## supplementary tables and figures of the antigen-antibody binding affinity landscapes for "Scalable Antigen-Antibody Binding Affinity Landscape: A Case Study with ENHERTU"

KIWI
© KIWI Press

### Scalable Antigen-Antibody Binding Affinity Landscape: A Case Study with ENHERTU: Supplementary Information

Wei Li

*Contrebola Institute of Computational Interstructural Biophysics, No. 88, Renaissance East Road, Nantong City 226000, Jiangsu Province, People's Republic of China*

**

Received (Day Month Year)
Revised (Day Month Year)
Accepted (Day Month Year)

#### Supplementary information

| **Chain ID** | **Molecular entity** | **Length** |
| --- | --- | --- |
| A | Receptor tyrosine-protein kinase erbB-2 (Homo sapiens, 9606) | 622 |
| B | Pertuzumab FAB LIGHT CHAIN (Homo sapiens, 9606) | 214 |
| C | Pertuzumab FAB HEAVY CHAIN (Homo sapiens, 9606) | 222 |
| D | Trastuzumab FAB LIGHT CHAIN (Homo sapiens, 9606) | 214 |
| E | Trastuzumab FAB HEAVY CHAIN (Homo sapiens, 9606) | 220 |
| BC | Pertuzumab FAB (Homo sapiens, 9606) | 436 |
| DE | Trastuzumab FAB (Homo sapiens, 9606) | 434 |
| Total | PDB entry 6OGE [1] | 1492 |

Table 1: Relationships between chain IDs and molecular entities of PDB entry 6OGE. In this table, length represents the number of amino acid residues, total means A+B+C+D+E.

| **Molecule pair** | **Scalable $K_d$ landscape** | **Histogram of $K_d$ distribution** |
|---|---|---|
| A + B | Table 3 (this file) | Figure 1 (this file) |
| A + C | Table 4 (this file) | Figure 2 (this file) |
| A + D | Table 5 (this file) | Figure 3 (this file) |
| A + E | Table 6 (this file) | Figure 4 (this file) |
| A + BC | Table 7 (this file) | Figure 5 (this file) |
| A + DE | Table 8 (this file) | Figure 6 (this file) |

Table 2: Scalable Her2-Trastuzumab-Pertuzumab binding affinity landscapes and their distribution patterns. In this table, the codes (A, B, C, etc) for binding partners are defined in Table 1.

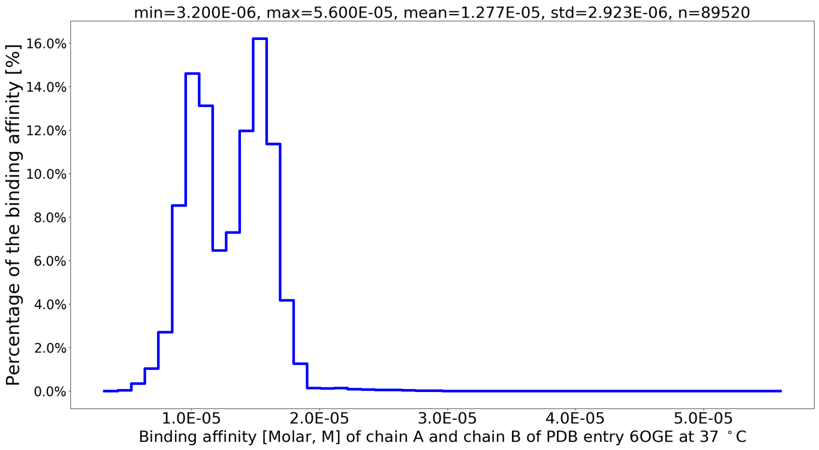


Figure 1: A histogram of the distribution pattern of the Her2-Trastuzumab-Pertuzumab binding affinities between chain A and chain B (listed in Table 1) of PDB entry 6OGE with one site-specific missense mutation.

| No. | ResID | Muta | ChainID | $K_{d1}$ (M) | $K_{d2}$ (M) | $K_{d3}$ (M) | Mean (M) | Std (M) | Min (M) | Max (M) |
|---|---|---|---|---|---|---|---|---|---|---|
| 1 | 1 | T1A | A | 1.5e-05 | 1.5e-05 | 1.5e-05 | 1.500E-05 | 0.000E+00 | 1.500E-05 | 1.500E-05 |
| 2 | 1 | T1R | A | 1.7e-05 | 1.7e-05 | 1.7e-05 | 1.700E-05 | 0.000E+00 | 1.700E-05 | 1.700E-05 |
| 3 | 1 | T1N | A | 1e-05 | 1e-05 | 1e-05 | 1.000E-05 | 0.000E+00 | 1.000E-05 | 1.000E-05 |
| 4 | 1 | T1D | A | 1.7e-05 | 1.7e-05 | 1.7e-05 | 1.700E-05 | 0.000E+00 | 1.700E-05 | 1.700E-05 |
| 5 | 1 | T1C | A | 1.1e-05 | 1.1e-05 | 1.1e-05 | 1.100E-05 | 2.075E-21 | 1.100E-05 | 1.100E-05 |
| 6 | 1 | T1E | A | 1.5e-05 | 1.5e-05 | 1.5e-05 | 1.500E-05 | 0.000E+00 | 1.500E-05 | 1.500E-05 |
| 7 | 1 | T1Q | A | 1.5e-05 | 1.5e-05 | 1.5e-05 | 1.500E-05 | 0.000E+00 | 1.500E-05 | 1.500E-05 |
| 8 | 1 | T1G | A | 1.7e-05 | 1.7e-05 | 1.7e-05 | 1.700E-05 | 0.000E+00 | 1.700E-05 | 1.700E-05 |
| 9 | 1 | T1H | A | 8.2e-06 | 8.2e-06 | 8.2e-06 | 8.200E-06 | 0.000E+00 | 8.200E-06 | 8.200E-06 |
| 10 | 1 | T1I | A | 1.6e-05 | 1.6e-05 | 1.6e-05 | 1.600E-05 | 0.000E+00 | 1.600E-05 | 1.600E-05 |
| 11 | 1 | T1L | A | 1.8e-05 | 1.8e-05 | 1.8e-05 | 1.800E-05 | 0.000E+00 | 1.800E-05 | 1.800E-05 |
| 12 | 1 | T1K | A | 9e-06 | 9e-06 | 9e-06 | 9.000E-06 | 0.000E+00 | 9.000E-06 | 9.000E-06 |
| 13 | 1 | T1M | A | 9.2e-06 | 9.2e-06 | 9.2e-06 | 9.200E-06 | 0.000E+00 | 9.200E-06 | 9.200E-06 |
| 14 | 1 | T1F | A | 1.1e-05 | 1.1e-05 | 1.1e-05 | 1.100E-05 | 2.075E-21 | 1.100E-05 | 1.100E-05 |
| 15 | 1 | T1P | A | 1.2e-05 | 1.2e-05 | 1.2e-05 | 1.200E-05 | 0.000E+00 | 1.200E-05 | 1.200E-05 |
| 16 | 1 | T1S | A | 1.1e-05 | 1.1e-05 | 1.1e-05 | 1.100E-05 | 2.075E-21 | 1.100E-05 | 1.100E-05 |
| 17 | 1 | T1T | A | 1.5e-05 | 1.5e-05 | 1.5e-05 | 1.500E-05 | 0.000E+00 | 1.500E-05 | 1.500E-05 |
| 18 | 1 | T1W | A | 9.5e-06 | 9.5e-06 | 9.5e-06 | 9.500E-06 | 0.000E+00 | 9.500E-06 | 9.500E-06 |
| 19 | 1 | T1Y | A | 1.1e-05 | 1.1e-05 | 1.1e-05 | 1.100E-05 | 2.075E-21 | 1.100E-05 | 1.100E-05 |
| 20 | 1 | T1V | A | 1.1e-05 | 1.1e-05 | 1.1e-05 | 1.100E-05 | 2.075E-21 | 1.100E-05 | 1.100E-05 |
| 21 | 2 | Q2A | A | 1.8e-05 | 1.8e-05 | 1.8e-05 | 1.800E-05 | 0.000E+00 | 1.800E-05 | 1.800E-05 |
| 22 | 2 | Q2R | A | 2.6e-05 | 2.6e-05 | 2.6e-05 | 2.600E-05 | 0.000E+00 | 2.600E-05 | 2.600E-05 |
| 23 | 2 | Q2N | A | 6e-06 | 6e-06 | 6e-06 | 6.000E-06 | 0.000E+00 | 6.000E-06 | 6.000E-06 |
| 24 | 2 | Q2D | A | 1.2e-05 | 1.2e-05 | 1.2e-05 | 1.200E-05 | 0.000E+00 | 1.200E-05 | 1.200E-05 |
| 25 | 2 | Q2C | A | 1.1e-05 | 1.6e-05 | 1.6e-05 | 1.433E-05 | 2.887E-06 | 1.100E-05 | 1.600E-05 |
| 26 | 2 | Q2E | A | 1.2e-05 | 1.3e-05 | 1.5e-05 | 1.333E-05 | 1.528E-06 | 1.200E-05 | 1.500E-05 |
| 27 | 2 | Q2Q | A | 1.3e-05 | 9.8e-06 | 1.5e-05 | 1.260E-05 | 2.623E-06 | 9.800E-06 | 1.500E-05 |
| 28 | 2 | Q2G | A | 1.6e-05 | 1.5e-05 | 9.1e-06 | 1.337E-05 | 3.729E-06 | 9.100E-06 | 1.600E-05 |
| 29 | 2 | Q2H | A | 1.7e-05 | 9.4e-06 | 9.4e-06 | 1.193E-05 | 4.388E-06 | 9.400E-06 | 1.700E-05 |
| 30 | 2 | Q2I | A | 1.1e-05 | 1.6e-05 | 1.1e-05 | 1.267E-05 | 2.887E-06 | 1.100E-05 | 1.600E-05 |
| 31 | 2 | Q2L | A | 1e-05 | 1.5e-05 | 1e-05 | 1.167E-05 | 2.887E-06 | 1.000E-05 | 1.500E-05 |
| 32 | 2 | Q2K | A | 1.7e-05 | 1.7e-05 | 1.2e-05 | 1.533E-05 | 2.887E-06 | 1.200E-05 | 1.700E-05 |
| 33 | 2 | Q2M | A | 1.8e-05 | 1.8e-05 | 1.5e-05 | 1.700E-05 | 1.732E-06 | 1.500E-05 | 1.800E-05 |
| 34 | 2 | Q2F | A | 1.5e-05 | 1.7e-05 | 1.5e-05 | 1.567E-05 | 1.155E-06 | 1.500E-05 | 1.700E-05 |
| 35 | 2 | Q2P | A | 1.4e-05 | 1.7e-05 | 1.4e-05 | 1.500E-05 | 1.732E-06 | 1.400E-05 | 1.700E-05 |
| 36 | 2 | Q2S | A | 1.2e-05 | 1.2e-05 | 1.2e-05 | 1.200E-05 | 0.000E+00 | 1.200E-05 | 1.200E-05 |
| 37 | 2 | Q2T | A | 9.2e-06 | 1.3e-05 | 1.5e-05 | 1.240E-05 | 2.946E-06 | 9.200E-06 | 1.500E-05 |
| 38 | 2 | Q2W | A | 1.4e-05 | 1.6e-05 | 1.3e-05 | 1.433E-05 | 1.528E-06 | 1.300E-05 | 1.600E-05 |
| 39 | 2 | Q2Y | A | 9.1e-06 | 1.3e-05 | 1.1e-05 | 1.103E-05 | 1.950E-06 | 9.100E-06 | 1.300E-05 |
| 40 | 2 | Q2V | A | 1.5e-05 | 8e-06 | 8e-06 | 1.033E-05 | 4.041E-06 | 8.000E-06 | 1.500E-05 |
| 41 | 3 | V3A | A | 1e-05 | 1e-05 | 1e-05 | 1.000E-05 | 0.000E+00 | 1.000E-05 | 1.000E-05 |
| 42 | 3 | V3R | A | 8.9e-06 | 1.4e-05 | 1.4e-05 | 1.230E-05 | 2.944E-06 | 8.900E-06 | 1.400E-05 |
| 43 | 3 | V3N | A | 8e-06 | 8e-06 | 8e-06 | 8.000E-06 | 0.000E+00 | 8.000E-06 | 8.000E-06 |
| 44 | 3 | V3D | A | 1e-05 | 1e-05 | 1e-05 | 1.000E-05 | 0.000E+00 | 1.000E-05 | 1.000E-05 |
| 45 | 3 | V3C | A | 1.7e-05 | 1.7e-05 | 1.3e-05 | 1.567E-05 | 2.309E-06 | 1.300E-05 | 1.700E-05 |
| 46 | 3 | V3E | A | 9.2e-06 | 9.2e-06 | 1.6e-05 | 1.147E-05 | 3.926E-06 | 9.200E-06 | 1.600E-05 |
| 47 | 3 | V3Q | A | 1.7e-05 | 1.7e-05 | 1.1e-05 | 1.500E-05 | 3.464E-06 | 1.100E-05 | 1.700E-05 |
| 48 | 3 | V3G | A | 1.4e-05 | 1.4e-05 | 1.4e-05 | 1.400E-05 | 0.000E+00 | 1.400E-05 | 1.400E-05 |
| 49 | 3 | V3H | A | 1.5e-05 | 9e-06 | 1.4e-05 | 1.267E-05 | 3.215E-06 | 9.000E-06 | 1.500E-05 |
| 50 | 3 | V3I | A | 1.2e-05 | 1e-05 | 1.2e-05 | 1.133E-05 | 1.155E-06 | 1.000E-05 | 1.200E-05 |
| 51 | 3 | V3L | A | 1.5e-05 | 9.7e-06 | 1.4e-05 | 1.290E-05 | 2.816E-06 | 9.700E-06 | 1.500E-05 |
| 52 | 3 | V3K | A | 1.3e-05 | 1.5e-05 | 1e-05 | 1.267E-05 | 2.517E-06 | 1.000E-05 | 1.500E-05 |
| 53 | 3 | V3M | A | 1.1e-05 | 1e-05 | 1.1e-05 | 1.067E-05 | 5.774E-07 | 1.000E-05 | 1.100E-05 |
| 54 | 3 | V3F | A | 1.5e-05 | 9.5e-06 | 1.6e-05 | 1.350E-05 | 3.500E-06 | 9.500E-06 | 1.600E-05 |
| 55 | 3 | V3P | A | 9.7e-06 | 1.5e-05 | 1.5e-05 | 1.323E-05 | 3.060E-06 | 9.700E-06 | 1.500E-05 |
| 56 | 3 | V3S | A | 1.3e-05 | 1.3e-05 | 1.5e-05 | 1.367E-05 | 1.155E-06 | 1.300E-05 | 1.500E-05 |
| 57 | 3 | V3T | A | 1.2e-05 | 7.7e-06 | 9.2e-06 | 9.633E-06 | 2.183E-06 | 7.700E-06 | 1.200E-05 |
| 58 | 3 | V3W | A | 1.7e-05 | 1.7e-05 | 1e-05 | 1.467E-05 | 4.041E-06 | 1.000E-05 | 1.700E-05 |
| 59 | 3 | V3Y | A | 1.3e-05 | 1.5e-05 | 1.3e-05 | 1.367E-05 | 1.155E-06 | 1.300E-05 | 1.500E-05 |
| 60 | 3 | V3V | A | 1.7e-05 | 1.3e-05 | 1.5e-05 | 1.500E-05 | 2.000E-06 | 1.300E-05 | 1.700E-05 |
| 61 | 4 | C4A | A | 8.9e-06 | 1e-05 | 1.2e-05 | 1.030E-05 | 1.572E-06 | 8.900E-06 | 1.200E-05 |
| 62 | 4 | C4R | A | 8e-06 | 8.1e-06 | 1.5e-05 | 1.037E-05 | 4.013E-06 | 8.000E-06 | 1.500E-05 |
| 63 | 4 | C4N | A | 1.5e-05 | 1.7e-05 | 1.4e-05 | 1.533E-05 | 1.528E-06 | 1.400E-05 | 1.700E-05 |
| 64 | 4 | C4D | A | 1.4e-05 | 1.6e-05 | 1.2e-05 | 1.400E-05 | 2.000E-06 | 1.200E-05 | 1.600E-05 |
| 65 | 4 | C4C | A | 1.7e-05 | 1.7e-05 | 1.7e-05 | 1.700E-05 | 0.000E+00 | 1.700E-05 | 1.700E-05 |
| 66 | 4 | C4E | A | 1.5e-05 | 1.6e-05 | 1e-05 | 1.367E-05 | 3.215E-06 | 1.000E-05 | 1.600E-05 |
| 67 | 4 | C4Q | A | 8.4e-06 | 1.5e-05 | 1.1e-05 | 1.147E-05 | 3.325E-06 | 8.400E-06 | 1.500E-05 |
| 68 | 4 | C4G | A | 1.5e-05 | 1.7e-05 | 1.3e-05 | 1.500E-05 | 2.000E-06 | 1.300E-05 | 1.700E-05 |
| 69 | 4 | C4H | A | 1.1e-05 | 9.2e-06 | 1.7e-05 | 1.240E-05 | 4.084E-06 | 9.200E-06 | 1.700E-05 |
| 70 | 4 | C4I | A | 1.3e-05 | 1.3e-05 | 1.7e-05 | 1.433E-05 | 2.309E-06 | 1.300E-05 | 1.700E-05 |
| 71 | 4 | C4L | A | 1.2e-05 | 1.5e-05 | 1.3e-05 | 1.333E-05 | 1.528E-06 | 1.200E-05 | 1.500E-05 |
| 72 | 4 | C4K | A | 1.7e-05 | 1.7e-05 | 9e-06 | 1.433E-05 | 4.619E-06 | 9.000E-06 | 1.700E-05 |
| 73 | 4 | C4M | A | 1e-05 | 1.8e-05 | 1.1e-05 | 1.300E-05 | 4.359E-06 | 1.000E-05 | 1.800E-05 |
| 74 | 4 | C4F | A | 9.6e-06 | 1.1e-05 | 1.2e-05 | 1.087E-05 | 1.206E-06 | 9.600E-06 | 1.200E-05 |
| 75 | 4 | C4P | A | 9.1e-06 | 1.6e-05 | 1.3e-05 | 1.270E-05 | 3.460E-06 | 9.100E-06 | 1.600E-05 |
| 76 | 4 | C4S | A | 1.6e-05 | 1.5e-05 | 1.5e-05 | 1.533E-05 | 5.774E-07 | 1.500E-05 | 1.600E-05 |
| 77 | 4 | C4T | A | 1.5e-05 | 1.4e-05 | 1.3e-05 | 1.400E-05 | 1.000E-06 | 1.300E-05 | 1.500E-05 |
| 78 | 4 | C4W | A | 1.5e-05 | 9.2e-06 | 1.5e-05 | 1.307E-05 | 3.349E-06 | 9.200E-06 | 1.500E-05 |
| 79 | 4 | C4Y | A | 9.8e-06 | 1.1e-05 | 1.2e-05 | 1.093E-05 | 1.102E-06 | 9.800E-06 | 1.200E-05 |
| 80 | 4 | C4V | A | 8.7e-06 | 1e-05 | 1.5e-05 | 1.123E-05 | 3.326E-06 | 8.700E-06 | 1.500E-05 |
| 81 | 5 | T5A | A | 1.5e-05 | 1.5e-05 | 1.6e-05 | 1.533E-05 | 5.774E-07 | 1.500E-05 | 1.600E-05 |
| 82 | 5 | T5R | A | 1.2e-05 | 1.2e-05 | 1.3e-05 | 1.233E-05 | 5.774E-07 | 1.200E-05 | 1.300E-05 |
| 83 | 5 | T5N | A | 8.6e-06 | 1.6e-05 | 1.6e-05 | 1.353E-05 | 4.272E-06 | 8.600E-06 | 1.600E-05 |
| 84 | 5 | T5D | A | 1.1e-05 | 1.2e-05 | 1.2e-05 | 1.167E-05 | 5.774E-07 | 1.100E-05 | 1.200E-05 |

| | | | | | | | | | | |
|---|---|---|---|---|---|---|---|---|---|---|
| 85 | 5 | T5C | A | 8.5e-06 | 1.3e-05 | 9.8e-06 | 1.043E-05 | 2.316E-06 | 8.500E-06 | 1.300E-05 |
| 86 | 5 | T5E | A | 1.1e-05 | 1.6e-05 | 1.3e-05 | 1.333E-05 | 2.517E-06 | 1.100E-05 | 1.600E-05 |
| 87 | 5 | T5Q | A | 1e-05 | 1.3e-05 | 1.2e-05 | 1.167E-05 | 1.528E-06 | 1.000E-05 | 1.300E-05 |
| 88 | 5 | T5G | A | 1.2e-05 | 1.4e-05 | 1.7e-05 | 1.433E-05 | 2.517E-06 | 1.200E-05 | 1.700E-05 |
| 89 | 5 | T5H | A | 1.2e-05 | 1.5e-05 | 1e-05 | 1.233E-05 | 2.517E-06 | 1.000E-05 | 1.500E-05 |
| 90 | 5 | T5I | A | 1.6e-05 | 1.1e-05 | 1e-05 | 1.233E-05 | 3.215E-06 | 1.000E-05 | 1.600E-05 |
| 91 | 5 | T5L | A | 8e-06 | 8e-06 | 1.2e-05 | 9.333E-06 | 2.309E-06 | 8.000E-06 | 1.200E-05 |
| 92 | 5 | T5K | A | 1.2e-05 | 9.1e-06 | 1.6e-05 | 1.237E-05 | 3.465E-06 | 9.100E-06 | 1.600E-05 |
| 93 | 5 | T5M | A | 1.7e-05 | 1.6e-05 | 1.6e-05 | 1.633E-05 | 5.774E-07 | 1.600E-05 | 1.700E-05 |
| 94 | 5 | T5F | A | 1.1e-05 | 1.6e-05 | 1.6e-05 | 1.433E-05 | 2.887E-06 | 1.100E-05 | 1.600E-05 |
| 95 | 5 | T5P | A | 1.6e-05 | 1.6e-05 | 1.1e-05 | 1.433E-05 | 2.887E-06 | 1.100E-05 | 1.600E-05 |
| 96 | 5 | T5S | A | 1e-05 | 8.5e-06 | 1.3e-05 | 1.050E-05 | 2.291E-06 | 8.500E-06 | 1.300E-05 |
| 97 | 5 | T5T | A | 1.5e-05 | 1.7e-05 | 1.6e-05 | 1.600E-05 | 1.000E-06 | 1.500E-05 | 1.700E-05 |
| 98 | 5 | T5W | A | 1.2e-05 | 1.4e-05 | 1e-05 | 1.200E-05 | 2.000E-06 | 1.000E-05 | 1.400E-05 |
| 99 | 5 | T5Y | A | 1.2e-05 | 1.2e-05 | 9.2e-06 | 1.107E-05 | 1.617E-06 | 9.200E-06 | 1.200E-05 |
| 100 | 5 | T5V | A | 1.2e-05 | 1.4e-05 | 1.6e-05 | 1.400E-05 | 2.000E-06 | 1.200E-05 | 1.600E-05 |
| 101 | 6 | G6A | A | 1.1e-05 | 1.3e-05 | 1.4e-05 | 1.267E-05 | 1.528E-06 | 1.100E-05 | 1.400E-05 |
| 102 | 6 | G6R | A | 9.3e-06 | 1e-05 | 8.5e-06 | 9.267E-06 | 7.506E-07 | 8.500E-06 | 1.000E-05 |
| 103 | 6 | G6N | A | 1e-05 | 1e-05 | 1.5e-05 | 1.167E-05 | 2.887E-06 | 1.000E-05 | 1.500E-05 |
| 104 | 6 | G6D | A | 1.1e-05 | 1.4e-05 | 1.1e-05 | 1.200E-05 | 1.732E-06 | 1.100E-05 | 1.400E-05 |
| 105 | 6 | G6C | A | 1.5e-05 | 1.4e-05 | 1.3e-05 | 1.400E-05 | 1.000E-06 | 1.300E-05 | 1.500E-05 |
| 106 | 6 | G6E | A | 1.1e-05 | 1.6e-05 | 1e-05 | 1.233E-05 | 3.215E-06 | 1.000E-05 | 1.600E-05 |
| 107 | 6 | G6Q | A | 1.3e-05 | 1.5e-05 | 6.7e-06 | 1.157E-05 | 4.332E-06 | 6.700E-06 | 1.500E-05 |
| 108 | 6 | G6G | A | 1.6e-05 | 1.1e-05 | 1.6e-05 | 1.433E-05 | 2.887E-06 | 1.100E-05 | 1.600E-05 |
| 109 | 6 | G6H | A | 1.5e-05 | 1.4e-05 | 1.2e-05 | 1.367E-05 | 1.528E-06 | 1.200E-05 | 1.500E-05 |
| 110 | 6 | G6I | A | 1.6e-05 | 7.7e-06 | 1e-05 | 1.123E-05 | 4.285E-06 | 7.700E-06 | 1.600E-05 |
| 111 | 6 | G6L | A | 1.1e-05 | 1.2e-05 | 1.2e-05 | 1.167E-05 | 5.774E-07 | 1.100E-05 | 1.200E-05 |
| 112 | 6 | G6K | A | 1.4e-05 | 1.5e-05 | 1.1e-05 | 1.333E-05 | 2.082E-06 | 1.100E-05 | 1.500E-05 |
| 113 | 6 | G6M | A | 1.8e-05 | 1.5e-05 | 1.6e-05 | 1.633E-05 | 1.528E-06 | 1.500E-05 | 1.800E-05 |
| 114 | 6 | G6F | A | 1.3e-05 | 1e-05 | 1e-05 | 1.100E-05 | 1.732E-06 | 1.000E-05 | 1.300E-05 |
| 115 | 6 | G6P | A | 9.2e-06 | 9.6e-06 | 8.6e-06 | 9.133E-06 | 5.033E-07 | 8.600E-06 | 9.600E-06 |
| 116 | 6 | G6S | A | 1e-05 | 1.7e-05 | 1.6e-05 | 1.433E-05 | 3.786E-06 | 1.000E-05 | 1.700E-05 |
| 117 | 6 | G6T | A | 1.5e-05 | 1.2e-05 | 1.6e-05 | 1.433E-05 | 2.082E-06 | 1.200E-05 | 1.600E-05 |
| 118 | 6 | G6W | A | 9.8e-06 | 1.4e-05 | 1e-05 | 1.127E-05 | 2.369E-06 | 9.800E-06 | 1.400E-05 |
| 119 | 6 | G6Y | A | 5.6e-06 | 8.7e-06 | 1.5e-05 | 9.767E-06 | 4.790E-06 | 5.600E-06 | 1.500E-05 |
| 120 | 6 | G6V | A | 1.3e-05 | 1e-05 | 1.3e-05 | 1.200E-05 | 1.732E-06 | 1.000E-05 | 1.300E-05 |
| 121 | 7 | T7A | A | 1.3e-05 | 8.4e-06 | 1.1e-05 | 1.080E-05 | 2.307E-06 | 8.400E-06 | 1.300E-05 |
| 122 | 7 | T7R | A | 1.4e-05 | 1.5e-05 | 1e-05 | 1.300E-05 | 2.646E-06 | 1.000E-05 | 1.500E-05 |
| 123 | 7 | T7N | A | 1.1e-05 | 1.3e-05 | 1.2e-05 | 1.200E-05 | 1.000E-06 | 1.100E-05 | 1.300E-05 |
| 124 | 7 | T7D | A | 1.7e-05 | 8.4e-06 | 9.9e-06 | 1.177E-05 | 4.594E-06 | 8.400E-06 | 1.700E-05 |
| 125 | 7 | T7C | A | 1.1e-05 | 1.1e-05 | 1.4e-05 | 1.200E-05 | 1.732E-06 | 1.100E-05 | 1.400E-05 |
| 126 | 7 | T7E | A | 9.9e-06 | 1.6e-05 | 1.4e-05 | 1.330E-05 | 3.110E-06 | 9.900E-06 | 1.600E-05 |
| 127 | 7 | T7Q | A | 1.5e-05 | 1.6e-05 | 9.1e-06 | 1.337E-05 | 3.729E-06 | 9.100E-06 | 1.600E-05 |
| 128 | 7 | T7G | A | 1.5e-05 | 1.2e-05 | 1e-05 | 1.233E-05 | 2.517E-06 | 1.000E-05 | 1.500E-05 |
| 129 | 7 | T7H | A | 1.1e-05 | 1.4e-05 | 1.4e-05 | 1.300E-05 | 1.732E-06 | 1.100E-05 | 1.400E-05 |
| 130 | 7 | T7I | A | 9.2e-06 | 1.2e-05 | 1.1e-05 | 1.073E-05 | 1.419E-06 | 9.200E-06 | 1.200E-05 |
| 131 | 7 | T7L | A | 1.4e-05 | 1e-05 | 1.3e-05 | 1.233E-05 | 2.082E-06 | 1.000E-05 | 1.400E-05 |
| 132 | 7 | T7K | A | 1.7e-05 | 1.6e-05 | 1.4e-05 | 1.567E-05 | 1.528E-06 | 1.400E-05 | 1.700E-05 |
| 133 | 7 | T7M | A | 1.2e-05 | 1.5e-05 | 1.1e-05 | 1.267E-05 | 2.082E-06 | 1.100E-05 | 1.500E-05 |
| 134 | 7 | T7F | A | 1.1e-05 | 1.3e-05 | 1.6e-05 | 1.333E-05 | 2.517E-06 | 1.100E-05 | 1.600E-05 |
| 135 | 7 | T7P | A | 1.7e-05 | 1.7e-05 | 1.3e-05 | 1.567E-05 | 2.309E-06 | 1.300E-05 | 1.700E-05 |
| 136 | 7 | T7S | A | 1.3e-05 | 1.3e-05 | 1.1e-05 | 1.233E-05 | 1.155E-06 | 1.100E-05 | 1.300E-05 |
| 137 | 7 | T7T | A | 1.5e-05 | 8e-06 | 1.5e-05 | 1.267E-05 | 4.041E-06 | 8.000E-06 | 1.500E-05 |
| 138 | 7 | T7W | A | 2.6e-05 | 1.6e-05 | 1.1e-05 | 1.767E-05 | 7.638E-06 | 1.100E-05 | 2.600E-05 |
| 139 | 7 | T7Y | A | 8.3e-06 | 7.9e-06 | 1.5e-05 | 1.040E-05 | 3.989E-06 | 7.900E-06 | 1.500E-05 |
| 140 | 7 | T7V | A | 1.2e-05 | 1.1e-05 | 9.9e-06 | 1.097E-05 | 1.050E-06 | 9.900E-06 | 1.200E-05 |
| 141 | 8 | D8A | A | 1.1e-05 | 1e-05 | 1.5e-05 | 1.200E-05 | 2.646E-06 | 1.000E-05 | 1.500E-05 |
| 142 | 8 | D8R | A | 1e-05 | 1e-05 | 1e-05 | 1.000E-05 | 0.000E+00 | 1.000E-05 | 1.000E-05 |
| 143 | 8 | D8N | A | 1.1e-05 | 1.6e-05 | 1.4e-05 | 1.367E-05 | 2.517E-06 | 1.100E-05 | 1.600E-05 |
| 144 | 8 | D8D | A | 8.4e-06 | 1.1e-05 | 1.5e-05 | 1.147E-05 | 3.325E-06 | 8.400E-06 | 1.500E-05 |
| 145 | 8 | D8C | A | 1.6e-05 | 1.6e-05 | 1.5e-05 | 1.567E-05 | 5.774E-07 | 1.500E-05 | 1.600E-05 |
| 146 | 8 | D8E | A | 1e-05 | 1.6e-05 | 1e-05 | 1.200E-05 | 3.464E-06 | 1.000E-05 | 1.600E-05 |
| 147 | 8 | D8Q | A | 8.5e-06 | 1.6e-05 | 1.6e-05 | 1.350E-05 | 4.330E-06 | 8.500E-06 | 1.600E-05 |
| 148 | 8 | D8G | A | 1e-05 | 1.5e-05 | 1.1e-05 | 1.200E-05 | 2.646E-06 | 1.000E-05 | 1.500E-05 |
| 149 | 8 | D8H | A | 1.4e-05 | 1.1e-05 | 1.4e-05 | 1.300E-05 | 1.732E-06 | 1.100E-05 | 1.400E-05 |
| 150 | 8 | D8I | A | 1.5e-05 | 1.6e-05 | 5.6e-06 | 1.220E-05 | 5.738E-06 | 5.600E-06 | 1.600E-05 |
| 151 | 8 | D8L | A | 1.4e-05 | 1.1e-05 | 1.5e-05 | 1.333E-05 | 2.082E-06 | 1.100E-05 | 1.500E-05 |
| 152 | 8 | D8K | A | 1.6e-05 | 1.5e-05 | 1.6e-05 | 1.567E-05 | 5.774E-07 | 1.500E-05 | 1.600E-05 |
| 153 | 8 | D8M | A | 1e-05 | 1.5e-05 | 1.4e-05 | 1.300E-05 | 2.646E-06 | 1.000E-05 | 1.500E-05 |
| 154 | 8 | D8F | A | 1.5e-05 | 1.2e-05 | 1.5e-05 | 1.400E-05 | 1.732E-06 | 1.200E-05 | 1.500E-05 |
| 155 | 8 | D8P | A | 1.6e-05 | 1e-05 | 1.6e-05 | 1.400E-05 | 3.464E-06 | 1.000E-05 | 1.600E-05 |
| 156 | 8 | D8S | A | 1.4e-05 | 1.4e-05 | 1e-05 | 1.267E-05 | 2.309E-06 | 1.000E-05 | 1.400E-05 |
| 157 | 8 | D8T | A | 1.1e-05 | 1e-05 | 1e-05 | 1.033E-05 | 5.774E-07 | 1.000E-05 | 1.100E-05 |
| 158 | 8 | D8W | A | 1e-05 | 1.6e-05 | 1e-05 | 1.200E-05 | 3.464E-06 | 1.000E-05 | 1.600E-05 |
| 159 | 8 | D8Y | A | 1.5e-05 | 1.4e-05 | 1e-05 | 1.300E-05 | 2.646E-06 | 1.000E-05 | 1.500E-05 |
| 160 | 8 | D8V | A | 1.5e-05 | 1.4e-05 | 1.1e-05 | 1.333E-05 | 2.082E-06 | 1.100E-05 | 1.500E-05 |
| 161 | 9 | M9A | A | 1.1e-05 | 9e-06 | 1.1e-05 | 1.033E-05 | 1.155E-06 | 9.000E-06 | 1.100E-05 |
| 162 | 9 | M9R | A | 1.4e-05 | 9.9e-06 | 1.5e-05 | 1.297E-05 | 2.702E-06 | 9.900E-06 | 1.500E-05 |
| 163 | 9 | M9N | A | 1.1e-05 | 1e-05 | 1e-05 | 1.033E-05 | 5.774E-07 | 1.000E-05 | 1.100E-05 |
| 164 | 9 | M9D | A | 9.4e-06 | 1.3e-05 | 1.5e-05 | 1.247E-05 | 2.838E-06 | 9.400E-06 | 1.500E-05 |
| 165 | 9 | M9C | A | 1e-05 | 9.3e-06 | 9.9e-06 | 9.733E-06 | 3.786E-07 | 9.300E-06 | 1.000E-05 |
| 166 | 9 | M9E | A | 9.8e-06 | 1.1e-05 | 1e-05 | 1.027E-05 | 6.429E-07 | 9.800E-06 | 1.100E-05 |
| 167 | 9 | M9Q | A | 1.4e-05 | 1.6e-05 | 1.1e-05 | 1.367E-05 | 2.517E-06 | 1.100E-05 | 1.600E-05 |
| 168 | 9 | M9G | A | 1.5e-05 | 1.2e-05 | 1e-05 | 1.233E-05 | 2.517E-06 | 1.000E-05 | 1.500E-05 |
| 169 | 9 | M9H | A | 1.2e-05 | 9.3e-06 | 1e-05 | 1.043E-05 | 1.401E-06 | 9.300E-06 | 1.200E-05 |

| | | | | | | | | | | |
|---|---|---|---|---|---|---|---|---|---|---|
| 170 | 9 | M9I | A | 1.6e-05 | 1.2e-05 | 1.5e-05 | 1.433E-05 | 2.082E-06 | 1.200E-05 | 1.600E-05 |
| 171 | 9 | M9L | A | 1.5e-05 | 9.8e-06 | 8.5e-06 | 1.110E-05 | 3.439E-06 | 8.500E-06 | 1.500E-05 |
| 172 | 9 | M9K | A | 1.6e-05 | 1.3e-05 | 6e-06 | 1.167E-05 | 5.132E-06 | 6.000E-06 | 1.600E-05 |
| 173 | 9 | M9M | A | 1.6e-05 | 1.1e-05 | 1.1e-05 | 1.267E-05 | 2.887E-06 | 1.100E-05 | 1.600E-05 |
| 174 | 9 | M9F | A | 9.2e-06 | 1.1e-05 | 9.7e-06 | 9.967E-06 | 9.292E-07 | 9.200E-06 | 1.100E-05 |
| 175 | 9 | M9P | A | 1e-05 | 1.1e-05 | 8.2e-06 | 9.733E-06 | 1.419E-06 | 8.200E-06 | 1.100E-05 |
| 176 | 9 | M9S | A | 1.5e-05 | 1.3e-05 | 9.5e-06 | 1.250E-05 | 2.784E-06 | 9.500E-06 | 1.500E-05 |
| 177 | 9 | M9T | A | 1.6e-05 | 1.1e-05 | 1.1e-05 | 1.267E-05 | 2.887E-06 | 1.100E-05 | 1.600E-05 |
| 178 | 9 | M9W | A | 1.4e-05 | 1.5e-05 | 1.5e-05 | 1.467E-05 | 5.774E-07 | 1.400E-05 | 1.500E-05 |
| 179 | 9 | M9Y | A | 1e-05 | 1.5e-05 | 1.1e-05 | 1.200E-05 | 2.646E-06 | 1.000E-05 | 1.500E-05 |
| 180 | 9 | M9V | A | 1.3e-05 | 1.1e-05 | 1.6e-05 | 1.333E-05 | 2.517E-06 | 1.100E-05 | 1.600E-05 |
| 181 | 10 | K10A | A | 9.4e-06 | 1.1e-05 | 1.1e-05 | 1.047E-05 | 9.238E-07 | 9.400E-06 | 1.100E-05 |
| 182 | 10 | K10R | A | 1e-05 | 1.2e-05 | 1.5e-05 | 1.233E-05 | 2.517E-06 | 1.000E-05 | 1.500E-05 |
| 183 | 10 | K10N | A | 1.4e-05 | 1.2e-05 | 1.6e-05 | 1.400E-05 | 2.000E-06 | 1.200E-05 | 1.600E-05 |
| 184 | 10 | K10D | A | 1.4e-05 | 1.9e-05 | 1.6e-05 | 1.633E-05 | 2.517E-06 | 1.400E-05 | 1.900E-05 |
| 185 | 10 | K10C | A | 9.9e-06 | 1.3e-05 | 1.4e-05 | 1.230E-05 | 2.138E-06 | 9.900E-06 | 1.400E-05 |
| 186 | 10 | K10E | A | 1.6e-05 | 1.4e-05 | 1.3e-05 | 1.433E-05 | 1.528E-06 | 1.300E-05 | 1.600E-05 |
| 187 | 10 | K10Q | A | 1.1e-05 | 9.8e-06 | 1.1e-05 | 1.060E-05 | 6.928E-07 | 9.800E-06 | 1.100E-05 |
| 188 | 10 | K10G | A | 1.4e-05 | 1.6e-05 | 9.6e-06 | 1.320E-05 | 3.274E-06 | 9.600E-06 | 1.600E-05 |
| 189 | 10 | K10H | A | 1.5e-05 | 1e-05 | 1.7e-05 | 1.400E-05 | 3.606E-06 | 1.000E-05 | 1.700E-05 |
| 190 | 10 | K10I | A | 1.6e-05 | 1.1e-05 | 1.1e-05 | 1.267E-05 | 2.887E-06 | 1.100E-05 | 1.600E-05 |
| 191 | 10 | K10L | A | 1.6e-05 | 1e-05 | 1.5e-05 | 1.367E-05 | 3.215E-06 | 1.000E-05 | 1.600E-05 |
| 192 | 10 | K10K | A | 1.6e-05 | 9.9e-06 | 9.3e-06 | 1.173E-05 | 3.707E-06 | 9.300E-06 | 1.600E-05 |
| 193 | 10 | K10M | A | 8.5e-06 | 9.1e-06 | 1.7e-05 | 1.153E-05 | 4.744E-06 | 8.500E-06 | 1.700E-05 |
| 194 | 10 | K10F | A | 1.4e-05 | 1.1e-05 | 1e-05 | 1.167E-05 | 2.082E-06 | 1.000E-05 | 1.400E-05 |
| 195 | 10 | K10P | A | 1.5e-05 | 1.2e-05 | 1.6e-05 | 1.433E-05 | 2.082E-06 | 1.200E-05 | 1.600E-05 |
| 196 | 10 | K10S | A | 1.4e-05 | 1.2e-05 | 8.1e-06 | 1.137E-05 | 3.001E-06 | 8.100E-06 | 1.400E-05 |
| 197 | 10 | K10T | A | 8.7e-06 | 1.4e-05 | 1.5e-05 | 1.257E-05 | 3.386E-06 | 8.700E-06 | 1.500E-05 |
| 198 | 10 | K10W | A | 8.9e-06 | 1.3e-05 | 1e-05 | 1.063E-05 | 2.122E-06 | 8.900E-06 | 1.300E-05 |
| 199 | 10 | K10Y | A | 1.4e-05 | 1.5e-05 | 1.3e-05 | 1.400E-05 | 1.000E-06 | 1.300E-05 | 1.500E-05 |
| 200 | 10 | K10V | A | 1.1e-05 | 1e-05 | 1e-05 | 1.033E-05 | 5.774E-07 | 1.000E-05 | 1.100E-05 |
| 201 | 11 | L11A | A | 1.8e-05 | 8.9e-06 | 8.1e-06 | 1.167E-05 | 5.499E-06 | 8.100E-06 | 1.800E-05 |
| 202 | 11 | L11R | A | 2.1e-05 | 1.4e-05 | 1.5e-05 | 1.667E-05 | 3.786E-06 | 1.400E-05 | 2.100E-05 |
| 203 | 11 | L11N | A | 6.9e-06 | 1e-05 | 8.6e-06 | 8.500E-06 | 1.552E-06 | 6.900E-06 | 1.000E-05 |
| 204 | 11 | L11D | A | 1.5e-05 | 1.1e-05 | 1.4e-05 | 1.333E-05 | 2.082E-06 | 1.100E-05 | 1.500E-05 |
| 205 | 11 | L11C | A | 1.3e-05 | 8.4e-06 | 1.6e-05 | 1.247E-05 | 3.828E-06 | 8.400E-06 | 1.600E-05 |
| 206 | 11 | L11E | A | 1.6e-05 | 1.5e-05 | 9.7e-06 | 1.357E-05 | 3.386E-06 | 9.700E-06 | 1.600E-05 |
| 207 | 11 | L11Q | A | 9.6e-06 | 1e-05 | 1.1e-05 | 1.020E-05 | 7.211E-07 | 9.600E-06 | 1.100E-05 |
| 208 | 11 | L11G | A | 1.4e-05 | 1.5e-05 | 1.1e-05 | 1.333E-05 | 2.082E-06 | 1.100E-05 | 1.500E-05 |
| 209 | 11 | L11H | A | 1.4e-05 | 1.1e-05 | 8.3e-06 | 1.110E-05 | 2.851E-06 | 8.300E-06 | 1.400E-05 |
| 210 | 11 | L11I | A | 1.5e-05 | 1.2e-05 | 9.8e-06 | 1.227E-05 | 2.610E-06 | 9.800E-06 | 1.500E-05 |
| 211 | 11 | L11L | A | 9.5e-06 | 1.2e-05 | 9.3e-06 | 1.027E-05 | 1.504E-06 | 9.300E-06 | 1.200E-05 |
| 212 | 11 | L11K | A | 1.2e-05 | 1.6e-05 | 9.5e-06 | 1.250E-05 | 3.279E-06 | 9.500E-06 | 1.600E-05 |
| 213 | 11 | L11M | A | 1.5e-05 | 8.1e-06 | 1.6e-05 | 1.303E-05 | 4.302E-06 | 8.100E-06 | 1.600E-05 |
| 214 | 11 | L11F | A | 1.6e-05 | 1.7e-05 | 1e-05 | 1.433E-05 | 3.786E-06 | 1.000E-05 | 1.700E-05 |
| 215 | 11 | L11P | A | 1.5e-05 | 1.5e-05 | 1.3e-05 | 1.433E-05 | 1.155E-06 | 1.300E-05 | 1.500E-05 |
| 216 | 11 | L11S | A | 1.3e-05 | 9.5e-06 | 1.6e-05 | 1.283E-05 | 3.253E-06 | 9.500E-06 | 1.600E-05 |
| 217 | 11 | L11T | A | 1.6e-05 | 1.6e-05 | 1.1e-05 | 1.433E-05 | 2.887E-06 | 1.100E-05 | 1.600E-05 |
| 218 | 11 | L11W | A | 1.1e-05 | 9.4e-06 | 1.3e-05 | 1.113E-05 | 1.804E-06 | 9.400E-06 | 1.300E-05 |
| 219 | 11 | L11Y | A | 1.1e-05 | 1e-05 | 1e-05 | 1.033E-05 | 5.774E-07 | 1.000E-05 | 1.100E-05 |
| 220 | 11 | L11V | A | 1.4e-05 | 1.1e-05 | 1.6e-05 | 1.367E-05 | 2.517E-06 | 1.100E-05 | 1.600E-05 |
| 221 | 12 | R12A | A | 8.4e-06 | 8.7e-06 | 1.3e-05 | 1.003E-05 | 2.574E-06 | 8.400E-06 | 1.300E-05 |
| 222 | 12 | R12R | A | 8.9e-06 | 1.1e-05 | 1.5e-05 | 1.163E-05 | 3.099E-06 | 8.900E-06 | 1.500E-05 |
| 223 | 12 | R12N | A | 1.5e-05 | 1.4e-05 | 1.5e-05 | 1.467E-05 | 5.774E-07 | 1.400E-05 | 1.500E-05 |
| 224 | 12 | R12D | A | 1.3e-05 | 1.4e-05 | 1.1e-05 | 1.267E-05 | 1.528E-06 | 1.100E-05 | 1.400E-05 |
| 225 | 12 | R12C | A | 1.6e-05 | 1.6e-05 | 1.4e-05 | 1.533E-05 | 1.155E-06 | 1.400E-05 | 1.600E-05 |
| 226 | 12 | R12E | A | 1.2e-05 | 1.5e-05 | 1.1e-05 | 1.267E-05 | 2.082E-06 | 1.100E-05 | 1.500E-05 |
| 227 | 12 | R12Q | A | 1e-05 | 1.7e-05 | 1e-05 | 1.233E-05 | 4.041E-06 | 1.000E-05 | 1.700E-05 |
| 228 | 12 | R12G | A | 9.8e-06 | 9.9e-06 | 1e-05 | 9.900E-06 | 1.000E-07 | 9.800E-06 | 1.000E-05 |
| 229 | 12 | R12H | A | 8.4e-06 | 1.6e-05 | 1.4e-05 | 1.280E-05 | 3.940E-06 | 8.400E-06 | 1.600E-05 |
| 230 | 12 | R12I | A | 1.4e-05 | 1.1e-05 | 1.7e-05 | 1.400E-05 | 3.000E-06 | 1.100E-05 | 1.700E-05 |
| 231 | 12 | R12L | A | 1e-05 | 1.5e-05 | 1.2e-05 | 1.233E-05 | 2.517E-06 | 1.000E-05 | 1.500E-05 |
| 232 | 12 | R12K | A | 1.6e-05 | 1e-05 | 1.1e-05 | 1.233E-05 | 3.215E-06 | 1.000E-05 | 1.600E-05 |
| 233 | 12 | R12M | A | 1.5e-05 | 1.6e-05 | 8e-06 | 1.300E-05 | 4.359E-06 | 8.000E-06 | 1.600E-05 |
| 234 | 12 | R12F | A | 6.3e-06 | 9.5e-06 | 1.6e-05 | 1.060E-05 | 4.943E-06 | 6.300E-06 | 1.600E-05 |
| 235 | 12 | R12P | A | 1.4e-05 | 1.3e-05 | 1.1e-05 | 1.267E-05 | 1.528E-06 | 1.100E-05 | 1.400E-05 |
| 236 | 12 | R12S | A | 9.6e-06 | 1.2e-05 | 1.7e-05 | 1.287E-05 | 3.775E-06 | 9.600E-06 | 1.700E-05 |
| 237 | 12 | R12T | A | 1e-05 | 1.6e-05 | 9.6e-06 | 1.187E-05 | 3.585E-06 | 9.600E-06 | 1.600E-05 |
| 238 | 12 | R12W | A | 1.3e-05 | 1.5e-05 | 1e-05 | 1.267E-05 | 2.517E-06 | 1.000E-05 | 1.500E-05 |
| 239 | 12 | R12Y | A | 1.5e-05 | 8.1e-06 | 1.4e-05 | 1.237E-05 | 3.729E-06 | 8.100E-06 | 1.500E-05 |
| 240 | 12 | R12V | A | 1e-05 | 1.1e-05 | 1e-05 | 1.033E-05 | 5.774E-07 | 1.000E-05 | 1.100E-05 |
| 241 | 13 | L13A | A | 1.6e-05 | 1.1e-05 | 1.3e-05 | 1.333E-05 | 2.517E-06 | 1.100E-05 | 1.600E-05 |
| 242 | 13 | L13R | A | 1.4e-05 | 1.4e-05 | 9.4e-06 | 1.247E-05 | 2.656E-06 | 9.400E-06 | 1.400E-05 |
| 243 | 13 | L13N | A | 1.4e-05 | 1.5e-05 | 1.4e-05 | 1.433E-05 | 5.774E-07 | 1.400E-05 | 1.500E-05 |
| 244 | 13 | L13D | A | 1e-05 | 1.1e-05 | 1.5e-05 | 1.200E-05 | 2.646E-06 | 1.000E-05 | 1.500E-05 |
| 245 | 13 | L13C | A | 8e-06 | 7.6e-06 | 1.5e-05 | 1.020E-05 | 4.162E-06 | 7.600E-06 | 1.500E-05 |
| 246 | 13 | L13E | A | 1.6e-05 | 8.5e-06 | 1.4e-05 | 1.283E-05 | 3.884E-06 | 8.500E-06 | 1.600E-05 |
| 247 | 13 | L13Q | A | 1.3e-05 | 1.2e-05 | 1.2e-05 | 1.233E-05 | 5.774E-07 | 1.200E-05 | 1.300E-05 |
| 248 | 13 | L13G | A | 1.1e-05 | 1.4e-05 | 1.6e-05 | 1.367E-05 | 2.517E-06 | 1.100E-05 | 1.600E-05 |
| 249 | 13 | L13H | A | 2.4e-05 | 1.1e-05 | 8e-06 | 1.433E-05 | 8.505E-06 | 8.000E-06 | 2.400E-05 |
| 250 | 13 | L13I | A | 1.5e-05 | 1.2e-05 | 1.4e-05 | 1.367E-05 | 1.528E-06 | 1.200E-05 | 1.500E-05 |
| 251 | 13 | L13L | A | 1.1e-05 | 1.6e-05 | 1.1e-05 | 1.267E-05 | 2.887E-06 | 1.100E-05 | 1.600E-05 |
| 252 | 13 | L13K | A | 1.5e-05 | 9e-06 | 1.5e-05 | 1.300E-05 | 3.464E-06 | 9.000E-06 | 1.500E-05 |
| 253 | 13 | L13M | A | 1.3e-05 | 1.4e-05 | 1.2e-05 | 1.300E-05 | 1.000E-06 | 1.200E-05 | 1.400E-05 |
| 254 | 13 | L13F | A | 1e-05 | 1.1e-05 | 1.5e-05 | 1.200E-05 | 2.646E-06 | 1.000E-05 | 1.500E-05 |

| | | | | | | | | | | |
|---|---|---|---|---|---|---|---|---|---|---|
| 255 | 13 | L13P | A | 9.9e-06 | 1.6e-05 | 9.7e-06 | 1.187E-05 | 3.581E-06 | 9.700E-06 | 1.600E-05 |
| 256 | 13 | L13S | A | 1.6e-05 | 1.5e-05 | 1.6e-05 | 1.567E-05 | 5.774E-07 | 1.500E-05 | 1.600E-05 |
| 257 | 13 | L13T | A | 1.4e-05 | 1.2e-05 | 1.2e-05 | 1.267E-05 | 1.155E-06 | 1.200E-05 | 1.400E-05 |
| 258 | 13 | L13W | A | 1.2e-05 | 1.5e-05 | 1.7e-05 | 1.467E-05 | 2.517E-06 | 1.200E-05 | 1.700E-05 |
| 259 | 13 | L13Y | A | 1.5e-05 | 1.2e-05 | 1e-05 | 1.233E-05 | 2.517E-06 | 1.000E-05 | 1.500E-05 |
| 260 | 13 | L13V | A | 9.7e-06 | 1.7e-05 | 1.6e-05 | 1.423E-05 | 3.958E-06 | 9.700E-06 | 1.700E-05 |
| 261 | 14 | P14A | A | 1.1e-05 | 1.6e-05 | 9.7e-06 | 1.223E-05 | 3.326E-06 | 9.700E-06 | 1.600E-05 |
| 262 | 14 | P14R | A | 1.6e-05 | 8.8e-06 | 8.7e-06 | 1.117E-05 | 4.186E-06 | 8.700E-06 | 1.600E-05 |
| 263 | 14 | P14N | A | 1.5e-05 | 1e-05 | 1.4e-05 | 1.300E-05 | 2.646E-06 | 1.000E-05 | 1.500E-05 |
| 264 | 14 | P14D | A | 1.1e-05 | 1.1e-05 | 8.1e-06 | 1.003E-05 | 1.674E-06 | 8.100E-06 | 1.100E-05 |
| 265 | 14 | P14C | A | 8.8e-06 | 1.4e-05 | 1.5e-05 | 1.260E-05 | 3.329E-06 | 8.800E-06 | 1.500E-05 |
| 266 | 14 | P14E | A | 6.1e-06 | 1.2e-05 | 1.5e-05 | 1.103E-05 | 4.528E-06 | 6.100E-06 | 1.500E-05 |
| 267 | 14 | P14Q | A | 1e-05 | 1.4e-05 | 1.1e-05 | 1.167E-05 | 2.082E-06 | 1.000E-05 | 1.400E-05 |
| 268 | 14 | P14G | A | 1.4e-05 | 9.9e-06 | 7.6e-06 | 1.050E-05 | 3.242E-06 | 7.600E-06 | 1.400E-05 |
| 269 | 14 | P14H | A | 1e-05 | 1.5e-05 | 1.6e-05 | 1.367E-05 | 3.215E-06 | 1.000E-05 | 1.600E-05 |
| 270 | 14 | P14I | A | 1.1e-05 | 1.6e-05 | 1.6e-05 | 1.433E-05 | 2.887E-06 | 1.100E-05 | 1.600E-05 |
| 271 | 14 | P14L | A | 1.4e-05 | 1e-05 | 1.2e-05 | 1.200E-05 | 2.000E-06 | 1.000E-05 | 1.400E-05 |
| 272 | 14 | P14K | A | 8.8e-06 | 1.5e-05 | 1.6e-05 | 1.327E-05 | 3.900E-06 | 8.800E-06 | 1.600E-05 |
| 273 | 14 | P14M | A | 8.4e-06 | 9.4e-06 | 8e-06 | 8.600E-06 | 7.211E-07 | 8.000E-06 | 9.400E-06 |
| 274 | 14 | P14F | A | 7.7e-06 | 8.8e-06 | 1.5e-05 | 1.050E-05 | 3.936E-06 | 7.700E-06 | 1.500E-05 |
| 275 | 14 | P14P | A | 2.2e-05 | 1.3e-05 | 1.4e-05 | 1.633E-05 | 4.933E-06 | 1.300E-05 | 2.200E-05 |
| 276 | 14 | P14S | A | 1.4e-05 | 1.6e-05 | 1.5e-05 | 1.500E-05 | 1.000E-06 | 1.400E-05 | 1.600E-05 |
| 277 | 14 | P14T | A | 8.9e-06 | 1.6e-05 | 8.2e-06 | 1.103E-05 | 4.315E-06 | 8.200E-06 | 1.600E-05 |
| 278 | 14 | P14W | A | 1e-05 | 1e-05 | 1.2e-05 | 1.067E-05 | 1.155E-06 | 1.000E-05 | 1.200E-05 |
| 279 | 14 | P14Y | A | 1.5e-05 | 1.4e-05 | 1e-05 | 1.300E-05 | 2.646E-06 | 1.000E-05 | 1.500E-05 |
| 280 | 14 | P14V | A | 1.4e-05 | 1.5e-05 | 1.7e-05 | 1.533E-05 | 1.528E-06 | 1.400E-05 | 1.700E-05 |
| 281 | 15 | A15A | A | 1.5e-05 | 9e-06 | 9.7e-06 | 1.123E-05 | 3.281E-06 | 9.000E-06 | 1.500E-05 |
| 282 | 15 | A15R | A | 8.5e-06 | 1.5e-05 | 1e-05 | 1.117E-05 | 3.403E-06 | 8.500E-06 | 1.500E-05 |
| 283 | 15 | A15N | A | 1.4e-05 | 1e-05 | 1.4e-05 | 1.267E-05 | 2.309E-06 | 1.000E-05 | 1.400E-05 |
| 284 | 15 | A15D | A | 2.2e-05 | 1.6e-05 | 9.9e-06 | 1.597E-05 | 6.050E-06 | 9.900E-06 | 2.200E-05 |
| 285 | 15 | A15C | A | 1.2e-05 | 1.1e-05 | 1.1e-05 | 1.133E-05 | 5.774E-07 | 1.100E-05 | 1.200E-05 |
| 286 | 15 | A15E | A | 1.7e-05 | 9.5e-06 | 1.7e-05 | 1.450E-05 | 4.330E-06 | 9.500E-06 | 1.700E-05 |
| 287 | 15 | A15Q | A | 1.6e-05 | 1.5e-05 | 1.7e-05 | 1.600E-05 | 1.000E-06 | 1.500E-05 | 1.700E-05 |
| 288 | 15 | A15G | A | 8.7e-06 | 1.1e-05 | 1.7e-05 | 1.223E-05 | 4.285E-06 | 8.700E-06 | 1.700E-05 |
| 289 | 15 | A15H | A | 9.1e-06 | 1.7e-05 | 1.4e-05 | 1.337E-05 | 3.988E-06 | 9.100E-06 | 1.700E-05 |
| 290 | 15 | A15I | A | 1.3e-05 | 1.1e-05 | 1.1e-05 | 1.167E-05 | 1.155E-06 | 1.100E-05 | 1.300E-05 |
| 291 | 15 | A15L | A | 1.6e-05 | 1.4e-05 | 1.3e-05 | 1.433E-05 | 1.528E-06 | 1.300E-05 | 1.600E-05 |
| 292 | 15 | A15K | A | 1.4e-05 | 1.5e-05 | 1.4e-05 | 1.433E-05 | 5.774E-07 | 1.400E-05 | 1.500E-05 |
| 293 | 15 | A15M | A | 1.1e-05 | 9.6e-06 | 9.7e-06 | 1.010E-05 | 7.810E-07 | 9.600E-06 | 1.100E-05 |
| 294 | 15 | A15F | A | 1.8e-05 | 1.5e-05 | 1.5e-05 | 1.600E-05 | 1.732E-06 | 1.500E-05 | 1.800E-05 |
| 295 | 15 | A15P | A | 1.4e-05 | 1.3e-05 | 8e-06 | 1.167E-05 | 3.215E-06 | 8.000E-06 | 1.400E-05 |
| 296 | 15 | A15S | A | 1.3e-05 | 8.7e-06 | 1.4e-05 | 1.190E-05 | 2.816E-06 | 8.700E-06 | 1.400E-05 |
| 297 | 15 | A15T | A | 1e-05 | 1.5e-05 | 9.4e-06 | 1.147E-05 | 3.075E-06 | 9.400E-06 | 1.500E-05 |
| 298 | 15 | A15W | A | 1.2e-05 | 1e-05 | 1.6e-05 | 1.267E-05 | 3.055E-06 | 1.000E-05 | 1.600E-05 |
| 299 | 15 | A15Y | A | 9.3e-06 | 1.1e-05 | 1.7e-05 | 1.243E-05 | 4.045E-06 | 9.300E-06 | 1.700E-05 |
| 300 | 15 | A15V | A | 9.5e-06 | 1.7e-05 | 8.1e-06 | 1.153E-05 | 4.786E-06 | 8.100E-06 | 1.700E-05 |
| 301 | 16 | S16A | A | 1.6e-05 | 9.9e-06 | 1.1e-05 | 1.230E-05 | 3.251E-06 | 9.900E-06 | 1.600E-05 |
| 302 | 16 | S16R | A | 1.3e-05 | 1.4e-05 | 1.1e-05 | 1.267E-05 | 1.528E-06 | 1.100E-05 | 1.400E-05 |
| 303 | 16 | S16N | A | 1.4e-05 | 1.6e-05 | 1.4e-05 | 1.467E-05 | 1.155E-06 | 1.400E-05 | 1.600E-05 |
| 304 | 16 | S16D | A | 1.5e-05 | 1.4e-05 | 1.4e-05 | 1.433E-05 | 5.774E-07 | 1.400E-05 | 1.500E-05 |
| 305 | 16 | S16C | A | 1.5e-05 | 1.1e-05 | 1.6e-05 | 1.400E-05 | 2.646E-06 | 1.100E-05 | 1.600E-05 |
| 306 | 16 | S16E | A | 1e-05 | 1.1e-05 | 1.4e-05 | 1.167E-05 | 2.082E-06 | 1.000E-05 | 1.400E-05 |
| 307 | 16 | S16Q | A | 1.6e-05 | 1.2e-05 | 1.1e-05 | 1.300E-05 | 2.646E-06 | 1.100E-05 | 1.600E-05 |
| 308 | 16 | S16G | A | 1e-05 | 1.5e-05 | 1.6e-05 | 1.367E-05 | 3.215E-06 | 1.000E-05 | 1.600E-05 |
| 309 | 16 | S16H | A | 1.6e-05 | 1.6e-05 | 1e-05 | 1.400E-05 | 3.464E-06 | 1.000E-05 | 1.600E-05 |
| 310 | 16 | S16I | A | 9.1e-06 | 1.4e-05 | 8.2e-06 | 1.043E-05 | 3.121E-06 | 8.200E-06 | 1.400E-05 |
| 311 | 16 | S16L | A | 9e-06 | 9.7e-06 | 1.6e-05 | 1.157E-05 | 3.855E-06 | 9.000E-06 | 1.600E-05 |
| 312 | 16 | S16K | A | 9e-06 | 1.5e-05 | 8.2e-06 | 1.073E-05 | 3.717E-06 | 8.200E-06 | 1.500E-05 |
| 313 | 16 | S16M | A | 1.4e-05 | 1.6e-05 | 1.1e-05 | 1.367E-05 | 2.517E-06 | 1.100E-05 | 1.600E-05 |
| 314 | 16 | S16F | A | 1.4e-05 | 1.4e-05 | 1e-05 | 1.267E-05 | 2.309E-06 | 1.000E-05 | 1.400E-05 |
| 315 | 16 | S16P | A | 1.3e-05 | 1e-05 | 8.7e-06 | 1.057E-05 | 2.205E-06 | 8.700E-06 | 1.300E-05 |
| 316 | 16 | S16S | A | 1.6e-05 | 1.2e-05 | 1.6e-05 | 1.467E-05 | 2.309E-06 | 1.200E-05 | 1.600E-05 |
| 317 | 16 | S16T | A | 1.5e-05 | 1.1e-05 | 1.4e-05 | 1.333E-05 | 2.082E-06 | 1.100E-05 | 1.500E-05 |
| 318 | 16 | S16W | A | 9e-06 | 1.1e-05 | 1.4e-05 | 1.133E-05 | 2.517E-06 | 9.000E-06 | 1.400E-05 |
| 319 | 16 | S16Y | A | 1.5e-05 | 1.1e-05 | 1.5e-05 | 1.367E-05 | 2.309E-06 | 1.100E-05 | 1.500E-05 |
| 320 | 16 | S16V | A | 1.1e-05 | 1.7e-05 | 1.1e-05 | 1.300E-05 | 3.464E-06 | 1.100E-05 | 1.700E-05 |
| 321 | 17 | P17A | A | 1.1e-05 | 1.6e-05 | 1e-05 | 1.233E-05 | 3.215E-06 | 1.000E-05 | 1.600E-05 |
| 322 | 17 | P17R | A | 1e-05 | 1.2e-05 | 1.1e-05 | 1.100E-05 | 1.000E-06 | 1.000E-05 | 1.200E-05 |
| 323 | 17 | P17N | A | 1.5e-05 | 1.5e-05 | 8.6e-06 | 1.287E-05 | 3.695E-06 | 8.600E-06 | 1.500E-05 |
| 324 | 17 | P17D | A | 1.6e-05 | 1.5e-05 | 1.5e-05 | 1.533E-05 | 5.774E-07 | 1.500E-05 | 1.600E-05 |
| 325 | 17 | P17C | A | 1e-05 | 1e-05 | 1.3e-05 | 1.100E-05 | 1.732E-06 | 1.000E-05 | 1.300E-05 |
| 326 | 17 | P17E | A | 8.4e-06 | 1.5e-05 | 1.4e-05 | 1.247E-05 | 3.557E-06 | 8.400E-06 | 1.500E-05 |
| 327 | 17 | P17Q | A | 1e-05 | 1.6e-05 | 1.1e-05 | 1.233E-05 | 3.215E-06 | 1.000E-05 | 1.600E-05 |
| 328 | 17 | P17G | A | 1.4e-05 | 1.4e-05 | 1e-05 | 1.267E-05 | 2.309E-06 | 1.000E-05 | 1.400E-05 |
| 329 | 17 | P17H | A | 1.5e-05 | 1.3e-05 | 9.4e-06 | 1.247E-05 | 2.838E-06 | 9.400E-06 | 1.500E-05 |
| 330 | 17 | P17I | A | 9e-06 | 1.3e-05 | 1.6e-05 | 1.267E-05 | 3.512E-06 | 9.000E-06 | 1.600E-05 |
| 331 | 17 | P17L | A | 1.5e-05 | 1.5e-05 | 1.2e-05 | 1.400E-05 | 1.732E-06 | 1.200E-05 | 1.500E-05 |
| 332 | 17 | P17K | A | 1.1e-05 | 1.4e-05 | 1.4e-05 | 1.300E-05 | 1.732E-06 | 1.100E-05 | 1.400E-05 |
| 333 | 17 | P17M | A | 1.5e-05 | 1.4e-05 | 1.5e-05 | 1.467E-05 | 5.774E-07 | 1.400E-05 | 1.500E-05 |
| 334 | 17 | P17F | A | 1.1e-05 | 1.3e-05 | 6.3e-06 | 1.010E-05 | 3.439E-06 | 6.300E-06 | 1.300E-05 |
| 335 | 17 | P17P | A | 1.5e-05 | 1.1e-05 | 9.5e-06 | 1.183E-05 | 2.843E-06 | 9.500E-06 | 1.500E-05 |
| 336 | 17 | P17S | A | 1.5e-05 | 1.5e-05 | 8e-06 | 1.267E-05 | 4.041E-06 | 8.000E-06 | 1.500E-05 |
| 337 | 17 | P17T | A | 1.6e-05 | 1.5e-05 | 9.8e-06 | 1.360E-05 | 3.329E-06 | 9.800E-06 | 1.600E-05 |
| 338 | 17 | P17W | A | 9.1e-06 | 1e-05 | 1.1e-05 | 1.003E-05 | 9.504E-07 | 9.100E-06 | 1.100E-05 |
| 339 | 17 | P17Y | A | 5.8e-06 | 1.4e-05 | 1e-05 | 9.933E-06 | 4.100E-06 | 5.800E-06 | 1.400E-05 |

| | | | | | | | | | | |
|---|---|---|---|---|---|---|---|---|---|---|
| 340 | 17 | P17V | A | 1.3e-05 | 1e-05 | 1.3e-05 | 1.200E-05 | 1.732E-06 | 1.000E-05 | 1.300E-05 |
| 341 | 18 | E18A | A | 1.4e-05 | 1.1e-05 | 1.4e-05 | 1.300E-05 | 1.732E-06 | 1.100E-05 | 1.400E-05 |
| 342 | 18 | E18R | A | 1.4e-05 | 1.1e-05 | 1.7e-05 | 1.400E-05 | 3.000E-06 | 1.100E-05 | 1.700E-05 |
| 343 | 18 | E18N | A | 1.6e-05 | 1.4e-05 | 1.2e-05 | 1.400E-05 | 2.000E-06 | 1.200E-05 | 1.600E-05 |
| 344 | 18 | E18D | A | 8.5e-06 | 1.1e-05 | 9.3e-06 | 9.600E-06 | 1.277E-06 | 8.500E-06 | 1.100E-05 |
| 345 | 18 | E18C | A | 9.8e-06 | 1.6e-05 | 1.1e-05 | 1.227E-05 | 3.288E-06 | 9.800E-06 | 1.600E-05 |
| 346 | 18 | E18E | A | 1.2e-05 | 1.5e-05 | 8.3e-06 | 1.177E-05 | 3.356E-06 | 8.300E-06 | 1.500E-05 |
| 347 | 18 | E18Q | A | 8.3e-06 | 1.4e-05 | 1.5e-05 | 1.243E-05 | 3.614E-06 | 8.300E-06 | 1.500E-05 |
| 348 | 18 | E18G | A | 1.5e-05 | 1.6e-05 | 1.4e-05 | 1.500E-05 | 1.000E-06 | 1.400E-05 | 1.600E-05 |
| 349 | 18 | E18H | A | 9.8e-06 | 1.5e-05 | 1.1e-05 | 1.193E-05 | 2.723E-06 | 9.800E-06 | 1.500E-05 |
| 350 | 18 | E18I | A | 9.1e-06 | 1.6e-05 | 1.5e-05 | 1.337E-05 | 3.729E-06 | 9.100E-06 | 1.600E-05 |
| 351 | 18 | E18L | A | 1e-05 | 1.7e-05 | 1.5e-05 | 1.400E-05 | 3.606E-06 | 1.000E-05 | 1.700E-05 |
| 352 | 18 | E18K | A | 1.5e-05 | 1.2e-05 | 1.4e-05 | 1.367E-05 | 1.528E-06 | 1.200E-05 | 1.500E-05 |
| 353 | 18 | E18M | A | 1.1e-05 | 1.1e-05 | 1.6e-05 | 1.267E-05 | 2.887E-06 | 1.100E-05 | 1.600E-05 |
| 354 | 18 | E18F | A | 8.7e-06 | 1.1e-05 | 1e-05 | 9.900E-06 | 1.153E-06 | 8.700E-06 | 1.100E-05 |
| 355 | 18 | E18P | A | 9.4e-06 | 1.5e-05 | 1.6e-05 | 1.347E-05 | 3.557E-06 | 9.400E-06 | 1.600E-05 |
| 356 | 18 | E18S | A | 8.9e-06 | 1.1e-05 | 1.4e-05 | 1.130E-05 | 2.563E-06 | 8.900E-06 | 1.400E-05 |
| 357 | 18 | E18T | A | 9.7e-06 | 1.4e-05 | 1.2e-05 | 1.190E-05 | 2.152E-06 | 9.700E-06 | 1.400E-05 |
| 358 | 18 | E18W | A | 1.5e-05 | 1.1e-05 | 7.9e-06 | 1.130E-05 | 3.559E-06 | 7.900E-06 | 1.500E-05 |
| 359 | 18 | E18Y | A | 1.4e-05 | 1.7e-05 | 1.4e-05 | 1.500E-05 | 1.732E-06 | 1.400E-05 | 1.700E-05 |
| 360 | 18 | E18V | A | 1.4e-05 | 1.5e-05 | 1.3e-05 | 1.400E-05 | 1.000E-06 | 1.300E-05 | 1.500E-05 |
| 361 | 19 | T19A | A | 1.1e-05 | 7.9e-06 | 2.9e-05 | 1.597E-05 | 1.139E-05 | 7.900E-06 | 2.900E-05 |
| 362 | 19 | T19R | A | 1.7e-05 | 1.1e-05 | 1.7e-05 | 1.500E-05 | 3.464E-06 | 1.100E-05 | 1.700E-05 |
| 363 | 19 | T19N | A | 9e-06 | 9.7e-06 | 1.5e-05 | 1.123E-05 | 3.281E-06 | 9.000E-06 | 1.500E-05 |
| 364 | 19 | T19D | A | 1.5e-05 | 1.5e-05 | 1.4e-05 | 1.467E-05 | 5.774E-07 | 1.400E-05 | 1.500E-05 |
| 365 | 19 | T19C | A | 9.6e-06 | 1.4e-05 | 1.5e-05 | 1.287E-05 | 2.873E-06 | 9.600E-06 | 1.500E-05 |
| 366 | 19 | T19E | A | 9.8e-06 | 1.6e-05 | 1.7e-05 | 1.427E-05 | 3.900E-06 | 9.800E-06 | 1.700E-05 |
| 367 | 19 | T19Q | A | 1e-05 | 9e-06 | 1e-05 | 9.667E-06 | 5.774E-07 | 9.000E-06 | 1.000E-05 |
| 368 | 19 | T19G | A | 1e-05 | 1.5e-05 | 1.1e-05 | 1.200E-05 | 2.646E-06 | 1.000E-05 | 1.500E-05 |
| 369 | 19 | T19H | A | 1.4e-05 | 1.6e-05 | 1.3e-05 | 1.433E-05 | 1.528E-06 | 1.300E-05 | 1.600E-05 |
| 370 | 19 | T19I | A | 1.3e-05 | 1.3e-05 | 1.3e-05 | 1.300E-05 | 0.000E+00 | 1.300E-05 | 1.300E-05 |
| 371 | 19 | T19L | A | 1.2e-05 | 1e-05 | 1.6e-05 | 1.267E-05 | 3.055E-06 | 1.000E-05 | 1.600E-05 |
| 372 | 19 | T19K | A | 1.6e-05 | 1e-05 | 1.4e-05 | 1.333E-05 | 3.055E-06 | 1.000E-05 | 1.600E-05 |
| 373 | 19 | T19M | A | 9.6e-06 | 1.5e-05 | 1.2e-05 | 1.220E-05 | 2.706E-06 | 9.600E-06 | 1.500E-05 |
| 374 | 19 | T19F | A | 1.4e-05 | 1e-05 | 1.4e-05 | 1.267E-05 | 2.309E-06 | 1.000E-05 | 1.400E-05 |
| 375 | 19 | T19P | A | 1e-05 | 1.7e-05 | 1.5e-05 | 1.400E-05 | 3.606E-06 | 1.000E-05 | 1.700E-05 |
| 376 | 19 | T19S | A | 1.3e-05 | 1.3e-05 | 9.6e-06 | 1.187E-05 | 1.963E-06 | 9.600E-06 | 1.300E-05 |
| 377 | 19 | T19T | A | 1e-05 | 1.4e-05 | 1.5e-05 | 1.300E-05 | 2.646E-06 | 1.000E-05 | 1.500E-05 |
| 378 | 19 | T19W | A | 1e-05 | 1.5e-05 | 1.6e-05 | 1.367E-05 | 3.215E-06 | 1.000E-05 | 1.600E-05 |
| 379 | 19 | T19Y | A | 1.5e-05 | 1e-05 | 1e-05 | 1.167E-05 | 2.887E-06 | 1.000E-05 | 1.500E-05 |
| 380 | 19 | T19V | A | 1.3e-05 | 1.1e-05 | 1.7e-05 | 1.367E-05 | 3.055E-06 | 1.100E-05 | 1.700E-05 |
| 381 | 20 | H20A | A | 1e-05 | 1.6e-05 | 1.2e-05 | 1.267E-05 | 3.055E-06 | 1.000E-05 | 1.600E-05 |
| 382 | 20 | H20R | A | 1.1e-05 | 1.4e-05 | 1.4e-05 | 1.300E-05 | 1.732E-06 | 1.100E-05 | 1.400E-05 |
| 383 | 20 | H20N | A | 1e-05 | 1.6e-05 | 1.2e-05 | 1.267E-05 | 3.055E-06 | 1.000E-05 | 1.600E-05 |
| 384 | 20 | H20D | A | 1e-05 | 8.2e-06 | 9e-06 | 9.067E-06 | 9.018E-07 | 8.200E-06 | 1.000E-05 |
| 385 | 20 | H20C | A | 1.5e-05 | 1.4e-05 | 1.4e-05 | 1.433E-05 | 5.774E-07 | 1.400E-05 | 1.500E-05 |
| 386 | 20 | H20E | A | 1.4e-05 | 9.6e-06 | 1.6e-05 | 1.320E-05 | 3.274E-06 | 9.600E-06 | 1.600E-05 |
| 387 | 20 | H20Q | A | 1.5e-05 | 1.1e-05 | 1.4e-05 | 1.333E-05 | 2.082E-06 | 1.100E-05 | 1.500E-05 |
| 388 | 20 | H20G | A | 1.3e-05 | 1.3e-05 | 1.3e-05 | 1.300E-05 | 0.000E+00 | 1.300E-05 | 1.300E-05 |
| 389 | 20 | H20H | A | 1.5e-05 | 1.7e-05 | 1.3e-05 | 1.500E-05 | 2.000E-06 | 1.300E-05 | 1.700E-05 |
| 390 | 20 | H20I | A | 1.7e-05 | 1e-05 | 1.7e-05 | 1.467E-05 | 4.041E-06 | 1.000E-05 | 1.700E-05 |
| 391 | 20 | H20L | A | 1.8e-05 | 1.6e-05 | 1.7e-05 | 1.700E-05 | 1.000E-06 | 1.600E-05 | 1.800E-05 |
| 392 | 20 | H20K | A | 1.7e-05 | 1.5e-05 | 1.5e-05 | 1.567E-05 | 1.155E-06 | 1.500E-05 | 1.700E-05 |
| 393 | 20 | H20M | A | 1.4e-05 | 1.4e-05 | 1.4e-05 | 1.400E-05 | 0.000E+00 | 1.400E-05 | 1.400E-05 |
| 394 | 20 | H20F | A | 1.2e-05 | 1.1e-05 | 1.4e-05 | 1.233E-05 | 1.528E-06 | 1.100E-05 | 1.400E-05 |
| 395 | 20 | H20P | A | 9.6e-06 | 1.4e-05 | 1.4e-05 | 1.253E-05 | 2.540E-06 | 9.600E-06 | 1.400E-05 |
| 396 | 20 | H20S | A | 1.6e-05 | 1.2e-05 | 1.8e-05 | 1.533E-05 | 3.055E-06 | 1.200E-05 | 1.800E-05 |
| 397 | 20 | H20T | A | 1e-05 | 9.7e-06 | 9.5e-06 | 9.733E-06 | 2.517E-07 | 9.500E-06 | 1.000E-05 |
| 398 | 20 | H20W | A | 1.6e-05 | 9.7e-06 | 1.1e-05 | 1.223E-05 | 3.326E-06 | 9.700E-06 | 1.600E-05 |
| 399 | 20 | H20Y | A | 1.5e-05 | 1.1e-05 | 1.1e-05 | 1.233E-05 | 2.309E-06 | 1.100E-05 | 1.500E-05 |
| 400 | 20 | H20V | A | 1.2e-05 | 1.1e-05 | 1.1e-05 | 1.133E-05 | 5.774E-07 | 1.100E-05 | 1.200E-05 |
| 401 | 21 | L21A | A | 1.2e-05 | 1.1e-05 | 1.6e-05 | 1.300E-05 | 2.646E-06 | 1.100E-05 | 1.600E-05 |
| 402 | 21 | L21R | A | 1.5e-05 | 1.3e-05 | 1.5e-05 | 1.433E-05 | 1.155E-06 | 1.300E-05 | 1.500E-05 |
| 403 | 21 | L21N | A | 1.2e-05 | 1.3e-05 | 1.5e-05 | 1.333E-05 | 1.528E-06 | 1.200E-05 | 1.500E-05 |
| 404 | 21 | L21D | A | 1.5e-05 | 1.5e-05 | 1e-05 | 1.333E-05 | 2.887E-06 | 1.000E-05 | 1.500E-05 |
| 405 | 21 | L21C | A | 1.5e-05 | 1.5e-05 | 1.5e-05 | 1.500E-05 | 0.000E+00 | 1.500E-05 | 1.500E-05 |
| 406 | 21 | L21E | A | 1.5e-05 | 1.5e-05 | 1.6e-05 | 1.533E-05 | 5.774E-07 | 1.500E-05 | 1.600E-05 |
| 407 | 21 | L21Q | A | 9.2e-06 | 1.5e-05 | 1.7e-05 | 1.373E-05 | 4.051E-06 | 9.200E-06 | 1.700E-05 |
| 408 | 21 | L21G | A | 9.9e-06 | 1.5e-05 | 8.8e-06 | 1.123E-05 | 3.308E-06 | 8.800E-06 | 1.500E-05 |
| 409 | 21 | L21H | A | 1.4e-05 | 1.5e-05 | 1e-05 | 1.300E-05 | 2.646E-06 | 1.000E-05 | 1.500E-05 |
| 410 | 21 | L21I | A | 1.6e-05 | 1.4e-05 | 1.1e-05 | 1.367E-05 | 2.517E-06 | 1.100E-05 | 1.600E-05 |
| 411 | 21 | L21L | A | 9.2e-06 | 1.7e-05 | 1e-05 | 1.207E-05 | 4.291E-06 | 9.200E-06 | 1.700E-05 |
| 412 | 21 | L21K | A | 1.5e-05 | 1.1e-05 | 1e-05 | 1.200E-05 | 2.646E-06 | 1.000E-05 | 1.500E-05 |
| 413 | 21 | L21M | A | 1.3e-05 | 1.1e-05 | 1.5e-05 | 1.300E-05 | 2.000E-06 | 1.100E-05 | 1.500E-05 |
| 414 | 21 | L21F | A | 3.8e-05 | 9.8e-06 | 1.6e-05 | 2.127E-05 | 1.482E-05 | 9.800E-06 | 3.800E-05 |
| 415 | 21 | L21P | A | 8.8e-06 | 1.5e-05 | 1.5e-05 | 1.293E-05 | 3.580E-06 | 8.800E-06 | 1.500E-05 |
| 416 | 21 | L21S | A | 9.7e-06 | 1e-05 | 1.3e-05 | 1.090E-05 | 1.825E-06 | 9.700E-06 | 1.300E-05 |
| 417 | 21 | L21T | A | 1.3e-05 | 9.9e-06 | 1.5e-05 | 1.263E-05 | 2.570E-06 | 9.900E-06 | 1.500E-05 |
| 418 | 21 | L21W | A | 1.4e-05 | 1.1e-05 | 1.4e-05 | 1.300E-05 | 1.732E-06 | 1.100E-05 | 1.400E-05 |
| 419 | 21 | L21Y | A | 1.5e-05 | 1e-05 | 1.3e-05 | 1.267E-05 | 2.517E-06 | 1.000E-05 | 1.500E-05 |
| 420 | 21 | L21V | A | 1.5e-05 | 1.5e-05 | 9.3e-06 | 1.310E-05 | 3.291E-06 | 9.300E-06 | 1.500E-05 |
| 421 | 22 | D22A | A | 1.6e-05 | 1.6e-05 | 1.1e-05 | 1.433E-05 | 2.887E-06 | 1.100E-05 | 1.600E-05 |
| 422 | 22 | D22R | A | 1.8e-05 | 1.4e-05 | 1.4e-05 | 1.533E-05 | 2.309E-06 | 1.400E-05 | 1.800E-05 |
| 423 | 22 | D22N | A | 1.5e-05 | 1.1e-05 | 1.1e-05 | 1.233E-05 | 2.309E-06 | 1.100E-05 | 1.500E-05 |
| 424 | 22 | D22D | A | 8.6e-06 | 1.5e-05 | 1.3e-05 | 1.220E-05 | 3.274E-06 | 8.600E-06 | 1.500E-05 |

| | | | | | | | | | | |
|---|---|---|---|---|---|---|---|---|---|---|
| 425 | 22 | D22C | A | 1e-05 | 1.4e-05 | 1.4e-05 | 1.267E-05 | 2.309E-06 | 1.000E-05 | 1.400E-05 |
| 426 | 22 | D22E | A | 1.2e-05 | 1.4e-05 | 1.5e-05 | 1.367E-05 | 1.528E-06 | 1.200E-05 | 1.500E-05 |
| 427 | 22 | D22Q | A | 1.4e-05 | 9.1e-06 | 8.7e-06 | 1.060E-05 | 2.951E-06 | 8.700E-06 | 1.400E-05 |
| 428 | 22 | D22G | A | 9.2e-06 | 9.9e-06 | 1.6e-05 | 1.170E-05 | 3.740E-06 | 9.200E-06 | 1.600E-05 |
| 429 | 22 | D22H | A | 9.7e-06 | 1e-05 | 1e-05 | 9.900E-06 | 1.732E-07 | 9.700E-06 | 1.000E-05 |
| 430 | 22 | D22I | A | 1.6e-05 | 1.5e-05 | 1.6e-05 | 1.567E-05 | 5.774E-07 | 1.500E-05 | 1.600E-05 |
| 431 | 22 | D22L | A | 1.5e-05 | 1.4e-05 | 8.8e-06 | 1.260E-05 | 3.329E-06 | 8.800E-06 | 1.500E-05 |
| 432 | 22 | D22K | A | 9.4e-06 | 1.6e-05 | 1.4e-05 | 1.313E-05 | 3.384E-06 | 9.400E-06 | 1.600E-05 |
| 433 | 22 | D22M | A | 1.7e-05 | 1.6e-05 | 9.2e-06 | 1.407E-05 | 4.244E-06 | 9.200E-06 | 1.700E-05 |
| 434 | 22 | D22F | A | 9.8e-06 | 1.4e-05 | 1.4e-05 | 1.260E-05 | 2.425E-06 | 9.800E-06 | 1.400E-05 |
| 435 | 22 | D22P | A | 1.7e-05 | 1.3e-05 | 9.1e-06 | 1.303E-05 | 3.950E-06 | 9.100E-06 | 1.700E-05 |
| 436 | 22 | D22S | A | 8.2e-06 | 1.5e-05 | 1.8e-05 | 1.373E-05 | 5.021E-06 | 8.200E-06 | 1.800E-05 |
| 437 | 22 | D22T | A | 1.5e-05 | 1.2e-05 | 1.6e-05 | 1.433E-05 | 2.082E-06 | 1.200E-05 | 1.600E-05 |
| 438 | 22 | D22W | A | 1.4e-05 | 1.5e-05 | 1.6e-05 | 1.500E-05 | 1.000E-06 | 1.400E-05 | 1.600E-05 |
| 439 | 22 | D22Y | A | 1.4e-05 | 8.6e-06 | 1.5e-05 | 1.253E-05 | 3.443E-06 | 8.600E-06 | 1.500E-05 |
| 440 | 22 | D22V | A | 1.3e-05 | 1.1e-05 | 1.4e-05 | 1.267E-05 | 1.528E-06 | 1.100E-05 | 1.400E-05 |
| 441 | 23 | M23A | A | 1e-05 | 1.5e-05 | 1.2e-05 | 1.233E-05 | 2.517E-06 | 1.000E-05 | 1.500E-05 |
| 442 | 23 | M23R | A | 1.4e-05 | 1.1e-05 | 1.5e-05 | 1.333E-05 | 2.082E-06 | 1.100E-05 | 1.500E-05 |
| 443 | 23 | M23N | A | 1e-05 | 1.2e-05 | 9.9e-06 | 1.063E-05 | 1.185E-06 | 9.900E-06 | 1.200E-05 |
| 444 | 23 | M23D | A | 1.4e-05 | 1.4e-05 | 8.8e-06 | 1.227E-05 | 3.002E-06 | 8.800E-06 | 1.400E-05 |
| 445 | 23 | M23C | A | 1.3e-05 | 9.8e-06 | 1.2e-05 | 1.160E-05 | 1.637E-06 | 9.800E-06 | 1.300E-05 |
| 446 | 23 | M23E | A | 1.4e-05 | 8.9e-06 | 1.5e-05 | 1.263E-05 | 3.272E-06 | 8.900E-06 | 1.500E-05 |
| 447 | 23 | M23Q | A | 1e-05 | 1.5e-05 | 9.7e-06 | 1.157E-05 | 2.977E-06 | 9.700E-06 | 1.500E-05 |
| 448 | 23 | M23G | A | 1.5e-05 | 1.1e-05 | 1.6e-05 | 1.400E-05 | 2.646E-06 | 1.100E-05 | 1.600E-05 |
| 449 | 23 | M23H | A | 1.7e-05 | 1.7e-05 | 1.5e-05 | 1.633E-05 | 1.155E-06 | 1.500E-05 | 1.700E-05 |
| 450 | 23 | M23I | A | 1.4e-05 | 1.1e-05 | 1.5e-05 | 1.333E-05 | 2.082E-06 | 1.100E-05 | 1.500E-05 |
| 451 | 23 | M23L | A | 1e-05 | 1e-05 | 1.1e-05 | 1.033E-05 | 5.774E-07 | 1.000E-05 | 1.100E-05 |
| 452 | 23 | M23K | A | 1.2e-05 | 1.3e-05 | 1.3e-05 | 1.267E-05 | 5.774E-07 | 1.200E-05 | 1.300E-05 |
| 453 | 23 | M23M | A | 8.5e-06 | 1.4e-05 | 7.9e-06 | 1.013E-05 | 3.362E-06 | 7.900E-06 | 1.400E-05 |
| 454 | 23 | M23F | A | 1.6e-05 | 1.5e-05 | 1.1e-05 | 1.400E-05 | 2.646E-06 | 1.100E-05 | 1.600E-05 |
| 455 | 23 | M23P | A | 1.4e-05 | 1.6e-05 | 1.4e-05 | 1.467E-05 | 1.155E-06 | 1.400E-05 | 1.600E-05 |
| 456 | 23 | M23S | A | 9.8e-06 | 1.5e-05 | 1.6e-05 | 1.360E-05 | 3.329E-06 | 9.800E-06 | 1.600E-05 |
| 457 | 23 | M23T | A | 9.3e-06 | 1.1e-05 | 1.3e-05 | 1.110E-05 | 1.852E-06 | 9.300E-06 | 1.300E-05 |
| 458 | 23 | M23W | A | 1.7e-05 | 9.4e-06 | 1.5e-05 | 1.380E-05 | 3.940E-06 | 9.400E-06 | 1.700E-05 |
| 459 | 23 | M23Y | A | 1.5e-05 | 1.3e-05 | 9.7e-06 | 1.257E-05 | 2.676E-06 | 9.700E-06 | 1.500E-05 |
| 460 | 23 | M23V | A | 1.3e-05 | 1.6e-05 | 1.6e-05 | 1.500E-05 | 1.732E-06 | 1.300E-05 | 1.600E-05 |
| 461 | 24 | L24A | A | 1.1e-05 | 8.4e-06 | 1.3e-05 | 1.080E-05 | 2.307E-06 | 8.400E-06 | 1.300E-05 |
| 462 | 24 | L24R | A | 1e-05 | 1.7e-05 | 1.1e-05 | 1.267E-05 | 3.786E-06 | 1.000E-05 | 1.700E-05 |
| 463 | 24 | L24N | A | 1.1e-05 | 1.1e-05 | 1.1e-05 | 1.100E-05 | 2.075E-21 | 1.100E-05 | 1.100E-05 |
| 464 | 24 | L24D | A | 1e-05 | 1.1e-05 | 1.6e-05 | 1.233E-05 | 3.215E-06 | 1.000E-05 | 1.600E-05 |
| 465 | 24 | L24C | A | 1e-05 | 1.3e-05 | 9.1e-06 | 1.070E-05 | 2.042E-06 | 9.100E-06 | 1.300E-05 |
| 466 | 24 | L24E | A | 1.5e-05 | 1.2e-05 | 1.4e-05 | 1.367E-05 | 1.528E-06 | 1.200E-05 | 1.500E-05 |
| 467 | 24 | L24Q | A | 1.3e-05 | 9.4e-06 | 1.4e-05 | 1.213E-05 | 2.419E-06 | 9.400E-06 | 1.400E-05 |
| 468 | 24 | L24G | A | 1e-05 | 9.7e-06 | 1.6e-05 | 1.190E-05 | 3.554E-06 | 9.700E-06 | 1.600E-05 |
| 469 | 24 | L24H | A | 1.7e-05 | 1.7e-05 | 1.4e-05 | 1.600E-05 | 1.732E-06 | 1.400E-05 | 1.700E-05 |
| 470 | 24 | L24I | A | 1.5e-05 | 1.2e-05 | 1.1e-05 | 1.267E-05 | 2.082E-06 | 1.100E-05 | 1.500E-05 |
| 471 | 24 | L24L | A | 1.2e-05 | 7.4e-06 | 7.2e-06 | 8.867E-06 | 2.715E-06 | 7.200E-06 | 1.200E-05 |
| 472 | 24 | L24K | A | 1.5e-05 | 9.9e-06 | 8.6e-06 | 1.117E-05 | 3.383E-06 | 8.600E-06 | 1.500E-05 |
| 473 | 24 | L24M | A | 1.5e-05 | 1.1e-05 | 1.6e-05 | 1.400E-05 | 2.646E-06 | 1.100E-05 | 1.600E-05 |
| 474 | 24 | L24F | A | 1.4e-05 | 9.2e-06 | 1.4e-05 | 1.240E-05 | 2.771E-06 | 9.200E-06 | 1.400E-05 |
| 475 | 24 | L24P | A | 1.4e-05 | 1.1e-05 | 1.4e-05 | 1.300E-05 | 1.732E-06 | 1.100E-05 | 1.400E-05 |
| 476 | 24 | L24S | A | 1.5e-05 | 1.5e-05 | 1e-05 | 1.333E-05 | 2.887E-06 | 1.000E-05 | 1.500E-05 |
| 477 | 24 | L24T | A | 1e-05 | 1.4e-05 | 1.3e-05 | 1.233E-05 | 2.082E-06 | 1.000E-05 | 1.400E-05 |
| 478 | 24 | L24W | A | 1.3e-05 | 1e-05 | 9.9e-06 | 1.097E-05 | 1.762E-06 | 9.900E-06 | 1.300E-05 |
| 479 | 24 | L24Y | A | 1.5e-05 | 9.6e-06 | 1.1e-05 | 1.187E-05 | 2.802E-06 | 9.600E-06 | 1.500E-05 |
| 480 | 24 | L24V | A | 1e-05 | 1e-05 | 1.5e-05 | 1.167E-05 | 2.887E-06 | 1.000E-05 | 1.500E-05 |
| 481 | 25 | R25A | A | 1.1e-05 | 1.2e-05 | 1.3e-05 | 1.200E-05 | 1.000E-06 | 1.100E-05 | 1.300E-05 |
| 482 | 25 | R25R | A | 9.1e-06 | 1.2e-05 | 1.5e-05 | 1.203E-05 | 2.950E-06 | 9.100E-06 | 1.500E-05 |
| 483 | 25 | R25N | A | 1.1e-05 | 1.4e-05 | 7.9e-06 | 1.097E-05 | 3.050E-06 | 7.900E-06 | 1.400E-05 |
| 484 | 25 | R25D | A | 8.1e-06 | 8.6e-06 | 1.5e-05 | 1.057E-05 | 3.848E-06 | 8.100E-06 | 1.500E-05 |
| 485 | 25 | R25C | A | 8.3e-06 | 8.3e-06 | 1.5e-05 | 1.053E-05 | 3.868E-06 | 8.300E-06 | 1.500E-05 |
| 486 | 25 | R25E | A | 1.1e-05 | 9.9e-06 | 1.6e-05 | 1.230E-05 | 3.251E-06 | 9.900E-06 | 1.600E-05 |
| 487 | 25 | R25Q | A | 1.5e-05 | 8.8e-06 | 1.4e-05 | 1.260E-05 | 3.329E-06 | 8.800E-06 | 1.500E-05 |
| 488 | 25 | R25G | A | 1.1e-05 | 1.1e-05 | 1.6e-05 | 1.267E-05 | 2.887E-06 | 1.100E-05 | 1.600E-05 |
| 489 | 25 | R25H | A | 1e-05 | 1.4e-05 | 1e-05 | 1.133E-05 | 2.309E-06 | 1.000E-05 | 1.400E-05 |
| 490 | 25 | R25I | A | 8.7e-06 | 1.3e-05 | 1.6e-05 | 1.257E-05 | 3.669E-06 | 8.700E-06 | 1.600E-05 |
| 491 | 25 | R25L | A | 1.4e-05 | 1.4e-05 | 1e-05 | 1.267E-05 | 2.309E-06 | 1.000E-05 | 1.400E-05 |
| 492 | 25 | R25K | A | 1.5e-05 | 1.6e-05 | 1.2e-05 | 1.433E-05 | 2.082E-06 | 1.200E-05 | 1.600E-05 |
| 493 | 25 | R25M | A | 1.5e-05 | 1.2e-05 | 1.1e-05 | 1.267E-05 | 2.082E-06 | 1.100E-05 | 1.500E-05 |
| 494 | 25 | R25F | A | 1.2e-05 | 1.4e-05 | 1.1e-05 | 1.233E-05 | 1.528E-06 | 1.100E-05 | 1.400E-05 |
| 495 | 25 | R25P | A | 1.1e-05 | 1.1e-05 | 8.7e-06 | 1.023E-05 | 1.328E-06 | 8.700E-06 | 1.100E-05 |
| 496 | 25 | R25S | A | 1.4e-05 | 1.2e-05 | 1.1e-05 | 1.233E-05 | 1.528E-06 | 1.100E-05 | 1.400E-05 |
| 497 | 25 | R25T | A | 1.5e-05 | 1.5e-05 | 1.1e-05 | 1.367E-05 | 2.309E-06 | 1.100E-05 | 1.500E-05 |
| 498 | 25 | R25W | A | 1.2e-05 | 1.1e-05 | 1.5e-05 | 1.267E-05 | 2.082E-06 | 1.100E-05 | 1.500E-05 |
| 499 | 25 | R25Y | A | 9.7e-06 | 1.5e-05 | 1e-05 | 1.157E-05 | 2.977E-06 | 9.700E-06 | 1.500E-05 |
| 500 | 25 | R25V | A | 8.2e-06 | 8.3e-06 | 8.4e-06 | 8.300E-06 | 1.000E-07 | 8.200E-06 | 8.400E-06 |
| 501 | 26 | H26A | A | 1.5e-05 | 1e-05 | 1.6e-05 | 1.367E-05 | 3.215E-06 | 1.000E-05 | 1.600E-05 |
| 502 | 26 | H26R | A | 7e-06 | 1.3e-05 | 1.4e-05 | 1.133E-05 | 3.786E-06 | 7.000E-06 | 1.400E-05 |
| 503 | 26 | H26N | A | 9.3e-06 | 1.8e-05 | 1.1e-05 | 1.277E-05 | 4.611E-06 | 9.300E-06 | 1.800E-05 |
| 504 | 26 | H26D | A | 1.2e-05 | 1.5e-05 | 1.7e-05 | 1.467E-05 | 2.517E-06 | 1.200E-05 | 1.700E-05 |
| 505 | 26 | H26C | A | 2e-05 | 9.3e-06 | 1.7e-05 | 1.543E-05 | 5.519E-06 | 9.300E-06 | 2.000E-05 |
| 506 | 26 | H26E | A | 1.1e-05 | 1.6e-05 | 1.5e-05 | 1.400E-05 | 2.646E-06 | 1.100E-05 | 1.600E-05 |
| 507 | 26 | H26Q | A | 1.5e-05 | 1.1e-05 | 1.1e-05 | 1.233E-05 | 2.309E-06 | 1.100E-05 | 1.500E-05 |
| 508 | 26 | H26G | A | 1.4e-05 | 1.6e-05 | 1.6e-05 | 1.533E-05 | 1.155E-06 | 1.400E-05 | 1.600E-05 |
| 509 | 26 | H26H | A | 1.4e-05 | 1.3e-05 | 1.5e-05 | 1.400E-05 | 1.000E-06 | 1.300E-05 | 1.500E-05 |

| | | | | | | | | | | |
|---|---|---|---|---|---|---|---|---|---|---|
| 510 | 26 | H26I | A | 1.6e-05 | 1.6e-05 | 1.1e-05 | 1.433E-05 | 2.887E-06 | 1.100E-05 | 1.600E-05 |
| 511 | 26 | H26L | A | 8.4e-06 | 1.5e-05 | 1.1e-05 | 1.147E-05 | 3.325E-06 | 8.400E-06 | 1.500E-05 |
| 512 | 26 | H26K | A | 1.6e-05 | 1.1e-05 | 1.5e-05 | 1.400E-05 | 2.646E-06 | 1.100E-05 | 1.600E-05 |
| 513 | 26 | H26M | A | 1.5e-05 | 1.2e-05 | 1.4e-05 | 1.367E-05 | 1.528E-06 | 1.200E-05 | 1.500E-05 |
| 514 | 26 | H26F | A | 1.6e-05 | 1.5e-05 | 9e-06 | 1.333E-05 | 3.786E-06 | 9.000E-06 | 1.600E-05 |
| 515 | 26 | H26P | A | 1e-05 | 1.5e-05 | 1.5e-05 | 1.333E-05 | 2.887E-06 | 1.000E-05 | 1.500E-05 |
| 516 | 26 | H26S | A | 1.2e-05 | 1.1e-05 | 9.8e-06 | 1.093E-05 | 1.102E-06 | 9.800E-06 | 1.200E-05 |
| 517 | 26 | H26T | A | 9.9e-06 | 1.4e-05 | 1.1e-05 | 1.163E-05 | 2.122E-06 | 9.900E-06 | 1.400E-05 |
| 518 | 26 | H26W | A | 1.7e-05 | 1.7e-05 | 1.3e-05 | 1.567E-05 | 2.309E-06 | 1.300E-05 | 1.700E-05 |
| 519 | 26 | H26Y | A | 2e-05 | 6.9e-06 | 1.1e-05 | 1.263E-05 | 6.701E-06 | 6.900E-06 | 2.000E-05 |
| 520 | 26 | H26V | A | 1.1e-05 | 1.5e-05 | 9.6e-06 | 1.187E-05 | 2.802E-06 | 9.600E-06 | 1.500E-05 |
| 521 | 27 | L27A | A | 1.5e-05 | 1e-05 | 1.6e-05 | 1.367E-05 | 3.215E-06 | 1.000E-05 | 1.600E-05 |
| 522 | 27 | L27R | A | 1.6e-05 | 9e-06 | 1.8e-05 | 1.433E-05 | 4.726E-06 | 9.000E-06 | 1.800E-05 |
| 523 | 27 | L27N | A | 1.5e-05 | 1.4e-05 | 1.5e-05 | 1.467E-05 | 5.774E-07 | 1.400E-05 | 1.500E-05 |
| 524 | 27 | L27D | A | 1.6e-05 | 9.3e-06 | 1.2e-05 | 1.243E-05 | 3.371E-06 | 9.300E-06 | 1.600E-05 |
| 525 | 27 | L27C | A | 1.4e-05 | 9.3e-06 | 1e-05 | 1.110E-05 | 2.536E-06 | 9.300E-06 | 1.400E-05 |
| 526 | 27 | L27E | A | 1.4e-05 | 1.1e-05 | 1.3e-05 | 1.267E-05 | 1.528E-06 | 1.100E-05 | 1.400E-05 |
| 527 | 27 | L27Q | A | 7.9e-06 | 1.5e-05 | 1.6e-05 | 1.297E-05 | 4.416E-06 | 7.900E-06 | 1.600E-05 |
| 528 | 27 | L27G | A | 7.9e-06 | 1.4e-05 | 1.3e-05 | 1.163E-05 | 3.272E-06 | 7.900E-06 | 1.400E-05 |
| 529 | 27 | L27H | A | 1.4e-05 | 1.2e-05 | 1.2e-05 | 1.267E-05 | 1.155E-06 | 1.200E-05 | 1.400E-05 |
| 530 | 27 | L27I | A | 1.4e-05 | 1.1e-05 | 1.5e-05 | 1.333E-05 | 2.082E-06 | 1.100E-05 | 1.500E-05 |
| 531 | 27 | L27L | A | 1.7e-05 | 1.6e-05 | 8.6e-06 | 1.387E-05 | 4.588E-06 | 8.600E-06 | 1.700E-05 |
| 532 | 27 | L27K | A | 8.6e-06 | 1.4e-05 | 9.2e-06 | 1.060E-05 | 2.960E-06 | 8.600E-06 | 1.400E-05 |
| 533 | 27 | L27M | A | 1e-05 | 1.3e-05 | 1e-05 | 1.100E-05 | 1.732E-06 | 1.000E-05 | 1.300E-05 |
| 534 | 27 | L27F | A | 1e-05 | 1.6e-05 | 1e-05 | 1.200E-05 | 3.464E-06 | 1.000E-05 | 1.600E-05 |
| 535 | 27 | L27P | A | 1e-05 | 6.7e-06 | 1.4e-05 | 1.023E-05 | 3.656E-06 | 6.700E-06 | 1.400E-05 |
| 536 | 27 | L27S | A | 1e-05 | 1e-05 | 1.6e-05 | 1.200E-05 | 3.464E-06 | 1.000E-05 | 1.600E-05 |
| 537 | 27 | L27T | A | 1e-05 | 1.4e-05 | 1.3e-05 | 1.233E-05 | 2.082E-06 | 1.000E-05 | 1.400E-05 |
| 538 | 27 | L27W | A | 1.1e-05 | 1.4e-05 | 9.6e-06 | 1.153E-05 | 2.248E-06 | 9.600E-06 | 1.400E-05 |
| 539 | 27 | L27Y | A | 1.3e-05 | 9.5e-06 | 1e-05 | 1.083E-05 | 1.893E-06 | 9.500E-06 | 1.300E-05 |
| 540 | 27 | L27V | A | 1.4e-05 | 9.5e-06 | 1.4e-05 | 1.250E-05 | 2.598E-06 | 9.500E-06 | 1.400E-05 |
| 541 | 28 | Y28A | A | 1.6e-05 | 1.2e-05 | 8.6e-06 | 1.220E-05 | 3.704E-06 | 8.600E-06 | 1.600E-05 |
| 542 | 28 | Y28R | A | 9.2e-06 | 1.6e-05 | 2e-05 | 1.507E-05 | 5.460E-06 | 9.200E-06 | 2.000E-05 |
| 543 | 28 | Y28N | A | 1.5e-05 | 9.7e-06 | 1.6e-05 | 1.357E-05 | 3.386E-06 | 9.700E-06 | 1.600E-05 |
| 544 | 28 | Y28D | A | 1.2e-05 | 1.7e-05 | 6.3e-06 | 1.177E-05 | 5.354E-06 | 6.300E-06 | 1.700E-05 |
| 545 | 28 | Y28C | A | 1.5e-05 | 1.6e-05 | 1.3e-05 | 1.467E-05 | 1.528E-06 | 1.300E-05 | 1.600E-05 |
| 546 | 28 | Y28E | A | 1.4e-05 | 1.3e-05 | 1.2e-05 | 1.300E-05 | 1.000E-06 | 1.200E-05 | 1.400E-05 |
| 547 | 28 | Y28Q | A | 1.1e-05 | 1.7e-05 | 7.7e-06 | 1.190E-05 | 4.715E-06 | 7.700E-06 | 1.700E-05 |
| 548 | 28 | Y28G | A | 5.9e-06 | 1.4e-05 | 1.3e-05 | 1.097E-05 | 4.416E-06 | 5.900E-06 | 1.400E-05 |
| 549 | 28 | Y28H | A | 1.5e-05 | 1.1e-05 | 1.6e-05 | 1.400E-05 | 2.646E-06 | 1.100E-05 | 1.600E-05 |
| 550 | 28 | Y28I | A | 1.3e-05 | 8.6e-06 | 1e-05 | 1.053E-05 | 2.248E-06 | 8.600E-06 | 1.300E-05 |
| 551 | 28 | Y28L | A | 1.6e-05 | 1e-05 | 1.5e-05 | 1.367E-05 | 3.215E-06 | 1.000E-05 | 1.600E-05 |
| 552 | 28 | Y28K | A | 1.5e-05 | 1.7e-05 | 1.1e-05 | 1.433E-05 | 3.055E-06 | 1.100E-05 | 1.700E-05 |
| 553 | 28 | Y28M | A | 1.4e-05 | 1.5e-05 | 1.2e-05 | 1.367E-05 | 1.528E-06 | 1.200E-05 | 1.500E-05 |
| 554 | 28 | Y28F | A | 1.5e-05 | 1.6e-05 | 1.7e-05 | 1.600E-05 | 1.000E-06 | 1.500E-05 | 1.700E-05 |
| 555 | 28 | Y28P | A | 1.7e-05 | 1e-05 | 1e-05 | 1.233E-05 | 4.041E-06 | 1.000E-05 | 1.700E-05 |
| 556 | 28 | Y28S | A | 1.6e-05 | 1.1e-05 | 1e-05 | 1.233E-05 | 3.215E-06 | 1.000E-05 | 1.600E-05 |
| 557 | 28 | Y28T | A | 1.2e-05 | 1.6e-05 | 8.1e-06 | 1.203E-05 | 3.950E-06 | 8.100E-06 | 1.600E-05 |
| 558 | 28 | Y28W | A | 8.5e-06 | 9.8e-06 | 1.7e-05 | 1.177E-05 | 4.579E-06 | 8.500E-06 | 1.700E-05 |
| 559 | 28 | Y28Y | A | 1e-05 | 1.5e-05 | 1.6e-05 | 1.367E-05 | 3.215E-06 | 1.000E-05 | 1.600E-05 |
| 560 | 28 | Y28V | A | 1.3e-05 | 9.3e-06 | 7.9e-06 | 1.007E-05 | 2.635E-06 | 7.900E-06 | 1.300E-05 |
| 561 | 29 | Q29A | A | 9.8e-06 | 1.1e-05 | 1.6e-05 | 1.227E-05 | 3.288E-06 | 9.800E-06 | 1.600E-05 |
| 562 | 29 | Q29R | A | 1.1e-05 | 1.5e-05 | 1.5e-05 | 1.367E-05 | 2.309E-06 | 1.100E-05 | 1.500E-05 |
| 563 | 29 | Q29N | A | 1.7e-05 | 1.5e-05 | 1.3e-05 | 1.500E-05 | 2.000E-06 | 1.300E-05 | 1.700E-05 |
| 564 | 29 | Q29D | A | 1.4e-05 | 2.1e-05 | 1e-05 | 1.500E-05 | 5.568E-06 | 1.000E-05 | 2.100E-05 |
| 565 | 29 | Q29C | A | 1e-05 | 1.7e-05 | 9.6e-06 | 1.220E-05 | 4.162E-06 | 9.600E-06 | 1.700E-05 |
| 566 | 29 | Q29E | A | 1.5e-05 | 1.5e-05 | 1.2e-05 | 1.400E-05 | 1.732E-06 | 1.200E-05 | 1.500E-05 |
| 567 | 29 | Q29Q | A | 9.8e-06 | 1.6e-05 | 9.5e-06 | 1.177E-05 | 3.669E-06 | 9.500E-06 | 1.600E-05 |
| 568 | 29 | Q29G | A | 1.4e-05 | 1.2e-05 | 1.5e-05 | 1.367E-05 | 1.528E-06 | 1.200E-05 | 1.500E-05 |
| 569 | 29 | Q29H | A | 1.5e-05 | 1.6e-05 | 8.4e-06 | 1.313E-05 | 4.130E-06 | 8.400E-06 | 1.600E-05 |
| 570 | 29 | Q29I | A | 1.3e-05 | 9.2e-06 | 1.3e-05 | 1.173E-05 | 2.194E-06 | 9.200E-06 | 1.300E-05 |
| 571 | 29 | Q29L | A | 9.4e-06 | 1.5e-05 | 1.1e-05 | 1.180E-05 | 2.884E-06 | 9.400E-06 | 1.500E-05 |
| 572 | 29 | Q29K | A | 1.2e-05 | 1.4e-05 | 1.5e-05 | 1.367E-05 | 1.528E-06 | 1.200E-05 | 1.500E-05 |
| 573 | 29 | Q29M | A | 1.6e-05 | 1.1e-05 | 1.1e-05 | 1.267E-05 | 2.887E-06 | 1.100E-05 | 1.600E-05 |
| 574 | 29 | Q29F | A | 9.4e-06 | 8.8e-06 | 1.1e-05 | 9.733E-06 | 1.137E-06 | 8.800E-06 | 1.100E-05 |
| 575 | 29 | Q29P | A | 1.6e-05 | 1.6e-05 | 1.5e-05 | 1.567E-05 | 5.774E-07 | 1.500E-05 | 1.600E-05 |
| 576 | 29 | Q29S | A | 1.2e-05 | 1.1e-05 | 1e-05 | 1.100E-05 | 1.000E-06 | 1.000E-05 | 1.200E-05 |
| 577 | 29 | Q29T | A | 1.3e-05 | 9.7e-06 | 8.9e-06 | 1.053E-05 | 2.173E-06 | 8.900E-06 | 1.300E-05 |
| 578 | 29 | Q29W | A | 1.6e-05 | 8.7e-06 | 1.4e-05 | 1.290E-05 | 3.772E-06 | 8.700E-06 | 1.600E-05 |
| 579 | 29 | Q29Y | A | 1.3e-05 | 7.1e-06 | 9.4e-06 | 9.833E-06 | 2.974E-06 | 7.100E-06 | 1.300E-05 |
| 580 | 29 | Q29V | A | 1.5e-05 | 1.2e-05 | 1.3e-05 | 1.333E-05 | 1.528E-06 | 1.200E-05 | 1.500E-05 |
| 581 | 30 | G30A | A | 1.5e-05 | 1e-05 | 1e-05 | 1.167E-05 | 2.887E-06 | 1.000E-05 | 1.500E-05 |
| 582 | 30 | G30R | A | 9.7e-06 | 1.4e-05 | 1.1e-05 | 1.157E-05 | 2.205E-06 | 9.700E-06 | 1.400E-05 |
| 583 | 30 | G30N | A | 9.8e-06 | 9.2e-06 | 1.3e-05 | 1.067E-05 | 2.043E-06 | 9.200E-06 | 1.300E-05 |
| 584 | 30 | G30D | A | 1.3e-05 | 8.6e-06 | 8.1e-06 | 9.900E-06 | 2.696E-06 | 8.100E-06 | 1.300E-05 |
| 585 | 30 | G30C | A | 1.2e-05 | 1e-05 | 9.1e-06 | 1.037E-05 | 1.484E-06 | 9.100E-06 | 1.200E-05 |
| 586 | 30 | G30E | A | 1.2e-05 | 1.6e-05 | 1.2e-05 | 1.333E-05 | 2.309E-06 | 1.200E-05 | 1.600E-05 |
| 587 | 30 | G30Q | A | 8.9e-06 | 1.2e-05 | 1.6e-05 | 1.230E-05 | 3.559E-06 | 8.900E-06 | 1.600E-05 |
| 588 | 30 | G30G | A | 7.3e-06 | 8.4e-06 | 1.6e-05 | 1.057E-05 | 4.737E-06 | 7.300E-06 | 1.600E-05 |
| 589 | 30 | G30H | A | 1.3e-05 | 1.2e-05 | 1.5e-05 | 1.333E-05 | 1.528E-06 | 1.200E-05 | 1.500E-05 |
| 590 | 30 | G30I | A | 8.9e-06 | 1.4e-05 | 1.6e-05 | 1.297E-05 | 3.661E-06 | 8.900E-06 | 1.600E-05 |
| 591 | 30 | G30L | A | 1.1e-05 | 1.5e-05 | 1.5e-05 | 1.367E-05 | 2.309E-06 | 1.100E-05 | 1.500E-05 |
| 592 | 30 | G30K | A | 1e-05 | 1e-05 | 1.2e-05 | 1.067E-05 | 1.155E-06 | 1.000E-05 | 1.200E-05 |
| 593 | 30 | G30M | A | 1.4e-05 | 1.4e-05 | 1e-05 | 1.267E-05 | 2.309E-06 | 1.000E-05 | 1.400E-05 |
| 594 | 30 | G30F | A | 2.7e-05 | 1.6e-05 | 1.6e-05 | 1.967E-05 | 6.351E-06 | 1.600E-05 | 2.700E-05 |

| | | | | | | | | | | |
|---|---|---|---|---|---|---|---|---|---|---|
| 595 | 30 | G30P | A | 1.6e-05 | 1.2e-05 | 1.8e-05 | 1.533E-05 | 3.055E-06 | 1.200E-05 | 1.800E-05 |
| 596 | 30 | G30S | A | 1.5e-05 | 8.8e-06 | 1.7e-05 | 1.360E-05 | 4.276E-06 | 8.800E-06 | 1.700E-05 |
| 597 | 30 | G30T | A | 8.8e-06 | 1.5e-05 | 1.2e-05 | 1.193E-05 | 3.101E-06 | 8.800E-06 | 1.500E-05 |
| 598 | 30 | G30W | A | 1.1e-05 | 1.6e-05 | 1e-05 | 1.233E-05 | 3.215E-06 | 1.000E-05 | 1.600E-05 |
| 599 | 30 | G30Y | A | 1.5e-05 | 1.6e-05 | 9e-06 | 1.333E-05 | 3.786E-06 | 9.000E-06 | 1.600E-05 |
| 600 | 30 | G30V | A | 1.4e-05 | 1.6e-05 | 9.9e-06 | 1.330E-05 | 3.110E-06 | 9.900E-06 | 1.600E-05 |
| 601 | 31 | C31A | A | 1.5e-05 | 1.5e-05 | 1.5e-05 | 1.500E-05 | 0.000E+00 | 1.500E-05 | 1.500E-05 |
| 602 | 31 | C31R | A | 1e-05 | 1.5e-05 | 1.5e-05 | 1.333E-05 | 2.887E-06 | 1.000E-05 | 1.500E-05 |
| 603 | 31 | C31N | A | 1.7e-05 | 1.3e-05 | 1e-05 | 1.333E-05 | 3.512E-06 | 1.000E-05 | 1.700E-05 |
| 604 | 31 | C31D | A | 1.1e-05 | 1.2e-05 | 1.2e-05 | 1.167E-05 | 5.774E-07 | 1.100E-05 | 1.200E-05 |
| 605 | 31 | C31C | A | 1.2e-05 | 1.8e-05 | 9.7e-06 | 1.323E-05 | 4.285E-06 | 9.700E-06 | 1.800E-05 |
| 606 | 31 | C31E | A | 1.5e-05 | 1e-05 | 1.1e-05 | 1.200E-05 | 2.646E-06 | 1.000E-05 | 1.500E-05 |
| 607 | 31 | C31Q | A | 1.1e-05 | 6.3e-06 | 1.3e-05 | 1.010E-05 | 3.439E-06 | 6.300E-06 | 1.300E-05 |
| 608 | 31 | C31G | A | 9.2e-06 | 1.4e-05 | 1.6e-05 | 1.307E-05 | 3.495E-06 | 9.200E-06 | 1.600E-05 |
| 609 | 31 | C31H | A | 1.5e-05 | 9.7e-06 | 1.4e-05 | 1.290E-05 | 2.816E-06 | 9.700E-06 | 1.500E-05 |
| 610 | 31 | C31I | A | 1.1e-05 | 1e-05 | 1.6e-05 | 1.233E-05 | 3.215E-06 | 1.000E-05 | 1.600E-05 |
| 611 | 31 | C31L | A | 9.6e-06 | 1.5e-05 | 1e-05 | 1.153E-05 | 3.009E-06 | 9.600E-06 | 1.500E-05 |
| 612 | 31 | C31K | A | 1.4e-05 | 1.8e-05 | 9.9e-06 | 1.397E-05 | 4.050E-06 | 9.900E-06 | 1.800E-05 |
| 613 | 31 | C31M | A | 1.5e-05 | 1.1e-05 | 1.2e-05 | 1.267E-05 | 2.082E-06 | 1.100E-05 | 1.500E-05 |
| 614 | 31 | C31F | A | 1.5e-05 | 8.9e-06 | 1.5e-05 | 1.297E-05 | 3.522E-06 | 8.900E-06 | 1.500E-05 |
| 615 | 31 | C31P | A | 1.4e-05 | 1.3e-05 | 1e-05 | 1.233E-05 | 2.082E-06 | 1.000E-05 | 1.400E-05 |
| 616 | 31 | C31S | A | 1.2e-05 | 1.1e-05 | 1.5e-05 | 1.267E-05 | 2.082E-06 | 1.100E-05 | 1.500E-05 |
| 617 | 31 | C31T | A | 9.6e-06 | 1.1e-05 | 1.7e-05 | 1.253E-05 | 3.931E-06 | 9.600E-06 | 1.700E-05 |
| 618 | 31 | C31W | A | 8.1e-06 | 1.6e-05 | 9.4e-06 | 1.117E-05 | 4.236E-06 | 8.100E-06 | 1.600E-05 |
| 619 | 31 | C31Y | A | 1.6e-05 | 1.5e-05 | 1.4e-05 | 1.500E-05 | 1.000E-06 | 1.400E-05 | 1.600E-05 |
| 620 | 31 | C31V | A | 1.6e-05 | 1.4e-05 | 1.4e-05 | 1.467E-05 | 1.155E-06 | 1.400E-05 | 1.600E-05 |
| 621 | 32 | Q32A | A | 1.5e-05 | 1.6e-05 | 1.6e-05 | 1.567E-05 | 5.774E-07 | 1.500E-05 | 1.600E-05 |
| 622 | 32 | Q32R | A | 8.4e-06 | 8.9e-06 | 1.5e-05 | 1.077E-05 | 3.675E-06 | 8.400E-06 | 1.500E-05 |
| 623 | 32 | Q32N | A | 1.5e-05 | 9.9e-06 | 1.5e-05 | 1.330E-05 | 2.944E-06 | 9.900E-06 | 1.500E-05 |
| 624 | 32 | Q32D | A | 1.7e-05 | 1.1e-05 | 1.7e-05 | 1.500E-05 | 3.464E-06 | 1.100E-05 | 1.700E-05 |
| 625 | 32 | Q32C | A | 8.2e-06 | 9.7e-06 | 1.5e-05 | 1.097E-05 | 3.573E-06 | 8.200E-06 | 1.500E-05 |
| 626 | 32 | Q32E | A | 8.8e-06 | 1.6e-05 | 1.4e-05 | 1.293E-05 | 3.717E-06 | 8.800E-06 | 1.600E-05 |
| 627 | 32 | Q32Q | A | 1e-05 | 1.3e-05 | 1.5e-05 | 1.267E-05 | 2.517E-06 | 1.000E-05 | 1.500E-05 |
| 628 | 32 | Q32G | A | 1.5e-05 | 1.7e-05 | 1.1e-05 | 1.433E-05 | 3.055E-06 | 1.100E-05 | 1.700E-05 |
| 629 | 32 | Q32H | A | 1.1e-05 | 1.1e-05 | 1.7e-05 | 1.300E-05 | 3.464E-06 | 1.100E-05 | 1.700E-05 |
| 630 | 32 | Q32I | A | 8.2e-06 | 1.5e-05 | 1.6e-05 | 1.307E-05 | 4.244E-06 | 8.200E-06 | 1.600E-05 |
| 631 | 32 | Q32L | A | 9.7e-06 | 1.5e-05 | 1.6e-05 | 1.357E-05 | 3.386E-06 | 9.700E-06 | 1.600E-05 |
| 632 | 32 | Q32K | A | 1.5e-05 | 9.5e-06 | 1.5e-05 | 1.317E-05 | 3.175E-06 | 9.500E-06 | 1.500E-05 |
| 633 | 32 | Q32M | A | 1.2e-05 | 1.1e-05 | 1e-05 | 1.100E-05 | 1.000E-06 | 1.000E-05 | 1.200E-05 |
| 634 | 32 | Q32F | A | 1.5e-05 | 1.5e-05 | 1.4e-05 | 1.467E-05 | 5.774E-07 | 1.400E-05 | 1.500E-05 |
| 635 | 32 | Q32P | A | 1.1e-05 | 9.8e-06 | 1.1e-05 | 1.060E-05 | 6.928E-07 | 9.800E-06 | 1.100E-05 |
| 636 | 32 | Q32S | A | 1.3e-05 | 1.5e-05 | 9.1e-06 | 1.237E-05 | 3.001E-06 | 9.100E-06 | 1.500E-05 |
| 637 | 32 | Q32T | A | 1.5e-05 | 7e-06 | 1e-05 | 1.067E-05 | 4.041E-06 | 7.000E-06 | 1.500E-05 |
| 638 | 32 | Q32W | A | 1e-05 | 1.7e-05 | 1.6e-05 | 1.433E-05 | 3.786E-06 | 1.000E-05 | 1.700E-05 |
| 639 | 32 | Q32Y | A | 1.4e-05 | 8.6e-06 | 1e-05 | 1.087E-05 | 2.802E-06 | 8.600E-06 | 1.400E-05 |
| 640 | 32 | Q32V | A | 1.4e-05 | 1.1e-05 | 1.6e-05 | 1.367E-05 | 2.517E-06 | 1.100E-05 | 1.600E-05 |
| 641 | 33 | V33A | A | 1.4e-05 | 1.1e-05 | 1e-05 | 1.167E-05 | 2.082E-06 | 1.000E-05 | 1.400E-05 |
| 642 | 33 | V33R | A | 9.4e-06 | 1.7e-05 | 1.3e-05 | 1.313E-05 | 3.802E-06 | 9.400E-06 | 1.700E-05 |
| 643 | 33 | V33N | A | 1.4e-05 | 1.2e-05 | 8.9e-06 | 1.163E-05 | 2.570E-06 | 8.900E-06 | 1.400E-05 |
| 644 | 33 | V33D | A | 1.4e-05 | 8.4e-06 | 6.3e-06 | 9.567E-06 | 3.980E-06 | 6.300E-06 | 1.400E-05 |
| 645 | 33 | V33C | A | 1.4e-05 | 9.6e-06 | 1.6e-05 | 1.320E-05 | 3.274E-06 | 9.600E-06 | 1.600E-05 |
| 646 | 33 | V33E | A | 8.6e-06 | 1.5e-05 | 1.5e-05 | 1.287E-05 | 3.695E-06 | 8.600E-06 | 1.500E-05 |
| 647 | 33 | V33Q | A | 9e-06 | 1.4e-05 | 1.1e-05 | 1.133E-05 | 2.517E-06 | 9.000E-06 | 1.400E-05 |
| 648 | 33 | V33G | A | 1.4e-05 | 1e-05 | 1e-05 | 1.133E-05 | 2.309E-06 | 1.000E-05 | 1.400E-05 |
| 649 | 33 | V33H | A | 1.4e-05 | 1e-05 | 1e-05 | 1.133E-05 | 2.309E-06 | 1.000E-05 | 1.400E-05 |
| 650 | 33 | V33I | A | 1.6e-05 | 1.1e-05 | 1.1e-05 | 1.267E-05 | 2.887E-06 | 1.100E-05 | 1.600E-05 |
| 651 | 33 | V33L | A | 9.9e-06 | 1e-05 | 1.5e-05 | 1.163E-05 | 2.916E-06 | 9.900E-06 | 1.500E-05 |
| 652 | 33 | V33K | A | 9.7e-06 | 1.6e-05 | 1.4e-05 | 1.323E-05 | 3.219E-06 | 9.700E-06 | 1.600E-05 |
| 653 | 33 | V33M | A | 1.1e-05 | 7.6e-06 | 9.2e-06 | 9.267E-06 | 1.701E-06 | 7.600E-06 | 1.100E-05 |
| 654 | 33 | V33F | A | 1.4e-05 | 1.5e-05 | 1.5e-05 | 1.467E-05 | 5.774E-07 | 1.400E-05 | 1.500E-05 |
| 655 | 33 | V33P | A | 1e-05 | 1.4e-05 | 8.2e-06 | 1.073E-05 | 2.969E-06 | 8.200E-06 | 1.400E-05 |
| 656 | 33 | V33S | A | 1.3e-05 | 1.6e-05 | 1.4e-05 | 1.433E-05 | 1.528E-06 | 1.300E-05 | 1.600E-05 |
| 657 | 33 | V33T | A | 1e-05 | 1.4e-05 | 9.7e-06 | 1.123E-05 | 2.401E-06 | 9.700E-06 | 1.400E-05 |
| 658 | 33 | V33W | A | 8.5e-06 | 9.9e-06 | 1.6e-05 | 1.147E-05 | 3.988E-06 | 8.500E-06 | 1.600E-05 |
| 659 | 33 | V33Y | A | 1.4e-05 | 1e-05 | 1.7e-05 | 1.367E-05 | 3.512E-06 | 1.000E-05 | 1.700E-05 |
| 660 | 33 | V33V | A | 1.1e-05 | 9.9e-06 | 1.3e-05 | 1.130E-05 | 1.572E-06 | 9.900E-06 | 1.300E-05 |
| 661 | 34 | V34A | A | 1e-05 | 1.6e-05 | 1.5e-05 | 1.367E-05 | 3.215E-06 | 1.000E-05 | 1.600E-05 |
| 662 | 34 | V34R | A | 1.5e-05 | 1.6e-05 | 8.7e-06 | 1.323E-05 | 3.958E-06 | 8.700E-06 | 1.600E-05 |
| 663 | 34 | V34N | A | 1e-05 | 1.6e-05 | 1.2e-05 | 1.267E-05 | 3.055E-06 | 1.000E-05 | 1.600E-05 |
| 664 | 34 | V34D | A | 1.3e-05 | 9.9e-06 | 1.4e-05 | 1.230E-05 | 2.138E-06 | 9.900E-06 | 1.400E-05 |
| 665 | 34 | V34C | A | 1.1e-05 | 8.1e-06 | 8.9e-06 | 9.333E-06 | 1.498E-06 | 8.100E-06 | 1.100E-05 |
| 666 | 34 | V34E | A | 1.1e-05 | 1.1e-05 | 1e-05 | 1.067E-05 | 5.774E-07 | 1.000E-05 | 1.100E-05 |
| 667 | 34 | V34Q | A | 9.4e-06 | 1.6e-05 | 1.7e-05 | 1.413E-05 | 4.130E-06 | 9.400E-06 | 1.700E-05 |
| 668 | 34 | V34G | A | 1.1e-05 | 1.5e-05 | 2.6e-05 | 1.733E-05 | 7.767E-06 | 1.100E-05 | 2.600E-05 |
| 669 | 34 | V34H | A | 1.6e-05 | 1.6e-05 | 1.8e-05 | 1.667E-05 | 1.155E-06 | 1.600E-05 | 1.800E-05 |
| 670 | 34 | V34I | A | 1.2e-05 | 1.2e-05 | 1.7e-05 | 1.367E-05 | 2.887E-06 | 1.200E-05 | 1.700E-05 |
| 671 | 34 | V34L | A | 1e-05 | 1.2e-05 | 1.6e-05 | 1.267E-05 | 3.055E-06 | 1.000E-05 | 1.600E-05 |
| 672 | 34 | V34K | A | 1.3e-05 | 1.1e-05 | 1.2e-05 | 1.200E-05 | 1.000E-06 | 1.100E-05 | 1.300E-05 |
| 673 | 34 | V34M | A | 1e-05 | 1.7e-05 | 1e-05 | 1.233E-05 | 4.041E-06 | 1.000E-05 | 1.700E-05 |
| 674 | 34 | V34F | A | 1.4e-05 | 8.2e-06 | 1.2e-05 | 1.140E-05 | 2.946E-06 | 8.200E-06 | 1.400E-05 |
| 675 | 34 | V34P | A | 1.1e-05 | 8.7e-06 | 1.5e-05 | 1.157E-05 | 3.188E-06 | 8.700E-06 | 1.500E-05 |
| 676 | 34 | V34S | A | 9.2e-06 | 1.1e-05 | 1.4e-05 | 1.140E-05 | 2.425E-06 | 9.200E-06 | 1.400E-05 |
| 677 | 34 | V34T | A | 9e-06 | 9.9e-06 | 1.1e-05 | 9.967E-06 | 1.002E-06 | 9.000E-06 | 1.100E-05 |
| 678 | 34 | V34W | A | 1.4e-05 | 8.9e-06 | 1.1e-05 | 1.130E-05 | 2.563E-06 | 8.900E-06 | 1.400E-05 |
| 679 | 34 | V34Y | A | 1.3e-05 | 1.2e-05 | 1e-05 | 1.167E-05 | 1.528E-06 | 1.000E-05 | 1.300E-05 |

| | | | | | | | | | | |
|---|---|---|---|---|---|---|---|---|---|---|
| 680 | 34 | V34V | A | 9.8e-06 | 1.5e-05 | 1.7e-05 | 1.393E-05 | 3.717E-06 | 9.800E-06 | 1.700E-05 |
| 681 | 35 | Q35A | A | 1.4e-05 | 1.3e-05 | 1.5e-05 | 1.400E-05 | 1.000E-06 | 1.300E-05 | 1.500E-05 |
| 682 | 35 | Q35R | A | 1.5e-05 | 1.4e-05 | 1.1e-05 | 1.333E-05 | 2.082E-06 | 1.100E-05 | 1.500E-05 |
| 683 | 35 | Q35N | A | 1.1e-05 | 1.6e-05 | 6.3e-06 | 1.110E-05 | 4.851E-06 | 6.300E-06 | 1.600E-05 |
| 684 | 35 | Q35D | A | 1.5e-05 | 1.2e-05 | 1.2e-05 | 1.300E-05 | 1.732E-06 | 1.200E-05 | 1.500E-05 |
| 685 | 35 | Q35C | A | 1.5e-05 | 1.5e-05 | 1.6e-05 | 1.533E-05 | 5.774E-07 | 1.500E-05 | 1.600E-05 |
| 686 | 35 | Q35E | A | 8.1e-06 | 1.3e-05 | 1.5e-05 | 1.203E-05 | 3.550E-06 | 8.100E-06 | 1.500E-05 |
| 687 | 35 | Q35Q | A | 1.2e-05 | 1.7e-05 | 9.7e-06 | 1.290E-05 | 3.732E-06 | 9.700E-06 | 1.700E-05 |
| 688 | 35 | Q35G | A | 1.2e-05 | 9.3e-06 | 1.5e-05 | 1.210E-05 | 2.851E-06 | 9.300E-06 | 1.500E-05 |
| 689 | 35 | Q35H | A | 1.6e-05 | 1.4e-05 | 1.6e-05 | 1.533E-05 | 1.155E-06 | 1.400E-05 | 1.600E-05 |
| 690 | 35 | Q35I | A | 6.5e-06 | 1.6e-05 | 8.7e-06 | 1.040E-05 | 4.973E-06 | 6.500E-06 | 1.600E-05 |
| 691 | 35 | Q35L | A | 1.6e-05 | 1.5e-05 | 9.7e-06 | 1.357E-05 | 3.386E-06 | 9.700E-06 | 1.600E-05 |
| 692 | 35 | Q35K | A | 1.4e-05 | 9.2e-06 | 8.9e-06 | 1.070E-05 | 2.862E-06 | 8.900E-06 | 1.400E-05 |
| 693 | 35 | Q35M | A | 1.4e-05 | 1.5e-05 | 8.7e-06 | 1.257E-05 | 3.386E-06 | 8.700E-06 | 1.500E-05 |
| 694 | 35 | Q35F | A | 1.1e-05 | 1.2e-05 | 9.7e-06 | 1.090E-05 | 1.153E-06 | 9.700E-06 | 1.200E-05 |
| 695 | 35 | Q35P | A | 1.2e-05 | 1.5e-05 | 1.4e-05 | 1.367E-05 | 1.528E-06 | 1.200E-05 | 1.500E-05 |
| 696 | 35 | Q35S | A | 1.6e-05 | 1.5e-05 | 1.5e-05 | 1.533E-05 | 5.774E-07 | 1.500E-05 | 1.600E-05 |
| 697 | 35 | Q35T | A | 1.1e-05 | 1e-05 | 1e-05 | 1.033E-05 | 5.774E-07 | 1.000E-05 | 1.100E-05 |
| 698 | 35 | Q35W | A | 1.6e-05 | 1.6e-05 | 8.7e-06 | 1.357E-05 | 4.215E-06 | 8.700E-06 | 1.600E-05 |
| 699 | 35 | Q35Y | A | 1.7e-05 | 1.8e-05 | 1.6e-05 | 1.700E-05 | 1.000E-06 | 1.600E-05 | 1.800E-05 |
| 700 | 35 | Q35V | A | 1.1e-05 | 1.4e-05 | 9.8e-06 | 1.160E-05 | 2.163E-06 | 9.800E-06 | 1.400E-05 |
| 701 | 36 | G36A | A | 1e-05 | 1.5e-05 | 9.6e-06 | 1.153E-05 | 3.009E-06 | 9.600E-06 | 1.500E-05 |
| 702 | 36 | G36R | A | 1.5e-05 | 1.1e-05 | 1.4e-05 | 1.333E-05 | 2.082E-06 | 1.100E-05 | 1.500E-05 |
| 703 | 36 | G36N | A | 1.5e-05 | 1.1e-05 | 1e-05 | 1.200E-05 | 2.646E-06 | 1.000E-05 | 1.500E-05 |
| 704 | 36 | G36D | A | 1.1e-05 | 1e-05 | 1.2e-05 | 1.100E-05 | 1.000E-06 | 1.000E-05 | 1.200E-05 |
| 705 | 36 | G36C | A | 1.4e-05 | 1.8e-05 | 1.1e-05 | 1.433E-05 | 3.512E-06 | 1.100E-05 | 1.800E-05 |
| 706 | 36 | G36E | A | 1.1e-05 | 1.7e-05 | 1.4e-05 | 1.400E-05 | 3.000E-06 | 1.100E-05 | 1.700E-05 |
| 707 | 36 | G36Q | A | 1.4e-05 | 9.1e-06 | 1.4e-05 | 1.237E-05 | 2.829E-06 | 9.100E-06 | 1.400E-05 |
| 708 | 36 | G36G | A | 1.4e-05 | 1.5e-05 | 1.5e-05 | 1.467E-05 | 5.774E-07 | 1.400E-05 | 1.500E-05 |
| 709 | 36 | G36H | A | 1.6e-05 | 9.8e-06 | 1.4e-05 | 1.327E-05 | 3.164E-06 | 9.800E-06 | 1.600E-05 |
| 710 | 36 | G36I | A | 1e-05 | 1e-05 | 1.2e-05 | 1.067E-05 | 1.155E-06 | 1.000E-05 | 1.200E-05 |
| 711 | 36 | G36L | A | 1.7e-05 | 1.1e-05 | 1.3e-05 | 1.367E-05 | 3.055E-06 | 1.100E-05 | 1.700E-05 |
| 712 | 36 | G36K | A | 1.8e-05 | 1.1e-05 | 7.4e-06 | 1.213E-05 | 5.390E-06 | 7.400E-06 | 1.800E-05 |
| 713 | 36 | G36M | A | 1.5e-05 | 1.4e-05 | 1.6e-05 | 1.500E-05 | 1.000E-06 | 1.400E-05 | 1.600E-05 |
| 714 | 36 | G36F | A | 9e-06 | 1.2e-05 | 8.9e-06 | 9.967E-06 | 1.762E-06 | 8.900E-06 | 1.200E-05 |
| 715 | 36 | G36P | A | 1.1e-05 | 1.5e-05 | 1.5e-05 | 1.367E-05 | 2.309E-06 | 1.100E-05 | 1.500E-05 |
| 716 | 36 | G36S | A | 8.3e-06 | 1.3e-05 | 1.5e-05 | 1.210E-05 | 3.439E-06 | 8.300E-06 | 1.500E-05 |
| 717 | 36 | G36T | A | 9.6e-06 | 1.5e-05 | 1.3e-05 | 1.253E-05 | 2.730E-06 | 9.600E-06 | 1.500E-05 |
| 718 | 36 | G36W | A | 1.3e-05 | 1.5e-05 | 1.1e-05 | 1.300E-05 | 2.000E-06 | 1.100E-05 | 1.500E-05 |
| 719 | 36 | G36Y | A | 7.5e-06 | 1.1e-05 | 1e-05 | 9.500E-06 | 1.803E-06 | 7.500E-06 | 1.100E-05 |
| 720 | 36 | G36V | A | 1.5e-05 | 9.4e-06 | 1e-05 | 1.147E-05 | 3.075E-06 | 9.400E-06 | 1.500E-05 |
| 721 | 37 | N37A | A | 1.1e-05 | 1.5e-05 | 1.5e-05 | 1.367E-05 | 2.309E-06 | 1.100E-05 | 1.500E-05 |
| 722 | 37 | N37R | A | 7.3e-06 | 1.6e-05 | 1.5e-05 | 1.277E-05 | 4.761E-06 | 7.300E-06 | 1.600E-05 |
| 723 | 37 | N37N | A | 1.5e-05 | 9.7e-06 | 1e-05 | 1.157E-05 | 2.977E-06 | 9.700E-06 | 1.500E-05 |
| 724 | 37 | N37D | A | 1.7e-05 | 1e-05 | 1.1e-05 | 1.267E-05 | 3.786E-06 | 1.000E-05 | 1.700E-05 |
| 725 | 37 | N37C | A | 1e-05 | 9.6e-06 | 1.8e-05 | 1.253E-05 | 4.738E-06 | 9.600E-06 | 1.800E-05 |
| 726 | 37 | N37E | A | 1.5e-05 | 1.1e-05 | 1.5e-05 | 1.367E-05 | 2.309E-06 | 1.100E-05 | 1.500E-05 |
| 727 | 37 | N37Q | A | 9.4e-06 | 1.5e-05 | 1e-05 | 1.147E-05 | 3.075E-06 | 9.400E-06 | 1.500E-05 |
| 728 | 37 | N37G | A | 1.4e-05 | 9.9e-06 | 1.3e-05 | 1.230E-05 | 2.138E-06 | 9.900E-06 | 1.400E-05 |
| 729 | 37 | N37H | A | 1.2e-05 | 8.8e-06 | 1.2e-05 | 1.093E-05 | 1.848E-06 | 8.800E-06 | 1.200E-05 |
| 730 | 37 | N37I | A | 6.9e-06 | 1.3e-05 | 7.4e-06 | 9.100E-06 | 3.387E-06 | 6.900E-06 | 1.300E-05 |
| 731 | 37 | N37L | A | 1.5e-05 | 9.9e-06 | 1.6e-05 | 1.363E-05 | 3.272E-06 | 9.900E-06 | 1.600E-05 |
| 732 | 37 | N37K | A | 1e-05 | 9.8e-06 | 1.5e-05 | 1.160E-05 | 2.946E-06 | 9.800E-06 | 1.500E-05 |
| 733 | 37 | N37M | A | 1.3e-05 | 1.5e-05 | 1.3e-05 | 1.367E-05 | 1.155E-06 | 1.300E-05 | 1.500E-05 |
| 734 | 37 | N37F | A | 8.7e-06 | 8.7e-06 | 1e-05 | 9.133E-06 | 7.506E-07 | 8.700E-06 | 1.000E-05 |
| 735 | 37 | N37P | A | 1.1e-05 | 9.8e-06 | 8.6e-06 | 9.800E-06 | 1.200E-06 | 8.600E-06 | 1.100E-05 |
| 736 | 37 | N37S | A | 5.7e-06 | 9.8e-06 | 1.6e-05 | 1.050E-05 | 5.186E-06 | 5.700E-06 | 1.600E-05 |
| 737 | 37 | N37T | A | 1.3e-05 | 1e-05 | 1.3e-05 | 1.200E-05 | 1.732E-06 | 1.000E-05 | 1.300E-05 |
| 738 | 37 | N37W | A | 1e-05 | 1.4e-05 | 9.8e-06 | 1.127E-05 | 2.369E-06 | 9.800E-06 | 1.400E-05 |
| 739 | 37 | N37Y | A | 1.5e-05 | 1.2e-05 | 1.5e-05 | 1.400E-05 | 1.732E-06 | 1.200E-05 | 1.500E-05 |
| 740 | 37 | N37V | A | 9.2e-06 | 1.6e-05 | 1.6e-05 | 1.373E-05 | 3.926E-06 | 9.200E-06 | 1.600E-05 |
| 741 | 38 | L38A | A | 9.4e-06 | 1.5e-05 | 1.6e-05 | 1.347E-05 | 3.557E-06 | 9.400E-06 | 1.600E-05 |
| 742 | 38 | L38R | A | 1.2e-05 | 1.4e-05 | 1e-05 | 1.200E-05 | 2.000E-06 | 1.000E-05 | 1.400E-05 |
| 743 | 38 | L38N | A | 1.3e-05 | 1.1e-05 | 1.4e-05 | 1.267E-05 | 1.528E-06 | 1.100E-05 | 1.400E-05 |
| 744 | 38 | L38D | A | 1e-05 | 1.5e-05 | 1.8e-05 | 1.433E-05 | 4.041E-06 | 1.000E-05 | 1.800E-05 |
| 745 | 38 | L38C | A | 1e-05 | 1.2e-05 | 1.5e-05 | 1.233E-05 | 2.517E-06 | 1.000E-05 | 1.500E-05 |
| 746 | 38 | L38E | A | 8.6e-06 | 1.4e-05 | 1.6e-05 | 1.287E-05 | 3.828E-06 | 8.600E-06 | 1.600E-05 |
| 747 | 38 | L38Q | A | 1e-05 | 1.1e-05 | 9.9e-06 | 1.030E-05 | 6.083E-07 | 9.900E-06 | 1.100E-05 |
| 748 | 38 | L38G | A | 8e-06 | 1.1e-05 | 1.5e-05 | 1.133E-05 | 3.512E-06 | 8.000E-06 | 1.500E-05 |
| 749 | 38 | L38H | A | 1.1e-05 | 1.5e-05 | 9.5e-06 | 1.183E-05 | 2.843E-06 | 9.500E-06 | 1.500E-05 |
| 750 | 38 | L38I | A | 1.4e-05 | 9.7e-06 | 1.7e-05 | 1.357E-05 | 3.669E-06 | 9.700E-06 | 1.700E-05 |
| 751 | 38 | L38L | A | 1.6e-05 | 1.1e-05 | 1.1e-05 | 1.267E-05 | 2.887E-06 | 1.100E-05 | 1.600E-05 |
| 752 | 38 | L38K | A | 1.3e-05 | 1e-05 | 1.5e-05 | 1.267E-05 | 2.517E-06 | 1.000E-05 | 1.500E-05 |
| 753 | 38 | L38M | A | 1.2e-05 | 1.5e-05 | 1.7e-05 | 1.467E-05 | 2.517E-06 | 1.200E-05 | 1.700E-05 |
| 754 | 38 | L38F | A | 1.6e-05 | 9.7e-06 | 1.5e-05 | 1.357E-05 | 3.386E-06 | 9.700E-06 | 1.600E-05 |
| 755 | 38 | L38P | A | 1.2e-05 | 1.4e-05 | 9.9e-06 | 1.197E-05 | 2.050E-06 | 9.900E-06 | 1.400E-05 |
| 756 | 38 | L38S | A | 1.3e-05 | 1.2e-05 | 1.4e-05 | 1.300E-05 | 1.000E-06 | 1.200E-05 | 1.400E-05 |
| 757 | 38 | L38T | A | 1.5e-05 | 1.4e-05 | 1.5e-05 | 1.467E-05 | 5.774E-07 | 1.400E-05 | 1.500E-05 |
| 758 | 38 | L38W | A | 1.8e-05 | 9.4e-06 | 1.2e-05 | 1.313E-05 | 4.411E-06 | 9.400E-06 | 1.800E-05 |
| 759 | 38 | L38Y | A | 1.3e-05 | 1.7e-05 | 1.3e-05 | 1.433E-05 | 2.309E-06 | 1.300E-05 | 1.700E-05 |
| 760 | 38 | L38V | A | 1.4e-05 | 1.7e-05 | 6.1e-06 | 1.237E-05 | 5.631E-06 | 6.100E-06 | 1.700E-05 |
| 761 | 39 | E39A | A | 1e-05 | 9.9e-06 | 1.6e-05 | 1.197E-05 | 3.493E-06 | 9.900E-06 | 1.600E-05 |
| 762 | 39 | E39R | A | 1e-05 | 9.6e-06 | 1.1e-05 | 1.020E-05 | 7.211E-07 | 9.600E-06 | 1.100E-05 |
| 763 | 39 | E39N | A | 9.6e-06 | 1.5e-05 | 1.2e-05 | 1.220E-05 | 2.706E-06 | 9.600E-06 | 1.500E-05 |
| 764 | 39 | E39D | A | 1.7e-05 | 1.4e-05 | 1.6e-05 | 1.567E-05 | 1.528E-06 | 1.400E-05 | 1.700E-05 |

| | | | | | | | | | | |
|---|---|---|---|---|---|---|---|---|---|---|
| 765 | 39 | E39C | A | 8.9e-06 | 1.4e-05 | 1.7e-05 | 1.330E-05 | 4.095E-06 | 8.900E-06 | 1.700E-05 |
| 766 | 39 | E39E | A | 8.2e-06 | 1.1e-05 | 1.1e-05 | 1.007E-05 | 1.617E-06 | 8.200E-06 | 1.100E-05 |
| 767 | 39 | E39Q | A | 9e-06 | 1.4e-05 | 1.5e-05 | 1.267E-05 | 3.215E-06 | 9.000E-06 | 1.500E-05 |
| 768 | 39 | E39G | A | 1.6e-05 | 1.1e-05 | 1e-05 | 1.233E-05 | 3.215E-06 | 1.000E-05 | 1.600E-05 |
| 769 | 39 | E39H | A | 2.4e-05 | 1.4e-05 | 1.2e-05 | 1.667E-05 | 6.429E-06 | 1.200E-05 | 2.400E-05 |
| 770 | 39 | E39I | A | 1.5e-05 | 1.3e-05 | 9.8e-06 | 1.260E-05 | 2.623E-06 | 9.800E-06 | 1.500E-05 |
| 771 | 39 | E39L | A | 1.5e-05 | 1.5e-05 | 4.4e-06 | 1.147E-05 | 6.120E-06 | 4.400E-06 | 1.500E-05 |
| 772 | 39 | E39K | A | 1.6e-05 | 1.4e-05 | 1.3e-05 | 1.433E-05 | 1.528E-06 | 1.300E-05 | 1.600E-05 |
| 773 | 39 | E39M | A | 1e-05 | 1.4e-05 | 9.6e-06 | 1.120E-05 | 2.433E-06 | 9.600E-06 | 1.400E-05 |
| 774 | 39 | E39F | A | 1.4e-05 | 1.2e-05 | 1.1e-05 | 1.233E-05 | 1.528E-06 | 1.100E-05 | 1.400E-05 |
| 775 | 39 | E39P | A | 1.6e-05 | 1.1e-05 | 1.6e-05 | 1.433E-05 | 2.887E-06 | 1.100E-05 | 1.600E-05 |
| 776 | 39 | E39S | A | 1e-05 | 1.4e-05 | 8.3e-06 | 1.077E-05 | 2.926E-06 | 8.300E-06 | 1.400E-05 |
| 777 | 39 | E39T | A | 1e-05 | 9.7e-06 | 1.2e-05 | 1.057E-05 | 1.250E-06 | 9.700E-06 | 1.200E-05 |
| 778 | 39 | E39W | A | 1.6e-05 | 9.9e-06 | 8.3e-06 | 1.140E-05 | 4.063E-06 | 8.300E-06 | 1.600E-05 |
| 779 | 39 | E39Y | A | 1.4e-05 | 1.6e-05 | 1.4e-05 | 1.467E-05 | 1.155E-06 | 1.400E-05 | 1.600E-05 |
| 780 | 39 | E39V | A | 1.4e-05 | 1e-05 | 1e-05 | 1.133E-05 | 2.309E-06 | 1.000E-05 | 1.400E-05 |
| 781 | 40 | L40A | A | 1.6e-05 | 7.8e-06 | 1.1e-05 | 1.160E-05 | 4.133E-06 | 7.800E-06 | 1.600E-05 |
| 782 | 40 | L40R | A | 1.4e-05 | 1.3e-05 | 1.1e-05 | 1.267E-05 | 1.528E-06 | 1.100E-05 | 1.400E-05 |
| 783 | 40 | L40N | A | 8.3e-06 | 1.5e-05 | 1.4e-05 | 1.243E-05 | 3.614E-06 | 8.300E-06 | 1.500E-05 |
| 784 | 40 | L40D | A | 1.2e-05 | 1.5e-05 | 1.6e-05 | 1.433E-05 | 2.082E-06 | 1.200E-05 | 1.600E-05 |
| 785 | 40 | L40C | A | 1e-05 | 8.7e-06 | 1.4e-05 | 1.090E-05 | 2.762E-06 | 8.700E-06 | 1.400E-05 |
| 786 | 40 | L40E | A | 1.6e-05 | 1.5e-05 | 1.2e-05 | 1.433E-05 | 2.082E-06 | 1.200E-05 | 1.600E-05 |
| 787 | 40 | L40Q | A | 9.6e-06 | 1.5e-05 | 1.3e-05 | 1.253E-05 | 2.730E-06 | 9.600E-06 | 1.500E-05 |
| 788 | 40 | L40G | A | 1.6e-05 | 8.2e-06 | 1.5e-05 | 1.307E-05 | 4.244E-06 | 8.200E-06 | 1.600E-05 |
| 789 | 40 | L40H | A | 1.7e-05 | 1.7e-05 | 1.2e-05 | 1.533E-05 | 2.887E-06 | 1.200E-05 | 1.700E-05 |
| 790 | 40 | L40I | A | 1.2e-05 | 1.1e-05 | 9.6e-06 | 1.087E-05 | 1.206E-06 | 9.600E-06 | 1.200E-05 |
| 791 | 40 | L40L | A | 9.8e-06 | 1e-05 | 1.5e-05 | 1.160E-05 | 2.946E-06 | 9.800E-06 | 1.500E-05 |
| 792 | 40 | L40K | A | 1.4e-05 | 1.8e-05 | 9.5e-06 | 1.383E-05 | 4.252E-06 | 9.500E-06 | 1.800E-05 |
| 793 | 40 | L40M | A | 1.4e-05 | 1.2e-05 | 1e-05 | 1.200E-05 | 2.000E-06 | 1.000E-05 | 1.400E-05 |
| 794 | 40 | L40F | A | 1.4e-05 | 1.3e-05 | 1.6e-05 | 1.433E-05 | 1.528E-06 | 1.300E-05 | 1.600E-05 |
| 795 | 40 | L40P | A | 1.6e-05 | 1.3e-05 | 1e-05 | 1.300E-05 | 3.000E-06 | 1.000E-05 | 1.600E-05 |
| 796 | 40 | L40S | A | 1.5e-05 | 1.1e-05 | 1.5e-05 | 1.367E-05 | 2.309E-06 | 1.100E-05 | 1.500E-05 |
| 797 | 40 | L40T | A | 9.7e-06 | 1.3e-05 | 1.6e-05 | 1.290E-05 | 3.151E-06 | 9.700E-06 | 1.600E-05 |
| 798 | 40 | L40W | A | 1.3e-05 | 2.7e-05 | 1.1e-05 | 1.700E-05 | 8.718E-06 | 1.100E-05 | 2.700E-05 |
| 799 | 40 | L40Y | A | 1.5e-05 | 1.3e-05 | 1.5e-05 | 1.433E-05 | 1.155E-06 | 1.300E-05 | 1.500E-05 |
| 800 | 40 | L40V | A | 1.3e-05 | 1.4e-05 | 1.3e-05 | 1.333E-05 | 5.774E-07 | 1.300E-05 | 1.400E-05 |
| 801 | 41 | T41A | A | 9.2e-06 | 8.7e-06 | 1.1e-05 | 9.633E-06 | 1.210E-06 | 8.700E-06 | 1.100E-05 |
| 802 | 41 | T41R | A | 1.2e-05 | 1.6e-05 | 1.1e-05 | 1.300E-05 | 2.646E-06 | 1.100E-05 | 1.600E-05 |
| 803 | 41 | T41N | A | 1.4e-05 | 1.1e-05 | 1.1e-05 | 1.200E-05 | 1.732E-06 | 1.100E-05 | 1.400E-05 |
| 804 | 41 | T41D | A | 1.1e-05 | 1.1e-05 | 1.3e-05 | 1.167E-05 | 1.155E-06 | 1.100E-05 | 1.300E-05 |
| 805 | 41 | T41C | A | 1.1e-05 | 1.4e-05 | 1.1e-05 | 1.200E-05 | 1.732E-06 | 1.100E-05 | 1.400E-05 |
| 806 | 41 | T41E | A | 8.3e-06 | 1.5e-05 | 1.4e-05 | 1.243E-05 | 3.614E-06 | 8.300E-06 | 1.500E-05 |
| 807 | 41 | T41Q | A | 1.6e-05 | 1.6e-05 | 1.5e-05 | 1.567E-05 | 5.774E-07 | 1.500E-05 | 1.600E-05 |
| 808 | 41 | T41G | A | 6.5e-06 | 1.6e-05 | 1.5e-05 | 1.250E-05 | 5.220E-06 | 6.500E-06 | 1.600E-05 |
| 809 | 41 | T41H | A | 1.4e-05 | 1.6e-05 | 1.4e-05 | 1.467E-05 | 1.155E-06 | 1.400E-05 | 1.600E-05 |
| 810 | 41 | T41I | A | 7.4e-06 | 1e-05 | 1.3e-05 | 1.013E-05 | 2.802E-06 | 7.400E-06 | 1.300E-05 |
| 811 | 41 | T41L | A | 1.2e-05 | 1.6e-05 | 9.8e-06 | 1.260E-05 | 3.143E-06 | 9.800E-06 | 1.600E-05 |
| 812 | 41 | T41K | A | 1.6e-05 | 1.5e-05 | 1.2e-05 | 1.433E-05 | 2.082E-06 | 1.200E-05 | 1.600E-05 |
| 813 | 41 | T41M | A | 9.9e-06 | 1.6e-05 | 1e-05 | 1.197E-05 | 3.493E-06 | 9.900E-06 | 1.600E-05 |
| 814 | 41 | T41F | A | 6.8e-06 | 1.1e-05 | 8.5e-06 | 8.767E-06 | 2.113E-06 | 6.800E-06 | 1.100E-05 |
| 815 | 41 | T41P | A | 1.1e-05 | 1.5e-05 | 1.1e-05 | 1.233E-05 | 2.309E-06 | 1.100E-05 | 1.500E-05 |
| 816 | 41 | T41S | A | 1.1e-05 | 1.4e-05 | 1.1e-05 | 1.200E-05 | 1.732E-06 | 1.100E-05 | 1.400E-05 |
| 817 | 41 | T41T | A | 1.3e-05 | 1.4e-05 | 1.5e-05 | 1.400E-05 | 1.000E-06 | 1.300E-05 | 1.500E-05 |
| 818 | 41 | T41W | A | 1.5e-05 | 1.7e-05 | 1e-05 | 1.400E-05 | 3.606E-06 | 1.000E-05 | 1.700E-05 |
| 819 | 41 | T41Y | A | 1.6e-05 | 1.7e-05 | 1.4e-05 | 1.567E-05 | 1.528E-06 | 1.400E-05 | 1.700E-05 |
| 820 | 41 | T41V | A | 1.1e-05 | 1e-05 | 1.5e-05 | 1.200E-05 | 2.646E-06 | 1.000E-05 | 1.500E-05 |
| 821 | 42 | Y42A | A | 1.7e-05 | 1.6e-05 | 1.7e-05 | 1.667E-05 | 5.774E-07 | 1.600E-05 | 1.700E-05 |
| 822 | 42 | Y42R | A | 9.2e-06 | 1.7e-05 | 1.2e-05 | 1.273E-05 | 3.951E-06 | 9.200E-06 | 1.700E-05 |
| 823 | 42 | Y42N | A | 1.4e-05 | 1.1e-05 | 9.7e-06 | 1.157E-05 | 2.205E-06 | 9.700E-06 | 1.400E-05 |
| 824 | 42 | Y42D | A | 1.1e-05 | 1.5e-05 | 1.3e-05 | 1.300E-05 | 2.000E-06 | 1.100E-05 | 1.500E-05 |
| 825 | 42 | Y42C | A | 1.4e-05 | 1.2e-05 | 1.1e-05 | 1.233E-05 | 1.528E-06 | 1.100E-05 | 1.400E-05 |
| 826 | 42 | Y42E | A | 1.7e-05 | 9.1e-06 | 1.6e-05 | 1.403E-05 | 4.302E-06 | 9.100E-06 | 1.700E-05 |
| 827 | 42 | Y42Q | A | 1e-05 | 1.6e-05 | 1.1e-05 | 1.233E-05 | 3.215E-06 | 1.000E-05 | 1.600E-05 |
| 828 | 42 | Y42G | A | 1.6e-05 | 1.5e-05 | 1.8e-05 | 1.633E-05 | 1.528E-06 | 1.500E-05 | 1.800E-05 |
| 829 | 42 | Y42H | A | 1e-05 | 1.4e-05 | 6.5e-06 | 1.017E-05 | 3.753E-06 | 6.500E-06 | 1.400E-05 |
| 830 | 42 | Y42I | A | 1.7e-05 | 1.2e-05 | 1.3e-05 | 1.400E-05 | 2.646E-06 | 1.200E-05 | 1.700E-05 |
| 831 | 42 | Y42L | A | 1.7e-05 | 1e-05 | 1.2e-05 | 1.300E-05 | 3.606E-06 | 1.000E-05 | 1.700E-05 |
| 832 | 42 | Y42K | A | 1.4e-05 | 1e-05 | 1.3e-05 | 1.233E-05 | 2.082E-06 | 1.000E-05 | 1.400E-05 |
| 833 | 42 | Y42M | A | 1.8e-05 | 1.5e-05 | 1.6e-05 | 1.633E-05 | 1.528E-06 | 1.500E-05 | 1.800E-05 |
| 834 | 42 | Y42F | A | 1.2e-05 | 1.2e-05 | 1e-05 | 1.133E-05 | 1.155E-06 | 1.000E-05 | 1.200E-05 |
| 835 | 42 | Y42P | A | 1.5e-05 | 8.1e-06 | 1.1e-05 | 1.137E-05 | 3.465E-06 | 8.100E-06 | 1.500E-05 |
| 836 | 42 | Y42S | A | 1.1e-05 | 1.5e-05 | 1.6e-05 | 1.400E-05 | 2.646E-06 | 1.100E-05 | 1.600E-05 |
| 837 | 42 | Y42T | A | 1.4e-05 | 1e-05 | 1.4e-05 | 1.267E-05 | 2.309E-06 | 1.000E-05 | 1.400E-05 |
| 838 | 42 | Y42W | A | 1.4e-05 | 1.5e-05 | 1.2e-05 | 1.367E-05 | 1.528E-06 | 1.200E-05 | 1.500E-05 |
| 839 | 42 | Y42Y | A | 9.8e-06 | 1e-05 | 1.1e-05 | 1.027E-05 | 6.429E-07 | 9.800E-06 | 1.100E-05 |
| 840 | 42 | Y42V | A | 8e-06 | 1.4e-05 | 1.6e-05 | 1.267E-05 | 4.163E-06 | 8.000E-06 | 1.600E-05 |
| 841 | 43 | L43A | A | 1.4e-05 | 1.1e-05 | 1.4e-05 | 1.300E-05 | 1.732E-06 | 1.100E-05 | 1.400E-05 |
| 842 | 43 | L43R | A | 1.9e-05 | 1.5e-05 | 1e-05 | 1.467E-05 | 4.509E-06 | 1.000E-05 | 1.900E-05 |
| 843 | 43 | L43N | A | 1.7e-05 | 8.7e-06 | 9.4e-06 | 1.170E-05 | 4.603E-06 | 8.700E-06 | 1.700E-05 |
| 844 | 43 | L43D | A | 7.2e-06 | 1.7e-05 | 7.3e-06 | 1.050E-05 | 5.629E-06 | 7.200E-06 | 1.700E-05 |
| 845 | 43 | L43C | A | 1.4e-05 | 1.7e-05 | 1.1e-05 | 1.400E-05 | 3.000E-06 | 1.100E-05 | 1.700E-05 |
| 846 | 43 | L43E | A | 9.6e-06 | 9.1e-06 | 1e-05 | 9.567E-06 | 4.509E-07 | 9.100E-06 | 1.000E-05 |
| 847 | 43 | L43Q | A | 1.1e-05 | 1.2e-05 | 1.5e-05 | 1.267E-05 | 2.082E-06 | 1.100E-05 | 1.500E-05 |
| 848 | 43 | L43G | A | 1.6e-05 | 1.5e-05 | 1.1e-05 | 1.400E-05 | 2.646E-06 | 1.100E-05 | 1.600E-05 |
| 849 | 43 | L43H | A | 1.1e-05 | 1.3e-05 | 1.8e-05 | 1.400E-05 | 3.606E-06 | 1.100E-05 | 1.800E-05 |

| | | | | | | | | | | |
|---|---|---|---|---|---|---|---|---|---|---|
| 850 | 43 | L43I | A | 1.6e-05 | 7.6e-06 | 1.1e-05 | 1.153E-05 | 4.225E-06 | 7.600E-06 | 1.600E-05 |
| 851 | 43 | L43L | A | 1.3e-05 | 1.2e-05 | 1.3e-05 | 1.267E-05 | 5.774E-07 | 1.200E-05 | 1.300E-05 |
| 852 | 43 | L43K | A | 1.2e-05 | 1.5e-05 | 1.1e-05 | 1.267E-05 | 2.082E-06 | 1.100E-05 | 1.500E-05 |
| 853 | 43 | L43M | A | 1.5e-05 | 1.1e-05 | 1.6e-05 | 1.400E-05 | 2.646E-06 | 1.100E-05 | 1.600E-05 |
| 854 | 43 | L43F | A | 1.5e-05 | 1.6e-05 | 1.2e-05 | 1.433E-05 | 2.082E-06 | 1.200E-05 | 1.600E-05 |
| 855 | 43 | L43P | A | 1.7e-05 | 1.4e-05 | 1.2e-05 | 1.433E-05 | 2.517E-06 | 1.200E-05 | 1.700E-05 |
| 856 | 43 | L43S | A | 1.3e-05 | 8.5e-06 | 1.4e-05 | 1.183E-05 | 2.930E-06 | 8.500E-06 | 1.400E-05 |
| 857 | 43 | L43T | A | 1.6e-05 | 1.6e-05 | 1.6e-05 | 1.600E-05 | 0.000E+00 | 1.600E-05 | 1.600E-05 |
| 858 | 43 | L43W | A | 1.4e-05 | 1.5e-05 | 9.2e-06 | 1.273E-05 | 3.101E-06 | 9.200E-06 | 1.500E-05 |
| 859 | 43 | L43Y | A | 1.6e-05 | 8.6e-06 | 1.1e-05 | 1.187E-05 | 3.775E-06 | 8.600E-06 | 1.600E-05 |
| 860 | 43 | L43V | A | 9.1e-06 | 1.1e-05 | 1.2e-05 | 1.070E-05 | 1.473E-06 | 9.100E-06 | 1.200E-05 |
| 861 | 44 | P44A | A | 1.2e-05 | 1.2e-05 | 9.5e-06 | 1.117E-05 | 1.443E-06 | 9.500E-06 | 1.200E-05 |
| 862 | 44 | P44R | A | 9.5e-06 | 1e-05 | 1.7e-05 | 1.217E-05 | 4.193E-06 | 9.500E-06 | 1.700E-05 |
| 863 | 44 | P44N | A | 1.1e-05 | 1e-05 | 1e-05 | 1.033E-05 | 5.774E-07 | 1.000E-05 | 1.100E-05 |
| 864 | 44 | P44D | A | 1.5e-05 | 1.4e-05 | 1.6e-05 | 1.500E-05 | 1.000E-06 | 1.400E-05 | 1.600E-05 |
| 865 | 44 | P44C | A | 1.3e-05 | 1.5e-05 | 1.5e-05 | 1.433E-05 | 1.155E-06 | 1.300E-05 | 1.500E-05 |
| 866 | 44 | P44E | A | 1.4e-05 | 1.1e-05 | 1.2e-05 | 1.233E-05 | 1.528E-06 | 1.100E-05 | 1.400E-05 |
| 867 | 44 | P44Q | A | 7.9e-06 | 1.8e-05 | 1.3e-05 | 1.297E-05 | 5.050E-06 | 7.900E-06 | 1.800E-05 |
| 868 | 44 | P44G | A | 1.5e-05 | 1.4e-05 | 1.5e-05 | 1.467E-05 | 5.774E-07 | 1.400E-05 | 1.500E-05 |
| 869 | 44 | P44H | A | 1.4e-05 | 1.4e-05 | 1.5e-05 | 1.433E-05 | 5.774E-07 | 1.400E-05 | 1.500E-05 |
| 870 | 44 | P44I | A | 8.9e-06 | 9.5e-06 | 1.5e-05 | 1.113E-05 | 3.362E-06 | 8.900E-06 | 1.500E-05 |
| 871 | 44 | P44L | A | 7.9e-06 | 1.5e-05 | 1.4e-05 | 1.230E-05 | 3.843E-06 | 7.900E-06 | 1.500E-05 |
| 872 | 44 | P44K | A | 1.5e-05 | 1.7e-05 | 1.3e-05 | 1.500E-05 | 2.000E-06 | 1.300E-05 | 1.700E-05 |
| 873 | 44 | P44M | A | 1.4e-05 | 1.2e-05 | 1.1e-05 | 1.233E-05 | 1.528E-06 | 1.100E-05 | 1.400E-05 |
| 874 | 44 | P44F | A | 8.4e-06 | 1.4e-05 | 1.1e-05 | 1.113E-05 | 2.802E-06 | 8.400E-06 | 1.400E-05 |
| 875 | 44 | P44P | A | 1.4e-05 | 9.7e-06 | 1e-05 | 1.123E-05 | 2.401E-06 | 9.700E-06 | 1.400E-05 |
| 876 | 44 | P44S | A | 1.4e-05 | 1.4e-05 | 1.6e-05 | 1.467E-05 | 1.155E-06 | 1.400E-05 | 1.600E-05 |
| 877 | 44 | P44T | A | 1.3e-05 | 1.3e-05 | 1.1e-05 | 1.233E-05 | 1.155E-06 | 1.100E-05 | 1.300E-05 |
| 878 | 44 | P44W | A | 9.5e-06 | 1.5e-05 | 1.5e-05 | 1.317E-05 | 3.175E-06 | 9.500E-06 | 1.500E-05 |
| 879 | 44 | P44Y | A | 1.3e-05 | 1.6e-05 | 1.6e-05 | 1.500E-05 | 1.732E-06 | 1.300E-05 | 1.600E-05 |
| 880 | 44 | P44V | A | 1.1e-05 | 1.3e-05 | 9.9e-06 | 1.130E-05 | 1.572E-06 | 9.900E-06 | 1.300E-05 |
| 881 | 45 | T45A | A | 1.6e-05 | 1.1e-05 | 1.1e-05 | 1.267E-05 | 2.887E-06 | 1.100E-05 | 1.600E-05 |
| 882 | 45 | T45R | A | 1.3e-05 | 1.3e-05 | 9.8e-06 | 1.193E-05 | 1.848E-06 | 9.800E-06 | 1.300E-05 |
| 883 | 45 | T45N | A | 1.1e-05 | 8.8e-06 | 9.4e-06 | 9.733E-06 | 1.137E-06 | 8.800E-06 | 1.100E-05 |
| 884 | 45 | T45D | A | 1e-05 | 1.2e-05 | 1.5e-05 | 1.233E-05 | 2.517E-06 | 1.000E-05 | 1.500E-05 |
| 885 | 45 | T45C | A | 9e-06 | 9.2e-06 | 1e-05 | 9.400E-06 | 5.292E-07 | 9.000E-06 | 1.000E-05 |
| 886 | 45 | T45E | A | 1.4e-05 | 1.3e-05 | 1.7e-05 | 1.467E-05 | 2.082E-06 | 1.300E-05 | 1.700E-05 |
| 887 | 45 | T45Q | A | 1.7e-05 | 1.2e-05 | 1.8e-05 | 1.567E-05 | 3.215E-06 | 1.200E-05 | 1.800E-05 |
| 888 | 45 | T45G | A | 1.1e-05 | 1.6e-05 | 1.6e-05 | 1.433E-05 | 2.887E-06 | 1.100E-05 | 1.600E-05 |
| 889 | 45 | T45H | A | 8.9e-06 | 1e-05 | 1e-05 | 9.633E-06 | 6.351E-07 | 8.900E-06 | 1.000E-05 |
| 890 | 45 | T45I | A | 1.5e-05 | 1.4e-05 | 1.7e-05 | 1.533E-05 | 1.528E-06 | 1.400E-05 | 1.700E-05 |
| 891 | 45 | T45L | A | 1.2e-05 | 1.5e-05 | 1.4e-05 | 1.367E-05 | 1.528E-06 | 1.200E-05 | 1.500E-05 |
| 892 | 45 | T45K | A | 1.1e-05 | 1.7e-05 | 1.6e-05 | 1.467E-05 | 3.215E-06 | 1.100E-05 | 1.700E-05 |
| 893 | 45 | T45M | A | 1.3e-05 | 1e-05 | 1.5e-05 | 1.267E-05 | 2.517E-06 | 1.000E-05 | 1.500E-05 |
| 894 | 45 | T45F | A | 1.7e-05 | 1.7e-05 | 1.4e-05 | 1.600E-05 | 1.732E-06 | 1.400E-05 | 1.700E-05 |
| 895 | 45 | T45P | A | 1.5e-05 | 1.5e-05 | 1.6e-05 | 1.533E-05 | 5.774E-07 | 1.500E-05 | 1.600E-05 |
| 896 | 45 | T45S | A | 1.6e-05 | 1.7e-05 | 9.9e-06 | 1.430E-05 | 3.843E-06 | 9.900E-06 | 1.700E-05 |
| 897 | 45 | T45T | A | 1.2e-05 | 1.6e-05 | 1.3e-05 | 1.367E-05 | 2.082E-06 | 1.200E-05 | 1.600E-05 |
| 898 | 45 | T45W | A | 1.4e-05 | 1.2e-05 | 9.7e-06 | 1.190E-05 | 2.152E-06 | 9.700E-06 | 1.400E-05 |
| 899 | 45 | T45Y | A | 1e-05 | 1.1e-05 | 1.3e-05 | 1.133E-05 | 1.528E-06 | 1.000E-05 | 1.300E-05 |
| 900 | 45 | T45V | A | 9.6e-06 | 1.1e-05 | 1.2e-05 | 1.087E-05 | 1.206E-06 | 9.600E-06 | 1.200E-05 |
| 901 | 46 | N46A | A | 8.6e-06 | 1.5e-05 | 1.1e-05 | 1.153E-05 | 3.233E-06 | 8.600E-06 | 1.500E-05 |
| 902 | 46 | N46R | A | 1.6e-05 | 1.3e-05 | 1.5e-05 | 1.467E-05 | 1.528E-06 | 1.300E-05 | 1.600E-05 |
| 903 | 46 | N46N | A | 1.7e-05 | 1.6e-05 | 1.4e-05 | 1.567E-05 | 1.528E-06 | 1.400E-05 | 1.700E-05 |
| 904 | 46 | N46D | A | 1e-05 | 1.1e-05 | 1.2e-05 | 1.100E-05 | 1.000E-06 | 1.000E-05 | 1.200E-05 |
| 905 | 46 | N46C | A | 1.3e-05 | 1.6e-05 | 1.5e-05 | 1.467E-05 | 1.528E-06 | 1.300E-05 | 1.600E-05 |
| 906 | 46 | N46E | A | 1.6e-05 | 1.3e-05 | 1.5e-05 | 1.467E-05 | 1.528E-06 | 1.300E-05 | 1.600E-05 |
| 907 | 46 | N46Q | A | 2e-05 | 1e-05 | 1.6e-05 | 1.533E-05 | 5.033E-06 | 1.000E-05 | 2.000E-05 |
| 908 | 46 | N46G | A | 1.1e-05 | 1.6e-05 | 1.5e-05 | 1.400E-05 | 2.646E-06 | 1.100E-05 | 1.600E-05 |
| 909 | 46 | N46H | A | 1.2e-05 | 1.5e-05 | 1.1e-05 | 1.267E-05 | 2.082E-06 | 1.100E-05 | 1.500E-05 |
| 910 | 46 | N46I | A | 1.5e-05 | 1.6e-05 | 1.1e-05 | 1.400E-05 | 2.646E-06 | 1.100E-05 | 1.600E-05 |
| 911 | 46 | N46L | A | 1e-05 | 1.5e-05 | 1.4e-05 | 1.300E-05 | 2.646E-06 | 1.000E-05 | 1.500E-05 |
| 912 | 46 | N46K | A | 1.5e-05 | 1.6e-05 | 1.6e-05 | 1.567E-05 | 5.774E-07 | 1.500E-05 | 1.600E-05 |
| 913 | 46 | N46M | A | 8.6e-06 | 1.5e-05 | 1.5e-05 | 1.287E-05 | 3.695E-06 | 8.600E-06 | 1.500E-05 |
| 914 | 46 | N46F | A | 9.8e-06 | 1.3e-05 | 1.3e-05 | 1.193E-05 | 1.848E-06 | 9.800E-06 | 1.300E-05 |
| 915 | 46 | N46P | A | 1.4e-05 | 1.2e-05 | 1.3e-05 | 1.300E-05 | 1.000E-06 | 1.200E-05 | 1.400E-05 |
| 916 | 46 | N46S | A | 1.6e-05 | 1.5e-05 | 1.7e-05 | 1.600E-05 | 1.000E-06 | 1.500E-05 | 1.700E-05 |
| 917 | 46 | N46T | A | 1.4e-05 | 1e-05 | 8.3e-06 | 1.077E-05 | 2.926E-06 | 8.300E-06 | 1.400E-05 |
| 918 | 46 | N46W | A | 1.2e-05 | 1.5e-05 | 1.4e-05 | 1.367E-05 | 1.528E-06 | 1.200E-05 | 1.500E-05 |
| 919 | 46 | N46Y | A | 8.7e-06 | 1.4e-05 | 1.5e-05 | 1.257E-05 | 3.386E-06 | 8.700E-06 | 1.500E-05 |
| 920 | 46 | N46V | A | 1.5e-05 | 9.9e-06 | 1.5e-05 | 1.330E-05 | 2.944E-06 | 9.900E-06 | 1.500E-05 |
| 921 | 47 | A47A | A | 1.6e-05 | 7e-06 | 1.2e-05 | 1.167E-05 | 4.509E-06 | 7.000E-06 | 1.600E-05 |
| 922 | 47 | A47R | A | 8.9e-06 | 1e-05 | 1.3e-05 | 1.063E-05 | 2.122E-06 | 8.900E-06 | 1.300E-05 |
| 923 | 47 | A47N | A | 1.3e-05 | 1.4e-05 | 2.1e-05 | 1.600E-05 | 4.359E-06 | 1.300E-05 | 2.100E-05 |
| 924 | 47 | A47D | A | 1e-05 | 1.1e-05 | 1.4e-05 | 1.167E-05 | 2.082E-06 | 1.000E-05 | 1.400E-05 |
| 925 | 47 | A47C | A | 1.5e-05 | 1.5e-05 | 1.4e-05 | 1.467E-05 | 5.774E-07 | 1.400E-05 | 1.500E-05 |
| 926 | 47 | A47E | A | 1.6e-05 | 1.4e-05 | 1.6e-05 | 1.533E-05 | 1.155E-06 | 1.400E-05 | 1.600E-05 |
| 927 | 47 | A47Q | A | 9.3e-06 | 9.3e-06 | 1.6e-05 | 1.153E-05 | 3.868E-06 | 9.300E-06 | 1.600E-05 |
| 928 | 47 | A47G | A | 1e-05 | 1.1e-05 | 9.9e-06 | 1.030E-05 | 6.083E-07 | 9.900E-06 | 1.100E-05 |
| 929 | 47 | A47H | A | 7.7e-06 | 1.3e-05 | 1.1e-05 | 1.057E-05 | 2.676E-06 | 7.700E-06 | 1.300E-05 |
| 930 | 47 | A47I | A | 1.3e-05 | 1e-05 | 1.5e-05 | 1.267E-05 | 2.517E-06 | 1.000E-05 | 1.500E-05 |
| 931 | 47 | A47L | A | 1.7e-05 | 1.3e-05 | 1.4e-05 | 1.467E-05 | 2.082E-06 | 1.300E-05 | 1.700E-05 |
| 932 | 47 | A47K | A | 8.7e-06 | 1.5e-05 | 1.6e-05 | 1.323E-05 | 3.958E-06 | 8.700E-06 | 1.600E-05 |
| 933 | 47 | A47M | A | 1.3e-05 | 1.5e-05 | 1.1e-05 | 1.300E-05 | 2.000E-06 | 1.100E-05 | 1.500E-05 |
| 934 | 47 | A47F | A | 8.8e-06 | 1.2e-05 | 9.8e-06 | 1.020E-05 | 1.637E-06 | 8.800E-06 | 1.200E-05 |

| | | | | | | | | | | |
|---|---|---|---|---|---|---|---|---|---|---|
| 935 | 47 | A47P | A | 1.1e-05 | 1.6e-05 | 1.7e-05 | 1.467E-05 | 3.215E-06 | 1.100E-05 | 1.700E-05 |
| 936 | 47 | A47S | A | 1.5e-05 | 1e-05 | 1.1e-05 | 1.200E-05 | 2.646E-06 | 1.000E-05 | 1.500E-05 |
| 937 | 47 | A47T | A | 1.1e-05 | 9.8e-06 | 1.1e-05 | 1.060E-05 | 6.928E-07 | 9.800E-06 | 1.100E-05 |
| 938 | 47 | A47W | A | 1.6e-05 | 2.2e-05 | 1.5e-05 | 1.767E-05 | 3.786E-06 | 1.500E-05 | 2.200E-05 |
| 939 | 47 | A47Y | A | 1e-05 | 1e-05 | 1.3e-05 | 1.100E-05 | 1.732E-06 | 1.000E-05 | 1.300E-05 |
| 940 | 47 | A47V | A | 1.3e-05 | 1.4e-05 | 1.5e-05 | 1.400E-05 | 1.000E-06 | 1.300E-05 | 1.500E-05 |
| 941 | 48 | S48A | A | 8.5e-06 | 1.5e-05 | 1.7e-05 | 1.350E-05 | 4.444E-06 | 8.500E-06 | 1.700E-05 |
| 942 | 48 | S48R | A | 9.4e-06 | 1.5e-05 | 1.4e-05 | 1.280E-05 | 2.987E-06 | 9.400E-06 | 1.500E-05 |
| 943 | 48 | S48N | A | 1e-05 | 1.1e-05 | 1.5e-05 | 1.200E-05 | 2.646E-06 | 1.000E-05 | 1.500E-05 |
| 944 | 48 | S48D | A | 1.1e-05 | 1.5e-05 | 1.5e-05 | 1.367E-05 | 2.309E-06 | 1.100E-05 | 1.500E-05 |
| 945 | 48 | S48C | A | 1.5e-05 | 1.1e-05 | 1.1e-05 | 1.233E-05 | 2.309E-06 | 1.100E-05 | 1.500E-05 |
| 946 | 48 | S48E | A | 1.1e-05 | 1.2e-05 | 1.1e-05 | 1.133E-05 | 5.774E-07 | 1.100E-05 | 1.200E-05 |
| 947 | 48 | S48Q | A | 1.2e-05 | 1.3e-05 | 1.3e-05 | 1.267E-05 | 5.774E-07 | 1.200E-05 | 1.300E-05 |
| 948 | 48 | S48G | A | 1.6e-05 | 1.4e-05 | 8.9e-06 | 1.297E-05 | 3.661E-06 | 8.900E-06 | 1.600E-05 |
| 949 | 48 | S48H | A | 1.3e-05 | 9.2e-06 | 7.4e-06 | 9.867E-06 | 2.859E-06 | 7.400E-06 | 1.300E-05 |
| 950 | 48 | S48I | A | 1.4e-05 | 1.1e-05 | 1.9e-05 | 1.467E-05 | 4.041E-06 | 1.100E-05 | 1.900E-05 |
| 951 | 48 | S48L | A | 1.7e-05 | 9.3e-06 | 9.1e-06 | 1.180E-05 | 4.504E-06 | 9.100E-06 | 1.700E-05 |
| 952 | 48 | S48K | A | 1.2e-05 | 1e-05 | 1.5e-05 | 1.233E-05 | 2.517E-06 | 1.000E-05 | 1.500E-05 |
| 953 | 48 | S48M | A | 1.6e-05 | 1.7e-05 | 1.1e-05 | 1.467E-05 | 3.215E-06 | 1.100E-05 | 1.700E-05 |
| 954 | 48 | S48F | A | 1.5e-05 | 9.5e-06 | 1.7e-05 | 1.383E-05 | 3.884E-06 | 9.500E-06 | 1.700E-05 |
| 955 | 48 | S48P | A | 1.7e-05 | 1.2e-05 | 1.1e-05 | 1.333E-05 | 3.215E-06 | 1.100E-05 | 1.700E-05 |
| 956 | 48 | S48S | A | 9.1e-06 | 9.5e-06 | 2.2e-05 | 1.353E-05 | 7.335E-06 | 9.100E-06 | 2.200E-05 |
| 957 | 48 | S48T | A | 1.4e-05 | 9.7e-06 | 1.1e-05 | 1.157E-05 | 2.205E-06 | 9.700E-06 | 1.400E-05 |
| 958 | 48 | S48W | A | 1e-05 | 1e-05 | 1.7e-05 | 1.233E-05 | 4.041E-06 | 1.000E-05 | 1.700E-05 |
| 959 | 48 | S48Y | A | 1.1e-05 | 1.2e-05 | 1.5e-05 | 1.267E-05 | 2.082E-06 | 1.100E-05 | 1.500E-05 |
| 960 | 48 | S48V | A | 1.1e-05 | 8.6e-06 | 1.4e-05 | 1.120E-05 | 2.706E-06 | 8.600E-06 | 1.400E-05 |
| 961 | 49 | L49A | A | 1.5e-05 | 1.4e-05 | 1.5e-05 | 1.467E-05 | 5.774E-07 | 1.400E-05 | 1.500E-05 |
| 962 | 49 | L49R | A | 1.1e-05 | 9.7e-06 | 1.2e-05 | 1.090E-05 | 1.153E-06 | 9.700E-06 | 1.200E-05 |
| 963 | 49 | L49N | A | 1.6e-05 | 1.1e-05 | 1.2e-05 | 1.300E-05 | 2.646E-06 | 1.100E-05 | 1.600E-05 |
| 964 | 49 | L49D | A | 5.9e-06 | 1e-05 | 1.5e-05 | 1.030E-05 | 4.557E-06 | 5.900E-06 | 1.500E-05 |
| 965 | 49 | L49C | A | 1.6e-05 | 8e-06 | 6.8e-06 | 1.027E-05 | 5.001E-06 | 6.800E-06 | 1.600E-05 |
| 966 | 49 | L49E | A | 9.7e-06 | 9.9e-06 | 8.8e-06 | 9.467E-06 | 5.859E-07 | 8.800E-06 | 9.900E-06 |
| 967 | 49 | L49Q | A | 1.4e-05 | 1.5e-05 | 1.6e-05 | 1.500E-05 | 1.000E-06 | 1.400E-05 | 1.600E-05 |
| 968 | 49 | L49G | A | 1e-05 | 1.5e-05 | 1.1e-05 | 1.200E-05 | 2.646E-06 | 1.000E-05 | 1.500E-05 |
| 969 | 49 | L49H | A | 1.4e-05 | 8.7e-06 | 1e-05 | 1.090E-05 | 2.762E-06 | 8.700E-06 | 1.400E-05 |
| 970 | 49 | L49I | A | 1.1e-05 | 7.8e-06 | 1.2e-05 | 1.027E-05 | 2.194E-06 | 7.800E-06 | 1.200E-05 |
| 971 | 49 | L49L | A | 1.1e-05 | 1.6e-05 | 1.5e-05 | 1.400E-05 | 2.646E-06 | 1.100E-05 | 1.600E-05 |
| 972 | 49 | L49K | A | 1.6e-05 | 1.6e-05 | 1.6e-05 | 1.600E-05 | 0.000E+00 | 1.600E-05 | 1.600E-05 |
| 973 | 49 | L49M | A | 1.5e-05 | 1.4e-05 | 1.5e-05 | 1.467E-05 | 5.774E-07 | 1.400E-05 | 1.500E-05 |
| 974 | 49 | L49F | A | 1.2e-05 | 8.9e-06 | 1.2e-05 | 1.097E-05 | 1.790E-06 | 8.900E-06 | 1.200E-05 |
| 975 | 49 | L49P | A | 1.4e-05 | 1.1e-05 | 1.2e-05 | 1.233E-05 | 1.528E-06 | 1.100E-05 | 1.400E-05 |
| 976 | 49 | L49S | A | 1.1e-05 | 1.5e-05 | 9.6e-06 | 1.187E-05 | 2.802E-06 | 9.600E-06 | 1.500E-05 |
| 977 | 49 | L49T | A | 1.4e-05 | 1.4e-05 | 1.3e-05 | 1.367E-05 | 5.774E-07 | 1.300E-05 | 1.400E-05 |
| 978 | 49 | L49W | A | 1.4e-05 | 1.5e-05 | 1.1e-05 | 1.333E-05 | 2.082E-06 | 1.100E-05 | 1.500E-05 |
| 979 | 49 | L49Y | A | 1.5e-05 | 1.3e-05 | 1.7e-05 | 1.500E-05 | 2.000E-06 | 1.300E-05 | 1.700E-05 |
| 980 | 49 | L49V | A | 1.5e-05 | 1.3e-05 | 1.4e-05 | 1.400E-05 | 1.000E-06 | 1.300E-05 | 1.500E-05 |
| 981 | 50 | S50A | A | 1.2e-05 | 1e-05 | 1.2e-05 | 1.133E-05 | 1.155E-06 | 1.000E-05 | 1.200E-05 |
| 982 | 50 | S50R | A | 1.5e-05 | 1.6e-05 | 1e-05 | 1.367E-05 | 3.215E-06 | 1.000E-05 | 1.600E-05 |
| 983 | 50 | S50N | A | 1.4e-05 | 1.5e-05 | 1.1e-05 | 1.333E-05 | 2.082E-06 | 1.100E-05 | 1.500E-05 |
| 984 | 50 | S50D | A | 7.5e-06 | 1.7e-05 | 9.9e-06 | 1.147E-05 | 4.940E-06 | 7.500E-06 | 1.700E-05 |
| 985 | 50 | S50C | A | 1e-05 | 1.4e-05 | 9e-06 | 1.100E-05 | 2.646E-06 | 9.000E-06 | 1.400E-05 |
| 986 | 50 | S50E | A | 1.1e-05 | 1e-05 | 1.4e-05 | 1.167E-05 | 2.082E-06 | 1.000E-05 | 1.400E-05 |
| 987 | 50 | S50Q | A | 9.7e-06 | 8.5e-06 | 1.1e-05 | 9.733E-06 | 1.250E-06 | 8.500E-06 | 1.100E-05 |
| 988 | 50 | S50G | A | 1.8e-05 | 1.6e-05 | 1.2e-05 | 1.533E-05 | 3.055E-06 | 1.200E-05 | 1.800E-05 |
| 989 | 50 | S50H | A | 1.3e-05 | 1e-05 | 1.6e-05 | 1.300E-05 | 3.000E-06 | 1.000E-05 | 1.600E-05 |
| 990 | 50 | S50I | A | 1.7e-05 | 1.4e-05 | 1.6e-05 | 1.567E-05 | 1.528E-06 | 1.400E-05 | 1.700E-05 |
| 991 | 50 | S50L | A | 1.1e-05 | 1.1e-05 | 1.6e-05 | 1.267E-05 | 2.887E-06 | 1.100E-05 | 1.600E-05 |
| 992 | 50 | S50K | A | 1.7e-05 | 1.5e-05 | 1.5e-05 | 1.567E-05 | 1.155E-06 | 1.500E-05 | 1.700E-05 |
| 993 | 50 | S50M | A | 1.6e-05 | 1.5e-05 | 1.2e-05 | 1.433E-05 | 2.082E-06 | 1.200E-05 | 1.600E-05 |
| 994 | 50 | S50F | A | 1.6e-05 | 1.8e-05 | 1.6e-05 | 1.667E-05 | 1.155E-06 | 1.600E-05 | 1.800E-05 |
| 995 | 50 | S50P | A | 1.5e-05 | 1.4e-05 | 1.1e-05 | 1.333E-05 | 2.082E-06 | 1.100E-05 | 1.500E-05 |
| 996 | 50 | S50S | A | 1.2e-05 | 9.2e-06 | 1.4e-05 | 1.173E-05 | 2.411E-06 | 9.200E-06 | 1.400E-05 |
| 997 | 50 | S50T | A | 1.6e-05 | 1.1e-05 | 1.4e-05 | 1.367E-05 | 2.517E-06 | 1.100E-05 | 1.600E-05 |
| 998 | 50 | S50W | A | 1.4e-05 | 1e-05 | 1.7e-05 | 1.367E-05 | 3.512E-06 | 1.000E-05 | 1.700E-05 |
| 999 | 50 | S50Y | A | 1.4e-05 | 1.5e-05 | 1.4e-05 | 1.433E-05 | 5.774E-07 | 1.400E-05 | 1.500E-05 |
| 1000 | 50 | S50V | A | 1.5e-05 | 7.1e-06 | 1.5e-05 | 1.237E-05 | 4.561E-06 | 7.100E-06 | 1.500E-05 |
| 1001 | 51 | F51A | A | 1.5e-05 | 1.4e-05 | 1e-05 | 1.300E-05 | 2.646E-06 | 1.000E-05 | 1.500E-05 |
| 1002 | 51 | F51R | A | 1.6e-05 | 8e-06 | 1e-05 | 1.133E-05 | 4.163E-06 | 8.000E-06 | 1.600E-05 |
| 1003 | 51 | F51N | A | 1.2e-05 | 1e-05 | 9.2e-06 | 1.040E-05 | 1.442E-06 | 9.200E-06 | 1.200E-05 |
| 1004 | 51 | F51D | A | 1.5e-05 | 8.6e-06 | 1.4e-05 | 1.253E-05 | 3.443E-06 | 8.600E-06 | 1.500E-05 |
| 1005 | 51 | F51C | A | 1.4e-05 | 1.1e-05 | 1.4e-05 | 1.300E-05 | 1.732E-06 | 1.100E-05 | 1.400E-05 |
| 1006 | 51 | F51E | A | 1.5e-05 | 1.7e-05 | 1.3e-05 | 1.500E-05 | 2.000E-06 | 1.300E-05 | 1.700E-05 |
| 1007 | 51 | F51Q | A | 1.2e-05 | 1.5e-05 | 1.1e-05 | 1.267E-05 | 2.082E-06 | 1.100E-05 | 1.500E-05 |
| 1008 | 51 | F51G | A | 1.5e-05 | 1e-05 | 1.4e-05 | 1.300E-05 | 2.646E-06 | 1.000E-05 | 1.500E-05 |
| 1009 | 51 | F51H | A | 1.1e-05 | 8.6e-06 | 1.3e-05 | 1.087E-05 | 2.203E-06 | 8.600E-06 | 1.300E-05 |
| 1010 | 51 | F51I | A | 1.3e-05 | 1.1e-05 | 1.1e-05 | 1.167E-05 | 1.155E-06 | 1.100E-05 | 1.300E-05 |
| 1011 | 51 | F51L | A | 1.4e-05 | 1.2e-05 | 1.6e-05 | 1.400E-05 | 2.000E-06 | 1.200E-05 | 1.600E-05 |
| 1012 | 51 | F51K | A | 1.3e-05 | 1.7e-05 | 1.6e-05 | 1.533E-05 | 2.082E-06 | 1.300E-05 | 1.700E-05 |
| 1013 | 51 | F51M | A | 1.7e-05 | 1.1e-05 | 1.7e-05 | 1.500E-05 | 3.464E-06 | 1.100E-05 | 1.700E-05 |
| 1014 | 51 | F51F | A | 1.3e-05 | 8.4e-06 | 1e-05 | 1.047E-05 | 2.335E-06 | 8.400E-06 | 1.300E-05 |
| 1015 | 51 | F51P | A | 9.9e-06 | 1.2e-05 | 1.5e-05 | 1.230E-05 | 2.563E-06 | 9.900E-06 | 1.500E-05 |
| 1016 | 51 | F51S | A | 1.3e-05 | 1.4e-05 | 1.2e-05 | 1.300E-05 | 1.000E-06 | 1.200E-05 | 1.400E-05 |
| 1017 | 51 | F51T | A | 1.1e-05 | 9.9e-06 | 1.7e-05 | 1.263E-05 | 3.821E-06 | 9.900E-06 | 1.700E-05 |
| 1018 | 51 | F51W | A | 1.4e-05 | 1.2e-05 | 1.1e-05 | 1.233E-05 | 1.528E-06 | 1.100E-05 | 1.400E-05 |
| 1019 | 51 | F51Y | A | 1.3e-05 | 9.9e-06 | 1.5e-05 | 1.263E-05 | 2.570E-06 | 9.900E-06 | 1.500E-05 |

| | | | | | | | | | | |
|---|---|---|---|---|---|---|---|---|---|---|
| 1020 | 51 | F51V | A | 9e-06 | 1.3e-05 | 1.2e-05 | 1.133E-05 | 2.082E-06 | 9.000E-06 | 1.300E-05 |
| 1021 | 52 | L52A | A | 1.2e-05 | 1.5e-05 | 1.6e-05 | 1.433E-05 | 2.082E-06 | 1.200E-05 | 1.600E-05 |
| 1022 | 52 | L52R | A | 1.5e-05 | 1.5e-05 | 1.5e-05 | 1.500E-05 | 0.000E+00 | 1.500E-05 | 1.500E-05 |
| 1023 | 52 | L52N | A | 1.6e-05 | 1.6e-05 | 1.5e-05 | 1.567E-05 | 5.774E-07 | 1.500E-05 | 1.600E-05 |
| 1024 | 52 | L52D | A | 1e-05 | 8.9e-06 | 1.5e-05 | 1.130E-05 | 3.251E-06 | 8.900E-06 | 1.500E-05 |
| 1025 | 52 | L52C | A | 1.7e-05 | 8.8e-06 | 1.2e-05 | 1.260E-05 | 4.133E-06 | 8.800E-06 | 1.700E-05 |
| 1026 | 52 | L52E | A | 1.2e-05 | 1.4e-05 | 1.6e-05 | 1.400E-05 | 2.000E-06 | 1.200E-05 | 1.600E-05 |
| 1027 | 52 | L52Q | A | 1.3e-05 | 9e-06 | 1.4e-05 | 1.200E-05 | 2.646E-06 | 9.000E-06 | 1.400E-05 |
| 1028 | 52 | L52G | A | 1.5e-05 | 1.5e-05 | 1.4e-05 | 1.467E-05 | 5.774E-07 | 1.400E-05 | 1.500E-05 |
| 1029 | 52 | L52H | A | 1.4e-05 | 1.5e-05 | 1.1e-05 | 1.333E-05 | 2.082E-06 | 1.100E-05 | 1.500E-05 |
| 1030 | 52 | L52I | A | 1.7e-05 | 7.2e-06 | 9.8e-06 | 1.133E-05 | 5.077E-06 | 7.200E-06 | 1.700E-05 |
| 1031 | 52 | L52L | A | 1.7e-05 | 1.1e-05 | 1.6e-05 | 1.467E-05 | 3.215E-06 | 1.100E-05 | 1.700E-05 |
| 1032 | 52 | L52K | A | 1.6e-05 | 1.7e-05 | 1.3e-05 | 1.533E-05 | 2.082E-06 | 1.300E-05 | 1.700E-05 |
| 1033 | 52 | L52M | A | 1.4e-05 | 1.3e-05 | 7.5e-06 | 1.150E-05 | 3.500E-06 | 7.500E-06 | 1.400E-05 |
| 1034 | 52 | L52F | A | 1.6e-05 | 1.6e-05 | 1.4e-05 | 1.533E-05 | 1.155E-06 | 1.400E-05 | 1.600E-05 |
| 1035 | 52 | L52P | A | 1.1e-05 | 1e-05 | 1.5e-05 | 1.200E-05 | 2.646E-06 | 1.000E-05 | 1.500E-05 |
| 1036 | 52 | L52S | A | 1.5e-05 | 1.4e-05 | 9.9e-06 | 1.297E-05 | 2.702E-06 | 9.900E-06 | 1.500E-05 |
| 1037 | 52 | L52T | A | 1.1e-05 | 1.6e-05 | 1.4e-05 | 1.367E-05 | 2.517E-06 | 1.100E-05 | 1.600E-05 |
| 1038 | 52 | L52W | A | 1e-05 | 1.6e-05 | 9.3e-06 | 1.177E-05 | 3.683E-06 | 9.300E-06 | 1.600E-05 |
| 1039 | 52 | L52Y | A | 1.5e-05 | 1.4e-05 | 8e-06 | 1.233E-05 | 3.786E-06 | 8.000E-06 | 1.500E-05 |
| 1040 | 52 | L52V | A | 1.4e-05 | 1.7e-05 | 1.4e-05 | 1.500E-05 | 1.732E-06 | 1.400E-05 | 1.700E-05 |
| 1041 | 53 | Q53A | A | 1.7e-05 | 1.1e-05 | 1.6e-05 | 1.467E-05 | 3.215E-06 | 1.100E-05 | 1.700E-05 |
| 1042 | 53 | Q53R | A | 1.5e-05 | 9.6e-06 | 1.2e-05 | 1.220E-05 | 2.706E-06 | 9.600E-06 | 1.500E-05 |
| 1043 | 53 | Q53N | A | 1.1e-05 | 9.7e-06 | 9e-06 | 9.900E-06 | 1.015E-06 | 9.000E-06 | 1.100E-05 |
| 1044 | 53 | Q53D | A | 1.7e-05 | 9.1e-06 | 1.4e-05 | 1.337E-05 | 3.988E-06 | 9.100E-06 | 1.700E-05 |
| 1045 | 53 | Q53C | A | 1.4e-05 | 1.7e-05 | 1.4e-05 | 1.500E-05 | 1.732E-06 | 1.400E-05 | 1.700E-05 |
| 1046 | 53 | Q53E | A | 1.1e-05 | 7.8e-06 | 1.4e-05 | 1.093E-05 | 3.101E-06 | 7.800E-06 | 1.400E-05 |
| 1047 | 53 | Q53Q | A | 1.7e-05 | 1.2e-05 | 2.7e-05 | 1.867E-05 | 7.638E-06 | 1.200E-05 | 2.700E-05 |
| 1048 | 53 | Q53G | A | 1e-05 | 1.6e-05 | 1.6e-05 | 1.400E-05 | 3.464E-06 | 1.000E-05 | 1.600E-05 |
| 1049 | 53 | Q53H | A | 1.1e-05 | 1.1e-05 | 1.1e-05 | 1.100E-05 | 2.075E-21 | 1.100E-05 | 1.100E-05 |
| 1050 | 53 | Q53I | A | 1.6e-05 | 1.5e-05 | 9.4e-06 | 1.347E-05 | 3.557E-06 | 9.400E-06 | 1.600E-05 |
| 1051 | 53 | Q53L | A | 1.1e-05 | 1.3e-05 | 1.2e-05 | 1.200E-05 | 1.000E-06 | 1.100E-05 | 1.300E-05 |
| 1052 | 53 | Q53K | A | 7.8e-06 | 1.6e-05 | 1.6e-05 | 1.327E-05 | 4.734E-06 | 7.800E-06 | 1.600E-05 |
| 1053 | 53 | Q53M | A | 1.5e-05 | 1.3e-05 | 1.4e-05 | 1.400E-05 | 1.000E-06 | 1.300E-05 | 1.500E-05 |
| 1054 | 53 | Q53F | A | 1.1e-05 | 1.6e-05 | 1.1e-05 | 1.267E-05 | 2.887E-06 | 1.100E-05 | 1.600E-05 |
| 1055 | 53 | Q53P | A | 1.3e-05 | 1.1e-05 | 9.2e-06 | 1.107E-05 | 1.901E-06 | 9.200E-06 | 1.300E-05 |
| 1056 | 53 | Q53S | A | 1.1e-05 | 1.7e-05 | 1.1e-05 | 1.300E-05 | 3.464E-06 | 1.100E-05 | 1.700E-05 |
| 1057 | 53 | Q53T | A | 1.3e-05 | 1.5e-05 | 1.2e-05 | 1.333E-05 | 1.528E-06 | 1.200E-05 | 1.500E-05 |
| 1058 | 53 | Q53W | A | 1.4e-05 | 8.4e-06 | 1.5e-05 | 1.247E-05 | 3.557E-06 | 8.400E-06 | 1.500E-05 |
| 1059 | 53 | Q53Y | A | 1.4e-05 | 1.1e-05 | 1.6e-05 | 1.367E-05 | 2.517E-06 | 1.100E-05 | 1.600E-05 |
| 1060 | 53 | Q53V | A | 9.8e-06 | 1.1e-05 | 1.6e-05 | 1.227E-05 | 3.288E-06 | 9.800E-06 | 1.600E-05 |
| 1061 | 54 | D54A | A | 1.2e-05 | 1.5e-05 | 1e-05 | 1.233E-05 | 2.517E-06 | 1.000E-05 | 1.500E-05 |
| 1062 | 54 | D54R | A | 1.8e-05 | 9.9e-06 | 1e-05 | 1.263E-05 | 4.648E-06 | 9.900E-06 | 1.800E-05 |
| 1063 | 54 | D54N | A | 1.1e-05 | 1e-05 | 1e-05 | 1.033E-05 | 5.774E-07 | 1.000E-05 | 1.100E-05 |
| 1064 | 54 | D54D | A | 1.4e-05 | 1.1e-05 | 1.3e-05 | 1.267E-05 | 1.528E-06 | 1.100E-05 | 1.400E-05 |
| 1065 | 54 | D54C | A | 1e-05 | 1.5e-05 | 1.4e-05 | 1.300E-05 | 2.646E-06 | 1.000E-05 | 1.500E-05 |
| 1066 | 54 | D54E | A | 1e-05 | 1.1e-05 | 1.2e-05 | 1.100E-05 | 1.000E-06 | 1.000E-05 | 1.200E-05 |
| 1067 | 54 | D54Q | A | 1.8e-05 | 1.5e-05 | 4.7e-06 | 1.257E-05 | 6.976E-06 | 4.700E-06 | 1.800E-05 |
| 1068 | 54 | D54G | A | 1e-05 | 1e-05 | 1e-05 | 1.000E-05 | 0.000E+00 | 1.000E-05 | 1.000E-05 |
| 1069 | 54 | D54H | A | 1.1e-05 | 9.2e-06 | 1.3e-05 | 1.107E-05 | 1.901E-06 | 9.200E-06 | 1.300E-05 |
| 1070 | 54 | D54I | A | 9.9e-06 | 1.6e-05 | 1e-05 | 1.197E-05 | 3.493E-06 | 9.900E-06 | 1.600E-05 |
| 1071 | 54 | D54L | A | 1.5e-05 | 1.6e-05 | 1.5e-05 | 1.533E-05 | 5.774E-07 | 1.500E-05 | 1.600E-05 |
| 1072 | 54 | D54K | A | 1.1e-05 | 1.2e-05 | 1.5e-05 | 1.267E-05 | 2.082E-06 | 1.100E-05 | 1.500E-05 |
| 1073 | 54 | D54M | A | 1.7e-05 | 1.5e-05 | 1.6e-05 | 1.600E-05 | 1.000E-06 | 1.500E-05 | 1.700E-05 |
| 1074 | 54 | D54F | A | 1.6e-05 | 2.1e-05 | 8.4e-06 | 1.513E-05 | 6.345E-06 | 8.400E-06 | 2.100E-05 |
| 1075 | 54 | D54P | A | 1.1e-05 | 9.7e-06 | 1.1e-05 | 1.057E-05 | 7.506E-07 | 9.700E-06 | 1.100E-05 |
| 1076 | 54 | D54S | A | 1.2e-05 | 1.4e-05 | 1.5e-05 | 1.367E-05 | 1.528E-06 | 1.200E-05 | 1.500E-05 |
| 1077 | 54 | D54T | A | 1e-05 | 1.6e-05 | 1.1e-05 | 1.233E-05 | 3.215E-06 | 1.000E-05 | 1.600E-05 |
| 1078 | 54 | D54W | A | 1.1e-05 | 1.1e-05 | 1.5e-05 | 1.233E-05 | 2.309E-06 | 1.100E-05 | 1.500E-05 |
| 1079 | 54 | D54Y | A | 1.4e-05 | 1.5e-05 | 1.2e-05 | 1.367E-05 | 1.528E-06 | 1.200E-05 | 1.500E-05 |
| 1080 | 54 | D54V | A | 1.1e-05 | 9.9e-06 | 1.4e-05 | 1.163E-05 | 2.122E-06 | 9.900E-06 | 1.400E-05 |
| 1081 | 55 | I55A | A | 1.1e-05 | 1.1e-05 | 1.3e-05 | 1.167E-05 | 1.155E-06 | 1.100E-05 | 1.300E-05 |
| 1082 | 55 | I55R | A | 1.3e-05 | 1.1e-05 | 1.5e-05 | 1.300E-05 | 2.000E-06 | 1.100E-05 | 1.500E-05 |
| 1083 | 55 | I55N | A | 1.3e-05 | 1.5e-05 | 1.5e-05 | 1.433E-05 | 1.155E-06 | 1.300E-05 | 1.500E-05 |
| 1084 | 55 | I55D | A | 8.6e-06 | 1.1e-05 | 1.2e-05 | 1.053E-05 | 1.747E-06 | 8.600E-06 | 1.200E-05 |
| 1085 | 55 | I55C | A | 1e-05 | 9.7e-06 | 7.8e-06 | 9.167E-06 | 1.193E-06 | 7.800E-06 | 1.000E-05 |
| 1086 | 55 | I55E | A | 1e-05 | 1.5e-05 | 1.7e-05 | 1.400E-05 | 3.606E-06 | 1.000E-05 | 1.700E-05 |
| 1087 | 55 | I55Q | A | 1.2e-05 | 1.5e-05 | 1.3e-05 | 1.333E-05 | 1.528E-06 | 1.200E-05 | 1.500E-05 |
| 1088 | 55 | I55G | A | 1.3e-05 | 1.2e-05 | 1.7e-05 | 1.400E-05 | 2.646E-06 | 1.200E-05 | 1.700E-05 |
| 1089 | 55 | I55H | A | 1.7e-05 | 1.4e-05 | 1e-05 | 1.367E-05 | 3.512E-06 | 1.000E-05 | 1.700E-05 |
| 1090 | 55 | I55I | A | 1.5e-05 | 1.1e-05 | 1e-05 | 1.200E-05 | 2.646E-06 | 1.000E-05 | 1.500E-05 |
| 1091 | 55 | I55L | A | 1e-05 | 1.3e-05 | 1e-05 | 1.100E-05 | 1.732E-06 | 1.000E-05 | 1.300E-05 |
| 1092 | 55 | I55K | A | 8.7e-06 | 1.4e-05 | 9.1e-06 | 1.060E-05 | 2.951E-06 | 8.700E-06 | 1.400E-05 |
| 1093 | 55 | I55M | A | 1.3e-05 | 1e-05 | 1.1e-05 | 1.133E-05 | 1.528E-06 | 1.000E-05 | 1.300E-05 |
| 1094 | 55 | I55F | A | 9.8e-06 | 1.2e-05 | 1.5e-05 | 1.227E-05 | 2.610E-06 | 9.800E-06 | 1.500E-05 |
| 1095 | 55 | I55P | A | 1.7e-05 | 1.3e-05 | 1.4e-05 | 1.467E-05 | 2.082E-06 | 1.300E-05 | 1.700E-05 |
| 1096 | 55 | I55S | A | 1.1e-05 | 1e-05 | 1.3e-05 | 1.133E-05 | 1.528E-06 | 1.000E-05 | 1.300E-05 |
| 1097 | 55 | I55T | A | 1.5e-05 | 1.7e-05 | 1e-05 | 1.400E-05 | 3.606E-06 | 1.000E-05 | 1.700E-05 |
| 1098 | 55 | I55W | A | 1.1e-05 | 1.1e-05 | 1.3e-05 | 1.167E-05 | 1.155E-06 | 1.100E-05 | 1.300E-05 |
| 1099 | 55 | I55Y | A | 1.2e-05 | 9.4e-06 | 1.6e-05 | 1.247E-05 | 3.325E-06 | 9.400E-06 | 1.600E-05 |
| 1100 | 55 | I55V | A | 9.6e-06 | 1.6e-05 | 1.5e-05 | 1.353E-05 | 3.443E-06 | 9.600E-06 | 1.600E-05 |
| 1101 | 56 | Q56A | A | 1.1e-05 | 1.2e-05 | 7.8e-06 | 1.027E-05 | 2.194E-06 | 7.800E-06 | 1.200E-05 |
| 1102 | 56 | Q56R | A | 1e-05 | 1.1e-05 | 1.6e-05 | 1.233E-05 | 3.215E-06 | 1.000E-05 | 1.600E-05 |
| 1103 | 56 | Q56N | A | 1.4e-05 | 1.1e-05 | 7.8e-06 | 1.093E-05 | 3.101E-06 | 7.800E-06 | 1.400E-05 |
| 1104 | 56 | Q56D | A | 1.6e-05 | 8e-06 | 1.3e-05 | 1.233E-05 | 4.041E-06 | 8.000E-06 | 1.600E-05 |

| | | | | | | | | | | |
|---|---|---|---|---|---|---|---|---|---|---|
| 1105 | 56 | Q56C | A | 1.6e-05 | 1.1e-05 | 1.4e-05 | 1.367E-05 | 2.517E-06 | 1.100E-05 | 1.600E-05 |
| 1106 | 56 | Q56E | A | 1.5e-05 | 1.3e-05 | 1.4e-05 | 1.400E-05 | 1.000E-06 | 1.300E-05 | 1.500E-05 |
| 1107 | 56 | Q56Q | A | 1.5e-05 | 1.4e-05 | 1.5e-05 | 1.467E-05 | 5.774E-07 | 1.400E-05 | 1.500E-05 |
| 1108 | 56 | Q56G | A | 1.3e-05 | 1.1e-05 | 1.6e-05 | 1.333E-05 | 2.517E-06 | 1.100E-05 | 1.600E-05 |
| 1109 | 56 | Q56H | A | 1.2e-05 | 9.4e-06 | 1.4e-05 | 1.180E-05 | 2.307E-06 | 9.400E-06 | 1.400E-05 |
| 1110 | 56 | Q56I | A | 1e-05 | 9.4e-06 | 1.1e-05 | 1.013E-05 | 8.083E-07 | 9.400E-06 | 1.100E-05 |
| 1111 | 56 | Q56L | A | 1.1e-05 | 1.3e-05 | 1.8e-05 | 1.400E-05 | 3.606E-06 | 1.100E-05 | 1.800E-05 |
| 1112 | 56 | Q56K | A | 1.6e-05 | 1.5e-05 | 1.5e-05 | 1.533E-05 | 5.774E-07 | 1.500E-05 | 1.600E-05 |
| 1113 | 56 | Q56M | A | 1.7e-05 | 1.5e-05 | 1.6e-05 | 1.600E-05 | 1.000E-06 | 1.500E-05 | 1.700E-05 |
| 1114 | 56 | Q56F | A | 1.6e-05 | 1.6e-05 | 1.1e-05 | 1.433E-05 | 2.887E-06 | 1.100E-05 | 1.600E-05 |
| 1115 | 56 | Q56P | A | 1.5e-05 | 1.3e-05 | 1.1e-05 | 1.300E-05 | 2.000E-06 | 1.100E-05 | 1.500E-05 |
| 1116 | 56 | Q56S | A | 1.4e-05 | 1.2e-05 | 1.1e-05 | 1.233E-05 | 1.528E-06 | 1.100E-05 | 1.400E-05 |
| 1117 | 56 | Q56T | A | 1.2e-05 | 1.5e-05 | 1.4e-05 | 1.367E-05 | 1.528E-06 | 1.200E-05 | 1.500E-05 |
| 1118 | 56 | Q56W | A | 1.5e-05 | 9.6e-06 | 1.6e-05 | 1.353E-05 | 3.443E-06 | 9.600E-06 | 1.600E-05 |
| 1119 | 56 | Q56Y | A | 1.1e-05 | 8.6e-06 | 1.5e-05 | 1.153E-05 | 3.233E-06 | 8.600E-06 | 1.500E-05 |
| 1120 | 56 | Q56V | A | 7.3e-06 | 1e-05 | 1.3e-05 | 1.010E-05 | 2.851E-06 | 7.300E-06 | 1.300E-05 |
| 1121 | 57 | E57A | A | 1.1e-05 | 1.2e-05 | 8.9e-06 | 1.063E-05 | 1.582E-06 | 8.900E-06 | 1.200E-05 |
| 1122 | 57 | E57R | A | 9.3e-06 | 1.4e-05 | 1.1e-05 | 1.143E-05 | 2.380E-06 | 9.300E-06 | 1.400E-05 |
| 1123 | 57 | E57N | A | 8.3e-06 | 9.2e-06 | 1.4e-05 | 1.050E-05 | 3.064E-06 | 8.300E-06 | 1.400E-05 |
| 1124 | 57 | E57D | A | 1.2e-05 | 1.1e-05 | 1.4e-05 | 1.233E-05 | 1.528E-06 | 1.100E-05 | 1.400E-05 |
| 1125 | 57 | E57C | A | 9.8e-06 | 9.1e-06 | 1.1e-05 | 9.967E-06 | 9.609E-07 | 9.100E-06 | 1.100E-05 |
| 1126 | 57 | E57E | A | 1.7e-05 | 1.6e-05 | 1e-05 | 1.433E-05 | 3.786E-06 | 1.000E-05 | 1.700E-05 |
| 1127 | 57 | E57Q | A | 1.5e-05 | 9e-06 | 1.1e-05 | 1.167E-05 | 3.055E-06 | 9.000E-06 | 1.500E-05 |
| 1128 | 57 | E57G | A | 1.1e-05 | 1.6e-05 | 8.1e-06 | 1.170E-05 | 3.996E-06 | 8.100E-06 | 1.600E-05 |
| 1129 | 57 | E57H | A | 1.4e-05 | 9.5e-06 | 1e-05 | 1.117E-05 | 2.466E-06 | 9.500E-06 | 1.400E-05 |
| 1130 | 57 | E57I | A | 1.5e-05 | 1.1e-05 | 1.2e-05 | 1.267E-05 | 2.082E-06 | 1.100E-05 | 1.500E-05 |
| 1131 | 57 | E57L | A | 1.5e-05 | 1.6e-05 | 1.4e-05 | 1.500E-05 | 1.000E-06 | 1.400E-05 | 1.600E-05 |
| 1132 | 57 | E57K | A | 1.4e-05 | 1.8e-05 | 1.4e-05 | 1.533E-05 | 2.309E-06 | 1.400E-05 | 1.800E-05 |
| 1133 | 57 | E57M | A | 1.2e-05 | 1.1e-05 | 1.1e-05 | 1.133E-05 | 5.774E-07 | 1.100E-05 | 1.200E-05 |
| 1134 | 57 | E57F | A | 1.5e-05 | 1e-05 | 1.5e-05 | 1.333E-05 | 2.887E-06 | 1.000E-05 | 1.500E-05 |
| 1135 | 57 | E57P | A | 1.6e-05 | 1.1e-05 | 1.5e-05 | 1.400E-05 | 2.646E-06 | 1.100E-05 | 1.600E-05 |
| 1136 | 57 | E57S | A | 9.2e-06 | 1.4e-05 | 9.5e-06 | 1.090E-05 | 2.689E-06 | 9.200E-06 | 1.400E-05 |
| 1137 | 57 | E57T | A | 1.3e-05 | 1.4e-05 | 1.6e-05 | 1.433E-05 | 1.528E-06 | 1.300E-05 | 1.600E-05 |
| 1138 | 57 | E57W | A | 1.4e-05 | 1.1e-05 | 8.3e-06 | 1.110E-05 | 2.851E-06 | 8.300E-06 | 1.400E-05 |
| 1139 | 57 | E57Y | A | 1.3e-05 | 9.7e-06 | 1.1e-05 | 1.123E-05 | 1.662E-06 | 9.700E-06 | 1.300E-05 |
| 1140 | 57 | E57V | A | 1e-05 | 1e-05 | 1.1e-05 | 1.033E-05 | 5.774E-07 | 1.000E-05 | 1.100E-05 |
| 1141 | 58 | V58A | A | 9e-06 | 1.4e-05 | 1.2e-05 | 1.167E-05 | 2.517E-06 | 9.000E-06 | 1.400E-05 |
| 1142 | 58 | V58R | A | 1.5e-05 | 1.1e-05 | 1.5e-05 | 1.367E-05 | 2.309E-06 | 1.100E-05 | 1.500E-05 |
| 1143 | 58 | V58N | A | 1.2e-05 | 1.6e-05 | 1.6e-05 | 1.467E-05 | 2.309E-06 | 1.200E-05 | 1.600E-05 |
| 1144 | 58 | V58D | A | 1.1e-05 | 8.6e-06 | 9.8e-06 | 9.800E-06 | 1.200E-06 | 8.600E-06 | 1.100E-05 |
| 1145 | 58 | V58C | A | 1.5e-05 | 9.6e-06 | 1.4e-05 | 1.287E-05 | 2.873E-06 | 9.600E-06 | 1.500E-05 |
| 1146 | 58 | V58E | A | 1.5e-05 | 1.6e-05 | 1.1e-05 | 1.400E-05 | 2.646E-06 | 1.100E-05 | 1.600E-05 |
| 1147 | 58 | V58Q | A | 1.1e-05 | 1.5e-05 | 1.4e-05 | 1.333E-05 | 2.082E-06 | 1.100E-05 | 1.500E-05 |
| 1148 | 58 | V58G | A | 1.6e-05 | 1.6e-05 | 1.5e-05 | 1.567E-05 | 5.774E-07 | 1.500E-05 | 1.600E-05 |
| 1149 | 58 | V58H | A | 1e-05 | 1e-05 | 9.6e-06 | 9.867E-06 | 2.309E-07 | 9.600E-06 | 1.000E-05 |
| 1150 | 58 | V58I | A | 1.4e-05 | 1e-05 | 5.9e-06 | 9.967E-06 | 4.050E-06 | 5.900E-06 | 1.400E-05 |
| 1151 | 58 | V58L | A | 1.1e-05 | 1.1e-05 | 1.3e-05 | 1.167E-05 | 1.155E-06 | 1.100E-05 | 1.300E-05 |
| 1152 | 58 | V58K | A | 1.1e-05 | 1.4e-05 | 1.4e-05 | 1.300E-05 | 1.732E-06 | 1.100E-05 | 1.400E-05 |
| 1153 | 58 | V58M | A | 9.2e-06 | 1e-05 | 1.4e-05 | 1.107E-05 | 2.572E-06 | 9.200E-06 | 1.400E-05 |
| 1154 | 58 | V58F | A | 1.6e-05 | 1.5e-05 | 9.9e-06 | 1.363E-05 | 3.272E-06 | 9.900E-06 | 1.600E-05 |
| 1155 | 58 | V58P | A | 1.1e-05 | 1.4e-05 | 1.3e-05 | 1.267E-05 | 1.528E-06 | 1.100E-05 | 1.400E-05 |
| 1156 | 58 | V58S | A | 1e-05 | 1.7e-05 | 1.6e-05 | 1.433E-05 | 3.786E-06 | 1.000E-05 | 1.700E-05 |
| 1157 | 58 | V58T | A | 1.5e-05 | 1.5e-05 | 1e-05 | 1.333E-05 | 2.887E-06 | 1.000E-05 | 1.500E-05 |
| 1158 | 58 | V58W | A | 9.1e-06 | 1.5e-05 | 1.4e-05 | 1.270E-05 | 3.158E-06 | 9.100E-06 | 1.500E-05 |
| 1159 | 58 | V58Y | A | 1e-05 | 1e-05 | 9.9e-06 | 9.967E-06 | 5.774E-08 | 9.900E-06 | 1.000E-05 |
| 1160 | 58 | V58V | A | 1.5e-05 | 1.3e-05 | 1e-05 | 1.267E-05 | 2.517E-06 | 1.000E-05 | 1.500E-05 |
| 1161 | 59 | Q59A | A | 1.4e-05 | 1.1e-05 | 1.5e-05 | 1.333E-05 | 2.082E-06 | 1.100E-05 | 1.500E-05 |
| 1162 | 59 | Q59R | A | 1.3e-05 | 1.4e-05 | 1.5e-05 | 1.400E-05 | 1.000E-06 | 1.300E-05 | 1.500E-05 |
| 1163 | 59 | Q59N | A | 9.7e-06 | 1.3e-05 | 1.7e-05 | 1.323E-05 | 3.656E-06 | 9.700E-06 | 1.700E-05 |
| 1164 | 59 | Q59D | A | 1.6e-05 | 1e-05 | 1.5e-05 | 1.367E-05 | 3.215E-06 | 1.000E-05 | 1.600E-05 |
| 1165 | 59 | Q59C | A | 1.5e-05 | 1.5e-05 | 1.3e-05 | 1.433E-05 | 1.155E-06 | 1.300E-05 | 1.500E-05 |
| 1166 | 59 | Q59E | A | 1.5e-05 | 1e-05 | 1.5e-05 | 1.333E-05 | 2.887E-06 | 1.000E-05 | 1.500E-05 |
| 1167 | 59 | Q59Q | A | 1.6e-05 | 1.1e-05 | 1.3e-05 | 1.333E-05 | 2.517E-06 | 1.100E-05 | 1.600E-05 |
| 1168 | 59 | Q59G | A | 1.5e-05 | 1.4e-05 | 1e-05 | 1.300E-05 | 2.646E-06 | 1.000E-05 | 1.500E-05 |
| 1169 | 59 | Q59H | A | 1.6e-05 | 1.5e-05 | 1.5e-05 | 1.533E-05 | 5.774E-07 | 1.500E-05 | 1.600E-05 |
| 1170 | 59 | Q59I | A | 1.1e-05 | 1.5e-05 | 1.4e-05 | 1.333E-05 | 2.082E-06 | 1.100E-05 | 1.500E-05 |
| 1171 | 59 | Q59L | A | 9.7e-06 | 1.3e-05 | 1.4e-05 | 1.223E-05 | 2.250E-06 | 9.700E-06 | 1.400E-05 |
| 1172 | 59 | Q59K | A | 1e-05 | 1.5e-05 | 9.2e-06 | 1.140E-05 | 3.143E-06 | 9.200E-06 | 1.500E-05 |
| 1173 | 59 | Q59M | A | 1.1e-05 | 1.7e-05 | 1.6e-05 | 1.467E-05 | 3.215E-06 | 1.100E-05 | 1.700E-05 |
| 1174 | 59 | Q59F | A | 1.3e-05 | 1.6e-05 | 1.6e-05 | 1.500E-05 | 1.732E-06 | 1.300E-05 | 1.600E-05 |
| 1175 | 59 | Q59P | A | 1.1e-05 | 1.6e-05 | 1.1e-05 | 1.267E-05 | 2.887E-06 | 1.100E-05 | 1.600E-05 |
| 1176 | 59 | Q59S | A | 1e-05 | 1e-05 | 9.9e-06 | 9.967E-06 | 5.774E-08 | 9.900E-06 | 1.000E-05 |
| 1177 | 59 | Q59T | A | 9.6e-06 | 1.7e-05 | 1.1e-05 | 1.253E-05 | 3.931E-06 | 9.600E-06 | 1.700E-05 |
| 1178 | 59 | Q59W | A | 1.7e-05 | 9.6e-06 | 1.2e-05 | 1.287E-05 | 3.775E-06 | 9.600E-06 | 1.700E-05 |
| 1179 | 59 | Q59Y | A | 1.5e-05 | 1.6e-05 | 1.5e-05 | 1.533E-05 | 5.774E-07 | 1.500E-05 | 1.600E-05 |
| 1180 | 59 | Q59V | A | 1.2e-05 | 1.8e-05 | 1.6e-05 | 1.533E-05 | 3.055E-06 | 1.200E-05 | 1.800E-05 |
| 1181 | 60 | G60A | A | 1.5e-05 | 1.4e-05 | 9.7e-06 | 1.290E-05 | 2.816E-06 | 9.700E-06 | 1.500E-05 |
| 1182 | 60 | G60R | A | 9.7e-06 | 9.3e-06 | 9.7e-06 | 9.567E-06 | 2.309E-07 | 9.300E-06 | 9.700E-06 |
| 1183 | 60 | G60N | A | 1.1e-05 | 1.3e-05 | 1.1e-05 | 1.167E-05 | 1.155E-06 | 1.100E-05 | 1.300E-05 |
| 1184 | 60 | G60D | A | 1.1e-05 | 1.1e-05 | 1.1e-05 | 1.100E-05 | 2.075E-21 | 1.100E-05 | 1.100E-05 |
| 1185 | 60 | G60C | A | 1.1e-05 | 1.6e-05 | 1.3e-05 | 1.333E-05 | 2.517E-06 | 1.100E-05 | 1.600E-05 |
| 1186 | 60 | G60E | A | 1.3e-05 | 1.1e-05 | 1.6e-05 | 1.333E-05 | 2.517E-06 | 1.100E-05 | 1.600E-05 |
| 1187 | 60 | G60Q | A | 1.3e-05 | 1.6e-05 | 9.4e-06 | 1.280E-05 | 3.305E-06 | 9.400E-06 | 1.600E-05 |
| 1188 | 60 | G60G | A | 9.4e-06 | 1.4e-05 | 1.4e-05 | 1.247E-05 | 2.656E-06 | 9.400E-06 | 1.400E-05 |
| 1189 | 60 | G60H | A | 1.1e-05 | 1.6e-05 | 1.4e-05 | 1.367E-05 | 2.517E-06 | 1.100E-05 | 1.600E-05 |

| | | | | | | | | | | |
|---|---|---|---|---|---|---|---|---|---|---|
| 1190 | 60 | G60I | A | 8.2e-06 | 1e-05 | 9.4e-06 | 9.200E-06 | 9.165E-07 | 8.200E-06 | 1.000E-05 |
| 1191 | 60 | G60L | A | 9.2e-06 | 1.5e-05 | 1.6e-05 | 1.340E-05 | 3.672E-06 | 9.200E-06 | 1.600E-05 |
| 1192 | 60 | G60K | A | 1e-05 | 1.5e-05 | 1e-05 | 1.167E-05 | 2.887E-06 | 1.000E-05 | 1.500E-05 |
| 1193 | 60 | G60M | A | 1.6e-05 | 1.8e-05 | 1.1e-05 | 1.500E-05 | 3.606E-06 | 1.100E-05 | 1.800E-05 |
| 1194 | 60 | G60F | A | 1.9e-05 | 1.6e-05 | 1.4e-05 | 1.633E-05 | 2.517E-06 | 1.400E-05 | 1.900E-05 |
| 1195 | 60 | G60P | A | 1.1e-05 | 1.1e-05 | 9.4e-06 | 1.047E-05 | 9.238E-07 | 9.400E-06 | 1.100E-05 |
| 1196 | 60 | G60S | A | 1e-05 | 1.4e-05 | 1.3e-05 | 1.233E-05 | 2.082E-06 | 1.000E-05 | 1.400E-05 |
| 1197 | 60 | G60T | A | 1.5e-05 | 1.3e-05 | 1.5e-05 | 1.433E-05 | 1.155E-06 | 1.300E-05 | 1.500E-05 |
| 1198 | 60 | G60W | A | 1.7e-05 | 9.4e-06 | 9.8e-06 | 1.207E-05 | 4.277E-06 | 9.400E-06 | 1.700E-05 |
| 1199 | 60 | G60Y | A | 9.2e-06 | 1.4e-05 | 1.5e-05 | 1.273E-05 | 3.101E-06 | 9.200E-06 | 1.500E-05 |
| 1200 | 60 | G60V | A | 1.6e-05 | 1.5e-05 | 1.5e-05 | 1.533E-05 | 5.774E-07 | 1.500E-05 | 1.600E-05 |
| 1201 | 61 | Y61A | A | 1.5e-05 | 1e-05 | 9.9e-06 | 1.163E-05 | 2.916E-06 | 9.900E-06 | 1.500E-05 |
| 1202 | 61 | Y61R | A | 1.4e-05 | 1.1e-05 | 1.1e-05 | 1.200E-05 | 1.732E-06 | 1.100E-05 | 1.400E-05 |
| 1203 | 61 | Y61N | A | 1.6e-05 | 1e-05 | 1.5e-05 | 1.367E-05 | 3.215E-06 | 1.000E-05 | 1.600E-05 |
| 1204 | 61 | Y61D | A | 1e-05 | 1.3e-05 | 8.6e-06 | 1.053E-05 | 2.248E-06 | 8.600E-06 | 1.300E-05 |
| 1205 | 61 | Y61C | A | 8.6e-06 | 9.1e-06 | 1.6e-05 | 1.123E-05 | 4.136E-06 | 8.600E-06 | 1.600E-05 |
| 1206 | 61 | Y61E | A | 1.1e-05 | 1.1e-05 | 1.1e-05 | 1.100E-05 | 2.075E-21 | 1.100E-05 | 1.100E-05 |
| 1207 | 61 | Y61Q | A | 1.3e-05 | 1.4e-05 | 1.5e-05 | 1.400E-05 | 1.000E-06 | 1.300E-05 | 1.500E-05 |
| 1208 | 61 | Y61G | A | 1.4e-05 | 1.5e-05 | 1.1e-05 | 1.333E-05 | 2.082E-06 | 1.100E-05 | 1.500E-05 |
| 1209 | 61 | Y61H | A | 9.6e-06 | 1.4e-05 | 1.3e-05 | 1.220E-05 | 2.307E-06 | 9.600E-06 | 1.400E-05 |
| 1210 | 61 | Y61I | A | 1.2e-05 | 1.6e-05 | 1.4e-05 | 1.400E-05 | 2.000E-06 | 1.200E-05 | 1.600E-05 |
| 1211 | 61 | Y61L | A | 1.7e-05 | 1.6e-05 | 1.5e-05 | 1.600E-05 | 1.000E-06 | 1.500E-05 | 1.700E-05 |
| 1212 | 61 | Y61K | A | 1e-05 | 1.6e-05 | 1.5e-05 | 1.367E-05 | 3.215E-06 | 1.000E-05 | 1.600E-05 |
| 1213 | 61 | Y61M | A | 1e-05 | 1.6e-05 | 1.3e-05 | 1.300E-05 | 3.000E-06 | 1.000E-05 | 1.600E-05 |
| 1214 | 61 | Y61F | A | 1.4e-05 | 1.2e-05 | 1.7e-05 | 1.433E-05 | 2.517E-06 | 1.200E-05 | 1.700E-05 |
| 1215 | 61 | Y61P | A | 9.2e-06 | 1e-05 | 1.6e-05 | 1.173E-05 | 3.717E-06 | 9.200E-06 | 1.600E-05 |
| 1216 | 61 | Y61S | A | 1e-05 | 1.5e-05 | 1.1e-05 | 1.200E-05 | 2.646E-06 | 1.000E-05 | 1.500E-05 |
| 1217 | 61 | Y61T | A | 1.1e-05 | 1.4e-05 | 1.4e-05 | 1.300E-05 | 1.732E-06 | 1.100E-05 | 1.400E-05 |
| 1218 | 61 | Y61W | A | 1.7e-05 | 1.4e-05 | 6.1e-06 | 1.237E-05 | 5.631E-06 | 6.100E-06 | 1.700E-05 |
| 1219 | 61 | Y61Y | A | 1.5e-05 | 1.5e-05 | 1.1e-05 | 1.367E-05 | 2.309E-06 | 1.100E-05 | 1.500E-05 |
| 1220 | 61 | Y61V | A | 9.2e-06 | 1e-05 | 8.6e-06 | 9.267E-06 | 7.024E-07 | 8.600E-06 | 1.000E-05 |
| 1221 | 62 | V62A | A | 1.5e-05 | 1.4e-05 | 1e-05 | 1.300E-05 | 2.646E-06 | 1.000E-05 | 1.500E-05 |
| 1222 | 62 | V62R | A | 1.7e-05 | 1.3e-05 | 1.1e-05 | 1.367E-05 | 3.055E-06 | 1.100E-05 | 1.700E-05 |
| 1223 | 62 | V62N | A | 1.6e-05 | 1.3e-05 | 1.1e-05 | 1.333E-05 | 2.517E-06 | 1.100E-05 | 1.600E-05 |
| 1224 | 62 | V62D | A | 1.6e-05 | 7.9e-06 | 1.1e-05 | 1.163E-05 | 4.087E-06 | 7.900E-06 | 1.600E-05 |
| 1225 | 62 | V62C | A | 9.3e-06 | 1.4e-05 | 7.5e-06 | 1.027E-05 | 3.356E-06 | 7.500E-06 | 1.400E-05 |
| 1226 | 62 | V62E | A | 1.4e-05 | 9.1e-06 | 1.3e-05 | 1.203E-05 | 2.589E-06 | 9.100E-06 | 1.400E-05 |
| 1227 | 62 | V62Q | A | 1.1e-05 | 1.3e-05 | 1.6e-05 | 1.333E-05 | 2.517E-06 | 1.100E-05 | 1.600E-05 |
| 1228 | 62 | V62G | A | 8.9e-06 | 1.1e-05 | 1.6e-05 | 1.197E-05 | 3.647E-06 | 8.900E-06 | 1.600E-05 |
| 1229 | 62 | V62H | A | 1.4e-05 | 1.7e-05 | 1.5e-05 | 1.533E-05 | 1.528E-06 | 1.400E-05 | 1.700E-05 |
| 1230 | 62 | V62I | A | 1.7e-05 | 1.6e-05 | 1.4e-05 | 1.567E-05 | 1.528E-06 | 1.400E-05 | 1.700E-05 |
| 1231 | 62 | V62L | A | 1.5e-05 | 1.2e-05 | 1.1e-05 | 1.267E-05 | 2.082E-06 | 1.100E-05 | 1.500E-05 |
| 1232 | 62 | V62K | A | 8.6e-06 | 9.3e-06 | 1.4e-05 | 1.063E-05 | 2.937E-06 | 8.600E-06 | 1.400E-05 |
| 1233 | 62 | V62M | A | 9.5e-06 | 1.1e-05 | 1.4e-05 | 1.150E-05 | 2.291E-06 | 9.500E-06 | 1.400E-05 |
| 1234 | 62 | V62F | A | 2.3e-05 | 1.5e-05 | 9.4e-06 | 1.580E-05 | 6.835E-06 | 9.400E-06 | 2.300E-05 |
| 1235 | 62 | V62P | A | 1.1e-05 | 1e-05 | 1.1e-05 | 1.067E-05 | 5.774E-07 | 1.000E-05 | 1.100E-05 |
| 1236 | 62 | V62S | A | 1.2e-05 | 1.5e-05 | 1.4e-05 | 1.367E-05 | 1.528E-06 | 1.200E-05 | 1.500E-05 |
| 1237 | 62 | V62T | A | 1.4e-05 | 1.6e-05 | 1.1e-05 | 1.367E-05 | 2.517E-06 | 1.100E-05 | 1.600E-05 |
| 1238 | 62 | V62W | A | 1.4e-05 | 1e-05 | 1.1e-05 | 1.167E-05 | 2.082E-06 | 1.000E-05 | 1.400E-05 |
| 1239 | 62 | V62Y | A | 1.7e-05 | 6.1e-06 | 1.1e-05 | 1.137E-05 | 5.459E-06 | 6.100E-06 | 1.700E-05 |
| 1240 | 62 | V62V | A | 1.2e-05 | 1.1e-05 | 8.8e-06 | 1.060E-05 | 1.637E-06 | 8.800E-06 | 1.200E-05 |
| 1241 | 63 | L63A | A | 8.4e-06 | 1.3e-05 | 1.1e-05 | 1.080E-05 | 2.307E-06 | 8.400E-06 | 1.300E-05 |
| 1242 | 63 | L63R | A | 1.5e-05 | 8.9e-06 | 6.7e-06 | 1.020E-05 | 4.300E-06 | 6.700E-06 | 1.500E-05 |
| 1243 | 63 | L63N | A | 1.1e-05 | 1.5e-05 | 1.3e-05 | 1.300E-05 | 2.000E-06 | 1.100E-05 | 1.500E-05 |
| 1244 | 63 | L63D | A | 1.4e-05 | 1.6e-05 | 1.1e-05 | 1.367E-05 | 2.517E-06 | 1.100E-05 | 1.600E-05 |
| 1245 | 63 | L63C | A | 1.4e-05 | 1e-05 | 1.3e-05 | 1.233E-05 | 2.082E-06 | 1.000E-05 | 1.400E-05 |
| 1246 | 63 | L63E | A | 1e-05 | 6e-06 | 1.4e-05 | 1.000E-05 | 4.000E-06 | 6.000E-06 | 1.400E-05 |
| 1247 | 63 | L63Q | A | 1.6e-05 | 1.6e-05 | 1e-05 | 1.400E-05 | 3.464E-06 | 1.000E-05 | 1.600E-05 |
| 1248 | 63 | L63G | A | 1.6e-05 | 1e-05 | 1.5e-05 | 1.367E-05 | 3.215E-06 | 1.000E-05 | 1.600E-05 |
| 1249 | 63 | L63H | A | 1e-05 | 1e-05 | 1.5e-05 | 1.167E-05 | 2.887E-06 | 1.000E-05 | 1.500E-05 |
| 1250 | 63 | L63I | A | 1.5e-05 | 1.1e-05 | 1e-05 | 1.200E-05 | 2.646E-06 | 1.000E-05 | 1.500E-05 |
| 1251 | 63 | L63L | A | 1.2e-05 | 1.5e-05 | 1.1e-05 | 1.267E-05 | 2.082E-06 | 1.100E-05 | 1.500E-05 |
| 1252 | 63 | L63K | A | 1.5e-05 | 1e-05 | 1.6e-05 | 1.367E-05 | 3.215E-06 | 1.000E-05 | 1.600E-05 |
| 1253 | 63 | L63M | A | 1.6e-05 | 1.1e-05 | 1.2e-05 | 1.300E-05 | 2.646E-06 | 1.100E-05 | 1.600E-05 |
| 1254 | 63 | L63F | A | 1.2e-05 | 1.5e-05 | 9.7e-06 | 1.223E-05 | 2.658E-06 | 9.700E-06 | 1.500E-05 |
| 1255 | 63 | L63P | A | 1.2e-05 | 1.7e-05 | 1.5e-05 | 1.467E-05 | 2.517E-06 | 1.200E-05 | 1.700E-05 |
| 1256 | 63 | L63S | A | 1e-05 | 1.2e-05 | 1.3e-05 | 1.167E-05 | 1.528E-06 | 1.000E-05 | 1.300E-05 |
| 1257 | 63 | L63T | A | 1.5e-05 | 1.4e-05 | 1.5e-05 | 1.467E-05 | 5.774E-07 | 1.400E-05 | 1.500E-05 |
| 1258 | 63 | L63W | A | 1.6e-05 | 1.5e-05 | 1.1e-05 | 1.400E-05 | 2.646E-06 | 1.100E-05 | 1.600E-05 |
| 1259 | 63 | L63Y | A | 1.4e-05 | 9.7e-06 | 1.3e-05 | 1.223E-05 | 2.250E-06 | 9.700E-06 | 1.400E-05 |
| 1260 | 63 | L63V | A | 1.2e-05 | 1.6e-05 | 1e-05 | 1.267E-05 | 3.055E-06 | 1.000E-05 | 1.600E-05 |
| 1261 | 64 | I64A | A | 1.4e-05 | 1.1e-05 | 9.6e-06 | 1.153E-05 | 2.248E-06 | 9.600E-06 | 1.400E-05 |
| 1262 | 64 | I64R | A | 9.6e-06 | 1.1e-05 | 1.2e-05 | 1.087E-05 | 1.206E-06 | 9.600E-06 | 1.200E-05 |
| 1263 | 64 | I64N | A | 1.5e-05 | 8.6e-06 | 1.2e-05 | 1.187E-05 | 3.202E-06 | 8.600E-06 | 1.500E-05 |
| 1264 | 64 | I64D | A | 1.6e-05 | 1.1e-05 | 1.6e-05 | 1.433E-05 | 2.887E-06 | 1.100E-05 | 1.600E-05 |
| 1265 | 64 | I64C | A | 1.5e-05 | 1.1e-05 | 1e-05 | 1.200E-05 | 2.646E-06 | 1.000E-05 | 1.500E-05 |
| 1266 | 64 | I64E | A | 1.4e-05 | 1e-05 | 1.6e-05 | 1.333E-05 | 3.055E-06 | 1.000E-05 | 1.600E-05 |
| 1267 | 64 | I64Q | A | 1.3e-05 | 1.5e-05 | 1.4e-05 | 1.400E-05 | 1.000E-06 | 1.300E-05 | 1.500E-05 |
| 1268 | 64 | I64G | A | 1.5e-05 | 9e-06 | 9.9e-06 | 1.130E-05 | 3.236E-06 | 9.000E-06 | 1.500E-05 |
| 1269 | 64 | I64H | A | 1.1e-05 | 1.1e-05 | 1.5e-05 | 1.233E-05 | 2.309E-06 | 1.100E-05 | 1.500E-05 |
| 1270 | 64 | I64I | A | 1.3e-05 | 1.5e-05 | 1.4e-05 | 1.400E-05 | 1.000E-06 | 1.300E-05 | 1.500E-05 |
| 1271 | 64 | I64L | A | 1.4e-05 | 1.2e-05 | 7.9e-06 | 1.130E-05 | 3.110E-06 | 7.900E-06 | 1.400E-05 |
| 1272 | 64 | I64K | A | 1.4e-05 | 9.8e-06 | 1.3e-05 | 1.227E-05 | 2.194E-06 | 9.800E-06 | 1.400E-05 |
| 1273 | 64 | I64M | A | 9.9e-06 | 1.3e-05 | 1.1e-05 | 1.130E-05 | 1.572E-06 | 9.900E-06 | 1.300E-05 |
| 1274 | 64 | I64F | A | 9.7e-06 | 8.5e-06 | 1.4e-05 | 1.073E-05 | 2.892E-06 | 8.500E-06 | 1.400E-05 |

| | | | | | | | | | | |
|---|---|---|---|---|---|---|---|---|---|---|
| 1275 | 64 | I64P | A | 1.1e-05 | 1.2e-05 | 1.6e-05 | 1.300E-05 | 2.646E-06 | 1.100E-05 | 1.600E-05 |
| 1276 | 64 | I64S | A | 1.2e-05 | 1.4e-05 | 9.4e-06 | 1.180E-05 | 2.307E-06 | 9.400E-06 | 1.400E-05 |
| 1277 | 64 | I64T | A | 1.1e-05 | 1.1e-05 | 1.8e-05 | 1.333E-05 | 4.041E-06 | 1.100E-05 | 1.800E-05 |
| 1278 | 64 | I64W | A | 1.6e-05 | 1e-05 | 9.6e-06 | 1.187E-05 | 3.585E-06 | 9.600E-06 | 1.600E-05 |
| 1279 | 64 | I64Y | A | 1.4e-05 | 1e-05 | 1.3e-05 | 1.233E-05 | 2.082E-06 | 1.000E-05 | 1.400E-05 |
| 1280 | 64 | I64V | A | 1.6e-05 | 1.3e-05 | 1.6e-05 | 1.500E-05 | 1.732E-06 | 1.300E-05 | 1.600E-05 |
| 1281 | 65 | A65A | A | 1e-05 | 1.4e-05 | 1.6e-05 | 1.333E-05 | 3.055E-06 | 1.000E-05 | 1.600E-05 |
| 1282 | 65 | A65R | A | 1.3e-05 | 1.4e-05 | 1.7e-05 | 1.467E-05 | 2.082E-06 | 1.300E-05 | 1.700E-05 |
| 1283 | 65 | A65N | A | 1.6e-05 | 1.4e-05 | 1.1e-05 | 1.367E-05 | 2.517E-06 | 1.100E-05 | 1.600E-05 |
| 1284 | 65 | A65D | A | 9.7e-06 | 1.2e-05 | 1.7e-05 | 1.290E-05 | 3.732E-06 | 9.700E-06 | 1.700E-05 |
| 1285 | 65 | A65C | A | 1e-05 | 8.6e-06 | 1e-05 | 9.533E-06 | 8.083E-07 | 8.600E-06 | 1.000E-05 |
| 1286 | 65 | A65E | A | 1e-05 | 1.5e-05 | 1e-05 | 1.167E-05 | 2.887E-06 | 1.000E-05 | 1.500E-05 |
| 1287 | 65 | A65Q | A | 9.7e-06 | 9.6e-06 | 9.6e-06 | 9.633E-06 | 5.774E-08 | 9.600E-06 | 9.700E-06 |
| 1288 | 65 | A65G | A | 1.5e-05 | 1.6e-05 | 1.5e-05 | 1.533E-05 | 5.774E-07 | 1.500E-05 | 1.600E-05 |
| 1289 | 65 | A65H | A | 1e-05 | 1.3e-05 | 8e-06 | 1.033E-05 | 2.517E-06 | 8.000E-06 | 1.300E-05 |
| 1290 | 65 | A65I | A | 9.1e-06 | 1e-05 | 1.1e-05 | 1.003E-05 | 9.504E-07 | 9.100E-06 | 1.100E-05 |
| 1291 | 65 | A65L | A | 1.6e-05 | 1e-05 | 6.3e-06 | 1.077E-05 | 4.895E-06 | 6.300E-06 | 1.600E-05 |
| 1292 | 65 | A65K | A | 1.4e-05 | 1.1e-05 | 1.3e-05 | 1.267E-05 | 1.528E-06 | 1.100E-05 | 1.400E-05 |
| 1293 | 65 | A65M | A | 1.5e-05 | 1.2e-05 | 1.4e-05 | 1.367E-05 | 1.528E-06 | 1.200E-05 | 1.500E-05 |
| 1294 | 65 | A65F | A | 1.5e-05 | 1e-05 | 1.5e-05 | 1.333E-05 | 2.887E-06 | 1.000E-05 | 1.500E-05 |
| 1295 | 65 | A65P | A | 1.5e-05 | 9.8e-06 | 1.5e-05 | 1.327E-05 | 3.002E-06 | 9.800E-06 | 1.500E-05 |
| 1296 | 65 | A65S | A | 1.4e-05 | 2.3e-05 | 1.3e-05 | 1.667E-05 | 5.508E-06 | 1.300E-05 | 2.300E-05 |
| 1297 | 65 | A65T | A | 1.2e-05 | 9.7e-06 | 1.1e-05 | 1.090E-05 | 1.153E-06 | 9.700E-06 | 1.200E-05 |
| 1298 | 65 | A65W | A | 1.6e-05 | 1.6e-05 | 1.5e-05 | 1.567E-05 | 5.774E-07 | 1.500E-05 | 1.600E-05 |
| 1299 | 65 | A65Y | A | 9.5e-06 | 1.7e-05 | 1.2e-05 | 1.283E-05 | 3.819E-06 | 9.500E-06 | 1.700E-05 |
| 1300 | 65 | A65V | A | 1.4e-05 | 1.3e-05 | 8e-06 | 1.167E-05 | 3.215E-06 | 8.000E-06 | 1.400E-05 |
| 1301 | 66 | H66A | A | 1.1e-05 | 1.6e-05 | 7.1e-06 | 1.137E-05 | 4.461E-06 | 7.100E-06 | 1.600E-05 |
| 1302 | 66 | H66R | A | 1.1e-05 | 9.6e-06 | 1e-05 | 1.020E-05 | 7.211E-07 | 9.600E-06 | 1.100E-05 |
| 1303 | 66 | H66N | A | 1.6e-05 | 1.3e-05 | 1.5e-05 | 1.467E-05 | 1.528E-06 | 1.300E-05 | 1.600E-05 |
| 1304 | 66 | H66D | A | 1.3e-05 | 9e-06 | 1.1e-05 | 1.100E-05 | 2.000E-06 | 9.000E-06 | 1.300E-05 |
| 1305 | 66 | H66C | A | 1e-05 | 1.1e-05 | 1.4e-05 | 1.167E-05 | 2.082E-06 | 1.000E-05 | 1.400E-05 |
| 1306 | 66 | H66E | A | 1.1e-05 | 1.3e-05 | 1.2e-05 | 1.200E-05 | 1.000E-06 | 1.100E-05 | 1.300E-05 |
| 1307 | 66 | H66Q | A | 8.9e-06 | 1.5e-05 | 1.5e-05 | 1.297E-05 | 3.522E-06 | 8.900E-06 | 1.500E-05 |
| 1308 | 66 | H66G | A | 9.2e-06 | 1.5e-05 | 1.3e-05 | 1.240E-05 | 2.946E-06 | 9.200E-06 | 1.500E-05 |
| 1309 | 66 | H66H | A | 1.6e-05 | 1.5e-05 | 1.7e-05 | 1.600E-05 | 1.000E-06 | 1.500E-05 | 1.700E-05 |
| 1310 | 66 | H66I | A | 1.6e-05 | 1.8e-05 | 1.5e-05 | 1.633E-05 | 1.528E-06 | 1.500E-05 | 1.800E-05 |
| 1311 | 66 | H66L | A | 1.2e-05 | 1.4e-05 | 1.2e-05 | 1.267E-05 | 1.155E-06 | 1.200E-05 | 1.400E-05 |
| 1312 | 66 | H66K | A | 1.1e-05 | 1.1e-05 | 1.6e-05 | 1.267E-05 | 2.887E-06 | 1.100E-05 | 1.600E-05 |
| 1313 | 66 | H66M | A | 1.1e-05 | 1e-05 | 1.1e-05 | 1.067E-05 | 5.774E-07 | 1.000E-05 | 1.100E-05 |
| 1314 | 66 | H66F | A | 1.1e-05 | 1.4e-05 | 1.4e-05 | 1.300E-05 | 1.732E-06 | 1.100E-05 | 1.400E-05 |
| 1315 | 66 | H66P | A | 1.5e-05 | 9.7e-06 | 9.1e-06 | 1.127E-05 | 3.247E-06 | 9.100E-06 | 1.500E-05 |
| 1316 | 66 | H66S | A | 1.6e-05 | 1.5e-05 | 1.1e-05 | 1.400E-05 | 2.646E-06 | 1.100E-05 | 1.600E-05 |
| 1317 | 66 | H66T | A | 1e-05 | 1.6e-05 | 1.1e-05 | 1.233E-05 | 3.215E-06 | 1.000E-05 | 1.600E-05 |
| 1318 | 66 | H66W | A | 5.8e-06 | 9.3e-06 | 1.6e-05 | 1.037E-05 | 5.183E-06 | 5.800E-06 | 1.600E-05 |
| 1319 | 66 | H66Y | A | 8e-06 | 1.1e-05 | 1e-05 | 9.667E-06 | 1.528E-06 | 8.000E-06 | 1.100E-05 |
| 1320 | 66 | H66V | A | 1.5e-05 | 1.5e-05 | 1.6e-05 | 1.533E-05 | 5.774E-07 | 1.500E-05 | 1.600E-05 |
| 1321 | 67 | N67A | A | 1.1e-05 | 1.1e-05 | 1.7e-05 | 1.300E-05 | 3.464E-06 | 1.100E-05 | 1.700E-05 |
| 1322 | 67 | N67R | A | 1.4e-05 | 1.3e-05 | 7.4e-06 | 1.147E-05 | 3.557E-06 | 7.400E-06 | 1.400E-05 |
| 1323 | 67 | N67N | A | 1.5e-05 | 1.1e-05 | 1.5e-05 | 1.367E-05 | 2.309E-06 | 1.100E-05 | 1.500E-05 |
| 1324 | 67 | N67D | A | 1.3e-05 | 1.6e-05 | 1.4e-05 | 1.433E-05 | 1.528E-06 | 1.300E-05 | 1.600E-05 |
| 1325 | 67 | N67C | A | 1.5e-05 | 1.1e-05 | 1.6e-05 | 1.400E-05 | 2.646E-06 | 1.100E-05 | 1.600E-05 |
| 1326 | 67 | N67E | A | 9.9e-06 | 1.4e-05 | 1.1e-05 | 1.163E-05 | 2.122E-06 | 9.900E-06 | 1.400E-05 |
| 1327 | 67 | N67Q | A | 1.5e-05 | 1.5e-05 | 1.3e-05 | 1.433E-05 | 1.155E-06 | 1.300E-05 | 1.500E-05 |
| 1328 | 67 | N67G | A | 9.2e-06 | 1e-05 | 1.6e-05 | 1.173E-05 | 3.717E-06 | 9.200E-06 | 1.600E-05 |
| 1329 | 67 | N67H | A | 1e-05 | 1.2e-05 | 1.1e-05 | 1.100E-05 | 1.000E-06 | 1.000E-05 | 1.200E-05 |
| 1330 | 67 | N67I | A | 1.5e-05 | 1.5e-05 | 8e-06 | 1.267E-05 | 4.041E-06 | 8.000E-06 | 1.500E-05 |
| 1331 | 67 | N67L | A | 1.5e-05 | 1e-05 | 1.3e-05 | 1.267E-05 | 2.517E-06 | 1.000E-05 | 1.500E-05 |
| 1332 | 67 | N67K | A | 1.1e-05 | 1.2e-05 | 1.6e-05 | 1.300E-05 | 2.646E-06 | 1.100E-05 | 1.600E-05 |
| 1333 | 67 | N67M | A | 1.7e-05 | 7.7e-06 | 1.4e-05 | 1.290E-05 | 4.747E-06 | 7.700E-06 | 1.700E-05 |
| 1334 | 67 | N67F | A | 9e-06 | 1e-05 | 1e-05 | 9.667E-06 | 5.774E-07 | 9.000E-06 | 1.000E-05 |
| 1335 | 67 | N67P | A | 1.5e-05 | 1.1e-05 | 1.4e-05 | 1.333E-05 | 2.082E-06 | 1.100E-05 | 1.500E-05 |
| 1336 | 67 | N67S | A | 1.4e-05 | 1.7e-05 | 1.1e-05 | 1.400E-05 | 3.000E-06 | 1.100E-05 | 1.700E-05 |
| 1337 | 67 | N67T | A | 8.1e-06 | 1.6e-05 | 1.6e-05 | 1.337E-05 | 4.561E-06 | 8.100E-06 | 1.600E-05 |
| 1338 | 67 | N67W | A | 1.3e-05 | 1.7e-05 | 1.3e-05 | 1.433E-05 | 2.309E-06 | 1.300E-05 | 1.700E-05 |
| 1339 | 67 | N67Y | A | 1.6e-05 | 1.2e-05 | 8.5e-06 | 1.217E-05 | 3.753E-06 | 8.500E-06 | 1.600E-05 |
| 1340 | 67 | N67V | A | 1.6e-05 | 1.4e-05 | 8.8e-06 | 1.293E-05 | 3.717E-06 | 8.800E-06 | 1.600E-05 |
| 1341 | 68 | Q68A | A | 1.6e-05 | 1.4e-05 | 1.4e-05 | 1.467E-05 | 1.155E-06 | 1.400E-05 | 1.600E-05 |
| 1342 | 68 | Q68R | A | 1.1e-05 | 1.6e-05 | 1.5e-05 | 1.400E-05 | 2.646E-06 | 1.100E-05 | 1.600E-05 |
| 1343 | 68 | Q68N | A | 1.4e-05 | 1.3e-05 | 1.3e-05 | 1.333E-05 | 5.774E-07 | 1.300E-05 | 1.400E-05 |
| 1344 | 68 | Q68D | A | 1.5e-05 | 1.5e-05 | 1.7e-05 | 1.567E-05 | 1.155E-06 | 1.500E-05 | 1.700E-05 |
| 1345 | 68 | Q68C | A | 1e-05 | 1.4e-05 | 1.5e-05 | 1.300E-05 | 2.646E-06 | 1.000E-05 | 1.500E-05 |
| 1346 | 68 | Q68E | A | 9e-06 | 1.1e-05 | 1.1e-05 | 1.033E-05 | 1.155E-06 | 9.000E-06 | 1.100E-05 |
| 1347 | 68 | Q68Q | A | 9.8e-06 | 1.4e-05 | 1.3e-05 | 1.227E-05 | 2.194E-06 | 9.800E-06 | 1.400E-05 |
| 1348 | 68 | Q68G | A | 1.6e-05 | 1.1e-05 | 1.5e-05 | 1.400E-05 | 2.646E-06 | 1.100E-05 | 1.600E-05 |
| 1349 | 68 | Q68H | A | 1.6e-05 | 1.1e-05 | 1.3e-05 | 1.333E-05 | 2.517E-06 | 1.100E-05 | 1.600E-05 |
| 1350 | 68 | Q68I | A | 8.9e-06 | 1.5e-05 | 9.5e-06 | 1.113E-05 | 3.362E-06 | 8.900E-06 | 1.500E-05 |
| 1351 | 68 | Q68L | A | 1e-05 | 1.2e-05 | 1.7e-05 | 1.300E-05 | 3.606E-06 | 1.000E-05 | 1.700E-05 |
| 1352 | 68 | Q68K | A | 1.4e-05 | 1.6e-05 | 1.1e-05 | 1.367E-05 | 2.517E-06 | 1.100E-05 | 1.600E-05 |
| 1353 | 68 | Q68M | A | 9.4e-06 | 1e-05 | 1.6e-05 | 1.180E-05 | 3.650E-06 | 9.400E-06 | 1.600E-05 |
| 1354 | 68 | Q68F | A | 1.2e-05 | 1.2e-05 | 1.6e-05 | 1.333E-05 | 2.309E-06 | 1.200E-05 | 1.600E-05 |
| 1355 | 68 | Q68P | A | 1.1e-05 | 9.5e-06 | 1.5e-05 | 1.183E-05 | 2.843E-06 | 9.500E-06 | 1.500E-05 |
| 1356 | 68 | Q68S | A | 1.4e-05 | 1.7e-05 | 1.7e-05 | 1.600E-05 | 1.732E-06 | 1.400E-05 | 1.700E-05 |
| 1357 | 68 | Q68T | A | 1e-05 | 6e-06 | 9.8e-06 | 8.600E-06 | 2.254E-06 | 6.000E-06 | 1.000E-05 |
| 1358 | 68 | Q68W | A | 1e-05 | 1.6e-05 | 1.1e-05 | 1.233E-05 | 3.215E-06 | 1.000E-05 | 1.600E-05 |
| 1359 | 68 | Q68Y | A | 1.7e-05 | 8.2e-06 | 8.9e-06 | 1.137E-05 | 4.891E-06 | 8.200E-06 | 1.700E-05 |

| | | | | | | | | | | |
|---|---|---|---|---|---|---|---|---|---|---|
| 1360 | 68 | Q68V | A | 1.6e-05 | 1.7e-05 | 1.4e-05 | 1.567E-05 | 1.528E-06 | 1.400E-05 | 1.700E-05 |
| 1361 | 69 | V69A | A | 1.2e-05 | 1.5e-05 | 1.6e-05 | 1.433E-05 | 2.082E-06 | 1.200E-05 | 1.600E-05 |
| 1362 | 69 | V69R | A | 1.4e-05 | 1.3e-05 | 1e-05 | 1.233E-05 | 2.082E-06 | 1.000E-05 | 1.400E-05 |
| 1363 | 69 | V69N | A | 1e-05 | 1.6e-05 | 1.4e-05 | 1.333E-05 | 3.055E-06 | 1.000E-05 | 1.600E-05 |
| 1364 | 69 | V69D | A | 9.9e-06 | 1.7e-05 | 1.5e-05 | 1.397E-05 | 3.661E-06 | 9.900E-06 | 1.700E-05 |
| 1365 | 69 | V69C | A | 1.4e-05 | 9.8e-06 | 1.4e-05 | 1.260E-05 | 2.425E-06 | 9.800E-06 | 1.400E-05 |
| 1366 | 69 | V69E | A | 1.2e-05 | 1.1e-05 | 1.5e-05 | 1.267E-05 | 2.082E-06 | 1.100E-05 | 1.500E-05 |
| 1367 | 69 | V69Q | A | 1.4e-05 | 8.3e-06 | 1.4e-05 | 1.210E-05 | 3.291E-06 | 8.300E-06 | 1.400E-05 |
| 1368 | 69 | V69G | A | 1.4e-05 | 1.6e-05 | 1.6e-05 | 1.533E-05 | 1.155E-06 | 1.400E-05 | 1.600E-05 |
| 1369 | 69 | V69H | A | 1.5e-05 | 1.5e-05 | 1.4e-05 | 1.467E-05 | 5.774E-07 | 1.400E-05 | 1.500E-05 |
| 1370 | 69 | V69I | A | 1.6e-05 | 1.5e-05 | 8.8e-06 | 1.327E-05 | 3.900E-06 | 8.800E-06 | 1.600E-05 |
| 1371 | 69 | V69L | A | 1.1e-05 | 1.4e-05 | 1.1e-05 | 1.200E-05 | 1.732E-06 | 1.100E-05 | 1.400E-05 |
| 1372 | 69 | V69K | A | 1.4e-05 | 9.4e-06 | 1.1e-05 | 1.147E-05 | 2.335E-06 | 9.400E-06 | 1.400E-05 |
| 1373 | 69 | V69M | A | 1.2e-05 | 1.6e-05 | 1.1e-05 | 1.300E-05 | 2.646E-06 | 1.100E-05 | 1.600E-05 |
| 1374 | 69 | V69F | A | 1.2e-05 | 1.5e-05 | 1.7e-05 | 1.467E-05 | 2.517E-06 | 1.200E-05 | 1.700E-05 |
| 1375 | 69 | V69P | A | 9.9e-06 | 1.1e-05 | 1.8e-05 | 1.297E-05 | 4.394E-06 | 9.900E-06 | 1.800E-05 |
| 1376 | 69 | V69S | A | 1.5e-05 | 1.4e-05 | 1.6e-05 | 1.500E-05 | 1.000E-06 | 1.400E-05 | 1.600E-05 |
| 1377 | 69 | V69T | A | 1e-05 | 1.4e-05 | 1e-05 | 1.133E-05 | 2.309E-06 | 1.000E-05 | 1.400E-05 |
| 1378 | 69 | V69W | A | 1.5e-05 | 8.8e-06 | 1.4e-05 | 1.260E-05 | 3.329E-06 | 8.800E-06 | 1.500E-05 |
| 1379 | 69 | V69Y | A | 1.4e-05 | 1.4e-05 | 8.4e-06 | 1.213E-05 | 3.233E-06 | 8.400E-06 | 1.400E-05 |
| 1380 | 69 | V69V | A | 1.1e-05 | 1.5e-05 | 1.5e-05 | 1.367E-05 | 2.309E-06 | 1.100E-05 | 1.500E-05 |
| 1381 | 70 | R70A | A | 1.2e-05 | 1.7e-05 | 1e-05 | 1.300E-05 | 3.606E-06 | 1.000E-05 | 1.700E-05 |
| 1382 | 70 | R70R | A | 9.2e-06 | 1.3e-05 | 1.3e-05 | 1.173E-05 | 2.194E-06 | 9.200E-06 | 1.300E-05 |
| 1383 | 70 | R70N | A | 1.6e-05 | 1.2e-05 | 1.6e-05 | 1.467E-05 | 2.309E-06 | 1.200E-05 | 1.600E-05 |
| 1384 | 70 | R70D | A | 1.8e-05 | 1.6e-05 | 1.5e-05 | 1.633E-05 | 1.528E-06 | 1.500E-05 | 1.800E-05 |
| 1385 | 70 | R70C | A | 1.6e-05 | 1.5e-05 | 1.5e-05 | 1.533E-05 | 5.774E-07 | 1.500E-05 | 1.600E-05 |
| 1386 | 70 | R70E | A | 1.5e-05 | 1.6e-05 | 9.7e-06 | 1.357E-05 | 3.386E-06 | 9.700E-06 | 1.600E-05 |
| 1387 | 70 | R70Q | A | 1.2e-05 | 1.4e-05 | 1.5e-05 | 1.367E-05 | 1.528E-06 | 1.200E-05 | 1.500E-05 |
| 1388 | 70 | R70G | A | 1.6e-05 | 1.5e-05 | 1.4e-05 | 1.500E-05 | 1.000E-06 | 1.400E-05 | 1.600E-05 |
| 1389 | 70 | R70H | A | 9.5e-06 | 1.5e-05 | 1.6e-05 | 1.350E-05 | 3.500E-06 | 9.500E-06 | 1.600E-05 |
| 1390 | 70 | R70I | A | 1.7e-05 | 1.2e-05 | 1.3e-05 | 1.400E-05 | 2.646E-06 | 1.200E-05 | 1.700E-05 |
| 1391 | 70 | R70L | A | 7.3e-06 | 1.2e-05 | 1.4e-05 | 1.110E-05 | 3.439E-06 | 7.300E-06 | 1.400E-05 |
| 1392 | 70 | R70K | A | 1.6e-05 | 1.1e-05 | 1.6e-05 | 1.433E-05 | 2.887E-06 | 1.100E-05 | 1.600E-05 |
| 1393 | 70 | R70M | A | 9e-06 | 1.6e-05 | 1.5e-05 | 1.333E-05 | 3.786E-06 | 9.000E-06 | 1.600E-05 |
| 1394 | 70 | R70F | A | 1e-05 | 1.5e-05 | 1.4e-05 | 1.300E-05 | 2.646E-06 | 1.000E-05 | 1.500E-05 |
| 1395 | 70 | R70P | A | 1.4e-05 | 1.5e-05 | 8.4e-06 | 1.247E-05 | 3.557E-06 | 8.400E-06 | 1.500E-05 |
| 1396 | 70 | R70S | A | 9.4e-06 | 1.4e-05 | 1.3e-05 | 1.213E-05 | 2.419E-06 | 9.400E-06 | 1.400E-05 |
| 1397 | 70 | R70T | A | 9e-06 | 8.9e-06 | 1.2e-05 | 9.967E-06 | 1.762E-06 | 8.900E-06 | 1.200E-05 |
| 1398 | 70 | R70W | A | 1.2e-05 | 8.2e-06 | 1e-05 | 1.007E-05 | 1.901E-06 | 8.200E-06 | 1.200E-05 |
| 1399 | 70 | R70Y | A | 1.5e-05 | 1.3e-05 | 1.6e-05 | 1.467E-05 | 1.528E-06 | 1.300E-05 | 1.600E-05 |
| 1400 | 70 | R70V | A | 1.5e-05 | 1e-05 | 1.5e-05 | 1.333E-05 | 2.887E-06 | 1.000E-05 | 1.500E-05 |
| 1401 | 71 | Q71A | A | 1.5e-05 | 1.1e-05 | 1.6e-05 | 1.400E-05 | 2.646E-06 | 1.100E-05 | 1.600E-05 |
| 1402 | 71 | Q71R | A | 1.3e-05 | 1.5e-05 | 1.5e-05 | 1.433E-05 | 1.155E-06 | 1.300E-05 | 1.500E-05 |
| 1403 | 71 | Q71N | A | 1.8e-05 | 1.7e-05 | 9.3e-06 | 1.477E-05 | 4.761E-06 | 9.300E-06 | 1.800E-05 |
| 1404 | 71 | Q71D | A | 7.6e-06 | 1.1e-05 | 8.8e-06 | 9.133E-06 | 1.724E-06 | 7.600E-06 | 1.100E-05 |
| 1405 | 71 | Q71C | A | 1.3e-05 | 8.9e-06 | 1.4e-05 | 1.197E-05 | 2.702E-06 | 8.900E-06 | 1.400E-05 |
| 1406 | 71 | Q71E | A | 1.3e-05 | 1.5e-05 | 1.1e-05 | 1.300E-05 | 2.000E-06 | 1.100E-05 | 1.500E-05 |
| 1407 | 71 | Q71Q | A | 1.5e-05 | 9.9e-06 | 8.9e-06 | 1.127E-05 | 3.272E-06 | 8.900E-06 | 1.500E-05 |
| 1408 | 71 | Q71G | A | 1.3e-05 | 1.5e-05 | 1.8e-05 | 1.533E-05 | 2.517E-06 | 1.300E-05 | 1.800E-05 |
| 1409 | 71 | Q71H | A | 1.6e-05 | 1.6e-05 | 1e-05 | 1.400E-05 | 3.464E-06 | 1.000E-05 | 1.600E-05 |
| 1410 | 71 | Q71I | A | 1.4e-05 | 1.1e-05 | 1.4e-05 | 1.300E-05 | 1.732E-06 | 1.100E-05 | 1.400E-05 |
| 1411 | 71 | Q71L | A | 1.5e-05 | 1e-05 | 1.6e-05 | 1.367E-05 | 3.215E-06 | 1.000E-05 | 1.600E-05 |
| 1412 | 71 | Q71K | A | 1.4e-05 | 1.2e-05 | 9.8e-06 | 1.193E-05 | 2.101E-06 | 9.800E-06 | 1.400E-05 |
| 1413 | 71 | Q71M | A | 1.4e-05 | 1.1e-05 | 9.1e-06 | 1.137E-05 | 2.470E-06 | 9.100E-06 | 1.400E-05 |
| 1414 | 71 | Q71F | A | 1.4e-05 | 1.2e-05 | 1.2e-05 | 1.267E-05 | 1.155E-06 | 1.200E-05 | 1.400E-05 |
| 1415 | 71 | Q71P | A | 1.7e-05 | 1e-05 | 1.6e-05 | 1.433E-05 | 3.786E-06 | 1.000E-05 | 1.700E-05 |
| 1416 | 71 | Q71S | A | 1.5e-05 | 1.5e-05 | 1.7e-05 | 1.567E-05 | 1.155E-06 | 1.500E-05 | 1.700E-05 |
| 1417 | 71 | Q71T | A | 1.3e-05 | 1.3e-05 | 1.5e-05 | 1.367E-05 | 1.155E-06 | 1.300E-05 | 1.500E-05 |
| 1418 | 71 | Q71W | A | 1.1e-05 | 1.5e-05 | 1.2e-05 | 1.267E-05 | 2.082E-06 | 1.100E-05 | 1.500E-05 |
| 1419 | 71 | Q71Y | A | 1.2e-05 | 1.6e-05 | 1e-05 | 1.267E-05 | 3.055E-06 | 1.000E-05 | 1.600E-05 |
| 1420 | 71 | Q71V | A | 1.7e-05 | 8.6e-06 | 1.6e-05 | 1.387E-05 | 4.588E-06 | 8.600E-06 | 1.700E-05 |
| 1421 | 72 | V72A | A | 1.3e-05 | 1.4e-05 | 1e-05 | 1.233E-05 | 2.082E-06 | 1.000E-05 | 1.400E-05 |
| 1422 | 72 | V72R | A | 1.3e-05 | 1.3e-05 | 9.7e-06 | 1.190E-05 | 1.905E-06 | 9.700E-06 | 1.300E-05 |
| 1423 | 72 | V72N | A | 8.7e-06 | 1.2e-05 | 1.5e-05 | 1.190E-05 | 3.151E-06 | 8.700E-06 | 1.500E-05 |
| 1424 | 72 | V72D | A | 1.6e-05 | 1.2e-05 | 9.8e-06 | 1.260E-05 | 3.143E-06 | 9.800E-06 | 1.600E-05 |
| 1425 | 72 | V72C | A | 1.2e-05 | 8.1e-06 | 1.5e-05 | 1.170E-05 | 3.460E-06 | 8.100E-06 | 1.500E-05 |
| 1426 | 72 | V72E | A | 6e-06 | 1.4e-05 | 1.1e-05 | 1.033E-05 | 4.041E-06 | 6.000E-06 | 1.400E-05 |
| 1427 | 72 | V72Q | A | 1.2e-05 | 1.3e-05 | 1.6e-05 | 1.367E-05 | 2.082E-06 | 1.200E-05 | 1.600E-05 |
| 1428 | 72 | V72G | A | 1.6e-05 | 2e-05 | 1.7e-05 | 1.767E-05 | 2.082E-06 | 1.600E-05 | 2.000E-05 |
| 1429 | 72 | V72H | A | 1.3e-05 | 9.1e-06 | 1.5e-05 | 1.237E-05 | 3.001E-06 | 9.100E-06 | 1.500E-05 |
| 1430 | 72 | V72I | A | 1.2e-05 | 1.4e-05 | 1.3e-05 | 1.300E-05 | 1.000E-06 | 1.200E-05 | 1.400E-05 |
| 1431 | 72 | V72L | A | 1.1e-05 | 1.3e-05 | 1.5e-05 | 1.300E-05 | 2.000E-06 | 1.100E-05 | 1.500E-05 |
| 1432 | 72 | V72K | A | 1e-05 | 1.7e-05 | 1.7e-05 | 1.467E-05 | 4.041E-06 | 1.000E-05 | 1.700E-05 |
| 1433 | 72 | V72M | A | 7e-06 | 1.5e-05 | 1.4e-05 | 1.200E-05 | 4.359E-06 | 7.000E-06 | 1.500E-05 |
| 1434 | 72 | V72F | A | 1.1e-05 | 9.3e-06 | 9e-06 | 9.767E-06 | 1.079E-06 | 9.000E-06 | 1.100E-05 |
| 1435 | 72 | V72P | A | 1.6e-05 | 1.6e-05 | 1.3e-05 | 1.500E-05 | 1.732E-06 | 1.300E-05 | 1.600E-05 |
| 1436 | 72 | V72S | A | 1.6e-05 | 1.3e-05 | 1.4e-05 | 1.433E-05 | 1.528E-06 | 1.300E-05 | 1.600E-05 |
| 1437 | 72 | V72T | A | 1.4e-05 | 1.5e-05 | 1.5e-05 | 1.467E-05 | 5.774E-07 | 1.400E-05 | 1.500E-05 |
| 1438 | 72 | V72W | A | 1.4e-05 | 1.1e-05 | 1.4e-05 | 1.300E-05 | 1.732E-06 | 1.100E-05 | 1.400E-05 |
| 1439 | 72 | V72Y | A | 1.6e-05 | 1.6e-05 | 9.6e-06 | 1.387E-05 | 3.695E-06 | 9.600E-06 | 1.600E-05 |
| 1440 | 72 | V72V | A | 1.3e-05 | 9.7e-06 | 1.6e-05 | 1.290E-05 | 3.151E-06 | 9.700E-06 | 1.600E-05 |
| 1441 | 73 | P73A | A | 9.9e-06 | 1.5e-05 | 1.2e-05 | 1.230E-05 | 2.563E-06 | 9.900E-06 | 1.500E-05 |
| 1442 | 73 | P73R | A | 1.5e-05 | 1.7e-05 | 1.4e-05 | 1.533E-05 | 1.528E-06 | 1.400E-05 | 1.700E-05 |
| 1443 | 73 | P73N | A | 1.5e-05 | 1.2e-05 | 9.2e-06 | 1.207E-05 | 2.901E-06 | 9.200E-06 | 1.500E-05 |
| 1444 | 73 | P73D | A | 1.4e-05 | 1.3e-05 | 1.3e-05 | 1.333E-05 | 5.774E-07 | 1.300E-05 | 1.400E-05 |

| | | | | | | | | | | |
|---|---|---|---|---|---|---|---|---|---|---|
| 1445 | 73 | P73C | A | 1.4e-05 | 1.6e-05 | 1.1e-05 | 1.367E-05 | 2.517E-06 | 1.100E-05 | 1.600E-05 |
| 1446 | 73 | P73E | A | 1.6e-05 | 8.5e-06 | 1.4e-05 | 1.283E-05 | 3.884E-06 | 8.500E-06 | 1.600E-05 |
| 1447 | 73 | P73Q | A | 8.1e-06 | 1.1e-05 | 1.4e-05 | 1.103E-05 | 2.950E-06 | 8.100E-06 | 1.400E-05 |
| 1448 | 73 | P73G | A | 1.5e-05 | 9.9e-06 | 9.4e-06 | 1.143E-05 | 3.099E-06 | 9.400E-06 | 1.500E-05 |
| 1449 | 73 | P73H | A | 1.2e-05 | 1.1e-05 | 1.5e-05 | 1.267E-05 | 2.082E-06 | 1.100E-05 | 1.500E-05 |
| 1450 | 73 | P73I | A | 1.4e-05 | 1.2e-05 | 1.4e-05 | 1.333E-05 | 1.155E-06 | 1.200E-05 | 1.400E-05 |
| 1451 | 73 | P73L | A | 1.4e-05 | 1.2e-05 | 1.5e-05 | 1.367E-05 | 1.528E-06 | 1.200E-05 | 1.500E-05 |
| 1452 | 73 | P73K | A | 1.3e-05 | 1.4e-05 | 1.1e-05 | 1.267E-05 | 1.528E-06 | 1.100E-05 | 1.400E-05 |
| 1453 | 73 | P73M | A | 1.1e-05 | 1.7e-05 | 1e-05 | 1.267E-05 | 3.786E-06 | 1.000E-05 | 1.700E-05 |
| 1454 | 73 | P73F | A | 9.5e-06 | 1.4e-05 | 1.1e-05 | 1.150E-05 | 2.291E-06 | 9.500E-06 | 1.400E-05 |
| 1455 | 73 | P73P | A | 1.5e-05 | 1e-05 | 7.6e-06 | 1.087E-05 | 3.775E-06 | 7.600E-06 | 1.500E-05 |
| 1456 | 73 | P73S | A | 7.9e-06 | 1e-05 | 1.2e-05 | 9.967E-06 | 2.050E-06 | 7.900E-06 | 1.200E-05 |
| 1457 | 73 | P73T | A | 1.4e-05 | 1e-05 | 1.4e-05 | 1.267E-05 | 2.309E-06 | 1.000E-05 | 1.400E-05 |
| 1458 | 73 | P73W | A | 1.6e-05 | 1.5e-05 | 1.5e-05 | 1.533E-05 | 5.774E-07 | 1.500E-05 | 1.600E-05 |
| 1459 | 73 | P73Y | A | 1.6e-05 | 1.5e-05 | 1.5e-05 | 1.533E-05 | 5.774E-07 | 1.500E-05 | 1.600E-05 |
| 1460 | 73 | P73V | A | 1.1e-05 | 9.5e-06 | 1.3e-05 | 1.117E-05 | 1.756E-06 | 9.500E-06 | 1.300E-05 |
| 1461 | 74 | L74A | A | 1.5e-05 | 1.1e-05 | 1.7e-05 | 1.433E-05 | 3.055E-06 | 1.100E-05 | 1.700E-05 |
| 1462 | 74 | L74R | A | 1.4e-05 | 1e-05 | 9.2e-06 | 1.107E-05 | 2.572E-06 | 9.200E-06 | 1.400E-05 |
| 1463 | 74 | L74N | A | 9.8e-06 | 1.6e-05 | 9.5e-06 | 1.177E-05 | 3.669E-06 | 9.500E-06 | 1.600E-05 |
| 1464 | 74 | L74D | A | 1.1e-05 | 1.2e-05 | 1.4e-05 | 1.233E-05 | 1.528E-06 | 1.100E-05 | 1.400E-05 |
| 1465 | 74 | L74C | A | 1.1e-05 | 1.5e-05 | 1.3e-05 | 1.300E-05 | 2.000E-06 | 1.100E-05 | 1.500E-05 |
| 1466 | 74 | L74E | A | 1.5e-05 | 1.6e-05 | 1e-05 | 1.367E-05 | 3.215E-06 | 1.000E-05 | 1.600E-05 |
| 1467 | 74 | L74Q | A | 1.6e-05 | 1.7e-05 | 1.3e-05 | 1.533E-05 | 2.082E-06 | 1.300E-05 | 1.700E-05 |
| 1468 | 74 | L74G | A | 1e-05 | 1.5e-05 | 1.7e-05 | 1.400E-05 | 3.606E-06 | 1.000E-05 | 1.700E-05 |
| 1469 | 74 | L74H | A | 1.1e-05 | 1.3e-05 | 1.5e-05 | 1.300E-05 | 2.000E-06 | 1.100E-05 | 1.500E-05 |
| 1470 | 74 | L74I | A | 1e-05 | 1.6e-05 | 1e-05 | 1.200E-05 | 3.464E-06 | 1.000E-05 | 1.600E-05 |
| 1471 | 74 | L74L | A | 1.3e-05 | 9.5e-06 | 9.5e-06 | 1.067E-05 | 2.021E-06 | 9.500E-06 | 1.300E-05 |
| 1472 | 74 | L74K | A | 1.5e-05 | 1.5e-05 | 9.9e-06 | 1.330E-05 | 2.944E-06 | 9.900E-06 | 1.500E-05 |
| 1473 | 74 | L74M | A | 1.3e-05 | 1.1e-05 | 1.5e-05 | 1.300E-05 | 2.000E-06 | 1.100E-05 | 1.500E-05 |
| 1474 | 74 | L74F | A | 1.6e-05 | 1.1e-05 | 1.5e-05 | 1.400E-05 | 2.646E-06 | 1.100E-05 | 1.600E-05 |
| 1475 | 74 | L74P | A | 1.4e-05 | 1.1e-05 | 1.1e-05 | 1.200E-05 | 1.732E-06 | 1.100E-05 | 1.400E-05 |
| 1476 | 74 | L74S | A | 1.5e-05 | 1.6e-05 | 1e-05 | 1.367E-05 | 3.215E-06 | 1.000E-05 | 1.600E-05 |
| 1477 | 74 | L74T | A | 6.6e-06 | 1.6e-05 | 1.5e-05 | 1.253E-05 | 5.163E-06 | 6.600E-06 | 1.600E-05 |
| 1478 | 74 | L74W | A | 9.5e-06 | 1.6e-05 | 6e-06 | 1.050E-05 | 5.074E-06 | 6.000E-06 | 1.600E-05 |
| 1479 | 74 | L74Y | A | 8.9e-06 | 9.7e-06 | 1e-05 | 9.533E-06 | 5.686E-07 | 8.900E-06 | 1.000E-05 |
| 1480 | 74 | L74V | A | 8.9e-06 | 1.6e-05 | 1.7e-05 | 1.397E-05 | 4.416E-06 | 8.900E-06 | 1.700E-05 |
| 1481 | 75 | Q75A | A | 1.1e-05 | 1.3e-05 | 1.1e-05 | 1.167E-05 | 1.155E-06 | 1.100E-05 | 1.300E-05 |
| 1482 | 75 | Q75R | A | 1.5e-05 | 1.1e-05 | 1.6e-05 | 1.400E-05 | 2.646E-06 | 1.100E-05 | 1.600E-05 |
| 1483 | 75 | Q75N | A | 1.2e-05 | 1.7e-05 | 1.5e-05 | 1.467E-05 | 2.517E-06 | 1.200E-05 | 1.700E-05 |
| 1484 | 75 | Q75D | A | 1.4e-05 | 1.8e-05 | 1.2e-05 | 1.467E-05 | 3.055E-06 | 1.200E-05 | 1.800E-05 |
| 1485 | 75 | Q75C | A | 1e-05 | 1.2e-05 | 9.2e-06 | 1.040E-05 | 1.442E-06 | 9.200E-06 | 1.200E-05 |
| 1486 | 75 | Q75E | A | 1.8e-05 | 1.7e-05 | 1e-05 | 1.500E-05 | 4.359E-06 | 1.000E-05 | 1.800E-05 |
| 1487 | 75 | Q75Q | A | 1.2e-05 | 1.4e-05 | 1.6e-05 | 1.400E-05 | 2.000E-06 | 1.200E-05 | 1.600E-05 |
| 1488 | 75 | Q75G | A | 1.8e-05 | 7.5e-06 | 1.1e-05 | 1.217E-05 | 5.346E-06 | 7.500E-06 | 1.800E-05 |
| 1489 | 75 | Q75H | A | 9.2e-06 | 1.5e-05 | 1e-05 | 1.140E-05 | 3.143E-06 | 9.200E-06 | 1.500E-05 |
| 1490 | 75 | Q75I | A | 1.1e-05 | 1.4e-05 | 1e-05 | 1.167E-05 | 2.082E-06 | 1.000E-05 | 1.400E-05 |
| 1491 | 75 | Q75L | A | 1.7e-05 | 1.1e-05 | 1.6e-05 | 1.467E-05 | 3.215E-06 | 1.100E-05 | 1.700E-05 |
| 1492 | 75 | Q75K | A | 1.1e-05 | 1.4e-05 | 1.1e-05 | 1.200E-05 | 1.732E-06 | 1.100E-05 | 1.400E-05 |
| 1493 | 75 | Q75M | A | 1.6e-05 | 1.3e-05 | 1.4e-05 | 1.433E-05 | 1.528E-06 | 1.300E-05 | 1.600E-05 |
| 1494 | 75 | Q75F | A | 7.6e-06 | 1.6e-05 | 1.3e-05 | 1.220E-05 | 4.257E-06 | 7.600E-06 | 1.600E-05 |
| 1495 | 75 | Q75P | A | 1.6e-05 | 1.4e-05 | 8.2e-06 | 1.273E-05 | 4.051E-06 | 8.200E-06 | 1.600E-05 |
| 1496 | 75 | Q75S | A | 1.5e-05 | 1.5e-05 | 1.5e-05 | 1.500E-05 | 0.000E+00 | 1.500E-05 | 1.500E-05 |
| 1497 | 75 | Q75T | A | 1.4e-05 | 1.1e-05 | 1.5e-05 | 1.333E-05 | 2.082E-06 | 1.100E-05 | 1.500E-05 |
| 1498 | 75 | Q75W | A | 1e-05 | 1.5e-05 | 1.5e-05 | 1.333E-05 | 2.887E-06 | 1.000E-05 | 1.500E-05 |
| 1499 | 75 | Q75Y | A | 9.9e-06 | 9.8e-06 | 1.4e-05 | 1.123E-05 | 2.397E-06 | 9.800E-06 | 1.400E-05 |
| 1500 | 75 | Q75V | A | 1.3e-05 | 1.6e-05 | 9.6e-06 | 1.287E-05 | 3.202E-06 | 9.600E-06 | 1.600E-05 |
| 1501 | 76 | R76A | A | 1.1e-05 | 9.4e-06 | 1.1e-05 | 1.047E-05 | 9.238E-07 | 9.400E-06 | 1.100E-05 |
| 1502 | 76 | R76R | A | 1.5e-05 | 9e-06 | 1.5e-05 | 1.300E-05 | 3.464E-06 | 9.000E-06 | 1.500E-05 |
| 1503 | 76 | R76N | A | 1.1e-05 | 1.7e-05 | 1.5e-05 | 1.433E-05 | 3.055E-06 | 1.100E-05 | 1.700E-05 |
| 1504 | 76 | R76D | A | 8.4e-06 | 1.1e-05 | 1.1e-05 | 1.013E-05 | 1.501E-06 | 8.400E-06 | 1.100E-05 |
| 1505 | 76 | R76C | A | 9.3e-06 | 1.6e-05 | 8.7e-06 | 1.133E-05 | 4.053E-06 | 8.700E-06 | 1.600E-05 |
| 1506 | 76 | R76E | A | 1.6e-05 | 1.5e-05 | 1.2e-05 | 1.433E-05 | 2.082E-06 | 1.200E-05 | 1.600E-05 |
| 1507 | 76 | R76Q | A | 9.6e-06 | 1.5e-05 | 1.3e-05 | 1.253E-05 | 2.730E-06 | 9.600E-06 | 1.500E-05 |
| 1508 | 76 | R76G | A | 1.5e-05 | 1.6e-05 | 7.8e-06 | 1.293E-05 | 4.474E-06 | 7.800E-06 | 1.600E-05 |
| 1509 | 76 | R76H | A | 1.1e-05 | 1.1e-05 | 6.4e-06 | 9.467E-06 | 2.656E-06 | 6.400E-06 | 1.100E-05 |
| 1510 | 76 | R76I | A | 1.5e-05 | 1e-05 | 1.5e-05 | 1.333E-05 | 2.887E-06 | 1.000E-05 | 1.500E-05 |
| 1511 | 76 | R76L | A | 1.3e-05 | 1.5e-05 | 1.2e-05 | 1.333E-05 | 1.528E-06 | 1.200E-05 | 1.500E-05 |
| 1512 | 76 | R76K | A | 1.1e-05 | 1.6e-05 | 1.6e-05 | 1.433E-05 | 2.887E-06 | 1.100E-05 | 1.600E-05 |
| 1513 | 76 | R76M | A | 1.2e-05 | 8.8e-06 | 1.2e-05 | 1.093E-05 | 1.848E-06 | 8.800E-06 | 1.200E-05 |
| 1514 | 76 | R76F | A | 1.5e-05 | 1.6e-05 | 1.5e-05 | 1.533E-05 | 5.774E-07 | 1.500E-05 | 1.600E-05 |
| 1515 | 76 | R76P | A | 1.3e-05 | 9.6e-06 | 1.5e-05 | 1.253E-05 | 2.730E-06 | 9.600E-06 | 1.500E-05 |
| 1516 | 76 | R76S | A | 1.4e-05 | 1.5e-05 | 1.5e-05 | 1.467E-05 | 5.774E-07 | 1.400E-05 | 1.500E-05 |
| 1517 | 76 | R76T | A | 1e-05 | 1.1e-05 | 1.1e-05 | 1.067E-05 | 5.774E-07 | 1.000E-05 | 1.100E-05 |
| 1518 | 76 | R76W | A | 1.6e-05 | 1.4e-05 | 1.6e-05 | 1.533E-05 | 1.155E-06 | 1.400E-05 | 1.600E-05 |
| 1519 | 76 | R76Y | A | 1.5e-05 | 1e-05 | 1.4e-05 | 1.300E-05 | 2.646E-06 | 1.000E-05 | 1.500E-05 |
| 1520 | 76 | R76V | A | 1.1e-05 | 1.2e-05 | 1e-05 | 1.100E-05 | 1.000E-06 | 1.000E-05 | 1.200E-05 |
| 1521 | 77 | L77A | A | 1.1e-05 | 1.6e-05 | 1e-05 | 1.233E-05 | 3.215E-06 | 1.000E-05 | 1.600E-05 |
| 1522 | 77 | L77R | A | 1.1e-05 | 1e-05 | 1.1e-05 | 1.067E-05 | 5.774E-07 | 1.000E-05 | 1.100E-05 |
| 1523 | 77 | L77N | A | 1.8e-05 | 1.5e-05 | 1.5e-05 | 1.600E-05 | 1.732E-06 | 1.500E-05 | 1.800E-05 |
| 1524 | 77 | L77D | A | 1.4e-05 | 9e-06 | 1.5e-05 | 1.267E-05 | 3.215E-06 | 9.000E-06 | 1.500E-05 |
| 1525 | 77 | L77C | A | 1.2e-05 | 1.5e-05 | 1.1e-05 | 1.267E-05 | 2.082E-06 | 1.100E-05 | 1.500E-05 |
| 1526 | 77 | L77E | A | 1e-05 | 1.5e-05 | 9.8e-06 | 1.160E-05 | 2.946E-06 | 9.800E-06 | 1.500E-05 |
| 1527 | 77 | L77Q | A | 1.5e-05 | 1.1e-05 | 9.6e-06 | 1.187E-05 | 2.802E-06 | 9.600E-06 | 1.500E-05 |
| 1528 | 77 | L77G | A | 1.3e-05 | 1.4e-05 | 1e-05 | 1.233E-05 | 2.082E-06 | 1.000E-05 | 1.400E-05 |
| 1529 | 77 | L77H | A | 1.3e-05 | 1.6e-05 | 1e-05 | 1.300E-05 | 3.000E-06 | 1.000E-05 | 1.600E-05 |

| | | | | | | | | | | |
|---|---|---|---|---|---|---|---|---|---|---|
| 1530 | 77 | L77I | A | 1.2e-05 | 8.3e-06 | 1.3e-05 | 1.110E-05 | 2.476E-06 | 8.300E-06 | 1.300E-05 |
| 1531 | 77 | L77L | A | 1.5e-05 | 1.5e-05 | 9.1e-06 | 1.303E-05 | 3.406E-06 | 9.100E-06 | 1.500E-05 |
| 1532 | 77 | L77K | A | 1.3e-05 | 9.6e-06 | 1e-05 | 1.087E-05 | 1.858E-06 | 9.600E-06 | 1.300E-05 |
| 1533 | 77 | L77M | A | 1e-05 | 1.4e-05 | 9.4e-06 | 1.113E-05 | 2.501E-06 | 9.400E-06 | 1.400E-05 |
| 1534 | 77 | L77F | A | 9.3e-06 | 1.4e-05 | 1.1e-05 | 1.143E-05 | 2.380E-06 | 9.300E-06 | 1.400E-05 |
| 1535 | 77 | L77P | A | 9.5e-06 | 1e-05 | 1.7e-05 | 1.217E-05 | 4.193E-06 | 9.500E-06 | 1.700E-05 |
| 1536 | 77 | L77S | A | 1.1e-05 | 1.1e-05 | 9.3e-06 | 1.043E-05 | 9.815E-07 | 9.300E-06 | 1.100E-05 |
| 1537 | 77 | L77T | A | 1.1e-05 | 1.6e-05 | 1.4e-05 | 1.367E-05 | 2.517E-06 | 1.100E-05 | 1.600E-05 |
| 1538 | 77 | L77W | A | 1.4e-05 | 1.2e-05 | 1e-05 | 1.200E-05 | 2.000E-06 | 1.000E-05 | 1.400E-05 |
| 1539 | 77 | L77Y | A | 1.4e-05 | 1.5e-05 | 1.6e-05 | 1.500E-05 | 1.000E-06 | 1.400E-05 | 1.600E-05 |
| 1540 | 77 | L77V | A | 9.4e-06 | 1e-05 | 1.5e-05 | 1.147E-05 | 3.075E-06 | 9.400E-06 | 1.500E-05 |
| 1541 | 78 | R78A | A | 1.8e-05 | 9.8e-06 | 1.2e-05 | 1.327E-05 | 4.244E-06 | 9.800E-06 | 1.800E-05 |
| 1542 | 78 | R78R | A | 1.2e-05 | 1.5e-05 | 1.4e-05 | 1.367E-05 | 1.528E-06 | 1.200E-05 | 1.500E-05 |
| 1543 | 78 | R78N | A | 1.5e-05 | 1.6e-05 | 1.4e-05 | 1.500E-05 | 1.000E-06 | 1.400E-05 | 1.600E-05 |
| 1544 | 78 | R78D | A | 1.4e-05 | 7.8e-06 | 1.6e-05 | 1.260E-05 | 4.276E-06 | 7.800E-06 | 1.600E-05 |
| 1545 | 78 | R78C | A | 1e-05 | 8.6e-06 | 1e-05 | 9.533E-06 | 8.083E-07 | 8.600E-06 | 1.000E-05 |
| 1546 | 78 | R78E | A | 1.4e-05 | 1.7e-05 | 1.1e-05 | 1.400E-05 | 3.000E-06 | 1.100E-05 | 1.700E-05 |
| 1547 | 78 | R78Q | A | 1.6e-05 | 9e-06 | 1.3e-05 | 1.267E-05 | 3.512E-06 | 9.000E-06 | 1.600E-05 |
| 1548 | 78 | R78G | A | 1.2e-05 | 9.8e-06 | 1.6e-05 | 1.260E-05 | 3.143E-06 | 9.800E-06 | 1.600E-05 |
| 1549 | 78 | R78H | A | 1.1e-05 | 1.6e-05 | 9.9e-06 | 1.230E-05 | 3.251E-06 | 9.900E-06 | 1.600E-05 |
| 1550 | 78 | R78I | A | 1.6e-05 | 1.8e-05 | 1.5e-05 | 1.633E-05 | 1.528E-06 | 1.500E-05 | 1.800E-05 |
| 1551 | 78 | R78L | A | 1e-05 | 8.8e-06 | 1.4e-05 | 1.093E-05 | 2.723E-06 | 8.800E-06 | 1.400E-05 |
| 1552 | 78 | R78K | A | 8.5e-06 | 1.4e-05 | 9.5e-06 | 1.067E-05 | 2.930E-06 | 8.500E-06 | 1.400E-05 |
| 1553 | 78 | R78M | A | 1.1e-05 | 1.1e-05 | 1.1e-05 | 1.100E-05 | 2.075E-21 | 1.100E-05 | 1.100E-05 |
| 1554 | 78 | R78F | A | 9e-06 | 1.5e-05 | 1.3e-05 | 1.233E-05 | 3.055E-06 | 9.000E-06 | 1.500E-05 |
| 1555 | 78 | R78P | A | 1.3e-05 | 1.3e-05 | 1.6e-05 | 1.400E-05 | 1.732E-06 | 1.300E-05 | 1.600E-05 |
| 1556 | 78 | R78S | A | 1.2e-05 | 1e-05 | 1e-05 | 1.067E-05 | 1.155E-06 | 1.000E-05 | 1.200E-05 |
| 1557 | 78 | R78T | A | 9.3e-06 | 1.5e-05 | 1.5e-05 | 1.310E-05 | 3.291E-06 | 9.300E-06 | 1.500E-05 |
| 1558 | 78 | R78W | A | 9.9e-06 | 1.2e-05 | 1.5e-05 | 1.230E-05 | 2.563E-06 | 9.900E-06 | 1.500E-05 |
| 1559 | 78 | R78Y | A | 1.1e-05 | 8.8e-06 | 1.6e-05 | 1.193E-05 | 3.690E-06 | 8.800E-06 | 1.600E-05 |
| 1560 | 78 | R78V | A | 1.1e-05 | 1.5e-05 | 1.6e-05 | 1.400E-05 | 2.646E-06 | 1.100E-05 | 1.600E-05 |
| 1561 | 79 | I79A | A | 1.8e-05 | 1.3e-05 | 8.8e-06 | 1.327E-05 | 4.606E-06 | 8.800E-06 | 1.800E-05 |
| 1562 | 79 | I79R | A | 1.2e-05 | 9.7e-06 | 9.7e-06 | 1.047E-05 | 1.328E-06 | 9.700E-06 | 1.200E-05 |
| 1563 | 79 | I79N | A | 1.1e-05 | 9.6e-06 | 1.8e-05 | 1.287E-05 | 4.500E-06 | 9.600E-06 | 1.800E-05 |
| 1564 | 79 | I79D | A | 1.6e-05 | 1.4e-05 | 1.4e-05 | 1.467E-05 | 1.155E-06 | 1.400E-05 | 1.600E-05 |
| 1565 | 79 | I79C | A | 1.7e-05 | 9.6e-06 | 1.6e-05 | 1.420E-05 | 4.015E-06 | 9.600E-06 | 1.700E-05 |
| 1566 | 79 | I79E | A | 6.9e-06 | 1.4e-05 | 1.5e-05 | 1.197E-05 | 4.416E-06 | 6.900E-06 | 1.500E-05 |
| 1567 | 79 | I79Q | A | 1e-05 | 1.4e-05 | 1.3e-05 | 1.233E-05 | 2.082E-06 | 1.000E-05 | 1.400E-05 |
| 1568 | 79 | I79G | A | 1.6e-05 | 1.5e-05 | 1.5e-05 | 1.533E-05 | 5.774E-07 | 1.500E-05 | 1.600E-05 |
| 1569 | 79 | I79H | A | 1.3e-05 | 1.4e-05 | 1.1e-05 | 1.267E-05 | 1.528E-06 | 1.100E-05 | 1.400E-05 |
| 1570 | 79 | I79I | A | 7.2e-06 | 1.3e-05 | 1e-05 | 1.007E-05 | 2.901E-06 | 7.200E-06 | 1.300E-05 |
| 1571 | 79 | I79L | A | 1.4e-05 | 1.3e-05 | 9.8e-06 | 1.227E-05 | 2.194E-06 | 9.800E-06 | 1.400E-05 |
| 1572 | 79 | I79K | A | 1e-05 | 1.1e-05 | 7.8e-06 | 9.600E-06 | 1.637E-06 | 7.800E-06 | 1.100E-05 |
| 1573 | 79 | I79M | A | 1.5e-05 | 9.8e-06 | 1.5e-05 | 1.327E-05 | 3.002E-06 | 9.800E-06 | 1.500E-05 |
| 1574 | 79 | I79F | A | 1.1e-05 | 1.5e-05 | 9e-06 | 1.167E-05 | 3.055E-06 | 9.000E-06 | 1.500E-05 |
| 1575 | 79 | I79P | A | 1.7e-05 | 1e-05 | 1.1e-05 | 1.267E-05 | 3.786E-06 | 1.000E-05 | 1.700E-05 |
| 1576 | 79 | I79S | A | 1.6e-05 | 1.4e-05 | 1.1e-05 | 1.367E-05 | 2.517E-06 | 1.100E-05 | 1.600E-05 |
| 1577 | 79 | I79T | A | 1.6e-05 | 6e-06 | 1.4e-05 | 1.200E-05 | 5.292E-06 | 6.000E-06 | 1.600E-05 |
| 1578 | 79 | I79W | A | 1.1e-05 | 1.3e-05 | 9.1e-06 | 1.103E-05 | 1.950E-06 | 9.100E-06 | 1.300E-05 |
| 1579 | 79 | I79Y | A | 1.6e-05 | 8.7e-06 | 1e-05 | 1.157E-05 | 3.894E-06 | 8.700E-06 | 1.600E-05 |
| 1580 | 79 | I79V | A | 1.3e-05 | 1.1e-05 | 1.5e-05 | 1.300E-05 | 2.000E-06 | 1.100E-05 | 1.500E-05 |
| 1581 | 80 | V80A | A | 1.5e-05 | 1.6e-05 | 1.3e-05 | 1.467E-05 | 1.528E-06 | 1.300E-05 | 1.600E-05 |
| 1582 | 80 | V80R | A | 1.1e-05 | 9.9e-06 | 1.5e-05 | 1.197E-05 | 2.684E-06 | 9.900E-06 | 1.500E-05 |
| 1583 | 80 | V80N | A | 1.2e-05 | 2.1e-05 | 9.4e-06 | 1.413E-05 | 6.087E-06 | 9.400E-06 | 2.100E-05 |
| 1584 | 80 | V80D | A | 9.5e-06 | 1.4e-05 | 1.2e-05 | 1.183E-05 | 2.255E-06 | 9.500E-06 | 1.400E-05 |
| 1585 | 80 | V80C | A | 9.3e-06 | 1.2e-05 | 9.5e-06 | 1.027E-05 | 1.504E-06 | 9.300E-06 | 1.200E-05 |
| 1586 | 80 | V80E | A | 8.9e-06 | 6.3e-06 | 1.4e-05 | 9.733E-06 | 3.917E-06 | 6.300E-06 | 1.400E-05 |
| 1587 | 80 | V80Q | A | 1.8e-05 | 1.3e-05 | 1.5e-05 | 1.533E-05 | 2.517E-06 | 1.300E-05 | 1.800E-05 |
| 1588 | 80 | V80G | A | 1.5e-05 | 1.5e-05 | 1.5e-05 | 1.500E-05 | 0.000E+00 | 1.500E-05 | 1.500E-05 |
| 1589 | 80 | V80H | A | 1.4e-05 | 1.3e-05 | 1.3e-05 | 1.333E-05 | 5.774E-07 | 1.300E-05 | 1.400E-05 |
| 1590 | 80 | V80I | A | 1.5e-05 | 1.1e-05 | 1.3e-05 | 1.300E-05 | 2.000E-06 | 1.100E-05 | 1.500E-05 |
| 1591 | 80 | V80L | A | 1.6e-05 | 1.1e-05 | 1.1e-05 | 1.267E-05 | 2.887E-06 | 1.100E-05 | 1.600E-05 |
| 1592 | 80 | V80K | A | 1.1e-05 | 1.5e-05 | 1.5e-05 | 1.367E-05 | 2.309E-06 | 1.100E-05 | 1.500E-05 |
| 1593 | 80 | V80M | A | 1.5e-05 | 1.3e-05 | 1.5e-05 | 1.433E-05 | 1.155E-06 | 1.300E-05 | 1.500E-05 |
| 1594 | 80 | V80F | A | 8.6e-06 | 7.6e-06 | 1.1e-05 | 9.067E-06 | 1.747E-06 | 7.600E-06 | 1.100E-05 |
| 1595 | 80 | V80P | A | 1.1e-05 | 1.6e-05 | 9.1e-06 | 1.203E-05 | 3.564E-06 | 9.100E-06 | 1.600E-05 |
| 1596 | 80 | V80S | A | 1.3e-05 | 1.2e-05 | 1.1e-05 | 1.200E-05 | 1.000E-06 | 1.100E-05 | 1.300E-05 |
| 1597 | 80 | V80T | A | 1.6e-05 | 1.6e-05 | 1.2e-05 | 1.467E-05 | 2.309E-06 | 1.200E-05 | 1.600E-05 |
| 1598 | 80 | V80W | A | 1.5e-05 | 1.1e-05 | 1e-05 | 1.200E-05 | 2.646E-06 | 1.000E-05 | 1.500E-05 |
| 1599 | 80 | V80Y | A | 1.1e-05 | 1.3e-05 | 1.4e-05 | 1.267E-05 | 1.528E-06 | 1.100E-05 | 1.400E-05 |
| 1600 | 80 | V80V | A | 1.4e-05 | 1.1e-05 | 1.5e-05 | 1.333E-05 | 2.082E-06 | 1.100E-05 | 1.500E-05 |
| 1601 | 81 | R81A | A | 9.1e-06 | 1e-05 | 1.3e-05 | 1.070E-05 | 2.042E-06 | 9.100E-06 | 1.300E-05 |
| 1602 | 81 | R81R | A | 9.5e-06 | 1.3e-05 | 1e-05 | 1.083E-05 | 1.893E-06 | 9.500E-06 | 1.300E-05 |
| 1603 | 81 | R81N | A | 1.5e-05 | 1.5e-05 | 1.5e-05 | 1.500E-05 | 0.000E+00 | 1.500E-05 | 1.500E-05 |
| 1604 | 81 | R81D | A | 1.5e-05 | 1.1e-05 | 1e-05 | 1.200E-05 | 2.646E-06 | 1.000E-05 | 1.500E-05 |
| 1605 | 81 | R81C | A | 1.6e-05 | 9.8e-06 | 1e-05 | 1.193E-05 | 3.523E-06 | 9.800E-06 | 1.600E-05 |
| 1606 | 81 | R81E | A | 1.7e-05 | 1.6e-05 | 1e-05 | 1.433E-05 | 3.786E-06 | 1.000E-05 | 1.700E-05 |
| 1607 | 81 | R81Q | A | 1.4e-05 | 1.6e-05 | 8.8e-06 | 1.293E-05 | 3.717E-06 | 8.800E-06 | 1.600E-05 |
| 1608 | 81 | R81G | A | 1.4e-05 | 9.3e-06 | 1e-05 | 1.110E-05 | 2.536E-06 | 9.300E-06 | 1.400E-05 |
| 1609 | 81 | R81H | A | 1.2e-05 | 1e-05 | 9.5e-06 | 1.050E-05 | 1.323E-06 | 9.500E-06 | 1.200E-05 |
| 1610 | 81 | R81I | A | 1.5e-05 | 1e-05 | 1.2e-05 | 1.233E-05 | 2.517E-06 | 1.000E-05 | 1.500E-05 |
| 1611 | 81 | R81L | A | 1.6e-05 | 9.2e-06 | 1.1e-05 | 1.207E-05 | 3.523E-06 | 9.200E-06 | 1.600E-05 |
| 1612 | 81 | R81K | A | 1e-05 | 9.5e-06 | 7.7e-06 | 9.067E-06 | 1.210E-06 | 7.700E-06 | 1.000E-05 |
| 1613 | 81 | R81M | A | 1.1e-05 | 1.1e-05 | 1.4e-05 | 1.200E-05 | 1.732E-06 | 1.100E-05 | 1.400E-05 |
| 1614 | 81 | R81F | A | 1.5e-05 | 1.6e-05 | 1.5e-05 | 1.533E-05 | 5.774E-07 | 1.500E-05 | 1.600E-05 |

| | | | | | | | | | | |
|---|---|---|---|---|---|---|---|---|---|---|
| 1615 | 81 | R81P | A | 1.5e-05 | 1.6e-05 | 1.2e-05 | 1.433E-05 | 2.082E-06 | 1.200E-05 | 1.600E-05 |
| 1616 | 81 | R81S | A | 1.3e-05 | 1.6e-05 | 1.5e-05 | 1.467E-05 | 1.528E-06 | 1.300E-05 | 1.600E-05 |
| 1617 | 81 | R81T | A | 2.1e-05 | 1.6e-05 | 1.3e-05 | 1.667E-05 | 4.041E-06 | 1.300E-05 | 2.100E-05 |
| 1618 | 81 | R81W | A | 1.1e-05 | 1.4e-05 | 1.5e-05 | 1.333E-05 | 2.082E-06 | 1.100E-05 | 1.500E-05 |
| 1619 | 81 | R81Y | A | 1.2e-05 | 1.6e-05 | 1e-05 | 1.267E-05 | 3.055E-06 | 1.000E-05 | 1.600E-05 |
| 1620 | 81 | R81V | A | 1.6e-05 | 1.2e-05 | 1.5e-05 | 1.433E-05 | 2.082E-06 | 1.200E-05 | 1.600E-05 |
| 1621 | 82 | G82A | A | 1.7e-05 | 1.6e-05 | 1.2e-05 | 1.500E-05 | 2.646E-06 | 1.200E-05 | 1.700E-05 |
| 1622 | 82 | G82R | A | 1e-05 | 1.4e-05 | 1.3e-05 | 1.233E-05 | 2.082E-06 | 1.000E-05 | 1.400E-05 |
| 1623 | 82 | G82N | A | 1.5e-05 | 1.4e-05 | 1.7e-05 | 1.533E-05 | 1.528E-06 | 1.400E-05 | 1.700E-05 |
| 1624 | 82 | G82D | A | 1.4e-05 | 9.2e-06 | 1.1e-05 | 1.140E-05 | 2.425E-06 | 9.200E-06 | 1.400E-05 |
| 1625 | 82 | G82C | A | 1.8e-05 | 1.3e-05 | 2.1e-05 | 1.733E-05 | 4.041E-06 | 1.300E-05 | 2.100E-05 |
| 1626 | 82 | G82E | A | 9.6e-06 | 1e-05 | 1e-05 | 9.867E-06 | 2.309E-07 | 9.600E-06 | 1.000E-05 |
| 1627 | 82 | G82Q | A | 1.3e-05 | 7.7e-06 | 1.2e-05 | 1.090E-05 | 2.816E-06 | 7.700E-06 | 1.300E-05 |
| 1628 | 82 | G82G | A | 1.1e-05 | 1e-05 | 1.7e-05 | 1.267E-05 | 3.786E-06 | 1.000E-05 | 1.700E-05 |
| 1629 | 82 | G82H | A | 8.7e-06 | 9.8e-06 | 1.8e-05 | 1.217E-05 | 5.082E-06 | 8.700E-06 | 1.800E-05 |
| 1630 | 82 | G82I | A | 8.9e-06 | 9.8e-06 | 9.8e-06 | 9.500E-06 | 5.196E-07 | 8.900E-06 | 9.800E-06 |
| 1631 | 82 | G82L | A | 9.1e-06 | 1.5e-05 | 9e-06 | 1.103E-05 | 3.436E-06 | 9.000E-06 | 1.500E-05 |
| 1632 | 82 | G82K | A | 1.1e-05 | 1e-05 | 1.6e-05 | 1.233E-05 | 3.215E-06 | 1.000E-05 | 1.600E-05 |
| 1633 | 82 | G82M | A | 1.4e-05 | 9.7e-06 | 1.5e-05 | 1.290E-05 | 2.816E-06 | 9.700E-06 | 1.500E-05 |
| 1634 | 82 | G82F | A | 9.5e-06 | 1.1e-05 | 1.1e-05 | 1.050E-05 | 8.660E-07 | 9.500E-06 | 1.100E-05 |
| 1635 | 82 | G82P | A | 1.4e-05 | 1.5e-05 | 1.7e-05 | 1.533E-05 | 1.528E-06 | 1.400E-05 | 1.700E-05 |
| 1636 | 82 | G82S | A | 1.6e-05 | 1.4e-05 | 1.4e-05 | 1.467E-05 | 1.155E-06 | 1.400E-05 | 1.600E-05 |
| 1637 | 82 | G82T | A | 9.5e-06 | 9.6e-06 | 1.5e-05 | 1.137E-05 | 3.147E-06 | 9.500E-06 | 1.500E-05 |
| 1638 | 82 | G82W | A | 1e-05 | 1.2e-05 | 1.4e-05 | 1.200E-05 | 2.000E-06 | 1.000E-05 | 1.400E-05 |
| 1639 | 82 | G82Y | A | 9.9e-06 | 1.6e-05 | 1.2e-05 | 1.263E-05 | 3.099E-06 | 9.900E-06 | 1.600E-05 |
| 1640 | 82 | G82V | A | 1.5e-05 | 1.3e-05 | 1.4e-05 | 1.400E-05 | 1.000E-06 | 1.300E-05 | 1.500E-05 |
| 1641 | 83 | T83A | A | 1.3e-05 | 9.2e-06 | 1.1e-05 | 1.107E-05 | 1.901E-06 | 9.200E-06 | 1.300E-05 |
| 1642 | 83 | T83R | A | 1e-05 | 1.1e-05 | 8.2e-06 | 9.733E-06 | 1.419E-06 | 8.200E-06 | 1.100E-05 |
| 1643 | 83 | T83N | A | 1.1e-05 | 1.7e-05 | 1.1e-05 | 1.300E-05 | 3.464E-06 | 1.100E-05 | 1.700E-05 |
| 1644 | 83 | T83D | A | 1.1e-05 | 9e-06 | 1.6e-05 | 1.200E-05 | 3.606E-06 | 9.000E-06 | 1.600E-05 |
| 1645 | 83 | T83C | A | 1.7e-05 | 1.5e-05 | 1.1e-05 | 1.433E-05 | 3.055E-06 | 1.100E-05 | 1.700E-05 |
| 1646 | 83 | T83E | A | 1.6e-05 | 1.2e-05 | 1.6e-05 | 1.467E-05 | 2.309E-06 | 1.200E-05 | 1.600E-05 |
| 1647 | 83 | T83Q | A | 1.7e-05 | 1.4e-05 | 1.4e-05 | 1.500E-05 | 1.732E-06 | 1.400E-05 | 1.700E-05 |
| 1648 | 83 | T83G | A | 1.5e-05 | 7e-06 | 9.5e-06 | 1.050E-05 | 4.093E-06 | 7.000E-06 | 1.500E-05 |
| 1649 | 83 | T83H | A | 1.2e-05 | 1.6e-05 | 1.6e-05 | 1.467E-05 | 2.309E-06 | 1.200E-05 | 1.600E-05 |
| 1650 | 83 | T83I | A | 1.3e-05 | 1.2e-05 | 1.2e-05 | 1.233E-05 | 5.774E-07 | 1.200E-05 | 1.300E-05 |
| 1651 | 83 | T83L | A | 1.1e-05 | 9.2e-06 | 9.8e-06 | 1.000E-05 | 9.165E-07 | 9.200E-06 | 1.100E-05 |
| 1652 | 83 | T83K | A | 1.6e-05 | 1.6e-05 | 1e-05 | 1.400E-05 | 3.464E-06 | 1.000E-05 | 1.600E-05 |
| 1653 | 83 | T83M | A | 1.1e-05 | 5.9e-06 | 7.7e-06 | 8.200E-06 | 2.587E-06 | 5.900E-06 | 1.100E-05 |
| 1654 | 83 | T83F | A | 1.4e-05 | 1.1e-05 | 1.8e-05 | 1.433E-05 | 3.512E-06 | 1.100E-05 | 1.800E-05 |
| 1655 | 83 | T83P | A | 1.5e-05 | 1.2e-05 | 1e-05 | 1.233E-05 | 2.517E-06 | 1.000E-05 | 1.500E-05 |
| 1656 | 83 | T83S | A | 1.2e-05 | 8.8e-06 | 1.4e-05 | 1.160E-05 | 2.623E-06 | 8.800E-06 | 1.400E-05 |
| 1657 | 83 | T83T | A | 1.2e-05 | 1.7e-05 | 1.4e-05 | 1.433E-05 | 2.517E-06 | 1.200E-05 | 1.700E-05 |
| 1658 | 83 | T83W | A | 1.1e-05 | 1.6e-05 | 1.4e-05 | 1.367E-05 | 2.517E-06 | 1.100E-05 | 1.600E-05 |
| 1659 | 83 | T83Y | A | 1e-05 | 1.5e-05 | 1.6e-05 | 1.367E-05 | 3.215E-06 | 1.000E-05 | 1.600E-05 |
| 1660 | 83 | T83V | A | 9.5e-06 | 1.4e-05 | 1e-05 | 1.117E-05 | 2.466E-06 | 9.500E-06 | 1.400E-05 |
| 1661 | 84 | Q84A | A | 1.4e-05 | 1.3e-05 | 1.4e-05 | 1.367E-05 | 5.774E-07 | 1.300E-05 | 1.400E-05 |
| 1662 | 84 | Q84R | A | 1.5e-05 | 1.1e-05 | 1.6e-05 | 1.400E-05 | 2.646E-06 | 1.100E-05 | 1.600E-05 |
| 1663 | 84 | Q84N | A | 1.3e-05 | 1.5e-05 | 1.4e-05 | 1.400E-05 | 1.000E-06 | 1.300E-05 | 1.500E-05 |
| 1664 | 84 | Q84D | A | 1.6e-05 | 1.2e-05 | 1.1e-05 | 1.300E-05 | 2.646E-06 | 1.100E-05 | 1.600E-05 |
| 1665 | 84 | Q84C | A | 1.3e-05 | 1.4e-05 | 1.5e-05 | 1.400E-05 | 1.000E-06 | 1.300E-05 | 1.500E-05 |
| 1666 | 84 | Q84E | A | 9e-06 | 1.6e-05 | 1.2e-05 | 1.233E-05 | 3.512E-06 | 9.000E-06 | 1.600E-05 |
| 1667 | 84 | Q84Q | A | 1.6e-05 | 2.6e-05 | 8.3e-06 | 1.677E-05 | 8.875E-06 | 8.300E-06 | 2.600E-05 |
| 1668 | 84 | Q84G | A | 1.1e-05 | 1.1e-05 | 1.1e-05 | 1.100E-05 | 2.075E-21 | 1.100E-05 | 1.100E-05 |
| 1669 | 84 | Q84H | A | 9.8e-06 | 1.6e-05 | 1.2e-05 | 1.260E-05 | 3.143E-06 | 9.800E-06 | 1.600E-05 |
| 1670 | 84 | Q84I | A | 1.4e-05 | 1.6e-05 | 1.1e-05 | 1.367E-05 | 2.517E-06 | 1.100E-05 | 1.600E-05 |
| 1671 | 84 | Q84L | A | 1.1e-05 | 1.6e-05 | 9.6e-06 | 1.220E-05 | 3.365E-06 | 9.600E-06 | 1.600E-05 |
| 1672 | 84 | Q84K | A | 1.2e-05 | 1e-05 | 8.5e-06 | 1.017E-05 | 1.756E-06 | 8.500E-06 | 1.200E-05 |
| 1673 | 84 | Q84M | A | 1.3e-05 | 1.5e-05 | 1.2e-05 | 1.333E-05 | 1.528E-06 | 1.200E-05 | 1.500E-05 |
| 1674 | 84 | Q84F | A | 1.1e-05 | 1.4e-05 | 9.5e-06 | 1.150E-05 | 2.291E-06 | 9.500E-06 | 1.400E-05 |
| 1675 | 84 | Q84P | A | 1.6e-05 | 1.3e-05 | 8.9e-06 | 1.263E-05 | 3.564E-06 | 8.900E-06 | 1.600E-05 |
| 1676 | 84 | Q84S | A | 8.8e-06 | 9.6e-06 | 1.1e-05 | 9.800E-06 | 1.114E-06 | 8.800E-06 | 1.100E-05 |
| 1677 | 84 | Q84T | A | 1.5e-05 | 1.3e-05 | 1.6e-05 | 1.467E-05 | 1.528E-06 | 1.300E-05 | 1.600E-05 |
| 1678 | 84 | Q84W | A | 1.5e-05 | 4.1e-05 | 1.2e-05 | 2.267E-05 | 1.595E-05 | 1.200E-05 | 4.100E-05 |
| 1679 | 84 | Q84Y | A | 1.5e-05 | 1e-05 | 1.1e-05 | 1.200E-05 | 2.646E-06 | 1.000E-05 | 1.500E-05 |
| 1680 | 84 | Q84V | A | 1.5e-05 | 1.4e-05 | 1e-05 | 1.300E-05 | 2.646E-06 | 1.000E-05 | 1.500E-05 |
| 1681 | 85 | L85A | A | 9.3e-06 | 1.1e-05 | 1.5e-05 | 1.177E-05 | 2.926E-06 | 9.300E-06 | 1.500E-05 |
| 1682 | 85 | L85R | A | 1.6e-05 | 1.5e-05 | 1.1e-05 | 1.400E-05 | 2.646E-06 | 1.100E-05 | 1.600E-05 |
| 1683 | 85 | L85N | A | 1.5e-05 | 1e-05 | 9.7e-06 | 1.157E-05 | 2.977E-06 | 9.700E-06 | 1.500E-05 |
| 1684 | 85 | L85D | A | 1e-05 | 1.5e-05 | 1.4e-05 | 1.300E-05 | 2.646E-06 | 1.000E-05 | 1.500E-05 |
| 1685 | 85 | L85C | A | 8.4e-06 | 9.2e-06 | 1.4e-05 | 1.053E-05 | 3.029E-06 | 8.400E-06 | 1.400E-05 |
| 1686 | 85 | L85E | A | 1.4e-05 | 1.5e-05 | 9.3e-06 | 1.277E-05 | 3.044E-06 | 9.300E-06 | 1.500E-05 |
| 1687 | 85 | L85Q | A | 1e-05 | 9.2e-06 | 9.3e-06 | 9.500E-06 | 4.359E-07 | 9.200E-06 | 1.000E-05 |
| 1688 | 85 | L85G | A | 1.5e-05 | 9.3e-06 | 9.4e-06 | 1.123E-05 | 3.262E-06 | 9.300E-06 | 1.500E-05 |
| 1689 | 85 | L85H | A | 1.3e-05 | 1e-05 | 1.3e-05 | 1.200E-05 | 1.732E-06 | 1.000E-05 | 1.300E-05 |
| 1690 | 85 | L85I | A | 1.6e-05 | 1.1e-05 | 1e-05 | 1.233E-05 | 3.215E-06 | 1.000E-05 | 1.600E-05 |
| 1691 | 85 | L85L | A | 1.5e-05 | 1.1e-05 | 1.5e-05 | 1.367E-05 | 2.309E-06 | 1.100E-05 | 1.500E-05 |
| 1692 | 85 | L85K | A | 1.4e-05 | 9.8e-06 | 9.1e-06 | 1.097E-05 | 2.650E-06 | 9.100E-06 | 1.400E-05 |
| 1693 | 85 | L85M | A | 1.5e-05 | 1.1e-05 | 9.6e-06 | 1.187E-05 | 2.802E-06 | 9.600E-06 | 1.500E-05 |
| 1694 | 85 | L85F | A | 1.1e-05 | 1.4e-05 | 9e-06 | 1.133E-05 | 2.517E-06 | 9.000E-06 | 1.400E-05 |
| 1695 | 85 | L85P | A | 9.4e-06 | 1.1e-05 | 9.5e-06 | 9.967E-06 | 8.963E-07 | 9.400E-06 | 1.100E-05 |
| 1696 | 85 | L85S | A | 9.7e-06 | 9.7e-06 | 1.6e-05 | 1.180E-05 | 3.637E-06 | 9.700E-06 | 1.600E-05 |
| 1697 | 85 | L85T | A | 1.1e-05 | 9.8e-06 | 1.5e-05 | 1.193E-05 | 2.723E-06 | 9.800E-06 | 1.500E-05 |
| 1698 | 85 | L85W | A | 1.2e-05 | 1.4e-05 | 1e-05 | 1.200E-05 | 2.000E-06 | 1.000E-05 | 1.400E-05 |
| 1699 | 85 | L85Y | A | 1.2e-05 | 1.4e-05 | 1.4e-05 | 1.333E-05 | 1.155E-06 | 1.200E-05 | 1.400E-05 |

| | | | | | | | | | | |
|---|---|---|---|---|---|---|---|---|---|---|
| 1700 | 85 | L85V | A | 1.6e-05 | 1.5e-05 | 1.6e-05 | 1.567E-05 | 5.774E-07 | 1.500E-05 | 1.600E-05 |
| 1701 | 86 | F86A | A | 1e-05 | 1.5e-05 | 1.7e-05 | 1.400E-05 | 3.606E-06 | 1.000E-05 | 1.700E-05 |
| 1702 | 86 | F86R | A | 1.5e-05 | 9.5e-06 | 1.6e-05 | 1.350E-05 | 3.500E-06 | 9.500E-06 | 1.600E-05 |
| 1703 | 86 | F86N | A | 9.4e-06 | 1.4e-05 | 1.1e-05 | 1.147E-05 | 2.335E-06 | 9.400E-06 | 1.400E-05 |
| 1704 | 86 | F86D | A | 9.8e-06 | 9.5e-06 | 1e-05 | 9.767E-06 | 2.517E-07 | 9.500E-06 | 1.000E-05 |
| 1705 | 86 | F86C | A | 1.6e-05 | 8.3e-06 | 1.4e-05 | 1.277E-05 | 3.995E-06 | 8.300E-06 | 1.600E-05 |
| 1706 | 86 | F86E | A | 1.5e-05 | 9.7e-06 | 1.3e-05 | 1.257E-05 | 2.676E-06 | 9.700E-06 | 1.500E-05 |
| 1707 | 86 | F86Q | A | 9e-06 | 9.5e-06 | 1.4e-05 | 1.083E-05 | 2.754E-06 | 9.000E-06 | 1.400E-05 |
| 1708 | 86 | F86G | A | 1.7e-05 | 1.1e-05 | 1e-05 | 1.267E-05 | 3.786E-06 | 1.000E-05 | 1.700E-05 |
| 1709 | 86 | F86H | A | 1.1e-05 | 1.6e-05 | 1.1e-05 | 1.267E-05 | 2.887E-06 | 1.100E-05 | 1.600E-05 |
| 1710 | 86 | F86I | A | 9.9e-06 | 1.7e-05 | 1.2e-05 | 1.297E-05 | 3.647E-06 | 9.900E-06 | 1.700E-05 |
| 1711 | 86 | F86L | A | 1.3e-05 | 1.5e-05 | 1.4e-05 | 1.400E-05 | 1.000E-06 | 1.300E-05 | 1.500E-05 |
| 1712 | 86 | F86K | A | 9.9e-06 | 1.1e-05 | 1.3e-05 | 1.130E-05 | 1.572E-06 | 9.900E-06 | 1.300E-05 |
| 1713 | 86 | F86M | A | 1.6e-05 | 1e-05 | 1.9e-05 | 1.500E-05 | 4.583E-06 | 1.000E-05 | 1.900E-05 |
| 1714 | 86 | F86F | A | 1.1e-05 | 1.1e-05 | 1.2e-05 | 1.133E-05 | 5.774E-07 | 1.100E-05 | 1.200E-05 |
| 1715 | 86 | F86P | A | 1e-05 | 1.8e-05 | 9.3e-06 | 1.243E-05 | 4.834E-06 | 9.300E-06 | 1.800E-05 |
| 1716 | 86 | F86S | A | 1.4e-05 | 1.1e-05 | 1.6e-05 | 1.367E-05 | 2.517E-06 | 1.100E-05 | 1.600E-05 |
| 1717 | 86 | F86T | A | 9.9e-06 | 1.5e-05 | 1.4e-05 | 1.297E-05 | 2.702E-06 | 9.900E-06 | 1.500E-05 |
| 1718 | 86 | F86W | A | 1.4e-05 | 1.5e-05 | 1.4e-05 | 1.433E-05 | 5.774E-07 | 1.400E-05 | 1.500E-05 |
| 1719 | 86 | F86Y | A | 1.4e-05 | 9.1e-06 | 1.1e-05 | 1.137E-05 | 2.470E-06 | 9.100E-06 | 1.400E-05 |
| 1720 | 86 | F86V | A | 1.1e-05 | 1.2e-05 | 1e-05 | 1.100E-05 | 1.000E-06 | 1.000E-05 | 1.200E-05 |
| 1721 | 87 | E87A | A | 1.7e-05 | 1.1e-05 | 1.5e-05 | 1.433E-05 | 3.055E-06 | 1.100E-05 | 1.700E-05 |
| 1722 | 87 | E87R | A | 1.1e-05 | 1e-05 | 1.4e-05 | 1.167E-05 | 2.082E-06 | 1.000E-05 | 1.400E-05 |
| 1723 | 87 | E87N | A | 1.1e-05 | 8.6e-06 | 9.3e-06 | 9.633E-06 | 1.234E-06 | 8.600E-06 | 1.100E-05 |
| 1724 | 87 | E87D | A | 9.2e-06 | 9.2e-06 | 9.5e-06 | 9.300E-06 | 1.732E-07 | 9.200E-06 | 9.500E-06 |
| 1725 | 87 | E87C | A | 1.2e-05 | 1e-05 | 1.1e-05 | 1.100E-05 | 1.000E-06 | 1.000E-05 | 1.200E-05 |
| 1726 | 87 | E87E | A | 1.5e-05 | 1.1e-05 | 1.5e-05 | 1.367E-05 | 2.309E-06 | 1.100E-05 | 1.500E-05 |
| 1727 | 87 | E87Q | A | 9.7e-06 | 1.4e-05 | 1.5e-05 | 1.290E-05 | 2.816E-06 | 9.700E-06 | 1.500E-05 |
| 1728 | 87 | E87G | A | 1.1e-05 | 1.8e-05 | 1.6e-05 | 1.500E-05 | 3.606E-06 | 1.100E-05 | 1.800E-05 |
| 1729 | 87 | E87H | A | 1.6e-05 | 1.5e-05 | 1.4e-05 | 1.500E-05 | 1.000E-06 | 1.400E-05 | 1.600E-05 |
| 1730 | 87 | E87I | A | 9.8e-06 | 1.6e-05 | 9.1e-06 | 1.163E-05 | 3.798E-06 | 9.100E-06 | 1.600E-05 |
| 1731 | 87 | E87L | A | 1.2e-05 | 1e-05 | 1.5e-05 | 1.233E-05 | 2.517E-06 | 1.000E-05 | 1.500E-05 |
| 1732 | 87 | E87K | A | 1.1e-05 | 9e-06 | 1.6e-05 | 1.200E-05 | 3.606E-06 | 9.000E-06 | 1.600E-05 |
| 1733 | 87 | E87M | A | 1.2e-05 | 1.5e-05 | 1.3e-05 | 1.333E-05 | 1.528E-06 | 1.200E-05 | 1.500E-05 |
| 1734 | 87 | E87F | A | 1.6e-05 | 1.1e-05 | 1.2e-05 | 1.300E-05 | 2.646E-06 | 1.100E-05 | 1.600E-05 |
| 1735 | 87 | E87P | A | 1e-05 | 1.5e-05 | 9.8e-06 | 1.160E-05 | 2.946E-06 | 9.800E-06 | 1.500E-05 |
| 1736 | 87 | E87S | A | 1.5e-05 | 1.5e-05 | 9.3e-06 | 1.310E-05 | 3.291E-06 | 9.300E-06 | 1.500E-05 |
| 1737 | 87 | E87T | A | 9.7e-06 | 1.5e-05 | 1.3e-05 | 1.257E-05 | 2.676E-06 | 9.700E-06 | 1.500E-05 |
| 1738 | 87 | E87W | A | 8.5e-06 | 1.5e-05 | 1.4e-05 | 1.250E-05 | 3.500E-06 | 8.500E-06 | 1.500E-05 |
| 1739 | 87 | E87Y | A | 1e-05 | 1.2e-05 | 2.3e-05 | 1.500E-05 | 7.000E-06 | 1.000E-05 | 2.300E-05 |
| 1740 | 87 | E87V | A | 7.7e-06 | 1.1e-05 | 1.5e-05 | 1.123E-05 | 3.656E-06 | 7.700E-06 | 1.500E-05 |
| 1741 | 88 | D88A | A | 1.6e-05 | 1e-05 | 1.7e-05 | 1.433E-05 | 3.786E-06 | 1.000E-05 | 1.700E-05 |
| 1742 | 88 | D88R | A | 1.1e-05 | 1.3e-05 | 1.1e-05 | 1.167E-05 | 1.155E-06 | 1.100E-05 | 1.300E-05 |
| 1743 | 88 | D88N | A | 1.1e-05 | 1.5e-05 | 8.8e-06 | 1.160E-05 | 3.143E-06 | 8.800E-06 | 1.500E-05 |
| 1744 | 88 | D88D | A | 1.3e-05 | 9.2e-06 | 1e-05 | 1.073E-05 | 2.003E-06 | 9.200E-06 | 1.300E-05 |
| 1745 | 88 | D88C | A | 1.4e-05 | 9.6e-06 | 8.5e-06 | 1.070E-05 | 2.910E-06 | 8.500E-06 | 1.400E-05 |
| 1746 | 88 | D88E | A | 1.5e-05 | 1.1e-05 | 1.6e-05 | 1.400E-05 | 2.646E-06 | 1.100E-05 | 1.600E-05 |
| 1747 | 88 | D88Q | A | 9.6e-06 | 1e-05 | 1e-05 | 9.867E-06 | 2.309E-07 | 9.600E-06 | 1.000E-05 |
| 1748 | 88 | D88G | A | 1.4e-05 | 1e-05 | 1.1e-05 | 1.167E-05 | 2.082E-06 | 1.000E-05 | 1.400E-05 |
| 1749 | 88 | D88H | A | 9.2e-06 | 9.5e-06 | 1.4e-05 | 1.090E-05 | 2.689E-06 | 9.200E-06 | 1.400E-05 |
| 1750 | 88 | D88I | A | 1e-05 | 1e-05 | 9.3e-06 | 9.767E-06 | 4.041E-07 | 9.300E-06 | 1.000E-05 |
| 1751 | 88 | D88L | A | 1e-05 | 1.2e-05 | 2.6e-05 | 1.600E-05 | 8.718E-06 | 1.000E-05 | 2.600E-05 |
| 1752 | 88 | D88K | A | 1.5e-05 | 1.5e-05 | 9.2e-06 | 1.307E-05 | 3.349E-06 | 9.200E-06 | 1.500E-05 |
| 1753 | 88 | D88M | A | 1.6e-05 | 1.4e-05 | 1.6e-05 | 1.533E-05 | 1.155E-06 | 1.400E-05 | 1.600E-05 |
| 1754 | 88 | D88F | A | 1.4e-05 | 9.6e-06 | 1.6e-05 | 1.320E-05 | 3.274E-06 | 9.600E-06 | 1.600E-05 |
| 1755 | 88 | D88P | A | 1.1e-05 | 1.1e-05 | 1.7e-05 | 1.300E-05 | 3.464E-06 | 1.100E-05 | 1.700E-05 |
| 1756 | 88 | D88S | A | 1.5e-05 | 9.8e-06 | 1.5e-05 | 1.327E-05 | 3.002E-06 | 9.800E-06 | 1.500E-05 |
| 1757 | 88 | D88T | A | 1.5e-05 | 1.5e-05 | 1.6e-05 | 1.533E-05 | 5.774E-07 | 1.500E-05 | 1.600E-05 |
| 1758 | 88 | D88W | A | 1.4e-05 | 9.9e-06 | 1e-05 | 1.130E-05 | 2.339E-06 | 9.900E-06 | 1.400E-05 |
| 1759 | 88 | D88Y | A | 1.7e-05 | 1.5e-05 | 1.4e-05 | 1.533E-05 | 1.528E-06 | 1.400E-05 | 1.700E-05 |
| 1760 | 88 | D88V | A | 8.9e-06 | 1.6e-05 | 1.4e-05 | 1.297E-05 | 3.661E-06 | 8.900E-06 | 1.600E-05 |
| 1761 | 89 | N89A | A | 1.1e-05 | 9e-06 | 1e-05 | 1.000E-05 | 1.000E-06 | 9.000E-06 | 1.100E-05 |
| 1762 | 89 | N89R | A | 1.4e-05 | 1e-05 | 8.5e-06 | 1.083E-05 | 2.843E-06 | 8.500E-06 | 1.400E-05 |
| 1763 | 89 | N89N | A | 1e-05 | 1.6e-05 | 1.1e-05 | 1.233E-05 | 3.215E-06 | 1.000E-05 | 1.600E-05 |
| 1764 | 89 | N89D | A | 9.9e-06 | 1.2e-05 | 1.2e-05 | 1.130E-05 | 1.212E-06 | 9.900E-06 | 1.200E-05 |
| 1765 | 89 | N89C | A | 8.6e-06 | 1.5e-05 | 1.5e-05 | 1.287E-05 | 3.695E-06 | 8.600E-06 | 1.500E-05 |
| 1766 | 89 | N89E | A | 1.7e-05 | 1.1e-05 | 1.3e-05 | 1.367E-05 | 3.055E-06 | 1.100E-05 | 1.700E-05 |
| 1767 | 89 | N89Q | A | 1.6e-05 | 1.3e-05 | 1.3e-05 | 1.400E-05 | 1.732E-06 | 1.300E-05 | 1.600E-05 |
| 1768 | 89 | N89G | A | 1.1e-05 | 1e-05 | 1.2e-05 | 1.100E-05 | 1.000E-06 | 1.000E-05 | 1.200E-05 |
| 1769 | 89 | N89H | A | 1.1e-05 | 1.7e-05 | 1.6e-05 | 1.467E-05 | 3.215E-06 | 1.100E-05 | 1.700E-05 |
| 1770 | 89 | N89I | A | 1.7e-05 | 1.5e-05 | 1.1e-05 | 1.433E-05 | 3.055E-06 | 1.100E-05 | 1.700E-05 |
| 1771 | 89 | N89L | A | 1.7e-05 | 1.5e-05 | 1.5e-05 | 1.567E-05 | 1.155E-06 | 1.500E-05 | 1.700E-05 |
| 1772 | 89 | N89K | A | 1.6e-05 | 1.1e-05 | 9.8e-06 | 1.227E-05 | 3.288E-06 | 9.800E-06 | 1.600E-05 |
| 1773 | 89 | N89M | A | 9.9e-06 | 1e-05 | 9.7e-06 | 9.867E-06 | 1.528E-07 | 9.700E-06 | 1.000E-05 |
| 1774 | 89 | N89F | A | 1.5e-05 | 1.1e-05 | 1.6e-05 | 1.400E-05 | 2.646E-06 | 1.100E-05 | 1.600E-05 |
| 1775 | 89 | N89P | A | 1.5e-05 | 1.4e-05 | 1e-05 | 1.300E-05 | 2.646E-06 | 1.000E-05 | 1.500E-05 |
| 1776 | 89 | N89S | A | 1e-05 | 9.8e-06 | 1.5e-05 | 1.160E-05 | 2.946E-06 | 9.800E-06 | 1.500E-05 |
| 1777 | 89 | N89T | A | 9.9e-06 | 1.6e-05 | 1.2e-05 | 1.263E-05 | 3.099E-06 | 9.900E-06 | 1.600E-05 |
| 1778 | 89 | N89W | A | 1.4e-05 | 1.6e-05 | 1.7e-05 | 1.567E-05 | 1.528E-06 | 1.400E-05 | 1.700E-05 |
| 1779 | 89 | N89Y | A | 1.5e-05 | 1.2e-05 | 1e-05 | 1.233E-05 | 2.517E-06 | 1.000E-05 | 1.500E-05 |
| 1780 | 89 | N89V | A | 1.4e-05 | 1.7e-05 | 1.4e-05 | 1.500E-05 | 1.732E-06 | 1.400E-05 | 1.700E-05 |
| 1781 | 90 | Y90A | A | 1.6e-05 | 1.1e-05 | 1.4e-05 | 1.367E-05 | 2.517E-06 | 1.100E-05 | 1.600E-05 |
| 1782 | 90 | Y90R | A | 9.4e-06 | 1.4e-05 | 1.4e-05 | 1.247E-05 | 2.656E-06 | 9.400E-06 | 1.400E-05 |
| 1783 | 90 | Y90N | A | 9.4e-06 | 1.1e-05 | 1.6e-05 | 1.213E-05 | 3.443E-06 | 9.400E-06 | 1.600E-05 |
| 1784 | 90 | Y90D | A | 9e-06 | 1.1e-05 | 1.6e-05 | 1.200E-05 | 3.606E-06 | 9.000E-06 | 1.600E-05 |

| | | | | | | | | | | |
|---|---|---|---|---|---|---|---|---|---|---|
| 1785 | 90 | Y90C | A | 7.3e-06 | 1e-05 | 9.9e-06 | 9.067E-06 | 1.531E-06 | 7.300E-06 | 1.000E-05 |
| 1786 | 90 | Y90E | A | 1.6e-05 | 1.1e-05 | 9.3e-06 | 1.210E-05 | 3.483E-06 | 9.300E-06 | 1.600E-05 |
| 1787 | 90 | Y90Q | A | 9.3e-06 | 1.4e-05 | 1.6e-05 | 1.310E-05 | 3.439E-06 | 9.300E-06 | 1.600E-05 |
| 1788 | 90 | Y90G | A | 1.1e-05 | 9.3e-06 | 9.2e-06 | 9.833E-06 | 1.012E-06 | 9.200E-06 | 1.100E-05 |
| 1789 | 90 | Y90H | A | 1.5e-05 | 1.4e-05 | 1.4e-05 | 1.433E-05 | 5.774E-07 | 1.400E-05 | 1.500E-05 |
| 1790 | 90 | Y90I | A | 1.8e-05 | 1.1e-05 | 8.8e-06 | 1.260E-05 | 4.804E-06 | 8.800E-06 | 1.800E-05 |
| 1791 | 90 | Y90L | A | 1.1e-05 | 1.6e-05 | 1.4e-05 | 1.367E-05 | 2.517E-06 | 1.100E-05 | 1.600E-05 |
| 1792 | 90 | Y90K | A | 1.3e-05 | 1.6e-05 | 1.2e-05 | 1.367E-05 | 2.082E-06 | 1.200E-05 | 1.600E-05 |
| 1793 | 90 | Y90M | A | 1.1e-05 | 1.4e-05 | 1.5e-05 | 1.333E-05 | 2.082E-06 | 1.100E-05 | 1.500E-05 |
| 1794 | 90 | Y90F | A | 1.4e-05 | 1.1e-05 | 9.7e-06 | 1.157E-05 | 2.205E-06 | 9.700E-06 | 1.400E-05 |
| 1795 | 90 | Y90P | A | 1.5e-05 | 1.1e-05 | 1.5e-05 | 1.367E-05 | 2.309E-06 | 1.100E-05 | 1.500E-05 |
| 1796 | 90 | Y90S | A | 1.4e-05 | 1.4e-05 | 1e-05 | 1.267E-05 | 2.309E-06 | 1.000E-05 | 1.400E-05 |
| 1797 | 90 | Y90T | A | 9.6e-06 | 1.4e-05 | 1.1e-05 | 1.153E-05 | 2.248E-06 | 9.600E-06 | 1.400E-05 |
| 1798 | 90 | Y90W | A | 9.8e-06 | 1.2e-05 | 8.7e-06 | 1.017E-05 | 1.680E-06 | 8.700E-06 | 1.200E-05 |
| 1799 | 90 | Y90Y | A | 1.5e-05 | 1.3e-05 | 1.4e-05 | 1.400E-05 | 1.000E-06 | 1.300E-05 | 1.500E-05 |
| 1800 | 90 | Y90V | A | 1.1e-05 | 1.6e-05 | 1.5e-05 | 1.400E-05 | 2.646E-06 | 1.100E-05 | 1.600E-05 |
| 1801 | 91 | A91A | A | 1.3e-05 | 1.8e-05 | 1.1e-05 | 1.400E-05 | 3.606E-06 | 1.100E-05 | 1.800E-05 |
| 1802 | 91 | A91R | A | 1.1e-05 | 1.8e-05 | 1.3e-05 | 1.400E-05 | 3.606E-06 | 1.100E-05 | 1.800E-05 |
| 1803 | 91 | A91N | A | 1.4e-05 | 1.6e-05 | 9.8e-06 | 1.327E-05 | 3.164E-06 | 9.800E-06 | 1.600E-05 |
| 1804 | 91 | A91D | A | 1.1e-05 | 1.4e-05 | 1.4e-05 | 1.300E-05 | 1.732E-06 | 1.100E-05 | 1.400E-05 |
| 1805 | 91 | A91C | A | 9.3e-06 | 1.5e-05 | 1e-05 | 1.143E-05 | 3.109E-06 | 9.300E-06 | 1.500E-05 |
| 1806 | 91 | A91E | A | 1.6e-05 | 1.1e-05 | 1.5e-05 | 1.400E-05 | 2.646E-06 | 1.100E-05 | 1.600E-05 |
| 1807 | 91 | A91Q | A | 1e-05 | 9.4e-06 | 1.6e-05 | 1.180E-05 | 3.650E-06 | 9.400E-06 | 1.600E-05 |
| 1808 | 91 | A91G | A | 2.2e-05 | 1.7e-05 | 9.3e-06 | 1.610E-05 | 6.398E-06 | 9.300E-06 | 2.200E-05 |
| 1809 | 91 | A91H | A | 1e-05 | 1.4e-05 | 1.5e-05 | 1.300E-05 | 2.646E-06 | 1.000E-05 | 1.500E-05 |
| 1810 | 91 | A91I | A | 1.4e-05 | 1.4e-05 | 8.4e-06 | 1.213E-05 | 3.233E-06 | 8.400E-06 | 1.400E-05 |
| 1811 | 91 | A91L | A | 9.6e-06 | 1.4e-05 | 1.3e-05 | 1.220E-05 | 2.307E-06 | 9.600E-06 | 1.400E-05 |
| 1812 | 91 | A91K | A | 1e-05 | 1.1e-05 | 1.1e-05 | 1.067E-05 | 5.774E-07 | 1.000E-05 | 1.100E-05 |
| 1813 | 91 | A91M | A | 1.1e-05 | 1.2e-05 | 9.6e-06 | 1.087E-05 | 1.206E-06 | 9.600E-06 | 1.200E-05 |
| 1814 | 91 | A91F | A | 1.1e-05 | 1.4e-05 | 1.2e-05 | 1.233E-05 | 1.528E-06 | 1.100E-05 | 1.400E-05 |
| 1815 | 91 | A91P | A | 9.7e-06 | 1.1e-05 | 1.3e-05 | 1.123E-05 | 1.662E-06 | 9.700E-06 | 1.300E-05 |
| 1816 | 91 | A91S | A | 1.4e-05 | 1.5e-05 | 1.4e-05 | 1.433E-05 | 5.774E-07 | 1.400E-05 | 1.500E-05 |
| 1817 | 91 | A91T | A | 1.3e-05 | 1.2e-05 | 1.1e-05 | 1.200E-05 | 1.000E-06 | 1.100E-05 | 1.300E-05 |
| 1818 | 91 | A91W | A | 1.6e-05 | 1e-05 | 1.5e-05 | 1.367E-05 | 3.215E-06 | 1.000E-05 | 1.600E-05 |
| 1819 | 91 | A91Y | A | 1.7e-05 | 9.7e-06 | 1.5e-05 | 1.390E-05 | 3.772E-06 | 9.700E-06 | 1.700E-05 |
| 1820 | 91 | A91V | A | 1.1e-05 | 1.1e-05 | 1.1e-05 | 1.100E-05 | 2.075E-21 | 1.100E-05 | 1.100E-05 |
| 1821 | 92 | L92A | A | 1.2e-05 | 1.6e-05 | 1.7e-05 | 1.500E-05 | 2.646E-06 | 1.200E-05 | 1.700E-05 |
| 1822 | 92 | L92R | A | 1.5e-05 | 1.3e-05 | 1.1e-05 | 1.300E-05 | 2.000E-06 | 1.100E-05 | 1.500E-05 |
| 1823 | 92 | L92N | A | 1.6e-05 | 1.1e-05 | 8.9e-06 | 1.197E-05 | 3.647E-06 | 8.900E-06 | 1.600E-05 |
| 1824 | 92 | L92D | A | 1.5e-05 | 1.7e-05 | 1.6e-05 | 1.600E-05 | 1.000E-06 | 1.500E-05 | 1.700E-05 |
| 1825 | 92 | L92C | A | 9.1e-06 | 1.3e-05 | 1.6e-05 | 1.270E-05 | 3.460E-06 | 9.100E-06 | 1.600E-05 |
| 1826 | 92 | L92E | A | 1.4e-05 | 1.4e-05 | 1.6e-05 | 1.467E-05 | 1.155E-06 | 1.400E-05 | 1.600E-05 |
| 1827 | 92 | L92Q | A | 1e-05 | 7.9e-06 | 1e-05 | 9.300E-06 | 1.212E-06 | 7.900E-06 | 1.000E-05 |
| 1828 | 92 | L92G | A | 1.6e-05 | 1.1e-05 | 1.5e-05 | 1.400E-05 | 2.646E-06 | 1.100E-05 | 1.600E-05 |
| 1829 | 92 | L92H | A | 8.3e-06 | 9.1e-06 | 1.4e-05 | 1.047E-05 | 3.086E-06 | 8.300E-06 | 1.400E-05 |
| 1830 | 92 | L92I | A | 9.5e-06 | 1.4e-05 | 1.1e-05 | 1.150E-05 | 2.291E-06 | 9.500E-06 | 1.400E-05 |
| 1831 | 92 | L92L | A | 1.1e-05 | 1.1e-05 | 1.6e-05 | 1.267E-05 | 2.887E-06 | 1.100E-05 | 1.600E-05 |
| 1832 | 92 | L92K | A | 1.3e-05 | 1.4e-05 | 1.5e-05 | 1.400E-05 | 1.000E-06 | 1.300E-05 | 1.500E-05 |
| 1833 | 92 | L92M | A | 1.5e-05 | 1e-05 | 1.4e-05 | 1.300E-05 | 2.646E-06 | 1.000E-05 | 1.500E-05 |
| 1834 | 92 | L92F | A | 1.5e-05 | 7.4e-06 | 1.1e-05 | 1.113E-05 | 3.802E-06 | 7.400E-06 | 1.500E-05 |
| 1835 | 92 | L92P | A | 1e-05 | 1.3e-05 | 1.1e-05 | 1.133E-05 | 1.528E-06 | 1.000E-05 | 1.300E-05 |
| 1836 | 92 | L92S | A | 1.4e-05 | 1.5e-05 | 1.4e-05 | 1.433E-05 | 5.774E-07 | 1.400E-05 | 1.500E-05 |
| 1837 | 92 | L92T | A | 1.5e-05 | 9.6e-06 | 1.5e-05 | 1.320E-05 | 3.118E-06 | 9.600E-06 | 1.500E-05 |
| 1838 | 92 | L92W | A | 1.2e-05 | 1.1e-05 | 1.4e-05 | 1.233E-05 | 1.528E-06 | 1.100E-05 | 1.400E-05 |
| 1839 | 92 | L92Y | A | 9.9e-06 | 1.3e-05 | 9.8e-06 | 1.090E-05 | 1.819E-06 | 9.800E-06 | 1.300E-05 |
| 1840 | 92 | L92V | A | 9e-06 | 1.6e-05 | 1.4e-05 | 1.300E-05 | 3.606E-06 | 9.000E-06 | 1.600E-05 |
| 1841 | 93 | A93A | A | 1.3e-05 | 1.4e-05 | 9.9e-06 | 1.230E-05 | 2.138E-06 | 9.900E-06 | 1.400E-05 |
| 1842 | 93 | A93R | A | 1.1e-05 | 1.6e-05 | 1.3e-05 | 1.333E-05 | 2.517E-06 | 1.100E-05 | 1.600E-05 |
| 1843 | 93 | A93N | A | 1.2e-05 | 1.4e-05 | 1e-05 | 1.200E-05 | 2.000E-06 | 1.000E-05 | 1.400E-05 |
| 1844 | 93 | A93D | A | 9.7e-06 | 1.1e-05 | 1.4e-05 | 1.157E-05 | 2.205E-06 | 9.700E-06 | 1.400E-05 |
| 1845 | 93 | A93C | A | 1.4e-05 | 1.4e-05 | 8.8e-06 | 1.227E-05 | 3.002E-06 | 8.800E-06 | 1.400E-05 |
| 1846 | 93 | A93E | A | 8.3e-06 | 1.5e-05 | 1.6e-05 | 1.310E-05 | 4.187E-06 | 8.300E-06 | 1.600E-05 |
| 1847 | 93 | A93Q | A | 1.4e-05 | 1.5e-05 | 1.1e-05 | 1.333E-05 | 2.082E-06 | 1.100E-05 | 1.500E-05 |
| 1848 | 93 | A93G | A | 1.6e-05 | 1.6e-05 | 1.7e-05 | 1.633E-05 | 5.774E-07 | 1.600E-05 | 1.700E-05 |
| 1849 | 93 | A93H | A | 1.6e-05 | 1.6e-05 | 1.5e-05 | 1.567E-05 | 5.774E-07 | 1.500E-05 | 1.600E-05 |
| 1850 | 93 | A93I | A | 1.1e-05 | 1.3e-05 | 1.5e-05 | 1.300E-05 | 2.000E-06 | 1.100E-05 | 1.500E-05 |
| 1851 | 93 | A93L | A | 9.7e-06 | 1.5e-05 | 1.4e-05 | 1.290E-05 | 2.816E-06 | 9.700E-06 | 1.500E-05 |
| 1852 | 93 | A93K | A | 1.6e-05 | 1.3e-05 | 1e-05 | 1.300E-05 | 3.000E-06 | 1.000E-05 | 1.600E-05 |
| 1853 | 93 | A93M | A | 1.1e-05 | 8.3e-06 | 1.5e-05 | 1.143E-05 | 3.371E-06 | 8.300E-06 | 1.500E-05 |
| 1854 | 93 | A93F | A | 9.7e-06 | 1.1e-05 | 1.7e-05 | 1.257E-05 | 3.894E-06 | 9.700E-06 | 1.700E-05 |
| 1855 | 93 | A93P | A | 8.9e-06 | 7.7e-06 | 1.4e-05 | 1.020E-05 | 3.345E-06 | 7.700E-06 | 1.400E-05 |
| 1856 | 93 | A93S | A | 9.4e-06 | 1.3e-05 | 1.4e-05 | 1.213E-05 | 2.419E-06 | 9.400E-06 | 1.400E-05 |
| 1857 | 93 | A93T | A | 1.1e-05 | 1.6e-05 | 1.4e-05 | 1.367E-05 | 2.517E-06 | 1.100E-05 | 1.600E-05 |
| 1858 | 93 | A93W | A | 3.2e-05 | 8.5e-06 | 1.5e-05 | 1.850E-05 | 1.213E-05 | 8.500E-06 | 3.200E-05 |
| 1859 | 93 | A93Y | A | 1e-05 | 9.7e-06 | 1.2e-05 | 1.057E-05 | 1.250E-06 | 9.700E-06 | 1.200E-05 |
| 1860 | 93 | A93V | A | 1e-05 | 1.3e-05 | 1e-05 | 1.100E-05 | 1.732E-06 | 1.000E-05 | 1.300E-05 |
| 1861 | 94 | V94A | A | 1.2e-05 | 1.6e-05 | 1.6e-05 | 1.467E-05 | 2.309E-06 | 1.200E-05 | 1.600E-05 |
| 1862 | 94 | V94R | A | 8.4e-06 | 1.4e-05 | 1.1e-05 | 1.113E-05 | 2.802E-06 | 8.400E-06 | 1.400E-05 |
| 1863 | 94 | V94N | A | 1.4e-05 | 1.4e-05 | 1.5e-05 | 1.433E-05 | 5.774E-07 | 1.400E-05 | 1.500E-05 |
| 1864 | 94 | V94D | A | 1.6e-05 | 1.6e-05 | 1.5e-05 | 1.567E-05 | 5.774E-07 | 1.500E-05 | 1.600E-05 |
| 1865 | 94 | V94C | A | 1e-05 | 1.1e-05 | 9e-06 | 1.000E-05 | 1.000E-06 | 9.000E-06 | 1.100E-05 |
| 1866 | 94 | V94E | A | 9.8e-06 | 1.5e-05 | 1.4e-05 | 1.293E-05 | 2.759E-06 | 9.800E-06 | 1.500E-05 |
| 1867 | 94 | V94Q | A | 1.4e-05 | 1e-05 | 1.6e-05 | 1.333E-05 | 3.055E-06 | 1.000E-05 | 1.600E-05 |
| 1868 | 94 | V94G | A | 1.7e-05 | 1e-05 | 1.3e-05 | 1.333E-05 | 3.512E-06 | 1.000E-05 | 1.700E-05 |
| 1869 | 94 | V94H | A | 1.5e-05 | 1.3e-05 | 1e-05 | 1.267E-05 | 2.517E-06 | 1.000E-05 | 1.500E-05 |

| | | | | | | | | | | |
|---|---|---|---|---|---|---|---|---|---|---|
| 1870 | 94 | V94I | A | 1.5e-05 | 1e-05 | 1.8e-05 | 1.433E-05 | 4.041E-06 | 1.000E-05 | 1.800E-05 |
| 1871 | 94 | V94L | A | 9.8e-06 | 8.7e-06 | 1.6e-05 | 1.150E-05 | 3.936E-06 | 8.700E-06 | 1.600E-05 |
| 1872 | 94 | V94K | A | 8.2e-06 | 1.5e-05 | 1.1e-05 | 1.140E-05 | 3.418E-06 | 8.200E-06 | 1.500E-05 |
| 1873 | 94 | V94M | A | 1.3e-05 | 9.4e-06 | 9.3e-06 | 1.057E-05 | 2.108E-06 | 9.300E-06 | 1.300E-05 |
| 1874 | 94 | V94F | A | 1.1e-05 | 1.1e-05 | 1.6e-05 | 1.267E-05 | 2.887E-06 | 1.100E-05 | 1.600E-05 |
| 1875 | 94 | V94P | A | 1.3e-05 | 1.1e-05 | 1.1e-05 | 1.167E-05 | 1.155E-06 | 1.100E-05 | 1.300E-05 |
| 1876 | 94 | V94S | A | 9.1e-06 | 1.5e-05 | 8.4e-06 | 1.083E-05 | 3.625E-06 | 8.400E-06 | 1.500E-05 |
| 1877 | 94 | V94T | A | 1.5e-05 | 1.4e-05 | 9.9e-06 | 1.297E-05 | 2.702E-06 | 9.900E-06 | 1.500E-05 |
| 1878 | 94 | V94W | A | 1.5e-05 | 1e-05 | 1.4e-05 | 1.300E-05 | 2.646E-06 | 1.000E-05 | 1.500E-05 |
| 1879 | 94 | V94Y | A | 1.4e-05 | 8.1e-06 | 1.4e-05 | 1.203E-05 | 3.406E-06 | 8.100E-06 | 1.400E-05 |
| 1880 | 94 | V94V | A | 1.3e-05 | 9.8e-06 | 8.1e-06 | 1.030E-05 | 2.488E-06 | 8.100E-06 | 1.300E-05 |
| 1881 | 95 | L95A | A | 1.5e-05 | 1.1e-05 | 1.1e-05 | 1.233E-05 | 2.309E-06 | 1.100E-05 | 1.500E-05 |
| 1882 | 95 | L95R | A | 1.5e-05 | 1.1e-05 | 1.1e-05 | 1.233E-05 | 2.309E-06 | 1.100E-05 | 1.500E-05 |
| 1883 | 95 | L95N | A | 9.8e-06 | 1.6e-05 | 9.1e-06 | 1.163E-05 | 3.798E-06 | 9.100E-06 | 1.600E-05 |
| 1884 | 95 | L95D | A | 1.5e-05 | 1.4e-05 | 1.5e-05 | 1.467E-05 | 5.774E-07 | 1.400E-05 | 1.500E-05 |
| 1885 | 95 | L95C | A | 1.1e-05 | 1e-05 | 1e-05 | 1.033E-05 | 5.774E-07 | 1.000E-05 | 1.100E-05 |
| 1886 | 95 | L95E | A | 1.3e-05 | 1.3e-05 | 1.4e-05 | 1.333E-05 | 5.774E-07 | 1.300E-05 | 1.400E-05 |
| 1887 | 95 | L95Q | A | 8.8e-06 | 1.4e-05 | 9.2e-06 | 1.067E-05 | 2.894E-06 | 8.800E-06 | 1.400E-05 |
| 1888 | 95 | L95G | A | 1.7e-05 | 1.2e-05 | 1.3e-05 | 1.400E-05 | 2.646E-06 | 1.200E-05 | 1.700E-05 |
| 1889 | 95 | L95H | A | 1.6e-05 | 1.1e-05 | 1.3e-05 | 1.333E-05 | 2.517E-06 | 1.100E-05 | 1.600E-05 |
| 1890 | 95 | L95I | A | 1.1e-05 | 8.9e-06 | 1.5e-05 | 1.163E-05 | 3.099E-06 | 8.900E-06 | 1.500E-05 |
| 1891 | 95 | L95L | A | 1.4e-05 | 1.1e-05 | 1.1e-05 | 1.200E-05 | 1.732E-06 | 1.100E-05 | 1.400E-05 |
| 1892 | 95 | L95K | A | 7.9e-06 | 1.6e-05 | 1.5e-05 | 1.297E-05 | 4.416E-06 | 7.900E-06 | 1.600E-05 |
| 1893 | 95 | L95M | A | 1.6e-05 | 1.4e-05 | 1.1e-05 | 1.367E-05 | 2.517E-06 | 1.100E-05 | 1.600E-05 |
| 1894 | 95 | L95F | A | 9.1e-06 | 1.1e-05 | 9.5e-06 | 9.867E-06 | 1.002E-06 | 9.100E-06 | 1.100E-05 |
| 1895 | 95 | L95P | A | 1.1e-05 | 1.3e-05 | 1e-05 | 1.133E-05 | 1.528E-06 | 1.000E-05 | 1.300E-05 |
| 1896 | 95 | L95S | A | 1.4e-05 | 1e-05 | 9.5e-06 | 1.117E-05 | 2.466E-06 | 9.500E-06 | 1.400E-05 |
| 1897 | 95 | L95T | A | 9.8e-06 | 1.5e-05 | 8e-06 | 1.093E-05 | 3.635E-06 | 8.000E-06 | 1.500E-05 |
| 1898 | 95 | L95W | A | 8.2e-06 | 9.4e-06 | 1e-05 | 9.200E-06 | 9.165E-07 | 8.200E-06 | 1.000E-05 |
| 1899 | 95 | L95Y | A | 1.3e-05 | 1.3e-05 | 1.2e-05 | 1.267E-05 | 5.774E-07 | 1.200E-05 | 1.300E-05 |
| 1900 | 95 | L95V | A | 1e-05 | 1.6e-05 | 1.2e-05 | 1.267E-05 | 3.055E-06 | 1.000E-05 | 1.600E-05 |
| 1901 | 96 | D96A | A | 1.5e-05 | 8.8e-06 | 1e-05 | 1.127E-05 | 3.288E-06 | 8.800E-06 | 1.500E-05 |
| 1902 | 96 | D96R | A | 1.3e-05 | 1.6e-05 | 8.7e-06 | 1.257E-05 | 3.669E-06 | 8.700E-06 | 1.600E-05 |
| 1903 | 96 | D96N | A | 1.1e-05 | 9e-06 | 8.4e-06 | 9.467E-06 | 1.361E-06 | 8.400E-06 | 1.100E-05 |
| 1904 | 96 | D96D | A | 1.5e-05 | 1.5e-05 | 1e-05 | 1.333E-05 | 2.887E-06 | 1.000E-05 | 1.500E-05 |
| 1905 | 96 | D96C | A | 1.4e-05 | 1.2e-05 | 1.5e-05 | 1.367E-05 | 1.528E-06 | 1.200E-05 | 1.500E-05 |
| 1906 | 96 | D96E | A | 1.5e-05 | 1.5e-05 | 1.6e-05 | 1.533E-05 | 5.774E-07 | 1.500E-05 | 1.600E-05 |
| 1907 | 96 | D96Q | A | 9.9e-06 | 1.8e-05 | 9.8e-06 | 1.257E-05 | 4.706E-06 | 9.800E-06 | 1.800E-05 |
| 1908 | 96 | D96G | A | 1.7e-05 | 9.7e-06 | 1.4e-05 | 1.357E-05 | 3.669E-06 | 9.700E-06 | 1.700E-05 |
| 1909 | 96 | D96H | A | 1e-05 | 1.5e-05 | 1e-05 | 1.167E-05 | 2.887E-06 | 1.000E-05 | 1.500E-05 |
| 1910 | 96 | D96I | A | 1.6e-05 | 1.1e-05 | 1.6e-05 | 1.433E-05 | 2.887E-06 | 1.100E-05 | 1.600E-05 |
| 1911 | 96 | D96L | A | 1.1e-05 | 1.1e-05 | 1e-05 | 1.067E-05 | 5.774E-07 | 1.000E-05 | 1.100E-05 |
| 1912 | 96 | D96K | A | 1.4e-05 | 1.6e-05 | 1e-05 | 1.333E-05 | 3.055E-06 | 1.000E-05 | 1.600E-05 |
| 1913 | 96 | D96M | A | 1e-05 | 1.7e-05 | 1.4e-05 | 1.367E-05 | 3.512E-06 | 1.000E-05 | 1.700E-05 |
| 1914 | 96 | D96F | A | 1.5e-05 | 1e-05 | 1.6e-05 | 1.367E-05 | 3.215E-06 | 1.000E-05 | 1.600E-05 |
| 1915 | 96 | D96P | A | 1e-05 | 1.6e-05 | 1.4e-05 | 1.333E-05 | 3.055E-06 | 1.000E-05 | 1.600E-05 |
| 1916 | 96 | D96S | A | 1e-05 | 6.9e-06 | 1.1e-05 | 9.300E-06 | 2.138E-06 | 6.900E-06 | 1.100E-05 |
| 1917 | 96 | D96T | A | 1.4e-05 | 9.8e-06 | 1e-05 | 1.127E-05 | 2.369E-06 | 9.800E-06 | 1.400E-05 |
| 1918 | 96 | D96W | A | 1.3e-05 | 8.7e-06 | 1.4e-05 | 1.190E-05 | 2.816E-06 | 8.700E-06 | 1.400E-05 |
| 1919 | 96 | D96Y | A | 1.4e-05 | 1.4e-05 | 1.4e-05 | 1.400E-05 | 0.000E+00 | 1.400E-05 | 1.400E-05 |
| 1920 | 96 | D96V | A | 9.7e-06 | 1.3e-05 | 7.8e-06 | 1.017E-05 | 2.631E-06 | 7.800E-06 | 1.300E-05 |
| 1921 | 97 | N97A | A | 1.6e-05 | 1.1e-05 | 1.7e-05 | 1.467E-05 | 3.215E-06 | 1.100E-05 | 1.700E-05 |
| 1922 | 97 | N97R | A | 1e-05 | 1.1e-05 | 1.4e-05 | 1.167E-05 | 2.082E-06 | 1.000E-05 | 1.400E-05 |
| 1923 | 97 | N97N | A | 9.9e-06 | 1.3e-05 | 1.3e-05 | 1.197E-05 | 1.790E-06 | 9.900E-06 | 1.300E-05 |
| 1924 | 97 | N97D | A | 1.2e-05 | 8.6e-06 | 1.5e-05 | 1.187E-05 | 3.202E-06 | 8.600E-06 | 1.500E-05 |
| 1925 | 97 | N97C | A | 1.8e-05 | 1e-05 | 1.5e-05 | 1.433E-05 | 4.041E-06 | 1.000E-05 | 1.800E-05 |
| 1926 | 97 | N97E | A | 1.4e-05 | 1.5e-05 | 1.6e-05 | 1.500E-05 | 1.000E-06 | 1.400E-05 | 1.600E-05 |
| 1927 | 97 | N97Q | A | 8.3e-06 | 1.1e-05 | 9.2e-06 | 9.500E-06 | 1.375E-06 | 8.300E-06 | 1.100E-05 |
| 1928 | 97 | N97G | A | 1e-05 | 1.4e-05 | 1.5e-05 | 1.300E-05 | 2.646E-06 | 1.000E-05 | 1.500E-05 |
| 1929 | 97 | N97H | A | 1.4e-05 | 1.5e-05 | 1.9e-05 | 1.600E-05 | 2.646E-06 | 1.400E-05 | 1.900E-05 |
| 1930 | 97 | N97I | A | 1.7e-05 | 7.9e-06 | 9.8e-06 | 1.157E-05 | 4.800E-06 | 7.900E-06 | 1.700E-05 |
| 1931 | 97 | N97L | A | 1.7e-05 | 1.3e-05 | 7.8e-06 | 1.260E-05 | 4.613E-06 | 7.800E-06 | 1.700E-05 |
| 1932 | 97 | N97K | A | 1.6e-05 | 1e-05 | 9.2e-06 | 1.173E-05 | 3.717E-06 | 9.200E-06 | 1.600E-05 |
| 1933 | 97 | N97M | A | 1.2e-05 | 9.8e-06 | 1.7e-05 | 1.293E-05 | 3.690E-06 | 9.800E-06 | 1.700E-05 |
| 1934 | 97 | N97F | A | 1.2e-05 | 1.6e-05 | 1.4e-05 | 1.400E-05 | 2.000E-06 | 1.200E-05 | 1.600E-05 |
| 1935 | 97 | N97P | A | 8.5e-06 | 1.6e-05 | 1.6e-05 | 1.350E-05 | 4.330E-06 | 8.500E-06 | 1.600E-05 |
| 1936 | 97 | N97S | A | 1.1e-05 | 8.4e-06 | 1.1e-05 | 1.013E-05 | 1.501E-06 | 8.400E-06 | 1.100E-05 |
| 1937 | 97 | N97T | A | 1.1e-05 | 1e-05 | 1.1e-05 | 1.067E-05 | 5.774E-07 | 1.000E-05 | 1.100E-05 |
| 1938 | 97 | N97W | A | 1.5e-05 | 8.7e-06 | 9.4e-06 | 1.103E-05 | 3.453E-06 | 8.700E-06 | 1.500E-05 |
| 1939 | 97 | N97Y | A | 1.2e-05 | 1.6e-05 | 1.8e-05 | 1.533E-05 | 3.055E-06 | 1.200E-05 | 1.800E-05 |
| 1940 | 97 | N97V | A | 1.1e-05 | 1.5e-05 | 1.3e-05 | 1.300E-05 | 2.000E-06 | 1.100E-05 | 1.500E-05 |
| 1941 | 98 | G98A | A | 1.3e-05 | 1.5e-05 | 1.1e-05 | 1.300E-05 | 2.000E-06 | 1.100E-05 | 1.500E-05 |
| 1942 | 98 | G98R | A | 1e-05 | 1.4e-05 | 1.6e-05 | 1.333E-05 | 3.055E-06 | 1.000E-05 | 1.600E-05 |
| 1943 | 98 | G98N | A | 9.9e-06 | 1e-05 | 1.4e-05 | 1.130E-05 | 2.339E-06 | 9.900E-06 | 1.400E-05 |
| 1944 | 98 | G98D | A | 1.4e-05 | 1.7e-05 | 1e-05 | 1.367E-05 | 3.512E-06 | 1.000E-05 | 1.700E-05 |
| 1945 | 98 | G98C | A | 1.1e-05 | 8.3e-06 | 1.4e-05 | 1.110E-05 | 2.851E-06 | 8.300E-06 | 1.400E-05 |
| 1946 | 98 | G98E | A | 1.5e-05 | 1.3e-05 | 9.1e-06 | 1.237E-05 | 3.001E-06 | 9.100E-06 | 1.500E-05 |
| 1947 | 98 | G98Q | A | 1e-05 | 1.3e-05 | 1.4e-05 | 1.233E-05 | 2.082E-06 | 1.000E-05 | 1.400E-05 |
| 1948 | 98 | G98G | A | 1.6e-05 | 1e-05 | 1.7e-05 | 1.433E-05 | 3.786E-06 | 1.000E-05 | 1.700E-05 |
| 1949 | 98 | G98H | A | 9.6e-06 | 8.9e-06 | 1e-05 | 9.500E-06 | 5.568E-07 | 8.900E-06 | 1.000E-05 |
| 1950 | 98 | G98I | A | 1.1e-05 | 1.7e-05 | 9.9e-06 | 1.263E-05 | 3.821E-06 | 9.900E-06 | 1.700E-05 |
| 1951 | 98 | G98L | A | 9.1e-06 | 1.3e-05 | 1.8e-05 | 1.337E-05 | 4.461E-06 | 9.100E-06 | 1.800E-05 |
| 1952 | 98 | G98K | A | 9.9e-06 | 9.3e-06 | 1.6e-05 | 1.173E-05 | 3.707E-06 | 9.300E-06 | 1.600E-05 |
| 1953 | 98 | G98M | A | 1.6e-05 | 1.2e-05 | 1e-05 | 1.267E-05 | 3.055E-06 | 1.000E-05 | 1.600E-05 |
| 1954 | 98 | G98F | A | 1.6e-05 | 9.3e-06 | 1.1e-05 | 1.210E-05 | 3.483E-06 | 9.300E-06 | 1.600E-05 |

| | | | | | | | | | | |
|---|---|---|---|---|---|---|---|---|---|---|
| 1955 | 98 | G98P | A | 1e-05 | 1.5e-05 | 5.8e-06 | 1.027E-05 | 4.606E-06 | 5.800E-06 | 1.500E-05 |
| 1956 | 98 | G98S | A | 1.5e-05 | 8.4e-06 | 9.5e-06 | 1.097E-05 | 3.536E-06 | 8.400E-06 | 1.500E-05 |
| 1957 | 98 | G98T | A | 1.3e-05 | 1.3e-05 | 1.2e-05 | 1.267E-05 | 5.774E-07 | 1.200E-05 | 1.300E-05 |
| 1958 | 98 | G98W | A | 1.6e-05 | 1.1e-05 | 1.1e-05 | 1.267E-05 | 2.887E-06 | 1.100E-05 | 1.600E-05 |
| 1959 | 98 | G98Y | A | 8.3e-06 | 8.5e-06 | 1.4e-05 | 1.027E-05 | 3.235E-06 | 8.300E-06 | 1.400E-05 |
| 1960 | 98 | G98V | A | 8.8e-06 | 9.6e-06 | 8.2e-06 | 8.867E-06 | 7.024E-07 | 8.200E-06 | 9.600E-06 |
| 1961 | 99 | D99A | A | 8.1e-06 | 1.2e-05 | 9.7e-06 | 9.933E-06 | 1.960E-06 | 8.100E-06 | 1.200E-05 |
| 1962 | 99 | D99R | A | 7.1e-06 | 1.1e-05 | 9e-06 | 9.033E-06 | 1.950E-06 | 7.100E-06 | 1.100E-05 |
| 1963 | 99 | D99N | A | 1e-05 | 1.6e-05 | 1.7e-05 | 1.433E-05 | 3.786E-06 | 1.000E-05 | 1.700E-05 |
| 1964 | 99 | D99D | A | 1.1e-05 | 1.3e-05 | 1.1e-05 | 1.167E-05 | 1.155E-06 | 1.100E-05 | 1.300E-05 |
| 1965 | 99 | D99C | A | 1.1e-05 | 9.5e-06 | 1.5e-05 | 1.183E-05 | 2.843E-06 | 9.500E-06 | 1.500E-05 |
| 1966 | 99 | D99E | A | 9.2e-06 | 1.4e-05 | 1.4e-05 | 1.240E-05 | 2.771E-06 | 9.200E-06 | 1.400E-05 |
| 1967 | 99 | D99Q | A | 1.1e-05 | 1.4e-05 | 1.2e-05 | 1.233E-05 | 1.528E-06 | 1.100E-05 | 1.400E-05 |
| 1968 | 99 | D99G | A | 1.6e-05 | 9.4e-06 | 1.5e-05 | 1.347E-05 | 3.557E-06 | 9.400E-06 | 1.600E-05 |
| 1969 | 99 | D99H | A | 1.6e-05 | 1.1e-05 | 1.6e-05 | 1.433E-05 | 2.887E-06 | 1.100E-05 | 1.600E-05 |
| 1970 | 99 | D99I | A | 1.1e-05 | 1.5e-05 | 1.2e-05 | 1.267E-05 | 2.082E-06 | 1.100E-05 | 1.500E-05 |
| 1971 | 99 | D99L | A | 9.8e-06 | 1.1e-05 | 9e-06 | 9.933E-06 | 1.007E-06 | 9.000E-06 | 1.100E-05 |
| 1972 | 99 | D99K | A | 9.4e-06 | 1.4e-05 | 1.2e-05 | 1.180E-05 | 2.307E-06 | 9.400E-06 | 1.400E-05 |
| 1973 | 99 | D99M | A | 1.5e-05 | 9.5e-06 | 1.6e-05 | 1.350E-05 | 3.500E-06 | 9.500E-06 | 1.600E-05 |
| 1974 | 99 | D99F | A | 1.5e-05 | 1.5e-05 | 1.1e-05 | 1.367E-05 | 2.309E-06 | 1.100E-05 | 1.500E-05 |
| 1975 | 99 | D99P | A | 1.3e-05 | 1.6e-05 | 1.6e-05 | 1.500E-05 | 1.732E-06 | 1.300E-05 | 1.600E-05 |
| 1976 | 99 | D99S | A | 1.2e-05 | 1.5e-05 | 1.4e-05 | 1.367E-05 | 1.528E-06 | 1.200E-05 | 1.500E-05 |
| 1977 | 99 | D99T | A | 1.3e-05 | 1.4e-05 | 1.3e-05 | 1.333E-05 | 5.774E-07 | 1.300E-05 | 1.400E-05 |
| 1978 | 99 | D99W | A | 1.5e-05 | 7.8e-06 | 1e-05 | 1.093E-05 | 3.690E-06 | 7.800E-06 | 1.500E-05 |
| 1979 | 99 | D99Y | A | 1.3e-05 | 1.4e-05 | 1e-05 | 1.233E-05 | 2.082E-06 | 1.000E-05 | 1.400E-05 |
| 1980 | 99 | D99V | A | 1.6e-05 | 1.1e-05 | 1.4e-05 | 1.367E-05 | 2.517E-06 | 1.100E-05 | 1.600E-05 |
| 1981 | 100 | P100A | A | 1.1e-05 | 1.6e-05 | 1e-05 | 1.233E-05 | 3.215E-06 | 1.000E-05 | 1.600E-05 |
| 1982 | 100 | P100R | A | 1.3e-05 | 1.4e-05 | 1.5e-05 | 1.400E-05 | 1.000E-06 | 1.300E-05 | 1.500E-05 |
| 1983 | 100 | P100N | A | 1.6e-05 | 1.1e-05 | 1.2e-05 | 1.300E-05 | 2.646E-06 | 1.100E-05 | 1.600E-05 |
| 1984 | 100 | P100D | A | 1.1e-05 | 9.6e-06 | 1.2e-05 | 1.087E-05 | 1.206E-06 | 9.600E-06 | 1.200E-05 |
| 1985 | 100 | P100C | A | 1.1e-05 | 1.2e-05 | 1.4e-05 | 1.233E-05 | 1.528E-06 | 1.100E-05 | 1.400E-05 |
| 1986 | 100 | P100E | A | 1.8e-05 | 2.1e-05 | 1.6e-05 | 1.833E-05 | 2.517E-06 | 1.600E-05 | 2.100E-05 |
| 1987 | 100 | P100Q | A | 1.5e-05 | 9.9e-06 | 1.1e-05 | 1.197E-05 | 2.684E-06 | 9.900E-06 | 1.500E-05 |
| 1988 | 100 | P100G | A | 1.1e-05 | 1.1e-05 | 1.5e-05 | 1.233E-05 | 2.309E-06 | 1.100E-05 | 1.500E-05 |
| 1989 | 100 | P100H | A | 1e-05 | 1.5e-05 | 1.5e-05 | 1.333E-05 | 2.887E-06 | 1.000E-05 | 1.500E-05 |
| 1990 | 100 | P100I | A | 1.1e-05 | 1.3e-05 | 1.7e-05 | 1.367E-05 | 3.055E-06 | 1.100E-05 | 1.700E-05 |
| 1991 | 100 | P100L | A | 1.1e-05 | 1.5e-05 | 9.8e-06 | 1.193E-05 | 2.723E-06 | 9.800E-06 | 1.500E-05 |
| 1992 | 100 | P100K | A | 1.1e-05 | 1e-05 | 1.3e-05 | 1.133E-05 | 1.528E-06 | 1.000E-05 | 1.300E-05 |
| 1993 | 100 | P100M | A | 1e-05 | 1e-05 | 1.5e-05 | 1.167E-05 | 2.887E-06 | 1.000E-05 | 1.500E-05 |
| 1994 | 100 | P100F | A | 1e-05 | 1.1e-05 | 9.2e-06 | 1.007E-05 | 9.018E-07 | 9.200E-06 | 1.100E-05 |
| 1995 | 100 | P100P | A | 1.1e-05 | 8.3e-06 | 1.3e-05 | 1.077E-05 | 2.359E-06 | 8.300E-06 | 1.300E-05 |
| 1996 | 100 | P100S | A | 1.1e-05 | 9.3e-06 | 1.4e-05 | 1.143E-05 | 2.380E-06 | 9.300E-06 | 1.400E-05 |
| 1997 | 100 | P100T | A | 1.1e-05 | 1.2e-05 | 1.2e-05 | 1.167E-05 | 5.774E-07 | 1.100E-05 | 1.200E-05 |
| 1998 | 100 | P100W | A | 1.5e-05 | 1.1e-05 | 1.5e-05 | 1.367E-05 | 2.309E-06 | 1.100E-05 | 1.500E-05 |
| 1999 | 100 | P100Y | A | 1.5e-05 | 1.3e-05 | 1.4e-05 | 1.400E-05 | 1.000E-06 | 1.300E-05 | 1.500E-05 |
| 2000 | 100 | P100V | A | 6.7e-06 | 1e-05 | 1.5e-05 | 1.057E-05 | 4.179E-06 | 6.700E-06 | 1.500E-05 |
| 2001 | 101 | L101A | A | 9.6e-06 | 1.6e-05 | 1.5e-05 | 1.353E-05 | 3.443E-06 | 9.600E-06 | 1.600E-05 |
| 2002 | 101 | L101R | A | 1.7e-05 | 9.7e-06 | 9.5e-06 | 1.207E-05 | 4.274E-06 | 9.500E-06 | 1.700E-05 |
| 2003 | 101 | L101N | A | 1.6e-05 | 1.5e-05 | 1.6e-05 | 1.567E-05 | 5.774E-07 | 1.500E-05 | 1.600E-05 |
| 2004 | 101 | L101D | A | 1.3e-05 | 5.8e-06 | 1.6e-05 | 1.160E-05 | 5.242E-06 | 5.800E-06 | 1.600E-05 |
| 2005 | 101 | L101C | A | 1.5e-05 | 1.2e-05 | 8.3e-06 | 1.177E-05 | 3.356E-06 | 8.300E-06 | 1.500E-05 |
| 2006 | 101 | L101E | A | 9.7e-06 | 1.1e-05 | 1.4e-05 | 1.157E-05 | 2.205E-06 | 9.700E-06 | 1.400E-05 |
| 2007 | 101 | L101Q | A | 9.8e-06 | 1.2e-05 | 1.5e-05 | 1.227E-05 | 2.610E-06 | 9.800E-06 | 1.500E-05 |
| 2008 | 101 | L101G | A | 1.7e-05 | 1e-05 | 6.1e-06 | 1.103E-05 | 5.523E-06 | 6.100E-06 | 1.700E-05 |
| 2009 | 101 | L101H | A | 1.4e-05 | 9.7e-06 | 6.2e-06 | 9.967E-06 | 3.907E-06 | 6.200E-06 | 1.400E-05 |
| 2010 | 101 | L101I | A | 9.9e-06 | 1.1e-05 | 1.5e-05 | 1.197E-05 | 2.684E-06 | 9.900E-06 | 1.500E-05 |
| 2011 | 101 | L101L | A | 1.5e-05 | 1.7e-05 | 7.3e-06 | 1.310E-05 | 5.122E-06 | 7.300E-06 | 1.700E-05 |
| 2012 | 101 | L101K | A | 1.3e-05 | 9.6e-06 | 1e-05 | 1.087E-05 | 1.858E-06 | 9.600E-06 | 1.300E-05 |
| 2013 | 101 | L101M | A | 8.7e-06 | 1.3e-05 | 1.3e-05 | 1.157E-05 | 2.483E-06 | 8.700E-06 | 1.300E-05 |
| 2014 | 101 | L101F | A | 1.1e-05 | 1.4e-05 | 1.2e-05 | 1.233E-05 | 1.528E-06 | 1.100E-05 | 1.400E-05 |
| 2015 | 101 | L101P | A | 1.1e-05 | 1.3e-05 | 1.4e-05 | 1.267E-05 | 1.528E-06 | 1.100E-05 | 1.400E-05 |
| 2016 | 101 | L101S | A | 9.4e-06 | 1.5e-05 | 1.6e-05 | 1.347E-05 | 3.557E-06 | 9.400E-06 | 1.600E-05 |
| 2017 | 101 | L101T | A | 8.4e-06 | 1.3e-05 | 1.4e-05 | 1.180E-05 | 2.987E-06 | 8.400E-06 | 1.400E-05 |
| 2018 | 101 | L101W | A | 1e-05 | 1.3e-05 | 9.4e-06 | 1.080E-05 | 1.929E-06 | 9.400E-06 | 1.300E-05 |
| 2019 | 101 | L101Y | A | 1.3e-05 | 1.6e-05 | 9.1e-06 | 1.270E-05 | 3.460E-06 | 9.100E-06 | 1.600E-05 |
| 2020 | 101 | L101V | A | 8.6e-06 | 1.1e-05 | 1.2e-05 | 1.053E-05 | 1.747E-06 | 8.600E-06 | 1.200E-05 |
| 2021 | 102 | N102A | A | 1.6e-05 | 1.3e-05 | 1.6e-05 | 1.500E-05 | 1.732E-06 | 1.300E-05 | 1.600E-05 |
| 2022 | 102 | N102R | A | 1.4e-05 | 1e-05 | 9.3e-06 | 1.110E-05 | 2.536E-06 | 9.300E-06 | 1.400E-05 |
| 2023 | 102 | N102N | A | 1.1e-05 | 1.5e-05 | 1.6e-05 | 1.400E-05 | 2.646E-06 | 1.100E-05 | 1.600E-05 |
| 2024 | 102 | N102D | A | 1.1e-05 | 1.4e-05 | 1e-05 | 1.167E-05 | 2.082E-06 | 1.000E-05 | 1.400E-05 |
| 2025 | 102 | N102C | A | 9.6e-06 | 1.4e-05 | 1.1e-05 | 1.153E-05 | 2.248E-06 | 9.600E-06 | 1.400E-05 |
| 2026 | 102 | N102E | A | 1.1e-05 | 1.5e-05 | 9.2e-06 | 1.173E-05 | 2.969E-06 | 9.200E-06 | 1.500E-05 |
| 2027 | 102 | N102Q | A | 8.8e-06 | 1.5e-05 | 1.6e-05 | 1.327E-05 | 3.900E-06 | 8.800E-06 | 1.600E-05 |
| 2028 | 102 | N102G | A | 1.8e-05 | 1.3e-05 | 1e-05 | 1.367E-05 | 4.041E-06 | 1.000E-05 | 1.800E-05 |
| 2029 | 102 | N102H | A | 1.5e-05 | 1.3e-05 | 1e-05 | 1.267E-05 | 2.517E-06 | 1.000E-05 | 1.500E-05 |
| 2030 | 102 | N102I | A | 9.3e-06 | 1.5e-05 | 1e-05 | 1.143E-05 | 3.109E-06 | 9.300E-06 | 1.500E-05 |
| 2031 | 102 | N102L | A | 1.8e-05 | 1.2e-05 | 1.2e-05 | 1.400E-05 | 3.464E-06 | 1.200E-05 | 1.800E-05 |
| 2032 | 102 | N102K | A | 1.1e-05 | 1.4e-05 | 1.4e-05 | 1.300E-05 | 1.732E-06 | 1.100E-05 | 1.400E-05 |
| 2033 | 102 | N102M | A | 1.3e-05 | 9.3e-06 | 1.5e-05 | 1.243E-05 | 2.892E-06 | 9.300E-06 | 1.500E-05 |
| 2034 | 102 | N102F | A | 1.1e-05 | 1.4e-05 | 1.5e-05 | 1.333E-05 | 2.082E-06 | 1.100E-05 | 1.500E-05 |
| 2035 | 102 | N102P | A | 1.5e-05 | 1.6e-05 | 1.7e-05 | 1.600E-05 | 1.000E-06 | 1.500E-05 | 1.700E-05 |
| 2036 | 102 | N102S | A | 1.1e-05 | 1.2e-05 | 1.6e-05 | 1.300E-05 | 2.646E-06 | 1.100E-05 | 1.600E-05 |
| 2037 | 102 | N102T | A | 1e-05 | 1.6e-05 | 1.5e-05 | 1.367E-05 | 3.215E-06 | 1.000E-05 | 1.600E-05 |
| 2038 | 102 | N102W | A | 1.5e-05 | 1.1e-05 | 1.5e-05 | 1.367E-05 | 2.309E-06 | 1.100E-05 | 1.500E-05 |
| 2039 | 102 | N102Y | A | 1.1e-05 | 1e-05 | 1.7e-05 | 1.267E-05 | 3.786E-06 | 1.000E-05 | 1.700E-05 |

| | | | | | | | | | | |
|---|---|---|---|---|---|---|---|---|---|---|
| 2040 | 102 | N102V | A | 9.8e-06 | 1.5e-05 | 1.1e-05 | 1.193E-05 | 2.723E-06 | 9.800E-06 | 1.500E-05 |
| 2041 | 103 | N103A | A | 7.2e-06 | 1.7e-05 | 1.4e-05 | 1.273E-05 | 5.021E-06 | 7.200E-06 | 1.700E-05 |
| 2042 | 103 | N103R | A | 9.6e-06 | 7.5e-06 | 1.4e-05 | 1.037E-05 | 3.317E-06 | 7.500E-06 | 1.400E-05 |
| 2043 | 103 | N103N | A | 1.5e-05 | 1.7e-05 | 8.6e-06 | 1.353E-05 | 4.388E-06 | 8.600E-06 | 1.700E-05 |
| 2044 | 103 | N103D | A | 1.5e-05 | 1.1e-05 | 1.5e-05 | 1.367E-05 | 2.309E-06 | 1.100E-05 | 1.500E-05 |
| 2045 | 103 | N103C | A | 1e-05 | 1.4e-05 | 1.1e-05 | 1.167E-05 | 2.082E-06 | 1.000E-05 | 1.400E-05 |
| 2046 | 103 | N103E | A | 1.6e-05 | 1.5e-05 | 1.1e-05 | 1.400E-05 | 2.646E-06 | 1.100E-05 | 1.600E-05 |
| 2047 | 103 | N103Q | A | 1.5e-05 | 8.9e-06 | 1.6e-05 | 1.330E-05 | 3.843E-06 | 8.900E-06 | 1.600E-05 |
| 2048 | 103 | N103G | A | 1e-05 | 1.1e-05 | 1.7e-05 | 1.267E-05 | 3.786E-06 | 1.000E-05 | 1.700E-05 |
| 2049 | 103 | N103H | A | 1.1e-05 | 9e-06 | 1e-05 | 1.000E-05 | 1.000E-06 | 9.000E-06 | 1.100E-05 |
| 2050 | 103 | N103I | A | 1.2e-05 | 1.6e-05 | 1.1e-05 | 1.300E-05 | 2.646E-06 | 1.100E-05 | 1.600E-05 |
| 2051 | 103 | N103L | A | 1.4e-05 | 1.1e-05 | 1.4e-05 | 1.300E-05 | 1.732E-06 | 1.100E-05 | 1.400E-05 |
| 2052 | 103 | N103K | A | 8.5e-06 | 9.1e-06 | 1.6e-05 | 1.120E-05 | 4.168E-06 | 8.500E-06 | 1.600E-05 |
| 2053 | 103 | N103M | A | 1.1e-05 | 1.5e-05 | 1.6e-05 | 1.400E-05 | 2.646E-06 | 1.100E-05 | 1.600E-05 |
| 2054 | 103 | N103F | A | 9.3e-06 | 1.4e-05 | 1.1e-05 | 1.143E-05 | 2.380E-06 | 9.300E-06 | 1.400E-05 |
| 2055 | 103 | N103P | A | 1.7e-05 | 1e-05 | 1e-05 | 1.233E-05 | 4.041E-06 | 1.000E-05 | 1.700E-05 |
| 2056 | 103 | N103S | A | 9.6e-06 | 6.7e-06 | 1.4e-05 | 1.010E-05 | 3.676E-06 | 6.700E-06 | 1.400E-05 |
| 2057 | 103 | N103T | A | 1.3e-05 | 1e-05 | 9.7e-06 | 1.090E-05 | 1.825E-06 | 9.700E-06 | 1.300E-05 |
| 2058 | 103 | N103W | A | 1.6e-05 | 1.5e-05 | 1.5e-05 | 1.533E-05 | 5.774E-07 | 1.500E-05 | 1.600E-05 |
| 2059 | 103 | N103Y | A | 1e-05 | 1.5e-05 | 1.9e-05 | 1.467E-05 | 4.509E-06 | 1.000E-05 | 1.900E-05 |
| 2060 | 103 | N103V | A | 1.2e-05 | 1.1e-05 | 1e-05 | 1.100E-05 | 1.000E-06 | 1.000E-05 | 1.200E-05 |
| 2061 | 104 | T104A | A | 1e-05 | 1.1e-05 | 1e-05 | 1.033E-05 | 5.774E-07 | 1.000E-05 | 1.100E-05 |
| 2062 | 104 | T104R | A | 1.1e-05 | 1.5e-05 | 1.1e-05 | 1.233E-05 | 2.309E-06 | 1.100E-05 | 1.500E-05 |
| 2063 | 104 | T104N | A | 1.2e-05 | 1.5e-05 | 1e-05 | 1.233E-05 | 2.517E-06 | 1.000E-05 | 1.500E-05 |
| 2064 | 104 | T104D | A | 1.7e-05 | 1.5e-05 | 1.3e-05 | 1.500E-05 | 2.000E-06 | 1.300E-05 | 1.700E-05 |
| 2065 | 104 | T104C | A | 1.4e-05 | 1e-05 | 1.5e-05 | 1.300E-05 | 2.646E-06 | 1.000E-05 | 1.500E-05 |
| 2066 | 104 | T104E | A | 9.9e-06 | 9.5e-06 | 1e-05 | 9.800E-06 | 2.646E-07 | 9.500E-06 | 1.000E-05 |
| 2067 | 104 | T104Q | A | 9.9e-06 | 1.6e-05 | 1e-05 | 1.197E-05 | 3.493E-06 | 9.900E-06 | 1.600E-05 |
| 2068 | 104 | T104G | A | 1e-05 | 1.1e-05 | 1.5e-05 | 1.200E-05 | 2.646E-06 | 1.000E-05 | 1.500E-05 |
| 2069 | 104 | T104H | A | 1.2e-05 | 1.6e-05 | 1.3e-05 | 1.367E-05 | 2.082E-06 | 1.200E-05 | 1.600E-05 |
| 2070 | 104 | T104I | A | 1.4e-05 | 1.4e-05 | 1.5e-05 | 1.433E-05 | 5.774E-07 | 1.400E-05 | 1.500E-05 |
| 2071 | 104 | T104L | A | 7.3e-06 | 1.7e-05 | 7.7e-06 | 1.067E-05 | 5.488E-06 | 7.300E-06 | 1.700E-05 |
| 2072 | 104 | T104K | A | 1.7e-05 | 1.8e-05 | 1.1e-05 | 1.533E-05 | 3.786E-06 | 1.100E-05 | 1.800E-05 |
| 2073 | 104 | T104M | A | 1.1e-05 | 1.4e-05 | 9.3e-06 | 1.143E-05 | 2.380E-06 | 9.300E-06 | 1.400E-05 |
| 2074 | 104 | T104F | A | 1.3e-05 | 8.4e-06 | 1.4e-05 | 1.180E-05 | 2.987E-06 | 8.400E-06 | 1.400E-05 |
| 2075 | 104 | T104P | A | 1.1e-05 | 2.4e-05 | 8.5e-06 | 1.450E-05 | 8.322E-06 | 8.500E-06 | 2.400E-05 |
| 2076 | 104 | T104S | A | 1.6e-05 | 1.7e-05 | 1.3e-05 | 1.533E-05 | 2.082E-06 | 1.300E-05 | 1.700E-05 |
| 2077 | 104 | T104T | A | 9.3e-06 | 1.6e-05 | 1.3e-05 | 1.277E-05 | 3.356E-06 | 9.300E-06 | 1.600E-05 |
| 2078 | 104 | T104W | A | 1.5e-05 | 1.6e-05 | 1.6e-05 | 1.567E-05 | 5.774E-07 | 1.500E-05 | 1.600E-05 |
| 2079 | 104 | T104Y | A | 1.5e-05 | 1e-05 | 7.3e-06 | 1.077E-05 | 3.907E-06 | 7.300E-06 | 1.500E-05 |
| 2080 | 104 | T104V | A | 1.5e-05 | 1e-05 | 1.9e-05 | 1.467E-05 | 4.509E-06 | 1.000E-05 | 1.900E-05 |
| 2081 | 105 | T105A | A | 1.4e-05 | 1.4e-05 | 1.1e-05 | 1.300E-05 | 1.732E-06 | 1.100E-05 | 1.400E-05 |
| 2082 | 105 | T105R | A | 1.5e-05 | 1.3e-05 | 8.7e-06 | 1.223E-05 | 3.219E-06 | 8.700E-06 | 1.500E-05 |
| 2083 | 105 | T105N | A | 1.6e-05 | 1e-05 | 1.6e-05 | 1.400E-05 | 3.464E-06 | 1.000E-05 | 1.600E-05 |
| 2084 | 105 | T105D | A | 1.4e-05 | 1.5e-05 | 9.9e-06 | 1.297E-05 | 2.702E-06 | 9.900E-06 | 1.500E-05 |
| 2085 | 105 | T105C | A | 1e-05 | 1.7e-05 | 1.1e-05 | 1.267E-05 | 3.786E-06 | 1.000E-05 | 1.700E-05 |
| 2086 | 105 | T105E | A | 1.4e-05 | 1.7e-05 | 1.8e-05 | 1.633E-05 | 2.082E-06 | 1.400E-05 | 1.800E-05 |
| 2087 | 105 | T105Q | A | 1.5e-05 | 9.4e-06 | 1.2e-05 | 1.213E-05 | 2.802E-06 | 9.400E-06 | 1.500E-05 |
| 2088 | 105 | T105G | A | 1.3e-05 | 1e-05 | 8.8e-06 | 1.060E-05 | 2.163E-06 | 8.800E-06 | 1.300E-05 |
| 2089 | 105 | T105H | A | 1e-05 | 7.8e-06 | 1.5e-05 | 1.093E-05 | 3.690E-06 | 7.800E-06 | 1.500E-05 |
| 2090 | 105 | T105I | A | 1.4e-05 | 1.8e-05 | 7.2e-06 | 1.307E-05 | 5.460E-06 | 7.200E-06 | 1.800E-05 |
| 2091 | 105 | T105L | A | 1.1e-05 | 1.5e-05 | 1.1e-05 | 1.233E-05 | 2.309E-06 | 1.100E-05 | 1.500E-05 |
| 2092 | 105 | T105K | A | 1.4e-05 | 1.1e-05 | 1.4e-05 | 1.300E-05 | 1.732E-06 | 1.100E-05 | 1.400E-05 |
| 2093 | 105 | T105M | A | 1.2e-05 | 1.1e-05 | 9.4e-06 | 1.080E-05 | 1.311E-06 | 9.400E-06 | 1.200E-05 |
| 2094 | 105 | T105F | A | 1e-05 | 1.8e-05 | 7e-06 | 1.167E-05 | 5.686E-06 | 7.000E-06 | 1.800E-05 |
| 2095 | 105 | T105P | A | 1.6e-05 | 1.2e-05 | 4.7e-06 | 1.090E-05 | 5.730E-06 | 4.700E-06 | 1.600E-05 |
| 2096 | 105 | T105S | A | 9.4e-06 | 1.4e-05 | 6.8e-06 | 1.007E-05 | 3.646E-06 | 6.800E-06 | 1.400E-05 |
| 2097 | 105 | T105T | A | 1.3e-05 | 9.8e-06 | 1.3e-05 | 1.193E-05 | 1.848E-06 | 9.800E-06 | 1.300E-05 |
| 2098 | 105 | T105W | A | 9.6e-06 | 1.6e-05 | 1.1e-05 | 1.220E-05 | 3.365E-06 | 9.600E-06 | 1.600E-05 |
| 2099 | 105 | T105Y | A | 1.8e-05 | 1e-05 | 1.1e-05 | 1.300E-05 | 4.359E-06 | 1.000E-05 | 1.800E-05 |
| 2100 | 105 | T105V | A | 1.8e-05 | 1.3e-05 | 1.4e-05 | 1.500E-05 | 2.646E-06 | 1.300E-05 | 1.800E-05 |
| 2101 | 106 | P106A | A | 9.7e-06 | 9.5e-06 | 1.5e-05 | 1.140E-05 | 3.119E-06 | 9.500E-06 | 1.500E-05 |
| 2102 | 106 | P106R | A | 9.9e-06 | 8.4e-06 | 1e-05 | 9.433E-06 | 8.963E-07 | 8.400E-06 | 1.000E-05 |
| 2103 | 106 | P106N | A | 1.2e-05 | 1.3e-05 | 1.6e-05 | 1.367E-05 | 2.082E-06 | 1.200E-05 | 1.600E-05 |
| 2104 | 106 | P106D | A | 1.6e-05 | 1.4e-05 | 1.5e-05 | 1.500E-05 | 1.000E-06 | 1.400E-05 | 1.600E-05 |
| 2105 | 106 | P106C | A | 1.5e-05 | 1.5e-05 | 1e-05 | 1.333E-05 | 2.887E-06 | 1.000E-05 | 1.500E-05 |
| 2106 | 106 | P106E | A | 1.4e-05 | 1.6e-05 | 1e-05 | 1.333E-05 | 3.055E-06 | 1.000E-05 | 1.600E-05 |
| 2107 | 106 | P106Q | A | 1.6e-05 | 1.9e-05 | 8.9e-06 | 1.463E-05 | 5.187E-06 | 8.900E-06 | 1.900E-05 |
| 2108 | 106 | P106G | A | 1e-05 | 1.4e-05 | 1.5e-05 | 1.300E-05 | 2.646E-06 | 1.000E-05 | 1.500E-05 |
| 2109 | 106 | P106H | A | 9.5e-06 | 1.1e-05 | 1.6e-05 | 1.217E-05 | 3.403E-06 | 9.500E-06 | 1.600E-05 |
| 2110 | 106 | P106I | A | 1.5e-05 | 9.9e-06 | 1.2e-05 | 1.230E-05 | 2.563E-06 | 9.900E-06 | 1.500E-05 |
| 2111 | 106 | P106L | A | 1.5e-05 | 1e-05 | 1.5e-05 | 1.333E-05 | 2.887E-06 | 1.000E-05 | 1.500E-05 |
| 2112 | 106 | P106K | A | 9.8e-06 | 1.6e-05 | 1.6e-05 | 1.393E-05 | 3.580E-06 | 9.800E-06 | 1.600E-05 |
| 2113 | 106 | P106M | A | 1.5e-05 | 1.1e-05 | 9.8e-06 | 1.193E-05 | 2.723E-06 | 9.800E-06 | 1.500E-05 |
| 2114 | 106 | P106F | A | 1.7e-05 | 1.6e-05 | 1e-05 | 1.433E-05 | 3.786E-06 | 1.000E-05 | 1.700E-05 |
| 2115 | 106 | P106P | A | 1.2e-05 | 1.6e-05 | 1e-05 | 1.267E-05 | 3.055E-06 | 1.000E-05 | 1.600E-05 |
| 2116 | 106 | P106S | A | 9e-06 | 1e-05 | 1.4e-05 | 1.100E-05 | 2.646E-06 | 9.000E-06 | 1.400E-05 |
| 2117 | 106 | P106T | A | 1.2e-05 | 9.7e-06 | 1.1e-05 | 1.090E-05 | 1.153E-06 | 9.700E-06 | 1.200E-05 |
| 2118 | 106 | P106W | A | 1.1e-05 | 1.2e-05 | 1.5e-05 | 1.267E-05 | 2.082E-06 | 1.100E-05 | 1.500E-05 |
| 2119 | 106 | P106Y | A | 1.6e-05 | 1.4e-05 | 9.2e-06 | 1.307E-05 | 3.495E-06 | 9.200E-06 | 1.600E-05 |
| 2120 | 106 | P106V | A | 9.3e-06 | 1.5e-05 | 1.6e-05 | 1.343E-05 | 3.614E-06 | 9.300E-06 | 1.600E-05 |
| 2121 | 107 | V107A | A | 1.1e-05 | 1.6e-05 | 1.6e-05 | 1.433E-05 | 2.887E-06 | 1.100E-05 | 1.600E-05 |
| 2122 | 107 | V107R | A | 9.7e-06 | 1.6e-05 | 1.1e-05 | 1.223E-05 | 3.326E-06 | 9.700E-06 | 1.600E-05 |
| 2123 | 107 | V107N | A | 1.3e-05 | 1.5e-05 | 1.5e-05 | 1.433E-05 | 1.155E-06 | 1.300E-05 | 1.500E-05 |
| 2124 | 107 | V107D | A | 1.5e-05 | 1.1e-05 | 1.1e-05 | 1.233E-05 | 2.309E-06 | 1.100E-05 | 1.500E-05 |

| | | | | | | | | | | |
|---|---|---|---|---|---|---|---|---|---|---|
| 2125 | 107 | V107C | A | 1.4e-05 | 8.1e-06 | 9.9e-06 | 1.067E-05 | 3.024E-06 | 8.100E-06 | 1.400E-05 |
| 2126 | 107 | V107E | A | 1.1e-05 | 1.5e-05 | 9.6e-06 | 1.187E-05 | 2.802E-06 | 9.600E-06 | 1.500E-05 |
| 2127 | 107 | V107Q | A | 1.5e-05 | 1.3e-05 | 1.5e-05 | 1.433E-05 | 1.155E-06 | 1.300E-05 | 1.500E-05 |
| 2128 | 107 | V107G | A | 9.1e-06 | 1.4e-05 | 1.6e-05 | 1.303E-05 | 3.550E-06 | 9.100E-06 | 1.600E-05 |
| 2129 | 107 | V107H | A | 1.5e-05 | 1.2e-05 | 1.4e-05 | 1.367E-05 | 1.528E-06 | 1.200E-05 | 1.500E-05 |
| 2130 | 107 | V107I | A | 1.1e-05 | 1.4e-05 | 9.3e-06 | 1.143E-05 | 2.380E-06 | 9.300E-06 | 1.400E-05 |
| 2131 | 107 | V107L | A | 1.1e-05 | 1.3e-05 | 1.5e-05 | 1.300E-05 | 2.000E-06 | 1.100E-05 | 1.500E-05 |
| 2132 | 107 | V107K | A | 1.5e-05 | 1.4e-05 | 1.7e-05 | 1.533E-05 | 1.528E-06 | 1.400E-05 | 1.700E-05 |
| 2133 | 107 | V107M | A | 8.5e-06 | 1e-05 | 1.5e-05 | 1.117E-05 | 3.403E-06 | 8.500E-06 | 1.500E-05 |
| 2134 | 107 | V107F | A | 1.7e-05 | 8.9e-06 | 1.4e-05 | 1.330E-05 | 4.095E-06 | 8.900E-06 | 1.700E-05 |
| 2135 | 107 | V107P | A | 1.5e-05 | 1.8e-05 | 1.2e-05 | 1.500E-05 | 3.000E-06 | 1.200E-05 | 1.800E-05 |
| 2136 | 107 | V107S | A | 1.4e-05 | 1.5e-05 | 1.4e-05 | 1.433E-05 | 5.774E-07 | 1.400E-05 | 1.500E-05 |
| 2137 | 107 | V107T | A | 1.4e-05 | 1.4e-05 | 9.2e-06 | 1.240E-05 | 2.771E-06 | 9.200E-06 | 1.400E-05 |
| 2138 | 107 | V107W | A | 8.2e-06 | 1.5e-05 | 1.3e-05 | 1.207E-05 | 3.495E-06 | 8.200E-06 | 1.500E-05 |
| 2139 | 107 | V107Y | A | 1.4e-05 | 1.5e-05 | 1.2e-05 | 1.367E-05 | 1.528E-06 | 1.200E-05 | 1.500E-05 |
| 2140 | 107 | V107V | A | 1.5e-05 | 1.5e-05 | 1.4e-05 | 1.467E-05 | 5.774E-07 | 1.400E-05 | 1.500E-05 |
| 2141 | 108 | T108A | A | 1e-05 | 1.7e-05 | 1.5e-05 | 1.400E-05 | 3.606E-06 | 1.000E-05 | 1.700E-05 |
| 2142 | 108 | T108R | A | 1.4e-05 | 1.6e-05 | 1.6e-05 | 1.533E-05 | 1.155E-06 | 1.400E-05 | 1.600E-05 |
| 2143 | 108 | T108N | A | 5.6e-06 | 1.6e-05 | 1.4e-05 | 1.187E-05 | 5.518E-06 | 5.600E-06 | 1.600E-05 |
| 2144 | 108 | T108D | A | 1.2e-05 | 1.4e-05 | 1.4e-05 | 1.333E-05 | 1.155E-06 | 1.200E-05 | 1.400E-05 |
| 2145 | 108 | T108C | A | 1.5e-05 | 8.4e-06 | 9.5e-06 | 1.097E-05 | 3.536E-06 | 8.400E-06 | 1.500E-05 |
| 2146 | 108 | T108E | A | 1.1e-05 | 1.4e-05 | 1.5e-05 | 1.333E-05 | 2.082E-06 | 1.100E-05 | 1.500E-05 |
| 2147 | 108 | T108Q | A | 1.6e-05 | 1.5e-05 | 1.5e-05 | 1.533E-05 | 5.774E-07 | 1.500E-05 | 1.600E-05 |
| 2148 | 108 | T108G | A | 1.6e-05 | 8.8e-06 | 1.6e-05 | 1.360E-05 | 4.157E-06 | 8.800E-06 | 1.600E-05 |
| 2149 | 108 | T108H | A | 1.6e-05 | 1.6e-05 | 1e-05 | 1.400E-05 | 3.464E-06 | 1.000E-05 | 1.600E-05 |
| 2150 | 108 | T108I | A | 1.6e-05 | 1.4e-05 | 1.5e-05 | 1.500E-05 | 1.000E-06 | 1.400E-05 | 1.600E-05 |
| 2151 | 108 | T108L | A | 9.2e-06 | 1.2e-05 | 1.1e-05 | 1.073E-05 | 1.419E-06 | 9.200E-06 | 1.200E-05 |
| 2152 | 108 | T108K | A | 1.1e-05 | 9.6e-06 | 1.1e-05 | 1.053E-05 | 8.083E-07 | 9.600E-06 | 1.100E-05 |
| 2153 | 108 | T108M | A | 1.5e-05 | 1.6e-05 | 1.1e-05 | 1.400E-05 | 2.646E-06 | 1.100E-05 | 1.600E-05 |
| 2154 | 108 | T108F | A | 1.2e-05 | 1.4e-05 | 1.5e-05 | 1.367E-05 | 1.528E-06 | 1.200E-05 | 1.500E-05 |
| 2155 | 108 | T108P | A | 1.5e-05 | 1.2e-05 | 1.2e-05 | 1.300E-05 | 1.732E-06 | 1.200E-05 | 1.500E-05 |
| 2156 | 108 | T108S | A | 1.6e-05 | 1.5e-05 | 1.2e-05 | 1.433E-05 | 2.082E-06 | 1.200E-05 | 1.600E-05 |
| 2157 | 108 | T108T | A | 1.4e-05 | 1.1e-05 | 1.1e-05 | 1.200E-05 | 1.732E-06 | 1.100E-05 | 1.400E-05 |
| 2158 | 108 | T108W | A | 1.5e-05 | 1.5e-05 | 1.4e-05 | 1.467E-05 | 5.774E-07 | 1.400E-05 | 1.500E-05 |
| 2159 | 108 | T108Y | A | 8.1e-06 | 8.2e-06 | 1e-05 | 8.767E-06 | 1.069E-06 | 8.100E-06 | 1.000E-05 |
| 2160 | 108 | T108V | A | 1.1e-05 | 1e-05 | 1.1e-05 | 1.067E-05 | 5.774E-07 | 1.000E-05 | 1.100E-05 |
| 2161 | 109 | G109A | A | 1.5e-05 | 8.4e-06 | 1.3e-05 | 1.213E-05 | 3.384E-06 | 8.400E-06 | 1.500E-05 |
| 2162 | 109 | G109R | A | 9.9e-06 | 1.2e-05 | 9.3e-06 | 1.040E-05 | 1.418E-06 | 9.300E-06 | 1.200E-05 |
| 2163 | 109 | G109N | A | 1.6e-05 | 1.1e-05 | 9e-06 | 1.200E-05 | 3.606E-06 | 9.000E-06 | 1.600E-05 |
| 2164 | 109 | G109D | A | 8.9e-06 | 1.1e-05 | 1e-05 | 9.967E-06 | 1.050E-06 | 8.900E-06 | 1.100E-05 |
| 2165 | 109 | G109C | A | 1.5e-05 | 1.6e-05 | 8.5e-06 | 1.317E-05 | 4.072E-06 | 8.500E-06 | 1.600E-05 |
| 2166 | 109 | G109E | A | 1.6e-05 | 1e-05 | 1.2e-05 | 1.267E-05 | 3.055E-06 | 1.000E-05 | 1.600E-05 |
| 2167 | 109 | G109Q | A | 1.4e-05 | 1.4e-05 | 1.7e-05 | 1.500E-05 | 1.732E-06 | 1.400E-05 | 1.700E-05 |
| 2168 | 109 | G109G | A | 8.6e-06 | 9.8e-06 | 9.8e-06 | 9.400E-06 | 6.928E-07 | 8.600E-06 | 9.800E-06 |
| 2169 | 109 | G109H | A | 1.1e-05 | 1e-05 | 1.5e-05 | 1.200E-05 | 2.646E-06 | 1.000E-05 | 1.500E-05 |
| 2170 | 109 | G109I | A | 1.2e-05 | 1.4e-05 | 1.5e-05 | 1.367E-05 | 1.528E-06 | 1.200E-05 | 1.500E-05 |
| 2171 | 109 | G109L | A | 1.2e-05 | 1.4e-05 | 1.6e-05 | 1.400E-05 | 2.000E-06 | 1.200E-05 | 1.600E-05 |
| 2172 | 109 | G109K | A | 1.3e-05 | 8.9e-06 | 1.6e-05 | 1.263E-05 | 3.564E-06 | 8.900E-06 | 1.600E-05 |
| 2173 | 109 | G109M | A | 2e-05 | 1.3e-05 | 1.5e-05 | 1.600E-05 | 3.606E-06 | 1.300E-05 | 2.000E-05 |
| 2174 | 109 | G109F | A | 1.5e-05 | 7e-06 | 1e-05 | 1.067E-05 | 4.041E-06 | 7.000E-06 | 1.500E-05 |
| 2175 | 109 | G109P | A | 1.4e-05 | 9.8e-06 | 1.6e-05 | 1.327E-05 | 3.164E-06 | 9.800E-06 | 1.600E-05 |
| 2176 | 109 | G109S | A | 1.5e-05 | 1e-05 | 1e-05 | 1.167E-05 | 2.887E-06 | 1.000E-05 | 1.500E-05 |
| 2177 | 109 | G109T | A | 1.1e-05 | 1e-05 | 9.2e-06 | 1.007E-05 | 9.018E-07 | 9.200E-06 | 1.100E-05 |
| 2178 | 109 | G109W | A | 1.1e-05 | 1.5e-05 | 1.1e-05 | 1.233E-05 | 2.309E-06 | 1.100E-05 | 1.500E-05 |
| 2179 | 109 | G109Y | A | 7.6e-06 | 9.4e-06 | 1.5e-05 | 1.067E-05 | 3.859E-06 | 7.600E-06 | 1.500E-05 |
| 2180 | 109 | G109V | A | 1.1e-05 | 1.1e-05 | 1.6e-05 | 1.267E-05 | 2.887E-06 | 1.100E-05 | 1.600E-05 |
| 2181 | 110 | A110A | A | 1.2e-05 | 9.5e-06 | 1e-05 | 1.050E-05 | 1.323E-06 | 9.500E-06 | 1.200E-05 |
| 2182 | 110 | A110R | A | 1e-05 | 1.3e-05 | 1.6e-05 | 1.300E-05 | 3.000E-06 | 1.000E-05 | 1.600E-05 |
| 2183 | 110 | A110N | A | 9.8e-06 | 1e-05 | 1e-05 | 9.933E-06 | 1.155E-07 | 9.800E-06 | 1.000E-05 |
| 2184 | 110 | A110D | A | 1.4e-05 | 1.1e-05 | 9e-06 | 1.133E-05 | 2.517E-06 | 9.000E-06 | 1.400E-05 |
| 2185 | 110 | A110C | A | 1.3e-05 | 8e-06 | 1.5e-05 | 1.200E-05 | 3.606E-06 | 8.000E-06 | 1.500E-05 |
| 2186 | 110 | A110E | A | 1.2e-05 | 9.6e-06 | 9.4e-06 | 1.033E-05 | 1.447E-06 | 9.400E-06 | 1.200E-05 |
| 2187 | 110 | A110Q | A | 7.1e-06 | 9.8e-06 | 9.9e-06 | 8.933E-06 | 1.589E-06 | 7.100E-06 | 9.900E-06 |
| 2188 | 110 | A110G | A | 1.6e-05 | 1.2e-05 | 1.3e-05 | 1.367E-05 | 2.082E-06 | 1.200E-05 | 1.600E-05 |
| 2189 | 110 | A110H | A | 1.1e-05 | 1.7e-05 | 1.5e-05 | 1.433E-05 | 3.055E-06 | 1.100E-05 | 1.700E-05 |
| 2190 | 110 | A110I | A | 1.3e-05 | 7.2e-06 | 1.4e-05 | 1.140E-05 | 3.672E-06 | 7.200E-06 | 1.400E-05 |
| 2191 | 110 | A110L | A | 1.4e-05 | 8.2e-06 | 1.5e-05 | 1.240E-05 | 3.672E-06 | 8.200E-06 | 1.500E-05 |
| 2192 | 110 | A110K | A | 1.2e-05 | 9.7e-06 | 1.5e-05 | 1.223E-05 | 2.658E-06 | 9.700E-06 | 1.500E-05 |
| 2193 | 110 | A110M | A | 1.6e-05 | 1.2e-05 | 1.4e-05 | 1.400E-05 | 2.000E-06 | 1.200E-05 | 1.600E-05 |
| 2194 | 110 | A110F | A | 1.3e-05 | 1.5e-05 | 1.5e-05 | 1.433E-05 | 1.155E-06 | 1.300E-05 | 1.500E-05 |
| 2195 | 110 | A110P | A | 1e-05 | 1.7e-05 | 1.1e-05 | 1.267E-05 | 3.786E-06 | 1.000E-05 | 1.700E-05 |
| 2196 | 110 | A110S | A | 1.1e-05 | 1.3e-05 | 6.2e-06 | 1.007E-05 | 3.495E-06 | 6.200E-06 | 1.300E-05 |
| 2197 | 110 | A110T | A | 1.4e-05 | 9.6e-06 | 1.3e-05 | 1.220E-05 | 2.307E-06 | 9.600E-06 | 1.400E-05 |
| 2198 | 110 | A110W | A | 1.6e-05 | 1.2e-05 | 1e-05 | 1.267E-05 | 3.055E-06 | 1.000E-05 | 1.600E-05 |
| 2199 | 110 | A110Y | A | 1e-05 | 1.1e-05 | 1.6e-05 | 1.233E-05 | 3.215E-06 | 1.000E-05 | 1.600E-05 |
| 2200 | 110 | A110V | A | 9.9e-06 | 1.1e-05 | 1.5e-05 | 1.197E-05 | 2.684E-06 | 9.900E-06 | 1.500E-05 |
| 2201 | 111 | S111A | A | 7e-06 | 1.1e-05 | 1.5e-05 | 1.100E-05 | 4.000E-06 | 7.000E-06 | 1.500E-05 |
| 2202 | 111 | S111R | A | 1e-05 | 9.1e-06 | 1.5e-05 | 1.137E-05 | 3.179E-06 | 9.100E-06 | 1.500E-05 |
| 2203 | 111 | S111N | A | 1.5e-05 | 1.5e-05 | 6.2e-06 | 1.207E-05 | 5.081E-06 | 6.200E-06 | 1.500E-05 |
| 2204 | 111 | S111D | A | 1.4e-05 | 1.1e-05 | 1.1e-05 | 1.200E-05 | 1.732E-06 | 1.100E-05 | 1.400E-05 |
| 2205 | 111 | S111C | A | 1.5e-05 | 1.6e-05 | 1.7e-05 | 1.600E-05 | 1.000E-06 | 1.500E-05 | 1.700E-05 |
| 2206 | 111 | S111E | A | 1.5e-05 | 1.5e-05 | 1.5e-05 | 1.500E-05 | 0.000E+00 | 1.500E-05 | 1.500E-05 |
| 2207 | 111 | S111Q | A | 1.4e-05 | 7.8e-06 | 1.2e-05 | 1.127E-05 | 3.164E-06 | 7.800E-06 | 1.400E-05 |
| 2208 | 111 | S111G | A | 1.4e-05 | 1.5e-05 | 1.6e-05 | 1.500E-05 | 1.000E-06 | 1.400E-05 | 1.600E-05 |
| 2209 | 111 | S111H | A | 1.1e-05 | 1.1e-05 | 1.5e-05 | 1.233E-05 | 2.309E-06 | 1.100E-05 | 1.500E-05 |

| | | | | | | | | | | |
|---|---|---|---|---|---|---|---|---|---|---|
| 2210 | 111 | S111I | A | 1e-05 | 1.4e-05 | 1.7e-05 | 1.367E-05 | 3.512E-06 | 1.000E-05 | 1.700E-05 |
| 2211 | 111 | S111L | A | 1.1e-05 | 1.5e-05 | 9.9e-06 | 1.197E-05 | 2.684E-06 | 9.900E-06 | 1.500E-05 |
| 2212 | 111 | S111K | A | 1.4e-05 | 1.3e-05 | 9.5e-06 | 1.217E-05 | 2.363E-06 | 9.500E-06 | 1.400E-05 |
| 2213 | 111 | S111M | A | 1.1e-05 | 9.3e-06 | 1.4e-05 | 1.143E-05 | 2.380E-06 | 9.300E-06 | 1.400E-05 |
| 2214 | 111 | S111F | A | 1.7e-05 | 1.5e-05 | 5.6e-06 | 1.253E-05 | 6.087E-06 | 5.600E-06 | 1.700E-05 |
| 2215 | 111 | S111P | A | 1e-05 | 9.9e-06 | 1.1e-05 | 1.030E-05 | 6.083E-07 | 9.900E-06 | 1.100E-05 |
| 2216 | 111 | S111S | A | 1.1e-05 | 1.5e-05 | 9.8e-06 | 1.193E-05 | 2.723E-06 | 9.800E-06 | 1.500E-05 |
| 2217 | 111 | S111T | A | 9.4e-06 | 1.3e-05 | 1.5e-05 | 1.247E-05 | 2.838E-06 | 9.400E-06 | 1.500E-05 |
| 2218 | 111 | S111W | A | 9.7e-06 | 1.1e-05 | 9.5e-06 | 1.007E-05 | 8.145E-07 | 9.500E-06 | 1.100E-05 |
| 2219 | 111 | S111Y | A | 1.5e-05 | 1e-05 | 1.1e-05 | 1.200E-05 | 2.646E-06 | 1.000E-05 | 1.500E-05 |
| 2220 | 111 | S111V | A | 1.6e-05 | 1e-05 | 1.2e-05 | 1.267E-05 | 3.055E-06 | 1.000E-05 | 1.600E-05 |
| 2221 | 112 | P112A | A | 1.5e-05 | 8.8e-06 | 1.4e-05 | 1.260E-05 | 3.329E-06 | 8.800E-06 | 1.500E-05 |
| 2222 | 112 | P112R | A | 1.2e-05 | 8.9e-06 | 1e-05 | 1.030E-05 | 1.572E-06 | 8.900E-06 | 1.200E-05 |
| 2223 | 112 | P112N | A | 1.1e-05 | 1e-05 | 1.3e-05 | 1.133E-05 | 1.528E-06 | 1.000E-05 | 1.300E-05 |
| 2224 | 112 | P112D | A | 9.9e-06 | 1.3e-05 | 9.7e-06 | 1.087E-05 | 1.850E-06 | 9.700E-06 | 1.300E-05 |
| 2225 | 112 | P112C | A | 1.4e-05 | 1.4e-05 | 1.3e-05 | 1.367E-05 | 5.774E-07 | 1.300E-05 | 1.400E-05 |
| 2226 | 112 | P112E | A | 1.1e-05 | 1.3e-05 | 1.5e-05 | 1.300E-05 | 2.000E-06 | 1.100E-05 | 1.500E-05 |
| 2227 | 112 | P112Q | A | 1.4e-05 | 8.2e-06 | 1.7e-05 | 1.307E-05 | 4.474E-06 | 8.200E-06 | 1.700E-05 |
| 2228 | 112 | P112G | A | 1.4e-05 | 1.7e-05 | 1.5e-05 | 1.533E-05 | 1.528E-06 | 1.400E-05 | 1.700E-05 |
| 2229 | 112 | P112H | A | 1.6e-05 | 1.2e-05 | 1.5e-05 | 1.433E-05 | 2.082E-06 | 1.200E-05 | 1.600E-05 |
| 2230 | 112 | P112I | A | 1.2e-05 | 1.3e-05 | 9.9e-06 | 1.163E-05 | 1.582E-06 | 9.900E-06 | 1.300E-05 |
| 2231 | 112 | P112L | A | 1e-05 | 9e-06 | 1.6e-05 | 1.167E-05 | 3.786E-06 | 9.000E-06 | 1.600E-05 |
| 2232 | 112 | P112K | A | 8.6e-06 | 1.2e-05 | 1.1e-05 | 1.053E-05 | 1.747E-06 | 8.600E-06 | 1.200E-05 |
| 2233 | 112 | P112M | A | 1.6e-05 | 9.4e-06 | 1.5e-05 | 1.347E-05 | 3.557E-06 | 9.400E-06 | 1.600E-05 |
| 2234 | 112 | P112F | A | 1.1e-05 | 1.8e-05 | 9.7e-06 | 1.290E-05 | 4.464E-06 | 9.700E-06 | 1.800E-05 |
| 2235 | 112 | P112P | A | 1.3e-05 | 1.8e-05 | 9.1e-06 | 1.337E-05 | 4.461E-06 | 9.100E-06 | 1.800E-05 |
| 2236 | 112 | P112S | A | 1.6e-05 | 1.4e-05 | 1.5e-05 | 1.500E-05 | 1.000E-06 | 1.400E-05 | 1.600E-05 |
| 2237 | 112 | P112T | A | 9.6e-06 | 1.5e-05 | 1.1e-05 | 1.187E-05 | 2.802E-06 | 9.600E-06 | 1.500E-05 |
| 2238 | 112 | P112W | A | 1.4e-05 | 1e-05 | 1.1e-05 | 1.167E-05 | 2.082E-06 | 1.000E-05 | 1.400E-05 |
| 2239 | 112 | P112Y | A | 1.1e-05 | 1.1e-05 | 1.1e-05 | 1.100E-05 | 2.075E-21 | 1.100E-05 | 1.100E-05 |
| 2240 | 112 | P112V | A | 2e-05 | 1.1e-05 | 7.1e-06 | 1.270E-05 | 6.616E-06 | 7.100E-06 | 2.000E-05 |
| 2241 | 113 | G113A | A | 8.9e-06 | 1.6e-05 | 1.1e-05 | 1.197E-05 | 3.647E-06 | 8.900E-06 | 1.600E-05 |
| 2242 | 113 | G113R | A | 8.4e-06 | 1e-05 | 9.5e-06 | 9.300E-06 | 8.185E-07 | 8.400E-06 | 1.000E-05 |
| 2243 | 113 | G113N | A | 1.4e-05 | 1.5e-05 | 1.7e-05 | 1.533E-05 | 1.528E-06 | 1.400E-05 | 1.700E-05 |
| 2244 | 113 | G113D | A | 1.3e-05 | 1.2e-05 | 1.4e-05 | 1.300E-05 | 1.000E-06 | 1.200E-05 | 1.400E-05 |
| 2245 | 113 | G113C | A | 9.7e-06 | 1.6e-05 | 1e-05 | 1.190E-05 | 3.554E-06 | 9.700E-06 | 1.600E-05 |
| 2246 | 113 | G113E | A | 1.5e-05 | 9.1e-06 | 1e-05 | 1.137E-05 | 3.179E-06 | 9.100E-06 | 1.500E-05 |
| 2247 | 113 | G113Q | A | 1.5e-05 | 1.5e-05 | 1.4e-05 | 1.467E-05 | 5.774E-07 | 1.400E-05 | 1.500E-05 |
| 2248 | 113 | G113G | A | 1.4e-05 | 1.5e-05 | 1.6e-05 | 1.500E-05 | 1.000E-06 | 1.400E-05 | 1.600E-05 |
| 2249 | 113 | G113H | A | 1e-05 | 1.1e-05 | 9.8e-06 | 1.027E-05 | 6.429E-07 | 9.800E-06 | 1.100E-05 |
| 2250 | 113 | G113I | A | 1e-05 | 1e-05 | 1e-05 | 1.000E-05 | 0.000E+00 | 1.000E-05 | 1.000E-05 |
| 2251 | 113 | G113L | A | 1.2e-05 | 1.6e-05 | 1e-05 | 1.267E-05 | 3.055E-06 | 1.000E-05 | 1.600E-05 |
| 2252 | 113 | G113K | A | 1.5e-05 | 1.5e-05 | 1.5e-05 | 1.500E-05 | 0.000E+00 | 1.500E-05 | 1.500E-05 |
| 2253 | 113 | G113M | A | 1.6e-05 | 1.7e-05 | 1e-05 | 1.433E-05 | 3.786E-06 | 1.000E-05 | 1.700E-05 |
| 2254 | 113 | G113F | A | 1.8e-05 | 8.6e-06 | 1.3e-05 | 1.320E-05 | 4.703E-06 | 8.600E-06 | 1.800E-05 |
| 2255 | 113 | G113P | A | 9.1e-06 | 8.2e-06 | 9.4e-06 | 8.900E-06 | 6.245E-07 | 8.200E-06 | 9.400E-06 |
| 2256 | 113 | G113S | A | 1.4e-05 | 1.5e-05 | 1.6e-05 | 1.500E-05 | 1.000E-06 | 1.400E-05 | 1.600E-05 |
| 2257 | 113 | G113T | A | 1.4e-05 | 5.3e-06 | 1.1e-05 | 1.010E-05 | 4.419E-06 | 5.300E-06 | 1.400E-05 |
| 2258 | 113 | G113W | A | 1.6e-05 | 1.3e-05 | 1e-05 | 1.300E-05 | 3.000E-06 | 1.000E-05 | 1.600E-05 |
| 2259 | 113 | G113Y | A | 1.5e-05 | 1.3e-05 | 1.7e-05 | 1.500E-05 | 2.000E-06 | 1.300E-05 | 1.700E-05 |
| 2260 | 113 | G113V | A | 1e-05 | 1.6e-05 | 1e-05 | 1.200E-05 | 3.464E-06 | 1.000E-05 | 1.600E-05 |
| 2261 | 114 | G114A | A | 9.3e-06 | 1e-05 | 1.1e-05 | 1.010E-05 | 8.544E-07 | 9.300E-06 | 1.100E-05 |
| 2262 | 114 | G114R | A | 1.5e-05 | 1.3e-05 | 1.6e-05 | 1.467E-05 | 1.528E-06 | 1.300E-05 | 1.600E-05 |
| 2263 | 114 | G114N | A | 1.6e-05 | 1.5e-05 | 1.6e-05 | 1.567E-05 | 5.774E-07 | 1.500E-05 | 1.600E-05 |
| 2264 | 114 | G114D | A | 1.2e-05 | 1.1e-05 | 1.1e-05 | 1.133E-05 | 5.774E-07 | 1.100E-05 | 1.200E-05 |
| 2265 | 114 | G114C | A | 1.6e-05 | 1.1e-05 | 1.5e-05 | 1.400E-05 | 2.646E-06 | 1.100E-05 | 1.600E-05 |
| 2266 | 114 | G114E | A | 1.5e-05 | 1.3e-05 | 1.3e-05 | 1.367E-05 | 1.155E-06 | 1.300E-05 | 1.500E-05 |
| 2267 | 114 | G114Q | A | 8.5e-06 | 1e-05 | 1.5e-05 | 1.117E-05 | 3.403E-06 | 8.500E-06 | 1.500E-05 |
| 2268 | 114 | G114G | A | 1.7e-05 | 1.5e-05 | 1.5e-05 | 1.567E-05 | 1.155E-06 | 1.500E-05 | 1.700E-05 |
| 2269 | 114 | G114H | A | 1.4e-05 | 1.2e-05 | 1.4e-05 | 1.333E-05 | 1.155E-06 | 1.200E-05 | 1.400E-05 |
| 2270 | 114 | G114I | A | 1e-05 | 1.3e-05 | 1.6e-05 | 1.300E-05 | 3.000E-06 | 1.000E-05 | 1.600E-05 |
| 2271 | 114 | G114L | A | 9.7e-06 | 1.2e-05 | 1.3e-05 | 1.157E-05 | 1.692E-06 | 9.700E-06 | 1.300E-05 |
| 2272 | 114 | G114K | A | 1e-05 | 1.6e-05 | 1.3e-05 | 1.300E-05 | 3.000E-06 | 1.000E-05 | 1.600E-05 |
| 2273 | 114 | G114M | A | 1.3e-05 | 9.8e-06 | 9.6e-06 | 1.080E-05 | 1.908E-06 | 9.600E-06 | 1.300E-05 |
| 2274 | 114 | G114F | A | 1.5e-05 | 1.5e-05 | 1.1e-05 | 1.367E-05 | 2.309E-06 | 1.100E-05 | 1.500E-05 |
| 2275 | 114 | G114P | A | 1.5e-05 | 9.3e-06 | 1.4e-05 | 1.277E-05 | 3.044E-06 | 9.300E-06 | 1.500E-05 |
| 2276 | 114 | G114S | A | 1.3e-05 | 1.4e-05 | 1e-05 | 1.233E-05 | 2.082E-06 | 1.000E-05 | 1.400E-05 |
| 2277 | 114 | G114T | A | 1.6e-05 | 1.1e-05 | 7.8e-06 | 1.160E-05 | 4.133E-06 | 7.800E-06 | 1.600E-05 |
| 2278 | 114 | G114W | A | 1.1e-05 | 1.1e-05 | 1e-05 | 1.067E-05 | 5.774E-07 | 1.000E-05 | 1.100E-05 |
| 2279 | 114 | G114Y | A | 1.6e-05 | 1.5e-05 | 1.6e-05 | 1.567E-05 | 5.774E-07 | 1.500E-05 | 1.600E-05 |
| 2280 | 114 | G114V | A | 1.6e-05 | 1.6e-05 | 1.5e-05 | 1.567E-05 | 5.774E-07 | 1.500E-05 | 1.600E-05 |
| 2281 | 115 | L115A | A | 1.2e-05 | 1.4e-05 | 1.4e-05 | 1.333E-05 | 1.155E-06 | 1.200E-05 | 1.400E-05 |
| 2282 | 115 | L115R | A | 1.2e-05 | 9.1e-06 | 1.4e-05 | 1.170E-05 | 2.464E-06 | 9.100E-06 | 1.400E-05 |
| 2283 | 115 | L115N | A | 1.5e-05 | 1.2e-05 | 1e-05 | 1.233E-05 | 2.517E-06 | 1.000E-05 | 1.500E-05 |
| 2284 | 115 | L115D | A | 1.4e-05 | 1.2e-05 | 1.4e-05 | 1.333E-05 | 1.155E-06 | 1.200E-05 | 1.400E-05 |
| 2285 | 115 | L115C | A | 1.4e-05 | 1.7e-05 | 1.6e-05 | 1.567E-05 | 1.528E-06 | 1.400E-05 | 1.700E-05 |
| 2286 | 115 | L115E | A | 1.5e-05 | 1.5e-05 | 1.1e-05 | 1.367E-05 | 2.309E-06 | 1.100E-05 | 1.500E-05 |
| 2287 | 115 | L115Q | A | 1.4e-05 | 9.4e-06 | 1.4e-05 | 1.247E-05 | 2.656E-06 | 9.400E-06 | 1.400E-05 |
| 2288 | 115 | L115G | A | 1.2e-05 | 1.6e-05 | 9.2e-06 | 1.240E-05 | 3.418E-06 | 9.200E-06 | 1.600E-05 |
| 2289 | 115 | L115H | A | 9.9e-06 | 1e-05 | 1.5e-05 | 1.163E-05 | 2.916E-06 | 9.900E-06 | 1.500E-05 |
| 2290 | 115 | L115I | A | 1.4e-05 | 1.5e-05 | 1.1e-05 | 1.333E-05 | 2.082E-06 | 1.100E-05 | 1.500E-05 |
| 2291 | 115 | L115L | A | 1.6e-05 | 1.3e-05 | 9.6e-06 | 1.287E-05 | 3.202E-06 | 9.600E-06 | 1.600E-05 |
| 2292 | 115 | L115K | A | 8.4e-06 | 8.5e-06 | 9e-06 | 8.633E-06 | 3.215E-07 | 8.400E-06 | 9.000E-06 |
| 2293 | 115 | L115M | A | 1e-05 | 1e-05 | 1.6e-05 | 1.200E-05 | 3.464E-06 | 1.000E-05 | 1.600E-05 |
| 2294 | 115 | L115F | A | 9e-06 | 1.1e-05 | 1.1e-05 | 1.033E-05 | 1.155E-06 | 9.000E-06 | 1.100E-05 |

| | | | | | | | | | | |
|---|---|---|---|---|---|---|---|---|---|---|
| 2295 | 115 | L115P | A | 1e-05 | 1.5e-05 | 1.6e-05 | 1.367E-05 | 3.215E-06 | 1.000E-05 | 1.600E-05 |
| 2296 | 115 | L115S | A | 1.5e-05 | 1.2e-05 | 1.5e-05 | 1.400E-05 | 1.732E-06 | 1.200E-05 | 1.500E-05 |
| 2297 | 115 | L115T | A | 9.7e-06 | 1e-05 | 1.5e-05 | 1.157E-05 | 2.977E-06 | 9.700E-06 | 1.500E-05 |
| 2298 | 115 | L115W | A | 1.5e-05 | 1.3e-05 | 1.1e-05 | 1.300E-05 | 2.000E-06 | 1.100E-05 | 1.500E-05 |
| 2299 | 115 | L115Y | A | 1.3e-05 | 8.6e-06 | 1e-05 | 1.053E-05 | 2.248E-06 | 8.600E-06 | 1.300E-05 |
| 2300 | 115 | L115V | A | 1e-05 | 1.4e-05 | 1.1e-05 | 1.167E-05 | 2.082E-06 | 1.000E-05 | 1.400E-05 |
| 2301 | 116 | R116A | A | 1.5e-05 | 1.1e-05 | 1.4e-05 | 1.333E-05 | 2.082E-06 | 1.100E-05 | 1.500E-05 |
| 2302 | 116 | R116R | A | 1.4e-05 | 1.2e-05 | 1.3e-05 | 1.300E-05 | 1.000E-06 | 1.200E-05 | 1.400E-05 |
| 2303 | 116 | R116N | A | 8.6e-06 | 1e-05 | 2.7e-05 | 1.520E-05 | 1.024E-05 | 8.600E-06 | 2.700E-05 |
| 2304 | 116 | R116D | A | 1.4e-05 | 1e-05 | 8.9e-06 | 1.097E-05 | 2.684E-06 | 8.900E-06 | 1.400E-05 |
| 2305 | 116 | R116C | A | 1.3e-05 | 1.2e-05 | 1.7e-05 | 1.400E-05 | 2.646E-06 | 1.200E-05 | 1.700E-05 |
| 2306 | 116 | R116E | A | 1.2e-05 | 1.6e-05 | 1.3e-05 | 1.367E-05 | 2.082E-06 | 1.200E-05 | 1.600E-05 |
| 2307 | 116 | R116Q | A | 9.6e-06 | 1.1e-05 | 9.9e-06 | 1.017E-05 | 7.371E-07 | 9.600E-06 | 1.100E-05 |
| 2308 | 116 | R116G | A | 1.2e-05 | 1e-05 | 9e-06 | 1.033E-05 | 1.528E-06 | 9.000E-06 | 1.200E-05 |
| 2309 | 116 | R116H | A | 1.1e-05 | 1.5e-05 | 1.6e-05 | 1.400E-05 | 2.646E-06 | 1.100E-05 | 1.600E-05 |
| 2310 | 116 | R116I | A | 1.5e-05 | 2.6e-05 | 1.6e-05 | 1.900E-05 | 6.083E-06 | 1.500E-05 | 2.600E-05 |
| 2311 | 116 | R116L | A | 1e-05 | 1.3e-05 | 1.6e-05 | 1.300E-05 | 3.000E-06 | 1.000E-05 | 1.600E-05 |
| 2312 | 116 | R116K | A | 1e-05 | 1e-05 | 1.1e-05 | 1.033E-05 | 5.774E-07 | 1.000E-05 | 1.100E-05 |
| 2313 | 116 | R116M | A | 1.4e-05 | 1.1e-05 | 1.5e-05 | 1.333E-05 | 2.082E-06 | 1.100E-05 | 1.500E-05 |
| 2314 | 116 | R116F | A | 1.2e-05 | 1.4e-05 | 1.4e-05 | 1.333E-05 | 1.155E-06 | 1.200E-05 | 1.400E-05 |
| 2315 | 116 | R116P | A | 1.3e-05 | 1.3e-05 | 9.9e-06 | 1.197E-05 | 1.790E-06 | 9.900E-06 | 1.300E-05 |
| 2316 | 116 | R116S | A | 9.9e-06 | 1.3e-05 | 7.5e-06 | 1.013E-05 | 2.757E-06 | 7.500E-06 | 1.300E-05 |
| 2317 | 116 | R116T | A | 1.4e-05 | 1.4e-05 | 1.2e-05 | 1.333E-05 | 1.155E-06 | 1.200E-05 | 1.400E-05 |
| 2318 | 116 | R116W | A | 1.5e-05 | 1.2e-05 | 7.6e-06 | 1.153E-05 | 3.722E-06 | 7.600E-06 | 1.500E-05 |
| 2319 | 116 | R116Y | A | 1.4e-05 | 8.3e-06 | 1.5e-05 | 1.243E-05 | 3.614E-06 | 8.300E-06 | 1.500E-05 |
| 2320 | 116 | R116V | A | 1.2e-05 | 1.1e-05 | 1.7e-05 | 1.333E-05 | 3.215E-06 | 1.100E-05 | 1.700E-05 |
| 2321 | 117 | E117A | A | 1.6e-05 | 1.6e-05 | 9.4e-06 | 1.380E-05 | 3.811E-06 | 9.400E-06 | 1.600E-05 |
| 2322 | 117 | E117R | A | 1.5e-05 | 1.6e-05 | 1.3e-05 | 1.467E-05 | 1.528E-06 | 1.300E-05 | 1.600E-05 |
| 2323 | 117 | E117N | A | 9.6e-06 | 1e-05 | 1.5e-05 | 1.153E-05 | 3.009E-06 | 9.600E-06 | 1.500E-05 |
| 2324 | 117 | E117D | A | 1.5e-05 | 1.5e-05 | 1.7e-05 | 1.567E-05 | 1.155E-06 | 1.500E-05 | 1.700E-05 |
| 2325 | 117 | E117C | A | 1.1e-05 | 1.3e-05 | 1.3e-05 | 1.233E-05 | 1.155E-06 | 1.100E-05 | 1.300E-05 |
| 2326 | 117 | E117E | A | 1.4e-05 | 1.5e-05 | 1.6e-05 | 1.500E-05 | 1.000E-06 | 1.400E-05 | 1.600E-05 |
| 2327 | 117 | E117Q | A | 1.4e-05 | 1.5e-05 | 1.5e-05 | 1.467E-05 | 5.774E-07 | 1.400E-05 | 1.500E-05 |
| 2328 | 117 | E117G | A | 1e-05 | 1.6e-05 | 1.1e-05 | 1.233E-05 | 3.215E-06 | 1.000E-05 | 1.600E-05 |
| 2329 | 117 | E117H | A | 8.8e-06 | 1.4e-05 | 1e-05 | 1.093E-05 | 2.723E-06 | 8.800E-06 | 1.400E-05 |
| 2330 | 117 | E117I | A | 1.5e-05 | 1.5e-05 | 1.7e-05 | 1.567E-05 | 1.155E-06 | 1.500E-05 | 1.700E-05 |
| 2331 | 117 | E117L | A | 1e-05 | 1.5e-05 | 8.9e-06 | 1.130E-05 | 3.251E-06 | 8.900E-06 | 1.500E-05 |
| 2332 | 117 | E117K | A | 9.1e-06 | 7.9e-06 | 6.5e-06 | 7.833E-06 | 1.301E-06 | 6.500E-06 | 9.100E-06 |
| 2333 | 117 | E117M | A | 8.9e-06 | 1.5e-05 | 1.1e-05 | 1.163E-05 | 3.099E-06 | 8.900E-06 | 1.500E-05 |
| 2334 | 117 | E117F | A | 1.3e-05 | 1.4e-05 | 1.5e-05 | 1.400E-05 | 1.000E-06 | 1.300E-05 | 1.500E-05 |
| 2335 | 117 | E117P | A | 1e-05 | 1.4e-05 | 1.3e-05 | 1.233E-05 | 2.082E-06 | 1.000E-05 | 1.400E-05 |
| 2336 | 117 | E117S | A | 1e-05 | 1.2e-05 | 1.1e-05 | 1.100E-05 | 1.000E-06 | 1.000E-05 | 1.200E-05 |
| 2337 | 117 | E117T | A | 1.6e-05 | 1.2e-05 | 1.2e-05 | 1.333E-05 | 2.309E-06 | 1.200E-05 | 1.600E-05 |
| 2338 | 117 | E117W | A | 1.3e-05 | 8.3e-06 | 1.2e-05 | 1.110E-05 | 2.476E-06 | 8.300E-06 | 1.300E-05 |
| 2339 | 117 | E117Y | A | 1.1e-05 | 9.3e-06 | 9.5e-06 | 9.933E-06 | 9.292E-07 | 9.300E-06 | 1.100E-05 |
| 2340 | 117 | E117V | A | 1.7e-05 | 1.6e-05 | 1.1e-05 | 1.467E-05 | 3.215E-06 | 1.100E-05 | 1.700E-05 |
| 2341 | 118 | L118A | A | 1.2e-05 | 2.4e-05 | 8.2e-06 | 1.473E-05 | 8.247E-06 | 8.200E-06 | 2.400E-05 |
| 2342 | 118 | L118R | A | 1e-05 | 1.4e-05 | 1.4e-05 | 1.267E-05 | 2.309E-06 | 1.000E-05 | 1.400E-05 |
| 2343 | 118 | L118N | A | 1.4e-05 | 1.5e-05 | 1.3e-05 | 1.400E-05 | 1.000E-06 | 1.300E-05 | 1.500E-05 |
| 2344 | 118 | L118D | A | 1.5e-05 | 1.3e-05 | 1.4e-05 | 1.400E-05 | 1.000E-06 | 1.300E-05 | 1.500E-05 |
| 2345 | 118 | L118C | A | 1.4e-05 | 1.5e-05 | 1.7e-05 | 1.533E-05 | 1.528E-06 | 1.400E-05 | 1.700E-05 |
| 2346 | 118 | L118E | A | 1.3e-05 | 1.6e-05 | 1.1e-05 | 1.333E-05 | 2.517E-06 | 1.100E-05 | 1.600E-05 |
| 2347 | 118 | L118Q | A | 9.7e-06 | 8.1e-06 | 1e-05 | 9.267E-06 | 1.021E-06 | 8.100E-06 | 1.000E-05 |
| 2348 | 118 | L118G | A | 9.1e-06 | 1.5e-05 | 9.1e-06 | 1.107E-05 | 3.406E-06 | 9.100E-06 | 1.500E-05 |
| 2349 | 118 | L118H | A | 1.6e-05 | 8.9e-06 | 9.2e-06 | 1.137E-05 | 4.015E-06 | 8.900E-06 | 1.600E-05 |
| 2350 | 118 | L118I | A | 9e-06 | 8.8e-06 | 1.3e-05 | 1.027E-05 | 2.369E-06 | 8.800E-06 | 1.300E-05 |
| 2351 | 118 | L118L | A | 1.7e-05 | 1.4e-05 | 1.3e-05 | 1.467E-05 | 2.082E-06 | 1.300E-05 | 1.700E-05 |
| 2352 | 118 | L118K | A | 1.7e-05 | 1e-05 | 1.4e-05 | 1.367E-05 | 3.512E-06 | 1.000E-05 | 1.700E-05 |
| 2353 | 118 | L118M | A | 1.1e-05 | 1.3e-05 | 9e-06 | 1.100E-05 | 2.000E-06 | 9.000E-06 | 1.300E-05 |
| 2354 | 118 | L118F | A | 1e-05 | 1.4e-05 | 1.2e-05 | 1.200E-05 | 2.000E-06 | 1.000E-05 | 1.400E-05 |
| 2355 | 118 | L118P | A | 1e-05 | 1.7e-05 | 1.4e-05 | 1.367E-05 | 3.512E-06 | 1.000E-05 | 1.700E-05 |
| 2356 | 118 | L118S | A | 1.6e-05 | 1.4e-05 | 1.4e-05 | 1.467E-05 | 1.155E-06 | 1.400E-05 | 1.600E-05 |
| 2357 | 118 | L118T | A | 1.5e-05 | 1.4e-05 | 1.1e-05 | 1.333E-05 | 2.082E-06 | 1.100E-05 | 1.500E-05 |
| 2358 | 118 | L118W | A | 8.6e-06 | 9.4e-06 | 1.4e-05 | 1.067E-05 | 2.914E-06 | 8.600E-06 | 1.400E-05 |
| 2359 | 118 | L118Y | A | 9.8e-06 | 1.6e-05 | 9.6e-06 | 1.180E-05 | 3.639E-06 | 9.600E-06 | 1.600E-05 |
| 2360 | 118 | L118V | A | 5.5e-06 | 1e-05 | 7.6e-06 | 7.700E-06 | 2.252E-06 | 5.500E-06 | 1.000E-05 |
| 2361 | 119 | Q119A | A | 1.2e-05 | 1.1e-05 | 1.2e-05 | 1.167E-05 | 5.774E-07 | 1.100E-05 | 1.200E-05 |
| 2362 | 119 | Q119R | A | 1.1e-05 | 1.8e-05 | 1.6e-05 | 1.500E-05 | 3.606E-06 | 1.100E-05 | 1.800E-05 |
| 2363 | 119 | Q119N | A | 1.3e-05 | 1.5e-05 | 2.2e-05 | 1.667E-05 | 4.726E-06 | 1.300E-05 | 2.200E-05 |
| 2364 | 119 | Q119D | A | 9e-06 | 1e-05 | 1.5e-05 | 1.133E-05 | 3.215E-06 | 9.000E-06 | 1.500E-05 |
| 2365 | 119 | Q119C | A | 1.5e-05 | 8.8e-06 | 9.6e-06 | 1.113E-05 | 3.372E-06 | 8.800E-06 | 1.500E-05 |
| 2366 | 119 | Q119E | A | 1.4e-05 | 1.4e-05 | 7.4e-06 | 1.180E-05 | 3.811E-06 | 7.400E-06 | 1.400E-05 |
| 2367 | 119 | Q119Q | A | 1e-05 | 1.5e-05 | 1.4e-05 | 1.300E-05 | 2.646E-06 | 1.000E-05 | 1.500E-05 |
| 2368 | 119 | Q119G | A | 1.6e-05 | 1.2e-05 | 1.1e-05 | 1.300E-05 | 2.646E-06 | 1.100E-05 | 1.600E-05 |
| 2369 | 119 | Q119H | A | 1e-05 | 1.4e-05 | 1.4e-05 | 1.267E-05 | 2.309E-06 | 1.000E-05 | 1.400E-05 |
| 2370 | 119 | Q119I | A | 1.4e-05 | 1e-05 | 1.1e-05 | 1.167E-05 | 2.082E-06 | 1.000E-05 | 1.400E-05 |
| 2371 | 119 | Q119L | A | 1e-05 | 1.4e-05 | 9.5e-06 | 1.117E-05 | 2.466E-06 | 9.500E-06 | 1.400E-05 |
| 2372 | 119 | Q119K | A | 9.4e-06 | 1.5e-05 | 1.3e-05 | 1.247E-05 | 2.838E-06 | 9.400E-06 | 1.500E-05 |
| 2373 | 119 | Q119M | A | 1.6e-05 | 1.5e-05 | 9.9e-06 | 1.363E-05 | 3.272E-06 | 9.900E-06 | 1.600E-05 |
| 2374 | 119 | Q119F | A | 1.4e-05 | 1.7e-05 | 1.4e-05 | 1.500E-05 | 1.732E-06 | 1.400E-05 | 1.700E-05 |
| 2375 | 119 | Q119P | A | 1.1e-05 | 1.6e-05 | 1.4e-05 | 1.367E-05 | 2.517E-06 | 1.100E-05 | 1.600E-05 |
| 2376 | 119 | Q119S | A | 1.4e-05 | 1.5e-05 | 1.5e-05 | 1.467E-05 | 5.774E-07 | 1.400E-05 | 1.500E-05 |
| 2377 | 119 | Q119T | A | 1.5e-05 | 1.8e-05 | 1.6e-05 | 1.633E-05 | 1.528E-06 | 1.500E-05 | 1.800E-05 |
| 2378 | 119 | Q119W | A | 1.2e-05 | 1e-05 | 1.7e-05 | 1.300E-05 | 3.606E-06 | 1.000E-05 | 1.700E-05 |
| 2379 | 119 | Q119Y | A | 1.3e-05 | 1.6e-05 | 1.6e-05 | 1.500E-05 | 1.732E-06 | 1.300E-05 | 1.600E-05 |

| | | | | | | | | | | |
|---|---|---|---|---|---|---|---|---|---|---|
| 2380 | 119 | Q119V | A | 1.6e-05 | 1.1e-05 | 9.7e-06 | 1.223E-05 | 3.326E-06 | 9.700E-06 | 1.600E-05 |
| 2381 | 120 | L120A | A | 1.6e-05 | 1.3e-05 | 2.9e-05 | 1.933E-05 | 8.505E-06 | 1.300E-05 | 2.900E-05 |
| 2382 | 120 | L120R | A | 9.2e-06 | 1.4e-05 | 1.2e-05 | 1.173E-05 | 2.411E-06 | 9.200E-06 | 1.400E-05 |
| 2383 | 120 | L120N | A | 1e-05 | 1.2e-05 | 1.5e-05 | 1.233E-05 | 2.517E-06 | 1.000E-05 | 1.500E-05 |
| 2384 | 120 | L120D | A | 1.4e-05 | 9.4e-06 | 1.5e-05 | 1.280E-05 | 2.987E-06 | 9.400E-06 | 1.500E-05 |
| 2385 | 120 | L120C | A | 1.8e-05 | 1.4e-05 | 1.4e-05 | 1.533E-05 | 2.309E-06 | 1.400E-05 | 1.800E-05 |
| 2386 | 120 | L120E | A | 1.3e-05 | 1.4e-05 | 1.5e-05 | 1.400E-05 | 1.000E-06 | 1.300E-05 | 1.500E-05 |
| 2387 | 120 | L120Q | A | 1.7e-05 | 1.5e-05 | 9.4e-06 | 1.380E-05 | 3.940E-06 | 9.400E-06 | 1.700E-05 |
| 2388 | 120 | L120G | A | 1.6e-05 | 1.6e-05 | 1.4e-05 | 1.533E-05 | 1.155E-06 | 1.400E-05 | 1.600E-05 |
| 2389 | 120 | L120H | A | 1.4e-05 | 1.1e-05 | 1.4e-05 | 1.300E-05 | 1.732E-06 | 1.100E-05 | 1.400E-05 |
| 2390 | 120 | L120I | A | 1.1e-05 | 1.3e-05 | 8.4e-06 | 1.080E-05 | 2.307E-06 | 8.400E-06 | 1.300E-05 |
| 2391 | 120 | L120L | A | 1.6e-05 | 1.4e-05 | 2.7e-05 | 1.900E-05 | 7.000E-06 | 1.400E-05 | 2.700E-05 |
| 2392 | 120 | L120K | A | 1.4e-05 | 1.6e-05 | 1.6e-05 | 1.533E-05 | 1.155E-06 | 1.400E-05 | 1.600E-05 |
| 2393 | 120 | L120M | A | 1.4e-05 | 9.7e-06 | 1e-05 | 1.123E-05 | 2.401E-06 | 9.700E-06 | 1.400E-05 |
| 2394 | 120 | L120F | A | 1.1e-05 | 1.1e-05 | 1.7e-05 | 1.300E-05 | 3.464E-06 | 1.100E-05 | 1.700E-05 |
| 2395 | 120 | L120P | A | 1e-05 | 1.6e-05 | 7.1e-06 | 1.103E-05 | 4.539E-06 | 7.100E-06 | 1.600E-05 |
| 2396 | 120 | L120S | A | 9.7e-06 | 1.3e-05 | 1.1e-05 | 1.123E-05 | 1.662E-06 | 9.700E-06 | 1.300E-05 |
| 2397 | 120 | L120T | A | 1e-05 | 1.5e-05 | 1.1e-05 | 1.200E-05 | 2.646E-06 | 1.000E-05 | 1.500E-05 |
| 2398 | 120 | L120W | A | 9.9e-06 | 1e-05 | 1.7e-05 | 1.230E-05 | 4.071E-06 | 9.900E-06 | 1.700E-05 |
| 2399 | 120 | L120Y | A | 1.4e-05 | 1.7e-05 | 6.5e-06 | 1.250E-05 | 5.408E-06 | 6.500E-06 | 1.700E-05 |
| 2400 | 120 | L120V | A | 1.1e-05 | 1.6e-05 | 1.4e-05 | 1.367E-05 | 2.517E-06 | 1.100E-05 | 1.600E-05 |
| 2401 | 121 | R121A | A | 9.4e-06 | 1.4e-05 | 1.1e-05 | 1.147E-05 | 2.335E-06 | 9.400E-06 | 1.400E-05 |
| 2402 | 121 | R121R | A | 1.1e-05 | 1.1e-05 | 1.2e-05 | 1.133E-05 | 5.774E-07 | 1.100E-05 | 1.200E-05 |
| 2403 | 121 | R121N | A | 9.6e-06 | 1.5e-05 | 1e-05 | 1.153E-05 | 3.009E-06 | 9.600E-06 | 1.500E-05 |
| 2404 | 121 | R121D | A | 1.3e-05 | 1.4e-05 | 1.5e-05 | 1.400E-05 | 1.000E-06 | 1.300E-05 | 1.500E-05 |
| 2405 | 121 | R121C | A | 1.5e-05 | 1.5e-05 | 1.1e-05 | 1.367E-05 | 2.309E-06 | 1.100E-05 | 1.500E-05 |
| 2406 | 121 | R121E | A | 1.5e-05 | 1.4e-05 | 1.4e-05 | 1.433E-05 | 5.774E-07 | 1.400E-05 | 1.500E-05 |
| 2407 | 121 | R121Q | A | 1.2e-05 | 9.9e-06 | 1.5e-05 | 1.230E-05 | 2.563E-06 | 9.900E-06 | 1.500E-05 |
| 2408 | 121 | R121G | A | 1.6e-05 | 9.8e-06 | 1.6e-05 | 1.393E-05 | 3.580E-06 | 9.800E-06 | 1.600E-05 |
| 2409 | 121 | R121H | A | 8.5e-06 | 1.3e-05 | 1.4e-05 | 1.183E-05 | 2.930E-06 | 8.500E-06 | 1.400E-05 |
| 2410 | 121 | R121I | A | 7.3e-06 | 1.6e-05 | 1.5e-05 | 1.277E-05 | 4.761E-06 | 7.300E-06 | 1.600E-05 |
| 2411 | 121 | R121L | A | 1.2e-05 | 1.4e-05 | 1.6e-05 | 1.400E-05 | 2.000E-06 | 1.200E-05 | 1.600E-05 |
| 2412 | 121 | R121K | A | 1.2e-05 | 1e-05 | 9.4e-06 | 1.047E-05 | 1.361E-06 | 9.400E-06 | 1.200E-05 |
| 2413 | 121 | R121M | A | 1.8e-05 | 1e-05 | 1.6e-05 | 1.467E-05 | 4.163E-06 | 1.000E-05 | 1.800E-05 |
| 2414 | 121 | R121F | A | 8e-06 | 1.2e-05 | 1.3e-05 | 1.100E-05 | 2.646E-06 | 8.000E-06 | 1.300E-05 |
| 2415 | 121 | R121P | A | 1.3e-05 | 1.5e-05 | 1.5e-05 | 1.433E-05 | 1.155E-06 | 1.300E-05 | 1.500E-05 |
| 2416 | 121 | R121S | A | 7.4e-06 | 1e-05 | 1.6e-05 | 1.113E-05 | 4.411E-06 | 7.400E-06 | 1.600E-05 |
| 2417 | 121 | R121T | A | 1.1e-05 | 1.4e-05 | 9.7e-06 | 1.157E-05 | 2.205E-06 | 9.700E-06 | 1.400E-05 |
| 2418 | 121 | R121W | A | 1.4e-05 | 1.5e-05 | 1.1e-05 | 1.333E-05 | 2.082E-06 | 1.100E-05 | 1.500E-05 |
| 2419 | 121 | R121Y | A | 6.9e-06 | 1.1e-05 | 1.1e-05 | 9.633E-06 | 2.367E-06 | 6.900E-06 | 1.100E-05 |
| 2420 | 121 | R121V | A | 9e-06 | 1.5e-05 | 1.4e-05 | 1.267E-05 | 3.215E-06 | 9.000E-06 | 1.500E-05 |
| 2421 | 122 | S122A | A | 1.1e-05 | 1.1e-05 | 1.4e-05 | 1.200E-05 | 1.732E-06 | 1.100E-05 | 1.400E-05 |
| 2422 | 122 | S122R | A | 1.4e-05 | 1.4e-05 | 9.7e-06 | 1.257E-05 | 2.483E-06 | 9.700E-06 | 1.400E-05 |
| 2423 | 122 | S122N | A | 1.1e-05 | 1e-05 | 1.6e-05 | 1.233E-05 | 3.215E-06 | 1.000E-05 | 1.600E-05 |
| 2424 | 122 | S122D | A | 1.1e-05 | 1.7e-05 | 1.1e-05 | 1.300E-05 | 3.464E-06 | 1.100E-05 | 1.700E-05 |
| 2425 | 122 | S122C | A | 1.4e-05 | 1.5e-05 | 1e-05 | 1.300E-05 | 2.646E-06 | 1.000E-05 | 1.500E-05 |
| 2426 | 122 | S122E | A | 1.5e-05 | 1e-05 | 1.1e-05 | 1.200E-05 | 2.646E-06 | 1.000E-05 | 1.500E-05 |
| 2427 | 122 | S122Q | A | 1.3e-05 | 1.4e-05 | 1.7e-05 | 1.467E-05 | 2.082E-06 | 1.300E-05 | 1.700E-05 |
| 2428 | 122 | S122G | A | 1.6e-05 | 1.5e-05 | 9.6e-06 | 1.353E-05 | 3.443E-06 | 9.600E-06 | 1.600E-05 |
| 2429 | 122 | S122H | A | 1.9e-05 | 1.3e-05 | 1.6e-05 | 1.600E-05 | 3.000E-06 | 1.300E-05 | 1.900E-05 |
| 2430 | 122 | S122I | A | 1.3e-05 | 1.3e-05 | 1.5e-05 | 1.367E-05 | 1.155E-06 | 1.300E-05 | 1.500E-05 |
| 2431 | 122 | S122L | A | 1.6e-05 | 1.5e-05 | 1.3e-05 | 1.467E-05 | 1.528E-06 | 1.300E-05 | 1.600E-05 |
| 2432 | 122 | S122K | A | 1.1e-05 | 1.7e-05 | 1.5e-05 | 1.433E-05 | 3.055E-06 | 1.100E-05 | 1.700E-05 |
| 2433 | 122 | S122M | A | 1.2e-05 | 1e-05 | 1.1e-05 | 1.100E-05 | 1.000E-06 | 1.000E-05 | 1.200E-05 |
| 2434 | 122 | S122F | A | 1.2e-05 | 1.9e-05 | 1.1e-05 | 1.400E-05 | 4.359E-06 | 1.100E-05 | 1.900E-05 |
| 2435 | 122 | S122P | A | 9.2e-06 | 1.2e-05 | 9.1e-06 | 1.010E-05 | 1.646E-06 | 9.100E-06 | 1.200E-05 |
| 2436 | 122 | S122S | A | 1.2e-05 | 1.2e-05 | 1.4e-05 | 1.267E-05 | 1.155E-06 | 1.200E-05 | 1.400E-05 |
| 2437 | 122 | S122T | A | 1.6e-05 | 1.5e-05 | 1.4e-05 | 1.500E-05 | 1.000E-06 | 1.400E-05 | 1.600E-05 |
| 2438 | 122 | S122W | A | 1.2e-05 | 1.4e-05 | 1.1e-05 | 1.233E-05 | 1.528E-06 | 1.100E-05 | 1.400E-05 |
| 2439 | 122 | S122Y | A | 8.5e-06 | 1.5e-05 | 1.1e-05 | 1.150E-05 | 3.279E-06 | 8.500E-06 | 1.500E-05 |
| 2440 | 122 | S122V | A | 1.7e-05 | 8.6e-06 | 1.4e-05 | 1.320E-05 | 4.257E-06 | 8.600E-06 | 1.700E-05 |
| 2441 | 123 | L123A | A | 1.3e-05 | 1.7e-05 | 1.6e-05 | 1.533E-05 | 2.082E-06 | 1.300E-05 | 1.700E-05 |
| 2442 | 123 | L123R | A | 1.3e-05 | 1.4e-05 | 1.5e-05 | 1.400E-05 | 1.000E-06 | 1.300E-05 | 1.500E-05 |
| 2443 | 123 | L123N | A | 1.4e-05 | 1.5e-05 | 1.1e-05 | 1.333E-05 | 2.082E-06 | 1.100E-05 | 1.500E-05 |
| 2444 | 123 | L123D | A | 1.4e-05 | 1.6e-05 | 1.5e-05 | 1.500E-05 | 1.000E-06 | 1.400E-05 | 1.600E-05 |
| 2445 | 123 | L123C | A | 8.2e-06 | 8.6e-06 | 1.6e-05 | 1.093E-05 | 4.392E-06 | 8.200E-06 | 1.600E-05 |
| 2446 | 123 | L123E | A | 1.4e-05 | 1e-05 | 1.1e-05 | 1.167E-05 | 2.082E-06 | 1.000E-05 | 1.400E-05 |
| 2447 | 123 | L123Q | A | 1.1e-05 | 1.5e-05 | 1.3e-05 | 1.300E-05 | 2.000E-06 | 1.100E-05 | 1.500E-05 |
| 2448 | 123 | L123G | A | 1.5e-05 | 1.2e-05 | 1.6e-05 | 1.433E-05 | 2.082E-06 | 1.200E-05 | 1.600E-05 |
| 2449 | 123 | L123H | A | 1.5e-05 | 1.6e-05 | 1.7e-05 | 1.600E-05 | 1.000E-06 | 1.500E-05 | 1.700E-05 |
| 2450 | 123 | L123I | A | 9.5e-06 | 1.5e-05 | 1e-05 | 1.150E-05 | 3.041E-06 | 9.500E-06 | 1.500E-05 |
| 2451 | 123 | L123L | A | 1e-05 | 1.3e-05 | 1e-05 | 1.100E-05 | 1.732E-06 | 1.000E-05 | 1.300E-05 |
| 2452 | 123 | L123K | A | 8.3e-06 | 1.5e-05 | 1.5e-05 | 1.277E-05 | 3.868E-06 | 8.300E-06 | 1.500E-05 |
| 2453 | 123 | L123M | A | 1.4e-05 | 1.4e-05 | 1e-05 | 1.267E-05 | 2.309E-06 | 1.000E-05 | 1.400E-05 |
| 2454 | 123 | L123F | A | 1.1e-05 | 1.5e-05 | 1.2e-05 | 1.267E-05 | 2.082E-06 | 1.100E-05 | 1.500E-05 |
| 2455 | 123 | L123P | A | 8.6e-06 | 1.8e-05 | 1.1e-05 | 1.253E-05 | 4.884E-06 | 8.600E-06 | 1.800E-05 |
| 2456 | 123 | L123S | A | 9e-06 | 7.3e-06 | 1e-05 | 8.767E-06 | 1.365E-06 | 7.300E-06 | 1.000E-05 |
| 2457 | 123 | L123T | A | 1.7e-05 | 1.5e-05 | 1.4e-05 | 1.533E-05 | 1.528E-06 | 1.400E-05 | 1.700E-05 |
| 2458 | 123 | L123W | A | 1.4e-05 | 1e-05 | 1.1e-05 | 1.167E-05 | 2.082E-06 | 1.000E-05 | 1.400E-05 |
| 2459 | 123 | L123Y | A | 1e-05 | 2.1e-05 | 1.4e-05 | 1.500E-05 | 5.568E-06 | 1.000E-05 | 2.100E-05 |
| 2460 | 123 | L123V | A | 7.5e-06 | 1.8e-05 | 8.7e-06 | 1.140E-05 | 5.747E-06 | 7.500E-06 | 1.800E-05 |
| 2461 | 124 | T124A | A | 1.5e-05 | 1.5e-05 | 1.5e-05 | 1.500E-05 | 0.000E+00 | 1.500E-05 | 1.500E-05 |
| 2462 | 124 | T124R | A | 1.6e-05 | 1.2e-05 | 1.8e-05 | 1.533E-05 | 3.055E-06 | 1.200E-05 | 1.800E-05 |
| 2463 | 124 | T124N | A | 1e-05 | 1.6e-05 | 1.6e-05 | 1.400E-05 | 3.464E-06 | 1.000E-05 | 1.600E-05 |
| 2464 | 124 | T124D | A | 1.6e-05 | 1.6e-05 | 1.2e-05 | 1.467E-05 | 2.309E-06 | 1.200E-05 | 1.600E-05 |

| | | | | | | | | | | |
|---|---|---|---|---|---|---|---|---|---|---|
| 2465 | 124 | T124C | A | 1.4e-05 | 1e-05 | 9.7e-06 | 1.123E-05 | 2.401E-06 | 9.700E-06 | 1.400E-05 |
| 2466 | 124 | T124E | A | 1.1e-05 | 1.6e-05 | 1.4e-05 | 1.367E-05 | 2.517E-06 | 1.100E-05 | 1.600E-05 |
| 2467 | 124 | T124Q | A | 1.2e-05 | 1.1e-05 | 1.6e-05 | 1.300E-05 | 2.646E-06 | 1.100E-05 | 1.600E-05 |
| 2468 | 124 | T124G | A | 1.5e-05 | 6.4e-06 | 9.9e-06 | 1.043E-05 | 4.325E-06 | 6.400E-06 | 1.500E-05 |
| 2469 | 124 | T124H | A | 1.1e-05 | 9.2e-06 | 1.1e-05 | 1.040E-05 | 1.039E-06 | 9.200E-06 | 1.100E-05 |
| 2470 | 124 | T124I | A | 1.1e-05 | 1.4e-05 | 1.2e-05 | 1.233E-05 | 1.528E-06 | 1.100E-05 | 1.400E-05 |
| 2471 | 124 | T124L | A | 9.7e-06 | 1.5e-05 | 1.6e-05 | 1.357E-05 | 3.386E-06 | 9.700E-06 | 1.600E-05 |
| 2472 | 124 | T124K | A | 9.4e-06 | 1.2e-05 | 1.7e-05 | 1.280E-05 | 3.863E-06 | 9.400E-06 | 1.700E-05 |
| 2473 | 124 | T124M | A | 1.4e-05 | 1.2e-05 | 6.8e-06 | 1.093E-05 | 3.717E-06 | 6.800E-06 | 1.400E-05 |
| 2474 | 124 | T124F | A | 1.5e-05 | 9.9e-06 | 1.5e-05 | 1.330E-05 | 2.944E-06 | 9.900E-06 | 1.500E-05 |
| 2475 | 124 | T124P | A | 1e-05 | 1.1e-05 | 6.6e-06 | 9.200E-06 | 2.307E-06 | 6.600E-06 | 1.100E-05 |
| 2476 | 124 | T124S | A | 1e-05 | 8.9e-06 | 1.1e-05 | 9.967E-06 | 1.050E-06 | 8.900E-06 | 1.100E-05 |
| 2477 | 124 | T124T | A | 6e-06 | 9.8e-06 | 9.8e-06 | 8.533E-06 | 2.194E-06 | 6.000E-06 | 9.800E-06 |
| 2478 | 124 | T124W | A | 1.2e-05 | 1.4e-05 | 1.5e-05 | 1.367E-05 | 1.528E-06 | 1.200E-05 | 1.500E-05 |
| 2479 | 124 | T124Y | A | 1e-05 | 1.5e-05 | 1.2e-05 | 1.233E-05 | 2.517E-06 | 1.000E-05 | 1.500E-05 |
| 2480 | 124 | T124V | A | 1.2e-05 | 1.5e-05 | 1.2e-05 | 1.300E-05 | 1.732E-06 | 1.200E-05 | 1.500E-05 |
| 2481 | 125 | E125A | A | 1.5e-05 | 1.4e-05 | 1.2e-05 | 1.367E-05 | 1.528E-06 | 1.200E-05 | 1.500E-05 |
| 2482 | 125 | E125R | A | 1.4e-05 | 1.5e-05 | 1.4e-05 | 1.433E-05 | 5.774E-07 | 1.400E-05 | 1.500E-05 |
| 2483 | 125 | E125N | A | 1e-05 | 8.8e-06 | 9e-06 | 9.267E-06 | 6.429E-07 | 8.800E-06 | 1.000E-05 |
| 2484 | 125 | E125D | A | 9.7e-06 | 8e-06 | 1.5e-05 | 1.090E-05 | 3.651E-06 | 8.000E-06 | 1.500E-05 |
| 2485 | 125 | E125C | A | 9.5e-06 | 1.7e-05 | 1.6e-05 | 1.417E-05 | 4.072E-06 | 9.500E-06 | 1.700E-05 |
| 2486 | 125 | E125E | A | 1.5e-05 | 1.5e-05 | 1.6e-05 | 1.533E-05 | 5.774E-07 | 1.500E-05 | 1.600E-05 |
| 2487 | 125 | E125Q | A | 1.4e-05 | 1e-05 | 1e-05 | 1.133E-05 | 2.309E-06 | 1.000E-05 | 1.400E-05 |
| 2488 | 125 | E125G | A | 1.6e-05 | 1.5e-05 | 1.2e-05 | 1.433E-05 | 2.082E-06 | 1.200E-05 | 1.600E-05 |
| 2489 | 125 | E125H | A | 8.9e-06 | 1.4e-05 | 1e-05 | 1.097E-05 | 2.684E-06 | 8.900E-06 | 1.400E-05 |
| 2490 | 125 | E125I | A | 1.5e-05 | 7.8e-06 | 1.1e-05 | 1.127E-05 | 3.607E-06 | 7.800E-06 | 1.500E-05 |
| 2491 | 125 | E125L | A | 1.1e-05 | 1.4e-05 | 8.9e-06 | 1.130E-05 | 2.563E-06 | 8.900E-06 | 1.400E-05 |
| 2492 | 125 | E125K | A | 1.2e-05 | 1.6e-05 | 1.4e-05 | 1.400E-05 | 2.000E-06 | 1.200E-05 | 1.600E-05 |
| 2493 | 125 | E125M | A | 9.6e-06 | 1.5e-05 | 1.3e-05 | 1.253E-05 | 2.730E-06 | 9.600E-06 | 1.500E-05 |
| 2494 | 125 | E125F | A | 1.6e-05 | 1.2e-05 | 9.2e-06 | 1.240E-05 | 3.418E-06 | 9.200E-06 | 1.600E-05 |
| 2495 | 125 | E125P | A | 6.2e-06 | 1e-05 | 1.3e-05 | 9.733E-06 | 3.408E-06 | 6.200E-06 | 1.300E-05 |
| 2496 | 125 | E125S | A | 1.1e-05 | 1.5e-05 | 1.5e-05 | 1.367E-05 | 2.309E-06 | 1.100E-05 | 1.500E-05 |
| 2497 | 125 | E125T | A | 1.4e-05 | 1.2e-05 | 1.5e-05 | 1.367E-05 | 1.528E-06 | 1.200E-05 | 1.500E-05 |
| 2498 | 125 | E125W | A | 1.6e-05 | 1.4e-05 | 9.9e-06 | 1.330E-05 | 3.110E-06 | 9.900E-06 | 1.600E-05 |
| 2499 | 125 | E125Y | A | 1.6e-05 | 8.2e-06 | 1.3e-05 | 1.240E-05 | 3.934E-06 | 8.200E-06 | 1.600E-05 |
| 2500 | 125 | E125V | A | 1.7e-05 | 1.4e-05 | 1.1e-05 | 1.400E-05 | 3.000E-06 | 1.100E-05 | 1.700E-05 |
| 2501 | 126 | I126A | A | 8.9e-06 | 1.5e-05 | 1.6e-05 | 1.330E-05 | 3.843E-06 | 8.900E-06 | 1.600E-05 |
| 2502 | 126 | I126R | A | 7.2e-06 | 1.5e-05 | 8e-06 | 1.007E-05 | 4.291E-06 | 7.200E-06 | 1.500E-05 |
| 2503 | 126 | I126N | A | 1.5e-05 | 1.1e-05 | 1.5e-05 | 1.367E-05 | 2.309E-06 | 1.100E-05 | 1.500E-05 |
| 2504 | 126 | I126D | A | 1.6e-05 | 1.5e-05 | 1e-05 | 1.367E-05 | 3.215E-06 | 1.000E-05 | 1.600E-05 |
| 2505 | 126 | I126C | A | 1.6e-05 | 1.4e-05 | 1.6e-05 | 1.533E-05 | 1.155E-06 | 1.400E-05 | 1.600E-05 |
| 2506 | 126 | I126E | A | 8.3e-06 | 1.6e-05 | 6.5e-06 | 1.027E-05 | 5.046E-06 | 6.500E-06 | 1.600E-05 |
| 2507 | 126 | I126Q | A | 1.4e-05 | 1.4e-05 | 1.6e-05 | 1.467E-05 | 1.155E-06 | 1.400E-05 | 1.600E-05 |
| 2508 | 126 | I126G | A | 1.1e-05 | 1.6e-05 | 1.1e-05 | 1.267E-05 | 2.887E-06 | 1.100E-05 | 1.600E-05 |
| 2509 | 126 | I126H | A | 1.4e-05 | 8.6e-06 | 8.1e-06 | 1.023E-05 | 3.272E-06 | 8.100E-06 | 1.400E-05 |
| 2510 | 126 | I126I | A | 1e-05 | 1.5e-05 | 1.5e-05 | 1.333E-05 | 2.887E-06 | 1.000E-05 | 1.500E-05 |
| 2511 | 126 | I126L | A | 1.6e-05 | 1.5e-05 | 1e-05 | 1.367E-05 | 3.215E-06 | 1.000E-05 | 1.600E-05 |
| 2512 | 126 | I126K | A | 1e-05 | 1.6e-05 | 1.1e-05 | 1.233E-05 | 3.215E-06 | 1.000E-05 | 1.600E-05 |
| 2513 | 126 | I126M | A | 1.1e-05 | 9.5e-06 | 2.2e-05 | 1.417E-05 | 6.825E-06 | 9.500E-06 | 2.200E-05 |
| 2514 | 126 | I126F | A | 1.1e-05 | 1.5e-05 | 9.6e-06 | 1.187E-05 | 2.802E-06 | 9.600E-06 | 1.500E-05 |
| 2515 | 126 | I126P | A | 1.4e-05 | 9.8e-06 | 1.5e-05 | 1.293E-05 | 2.759E-06 | 9.800E-06 | 1.500E-05 |
| 2516 | 126 | I126S | A | 1.2e-05 | 7.2e-06 | 1.6e-05 | 1.173E-05 | 4.406E-06 | 7.200E-06 | 1.600E-05 |
| 2517 | 126 | I126T | A | 1.2e-05 | 1.5e-05 | 1.5e-05 | 1.400E-05 | 1.732E-06 | 1.200E-05 | 1.500E-05 |
| 2518 | 126 | I126W | A | 1e-05 | 6.9e-06 | 1.4e-05 | 1.030E-05 | 3.559E-06 | 6.900E-06 | 1.400E-05 |
| 2519 | 126 | I126Y | A | 9.6e-06 | 1.7e-05 | 1.4e-05 | 1.353E-05 | 3.722E-06 | 9.600E-06 | 1.700E-05 |
| 2520 | 126 | I126V | A | 2.9e-05 | 1.2e-05 | 9.3e-06 | 1.677E-05 | 1.068E-05 | 9.300E-06 | 2.900E-05 |
| 2521 | 127 | L127A | A | 1.6e-05 | 1.6e-05 | 1.6e-05 | 1.600E-05 | 0.000E+00 | 1.600E-05 | 1.600E-05 |
| 2522 | 127 | L127R | A | 1.8e-05 | 9e-06 | 1.3e-05 | 1.333E-05 | 4.509E-06 | 9.000E-06 | 1.800E-05 |
| 2523 | 127 | L127N | A | 1.1e-05 | 1.1e-05 | 1.1e-05 | 1.100E-05 | 2.075E-21 | 1.100E-05 | 1.100E-05 |
| 2524 | 127 | L127D | A | 1.2e-05 | 1.5e-05 | 1.5e-05 | 1.400E-05 | 1.732E-06 | 1.200E-05 | 1.500E-05 |
| 2525 | 127 | L127C | A | 1.6e-05 | 1e-05 | 1.2e-05 | 1.267E-05 | 3.055E-06 | 1.000E-05 | 1.600E-05 |
| 2526 | 127 | L127E | A | 9.7e-06 | 1.5e-05 | 1.3e-05 | 1.257E-05 | 2.676E-06 | 9.700E-06 | 1.500E-05 |
| 2527 | 127 | L127Q | A | 1.4e-05 | 1.3e-05 | 1.5e-05 | 1.400E-05 | 1.000E-06 | 1.300E-05 | 1.500E-05 |
| 2528 | 127 | L127G | A | 1.5e-05 | 9.2e-06 | 1.4e-05 | 1.273E-05 | 3.101E-06 | 9.200E-06 | 1.500E-05 |
| 2529 | 127 | L127H | A | 9.9e-06 | 1.6e-05 | 1.4e-05 | 1.330E-05 | 3.110E-06 | 9.900E-06 | 1.600E-05 |
| 2530 | 127 | L127I | A | 1.3e-05 | 1.6e-05 | 1.4e-05 | 1.433E-05 | 1.528E-06 | 1.300E-05 | 1.600E-05 |
| 2531 | 127 | L127L | A | 1e-05 | 1.1e-05 | 1.1e-05 | 1.067E-05 | 5.774E-07 | 1.000E-05 | 1.100E-05 |
| 2532 | 127 | L127K | A | 9.6e-06 | 1.6e-05 | 9.2e-06 | 1.160E-05 | 3.816E-06 | 9.200E-06 | 1.600E-05 |
| 2533 | 127 | L127M | A | 1.6e-05 | 8.1e-06 | 1.2e-05 | 1.203E-05 | 3.950E-06 | 8.100E-06 | 1.600E-05 |
| 2534 | 127 | L127F | A | 1.6e-05 | 1.3e-05 | 1.2e-05 | 1.367E-05 | 2.082E-06 | 1.200E-05 | 1.600E-05 |
| 2535 | 127 | L127P | A | 1e-05 | 9.3e-06 | 1.1e-05 | 1.010E-05 | 8.544E-07 | 9.300E-06 | 1.100E-05 |
| 2536 | 127 | L127S | A | 1.4e-05 | 1.4e-05 | 1.2e-05 | 1.333E-05 | 1.155E-06 | 1.200E-05 | 1.400E-05 |
| 2537 | 127 | L127T | A | 9.8e-06 | 1.4e-05 | 9.8e-06 | 1.120E-05 | 2.425E-06 | 9.800E-06 | 1.400E-05 |
| 2538 | 127 | L127W | A | 1e-05 | 7.9e-06 | 1.1e-05 | 9.633E-06 | 1.582E-06 | 7.900E-06 | 1.100E-05 |
| 2539 | 127 | L127Y | A | 1.4e-05 | 9.6e-06 | 1.6e-05 | 1.320E-05 | 3.274E-06 | 9.600E-06 | 1.600E-05 |
| 2540 | 127 | L127V | A | 6.8e-06 | 9.8e-06 | 1e-05 | 8.867E-06 | 1.793E-06 | 6.800E-06 | 1.000E-05 |
| 2541 | 128 | K128A | A | 1.4e-05 | 1.6e-05 | 1.3e-05 | 1.433E-05 | 1.528E-06 | 1.300E-05 | 1.600E-05 |
| 2542 | 128 | K128R | A | 1.5e-05 | 9.7e-06 | 1.5e-05 | 1.323E-05 | 3.060E-06 | 9.700E-06 | 1.500E-05 |
| 2543 | 128 | K128N | A | 9.7e-06 | 9.7e-06 | 1.5e-05 | 1.147E-05 | 3.060E-06 | 9.700E-06 | 1.500E-05 |
| 2544 | 128 | K128D | A | 1.1e-05 | 2.3e-05 | 1.4e-05 | 1.600E-05 | 6.245E-06 | 1.100E-05 | 2.300E-05 |
| 2545 | 128 | K128C | A | 1.4e-05 | 1.5e-05 | 1.2e-05 | 1.367E-05 | 1.528E-06 | 1.200E-05 | 1.500E-05 |
| 2546 | 128 | K128E | A | 1e-05 | 1.3e-05 | 1.5e-05 | 1.267E-05 | 2.517E-06 | 1.000E-05 | 1.500E-05 |
| 2547 | 128 | K128Q | A | 1.3e-05 | 1.4e-05 | 7.6e-06 | 1.153E-05 | 3.443E-06 | 7.600E-06 | 1.400E-05 |
| 2548 | 128 | K128G | A | 1.2e-05 | 1.5e-05 | 8e-06 | 1.167E-05 | 3.512E-06 | 8.000E-06 | 1.500E-05 |
| 2549 | 128 | K128H | A | 1e-05 | 1e-05 | 1.4e-05 | 1.133E-05 | 2.309E-06 | 1.000E-05 | 1.400E-05 |

| | | | | | | | | | | |
|---|---|---|---|---|---|---|---|---|---|---|
| 2550 | 128 | K128I | A | 1.6e-05 | 1.2e-05 | 1e-05 | 1.267E-05 | 3.055E-06 | 1.000E-05 | 1.600E-05 |
| 2551 | 128 | K128L | A | 1.8e-05 | 1.3e-05 | 1.1e-05 | 1.400E-05 | 3.606E-06 | 1.100E-05 | 1.800E-05 |
| 2552 | 128 | K128K | A | 1.3e-05 | 1.6e-05 | 1.6e-05 | 1.500E-05 | 1.732E-06 | 1.300E-05 | 1.600E-05 |
| 2553 | 128 | K128M | A | 1.1e-05 | 1.1e-05 | 1.2e-05 | 1.133E-05 | 5.774E-07 | 1.100E-05 | 1.200E-05 |
| 2554 | 128 | K128F | A | 9.4e-06 | 1.5e-05 | 1.5e-05 | 1.313E-05 | 3.233E-06 | 9.400E-06 | 1.500E-05 |
| 2555 | 128 | K128P | A | 9.9e-06 | 1.5e-05 | 1.1e-05 | 1.197E-05 | 2.684E-06 | 9.900E-06 | 1.500E-05 |
| 2556 | 128 | K128S | A | 1.1e-05 | 1.5e-05 | 1.3e-05 | 1.300E-05 | 2.000E-06 | 1.100E-05 | 1.500E-05 |
| 2557 | 128 | K128T | A | 1.5e-05 | 1.5e-05 | 1.4e-05 | 1.467E-05 | 5.774E-07 | 1.400E-05 | 1.500E-05 |
| 2558 | 128 | K128W | A | 1.6e-05 | 1.5e-05 | 1.2e-05 | 1.433E-05 | 2.082E-06 | 1.200E-05 | 1.600E-05 |
| 2559 | 128 | K128Y | A | 7.3e-06 | 6.7e-06 | 1.2e-05 | 8.667E-06 | 2.902E-06 | 6.700E-06 | 1.200E-05 |
| 2560 | 128 | K128V | A | 1.3e-05 | 1.4e-05 | 1.6e-05 | 1.433E-05 | 1.528E-06 | 1.300E-05 | 1.600E-05 |
| 2561 | 129 | G129A | A | 9.1e-06 | 1.3e-05 | 1.4e-05 | 1.203E-05 | 2.589E-06 | 9.100E-06 | 1.400E-05 |
| 2562 | 129 | G129R | A | 9.7e-06 | 1.5e-05 | 1.5e-05 | 1.323E-05 | 3.060E-06 | 9.700E-06 | 1.500E-05 |
| 2563 | 129 | G129N | A | 8.6e-06 | 1.7e-05 | 1e-05 | 1.187E-05 | 4.500E-06 | 8.600E-06 | 1.700E-05 |
| 2564 | 129 | G129D | A | 1.4e-05 | 1.5e-05 | 1.6e-05 | 1.500E-05 | 1.000E-06 | 1.400E-05 | 1.600E-05 |
| 2565 | 129 | G129C | A | 1.6e-05 | 1.4e-05 | 9.1e-06 | 1.303E-05 | 3.550E-06 | 9.100E-06 | 1.600E-05 |
| 2566 | 129 | G129E | A | 9.6e-06 | 1.2e-05 | 8.7e-06 | 1.010E-05 | 1.706E-06 | 8.700E-06 | 1.200E-05 |
| 2567 | 129 | G129Q | A | 1.1e-05 | 6.5e-06 | 1.2e-05 | 9.833E-06 | 2.930E-06 | 6.500E-06 | 1.200E-05 |
| 2568 | 129 | G129G | A | 1.2e-05 | 1.2e-05 | 1.5e-05 | 1.300E-05 | 1.732E-06 | 1.200E-05 | 1.500E-05 |
| 2569 | 129 | G129H | A | 1.6e-05 | 9.9e-06 | 7.4e-06 | 1.110E-05 | 4.424E-06 | 7.400E-06 | 1.600E-05 |
| 2570 | 129 | G129I | A | 7.3e-06 | 1.7e-05 | 1e-05 | 1.143E-05 | 5.006E-06 | 7.300E-06 | 1.700E-05 |
| 2571 | 129 | G129L | A | 1.3e-05 | 1.6e-05 | 1.5e-05 | 1.467E-05 | 1.528E-06 | 1.300E-05 | 1.600E-05 |
| 2572 | 129 | G129K | A | 1e-05 | 1.2e-05 | 1.2e-05 | 1.133E-05 | 1.155E-06 | 1.000E-05 | 1.200E-05 |
| 2573 | 129 | G129M | A | 1.2e-05 | 1.4e-05 | 1.5e-05 | 1.367E-05 | 1.528E-06 | 1.200E-05 | 1.500E-05 |
| 2574 | 129 | G129F | A | 1.6e-05 | 1.6e-05 | 1.3e-05 | 1.500E-05 | 1.732E-06 | 1.300E-05 | 1.600E-05 |
| 2575 | 129 | G129P | A | 1.4e-05 | 1.3e-05 | 1.5e-05 | 1.400E-05 | 1.000E-06 | 1.300E-05 | 1.500E-05 |
| 2576 | 129 | G129S | A | 1.4e-05 | 1.3e-05 | 6.1e-06 | 1.103E-05 | 4.302E-06 | 6.100E-06 | 1.400E-05 |
| 2577 | 129 | G129T | A | 1.6e-05 | 1.4e-05 | 1.2e-05 | 1.400E-05 | 2.000E-06 | 1.200E-05 | 1.600E-05 |
| 2578 | 129 | G129W | A | 1.1e-05 | 1.1e-05 | 1.2e-05 | 1.133E-05 | 5.774E-07 | 1.100E-05 | 1.200E-05 |
| 2579 | 129 | G129Y | A | 1.4e-05 | 1.5e-05 | 1.3e-05 | 1.400E-05 | 1.000E-06 | 1.300E-05 | 1.500E-05 |
| 2580 | 129 | G129V | A | 1.1e-05 | 8.4e-06 | 1.6e-05 | 1.180E-05 | 3.863E-06 | 8.400E-06 | 1.600E-05 |
| 2581 | 130 | G130A | A | 1.7e-05 | 1.6e-05 | 1.5e-05 | 1.600E-05 | 1.000E-06 | 1.500E-05 | 1.700E-05 |
| 2582 | 130 | G130R | A | 1.1e-05 | 9.7e-06 | 1.1e-05 | 1.057E-05 | 7.506E-07 | 9.700E-06 | 1.100E-05 |
| 2583 | 130 | G130N | A | 1.7e-05 | 1.1e-05 | 1.2e-05 | 1.333E-05 | 3.215E-06 | 1.100E-05 | 1.700E-05 |
| 2584 | 130 | G130D | A | 1.1e-05 | 1e-05 | 1.1e-05 | 1.067E-05 | 5.774E-07 | 1.000E-05 | 1.100E-05 |
| 2585 | 130 | G130C | A | 1.3e-05 | 1.5e-05 | 1.5e-05 | 1.433E-05 | 1.155E-06 | 1.300E-05 | 1.500E-05 |
| 2586 | 130 | G130E | A | 1.4e-05 | 9.6e-06 | 1e-05 | 1.120E-05 | 2.433E-06 | 9.600E-06 | 1.400E-05 |
| 2587 | 130 | G130Q | A | 1.3e-05 | 1.7e-05 | 1.1e-05 | 1.367E-05 | 3.055E-06 | 1.100E-05 | 1.700E-05 |
| 2588 | 130 | G130G | A | 9.8e-06 | 1e-05 | 1.6e-05 | 1.193E-05 | 3.523E-06 | 9.800E-06 | 1.600E-05 |
| 2589 | 130 | G130H | A | 1.6e-05 | 1.4e-05 | 1.4e-05 | 1.467E-05 | 1.155E-06 | 1.400E-05 | 1.600E-05 |
| 2590 | 130 | G130I | A | 1.5e-05 | 1.4e-05 | 9.2e-06 | 1.273E-05 | 3.101E-06 | 9.200E-06 | 1.500E-05 |
| 2591 | 130 | G130L | A | 1.6e-05 | 1.3e-05 | 1e-05 | 1.300E-05 | 3.000E-06 | 1.000E-05 | 1.600E-05 |
| 2592 | 130 | G130K | A | 1e-05 | 1e-05 | 1.1e-05 | 1.033E-05 | 5.774E-07 | 1.000E-05 | 1.100E-05 |
| 2593 | 130 | G130M | A | 1.5e-05 | 5.9e-06 | 1.3e-05 | 1.130E-05 | 4.782E-06 | 5.900E-06 | 1.500E-05 |
| 2594 | 130 | G130F | A | 7e-06 | 1.4e-05 | 1.1e-05 | 1.067E-05 | 3.512E-06 | 7.000E-06 | 1.400E-05 |
| 2595 | 130 | G130P | A | 1.2e-05 | 1.6e-05 | 1.4e-05 | 1.400E-05 | 2.000E-06 | 1.200E-05 | 1.600E-05 |
| 2596 | 130 | G130S | A | 1.4e-05 | 1.1e-05 | 1.4e-05 | 1.300E-05 | 1.732E-06 | 1.100E-05 | 1.400E-05 |
| 2597 | 130 | G130T | A | 1.3e-05 | 1.4e-05 | 1.4e-05 | 1.367E-05 | 5.774E-07 | 1.300E-05 | 1.400E-05 |
| 2598 | 130 | G130W | A | 1.5e-05 | 1.5e-05 | 1.4e-05 | 1.467E-05 | 5.774E-07 | 1.400E-05 | 1.500E-05 |
| 2599 | 130 | G130Y | A | 1e-05 | 1.7e-05 | 8.9e-06 | 1.197E-05 | 4.394E-06 | 8.900E-06 | 1.700E-05 |
| 2600 | 130 | G130V | A | 1.5e-05 | 1e-05 | 1.1e-05 | 1.200E-05 | 2.646E-06 | 1.000E-05 | 1.500E-05 |
| 2601 | 131 | V131A | A | 1.7e-05 | 1.2e-05 | 1.5e-05 | 1.467E-05 | 2.517E-06 | 1.200E-05 | 1.700E-05 |
| 2602 | 131 | V131R | A | 1.3e-05 | 1.5e-05 | 1.7e-05 | 1.500E-05 | 2.000E-06 | 1.300E-05 | 1.700E-05 |
| 2603 | 131 | V131N | A | 1.3e-05 | 1.5e-05 | 9e-06 | 1.233E-05 | 3.055E-06 | 9.000E-06 | 1.500E-05 |
| 2604 | 131 | V131D | A | 1.8e-05 | 9.2e-06 | 1.4e-05 | 1.373E-05 | 4.406E-06 | 9.200E-06 | 1.800E-05 |
| 2605 | 131 | V131C | A | 9.5e-06 | 1.1e-05 | 1.4e-05 | 1.150E-05 | 2.291E-06 | 9.500E-06 | 1.400E-05 |
| 2606 | 131 | V131E | A | 1.5e-05 | 6.9e-06 | 1.4e-05 | 1.197E-05 | 4.416E-06 | 6.900E-06 | 1.500E-05 |
| 2607 | 131 | V131Q | A | 1.3e-05 | 1.5e-05 | 1.4e-05 | 1.400E-05 | 1.000E-06 | 1.300E-05 | 1.500E-05 |
| 2608 | 131 | V131G | A | 1e-05 | 1.7e-05 | 1.1e-05 | 1.267E-05 | 3.786E-06 | 1.000E-05 | 1.700E-05 |
| 2609 | 131 | V131H | A | 1.3e-05 | 1.5e-05 | 1.6e-05 | 1.467E-05 | 1.528E-06 | 1.300E-05 | 1.600E-05 |
| 2610 | 131 | V131I | A | 1.7e-05 | 1.2e-05 | 1.7e-05 | 1.533E-05 | 2.887E-06 | 1.200E-05 | 1.700E-05 |
| 2611 | 131 | V131L | A | 1.4e-05 | 1.6e-05 | 1.5e-05 | 1.500E-05 | 1.000E-06 | 1.400E-05 | 1.600E-05 |
| 2612 | 131 | V131K | A | 1.6e-05 | 1.5e-05 | 1.5e-05 | 1.533E-05 | 5.774E-07 | 1.500E-05 | 1.600E-05 |
| 2613 | 131 | V131M | A | 1.3e-05 | 1e-05 | 1.7e-05 | 1.333E-05 | 3.512E-06 | 1.000E-05 | 1.700E-05 |
| 2614 | 131 | V131F | A | 1.5e-05 | 1e-05 | 1.5e-05 | 1.333E-05 | 2.887E-06 | 1.000E-05 | 1.500E-05 |
| 2615 | 131 | V131P | A | 1.5e-05 | 9.5e-06 | 1.1e-05 | 1.183E-05 | 2.843E-06 | 9.500E-06 | 1.500E-05 |
| 2616 | 131 | V131S | A | 1.3e-05 | 1.2e-05 | 1.1e-05 | 1.200E-05 | 1.000E-06 | 1.100E-05 | 1.300E-05 |
| 2617 | 131 | V131T | A | 1.2e-05 | 1.3e-05 | 9.8e-06 | 1.160E-05 | 1.637E-06 | 9.800E-06 | 1.300E-05 |
| 2618 | 131 | V131W | A | 9.1e-06 | 1.6e-05 | 1.4e-05 | 1.303E-05 | 3.550E-06 | 9.100E-06 | 1.600E-05 |
| 2619 | 131 | V131Y | A | 1.5e-05 | 1.3e-05 | 1.4e-05 | 1.400E-05 | 1.000E-06 | 1.300E-05 | 1.500E-05 |
| 2620 | 131 | V131V | A | 1.2e-05 | 1.4e-05 | 9e-06 | 1.167E-05 | 2.517E-06 | 9.000E-06 | 1.400E-05 |
| 2621 | 132 | L132A | A | 1.1e-05 | 1.3e-05 | 1.7e-05 | 1.367E-05 | 3.055E-06 | 1.100E-05 | 1.700E-05 |
| 2622 | 132 | L132R | A | 8.1e-06 | 9e-06 | 1.3e-05 | 1.003E-05 | 2.608E-06 | 8.100E-06 | 1.300E-05 |
| 2623 | 132 | L132N | A | 7.1e-06 | 1.5e-05 | 1e-05 | 1.070E-05 | 3.996E-06 | 7.100E-06 | 1.500E-05 |
| 2624 | 132 | L132D | A | 1e-05 | 9.2e-06 | 1.5e-05 | 1.140E-05 | 3.143E-06 | 9.200E-06 | 1.500E-05 |
| 2625 | 132 | L132C | A | 1.6e-05 | 1.3e-05 | 1.6e-05 | 1.500E-05 | 1.732E-06 | 1.300E-05 | 1.600E-05 |
| 2626 | 132 | L132E | A | 1.6e-05 | 1.4e-05 | 1.4e-05 | 1.467E-05 | 1.155E-06 | 1.400E-05 | 1.600E-05 |
| 2627 | 132 | L132Q | A | 1.5e-05 | 1.6e-05 | 1.4e-05 | 1.500E-05 | 1.000E-06 | 1.400E-05 | 1.600E-05 |
| 2628 | 132 | L132G | A | 1.5e-05 | 6.8e-06 | 1.3e-05 | 1.160E-05 | 4.276E-06 | 6.800E-06 | 1.500E-05 |
| 2629 | 132 | L132H | A | 1e-05 | 1e-05 | 1.6e-05 | 1.200E-05 | 3.464E-06 | 1.000E-05 | 1.600E-05 |
| 2630 | 132 | L132I | A | 1e-05 | 1.4e-05 | 1e-05 | 1.133E-05 | 2.309E-06 | 1.000E-05 | 1.400E-05 |
| 2631 | 132 | L132L | A | 8e-06 | 1.7e-05 | 9e-06 | 1.133E-05 | 4.933E-06 | 8.000E-06 | 1.700E-05 |
| 2632 | 132 | L132K | A | 1.6e-05 | 1.8e-05 | 1.3e-05 | 1.567E-05 | 2.517E-06 | 1.300E-05 | 1.800E-05 |
| 2633 | 132 | L132M | A | 1.4e-05 | 1.3e-05 | 1e-05 | 1.233E-05 | 2.082E-06 | 1.000E-05 | 1.400E-05 |
| 2634 | 132 | L132F | A | 8e-06 | 1.2e-05 | 1.6e-05 | 1.200E-05 | 4.000E-06 | 8.000E-06 | 1.600E-05 |

| | | | | | | | | | | |
|---|---|---|---|---|---|---|---|---|---|---|
| 2635 | 132 | L132P | A | 1.5e-05 | 1.3e-05 | 1.4e-05 | 1.400E-05 | 1.000E-06 | 1.300E-05 | 1.500E-05 |
| 2636 | 132 | L132S | A | 1.5e-05 | 8.3e-06 | 9.4e-06 | 1.090E-05 | 3.593E-06 | 8.300E-06 | 1.500E-05 |
| 2637 | 132 | L132T | A | 1.7e-05 | 9.9e-06 | 8.2e-06 | 1.170E-05 | 4.668E-06 | 8.200E-06 | 1.700E-05 |
| 2638 | 132 | L132W | A | 1.2e-05 | 1e-05 | 1.3e-05 | 1.167E-05 | 1.528E-06 | 1.000E-05 | 1.300E-05 |
| 2639 | 132 | L132Y | A | 1.3e-05 | 1e-05 | 1.6e-05 | 1.300E-05 | 3.000E-06 | 1.000E-05 | 1.600E-05 |
| 2640 | 132 | L132V | A | 1.4e-05 | 9.8e-06 | 1.5e-05 | 1.293E-05 | 2.759E-06 | 9.800E-06 | 1.500E-05 |
| 2641 | 133 | I133A | A | 1.7e-05 | 1.1e-05 | 1.6e-05 | 1.467E-05 | 3.215E-06 | 1.100E-05 | 1.700E-05 |
| 2642 | 133 | I133R | A | 9.8e-06 | 1.6e-05 | 1.1e-05 | 1.227E-05 | 3.288E-06 | 9.800E-06 | 1.600E-05 |
| 2643 | 133 | I133N | A | 1.6e-05 | 8.8e-06 | 1.4e-05 | 1.293E-05 | 3.717E-06 | 8.800E-06 | 1.600E-05 |
| 2644 | 133 | I133D | A | 1.1e-05 | 1.6e-05 | 1.5e-05 | 1.400E-05 | 2.646E-06 | 1.100E-05 | 1.600E-05 |
| 2645 | 133 | I133C | A | 1.1e-05 | 1.6e-05 | 1.4e-05 | 1.367E-05 | 2.517E-06 | 1.100E-05 | 1.600E-05 |
| 2646 | 133 | I133E | A | 1.5e-05 | 2.2e-05 | 1.4e-05 | 1.700E-05 | 4.359E-06 | 1.400E-05 | 2.200E-05 |
| 2647 | 133 | I133Q | A | 1.5e-05 | 8.6e-06 | 1.6e-05 | 1.320E-05 | 4.015E-06 | 8.600E-06 | 1.600E-05 |
| 2648 | 133 | I133G | A | 1.6e-05 | 1.2e-05 | 8.8e-06 | 1.227E-05 | 3.607E-06 | 8.800E-06 | 1.600E-05 |
| 2649 | 133 | I133H | A | 9.2e-06 | 1.3e-05 | 1.4e-05 | 1.207E-05 | 2.532E-06 | 9.200E-06 | 1.400E-05 |
| 2650 | 133 | I133I | A | 1.5e-05 | 1.1e-05 | 1.5e-05 | 1.367E-05 | 2.309E-06 | 1.100E-05 | 1.500E-05 |
| 2651 | 133 | I133L | A | 1.4e-05 | 1.1e-05 | 1.6e-05 | 1.367E-05 | 2.517E-06 | 1.100E-05 | 1.600E-05 |
| 2652 | 133 | I133K | A | 1.2e-05 | 1.1e-05 | 1.1e-05 | 1.133E-05 | 5.774E-07 | 1.100E-05 | 1.200E-05 |
| 2653 | 133 | I133M | A | 1.4e-05 | 1.1e-05 | 9.9e-06 | 1.163E-05 | 2.122E-06 | 9.900E-06 | 1.400E-05 |
| 2654 | 133 | I133F | A | 1.5e-05 | 1.1e-05 | 1.3e-05 | 1.300E-05 | 2.000E-06 | 1.100E-05 | 1.500E-05 |
| 2655 | 133 | I133P | A | 1.1e-05 | 1.7e-05 | 1.3e-05 | 1.367E-05 | 3.055E-06 | 1.100E-05 | 1.700E-05 |
| 2656 | 133 | I133S | A | 1.8e-05 | 9.7e-06 | 1.5e-05 | 1.423E-05 | 4.203E-06 | 9.700E-06 | 1.800E-05 |
| 2657 | 133 | I133T | A | 6.8e-06 | 1.4e-05 | 1.5e-05 | 1.193E-05 | 4.474E-06 | 6.800E-06 | 1.500E-05 |
| 2658 | 133 | I133W | A | 1e-05 | 1.6e-05 | 1e-05 | 1.200E-05 | 3.464E-06 | 1.000E-05 | 1.600E-05 |
| 2659 | 133 | I133Y | A | 1e-05 | 1e-05 | 1.3e-05 | 1.100E-05 | 1.732E-06 | 1.000E-05 | 1.300E-05 |
| 2660 | 133 | I133V | A | 8e-06 | 1.1e-05 | 9e-06 | 9.333E-06 | 1.528E-06 | 8.000E-06 | 1.100E-05 |
| 2661 | 134 | Q134A | A | 1.4e-05 | 1.4e-05 | 1.6e-05 | 1.467E-05 | 1.155E-06 | 1.400E-05 | 1.600E-05 |
| 2662 | 134 | Q134R | A | 1.1e-05 | 1.5e-05 | 1.6e-05 | 1.400E-05 | 2.646E-06 | 1.100E-05 | 1.600E-05 |
| 2663 | 134 | Q134N | A | 1.1e-05 | 1.5e-05 | 1.2e-05 | 1.267E-05 | 2.082E-06 | 1.100E-05 | 1.500E-05 |
| 2664 | 134 | Q134D | A | 9.9e-06 | 1.2e-05 | 1.3e-05 | 1.163E-05 | 1.582E-06 | 9.900E-06 | 1.300E-05 |
| 2665 | 134 | Q134C | A | 1e-05 | 1e-05 | 1.3e-05 | 1.100E-05 | 1.732E-06 | 1.000E-05 | 1.300E-05 |
| 2666 | 134 | Q134E | A | 9.3e-06 | 1.4e-05 | 1.3e-05 | 1.210E-05 | 2.476E-06 | 9.300E-06 | 1.400E-05 |
| 2667 | 134 | Q134Q | A | 1.4e-05 | 1.2e-05 | 1e-05 | 1.200E-05 | 2.000E-06 | 1.000E-05 | 1.400E-05 |
| 2668 | 134 | Q134G | A | 1.5e-05 | 1.1e-05 | 1.5e-05 | 1.367E-05 | 2.309E-06 | 1.100E-05 | 1.500E-05 |
| 2669 | 134 | Q134H | A | 1.1e-05 | 1.4e-05 | 1.7e-05 | 1.400E-05 | 3.000E-06 | 1.100E-05 | 1.700E-05 |
| 2670 | 134 | Q134I | A | 1.5e-05 | 1.1e-05 | 1.3e-05 | 1.300E-05 | 2.000E-06 | 1.100E-05 | 1.500E-05 |
| 2671 | 134 | Q134L | A | 1e-05 | 1.5e-05 | 1.3e-05 | 1.267E-05 | 2.517E-06 | 1.000E-05 | 1.500E-05 |
| 2672 | 134 | Q134K | A | 1.6e-05 | 9.7e-06 | 9.4e-06 | 1.170E-05 | 3.727E-06 | 9.400E-06 | 1.600E-05 |
| 2673 | 134 | Q134M | A | 1.4e-05 | 1.5e-05 | 1.5e-05 | 1.467E-05 | 5.774E-07 | 1.400E-05 | 1.500E-05 |
| 2674 | 134 | Q134F | A | 1.6e-05 | 1.6e-05 | 1.6e-05 | 1.600E-05 | 0.000E+00 | 1.600E-05 | 1.600E-05 |
| 2675 | 134 | Q134P | A | 1.3e-05 | 1.4e-05 | 1.1e-05 | 1.267E-05 | 1.528E-06 | 1.100E-05 | 1.400E-05 |
| 2676 | 134 | Q134S | A | 9.1e-06 | 1.2e-05 | 1.3e-05 | 1.137E-05 | 2.026E-06 | 9.100E-06 | 1.300E-05 |
| 2677 | 134 | Q134T | A | 1.7e-05 | 1.6e-05 | 1.5e-05 | 1.600E-05 | 1.000E-06 | 1.500E-05 | 1.700E-05 |
| 2678 | 134 | Q134W | A | 9.3e-06 | 1.5e-05 | 1.2e-05 | 1.210E-05 | 2.851E-06 | 9.300E-06 | 1.500E-05 |
| 2679 | 134 | Q134Y | A | 1.6e-05 | 1.1e-05 | 1.5e-05 | 1.400E-05 | 2.646E-06 | 1.100E-05 | 1.600E-05 |
| 2680 | 134 | Q134V | A | 1.5e-05 | 1.6e-05 | 9.1e-06 | 1.337E-05 | 3.729E-06 | 9.100E-06 | 1.600E-05 |
| 2681 | 135 | R135A | A | 1e-05 | 1.5e-05 | 1.4e-05 | 1.300E-05 | 2.646E-06 | 1.000E-05 | 1.500E-05 |
| 2682 | 135 | R135R | A | 1.1e-05 | 9.4e-06 | 1.5e-05 | 1.180E-05 | 2.884E-06 | 9.400E-06 | 1.500E-05 |
| 2683 | 135 | R135N | A | 9.3e-06 | 9.7e-06 | 1.5e-05 | 1.133E-05 | 3.182E-06 | 9.300E-06 | 1.500E-05 |
| 2684 | 135 | R135D | A | 1.6e-05 | 1.4e-05 | 1.6e-05 | 1.533E-05 | 1.155E-06 | 1.400E-05 | 1.600E-05 |
| 2685 | 135 | R135C | A | 1.5e-05 | 1.4e-05 | 1.3e-05 | 1.400E-05 | 1.000E-06 | 1.300E-05 | 1.500E-05 |
| 2686 | 135 | R135E | A | 1.3e-05 | 1.3e-05 | 9.7e-06 | 1.190E-05 | 1.905E-06 | 9.700E-06 | 1.300E-05 |
| 2687 | 135 | R135Q | A | 1.1e-05 | 6.6e-06 | 1.3e-05 | 1.020E-05 | 3.274E-06 | 6.600E-06 | 1.300E-05 |
| 2688 | 135 | R135G | A | 2.5e-05 | 1.6e-05 | 1e-05 | 1.700E-05 | 7.550E-06 | 1.000E-05 | 2.500E-05 |
| 2689 | 135 | R135H | A | 1.5e-05 | 1.1e-05 | 1.3e-05 | 1.300E-05 | 2.000E-06 | 1.100E-05 | 1.500E-05 |
| 2690 | 135 | R135I | A | 1.3e-05 | 1e-05 | 1e-05 | 1.100E-05 | 1.732E-06 | 1.000E-05 | 1.300E-05 |
| 2691 | 135 | R135L | A | 1.6e-05 | 1.1e-05 | 8.2e-06 | 1.173E-05 | 3.951E-06 | 8.200E-06 | 1.600E-05 |
| 2692 | 135 | R135K | A | 1.2e-05 | 8.4e-06 | 6.1e-06 | 8.833E-06 | 2.974E-06 | 6.100E-06 | 1.200E-05 |
| 2693 | 135 | R135M | A | 1.4e-05 | 1.3e-05 | 1.1e-05 | 1.267E-05 | 1.528E-06 | 1.100E-05 | 1.400E-05 |
| 2694 | 135 | R135F | A | 1.1e-05 | 1.1e-05 | 1e-05 | 1.067E-05 | 5.774E-07 | 1.000E-05 | 1.100E-05 |
| 2695 | 135 | R135P | A | 1.5e-05 | 1.3e-05 | 1.5e-05 | 1.433E-05 | 1.155E-06 | 1.300E-05 | 1.500E-05 |
| 2696 | 135 | R135S | A | 1.3e-05 | 1e-05 | 8.9e-06 | 1.063E-05 | 2.122E-06 | 8.900E-06 | 1.300E-05 |
| 2697 | 135 | R135T | A | 1.4e-05 | 6.8e-06 | 9.5e-06 | 1.010E-05 | 3.637E-06 | 6.800E-06 | 1.400E-05 |
| 2698 | 135 | R135W | A | 9.4e-06 | 1.4e-05 | 1.1e-05 | 1.147E-05 | 2.335E-06 | 9.400E-06 | 1.400E-05 |
| 2699 | 135 | R135Y | A | 1.3e-05 | 1.1e-05 | 8.8e-06 | 1.093E-05 | 2.101E-06 | 8.800E-06 | 1.300E-05 |
| 2700 | 135 | R135V | A | 1.4e-05 | 1.7e-05 | 9.9e-06 | 1.363E-05 | 3.564E-06 | 9.900E-06 | 1.700E-05 |
| 2701 | 136 | N136A | A | 1.5e-05 | 1.3e-05 | 9.2e-06 | 1.240E-05 | 2.946E-06 | 9.200E-06 | 1.500E-05 |
| 2702 | 136 | N136R | A | 1.5e-05 | 1.4e-05 | 1.6e-05 | 1.500E-05 | 1.000E-06 | 1.400E-05 | 1.600E-05 |
| 2703 | 136 | N136N | A | 1.4e-05 | 1.5e-05 | 1e-05 | 1.300E-05 | 2.646E-06 | 1.000E-05 | 1.500E-05 |
| 2704 | 136 | N136D | A | 1.7e-05 | 1.6e-05 | 9.5e-06 | 1.417E-05 | 4.072E-06 | 9.500E-06 | 1.700E-05 |
| 2705 | 136 | N136C | A | 1.6e-05 | 1.3e-05 | 9.8e-06 | 1.293E-05 | 3.101E-06 | 9.800E-06 | 1.600E-05 |
| 2706 | 136 | N136E | A | 1e-05 | 1.4e-05 | 8.6e-06 | 1.087E-05 | 2.802E-06 | 8.600E-06 | 1.400E-05 |
| 2707 | 136 | N136Q | A | 1.4e-05 | 1.1e-05 | 1.1e-05 | 1.200E-05 | 1.732E-06 | 1.100E-05 | 1.400E-05 |
| 2708 | 136 | N136G | A | 9.9e-06 | 1.6e-05 | 1.6e-05 | 1.397E-05 | 3.522E-06 | 9.900E-06 | 1.600E-05 |
| 2709 | 136 | N136H | A | 1.1e-05 | 1.6e-05 | 1.2e-05 | 1.300E-05 | 2.646E-06 | 1.100E-05 | 1.600E-05 |
| 2710 | 136 | N136I | A | 1.3e-05 | 1e-05 | 1.7e-05 | 1.333E-05 | 3.512E-06 | 1.000E-05 | 1.700E-05 |
| 2711 | 136 | N136L | A | 1.4e-05 | 1.3e-05 | 9.9e-06 | 1.230E-05 | 2.138E-06 | 9.900E-06 | 1.400E-05 |
| 2712 | 136 | N136K | A | 1.8e-05 | 1.3e-05 | 1.6e-05 | 1.567E-05 | 2.517E-06 | 1.300E-05 | 1.800E-05 |
| 2713 | 136 | N136M | A | 1.4e-05 | 1.5e-05 | 1.1e-05 | 1.333E-05 | 2.082E-06 | 1.100E-05 | 1.500E-05 |
| 2714 | 136 | N136F | A | 1.1e-05 | 1.5e-05 | 9.1e-06 | 1.170E-05 | 3.012E-06 | 9.100E-06 | 1.500E-05 |
| 2715 | 136 | N136P | A | 9.7e-06 | 1.2e-05 | 1e-05 | 1.057E-05 | 1.250E-06 | 9.700E-06 | 1.200E-05 |
| 2716 | 136 | N136S | A | 1.1e-05 | 1.6e-05 | 8.2e-06 | 1.173E-05 | 3.951E-06 | 8.200E-06 | 1.600E-05 |
| 2717 | 136 | N136T | A | 1.1e-05 | 1.4e-05 | 1.5e-05 | 1.333E-05 | 2.082E-06 | 1.100E-05 | 1.500E-05 |
| 2718 | 136 | N136W | A | 1.1e-05 | 1.5e-05 | 1.7e-05 | 1.433E-05 | 3.055E-06 | 1.100E-05 | 1.700E-05 |
| 2719 | 136 | N136Y | A | 1e-05 | 1.4e-05 | 1.6e-05 | 1.333E-05 | 3.055E-06 | 1.000E-05 | 1.600E-05 |

| | | | | | | | | | | |
|---|---|---|---|---|---|---|---|---|---|---|
| 2720 | 136 | N136V | A | 1.6e-05 | 1.5e-05 | 1.7e-05 | 1.600E-05 | 1.000E-06 | 1.500E-05 | 1.700E-05 |
| 2721 | 137 | P137A | A | 9.6e-06 | 2.4e-05 | 1.4e-05 | 1.587E-05 | 7.379E-06 | 9.600E-06 | 2.400E-05 |
| 2722 | 137 | P137R | A | 1.4e-05 | 9.9e-06 | 1.2e-05 | 1.197E-05 | 2.050E-06 | 9.900E-06 | 1.400E-05 |
| 2723 | 137 | P137N | A | 1.6e-05 | 9.1e-06 | 1.2e-05 | 1.237E-05 | 3.465E-06 | 9.100E-06 | 1.600E-05 |
| 2724 | 137 | P137D | A | 1.4e-05 | 1e-05 | 1e-05 | 1.133E-05 | 2.309E-06 | 1.000E-05 | 1.400E-05 |
| 2725 | 137 | P137C | A | 1.6e-05 | 1.4e-05 | 1e-05 | 1.333E-05 | 3.055E-06 | 1.000E-05 | 1.600E-05 |
| 2726 | 137 | P137E | A | 1.2e-05 | 1.7e-05 | 1e-05 | 1.300E-05 | 3.606E-06 | 1.000E-05 | 1.700E-05 |
| 2727 | 137 | P137Q | A | 1.1e-05 | 1.6e-05 | 1.4e-05 | 1.367E-05 | 2.517E-06 | 1.100E-05 | 1.600E-05 |
| 2728 | 137 | P137G | A | 1e-05 | 1.5e-05 | 1e-05 | 1.167E-05 | 2.887E-06 | 1.000E-05 | 1.500E-05 |
| 2729 | 137 | P137H | A | 8.4e-06 | 7.7e-06 | 1.5e-05 | 1.037E-05 | 4.028E-06 | 7.700E-06 | 1.500E-05 |
| 2730 | 137 | P137I | A | 1.4e-05 | 1.1e-05 | 1.1e-05 | 1.200E-05 | 1.732E-06 | 1.100E-05 | 1.400E-05 |
| 2731 | 137 | P137L | A | 1.5e-05 | 9.9e-06 | 1.5e-05 | 1.330E-05 | 2.944E-06 | 9.900E-06 | 1.500E-05 |
| 2732 | 137 | P137K | A | 9.3e-06 | 1e-05 | 1.4e-05 | 1.110E-05 | 2.536E-06 | 9.300E-06 | 1.400E-05 |
| 2733 | 137 | P137M | A | 1.1e-05 | 1.1e-05 | 1.6e-05 | 1.267E-05 | 2.887E-06 | 1.100E-05 | 1.600E-05 |
| 2734 | 137 | P137F | A | 1.7e-05 | 6.5e-06 | 1.4e-05 | 1.250E-05 | 5.408E-06 | 6.500E-06 | 1.700E-05 |
| 2735 | 137 | P137P | A | 1e-05 | 1.3e-05 | 1.3e-05 | 1.200E-05 | 1.732E-06 | 1.000E-05 | 1.300E-05 |
| 2736 | 137 | P137S | A | 1.7e-05 | 1.4e-05 | 1.3e-05 | 1.467E-05 | 2.082E-06 | 1.300E-05 | 1.700E-05 |
| 2737 | 137 | P137T | A | 9.9e-06 | 1.2e-05 | 1.4e-05 | 1.197E-05 | 2.050E-06 | 9.900E-06 | 1.400E-05 |
| 2738 | 137 | P137W | A | 9.9e-06 | 1.4e-05 | 1.1e-05 | 1.163E-05 | 2.122E-06 | 9.900E-06 | 1.400E-05 |
| 2739 | 137 | P137Y | A | 1.5e-05 | 1.5e-05 | 8.7e-06 | 1.290E-05 | 3.637E-06 | 8.700E-06 | 1.500E-05 |
| 2740 | 137 | P137V | A | 1e-05 | 1e-05 | 1.5e-05 | 1.167E-05 | 2.887E-06 | 1.000E-05 | 1.500E-05 |
| 2741 | 138 | Q138A | A | 1.5e-05 | 1.4e-05 | 1.5e-05 | 1.467E-05 | 5.774E-07 | 1.400E-05 | 1.500E-05 |
| 2742 | 138 | Q138R | A | 1.6e-05 | 9.5e-06 | 1.5e-05 | 1.350E-05 | 3.500E-06 | 9.500E-06 | 1.600E-05 |
| 2743 | 138 | Q138N | A | 1.1e-05 | 1.2e-05 | 1.2e-05 | 1.167E-05 | 5.774E-07 | 1.100E-05 | 1.200E-05 |
| 2744 | 138 | Q138D | A | 1.2e-05 | 1.5e-05 | 1.4e-05 | 1.367E-05 | 1.528E-06 | 1.200E-05 | 1.500E-05 |
| 2745 | 138 | Q138C | A | 1.2e-05 | 8.8e-06 | 9.8e-06 | 1.020E-05 | 1.637E-06 | 8.800E-06 | 1.200E-05 |
| 2746 | 138 | Q138E | A | 9.1e-06 | 1.1e-05 | 9.1e-06 | 9.733E-06 | 1.097E-06 | 9.100E-06 | 1.100E-05 |
| 2747 | 138 | Q138Q | A | 1.2e-05 | 1.6e-05 | 1.1e-05 | 1.300E-05 | 2.646E-06 | 1.100E-05 | 1.600E-05 |
| 2748 | 138 | Q138G | A | 8.4e-06 | 1.7e-05 | 1.6e-05 | 1.380E-05 | 4.703E-06 | 8.400E-06 | 1.700E-05 |
| 2749 | 138 | Q138H | A | 9.2e-06 | 1.1e-05 | 9.3e-06 | 9.833E-06 | 1.012E-06 | 9.200E-06 | 1.100E-05 |
| 2750 | 138 | Q138I | A | 1.4e-05 | 1.5e-05 | 1.1e-05 | 1.333E-05 | 2.082E-06 | 1.100E-05 | 1.500E-05 |
| 2751 | 138 | Q138L | A | 1.1e-05 | 1.1e-05 | 9.3e-06 | 1.043E-05 | 9.815E-07 | 9.300E-06 | 1.100E-05 |
| 2752 | 138 | Q138K | A | 1.5e-05 | 1.2e-05 | 9.7e-06 | 1.223E-05 | 2.658E-06 | 9.700E-06 | 1.500E-05 |
| 2753 | 138 | Q138M | A | 1.1e-05 | 1.2e-05 | 1.6e-05 | 1.300E-05 | 2.646E-06 | 1.100E-05 | 1.600E-05 |
| 2754 | 138 | Q138F | A | 1.1e-05 | 1.5e-05 | 9.8e-06 | 1.193E-05 | 2.723E-06 | 9.800E-06 | 1.500E-05 |
| 2755 | 138 | Q138P | A | 1.3e-05 | 1.5e-05 | 1.2e-05 | 1.333E-05 | 1.528E-06 | 1.200E-05 | 1.500E-05 |
| 2756 | 138 | Q138S | A | 1.3e-05 | 1.6e-05 | 1.1e-05 | 1.333E-05 | 2.517E-06 | 1.100E-05 | 1.600E-05 |
| 2757 | 138 | Q138T | A | 1.3e-05 | 1.2e-05 | 1.6e-05 | 1.367E-05 | 2.082E-06 | 1.200E-05 | 1.600E-05 |
| 2758 | 138 | Q138W | A | 1.2e-05 | 6.1e-06 | 9e-06 | 9.033E-06 | 2.950E-06 | 6.100E-06 | 1.200E-05 |
| 2759 | 138 | Q138Y | A | 1e-05 | 1.1e-05 | 1.4e-05 | 1.167E-05 | 2.082E-06 | 1.000E-05 | 1.400E-05 |
| 2760 | 138 | Q138V | A | 1.1e-05 | 1.3e-05 | 1.4e-05 | 1.267E-05 | 1.528E-06 | 1.100E-05 | 1.400E-05 |
| 2761 | 139 | L139A | A | 1.4e-05 | 1.2e-05 | 1.5e-05 | 1.367E-05 | 1.528E-06 | 1.200E-05 | 1.500E-05 |
| 2762 | 139 | L139R | A | 9.6e-06 | 1.3e-05 | 9.3e-06 | 1.063E-05 | 2.055E-06 | 9.300E-06 | 1.300E-05 |
| 2763 | 139 | L139N | A | 1.2e-05 | 1.1e-05 | 1.5e-05 | 1.267E-05 | 2.082E-06 | 1.100E-05 | 1.500E-05 |
| 2764 | 139 | L139D | A | 1.6e-05 | 8.7e-06 | 9e-06 | 1.123E-05 | 4.131E-06 | 8.700E-06 | 1.600E-05 |
| 2765 | 139 | L139C | A | 1.4e-05 | 9.9e-06 | 9.7e-06 | 1.120E-05 | 2.427E-06 | 9.700E-06 | 1.400E-05 |
| 2766 | 139 | L139E | A | 9.3e-06 | 1.4e-05 | 1.1e-05 | 1.143E-05 | 2.380E-06 | 9.300E-06 | 1.400E-05 |
| 2767 | 139 | L139Q | A | 9.4e-06 | 8.5e-06 | 1.3e-05 | 1.030E-05 | 2.381E-06 | 8.500E-06 | 1.300E-05 |
| 2768 | 139 | L139G | A | 1e-05 | 1.1e-05 | 1.1e-05 | 1.067E-05 | 5.774E-07 | 1.000E-05 | 1.100E-05 |
| 2769 | 139 | L139H | A | 1.1e-05 | 1.6e-05 | 1.1e-05 | 1.267E-05 | 2.887E-06 | 1.100E-05 | 1.600E-05 |
| 2770 | 139 | L139I | A | 1.5e-05 | 1e-05 | 1.2e-05 | 1.233E-05 | 2.517E-06 | 1.000E-05 | 1.500E-05 |
| 2771 | 139 | L139L | A | 1.7e-05 | 1.6e-05 | 1.3e-05 | 1.533E-05 | 2.082E-06 | 1.300E-05 | 1.700E-05 |
| 2772 | 139 | L139K | A | 1.1e-05 | 1e-05 | 1.3e-05 | 1.133E-05 | 1.528E-06 | 1.000E-05 | 1.300E-05 |
| 2773 | 139 | L139M | A | 1.5e-05 | 1.6e-05 | 1.3e-05 | 1.467E-05 | 1.528E-06 | 1.300E-05 | 1.600E-05 |
| 2774 | 139 | L139F | A | 1.5e-05 | 1.6e-05 | 1.6e-05 | 1.567E-05 | 5.774E-07 | 1.500E-05 | 1.600E-05 |
| 2775 | 139 | L139P | A | 9e-06 | 9.9e-06 | 1.2e-05 | 1.030E-05 | 1.539E-06 | 9.000E-06 | 1.200E-05 |
| 2776 | 139 | L139S | A | 1.1e-05 | 1.5e-05 | 9.7e-06 | 1.190E-05 | 2.762E-06 | 9.700E-06 | 1.500E-05 |
| 2777 | 139 | L139T | A | 1.1e-05 | 1.2e-05 | 1.3e-05 | 1.200E-05 | 1.000E-06 | 1.100E-05 | 1.300E-05 |
| 2778 | 139 | L139W | A | 9.8e-06 | 9.5e-06 | 1.4e-05 | 1.110E-05 | 2.516E-06 | 9.500E-06 | 1.400E-05 |
| 2779 | 139 | L139Y | A | 9.9e-06 | 1.6e-05 | 8.6e-06 | 1.150E-05 | 3.951E-06 | 8.600E-06 | 1.600E-05 |
| 2780 | 139 | L139V | A | 1e-05 | 1e-05 | 1.1e-05 | 1.033E-05 | 5.774E-07 | 1.000E-05 | 1.100E-05 |
| 2781 | 140 | C140A | A | 1.4e-05 | 1.3e-05 | 6.9e-06 | 1.130E-05 | 3.843E-06 | 6.900E-06 | 1.400E-05 |
| 2782 | 140 | C140R | A | 1.1e-05 | 1.1e-05 | 1.2e-05 | 1.133E-05 | 5.774E-07 | 1.100E-05 | 1.200E-05 |
| 2783 | 140 | C140N | A | 1.6e-05 | 1.3e-05 | 1.4e-05 | 1.433E-05 | 1.528E-06 | 1.300E-05 | 1.600E-05 |
| 2784 | 140 | C140D | A | 1.5e-05 | 8.9e-06 | 1.2e-05 | 1.197E-05 | 3.050E-06 | 8.900E-06 | 1.500E-05 |
| 2785 | 140 | C140C | A | 9.6e-06 | 1.1e-05 | 1e-05 | 1.020E-05 | 7.211E-07 | 9.600E-06 | 1.100E-05 |
| 2786 | 140 | C140E | A | 1.1e-05 | 1.7e-05 | 1.4e-05 | 1.400E-05 | 3.000E-06 | 1.100E-05 | 1.700E-05 |
| 2787 | 140 | C140Q | A | 1e-05 | 1.1e-05 | 1.4e-05 | 1.167E-05 | 2.082E-06 | 1.000E-05 | 1.400E-05 |
| 2788 | 140 | C140G | A | 1.1e-05 | 1.5e-05 | 1.2e-05 | 1.267E-05 | 2.082E-06 | 1.100E-05 | 1.500E-05 |
| 2789 | 140 | C140H | A | 1.5e-05 | 1.4e-05 | 1.5e-05 | 1.467E-05 | 5.774E-07 | 1.400E-05 | 1.500E-05 |
| 2790 | 140 | C140I | A | 1.5e-05 | 7.9e-06 | 1.3e-05 | 1.197E-05 | 3.661E-06 | 7.900E-06 | 1.500E-05 |
| 2791 | 140 | C140L | A | 1.2e-05 | 1.6e-05 | 1.4e-05 | 1.400E-05 | 2.000E-06 | 1.200E-05 | 1.600E-05 |
| 2792 | 140 | C140K | A | 1.4e-05 | 1.4e-05 | 1.3e-05 | 1.367E-05 | 5.774E-07 | 1.300E-05 | 1.400E-05 |
| 2793 | 140 | C140M | A | 1.2e-05 | 1e-05 | 1e-05 | 1.067E-05 | 1.155E-06 | 1.000E-05 | 1.200E-05 |
| 2794 | 140 | C140F | A | 1.6e-05 | 1.3e-05 | 1.1e-05 | 1.333E-05 | 2.517E-06 | 1.100E-05 | 1.600E-05 |
| 2795 | 140 | C140P | A | 1.2e-05 | 1.5e-05 | 1.4e-05 | 1.367E-05 | 1.528E-06 | 1.200E-05 | 1.500E-05 |
| 2796 | 140 | C140S | A | 1.5e-05 | 1.4e-05 | 1.1e-05 | 1.333E-05 | 2.082E-06 | 1.100E-05 | 1.500E-05 |
| 2797 | 140 | C140T | A | 1e-05 | 1.3e-05 | 1.7e-05 | 1.333E-05 | 3.512E-06 | 1.000E-05 | 1.700E-05 |
| 2798 | 140 | C140W | A | 1.1e-05 | 1.4e-05 | 1.3e-05 | 1.267E-05 | 1.528E-06 | 1.100E-05 | 1.400E-05 |
| 2799 | 140 | C140Y | A | 1.2e-05 | 1.6e-05 | 1.4e-05 | 1.400E-05 | 2.000E-06 | 1.200E-05 | 1.600E-05 |
| 2800 | 140 | C140V | A | 1.2e-05 | 1.6e-05 | 1.1e-05 | 1.300E-05 | 2.646E-06 | 1.100E-05 | 1.600E-05 |
| 2801 | 141 | Y141A | A | 9.8e-06 | 1.5e-05 | 2.7e-05 | 1.727E-05 | 8.821E-06 | 9.800E-06 | 2.700E-05 |
| 2802 | 141 | Y141R | A | 1.3e-05 | 1.1e-05 | 1.1e-05 | 1.167E-05 | 1.155E-06 | 1.100E-05 | 1.300E-05 |
| 2803 | 141 | Y141N | A | 1.3e-05 | 1.5e-05 | 9.1e-06 | 1.237E-05 | 3.001E-06 | 9.100E-06 | 1.500E-05 |
| 2804 | 141 | Y141D | A | 1.8e-05 | 1.6e-05 | 1.3e-05 | 1.567E-05 | 2.517E-06 | 1.300E-05 | 1.800E-05 |

| | | | | | | | | | | |
|---|---|---|---|---|---|---|---|---|---|---|
| 2805 | 141 | Y141C | A | 1e-05 | 1.5e-05 | 8.5e-06 | 1.117E-05 | 3.403E-06 | 8.500E-06 | 1.500E-05 |
| 2806 | 141 | Y141E | A | 1.1e-05 | 1.5e-05 | 1.6e-05 | 1.400E-05 | 2.646E-06 | 1.100E-05 | 1.600E-05 |
| 2807 | 141 | Y141Q | A | 1.6e-05 | 9.5e-06 | 1e-05 | 1.183E-05 | 3.617E-06 | 9.500E-06 | 1.600E-05 |
| 2808 | 141 | Y141G | A | 9e-06 | 1.2e-05 | 9.5e-06 | 1.017E-05 | 1.607E-06 | 9.000E-06 | 1.200E-05 |
| 2809 | 141 | Y141H | A | 1.6e-05 | 1.6e-05 | 1e-05 | 1.400E-05 | 3.464E-06 | 1.000E-05 | 1.600E-05 |
| 2810 | 141 | Y141I | A | 1.1e-05 | 1.1e-05 | 1.6e-05 | 1.267E-05 | 2.887E-06 | 1.100E-05 | 1.600E-05 |
| 2811 | 141 | Y141L | A | 1.8e-05 | 1.4e-05 | 1e-05 | 1.400E-05 | 4.000E-06 | 1.000E-05 | 1.800E-05 |
| 2812 | 141 | Y141K | A | 1.5e-05 | 1.5e-05 | 1.6e-05 | 1.533E-05 | 5.774E-07 | 1.500E-05 | 1.600E-05 |
| 2813 | 141 | Y141M | A | 1.4e-05 | 1.6e-05 | 1.7e-05 | 1.567E-05 | 1.528E-06 | 1.400E-05 | 1.700E-05 |
| 2814 | 141 | Y141F | A | 1.6e-05 | 1.6e-05 | 1.4e-05 | 1.533E-05 | 1.155E-06 | 1.400E-05 | 1.600E-05 |
| 2815 | 141 | Y141P | A | 1.1e-05 | 1e-05 | 1.1e-05 | 1.067E-05 | 5.774E-07 | 1.000E-05 | 1.100E-05 |
| 2816 | 141 | Y141S | A | 9.4e-06 | 9.7e-06 | 1e-05 | 9.700E-06 | 3.000E-07 | 9.400E-06 | 1.000E-05 |
| 2817 | 141 | Y141T | A | 1e-05 | 1.6e-05 | 1.4e-05 | 1.333E-05 | 3.055E-06 | 1.000E-05 | 1.600E-05 |
| 2818 | 141 | Y141W | A | 1e-05 | 8.1e-06 | 1.3e-05 | 1.037E-05 | 2.470E-06 | 8.100E-06 | 1.300E-05 |
| 2819 | 141 | Y141Y | A | 9.4e-06 | 1.6e-05 | 9.8e-06 | 1.173E-05 | 3.700E-06 | 9.400E-06 | 1.600E-05 |
| 2820 | 141 | Y141V | A | 1.4e-05 | 1.4e-05 | 1.3e-05 | 1.367E-05 | 5.774E-07 | 1.300E-05 | 1.400E-05 |
| 2821 | 142 | Q142A | A | 8.6e-06 | 1.7e-05 | 1.1e-05 | 1.220E-05 | 4.327E-06 | 8.600E-06 | 1.700E-05 |
| 2822 | 142 | Q142R | A | 8.8e-06 | 1.5e-05 | 1.5e-05 | 1.293E-05 | 3.580E-06 | 8.800E-06 | 1.500E-05 |
| 2823 | 142 | Q142N | A | 1.2e-05 | 1.2e-05 | 1.2e-05 | 1.200E-05 | 0.000E+00 | 1.200E-05 | 1.200E-05 |
| 2824 | 142 | Q142D | A | 1.1e-05 | 1.3e-05 | 1.5e-05 | 1.300E-05 | 2.000E-06 | 1.100E-05 | 1.500E-05 |
| 2825 | 142 | Q142C | A | 1.1e-05 | 7.3e-06 | 1.6e-05 | 1.143E-05 | 4.366E-06 | 7.300E-06 | 1.600E-05 |
| 2826 | 142 | Q142E | A | 1e-05 | 1.3e-05 | 1.3e-05 | 1.200E-05 | 1.732E-06 | 1.000E-05 | 1.300E-05 |
| 2827 | 142 | Q142Q | A | 9e-06 | 1.4e-05 | 1.6e-05 | 1.300E-05 | 3.606E-06 | 9.000E-06 | 1.600E-05 |
| 2828 | 142 | Q142G | A | 1.6e-05 | 1.5e-05 | 1.1e-05 | 1.400E-05 | 2.646E-06 | 1.100E-05 | 1.600E-05 |
| 2829 | 142 | Q142H | A | 1.5e-05 | 1.7e-05 | 6.2e-06 | 1.273E-05 | 5.746E-06 | 6.200E-06 | 1.700E-05 |
| 2830 | 142 | Q142I | A | 8.4e-06 | 1.1e-05 | 1.5e-05 | 1.147E-05 | 3.325E-06 | 8.400E-06 | 1.500E-05 |
| 2831 | 142 | Q142L | A | 1.3e-05 | 9.7e-06 | 1.5e-05 | 1.257E-05 | 2.676E-06 | 9.700E-06 | 1.500E-05 |
| 2832 | 142 | Q142K | A | 1.1e-05 | 7.6e-06 | 8.8e-06 | 9.133E-06 | 1.724E-06 | 7.600E-06 | 1.100E-05 |
| 2833 | 142 | Q142M | A | 1.4e-05 | 1.5e-05 | 1.5e-05 | 1.467E-05 | 5.774E-07 | 1.400E-05 | 1.500E-05 |
| 2834 | 142 | Q142F | A | 1.4e-05 | 8.8e-06 | 1.1e-05 | 1.127E-05 | 2.610E-06 | 8.800E-06 | 1.400E-05 |
| 2835 | 142 | Q142P | A | 8.8e-06 | 1.7e-05 | 1.1e-05 | 1.227E-05 | 4.244E-06 | 8.800E-06 | 1.700E-05 |
| 2836 | 142 | Q142S | A | 1.1e-05 | 6.4e-06 | 1.2e-05 | 9.800E-06 | 2.987E-06 | 6.400E-06 | 1.200E-05 |
| 2837 | 142 | Q142T | A | 1.1e-05 | 1.1e-05 | 2e-05 | 1.400E-05 | 5.196E-06 | 1.100E-05 | 2.000E-05 |
| 2838 | 142 | Q142W | A | 1.6e-05 | 1.2e-05 | 1.4e-05 | 1.400E-05 | 2.000E-06 | 1.200E-05 | 1.600E-05 |
| 2839 | 142 | Q142Y | A | 9.7e-06 | 1.6e-05 | 1e-05 | 1.190E-05 | 3.554E-06 | 9.700E-06 | 1.600E-05 |
| 2840 | 142 | Q142V | A | 1e-05 | 9.4e-06 | 1.6e-05 | 1.180E-05 | 3.650E-06 | 9.400E-06 | 1.600E-05 |
| 2841 | 143 | D143A | A | 1.5e-05 | 1.4e-05 | 9.5e-06 | 1.283E-05 | 2.930E-06 | 9.500E-06 | 1.500E-05 |
| 2842 | 143 | D143R | A | 1.5e-05 | 9.1e-06 | 1e-05 | 1.137E-05 | 3.179E-06 | 9.100E-06 | 1.500E-05 |
| 2843 | 143 | D143N | A | 1.6e-05 | 1.3e-05 | 1e-05 | 1.300E-05 | 3.000E-06 | 1.000E-05 | 1.600E-05 |
| 2844 | 143 | D143D | A | 1.1e-05 | 1.3e-05 | 1.3e-05 | 1.233E-05 | 1.155E-06 | 1.100E-05 | 1.300E-05 |
| 2845 | 143 | D143C | A | 1.4e-05 | 1.3e-05 | 1.1e-05 | 1.267E-05 | 1.528E-06 | 1.100E-05 | 1.400E-05 |
| 2846 | 143 | D143E | A | 1.8e-05 | 9.6e-06 | 1.6e-05 | 1.453E-05 | 4.388E-06 | 9.600E-06 | 1.800E-05 |
| 2847 | 143 | D143Q | A | 1.2e-05 | 6.3e-06 | 1.4e-05 | 1.077E-05 | 3.995E-06 | 6.300E-06 | 1.400E-05 |
| 2848 | 143 | D143G | A | 1.6e-05 | 8.7e-06 | 1.3e-05 | 1.257E-05 | 3.669E-06 | 8.700E-06 | 1.600E-05 |
| 2849 | 143 | D143H | A | 1.4e-05 | 1.5e-05 | 1.2e-05 | 1.367E-05 | 1.528E-06 | 1.200E-05 | 1.500E-05 |
| 2850 | 143 | D143I | A | 1.2e-05 | 9.6e-06 | 1.1e-05 | 1.087E-05 | 1.206E-06 | 9.600E-06 | 1.200E-05 |
| 2851 | 143 | D143L | A | 1e-05 | 1.5e-05 | 1.4e-05 | 1.300E-05 | 2.646E-06 | 1.000E-05 | 1.500E-05 |
| 2852 | 143 | D143K | A | 1.5e-05 | 1.3e-05 | 1e-05 | 1.267E-05 | 2.517E-06 | 1.000E-05 | 1.500E-05 |
| 2853 | 143 | D143M | A | 9.8e-06 | 1e-05 | 1.3e-05 | 1.093E-05 | 1.793E-06 | 9.800E-06 | 1.300E-05 |
| 2854 | 143 | D143F | A | 1e-05 | 9.8e-06 | 1.6e-05 | 1.193E-05 | 3.523E-06 | 9.800E-06 | 1.600E-05 |
| 2855 | 143 | D143P | A | 8.6e-06 | 1.5e-05 | 1.1e-05 | 1.153E-05 | 3.233E-06 | 8.600E-06 | 1.500E-05 |
| 2856 | 143 | D143S | A | 1.6e-05 | 1.5e-05 | 8.9e-06 | 1.330E-05 | 3.843E-06 | 8.900E-06 | 1.600E-05 |
| 2857 | 143 | D143T | A | 1e-05 | 1.1e-05 | 1.1e-05 | 1.067E-05 | 5.774E-07 | 1.000E-05 | 1.100E-05 |
| 2858 | 143 | D143W | A | 9.8e-06 | 9.3e-06 | 1.1e-05 | 1.003E-05 | 8.737E-07 | 9.300E-06 | 1.100E-05 |
| 2859 | 143 | D143Y | A | 9.5e-06 | 1.6e-05 | 1.5e-05 | 1.350E-05 | 3.500E-06 | 9.500E-06 | 1.600E-05 |
| 2860 | 143 | D143V | A | 1.1e-05 | 1.6e-05 | 1.5e-05 | 1.400E-05 | 2.646E-06 | 1.100E-05 | 1.600E-05 |
| 2861 | 144 | T144A | A | 9.3e-06 | 9.9e-06 | 1.4e-05 | 1.107E-05 | 2.558E-06 | 9.300E-06 | 1.400E-05 |
| 2862 | 144 | T144R | A | 1.6e-05 | 1.4e-05 | 1e-05 | 1.333E-05 | 3.055E-06 | 1.000E-05 | 1.600E-05 |
| 2863 | 144 | T144N | A | 1.3e-05 | 1.3e-05 | 1.6e-05 | 1.400E-05 | 1.732E-06 | 1.300E-05 | 1.600E-05 |
| 2864 | 144 | T144D | A | 1.8e-05 | 1.5e-05 | 1.6e-05 | 1.633E-05 | 1.528E-06 | 1.500E-05 | 1.800E-05 |
| 2865 | 144 | T144C | A | 1.6e-05 | 1.1e-05 | 6.8e-06 | 1.127E-05 | 4.606E-06 | 6.800E-06 | 1.600E-05 |
| 2866 | 144 | T144E | A | 1.2e-05 | 1.6e-05 | 1.8e-05 | 1.533E-05 | 3.055E-06 | 1.200E-05 | 1.800E-05 |
| 2867 | 144 | T144Q | A | 1.6e-05 | 8.8e-06 | 8.5e-06 | 1.110E-05 | 4.246E-06 | 8.500E-06 | 1.600E-05 |
| 2868 | 144 | T144G | A | 1e-05 | 1.2e-05 | 1.2e-05 | 1.133E-05 | 1.155E-06 | 1.000E-05 | 1.200E-05 |
| 2869 | 144 | T144H | A | 1.6e-05 | 2e-05 | 1.1e-05 | 1.567E-05 | 4.509E-06 | 1.100E-05 | 2.000E-05 |
| 2870 | 144 | T144I | A | 1.6e-05 | 8.5e-06 | 1e-05 | 1.150E-05 | 3.969E-06 | 8.500E-06 | 1.600E-05 |
| 2871 | 144 | T144L | A | 1.5e-05 | 1.5e-05 | 1.1e-05 | 1.367E-05 | 2.309E-06 | 1.100E-05 | 1.500E-05 |
| 2872 | 144 | T144K | A | 9.5e-06 | 1e-05 | 8.6e-06 | 9.367E-06 | 7.095E-07 | 8.600E-06 | 1.000E-05 |
| 2873 | 144 | T144M | A | 1.6e-05 | 1.1e-05 | 1.1e-05 | 1.267E-05 | 2.887E-06 | 1.100E-05 | 1.600E-05 |
| 2874 | 144 | T144F | A | 1e-05 | 8.6e-06 | 9.5e-06 | 9.367E-06 | 7.095E-07 | 8.600E-06 | 1.000E-05 |
| 2875 | 144 | T144P | A | 1.4e-05 | 1.1e-05 | 1.1e-05 | 1.200E-05 | 1.732E-06 | 1.100E-05 | 1.400E-05 |
| 2876 | 144 | T144S | A | 8.3e-06 | 1.6e-05 | 9.5e-06 | 1.127E-05 | 4.143E-06 | 8.300E-06 | 1.600E-05 |
| 2877 | 144 | T144T | A | 1.1e-05 | 1.3e-05 | 1e-05 | 1.133E-05 | 1.528E-06 | 1.000E-05 | 1.300E-05 |
| 2878 | 144 | T144W | A | 1.4e-05 | 1.4e-05 | 1.7e-05 | 1.500E-05 | 1.732E-06 | 1.400E-05 | 1.700E-05 |
| 2879 | 144 | T144Y | A | 1e-05 | 1.2e-05 | 1.3e-05 | 1.167E-05 | 1.528E-06 | 1.000E-05 | 1.300E-05 |
| 2880 | 144 | T144V | A | 1.6e-05 | 1.6e-05 | 1.3e-05 | 1.500E-05 | 1.732E-06 | 1.300E-05 | 1.600E-05 |
| 2881 | 145 | I145A | A | 1.1e-05 | 1.1e-05 | 1e-05 | 1.067E-05 | 5.774E-07 | 1.000E-05 | 1.100E-05 |
| 2882 | 145 | I145R | A | 1.1e-05 | 8.4e-06 | 8.7e-06 | 9.367E-06 | 1.422E-06 | 8.400E-06 | 1.100E-05 |
| 2883 | 145 | I145N | A | 1.1e-05 | 1.1e-05 | 1.6e-05 | 1.267E-05 | 2.887E-06 | 1.100E-05 | 1.600E-05 |
| 2884 | 145 | I145D | A | 1.2e-05 | 1.1e-05 | 1.6e-05 | 1.300E-05 | 2.646E-06 | 1.100E-05 | 1.600E-05 |
| 2885 | 145 | I145C | A | 1.2e-05 | 1.3e-05 | 1.7e-05 | 1.400E-05 | 2.646E-06 | 1.200E-05 | 1.700E-05 |
| 2886 | 145 | I145E | A | 1.3e-05 | 1.2e-05 | 1e-05 | 1.167E-05 | 1.528E-06 | 1.000E-05 | 1.300E-05 |
| 2887 | 145 | I145Q | A | 9.8e-06 | 7.9e-06 | 9.4e-06 | 9.033E-06 | 1.002E-06 | 7.900E-06 | 9.800E-06 |
| 2888 | 145 | I145G | A | 1e-05 | 9.7e-06 | 1.7e-05 | 1.223E-05 | 4.131E-06 | 9.700E-06 | 1.700E-05 |
| 2889 | 145 | I145H | A | 1.2e-05 | 1e-05 | 6.2e-06 | 9.400E-06 | 2.946E-06 | 6.200E-06 | 1.200E-05 |

| | | | | | | | | | | |
|---|---|---|---|---|---|---|---|---|---|---|
| 2890 | 145 | I145I | A | 1.2e-05 | 1.6e-05 | 8.5e-06 | 1.217E-05 | 3.753E-06 | 8.500E-06 | 1.600E-05 |
| 2891 | 145 | I145L | A | 1.4e-05 | 1e-05 | 1.7e-05 | 1.367E-05 | 3.512E-06 | 1.000E-05 | 1.700E-05 |
| 2892 | 145 | I145K | A | 1.6e-05 | 8.9e-06 | 8.3e-06 | 1.107E-05 | 4.283E-06 | 8.300E-06 | 1.600E-05 |
| 2893 | 145 | I145M | A | 1.4e-05 | 9.9e-06 | 1.6e-05 | 1.330E-05 | 3.110E-06 | 9.900E-06 | 1.600E-05 |
| 2894 | 145 | I145F | A | 1.6e-05 | 1.4e-05 | 2e-05 | 1.667E-05 | 3.055E-06 | 1.400E-05 | 2.000E-05 |
| 2895 | 145 | I145P | A | 1e-05 | 1.6e-05 | 1.5e-05 | 1.367E-05 | 3.215E-06 | 1.000E-05 | 1.600E-05 |
| 2896 | 145 | I145S | A | 1e-05 | 1.4e-05 | 1.1e-05 | 1.167E-05 | 2.082E-06 | 1.000E-05 | 1.400E-05 |
| 2897 | 145 | I145T | A | 1.1e-05 | 9.5e-06 | 1.1e-05 | 1.050E-05 | 8.660E-07 | 9.500E-06 | 1.100E-05 |
| 2898 | 145 | I145W | A | 1.3e-05 | 1.6e-05 | 1.1e-05 | 1.333E-05 | 2.517E-06 | 1.100E-05 | 1.600E-05 |
| 2899 | 145 | I145Y | A | 1.3e-05 | 1.4e-05 | 1.2e-05 | 1.300E-05 | 1.000E-06 | 1.200E-05 | 1.400E-05 |
| 2900 | 145 | I145V | A | 1.3e-05 | 1.4e-05 | 9.7e-06 | 1.223E-05 | 2.250E-06 | 9.700E-06 | 1.400E-05 |
| 2901 | 146 | L146A | A | 1.1e-05 | 1.7e-05 | 1e-05 | 1.267E-05 | 3.786E-06 | 1.000E-05 | 1.700E-05 |
| 2902 | 146 | L146R | A | 1.6e-05 | 1e-05 | 1.5e-05 | 1.367E-05 | 3.215E-06 | 1.000E-05 | 1.600E-05 |
| 2903 | 146 | L146N | A | 1.5e-05 | 1.4e-05 | 1.4e-05 | 1.433E-05 | 5.774E-07 | 1.400E-05 | 1.500E-05 |
| 2904 | 146 | L146D | A | 1.3e-05 | 7.2e-06 | 1.5e-05 | 1.173E-05 | 4.051E-06 | 7.200E-06 | 1.500E-05 |
| 2905 | 146 | L146C | A | 1.6e-05 | 1.1e-05 | 1e-05 | 1.233E-05 | 3.215E-06 | 1.000E-05 | 1.600E-05 |
| 2906 | 146 | L146E | A | 1.4e-05 | 1.7e-05 | 1.4e-05 | 1.500E-05 | 1.732E-06 | 1.400E-05 | 1.700E-05 |
| 2907 | 146 | L146Q | A | 1e-05 | 1.2e-05 | 1.1e-05 | 1.100E-05 | 1.000E-06 | 1.000E-05 | 1.200E-05 |
| 2908 | 146 | L146G | A | 1.1e-05 | 1.2e-05 | 1.5e-05 | 1.267E-05 | 2.082E-06 | 1.100E-05 | 1.500E-05 |
| 2909 | 146 | L146H | A | 1.5e-05 | 1e-05 | 1.1e-05 | 1.200E-05 | 2.646E-06 | 1.000E-05 | 1.500E-05 |
| 2910 | 146 | L146I | A | 1.6e-05 | 1.4e-05 | 1.3e-05 | 1.433E-05 | 1.528E-06 | 1.300E-05 | 1.600E-05 |
| 2911 | 146 | L146L | A | 1.5e-05 | 1.4e-05 | 1.1e-05 | 1.333E-05 | 2.082E-06 | 1.100E-05 | 1.500E-05 |
| 2912 | 146 | L146K | A | 8.9e-06 | 1.4e-05 | 9.1e-06 | 1.067E-05 | 2.888E-06 | 8.900E-06 | 1.400E-05 |
| 2913 | 146 | L146M | A | 1.5e-05 | 1.1e-05 | 1.2e-05 | 1.267E-05 | 2.082E-06 | 1.100E-05 | 1.500E-05 |
| 2914 | 146 | L146F | A | 1e-05 | 1.1e-05 | 1.4e-05 | 1.167E-05 | 2.082E-06 | 1.000E-05 | 1.400E-05 |
| 2915 | 146 | L146P | A | 9e-06 | 1.3e-05 | 1.6e-05 | 1.267E-05 | 3.512E-06 | 9.000E-06 | 1.600E-05 |
| 2916 | 146 | L146S | A | 9.3e-06 | 1.5e-05 | 1.4e-05 | 1.277E-05 | 3.044E-06 | 9.300E-06 | 1.500E-05 |
| 2917 | 146 | L146T | A | 8.2e-06 | 1.5e-05 | 1e-05 | 1.107E-05 | 3.523E-06 | 8.200E-06 | 1.500E-05 |
| 2918 | 146 | L146W | A | 1.2e-05 | 1.5e-05 | 1e-05 | 1.233E-05 | 2.517E-06 | 1.000E-05 | 1.500E-05 |
| 2919 | 146 | L146Y | A | 1.5e-05 | 1.8e-05 | 1.4e-05 | 1.567E-05 | 2.082E-06 | 1.400E-05 | 1.800E-05 |
| 2920 | 146 | L146V | A | 1.6e-05 | 9.9e-06 | 1.4e-05 | 1.330E-05 | 3.110E-06 | 9.900E-06 | 1.600E-05 |
| 2921 | 147 | W147A | A | 1.1e-05 | 1.1e-05 | 1.6e-05 | 1.267E-05 | 2.887E-06 | 1.100E-05 | 1.600E-05 |
| 2922 | 147 | W147R | A | 1e-05 | 1.7e-05 | 1.7e-05 | 1.467E-05 | 4.041E-06 | 1.000E-05 | 1.700E-05 |
| 2923 | 147 | W147N | A | 8.5e-06 | 9.7e-06 | 1.5e-05 | 1.107E-05 | 3.459E-06 | 8.500E-06 | 1.500E-05 |
| 2924 | 147 | W147D | A | 8.4e-06 | 1.6e-05 | 1.4e-05 | 1.280E-05 | 3.940E-06 | 8.400E-06 | 1.600E-05 |
| 2925 | 147 | W147C | A | 9.7e-06 | 1.3e-05 | 1.2e-05 | 1.157E-05 | 1.692E-06 | 9.700E-06 | 1.300E-05 |
| 2926 | 147 | W147E | A | 1.5e-05 | 2.3e-05 | 1.7e-05 | 1.833E-05 | 4.163E-06 | 1.500E-05 | 2.300E-05 |
| 2927 | 147 | W147Q | A | 1.2e-05 | 1.1e-05 | 1.3e-05 | 1.200E-05 | 1.000E-06 | 1.100E-05 | 1.300E-05 |
| 2928 | 147 | W147G | A | 1.5e-05 | 1.6e-05 | 1.7e-05 | 1.600E-05 | 1.000E-06 | 1.500E-05 | 1.700E-05 |
| 2929 | 147 | W147H | A | 1.1e-05 | 1.5e-05 | 1.1e-05 | 1.233E-05 | 2.309E-06 | 1.100E-05 | 1.500E-05 |
| 2930 | 147 | W147I | A | 1.4e-05 | 1.6e-05 | 1.7e-05 | 1.567E-05 | 1.528E-06 | 1.400E-05 | 1.700E-05 |
| 2931 | 147 | W147L | A | 1.1e-05 | 1.1e-05 | 7.7e-06 | 9.900E-06 | 1.905E-06 | 7.700E-06 | 1.100E-05 |
| 2932 | 147 | W147K | A | 1.7e-05 | 1e-05 | 1.4e-05 | 1.367E-05 | 3.512E-06 | 1.000E-05 | 1.700E-05 |
| 2933 | 147 | W147M | A | 1.4e-05 | 1.6e-05 | 1.1e-05 | 1.367E-05 | 2.517E-06 | 1.100E-05 | 1.600E-05 |
| 2934 | 147 | W147F | A | 1.2e-05 | 1.4e-05 | 9.6e-06 | 1.187E-05 | 2.203E-06 | 9.600E-06 | 1.400E-05 |
| 2935 | 147 | W147P | A | 1.4e-05 | 1e-05 | 1.2e-05 | 1.200E-05 | 2.000E-06 | 1.000E-05 | 1.400E-05 |
| 2936 | 147 | W147S | A | 1.1e-05 | 1.6e-05 | 1.1e-05 | 1.267E-05 | 2.887E-06 | 1.100E-05 | 1.600E-05 |
| 2937 | 147 | W147T | A | 1.6e-05 | 8.5e-06 | 1.3e-05 | 1.250E-05 | 3.775E-06 | 8.500E-06 | 1.600E-05 |
| 2938 | 147 | W147W | A | 1.4e-05 | 1.4e-05 | 1.7e-05 | 1.500E-05 | 1.732E-06 | 1.400E-05 | 1.700E-05 |
| 2939 | 147 | W147Y | A | 1.6e-05 | 1.1e-05 | 1.4e-05 | 1.367E-05 | 2.517E-06 | 1.100E-05 | 1.600E-05 |
| 2940 | 147 | W147V | A | 1.4e-05 | 1.8e-05 | 1.4e-05 | 1.533E-05 | 2.309E-06 | 1.400E-05 | 1.800E-05 |
| 2941 | 148 | K148A | A | 1.1e-05 | 1.1e-05 | 1.6e-05 | 1.267E-05 | 2.887E-06 | 1.100E-05 | 1.600E-05 |
| 2942 | 148 | K148R | A | 1.6e-05 | 1e-05 | 8e-06 | 1.133E-05 | 4.163E-06 | 8.000E-06 | 1.600E-05 |
| 2943 | 148 | K148N | A | 1.4e-05 | 1.6e-05 | 1e-05 | 1.333E-05 | 3.055E-06 | 1.000E-05 | 1.600E-05 |
| 2944 | 148 | K148D | A | 1e-05 | 9.4e-06 | 1.3e-05 | 1.080E-05 | 1.929E-06 | 9.400E-06 | 1.300E-05 |
| 2945 | 148 | K148C | A | 1.5e-05 | 1e-05 | 1.3e-05 | 1.267E-05 | 2.517E-06 | 1.000E-05 | 1.500E-05 |
| 2946 | 148 | K148E | A | 1e-05 | 8.6e-06 | 1.7e-05 | 1.187E-05 | 4.500E-06 | 8.600E-06 | 1.700E-05 |
| 2947 | 148 | K148Q | A | 1.5e-05 | 1.5e-05 | 1.5e-05 | 1.500E-05 | 0.000E+00 | 1.500E-05 | 1.500E-05 |
| 2948 | 148 | K148G | A | 1.4e-05 | 1.3e-05 | 1.3e-05 | 1.333E-05 | 5.774E-07 | 1.300E-05 | 1.400E-05 |
| 2949 | 148 | K148H | A | 1.6e-05 | 1.5e-05 | 1.3e-05 | 1.467E-05 | 1.528E-06 | 1.300E-05 | 1.600E-05 |
| 2950 | 148 | K148I | A | 1.7e-05 | 7.8e-06 | 1.5e-05 | 1.327E-05 | 4.839E-06 | 7.800E-06 | 1.700E-05 |
| 2951 | 148 | K148L | A | 1.1e-05 | 1.5e-05 | 8.3e-06 | 1.143E-05 | 3.371E-06 | 8.300E-06 | 1.500E-05 |
| 2952 | 148 | K148K | A | 1.3e-05 | 1.5e-05 | 1.7e-05 | 1.500E-05 | 2.000E-06 | 1.300E-05 | 1.700E-05 |
| 2953 | 148 | K148M | A | 1.3e-05 | 1.1e-05 | 1e-05 | 1.133E-05 | 1.528E-06 | 1.000E-05 | 1.300E-05 |
| 2954 | 148 | K148F | A | 1.3e-05 | 1.6e-05 | 1.6e-05 | 1.500E-05 | 1.732E-06 | 1.300E-05 | 1.600E-05 |
| 2955 | 148 | K148P | A | 1.7e-05 | 1.6e-05 | 1.1e-05 | 1.467E-05 | 3.215E-06 | 1.100E-05 | 1.700E-05 |
| 2956 | 148 | K148S | A | 1.6e-05 | 9.9e-06 | 1.4e-05 | 1.330E-05 | 3.110E-06 | 9.900E-06 | 1.600E-05 |
| 2957 | 148 | K148T | A | 1e-05 | 9.5e-06 | 9.1e-06 | 9.533E-06 | 4.509E-07 | 9.100E-06 | 1.000E-05 |
| 2958 | 148 | K148W | A | 1.4e-05 | 1.6e-05 | 1e-05 | 1.333E-05 | 3.055E-06 | 1.000E-05 | 1.600E-05 |
| 2959 | 148 | K148Y | A | 1.5e-05 | 1.8e-05 | 7.2e-06 | 1.340E-05 | 5.575E-06 | 7.200E-06 | 1.800E-05 |
| 2960 | 148 | K148V | A | 1.5e-05 | 1e-05 | 1.3e-05 | 1.267E-05 | 2.517E-06 | 1.000E-05 | 1.500E-05 |
| 2961 | 149 | D149A | A | 1.3e-05 | 1.4e-05 | 1.1e-05 | 1.267E-05 | 1.528E-06 | 1.100E-05 | 1.400E-05 |
| 2962 | 149 | D149R | A | 1.4e-05 | 1.1e-05 | 1.1e-05 | 1.200E-05 | 1.732E-06 | 1.100E-05 | 1.400E-05 |
| 2963 | 149 | D149N | A | 8.1e-06 | 1.4e-05 | 1.4e-05 | 1.203E-05 | 3.406E-06 | 8.100E-06 | 1.400E-05 |
| 2964 | 149 | D149D | A | 1.2e-05 | 8.9e-06 | 1e-05 | 1.030E-05 | 1.572E-06 | 8.900E-06 | 1.200E-05 |
| 2965 | 149 | D149C | A | 1.3e-05 | 1.1e-05 | 1.5e-05 | 1.300E-05 | 2.000E-06 | 1.100E-05 | 1.500E-05 |
| 2966 | 149 | D149E | A | 1e-05 | 1.1e-05 | 8.1e-06 | 9.700E-06 | 1.473E-06 | 8.100E-06 | 1.100E-05 |
| 2967 | 149 | D149Q | A | 7.9e-06 | 1e-05 | 1.5e-05 | 1.097E-05 | 3.647E-06 | 7.900E-06 | 1.500E-05 |
| 2968 | 149 | D149G | A | 1.4e-05 | 6.5e-06 | 1.8e-05 | 1.283E-05 | 5.838E-06 | 6.500E-06 | 1.800E-05 |
| 2969 | 149 | D149H | A | 1.6e-05 | 1.5e-05 | 1.2e-05 | 1.433E-05 | 2.082E-06 | 1.200E-05 | 1.600E-05 |
| 2970 | 149 | D149I | A | 1.3e-05 | 1.1e-05 | 8.4e-06 | 1.080E-05 | 2.307E-06 | 8.400E-06 | 1.300E-05 |
| 2971 | 149 | D149L | A | 1.3e-05 | 1.1e-05 | 1.5e-05 | 1.300E-05 | 2.000E-06 | 1.100E-05 | 1.500E-05 |
| 2972 | 149 | D149K | A | 1.2e-05 | 1.5e-05 | 1.3e-05 | 1.333E-05 | 1.528E-06 | 1.200E-05 | 1.500E-05 |
| 2973 | 149 | D149M | A | 1.6e-05 | 8.4e-06 | 1.4e-05 | 1.280E-05 | 3.940E-06 | 8.400E-06 | 1.600E-05 |
| 2974 | 149 | D149F | A | 1.6e-05 | 1.4e-05 | 1e-05 | 1.333E-05 | 3.055E-06 | 1.000E-05 | 1.600E-05 |

| | | | | | | | | | | |
|---|---|---|---|---|---|---|---|---|---|---|
| 2975 | 149 | D149P | A | 1.4e-05 | 1.1e-05 | 1.4e-05 | 1.300E-05 | 1.732E-06 | 1.100E-05 | 1.400E-05 |
| 2976 | 149 | D149S | A | 1.2e-05 | 1.4e-05 | 1.6e-05 | 1.400E-05 | 2.000E-06 | 1.200E-05 | 1.600E-05 |
| 2977 | 149 | D149T | A | 7.4e-06 | 1.4e-05 | 1.5e-05 | 1.213E-05 | 4.130E-06 | 7.400E-06 | 1.500E-05 |
| 2978 | 149 | D149W | A | 9.2e-06 | 1.1e-05 | 9e-06 | 9.733E-06 | 1.102E-06 | 9.000E-06 | 1.100E-05 |
| 2979 | 149 | D149Y | A | 1.6e-05 | 1.4e-05 | 1.1e-05 | 1.367E-05 | 2.517E-06 | 1.100E-05 | 1.600E-05 |
| 2980 | 149 | D149V | A | 1.5e-05 | 1.1e-05 | 1.1e-05 | 1.233E-05 | 2.309E-06 | 1.100E-05 | 1.500E-05 |
| 2981 | 150 | I150A | A | 1.3e-05 | 1.1e-05 | 1.1e-05 | 1.167E-05 | 1.155E-06 | 1.100E-05 | 1.300E-05 |
| 2982 | 150 | I150R | A | 1.4e-05 | 1.6e-05 | 1e-05 | 1.333E-05 | 3.055E-06 | 1.000E-05 | 1.600E-05 |
| 2983 | 150 | I150N | A | 1.3e-05 | 1.5e-05 | 1.4e-05 | 1.400E-05 | 1.000E-06 | 1.300E-05 | 1.500E-05 |
| 2984 | 150 | I150D | A | 1.6e-05 | 1e-05 | 1.5e-05 | 1.367E-05 | 3.215E-06 | 1.000E-05 | 1.600E-05 |
| 2985 | 150 | I150C | A | 8.6e-06 | 1.5e-05 | 1.5e-05 | 1.287E-05 | 3.695E-06 | 8.600E-06 | 1.500E-05 |
| 2986 | 150 | I150E | A | 9.5e-06 | 1.5e-05 | 1e-05 | 1.150E-05 | 3.041E-06 | 9.500E-06 | 1.500E-05 |
| 2987 | 150 | I150Q | A | 1.3e-05 | 1.1e-05 | 1.3e-05 | 1.233E-05 | 1.155E-06 | 1.100E-05 | 1.300E-05 |
| 2988 | 150 | I150G | A | 1.1e-05 | 1.6e-05 | 1e-05 | 1.233E-05 | 3.215E-06 | 1.000E-05 | 1.600E-05 |
| 2989 | 150 | I150H | A | 1.6e-05 | 9.7e-06 | 1.2e-05 | 1.257E-05 | 3.188E-06 | 9.700E-06 | 1.600E-05 |
| 2990 | 150 | I150I | A | 1e-05 | 1.5e-05 | 1.2e-05 | 1.233E-05 | 2.517E-06 | 1.000E-05 | 1.500E-05 |
| 2991 | 150 | I150L | A | 1.1e-05 | 9.9e-06 | 1.6e-05 | 1.230E-05 | 3.251E-06 | 9.900E-06 | 1.600E-05 |
| 2992 | 150 | I150K | A | 1.1e-05 | 1e-05 | 2.2e-05 | 1.433E-05 | 6.658E-06 | 1.000E-05 | 2.200E-05 |
| 2993 | 150 | I150M | A | 1.5e-05 | 1.1e-05 | 1.1e-05 | 1.233E-05 | 2.309E-06 | 1.100E-05 | 1.500E-05 |
| 2994 | 150 | I150F | A | 1.1e-05 | 1.4e-05 | 1e-05 | 1.167E-05 | 2.082E-06 | 1.000E-05 | 1.400E-05 |
| 2995 | 150 | I150P | A | 1.5e-05 | 1.1e-05 | 1.5e-05 | 1.367E-05 | 2.309E-06 | 1.100E-05 | 1.500E-05 |
| 2996 | 150 | I150S | A | 1.5e-05 | 1.5e-05 | 1.5e-05 | 1.500E-05 | 0.000E+00 | 1.500E-05 | 1.500E-05 |
| 2997 | 150 | I150T | A | 1.6e-05 | 1.5e-05 | 9.1e-06 | 1.337E-05 | 3.729E-06 | 9.100E-06 | 1.600E-05 |
| 2998 | 150 | I150W | A | 1.6e-05 | 1.4e-05 | 1.1e-05 | 1.367E-05 | 2.517E-06 | 1.100E-05 | 1.600E-05 |
| 2999 | 150 | I150Y | A | 1.5e-05 | 1.1e-05 | 1.7e-05 | 1.433E-05 | 3.055E-06 | 1.100E-05 | 1.700E-05 |
| 3000 | 150 | I150V | A | 1.1e-05 | 1.5e-05 | 1.4e-05 | 1.333E-05 | 2.082E-06 | 1.100E-05 | 1.500E-05 |
| 3001 | 151 | F151A | A | 8.8e-06 | 8.4e-06 | 1.1e-05 | 9.400E-06 | 1.400E-06 | 8.400E-06 | 1.100E-05 |
| 3002 | 151 | F151R | A | 1.5e-05 | 1.5e-05 | 1.4e-05 | 1.467E-05 | 5.774E-07 | 1.400E-05 | 1.500E-05 |
| 3003 | 151 | F151N | A | 1.1e-05 | 8.9e-06 | 1.1e-05 | 1.030E-05 | 1.212E-06 | 8.900E-06 | 1.100E-05 |
| 3004 | 151 | F151D | A | 1.1e-05 | 1.6e-05 | 1e-05 | 1.233E-05 | 3.215E-06 | 1.000E-05 | 1.600E-05 |
| 3005 | 151 | F151C | A | 9.4e-06 | 8.3e-06 | 1.1e-05 | 9.567E-06 | 1.358E-06 | 8.300E-06 | 1.100E-05 |
| 3006 | 151 | F151E | A | 1.8e-05 | 1.1e-05 | 1.1e-05 | 1.333E-05 | 4.041E-06 | 1.100E-05 | 1.800E-05 |
| 3007 | 151 | F151Q | A | 1.7e-05 | 1.1e-05 | 1.5e-05 | 1.433E-05 | 3.055E-06 | 1.100E-05 | 1.700E-05 |
| 3008 | 151 | F151G | A | 1.5e-05 | 1.6e-05 | 1e-05 | 1.367E-05 | 3.215E-06 | 1.000E-05 | 1.600E-05 |
| 3009 | 151 | F151H | A | 1.3e-05 | 1e-05 | 1.1e-05 | 1.133E-05 | 1.528E-06 | 1.000E-05 | 1.300E-05 |
| 3010 | 151 | F151I | A | 8.5e-06 | 1.1e-05 | 1.1e-05 | 1.017E-05 | 1.443E-06 | 8.500E-06 | 1.100E-05 |
| 3011 | 151 | F151L | A | 1.1e-05 | 1.1e-05 | 1.5e-05 | 1.233E-05 | 2.309E-06 | 1.100E-05 | 1.500E-05 |
| 3012 | 151 | F151K | A | 9.8e-06 | 1.2e-05 | 1.8e-05 | 1.327E-05 | 4.244E-06 | 9.800E-06 | 1.800E-05 |
| 3013 | 151 | F151M | A | 1.2e-05 | 9.5e-06 | 7.3e-06 | 9.600E-06 | 2.352E-06 | 7.300E-06 | 1.200E-05 |
| 3014 | 151 | F151F | A | 9.9e-06 | 1.5e-05 | 1.1e-05 | 1.197E-05 | 2.684E-06 | 9.900E-06 | 1.500E-05 |
| 3015 | 151 | F151P | A | 1.4e-05 | 1e-05 | 1.5e-05 | 1.300E-05 | 2.646E-06 | 1.000E-05 | 1.500E-05 |
| 3016 | 151 | F151S | A | 1.5e-05 | 1.6e-05 | 9.1e-06 | 1.337E-05 | 3.729E-06 | 9.100E-06 | 1.600E-05 |
| 3017 | 151 | F151T | A | 1.3e-05 | 1.3e-05 | 1e-05 | 1.200E-05 | 1.732E-06 | 1.000E-05 | 1.300E-05 |
| 3018 | 151 | F151W | A | 1.3e-05 | 1e-05 | 1.5e-05 | 1.267E-05 | 2.517E-06 | 1.000E-05 | 1.500E-05 |
| 3019 | 151 | F151Y | A | 1.6e-05 | 9.7e-06 | 8.5e-06 | 1.140E-05 | 4.029E-06 | 8.500E-06 | 1.600E-05 |
| 3020 | 151 | F151V | A | 8.7e-06 | 1.9e-05 | 1.1e-05 | 1.290E-05 | 5.406E-06 | 8.700E-06 | 1.900E-05 |
| 3021 | 152 | H152A | A | 1.1e-05 | 1e-05 | 1.6e-05 | 1.233E-05 | 3.215E-06 | 1.000E-05 | 1.600E-05 |
| 3022 | 152 | H152R | A | 1e-05 | 1.5e-05 | 1e-05 | 1.167E-05 | 2.887E-06 | 1.000E-05 | 1.500E-05 |
| 3023 | 152 | H152N | A | 1.1e-05 | 1e-05 | 1.8e-05 | 1.300E-05 | 4.359E-06 | 1.000E-05 | 1.800E-05 |
| 3024 | 152 | H152D | A | 1e-05 | 1.2e-05 | 1e-05 | 1.067E-05 | 1.155E-06 | 1.000E-05 | 1.200E-05 |
| 3025 | 152 | H152C | A | 1.5e-05 | 1.5e-05 | 1.2e-05 | 1.400E-05 | 1.732E-06 | 1.200E-05 | 1.500E-05 |
| 3026 | 152 | H152E | A | 1.4e-05 | 9.2e-06 | 1.7e-05 | 1.340E-05 | 3.934E-06 | 9.200E-06 | 1.700E-05 |
| 3027 | 152 | H152Q | A | 9.5e-06 | 1e-05 | 1.4e-05 | 1.117E-05 | 2.466E-06 | 9.500E-06 | 1.400E-05 |
| 3028 | 152 | H152G | A | 1e-05 | 7.3e-06 | 9e-06 | 8.767E-06 | 1.365E-06 | 7.300E-06 | 1.000E-05 |
| 3029 | 152 | H152H | A | 1.4e-05 | 1.8e-05 | 1e-05 | 1.400E-05 | 4.000E-06 | 1.000E-05 | 1.800E-05 |
| 3030 | 152 | H152I | A | 1.5e-05 | 1.1e-05 | 1.3e-05 | 1.300E-05 | 2.000E-06 | 1.100E-05 | 1.500E-05 |
| 3031 | 152 | H152L | A | 9.4e-06 | 1.4e-05 | 1e-05 | 1.113E-05 | 2.501E-06 | 9.400E-06 | 1.400E-05 |
| 3032 | 152 | H152K | A | 1.6e-05 | 9.5e-06 | 1.6e-05 | 1.383E-05 | 3.753E-06 | 9.500E-06 | 1.600E-05 |
| 3033 | 152 | H152M | A | 1.6e-05 | 1.1e-05 | 1.6e-05 | 1.433E-05 | 2.887E-06 | 1.100E-05 | 1.600E-05 |
| 3034 | 152 | H152F | A | 1.1e-05 | 1.4e-05 | 1.7e-05 | 1.400E-05 | 3.000E-06 | 1.100E-05 | 1.700E-05 |
| 3035 | 152 | H152P | A | 1.1e-05 | 1.4e-05 | 1.5e-05 | 1.333E-05 | 2.082E-06 | 1.100E-05 | 1.500E-05 |
| 3036 | 152 | H152S | A | 1.2e-05 | 1.1e-05 | 7.7e-06 | 1.023E-05 | 2.250E-06 | 7.700E-06 | 1.200E-05 |
| 3037 | 152 | H152T | A | 1.4e-05 | 1.1e-05 | 9.5e-06 | 1.150E-05 | 2.291E-06 | 9.500E-06 | 1.400E-05 |
| 3038 | 152 | H152W | A | 1.1e-05 | 1e-05 | 9.9e-06 | 1.030E-05 | 6.083E-07 | 9.900E-06 | 1.100E-05 |
| 3039 | 152 | H152Y | A | 1.1e-05 | 1.5e-05 | 1.5e-05 | 1.367E-05 | 2.309E-06 | 1.100E-05 | 1.500E-05 |
| 3040 | 152 | H152V | A | 1.2e-05 | 1e-05 | 7.7e-06 | 9.900E-06 | 2.152E-06 | 7.700E-06 | 1.200E-05 |
| 3041 | 153 | K153A | A | 1.1e-05 | 1e-05 | 1.3e-05 | 1.133E-05 | 1.528E-06 | 1.000E-05 | 1.300E-05 |
| 3042 | 153 | K153R | A | 1.4e-05 | 1.4e-05 | 1e-05 | 1.267E-05 | 2.309E-06 | 1.000E-05 | 1.400E-05 |
| 3043 | 153 | K153N | A | 1.4e-05 | 1.1e-05 | 8.2e-06 | 1.107E-05 | 2.901E-06 | 8.200E-06 | 1.400E-05 |
| 3044 | 153 | K153D | A | 1.5e-05 | 1.4e-05 | 8.3e-06 | 1.243E-05 | 3.614E-06 | 8.300E-06 | 1.500E-05 |
| 3045 | 153 | K153C | A | 1.4e-05 | 1.3e-05 | 1.6e-05 | 1.433E-05 | 1.528E-06 | 1.300E-05 | 1.600E-05 |
| 3046 | 153 | K153E | A | 1.5e-05 | 1e-05 | 1.5e-05 | 1.333E-05 | 2.887E-06 | 1.000E-05 | 1.500E-05 |
| 3047 | 153 | K153Q | A | 1.1e-05 | 1.3e-05 | 7.2e-06 | 1.040E-05 | 2.946E-06 | 7.200E-06 | 1.300E-05 |
| 3048 | 153 | K153G | A | 1.6e-05 | 1.4e-05 | 1e-05 | 1.333E-05 | 3.055E-06 | 1.000E-05 | 1.600E-05 |
| 3049 | 153 | K153H | A | 1.5e-05 | 1.5e-05 | 8.7e-06 | 1.290E-05 | 3.637E-06 | 8.700E-06 | 1.500E-05 |
| 3050 | 153 | K153I | A | 1.4e-05 | 1.4e-05 | 1.6e-05 | 1.467E-05 | 1.155E-06 | 1.400E-05 | 1.600E-05 |
| 3051 | 153 | K153L | A | 1.1e-05 | 9.9e-06 | 1.5e-05 | 1.197E-05 | 2.684E-06 | 9.900E-06 | 1.500E-05 |
| 3052 | 153 | K153K | A | 9.8e-06 | 1.6e-05 | 1.6e-05 | 1.393E-05 | 3.580E-06 | 9.800E-06 | 1.600E-05 |
| 3053 | 153 | K153M | A | 1.1e-05 | 1.1e-05 | 1.3e-05 | 1.167E-05 | 1.155E-06 | 1.100E-05 | 1.300E-05 |
| 3054 | 153 | K153F | A | 1.3e-05 | 1.2e-05 | 1e-05 | 1.167E-05 | 1.528E-06 | 1.000E-05 | 1.300E-05 |
| 3055 | 153 | K153P | A | 1.4e-05 | 1.1e-05 | 1.1e-05 | 1.200E-05 | 1.732E-06 | 1.100E-05 | 1.400E-05 |
| 3056 | 153 | K153S | A | 1e-05 | 1.8e-05 | 1.6e-05 | 1.467E-05 | 4.163E-06 | 1.000E-05 | 1.800E-05 |
| 3057 | 153 | K153T | A | 1.4e-05 | 1.2e-05 | 1.1e-05 | 1.233E-05 | 1.528E-06 | 1.100E-05 | 1.400E-05 |
| 3058 | 153 | K153W | A | 1.4e-05 | 8.3e-06 | 9.6e-06 | 1.063E-05 | 2.987E-06 | 8.300E-06 | 1.400E-05 |
| 3059 | 153 | K153Y | A | 9.2e-06 | 1.6e-05 | 1.3e-05 | 1.273E-05 | 3.408E-06 | 9.200E-06 | 1.600E-05 |

| | | | | | | | | | | |
|---|---|---|---|---|---|---|---|---|---|---|
| 3060 | 153 | K153V | A | 1.1e-05 | 1.5e-05 | 1.1e-05 | 1.233E-05 | 2.309E-06 | 1.100E-05 | 1.500E-05 |
| 3061 | 154 | N154A | A | 1.6e-05 | 1.1e-05 | 1.7e-05 | 1.467E-05 | 3.215E-06 | 1.100E-05 | 1.700E-05 |
| 3062 | 154 | N154R | A | 1.2e-05 | 1.2e-05 | 8.5e-06 | 1.083E-05 | 2.021E-06 | 8.500E-06 | 1.200E-05 |
| 3063 | 154 | N154N | A | 8.7e-06 | 1.4e-05 | 1.4e-05 | 1.223E-05 | 3.060E-06 | 8.700E-06 | 1.400E-05 |
| 3064 | 154 | N154D | A | 9e-06 | 1.5e-05 | 1.7e-05 | 1.367E-05 | 4.163E-06 | 9.000E-06 | 1.700E-05 |
| 3065 | 154 | N154C | A | 1.3e-05 | 1.3e-05 | 1.1e-05 | 1.233E-05 | 1.155E-06 | 1.100E-05 | 1.300E-05 |
| 3066 | 154 | N154E | A | 1.3e-05 | 1.5e-05 | 1.5e-05 | 1.433E-05 | 1.155E-06 | 1.300E-05 | 1.500E-05 |
| 3067 | 154 | N154Q | A | 1.6e-05 | 1.5e-05 | 1e-05 | 1.367E-05 | 3.215E-06 | 1.000E-05 | 1.600E-05 |
| 3068 | 154 | N154G | A | 1e-05 | 9.1e-06 | 1.1e-05 | 1.003E-05 | 9.504E-07 | 9.100E-06 | 1.100E-05 |
| 3069 | 154 | N154H | A | 1.5e-05 | 1.1e-05 | 1.6e-05 | 1.400E-05 | 2.646E-06 | 1.100E-05 | 1.600E-05 |
| 3070 | 154 | N154I | A | 1e-05 | 1.2e-05 | 1.7e-05 | 1.300E-05 | 3.606E-06 | 1.000E-05 | 1.700E-05 |
| 3071 | 154 | N154L | A | 1.3e-05 | 9e-06 | 1.5e-05 | 1.233E-05 | 3.055E-06 | 9.000E-06 | 1.500E-05 |
| 3072 | 154 | N154K | A | 1.2e-05 | 1.1e-05 | 1.1e-05 | 1.133E-05 | 5.774E-07 | 1.100E-05 | 1.200E-05 |
| 3073 | 154 | N154M | A | 9.6e-06 | 1.3e-05 | 8.6e-06 | 1.040E-05 | 2.307E-06 | 8.600E-06 | 1.300E-05 |
| 3074 | 154 | N154F | A | 1.7e-05 | 1.5e-05 | 1.2e-05 | 1.467E-05 | 2.517E-06 | 1.200E-05 | 1.700E-05 |
| 3075 | 154 | N154P | A | 1.5e-05 | 1.6e-05 | 1.1e-05 | 1.400E-05 | 2.646E-06 | 1.100E-05 | 1.600E-05 |
| 3076 | 154 | N154S | A | 1.5e-05 | 1.1e-05 | 1e-05 | 1.200E-05 | 2.646E-06 | 1.000E-05 | 1.500E-05 |
| 3077 | 154 | N154T | A | 1.6e-05 | 1.2e-05 | 1.4e-05 | 1.400E-05 | 2.000E-06 | 1.200E-05 | 1.600E-05 |
| 3078 | 154 | N154W | A | 1.6e-05 | 1.4e-05 | 1.4e-05 | 1.467E-05 | 1.155E-06 | 1.400E-05 | 1.600E-05 |
| 3079 | 154 | N154Y | A | 1.4e-05 | 1.4e-05 | 1.4e-05 | 1.400E-05 | 0.000E+00 | 1.400E-05 | 1.400E-05 |
| 3080 | 154 | N154V | A | 1.1e-05 | 1.6e-05 | 1.2e-05 | 1.300E-05 | 2.646E-06 | 1.100E-05 | 1.600E-05 |
| 3081 | 155 | N155A | A | 1.2e-05 | 1e-05 | 1.6e-05 | 1.267E-05 | 3.055E-06 | 1.000E-05 | 1.600E-05 |
| 3082 | 155 | N155R | A | 1.3e-05 | 1.6e-05 | 1.5e-05 | 1.467E-05 | 1.528E-06 | 1.300E-05 | 1.600E-05 |
| 3083 | 155 | N155N | A | 8.6e-06 | 1.6e-05 | 1.1e-05 | 1.187E-05 | 3.775E-06 | 8.600E-06 | 1.600E-05 |
| 3084 | 155 | N155D | A | 1.6e-05 | 1.5e-05 | 1e-05 | 1.367E-05 | 3.215E-06 | 1.000E-05 | 1.600E-05 |
| 3085 | 155 | N155C | A | 1.7e-05 | 1.4e-05 | 1.5e-05 | 1.533E-05 | 1.528E-06 | 1.400E-05 | 1.700E-05 |
| 3086 | 155 | N155E | A | 1.1e-05 | 7.6e-06 | 1.2e-05 | 1.020E-05 | 2.307E-06 | 7.600E-06 | 1.200E-05 |
| 3087 | 155 | N155Q | A | 1.4e-05 | 8.9e-06 | 1.4e-05 | 1.230E-05 | 2.944E-06 | 8.900E-06 | 1.400E-05 |
| 3088 | 155 | N155G | A | 1.5e-05 | 1.3e-05 | 1.5e-05 | 1.433E-05 | 1.155E-06 | 1.300E-05 | 1.500E-05 |
| 3089 | 155 | N155H | A | 1.2e-05 | 1e-05 | 1.6e-05 | 1.267E-05 | 3.055E-06 | 1.000E-05 | 1.600E-05 |
| 3090 | 155 | N155I | A | 1.5e-05 | 1.6e-05 | 9.9e-06 | 1.363E-05 | 3.272E-06 | 9.900E-06 | 1.600E-05 |
| 3091 | 155 | N155L | A | 8e-06 | 1e-05 | 1e-05 | 9.333E-06 | 1.155E-06 | 8.000E-06 | 1.000E-05 |
| 3092 | 155 | N155K | A | 1.1e-05 | 8.9e-06 | 1e-05 | 9.967E-06 | 1.050E-06 | 8.900E-06 | 1.100E-05 |
| 3093 | 155 | N155M | A | 1.5e-05 | 1.6e-05 | 6.5e-06 | 1.250E-05 | 5.220E-06 | 6.500E-06 | 1.600E-05 |
| 3094 | 155 | N155F | A | 9.9e-06 | 1.4e-05 | 9.4e-06 | 1.110E-05 | 2.524E-06 | 9.400E-06 | 1.400E-05 |
| 3095 | 155 | N155P | A | 1.5e-05 | 1.4e-05 | 1.5e-05 | 1.467E-05 | 5.774E-07 | 1.400E-05 | 1.500E-05 |
| 3096 | 155 | N155S | A | 1.5e-05 | 1.4e-05 | 1.5e-05 | 1.467E-05 | 5.774E-07 | 1.400E-05 | 1.500E-05 |
| 3097 | 155 | N155T | A | 1.6e-05 | 1.5e-05 | 1.1e-05 | 1.400E-05 | 2.646E-06 | 1.100E-05 | 1.600E-05 |
| 3098 | 155 | N155W | A | 9e-06 | 1e-05 | 1.2e-05 | 1.033E-05 | 1.528E-06 | 9.000E-06 | 1.200E-05 |
| 3099 | 155 | N155Y | A | 1.1e-05 | 1.1e-05 | 1.1e-05 | 1.100E-05 | 2.075E-21 | 1.100E-05 | 1.100E-05 |
| 3100 | 155 | N155V | A | 1.5e-05 | 1.6e-05 | 1.1e-05 | 1.400E-05 | 2.646E-06 | 1.100E-05 | 1.600E-05 |
| 3101 | 156 | Q156A | A | 1.6e-05 | 1.3e-05 | 1.1e-05 | 1.333E-05 | 2.517E-06 | 1.100E-05 | 1.600E-05 |
| 3102 | 156 | Q156R | A | 1.1e-05 | 1.5e-05 | 8.9e-06 | 1.163E-05 | 3.099E-06 | 8.900E-06 | 1.500E-05 |
| 3103 | 156 | Q156N | A | 1.1e-05 | 9.2e-06 | 1.5e-05 | 1.173E-05 | 2.969E-06 | 9.200E-06 | 1.500E-05 |
| 3104 | 156 | Q156D | A | 1.5e-05 | 1.4e-05 | 9e-06 | 1.267E-05 | 3.215E-06 | 9.000E-06 | 1.500E-05 |
| 3105 | 156 | Q156C | A | 1.1e-05 | 1.3e-05 | 1.1e-05 | 1.167E-05 | 1.155E-06 | 1.100E-05 | 1.300E-05 |
| 3106 | 156 | Q156E | A | 1.1e-05 | 1.2e-05 | 1.5e-05 | 1.267E-05 | 2.082E-06 | 1.100E-05 | 1.500E-05 |
| 3107 | 156 | Q156Q | A | 9.6e-06 | 1.1e-05 | 1.3e-05 | 1.120E-05 | 1.709E-06 | 9.600E-06 | 1.300E-05 |
| 3108 | 156 | Q156G | A | 1.1e-05 | 1.1e-05 | 9.5e-06 | 1.050E-05 | 8.660E-07 | 9.500E-06 | 1.100E-05 |
| 3109 | 156 | Q156H | A | 1.8e-05 | 1.7e-05 | 1.5e-05 | 1.667E-05 | 1.528E-06 | 1.500E-05 | 1.800E-05 |
| 3110 | 156 | Q156I | A | 9.6e-06 | 1.2e-05 | 1.2e-05 | 1.120E-05 | 1.386E-06 | 9.600E-06 | 1.200E-05 |
| 3111 | 156 | Q156L | A | 1.1e-05 | 1.5e-05 | 1.3e-05 | 1.300E-05 | 2.000E-06 | 1.100E-05 | 1.500E-05 |
| 3112 | 156 | Q156K | A | 1.1e-05 | 1e-05 | 8.4e-06 | 9.800E-06 | 1.311E-06 | 8.400E-06 | 1.100E-05 |
| 3113 | 156 | Q156M | A | 1.4e-05 | 1.2e-05 | 1.3e-05 | 1.300E-05 | 1.000E-06 | 1.200E-05 | 1.400E-05 |
| 3114 | 156 | Q156F | A | 1.2e-05 | 1.5e-05 | 1.3e-05 | 1.333E-05 | 1.528E-06 | 1.200E-05 | 1.500E-05 |
| 3115 | 156 | Q156P | A | 8.9e-06 | 1.5e-05 | 1.3e-05 | 1.230E-05 | 3.110E-06 | 8.900E-06 | 1.500E-05 |
| 3116 | 156 | Q156S | A | 1.4e-05 | 1.2e-05 | 1e-05 | 1.200E-05 | 2.000E-06 | 1.000E-05 | 1.400E-05 |
| 3117 | 156 | Q156T | A | 1e-05 | 1.7e-05 | 1.4e-05 | 1.367E-05 | 3.512E-06 | 1.000E-05 | 1.700E-05 |
| 3118 | 156 | Q156W | A | 1.1e-05 | 1.1e-05 | 1.7e-05 | 1.300E-05 | 3.464E-06 | 1.100E-05 | 1.700E-05 |
| 3119 | 156 | Q156Y | A | 1.5e-05 | 1.7e-05 | 8.9e-06 | 1.363E-05 | 4.219E-06 | 8.900E-06 | 1.700E-05 |
| 3120 | 156 | Q156V | A | 1e-05 | 1e-05 | 1e-05 | 1.000E-05 | 0.000E+00 | 1.000E-05 | 1.000E-05 |
| 3121 | 157 | L157A | A | 1.3e-05 | 9.7e-06 | 1e-05 | 1.090E-05 | 1.825E-06 | 9.700E-06 | 1.300E-05 |
| 3122 | 157 | L157R | A | 1.5e-05 | 1.4e-05 | 1.4e-05 | 1.433E-05 | 5.774E-07 | 1.400E-05 | 1.500E-05 |
| 3123 | 157 | L157N | A | 9.8e-06 | 9.2e-06 | 1.3e-05 | 1.067E-05 | 2.043E-06 | 9.200E-06 | 1.300E-05 |
| 3124 | 157 | L157D | A | 1.1e-05 | 1.2e-05 | 1.6e-05 | 1.300E-05 | 2.646E-06 | 1.100E-05 | 1.600E-05 |
| 3125 | 157 | L157C | A | 9.4e-06 | 1.5e-05 | 1.3e-05 | 1.247E-05 | 2.838E-06 | 9.400E-06 | 1.500E-05 |
| 3126 | 157 | L157E | A | 1e-05 | 9.1e-06 | 7.4e-06 | 8.833E-06 | 1.320E-06 | 7.400E-06 | 1.000E-05 |
| 3127 | 157 | L157Q | A | 8.5e-06 | 1.1e-05 | 1e-05 | 9.833E-06 | 1.258E-06 | 8.500E-06 | 1.100E-05 |
| 3128 | 157 | L157G | A | 9.5e-06 | 1.5e-05 | 1.1e-05 | 1.183E-05 | 2.843E-06 | 9.500E-06 | 1.500E-05 |
| 3129 | 157 | L157H | A | 1.5e-05 | 1.3e-05 | 1e-05 | 1.267E-05 | 2.517E-06 | 1.000E-05 | 1.500E-05 |
| 3130 | 157 | L157I | A | 1.6e-05 | 1.6e-05 | 1.4e-05 | 1.533E-05 | 1.155E-06 | 1.400E-05 | 1.600E-05 |
| 3131 | 157 | L157L | A | 1.8e-05 | 1.5e-05 | 1.6e-05 | 1.633E-05 | 1.528E-06 | 1.500E-05 | 1.800E-05 |
| 3132 | 157 | L157K | A | 1.7e-05 | 1.4e-05 | 1.5e-05 | 1.533E-05 | 1.528E-06 | 1.400E-05 | 1.700E-05 |
| 3133 | 157 | L157M | A | 1e-05 | 1e-05 | 1.5e-05 | 1.167E-05 | 2.887E-06 | 1.000E-05 | 1.500E-05 |
| 3134 | 157 | L157F | A | 1.2e-05 | 1.5e-05 | 1.4e-05 | 1.367E-05 | 1.528E-06 | 1.200E-05 | 1.500E-05 |
| 3135 | 157 | L157P | A | 1.3e-05 | 9.9e-06 | 1.1e-05 | 1.130E-05 | 1.572E-06 | 9.900E-06 | 1.300E-05 |
| 3136 | 157 | L157S | A | 1.3e-05 | 1.4e-05 | 1e-05 | 1.233E-05 | 2.082E-06 | 1.000E-05 | 1.400E-05 |
| 3137 | 157 | L157T | A | 1.4e-05 | 1.1e-05 | 1.2e-05 | 1.233E-05 | 1.528E-06 | 1.100E-05 | 1.400E-05 |
| 3138 | 157 | L157W | A | 1e-05 | 1.2e-05 | 9.3e-06 | 1.043E-05 | 1.401E-06 | 9.300E-06 | 1.200E-05 |
| 3139 | 157 | L157Y | A | 1.2e-05 | 1.1e-05 | 1e-05 | 1.100E-05 | 1.000E-06 | 1.000E-05 | 1.200E-05 |
| 3140 | 157 | L157V | A | 8.8e-06 | 9.4e-06 | 1.4e-05 | 1.073E-05 | 2.845E-06 | 8.800E-06 | 1.400E-05 |
| 3141 | 158 | A158A | A | 1e-05 | 1.3e-05 | 1.1e-05 | 1.133E-05 | 1.528E-06 | 1.000E-05 | 1.300E-05 |
| 3142 | 158 | A158R | A | 8.7e-06 | 1.3e-05 | 1.5e-05 | 1.223E-05 | 3.219E-06 | 8.700E-06 | 1.500E-05 |
| 3143 | 158 | A158N | A | 1.5e-05 | 9.5e-06 | 1.4e-05 | 1.283E-05 | 2.930E-06 | 9.500E-06 | 1.500E-05 |
| 3144 | 158 | A158D | A | 1.5e-05 | 1.1e-05 | 9.7e-06 | 1.190E-05 | 2.762E-06 | 9.700E-06 | 1.500E-05 |

| | | | | | | | | | | |
|---|---|---|---|---|---|---|---|---|---|---|
| 3145 | 158 | A158C | A | 8.7e-06 | 1.1e-05 | 1e-05 | 9.900E-06 | 1.153E-06 | 8.700E-06 | 1.100E-05 |
| 3146 | 158 | A158E | A | 1.5e-05 | 8.5e-06 | 1.6e-05 | 1.317E-05 | 4.072E-06 | 8.500E-06 | 1.600E-05 |
| 3147 | 158 | A158Q | A | 1.1e-05 | 9.2e-06 | 1.5e-05 | 1.173E-05 | 2.969E-06 | 9.200E-06 | 1.500E-05 |
| 3148 | 158 | A158G | A | 8.6e-06 | 1.6e-05 | 1.6e-05 | 1.353E-05 | 4.272E-06 | 8.600E-06 | 1.600E-05 |
| 3149 | 158 | A158H | A | 9.4e-06 | 1.4e-05 | 1e-05 | 1.113E-05 | 2.501E-06 | 9.400E-06 | 1.400E-05 |
| 3150 | 158 | A158I | A | 1.4e-05 | 9.1e-06 | 1.4e-05 | 1.237E-05 | 2.829E-06 | 9.100E-06 | 1.400E-05 |
| 3151 | 158 | A158L | A | 1.5e-05 | 1.5e-05 | 1.4e-05 | 1.467E-05 | 5.774E-07 | 1.400E-05 | 1.500E-05 |
| 3152 | 158 | A158K | A | 1.3e-05 | 1.5e-05 | 1e-05 | 1.267E-05 | 2.517E-06 | 1.000E-05 | 1.500E-05 |
| 3153 | 158 | A158M | A | 1.3e-05 | 1.7e-05 | 1.6e-05 | 1.533E-05 | 2.082E-06 | 1.300E-05 | 1.700E-05 |
| 3154 | 158 | A158F | A | 1.5e-05 | 9.9e-06 | 9.4e-06 | 1.143E-05 | 3.099E-06 | 9.400E-06 | 1.500E-05 |
| 3155 | 158 | A158P | A | 1.4e-05 | 1e-05 | 1.1e-05 | 1.167E-05 | 2.082E-06 | 1.000E-05 | 1.400E-05 |
| 3156 | 158 | A158S | A | 1.4e-05 | 7e-06 | 8.9e-06 | 9.967E-06 | 3.620E-06 | 7.000E-06 | 1.400E-05 |
| 3157 | 158 | A158T | A | 1e-05 | 1.1e-05 | 8e-06 | 9.667E-06 | 1.528E-06 | 8.000E-06 | 1.100E-05 |
| 3158 | 158 | A158W | A | 1.5e-05 | 1.4e-05 | 1.4e-05 | 1.433E-05 | 5.774E-07 | 1.400E-05 | 1.500E-05 |
| 3159 | 158 | A158Y | A | 1.4e-05 | 1.3e-05 | 1.3e-05 | 1.333E-05 | 5.774E-07 | 1.300E-05 | 1.400E-05 |
| 3160 | 158 | A158V | A | 1.4e-05 | 1.3e-05 | 1e-05 | 1.233E-05 | 2.082E-06 | 1.000E-05 | 1.400E-05 |
| 3161 | 159 | L159A | A | 1.5e-05 | 1.1e-05 | 1e-05 | 1.200E-05 | 2.646E-06 | 1.000E-05 | 1.500E-05 |
| 3162 | 159 | L159R | A | 1.7e-05 | 1.4e-05 | 7.7e-06 | 1.290E-05 | 4.747E-06 | 7.700E-06 | 1.700E-05 |
| 3163 | 159 | L159N | A | 1.4e-05 | 9.9e-06 | 1.6e-05 | 1.330E-05 | 3.110E-06 | 9.900E-06 | 1.600E-05 |
| 3164 | 159 | L159D | A | 1.3e-05 | 1.5e-05 | 1.3e-05 | 1.367E-05 | 1.155E-06 | 1.300E-05 | 1.500E-05 |
| 3165 | 159 | L159C | A | 1.1e-05 | 9.7e-06 | 1.3e-05 | 1.123E-05 | 1.662E-06 | 9.700E-06 | 1.300E-05 |
| 3166 | 159 | L159E | A | 1.1e-05 | 1.6e-05 | 1.7e-05 | 1.467E-05 | 3.215E-06 | 1.100E-05 | 1.700E-05 |
| 3167 | 159 | L159Q | A | 1e-05 | 8.8e-06 | 1e-05 | 9.600E-06 | 6.928E-07 | 8.800E-06 | 1.000E-05 |
| 3168 | 159 | L159G | A | 1.5e-05 | 1.5e-05 | 1.1e-05 | 1.367E-05 | 2.309E-06 | 1.100E-05 | 1.500E-05 |
| 3169 | 159 | L159H | A | 1.4e-05 | 1.4e-05 | 8.4e-06 | 1.213E-05 | 3.233E-06 | 8.400E-06 | 1.400E-05 |
| 3170 | 159 | L159I | A | 1.1e-05 | 1e-05 | 1.1e-05 | 1.067E-05 | 5.774E-07 | 1.000E-05 | 1.100E-05 |
| 3171 | 159 | L159L | A | 1.4e-05 | 1.1e-05 | 1.3e-05 | 1.267E-05 | 1.528E-06 | 1.100E-05 | 1.400E-05 |
| 3172 | 159 | L159K | A | 9.2e-06 | 1.5e-05 | 1.4e-05 | 1.273E-05 | 3.101E-06 | 9.200E-06 | 1.500E-05 |
| 3173 | 159 | L159M | A | 1.5e-05 | 1e-05 | 1.6e-05 | 1.367E-05 | 3.215E-06 | 1.000E-05 | 1.600E-05 |
| 3174 | 159 | L159F | A | 1.6e-05 | 1.5e-05 | 1.6e-05 | 1.567E-05 | 5.774E-07 | 1.500E-05 | 1.600E-05 |
| 3175 | 159 | L159P | A | 1.5e-05 | 1.5e-05 | 9.4e-06 | 1.313E-05 | 3.233E-06 | 9.400E-06 | 1.500E-05 |
| 3176 | 159 | L159S | A | 1.5e-05 | 1.1e-05 | 7.3e-06 | 1.110E-05 | 3.851E-06 | 7.300E-06 | 1.500E-05 |
| 3177 | 159 | L159T | A | 1.4e-05 | 8.8e-06 | 1.3e-05 | 1.193E-05 | 2.759E-06 | 8.800E-06 | 1.400E-05 |
| 3178 | 159 | L159W | A | 1.1e-05 | 1.1e-05 | 1.4e-05 | 1.200E-05 | 1.732E-06 | 1.100E-05 | 1.400E-05 |
| 3179 | 159 | L159Y | A | 1.5e-05 | 9.4e-06 | 1.5e-05 | 1.313E-05 | 3.233E-06 | 9.400E-06 | 1.500E-05 |
| 3180 | 159 | L159V | A | 9.6e-06 | 1.8e-05 | 1.1e-05 | 1.287E-05 | 4.500E-06 | 9.600E-06 | 1.800E-05 |
| 3181 | 160 | T160A | A | 9.5e-06 | 1.2e-05 | 1.4e-05 | 1.183E-05 | 2.255E-06 | 9.500E-06 | 1.400E-05 |
| 3182 | 160 | T160R | A | 1.6e-05 | 2.9e-05 | 1.6e-05 | 2.033E-05 | 7.506E-06 | 1.600E-05 | 2.900E-05 |
| 3183 | 160 | T160N | A | 9.4e-06 | 1.2e-05 | 1.4e-05 | 1.180E-05 | 2.307E-06 | 9.400E-06 | 1.400E-05 |
| 3184 | 160 | T160D | A | 1.5e-05 | 1.2e-05 | 1.6e-05 | 1.433E-05 | 2.082E-06 | 1.200E-05 | 1.600E-05 |
| 3185 | 160 | T160C | A | 1.5e-05 | 1.5e-05 | 1.4e-05 | 1.467E-05 | 5.774E-07 | 1.400E-05 | 1.500E-05 |
| 3186 | 160 | T160E | A | 1.4e-05 | 1.5e-05 | 1e-05 | 1.300E-05 | 2.646E-06 | 1.000E-05 | 1.500E-05 |
| 3187 | 160 | T160Q | A | 1.4e-05 | 1e-05 | 1.5e-05 | 1.300E-05 | 2.646E-06 | 1.000E-05 | 1.500E-05 |
| 3188 | 160 | T160G | A | 1.4e-05 | 9.8e-06 | 1.5e-05 | 1.293E-05 | 2.759E-06 | 9.800E-06 | 1.500E-05 |
| 3189 | 160 | T160H | A | 1.5e-05 | 9.4e-06 | 8.9e-06 | 1.110E-05 | 3.387E-06 | 8.900E-06 | 1.500E-05 |
| 3190 | 160 | T160I | A | 8.5e-06 | 1.5e-05 | 1.3e-05 | 1.217E-05 | 3.329E-06 | 8.500E-06 | 1.500E-05 |
| 3191 | 160 | T160L | A | 8.5e-06 | 9.4e-06 | 1.5e-05 | 1.097E-05 | 3.522E-06 | 8.500E-06 | 1.500E-05 |
| 3192 | 160 | T160K | A | 9.1e-06 | 1.6e-05 | 1.5e-05 | 1.337E-05 | 3.729E-06 | 9.100E-06 | 1.600E-05 |
| 3193 | 160 | T160M | A | 1.1e-05 | 1.1e-05 | 9.7e-06 | 1.057E-05 | 7.506E-07 | 9.700E-06 | 1.100E-05 |
| 3194 | 160 | T160F | A | 1.3e-05 | 1e-05 | 1.1e-05 | 1.133E-05 | 1.528E-06 | 1.000E-05 | 1.300E-05 |
| 3195 | 160 | T160P | A | 1.1e-05 | 1.6e-05 | 1.3e-05 | 1.333E-05 | 2.517E-06 | 1.100E-05 | 1.600E-05 |
| 3196 | 160 | T160S | A | 1.7e-05 | 9.8e-06 | 1.6e-05 | 1.427E-05 | 3.900E-06 | 9.800E-06 | 1.700E-05 |
| 3197 | 160 | T160T | A | 1.3e-05 | 1.6e-05 | 9.2e-06 | 1.273E-05 | 3.408E-06 | 9.200E-06 | 1.600E-05 |
| 3198 | 160 | T160W | A | 1.5e-05 | 9.5e-06 | 1.3e-05 | 1.250E-05 | 2.784E-06 | 9.500E-06 | 1.500E-05 |
| 3199 | 160 | T160Y | A | 1.4e-05 | 1.1e-05 | 9.8e-06 | 1.160E-05 | 2.163E-06 | 9.800E-06 | 1.400E-05 |
| 3200 | 160 | T160V | A | 8.6e-06 | 1.1e-05 | 1.1e-05 | 1.020E-05 | 1.386E-06 | 8.600E-06 | 1.100E-05 |
| 3201 | 161 | L161A | A | 1.6e-05 | 7e-06 | 6e-06 | 9.667E-06 | 5.508E-06 | 6.000E-06 | 1.600E-05 |
| 3202 | 161 | L161R | A | 9.1e-06 | 1.1e-05 | 1.1e-05 | 1.037E-05 | 1.097E-06 | 9.100E-06 | 1.100E-05 |
| 3203 | 161 | L161N | A | 1.5e-05 | 1.4e-05 | 1.4e-05 | 1.433E-05 | 5.774E-07 | 1.400E-05 | 1.500E-05 |
| 3204 | 161 | L161D | A | 1.5e-05 | 1.6e-05 | 1.3e-05 | 1.467E-05 | 1.528E-06 | 1.300E-05 | 1.600E-05 |
| 3205 | 161 | L161C | A | 8.7e-06 | 1.5e-05 | 1.5e-05 | 1.290E-05 | 3.637E-06 | 8.700E-06 | 1.500E-05 |
| 3206 | 161 | L161E | A | 1.5e-05 | 1.4e-05 | 9.5e-06 | 1.283E-05 | 2.930E-06 | 9.500E-06 | 1.500E-05 |
| 3207 | 161 | L161Q | A | 9.8e-06 | 1.1e-05 | 7.8e-06 | 9.533E-06 | 1.617E-06 | 7.800E-06 | 1.100E-05 |
| 3208 | 161 | L161G | A | 1.1e-05 | 1.3e-05 | 1.1e-05 | 1.167E-05 | 1.155E-06 | 1.100E-05 | 1.300E-05 |
| 3209 | 161 | L161H | A | 1.5e-05 | 1.5e-05 | 8.6e-06 | 1.287E-05 | 3.695E-06 | 8.600E-06 | 1.500E-05 |
| 3210 | 161 | L161I | A | 1.2e-05 | 1.5e-05 | 1.5e-05 | 1.400E-05 | 1.732E-06 | 1.200E-05 | 1.500E-05 |
| 3211 | 161 | L161L | A | 1.3e-05 | 1.4e-05 | 1e-05 | 1.233E-05 | 2.082E-06 | 1.000E-05 | 1.400E-05 |
| 3212 | 161 | L161K | A | 1.7e-05 | 1.5e-05 | 7.8e-06 | 1.327E-05 | 4.839E-06 | 7.800E-06 | 1.700E-05 |
| 3213 | 161 | L161M | A | 9.6e-06 | 1.5e-05 | 1.5e-05 | 1.320E-05 | 3.118E-06 | 9.600E-06 | 1.500E-05 |
| 3214 | 161 | L161F | A | 1.1e-05 | 1.4e-05 | 1.6e-05 | 1.367E-05 | 2.517E-06 | 1.100E-05 | 1.600E-05 |
| 3215 | 161 | L161P | A | 1.5e-05 | 1.2e-05 | 1.2e-05 | 1.300E-05 | 1.732E-06 | 1.200E-05 | 1.500E-05 |
| 3216 | 161 | L161S | A | 1e-05 | 1.5e-05 | 1.5e-05 | 1.333E-05 | 2.887E-06 | 1.000E-05 | 1.500E-05 |
| 3217 | 161 | L161T | A | 1.3e-05 | 9.8e-06 | 2.1e-05 | 1.460E-05 | 5.769E-06 | 9.800E-06 | 2.100E-05 |
| 3218 | 161 | L161W | A | 1e-05 | 1.3e-05 | 1.4e-05 | 1.233E-05 | 2.082E-06 | 1.000E-05 | 1.400E-05 |
| 3219 | 161 | L161Y | A | 1.4e-05 | 9.6e-06 | 1.4e-05 | 1.253E-05 | 2.540E-06 | 9.600E-06 | 1.400E-05 |
| 3220 | 161 | L161V | A | 1.2e-05 | 1.2e-05 | 1.3e-05 | 1.233E-05 | 5.774E-07 | 1.200E-05 | 1.300E-05 |
| 3221 | 162 | I162A | A | 1.3e-05 | 1.5e-05 | 8.7e-06 | 1.223E-05 | 3.219E-06 | 8.700E-06 | 1.500E-05 |
| 3222 | 162 | I162R | A | 1.6e-05 | 9.4e-06 | 1.4e-05 | 1.313E-05 | 3.384E-06 | 9.400E-06 | 1.600E-05 |
| 3223 | 162 | I162N | A | 1.3e-05 | 1.2e-05 | 7.8e-06 | 1.093E-05 | 2.759E-06 | 7.800E-06 | 1.300E-05 |
| 3224 | 162 | I162D | A | 1.5e-05 | 8.2e-06 | 1.8e-05 | 1.373E-05 | 5.021E-06 | 8.200E-06 | 1.800E-05 |
| 3225 | 162 | I162C | A | 9.1e-06 | 1e-05 | 8.3e-06 | 9.133E-06 | 8.505E-07 | 8.300E-06 | 1.000E-05 |
| 3226 | 162 | I162E | A | 1.3e-05 | 1.1e-05 | 9.5e-06 | 1.117E-05 | 1.756E-06 | 9.500E-06 | 1.300E-05 |
| 3227 | 162 | I162Q | A | 9.2e-06 | 3.7e-05 | 1.1e-05 | 1.907E-05 | 1.556E-05 | 9.200E-06 | 3.700E-05 |
| 3228 | 162 | I162G | A | 1.5e-05 | 1e-05 | 6.5e-06 | 1.050E-05 | 4.272E-06 | 6.500E-06 | 1.500E-05 |
| 3229 | 162 | I162H | A | 1.3e-05 | 1.3e-05 | 1.5e-05 | 1.367E-05 | 1.155E-06 | 1.300E-05 | 1.500E-05 |

| | | | | | | | | | | |
|---|---|---|---|---|---|---|---|---|---|---|
| 3230 | 162 | I162I | A | 9.9e-06 | 1.1e-05 | 1.1e-05 | 1.063E-05 | 6.351E-07 | 9.900E-06 | 1.100E-05 |
| 3231 | 162 | I162L | A | 1.3e-05 | 1.1e-05 | 1e-05 | 1.133E-05 | 1.528E-06 | 1.000E-05 | 1.300E-05 |
| 3232 | 162 | I162K | A | 1.8e-05 | 1.8e-05 | 8.2e-06 | 1.473E-05 | 5.658E-06 | 8.200E-06 | 1.800E-05 |
| 3233 | 162 | I162M | A | 1.4e-05 | 8.6e-06 | 1.6e-05 | 1.287E-05 | 3.828E-06 | 8.600E-06 | 1.600E-05 |
| 3234 | 162 | I162F | A | 9.7e-06 | 1.4e-05 | 1.5e-05 | 1.290E-05 | 2.816E-06 | 9.700E-06 | 1.500E-05 |
| 3235 | 162 | I162P | A | 1.1e-05 | 1.1e-05 | 1e-05 | 1.067E-05 | 5.774E-07 | 1.000E-05 | 1.100E-05 |
| 3236 | 162 | I162S | A | 9.8e-06 | 1.5e-05 | 1.6e-05 | 1.360E-05 | 3.329E-06 | 9.800E-06 | 1.600E-05 |
| 3237 | 162 | I162T | A | 1.1e-05 | 9.5e-06 | 1.6e-05 | 1.217E-05 | 3.403E-06 | 9.500E-06 | 1.600E-05 |
| 3238 | 162 | I162W | A | 1e-05 | 1.5e-05 | 9.6e-06 | 1.153E-05 | 3.009E-06 | 9.600E-06 | 1.500E-05 |
| 3239 | 162 | I162Y | A | 1e-05 | 8.9e-06 | 1.4e-05 | 1.097E-05 | 2.684E-06 | 8.900E-06 | 1.400E-05 |
| 3240 | 162 | I162V | A | 1.2e-05 | 1.2e-05 | 1.5e-05 | 1.300E-05 | 1.732E-06 | 1.200E-05 | 1.500E-05 |
| 3241 | 163 | D163A | A | 1.1e-05 | 1e-05 | 1.7e-05 | 1.267E-05 | 3.786E-06 | 1.000E-05 | 1.700E-05 |
| 3242 | 163 | D163R | A | 1e-05 | 1.7e-05 | 1.4e-05 | 1.367E-05 | 3.512E-06 | 1.000E-05 | 1.700E-05 |
| 3243 | 163 | D163N | A | 1.1e-05 | 1.3e-05 | 1.4e-05 | 1.267E-05 | 1.528E-06 | 1.100E-05 | 1.400E-05 |
| 3244 | 163 | D163D | A | 1.5e-05 | 1.3e-05 | 1.8e-05 | 1.533E-05 | 2.517E-06 | 1.300E-05 | 1.800E-05 |
| 3245 | 163 | D163C | A | 1.3e-05 | 8.8e-06 | 1.5e-05 | 1.227E-05 | 3.164E-06 | 8.800E-06 | 1.500E-05 |
| 3246 | 163 | D163E | A | 1e-05 | 1.7e-05 | 1.6e-05 | 1.433E-05 | 3.786E-06 | 1.000E-05 | 1.700E-05 |
| 3247 | 163 | D163Q | A | 1.5e-05 | 1.7e-05 | 1.2e-05 | 1.467E-05 | 2.517E-06 | 1.200E-05 | 1.700E-05 |
| 3248 | 163 | D163G | A | 1.3e-05 | 1.5e-05 | 1.5e-05 | 1.433E-05 | 1.155E-06 | 1.300E-05 | 1.500E-05 |
| 3249 | 163 | D163H | A | 1.6e-05 | 1.5e-05 | 1.8e-05 | 1.633E-05 | 1.528E-06 | 1.500E-05 | 1.800E-05 |
| 3250 | 163 | D163I | A | 9.5e-06 | 1.5e-05 | 1.5e-05 | 1.317E-05 | 3.175E-06 | 9.500E-06 | 1.500E-05 |
| 3251 | 163 | D163L | A | 1.4e-05 | 1e-05 | 8.4e-06 | 1.080E-05 | 2.884E-06 | 8.400E-06 | 1.400E-05 |
| 3252 | 163 | D163K | A | 1.4e-05 | 1.1e-05 | 1.4e-05 | 1.300E-05 | 1.732E-06 | 1.100E-05 | 1.400E-05 |
| 3253 | 163 | D163M | A | 1.5e-05 | 1.1e-05 | 1.5e-05 | 1.367E-05 | 2.309E-06 | 1.100E-05 | 1.500E-05 |
| 3254 | 163 | D163F | A | 1.3e-05 | 1.5e-05 | 1.3e-05 | 1.367E-05 | 1.155E-06 | 1.300E-05 | 1.500E-05 |
| 3255 | 163 | D163P | A | 1.3e-05 | 1e-05 | 1.6e-05 | 1.300E-05 | 3.000E-06 | 1.000E-05 | 1.600E-05 |
| 3256 | 163 | D163S | A | 8.4e-06 | 8.4e-06 | 1.2e-05 | 9.600E-06 | 2.078E-06 | 8.400E-06 | 1.200E-05 |
| 3257 | 163 | D163T | A | 1.7e-05 | 1.4e-05 | 8e-06 | 1.300E-05 | 4.583E-06 | 8.000E-06 | 1.700E-05 |
| 3258 | 163 | D163W | A | 1e-05 | 9.9e-06 | 1.1e-05 | 1.030E-05 | 6.083E-07 | 9.900E-06 | 1.100E-05 |
| 3259 | 163 | D163Y | A | 1.4e-05 | 1.5e-05 | 1.5e-05 | 1.467E-05 | 5.774E-07 | 1.400E-05 | 1.500E-05 |
| 3260 | 163 | D163V | A | 1.6e-05 | 1.2e-05 | 1.5e-05 | 1.433E-05 | 2.082E-06 | 1.200E-05 | 1.600E-05 |
| 3261 | 164 | T164A | A | 1.4e-05 | 8.5e-06 | 1.2e-05 | 1.150E-05 | 2.784E-06 | 8.500E-06 | 1.400E-05 |
| 3262 | 164 | T164R | A | 9.9e-06 | 1.2e-05 | 1.1e-05 | 1.097E-05 | 1.050E-06 | 9.900E-06 | 1.200E-05 |
| 3263 | 164 | T164N | A | 9.5e-06 | 8.4e-06 | 1.4e-05 | 1.063E-05 | 2.967E-06 | 8.400E-06 | 1.400E-05 |
| 3264 | 164 | T164D | A | 9.4e-06 | 8.4e-06 | 1e-05 | 9.267E-06 | 8.083E-07 | 8.400E-06 | 1.000E-05 |
| 3265 | 164 | T164C | A | 1.1e-05 | 1e-05 | 1.2e-05 | 1.100E-05 | 1.000E-06 | 1.000E-05 | 1.200E-05 |
| 3266 | 164 | T164E | A | 1.1e-05 | 1e-05 | 1.1e-05 | 1.067E-05 | 5.774E-07 | 1.000E-05 | 1.100E-05 |
| 3267 | 164 | T164Q | A | 1e-05 | 1.6e-05 | 1.5e-05 | 1.367E-05 | 3.215E-06 | 1.000E-05 | 1.600E-05 |
| 3268 | 164 | T164G | A | 1.4e-05 | 1.6e-05 | 1.1e-05 | 1.367E-05 | 2.517E-06 | 1.100E-05 | 1.600E-05 |
| 3269 | 164 | T164H | A | 1.4e-05 | 1e-05 | 1e-05 | 1.133E-05 | 2.309E-06 | 1.000E-05 | 1.400E-05 |
| 3270 | 164 | T164I | A | 1.3e-05 | 1.5e-05 | 1.6e-05 | 1.467E-05 | 1.528E-06 | 1.300E-05 | 1.600E-05 |
| 3271 | 164 | T164L | A | 1.6e-05 | 1.1e-05 | 1.4e-05 | 1.367E-05 | 2.517E-06 | 1.100E-05 | 1.600E-05 |
| 3272 | 164 | T164K | A | 1.1e-05 | 9.7e-06 | 1.1e-05 | 1.057E-05 | 7.506E-07 | 9.700E-06 | 1.100E-05 |
| 3273 | 164 | T164M | A | 1e-05 | 1.5e-05 | 1e-05 | 1.167E-05 | 2.887E-06 | 1.000E-05 | 1.500E-05 |
| 3274 | 164 | T164F | A | 1.6e-05 | 1.1e-05 | 1.6e-05 | 1.433E-05 | 2.887E-06 | 1.100E-05 | 1.600E-05 |
| 3275 | 164 | T164P | A | 8.3e-06 | 1.1e-05 | 7.9e-06 | 9.067E-06 | 1.686E-06 | 7.900E-06 | 1.100E-05 |
| 3276 | 164 | T164S | A | 1.3e-05 | 1e-05 | 9.7e-06 | 1.090E-05 | 1.825E-06 | 9.700E-06 | 1.300E-05 |
| 3277 | 164 | T164T | A | 1.1e-05 | 8.5e-06 | 1e-05 | 9.833E-06 | 1.258E-06 | 8.500E-06 | 1.100E-05 |
| 3278 | 164 | T164W | A | 1.7e-05 | 1.1e-05 | 1.5e-05 | 1.433E-05 | 3.055E-06 | 1.100E-05 | 1.700E-05 |
| 3279 | 164 | T164Y | A | 1.7e-05 | 1e-05 | 1.4e-05 | 1.367E-05 | 3.512E-06 | 1.000E-05 | 1.700E-05 |
| 3280 | 164 | T164V | A | 1.4e-05 | 1.1e-05 | 1.1e-05 | 1.200E-05 | 1.732E-06 | 1.100E-05 | 1.400E-05 |
| 3281 | 165 | N165A | A | 1.6e-05 | 1.1e-05 | 1.2e-05 | 1.300E-05 | 2.646E-06 | 1.100E-05 | 1.600E-05 |
| 3282 | 165 | N165R | A | 1.3e-05 | 1.3e-05 | 1.5e-05 | 1.367E-05 | 1.155E-06 | 1.300E-05 | 1.500E-05 |
| 3283 | 165 | N165N | A | 1e-05 | 1.4e-05 | 1.5e-05 | 1.300E-05 | 2.646E-06 | 1.000E-05 | 1.500E-05 |
| 3284 | 165 | N165D | A | 1.7e-05 | 1.4e-05 | 1.4e-05 | 1.500E-05 | 1.732E-06 | 1.400E-05 | 1.700E-05 |
| 3285 | 165 | N165C | A | 1.7e-05 | 8.3e-06 | 1.6e-05 | 1.377E-05 | 4.761E-06 | 8.300E-06 | 1.700E-05 |
| 3286 | 165 | N165E | A | 1.6e-05 | 1.1e-05 | 1.4e-05 | 1.367E-05 | 2.517E-06 | 1.100E-05 | 1.600E-05 |
| 3287 | 165 | N165Q | A | 1.2e-05 | 1.4e-05 | 1.4e-05 | 1.333E-05 | 1.155E-06 | 1.200E-05 | 1.400E-05 |
| 3288 | 165 | N165G | A | 8.3e-06 | 9.2e-06 | 1.9e-05 | 1.217E-05 | 5.935E-06 | 8.300E-06 | 1.900E-05 |
| 3289 | 165 | N165H | A | 1.2e-05 | 1.4e-05 | 1e-05 | 1.200E-05 | 2.000E-06 | 1.000E-05 | 1.400E-05 |
| 3290 | 165 | N165I | A | 1e-05 | 1.1e-05 | 9.8e-06 | 1.027E-05 | 6.429E-07 | 9.800E-06 | 1.100E-05 |
| 3291 | 165 | N165L | A | 1.5e-05 | 9.9e-06 | 6.8e-06 | 1.057E-05 | 4.140E-06 | 6.800E-06 | 1.500E-05 |
| 3292 | 165 | N165K | A | 1.4e-05 | 1.2e-05 | 1.6e-05 | 1.400E-05 | 2.000E-06 | 1.200E-05 | 1.600E-05 |
| 3293 | 165 | N165M | A | 1.6e-05 | 1.9e-05 | 1e-05 | 1.500E-05 | 4.583E-06 | 1.000E-05 | 1.900E-05 |
| 3294 | 165 | N165F | A | 1.3e-05 | 1.5e-05 | 1.5e-05 | 1.433E-05 | 1.155E-06 | 1.300E-05 | 1.500E-05 |
| 3295 | 165 | N165P | A | 1.6e-05 | 1.6e-05 | 1.5e-05 | 1.567E-05 | 5.774E-07 | 1.500E-05 | 1.600E-05 |
| 3296 | 165 | N165S | A | 1.7e-05 | 9.7e-06 | 1.5e-05 | 1.390E-05 | 3.772E-06 | 9.700E-06 | 1.700E-05 |
| 3297 | 165 | N165T | A | 1.6e-05 | 1.3e-05 | 9.1e-06 | 1.270E-05 | 3.460E-06 | 9.100E-06 | 1.600E-05 |
| 3298 | 165 | N165W | A | 1e-05 | 1.3e-05 | 1e-05 | 1.100E-05 | 1.732E-06 | 1.000E-05 | 1.300E-05 |
| 3299 | 165 | N165Y | A | 1.5e-05 | 1.3e-05 | 1.5e-05 | 1.433E-05 | 1.155E-06 | 1.300E-05 | 1.500E-05 |
| 3300 | 165 | N165V | A | 1.5e-05 | 9.7e-06 | 1.3e-05 | 1.257E-05 | 2.676E-06 | 9.700E-06 | 1.500E-05 |
| 3301 | 166 | R166A | A | 9.7e-06 | 1.6e-05 | 1.4e-05 | 1.323E-05 | 3.219E-06 | 9.700E-06 | 1.600E-05 |
| 3302 | 166 | R166R | A | 1e-05 | 1.5e-05 | 8.3e-06 | 1.110E-05 | 3.483E-06 | 8.300E-06 | 1.500E-05 |
| 3303 | 166 | R166N | A | 1e-05 | 1.6e-05 | 1.2e-05 | 1.267E-05 | 3.055E-06 | 1.000E-05 | 1.600E-05 |
| 3304 | 166 | R166D | A | 1.6e-05 | 1.5e-05 | 9e-06 | 1.333E-05 | 3.786E-06 | 9.000E-06 | 1.600E-05 |
| 3305 | 166 | R166C | A | 1.3e-05 | 9.2e-06 | 1.4e-05 | 1.207E-05 | 2.532E-06 | 9.200E-06 | 1.400E-05 |
| 3306 | 166 | R166E | A | 9.2e-06 | 1.1e-05 | 1.2e-05 | 1.073E-05 | 1.419E-06 | 9.200E-06 | 1.200E-05 |
| 3307 | 166 | R166Q | A | 1.4e-05 | 7.9e-06 | 1.4e-05 | 1.197E-05 | 3.522E-06 | 7.900E-06 | 1.400E-05 |
| 3308 | 166 | R166G | A | 9.6e-06 | 1.2e-05 | 1.5e-05 | 1.220E-05 | 2.706E-06 | 9.600E-06 | 1.500E-05 |
| 3309 | 166 | R166H | A | 1.2e-05 | 1.2e-05 | 7.1e-06 | 1.037E-05 | 2.829E-06 | 7.100E-06 | 1.200E-05 |
| 3310 | 166 | R166I | A | 1.1e-05 | 1.6e-05 | 7.1e-06 | 1.137E-05 | 4.461E-06 | 7.100E-06 | 1.600E-05 |
| 3311 | 166 | R166L | A | 9.2e-06 | 1.5e-05 | 1.5e-05 | 1.307E-05 | 3.349E-06 | 9.200E-06 | 1.500E-05 |
| 3312 | 166 | R166K | A | 1.6e-05 | 6.6e-06 | 1.2e-05 | 1.153E-05 | 4.717E-06 | 6.600E-06 | 1.600E-05 |
| 3313 | 166 | R166M | A | 1.5e-05 | 1.6e-05 | 1.7e-05 | 1.600E-05 | 1.000E-06 | 1.500E-05 | 1.700E-05 |
| 3314 | 166 | R166F | A | 1.9e-05 | 1.6e-05 | 1e-05 | 1.500E-05 | 4.583E-06 | 1.000E-05 | 1.900E-05 |

| | | | | | | | | | | |
|---|---|---|---|---|---|---|---|---|---|---|
| 3315 | 166 | R166P | A | 1.2e-05 | 1.5e-05 | 1.5e-05 | 1.400E-05 | 1.732E-06 | 1.200E-05 | 1.500E-05 |
| 3316 | 166 | R166S | A | 1.5e-05 | 1.7e-05 | 1.4e-05 | 1.533E-05 | 1.528E-06 | 1.400E-05 | 1.700E-05 |
| 3317 | 166 | R166T | A | 1.4e-05 | 9.2e-06 | 1.5e-05 | 1.273E-05 | 3.101E-06 | 9.200E-06 | 1.500E-05 |
| 3318 | 166 | R166W | A | 1.1e-05 | 1.5e-05 | 1.4e-05 | 1.333E-05 | 2.082E-06 | 1.100E-05 | 1.500E-05 |
| 3319 | 166 | R166Y | A | 1.5e-05 | 1.6e-05 | 1e-05 | 1.367E-05 | 3.215E-06 | 1.000E-05 | 1.600E-05 |
| 3320 | 166 | R166V | A | 1.8e-05 | 1.7e-05 | 1.4e-05 | 1.633E-05 | 2.082E-06 | 1.400E-05 | 1.800E-05 |
| 3321 | 167 | S167A | A | 1.7e-05 | 9.8e-06 | 1.4e-05 | 1.360E-05 | 3.617E-06 | 9.800E-06 | 1.700E-05 |
| 3322 | 167 | S167R | A | 1.6e-05 | 1.4e-05 | 1.1e-05 | 1.367E-05 | 2.517E-06 | 1.100E-05 | 1.600E-05 |
| 3323 | 167 | S167N | A | 1.2e-05 | 1.1e-05 | 9.6e-06 | 1.087E-05 | 1.206E-06 | 9.600E-06 | 1.200E-05 |
| 3324 | 167 | S167D | A | 1.1e-05 | 1.6e-05 | 1.1e-05 | 1.267E-05 | 2.887E-06 | 1.100E-05 | 1.600E-05 |
| 3325 | 167 | S167C | A | 9.9e-06 | 1.1e-05 | 1.1e-05 | 1.063E-05 | 6.351E-07 | 9.900E-06 | 1.100E-05 |
| 3326 | 167 | S167E | A | 1.5e-05 | 1.6e-05 | 9.8e-06 | 1.360E-05 | 3.329E-06 | 9.800E-06 | 1.600E-05 |
| 3327 | 167 | S167Q | A | 1.1e-05 | 1.3e-05 | 1e-05 | 1.133E-05 | 1.528E-06 | 1.000E-05 | 1.300E-05 |
| 3328 | 167 | S167G | A | 1.3e-05 | 1.5e-05 | 1.1e-05 | 1.300E-05 | 2.000E-06 | 1.100E-05 | 1.500E-05 |
| 3329 | 167 | S167H | A | 1.5e-05 | 9.9e-06 | 8.6e-06 | 1.117E-05 | 3.383E-06 | 8.600E-06 | 1.500E-05 |
| 3330 | 167 | S167I | A | 1.1e-05 | 1.5e-05 | 1e-05 | 1.200E-05 | 2.646E-06 | 1.000E-05 | 1.500E-05 |
| 3331 | 167 | S167L | A | 1.6e-05 | 1.6e-05 | 1.5e-05 | 1.567E-05 | 5.774E-07 | 1.500E-05 | 1.600E-05 |
| 3332 | 167 | S167K | A | 1e-05 | 1.5e-05 | 1e-05 | 1.167E-05 | 2.887E-06 | 1.000E-05 | 1.500E-05 |
| 3333 | 167 | S167M | A | 1.6e-05 | 9.6e-06 | 1.6e-05 | 1.387E-05 | 3.695E-06 | 9.600E-06 | 1.600E-05 |
| 3334 | 167 | S167F | A | 1.7e-05 | 1.6e-05 | 1e-05 | 1.433E-05 | 3.786E-06 | 1.000E-05 | 1.700E-05 |
| 3335 | 167 | S167P | A | 1.4e-05 | 1e-05 | 1.5e-05 | 1.300E-05 | 2.646E-06 | 1.000E-05 | 1.500E-05 |
| 3336 | 167 | S167S | A | 9.7e-06 | 1.3e-05 | 1.6e-05 | 1.290E-05 | 3.151E-06 | 9.700E-06 | 1.600E-05 |
| 3337 | 167 | S167T | A | 8.6e-06 | 1.5e-05 | 7.5e-06 | 1.037E-05 | 4.050E-06 | 7.500E-06 | 1.500E-05 |
| 3338 | 167 | S167W | A | 9.5e-06 | 9.8e-06 | 1.2e-05 | 1.043E-05 | 1.365E-06 | 9.500E-06 | 1.200E-05 |
| 3339 | 167 | S167Y | A | 9e-06 | 1.5e-05 | 8.6e-06 | 1.087E-05 | 3.585E-06 | 8.600E-06 | 1.500E-05 |
| 3340 | 167 | S167V | A | 1.1e-05 | 1e-05 | 1.2e-05 | 1.100E-05 | 1.000E-06 | 1.000E-05 | 1.200E-05 |
| 3341 | 168 | R168A | A | 1.3e-05 | 1.5e-05 | 1.1e-05 | 1.300E-05 | 2.000E-06 | 1.100E-05 | 1.500E-05 |
| 3342 | 168 | R168R | A | 1.3e-05 | 1.4e-05 | 1.1e-05 | 1.267E-05 | 1.528E-06 | 1.100E-05 | 1.400E-05 |
| 3343 | 168 | R168N | A | 1.4e-05 | 1.1e-05 | 1e-05 | 1.167E-05 | 2.082E-06 | 1.000E-05 | 1.400E-05 |
| 3344 | 168 | R168D | A | 8.9e-06 | 1.5e-05 | 1.7e-05 | 1.363E-05 | 4.219E-06 | 8.900E-06 | 1.700E-05 |
| 3345 | 168 | R168C | A | 9.8e-06 | 9.8e-06 | 1e-05 | 9.867E-06 | 1.155E-07 | 9.800E-06 | 1.000E-05 |
| 3346 | 168 | R168E | A | 1.5e-05 | 1.5e-05 | 1.3e-05 | 1.433E-05 | 1.155E-06 | 1.300E-05 | 1.500E-05 |
| 3347 | 168 | R168Q | A | 1.3e-05 | 9.4e-06 | 1.5e-05 | 1.247E-05 | 2.838E-06 | 9.400E-06 | 1.500E-05 |
| 3348 | 168 | R168G | A | 1e-05 | 1.2e-05 | 1e-05 | 1.067E-05 | 1.155E-06 | 1.000E-05 | 1.200E-05 |
| 3349 | 168 | R168H | A | 9.9e-06 | 1.4e-05 | 1.5e-05 | 1.297E-05 | 2.702E-06 | 9.900E-06 | 1.500E-05 |
| 3350 | 168 | R168I | A | 1.7e-05 | 1e-05 | 1.1e-05 | 1.267E-05 | 3.786E-06 | 1.000E-05 | 1.700E-05 |
| 3351 | 168 | R168L | A | 1.6e-05 | 1.5e-05 | 9.4e-06 | 1.347E-05 | 3.557E-06 | 9.400E-06 | 1.600E-05 |
| 3352 | 168 | R168K | A | 1.6e-05 | 1.4e-05 | 1.4e-05 | 1.467E-05 | 1.155E-06 | 1.400E-05 | 1.600E-05 |
| 3353 | 168 | R168M | A | 1.1e-05 | 1.4e-05 | 1.5e-05 | 1.333E-05 | 2.082E-06 | 1.100E-05 | 1.500E-05 |
| 3354 | 168 | R168F | A | 1.6e-05 | 9.9e-06 | 1.1e-05 | 1.230E-05 | 3.251E-06 | 9.900E-06 | 1.600E-05 |
| 3355 | 168 | R168P | A | 1.7e-05 | 1.1e-05 | 6.7e-06 | 1.157E-05 | 5.173E-06 | 6.700E-06 | 1.700E-05 |
| 3356 | 168 | R168S | A | 1e-05 | 8.2e-06 | 1.1e-05 | 9.733E-06 | 1.419E-06 | 8.200E-06 | 1.100E-05 |
| 3357 | 168 | R168T | A | 1.4e-05 | 1.5e-05 | 1.1e-05 | 1.333E-05 | 2.082E-06 | 1.100E-05 | 1.500E-05 |
| 3358 | 168 | R168W | A | 1.5e-05 | 1.1e-05 | 1.7e-05 | 1.433E-05 | 3.055E-06 | 1.100E-05 | 1.700E-05 |
| 3359 | 168 | R168Y | A | 1.6e-05 | 1e-05 | 8.2e-06 | 1.140E-05 | 4.084E-06 | 8.200E-06 | 1.600E-05 |
| 3360 | 168 | R168V | A | 1e-05 | 1.5e-05 | 1.7e-05 | 1.400E-05 | 3.606E-06 | 1.000E-05 | 1.700E-05 |
| 3361 | 169 | A169A | A | 1e-05 | 1.1e-05 | 1.5e-05 | 1.200E-05 | 2.646E-06 | 1.000E-05 | 1.500E-05 |
| 3362 | 169 | A169R | A | 9.5e-06 | 1.3e-05 | 1.4e-05 | 1.217E-05 | 2.363E-06 | 9.500E-06 | 1.400E-05 |
| 3363 | 169 | A169N | A | 1e-05 | 1.2e-05 | 1.7e-05 | 1.300E-05 | 3.606E-06 | 1.000E-05 | 1.700E-05 |
| 3364 | 169 | A169D | A | 1e-05 | 1.6e-05 | 1e-05 | 1.200E-05 | 3.464E-06 | 1.000E-05 | 1.600E-05 |
| 3365 | 169 | A169C | A | 1.4e-05 | 1.1e-05 | 9.7e-06 | 1.157E-05 | 2.205E-06 | 9.700E-06 | 1.400E-05 |
| 3366 | 169 | A169E | A | 1.1e-05 | 1.6e-05 | 1.4e-05 | 1.367E-05 | 2.517E-06 | 1.100E-05 | 1.600E-05 |
| 3367 | 169 | A169Q | A | 9.9e-06 | 1.5e-05 | 1.6e-05 | 1.363E-05 | 3.272E-06 | 9.900E-06 | 1.600E-05 |
| 3368 | 169 | A169G | A | 1.4e-05 | 9.7e-06 | 8.6e-06 | 1.077E-05 | 2.854E-06 | 8.600E-06 | 1.400E-05 |
| 3369 | 169 | A169H | A | 1.5e-05 | 1.4e-05 | 9.7e-06 | 1.290E-05 | 2.816E-06 | 9.700E-06 | 1.500E-05 |
| 3370 | 169 | A169I | A | 9.5e-06 | 1e-05 | 1.1e-05 | 1.017E-05 | 7.638E-07 | 9.500E-06 | 1.100E-05 |
| 3371 | 169 | A169L | A | 1.5e-05 | 1.2e-05 | 1.3e-05 | 1.333E-05 | 1.528E-06 | 1.200E-05 | 1.500E-05 |
| 3372 | 169 | A169K | A | 9.3e-06 | 1.4e-05 | 1.5e-05 | 1.277E-05 | 3.044E-06 | 9.300E-06 | 1.500E-05 |
| 3373 | 169 | A169M | A | 1.4e-05 | 1.3e-05 | 1.7e-05 | 1.467E-05 | 2.082E-06 | 1.300E-05 | 1.700E-05 |
| 3374 | 169 | A169F | A | 1.4e-05 | 1.1e-05 | 9.5e-06 | 1.150E-05 | 2.291E-06 | 9.500E-06 | 1.400E-05 |
| 3375 | 169 | A169P | A | 9.5e-06 | 9.9e-06 | 1.6e-05 | 1.180E-05 | 3.643E-06 | 9.500E-06 | 1.600E-05 |
| 3376 | 169 | A169S | A | 1.5e-05 | 1.1e-05 | 1.4e-05 | 1.333E-05 | 2.082E-06 | 1.100E-05 | 1.500E-05 |
| 3377 | 169 | A169T | A | 1.5e-05 | 1.1e-05 | 1.5e-05 | 1.367E-05 | 2.309E-06 | 1.100E-05 | 1.500E-05 |
| 3378 | 169 | A169W | A | 1.4e-05 | 1.1e-05 | 1.2e-05 | 1.233E-05 | 1.528E-06 | 1.100E-05 | 1.400E-05 |
| 3379 | 169 | A169Y | A | 1.4e-05 | 1.2e-05 | 1.3e-05 | 1.300E-05 | 1.000E-06 | 1.200E-05 | 1.400E-05 |
| 3380 | 169 | A169V | A | 1.4e-05 | 1.5e-05 | 1.5e-05 | 1.467E-05 | 5.774E-07 | 1.400E-05 | 1.500E-05 |
| 3381 | 170 | C170A | A | 1.6e-05 | 1e-05 | 9.8e-06 | 1.193E-05 | 3.523E-06 | 9.800E-06 | 1.600E-05 |
| 3382 | 170 | C170R | A | 1.9e-05 | 1.5e-05 | 8.6e-06 | 1.420E-05 | 5.246E-06 | 8.600E-06 | 1.900E-05 |
| 3383 | 170 | C170N | A | 1.5e-05 | 1.5e-05 | 8.5e-06 | 1.283E-05 | 3.753E-06 | 8.500E-06 | 1.500E-05 |
| 3384 | 170 | C170D | A | 2.5e-05 | 8.8e-06 | 1.6e-05 | 1.660E-05 | 8.117E-06 | 8.800E-06 | 2.500E-05 |
| 3385 | 170 | C170C | A | 1.1e-05 | 9e-06 | 1.1e-05 | 1.033E-05 | 1.155E-06 | 9.000E-06 | 1.100E-05 |
| 3386 | 170 | C170E | A | 1.2e-05 | 1.4e-05 | 1.4e-05 | 1.333E-05 | 1.155E-06 | 1.200E-05 | 1.400E-05 |
| 3387 | 170 | C170Q | A | 1.4e-05 | 1.5e-05 | 8.4e-06 | 1.247E-05 | 3.557E-06 | 8.400E-06 | 1.500E-05 |
| 3388 | 170 | C170G | A | 1.4e-05 | 8.7e-06 | 1.4e-05 | 1.223E-05 | 3.060E-06 | 8.700E-06 | 1.400E-05 |
| 3389 | 170 | C170H | A | 9.8e-06 | 1e-05 | 1.5e-05 | 1.160E-05 | 2.946E-06 | 9.800E-06 | 1.500E-05 |
| 3390 | 170 | C170I | A | 9.7e-06 | 9.2e-06 | 1.2e-05 | 1.030E-05 | 1.493E-06 | 9.200E-06 | 1.200E-05 |
| 3391 | 170 | C170L | A | 1.5e-05 | 1e-05 | 1.7e-05 | 1.400E-05 | 3.606E-06 | 1.000E-05 | 1.700E-05 |
| 3392 | 170 | C170K | A | 9.3e-06 | 1.1e-05 | 1.5e-05 | 1.177E-05 | 2.926E-06 | 9.300E-06 | 1.500E-05 |
| 3393 | 170 | C170M | A | 1e-05 | 1.6e-05 | 1.4e-05 | 1.333E-05 | 3.055E-06 | 1.000E-05 | 1.600E-05 |
| 3394 | 170 | C170F | A | 8.6e-06 | 1.7e-05 | 9.4e-06 | 1.167E-05 | 4.636E-06 | 8.600E-06 | 1.700E-05 |
| 3395 | 170 | C170P | A | 1.4e-05 | 1.2e-05 | 1e-05 | 1.200E-05 | 2.000E-06 | 1.000E-05 | 1.400E-05 |
| 3396 | 170 | C170S | A | 1.2e-05 | 1.5e-05 | 1e-05 | 1.233E-05 | 2.517E-06 | 1.000E-05 | 1.500E-05 |
| 3397 | 170 | C170T | A | 1.1e-05 | 1e-05 | 1.3e-05 | 1.133E-05 | 1.528E-06 | 1.000E-05 | 1.300E-05 |
| 3398 | 170 | C170W | A | 1.5e-05 | 1.3e-05 | 1.7e-05 | 1.500E-05 | 2.000E-06 | 1.300E-05 | 1.700E-05 |
| 3399 | 170 | C170Y | A | 1.5e-05 | 9.5e-06 | 9.9e-06 | 1.147E-05 | 3.066E-06 | 9.500E-06 | 1.500E-05 |

| | | | | | | | | | | |
|---|---|---|---|---|---|---|---|---|---|---|
| 3400 | 170 | C170V | A | 8e-06 | 1.7e-05 | 1e-05 | 1.167E-05 | 4.726E-06 | 8.000E-06 | 1.700E-05 |
| 3401 | 171 | H171A | A | 1.4e-05 | 9.5e-06 | 1.1e-05 | 1.150E-05 | 2.291E-06 | 9.500E-06 | 1.400E-05 |
| 3402 | 171 | H171R | A | 1e-05 | 7.8e-06 | 1.1e-05 | 9.600E-06 | 1.637E-06 | 7.800E-06 | 1.100E-05 |
| 3403 | 171 | H171N | A | 8.3e-06 | 1.1e-05 | 1.4e-05 | 1.110E-05 | 2.851E-06 | 8.300E-06 | 1.400E-05 |
| 3404 | 171 | H171D | A | 1.5e-05 | 1.3e-05 | 1.5e-05 | 1.433E-05 | 1.155E-06 | 1.300E-05 | 1.500E-05 |
| 3405 | 171 | H171C | A | 1.3e-05 | 1.1e-05 | 1.5e-05 | 1.300E-05 | 2.000E-06 | 1.100E-05 | 1.500E-05 |
| 3406 | 171 | H171E | A | 1.1e-05 | 1.6e-05 | 9.9e-06 | 1.230E-05 | 3.251E-06 | 9.900E-06 | 1.600E-05 |
| 3407 | 171 | H171Q | A | 1.5e-05 | 1.1e-05 | 1.5e-05 | 1.367E-05 | 2.309E-06 | 1.100E-05 | 1.500E-05 |
| 3408 | 171 | H171G | A | 1.5e-05 | 1.6e-05 | 1.1e-05 | 1.400E-05 | 2.646E-06 | 1.100E-05 | 1.600E-05 |
| 3409 | 171 | H171H | A | 1.1e-05 | 1e-05 | 1e-05 | 1.033E-05 | 5.774E-07 | 1.000E-05 | 1.100E-05 |
| 3410 | 171 | H171I | A | 1.3e-05 | 1.5e-05 | 1.5e-05 | 1.433E-05 | 1.155E-06 | 1.300E-05 | 1.500E-05 |
| 3411 | 171 | H171L | A | 9.7e-06 | 2.1e-05 | 1.5e-05 | 1.523E-05 | 5.654E-06 | 9.700E-06 | 2.100E-05 |
| 3412 | 171 | H171K | A | 9.3e-06 | 1.1e-05 | 1.4e-05 | 1.143E-05 | 2.380E-06 | 9.300E-06 | 1.400E-05 |
| 3413 | 171 | H171M | A | 1.4e-05 | 1.1e-05 | 1.5e-05 | 1.333E-05 | 2.082E-06 | 1.100E-05 | 1.500E-05 |
| 3414 | 171 | H171F | A | 1.2e-05 | 1.5e-05 | 8.8e-06 | 1.193E-05 | 3.101E-06 | 8.800E-06 | 1.500E-05 |
| 3415 | 171 | H171P | A | 1.2e-05 | 1.4e-05 | 1e-05 | 1.200E-05 | 2.000E-06 | 1.000E-05 | 1.400E-05 |
| 3416 | 171 | H171S | A | 1.5e-05 | 1.7e-05 | 6.9e-06 | 1.297E-05 | 5.348E-06 | 6.900E-06 | 1.700E-05 |
| 3417 | 171 | H171T | A | 1.5e-05 | 1.4e-05 | 1.6e-05 | 1.500E-05 | 1.000E-06 | 1.400E-05 | 1.600E-05 |
| 3418 | 171 | H171W | A | 1.2e-05 | 1.7e-05 | 1e-05 | 1.300E-05 | 3.606E-06 | 1.000E-05 | 1.700E-05 |
| 3419 | 171 | H171Y | A | 1.2e-05 | 1.4e-05 | 9.7e-06 | 1.190E-05 | 2.152E-06 | 9.700E-06 | 1.400E-05 |
| 3420 | 171 | H171V | A | 1e-05 | 6.3e-06 | 1.1e-05 | 9.100E-06 | 2.476E-06 | 6.300E-06 | 1.100E-05 |
| 3421 | 172 | P172A | A | 1.1e-05 | 1.6e-05 | 9.8e-06 | 1.227E-05 | 3.288E-06 | 9.800E-06 | 1.600E-05 |
| 3422 | 172 | P172R | A | 1.5e-05 | 1.4e-05 | 1.3e-05 | 1.400E-05 | 1.000E-06 | 1.300E-05 | 1.500E-05 |
| 3423 | 172 | P172N | A | 1.1e-05 | 1.3e-05 | 1.4e-05 | 1.267E-05 | 1.528E-06 | 1.100E-05 | 1.400E-05 |
| 3424 | 172 | P172D | A | 1e-05 | 9.7e-06 | 1.4e-05 | 1.123E-05 | 2.401E-06 | 9.700E-06 | 1.400E-05 |
| 3425 | 172 | P172C | A | 1.1e-05 | 1.1e-05 | 1e-05 | 1.067E-05 | 5.774E-07 | 1.000E-05 | 1.100E-05 |
| 3426 | 172 | P172E | A | 1.2e-05 | 1.1e-05 | 1.5e-05 | 1.267E-05 | 2.082E-06 | 1.100E-05 | 1.500E-05 |
| 3427 | 172 | P172Q | A | 1e-05 | 9.4e-06 | 1e-05 | 9.800E-06 | 3.464E-07 | 9.400E-06 | 1.000E-05 |
| 3428 | 172 | P172G | A | 9.9e-06 | 1.5e-05 | 1.6e-05 | 1.363E-05 | 3.272E-06 | 9.900E-06 | 1.600E-05 |
| 3429 | 172 | P172H | A | 1.2e-05 | 9.1e-06 | 1.7e-05 | 1.270E-05 | 3.996E-06 | 9.100E-06 | 1.700E-05 |
| 3430 | 172 | P172I | A | 1.5e-05 | 1e-05 | 1.4e-05 | 1.300E-05 | 2.646E-06 | 1.000E-05 | 1.500E-05 |
| 3431 | 172 | P172L | A | 1.4e-05 | 1.4e-05 | 1.2e-05 | 1.333E-05 | 1.155E-06 | 1.200E-05 | 1.400E-05 |
| 3432 | 172 | P172K | A | 9.1e-06 | 1.1e-05 | 1.6e-05 | 1.203E-05 | 3.564E-06 | 9.100E-06 | 1.600E-05 |
| 3433 | 172 | P172M | A | 1.1e-05 | 1.5e-05 | 1.6e-05 | 1.400E-05 | 2.646E-06 | 1.100E-05 | 1.600E-05 |
| 3434 | 172 | P172F | A | 1.6e-05 | 2.1e-05 | 1e-05 | 1.567E-05 | 5.508E-06 | 1.000E-05 | 2.100E-05 |
| 3435 | 172 | P172P | A | 1.7e-05 | 1.6e-05 | 1.1e-05 | 1.467E-05 | 3.215E-06 | 1.100E-05 | 1.700E-05 |
| 3436 | 172 | P172S | A | 1.5e-05 | 1.4e-05 | 1e-05 | 1.300E-05 | 2.646E-06 | 1.000E-05 | 1.500E-05 |
| 3437 | 172 | P172T | A | 1e-05 | 1e-05 | 1e-05 | 1.000E-05 | 0.000E+00 | 1.000E-05 | 1.000E-05 |
| 3438 | 172 | P172W | A | 1.4e-05 | 1.6e-05 | 1.6e-05 | 1.533E-05 | 1.155E-06 | 1.400E-05 | 1.600E-05 |
| 3439 | 172 | P172Y | A | 1.5e-05 | 1.1e-05 | 8.6e-06 | 1.153E-05 | 3.233E-06 | 8.600E-06 | 1.500E-05 |
| 3440 | 172 | P172V | A | 1.5e-05 | 7.7e-06 | 9.8e-06 | 1.083E-05 | 3.758E-06 | 7.700E-06 | 1.500E-05 |
| 3441 | 173 | C173A | A | 1.4e-05 | 1e-05 | 1.6e-05 | 1.333E-05 | 3.055E-06 | 1.000E-05 | 1.600E-05 |
| 3442 | 173 | C173R | A | 1.1e-05 | 1.1e-05 | 1.4e-05 | 1.200E-05 | 1.732E-06 | 1.100E-05 | 1.400E-05 |
| 3443 | 173 | C173N | A | 1.2e-05 | 1.2e-05 | 1.4e-05 | 1.267E-05 | 1.155E-06 | 1.200E-05 | 1.400E-05 |
| 3444 | 173 | C173D | A | 1.6e-05 | 1.5e-05 | 1.2e-05 | 1.433E-05 | 2.082E-06 | 1.200E-05 | 1.600E-05 |
| 3445 | 173 | C173C | A | 1.4e-05 | 1.2e-05 | 6.9e-06 | 1.097E-05 | 3.661E-06 | 6.900E-06 | 1.400E-05 |
| 3446 | 173 | C173E | A | 1.4e-05 | 1.5e-05 | 1.3e-05 | 1.400E-05 | 1.000E-06 | 1.300E-05 | 1.500E-05 |
| 3447 | 173 | C173Q | A | 1.1e-05 | 1.1e-05 | 1e-05 | 1.067E-05 | 5.774E-07 | 1.000E-05 | 1.100E-05 |
| 3448 | 173 | C173G | A | 1.6e-05 | 9.7e-06 | 1.6e-05 | 1.390E-05 | 3.637E-06 | 9.700E-06 | 1.600E-05 |
| 3449 | 173 | C173H | A | 1.1e-05 | 9.7e-06 | 1.2e-05 | 1.090E-05 | 1.153E-06 | 9.700E-06 | 1.200E-05 |
| 3450 | 173 | C173I | A | 1.4e-05 | 1.1e-05 | 1.6e-05 | 1.367E-05 | 2.517E-06 | 1.100E-05 | 1.600E-05 |
| 3451 | 173 | C173L | A | 1.3e-05 | 1.5e-05 | 1.3e-05 | 1.367E-05 | 1.155E-06 | 1.300E-05 | 1.500E-05 |
| 3452 | 173 | C173K | A | 1.2e-05 | 1.5e-05 | 1.5e-05 | 1.400E-05 | 1.732E-06 | 1.200E-05 | 1.500E-05 |
| 3453 | 173 | C173M | A | 9.7e-06 | 1.2e-05 | 8.7e-06 | 1.013E-05 | 1.692E-06 | 8.700E-06 | 1.200E-05 |
| 3454 | 173 | C173F | A | 1.3e-05 | 1.4e-05 | 7.6e-06 | 1.153E-05 | 3.443E-06 | 7.600E-06 | 1.400E-05 |
| 3455 | 173 | C173P | A | 1.6e-05 | 9.5e-06 | 9.2e-06 | 1.157E-05 | 3.842E-06 | 9.200E-06 | 1.600E-05 |
| 3456 | 173 | C173S | A | 9.7e-06 | 1.3e-05 | 1.6e-05 | 1.290E-05 | 3.151E-06 | 9.700E-06 | 1.600E-05 |
| 3457 | 173 | C173T | A | 1.5e-05 | 1e-05 | 1.5e-05 | 1.333E-05 | 2.887E-06 | 1.000E-05 | 1.500E-05 |
| 3458 | 173 | C173W | A | 1.1e-05 | 1.6e-05 | 1.1e-05 | 1.267E-05 | 2.887E-06 | 1.100E-05 | 1.600E-05 |
| 3459 | 173 | C173Y | A | 1.2e-05 | 1.6e-05 | 1.4e-05 | 1.400E-05 | 2.000E-06 | 1.200E-05 | 1.600E-05 |
| 3460 | 173 | C173V | A | 9.6e-06 | 1.8e-05 | 1e-05 | 1.253E-05 | 4.738E-06 | 9.600E-06 | 1.800E-05 |
| 3461 | 174 | S174A | A | 1.5e-05 | 1.1e-05 | 1e-05 | 1.200E-05 | 2.646E-06 | 1.000E-05 | 1.500E-05 |
| 3462 | 174 | S174R | A | 1.1e-05 | 1.4e-05 | 1.4e-05 | 1.300E-05 | 1.732E-06 | 1.100E-05 | 1.400E-05 |
| 3463 | 174 | S174N | A | 1.1e-05 | 1.5e-05 | 1e-05 | 1.200E-05 | 2.646E-06 | 1.000E-05 | 1.500E-05 |
| 3464 | 174 | S174D | A | 1.1e-05 | 1.6e-05 | 1.5e-05 | 1.400E-05 | 2.646E-06 | 1.100E-05 | 1.600E-05 |
| 3465 | 174 | S174C | A | 1.5e-05 | 1e-05 | 1.6e-05 | 1.367E-05 | 3.215E-06 | 1.000E-05 | 1.600E-05 |
| 3466 | 174 | S174E | A | 1.6e-05 | 8.5e-06 | 1.2e-05 | 1.217E-05 | 3.753E-06 | 8.500E-06 | 1.600E-05 |
| 3467 | 174 | S174Q | A | 9.8e-06 | 1.5e-05 | 1.4e-05 | 1.293E-05 | 2.759E-06 | 9.800E-06 | 1.500E-05 |
| 3468 | 174 | S174G | A | 1.7e-05 | 1.5e-05 | 1e-05 | 1.400E-05 | 3.606E-06 | 1.000E-05 | 1.700E-05 |
| 3469 | 174 | S174H | A | 1.6e-05 | 1.4e-05 | 1.1e-05 | 1.367E-05 | 2.517E-06 | 1.100E-05 | 1.600E-05 |
| 3470 | 174 | S174I | A | 1.6e-05 | 1.6e-05 | 1.6e-05 | 1.600E-05 | 0.000E+00 | 1.600E-05 | 1.600E-05 |
| 3471 | 174 | S174L | A | 1.5e-05 | 1e-05 | 1.4e-05 | 1.300E-05 | 2.646E-06 | 1.000E-05 | 1.500E-05 |
| 3472 | 174 | S174K | A | 1.5e-05 | 1.1e-05 | 1.4e-05 | 1.333E-05 | 2.082E-06 | 1.100E-05 | 1.500E-05 |
| 3473 | 174 | S174M | A | 1.3e-05 | 1.5e-05 | 1e-05 | 1.267E-05 | 2.517E-06 | 1.000E-05 | 1.500E-05 |
| 3474 | 174 | S174F | A | 1.4e-05 | 1.1e-05 | 1.9e-05 | 1.467E-05 | 4.041E-06 | 1.100E-05 | 1.900E-05 |
| 3475 | 174 | S174P | A | 1.1e-05 | 1.1e-05 | 1.7e-05 | 1.300E-05 | 3.464E-06 | 1.100E-05 | 1.700E-05 |
| 3476 | 174 | S174S | A | 1.1e-05 | 1.3e-05 | 1e-05 | 1.133E-05 | 1.528E-06 | 1.000E-05 | 1.300E-05 |
| 3477 | 174 | S174T | A | 1e-05 | 1.3e-05 | 1.4e-05 | 1.233E-05 | 2.082E-06 | 1.000E-05 | 1.400E-05 |
| 3478 | 174 | S174W | A | 1e-05 | 1.6e-05 | 1.7e-05 | 1.433E-05 | 3.786E-06 | 1.000E-05 | 1.700E-05 |
| 3479 | 174 | S174Y | A | 1.4e-05 | 1.1e-05 | 8.7e-06 | 1.123E-05 | 2.658E-06 | 8.700E-06 | 1.400E-05 |
| 3480 | 174 | S174V | A | 1.1e-05 | 1.2e-05 | 1.2e-05 | 1.167E-05 | 5.774E-07 | 1.100E-05 | 1.200E-05 |
| 3481 | 175 | P175A | A | 1.2e-05 | 1.9e-05 | 1.1e-05 | 1.400E-05 | 4.359E-06 | 1.100E-05 | 1.900E-05 |
| 3482 | 175 | P175R | A | 1.1e-05 | 1.2e-05 | 7.8e-06 | 1.027E-05 | 2.194E-06 | 7.800E-06 | 1.200E-05 |
| 3483 | 175 | P175N | A | 9.7e-06 | 1e-05 | 1.4e-05 | 1.123E-05 | 2.401E-06 | 9.700E-06 | 1.400E-05 |
| 3484 | 175 | P175D | A | 2.2e-05 | 1.4e-05 | 1.1e-05 | 1.567E-05 | 5.686E-06 | 1.100E-05 | 2.200E-05 |

| | | | | | | | | | | |
|---|---|---|---|---|---|---|---|---|---|---|
| 3485 | 175 | P175C | A | 1.3e-05 | 1.1e-05 | 1.1e-05 | 1.167E-05 | 1.155E-06 | 1.100E-05 | 1.300E-05 |
| 3486 | 175 | P175E | A | 1.6e-05 | 1.4e-05 | 1.6e-05 | 1.533E-05 | 1.155E-06 | 1.400E-05 | 1.600E-05 |
| 3487 | 175 | P175Q | A | 1.4e-05 | 1.3e-05 | 1.5e-05 | 1.400E-05 | 1.000E-06 | 1.300E-05 | 1.500E-05 |
| 3488 | 175 | P175G | A | 1.2e-05 | 1.1e-05 | 1.1e-05 | 1.133E-05 | 5.774E-07 | 1.100E-05 | 1.200E-05 |
| 3489 | 175 | P175H | A | 1.5e-05 | 1.6e-05 | 1.5e-05 | 1.533E-05 | 5.774E-07 | 1.500E-05 | 1.600E-05 |
| 3490 | 175 | P175I | A | 1.4e-05 | 1.1e-05 | 9.6e-06 | 1.153E-05 | 2.248E-06 | 9.600E-06 | 1.400E-05 |
| 3491 | 175 | P175L | A | 1e-05 | 1.5e-05 | 1.4e-05 | 1.300E-05 | 2.646E-06 | 1.000E-05 | 1.500E-05 |
| 3492 | 175 | P175K | A | 9.4e-06 | 1.6e-05 | 1.4e-05 | 1.313E-05 | 3.384E-06 | 9.400E-06 | 1.600E-05 |
| 3493 | 175 | P175M | A | 1.4e-05 | 1.5e-05 | 1e-05 | 1.300E-05 | 2.646E-06 | 1.000E-05 | 1.500E-05 |
| 3494 | 175 | P175F | A | 2.8e-05 | 1.5e-05 | 1.3e-05 | 1.867E-05 | 8.145E-06 | 1.300E-05 | 2.800E-05 |
| 3495 | 175 | P175P | A | 1e-05 | 8.9e-06 | 1.6e-05 | 1.163E-05 | 3.821E-06 | 8.900E-06 | 1.600E-05 |
| 3496 | 175 | P175S | A | 9.2e-06 | 1.5e-05 | 9e-06 | 1.107E-05 | 3.408E-06 | 9.000E-06 | 1.500E-05 |
| 3497 | 175 | P175T | A | 1.1e-05 | 1e-05 | 1.5e-05 | 1.200E-05 | 2.646E-06 | 1.000E-05 | 1.500E-05 |
| 3498 | 175 | P175W | A | 1.6e-05 | 1.6e-05 | 1.4e-05 | 1.533E-05 | 1.155E-06 | 1.400E-05 | 1.600E-05 |
| 3499 | 175 | P175Y | A | 9.6e-06 | 1.6e-05 | 1e-05 | 1.187E-05 | 3.585E-06 | 9.600E-06 | 1.600E-05 |
| 3500 | 175 | P175V | A | 9.6e-06 | 1.1e-05 | 9.3e-06 | 9.967E-06 | 9.074E-07 | 9.300E-06 | 1.100E-05 |
| 3501 | 176 | M176A | A | 1.4e-05 | 1.5e-05 | 1e-05 | 1.300E-05 | 2.646E-06 | 1.000E-05 | 1.500E-05 |
| 3502 | 176 | M176R | A | 1.7e-05 | 1.4e-05 | 1.5e-05 | 1.533E-05 | 1.528E-06 | 1.400E-05 | 1.700E-05 |
| 3503 | 176 | M176N | A | 1.5e-05 | 1.4e-05 | 6.5e-06 | 1.183E-05 | 4.646E-06 | 6.500E-06 | 1.500E-05 |
| 3504 | 176 | M176D | A | 1.6e-05 | 7e-06 | 9.6e-06 | 1.087E-05 | 4.632E-06 | 7.000E-06 | 1.600E-05 |
| 3505 | 176 | M176C | A | 1.4e-05 | 1.6e-05 | 1.5e-05 | 1.500E-05 | 1.000E-06 | 1.400E-05 | 1.600E-05 |
| 3506 | 176 | M176E | A | 1.4e-05 | 1.1e-05 | 1.6e-05 | 1.367E-05 | 2.517E-06 | 1.100E-05 | 1.600E-05 |
| 3507 | 176 | M176Q | A | 1.1e-05 | 1.6e-05 | 1.6e-05 | 1.433E-05 | 2.887E-06 | 1.100E-05 | 1.600E-05 |
| 3508 | 176 | M176G | A | 1.6e-05 | 1.5e-05 | 1e-05 | 1.367E-05 | 3.215E-06 | 1.000E-05 | 1.600E-05 |
| 3509 | 176 | M176H | A | 8.6e-06 | 1.5e-05 | 1.3e-05 | 1.220E-05 | 3.274E-06 | 8.600E-06 | 1.500E-05 |
| 3510 | 176 | M176I | A | 1.1e-05 | 9.2e-06 | 1.1e-05 | 1.040E-05 | 1.039E-06 | 9.200E-06 | 1.100E-05 |
| 3511 | 176 | M176L | A | 1.4e-05 | 1.3e-05 | 8.9e-06 | 1.197E-05 | 2.702E-06 | 8.900E-06 | 1.400E-05 |
| 3512 | 176 | M176K | A | 1.7e-05 | 1e-05 | 1.5e-05 | 1.400E-05 | 3.606E-06 | 1.000E-05 | 1.700E-05 |
| 3513 | 176 | M176M | A | 1e-05 | 1e-05 | 9.8e-06 | 9.933E-06 | 1.155E-07 | 9.800E-06 | 1.000E-05 |
| 3514 | 176 | M176F | A | 1.6e-05 | 1.5e-05 | 9.3e-06 | 1.343E-05 | 3.614E-06 | 9.300E-06 | 1.600E-05 |
| 3515 | 176 | M176P | A | 1.3e-05 | 1.1e-05 | 1.5e-05 | 1.300E-05 | 2.000E-06 | 1.100E-05 | 1.500E-05 |
| 3516 | 176 | M176S | A | 1.3e-05 | 1.4e-05 | 8.6e-06 | 1.187E-05 | 2.873E-06 | 8.600E-06 | 1.400E-05 |
| 3517 | 176 | M176T | A | 9.6e-06 | 1.4e-05 | 1.4e-05 | 1.253E-05 | 2.540E-06 | 9.600E-06 | 1.400E-05 |
| 3518 | 176 | M176W | A | 1.5e-05 | 1.4e-05 | 1.3e-05 | 1.400E-05 | 1.000E-06 | 1.300E-05 | 1.500E-05 |
| 3519 | 176 | M176Y | A | 1.5e-05 | 1.1e-05 | 9.3e-06 | 1.177E-05 | 2.926E-06 | 9.300E-06 | 1.500E-05 |
| 3520 | 176 | M176V | A | 1.4e-05 | 1.4e-05 | 1.6e-05 | 1.467E-05 | 1.155E-06 | 1.400E-05 | 1.600E-05 |
| 3521 | 177 | C177A | A | 1.4e-05 | 1.6e-05 | 1.7e-05 | 1.567E-05 | 1.528E-06 | 1.400E-05 | 1.700E-05 |
| 3522 | 177 | C177R | A | 1.4e-05 | 1.5e-05 | 1e-05 | 1.300E-05 | 2.646E-06 | 1.000E-05 | 1.500E-05 |
| 3523 | 177 | C177N | A | 1.4e-05 | 1.1e-05 | 1.5e-05 | 1.333E-05 | 2.082E-06 | 1.100E-05 | 1.500E-05 |
| 3524 | 177 | C177D | A | 1.7e-05 | 1e-05 | 1e-05 | 1.233E-05 | 4.041E-06 | 1.000E-05 | 1.700E-05 |
| 3525 | 177 | C177C | A | 7.2e-06 | 1.4e-05 | 1.3e-05 | 1.140E-05 | 3.672E-06 | 7.200E-06 | 1.400E-05 |
| 3526 | 177 | C177E | A | 1.1e-05 | 1.4e-05 | 1.5e-05 | 1.333E-05 | 2.082E-06 | 1.100E-05 | 1.500E-05 |
| 3527 | 177 | C177Q | A | 1.6e-05 | 1.1e-05 | 9.2e-06 | 1.207E-05 | 3.523E-06 | 9.200E-06 | 1.600E-05 |
| 3528 | 177 | C177G | A | 8.5e-06 | 9.4e-06 | 6.8e-06 | 8.233E-06 | 1.320E-06 | 6.800E-06 | 9.400E-06 |
| 3529 | 177 | C177H | A | 1.6e-05 | 1.5e-05 | 1.6e-05 | 1.567E-05 | 5.774E-07 | 1.500E-05 | 1.600E-05 |
| 3530 | 177 | C177I | A | 1.5e-05 | 1.1e-05 | 1.7e-05 | 1.433E-05 | 3.055E-06 | 1.100E-05 | 1.700E-05 |
| 3531 | 177 | C177L | A | 1.6e-05 | 1.7e-05 | 9.6e-06 | 1.420E-05 | 4.015E-06 | 9.600E-06 | 1.700E-05 |
| 3532 | 177 | C177K | A | 1.1e-05 | 1e-05 | 1.1e-05 | 1.067E-05 | 5.774E-07 | 1.000E-05 | 1.100E-05 |
| 3533 | 177 | C177M | A | 1.5e-05 | 1.6e-05 | 1.4e-05 | 1.500E-05 | 1.000E-06 | 1.400E-05 | 1.600E-05 |
| 3534 | 177 | C177F | A | 1.9e-05 | 1.4e-05 | 1.6e-05 | 1.633E-05 | 2.517E-06 | 1.400E-05 | 1.900E-05 |
| 3535 | 177 | C177P | A | 1.8e-05 | 9.5e-06 | 1.1e-05 | 1.283E-05 | 4.537E-06 | 9.500E-06 | 1.800E-05 |
| 3536 | 177 | C177S | A | 1.2e-05 | 1.6e-05 | 1.2e-05 | 1.333E-05 | 2.309E-06 | 1.200E-05 | 1.600E-05 |
| 3537 | 177 | C177T | A | 1.6e-05 | 1.8e-05 | 1.4e-05 | 1.600E-05 | 2.000E-06 | 1.400E-05 | 1.800E-05 |
| 3538 | 177 | C177W | A | 1.1e-05 | 1.1e-05 | 1.6e-05 | 1.267E-05 | 2.887E-06 | 1.100E-05 | 1.600E-05 |
| 3539 | 177 | C177Y | A | 1.5e-05 | 1.2e-05 | 1.4e-05 | 1.367E-05 | 1.528E-06 | 1.200E-05 | 1.500E-05 |
| 3540 | 177 | C177V | A | 1.6e-05 | 7.2e-06 | 1.3e-05 | 1.207E-05 | 4.474E-06 | 7.200E-06 | 1.600E-05 |
| 3541 | 178 | K178A | A | 1.1e-05 | 1.5e-05 | 1.5e-05 | 1.367E-05 | 2.309E-06 | 1.100E-05 | 1.500E-05 |
| 3542 | 178 | K178R | A | 1.3e-05 | 1.6e-05 | 1e-05 | 1.300E-05 | 3.000E-06 | 1.000E-05 | 1.600E-05 |
| 3543 | 178 | K178N | A | 1.3e-05 | 1.5e-05 | 9.5e-06 | 1.250E-05 | 2.784E-06 | 9.500E-06 | 1.500E-05 |
| 3544 | 178 | K178D | A | 1.1e-05 | 9.6e-06 | 1.1e-05 | 1.053E-05 | 8.083E-07 | 9.600E-06 | 1.100E-05 |
| 3545 | 178 | K178C | A | 1.2e-05 | 1.5e-05 | 1.1e-05 | 1.267E-05 | 2.082E-06 | 1.100E-05 | 1.500E-05 |
| 3546 | 178 | K178E | A | 9.9e-06 | 1.2e-05 | 1.1e-05 | 1.097E-05 | 1.050E-06 | 9.900E-06 | 1.200E-05 |
| 3547 | 178 | K178Q | A | 1.3e-05 | 1.4e-05 | 1.5e-05 | 1.400E-05 | 1.000E-06 | 1.300E-05 | 1.500E-05 |
| 3548 | 178 | K178G | A | 1.2e-05 | 1.1e-05 | 8.7e-06 | 1.057E-05 | 1.692E-06 | 8.700E-06 | 1.200E-05 |
| 3549 | 178 | K178H | A | 1e-05 | 7.5e-06 | 1e-05 | 9.167E-06 | 1.443E-06 | 7.500E-06 | 1.000E-05 |
| 3550 | 178 | K178I | A | 1.5e-05 | 1.3e-05 | 1.8e-05 | 1.533E-05 | 2.517E-06 | 1.300E-05 | 1.800E-05 |
| 3551 | 178 | K178L | A | 1.3e-05 | 1.4e-05 | 1.1e-05 | 1.267E-05 | 1.528E-06 | 1.100E-05 | 1.400E-05 |
| 3552 | 178 | K178K | A | 1.3e-05 | 1.2e-05 | 1.5e-05 | 1.333E-05 | 1.528E-06 | 1.200E-05 | 1.500E-05 |
| 3553 | 178 | K178M | A | 1.4e-05 | 8.6e-06 | 9.6e-06 | 1.073E-05 | 2.873E-06 | 8.600E-06 | 1.400E-05 |
| 3554 | 178 | K178F | A | 9e-06 | 3.1e-05 | 9.5e-06 | 1.650E-05 | 1.256E-05 | 9.000E-06 | 3.100E-05 |
| 3555 | 178 | K178P | A | 9.8e-06 | 1.1e-05 | 1.1e-05 | 1.060E-05 | 6.928E-07 | 9.800E-06 | 1.100E-05 |
| 3556 | 178 | K178S | A | 1.1e-05 | 1.3e-05 | 9e-06 | 1.100E-05 | 2.000E-06 | 9.000E-06 | 1.300E-05 |
| 3557 | 178 | K178T | A | 8.3e-06 | 1.6e-05 | 9.3e-06 | 1.120E-05 | 4.187E-06 | 8.300E-06 | 1.600E-05 |
| 3558 | 178 | K178W | A | 1.1e-05 | 1.6e-05 | 1.2e-05 | 1.300E-05 | 2.646E-06 | 1.100E-05 | 1.600E-05 |
| 3559 | 178 | K178Y | A | 1.3e-05 | 1.3e-05 | 8e-06 | 1.133E-05 | 2.887E-06 | 8.000E-06 | 1.300E-05 |
| 3560 | 178 | K178V | A | 1.7e-05 | 1.4e-05 | 1e-05 | 1.367E-05 | 3.512E-06 | 1.000E-05 | 1.700E-05 |
| 3561 | 179 | G179A | A | 1.3e-05 | 1.1e-05 | 1.6e-05 | 1.333E-05 | 2.517E-06 | 1.100E-05 | 1.600E-05 |
| 3562 | 179 | G179R | A | 9.7e-06 | 8.7e-06 | 1.1e-05 | 9.800E-06 | 1.153E-06 | 8.700E-06 | 1.100E-05 |
| 3563 | 179 | G179N | A | 1.2e-05 | 6.4e-06 | 9.5e-06 | 9.300E-06 | 2.805E-06 | 6.400E-06 | 1.200E-05 |
| 3564 | 179 | G179D | A | 8e-06 | 1.5e-05 | 1.3e-05 | 1.200E-05 | 3.606E-06 | 8.000E-06 | 1.500E-05 |
| 3565 | 179 | G179C | A | 1.3e-05 | 1.3e-05 | 1.6e-05 | 1.400E-05 | 1.732E-06 | 1.300E-05 | 1.600E-05 |
| 3566 | 179 | G179E | A | 1.2e-05 | 1.1e-05 | 1.1e-05 | 1.133E-05 | 5.774E-07 | 1.100E-05 | 1.200E-05 |
| 3567 | 179 | G179Q | A | 1.5e-05 | 2e-05 | 1.5e-05 | 1.667E-05 | 2.887E-06 | 1.500E-05 | 2.000E-05 |
| 3568 | 179 | G179G | A | 1.4e-05 | 1.4e-05 | 1.5e-05 | 1.433E-05 | 5.774E-07 | 1.400E-05 | 1.500E-05 |
| 3569 | 179 | G179H | A | 1.5e-05 | 1.4e-05 | 1.6e-05 | 1.500E-05 | 1.000E-06 | 1.400E-05 | 1.600E-05 |

| | | | | | | | | | | |
|---|---|---|---|---|---|---|---|---|---|---|
| 3570 | 179 | G179I | A | 9.8e-06 | 1.4e-05 | 1e-05 | 1.127E-05 | 2.369E-06 | 9.800E-06 | 1.400E-05 |
| 3571 | 179 | G179L | A | 1e-05 | 1.5e-05 | 1.3e-05 | 1.267E-05 | 2.517E-06 | 1.000E-05 | 1.500E-05 |
| 3572 | 179 | G179K | A | 1.4e-05 | 1.3e-05 | 9.9e-06 | 1.230E-05 | 2.138E-06 | 9.900E-06 | 1.400E-05 |
| 3573 | 179 | G179M | A | 1.5e-05 | 9.5e-06 | 9.7e-06 | 1.140E-05 | 3.119E-06 | 9.500E-06 | 1.500E-05 |
| 3574 | 179 | G179F | A | 1.1e-05 | 1e-05 | 1.4e-05 | 1.167E-05 | 2.082E-06 | 1.000E-05 | 1.400E-05 |
| 3575 | 179 | G179P | A | 9e-06 | 1e-05 | 1.2e-05 | 1.033E-05 | 1.528E-06 | 9.000E-06 | 1.200E-05 |
| 3576 | 179 | G179S | A | 1.1e-05 | 1.5e-05 | 1.6e-05 | 1.400E-05 | 2.646E-06 | 1.100E-05 | 1.600E-05 |
| 3577 | 179 | G179T | A | 1.4e-05 | 9.9e-06 | 1.4e-05 | 1.263E-05 | 2.367E-06 | 9.900E-06 | 1.400E-05 |
| 3578 | 179 | G179W | A | 1e-05 | 1.2e-05 | 1e-05 | 1.067E-05 | 1.155E-06 | 1.000E-05 | 1.200E-05 |
| 3579 | 179 | G179Y | A | 9.8e-06 | 1.3e-05 | 9e-06 | 1.060E-05 | 2.117E-06 | 9.000E-06 | 1.300E-05 |
| 3580 | 179 | G179V | A | 1e-05 | 1.4e-05 | 1.5e-05 | 1.300E-05 | 2.646E-06 | 1.000E-05 | 1.500E-05 |
| 3581 | 180 | S180A | A | 1.4e-05 | 9.2e-06 | 1.1e-05 | 1.140E-05 | 2.425E-06 | 9.200E-06 | 1.400E-05 |
| 3582 | 180 | S180R | A | 9.7e-06 | 1e-05 | 1.1e-05 | 1.023E-05 | 6.807E-07 | 9.700E-06 | 1.100E-05 |
| 3583 | 180 | S180N | A | 1.5e-05 | 1e-05 | 1.1e-05 | 1.200E-05 | 2.646E-06 | 1.000E-05 | 1.500E-05 |
| 3584 | 180 | S180D | A | 1e-05 | 1.1e-05 | 1.1e-05 | 1.067E-05 | 5.774E-07 | 1.000E-05 | 1.100E-05 |
| 3585 | 180 | S180C | A | 1.1e-05 | 1.4e-05 | 1.4e-05 | 1.300E-05 | 1.732E-06 | 1.100E-05 | 1.400E-05 |
| 3586 | 180 | S180E | A | 1e-05 | 1.1e-05 | 8.8e-06 | 9.933E-06 | 1.102E-06 | 8.800E-06 | 1.100E-05 |
| 3587 | 180 | S180Q | A | 9.2e-06 | 1.5e-05 | 1.5e-05 | 1.307E-05 | 3.349E-06 | 9.200E-06 | 1.500E-05 |
| 3588 | 180 | S180G | A | 1e-05 | 1.4e-05 | 6.2e-06 | 1.007E-05 | 3.900E-06 | 6.200E-06 | 1.400E-05 |
| 3589 | 180 | S180H | A | 1.4e-05 | 1.7e-05 | 1.2e-05 | 1.433E-05 | 2.517E-06 | 1.200E-05 | 1.700E-05 |
| 3590 | 180 | S180I | A | 1.1e-05 | 1.5e-05 | 1.2e-05 | 1.267E-05 | 2.082E-06 | 1.100E-05 | 1.500E-05 |
| 3591 | 180 | S180L | A | 1.2e-05 | 1.7e-05 | 1.7e-05 | 1.533E-05 | 2.887E-06 | 1.200E-05 | 1.700E-05 |
| 3592 | 180 | S180K | A | 1.2e-05 | 1.1e-05 | 1e-05 | 1.100E-05 | 1.000E-06 | 1.000E-05 | 1.200E-05 |
| 3593 | 180 | S180M | A | 1e-05 | 1.4e-05 | 1.6e-05 | 1.333E-05 | 3.055E-06 | 1.000E-05 | 1.600E-05 |
| 3594 | 180 | S180F | A | 1.4e-05 | 1.4e-05 | 1.5e-05 | 1.433E-05 | 5.774E-07 | 1.400E-05 | 1.500E-05 |
| 3595 | 180 | S180P | A | 1.1e-05 | 1.6e-05 | 9.7e-06 | 1.223E-05 | 3.326E-06 | 9.700E-06 | 1.600E-05 |
| 3596 | 180 | S180S | A | 1.3e-05 | 1.5e-05 | 1.6e-05 | 1.467E-05 | 1.528E-06 | 1.300E-05 | 1.600E-05 |
| 3597 | 180 | S180T | A | 1.5e-05 | 9.5e-06 | 1e-05 | 1.150E-05 | 3.041E-06 | 9.500E-06 | 1.500E-05 |
| 3598 | 180 | S180W | A | 8.7e-06 | 1.6e-05 | 1.4e-05 | 1.290E-05 | 3.772E-06 | 8.700E-06 | 1.600E-05 |
| 3599 | 180 | S180Y | A | 1.7e-05 | 1.3e-05 | 8.9e-06 | 1.297E-05 | 4.050E-06 | 8.900E-06 | 1.700E-05 |
| 3600 | 180 | S180V | A | 1.5e-05 | 1.6e-05 | 1.3e-05 | 1.467E-05 | 1.528E-06 | 1.300E-05 | 1.600E-05 |
| 3601 | 181 | R181A | A | 1.4e-05 | 1.1e-05 | 1.5e-05 | 1.333E-05 | 2.082E-06 | 1.100E-05 | 1.500E-05 |
| 3602 | 181 | R181R | A | 1.6e-05 | 1.2e-05 | 1.4e-05 | 1.400E-05 | 2.000E-06 | 1.200E-05 | 1.600E-05 |
| 3603 | 181 | R181N | A | 1e-05 | 1.3e-05 | 1e-05 | 1.100E-05 | 1.732E-06 | 1.000E-05 | 1.300E-05 |
| 3604 | 181 | R181D | A | 1.6e-05 | 1.4e-05 | 1.9e-05 | 1.633E-05 | 2.517E-06 | 1.400E-05 | 1.900E-05 |
| 3605 | 181 | R181C | A | 1.4e-05 | 8.6e-06 | 1.6e-05 | 1.287E-05 | 3.828E-06 | 8.600E-06 | 1.600E-05 |
| 3606 | 181 | R181E | A | 1.4e-05 | 8.8e-06 | 1e-05 | 1.093E-05 | 2.723E-06 | 8.800E-06 | 1.400E-05 |
| 3607 | 181 | R181Q | A | 8.3e-06 | 1.7e-05 | 1e-05 | 1.177E-05 | 4.611E-06 | 8.300E-06 | 1.700E-05 |
| 3608 | 181 | R181G | A | 1.3e-05 | 1.1e-05 | 8.5e-06 | 1.083E-05 | 2.255E-06 | 8.500E-06 | 1.300E-05 |
| 3609 | 181 | R181H | A | 1.4e-05 | 1.5e-05 | 1.6e-05 | 1.500E-05 | 1.000E-06 | 1.400E-05 | 1.600E-05 |
| 3610 | 181 | R181I | A | 1.1e-05 | 1.5e-05 | 1.5e-05 | 1.367E-05 | 2.309E-06 | 1.100E-05 | 1.500E-05 |
| 3611 | 181 | R181L | A | 1.1e-05 | 1.1e-05 | 1.4e-05 | 1.200E-05 | 1.732E-06 | 1.100E-05 | 1.400E-05 |
| 3612 | 181 | R181K | A | 1.4e-05 | 1.7e-05 | 1.1e-05 | 1.400E-05 | 3.000E-06 | 1.100E-05 | 1.700E-05 |
| 3613 | 181 | R181M | A | 1.5e-05 | 1.5e-05 | 1.2e-05 | 1.400E-05 | 1.732E-06 | 1.200E-05 | 1.500E-05 |
| 3614 | 181 | R181F | A | 1.5e-05 | 1.4e-05 | 1.3e-05 | 1.400E-05 | 1.000E-06 | 1.300E-05 | 1.500E-05 |
| 3615 | 181 | R181P | A | 9.3e-06 | 1.3e-05 | 9.9e-06 | 1.073E-05 | 1.986E-06 | 9.300E-06 | 1.300E-05 |
| 3616 | 181 | R181S | A | 1.5e-05 | 1.5e-05 | 8.3e-06 | 1.277E-05 | 3.868E-06 | 8.300E-06 | 1.500E-05 |
| 3617 | 181 | R181T | A | 1.1e-05 | 1.4e-05 | 1.4e-05 | 1.300E-05 | 1.732E-06 | 1.100E-05 | 1.400E-05 |
| 3618 | 181 | R181W | A | 9.8e-06 | 8.6e-06 | 1.5e-05 | 1.113E-05 | 3.402E-06 | 8.600E-06 | 1.500E-05 |
| 3619 | 181 | R181Y | A | 1.5e-05 | 1.3e-05 | 1.1e-05 | 1.300E-05 | 2.000E-06 | 1.100E-05 | 1.500E-05 |
| 3620 | 181 | R181V | A | 1.5e-05 | 9.8e-06 | 1.1e-05 | 1.193E-05 | 2.723E-06 | 9.800E-06 | 1.500E-05 |
| 3621 | 182 | C182A | A | 1.6e-05 | 1.4e-05 | 8.5e-06 | 1.283E-05 | 3.884E-06 | 8.500E-06 | 1.600E-05 |
| 3622 | 182 | C182R | A | 1.7e-05 | 1.6e-05 | 1.6e-05 | 1.633E-05 | 5.774E-07 | 1.600E-05 | 1.700E-05 |
| 3623 | 182 | C182N | A | 1.4e-05 | 1.5e-05 | 1.2e-05 | 1.367E-05 | 1.528E-06 | 1.200E-05 | 1.500E-05 |
| 3624 | 182 | C182D | A | 1.3e-05 | 1.2e-05 | 1e-05 | 1.167E-05 | 1.528E-06 | 1.000E-05 | 1.300E-05 |
| 3625 | 182 | C182C | A | 1.6e-05 | 1.6e-05 | 1.3e-05 | 1.500E-05 | 1.732E-06 | 1.300E-05 | 1.600E-05 |
| 3626 | 182 | C182E | A | 9e-06 | 1.6e-05 | 9.1e-06 | 1.137E-05 | 4.013E-06 | 9.000E-06 | 1.600E-05 |
| 3627 | 182 | C182Q | A | 1.2e-05 | 1.3e-05 | 9.5e-06 | 1.150E-05 | 1.803E-06 | 9.500E-06 | 1.300E-05 |
| 3628 | 182 | C182G | A | 1e-05 | 8.7e-06 | 1.5e-05 | 1.123E-05 | 3.326E-06 | 8.700E-06 | 1.500E-05 |
| 3629 | 182 | C182H | A | 1.5e-05 | 1.5e-05 | 1.1e-05 | 1.367E-05 | 2.309E-06 | 1.100E-05 | 1.500E-05 |
| 3630 | 182 | C182I | A | 1e-05 | 1.5e-05 | 1.1e-05 | 1.200E-05 | 2.646E-06 | 1.000E-05 | 1.500E-05 |
| 3631 | 182 | C182L | A | 1.3e-05 | 1.6e-05 | 1.5e-05 | 1.467E-05 | 1.528E-06 | 1.300E-05 | 1.600E-05 |
| 3632 | 182 | C182K | A | 1e-05 | 1.5e-05 | 1e-05 | 1.167E-05 | 2.887E-06 | 1.000E-05 | 1.500E-05 |
| 3633 | 182 | C182M | A | 1.5e-05 | 1.5e-05 | 1.1e-05 | 1.367E-05 | 2.309E-06 | 1.100E-05 | 1.500E-05 |
| 3634 | 182 | C182F | A | 1.3e-05 | 1.3e-05 | 1.1e-05 | 1.233E-05 | 1.155E-06 | 1.100E-05 | 1.300E-05 |
| 3635 | 182 | C182P | A | 1.4e-05 | 1e-05 | 1.4e-05 | 1.267E-05 | 2.309E-06 | 1.000E-05 | 1.400E-05 |
| 3636 | 182 | C182S | A | 1.7e-05 | 1.2e-05 | 1.3e-05 | 1.400E-05 | 2.646E-06 | 1.200E-05 | 1.700E-05 |
| 3637 | 182 | C182T | A | 1.2e-05 | 1.6e-05 | 1.5e-05 | 1.433E-05 | 2.082E-06 | 1.200E-05 | 1.600E-05 |
| 3638 | 182 | C182W | A | 1.4e-05 | 1.2e-05 | 1e-05 | 1.200E-05 | 2.000E-06 | 1.000E-05 | 1.400E-05 |
| 3639 | 182 | C182Y | A | 1.6e-05 | 1.5e-05 | 1.5e-05 | 1.533E-05 | 5.774E-07 | 1.500E-05 | 1.600E-05 |
| 3640 | 182 | C182V | A | 1.2e-05 | 1.1e-05 | 1.3e-05 | 1.200E-05 | 1.000E-06 | 1.100E-05 | 1.300E-05 |
| 3641 | 183 | W183A | A | 1.1e-05 | 1.6e-05 | 1.5e-05 | 1.400E-05 | 2.646E-06 | 1.100E-05 | 1.600E-05 |
| 3642 | 183 | W183R | A | 1.5e-05 | 1.5e-05 | 1.6e-05 | 1.533E-05 | 5.774E-07 | 1.500E-05 | 1.600E-05 |
| 3643 | 183 | W183N | A | 2.1e-05 | 1.1e-05 | 1.5e-05 | 1.567E-05 | 5.033E-06 | 1.100E-05 | 2.100E-05 |
| 3644 | 183 | W183D | A | 1.3e-05 | 1.5e-05 | 1.1e-05 | 1.300E-05 | 2.000E-06 | 1.100E-05 | 1.500E-05 |
| 3645 | 183 | W183C | A | 1e-05 | 1.4e-05 | 1.4e-05 | 1.267E-05 | 2.309E-06 | 1.000E-05 | 1.400E-05 |
| 3646 | 183 | W183E | A | 1.1e-05 | 1.6e-05 | 1.1e-05 | 1.267E-05 | 2.887E-06 | 1.100E-05 | 1.600E-05 |
| 3647 | 183 | W183Q | A | 1.2e-05 | 1.6e-05 | 1e-05 | 1.267E-05 | 3.055E-06 | 1.000E-05 | 1.600E-05 |
| 3648 | 183 | W183G | A | 1.7e-05 | 1.4e-05 | 1.5e-05 | 1.533E-05 | 1.528E-06 | 1.400E-05 | 1.700E-05 |
| 3649 | 183 | W183H | A | 1.7e-05 | 8.5e-06 | 1e-05 | 1.183E-05 | 4.537E-06 | 8.500E-06 | 1.700E-05 |
| 3650 | 183 | W183I | A | 1.3e-05 | 1.5e-05 | 1.3e-05 | 1.367E-05 | 1.155E-06 | 1.300E-05 | 1.500E-05 |
| 3651 | 183 | W183L | A | 1.4e-05 | 1.5e-05 | 1.1e-05 | 1.333E-05 | 2.082E-06 | 1.100E-05 | 1.500E-05 |
| 3652 | 183 | W183K | A | 7.9e-06 | 1.4e-05 | 1.6e-05 | 1.263E-05 | 4.219E-06 | 7.900E-06 | 1.600E-05 |
| 3653 | 183 | W183M | A | 1.7e-05 | 1.5e-05 | 1.5e-05 | 1.567E-05 | 1.155E-06 | 1.500E-05 | 1.700E-05 |
| 3654 | 183 | W183F | A | 1.1e-05 | 8.3e-06 | 1.7e-05 | 1.210E-05 | 4.453E-06 | 8.300E-06 | 1.700E-05 |

| | | | | | | | | | | |
|---|---|---|---|---|---|---|---|---|---|---|
| 3655 | 183 | W183P | A | 1.1e-05 | 1e-05 | 1.5e-05 | 1.200E-05 | 2.646E-06 | 1.000E-05 | 1.500E-05 |
| 3656 | 183 | W183S | A | 9.7e-06 | 1.5e-05 | 1.7e-05 | 1.390E-05 | 3.772E-06 | 9.700E-06 | 1.700E-05 |
| 3657 | 183 | W183T | A | 1.6e-05 | 1.1e-05 | 1.2e-05 | 1.300E-05 | 2.646E-06 | 1.100E-05 | 1.600E-05 |
| 3658 | 183 | W183W | A | 9.2e-06 | 1.6e-05 | 1.6e-05 | 1.373E-05 | 3.926E-06 | 9.200E-06 | 1.600E-05 |
| 3659 | 183 | W183Y | A | 1.3e-05 | 9e-06 | 1e-05 | 1.067E-05 | 2.082E-06 | 9.000E-06 | 1.300E-05 |
| 3660 | 183 | W183V | A | 9.4e-06 | 1.6e-05 | 1.6e-05 | 1.380E-05 | 3.811E-06 | 9.400E-06 | 1.600E-05 |
| 3661 | 184 | G184A | A | 1.4e-05 | 1.5e-05 | 1.4e-05 | 1.433E-05 | 5.774E-07 | 1.400E-05 | 1.500E-05 |
| 3662 | 184 | G184R | A | 1.5e-05 | 1.2e-05 | 2.1e-05 | 1.600E-05 | 4.583E-06 | 1.200E-05 | 2.100E-05 |
| 3663 | 184 | G184N | A | 1.6e-05 | 1.8e-05 | 1.4e-05 | 1.600E-05 | 2.000E-06 | 1.400E-05 | 1.800E-05 |
| 3664 | 184 | G184D | A | 1.2e-05 | 9.9e-06 | 1.7e-05 | 1.297E-05 | 3.647E-06 | 9.900E-06 | 1.700E-05 |
| 3665 | 184 | G184C | A | 1.7e-05 | 1.2e-05 | 1e-05 | 1.300E-05 | 3.606E-06 | 1.000E-05 | 1.700E-05 |
| 3666 | 184 | G184E | A | 1.5e-05 | 1.7e-05 | 1.1e-05 | 1.433E-05 | 3.055E-06 | 1.100E-05 | 1.700E-05 |
| 3667 | 184 | G184Q | A | 1e-05 | 1.2e-05 | 1.4e-05 | 1.200E-05 | 2.000E-06 | 1.000E-05 | 1.400E-05 |
| 3668 | 184 | G184G | A | 1.3e-05 | 1.2e-05 | 1.4e-05 | 1.300E-05 | 1.000E-06 | 1.200E-05 | 1.400E-05 |
| 3669 | 184 | G184H | A | 1.4e-05 | 1.7e-05 | 1e-05 | 1.367E-05 | 3.512E-06 | 1.000E-05 | 1.700E-05 |
| 3670 | 184 | G184I | A | 1.1e-05 | 1e-05 | 1e-05 | 1.033E-05 | 5.774E-07 | 1.000E-05 | 1.100E-05 |
| 3671 | 184 | G184L | A | 1.2e-05 | 9.4e-06 | 1.1e-05 | 1.080E-05 | 1.311E-06 | 9.400E-06 | 1.200E-05 |
| 3672 | 184 | G184K | A | 8.9e-06 | 1.2e-05 | 9.7e-06 | 1.020E-05 | 1.609E-06 | 8.900E-06 | 1.200E-05 |
| 3673 | 184 | G184M | A | 1.6e-05 | 1.5e-05 | 1.5e-05 | 1.533E-05 | 5.774E-07 | 1.500E-05 | 1.600E-05 |
| 3674 | 184 | G184F | A | 1.6e-05 | 1e-05 | 1.6e-05 | 1.400E-05 | 3.464E-06 | 1.000E-05 | 1.600E-05 |
| 3675 | 184 | G184P | A | 1.5e-05 | 1.6e-05 | 1.5e-05 | 1.533E-05 | 5.774E-07 | 1.500E-05 | 1.600E-05 |
| 3676 | 184 | G184S | A | 1.1e-05 | 8.7e-06 | 1.6e-05 | 1.190E-05 | 3.732E-06 | 8.700E-06 | 1.600E-05 |
| 3677 | 184 | G184T | A | 1.6e-05 | 1.5e-05 | 1.1e-05 | 1.400E-05 | 2.646E-06 | 1.100E-05 | 1.600E-05 |
| 3678 | 184 | G184W | A | 1.4e-05 | 1.3e-05 | 1.7e-05 | 1.467E-05 | 2.082E-06 | 1.300E-05 | 1.700E-05 |
| 3679 | 184 | G184Y | A | 8.8e-06 | 1.3e-05 | 1.5e-05 | 1.227E-05 | 3.164E-06 | 8.800E-06 | 1.500E-05 |
| 3680 | 184 | G184V | A | 1e-05 | 1e-05 | 1.3e-05 | 1.100E-05 | 1.732E-06 | 1.000E-05 | 1.300E-05 |
| 3681 | 185 | E185A | A | 1.6e-05 | 1.7e-05 | 1.2e-05 | 1.500E-05 | 2.646E-06 | 1.200E-05 | 1.700E-05 |
| 3682 | 185 | E185R | A | 1.2e-05 | 1e-05 | 8.6e-06 | 1.020E-05 | 1.709E-06 | 8.600E-06 | 1.200E-05 |
| 3683 | 185 | E185N | A | 1e-05 | 1.5e-05 | 8.6e-06 | 1.120E-05 | 3.365E-06 | 8.600E-06 | 1.500E-05 |
| 3684 | 185 | E185D | A | 1e-05 | 1.6e-05 | 1.5e-05 | 1.367E-05 | 3.215E-06 | 1.000E-05 | 1.600E-05 |
| 3685 | 185 | E185C | A | 1.3e-05 | 1e-05 | 8.4e-06 | 1.047E-05 | 2.335E-06 | 8.400E-06 | 1.300E-05 |
| 3686 | 185 | E185E | A | 1.5e-05 | 1e-05 | 1.1e-05 | 1.200E-05 | 2.646E-06 | 1.000E-05 | 1.500E-05 |
| 3687 | 185 | E185Q | A | 1.4e-05 | 8.3e-06 | 1.5e-05 | 1.243E-05 | 3.614E-06 | 8.300E-06 | 1.500E-05 |
| 3688 | 185 | E185G | A | 1.5e-05 | 1.7e-05 | 1.4e-05 | 1.533E-05 | 1.528E-06 | 1.400E-05 | 1.700E-05 |
| 3689 | 185 | E185H | A | 1.5e-05 | 1.4e-05 | 1.3e-05 | 1.400E-05 | 1.000E-06 | 1.300E-05 | 1.500E-05 |
| 3690 | 185 | E185I | A | 8.5e-06 | 1.4e-05 | 1.6e-05 | 1.283E-05 | 3.884E-06 | 8.500E-06 | 1.600E-05 |
| 3691 | 185 | E185L | A | 1.3e-05 | 1.5e-05 | 1.5e-05 | 1.433E-05 | 1.155E-06 | 1.300E-05 | 1.500E-05 |
| 3692 | 185 | E185K | A | 1.6e-05 | 1.4e-05 | 1.7e-05 | 1.567E-05 | 1.528E-06 | 1.400E-05 | 1.700E-05 |
| 3693 | 185 | E185M | A | 9.2e-06 | 8.5e-06 | 1.5e-05 | 1.090E-05 | 3.568E-06 | 8.500E-06 | 1.500E-05 |
| 3694 | 185 | E185F | A | 1.2e-05 | 9.3e-06 | 1.1e-05 | 1.077E-05 | 1.365E-06 | 9.300E-06 | 1.200E-05 |
| 3695 | 185 | E185P | A | 1.6e-05 | 1e-05 | 1.1e-05 | 1.233E-05 | 3.215E-06 | 1.000E-05 | 1.600E-05 |
| 3696 | 185 | E185S | A | 1e-05 | 9.2e-06 | 9.7e-06 | 9.633E-06 | 4.041E-07 | 9.200E-06 | 1.000E-05 |
| 3697 | 185 | E185T | A | 1.1e-05 | 1.1e-05 | 1.4e-05 | 1.200E-05 | 1.732E-06 | 1.100E-05 | 1.400E-05 |
| 3698 | 185 | E185W | A | 1.6e-05 | 1.1e-05 | 1.1e-05 | 1.267E-05 | 2.887E-06 | 1.100E-05 | 1.600E-05 |
| 3699 | 185 | E185Y | A | 9.2e-06 | 8.7e-06 | 8.9e-06 | 8.933E-06 | 2.517E-07 | 8.700E-06 | 9.200E-06 |
| 3700 | 185 | E185V | A | 1.1e-05 | 9.1e-06 | 1.6e-05 | 1.203E-05 | 3.564E-06 | 9.100E-06 | 1.600E-05 |
| 3701 | 186 | S186A | A | 1.3e-05 | 1e-05 | 1.3e-05 | 1.200E-05 | 1.732E-06 | 1.000E-05 | 1.300E-05 |
| 3702 | 186 | S186R | A | 1.7e-05 | 1.1e-05 | 1e-05 | 1.267E-05 | 3.786E-06 | 1.000E-05 | 1.700E-05 |
| 3703 | 186 | S186N | A | 1e-05 | 1.1e-05 | 1.1e-05 | 1.067E-05 | 5.774E-07 | 1.000E-05 | 1.100E-05 |
| 3704 | 186 | S186D | A | 1.6e-05 | 9.2e-06 | 1.6e-05 | 1.373E-05 | 3.926E-06 | 9.200E-06 | 1.600E-05 |
| 3705 | 186 | S186C | A | 1.3e-05 | 1.6e-05 | 1.5e-05 | 1.467E-05 | 1.528E-06 | 1.300E-05 | 1.600E-05 |
| 3706 | 186 | S186E | A | 1.4e-05 | 1.1e-05 | 1.6e-05 | 1.367E-05 | 2.517E-06 | 1.100E-05 | 1.600E-05 |
| 3707 | 186 | S186Q | A | 9.3e-06 | 1.8e-05 | 8.8e-06 | 1.203E-05 | 5.173E-06 | 8.800E-06 | 1.800E-05 |
| 3708 | 186 | S186G | A | 1.5e-05 | 1.6e-05 | 1.5e-05 | 1.533E-05 | 5.774E-07 | 1.500E-05 | 1.600E-05 |
| 3709 | 186 | S186H | A | 1.4e-05 | 1.5e-05 | 1e-05 | 1.300E-05 | 2.646E-06 | 1.000E-05 | 1.500E-05 |
| 3710 | 186 | S186I | A | 1e-05 | 8.8e-06 | 1.4e-05 | 1.093E-05 | 2.723E-06 | 8.800E-06 | 1.400E-05 |
| 3711 | 186 | S186L | A | 1.1e-05 | 9.8e-06 | 1.1e-05 | 1.060E-05 | 6.928E-07 | 9.800E-06 | 1.100E-05 |
| 3712 | 186 | S186K | A | 1.1e-05 | 1.3e-05 | 1.3e-05 | 1.233E-05 | 1.155E-06 | 1.100E-05 | 1.300E-05 |
| 3713 | 186 | S186M | A | 1.1e-05 | 1.5e-05 | 1.1e-05 | 1.233E-05 | 2.309E-06 | 1.100E-05 | 1.500E-05 |
| 3714 | 186 | S186F | A | 1.2e-05 | 1.8e-05 | 1.6e-05 | 1.533E-05 | 3.055E-06 | 1.200E-05 | 1.800E-05 |
| 3715 | 186 | S186P | A | 1.4e-05 | 1.2e-05 | 1e-05 | 1.200E-05 | 2.000E-06 | 1.000E-05 | 1.400E-05 |
| 3716 | 186 | S186S | A | 1.1e-05 | 1.5e-05 | 1.2e-05 | 1.267E-05 | 2.082E-06 | 1.100E-05 | 1.500E-05 |
| 3717 | 186 | S186T | A | 1.2e-05 | 1.1e-05 | 8.7e-06 | 1.057E-05 | 1.692E-06 | 8.700E-06 | 1.200E-05 |
| 3718 | 186 | S186W | A | 1.4e-05 | 1.4e-05 | 8.4e-06 | 1.213E-05 | 3.233E-06 | 8.400E-06 | 1.400E-05 |
| 3719 | 186 | S186Y | A | 1e-05 | 9.4e-06 | 1.6e-05 | 1.180E-05 | 3.650E-06 | 9.400E-06 | 1.600E-05 |
| 3720 | 186 | S186V | A | 1.7e-05 | 1.5e-05 | 1e-05 | 1.400E-05 | 3.606E-06 | 1.000E-05 | 1.700E-05 |
| 3721 | 187 | S187A | A | 1.3e-05 | 1.4e-05 | 7.5e-06 | 1.150E-05 | 3.500E-06 | 7.500E-06 | 1.400E-05 |
| 3722 | 187 | S187R | A | 1.9e-05 | 1.4e-05 | 7.9e-06 | 1.363E-05 | 5.559E-06 | 7.900E-06 | 1.900E-05 |
| 3723 | 187 | S187N | A | 1.4e-05 | 1.1e-05 | 1.4e-05 | 1.300E-05 | 1.732E-06 | 1.100E-05 | 1.400E-05 |
| 3724 | 187 | S187D | A | 1e-05 | 1.6e-05 | 9.9e-06 | 1.197E-05 | 3.493E-06 | 9.900E-06 | 1.600E-05 |
| 3725 | 187 | S187C | A | 1.4e-05 | 1.1e-05 | 1.2e-05 | 1.233E-05 | 1.528E-06 | 1.100E-05 | 1.400E-05 |
| 3726 | 187 | S187E | A | 1.4e-05 | 1.5e-05 | 6.6e-06 | 1.187E-05 | 4.588E-06 | 6.600E-06 | 1.500E-05 |
| 3727 | 187 | S187Q | A | 2.5e-05 | 1.5e-05 | 1e-05 | 1.667E-05 | 7.638E-06 | 1.000E-05 | 2.500E-05 |
| 3728 | 187 | S187G | A | 1.5e-05 | 1.4e-05 | 1.3e-05 | 1.400E-05 | 1.000E-06 | 1.300E-05 | 1.500E-05 |
| 3729 | 187 | S187H | A | 1.6e-05 | 1.6e-05 | 1e-05 | 1.400E-05 | 3.464E-06 | 1.000E-05 | 1.600E-05 |
| 3730 | 187 | S187I | A | 1.2e-05 | 1.1e-05 | 1.1e-05 | 1.133E-05 | 5.774E-07 | 1.100E-05 | 1.200E-05 |
| 3731 | 187 | S187L | A | 9e-06 | 1.3e-05 | 1.6e-05 | 1.267E-05 | 3.512E-06 | 9.000E-06 | 1.600E-05 |
| 3732 | 187 | S187K | A | 1.4e-05 | 1.2e-05 | 1.4e-05 | 1.333E-05 | 1.155E-06 | 1.200E-05 | 1.400E-05 |
| 3733 | 187 | S187M | A | 1.3e-05 | 1.4e-05 | 1.6e-05 | 1.433E-05 | 1.528E-06 | 1.300E-05 | 1.600E-05 |
| 3734 | 187 | S187F | A | 1.5e-05 | 1.9e-05 | 1.3e-05 | 1.567E-05 | 3.055E-06 | 1.300E-05 | 1.900E-05 |
| 3735 | 187 | S187P | A | 1.1e-05 | 1e-05 | 9.2e-06 | 1.007E-05 | 9.018E-07 | 9.200E-06 | 1.100E-05 |
| 3736 | 187 | S187S | A | 8.1e-06 | 1.5e-05 | 1e-05 | 1.103E-05 | 3.564E-06 | 8.100E-06 | 1.500E-05 |
| 3737 | 187 | S187T | A | 1.8e-05 | 1.4e-05 | 1.4e-05 | 1.533E-05 | 2.309E-06 | 1.400E-05 | 1.800E-05 |
| 3738 | 187 | S187W | A | 1.4e-05 | 1.6e-05 | 1.1e-05 | 1.367E-05 | 2.517E-06 | 1.100E-05 | 1.600E-05 |
| 3739 | 187 | S187Y | A | 1.3e-05 | 9e-06 | 1e-05 | 1.067E-05 | 2.082E-06 | 9.000E-06 | 1.300E-05 |

| | | | | | | | | | | |
|---|---|---|---|---|---|---|---|---|---|---|
| 3740 | 187 | S187V | A | 1.6e-05 | 1.5e-05 | 1.2e-05 | 1.433E-05 | 2.082E-06 | 1.200E-05 | 1.600E-05 |
| 3741 | 188 | E188A | A | 9.9e-06 | 1.3e-05 | 1.7e-05 | 1.330E-05 | 3.559E-06 | 9.900E-06 | 1.700E-05 |
| 3742 | 188 | E188R | A | 1.4e-05 | 1.5e-05 | 1.5e-05 | 1.467E-05 | 5.774E-07 | 1.400E-05 | 1.500E-05 |
| 3743 | 188 | E188N | A | 1.8e-05 | 1.5e-05 | 1.5e-05 | 1.600E-05 | 1.732E-06 | 1.500E-05 | 1.800E-05 |
| 3744 | 188 | E188D | A | 1e-05 | 1.3e-05 | 1.6e-05 | 1.300E-05 | 3.000E-06 | 1.000E-05 | 1.600E-05 |
| 3745 | 188 | E188C | A | 1.6e-05 | 1.6e-05 | 1e-05 | 1.400E-05 | 3.464E-06 | 1.000E-05 | 1.600E-05 |
| 3746 | 188 | E188E | A | 1.5e-05 | 1.1e-05 | 1.5e-05 | 1.367E-05 | 2.309E-06 | 1.100E-05 | 1.500E-05 |
| 3747 | 188 | E188Q | A | 1.4e-05 | 1.5e-05 | 1.6e-05 | 1.500E-05 | 1.000E-06 | 1.400E-05 | 1.600E-05 |
| 3748 | 188 | E188G | A | 1.1e-05 | 1.2e-05 | 1.1e-05 | 1.133E-05 | 5.774E-07 | 1.100E-05 | 1.200E-05 |
| 3749 | 188 | E188H | A | 1.1e-05 | 1.5e-05 | 1e-05 | 1.200E-05 | 2.646E-06 | 1.000E-05 | 1.500E-05 |
| 3750 | 188 | E188I | A | 1.7e-05 | 1.1e-05 | 1.2e-05 | 1.333E-05 | 3.215E-06 | 1.100E-05 | 1.700E-05 |
| 3751 | 188 | E188L | A | 1.6e-05 | 1.1e-05 | 1.5e-05 | 1.400E-05 | 2.646E-06 | 1.100E-05 | 1.600E-05 |
| 3752 | 188 | E188K | A | 1.1e-05 | 1.2e-05 | 1.7e-05 | 1.333E-05 | 3.215E-06 | 1.100E-05 | 1.700E-05 |
| 3753 | 188 | E188M | A | 1.5e-05 | 1.4e-05 | 1.5e-05 | 1.467E-05 | 5.774E-07 | 1.400E-05 | 1.500E-05 |
| 3754 | 188 | E188F | A | 8.3e-06 | 1.5e-05 | 1.5e-05 | 1.277E-05 | 3.868E-06 | 8.300E-06 | 1.500E-05 |
| 3755 | 188 | E188P | A | 1.6e-05 | 1.4e-05 | 1.1e-05 | 1.367E-05 | 2.517E-06 | 1.100E-05 | 1.600E-05 |
| 3756 | 188 | E188S | A | 1.4e-05 | 1.6e-05 | 1.5e-05 | 1.500E-05 | 1.000E-06 | 1.400E-05 | 1.600E-05 |
| 3757 | 188 | E188T | A | 1.3e-05 | 9e-06 | 1.4e-05 | 1.200E-05 | 2.646E-06 | 9.000E-06 | 1.400E-05 |
| 3758 | 188 | E188W | A | 1e-05 | 1.1e-05 | 1.2e-05 | 1.100E-05 | 1.000E-06 | 1.000E-05 | 1.200E-05 |
| 3759 | 188 | E188Y | A | 1e-05 | 1.1e-05 | 1.5e-05 | 1.200E-05 | 2.646E-06 | 1.000E-05 | 1.500E-05 |
| 3760 | 188 | E188V | A | 1.5e-05 | 1.4e-05 | 1.3e-05 | 1.400E-05 | 1.000E-06 | 1.300E-05 | 1.500E-05 |
| 3761 | 189 | D189A | A | 1.5e-05 | 9.1e-06 | 1.4e-05 | 1.270E-05 | 3.158E-06 | 9.100E-06 | 1.500E-05 |
| 3762 | 189 | D189R | A | 1.1e-05 | 1e-05 | 1.6e-05 | 1.233E-05 | 3.215E-06 | 1.000E-05 | 1.600E-05 |
| 3763 | 189 | D189N | A | 1.4e-05 | 1.1e-05 | 1.5e-05 | 1.333E-05 | 2.082E-06 | 1.100E-05 | 1.500E-05 |
| 3764 | 189 | D189D | A | 1.2e-05 | 1.4e-05 | 9.2e-06 | 1.173E-05 | 2.411E-06 | 9.200E-06 | 1.400E-05 |
| 3765 | 189 | D189C | A | 9.3e-06 | 1.6e-05 | 1.5e-05 | 1.343E-05 | 3.614E-06 | 9.300E-06 | 1.600E-05 |
| 3766 | 189 | D189E | A | 1.5e-05 | 1.6e-05 | 1.9e-05 | 1.667E-05 | 2.082E-06 | 1.500E-05 | 1.900E-05 |
| 3767 | 189 | D189Q | A | 1.2e-05 | 9e-06 | 9.1e-06 | 1.003E-05 | 1.704E-06 | 9.000E-06 | 1.200E-05 |
| 3768 | 189 | D189G | A | 1.7e-05 | 1.5e-05 | 1.6e-05 | 1.600E-05 | 1.000E-06 | 1.500E-05 | 1.700E-05 |
| 3769 | 189 | D189H | A | 1.4e-05 | 1.6e-05 | 9e-06 | 1.300E-05 | 3.606E-06 | 9.000E-06 | 1.600E-05 |
| 3770 | 189 | D189I | A | 1e-05 | 1.6e-05 | 1.6e-05 | 1.400E-05 | 3.464E-06 | 1.000E-05 | 1.600E-05 |
| 3771 | 189 | D189L | A | 1.6e-05 | 1.8e-05 | 9.5e-06 | 1.450E-05 | 4.444E-06 | 9.500E-06 | 1.800E-05 |
| 3772 | 189 | D189K | A | 1.3e-05 | 1.1e-05 | 1.6e-05 | 1.333E-05 | 2.517E-06 | 1.100E-05 | 1.600E-05 |
| 3773 | 189 | D189M | A | 1.6e-05 | 8.2e-06 | 1.5e-05 | 1.307E-05 | 4.244E-06 | 8.200E-06 | 1.600E-05 |
| 3774 | 189 | D189F | A | 1.1e-05 | 1.1e-05 | 1.6e-05 | 1.267E-05 | 2.887E-06 | 1.100E-05 | 1.600E-05 |
| 3775 | 189 | D189P | A | 1.6e-05 | 1.4e-05 | 1.7e-05 | 1.567E-05 | 1.528E-06 | 1.400E-05 | 1.700E-05 |
| 3776 | 189 | D189S | A | 1.1e-05 | 9e-06 | 1.7e-05 | 1.233E-05 | 4.163E-06 | 9.000E-06 | 1.700E-05 |
| 3777 | 189 | D189T | A | 1.5e-05 | 8.4e-06 | 1.5e-05 | 1.280E-05 | 3.811E-06 | 8.400E-06 | 1.500E-05 |
| 3778 | 189 | D189W | A | 1.5e-05 | 1.6e-05 | 1.6e-05 | 1.567E-05 | 5.774E-07 | 1.500E-05 | 1.600E-05 |
| 3779 | 189 | D189Y | A | 1.5e-05 | 8.6e-06 | 1.5e-05 | 1.287E-05 | 3.695E-06 | 8.600E-06 | 1.500E-05 |
| 3780 | 189 | D189V | A | 1.6e-05 | 1.4e-05 | 1.4e-05 | 1.467E-05 | 1.155E-06 | 1.400E-05 | 1.600E-05 |
| 3781 | 190 | C190A | A | 1.4e-05 | 9.5e-06 | 1.8e-05 | 1.383E-05 | 4.252E-06 | 9.500E-06 | 1.800E-05 |
| 3782 | 190 | C190R | A | 1.9e-05 | 1.1e-05 | 1e-05 | 1.333E-05 | 4.933E-06 | 1.000E-05 | 1.900E-05 |
| 3783 | 190 | C190N | A | 1.5e-05 | 1.1e-05 | 1.4e-05 | 1.333E-05 | 2.082E-06 | 1.100E-05 | 1.500E-05 |
| 3784 | 190 | C190D | A | 1.2e-05 | 1.4e-05 | 1.6e-05 | 1.400E-05 | 2.000E-06 | 1.200E-05 | 1.600E-05 |
| 3785 | 190 | C190C | A | 8.5e-06 | 9.7e-06 | 1.8e-05 | 1.207E-05 | 5.173E-06 | 8.500E-06 | 1.800E-05 |
| 3786 | 190 | C190E | A | 1.9e-05 | 1.7e-05 | 1.6e-05 | 1.733E-05 | 1.528E-06 | 1.600E-05 | 1.900E-05 |
| 3787 | 190 | C190Q | A | 9.4e-06 | 1e-05 | 1.1e-05 | 1.013E-05 | 8.083E-07 | 9.400E-06 | 1.100E-05 |
| 3788 | 190 | C190G | A | 1.2e-05 | 1.4e-05 | 1.3e-05 | 1.300E-05 | 1.000E-06 | 1.200E-05 | 1.400E-05 |
| 3789 | 190 | C190H | A | 1.4e-05 | 1.5e-05 | 1.5e-05 | 1.467E-05 | 5.774E-07 | 1.400E-05 | 1.500E-05 |
| 3790 | 190 | C190I | A | 1.6e-05 | 1.5e-05 | 1.6e-05 | 1.567E-05 | 5.774E-07 | 1.500E-05 | 1.600E-05 |
| 3791 | 190 | C190L | A | 1.5e-05 | 1.4e-05 | 1.4e-05 | 1.433E-05 | 5.774E-07 | 1.400E-05 | 1.500E-05 |
| 3792 | 190 | C190K | A | 1.4e-05 | 1.1e-05 | 1.5e-05 | 1.333E-05 | 2.082E-06 | 1.100E-05 | 1.500E-05 |
| 3793 | 190 | C190M | A | 1.6e-05 | 1.4e-05 | 9.9e-06 | 1.330E-05 | 3.110E-06 | 9.900E-06 | 1.600E-05 |
| 3794 | 190 | C190F | A | 1.1e-05 | 1e-05 | 1.4e-05 | 1.167E-05 | 2.082E-06 | 1.000E-05 | 1.400E-05 |
| 3795 | 190 | C190P | A | 1.4e-05 | 1.6e-05 | 1.6e-05 | 1.533E-05 | 1.155E-06 | 1.400E-05 | 1.600E-05 |
| 3796 | 190 | C190S | A | 1.1e-05 | 1.4e-05 | 1.6e-05 | 1.367E-05 | 2.517E-06 | 1.100E-05 | 1.600E-05 |
| 3797 | 190 | C190T | A | 1.5e-05 | 2.4e-05 | 1.1e-05 | 1.667E-05 | 6.658E-06 | 1.100E-05 | 2.400E-05 |
| 3798 | 190 | C190W | A | 1.5e-05 | 1.4e-05 | 1.1e-05 | 1.333E-05 | 2.082E-06 | 1.100E-05 | 1.500E-05 |
| 3799 | 190 | C190Y | A | 9.5e-06 | 1.4e-05 | 1.5e-05 | 1.283E-05 | 2.930E-06 | 9.500E-06 | 1.500E-05 |
| 3800 | 190 | C190V | A | 1.6e-05 | 1e-05 | 1.2e-05 | 1.267E-05 | 3.055E-06 | 1.000E-05 | 1.600E-05 |
| 3801 | 191 | Q191A | A | 1.6e-05 | 1.1e-05 | 1e-05 | 1.233E-05 | 3.215E-06 | 1.000E-05 | 1.600E-05 |
| 3802 | 191 | Q191R | A | 1.5e-05 | 1.3e-05 | 1.5e-05 | 1.433E-05 | 1.155E-06 | 1.300E-05 | 1.500E-05 |
| 3803 | 191 | Q191N | A | 1.7e-05 | 1.4e-05 | 1.5e-05 | 1.533E-05 | 1.528E-06 | 1.400E-05 | 1.700E-05 |
| 3804 | 191 | Q191D | A | 1.2e-05 | 9.9e-06 | 1e-05 | 1.063E-05 | 1.185E-06 | 9.900E-06 | 1.200E-05 |
| 3805 | 191 | Q191C | A | 1.5e-05 | 1e-05 | 8.4e-06 | 1.113E-05 | 3.443E-06 | 8.400E-06 | 1.500E-05 |
| 3806 | 191 | Q191E | A | 1.4e-05 | 1.6e-05 | 1.1e-05 | 1.367E-05 | 2.517E-06 | 1.100E-05 | 1.600E-05 |
| 3807 | 191 | Q191Q | A | 1.1e-05 | 1e-05 | 1.2e-05 | 1.100E-05 | 1.000E-06 | 1.000E-05 | 1.200E-05 |
| 3808 | 191 | Q191G | A | 1.6e-05 | 1.5e-05 | 1.1e-05 | 1.400E-05 | 2.646E-06 | 1.100E-05 | 1.600E-05 |
| 3809 | 191 | Q191H | A | 1.6e-05 | 6.7e-06 | 1.1e-05 | 1.123E-05 | 4.654E-06 | 6.700E-06 | 1.600E-05 |
| 3810 | 191 | Q191I | A | 1.8e-05 | 1.3e-05 | 1e-05 | 1.367E-05 | 4.041E-06 | 1.000E-05 | 1.800E-05 |
| 3811 | 191 | Q191L | A | 1.1e-05 | 1.4e-05 | 1.5e-05 | 1.333E-05 | 2.082E-06 | 1.100E-05 | 1.500E-05 |
| 3812 | 191 | Q191K | A | 1.2e-05 | 1.7e-05 | 1.1e-05 | 1.333E-05 | 3.215E-06 | 1.100E-05 | 1.700E-05 |
| 3813 | 191 | Q191M | A | 1.2e-05 | 1.1e-05 | 1.6e-05 | 1.300E-05 | 2.646E-06 | 1.100E-05 | 1.600E-05 |
| 3814 | 191 | Q191F | A | 9.8e-06 | 1.1e-05 | 1.4e-05 | 1.160E-05 | 2.163E-06 | 9.800E-06 | 1.400E-05 |
| 3815 | 191 | Q191P | A | 1.6e-05 | 1.2e-05 | 9.6e-06 | 1.253E-05 | 3.233E-06 | 9.600E-06 | 1.600E-05 |
| 3816 | 191 | Q191S | A | 1.5e-05 | 1.6e-05 | 1.1e-05 | 1.400E-05 | 2.646E-06 | 1.100E-05 | 1.600E-05 |
| 3817 | 191 | Q191T | A | 1.6e-05 | 1.4e-05 | 1.6e-05 | 1.533E-05 | 1.155E-06 | 1.400E-05 | 1.600E-05 |
| 3818 | 191 | Q191W | A | 1.3e-05 | 1.5e-05 | 9.8e-06 | 1.260E-05 | 2.623E-06 | 9.800E-06 | 1.500E-05 |
| 3819 | 191 | Q191Y | A | 1e-05 | 1.4e-05 | 1.6e-05 | 1.333E-05 | 3.055E-06 | 1.000E-05 | 1.600E-05 |
| 3820 | 191 | Q191V | A | 1.5e-05 | 1e-05 | 1.7e-05 | 1.400E-05 | 3.606E-06 | 1.000E-05 | 1.700E-05 |
| 3821 | 192 | S192A | A | 1.2e-05 | 1.5e-05 | 9.5e-06 | 1.217E-05 | 2.754E-06 | 9.500E-06 | 1.500E-05 |
| 3822 | 192 | S192R | A | 9.9e-06 | 1e-05 | 1.1e-05 | 1.030E-05 | 6.083E-07 | 9.900E-06 | 1.100E-05 |
| 3823 | 192 | S192N | A | 1.6e-05 | 1e-05 | 9.4e-06 | 1.180E-05 | 3.650E-06 | 9.400E-06 | 1.600E-05 |
| 3824 | 192 | S192D | A | 1.6e-05 | 1e-05 | 1.1e-05 | 1.233E-05 | 3.215E-06 | 1.000E-05 | 1.600E-05 |

| | | | | | | | | | | |
|---|---|---|---|---|---|---|---|---|---|---|
| 3825 | 192 | S192C | A | 1.1e-05 | 1.4e-05 | 1.3e-05 | 1.267E-05 | 1.528E-06 | 1.100E-05 | 1.400E-05 |
| 3826 | 192 | S192E | A | 1.6e-05 | 1.1e-05 | 1.5e-05 | 1.400E-05 | 2.646E-06 | 1.100E-05 | 1.600E-05 |
| 3827 | 192 | S192Q | A | 1e-05 | 1e-05 | 1.4e-05 | 1.133E-05 | 2.309E-06 | 1.000E-05 | 1.400E-05 |
| 3828 | 192 | S192G | A | 1.4e-05 | 1.5e-05 | 1.5e-05 | 1.467E-05 | 5.774E-07 | 1.400E-05 | 1.500E-05 |
| 3829 | 192 | S192H | A | 9.3e-06 | 1e-05 | 1.4e-05 | 1.110E-05 | 2.536E-06 | 9.300E-06 | 1.400E-05 |
| 3830 | 192 | S192I | A | 1.4e-05 | 1.1e-05 | 1.7e-05 | 1.400E-05 | 3.000E-06 | 1.100E-05 | 1.700E-05 |
| 3831 | 192 | S192L | A | 8.9e-06 | 1.4e-05 | 1.5e-05 | 1.263E-05 | 3.272E-06 | 8.900E-06 | 1.500E-05 |
| 3832 | 192 | S192K | A | 1.1e-05 | 1.5e-05 | 1.4e-05 | 1.333E-05 | 2.082E-06 | 1.100E-05 | 1.500E-05 |
| 3833 | 192 | S192M | A | 1e-05 | 1.2e-05 | 1.1e-05 | 1.100E-05 | 1.000E-06 | 1.000E-05 | 1.200E-05 |
| 3834 | 192 | S192F | A | 1.1e-05 | 1.6e-05 | 1.7e-05 | 1.467E-05 | 3.215E-06 | 1.100E-05 | 1.700E-05 |
| 3835 | 192 | S192P | A | 1.8e-05 | 1e-05 | 1e-05 | 1.267E-05 | 4.619E-06 | 1.000E-05 | 1.800E-05 |
| 3836 | 192 | S192S | A | 1.1e-05 | 1e-05 | 1.2e-05 | 1.100E-05 | 1.000E-06 | 1.000E-05 | 1.200E-05 |
| 3837 | 192 | S192T | A | 5.8e-06 | 1e-05 | 1.5e-05 | 1.027E-05 | 4.606E-06 | 5.800E-06 | 1.500E-05 |
| 3838 | 192 | S192W | A | 1.6e-05 | 1.4e-05 | 1.7e-05 | 1.567E-05 | 1.528E-06 | 1.400E-05 | 1.700E-05 |
| 3839 | 192 | S192Y | A | 1.4e-05 | 1e-05 | 1.6e-05 | 1.333E-05 | 3.055E-06 | 1.000E-05 | 1.600E-05 |
| 3840 | 192 | S192V | A | 1.5e-05 | 1.3e-05 | 1.5e-05 | 1.433E-05 | 1.155E-06 | 1.300E-05 | 1.500E-05 |
| 3841 | 193 | L193A | A | 1.2e-05 | 1.5e-05 | 9.3e-06 | 1.210E-05 | 2.851E-06 | 9.300E-06 | 1.500E-05 |
| 3842 | 193 | L193R | A | 1.4e-05 | 1.5e-05 | 1e-05 | 1.300E-05 | 2.646E-06 | 1.000E-05 | 1.500E-05 |
| 3843 | 193 | L193N | A | 9.3e-06 | 8.1e-06 | 1e-05 | 9.133E-06 | 9.609E-07 | 8.100E-06 | 1.000E-05 |
| 3844 | 193 | L193D | A | 1e-05 | 9.2e-06 | 9.8e-06 | 9.667E-06 | 4.163E-07 | 9.200E-06 | 1.000E-05 |
| 3845 | 193 | L193C | A | 1.4e-05 | 9e-06 | 1e-05 | 1.100E-05 | 2.646E-06 | 9.000E-06 | 1.400E-05 |
| 3846 | 193 | L193E | A | 7.2e-06 | 1.2e-05 | 1.5e-05 | 1.140E-05 | 3.934E-06 | 7.200E-06 | 1.500E-05 |
| 3847 | 193 | L193Q | A | 8.7e-06 | 1.6e-05 | 1.3e-05 | 1.257E-05 | 3.669E-06 | 8.700E-06 | 1.600E-05 |
| 3848 | 193 | L193G | A | 1.4e-05 | 1.5e-05 | 9.1e-06 | 1.270E-05 | 3.158E-06 | 9.100E-06 | 1.500E-05 |
| 3849 | 193 | L193H | A | 1.1e-05 | 1.3e-05 | 1.3e-05 | 1.233E-05 | 1.155E-06 | 1.100E-05 | 1.300E-05 |
| 3850 | 193 | L193I | A | 1.6e-05 | 1.3e-05 | 1.5e-05 | 1.467E-05 | 1.528E-06 | 1.300E-05 | 1.600E-05 |
| 3851 | 193 | L193L | A | 1.1e-05 | 2.2e-05 | 9.5e-06 | 1.417E-05 | 6.825E-06 | 9.500E-06 | 2.200E-05 |
| 3852 | 193 | L193K | A | 1.5e-05 | 1.5e-05 | 1.4e-05 | 1.467E-05 | 5.774E-07 | 1.400E-05 | 1.500E-05 |
| 3853 | 193 | L193M | A | 7.7e-06 | 1.6e-05 | 1.5e-05 | 1.290E-05 | 4.531E-06 | 7.700E-06 | 1.600E-05 |
| 3854 | 193 | L193F | A | 9.3e-06 | 1.1e-05 | 1.5e-05 | 1.177E-05 | 2.926E-06 | 9.300E-06 | 1.500E-05 |
| 3855 | 193 | L193P | A | 1e-05 | 1e-05 | 1.3e-05 | 1.100E-05 | 1.732E-06 | 1.000E-05 | 1.300E-05 |
| 3856 | 193 | L193S | A | 1.2e-05 | 1.5e-05 | 9e-06 | 1.200E-05 | 3.000E-06 | 9.000E-06 | 1.500E-05 |
| 3857 | 193 | L193T | A | 1.6e-05 | 1.6e-05 | 9.9e-06 | 1.397E-05 | 3.522E-06 | 9.900E-06 | 1.600E-05 |
| 3858 | 193 | L193W | A | 1.4e-05 | 1.5e-05 | 1.5e-05 | 1.467E-05 | 5.774E-07 | 1.400E-05 | 1.500E-05 |
| 3859 | 193 | L193Y | A | 1.1e-05 | 1.4e-05 | 1.7e-05 | 1.400E-05 | 3.000E-06 | 1.100E-05 | 1.700E-05 |
| 3860 | 193 | L193V | A | 1.6e-05 | 1.2e-05 | 9.5e-06 | 1.250E-05 | 3.279E-06 | 9.500E-06 | 1.600E-05 |
| 3861 | 194 | T194A | A | 9.9e-06 | 1.5e-05 | 8.6e-06 | 1.117E-05 | 3.383E-06 | 8.600E-06 | 1.500E-05 |
| 3862 | 194 | T194R | A | 1.3e-05 | 9.9e-06 | 1e-05 | 1.097E-05 | 1.762E-06 | 9.900E-06 | 1.300E-05 |
| 3863 | 194 | T194N | A | 1.5e-05 | 1.1e-05 | 8.5e-06 | 1.150E-05 | 3.279E-06 | 8.500E-06 | 1.500E-05 |
| 3864 | 194 | T194D | A | 1.1e-05 | 8.2e-06 | 1e-05 | 9.733E-06 | 1.419E-06 | 8.200E-06 | 1.100E-05 |
| 3865 | 194 | T194C | A | 1.6e-05 | 9.2e-06 | 1.2e-05 | 1.240E-05 | 3.418E-06 | 9.200E-06 | 1.600E-05 |
| 3866 | 194 | T194E | A | 1.4e-05 | 1.2e-05 | 1.1e-05 | 1.233E-05 | 1.528E-06 | 1.100E-05 | 1.400E-05 |
| 3867 | 194 | T194Q | A | 1.4e-05 | 1.6e-05 | 1e-05 | 1.333E-05 | 3.055E-06 | 1.000E-05 | 1.600E-05 |
| 3868 | 194 | T194G | A | 1.5e-05 | 1.1e-05 | 1.6e-05 | 1.400E-05 | 2.646E-06 | 1.100E-05 | 1.600E-05 |
| 3869 | 194 | T194H | A | 1.6e-05 | 1.4e-05 | 1.6e-05 | 1.533E-05 | 1.155E-06 | 1.400E-05 | 1.600E-05 |
| 3870 | 194 | T194I | A | 8.9e-06 | 9.3e-06 | 1.5e-05 | 1.107E-05 | 3.412E-06 | 8.900E-06 | 1.500E-05 |
| 3871 | 194 | T194L | A | 1.5e-05 | 1.5e-05 | 9.3e-06 | 1.310E-05 | 3.291E-06 | 9.300E-06 | 1.500E-05 |
| 3872 | 194 | T194K | A | 1.6e-05 | 1.1e-05 | 1.1e-05 | 1.267E-05 | 2.887E-06 | 1.100E-05 | 1.600E-05 |
| 3873 | 194 | T194M | A | 1e-05 | 1.5e-05 | 1.7e-05 | 1.400E-05 | 3.606E-06 | 1.000E-05 | 1.700E-05 |
| 3874 | 194 | T194F | A | 1.4e-05 | 1.1e-05 | 1e-05 | 1.167E-05 | 2.082E-06 | 1.000E-05 | 1.400E-05 |
| 3875 | 194 | T194P | A | 1.6e-05 | 1.6e-05 | 1e-05 | 1.400E-05 | 3.464E-06 | 1.000E-05 | 1.600E-05 |
| 3876 | 194 | T194S | A | 8.5e-06 | 1.5e-05 | 1e-05 | 1.117E-05 | 3.403E-06 | 8.500E-06 | 1.500E-05 |
| 3877 | 194 | T194T | A | 1.5e-05 | 9.5e-06 | 1.6e-05 | 1.350E-05 | 3.500E-06 | 9.500E-06 | 1.600E-05 |
| 3878 | 194 | T194W | A | 1.1e-05 | 9.6e-06 | 9e-06 | 9.867E-06 | 1.026E-06 | 9.000E-06 | 1.100E-05 |
| 3879 | 194 | T194Y | A | 1.3e-05 | 8.9e-06 | 8.8e-06 | 1.023E-05 | 2.397E-06 | 8.800E-06 | 1.300E-05 |
| 3880 | 194 | T194V | A | 1.3e-05 | 1.5e-05 | 9.5e-06 | 1.250E-05 | 2.784E-06 | 9.500E-06 | 1.500E-05 |
| 3881 | 195 | R195A | A | 1.2e-05 | 1.1e-05 | 1.6e-05 | 1.300E-05 | 2.646E-06 | 1.100E-05 | 1.600E-05 |
| 3882 | 195 | R195R | A | 1e-05 | 8.9e-06 | 1.1e-05 | 9.967E-06 | 1.050E-06 | 8.900E-06 | 1.100E-05 |
| 3883 | 195 | R195N | A | 8.7e-06 | 1e-05 | 1.3e-05 | 1.057E-05 | 2.205E-06 | 8.700E-06 | 1.300E-05 |
| 3884 | 195 | R195D | A | 8.9e-06 | 1.4e-05 | 1.1e-05 | 1.130E-05 | 2.563E-06 | 8.900E-06 | 1.400E-05 |
| 3885 | 195 | R195C | A | 8.2e-06 | 1.5e-05 | 1.4e-05 | 1.240E-05 | 3.672E-06 | 8.200E-06 | 1.500E-05 |
| 3886 | 195 | R195E | A | 9.9e-06 | 1.6e-05 | 1.5e-05 | 1.363E-05 | 3.272E-06 | 9.900E-06 | 1.600E-05 |
| 3887 | 195 | R195Q | A | 1.3e-05 | 1.6e-05 | 1.4e-05 | 1.433E-05 | 1.528E-06 | 1.300E-05 | 1.600E-05 |
| 3888 | 195 | R195G | A | 9.6e-06 | 1.5e-05 | 1.5e-05 | 1.320E-05 | 3.118E-06 | 9.600E-06 | 1.500E-05 |
| 3889 | 195 | R195H | A | 8.6e-06 | 1.1e-05 | 1.4e-05 | 1.120E-05 | 2.706E-06 | 8.600E-06 | 1.400E-05 |
| 3890 | 195 | R195I | A | 1.6e-05 | 1.3e-05 | 9.9e-06 | 1.297E-05 | 3.050E-06 | 9.900E-06 | 1.600E-05 |
| 3891 | 195 | R195L | A | 8.8e-06 | 7.9e-06 | 1.5e-05 | 1.057E-05 | 3.866E-06 | 7.900E-06 | 1.500E-05 |
| 3892 | 195 | R195K | A | 1e-05 | 1.1e-05 | 1e-05 | 1.033E-05 | 5.774E-07 | 1.000E-05 | 1.100E-05 |
| 3893 | 195 | R195M | A | 1e-05 | 1.6e-05 | 9.8e-06 | 1.193E-05 | 3.523E-06 | 9.800E-06 | 1.600E-05 |
| 3894 | 195 | R195F | A | 1.2e-05 | 1.6e-05 | 1.6e-05 | 1.467E-05 | 2.309E-06 | 1.200E-05 | 1.600E-05 |
| 3895 | 195 | R195P | A | 1e-05 | 1.5e-05 | 1.5e-05 | 1.333E-05 | 2.887E-06 | 1.000E-05 | 1.500E-05 |
| 3896 | 195 | R195S | A | 9.5e-06 | 9.7e-06 | 1.4e-05 | 1.107E-05 | 2.542E-06 | 9.500E-06 | 1.400E-05 |
| 3897 | 195 | R195T | A | 1e-05 | 1.5e-05 | 1.6e-05 | 1.367E-05 | 3.215E-06 | 1.000E-05 | 1.600E-05 |
| 3898 | 195 | R195W | A | 1.1e-05 | 8.7e-06 | 1.1e-05 | 1.023E-05 | 1.328E-06 | 8.700E-06 | 1.100E-05 |
| 3899 | 195 | R195Y | A | 1.4e-05 | 1.5e-05 | 1.1e-05 | 1.333E-05 | 2.082E-06 | 1.100E-05 | 1.500E-05 |
| 3900 | 195 | R195V | A | 1.2e-05 | 1.5e-05 | 1.1e-05 | 1.267E-05 | 2.082E-06 | 1.100E-05 | 1.500E-05 |
| 3901 | 196 | T196A | A | 1.5e-05 | 1.8e-05 | 1.1e-05 | 1.467E-05 | 3.512E-06 | 1.100E-05 | 1.800E-05 |
| 3902 | 196 | T196R | A | 1.9e-05 | 1.7e-05 | 1e-05 | 1.533E-05 | 4.726E-06 | 1.000E-05 | 1.900E-05 |
| 3903 | 196 | T196N | A | 9e-06 | 1.3e-05 | 9.8e-06 | 1.060E-05 | 2.117E-06 | 9.000E-06 | 1.300E-05 |
| 3904 | 196 | T196D | A | 1.7e-05 | 2.5e-05 | 1.6e-05 | 1.933E-05 | 4.933E-06 | 1.600E-05 | 2.500E-05 |
| 3905 | 196 | T196C | A | 1.6e-05 | 9.8e-06 | 8.8e-06 | 1.153E-05 | 3.900E-06 | 8.800E-06 | 1.600E-05 |
| 3906 | 196 | T196E | A | 1.1e-05 | 1.5e-05 | 1.1e-05 | 1.233E-05 | 2.309E-06 | 1.100E-05 | 1.500E-05 |
| 3907 | 196 | T196Q | A | 1.5e-05 | 1.1e-05 | 1.3e-05 | 1.300E-05 | 2.000E-06 | 1.100E-05 | 1.500E-05 |
| 3908 | 196 | T196G | A | 9.6e-06 | 1.2e-05 | 1.6e-05 | 1.253E-05 | 3.233E-06 | 9.600E-06 | 1.600E-05 |
| 3909 | 196 | T196H | A | 1.4e-05 | 1.6e-05 | 9.8e-06 | 1.327E-05 | 3.164E-06 | 9.800E-06 | 1.600E-05 |

| | | | | | | | | | | |
|---|---|---|---|---|---|---|---|---|---|---|
| 3910 | 196 | T196I | A | 1.6e-05 | 2.5e-05 | 1.1e-05 | 1.733E-05 | 7.095E-06 | 1.100E-05 | 2.500E-05 |
| 3911 | 196 | T196L | A | 1.5e-05 | 1.6e-05 | 1.2e-05 | 1.433E-05 | 2.082E-06 | 1.200E-05 | 1.600E-05 |
| 3912 | 196 | T196K | A | 1e-05 | 1e-05 | 1.1e-05 | 1.033E-05 | 5.774E-07 | 1.000E-05 | 1.100E-05 |
| 3913 | 196 | T196M | A | 1e-05 | 1.5e-05 | 1.6e-05 | 1.367E-05 | 3.215E-06 | 1.000E-05 | 1.600E-05 |
| 3914 | 196 | T196F | A | 1.1e-05 | 1.6e-05 | 1.3e-05 | 1.333E-05 | 2.517E-06 | 1.100E-05 | 1.600E-05 |
| 3915 | 196 | T196P | A | 1.5e-05 | 1.5e-05 | 1e-05 | 1.333E-05 | 2.887E-06 | 1.000E-05 | 1.500E-05 |
| 3916 | 196 | T196S | A | 1e-05 | 1e-05 | 1.3e-05 | 1.100E-05 | 1.732E-06 | 1.000E-05 | 1.300E-05 |
| 3917 | 196 | T196T | A | 9.7e-06 | 1e-05 | 1.1e-05 | 1.023E-05 | 6.807E-07 | 9.700E-06 | 1.100E-05 |
| 3918 | 196 | T196W | A | 9.3e-06 | 1.1e-05 | 1.1e-05 | 1.043E-05 | 9.815E-07 | 9.300E-06 | 1.100E-05 |
| 3919 | 196 | T196Y | A | 1.6e-05 | 9.2e-06 | 9.4e-06 | 1.153E-05 | 3.870E-06 | 9.200E-06 | 1.600E-05 |
| 3920 | 196 | T196V | A | 1.5e-05 | 1.5e-05 | 1e-05 | 1.333E-05 | 2.887E-06 | 1.000E-05 | 1.500E-05 |
| 3921 | 197 | V197A | A | 1.7e-05 | 1e-05 | 1.1e-05 | 1.267E-05 | 3.786E-06 | 1.000E-05 | 1.700E-05 |
| 3922 | 197 | V197R | A | 7.5e-06 | 1.1e-05 | 1.4e-05 | 1.083E-05 | 3.253E-06 | 7.500E-06 | 1.400E-05 |
| 3923 | 197 | V197N | A | 1.2e-05 | 1.5e-05 | 1.1e-05 | 1.267E-05 | 2.082E-06 | 1.100E-05 | 1.500E-05 |
| 3924 | 197 | V197D | A | 1.5e-05 | 1.6e-05 | 1.1e-05 | 1.400E-05 | 2.646E-06 | 1.100E-05 | 1.600E-05 |
| 3925 | 197 | V197C | A | 1e-05 | 1.1e-05 | 9.4e-06 | 1.013E-05 | 8.083E-07 | 9.400E-06 | 1.100E-05 |
| 3926 | 197 | V197E | A | 1.1e-05 | 1.2e-05 | 9.3e-06 | 1.077E-05 | 1.365E-06 | 9.300E-06 | 1.200E-05 |
| 3927 | 197 | V197Q | A | 1.1e-05 | 1.7e-05 | 1e-05 | 1.267E-05 | 3.786E-06 | 1.000E-05 | 1.700E-05 |
| 3928 | 197 | V197G | A | 1e-05 | 1.7e-05 | 9e-06 | 1.200E-05 | 4.359E-06 | 9.000E-06 | 1.700E-05 |
| 3929 | 197 | V197H | A | 1.6e-05 | 1.7e-05 | 8.7e-06 | 1.390E-05 | 4.531E-06 | 8.700E-06 | 1.700E-05 |
| 3930 | 197 | V197I | A | 1.5e-05 | 1e-05 | 8.4e-06 | 1.113E-05 | 3.443E-06 | 8.400E-06 | 1.500E-05 |
| 3931 | 197 | V197L | A | 1.5e-05 | 7.8e-06 | 1.5e-05 | 1.260E-05 | 4.157E-06 | 7.800E-06 | 1.500E-05 |
| 3932 | 197 | V197K | A | 8.5e-06 | 1.4e-05 | 1e-05 | 1.083E-05 | 2.843E-06 | 8.500E-06 | 1.400E-05 |
| 3933 | 197 | V197M | A | 1.6e-05 | 1.5e-05 | 1.6e-05 | 1.567E-05 | 5.774E-07 | 1.500E-05 | 1.600E-05 |
| 3934 | 197 | V197F | A | 1.1e-05 | 1e-05 | 1.1e-05 | 1.067E-05 | 5.774E-07 | 1.000E-05 | 1.100E-05 |
| 3935 | 197 | V197P | A | 2.3e-05 | 1.5e-05 | 1e-05 | 1.600E-05 | 6.557E-06 | 1.000E-05 | 2.300E-05 |
| 3936 | 197 | V197S | A | 1.6e-05 | 7.7e-06 | 1e-05 | 1.123E-05 | 4.285E-06 | 7.700E-06 | 1.600E-05 |
| 3937 | 197 | V197T | A | 9.5e-06 | 8.5e-06 | 1.5e-05 | 1.100E-05 | 3.500E-06 | 8.500E-06 | 1.500E-05 |
| 3938 | 197 | V197W | A | 9.5e-06 | 1.4e-05 | 1.6e-05 | 1.317E-05 | 3.329E-06 | 9.500E-06 | 1.600E-05 |
| 3939 | 197 | V197Y | A | 1.7e-05 | 1e-05 | 1.6e-05 | 1.433E-05 | 3.786E-06 | 1.000E-05 | 1.700E-05 |
| 3940 | 197 | V197V | A | 1.6e-05 | 1.6e-05 | 1.4e-05 | 1.533E-05 | 1.155E-06 | 1.400E-05 | 1.600E-05 |
| 3941 | 198 | C198A | A | 1.6e-05 | 1.8e-05 | 1e-05 | 1.467E-05 | 4.163E-06 | 1.000E-05 | 1.800E-05 |
| 3942 | 198 | C198R | A | 1.5e-05 | 1.7e-05 | 1.4e-05 | 1.533E-05 | 1.528E-06 | 1.400E-05 | 1.700E-05 |
| 3943 | 198 | C198N | A | 1.7e-05 | 1e-05 | 1e-05 | 1.233E-05 | 4.041E-06 | 1.000E-05 | 1.700E-05 |
| 3944 | 198 | C198D | A | 1.5e-05 | 1.6e-05 | 1.5e-05 | 1.533E-05 | 5.774E-07 | 1.500E-05 | 1.600E-05 |
| 3945 | 198 | C198C | A | 1.2e-05 | 1.4e-05 | 1.5e-05 | 1.367E-05 | 1.528E-06 | 1.200E-05 | 1.500E-05 |
| 3946 | 198 | C198E | A | 1.1e-05 | 1.1e-05 | 1.7e-05 | 1.300E-05 | 3.464E-06 | 1.100E-05 | 1.700E-05 |
| 3947 | 198 | C198Q | A | 1e-05 | 9e-06 | 1.1e-05 | 1.000E-05 | 1.000E-06 | 9.000E-06 | 1.100E-05 |
| 3948 | 198 | C198G | A | 1.1e-05 | 1.1e-05 | 1.6e-05 | 1.267E-05 | 2.887E-06 | 1.100E-05 | 1.600E-05 |
| 3949 | 198 | C198H | A | 1.3e-05 | 1.4e-05 | 9.6e-06 | 1.220E-05 | 2.307E-06 | 9.600E-06 | 1.400E-05 |
| 3950 | 198 | C198I | A | 1e-05 | 1.6e-05 | 1.2e-05 | 1.267E-05 | 3.055E-06 | 1.000E-05 | 1.600E-05 |
| 3951 | 198 | C198L | A | 1.1e-05 | 1e-05 | 1.6e-05 | 1.233E-05 | 3.215E-06 | 1.000E-05 | 1.600E-05 |
| 3952 | 198 | C198K | A | 1e-05 | 1.5e-05 | 1.7e-05 | 1.400E-05 | 3.606E-06 | 1.000E-05 | 1.700E-05 |
| 3953 | 198 | C198M | A | 1.5e-05 | 1e-05 | 9.5e-06 | 1.150E-05 | 3.041E-06 | 9.500E-06 | 1.500E-05 |
| 3954 | 198 | C198F | A | 1.7e-05 | 1.7e-05 | 1.5e-05 | 1.633E-05 | 1.155E-06 | 1.500E-05 | 1.700E-05 |
| 3955 | 198 | C198P | A | 9e-06 | 1.5e-05 | 1.5e-05 | 1.300E-05 | 3.464E-06 | 9.000E-06 | 1.500E-05 |
| 3956 | 198 | C198S | A | 1.3e-05 | 1.5e-05 | 1.6e-05 | 1.467E-05 | 1.528E-06 | 1.300E-05 | 1.600E-05 |
| 3957 | 198 | C198T | A | 1.5e-05 | 1.6e-05 | 1.5e-05 | 1.533E-05 | 5.774E-07 | 1.500E-05 | 1.600E-05 |
| 3958 | 198 | C198W | A | 9.3e-06 | 1.6e-05 | 1.5e-05 | 1.343E-05 | 3.614E-06 | 9.300E-06 | 1.600E-05 |
| 3959 | 198 | C198Y | A | 1.1e-05 | 1.3e-05 | 1.1e-05 | 1.167E-05 | 1.155E-06 | 1.100E-05 | 1.300E-05 |
| 3960 | 198 | C198V | A | 1.1e-05 | 9e-06 | 1.6e-05 | 1.200E-05 | 3.606E-06 | 9.000E-06 | 1.600E-05 |
| 3961 | 199 | A199A | A | 1.6e-05 | 1.5e-05 | 8.4e-06 | 1.313E-05 | 4.130E-06 | 8.400E-06 | 1.600E-05 |
| 3962 | 199 | A199R | A | 1.6e-05 | 1.3e-05 | 8.6e-06 | 1.253E-05 | 3.722E-06 | 8.600E-06 | 1.600E-05 |
| 3963 | 199 | A199N | A | 1.4e-05 | 1.6e-05 | 9e-06 | 1.300E-05 | 3.606E-06 | 9.000E-06 | 1.600E-05 |
| 3964 | 199 | A199D | A | 9e-06 | 1.1e-05 | 1.4e-05 | 1.133E-05 | 2.517E-06 | 9.000E-06 | 1.400E-05 |
| 3965 | 199 | A199C | A | 1.6e-05 | 9.9e-06 | 1.2e-05 | 1.263E-05 | 3.099E-06 | 9.900E-06 | 1.600E-05 |
| 3966 | 199 | A199E | A | 1.3e-05 | 1.3e-05 | 1e-05 | 1.200E-05 | 1.732E-06 | 1.000E-05 | 1.300E-05 |
| 3967 | 199 | A199Q | A | 1.5e-05 | 1.4e-05 | 1.1e-05 | 1.333E-05 | 2.082E-06 | 1.100E-05 | 1.500E-05 |
| 3968 | 199 | A199G | A | 1.6e-05 | 1.1e-05 | 9.8e-06 | 1.227E-05 | 3.288E-06 | 9.800E-06 | 1.600E-05 |
| 3969 | 199 | A199H | A | 9.3e-06 | 1.1e-05 | 7.4e-06 | 9.233E-06 | 1.801E-06 | 7.400E-06 | 1.100E-05 |
| 3970 | 199 | A199I | A | 1.3e-05 | 9.2e-06 | 1.6e-05 | 1.273E-05 | 3.408E-06 | 9.200E-06 | 1.600E-05 |
| 3971 | 199 | A199L | A | 8.9e-06 | 1.6e-05 | 1e-05 | 1.163E-05 | 3.821E-06 | 8.900E-06 | 1.600E-05 |
| 3972 | 199 | A199K | A | 1.1e-05 | 9.2e-06 | 1.6e-05 | 1.207E-05 | 3.523E-06 | 9.200E-06 | 1.600E-05 |
| 3973 | 199 | A199M | A | 2.6e-05 | 1.2e-05 | 1.6e-05 | 1.800E-05 | 7.211E-06 | 1.200E-05 | 2.600E-05 |
| 3974 | 199 | A199F | A | 1.5e-05 | 1.2e-05 | 1.3e-05 | 1.333E-05 | 1.528E-06 | 1.200E-05 | 1.500E-05 |
| 3975 | 199 | A199P | A | 9.9e-06 | 1.5e-05 | 8.8e-06 | 1.123E-05 | 3.308E-06 | 8.800E-06 | 1.500E-05 |
| 3976 | 199 | A199S | A | 1.1e-05 | 1.1e-05 | 1.4e-05 | 1.200E-05 | 1.732E-06 | 1.100E-05 | 1.400E-05 |
| 3977 | 199 | A199T | A | 1.3e-05 | 9.4e-06 | 1.4e-05 | 1.213E-05 | 2.419E-06 | 9.400E-06 | 1.400E-05 |
| 3978 | 199 | A199W | A | 1.1e-05 | 1.3e-05 | 1.2e-05 | 1.200E-05 | 1.000E-06 | 1.100E-05 | 1.300E-05 |
| 3979 | 199 | A199Y | A | 1.5e-05 | 1.6e-05 | 1.5e-05 | 1.533E-05 | 5.774E-07 | 1.500E-05 | 1.600E-05 |
| 3980 | 199 | A199V | A | 1.7e-05 | 1.7e-05 | 1.5e-05 | 1.633E-05 | 1.155E-06 | 1.500E-05 | 1.700E-05 |
| 3981 | 200 | G200A | A | 9.7e-06 | 1.4e-05 | 9.9e-06 | 1.120E-05 | 2.427E-06 | 9.700E-06 | 1.400E-05 |
| 3982 | 200 | G200R | A | 1.4e-05 | 1.1e-05 | 1.6e-05 | 1.367E-05 | 2.517E-06 | 1.100E-05 | 1.600E-05 |
| 3983 | 200 | G200N | A | 1e-05 | 6.7e-06 | 1.5e-05 | 1.057E-05 | 4.179E-06 | 6.700E-06 | 1.500E-05 |
| 3984 | 200 | G200D | A | 1.5e-05 | 9.8e-06 | 1.6e-05 | 1.360E-05 | 3.329E-06 | 9.800E-06 | 1.600E-05 |
| 3985 | 200 | G200C | A | 7.5e-06 | 9.6e-06 | 1e-05 | 9.033E-06 | 1.343E-06 | 7.500E-06 | 1.000E-05 |
| 3986 | 200 | G200E | A | 1.4e-05 | 1.4e-05 | 6.3e-06 | 1.143E-05 | 4.446E-06 | 6.300E-06 | 1.400E-05 |
| 3987 | 200 | G200Q | A | 1.5e-05 | 1.4e-05 | 1.6e-05 | 1.500E-05 | 1.000E-06 | 1.400E-05 | 1.600E-05 |
| 3988 | 200 | G200G | A | 1e-05 | 1.5e-05 | 1.5e-05 | 1.333E-05 | 2.887E-06 | 1.000E-05 | 1.500E-05 |
| 3989 | 200 | G200H | A | 9.2e-06 | 1.4e-05 | 1.5e-05 | 1.273E-05 | 3.101E-06 | 9.200E-06 | 1.500E-05 |
| 3990 | 200 | G200I | A | 1e-05 | 1e-05 | 8.7e-06 | 9.567E-06 | 7.506E-07 | 8.700E-06 | 1.000E-05 |
| 3991 | 200 | G200L | A | 1.5e-05 | 1.7e-05 | 1.1e-05 | 1.433E-05 | 3.055E-06 | 1.100E-05 | 1.700E-05 |
| 3992 | 200 | G200K | A | 1.6e-05 | 1.4e-05 | 9.7e-06 | 1.323E-05 | 3.219E-06 | 9.700E-06 | 1.600E-05 |
| 3993 | 200 | G200M | A | 8e-06 | 1.7e-05 | 1.4e-05 | 1.300E-05 | 4.583E-06 | 8.000E-06 | 1.700E-05 |
| 3994 | 200 | G200F | A | 8.9e-06 | 9.5e-06 | 1.6e-05 | 1.147E-05 | 3.937E-06 | 8.900E-06 | 1.600E-05 |

| | | | | | | | | | | |
|---|---|---|---|---|---|---|---|---|---|---|
| 3995 | 200 | G200P | A | 1.6e-05 | 1.1e-05 | 1.4e-05 | 1.367E-05 | 2.517E-06 | 1.100E-05 | 1.600E-05 |
| 3996 | 200 | G200S | A | 1.4e-05 | 1.2e-05 | 1.1e-05 | 1.233E-05 | 1.528E-06 | 1.100E-05 | 1.400E-05 |
| 3997 | 200 | G200T | A | 1.1e-05 | 9e-06 | 1e-05 | 1.000E-05 | 1.000E-06 | 9.000E-06 | 1.100E-05 |
| 3998 | 200 | G200W | A | 1.5e-05 | 1e-05 | 1.3e-05 | 1.267E-05 | 2.517E-06 | 1.000E-05 | 1.500E-05 |
| 3999 | 200 | G200Y | A | 1.3e-05 | 1.4e-05 | 1.5e-05 | 1.400E-05 | 1.000E-06 | 1.300E-05 | 1.500E-05 |
| 4000 | 200 | G200V | A | 1.5e-05 | 1e-05 | 1e-05 | 1.167E-05 | 2.887E-06 | 1.000E-05 | 1.500E-05 |
| 4001 | 201 | G201A | A | 1.2e-05 | 1.5e-05 | 1.1e-05 | 1.267E-05 | 2.082E-06 | 1.100E-05 | 1.500E-05 |
| 4002 | 201 | G201R | A | 1.4e-05 | 1.5e-05 | 1e-05 | 1.300E-05 | 2.646E-06 | 1.000E-05 | 1.500E-05 |
| 4003 | 201 | G201N | A | 1.3e-05 | 1.1e-05 | 1.3e-05 | 1.233E-05 | 1.155E-06 | 1.100E-05 | 1.300E-05 |
| 4004 | 201 | G201D | A | 1.1e-05 | 1.3e-05 | 1.3e-05 | 1.233E-05 | 1.155E-06 | 1.100E-05 | 1.300E-05 |
| 4005 | 201 | G201C | A | 1.4e-05 | 1.5e-05 | 1.3e-05 | 1.400E-05 | 1.000E-06 | 1.300E-05 | 1.500E-05 |
| 4006 | 201 | G201E | A | 1.4e-05 | 9.9e-06 | 1.6e-05 | 1.330E-05 | 3.110E-06 | 9.900E-06 | 1.600E-05 |
| 4007 | 201 | G201Q | A | 1.5e-05 | 9.3e-06 | 1.4e-05 | 1.277E-05 | 3.044E-06 | 9.300E-06 | 1.500E-05 |
| 4008 | 201 | G201G | A | 1.5e-05 | 1.7e-05 | 1.5e-05 | 1.567E-05 | 1.155E-06 | 1.500E-05 | 1.700E-05 |
| 4009 | 201 | G201H | A | 1.4e-05 | 9.7e-06 | 1.5e-05 | 1.290E-05 | 2.816E-06 | 9.700E-06 | 1.500E-05 |
| 4010 | 201 | G201I | A | 9.8e-06 | 1.3e-05 | 1e-05 | 1.093E-05 | 1.793E-06 | 9.800E-06 | 1.300E-05 |
| 4011 | 201 | G201L | A | 9e-06 | 1.2e-05 | 1.5e-05 | 1.200E-05 | 3.000E-06 | 9.000E-06 | 1.500E-05 |
| 4012 | 201 | G201K | A | 1.8e-05 | 8.7e-06 | 8.3e-06 | 1.167E-05 | 5.488E-06 | 8.300E-06 | 1.800E-05 |
| 4013 | 201 | G201M | A | 9.5e-06 | 1.2e-05 | 1.3e-05 | 1.150E-05 | 1.803E-06 | 9.500E-06 | 1.300E-05 |
| 4014 | 201 | G201F | A | 1.5e-05 | 1.9e-05 | 9.7e-06 | 1.457E-05 | 4.665E-06 | 9.700E-06 | 1.900E-05 |
| 4015 | 201 | G201P | A | 1.4e-05 | 1.4e-05 | 1.3e-05 | 1.367E-05 | 5.774E-07 | 1.300E-05 | 1.400E-05 |
| 4016 | 201 | G201S | A | 1.1e-05 | 2e-05 | 1.3e-05 | 1.467E-05 | 4.726E-06 | 1.100E-05 | 2.000E-05 |
| 4017 | 201 | G201T | A | 1.7e-05 | 1.4e-05 | 9e-06 | 1.333E-05 | 4.041E-06 | 9.000E-06 | 1.700E-05 |
| 4018 | 201 | G201W | A | 1.5e-05 | 2.2e-05 | 1.4e-05 | 1.700E-05 | 4.359E-06 | 1.400E-05 | 2.200E-05 |
| 4019 | 201 | G201Y | A | 7.4e-06 | 1.5e-05 | 9.7e-06 | 1.070E-05 | 3.897E-06 | 7.400E-06 | 1.500E-05 |
| 4020 | 201 | G201V | A | 1e-05 | 1.3e-05 | 1.1e-05 | 1.133E-05 | 1.528E-06 | 1.000E-05 | 1.300E-05 |
| 4021 | 202 | C202A | A | 1.1e-05 | 1.4e-05 | 1.5e-05 | 1.333E-05 | 2.082E-06 | 1.100E-05 | 1.500E-05 |
| 4022 | 202 | C202R | A | 1e-05 | 1.5e-05 | 9.4e-06 | 1.147E-05 | 3.075E-06 | 9.400E-06 | 1.500E-05 |
| 4023 | 202 | C202N | A | 1.5e-05 | 1.6e-05 | 1.5e-05 | 1.533E-05 | 5.774E-07 | 1.500E-05 | 1.600E-05 |
| 4024 | 202 | C202D | A | 1.4e-05 | 1.7e-05 | 1.1e-05 | 1.400E-05 | 3.000E-06 | 1.100E-05 | 1.700E-05 |
| 4025 | 202 | C202C | A | 1.1e-05 | 1.6e-05 | 1.1e-05 | 1.267E-05 | 2.887E-06 | 1.100E-05 | 1.600E-05 |
| 4026 | 202 | C202E | A | 5.9e-06 | 1.4e-05 | 1.3e-05 | 1.097E-05 | 4.416E-06 | 5.900E-06 | 1.400E-05 |
| 4027 | 202 | C202Q | A | 1.6e-05 | 1.4e-05 | 1.1e-05 | 1.367E-05 | 2.517E-06 | 1.100E-05 | 1.600E-05 |
| 4028 | 202 | C202G | A | 1.5e-05 | 1.5e-05 | 9.7e-06 | 1.323E-05 | 3.060E-06 | 9.700E-06 | 1.500E-05 |
| 4029 | 202 | C202H | A | 1e-05 | 1.5e-05 | 1.7e-05 | 1.400E-05 | 3.606E-06 | 1.000E-05 | 1.700E-05 |
| 4030 | 202 | C202I | A | 1.1e-05 | 1.4e-05 | 1.3e-05 | 1.267E-05 | 1.528E-06 | 1.100E-05 | 1.400E-05 |
| 4031 | 202 | C202L | A | 1.4e-05 | 1.6e-05 | 1.3e-05 | 1.433E-05 | 1.528E-06 | 1.300E-05 | 1.600E-05 |
| 4032 | 202 | C202K | A | 1.2e-05 | 1e-05 | 1e-05 | 1.067E-05 | 1.155E-06 | 1.000E-05 | 1.200E-05 |
| 4033 | 202 | C202M | A | 1.1e-05 | 1e-05 | 1.1e-05 | 1.067E-05 | 5.774E-07 | 1.000E-05 | 1.100E-05 |
| 4034 | 202 | C202F | A | 8.9e-06 | 9.5e-06 | 1.4e-05 | 1.080E-05 | 2.787E-06 | 8.900E-06 | 1.400E-05 |
| 4035 | 202 | C202P | A | 1.3e-05 | 1.4e-05 | 1.3e-05 | 1.333E-05 | 5.774E-07 | 1.300E-05 | 1.400E-05 |
| 4036 | 202 | C202S | A | 1.4e-05 | 1.5e-05 | 1.4e-05 | 1.433E-05 | 5.774E-07 | 1.400E-05 | 1.500E-05 |
| 4037 | 202 | C202T | A | 1.4e-05 | 1.1e-05 | 7.8e-06 | 1.093E-05 | 3.101E-06 | 7.800E-06 | 1.400E-05 |
| 4038 | 202 | C202W | A | 1.4e-05 | 1.4e-05 | 1.6e-05 | 1.467E-05 | 1.155E-06 | 1.400E-05 | 1.600E-05 |
| 4039 | 202 | C202Y | A | 1.7e-05 | 9.1e-06 | 1.6e-05 | 1.403E-05 | 4.302E-06 | 9.100E-06 | 1.700E-05 |
| 4040 | 202 | C202V | A | 1.5e-05 | 1.2e-05 | 6.3e-06 | 1.110E-05 | 4.419E-06 | 6.300E-06 | 1.500E-05 |
| 4041 | 203 | A203A | A | 1.3e-05 | 1.1e-05 | 1.2e-05 | 1.200E-05 | 1.000E-06 | 1.100E-05 | 1.300E-05 |
| 4042 | 203 | A203R | A | 1.5e-05 | 1.2e-05 | 7e-06 | 1.133E-05 | 4.041E-06 | 7.000E-06 | 1.500E-05 |
| 4043 | 203 | A203N | A | 1e-05 | 1.5e-05 | 1e-05 | 1.167E-05 | 2.887E-06 | 1.000E-05 | 1.500E-05 |
| 4044 | 203 | A203D | A | 1.5e-05 | 1.3e-05 | 1.2e-05 | 1.333E-05 | 1.528E-06 | 1.200E-05 | 1.500E-05 |
| 4045 | 203 | A203C | A | 1.4e-05 | 1.4e-05 | 8e-06 | 1.200E-05 | 3.464E-06 | 8.000E-06 | 1.400E-05 |
| 4046 | 203 | A203E | A | 9.7e-06 | 1.4e-05 | 1.4e-05 | 1.257E-05 | 2.483E-06 | 9.700E-06 | 1.400E-05 |
| 4047 | 203 | A203Q | A | 9.4e-06 | 1.5e-05 | 1.1e-05 | 1.180E-05 | 2.884E-06 | 9.400E-06 | 1.500E-05 |
| 4048 | 203 | A203G | A | 1.1e-05 | 1.5e-05 | 1.2e-05 | 1.267E-05 | 2.082E-06 | 1.100E-05 | 1.500E-05 |
| 4049 | 203 | A203H | A | 1e-05 | 9e-06 | 1.2e-05 | 1.033E-05 | 1.528E-06 | 9.000E-06 | 1.200E-05 |
| 4050 | 203 | A203I | A | 6.9e-06 | 1.5e-05 | 2e-05 | 1.397E-05 | 6.611E-06 | 6.900E-06 | 2.000E-05 |
| 4051 | 203 | A203L | A | 1.5e-05 | 1.4e-05 | 1.4e-05 | 1.433E-05 | 5.774E-07 | 1.400E-05 | 1.500E-05 |
| 4052 | 203 | A203K | A | 1.7e-05 | 9.6e-06 | 1.1e-05 | 1.253E-05 | 3.931E-06 | 9.600E-06 | 1.700E-05 |
| 4053 | 203 | A203M | A | 1.5e-05 | 1.6e-05 | 1.5e-05 | 1.533E-05 | 5.774E-07 | 1.500E-05 | 1.600E-05 |
| 4054 | 203 | A203F | A | 9.3e-06 | 1.6e-05 | 1.6e-05 | 1.377E-05 | 3.868E-06 | 9.300E-06 | 1.600E-05 |
| 4055 | 203 | A203P | A | 1.4e-05 | 1.4e-05 | 9.8e-06 | 1.260E-05 | 2.425E-06 | 9.800E-06 | 1.400E-05 |
| 4056 | 203 | A203S | A | 8.6e-06 | 6.7e-06 | 1.1e-05 | 8.767E-06 | 2.155E-06 | 6.700E-06 | 1.100E-05 |
| 4057 | 203 | A203T | A | 8.4e-06 | 1.4e-05 | 7.9e-06 | 1.010E-05 | 3.387E-06 | 7.900E-06 | 1.400E-05 |
| 4058 | 203 | A203W | A | 9.5e-06 | 1.1e-05 | 8.7e-06 | 9.733E-06 | 1.168E-06 | 8.700E-06 | 1.100E-05 |
| 4059 | 203 | A203Y | A | 1.3e-05 | 1.7e-05 | 1.4e-05 | 1.467E-05 | 2.082E-06 | 1.300E-05 | 1.700E-05 |
| 4060 | 203 | A203V | A | 1.1e-05 | 1.3e-05 | 1e-05 | 1.133E-05 | 1.528E-06 | 1.000E-05 | 1.300E-05 |
| 4061 | 204 | R204A | A | 1.3e-05 | 1.5e-05 | 1.4e-05 | 1.400E-05 | 1.000E-06 | 1.300E-05 | 1.500E-05 |
| 4062 | 204 | R204R | A | 1.6e-05 | 1.5e-05 | 1.4e-05 | 1.500E-05 | 1.000E-06 | 1.400E-05 | 1.600E-05 |
| 4063 | 204 | R204N | A | 1.6e-05 | 1.1e-05 | 1.1e-05 | 1.267E-05 | 2.887E-06 | 1.100E-05 | 1.600E-05 |
| 4064 | 204 | R204D | A | 1.3e-05 | 1.4e-05 | 1.7e-05 | 1.467E-05 | 2.082E-06 | 1.300E-05 | 1.700E-05 |
| 4065 | 204 | R204C | A | 1e-05 | 1.3e-05 | 1.5e-05 | 1.267E-05 | 2.517E-06 | 1.000E-05 | 1.500E-05 |
| 4066 | 204 | R204E | A | 1.2e-05 | 9.5e-06 | 9.5e-06 | 1.033E-05 | 1.443E-06 | 9.500E-06 | 1.200E-05 |
| 4067 | 204 | R204Q | A | 1.4e-05 | 9.7e-06 | 1.7e-05 | 1.357E-05 | 3.669E-06 | 9.700E-06 | 1.700E-05 |
| 4068 | 204 | R204G | A | 1.3e-05 | 1.3e-05 | 8.8e-06 | 1.160E-05 | 2.425E-06 | 8.800E-06 | 1.300E-05 |
| 4069 | 204 | R204H | A | 1.5e-05 | 1.3e-05 | 1.4e-05 | 1.400E-05 | 1.000E-06 | 1.300E-05 | 1.500E-05 |
| 4070 | 204 | R204I | A | 1.5e-05 | 1.6e-05 | 1.4e-05 | 1.500E-05 | 1.000E-06 | 1.400E-05 | 1.600E-05 |
| 4071 | 204 | R204L | A | 1.3e-05 | 1.1e-05 | 1.4e-05 | 1.267E-05 | 1.528E-06 | 1.100E-05 | 1.400E-05 |
| 4072 | 204 | R204K | A | 1.4e-05 | 1.5e-05 | 9.2e-06 | 1.273E-05 | 3.101E-06 | 9.200E-06 | 1.500E-05 |
| 4073 | 204 | R204M | A | 1.4e-05 | 1.5e-05 | 1.6e-05 | 1.500E-05 | 1.000E-06 | 1.400E-05 | 1.600E-05 |
| 4074 | 204 | R204F | A | 8e-06 | 1.4e-05 | 1.5e-05 | 1.233E-05 | 3.786E-06 | 8.000E-06 | 1.500E-05 |
| 4075 | 204 | R204P | A | 1.5e-05 | 1.4e-05 | 1.1e-05 | 1.333E-05 | 2.082E-06 | 1.100E-05 | 1.500E-05 |
| 4076 | 204 | R204S | A | 8.3e-06 | 1.1e-05 | 1.5e-05 | 1.143E-05 | 3.371E-06 | 8.300E-06 | 1.500E-05 |
| 4077 | 204 | R204T | A | 1.5e-05 | 1.5e-05 | 1.2e-05 | 1.400E-05 | 1.732E-06 | 1.200E-05 | 1.500E-05 |
| 4078 | 204 | R204W | A | 1.5e-05 | 1.5e-05 | 1.2e-05 | 1.400E-05 | 1.732E-06 | 1.200E-05 | 1.500E-05 |
| 4079 | 204 | R204Y | A | 1.4e-05 | 1.5e-05 | 1.2e-05 | 1.367E-05 | 1.528E-06 | 1.200E-05 | 1.500E-05 |

| | | | | | | | | | | |
|---|---|---|---|---|---|---|---|---|---|---|
| 4080 | 204 | R204V | A | 1.4e-05 | 1e-05 | 1.5e-05 | 1.300E-05 | 2.646E-06 | 1.000E-05 | 1.500E-05 |
| 4081 | 205 | C205A | A | 1.1e-05 | 1.2e-05 | 1.1e-05 | 1.133E-05 | 5.774E-07 | 1.100E-05 | 1.200E-05 |
| 4082 | 205 | C205R | A | 1.4e-05 | 1.6e-05 | 1.1e-05 | 1.367E-05 | 2.517E-06 | 1.100E-05 | 1.600E-05 |
| 4083 | 205 | C205N | A | 1.8e-05 | 1e-05 | 1e-05 | 1.267E-05 | 4.619E-06 | 1.000E-05 | 1.800E-05 |
| 4084 | 205 | C205D | A | 1.6e-05 | 1.1e-05 | 1e-05 | 1.233E-05 | 3.215E-06 | 1.000E-05 | 1.600E-05 |
| 4085 | 205 | C205C | A | 1.6e-05 | 1.1e-05 | 1.1e-05 | 1.267E-05 | 2.887E-06 | 1.100E-05 | 1.600E-05 |
| 4086 | 205 | C205E | A | 1.6e-05 | 1.2e-05 | 1.6e-05 | 1.467E-05 | 2.309E-06 | 1.200E-05 | 1.600E-05 |
| 4087 | 205 | C205Q | A | 1.3e-05 | 9.2e-06 | 1.1e-05 | 1.107E-05 | 1.901E-06 | 9.200E-06 | 1.300E-05 |
| 4088 | 205 | C205G | A | 1.5e-05 | 1.1e-05 | 1e-05 | 1.200E-05 | 2.646E-06 | 1.000E-05 | 1.500E-05 |
| 4089 | 205 | C205H | A | 1.6e-05 | 1.1e-05 | 1.4e-05 | 1.367E-05 | 2.517E-06 | 1.100E-05 | 1.600E-05 |
| 4090 | 205 | C205I | A | 1.6e-05 | 1.1e-05 | 1e-05 | 1.233E-05 | 3.215E-06 | 1.000E-05 | 1.600E-05 |
| 4091 | 205 | C205L | A | 1.3e-05 | 9.7e-06 | 1.5e-05 | 1.257E-05 | 2.676E-06 | 9.700E-06 | 1.500E-05 |
| 4092 | 205 | C205K | A | 1.1e-05 | 1.5e-05 | 1.5e-05 | 1.367E-05 | 2.309E-06 | 1.100E-05 | 1.500E-05 |
| 4093 | 205 | C205M | A | 1.4e-05 | 1.1e-05 | 1e-05 | 1.167E-05 | 2.082E-06 | 1.000E-05 | 1.400E-05 |
| 4094 | 205 | C205F | A | 1.3e-05 | 1.6e-05 | 1.7e-05 | 1.533E-05 | 2.082E-06 | 1.300E-05 | 1.700E-05 |
| 4095 | 205 | C205P | A | 1.6e-05 | 9.8e-06 | 1e-05 | 1.193E-05 | 3.523E-06 | 9.800E-06 | 1.600E-05 |
| 4096 | 205 | C205S | A | 1.3e-05 | 9.7e-06 | 1.5e-05 | 1.257E-05 | 2.676E-06 | 9.700E-06 | 1.500E-05 |
| 4097 | 205 | C205T | A | 1e-05 | 1.6e-05 | 1.1e-05 | 1.233E-05 | 3.215E-06 | 1.000E-05 | 1.600E-05 |
| 4098 | 205 | C205W | A | 1.1e-05 | 1.5e-05 | 1.6e-05 | 1.400E-05 | 2.646E-06 | 1.100E-05 | 1.600E-05 |
| 4099 | 205 | C205Y | A | 9.5e-06 | 1.2e-05 | 1.6e-05 | 1.250E-05 | 3.279E-06 | 9.500E-06 | 1.600E-05 |
| 4100 | 205 | C205V | A | 1.6e-05 | 1.4e-05 | 1.4e-05 | 1.467E-05 | 1.155E-06 | 1.400E-05 | 1.600E-05 |
| 4101 | 206 | K206A | A | 9e-06 | 9.9e-06 | 7.7e-06 | 8.867E-06 | 1.106E-06 | 7.700E-06 | 9.900E-06 |
| 4102 | 206 | K206R | A | 9.7e-06 | 9.9e-06 | 9e-06 | 9.533E-06 | 4.726E-07 | 9.000E-06 | 9.900E-06 |
| 4103 | 206 | K206N | A | 1.1e-05 | 1.5e-05 | 1.6e-05 | 1.400E-05 | 2.646E-06 | 1.100E-05 | 1.600E-05 |
| 4104 | 206 | K206D | A | 1.5e-05 | 1.1e-05 | 1.7e-05 | 1.433E-05 | 3.055E-06 | 1.100E-05 | 1.700E-05 |
| 4105 | 206 | K206C | A | 1.4e-05 | 1.5e-05 | 1.2e-05 | 1.367E-05 | 1.528E-06 | 1.200E-05 | 1.500E-05 |
| 4106 | 206 | K206E | A | 1.6e-05 | 1.5e-05 | 2.6e-05 | 1.900E-05 | 6.083E-06 | 1.500E-05 | 2.600E-05 |
| 4107 | 206 | K206Q | A | 1.1e-05 | 9.8e-06 | 9.8e-06 | 1.020E-05 | 6.928E-07 | 9.800E-06 | 1.100E-05 |
| 4108 | 206 | K206G | A | 1.7e-05 | 1.5e-05 | 1.5e-05 | 1.567E-05 | 1.155E-06 | 1.500E-05 | 1.700E-05 |
| 4109 | 206 | K206H | A | 1.7e-05 | 1.6e-05 | 1.4e-05 | 1.567E-05 | 1.528E-06 | 1.400E-05 | 1.700E-05 |
| 4110 | 206 | K206I | A | 1.5e-05 | 1e-05 | 1.8e-05 | 1.433E-05 | 4.041E-06 | 1.000E-05 | 1.800E-05 |
| 4111 | 206 | K206L | A | 1.5e-05 | 1.4e-05 | 1.1e-05 | 1.333E-05 | 2.082E-06 | 1.100E-05 | 1.500E-05 |
| 4112 | 206 | K206K | A | 1.4e-05 | 1.5e-05 | 1e-05 | 1.300E-05 | 2.646E-06 | 1.000E-05 | 1.500E-05 |
| 4113 | 206 | K206M | A | 9e-06 | 1.5e-05 | 2.2e-05 | 1.533E-05 | 6.506E-06 | 9.000E-06 | 2.200E-05 |
| 4114 | 206 | K206F | A | 1.5e-05 | 1.7e-05 | 1.3e-05 | 1.500E-05 | 2.000E-06 | 1.300E-05 | 1.700E-05 |
| 4115 | 206 | K206P | A | 9.9e-06 | 1.1e-05 | 1.6e-05 | 1.230E-05 | 3.251E-06 | 9.900E-06 | 1.600E-05 |
| 4116 | 206 | K206S | A | 8.6e-06 | 1.1e-05 | 1.4e-05 | 1.120E-05 | 2.706E-06 | 8.600E-06 | 1.400E-05 |
| 4117 | 206 | K206T | A | 1e-05 | 9.4e-06 | 1.6e-05 | 1.180E-05 | 3.650E-06 | 9.400E-06 | 1.600E-05 |
| 4118 | 206 | K206W | A | 1.1e-05 | 9.8e-06 | 1e-05 | 1.027E-05 | 6.429E-07 | 9.800E-06 | 1.100E-05 |
| 4119 | 206 | K206Y | A | 1.5e-05 | 1.5e-05 | 1.5e-05 | 1.500E-05 | 0.000E+00 | 1.500E-05 | 1.500E-05 |
| 4120 | 206 | K206V | A | 1.4e-05 | 1e-05 | 1.1e-05 | 1.167E-05 | 2.082E-06 | 1.000E-05 | 1.400E-05 |
| 4121 | 207 | G207A | A | 1.4e-05 | 1.5e-05 | 1.1e-05 | 1.333E-05 | 2.082E-06 | 1.100E-05 | 1.500E-05 |
| 4122 | 207 | G207R | A | 1.6e-05 | 1e-05 | 1.5e-05 | 1.367E-05 | 3.215E-06 | 1.000E-05 | 1.600E-05 |
| 4123 | 207 | G207N | A | 1.5e-05 | 1.1e-05 | 1.3e-05 | 1.300E-05 | 2.000E-06 | 1.100E-05 | 1.500E-05 |
| 4124 | 207 | G207D | A | 1.5e-05 | 1.3e-05 | 1.4e-05 | 1.400E-05 | 1.000E-06 | 1.300E-05 | 1.500E-05 |
| 4125 | 207 | G207C | A | 1e-05 | 9.5e-06 | 1.2e-05 | 1.050E-05 | 1.323E-06 | 9.500E-06 | 1.200E-05 |
| 4126 | 207 | G207E | A | 1.5e-05 | 1.6e-05 | 1.5e-05 | 1.533E-05 | 5.774E-07 | 1.500E-05 | 1.600E-05 |
| 4127 | 207 | G207Q | A | 9.1e-06 | 1.7e-05 | 7.9e-06 | 1.133E-05 | 4.944E-06 | 7.900E-06 | 1.700E-05 |
| 4128 | 207 | G207G | A | 9.9e-06 | 8.6e-06 | 1.5e-05 | 1.117E-05 | 3.383E-06 | 8.600E-06 | 1.500E-05 |
| 4129 | 207 | G207H | A | 9.8e-06 | 1.6e-05 | 1.4e-05 | 1.327E-05 | 3.164E-06 | 9.800E-06 | 1.600E-05 |
| 4130 | 207 | G207I | A | 9.3e-06 | 9.1e-06 | 1.6e-05 | 1.147E-05 | 3.927E-06 | 9.100E-06 | 1.600E-05 |
| 4131 | 207 | G207L | A | 1.4e-05 | 1.6e-05 | 1.6e-05 | 1.533E-05 | 1.155E-06 | 1.400E-05 | 1.600E-05 |
| 4132 | 207 | G207K | A | 9.5e-06 | 1.6e-05 | 1e-05 | 1.183E-05 | 3.617E-06 | 9.500E-06 | 1.600E-05 |
| 4133 | 207 | G207M | A | 9.8e-06 | 1.6e-05 | 1.6e-05 | 1.393E-05 | 3.580E-06 | 9.800E-06 | 1.600E-05 |
| 4134 | 207 | G207F | A | 1e-05 | 1.6e-05 | 1.6e-05 | 1.400E-05 | 3.464E-06 | 1.000E-05 | 1.600E-05 |
| 4135 | 207 | G207P | A | 9.7e-06 | 1.5e-05 | 9.5e-06 | 1.140E-05 | 3.119E-06 | 9.500E-06 | 1.500E-05 |
| 4136 | 207 | G207S | A | 1.3e-05 | 1.1e-05 | 1.7e-05 | 1.367E-05 | 3.055E-06 | 1.100E-05 | 1.700E-05 |
| 4137 | 207 | G207T | A | 1.1e-05 | 1.1e-05 | 1e-05 | 1.067E-05 | 5.774E-07 | 1.000E-05 | 1.100E-05 |
| 4138 | 207 | G207W | A | 1.6e-05 | 9e-06 | 9.7e-06 | 1.157E-05 | 3.855E-06 | 9.000E-06 | 1.600E-05 |
| 4139 | 207 | G207Y | A | 1e-05 | 1.1e-05 | 1.2e-05 | 1.100E-05 | 1.000E-06 | 1.000E-05 | 1.200E-05 |
| 4140 | 207 | G207V | A | 1.2e-05 | 1e-05 | 1.5e-05 | 1.233E-05 | 2.517E-06 | 1.000E-05 | 1.500E-05 |
| 4141 | 208 | P208A | A | 1.6e-05 | 1.1e-05 | 1.2e-05 | 1.300E-05 | 2.646E-06 | 1.100E-05 | 1.600E-05 |
| 4142 | 208 | P208R | A | 9.2e-06 | 9.9e-06 | 8.6e-06 | 9.233E-06 | 6.506E-07 | 8.600E-06 | 9.900E-06 |
| 4143 | 208 | P208N | A | 8.7e-06 | 1.7e-05 | 1.2e-05 | 1.257E-05 | 4.179E-06 | 8.700E-06 | 1.700E-05 |
| 4144 | 208 | P208D | A | 1.2e-05 | 1.3e-05 | 1.2e-05 | 1.233E-05 | 5.774E-07 | 1.200E-05 | 1.300E-05 |
| 4145 | 208 | P208C | A | 1.6e-05 | 1.5e-05 | 1.3e-05 | 1.467E-05 | 1.528E-06 | 1.300E-05 | 1.600E-05 |
| 4146 | 208 | P208E | A | 1.1e-05 | 1.5e-05 | 1.1e-05 | 1.233E-05 | 2.309E-06 | 1.100E-05 | 1.500E-05 |
| 4147 | 208 | P208Q | A | 1.2e-05 | 1.3e-05 | 1.5e-05 | 1.333E-05 | 1.528E-06 | 1.200E-05 | 1.500E-05 |
| 4148 | 208 | P208G | A | 1.6e-05 | 9.1e-06 | 1.6e-05 | 1.370E-05 | 3.984E-06 | 9.100E-06 | 1.600E-05 |
| 4149 | 208 | P208H | A | 1.5e-05 | 1.5e-05 | 1.4e-05 | 1.467E-05 | 5.774E-07 | 1.400E-05 | 1.500E-05 |
| 4150 | 208 | P208I | A | 1.5e-05 | 9e-06 | 1.6e-05 | 1.333E-05 | 3.786E-06 | 9.000E-06 | 1.600E-05 |
| 4151 | 208 | P208L | A | 9.9e-06 | 9.7e-06 | 1.5e-05 | 1.153E-05 | 3.004E-06 | 9.700E-06 | 1.500E-05 |
| 4152 | 208 | P208K | A | 1.1e-05 | 1.5e-05 | 1.1e-05 | 1.233E-05 | 2.309E-06 | 1.100E-05 | 1.500E-05 |
| 4153 | 208 | P208M | A | 1e-05 | 1e-05 | 9.3e-06 | 9.767E-06 | 4.041E-07 | 9.300E-06 | 1.000E-05 |
| 4154 | 208 | P208F | A | 1.6e-05 | 1.1e-05 | 1.5e-05 | 1.400E-05 | 2.646E-06 | 1.100E-05 | 1.600E-05 |
| 4155 | 208 | P208P | A | 1.1e-05 | 1e-05 | 1.1e-05 | 1.067E-05 | 5.774E-07 | 1.000E-05 | 1.100E-05 |
| 4156 | 208 | P208S | A | 9.4e-06 | 9.7e-06 | 9.5e-06 | 9.533E-06 | 1.528E-07 | 9.400E-06 | 9.700E-06 |
| 4157 | 208 | P208T | A | 1.2e-05 | 1.6e-05 | 9.8e-06 | 1.260E-05 | 3.143E-06 | 9.800E-06 | 1.600E-05 |
| 4158 | 208 | P208W | A | 1e-05 | 1.4e-05 | 1.4e-05 | 1.267E-05 | 2.309E-06 | 1.000E-05 | 1.400E-05 |
| 4159 | 208 | P208Y | A | 1.5e-05 | 1e-05 | 1.5e-05 | 1.333E-05 | 2.887E-06 | 1.000E-05 | 1.500E-05 |
| 4160 | 208 | P208V | A | 1.5e-05 | 1.6e-05 | 1.1e-05 | 1.400E-05 | 2.646E-06 | 1.100E-05 | 1.600E-05 |
| 4161 | 209 | L209A | A | 9.9e-06 | 1.5e-05 | 1.1e-05 | 1.197E-05 | 2.684E-06 | 9.900E-06 | 1.500E-05 |
| 4162 | 209 | L209R | A | 1.4e-05 | 1.2e-05 | 1.5e-05 | 1.367E-05 | 1.528E-06 | 1.200E-05 | 1.500E-05 |
| 4163 | 209 | L209N | A | 1.3e-05 | 1.6e-05 | 1.4e-05 | 1.433E-05 | 1.528E-06 | 1.300E-05 | 1.600E-05 |
| 4164 | 209 | L209D | A | 1.1e-05 | 9.4e-06 | 1.1e-05 | 1.047E-05 | 9.238E-07 | 9.400E-06 | 1.100E-05 |

| | | | | | | | | | | |
|---|---|---|---|---|---|---|---|---|---|---|
| 4165 | 209 | L209C | A | 1.7e-05 | 9.5e-06 | 1.7e-05 | 1.450E-05 | 4.330E-06 | 9.500E-06 | 1.700E-05 |
| 4166 | 209 | L209E | A | 1.6e-05 | 1.5e-05 | 1.4e-05 | 1.500E-05 | 1.000E-06 | 1.400E-05 | 1.600E-05 |
| 4167 | 209 | L209Q | A | 1e-05 | 1.4e-05 | 1.4e-05 | 1.267E-05 | 2.309E-06 | 1.000E-05 | 1.400E-05 |
| 4168 | 209 | L209G | A | 1.1e-05 | 1.4e-05 | 1.3e-05 | 1.267E-05 | 1.528E-06 | 1.100E-05 | 1.400E-05 |
| 4169 | 209 | L209H | A | 1.4e-05 | 1.4e-05 | 1.3e-05 | 1.367E-05 | 5.774E-07 | 1.300E-05 | 1.400E-05 |
| 4170 | 209 | L209I | A | 1.5e-05 | 1.3e-05 | 1.1e-05 | 1.300E-05 | 2.000E-06 | 1.100E-05 | 1.500E-05 |
| 4171 | 209 | L209L | A | 1.4e-05 | 1.4e-05 | 1e-05 | 1.267E-05 | 2.309E-06 | 1.000E-05 | 1.400E-05 |
| 4172 | 209 | L209K | A | 1.6e-05 | 1.4e-05 | 1.1e-05 | 1.367E-05 | 2.517E-06 | 1.100E-05 | 1.600E-05 |
| 4173 | 209 | L209M | A | 8.4e-06 | 1.4e-05 | 1.4e-05 | 1.213E-05 | 3.233E-06 | 8.400E-06 | 1.400E-05 |
| 4174 | 209 | L209F | A | 1e-05 | 1.6e-05 | 1.1e-05 | 1.233E-05 | 3.215E-06 | 1.000E-05 | 1.600E-05 |
| 4175 | 209 | L209P | A | 2.5e-05 | 1.5e-05 | 9.6e-06 | 1.653E-05 | 7.814E-06 | 9.600E-06 | 2.500E-05 |
| 4176 | 209 | L209S | A | 1.3e-05 | 1.6e-05 | 1.1e-05 | 1.333E-05 | 2.517E-06 | 1.100E-05 | 1.600E-05 |
| 4177 | 209 | L209T | A | 1.4e-05 | 1.2e-05 | 7.7e-06 | 1.123E-05 | 3.219E-06 | 7.700E-06 | 1.400E-05 |
| 4178 | 209 | L209W | A | 1.7e-05 | 1.1e-05 | 1.5e-05 | 1.433E-05 | 3.055E-06 | 1.100E-05 | 1.700E-05 |
| 4179 | 209 | L209Y | A | 9e-06 | 8.6e-06 | 9e-06 | 8.867E-06 | 2.309E-07 | 8.600E-06 | 9.000E-06 |
| 4180 | 209 | L209V | A | 1.1e-05 | 1.6e-05 | 1.4e-05 | 1.367E-05 | 2.517E-06 | 1.100E-05 | 1.600E-05 |
| 4181 | 210 | P210A | A | 7.3e-06 | 8.1e-06 | 1.6e-05 | 1.047E-05 | 4.809E-06 | 7.300E-06 | 1.600E-05 |
| 4182 | 210 | P210R | A | 1.4e-05 | 1.4e-05 | 1.4e-05 | 1.400E-05 | 0.000E+00 | 1.400E-05 | 1.400E-05 |
| 4183 | 210 | P210N | A | 1.1e-05 | 9.3e-06 | 1.4e-05 | 1.143E-05 | 2.380E-06 | 9.300E-06 | 1.400E-05 |
| 4184 | 210 | P210D | A | 8.2e-06 | 1.1e-05 | 9.7e-06 | 9.633E-06 | 1.401E-06 | 8.200E-06 | 1.100E-05 |
| 4185 | 210 | P210C | A | 8.2e-06 | 1.4e-05 | 1.6e-05 | 1.273E-05 | 4.051E-06 | 8.200E-06 | 1.600E-05 |
| 4186 | 210 | P210E | A | 1.6e-05 | 1.6e-05 | 1.6e-05 | 1.600E-05 | 0.000E+00 | 1.600E-05 | 1.600E-05 |
| 4187 | 210 | P210Q | A | 1.6e-05 | 9.5e-06 | 1.3e-05 | 1.283E-05 | 3.253E-06 | 9.500E-06 | 1.600E-05 |
| 4188 | 210 | P210G | A | 1e-05 | 1.3e-05 | 1e-05 | 1.100E-05 | 1.732E-06 | 1.000E-05 | 1.300E-05 |
| 4189 | 210 | P210H | A | 1.7e-05 | 1.6e-05 | 9.3e-06 | 1.410E-05 | 4.187E-06 | 9.300E-06 | 1.700E-05 |
| 4190 | 210 | P210I | A | 9.9e-06 | 9.8e-06 | 1.2e-05 | 1.057E-05 | 1.242E-06 | 9.800E-06 | 1.200E-05 |
| 4191 | 210 | P210L | A | 1.4e-05 | 1.7e-05 | 1.5e-05 | 1.533E-05 | 1.528E-06 | 1.400E-05 | 1.700E-05 |
| 4192 | 210 | P210K | A | 1.6e-05 | 1.6e-05 | 1.4e-05 | 1.533E-05 | 1.155E-06 | 1.400E-05 | 1.600E-05 |
| 4193 | 210 | P210M | A | 6.9e-06 | 7.7e-06 | 1.5e-05 | 9.867E-06 | 4.464E-06 | 6.900E-06 | 1.500E-05 |
| 4194 | 210 | P210F | A | 1e-05 | 1.5e-05 | 1.6e-05 | 1.367E-05 | 3.215E-06 | 1.000E-05 | 1.600E-05 |
| 4195 | 210 | P210P | A | 1.5e-05 | 1.7e-05 | 1.6e-05 | 1.600E-05 | 1.000E-06 | 1.500E-05 | 1.700E-05 |
| 4196 | 210 | P210S | A | 8.6e-06 | 9.9e-06 | 9.8e-06 | 9.433E-06 | 7.234E-07 | 8.600E-06 | 9.900E-06 |
| 4197 | 210 | P210T | A | 1.4e-05 | 6.6e-06 | 1.2e-05 | 1.087E-05 | 3.828E-06 | 6.600E-06 | 1.400E-05 |
| 4198 | 210 | P210W | A | 8.8e-06 | 1.4e-05 | 1.5e-05 | 1.260E-05 | 3.329E-06 | 8.800E-06 | 1.500E-05 |
| 4199 | 210 | P210Y | A | 1.4e-05 | 9.4e-06 | 1.7e-05 | 1.347E-05 | 3.828E-06 | 9.400E-06 | 1.700E-05 |
| 4200 | 210 | P210V | A | 1.5e-05 | 1.1e-05 | 9.8e-06 | 1.193E-05 | 2.723E-06 | 9.800E-06 | 1.500E-05 |
| 4201 | 211 | T211A | A | 1.5e-05 | 1.1e-05 | 1.4e-05 | 1.333E-05 | 2.082E-06 | 1.100E-05 | 1.500E-05 |
| 4202 | 211 | T211R | A | 1.4e-05 | 1.5e-05 | 1.1e-05 | 1.333E-05 | 2.082E-06 | 1.100E-05 | 1.500E-05 |
| 4203 | 211 | T211N | A | 9.9e-06 | 1.2e-05 | 1.7e-05 | 1.297E-05 | 3.647E-06 | 9.900E-06 | 1.700E-05 |
| 4204 | 211 | T211D | A | 1.3e-05 | 1.2e-05 | 1.4e-05 | 1.300E-05 | 1.000E-06 | 1.200E-05 | 1.400E-05 |
| 4205 | 211 | T211C | A | 1.5e-05 | 1.2e-05 | 1.3e-05 | 1.333E-05 | 1.528E-06 | 1.200E-05 | 1.500E-05 |
| 4206 | 211 | T211E | A | 1e-05 | 7.4e-06 | 1.1e-05 | 9.467E-06 | 1.858E-06 | 7.400E-06 | 1.100E-05 |
| 4207 | 211 | T211Q | A | 1.5e-05 | 9.4e-06 | 9.8e-06 | 1.140E-05 | 3.124E-06 | 9.400E-06 | 1.500E-05 |
| 4208 | 211 | T211G | A | 1.1e-05 | 1.5e-05 | 1.4e-05 | 1.333E-05 | 2.082E-06 | 1.100E-05 | 1.500E-05 |
| 4209 | 211 | T211H | A | 1.4e-05 | 9.2e-06 | 1.1e-05 | 1.140E-05 | 2.425E-06 | 9.200E-06 | 1.400E-05 |
| 4210 | 211 | T211I | A | 1.5e-05 | 1e-05 | 1.5e-05 | 1.333E-05 | 2.887E-06 | 1.000E-05 | 1.500E-05 |
| 4211 | 211 | T211L | A | 1.4e-05 | 1.2e-05 | 1.1e-05 | 1.233E-05 | 1.528E-06 | 1.100E-05 | 1.400E-05 |
| 4212 | 211 | T211K | A | 1.4e-05 | 1.5e-05 | 1e-05 | 1.300E-05 | 2.646E-06 | 1.000E-05 | 1.500E-05 |
| 4213 | 211 | T211M | A | 1.7e-05 | 1.1e-05 | 1e-05 | 1.267E-05 | 3.786E-06 | 1.000E-05 | 1.700E-05 |
| 4214 | 211 | T211F | A | 1.5e-05 | 1.1e-05 | 9e-06 | 1.167E-05 | 3.055E-06 | 9.000E-06 | 1.500E-05 |
| 4215 | 211 | T211P | A | 1.5e-05 | 1.5e-05 | 1.1e-05 | 1.367E-05 | 2.309E-06 | 1.100E-05 | 1.500E-05 |
| 4216 | 211 | T211S | A | 1.5e-05 | 1.4e-05 | 1.6e-05 | 1.500E-05 | 1.000E-06 | 1.400E-05 | 1.600E-05 |
| 4217 | 211 | T211T | A | 1.4e-05 | 1.8e-05 | 1e-05 | 1.400E-05 | 4.000E-06 | 1.000E-05 | 1.800E-05 |
| 4218 | 211 | T211W | A | 1.4e-05 | 1.6e-05 | 1.7e-05 | 1.567E-05 | 1.528E-06 | 1.400E-05 | 1.700E-05 |
| 4219 | 211 | T211Y | A | 1.5e-05 | 1e-05 | 1.6e-05 | 1.367E-05 | 3.215E-06 | 1.000E-05 | 1.600E-05 |
| 4220 | 211 | T211V | A | 1.5e-05 | 1.4e-05 | 1.2e-05 | 1.367E-05 | 1.528E-06 | 1.200E-05 | 1.500E-05 |
| 4221 | 212 | D212A | A | 1.6e-05 | 1e-05 | 9.9e-06 | 1.197E-05 | 3.493E-06 | 9.900E-06 | 1.600E-05 |
| 4222 | 212 | D212R | A | 8.6e-06 | 1.6e-05 | 1.5e-05 | 1.320E-05 | 4.015E-06 | 8.600E-06 | 1.600E-05 |
| 4223 | 212 | D212N | A | 1.4e-05 | 9e-06 | 1.5e-05 | 1.267E-05 | 3.215E-06 | 9.000E-06 | 1.500E-05 |
| 4224 | 212 | D212D | A | 1e-05 | 1e-05 | 1.5e-05 | 1.167E-05 | 2.887E-06 | 1.000E-05 | 1.500E-05 |
| 4225 | 212 | D212C | A | 1.2e-05 | 1.1e-05 | 1.4e-05 | 1.233E-05 | 1.528E-06 | 1.100E-05 | 1.400E-05 |
| 4226 | 212 | D212E | A | 1.5e-05 | 1.3e-05 | 6.7e-06 | 1.157E-05 | 4.332E-06 | 6.700E-06 | 1.500E-05 |
| 4227 | 212 | D212Q | A | 1.5e-05 | 1.5e-05 | 6.9e-06 | 1.230E-05 | 4.677E-06 | 6.900E-06 | 1.500E-05 |
| 4228 | 212 | D212G | A | 9.8e-06 | 1.2e-05 | 8.6e-06 | 1.013E-05 | 1.724E-06 | 8.600E-06 | 1.200E-05 |
| 4229 | 212 | D212H | A | 1.5e-05 | 1e-05 | 1.4e-05 | 1.300E-05 | 2.646E-06 | 1.000E-05 | 1.500E-05 |
| 4230 | 212 | D212I | A | 9.7e-06 | 1.7e-05 | 1e-05 | 1.223E-05 | 4.131E-06 | 9.700E-06 | 1.700E-05 |
| 4231 | 212 | D212L | A | 1.3e-05 | 1.2e-05 | 1.7e-05 | 1.400E-05 | 2.646E-06 | 1.200E-05 | 1.700E-05 |
| 4232 | 212 | D212K | A | 1.2e-05 | 1.5e-05 | 2.5e-05 | 1.733E-05 | 6.807E-06 | 1.200E-05 | 2.500E-05 |
| 4233 | 212 | D212M | A | 1.1e-05 | 1.6e-05 | 1.5e-05 | 1.400E-05 | 2.646E-06 | 1.100E-05 | 1.600E-05 |
| 4234 | 212 | D212F | A | 1e-05 | 9.8e-06 | 1.6e-05 | 1.193E-05 | 3.523E-06 | 9.800E-06 | 1.600E-05 |
| 4235 | 212 | D212P | A | 2.2e-05 | 1.1e-05 | 9.8e-06 | 1.427E-05 | 6.724E-06 | 9.800E-06 | 2.200E-05 |
| 4236 | 212 | D212S | A | 1.4e-05 | 1.5e-05 | 1.6e-05 | 1.500E-05 | 1.000E-06 | 1.400E-05 | 1.600E-05 |
| 4237 | 212 | D212T | A | 9.9e-06 | 1.3e-05 | 1.3e-05 | 1.197E-05 | 1.790E-06 | 9.900E-06 | 1.300E-05 |
| 4238 | 212 | D212W | A | 9e-06 | 7e-06 | 1.4e-05 | 1.000E-05 | 3.606E-06 | 7.000E-06 | 1.400E-05 |
| 4239 | 212 | D212Y | A | 1.3e-05 | 6e-06 | 1.1e-05 | 1.000E-05 | 3.606E-06 | 6.000E-06 | 1.300E-05 |
| 4240 | 212 | D212V | A | 9.8e-06 | 1.5e-05 | 1.4e-05 | 1.293E-05 | 2.759E-06 | 9.800E-06 | 1.500E-05 |
| 4241 | 213 | C213A | A | 8.7e-06 | 8.6e-06 | 1.1e-05 | 9.433E-06 | 1.358E-06 | 8.600E-06 | 1.100E-05 |
| 4242 | 213 | C213R | A | 8.9e-06 | 1.1e-05 | 1.3e-05 | 1.097E-05 | 2.050E-06 | 8.900E-06 | 1.300E-05 |
| 4243 | 213 | C213N | A | 1.3e-05 | 1.5e-05 | 1.5e-05 | 1.433E-05 | 1.155E-06 | 1.300E-05 | 1.500E-05 |
| 4244 | 213 | C213D | A | 9.9e-06 | 1.3e-05 | 1.1e-05 | 1.130E-05 | 1.572E-06 | 9.900E-06 | 1.300E-05 |
| 4245 | 213 | C213C | A | 1.2e-05 | 1.5e-05 | 1e-05 | 1.233E-05 | 2.517E-06 | 1.000E-05 | 1.500E-05 |
| 4246 | 213 | C213E | A | 9.4e-06 | 1.6e-05 | 1.7e-05 | 1.413E-05 | 4.130E-06 | 9.400E-06 | 1.700E-05 |
| 4247 | 213 | C213Q | A | 1.4e-05 | 1.4e-05 | 1.5e-05 | 1.433E-05 | 5.774E-07 | 1.400E-05 | 1.500E-05 |
| 4248 | 213 | C213G | A | 1.6e-05 | 1.2e-05 | 1.5e-05 | 1.433E-05 | 2.082E-06 | 1.200E-05 | 1.600E-05 |
| 4249 | 213 | C213H | A | 1e-05 | 1.4e-05 | 8.7e-06 | 1.090E-05 | 2.762E-06 | 8.700E-06 | 1.400E-05 |

| | | | | | | | | | | |
|---|---|---|---|---|---|---|---|---|---|---|
| 4250 | 213 | C213I | A | 1.6e-05 | 1e-05 | 1.1e-05 | 1.233E-05 | 3.215E-06 | 1.000E-05 | 1.600E-05 |
| 4251 | 213 | C213L | A | 9.4e-06 | 8.3e-06 | 1e-05 | 9.233E-06 | 8.622E-07 | 8.300E-06 | 1.000E-05 |
| 4252 | 213 | C213K | A | 7.1e-06 | 1e-05 | 1.6e-05 | 1.103E-05 | 4.539E-06 | 7.100E-06 | 1.600E-05 |
| 4253 | 213 | C213M | A | 1.1e-05 | 1.7e-05 | 9.5e-06 | 1.250E-05 | 3.969E-06 | 9.500E-06 | 1.700E-05 |
| 4254 | 213 | C213F | A | 9.5e-06 | 1.9e-05 | 1.1e-05 | 1.317E-05 | 5.107E-06 | 9.500E-06 | 1.900E-05 |
| 4255 | 213 | C213P | A | 1.4e-05 | 8.9e-06 | 1.6e-05 | 1.297E-05 | 3.661E-06 | 8.900E-06 | 1.600E-05 |
| 4256 | 213 | C213S | A | 1.6e-05 | 1e-05 | 1.6e-05 | 1.400E-05 | 3.464E-06 | 1.000E-05 | 1.600E-05 |
| 4257 | 213 | C213T | A | 8.5e-06 | 1.4e-05 | 1.2e-05 | 1.150E-05 | 2.784E-06 | 8.500E-06 | 1.400E-05 |
| 4258 | 213 | C213W | A | 1.5e-05 | 1.4e-05 | 1.4e-05 | 1.433E-05 | 5.774E-07 | 1.400E-05 | 1.500E-05 |
| 4259 | 213 | C213Y | A | 1.5e-05 | 1.1e-05 | 9.8e-06 | 1.193E-05 | 2.723E-06 | 9.800E-06 | 1.500E-05 |
| 4260 | 213 | C213V | A | 1e-05 | 1.1e-05 | 1.1e-05 | 1.067E-05 | 5.774E-07 | 1.000E-05 | 1.100E-05 |
| 4261 | 214 | C214A | A | 1.5e-05 | 1.6e-05 | 1e-05 | 1.367E-05 | 3.215E-06 | 1.000E-05 | 1.600E-05 |
| 4262 | 214 | C214R | A | 1.6e-05 | 1.6e-05 | 1.2e-05 | 1.467E-05 | 2.309E-06 | 1.200E-05 | 1.600E-05 |
| 4263 | 214 | C214N | A | 1.5e-05 | 1.4e-05 | 1.4e-05 | 1.433E-05 | 5.774E-07 | 1.400E-05 | 1.500E-05 |
| 4264 | 214 | C214D | A | 1.1e-05 | 1.6e-05 | 1.1e-05 | 1.267E-05 | 2.887E-06 | 1.100E-05 | 1.600E-05 |
| 4265 | 214 | C214C | A | 1e-05 | 1.2e-05 | 1.3e-05 | 1.167E-05 | 1.528E-06 | 1.000E-05 | 1.300E-05 |
| 4266 | 214 | C214E | A | 1.4e-05 | 1.8e-05 | 1.7e-05 | 1.633E-05 | 2.082E-06 | 1.400E-05 | 1.800E-05 |
| 4267 | 214 | C214Q | A | 1.3e-05 | 1.3e-05 | 1.6e-05 | 1.400E-05 | 1.732E-06 | 1.300E-05 | 1.600E-05 |
| 4268 | 214 | C214G | A | 1.1e-05 | 1.2e-05 | 1e-05 | 1.100E-05 | 1.000E-06 | 1.000E-05 | 1.200E-05 |
| 4269 | 214 | C214H | A | 9.8e-06 | 9.7e-06 | 1.5e-05 | 1.150E-05 | 3.032E-06 | 9.700E-06 | 1.500E-05 |
| 4270 | 214 | C214I | A | 1.5e-05 | 1.1e-05 | 1.5e-05 | 1.367E-05 | 2.309E-06 | 1.100E-05 | 1.500E-05 |
| 4271 | 214 | C214L | A | 1.1e-05 | 1.5e-05 | 1.4e-05 | 1.333E-05 | 2.082E-06 | 1.100E-05 | 1.500E-05 |
| 4272 | 214 | C214K | A | 1.5e-05 | 1.4e-05 | 1.1e-05 | 1.333E-05 | 2.082E-06 | 1.100E-05 | 1.500E-05 |
| 4273 | 214 | C214M | A | 1.7e-05 | 9.2e-06 | 1.7e-05 | 1.440E-05 | 4.503E-06 | 9.200E-06 | 1.700E-05 |
| 4274 | 214 | C214F | A | 1.1e-05 | 1.2e-05 | 9.6e-06 | 1.087E-05 | 1.206E-06 | 9.600E-06 | 1.200E-05 |
| 4275 | 214 | C214P | A | 8.9e-06 | 1.4e-05 | 1.5e-05 | 1.263E-05 | 3.272E-06 | 8.900E-06 | 1.500E-05 |
| 4276 | 214 | C214S | A | 1.3e-05 | 1.5e-05 | 1.8e-05 | 1.533E-05 | 2.517E-06 | 1.300E-05 | 1.800E-05 |
| 4277 | 214 | C214T | A | 1.3e-05 | 1.4e-05 | 1.6e-05 | 1.433E-05 | 1.528E-06 | 1.300E-05 | 1.600E-05 |
| 4278 | 214 | C214W | A | 1.4e-05 | 1.2e-05 | 1.4e-05 | 1.333E-05 | 1.155E-06 | 1.200E-05 | 1.400E-05 |
| 4279 | 214 | C214Y | A | 1.3e-05 | 1.6e-05 | 1.6e-05 | 1.500E-05 | 1.732E-06 | 1.300E-05 | 1.600E-05 |
| 4280 | 214 | C214V | A | 1.4e-05 | 1.7e-05 | 1e-05 | 1.367E-05 | 3.512E-06 | 1.000E-05 | 1.700E-05 |
| 4281 | 215 | H215A | A | 1.4e-05 | 1.3e-05 | 1.1e-05 | 1.267E-05 | 1.528E-06 | 1.100E-05 | 1.400E-05 |
| 4282 | 215 | H215R | A | 1e-05 | 7.2e-06 | 1.3e-05 | 1.007E-05 | 2.901E-06 | 7.200E-06 | 1.300E-05 |
| 4283 | 215 | H215N | A | 1.4e-05 | 9.8e-06 | 1.6e-05 | 1.327E-05 | 3.164E-06 | 9.800E-06 | 1.600E-05 |
| 4284 | 215 | H215D | A | 9.8e-06 | 1.3e-05 | 1.6e-05 | 1.293E-05 | 3.101E-06 | 9.800E-06 | 1.600E-05 |
| 4285 | 215 | H215C | A | 1.6e-05 | 1.5e-05 | 1.1e-05 | 1.400E-05 | 2.646E-06 | 1.100E-05 | 1.600E-05 |
| 4286 | 215 | H215E | A | 1.4e-05 | 1.2e-05 | 1.6e-05 | 1.400E-05 | 2.000E-06 | 1.200E-05 | 1.600E-05 |
| 4287 | 215 | H215Q | A | 8.5e-06 | 1.1e-05 | 9.6e-06 | 9.700E-06 | 1.253E-06 | 8.500E-06 | 1.100E-05 |
| 4288 | 215 | H215G | A | 1.5e-05 | 1.5e-05 | 1.7e-05 | 1.567E-05 | 1.155E-06 | 1.500E-05 | 1.700E-05 |
| 4289 | 215 | H215H | A | 1.6e-05 | 1.4e-05 | 1.3e-05 | 1.433E-05 | 1.528E-06 | 1.300E-05 | 1.600E-05 |
| 4290 | 215 | H215I | A | 1.6e-05 | 1.6e-05 | 1.6e-05 | 1.600E-05 | 0.000E+00 | 1.600E-05 | 1.600E-05 |
| 4291 | 215 | H215L | A | 1.3e-05 | 1.5e-05 | 1.2e-05 | 1.333E-05 | 1.528E-06 | 1.200E-05 | 1.500E-05 |
| 4292 | 215 | H215K | A | 1.5e-05 | 9.1e-06 | 1.6e-05 | 1.337E-05 | 3.729E-06 | 9.100E-06 | 1.600E-05 |
| 4293 | 215 | H215M | A | 9.2e-06 | 1.1e-05 | 1.3e-05 | 1.107E-05 | 1.901E-06 | 9.200E-06 | 1.300E-05 |
| 4294 | 215 | H215F | A | 1.1e-05 | 9.6e-06 | 7.7e-06 | 9.433E-06 | 1.656E-06 | 7.700E-06 | 1.100E-05 |
| 4295 | 215 | H215P | A | 9.1e-06 | 1.2e-05 | 1.2e-05 | 1.103E-05 | 1.674E-06 | 9.100E-06 | 1.200E-05 |
| 4296 | 215 | H215S | A | 1.5e-05 | 7.9e-06 | 1.3e-05 | 1.197E-05 | 3.661E-06 | 7.900E-06 | 1.500E-05 |
| 4297 | 215 | H215T | A | 1.1e-05 | 1.4e-05 | 1.3e-05 | 1.267E-05 | 1.528E-06 | 1.100E-05 | 1.400E-05 |
| 4298 | 215 | H215W | A | 1.1e-05 | 1e-05 | 9.2e-06 | 1.007E-05 | 9.018E-07 | 9.200E-06 | 1.100E-05 |
| 4299 | 215 | H215Y | A | 1.6e-05 | 9e-06 | 1.5e-05 | 1.333E-05 | 3.786E-06 | 9.000E-06 | 1.600E-05 |
| 4300 | 215 | H215V | A | 1.3e-05 | 1.1e-05 | 1.3e-05 | 1.233E-05 | 1.155E-06 | 1.100E-05 | 1.300E-05 |
| 4301 | 216 | E216A | A | 1.1e-05 | 1.1e-05 | 7.8e-06 | 9.933E-06 | 1.848E-06 | 7.800E-06 | 1.100E-05 |
| 4302 | 216 | E216R | A | 1.6e-05 | 1e-05 | 1.5e-05 | 1.367E-05 | 3.215E-06 | 1.000E-05 | 1.600E-05 |
| 4303 | 216 | E216N | A | 1.4e-05 | 9.5e-06 | 7.8e-06 | 1.043E-05 | 3.204E-06 | 7.800E-06 | 1.400E-05 |
| 4304 | 216 | E216D | A | 9e-06 | 2.2e-05 | 1.1e-05 | 1.400E-05 | 7.000E-06 | 9.000E-06 | 2.200E-05 |
| 4305 | 216 | E216C | A | 8.6e-06 | 7.7e-06 | 1.5e-05 | 1.043E-05 | 3.980E-06 | 7.700E-06 | 1.500E-05 |
| 4306 | 216 | E216E | A | 1.2e-05 | 1.5e-05 | 1.4e-05 | 1.367E-05 | 1.528E-06 | 1.200E-05 | 1.500E-05 |
| 4307 | 216 | E216Q | A | 1.7e-05 | 1.6e-05 | 1.1e-05 | 1.467E-05 | 3.215E-06 | 1.100E-05 | 1.700E-05 |
| 4308 | 216 | E216G | A | 1e-05 | 1.5e-05 | 1.4e-05 | 1.300E-05 | 2.646E-06 | 1.000E-05 | 1.500E-05 |
| 4309 | 216 | E216H | A | 1.4e-05 | 2e-05 | 1.5e-05 | 1.633E-05 | 3.215E-06 | 1.400E-05 | 2.000E-05 |
| 4310 | 216 | E216I | A | 1.1e-05 | 1.1e-05 | 1.7e-05 | 1.300E-05 | 3.464E-06 | 1.100E-05 | 1.700E-05 |
| 4311 | 216 | E216L | A | 1.1e-05 | 1.1e-05 | 6.9e-06 | 9.633E-06 | 2.367E-06 | 6.900E-06 | 1.100E-05 |
| 4312 | 216 | E216K | A | 1e-05 | 8.4e-06 | 9.4e-06 | 9.267E-06 | 8.083E-07 | 8.400E-06 | 1.000E-05 |
| 4313 | 216 | E216M | A | 1.5e-05 | 1.3e-05 | 9.3e-06 | 1.243E-05 | 2.892E-06 | 9.300E-06 | 1.500E-05 |
| 4314 | 216 | E216F | A | 1.1e-05 | 1e-05 | 1.3e-05 | 1.133E-05 | 1.528E-06 | 1.000E-05 | 1.300E-05 |
| 4315 | 216 | E216P | A | 1.2e-05 | 8.8e-06 | 1.7e-05 | 1.260E-05 | 4.133E-06 | 8.800E-06 | 1.700E-05 |
| 4316 | 216 | E216S | A | 1.6e-05 | 9e-06 | 1.3e-05 | 1.267E-05 | 3.512E-06 | 9.000E-06 | 1.600E-05 |
| 4317 | 216 | E216T | A | 1e-05 | 1.1e-05 | 1.5e-05 | 1.200E-05 | 2.646E-06 | 1.000E-05 | 1.500E-05 |
| 4318 | 216 | E216W | A | 1.2e-05 | 8.9e-06 | 1.5e-05 | 1.197E-05 | 3.050E-06 | 8.900E-06 | 1.500E-05 |
| 4319 | 216 | E216Y | A | 1.2e-05 | 8.4e-06 | 1.3e-05 | 1.113E-05 | 2.419E-06 | 8.400E-06 | 1.300E-05 |
| 4320 | 216 | E216V | A | 1.7e-05 | 1e-05 | 1.6e-05 | 1.433E-05 | 3.786E-06 | 1.000E-05 | 1.700E-05 |
| 4321 | 217 | Q217A | A | 1.1e-05 | 1.2e-05 | 1.2e-05 | 1.167E-05 | 5.774E-07 | 1.100E-05 | 1.200E-05 |
| 4322 | 217 | Q217R | A | 8e-06 | 9.7e-06 | 1e-05 | 9.233E-06 | 1.079E-06 | 8.000E-06 | 1.000E-05 |
| 4323 | 217 | Q217N | A | 1e-05 | 1.6e-05 | 1.2e-05 | 1.267E-05 | 3.055E-06 | 1.000E-05 | 1.600E-05 |
| 4324 | 217 | Q217D | A | 1.1e-05 | 1.2e-05 | 1.7e-05 | 1.333E-05 | 3.215E-06 | 1.100E-05 | 1.700E-05 |
| 4325 | 217 | Q217C | A | 1.2e-05 | 9.6e-06 | 9.9e-06 | 1.050E-05 | 1.308E-06 | 9.600E-06 | 1.200E-05 |
| 4326 | 217 | Q217E | A | 2.1e-05 | 9.4e-06 | 8.5e-06 | 1.297E-05 | 6.972E-06 | 8.500E-06 | 2.100E-05 |
| 4327 | 217 | Q217Q | A | 1e-05 | 1.3e-05 | 1.4e-05 | 1.233E-05 | 2.082E-06 | 1.000E-05 | 1.400E-05 |
| 4328 | 217 | Q217G | A | 1.5e-05 | 1.4e-05 | 1.4e-05 | 1.433E-05 | 5.774E-07 | 1.400E-05 | 1.500E-05 |
| 4329 | 217 | Q217H | A | 9e-06 | 1e-05 | 9.5e-06 | 9.500E-06 | 5.000E-07 | 9.000E-06 | 1.000E-05 |
| 4330 | 217 | Q217I | A | 1.6e-05 | 1.2e-05 | 1.5e-05 | 1.433E-05 | 2.082E-06 | 1.200E-05 | 1.600E-05 |
| 4331 | 217 | Q217L | A | 1.1e-05 | 1.7e-05 | 1.2e-05 | 1.333E-05 | 3.215E-06 | 1.100E-05 | 1.700E-05 |
| 4332 | 217 | Q217K | A | 1.5e-05 | 9.7e-06 | 1.7e-05 | 1.390E-05 | 3.772E-06 | 9.700E-06 | 1.700E-05 |
| 4333 | 217 | Q217M | A | 1.5e-05 | 2.8e-05 | 1.5e-05 | 1.933E-05 | 7.506E-06 | 1.500E-05 | 2.800E-05 |
| 4334 | 217 | Q217F | A | 1.6e-05 | 1.5e-05 | 8.5e-06 | 1.317E-05 | 4.072E-06 | 8.500E-06 | 1.600E-05 |

| | | | | | | | | | | |
|---|---|---|---|---|---|---|---|---|---|---|
| 4335 | 217 | Q217P | A | 1.3e-05 | 1.1e-05 | 1e-05 | 1.133E-05 | 1.528E-06 | 1.000E-05 | 1.300E-05 |
| 4336 | 217 | Q217S | A | 8.9e-06 | 1e-05 | 1e-05 | 9.633E-06 | 6.351E-07 | 8.900E-06 | 1.000E-05 |
| 4337 | 217 | Q217T | A | 1.2e-05 | 6.9e-06 | 1.1e-05 | 9.967E-06 | 2.702E-06 | 6.900E-06 | 1.200E-05 |
| 4338 | 217 | Q217W | A | 1.2e-05 | 1.6e-05 | 1.5e-05 | 1.433E-05 | 2.082E-06 | 1.200E-05 | 1.600E-05 |
| 4339 | 217 | Q217Y | A | 1.6e-05 | 1.6e-05 | 9.2e-06 | 1.373E-05 | 3.926E-06 | 9.200E-06 | 1.600E-05 |
| 4340 | 217 | Q217V | A | 1.7e-05 | 1.7e-05 | 1.2e-05 | 1.533E-05 | 2.887E-06 | 1.200E-05 | 1.700E-05 |
| 4341 | 218 | C218A | A | 1e-05 | 1.7e-05 | 1.1e-05 | 1.267E-05 | 3.786E-06 | 1.000E-05 | 1.700E-05 |
| 4342 | 218 | C218R | A | 1.3e-05 | 9.4e-06 | 1.5e-05 | 1.247E-05 | 2.838E-06 | 9.400E-06 | 1.500E-05 |
| 4343 | 218 | C218N | A | 9.9e-06 | 1.5e-05 | 1e-05 | 1.163E-05 | 2.916E-06 | 9.900E-06 | 1.500E-05 |
| 4344 | 218 | C218D | A | 1.4e-05 | 8.7e-06 | 8.3e-06 | 1.033E-05 | 3.182E-06 | 8.300E-06 | 1.400E-05 |
| 4345 | 218 | C218C | A | 1.5e-05 | 1.5e-05 | 1.6e-05 | 1.533E-05 | 5.774E-07 | 1.500E-05 | 1.600E-05 |
| 4346 | 218 | C218E | A | 8e-06 | 8.9e-06 | 9.9e-06 | 8.933E-06 | 9.504E-07 | 8.000E-06 | 9.900E-06 |
| 4347 | 218 | C218Q | A | 1.6e-05 | 1.3e-05 | 1.2e-05 | 1.367E-05 | 2.082E-06 | 1.200E-05 | 1.600E-05 |
| 4348 | 218 | C218G | A | 1.7e-05 | 1.3e-05 | 1e-05 | 1.333E-05 | 3.512E-06 | 1.000E-05 | 1.700E-05 |
| 4349 | 218 | C218H | A | 1.6e-05 | 2.2e-05 | 1.4e-05 | 1.733E-05 | 4.163E-06 | 1.400E-05 | 2.200E-05 |
| 4350 | 218 | C218I | A | 1.6e-05 | 1.5e-05 | 1.1e-05 | 1.400E-05 | 2.646E-06 | 1.100E-05 | 1.600E-05 |
| 4351 | 218 | C218L | A | 1.5e-05 | 1.1e-05 | 9.3e-06 | 1.177E-05 | 2.926E-06 | 9.300E-06 | 1.500E-05 |
| 4352 | 218 | C218K | A | 1.3e-05 | 1e-05 | 1.1e-05 | 1.133E-05 | 1.528E-06 | 1.000E-05 | 1.300E-05 |
| 4353 | 218 | C218M | A | 1.1e-05 | 1.3e-05 | 1.7e-05 | 1.367E-05 | 3.055E-06 | 1.100E-05 | 1.700E-05 |
| 4354 | 218 | C218F | A | 1.4e-05 | 1.1e-05 | 1e-05 | 1.167E-05 | 2.082E-06 | 1.000E-05 | 1.400E-05 |
| 4355 | 218 | C218P | A | 1.2e-05 | 1.6e-05 | 9.1e-06 | 1.237E-05 | 3.465E-06 | 9.100E-06 | 1.600E-05 |
| 4356 | 218 | C218S | A | 1.5e-05 | 9.1e-06 | 1.7e-05 | 1.370E-05 | 4.107E-06 | 9.100E-06 | 1.700E-05 |
| 4357 | 218 | C218T | A | 1.5e-05 | 1e-05 | 1.7e-05 | 1.400E-05 | 3.606E-06 | 1.000E-05 | 1.700E-05 |
| 4358 | 218 | C218W | A | 1e-05 | 1.5e-05 | 9.8e-06 | 1.160E-05 | 2.946E-06 | 9.800E-06 | 1.500E-05 |
| 4359 | 218 | C218Y | A | 1.5e-05 | 1.4e-05 | 1.6e-05 | 1.500E-05 | 1.000E-06 | 1.400E-05 | 1.600E-05 |
| 4360 | 218 | C218V | A | 1.1e-05 | 1.6e-05 | 1.2e-05 | 1.300E-05 | 2.646E-06 | 1.100E-05 | 1.600E-05 |
| 4361 | 219 | A219A | A | 1.5e-05 | 1.4e-05 | 1.2e-05 | 1.367E-05 | 1.528E-06 | 1.200E-05 | 1.500E-05 |
| 4362 | 219 | A219R | A | 1.4e-05 | 1.1e-05 | 9.9e-06 | 1.163E-05 | 2.122E-06 | 9.900E-06 | 1.400E-05 |
| 4363 | 219 | A219N | A | 1.2e-05 | 1.2e-05 | 1e-05 | 1.133E-05 | 1.155E-06 | 1.000E-05 | 1.200E-05 |
| 4364 | 219 | A219D | A | 1.5e-05 | 1.6e-05 | 1.6e-05 | 1.567E-05 | 5.774E-07 | 1.500E-05 | 1.600E-05 |
| 4365 | 219 | A219C | A | 1.7e-05 | 1.5e-05 | 1.4e-05 | 1.533E-05 | 1.528E-06 | 1.400E-05 | 1.700E-05 |
| 4366 | 219 | A219E | A | 1e-05 | 1.6e-05 | 1.5e-05 | 1.367E-05 | 3.215E-06 | 1.000E-05 | 1.600E-05 |
| 4367 | 219 | A219Q | A | 1.1e-05 | 1.3e-05 | 1.4e-05 | 1.267E-05 | 1.528E-06 | 1.100E-05 | 1.400E-05 |
| 4368 | 219 | A219G | A | 1e-05 | 1.3e-05 | 9.6e-06 | 1.087E-05 | 1.858E-06 | 9.600E-06 | 1.300E-05 |
| 4369 | 219 | A219H | A | 9.8e-06 | 1.3e-05 | 1.1e-05 | 1.127E-05 | 1.617E-06 | 9.800E-06 | 1.300E-05 |
| 4370 | 219 | A219I | A | 1.1e-05 | 1.1e-05 | 1.4e-05 | 1.200E-05 | 1.732E-06 | 1.100E-05 | 1.400E-05 |
| 4371 | 219 | A219L | A | 1e-05 | 1.6e-05 | 9.3e-06 | 1.177E-05 | 3.683E-06 | 9.300E-06 | 1.600E-05 |
| 4372 | 219 | A219K | A | 1.4e-05 | 1.4e-05 | 1.4e-05 | 1.400E-05 | 0.000E+00 | 1.400E-05 | 1.400E-05 |
| 4373 | 219 | A219M | A | 1.3e-05 | 8e-06 | 7.8e-06 | 9.600E-06 | 2.946E-06 | 7.800E-06 | 1.300E-05 |
| 4374 | 219 | A219F | A | 6.5e-06 | 1e-05 | 9.5e-06 | 8.667E-06 | 1.893E-06 | 6.500E-06 | 1.000E-05 |
| 4375 | 219 | A219P | A | 1.5e-05 | 1.5e-05 | 1.3e-05 | 1.433E-05 | 1.155E-06 | 1.300E-05 | 1.500E-05 |
| 4376 | 219 | A219S | A | 1e-05 | 9.2e-06 | 1.4e-05 | 1.107E-05 | 2.572E-06 | 9.200E-06 | 1.400E-05 |
| 4377 | 219 | A219T | A | 1.2e-05 | 1.2e-05 | 9e-06 | 1.100E-05 | 1.732E-06 | 9.000E-06 | 1.200E-05 |
| 4378 | 219 | A219W | A | 1.7e-05 | 1.6e-05 | 1.4e-05 | 1.567E-05 | 1.528E-06 | 1.400E-05 | 1.700E-05 |
| 4379 | 219 | A219Y | A | 1.5e-05 | 1e-05 | 1.2e-05 | 1.233E-05 | 2.517E-06 | 1.000E-05 | 1.500E-05 |
| 4380 | 219 | A219V | A | 1.6e-05 | 1.5e-05 | 1.3e-05 | 1.467E-05 | 1.528E-06 | 1.300E-05 | 1.600E-05 |
| 4381 | 220 | A220A | A | 1.5e-05 | 1.2e-05 | 1e-05 | 1.233E-05 | 2.517E-06 | 1.000E-05 | 1.500E-05 |
| 4382 | 220 | A220R | A | 9.6e-06 | 9.9e-06 | 1.5e-05 | 1.150E-05 | 3.035E-06 | 9.600E-06 | 1.500E-05 |
| 4383 | 220 | A220N | A | 9.3e-06 | 8e-06 | 1.3e-05 | 1.010E-05 | 2.594E-06 | 8.000E-06 | 1.300E-05 |
| 4384 | 220 | A220D | A | 1.2e-05 | 1.3e-05 | 1.4e-05 | 1.300E-05 | 1.000E-06 | 1.200E-05 | 1.400E-05 |
| 4385 | 220 | A220C | A | 1.6e-05 | 1.5e-05 | 1.1e-05 | 1.400E-05 | 2.646E-06 | 1.100E-05 | 1.600E-05 |
| 4386 | 220 | A220E | A | 9.6e-06 | 1.3e-05 | 1.5e-05 | 1.253E-05 | 2.730E-06 | 9.600E-06 | 1.500E-05 |
| 4387 | 220 | A220Q | A | 1e-05 | 1e-05 | 1.5e-05 | 1.167E-05 | 2.887E-06 | 1.000E-05 | 1.500E-05 |
| 4388 | 220 | A220G | A | 7.9e-06 | 1.1e-05 | 1.6e-05 | 1.163E-05 | 4.087E-06 | 7.900E-06 | 1.600E-05 |
| 4389 | 220 | A220H | A | 1.1e-05 | 1.5e-05 | 1.7e-05 | 1.433E-05 | 3.055E-06 | 1.100E-05 | 1.700E-05 |
| 4390 | 220 | A220I | A | 8.9e-06 | 1.4e-05 | 1.6e-05 | 1.297E-05 | 3.661E-06 | 8.900E-06 | 1.600E-05 |
| 4391 | 220 | A220L | A | 9.1e-06 | 1.1e-05 | 1e-05 | 1.003E-05 | 9.504E-07 | 9.100E-06 | 1.100E-05 |
| 4392 | 220 | A220K | A | 1e-05 | 1.5e-05 | 8e-06 | 1.100E-05 | 3.606E-06 | 8.000E-06 | 1.500E-05 |
| 4393 | 220 | A220M | A | 1.5e-05 | 1.1e-05 | 9.2e-06 | 1.173E-05 | 2.969E-06 | 9.200E-06 | 1.500E-05 |
| 4394 | 220 | A220F | A | 1.2e-05 | 1.1e-05 | 1.1e-05 | 1.133E-05 | 5.774E-07 | 1.100E-05 | 1.200E-05 |
| 4395 | 220 | A220P | A | 1.5e-05 | 8.3e-06 | 1.1e-05 | 1.143E-05 | 3.371E-06 | 8.300E-06 | 1.500E-05 |
| 4396 | 220 | A220S | A | 1.2e-05 | 1.1e-05 | 1.9e-05 | 1.400E-05 | 4.359E-06 | 1.100E-05 | 1.900E-05 |
| 4397 | 220 | A220T | A | 8.5e-06 | 9.1e-06 | 1e-05 | 9.200E-06 | 7.550E-07 | 8.500E-06 | 1.000E-05 |
| 4398 | 220 | A220W | A | 1.6e-05 | 1.6e-05 | 1.2e-05 | 1.467E-05 | 2.309E-06 | 1.200E-05 | 1.600E-05 |
| 4399 | 220 | A220Y | A | 1.6e-05 | 1.1e-05 | 1.5e-05 | 1.400E-05 | 2.646E-06 | 1.100E-05 | 1.600E-05 |
| 4400 | 220 | A220V | A | 1.2e-05 | 1.6e-05 | 9.1e-06 | 1.237E-05 | 3.465E-06 | 9.100E-06 | 1.600E-05 |
| 4401 | 221 | G221A | A | 1e-05 | 1.6e-05 | 1.6e-05 | 1.400E-05 | 3.464E-06 | 1.000E-05 | 1.600E-05 |
| 4402 | 221 | G221R | A | 1.3e-05 | 1.2e-05 | 7.9e-06 | 1.097E-05 | 2.702E-06 | 7.900E-06 | 1.300E-05 |
| 4403 | 221 | G221N | A | 1.4e-05 | 1e-05 | 8.9e-06 | 1.097E-05 | 2.684E-06 | 8.900E-06 | 1.400E-05 |
| 4404 | 221 | G221D | A | 1.2e-05 | 1.1e-05 | 1.5e-05 | 1.267E-05 | 2.082E-06 | 1.100E-05 | 1.500E-05 |
| 4405 | 221 | G221C | A | 1e-05 | 1e-05 | 1.3e-05 | 1.100E-05 | 1.732E-06 | 1.000E-05 | 1.300E-05 |
| 4406 | 221 | G221E | A | 9.6e-06 | 1.1e-05 | 1.1e-05 | 1.053E-05 | 8.083E-07 | 9.600E-06 | 1.100E-05 |
| 4407 | 221 | G221Q | A | 9.8e-06 | 1.5e-05 | 1e-05 | 1.160E-05 | 2.946E-06 | 9.800E-06 | 1.500E-05 |
| 4408 | 221 | G221G | A | 8.4e-06 | 1.7e-05 | 1e-05 | 1.180E-05 | 4.574E-06 | 8.400E-06 | 1.700E-05 |
| 4409 | 221 | G221H | A | 1.4e-05 | 1.5e-05 | 1.6e-05 | 1.500E-05 | 1.000E-06 | 1.400E-05 | 1.600E-05 |
| 4410 | 221 | G221I | A | 1.6e-05 | 1.6e-05 | 1.7e-05 | 1.633E-05 | 5.774E-07 | 1.600E-05 | 1.700E-05 |
| 4411 | 221 | G221L | A | 1.1e-05 | 9.8e-06 | 1.3e-05 | 1.127E-05 | 1.617E-06 | 9.800E-06 | 1.300E-05 |
| 4412 | 221 | G221K | A | 2e-05 | 1.1e-05 | 1.5e-05 | 1.533E-05 | 4.509E-06 | 1.100E-05 | 2.000E-05 |
| 4413 | 221 | G221M | A | 1.5e-05 | 1.6e-05 | 1.5e-05 | 1.533E-05 | 5.774E-07 | 1.500E-05 | 1.600E-05 |
| 4414 | 221 | G221F | A | 1.1e-05 | 1.3e-05 | 1.1e-05 | 1.167E-05 | 1.155E-06 | 1.100E-05 | 1.300E-05 |
| 4415 | 221 | G221P | A | 8.7e-06 | 1.7e-05 | 1.3e-05 | 1.290E-05 | 4.151E-06 | 8.700E-06 | 1.700E-05 |
| 4416 | 221 | G221S | A | 1.5e-05 | 2.1e-05 | 1.4e-05 | 1.667E-05 | 3.786E-06 | 1.400E-05 | 2.100E-05 |
| 4417 | 221 | G221T | A | 1.1e-05 | 1.2e-05 | 1.3e-05 | 1.200E-05 | 1.000E-06 | 1.100E-05 | 1.300E-05 |
| 4418 | 221 | G221W | A | 1.1e-05 | 1.5e-05 | 1e-05 | 1.200E-05 | 2.646E-06 | 1.000E-05 | 1.500E-05 |
| 4419 | 221 | G221Y | A | 1.6e-05 | 1.5e-05 | 1.6e-05 | 1.567E-05 | 5.774E-07 | 1.500E-05 | 1.600E-05 |

| | | | | | | | | | | |
|---|---|---|---|---|---|---|---|---|---|---|
| 4420 | 221 | G221V | A | 1.1e-05 | 1e-05 | 1.6e-05 | 1.233E-05 | 3.215E-06 | 1.000E-05 | 1.600E-05 |
| 4421 | 222 | C222A | A | 3.7e-05 | 1.4e-05 | 8.2e-06 | 1.973E-05 | 1.523E-05 | 8.200E-06 | 3.700E-05 |
| 4422 | 222 | C222R | A | 9.8e-06 | 1.1e-05 | 1.5e-05 | 1.193E-05 | 2.723E-06 | 9.800E-06 | 1.500E-05 |
| 4423 | 222 | C222N | A | 1.2e-05 | 9.2e-06 | 1e-05 | 1.040E-05 | 1.442E-06 | 9.200E-06 | 1.200E-05 |
| 4424 | 222 | C222D | A | 9.9e-06 | 1.5e-05 | 1.5e-05 | 1.330E-05 | 2.944E-06 | 9.900E-06 | 1.500E-05 |
| 4425 | 222 | C222C | A | 1.4e-05 | 1.4e-05 | 1.1e-05 | 1.300E-05 | 1.732E-06 | 1.100E-05 | 1.400E-05 |
| 4426 | 222 | C222E | A | 8.7e-06 | 9.3e-06 | 1.2e-05 | 1.000E-05 | 1.758E-06 | 8.700E-06 | 1.200E-05 |
| 4427 | 222 | C222Q | A | 1e-05 | 7.8e-06 | 1e-05 | 9.267E-06 | 1.270E-06 | 7.800E-06 | 1.000E-05 |
| 4428 | 222 | C222G | A | 1.5e-05 | 1.1e-05 | 1.3e-05 | 1.300E-05 | 2.000E-06 | 1.100E-05 | 1.500E-05 |
| 4429 | 222 | C222H | A | 1.5e-05 | 1.1e-05 | 9.9e-06 | 1.197E-05 | 2.684E-06 | 9.900E-06 | 1.500E-05 |
| 4430 | 222 | C222I | A | 1.8e-05 | 1.3e-05 | 1.4e-05 | 1.500E-05 | 2.646E-06 | 1.300E-05 | 1.800E-05 |
| 4431 | 222 | C222L | A | 1.3e-05 | 1e-05 | 1.2e-05 | 1.167E-05 | 1.528E-06 | 1.000E-05 | 1.300E-05 |
| 4432 | 222 | C222K | A | 1.3e-05 | 1.7e-05 | 1e-05 | 1.333E-05 | 3.512E-06 | 1.000E-05 | 1.700E-05 |
| 4433 | 222 | C222M | A | 1.1e-05 | 1.1e-05 | 1.5e-05 | 1.233E-05 | 2.309E-06 | 1.100E-05 | 1.500E-05 |
| 4434 | 222 | C222F | A | 1.5e-05 | 1.2e-05 | 1.5e-05 | 1.400E-05 | 1.732E-06 | 1.200E-05 | 1.500E-05 |
| 4435 | 222 | C222P | A | 1.7e-05 | 1.5e-05 | 1.7e-05 | 1.633E-05 | 1.155E-06 | 1.500E-05 | 1.700E-05 |
| 4436 | 222 | C222S | A | 1.6e-05 | 1.5e-05 | 6.3e-06 | 1.243E-05 | 5.335E-06 | 6.300E-06 | 1.600E-05 |
| 4437 | 222 | C222T | A | 1e-05 | 9.8e-06 | 1.1e-05 | 1.027E-05 | 6.429E-07 | 9.800E-06 | 1.100E-05 |
| 4438 | 222 | C222W | A | 1.5e-05 | 1.2e-05 | 1.5e-05 | 1.400E-05 | 1.732E-06 | 1.200E-05 | 1.500E-05 |
| 4439 | 222 | C222Y | A | 1.5e-05 | 1.3e-05 | 1.1e-05 | 1.300E-05 | 2.000E-06 | 1.100E-05 | 1.500E-05 |
| 4440 | 222 | C222V | A | 1.3e-05 | 8.6e-06 | 1.4e-05 | 1.187E-05 | 2.873E-06 | 8.600E-06 | 1.400E-05 |
| 4441 | 223 | T223A | A | 1.2e-05 | 1.1e-05 | 1.4e-05 | 1.233E-05 | 1.528E-06 | 1.100E-05 | 1.400E-05 |
| 4442 | 223 | T223R | A | 1.6e-05 | 1.4e-05 | 1.5e-05 | 1.500E-05 | 1.000E-06 | 1.400E-05 | 1.600E-05 |
| 4443 | 223 | T223N | A | 1.1e-05 | 9.9e-06 | 1.1e-05 | 1.063E-05 | 6.351E-07 | 9.900E-06 | 1.100E-05 |
| 4444 | 223 | T223D | A | 1e-05 | 1.1e-05 | 1.7e-05 | 1.267E-05 | 3.786E-06 | 1.000E-05 | 1.700E-05 |
| 4445 | 223 | T223C | A | 1.3e-05 | 1e-05 | 1.5e-05 | 1.267E-05 | 2.517E-06 | 1.000E-05 | 1.500E-05 |
| 4446 | 223 | T223E | A | 8.7e-06 | 1.2e-05 | 1.5e-05 | 1.190E-05 | 3.151E-06 | 8.700E-06 | 1.500E-05 |
| 4447 | 223 | T223Q | A | 1.4e-05 | 1.4e-05 | 1.1e-05 | 1.300E-05 | 1.732E-06 | 1.100E-05 | 1.400E-05 |
| 4448 | 223 | T223G | A | 9.6e-06 | 1.2e-05 | 1.4e-05 | 1.187E-05 | 2.203E-06 | 9.600E-06 | 1.400E-05 |
| 4449 | 223 | T223H | A | 8.7e-06 | 1.4e-05 | 1.1e-05 | 1.123E-05 | 2.658E-06 | 8.700E-06 | 1.400E-05 |
| 4450 | 223 | T223I | A | 1.4e-05 | 1.4e-05 | 1.6e-05 | 1.467E-05 | 1.155E-06 | 1.400E-05 | 1.600E-05 |
| 4451 | 223 | T223L | A | 1.5e-05 | 1e-05 | 1.1e-05 | 1.200E-05 | 2.646E-06 | 1.000E-05 | 1.500E-05 |
| 4452 | 223 | T223K | A | 1.1e-05 | 1.1e-05 | 1.6e-05 | 1.267E-05 | 2.887E-06 | 1.100E-05 | 1.600E-05 |
| 4453 | 223 | T223M | A | 1.5e-05 | 1.4e-05 | 1.1e-05 | 1.333E-05 | 2.082E-06 | 1.100E-05 | 1.500E-05 |
| 4454 | 223 | T223F | A | 1.2e-05 | 1.4e-05 | 1e-05 | 1.200E-05 | 2.000E-06 | 1.000E-05 | 1.400E-05 |
| 4455 | 223 | T223P | A | 1e-05 | 1.6e-05 | 1.5e-05 | 1.367E-05 | 3.215E-06 | 1.000E-05 | 1.600E-05 |
| 4456 | 223 | T223S | A | 1.6e-05 | 1.4e-05 | 1.5e-05 | 1.500E-05 | 1.000E-06 | 1.400E-05 | 1.600E-05 |
| 4457 | 223 | T223T | A | 1.6e-05 | 1.6e-05 | 1.1e-05 | 1.433E-05 | 2.887E-06 | 1.100E-05 | 1.600E-05 |
| 4458 | 223 | T223W | A | 1.7e-05 | 8.7e-06 | 9.7e-06 | 1.180E-05 | 4.531E-06 | 8.700E-06 | 1.700E-05 |
| 4459 | 223 | T223Y | A | 1.1e-05 | 1e-05 | 9.9e-06 | 1.030E-05 | 6.083E-07 | 9.900E-06 | 1.100E-05 |
| 4460 | 223 | T223V | A | 1.5e-05 | 1.5e-05 | 1.5e-05 | 1.500E-05 | 0.000E+00 | 1.500E-05 | 1.500E-05 |
| 4461 | 224 | G224A | A | 1.6e-05 | 1.4e-05 | 1.4e-05 | 1.467E-05 | 1.155E-06 | 1.400E-05 | 1.600E-05 |
| 4462 | 224 | G224R | A | 1.5e-05 | 1.2e-05 | 1.3e-05 | 1.333E-05 | 1.528E-06 | 1.200E-05 | 1.500E-05 |
| 4463 | 224 | G224N | A | 1.6e-05 | 9.8e-06 | 1.1e-05 | 1.227E-05 | 3.288E-06 | 9.800E-06 | 1.600E-05 |
| 4464 | 224 | G224D | A | 1e-05 | 8.8e-06 | 1.5e-05 | 1.127E-05 | 3.288E-06 | 8.800E-06 | 1.500E-05 |
| 4465 | 224 | G224C | A | 1.4e-05 | 1.3e-05 | 9.4e-06 | 1.213E-05 | 2.419E-06 | 9.400E-06 | 1.400E-05 |
| 4466 | 224 | G224E | A | 1.4e-05 | 1.1e-05 | 1.4e-05 | 1.300E-05 | 1.732E-06 | 1.100E-05 | 1.400E-05 |
| 4467 | 224 | G224Q | A | 1.6e-05 | 1.5e-05 | 1.1e-05 | 1.400E-05 | 2.646E-06 | 1.100E-05 | 1.600E-05 |
| 4468 | 224 | G224G | A | 1.4e-05 | 1.1e-05 | 1.6e-05 | 1.367E-05 | 2.517E-06 | 1.100E-05 | 1.600E-05 |
| 4469 | 224 | G224H | A | 9.6e-06 | 1.6e-05 | 1.1e-05 | 1.220E-05 | 3.365E-06 | 9.600E-06 | 1.600E-05 |
| 4470 | 224 | G224I | A | 1e-05 | 1.4e-05 | 8.6e-06 | 1.087E-05 | 2.802E-06 | 8.600E-06 | 1.400E-05 |
| 4471 | 224 | G224L | A | 9.4e-06 | 1.5e-05 | 1.5e-05 | 1.313E-05 | 3.233E-06 | 9.400E-06 | 1.500E-05 |
| 4472 | 224 | G224K | A | 1.1e-05 | 1e-05 | 9.6e-06 | 1.020E-05 | 7.211E-07 | 9.600E-06 | 1.100E-05 |
| 4473 | 224 | G224M | A | 1.6e-05 | 1.5e-05 | 1e-05 | 1.367E-05 | 3.215E-06 | 1.000E-05 | 1.600E-05 |
| 4474 | 224 | G224F | A | 9.6e-06 | 1.8e-05 | 8.2e-06 | 1.193E-05 | 5.300E-06 | 8.200E-06 | 1.800E-05 |
| 4475 | 224 | G224P | A | 1.6e-05 | 1.5e-05 | 1.1e-05 | 1.400E-05 | 2.646E-06 | 1.100E-05 | 1.600E-05 |
| 4476 | 224 | G224S | A | 1e-05 | 1.2e-05 | 1.4e-05 | 1.200E-05 | 2.000E-06 | 1.000E-05 | 1.400E-05 |
| 4477 | 224 | G224T | A | 1.4e-05 | 1.1e-05 | 1.3e-05 | 1.267E-05 | 1.528E-06 | 1.100E-05 | 1.400E-05 |
| 4478 | 224 | G224W | A | 6.7e-06 | 1.1e-05 | 1.6e-05 | 1.123E-05 | 4.654E-06 | 6.700E-06 | 1.600E-05 |
| 4479 | 224 | G224Y | A | 9.6e-06 | 1.4e-05 | 1.4e-05 | 1.253E-05 | 2.540E-06 | 9.600E-06 | 1.400E-05 |
| 4480 | 224 | G224V | A | 1.4e-05 | 1.2e-05 | 1.6e-05 | 1.400E-05 | 2.000E-06 | 1.200E-05 | 1.600E-05 |
| 4481 | 225 | P225A | A | 1.4e-05 | 1.3e-05 | 1.5e-05 | 1.400E-05 | 1.000E-06 | 1.300E-05 | 1.500E-05 |
| 4482 | 225 | P225R | A | 9.4e-06 | 1.1e-05 | 1.4e-05 | 1.147E-05 | 2.335E-06 | 9.400E-06 | 1.400E-05 |
| 4483 | 225 | P225N | A | 1.6e-05 | 1.4e-05 | 1.4e-05 | 1.467E-05 | 1.155E-06 | 1.400E-05 | 1.600E-05 |
| 4484 | 225 | P225D | A | 1.5e-05 | 1.5e-05 | 1.5e-05 | 1.500E-05 | 0.000E+00 | 1.500E-05 | 1.500E-05 |
| 4485 | 225 | P225C | A | 1e-05 | 6.7e-06 | 1e-05 | 8.900E-06 | 1.905E-06 | 6.700E-06 | 1.000E-05 |
| 4486 | 225 | P225E | A | 1.1e-05 | 1.3e-05 | 9.3e-06 | 1.110E-05 | 1.852E-06 | 9.300E-06 | 1.300E-05 |
| 4487 | 225 | P225Q | A | 1.5e-05 | 1.3e-05 | 1.5e-05 | 1.433E-05 | 1.155E-06 | 1.300E-05 | 1.500E-05 |
| 4488 | 225 | P225G | A | 1.5e-05 | 1.2e-05 | 8.5e-06 | 1.183E-05 | 3.253E-06 | 8.500E-06 | 1.500E-05 |
| 4489 | 225 | P225H | A | 1e-05 | 1e-05 | 9.4e-06 | 9.800E-06 | 3.464E-07 | 9.400E-06 | 1.000E-05 |
| 4490 | 225 | P225I | A | 1.1e-05 | 1.5e-05 | 1.5e-05 | 1.367E-05 | 2.309E-06 | 1.100E-05 | 1.500E-05 |
| 4491 | 225 | P225L | A | 1e-05 | 8.4e-06 | 1.4e-05 | 1.080E-05 | 2.884E-06 | 8.400E-06 | 1.400E-05 |
| 4492 | 225 | P225K | A | 1e-05 | 1.7e-05 | 1e-05 | 1.233E-05 | 4.041E-06 | 1.000E-05 | 1.700E-05 |
| 4493 | 225 | P225M | A | 1.1e-05 | 5.9e-06 | 1.5e-05 | 1.063E-05 | 4.561E-06 | 5.900E-06 | 1.500E-05 |
| 4494 | 225 | P225F | A | 1.1e-05 | 1e-05 | 1.3e-05 | 1.133E-05 | 1.528E-06 | 1.000E-05 | 1.300E-05 |
| 4495 | 225 | P225P | A | 1.6e-05 | 1.3e-05 | 1.5e-05 | 1.467E-05 | 1.528E-06 | 1.300E-05 | 1.600E-05 |
| 4496 | 225 | P225S | A | 6.7e-06 | 1.3e-05 | 8.6e-06 | 9.433E-06 | 3.232E-06 | 6.700E-06 | 1.300E-05 |
| 4497 | 225 | P225T | A | 1.5e-05 | 1.1e-05 | 9.6e-06 | 1.187E-05 | 2.802E-06 | 9.600E-06 | 1.500E-05 |
| 4498 | 225 | P225W | A | 1.4e-05 | 8.6e-06 | 1.5e-05 | 1.253E-05 | 3.443E-06 | 8.600E-06 | 1.500E-05 |
| 4499 | 225 | P225Y | A | 1.2e-05 | 1.1e-05 | 1.1e-05 | 1.133E-05 | 5.774E-07 | 1.100E-05 | 1.200E-05 |
| 4500 | 225 | P225V | A | 1.2e-05 | 1.1e-05 | 1.6e-05 | 1.300E-05 | 2.646E-06 | 1.100E-05 | 1.600E-05 |
| 4501 | 226 | K226A | A | 9.3e-06 | 1e-05 | 1.1e-05 | 1.010E-05 | 8.544E-07 | 9.300E-06 | 1.100E-05 |
| 4502 | 226 | K226R | A | 1.4e-05 | 9.8e-06 | 1.1e-05 | 1.160E-05 | 2.163E-06 | 9.800E-06 | 1.400E-05 |
| 4503 | 226 | K226N | A | 8.6e-06 | 9.3e-06 | 8.8e-06 | 8.900E-06 | 3.606E-07 | 8.600E-06 | 9.300E-06 |
| 4504 | 226 | K226D | A | 1.1e-05 | 1.3e-05 | 1e-05 | 1.133E-05 | 1.528E-06 | 1.000E-05 | 1.300E-05 |

| | | | | | | | | | | |
|---|---|---|---|---|---|---|---|---|---|---|
| 4505 | 226 | K226C | A | 1.4e-05 | 1.6e-05 | 9.9e-06 | 1.330E-05 | 3.110E-06 | 9.900E-06 | 1.600E-05 |
| 4506 | 226 | K226E | A | 1.1e-05 | 1.3e-05 | 1.1e-05 | 1.167E-05 | 1.155E-06 | 1.100E-05 | 1.300E-05 |
| 4507 | 226 | K226Q | A | 1.5e-05 | 1.5e-05 | 1.6e-05 | 1.533E-05 | 5.774E-07 | 1.500E-05 | 1.600E-05 |
| 4508 | 226 | K226G | A | 1.1e-05 | 1.5e-05 | 1.4e-05 | 1.333E-05 | 2.082E-06 | 1.100E-05 | 1.500E-05 |
| 4509 | 226 | K226H | A | 1.3e-05 | 1.6e-05 | 2.1e-05 | 1.667E-05 | 4.041E-06 | 1.300E-05 | 2.100E-05 |
| 4510 | 226 | K226I | A | 1.1e-05 | 9.9e-06 | 1.4e-05 | 1.163E-05 | 2.122E-06 | 9.900E-06 | 1.400E-05 |
| 4511 | 226 | K226L | A | 1.6e-05 | 1.5e-05 | 9.8e-06 | 1.360E-05 | 3.329E-06 | 9.800E-06 | 1.600E-05 |
| 4512 | 226 | K226K | A | 1.4e-05 | 9.8e-06 | 1.6e-05 | 1.327E-05 | 3.164E-06 | 9.800E-06 | 1.600E-05 |
| 4513 | 226 | K226M | A | 9.9e-06 | 1e-05 | 1.1e-05 | 1.030E-05 | 6.083E-07 | 9.900E-06 | 1.100E-05 |
| 4514 | 226 | K226F | A | 9.7e-06 | 1.7e-05 | 1.5e-05 | 1.390E-05 | 3.772E-06 | 9.700E-06 | 1.700E-05 |
| 4515 | 226 | K226P | A | 1.1e-05 | 1.6e-05 | 9.2e-06 | 1.207E-05 | 3.523E-06 | 9.200E-06 | 1.600E-05 |
| 4516 | 226 | K226S | A | 6.8e-06 | 1.6e-05 | 1.5e-05 | 1.260E-05 | 5.048E-06 | 6.800E-06 | 1.600E-05 |
| 4517 | 226 | K226T | A | 1.4e-05 | 1.5e-05 | 1.5e-05 | 1.467E-05 | 5.774E-07 | 1.400E-05 | 1.500E-05 |
| 4518 | 226 | K226W | A | 1.5e-05 | 9.6e-06 | 1.5e-05 | 1.320E-05 | 3.118E-06 | 9.600E-06 | 1.500E-05 |
| 4519 | 226 | K226Y | A | 8.9e-06 | 1.5e-05 | 1.6e-05 | 1.330E-05 | 3.843E-06 | 8.900E-06 | 1.600E-05 |
| 4520 | 226 | K226V | A | 9.5e-06 | 1e-05 | 1.5e-05 | 1.150E-05 | 3.041E-06 | 9.500E-06 | 1.500E-05 |
| 4521 | 227 | H227A | A | 1.3e-05 | 1.3e-05 | 1.1e-05 | 1.233E-05 | 1.155E-06 | 1.100E-05 | 1.300E-05 |
| 4522 | 227 | H227R | A | 1.4e-05 | 1.1e-05 | 1.7e-05 | 1.400E-05 | 3.000E-06 | 1.100E-05 | 1.700E-05 |
| 4523 | 227 | H227N | A | 1.1e-05 | 9.5e-06 | 1e-05 | 1.017E-05 | 7.638E-07 | 9.500E-06 | 1.100E-05 |
| 4524 | 227 | H227D | A | 1.4e-05 | 1.6e-05 | 1.3e-05 | 1.433E-05 | 1.528E-06 | 1.300E-05 | 1.600E-05 |
| 4525 | 227 | H227C | A | 1.6e-05 | 9.7e-06 | 1.1e-05 | 1.223E-05 | 3.326E-06 | 9.700E-06 | 1.600E-05 |
| 4526 | 227 | H227E | A | 1.2e-05 | 1.7e-05 | 1.4e-05 | 1.433E-05 | 2.517E-06 | 1.200E-05 | 1.700E-05 |
| 4527 | 227 | H227Q | A | 1.3e-05 | 1e-05 | 1.1e-05 | 1.133E-05 | 1.528E-06 | 1.000E-05 | 1.300E-05 |
| 4528 | 227 | H227G | A | 8.9e-06 | 7.7e-06 | 1.1e-05 | 9.200E-06 | 1.670E-06 | 7.700E-06 | 1.100E-05 |
| 4529 | 227 | H227H | A | 9.7e-06 | 1.7e-05 | 9.2e-06 | 1.197E-05 | 4.366E-06 | 9.200E-06 | 1.700E-05 |
| 4530 | 227 | H227I | A | 1.6e-05 | 1.3e-05 | 1.5e-05 | 1.467E-05 | 1.528E-06 | 1.300E-05 | 1.600E-05 |
| 4531 | 227 | H227L | A | 1.4e-05 | 1.6e-05 | 1.5e-05 | 1.500E-05 | 1.000E-06 | 1.400E-05 | 1.600E-05 |
| 4532 | 227 | H227K | A | 1.7e-05 | 1.5e-05 | 1.2e-05 | 1.467E-05 | 2.517E-06 | 1.200E-05 | 1.700E-05 |
| 4533 | 227 | H227M | A | 1e-05 | 1.1e-05 | 9.8e-06 | 1.027E-05 | 6.429E-07 | 9.800E-06 | 1.100E-05 |
| 4534 | 227 | H227F | A | 1.4e-05 | 1.6e-05 | 8.9e-06 | 1.297E-05 | 3.661E-06 | 8.900E-06 | 1.600E-05 |
| 4535 | 227 | H227P | A | 1.6e-05 | 1.2e-05 | 1.5e-05 | 1.433E-05 | 2.082E-06 | 1.200E-05 | 1.600E-05 |
| 4536 | 227 | H227S | A | 1.5e-05 | 1.4e-05 | 8.6e-06 | 1.253E-05 | 3.443E-06 | 8.600E-06 | 1.500E-05 |
| 4537 | 227 | H227T | A | 1.1e-05 | 8.7e-06 | 1.7e-05 | 1.223E-05 | 4.285E-06 | 8.700E-06 | 1.700E-05 |
| 4538 | 227 | H227W | A | 1.2e-05 | 1.4e-05 | 1.6e-05 | 1.400E-05 | 2.000E-06 | 1.200E-05 | 1.600E-05 |
| 4539 | 227 | H227Y | A | 1.4e-05 | 1.2e-05 | 1.8e-05 | 1.467E-05 | 3.055E-06 | 1.200E-05 | 1.800E-05 |
| 4540 | 227 | H227V | A | 1.5e-05 | 1.6e-05 | 1.1e-05 | 1.400E-05 | 2.646E-06 | 1.100E-05 | 1.600E-05 |
| 4541 | 228 | S228A | A | 1e-05 | 1e-05 | 1.7e-05 | 1.233E-05 | 4.041E-06 | 1.000E-05 | 1.700E-05 |
| 4542 | 228 | S228R | A | 1.6e-05 | 1.5e-05 | 1.5e-05 | 1.533E-05 | 5.774E-07 | 1.500E-05 | 1.600E-05 |
| 4543 | 228 | S228N | A | 1.1e-05 | 1.3e-05 | 1.6e-05 | 1.333E-05 | 2.517E-06 | 1.100E-05 | 1.600E-05 |
| 4544 | 228 | S228D | A | 1.1e-05 | 1.1e-05 | 1.1e-05 | 1.100E-05 | 2.075E-21 | 1.100E-05 | 1.100E-05 |
| 4545 | 228 | S228C | A | 8.4e-06 | 1.7e-05 | 1.1e-05 | 1.213E-05 | 4.411E-06 | 8.400E-06 | 1.700E-05 |
| 4546 | 228 | S228E | A | 9.7e-06 | 1.4e-05 | 1.5e-05 | 1.290E-05 | 2.816E-06 | 9.700E-06 | 1.500E-05 |
| 4547 | 228 | S228Q | A | 8.8e-06 | 1.1e-05 | 9.9e-06 | 9.900E-06 | 1.100E-06 | 8.800E-06 | 1.100E-05 |
| 4548 | 228 | S228G | A | 1.7e-05 | 1.1e-05 | 9.4e-06 | 1.247E-05 | 4.007E-06 | 9.400E-06 | 1.700E-05 |
| 4549 | 228 | S228H | A | 9.1e-06 | 1.6e-05 | 1.5e-05 | 1.337E-05 | 3.729E-06 | 9.100E-06 | 1.600E-05 |
| 4550 | 228 | S228I | A | 1e-05 | 1.4e-05 | 1.7e-05 | 1.367E-05 | 3.512E-06 | 1.000E-05 | 1.700E-05 |
| 4551 | 228 | S228L | A | 1.3e-05 | 1.5e-05 | 1.6e-05 | 1.467E-05 | 1.528E-06 | 1.300E-05 | 1.600E-05 |
| 4552 | 228 | S228K | A | 1.5e-05 | 1.1e-05 | 1e-05 | 1.200E-05 | 2.646E-06 | 1.000E-05 | 1.500E-05 |
| 4553 | 228 | S228M | A | 1.6e-05 | 1.3e-05 | 1.5e-05 | 1.467E-05 | 1.528E-06 | 1.300E-05 | 1.600E-05 |
| 4554 | 228 | S228F | A | 1.7e-05 | 1.5e-05 | 1.6e-05 | 1.600E-05 | 1.000E-06 | 1.500E-05 | 1.700E-05 |
| 4555 | 228 | S228P | A | 9.8e-06 | 1.1e-05 | 1.5e-05 | 1.193E-05 | 2.723E-06 | 9.800E-06 | 1.500E-05 |
| 4556 | 228 | S228S | A | 1.7e-05 | 1e-05 | 1.5e-05 | 1.400E-05 | 3.606E-06 | 1.000E-05 | 1.700E-05 |
| 4557 | 228 | S228T | A | 1.5e-05 | 1.6e-05 | 1.2e-05 | 1.433E-05 | 2.082E-06 | 1.200E-05 | 1.600E-05 |
| 4558 | 228 | S228W | A | 1.6e-05 | 1.7e-05 | 1.3e-05 | 1.533E-05 | 2.082E-06 | 1.300E-05 | 1.700E-05 |
| 4559 | 228 | S228Y | A | 1.3e-05 | 7.1e-06 | 9.3e-06 | 9.800E-06 | 2.982E-06 | 7.100E-06 | 1.300E-05 |
| 4560 | 228 | S228V | A | 1.3e-05 | 1.6e-05 | 1.6e-05 | 1.500E-05 | 1.732E-06 | 1.300E-05 | 1.600E-05 |
| 4561 | 229 | D229A | A | 1.6e-05 | 1.2e-05 | 1.7e-05 | 1.500E-05 | 2.646E-06 | 1.200E-05 | 1.700E-05 |
| 4562 | 229 | D229R | A | 1e-05 | 9.4e-06 | 1.1e-05 | 1.013E-05 | 8.083E-07 | 9.400E-06 | 1.100E-05 |
| 4563 | 229 | D229N | A | 9.5e-06 | 1.5e-05 | 1.4e-05 | 1.283E-05 | 2.930E-06 | 9.500E-06 | 1.500E-05 |
| 4564 | 229 | D229D | A | 1.6e-05 | 1.1e-05 | 1.5e-05 | 1.400E-05 | 2.646E-06 | 1.100E-05 | 1.600E-05 |
| 4565 | 229 | D229C | A | 1.1e-05 | 1.5e-05 | 1.5e-05 | 1.367E-05 | 2.309E-06 | 1.100E-05 | 1.500E-05 |
| 4566 | 229 | D229E | A | 1.6e-05 | 1.7e-05 | 1.1e-05 | 1.467E-05 | 3.215E-06 | 1.100E-05 | 1.700E-05 |
| 4567 | 229 | D229Q | A | 1.3e-05 | 1.5e-05 | 1e-05 | 1.267E-05 | 2.517E-06 | 1.000E-05 | 1.500E-05 |
| 4568 | 229 | D229G | A | 6.7e-06 | 1.4e-05 | 1.2e-05 | 1.090E-05 | 3.772E-06 | 6.700E-06 | 1.400E-05 |
| 4569 | 229 | D229H | A | 1.1e-05 | 1.4e-05 | 1.2e-05 | 1.233E-05 | 1.528E-06 | 1.100E-05 | 1.400E-05 |
| 4570 | 229 | D229I | A | 9.7e-06 | 1e-05 | 1.3e-05 | 1.090E-05 | 1.825E-06 | 9.700E-06 | 1.300E-05 |
| 4571 | 229 | D229L | A | 1.6e-05 | 1.1e-05 | 1e-05 | 1.233E-05 | 3.215E-06 | 1.000E-05 | 1.600E-05 |
| 4572 | 229 | D229K | A | 1.1e-05 | 9e-06 | 1.7e-05 | 1.233E-05 | 4.163E-06 | 9.000E-06 | 1.700E-05 |
| 4573 | 229 | D229M | A | 1.5e-05 | 1.7e-05 | 1.5e-05 | 1.567E-05 | 1.155E-06 | 1.500E-05 | 1.700E-05 |
| 4574 | 229 | D229F | A | 9.9e-06 | 1.1e-05 | 1.4e-05 | 1.163E-05 | 2.122E-06 | 9.900E-06 | 1.400E-05 |
| 4575 | 229 | D229P | A | 9.2e-06 | 1.5e-05 | 8e-06 | 1.073E-05 | 3.743E-06 | 8.000E-06 | 1.500E-05 |
| 4576 | 229 | D229S | A | 1.3e-05 | 9e-06 | 1.1e-05 | 1.100E-05 | 2.000E-06 | 9.000E-06 | 1.300E-05 |
| 4577 | 229 | D229T | A | 1e-05 | 9.2e-06 | 1.6e-05 | 1.173E-05 | 3.717E-06 | 9.200E-06 | 1.600E-05 |
| 4578 | 229 | D229W | A | 1.6e-05 | 1.3e-05 | 8.1e-06 | 1.237E-05 | 3.988E-06 | 8.100E-06 | 1.600E-05 |
| 4579 | 229 | D229Y | A | 9.2e-06 | 1.4e-05 | 1.6e-05 | 1.307E-05 | 3.495E-06 | 9.200E-06 | 1.600E-05 |
| 4580 | 229 | D229V | A | 1.1e-05 | 1.6e-05 | 1.1e-05 | 1.267E-05 | 2.887E-06 | 1.100E-05 | 1.600E-05 |
| 4581 | 230 | C230A | A | 1.7e-05 | 1.1e-05 | 1.2e-05 | 1.333E-05 | 3.215E-06 | 1.100E-05 | 1.700E-05 |
| 4582 | 230 | C230R | A | 9.2e-06 | 1.6e-05 | 8.4e-06 | 1.120E-05 | 4.176E-06 | 8.400E-06 | 1.600E-05 |
| 4583 | 230 | C230N | A | 1.7e-05 | 1.5e-05 | 9.2e-06 | 1.373E-05 | 4.051E-06 | 9.200E-06 | 1.700E-05 |
| 4584 | 230 | C230D | A | 1.1e-05 | 1e-05 | 1.1e-05 | 1.067E-05 | 5.774E-07 | 1.000E-05 | 1.100E-05 |
| 4585 | 230 | C230C | A | 7.8e-06 | 1e-05 | 1.6e-05 | 1.127E-05 | 4.244E-06 | 7.800E-06 | 1.600E-05 |
| 4586 | 230 | C230E | A | 9.7e-06 | 1.6e-05 | 8.6e-06 | 1.143E-05 | 3.993E-06 | 8.600E-06 | 1.600E-05 |
| 4587 | 230 | C230Q | A | 8.3e-06 | 1e-05 | 1.7e-05 | 1.177E-05 | 4.611E-06 | 8.300E-06 | 1.700E-05 |
| 4588 | 230 | C230G | A | 1.4e-05 | 1.2e-05 | 1.5e-05 | 1.367E-05 | 1.528E-06 | 1.200E-05 | 1.500E-05 |
| 4589 | 230 | C230H | A | 1.6e-05 | 1.1e-05 | 1.6e-05 | 1.433E-05 | 2.887E-06 | 1.100E-05 | 1.600E-05 |

| | | | | | | | | | | |
|---|---|---|---|---|---|---|---|---|---|---|
| 4590 | 230 | C230I | A | 1.1e-05 | 9.4e-06 | 1.6e-05 | 1.213E-05 | 3.443E-06 | 9.400E-06 | 1.600E-05 |
| 4591 | 230 | C230L | A | 1.2e-05 | 1.6e-05 | 1.7e-05 | 1.500E-05 | 2.646E-06 | 1.200E-05 | 1.700E-05 |
| 4592 | 230 | C230K | A | 8.9e-06 | 8.8e-06 | 1.6e-05 | 1.123E-05 | 4.128E-06 | 8.800E-06 | 1.600E-05 |
| 4593 | 230 | C230M | A | 1.6e-05 | 1.4e-05 | 1.1e-05 | 1.367E-05 | 2.517E-06 | 1.100E-05 | 1.600E-05 |
| 4594 | 230 | C230F | A | 1.6e-05 | 1.3e-05 | 1.6e-05 | 1.500E-05 | 1.732E-06 | 1.300E-05 | 1.600E-05 |
| 4595 | 230 | C230P | A | 9.6e-06 | 1.3e-05 | 1.1e-05 | 1.120E-05 | 1.709E-06 | 9.600E-06 | 1.300E-05 |
| 4596 | 230 | C230S | A | 1e-05 | 1e-05 | 1.1e-05 | 1.033E-05 | 5.774E-07 | 1.000E-05 | 1.100E-05 |
| 4597 | 230 | C230T | A | 1.2e-05 | 1.5e-05 | 9.8e-06 | 1.227E-05 | 2.610E-06 | 9.800E-06 | 1.500E-05 |
| 4598 | 230 | C230W | A | 1.2e-05 | 1.5e-05 | 1.4e-05 | 1.367E-05 | 1.528E-06 | 1.200E-05 | 1.500E-05 |
| 4599 | 230 | C230Y | A | 1.4e-05 | 1.1e-05 | 1e-05 | 1.167E-05 | 2.082E-06 | 1.000E-05 | 1.400E-05 |
| 4600 | 230 | C230V | A | 1.2e-05 | 1.1e-05 | 1.1e-05 | 1.133E-05 | 5.774E-07 | 1.100E-05 | 1.200E-05 |
| 4601 | 231 | L231A | A | 9.8e-06 | 1.3e-05 | 1.3e-05 | 1.193E-05 | 1.848E-06 | 9.800E-06 | 1.300E-05 |
| 4602 | 231 | L231R | A | 1.5e-05 | 9.7e-06 | 1e-05 | 1.157E-05 | 2.977E-06 | 9.700E-06 | 1.500E-05 |
| 4603 | 231 | L231N | A | 1.6e-05 | 1.5e-05 | 1.4e-05 | 1.500E-05 | 1.000E-06 | 1.400E-05 | 1.600E-05 |
| 4604 | 231 | L231D | A | 1.4e-05 | 1.3e-05 | 1e-05 | 1.233E-05 | 2.082E-06 | 1.000E-05 | 1.400E-05 |
| 4605 | 231 | L231C | A | 1.1e-05 | 1.3e-05 | 1.3e-05 | 1.233E-05 | 1.155E-06 | 1.100E-05 | 1.300E-05 |
| 4606 | 231 | L231E | A | 1.6e-05 | 1.1e-05 | 1e-05 | 1.233E-05 | 3.215E-06 | 1.000E-05 | 1.600E-05 |
| 4607 | 231 | L231Q | A | 1.1e-05 | 1e-05 | 1.5e-05 | 1.200E-05 | 2.646E-06 | 1.000E-05 | 1.500E-05 |
| 4608 | 231 | L231G | A | 9.9e-06 | 1.8e-05 | 1.6e-05 | 1.463E-05 | 4.219E-06 | 9.900E-06 | 1.800E-05 |
| 4609 | 231 | L231H | A | 1e-05 | 1.6e-05 | 1.7e-05 | 1.433E-05 | 3.786E-06 | 1.000E-05 | 1.700E-05 |
| 4610 | 231 | L231I | A | 1.4e-05 | 1.5e-05 | 1.6e-05 | 1.500E-05 | 1.000E-06 | 1.400E-05 | 1.600E-05 |
| 4611 | 231 | L231L | A | 8.9e-06 | 1.1e-05 | 1.1e-05 | 1.030E-05 | 1.212E-06 | 8.900E-06 | 1.100E-05 |
| 4612 | 231 | L231K | A | 8.2e-06 | 1.1e-05 | 1.5e-05 | 1.140E-05 | 3.418E-06 | 8.200E-06 | 1.500E-05 |
| 4613 | 231 | L231M | A | 1.1e-05 | 9.1e-06 | 1.5e-05 | 1.170E-05 | 3.012E-06 | 9.100E-06 | 1.500E-05 |
| 4614 | 231 | L231F | A | 1.1e-05 | 1.4e-05 | 1.5e-05 | 1.333E-05 | 2.082E-06 | 1.100E-05 | 1.500E-05 |
| 4615 | 231 | L231P | A | 8.4e-06 | 1.5e-05 | 1.2e-05 | 1.180E-05 | 3.305E-06 | 8.400E-06 | 1.500E-05 |
| 4616 | 231 | L231S | A | 1.4e-05 | 1.4e-05 | 1.5e-05 | 1.433E-05 | 5.774E-07 | 1.400E-05 | 1.500E-05 |
| 4617 | 231 | L231T | A | 1.1e-05 | 1e-05 | 1.1e-05 | 1.067E-05 | 5.774E-07 | 1.000E-05 | 1.100E-05 |
| 4618 | 231 | L231W | A | 1.4e-05 | 8.8e-06 | 1e-05 | 1.093E-05 | 2.723E-06 | 8.800E-06 | 1.400E-05 |
| 4619 | 231 | L231Y | A | 8.3e-06 | 1e-05 | 1.4e-05 | 1.077E-05 | 2.926E-06 | 8.300E-06 | 1.400E-05 |
| 4620 | 231 | L231V | A | 1.5e-05 | 1.3e-05 | 1.6e-05 | 1.467E-05 | 1.528E-06 | 1.300E-05 | 1.600E-05 |
| 4621 | 232 | A232A | A | 1.6e-05 | 1e-05 | 1.5e-05 | 1.367E-05 | 3.215E-06 | 1.000E-05 | 1.600E-05 |
| 4622 | 232 | A232R | A | 1.5e-05 | 1e-05 | 1.1e-05 | 1.200E-05 | 2.646E-06 | 1.000E-05 | 1.500E-05 |
| 4623 | 232 | A232N | A | 1.6e-05 | 1.2e-05 | 9.1e-06 | 1.237E-05 | 3.465E-06 | 9.100E-06 | 1.600E-05 |
| 4624 | 232 | A232D | A | 1.6e-05 | 1.4e-05 | 1.4e-05 | 1.467E-05 | 1.155E-06 | 1.400E-05 | 1.600E-05 |
| 4625 | 232 | A232C | A | 1.6e-05 | 1.5e-05 | 1.2e-05 | 1.433E-05 | 2.082E-06 | 1.200E-05 | 1.600E-05 |
| 4626 | 232 | A232E | A | 1.5e-05 | 1.6e-05 | 1.1e-05 | 1.400E-05 | 2.646E-06 | 1.100E-05 | 1.600E-05 |
| 4627 | 232 | A232Q | A | 1.2e-05 | 9.8e-06 | 1.3e-05 | 1.160E-05 | 1.637E-06 | 9.800E-06 | 1.300E-05 |
| 4628 | 232 | A232G | A | 9.5e-06 | 8.7e-06 | 1.6e-05 | 1.140E-05 | 4.004E-06 | 8.700E-06 | 1.600E-05 |
| 4629 | 232 | A232H | A | 1.1e-05 | 1e-05 | 8.6e-06 | 9.867E-06 | 1.206E-06 | 8.600E-06 | 1.100E-05 |
| 4630 | 232 | A232I | A | 1.5e-05 | 8.4e-06 | 1.6e-05 | 1.313E-05 | 4.130E-06 | 8.400E-06 | 1.600E-05 |
| 4631 | 232 | A232L | A | 9.9e-06 | 1.2e-05 | 9.5e-06 | 1.047E-05 | 1.343E-06 | 9.500E-06 | 1.200E-05 |
| 4632 | 232 | A232K | A | 1.4e-05 | 1.2e-05 | 9.7e-06 | 1.190E-05 | 2.152E-06 | 9.700E-06 | 1.400E-05 |
| 4633 | 232 | A232M | A | 1.1e-05 | 1.7e-05 | 1.8e-05 | 1.533E-05 | 3.786E-06 | 1.100E-05 | 1.800E-05 |
| 4634 | 232 | A232F | A | 1e-05 | 1.5e-05 | 1.1e-05 | 1.200E-05 | 2.646E-06 | 1.000E-05 | 1.500E-05 |
| 4635 | 232 | A232P | A | 1.3e-05 | 1.1e-05 | 1.6e-05 | 1.333E-05 | 2.517E-06 | 1.100E-05 | 1.600E-05 |
| 4636 | 232 | A232S | A | 1.4e-05 | 1.4e-05 | 9.9e-06 | 1.263E-05 | 2.367E-06 | 9.900E-06 | 1.400E-05 |
| 4637 | 232 | A232T | A | 1.3e-05 | 9.9e-06 | 1e-05 | 1.097E-05 | 1.762E-06 | 9.900E-06 | 1.300E-05 |
| 4638 | 232 | A232W | A | 1e-05 | 1.2e-05 | 1.1e-05 | 1.100E-05 | 1.000E-06 | 1.000E-05 | 1.200E-05 |
| 4639 | 232 | A232Y | A | 1.2e-05 | 1.2e-05 | 1.4e-05 | 1.267E-05 | 1.155E-06 | 1.200E-05 | 1.400E-05 |
| 4640 | 232 | A232V | A | 1.1e-05 | 1.1e-05 | 1.1e-05 | 1.100E-05 | 2.075E-21 | 1.100E-05 | 1.100E-05 |
| 4641 | 233 | C233A | A | 1.2e-05 | 1.5e-05 | 8.5e-06 | 1.183E-05 | 3.253E-06 | 8.500E-06 | 1.500E-05 |
| 4642 | 233 | C233R | A | 7.7e-06 | 1.6e-05 | 1.7e-05 | 1.357E-05 | 5.105E-06 | 7.700E-06 | 1.700E-05 |
| 4643 | 233 | C233N | A | 1.5e-05 | 1.4e-05 | 7e-06 | 1.200E-05 | 4.359E-06 | 7.000E-06 | 1.500E-05 |
| 4644 | 233 | C233D | A | 1.4e-05 | 1.4e-05 | 1.5e-05 | 1.433E-05 | 5.774E-07 | 1.400E-05 | 1.500E-05 |
| 4645 | 233 | C233C | A | 1.9e-05 | 1.5e-05 | 1.1e-05 | 1.500E-05 | 4.000E-06 | 1.100E-05 | 1.900E-05 |
| 4646 | 233 | C233E | A | 1.1e-05 | 1.1e-05 | 1.5e-05 | 1.233E-05 | 2.309E-06 | 1.100E-05 | 1.500E-05 |
| 4647 | 233 | C233Q | A | 1e-05 | 1.5e-05 | 1.5e-05 | 1.333E-05 | 2.887E-06 | 1.000E-05 | 1.500E-05 |
| 4648 | 233 | C233G | A | 1.5e-05 | 8.9e-06 | 1.5e-05 | 1.297E-05 | 3.522E-06 | 8.900E-06 | 1.500E-05 |
| 4649 | 233 | C233H | A | 1.1e-05 | 1.3e-05 | 1e-05 | 1.133E-05 | 1.528E-06 | 1.000E-05 | 1.300E-05 |
| 4650 | 233 | C233I | A | 1.4e-05 | 1.5e-05 | 1.4e-05 | 1.433E-05 | 5.774E-07 | 1.400E-05 | 1.500E-05 |
| 4651 | 233 | C233L | A | 9e-06 | 1.8e-05 | 1e-05 | 1.233E-05 | 4.933E-06 | 9.000E-06 | 1.800E-05 |
| 4652 | 233 | C233K | A | 1.1e-05 | 1.4e-05 | 1.7e-05 | 1.400E-05 | 3.000E-06 | 1.100E-05 | 1.700E-05 |
| 4653 | 233 | C233M | A | 1.2e-05 | 1.4e-05 | 9.4e-06 | 1.180E-05 | 2.307E-06 | 9.400E-06 | 1.400E-05 |
| 4654 | 233 | C233F | A | 2.5e-05 | 1.6e-05 | 1.6e-05 | 1.900E-05 | 5.196E-06 | 1.600E-05 | 2.500E-05 |
| 4655 | 233 | C233P | A | 1e-05 | 1.8e-05 | 1.6e-05 | 1.467E-05 | 4.163E-06 | 1.000E-05 | 1.800E-05 |
| 4656 | 233 | C233S | A | 1e-05 | 1.1e-05 | 1.5e-05 | 1.200E-05 | 2.646E-06 | 1.000E-05 | 1.500E-05 |
| 4657 | 233 | C233T | A | 1.6e-05 | 1.5e-05 | 1.8e-05 | 1.633E-05 | 1.528E-06 | 1.500E-05 | 1.800E-05 |
| 4658 | 233 | C233W | A | 1.1e-05 | 1.1e-05 | 1.4e-05 | 1.200E-05 | 1.732E-06 | 1.100E-05 | 1.400E-05 |
| 4659 | 233 | C233Y | A | 8.3e-06 | 1.5e-05 | 1.3e-05 | 1.210E-05 | 3.439E-06 | 8.300E-06 | 1.500E-05 |
| 4660 | 233 | C233V | A | 1.1e-05 | 8.7e-06 | 1.5e-05 | 1.157E-05 | 3.188E-06 | 8.700E-06 | 1.500E-05 |
| 4661 | 234 | L234A | A | 1.5e-05 | 1.4e-05 | 1.5e-05 | 1.467E-05 | 5.774E-07 | 1.400E-05 | 1.500E-05 |
| 4662 | 234 | L234R | A | 1.5e-05 | 1.5e-05 | 1e-05 | 1.333E-05 | 2.887E-06 | 1.000E-05 | 1.500E-05 |
| 4663 | 234 | L234N | A | 1.1e-05 | 1.7e-05 | 1.3e-05 | 1.367E-05 | 3.055E-06 | 1.100E-05 | 1.700E-05 |
| 4664 | 234 | L234D | A | 1.4e-05 | 1.4e-05 | 1.4e-05 | 1.400E-05 | 0.000E+00 | 1.400E-05 | 1.400E-05 |
| 4665 | 234 | L234C | A | 1.7e-05 | 1.7e-05 | 8.7e-06 | 1.423E-05 | 4.792E-06 | 8.700E-06 | 1.700E-05 |
| 4666 | 234 | L234E | A | 1.4e-05 | 1.6e-05 | 1.1e-05 | 1.367E-05 | 2.517E-06 | 1.100E-05 | 1.600E-05 |
| 4667 | 234 | L234Q | A | 1.3e-05 | 1.1e-05 | 1.3e-05 | 1.233E-05 | 1.155E-06 | 1.100E-05 | 1.300E-05 |
| 4668 | 234 | L234G | A | 8.6e-06 | 1.1e-05 | 9.1e-06 | 9.567E-06 | 1.266E-06 | 8.600E-06 | 1.100E-05 |
| 4669 | 234 | L234H | A | 1.3e-05 | 1.6e-05 | 2.8e-05 | 1.900E-05 | 7.937E-06 | 1.300E-05 | 2.800E-05 |
| 4670 | 234 | L234I | A | 9.8e-06 | 8.4e-06 | 1.4e-05 | 1.073E-05 | 2.914E-06 | 8.400E-06 | 1.400E-05 |
| 4671 | 234 | L234L | A | 1.7e-05 | 6.8e-06 | 3.8e-05 | 2.060E-05 | 1.591E-05 | 6.800E-06 | 3.800E-05 |
| 4672 | 234 | L234K | A | 1.5e-05 | 1.4e-05 | 1.3e-05 | 1.400E-05 | 1.000E-06 | 1.300E-05 | 1.500E-05 |
| 4673 | 234 | L234M | A | 1.2e-05 | 1.3e-05 | 6.5e-06 | 1.050E-05 | 3.500E-06 | 6.500E-06 | 1.300E-05 |
| 4674 | 234 | L234F | A | 1.6e-05 | 1e-05 | 1.6e-05 | 1.400E-05 | 3.464E-06 | 1.000E-05 | 1.600E-05 |

| | | | | | | | | | | |
|---|---|---|---|---|---|---|---|---|---|---|
| 4675 | 234 | L234P | A | 1.1e-05 | 1.6e-05 | 1.4e-05 | 1.367E-05 | 2.517E-06 | 1.100E-05 | 1.600E-05 |
| 4676 | 234 | L234S | A | 9.7e-06 | 1e-05 | 9.3e-06 | 9.667E-06 | 3.512E-07 | 9.300E-06 | 1.000E-05 |
| 4677 | 234 | L234T | A | 1.1e-05 | 1e-05 | 1.3e-05 | 1.133E-05 | 1.528E-06 | 1.000E-05 | 1.300E-05 |
| 4678 | 234 | L234W | A | 1.4e-05 | 9.4e-06 | 1.4e-05 | 1.247E-05 | 2.656E-06 | 9.400E-06 | 1.400E-05 |
| 4679 | 234 | L234Y | A | 1.1e-05 | 1.5e-05 | 1.1e-05 | 1.233E-05 | 2.309E-06 | 1.100E-05 | 1.500E-05 |
| 4680 | 234 | L234V | A | 1.1e-05 | 1.7e-05 | 1e-05 | 1.267E-05 | 3.786E-06 | 1.000E-05 | 1.700E-05 |
| 4681 | 235 | H235A | A | 1.1e-05 | 1.5e-05 | 9.8e-06 | 1.193E-05 | 2.723E-06 | 9.800E-06 | 1.500E-05 |
| 4682 | 235 | H235R | A | 1e-05 | 9.5e-06 | 1.4e-05 | 1.117E-05 | 2.466E-06 | 9.500E-06 | 1.400E-05 |
| 4683 | 235 | H235N | A | 9.9e-06 | 1.6e-05 | 1.6e-05 | 1.397E-05 | 3.522E-06 | 9.900E-06 | 1.600E-05 |
| 4684 | 235 | H235D | A | 6.1e-06 | 1.5e-05 | 1.1e-05 | 1.070E-05 | 4.458E-06 | 6.100E-06 | 1.500E-05 |
| 4685 | 235 | H235C | A | 1.1e-05 | 8.5e-06 | 1.4e-05 | 1.117E-05 | 2.754E-06 | 8.500E-06 | 1.400E-05 |
| 4686 | 235 | H235E | A | 1.5e-05 | 1.8e-05 | 1.5e-05 | 1.600E-05 | 1.732E-06 | 1.500E-05 | 1.800E-05 |
| 4687 | 235 | H235Q | A | 7.6e-06 | 1.5e-05 | 1.4e-05 | 1.220E-05 | 4.015E-06 | 7.600E-06 | 1.500E-05 |
| 4688 | 235 | H235G | A | 9.7e-06 | 1.5e-05 | 1.3e-05 | 1.257E-05 | 2.676E-06 | 9.700E-06 | 1.500E-05 |
| 4689 | 235 | H235H | A | 1.6e-05 | 1e-05 | 1.1e-05 | 1.233E-05 | 3.215E-06 | 1.000E-05 | 1.600E-05 |
| 4690 | 235 | H235I | A | 9.2e-06 | 1.5e-05 | 1e-05 | 1.140E-05 | 3.143E-06 | 9.200E-06 | 1.500E-05 |
| 4691 | 235 | H235L | A | 1.5e-05 | 1.1e-05 | 1e-05 | 1.200E-05 | 2.646E-06 | 1.000E-05 | 1.500E-05 |
| 4692 | 235 | H235K | A | 1.6e-05 | 1e-05 | 9.6e-06 | 1.187E-05 | 3.585E-06 | 9.600E-06 | 1.600E-05 |
| 4693 | 235 | H235M | A | 1.5e-05 | 9.8e-06 | 1e-05 | 1.160E-05 | 2.946E-06 | 9.800E-06 | 1.500E-05 |
| 4694 | 235 | H235F | A | 9.1e-06 | 1.5e-05 | 8.8e-06 | 1.097E-05 | 3.496E-06 | 8.800E-06 | 1.500E-05 |
| 4695 | 235 | H235P | A | 1.1e-05 | 1.3e-05 | 1.3e-05 | 1.233E-05 | 1.155E-06 | 1.100E-05 | 1.300E-05 |
| 4696 | 235 | H235S | A | 1.5e-05 | 1.1e-05 | 1.5e-05 | 1.367E-05 | 2.309E-06 | 1.100E-05 | 1.500E-05 |
| 4697 | 235 | H235T | A | 8.5e-06 | 2.3e-05 | 1.4e-05 | 1.517E-05 | 7.320E-06 | 8.500E-06 | 2.300E-05 |
| 4698 | 235 | H235W | A | 1.1e-05 | 1.1e-05 | 1.2e-05 | 1.133E-05 | 5.774E-07 | 1.100E-05 | 1.200E-05 |
| 4699 | 235 | H235Y | A | 8.6e-06 | 3.1e-05 | 1.4e-05 | 1.787E-05 | 1.169E-05 | 8.600E-06 | 3.100E-05 |
| 4700 | 235 | H235V | A | 1.5e-05 | 1e-05 | 1.1e-05 | 1.200E-05 | 2.646E-06 | 1.000E-05 | 1.500E-05 |
| 4701 | 236 | F236A | A | 9.9e-06 | 1.1e-05 | 1e-05 | 1.030E-05 | 6.083E-07 | 9.900E-06 | 1.100E-05 |
| 4702 | 236 | F236R | A | 1.5e-05 | 9.6e-06 | 1.2e-05 | 1.220E-05 | 2.706E-06 | 9.600E-06 | 1.500E-05 |
| 4703 | 236 | F236N | A | 1.2e-05 | 1.2e-05 | 1.6e-05 | 1.333E-05 | 2.309E-06 | 1.200E-05 | 1.600E-05 |
| 4704 | 236 | F236D | A | 1.4e-05 | 1.5e-05 | 1e-05 | 1.300E-05 | 2.646E-06 | 1.000E-05 | 1.500E-05 |
| 4705 | 236 | F236C | A | 9.9e-06 | 1.4e-05 | 1.3e-05 | 1.230E-05 | 2.138E-06 | 9.900E-06 | 1.400E-05 |
| 4706 | 236 | F236E | A | 1e-05 | 1.1e-05 | 9.9e-06 | 1.030E-05 | 6.083E-07 | 9.900E-06 | 1.100E-05 |
| 4707 | 236 | F236Q | A | 1.5e-05 | 1.3e-05 | 1.1e-05 | 1.300E-05 | 2.000E-06 | 1.100E-05 | 1.500E-05 |
| 4708 | 236 | F236G | A | 9.6e-06 | 9.9e-06 | 9.3e-06 | 9.600E-06 | 3.000E-07 | 9.300E-06 | 9.900E-06 |
| 4709 | 236 | F236H | A | 1.5e-05 | 1e-05 | 1.2e-05 | 1.233E-05 | 2.517E-06 | 1.000E-05 | 1.500E-05 |
| 4710 | 236 | F236I | A | 1e-05 | 1.6e-05 | 1.3e-05 | 1.300E-05 | 3.000E-06 | 1.000E-05 | 1.600E-05 |
| 4711 | 236 | F236L | A | 1e-05 | 1.5e-05 | 1.5e-05 | 1.333E-05 | 2.887E-06 | 1.000E-05 | 1.500E-05 |
| 4712 | 236 | F236K | A | 1.5e-05 | 9.3e-06 | 1.5e-05 | 1.310E-05 | 3.291E-06 | 9.300E-06 | 1.500E-05 |
| 4713 | 236 | F236M | A | 1.7e-05 | 1.1e-05 | 1.5e-05 | 1.433E-05 | 3.055E-06 | 1.100E-05 | 1.700E-05 |
| 4714 | 236 | F236F | A | 9.5e-06 | 1.6e-05 | 8.8e-06 | 1.143E-05 | 3.970E-06 | 8.800E-06 | 1.600E-05 |
| 4715 | 236 | F236P | A | 9.8e-06 | 1.7e-05 | 1.5e-05 | 1.393E-05 | 3.717E-06 | 9.800E-06 | 1.700E-05 |
| 4716 | 236 | F236S | A | 1.1e-05 | 9.4e-06 | 1.4e-05 | 1.147E-05 | 2.335E-06 | 9.400E-06 | 1.400E-05 |
| 4717 | 236 | F236T | A | 1.3e-05 | 9.7e-06 | 1.3e-05 | 1.190E-05 | 1.905E-06 | 9.700E-06 | 1.300E-05 |
| 4718 | 236 | F236W | A | 1.4e-05 | 1.5e-05 | 1.2e-05 | 1.367E-05 | 1.528E-06 | 1.200E-05 | 1.500E-05 |
| 4719 | 236 | F236Y | A | 1e-05 | 1.5e-05 | 1.5e-05 | 1.333E-05 | 2.887E-06 | 1.000E-05 | 1.500E-05 |
| 4720 | 236 | F236V | A | 9.2e-06 | 1.1e-05 | 1.4e-05 | 1.140E-05 | 2.425E-06 | 9.200E-06 | 1.400E-05 |
| 4721 | 237 | N237A | A | 1.6e-05 | 1.6e-05 | 1e-05 | 1.400E-05 | 3.464E-06 | 1.000E-05 | 1.600E-05 |
| 4722 | 237 | N237R | A | 9.2e-06 | 1.5e-05 | 1.6e-05 | 1.340E-05 | 3.672E-06 | 9.200E-06 | 1.600E-05 |
| 4723 | 237 | N237N | A | 1.5e-05 | 1.4e-05 | 1.3e-05 | 1.400E-05 | 1.000E-06 | 1.300E-05 | 1.500E-05 |
| 4724 | 237 | N237D | A | 1.6e-05 | 1.8e-05 | 1.2e-05 | 1.533E-05 | 3.055E-06 | 1.200E-05 | 1.800E-05 |
| 4725 | 237 | N237C | A | 1.1e-05 | 1.6e-05 | 1.3e-05 | 1.333E-05 | 2.517E-06 | 1.100E-05 | 1.600E-05 |
| 4726 | 237 | N237E | A | 1.5e-05 | 1.6e-05 | 1.6e-05 | 1.567E-05 | 5.774E-07 | 1.500E-05 | 1.600E-05 |
| 4727 | 237 | N237Q | A | 1.1e-05 | 1.4e-05 | 8.9e-06 | 1.130E-05 | 2.563E-06 | 8.900E-06 | 1.400E-05 |
| 4728 | 237 | N237G | A | 1.2e-05 | 1.5e-05 | 1.1e-05 | 1.267E-05 | 2.082E-06 | 1.100E-05 | 1.500E-05 |
| 4729 | 237 | N237H | A | 1.3e-05 | 8.2e-06 | 9.8e-06 | 1.033E-05 | 2.444E-06 | 8.200E-06 | 1.300E-05 |
| 4730 | 237 | N237I | A | 9.2e-06 | 1.3e-05 | 1.5e-05 | 1.240E-05 | 2.946E-06 | 9.200E-06 | 1.500E-05 |
| 4731 | 237 | N237L | A | 1.4e-05 | 1.8e-05 | 8.9e-06 | 1.363E-05 | 4.561E-06 | 8.900E-06 | 1.800E-05 |
| 4732 | 237 | N237K | A | 1.1e-05 | 1.6e-05 | 1.4e-05 | 1.367E-05 | 2.517E-06 | 1.100E-05 | 1.600E-05 |
| 4733 | 237 | N237M | A | 5.8e-06 | 1.6e-05 | 1.6e-05 | 1.260E-05 | 5.889E-06 | 5.800E-06 | 1.600E-05 |
| 4734 | 237 | N237F | A | 1.3e-05 | 1.3e-05 | 1.6e-05 | 1.400E-05 | 1.732E-06 | 1.300E-05 | 1.600E-05 |
| 4735 | 237 | N237P | A | 1.8e-05 | 1.1e-05 | 1.5e-05 | 1.467E-05 | 3.512E-06 | 1.100E-05 | 1.800E-05 |
| 4736 | 237 | N237S | A | 1.1e-05 | 1.7e-05 | 1.6e-05 | 1.467E-05 | 3.215E-06 | 1.100E-05 | 1.700E-05 |
| 4737 | 237 | N237T | A | 8.8e-06 | 1.1e-05 | 1.6e-05 | 1.193E-05 | 3.690E-06 | 8.800E-06 | 1.600E-05 |
| 4738 | 237 | N237W | A | 8.1e-06 | 1.2e-05 | 9.4e-06 | 9.833E-06 | 1.986E-06 | 8.100E-06 | 1.200E-05 |
| 4739 | 237 | N237Y | A | 1e-05 | 1.5e-05 | 1.5e-05 | 1.333E-05 | 2.887E-06 | 1.000E-05 | 1.500E-05 |
| 4740 | 237 | N237V | A | 1.2e-05 | 1e-05 | 1.1e-05 | 1.100E-05 | 1.000E-06 | 1.000E-05 | 1.200E-05 |
| 4741 | 238 | H238A | A | 1.5e-05 | 1e-05 | 1.5e-05 | 1.333E-05 | 2.887E-06 | 1.000E-05 | 1.500E-05 |
| 4742 | 238 | H238R | A | 1.5e-05 | 1e-05 | 1.4e-05 | 1.300E-05 | 2.646E-06 | 1.000E-05 | 1.500E-05 |
| 4743 | 238 | H238N | A | 9.8e-06 | 1.5e-05 | 1.4e-05 | 1.293E-05 | 2.759E-06 | 9.800E-06 | 1.500E-05 |
| 4744 | 238 | H238D | A | 9.2e-06 | 1.6e-05 | 1.1e-05 | 1.207E-05 | 3.523E-06 | 9.200E-06 | 1.600E-05 |
| 4745 | 238 | H238C | A | 1.5e-05 | 9.7e-06 | 1.7e-05 | 1.390E-05 | 3.772E-06 | 9.700E-06 | 1.700E-05 |
| 4746 | 238 | H238E | A | 1.5e-05 | 6.8e-06 | 1.2e-05 | 1.127E-05 | 4.149E-06 | 6.800E-06 | 1.500E-05 |
| 4747 | 238 | H238Q | A | 9.3e-06 | 1.5e-05 | 1.1e-05 | 1.177E-05 | 2.926E-06 | 9.300E-06 | 1.500E-05 |
| 4748 | 238 | H238G | A | 1.7e-05 | 1.4e-05 | 1.2e-05 | 1.433E-05 | 2.517E-06 | 1.200E-05 | 1.700E-05 |
| 4749 | 238 | H238H | A | 1.4e-05 | 1.2e-05 | 1.2e-05 | 1.267E-05 | 1.155E-06 | 1.200E-05 | 1.400E-05 |
| 4750 | 238 | H238I | A | 1.6e-05 | 1.4e-05 | 1.7e-05 | 1.567E-05 | 1.528E-06 | 1.400E-05 | 1.700E-05 |
| 4751 | 238 | H238L | A | 1.4e-05 | 9.9e-06 | 9.2e-06 | 1.103E-05 | 2.593E-06 | 9.200E-06 | 1.400E-05 |
| 4752 | 238 | H238K | A | 1.5e-05 | 1.5e-05 | 1e-05 | 1.333E-05 | 2.887E-06 | 1.000E-05 | 1.500E-05 |
| 4753 | 238 | H238M | A | 1.2e-05 | 1.5e-05 | 1.2e-05 | 1.300E-05 | 1.732E-06 | 1.200E-05 | 1.500E-05 |
| 4754 | 238 | H238F | A | 1.5e-05 | 1.8e-05 | 1.8e-05 | 1.700E-05 | 1.732E-06 | 1.500E-05 | 1.800E-05 |
| 4755 | 238 | H238P | A | 1.4e-05 | 1.5e-05 | 9.3e-06 | 1.277E-05 | 3.044E-06 | 9.300E-06 | 1.500E-05 |
| 4756 | 238 | H238S | A | 1.5e-05 | 1.5e-05 | 1.3e-05 | 1.433E-05 | 1.155E-06 | 1.300E-05 | 1.500E-05 |
| 4757 | 238 | H238T | A | 1.1e-05 | 1.4e-05 | 1.1e-05 | 1.200E-05 | 1.732E-06 | 1.100E-05 | 1.400E-05 |
| 4758 | 238 | H238W | A | 1.5e-05 | 1e-05 | 1.7e-05 | 1.400E-05 | 3.606E-06 | 1.000E-05 | 1.700E-05 |
| 4759 | 238 | H238Y | A | 1.3e-05 | 1.1e-05 | 1.6e-05 | 1.333E-05 | 2.517E-06 | 1.100E-05 | 1.600E-05 |

| | | | | | | | | | | |
|---|---|---|---|---|---|---|---|---|---|---|
| 4760 | 238 | H238V | A | 1.6e-05 | 1.9e-05 | 1.4e-05 | 1.633E-05 | 2.517E-06 | 1.400E-05 | 1.900E-05 |
| 4761 | 239 | S239A | A | 1.1e-05 | 1.2e-05 | 1.7e-05 | 1.333E-05 | 3.215E-06 | 1.100E-05 | 1.700E-05 |
| 4762 | 239 | S239R | A | 1.7e-05 | 1.1e-05 | 1.8e-05 | 1.533E-05 | 3.786E-06 | 1.100E-05 | 1.800E-05 |
| 4763 | 239 | S239N | A | 1.5e-05 | 1.1e-05 | 1.4e-05 | 1.333E-05 | 2.082E-06 | 1.100E-05 | 1.500E-05 |
| 4764 | 239 | S239D | A | 1.6e-05 | 1.4e-05 | 1.2e-05 | 1.400E-05 | 2.000E-06 | 1.200E-05 | 1.600E-05 |
| 4765 | 239 | S239C | A | 1.4e-05 | 1.5e-05 | 1.2e-05 | 1.367E-05 | 1.528E-06 | 1.200E-05 | 1.500E-05 |
| 4766 | 239 | S239E | A | 1.1e-05 | 1.3e-05 | 1.6e-05 | 1.333E-05 | 2.517E-06 | 1.100E-05 | 1.600E-05 |
| 4767 | 239 | S239Q | A | 1.6e-05 | 1.2e-05 | 6.8e-06 | 1.160E-05 | 4.613E-06 | 6.800E-06 | 1.600E-05 |
| 4768 | 239 | S239G | A | 1.2e-05 | 7.4e-06 | 1.5e-05 | 1.147E-05 | 3.828E-06 | 7.400E-06 | 1.500E-05 |
| 4769 | 239 | S239H | A | 1.5e-05 | 1e-05 | 1.3e-05 | 1.267E-05 | 2.517E-06 | 1.000E-05 | 1.500E-05 |
| 4770 | 239 | S239I | A | 1.6e-05 | 1.3e-05 | 1.1e-05 | 1.333E-05 | 2.517E-06 | 1.100E-05 | 1.600E-05 |
| 4771 | 239 | S239L | A | 9.8e-06 | 9.1e-06 | 8.9e-06 | 9.267E-06 | 4.726E-07 | 8.900E-06 | 9.800E-06 |
| 4772 | 239 | S239K | A | 1.3e-05 | 1.5e-05 | 1.4e-05 | 1.400E-05 | 1.000E-06 | 1.300E-05 | 1.500E-05 |
| 4773 | 239 | S239M | A | 9.9e-06 | 1.1e-05 | 1.6e-05 | 1.230E-05 | 3.251E-06 | 9.900E-06 | 1.600E-05 |
| 4774 | 239 | S239F | A | 6.4e-06 | 8.4e-06 | 1e-05 | 8.267E-06 | 1.804E-06 | 6.400E-06 | 1.000E-05 |
| 4775 | 239 | S239P | A | 1.1e-05 | 1.6e-05 | 1.3e-05 | 1.333E-05 | 2.517E-06 | 1.100E-05 | 1.600E-05 |
| 4776 | 239 | S239S | A | 1.7e-05 | 1.2e-05 | 1.6e-05 | 1.500E-05 | 2.646E-06 | 1.200E-05 | 1.700E-05 |
| 4777 | 239 | S239T | A | 1e-05 | 1.1e-05 | 1.7e-05 | 1.267E-05 | 3.786E-06 | 1.000E-05 | 1.700E-05 |
| 4778 | 239 | S239W | A | 1.2e-05 | 1.6e-05 | 1.7e-05 | 1.500E-05 | 2.646E-06 | 1.200E-05 | 1.700E-05 |
| 4779 | 239 | S239Y | A | 1.6e-05 | 1.5e-05 | 1.1e-05 | 1.400E-05 | 2.646E-06 | 1.100E-05 | 1.600E-05 |
| 4780 | 239 | S239V | A | 1.1e-05 | 1.6e-05 | 1.3e-05 | 1.333E-05 | 2.517E-06 | 1.100E-05 | 1.600E-05 |
| 4781 | 240 | G240A | A | 1.3e-05 | 1.6e-05 | 1.3e-05 | 1.400E-05 | 1.732E-06 | 1.300E-05 | 1.600E-05 |
| 4782 | 240 | G240R | A | 1.6e-05 | 1e-05 | 1.3e-05 | 1.300E-05 | 3.000E-06 | 1.000E-05 | 1.600E-05 |
| 4783 | 240 | G240N | A | 1.2e-05 | 7.9e-06 | 8.9e-06 | 9.600E-06 | 2.138E-06 | 7.900E-06 | 1.200E-05 |
| 4784 | 240 | G240D | A | 1.5e-05 | 1.4e-05 | 1.6e-05 | 1.500E-05 | 1.000E-06 | 1.400E-05 | 1.600E-05 |
| 4785 | 240 | G240C | A | 9.6e-06 | 1.3e-05 | 1.2e-05 | 1.153E-05 | 1.747E-06 | 9.600E-06 | 1.300E-05 |
| 4786 | 240 | G240E | A | 1e-05 | 1.6e-05 | 1.3e-05 | 1.300E-05 | 3.000E-06 | 1.000E-05 | 1.600E-05 |
| 4787 | 240 | G240Q | A | 1.4e-05 | 1.2e-05 | 1.2e-05 | 1.267E-05 | 1.155E-06 | 1.200E-05 | 1.400E-05 |
| 4788 | 240 | G240G | A | 1.5e-05 | 1.5e-05 | 1.4e-05 | 1.467E-05 | 5.774E-07 | 1.400E-05 | 1.500E-05 |
| 4789 | 240 | G240H | A | 2.1e-05 | 1.4e-05 | 1.5e-05 | 1.667E-05 | 3.786E-06 | 1.400E-05 | 2.100E-05 |
| 4790 | 240 | G240I | A | 1.6e-05 | 8.1e-06 | 1.1e-05 | 1.170E-05 | 3.996E-06 | 8.100E-06 | 1.600E-05 |
| 4791 | 240 | G240L | A | 1.4e-05 | 1.7e-05 | 1.1e-05 | 1.400E-05 | 3.000E-06 | 1.100E-05 | 1.700E-05 |
| 4792 | 240 | G240K | A | 1.5e-05 | 1e-05 | 1.1e-05 | 1.200E-05 | 2.646E-06 | 1.000E-05 | 1.500E-05 |
| 4793 | 240 | G240M | A | 1.6e-05 | 1e-05 | 1.5e-05 | 1.367E-05 | 3.215E-06 | 1.000E-05 | 1.600E-05 |
| 4794 | 240 | G240F | A | 1.4e-05 | 9.1e-06 | 1.3e-05 | 1.203E-05 | 2.589E-06 | 9.100E-06 | 1.400E-05 |
| 4795 | 240 | G240P | A | 1.6e-05 | 8.8e-06 | 6.6e-06 | 1.047E-05 | 4.917E-06 | 6.600E-06 | 1.600E-05 |
| 4796 | 240 | G240S | A | 8.5e-06 | 1e-05 | 1.3e-05 | 1.050E-05 | 2.291E-06 | 8.500E-06 | 1.300E-05 |
| 4797 | 240 | G240T | A | 1.4e-05 | 1.5e-05 | 1.6e-05 | 1.500E-05 | 1.000E-06 | 1.400E-05 | 1.600E-05 |
| 4798 | 240 | G240W | A | 9.2e-06 | 1.5e-05 | 1.1e-05 | 1.173E-05 | 2.969E-06 | 9.200E-06 | 1.500E-05 |
| 4799 | 240 | G240Y | A | 1.5e-05 | 8.6e-06 | 2.4e-05 | 1.587E-05 | 7.736E-06 | 8.600E-06 | 2.400E-05 |
| 4800 | 240 | G240V | A | 1.1e-05 | 1.1e-05 | 9.8e-06 | 1.060E-05 | 6.928E-07 | 9.800E-06 | 1.100E-05 |
| 4801 | 241 | I241A | A | 1.1e-05 | 9.2e-06 | 1.1e-05 | 1.040E-05 | 1.039E-06 | 9.200E-06 | 1.100E-05 |
| 4802 | 241 | I241R | A | 8e-06 | 9.5e-06 | 1.5e-05 | 1.083E-05 | 3.686E-06 | 8.000E-06 | 1.500E-05 |
| 4803 | 241 | I241N | A | 1e-05 | 1.5e-05 | 1.4e-05 | 1.300E-05 | 2.646E-06 | 1.000E-05 | 1.500E-05 |
| 4804 | 241 | I241D | A | 8.9e-06 | 1.5e-05 | 1.1e-05 | 1.163E-05 | 3.099E-06 | 8.900E-06 | 1.500E-05 |
| 4805 | 241 | I241C | A | 1.7e-05 | 9.7e-06 | 1.7e-05 | 1.457E-05 | 4.215E-06 | 9.700E-06 | 1.700E-05 |
| 4806 | 241 | I241E | A | 1.6e-05 | 1.5e-05 | 9.1e-06 | 1.337E-05 | 3.729E-06 | 9.100E-06 | 1.600E-05 |
| 4807 | 241 | I241Q | A | 1e-05 | 1.6e-05 | 1e-05 | 1.200E-05 | 3.464E-06 | 1.000E-05 | 1.600E-05 |
| 4808 | 241 | I241G | A | 1.5e-05 | 1.2e-05 | 1.1e-05 | 1.267E-05 | 2.082E-06 | 1.100E-05 | 1.500E-05 |
| 4809 | 241 | I241H | A | 1.4e-05 | 1.5e-05 | 1e-05 | 1.300E-05 | 2.646E-06 | 1.000E-05 | 1.500E-05 |
| 4810 | 241 | I241I | A | 1.2e-05 | 1.4e-05 | 1.6e-05 | 1.400E-05 | 2.000E-06 | 1.200E-05 | 1.600E-05 |
| 4811 | 241 | I241L | A | 9.8e-06 | 7.7e-06 | 8.8e-06 | 8.767E-06 | 1.050E-06 | 7.700E-06 | 9.800E-06 |
| 4812 | 241 | I241K | A | 1.6e-05 | 1.3e-05 | 1.5e-05 | 1.467E-05 | 1.528E-06 | 1.300E-05 | 1.600E-05 |
| 4813 | 241 | I241M | A | 9.4e-06 | 1.7e-05 | 9.2e-06 | 1.187E-05 | 4.447E-06 | 9.200E-06 | 1.700E-05 |
| 4814 | 241 | I241F | A | 1.4e-05 | 1.6e-05 | 1.5e-05 | 1.500E-05 | 1.000E-06 | 1.400E-05 | 1.600E-05 |
| 4815 | 241 | I241P | A | 1.1e-05 | 1.6e-05 | 1.5e-05 | 1.400E-05 | 2.646E-06 | 1.100E-05 | 1.600E-05 |
| 4816 | 241 | I241S | A | 1.6e-05 | 1.5e-05 | 1.4e-05 | 1.500E-05 | 1.000E-06 | 1.400E-05 | 1.600E-05 |
| 4817 | 241 | I241T | A | 1.2e-05 | 1.2e-05 | 1.4e-05 | 1.267E-05 | 1.155E-06 | 1.200E-05 | 1.400E-05 |
| 4818 | 241 | I241W | A | 1.3e-05 | 9.6e-06 | 6.7e-06 | 9.767E-06 | 3.153E-06 | 6.700E-06 | 1.300E-05 |
| 4819 | 241 | I241Y | A | 1.7e-05 | 1.6e-05 | 1e-05 | 1.433E-05 | 3.786E-06 | 1.000E-05 | 1.700E-05 |
| 4820 | 241 | I241V | A | 1.6e-05 | 1.4e-05 | 1.5e-05 | 1.500E-05 | 1.000E-06 | 1.400E-05 | 1.600E-05 |
| 4821 | 242 | C242A | A | 1.1e-05 | 1.5e-05 | 1.1e-05 | 1.233E-05 | 2.309E-06 | 1.100E-05 | 1.500E-05 |
| 4822 | 242 | C242R | A | 1e-05 | 1.1e-05 | 1e-05 | 1.033E-05 | 5.774E-07 | 1.000E-05 | 1.100E-05 |
| 4823 | 242 | C242N | A | 1.3e-05 | 1.6e-05 | 9.8e-06 | 1.293E-05 | 3.101E-06 | 9.800E-06 | 1.600E-05 |
| 4824 | 242 | C242D | A | 1.4e-05 | 1.1e-05 | 1.5e-05 | 1.333E-05 | 2.082E-06 | 1.100E-05 | 1.500E-05 |
| 4825 | 242 | C242C | A | 1.1e-05 | 1.2e-05 | 1.6e-05 | 1.300E-05 | 2.646E-06 | 1.100E-05 | 1.600E-05 |
| 4826 | 242 | C242E | A | 1.6e-05 | 1.5e-05 | 1.1e-05 | 1.400E-05 | 2.646E-06 | 1.100E-05 | 1.600E-05 |
| 4827 | 242 | C242Q | A | 9.3e-06 | 1e-05 | 1.1e-05 | 1.010E-05 | 8.544E-07 | 9.300E-06 | 1.100E-05 |
| 4828 | 242 | C242G | A | 1.5e-05 | 9.7e-06 | 1e-05 | 1.157E-05 | 2.977E-06 | 9.700E-06 | 1.500E-05 |
| 4829 | 242 | C242H | A | 8.7e-06 | 1e-05 | 1.8e-05 | 1.223E-05 | 5.036E-06 | 8.700E-06 | 1.800E-05 |
| 4830 | 242 | C242I | A | 1.6e-05 | 1.6e-05 | 1.3e-05 | 1.500E-05 | 1.732E-06 | 1.300E-05 | 1.600E-05 |
| 4831 | 242 | C242L | A | 1.4e-05 | 1.6e-05 | 1.4e-05 | 1.467E-05 | 1.155E-06 | 1.400E-05 | 1.600E-05 |
| 4832 | 242 | C242K | A | 1.2e-05 | 1.1e-05 | 1.7e-05 | 1.333E-05 | 3.215E-06 | 1.100E-05 | 1.700E-05 |
| 4833 | 242 | C242M | A | 1.4e-05 | 1.4e-05 | 1.5e-05 | 1.433E-05 | 5.774E-07 | 1.400E-05 | 1.500E-05 |
| 4834 | 242 | C242F | A | 1.1e-05 | 1e-05 | 1.5e-05 | 1.200E-05 | 2.646E-06 | 1.000E-05 | 1.500E-05 |
| 4835 | 242 | C242P | A | 1.2e-05 | 9.7e-06 | 1.6e-05 | 1.257E-05 | 3.188E-06 | 9.700E-06 | 1.600E-05 |
| 4836 | 242 | C242S | A | 1e-05 | 1e-05 | 9e-06 | 9.667E-06 | 5.774E-07 | 9.000E-06 | 1.000E-05 |
| 4837 | 242 | C242T | A | 1.5e-05 | 1.4e-05 | 1e-05 | 1.300E-05 | 2.646E-06 | 1.000E-05 | 1.500E-05 |
| 4838 | 242 | C242W | A | 1e-05 | 1.2e-05 | 1.4e-05 | 1.200E-05 | 2.000E-06 | 1.000E-05 | 1.400E-05 |
| 4839 | 242 | C242Y | A | 1.6e-05 | 1.5e-05 | 1.7e-05 | 1.600E-05 | 1.000E-06 | 1.500E-05 | 1.700E-05 |
| 4840 | 242 | C242V | A | 1.4e-05 | 8.8e-06 | 1.6e-05 | 1.293E-05 | 3.717E-06 | 8.800E-06 | 1.600E-05 |
| 4841 | 243 | E243A | A | 1.5e-05 | 1.6e-05 | 1e-05 | 1.367E-05 | 3.215E-06 | 1.000E-05 | 1.600E-05 |
| 4842 | 243 | E243R | A | 7e-06 | 1.3e-05 | 1.1e-05 | 1.033E-05 | 3.055E-06 | 7.000E-06 | 1.300E-05 |
| 4843 | 243 | E243N | A | 1.5e-05 | 1.2e-05 | 1.1e-05 | 1.267E-05 | 2.082E-06 | 1.100E-05 | 1.500E-05 |
| 4844 | 243 | E243D | A | 1.6e-05 | 1.5e-05 | 1.1e-05 | 1.400E-05 | 2.646E-06 | 1.100E-05 | 1.600E-05 |

| | | | | | | | | | | |
|---|---|---|---|---|---|---|---|---|---|---|
| 4845 | 243 | E243C | A | 1.3e-05 | 1.5e-05 | 1.1e-05 | 1.300E-05 | 2.000E-06 | 1.100E-05 | 1.500E-05 |
| 4846 | 243 | E243E | A | 9.7e-06 | 1.7e-05 | 1.1e-05 | 1.257E-05 | 3.894E-06 | 9.700E-06 | 1.700E-05 |
| 4847 | 243 | E243Q | A | 1.6e-05 | 1e-05 | 9.4e-06 | 1.180E-05 | 3.650E-06 | 9.400E-06 | 1.600E-05 |
| 4848 | 243 | E243G | A | 1.5e-05 | 1.6e-05 | 1.5e-05 | 1.533E-05 | 5.774E-07 | 1.500E-05 | 1.600E-05 |
| 4849 | 243 | E243H | A | 9.8e-06 | 1.6e-05 | 1.5e-05 | 1.360E-05 | 3.329E-06 | 9.800E-06 | 1.600E-05 |
| 4850 | 243 | E243I | A | 9.4e-06 | 9.9e-06 | 1.3e-05 | 1.077E-05 | 1.950E-06 | 9.400E-06 | 1.300E-05 |
| 4851 | 243 | E243L | A | 9.4e-06 | 1.5e-05 | 1.4e-05 | 1.280E-05 | 2.987E-06 | 9.400E-06 | 1.500E-05 |
| 4852 | 243 | E243K | A | 1.1e-05 | 1.3e-05 | 1.6e-05 | 1.333E-05 | 2.517E-06 | 1.100E-05 | 1.600E-05 |
| 4853 | 243 | E243M | A | 1.6e-05 | 1e-05 | 1.5e-05 | 1.367E-05 | 3.215E-06 | 1.000E-05 | 1.600E-05 |
| 4854 | 243 | E243F | A | 1.2e-05 | 1.5e-05 | 8.9e-06 | 1.197E-05 | 3.050E-06 | 8.900E-06 | 1.500E-05 |
| 4855 | 243 | E243P | A | 1.1e-05 | 1.4e-05 | 1.5e-05 | 1.333E-05 | 2.082E-06 | 1.100E-05 | 1.500E-05 |
| 4856 | 243 | E243S | A | 1.5e-05 | 1.2e-05 | 8.5e-06 | 1.183E-05 | 3.253E-06 | 8.500E-06 | 1.500E-05 |
| 4857 | 243 | E243T | A | 1.1e-05 | 1.6e-05 | 1.6e-05 | 1.433E-05 | 2.887E-06 | 1.100E-05 | 1.600E-05 |
| 4858 | 243 | E243W | A | 1.6e-05 | 1.4e-05 | 1.4e-05 | 1.467E-05 | 1.155E-06 | 1.400E-05 | 1.600E-05 |
| 4859 | 243 | E243Y | A | 9.9e-06 | 9.4e-06 | 1.3e-05 | 1.077E-05 | 1.950E-06 | 9.400E-06 | 1.300E-05 |
| 4860 | 243 | E243V | A | 1.4e-05 | 1.5e-05 | 1.4e-05 | 1.433E-05 | 5.774E-07 | 1.400E-05 | 1.500E-05 |
| 4861 | 244 | L244A | A | 1.4e-05 | 1.4e-05 | 1.3e-05 | 1.367E-05 | 5.774E-07 | 1.300E-05 | 1.400E-05 |
| 4862 | 244 | L244R | A | 1e-05 | 1.2e-05 | 1.6e-05 | 1.267E-05 | 3.055E-06 | 1.000E-05 | 1.600E-05 |
| 4863 | 244 | L244N | A | 1.3e-05 | 1.3e-05 | 2.5e-05 | 1.700E-05 | 6.928E-06 | 1.300E-05 | 2.500E-05 |
| 4864 | 244 | L244D | A | 1.5e-05 | 7.9e-06 | 1.1e-05 | 1.130E-05 | 3.559E-06 | 7.900E-06 | 1.500E-05 |
| 4865 | 244 | L244C | A | 9.2e-06 | 1.1e-05 | 9.5e-06 | 9.900E-06 | 9.644E-07 | 9.200E-06 | 1.100E-05 |
| 4866 | 244 | L244E | A | 1.6e-05 | 1.4e-05 | 9.6e-06 | 1.320E-05 | 3.274E-06 | 9.600E-06 | 1.600E-05 |
| 4867 | 244 | L244Q | A | 1.1e-05 | 1.4e-05 | 1.2e-05 | 1.233E-05 | 1.528E-06 | 1.100E-05 | 1.400E-05 |
| 4868 | 244 | L244G | A | 1.5e-05 | 9.2e-06 | 1e-05 | 1.140E-05 | 3.143E-06 | 9.200E-06 | 1.500E-05 |
| 4869 | 244 | L244H | A | 1.5e-05 | 9.6e-06 | 1.2e-05 | 1.220E-05 | 2.706E-06 | 9.600E-06 | 1.500E-05 |
| 4870 | 244 | L244I | A | 1.6e-05 | 1.4e-05 | 1.1e-05 | 1.367E-05 | 2.517E-06 | 1.100E-05 | 1.600E-05 |
| 4871 | 244 | L244L | A | 1.7e-05 | 1.7e-05 | 1.6e-05 | 1.667E-05 | 5.774E-07 | 1.600E-05 | 1.700E-05 |
| 4872 | 244 | L244K | A | 1.6e-05 | 1.5e-05 | 1.2e-05 | 1.433E-05 | 2.082E-06 | 1.200E-05 | 1.600E-05 |
| 4873 | 244 | L244M | A | 1.6e-05 | 1e-05 | 8.7e-06 | 1.157E-05 | 3.894E-06 | 8.700E-06 | 1.600E-05 |
| 4874 | 244 | L244F | A | 1e-05 | 1.5e-05 | 9.7e-06 | 1.157E-05 | 2.977E-06 | 9.700E-06 | 1.500E-05 |
| 4875 | 244 | L244P | A | 9.6e-06 | 9.7e-06 | 1.2e-05 | 1.043E-05 | 1.358E-06 | 9.600E-06 | 1.200E-05 |
| 4876 | 244 | L244S | A | 1.5e-05 | 1e-05 | 1e-05 | 1.167E-05 | 2.887E-06 | 1.000E-05 | 1.500E-05 |
| 4877 | 244 | L244T | A | 1.5e-05 | 1e-05 | 1.5e-05 | 1.333E-05 | 2.887E-06 | 1.000E-05 | 1.500E-05 |
| 4878 | 244 | L244W | A | 1.3e-05 | 1.5e-05 | 1.4e-05 | 1.400E-05 | 1.000E-06 | 1.300E-05 | 1.500E-05 |
| 4879 | 244 | L244Y | A | 1.4e-05 | 1.4e-05 | 1.2e-05 | 1.333E-05 | 1.155E-06 | 1.200E-05 | 1.400E-05 |
| 4880 | 244 | L244V | A | 1.4e-05 | 1.1e-05 | 1.5e-05 | 1.333E-05 | 2.082E-06 | 1.100E-05 | 1.500E-05 |
| 4881 | 245 | H245A | A | 1.7e-05 | 1.1e-05 | 1.6e-05 | 1.467E-05 | 3.215E-06 | 1.100E-05 | 1.700E-05 |
| 4882 | 245 | H245R | A | 1.5e-05 | 1.6e-05 | 1.2e-05 | 1.433E-05 | 2.082E-06 | 1.200E-05 | 1.600E-05 |
| 4883 | 245 | H245N | A | 1.6e-05 | 1e-05 | 1.6e-05 | 1.400E-05 | 3.464E-06 | 1.000E-05 | 1.600E-05 |
| 4884 | 245 | H245D | A | 1.5e-05 | 1.7e-05 | 1.6e-05 | 1.600E-05 | 1.000E-06 | 1.500E-05 | 1.700E-05 |
| 4885 | 245 | H245C | A | 1.8e-05 | 1.2e-05 | 1.5e-05 | 1.500E-05 | 3.000E-06 | 1.200E-05 | 1.800E-05 |
| 4886 | 245 | H245E | A | 1.2e-05 | 1.2e-05 | 1.2e-05 | 1.200E-05 | 0.000E+00 | 1.200E-05 | 1.200E-05 |
| 4887 | 245 | H245Q | A | 1.2e-05 | 9.6e-06 | 6.4e-06 | 9.333E-06 | 2.810E-06 | 6.400E-06 | 1.200E-05 |
| 4888 | 245 | H245G | A | 7.8e-06 | 9.6e-06 | 1.3e-05 | 1.013E-05 | 2.641E-06 | 7.800E-06 | 1.300E-05 |
| 4889 | 245 | H245H | A | 1.6e-05 | 9.7e-06 | 1.5e-05 | 1.357E-05 | 3.386E-06 | 9.700E-06 | 1.600E-05 |
| 4890 | 245 | H245I | A | 1.1e-05 | 1.2e-05 | 1.6e-05 | 1.300E-05 | 2.646E-06 | 1.100E-05 | 1.600E-05 |
| 4891 | 245 | H245L | A | 9.8e-06 | 1.6e-05 | 1.3e-05 | 1.293E-05 | 3.101E-06 | 9.800E-06 | 1.600E-05 |
| 4892 | 245 | H245K | A | 1.1e-05 | 1.3e-05 | 1.5e-05 | 1.300E-05 | 2.000E-06 | 1.100E-05 | 1.500E-05 |
| 4893 | 245 | H245M | A | 1.5e-05 | 1.1e-05 | 1.5e-05 | 1.367E-05 | 2.309E-06 | 1.100E-05 | 1.500E-05 |
| 4894 | 245 | H245F | A | 1.5e-05 | 1e-05 | 1.5e-05 | 1.333E-05 | 2.887E-06 | 1.000E-05 | 1.500E-05 |
| 4895 | 245 | H245P | A | 1.2e-05 | 1.4e-05 | 1.6e-05 | 1.400E-05 | 2.000E-06 | 1.200E-05 | 1.600E-05 |
| 4896 | 245 | H245S | A | 1.4e-05 | 1e-05 | 1.1e-05 | 1.167E-05 | 2.082E-06 | 1.000E-05 | 1.400E-05 |
| 4897 | 245 | H245T | A | 1.1e-05 | 1.6e-05 | 1.3e-05 | 1.333E-05 | 2.517E-06 | 1.100E-05 | 1.600E-05 |
| 4898 | 245 | H245W | A | 1.5e-05 | 1.1e-05 | 1.7e-05 | 1.433E-05 | 3.055E-06 | 1.100E-05 | 1.700E-05 |
| 4899 | 245 | H245Y | A | 9.9e-06 | 1.1e-05 | 1.1e-05 | 1.063E-05 | 6.351E-07 | 9.900E-06 | 1.100E-05 |
| 4900 | 245 | H245V | A | 1.2e-05 | 1.4e-05 | 9.3e-06 | 1.177E-05 | 2.359E-06 | 9.300E-06 | 1.400E-05 |
| 4901 | 246 | C246A | A | 1.5e-05 | 1.1e-05 | 1.1e-05 | 1.233E-05 | 2.309E-06 | 1.100E-05 | 1.500E-05 |
| 4902 | 246 | C246R | A | 1e-05 | 8.4e-06 | 9.7e-06 | 9.367E-06 | 8.505E-07 | 8.400E-06 | 1.000E-05 |
| 4903 | 246 | C246N | A | 8.6e-06 | 1.2e-05 | 1.5e-05 | 1.187E-05 | 3.202E-06 | 8.600E-06 | 1.500E-05 |
| 4904 | 246 | C246D | A | 1.3e-05 | 1.5e-05 | 1.1e-05 | 1.300E-05 | 2.000E-06 | 1.100E-05 | 1.500E-05 |
| 4905 | 246 | C246C | A | 1.4e-05 | 1.5e-05 | 1.1e-05 | 1.333E-05 | 2.082E-06 | 1.100E-05 | 1.500E-05 |
| 4906 | 246 | C246E | A | 1e-05 | 1.5e-05 | 1.4e-05 | 1.300E-05 | 2.646E-06 | 1.000E-05 | 1.500E-05 |
| 4907 | 246 | C246Q | A | 1.3e-05 | 1.1e-05 | 1.1e-05 | 1.167E-05 | 1.155E-06 | 1.100E-05 | 1.300E-05 |
| 4908 | 246 | C246G | A | 1.1e-05 | 9.6e-06 | 1.5e-05 | 1.187E-05 | 2.802E-06 | 9.600E-06 | 1.500E-05 |
| 4909 | 246 | C246H | A | 9.2e-06 | 1.6e-05 | 1.5e-05 | 1.340E-05 | 3.672E-06 | 9.200E-06 | 1.600E-05 |
| 4910 | 246 | C246I | A | 1.1e-05 | 1.6e-05 | 1.7e-05 | 1.467E-05 | 3.215E-06 | 1.100E-05 | 1.700E-05 |
| 4911 | 246 | C246L | A | 1e-05 | 1.2e-05 | 1.5e-05 | 1.233E-05 | 2.517E-06 | 1.000E-05 | 1.500E-05 |
| 4912 | 246 | C246K | A | 1.4e-05 | 1.6e-05 | 1.5e-05 | 1.500E-05 | 1.000E-06 | 1.400E-05 | 1.600E-05 |
| 4913 | 246 | C246M | A | 1.3e-05 | 1.7e-05 | 8.8e-06 | 1.293E-05 | 4.100E-06 | 8.800E-06 | 1.700E-05 |
| 4914 | 246 | C246F | A | 1.5e-05 | 1.1e-05 | 1.4e-05 | 1.333E-05 | 2.082E-06 | 1.100E-05 | 1.500E-05 |
| 4915 | 246 | C246P | A | 1.2e-05 | 1.5e-05 | 1.3e-05 | 1.333E-05 | 1.528E-06 | 1.200E-05 | 1.500E-05 |
| 4916 | 246 | C246S | A | 3.2e-05 | 1.6e-05 | 1e-05 | 1.933E-05 | 1.137E-05 | 1.000E-05 | 3.200E-05 |
| 4917 | 246 | C246T | A | 1.1e-05 | 1.5e-05 | 1.1e-05 | 1.233E-05 | 2.309E-06 | 1.100E-05 | 1.500E-05 |
| 4918 | 246 | C246W | A | 1.6e-05 | 1.3e-05 | 1.1e-05 | 1.333E-05 | 2.517E-06 | 1.100E-05 | 1.600E-05 |
| 4919 | 246 | C246Y | A | 1e-05 | 1.6e-05 | 1.1e-05 | 1.233E-05 | 3.215E-06 | 1.000E-05 | 1.600E-05 |
| 4920 | 246 | C246V | A | 1.4e-05 | 1.5e-05 | 1.2e-05 | 1.367E-05 | 1.528E-06 | 1.200E-05 | 1.500E-05 |
| 4921 | 247 | P247A | A | 7.1e-06 | 1.1e-05 | 1e-05 | 9.367E-06 | 2.026E-06 | 7.100E-06 | 1.100E-05 |
| 4922 | 247 | P247R | A | 1.5e-05 | 1.4e-05 | 8.6e-06 | 1.253E-05 | 3.443E-06 | 8.600E-06 | 1.500E-05 |
| 4923 | 247 | P247N | A | 1.2e-05 | 1.5e-05 | 1.5e-05 | 1.400E-05 | 1.732E-06 | 1.200E-05 | 1.500E-05 |
| 4924 | 247 | P247D | A | 1e-05 | 1.1e-05 | 9.4e-06 | 1.013E-05 | 8.083E-07 | 9.400E-06 | 1.100E-05 |
| 4925 | 247 | P247C | A | 1.5e-05 | 8.1e-06 | 1.1e-05 | 1.137E-05 | 3.465E-06 | 8.100E-06 | 1.500E-05 |
| 4926 | 247 | P247E | A | 1.6e-05 | 1.1e-05 | 9.6e-06 | 1.220E-05 | 3.365E-06 | 9.600E-06 | 1.600E-05 |
| 4927 | 247 | P247Q | A | 1.5e-05 | 9.9e-06 | 1e-05 | 1.163E-05 | 2.916E-06 | 9.900E-06 | 1.500E-05 |
| 4928 | 247 | P247G | A | 7e-06 | 1.2e-05 | 1.5e-05 | 1.133E-05 | 4.041E-06 | 7.000E-06 | 1.500E-05 |
| 4929 | 247 | P247H | A | 1.4e-05 | 1.5e-05 | 1.6e-05 | 1.500E-05 | 1.000E-06 | 1.400E-05 | 1.600E-05 |

| | | | | | | | | | | |
|---|---|---|---|---|---|---|---|---|---|---|
| 4930 | 247 | P247I | A | 9.5e-06 | 1.7e-05 | 1.1e-05 | 1.250E-05 | 3.969E-06 | 9.500E-06 | 1.700E-05 |
| 4931 | 247 | P247L | A | 9.3e-06 | 1.4e-05 | 9.9e-06 | 1.107E-05 | 2.558E-06 | 9.300E-06 | 1.400E-05 |
| 4932 | 247 | P247K | A | 1.5e-05 | 1.5e-05 | 1.7e-05 | 1.567E-05 | 1.155E-06 | 1.500E-05 | 1.700E-05 |
| 4933 | 247 | P247M | A | 1e-05 | 9.9e-06 | 1.1e-05 | 1.030E-05 | 6.083E-07 | 9.900E-06 | 1.100E-05 |
| 4934 | 247 | P247F | A | 1e-05 | 9.2e-06 | 1.1e-05 | 1.007E-05 | 9.018E-07 | 9.200E-06 | 1.100E-05 |
| 4935 | 247 | P247P | A | 1.6e-05 | 1e-05 | 1.6e-05 | 1.400E-05 | 3.464E-06 | 1.000E-05 | 1.600E-05 |
| 4936 | 247 | P247S | A | 1.6e-05 | 1.3e-05 | 1.2e-05 | 1.367E-05 | 2.082E-06 | 1.200E-05 | 1.600E-05 |
| 4937 | 247 | P247T | A | 1.5e-05 | 1e-05 | 1e-05 | 1.167E-05 | 2.887E-06 | 1.000E-05 | 1.500E-05 |
| 4938 | 247 | P247W | A | 8.5e-06 | 1.5e-05 | 9e-06 | 1.083E-05 | 3.617E-06 | 8.500E-06 | 1.500E-05 |
| 4939 | 247 | P247Y | A | 1.1e-05 | 1.4e-05 | 9.2e-06 | 1.140E-05 | 2.425E-06 | 9.200E-06 | 1.400E-05 |
| 4940 | 247 | P247V | A | 1.4e-05 | 1.4e-05 | 1.4e-05 | 1.400E-05 | 0.000E+00 | 1.400E-05 | 1.400E-05 |
| 4941 | 248 | A248A | A | 1.7e-05 | 8.4e-06 | 1.5e-05 | 1.347E-05 | 4.500E-06 | 8.400E-06 | 1.700E-05 |
| 4942 | 248 | A248R | A | 1.5e-05 | 1.3e-05 | 1.6e-05 | 1.467E-05 | 1.528E-06 | 1.300E-05 | 1.600E-05 |
| 4943 | 248 | A248N | A | 1.1e-05 | 1.5e-05 | 1.5e-05 | 1.367E-05 | 2.309E-06 | 1.100E-05 | 1.500E-05 |
| 4944 | 248 | A248D | A | 1.5e-05 | 1.6e-05 | 1.1e-05 | 1.400E-05 | 2.646E-06 | 1.100E-05 | 1.600E-05 |
| 4945 | 248 | A248C | A | 1.5e-05 | 9.5e-06 | 8.2e-06 | 1.090E-05 | 3.610E-06 | 8.200E-06 | 1.500E-05 |
| 4946 | 248 | A248E | A | 8.1e-06 | 1.6e-05 | 1.5e-05 | 1.303E-05 | 4.302E-06 | 8.100E-06 | 1.600E-05 |
| 4947 | 248 | A248Q | A | 1.4e-05 | 1.5e-05 | 9.1e-06 | 1.270E-05 | 3.158E-06 | 9.100E-06 | 1.500E-05 |
| 4948 | 248 | A248G | A | 1e-05 | 1.1e-05 | 1.6e-05 | 1.233E-05 | 3.215E-06 | 1.000E-05 | 1.600E-05 |
| 4949 | 248 | A248H | A | 1.6e-05 | 9.6e-06 | 1.5e-05 | 1.353E-05 | 3.443E-06 | 9.600E-06 | 1.600E-05 |
| 4950 | 248 | A248I | A | 1.1e-05 | 1e-05 | 1.5e-05 | 1.200E-05 | 2.646E-06 | 1.000E-05 | 1.500E-05 |
| 4951 | 248 | A248L | A | 9.9e-06 | 8.4e-06 | 1.1e-05 | 9.767E-06 | 1.305E-06 | 8.400E-06 | 1.100E-05 |
| 4952 | 248 | A248K | A | 1.4e-05 | 1.5e-05 | 1.4e-05 | 1.433E-05 | 5.774E-07 | 1.400E-05 | 1.500E-05 |
| 4953 | 248 | A248M | A | 1.6e-05 | 1.1e-05 | 1.6e-05 | 1.433E-05 | 2.887E-06 | 1.100E-05 | 1.600E-05 |
| 4954 | 248 | A248F | A | 1.6e-05 | 1.5e-05 | 1.4e-05 | 1.500E-05 | 1.000E-06 | 1.400E-05 | 1.600E-05 |
| 4955 | 248 | A248P | A | 1.5e-05 | 1.6e-05 | 1.1e-05 | 1.400E-05 | 2.646E-06 | 1.100E-05 | 1.600E-05 |
| 4956 | 248 | A248S | A | 1.4e-05 | 1.5e-05 | 9.7e-06 | 1.290E-05 | 2.816E-06 | 9.700E-06 | 1.500E-05 |
| 4957 | 248 | A248T | A | 1.3e-05 | 1.6e-05 | 1.3e-05 | 1.400E-05 | 1.732E-06 | 1.300E-05 | 1.600E-05 |
| 4958 | 248 | A248W | A | 1.2e-05 | 1.5e-05 | 1.5e-05 | 1.400E-05 | 1.732E-06 | 1.200E-05 | 1.500E-05 |
| 4959 | 248 | A248Y | A | 1.1e-05 | 9.7e-06 | 1.5e-05 | 1.190E-05 | 2.762E-06 | 9.700E-06 | 1.500E-05 |
| 4960 | 248 | A248V | A | 7.4e-06 | 1.5e-05 | 1.5e-05 | 1.247E-05 | 4.388E-06 | 7.400E-06 | 1.500E-05 |
| 4961 | 249 | L249A | A | 1.6e-05 | 1.1e-05 | 1e-05 | 1.233E-05 | 3.215E-06 | 1.000E-05 | 1.600E-05 |
| 4962 | 249 | L249R | A | 1.4e-05 | 1e-05 | 1.4e-05 | 1.267E-05 | 2.309E-06 | 1.000E-05 | 1.400E-05 |
| 4963 | 249 | L249N | A | 1.6e-05 | 1.4e-05 | 1.5e-05 | 1.500E-05 | 1.000E-06 | 1.400E-05 | 1.600E-05 |
| 4964 | 249 | L249D | A | 7.2e-06 | 1.4e-05 | 1.2e-05 | 1.107E-05 | 3.495E-06 | 7.200E-06 | 1.400E-05 |
| 4965 | 249 | L249C | A | 1e-05 | 6e-06 | 8.5e-06 | 8.167E-06 | 2.021E-06 | 6.000E-06 | 1.000E-05 |
| 4966 | 249 | L249E | A | 1.6e-05 | 1.5e-05 | 1e-05 | 1.367E-05 | 3.215E-06 | 1.000E-05 | 1.600E-05 |
| 4967 | 249 | L249Q | A | 1.3e-05 | 1.5e-05 | 1.2e-05 | 1.333E-05 | 1.528E-06 | 1.200E-05 | 1.500E-05 |
| 4968 | 249 | L249G | A | 1.7e-05 | 9.5e-06 | 1.1e-05 | 1.250E-05 | 3.969E-06 | 9.500E-06 | 1.700E-05 |
| 4969 | 249 | L249H | A | 1.5e-05 | 9.2e-06 | 9.3e-06 | 1.117E-05 | 3.320E-06 | 9.200E-06 | 1.500E-05 |
| 4970 | 249 | L249I | A | 1.4e-05 | 1.4e-05 | 1.5e-05 | 1.433E-05 | 5.774E-07 | 1.400E-05 | 1.500E-05 |
| 4971 | 249 | L249L | A | 1.6e-05 | 1.5e-05 | 1.2e-05 | 1.433E-05 | 2.082E-06 | 1.200E-05 | 1.600E-05 |
| 4972 | 249 | L249K | A | 1.1e-05 | 1e-05 | 1.1e-05 | 1.067E-05 | 5.774E-07 | 1.000E-05 | 1.100E-05 |
| 4973 | 249 | L249M | A | 1.1e-05 | 1.5e-05 | 9.3e-06 | 1.177E-05 | 2.926E-06 | 9.300E-06 | 1.500E-05 |
| 4974 | 249 | L249F | A | 1.5e-05 | 1.1e-05 | 1.6e-05 | 1.400E-05 | 2.646E-06 | 1.100E-05 | 1.600E-05 |
| 4975 | 249 | L249P | A | 1e-05 | 1.6e-05 | 1.3e-05 | 1.300E-05 | 3.000E-06 | 1.000E-05 | 1.600E-05 |
| 4976 | 249 | L249S | A | 1.6e-05 | 1.3e-05 | 1.6e-05 | 1.500E-05 | 1.732E-06 | 1.300E-05 | 1.600E-05 |
| 4977 | 249 | L249T | A | 1e-05 | 9.5e-06 | 1.4e-05 | 1.117E-05 | 2.466E-06 | 9.500E-06 | 1.400E-05 |
| 4978 | 249 | L249W | A | 1.1e-05 | 1.5e-05 | 8.2e-06 | 1.140E-05 | 3.418E-06 | 8.200E-06 | 1.500E-05 |
| 4979 | 249 | L249Y | A | 1.4e-05 | 1.5e-05 | 9.8e-06 | 1.293E-05 | 2.759E-06 | 9.800E-06 | 1.500E-05 |
| 4980 | 249 | L249V | A | 1e-05 | 1.4e-05 | 1.5e-05 | 1.300E-05 | 2.646E-06 | 1.000E-05 | 1.500E-05 |
| 4981 | 250 | V250A | A | 1.7e-05 | 1.2e-05 | 9.5e-06 | 1.283E-05 | 3.819E-06 | 9.500E-06 | 1.700E-05 |
| 4982 | 250 | V250R | A | 1.1e-05 | 1.6e-05 | 9.9e-06 | 1.230E-05 | 3.251E-06 | 9.900E-06 | 1.600E-05 |
| 4983 | 250 | V250N | A | 1.8e-05 | 1.5e-05 | 1.5e-05 | 1.600E-05 | 1.732E-06 | 1.500E-05 | 1.800E-05 |
| 4984 | 250 | V250D | A | 1.2e-05 | 1.5e-05 | 1.5e-05 | 1.400E-05 | 1.732E-06 | 1.200E-05 | 1.500E-05 |
| 4985 | 250 | V250C | A | 1.7e-05 | 1.1e-05 | 1.6e-05 | 1.467E-05 | 3.215E-06 | 1.100E-05 | 1.700E-05 |
| 4986 | 250 | V250E | A | 1.6e-05 | 1.6e-05 | 8.8e-06 | 1.360E-05 | 4.157E-06 | 8.800E-06 | 1.600E-05 |
| 4987 | 250 | V250Q | A | 1.3e-05 | 1.5e-05 | 1e-05 | 1.267E-05 | 2.517E-06 | 1.000E-05 | 1.500E-05 |
| 4988 | 250 | V250G | A | 1.1e-05 | 1.2e-05 | 1.5e-05 | 1.267E-05 | 2.082E-06 | 1.100E-05 | 1.500E-05 |
| 4989 | 250 | V250H | A | 9.7e-06 | 1.3e-05 | 9.7e-06 | 1.080E-05 | 1.905E-06 | 9.700E-06 | 1.300E-05 |
| 4990 | 250 | V250I | A | 1.6e-05 | 1.3e-05 | 9.9e-06 | 1.297E-05 | 3.050E-06 | 9.900E-06 | 1.600E-05 |
| 4991 | 250 | V250L | A | 1.3e-05 | 1e-05 | 7.7e-06 | 1.023E-05 | 2.658E-06 | 7.700E-06 | 1.300E-05 |
| 4992 | 250 | V250K | A | 1.3e-05 | 1.6e-05 | 1.6e-05 | 1.500E-05 | 1.732E-06 | 1.300E-05 | 1.600E-05 |
| 4993 | 250 | V250M | A | 1.6e-05 | 1.3e-05 | 1.2e-05 | 1.367E-05 | 2.082E-06 | 1.200E-05 | 1.600E-05 |
| 4994 | 250 | V250F | A | 2.1e-05 | 1.1e-05 | 1.6e-05 | 1.600E-05 | 5.000E-06 | 1.100E-05 | 2.100E-05 |
| 4995 | 250 | V250P | A | 2.6e-05 | 1.2e-05 | 1.6e-05 | 1.800E-05 | 7.211E-06 | 1.200E-05 | 2.600E-05 |
| 4996 | 250 | V250S | A | 1.1e-05 | 9.7e-06 | 1.6e-05 | 1.223E-05 | 3.326E-06 | 9.700E-06 | 1.600E-05 |
| 4997 | 250 | V250T | A | 1.4e-05 | 9.3e-06 | 1.4e-05 | 1.243E-05 | 2.714E-06 | 9.300E-06 | 1.400E-05 |
| 4998 | 250 | V250W | A | 8.9e-06 | 8.9e-06 | 1.5e-05 | 1.093E-05 | 3.522E-06 | 8.900E-06 | 1.500E-05 |
| 4999 | 250 | V250Y | A | 1.1e-05 | 1e-05 | 9.7e-06 | 1.023E-05 | 6.807E-07 | 9.700E-06 | 1.100E-05 |
| 5000 | 250 | V250V | A | 1.6e-05 | 1e-05 | 1.6e-05 | 1.400E-05 | 3.464E-06 | 1.000E-05 | 1.600E-05 |
| 5001 | 251 | T251A | A | 1.1e-05 | 9.8e-06 | 1.4e-05 | 1.160E-05 | 2.163E-06 | 9.800E-06 | 1.400E-05 |
| 5002 | 251 | T251R | A | 7.5e-06 | 1.6e-05 | 1.2e-05 | 1.183E-05 | 4.252E-06 | 7.500E-06 | 1.600E-05 |
| 5003 | 251 | T251N | A | 1.3e-05 | 1.4e-05 | 1.1e-05 | 1.267E-05 | 1.528E-06 | 1.100E-05 | 1.400E-05 |
| 5004 | 251 | T251D | A | 1e-05 | 1.5e-05 | 1.2e-05 | 1.233E-05 | 2.517E-06 | 1.000E-05 | 1.500E-05 |
| 5005 | 251 | T251C | A | 1.6e-05 | 1.1e-05 | 1.2e-05 | 1.300E-05 | 2.646E-06 | 1.100E-05 | 1.600E-05 |
| 5006 | 251 | T251E | A | 1.1e-05 | 1.4e-05 | 8.6e-06 | 1.120E-05 | 2.706E-06 | 8.600E-06 | 1.400E-05 |
| 5007 | 251 | T251Q | A | 1.1e-05 | 1.3e-05 | 1.5e-05 | 1.300E-05 | 2.000E-06 | 1.100E-05 | 1.500E-05 |
| 5008 | 251 | T251G | A | 1e-05 | 1.1e-05 | 7.5e-06 | 9.500E-06 | 1.803E-06 | 7.500E-06 | 1.100E-05 |
| 5009 | 251 | T251H | A | 1.1e-05 | 1e-05 | 1.1e-05 | 1.067E-05 | 5.774E-07 | 1.000E-05 | 1.100E-05 |
| 5010 | 251 | T251I | A | 1e-05 | 1.7e-05 | 1.1e-05 | 1.267E-05 | 3.786E-06 | 1.000E-05 | 1.700E-05 |
| 5011 | 251 | T251L | A | 1.7e-05 | 1.9e-05 | 1.1e-05 | 1.567E-05 | 4.163E-06 | 1.100E-05 | 1.900E-05 |
| 5012 | 251 | T251K | A | 6.8e-06 | 1.1e-05 | 1.7e-05 | 1.160E-05 | 5.126E-06 | 6.800E-06 | 1.700E-05 |
| 5013 | 251 | T251M | A | 1e-05 | 1.5e-05 | 1.5e-05 | 1.333E-05 | 2.887E-06 | 1.000E-05 | 1.500E-05 |
| 5014 | 251 | T251F | A | 1.7e-05 | 1.1e-05 | 1.5e-05 | 1.433E-05 | 3.055E-06 | 1.100E-05 | 1.700E-05 |

| | | | | | | | | | | |
|---|---|---|---|---|---|---|---|---|---|---|
| 5015 | 251 | T251P | A | 1.5e-05 | 1.5e-05 | 9.7e-06 | 1.323E-05 | 3.060E-06 | 9.700E-06 | 1.500E-05 |
| 5016 | 251 | T251S | A | 1.4e-05 | 1.3e-05 | 1.9e-05 | 1.533E-05 | 3.215E-06 | 1.300E-05 | 1.900E-05 |
| 5017 | 251 | T251T | A | 1e-05 | 1.1e-05 | 1.6e-05 | 1.233E-05 | 3.215E-06 | 1.000E-05 | 1.600E-05 |
| 5018 | 251 | T251W | A | 1.2e-05 | 1.2e-05 | 1.5e-05 | 1.300E-05 | 1.732E-06 | 1.200E-05 | 1.500E-05 |
| 5019 | 251 | T251Y | A | 1.6e-05 | 9.4e-06 | 1.1e-05 | 1.213E-05 | 3.443E-06 | 9.400E-06 | 1.600E-05 |
| 5020 | 251 | T251V | A | 1.1e-05 | 1.1e-05 | 1.1e-05 | 1.100E-05 | 2.075E-21 | 1.100E-05 | 1.100E-05 |
| 5021 | 252 | Y252A | A | 1.4e-05 | 1.6e-05 | 1.4e-05 | 1.467E-05 | 1.155E-06 | 1.400E-05 | 1.600E-05 |
| 5022 | 252 | Y252R | A | 1.6e-05 | 1.2e-05 | 1.5e-05 | 1.433E-05 | 2.082E-06 | 1.200E-05 | 1.600E-05 |
| 5023 | 252 | Y252N | A | 1.4e-05 | 1.3e-05 | 1.2e-05 | 1.300E-05 | 1.000E-06 | 1.200E-05 | 1.400E-05 |
| 5024 | 252 | Y252D | A | 1.5e-05 | 1.6e-05 | 9.7e-06 | 1.357E-05 | 3.386E-06 | 9.700E-06 | 1.600E-05 |
| 5025 | 252 | Y252C | A | 1.5e-05 | 7.3e-06 | 1.5e-05 | 1.243E-05 | 4.446E-06 | 7.300E-06 | 1.500E-05 |
| 5026 | 252 | Y252E | A | 1.5e-05 | 1.4e-05 | 1e-05 | 1.300E-05 | 2.646E-06 | 1.000E-05 | 1.500E-05 |
| 5027 | 252 | Y252Q | A | 1.3e-05 | 1.4e-05 | 8.2e-06 | 1.173E-05 | 3.101E-06 | 8.200E-06 | 1.400E-05 |
| 5028 | 252 | Y252G | A | 1e-05 | 8.7e-06 | 1.6e-05 | 1.157E-05 | 3.894E-06 | 8.700E-06 | 1.600E-05 |
| 5029 | 252 | Y252H | A | 1.6e-05 | 1.1e-05 | 1.6e-05 | 1.433E-05 | 2.887E-06 | 1.100E-05 | 1.600E-05 |
| 5030 | 252 | Y252I | A | 1.7e-05 | 1.6e-05 | 1.4e-05 | 1.567E-05 | 1.528E-06 | 1.400E-05 | 1.700E-05 |
| 5031 | 252 | Y252L | A | 1.1e-05 | 1.6e-05 | 1.1e-05 | 1.267E-05 | 2.887E-06 | 1.100E-05 | 1.600E-05 |
| 5032 | 252 | Y252K | A | 1.1e-05 | 1.4e-05 | 1.4e-05 | 1.300E-05 | 1.732E-06 | 1.100E-05 | 1.400E-05 |
| 5033 | 252 | Y252M | A | 1.1e-05 | 1.7e-05 | 1.5e-05 | 1.433E-05 | 3.055E-06 | 1.100E-05 | 1.700E-05 |
| 5034 | 252 | Y252F | A | 1.1e-05 | 1.5e-05 | 1.1e-05 | 1.233E-05 | 2.309E-06 | 1.100E-05 | 1.500E-05 |
| 5035 | 252 | Y252P | A | 1.7e-05 | 7.7e-06 | 1.5e-05 | 1.323E-05 | 4.895E-06 | 7.700E-06 | 1.700E-05 |
| 5036 | 252 | Y252S | A | 1.6e-05 | 1e-05 | 1.1e-05 | 1.233E-05 | 3.215E-06 | 1.000E-05 | 1.600E-05 |
| 5037 | 252 | Y252T | A | 1e-05 | 1.4e-05 | 7.1e-06 | 1.037E-05 | 3.465E-06 | 7.100E-06 | 1.400E-05 |
| 5038 | 252 | Y252W | A | 1.4e-05 | 8.8e-06 | 1.7e-05 | 1.327E-05 | 4.149E-06 | 8.800E-06 | 1.700E-05 |
| 5039 | 252 | Y252Y | A | 9.5e-06 | 1.6e-05 | 1e-05 | 1.183E-05 | 3.617E-06 | 9.500E-06 | 1.600E-05 |
| 5040 | 252 | Y252V | A | 1.5e-05 | 1.5e-05 | 1.6e-05 | 1.533E-05 | 5.774E-07 | 1.500E-05 | 1.600E-05 |
| 5041 | 253 | N253A | A | 1.7e-05 | 1.1e-05 | 8.9e-06 | 1.230E-05 | 4.204E-06 | 8.900E-06 | 1.700E-05 |
| 5042 | 253 | N253R | A | 1.5e-05 | 8.8e-06 | 1.6e-05 | 1.327E-05 | 3.900E-06 | 8.800E-06 | 1.600E-05 |
| 5043 | 253 | N253N | A | 8.4e-06 | 1.4e-05 | 9.1e-06 | 1.050E-05 | 3.051E-06 | 8.400E-06 | 1.400E-05 |
| 5044 | 253 | N253D | A | 1.6e-05 | 1.1e-05 | 1.5e-05 | 1.400E-05 | 2.646E-06 | 1.100E-05 | 1.600E-05 |
| 5045 | 253 | N253C | A | 1.3e-05 | 1.4e-05 | 1.5e-05 | 1.400E-05 | 1.000E-06 | 1.300E-05 | 1.500E-05 |
| 5046 | 253 | N253E | A | 1.5e-05 | 1.2e-05 | 1.6e-05 | 1.433E-05 | 2.082E-06 | 1.200E-05 | 1.600E-05 |
| 5047 | 253 | N253Q | A | 1.2e-05 | 1.5e-05 | 1.6e-05 | 1.433E-05 | 2.082E-06 | 1.200E-05 | 1.600E-05 |
| 5048 | 253 | N253G | A | 1.5e-05 | 1.2e-05 | 1.5e-05 | 1.400E-05 | 1.732E-06 | 1.200E-05 | 1.500E-05 |
| 5049 | 253 | N253H | A | 1.6e-05 | 1.5e-05 | 9.9e-06 | 1.363E-05 | 3.272E-06 | 9.900E-06 | 1.600E-05 |
| 5050 | 253 | N253I | A | 1.1e-05 | 1.7e-05 | 2e-05 | 1.600E-05 | 4.583E-06 | 1.100E-05 | 2.000E-05 |
| 5051 | 253 | N253L | A | 1.3e-05 | 1.6e-05 | 9.3e-06 | 1.277E-05 | 3.356E-06 | 9.300E-06 | 1.600E-05 |
| 5052 | 253 | N253K | A | 1.6e-05 | 1.7e-05 | 1.4e-05 | 1.567E-05 | 1.528E-06 | 1.400E-05 | 1.700E-05 |
| 5053 | 253 | N253M | A | 7.2e-06 | 1.3e-05 | 1.7e-05 | 1.240E-05 | 4.927E-06 | 7.200E-06 | 1.700E-05 |
| 5054 | 253 | N253F | A | 1.1e-05 | 1.1e-05 | 1.5e-05 | 1.233E-05 | 2.309E-06 | 1.100E-05 | 1.500E-05 |
| 5055 | 253 | N253P | A | 1.6e-05 | 1.1e-05 | 1.1e-05 | 1.267E-05 | 2.887E-06 | 1.100E-05 | 1.600E-05 |
| 5056 | 253 | N253S | A | 1.7e-05 | 1.1e-05 | 1.1e-05 | 1.300E-05 | 3.464E-06 | 1.100E-05 | 1.700E-05 |
| 5057 | 253 | N253T | A | 1.3e-05 | 1.5e-05 | 1.6e-05 | 1.467E-05 | 1.528E-06 | 1.300E-05 | 1.600E-05 |
| 5058 | 253 | N253W | A | 1.6e-05 | 9.3e-06 | 1.5e-05 | 1.343E-05 | 3.614E-06 | 9.300E-06 | 1.600E-05 |
| 5059 | 253 | N253Y | A | 1.5e-05 | 1.2e-05 | 9.3e-06 | 1.210E-05 | 2.851E-06 | 9.300E-06 | 1.500E-05 |
| 5060 | 253 | N253V | A | 1.1e-05 | 1.3e-05 | 1.1e-05 | 1.167E-05 | 1.155E-06 | 1.100E-05 | 1.300E-05 |
| 5061 | 254 | T254A | A | 1.6e-05 | 1.7e-05 | 1.5e-05 | 1.600E-05 | 1.000E-06 | 1.500E-05 | 1.700E-05 |
| 5062 | 254 | T254R | A | 9.1e-06 | 7.6e-06 | 1.5e-05 | 1.057E-05 | 3.912E-06 | 7.600E-06 | 1.500E-05 |
| 5063 | 254 | T254N | A | 1.2e-05 | 8.8e-06 | 1e-05 | 1.027E-05 | 1.617E-06 | 8.800E-06 | 1.200E-05 |
| 5064 | 254 | T254D | A | 1.4e-05 | 1.6e-05 | 1.6e-05 | 1.533E-05 | 1.155E-06 | 1.400E-05 | 1.600E-05 |
| 5065 | 254 | T254C | A | 1.5e-05 | 1.1e-05 | 1.4e-05 | 1.333E-05 | 2.082E-06 | 1.100E-05 | 1.500E-05 |
| 5066 | 254 | T254E | A | 1.1e-05 | 1.8e-05 | 1.7e-05 | 1.533E-05 | 3.786E-06 | 1.100E-05 | 1.800E-05 |
| 5067 | 254 | T254Q | A | 1.6e-05 | 1.4e-05 | 1.3e-05 | 1.433E-05 | 1.528E-06 | 1.300E-05 | 1.600E-05 |
| 5068 | 254 | T254G | A | 1.4e-05 | 8.8e-06 | 1.6e-05 | 1.293E-05 | 3.717E-06 | 8.800E-06 | 1.600E-05 |
| 5069 | 254 | T254H | A | 1.4e-05 | 1.2e-05 | 9.6e-06 | 1.187E-05 | 2.203E-06 | 9.600E-06 | 1.400E-05 |
| 5070 | 254 | T254I | A | 1.3e-05 | 1.4e-05 | 1.7e-05 | 1.467E-05 | 2.082E-06 | 1.300E-05 | 1.700E-05 |
| 5071 | 254 | T254L | A | 1.3e-05 | 1.1e-05 | 1.2e-05 | 1.200E-05 | 1.000E-06 | 1.100E-05 | 1.300E-05 |
| 5072 | 254 | T254K | A | 1.1e-05 | 1.7e-05 | 1.6e-05 | 1.467E-05 | 3.215E-06 | 1.100E-05 | 1.700E-05 |
| 5073 | 254 | T254M | A | 1.5e-05 | 1.8e-05 | 1.7e-05 | 1.667E-05 | 1.528E-06 | 1.500E-05 | 1.800E-05 |
| 5074 | 254 | T254F | A | 1.3e-05 | 1.6e-05 | 1.3e-05 | 1.400E-05 | 1.732E-06 | 1.300E-05 | 1.600E-05 |
| 5075 | 254 | T254P | A | 1.6e-05 | 1.4e-05 | 1.3e-05 | 1.433E-05 | 1.528E-06 | 1.300E-05 | 1.600E-05 |
| 5076 | 254 | T254S | A | 1.3e-05 | 1.5e-05 | 1.4e-05 | 1.400E-05 | 1.000E-06 | 1.300E-05 | 1.500E-05 |
| 5077 | 254 | T254T | A | 1e-05 | 1e-05 | 1.6e-05 | 1.200E-05 | 3.464E-06 | 1.000E-05 | 1.600E-05 |
| 5078 | 254 | T254W | A | 1.3e-05 | 1.7e-05 | 1.5e-05 | 1.500E-05 | 2.000E-06 | 1.300E-05 | 1.700E-05 |
| 5079 | 254 | T254Y | A | 1.5e-05 | 1.6e-05 | 2.8e-05 | 1.967E-05 | 7.234E-06 | 1.500E-05 | 2.800E-05 |
| 5080 | 254 | T254V | A | 1.4e-05 | 1.2e-05 | 1.5e-05 | 1.367E-05 | 1.528E-06 | 1.200E-05 | 1.500E-05 |
| 5081 | 255 | D255A | A | 1.7e-05 | 1.4e-05 | 1.9e-05 | 1.667E-05 | 2.517E-06 | 1.400E-05 | 1.900E-05 |
| 5082 | 255 | D255R | A | 1.4e-05 | 9.1e-06 | 1.5e-05 | 1.270E-05 | 3.158E-06 | 9.100E-06 | 1.500E-05 |
| 5083 | 255 | D255N | A | 1.3e-05 | 1.4e-05 | 1.2e-05 | 1.300E-05 | 1.000E-06 | 1.200E-05 | 1.400E-05 |
| 5084 | 255 | D255D | A | 1.3e-05 | 1.2e-05 | 1.5e-05 | 1.333E-05 | 1.528E-06 | 1.200E-05 | 1.500E-05 |
| 5085 | 255 | D255C | A | 1e-05 | 1.2e-05 | 1.5e-05 | 1.233E-05 | 2.517E-06 | 1.000E-05 | 1.500E-05 |
| 5086 | 255 | D255E | A | 8.7e-06 | 1.4e-05 | 1.5e-05 | 1.257E-05 | 3.386E-06 | 8.700E-06 | 1.500E-05 |
| 5087 | 255 | D255Q | A | 8.5e-06 | 8.3e-06 | 1.3e-05 | 9.933E-06 | 2.658E-06 | 8.300E-06 | 1.300E-05 |
| 5088 | 255 | D255G | A | 1.7e-05 | 2e-05 | 1.7e-05 | 1.800E-05 | 1.732E-06 | 1.700E-05 | 2.000E-05 |
| 5089 | 255 | D255H | A | 1.2e-05 | 1e-05 | 1e-05 | 1.067E-05 | 1.155E-06 | 1.000E-05 | 1.200E-05 |
| 5090 | 255 | D255I | A | 1.8e-05 | 1.1e-05 | 1.3e-05 | 1.400E-05 | 3.606E-06 | 1.100E-05 | 1.800E-05 |
| 5091 | 255 | D255L | A | 3e-05 | 1.6e-05 | 1.7e-05 | 2.100E-05 | 7.810E-06 | 1.600E-05 | 3.000E-05 |
| 5092 | 255 | D255K | A | 9.8e-06 | 1.4e-05 | 9.8e-06 | 1.120E-05 | 2.425E-06 | 9.800E-06 | 1.400E-05 |
| 5093 | 255 | D255M | A | 1.3e-05 | 1.7e-05 | 2e-05 | 1.667E-05 | 3.512E-06 | 1.300E-05 | 2.000E-05 |
| 5094 | 255 | D255F | A | 1.8e-05 | 1.7e-05 | 9.7e-06 | 1.490E-05 | 4.531E-06 | 9.700E-06 | 1.800E-05 |
| 5095 | 255 | D255P | A | 1.1e-05 | 1.7e-05 | 1.6e-05 | 1.467E-05 | 3.215E-06 | 1.100E-05 | 1.700E-05 |
| 5096 | 255 | D255S | A | 1.3e-05 | 7.7e-06 | 1.1e-05 | 1.057E-05 | 2.676E-06 | 7.700E-06 | 1.300E-05 |
| 5097 | 255 | D255T | A | 1.2e-05 | 9.2e-06 | 1.2e-05 | 1.107E-05 | 1.617E-06 | 9.200E-06 | 1.200E-05 |
| 5098 | 255 | D255W | A | 1.7e-05 | 1.7e-05 | 1.3e-05 | 1.567E-05 | 2.309E-06 | 1.300E-05 | 1.700E-05 |
| 5099 | 255 | D255Y | A | 1.3e-05 | 1.3e-05 | 1.1e-05 | 1.233E-05 | 1.155E-06 | 1.100E-05 | 1.300E-05 |

| | | | | | | | | | | |
|---|---|---|---|---|---|---|---|---|---|---|
| 5100 | 255 | D255V | A | 1.8e-05 | 1.5e-05 | 1.7e-05 | 1.667E-05 | 1.528E-06 | 1.500E-05 | 1.800E-05 |
| 5101 | 256 | T256A | A | 2.2e-05 | 1.6e-05 | 2e-05 | 1.933E-05 | 3.055E-06 | 1.600E-05 | 2.200E-05 |
| 5102 | 256 | T256R | A | 1.8e-05 | 2.1e-05 | 1.8e-05 | 1.900E-05 | 1.732E-06 | 1.800E-05 | 2.100E-05 |
| 5103 | 256 | T256N | A | 1.1e-05 | 1.6e-05 | 1.3e-05 | 1.333E-05 | 2.517E-06 | 1.100E-05 | 1.600E-05 |
| 5104 | 256 | T256D | A | 1.7e-05 | 1.7e-05 | 1.9e-05 | 1.767E-05 | 1.155E-06 | 1.700E-05 | 1.900E-05 |
| 5105 | 256 | T256C | A | 1.5e-05 | 1.5e-05 | 2.2e-05 | 1.733E-05 | 4.041E-06 | 1.500E-05 | 2.200E-05 |
| 5106 | 256 | T256E | A | 1.2e-05 | 1.6e-05 | 1.9e-05 | 1.567E-05 | 3.512E-06 | 1.200E-05 | 1.900E-05 |
| 5107 | 256 | T256Q | A | 1.1e-05 | 1.7e-05 | 3.1e-05 | 1.967E-05 | 1.026E-05 | 1.100E-05 | 3.100E-05 |
| 5108 | 256 | T256G | A | 2.1e-05 | 2.1e-05 | 1.9e-05 | 2.033E-05 | 1.155E-06 | 1.900E-05 | 2.100E-05 |
| 5109 | 256 | T256H | A | 1.5e-05 | 1.1e-05 | 1.2e-05 | 1.267E-05 | 2.082E-06 | 1.100E-05 | 1.500E-05 |
| 5110 | 256 | T256I | A | 9.6e-06 | 3.1e-05 | 1.3e-05 | 1.787E-05 | 1.150E-05 | 9.600E-06 | 3.100E-05 |
| 5111 | 256 | T256L | A | 1.4e-05 | 2.1e-05 | 2.3e-05 | 1.933E-05 | 4.726E-06 | 1.400E-05 | 2.300E-05 |
| 5112 | 256 | T256K | A | 1.8e-05 | 1.3e-05 | 2.1e-05 | 1.733E-05 | 4.041E-06 | 1.300E-05 | 2.100E-05 |
| 5113 | 256 | T256M | A | 2.4e-05 | 2.1e-05 | 2.2e-05 | 2.233E-05 | 1.528E-06 | 2.100E-05 | 2.400E-05 |
| 5114 | 256 | T256F | A | 1.6e-05 | 2.3e-05 | 1.4e-05 | 1.767E-05 | 4.726E-06 | 1.400E-05 | 2.300E-05 |
| 5115 | 256 | T256P | A | 2.1e-05 | 2.4e-05 | 1.2e-05 | 1.900E-05 | 6.245E-06 | 1.200E-05 | 2.400E-05 |
| 5116 | 256 | T256S | A | 1.6e-05 | 1.1e-05 | 9.8e-06 | 1.227E-05 | 3.288E-06 | 9.800E-06 | 1.600E-05 |
| 5117 | 256 | T256T | A | 1.2e-05 | 1.5e-05 | 1.6e-05 | 1.433E-05 | 2.082E-06 | 1.200E-05 | 1.600E-05 |
| 5118 | 256 | T256W | A | 1.2e-05 | 1.7e-05 | 1.5e-05 | 1.467E-05 | 2.517E-06 | 1.200E-05 | 1.700E-05 |
| 5119 | 256 | T256Y | A | 1.2e-05 | 1.4e-05 | 5.6e-05 | 2.733E-05 | 2.485E-05 | 1.200E-05 | 5.600E-05 |
| 5120 | 256 | T256V | A | 1.3e-05 | 1.3e-05 | 1.4e-05 | 1.333E-05 | 5.774E-07 | 1.300E-05 | 1.400E-05 |
| 5121 | 257 | F257A | A | 1.4e-05 | 1.1e-05 | 1.5e-05 | 1.333E-05 | 2.082E-06 | 1.100E-05 | 1.500E-05 |
| 5122 | 257 | F257R | A | 1.1e-05 | 9.3e-06 | 7.7e-06 | 9.333E-06 | 1.650E-06 | 7.700E-06 | 1.100E-05 |
| 5123 | 257 | F257N | A | 9.1e-06 | 6.2e-06 | 7.6e-06 | 7.633E-06 | 1.450E-06 | 6.200E-06 | 9.100E-06 |
| 5124 | 257 | F257D | A | 1.3e-05 | 8.2e-06 | 1.1e-05 | 1.073E-05 | 2.411E-06 | 8.200E-06 | 1.300E-05 |
| 5125 | 257 | F257C | A | 1.1e-05 | 1.4e-05 | 1.1e-05 | 1.200E-05 | 1.732E-06 | 1.100E-05 | 1.400E-05 |
| 5126 | 257 | F257E | A | 1.1e-05 | 9.6e-06 | 8.6e-06 | 9.733E-06 | 1.206E-06 | 8.600E-06 | 1.100E-05 |
| 5127 | 257 | F257Q | A | 7.8e-06 | 5.8e-06 | 8.1e-06 | 7.233E-06 | 1.250E-06 | 5.800E-06 | 8.100E-06 |
| 5128 | 257 | F257G | A | 1.6e-05 | 1.7e-05 | 1.3e-05 | 1.533E-05 | 2.082E-06 | 1.300E-05 | 1.700E-05 |
| 5129 | 257 | F257H | A | 1e-05 | 8.1e-06 | 9.2e-06 | 9.100E-06 | 9.539E-07 | 8.100E-06 | 1.000E-05 |
| 5130 | 257 | F257I | A | 8.9e-06 | 1.2e-05 | 7.9e-06 | 9.600E-06 | 2.138E-06 | 7.900E-06 | 1.200E-05 |
| 5131 | 257 | F257L | A | 1.4e-05 | 1.5e-05 | 1.1e-05 | 1.333E-05 | 2.082E-06 | 1.100E-05 | 1.500E-05 |
| 5132 | 257 | F257K | A | 9.4e-06 | 8.9e-06 | 9e-06 | 9.100E-06 | 2.646E-07 | 8.900E-06 | 9.400E-06 |
| 5133 | 257 | F257M | A | 9.3e-06 | 9.7e-06 | 1.6e-05 | 1.167E-05 | 3.758E-06 | 9.300E-06 | 1.600E-05 |
| 5134 | 257 | F257F | A | 1.3e-05 | 1.5e-05 | 1.1e-05 | 1.300E-05 | 2.000E-06 | 1.100E-05 | 1.500E-05 |
| 5135 | 257 | F257P | A | 1.3e-05 | 1.4e-05 | 8.8e-06 | 1.193E-05 | 2.759E-06 | 8.800E-06 | 1.400E-05 |
| 5136 | 257 | F257S | A | 1.1e-05 | 7.5e-06 | 8.7e-06 | 9.067E-06 | 1.779E-06 | 7.500E-06 | 1.100E-05 |
| 5137 | 257 | F257T | A | 1e-05 | 7e-06 | 7.7e-06 | 8.233E-06 | 1.570E-06 | 7.000E-06 | 1.000E-05 |
| 5138 | 257 | F257W | A | 1.5e-05 | 1.1e-05 | 1.6e-05 | 1.400E-05 | 2.646E-06 | 1.100E-05 | 1.600E-05 |
| 5139 | 257 | F257Y | A | 9.3e-06 | 1.2e-05 | 1.1e-05 | 1.077E-05 | 1.365E-06 | 9.300E-06 | 1.200E-05 |
| 5140 | 257 | F257V | A | 1.1e-05 | 1.2e-05 | 8.8e-06 | 1.060E-05 | 1.637E-06 | 8.800E-06 | 1.200E-05 |
| 5141 | 258 | E258A | A | 1.7e-05 | 1.4e-05 | 1.1e-05 | 1.400E-05 | 3.000E-06 | 1.100E-05 | 1.700E-05 |
| 5142 | 258 | E258R | A | 1.3e-05 | 1.5e-05 | 1.5e-05 | 1.433E-05 | 1.155E-06 | 1.300E-05 | 1.500E-05 |
| 5143 | 258 | E258N | A | 9.5e-06 | 1.1e-05 | 9.2e-06 | 9.900E-06 | 9.644E-07 | 9.200E-06 | 1.100E-05 |
| 5144 | 258 | E258D | A | 1.6e-05 | 1.1e-05 | 1.5e-05 | 1.400E-05 | 2.646E-06 | 1.100E-05 | 1.600E-05 |
| 5145 | 258 | E258C | A | 1.2e-05 | 1e-05 | 1.5e-05 | 1.233E-05 | 2.517E-06 | 1.000E-05 | 1.500E-05 |
| 5146 | 258 | E258E | A | 8.9e-06 | 1.8e-05 | 1.1e-05 | 1.263E-05 | 4.765E-06 | 8.900E-06 | 1.800E-05 |
| 5147 | 258 | E258Q | A | 1.6e-05 | 1.4e-05 | 1.6e-05 | 1.533E-05 | 1.155E-06 | 1.400E-05 | 1.600E-05 |
| 5148 | 258 | E258G | A | 1.8e-05 | 1e-05 | 1e-05 | 1.267E-05 | 4.619E-06 | 1.000E-05 | 1.800E-05 |
| 5149 | 258 | E258H | A | 1.1e-05 | 1.1e-05 | 8.1e-06 | 1.003E-05 | 1.674E-06 | 8.100E-06 | 1.100E-05 |
| 5150 | 258 | E258I | A | 1.1e-05 | 1.1e-05 | 1.1e-05 | 1.100E-05 | 2.075E-21 | 1.100E-05 | 1.100E-05 |
| 5151 | 258 | E258L | A | 1.7e-05 | 8.2e-06 | 1.4e-05 | 1.307E-05 | 4.474E-06 | 8.200E-06 | 1.700E-05 |
| 5152 | 258 | E258K | A | 1.7e-05 | 1.6e-05 | 1.7e-05 | 1.667E-05 | 5.774E-07 | 1.600E-05 | 1.700E-05 |
| 5153 | 258 | E258M | A | 1.4e-05 | 1.6e-05 | 1.5e-05 | 1.500E-05 | 1.000E-06 | 1.400E-05 | 1.600E-05 |
| 5154 | 258 | E258F | A | 8.4e-06 | 9.4e-06 | 1.1e-05 | 9.600E-06 | 1.311E-06 | 8.400E-06 | 1.100E-05 |
| 5155 | 258 | E258P | A | 1.7e-05 | 1.1e-05 | 1.4e-05 | 1.400E-05 | 3.000E-06 | 1.100E-05 | 1.700E-05 |
| 5156 | 258 | E258S | A | 1.4e-05 | 1e-05 | 1.4e-05 | 1.267E-05 | 2.309E-06 | 1.000E-05 | 1.400E-05 |
| 5157 | 258 | E258T | A | 9.5e-06 | 1.3e-05 | 1.5e-05 | 1.250E-05 | 2.784E-06 | 9.500E-06 | 1.500E-05 |
| 5158 | 258 | E258W | A | 1.1e-05 | 1.3e-05 | 8.9e-06 | 1.097E-05 | 2.050E-06 | 8.900E-06 | 1.300E-05 |
| 5159 | 258 | E258Y | A | 1.5e-05 | 1e-05 | 1.4e-05 | 1.300E-05 | 2.646E-06 | 1.000E-05 | 1.500E-05 |
| 5160 | 258 | E258V | A | 1.5e-05 | 8.7e-06 | 1.6e-05 | 1.323E-05 | 3.958E-06 | 8.700E-06 | 1.600E-05 |
| 5161 | 259 | S259A | A | 1.8e-05 | 1.4e-05 | 1e-05 | 1.400E-05 | 4.000E-06 | 1.000E-05 | 1.800E-05 |
| 5162 | 259 | S259R | A | 1.2e-05 | 1.6e-05 | 1e-05 | 1.267E-05 | 3.055E-06 | 1.000E-05 | 1.600E-05 |
| 5163 | 259 | S259N | A | 1.5e-05 | 1.6e-05 | 1.1e-05 | 1.400E-05 | 2.646E-06 | 1.100E-05 | 1.600E-05 |
| 5164 | 259 | S259D | A | 1.5e-05 | 1.1e-05 | 1.4e-05 | 1.333E-05 | 2.082E-06 | 1.100E-05 | 1.500E-05 |
| 5165 | 259 | S259C | A | 1e-05 | 9.5e-06 | 1.1e-05 | 1.017E-05 | 7.638E-07 | 9.500E-06 | 1.100E-05 |
| 5166 | 259 | S259E | A | 1.8e-05 | 1.8e-05 | 1.2e-05 | 1.600E-05 | 3.464E-06 | 1.200E-05 | 1.800E-05 |
| 5167 | 259 | S259Q | A | 1e-05 | 1.1e-05 | 1.4e-05 | 1.167E-05 | 2.082E-06 | 1.000E-05 | 1.400E-05 |
| 5168 | 259 | S259G | A | 1.2e-05 | 1.6e-05 | 1.5e-05 | 1.433E-05 | 2.082E-06 | 1.200E-05 | 1.600E-05 |
| 5169 | 259 | S259H | A | 1.6e-05 | 9.9e-06 | 1.6e-05 | 1.397E-05 | 3.522E-06 | 9.900E-06 | 1.600E-05 |
| 5170 | 259 | S259I | A | 1e-05 | 1.5e-05 | 8.8e-06 | 1.127E-05 | 3.288E-06 | 8.800E-06 | 1.500E-05 |
| 5171 | 259 | S259L | A | 9.7e-06 | 1.3e-05 | 1.2e-05 | 1.157E-05 | 1.692E-06 | 9.700E-06 | 1.300E-05 |
| 5172 | 259 | S259K | A | 1e-05 | 6.7e-06 | 1.5e-05 | 1.057E-05 | 4.179E-06 | 6.700E-06 | 1.500E-05 |
| 5173 | 259 | S259M | A | 1.5e-05 | 1.3e-05 | 1.5e-05 | 1.433E-05 | 1.155E-06 | 1.300E-05 | 1.500E-05 |
| 5174 | 259 | S259F | A | 1.7e-05 | 1.3e-05 | 9.9e-06 | 1.330E-05 | 3.559E-06 | 9.900E-06 | 1.700E-05 |
| 5175 | 259 | S259P | A | 1.6e-05 | 1.2e-05 | 9.1e-06 | 1.237E-05 | 3.465E-06 | 9.100E-06 | 1.600E-05 |
| 5176 | 259 | S259S | A | 1.5e-05 | 1e-05 | 1.6e-05 | 1.367E-05 | 3.215E-06 | 1.000E-05 | 1.600E-05 |
| 5177 | 259 | S259T | A | 1.2e-05 | 1.1e-05 | 1.1e-05 | 1.133E-05 | 5.774E-07 | 1.100E-05 | 1.200E-05 |
| 5178 | 259 | S259W | A | 1.5e-05 | 1.1e-05 | 1.4e-05 | 1.333E-05 | 2.082E-06 | 1.100E-05 | 1.500E-05 |
| 5179 | 259 | S259Y | A | 8.3e-06 | 1.5e-05 | 1.7e-05 | 1.343E-05 | 4.557E-06 | 8.300E-06 | 1.700E-05 |
| 5180 | 259 | S259V | A | 1.7e-05 | 1.5e-05 | 1.4e-05 | 1.533E-05 | 1.528E-06 | 1.400E-05 | 1.700E-05 |
| 5181 | 260 | M260A | A | 9.3e-06 | 9.5e-06 | 1.1e-05 | 9.933E-06 | 9.292E-07 | 9.300E-06 | 1.100E-05 |
| 5182 | 260 | M260R | A | 1.3e-05 | 8.1e-06 | 1.5e-05 | 1.203E-05 | 3.550E-06 | 8.100E-06 | 1.500E-05 |
| 5183 | 260 | M260N | A | 1.4e-05 | 1.4e-05 | 9.5e-06 | 1.250E-05 | 2.598E-06 | 9.500E-06 | 1.400E-05 |
| 5184 | 260 | M260D | A | 1.5e-05 | 1.1e-05 | 8.5e-06 | 1.150E-05 | 3.279E-06 | 8.500E-06 | 1.500E-05 |

| | | | | | | | | | | |
|---|---|---|---|---|---|---|---|---|---|---|
| 5185 | 260 | M260C | A | 1.7e-05 | 9e-06 | 9.4e-06 | 1.180E-05 | 4.508E-06 | 9.000E-06 | 1.700E-05 |
| 5186 | 260 | M260E | A | 9.7e-06 | 1.1e-05 | 1e-05 | 1.023E-05 | 6.807E-07 | 9.700E-06 | 1.100E-05 |
| 5187 | 260 | M260Q | A | 1.1e-05 | 1.1e-05 | 1e-05 | 1.067E-05 | 5.774E-07 | 1.000E-05 | 1.100E-05 |
| 5188 | 260 | M260G | A | 1.5e-05 | 1.4e-05 | 1.4e-05 | 1.433E-05 | 5.774E-07 | 1.400E-05 | 1.500E-05 |
| 5189 | 260 | M260H | A | 1.4e-05 | 1e-05 | 1.3e-05 | 1.233E-05 | 2.082E-06 | 1.000E-05 | 1.400E-05 |
| 5190 | 260 | M260I | A | 1.3e-05 | 9e-06 | 1.6e-05 | 1.267E-05 | 3.512E-06 | 9.000E-06 | 1.600E-05 |
| 5191 | 260 | M260L | A | 1.4e-05 | 1.5e-05 | 6.8e-06 | 1.193E-05 | 4.474E-06 | 6.800E-06 | 1.500E-05 |
| 5192 | 260 | M260K | A | 1.1e-05 | 1.6e-05 | 8.5e-06 | 1.183E-05 | 3.819E-06 | 8.500E-06 | 1.600E-05 |
| 5193 | 260 | M260M | A | 1.1e-05 | 1.3e-05 | 7e-06 | 1.033E-05 | 3.055E-06 | 7.000E-06 | 1.300E-05 |
| 5194 | 260 | M260F | A | 1.4e-05 | 1e-05 | 1.3e-05 | 1.233E-05 | 2.082E-06 | 1.000E-05 | 1.400E-05 |
| 5195 | 260 | M260P | A | 8.6e-06 | 7.3e-06 | 1.7e-05 | 1.097E-05 | 5.265E-06 | 7.300E-06 | 1.700E-05 |
| 5196 | 260 | M260S | A | 9.3e-06 | 1.6e-05 | 1.2e-05 | 1.243E-05 | 3.371E-06 | 9.300E-06 | 1.600E-05 |
| 5197 | 260 | M260T | A | 1.6e-05 | 1.1e-05 | 1e-05 | 1.233E-05 | 3.215E-06 | 1.000E-05 | 1.600E-05 |
| 5198 | 260 | M260W | A | 1.5e-05 | 9.1e-06 | 1.4e-05 | 1.270E-05 | 3.158E-06 | 9.100E-06 | 1.500E-05 |
| 5199 | 260 | M260Y | A | 8.3e-06 | 1.1e-05 | 1.5e-05 | 1.143E-05 | 3.371E-06 | 8.300E-06 | 1.500E-05 |
| 5200 | 260 | M260V | A | 1.5e-05 | 1.6e-05 | 1.6e-05 | 1.567E-05 | 5.774E-07 | 1.500E-05 | 1.600E-05 |
| 5201 | 261 | P261A | A | 1.7e-05 | 9.6e-06 | 1.2e-05 | 1.287E-05 | 3.775E-06 | 9.600E-06 | 1.700E-05 |
| 5202 | 261 | P261R | A | 8.1e-06 | 9.4e-06 | 1.2e-05 | 9.833E-06 | 1.986E-06 | 8.100E-06 | 1.200E-05 |
| 5203 | 261 | P261N | A | 9.7e-06 | 2.6e-05 | 9.7e-06 | 1.513E-05 | 9.411E-06 | 9.700E-06 | 2.600E-05 |
| 5204 | 261 | P261D | A | 9.3e-06 | 1.1e-05 | 1.5e-05 | 1.177E-05 | 2.926E-06 | 9.300E-06 | 1.500E-05 |
| 5205 | 261 | P261C | A | 9.4e-06 | 1.3e-05 | 1.6e-05 | 1.280E-05 | 3.305E-06 | 9.400E-06 | 1.600E-05 |
| 5206 | 261 | P261E | A | 1.1e-05 | 8.6e-06 | 1.5e-05 | 1.153E-05 | 3.233E-06 | 8.600E-06 | 1.500E-05 |
| 5207 | 261 | P261Q | A | 9.6e-06 | 1.5e-05 | 1.4e-05 | 1.287E-05 | 2.873E-06 | 9.600E-06 | 1.500E-05 |
| 5208 | 261 | P261G | A | 1.5e-05 | 9.1e-06 | 9.1e-06 | 1.107E-05 | 3.406E-06 | 9.100E-06 | 1.500E-05 |
| 5209 | 261 | P261H | A | 7.6e-06 | 1e-05 | 1.5e-05 | 1.087E-05 | 3.775E-06 | 7.600E-06 | 1.500E-05 |
| 5210 | 261 | P261I | A | 7.4e-06 | 1.6e-05 | 1.3e-05 | 1.213E-05 | 4.365E-06 | 7.400E-06 | 1.600E-05 |
| 5211 | 261 | P261L | A | 1.4e-05 | 1.5e-05 | 1e-05 | 1.300E-05 | 2.646E-06 | 1.000E-05 | 1.500E-05 |
| 5212 | 261 | P261K | A | 1e-05 | 1.7e-05 | 1.2e-05 | 1.300E-05 | 3.606E-06 | 1.000E-05 | 1.700E-05 |
| 5213 | 261 | P261M | A | 9.9e-06 | 1.1e-05 | 1.2e-05 | 1.097E-05 | 1.050E-06 | 9.900E-06 | 1.200E-05 |
| 5214 | 261 | P261F | A | 9.6e-06 | 1.4e-05 | 1.2e-05 | 1.187E-05 | 2.203E-06 | 9.600E-06 | 1.400E-05 |
| 5215 | 261 | P261P | A | 1.5e-05 | 6.1e-06 | 1.6e-05 | 1.237E-05 | 5.450E-06 | 6.100E-06 | 1.600E-05 |
| 5216 | 261 | P261S | A | 1.3e-05 | 1.4e-05 | 1.1e-05 | 1.267E-05 | 1.528E-06 | 1.100E-05 | 1.400E-05 |
| 5217 | 261 | P261T | A | 9.9e-06 | 1.1e-05 | 1.4e-05 | 1.163E-05 | 2.122E-06 | 9.900E-06 | 1.400E-05 |
| 5218 | 261 | P261W | A | 1e-05 | 1.5e-05 | 1.5e-05 | 1.333E-05 | 2.887E-06 | 1.000E-05 | 1.500E-05 |
| 5219 | 261 | P261Y | A | 9.9e-06 | 9.8e-06 | 7.3e-06 | 9.000E-06 | 1.473E-06 | 7.300E-06 | 9.900E-06 |
| 5220 | 261 | P261V | A | 1.2e-05 | 1.5e-05 | 1.4e-05 | 1.367E-05 | 1.528E-06 | 1.200E-05 | 1.500E-05 |
| 5221 | 262 | N262A | A | 1.2e-05 | 1.1e-05 | 1.5e-05 | 1.267E-05 | 2.082E-06 | 1.100E-05 | 1.500E-05 |
| 5222 | 262 | N262R | A | 1.2e-05 | 1.6e-05 | 1.4e-05 | 1.400E-05 | 2.000E-06 | 1.200E-05 | 1.600E-05 |
| 5223 | 262 | N262N | A | 1.7e-05 | 1.5e-05 | 9.1e-06 | 1.370E-05 | 4.107E-06 | 9.100E-06 | 1.700E-05 |
| 5224 | 262 | N262D | A | 1.6e-05 | 1.4e-05 | 1.6e-05 | 1.533E-05 | 1.155E-06 | 1.400E-05 | 1.600E-05 |
| 5225 | 262 | N262C | A | 1.1e-05 | 1.4e-05 | 1.5e-05 | 1.333E-05 | 2.082E-06 | 1.100E-05 | 1.500E-05 |
| 5226 | 262 | N262E | A | 1.1e-05 | 1.1e-05 | 9.7e-06 | 1.057E-05 | 7.506E-07 | 9.700E-06 | 1.100E-05 |
| 5227 | 262 | N262Q | A | 1.1e-05 | 1.5e-05 | 9.3e-06 | 1.177E-05 | 2.926E-06 | 9.300E-06 | 1.500E-05 |
| 5228 | 262 | N262G | A | 8.8e-06 | 1.4e-05 | 1.5e-05 | 1.260E-05 | 3.329E-06 | 8.800E-06 | 1.500E-05 |
| 5229 | 262 | N262H | A | 1.3e-05 | 9.2e-06 | 1.2e-05 | 1.140E-05 | 1.970E-06 | 9.200E-06 | 1.300E-05 |
| 5230 | 262 | N262I | A | 1.5e-05 | 1.1e-05 | 1.6e-05 | 1.400E-05 | 2.646E-06 | 1.100E-05 | 1.600E-05 |
| 5231 | 262 | N262L | A | 1.4e-05 | 1.4e-05 | 1.5e-05 | 1.433E-05 | 5.774E-07 | 1.400E-05 | 1.500E-05 |
| 5232 | 262 | N262K | A | 8.9e-06 | 1e-05 | 1.4e-05 | 1.097E-05 | 2.684E-06 | 8.900E-06 | 1.400E-05 |
| 5233 | 262 | N262M | A | 1e-05 | 1.6e-05 | 1e-05 | 1.200E-05 | 3.464E-06 | 1.000E-05 | 1.600E-05 |
| 5234 | 262 | N262F | A | 1.5e-05 | 9.7e-06 | 1.3e-05 | 1.257E-05 | 2.676E-06 | 9.700E-06 | 1.500E-05 |
| 5235 | 262 | N262P | A | 2.1e-05 | 1e-05 | 1.7e-05 | 1.600E-05 | 5.568E-06 | 1.000E-05 | 2.100E-05 |
| 5236 | 262 | N262S | A | 1.7e-05 | 9.9e-06 | 9.9e-06 | 1.227E-05 | 4.099E-06 | 9.900E-06 | 1.700E-05 |
| 5237 | 262 | N262T | A | 1.1e-05 | 1.2e-05 | 1.4e-05 | 1.233E-05 | 1.528E-06 | 1.100E-05 | 1.400E-05 |
| 5238 | 262 | N262W | A | 1.3e-05 | 9.1e-06 | 1.4e-05 | 1.203E-05 | 2.589E-06 | 9.100E-06 | 1.400E-05 |
| 5239 | 262 | N262Y | A | 1e-05 | 1.4e-05 | 9.7e-06 | 1.123E-05 | 2.401E-06 | 9.700E-06 | 1.400E-05 |
| 5240 | 262 | N262V | A | 1.5e-05 | 1.5e-05 | 1.5e-05 | 1.500E-05 | 0.000E+00 | 1.500E-05 | 1.500E-05 |
| 5241 | 263 | P263A | A | 1.4e-05 | 1.4e-05 | 1.6e-05 | 1.467E-05 | 1.155E-06 | 1.400E-05 | 1.600E-05 |
| 5242 | 263 | P263R | A | 1.5e-05 | 1e-05 | 1e-05 | 1.167E-05 | 2.887E-06 | 1.000E-05 | 1.500E-05 |
| 5243 | 263 | P263N | A | 1.5e-05 | 9.5e-06 | 1.6e-05 | 1.350E-05 | 3.500E-06 | 9.500E-06 | 1.600E-05 |
| 5244 | 263 | P263D | A | 1.3e-05 | 1.5e-05 | 9e-06 | 1.233E-05 | 3.055E-06 | 9.000E-06 | 1.500E-05 |
| 5245 | 263 | P263C | A | 9.5e-06 | 1.1e-05 | 1.5e-05 | 1.183E-05 | 2.843E-06 | 9.500E-06 | 1.500E-05 |
| 5246 | 263 | P263E | A | 1.5e-05 | 1.5e-05 | 1e-05 | 1.333E-05 | 2.887E-06 | 1.000E-05 | 1.500E-05 |
| 5247 | 263 | P263Q | A | 1.5e-05 | 1e-05 | 1.9e-05 | 1.467E-05 | 4.509E-06 | 1.000E-05 | 1.900E-05 |
| 5248 | 263 | P263G | A | 1.5e-05 | 8.7e-06 | 1.2e-05 | 1.190E-05 | 3.151E-06 | 8.700E-06 | 1.500E-05 |
| 5249 | 263 | P263H | A | 1.1e-05 | 1.5e-05 | 1.4e-05 | 1.333E-05 | 2.082E-06 | 1.100E-05 | 1.500E-05 |
| 5250 | 263 | P263I | A | 1.6e-05 | 1.6e-05 | 1.5e-05 | 1.567E-05 | 5.774E-07 | 1.500E-05 | 1.600E-05 |
| 5251 | 263 | P263L | A | 1e-05 | 7.5e-06 | 1.3e-05 | 1.017E-05 | 2.754E-06 | 7.500E-06 | 1.300E-05 |
| 5252 | 263 | P263K | A | 8.2e-06 | 9.5e-06 | 1.3e-05 | 1.023E-05 | 2.483E-06 | 8.200E-06 | 1.300E-05 |
| 5253 | 263 | P263M | A | 1.4e-05 | 1.5e-05 | 1.7e-05 | 1.533E-05 | 1.528E-06 | 1.400E-05 | 1.700E-05 |
| 5254 | 263 | P263F | A | 1.6e-05 | 1.5e-05 | 1e-05 | 1.367E-05 | 3.215E-06 | 1.000E-05 | 1.600E-05 |
| 5255 | 263 | P263P | A | 1.4e-05 | 8.8e-06 | 1.1e-05 | 1.127E-05 | 2.610E-06 | 8.800E-06 | 1.400E-05 |
| 5256 | 263 | P263S | A | 9.7e-06 | 1.4e-05 | 1.5e-05 | 1.290E-05 | 2.816E-06 | 9.700E-06 | 1.500E-05 |
| 5257 | 263 | P263T | A | 1.1e-05 | 1.1e-05 | 9.7e-06 | 1.057E-05 | 7.506E-07 | 9.700E-06 | 1.100E-05 |
| 5258 | 263 | P263W | A | 1.6e-05 | 1.3e-05 | 1.4e-05 | 1.433E-05 | 1.528E-06 | 1.300E-05 | 1.600E-05 |
| 5259 | 263 | P263Y | A | 1.4e-05 | 1.1e-05 | 1.6e-05 | 1.367E-05 | 2.517E-06 | 1.100E-05 | 1.600E-05 |
| 5260 | 263 | P263V | A | 1.6e-05 | 1.7e-05 | 8.9e-06 | 1.397E-05 | 4.416E-06 | 8.900E-06 | 1.700E-05 |
| 5261 | 264 | E264A | A | 2e-05 | 9.6e-06 | 1.5e-05 | 1.487E-05 | 5.201E-06 | 9.600E-06 | 2.000E-05 |
| 5262 | 264 | E264R | A | 9.5e-06 | 1.5e-05 | 1.1e-05 | 1.183E-05 | 2.843E-06 | 9.500E-06 | 1.500E-05 |
| 5263 | 264 | E264N | A | 1.3e-05 | 1.4e-05 | 1e-05 | 1.233E-05 | 2.082E-06 | 1.000E-05 | 1.400E-05 |
| 5264 | 264 | E264D | A | 1.4e-05 | 9e-06 | 1.2e-05 | 1.167E-05 | 2.517E-06 | 9.000E-06 | 1.400E-05 |
| 5265 | 264 | E264C | A | 1.6e-05 | 1.3e-05 | 1.1e-05 | 1.333E-05 | 2.517E-06 | 1.100E-05 | 1.600E-05 |
| 5266 | 264 | E264E | A | 1.2e-05 | 1.5e-05 | 1.5e-05 | 1.400E-05 | 1.732E-06 | 1.200E-05 | 1.500E-05 |
| 5267 | 264 | E264Q | A | 1.3e-05 | 1e-05 | 1e-05 | 1.100E-05 | 1.732E-06 | 1.000E-05 | 1.300E-05 |
| 5268 | 264 | E264G | A | 1.1e-05 | 1.1e-05 | 1.6e-05 | 1.267E-05 | 2.887E-06 | 1.100E-05 | 1.600E-05 |
| 5269 | 264 | E264H | A | 1.4e-05 | 1e-05 | 1.2e-05 | 1.200E-05 | 2.000E-06 | 1.000E-05 | 1.400E-05 |

| | | | | | | | | | | |
|---|---|---|---|---|---|---|---|---|---|---|
| 5270 | 264 | E264I | A | 1e-05 | 1.5e-05 | 9.8e-06 | 1.160E-05 | 2.946E-06 | 9.800E-06 | 1.500E-05 |
| 5271 | 264 | E264L | A | 1.4e-05 | 1.1e-05 | 1.6e-05 | 1.367E-05 | 2.517E-06 | 1.100E-05 | 1.600E-05 |
| 5272 | 264 | E264K | A | 1e-05 | 1.4e-05 | 1.1e-05 | 1.167E-05 | 2.082E-06 | 1.000E-05 | 1.400E-05 |
| 5273 | 264 | E264M | A | 1.1e-05 | 8.1e-06 | 1e-05 | 9.700E-06 | 1.473E-06 | 8.100E-06 | 1.100E-05 |
| 5274 | 264 | E264F | A | 1.1e-05 | 1.5e-05 | 9.8e-06 | 1.193E-05 | 2.723E-06 | 9.800E-06 | 1.500E-05 |
| 5275 | 264 | E264P | A | 1.1e-05 | 1.6e-05 | 1.3e-05 | 1.333E-05 | 2.517E-06 | 1.100E-05 | 1.600E-05 |
| 5276 | 264 | E264S | A | 1.6e-05 | 1.3e-05 | 1.2e-05 | 1.367E-05 | 2.082E-06 | 1.200E-05 | 1.600E-05 |
| 5277 | 264 | E264T | A | 1.5e-05 | 1.6e-05 | 1e-05 | 1.367E-05 | 3.215E-06 | 1.000E-05 | 1.600E-05 |
| 5278 | 264 | E264W | A | 1.4e-05 | 6.3e-06 | 1e-05 | 1.010E-05 | 3.851E-06 | 6.300E-06 | 1.400E-05 |
| 5279 | 264 | E264Y | A | 1.2e-05 | 1.5e-05 | 1.4e-05 | 1.367E-05 | 1.528E-06 | 1.200E-05 | 1.500E-05 |
| 5280 | 264 | E264V | A | 1.5e-05 | 1.6e-05 | 1.5e-05 | 1.533E-05 | 5.774E-07 | 1.500E-05 | 1.600E-05 |
| 5281 | 265 | G265A | A | 9.1e-06 | 1.5e-05 | 1.2e-05 | 1.203E-05 | 2.950E-06 | 9.100E-06 | 1.500E-05 |
| 5282 | 265 | G265R | A | 1e-05 | 1.1e-05 | 1.3e-05 | 1.133E-05 | 1.528E-06 | 1.000E-05 | 1.300E-05 |
| 5283 | 265 | G265N | A | 1.4e-05 | 1.4e-05 | 8.3e-06 | 1.210E-05 | 3.291E-06 | 8.300E-06 | 1.400E-05 |
| 5284 | 265 | G265D | A | 1.1e-05 | 1.4e-05 | 1.6e-05 | 1.367E-05 | 2.517E-06 | 1.100E-05 | 1.600E-05 |
| 5285 | 265 | G265C | A | 1.4e-05 | 1.2e-05 | 8.7e-06 | 1.157E-05 | 2.676E-06 | 8.700E-06 | 1.400E-05 |
| 5286 | 265 | G265E | A | 9.6e-06 | 1e-05 | 1.4e-05 | 1.120E-05 | 2.433E-06 | 9.600E-06 | 1.400E-05 |
| 5287 | 265 | G265Q | A | 1.4e-05 | 1.2e-05 | 9.8e-06 | 1.193E-05 | 2.101E-06 | 9.800E-06 | 1.400E-05 |
| 5288 | 265 | G265G | A | 1.1e-05 | 9.4e-06 | 1.7e-05 | 1.247E-05 | 4.007E-06 | 9.400E-06 | 1.700E-05 |
| 5289 | 265 | G265H | A | 1.6e-05 | 1.3e-05 | 1e-05 | 1.300E-05 | 3.000E-06 | 1.000E-05 | 1.600E-05 |
| 5290 | 265 | G265I | A | 1.4e-05 | 1.5e-05 | 1.1e-05 | 1.333E-05 | 2.082E-06 | 1.100E-05 | 1.500E-05 |
| 5291 | 265 | G265L | A | 1e-05 | 1.1e-05 | 8.4e-06 | 9.800E-06 | 1.311E-06 | 8.400E-06 | 1.100E-05 |
| 5292 | 265 | G265K | A | 1.4e-05 | 1.6e-05 | 9.7e-06 | 1.323E-05 | 3.219E-06 | 9.700E-06 | 1.600E-05 |
| 5293 | 265 | G265M | A | 1.7e-05 | 1.3e-05 | 1.7e-05 | 1.567E-05 | 2.309E-06 | 1.300E-05 | 1.700E-05 |
| 5294 | 265 | G265F | A | 1.4e-05 | 1.2e-05 | 1.3e-05 | 1.300E-05 | 1.000E-06 | 1.200E-05 | 1.400E-05 |
| 5295 | 265 | G265P | A | 1.1e-05 | 1.1e-05 | 6.4e-06 | 9.467E-06 | 2.656E-06 | 6.400E-06 | 1.100E-05 |
| 5296 | 265 | G265S | A | 1.7e-05 | 1.3e-05 | 1.4e-05 | 1.467E-05 | 2.082E-06 | 1.300E-05 | 1.700E-05 |
| 5297 | 265 | G265T | A | 1.1e-05 | 1.4e-05 | 1.1e-05 | 1.200E-05 | 1.732E-06 | 1.100E-05 | 1.400E-05 |
| 5298 | 265 | G265W | A | 9.9e-06 | 1.3e-05 | 9.7e-06 | 1.087E-05 | 1.850E-06 | 9.700E-06 | 1.300E-05 |
| 5299 | 265 | G265Y | A | 1.2e-05 | 1.3e-05 | 9.8e-06 | 1.160E-05 | 1.637E-06 | 9.800E-06 | 1.300E-05 |
| 5300 | 265 | G265V | A | 1e-05 | 1.2e-05 | 9.7e-06 | 1.057E-05 | 1.250E-06 | 9.700E-06 | 1.200E-05 |
| 5301 | 266 | R266A | A | 1.4e-05 | 1.5e-05 | 1.4e-05 | 1.433E-05 | 5.774E-07 | 1.400E-05 | 1.500E-05 |
| 5302 | 266 | R266R | A | 1.5e-05 | 1.3e-05 | 1e-05 | 1.267E-05 | 2.517E-06 | 1.000E-05 | 1.500E-05 |
| 5303 | 266 | R266N | A | 1e-05 | 1.4e-05 | 1e-05 | 1.133E-05 | 2.309E-06 | 1.000E-05 | 1.400E-05 |
| 5304 | 266 | R266D | A | 2.3e-05 | 1.6e-05 | 1.4e-05 | 1.767E-05 | 4.726E-06 | 1.400E-05 | 2.300E-05 |
| 5305 | 266 | R266C | A | 1.3e-05 | 1.6e-05 | 1.4e-05 | 1.433E-05 | 1.528E-06 | 1.300E-05 | 1.600E-05 |
| 5306 | 266 | R266E | A | 1.3e-05 | 1.2e-05 | 1.7e-05 | 1.400E-05 | 2.646E-06 | 1.200E-05 | 1.700E-05 |
| 5307 | 266 | R266Q | A | 1.2e-05 | 1.4e-05 | 1.4e-05 | 1.333E-05 | 1.155E-06 | 1.200E-05 | 1.400E-05 |
| 5308 | 266 | R266G | A | 1.6e-05 | 1.6e-05 | 1e-05 | 1.400E-05 | 3.464E-06 | 1.000E-05 | 1.600E-05 |
| 5309 | 266 | R266H | A | 1.2e-05 | 1e-05 | 8.7e-06 | 1.023E-05 | 1.662E-06 | 8.700E-06 | 1.200E-05 |
| 5310 | 266 | R266I | A | 1.2e-05 | 1e-05 | 1.3e-05 | 1.167E-05 | 1.528E-06 | 1.000E-05 | 1.300E-05 |
| 5311 | 266 | R266L | A | 8.7e-06 | 1.5e-05 | 1.5e-05 | 1.290E-05 | 3.637E-06 | 8.700E-06 | 1.500E-05 |
| 5312 | 266 | R266K | A | 1e-05 | 1.5e-05 | 9e-06 | 1.133E-05 | 3.215E-06 | 9.000E-06 | 1.500E-05 |
| 5313 | 266 | R266M | A | 1.5e-05 | 9.5e-06 | 1.4e-05 | 1.283E-05 | 2.930E-06 | 9.500E-06 | 1.500E-05 |
| 5314 | 266 | R266F | A | 1.5e-05 | 1.1e-05 | 1.6e-05 | 1.400E-05 | 2.646E-06 | 1.100E-05 | 1.600E-05 |
| 5315 | 266 | R266P | A | 9.2e-06 | 9.8e-06 | 1e-05 | 9.667E-06 | 4.163E-07 | 9.200E-06 | 1.000E-05 |
| 5316 | 266 | R266S | A | 1.3e-05 | 1.6e-05 | 1.5e-05 | 1.467E-05 | 1.528E-06 | 1.300E-05 | 1.600E-05 |
| 5317 | 266 | R266T | A | 1.1e-05 | 1e-05 | 1.4e-05 | 1.167E-05 | 2.082E-06 | 1.000E-05 | 1.400E-05 |
| 5318 | 266 | R266W | A | 1.6e-05 | 1.4e-05 | 9e-06 | 1.300E-05 | 3.606E-06 | 9.000E-06 | 1.600E-05 |
| 5319 | 266 | R266Y | A | 1.5e-05 | 8.6e-06 | 1.4e-05 | 1.253E-05 | 3.443E-06 | 8.600E-06 | 1.500E-05 |
| 5320 | 266 | R266V | A | 1.5e-05 | 1.4e-05 | 1.4e-05 | 1.433E-05 | 5.774E-07 | 1.400E-05 | 1.500E-05 |
| 5321 | 267 | Y267A | A | 1.1e-05 | 1.6e-05 | 1.6e-05 | 1.433E-05 | 2.887E-06 | 1.100E-05 | 1.600E-05 |
| 5322 | 267 | Y267R | A | 7e-06 | 9.6e-06 | 1.1e-05 | 9.200E-06 | 2.030E-06 | 7.000E-06 | 1.100E-05 |
| 5323 | 267 | Y267N | A | 1.4e-05 | 1.4e-05 | 1.7e-05 | 1.500E-05 | 1.732E-06 | 1.400E-05 | 1.700E-05 |
| 5324 | 267 | Y267D | A | 1.7e-05 | 1.5e-05 | 1.2e-05 | 1.467E-05 | 2.517E-06 | 1.200E-05 | 1.700E-05 |
| 5325 | 267 | Y267C | A | 1.5e-05 | 9.8e-06 | 7.5e-06 | 1.077E-05 | 3.842E-06 | 7.500E-06 | 1.500E-05 |
| 5326 | 267 | Y267E | A | 1.5e-05 | 1e-05 | 9.5e-06 | 1.150E-05 | 3.041E-06 | 9.500E-06 | 1.500E-05 |
| 5327 | 267 | Y267Q | A | 1.4e-05 | 1e-05 | 1.7e-05 | 1.367E-05 | 3.512E-06 | 1.000E-05 | 1.700E-05 |
| 5328 | 267 | Y267G | A | 1.1e-05 | 1.1e-05 | 1.5e-05 | 1.233E-05 | 2.309E-06 | 1.100E-05 | 1.500E-05 |
| 5329 | 267 | Y267H | A | 1.5e-05 | 7.2e-06 | 1.3e-05 | 1.173E-05 | 4.051E-06 | 7.200E-06 | 1.500E-05 |
| 5330 | 267 | Y267I | A | 1.1e-05 | 1.5e-05 | 1.4e-05 | 1.333E-05 | 2.082E-06 | 1.100E-05 | 1.500E-05 |
| 5331 | 267 | Y267L | A | 1.5e-05 | 1.1e-05 | 1.5e-05 | 1.367E-05 | 2.309E-06 | 1.100E-05 | 1.500E-05 |
| 5332 | 267 | Y267K | A | 1.6e-05 | 9.5e-06 | 1.5e-05 | 1.350E-05 | 3.500E-06 | 9.500E-06 | 1.600E-05 |
| 5333 | 267 | Y267M | A | 1.1e-05 | 1.7e-05 | 9.5e-06 | 1.250E-05 | 3.969E-06 | 9.500E-06 | 1.700E-05 |
| 5334 | 267 | Y267F | A | 1.7e-05 | 1e-05 | 1.6e-05 | 1.433E-05 | 3.786E-06 | 1.000E-05 | 1.700E-05 |
| 5335 | 267 | Y267P | A | 1.1e-05 | 1.5e-05 | 1.6e-05 | 1.400E-05 | 2.646E-06 | 1.100E-05 | 1.600E-05 |
| 5336 | 267 | Y267S | A | 1.6e-05 | 9.4e-06 | 1.1e-05 | 1.213E-05 | 3.443E-06 | 9.400E-06 | 1.600E-05 |
| 5337 | 267 | Y267T | A | 1.4e-05 | 1.5e-05 | 1.1e-05 | 1.333E-05 | 2.082E-06 | 1.100E-05 | 1.500E-05 |
| 5338 | 267 | Y267W | A | 6.9e-06 | 1.1e-05 | 1.4e-05 | 1.063E-05 | 3.564E-06 | 6.900E-06 | 1.400E-05 |
| 5339 | 267 | Y267Y | A | 1.1e-05 | 9.8e-06 | 1.3e-05 | 1.127E-05 | 1.617E-06 | 9.800E-06 | 1.300E-05 |
| 5340 | 267 | Y267V | A | 1.6e-05 | 1.5e-05 | 1.5e-05 | 1.533E-05 | 5.774E-07 | 1.500E-05 | 1.600E-05 |
| 5341 | 268 | T268A | A | 9.9e-06 | 1.3e-05 | 9.4e-06 | 1.077E-05 | 1.950E-06 | 9.400E-06 | 1.300E-05 |
| 5342 | 268 | T268R | A | 6.8e-06 | 1.1e-05 | 1.1e-05 | 9.600E-06 | 2.425E-06 | 6.800E-06 | 1.100E-05 |
| 5343 | 268 | T268N | A | 1.5e-05 | 9.3e-06 | 8.3e-06 | 1.087E-05 | 3.614E-06 | 8.300E-06 | 1.500E-05 |
| 5344 | 268 | T268D | A | 1.6e-05 | 1.6e-05 | 1.3e-05 | 1.500E-05 | 1.732E-06 | 1.300E-05 | 1.600E-05 |
| 5345 | 268 | T268C | A | 9.3e-06 | 1.4e-05 | 1.7e-05 | 1.343E-05 | 3.881E-06 | 9.300E-06 | 1.700E-05 |
| 5346 | 268 | T268E | A | 1.7e-05 | 7.4e-06 | 1.4e-05 | 1.280E-05 | 4.911E-06 | 7.400E-06 | 1.700E-05 |
| 5347 | 268 | T268Q | A | 1.2e-05 | 1.1e-05 | 9.5e-06 | 1.083E-05 | 1.258E-06 | 9.500E-06 | 1.200E-05 |
| 5348 | 268 | T268G | A | 1.7e-05 | 1.5e-05 | 1.6e-05 | 1.600E-05 | 1.000E-06 | 1.500E-05 | 1.700E-05 |
| 5349 | 268 | T268H | A | 1.7e-05 | 1.6e-05 | 1.5e-05 | 1.600E-05 | 1.000E-06 | 1.500E-05 | 1.700E-05 |
| 5350 | 268 | T268I | A | 9.4e-06 | 1.6e-05 | 1.1e-05 | 1.213E-05 | 3.443E-06 | 9.400E-06 | 1.600E-05 |
| 5351 | 268 | T268L | A | 1.4e-05 | 9.3e-06 | 1.5e-05 | 1.277E-05 | 3.044E-06 | 9.300E-06 | 1.500E-05 |
| 5352 | 268 | T268K | A | 1e-05 | 1.6e-05 | 9.1e-06 | 1.170E-05 | 3.751E-06 | 9.100E-06 | 1.600E-05 |
| 5353 | 268 | T268M | A | 1.2e-05 | 1e-05 | 1.4e-05 | 1.200E-05 | 2.000E-06 | 1.000E-05 | 1.400E-05 |
| 5354 | 268 | T268F | A | 1e-05 | 1.1e-05 | 1.3e-05 | 1.133E-05 | 1.528E-06 | 1.000E-05 | 1.300E-05 |

| | | | | | | | | | | |
|---|---|---|---|---|---|---|---|---|---|---|
| 5355 | 268 | T268P | A | 1.4e-05 | 1.4e-05 | 1.5e-05 | 1.433E-05 | 5.774E-07 | 1.400E-05 | 1.500E-05 |
| 5356 | 268 | T268S | A | 9.7e-06 | 1.5e-05 | 1e-05 | 1.157E-05 | 2.977E-06 | 9.700E-06 | 1.500E-05 |
| 5357 | 268 | T268T | A | 9.7e-06 | 1.1e-05 | 1.1e-05 | 1.057E-05 | 7.506E-07 | 9.700E-06 | 1.100E-05 |
| 5358 | 268 | T268W | A | 8.6e-06 | 1e-05 | 6.9e-06 | 8.500E-06 | 1.552E-06 | 6.900E-06 | 1.000E-05 |
| 5359 | 268 | T268Y | A | 1.1e-05 | 1.6e-05 | 1e-05 | 1.233E-05 | 3.215E-06 | 1.000E-05 | 1.600E-05 |
| 5360 | 268 | T268V | A | 1.8e-05 | 1.5e-05 | 1.3e-05 | 1.533E-05 | 2.517E-06 | 1.300E-05 | 1.800E-05 |
| 5361 | 269 | F269A | A | 1.3e-05 | 1.2e-05 | 8.4e-06 | 1.113E-05 | 2.419E-06 | 8.400E-06 | 1.300E-05 |
| 5362 | 269 | F269R | A | 9.5e-06 | 1.3e-05 | 1.3e-05 | 1.183E-05 | 2.021E-06 | 9.500E-06 | 1.300E-05 |
| 5363 | 269 | F269N | A | 8.6e-06 | 1.4e-05 | 1.4e-05 | 1.220E-05 | 3.118E-06 | 8.600E-06 | 1.400E-05 |
| 5364 | 269 | F269D | A | 1.3e-05 | 1.1e-05 | 1.2e-05 | 1.200E-05 | 1.000E-06 | 1.100E-05 | 1.300E-05 |
| 5365 | 269 | F269C | A | 1.6e-05 | 1.4e-05 | 1.4e-05 | 1.467E-05 | 1.155E-06 | 1.400E-05 | 1.600E-05 |
| 5366 | 269 | F269E | A | 1.1e-05 | 1.6e-05 | 1.6e-05 | 1.433E-05 | 2.887E-06 | 1.100E-05 | 1.600E-05 |
| 5367 | 269 | F269Q | A | 1.7e-05 | 1.5e-05 | 1e-05 | 1.400E-05 | 3.606E-06 | 1.000E-05 | 1.700E-05 |
| 5368 | 269 | F269G | A | 1.1e-05 | 1.5e-05 | 1.4e-05 | 1.333E-05 | 2.082E-06 | 1.100E-05 | 1.500E-05 |
| 5369 | 269 | F269H | A | 1.1e-05 | 1e-05 | 1.6e-05 | 1.233E-05 | 3.215E-06 | 1.000E-05 | 1.600E-05 |
| 5370 | 269 | F269I | A | 1.5e-05 | 1.4e-05 | 1.5e-05 | 1.467E-05 | 5.774E-07 | 1.400E-05 | 1.500E-05 |
| 5371 | 269 | F269L | A | 9.2e-06 | 1.4e-05 | 9.7e-06 | 1.097E-05 | 2.639E-06 | 9.200E-06 | 1.400E-05 |
| 5372 | 269 | F269K | A | 1e-05 | 1.1e-05 | 1.1e-05 | 1.067E-05 | 5.774E-07 | 1.000E-05 | 1.100E-05 |
| 5373 | 269 | F269M | A | 1.6e-05 | 8.2e-06 | 1.7e-05 | 1.373E-05 | 4.818E-06 | 8.200E-06 | 1.700E-05 |
| 5374 | 269 | F269F | A | 1.1e-05 | 1.2e-05 | 1.3e-05 | 1.200E-05 | 1.000E-06 | 1.100E-05 | 1.300E-05 |
| 5375 | 269 | F269P | A | 1.5e-05 | 1.1e-05 | 1.6e-05 | 1.400E-05 | 2.646E-06 | 1.100E-05 | 1.600E-05 |
| 5376 | 269 | F269S | A | 9.2e-06 | 1e-05 | 8e-06 | 9.067E-06 | 1.007E-06 | 8.000E-06 | 1.000E-05 |
| 5377 | 269 | F269T | A | 1e-05 | 1.5e-05 | 8.7e-06 | 1.123E-05 | 3.326E-06 | 8.700E-06 | 1.500E-05 |
| 5378 | 269 | F269W | A | 1.2e-05 | 1.6e-05 | 1.5e-05 | 1.433E-05 | 2.082E-06 | 1.200E-05 | 1.600E-05 |
| 5379 | 269 | F269Y | A | 1.2e-05 | 1.1e-05 | 1.6e-05 | 1.300E-05 | 2.646E-06 | 1.100E-05 | 1.600E-05 |
| 5380 | 269 | F269V | A | 1.1e-05 | 9.9e-06 | 1.5e-05 | 1.197E-05 | 2.684E-06 | 9.900E-06 | 1.500E-05 |
| 5381 | 270 | G270A | A | 1.5e-05 | 9.4e-06 | 9.9e-06 | 1.143E-05 | 3.099E-06 | 9.400E-06 | 1.500E-05 |
| 5382 | 270 | G270R | A | 1.5e-05 | 1.4e-05 | 1e-05 | 1.300E-05 | 2.646E-06 | 1.000E-05 | 1.500E-05 |
| 5383 | 270 | G270N | A | 9e-06 | 1.1e-05 | 1.6e-05 | 1.200E-05 | 3.606E-06 | 9.000E-06 | 1.600E-05 |
| 5384 | 270 | G270D | A | 1.7e-05 | 1.8e-05 | 2.5e-05 | 2.000E-05 | 4.359E-06 | 1.700E-05 | 2.500E-05 |
| 5385 | 270 | G270C | A | 1.2e-05 | 1.3e-05 | 1.3e-05 | 1.267E-05 | 5.774E-07 | 1.200E-05 | 1.300E-05 |
| 5386 | 270 | G270E | A | 1e-05 | 8.9e-06 | 1.7e-05 | 1.197E-05 | 4.394E-06 | 8.900E-06 | 1.700E-05 |
| 5387 | 270 | G270Q | A | 1.2e-05 | 1.5e-05 | 1.1e-05 | 1.267E-05 | 2.082E-06 | 1.100E-05 | 1.500E-05 |
| 5388 | 270 | G270G | A | 8.3e-06 | 1.3e-05 | 1.1e-05 | 1.077E-05 | 2.359E-06 | 8.300E-06 | 1.300E-05 |
| 5389 | 270 | G270H | A | 9.4e-06 | 2e-05 | 1e-05 | 1.313E-05 | 5.954E-06 | 9.400E-06 | 2.000E-05 |
| 5390 | 270 | G270I | A | 9.7e-06 | 1.1e-05 | 1.4e-05 | 1.157E-05 | 2.205E-06 | 9.700E-06 | 1.400E-05 |
| 5391 | 270 | G270L | A | 1e-05 | 8e-06 | 1.4e-05 | 1.067E-05 | 3.055E-06 | 8.000E-06 | 1.400E-05 |
| 5392 | 270 | G270K | A | 1.4e-05 | 1.1e-05 | 6.2e-06 | 1.040E-05 | 3.934E-06 | 6.200E-06 | 1.400E-05 |
| 5393 | 270 | G270M | A | 1.4e-05 | 1e-05 | 1.4e-05 | 1.267E-05 | 2.309E-06 | 1.000E-05 | 1.400E-05 |
| 5394 | 270 | G270F | A | 1.3e-05 | 1.6e-05 | 8.5e-06 | 1.250E-05 | 3.775E-06 | 8.500E-06 | 1.600E-05 |
| 5395 | 270 | G270P | A | 1.1e-05 | 1.5e-05 | 1.6e-05 | 1.400E-05 | 2.646E-06 | 1.100E-05 | 1.600E-05 |
| 5396 | 270 | G270S | A | 1.2e-05 | 1e-05 | 1.4e-05 | 1.200E-05 | 2.000E-06 | 1.000E-05 | 1.400E-05 |
| 5397 | 270 | G270T | A | 1.1e-05 | 1.2e-05 | 1.6e-05 | 1.300E-05 | 2.646E-06 | 1.100E-05 | 1.600E-05 |
| 5398 | 270 | G270W | A | 9.6e-06 | 9.3e-06 | 9.3e-06 | 9.400E-06 | 1.732E-07 | 9.300E-06 | 9.600E-06 |
| 5399 | 270 | G270Y | A | 1.1e-05 | 1.5e-05 | 1.3e-05 | 1.300E-05 | 2.000E-06 | 1.100E-05 | 1.500E-05 |
| 5400 | 270 | G270V | A | 9.1e-06 | 9.6e-06 | 1.1e-05 | 9.900E-06 | 9.849E-07 | 9.100E-06 | 1.100E-05 |
| 5401 | 271 | A271A | A | 1.1e-05 | 1.8e-05 | 9.8e-06 | 1.293E-05 | 4.429E-06 | 9.800E-06 | 1.800E-05 |
| 5402 | 271 | A271R | A | 1.6e-05 | 1.5e-05 | 9.6e-06 | 1.353E-05 | 3.443E-06 | 9.600E-06 | 1.600E-05 |
| 5403 | 271 | A271N | A | 8.5e-06 | 1.3e-05 | 1.6e-05 | 1.250E-05 | 3.775E-06 | 8.500E-06 | 1.600E-05 |
| 5404 | 271 | A271D | A | 1.1e-05 | 1.4e-05 | 1.1e-05 | 1.200E-05 | 1.732E-06 | 1.100E-05 | 1.400E-05 |
| 5405 | 271 | A271C | A | 1.5e-05 | 1.7e-05 | 1.5e-05 | 1.567E-05 | 1.155E-06 | 1.500E-05 | 1.700E-05 |
| 5406 | 271 | A271E | A | 1e-05 | 1.3e-05 | 1e-05 | 1.100E-05 | 1.732E-06 | 1.000E-05 | 1.300E-05 |
| 5407 | 271 | A271Q | A | 1.1e-05 | 1.5e-05 | 9.7e-06 | 1.190E-05 | 2.762E-06 | 9.700E-06 | 1.500E-05 |
| 5408 | 271 | A271G | A | 1.6e-05 | 1.2e-05 | 1.3e-05 | 1.367E-05 | 2.082E-06 | 1.200E-05 | 1.600E-05 |
| 5409 | 271 | A271H | A | 9.4e-06 | 1.4e-05 | 1.1e-05 | 1.147E-05 | 2.335E-06 | 9.400E-06 | 1.400E-05 |
| 5410 | 271 | A271I | A | 9.2e-06 | 9.2e-06 | 1.4e-05 | 1.080E-05 | 2.771E-06 | 9.200E-06 | 1.400E-05 |
| 5411 | 271 | A271L | A | 9.6e-06 | 7.5e-06 | 1.5e-05 | 1.070E-05 | 3.869E-06 | 7.500E-06 | 1.500E-05 |
| 5412 | 271 | A271K | A | 1.1e-05 | 1.5e-05 | 1.5e-05 | 1.367E-05 | 2.309E-06 | 1.100E-05 | 1.500E-05 |
| 5413 | 271 | A271M | A | 1.2e-05 | 1.5e-05 | 1.4e-05 | 1.367E-05 | 1.528E-06 | 1.200E-05 | 1.500E-05 |
| 5414 | 271 | A271F | A | 1e-05 | 1.5e-05 | 1.4e-05 | 1.300E-05 | 2.646E-06 | 1.000E-05 | 1.500E-05 |
| 5415 | 271 | A271P | A | 1.5e-05 | 5.8e-06 | 1.7e-05 | 1.260E-05 | 5.973E-06 | 5.800E-06 | 1.700E-05 |
| 5416 | 271 | A271S | A | 1e-05 | 8.9e-06 | 1.4e-05 | 1.097E-05 | 2.684E-06 | 8.900E-06 | 1.400E-05 |
| 5417 | 271 | A271T | A | 1.5e-05 | 1.1e-05 | 8.3e-06 | 1.143E-05 | 3.371E-06 | 8.300E-06 | 1.500E-05 |
| 5418 | 271 | A271W | A | 2e-05 | 1.4e-05 | 1.7e-05 | 1.700E-05 | 3.000E-06 | 1.400E-05 | 2.000E-05 |
| 5419 | 271 | A271Y | A | 9.7e-06 | 1.4e-05 | 1.6e-05 | 1.323E-05 | 3.219E-06 | 9.700E-06 | 1.600E-05 |
| 5420 | 271 | A271V | A | 1.6e-05 | 1.2e-05 | 8.3e-06 | 1.210E-05 | 3.851E-06 | 8.300E-06 | 1.600E-05 |
| 5421 | 272 | S272A | A | 1.7e-05 | 1.4e-05 | 1.4e-05 | 1.500E-05 | 1.732E-06 | 1.400E-05 | 1.700E-05 |
| 5422 | 272 | S272R | A | 5.6e-06 | 1.1e-05 | 1.6e-05 | 1.087E-05 | 5.201E-06 | 5.600E-06 | 1.600E-05 |
| 5423 | 272 | S272N | A | 9.9e-06 | 1.3e-05 | 1.6e-05 | 1.297E-05 | 3.050E-06 | 9.900E-06 | 1.600E-05 |
| 5424 | 272 | S272D | A | 1.6e-05 | 8.8e-06 | 9.3e-06 | 1.137E-05 | 4.020E-06 | 8.800E-06 | 1.600E-05 |
| 5425 | 272 | S272C | A | 1.1e-05 | 1.3e-05 | 1e-05 | 1.133E-05 | 1.528E-06 | 1.000E-05 | 1.300E-05 |
| 5426 | 272 | S272E | A | 1.1e-05 | 1.4e-05 | 1.6e-05 | 1.367E-05 | 2.517E-06 | 1.100E-05 | 1.600E-05 |
| 5427 | 272 | S272Q | A | 1.5e-05 | 9.9e-06 | 1.3e-05 | 1.263E-05 | 2.570E-06 | 9.900E-06 | 1.500E-05 |
| 5428 | 272 | S272G | A | 1.5e-05 | 1.4e-05 | 1.5e-05 | 1.467E-05 | 5.774E-07 | 1.400E-05 | 1.500E-05 |
| 5429 | 272 | S272H | A | 1.6e-05 | 1.6e-05 | 1.1e-05 | 1.433E-05 | 2.887E-06 | 1.100E-05 | 1.600E-05 |
| 5430 | 272 | S272I | A | 1.5e-05 | 1.6e-05 | 7.5e-06 | 1.283E-05 | 4.646E-06 | 7.500E-06 | 1.600E-05 |
| 5431 | 272 | S272L | A | 9.5e-06 | 1.6e-05 | 1.4e-05 | 1.317E-05 | 3.329E-06 | 9.500E-06 | 1.600E-05 |
| 5432 | 272 | S272K | A | 1.4e-05 | 1.1e-05 | 1.7e-05 | 1.400E-05 | 3.000E-06 | 1.100E-05 | 1.700E-05 |
| 5433 | 272 | S272M | A | 1.5e-05 | 1.4e-05 | 1.7e-05 | 1.533E-05 | 1.528E-06 | 1.400E-05 | 1.700E-05 |
| 5434 | 272 | S272F | A | 1.5e-05 | 1.7e-05 | 9.4e-06 | 1.380E-05 | 3.940E-06 | 9.400E-06 | 1.700E-05 |
| 5435 | 272 | S272P | A | 1e-05 | 1.6e-05 | 1.6e-05 | 1.400E-05 | 3.464E-06 | 1.000E-05 | 1.600E-05 |
| 5436 | 272 | S272S | A | 1.5e-05 | 1.5e-05 | 1.4e-05 | 1.467E-05 | 5.774E-07 | 1.400E-05 | 1.500E-05 |
| 5437 | 272 | S272T | A | 1.4e-05 | 1.4e-05 | 1.5e-05 | 1.433E-05 | 5.774E-07 | 1.400E-05 | 1.500E-05 |
| 5438 | 272 | S272W | A | 9e-06 | 7.2e-06 | 1.1e-05 | 9.067E-06 | 1.901E-06 | 7.200E-06 | 1.100E-05 |
| 5439 | 272 | S272Y | A | 1.4e-05 | 1.5e-05 | 1.1e-05 | 1.333E-05 | 2.082E-06 | 1.100E-05 | 1.500E-05 |

| | | | | | | | | | | |
|---|---|---|---|---|---|---|---|---|---|---|
| 5440 | 272 | S272V | A | 1.5e-05 | 1.5e-05 | 9.8e-06 | 1.327E-05 | 3.002E-06 | 9.800E-06 | 1.500E-05 |
| 5441 | 273 | C273A | A | 1.7e-05 | 8e-06 | 1.3e-05 | 1.267E-05 | 4.509E-06 | 8.000E-06 | 1.700E-05 |
| 5442 | 273 | C273R | A | 1.5e-05 | 1e-05 | 1.7e-05 | 1.400E-05 | 3.606E-06 | 1.000E-05 | 1.700E-05 |
| 5443 | 273 | C273N | A | 1.5e-05 | 1.4e-05 | 1.5e-05 | 1.467E-05 | 5.774E-07 | 1.400E-05 | 1.500E-05 |
| 5444 | 273 | C273D | A | 1.5e-05 | 1.1e-05 | 1.5e-05 | 1.367E-05 | 2.309E-06 | 1.100E-05 | 1.500E-05 |
| 5445 | 273 | C273C | A | 1e-05 | 1.5e-05 | 1.4e-05 | 1.300E-05 | 2.646E-06 | 1.000E-05 | 1.500E-05 |
| 5446 | 273 | C273E | A | 1.8e-05 | 2.7e-05 | 1.3e-05 | 1.933E-05 | 7.095E-06 | 1.300E-05 | 2.700E-05 |
| 5447 | 273 | C273Q | A | 1.1e-05 | 1e-05 | 1.4e-05 | 1.167E-05 | 2.082E-06 | 1.000E-05 | 1.400E-05 |
| 5448 | 273 | C273G | A | 1.1e-05 | 1.2e-05 | 1.5e-05 | 1.267E-05 | 2.082E-06 | 1.100E-05 | 1.500E-05 |
| 5449 | 273 | C273H | A | 7.1e-06 | 9.1e-06 | 1.5e-05 | 1.040E-05 | 4.107E-06 | 7.100E-06 | 1.500E-05 |
| 5450 | 273 | C273I | A | 1.5e-05 | 9.6e-06 | 1.2e-05 | 1.220E-05 | 2.706E-06 | 9.600E-06 | 1.500E-05 |
| 5451 | 273 | C273L | A | 1.5e-05 | 1.7e-05 | 1.7e-05 | 1.633E-05 | 1.155E-06 | 1.500E-05 | 1.700E-05 |
| 5452 | 273 | C273K | A | 1.6e-05 | 1e-05 | 1.6e-05 | 1.400E-05 | 3.464E-06 | 1.000E-05 | 1.600E-05 |
| 5453 | 273 | C273M | A | 1e-05 | 1.5e-05 | 1.2e-05 | 1.233E-05 | 2.517E-06 | 1.000E-05 | 1.500E-05 |
| 5454 | 273 | C273F | A | 1.1e-05 | 1.5e-05 | 1.7e-05 | 1.433E-05 | 3.055E-06 | 1.100E-05 | 1.700E-05 |
| 5455 | 273 | C273P | A | 1e-05 | 9.9e-06 | 9.6e-06 | 9.833E-06 | 2.082E-07 | 9.600E-06 | 1.000E-05 |
| 5456 | 273 | C273S | A | 9.7e-06 | 9e-06 | 1.7e-05 | 1.190E-05 | 4.431E-06 | 9.000E-06 | 1.700E-05 |
| 5457 | 273 | C273T | A | 1.4e-05 | 1.1e-05 | 9.4e-06 | 1.147E-05 | 2.335E-06 | 9.400E-06 | 1.400E-05 |
| 5458 | 273 | C273W | A | 1e-05 | 1.1e-05 | 1.4e-05 | 1.167E-05 | 2.082E-06 | 1.000E-05 | 1.400E-05 |
| 5459 | 273 | C273Y | A | 1.6e-05 | 7.1e-06 | 1.6e-05 | 1.303E-05 | 5.138E-06 | 7.100E-06 | 1.600E-05 |
| 5460 | 273 | C273V | A | 1e-05 | 1.3e-05 | 1.4e-05 | 1.233E-05 | 2.082E-06 | 1.000E-05 | 1.400E-05 |
| 5461 | 274 | V274A | A | 1.3e-05 | 1.5e-05 | 1.1e-05 | 1.300E-05 | 2.000E-06 | 1.100E-05 | 1.500E-05 |
| 5462 | 274 | V274R | A | 1.5e-05 | 1.6e-05 | 1.4e-05 | 1.500E-05 | 1.000E-06 | 1.400E-05 | 1.600E-05 |
| 5463 | 274 | V274N | A | 9.3e-06 | 8.8e-06 | 1.3e-05 | 1.037E-05 | 2.294E-06 | 8.800E-06 | 1.300E-05 |
| 5464 | 274 | V274D | A | 1.2e-05 | 1.2e-05 | 1.1e-05 | 1.167E-05 | 5.774E-07 | 1.100E-05 | 1.200E-05 |
| 5465 | 274 | V274C | A | 9.4e-06 | 9.7e-06 | 1.4e-05 | 1.103E-05 | 2.574E-06 | 9.400E-06 | 1.400E-05 |
| 5466 | 274 | V274E | A | 9.3e-06 | 7e-06 | 1.6e-05 | 1.077E-05 | 4.676E-06 | 7.000E-06 | 1.600E-05 |
| 5467 | 274 | V274Q | A | 1.4e-05 | 1.1e-05 | 1.1e-05 | 1.200E-05 | 1.732E-06 | 1.100E-05 | 1.400E-05 |
| 5468 | 274 | V274G | A | 9.5e-06 | 1.1e-05 | 9.8e-06 | 1.010E-05 | 7.937E-07 | 9.500E-06 | 1.100E-05 |
| 5469 | 274 | V274H | A | 5.9e-06 | 1.3e-05 | 1e-05 | 9.633E-06 | 3.564E-06 | 5.900E-06 | 1.300E-05 |
| 5470 | 274 | V274I | A | 1.6e-05 | 1.1e-05 | 1e-05 | 1.233E-05 | 3.215E-06 | 1.000E-05 | 1.600E-05 |
| 5471 | 274 | V274L | A | 1.1e-05 | 1.1e-05 | 1.3e-05 | 1.167E-05 | 1.155E-06 | 1.100E-05 | 1.300E-05 |
| 5472 | 274 | V274K | A | 1.3e-05 | 1.6e-05 | 1.4e-05 | 1.433E-05 | 1.528E-06 | 1.300E-05 | 1.600E-05 |
| 5473 | 274 | V274M | A | 1.5e-05 | 1.1e-05 | 1.6e-05 | 1.400E-05 | 2.646E-06 | 1.100E-05 | 1.600E-05 |
| 5474 | 274 | V274F | A | 9.6e-06 | 1.1e-05 | 1.4e-05 | 1.153E-05 | 2.248E-06 | 9.600E-06 | 1.400E-05 |
| 5475 | 274 | V274P | A | 1.5e-05 | 1.6e-05 | 1.6e-05 | 1.567E-05 | 5.774E-07 | 1.500E-05 | 1.600E-05 |
| 5476 | 274 | V274S | A | 1.8e-05 | 1.5e-05 | 1.7e-05 | 1.667E-05 | 1.528E-06 | 1.500E-05 | 1.800E-05 |
| 5477 | 274 | V274T | A | 1e-05 | 1.6e-05 | 9.8e-06 | 1.193E-05 | 3.523E-06 | 9.800E-06 | 1.600E-05 |
| 5478 | 274 | V274W | A | 1.4e-05 | 1.6e-05 | 1.4e-05 | 1.467E-05 | 1.155E-06 | 1.400E-05 | 1.600E-05 |
| 5479 | 274 | V274Y | A | 1.5e-05 | 1.1e-05 | 1.2e-05 | 1.267E-05 | 2.082E-06 | 1.100E-05 | 1.500E-05 |
| 5480 | 274 | V274V | A | 1.6e-05 | 1.7e-05 | 1.4e-05 | 1.567E-05 | 1.528E-06 | 1.400E-05 | 1.700E-05 |
| 5481 | 275 | T275A | A | 1.7e-05 | 1.6e-05 | 1.5e-05 | 1.600E-05 | 1.000E-06 | 1.500E-05 | 1.700E-05 |
| 5482 | 275 | T275R | A | 1.4e-05 | 1.6e-05 | 1.4e-05 | 1.467E-05 | 1.155E-06 | 1.400E-05 | 1.600E-05 |
| 5483 | 275 | T275N | A | 1.2e-05 | 9e-06 | 1.4e-05 | 1.167E-05 | 2.517E-06 | 9.000E-06 | 1.400E-05 |
| 5484 | 275 | T275D | A | 8.7e-06 | 1.3e-05 | 1.7e-05 | 1.290E-05 | 4.151E-06 | 8.700E-06 | 1.700E-05 |
| 5485 | 275 | T275C | A | 1.5e-05 | 1.3e-05 | 1.1e-05 | 1.300E-05 | 2.000E-06 | 1.100E-05 | 1.500E-05 |
| 5486 | 275 | T275E | A | 1.1e-05 | 1.2e-05 | 1.3e-05 | 1.200E-05 | 1.000E-06 | 1.100E-05 | 1.300E-05 |
| 5487 | 275 | T275Q | A | 9.7e-06 | 1.5e-05 | 1.5e-05 | 1.323E-05 | 3.060E-06 | 9.700E-06 | 1.500E-05 |
| 5488 | 275 | T275G | A | 1e-05 | 1e-05 | 1.6e-05 | 1.200E-05 | 3.464E-06 | 1.000E-05 | 1.600E-05 |
| 5489 | 275 | T275H | A | 9.9e-06 | 1.6e-05 | 1.1e-05 | 1.230E-05 | 3.251E-06 | 9.900E-06 | 1.600E-05 |
| 5490 | 275 | T275I | A | 1e-05 | 9.3e-06 | 1e-05 | 9.767E-06 | 4.041E-07 | 9.300E-06 | 1.000E-05 |
| 5491 | 275 | T275L | A | 1.6e-05 | 1.5e-05 | 1.1e-05 | 1.400E-05 | 2.646E-06 | 1.100E-05 | 1.600E-05 |
| 5492 | 275 | T275K | A | 9.7e-06 | 1.5e-05 | 1.6e-05 | 1.357E-05 | 3.386E-06 | 9.700E-06 | 1.600E-05 |
| 5493 | 275 | T275M | A | 1.5e-05 | 1.5e-05 | 1.1e-05 | 1.367E-05 | 2.309E-06 | 1.100E-05 | 1.500E-05 |
| 5494 | 275 | T275F | A | 1.4e-05 | 1e-05 | 1.6e-05 | 1.333E-05 | 3.055E-06 | 1.000E-05 | 1.600E-05 |
| 5495 | 275 | T275P | A | 6.5e-06 | 1.3e-05 | 1.7e-05 | 1.217E-05 | 5.299E-06 | 6.500E-06 | 1.700E-05 |
| 5496 | 275 | T275S | A | 1.6e-05 | 9.2e-06 | 6.6e-06 | 1.060E-05 | 4.854E-06 | 6.600E-06 | 1.600E-05 |
| 5497 | 275 | T275T | A | 1e-05 | 1.5e-05 | 1.5e-05 | 1.333E-05 | 2.887E-06 | 1.000E-05 | 1.500E-05 |
| 5498 | 275 | T275W | A | 8.4e-06 | 8.1e-06 | 1.1e-05 | 9.167E-06 | 1.595E-06 | 8.100E-06 | 1.100E-05 |
| 5499 | 275 | T275Y | A | 9.9e-06 | 1.6e-05 | 1e-05 | 1.197E-05 | 3.493E-06 | 9.900E-06 | 1.600E-05 |
| 5500 | 275 | T275V | A | 1.1e-05 | 1.4e-05 | 1.3e-05 | 1.267E-05 | 1.528E-06 | 1.100E-05 | 1.400E-05 |
| 5501 | 276 | A276A | A | 7.1e-06 | 1.4e-05 | 1.5e-05 | 1.203E-05 | 4.302E-06 | 7.100E-06 | 1.500E-05 |
| 5502 | 276 | A276R | A | 1e-05 | 9.5e-06 | 1e-05 | 9.833E-06 | 2.887E-07 | 9.500E-06 | 1.000E-05 |
| 5503 | 276 | A276N | A | 8.6e-06 | 1.1e-05 | 1.5e-05 | 1.153E-05 | 3.233E-06 | 8.600E-06 | 1.500E-05 |
| 5504 | 276 | A276D | A | 1.4e-05 | 1e-05 | 1.6e-05 | 1.333E-05 | 3.055E-06 | 1.000E-05 | 1.600E-05 |
| 5505 | 276 | A276C | A | 8.7e-06 | 1.4e-05 | 1.3e-05 | 1.190E-05 | 2.816E-06 | 8.700E-06 | 1.400E-05 |
| 5506 | 276 | A276E | A | 1.1e-05 | 1.1e-05 | 1e-05 | 1.067E-05 | 5.774E-07 | 1.000E-05 | 1.100E-05 |
| 5507 | 276 | A276Q | A | 1.5e-05 | 1.1e-05 | 1.4e-05 | 1.333E-05 | 2.082E-06 | 1.100E-05 | 1.500E-05 |
| 5508 | 276 | A276G | A | 1.1e-05 | 1.2e-05 | 9.8e-06 | 1.093E-05 | 1.102E-06 | 9.800E-06 | 1.200E-05 |
| 5509 | 276 | A276H | A | 1.3e-05 | 1.5e-05 | 1.5e-05 | 1.433E-05 | 1.155E-06 | 1.300E-05 | 1.500E-05 |
| 5510 | 276 | A276I | A | 8.9e-06 | 9.6e-06 | 1.4e-05 | 1.083E-05 | 2.765E-06 | 8.900E-06 | 1.400E-05 |
| 5511 | 276 | A276L | A | 1.5e-05 | 6.4e-06 | 1.3e-05 | 1.147E-05 | 4.500E-06 | 6.400E-06 | 1.500E-05 |
| 5512 | 276 | A276K | A | 1.6e-05 | 1.1e-05 | 1.7e-05 | 1.467E-05 | 3.215E-06 | 1.100E-05 | 1.700E-05 |
| 5513 | 276 | A276M | A | 1.1e-05 | 1.5e-05 | 8.7e-06 | 1.157E-05 | 3.188E-06 | 8.700E-06 | 1.500E-05 |
| 5514 | 276 | A276F | A | 1e-05 | 1.6e-05 | 1.6e-05 | 1.400E-05 | 3.464E-06 | 1.000E-05 | 1.600E-05 |
| 5515 | 276 | A276P | A | 1.6e-05 | 1.2e-05 | 1.6e-05 | 1.467E-05 | 2.309E-06 | 1.200E-05 | 1.600E-05 |
| 5516 | 276 | A276S | A | 1.1e-05 | 9.3e-06 | 1.4e-05 | 1.143E-05 | 2.380E-06 | 9.300E-06 | 1.400E-05 |
| 5517 | 276 | A276T | A | 7.8e-06 | 8.1e-06 | 1.4e-05 | 9.967E-06 | 3.496E-06 | 7.800E-06 | 1.400E-05 |
| 5518 | 276 | A276W | A | 1.3e-05 | 1.2e-05 | 1e-05 | 1.167E-05 | 1.528E-06 | 1.000E-05 | 1.300E-05 |
| 5519 | 276 | A276Y | A | 1.4e-05 | 1.5e-05 | 1.5e-05 | 1.467E-05 | 5.774E-07 | 1.400E-05 | 1.500E-05 |
| 5520 | 276 | A276V | A | 1.5e-05 | 1.1e-05 | 9.6e-06 | 1.187E-05 | 2.802E-06 | 9.600E-06 | 1.500E-05 |
| 5521 | 277 | C277A | A | 1.4e-05 | 1.7e-05 | 9.8e-06 | 1.360E-05 | 3.617E-06 | 9.800E-06 | 1.700E-05 |
| 5522 | 277 | C277R | A | 9.6e-06 | 1.1e-05 | 1.8e-05 | 1.287E-05 | 4.500E-06 | 9.600E-06 | 1.800E-05 |
| 5523 | 277 | C277N | A | 1.1e-05 | 1.1e-05 | 1.6e-05 | 1.267E-05 | 2.887E-06 | 1.100E-05 | 1.600E-05 |
| 5524 | 277 | C277D | A | 1e-05 | 1.2e-05 | 1.3e-05 | 1.167E-05 | 1.528E-06 | 1.000E-05 | 1.300E-05 |

| | | | | | | | | | | |
|---|---|---|---|---|---|---|---|---|---|---|
| 5525 | 277 | C277C | A | 6.1e-06 | 1.5e-05 | 1.1e-05 | 1.070E-05 | 4.458E-06 | 6.100E-06 | 1.500E-05 |
| 5526 | 277 | C277E | A | 1.4e-05 | 1.8e-05 | 1.2e-05 | 1.467E-05 | 3.055E-06 | 1.200E-05 | 1.800E-05 |
| 5527 | 277 | C277Q | A | 8.6e-06 | 1.1e-05 | 1.4e-05 | 1.120E-05 | 2.706E-06 | 8.600E-06 | 1.400E-05 |
| 5528 | 277 | C277G | A | 1.6e-05 | 1.2e-05 | 1.5e-05 | 1.433E-05 | 2.082E-06 | 1.200E-05 | 1.600E-05 |
| 5529 | 277 | C277H | A | 1.1e-05 | 1.6e-05 | 8.3e-06 | 1.177E-05 | 3.907E-06 | 8.300E-06 | 1.600E-05 |
| 5530 | 277 | C277I | A | 1.2e-05 | 1.6e-05 | 1.6e-05 | 1.467E-05 | 2.309E-06 | 1.200E-05 | 1.600E-05 |
| 5531 | 277 | C277L | A | 1.7e-05 | 1.7e-05 | 8.6e-06 | 1.420E-05 | 4.850E-06 | 8.600E-06 | 1.700E-05 |
| 5532 | 277 | C277K | A | 1.3e-05 | 1.4e-05 | 9.6e-06 | 1.220E-05 | 2.307E-06 | 9.600E-06 | 1.400E-05 |
| 5533 | 277 | C277M | A | 1.6e-05 | 1.5e-05 | 1.6e-05 | 1.567E-05 | 5.774E-07 | 1.500E-05 | 1.600E-05 |
| 5534 | 277 | C277F | A | 1.4e-05 | 1.6e-05 | 1e-05 | 1.333E-05 | 3.055E-06 | 1.000E-05 | 1.600E-05 |
| 5535 | 277 | C277P | A | 9.5e-06 | 1.1e-05 | 7.1e-06 | 9.200E-06 | 1.967E-06 | 7.100E-06 | 1.100E-05 |
| 5536 | 277 | C277S | A | 1.2e-05 | 1.2e-05 | 1.1e-05 | 1.167E-05 | 5.774E-07 | 1.100E-05 | 1.200E-05 |
| 5537 | 277 | C277T | A | 1e-05 | 1.1e-05 | 1.5e-05 | 1.200E-05 | 2.646E-06 | 1.000E-05 | 1.500E-05 |
| 5538 | 277 | C277W | A | 1.5e-05 | 9.1e-06 | 1.5e-05 | 1.303E-05 | 3.406E-06 | 9.100E-06 | 1.500E-05 |
| 5539 | 277 | C277Y | A | 1.5e-05 | 1.6e-05 | 1.4e-05 | 1.500E-05 | 1.000E-06 | 1.400E-05 | 1.600E-05 |
| 5540 | 277 | C277V | A | 1.5e-05 | 1.1e-05 | 1.2e-05 | 1.267E-05 | 2.082E-06 | 1.100E-05 | 1.500E-05 |
| 5541 | 278 | P278A | A | 1.2e-05 | 1.1e-05 | 9.4e-06 | 1.080E-05 | 1.311E-06 | 9.400E-06 | 1.200E-05 |
| 5542 | 278 | P278R | A | 9.9e-06 | 1.2e-05 | 8.1e-06 | 1.000E-05 | 1.952E-06 | 8.100E-06 | 1.200E-05 |
| 5543 | 278 | P278N | A | 1.6e-05 | 1.6e-05 | 1e-05 | 1.400E-05 | 3.464E-06 | 1.000E-05 | 1.600E-05 |
| 5544 | 278 | P278D | A | 1.6e-05 | 1.6e-05 | 9.7e-06 | 1.390E-05 | 3.637E-06 | 9.700E-06 | 1.600E-05 |
| 5545 | 278 | P278C | A | 1.4e-05 | 1.8e-05 | 1.5e-05 | 1.567E-05 | 2.082E-06 | 1.400E-05 | 1.800E-05 |
| 5546 | 278 | P278E | A | 1.6e-05 | 1.1e-05 | 1.6e-05 | 1.433E-05 | 2.887E-06 | 1.100E-05 | 1.600E-05 |
| 5547 | 278 | P278Q | A | 1.2e-05 | 1.4e-05 | 1.1e-05 | 1.233E-05 | 1.528E-06 | 1.100E-05 | 1.400E-05 |
| 5548 | 278 | P278G | A | 1.6e-05 | 1.2e-05 | 1.3e-05 | 1.367E-05 | 2.082E-06 | 1.200E-05 | 1.600E-05 |
| 5549 | 278 | P278H | A | 1.5e-05 | 1e-05 | 1.7e-05 | 1.400E-05 | 3.606E-06 | 1.000E-05 | 1.700E-05 |
| 5550 | 278 | P278I | A | 1.3e-05 | 1.4e-05 | 1.6e-05 | 1.433E-05 | 1.528E-06 | 1.300E-05 | 1.600E-05 |
| 5551 | 278 | P278L | A | 1.3e-05 | 1.7e-05 | 1.1e-05 | 1.367E-05 | 3.055E-06 | 1.100E-05 | 1.700E-05 |
| 5552 | 278 | P278K | A | 1.5e-05 | 9.6e-06 | 1e-05 | 1.153E-05 | 3.009E-06 | 9.600E-06 | 1.500E-05 |
| 5553 | 278 | P278M | A | 1.1e-05 | 1.5e-05 | 1.7e-05 | 1.433E-05 | 3.055E-06 | 1.100E-05 | 1.700E-05 |
| 5554 | 278 | P278F | A | 1.3e-05 | 1.3e-05 | 1.1e-05 | 1.233E-05 | 1.155E-06 | 1.100E-05 | 1.300E-05 |
| 5555 | 278 | P278P | A | 1.3e-05 | 1.6e-05 | 1.5e-05 | 1.467E-05 | 1.528E-06 | 1.300E-05 | 1.600E-05 |
| 5556 | 278 | P278S | A | 9.1e-06 | 8.3e-06 | 1.1e-05 | 9.467E-06 | 1.387E-06 | 8.300E-06 | 1.100E-05 |
| 5557 | 278 | P278T | A | 1.4e-05 | 9e-06 | 1.5e-05 | 1.267E-05 | 3.215E-06 | 9.000E-06 | 1.500E-05 |
| 5558 | 278 | P278W | A | 1.3e-05 | 9.8e-06 | 8.4e-06 | 1.040E-05 | 2.358E-06 | 8.400E-06 | 1.300E-05 |
| 5559 | 278 | P278Y | A | 1.5e-05 | 1.1e-05 | 1.5e-05 | 1.367E-05 | 2.309E-06 | 1.100E-05 | 1.500E-05 |
| 5560 | 278 | P278V | A | 9.4e-06 | 8.5e-06 | 1.1e-05 | 9.633E-06 | 1.266E-06 | 8.500E-06 | 1.100E-05 |
| 5561 | 279 | Y279A | A | 1.2e-05 | 1.1e-05 | 9.4e-06 | 1.080E-05 | 1.311E-06 | 9.400E-06 | 1.200E-05 |
| 5562 | 279 | Y279R | A | 1.1e-05 | 1.6e-05 | 8.6e-06 | 1.187E-05 | 3.775E-06 | 8.600E-06 | 1.600E-05 |
| 5563 | 279 | Y279N | A | 1.5e-05 | 7.5e-06 | 1.2e-05 | 1.150E-05 | 3.775E-06 | 7.500E-06 | 1.500E-05 |
| 5564 | 279 | Y279D | A | 1.6e-05 | 1.1e-05 | 8.3e-06 | 1.177E-05 | 3.907E-06 | 8.300E-06 | 1.600E-05 |
| 5565 | 279 | Y279C | A | 1.6e-05 | 1.6e-05 | 1e-05 | 1.400E-05 | 3.464E-06 | 1.000E-05 | 1.600E-05 |
| 5566 | 279 | Y279E | A | 2.1e-05 | 1.6e-05 | 1.2e-05 | 1.633E-05 | 4.509E-06 | 1.200E-05 | 2.100E-05 |
| 5567 | 279 | Y279Q | A | 1.3e-05 | 1.2e-05 | 1.1e-05 | 1.200E-05 | 1.000E-06 | 1.100E-05 | 1.300E-05 |
| 5568 | 279 | Y279G | A | 1.2e-05 | 1.2e-05 | 1.5e-05 | 1.300E-05 | 1.732E-06 | 1.200E-05 | 1.500E-05 |
| 5569 | 279 | Y279H | A | 1.5e-05 | 9.7e-06 | 1.4e-05 | 1.290E-05 | 2.816E-06 | 9.700E-06 | 1.500E-05 |
| 5570 | 279 | Y279I | A | 1.6e-05 | 1.8e-05 | 1.5e-05 | 1.633E-05 | 1.528E-06 | 1.500E-05 | 1.800E-05 |
| 5571 | 279 | Y279L | A | 1.2e-05 | 1.6e-05 | 1.4e-05 | 1.400E-05 | 2.000E-06 | 1.200E-05 | 1.600E-05 |
| 5572 | 279 | Y279K | A | 1.3e-05 | 1.4e-05 | 1.1e-05 | 1.267E-05 | 1.528E-06 | 1.100E-05 | 1.400E-05 |
| 5573 | 279 | Y279M | A | 1.7e-05 | 1.2e-05 | 1.7e-05 | 1.533E-05 | 2.887E-06 | 1.200E-05 | 1.700E-05 |
| 5574 | 279 | Y279F | A | 1.5e-05 | 9.9e-06 | 9.1e-06 | 1.133E-05 | 3.201E-06 | 9.100E-06 | 1.500E-05 |
| 5575 | 279 | Y279P | A | 1.2e-05 | 1.3e-05 | 1.1e-05 | 1.200E-05 | 1.000E-06 | 1.100E-05 | 1.300E-05 |
| 5576 | 279 | Y279S | A | 1.1e-05 | 1.6e-05 | 1.8e-05 | 1.500E-05 | 3.606E-06 | 1.100E-05 | 1.800E-05 |
| 5577 | 279 | Y279T | A | 1.1e-05 | 8.6e-06 | 1.2e-05 | 1.053E-05 | 1.747E-06 | 8.600E-06 | 1.200E-05 |
| 5578 | 279 | Y279W | A | 1.5e-05 | 9.8e-06 | 7.8e-06 | 1.087E-05 | 3.717E-06 | 7.800E-06 | 1.500E-05 |
| 5579 | 279 | Y279Y | A | 1.5e-05 | 1.7e-05 | 1.1e-05 | 1.433E-05 | 3.055E-06 | 1.100E-05 | 1.700E-05 |
| 5580 | 279 | Y279V | A | 1.5e-05 | 1.5e-05 | 1e-05 | 1.333E-05 | 2.887E-06 | 1.000E-05 | 1.500E-05 |
| 5581 | 280 | N280A | A | 1.5e-05 | 1e-05 | 1.3e-05 | 1.267E-05 | 2.517E-06 | 1.000E-05 | 1.500E-05 |
| 5582 | 280 | N280R | A | 1.4e-05 | 1.6e-05 | 1.7e-05 | 1.567E-05 | 1.528E-06 | 1.400E-05 | 1.700E-05 |
| 5583 | 280 | N280N | A | 9.7e-06 | 8.5e-06 | 1.5e-05 | 1.107E-05 | 3.459E-06 | 8.500E-06 | 1.500E-05 |
| 5584 | 280 | N280D | A | 1.1e-05 | 7.2e-06 | 1.4e-05 | 1.073E-05 | 3.408E-06 | 7.200E-06 | 1.400E-05 |
| 5585 | 280 | N280C | A | 1.1e-05 | 1.4e-05 | 1.7e-05 | 1.400E-05 | 3.000E-06 | 1.100E-05 | 1.700E-05 |
| 5586 | 280 | N280E | A | 1.4e-05 | 8.8e-06 | 1.2e-05 | 1.160E-05 | 2.623E-06 | 8.800E-06 | 1.400E-05 |
| 5587 | 280 | N280Q | A | 1.5e-05 | 9.7e-06 | 8.2e-06 | 1.097E-05 | 3.573E-06 | 8.200E-06 | 1.500E-05 |
| 5588 | 280 | N280G | A | 1.4e-05 | 1.3e-05 | 1.5e-05 | 1.400E-05 | 1.000E-06 | 1.300E-05 | 1.500E-05 |
| 5589 | 280 | N280H | A | 9.9e-06 | 1.7e-05 | 1e-05 | 1.230E-05 | 4.071E-06 | 9.900E-06 | 1.700E-05 |
| 5590 | 280 | N280I | A | 1.3e-05 | 9.9e-06 | 9.9e-06 | 1.093E-05 | 1.790E-06 | 9.900E-06 | 1.300E-05 |
| 5591 | 280 | N280L | A | 1.5e-05 | 1.5e-05 | 7.2e-06 | 1.240E-05 | 4.503E-06 | 7.200E-06 | 1.500E-05 |
| 5592 | 280 | N280K | A | 1.1e-05 | 1.2e-05 | 9.5e-06 | 1.083E-05 | 1.258E-06 | 9.500E-06 | 1.200E-05 |
| 5593 | 280 | N280M | A | 1.5e-05 | 1.5e-05 | 1.1e-05 | 1.367E-05 | 2.309E-06 | 1.100E-05 | 1.500E-05 |
| 5594 | 280 | N280F | A | 1.3e-05 | 1.5e-05 | 1.6e-05 | 1.467E-05 | 1.528E-06 | 1.300E-05 | 1.600E-05 |
| 5595 | 280 | N280P | A | 8.5e-06 | 1.1e-05 | 1.6e-05 | 1.183E-05 | 3.819E-06 | 8.500E-06 | 1.600E-05 |
| 5596 | 280 | N280S | A | 1.1e-05 | 1.2e-05 | 1.4e-05 | 1.233E-05 | 1.528E-06 | 1.100E-05 | 1.400E-05 |
| 5597 | 280 | N280T | A | 1.4e-05 | 1.5e-05 | 1.1e-05 | 1.333E-05 | 2.082E-06 | 1.100E-05 | 1.500E-05 |
| 5598 | 280 | N280W | A | 1.1e-05 | 1.2e-05 | 1.5e-05 | 1.267E-05 | 2.082E-06 | 1.100E-05 | 1.500E-05 |
| 5599 | 280 | N280Y | A | 1.5e-05 | 1.5e-05 | 1.6e-05 | 1.533E-05 | 5.774E-07 | 1.500E-05 | 1.600E-05 |
| 5600 | 280 | N280V | A | 1.7e-05 | 7.8e-06 | 1.2e-05 | 1.227E-05 | 4.606E-06 | 7.800E-06 | 1.700E-05 |
| 5601 | 281 | Y281A | A | 1.1e-05 | 1e-05 | 1.5e-05 | 1.200E-05 | 2.646E-06 | 1.000E-05 | 1.500E-05 |
| 5602 | 281 | Y281R | A | 9.7e-06 | 1.8e-05 | 1.1e-05 | 1.290E-05 | 4.464E-06 | 9.700E-06 | 1.800E-05 |
| 5603 | 281 | Y281N | A | 1.4e-05 | 1.5e-05 | 1e-05 | 1.300E-05 | 2.646E-06 | 1.000E-05 | 1.500E-05 |
| 5604 | 281 | Y281D | A | 9.5e-06 | 1.2e-05 | 1.6e-05 | 1.250E-05 | 3.279E-06 | 9.500E-06 | 1.600E-05 |
| 5605 | 281 | Y281C | A | 1.4e-05 | 1.5e-05 | 8e-06 | 1.233E-05 | 3.786E-06 | 8.000E-06 | 1.500E-05 |
| 5606 | 281 | Y281E | A | 1.5e-05 | 1.1e-05 | 1.5e-05 | 1.367E-05 | 2.309E-06 | 1.100E-05 | 1.500E-05 |
| 5607 | 281 | Y281Q | A | 1.5e-05 | 9.8e-06 | 9.1e-06 | 1.130E-05 | 3.223E-06 | 9.100E-06 | 1.500E-05 |
| 5608 | 281 | Y281G | A | 1.6e-05 | 1.1e-05 | 1.6e-05 | 1.433E-05 | 2.887E-06 | 1.100E-05 | 1.600E-05 |
| 5609 | 281 | Y281H | A | 1e-05 | 1.1e-05 | 9.9e-06 | 1.030E-05 | 6.083E-07 | 9.900E-06 | 1.100E-05 |

| | | | | | | | | | | |
|---|---|---|---|---|---|---|---|---|---|---|
| 5610 | 281 | Y281I | A | 1.4e-05 | 1.1e-05 | 1.2e-05 | 1.233E-05 | 1.528E-06 | 1.100E-05 | 1.400E-05 |
| 5611 | 281 | Y281L | A | 1.6e-05 | 1.6e-05 | 8.6e-06 | 1.353E-05 | 4.272E-06 | 8.600E-06 | 1.600E-05 |
| 5612 | 281 | Y281K | A | 1.4e-05 | 9.9e-06 | 1.6e-05 | 1.330E-05 | 3.110E-06 | 9.900E-06 | 1.600E-05 |
| 5613 | 281 | Y281M | A | 1.4e-05 | 1.4e-05 | 1.5e-05 | 1.433E-05 | 5.774E-07 | 1.400E-05 | 1.500E-05 |
| 5614 | 281 | Y281F | A | 1.6e-05 | 1.2e-05 | 1.3e-05 | 1.367E-05 | 2.082E-06 | 1.200E-05 | 1.600E-05 |
| 5615 | 281 | Y281P | A | 1.1e-05 | 8.1e-06 | 1.1e-05 | 1.003E-05 | 1.674E-06 | 8.100E-06 | 1.100E-05 |
| 5616 | 281 | Y281S | A | 1.6e-05 | 8.9e-06 | 1.1e-05 | 1.197E-05 | 3.647E-06 | 8.900E-06 | 1.600E-05 |
| 5617 | 281 | Y281T | A | 1.5e-05 | 1.6e-05 | 1.4e-05 | 1.500E-05 | 1.000E-06 | 1.400E-05 | 1.600E-05 |
| 5618 | 281 | Y281W | A | 1.5e-05 | 9.6e-06 | 1.6e-05 | 1.353E-05 | 3.443E-06 | 9.600E-06 | 1.600E-05 |
| 5619 | 281 | Y281Y | A | 1e-05 | 1.3e-05 | 8.6e-06 | 1.053E-05 | 2.248E-06 | 8.600E-06 | 1.300E-05 |
| 5620 | 281 | Y281V | A | 1.7e-05 | 1.5e-05 | 9.6e-06 | 1.387E-05 | 3.828E-06 | 9.600E-06 | 1.700E-05 |
| 5621 | 282 | L282A | A | 1.5e-05 | 9.6e-06 | 9.6e-06 | 1.140E-05 | 3.118E-06 | 9.600E-06 | 1.500E-05 |
| 5622 | 282 | L282R | A | 1.2e-05 | 1.4e-05 | 8.5e-06 | 1.150E-05 | 2.784E-06 | 8.500E-06 | 1.400E-05 |
| 5623 | 282 | L282N | A | 1.4e-05 | 1.3e-05 | 8.1e-06 | 1.170E-05 | 3.158E-06 | 8.100E-06 | 1.400E-05 |
| 5624 | 282 | L282D | A | 9.6e-06 | 1.5e-05 | 1.5e-05 | 1.320E-05 | 3.118E-06 | 9.600E-06 | 1.500E-05 |
| 5625 | 282 | L282C | A | 1.5e-05 | 1.1e-05 | 1.1e-05 | 1.233E-05 | 2.309E-06 | 1.100E-05 | 1.500E-05 |
| 5626 | 282 | L282E | A | 1.4e-05 | 1.6e-05 | 1.1e-05 | 1.367E-05 | 2.517E-06 | 1.100E-05 | 1.600E-05 |
| 5627 | 282 | L282Q | A | 1.3e-05 | 8.2e-06 | 1.4e-05 | 1.173E-05 | 3.101E-06 | 8.200E-06 | 1.400E-05 |
| 5628 | 282 | L282G | A | 1.6e-05 | 1.6e-05 | 1.6e-05 | 1.600E-05 | 0.000E+00 | 1.600E-05 | 1.600E-05 |
| 5629 | 282 | L282H | A | 1e-05 | 1.5e-05 | 1e-05 | 1.167E-05 | 2.887E-06 | 1.000E-05 | 1.500E-05 |
| 5630 | 282 | L282I | A | 1.6e-05 | 1.2e-05 | 1.3e-05 | 1.367E-05 | 2.082E-06 | 1.200E-05 | 1.600E-05 |
| 5631 | 282 | L282L | A | 1.9e-05 | 1.4e-05 | 1.1e-05 | 1.467E-05 | 4.041E-06 | 1.100E-05 | 1.900E-05 |
| 5632 | 282 | L282K | A | 1.5e-05 | 1.2e-05 | 1.5e-05 | 1.400E-05 | 1.732E-06 | 1.200E-05 | 1.500E-05 |
| 5633 | 282 | L282M | A | 9.6e-06 | 1e-05 | 1.1e-05 | 1.020E-05 | 7.211E-07 | 9.600E-06 | 1.100E-05 |
| 5634 | 282 | L282F | A | 9.6e-06 | 1.6e-05 | 1.6e-05 | 1.387E-05 | 3.695E-06 | 9.600E-06 | 1.600E-05 |
| 5635 | 282 | L282P | A | 1.4e-05 | 1.3e-05 | 1.6e-05 | 1.433E-05 | 1.528E-06 | 1.300E-05 | 1.600E-05 |
| 5636 | 282 | L282S | A | 1.3e-05 | 1.5e-05 | 1.6e-05 | 1.467E-05 | 1.528E-06 | 1.300E-05 | 1.600E-05 |
| 5637 | 282 | L282T | A | 1.3e-05 | 1.5e-05 | 1.1e-05 | 1.300E-05 | 2.000E-06 | 1.100E-05 | 1.500E-05 |
| 5638 | 282 | L282W | A | 1.2e-05 | 1e-05 | 1.5e-05 | 1.233E-05 | 2.517E-06 | 1.000E-05 | 1.500E-05 |
| 5639 | 282 | L282Y | A | 9.3e-06 | 1.1e-05 | 1.4e-05 | 1.143E-05 | 2.380E-06 | 9.300E-06 | 1.400E-05 |
| 5640 | 282 | L282V | A | 1.1e-05 | 9.4e-06 | 9.2e-06 | 9.867E-06 | 9.866E-07 | 9.200E-06 | 1.100E-05 |
| 5641 | 283 | S283A | A | 1.6e-05 | 1.2e-05 | 1.1e-05 | 1.300E-05 | 2.646E-06 | 1.100E-05 | 1.600E-05 |
| 5642 | 283 | S283R | A | 1.6e-05 | 1.1e-05 | 1.5e-05 | 1.400E-05 | 2.646E-06 | 1.100E-05 | 1.600E-05 |
| 5643 | 283 | S283N | A | 1.4e-05 | 9.9e-06 | 1.6e-05 | 1.330E-05 | 3.110E-06 | 9.900E-06 | 1.600E-05 |
| 5644 | 283 | S283D | A | 1.1e-05 | 1.5e-05 | 1.5e-05 | 1.367E-05 | 2.309E-06 | 1.100E-05 | 1.500E-05 |
| 5645 | 283 | S283C | A | 1e-05 | 8.2e-06 | 1.2e-05 | 1.007E-05 | 1.901E-06 | 8.200E-06 | 1.200E-05 |
| 5646 | 283 | S283E | A | 1.7e-05 | 1.3e-05 | 1.1e-05 | 1.367E-05 | 3.055E-06 | 1.100E-05 | 1.700E-05 |
| 5647 | 283 | S283Q | A | 1.1e-05 | 1.6e-05 | 1e-05 | 1.233E-05 | 3.215E-06 | 1.000E-05 | 1.600E-05 |
| 5648 | 283 | S283G | A | 1e-05 | 1.4e-05 | 1e-05 | 1.133E-05 | 2.309E-06 | 1.000E-05 | 1.400E-05 |
| 5649 | 283 | S283H | A | 9.1e-06 | 1.4e-05 | 1.3e-05 | 1.203E-05 | 2.589E-06 | 9.100E-06 | 1.400E-05 |
| 5650 | 283 | S283I | A | 1.1e-05 | 1.7e-05 | 1e-05 | 1.267E-05 | 3.786E-06 | 1.000E-05 | 1.700E-05 |
| 5651 | 283 | S283L | A | 1.3e-05 | 1.2e-05 | 1.5e-05 | 1.333E-05 | 1.528E-06 | 1.200E-05 | 1.500E-05 |
| 5652 | 283 | S283K | A | 1e-05 | 9.9e-06 | 1e-05 | 9.967E-06 | 5.774E-08 | 9.900E-06 | 1.000E-05 |
| 5653 | 283 | S283M | A | 1.6e-05 | 1.4e-05 | 1.4e-05 | 1.467E-05 | 1.155E-06 | 1.400E-05 | 1.600E-05 |
| 5654 | 283 | S283F | A | 1.1e-05 | 1.6e-05 | 1.8e-05 | 1.500E-05 | 3.606E-06 | 1.100E-05 | 1.800E-05 |
| 5655 | 283 | S283P | A | 1e-05 | 1.3e-05 | 1.1e-05 | 1.133E-05 | 1.528E-06 | 1.000E-05 | 1.300E-05 |
| 5656 | 283 | S283S | A | 9.9e-06 | 1.6e-05 | 1.3e-05 | 1.297E-05 | 3.050E-06 | 9.900E-06 | 1.600E-05 |
| 5657 | 283 | S283T | A | 1.6e-05 | 1.1e-05 | 9.7e-06 | 1.223E-05 | 3.326E-06 | 9.700E-06 | 1.600E-05 |
| 5658 | 283 | S283W | A | 9.3e-06 | 1.6e-05 | 2.1e-05 | 1.543E-05 | 5.871E-06 | 9.300E-06 | 2.100E-05 |
| 5659 | 283 | S283Y | A | 1.5e-05 | 1.6e-05 | 1.6e-05 | 1.567E-05 | 5.774E-07 | 1.500E-05 | 1.600E-05 |
| 5660 | 283 | S283V | A | 1.1e-05 | 1.4e-05 | 1.4e-05 | 1.300E-05 | 1.732E-06 | 1.100E-05 | 1.400E-05 |
| 5661 | 284 | T284A | A | 1.6e-05 | 1.7e-05 | 1.4e-05 | 1.567E-05 | 1.528E-06 | 1.400E-05 | 1.700E-05 |
| 5662 | 284 | T284R | A | 9.9e-06 | 1.4e-05 | 3.2e-05 | 1.863E-05 | 1.176E-05 | 9.900E-06 | 3.200E-05 |
| 5663 | 284 | T284N | A | 1.4e-05 | 1.1e-05 | 9.7e-06 | 1.157E-05 | 2.205E-06 | 9.700E-06 | 1.400E-05 |
| 5664 | 284 | T284D | A | 1e-05 | 1.5e-05 | 1.1e-05 | 1.200E-05 | 2.646E-06 | 1.000E-05 | 1.500E-05 |
| 5665 | 284 | T284C | A | 1.1e-05 | 1.2e-05 | 1e-05 | 1.100E-05 | 1.000E-06 | 1.000E-05 | 1.200E-05 |
| 5666 | 284 | T284E | A | 1.2e-05 | 1.4e-05 | 1.3e-05 | 1.300E-05 | 1.000E-06 | 1.200E-05 | 1.400E-05 |
| 5667 | 284 | T284Q | A | 9.5e-06 | 9.9e-06 | 1.4e-05 | 1.113E-05 | 2.491E-06 | 9.500E-06 | 1.400E-05 |
| 5668 | 284 | T284G | A | 1e-05 | 7.6e-06 | 1e-05 | 9.200E-06 | 1.386E-06 | 7.600E-06 | 1.000E-05 |
| 5669 | 284 | T284H | A | 1.5e-05 | 1e-05 | 1.6e-05 | 1.367E-05 | 3.215E-06 | 1.000E-05 | 1.600E-05 |
| 5670 | 284 | T284I | A | 1.8e-05 | 1.5e-05 | 1.3e-05 | 1.533E-05 | 2.517E-06 | 1.300E-05 | 1.800E-05 |
| 5671 | 284 | T284L | A | 1.4e-05 | 1.4e-05 | 1e-05 | 1.267E-05 | 2.309E-06 | 1.000E-05 | 1.400E-05 |
| 5672 | 284 | T284K | A | 1.2e-05 | 1.4e-05 | 1.4e-05 | 1.333E-05 | 1.155E-06 | 1.200E-05 | 1.400E-05 |
| 5673 | 284 | T284M | A | 1.5e-05 | 1.4e-05 | 1e-05 | 1.300E-05 | 2.646E-06 | 1.000E-05 | 1.500E-05 |
| 5674 | 284 | T284F | A | 1e-05 | 9.6e-06 | 1e-05 | 9.867E-06 | 2.309E-07 | 9.600E-06 | 1.000E-05 |
| 5675 | 284 | T284P | A | 1.1e-05 | 1.1e-05 | 1.7e-05 | 1.300E-05 | 3.464E-06 | 1.100E-05 | 1.700E-05 |
| 5676 | 284 | T284S | A | 9.4e-06 | 1.1e-05 | 1.5e-05 | 1.180E-05 | 2.884E-06 | 9.400E-06 | 1.500E-05 |
| 5677 | 284 | T284T | A | 1.2e-05 | 1.4e-05 | 9.3e-06 | 1.177E-05 | 2.359E-06 | 9.300E-06 | 1.400E-05 |
| 5678 | 284 | T284W | A | 1.6e-05 | 1.5e-05 | 9.5e-06 | 1.350E-05 | 3.500E-06 | 9.500E-06 | 1.600E-05 |
| 5679 | 284 | T284Y | A | 1.2e-05 | 1.4e-05 | 1.6e-05 | 1.400E-05 | 2.000E-06 | 1.200E-05 | 1.600E-05 |
| 5680 | 284 | T284V | A | 1.1e-05 | 1.5e-05 | 1.4e-05 | 1.333E-05 | 2.082E-06 | 1.100E-05 | 1.500E-05 |
| 5681 | 285 | D285A | A | 1.3e-05 | 1.6e-05 | 1e-05 | 1.300E-05 | 3.000E-06 | 1.000E-05 | 1.600E-05 |
| 5682 | 285 | D285R | A | 1.7e-05 | 1.5e-05 | 9.6e-06 | 1.387E-05 | 3.828E-06 | 9.600E-06 | 1.700E-05 |
| 5683 | 285 | D285N | A | 1e-05 | 5.9e-06 | 1.4e-05 | 9.967E-06 | 4.050E-06 | 5.900E-06 | 1.400E-05 |
| 5684 | 285 | D285D | A | 8e-06 | 1.2e-05 | 1.4e-05 | 1.133E-05 | 3.055E-06 | 8.000E-06 | 1.400E-05 |
| 5685 | 285 | D285C | A | 1.2e-05 | 8.4e-06 | 8.7e-06 | 9.700E-06 | 1.997E-06 | 8.400E-06 | 1.200E-05 |
| 5686 | 285 | D285E | A | 1.6e-05 | 1.7e-05 | 7.3e-06 | 1.343E-05 | 5.335E-06 | 7.300E-06 | 1.700E-05 |
| 5687 | 285 | D285Q | A | 1.2e-05 | 9.4e-06 | 1.6e-05 | 1.247E-05 | 3.325E-06 | 9.400E-06 | 1.600E-05 |
| 5688 | 285 | D285G | A | 8.9e-06 | 1.4e-05 | 1.4e-05 | 1.230E-05 | 2.944E-06 | 8.900E-06 | 1.400E-05 |
| 5689 | 285 | D285H | A | 1.5e-05 | 1.3e-05 | 1.1e-05 | 1.300E-05 | 2.000E-06 | 1.100E-05 | 1.500E-05 |
| 5690 | 285 | D285I | A | 1.5e-05 | 1.5e-05 | 1.4e-05 | 1.467E-05 | 5.774E-07 | 1.400E-05 | 1.500E-05 |
| 5691 | 285 | D285L | A | 1.6e-05 | 1.2e-05 | 1.3e-05 | 1.367E-05 | 2.082E-06 | 1.200E-05 | 1.600E-05 |
| 5692 | 285 | D285K | A | 8.7e-06 | 1.1e-05 | 1.3e-05 | 1.090E-05 | 2.152E-06 | 8.700E-06 | 1.300E-05 |
| 5693 | 285 | D285M | A | 1.6e-05 | 7.4e-06 | 1.5e-05 | 1.280E-05 | 4.703E-06 | 7.400E-06 | 1.600E-05 |
| 5694 | 285 | D285F | A | 1.5e-05 | 9.8e-06 | 9.6e-06 | 1.147E-05 | 3.062E-06 | 9.600E-06 | 1.500E-05 |

| | | | | | | | | | | |
|---|---|---|---|---|---|---|---|---|---|---|
| 5695 | 285 | D285P | A | 1.9e-05 | 8.4e-06 | 1e-05 | 1.247E-05 | 5.714E-06 | 8.400E-06 | 1.900E-05 |
| 5696 | 285 | D285S | A | 1.1e-05 | 1.4e-05 | 1.1e-05 | 1.200E-05 | 1.732E-06 | 1.100E-05 | 1.400E-05 |
| 5697 | 285 | D285T | A | 1.1e-05 | 9.2e-06 | 1.1e-05 | 1.040E-05 | 1.039E-06 | 9.200E-06 | 1.100E-05 |
| 5698 | 285 | D285W | A | 1.5e-05 | 9.8e-06 | 9.5e-06 | 1.143E-05 | 3.092E-06 | 9.500E-06 | 1.500E-05 |
| 5699 | 285 | D285Y | A | 1.5e-05 | 1.7e-05 | 1.4e-05 | 1.533E-05 | 1.528E-06 | 1.400E-05 | 1.700E-05 |
| 5700 | 285 | D285V | A | 1.4e-05 | 1.5e-05 | 9.5e-06 | 1.283E-05 | 2.930E-06 | 9.500E-06 | 1.500E-05 |
| 5701 | 286 | V286A | A | 1.5e-05 | 8.1e-06 | 1.7e-05 | 1.337E-05 | 4.669E-06 | 8.100E-06 | 1.700E-05 |
| 5702 | 286 | V286R | A | 1.5e-05 | 1.6e-05 | 8.8e-06 | 1.327E-05 | 3.900E-06 | 8.800E-06 | 1.600E-05 |
| 5703 | 286 | V286N | A | 1.4e-05 | 1.3e-05 | 1.1e-05 | 1.267E-05 | 1.528E-06 | 1.100E-05 | 1.400E-05 |
| 5704 | 286 | V286D | A | 1e-05 | 1.3e-05 | 1.3e-05 | 1.200E-05 | 1.732E-06 | 1.000E-05 | 1.300E-05 |
| 5705 | 286 | V286C | A | 1.1e-05 | 1e-05 | 9.8e-06 | 1.027E-05 | 6.429E-07 | 9.800E-06 | 1.100E-05 |
| 5706 | 286 | V286E | A | 1.5e-05 | 1.5e-05 | 1.5e-05 | 1.500E-05 | 0.000E+00 | 1.500E-05 | 1.500E-05 |
| 5707 | 286 | V286Q | A | 9.7e-06 | 1.4e-05 | 1.6e-05 | 1.323E-05 | 3.219E-06 | 9.700E-06 | 1.600E-05 |
| 5708 | 286 | V286G | A | 8.1e-06 | 9.5e-06 | 1.5e-05 | 1.087E-05 | 3.647E-06 | 8.100E-06 | 1.500E-05 |
| 5709 | 286 | V286H | A | 1.6e-05 | 1.1e-05 | 1.1e-05 | 1.267E-05 | 2.887E-06 | 1.100E-05 | 1.600E-05 |
| 5710 | 286 | V286I | A | 1.3e-05 | 9.7e-06 | 1.5e-05 | 1.257E-05 | 2.676E-06 | 9.700E-06 | 1.500E-05 |
| 5711 | 286 | V286L | A | 1.3e-05 | 1.4e-05 | 1.3e-05 | 1.333E-05 | 5.774E-07 | 1.300E-05 | 1.400E-05 |
| 5712 | 286 | V286K | A | 1.6e-05 | 9.6e-06 | 1.1e-05 | 1.220E-05 | 3.365E-06 | 9.600E-06 | 1.600E-05 |
| 5713 | 286 | V286M | A | 1.2e-05 | 9.6e-06 | 1.5e-05 | 1.220E-05 | 2.706E-06 | 9.600E-06 | 1.500E-05 |
| 5714 | 286 | V286F | A | 1.6e-05 | 1e-05 | 1.1e-05 | 1.233E-05 | 3.215E-06 | 1.000E-05 | 1.600E-05 |
| 5715 | 286 | V286P | A | 1.6e-05 | 1e-05 | 1.2e-05 | 1.267E-05 | 3.055E-06 | 1.000E-05 | 1.600E-05 |
| 5716 | 286 | V286S | A | 1.5e-05 | 1.3e-05 | 5.4e-06 | 1.113E-05 | 5.065E-06 | 5.400E-06 | 1.500E-05 |
| 5717 | 286 | V286T | A | 1.6e-05 | 1.3e-05 | 1e-05 | 1.300E-05 | 3.000E-06 | 1.000E-05 | 1.600E-05 |
| 5718 | 286 | V286W | A | 1.1e-05 | 1.4e-05 | 1.6e-05 | 1.367E-05 | 2.517E-06 | 1.100E-05 | 1.600E-05 |
| 5719 | 286 | V286Y | A | 1.4e-05 | 9.8e-06 | 1.5e-05 | 1.293E-05 | 2.759E-06 | 9.800E-06 | 1.500E-05 |
| 5720 | 286 | V286V | A | 1.2e-05 | 6.8e-06 | 1.1e-05 | 9.933E-06 | 2.759E-06 | 6.800E-06 | 1.200E-05 |
| 5721 | 287 | G287A | A | 1.6e-05 | 1.1e-05 | 1.7e-05 | 1.467E-05 | 3.215E-06 | 1.100E-05 | 1.700E-05 |
| 5722 | 287 | G287R | A | 1e-05 | 1.3e-05 | 1.5e-05 | 1.267E-05 | 2.517E-06 | 1.000E-05 | 1.500E-05 |
| 5723 | 287 | G287N | A | 1.1e-05 | 1.2e-05 | 9.7e-06 | 1.090E-05 | 1.153E-06 | 9.700E-06 | 1.200E-05 |
| 5724 | 287 | G287D | A | 1.5e-05 | 1e-05 | 1.1e-05 | 1.200E-05 | 2.646E-06 | 1.000E-05 | 1.500E-05 |
| 5725 | 287 | G287C | A | 1.1e-05 | 2e-05 | 9.9e-06 | 1.363E-05 | 5.541E-06 | 9.900E-06 | 2.000E-05 |
| 5726 | 287 | G287E | A | 1.3e-05 | 1.1e-05 | 1.7e-05 | 1.367E-05 | 3.055E-06 | 1.100E-05 | 1.700E-05 |
| 5727 | 287 | G287Q | A | 1.1e-05 | 1.1e-05 | 1.5e-05 | 1.233E-05 | 2.309E-06 | 1.100E-05 | 1.500E-05 |
| 5728 | 287 | G287G | A | 1.5e-05 | 1.5e-05 | 9.8e-06 | 1.327E-05 | 3.002E-06 | 9.800E-06 | 1.500E-05 |
| 5729 | 287 | G287H | A | 1.4e-05 | 1.3e-05 | 1.5e-05 | 1.400E-05 | 1.000E-06 | 1.300E-05 | 1.500E-05 |
| 5730 | 287 | G287I | A | 9.4e-06 | 1.7e-05 | 1.2e-05 | 1.280E-05 | 3.863E-06 | 9.400E-06 | 1.700E-05 |
| 5731 | 287 | G287L | A | 1.5e-05 | 1.6e-05 | 1.3e-05 | 1.467E-05 | 1.528E-06 | 1.300E-05 | 1.600E-05 |
| 5732 | 287 | G287K | A | 1.4e-05 | 1.4e-05 | 8.6e-06 | 1.220E-05 | 3.118E-06 | 8.600E-06 | 1.400E-05 |
| 5733 | 287 | G287M | A | 1.3e-05 | 1.1e-05 | 1.4e-05 | 1.267E-05 | 1.528E-06 | 1.100E-05 | 1.400E-05 |
| 5734 | 287 | G287F | A | 1.4e-05 | 1.4e-05 | 1.1e-05 | 1.300E-05 | 1.732E-06 | 1.100E-05 | 1.400E-05 |
| 5735 | 287 | G287P | A | 1.4e-05 | 1e-05 | 1e-05 | 1.133E-05 | 2.309E-06 | 1.000E-05 | 1.400E-05 |
| 5736 | 287 | G287S | A | 1.1e-05 | 1.4e-05 | 1.6e-05 | 1.367E-05 | 2.517E-06 | 1.100E-05 | 1.600E-05 |
| 5737 | 287 | G287T | A | 1.7e-05 | 1e-05 | 1e-05 | 1.233E-05 | 4.041E-06 | 1.000E-05 | 1.700E-05 |
| 5738 | 287 | G287W | A | 1.5e-05 | 1e-05 | 1.5e-05 | 1.333E-05 | 2.887E-06 | 1.000E-05 | 1.500E-05 |
| 5739 | 287 | G287Y | A | 1.3e-05 | 1.1e-05 | 9.7e-06 | 1.123E-05 | 1.662E-06 | 9.700E-06 | 1.300E-05 |
| 5740 | 287 | G287V | A | 1e-05 | 9.3e-06 | 1.5e-05 | 1.143E-05 | 3.109E-06 | 9.300E-06 | 1.500E-05 |
| 5741 | 288 | S288A | A | 9.6e-06 | 1.2e-05 | 1e-05 | 1.053E-05 | 1.286E-06 | 9.600E-06 | 1.200E-05 |
| 5742 | 288 | S288R | A | 1.5e-05 | 1.5e-05 | 8.6e-06 | 1.287E-05 | 3.695E-06 | 8.600E-06 | 1.500E-05 |
| 5743 | 288 | S288N | A | 1.1e-05 | 9.8e-06 | 9.8e-06 | 1.020E-05 | 6.928E-07 | 9.800E-06 | 1.100E-05 |
| 5744 | 288 | S288D | A | 1e-05 | 1.2e-05 | 1.1e-05 | 1.100E-05 | 1.000E-06 | 1.000E-05 | 1.200E-05 |
| 5745 | 288 | S288C | A | 9.8e-06 | 1.3e-05 | 1.5e-05 | 1.260E-05 | 2.623E-06 | 9.800E-06 | 1.500E-05 |
| 5746 | 288 | S288E | A | 1.1e-05 | 1.2e-05 | 1.5e-05 | 1.267E-05 | 2.082E-06 | 1.100E-05 | 1.500E-05 |
| 5747 | 288 | S288Q | A | 1.3e-05 | 1.6e-05 | 9.9e-06 | 1.297E-05 | 3.050E-06 | 9.900E-06 | 1.600E-05 |
| 5748 | 288 | S288G | A | 9.9e-06 | 1.2e-05 | 1.5e-05 | 1.230E-05 | 2.563E-06 | 9.900E-06 | 1.500E-05 |
| 5749 | 288 | S288H | A | 1.7e-05 | 1.1e-05 | 1.6e-05 | 1.467E-05 | 3.215E-06 | 1.100E-05 | 1.700E-05 |
| 5750 | 288 | S288I | A | 1.1e-05 | 1.6e-05 | 1.6e-05 | 1.433E-05 | 2.887E-06 | 1.100E-05 | 1.600E-05 |
| 5751 | 288 | S288L | A | 1.1e-05 | 8.7e-06 | 1.4e-05 | 1.123E-05 | 2.658E-06 | 8.700E-06 | 1.400E-05 |
| 5752 | 288 | S288K | A | 1.5e-05 | 1.3e-05 | 9.3e-06 | 1.243E-05 | 2.892E-06 | 9.300E-06 | 1.500E-05 |
| 5753 | 288 | S288M | A | 1.5e-05 | 1.5e-05 | 1.2e-05 | 1.400E-05 | 1.732E-06 | 1.200E-05 | 1.500E-05 |
| 5754 | 288 | S288F | A | 1.7e-05 | 1.8e-05 | 1.6e-05 | 1.700E-05 | 1.000E-06 | 1.600E-05 | 1.800E-05 |
| 5755 | 288 | S288P | A | 1.2e-05 | 1e-05 | 6.3e-06 | 9.433E-06 | 2.892E-06 | 6.300E-06 | 1.200E-05 |
| 5756 | 288 | S288S | A | 1.6e-05 | 1.5e-05 | 1.7e-05 | 1.600E-05 | 1.000E-06 | 1.500E-05 | 1.700E-05 |
| 5757 | 288 | S288T | A | 8.9e-06 | 1.5e-05 | 1.4e-05 | 1.263E-05 | 3.272E-06 | 8.900E-06 | 1.500E-05 |
| 5758 | 288 | S288W | A | 1.4e-05 | 1.6e-05 | 1.3e-05 | 1.433E-05 | 1.528E-06 | 1.300E-05 | 1.600E-05 |
| 5759 | 288 | S288Y | A | 1.4e-05 | 1.2e-05 | 1.1e-05 | 1.233E-05 | 1.528E-06 | 1.100E-05 | 1.400E-05 |
| 5760 | 288 | S288V | A | 1.1e-05 | 1.6e-05 | 1.5e-05 | 1.400E-05 | 2.646E-06 | 1.100E-05 | 1.600E-05 |
| 5761 | 289 | C289A | A | 1.1e-05 | 1.2e-05 | 1.7e-05 | 1.333E-05 | 3.215E-06 | 1.100E-05 | 1.700E-05 |
| 5762 | 289 | C289R | A | 1.5e-05 | 1.5e-05 | 1e-05 | 1.333E-05 | 2.887E-06 | 1.000E-05 | 1.500E-05 |
| 5763 | 289 | C289N | A | 1.2e-05 | 1.4e-05 | 1.4e-05 | 1.333E-05 | 1.155E-06 | 1.200E-05 | 1.400E-05 |
| 5764 | 289 | C289D | A | 1.6e-05 | 1e-05 | 1.7e-05 | 1.433E-05 | 3.786E-06 | 1.000E-05 | 1.700E-05 |
| 5765 | 289 | C289C | A | 1.7e-05 | 1.1e-05 | 1e-05 | 1.267E-05 | 3.786E-06 | 1.000E-05 | 1.700E-05 |
| 5766 | 289 | C289E | A | 1.4e-05 | 1e-05 | 1.6e-05 | 1.333E-05 | 3.055E-06 | 1.000E-05 | 1.600E-05 |
| 5767 | 289 | C289Q | A | 1.6e-05 | 1.4e-05 | 1.1e-05 | 1.367E-05 | 2.517E-06 | 1.100E-05 | 1.600E-05 |
| 5768 | 289 | C289G | A | 1.5e-05 | 1.4e-05 | 1.2e-05 | 1.367E-05 | 1.528E-06 | 1.200E-05 | 1.500E-05 |
| 5769 | 289 | C289H | A | 1.4e-05 | 1.7e-05 | 1.5e-05 | 1.533E-05 | 1.528E-06 | 1.400E-05 | 1.700E-05 |
| 5770 | 289 | C289I | A | 1e-05 | 1.4e-05 | 1.6e-05 | 1.333E-05 | 3.055E-06 | 1.000E-05 | 1.600E-05 |
| 5771 | 289 | C289L | A | 1.8e-05 | 1.5e-05 | 1.1e-05 | 1.467E-05 | 3.512E-06 | 1.100E-05 | 1.800E-05 |
| 5772 | 289 | C289K | A | 1.6e-05 | 9.8e-06 | 1.1e-05 | 1.227E-05 | 3.288E-06 | 9.800E-06 | 1.600E-05 |
| 5773 | 289 | C289M | A | 1.1e-05 | 1.7e-05 | 1.6e-05 | 1.467E-05 | 3.215E-06 | 1.100E-05 | 1.700E-05 |
| 5774 | 289 | C289F | A | 9.8e-06 | 9.9e-06 | 1.1e-05 | 1.023E-05 | 6.658E-07 | 9.800E-06 | 1.100E-05 |
| 5775 | 289 | C289P | A | 1.1e-05 | 1.2e-05 | 1.2e-05 | 1.167E-05 | 5.774E-07 | 1.100E-05 | 1.200E-05 |
| 5776 | 289 | C289S | A | 1.2e-05 | 1.5e-05 | 1e-05 | 1.233E-05 | 2.517E-06 | 1.000E-05 | 1.500E-05 |
| 5777 | 289 | C289T | A | 1.1e-05 | 9.3e-06 | 1.4e-05 | 1.143E-05 | 2.380E-06 | 9.300E-06 | 1.400E-05 |
| 5778 | 289 | C289W | A | 1.1e-05 | 1.2e-05 | 1.5e-05 | 1.267E-05 | 2.082E-06 | 1.100E-05 | 1.500E-05 |
| 5779 | 289 | C289Y | A | 8.3e-06 | 9.9e-06 | 1.1e-05 | 9.733E-06 | 1.358E-06 | 8.300E-06 | 1.100E-05 |

| | | | | | | | | | | |
|---|---|---|---|---|---|---|---|---|---|---|
| 5780 | 289 | C289V | A | 1.6e-05 | 1e-05 | 1.5e-05 | 1.367E-05 | 3.215E-06 | 1.000E-05 | 1.600E-05 |
| 5781 | 290 | T290A | A | 1.5e-05 | 1.1e-05 | 1.1e-05 | 1.233E-05 | 2.309E-06 | 1.100E-05 | 1.500E-05 |
| 5782 | 290 | T290R | A | 1.1e-05 | 1.6e-05 | 1.4e-05 | 1.367E-05 | 2.517E-06 | 1.100E-05 | 1.600E-05 |
| 5783 | 290 | T290N | A | 7.6e-06 | 1.3e-05 | 1.5e-05 | 1.187E-05 | 3.828E-06 | 7.600E-06 | 1.500E-05 |
| 5784 | 290 | T290D | A | 1.2e-05 | 1e-05 | 1.3e-05 | 1.167E-05 | 1.528E-06 | 1.000E-05 | 1.300E-05 |
| 5785 | 290 | T290C | A | 1.6e-05 | 1.5e-05 | 1.4e-05 | 1.500E-05 | 1.000E-06 | 1.400E-05 | 1.600E-05 |
| 5786 | 290 | T290E | A | 1.6e-05 | 1e-05 | 1.7e-05 | 1.433E-05 | 3.786E-06 | 1.000E-05 | 1.700E-05 |
| 5787 | 290 | T290Q | A | 1.8e-05 | 1.5e-05 | 1.5e-05 | 1.600E-05 | 1.732E-06 | 1.500E-05 | 1.800E-05 |
| 5788 | 290 | T290G | A | 1.7e-05 | 7e-06 | 1.6e-05 | 1.333E-05 | 5.508E-06 | 7.000E-06 | 1.700E-05 |
| 5789 | 290 | T290H | A | 8e-06 | 9.4e-06 | 1.6e-05 | 1.113E-05 | 4.272E-06 | 8.000E-06 | 1.600E-05 |
| 5790 | 290 | T290I | A | 1.4e-05 | 1.1e-05 | 1.2e-05 | 1.233E-05 | 1.528E-06 | 1.100E-05 | 1.400E-05 |
| 5791 | 290 | T290L | A | 1e-05 | 1.6e-05 | 6.9e-06 | 1.097E-05 | 4.626E-06 | 6.900E-06 | 1.600E-05 |
| 5792 | 290 | T290K | A | 1.1e-05 | 1.5e-05 | 1.1e-05 | 1.233E-05 | 2.309E-06 | 1.100E-05 | 1.500E-05 |
| 5793 | 290 | T290M | A | 1.6e-05 | 1.6e-05 | 1.5e-05 | 1.567E-05 | 5.774E-07 | 1.500E-05 | 1.600E-05 |
| 5794 | 290 | T290F | A | 1e-05 | 1.4e-05 | 1.7e-05 | 1.367E-05 | 3.512E-06 | 1.000E-05 | 1.700E-05 |
| 5795 | 290 | T290P | A | 1.4e-05 | 1.1e-05 | 1.4e-05 | 1.300E-05 | 1.732E-06 | 1.100E-05 | 1.400E-05 |
| 5796 | 290 | T290S | A | 1.1e-05 | 1.3e-05 | 1.5e-05 | 1.300E-05 | 2.000E-06 | 1.100E-05 | 1.500E-05 |
| 5797 | 290 | T290T | A | 1.5e-05 | 1.6e-05 | 1.6e-05 | 1.567E-05 | 5.774E-07 | 1.500E-05 | 1.600E-05 |
| 5798 | 290 | T290W | A | 9.8e-06 | 1.5e-05 | 1e-05 | 1.160E-05 | 2.946E-06 | 9.800E-06 | 1.500E-05 |
| 5799 | 290 | T290Y | A | 1.6e-05 | 1.6e-05 | 8.4e-06 | 1.347E-05 | 4.388E-06 | 8.400E-06 | 1.600E-05 |
| 5800 | 290 | T290V | A | 1.1e-05 | 8.4e-06 | 1.5e-05 | 1.147E-05 | 3.325E-06 | 8.400E-06 | 1.500E-05 |
| 5801 | 291 | L291A | A | 1.7e-05 | 1.7e-05 | 9.2e-06 | 1.440E-05 | 4.503E-06 | 9.200E-06 | 1.700E-05 |
| 5802 | 291 | L291R | A | 1.2e-05 | 1.4e-05 | 1.5e-05 | 1.367E-05 | 1.528E-06 | 1.200E-05 | 1.500E-05 |
| 5803 | 291 | L291N | A | 1.1e-05 | 1.4e-05 | 8.7e-06 | 1.123E-05 | 2.658E-06 | 8.700E-06 | 1.400E-05 |
| 5804 | 291 | L291D | A | 1.1e-05 | 1.1e-05 | 1e-05 | 1.067E-05 | 5.774E-07 | 1.000E-05 | 1.100E-05 |
| 5805 | 291 | L291C | A | 1.4e-05 | 1.3e-05 | 1.4e-05 | 1.367E-05 | 5.774E-07 | 1.300E-05 | 1.400E-05 |
| 5806 | 291 | L291E | A | 1.2e-05 | 1.4e-05 | 1e-05 | 1.200E-05 | 2.000E-06 | 1.000E-05 | 1.400E-05 |
| 5807 | 291 | L291Q | A | 9.4e-06 | 1.6e-05 | 1.6e-05 | 1.380E-05 | 3.811E-06 | 9.400E-06 | 1.600E-05 |
| 5808 | 291 | L291G | A | 1.4e-05 | 1.4e-05 | 1.6e-05 | 1.467E-05 | 1.155E-06 | 1.400E-05 | 1.600E-05 |
| 5809 | 291 | L291H | A | 1.3e-05 | 1e-05 | 1.3e-05 | 1.200E-05 | 1.732E-06 | 1.000E-05 | 1.300E-05 |
| 5810 | 291 | L291I | A | 9.7e-06 | 1.1e-05 | 1e-05 | 1.023E-05 | 6.807E-07 | 9.700E-06 | 1.100E-05 |
| 5811 | 291 | L291L | A | 1.5e-05 | 1.4e-05 | 9.6e-06 | 1.287E-05 | 2.873E-06 | 9.600E-06 | 1.500E-05 |
| 5812 | 291 | L291K | A | 1.1e-05 | 8.4e-06 | 1.5e-05 | 1.147E-05 | 3.325E-06 | 8.400E-06 | 1.500E-05 |
| 5813 | 291 | L291M | A | 1.3e-05 | 1.1e-05 | 1.5e-05 | 1.300E-05 | 2.000E-06 | 1.100E-05 | 1.500E-05 |
| 5814 | 291 | L291F | A | 1.4e-05 | 1e-05 | 8.5e-06 | 1.083E-05 | 2.843E-06 | 8.500E-06 | 1.400E-05 |
| 5815 | 291 | L291P | A | 1e-05 | 1.2e-05 | 9.4e-06 | 1.047E-05 | 1.361E-06 | 9.400E-06 | 1.200E-05 |
| 5816 | 291 | L291S | A | 1e-05 | 1.6e-05 | 1.1e-05 | 1.233E-05 | 3.215E-06 | 1.000E-05 | 1.600E-05 |
| 5817 | 291 | L291T | A | 8.8e-06 | 1.1e-05 | 9.2e-06 | 9.667E-06 | 1.172E-06 | 8.800E-06 | 1.100E-05 |
| 5818 | 291 | L291W | A | 1.6e-05 | 9.9e-06 | 9.2e-06 | 1.170E-05 | 3.740E-06 | 9.200E-06 | 1.600E-05 |
| 5819 | 291 | L291Y | A | 1.4e-05 | 1.5e-05 | 1.1e-05 | 1.333E-05 | 2.082E-06 | 1.100E-05 | 1.500E-05 |
| 5820 | 291 | L291V | A | 8.2e-06 | 9.4e-06 | 9.3e-06 | 8.967E-06 | 6.658E-07 | 8.200E-06 | 9.400E-06 |
| 5821 | 292 | V292A | A | 1.6e-05 | 8.9e-06 | 8.5e-06 | 1.113E-05 | 4.219E-06 | 8.500E-06 | 1.600E-05 |
| 5822 | 292 | V292R | A | 1.3e-05 | 2e-05 | 1.4e-05 | 1.567E-05 | 3.786E-06 | 1.300E-05 | 2.000E-05 |
| 5823 | 292 | V292N | A | 1.5e-05 | 1.3e-05 | 1.3e-05 | 1.367E-05 | 1.155E-06 | 1.300E-05 | 1.500E-05 |
| 5824 | 292 | V292D | A | 1.2e-05 | 1.5e-05 | 8.9e-06 | 1.197E-05 | 3.050E-06 | 8.900E-06 | 1.500E-05 |
| 5825 | 292 | V292C | A | 1.4e-05 | 7.7e-06 | 1.4e-05 | 1.190E-05 | 3.637E-06 | 7.700E-06 | 1.400E-05 |
| 5826 | 292 | V292E | A | 1.6e-05 | 1e-05 | 1.3e-05 | 1.300E-05 | 3.000E-06 | 1.000E-05 | 1.600E-05 |
| 5827 | 292 | V292Q | A | 1.1e-05 | 8.7e-06 | 9.1e-06 | 9.600E-06 | 1.229E-06 | 8.700E-06 | 1.100E-05 |
| 5828 | 292 | V292G | A | 1.4e-05 | 9.8e-06 | 1.5e-05 | 1.293E-05 | 2.759E-06 | 9.800E-06 | 1.500E-05 |
| 5829 | 292 | V292H | A | 1.4e-05 | 9.9e-06 | 1.6e-05 | 1.330E-05 | 3.110E-06 | 9.900E-06 | 1.600E-05 |
| 5830 | 292 | V292I | A | 1.6e-05 | 1.1e-05 | 1.5e-05 | 1.400E-05 | 2.646E-06 | 1.100E-05 | 1.600E-05 |
| 5831 | 292 | V292L | A | 7.6e-06 | 1.4e-05 | 1e-05 | 1.053E-05 | 3.233E-06 | 7.600E-06 | 1.400E-05 |
| 5832 | 292 | V292K | A | 1.6e-05 | 9.9e-06 | 7.8e-06 | 1.123E-05 | 4.259E-06 | 7.800E-06 | 1.600E-05 |
| 5833 | 292 | V292M | A | 9.6e-06 | 1.6e-05 | 1.8e-05 | 1.453E-05 | 4.388E-06 | 9.600E-06 | 1.800E-05 |
| 5834 | 292 | V292F | A | 1.1e-05 | 1.6e-05 | 1e-05 | 1.233E-05 | 3.215E-06 | 1.000E-05 | 1.600E-05 |
| 5835 | 292 | V292P | A | 1.4e-05 | 9.8e-06 | 1.5e-05 | 1.293E-05 | 2.759E-06 | 9.800E-06 | 1.500E-05 |
| 5836 | 292 | V292S | A | 1.1e-05 | 1.1e-05 | 1.8e-05 | 1.333E-05 | 4.041E-06 | 1.100E-05 | 1.800E-05 |
| 5837 | 292 | V292T | A | 1.3e-05 | 1.4e-05 | 1.3e-05 | 1.333E-05 | 5.774E-07 | 1.300E-05 | 1.400E-05 |
| 5838 | 292 | V292W | A | 7.8e-06 | 1.2e-05 | 1.7e-05 | 1.227E-05 | 4.606E-06 | 7.800E-06 | 1.700E-05 |
| 5839 | 292 | V292Y | A | 1.7e-05 | 1.5e-05 | 1e-05 | 1.400E-05 | 3.606E-06 | 1.000E-05 | 1.700E-05 |
| 5840 | 292 | V292V | A | 1.4e-05 | 9.2e-06 | 1.1e-05 | 1.140E-05 | 2.425E-06 | 9.200E-06 | 1.400E-05 |
| 5841 | 293 | C293A | A | 1.1e-05 | 1.5e-05 | 1.6e-05 | 1.400E-05 | 2.646E-06 | 1.100E-05 | 1.600E-05 |
| 5842 | 293 | C293R | A | 1.5e-05 | 1.2e-05 | 1.5e-05 | 1.400E-05 | 1.732E-06 | 1.200E-05 | 1.500E-05 |
| 5843 | 293 | C293N | A | 1.4e-05 | 2e-05 | 1.4e-05 | 1.600E-05 | 3.464E-06 | 1.400E-05 | 2.000E-05 |
| 5844 | 293 | C293D | A | 1.3e-05 | 1.5e-05 | 1.4e-05 | 1.400E-05 | 1.000E-06 | 1.300E-05 | 1.500E-05 |
| 5845 | 293 | C293C | A | 1.5e-05 | 1.5e-05 | 1.4e-05 | 1.467E-05 | 5.774E-07 | 1.400E-05 | 1.500E-05 |
| 5846 | 293 | C293E | A | 1.5e-05 | 1.7e-05 | 1.1e-05 | 1.433E-05 | 3.055E-06 | 1.100E-05 | 1.700E-05 |
| 5847 | 293 | C293Q | A | 1.4e-05 | 1.3e-05 | 1.8e-05 | 1.500E-05 | 2.646E-06 | 1.300E-05 | 1.800E-05 |
| 5848 | 293 | C293G | A | 1.6e-05 | 1.6e-05 | 9.3e-06 | 1.377E-05 | 3.868E-06 | 9.300E-06 | 1.600E-05 |
| 5849 | 293 | C293H | A | 1e-05 | 1.1e-05 | 1.4e-05 | 1.167E-05 | 2.082E-06 | 1.000E-05 | 1.400E-05 |
| 5850 | 293 | C293I | A | 9.3e-06 | 1.2e-05 | 1.5e-05 | 1.210E-05 | 2.851E-06 | 9.300E-06 | 1.500E-05 |
| 5851 | 293 | C293L | A | 9e-06 | 1.5e-05 | 1.1e-05 | 1.167E-05 | 3.055E-06 | 9.000E-06 | 1.500E-05 |
| 5852 | 293 | C293K | A | 1.6e-05 | 1.7e-05 | 1.8e-05 | 1.700E-05 | 1.000E-06 | 1.600E-05 | 1.800E-05 |
| 5853 | 293 | C293M | A | 1.4e-05 | 9.3e-06 | 1.5e-05 | 1.277E-05 | 3.044E-06 | 9.300E-06 | 1.500E-05 |
| 5854 | 293 | C293F | A | 1.1e-05 | 1.6e-05 | 9.5e-06 | 1.217E-05 | 3.403E-06 | 9.500E-06 | 1.600E-05 |
| 5855 | 293 | C293P | A | 1.6e-05 | 1.7e-05 | 1.1e-05 | 1.467E-05 | 3.215E-06 | 1.100E-05 | 1.700E-05 |
| 5856 | 293 | C293S | A | 1e-05 | 1.1e-05 | 1.3e-05 | 1.133E-05 | 1.528E-06 | 1.000E-05 | 1.300E-05 |
| 5857 | 293 | C293T | A | 1.6e-05 | 9.7e-06 | 1.4e-05 | 1.323E-05 | 3.219E-06 | 9.700E-06 | 1.600E-05 |
| 5858 | 293 | C293W | A | 1e-05 | 1.4e-05 | 1e-05 | 1.133E-05 | 2.309E-06 | 1.000E-05 | 1.400E-05 |
| 5859 | 293 | C293Y | A | 1.2e-05 | 8.2e-06 | 1.1e-05 | 1.040E-05 | 1.970E-06 | 8.200E-06 | 1.200E-05 |
| 5860 | 293 | C293V | A | 1.1e-05 | 1.5e-05 | 1.1e-05 | 1.233E-05 | 2.309E-06 | 1.100E-05 | 1.500E-05 |
| 5861 | 294 | P294A | A | 9.1e-06 | 1.7e-05 | 1.4e-05 | 1.337E-05 | 3.988E-06 | 9.100E-06 | 1.700E-05 |
| 5862 | 294 | P294R | A | 1.1e-05 | 1.4e-05 | 1.6e-05 | 1.367E-05 | 2.517E-06 | 1.100E-05 | 1.600E-05 |
| 5863 | 294 | P294N | A | 1.4e-05 | 1.2e-05 | 9.4e-06 | 1.180E-05 | 2.307E-06 | 9.400E-06 | 1.400E-05 |
| 5864 | 294 | P294D | A | 1.5e-05 | 1e-05 | 1.4e-05 | 1.300E-05 | 2.646E-06 | 1.000E-05 | 1.500E-05 |

| | | | | | | | | | | |
|---|---|---|---|---|---|---|---|---|---|---|
| 5865 | 294 | P294C | A | 1.7e-05 | 1.1e-05 | 1e-05 | 1.267E-05 | 3.786E-06 | 1.000E-05 | 1.700E-05 |
| 5866 | 294 | P294E | A | 8.8e-06 | 1.5e-05 | 1.2e-05 | 1.193E-05 | 3.101E-06 | 8.800E-06 | 1.500E-05 |
| 5867 | 294 | P294Q | A | 1.3e-05 | 9.8e-06 | 1.1e-05 | 1.127E-05 | 1.617E-06 | 9.800E-06 | 1.300E-05 |
| 5868 | 294 | P294G | A | 1.4e-05 | 1.6e-05 | 8.9e-06 | 1.297E-05 | 3.661E-06 | 8.900E-06 | 1.600E-05 |
| 5869 | 294 | P294H | A | 1.3e-05 | 1.7e-05 | 1e-05 | 1.333E-05 | 3.512E-06 | 1.000E-05 | 1.700E-05 |
| 5870 | 294 | P294I | A | 1.4e-05 | 1.4e-05 | 1.6e-05 | 1.467E-05 | 1.155E-06 | 1.400E-05 | 1.600E-05 |
| 5871 | 294 | P294L | A | 1.1e-05 | 1.3e-05 | 1.5e-05 | 1.300E-05 | 2.000E-06 | 1.100E-05 | 1.500E-05 |
| 5872 | 294 | P294K | A | 1.4e-05 | 1.5e-05 | 9.3e-06 | 1.277E-05 | 3.044E-06 | 9.300E-06 | 1.500E-05 |
| 5873 | 294 | P294M | A | 1.7e-05 | 1.6e-05 | 1.5e-05 | 1.600E-05 | 1.000E-06 | 1.500E-05 | 1.700E-05 |
| 5874 | 294 | P294F | A | 1.2e-05 | 1.5e-05 | 1e-05 | 1.233E-05 | 2.517E-06 | 1.000E-05 | 1.500E-05 |
| 5875 | 294 | P294P | A | 1.6e-05 | 1.6e-05 | 1.1e-05 | 1.433E-05 | 2.887E-06 | 1.100E-05 | 1.600E-05 |
| 5876 | 294 | P294S | A | 1.2e-05 | 1.4e-05 | 1.5e-05 | 1.367E-05 | 1.528E-06 | 1.200E-05 | 1.500E-05 |
| 5877 | 294 | P294T | A | 1.3e-05 | 1.5e-05 | 1e-05 | 1.267E-05 | 2.517E-06 | 1.000E-05 | 1.500E-05 |
| 5878 | 294 | P294W | A | 1.1e-05 | 1.2e-05 | 8.6e-06 | 1.053E-05 | 1.747E-06 | 8.600E-06 | 1.200E-05 |
| 5879 | 294 | P294Y | A | 8.2e-06 | 1.5e-05 | 1.5e-05 | 1.273E-05 | 3.926E-06 | 8.200E-06 | 1.500E-05 |
| 5880 | 294 | P294V | A | 9.8e-06 | 1.5e-05 | 1.4e-05 | 1.293E-05 | 2.759E-06 | 9.800E-06 | 1.500E-05 |
| 5881 | 295 | L295A | A | 1.5e-05 | 1.4e-05 | 1.3e-05 | 1.400E-05 | 1.000E-06 | 1.300E-05 | 1.500E-05 |
| 5882 | 295 | L295R | A | 1.2e-05 | 1.4e-05 | 6.9e-06 | 1.097E-05 | 3.661E-06 | 6.900E-06 | 1.400E-05 |
| 5883 | 295 | L295N | A | 5.5e-06 | 1.1e-05 | 1.3e-05 | 9.833E-06 | 3.884E-06 | 5.500E-06 | 1.300E-05 |
| 5884 | 295 | L295D | A | 1.1e-05 | 1.3e-05 | 6.7e-06 | 1.023E-05 | 3.219E-06 | 6.700E-06 | 1.300E-05 |
| 5885 | 295 | L295C | A | 1.3e-05 | 1.1e-05 | 1.4e-05 | 1.267E-05 | 1.528E-06 | 1.100E-05 | 1.400E-05 |
| 5886 | 295 | L295E | A | 8.7e-06 | 1.2e-05 | 9.4e-06 | 1.003E-05 | 1.739E-06 | 8.700E-06 | 1.200E-05 |
| 5887 | 295 | L295Q | A | 9.7e-06 | 7.8e-06 | 1.5e-05 | 1.083E-05 | 3.731E-06 | 7.800E-06 | 1.500E-05 |
| 5888 | 295 | L295G | A | 1.4e-05 | 1.4e-05 | 1.5e-05 | 1.433E-05 | 5.774E-07 | 1.400E-05 | 1.500E-05 |
| 5889 | 295 | L295H | A | 1e-05 | 8.8e-06 | 1.3e-05 | 1.060E-05 | 2.163E-06 | 8.800E-06 | 1.300E-05 |
| 5890 | 295 | L295I | A | 1.1e-05 | 1.5e-05 | 1.4e-05 | 1.333E-05 | 2.082E-06 | 1.100E-05 | 1.500E-05 |
| 5891 | 295 | L295L | A | 9.3e-06 | 9.4e-06 | 1.4e-05 | 1.090E-05 | 2.685E-06 | 9.300E-06 | 1.400E-05 |
| 5892 | 295 | L295K | A | 7.6e-06 | 1.2e-05 | 1.7e-05 | 1.220E-05 | 4.703E-06 | 7.600E-06 | 1.700E-05 |
| 5893 | 295 | L295M | A | 1.9e-05 | 1.6e-05 | 1.2e-05 | 1.567E-05 | 3.512E-06 | 1.200E-05 | 1.900E-05 |
| 5894 | 295 | L295F | A | 1e-05 | 1e-05 | 1e-05 | 1.000E-05 | 0.000E+00 | 1.000E-05 | 1.000E-05 |
| 5895 | 295 | L295P | A | 1.5e-05 | 9.3e-06 | 1.1e-05 | 1.177E-05 | 2.926E-06 | 9.300E-06 | 1.500E-05 |
| 5896 | 295 | L295S | A | 1.1e-05 | 7.7e-06 | 8.4e-06 | 9.033E-06 | 1.739E-06 | 7.700E-06 | 1.100E-05 |
| 5897 | 295 | L295T | A | 1.1e-05 | 8.7e-06 | 8.1e-06 | 9.267E-06 | 1.531E-06 | 8.100E-06 | 1.100E-05 |
| 5898 | 295 | L295W | A | 6.2e-06 | 1.6e-05 | 1.1e-05 | 1.107E-05 | 4.900E-06 | 6.200E-06 | 1.600E-05 |
| 5899 | 295 | L295Y | A | 7.5e-06 | 1.5e-05 | 7.8e-06 | 1.010E-05 | 4.246E-06 | 7.500E-06 | 1.500E-05 |
| 5900 | 295 | L295V | A | 1.6e-05 | 9.7e-06 | 1.5e-05 | 1.357E-05 | 3.386E-06 | 9.700E-06 | 1.600E-05 |
| 5901 | 296 | H296A | A | 2.3e-05 | 1.5e-05 | 1.7e-05 | 1.833E-05 | 4.163E-06 | 1.500E-05 | 2.300E-05 |
| 5902 | 296 | H296R | A | 1.7e-05 | 1.2e-05 | 1.1e-05 | 1.333E-05 | 3.215E-06 | 1.100E-05 | 1.700E-05 |
| 5903 | 296 | H296N | A | 1e-05 | 7e-06 | 7.1e-06 | 8.033E-06 | 1.704E-06 | 7.000E-06 | 1.000E-05 |
| 5904 | 296 | H296D | A | 1.1e-05 | 1.5e-05 | 1.5e-05 | 1.367E-05 | 2.309E-06 | 1.100E-05 | 1.500E-05 |
| 5905 | 296 | H296C | A | 2.1e-05 | 1.3e-05 | 1.3e-05 | 1.567E-05 | 4.619E-06 | 1.300E-05 | 2.100E-05 |
| 5906 | 296 | H296E | A | 1.7e-05 | 1.2e-05 | 1.1e-05 | 1.333E-05 | 3.215E-06 | 1.100E-05 | 1.700E-05 |
| 5907 | 296 | H296Q | A | 1e-05 | 9e-06 | 7.4e-06 | 8.800E-06 | 1.311E-06 | 7.400E-06 | 1.000E-05 |
| 5908 | 296 | H296G | A | 1.9e-05 | 2.2e-05 | 2.1e-05 | 2.067E-05 | 1.528E-06 | 1.900E-05 | 2.200E-05 |
| 5909 | 296 | H296H | A | 1.5e-05 | 1.5e-05 | 8.9e-06 | 1.297E-05 | 3.522E-06 | 8.900E-06 | 1.500E-05 |
| 5910 | 296 | H296I | A | 1.5e-05 | 2.2e-05 | 2.1e-05 | 1.933E-05 | 3.786E-06 | 1.500E-05 | 2.200E-05 |
| 5911 | 296 | H296L | A | 2.1e-05 | 2.3e-05 | 1.4e-05 | 1.933E-05 | 4.726E-06 | 1.400E-05 | 2.300E-05 |
| 5912 | 296 | H296K | A | 1.3e-05 | 9.1e-06 | 7.7e-06 | 9.933E-06 | 2.747E-06 | 7.700E-06 | 1.300E-05 |
| 5913 | 296 | H296M | A | 2.1e-05 | 1.6e-05 | 2.1e-05 | 1.933E-05 | 2.887E-06 | 1.600E-05 | 2.100E-05 |
| 5914 | 296 | H296F | A | 1.5e-05 | 2.2e-05 | 1.4e-05 | 1.700E-05 | 4.359E-06 | 1.400E-05 | 2.200E-05 |
| 5915 | 296 | H296P | A | 2.2e-05 | 2.4e-05 | 2e-05 | 2.200E-05 | 2.000E-06 | 2.000E-05 | 2.400E-05 |
| 5916 | 296 | H296S | A | 1.4e-05 | 9.7e-06 | 1.8e-05 | 1.390E-05 | 4.151E-06 | 9.700E-06 | 1.800E-05 |
| 5917 | 296 | H296T | A | 9.3e-06 | 1e-05 | 2e-05 | 1.310E-05 | 5.986E-06 | 9.300E-06 | 2.000E-05 |
| 5918 | 296 | H296W | A | 1.3e-05 | 1.3e-05 | 1.4e-05 | 1.333E-05 | 5.774E-07 | 1.300E-05 | 1.400E-05 |
| 5919 | 296 | H296Y | A | 1.5e-05 | 1.5e-05 | 1.6e-05 | 1.533E-05 | 5.774E-07 | 1.500E-05 | 1.600E-05 |
| 5920 | 296 | H296V | A | 1.6e-05 | 1.3e-05 | 2.3e-05 | 1.733E-05 | 5.132E-06 | 1.300E-05 | 2.300E-05 |
| 5921 | 297 | N297A | A | 1.5e-05 | 1.6e-05 | 9.5e-06 | 1.350E-05 | 3.500E-06 | 9.500E-06 | 1.600E-05 |
| 5922 | 297 | N297R | A | 1.5e-05 | 1.5e-05 | 1.4e-05 | 1.467E-05 | 5.774E-07 | 1.400E-05 | 1.500E-05 |
| 5923 | 297 | N297N | A | 1.5e-05 | 9.6e-06 | 1.5e-05 | 1.320E-05 | 3.118E-06 | 9.600E-06 | 1.500E-05 |
| 5924 | 297 | N297D | A | 1.7e-05 | 1.7e-05 | 1.5e-05 | 1.633E-05 | 1.155E-06 | 1.500E-05 | 1.700E-05 |
| 5925 | 297 | N297C | A | 1.2e-05 | 1.5e-05 | 1.5e-05 | 1.400E-05 | 1.732E-06 | 1.200E-05 | 1.500E-05 |
| 5926 | 297 | N297E | A | 1e-05 | 1.1e-05 | 8.7e-06 | 9.900E-06 | 1.153E-06 | 8.700E-06 | 1.100E-05 |
| 5927 | 297 | N297Q | A | 6.4e-06 | 6.7e-06 | 1.5e-05 | 9.367E-06 | 4.881E-06 | 6.400E-06 | 1.500E-05 |
| 5928 | 297 | N297G | A | 1.1e-05 | 1.8e-05 | 1.4e-05 | 1.433E-05 | 3.512E-06 | 1.100E-05 | 1.800E-05 |
| 5929 | 297 | N297H | A | 1.3e-05 | 1.8e-05 | 1.4e-05 | 1.500E-05 | 2.646E-06 | 1.300E-05 | 1.800E-05 |
| 5930 | 297 | N297I | A | 1.2e-05 | 1.6e-05 | 1.4e-05 | 1.400E-05 | 2.000E-06 | 1.200E-05 | 1.600E-05 |
| 5931 | 297 | N297L | A | 1.5e-05 | 1.6e-05 | 1e-05 | 1.367E-05 | 3.215E-06 | 1.000E-05 | 1.600E-05 |
| 5932 | 297 | N297K | A | 1.6e-05 | 1.7e-05 | 1.2e-05 | 1.500E-05 | 2.646E-06 | 1.200E-05 | 1.700E-05 |
| 5933 | 297 | N297M | A | 1.1e-05 | 1.2e-05 | 9.6e-06 | 1.087E-05 | 1.206E-06 | 9.600E-06 | 1.200E-05 |
| 5934 | 297 | N297F | A | 1.7e-05 | 1.6e-05 | 1.9e-05 | 1.733E-05 | 1.528E-06 | 1.600E-05 | 1.900E-05 |
| 5935 | 297 | N297P | A | 1.6e-05 | 1.2e-05 | 1.4e-05 | 1.400E-05 | 2.000E-06 | 1.200E-05 | 1.600E-05 |
| 5936 | 297 | N297S | A | 1.5e-05 | 8.7e-06 | 1e-05 | 1.123E-05 | 3.326E-06 | 8.700E-06 | 1.500E-05 |
| 5937 | 297 | N297T | A | 1e-05 | 1.6e-05 | 1.4e-05 | 1.333E-05 | 3.055E-06 | 1.000E-05 | 1.600E-05 |
| 5938 | 297 | N297W | A | 8.3e-06 | 1.5e-05 | 1e-05 | 1.110E-05 | 3.483E-06 | 8.300E-06 | 1.500E-05 |
| 5939 | 297 | N297Y | A | 1.7e-05 | 1.5e-05 | 1.2e-05 | 1.467E-05 | 2.517E-06 | 1.200E-05 | 1.700E-05 |
| 5940 | 297 | N297V | A | 9.7e-06 | 1.6e-05 | 1.6e-05 | 1.390E-05 | 3.637E-06 | 9.700E-06 | 1.600E-05 |
| 5941 | 298 | Q298A | A | 1.2e-05 | 1.1e-05 | 1.6e-05 | 1.300E-05 | 2.646E-06 | 1.100E-05 | 1.600E-05 |
| 5942 | 298 | Q298R | A | 1.8e-05 | 1.2e-05 | 1.1e-05 | 1.367E-05 | 3.786E-06 | 1.100E-05 | 1.800E-05 |
| 5943 | 298 | Q298N | A | 1.6e-05 | 1.3e-05 | 1.3e-05 | 1.400E-05 | 1.732E-06 | 1.300E-05 | 1.600E-05 |
| 5944 | 298 | Q298D | A | 1.6e-05 | 1.3e-05 | 1.4e-05 | 1.433E-05 | 1.528E-06 | 1.300E-05 | 1.600E-05 |
| 5945 | 298 | Q298C | A | 1.6e-05 | 1.6e-05 | 1.1e-05 | 1.433E-05 | 2.887E-06 | 1.100E-05 | 1.600E-05 |
| 5946 | 298 | Q298E | A | 2.5e-05 | 1.7e-05 | 1.6e-05 | 1.933E-05 | 4.933E-06 | 1.600E-05 | 2.500E-05 |
| 5947 | 298 | Q298Q | A | 9.5e-06 | 1.4e-05 | 1.7e-05 | 1.350E-05 | 3.775E-06 | 9.500E-06 | 1.700E-05 |
| 5948 | 298 | Q298G | A | 1.8e-05 | 1.5e-05 | 1.6e-05 | 1.633E-05 | 1.528E-06 | 1.500E-05 | 1.800E-05 |
| 5949 | 298 | Q298H | A | 2e-05 | 1.1e-05 | 8.6e-06 | 1.320E-05 | 6.010E-06 | 8.600E-06 | 2.000E-05 |

| | | | | | | | | | | |
|---|---|---|---|---|---|---|---|---|---|---|
| 5950 | 298 | Q298I | A | 1.4e-05 | 1.4e-05 | 1.5e-05 | 1.433E-05 | 5.774E-07 | 1.400E-05 | 1.500E-05 |
| 5951 | 298 | Q298L | A | 1.7e-05 | 9.8e-06 | 1.6e-05 | 1.427E-05 | 3.900E-06 | 9.800E-06 | 1.700E-05 |
| 5952 | 298 | Q298K | A | 1.3e-05 | 1e-05 | 1e-05 | 1.100E-05 | 1.732E-06 | 1.000E-05 | 1.300E-05 |
| 5953 | 298 | Q298M | A | 1.1e-05 | 1e-05 | 1.3e-05 | 1.133E-05 | 1.528E-06 | 1.000E-05 | 1.300E-05 |
| 5954 | 298 | Q298F | A | 1.8e-05 | 1.4e-05 | 1.6e-05 | 1.600E-05 | 2.000E-06 | 1.400E-05 | 1.800E-05 |
| 5955 | 298 | Q298P | A | 1.1e-05 | 1.1e-05 | 1.7e-05 | 1.300E-05 | 3.464E-06 | 1.100E-05 | 1.700E-05 |
| 5956 | 298 | Q298S | A | 1.1e-05 | 9.3e-06 | 8.2e-06 | 9.500E-06 | 1.411E-06 | 8.200E-06 | 1.100E-05 |
| 5957 | 298 | Q298T | A | 1.1e-05 | 1.5e-05 | 1.5e-05 | 1.367E-05 | 2.309E-06 | 1.100E-05 | 1.500E-05 |
| 5958 | 298 | Q298W | A | 1.8e-05 | 1.5e-05 | 1.5e-05 | 1.600E-05 | 1.732E-06 | 1.500E-05 | 1.800E-05 |
| 5959 | 298 | Q298Y | A | 1.5e-05 | 1.1e-05 | 1.2e-05 | 1.267E-05 | 2.082E-06 | 1.100E-05 | 1.500E-05 |
| 5960 | 298 | Q298V | A | 1.1e-05 | 1.5e-05 | 9.6e-06 | 1.187E-05 | 2.802E-06 | 9.600E-06 | 1.500E-05 |
| 5961 | 299 | E299A | A | 1.5e-05 | 1.2e-05 | 1.6e-05 | 1.433E-05 | 2.082E-06 | 1.200E-05 | 1.600E-05 |
| 5962 | 299 | E299R | A | 1.6e-05 | 1.3e-05 | 1.4e-05 | 1.433E-05 | 1.528E-06 | 1.300E-05 | 1.600E-05 |
| 5963 | 299 | E299N | A | 1e-05 | 9.5e-06 | 1e-05 | 9.833E-06 | 2.887E-07 | 9.500E-06 | 1.000E-05 |
| 5964 | 299 | E299D | A | 1.4e-05 | 6.5e-06 | 1.4e-05 | 1.150E-05 | 4.330E-06 | 6.500E-06 | 1.400E-05 |
| 5965 | 299 | E299C | A | 1.2e-05 | 1.5e-05 | 1.4e-05 | 1.367E-05 | 1.528E-06 | 1.200E-05 | 1.500E-05 |
| 5966 | 299 | E299E | A | 1e-05 | 1.7e-05 | 1.6e-05 | 1.433E-05 | 3.786E-06 | 1.000E-05 | 1.700E-05 |
| 5967 | 299 | E299Q | A | 1.6e-05 | 8.9e-06 | 9.9e-06 | 1.160E-05 | 3.843E-06 | 8.900E-06 | 1.600E-05 |
| 5968 | 299 | E299G | A | 1.1e-05 | 1.1e-05 | 9e-06 | 1.033E-05 | 1.155E-06 | 9.000E-06 | 1.100E-05 |
| 5969 | 299 | E299H | A | 1.2e-05 | 1.5e-05 | 1.1e-05 | 1.267E-05 | 2.082E-06 | 1.100E-05 | 1.500E-05 |
| 5970 | 299 | E299I | A | 1.8e-05 | 1.3e-05 | 1.2e-05 | 1.433E-05 | 3.215E-06 | 1.200E-05 | 1.800E-05 |
| 5971 | 299 | E299L | A | 1.6e-05 | 1.8e-05 | 9.6e-06 | 1.453E-05 | 4.388E-06 | 9.600E-06 | 1.800E-05 |
| 5972 | 299 | E299K | A | 1.2e-05 | 1.6e-05 | 1.4e-05 | 1.400E-05 | 2.000E-06 | 1.200E-05 | 1.600E-05 |
| 5973 | 299 | E299M | A | 1.4e-05 | 1.4e-05 | 9.7e-06 | 1.257E-05 | 2.483E-06 | 9.700E-06 | 1.400E-05 |
| 5974 | 299 | E299F | A | 1.6e-05 | 1.1e-05 | 1.4e-05 | 1.367E-05 | 2.517E-06 | 1.100E-05 | 1.600E-05 |
| 5975 | 299 | E299P | A | 1.1e-05 | 1.7e-05 | 1.3e-05 | 1.367E-05 | 3.055E-06 | 1.100E-05 | 1.700E-05 |
| 5976 | 299 | E299S | A | 8.7e-06 | 8.3e-06 | 1.3e-05 | 1.000E-05 | 2.606E-06 | 8.300E-06 | 1.300E-05 |
| 5977 | 299 | E299T | A | 1e-05 | 1.6e-05 | 8.7e-06 | 1.157E-05 | 3.894E-06 | 8.700E-06 | 1.600E-05 |
| 5978 | 299 | E299W | A | 1.5e-05 | 1.3e-05 | 1.4e-05 | 1.400E-05 | 1.000E-06 | 1.300E-05 | 1.500E-05 |
| 5979 | 299 | E299Y | A | 9.2e-06 | 1.5e-05 | 1e-05 | 1.140E-05 | 3.143E-06 | 9.200E-06 | 1.500E-05 |
| 5980 | 299 | E299V | A | 1e-05 | 1.4e-05 | 2.3e-05 | 1.567E-05 | 6.658E-06 | 1.000E-05 | 2.300E-05 |
| 5981 | 300 | V300A | A | 1.2e-05 | 1.2e-05 | 1.6e-05 | 1.333E-05 | 2.309E-06 | 1.200E-05 | 1.600E-05 |
| 5982 | 300 | V300R | A | 9.4e-06 | 1.6e-05 | 1e-05 | 1.180E-05 | 3.650E-06 | 9.400E-06 | 1.600E-05 |
| 5983 | 300 | V300N | A | 1.7e-05 | 9e-06 | 1.4e-05 | 1.333E-05 | 4.041E-06 | 9.000E-06 | 1.700E-05 |
| 5984 | 300 | V300D | A | 1.7e-05 | 1.4e-05 | 1.1e-05 | 1.400E-05 | 3.000E-06 | 1.100E-05 | 1.700E-05 |
| 5985 | 300 | V300C | A | 1e-05 | 8.4e-06 | 9e-06 | 9.133E-06 | 8.083E-07 | 8.400E-06 | 1.000E-05 |
| 5986 | 300 | V300E | A | 1.2e-05 | 1e-05 | 1.3e-05 | 1.167E-05 | 1.528E-06 | 1.000E-05 | 1.300E-05 |
| 5987 | 300 | V300Q | A | 1.6e-05 | 1.5e-05 | 1.4e-05 | 1.500E-05 | 1.000E-06 | 1.400E-05 | 1.600E-05 |
| 5988 | 300 | V300G | A | 1.2e-05 | 1e-05 | 1.5e-05 | 1.233E-05 | 2.517E-06 | 1.000E-05 | 1.500E-05 |
| 5989 | 300 | V300H | A | 1.6e-05 | 1.6e-05 | 9.8e-06 | 1.393E-05 | 3.580E-06 | 9.800E-06 | 1.600E-05 |
| 5990 | 300 | V300I | A | 1.5e-05 | 1e-05 | 1.4e-05 | 1.300E-05 | 2.646E-06 | 1.000E-05 | 1.500E-05 |
| 5991 | 300 | V300L | A | 1.1e-05 | 8.5e-06 | 1e-05 | 9.833E-06 | 1.258E-06 | 8.500E-06 | 1.100E-05 |
| 5992 | 300 | V300K | A | 1.4e-05 | 7.3e-06 | 1.6e-05 | 1.243E-05 | 4.557E-06 | 7.300E-06 | 1.600E-05 |
| 5993 | 300 | V300M | A | 1.5e-05 | 1e-05 | 7.7e-06 | 1.090E-05 | 3.732E-06 | 7.700E-06 | 1.500E-05 |
| 5994 | 300 | V300F | A | 1.5e-05 | 1.3e-05 | 1.5e-05 | 1.433E-05 | 1.155E-06 | 1.300E-05 | 1.500E-05 |
| 5995 | 300 | V300P | A | 1.5e-05 | 1.1e-05 | 1.1e-05 | 1.233E-05 | 2.309E-06 | 1.100E-05 | 1.500E-05 |
| 5996 | 300 | V300S | A | 1.1e-05 | 1.4e-05 | 1.5e-05 | 1.333E-05 | 2.082E-06 | 1.100E-05 | 1.500E-05 |
| 5997 | 300 | V300T | A | 1.1e-05 | 1.4e-05 | 8.5e-06 | 1.117E-05 | 2.754E-06 | 8.500E-06 | 1.400E-05 |
| 5998 | 300 | V300W | A | 1.4e-05 | 8.9e-06 | 1.4e-05 | 1.230E-05 | 2.944E-06 | 8.900E-06 | 1.400E-05 |
| 5999 | 300 | V300Y | A | 1.6e-05 | 1.7e-05 | 1.3e-05 | 1.533E-05 | 2.082E-06 | 1.300E-05 | 1.700E-05 |
| 6000 | 300 | V300V | A | 2.2e-05 | 1.6e-05 | 1.1e-05 | 1.633E-05 | 5.508E-06 | 1.100E-05 | 2.200E-05 |
| 6001 | 301 | T301A | A | 1.5e-05 | 1.5e-05 | 1e-05 | 1.333E-05 | 2.887E-06 | 1.000E-05 | 1.500E-05 |
| 6002 | 301 | T301R | A | 1.6e-05 | 1.1e-05 | 9.1e-06 | 1.203E-05 | 3.564E-06 | 9.100E-06 | 1.600E-05 |
| 6003 | 301 | T301N | A | 1.6e-05 | 1.1e-05 | 9.5e-06 | 1.217E-05 | 3.403E-06 | 9.500E-06 | 1.600E-05 |
| 6004 | 301 | T301D | A | 1.1e-05 | 8.2e-06 | 1.1e-05 | 1.007E-05 | 1.617E-06 | 8.200E-06 | 1.100E-05 |
| 6005 | 301 | T301C | A | 1.1e-05 | 1.5e-05 | 2.7e-05 | 1.767E-05 | 8.327E-06 | 1.100E-05 | 2.700E-05 |
| 6006 | 301 | T301E | A | 1.1e-05 | 1.5e-05 | 1.6e-05 | 1.400E-05 | 2.646E-06 | 1.100E-05 | 1.600E-05 |
| 6007 | 301 | T301Q | A | 1.2e-05 | 1.6e-05 | 1.1e-05 | 1.300E-05 | 2.646E-06 | 1.100E-05 | 1.600E-05 |
| 6008 | 301 | T301G | A | 1.6e-05 | 1.7e-05 | 1.5e-05 | 1.600E-05 | 1.000E-06 | 1.500E-05 | 1.700E-05 |
| 6009 | 301 | T301H | A | 1.1e-05 | 1.1e-05 | 9.4e-06 | 1.047E-05 | 9.238E-07 | 9.400E-06 | 1.100E-05 |
| 6010 | 301 | T301I | A | 1.2e-05 | 1.5e-05 | 9.9e-06 | 1.230E-05 | 2.563E-06 | 9.900E-06 | 1.500E-05 |
| 6011 | 301 | T301L | A | 9.5e-06 | 1.5e-05 | 1.4e-05 | 1.283E-05 | 2.930E-06 | 9.500E-06 | 1.500E-05 |
| 6012 | 301 | T301K | A | 1.8e-05 | 1.1e-05 | 7.9e-06 | 1.230E-05 | 5.174E-06 | 7.900E-06 | 1.800E-05 |
| 6013 | 301 | T301M | A | 1.1e-05 | 1.6e-05 | 1.5e-05 | 1.400E-05 | 2.646E-06 | 1.100E-05 | 1.600E-05 |
| 6014 | 301 | T301F | A | 9.3e-06 | 1.5e-05 | 1.6e-05 | 1.343E-05 | 3.614E-06 | 9.300E-06 | 1.600E-05 |
| 6015 | 301 | T301P | A | 1e-05 | 1.5e-05 | 1e-05 | 1.167E-05 | 2.887E-06 | 1.000E-05 | 1.500E-05 |
| 6016 | 301 | T301S | A | 3e-05 | 1e-05 | 1.7e-05 | 1.900E-05 | 1.015E-05 | 1.000E-05 | 3.000E-05 |
| 6017 | 301 | T301T | A | 1.5e-05 | 1e-05 | 1.5e-05 | 1.333E-05 | 2.887E-06 | 1.000E-05 | 1.500E-05 |
| 6018 | 301 | T301W | A | 1.1e-05 | 1.5e-05 | 1.5e-05 | 1.367E-05 | 2.309E-06 | 1.100E-05 | 1.500E-05 |
| 6019 | 301 | T301Y | A | 1.3e-05 | 1.5e-05 | 9e-06 | 1.233E-05 | 3.055E-06 | 9.000E-06 | 1.500E-05 |
| 6020 | 301 | T301V | A | 1.1e-05 | 1.1e-05 | 1.4e-05 | 1.200E-05 | 1.732E-06 | 1.100E-05 | 1.400E-05 |
| 6021 | 302 | A302A | A | 1.7e-05 | 9e-06 | 8.9e-06 | 1.163E-05 | 4.648E-06 | 8.900E-06 | 1.700E-05 |
| 6022 | 302 | A302R | A | 1.7e-05 | 1.4e-05 | 1.2e-05 | 1.433E-05 | 2.517E-06 | 1.200E-05 | 1.700E-05 |
| 6023 | 302 | A302N | A | 9.9e-06 | 1.1e-05 | 1.6e-05 | 1.230E-05 | 3.251E-06 | 9.900E-06 | 1.600E-05 |
| 6024 | 302 | A302D | A | 1.5e-05 | 1.1e-05 | 1.5e-05 | 1.367E-05 | 2.309E-06 | 1.100E-05 | 1.500E-05 |
| 6025 | 302 | A302C | A | 1.6e-05 | 1.1e-05 | 1.1e-05 | 1.267E-05 | 2.887E-06 | 1.100E-05 | 1.600E-05 |
| 6026 | 302 | A302E | A | 1.6e-05 | 1.2e-05 | 1.5e-05 | 1.433E-05 | 2.082E-06 | 1.200E-05 | 1.600E-05 |
| 6027 | 302 | A302Q | A | 1.4e-05 | 1.4e-05 | 9.9e-06 | 1.263E-05 | 2.367E-06 | 9.900E-06 | 1.400E-05 |
| 6028 | 302 | A302G | A | 1.6e-05 | 1.1e-05 | 1.4e-05 | 1.367E-05 | 2.517E-06 | 1.100E-05 | 1.600E-05 |
| 6029 | 302 | A302H | A | 9.8e-06 | 9.7e-06 | 1.3e-05 | 1.083E-05 | 1.877E-06 | 9.700E-06 | 1.300E-05 |
| 6030 | 302 | A302I | A | 1.4e-05 | 1.6e-05 | 1.5e-05 | 1.500E-05 | 1.000E-06 | 1.400E-05 | 1.600E-05 |
| 6031 | 302 | A302L | A | 1.5e-05 | 1.3e-05 | 1.3e-05 | 1.367E-05 | 1.155E-06 | 1.300E-05 | 1.500E-05 |
| 6032 | 302 | A302K | A | 1e-05 | 1.5e-05 | 1.6e-05 | 1.367E-05 | 3.215E-06 | 1.000E-05 | 1.600E-05 |
| 6033 | 302 | A302M | A | 9.1e-06 | 1.9e-05 | 1.1e-05 | 1.303E-05 | 5.254E-06 | 9.100E-06 | 1.900E-05 |
| 6034 | 302 | A302F | A | 1.6e-05 | 1.1e-05 | 9.9e-06 | 1.230E-05 | 3.251E-06 | 9.900E-06 | 1.600E-05 |

| | | | | | | | | | | |
|---|---|---|---|---|---|---|---|---|---|---|
| 6035 | 302 | A302P | A | 1.5e-05 | 1.4e-05 | 1.1e-05 | 1.333E-05 | 2.082E-06 | 1.100E-05 | 1.500E-05 |
| 6036 | 302 | A302S | A | 1.1e-05 | 1.1e-05 | 1.4e-05 | 1.200E-05 | 1.732E-06 | 1.100E-05 | 1.400E-05 |
| 6037 | 302 | A302T | A | 1.2e-05 | 1.4e-05 | 1.5e-05 | 1.367E-05 | 1.528E-06 | 1.200E-05 | 1.500E-05 |
| 6038 | 302 | A302W | A | 9.5e-06 | 1.3e-05 | 9.4e-06 | 1.063E-05 | 2.050E-06 | 9.400E-06 | 1.300E-05 |
| 6039 | 302 | A302Y | A | 9.3e-06 | 9.3e-06 | 1e-05 | 9.533E-06 | 4.041E-07 | 9.300E-06 | 1.000E-05 |
| 6040 | 302 | A302V | A | 1.1e-05 | 1.4e-05 | 1.3e-05 | 1.267E-05 | 1.528E-06 | 1.100E-05 | 1.400E-05 |
| 6041 | 303 | E303A | A | 1e-05 | 1.4e-05 | 1e-05 | 1.133E-05 | 2.309E-06 | 1.000E-05 | 1.400E-05 |
| 6042 | 303 | E303R | A | 1.2e-05 | 1.5e-05 | 9.2e-06 | 1.207E-05 | 2.901E-06 | 9.200E-06 | 1.500E-05 |
| 6043 | 303 | E303N | A | 1.4e-05 | 9.6e-06 | 1.1e-05 | 1.153E-05 | 2.248E-06 | 9.600E-06 | 1.400E-05 |
| 6044 | 303 | E303D | A | 9.4e-06 | 9.9e-06 | 1e-05 | 9.767E-06 | 3.215E-07 | 9.400E-06 | 1.000E-05 |
| 6045 | 303 | E303C | A | 1e-05 | 1.3e-05 | 1e-05 | 1.100E-05 | 1.732E-06 | 1.000E-05 | 1.300E-05 |
| 6046 | 303 | E303E | A | 1.5e-05 | 1.2e-05 | 8.6e-06 | 1.187E-05 | 3.202E-06 | 8.600E-06 | 1.500E-05 |
| 6047 | 303 | E303Q | A | 1.4e-05 | 1.5e-05 | 1.6e-05 | 1.500E-05 | 1.000E-06 | 1.400E-05 | 1.600E-05 |
| 6048 | 303 | E303G | A | 7.9e-06 | 1.4e-05 | 1.4e-05 | 1.197E-05 | 3.522E-06 | 7.900E-06 | 1.400E-05 |
| 6049 | 303 | E303H | A | 1.5e-05 | 9e-06 | 1.1e-05 | 1.167E-05 | 3.055E-06 | 9.000E-06 | 1.500E-05 |
| 6050 | 303 | E303I | A | 1.3e-05 | 1.1e-05 | 1.6e-05 | 1.333E-05 | 2.517E-06 | 1.100E-05 | 1.600E-05 |
| 6051 | 303 | E303L | A | 1.1e-05 | 1.3e-05 | 1.4e-05 | 1.267E-05 | 1.528E-06 | 1.100E-05 | 1.400E-05 |
| 6052 | 303 | E303K | A | 1.3e-05 | 1.5e-05 | 1.4e-05 | 1.400E-05 | 1.000E-06 | 1.300E-05 | 1.500E-05 |
| 6053 | 303 | E303M | A | 1.1e-05 | 1.4e-05 | 1e-05 | 1.167E-05 | 2.082E-06 | 1.000E-05 | 1.400E-05 |
| 6054 | 303 | E303F | A | 1.4e-05 | 1.1e-05 | 1.4e-05 | 1.300E-05 | 1.732E-06 | 1.100E-05 | 1.400E-05 |
| 6055 | 303 | E303P | A | 1.5e-05 | 1.5e-05 | 8.2e-06 | 1.273E-05 | 3.926E-06 | 8.200E-06 | 1.500E-05 |
| 6056 | 303 | E303S | A | 1.5e-05 | 1.4e-05 | 9e-06 | 1.267E-05 | 3.215E-06 | 9.000E-06 | 1.500E-05 |
| 6057 | 303 | E303T | A | 1.4e-05 | 1.2e-05 | 1.5e-05 | 1.367E-05 | 1.528E-06 | 1.200E-05 | 1.500E-05 |
| 6058 | 303 | E303W | A | 1.5e-05 | 8.4e-06 | 1e-05 | 1.113E-05 | 3.443E-06 | 8.400E-06 | 1.500E-05 |
| 6059 | 303 | E303Y | A | 1.4e-05 | 1.5e-05 | 8.7e-06 | 1.257E-05 | 3.386E-06 | 8.700E-06 | 1.500E-05 |
| 6060 | 303 | E303V | A | 1.7e-05 | 1.5e-05 | 1.6e-05 | 1.600E-05 | 1.000E-06 | 1.500E-05 | 1.700E-05 |
| 6061 | 304 | D304A | A | 1.6e-05 | 1.3e-05 | 1.7e-05 | 1.533E-05 | 2.082E-06 | 1.300E-05 | 1.700E-05 |
| 6062 | 304 | D304R | A | 1.3e-05 | 1.4e-05 | 1.1e-05 | 1.267E-05 | 1.528E-06 | 1.100E-05 | 1.400E-05 |
| 6063 | 304 | D304N | A | 1.2e-05 | 1e-05 | 1.1e-05 | 1.100E-05 | 1.000E-06 | 1.000E-05 | 1.200E-05 |
| 6064 | 304 | D304D | A | 1.4e-05 | 1.6e-05 | 1.4e-05 | 1.467E-05 | 1.155E-06 | 1.400E-05 | 1.600E-05 |
| 6065 | 304 | D304C | A | 1.5e-05 | 9.5e-06 | 1.6e-05 | 1.350E-05 | 3.500E-06 | 9.500E-06 | 1.600E-05 |
| 6066 | 304 | D304E | A | 1e-05 | 1.3e-05 | 1e-05 | 1.100E-05 | 1.732E-06 | 1.000E-05 | 1.300E-05 |
| 6067 | 304 | D304Q | A | 6.3e-06 | 1e-05 | 1e-05 | 8.767E-06 | 2.136E-06 | 6.300E-06 | 1.000E-05 |
| 6068 | 304 | D304G | A | 1.5e-05 | 1.1e-05 | 1e-05 | 1.200E-05 | 2.646E-06 | 1.000E-05 | 1.500E-05 |
| 6069 | 304 | D304H | A | 1.6e-05 | 1e-05 | 1.6e-05 | 1.400E-05 | 3.464E-06 | 1.000E-05 | 1.600E-05 |
| 6070 | 304 | D304I | A | 1.1e-05 | 1.5e-05 | 1.2e-05 | 1.267E-05 | 2.082E-06 | 1.100E-05 | 1.500E-05 |
| 6071 | 304 | D304L | A | 9.9e-06 | 1e-05 | 1.6e-05 | 1.197E-05 | 3.493E-06 | 9.900E-06 | 1.600E-05 |
| 6072 | 304 | D304K | A | 1.5e-05 | 1.1e-05 | 8.4e-06 | 1.147E-05 | 3.325E-06 | 8.400E-06 | 1.500E-05 |
| 6073 | 304 | D304M | A | 1.2e-05 | 1.4e-05 | 1e-05 | 1.200E-05 | 2.000E-06 | 1.000E-05 | 1.400E-05 |
| 6074 | 304 | D304F | A | 9.3e-06 | 1.5e-05 | 1.5e-05 | 1.310E-05 | 3.291E-06 | 9.300E-06 | 1.500E-05 |
| 6075 | 304 | D304P | A | 1.5e-05 | 1.4e-05 | 1.1e-05 | 1.333E-05 | 2.082E-06 | 1.100E-05 | 1.500E-05 |
| 6076 | 304 | D304S | A | 1.4e-05 | 1.5e-05 | 9e-06 | 1.267E-05 | 3.215E-06 | 9.000E-06 | 1.500E-05 |
| 6077 | 304 | D304T | A | 1.1e-05 | 1.6e-05 | 1.4e-05 | 1.367E-05 | 2.517E-06 | 1.100E-05 | 1.600E-05 |
| 6078 | 304 | D304W | A | 9.4e-06 | 1.3e-05 | 1.5e-05 | 1.247E-05 | 2.838E-06 | 9.400E-06 | 1.500E-05 |
| 6079 | 304 | D304Y | A | 1.5e-05 | 1.7e-05 | 1.5e-05 | 1.567E-05 | 1.155E-06 | 1.500E-05 | 1.700E-05 |
| 6080 | 304 | D304V | A | 1.5e-05 | 1.3e-05 | 1e-05 | 1.267E-05 | 2.517E-06 | 1.000E-05 | 1.500E-05 |
| 6081 | 305 | G305A | A | 1.1e-05 | 1.4e-05 | 1e-05 | 1.167E-05 | 2.082E-06 | 1.000E-05 | 1.400E-05 |
| 6082 | 305 | G305R | A | 1.6e-05 | 1.1e-05 | 1.7e-05 | 1.467E-05 | 3.215E-06 | 1.100E-05 | 1.700E-05 |
| 6083 | 305 | G305N | A | 1.3e-05 | 1.4e-05 | 1.4e-05 | 1.367E-05 | 5.774E-07 | 1.300E-05 | 1.400E-05 |
| 6084 | 305 | G305D | A | 9.4e-06 | 1.4e-05 | 9.8e-06 | 1.107E-05 | 2.548E-06 | 9.400E-06 | 1.400E-05 |
| 6085 | 305 | G305C | A | 9.6e-06 | 1.4e-05 | 2.2e-05 | 1.520E-05 | 6.286E-06 | 9.600E-06 | 2.200E-05 |
| 6086 | 305 | G305E | A | 1.1e-05 | 1e-05 | 1.4e-05 | 1.167E-05 | 2.082E-06 | 1.000E-05 | 1.400E-05 |
| 6087 | 305 | G305Q | A | 1e-05 | 1.5e-05 | 1.1e-05 | 1.200E-05 | 2.646E-06 | 1.000E-05 | 1.500E-05 |
| 6088 | 305 | G305G | A | 1.2e-05 | 1.2e-05 | 1.4e-05 | 1.267E-05 | 1.155E-06 | 1.200E-05 | 1.400E-05 |
| 6089 | 305 | G305H | A | 1.1e-05 | 1.6e-05 | 1.4e-05 | 1.367E-05 | 2.517E-06 | 1.100E-05 | 1.600E-05 |
| 6090 | 305 | G305I | A | 1.2e-05 | 1.6e-05 | 1.1e-05 | 1.300E-05 | 2.646E-06 | 1.100E-05 | 1.600E-05 |
| 6091 | 305 | G305L | A | 1.5e-05 | 1e-05 | 1.5e-05 | 1.333E-05 | 2.887E-06 | 1.000E-05 | 1.500E-05 |
| 6092 | 305 | G305K | A | 1.1e-05 | 1.6e-05 | 1e-05 | 1.233E-05 | 3.215E-06 | 1.000E-05 | 1.600E-05 |
| 6093 | 305 | G305M | A | 1.3e-05 | 9.2e-06 | 1.6e-05 | 1.273E-05 | 3.408E-06 | 9.200E-06 | 1.600E-05 |
| 6094 | 305 | G305F | A | 1.4e-05 | 1.2e-05 | 1.2e-05 | 1.267E-05 | 1.155E-06 | 1.200E-05 | 1.400E-05 |
| 6095 | 305 | G305P | A | 8.3e-06 | 1.1e-05 | 9.8e-06 | 9.700E-06 | 1.353E-06 | 8.300E-06 | 1.100E-05 |
| 6096 | 305 | G305S | A | 1.4e-05 | 8.8e-06 | 2.1e-05 | 1.460E-05 | 6.122E-06 | 8.800E-06 | 2.100E-05 |
| 6097 | 305 | G305T | A | 1e-05 | 1.1e-05 | 9.9e-06 | 1.030E-05 | 6.083E-07 | 9.900E-06 | 1.100E-05 |
| 6098 | 305 | G305W | A | 1.4e-05 | 9.6e-06 | 8.5e-06 | 1.070E-05 | 2.910E-06 | 8.500E-06 | 1.400E-05 |
| 6099 | 305 | G305Y | A | 1.6e-05 | 1.5e-05 | 1.2e-05 | 1.433E-05 | 2.082E-06 | 1.200E-05 | 1.600E-05 |
| 6100 | 305 | G305V | A | 8.3e-06 | 1.6e-05 | 1.5e-05 | 1.310E-05 | 4.187E-06 | 8.300E-06 | 1.600E-05 |
| 6101 | 306 | T306A | A | 1.6e-05 | 1.6e-05 | 1.4e-05 | 1.533E-05 | 1.155E-06 | 1.400E-05 | 1.600E-05 |
| 6102 | 306 | T306R | A | 1.7e-05 | 1.1e-05 | 1.1e-05 | 1.300E-05 | 3.464E-06 | 1.100E-05 | 1.700E-05 |
| 6103 | 306 | T306N | A | 1.5e-05 | 1.1e-05 | 1.5e-05 | 1.367E-05 | 2.309E-06 | 1.100E-05 | 1.500E-05 |
| 6104 | 306 | T306D | A | 1.2e-05 | 1.6e-05 | 1.4e-05 | 1.400E-05 | 2.000E-06 | 1.200E-05 | 1.600E-05 |
| 6105 | 306 | T306C | A | 1.4e-05 | 1.2e-05 | 1.5e-05 | 1.367E-05 | 1.528E-06 | 1.200E-05 | 1.500E-05 |
| 6106 | 306 | T306E | A | 1.7e-05 | 1.5e-05 | 1.6e-05 | 1.600E-05 | 1.000E-06 | 1.500E-05 | 1.700E-05 |
| 6107 | 306 | T306Q | A | 1.5e-05 | 1.6e-05 | 1.4e-05 | 1.500E-05 | 1.000E-06 | 1.400E-05 | 1.600E-05 |
| 6108 | 306 | T306G | A | 1.4e-05 | 1.1e-05 | 1.4e-05 | 1.300E-05 | 1.732E-06 | 1.100E-05 | 1.400E-05 |
| 6109 | 306 | T306H | A | 1.2e-05 | 1.4e-05 | 1.6e-05 | 1.400E-05 | 2.000E-06 | 1.200E-05 | 1.600E-05 |
| 6110 | 306 | T306I | A | 9.1e-06 | 1.5e-05 | 1.4e-05 | 1.270E-05 | 3.158E-06 | 9.100E-06 | 1.500E-05 |
| 6111 | 306 | T306L | A | 1.6e-05 | 9.6e-06 | 5.9e-06 | 1.050E-05 | 5.110E-06 | 5.900E-06 | 1.600E-05 |
| 6112 | 306 | T306K | A | 1.1e-05 | 1.5e-05 | 1.6e-05 | 1.400E-05 | 2.646E-06 | 1.100E-05 | 1.600E-05 |
| 6113 | 306 | T306M | A | 1.4e-05 | 9.7e-06 | 1.2e-05 | 1.190E-05 | 2.152E-06 | 9.700E-06 | 1.400E-05 |
| 6114 | 306 | T306F | A | 1.1e-05 | 1.4e-05 | 1.6e-05 | 1.367E-05 | 2.517E-06 | 1.100E-05 | 1.600E-05 |
| 6115 | 306 | T306P | A | 1.5e-05 | 1.1e-05 | 1.6e-05 | 1.400E-05 | 2.646E-06 | 1.100E-05 | 1.600E-05 |
| 6116 | 306 | T306S | A | 1.6e-05 | 1.1e-05 | 9.8e-06 | 1.227E-05 | 3.288E-06 | 9.800E-06 | 1.600E-05 |
| 6117 | 306 | T306T | A | 1.5e-05 | 1.5e-05 | 8.9e-06 | 1.297E-05 | 3.522E-06 | 8.900E-06 | 1.500E-05 |
| 6118 | 306 | T306W | A | 1.5e-05 | 1.1e-05 | 9.9e-06 | 1.197E-05 | 2.684E-06 | 9.900E-06 | 1.500E-05 |
| 6119 | 306 | T306Y | A | 1.5e-05 | 1.4e-05 | 1.1e-05 | 1.333E-05 | 2.082E-06 | 1.100E-05 | 1.500E-05 |

| | | | | | | | | | | |
|---|---|---|---|---|---|---|---|---|---|---|
| 6120 | 306 | T306V | A | 1.5e-05 | 1.4e-05 | 1.6e-05 | 1.500E-05 | 1.000E-06 | 1.400E-05 | 1.600E-05 |
| 6121 | 307 | Q307A | A | 1.5e-05 | 1.4e-05 | 1e-05 | 1.300E-05 | 2.646E-06 | 1.000E-05 | 1.500E-05 |
| 6122 | 307 | Q307R | A | 1e-05 | 1.1e-05 | 1.5e-05 | 1.200E-05 | 2.646E-06 | 1.000E-05 | 1.500E-05 |
| 6123 | 307 | Q307N | A | 1e-05 | 1.4e-05 | 1.6e-05 | 1.333E-05 | 3.055E-06 | 1.000E-05 | 1.600E-05 |
| 6124 | 307 | Q307D | A | 1.4e-05 | 1.8e-05 | 1.1e-05 | 1.433E-05 | 3.512E-06 | 1.100E-05 | 1.800E-05 |
| 6125 | 307 | Q307C | A | 1.7e-05 | 9.6e-06 | 1.6e-05 | 1.420E-05 | 4.015E-06 | 9.600E-06 | 1.700E-05 |
| 6126 | 307 | Q307E | A | 1.4e-05 | 1e-05 | 1.1e-05 | 1.167E-05 | 2.082E-06 | 1.000E-05 | 1.400E-05 |
| 6127 | 307 | Q307Q | A | 1.5e-05 | 9.3e-06 | 1.5e-05 | 1.310E-05 | 3.291E-06 | 9.300E-06 | 1.500E-05 |
| 6128 | 307 | Q307G | A | 1e-05 | 1.8e-05 | 1.6e-05 | 1.467E-05 | 4.163E-06 | 1.000E-05 | 1.800E-05 |
| 6129 | 307 | Q307H | A | 1.1e-05 | 1.4e-05 | 1.4e-05 | 1.300E-05 | 1.732E-06 | 1.100E-05 | 1.400E-05 |
| 6130 | 307 | Q307I | A | 1.6e-05 | 1.1e-05 | 1.1e-05 | 1.267E-05 | 2.887E-06 | 1.100E-05 | 1.600E-05 |
| 6131 | 307 | Q307L | A | 1.2e-05 | 1.5e-05 | 1.6e-05 | 1.433E-05 | 2.082E-06 | 1.200E-05 | 1.600E-05 |
| 6132 | 307 | Q307K | A | 1.1e-05 | 1.5e-05 | 1.1e-05 | 1.233E-05 | 2.309E-06 | 1.100E-05 | 1.500E-05 |
| 6133 | 307 | Q307M | A | 1.5e-05 | 1.2e-05 | 1.3e-05 | 1.333E-05 | 1.528E-06 | 1.200E-05 | 1.500E-05 |
| 6134 | 307 | Q307F | A | 1.3e-05 | 1.7e-05 | 9.8e-06 | 1.327E-05 | 3.607E-06 | 9.800E-06 | 1.700E-05 |
| 6135 | 307 | Q307P | A | 9.6e-06 | 1.1e-05 | 1.2e-05 | 1.087E-05 | 1.206E-06 | 9.600E-06 | 1.200E-05 |
| 6136 | 307 | Q307S | A | 1.5e-05 | 1.4e-05 | 1.6e-05 | 1.500E-05 | 1.000E-06 | 1.400E-05 | 1.600E-05 |
| 6137 | 307 | Q307T | A | 1e-05 | 1.2e-05 | 1.6e-05 | 1.267E-05 | 3.055E-06 | 1.000E-05 | 1.600E-05 |
| 6138 | 307 | Q307W | A | 7.3e-06 | 1.5e-05 | 1.2e-05 | 1.143E-05 | 3.881E-06 | 7.300E-06 | 1.500E-05 |
| 6139 | 307 | Q307Y | A | 9.8e-06 | 1.6e-05 | 1.1e-05 | 1.227E-05 | 3.288E-06 | 9.800E-06 | 1.600E-05 |
| 6140 | 307 | Q307V | A | 1.6e-05 | 1.1e-05 | 9.4e-06 | 1.213E-05 | 3.443E-06 | 9.400E-06 | 1.600E-05 |
| 6141 | 308 | R308A | A | 1.8e-05 | 1.1e-05 | 1.1e-05 | 1.333E-05 | 4.041E-06 | 1.100E-05 | 1.800E-05 |
| 6142 | 308 | R308R | A | 1e-05 | 9.6e-06 | 1e-05 | 9.867E-06 | 2.309E-07 | 9.600E-06 | 1.000E-05 |
| 6143 | 308 | R308N | A | 9.5e-06 | 1.5e-05 | 8.9e-06 | 1.113E-05 | 3.362E-06 | 8.900E-06 | 1.500E-05 |
| 6144 | 308 | R308D | A | 1.6e-05 | 7.9e-06 | 1.4e-05 | 1.263E-05 | 4.219E-06 | 7.900E-06 | 1.600E-05 |
| 6145 | 308 | R308C | A | 1.6e-05 | 1.5e-05 | 1.3e-05 | 1.467E-05 | 1.528E-06 | 1.300E-05 | 1.600E-05 |
| 6146 | 308 | R308E | A | 1.5e-05 | 9.9e-06 | 1.5e-05 | 1.330E-05 | 2.944E-06 | 9.900E-06 | 1.500E-05 |
| 6147 | 308 | R308Q | A | 1e-05 | 1.4e-05 | 1.3e-05 | 1.233E-05 | 2.082E-06 | 1.000E-05 | 1.400E-05 |
| 6148 | 308 | R308G | A | 1.2e-05 | 1.6e-05 | 1.5e-05 | 1.433E-05 | 2.082E-06 | 1.200E-05 | 1.600E-05 |
| 6149 | 308 | R308H | A | 1e-05 | 1.5e-05 | 1.3e-05 | 1.267E-05 | 2.517E-06 | 1.000E-05 | 1.500E-05 |
| 6150 | 308 | R308I | A | 1.5e-05 | 1.7e-05 | 1.7e-05 | 1.633E-05 | 1.155E-06 | 1.500E-05 | 1.700E-05 |
| 6151 | 308 | R308L | A | 6.7e-06 | 1.6e-05 | 1.4e-05 | 1.223E-05 | 4.895E-06 | 6.700E-06 | 1.600E-05 |
| 6152 | 308 | R308K | A | 1.3e-05 | 1e-05 | 1.6e-05 | 1.300E-05 | 3.000E-06 | 1.000E-05 | 1.600E-05 |
| 6153 | 308 | R308M | A | 1.2e-05 | 8.8e-06 | 1.4e-05 | 1.160E-05 | 2.623E-06 | 8.800E-06 | 1.400E-05 |
| 6154 | 308 | R308F | A | 1e-05 | 1e-05 | 1.2e-05 | 1.067E-05 | 1.155E-06 | 1.000E-05 | 1.200E-05 |
| 6155 | 308 | R308P | A | 1.4e-05 | 1.3e-05 | 1.1e-05 | 1.267E-05 | 1.528E-06 | 1.100E-05 | 1.400E-05 |
| 6156 | 308 | R308S | A | 1e-05 | 1.4e-05 | 1.2e-05 | 1.200E-05 | 2.000E-06 | 1.000E-05 | 1.400E-05 |
| 6157 | 308 | R308T | A | 1.4e-05 | 1e-05 | 8.3e-06 | 1.077E-05 | 2.926E-06 | 8.300E-06 | 1.400E-05 |
| 6158 | 308 | R308W | A | 1e-05 | 1.6e-05 | 1.6e-05 | 1.400E-05 | 3.464E-06 | 1.000E-05 | 1.600E-05 |
| 6159 | 308 | R308Y | A | 1.3e-05 | 1.8e-05 | 1.4e-05 | 1.500E-05 | 2.646E-06 | 1.300E-05 | 1.800E-05 |
| 6160 | 308 | R308V | A | 1.2e-05 | 1.1e-05 | 1.1e-05 | 1.133E-05 | 5.774E-07 | 1.100E-05 | 1.200E-05 |
| 6161 | 309 | C309A | A | 1.3e-05 | 1.4e-05 | 1.6e-05 | 1.433E-05 | 1.528E-06 | 1.300E-05 | 1.600E-05 |
| 6162 | 309 | C309R | A | 1.6e-05 | 1.6e-05 | 9.9e-06 | 1.397E-05 | 3.522E-06 | 9.900E-06 | 1.600E-05 |
| 6163 | 309 | C309N | A | 1.5e-05 | 9.6e-06 | 1.2e-05 | 1.220E-05 | 2.706E-06 | 9.600E-06 | 1.500E-05 |
| 6164 | 309 | C309D | A | 1e-05 | 1e-05 | 1e-05 | 1.000E-05 | 0.000E+00 | 1.000E-05 | 1.000E-05 |
| 6165 | 309 | C309C | A | 1.4e-05 | 8.9e-06 | 1e-05 | 1.097E-05 | 2.684E-06 | 8.900E-06 | 1.400E-05 |
| 6166 | 309 | C309E | A | 1.7e-05 | 1.4e-05 | 1e-05 | 1.367E-05 | 3.512E-06 | 1.000E-05 | 1.700E-05 |
| 6167 | 309 | C309Q | A | 1.5e-05 | 1.1e-05 | 1.1e-05 | 1.233E-05 | 2.309E-06 | 1.100E-05 | 1.500E-05 |
| 6168 | 309 | C309G | A | 1e-05 | 1.1e-05 | 1.5e-05 | 1.200E-05 | 2.646E-06 | 1.000E-05 | 1.500E-05 |
| 6169 | 309 | C309H | A | 1.6e-05 | 1.1e-05 | 1e-05 | 1.233E-05 | 3.215E-06 | 1.000E-05 | 1.600E-05 |
| 6170 | 309 | C309I | A | 1e-05 | 1.7e-05 | 8.7e-06 | 1.190E-05 | 4.464E-06 | 8.700E-06 | 1.700E-05 |
| 6171 | 309 | C309L | A | 1.2e-05 | 1.7e-05 | 9.5e-06 | 1.283E-05 | 3.819E-06 | 9.500E-06 | 1.700E-05 |
| 6172 | 309 | C309K | A | 1e-05 | 1.2e-05 | 9e-06 | 1.033E-05 | 1.528E-06 | 9.000E-06 | 1.200E-05 |
| 6173 | 309 | C309M | A | 2.2e-05 | 1e-05 | 7.5e-06 | 1.317E-05 | 7.751E-06 | 7.500E-06 | 2.200E-05 |
| 6174 | 309 | C309F | A | 1.4e-05 | 1.7e-05 | 1.2e-05 | 1.433E-05 | 2.517E-06 | 1.200E-05 | 1.700E-05 |
| 6175 | 309 | C309P | A | 1.5e-05 | 1.3e-05 | 1.5e-05 | 1.433E-05 | 1.155E-06 | 1.300E-05 | 1.500E-05 |
| 6176 | 309 | C309S | A | 1.6e-05 | 1.2e-05 | 1.4e-05 | 1.400E-05 | 2.000E-06 | 1.200E-05 | 1.600E-05 |
| 6177 | 309 | C309T | A | 1.1e-05 | 8e-06 | 8.7e-06 | 9.233E-06 | 1.570E-06 | 8.000E-06 | 1.100E-05 |
| 6178 | 309 | C309W | A | 1.2e-05 | 1.4e-05 | 1e-05 | 1.200E-05 | 2.000E-06 | 1.000E-05 | 1.400E-05 |
| 6179 | 309 | C309Y | A | 1e-05 | 1.5e-05 | 9e-06 | 1.133E-05 | 3.215E-06 | 9.000E-06 | 1.500E-05 |
| 6180 | 309 | C309V | A | 1.6e-05 | 1.2e-05 | 1.4e-05 | 1.400E-05 | 2.000E-06 | 1.200E-05 | 1.600E-05 |
| 6181 | 310 | E310A | A | 1.2e-05 | 1.1e-05 | 1.6e-05 | 1.300E-05 | 2.646E-06 | 1.100E-05 | 1.600E-05 |
| 6182 | 310 | E310R | A | 9e-06 | 1.6e-05 | 1.1e-05 | 1.200E-05 | 3.606E-06 | 9.000E-06 | 1.600E-05 |
| 6183 | 310 | E310N | A | 1.4e-05 | 1.4e-05 | 9.2e-06 | 1.240E-05 | 2.771E-06 | 9.200E-06 | 1.400E-05 |
| 6184 | 310 | E310D | A | 1.1e-05 | 1.6e-05 | 1.5e-05 | 1.400E-05 | 2.646E-06 | 1.100E-05 | 1.600E-05 |
| 6185 | 310 | E310C | A | 1.1e-05 | 1.2e-05 | 1.4e-05 | 1.233E-05 | 1.528E-06 | 1.100E-05 | 1.400E-05 |
| 6186 | 310 | E310E | A | 1.2e-05 | 1.4e-05 | 1.5e-05 | 1.367E-05 | 1.528E-06 | 1.200E-05 | 1.500E-05 |
| 6187 | 310 | E310Q | A | 8.3e-06 | 1e-05 | 1.1e-05 | 9.767E-06 | 1.365E-06 | 8.300E-06 | 1.100E-05 |
| 6188 | 310 | E310G | A | 1.6e-05 | 1.4e-05 | 8.9e-06 | 1.297E-05 | 3.661E-06 | 8.900E-06 | 1.600E-05 |
| 6189 | 310 | E310H | A | 1.3e-05 | 1e-05 | 1e-05 | 1.100E-05 | 1.732E-06 | 1.000E-05 | 1.300E-05 |
| 6190 | 310 | E310I | A | 8.6e-06 | 1e-05 | 1e-05 | 9.533E-06 | 8.083E-07 | 8.600E-06 | 1.000E-05 |
| 6191 | 310 | E310L | A | 1e-05 | 1.4e-05 | 9.8e-06 | 1.127E-05 | 2.369E-06 | 9.800E-06 | 1.400E-05 |
| 6192 | 310 | E310K | A | 1.4e-05 | 1.2e-05 | 8.8e-06 | 1.160E-05 | 2.623E-06 | 8.800E-06 | 1.400E-05 |
| 6193 | 310 | E310M | A | 1.1e-05 | 1.5e-05 | 1.3e-05 | 1.300E-05 | 2.000E-06 | 1.100E-05 | 1.500E-05 |
| 6194 | 310 | E310F | A | 1.4e-05 | 1.4e-05 | 1.6e-05 | 1.467E-05 | 1.155E-06 | 1.400E-05 | 1.600E-05 |
| 6195 | 310 | E310P | A | 1.3e-05 | 1.3e-05 | 1.5e-05 | 1.367E-05 | 1.155E-06 | 1.300E-05 | 1.500E-05 |
| 6196 | 310 | E310S | A | 1.5e-05 | 1e-05 | 1e-05 | 1.167E-05 | 2.887E-06 | 1.000E-05 | 1.500E-05 |
| 6197 | 310 | E310T | A | 1e-05 | 1.3e-05 | 1.4e-05 | 1.233E-05 | 2.082E-06 | 1.000E-05 | 1.400E-05 |
| 6198 | 310 | E310W | A | 1.5e-05 | 8.3e-06 | 1.3e-05 | 1.210E-05 | 3.439E-06 | 8.300E-06 | 1.500E-05 |
| 6199 | 310 | E310Y | A | 1.7e-05 | 9.8e-06 | 1.4e-05 | 1.360E-05 | 3.617E-06 | 9.800E-06 | 1.700E-05 |
| 6200 | 310 | E310V | A | 1.4e-05 | 1.5e-05 | 1e-05 | 1.300E-05 | 2.646E-06 | 1.000E-05 | 1.500E-05 |
| 6201 | 311 | K311A | A | 1.7e-05 | 1.6e-05 | 6.9e-06 | 1.330E-05 | 5.565E-06 | 6.900E-06 | 1.700E-05 |
| 6202 | 311 | K311R | A | 1.3e-05 | 1e-05 | 7.8e-06 | 1.027E-05 | 2.610E-06 | 7.800E-06 | 1.300E-05 |
| 6203 | 311 | K311N | A | 1.3e-05 | 1.1e-05 | 1.1e-05 | 1.167E-05 | 1.155E-06 | 1.100E-05 | 1.300E-05 |
| 6204 | 311 | K311D | A | 1.8e-05 | 1.5e-05 | 9.1e-06 | 1.403E-05 | 4.528E-06 | 9.100E-06 | 1.800E-05 |

| | | | | | | | | | | |
|---|---|---|---|---|---|---|---|---|---|---|
| 6205 | 311 | K311C | A | 1.6e-05 | 1.2e-05 | 1.5e-05 | 1.433E-05 | 2.082E-06 | 1.200E-05 | 1.600E-05 |
| 6206 | 311 | K311E | A | 8.6e-06 | 1.4e-05 | 1.3e-05 | 1.187E-05 | 2.873E-06 | 8.600E-06 | 1.400E-05 |
| 6207 | 311 | K311Q | A | 9.1e-06 | 1.1e-05 | 8.7e-06 | 9.600E-06 | 1.229E-06 | 8.700E-06 | 1.100E-05 |
| 6208 | 311 | K311G | A | 9.1e-06 | 1.5e-05 | 1.1e-05 | 1.170E-05 | 3.012E-06 | 9.100E-06 | 1.500E-05 |
| 6209 | 311 | K311H | A | 1.5e-05 | 9.4e-06 | 9.5e-06 | 1.130E-05 | 3.205E-06 | 9.400E-06 | 1.500E-05 |
| 6210 | 311 | K311I | A | 1.4e-05 | 1.7e-05 | 1.2e-05 | 1.433E-05 | 2.517E-06 | 1.200E-05 | 1.700E-05 |
| 6211 | 311 | K311L | A | 1.4e-05 | 1.5e-05 | 1.4e-05 | 1.433E-05 | 5.774E-07 | 1.400E-05 | 1.500E-05 |
| 6212 | 311 | K311K | A | 1.2e-05 | 1.4e-05 | 1.5e-05 | 1.367E-05 | 1.528E-06 | 1.200E-05 | 1.500E-05 |
| 6213 | 311 | K311M | A | 1.7e-05 | 9e-06 | 1.4e-05 | 1.333E-05 | 4.041E-06 | 9.000E-06 | 1.700E-05 |
| 6214 | 311 | K311F | A | 1.6e-05 | 9.8e-06 | 1.5e-05 | 1.360E-05 | 3.329E-06 | 9.800E-06 | 1.600E-05 |
| 6215 | 311 | K311P | A | 8.6e-06 | 1.4e-05 | 1.4e-05 | 1.220E-05 | 3.118E-06 | 8.600E-06 | 1.400E-05 |
| 6216 | 311 | K311S | A | 1.3e-05 | 9.3e-06 | 1.6e-05 | 1.277E-05 | 3.356E-06 | 9.300E-06 | 1.600E-05 |
| 6217 | 311 | K311T | A | 1.7e-05 | 1.5e-05 | 1.2e-05 | 1.467E-05 | 2.517E-06 | 1.200E-05 | 1.700E-05 |
| 6218 | 311 | K311W | A | 9.1e-06 | 1.1e-05 | 9.4e-06 | 9.833E-06 | 1.021E-06 | 9.100E-06 | 1.100E-05 |
| 6219 | 311 | K311Y | A | 1.5e-05 | 1.5e-05 | 8.5e-06 | 1.283E-05 | 3.753E-06 | 8.500E-06 | 1.500E-05 |
| 6220 | 311 | K311V | A | 1.4e-05 | 1.6e-05 | 9.6e-06 | 1.320E-05 | 3.274E-06 | 9.600E-06 | 1.600E-05 |
| 6221 | 312 | C312A | A | 1.2e-05 | 9.8e-06 | 9.1e-06 | 1.030E-05 | 1.513E-06 | 9.100E-06 | 1.200E-05 |
| 6222 | 312 | C312R | A | 9.7e-06 | 1.5e-05 | 6.7e-06 | 1.047E-05 | 4.203E-06 | 6.700E-06 | 1.500E-05 |
| 6223 | 312 | C312N | A | 1.4e-05 | 7.9e-06 | 1.9e-05 | 1.363E-05 | 5.559E-06 | 7.900E-06 | 1.900E-05 |
| 6224 | 312 | C312D | A | 9.4e-06 | 1.3e-05 | 1.3e-05 | 1.180E-05 | 2.078E-06 | 9.400E-06 | 1.300E-05 |
| 6225 | 312 | C312C | A | 1.3e-05 | 9.8e-06 | 1.5e-05 | 1.260E-05 | 2.623E-06 | 9.800E-06 | 1.500E-05 |
| 6226 | 312 | C312E | A | 1e-05 | 1.4e-05 | 8.5e-06 | 1.083E-05 | 2.843E-06 | 8.500E-06 | 1.400E-05 |
| 6227 | 312 | C312Q | A | 1.1e-05 | 9e-06 | 7.6e-06 | 9.200E-06 | 1.709E-06 | 7.600E-06 | 1.100E-05 |
| 6228 | 312 | C312G | A | 9.8e-06 | 1.3e-05 | 1.2e-05 | 1.160E-05 | 1.637E-06 | 9.800E-06 | 1.300E-05 |
| 6229 | 312 | C312H | A | 1.4e-05 | 9.5e-06 | 8.3e-06 | 1.060E-05 | 3.005E-06 | 8.300E-06 | 1.400E-05 |
| 6230 | 312 | C312I | A | 1.1e-05 | 1.4e-05 | 1.1e-05 | 1.200E-05 | 1.732E-06 | 1.100E-05 | 1.400E-05 |
| 6231 | 312 | C312L | A | 1.4e-05 | 1e-05 | 1.5e-05 | 1.300E-05 | 2.646E-06 | 1.000E-05 | 1.500E-05 |
| 6232 | 312 | C312K | A | 1.5e-05 | 1.2e-05 | 7e-06 | 1.133E-05 | 4.041E-06 | 7.000E-06 | 1.500E-05 |
| 6233 | 312 | C312M | A | 9.7e-06 | 1e-05 | 1.1e-05 | 1.023E-05 | 6.807E-07 | 9.700E-06 | 1.100E-05 |
| 6234 | 312 | C312F | A | 9.9e-06 | 8.9e-06 | 1.1e-05 | 9.933E-06 | 1.050E-06 | 8.900E-06 | 1.100E-05 |
| 6235 | 312 | C312P | A | 1.6e-05 | 1.5e-05 | 8.5e-06 | 1.317E-05 | 4.072E-06 | 8.500E-06 | 1.600E-05 |
| 6236 | 312 | C312S | A | 7.6e-06 | 1.4e-05 | 1.5e-05 | 1.220E-05 | 4.015E-06 | 7.600E-06 | 1.500E-05 |
| 6237 | 312 | C312T | A | 8.7e-06 | 1.4e-05 | 8.9e-06 | 1.053E-05 | 3.004E-06 | 8.700E-06 | 1.400E-05 |
| 6238 | 312 | C312W | A | 1.6e-05 | 1.1e-05 | 1.6e-05 | 1.433E-05 | 2.887E-06 | 1.100E-05 | 1.600E-05 |
| 6239 | 312 | C312Y | A | 1e-05 | 1.4e-05 | 1.3e-05 | 1.233E-05 | 2.082E-06 | 1.000E-05 | 1.400E-05 |
| 6240 | 312 | C312V | A | 1.4e-05 | 1.3e-05 | 1.2e-05 | 1.300E-05 | 1.000E-06 | 1.200E-05 | 1.400E-05 |
| 6241 | 313 | S313A | A | 1.7e-05 | 1.8e-05 | 1.2e-05 | 1.567E-05 | 3.215E-06 | 1.200E-05 | 1.800E-05 |
| 6242 | 313 | S313R | A | 9.1e-06 | 1.6e-05 | 1.5e-05 | 1.337E-05 | 3.729E-06 | 9.100E-06 | 1.600E-05 |
| 6243 | 313 | S313N | A | 1.1e-05 | 1.4e-05 | 1.4e-05 | 1.300E-05 | 1.732E-06 | 1.100E-05 | 1.400E-05 |
| 6244 | 313 | S313D | A | 1.7e-05 | 1.3e-05 | 1.2e-05 | 1.400E-05 | 2.646E-06 | 1.200E-05 | 1.700E-05 |
| 6245 | 313 | S313C | A | 1.5e-05 | 1.2e-05 | 1.9e-05 | 1.533E-05 | 3.512E-06 | 1.200E-05 | 1.900E-05 |
| 6246 | 313 | S313E | A | 9.6e-06 | 1.5e-05 | 1.7e-05 | 1.387E-05 | 3.828E-06 | 9.600E-06 | 1.700E-05 |
| 6247 | 313 | S313Q | A | 8.6e-06 | 9.6e-06 | 1.3e-05 | 1.040E-05 | 2.307E-06 | 8.600E-06 | 1.300E-05 |
| 6248 | 313 | S313G | A | 1.4e-05 | 9.6e-06 | 1.2e-05 | 1.187E-05 | 2.203E-06 | 9.600E-06 | 1.400E-05 |
| 6249 | 313 | S313H | A | 7.8e-06 | 7.4e-06 | 8e-06 | 7.733E-06 | 3.055E-07 | 7.400E-06 | 8.000E-06 |
| 6250 | 313 | S313I | A | 1.6e-05 | 1.2e-05 | 1.4e-05 | 1.400E-05 | 2.000E-06 | 1.200E-05 | 1.600E-05 |
| 6251 | 313 | S313L | A | 1.8e-05 | 1e-05 | 1.2e-05 | 1.333E-05 | 4.163E-06 | 1.000E-05 | 1.800E-05 |
| 6252 | 313 | S313K | A | 1.5e-05 | 9.7e-06 | 1.3e-05 | 1.257E-05 | 2.676E-06 | 9.700E-06 | 1.500E-05 |
| 6253 | 313 | S313M | A | 1.1e-05 | 1.7e-05 | 1.7e-05 | 1.500E-05 | 3.464E-06 | 1.100E-05 | 1.700E-05 |
| 6254 | 313 | S313F | A | 1.4e-05 | 1.2e-05 | 1.6e-05 | 1.400E-05 | 2.000E-06 | 1.200E-05 | 1.600E-05 |
| 6255 | 313 | S313P | A | 1.6e-05 | 1.2e-05 | 1.6e-05 | 1.467E-05 | 2.309E-06 | 1.200E-05 | 1.600E-05 |
| 6256 | 313 | S313S | A | 1.7e-05 | 1.4e-05 | 1.2e-05 | 1.433E-05 | 2.517E-06 | 1.200E-05 | 1.700E-05 |
| 6257 | 313 | S313T | A | 1.6e-05 | 7e-06 | 1.6e-05 | 1.300E-05 | 5.196E-06 | 7.000E-06 | 1.600E-05 |
| 6258 | 313 | S313W | A | 9e-06 | 1.2e-05 | 1.1e-05 | 1.067E-05 | 1.528E-06 | 9.000E-06 | 1.200E-05 |
| 6259 | 313 | S313Y | A | 1.8e-05 | 1.6e-05 | 1.5e-05 | 1.633E-05 | 1.528E-06 | 1.500E-05 | 1.800E-05 |
| 6260 | 313 | S313V | A | 1.1e-05 | 1.8e-05 | 1.4e-05 | 1.433E-05 | 3.512E-06 | 1.100E-05 | 1.800E-05 |
| 6261 | 314 | K314A | A | 2.1e-05 | 2.3e-05 | 9.3e-06 | 1.777E-05 | 7.400E-06 | 9.300E-06 | 2.300E-05 |
| 6262 | 314 | K314R | A | 1.6e-05 | 1e-05 | 1.3e-05 | 1.300E-05 | 3.000E-06 | 1.000E-05 | 1.600E-05 |
| 6263 | 314 | K314N | A | 6e-06 | 9.6e-06 | 1e-05 | 8.533E-06 | 2.203E-06 | 6.000E-06 | 1.000E-05 |
| 6264 | 314 | K314D | A | 1.1e-05 | 1.5e-05 | 1e-05 | 1.200E-05 | 2.646E-06 | 1.000E-05 | 1.500E-05 |
| 6265 | 314 | K314C | A | 2.2e-05 | 1.2e-05 | 2.1e-05 | 1.833E-05 | 5.508E-06 | 1.200E-05 | 2.200E-05 |
| 6266 | 314 | K314E | A | 1.5e-05 | 1.3e-05 | 1.6e-05 | 1.467E-05 | 1.528E-06 | 1.300E-05 | 1.600E-05 |
| 6267 | 314 | K314Q | A | 7.1e-06 | 7.2e-06 | 1e-05 | 8.100E-06 | 1.646E-06 | 7.100E-06 | 1.000E-05 |
| 6268 | 314 | K314G | A | 1.2e-05 | 1.5e-05 | 1.4e-05 | 1.367E-05 | 1.528E-06 | 1.200E-05 | 1.500E-05 |
| 6269 | 314 | K314H | A | 1.7e-05 | 1.1e-05 | 1.8e-05 | 1.533E-05 | 3.786E-06 | 1.100E-05 | 1.800E-05 |
| 6270 | 314 | K314I | A | 1.2e-05 | 1.2e-05 | 1.4e-05 | 1.267E-05 | 1.155E-06 | 1.200E-05 | 1.400E-05 |
| 6271 | 314 | K314L | A | 1.9e-05 | 1.4e-05 | 2.3e-05 | 1.867E-05 | 4.509E-06 | 1.400E-05 | 2.300E-05 |
| 6272 | 314 | K314K | A | 1.2e-05 | 1.5e-05 | 1.6e-05 | 1.433E-05 | 2.082E-06 | 1.200E-05 | 1.600E-05 |
| 6273 | 314 | K314M | A | 1.7e-05 | 1.8e-05 | 2e-05 | 1.833E-05 | 1.528E-06 | 1.700E-05 | 2.000E-05 |
| 6274 | 314 | K314F | A | 1.1e-05 | 1.6e-05 | 2e-05 | 1.567E-05 | 4.509E-06 | 1.100E-05 | 2.000E-05 |
| 6275 | 314 | K314P | A | 2e-05 | 1.2e-05 | 1.4e-05 | 1.533E-05 | 4.163E-06 | 1.200E-05 | 2.000E-05 |
| 6276 | 314 | K314S | A | 1.1e-05 | 5.7e-06 | 7.1e-06 | 7.933E-06 | 2.747E-06 | 5.700E-06 | 1.100E-05 |
| 6277 | 314 | K314T | A | 6.2e-06 | 7.5e-06 | 8.3e-06 | 7.333E-06 | 1.060E-06 | 6.200E-06 | 8.300E-06 |
| 6278 | 314 | K314W | A | 1.3e-05 | 1.3e-05 | 1.9e-05 | 1.500E-05 | 3.464E-06 | 1.300E-05 | 1.900E-05 |
| 6279 | 314 | K314Y | A | 1.4e-05 | 1.5e-05 | 2.1e-05 | 1.667E-05 | 3.786E-06 | 1.400E-05 | 2.100E-05 |
| 6280 | 314 | K314V | A | 1.4e-05 | 1.8e-05 | 1.3e-05 | 1.500E-05 | 2.646E-06 | 1.300E-05 | 1.800E-05 |
| 6281 | 315 | P315A | A | 1.5e-05 | 1.6e-05 | 1.6e-05 | 1.567E-05 | 5.774E-07 | 1.500E-05 | 1.600E-05 |
| 6282 | 315 | P315R | A | 5.8e-06 | 8.2e-06 | 8.2e-06 | 7.400E-06 | 1.386E-06 | 5.800E-06 | 8.200E-06 |
| 6283 | 315 | P315N | A | 7.2e-06 | 5.4e-06 | 4.5e-06 | 5.700E-06 | 1.375E-06 | 4.500E-06 | 7.200E-06 |
| 6284 | 315 | P315D | A | 8.9e-06 | 9.2e-06 | 1e-05 | 9.367E-06 | 5.686E-07 | 8.900E-06 | 1.000E-05 |
| 6285 | 315 | P315C | A | 9.4e-06 | 9.5e-06 | 1.2e-05 | 1.030E-05 | 1.473E-06 | 9.400E-06 | 1.200E-05 |
| 6286 | 315 | P315E | A | 9.9e-06 | 6.9e-06 | 7.9e-06 | 8.233E-06 | 1.528E-06 | 6.900E-06 | 9.900E-06 |
| 6287 | 315 | P315Q | A | 5.8e-06 | 3.2e-06 | 5.2e-06 | 4.733E-06 | 1.361E-06 | 3.200E-06 | 5.800E-06 |
| 6288 | 315 | P315G | A | 1.4e-05 | 1.4e-05 | 9.8e-06 | 1.260E-05 | 2.425E-06 | 9.800E-06 | 1.400E-05 |
| 6289 | 315 | P315H | A | 7.9e-06 | 5.5e-06 | 7.9e-06 | 7.100E-06 | 1.386E-06 | 5.500E-06 | 7.900E-06 |

| | | | | | | | | | | |
|---|---|---|---|---|---|---|---|---|---|---|
| 6290 | 315 | P315I | A | 1.4e-05 | 1.2e-05 | 8.4e-06 | 1.147E-05 | 2.838E-06 | 8.400E-06 | 1.400E-05 |
| 6291 | 315 | P315L | A | 1.3e-05 | 1e-05 | 1.2e-05 | 1.167E-05 | 1.528E-06 | 1.000E-05 | 1.300E-05 |
| 6292 | 315 | P315K | A | 1e-05 | 8.2e-06 | 8.1e-06 | 8.767E-06 | 1.069E-06 | 8.100E-06 | 1.000E-05 |
| 6293 | 315 | P315M | A | 9.7e-06 | 8.4e-06 | 1.2e-05 | 1.003E-05 | 1.823E-06 | 8.400E-06 | 1.200E-05 |
| 6294 | 315 | P315F | A | 8.5e-06 | 9.3e-06 | 8.8e-06 | 8.867E-06 | 4.041E-07 | 8.500E-06 | 9.300E-06 |
| 6295 | 315 | P315P | A | 1.1e-05 | 1.6e-05 | 1.3e-05 | 1.333E-05 | 2.517E-06 | 1.100E-05 | 1.600E-05 |
| 6296 | 315 | P315S | A | 7.6e-06 | 5.3e-06 | 6.5e-06 | 6.467E-06 | 1.150E-06 | 5.300E-06 | 7.600E-06 |
| 6297 | 315 | P315T | A | 6.8e-06 | 7.2e-06 | 4.4e-06 | 6.133E-06 | 1.514E-06 | 4.400E-06 | 7.200E-06 |
| 6298 | 315 | P315W | A | 7.5e-06 | 1.3e-05 | 9.5e-06 | 1.000E-05 | 2.784E-06 | 7.500E-06 | 1.300E-05 |
| 6299 | 315 | P315Y | A | 8.8e-06 | 8.4e-06 | 6e-06 | 7.733E-06 | 1.514E-06 | 6.000E-06 | 8.800E-06 |
| 6300 | 315 | P315V | A | 1.2e-05 | 1e-05 | 1.1e-05 | 1.100E-05 | 1.000E-06 | 1.000E-05 | 1.200E-05 |
| 6301 | 316 | C316A | A | 1.4e-05 | 1.7e-05 | 7.2e-06 | 1.273E-05 | 5.021E-06 | 7.200E-06 | 1.700E-05 |
| 6302 | 316 | C316R | A | 1.8e-05 | 1.4e-05 | 1.3e-05 | 1.500E-05 | 2.646E-06 | 1.300E-05 | 1.800E-05 |
| 6303 | 316 | C316N | A | 1.8e-05 | 1.1e-05 | 1.7e-05 | 1.533E-05 | 3.786E-06 | 1.100E-05 | 1.800E-05 |
| 6304 | 316 | C316D | A | 1.7e-05 | 1.2e-05 | 9.2e-06 | 1.273E-05 | 3.951E-06 | 9.200E-06 | 1.700E-05 |
| 6305 | 316 | C316C | A | 1.6e-05 | 1.4e-05 | 1.1e-05 | 1.367E-05 | 2.517E-06 | 1.100E-05 | 1.600E-05 |
| 6306 | 316 | C316E | A | 1.2e-05 | 7.7e-06 | 1.6e-05 | 1.190E-05 | 4.151E-06 | 7.700E-06 | 1.600E-05 |
| 6307 | 316 | C316Q | A | 7.6e-06 | 1.1e-05 | 9.8e-06 | 9.467E-06 | 1.724E-06 | 7.600E-06 | 1.100E-05 |
| 6308 | 316 | C316G | A | 9.9e-06 | 8.5e-06 | 1.4e-05 | 1.080E-05 | 2.858E-06 | 8.500E-06 | 1.400E-05 |
| 6309 | 316 | C316H | A | 8.1e-06 | 9.4e-06 | 1e-05 | 9.167E-06 | 9.713E-07 | 8.100E-06 | 1.000E-05 |
| 6310 | 316 | C316I | A | 1.7e-05 | 1.5e-05 | 1.2e-05 | 1.467E-05 | 2.517E-06 | 1.200E-05 | 1.700E-05 |
| 6311 | 316 | C316L | A | 8.7e-06 | 1.6e-05 | 1.3e-05 | 1.257E-05 | 3.669E-06 | 8.700E-06 | 1.600E-05 |
| 6312 | 316 | C316K | A | 9.1e-06 | 1.5e-05 | 1.7e-05 | 1.370E-05 | 4.107E-06 | 9.100E-06 | 1.700E-05 |
| 6313 | 316 | C316M | A | 1.6e-05 | 8.3e-06 | 1.1e-05 | 1.177E-05 | 3.907E-06 | 8.300E-06 | 1.600E-05 |
| 6314 | 316 | C316F | A | 1e-05 | 1.7e-05 | 2.1e-05 | 1.600E-05 | 5.568E-06 | 1.000E-05 | 2.100E-05 |
| 6315 | 316 | C316P | A | 9.1e-06 | 1.1e-05 | 1.3e-05 | 1.103E-05 | 1.950E-06 | 9.100E-06 | 1.300E-05 |
| 6316 | 316 | C316S | A | 1e-05 | 9.9e-06 | 1.4e-05 | 1.130E-05 | 2.339E-06 | 9.900E-06 | 1.400E-05 |
| 6317 | 316 | C316T | A | 1e-05 | 1.1e-05 | 1.3e-05 | 1.133E-05 | 1.528E-06 | 1.000E-05 | 1.300E-05 |
| 6318 | 316 | C316W | A | 1.5e-05 | 1.3e-05 | 1.1e-05 | 1.300E-05 | 2.000E-06 | 1.100E-05 | 1.500E-05 |
| 6319 | 316 | C316Y | A | 1.4e-05 | 1.6e-05 | 1.2e-05 | 1.400E-05 | 2.000E-06 | 1.200E-05 | 1.600E-05 |
| 6320 | 316 | C316V | A | 1e-05 | 1.1e-05 | 1.7e-05 | 1.267E-05 | 3.786E-06 | 1.000E-05 | 1.700E-05 |
| 6321 | 317 | A317A | A | 1e-05 | 1.4e-05 | 1.5e-05 | 1.300E-05 | 2.646E-06 | 1.000E-05 | 1.500E-05 |
| 6322 | 317 | A317R | A | 1.1e-05 | 1.5e-05 | 1.4e-05 | 1.333E-05 | 2.082E-06 | 1.100E-05 | 1.500E-05 |
| 6323 | 317 | A317N | A | 8.8e-06 | 1e-05 | 8e-06 | 8.933E-06 | 1.007E-06 | 8.000E-06 | 1.000E-05 |
| 6324 | 317 | A317D | A | 1.1e-05 | 9.8e-06 | 1.3e-05 | 1.127E-05 | 1.617E-06 | 9.800E-06 | 1.300E-05 |
| 6325 | 317 | A317C | A | 1.3e-05 | 1.7e-05 | 1.2e-05 | 1.400E-05 | 2.646E-06 | 1.200E-05 | 1.700E-05 |
| 6326 | 317 | A317E | A | 1e-05 | 1.6e-05 | 1.1e-05 | 1.233E-05 | 3.215E-06 | 1.000E-05 | 1.600E-05 |
| 6327 | 317 | A317Q | A | 1.1e-05 | 1.5e-05 | 1.4e-05 | 1.333E-05 | 2.082E-06 | 1.100E-05 | 1.500E-05 |
| 6328 | 317 | A317G | A | 1.5e-05 | 1.7e-05 | 9.7e-06 | 1.390E-05 | 3.772E-06 | 9.700E-06 | 1.700E-05 |
| 6329 | 317 | A317H | A | 9.1e-06 | 1.2e-05 | 8.7e-06 | 9.933E-06 | 1.801E-06 | 8.700E-06 | 1.200E-05 |
| 6330 | 317 | A317I | A | 1.2e-05 | 1.1e-05 | 1.5e-05 | 1.267E-05 | 2.082E-06 | 1.100E-05 | 1.500E-05 |
| 6331 | 317 | A317L | A | 1.6e-05 | 1.5e-05 | 1.5e-05 | 1.533E-05 | 5.774E-07 | 1.500E-05 | 1.600E-05 |
| 6332 | 317 | A317K | A | 1.1e-05 | 1.5e-05 | 1.5e-05 | 1.367E-05 | 2.309E-06 | 1.100E-05 | 1.500E-05 |
| 6333 | 317 | A317M | A | 1.3e-05 | 1.1e-05 | 5.9e-06 | 9.967E-06 | 3.661E-06 | 5.900E-06 | 1.300E-05 |
| 6334 | 317 | A317F | A | 8.6e-06 | 1.6e-05 | 1.4e-05 | 1.287E-05 | 3.828E-06 | 8.600E-06 | 1.600E-05 |
| 6335 | 317 | A317P | A | 1.1e-05 | 1.5e-05 | 9.5e-06 | 1.183E-05 | 2.843E-06 | 9.500E-06 | 1.500E-05 |
| 6336 | 317 | A317S | A | 1.6e-05 | 1.1e-05 | 1.1e-05 | 1.267E-05 | 2.887E-06 | 1.100E-05 | 1.600E-05 |
| 6337 | 317 | A317T | A | 1.6e-05 | 1.5e-05 | 1.4e-05 | 1.500E-05 | 1.000E-06 | 1.400E-05 | 1.600E-05 |
| 6338 | 317 | A317W | A | 1.5e-05 | 7.6e-06 | 1.4e-05 | 1.220E-05 | 4.015E-06 | 7.600E-06 | 1.500E-05 |
| 6339 | 317 | A317Y | A | 9.6e-06 | 1.5e-05 | 1.3e-05 | 1.253E-05 | 2.730E-06 | 9.600E-06 | 1.500E-05 |
| 6340 | 317 | A317V | A | 1.5e-05 | 1.4e-05 | 1.6e-05 | 1.500E-05 | 1.000E-06 | 1.400E-05 | 1.600E-05 |
| 6341 | 318 | R318A | A | 1.6e-05 | 1e-05 | 1.7e-05 | 1.433E-05 | 3.786E-06 | 1.000E-05 | 1.700E-05 |
| 6342 | 318 | R318R | A | 8.6e-06 | 1.4e-05 | 1.4e-05 | 1.220E-05 | 3.118E-06 | 8.600E-06 | 1.400E-05 |
| 6343 | 318 | R318N | A | 9.6e-06 | 1.7e-05 | 9.1e-06 | 1.190E-05 | 4.424E-06 | 9.100E-06 | 1.700E-05 |
| 6344 | 318 | R318D | A | 1.5e-05 | 1.6e-05 | 1.5e-05 | 1.533E-05 | 5.774E-07 | 1.500E-05 | 1.600E-05 |
| 6345 | 318 | R318C | A | 9.5e-06 | 1.5e-05 | 1.3e-05 | 1.250E-05 | 2.784E-06 | 9.500E-06 | 1.500E-05 |
| 6346 | 318 | R318E | A | 8.2e-06 | 1.6e-05 | 1.1e-05 | 1.173E-05 | 3.951E-06 | 8.200E-06 | 1.600E-05 |
| 6347 | 318 | R318Q | A | 1.9e-05 | 1.4e-05 | 1.6e-05 | 1.633E-05 | 2.517E-06 | 1.400E-05 | 1.900E-05 |
| 6348 | 318 | R318G | A | 1.2e-05 | 1.2e-05 | 9.5e-06 | 1.117E-05 | 1.443E-06 | 9.500E-06 | 1.200E-05 |
| 6349 | 318 | R318H | A | 9.8e-06 | 1.4e-05 | 1.7e-05 | 1.360E-05 | 3.617E-06 | 9.800E-06 | 1.700E-05 |
| 6350 | 318 | R318I | A | 1.7e-05 | 8.8e-06 | 1.6e-05 | 1.393E-05 | 4.474E-06 | 8.800E-06 | 1.700E-05 |
| 6351 | 318 | R318L | A | 1.4e-05 | 1.1e-05 | 9.5e-06 | 1.150E-05 | 2.291E-06 | 9.500E-06 | 1.400E-05 |
| 6352 | 318 | R318K | A | 1.5e-05 | 1.5e-05 | 1.7e-05 | 1.567E-05 | 1.155E-06 | 1.500E-05 | 1.700E-05 |
| 6353 | 318 | R318M | A | 1.4e-05 | 1.5e-05 | 1.2e-05 | 1.367E-05 | 1.528E-06 | 1.200E-05 | 1.500E-05 |
| 6354 | 318 | R318F | A | 1.5e-05 | 1.6e-05 | 1e-05 | 1.367E-05 | 3.215E-06 | 1.000E-05 | 1.600E-05 |
| 6355 | 318 | R318P | A | 1.4e-05 | 1.1e-05 | 1.6e-05 | 1.367E-05 | 2.517E-06 | 1.100E-05 | 1.600E-05 |
| 6356 | 318 | R318S | A | 9.7e-06 | 1.4e-05 | 1.1e-05 | 1.157E-05 | 2.205E-06 | 9.700E-06 | 1.400E-05 |
| 6357 | 318 | R318T | A | 1e-05 | 1.1e-05 | 1.5e-05 | 1.200E-05 | 2.646E-06 | 1.000E-05 | 1.500E-05 |
| 6358 | 318 | R318W | A | 1.6e-05 | 1.1e-05 | 1.1e-05 | 1.267E-05 | 2.887E-06 | 1.100E-05 | 1.600E-05 |
| 6359 | 318 | R318Y | A | 1.4e-05 | 1.4e-05 | 1.5e-05 | 1.433E-05 | 5.774E-07 | 1.400E-05 | 1.500E-05 |
| 6360 | 318 | R318V | A | 8.3e-06 | 1.3e-05 | 8.9e-06 | 1.007E-05 | 2.558E-06 | 8.300E-06 | 1.300E-05 |
| 6361 | 319 | V319A | A | 1e-05 | 1e-05 | 8.6e-06 | 9.533E-06 | 8.083E-07 | 8.600E-06 | 1.000E-05 |
| 6362 | 319 | V319R | A | 7.3e-06 | 9.3e-06 | 1.4e-05 | 1.020E-05 | 3.439E-06 | 7.300E-06 | 1.400E-05 |
| 6363 | 319 | V319N | A | 1.2e-05 | 1e-05 | 1.5e-05 | 1.233E-05 | 2.517E-06 | 1.000E-05 | 1.500E-05 |
| 6364 | 319 | V319D | A | 1.6e-05 | 1.5e-05 | 9.9e-06 | 1.363E-05 | 3.272E-06 | 9.900E-06 | 1.600E-05 |
| 6365 | 319 | V319C | A | 1.2e-05 | 1.5e-05 | 9.3e-06 | 1.210E-05 | 2.851E-06 | 9.300E-06 | 1.500E-05 |
| 6366 | 319 | V319E | A | 1.6e-05 | 9.1e-06 | 1e-05 | 1.170E-05 | 3.751E-06 | 9.100E-06 | 1.600E-05 |
| 6367 | 319 | V319Q | A | 1.1e-05 | 1.3e-05 | 9.6e-06 | 1.120E-05 | 1.709E-06 | 9.600E-06 | 1.300E-05 |
| 6368 | 319 | V319G | A | 1.5e-05 | 1.5e-05 | 1.5e-05 | 1.500E-05 | 0.000E+00 | 1.500E-05 | 1.500E-05 |
| 6369 | 319 | V319H | A | 1.6e-05 | 1.4e-05 | 1.3e-05 | 1.433E-05 | 1.528E-06 | 1.300E-05 | 1.600E-05 |
| 6370 | 319 | V319I | A | 1.7e-05 | 7.7e-06 | 1.1e-05 | 1.190E-05 | 4.715E-06 | 7.700E-06 | 1.700E-05 |
| 6371 | 319 | V319L | A | 1.3e-05 | 9.8e-06 | 1.5e-05 | 1.260E-05 | 2.623E-06 | 9.800E-06 | 1.500E-05 |
| 6372 | 319 | V319K | A | 1.3e-05 | 1e-05 | 1.1e-05 | 1.133E-05 | 1.528E-06 | 1.000E-05 | 1.300E-05 |
| 6373 | 319 | V319M | A | 8.2e-06 | 1.6e-05 | 1.7e-05 | 1.373E-05 | 4.818E-06 | 8.200E-06 | 1.700E-05 |
| 6374 | 319 | V319F | A | 1.2e-05 | 9.5e-06 | 1.2e-05 | 1.117E-05 | 1.443E-06 | 9.500E-06 | 1.200E-05 |

| | | | | | | | | | | |
|---|---|---|---|---|---|---|---|---|---|---|
| 6375 | 319 | V319P | A | 1.5e-05 | 1.5e-05 | 9.2e-06 | 1.307E-05 | 3.349E-06 | 9.200E-06 | 1.500E-05 |
| 6376 | 319 | V319S | A | 1.6e-05 | 1.5e-05 | 1.5e-05 | 1.533E-05 | 5.774E-07 | 1.500E-05 | 1.600E-05 |
| 6377 | 319 | V319T | A | 1.1e-05 | 8.7e-06 | 1.6e-05 | 1.190E-05 | 3.732E-06 | 8.700E-06 | 1.600E-05 |
| 6378 | 319 | V319W | A | 1.4e-05 | 1.1e-05 | 1.3e-05 | 1.267E-05 | 1.528E-06 | 1.100E-05 | 1.400E-05 |
| 6379 | 319 | V319Y | A | 1e-05 | 1.5e-05 | 8.7e-06 | 1.123E-05 | 3.326E-06 | 8.700E-06 | 1.500E-05 |
| 6380 | 319 | V319V | A | 8.4e-06 | 1.4e-05 | 1.1e-05 | 1.113E-05 | 2.802E-06 | 8.400E-06 | 1.400E-05 |
| 6381 | 320 | C320A | A | 1.4e-05 | 1.1e-05 | 1.3e-05 | 1.267E-05 | 1.528E-06 | 1.100E-05 | 1.400E-05 |
| 6382 | 320 | C320R | A | 1.5e-05 | 1.4e-05 | 1.5e-05 | 1.467E-05 | 5.774E-07 | 1.400E-05 | 1.500E-05 |
| 6383 | 320 | C320N | A | 1.7e-05 | 1.4e-05 | 1e-05 | 1.367E-05 | 3.512E-06 | 1.000E-05 | 1.700E-05 |
| 6384 | 320 | C320D | A | 1.5e-05 | 1.3e-05 | 9e-06 | 1.233E-05 | 3.055E-06 | 9.000E-06 | 1.500E-05 |
| 6385 | 320 | C320C | A | 1.5e-05 | 1e-05 | 1.7e-05 | 1.400E-05 | 3.606E-06 | 1.000E-05 | 1.700E-05 |
| 6386 | 320 | C320E | A | 1.4e-05 | 1.2e-05 | 1.5e-05 | 1.367E-05 | 1.528E-06 | 1.200E-05 | 1.500E-05 |
| 6387 | 320 | C320Q | A | 1.5e-05 | 1.1e-05 | 1.1e-05 | 1.233E-05 | 2.309E-06 | 1.100E-05 | 1.500E-05 |
| 6388 | 320 | C320G | A | 1.1e-05 | 1.2e-05 | 1.6e-05 | 1.300E-05 | 2.646E-06 | 1.100E-05 | 1.600E-05 |
| 6389 | 320 | C320H | A | 1.1e-05 | 1.6e-05 | 8.2e-06 | 1.173E-05 | 3.951E-06 | 8.200E-06 | 1.600E-05 |
| 6390 | 320 | C320I | A | 1.3e-05 | 1.5e-05 | 1.9e-05 | 1.567E-05 | 3.055E-06 | 1.300E-05 | 1.900E-05 |
| 6391 | 320 | C320L | A | 9.3e-06 | 1.6e-05 | 1.5e-05 | 1.343E-05 | 3.614E-06 | 9.300E-06 | 1.600E-05 |
| 6392 | 320 | C320K | A | 1.3e-05 | 1.5e-05 | 1.1e-05 | 1.300E-05 | 2.000E-06 | 1.100E-05 | 1.500E-05 |
| 6393 | 320 | C320M | A | 1.6e-05 | 9.2e-06 | 1.5e-05 | 1.340E-05 | 3.672E-06 | 9.200E-06 | 1.600E-05 |
| 6394 | 320 | C320F | A | 1.5e-05 | 1.7e-05 | 9.9e-06 | 1.397E-05 | 3.661E-06 | 9.900E-06 | 1.700E-05 |
| 6395 | 320 | C320P | A | 1.1e-05 | 1.5e-05 | 1.7e-05 | 1.433E-05 | 3.055E-06 | 1.100E-05 | 1.700E-05 |
| 6396 | 320 | C320S | A | 1e-05 | 1.7e-05 | 1.5e-05 | 1.400E-05 | 3.606E-06 | 1.000E-05 | 1.700E-05 |
| 6397 | 320 | C320T | A | 1.7e-05 | 9.8e-06 | 1.5e-05 | 1.393E-05 | 3.717E-06 | 9.800E-06 | 1.700E-05 |
| 6398 | 320 | C320W | A | 1.3e-05 | 1.4e-05 | 1.5e-05 | 1.400E-05 | 1.000E-06 | 1.300E-05 | 1.500E-05 |
| 6399 | 320 | C320Y | A | 1.3e-05 | 8.8e-06 | 1.5e-05 | 1.227E-05 | 3.164E-06 | 8.800E-06 | 1.500E-05 |
| 6400 | 320 | C320V | A | 1.4e-05 | 1.2e-05 | 1.4e-05 | 1.333E-05 | 1.155E-06 | 1.200E-05 | 1.400E-05 |
| 6401 | 321 | Y321A | A | 1e-05 | 1e-05 | 1.4e-05 | 1.133E-05 | 2.309E-06 | 1.000E-05 | 1.400E-05 |
| 6402 | 321 | Y321R | A | 1e-05 | 7.7e-06 | 1.1e-05 | 9.567E-06 | 1.692E-06 | 7.700E-06 | 1.100E-05 |
| 6403 | 321 | Y321N | A | 1.3e-05 | 1.4e-05 | 1.7e-05 | 1.467E-05 | 2.082E-06 | 1.300E-05 | 1.700E-05 |
| 6404 | 321 | Y321D | A | 1.7e-05 | 1.1e-05 | 1.1e-05 | 1.300E-05 | 3.464E-06 | 1.100E-05 | 1.700E-05 |
| 6405 | 321 | Y321C | A | 1.1e-05 | 1e-05 | 1.3e-05 | 1.133E-05 | 1.528E-06 | 1.000E-05 | 1.300E-05 |
| 6406 | 321 | Y321E | A | 9.8e-06 | 1.7e-05 | 1.5e-05 | 1.393E-05 | 3.717E-06 | 9.800E-06 | 1.700E-05 |
| 6407 | 321 | Y321Q | A | 9.8e-06 | 1e-05 | 8.6e-06 | 9.467E-06 | 7.572E-07 | 8.600E-06 | 1.000E-05 |
| 6408 | 321 | Y321G | A | 1.2e-05 | 1.3e-05 | 1.7e-05 | 1.400E-05 | 2.646E-06 | 1.200E-05 | 1.700E-05 |
| 6409 | 321 | Y321H | A | 1.5e-05 | 1.4e-05 | 7.9e-06 | 1.230E-05 | 3.843E-06 | 7.900E-06 | 1.500E-05 |
| 6410 | 321 | Y321I | A | 9.2e-06 | 1.5e-05 | 1.4e-05 | 1.273E-05 | 3.101E-06 | 9.200E-06 | 1.500E-05 |
| 6411 | 321 | Y321L | A | 6.5e-06 | 1.5e-05 | 1.6e-05 | 1.250E-05 | 5.220E-06 | 6.500E-06 | 1.600E-05 |
| 6412 | 321 | Y321K | A | 8.9e-06 | 1.6e-05 | 1.1e-05 | 1.197E-05 | 3.647E-06 | 8.900E-06 | 1.600E-05 |
| 6413 | 321 | Y321M | A | 1.1e-05 | 1.5e-05 | 1.3e-05 | 1.300E-05 | 2.000E-06 | 1.100E-05 | 1.500E-05 |
| 6414 | 321 | Y321F | A | 1.5e-05 | 9.3e-06 | 1e-05 | 1.143E-05 | 3.109E-06 | 9.300E-06 | 1.500E-05 |
| 6415 | 321 | Y321P | A | 1.7e-05 | 9.4e-06 | 9e-06 | 1.180E-05 | 4.508E-06 | 9.000E-06 | 1.700E-05 |
| 6416 | 321 | Y321S | A | 1.5e-05 | 1.5e-05 | 1.6e-05 | 1.533E-05 | 5.774E-07 | 1.500E-05 | 1.600E-05 |
| 6417 | 321 | Y321T | A | 8.6e-06 | 9.7e-06 | 9.4e-06 | 9.233E-06 | 5.686E-07 | 8.600E-06 | 9.700E-06 |
| 6418 | 321 | Y321W | A | 1.5e-05 | 8.9e-06 | 1.2e-05 | 1.197E-05 | 3.050E-06 | 8.900E-06 | 1.500E-05 |
| 6419 | 321 | Y321Y | A | 1e-05 | 1e-05 | 9.3e-06 | 9.767E-06 | 4.041E-07 | 9.300E-06 | 1.000E-05 |
| 6420 | 321 | Y321V | A | 1.2e-05 | 1.7e-05 | 1.6e-05 | 1.500E-05 | 2.646E-06 | 1.200E-05 | 1.700E-05 |
| 6421 | 322 | G322A | A | 1.1e-05 | 1e-05 | 1.8e-05 | 1.300E-05 | 4.359E-06 | 1.000E-05 | 1.800E-05 |
| 6422 | 322 | G322R | A | 1.4e-05 | 1.5e-05 | 9.1e-06 | 1.270E-05 | 3.158E-06 | 9.100E-06 | 1.500E-05 |
| 6423 | 322 | G322N | A | 9.9e-06 | 1.5e-05 | 1.5e-05 | 1.330E-05 | 2.944E-06 | 9.900E-06 | 1.500E-05 |
| 6424 | 322 | G322D | A | 1.5e-05 | 1.3e-05 | 1e-05 | 1.267E-05 | 2.517E-06 | 1.000E-05 | 1.500E-05 |
| 6425 | 322 | G322C | A | 1e-05 | 1.3e-05 | 1.4e-05 | 1.233E-05 | 2.082E-06 | 1.000E-05 | 1.400E-05 |
| 6426 | 322 | G322E | A | 1.4e-05 | 1.7e-05 | 1.5e-05 | 1.533E-05 | 1.528E-06 | 1.400E-05 | 1.700E-05 |
| 6427 | 322 | G322Q | A | 1.7e-05 | 9.5e-06 | 1.4e-05 | 1.350E-05 | 3.775E-06 | 9.500E-06 | 1.700E-05 |
| 6428 | 322 | G322G | A | 1.4e-05 | 1.1e-05 | 1.7e-05 | 1.400E-05 | 3.000E-06 | 1.100E-05 | 1.700E-05 |
| 6429 | 322 | G322H | A | 9e-06 | 1.6e-05 | 1.5e-05 | 1.333E-05 | 3.786E-06 | 9.000E-06 | 1.600E-05 |
| 6430 | 322 | G322I | A | 1e-05 | 8.8e-06 | 1.1e-05 | 9.933E-06 | 1.102E-06 | 8.800E-06 | 1.100E-05 |
| 6431 | 322 | G322L | A | 1e-05 | 1.5e-05 | 1.6e-05 | 1.367E-05 | 3.215E-06 | 1.000E-05 | 1.600E-05 |
| 6432 | 322 | G322K | A | 1.6e-05 | 1.2e-05 | 1.3e-05 | 1.367E-05 | 2.082E-06 | 1.200E-05 | 1.600E-05 |
| 6433 | 322 | G322M | A | 1.5e-05 | 1.5e-05 | 1.4e-05 | 1.467E-05 | 5.774E-07 | 1.400E-05 | 1.500E-05 |
| 6434 | 322 | G322F | A | 8.1e-06 | 1.5e-05 | 1.4e-05 | 1.237E-05 | 3.729E-06 | 8.100E-06 | 1.500E-05 |
| 6435 | 322 | G322P | A | 1.6e-05 | 1.4e-05 | 9.6e-06 | 1.320E-05 | 3.274E-06 | 9.600E-06 | 1.600E-05 |
| 6436 | 322 | G322S | A | 1.3e-05 | 1.6e-05 | 9.6e-06 | 1.287E-05 | 3.202E-06 | 9.600E-06 | 1.600E-05 |
| 6437 | 322 | G322T | A | 1.1e-05 | 1.2e-05 | 9.2e-06 | 1.073E-05 | 1.419E-06 | 9.200E-06 | 1.200E-05 |
| 6438 | 322 | G322W | A | 6.2e-06 | 1e-05 | 1.3e-05 | 9.733E-06 | 3.408E-06 | 6.200E-06 | 1.300E-05 |
| 6439 | 322 | G322Y | A | 9.8e-06 | 1.5e-05 | 1.1e-05 | 1.193E-05 | 2.723E-06 | 9.800E-06 | 1.500E-05 |
| 6440 | 322 | G322V | A | 1e-05 | 9.6e-06 | 1.5e-05 | 1.153E-05 | 3.009E-06 | 9.600E-06 | 1.500E-05 |
| 6441 | 323 | L323A | A | 1.4e-05 | 9.2e-06 | 1.4e-05 | 1.240E-05 | 2.771E-06 | 9.200E-06 | 1.400E-05 |
| 6442 | 323 | L323R | A | 9.4e-06 | 8.3e-06 | 1.6e-05 | 1.123E-05 | 4.165E-06 | 8.300E-06 | 1.600E-05 |
| 6443 | 323 | L323N | A | 8.2e-06 | 1.5e-05 | 1.4e-05 | 1.240E-05 | 3.672E-06 | 8.200E-06 | 1.500E-05 |
| 6444 | 323 | L323D | A | 1.7e-05 | 1.4e-05 | 1.2e-05 | 1.433E-05 | 2.517E-06 | 1.200E-05 | 1.700E-05 |
| 6445 | 323 | L323C | A | 1e-05 | 1.4e-05 | 2e-05 | 1.467E-05 | 5.033E-06 | 1.000E-05 | 2.000E-05 |
| 6446 | 323 | L323E | A | 1.4e-05 | 1.5e-05 | 1.3e-05 | 1.400E-05 | 1.000E-06 | 1.300E-05 | 1.500E-05 |
| 6447 | 323 | L323Q | A | 1.9e-05 | 1.4e-05 | 1.3e-05 | 1.533E-05 | 3.215E-06 | 1.300E-05 | 1.900E-05 |
| 6448 | 323 | L323G | A | 8.6e-06 | 1.4e-05 | 1.1e-05 | 1.120E-05 | 2.706E-06 | 8.600E-06 | 1.400E-05 |
| 6449 | 323 | L323H | A | 1e-05 | 8.5e-06 | 1.5e-05 | 1.117E-05 | 3.403E-06 | 8.500E-06 | 1.500E-05 |
| 6450 | 323 | L323I | A | 1e-05 | 9.8e-06 | 1.7e-05 | 1.227E-05 | 4.100E-06 | 9.800E-06 | 1.700E-05 |
| 6451 | 323 | L323L | A | 1.1e-05 | 1.4e-05 | 1.2e-05 | 1.233E-05 | 1.528E-06 | 1.100E-05 | 1.400E-05 |
| 6452 | 323 | L323K | A | 1.5e-05 | 1.6e-05 | 1.1e-05 | 1.400E-05 | 2.646E-06 | 1.100E-05 | 1.600E-05 |
| 6453 | 323 | L323M | A | 1e-05 | 1.2e-05 | 1.3e-05 | 1.167E-05 | 1.528E-06 | 1.000E-05 | 1.300E-05 |
| 6454 | 323 | L323F | A | 1.6e-05 | 1.5e-05 | 1.5e-05 | 1.533E-05 | 5.774E-07 | 1.500E-05 | 1.600E-05 |
| 6455 | 323 | L323P | A | 1e-05 | 1e-05 | 1.4e-05 | 1.133E-05 | 2.309E-06 | 1.000E-05 | 1.400E-05 |
| 6456 | 323 | L323S | A | 1.5e-05 | 9.7e-06 | 1.6e-05 | 1.357E-05 | 3.386E-06 | 9.700E-06 | 1.600E-05 |
| 6457 | 323 | L323T | A | 1e-05 | 1.4e-05 | 1.5e-05 | 1.300E-05 | 2.646E-06 | 1.000E-05 | 1.500E-05 |
| 6458 | 323 | L323W | A | 1.6e-05 | 9.9e-06 | 1.2e-05 | 1.263E-05 | 3.099E-06 | 9.900E-06 | 1.600E-05 |
| 6459 | 323 | L323Y | A | 9.3e-06 | 1.6e-05 | 1.1e-05 | 1.210E-05 | 3.483E-06 | 9.300E-06 | 1.600E-05 |

| | | | | | | | | | | |
|---|---|---|---|---|---|---|---|---|---|---|
| 6460 | 323 | L323V | A | 1.4e-05 | 1.4e-05 | 1.5e-05 | 1.433E-05 | 5.774E-07 | 1.400E-05 | 1.500E-05 |
| 6461 | 324 | G324A | A | 1.5e-05 | 1.3e-05 | 1.1e-05 | 1.300E-05 | 2.000E-06 | 1.100E-05 | 1.500E-05 |
| 6462 | 324 | G324R | A | 1.8e-05 | 1.6e-05 | 1.5e-05 | 1.633E-05 | 1.528E-06 | 1.500E-05 | 1.800E-05 |
| 6463 | 324 | G324N | A | 6.8e-06 | 1.5e-05 | 1e-05 | 1.060E-05 | 4.133E-06 | 6.800E-06 | 1.500E-05 |
| 6464 | 324 | G324D | A | 1.6e-05 | 1.7e-05 | 1.5e-05 | 1.600E-05 | 1.000E-06 | 1.500E-05 | 1.700E-05 |
| 6465 | 324 | G324C | A | 1.5e-05 | 1.1e-05 | 8.3e-06 | 1.143E-05 | 3.371E-06 | 8.300E-06 | 1.500E-05 |
| 6466 | 324 | G324E | A | 1.2e-05 | 1.4e-05 | 1.4e-05 | 1.333E-05 | 1.155E-06 | 1.200E-05 | 1.400E-05 |
| 6467 | 324 | G324Q | A | 1.5e-05 | 1.4e-05 | 1.6e-05 | 1.500E-05 | 1.000E-06 | 1.400E-05 | 1.600E-05 |
| 6468 | 324 | G324G | A | 1.1e-05 | 8e-06 | 1.4e-05 | 1.100E-05 | 3.000E-06 | 8.000E-06 | 1.400E-05 |
| 6469 | 324 | G324H | A | 8.4e-06 | 1.5e-05 | 1.1e-05 | 1.147E-05 | 3.325E-06 | 8.400E-06 | 1.500E-05 |
| 6470 | 324 | G324I | A | 1.1e-05 | 1.6e-05 | 1.3e-05 | 1.333E-05 | 2.517E-06 | 1.100E-05 | 1.600E-05 |
| 6471 | 324 | G324L | A | 1.3e-05 | 1.2e-05 | 1.5e-05 | 1.333E-05 | 1.528E-06 | 1.200E-05 | 1.500E-05 |
| 6472 | 324 | G324K | A | 9.1e-06 | 8.9e-06 | 1.6e-05 | 1.133E-05 | 4.043E-06 | 8.900E-06 | 1.600E-05 |
| 6473 | 324 | G324M | A | 1e-05 | 1.6e-05 | 1.6e-05 | 1.400E-05 | 3.464E-06 | 1.000E-05 | 1.600E-05 |
| 6474 | 324 | G324F | A | 1.3e-05 | 1.4e-05 | 1.7e-05 | 1.467E-05 | 2.082E-06 | 1.300E-05 | 1.700E-05 |
| 6475 | 324 | G324P | A | 1e-05 | 9.5e-06 | 1.5e-05 | 1.150E-05 | 3.041E-06 | 9.500E-06 | 1.500E-05 |
| 6476 | 324 | G324S | A | 1.1e-05 | 1.2e-05 | 1.4e-05 | 1.233E-05 | 1.528E-06 | 1.100E-05 | 1.400E-05 |
| 6477 | 324 | G324T | A | 1.1e-05 | 1e-05 | 9.1e-06 | 1.003E-05 | 9.504E-07 | 9.100E-06 | 1.100E-05 |
| 6478 | 324 | G324W | A | 2e-05 | 1.6e-05 | 9.8e-06 | 1.527E-05 | 5.139E-06 | 9.800E-06 | 2.000E-05 |
| 6479 | 324 | G324Y | A | 9.1e-06 | 1.5e-05 | 1.3e-05 | 1.237E-05 | 3.001E-06 | 9.100E-06 | 1.500E-05 |
| 6480 | 324 | G324V | A | 1.6e-05 | 8.6e-06 | 1.5e-05 | 1.320E-05 | 4.015E-06 | 8.600E-06 | 1.600E-05 |
| 6481 | 325 | M325A | A | 1.4e-05 | 6.5e-06 | 1.5e-05 | 1.183E-05 | 4.646E-06 | 6.500E-06 | 1.500E-05 |
| 6482 | 325 | M325R | A | 1.5e-05 | 1e-05 | 1.5e-05 | 1.333E-05 | 2.887E-06 | 1.000E-05 | 1.500E-05 |
| 6483 | 325 | M325N | A | 1.5e-05 | 1.1e-05 | 1.5e-05 | 1.367E-05 | 2.309E-06 | 1.100E-05 | 1.500E-05 |
| 6484 | 325 | M325D | A | 1.7e-05 | 1.4e-05 | 1.2e-05 | 1.433E-05 | 2.517E-06 | 1.200E-05 | 1.700E-05 |
| 6485 | 325 | M325C | A | 9.4e-06 | 1.4e-05 | 1.3e-05 | 1.213E-05 | 2.419E-06 | 9.400E-06 | 1.400E-05 |
| 6486 | 325 | M325E | A | 1.6e-05 | 1.7e-05 | 1.4e-05 | 1.567E-05 | 1.528E-06 | 1.400E-05 | 1.700E-05 |
| 6487 | 325 | M325Q | A | 1.4e-05 | 9.9e-06 | 1.3e-05 | 1.230E-05 | 2.138E-06 | 9.900E-06 | 1.400E-05 |
| 6488 | 325 | M325G | A | 9.4e-06 | 1.5e-05 | 1.4e-05 | 1.280E-05 | 2.987E-06 | 9.400E-06 | 1.500E-05 |
| 6489 | 325 | M325H | A | 9.9e-06 | 1.6e-05 | 1e-05 | 1.197E-05 | 3.493E-06 | 9.900E-06 | 1.600E-05 |
| 6490 | 325 | M325I | A | 1.5e-05 | 1.5e-05 | 1.1e-05 | 1.367E-05 | 2.309E-06 | 1.100E-05 | 1.500E-05 |
| 6491 | 325 | M325L | A | 1.3e-05 | 1.5e-05 | 1.4e-05 | 1.400E-05 | 1.000E-06 | 1.300E-05 | 1.500E-05 |
| 6492 | 325 | M325K | A | 1.5e-05 | 1.2e-05 | 1.3e-05 | 1.333E-05 | 1.528E-06 | 1.200E-05 | 1.500E-05 |
| 6493 | 325 | M325M | A | 1.6e-05 | 1.6e-05 | 1.6e-05 | 1.600E-05 | 0.000E+00 | 1.600E-05 | 1.600E-05 |
| 6494 | 325 | M325F | A | 1.5e-05 | 2.9e-05 | 8.8e-06 | 1.760E-05 | 1.035E-05 | 8.800E-06 | 2.900E-05 |
| 6495 | 325 | M325P | A | 8.4e-06 | 9.6e-06 | 1.2e-05 | 1.000E-05 | 1.833E-06 | 8.400E-06 | 1.200E-05 |
| 6496 | 325 | M325S | A | 9.9e-06 | 1.4e-05 | 1.1e-05 | 1.163E-05 | 2.122E-06 | 9.900E-06 | 1.400E-05 |
| 6497 | 325 | M325T | A | 1e-05 | 1.4e-05 | 1.2e-05 | 1.200E-05 | 2.000E-06 | 1.000E-05 | 1.400E-05 |
| 6498 | 325 | M325W | A | 8.7e-06 | 1.5e-05 | 1.5e-05 | 1.290E-05 | 3.637E-06 | 8.700E-06 | 1.500E-05 |
| 6499 | 325 | M325Y | A | 9.2e-06 | 1.7e-05 | 1.4e-05 | 1.340E-05 | 3.934E-06 | 9.200E-06 | 1.700E-05 |
| 6500 | 325 | M325V | A | 1.4e-05 | 1.6e-05 | 1.9e-05 | 1.633E-05 | 2.517E-06 | 1.400E-05 | 1.900E-05 |
| 6501 | 326 | E326A | A | 1.6e-05 | 1.5e-05 | 9.9e-06 | 1.363E-05 | 3.272E-06 | 9.900E-06 | 1.600E-05 |
| 6502 | 326 | E326R | A | 1.5e-05 | 1.4e-05 | 8.8e-06 | 1.260E-05 | 3.329E-06 | 8.800E-06 | 1.500E-05 |
| 6503 | 326 | E326N | A | 2.2e-05 | 1.1e-05 | 1.5e-05 | 1.600E-05 | 5.568E-06 | 1.100E-05 | 2.200E-05 |
| 6504 | 326 | E326D | A | 1.5e-05 | 1.4e-05 | 1.4e-05 | 1.433E-05 | 5.774E-07 | 1.400E-05 | 1.500E-05 |
| 6505 | 326 | E326C | A | 1.5e-05 | 1.2e-05 | 1.6e-05 | 1.433E-05 | 2.082E-06 | 1.200E-05 | 1.600E-05 |
| 6506 | 326 | E326E | A | 1.6e-05 | 9.1e-06 | 1.5e-05 | 1.337E-05 | 3.729E-06 | 9.100E-06 | 1.600E-05 |
| 6507 | 326 | E326Q | A | 8.3e-06 | 1.1e-05 | 1.4e-05 | 1.110E-05 | 2.851E-06 | 8.300E-06 | 1.400E-05 |
| 6508 | 326 | E326G | A | 1.5e-05 | 9.9e-06 | 8.2e-06 | 1.103E-05 | 3.539E-06 | 8.200E-06 | 1.500E-05 |
| 6509 | 326 | E326H | A | 1.4e-05 | 1.5e-05 | 1.6e-05 | 1.500E-05 | 1.000E-06 | 1.400E-05 | 1.600E-05 |
| 6510 | 326 | E326I | A | 1.7e-05 | 1.6e-05 | 1.5e-05 | 1.600E-05 | 1.000E-06 | 1.500E-05 | 1.700E-05 |
| 6511 | 326 | E326L | A | 1.7e-05 | 1e-05 | 1.4e-05 | 1.367E-05 | 3.512E-06 | 1.000E-05 | 1.700E-05 |
| 6512 | 326 | E326K | A | 1.1e-05 | 1.6e-05 | 1.1e-05 | 1.267E-05 | 2.887E-06 | 1.100E-05 | 1.600E-05 |
| 6513 | 326 | E326M | A | 1.5e-05 | 1.1e-05 | 1.3e-05 | 1.300E-05 | 2.000E-06 | 1.100E-05 | 1.500E-05 |
| 6514 | 326 | E326F | A | 1.5e-05 | 1.3e-05 | 1.5e-05 | 1.433E-05 | 1.155E-06 | 1.300E-05 | 1.500E-05 |
| 6515 | 326 | E326P | A | 1.2e-05 | 1.5e-05 | 1.2e-05 | 1.300E-05 | 1.732E-06 | 1.200E-05 | 1.500E-05 |
| 6516 | 326 | E326S | A | 1.5e-05 | 6.1e-06 | 1.1e-05 | 1.070E-05 | 4.458E-06 | 6.100E-06 | 1.500E-05 |
| 6517 | 326 | E326T | A | 1e-05 | 1.5e-05 | 1.6e-05 | 1.367E-05 | 3.215E-06 | 1.000E-05 | 1.600E-05 |
| 6518 | 326 | E326W | A | 1e-05 | 7.1e-06 | 7.3e-06 | 8.133E-06 | 1.620E-06 | 7.100E-06 | 1.000E-05 |
| 6519 | 326 | E326Y | A | 9.9e-06 | 1e-05 | 1.4e-05 | 1.130E-05 | 2.339E-06 | 9.900E-06 | 1.400E-05 |
| 6520 | 326 | E326V | A | 1.7e-05 | 1.3e-05 | 1.2e-05 | 1.400E-05 | 2.646E-06 | 1.200E-05 | 1.700E-05 |
| 6521 | 327 | H327A | A | 8e-06 | 1.4e-05 | 1.7e-05 | 1.300E-05 | 4.583E-06 | 8.000E-06 | 1.700E-05 |
| 6522 | 327 | H327R | A | 1e-05 | 1.4e-05 | 1.2e-05 | 1.200E-05 | 2.000E-06 | 1.000E-05 | 1.400E-05 |
| 6523 | 327 | H327N | A | 1.3e-05 | 1.6e-05 | 1.4e-05 | 1.433E-05 | 1.528E-06 | 1.300E-05 | 1.600E-05 |
| 6524 | 327 | H327D | A | 1e-05 | 1e-05 | 1.5e-05 | 1.167E-05 | 2.887E-06 | 1.000E-05 | 1.500E-05 |
| 6525 | 327 | H327C | A | 1.3e-05 | 1.1e-05 | 9.5e-06 | 1.117E-05 | 1.756E-06 | 9.500E-06 | 1.300E-05 |
| 6526 | 327 | H327E | A | 1e-05 | 1.1e-05 | 1.6e-05 | 1.233E-05 | 3.215E-06 | 1.000E-05 | 1.600E-05 |
| 6527 | 327 | H327Q | A | 1.5e-05 | 8.6e-06 | 1.1e-05 | 1.153E-05 | 3.233E-06 | 8.600E-06 | 1.500E-05 |
| 6528 | 327 | H327G | A | 1.6e-05 | 1e-05 | 8.9e-06 | 1.163E-05 | 3.821E-06 | 8.900E-06 | 1.600E-05 |
| 6529 | 327 | H327H | A | 1.6e-05 | 1.5e-05 | 9.6e-06 | 1.353E-05 | 3.443E-06 | 9.600E-06 | 1.600E-05 |
| 6530 | 327 | H327I | A | 9.3e-06 | 1e-05 | 1.7e-05 | 1.210E-05 | 4.258E-06 | 9.300E-06 | 1.700E-05 |
| 6531 | 327 | H327L | A | 1e-05 | 1.3e-05 | 1.6e-05 | 1.300E-05 | 3.000E-06 | 1.000E-05 | 1.600E-05 |
| 6532 | 327 | H327K | A | 1.7e-05 | 2.6e-05 | 1.6e-05 | 1.967E-05 | 5.508E-06 | 1.600E-05 | 2.600E-05 |
| 6533 | 327 | H327M | A | 1.5e-05 | 1.3e-05 | 1.7e-05 | 1.500E-05 | 2.000E-06 | 1.300E-05 | 1.700E-05 |
| 6534 | 327 | H327F | A | 1.5e-05 | 1e-05 | 1.7e-05 | 1.400E-05 | 3.606E-06 | 1.000E-05 | 1.700E-05 |
| 6535 | 327 | H327P | A | 9.3e-06 | 9.1e-06 | 9.7e-06 | 9.367E-06 | 3.055E-07 | 9.100E-06 | 9.700E-06 |
| 6536 | 327 | H327S | A | 1.4e-05 | 1.5e-05 | 1.2e-05 | 1.367E-05 | 1.528E-06 | 1.200E-05 | 1.500E-05 |
| 6537 | 327 | H327T | A | 1.6e-05 | 1.4e-05 | 1.7e-05 | 1.567E-05 | 1.528E-06 | 1.400E-05 | 1.700E-05 |
| 6538 | 327 | H327W | A | 1.4e-05 | 1e-05 | 1.6e-05 | 1.333E-05 | 3.055E-06 | 1.000E-05 | 1.600E-05 |
| 6539 | 327 | H327Y | A | 1.3e-05 | 1.1e-05 | 1.4e-05 | 1.267E-05 | 1.528E-06 | 1.100E-05 | 1.400E-05 |
| 6540 | 327 | H327V | A | 1.3e-05 | 1.6e-05 | 1.4e-05 | 1.433E-05 | 1.528E-06 | 1.300E-05 | 1.600E-05 |
| 6541 | 328 | L328A | A | 1.4e-05 | 9.9e-06 | 1.3e-05 | 1.230E-05 | 2.138E-06 | 9.900E-06 | 1.400E-05 |
| 6542 | 328 | L328R | A | 1.2e-05 | 9.9e-06 | 1.5e-05 | 1.230E-05 | 2.563E-06 | 9.900E-06 | 1.500E-05 |
| 6543 | 328 | L328N | A | 1.4e-05 | 1.3e-05 | 1.3e-05 | 1.333E-05 | 5.774E-07 | 1.300E-05 | 1.400E-05 |
| 6544 | 328 | L328D | A | 1.1e-05 | 1.1e-05 | 1.1e-05 | 1.100E-05 | 2.075E-21 | 1.100E-05 | 1.100E-05 |

| | | | | | | | | | | |
|---|---|---|---|---|---|---|---|---|---|---|
| 6545 | 328 | L328C | A | 1.5e-05 | 8.4e-06 | 9.3e-06 | 1.090E-05 | 3.579E-06 | 8.400E-06 | 1.500E-05 |
| 6546 | 328 | L328E | A | 1.1e-05 | 1e-05 | 8.9e-06 | 9.967E-06 | 1.050E-06 | 8.900E-06 | 1.100E-05 |
| 6547 | 328 | L328Q | A | 9.9e-06 | 1.2e-05 | 9.4e-06 | 1.043E-05 | 1.380E-06 | 9.400E-06 | 1.200E-05 |
| 6548 | 328 | L328G | A | 1.1e-05 | 1.5e-05 | 1.7e-05 | 1.433E-05 | 3.055E-06 | 1.100E-05 | 1.700E-05 |
| 6549 | 328 | L328H | A | 1.4e-05 | 1.6e-05 | 1.4e-05 | 1.467E-05 | 1.155E-06 | 1.400E-05 | 1.600E-05 |
| 6550 | 328 | L328I | A | 1e-05 | 1.4e-05 | 9.4e-06 | 1.113E-05 | 2.501E-06 | 9.400E-06 | 1.400E-05 |
| 6551 | 328 | L328L | A | 1.3e-05 | 1.5e-05 | 8.6e-06 | 1.220E-05 | 3.274E-06 | 8.600E-06 | 1.500E-05 |
| 6552 | 328 | L328K | A | 1.6e-05 | 1.4e-05 | 1e-05 | 1.333E-05 | 3.055E-06 | 1.000E-05 | 1.600E-05 |
| 6553 | 328 | L328M | A | 1.5e-05 | 1.6e-05 | 1e-05 | 1.367E-05 | 3.215E-06 | 1.000E-05 | 1.600E-05 |
| 6554 | 328 | L328F | A | 1.1e-05 | 8.9e-06 | 1.5e-05 | 1.163E-05 | 3.099E-06 | 8.900E-06 | 1.500E-05 |
| 6555 | 328 | L328P | A | 1.4e-05 | 1.7e-05 | 1.5e-05 | 1.533E-05 | 1.528E-06 | 1.400E-05 | 1.700E-05 |
| 6556 | 328 | L328S | A | 9.9e-06 | 1.5e-05 | 1.4e-05 | 1.297E-05 | 2.702E-06 | 9.900E-06 | 1.500E-05 |
| 6557 | 328 | L328T | A | 1e-05 | 1e-05 | 1.3e-05 | 1.100E-05 | 1.732E-06 | 1.000E-05 | 1.300E-05 |
| 6558 | 328 | L328W | A | 1.5e-05 | 1.4e-05 | 1.6e-05 | 1.500E-05 | 1.000E-06 | 1.400E-05 | 1.600E-05 |
| 6559 | 328 | L328Y | A | 9.2e-06 | 6.2e-06 | 1.5e-05 | 1.013E-05 | 4.474E-06 | 6.200E-06 | 1.500E-05 |
| 6560 | 328 | L328V | A | 1.5e-05 | 1.6e-05 | 1.3e-05 | 1.467E-05 | 1.528E-06 | 1.300E-05 | 1.600E-05 |
| 6561 | 329 | R329A | A | 1.7e-05 | 8.4e-06 | 1e-05 | 1.180E-05 | 4.574E-06 | 8.400E-06 | 1.700E-05 |
| 6562 | 329 | R329R | A | 1.5e-05 | 9.5e-06 | 9.9e-06 | 1.147E-05 | 3.066E-06 | 9.500E-06 | 1.500E-05 |
| 6563 | 329 | R329N | A | 1.2e-05 | 1e-05 | 1e-05 | 1.067E-05 | 1.155E-06 | 1.000E-05 | 1.200E-05 |
| 6564 | 329 | R329D | A | 1.6e-05 | 1.1e-05 | 1e-05 | 1.233E-05 | 3.215E-06 | 1.000E-05 | 1.600E-05 |
| 6565 | 329 | R329C | A | 1e-05 | 1.3e-05 | 9.6e-06 | 1.087E-05 | 1.858E-06 | 9.600E-06 | 1.300E-05 |
| 6566 | 329 | R329E | A | 1.1e-05 | 1.2e-05 | 1.1e-05 | 1.133E-05 | 5.774E-07 | 1.100E-05 | 1.200E-05 |
| 6567 | 329 | R329Q | A | 1.5e-05 | 9.8e-06 | 9.1e-06 | 1.130E-05 | 3.223E-06 | 9.100E-06 | 1.500E-05 |
| 6568 | 329 | R329G | A | 8.7e-06 | 1.4e-05 | 1.1e-05 | 1.123E-05 | 2.658E-06 | 8.700E-06 | 1.400E-05 |
| 6569 | 329 | R329H | A | 1.2e-05 | 1e-05 | 1.7e-05 | 1.300E-05 | 3.606E-06 | 1.000E-05 | 1.700E-05 |
| 6570 | 329 | R329I | A | 1.6e-05 | 1e-05 | 1.6e-05 | 1.400E-05 | 3.464E-06 | 1.000E-05 | 1.600E-05 |
| 6571 | 329 | R329L | A | 1.4e-05 | 9.9e-06 | 1.1e-05 | 1.163E-05 | 2.122E-06 | 9.900E-06 | 1.400E-05 |
| 6572 | 329 | R329K | A | 1.4e-05 | 1.5e-05 | 1.6e-05 | 1.500E-05 | 1.000E-06 | 1.400E-05 | 1.600E-05 |
| 6573 | 329 | R329M | A | 1.1e-05 | 9.1e-06 | 1.5e-05 | 1.170E-05 | 3.012E-06 | 9.100E-06 | 1.500E-05 |
| 6574 | 329 | R329F | A | 1e-05 | 1.2e-05 | 1e-05 | 1.067E-05 | 1.155E-06 | 1.000E-05 | 1.200E-05 |
| 6575 | 329 | R329P | A | 9.1e-06 | 1e-05 | 1.4e-05 | 1.103E-05 | 2.608E-06 | 9.100E-06 | 1.400E-05 |
| 6576 | 329 | R329S | A | 1.6e-05 | 9.4e-06 | 1.2e-05 | 1.247E-05 | 3.325E-06 | 9.400E-06 | 1.600E-05 |
| 6577 | 329 | R329T | A | 1.5e-05 | 1.5e-05 | 1.5e-05 | 1.500E-05 | 0.000E+00 | 1.500E-05 | 1.500E-05 |
| 6578 | 329 | R329W | A | 9.7e-06 | 1.1e-05 | 1.4e-05 | 1.157E-05 | 2.205E-06 | 9.700E-06 | 1.400E-05 |
| 6579 | 329 | R329Y | A | 1.7e-05 | 1.2e-05 | 1.5e-05 | 1.467E-05 | 2.517E-06 | 1.200E-05 | 1.700E-05 |
| 6580 | 329 | R329V | A | 7.6e-06 | 1.4e-05 | 1.2e-05 | 1.120E-05 | 3.274E-06 | 7.600E-06 | 1.400E-05 |
| 6581 | 330 | E330A | A | 1.6e-05 | 1.8e-05 | 1.6e-05 | 1.667E-05 | 1.155E-06 | 1.600E-05 | 1.800E-05 |
| 6582 | 330 | E330R | A | 1.5e-05 | 1.6e-05 | 8.3e-06 | 1.310E-05 | 4.187E-06 | 8.300E-06 | 1.600E-05 |
| 6583 | 330 | E330N | A | 7.3e-06 | 1.1e-05 | 1.6e-05 | 1.143E-05 | 4.366E-06 | 7.300E-06 | 1.600E-05 |
| 6584 | 330 | E330D | A | 1.2e-05 | 8.5e-06 | 1.5e-05 | 1.183E-05 | 3.253E-06 | 8.500E-06 | 1.500E-05 |
| 6585 | 330 | E330C | A | 1.3e-05 | 1.1e-05 | 1.5e-05 | 1.300E-05 | 2.000E-06 | 1.100E-05 | 1.500E-05 |
| 6586 | 330 | E330E | A | 1.6e-05 | 1.1e-05 | 1e-05 | 1.233E-05 | 3.215E-06 | 1.000E-05 | 1.600E-05 |
| 6587 | 330 | E330Q | A | 1.7e-05 | 1e-05 | 8.9e-06 | 1.197E-05 | 4.394E-06 | 8.900E-06 | 1.700E-05 |
| 6588 | 330 | E330G | A | 1.5e-05 | 1.4e-05 | 1.5e-05 | 1.467E-05 | 5.774E-07 | 1.400E-05 | 1.500E-05 |
| 6589 | 330 | E330H | A | 1.5e-05 | 8.9e-06 | 9.4e-06 | 1.110E-05 | 3.387E-06 | 8.900E-06 | 1.500E-05 |
| 6590 | 330 | E330I | A | 1.8e-05 | 1e-05 | 9.6e-06 | 1.253E-05 | 4.738E-06 | 9.600E-06 | 1.800E-05 |
| 6591 | 330 | E330L | A | 1.6e-05 | 9.9e-06 | 1.3e-05 | 1.297E-05 | 3.050E-06 | 9.900E-06 | 1.600E-05 |
| 6592 | 330 | E330K | A | 9.4e-06 | 1.6e-05 | 9.4e-06 | 1.160E-05 | 3.811E-06 | 9.400E-06 | 1.600E-05 |
| 6593 | 330 | E330M | A | 1.6e-05 | 1.6e-05 | 1.5e-05 | 1.567E-05 | 5.774E-07 | 1.500E-05 | 1.600E-05 |
| 6594 | 330 | E330F | A | 1.3e-05 | 1.3e-05 | 1.4e-05 | 1.333E-05 | 5.774E-07 | 1.300E-05 | 1.400E-05 |
| 6595 | 330 | E330P | A | 1.5e-05 | 1.6e-05 | 1.1e-05 | 1.400E-05 | 2.646E-06 | 1.100E-05 | 1.600E-05 |
| 6596 | 330 | E330S | A | 1.5e-05 | 1.5e-05 | 1.1e-05 | 1.367E-05 | 2.309E-06 | 1.100E-05 | 1.500E-05 |
| 6597 | 330 | E330T | A | 1.4e-05 | 1.3e-05 | 1.7e-05 | 1.467E-05 | 2.082E-06 | 1.300E-05 | 1.700E-05 |
| 6598 | 330 | E330W | A | 8.2e-06 | 1.1e-05 | 1.5e-05 | 1.140E-05 | 3.418E-06 | 8.200E-06 | 1.500E-05 |
| 6599 | 330 | E330Y | A | 9e-06 | 1.5e-05 | 1.3e-05 | 1.233E-05 | 3.055E-06 | 9.000E-06 | 1.500E-05 |
| 6600 | 330 | E330V | A | 1.4e-05 | 1.5e-05 | 9.8e-06 | 1.293E-05 | 2.759E-06 | 9.800E-06 | 1.500E-05 |
| 6601 | 331 | V331A | A | 1.3e-05 | 1.5e-05 | 1.6e-05 | 1.467E-05 | 1.528E-06 | 1.300E-05 | 1.600E-05 |
| 6602 | 331 | V331R | A | 9.4e-06 | 1.6e-05 | 9.8e-06 | 1.173E-05 | 3.700E-06 | 9.400E-06 | 1.600E-05 |
| 6603 | 331 | V331N | A | 8.6e-06 | 1.3e-05 | 1.5e-05 | 1.220E-05 | 3.274E-06 | 8.600E-06 | 1.500E-05 |
| 6604 | 331 | V331D | A | 1.1e-05 | 1.1e-05 | 1.4e-05 | 1.200E-05 | 1.732E-06 | 1.100E-05 | 1.400E-05 |
| 6605 | 331 | V331C | A | 9.5e-06 | 1.4e-05 | 1.6e-05 | 1.317E-05 | 3.329E-06 | 9.500E-06 | 1.600E-05 |
| 6606 | 331 | V331E | A | 1.2e-05 | 1.4e-05 | 1.7e-05 | 1.433E-05 | 2.517E-06 | 1.200E-05 | 1.700E-05 |
| 6607 | 331 | V331Q | A | 9e-06 | 9.6e-06 | 1.5e-05 | 1.120E-05 | 3.305E-06 | 9.000E-06 | 1.500E-05 |
| 6608 | 331 | V331G | A | 1.6e-05 | 8.5e-06 | 1.5e-05 | 1.317E-05 | 4.072E-06 | 8.500E-06 | 1.600E-05 |
| 6609 | 331 | V331H | A | 1.5e-05 | 9.9e-06 | 1e-05 | 1.163E-05 | 2.916E-06 | 9.900E-06 | 1.500E-05 |
| 6610 | 331 | V331I | A | 1.5e-05 | 1.1e-05 | 1.1e-05 | 1.233E-05 | 2.309E-06 | 1.100E-05 | 1.500E-05 |
| 6611 | 331 | V331L | A | 1.4e-05 | 1.6e-05 | 1.6e-05 | 1.533E-05 | 1.155E-06 | 1.400E-05 | 1.600E-05 |
| 6612 | 331 | V331K | A | 1.6e-05 | 1e-05 | 8.8e-06 | 1.160E-05 | 3.857E-06 | 8.800E-06 | 1.600E-05 |
| 6613 | 331 | V331M | A | 1.8e-05 | 1.2e-05 | 1.2e-05 | 1.400E-05 | 3.464E-06 | 1.200E-05 | 1.800E-05 |
| 6614 | 331 | V331F | A | 1.3e-05 | 1.6e-05 | 1.5e-05 | 1.467E-05 | 1.528E-06 | 1.300E-05 | 1.600E-05 |
| 6615 | 331 | V331P | A | 1.4e-05 | 1.4e-05 | 1.5e-05 | 1.433E-05 | 5.774E-07 | 1.400E-05 | 1.500E-05 |
| 6616 | 331 | V331S | A | 9.6e-06 | 1.1e-05 | 9.2e-06 | 9.933E-06 | 9.452E-07 | 9.200E-06 | 1.100E-05 |
| 6617 | 331 | V331T | A | 1.1e-05 | 8.6e-06 | 7.2e-06 | 8.933E-06 | 1.922E-06 | 7.200E-06 | 1.100E-05 |
| 6618 | 331 | V331W | A | 1.2e-05 | 1.5e-05 | 1e-05 | 1.233E-05 | 2.517E-06 | 1.000E-05 | 1.500E-05 |
| 6619 | 331 | V331Y | A | 8.7e-06 | 1.5e-05 | 9.4e-06 | 1.103E-05 | 3.453E-06 | 8.700E-06 | 1.500E-05 |
| 6620 | 331 | V331V | A | 1e-05 | 1.3e-05 | 1.5e-05 | 1.267E-05 | 2.517E-06 | 1.000E-05 | 1.500E-05 |
| 6621 | 332 | R332A | A | 1.2e-05 | 1.2e-05 | 1.1e-05 | 1.167E-05 | 5.774E-07 | 1.100E-05 | 1.200E-05 |
| 6622 | 332 | R332R | A | 1.4e-05 | 1.5e-05 | 1.7e-05 | 1.533E-05 | 1.528E-06 | 1.400E-05 | 1.700E-05 |
| 6623 | 332 | R332N | A | 1.6e-05 | 8.9e-06 | 1.5e-05 | 1.330E-05 | 3.843E-06 | 8.900E-06 | 1.600E-05 |
| 6624 | 332 | R332D | A | 1.1e-05 | 1.5e-05 | 1.5e-05 | 1.367E-05 | 2.309E-06 | 1.100E-05 | 1.500E-05 |
| 6625 | 332 | R332C | A | 9.1e-06 | 6.3e-06 | 8.4e-06 | 7.933E-06 | 1.457E-06 | 6.300E-06 | 9.100E-06 |
| 6626 | 332 | R332E | A | 1.2e-05 | 1.2e-05 | 1.1e-05 | 1.167E-05 | 5.774E-07 | 1.100E-05 | 1.200E-05 |
| 6627 | 332 | R332Q | A | 1.5e-05 | 1.3e-05 | 1.4e-05 | 1.400E-05 | 1.000E-06 | 1.300E-05 | 1.500E-05 |
| 6628 | 332 | R332G | A | 1.1e-05 | 1.5e-05 | 1.5e-05 | 1.367E-05 | 2.309E-06 | 1.100E-05 | 1.500E-05 |
| 6629 | 332 | R332H | A | 1.4e-05 | 9.8e-06 | 1.5e-05 | 1.293E-05 | 2.759E-06 | 9.800E-06 | 1.500E-05 |

| | | | | | | | | | | |
|---|---|---|---|---|---|---|---|---|---|---|
| 6630 | 332 | R332I | A | 1.1e-05 | 1.6e-05 | 1.4e-05 | 1.367E-05 | 2.517E-06 | 1.100E-05 | 1.600E-05 |
| 6631 | 332 | R332L | A | 9.6e-06 | 1.4e-05 | 9.5e-06 | 1.103E-05 | 2.570E-06 | 9.500E-06 | 1.400E-05 |
| 6632 | 332 | R332K | A | 1.5e-05 | 6.1e-06 | 1.6e-05 | 1.237E-05 | 5.450E-06 | 6.100E-06 | 1.600E-05 |
| 6633 | 332 | R332M | A | 9.1e-06 | 1.2e-05 | 1.7e-05 | 1.270E-05 | 3.996E-06 | 9.100E-06 | 1.700E-05 |
| 6634 | 332 | R332F | A | 1e-05 | 1.5e-05 | 1.7e-05 | 1.400E-05 | 3.606E-06 | 1.000E-05 | 1.700E-05 |
| 6635 | 332 | R332P | A | 1.4e-05 | 1.5e-05 | 9.8e-06 | 1.293E-05 | 2.759E-06 | 9.800E-06 | 1.500E-05 |
| 6636 | 332 | R332S | A | 5.8e-06 | 1e-05 | 1.6e-05 | 1.060E-05 | 5.126E-06 | 5.800E-06 | 1.600E-05 |
| 6637 | 332 | R332T | A | 1.6e-05 | 9.6e-06 | 1e-05 | 1.187E-05 | 3.585E-06 | 9.600E-06 | 1.600E-05 |
| 6638 | 332 | R332W | A | 1.6e-05 | 1.6e-05 | 1.6e-05 | 1.600E-05 | 0.000E+00 | 1.600E-05 | 1.600E-05 |
| 6639 | 332 | R332Y | A | 1.3e-05 | 9.5e-06 | 1.5e-05 | 1.250E-05 | 2.784E-06 | 9.500E-06 | 1.500E-05 |
| 6640 | 332 | R332V | A | 1.7e-05 | 1.1e-05 | 1.5e-05 | 1.433E-05 | 3.055E-06 | 1.100E-05 | 1.700E-05 |
| 6641 | 333 | A333A | A | 1.6e-05 | 9.9e-06 | 1.5e-05 | 1.363E-05 | 3.272E-06 | 9.900E-06 | 1.600E-05 |
| 6642 | 333 | A333R | A | 1e-05 | 9.9e-06 | 1.5e-05 | 1.163E-05 | 2.916E-06 | 9.900E-06 | 1.500E-05 |
| 6643 | 333 | A333N | A | 1.5e-05 | 1.7e-05 | 1.2e-05 | 1.467E-05 | 2.517E-06 | 1.200E-05 | 1.700E-05 |
| 6644 | 333 | A333D | A | 1.5e-05 | 1.4e-05 | 1.6e-05 | 1.500E-05 | 1.000E-06 | 1.400E-05 | 1.600E-05 |
| 6645 | 333 | A333C | A | 1.2e-05 | 1.4e-05 | 1.1e-05 | 1.233E-05 | 1.528E-06 | 1.100E-05 | 1.400E-05 |
| 6646 | 333 | A333E | A | 1.2e-05 | 1.7e-05 | 1.2e-05 | 1.367E-05 | 2.887E-06 | 1.200E-05 | 1.700E-05 |
| 6647 | 333 | A333Q | A | 1.2e-05 | 1.6e-05 | 8.4e-06 | 1.213E-05 | 3.802E-06 | 8.400E-06 | 1.600E-05 |
| 6648 | 333 | A333G | A | 1.4e-05 | 1.1e-05 | 8.9e-06 | 1.130E-05 | 2.563E-06 | 8.900E-06 | 1.400E-05 |
| 6649 | 333 | A333H | A | 1.4e-05 | 1.1e-05 | 1.2e-05 | 1.233E-05 | 1.528E-06 | 1.100E-05 | 1.400E-05 |
| 6650 | 333 | A333I | A | 9.7e-06 | 1e-05 | 1e-05 | 9.900E-06 | 1.732E-07 | 9.700E-06 | 1.000E-05 |
| 6651 | 333 | A333L | A | 1.6e-05 | 1.4e-05 | 9.3e-06 | 1.310E-05 | 3.439E-06 | 9.300E-06 | 1.600E-05 |
| 6652 | 333 | A333K | A | 9.6e-06 | 1.2e-05 | 9.2e-06 | 1.027E-05 | 1.514E-06 | 9.200E-06 | 1.200E-05 |
| 6653 | 333 | A333M | A | 9.9e-06 | 1.6e-05 | 1.3e-05 | 1.297E-05 | 3.050E-06 | 9.900E-06 | 1.600E-05 |
| 6654 | 333 | A333F | A | 1.7e-05 | 1.7e-05 | 1.1e-05 | 1.500E-05 | 3.464E-06 | 1.100E-05 | 1.700E-05 |
| 6655 | 333 | A333P | A | 1.4e-05 | 6.8e-06 | 1.6e-05 | 1.227E-05 | 4.839E-06 | 6.800E-06 | 1.600E-05 |
| 6656 | 333 | A333S | A | 9e-06 | 1.4e-05 | 1.5e-05 | 1.267E-05 | 3.215E-06 | 9.000E-06 | 1.500E-05 |
| 6657 | 333 | A333T | A | 1.1e-05 | 1.3e-05 | 9.7e-06 | 1.123E-05 | 1.662E-06 | 9.700E-06 | 1.300E-05 |
| 6658 | 333 | A333W | A | 1.3e-05 | 1.5e-05 | 9.6e-06 | 1.253E-05 | 2.730E-06 | 9.600E-06 | 1.500E-05 |
| 6659 | 333 | A333Y | A | 1.1e-05 | 1.7e-05 | 1.1e-05 | 1.300E-05 | 3.464E-06 | 1.100E-05 | 1.700E-05 |
| 6660 | 333 | A333V | A | 1.6e-05 | 1.6e-05 | 8.7e-06 | 1.357E-05 | 4.215E-06 | 8.700E-06 | 1.600E-05 |
| 6661 | 334 | V334A | A | 1.5e-05 | 1.1e-05 | 1.8e-05 | 1.467E-05 | 3.512E-06 | 1.100E-05 | 1.800E-05 |
| 6662 | 334 | V334R | A | 1.4e-05 | 9.9e-06 | 9.8e-06 | 1.123E-05 | 2.397E-06 | 9.800E-06 | 1.400E-05 |
| 6663 | 334 | V334N | A | 8.1e-06 | 9.9e-06 | 1.2e-05 | 1.000E-05 | 1.952E-06 | 8.100E-06 | 1.200E-05 |
| 6664 | 334 | V334D | A | 1.5e-05 | 1.5e-05 | 1.3e-05 | 1.433E-05 | 1.155E-06 | 1.300E-05 | 1.500E-05 |
| 6665 | 334 | V334C | A | 1.4e-05 | 1.4e-05 | 1.5e-05 | 1.433E-05 | 5.774E-07 | 1.400E-05 | 1.500E-05 |
| 6666 | 334 | V334E | A | 1.5e-05 | 1.2e-05 | 9.2e-06 | 1.207E-05 | 2.901E-06 | 9.200E-06 | 1.500E-05 |
| 6667 | 334 | V334Q | A | 1.5e-05 | 1.2e-05 | 1.1e-05 | 1.267E-05 | 2.082E-06 | 1.100E-05 | 1.500E-05 |
| 6668 | 334 | V334G | A | 1.6e-05 | 1.3e-05 | 1.4e-05 | 1.433E-05 | 1.528E-06 | 1.300E-05 | 1.600E-05 |
| 6669 | 334 | V334H | A | 9.9e-06 | 9.9e-06 | 1.6e-05 | 1.193E-05 | 3.522E-06 | 9.900E-06 | 1.600E-05 |
| 6670 | 334 | V334I | A | 1.6e-05 | 7.6e-06 | 1.3e-05 | 1.220E-05 | 4.257E-06 | 7.600E-06 | 1.600E-05 |
| 6671 | 334 | V334L | A | 1.5e-05 | 1.4e-05 | 1.2e-05 | 1.367E-05 | 1.528E-06 | 1.200E-05 | 1.500E-05 |
| 6672 | 334 | V334K | A | 1.1e-05 | 1.1e-05 | 1.4e-05 | 1.200E-05 | 1.732E-06 | 1.100E-05 | 1.400E-05 |
| 6673 | 334 | V334M | A | 1.6e-05 | 1.1e-05 | 1.2e-05 | 1.300E-05 | 2.646E-06 | 1.100E-05 | 1.600E-05 |
| 6674 | 334 | V334F | A | 8.7e-06 | 1.5e-05 | 1.4e-05 | 1.257E-05 | 3.386E-06 | 8.700E-06 | 1.500E-05 |
| 6675 | 334 | V334P | A | 1.6e-05 | 1.1e-05 | 1.4e-05 | 1.367E-05 | 2.517E-06 | 1.100E-05 | 1.600E-05 |
| 6676 | 334 | V334S | A | 1e-05 | 6.8e-06 | 1.5e-05 | 1.060E-05 | 4.133E-06 | 6.800E-06 | 1.500E-05 |
| 6677 | 334 | V334T | A | 1.1e-05 | 8.7e-06 | 1.7e-05 | 1.223E-05 | 4.285E-06 | 8.700E-06 | 1.700E-05 |
| 6678 | 334 | V334W | A | 1.1e-05 | 1.5e-05 | 8e-06 | 1.133E-05 | 3.512E-06 | 8.000E-06 | 1.500E-05 |
| 6679 | 334 | V334Y | A | 1e-05 | 9.3e-06 | 9.3e-06 | 9.533E-06 | 4.041E-07 | 9.300E-06 | 1.000E-05 |
| 6680 | 334 | V334V | A | 1.5e-05 | 1.2e-05 | 1.5e-05 | 1.400E-05 | 1.732E-06 | 1.200E-05 | 1.500E-05 |
| 6681 | 335 | T335A | A | 1.5e-05 | 1.2e-05 | 1.1e-05 | 1.267E-05 | 2.082E-06 | 1.100E-05 | 1.500E-05 |
| 6682 | 335 | T335R | A | 1e-05 | 1.6e-05 | 1.3e-05 | 1.300E-05 | 3.000E-06 | 1.000E-05 | 1.600E-05 |
| 6683 | 335 | T335N | A | 1e-05 | 1.5e-05 | 1.7e-05 | 1.400E-05 | 3.606E-06 | 1.000E-05 | 1.700E-05 |
| 6684 | 335 | T335D | A | 1.4e-05 | 1.1e-05 | 1.5e-05 | 1.333E-05 | 2.082E-06 | 1.100E-05 | 1.500E-05 |
| 6685 | 335 | T335C | A | 1e-05 | 1.2e-05 | 9.8e-06 | 1.060E-05 | 1.217E-06 | 9.800E-06 | 1.200E-05 |
| 6686 | 335 | T335E | A | 9.7e-06 | 1e-05 | 1e-05 | 9.900E-06 | 1.732E-07 | 9.700E-06 | 1.000E-05 |
| 6687 | 335 | T335Q | A | 1.4e-05 | 1.5e-05 | 9.6e-06 | 1.287E-05 | 2.873E-06 | 9.600E-06 | 1.500E-05 |
| 6688 | 335 | T335G | A | 1.6e-05 | 1e-05 | 1.8e-05 | 1.467E-05 | 4.163E-06 | 1.000E-05 | 1.800E-05 |
| 6689 | 335 | T335H | A | 1.5e-05 | 1e-05 | 1.4e-05 | 1.300E-05 | 2.646E-06 | 1.000E-05 | 1.500E-05 |
| 6690 | 335 | T335I | A | 1.1e-05 | 1.1e-05 | 1.6e-05 | 1.267E-05 | 2.887E-06 | 1.100E-05 | 1.600E-05 |
| 6691 | 335 | T335L | A | 1.3e-05 | 1e-05 | 1.5e-05 | 1.267E-05 | 2.517E-06 | 1.000E-05 | 1.500E-05 |
| 6692 | 335 | T335K | A | 1.2e-05 | 1.1e-05 | 1.3e-05 | 1.200E-05 | 1.000E-06 | 1.100E-05 | 1.300E-05 |
| 6693 | 335 | T335M | A | 9.2e-06 | 1.5e-05 | 1.1e-05 | 1.173E-05 | 2.969E-06 | 9.200E-06 | 1.500E-05 |
| 6694 | 335 | T335F | A | 1.1e-05 | 1.5e-05 | 1.5e-05 | 1.367E-05 | 2.309E-06 | 1.100E-05 | 1.500E-05 |
| 6695 | 335 | T335P | A | 1.5e-05 | 1.4e-05 | 1.3e-05 | 1.400E-05 | 1.000E-06 | 1.300E-05 | 1.500E-05 |
| 6696 | 335 | T335S | A | 1.2e-05 | 1.3e-05 | 1.5e-05 | 1.333E-05 | 1.528E-06 | 1.200E-05 | 1.500E-05 |
| 6697 | 335 | T335T | A | 1.1e-05 | 1.5e-05 | 1.5e-05 | 1.367E-05 | 2.309E-06 | 1.100E-05 | 1.500E-05 |
| 6698 | 335 | T335W | A | 9.7e-06 | 1.5e-05 | 1.6e-05 | 1.357E-05 | 3.386E-06 | 9.700E-06 | 1.600E-05 |
| 6699 | 335 | T335Y | A | 1.5e-05 | 1e-05 | 1.2e-05 | 1.233E-05 | 2.517E-06 | 1.000E-05 | 1.500E-05 |
| 6700 | 335 | T335V | A | 1.4e-05 | 8e-06 | 1.2e-05 | 1.133E-05 | 3.055E-06 | 8.000E-06 | 1.400E-05 |
| 6701 | 336 | S336A | A | 1.4e-05 | 1.1e-05 | 1.1e-05 | 1.200E-05 | 1.732E-06 | 1.100E-05 | 1.400E-05 |
| 6702 | 336 | S336R | A | 1.4e-05 | 1.3e-05 | 9.9e-06 | 1.230E-05 | 2.138E-06 | 9.900E-06 | 1.400E-05 |
| 6703 | 336 | S336N | A | 1.6e-05 | 8.8e-06 | 1.4e-05 | 1.293E-05 | 3.717E-06 | 8.800E-06 | 1.600E-05 |
| 6704 | 336 | S336D | A | 1.7e-05 | 1.5e-05 | 3.6e-05 | 2.267E-05 | 1.159E-05 | 1.500E-05 | 3.600E-05 |
| 6705 | 336 | S336C | A | 8.9e-06 | 1.6e-05 | 1.6e-05 | 1.363E-05 | 4.099E-06 | 8.900E-06 | 1.600E-05 |
| 6706 | 336 | S336E | A | 1.5e-05 | 1e-05 | 1.3e-05 | 1.267E-05 | 2.517E-06 | 1.000E-05 | 1.500E-05 |
| 6707 | 336 | S336Q | A | 1.1e-05 | 1.5e-05 | 1.5e-05 | 1.367E-05 | 2.309E-06 | 1.100E-05 | 1.500E-05 |
| 6708 | 336 | S336G | A | 9.2e-06 | 1.7e-05 | 1.5e-05 | 1.373E-05 | 4.051E-06 | 9.200E-06 | 1.700E-05 |
| 6709 | 336 | S336H | A | 1.5e-05 | 1.5e-05 | 9e-06 | 1.300E-05 | 3.464E-06 | 9.000E-06 | 1.500E-05 |
| 6710 | 336 | S336I | A | 1.6e-05 | 1.7e-05 | 1.1e-05 | 1.467E-05 | 3.215E-06 | 1.100E-05 | 1.700E-05 |
| 6711 | 336 | S336L | A | 1.4e-05 | 1.5e-05 | 1.4e-05 | 1.433E-05 | 5.774E-07 | 1.400E-05 | 1.500E-05 |
| 6712 | 336 | S336K | A | 1.1e-05 | 1.4e-05 | 1.3e-05 | 1.267E-05 | 1.528E-06 | 1.100E-05 | 1.400E-05 |
| 6713 | 336 | S336M | A | 1.1e-05 | 1.5e-05 | 1.1e-05 | 1.233E-05 | 2.309E-06 | 1.100E-05 | 1.500E-05 |
| 6714 | 336 | S336F | A | 1.1e-05 | 1.5e-05 | 1.2e-05 | 1.267E-05 | 2.082E-06 | 1.100E-05 | 1.500E-05 |

| | | | | | | | | | | |
|---|---|---|---|---|---|---|---|---|---|---|
| 6715 | 336 | S336P | A | 9.4e-06 | 1.6e-05 | 1.5e-05 | 1.347E-05 | 3.557E-06 | 9.400E-06 | 1.600E-05 |
| 6716 | 336 | S336S | A | 1.6e-05 | 1.1e-05 | 1.1e-05 | 1.267E-05 | 2.887E-06 | 1.100E-05 | 1.600E-05 |
| 6717 | 336 | S336T | A | 1.4e-05 | 7.9e-06 | 1.5e-05 | 1.230E-05 | 3.843E-06 | 7.900E-06 | 1.500E-05 |
| 6718 | 336 | S336W | A | 1.6e-05 | 1.6e-05 | 1.3e-05 | 1.500E-05 | 1.732E-06 | 1.300E-05 | 1.600E-05 |
| 6719 | 336 | S336Y | A | 9.3e-06 | 1e-05 | 9.3e-06 | 9.533E-06 | 4.041E-07 | 9.300E-06 | 1.000E-05 |
| 6720 | 336 | S336V | A | 1.1e-05 | 1.8e-05 | 1.2e-05 | 1.367E-05 | 3.786E-06 | 1.100E-05 | 1.800E-05 |
| 6721 | 337 | A337A | A | 8.8e-06 | 1.1e-05 | 1e-05 | 9.933E-06 | 1.102E-06 | 8.800E-06 | 1.100E-05 |
| 6722 | 337 | A337R | A | 1.3e-05 | 1.1e-05 | 1.3e-05 | 1.233E-05 | 1.155E-06 | 1.100E-05 | 1.300E-05 |
| 6723 | 337 | A337N | A | 1.6e-05 | 8.3e-06 | 2.3e-05 | 1.577E-05 | 7.353E-06 | 8.300E-06 | 2.300E-05 |
| 6724 | 337 | A337D | A | 1.1e-05 | 1.1e-05 | 1.5e-05 | 1.233E-05 | 2.309E-06 | 1.100E-05 | 1.500E-05 |
| 6725 | 337 | A337C | A | 9e-06 | 1.1e-05 | 9.2e-06 | 9.733E-06 | 1.102E-06 | 9.000E-06 | 1.100E-05 |
| 6726 | 337 | A337E | A | 1.5e-05 | 1.4e-05 | 1.5e-05 | 1.467E-05 | 5.774E-07 | 1.400E-05 | 1.500E-05 |
| 6727 | 337 | A337Q | A | 1.5e-05 | 1.5e-05 | 1.1e-05 | 1.367E-05 | 2.309E-06 | 1.100E-05 | 1.500E-05 |
| 6728 | 337 | A337G | A | 1.1e-05 | 1.5e-05 | 1.6e-05 | 1.400E-05 | 2.646E-06 | 1.100E-05 | 1.600E-05 |
| 6729 | 337 | A337H | A | 1.1e-05 | 1e-05 | 1.2e-05 | 1.100E-05 | 1.000E-06 | 1.000E-05 | 1.200E-05 |
| 6730 | 337 | A337I | A | 8.6e-06 | 1.5e-05 | 7.8e-06 | 1.047E-05 | 3.946E-06 | 7.800E-06 | 1.500E-05 |
| 6731 | 337 | A337L | A | 1.1e-05 | 1.5e-05 | 9.3e-06 | 1.177E-05 | 2.926E-06 | 9.300E-06 | 1.500E-05 |
| 6732 | 337 | A337K | A | 1.3e-05 | 1.1e-05 | 1.2e-05 | 1.200E-05 | 1.000E-06 | 1.100E-05 | 1.300E-05 |
| 6733 | 337 | A337M | A | 9.5e-06 | 1.1e-05 | 1.6e-05 | 1.217E-05 | 3.403E-06 | 9.500E-06 | 1.600E-05 |
| 6734 | 337 | A337F | A | 1.7e-05 | 1.2e-05 | 1.2e-05 | 1.367E-05 | 2.887E-06 | 1.200E-05 | 1.700E-05 |
| 6735 | 337 | A337P | A | 1.7e-05 | 1.3e-05 | 1.1e-05 | 1.367E-05 | 3.055E-06 | 1.100E-05 | 1.700E-05 |
| 6736 | 337 | A337S | A | 7.7e-06 | 1.4e-05 | 9.9e-06 | 1.053E-05 | 3.197E-06 | 7.700E-06 | 1.400E-05 |
| 6737 | 337 | A337T | A | 1.6e-05 | 1e-05 | 7.7e-06 | 1.123E-05 | 4.285E-06 | 7.700E-06 | 1.600E-05 |
| 6738 | 337 | A337W | A | 1.5e-05 | 8.7e-06 | 1.5e-05 | 1.290E-05 | 3.637E-06 | 8.700E-06 | 1.500E-05 |
| 6739 | 337 | A337Y | A | 1.5e-05 | 1.4e-05 | 1.6e-05 | 1.500E-05 | 1.000E-06 | 1.400E-05 | 1.600E-05 |
| 6740 | 337 | A337V | A | 1.5e-05 | 1.1e-05 | 1.1e-05 | 1.233E-05 | 2.309E-06 | 1.100E-05 | 1.500E-05 |
| 6741 | 338 | N338A | A | 1.3e-05 | 8.5e-06 | 1.6e-05 | 1.250E-05 | 3.775E-06 | 8.500E-06 | 1.600E-05 |
| 6742 | 338 | N338R | A | 1.4e-05 | 1.4e-05 | 1e-05 | 1.267E-05 | 2.309E-06 | 1.000E-05 | 1.400E-05 |
| 6743 | 338 | N338N | A | 1.5e-05 | 1.5e-05 | 1.1e-05 | 1.367E-05 | 2.309E-06 | 1.100E-05 | 1.500E-05 |
| 6744 | 338 | N338D | A | 1.5e-05 | 1.5e-05 | 9.8e-06 | 1.327E-05 | 3.002E-06 | 9.800E-06 | 1.500E-05 |
| 6745 | 338 | N338C | A | 1.5e-05 | 1.6e-05 | 1.2e-05 | 1.433E-05 | 2.082E-06 | 1.200E-05 | 1.600E-05 |
| 6746 | 338 | N338E | A | 1.1e-05 | 1.5e-05 | 9.3e-06 | 1.177E-05 | 2.926E-06 | 9.300E-06 | 1.500E-05 |
| 6747 | 338 | N338Q | A | 1.1e-05 | 1.1e-05 | 1.5e-05 | 1.233E-05 | 2.309E-06 | 1.100E-05 | 1.500E-05 |
| 6748 | 338 | N338G | A | 9.3e-06 | 7.1e-06 | 1.7e-05 | 1.113E-05 | 5.198E-06 | 7.100E-06 | 1.700E-05 |
| 6749 | 338 | N338H | A | 1.5e-05 | 1.4e-05 | 9.4e-06 | 1.280E-05 | 2.987E-06 | 9.400E-06 | 1.500E-05 |
| 6750 | 338 | N338I | A | 1.5e-05 | 1.5e-05 | 1.1e-05 | 1.367E-05 | 2.309E-06 | 1.100E-05 | 1.500E-05 |
| 6751 | 338 | N338L | A | 1.7e-05 | 1.3e-05 | 1.5e-05 | 1.500E-05 | 2.000E-06 | 1.300E-05 | 1.700E-05 |
| 6752 | 338 | N338K | A | 1.5e-05 | 1.5e-05 | 1.5e-05 | 1.500E-05 | 0.000E+00 | 1.500E-05 | 1.500E-05 |
| 6753 | 338 | N338M | A | 1.6e-05 | 1e-05 | 1.1e-05 | 1.233E-05 | 3.215E-06 | 1.000E-05 | 1.600E-05 |
| 6754 | 338 | N338F | A | 1.7e-05 | 1.5e-05 | 1.1e-05 | 1.433E-05 | 3.055E-06 | 1.100E-05 | 1.700E-05 |
| 6755 | 338 | N338P | A | 1.5e-05 | 1.5e-05 | 1e-05 | 1.333E-05 | 2.887E-06 | 1.000E-05 | 1.500E-05 |
| 6756 | 338 | N338S | A | 1.1e-05 | 1.5e-05 | 1.6e-05 | 1.400E-05 | 2.646E-06 | 1.100E-05 | 1.600E-05 |
| 6757 | 338 | N338T | A | 1.6e-05 | 1.6e-05 | 1.6e-05 | 1.600E-05 | 0.000E+00 | 1.600E-05 | 1.600E-05 |
| 6758 | 338 | N338W | A | 1.7e-05 | 1.6e-05 | 1.3e-05 | 1.533E-05 | 2.082E-06 | 1.300E-05 | 1.700E-05 |
| 6759 | 338 | N338Y | A | 1.6e-05 | 1.7e-05 | 1.5e-05 | 1.600E-05 | 1.000E-06 | 1.500E-05 | 1.700E-05 |
| 6760 | 338 | N338V | A | 1.5e-05 | 8.9e-06 | 1e-05 | 1.130E-05 | 3.251E-06 | 8.900E-06 | 1.500E-05 |
| 6761 | 339 | I339A | A | 9.8e-06 | 1.5e-05 | 1.1e-05 | 1.193E-05 | 2.723E-06 | 9.800E-06 | 1.500E-05 |
| 6762 | 339 | I339R | A | 1.6e-05 | 8.1e-06 | 1.6e-05 | 1.337E-05 | 4.561E-06 | 8.100E-06 | 1.600E-05 |
| 6763 | 339 | I339N | A | 1.4e-05 | 1.1e-05 | 7.9e-06 | 1.097E-05 | 3.050E-06 | 7.900E-06 | 1.400E-05 |
| 6764 | 339 | I339D | A | 1.4e-05 | 1.3e-05 | 1.4e-05 | 1.367E-05 | 5.774E-07 | 1.300E-05 | 1.400E-05 |
| 6765 | 339 | I339C | A | 1.1e-05 | 1.4e-05 | 1.1e-05 | 1.200E-05 | 1.732E-06 | 1.100E-05 | 1.400E-05 |
| 6766 | 339 | I339E | A | 1.1e-05 | 9.9e-06 | 1e-05 | 1.030E-05 | 6.083E-07 | 9.900E-06 | 1.100E-05 |
| 6767 | 339 | I339Q | A | 8.2e-06 | 1.6e-05 | 1.4e-05 | 1.273E-05 | 4.051E-06 | 8.200E-06 | 1.600E-05 |
| 6768 | 339 | I339G | A | 1.4e-05 | 9.9e-06 | 1.5e-05 | 1.297E-05 | 2.702E-06 | 9.900E-06 | 1.500E-05 |
| 6769 | 339 | I339H | A | 1.6e-05 | 1.5e-05 | 9.8e-06 | 1.360E-05 | 3.329E-06 | 9.800E-06 | 1.600E-05 |
| 6770 | 339 | I339I | A | 1e-05 | 1.1e-05 | 1.1e-05 | 1.067E-05 | 5.774E-07 | 1.000E-05 | 1.100E-05 |
| 6771 | 339 | I339L | A | 9.5e-06 | 9.7e-06 | 1.4e-05 | 1.107E-05 | 2.542E-06 | 9.500E-06 | 1.400E-05 |
| 6772 | 339 | I339K | A | 1.2e-05 | 1.1e-05 | 1.3e-05 | 1.200E-05 | 1.000E-06 | 1.100E-05 | 1.300E-05 |
| 6773 | 339 | I339M | A | 1.4e-05 | 1.5e-05 | 1.2e-05 | 1.367E-05 | 1.528E-06 | 1.200E-05 | 1.500E-05 |
| 6774 | 339 | I339F | A | 1.3e-05 | 8.1e-06 | 1.1e-05 | 1.070E-05 | 2.464E-06 | 8.100E-06 | 1.300E-05 |
| 6775 | 339 | I339P | A | 1.5e-05 | 1.7e-05 | 1.2e-05 | 1.467E-05 | 2.517E-06 | 1.200E-05 | 1.700E-05 |
| 6776 | 339 | I339S | A | 1.5e-05 | 9.3e-06 | 1e-05 | 1.143E-05 | 3.109E-06 | 9.300E-06 | 1.500E-05 |
| 6777 | 339 | I339T | A | 1.2e-05 | 1.5e-05 | 8.8e-06 | 1.193E-05 | 3.101E-06 | 8.800E-06 | 1.500E-05 |
| 6778 | 339 | I339W | A | 1.4e-05 | 1.4e-05 | 1e-05 | 1.267E-05 | 2.309E-06 | 1.000E-05 | 1.400E-05 |
| 6779 | 339 | I339Y | A | 1.5e-05 | 1.1e-05 | 9.4e-06 | 1.180E-05 | 2.884E-06 | 9.400E-06 | 1.500E-05 |
| 6780 | 339 | I339V | A | 1.6e-05 | 1.7e-05 | 1.6e-05 | 1.633E-05 | 5.774E-07 | 1.600E-05 | 1.700E-05 |
| 6781 | 340 | Q340A | A | 1.5e-05 | 1.5e-05 | 1.6e-05 | 1.533E-05 | 5.774E-07 | 1.500E-05 | 1.600E-05 |
| 6782 | 340 | Q340R | A | 1.3e-05 | 1.5e-05 | 1.6e-05 | 1.467E-05 | 1.528E-06 | 1.300E-05 | 1.600E-05 |
| 6783 | 340 | Q340N | A | 8.7e-06 | 1.1e-05 | 1.6e-05 | 1.190E-05 | 3.732E-06 | 8.700E-06 | 1.600E-05 |
| 6784 | 340 | Q340D | A | 1.5e-05 | 1.5e-05 | 9.9e-06 | 1.330E-05 | 2.944E-06 | 9.900E-06 | 1.500E-05 |
| 6785 | 340 | Q340C | A | 1.4e-05 | 1.1e-05 | 1.4e-05 | 1.300E-05 | 1.732E-06 | 1.100E-05 | 1.400E-05 |
| 6786 | 340 | Q340E | A | 1.8e-05 | 1.2e-05 | 1.2e-05 | 1.400E-05 | 3.464E-06 | 1.200E-05 | 1.800E-05 |
| 6787 | 340 | Q340Q | A | 1e-05 | 1e-05 | 1.4e-05 | 1.133E-05 | 2.309E-06 | 1.000E-05 | 1.400E-05 |
| 6788 | 340 | Q340G | A | 1.1e-05 | 1.5e-05 | 1e-05 | 1.200E-05 | 2.646E-06 | 1.000E-05 | 1.500E-05 |
| 6789 | 340 | Q340H | A | 1.5e-05 | 9.6e-06 | 9.9e-06 | 1.150E-05 | 3.035E-06 | 9.600E-06 | 1.500E-05 |
| 6790 | 340 | Q340I | A | 1.3e-05 | 1.8e-05 | 1.4e-05 | 1.500E-05 | 2.646E-06 | 1.300E-05 | 1.800E-05 |
| 6791 | 340 | Q340L | A | 1.2e-05 | 1e-05 | 1.4e-05 | 1.200E-05 | 2.000E-06 | 1.000E-05 | 1.400E-05 |
| 6792 | 340 | Q340K | A | 1.6e-05 | 1.1e-05 | 1.2e-05 | 1.300E-05 | 2.646E-06 | 1.100E-05 | 1.600E-05 |
| 6793 | 340 | Q340M | A | 1.6e-05 | 1.6e-05 | 1.5e-05 | 1.567E-05 | 5.774E-07 | 1.500E-05 | 1.600E-05 |
| 6794 | 340 | Q340F | A | 1.1e-05 | 1.6e-05 | 1e-05 | 1.233E-05 | 3.215E-06 | 1.000E-05 | 1.600E-05 |
| 6795 | 340 | Q340P | A | 9.3e-06 | 8.9e-06 | 1.7e-05 | 1.173E-05 | 4.565E-06 | 8.900E-06 | 1.700E-05 |
| 6796 | 340 | Q340S | A | 1.5e-05 | 1.7e-05 | 1e-05 | 1.400E-05 | 3.606E-06 | 1.000E-05 | 1.700E-05 |
| 6797 | 340 | Q340T | A | 1.2e-05 | 1.2e-05 | 1.2e-05 | 1.200E-05 | 0.000E+00 | 1.200E-05 | 1.200E-05 |
| 6798 | 340 | Q340W | A | 9.7e-06 | 1.1e-05 | 1.5e-05 | 1.190E-05 | 2.762E-06 | 9.700E-06 | 1.500E-05 |
| 6799 | 340 | Q340Y | A | 1.5e-05 | 8.3e-06 | 1.2e-05 | 1.177E-05 | 3.356E-06 | 8.300E-06 | 1.500E-05 |

| | | | | | | | | | | |
|---|---|---|---|---|---|---|---|---|---|---|
| 6800 | 340 | Q340V | A | 8.7e-06 | 1.1e-05 | 1.4e-05 | 1.123E-05 | 2.658E-06 | 8.700E-06 | 1.400E-05 |
| 6801 | 341 | E341A | A | 1.7e-05 | 1.7e-05 | 1.4e-05 | 1.600E-05 | 1.732E-06 | 1.400E-05 | 1.700E-05 |
| 6802 | 341 | E341R | A | 1.3e-05 | 1.6e-05 | 2.2e-05 | 1.700E-05 | 4.583E-06 | 1.300E-05 | 2.200E-05 |
| 6803 | 341 | E341N | A | 1.1e-05 | 1.3e-05 | 1.6e-05 | 1.333E-05 | 2.517E-06 | 1.100E-05 | 1.600E-05 |
| 6804 | 341 | E341D | A | 1.5e-05 | 1.6e-05 | 8.9e-06 | 1.330E-05 | 3.843E-06 | 8.900E-06 | 1.600E-05 |
| 6805 | 341 | E341C | A | 1.3e-05 | 1.1e-05 | 1.4e-05 | 1.267E-05 | 1.528E-06 | 1.100E-05 | 1.400E-05 |
| 6806 | 341 | E341E | A | 1.5e-05 | 1.3e-05 | 1.3e-05 | 1.367E-05 | 1.155E-06 | 1.300E-05 | 1.500E-05 |
| 6807 | 341 | E341Q | A | 8.9e-06 | 1.1e-05 | 1.5e-05 | 1.163E-05 | 3.099E-06 | 8.900E-06 | 1.500E-05 |
| 6808 | 341 | E341G | A | 1.2e-05 | 7.6e-06 | 1.4e-05 | 1.120E-05 | 3.274E-06 | 7.600E-06 | 1.400E-05 |
| 6809 | 341 | E341H | A | 1.5e-05 | 1.2e-05 | 1.7e-05 | 1.467E-05 | 2.517E-06 | 1.200E-05 | 1.700E-05 |
| 6810 | 341 | E341I | A | 9e-06 | 1.4e-05 | 1.5e-05 | 1.267E-05 | 3.215E-06 | 9.000E-06 | 1.500E-05 |
| 6811 | 341 | E341L | A | 1.5e-05 | 8.9e-06 | 1.4e-05 | 1.263E-05 | 3.272E-06 | 8.900E-06 | 1.500E-05 |
| 6812 | 341 | E341K | A | 9.7e-06 | 1.1e-05 | 1.6e-05 | 1.223E-05 | 3.326E-06 | 9.700E-06 | 1.600E-05 |
| 6813 | 341 | E341M | A | 1.4e-05 | 1.3e-05 | 1.4e-05 | 1.367E-05 | 5.774E-07 | 1.300E-05 | 1.400E-05 |
| 6814 | 341 | E341F | A | 8.7e-06 | 1.6e-05 | 1.5e-05 | 1.323E-05 | 3.958E-06 | 8.700E-06 | 1.600E-05 |
| 6815 | 341 | E341P | A | 1.7e-05 | 1e-05 | 9.9e-06 | 1.230E-05 | 4.071E-06 | 9.900E-06 | 1.700E-05 |
| 6816 | 341 | E341S | A | 9.5e-06 | 1.7e-05 | 1.1e-05 | 1.250E-05 | 3.969E-06 | 9.500E-06 | 1.700E-05 |
| 6817 | 341 | E341T | A | 1.7e-05 | 1.6e-05 | 1.5e-05 | 1.600E-05 | 1.000E-06 | 1.500E-05 | 1.700E-05 |
| 6818 | 341 | E341W | A | 9.7e-06 | 9e-06 | 1.1e-05 | 9.900E-06 | 1.015E-06 | 9.000E-06 | 1.100E-05 |
| 6819 | 341 | E341Y | A | 1.1e-05 | 8.9e-06 | 1.1e-05 | 1.030E-05 | 1.212E-06 | 8.900E-06 | 1.100E-05 |
| 6820 | 341 | E341V | A | 1.5e-05 | 1.4e-05 | 8.2e-06 | 1.240E-05 | 3.672E-06 | 8.200E-06 | 1.500E-05 |
| 6821 | 342 | F342A | A | 1.1e-05 | 1.3e-05 | 1.3e-05 | 1.233E-05 | 1.155E-06 | 1.100E-05 | 1.300E-05 |
| 6822 | 342 | F342R | A | 1.1e-05 | 9.6e-06 | 1e-05 | 1.020E-05 | 7.211E-07 | 9.600E-06 | 1.100E-05 |
| 6823 | 342 | F342N | A | 1.2e-05 | 1.4e-05 | 1.5e-05 | 1.367E-05 | 1.528E-06 | 1.200E-05 | 1.500E-05 |
| 6824 | 342 | F342D | A | 1.6e-05 | 1.3e-05 | 8.3e-06 | 1.243E-05 | 3.881E-06 | 8.300E-06 | 1.600E-05 |
| 6825 | 342 | F342C | A | 1.7e-05 | 1.2e-05 | 1.5e-05 | 1.467E-05 | 2.517E-06 | 1.200E-05 | 1.700E-05 |
| 6826 | 342 | F342E | A | 1e-05 | 8.8e-06 | 1.2e-05 | 1.027E-05 | 1.617E-06 | 8.800E-06 | 1.200E-05 |
| 6827 | 342 | F342Q | A | 9.5e-06 | 6.9e-06 | 1.6e-05 | 1.080E-05 | 4.687E-06 | 6.900E-06 | 1.600E-05 |
| 6828 | 342 | F342G | A | 1.6e-05 | 9.4e-06 | 1.1e-05 | 1.213E-05 | 3.443E-06 | 9.400E-06 | 1.600E-05 |
| 6829 | 342 | F342H | A | 1.1e-05 | 1.6e-05 | 9.1e-06 | 1.203E-05 | 3.564E-06 | 9.100E-06 | 1.600E-05 |
| 6830 | 342 | F342I | A | 8.7e-06 | 1.5e-05 | 1.6e-05 | 1.323E-05 | 3.958E-06 | 8.700E-06 | 1.600E-05 |
| 6831 | 342 | F342L | A | 1.1e-05 | 1.4e-05 | 9.6e-06 | 1.153E-05 | 2.248E-06 | 9.600E-06 | 1.400E-05 |
| 6832 | 342 | F342K | A | 1.2e-05 | 9.9e-06 | 8.3e-06 | 1.007E-05 | 1.856E-06 | 8.300E-06 | 1.200E-05 |
| 6833 | 342 | F342M | A | 1.4e-05 | 1.6e-05 | 1.6e-05 | 1.533E-05 | 1.155E-06 | 1.400E-05 | 1.600E-05 |
| 6834 | 342 | F342F | A | 9.7e-06 | 1.3e-05 | 1.5e-05 | 1.257E-05 | 2.676E-06 | 9.700E-06 | 1.500E-05 |
| 6835 | 342 | F342P | A | 1.4e-05 | 1.1e-05 | 1e-05 | 1.167E-05 | 2.082E-06 | 1.000E-05 | 1.400E-05 |
| 6836 | 342 | F342S | A | 1.5e-05 | 1.3e-05 | 1.6e-05 | 1.467E-05 | 1.528E-06 | 1.300E-05 | 1.600E-05 |
| 6837 | 342 | F342T | A | 1.5e-05 | 1e-05 | 1.6e-05 | 1.367E-05 | 3.215E-06 | 1.000E-05 | 1.600E-05 |
| 6838 | 342 | F342W | A | 1.4e-05 | 1e-05 | 1.4e-05 | 1.267E-05 | 2.309E-06 | 1.000E-05 | 1.400E-05 |
| 6839 | 342 | F342Y | A | 1.1e-05 | 1.1e-05 | 9.3e-06 | 1.043E-05 | 9.815E-07 | 9.300E-06 | 1.100E-05 |
| 6840 | 342 | F342V | A | 1.6e-05 | 1.1e-05 | 1.4e-05 | 1.367E-05 | 2.517E-06 | 1.100E-05 | 1.600E-05 |
| 6841 | 343 | A343A | A | 1.5e-05 | 1.5e-05 | 1.1e-05 | 1.367E-05 | 2.309E-06 | 1.100E-05 | 1.500E-05 |
| 6842 | 343 | A343R | A | 1e-05 | 1.2e-05 | 9.9e-06 | 1.063E-05 | 1.185E-06 | 9.900E-06 | 1.200E-05 |
| 6843 | 343 | A343N | A | 1e-05 | 7e-06 | 1.4e-05 | 1.033E-05 | 3.512E-06 | 7.000E-06 | 1.400E-05 |
| 6844 | 343 | A343D | A | 1.3e-05 | 1.4e-05 | 1.3e-05 | 1.333E-05 | 5.774E-07 | 1.300E-05 | 1.400E-05 |
| 6845 | 343 | A343C | A | 1.1e-05 | 1.1e-05 | 1.1e-05 | 1.100E-05 | 2.075E-21 | 1.100E-05 | 1.100E-05 |
| 6846 | 343 | A343E | A | 1.6e-05 | 1.5e-05 | 1.5e-05 | 1.533E-05 | 5.774E-07 | 1.500E-05 | 1.600E-05 |
| 6847 | 343 | A343Q | A | 1e-05 | 9.7e-06 | 1e-05 | 9.900E-06 | 1.732E-07 | 9.700E-06 | 1.000E-05 |
| 6848 | 343 | A343G | A | 1.1e-05 | 1.7e-05 | 1.5e-05 | 1.433E-05 | 3.055E-06 | 1.100E-05 | 1.700E-05 |
| 6849 | 343 | A343H | A | 1.7e-05 | 1.4e-05 | 1.4e-05 | 1.500E-05 | 1.732E-06 | 1.400E-05 | 1.700E-05 |
| 6850 | 343 | A343I | A | 1.1e-05 | 1.7e-05 | 1.7e-05 | 1.500E-05 | 3.464E-06 | 1.100E-05 | 1.700E-05 |
| 6851 | 343 | A343L | A | 1.4e-05 | 1.5e-05 | 1.4e-05 | 1.433E-05 | 5.774E-07 | 1.400E-05 | 1.500E-05 |
| 6852 | 343 | A343K | A | 6.3e-06 | 1.1e-05 | 1.1e-05 | 9.433E-06 | 2.714E-06 | 6.300E-06 | 1.100E-05 |
| 6853 | 343 | A343M | A | 1.6e-05 | 1.1e-05 | 1e-05 | 1.233E-05 | 3.215E-06 | 1.000E-05 | 1.600E-05 |
| 6854 | 343 | A343F | A | 1.1e-05 | 1.1e-05 | 1.4e-05 | 1.200E-05 | 1.732E-06 | 1.100E-05 | 1.400E-05 |
| 6855 | 343 | A343P | A | 1.5e-05 | 1.6e-05 | 1.4e-05 | 1.500E-05 | 1.000E-06 | 1.400E-05 | 1.600E-05 |
| 6856 | 343 | A343S | A | 1.3e-05 | 1.4e-05 | 1.4e-05 | 1.367E-05 | 5.774E-07 | 1.300E-05 | 1.400E-05 |
| 6857 | 343 | A343T | A | 1.1e-05 | 1.2e-05 | 1.5e-05 | 1.267E-05 | 2.082E-06 | 1.100E-05 | 1.500E-05 |
| 6858 | 343 | A343W | A | 1.5e-05 | 1.5e-05 | 1.6e-05 | 1.533E-05 | 5.774E-07 | 1.500E-05 | 1.600E-05 |
| 6859 | 343 | A343Y | A | 1.4e-05 | 1.1e-05 | 8.7e-06 | 1.123E-05 | 2.658E-06 | 8.700E-06 | 1.400E-05 |
| 6860 | 343 | A343V | A | 9.5e-06 | 1.5e-05 | 1.6e-05 | 1.350E-05 | 3.500E-06 | 9.500E-06 | 1.600E-05 |
| 6861 | 344 | G344A | A | 2.7e-05 | 1.1e-05 | 1.5e-05 | 1.767E-05 | 8.327E-06 | 1.100E-05 | 2.700E-05 |
| 6862 | 344 | G344R | A | 1.4e-05 | 1.5e-05 | 1.1e-05 | 1.333E-05 | 2.082E-06 | 1.100E-05 | 1.500E-05 |
| 6863 | 344 | G344N | A | 1.2e-05 | 1.5e-05 | 1.6e-05 | 1.433E-05 | 2.082E-06 | 1.200E-05 | 1.600E-05 |
| 6864 | 344 | G344D | A | 1.4e-05 | 1.7e-05 | 1.3e-05 | 1.467E-05 | 2.082E-06 | 1.300E-05 | 1.700E-05 |
| 6865 | 344 | G344C | A | 1.5e-05 | 9.5e-06 | 9.1e-06 | 1.120E-05 | 3.297E-06 | 9.100E-06 | 1.500E-05 |
| 6866 | 344 | G344E | A | 1.4e-05 | 1.6e-05 | 1.2e-05 | 1.400E-05 | 2.000E-06 | 1.200E-05 | 1.600E-05 |
| 6867 | 344 | G344Q | A | 9.6e-06 | 1.1e-05 | 8.8e-06 | 9.800E-06 | 1.114E-06 | 8.800E-06 | 1.100E-05 |
| 6868 | 344 | G344G | A | 1.1e-05 | 1.6e-05 | 1.3e-05 | 1.333E-05 | 2.517E-06 | 1.100E-05 | 1.600E-05 |
| 6869 | 344 | G344H | A | 1e-05 | 1.5e-05 | 1.1e-05 | 1.200E-05 | 2.646E-06 | 1.000E-05 | 1.500E-05 |
| 6870 | 344 | G344I | A | 1.3e-05 | 1.3e-05 | 1.7e-05 | 1.433E-05 | 2.309E-06 | 1.300E-05 | 1.700E-05 |
| 6871 | 344 | G344L | A | 1e-05 | 1.1e-05 | 1.6e-05 | 1.233E-05 | 3.215E-06 | 1.000E-05 | 1.600E-05 |
| 6872 | 344 | G344K | A | 1e-05 | 1.5e-05 | 1.5e-05 | 1.333E-05 | 2.887E-06 | 1.000E-05 | 1.500E-05 |
| 6873 | 344 | G344M | A | 1.3e-05 | 1.5e-05 | 1.4e-05 | 1.400E-05 | 1.000E-06 | 1.300E-05 | 1.500E-05 |
| 6874 | 344 | G344F | A | 1.7e-05 | 1.5e-05 | 1.1e-05 | 1.433E-05 | 3.055E-06 | 1.100E-05 | 1.700E-05 |
| 6875 | 344 | G344P | A | 1.1e-05 | 1.2e-05 | 1.4e-05 | 1.233E-05 | 1.528E-06 | 1.100E-05 | 1.400E-05 |
| 6876 | 344 | G344S | A | 1.6e-05 | 1.4e-05 | 9.7e-06 | 1.323E-05 | 3.219E-06 | 9.700E-06 | 1.600E-05 |
| 6877 | 344 | G344T | A | 1.6e-05 | 9.6e-06 | 1.6e-05 | 1.387E-05 | 3.695E-06 | 9.600E-06 | 1.600E-05 |
| 6878 | 344 | G344W | A | 1.1e-05 | 1.2e-05 | 1.5e-05 | 1.267E-05 | 2.082E-06 | 1.100E-05 | 1.500E-05 |
| 6879 | 344 | G344Y | A | 1e-05 | 9.8e-06 | 9.3e-06 | 9.700E-06 | 3.606E-07 | 9.300E-06 | 1.000E-05 |
| 6880 | 344 | G344V | A | 1.3e-05 | 9e-06 | 8.5e-06 | 1.017E-05 | 2.466E-06 | 8.500E-06 | 1.300E-05 |
| 6881 | 345 | C345A | A | 1.1e-05 | 1.8e-05 | 1.7e-05 | 1.533E-05 | 3.786E-06 | 1.100E-05 | 1.800E-05 |
| 6882 | 345 | C345R | A | 1.4e-05 | 1.1e-05 | 1.3e-05 | 1.267E-05 | 1.528E-06 | 1.100E-05 | 1.400E-05 |
| 6883 | 345 | C345N | A | 1e-05 | 1.1e-05 | 1e-05 | 1.033E-05 | 5.774E-07 | 1.000E-05 | 1.100E-05 |
| 6884 | 345 | C345D | A | 1e-05 | 1.5e-05 | 1.2e-05 | 1.233E-05 | 2.517E-06 | 1.000E-05 | 1.500E-05 |

| | | | | | | | | | | |
|---|---|---|---|---|---|---|---|---|---|---|
| 6885 | 345 | C345C | A | 1.6e-05 | 1.6e-05 | 1e-05 | 1.400E-05 | 3.464E-06 | 1.000E-05 | 1.600E-05 |
| 6886 | 345 | C345E | A | 9.3e-06 | 9.7e-06 | 1.1e-05 | 1.000E-05 | 8.888E-07 | 9.300E-06 | 1.100E-05 |
| 6887 | 345 | C345Q | A | 1.4e-05 | 9.4e-06 | 1.2e-05 | 1.180E-05 | 2.307E-06 | 9.400E-06 | 1.400E-05 |
| 6888 | 345 | C345G | A | 1.6e-05 | 1.6e-05 | 1.5e-05 | 1.567E-05 | 5.774E-07 | 1.500E-05 | 1.600E-05 |
| 6889 | 345 | C345H | A | 1e-05 | 9.8e-06 | 1.5e-05 | 1.160E-05 | 2.946E-06 | 9.800E-06 | 1.500E-05 |
| 6890 | 345 | C345I | A | 9.8e-06 | 1.6e-05 | 1.1e-05 | 1.227E-05 | 3.288E-06 | 9.800E-06 | 1.600E-05 |
| 6891 | 345 | C345L | A | 1.6e-05 | 1.6e-05 | 1.5e-05 | 1.567E-05 | 5.774E-07 | 1.500E-05 | 1.600E-05 |
| 6892 | 345 | C345K | A | 1.1e-05 | 8.5e-06 | 1.6e-05 | 1.183E-05 | 3.819E-06 | 8.500E-06 | 1.600E-05 |
| 6893 | 345 | C345M | A | 1.8e-05 | 1.1e-05 | 1.3e-05 | 1.400E-05 | 3.606E-06 | 1.100E-05 | 1.800E-05 |
| 6894 | 345 | C345F | A | 1.7e-05 | 1.2e-05 | 1.4e-05 | 1.433E-05 | 2.517E-06 | 1.200E-05 | 1.700E-05 |
| 6895 | 345 | C345P | A | 1.2e-05 | 1.2e-05 | 9.4e-06 | 1.113E-05 | 1.501E-06 | 9.400E-06 | 1.200E-05 |
| 6896 | 345 | C345S | A | 1.5e-05 | 1.1e-05 | 1.1e-05 | 1.233E-05 | 2.309E-06 | 1.100E-05 | 1.500E-05 |
| 6897 | 345 | C345T | A | 1.3e-05 | 1.6e-05 | 1.1e-05 | 1.333E-05 | 2.517E-06 | 1.100E-05 | 1.600E-05 |
| 6898 | 345 | C345W | A | 1.2e-05 | 1.3e-05 | 1.1e-05 | 1.200E-05 | 1.000E-06 | 1.100E-05 | 1.300E-05 |
| 6899 | 345 | C345Y | A | 1e-05 | 9.8e-06 | 1e-05 | 9.933E-06 | 1.155E-07 | 9.800E-06 | 1.000E-05 |
| 6900 | 345 | C345V | A | 1.4e-05 | 1e-05 | 1.5e-05 | 1.300E-05 | 2.646E-06 | 1.000E-05 | 1.500E-05 |
| 6901 | 346 | K346A | A | 1.1e-05 | 1.7e-05 | 1.5e-05 | 1.433E-05 | 3.055E-06 | 1.100E-05 | 1.700E-05 |
| 6902 | 346 | K346R | A | 1.5e-05 | 1.4e-05 | 8.9e-06 | 1.263E-05 | 3.272E-06 | 8.900E-06 | 1.500E-05 |
| 6903 | 346 | K346N | A | 1e-05 | 1.6e-05 | 1.5e-05 | 1.367E-05 | 3.215E-06 | 1.000E-05 | 1.600E-05 |
| 6904 | 346 | K346D | A | 1e-05 | 1.1e-05 | 1.6e-05 | 1.233E-05 | 3.215E-06 | 1.000E-05 | 1.600E-05 |
| 6905 | 346 | K346C | A | 9.8e-06 | 1.6e-05 | 1.3e-05 | 1.293E-05 | 3.101E-06 | 9.800E-06 | 1.600E-05 |
| 6906 | 346 | K346E | A | 1.3e-05 | 1.6e-05 | 1.3e-05 | 1.400E-05 | 1.732E-06 | 1.300E-05 | 1.600E-05 |
| 6907 | 346 | K346Q | A | 1.6e-05 | 1.3e-05 | 1e-05 | 1.300E-05 | 3.000E-06 | 1.000E-05 | 1.600E-05 |
| 6908 | 346 | K346G | A | 1.4e-05 | 9.5e-06 | 1.7e-05 | 1.350E-05 | 3.775E-06 | 9.500E-06 | 1.700E-05 |
| 6909 | 346 | K346H | A | 9.8e-06 | 1e-05 | 1.4e-05 | 1.127E-05 | 2.369E-06 | 9.800E-06 | 1.400E-05 |
| 6910 | 346 | K346I | A | 1e-05 | 1e-05 | 9.9e-06 | 9.967E-06 | 5.774E-08 | 9.900E-06 | 1.000E-05 |
| 6911 | 346 | K346L | A | 1.7e-05 | 1.5e-05 | 1e-05 | 1.400E-05 | 3.606E-06 | 1.000E-05 | 1.700E-05 |
| 6912 | 346 | K346K | A | 9.6e-06 | 8.7e-06 | 1e-05 | 9.433E-06 | 6.658E-07 | 8.700E-06 | 1.000E-05 |
| 6913 | 346 | K346M | A | 1.1e-05 | 1.3e-05 | 9.6e-06 | 1.120E-05 | 1.709E-06 | 9.600E-06 | 1.300E-05 |
| 6914 | 346 | K346F | A | 1.1e-05 | 1.2e-05 | 1.1e-05 | 1.133E-05 | 5.774E-07 | 1.100E-05 | 1.200E-05 |
| 6915 | 346 | K346P | A | 1.5e-05 | 1.5e-05 | 9.3e-06 | 1.310E-05 | 3.291E-06 | 9.300E-06 | 1.500E-05 |
| 6916 | 346 | K346S | A | 1.4e-05 | 1.1e-05 | 1.5e-05 | 1.333E-05 | 2.082E-06 | 1.100E-05 | 1.500E-05 |
| 6917 | 346 | K346T | A | 1.2e-05 | 1.1e-05 | 1e-05 | 1.100E-05 | 1.000E-06 | 1.000E-05 | 1.200E-05 |
| 6918 | 346 | K346W | A | 1.2e-05 | 1.2e-05 | 1e-05 | 1.133E-05 | 1.155E-06 | 1.000E-05 | 1.200E-05 |
| 6919 | 346 | K346Y | A | 1.1e-05 | 1e-05 | 1.6e-05 | 1.233E-05 | 3.215E-06 | 1.000E-05 | 1.600E-05 |
| 6920 | 346 | K346V | A | 8.7e-06 | 1.1e-05 | 1.1e-05 | 1.023E-05 | 1.328E-06 | 8.700E-06 | 1.100E-05 |
| 6921 | 347 | K347A | A | 1.5e-05 | 6.3e-06 | 1.3e-05 | 1.143E-05 | 4.557E-06 | 6.300E-06 | 1.500E-05 |
| 6922 | 347 | K347R | A | 1.1e-05 | 1.6e-05 | 1.6e-05 | 1.433E-05 | 2.887E-06 | 1.100E-05 | 1.600E-05 |
| 6923 | 347 | K347N | A | 1.5e-05 | 1.5e-05 | 1.1e-05 | 1.367E-05 | 2.309E-06 | 1.100E-05 | 1.500E-05 |
| 6924 | 347 | K347D | A | 1.4e-05 | 9.5e-06 | 1.4e-05 | 1.250E-05 | 2.598E-06 | 9.500E-06 | 1.400E-05 |
| 6925 | 347 | K347C | A | 8.9e-06 | 9.6e-06 | 1.1e-05 | 9.833E-06 | 1.069E-06 | 8.900E-06 | 1.100E-05 |
| 6926 | 347 | K347E | A | 9.4e-06 | 1.6e-05 | 1.6e-05 | 1.380E-05 | 3.811E-06 | 9.400E-06 | 1.600E-05 |
| 6927 | 347 | K347Q | A | 1.4e-05 | 1.4e-05 | 8.6e-06 | 1.220E-05 | 3.118E-06 | 8.600E-06 | 1.400E-05 |
| 6928 | 347 | K347G | A | 1e-05 | 1.7e-05 | 1e-05 | 1.233E-05 | 4.041E-06 | 1.000E-05 | 1.700E-05 |
| 6929 | 347 | K347H | A | 1.3e-05 | 1.3e-05 | 1.5e-05 | 1.367E-05 | 1.155E-06 | 1.300E-05 | 1.500E-05 |
| 6930 | 347 | K347I | A | 1e-05 | 9.1e-06 | 1e-05 | 9.700E-06 | 5.196E-07 | 9.100E-06 | 1.000E-05 |
| 6931 | 347 | K347L | A | 1.5e-05 | 1.6e-05 | 1.6e-05 | 1.567E-05 | 5.774E-07 | 1.500E-05 | 1.600E-05 |
| 6932 | 347 | K347K | A | 1.5e-05 | 1.1e-05 | 1.1e-05 | 1.233E-05 | 2.309E-06 | 1.100E-05 | 1.500E-05 |
| 6933 | 347 | K347M | A | 1.6e-05 | 1.5e-05 | 1.6e-05 | 1.567E-05 | 5.774E-07 | 1.500E-05 | 1.600E-05 |
| 6934 | 347 | K347F | A | 1.5e-05 | 8.5e-06 | 1e-05 | 1.117E-05 | 3.403E-06 | 8.500E-06 | 1.500E-05 |
| 6935 | 347 | K347P | A | 1.1e-05 | 1.1e-05 | 1.6e-05 | 1.267E-05 | 2.887E-06 | 1.100E-05 | 1.600E-05 |
| 6936 | 347 | K347S | A | 1.5e-05 | 8.4e-06 | 1.1e-05 | 1.147E-05 | 3.325E-06 | 8.400E-06 | 1.500E-05 |
| 6937 | 347 | K347T | A | 9.5e-06 | 1.7e-05 | 1.4e-05 | 1.350E-05 | 3.775E-06 | 9.500E-06 | 1.700E-05 |
| 6938 | 347 | K347W | A | 1.6e-05 | 1.3e-05 | 1.6e-05 | 1.500E-05 | 1.732E-06 | 1.300E-05 | 1.600E-05 |
| 6939 | 347 | K347Y | A | 1.1e-05 | 9.8e-06 | 1.2e-05 | 1.093E-05 | 1.102E-06 | 9.800E-06 | 1.200E-05 |
| 6940 | 347 | K347V | A | 1.6e-05 | 1.2e-05 | 1.3e-05 | 1.367E-05 | 2.082E-06 | 1.200E-05 | 1.600E-05 |
| 6941 | 348 | I348A | A | 1.4e-05 | 1.1e-05 | 1.2e-05 | 1.233E-05 | 1.528E-06 | 1.100E-05 | 1.400E-05 |
| 6942 | 348 | I348R | A | 9.1e-06 | 1.4e-05 | 8.6e-06 | 1.057E-05 | 2.984E-06 | 8.600E-06 | 1.400E-05 |
| 6943 | 348 | I348N | A | 1.4e-05 | 1.1e-05 | 1.5e-05 | 1.333E-05 | 2.082E-06 | 1.100E-05 | 1.500E-05 |
| 6944 | 348 | I348D | A | 1.1e-05 | 1.7e-05 | 1.6e-05 | 1.467E-05 | 3.215E-06 | 1.100E-05 | 1.700E-05 |
| 6945 | 348 | I348C | A | 1.4e-05 | 1.8e-05 | 1.7e-05 | 1.633E-05 | 2.082E-06 | 1.400E-05 | 1.800E-05 |
| 6946 | 348 | I348E | A | 1e-05 | 1.7e-05 | 1.4e-05 | 1.367E-05 | 3.512E-06 | 1.000E-05 | 1.700E-05 |
| 6947 | 348 | I348Q | A | 1.6e-05 | 1.3e-05 | 9.7e-06 | 1.290E-05 | 3.151E-06 | 9.700E-06 | 1.600E-05 |
| 6948 | 348 | I348G | A | 8.4e-06 | 1.1e-05 | 1.5e-05 | 1.147E-05 | 3.325E-06 | 8.400E-06 | 1.500E-05 |
| 6949 | 348 | I348H | A | 1e-05 | 1.8e-05 | 1.4e-05 | 1.400E-05 | 4.000E-06 | 1.000E-05 | 1.800E-05 |
| 6950 | 348 | I348I | A | 9.8e-06 | 1.5e-05 | 1e-05 | 1.160E-05 | 2.946E-06 | 9.800E-06 | 1.500E-05 |
| 6951 | 348 | I348L | A | 1.3e-05 | 9.8e-06 | 1.5e-05 | 1.260E-05 | 2.623E-06 | 9.800E-06 | 1.500E-05 |
| 6952 | 348 | I348K | A | 1.5e-05 | 8.2e-06 | 1.6e-05 | 1.307E-05 | 4.244E-06 | 8.200E-06 | 1.600E-05 |
| 6953 | 348 | I348M | A | 1.6e-05 | 1.7e-05 | 1.6e-05 | 1.633E-05 | 5.774E-07 | 1.600E-05 | 1.700E-05 |
| 6954 | 348 | I348F | A | 1.1e-05 | 1.5e-05 | 1e-05 | 1.200E-05 | 2.646E-06 | 1.000E-05 | 1.500E-05 |
| 6955 | 348 | I348P | A | 1.1e-05 | 1.3e-05 | 8.8e-06 | 1.093E-05 | 2.101E-06 | 8.800E-06 | 1.300E-05 |
| 6956 | 348 | I348S | A | 1.1e-05 | 1.5e-05 | 1.1e-05 | 1.233E-05 | 2.309E-06 | 1.100E-05 | 1.500E-05 |
| 6957 | 348 | I348T | A | 1.6e-05 | 1.3e-05 | 1.1e-05 | 1.333E-05 | 2.517E-06 | 1.100E-05 | 1.600E-05 |
| 6958 | 348 | I348W | A | 1.7e-05 | 1.4e-05 | 1.6e-05 | 1.567E-05 | 1.528E-06 | 1.400E-05 | 1.700E-05 |
| 6959 | 348 | I348Y | A | 1.4e-05 | 1.1e-05 | 1.7e-05 | 1.400E-05 | 3.000E-06 | 1.100E-05 | 1.700E-05 |
| 6960 | 348 | I348V | A | 1.4e-05 | 1.4e-05 | 1.8e-05 | 1.533E-05 | 2.309E-06 | 1.400E-05 | 1.800E-05 |
| 6961 | 349 | F349A | A | 8.5e-06 | 1.7e-05 | 8.4e-06 | 1.130E-05 | 4.937E-06 | 8.400E-06 | 1.700E-05 |
| 6962 | 349 | F349R | A | 2e-05 | 1.5e-05 | 9.3e-06 | 1.477E-05 | 5.354E-06 | 9.300E-06 | 2.000E-05 |
| 6963 | 349 | F349N | A | 1.4e-05 | 9.6e-06 | 1.3e-05 | 1.220E-05 | 2.307E-06 | 9.600E-06 | 1.400E-05 |
| 6964 | 349 | F349D | A | 1.5e-05 | 1.5e-05 | 1.1e-05 | 1.367E-05 | 2.309E-06 | 1.100E-05 | 1.500E-05 |
| 6965 | 349 | F349C | A | 1.1e-05 | 1.1e-05 | 1.1e-05 | 1.100E-05 | 2.075E-21 | 1.100E-05 | 1.100E-05 |
| 6966 | 349 | F349E | A | 1e-05 | 1.6e-05 | 8.2e-06 | 1.140E-05 | 4.084E-06 | 8.200E-06 | 1.600E-05 |
| 6967 | 349 | F349Q | A | 1.4e-05 | 9.5e-06 | 1.2e-05 | 1.183E-05 | 2.255E-06 | 9.500E-06 | 1.400E-05 |
| 6968 | 349 | F349G | A | 9.7e-06 | 1e-05 | 9.8e-06 | 9.833E-06 | 1.528E-07 | 9.700E-06 | 1.000E-05 |
| 6969 | 349 | F349H | A | 1.2e-05 | 1.6e-05 | 1.5e-05 | 1.433E-05 | 2.082E-06 | 1.200E-05 | 1.600E-05 |

| | | | | | | | | | | |
|---|---|---|---|---|---|---|---|---|---|---|
| 6970 | 349 | F349I | A | 1.1e-05 | 1.3e-05 | 1.1e-05 | 1.167E-05 | 1.155E-06 | 1.100E-05 | 1.300E-05 |
| 6971 | 349 | F349L | A | 1.2e-05 | 1.5e-05 | 1.5e-05 | 1.400E-05 | 1.732E-06 | 1.200E-05 | 1.500E-05 |
| 6972 | 349 | F349K | A | 8.6e-06 | 9.4e-06 | 1.6e-05 | 1.133E-05 | 4.061E-06 | 8.600E-06 | 1.600E-05 |
| 6973 | 349 | F349M | A | 1.5e-05 | 8.7e-06 | 1.6e-05 | 1.323E-05 | 3.958E-06 | 8.700E-06 | 1.600E-05 |
| 6974 | 349 | F349F | A | 6.7e-06 | 1.4e-05 | 1.1e-05 | 1.057E-05 | 3.669E-06 | 6.700E-06 | 1.400E-05 |
| 6975 | 349 | F349P | A | 9.4e-06 | 1.4e-05 | 1.1e-05 | 1.147E-05 | 2.335E-06 | 9.400E-06 | 1.400E-05 |
| 6976 | 349 | F349S | A | 1e-05 | 1.3e-05 | 1.6e-05 | 1.300E-05 | 3.000E-06 | 1.000E-05 | 1.600E-05 |
| 6977 | 349 | F349T | A | 1.4e-05 | 1.5e-05 | 1e-05 | 1.300E-05 | 2.646E-06 | 1.000E-05 | 1.500E-05 |
| 6978 | 349 | F349W | A | 8.8e-06 | 1.4e-05 | 9.3e-06 | 1.070E-05 | 2.869E-06 | 8.800E-06 | 1.400E-05 |
| 6979 | 349 | F349Y | A | 1.5e-05 | 1e-05 | 1.6e-05 | 1.367E-05 | 3.215E-06 | 1.000E-05 | 1.600E-05 |
| 6980 | 349 | F349V | A | 1.1e-05 | 1.5e-05 | 1.7e-05 | 1.433E-05 | 3.055E-06 | 1.100E-05 | 1.700E-05 |
| 6981 | 350 | G350A | A | 1.6e-05 | 1.5e-05 | 1e-05 | 1.367E-05 | 3.215E-06 | 1.000E-05 | 1.600E-05 |
| 6982 | 350 | G350R | A | 1.1e-05 | 1.4e-05 | 1e-05 | 1.167E-05 | 2.082E-06 | 1.000E-05 | 1.400E-05 |
| 6983 | 350 | G350N | A | 1.5e-05 | 1.1e-05 | 1.6e-05 | 1.400E-05 | 2.646E-06 | 1.100E-05 | 1.600E-05 |
| 6984 | 350 | G350D | A | 1.4e-05 | 1.4e-05 | 8.4e-06 | 1.213E-05 | 3.233E-06 | 8.400E-06 | 1.400E-05 |
| 6985 | 350 | G350C | A | 1.4e-05 | 1.7e-05 | 1.4e-05 | 1.500E-05 | 1.732E-06 | 1.400E-05 | 1.700E-05 |
| 6986 | 350 | G350E | A | 1.3e-05 | 1e-05 | 9.6e-06 | 1.087E-05 | 1.858E-06 | 9.600E-06 | 1.300E-05 |
| 6987 | 350 | G350Q | A | 8.4e-06 | 9.5e-06 | 8.1e-06 | 8.667E-06 | 7.371E-07 | 8.100E-06 | 9.500E-06 |
| 6988 | 350 | G350G | A | 1.1e-05 | 1.5e-05 | 9.4e-06 | 1.180E-05 | 2.884E-06 | 9.400E-06 | 1.500E-05 |
| 6989 | 350 | G350H | A | 1.5e-05 | 1.3e-05 | 1e-05 | 1.267E-05 | 2.517E-06 | 1.000E-05 | 1.500E-05 |
| 6990 | 350 | G350I | A | 1e-05 | 1.2e-05 | 1.7e-05 | 1.300E-05 | 3.606E-06 | 1.000E-05 | 1.700E-05 |
| 6991 | 350 | G350L | A | 1.1e-05 | 1.2e-05 | 1.2e-05 | 1.167E-05 | 5.774E-07 | 1.100E-05 | 1.200E-05 |
| 6992 | 350 | G350K | A | 1.6e-05 | 9.6e-06 | 1.1e-05 | 1.220E-05 | 3.365E-06 | 9.600E-06 | 1.600E-05 |
| 6993 | 350 | G350M | A | 1.8e-05 | 5.9e-06 | 1.6e-05 | 1.330E-05 | 6.486E-06 | 5.900E-06 | 1.800E-05 |
| 6994 | 350 | G350F | A | 1e-05 | 9.8e-06 | 1.4e-05 | 1.127E-05 | 2.369E-06 | 9.800E-06 | 1.400E-05 |
| 6995 | 350 | G350P | A | 1.2e-05 | 1.4e-05 | 1.8e-05 | 1.467E-05 | 3.055E-06 | 1.200E-05 | 1.800E-05 |
| 6996 | 350 | G350S | A | 1.4e-05 | 1.1e-05 | 1.7e-05 | 1.400E-05 | 3.000E-06 | 1.100E-05 | 1.700E-05 |
| 6997 | 350 | G350T | A | 8.9e-06 | 1.6e-05 | 1.6e-05 | 1.363E-05 | 4.099E-06 | 8.900E-06 | 1.600E-05 |
| 6998 | 350 | G350W | A | 1.4e-05 | 1.5e-05 | 1.5e-05 | 1.467E-05 | 5.774E-07 | 1.400E-05 | 1.500E-05 |
| 6999 | 350 | G350Y | A | 1.5e-05 | 1.3e-05 | 1.5e-05 | 1.433E-05 | 1.155E-06 | 1.300E-05 | 1.500E-05 |
| 7000 | 350 | G350V | A | 1.3e-05 | 1.2e-05 | 1.2e-05 | 1.233E-05 | 5.774E-07 | 1.200E-05 | 1.300E-05 |
| 7001 | 351 | S351A | A | 1.7e-05 | 1.6e-05 | 1e-05 | 1.433E-05 | 3.786E-06 | 1.000E-05 | 1.700E-05 |
| 7002 | 351 | S351R | A | 1.4e-05 | 1.3e-05 | 1.3e-05 | 1.333E-05 | 5.774E-07 | 1.300E-05 | 1.400E-05 |
| 7003 | 351 | S351N | A | 9.8e-06 | 1.1e-05 | 9.3e-06 | 1.003E-05 | 8.737E-07 | 9.300E-06 | 1.100E-05 |
| 7004 | 351 | S351D | A | 1.5e-05 | 2.4e-05 | 1.3e-05 | 1.733E-05 | 5.859E-06 | 1.300E-05 | 2.400E-05 |
| 7005 | 351 | S351C | A | 1.4e-05 | 1.8e-05 | 1e-05 | 1.400E-05 | 4.000E-06 | 1.000E-05 | 1.800E-05 |
| 7006 | 351 | S351E | A | 1.1e-05 | 1.3e-05 | 1.1e-05 | 1.167E-05 | 1.155E-06 | 1.100E-05 | 1.300E-05 |
| 7007 | 351 | S351Q | A | 8.4e-06 | 1.5e-05 | 7.6e-06 | 1.033E-05 | 4.061E-06 | 7.600E-06 | 1.500E-05 |
| 7008 | 351 | S351G | A | 1.5e-05 | 1.5e-05 | 1.4e-05 | 1.467E-05 | 5.774E-07 | 1.400E-05 | 1.500E-05 |
| 7009 | 351 | S351H | A | 1.1e-05 | 1.3e-05 | 1.6e-05 | 1.333E-05 | 2.517E-06 | 1.100E-05 | 1.600E-05 |
| 7010 | 351 | S351I | A | 1.5e-05 | 1.6e-05 | 9.9e-06 | 1.363E-05 | 3.272E-06 | 9.900E-06 | 1.600E-05 |
| 7011 | 351 | S351L | A | 1.6e-05 | 1.5e-05 | 1e-05 | 1.367E-05 | 3.215E-06 | 1.000E-05 | 1.600E-05 |
| 7012 | 351 | S351K | A | 1.3e-05 | 1.5e-05 | 1.6e-05 | 1.467E-05 | 1.528E-06 | 1.300E-05 | 1.600E-05 |
| 7013 | 351 | S351M | A | 1.5e-05 | 1.4e-05 | 1.6e-05 | 1.500E-05 | 1.000E-06 | 1.400E-05 | 1.600E-05 |
| 7014 | 351 | S351F | A | 1.6e-05 | 1.5e-05 | 1e-05 | 1.367E-05 | 3.215E-06 | 1.000E-05 | 1.600E-05 |
| 7015 | 351 | S351P | A | 1.3e-05 | 1.2e-05 | 1.4e-05 | 1.300E-05 | 1.000E-06 | 1.200E-05 | 1.400E-05 |
| 7016 | 351 | S351S | A | 1.5e-05 | 9.3e-06 | 1.4e-05 | 1.277E-05 | 3.044E-06 | 9.300E-06 | 1.500E-05 |
| 7017 | 351 | S351T | A | 9.6e-06 | 1.2e-05 | 1.2e-05 | 1.120E-05 | 1.386E-06 | 9.600E-06 | 1.200E-05 |
| 7018 | 351 | S351W | A | 1.4e-05 | 1.3e-05 | 1.3e-05 | 1.333E-05 | 5.774E-07 | 1.300E-05 | 1.400E-05 |
| 7019 | 351 | S351Y | A | 1.5e-05 | 1.5e-05 | 1.5e-05 | 1.500E-05 | 0.000E+00 | 1.500E-05 | 1.500E-05 |
| 7020 | 351 | S351V | A | 1.6e-05 | 9.8e-06 | 9.4e-06 | 1.173E-05 | 3.700E-06 | 9.400E-06 | 1.600E-05 |
| 7021 | 352 | L352A | A | 1.2e-05 | 1.5e-05 | 1.1e-05 | 1.267E-05 | 2.082E-06 | 1.100E-05 | 1.500E-05 |
| 7022 | 352 | L352R | A | 1.8e-05 | 1.4e-05 | 9.4e-06 | 1.380E-05 | 4.303E-06 | 9.400E-06 | 1.800E-05 |
| 7023 | 352 | L352N | A | 1.4e-05 | 1.5e-05 | 1.1e-05 | 1.333E-05 | 2.082E-06 | 1.100E-05 | 1.500E-05 |
| 7024 | 352 | L352D | A | 1.5e-05 | 1.7e-05 | 8.9e-06 | 1.363E-05 | 4.219E-06 | 8.900E-06 | 1.700E-05 |
| 7025 | 352 | L352C | A | 1.4e-05 | 1.3e-05 | 1.5e-05 | 1.400E-05 | 1.000E-06 | 1.300E-05 | 1.500E-05 |
| 7026 | 352 | L352E | A | 8.9e-06 | 1.3e-05 | 9.7e-06 | 1.053E-05 | 2.173E-06 | 8.900E-06 | 1.300E-05 |
| 7027 | 352 | L352Q | A | 1.5e-05 | 1.1e-05 | 1.4e-05 | 1.333E-05 | 2.082E-06 | 1.100E-05 | 1.500E-05 |
| 7028 | 352 | L352G | A | 8.4e-06 | 1.2e-05 | 1.7e-05 | 1.247E-05 | 4.319E-06 | 8.400E-06 | 1.700E-05 |
| 7029 | 352 | L352H | A | 1.1e-05 | 1e-05 | 8.7e-06 | 9.900E-06 | 1.153E-06 | 8.700E-06 | 1.100E-05 |
| 7030 | 352 | L352I | A | 1.1e-05 | 1.5e-05 | 1.7e-05 | 1.433E-05 | 3.055E-06 | 1.100E-05 | 1.700E-05 |
| 7031 | 352 | L352L | A | 1.4e-05 | 1.1e-05 | 1.5e-05 | 1.333E-05 | 2.082E-06 | 1.100E-05 | 1.500E-05 |
| 7032 | 352 | L352K | A | 1.2e-05 | 9.8e-06 | 1.3e-05 | 1.160E-05 | 1.637E-06 | 9.800E-06 | 1.300E-05 |
| 7033 | 352 | L352M | A | 1.3e-05 | 1.3e-05 | 9.4e-06 | 1.180E-05 | 2.078E-06 | 9.400E-06 | 1.300E-05 |
| 7034 | 352 | L352F | A | 1.5e-05 | 1.6e-05 | 1.4e-05 | 1.500E-05 | 1.000E-06 | 1.400E-05 | 1.600E-05 |
| 7035 | 352 | L352P | A | 1e-05 | 1.4e-05 | 1.3e-05 | 1.233E-05 | 2.082E-06 | 1.000E-05 | 1.400E-05 |
| 7036 | 352 | L352S | A | 1.6e-05 | 1.6e-05 | 1.3e-05 | 1.500E-05 | 1.732E-06 | 1.300E-05 | 1.600E-05 |
| 7037 | 352 | L352T | A | 9.7e-06 | 1e-05 | 1.5e-05 | 1.157E-05 | 2.977E-06 | 9.700E-06 | 1.500E-05 |
| 7038 | 352 | L352W | A | 1.5e-05 | 9.9e-06 | 1.6e-05 | 1.363E-05 | 3.272E-06 | 9.900E-06 | 1.600E-05 |
| 7039 | 352 | L352Y | A | 1.5e-05 | 1.1e-05 | 1e-05 | 1.200E-05 | 2.646E-06 | 1.000E-05 | 1.500E-05 |
| 7040 | 352 | L352V | A | 8.4e-06 | 1.6e-05 | 1.1e-05 | 1.180E-05 | 3.863E-06 | 8.400E-06 | 1.600E-05 |
| 7041 | 353 | A353A | A | 1.9e-05 | 1.3e-05 | 1.5e-05 | 1.567E-05 | 3.055E-06 | 1.300E-05 | 1.900E-05 |
| 7042 | 353 | A353R | A | 1.5e-05 | 1.4e-05 | 1.3e-05 | 1.400E-05 | 1.000E-06 | 1.300E-05 | 1.500E-05 |
| 7043 | 353 | A353N | A | 1e-05 | 1.3e-05 | 1.5e-05 | 1.267E-05 | 2.517E-06 | 1.000E-05 | 1.500E-05 |
| 7044 | 353 | A353D | A | 1e-05 | 1e-05 | 1.5e-05 | 1.167E-05 | 2.887E-06 | 1.000E-05 | 1.500E-05 |
| 7045 | 353 | A353C | A | 5.5e-06 | 1.5e-05 | 1.5e-05 | 1.183E-05 | 5.485E-06 | 5.500E-06 | 1.500E-05 |
| 7046 | 353 | A353E | A | 1.2e-05 | 1.6e-05 | 1.4e-05 | 1.400E-05 | 2.000E-06 | 1.200E-05 | 1.600E-05 |
| 7047 | 353 | A353Q | A | 1.2e-05 | 1.5e-05 | 1.1e-05 | 1.267E-05 | 2.082E-06 | 1.100E-05 | 1.500E-05 |
| 7048 | 353 | A353G | A | 8.9e-06 | 2.4e-05 | 1.7e-05 | 1.663E-05 | 7.557E-06 | 8.900E-06 | 2.400E-05 |
| 7049 | 353 | A353H | A | 1.2e-05 | 9.9e-06 | 9.5e-06 | 1.047E-05 | 1.343E-06 | 9.500E-06 | 1.200E-05 |
| 7050 | 353 | A353I | A | 1.1e-05 | 8.7e-06 | 1.5e-05 | 1.157E-05 | 3.188E-06 | 8.700E-06 | 1.500E-05 |
| 7051 | 353 | A353L | A | 1.6e-05 | 1.7e-05 | 1.5e-05 | 1.600E-05 | 1.000E-06 | 1.500E-05 | 1.700E-05 |
| 7052 | 353 | A353K | A | 1.6e-05 | 1e-05 | 1.4e-05 | 1.333E-05 | 3.055E-06 | 1.000E-05 | 1.600E-05 |
| 7053 | 353 | A353M | A | 1.3e-05 | 1.5e-05 | 9.8e-06 | 1.260E-05 | 2.623E-06 | 9.800E-06 | 1.500E-05 |
| 7054 | 353 | A353F | A | 1e-05 | 1.1e-05 | 9.1e-06 | 1.003E-05 | 9.504E-07 | 9.100E-06 | 1.100E-05 |

| | | | | | | | | | | |
|---|---|---|---|---|---|---|---|---|---|---|
| 7055 | 353 | A353P | A | 1.6e-05 | 1.6e-05 | 1.1e-05 | 1.433E-05 | 2.887E-06 | 1.100E-05 | 1.600E-05 |
| 7056 | 353 | A353S | A | 8.4e-06 | 1.6e-05 | 1.4e-05 | 1.280E-05 | 3.940E-06 | 8.400E-06 | 1.600E-05 |
| 7057 | 353 | A353T | A | 1.1e-05 | 1.5e-05 | 1.6e-05 | 1.400E-05 | 2.646E-06 | 1.100E-05 | 1.600E-05 |
| 7058 | 353 | A353W | A | 1.4e-05 | 7.9e-06 | 1.5e-05 | 1.230E-05 | 3.843E-06 | 7.900E-06 | 1.500E-05 |
| 7059 | 353 | A353Y | A | 9e-06 | 1.4e-05 | 8.2e-06 | 1.040E-05 | 3.143E-06 | 8.200E-06 | 1.400E-05 |
| 7060 | 353 | A353V | A | 1.5e-05 | 1.1e-05 | 1.5e-05 | 1.367E-05 | 2.309E-06 | 1.100E-05 | 1.500E-05 |
| 7061 | 354 | F354A | A | 1.6e-05 | 1.6e-05 | 1.6e-05 | 1.600E-05 | 0.000E+00 | 1.600E-05 | 1.600E-05 |
| 7062 | 354 | F354R | A | 1.1e-05 | 9.2e-06 | 1.5e-05 | 1.173E-05 | 2.969E-06 | 9.200E-06 | 1.500E-05 |
| 7063 | 354 | F354N | A | 1.1e-05 | 1.6e-05 | 1.1e-05 | 1.267E-05 | 2.887E-06 | 1.100E-05 | 1.600E-05 |
| 7064 | 354 | F354D | A | 1.5e-05 | 1.5e-05 | 1.6e-05 | 1.533E-05 | 5.774E-07 | 1.500E-05 | 1.600E-05 |
| 7065 | 354 | F354C | A | 1.6e-05 | 1.6e-05 | 1.6e-05 | 1.600E-05 | 0.000E+00 | 1.600E-05 | 1.600E-05 |
| 7066 | 354 | F354E | A | 1.2e-05 | 1e-05 | 1.6e-05 | 1.267E-05 | 3.055E-06 | 1.000E-05 | 1.600E-05 |
| 7067 | 354 | F354Q | A | 7e-06 | 1.6e-05 | 1.3e-05 | 1.200E-05 | 4.583E-06 | 7.000E-06 | 1.600E-05 |
| 7068 | 354 | F354G | A | 9.7e-06 | 1.5e-05 | 1.2e-05 | 1.223E-05 | 2.658E-06 | 9.700E-06 | 1.500E-05 |
| 7069 | 354 | F354H | A | 1.3e-05 | 1.6e-05 | 1e-05 | 1.300E-05 | 3.000E-06 | 1.000E-05 | 1.600E-05 |
| 7070 | 354 | F354I | A | 1.1e-05 | 1.8e-05 | 1.4e-05 | 1.433E-05 | 3.512E-06 | 1.100E-05 | 1.800E-05 |
| 7071 | 354 | F354L | A | 1.6e-05 | 1.4e-05 | 1.1e-05 | 1.367E-05 | 2.517E-06 | 1.100E-05 | 1.600E-05 |
| 7072 | 354 | F354K | A | 1e-05 | 1.5e-05 | 1.1e-05 | 1.200E-05 | 2.646E-06 | 1.000E-05 | 1.500E-05 |
| 7073 | 354 | F354M | A | 1.6e-05 | 9.7e-06 | 1.6e-05 | 1.390E-05 | 3.637E-06 | 9.700E-06 | 1.600E-05 |
| 7074 | 354 | F354F | A | 1.6e-05 | 1.3e-05 | 1.1e-05 | 1.333E-05 | 2.517E-06 | 1.100E-05 | 1.600E-05 |
| 7075 | 354 | F354P | A | 1.7e-05 | 2.7e-05 | 1.2e-05 | 1.867E-05 | 7.638E-06 | 1.200E-05 | 2.700E-05 |
| 7076 | 354 | F354S | A | 9.6e-06 | 1.1e-05 | 1.2e-05 | 1.087E-05 | 1.206E-06 | 9.600E-06 | 1.200E-05 |
| 7077 | 354 | F354T | A | 1.3e-05 | 1.4e-05 | 1.5e-05 | 1.400E-05 | 1.000E-06 | 1.300E-05 | 1.500E-05 |
| 7078 | 354 | F354W | A | 1e-05 | 8.9e-06 | 8.5e-06 | 9.133E-06 | 7.767E-07 | 8.500E-06 | 1.000E-05 |
| 7079 | 354 | F354Y | A | 1e-05 | 1.5e-05 | 8.1e-06 | 1.103E-05 | 3.564E-06 | 8.100E-06 | 1.500E-05 |
| 7080 | 354 | F354V | A | 1.4e-05 | 1.6e-05 | 1.1e-05 | 1.367E-05 | 2.517E-06 | 1.100E-05 | 1.600E-05 |
| 7081 | 355 | L355A | A | 1.4e-05 | 1.1e-05 | 9.9e-06 | 1.163E-05 | 2.122E-06 | 9.900E-06 | 1.400E-05 |
| 7082 | 355 | L355R | A | 1.7e-05 | 9.6e-06 | 1.4e-05 | 1.353E-05 | 3.722E-06 | 9.600E-06 | 1.700E-05 |
| 7083 | 355 | L355N | A | 1.4e-05 | 7.9e-06 | 1.4e-05 | 1.197E-05 | 3.522E-06 | 7.900E-06 | 1.400E-05 |
| 7084 | 355 | L355D | A | 1.5e-05 | 1.3e-05 | 9.6e-06 | 1.253E-05 | 2.730E-06 | 9.600E-06 | 1.500E-05 |
| 7085 | 355 | L355C | A | 1.4e-05 | 1.3e-05 | 9.6e-06 | 1.220E-05 | 2.307E-06 | 9.600E-06 | 1.400E-05 |
| 7086 | 355 | L355E | A | 9.2e-06 | 1.5e-05 | 1.4e-05 | 1.273E-05 | 3.101E-06 | 9.200E-06 | 1.500E-05 |
| 7087 | 355 | L355Q | A | 1.1e-05 | 1.4e-05 | 1.4e-05 | 1.300E-05 | 1.732E-06 | 1.100E-05 | 1.400E-05 |
| 7088 | 355 | L355G | A | 1.8e-05 | 1.5e-05 | 1.4e-05 | 1.567E-05 | 2.082E-06 | 1.400E-05 | 1.800E-05 |
| 7089 | 355 | L355H | A | 1.4e-05 | 9.8e-06 | 1.1e-05 | 1.160E-05 | 2.163E-06 | 9.800E-06 | 1.400E-05 |
| 7090 | 355 | L355I | A | 1.7e-05 | 1e-05 | 7.9e-06 | 1.163E-05 | 4.765E-06 | 7.900E-06 | 1.700E-05 |
| 7091 | 355 | L355L | A | 1.4e-05 | 1.2e-05 | 1.1e-05 | 1.233E-05 | 1.528E-06 | 1.100E-05 | 1.400E-05 |
| 7092 | 355 | L355K | A | 1.1e-05 | 1e-05 | 1.4e-05 | 1.167E-05 | 2.082E-06 | 1.000E-05 | 1.400E-05 |
| 7093 | 355 | L355M | A | 1.7e-05 | 1.1e-05 | 1.6e-05 | 1.467E-05 | 3.215E-06 | 1.100E-05 | 1.700E-05 |
| 7094 | 355 | L355F | A | 1e-05 | 1.7e-05 | 9.9e-06 | 1.230E-05 | 4.071E-06 | 9.900E-06 | 1.700E-05 |
| 7095 | 355 | L355P | A | 1.5e-05 | 1.6e-05 | 6.1e-06 | 1.237E-05 | 5.450E-06 | 6.100E-06 | 1.600E-05 |
| 7096 | 355 | L355S | A | 9.8e-06 | 1.1e-05 | 1.6e-05 | 1.227E-05 | 3.288E-06 | 9.800E-06 | 1.600E-05 |
| 7097 | 355 | L355T | A | 1.3e-05 | 1.5e-05 | 1.4e-05 | 1.400E-05 | 1.000E-06 | 1.300E-05 | 1.500E-05 |
| 7098 | 355 | L355W | A | 7.3e-06 | 9.9e-06 | 1.1e-05 | 9.400E-06 | 1.900E-06 | 7.300E-06 | 1.100E-05 |
| 7099 | 355 | L355Y | A | 1.6e-05 | 1e-05 | 1.4e-05 | 1.333E-05 | 3.055E-06 | 1.000E-05 | 1.600E-05 |
| 7100 | 355 | L355V | A | 1.4e-05 | 1.5e-05 | 1.5e-05 | 1.467E-05 | 5.774E-07 | 1.400E-05 | 1.500E-05 |
| 7101 | 356 | P356A | A | 1.6e-05 | 6.2e-06 | 1e-05 | 1.073E-05 | 4.941E-06 | 6.200E-06 | 1.600E-05 |
| 7102 | 356 | P356R | A | 1.6e-05 | 1.1e-05 | 9.3e-06 | 1.210E-05 | 3.483E-06 | 9.300E-06 | 1.600E-05 |
| 7103 | 356 | P356N | A | 1.1e-05 | 1e-05 | 1.5e-05 | 1.200E-05 | 2.646E-06 | 1.000E-05 | 1.500E-05 |
| 7104 | 356 | P356D | A | 9.9e-06 | 1.6e-05 | 1.6e-05 | 1.397E-05 | 3.522E-06 | 9.900E-06 | 1.600E-05 |
| 7105 | 356 | P356C | A | 1.2e-05 | 1.4e-05 | 9.8e-06 | 1.193E-05 | 2.101E-06 | 9.800E-06 | 1.400E-05 |
| 7106 | 356 | P356E | A | 1e-05 | 1.6e-05 | 1.3e-05 | 1.300E-05 | 3.000E-06 | 1.000E-05 | 1.600E-05 |
| 7107 | 356 | P356Q | A | 1.6e-05 | 1.1e-05 | 1.4e-05 | 1.367E-05 | 2.517E-06 | 1.100E-05 | 1.600E-05 |
| 7108 | 356 | P356G | A | 9.8e-06 | 1.4e-05 | 1.4e-05 | 1.260E-05 | 2.425E-06 | 9.800E-06 | 1.400E-05 |
| 7109 | 356 | P356H | A | 8.9e-06 | 1.5e-05 | 1e-05 | 1.130E-05 | 3.251E-06 | 8.900E-06 | 1.500E-05 |
| 7110 | 356 | P356I | A | 1.3e-05 | 1.5e-05 | 1.6e-05 | 1.467E-05 | 1.528E-06 | 1.300E-05 | 1.600E-05 |
| 7111 | 356 | P356L | A | 1.6e-05 | 1.3e-05 | 9.4e-06 | 1.280E-05 | 3.305E-06 | 9.400E-06 | 1.600E-05 |
| 7112 | 356 | P356K | A | 1e-05 | 1.4e-05 | 1.3e-05 | 1.233E-05 | 2.082E-06 | 1.000E-05 | 1.400E-05 |
| 7113 | 356 | P356M | A | 1.4e-05 | 1.5e-05 | 1.5e-05 | 1.467E-05 | 5.774E-07 | 1.400E-05 | 1.500E-05 |
| 7114 | 356 | P356F | A | 1.7e-05 | 1.6e-05 | 1.3e-05 | 1.533E-05 | 2.082E-06 | 1.300E-05 | 1.700E-05 |
| 7115 | 356 | P356P | A | 9.7e-06 | 1.4e-05 | 8e-06 | 1.057E-05 | 3.092E-06 | 8.000E-06 | 1.400E-05 |
| 7116 | 356 | P356S | A | 9.7e-06 | 1e-05 | 1.4e-05 | 1.123E-05 | 2.401E-06 | 9.700E-06 | 1.400E-05 |
| 7117 | 356 | P356T | A | 1e-05 | 1.5e-05 | 1.3e-05 | 1.267E-05 | 2.517E-06 | 1.000E-05 | 1.500E-05 |
| 7118 | 356 | P356W | A | 1e-05 | 1.1e-05 | 1.1e-05 | 1.067E-05 | 5.774E-07 | 1.000E-05 | 1.100E-05 |
| 7119 | 356 | P356Y | A | 1.1e-05 | 1.5e-05 | 1.6e-05 | 1.400E-05 | 2.646E-06 | 1.100E-05 | 1.600E-05 |
| 7120 | 356 | P356V | A | 1.4e-05 | 1.5e-05 | 1e-05 | 1.300E-05 | 2.646E-06 | 1.000E-05 | 1.500E-05 |
| 7121 | 357 | E357A | A | 1.6e-05 | 9.9e-06 | 1.2e-05 | 1.263E-05 | 3.099E-06 | 9.900E-06 | 1.600E-05 |
| 7122 | 357 | E357R | A | 1.6e-05 | 1.5e-05 | 1.2e-05 | 1.433E-05 | 2.082E-06 | 1.200E-05 | 1.600E-05 |
| 7123 | 357 | E357N | A | 9.3e-06 | 1.6e-05 | 1.4e-05 | 1.310E-05 | 3.439E-06 | 9.300E-06 | 1.600E-05 |
| 7124 | 357 | E357D | A | 1.4e-05 | 1.3e-05 | 9.5e-06 | 1.217E-05 | 2.363E-06 | 9.500E-06 | 1.400E-05 |
| 7125 | 357 | E357C | A | 1.4e-05 | 1.4e-05 | 1.7e-05 | 1.500E-05 | 1.732E-06 | 1.400E-05 | 1.700E-05 |
| 7126 | 357 | E357E | A | 1.4e-05 | 1.4e-05 | 1.1e-05 | 1.300E-05 | 1.732E-06 | 1.100E-05 | 1.400E-05 |
| 7127 | 357 | E357Q | A | 1e-05 | 1e-05 | 9.7e-06 | 9.900E-06 | 1.732E-07 | 9.700E-06 | 1.000E-05 |
| 7128 | 357 | E357G | A | 1.6e-05 | 1.5e-05 | 1.5e-05 | 1.533E-05 | 5.774E-07 | 1.500E-05 | 1.600E-05 |
| 7129 | 357 | E357H | A | 1.6e-05 | 1.6e-05 | 9.7e-06 | 1.390E-05 | 3.637E-06 | 9.700E-06 | 1.600E-05 |
| 7130 | 357 | E357I | A | 1.4e-05 | 1e-05 | 1.5e-05 | 1.300E-05 | 2.646E-06 | 1.000E-05 | 1.500E-05 |
| 7131 | 357 | E357L | A | 1.5e-05 | 1.6e-05 | 1.4e-05 | 1.500E-05 | 1.000E-06 | 1.400E-05 | 1.600E-05 |
| 7132 | 357 | E357K | A | 1.6e-05 | 8.8e-06 | 1.5e-05 | 1.327E-05 | 3.900E-06 | 8.800E-06 | 1.600E-05 |
| 7133 | 357 | E357M | A | 1e-05 | 1.7e-05 | 9.3e-06 | 1.210E-05 | 4.258E-06 | 9.300E-06 | 1.700E-05 |
| 7134 | 357 | E357F | A | 1.7e-05 | 1.1e-05 | 9.7e-06 | 1.257E-05 | 3.894E-06 | 9.700E-06 | 1.700E-05 |
| 7135 | 357 | E357P | A | 1.1e-05 | 1.2e-05 | 1.6e-05 | 1.300E-05 | 2.646E-06 | 1.100E-05 | 1.600E-05 |
| 7136 | 357 | E357S | A | 1.4e-05 | 1.5e-05 | 1.5e-05 | 1.467E-05 | 5.774E-07 | 1.400E-05 | 1.500E-05 |
| 7137 | 357 | E357T | A | 1.6e-05 | 1.1e-05 | 1.6e-05 | 1.433E-05 | 2.887E-06 | 1.100E-05 | 1.600E-05 |
| 7138 | 357 | E357W | A | 7.9e-06 | 9.7e-06 | 1.3e-05 | 1.020E-05 | 2.587E-06 | 7.900E-06 | 1.300E-05 |
| 7139 | 357 | E357Y | A | 1.4e-05 | 1.4e-05 | 9.5e-06 | 1.250E-05 | 2.598E-06 | 9.500E-06 | 1.400E-05 |

| | | | | | | | | | | |
|---|---|---|---|---|---|---|---|---|---|---|
| 7140 | 357 | E357V | A | 1.4e-05 | 1.2e-05 | 1.4e-05 | 1.333E-05 | 1.155E-06 | 1.200E-05 | 1.400E-05 |
| 7141 | 358 | S358A | A | 1.5e-05 | 1.4e-05 | 1.6e-05 | 1.500E-05 | 1.000E-06 | 1.400E-05 | 1.600E-05 |
| 7142 | 358 | S358R | A | 1.3e-05 | 1.2e-05 | 1.6e-05 | 1.367E-05 | 2.082E-06 | 1.200E-05 | 1.600E-05 |
| 7143 | 358 | S358N | A | 1.2e-05 | 1.1e-05 | 1.3e-05 | 1.200E-05 | 1.000E-06 | 1.100E-05 | 1.300E-05 |
| 7144 | 358 | S358D | A | 1.5e-05 | 9.6e-06 | 1.5e-05 | 1.320E-05 | 3.118E-06 | 9.600E-06 | 1.500E-05 |
| 7145 | 358 | S358C | A | 9.8e-06 | 9.3e-06 | 1.4e-05 | 1.103E-05 | 2.581E-06 | 9.300E-06 | 1.400E-05 |
| 7146 | 358 | S358E | A | 1.5e-05 | 1.2e-05 | 1.2e-05 | 1.300E-05 | 1.732E-06 | 1.200E-05 | 1.500E-05 |
| 7147 | 358 | S358Q | A | 1.1e-05 | 1.1e-05 | 1.1e-05 | 1.100E-05 | 2.075E-21 | 1.100E-05 | 1.100E-05 |
| 7148 | 358 | S358G | A | 9.6e-06 | 1.8e-05 | 8.9e-06 | 1.217E-05 | 5.064E-06 | 8.900E-06 | 1.800E-05 |
| 7149 | 358 | S358H | A | 1.4e-05 | 1.4e-05 | 1.6e-05 | 1.467E-05 | 1.155E-06 | 1.400E-05 | 1.600E-05 |
| 7150 | 358 | S358I | A | 1.6e-05 | 1.6e-05 | 1.1e-05 | 1.433E-05 | 2.887E-06 | 1.100E-05 | 1.600E-05 |
| 7151 | 358 | S358L | A | 1.5e-05 | 9.3e-06 | 1e-05 | 1.143E-05 | 3.109E-06 | 9.300E-06 | 1.500E-05 |
| 7152 | 358 | S358K | A | 1.5e-05 | 9.7e-06 | 1.3e-05 | 1.257E-05 | 2.676E-06 | 9.700E-06 | 1.500E-05 |
| 7153 | 358 | S358M | A | 8.9e-06 | 1e-05 | 1.7e-05 | 1.197E-05 | 4.394E-06 | 8.900E-06 | 1.700E-05 |
| 7154 | 358 | S358F | A | 1e-05 | 9.2e-06 | 9.4e-06 | 9.533E-06 | 4.163E-07 | 9.200E-06 | 1.000E-05 |
| 7155 | 358 | S358P | A | 1.6e-05 | 1.1e-05 | 1e-05 | 1.233E-05 | 3.215E-06 | 1.000E-05 | 1.600E-05 |
| 7156 | 358 | S358S | A | 9.9e-06 | 9.4e-06 | 1.3e-05 | 1.077E-05 | 1.950E-06 | 9.400E-06 | 1.300E-05 |
| 7157 | 358 | S358T | A | 1.5e-05 | 1.6e-05 | 9.5e-06 | 1.350E-05 | 3.500E-06 | 9.500E-06 | 1.600E-05 |
| 7158 | 358 | S358W | A | 1.4e-05 | 1.3e-05 | 1.7e-05 | 1.467E-05 | 2.082E-06 | 1.300E-05 | 1.700E-05 |
| 7159 | 358 | S358Y | A | 8.9e-06 | 1e-05 | 1.5e-05 | 1.130E-05 | 3.251E-06 | 8.900E-06 | 1.500E-05 |
| 7160 | 358 | S358V | A | 1.5e-05 | 1.3e-05 | 1.4e-05 | 1.400E-05 | 1.000E-06 | 1.300E-05 | 1.500E-05 |
| 7161 | 359 | F359A | A | 1.2e-05 | 1e-05 | 1.5e-05 | 1.233E-05 | 2.517E-06 | 1.000E-05 | 1.500E-05 |
| 7162 | 359 | F359R | A | 1.6e-05 | 1.1e-05 | 1e-05 | 1.233E-05 | 3.215E-06 | 1.000E-05 | 1.600E-05 |
| 7163 | 359 | F359N | A | 9.8e-06 | 8.7e-06 | 9.4e-06 | 9.300E-06 | 5.568E-07 | 8.700E-06 | 9.800E-06 |
| 7164 | 359 | F359D | A | 1.6e-05 | 8.1e-06 | 8.9e-06 | 1.100E-05 | 4.349E-06 | 8.100E-06 | 1.600E-05 |
| 7165 | 359 | F359C | A | 1.5e-05 | 1.1e-05 | 1.4e-05 | 1.333E-05 | 2.082E-06 | 1.100E-05 | 1.500E-05 |
| 7166 | 359 | F359E | A | 8.4e-06 | 1e-05 | 1e-05 | 9.467E-06 | 9.238E-07 | 8.400E-06 | 1.000E-05 |
| 7167 | 359 | F359Q | A | 1e-05 | 9.2e-06 | 1.4e-05 | 1.107E-05 | 2.572E-06 | 9.200E-06 | 1.400E-05 |
| 7168 | 359 | F359G | A | 1.3e-05 | 1e-05 | 1.3e-05 | 1.200E-05 | 1.732E-06 | 1.000E-05 | 1.300E-05 |
| 7169 | 359 | F359H | A | 1.3e-05 | 1.2e-05 | 9.2e-06 | 1.140E-05 | 1.970E-06 | 9.200E-06 | 1.300E-05 |
| 7170 | 359 | F359I | A | 1.4e-05 | 1.1e-05 | 1.4e-05 | 1.300E-05 | 1.732E-06 | 1.100E-05 | 1.400E-05 |
| 7171 | 359 | F359L | A | 1.5e-05 | 1.7e-05 | 1.6e-05 | 1.600E-05 | 1.000E-06 | 1.500E-05 | 1.700E-05 |
| 7172 | 359 | F359K | A | 1e-05 | 1.5e-05 | 1e-05 | 1.167E-05 | 2.887E-06 | 1.000E-05 | 1.500E-05 |
| 7173 | 359 | F359M | A | 1.5e-05 | 1.6e-05 | 1.5e-05 | 1.533E-05 | 5.774E-07 | 1.500E-05 | 1.600E-05 |
| 7174 | 359 | F359F | A | 1.6e-05 | 1.6e-05 | 1.2e-05 | 1.467E-05 | 2.309E-06 | 1.200E-05 | 1.600E-05 |
| 7175 | 359 | F359P | A | 1.2e-05 | 9.9e-06 | 1.1e-05 | 1.097E-05 | 1.050E-06 | 9.900E-06 | 1.200E-05 |
| 7176 | 359 | F359S | A | 1.5e-05 | 1.6e-05 | 1.6e-05 | 1.567E-05 | 5.774E-07 | 1.500E-05 | 1.600E-05 |
| 7177 | 359 | F359T | A | 1.6e-05 | 9e-06 | 1e-05 | 1.167E-05 | 3.786E-06 | 9.000E-06 | 1.600E-05 |
| 7178 | 359 | F359W | A | 1.5e-05 | 1e-05 | 1.1e-05 | 1.200E-05 | 2.646E-06 | 1.000E-05 | 1.500E-05 |
| 7179 | 359 | F359Y | A | 9.5e-06 | 1.5e-05 | 1.1e-05 | 1.183E-05 | 2.843E-06 | 9.500E-06 | 1.500E-05 |
| 7180 | 359 | F359V | A | 1.1e-05 | 1.6e-05 | 1.1e-05 | 1.267E-05 | 2.887E-06 | 1.100E-05 | 1.600E-05 |
| 7181 | 360 | D360A | A | 1.5e-05 | 1.4e-05 | 1.4e-05 | 1.433E-05 | 5.774E-07 | 1.400E-05 | 1.500E-05 |
| 7182 | 360 | D360R | A | 1.1e-05 | 1.5e-05 | 1.5e-05 | 1.367E-05 | 2.309E-06 | 1.100E-05 | 1.500E-05 |
| 7183 | 360 | D360N | A | 8.8e-06 | 1.3e-05 | 1.8e-05 | 1.327E-05 | 4.606E-06 | 8.800E-06 | 1.800E-05 |
| 7184 | 360 | D360D | A | 1.1e-05 | 7.6e-06 | 1.6e-05 | 1.153E-05 | 4.225E-06 | 7.600E-06 | 1.600E-05 |
| 7185 | 360 | D360C | A | 1.1e-05 | 1e-05 | 9.3e-06 | 1.010E-05 | 8.544E-07 | 9.300E-06 | 1.100E-05 |
| 7186 | 360 | D360E | A | 1e-05 | 1.3e-05 | 1.4e-05 | 1.233E-05 | 2.082E-06 | 1.000E-05 | 1.400E-05 |
| 7187 | 360 | D360Q | A | 9.4e-06 | 1.1e-05 | 1.4e-05 | 1.147E-05 | 2.335E-06 | 9.400E-06 | 1.400E-05 |
| 7188 | 360 | D360G | A | 9.6e-06 | 1.6e-05 | 1.5e-05 | 1.353E-05 | 3.443E-06 | 9.600E-06 | 1.600E-05 |
| 7189 | 360 | D360H | A | 9.9e-06 | 1.5e-05 | 1.1e-05 | 1.197E-05 | 2.684E-06 | 9.900E-06 | 1.500E-05 |
| 7190 | 360 | D360I | A | 1.7e-05 | 1.6e-05 | 9.1e-06 | 1.403E-05 | 4.302E-06 | 9.100E-06 | 1.700E-05 |
| 7191 | 360 | D360L | A | 1.4e-05 | 1.6e-05 | 1.1e-05 | 1.367E-05 | 2.517E-06 | 1.100E-05 | 1.600E-05 |
| 7192 | 360 | D360K | A | 1.5e-05 | 1.5e-05 | 1e-05 | 1.333E-05 | 2.887E-06 | 1.000E-05 | 1.500E-05 |
| 7193 | 360 | D360M | A | 9.7e-06 | 1.4e-05 | 1e-05 | 1.123E-05 | 2.401E-06 | 9.700E-06 | 1.400E-05 |
| 7194 | 360 | D360F | A | 1.3e-05 | 1.4e-05 | 1.6e-05 | 1.433E-05 | 1.528E-06 | 1.300E-05 | 1.600E-05 |
| 7195 | 360 | D360P | A | 1.6e-05 | 1.2e-05 | 1.8e-05 | 1.533E-05 | 3.055E-06 | 1.200E-05 | 1.800E-05 |
| 7196 | 360 | D360S | A | 1e-05 | 1.3e-05 | 1e-05 | 1.100E-05 | 1.732E-06 | 1.000E-05 | 1.300E-05 |
| 7197 | 360 | D360T | A | 1.4e-05 | 1.5e-05 | 1.5e-05 | 1.467E-05 | 5.774E-07 | 1.400E-05 | 1.500E-05 |
| 7198 | 360 | D360W | A | 1e-05 | 1.2e-05 | 8.9e-06 | 1.030E-05 | 1.572E-06 | 8.900E-06 | 1.200E-05 |
| 7199 | 360 | D360Y | A | 8.3e-06 | 1.1e-05 | 1.1e-05 | 1.010E-05 | 1.559E-06 | 8.300E-06 | 1.100E-05 |
| 7200 | 360 | D360V | A | 1.5e-05 | 1e-05 | 1.3e-05 | 1.267E-05 | 2.517E-06 | 1.000E-05 | 1.500E-05 |
| 7201 | 361 | G361A | A | 1.5e-05 | 1.6e-05 | 1.6e-05 | 1.567E-05 | 5.774E-07 | 1.500E-05 | 1.600E-05 |
| 7202 | 361 | G361R | A | 1e-05 | 1.1e-05 | 1.5e-05 | 1.200E-05 | 2.646E-06 | 1.000E-05 | 1.500E-05 |
| 7203 | 361 | G361N | A | 1.6e-05 | 1.5e-05 | 1.4e-05 | 1.500E-05 | 1.000E-06 | 1.400E-05 | 1.600E-05 |
| 7204 | 361 | G361D | A | 1e-05 | 1.5e-05 | 1.5e-05 | 1.333E-05 | 2.887E-06 | 1.000E-05 | 1.500E-05 |
| 7205 | 361 | G361C | A | 1.5e-05 | 1.6e-05 | 1.4e-05 | 1.500E-05 | 1.000E-06 | 1.400E-05 | 1.600E-05 |
| 7206 | 361 | G361E | A | 1.1e-05 | 1.6e-05 | 1.4e-05 | 1.367E-05 | 2.517E-06 | 1.100E-05 | 1.600E-05 |
| 7207 | 361 | G361Q | A | 9.6e-06 | 1e-05 | 2.5e-05 | 1.487E-05 | 8.778E-06 | 9.600E-06 | 2.500E-05 |
| 7208 | 361 | G361G | A | 9.5e-06 | 1.6e-05 | 1.4e-05 | 1.317E-05 | 3.329E-06 | 9.500E-06 | 1.600E-05 |
| 7209 | 361 | G361H | A | 1.5e-05 | 1e-05 | 1.5e-05 | 1.333E-05 | 2.887E-06 | 1.000E-05 | 1.500E-05 |
| 7210 | 361 | G361I | A | 1.3e-05 | 9.9e-06 | 9.8e-06 | 1.090E-05 | 1.819E-06 | 9.800E-06 | 1.300E-05 |
| 7211 | 361 | G361L | A | 1.5e-05 | 1.1e-05 | 1.2e-05 | 1.267E-05 | 2.082E-06 | 1.100E-05 | 1.500E-05 |
| 7212 | 361 | G361K | A | 1.5e-05 | 1.4e-05 | 1.5e-05 | 1.467E-05 | 5.774E-07 | 1.400E-05 | 1.500E-05 |
| 7213 | 361 | G361M | A | 1.6e-05 | 1.5e-05 | 1e-05 | 1.367E-05 | 3.215E-06 | 1.000E-05 | 1.600E-05 |
| 7214 | 361 | G361F | A | 1.5e-05 | 1.1e-05 | 1.3e-05 | 1.300E-05 | 2.000E-06 | 1.100E-05 | 1.500E-05 |
| 7215 | 361 | G361P | A | 1.1e-05 | 1.5e-05 | 9.6e-06 | 1.187E-05 | 2.802E-06 | 9.600E-06 | 1.500E-05 |
| 7216 | 361 | G361S | A | 1.5e-05 | 1.5e-05 | 1.5e-05 | 1.500E-05 | 0.000E+00 | 1.500E-05 | 1.500E-05 |
| 7217 | 361 | G361T | A | 2.5e-05 | 8.6e-06 | 1.2e-05 | 1.520E-05 | 8.656E-06 | 8.600E-06 | 2.500E-05 |
| 7218 | 361 | G361W | A | 1.5e-05 | 1.1e-05 | 1.5e-05 | 1.367E-05 | 2.309E-06 | 1.100E-05 | 1.500E-05 |
| 7219 | 361 | G361Y | A | 1.3e-05 | 9.5e-06 | 1.1e-05 | 1.117E-05 | 1.756E-06 | 9.500E-06 | 1.300E-05 |
| 7220 | 361 | G361V | A | 1.4e-05 | 9.1e-06 | 1.4e-05 | 1.237E-05 | 2.829E-06 | 9.100E-06 | 1.400E-05 |
| 7221 | 362 | D362A | A | 1e-05 | 1.5e-05 | 1e-05 | 1.167E-05 | 2.887E-06 | 1.000E-05 | 1.500E-05 |
| 7222 | 362 | D362R | A | 1.4e-05 | 1e-05 | 1.5e-05 | 1.300E-05 | 2.646E-06 | 1.000E-05 | 1.500E-05 |
| 7223 | 362 | D362N | A | 1.1e-05 | 1.5e-05 | 1.5e-05 | 1.367E-05 | 2.309E-06 | 1.100E-05 | 1.500E-05 |
| 7224 | 362 | D362D | A | 1.4e-05 | 1.1e-05 | 1.1e-05 | 1.200E-05 | 1.732E-06 | 1.100E-05 | 1.400E-05 |

| | | | | | | | | | | |
|---|---|---|---|---|---|---|---|---|---|---|
| 7225 | 362 | D362C | A | 1.6e-05 | 1.1e-05 | 1.5e-05 | 1.400E-05 | 2.646E-06 | 1.100E-05 | 1.600E-05 |
| 7226 | 362 | D362E | A | 1.9e-05 | 1.6e-05 | 1.4e-05 | 1.633E-05 | 2.517E-06 | 1.400E-05 | 1.900E-05 |
| 7227 | 362 | D362Q | A | 1.2e-05 | 1.4e-05 | 1.6e-05 | 1.400E-05 | 2.000E-06 | 1.200E-05 | 1.600E-05 |
| 7228 | 362 | D362G | A | 1.4e-05 | 1.4e-05 | 1.8e-05 | 1.533E-05 | 2.309E-06 | 1.400E-05 | 1.800E-05 |
| 7229 | 362 | D362H | A | 1.2e-05 | 9.6e-06 | 1.4e-05 | 1.187E-05 | 2.203E-06 | 9.600E-06 | 1.400E-05 |
| 7230 | 362 | D362I | A | 1.6e-05 | 1.2e-05 | 1.1e-05 | 1.300E-05 | 2.646E-06 | 1.100E-05 | 1.600E-05 |
| 7231 | 362 | D362L | A | 1.4e-05 | 1.4e-05 | 1e-05 | 1.267E-05 | 2.309E-06 | 1.000E-05 | 1.400E-05 |
| 7232 | 362 | D362K | A | 1.5e-05 | 1.6e-05 | 2.1e-05 | 1.733E-05 | 3.215E-06 | 1.500E-05 | 2.100E-05 |
| 7233 | 362 | D362M | A | 1.1e-05 | 1.5e-05 | 1.6e-05 | 1.400E-05 | 2.646E-06 | 1.100E-05 | 1.600E-05 |
| 7234 | 362 | D362F | A | 8.6e-06 | 1.5e-05 | 1.6e-05 | 1.320E-05 | 4.015E-06 | 8.600E-06 | 1.600E-05 |
| 7235 | 362 | D362P | A | 8.6e-06 | 2.3e-05 | 9.6e-06 | 1.373E-05 | 8.041E-06 | 8.600E-06 | 2.300E-05 |
| 7236 | 362 | D362S | A | 1e-05 | 1e-05 | 1.6e-05 | 1.200E-05 | 3.464E-06 | 1.000E-05 | 1.600E-05 |
| 7237 | 362 | D362T | A | 1.6e-05 | 1.6e-05 | 1.6e-05 | 1.600E-05 | 0.000E+00 | 1.600E-05 | 1.600E-05 |
| 7238 | 362 | D362W | A | 6.6e-06 | 1.6e-05 | 1.6e-05 | 1.287E-05 | 5.427E-06 | 6.600E-06 | 1.600E-05 |
| 7239 | 362 | D362Y | A | 1.2e-05 | 1.3e-05 | 1.7e-05 | 1.400E-05 | 2.646E-06 | 1.200E-05 | 1.700E-05 |
| 7240 | 362 | D362V | A | 1.5e-05 | 1.6e-05 | 8.9e-06 | 1.330E-05 | 3.843E-06 | 8.900E-06 | 1.600E-05 |
| 7241 | 363 | P363A | A | 1.2e-05 | 1.2e-05 | 1e-05 | 1.133E-05 | 1.155E-06 | 1.000E-05 | 1.200E-05 |
| 7242 | 363 | P363R | A | 1.1e-05 | 1.1e-05 | 1.3e-05 | 1.167E-05 | 1.155E-06 | 1.100E-05 | 1.300E-05 |
| 7243 | 363 | P363N | A | 1.4e-05 | 1.5e-05 | 1.5e-05 | 1.467E-05 | 5.774E-07 | 1.400E-05 | 1.500E-05 |
| 7244 | 363 | P363D | A | 1.5e-05 | 1.6e-05 | 1e-05 | 1.367E-05 | 3.215E-06 | 1.000E-05 | 1.600E-05 |
| 7245 | 363 | P363C | A | 1.5e-05 | 8.7e-06 | 8.1e-06 | 1.060E-05 | 3.822E-06 | 8.100E-06 | 1.500E-05 |
| 7246 | 363 | P363E | A | 1.1e-05 | 1.5e-05 | 9.7e-06 | 1.190E-05 | 2.762E-06 | 9.700E-06 | 1.500E-05 |
| 7247 | 363 | P363Q | A | 1.2e-05 | 1.4e-05 | 1.2e-05 | 1.267E-05 | 1.155E-06 | 1.200E-05 | 1.400E-05 |
| 7248 | 363 | P363G | A | 9.8e-06 | 1.1e-05 | 1.2e-05 | 1.093E-05 | 1.102E-06 | 9.800E-06 | 1.200E-05 |
| 7249 | 363 | P363H | A | 8.8e-06 | 1.5e-05 | 1.1e-05 | 1.160E-05 | 3.143E-06 | 8.800E-06 | 1.500E-05 |
| 7250 | 363 | P363I | A | 1.4e-05 | 8.7e-06 | 1.1e-05 | 1.123E-05 | 2.658E-06 | 8.700E-06 | 1.400E-05 |
| 7251 | 363 | P363L | A | 1.4e-05 | 1.5e-05 | 1.6e-05 | 1.500E-05 | 1.000E-06 | 1.400E-05 | 1.600E-05 |
| 7252 | 363 | P363K | A | 1.6e-05 | 1e-05 | 1.2e-05 | 1.267E-05 | 3.055E-06 | 1.000E-05 | 1.600E-05 |
| 7253 | 363 | P363M | A | 1.1e-05 | 1.3e-05 | 1.2e-05 | 1.200E-05 | 1.000E-06 | 1.100E-05 | 1.300E-05 |
| 7254 | 363 | P363F | A | 1.5e-05 | 1.4e-05 | 9.5e-06 | 1.283E-05 | 2.930E-06 | 9.500E-06 | 1.500E-05 |
| 7255 | 363 | P363P | A | 8.4e-06 | 1.1e-05 | 1.1e-05 | 1.013E-05 | 1.501E-06 | 8.400E-06 | 1.100E-05 |
| 7256 | 363 | P363S | A | 1.6e-05 | 1e-05 | 1.5e-05 | 1.367E-05 | 3.215E-06 | 1.000E-05 | 1.600E-05 |
| 7257 | 363 | P363T | A | 9.9e-06 | 1.1e-05 | 1.2e-05 | 1.097E-05 | 1.050E-06 | 9.900E-06 | 1.200E-05 |
| 7258 | 363 | P363W | A | 1.5e-05 | 1.2e-05 | 1.3e-05 | 1.333E-05 | 1.528E-06 | 1.200E-05 | 1.500E-05 |
| 7259 | 363 | P363Y | A | 1e-05 | 1.3e-05 | 9.9e-06 | 1.097E-05 | 1.762E-06 | 9.900E-06 | 1.300E-05 |
| 7260 | 363 | P363V | A | 1.1e-05 | 9.6e-06 | 1.1e-05 | 1.053E-05 | 8.083E-07 | 9.600E-06 | 1.100E-05 |
| 7261 | 364 | A364A | A | 1.1e-05 | 1.1e-05 | 9.5e-06 | 1.050E-05 | 8.660E-07 | 9.500E-06 | 1.100E-05 |
| 7262 | 364 | A364R | A | 9.6e-06 | 9.9e-06 | 9.5e-06 | 9.667E-06 | 2.082E-07 | 9.500E-06 | 9.900E-06 |
| 7263 | 364 | A364N | A | 7.9e-06 | 1.4e-05 | 1e-05 | 1.063E-05 | 3.099E-06 | 7.900E-06 | 1.400E-05 |
| 7264 | 364 | A364D | A | 1.1e-05 | 1.4e-05 | 8.4e-06 | 1.113E-05 | 2.802E-06 | 8.400E-06 | 1.400E-05 |
| 7265 | 364 | A364C | A | 1e-05 | 1.1e-05 | 6.4e-06 | 9.133E-06 | 2.419E-06 | 6.400E-06 | 1.100E-05 |
| 7266 | 364 | A364E | A | 1.6e-05 | 1.7e-05 | 1e-05 | 1.433E-05 | 3.786E-06 | 1.000E-05 | 1.700E-05 |
| 7267 | 364 | A364Q | A | 1.5e-05 | 1.4e-05 | 1.6e-05 | 1.500E-05 | 1.000E-06 | 1.400E-05 | 1.600E-05 |
| 7268 | 364 | A364G | A | 1.5e-05 | 1.5e-05 | 8.2e-06 | 1.273E-05 | 3.926E-06 | 8.200E-06 | 1.500E-05 |
| 7269 | 364 | A364H | A | 1.2e-05 | 1e-05 | 1.1e-05 | 1.100E-05 | 1.000E-06 | 1.000E-05 | 1.200E-05 |
| 7270 | 364 | A364I | A | 1.5e-05 | 1.5e-05 | 1.6e-05 | 1.533E-05 | 5.774E-07 | 1.500E-05 | 1.600E-05 |
| 7271 | 364 | A364L | A | 1e-05 | 1.1e-05 | 1.9e-05 | 1.333E-05 | 4.933E-06 | 1.000E-05 | 1.900E-05 |
| 7272 | 364 | A364K | A | 1.6e-05 | 1e-05 | 1.1e-05 | 1.233E-05 | 3.215E-06 | 1.000E-05 | 1.600E-05 |
| 7273 | 364 | A364M | A | 1e-05 | 1e-05 | 1.5e-05 | 1.167E-05 | 2.887E-06 | 1.000E-05 | 1.500E-05 |
| 7274 | 364 | A364F | A | 1.5e-05 | 1.5e-05 | 1e-05 | 1.333E-05 | 2.887E-06 | 1.000E-05 | 1.500E-05 |
| 7275 | 364 | A364P | A | 1.6e-05 | 1.3e-05 | 1.7e-05 | 1.533E-05 | 2.082E-06 | 1.300E-05 | 1.700E-05 |
| 7276 | 364 | A364S | A | 9.5e-06 | 1.3e-05 | 1e-05 | 1.083E-05 | 1.893E-06 | 9.500E-06 | 1.300E-05 |
| 7277 | 364 | A364T | A | 1.5e-05 | 8.7e-06 | 1.3e-05 | 1.223E-05 | 3.219E-06 | 8.700E-06 | 1.500E-05 |
| 7278 | 364 | A364W | A | 9.3e-06 | 1.5e-05 | 9.3e-06 | 1.120E-05 | 3.291E-06 | 9.300E-06 | 1.500E-05 |
| 7279 | 364 | A364Y | A | 1e-05 | 1.1e-05 | 1.5e-05 | 1.200E-05 | 2.646E-06 | 1.000E-05 | 1.500E-05 |
| 7280 | 364 | A364V | A | 1.3e-05 | 1.6e-05 | 1.4e-05 | 1.433E-05 | 1.528E-06 | 1.300E-05 | 1.600E-05 |
| 7281 | 365 | S365A | A | 1e-05 | 8.7e-06 | 1.4e-05 | 1.090E-05 | 2.762E-06 | 8.700E-06 | 1.400E-05 |
| 7282 | 365 | S365R | A | 9.5e-06 | 1.7e-05 | 1.5e-05 | 1.383E-05 | 3.884E-06 | 9.500E-06 | 1.700E-05 |
| 7283 | 365 | S365N | A | 1.5e-05 | 1.6e-05 | 1.2e-05 | 1.433E-05 | 2.082E-06 | 1.200E-05 | 1.600E-05 |
| 7284 | 365 | S365D | A | 9.2e-06 | 1e-05 | 1.4e-05 | 1.107E-05 | 2.572E-06 | 9.200E-06 | 1.400E-05 |
| 7285 | 365 | S365C | A | 1.5e-05 | 1.3e-05 | 1.4e-05 | 1.400E-05 | 1.000E-06 | 1.300E-05 | 1.500E-05 |
| 7286 | 365 | S365E | A | 1.4e-05 | 1.2e-05 | 1.5e-05 | 1.367E-05 | 1.528E-06 | 1.200E-05 | 1.500E-05 |
| 7287 | 365 | S365Q | A | 1.3e-05 | 1.5e-05 | 1.4e-05 | 1.400E-05 | 1.000E-06 | 1.300E-05 | 1.500E-05 |
| 7288 | 365 | S365G | A | 1.6e-05 | 1.6e-05 | 1e-05 | 1.400E-05 | 3.464E-06 | 1.000E-05 | 1.600E-05 |
| 7289 | 365 | S365H | A | 1.4e-05 | 1.4e-05 | 7.2e-06 | 1.173E-05 | 3.926E-06 | 7.200E-06 | 1.400E-05 |
| 7290 | 365 | S365I | A | 1.2e-05 | 1.2e-05 | 1.3e-05 | 1.233E-05 | 5.774E-07 | 1.200E-05 | 1.300E-05 |
| 7291 | 365 | S365L | A | 1.7e-05 | 9.3e-06 | 1e-05 | 1.210E-05 | 4.258E-06 | 9.300E-06 | 1.700E-05 |
| 7292 | 365 | S365K | A | 1.7e-05 | 9.5e-06 | 8.9e-06 | 1.180E-05 | 4.513E-06 | 8.900E-06 | 1.700E-05 |
| 7293 | 365 | S365M | A | 1.5e-05 | 1.7e-05 | 1e-05 | 1.400E-05 | 3.606E-06 | 1.000E-05 | 1.700E-05 |
| 7294 | 365 | S365F | A | 1.7e-05 | 9.6e-06 | 1.5e-05 | 1.387E-05 | 3.828E-06 | 9.600E-06 | 1.700E-05 |
| 7295 | 365 | S365P | A | 1.4e-05 | 1.6e-05 | 9.7e-06 | 1.323E-05 | 3.219E-06 | 9.700E-06 | 1.600E-05 |
| 7296 | 365 | S365S | A | 1.7e-05 | 1.4e-05 | 1.1e-05 | 1.400E-05 | 3.000E-06 | 1.100E-05 | 1.700E-05 |
| 7297 | 365 | S365T | A | 1.6e-05 | 1.7e-05 | 1.5e-05 | 1.600E-05 | 1.000E-06 | 1.500E-05 | 1.700E-05 |
| 7298 | 365 | S365W | A | 9.8e-06 | 9.8e-06 | 1.3e-05 | 1.087E-05 | 1.848E-06 | 9.800E-06 | 1.300E-05 |
| 7299 | 365 | S365Y | A | 1.6e-05 | 8.5e-06 | 1.5e-05 | 1.317E-05 | 4.072E-06 | 8.500E-06 | 1.600E-05 |
| 7300 | 365 | S365V | A | 1.1e-05 | 1.4e-05 | 1.2e-05 | 1.233E-05 | 1.528E-06 | 1.100E-05 | 1.400E-05 |
| 7301 | 366 | N366A | A | 1e-05 | 9.9e-06 | 1e-05 | 9.967E-06 | 5.774E-08 | 9.900E-06 | 1.000E-05 |
| 7302 | 366 | N366R | A | 1.5e-05 | 2.9e-05 | 1e-05 | 1.800E-05 | 9.849E-06 | 1.000E-05 | 2.900E-05 |
| 7303 | 366 | N366N | A | 1.5e-05 | 1.5e-05 | 1.6e-05 | 1.533E-05 | 5.774E-07 | 1.500E-05 | 1.600E-05 |
| 7304 | 366 | N366D | A | 1.1e-05 | 1e-05 | 1e-05 | 1.033E-05 | 5.774E-07 | 1.000E-05 | 1.100E-05 |
| 7305 | 366 | N366C | A | 1e-05 | 1.7e-05 | 1.2e-05 | 1.300E-05 | 3.606E-06 | 1.000E-05 | 1.700E-05 |
| 7306 | 366 | N366E | A | 8.5e-06 | 1.4e-05 | 1.2e-05 | 1.150E-05 | 2.784E-06 | 8.500E-06 | 1.400E-05 |
| 7307 | 366 | N366Q | A | 1.2e-05 | 8.9e-06 | 1.3e-05 | 1.130E-05 | 2.138E-06 | 8.900E-06 | 1.300E-05 |
| 7308 | 366 | N366G | A | 1.7e-05 | 1.7e-05 | 1.1e-05 | 1.500E-05 | 3.464E-06 | 1.100E-05 | 1.700E-05 |
| 7309 | 366 | N366H | A | 1.5e-05 | 1.4e-05 | 8.2e-06 | 1.240E-05 | 3.672E-06 | 8.200E-06 | 1.500E-05 |

| | | | | | | | | | | |
|---|---|---|---|---|---|---|---|---|---|---|
| 7310 | 366 | N366I | A | 1.5e-05 | 1.5e-05 | 8.4e-06 | 1.280E-05 | 3.811E-06 | 8.400E-06 | 1.500E-05 |
| 7311 | 366 | N366L | A | 1.4e-05 | 1.5e-05 | 1.7e-05 | 1.533E-05 | 1.528E-06 | 1.400E-05 | 1.700E-05 |
| 7312 | 366 | N366K | A | 1.1e-05 | 9.8e-06 | 1.6e-05 | 1.227E-05 | 3.288E-06 | 9.800E-06 | 1.600E-05 |
| 7313 | 366 | N366M | A | 7.7e-06 | 1.5e-05 | 1.1e-05 | 1.123E-05 | 3.656E-06 | 7.700E-06 | 1.500E-05 |
| 7314 | 366 | N366F | A | 1.2e-05 | 9.4e-06 | 1.2e-05 | 1.113E-05 | 1.501E-06 | 9.400E-06 | 1.200E-05 |
| 7315 | 366 | N366P | A | 1.3e-05 | 1.7e-05 | 1e-05 | 1.333E-05 | 3.512E-06 | 1.000E-05 | 1.700E-05 |
| 7316 | 366 | N366S | A | 1.5e-05 | 1.6e-05 | 1.5e-05 | 1.533E-05 | 5.774E-07 | 1.500E-05 | 1.600E-05 |
| 7317 | 366 | N366T | A | 1e-05 | 1e-05 | 1.6e-05 | 1.200E-05 | 3.464E-06 | 1.000E-05 | 1.600E-05 |
| 7318 | 366 | N366W | A | 1.1e-05 | 1e-05 | 1e-05 | 1.033E-05 | 5.774E-07 | 1.000E-05 | 1.100E-05 |
| 7319 | 366 | N366Y | A | 1.5e-05 | 1.5e-05 | 1.6e-05 | 1.533E-05 | 5.774E-07 | 1.500E-05 | 1.600E-05 |
| 7320 | 366 | N366V | A | 1.6e-05 | 1.3e-05 | 9.2e-06 | 1.273E-05 | 3.408E-06 | 9.200E-06 | 1.600E-05 |
| 7321 | 367 | T367A | A | 1.6e-05 | 1.3e-05 | 1.5e-05 | 1.467E-05 | 1.528E-06 | 1.300E-05 | 1.600E-05 |
| 7322 | 367 | T367R | A | 9.8e-06 | 1.1e-05 | 1.1e-05 | 1.060E-05 | 6.928E-07 | 9.800E-06 | 1.100E-05 |
| 7323 | 367 | T367N | A | 1e-05 | 9e-06 | 1.6e-05 | 1.167E-05 | 3.786E-06 | 9.000E-06 | 1.600E-05 |
| 7324 | 367 | T367D | A | 1e-05 | 9.9e-06 | 1.5e-05 | 1.163E-05 | 2.916E-06 | 9.900E-06 | 1.500E-05 |
| 7325 | 367 | T367C | A | 1.4e-05 | 7.1e-06 | 1.2e-05 | 1.103E-05 | 3.550E-06 | 7.100E-06 | 1.400E-05 |
| 7326 | 367 | T367E | A | 1.6e-05 | 1.5e-05 | 1.6e-05 | 1.567E-05 | 5.774E-07 | 1.500E-05 | 1.600E-05 |
| 7327 | 367 | T367Q | A | 1.4e-05 | 1.5e-05 | 9.9e-06 | 1.297E-05 | 2.702E-06 | 9.900E-06 | 1.500E-05 |
| 7328 | 367 | T367G | A | 1.5e-05 | 9.8e-06 | 1.8e-05 | 1.427E-05 | 4.149E-06 | 9.800E-06 | 1.800E-05 |
| 7329 | 367 | T367H | A | 1.5e-05 | 1.6e-05 | 1.3e-05 | 1.467E-05 | 1.528E-06 | 1.300E-05 | 1.600E-05 |
| 7330 | 367 | T367I | A | 9.6e-06 | 1.1e-05 | 1.3e-05 | 1.120E-05 | 1.709E-06 | 9.600E-06 | 1.300E-05 |
| 7331 | 367 | T367L | A | 1.6e-05 | 1.2e-05 | 1.5e-05 | 1.433E-05 | 2.082E-06 | 1.200E-05 | 1.600E-05 |
| 7332 | 367 | T367K | A | 1.7e-05 | 1.1e-05 | 1e-05 | 1.267E-05 | 3.786E-06 | 1.000E-05 | 1.700E-05 |
| 7333 | 367 | T367M | A | 1.8e-05 | 1e-05 | 1.6e-05 | 1.467E-05 | 4.163E-06 | 1.000E-05 | 1.800E-05 |
| 7334 | 367 | T367F | A | 1.4e-05 | 1.7e-05 | 1.4e-05 | 1.500E-05 | 1.732E-06 | 1.400E-05 | 1.700E-05 |
| 7335 | 367 | T367P | A | 1.7e-05 | 1.7e-05 | 1.5e-05 | 1.633E-05 | 1.155E-06 | 1.500E-05 | 1.700E-05 |
| 7336 | 367 | T367S | A | 6.7e-06 | 1e-05 | 8.8e-06 | 8.500E-06 | 1.670E-06 | 6.700E-06 | 1.000E-05 |
| 7337 | 367 | T367T | A | 1.5e-05 | 1.4e-05 | 1.5e-05 | 1.467E-05 | 5.774E-07 | 1.400E-05 | 1.500E-05 |
| 7338 | 367 | T367W | A | 1.6e-05 | 1.1e-05 | 9.9e-06 | 1.230E-05 | 3.251E-06 | 9.900E-06 | 1.600E-05 |
| 7339 | 367 | T367Y | A | 1.1e-05 | 1.4e-05 | 1.5e-05 | 1.333E-05 | 2.082E-06 | 1.100E-05 | 1.500E-05 |
| 7340 | 367 | T367V | A | 1.3e-05 | 1.1e-05 | 1.3e-05 | 1.233E-05 | 1.155E-06 | 1.100E-05 | 1.300E-05 |
| 7341 | 368 | A368A | A | 9.3e-06 | 1.1e-05 | 1.4e-05 | 1.143E-05 | 2.380E-06 | 9.300E-06 | 1.400E-05 |
| 7342 | 368 | A368R | A | 1.1e-05 | 5.9e-06 | 1.4e-05 | 1.030E-05 | 4.095E-06 | 5.900E-06 | 1.400E-05 |
| 7343 | 368 | A368N | A | 8.3e-06 | 1e-05 | 9.5e-06 | 9.267E-06 | 8.737E-07 | 8.300E-06 | 1.000E-05 |
| 7344 | 368 | A368D | A | 9.7e-06 | 1.2e-05 | 1e-05 | 1.057E-05 | 1.250E-06 | 9.700E-06 | 1.200E-05 |
| 7345 | 368 | A368C | A | 1.5e-05 | 9.5e-06 | 1.5e-05 | 1.317E-05 | 3.175E-06 | 9.500E-06 | 1.500E-05 |
| 7346 | 368 | A368E | A | 1.6e-05 | 9.9e-06 | 1.3e-05 | 1.297E-05 | 3.050E-06 | 9.900E-06 | 1.600E-05 |
| 7347 | 368 | A368Q | A | 1.5e-05 | 1.5e-05 | 1.3e-05 | 1.433E-05 | 1.155E-06 | 1.300E-05 | 1.500E-05 |
| 7348 | 368 | A368G | A | 9.6e-06 | 1.6e-05 | 1.4e-05 | 1.320E-05 | 3.274E-06 | 9.600E-06 | 1.600E-05 |
| 7349 | 368 | A368H | A | 1.1e-05 | 9.4e-06 | 1.5e-05 | 1.180E-05 | 2.884E-06 | 9.400E-06 | 1.500E-05 |
| 7350 | 368 | A368I | A | 1.6e-05 | 1.4e-05 | 1.3e-05 | 1.433E-05 | 1.528E-06 | 1.300E-05 | 1.600E-05 |
| 7351 | 368 | A368L | A | 1.5e-05 | 1.4e-05 | 1.4e-05 | 1.433E-05 | 5.774E-07 | 1.400E-05 | 1.500E-05 |
| 7352 | 368 | A368K | A | 1e-05 | 1.3e-05 | 1e-05 | 1.100E-05 | 1.732E-06 | 1.000E-05 | 1.300E-05 |
| 7353 | 368 | A368M | A | 1e-05 | 1.5e-05 | 1.1e-05 | 1.200E-05 | 2.646E-06 | 1.000E-05 | 1.500E-05 |
| 7354 | 368 | A368F | A | 1.5e-05 | 9.5e-06 | 1.2e-05 | 1.217E-05 | 2.754E-06 | 9.500E-06 | 1.500E-05 |
| 7355 | 368 | A368P | A | 1.1e-05 | 1.5e-05 | 1.2e-05 | 1.267E-05 | 2.082E-06 | 1.100E-05 | 1.500E-05 |
| 7356 | 368 | A368S | A | 8.4e-06 | 1.1e-05 | 1.3e-05 | 1.080E-05 | 2.307E-06 | 8.400E-06 | 1.300E-05 |
| 7357 | 368 | A368T | A | 9.9e-06 | 1.6e-05 | 1.2e-05 | 1.263E-05 | 3.099E-06 | 9.900E-06 | 1.600E-05 |
| 7358 | 368 | A368W | A | 1.3e-05 | 1.4e-05 | 1e-05 | 1.233E-05 | 2.082E-06 | 1.000E-05 | 1.400E-05 |
| 7359 | 368 | A368Y | A | 8.9e-06 | 1.4e-05 | 1.5e-05 | 1.263E-05 | 3.272E-06 | 8.900E-06 | 1.500E-05 |
| 7360 | 368 | A368V | A | 1.6e-05 | 8.8e-06 | 1e-05 | 1.160E-05 | 3.857E-06 | 8.800E-06 | 1.600E-05 |
| 7361 | 369 | P369A | A | 9.8e-06 | 1.5e-05 | 7.4e-06 | 1.073E-05 | 3.885E-06 | 7.400E-06 | 1.500E-05 |
| 7362 | 369 | P369R | A | 1.6e-05 | 1.1e-05 | 1.2e-05 | 1.300E-05 | 2.646E-06 | 1.100E-05 | 1.600E-05 |
| 7363 | 369 | P369N | A | 1e-05 | 1.6e-05 | 1.1e-05 | 1.233E-05 | 3.215E-06 | 1.000E-05 | 1.600E-05 |
| 7364 | 369 | P369D | A | 1.5e-05 | 1.4e-05 | 1e-05 | 1.300E-05 | 2.646E-06 | 1.000E-05 | 1.500E-05 |
| 7365 | 369 | P369C | A | 1.3e-05 | 1.4e-05 | 6.7e-06 | 1.123E-05 | 3.958E-06 | 6.700E-06 | 1.400E-05 |
| 7366 | 369 | P369E | A | 1.5e-05 | 1.5e-05 | 1.6e-05 | 1.533E-05 | 5.774E-07 | 1.500E-05 | 1.600E-05 |
| 7367 | 369 | P369Q | A | 1.6e-05 | 1e-05 | 1.2e-05 | 1.267E-05 | 3.055E-06 | 1.000E-05 | 1.600E-05 |
| 7368 | 369 | P369G | A | 1.4e-05 | 1e-05 | 1.7e-05 | 1.367E-05 | 3.512E-06 | 1.000E-05 | 1.700E-05 |
| 7369 | 369 | P369H | A | 1.3e-05 | 8.2e-06 | 8.8e-06 | 1.000E-05 | 2.615E-06 | 8.200E-06 | 1.300E-05 |
| 7370 | 369 | P369I | A | 1.4e-05 | 1.5e-05 | 1.7e-05 | 1.533E-05 | 1.528E-06 | 1.400E-05 | 1.700E-05 |
| 7371 | 369 | P369L | A | 1.4e-05 | 1.6e-05 | 1.8e-05 | 1.600E-05 | 2.000E-06 | 1.400E-05 | 1.800E-05 |
| 7372 | 369 | P369K | A | 1e-05 | 9.9e-06 | 1.1e-05 | 1.030E-05 | 6.083E-07 | 9.900E-06 | 1.100E-05 |
| 7373 | 369 | P369M | A | 1.5e-05 | 1.2e-05 | 9.8e-06 | 1.227E-05 | 2.610E-06 | 9.800E-06 | 1.500E-05 |
| 7374 | 369 | P369F | A | 1.4e-05 | 1.6e-05 | 9.2e-06 | 1.307E-05 | 3.495E-06 | 9.200E-06 | 1.600E-05 |
| 7375 | 369 | P369P | A | 9.7e-06 | 1.5e-05 | 1.5e-05 | 1.323E-05 | 3.060E-06 | 9.700E-06 | 1.500E-05 |
| 7376 | 369 | P369S | A | 1.4e-05 | 1.3e-05 | 1e-05 | 1.233E-05 | 2.082E-06 | 1.000E-05 | 1.400E-05 |
| 7377 | 369 | P369T | A | 1.5e-05 | 1.4e-05 | 1.2e-05 | 1.367E-05 | 1.528E-06 | 1.200E-05 | 1.500E-05 |
| 7378 | 369 | P369W | A | 1.6e-05 | 8.6e-06 | 1.4e-05 | 1.287E-05 | 3.828E-06 | 8.600E-06 | 1.600E-05 |
| 7379 | 369 | P369Y | A | 1.6e-05 | 8.6e-06 | 1e-05 | 1.153E-05 | 3.931E-06 | 8.600E-06 | 1.600E-05 |
| 7380 | 369 | P369V | A | 1.2e-05 | 1.5e-05 | 1.4e-05 | 1.367E-05 | 1.528E-06 | 1.200E-05 | 1.500E-05 |
| 7381 | 370 | L370A | A | 1.6e-05 | 1.5e-05 | 1e-05 | 1.367E-05 | 3.215E-06 | 1.000E-05 | 1.600E-05 |
| 7382 | 370 | L370R | A | 1.3e-05 | 1.1e-05 | 1.5e-05 | 1.300E-05 | 2.000E-06 | 1.100E-05 | 1.500E-05 |
| 7383 | 370 | L370N | A | 1.4e-05 | 1.1e-05 | 1.6e-05 | 1.367E-05 | 2.517E-06 | 1.100E-05 | 1.600E-05 |
| 7384 | 370 | L370D | A | 1.1e-05 | 1.5e-05 | 1e-05 | 1.200E-05 | 2.646E-06 | 1.000E-05 | 1.500E-05 |
| 7385 | 370 | L370C | A | 8.9e-06 | 8.6e-06 | 1.1e-05 | 9.500E-06 | 1.308E-06 | 8.600E-06 | 1.100E-05 |
| 7386 | 370 | L370E | A | 8.3e-06 | 1.6e-05 | 1.1e-05 | 1.177E-05 | 3.907E-06 | 8.300E-06 | 1.600E-05 |
| 7387 | 370 | L370Q | A | 1.2e-05 | 1.5e-05 | 6.9e-06 | 1.130E-05 | 4.095E-06 | 6.900E-06 | 1.500E-05 |
| 7388 | 370 | L370G | A | 1.5e-05 | 1e-05 | 1.5e-05 | 1.333E-05 | 2.887E-06 | 1.000E-05 | 1.500E-05 |
| 7389 | 370 | L370H | A | 1.2e-05 | 9.1e-06 | 7.9e-06 | 9.667E-06 | 2.108E-06 | 7.900E-06 | 1.200E-05 |
| 7390 | 370 | L370I | A | 1.5e-05 | 1.1e-05 | 1.5e-05 | 1.367E-05 | 2.309E-06 | 1.100E-05 | 1.500E-05 |
| 7391 | 370 | L370L | A | 4.6e-05 | 1.2e-05 | 1e-05 | 2.267E-05 | 2.023E-05 | 1.000E-05 | 4.600E-05 |
| 7392 | 370 | L370K | A | 1.5e-05 | 1.1e-05 | 8.6e-06 | 1.153E-05 | 3.233E-06 | 8.600E-06 | 1.500E-05 |
| 7393 | 370 | L370M | A | 1.3e-05 | 8.7e-06 | 1.6e-05 | 1.257E-05 | 3.669E-06 | 8.700E-06 | 1.600E-05 |
| 7394 | 370 | L370F | A | 1.1e-05 | 1.1e-05 | 8.9e-06 | 1.030E-05 | 1.212E-06 | 8.900E-06 | 1.100E-05 |

| | | | | | | | | | | |
|---|---|---|---|---|---|---|---|---|---|---|
| 7395 | 370 | L370P | A | 1.6e-05 | 1.1e-05 | 1.1e-05 | 1.267E-05 | 2.887E-06 | 1.100E-05 | 1.600E-05 |
| 7396 | 370 | L370S | A | 1.6e-05 | 9.8e-06 | 1.1e-05 | 1.227E-05 | 3.288E-06 | 9.800E-06 | 1.600E-05 |
| 7397 | 370 | L370T | A | 1.1e-05 | 1.2e-05 | 9.6e-06 | 1.087E-05 | 1.206E-06 | 9.600E-06 | 1.200E-05 |
| 7398 | 370 | L370W | A | 1e-05 | 1.2e-05 | 7.2e-06 | 9.733E-06 | 2.411E-06 | 7.200E-06 | 1.200E-05 |
| 7399 | 370 | L370Y | A | 1.5e-05 | 1.5e-05 | 9.8e-06 | 1.327E-05 | 3.002E-06 | 9.800E-06 | 1.500E-05 |
| 7400 | 370 | L370V | A | 1.1e-05 | 1.3e-05 | 1.5e-05 | 1.300E-05 | 2.000E-06 | 1.100E-05 | 1.500E-05 |
| 7401 | 371 | Q371A | A | 1.7e-05 | 1.7e-05 | 1.1e-05 | 1.500E-05 | 3.464E-06 | 1.100E-05 | 1.700E-05 |
| 7402 | 371 | Q371R | A | 1.4e-05 | 1.7e-05 | 1.7e-05 | 1.600E-05 | 1.732E-06 | 1.400E-05 | 1.700E-05 |
| 7403 | 371 | Q371N | A | 9.8e-06 | 1.7e-05 | 1.5e-05 | 1.393E-05 | 3.717E-06 | 9.800E-06 | 1.700E-05 |
| 7404 | 371 | Q371D | A | 1.4e-05 | 8.2e-06 | 9.8e-06 | 1.067E-05 | 2.996E-06 | 8.200E-06 | 1.400E-05 |
| 7405 | 371 | Q371C | A | 1e-05 | 1.1e-05 | 1e-05 | 1.033E-05 | 5.774E-07 | 1.000E-05 | 1.100E-05 |
| 7406 | 371 | Q371E | A | 1e-05 | 1.2e-05 | 1.6e-05 | 1.267E-05 | 3.055E-06 | 1.000E-05 | 1.600E-05 |
| 7407 | 371 | Q371Q | A | 9.7e-06 | 1e-05 | 1.6e-05 | 1.190E-05 | 3.554E-06 | 9.700E-06 | 1.600E-05 |
| 7408 | 371 | Q371G | A | 9.3e-06 | 1.5e-05 | 1.1e-05 | 1.177E-05 | 2.926E-06 | 9.300E-06 | 1.500E-05 |
| 7409 | 371 | Q371H | A | 1.2e-05 | 1.4e-05 | 1.5e-05 | 1.367E-05 | 1.528E-06 | 1.200E-05 | 1.500E-05 |
| 7410 | 371 | Q371I | A | 1.5e-05 | 1.5e-05 | 1.6e-05 | 1.533E-05 | 5.774E-07 | 1.500E-05 | 1.600E-05 |
| 7411 | 371 | Q371L | A | 1.1e-05 | 1.6e-05 | 1.3e-05 | 1.333E-05 | 2.517E-06 | 1.100E-05 | 1.600E-05 |
| 7412 | 371 | Q371K | A | 1.5e-05 | 2.5e-05 | 1.5e-05 | 1.833E-05 | 5.774E-06 | 1.500E-05 | 2.500E-05 |
| 7413 | 371 | Q371M | A | 1.1e-05 | 1.6e-05 | 9.8e-06 | 1.227E-05 | 3.288E-06 | 9.800E-06 | 1.600E-05 |
| 7414 | 371 | Q371F | A | 1.7e-05 | 1.5e-05 | 1.1e-05 | 1.433E-05 | 3.055E-06 | 1.100E-05 | 1.700E-05 |
| 7415 | 371 | Q371P | A | 1.5e-05 | 1.1e-05 | 1.4e-05 | 1.333E-05 | 2.082E-06 | 1.100E-05 | 1.500E-05 |
| 7416 | 371 | Q371S | A | 8.7e-06 | 9e-06 | 1.6e-05 | 1.123E-05 | 4.131E-06 | 8.700E-06 | 1.600E-05 |
| 7417 | 371 | Q371T | A | 1.6e-05 | 1.2e-05 | 1.5e-05 | 1.433E-05 | 2.082E-06 | 1.200E-05 | 1.600E-05 |
| 7418 | 371 | Q371W | A | 1.1e-05 | 1e-05 | 1.3e-05 | 1.133E-05 | 1.528E-06 | 1.000E-05 | 1.300E-05 |
| 7419 | 371 | Q371Y | A | 1.1e-05 | 1.1e-05 | 1.3e-05 | 1.167E-05 | 1.155E-06 | 1.100E-05 | 1.300E-05 |
| 7420 | 371 | Q371V | A | 1.4e-05 | 1.5e-05 | 1.6e-05 | 1.500E-05 | 1.000E-06 | 1.400E-05 | 1.600E-05 |
| 7421 | 372 | P372A | A | 1e-05 | 1.6e-05 | 1.3e-05 | 1.300E-05 | 3.000E-06 | 1.000E-05 | 1.600E-05 |
| 7422 | 372 | P372R | A | 1.3e-05 | 1.6e-05 | 1.4e-05 | 1.433E-05 | 1.528E-06 | 1.300E-05 | 1.600E-05 |
| 7423 | 372 | P372N | A | 1.3e-05 | 1.3e-05 | 1.4e-05 | 1.333E-05 | 5.774E-07 | 1.300E-05 | 1.400E-05 |
| 7424 | 372 | P372D | A | 8.4e-06 | 8.9e-06 | 9.9e-06 | 9.067E-06 | 7.638E-07 | 8.400E-06 | 9.900E-06 |
| 7425 | 372 | P372C | A | 1e-05 | 1.6e-05 | 1.4e-05 | 1.333E-05 | 3.055E-06 | 1.000E-05 | 1.600E-05 |
| 7426 | 372 | P372E | A | 1e-05 | 1.4e-05 | 9.8e-06 | 1.127E-05 | 2.369E-06 | 9.800E-06 | 1.400E-05 |
| 7427 | 372 | P372Q | A | 1.5e-05 | 1.5e-05 | 1.2e-05 | 1.400E-05 | 1.732E-06 | 1.200E-05 | 1.500E-05 |
| 7428 | 372 | P372G | A | 1.5e-05 | 9.7e-06 | 1.5e-05 | 1.323E-05 | 3.060E-06 | 9.700E-06 | 1.500E-05 |
| 7429 | 372 | P372H | A | 1.3e-05 | 1.6e-05 | 1.1e-05 | 1.333E-05 | 2.517E-06 | 1.100E-05 | 1.600E-05 |
| 7430 | 372 | P372I | A | 1.6e-05 | 1.7e-05 | 1.4e-05 | 1.567E-05 | 1.528E-06 | 1.400E-05 | 1.700E-05 |
| 7431 | 372 | P372L | A | 1.1e-05 | 1.4e-05 | 1.1e-05 | 1.200E-05 | 1.732E-06 | 1.100E-05 | 1.400E-05 |
| 7432 | 372 | P372K | A | 1.6e-05 | 1e-05 | 1.5e-05 | 1.367E-05 | 3.215E-06 | 1.000E-05 | 1.600E-05 |
| 7433 | 372 | P372M | A | 1.3e-05 | 1.7e-05 | 1.4e-05 | 1.467E-05 | 2.082E-06 | 1.300E-05 | 1.700E-05 |
| 7434 | 372 | P372F | A | 1.3e-05 | 1.4e-05 | 1.4e-05 | 1.367E-05 | 5.774E-07 | 1.300E-05 | 1.400E-05 |
| 7435 | 372 | P372P | A | 1.3e-05 | 1.6e-05 | 1.4e-05 | 1.433E-05 | 1.528E-06 | 1.300E-05 | 1.600E-05 |
| 7436 | 372 | P372S | A | 1.1e-05 | 1.5e-05 | 1.4e-05 | 1.333E-05 | 2.082E-06 | 1.100E-05 | 1.500E-05 |
| 7437 | 372 | P372T | A | 7.8e-06 | 8.4e-06 | 1.6e-05 | 1.073E-05 | 4.571E-06 | 7.800E-06 | 1.600E-05 |
| 7438 | 372 | P372W | A | 1.4e-05 | 1.3e-05 | 1.3e-05 | 1.333E-05 | 5.774E-07 | 1.300E-05 | 1.400E-05 |
| 7439 | 372 | P372Y | A | 1.4e-05 | 1.6e-05 | 8.3e-06 | 1.277E-05 | 3.995E-06 | 8.300E-06 | 1.600E-05 |
| 7440 | 372 | P372V | A | 1.1e-05 | 1.4e-05 | 1.8e-05 | 1.433E-05 | 3.512E-06 | 1.100E-05 | 1.800E-05 |
| 7441 | 373 | E373A | A | 1.5e-05 | 1.6e-05 | 9.8e-06 | 1.360E-05 | 3.329E-06 | 9.800E-06 | 1.600E-05 |
| 7442 | 373 | E373R | A | 1.2e-05 | 9.4e-06 | 1e-05 | 1.047E-05 | 1.361E-06 | 9.400E-06 | 1.200E-05 |
| 7443 | 373 | E373N | A | 2.7e-05 | 1.5e-05 | 1.5e-05 | 1.900E-05 | 6.928E-06 | 1.500E-05 | 2.700E-05 |
| 7444 | 373 | E373D | A | 1.7e-05 | 8.7e-06 | 1.5e-05 | 1.357E-05 | 4.332E-06 | 8.700E-06 | 1.700E-05 |
| 7445 | 373 | E373C | A | 9.9e-06 | 1.1e-05 | 1.1e-05 | 1.063E-05 | 6.351E-07 | 9.900E-06 | 1.100E-05 |
| 7446 | 373 | E373E | A | 1.4e-05 | 1.1e-05 | 1.6e-05 | 1.367E-05 | 2.517E-06 | 1.100E-05 | 1.600E-05 |
| 7447 | 373 | E373Q | A | 1.6e-05 | 9.9e-06 | 1.4e-05 | 1.330E-05 | 3.110E-06 | 9.900E-06 | 1.600E-05 |
| 7448 | 373 | E373G | A | 2.1e-05 | 1.5e-05 | 1.6e-05 | 1.733E-05 | 3.215E-06 | 1.500E-05 | 2.100E-05 |
| 7449 | 373 | E373H | A | 6.9e-06 | 1.4e-05 | 1.4e-05 | 1.163E-05 | 4.099E-06 | 6.900E-06 | 1.400E-05 |
| 7450 | 373 | E373I | A | 1.7e-05 | 1.6e-05 | 1e-05 | 1.433E-05 | 3.786E-06 | 1.000E-05 | 1.700E-05 |
| 7451 | 373 | E373L | A | 1.5e-05 | 1.2e-05 | 1.2e-05 | 1.300E-05 | 1.732E-06 | 1.200E-05 | 1.500E-05 |
| 7452 | 373 | E373K | A | 1.2e-05 | 5.5e-06 | 1.5e-05 | 1.083E-05 | 4.856E-06 | 5.500E-06 | 1.500E-05 |
| 7453 | 373 | E373M | A | 1.5e-05 | 9.1e-06 | 9.6e-06 | 1.123E-05 | 3.272E-06 | 9.100E-06 | 1.500E-05 |
| 7454 | 373 | E373F | A | 1.6e-05 | 1.4e-05 | 1.1e-05 | 1.367E-05 | 2.517E-06 | 1.100E-05 | 1.600E-05 |
| 7455 | 373 | E373P | A | 1.5e-05 | 1e-05 | 1.1e-05 | 1.200E-05 | 2.646E-06 | 1.000E-05 | 1.500E-05 |
| 7456 | 373 | E373S | A | 8.8e-06 | 1.3e-05 | 1.6e-05 | 1.260E-05 | 3.617E-06 | 8.800E-06 | 1.600E-05 |
| 7457 | 373 | E373T | A | 1.5e-05 | 1.6e-05 | 9.9e-06 | 1.363E-05 | 3.272E-06 | 9.900E-06 | 1.600E-05 |
| 7458 | 373 | E373W | A | 9.5e-06 | 7.9e-06 | 8.4e-06 | 8.600E-06 | 8.185E-07 | 7.900E-06 | 9.500E-06 |
| 7459 | 373 | E373Y | A | 1.4e-05 | 9.8e-06 | 1.3e-05 | 1.227E-05 | 2.194E-06 | 9.800E-06 | 1.400E-05 |
| 7460 | 373 | E373V | A | 1.5e-05 | 1.3e-05 | 1.5e-05 | 1.433E-05 | 1.155E-06 | 1.300E-05 | 1.500E-05 |
| 7461 | 374 | Q374A | A | 1.4e-05 | 1.2e-05 | 1.1e-05 | 1.233E-05 | 1.528E-06 | 1.100E-05 | 1.400E-05 |
| 7462 | 374 | Q374R | A | 1.5e-05 | 1.6e-05 | 1.5e-05 | 1.533E-05 | 5.774E-07 | 1.500E-05 | 1.600E-05 |
| 7463 | 374 | Q374N | A | 9e-06 | 1.4e-05 | 1.5e-05 | 1.267E-05 | 3.215E-06 | 9.000E-06 | 1.500E-05 |
| 7464 | 374 | Q374D | A | 1.5e-05 | 1.6e-05 | 1.4e-05 | 1.500E-05 | 1.000E-06 | 1.400E-05 | 1.600E-05 |
| 7465 | 374 | Q374C | A | 9.8e-06 | 1.5e-05 | 1.1e-05 | 1.193E-05 | 2.723E-06 | 9.800E-06 | 1.500E-05 |
| 7466 | 374 | Q374E | A | 1e-05 | 9.5e-06 | 1.6e-05 | 1.183E-05 | 3.617E-06 | 9.500E-06 | 1.600E-05 |
| 7467 | 374 | Q374Q | A | 1.8e-05 | 1e-05 | 1.6e-05 | 1.467E-05 | 4.163E-06 | 1.000E-05 | 1.800E-05 |
| 7468 | 374 | Q374G | A | 1.6e-05 | 1.5e-05 | 1.5e-05 | 1.533E-05 | 5.774E-07 | 1.500E-05 | 1.600E-05 |
| 7469 | 374 | Q374H | A | 9e-06 | 1.4e-05 | 1.5e-05 | 1.267E-05 | 3.215E-06 | 9.000E-06 | 1.500E-05 |
| 7470 | 374 | Q374I | A | 1.4e-05 | 1.6e-05 | 9.1e-06 | 1.303E-05 | 3.550E-06 | 9.100E-06 | 1.600E-05 |
| 7471 | 374 | Q374L | A | 1.5e-05 | 7.8e-06 | 1.5e-05 | 1.260E-05 | 4.157E-06 | 7.800E-06 | 1.500E-05 |
| 7472 | 374 | Q374K | A | 1.2e-05 | 9e-06 | 1e-05 | 1.033E-05 | 1.528E-06 | 9.000E-06 | 1.200E-05 |
| 7473 | 374 | Q374M | A | 1.1e-05 | 9.4e-06 | 1.5e-05 | 1.180E-05 | 2.884E-06 | 9.400E-06 | 1.500E-05 |
| 7474 | 374 | Q374F | A | 2.2e-05 | 1.2e-05 | 1.2e-05 | 1.533E-05 | 5.774E-06 | 1.200E-05 | 2.200E-05 |
| 7475 | 374 | Q374P | A | 1.5e-05 | 1.3e-05 | 1.6e-05 | 1.467E-05 | 1.528E-06 | 1.300E-05 | 1.600E-05 |
| 7476 | 374 | Q374S | A | 1.5e-05 | 1.6e-05 | 1.6e-05 | 1.567E-05 | 5.774E-07 | 1.500E-05 | 1.600E-05 |
| 7477 | 374 | Q374T | A | 1.4e-05 | 1.5e-05 | 1.3e-05 | 1.400E-05 | 1.000E-06 | 1.300E-05 | 1.500E-05 |
| 7478 | 374 | Q374W | A | 1.3e-05 | 1.1e-05 | 9.9e-06 | 1.130E-05 | 1.572E-06 | 9.900E-06 | 1.300E-05 |
| 7479 | 374 | Q374Y | A | 1.8e-05 | 1.6e-05 | 1.5e-05 | 1.633E-05 | 1.528E-06 | 1.500E-05 | 1.800E-05 |

| | | | | | | | | | | |
|---|---|---|---|---|---|---|---|---|---|---|
| 7480 | 374 | Q374V | A | 1.8e-05 | 1.4e-05 | 1.4e-05 | 1.533E-05 | 2.309E-06 | 1.400E-05 | 1.800E-05 |
| 7481 | 375 | L375A | A | 9.5e-06 | 1.5e-05 | 1.6e-05 | 1.350E-05 | 3.500E-06 | 9.500E-06 | 1.600E-05 |
| 7482 | 375 | L375R | A | 1.3e-05 | 1.4e-05 | 1.5e-05 | 1.400E-05 | 1.000E-06 | 1.300E-05 | 1.500E-05 |
| 7483 | 375 | L375N | A | 9.4e-06 | 1.3e-05 | 1.2e-05 | 1.147E-05 | 1.858E-06 | 9.400E-06 | 1.300E-05 |
| 7484 | 375 | L375D | A | 1.4e-05 | 1.2e-05 | 1e-05 | 1.200E-05 | 2.000E-06 | 1.000E-05 | 1.400E-05 |
| 7485 | 375 | L375C | A | 1.4e-05 | 1.2e-05 | 1.4e-05 | 1.333E-05 | 1.155E-06 | 1.200E-05 | 1.400E-05 |
| 7486 | 375 | L375E | A | 1.2e-05 | 1.4e-05 | 1.2e-05 | 1.267E-05 | 1.155E-06 | 1.200E-05 | 1.400E-05 |
| 7487 | 375 | L375Q | A | 1.4e-05 | 1.1e-05 | 1e-05 | 1.167E-05 | 2.082E-06 | 1.000E-05 | 1.400E-05 |
| 7488 | 375 | L375G | A | 1.2e-05 | 1.3e-05 | 1.1e-05 | 1.200E-05 | 1.000E-06 | 1.100E-05 | 1.300E-05 |
| 7489 | 375 | L375H | A | 1.5e-05 | 1.1e-05 | 1.7e-05 | 1.433E-05 | 3.055E-06 | 1.100E-05 | 1.700E-05 |
| 7490 | 375 | L375I | A | 1.6e-05 | 1.5e-05 | 1.4e-05 | 1.500E-05 | 1.000E-06 | 1.400E-05 | 1.600E-05 |
| 7491 | 375 | L375L | A | 1.5e-05 | 8e-06 | 1.8e-05 | 1.367E-05 | 5.132E-06 | 8.000E-06 | 1.800E-05 |
| 7492 | 375 | L375K | A | 8.9e-06 | 1.1e-05 | 1.7e-05 | 1.230E-05 | 4.204E-06 | 8.900E-06 | 1.700E-05 |
| 7493 | 375 | L375M | A | 1.3e-05 | 9.5e-06 | 1.5e-05 | 1.250E-05 | 2.784E-06 | 9.500E-06 | 1.500E-05 |
| 7494 | 375 | L375F | A | 1.2e-05 | 1.1e-05 | 1.1e-05 | 1.133E-05 | 5.774E-07 | 1.100E-05 | 1.200E-05 |
| 7495 | 375 | L375P | A | 1.4e-05 | 1.5e-05 | 1.1e-05 | 1.333E-05 | 2.082E-06 | 1.100E-05 | 1.500E-05 |
| 7496 | 375 | L375S | A | 1.4e-05 | 1e-05 | 1.6e-05 | 1.333E-05 | 3.055E-06 | 1.000E-05 | 1.600E-05 |
| 7497 | 375 | L375T | A | 1.5e-05 | 1.6e-05 | 1e-05 | 1.367E-05 | 3.215E-06 | 1.000E-05 | 1.600E-05 |
| 7498 | 375 | L375W | A | 1.2e-05 | 1.1e-05 | 1.6e-05 | 1.300E-05 | 2.646E-06 | 1.100E-05 | 1.600E-05 |
| 7499 | 375 | L375Y | A | 1.7e-05 | 1.8e-05 | 8.7e-06 | 1.457E-05 | 5.105E-06 | 8.700E-06 | 1.800E-05 |
| 7500 | 375 | L375V | A | 9.6e-06 | 1.5e-05 | 1.3e-05 | 1.253E-05 | 2.730E-06 | 9.600E-06 | 1.500E-05 |
| 7501 | 376 | Q376A | A | 6.6e-06 | 1.1e-05 | 1.1e-05 | 9.533E-06 | 2.540E-06 | 6.600E-06 | 1.100E-05 |
| 7502 | 376 | Q376R | A | 1.1e-05 | 1.4e-05 | 1.7e-05 | 1.400E-05 | 3.000E-06 | 1.100E-05 | 1.700E-05 |
| 7503 | 376 | Q376N | A | 9.6e-06 | 1.6e-05 | 9.1e-06 | 1.157E-05 | 3.848E-06 | 9.100E-06 | 1.600E-05 |
| 7504 | 376 | Q376D | A | 1.4e-05 | 1.5e-05 | 1.4e-05 | 1.433E-05 | 5.774E-07 | 1.400E-05 | 1.500E-05 |
| 7505 | 376 | Q376C | A | 1.5e-05 | 7.6e-06 | 1.5e-05 | 1.253E-05 | 4.272E-06 | 7.600E-06 | 1.500E-05 |
| 7506 | 376 | Q376E | A | 1.5e-05 | 1.3e-05 | 1.6e-05 | 1.467E-05 | 1.528E-06 | 1.300E-05 | 1.600E-05 |
| 7507 | 376 | Q376Q | A | 1.5e-05 | 1.1e-05 | 9.6e-06 | 1.187E-05 | 2.802E-06 | 9.600E-06 | 1.500E-05 |
| 7508 | 376 | Q376G | A | 9.6e-06 | 1.5e-05 | 1.8e-05 | 1.420E-05 | 4.257E-06 | 9.600E-06 | 1.800E-05 |
| 7509 | 376 | Q376H | A | 1.2e-05 | 1.2e-05 | 1.1e-05 | 1.167E-05 | 5.774E-07 | 1.100E-05 | 1.200E-05 |
| 7510 | 376 | Q376I | A | 1.6e-05 | 1.7e-05 | 1.5e-05 | 1.600E-05 | 1.000E-06 | 1.500E-05 | 1.700E-05 |
| 7511 | 376 | Q376L | A | 1e-05 | 1.1e-05 | 1.5e-05 | 1.200E-05 | 2.646E-06 | 1.000E-05 | 1.500E-05 |
| 7512 | 376 | Q376K | A | 1.8e-05 | 1.6e-05 | 1.6e-05 | 1.667E-05 | 1.155E-06 | 1.600E-05 | 1.800E-05 |
| 7513 | 376 | Q376M | A | 1.5e-05 | 1.6e-05 | 1.6e-05 | 1.567E-05 | 5.774E-07 | 1.500E-05 | 1.600E-05 |
| 7514 | 376 | Q376F | A | 1.6e-05 | 1.6e-05 | 1e-05 | 1.400E-05 | 3.464E-06 | 1.000E-05 | 1.600E-05 |
| 7515 | 376 | Q376P | A | 8.8e-06 | 1e-05 | 1e-05 | 9.600E-06 | 6.928E-07 | 8.800E-06 | 1.000E-05 |
| 7516 | 376 | Q376S | A | 9.7e-06 | 1e-05 | 1.5e-05 | 1.157E-05 | 2.977E-06 | 9.700E-06 | 1.500E-05 |
| 7517 | 376 | Q376T | A | 1.1e-05 | 1.4e-05 | 1.4e-05 | 1.300E-05 | 1.732E-06 | 1.100E-05 | 1.400E-05 |
| 7518 | 376 | Q376W | A | 1.4e-05 | 1e-05 | 7.8e-06 | 1.060E-05 | 3.143E-06 | 7.800E-06 | 1.400E-05 |
| 7519 | 376 | Q376Y | A | 1.1e-05 | 8.1e-06 | 1.2e-05 | 1.037E-05 | 2.026E-06 | 8.100E-06 | 1.200E-05 |
| 7520 | 376 | Q376V | A | 1.5e-05 | 1e-05 | 2.2e-05 | 1.567E-05 | 6.028E-06 | 1.000E-05 | 2.200E-05 |
| 7521 | 377 | V377A | A | 1.1e-05 | 9.7e-06 | 9.9e-06 | 1.020E-05 | 7.000E-07 | 9.700E-06 | 1.100E-05 |
| 7522 | 377 | V377R | A | 9.8e-06 | 9.6e-06 | 1.1e-05 | 1.013E-05 | 7.572E-07 | 9.600E-06 | 1.100E-05 |
| 7523 | 377 | V377N | A | 1.6e-05 | 1.2e-05 | 1e-05 | 1.267E-05 | 3.055E-06 | 1.000E-05 | 1.600E-05 |
| 7524 | 377 | V377D | A | 1e-05 | 1.1e-05 | 1.7e-05 | 1.267E-05 | 3.786E-06 | 1.000E-05 | 1.700E-05 |
| 7525 | 377 | V377C | A | 1.4e-05 | 1.1e-05 | 1e-05 | 1.167E-05 | 2.082E-06 | 1.000E-05 | 1.400E-05 |
| 7526 | 377 | V377E | A | 1.6e-05 | 1.3e-05 | 1.4e-05 | 1.433E-05 | 1.528E-06 | 1.300E-05 | 1.600E-05 |
| 7527 | 377 | V377Q | A | 1.3e-05 | 1.5e-05 | 1.6e-05 | 1.467E-05 | 1.528E-06 | 1.300E-05 | 1.600E-05 |
| 7528 | 377 | V377G | A | 9e-06 | 1.4e-05 | 1.2e-05 | 1.167E-05 | 2.517E-06 | 9.000E-06 | 1.400E-05 |
| 7529 | 377 | V377H | A | 1e-05 | 1e-05 | 1.5e-05 | 1.167E-05 | 2.887E-06 | 1.000E-05 | 1.500E-05 |
| 7530 | 377 | V377I | A | 6.4e-06 | 1.6e-05 | 1.2e-05 | 1.147E-05 | 4.822E-06 | 6.400E-06 | 1.600E-05 |
| 7531 | 377 | V377L | A | 1.2e-05 | 8.7e-06 | 1.3e-05 | 1.123E-05 | 2.250E-06 | 8.700E-06 | 1.300E-05 |
| 7532 | 377 | V377K | A | 1.1e-05 | 6.2e-06 | 9.5e-06 | 8.900E-06 | 2.456E-06 | 6.200E-06 | 1.100E-05 |
| 7533 | 377 | V377M | A | 1.5e-05 | 9.5e-06 | 6.6e-06 | 1.037E-05 | 4.267E-06 | 6.600E-06 | 1.500E-05 |
| 7534 | 377 | V377F | A | 8.8e-06 | 1.1e-05 | 9.5e-06 | 9.767E-06 | 1.124E-06 | 8.800E-06 | 1.100E-05 |
| 7535 | 377 | V377P | A | 1.3e-05 | 1.5e-05 | 1.6e-05 | 1.467E-05 | 1.528E-06 | 1.300E-05 | 1.600E-05 |
| 7536 | 377 | V377S | A | 9.6e-06 | 1.6e-05 | 1.5e-05 | 1.353E-05 | 3.443E-06 | 9.600E-06 | 1.600E-05 |
| 7537 | 377 | V377T | A | 1.4e-05 | 1.5e-05 | 8e-06 | 1.233E-05 | 3.786E-06 | 8.000E-06 | 1.500E-05 |
| 7538 | 377 | V377W | A | 1e-05 | 9.2e-06 | 1e-05 | 9.733E-06 | 4.619E-07 | 9.200E-06 | 1.000E-05 |
| 7539 | 377 | V377Y | A | 1.2e-05 | 1.4e-05 | 1.1e-05 | 1.233E-05 | 1.528E-06 | 1.100E-05 | 1.400E-05 |
| 7540 | 377 | V377V | A | 1.5e-05 | 1.7e-05 | 1.4e-05 | 1.533E-05 | 1.528E-06 | 1.400E-05 | 1.700E-05 |
| 7541 | 378 | F378A | A | 1.6e-05 | 2.4e-05 | 1.5e-05 | 1.833E-05 | 4.933E-06 | 1.500E-05 | 2.400E-05 |
| 7542 | 378 | F378R | A | 1.7e-05 | 1.5e-05 | 7e-06 | 1.300E-05 | 5.292E-06 | 7.000E-06 | 1.700E-05 |
| 7543 | 378 | F378N | A | 1.1e-05 | 9.6e-06 | 1.2e-05 | 1.087E-05 | 1.206E-06 | 9.600E-06 | 1.200E-05 |
| 7544 | 378 | F378D | A | 1.3e-05 | 1.5e-05 | 1.6e-05 | 1.467E-05 | 1.528E-06 | 1.300E-05 | 1.600E-05 |
| 7545 | 378 | F378C | A | 1.4e-05 | 9.4e-06 | 9.1e-06 | 1.083E-05 | 2.747E-06 | 9.100E-06 | 1.400E-05 |
| 7546 | 378 | F378E | A | 1.5e-05 | 1e-05 | 1.6e-05 | 1.367E-05 | 3.215E-06 | 1.000E-05 | 1.600E-05 |
| 7547 | 378 | F378Q | A | 1.1e-05 | 1.4e-05 | 1.5e-05 | 1.333E-05 | 2.082E-06 | 1.100E-05 | 1.500E-05 |
| 7548 | 378 | F378G | A | 1.5e-05 | 9.6e-06 | 1.6e-05 | 1.353E-05 | 3.443E-06 | 9.600E-06 | 1.600E-05 |
| 7549 | 378 | F378H | A | 1.4e-05 | 1.5e-05 | 1e-05 | 1.300E-05 | 2.646E-06 | 1.000E-05 | 1.500E-05 |
| 7550 | 378 | F378I | A | 1.1e-05 | 1.1e-05 | 1.2e-05 | 1.133E-05 | 5.774E-07 | 1.100E-05 | 1.200E-05 |
| 7551 | 378 | F378L | A | 1.6e-05 | 1e-05 | 1.2e-05 | 1.267E-05 | 3.055E-06 | 1.000E-05 | 1.600E-05 |
| 7552 | 378 | F378K | A | 1.2e-05 | 1.7e-05 | 8.7e-06 | 1.257E-05 | 4.179E-06 | 8.700E-06 | 1.700E-05 |
| 7553 | 378 | F378M | A | 1.5e-05 | 1.3e-05 | 1.6e-05 | 1.467E-05 | 1.528E-06 | 1.300E-05 | 1.600E-05 |
| 7554 | 378 | F378F | A | 1.3e-05 | 1.2e-05 | 1.1e-05 | 1.200E-05 | 1.000E-06 | 1.100E-05 | 1.300E-05 |
| 7555 | 378 | F378P | A | 1.5e-05 | 1.2e-05 | 9.9e-06 | 1.230E-05 | 2.563E-06 | 9.900E-06 | 1.500E-05 |
| 7556 | 378 | F378S | A | 1.4e-05 | 1.7e-05 | 1.1e-05 | 1.400E-05 | 3.000E-06 | 1.100E-05 | 1.700E-05 |
| 7557 | 378 | F378T | A | 1.4e-05 | 1.6e-05 | 1.3e-05 | 1.433E-05 | 1.528E-06 | 1.300E-05 | 1.600E-05 |
| 7558 | 378 | F378W | A | 1.5e-05 | 1.4e-05 | 1e-05 | 1.300E-05 | 2.646E-06 | 1.000E-05 | 1.500E-05 |
| 7559 | 378 | F378Y | A | 9.5e-06 | 9.4e-06 | 1e-05 | 9.633E-06 | 3.215E-07 | 9.400E-06 | 1.000E-05 |
| 7560 | 378 | F378V | A | 1.8e-05 | 1.3e-05 | 1.7e-05 | 1.600E-05 | 2.646E-06 | 1.300E-05 | 1.800E-05 |
| 7561 | 379 | E379A | A | 1.1e-05 | 1.6e-05 | 1e-05 | 1.233E-05 | 3.215E-06 | 1.000E-05 | 1.600E-05 |
| 7562 | 379 | E379R | A | 1e-05 | 1.1e-05 | 8.1e-06 | 9.700E-06 | 1.473E-06 | 8.100E-06 | 1.100E-05 |
| 7563 | 379 | E379N | A | 1.3e-05 | 8.9e-06 | 1.4e-05 | 1.197E-05 | 2.702E-06 | 8.900E-06 | 1.400E-05 |
| 7564 | 379 | E379D | A | 1.3e-05 | 1.6e-05 | 8.9e-06 | 1.263E-05 | 3.564E-06 | 8.900E-06 | 1.600E-05 |

| | | | | | | | | | | |
|---|---|---|---|---|---|---|---|---|---|---|
| 7565 | 379 | E379C | A | 1.2e-05 | 1e-05 | 1.7e-05 | 1.300E-05 | 3.606E-06 | 1.000E-05 | 1.700E-05 |
| 7566 | 379 | E379E | A | 1.6e-05 | 1.1e-05 | 1.1e-05 | 1.267E-05 | 2.887E-06 | 1.100E-05 | 1.600E-05 |
| 7567 | 379 | E379Q | A | 1.1e-05 | 1e-05 | 1e-05 | 1.033E-05 | 5.774E-07 | 1.000E-05 | 1.100E-05 |
| 7568 | 379 | E379G | A | 7.5e-06 | 9.5e-06 | 1.4e-05 | 1.033E-05 | 3.329E-06 | 7.500E-06 | 1.400E-05 |
| 7569 | 379 | E379H | A | 2.3e-05 | 1.4e-05 | 1e-05 | 1.567E-05 | 6.658E-06 | 1.000E-05 | 2.300E-05 |
| 7570 | 379 | E379I | A | 1.2e-05 | 9.6e-06 | 1.6e-05 | 1.253E-05 | 3.233E-06 | 9.600E-06 | 1.600E-05 |
| 7571 | 379 | E379L | A | 8.9e-06 | 1.6e-05 | 1.4e-05 | 1.297E-05 | 3.661E-06 | 8.900E-06 | 1.600E-05 |
| 7572 | 379 | E379K | A | 1e-05 | 1.5e-05 | 9.6e-06 | 1.153E-05 | 3.009E-06 | 9.600E-06 | 1.500E-05 |
| 7573 | 379 | E379M | A | 1.3e-05 | 1.5e-05 | 1.4e-05 | 1.400E-05 | 1.000E-06 | 1.300E-05 | 1.500E-05 |
| 7574 | 379 | E379F | A | 6.8e-06 | 1.1e-05 | 1e-05 | 9.267E-06 | 2.194E-06 | 6.800E-06 | 1.100E-05 |
| 7575 | 379 | E379P | A | 1e-05 | 9.5e-06 | 1.1e-05 | 1.017E-05 | 7.638E-07 | 9.500E-06 | 1.100E-05 |
| 7576 | 379 | E379S | A | 9e-06 | 1.5e-05 | 1.1e-05 | 1.167E-05 | 3.055E-06 | 9.000E-06 | 1.500E-05 |
| 7577 | 379 | E379T | A | 1e-05 | 1.1e-05 | 7.9e-06 | 9.633E-06 | 1.582E-06 | 7.900E-06 | 1.100E-05 |
| 7578 | 379 | E379W | A | 1.1e-05 | 1.4e-05 | 1.1e-05 | 1.200E-05 | 1.732E-06 | 1.100E-05 | 1.400E-05 |
| 7579 | 379 | E379Y | A | 1.6e-05 | 9.9e-06 | 1.6e-05 | 1.397E-05 | 3.522E-06 | 9.900E-06 | 1.600E-05 |
| 7580 | 379 | E379V | A | 1.5e-05 | 1.4e-05 | 1.5e-05 | 1.467E-05 | 5.774E-07 | 1.400E-05 | 1.500E-05 |
| 7581 | 380 | T380A | A | 1.1e-05 | 1e-05 | 1.5e-05 | 1.200E-05 | 2.646E-06 | 1.000E-05 | 1.500E-05 |
| 7582 | 380 | T380R | A | 1.3e-05 | 9.3e-06 | 1.1e-05 | 1.110E-05 | 1.852E-06 | 9.300E-06 | 1.300E-05 |
| 7583 | 380 | T380N | A | 1.8e-05 | 1.4e-05 | 1.7e-05 | 1.633E-05 | 2.082E-06 | 1.400E-05 | 1.800E-05 |
| 7584 | 380 | T380D | A | 1.5e-05 | 1.6e-05 | 1.3e-05 | 1.467E-05 | 1.528E-06 | 1.300E-05 | 1.600E-05 |
| 7585 | 380 | T380C | A | 1.1e-05 | 1e-05 | 1.4e-05 | 1.167E-05 | 2.082E-06 | 1.000E-05 | 1.400E-05 |
| 7586 | 380 | T380E | A | 9.1e-06 | 9.6e-06 | 1.4e-05 | 1.090E-05 | 2.696E-06 | 9.100E-06 | 1.400E-05 |
| 7587 | 380 | T380Q | A | 9.8e-06 | 1.4e-05 | 1.6e-05 | 1.327E-05 | 3.164E-06 | 9.800E-06 | 1.600E-05 |
| 7588 | 380 | T380G | A | 1e-05 | 1.5e-05 | 1.5e-05 | 1.333E-05 | 2.887E-06 | 1.000E-05 | 1.500E-05 |
| 7589 | 380 | T380H | A | 9.7e-06 | 1.3e-05 | 1.1e-05 | 1.123E-05 | 1.662E-06 | 9.700E-06 | 1.300E-05 |
| 7590 | 380 | T380I | A | 1.7e-05 | 1.1e-05 | 6.9e-06 | 1.163E-05 | 5.080E-06 | 6.900E-06 | 1.700E-05 |
| 7591 | 380 | T380L | A | 1e-05 | 1.8e-05 | 1.7e-05 | 1.500E-05 | 4.359E-06 | 1.000E-05 | 1.800E-05 |
| 7592 | 380 | T380K | A | 1.8e-05 | 1.7e-05 | 1.6e-05 | 1.700E-05 | 1.000E-06 | 1.600E-05 | 1.800E-05 |
| 7593 | 380 | T380M | A | 9.9e-06 | 1.3e-05 | 1.7e-05 | 1.330E-05 | 3.559E-06 | 9.900E-06 | 1.700E-05 |
| 7594 | 380 | T380F | A | 1.5e-05 | 1.7e-05 | 2.3e-05 | 1.833E-05 | 4.163E-06 | 1.500E-05 | 2.300E-05 |
| 7595 | 380 | T380P | A | 1.1e-05 | 1.6e-05 | 1.4e-05 | 1.367E-05 | 2.517E-06 | 1.100E-05 | 1.600E-05 |
| 7596 | 380 | T380S | A | 1.2e-05 | 1e-05 | 1.1e-05 | 1.100E-05 | 1.000E-06 | 1.000E-05 | 1.200E-05 |
| 7597 | 380 | T380T | A | 7.3e-06 | 1.7e-05 | 1.1e-05 | 1.177E-05 | 4.895E-06 | 7.300E-06 | 1.700E-05 |
| 7598 | 380 | T380W | A | 9.5e-06 | 1.5e-05 | 9.7e-06 | 1.140E-05 | 3.119E-06 | 9.500E-06 | 1.500E-05 |
| 7599 | 380 | T380Y | A | 1e-05 | 1.1e-05 | 1.4e-05 | 1.167E-05 | 2.082E-06 | 1.000E-05 | 1.400E-05 |
| 7600 | 380 | T380V | A | 1.5e-05 | 1.3e-05 | 1.3e-05 | 1.367E-05 | 1.155E-06 | 1.300E-05 | 1.500E-05 |
| 7601 | 381 | L381A | A | 1.1e-05 | 1e-05 | 1.1e-05 | 1.067E-05 | 5.774E-07 | 1.000E-05 | 1.100E-05 |
| 7602 | 381 | L381R | A | 1.3e-05 | 1e-05 | 8.4e-06 | 1.047E-05 | 2.335E-06 | 8.400E-06 | 1.300E-05 |
| 7603 | 381 | L381N | A | 8e-06 | 9.8e-06 | 9e-06 | 8.933E-06 | 9.018E-07 | 8.000E-06 | 9.800E-06 |
| 7604 | 381 | L381D | A | 1.6e-05 | 8.8e-06 | 1.4e-05 | 1.293E-05 | 3.717E-06 | 8.800E-06 | 1.600E-05 |
| 7605 | 381 | L381C | A | 8.6e-06 | 1.1e-05 | 1.4e-05 | 1.120E-05 | 2.706E-06 | 8.600E-06 | 1.400E-05 |
| 7606 | 381 | L381E | A | 1e-05 | 1.6e-05 | 1e-05 | 1.200E-05 | 3.464E-06 | 1.000E-05 | 1.600E-05 |
| 7607 | 381 | L381Q | A | 9e-06 | 1.6e-05 | 1.3e-05 | 1.267E-05 | 3.512E-06 | 9.000E-06 | 1.600E-05 |
| 7608 | 381 | L381G | A | 9e-06 | 1.5e-05 | 1.5e-05 | 1.300E-05 | 3.464E-06 | 9.000E-06 | 1.500E-05 |
| 7609 | 381 | L381H | A | 1e-05 | 1e-05 | 1.4e-05 | 1.133E-05 | 2.309E-06 | 1.000E-05 | 1.400E-05 |
| 7610 | 381 | L381I | A | 1.4e-05 | 9.2e-06 | 1.1e-05 | 1.140E-05 | 2.425E-06 | 9.200E-06 | 1.400E-05 |
| 7611 | 381 | L381L | A | 1.1e-05 | 1.6e-05 | 9.1e-06 | 1.203E-05 | 3.564E-06 | 9.100E-06 | 1.600E-05 |
| 7612 | 381 | L381K | A | 1.4e-05 | 1.5e-05 | 1.5e-05 | 1.467E-05 | 5.774E-07 | 1.400E-05 | 1.500E-05 |
| 7613 | 381 | L381M | A | 1e-05 | 9.3e-06 | 1.2e-05 | 1.043E-05 | 1.401E-06 | 9.300E-06 | 1.200E-05 |
| 7614 | 381 | L381F | A | 9.1e-06 | 1.1e-05 | 1.5e-05 | 1.170E-05 | 3.012E-06 | 9.100E-06 | 1.500E-05 |
| 7615 | 381 | L381P | A | 1.8e-05 | 1.6e-05 | 1.1e-05 | 1.500E-05 | 3.606E-06 | 1.100E-05 | 1.800E-05 |
| 7616 | 381 | L381S | A | 1.4e-05 | 9.3e-06 | 1.4e-05 | 1.243E-05 | 2.714E-06 | 9.300E-06 | 1.400E-05 |
| 7617 | 381 | L381T | A | 9.7e-06 | 7.6e-06 | 1.1e-05 | 9.433E-06 | 1.716E-06 | 7.600E-06 | 1.100E-05 |
| 7618 | 381 | L381W | A | 1.1e-05 | 1.6e-05 | 1.5e-05 | 1.400E-05 | 2.646E-06 | 1.100E-05 | 1.600E-05 |
| 7619 | 381 | L381Y | A | 1.6e-05 | 1.4e-05 | 1.1e-05 | 1.367E-05 | 2.517E-06 | 1.100E-05 | 1.600E-05 |
| 7620 | 381 | L381V | A | 9.3e-06 | 9.7e-06 | 9.8e-06 | 9.600E-06 | 2.646E-07 | 9.300E-06 | 9.800E-06 |
| 7621 | 382 | E382A | A | 1.1e-05 | 1e-05 | 9e-06 | 1.000E-05 | 1.000E-06 | 9.000E-06 | 1.100E-05 |
| 7622 | 382 | E382R | A | 1.4e-05 | 1e-05 | 1.5e-05 | 1.300E-05 | 2.646E-06 | 1.000E-05 | 1.500E-05 |
| 7623 | 382 | E382N | A | 8.6e-06 | 1.6e-05 | 1e-05 | 1.153E-05 | 3.931E-06 | 8.600E-06 | 1.600E-05 |
| 7624 | 382 | E382D | A | 8.2e-06 | 1e-05 | 1e-05 | 9.400E-06 | 1.039E-06 | 8.200E-06 | 1.000E-05 |
| 7625 | 382 | E382C | A | 1.4e-05 | 1.3e-05 | 1.4e-05 | 1.367E-05 | 5.774E-07 | 1.300E-05 | 1.400E-05 |
| 7626 | 382 | E382E | A | 1.6e-05 | 1.6e-05 | 1.5e-05 | 1.567E-05 | 5.774E-07 | 1.500E-05 | 1.600E-05 |
| 7627 | 382 | E382Q | A | 8.7e-06 | 9.3e-06 | 1.1e-05 | 9.667E-06 | 1.193E-06 | 8.700E-06 | 1.100E-05 |
| 7628 | 382 | E382G | A | 1.8e-05 | 1.5e-05 | 1.5e-05 | 1.600E-05 | 1.732E-06 | 1.500E-05 | 1.800E-05 |
| 7629 | 382 | E382H | A | 1.4e-05 | 1.4e-05 | 1.2e-05 | 1.333E-05 | 1.155E-06 | 1.200E-05 | 1.400E-05 |
| 7630 | 382 | E382I | A | 1.5e-05 | 1.5e-05 | 1.5e-05 | 1.500E-05 | 0.000E+00 | 1.500E-05 | 1.500E-05 |
| 7631 | 382 | E382L | A | 9.4e-06 | 1.2e-05 | 1.4e-05 | 1.180E-05 | 2.307E-06 | 9.400E-06 | 1.400E-05 |
| 7632 | 382 | E382K | A | 1.7e-05 | 1.4e-05 | 1.4e-05 | 1.500E-05 | 1.732E-06 | 1.400E-05 | 1.700E-05 |
| 7633 | 382 | E382M | A | 1.5e-05 | 1.4e-05 | 9e-06 | 1.267E-05 | 3.215E-06 | 9.000E-06 | 1.500E-05 |
| 7634 | 382 | E382F | A | 1.5e-05 | 8.6e-06 | 1.4e-05 | 1.253E-05 | 3.443E-06 | 8.600E-06 | 1.500E-05 |
| 7635 | 382 | E382P | A | 1.1e-05 | 1.5e-05 | 1.1e-05 | 1.233E-05 | 2.309E-06 | 1.100E-05 | 1.500E-05 |
| 7636 | 382 | E382S | A | 8.2e-06 | 1.5e-05 | 1.3e-05 | 1.207E-05 | 3.495E-06 | 8.200E-06 | 1.500E-05 |
| 7637 | 382 | E382T | A | 1.6e-05 | 1.4e-05 | 1.2e-05 | 1.400E-05 | 2.000E-06 | 1.200E-05 | 1.600E-05 |
| 7638 | 382 | E382W | A | 1e-05 | 1.6e-05 | 1.9e-05 | 1.500E-05 | 4.583E-06 | 1.000E-05 | 1.900E-05 |
| 7639 | 382 | E382Y | A | 1.2e-05 | 1e-05 | 1.1e-05 | 1.100E-05 | 1.000E-06 | 1.000E-05 | 1.200E-05 |
| 7640 | 382 | E382V | A | 1.1e-05 | 1e-05 | 6.8e-06 | 9.267E-06 | 2.194E-06 | 6.800E-06 | 1.100E-05 |
| 7641 | 383 | E383A | A | 1.1e-05 | 1.5e-05 | 1.1e-05 | 1.233E-05 | 2.309E-06 | 1.100E-05 | 1.500E-05 |
| 7642 | 383 | E383R | A | 9.7e-06 | 1.4e-05 | 1.3e-05 | 1.223E-05 | 2.250E-06 | 9.700E-06 | 1.400E-05 |
| 7643 | 383 | E383N | A | 1.5e-05 | 1.4e-05 | 8.4e-06 | 1.247E-05 | 3.557E-06 | 8.400E-06 | 1.500E-05 |
| 7644 | 383 | E383D | A | 1.4e-05 | 1.6e-05 | 1.4e-05 | 1.467E-05 | 1.155E-06 | 1.400E-05 | 1.600E-05 |
| 7645 | 383 | E383C | A | 1.5e-05 | 1.7e-05 | 1.3e-05 | 1.500E-05 | 2.000E-06 | 1.300E-05 | 1.700E-05 |
| 7646 | 383 | E383E | A | 1.2e-05 | 9.8e-06 | 1e-05 | 1.060E-05 | 1.217E-06 | 9.800E-06 | 1.200E-05 |
| 7647 | 383 | E383Q | A | 1.3e-05 | 1.3e-05 | 1.2e-05 | 1.267E-05 | 5.774E-07 | 1.200E-05 | 1.300E-05 |
| 7648 | 383 | E383G | A | 1.1e-05 | 1.1e-05 | 1.4e-05 | 1.200E-05 | 1.732E-06 | 1.100E-05 | 1.400E-05 |
| 7649 | 383 | E383H | A | 1.3e-05 | 9.4e-06 | 1.3e-05 | 1.180E-05 | 2.078E-06 | 9.400E-06 | 1.300E-05 |

| | | | | | | | | | | |
|---|---|---|---|---|---|---|---|---|---|---|
| 7650 | 383 | E383I | A | 1.6e-05 | 9.6e-06 | 1.5e-05 | 1.353E-05 | 3.443E-06 | 9.600E-06 | 1.600E-05 |
| 7651 | 383 | E383L | A | 1.5e-05 | 1.2e-05 | 1.6e-05 | 1.433E-05 | 2.082E-06 | 1.200E-05 | 1.600E-05 |
| 7652 | 383 | E383K | A | 1.4e-05 | 1.1e-05 | 9e-06 | 1.133E-05 | 2.517E-06 | 9.000E-06 | 1.400E-05 |
| 7653 | 383 | E383M | A | 1.6e-05 | 1.6e-05 | 1.7e-05 | 1.633E-05 | 5.774E-07 | 1.600E-05 | 1.700E-05 |
| 7654 | 383 | E383F | A | 1.5e-05 | 8.8e-06 | 1.5e-05 | 1.293E-05 | 3.580E-06 | 8.800E-06 | 1.500E-05 |
| 7655 | 383 | E383P | A | 1.1e-05 | 1.7e-05 | 8.2e-06 | 1.207E-05 | 4.496E-06 | 8.200E-06 | 1.700E-05 |
| 7656 | 383 | E383S | A | 1.5e-05 | 1.3e-05 | 9.3e-06 | 1.243E-05 | 2.892E-06 | 9.300E-06 | 1.500E-05 |
| 7657 | 383 | E383T | A | 1.3e-05 | 1e-05 | 1.4e-05 | 1.233E-05 | 2.082E-06 | 1.000E-05 | 1.400E-05 |
| 7658 | 383 | E383W | A | 1.4e-05 | 1.5e-05 | 1.5e-05 | 1.467E-05 | 5.774E-07 | 1.400E-05 | 1.500E-05 |
| 7659 | 383 | E383Y | A | 1.4e-05 | 1.5e-05 | 9.5e-06 | 1.283E-05 | 2.930E-06 | 9.500E-06 | 1.500E-05 |
| 7660 | 383 | E383V | A | 1.5e-05 | 1.4e-05 | 1.5e-05 | 1.467E-05 | 5.774E-07 | 1.400E-05 | 1.500E-05 |
| 7661 | 384 | I384A | A | 1e-05 | 8.2e-06 | 1.5e-05 | 1.107E-05 | 3.523E-06 | 8.200E-06 | 1.500E-05 |
| 7662 | 384 | I384R | A | 6.1e-06 | 1.5e-05 | 1.4e-05 | 1.170E-05 | 4.875E-06 | 6.100E-06 | 1.500E-05 |
| 7663 | 384 | I384N | A | 7.5e-06 | 1.1e-05 | 1.7e-05 | 1.183E-05 | 4.805E-06 | 7.500E-06 | 1.700E-05 |
| 7664 | 384 | I384D | A | 1.1e-05 | 1.2e-05 | 1.6e-05 | 1.300E-05 | 2.646E-06 | 1.100E-05 | 1.600E-05 |
| 7665 | 384 | I384C | A | 1.7e-05 | 1.5e-05 | 1e-05 | 1.400E-05 | 3.606E-06 | 1.000E-05 | 1.700E-05 |
| 7666 | 384 | I384E | A | 1.2e-05 | 1.7e-05 | 1.6e-05 | 1.500E-05 | 2.646E-06 | 1.200E-05 | 1.700E-05 |
| 7667 | 384 | I384Q | A | 9.8e-06 | 1.3e-05 | 1.1e-05 | 1.127E-05 | 1.617E-06 | 9.800E-06 | 1.300E-05 |
| 7668 | 384 | I384G | A | 8.9e-06 | 1.1e-05 | 1.6e-05 | 1.197E-05 | 3.647E-06 | 8.900E-06 | 1.600E-05 |
| 7669 | 384 | I384H | A | 1.4e-05 | 1.4e-05 | 9e-06 | 1.233E-05 | 2.887E-06 | 9.000E-06 | 1.400E-05 |
| 7670 | 384 | I384I | A | 1.6e-05 | 1.4e-05 | 1.1e-05 | 1.367E-05 | 2.517E-06 | 1.100E-05 | 1.600E-05 |
| 7671 | 384 | I384L | A | 1.1e-05 | 1.6e-05 | 1.5e-05 | 1.400E-05 | 2.646E-06 | 1.100E-05 | 1.600E-05 |
| 7672 | 384 | I384K | A | 1.4e-05 | 1.4e-05 | 1.6e-05 | 1.467E-05 | 1.155E-06 | 1.400E-05 | 1.600E-05 |
| 7673 | 384 | I384M | A | 9.1e-06 | 7.9e-06 | 1.4e-05 | 1.033E-05 | 3.232E-06 | 7.900E-06 | 1.400E-05 |
| 7674 | 384 | I384F | A | 1.3e-05 | 1.2e-05 | 1.1e-05 | 1.200E-05 | 1.000E-06 | 1.100E-05 | 1.300E-05 |
| 7675 | 384 | I384P | A | 1.4e-05 | 1.5e-05 | 1.1e-05 | 1.333E-05 | 2.082E-06 | 1.100E-05 | 1.500E-05 |
| 7676 | 384 | I384S | A | 1.7e-05 | 1.2e-05 | 1e-05 | 1.300E-05 | 3.606E-06 | 1.000E-05 | 1.700E-05 |
| 7677 | 384 | I384T | A | 1.1e-05 | 1.5e-05 | 7.2e-06 | 1.107E-05 | 3.900E-06 | 7.200E-06 | 1.500E-05 |
| 7678 | 384 | I384W | A | 9.8e-06 | 1.4e-05 | 1.6e-05 | 1.327E-05 | 3.164E-06 | 9.800E-06 | 1.600E-05 |
| 7679 | 384 | I384Y | A | 1.4e-05 | 1e-05 | 1.5e-05 | 1.300E-05 | 2.646E-06 | 1.000E-05 | 1.500E-05 |
| 7680 | 384 | I384V | A | 1.1e-05 | 1.5e-05 | 1.1e-05 | 1.233E-05 | 2.309E-06 | 1.100E-05 | 1.500E-05 |
| 7681 | 385 | T385A | A | 8.8e-06 | 1.5e-05 | 1e-05 | 1.127E-05 | 3.288E-06 | 8.800E-06 | 1.500E-05 |
| 7682 | 385 | T385R | A | 1e-05 | 1.1e-05 | 1.3e-05 | 1.133E-05 | 1.528E-06 | 1.000E-05 | 1.300E-05 |
| 7683 | 385 | T385N | A | 1.5e-05 | 1.5e-05 | 1.4e-05 | 1.467E-05 | 5.774E-07 | 1.400E-05 | 1.500E-05 |
| 7684 | 385 | T385D | A | 1e-05 | 1.4e-05 | 1.7e-05 | 1.367E-05 | 3.512E-06 | 1.000E-05 | 1.700E-05 |
| 7685 | 385 | T385C | A | 1.1e-05 | 1.1e-05 | 1.1e-05 | 1.100E-05 | 2.075E-21 | 1.100E-05 | 1.100E-05 |
| 7686 | 385 | T385E | A | 1.5e-05 | 1.5e-05 | 7.4e-06 | 1.247E-05 | 4.388E-06 | 7.400E-06 | 1.500E-05 |
| 7687 | 385 | T385Q | A | 9.1e-06 | 7.1e-06 | 1.4e-05 | 1.007E-05 | 3.550E-06 | 7.100E-06 | 1.400E-05 |
| 7688 | 385 | T385G | A | 1.1e-05 | 1.4e-05 | 1.4e-05 | 1.300E-05 | 1.732E-06 | 1.100E-05 | 1.400E-05 |
| 7689 | 385 | T385H | A | 1.4e-05 | 9.8e-06 | 1.1e-05 | 1.160E-05 | 2.163E-06 | 9.800E-06 | 1.400E-05 |
| 7690 | 385 | T385I | A | 1.1e-05 | 1.5e-05 | 9.6e-06 | 1.187E-05 | 2.802E-06 | 9.600E-06 | 1.500E-05 |
| 7691 | 385 | T385L | A | 1.1e-05 | 1.2e-05 | 1e-05 | 1.100E-05 | 1.000E-06 | 1.000E-05 | 1.200E-05 |
| 7692 | 385 | T385K | A | 1.2e-05 | 1.5e-05 | 1.5e-05 | 1.400E-05 | 1.732E-06 | 1.200E-05 | 1.500E-05 |
| 7693 | 385 | T385M | A | 1.6e-05 | 9.8e-06 | 1.1e-05 | 1.227E-05 | 3.288E-06 | 9.800E-06 | 1.600E-05 |
| 7694 | 385 | T385F | A | 1e-05 | 1.4e-05 | 1.1e-05 | 1.167E-05 | 2.082E-06 | 1.000E-05 | 1.400E-05 |
| 7695 | 385 | T385P | A | 1.5e-05 | 1.6e-05 | 1.7e-05 | 1.600E-05 | 1.000E-06 | 1.500E-05 | 1.700E-05 |
| 7696 | 385 | T385S | A | 1.4e-05 | 1e-05 | 1.5e-05 | 1.300E-05 | 2.646E-06 | 1.000E-05 | 1.500E-05 |
| 7697 | 385 | T385T | A | 9.5e-06 | 1.3e-05 | 1.4e-05 | 1.217E-05 | 2.363E-06 | 9.500E-06 | 1.400E-05 |
| 7698 | 385 | T385W | A | 9.1e-06 | 1.2e-05 | 1.3e-05 | 1.137E-05 | 2.026E-06 | 9.100E-06 | 1.300E-05 |
| 7699 | 385 | T385Y | A | 9.8e-06 | 1.2e-05 | 1.2e-05 | 1.127E-05 | 1.270E-06 | 9.800E-06 | 1.200E-05 |
| 7700 | 385 | T385V | A | 1.1e-05 | 1.5e-05 | 9.1e-06 | 1.170E-05 | 3.012E-06 | 9.100E-06 | 1.500E-05 |
| 7701 | 386 | G386A | A | 1.6e-05 | 9.6e-06 | 1.5e-05 | 1.353E-05 | 3.443E-06 | 9.600E-06 | 1.600E-05 |
| 7702 | 386 | G386R | A | 1.4e-05 | 1.2e-05 | 7.2e-06 | 1.107E-05 | 3.495E-06 | 7.200E-06 | 1.400E-05 |
| 7703 | 386 | G386N | A | 1.5e-05 | 9.5e-06 | 1.6e-05 | 1.350E-05 | 3.500E-06 | 9.500E-06 | 1.600E-05 |
| 7704 | 386 | G386D | A | 1e-05 | 9.7e-06 | 1.6e-05 | 1.190E-05 | 3.554E-06 | 9.700E-06 | 1.600E-05 |
| 7705 | 386 | G386C | A | 1.4e-05 | 9.4e-06 | 1.4e-05 | 1.247E-05 | 2.656E-06 | 9.400E-06 | 1.400E-05 |
| 7706 | 386 | G386E | A | 1.6e-05 | 1.7e-05 | 1.1e-05 | 1.467E-05 | 3.215E-06 | 1.100E-05 | 1.700E-05 |
| 7707 | 386 | G386Q | A | 2.1e-05 | 1e-05 | 9.2e-06 | 1.340E-05 | 6.594E-06 | 9.200E-06 | 2.100E-05 |
| 7708 | 386 | G386G | A | 1.3e-05 | 1.6e-05 | 1.6e-05 | 1.500E-05 | 1.732E-06 | 1.300E-05 | 1.600E-05 |
| 7709 | 386 | G386H | A | 2.1e-05 | 1.1e-05 | 1e-05 | 1.400E-05 | 6.083E-06 | 1.000E-05 | 2.100E-05 |
| 7710 | 386 | G386I | A | 1.1e-05 | 1.2e-05 | 1.4e-05 | 1.233E-05 | 1.528E-06 | 1.100E-05 | 1.400E-05 |
| 7711 | 386 | G386L | A | 1.5e-05 | 1.2e-05 | 1.4e-05 | 1.367E-05 | 1.528E-06 | 1.200E-05 | 1.500E-05 |
| 7712 | 386 | G386K | A | 1.2e-05 | 9.8e-06 | 9.8e-06 | 1.053E-05 | 1.270E-06 | 9.800E-06 | 1.200E-05 |
| 7713 | 386 | G386M | A | 8.8e-06 | 1.3e-05 | 1.5e-05 | 1.227E-05 | 3.164E-06 | 8.800E-06 | 1.500E-05 |
| 7714 | 386 | G386F | A | 1.3e-05 | 1.5e-05 | 1.5e-05 | 1.433E-05 | 1.155E-06 | 1.300E-05 | 1.500E-05 |
| 7715 | 386 | G386P | A | 1.5e-05 | 1.7e-05 | 9.9e-06 | 1.397E-05 | 3.661E-06 | 9.900E-06 | 1.700E-05 |
| 7716 | 386 | G386S | A | 1.5e-05 | 8.6e-06 | 1.6e-05 | 1.320E-05 | 4.015E-06 | 8.600E-06 | 1.600E-05 |
| 7717 | 386 | G386T | A | 1.4e-05 | 1.1e-05 | 1.4e-05 | 1.300E-05 | 1.732E-06 | 1.100E-05 | 1.400E-05 |
| 7718 | 386 | G386W | A | 1.4e-05 | 1.1e-05 | 1.7e-05 | 1.400E-05 | 3.000E-06 | 1.100E-05 | 1.700E-05 |
| 7719 | 386 | G386Y | A | 1.6e-05 | 1.4e-05 | 1.6e-05 | 1.533E-05 | 1.155E-06 | 1.400E-05 | 1.600E-05 |
| 7720 | 386 | G386V | A | 1.5e-05 | 1.1e-05 | 1.3e-05 | 1.300E-05 | 2.000E-06 | 1.100E-05 | 1.500E-05 |
| 7721 | 387 | Y387A | A | 1.5e-05 | 1.3e-05 | 1.5e-05 | 1.433E-05 | 1.155E-06 | 1.300E-05 | 1.500E-05 |
| 7722 | 387 | Y387R | A | 1.1e-05 | 1.4e-05 | 1e-05 | 1.167E-05 | 2.082E-06 | 1.000E-05 | 1.400E-05 |
| 7723 | 387 | Y387N | A | 1.5e-05 | 1.2e-05 | 1.1e-05 | 1.267E-05 | 2.082E-06 | 1.100E-05 | 1.500E-05 |
| 7724 | 387 | Y387D | A | 1.5e-05 | 9.6e-06 | 1.5e-05 | 1.320E-05 | 3.118E-06 | 9.600E-06 | 1.500E-05 |
| 7725 | 387 | Y387C | A | 7.8e-06 | 1.4e-05 | 1.7e-05 | 1.293E-05 | 4.692E-06 | 7.800E-06 | 1.700E-05 |
| 7726 | 387 | Y387E | A | 1.7e-05 | 1.6e-05 | 1.7e-05 | 1.667E-05 | 5.774E-07 | 1.600E-05 | 1.700E-05 |
| 7727 | 387 | Y387Q | A | 1.4e-05 | 1.4e-05 | 8.5e-06 | 1.217E-05 | 3.175E-06 | 8.500E-06 | 1.400E-05 |
| 7728 | 387 | Y387G | A | 1.4e-05 | 1.1e-05 | 1e-05 | 1.167E-05 | 2.082E-06 | 1.000E-05 | 1.400E-05 |
| 7729 | 387 | Y387H | A | 1.3e-05 | 1.6e-05 | 9.8e-06 | 1.293E-05 | 3.101E-06 | 9.800E-06 | 1.600E-05 |
| 7730 | 387 | Y387I | A | 6.4e-06 | 1.5e-05 | 1.7e-05 | 1.280E-05 | 5.632E-06 | 6.400E-06 | 1.700E-05 |
| 7731 | 387 | Y387L | A | 1.4e-05 | 1.5e-05 | 1.3e-05 | 1.400E-05 | 1.000E-06 | 1.300E-05 | 1.500E-05 |
| 7732 | 387 | Y387K | A | 1.4e-05 | 1.7e-05 | 1.4e-05 | 1.500E-05 | 1.732E-06 | 1.400E-05 | 1.700E-05 |
| 7733 | 387 | Y387M | A | 1.5e-05 | 1.4e-05 | 1.1e-05 | 1.333E-05 | 2.082E-06 | 1.100E-05 | 1.500E-05 |
| 7734 | 387 | Y387F | A | 1.2e-05 | 1.7e-05 | 1.1e-05 | 1.333E-05 | 3.215E-06 | 1.100E-05 | 1.700E-05 |

| | | | | | | | | | | |
|---|---|---|---|---|---|---|---|---|---|---|
| 7735 | 387 | Y387P | A | 1.2e-05 | 1.1e-05 | 1.6e-05 | 1.300E-05 | 2.646E-06 | 1.100E-05 | 1.600E-05 |
| 7736 | 387 | Y387S | A | 8.1e-06 | 1.1e-05 | 9.4e-06 | 9.500E-06 | 1.453E-06 | 8.100E-06 | 1.100E-05 |
| 7737 | 387 | Y387T | A | 1.3e-05 | 1.6e-05 | 1.5e-05 | 1.467E-05 | 1.528E-06 | 1.300E-05 | 1.600E-05 |
| 7738 | 387 | Y387W | A | 1.4e-05 | 1.4e-05 | 1.6e-05 | 1.467E-05 | 1.155E-06 | 1.400E-05 | 1.600E-05 |
| 7739 | 387 | Y387Y | A | 8.8e-06 | 1.5e-05 | 7.4e-06 | 1.040E-05 | 4.045E-06 | 7.400E-06 | 1.500E-05 |
| 7740 | 387 | Y387V | A | 1.5e-05 | 1.6e-05 | 1.1e-05 | 1.400E-05 | 2.646E-06 | 1.100E-05 | 1.600E-05 |
| 7741 | 388 | L388A | A | 1.5e-05 | 1.6e-05 | 1.1e-05 | 1.400E-05 | 2.646E-06 | 1.100E-05 | 1.600E-05 |
| 7742 | 388 | L388R | A | 7.7e-06 | 1e-05 | 1.4e-05 | 1.057E-05 | 3.188E-06 | 7.700E-06 | 1.400E-05 |
| 7743 | 388 | L388N | A | 1.2e-05 | 9.5e-06 | 1.2e-05 | 1.117E-05 | 1.443E-06 | 9.500E-06 | 1.200E-05 |
| 7744 | 388 | L388D | A | 1e-05 | 1.5e-05 | 1.3e-05 | 1.267E-05 | 2.517E-06 | 1.000E-05 | 1.500E-05 |
| 7745 | 388 | L388C | A | 1.6e-05 | 1e-05 | 1e-05 | 1.200E-05 | 3.464E-06 | 1.000E-05 | 1.600E-05 |
| 7746 | 388 | L388E | A | 1.2e-05 | 9.9e-06 | 1.4e-05 | 1.197E-05 | 2.050E-06 | 9.900E-06 | 1.400E-05 |
| 7747 | 388 | L388Q | A | 1e-05 | 9.9e-06 | 1.5e-05 | 1.163E-05 | 2.916E-06 | 9.900E-06 | 1.500E-05 |
| 7748 | 388 | L388G | A | 1.5e-05 | 1e-05 | 1.2e-05 | 1.233E-05 | 2.517E-06 | 1.000E-05 | 1.500E-05 |
| 7749 | 388 | L388H | A | 1.1e-05 | 1.2e-05 | 1.6e-05 | 1.300E-05 | 2.646E-06 | 1.100E-05 | 1.600E-05 |
| 7750 | 388 | L388I | A | 1.4e-05 | 8.5e-06 | 9.9e-06 | 1.080E-05 | 2.858E-06 | 8.500E-06 | 1.400E-05 |
| 7751 | 388 | L388L | A | 1e-05 | 1.5e-05 | 9.5e-06 | 1.150E-05 | 3.041E-06 | 9.500E-06 | 1.500E-05 |
| 7752 | 388 | L388K | A | 1.1e-05 | 1.6e-05 | 8.6e-06 | 1.187E-05 | 3.775E-06 | 8.600E-06 | 1.600E-05 |
| 7753 | 388 | L388M | A | 1e-05 | 1.3e-05 | 2.9e-05 | 1.733E-05 | 1.021E-05 | 1.000E-05 | 2.900E-05 |
| 7754 | 388 | L388F | A | 1.6e-05 | 1.4e-05 | 1.3e-05 | 1.433E-05 | 1.528E-06 | 1.300E-05 | 1.600E-05 |
| 7755 | 388 | L388P | A | 1.6e-05 | 1e-05 | 1.3e-05 | 1.300E-05 | 3.000E-06 | 1.000E-05 | 1.600E-05 |
| 7756 | 388 | L388S | A | 1.6e-05 | 1.4e-05 | 9.8e-06 | 1.327E-05 | 3.164E-06 | 9.800E-06 | 1.600E-05 |
| 7757 | 388 | L388T | A | 1.5e-05 | 1.4e-05 | 1.5e-05 | 1.467E-05 | 5.774E-07 | 1.400E-05 | 1.500E-05 |
| 7758 | 388 | L388W | A | 1.2e-05 | 8.4e-06 | 1.7e-05 | 1.247E-05 | 4.319E-06 | 8.400E-06 | 1.700E-05 |
| 7759 | 388 | L388Y | A | 1.3e-05 | 1.7e-05 | 1.6e-05 | 1.533E-05 | 2.082E-06 | 1.300E-05 | 1.700E-05 |
| 7760 | 388 | L388V | A | 9.7e-06 | 1.5e-05 | 1e-05 | 1.157E-05 | 2.977E-06 | 9.700E-06 | 1.500E-05 |
| 7761 | 389 | Y389A | A | 1.2e-05 | 1.1e-05 | 8.7e-06 | 1.057E-05 | 1.692E-06 | 8.700E-06 | 1.200E-05 |
| 7762 | 389 | Y389R | A | 1.3e-05 | 1e-05 | 1.2e-05 | 1.167E-05 | 1.528E-06 | 1.000E-05 | 1.300E-05 |
| 7763 | 389 | Y389N | A | 1.2e-05 | 1e-05 | 1.1e-05 | 1.100E-05 | 1.000E-06 | 1.000E-05 | 1.200E-05 |
| 7764 | 389 | Y389D | A | 1e-05 | 1.5e-05 | 1.6e-05 | 1.367E-05 | 3.215E-06 | 1.000E-05 | 1.600E-05 |
| 7765 | 389 | Y389C | A | 1.3e-05 | 1e-05 | 1.4e-05 | 1.233E-05 | 2.082E-06 | 1.000E-05 | 1.400E-05 |
| 7766 | 389 | Y389E | A | 1.3e-05 | 1e-05 | 1.2e-05 | 1.167E-05 | 1.528E-06 | 1.000E-05 | 1.300E-05 |
| 7767 | 389 | Y389Q | A | 1.5e-05 | 1.6e-05 | 1.1e-05 | 1.400E-05 | 2.646E-06 | 1.100E-05 | 1.600E-05 |
| 7768 | 389 | Y389G | A | 1.5e-05 | 1.2e-05 | 1.5e-05 | 1.400E-05 | 1.732E-06 | 1.200E-05 | 1.500E-05 |
| 7769 | 389 | Y389H | A | 1.5e-05 | 1.4e-05 | 1e-05 | 1.300E-05 | 2.646E-06 | 1.000E-05 | 1.500E-05 |
| 7770 | 389 | Y389I | A | 1.5e-05 | 1.1e-05 | 1.4e-05 | 1.333E-05 | 2.082E-06 | 1.100E-05 | 1.500E-05 |
| 7771 | 389 | Y389L | A | 1.5e-05 | 1.1e-05 | 9.3e-06 | 1.177E-05 | 2.926E-06 | 9.300E-06 | 1.500E-05 |
| 7772 | 389 | Y389K | A | 1.2e-05 | 1e-05 | 1.8e-05 | 1.333E-05 | 4.163E-06 | 1.000E-05 | 1.800E-05 |
| 7773 | 389 | Y389M | A | 1.3e-05 | 1.5e-05 | 1.1e-05 | 1.300E-05 | 2.000E-06 | 1.100E-05 | 1.500E-05 |
| 7774 | 389 | Y389F | A | 1e-05 | 8.9e-06 | 1.2e-05 | 1.030E-05 | 1.572E-06 | 8.900E-06 | 1.200E-05 |
| 7775 | 389 | Y389P | A | 9.7e-06 | 1.3e-05 | 1.6e-05 | 1.290E-05 | 3.151E-06 | 9.700E-06 | 1.600E-05 |
| 7776 | 389 | Y389S | A | 1.5e-05 | 1.1e-05 | 1.7e-05 | 1.433E-05 | 3.055E-06 | 1.100E-05 | 1.700E-05 |
| 7777 | 389 | Y389T | A | 1.1e-05 | 1.3e-05 | 1.5e-05 | 1.300E-05 | 2.000E-06 | 1.100E-05 | 1.500E-05 |
| 7778 | 389 | Y389W | A | 1e-05 | 1.2e-05 | 1e-05 | 1.067E-05 | 1.155E-06 | 1.000E-05 | 1.200E-05 |
| 7779 | 389 | Y389Y | A | 1e-05 | 1.5e-05 | 1.4e-05 | 1.300E-05 | 2.646E-06 | 1.000E-05 | 1.500E-05 |
| 7780 | 389 | Y389V | A | 1e-05 | 9.3e-06 | 9.1e-06 | 9.467E-06 | 4.726E-07 | 9.100E-06 | 1.000E-05 |
| 7781 | 390 | I390A | A | 1.1e-05 | 1.3e-05 | 1.5e-05 | 1.300E-05 | 2.000E-06 | 1.100E-05 | 1.500E-05 |
| 7782 | 390 | I390R | A | 1e-05 | 1.8e-05 | 1.2e-05 | 1.333E-05 | 4.163E-06 | 1.000E-05 | 1.800E-05 |
| 7783 | 390 | I390N | A | 1.2e-05 | 7e-06 | 1.1e-05 | 1.000E-05 | 2.646E-06 | 7.000E-06 | 1.200E-05 |
| 7784 | 390 | I390D | A | 1e-05 | 1.2e-05 | 1.2e-05 | 1.133E-05 | 1.155E-06 | 1.000E-05 | 1.200E-05 |
| 7785 | 390 | I390C | A | 1e-05 | 1.5e-05 | 9.9e-06 | 1.163E-05 | 2.916E-06 | 9.900E-06 | 1.500E-05 |
| 7786 | 390 | I390E | A | 1.6e-05 | 1.2e-05 | 1.1e-05 | 1.300E-05 | 2.646E-06 | 1.100E-05 | 1.600E-05 |
| 7787 | 390 | I390Q | A | 1.6e-05 | 1.5e-05 | 1.8e-05 | 1.633E-05 | 1.528E-06 | 1.500E-05 | 1.800E-05 |
| 7788 | 390 | I390G | A | 1e-05 | 1.6e-05 | 1.5e-05 | 1.367E-05 | 3.215E-06 | 1.000E-05 | 1.600E-05 |
| 7789 | 390 | I390H | A | 1.5e-05 | 1.6e-05 | 1.2e-05 | 1.433E-05 | 2.082E-06 | 1.200E-05 | 1.600E-05 |
| 7790 | 390 | I390I | A | 1.1e-05 | 1.5e-05 | 1.5e-05 | 1.367E-05 | 2.309E-06 | 1.100E-05 | 1.500E-05 |
| 7791 | 390 | I390L | A | 9.5e-06 | 9.5e-06 | 9.8e-06 | 9.600E-06 | 1.732E-07 | 9.500E-06 | 9.800E-06 |
| 7792 | 390 | I390K | A | 1.4e-05 | 1.6e-05 | 1.2e-05 | 1.400E-05 | 2.000E-06 | 1.200E-05 | 1.600E-05 |
| 7793 | 390 | I390M | A | 1e-05 | 9.5e-06 | 1.1e-05 | 1.017E-05 | 7.638E-07 | 9.500E-06 | 1.100E-05 |
| 7794 | 390 | I390F | A | 1.1e-05 | 1.1e-05 | 1.5e-05 | 1.233E-05 | 2.309E-06 | 1.100E-05 | 1.500E-05 |
| 7795 | 390 | I390P | A | 1.5e-05 | 1.5e-05 | 7.5e-06 | 1.250E-05 | 4.330E-06 | 7.500E-06 | 1.500E-05 |
| 7796 | 390 | I390S | A | 1.6e-05 | 1.1e-05 | 8.2e-06 | 1.173E-05 | 3.951E-06 | 8.200E-06 | 1.600E-05 |
| 7797 | 390 | I390T | A | 1e-05 | 9.8e-06 | 1.6e-05 | 1.193E-05 | 3.523E-06 | 9.800E-06 | 1.600E-05 |
| 7798 | 390 | I390W | A | 1.2e-05 | 1.1e-05 | 1.5e-05 | 1.267E-05 | 2.082E-06 | 1.100E-05 | 1.500E-05 |
| 7799 | 390 | I390Y | A | 9e-06 | 1.2e-05 | 1.1e-05 | 1.067E-05 | 1.528E-06 | 9.000E-06 | 1.200E-05 |
| 7800 | 390 | I390V | A | 1.3e-05 | 6.5e-06 | 1.4e-05 | 1.117E-05 | 4.072E-06 | 6.500E-06 | 1.400E-05 |
| 7801 | 391 | S391A | A | 1.2e-05 | 1.5e-05 | 1.7e-05 | 1.467E-05 | 2.517E-06 | 1.200E-05 | 1.700E-05 |
| 7802 | 391 | S391R | A | 1.1e-05 | 1e-05 | 1.5e-05 | 1.200E-05 | 2.646E-06 | 1.000E-05 | 1.500E-05 |
| 7803 | 391 | S391N | A | 1.6e-05 | 1.1e-05 | 1.3e-05 | 1.333E-05 | 2.517E-06 | 1.100E-05 | 1.600E-05 |
| 7804 | 391 | S391D | A | 1.1e-05 | 1.7e-05 | 1e-05 | 1.267E-05 | 3.786E-06 | 1.000E-05 | 1.700E-05 |
| 7805 | 391 | S391C | A | 1.1e-05 | 1.3e-05 | 8.3e-06 | 1.077E-05 | 2.359E-06 | 8.300E-06 | 1.300E-05 |
| 7806 | 391 | S391E | A | 8.5e-06 | 1.1e-05 | 1.4e-05 | 1.117E-05 | 2.754E-06 | 8.500E-06 | 1.400E-05 |
| 7807 | 391 | S391Q | A | 1.5e-05 | 1.5e-05 | 9.5e-06 | 1.317E-05 | 3.175E-06 | 9.500E-06 | 1.500E-05 |
| 7808 | 391 | S391G | A | 1.6e-05 | 1.6e-05 | 1.3e-05 | 1.500E-05 | 1.732E-06 | 1.300E-05 | 1.600E-05 |
| 7809 | 391 | S391H | A | 1.4e-05 | 1.1e-05 | 1e-05 | 1.167E-05 | 2.082E-06 | 1.000E-05 | 1.400E-05 |
| 7810 | 391 | S391I | A | 1.6e-05 | 1.6e-05 | 8.6e-06 | 1.353E-05 | 4.272E-06 | 8.600E-06 | 1.600E-05 |
| 7811 | 391 | S391L | A | 8.4e-06 | 1.5e-05 | 1.1e-05 | 1.147E-05 | 3.325E-06 | 8.400E-06 | 1.500E-05 |
| 7812 | 391 | S391K | A | 1.3e-05 | 1.1e-05 | 1.5e-05 | 1.300E-05 | 2.000E-06 | 1.100E-05 | 1.500E-05 |
| 7813 | 391 | S391M | A | 1.6e-05 | 1.3e-05 | 1.2e-05 | 1.367E-05 | 2.082E-06 | 1.200E-05 | 1.600E-05 |
| 7814 | 391 | S391F | A | 1.6e-05 | 1.8e-05 | 1e-05 | 1.467E-05 | 4.163E-06 | 1.000E-05 | 1.800E-05 |
| 7815 | 391 | S391P | A | 1.5e-05 | 1.5e-05 | 1.5e-05 | 1.500E-05 | 0.000E+00 | 1.500E-05 | 1.500E-05 |
| 7816 | 391 | S391S | A | 8.1e-06 | 9.8e-06 | 1.7e-05 | 1.163E-05 | 4.725E-06 | 8.100E-06 | 1.700E-05 |
| 7817 | 391 | S391T | A | 8.9e-06 | 8.6e-06 | 1.5e-05 | 1.083E-05 | 3.612E-06 | 8.600E-06 | 1.500E-05 |
| 7818 | 391 | S391W | A | 1.6e-05 | 1e-05 | 8.8e-06 | 1.160E-05 | 3.857E-06 | 8.800E-06 | 1.600E-05 |
| 7819 | 391 | S391Y | A | 9.2e-06 | 1e-05 | 1.7e-05 | 1.207E-05 | 4.291E-06 | 9.200E-06 | 1.700E-05 |

| | | | | | | | | | | |
|---|---|---|---|---|---|---|---|---|---|---|
| 7820 | 391 | S391V | A | 1.5e-05 | 1e-05 | 1e-05 | 1.167E-05 | 2.887E-06 | 1.000E-05 | 1.500E-05 |
| 7821 | 392 | A392A | A | 1.6e-05 | 1.1e-05 | 1.2e-05 | 1.300E-05 | 2.646E-06 | 1.100E-05 | 1.600E-05 |
| 7822 | 392 | A392R | A | 1.3e-05 | 1e-05 | 1.5e-05 | 1.267E-05 | 2.517E-06 | 1.000E-05 | 1.500E-05 |
| 7823 | 392 | A392N | A | 1.5e-05 | 6.5e-06 | 1.5e-05 | 1.217E-05 | 4.907E-06 | 6.500E-06 | 1.500E-05 |
| 7824 | 392 | A392D | A | 1.3e-05 | 1.6e-05 | 1.5e-05 | 1.467E-05 | 1.528E-06 | 1.300E-05 | 1.600E-05 |
| 7825 | 392 | A392C | A | 1.3e-05 | 1e-05 | 1.4e-05 | 1.233E-05 | 2.082E-06 | 1.000E-05 | 1.400E-05 |
| 7826 | 392 | A392E | A | 1.1e-05 | 1.5e-05 | 1.5e-05 | 1.367E-05 | 2.309E-06 | 1.100E-05 | 1.500E-05 |
| 7827 | 392 | A392Q | A | 1.5e-05 | 1.5e-05 | 1.5e-05 | 1.500E-05 | 0.000E+00 | 1.500E-05 | 1.500E-05 |
| 7828 | 392 | A392G | A | 1.5e-05 | 1.2e-05 | 1.2e-05 | 1.300E-05 | 1.732E-06 | 1.200E-05 | 1.500E-05 |
| 7829 | 392 | A392H | A | 9.7e-06 | 9.9e-06 | 1e-05 | 9.867E-06 | 1.528E-07 | 9.700E-06 | 1.000E-05 |
| 7830 | 392 | A392I | A | 1.6e-05 | 1.6e-05 | 9.8e-06 | 1.393E-05 | 3.580E-06 | 9.800E-06 | 1.600E-05 |
| 7831 | 392 | A392L | A | 1.6e-05 | 1.6e-05 | 1.1e-05 | 1.433E-05 | 2.887E-06 | 1.100E-05 | 1.600E-05 |
| 7832 | 392 | A392K | A | 1.1e-05 | 9.4e-06 | 1.1e-05 | 1.047E-05 | 9.238E-07 | 9.400E-06 | 1.100E-05 |
| 7833 | 392 | A392M | A | 1.4e-05 | 1e-05 | 1.5e-05 | 1.300E-05 | 2.646E-06 | 1.000E-05 | 1.500E-05 |
| 7834 | 392 | A392F | A | 1.5e-05 | 8.7e-06 | 1.3e-05 | 1.223E-05 | 3.219E-06 | 8.700E-06 | 1.500E-05 |
| 7835 | 392 | A392P | A | 1.2e-05 | 1.4e-05 | 1.6e-05 | 1.400E-05 | 2.000E-06 | 1.200E-05 | 1.600E-05 |
| 7836 | 392 | A392S | A | 1e-05 | 1.3e-05 | 1.5e-05 | 1.267E-05 | 2.517E-06 | 1.000E-05 | 1.500E-05 |
| 7837 | 392 | A392T | A | 9.8e-06 | 1.3e-05 | 1.4e-05 | 1.227E-05 | 2.194E-06 | 9.800E-06 | 1.400E-05 |
| 7838 | 392 | A392W | A | 1.5e-05 | 1e-05 | 1.4e-05 | 1.300E-05 | 2.646E-06 | 1.000E-05 | 1.500E-05 |
| 7839 | 392 | A392Y | A | 1.3e-05 | 2.1e-05 | 9.4e-06 | 1.447E-05 | 5.937E-06 | 9.400E-06 | 2.100E-05 |
| 7840 | 392 | A392V | A | 1.5e-05 | 1.4e-05 | 1.6e-05 | 1.500E-05 | 1.000E-06 | 1.400E-05 | 1.600E-05 |
| 7841 | 393 | W393A | A | 1.6e-05 | 1.5e-05 | 1.1e-05 | 1.400E-05 | 2.646E-06 | 1.100E-05 | 1.600E-05 |
| 7842 | 393 | W393R | A | 1.7e-05 | 1.8e-05 | 1.6e-05 | 1.700E-05 | 1.000E-06 | 1.600E-05 | 1.800E-05 |
| 7843 | 393 | W393N | A | 1.1e-05 | 1.4e-05 | 1.6e-05 | 1.367E-05 | 2.517E-06 | 1.100E-05 | 1.600E-05 |
| 7844 | 393 | W393D | A | 1e-05 | 1.1e-05 | 1.4e-05 | 1.167E-05 | 2.082E-06 | 1.000E-05 | 1.400E-05 |
| 7845 | 393 | W393C | A | 1.1e-05 | 1.6e-05 | 1.5e-05 | 1.400E-05 | 2.646E-06 | 1.100E-05 | 1.600E-05 |
| 7846 | 393 | W393E | A | 1.6e-05 | 1.5e-05 | 1.5e-05 | 1.533E-05 | 5.774E-07 | 1.500E-05 | 1.600E-05 |
| 7847 | 393 | W393Q | A | 1.3e-05 | 1.4e-05 | 9.8e-06 | 1.227E-05 | 2.194E-06 | 9.800E-06 | 1.400E-05 |
| 7848 | 393 | W393G | A | 1.6e-05 | 2e-05 | 1e-05 | 1.533E-05 | 5.033E-06 | 1.000E-05 | 2.000E-05 |
| 7849 | 393 | W393H | A | 1.5e-05 | 1.7e-05 | 1.7e-05 | 1.633E-05 | 1.155E-06 | 1.500E-05 | 1.700E-05 |
| 7850 | 393 | W393I | A | 1.6e-05 | 1.6e-05 | 1.5e-05 | 1.567E-05 | 5.774E-07 | 1.500E-05 | 1.600E-05 |
| 7851 | 393 | W393L | A | 1.1e-05 | 1.1e-05 | 1.6e-05 | 1.267E-05 | 2.887E-06 | 1.100E-05 | 1.600E-05 |
| 7852 | 393 | W393K | A | 1.4e-05 | 1.3e-05 | 1.4e-05 | 1.367E-05 | 5.774E-07 | 1.300E-05 | 1.400E-05 |
| 7853 | 393 | W393M | A | 1.2e-05 | 1.3e-05 | 1.6e-05 | 1.367E-05 | 2.082E-06 | 1.200E-05 | 1.600E-05 |
| 7854 | 393 | W393F | A | 1.1e-05 | 1.8e-05 | 1.5e-05 | 1.467E-05 | 3.512E-06 | 1.100E-05 | 1.800E-05 |
| 7855 | 393 | W393P | A | 1.4e-05 | 8.2e-06 | 1.1e-05 | 1.107E-05 | 2.901E-06 | 8.200E-06 | 1.400E-05 |
| 7856 | 393 | W393S | A | 2e-05 | 1.7e-05 | 1.7e-05 | 1.800E-05 | 1.732E-06 | 1.700E-05 | 2.000E-05 |
| 7857 | 393 | W393T | A | 9.3e-06 | 1.5e-05 | 1.6e-05 | 1.343E-05 | 3.614E-06 | 9.300E-06 | 1.600E-05 |
| 7858 | 393 | W393W | A | 8.8e-06 | 1.6e-05 | 1.4e-05 | 1.293E-05 | 3.717E-06 | 8.800E-06 | 1.600E-05 |
| 7859 | 393 | W393Y | A | 1.1e-05 | 1.3e-05 | 1.3e-05 | 1.233E-05 | 1.155E-06 | 1.100E-05 | 1.300E-05 |
| 7860 | 393 | W393V | A | 1.1e-05 | 1.5e-05 | 1.1e-05 | 1.233E-05 | 2.309E-06 | 1.100E-05 | 1.500E-05 |
| 7861 | 394 | P394A | A | 1.6e-05 | 1.4e-05 | 7.2e-06 | 1.240E-05 | 4.613E-06 | 7.200E-06 | 1.600E-05 |
| 7862 | 394 | P394R | A | 1.4e-05 | 1.5e-05 | 9.3e-06 | 1.277E-05 | 3.044E-06 | 9.300E-06 | 1.500E-05 |
| 7863 | 394 | P394N | A | 9.4e-06 | 1.7e-05 | 1.4e-05 | 1.347E-05 | 3.828E-06 | 9.400E-06 | 1.700E-05 |
| 7864 | 394 | P394D | A | 1.6e-05 | 1.7e-05 | 1.5e-05 | 1.600E-05 | 1.000E-06 | 1.500E-05 | 1.700E-05 |
| 7865 | 394 | P394C | A | 1e-05 | 1.5e-05 | 1.3e-05 | 1.267E-05 | 2.517E-06 | 1.000E-05 | 1.500E-05 |
| 7866 | 394 | P394E | A | 1.5e-05 | 1.4e-05 | 8.6e-06 | 1.253E-05 | 3.443E-06 | 8.600E-06 | 1.500E-05 |
| 7867 | 394 | P394Q | A | 1.3e-05 | 1.2e-05 | 7.5e-06 | 1.083E-05 | 2.930E-06 | 7.500E-06 | 1.300E-05 |
| 7868 | 394 | P394G | A | 1.5e-05 | 1.8e-05 | 1.6e-05 | 1.633E-05 | 1.528E-06 | 1.500E-05 | 1.800E-05 |
| 7869 | 394 | P394H | A | 1.5e-05 | 1.5e-05 | 1.5e-05 | 1.500E-05 | 0.000E+00 | 1.500E-05 | 1.500E-05 |
| 7870 | 394 | P394I | A | 1.2e-05 | 1.1e-05 | 1.5e-05 | 1.267E-05 | 2.082E-06 | 1.100E-05 | 1.500E-05 |
| 7871 | 394 | P394L | A | 1.1e-05 | 8.9e-06 | 1.1e-05 | 1.030E-05 | 1.212E-06 | 8.900E-06 | 1.100E-05 |
| 7872 | 394 | P394K | A | 1e-05 | 1.5e-05 | 1e-05 | 1.167E-05 | 2.887E-06 | 1.000E-05 | 1.500E-05 |
| 7873 | 394 | P394M | A | 8.4e-06 | 1.4e-05 | 1.6e-05 | 1.280E-05 | 3.940E-06 | 8.400E-06 | 1.600E-05 |
| 7874 | 394 | P394F | A | 9.2e-06 | 1.6e-05 | 9.8e-06 | 1.167E-05 | 3.765E-06 | 9.200E-06 | 1.600E-05 |
| 7875 | 394 | P394P | A | 1e-05 | 1e-05 | 1.8e-05 | 1.267E-05 | 4.619E-06 | 1.000E-05 | 1.800E-05 |
| 7876 | 394 | P394S | A | 8.4e-06 | 1.4e-05 | 1.5e-05 | 1.247E-05 | 3.557E-06 | 8.400E-06 | 1.500E-05 |
| 7877 | 394 | P394T | A | 1.5e-05 | 1.1e-05 | 1.5e-05 | 1.367E-05 | 2.309E-06 | 1.100E-05 | 1.500E-05 |
| 7878 | 394 | P394W | A | 1.1e-05 | 1e-05 | 1.4e-05 | 1.167E-05 | 2.082E-06 | 1.000E-05 | 1.400E-05 |
| 7879 | 394 | P394Y | A | 1.3e-05 | 1.6e-05 | 9.6e-06 | 1.287E-05 | 3.202E-06 | 9.600E-06 | 1.600E-05 |
| 7880 | 394 | P394V | A | 1.4e-05 | 1.5e-05 | 1.3e-05 | 1.400E-05 | 1.000E-06 | 1.300E-05 | 1.500E-05 |
| 7881 | 395 | D395A | A | 1.5e-05 | 7.1e-06 | 1.6e-05 | 1.270E-05 | 4.875E-06 | 7.100E-06 | 1.600E-05 |
| 7882 | 395 | D395R | A | 1.6e-05 | 1.4e-05 | 1.4e-05 | 1.467E-05 | 1.155E-06 | 1.400E-05 | 1.600E-05 |
| 7883 | 395 | D395N | A | 8.9e-06 | 1e-05 | 9.8e-06 | 9.567E-06 | 5.859E-07 | 8.900E-06 | 1.000E-05 |
| 7884 | 395 | D395D | A | 1.4e-05 | 1.2e-05 | 9.1e-06 | 1.170E-05 | 2.464E-06 | 9.100E-06 | 1.400E-05 |
| 7885 | 395 | D395C | A | 9.6e-06 | 8.5e-06 | 1.4e-05 | 1.070E-05 | 2.910E-06 | 8.500E-06 | 1.400E-05 |
| 7886 | 395 | D395E | A | 1.4e-05 | 1e-05 | 1.5e-05 | 1.300E-05 | 2.646E-06 | 1.000E-05 | 1.500E-05 |
| 7887 | 395 | D395Q | A | 1.5e-05 | 1.5e-05 | 1.2e-05 | 1.400E-05 | 1.732E-06 | 1.200E-05 | 1.500E-05 |
| 7888 | 395 | D395G | A | 1e-05 | 1.2e-05 | 1.5e-05 | 1.233E-05 | 2.517E-06 | 1.000E-05 | 1.500E-05 |
| 7889 | 395 | D395H | A | 1.5e-05 | 1e-05 | 1.2e-05 | 1.233E-05 | 2.517E-06 | 1.000E-05 | 1.500E-05 |
| 7890 | 395 | D395I | A | 1.6e-05 | 1.7e-05 | 1.1e-05 | 1.467E-05 | 3.215E-06 | 1.100E-05 | 1.700E-05 |
| 7891 | 395 | D395L | A | 1e-05 | 1.4e-05 | 1.6e-05 | 1.333E-05 | 3.055E-06 | 1.000E-05 | 1.600E-05 |
| 7892 | 395 | D395K | A | 9.9e-06 | 9.5e-06 | 9.3e-06 | 9.567E-06 | 3.055E-07 | 9.300E-06 | 9.900E-06 |
| 7893 | 395 | D395M | A | 1.6e-05 | 1.1e-05 | 8.8e-06 | 1.193E-05 | 3.690E-06 | 8.800E-06 | 1.600E-05 |
| 7894 | 395 | D395F | A | 1.1e-05 | 1.6e-05 | 7.7e-06 | 1.157E-05 | 4.179E-06 | 7.700E-06 | 1.600E-05 |
| 7895 | 395 | D395P | A | 1.5e-05 | 1.3e-05 | 9.7e-06 | 1.257E-05 | 2.676E-06 | 9.700E-06 | 1.500E-05 |
| 7896 | 395 | D395S | A | 1.5e-05 | 1.1e-05 | 1.4e-05 | 1.333E-05 | 2.082E-06 | 1.100E-05 | 1.500E-05 |
| 7897 | 395 | D395T | A | 1.6e-05 | 1.6e-05 | 1.4e-05 | 1.533E-05 | 1.155E-06 | 1.400E-05 | 1.600E-05 |
| 7898 | 395 | D395W | A | 9.8e-06 | 1.4e-05 | 1.5e-05 | 1.293E-05 | 2.759E-06 | 9.800E-06 | 1.500E-05 |
| 7899 | 395 | D395Y | A | 1.7e-05 | 9.4e-06 | 9.3e-06 | 1.190E-05 | 4.417E-06 | 9.300E-06 | 1.700E-05 |
| 7900 | 395 | D395V | A | 1.7e-05 | 1.6e-05 | 1.7e-05 | 1.667E-05 | 5.774E-07 | 1.600E-05 | 1.700E-05 |
| 7901 | 396 | S396A | A | 1.6e-05 | 8.8e-06 | 1.1e-05 | 1.193E-05 | 3.690E-06 | 8.800E-06 | 1.600E-05 |
| 7902 | 396 | S396R | A | 1.5e-05 | 1.6e-05 | 8.9e-06 | 1.330E-05 | 3.843E-06 | 8.900E-06 | 1.600E-05 |
| 7903 | 396 | S396N | A | 1.5e-05 | 1.4e-05 | 1.1e-05 | 1.333E-05 | 2.082E-06 | 1.100E-05 | 1.500E-05 |
| 7904 | 396 | S396D | A | 1.3e-05 | 1.4e-05 | 1.5e-05 | 1.400E-05 | 1.000E-06 | 1.300E-05 | 1.500E-05 |

| | | | | | | | | | | |
|---|---|---|---|---|---|---|---|---|---|---|
| 7905 | 396 | S396C | A | 1.5e-05 | 1.5e-05 | 1.2e-05 | 1.400E-05 | 1.732E-06 | 1.200E-05 | 1.500E-05 |
| 7906 | 396 | S396E | A | 9.2e-06 | 1.5e-05 | 9.3e-06 | 1.117E-05 | 3.320E-06 | 9.200E-06 | 1.500E-05 |
| 7907 | 396 | S396Q | A | 1.6e-05 | 1.1e-05 | 8.1e-06 | 1.170E-05 | 3.996E-06 | 8.100E-06 | 1.600E-05 |
| 7908 | 396 | S396G | A | 1.5e-05 | 1.1e-05 | 9.5e-06 | 1.183E-05 | 2.843E-06 | 9.500E-06 | 1.500E-05 |
| 7909 | 396 | S396H | A | 1.4e-05 | 1e-05 | 1.4e-05 | 1.267E-05 | 2.309E-06 | 1.000E-05 | 1.400E-05 |
| 7910 | 396 | S396I | A | 1e-05 | 1.5e-05 | 1.5e-05 | 1.333E-05 | 2.887E-06 | 1.000E-05 | 1.500E-05 |
| 7911 | 396 | S396L | A | 1.2e-05 | 1.6e-05 | 1.6e-05 | 1.467E-05 | 2.309E-06 | 1.200E-05 | 1.600E-05 |
| 7912 | 396 | S396K | A | 1.1e-05 | 1.4e-05 | 7.3e-06 | 1.077E-05 | 3.356E-06 | 7.300E-06 | 1.400E-05 |
| 7913 | 396 | S396M | A | 1.1e-05 | 1.6e-05 | 1.6e-05 | 1.433E-05 | 2.887E-06 | 1.100E-05 | 1.600E-05 |
| 7914 | 396 | S396F | A | 9.6e-06 | 8.5e-06 | 1e-05 | 9.367E-06 | 7.767E-07 | 8.500E-06 | 1.000E-05 |
| 7915 | 396 | S396P | A | 1.4e-05 | 1.1e-05 | 1.6e-05 | 1.367E-05 | 2.517E-06 | 1.100E-05 | 1.600E-05 |
| 7916 | 396 | S396S | A | 1.1e-05 | 1e-05 | 1.5e-05 | 1.200E-05 | 2.646E-06 | 1.000E-05 | 1.500E-05 |
| 7917 | 396 | S396T | A | 9.6e-06 | 1e-05 | 1.2e-05 | 1.053E-05 | 1.286E-06 | 9.600E-06 | 1.200E-05 |
| 7918 | 396 | S396W | A | 1.7e-05 | 1.4e-05 | 1.4e-05 | 1.500E-05 | 1.732E-06 | 1.400E-05 | 1.700E-05 |
| 7919 | 396 | S396Y | A | 1.5e-05 | 1.6e-05 | 1.6e-05 | 1.567E-05 | 5.774E-07 | 1.500E-05 | 1.600E-05 |
| 7920 | 396 | S396V | A | 1.6e-05 | 1.5e-05 | 1.4e-05 | 1.500E-05 | 1.000E-06 | 1.400E-05 | 1.600E-05 |
| 7921 | 397 | L397A | A | 9.8e-06 | 7.8e-06 | 7.9e-06 | 8.500E-06 | 1.127E-06 | 7.800E-06 | 9.800E-06 |
| 7922 | 397 | L397R | A | 3.4e-05 | 1.1e-05 | 1.8e-05 | 2.100E-05 | 1.179E-05 | 1.100E-05 | 3.400E-05 |
| 7923 | 397 | L397N | A | 1e-05 | 1.1e-05 | 1.1e-05 | 1.067E-05 | 5.774E-07 | 1.000E-05 | 1.100E-05 |
| 7924 | 397 | L397D | A | 1.4e-05 | 1.6e-05 | 1e-05 | 1.333E-05 | 3.055E-06 | 1.000E-05 | 1.600E-05 |
| 7925 | 397 | L397C | A | 1.3e-05 | 8.3e-06 | 1.6e-05 | 1.243E-05 | 3.881E-06 | 8.300E-06 | 1.600E-05 |
| 7926 | 397 | L397E | A | 1.4e-05 | 1e-05 | 1e-05 | 1.133E-05 | 2.309E-06 | 1.000E-05 | 1.400E-05 |
| 7927 | 397 | L397Q | A | 1.2e-05 | 1.5e-05 | 1.1e-05 | 1.267E-05 | 2.082E-06 | 1.100E-05 | 1.500E-05 |
| 7928 | 397 | L397G | A | 1.6e-05 | 1.4e-05 | 7.7e-06 | 1.257E-05 | 4.332E-06 | 7.700E-06 | 1.600E-05 |
| 7929 | 397 | L397H | A | 1e-05 | 9.9e-06 | 5.6e-06 | 8.500E-06 | 2.512E-06 | 5.600E-06 | 1.000E-05 |
| 7930 | 397 | L397I | A | 1.5e-05 | 1.1e-05 | 1e-05 | 1.200E-05 | 2.646E-06 | 1.000E-05 | 1.500E-05 |
| 7931 | 397 | L397L | A | 1.4e-05 | 1.5e-05 | 1.5e-05 | 1.467E-05 | 5.774E-07 | 1.400E-05 | 1.500E-05 |
| 7932 | 397 | L397K | A | 1.6e-05 | 1.1e-05 | 1e-05 | 1.233E-05 | 3.215E-06 | 1.000E-05 | 1.600E-05 |
| 7933 | 397 | L397M | A | 1.5e-05 | 1.4e-05 | 1.2e-05 | 1.367E-05 | 1.528E-06 | 1.200E-05 | 1.500E-05 |
| 7934 | 397 | L397F | A | 1.6e-05 | 1.7e-05 | 1.5e-05 | 1.600E-05 | 1.000E-06 | 1.500E-05 | 1.700E-05 |
| 7935 | 397 | L397P | A | 9.9e-06 | 1e-05 | 1.6e-05 | 1.197E-05 | 3.493E-06 | 9.900E-06 | 1.600E-05 |
| 7936 | 397 | L397S | A | 1e-05 | 1.5e-05 | 9.7e-06 | 1.157E-05 | 2.977E-06 | 9.700E-06 | 1.500E-05 |
| 7937 | 397 | L397T | A | 1.5e-05 | 1.7e-05 | 1.3e-05 | 1.500E-05 | 2.000E-06 | 1.300E-05 | 1.700E-05 |
| 7938 | 397 | L397W | A | 1.1e-05 | 8.4e-06 | 1e-05 | 9.800E-06 | 1.311E-06 | 8.400E-06 | 1.100E-05 |
| 7939 | 397 | L397Y | A | 1.7e-05 | 1.2e-05 | 1.7e-05 | 1.533E-05 | 2.887E-06 | 1.200E-05 | 1.700E-05 |
| 7940 | 397 | L397V | A | 1e-05 | 1.2e-05 | 1.1e-05 | 1.100E-05 | 1.000E-06 | 1.000E-05 | 1.200E-05 |
| 7941 | 398 | P398A | A | 1.5e-05 | 1.3e-05 | 1.4e-05 | 1.400E-05 | 1.000E-06 | 1.300E-05 | 1.500E-05 |
| 7942 | 398 | P398R | A | 1.1e-05 | 1e-05 | 8e-06 | 9.667E-06 | 1.528E-06 | 8.000E-06 | 1.100E-05 |
| 7943 | 398 | P398N | A | 1.6e-05 | 9.3e-06 | 1.4e-05 | 1.310E-05 | 3.439E-06 | 9.300E-06 | 1.600E-05 |
| 7944 | 398 | P398D | A | 1.4e-05 | 1.6e-05 | 1.1e-05 | 1.367E-05 | 2.517E-06 | 1.100E-05 | 1.600E-05 |
| 7945 | 398 | P398C | A | 1.5e-05 | 1.4e-05 | 1.5e-05 | 1.467E-05 | 5.774E-07 | 1.400E-05 | 1.500E-05 |
| 7946 | 398 | P398E | A | 1e-05 | 1.5e-05 | 9.9e-06 | 1.163E-05 | 2.916E-06 | 9.900E-06 | 1.500E-05 |
| 7947 | 398 | P398Q | A | 8e-06 | 1e-05 | 9.4e-06 | 9.133E-06 | 1.026E-06 | 8.000E-06 | 1.000E-05 |
| 7948 | 398 | P398G | A | 9.5e-06 | 1.6e-05 | 1.3e-05 | 1.283E-05 | 3.253E-06 | 9.500E-06 | 1.600E-05 |
| 7949 | 398 | P398H | A | 1e-05 | 9.3e-06 | 1.5e-05 | 1.143E-05 | 3.109E-06 | 9.300E-06 | 1.500E-05 |
| 7950 | 398 | P398I | A | 9.9e-06 | 1.2e-05 | 1.1e-05 | 1.097E-05 | 1.050E-06 | 9.900E-06 | 1.200E-05 |
| 7951 | 398 | P398L | A | 9.5e-06 | 1.2e-05 | 1.2e-05 | 1.117E-05 | 1.443E-06 | 9.500E-06 | 1.200E-05 |
| 7952 | 398 | P398K | A | 1.5e-05 | 1.7e-05 | 1.6e-05 | 1.600E-05 | 1.000E-06 | 1.500E-05 | 1.700E-05 |
| 7953 | 398 | P398M | A | 1.6e-05 | 1.4e-05 | 1.4e-05 | 1.467E-05 | 1.155E-06 | 1.400E-05 | 1.600E-05 |
| 7954 | 398 | P398F | A | 1.7e-05 | 1.5e-05 | 1.6e-05 | 1.600E-05 | 1.000E-06 | 1.500E-05 | 1.700E-05 |
| 7955 | 398 | P398P | A | 8.4e-06 | 1.5e-05 | 1.6e-05 | 1.313E-05 | 4.130E-06 | 8.400E-06 | 1.600E-05 |
| 7956 | 398 | P398S | A | 1.1e-05 | 1.4e-05 | 1.5e-05 | 1.333E-05 | 2.082E-06 | 1.100E-05 | 1.500E-05 |
| 7957 | 398 | P398T | A | 1.6e-05 | 1.1e-05 | 1e-05 | 1.233E-05 | 3.215E-06 | 1.000E-05 | 1.600E-05 |
| 7958 | 398 | P398W | A | 1.6e-05 | 1.2e-05 | 8.6e-06 | 1.220E-05 | 3.704E-06 | 8.600E-06 | 1.600E-05 |
| 7959 | 398 | P398Y | A | 6.3e-06 | 1.1e-05 | 1.4e-05 | 1.043E-05 | 3.881E-06 | 6.300E-06 | 1.400E-05 |
| 7960 | 398 | P398V | A | 1.5e-05 | 1.7e-05 | 1.7e-05 | 1.633E-05 | 1.155E-06 | 1.500E-05 | 1.700E-05 |
| 7961 | 399 | D399A | A | 1.3e-05 | 1.1e-05 | 1.5e-05 | 1.300E-05 | 2.000E-06 | 1.100E-05 | 1.500E-05 |
| 7962 | 399 | D399R | A | 1.6e-05 | 1e-05 | 1.3e-05 | 1.300E-05 | 3.000E-06 | 1.000E-05 | 1.600E-05 |
| 7963 | 399 | D399N | A | 9.2e-06 | 8.7e-06 | 9.7e-06 | 9.200E-06 | 5.000E-07 | 8.700E-06 | 9.700E-06 |
| 7964 | 399 | D399D | A | 2.4e-05 | 9.4e-06 | 1.1e-05 | 1.480E-05 | 8.007E-06 | 9.400E-06 | 2.400E-05 |
| 7965 | 399 | D399C | A | 1.5e-05 | 1.4e-05 | 9.7e-06 | 1.290E-05 | 2.816E-06 | 9.700E-06 | 1.500E-05 |
| 7966 | 399 | D399E | A | 1.7e-05 | 1.8e-05 | 1.4e-05 | 1.633E-05 | 2.082E-06 | 1.400E-05 | 1.800E-05 |
| 7967 | 399 | D399Q | A | 8.7e-06 | 1.2e-05 | 8.3e-06 | 9.667E-06 | 2.031E-06 | 8.300E-06 | 1.200E-05 |
| 7968 | 399 | D399G | A | 1.6e-05 | 8.5e-06 | 1.1e-05 | 1.183E-05 | 3.819E-06 | 8.500E-06 | 1.600E-05 |
| 7969 | 399 | D399H | A | 9.7e-06 | 1.1e-05 | 7.1e-06 | 9.267E-06 | 1.986E-06 | 7.100E-06 | 1.100E-05 |
| 7970 | 399 | D399I | A | 1.6e-05 | 8.6e-06 | 1.5e-05 | 1.320E-05 | 4.015E-06 | 8.600E-06 | 1.600E-05 |
| 7971 | 399 | D399L | A | 1.1e-05 | 1.1e-05 | 7e-06 | 9.667E-06 | 2.309E-06 | 7.000E-06 | 1.100E-05 |
| 7972 | 399 | D399K | A | 1.4e-05 | 1.1e-05 | 9e-06 | 1.133E-05 | 2.517E-06 | 9.000E-06 | 1.400E-05 |
| 7973 | 399 | D399M | A | 1.5e-05 | 1.5e-05 | 1.5e-05 | 1.500E-05 | 0.000E+00 | 1.500E-05 | 1.500E-05 |
| 7974 | 399 | D399F | A | 1e-05 | 1.5e-05 | 1.1e-05 | 1.200E-05 | 2.646E-06 | 1.000E-05 | 1.500E-05 |
| 7975 | 399 | D399P | A | 1.5e-05 | 1.4e-05 | 1.6e-05 | 1.500E-05 | 1.000E-06 | 1.400E-05 | 1.600E-05 |
| 7976 | 399 | D399S | A | 1.5e-05 | 1e-05 | 1.4e-05 | 1.300E-05 | 2.646E-06 | 1.000E-05 | 1.500E-05 |
| 7977 | 399 | D399T | A | 1.4e-05 | 8.9e-06 | 1.4e-05 | 1.230E-05 | 2.944E-06 | 8.900E-06 | 1.400E-05 |
| 7978 | 399 | D399W | A | 1.5e-05 | 1.5e-05 | 1.1e-05 | 1.367E-05 | 2.309E-06 | 1.100E-05 | 1.500E-05 |
| 7979 | 399 | D399Y | A | 1.3e-05 | 9.7e-06 | 1.3e-05 | 1.190E-05 | 1.905E-06 | 9.700E-06 | 1.300E-05 |
| 7980 | 399 | D399V | A | 8.6e-06 | 9.6e-06 | 1e-05 | 9.400E-06 | 7.211E-07 | 8.600E-06 | 1.000E-05 |
| 7981 | 400 | L400A | A | 8.5e-06 | 1.4e-05 | 1.1e-05 | 1.117E-05 | 2.754E-06 | 8.500E-06 | 1.400E-05 |
| 7982 | 400 | L400R | A | 1.3e-05 | 1.1e-05 | 1.3e-05 | 1.233E-05 | 1.155E-06 | 1.100E-05 | 1.300E-05 |
| 7983 | 400 | L400N | A | 1.5e-05 | 7.9e-06 | 9.5e-06 | 1.080E-05 | 3.724E-06 | 7.900E-06 | 1.500E-05 |
| 7984 | 400 | L400D | A | 1.5e-05 | 9.2e-06 | 1.1e-05 | 1.173E-05 | 2.969E-06 | 9.200E-06 | 1.500E-05 |
| 7985 | 400 | L400C | A | 9.5e-06 | 1.5e-05 | 1.6e-05 | 1.350E-05 | 3.500E-06 | 9.500E-06 | 1.600E-05 |
| 7986 | 400 | L400E | A | 1.1e-05 | 1.5e-05 | 9.9e-06 | 1.197E-05 | 2.684E-06 | 9.900E-06 | 1.500E-05 |
| 7987 | 400 | L400Q | A | 8.5e-06 | 1.5e-05 | 1.7e-05 | 1.350E-05 | 4.444E-06 | 8.500E-06 | 1.700E-05 |
| 7988 | 400 | L400G | A | 1.8e-05 | 8.9e-06 | 1.6e-05 | 1.430E-05 | 4.782E-06 | 8.900E-06 | 1.800E-05 |
| 7989 | 400 | L400H | A | 9.6e-06 | 1e-05 | 2.2e-05 | 1.387E-05 | 7.047E-06 | 9.600E-06 | 2.200E-05 |

| | | | | | | | | | | |
|---|---|---|---|---|---|---|---|---|---|---|
| 7990 | 400 | L400I | A | 1.7e-05 | 4.6e-05 | 9.3e-06 | 2.410E-05 | 1.935E-05 | 9.300E-06 | 4.600E-05 |
| 7991 | 400 | L400L | A | 1.3e-05 | 1.7e-05 | 1.6e-05 | 1.533E-05 | 2.082E-06 | 1.300E-05 | 1.700E-05 |
| 7992 | 400 | L400K | A | 1.5e-05 | 1.4e-05 | 1.5e-05 | 1.467E-05 | 5.774E-07 | 1.400E-05 | 1.500E-05 |
| 7993 | 400 | L400M | A | 1.6e-05 | 1e-05 | 1e-05 | 1.200E-05 | 3.464E-06 | 1.000E-05 | 1.600E-05 |
| 7994 | 400 | L400F | A | 8.7e-06 | 1.2e-05 | 9.5e-06 | 1.007E-05 | 1.721E-06 | 8.700E-06 | 1.200E-05 |
| 7995 | 400 | L400P | A | 1.5e-05 | 1.7e-05 | 7.1e-06 | 1.303E-05 | 5.235E-06 | 7.100E-06 | 1.700E-05 |
| 7996 | 400 | L400S | A | 1.7e-05 | 1.5e-05 | 1.7e-05 | 1.633E-05 | 1.155E-06 | 1.500E-05 | 1.700E-05 |
| 7997 | 400 | L400T | A | 9.7e-06 | 1.6e-05 | 1.2e-05 | 1.257E-05 | 3.188E-06 | 9.700E-06 | 1.600E-05 |
| 7998 | 400 | L400W | A | 1.4e-05 | 1.4e-05 | 1.1e-05 | 1.300E-05 | 1.732E-06 | 1.100E-05 | 1.400E-05 |
| 7999 | 400 | L400Y | A | 9.5e-06 | 1.5e-05 | 8.6e-06 | 1.103E-05 | 3.465E-06 | 8.600E-06 | 1.500E-05 |
| 8000 | 400 | L400V | A | 1.3e-05 | 1.1e-05 | 1.3e-05 | 1.233E-05 | 1.155E-06 | 1.100E-05 | 1.300E-05 |
| 8001 | 401 | S401A | A | 1.5e-05 | 1.6e-05 | 1.1e-05 | 1.400E-05 | 2.646E-06 | 1.100E-05 | 1.600E-05 |
| 8002 | 401 | S401R | A | 1.3e-05 | 1.1e-05 | 1.1e-05 | 1.167E-05 | 1.155E-06 | 1.100E-05 | 1.300E-05 |
| 8003 | 401 | S401N | A | 9.4e-06 | 1.4e-05 | 1.3e-05 | 1.213E-05 | 2.419E-06 | 9.400E-06 | 1.400E-05 |
| 8004 | 401 | S401D | A | 1.2e-05 | 1.5e-05 | 9.4e-06 | 1.213E-05 | 2.802E-06 | 9.400E-06 | 1.500E-05 |
| 8005 | 401 | S401C | A | 1.1e-05 | 1.1e-05 | 1.6e-05 | 1.267E-05 | 2.887E-06 | 1.100E-05 | 1.600E-05 |
| 8006 | 401 | S401E | A | 8.1e-06 | 9.2e-06 | 1.4e-05 | 1.043E-05 | 3.137E-06 | 8.100E-06 | 1.400E-05 |
| 8007 | 401 | S401Q | A | 1.4e-05 | 1.2e-05 | 1.1e-05 | 1.233E-05 | 1.528E-06 | 1.100E-05 | 1.400E-05 |
| 8008 | 401 | S401G | A | 1.6e-05 | 1.1e-05 | 1.1e-05 | 1.267E-05 | 2.887E-06 | 1.100E-05 | 1.600E-05 |
| 8009 | 401 | S401H | A | 1.7e-05 | 1.5e-05 | 1.4e-05 | 1.533E-05 | 1.528E-06 | 1.400E-05 | 1.700E-05 |
| 8010 | 401 | S401I | A | 1e-05 | 1e-05 | 1.6e-05 | 1.200E-05 | 3.464E-06 | 1.000E-05 | 1.600E-05 |
| 8011 | 401 | S401L | A | 1.1e-05 | 8.9e-06 | 1.6e-05 | 1.197E-05 | 3.647E-06 | 8.900E-06 | 1.600E-05 |
| 8012 | 401 | S401K | A | 1.6e-05 | 6.7e-06 | 1.2e-05 | 1.157E-05 | 4.665E-06 | 6.700E-06 | 1.600E-05 |
| 8013 | 401 | S401M | A | 9.4e-06 | 1e-05 | 1.5e-05 | 1.147E-05 | 3.075E-06 | 9.400E-06 | 1.500E-05 |
| 8014 | 401 | S401F | A | 1.6e-05 | 1.7e-05 | 1.5e-05 | 1.600E-05 | 1.000E-06 | 1.500E-05 | 1.700E-05 |
| 8015 | 401 | S401P | A | 1.1e-05 | 8.8e-06 | 9.6e-06 | 9.800E-06 | 1.114E-06 | 8.800E-06 | 1.100E-05 |
| 8016 | 401 | S401S | A | 1.5e-05 | 1.1e-05 | 1.6e-05 | 1.400E-05 | 2.646E-06 | 1.100E-05 | 1.600E-05 |
| 8017 | 401 | S401T | A | 1.1e-05 | 1.3e-05 | 1.1e-05 | 1.167E-05 | 1.155E-06 | 1.100E-05 | 1.300E-05 |
| 8018 | 401 | S401W | A | 1.1e-05 | 1.5e-05 | 1.5e-05 | 1.367E-05 | 2.309E-06 | 1.100E-05 | 1.500E-05 |
| 8019 | 401 | S401Y | A | 1.5e-05 | 1.1e-05 | 1.4e-05 | 1.333E-05 | 2.082E-06 | 1.100E-05 | 1.500E-05 |
| 8020 | 401 | S401V | A | 1.5e-05 | 8.3e-06 | 1.6e-05 | 1.310E-05 | 4.187E-06 | 8.300E-06 | 1.600E-05 |
| 8021 | 402 | V402A | A | 1.3e-05 | 1.5e-05 | 1.4e-05 | 1.400E-05 | 1.000E-06 | 1.300E-05 | 1.500E-05 |
| 8022 | 402 | V402R | A | 1.7e-05 | 1.3e-05 | 1.7e-05 | 1.567E-05 | 2.309E-06 | 1.300E-05 | 1.700E-05 |
| 8023 | 402 | V402N | A | 9.1e-06 | 1e-05 | 9.9e-06 | 9.667E-06 | 4.933E-07 | 9.100E-06 | 1.000E-05 |
| 8024 | 402 | V402D | A | 1.6e-05 | 1.2e-05 | 1e-05 | 1.267E-05 | 3.055E-06 | 1.000E-05 | 1.600E-05 |
| 8025 | 402 | V402C | A | 1.4e-05 | 1.5e-05 | 1.1e-05 | 1.333E-05 | 2.082E-06 | 1.100E-05 | 1.500E-05 |
| 8026 | 402 | V402E | A | 9.5e-06 | 1.5e-05 | 1.2e-05 | 1.217E-05 | 2.754E-06 | 9.500E-06 | 1.500E-05 |
| 8027 | 402 | V402Q | A | 1.4e-05 | 1.6e-05 | 1.6e-05 | 1.533E-05 | 1.155E-06 | 1.400E-05 | 1.600E-05 |
| 8028 | 402 | V402G | A | 1.1e-05 | 1.2e-05 | 1.6e-05 | 1.300E-05 | 2.646E-06 | 1.100E-05 | 1.600E-05 |
| 8029 | 402 | V402H | A | 1.6e-05 | 1.3e-05 | 7.4e-06 | 1.213E-05 | 4.365E-06 | 7.400E-06 | 1.600E-05 |
| 8030 | 402 | V402I | A | 1.5e-05 | 1.4e-05 | 9.5e-06 | 1.283E-05 | 2.930E-06 | 9.500E-06 | 1.500E-05 |
| 8031 | 402 | V402L | A | 9.1e-06 | 1.5e-05 | 1e-05 | 1.137E-05 | 3.179E-06 | 9.100E-06 | 1.500E-05 |
| 8032 | 402 | V402K | A | 1.1e-05 | 1.4e-05 | 1e-05 | 1.167E-05 | 2.082E-06 | 1.000E-05 | 1.400E-05 |
| 8033 | 402 | V402M | A | 1.6e-05 | 8.7e-06 | 1.5e-05 | 1.323E-05 | 3.958E-06 | 8.700E-06 | 1.600E-05 |
| 8034 | 402 | V402F | A | 9.6e-06 | 1.7e-05 | 9.2e-06 | 1.193E-05 | 4.392E-06 | 9.200E-06 | 1.700E-05 |
| 8035 | 402 | V402P | A | 1.1e-05 | 1.2e-05 | 1e-05 | 1.100E-05 | 1.000E-06 | 1.000E-05 | 1.200E-05 |
| 8036 | 402 | V402S | A | 9.6e-06 | 8.7e-06 | 8.3e-06 | 8.867E-06 | 6.658E-07 | 8.300E-06 | 9.600E-06 |
| 8037 | 402 | V402T | A | 1.4e-05 | 1.6e-05 | 1.4e-05 | 1.467E-05 | 1.155E-06 | 1.400E-05 | 1.600E-05 |
| 8038 | 402 | V402W | A | 9.2e-06 | 8.4e-06 | 1.1e-05 | 9.533E-06 | 1.332E-06 | 8.400E-06 | 1.100E-05 |
| 8039 | 402 | V402Y | A | 1.4e-05 | 1.5e-05 | 1.1e-05 | 1.333E-05 | 2.082E-06 | 1.100E-05 | 1.500E-05 |
| 8040 | 402 | V402V | A | 1.5e-05 | 9.1e-06 | 1.3e-05 | 1.237E-05 | 3.001E-06 | 9.100E-06 | 1.500E-05 |
| 8041 | 403 | F403A | A | 1.5e-05 | 1.2e-05 | 1.5e-05 | 1.400E-05 | 1.732E-06 | 1.200E-05 | 1.500E-05 |
| 8042 | 403 | F403R | A | 1e-05 | 1.1e-05 | 1.5e-05 | 1.200E-05 | 2.646E-06 | 1.000E-05 | 1.500E-05 |
| 8043 | 403 | F403N | A | 1.4e-05 | 8.2e-06 | 1.3e-05 | 1.173E-05 | 3.101E-06 | 8.200E-06 | 1.400E-05 |
| 8044 | 403 | F403D | A | 1.1e-05 | 1.4e-05 | 1.6e-05 | 1.367E-05 | 2.517E-06 | 1.100E-05 | 1.600E-05 |
| 8045 | 403 | F403C | A | 1.1e-05 | 1e-05 | 1.6e-05 | 1.233E-05 | 3.215E-06 | 1.000E-05 | 1.600E-05 |
| 8046 | 403 | F403E | A | 1.1e-05 | 1.1e-05 | 1.1e-05 | 1.100E-05 | 2.075E-21 | 1.100E-05 | 1.100E-05 |
| 8047 | 403 | F403Q | A | 8.3e-06 | 1.1e-05 | 1.4e-05 | 1.110E-05 | 2.851E-06 | 8.300E-06 | 1.400E-05 |
| 8048 | 403 | F403G | A | 1.7e-05 | 1.6e-05 | 1.2e-05 | 1.500E-05 | 2.646E-06 | 1.200E-05 | 1.700E-05 |
| 8049 | 403 | F403H | A | 1.3e-05 | 9.2e-06 | 1.1e-05 | 1.107E-05 | 1.901E-06 | 9.200E-06 | 1.300E-05 |
| 8050 | 403 | F403I | A | 1.4e-05 | 1.7e-05 | 1.2e-05 | 1.433E-05 | 2.517E-06 | 1.200E-05 | 1.700E-05 |
| 8051 | 403 | F403L | A | 1.5e-05 | 1.5e-05 | 1.6e-05 | 1.533E-05 | 5.774E-07 | 1.500E-05 | 1.600E-05 |
| 8052 | 403 | F403K | A | 1.8e-05 | 1.2e-05 | 9.2e-06 | 1.307E-05 | 4.496E-06 | 9.200E-06 | 1.800E-05 |
| 8053 | 403 | F403M | A | 8.3e-06 | 1.1e-05 | 1.2e-05 | 1.043E-05 | 1.914E-06 | 8.300E-06 | 1.200E-05 |
| 8054 | 403 | F403F | A | 1.5e-05 | 1.1e-05 | 1.2e-05 | 1.267E-05 | 2.082E-06 | 1.100E-05 | 1.500E-05 |
| 8055 | 403 | F403P | A | 8.6e-06 | 1.2e-05 | 1.6e-05 | 1.220E-05 | 3.704E-06 | 8.600E-06 | 1.600E-05 |
| 8056 | 403 | F403S | A | 1.5e-05 | 1e-05 | 1e-05 | 1.167E-05 | 2.887E-06 | 1.000E-05 | 1.500E-05 |
| 8057 | 403 | F403T | A | 1.7e-05 | 1.5e-05 | 1.5e-05 | 1.567E-05 | 1.155E-06 | 1.500E-05 | 1.700E-05 |
| 8058 | 403 | F403W | A | 1e-05 | 1.1e-05 | 1.7e-05 | 1.267E-05 | 3.786E-06 | 1.000E-05 | 1.700E-05 |
| 8059 | 403 | F403Y | A | 1.4e-05 | 1.5e-05 | 1.6e-05 | 1.500E-05 | 1.000E-06 | 1.400E-05 | 1.600E-05 |
| 8060 | 403 | F403V | A | 8.3e-06 | 1.6e-05 | 1.5e-05 | 1.310E-05 | 4.187E-06 | 8.300E-06 | 1.600E-05 |
| 8061 | 404 | Q404A | A | 1.3e-05 | 1.4e-05 | 9.4e-06 | 1.213E-05 | 2.419E-06 | 9.400E-06 | 1.400E-05 |
| 8062 | 404 | Q404R | A | 1.6e-05 | 1.6e-05 | 1.5e-05 | 1.567E-05 | 5.774E-07 | 1.500E-05 | 1.600E-05 |
| 8063 | 404 | Q404N | A | 1.5e-05 | 1.1e-05 | 1.5e-05 | 1.367E-05 | 2.309E-06 | 1.100E-05 | 1.500E-05 |
| 8064 | 404 | Q404D | A | 1.7e-05 | 1e-05 | 9.6e-06 | 1.220E-05 | 4.162E-06 | 9.600E-06 | 1.700E-05 |
| 8065 | 404 | Q404C | A | 1e-05 | 1.6e-05 | 1.5e-05 | 1.367E-05 | 3.215E-06 | 1.000E-05 | 1.600E-05 |
| 8066 | 404 | Q404E | A | 1.6e-05 | 1.4e-05 | 1.1e-05 | 1.367E-05 | 2.517E-06 | 1.100E-05 | 1.600E-05 |
| 8067 | 404 | Q404Q | A | 1e-05 | 1.1e-05 | 1.6e-05 | 1.233E-05 | 3.215E-06 | 1.000E-05 | 1.600E-05 |
| 8068 | 404 | Q404G | A | 9.2e-06 | 1.7e-05 | 1.6e-05 | 1.407E-05 | 4.244E-06 | 9.200E-06 | 1.700E-05 |
| 8069 | 404 | Q404H | A | 9.2e-06 | 1.7e-05 | 1.1e-05 | 1.240E-05 | 4.084E-06 | 9.200E-06 | 1.700E-05 |
| 8070 | 404 | Q404I | A | 1.5e-05 | 1.4e-05 | 9.1e-06 | 1.270E-05 | 3.158E-06 | 9.100E-06 | 1.500E-05 |
| 8071 | 404 | Q404L | A | 1.6e-05 | 1.7e-05 | 1.5e-05 | 1.600E-05 | 1.000E-06 | 1.500E-05 | 1.700E-05 |
| 8072 | 404 | Q404K | A | 8.1e-06 | 1.5e-05 | 1.1e-05 | 1.137E-05 | 3.465E-06 | 8.100E-06 | 1.500E-05 |
| 8073 | 404 | Q404M | A | 9.9e-06 | 1.8e-05 | 1.4e-05 | 1.397E-05 | 4.050E-06 | 9.900E-06 | 1.800E-05 |
| 8074 | 404 | Q404F | A | 1.5e-05 | 1.9e-05 | 1.5e-05 | 1.633E-05 | 2.309E-06 | 1.500E-05 | 1.900E-05 |

| | | | | | | | | | | |
|---|---|---|---|---|---|---|---|---|---|---|
| 8075 | 404 | Q404P | A | 1.5e-05 | 1.6e-05 | 7e-06 | 1.267E-05 | 4.933E-06 | 7.000E-06 | 1.600E-05 |
| 8076 | 404 | Q404S | A | 1.3e-05 | 1.1e-05 | 1.6e-05 | 1.333E-05 | 2.517E-06 | 1.100E-05 | 1.600E-05 |
| 8077 | 404 | Q404T | A | 8.1e-06 | 1.5e-05 | 9.5e-06 | 1.087E-05 | 3.647E-06 | 8.100E-06 | 1.500E-05 |
| 8078 | 404 | Q404W | A | 1.4e-05 | 9.8e-06 | 1.5e-05 | 1.293E-05 | 2.759E-06 | 9.800E-06 | 1.500E-05 |
| 8079 | 404 | Q404Y | A | 1.4e-05 | 1.7e-05 | 1.3e-05 | 1.467E-05 | 2.082E-06 | 1.300E-05 | 1.700E-05 |
| 8080 | 404 | Q404V | A | 1.3e-05 | 1.5e-05 | 9.3e-06 | 1.243E-05 | 2.892E-06 | 9.300E-06 | 1.500E-05 |
| 8081 | 405 | N405A | A | 1.1e-05 | 8.2e-06 | 1.6e-05 | 1.173E-05 | 3.951E-06 | 8.200E-06 | 1.600E-05 |
| 8082 | 405 | N405R | A | 1e-05 | 1.6e-05 | 1.5e-05 | 1.367E-05 | 3.215E-06 | 1.000E-05 | 1.600E-05 |
| 8083 | 405 | N405N | A | 7.7e-06 | 1.3e-05 | 1.5e-05 | 1.190E-05 | 3.772E-06 | 7.700E-06 | 1.500E-05 |
| 8084 | 405 | N405D | A | 9.9e-06 | 1.3e-05 | 1.6e-05 | 1.297E-05 | 3.050E-06 | 9.900E-06 | 1.600E-05 |
| 8085 | 405 | N405C | A | 1.6e-05 | 7.6e-06 | 1.6e-05 | 1.320E-05 | 4.850E-06 | 7.600E-06 | 1.600E-05 |
| 8086 | 405 | N405E | A | 7.3e-06 | 1.6e-05 | 1.1e-05 | 1.143E-05 | 4.366E-06 | 7.300E-06 | 1.600E-05 |
| 8087 | 405 | N405Q | A | 5.6e-06 | 1.5e-05 | 1.7e-05 | 1.253E-05 | 6.087E-06 | 5.600E-06 | 1.700E-05 |
| 8088 | 405 | N405G | A | 1.4e-05 | 1.5e-05 | 1.3e-05 | 1.400E-05 | 1.000E-06 | 1.300E-05 | 1.500E-05 |
| 8089 | 405 | N405H | A | 9.6e-06 | 1.1e-05 | 1.7e-05 | 1.253E-05 | 3.931E-06 | 9.600E-06 | 1.700E-05 |
| 8090 | 405 | N405I | A | 1e-05 | 1.3e-05 | 1.3e-05 | 1.200E-05 | 1.732E-06 | 1.000E-05 | 1.300E-05 |
| 8091 | 405 | N405L | A | 1.1e-05 | 9.2e-06 | 1.5e-05 | 1.173E-05 | 2.969E-06 | 9.200E-06 | 1.500E-05 |
| 8092 | 405 | N405K | A | 1.1e-05 | 1.6e-05 | 1e-05 | 1.233E-05 | 3.215E-06 | 1.000E-05 | 1.600E-05 |
| 8093 | 405 | N405M | A | 9.7e-06 | 1.4e-05 | 1.5e-05 | 1.290E-05 | 2.816E-06 | 9.700E-06 | 1.500E-05 |
| 8094 | 405 | N405F | A | 1.5e-05 | 1.7e-05 | 1e-05 | 1.400E-05 | 3.606E-06 | 1.000E-05 | 1.700E-05 |
| 8095 | 405 | N405P | A | 1.5e-05 | 9.1e-06 | 1.3e-05 | 1.237E-05 | 3.001E-06 | 9.100E-06 | 1.500E-05 |
| 8096 | 405 | N405S | A | 1.1e-05 | 1.1e-05 | 1e-05 | 1.067E-05 | 5.774E-07 | 1.000E-05 | 1.100E-05 |
| 8097 | 405 | N405T | A | 9.5e-06 | 1.6e-05 | 1e-05 | 1.183E-05 | 3.617E-06 | 9.500E-06 | 1.600E-05 |
| 8098 | 405 | N405W | A | 1e-05 | 1.5e-05 | 1e-05 | 1.167E-05 | 2.887E-06 | 1.000E-05 | 1.500E-05 |
| 8099 | 405 | N405Y | A | 1.4e-05 | 1.5e-05 | 1.2e-05 | 1.367E-05 | 1.528E-06 | 1.200E-05 | 1.500E-05 |
| 8100 | 405 | N405V | A | 1.7e-05 | 1.1e-05 | 9.8e-06 | 1.260E-05 | 3.857E-06 | 9.800E-06 | 1.700E-05 |
| 8101 | 406 | L406A | A | 1.1e-05 | 1.4e-05 | 1.5e-05 | 1.333E-05 | 2.082E-06 | 1.100E-05 | 1.500E-05 |
| 8102 | 406 | L406R | A | 8.6e-06 | 1.3e-05 | 1.5e-05 | 1.220E-05 | 3.274E-06 | 8.600E-06 | 1.500E-05 |
| 8103 | 406 | L406N | A | 1.5e-05 | 1.4e-05 | 1.1e-05 | 1.333E-05 | 2.082E-06 | 1.100E-05 | 1.500E-05 |
| 8104 | 406 | L406D | A | 1.7e-05 | 1.4e-05 | 1.1e-05 | 1.400E-05 | 3.000E-06 | 1.100E-05 | 1.700E-05 |
| 8105 | 406 | L406C | A | 1.4e-05 | 1.4e-05 | 1e-05 | 1.267E-05 | 2.309E-06 | 1.000E-05 | 1.400E-05 |
| 8106 | 406 | L406E | A | 1.1e-05 | 9.5e-06 | 1.5e-05 | 1.183E-05 | 2.843E-06 | 9.500E-06 | 1.500E-05 |
| 8107 | 406 | L406Q | A | 1e-05 | 1.6e-05 | 1.7e-05 | 1.433E-05 | 3.786E-06 | 1.000E-05 | 1.700E-05 |
| 8108 | 406 | L406G | A | 1.5e-05 | 1.8e-05 | 1.1e-05 | 1.467E-05 | 3.512E-06 | 1.100E-05 | 1.800E-05 |
| 8109 | 406 | L406H | A | 1.6e-05 | 9.7e-06 | 1.6e-05 | 1.390E-05 | 3.637E-06 | 9.700E-06 | 1.600E-05 |
| 8110 | 406 | L406I | A | 1e-05 | 9.3e-06 | 1.1e-05 | 1.010E-05 | 8.544E-07 | 9.300E-06 | 1.100E-05 |
| 8111 | 406 | L406L | A | 1.5e-05 | 1e-05 | 1.6e-05 | 1.367E-05 | 3.215E-06 | 1.000E-05 | 1.600E-05 |
| 8112 | 406 | L406K | A | 1e-05 | 8.8e-06 | 1.7e-05 | 1.193E-05 | 4.429E-06 | 8.800E-06 | 1.700E-05 |
| 8113 | 406 | L406M | A | 1.3e-05 | 1.4e-05 | 1.1e-05 | 1.267E-05 | 1.528E-06 | 1.100E-05 | 1.400E-05 |
| 8114 | 406 | L406F | A | 1.1e-05 | 1.5e-05 | 1.6e-05 | 1.400E-05 | 2.646E-06 | 1.100E-05 | 1.600E-05 |
| 8115 | 406 | L406P | A | 1.6e-05 | 1.7e-05 | 1.1e-05 | 1.467E-05 | 3.215E-06 | 1.100E-05 | 1.700E-05 |
| 8116 | 406 | L406S | A | 1.6e-05 | 1.4e-05 | 1.2e-05 | 1.400E-05 | 2.000E-06 | 1.200E-05 | 1.600E-05 |
| 8117 | 406 | L406T | A | 1.4e-05 | 1e-05 | 1.6e-05 | 1.333E-05 | 3.055E-06 | 1.000E-05 | 1.600E-05 |
| 8118 | 406 | L406W | A | 1.5e-05 | 1.3e-05 | 1.5e-05 | 1.433E-05 | 1.155E-06 | 1.300E-05 | 1.500E-05 |
| 8119 | 406 | L406Y | A | 1.7e-05 | 1.7e-05 | 1.6e-05 | 1.667E-05 | 5.774E-07 | 1.600E-05 | 1.700E-05 |
| 8120 | 406 | L406V | A | 1.6e-05 | 1.1e-05 | 1.6e-05 | 1.433E-05 | 2.887E-06 | 1.100E-05 | 1.600E-05 |
| 8121 | 407 | Q407A | A | 9.1e-06 | 5.8e-06 | 1.7e-05 | 1.063E-05 | 5.755E-06 | 5.800E-06 | 1.700E-05 |
| 8122 | 407 | Q407R | A | 8.9e-06 | 1.4e-05 | 1.6e-05 | 1.297E-05 | 3.661E-06 | 8.900E-06 | 1.600E-05 |
| 8123 | 407 | Q407N | A | 1.7e-05 | 1.4e-05 | 1.1e-05 | 1.400E-05 | 3.000E-06 | 1.100E-05 | 1.700E-05 |
| 8124 | 407 | Q407D | A | 1.1e-05 | 1.1e-05 | 9.9e-06 | 1.063E-05 | 6.351E-07 | 9.900E-06 | 1.100E-05 |
| 8125 | 407 | Q407C | A | 1.4e-05 | 1.4e-05 | 1.5e-05 | 1.433E-05 | 5.774E-07 | 1.400E-05 | 1.500E-05 |
| 8126 | 407 | Q407E | A | 8.2e-06 | 1.2e-05 | 1.6e-05 | 1.207E-05 | 3.900E-06 | 8.200E-06 | 1.600E-05 |
| 8127 | 407 | Q407Q | A | 9.8e-06 | 1.3e-05 | 7.8e-06 | 1.020E-05 | 2.623E-06 | 7.800E-06 | 1.300E-05 |
| 8128 | 407 | Q407G | A | 8.7e-06 | 8.3e-06 | 1.1e-05 | 9.333E-06 | 1.457E-06 | 8.300E-06 | 1.100E-05 |
| 8129 | 407 | Q407H | A | 1.3e-05 | 1e-05 | 1.3e-05 | 1.200E-05 | 1.732E-06 | 1.000E-05 | 1.300E-05 |
| 8130 | 407 | Q407I | A | 1.3e-05 | 2.1e-05 | 1.3e-05 | 1.567E-05 | 4.619E-06 | 1.300E-05 | 2.100E-05 |
| 8131 | 407 | Q407L | A | 9.6e-06 | 1.5e-05 | 1.1e-05 | 1.187E-05 | 2.802E-06 | 9.600E-06 | 1.500E-05 |
| 8132 | 407 | Q407K | A | 8.6e-06 | 1.5e-05 | 9.9e-06 | 1.117E-05 | 3.383E-06 | 8.600E-06 | 1.500E-05 |
| 8133 | 407 | Q407M | A | 1.1e-05 | 1.6e-05 | 1.6e-05 | 1.433E-05 | 2.887E-06 | 1.100E-05 | 1.600E-05 |
| 8134 | 407 | Q407F | A | 9.8e-06 | 1e-05 | 1.6e-05 | 1.193E-05 | 3.523E-06 | 9.800E-06 | 1.600E-05 |
| 8135 | 407 | Q407P | A | 9.5e-06 | 1e-05 | 1.2e-05 | 1.050E-05 | 1.323E-06 | 9.500E-06 | 1.200E-05 |
| 8136 | 407 | Q407S | A | 1.4e-05 | 1e-05 | 1.5e-05 | 1.300E-05 | 2.646E-06 | 1.000E-05 | 1.500E-05 |
| 8137 | 407 | Q407T | A | 1.5e-05 | 1.5e-05 | 1.1e-05 | 1.367E-05 | 2.309E-06 | 1.100E-05 | 1.500E-05 |
| 8138 | 407 | Q407W | A | 1.2e-05 | 1.5e-05 | 1.4e-05 | 1.367E-05 | 1.528E-06 | 1.200E-05 | 1.500E-05 |
| 8139 | 407 | Q407Y | A | 1.5e-05 | 1.1e-05 | 1.1e-05 | 1.233E-05 | 2.309E-06 | 1.100E-05 | 1.500E-05 |
| 8140 | 407 | Q407V | A | 1.5e-05 | 8.9e-06 | 1.1e-05 | 1.163E-05 | 3.099E-06 | 8.900E-06 | 1.500E-05 |
| 8141 | 408 | V408A | A | 1.1e-05 | 1.2e-05 | 8.4e-06 | 1.047E-05 | 1.858E-06 | 8.400E-06 | 1.200E-05 |
| 8142 | 408 | V408R | A | 1.4e-05 | 1.2e-05 | 1.4e-05 | 1.333E-05 | 1.155E-06 | 1.200E-05 | 1.400E-05 |
| 8143 | 408 | V408N | A | 1.4e-05 | 9e-06 | 1.4e-05 | 1.233E-05 | 2.887E-06 | 9.000E-06 | 1.400E-05 |
| 8144 | 408 | V408D | A | 1.5e-05 | 1.5e-05 | 1.1e-05 | 1.367E-05 | 2.309E-06 | 1.100E-05 | 1.500E-05 |
| 8145 | 408 | V408C | A | 1.4e-05 | 1.1e-05 | 9.3e-06 | 1.143E-05 | 2.380E-06 | 9.300E-06 | 1.400E-05 |
| 8146 | 408 | V408E | A | 1e-05 | 1.3e-05 | 1.1e-05 | 1.133E-05 | 1.528E-06 | 1.000E-05 | 1.300E-05 |
| 8147 | 408 | V408Q | A | 1.2e-05 | 1.3e-05 | 1.1e-05 | 1.200E-05 | 1.000E-06 | 1.100E-05 | 1.300E-05 |
| 8148 | 408 | V408G | A | 1.1e-05 | 1.1e-05 | 1.5e-05 | 1.233E-05 | 2.309E-06 | 1.100E-05 | 1.500E-05 |
| 8149 | 408 | V408H | A | 1.6e-05 | 1.4e-05 | 1.5e-05 | 1.500E-05 | 1.000E-06 | 1.400E-05 | 1.600E-05 |
| 8150 | 408 | V408I | A | 1.1e-05 | 1e-05 | 1.6e-05 | 1.233E-05 | 3.215E-06 | 1.000E-05 | 1.600E-05 |
| 8151 | 408 | V408L | A | 9.6e-06 | 1.1e-05 | 1.2e-05 | 1.087E-05 | 1.206E-06 | 9.600E-06 | 1.200E-05 |
| 8152 | 408 | V408K | A | 1.6e-05 | 1.4e-05 | 1.5e-05 | 1.500E-05 | 1.000E-06 | 1.400E-05 | 1.600E-05 |
| 8153 | 408 | V408M | A | 1.6e-05 | 1.6e-05 | 9.9e-06 | 1.397E-05 | 3.522E-06 | 9.900E-06 | 1.600E-05 |
| 8154 | 408 | V408F | A | 9.8e-06 | 1e-05 | 1.3e-05 | 1.093E-05 | 1.793E-06 | 9.800E-06 | 1.300E-05 |
| 8155 | 408 | V408P | A | 9.5e-06 | 2.7e-05 | 1e-05 | 1.550E-05 | 9.962E-06 | 9.500E-06 | 2.700E-05 |
| 8156 | 408 | V408S | A | 1.5e-05 | 1.4e-05 | 8.7e-06 | 1.257E-05 | 3.386E-06 | 8.700E-06 | 1.500E-05 |
| 8157 | 408 | V408T | A | 1.4e-05 | 1.1e-05 | 1.5e-05 | 1.333E-05 | 2.082E-06 | 1.100E-05 | 1.500E-05 |
| 8158 | 408 | V408W | A | 1e-05 | 1.5e-05 | 1.3e-05 | 1.267E-05 | 2.517E-06 | 1.000E-05 | 1.500E-05 |
| 8159 | 408 | V408Y | A | 9.6e-06 | 9.4e-06 | 1.1e-05 | 1.000E-05 | 8.718E-07 | 9.400E-06 | 1.100E-05 |

| | | | | | | | | | | |
|---|---|---|---|---|---|---|---|---|---|---|
| 8160 | 408 | V408V | A | 1.6e-05 | 1.1e-05 | 1.2e-05 | 1.300E-05 | 2.646E-06 | 1.100E-05 | 1.600E-05 |
| 8161 | 409 | I409A | A | 1.6e-05 | 1.3e-05 | 1.6e-05 | 1.500E-05 | 1.732E-06 | 1.300E-05 | 1.600E-05 |
| 8162 | 409 | I409R | A | 1.7e-05 | 1.5e-05 | 1.6e-05 | 1.600E-05 | 1.000E-06 | 1.500E-05 | 1.700E-05 |
| 8163 | 409 | I409N | A | 1.5e-05 | 1.5e-05 | 9.3e-06 | 1.310E-05 | 3.291E-06 | 9.300E-06 | 1.500E-05 |
| 8164 | 409 | I409D | A | 1.1e-05 | 8.8e-06 | 9.2e-06 | 9.667E-06 | 1.172E-06 | 8.800E-06 | 1.100E-05 |
| 8165 | 409 | I409C | A | 1e-05 | 1.4e-05 | 1.5e-05 | 1.300E-05 | 2.646E-06 | 1.000E-05 | 1.500E-05 |
| 8166 | 409 | I409E | A | 1.4e-05 | 1.6e-05 | 1.5e-05 | 1.500E-05 | 1.000E-06 | 1.400E-05 | 1.600E-05 |
| 8167 | 409 | I409Q | A | 1.7e-05 | 1.3e-05 | 1.9e-05 | 1.633E-05 | 3.055E-06 | 1.300E-05 | 1.900E-05 |
| 8168 | 409 | I409G | A | 1.2e-05 | 1e-05 | 1.6e-05 | 1.267E-05 | 3.055E-06 | 1.000E-05 | 1.600E-05 |
| 8169 | 409 | I409H | A | 1.4e-05 | 1.6e-05 | 1.5e-05 | 1.500E-05 | 1.000E-06 | 1.400E-05 | 1.600E-05 |
| 8170 | 409 | I409I | A | 1.7e-05 | 1.4e-05 | 1.4e-05 | 1.500E-05 | 1.732E-06 | 1.400E-05 | 1.700E-05 |
| 8171 | 409 | I409L | A | 1.5e-05 | 1.1e-05 | 8.7e-06 | 1.157E-05 | 3.188E-06 | 8.700E-06 | 1.500E-05 |
| 8172 | 409 | I409K | A | 1.1e-05 | 1.4e-05 | 1.7e-05 | 1.400E-05 | 3.000E-06 | 1.100E-05 | 1.700E-05 |
| 8173 | 409 | I409M | A | 1.1e-05 | 1.4e-05 | 1.1e-05 | 1.200E-05 | 1.732E-06 | 1.100E-05 | 1.400E-05 |
| 8174 | 409 | I409F | A | 9.4e-06 | 1.5e-05 | 1.7e-05 | 1.380E-05 | 3.940E-06 | 9.400E-06 | 1.700E-05 |
| 8175 | 409 | I409P | A | 1.6e-05 | 1.6e-05 | 1.5e-05 | 1.567E-05 | 5.774E-07 | 1.500E-05 | 1.600E-05 |
| 8176 | 409 | I409S | A | 1.7e-05 | 1.1e-05 | 1.1e-05 | 1.300E-05 | 3.464E-06 | 1.100E-05 | 1.700E-05 |
| 8177 | 409 | I409T | A | 1.5e-05 | 9.9e-06 | 1.4e-05 | 1.297E-05 | 2.702E-06 | 9.900E-06 | 1.500E-05 |
| 8178 | 409 | I409W | A | 9.9e-06 | 1.1e-05 | 7.3e-06 | 9.400E-06 | 1.900E-06 | 7.300E-06 | 1.100E-05 |
| 8179 | 409 | I409Y | A | 1.5e-05 | 1e-05 | 1.7e-05 | 1.400E-05 | 3.606E-06 | 1.000E-05 | 1.700E-05 |
| 8180 | 409 | I409V | A | 1.7e-05 | 1.1e-05 | 1.2e-05 | 1.333E-05 | 3.215E-06 | 1.100E-05 | 1.700E-05 |
| 8181 | 410 | R410A | A | 9e-06 | 1.7e-05 | 1.6e-05 | 1.400E-05 | 4.359E-06 | 9.000E-06 | 1.700E-05 |
| 8182 | 410 | R410R | A | 1.4e-05 | 1.3e-05 | 9.6e-06 | 1.220E-05 | 2.307E-06 | 9.600E-06 | 1.400E-05 |
| 8183 | 410 | R410N | A | 1.4e-05 | 1.2e-05 | 1.2e-05 | 1.267E-05 | 1.155E-06 | 1.200E-05 | 1.400E-05 |
| 8184 | 410 | R410D | A | 1.1e-05 | 1.4e-05 | 1.1e-05 | 1.200E-05 | 1.732E-06 | 1.100E-05 | 1.400E-05 |
| 8185 | 410 | R410C | A | 1.4e-05 | 1.7e-05 | 1.5e-05 | 1.533E-05 | 1.528E-06 | 1.400E-05 | 1.700E-05 |
| 8186 | 410 | R410E | A | 8.4e-06 | 1.5e-05 | 1.5e-05 | 1.280E-05 | 3.811E-06 | 8.400E-06 | 1.500E-05 |
| 8187 | 410 | R410Q | A | 9.3e-06 | 8.7e-06 | 1e-05 | 9.333E-06 | 6.506E-07 | 8.700E-06 | 1.000E-05 |
| 8188 | 410 | R410G | A | 1.4e-05 | 1.5e-05 | 1.5e-05 | 1.467E-05 | 5.774E-07 | 1.400E-05 | 1.500E-05 |
| 8189 | 410 | R410H | A | 1.5e-05 | 1.5e-05 | 1.6e-05 | 1.533E-05 | 5.774E-07 | 1.500E-05 | 1.600E-05 |
| 8190 | 410 | R410I | A | 1.1e-05 | 1.4e-05 | 9.7e-06 | 1.157E-05 | 2.205E-06 | 9.700E-06 | 1.400E-05 |
| 8191 | 410 | R410L | A | 1.1e-05 | 8.2e-06 | 8.8e-06 | 9.333E-06 | 1.474E-06 | 8.200E-06 | 1.100E-05 |
| 8192 | 410 | R410K | A | 1.6e-05 | 7.2e-06 | 1.1e-05 | 1.140E-05 | 4.414E-06 | 7.200E-06 | 1.600E-05 |
| 8193 | 410 | R410M | A | 1e-05 | 8.8e-06 | 1.5e-05 | 1.127E-05 | 3.288E-06 | 8.800E-06 | 1.500E-05 |
| 8194 | 410 | R410F | A | 1.5e-05 | 1.2e-05 | 1.2e-05 | 1.300E-05 | 1.732E-06 | 1.200E-05 | 1.500E-05 |
| 8195 | 410 | R410P | A | 1.3e-05 | 1e-05 | 1.4e-05 | 1.233E-05 | 2.082E-06 | 1.000E-05 | 1.400E-05 |
| 8196 | 410 | R410S | A | 1.1e-05 | 1.3e-05 | 1.6e-05 | 1.333E-05 | 2.517E-06 | 1.100E-05 | 1.600E-05 |
| 8197 | 410 | R410T | A | 1.3e-05 | 8.3e-06 | 1.6e-05 | 1.243E-05 | 3.881E-06 | 8.300E-06 | 1.600E-05 |
| 8198 | 410 | R410W | A | 1.4e-05 | 1.3e-05 | 9.9e-06 | 1.230E-05 | 2.138E-06 | 9.900E-06 | 1.400E-05 |
| 8199 | 410 | R410Y | A | 1.2e-05 | 1.2e-05 | 1e-05 | 1.133E-05 | 1.155E-06 | 1.000E-05 | 1.200E-05 |
| 8200 | 410 | R410V | A | 1.3e-05 | 9.6e-06 | 1.4e-05 | 1.220E-05 | 2.307E-06 | 9.600E-06 | 1.400E-05 |
| 8201 | 411 | G411A | A | 1.3e-05 | 1.1e-05 | 7.2e-06 | 1.040E-05 | 2.946E-06 | 7.200E-06 | 1.300E-05 |
| 8202 | 411 | G411R | A | 1.1e-05 | 1.3e-05 | 9.2e-06 | 1.107E-05 | 1.901E-06 | 9.200E-06 | 1.300E-05 |
| 8203 | 411 | G411N | A | 1.2e-05 | 1.1e-05 | 1.1e-05 | 1.133E-05 | 5.774E-07 | 1.100E-05 | 1.200E-05 |
| 8204 | 411 | G411D | A | 1.6e-05 | 9.9e-06 | 9.3e-06 | 1.173E-05 | 3.707E-06 | 9.300E-06 | 1.600E-05 |
| 8205 | 411 | G411C | A | 1.7e-05 | 1.6e-05 | 1.6e-05 | 1.633E-05 | 5.774E-07 | 1.600E-05 | 1.700E-05 |
| 8206 | 411 | G411E | A | 1.5e-05 | 1.1e-05 | 1.7e-05 | 1.433E-05 | 3.055E-06 | 1.100E-05 | 1.700E-05 |
| 8207 | 411 | G411Q | A | 8.2e-06 | 1.1e-05 | 1.1e-05 | 1.007E-05 | 1.617E-06 | 8.200E-06 | 1.100E-05 |
| 8208 | 411 | G411G | A | 1.3e-05 | 1.1e-05 | 1.6e-05 | 1.333E-05 | 2.517E-06 | 1.100E-05 | 1.600E-05 |
| 8209 | 411 | G411H | A | 1.4e-05 | 1.3e-05 | 1.1e-05 | 1.267E-05 | 1.528E-06 | 1.100E-05 | 1.400E-05 |
| 8210 | 411 | G411I | A | 9.7e-06 | 1e-05 | 1.5e-05 | 1.157E-05 | 2.977E-06 | 9.700E-06 | 1.500E-05 |
| 8211 | 411 | G411L | A | 1.1e-05 | 1.5e-05 | 1.1e-05 | 1.233E-05 | 2.309E-06 | 1.100E-05 | 1.500E-05 |
| 8212 | 411 | G411K | A | 1.6e-05 | 8.1e-06 | 1.6e-05 | 1.337E-05 | 4.561E-06 | 8.100E-06 | 1.600E-05 |
| 8213 | 411 | G411M | A | 1.5e-05 | 1.6e-05 | 1.6e-05 | 1.567E-05 | 5.774E-07 | 1.500E-05 | 1.600E-05 |
| 8214 | 411 | G411F | A | 1e-05 | 1.4e-05 | 9.8e-06 | 1.127E-05 | 2.369E-06 | 9.800E-06 | 1.400E-05 |
| 8215 | 411 | G411P | A | 1.1e-05 | 1.6e-05 | 1e-05 | 1.233E-05 | 3.215E-06 | 1.000E-05 | 1.600E-05 |
| 8216 | 411 | G411S | A | 9.4e-06 | 1.4e-05 | 1.6e-05 | 1.313E-05 | 3.384E-06 | 9.400E-06 | 1.600E-05 |
| 8217 | 411 | G411T | A | 1.4e-05 | 1.6e-05 | 1.6e-05 | 1.533E-05 | 1.155E-06 | 1.400E-05 | 1.600E-05 |
| 8218 | 411 | G411W | A | 1.6e-05 | 1.6e-05 | 1.6e-05 | 1.600E-05 | 0.000E+00 | 1.600E-05 | 1.600E-05 |
| 8219 | 411 | G411Y | A | 9.8e-06 | 1e-05 | 1.4e-05 | 1.127E-05 | 2.369E-06 | 9.800E-06 | 1.400E-05 |
| 8220 | 411 | G411V | A | 9e-06 | 1.5e-05 | 1.1e-05 | 1.167E-05 | 3.055E-06 | 9.000E-06 | 1.500E-05 |
| 8221 | 412 | R412A | A | 1.3e-05 | 1.5e-05 | 1e-05 | 1.267E-05 | 2.517E-06 | 1.000E-05 | 1.500E-05 |
| 8222 | 412 | R412R | A | 1.7e-05 | 1.4e-05 | 1e-05 | 1.367E-05 | 3.512E-06 | 1.000E-05 | 1.700E-05 |
| 8223 | 412 | R412N | A | 1.5e-05 | 1.2e-05 | 1.1e-05 | 1.267E-05 | 2.082E-06 | 1.100E-05 | 1.500E-05 |
| 8224 | 412 | R412D | A | 1.6e-05 | 1.2e-05 | 1.5e-05 | 1.433E-05 | 2.082E-06 | 1.200E-05 | 1.600E-05 |
| 8225 | 412 | R412C | A | 1.6e-05 | 1.5e-05 | 1.2e-05 | 1.433E-05 | 2.082E-06 | 1.200E-05 | 1.600E-05 |
| 8226 | 412 | R412E | A | 1.1e-05 | 1.1e-05 | 1.5e-05 | 1.233E-05 | 2.309E-06 | 1.100E-05 | 1.500E-05 |
| 8227 | 412 | R412Q | A | 9e-06 | 8.5e-06 | 1e-05 | 9.167E-06 | 7.638E-07 | 8.500E-06 | 1.000E-05 |
| 8228 | 412 | R412G | A | 1e-05 | 1.4e-05 | 1.6e-05 | 1.333E-05 | 3.055E-06 | 1.000E-05 | 1.600E-05 |
| 8229 | 412 | R412H | A | 1.4e-05 | 1.4e-05 | 9.8e-06 | 1.260E-05 | 2.425E-06 | 9.800E-06 | 1.400E-05 |
| 8230 | 412 | R412I | A | 1.7e-05 | 8.8e-06 | 1.7e-05 | 1.427E-05 | 4.734E-06 | 8.800E-06 | 1.700E-05 |
| 8231 | 412 | R412L | A | 9.3e-06 | 7.4e-06 | 1.2e-05 | 9.567E-06 | 2.312E-06 | 7.400E-06 | 1.200E-05 |
| 8232 | 412 | R412K | A | 1.5e-05 | 1.4e-05 | 1.1e-05 | 1.333E-05 | 2.082E-06 | 1.100E-05 | 1.500E-05 |
| 8233 | 412 | R412M | A | 1.7e-05 | 1.5e-05 | 1.6e-05 | 1.600E-05 | 1.000E-06 | 1.500E-05 | 1.700E-05 |
| 8234 | 412 | R412F | A | 1.3e-05 | 1.4e-05 | 1.5e-05 | 1.400E-05 | 1.000E-06 | 1.300E-05 | 1.500E-05 |
| 8235 | 412 | R412P | A | 1.5e-05 | 1.5e-05 | 1.6e-05 | 1.533E-05 | 5.774E-07 | 1.500E-05 | 1.600E-05 |
| 8236 | 412 | R412S | A | 1.3e-05 | 1.5e-05 | 1.2e-05 | 1.333E-05 | 1.528E-06 | 1.200E-05 | 1.500E-05 |
| 8237 | 412 | R412T | A | 1.6e-05 | 1.1e-05 | 9.8e-06 | 1.227E-05 | 3.288E-06 | 9.800E-06 | 1.600E-05 |
| 8238 | 412 | R412W | A | 1.1e-05 | 9.2e-06 | 8.9e-06 | 9.700E-06 | 1.136E-06 | 8.900E-06 | 1.100E-05 |
| 8239 | 412 | R412Y | A | 1.6e-05 | 1.5e-05 | 9.9e-06 | 1.363E-05 | 3.272E-06 | 9.900E-06 | 1.600E-05 |
| 8240 | 412 | R412V | A | 9.5e-06 | 1e-05 | 1.6e-05 | 1.183E-05 | 3.617E-06 | 9.500E-06 | 1.600E-05 |
| 8241 | 413 | I413A | A | 1.2e-05 | 1.6e-05 | 1.3e-05 | 1.367E-05 | 2.082E-06 | 1.200E-05 | 1.600E-05 |
| 8242 | 413 | I413R | A | 1.2e-05 | 1.7e-05 | 1.4e-05 | 1.433E-05 | 2.517E-06 | 1.200E-05 | 1.700E-05 |
| 8243 | 413 | I413N | A | 1e-05 | 1.3e-05 | 1.2e-05 | 1.167E-05 | 1.528E-06 | 1.000E-05 | 1.300E-05 |
| 8244 | 413 | I413D | A | 1.8e-05 | 1.2e-05 | 1.4e-05 | 1.467E-05 | 3.055E-06 | 1.200E-05 | 1.800E-05 |

| | | | | | | | | | | |
|---|---|---|---|---|---|---|---|---|---|---|
| 8245 | 413 | I413C | A | 1e-05 | 1.4e-05 | 1.5e-05 | 1.300E-05 | 2.646E-06 | 1.000E-05 | 1.500E-05 |
| 8246 | 413 | I413E | A | 1.6e-05 | 1.5e-05 | 1.6e-05 | 1.567E-05 | 5.774E-07 | 1.500E-05 | 1.600E-05 |
| 8247 | 413 | I413Q | A | 1.6e-05 | 1.7e-05 | 1.5e-05 | 1.600E-05 | 1.000E-06 | 1.500E-05 | 1.700E-05 |
| 8248 | 413 | I413G | A | 1.5e-05 | 1.2e-05 | 1.6e-05 | 1.433E-05 | 2.082E-06 | 1.200E-05 | 1.600E-05 |
| 8249 | 413 | I413H | A | 1.5e-05 | 1.4e-05 | 1e-05 | 1.300E-05 | 2.646E-06 | 1.000E-05 | 1.500E-05 |
| 8250 | 413 | I413I | A | 1.5e-05 | 9.9e-06 | 1.1e-05 | 1.197E-05 | 2.684E-06 | 9.900E-06 | 1.500E-05 |
| 8251 | 413 | I413L | A | 1.4e-05 | 1.5e-05 | 1.5e-05 | 1.467E-05 | 5.774E-07 | 1.400E-05 | 1.500E-05 |
| 8252 | 413 | I413K | A | 9.9e-06 | 1.6e-05 | 9.1e-06 | 1.167E-05 | 3.774E-06 | 9.100E-06 | 1.600E-05 |
| 8253 | 413 | I413M | A | 9.8e-06 | 1.6e-05 | 8.9e-06 | 1.157E-05 | 3.866E-06 | 8.900E-06 | 1.600E-05 |
| 8254 | 413 | I413F | A | 1.3e-05 | 1.4e-05 | 1.6e-05 | 1.433E-05 | 1.528E-06 | 1.300E-05 | 1.600E-05 |
| 8255 | 413 | I413P | A | 9.8e-06 | 1.4e-05 | 1.4e-05 | 1.260E-05 | 2.425E-06 | 9.800E-06 | 1.400E-05 |
| 8256 | 413 | I413S | A | 1.2e-05 | 9.7e-06 | 1.7e-05 | 1.290E-05 | 3.732E-06 | 9.700E-06 | 1.700E-05 |
| 8257 | 413 | I413T | A | 1.2e-05 | 1.4e-05 | 1e-05 | 1.200E-05 | 2.000E-06 | 1.000E-05 | 1.400E-05 |
| 8258 | 413 | I413W | A | 1.4e-05 | 7.8e-06 | 1.5e-05 | 1.227E-05 | 3.900E-06 | 7.800E-06 | 1.500E-05 |
| 8259 | 413 | I413Y | A | 1.1e-05 | 1e-05 | 1.3e-05 | 1.133E-05 | 1.528E-06 | 1.000E-05 | 1.300E-05 |
| 8260 | 413 | I413V | A | 8.7e-06 | 1.6e-05 | 6.8e-06 | 1.050E-05 | 4.857E-06 | 6.800E-06 | 1.600E-05 |
| 8261 | 414 | L414A | A | 1.5e-05 | 9.7e-06 | 9.6e-06 | 1.143E-05 | 3.089E-06 | 9.600E-06 | 1.500E-05 |
| 8262 | 414 | L414R | A | 7.6e-06 | 1e-05 | 1e-05 | 9.200E-06 | 1.386E-06 | 7.600E-06 | 1.000E-05 |
| 8263 | 414 | L414N | A | 2.2e-05 | 9.2e-06 | 1.5e-05 | 1.540E-05 | 6.409E-06 | 9.200E-06 | 2.200E-05 |
| 8264 | 414 | L414D | A | 1.5e-05 | 1.1e-05 | 1.5e-05 | 1.367E-05 | 2.309E-06 | 1.100E-05 | 1.500E-05 |
| 8265 | 414 | L414C | A | 1.6e-05 | 8.6e-06 | 9.6e-06 | 1.140E-05 | 4.015E-06 | 8.600E-06 | 1.600E-05 |
| 8266 | 414 | L414E | A | 1.5e-05 | 1.6e-05 | 8.7e-06 | 1.323E-05 | 3.958E-06 | 8.700E-06 | 1.600E-05 |
| 8267 | 414 | L414Q | A | 1.1e-05 | 9.8e-06 | 1.2e-05 | 1.093E-05 | 1.102E-06 | 9.800E-06 | 1.200E-05 |
| 8268 | 414 | L414G | A | 1e-05 | 1e-05 | 1.6e-05 | 1.200E-05 | 3.464E-06 | 1.000E-05 | 1.600E-05 |
| 8269 | 414 | L414H | A | 1.3e-05 | 1.5e-05 | 9e-06 | 1.233E-05 | 3.055E-06 | 9.000E-06 | 1.500E-05 |
| 8270 | 414 | L414I | A | 7.7e-06 | 1.5e-05 | 1.5e-05 | 1.257E-05 | 4.215E-06 | 7.700E-06 | 1.500E-05 |
| 8271 | 414 | L414L | A | 1.7e-05 | 1.2e-05 | 1.5e-05 | 1.467E-05 | 2.517E-06 | 1.200E-05 | 1.700E-05 |
| 8272 | 414 | L414K | A | 1.3e-05 | 9.6e-06 | 1.6e-05 | 1.287E-05 | 3.202E-06 | 9.600E-06 | 1.600E-05 |
| 8273 | 414 | L414M | A | 1.1e-05 | 1.1e-05 | 1.4e-05 | 1.200E-05 | 1.732E-06 | 1.100E-05 | 1.400E-05 |
| 8274 | 414 | L414F | A | 1e-05 | 1.3e-05 | 1.6e-05 | 1.300E-05 | 3.000E-06 | 1.000E-05 | 1.600E-05 |
| 8275 | 414 | L414P | A | 1.6e-05 | 8.2e-06 | 1.2e-05 | 1.207E-05 | 3.900E-06 | 8.200E-06 | 1.600E-05 |
| 8276 | 414 | L414S | A | 1.5e-05 | 1e-05 | 1e-05 | 1.167E-05 | 2.887E-06 | 1.000E-05 | 1.500E-05 |
| 8277 | 414 | L414T | A | 1.4e-05 | 9.1e-06 | 1.4e-05 | 1.237E-05 | 2.829E-06 | 9.100E-06 | 1.400E-05 |
| 8278 | 414 | L414W | A | 9.4e-06 | 8.3e-06 | 1.7e-05 | 1.157E-05 | 4.737E-06 | 8.300E-06 | 1.700E-05 |
| 8279 | 414 | L414Y | A | 1.4e-05 | 9.9e-06 | 1.1e-05 | 1.163E-05 | 2.122E-06 | 9.900E-06 | 1.400E-05 |
| 8280 | 414 | L414V | A | 1e-05 | 1.4e-05 | 1.1e-05 | 1.167E-05 | 2.082E-06 | 1.000E-05 | 1.400E-05 |
| 8281 | 415 | H415A | A | 1.3e-05 | 1.7e-05 | 1.3e-05 | 1.433E-05 | 2.309E-06 | 1.300E-05 | 1.700E-05 |
| 8282 | 415 | H415R | A | 1.6e-05 | 1.7e-05 | 1.1e-05 | 1.467E-05 | 3.215E-06 | 1.100E-05 | 1.700E-05 |
| 8283 | 415 | H415N | A | 1.1e-05 | 1.5e-05 | 1e-05 | 1.200E-05 | 2.646E-06 | 1.000E-05 | 1.500E-05 |
| 8284 | 415 | H415D | A | 1.6e-05 | 1.7e-05 | 1.3e-05 | 1.533E-05 | 2.082E-06 | 1.300E-05 | 1.700E-05 |
| 8285 | 415 | H415C | A | 1.7e-05 | 9.8e-06 | 1.2e-05 | 1.293E-05 | 3.690E-06 | 9.800E-06 | 1.700E-05 |
| 8286 | 415 | H415E | A | 1.3e-05 | 1.4e-05 | 8.7e-06 | 1.190E-05 | 2.816E-06 | 8.700E-06 | 1.400E-05 |
| 8287 | 415 | H415Q | A | 1.1e-05 | 1.6e-05 | 1.6e-05 | 1.433E-05 | 2.887E-06 | 1.100E-05 | 1.600E-05 |
| 8288 | 415 | H415G | A | 1.7e-05 | 6.8e-06 | 9.5e-06 | 1.110E-05 | 5.285E-06 | 6.800E-06 | 1.700E-05 |
| 8289 | 415 | H415H | A | 1e-05 | 1.8e-05 | 1.3e-05 | 1.367E-05 | 4.041E-06 | 1.000E-05 | 1.800E-05 |
| 8290 | 415 | H415I | A | 1.2e-05 | 1.5e-05 | 1.2e-05 | 1.300E-05 | 1.732E-06 | 1.200E-05 | 1.500E-05 |
| 8291 | 415 | H415L | A | 1.1e-05 | 1.5e-05 | 1e-05 | 1.200E-05 | 2.646E-06 | 1.000E-05 | 1.500E-05 |
| 8292 | 415 | H415K | A | 1.3e-05 | 1.2e-05 | 1.1e-05 | 1.200E-05 | 1.000E-06 | 1.100E-05 | 1.300E-05 |
| 8293 | 415 | H415M | A | 1.3e-05 | 1.5e-05 | 9.4e-06 | 1.247E-05 | 2.838E-06 | 9.400E-06 | 1.500E-05 |
| 8294 | 415 | H415F | A | 1e-05 | 1.7e-05 | 8.9e-06 | 1.197E-05 | 4.394E-06 | 8.900E-06 | 1.700E-05 |
| 8295 | 415 | H415P | A | 1.4e-05 | 1.4e-05 | 1.5e-05 | 1.433E-05 | 5.774E-07 | 1.400E-05 | 1.500E-05 |
| 8296 | 415 | H415S | A | 1.7e-05 | 1e-05 | 1.2e-05 | 1.300E-05 | 3.606E-06 | 1.000E-05 | 1.700E-05 |
| 8297 | 415 | H415T | A | 8.9e-06 | 9.5e-06 | 1.3e-05 | 1.047E-05 | 2.214E-06 | 8.900E-06 | 1.300E-05 |
| 8298 | 415 | H415W | A | 1.1e-05 | 1.1e-05 | 1.5e-05 | 1.233E-05 | 2.309E-06 | 1.100E-05 | 1.500E-05 |
| 8299 | 415 | H415Y | A | 1e-05 | 1.8e-05 | 9.9e-06 | 1.263E-05 | 4.648E-06 | 9.900E-06 | 1.800E-05 |
| 8300 | 415 | H415V | A | 7.7e-06 | 9.6e-06 | 1.7e-05 | 1.143E-05 | 4.914E-06 | 7.700E-06 | 1.700E-05 |
| 8301 | 416 | N416A | A | 1.4e-05 | 1.4e-05 | 1e-05 | 1.267E-05 | 2.309E-06 | 1.000E-05 | 1.400E-05 |
| 8302 | 416 | N416R | A | 1.1e-05 | 1.5e-05 | 1.3e-05 | 1.300E-05 | 2.000E-06 | 1.100E-05 | 1.500E-05 |
| 8303 | 416 | N416N | A | 1.6e-05 | 1.6e-05 | 1.2e-05 | 1.467E-05 | 2.309E-06 | 1.200E-05 | 1.600E-05 |
| 8304 | 416 | N416D | A | 1.5e-05 | 1.3e-05 | 1.4e-05 | 1.400E-05 | 1.000E-06 | 1.300E-05 | 1.500E-05 |
| 8305 | 416 | N416C | A | 1.2e-05 | 1.6e-05 | 6.5e-06 | 1.150E-05 | 4.770E-06 | 6.500E-06 | 1.600E-05 |
| 8306 | 416 | N416E | A | 1.5e-05 | 1.5e-05 | 1e-05 | 1.333E-05 | 2.887E-06 | 1.000E-05 | 1.500E-05 |
| 8307 | 416 | N416Q | A | 8.7e-06 | 9.6e-06 | 1.2e-05 | 1.010E-05 | 1.706E-06 | 8.700E-06 | 1.200E-05 |
| 8308 | 416 | N416G | A | 1.1e-05 | 1.1e-05 | 1.2e-05 | 1.133E-05 | 5.774E-07 | 1.100E-05 | 1.200E-05 |
| 8309 | 416 | N416H | A | 1.6e-05 | 1.4e-05 | 1.6e-05 | 1.533E-05 | 1.155E-06 | 1.400E-05 | 1.600E-05 |
| 8310 | 416 | N416I | A | 1.4e-05 | 1.2e-05 | 1.8e-05 | 1.467E-05 | 3.055E-06 | 1.200E-05 | 1.800E-05 |
| 8311 | 416 | N416L | A | 1.1e-05 | 1.7e-05 | 9.6e-06 | 1.253E-05 | 3.931E-06 | 9.600E-06 | 1.700E-05 |
| 8312 | 416 | N416K | A | 9.4e-06 | 9.7e-06 | 1.5e-05 | 1.137E-05 | 3.150E-06 | 9.400E-06 | 1.500E-05 |
| 8313 | 416 | N416M | A | 1.1e-05 | 1.3e-05 | 1.4e-05 | 1.267E-05 | 1.528E-06 | 1.100E-05 | 1.400E-05 |
| 8314 | 416 | N416F | A | 1.1e-05 | 1e-05 | 1.6e-05 | 1.233E-05 | 3.215E-06 | 1.000E-05 | 1.600E-05 |
| 8315 | 416 | N416P | A | 1.4e-05 | 1.6e-05 | 8.7e-06 | 1.290E-05 | 3.772E-06 | 8.700E-06 | 1.600E-05 |
| 8316 | 416 | N416S | A | 1.1e-05 | 8.5e-06 | 1.2e-05 | 1.050E-05 | 1.803E-06 | 8.500E-06 | 1.200E-05 |
| 8317 | 416 | N416T | A | 1.4e-05 | 1.3e-05 | 1.5e-05 | 1.400E-05 | 1.000E-06 | 1.300E-05 | 1.500E-05 |
| 8318 | 416 | N416W | A | 1.1e-05 | 1.4e-05 | 1.1e-05 | 1.200E-05 | 1.732E-06 | 1.100E-05 | 1.400E-05 |
| 8319 | 416 | N416Y | A | 1.4e-05 | 1.5e-05 | 1.4e-05 | 1.433E-05 | 5.774E-07 | 1.400E-05 | 1.500E-05 |
| 8320 | 416 | N416V | A | 1.3e-05 | 2.3e-05 | 1.5e-05 | 1.700E-05 | 5.292E-06 | 1.300E-05 | 2.300E-05 |
| 8321 | 417 | G417A | A | 1.5e-05 | 1.5e-05 | 1.1e-05 | 1.367E-05 | 2.309E-06 | 1.100E-05 | 1.500E-05 |
| 8322 | 417 | G417R | A | 1.6e-05 | 1e-05 | 1.4e-05 | 1.333E-05 | 3.055E-06 | 1.000E-05 | 1.600E-05 |
| 8323 | 417 | G417N | A | 1.7e-05 | 1.5e-05 | 8.5e-06 | 1.350E-05 | 4.444E-06 | 8.500E-06 | 1.700E-05 |
| 8324 | 417 | G417D | A | 1.4e-05 | 8.8e-06 | 1.7e-05 | 1.327E-05 | 4.149E-06 | 8.800E-06 | 1.700E-05 |
| 8325 | 417 | G417C | A | 1.5e-05 | 1.4e-05 | 1.5e-05 | 1.467E-05 | 5.774E-07 | 1.400E-05 | 1.500E-05 |
| 8326 | 417 | G417E | A | 1.3e-05 | 1e-05 | 9e-06 | 1.067E-05 | 2.082E-06 | 9.000E-06 | 1.300E-05 |
| 8327 | 417 | G417Q | A | 1.4e-05 | 1.5e-05 | 1.6e-05 | 1.500E-05 | 1.000E-06 | 1.400E-05 | 1.600E-05 |
| 8328 | 417 | G417G | A | 9.4e-06 | 1.2e-05 | 1.1e-05 | 1.080E-05 | 1.311E-06 | 9.400E-06 | 1.200E-05 |
| 8329 | 417 | G417H | A | 1.5e-05 | 1.6e-05 | 1.4e-05 | 1.500E-05 | 1.000E-06 | 1.400E-05 | 1.600E-05 |

| | | | | | | | | | | |
|---|---|---|---|---|---|---|---|---|---|---|
| 8330 | 417 | G417I | A | 1.6e-05 | 1e-05 | 1.1e-05 | 1.233E-05 | 3.215E-06 | 1.000E-05 | 1.600E-05 |
| 8331 | 417 | G417L | A | 1.4e-05 | 1.1e-05 | 1.1e-05 | 1.200E-05 | 1.732E-06 | 1.100E-05 | 1.400E-05 |
| 8332 | 417 | G417K | A | 1e-05 | 1.1e-05 | 9e-06 | 1.000E-05 | 1.000E-06 | 9.000E-06 | 1.100E-05 |
| 8333 | 417 | G417M | A | 1.4e-05 | 1.5e-05 | 9.8e-06 | 1.293E-05 | 2.759E-06 | 9.800E-06 | 1.500E-05 |
| 8334 | 417 | G417F | A | 1.1e-05 | 1.2e-05 | 1.3e-05 | 1.200E-05 | 1.000E-06 | 1.100E-05 | 1.300E-05 |
| 8335 | 417 | G417P | A | 1.1e-05 | 1.3e-05 | 9.5e-06 | 1.117E-05 | 1.756E-06 | 9.500E-06 | 1.300E-05 |
| 8336 | 417 | G417S | A | 1.3e-05 | 1.4e-05 | 9.9e-06 | 1.230E-05 | 2.138E-06 | 9.900E-06 | 1.400E-05 |
| 8337 | 417 | G417T | A | 1.1e-05 | 1.5e-05 | 1.5e-05 | 1.367E-05 | 2.309E-06 | 1.100E-05 | 1.500E-05 |
| 8338 | 417 | G417W | A | 1.4e-05 | 9.7e-06 | 1.2e-05 | 1.190E-05 | 2.152E-06 | 9.700E-06 | 1.400E-05 |
| 8339 | 417 | G417Y | A | 1.2e-05 | 1.5e-05 | 1.3e-05 | 1.333E-05 | 1.528E-06 | 1.200E-05 | 1.500E-05 |
| 8340 | 417 | G417V | A | 1.7e-05 | 1.4e-05 | 1e-05 | 1.367E-05 | 3.512E-06 | 1.000E-05 | 1.700E-05 |
| 8341 | 418 | A418A | A | 1e-05 | 1.3e-05 | 1.3e-05 | 1.200E-05 | 1.732E-06 | 1.000E-05 | 1.300E-05 |
| 8342 | 418 | A418R | A | 1.2e-05 | 1.4e-05 | 9.9e-06 | 1.197E-05 | 2.050E-06 | 9.900E-06 | 1.400E-05 |
| 8343 | 418 | A418N | A | 8.9e-06 | 9.8e-06 | 1.1e-05 | 9.900E-06 | 1.054E-06 | 8.900E-06 | 1.100E-05 |
| 8344 | 418 | A418D | A | 1.9e-05 | 1.2e-05 | 9.9e-06 | 1.363E-05 | 4.765E-06 | 9.900E-06 | 1.900E-05 |
| 8345 | 418 | A418C | A | 8.3e-06 | 1.1e-05 | 1e-05 | 9.767E-06 | 1.365E-06 | 8.300E-06 | 1.100E-05 |
| 8346 | 418 | A418E | A | 1.4e-05 | 1.5e-05 | 1.5e-05 | 1.467E-05 | 5.774E-07 | 1.400E-05 | 1.500E-05 |
| 8347 | 418 | A418Q | A | 8.2e-06 | 1.3e-05 | 1.5e-05 | 1.207E-05 | 3.495E-06 | 8.200E-06 | 1.500E-05 |
| 8348 | 418 | A418G | A | 1.3e-05 | 1.2e-05 | 8.9e-06 | 1.130E-05 | 2.138E-06 | 8.900E-06 | 1.300E-05 |
| 8349 | 418 | A418H | A | 1.1e-05 | 1.4e-05 | 1.6e-05 | 1.367E-05 | 2.517E-06 | 1.100E-05 | 1.600E-05 |
| 8350 | 418 | A418I | A | 1e-05 | 1.2e-05 | 1.1e-05 | 1.100E-05 | 1.000E-06 | 1.000E-05 | 1.200E-05 |
| 8351 | 418 | A418L | A | 1e-05 | 1e-05 | 1.3e-05 | 1.100E-05 | 1.732E-06 | 1.000E-05 | 1.300E-05 |
| 8352 | 418 | A418K | A | 1.3e-05 | 1.5e-05 | 1e-05 | 1.267E-05 | 2.517E-06 | 1.000E-05 | 1.500E-05 |
| 8353 | 418 | A418M | A | 9.2e-06 | 1.8e-05 | 1.5e-05 | 1.407E-05 | 4.474E-06 | 9.200E-06 | 1.800E-05 |
| 8354 | 418 | A418F | A | 1.5e-05 | 1.4e-05 | 1.6e-05 | 1.500E-05 | 1.000E-06 | 1.400E-05 | 1.600E-05 |
| 8355 | 418 | A418P | A | 1.5e-05 | 1.4e-05 | 1.4e-05 | 1.433E-05 | 5.774E-07 | 1.400E-05 | 1.500E-05 |
| 8356 | 418 | A418S | A | 1.5e-05 | 1.5e-05 | 1.1e-05 | 1.367E-05 | 2.309E-06 | 1.100E-05 | 1.500E-05 |
| 8357 | 418 | A418T | A | 1.6e-05 | 1.5e-05 | 1.5e-05 | 1.533E-05 | 5.774E-07 | 1.500E-05 | 1.600E-05 |
| 8358 | 418 | A418W | A | 1.4e-05 | 1.6e-05 | 1.1e-05 | 1.367E-05 | 2.517E-06 | 1.100E-05 | 1.600E-05 |
| 8359 | 418 | A418Y | A | 1.5e-05 | 1e-05 | 1.6e-05 | 1.367E-05 | 3.215E-06 | 1.000E-05 | 1.600E-05 |
| 8360 | 418 | A418V | A | 1e-05 | 1.4e-05 | 1.1e-05 | 1.167E-05 | 2.082E-06 | 1.000E-05 | 1.400E-05 |
| 8361 | 419 | Y419A | A | 1.1e-05 | 1.1e-05 | 1.6e-05 | 1.267E-05 | 2.887E-06 | 1.100E-05 | 1.600E-05 |
| 8362 | 419 | Y419R | A | 1.1e-05 | 1.5e-05 | 1.1e-05 | 1.233E-05 | 2.309E-06 | 1.100E-05 | 1.500E-05 |
| 8363 | 419 | Y419N | A | 1.8e-05 | 1.3e-05 | 1.2e-05 | 1.433E-05 | 3.215E-06 | 1.200E-05 | 1.800E-05 |
| 8364 | 419 | Y419D | A | 1.5e-05 | 8.7e-06 | 1.1e-05 | 1.157E-05 | 3.188E-06 | 8.700E-06 | 1.500E-05 |
| 8365 | 419 | Y419C | A | 9.3e-06 | 9.3e-06 | 1.6e-05 | 1.153E-05 | 3.868E-06 | 9.300E-06 | 1.600E-05 |
| 8366 | 419 | Y419E | A | 1.1e-05 | 1.8e-05 | 1.7e-05 | 1.533E-05 | 3.786E-06 | 1.100E-05 | 1.800E-05 |
| 8367 | 419 | Y419Q | A | 1.2e-05 | 1.5e-05 | 1.1e-05 | 1.267E-05 | 2.082E-06 | 1.100E-05 | 1.500E-05 |
| 8368 | 419 | Y419G | A | 1.2e-05 | 7.3e-06 | 1.4e-05 | 1.110E-05 | 3.439E-06 | 7.300E-06 | 1.400E-05 |
| 8369 | 419 | Y419H | A | 9.6e-06 | 1.3e-05 | 1.4e-05 | 1.220E-05 | 2.307E-06 | 9.600E-06 | 1.400E-05 |
| 8370 | 419 | Y419I | A | 2.3e-05 | 8.9e-06 | 1.6e-05 | 1.597E-05 | 7.050E-06 | 8.900E-06 | 2.300E-05 |
| 8371 | 419 | Y419L | A | 1.6e-05 | 1.1e-05 | 1.6e-05 | 1.433E-05 | 2.887E-06 | 1.100E-05 | 1.600E-05 |
| 8372 | 419 | Y419K | A | 1.5e-05 | 1.4e-05 | 1.5e-05 | 1.467E-05 | 5.774E-07 | 1.400E-05 | 1.500E-05 |
| 8373 | 419 | Y419M | A | 9.6e-06 | 1.1e-05 | 1.5e-05 | 1.187E-05 | 2.802E-06 | 9.600E-06 | 1.500E-05 |
| 8374 | 419 | Y419F | A | 1.1e-05 | 7.7e-06 | 1.5e-05 | 1.123E-05 | 3.656E-06 | 7.700E-06 | 1.500E-05 |
| 8375 | 419 | Y419P | A | 1.5e-05 | 1.4e-05 | 9.2e-06 | 1.273E-05 | 3.101E-06 | 9.200E-06 | 1.500E-05 |
| 8376 | 419 | Y419S | A | 1.6e-05 | 1.5e-05 | 1e-05 | 1.367E-05 | 3.215E-06 | 1.000E-05 | 1.600E-05 |
| 8377 | 419 | Y419T | A | 1.6e-05 | 1.4e-05 | 2.7e-05 | 1.900E-05 | 7.000E-06 | 1.400E-05 | 2.700E-05 |
| 8378 | 419 | Y419W | A | 1.5e-05 | 1.5e-05 | 1.4e-05 | 1.467E-05 | 5.774E-07 | 1.400E-05 | 1.500E-05 |
| 8379 | 419 | Y419Y | A | 1.3e-05 | 9.4e-06 | 9.1e-06 | 1.050E-05 | 2.170E-06 | 9.100E-06 | 1.300E-05 |
| 8380 | 419 | Y419V | A | 1.3e-05 | 1.5e-05 | 1.5e-05 | 1.433E-05 | 1.155E-06 | 1.300E-05 | 1.500E-05 |
| 8381 | 420 | S420A | A | 1.2e-05 | 1.4e-05 | 1.5e-05 | 1.367E-05 | 1.528E-06 | 1.200E-05 | 1.500E-05 |
| 8382 | 420 | S420R | A | 1e-05 | 1.5e-05 | 1.3e-05 | 1.267E-05 | 2.517E-06 | 1.000E-05 | 1.500E-05 |
| 8383 | 420 | S420N | A | 1.2e-05 | 1.1e-05 | 1.5e-05 | 1.267E-05 | 2.082E-06 | 1.100E-05 | 1.500E-05 |
| 8384 | 420 | S420D | A | 1.1e-05 | 1.4e-05 | 1.1e-05 | 1.200E-05 | 1.732E-06 | 1.100E-05 | 1.400E-05 |
| 8385 | 420 | S420C | A | 1.4e-05 | 1.1e-05 | 1.1e-05 | 1.200E-05 | 1.732E-06 | 1.100E-05 | 1.400E-05 |
| 8386 | 420 | S420E | A | 1.6e-05 | 1.1e-05 | 1.2e-05 | 1.300E-05 | 2.646E-06 | 1.100E-05 | 1.600E-05 |
| 8387 | 420 | S420Q | A | 1.5e-05 | 1.5e-05 | 1.1e-05 | 1.367E-05 | 2.309E-06 | 1.100E-05 | 1.500E-05 |
| 8388 | 420 | S420G | A | 9.7e-06 | 1.6e-05 | 1.1e-05 | 1.223E-05 | 3.326E-06 | 9.700E-06 | 1.600E-05 |
| 8389 | 420 | S420H | A | 1.5e-05 | 1.4e-05 | 1.2e-05 | 1.367E-05 | 1.528E-06 | 1.200E-05 | 1.500E-05 |
| 8390 | 420 | S420I | A | 1.5e-05 | 1.4e-05 | 1.1e-05 | 1.333E-05 | 2.082E-06 | 1.100E-05 | 1.500E-05 |
| 8391 | 420 | S420L | A | 1.3e-05 | 1.5e-05 | 1.4e-05 | 1.400E-05 | 1.000E-06 | 1.300E-05 | 1.500E-05 |
| 8392 | 420 | S420K | A | 1.4e-05 | 1.3e-05 | 1e-05 | 1.233E-05 | 2.082E-06 | 1.000E-05 | 1.400E-05 |
| 8393 | 420 | S420M | A | 8.7e-06 | 1.5e-05 | 1.4e-05 | 1.257E-05 | 3.386E-06 | 8.700E-06 | 1.500E-05 |
| 8394 | 420 | S420F | A | 1.6e-05 | 7.9e-06 | 1.1e-05 | 1.163E-05 | 4.087E-06 | 7.900E-06 | 1.600E-05 |
| 8395 | 420 | S420P | A | 1.5e-05 | 1.5e-05 | 1.1e-05 | 1.367E-05 | 2.309E-06 | 1.100E-05 | 1.500E-05 |
| 8396 | 420 | S420S | A | 7.8e-06 | 1.5e-05 | 1.4e-05 | 1.227E-05 | 3.900E-06 | 7.800E-06 | 1.500E-05 |
| 8397 | 420 | S420T | A | 8.4e-06 | 1.4e-05 | 1.6e-05 | 1.280E-05 | 3.940E-06 | 8.400E-06 | 1.600E-05 |
| 8398 | 420 | S420W | A | 6.6e-06 | 9.9e-06 | 1.4e-05 | 1.017E-05 | 3.707E-06 | 6.600E-06 | 1.400E-05 |
| 8399 | 420 | S420Y | A | 1.5e-05 | 1e-05 | 1.1e-05 | 1.200E-05 | 2.646E-06 | 1.000E-05 | 1.500E-05 |
| 8400 | 420 | S420V | A | 9.7e-06 | 1.4e-05 | 1.2e-05 | 1.190E-05 | 2.152E-06 | 9.700E-06 | 1.400E-05 |
| 8401 | 421 | L421A | A | 1.6e-05 | 1.5e-05 | 7.1e-06 | 1.270E-05 | 4.875E-06 | 7.100E-06 | 1.600E-05 |
| 8402 | 421 | L421R | A | 1.2e-05 | 9.7e-06 | 9.2e-06 | 1.030E-05 | 1.493E-06 | 9.200E-06 | 1.200E-05 |
| 8403 | 421 | L421N | A | 9.1e-06 | 1.3e-05 | 1.5e-05 | 1.237E-05 | 3.001E-06 | 9.100E-06 | 1.500E-05 |
| 8404 | 421 | L421D | A | 1.7e-05 | 1.1e-05 | 1.6e-05 | 1.467E-05 | 3.215E-06 | 1.100E-05 | 1.700E-05 |
| 8405 | 421 | L421C | A | 1.5e-05 | 1.5e-05 | 1.5e-05 | 1.500E-05 | 0.000E+00 | 1.500E-05 | 1.500E-05 |
| 8406 | 421 | L421E | A | 1.2e-05 | 1.5e-05 | 1.3e-05 | 1.333E-05 | 1.528E-06 | 1.200E-05 | 1.500E-05 |
| 8407 | 421 | L421Q | A | 1.5e-05 | 1.6e-05 | 1.6e-05 | 1.567E-05 | 5.774E-07 | 1.500E-05 | 1.600E-05 |
| 8408 | 421 | L421G | A | 1.1e-05 | 1.6e-05 | 1.4e-05 | 1.367E-05 | 2.517E-06 | 1.100E-05 | 1.600E-05 |
| 8409 | 421 | L421H | A | 9.8e-06 | 1e-05 | 1.2e-05 | 1.060E-05 | 1.217E-06 | 9.800E-06 | 1.200E-05 |
| 8410 | 421 | L421I | A | 1.4e-05 | 1.1e-05 | 1.7e-05 | 1.400E-05 | 3.000E-06 | 1.100E-05 | 1.700E-05 |
| 8411 | 421 | L421L | A | 1.5e-05 | 1.2e-05 | 1.3e-05 | 1.333E-05 | 1.528E-06 | 1.200E-05 | 1.500E-05 |
| 8412 | 421 | L421K | A | 1.6e-05 | 1.1e-05 | 1.7e-05 | 1.467E-05 | 3.215E-06 | 1.100E-05 | 1.700E-05 |
| 8413 | 421 | L421M | A | 1.6e-05 | 1e-05 | 1.5e-05 | 1.367E-05 | 3.215E-06 | 1.000E-05 | 1.600E-05 |
| 8414 | 421 | L421F | A | 1.8e-05 | 1e-05 | 9.9e-06 | 1.263E-05 | 4.648E-06 | 9.900E-06 | 1.800E-05 |

| | | | | | | | | | | |
|---|---|---|---|---|---|---|---|---|---|---|
| 8415 | 421 | L421P | A | 1.1e-05 | 1.5e-05 | 1.1e-05 | 1.233E-05 | 2.309E-06 | 1.100E-05 | 1.500E-05 |
| 8416 | 421 | L421S | A | 1.4e-05 | 1.1e-05 | 1.8e-05 | 1.433E-05 | 3.512E-06 | 1.100E-05 | 1.800E-05 |
| 8417 | 421 | L421T | A | 1.3e-05 | 1.1e-05 | 1.5e-05 | 1.300E-05 | 2.000E-06 | 1.100E-05 | 1.500E-05 |
| 8418 | 421 | L421W | A | 1.6e-05 | 1.4e-05 | 1.7e-05 | 1.567E-05 | 1.528E-06 | 1.400E-05 | 1.700E-05 |
| 8419 | 421 | L421Y | A | 7.2e-06 | 1.6e-05 | 1.5e-05 | 1.273E-05 | 4.818E-06 | 7.200E-06 | 1.600E-05 |
| 8420 | 421 | L421V | A | 8.5e-06 | 1e-05 | 1.5e-05 | 1.117E-05 | 3.403E-06 | 8.500E-06 | 1.500E-05 |
| 8421 | 422 | T422A | A | 1.5e-05 | 1.7e-05 | 9.9e-06 | 1.397E-05 | 3.661E-06 | 9.900E-06 | 1.700E-05 |
| 8422 | 422 | T422R | A | 1.4e-05 | 1.5e-05 | 1.6e-05 | 1.500E-05 | 1.000E-06 | 1.400E-05 | 1.600E-05 |
| 8423 | 422 | T422N | A | 8.3e-06 | 1e-05 | 1.7e-05 | 1.177E-05 | 4.611E-06 | 8.300E-06 | 1.700E-05 |
| 8424 | 422 | T422D | A | 1.2e-05 | 1e-05 | 1.1e-05 | 1.100E-05 | 1.000E-06 | 1.000E-05 | 1.200E-05 |
| 8425 | 422 | T422C | A | 1.5e-05 | 1e-05 | 1.5e-05 | 1.333E-05 | 2.887E-06 | 1.000E-05 | 1.500E-05 |
| 8426 | 422 | T422E | A | 1.2e-05 | 1.1e-05 | 1e-05 | 1.100E-05 | 1.000E-06 | 1.000E-05 | 1.200E-05 |
| 8427 | 422 | T422Q | A | 8e-06 | 1.3e-05 | 1e-05 | 1.033E-05 | 2.517E-06 | 8.000E-06 | 1.300E-05 |
| 8428 | 422 | T422G | A | 1.4e-05 | 1.2e-05 | 1.6e-05 | 1.400E-05 | 2.000E-06 | 1.200E-05 | 1.600E-05 |
| 8429 | 422 | T422H | A | 1.6e-05 | 1.5e-05 | 1.1e-05 | 1.400E-05 | 2.646E-06 | 1.100E-05 | 1.600E-05 |
| 8430 | 422 | T422I | A | 1.8e-05 | 1.3e-05 | 1.6e-05 | 1.567E-05 | 2.517E-06 | 1.300E-05 | 1.800E-05 |
| 8431 | 422 | T422L | A | 1.2e-05 | 1.5e-05 | 1.6e-05 | 1.433E-05 | 2.082E-06 | 1.200E-05 | 1.600E-05 |
| 8432 | 422 | T422K | A | 1.6e-05 | 1.6e-05 | 6.7e-06 | 1.290E-05 | 5.369E-06 | 6.700E-06 | 1.600E-05 |
| 8433 | 422 | T422M | A | 8.7e-06 | 1.1e-05 | 1.6e-05 | 1.190E-05 | 3.732E-06 | 8.700E-06 | 1.600E-05 |
| 8434 | 422 | T422F | A | 1.6e-05 | 1.4e-05 | 1.7e-05 | 1.567E-05 | 1.528E-06 | 1.400E-05 | 1.700E-05 |
| 8435 | 422 | T422P | A | 1.4e-05 | 1.6e-05 | 1e-05 | 1.333E-05 | 3.055E-06 | 1.000E-05 | 1.600E-05 |
| 8436 | 422 | T422S | A | 1.5e-05 | 1.2e-05 | 9.8e-06 | 1.227E-05 | 2.610E-06 | 9.800E-06 | 1.500E-05 |
| 8437 | 422 | T422T | A | 1.2e-05 | 1.2e-05 | 1.5e-05 | 1.300E-05 | 1.732E-06 | 1.200E-05 | 1.500E-05 |
| 8438 | 422 | T422W | A | 1.5e-05 | 1.5e-05 | 9.5e-06 | 1.317E-05 | 3.175E-06 | 9.500E-06 | 1.500E-05 |
| 8439 | 422 | T422Y | A | 1.2e-05 | 1.1e-05 | 1.5e-05 | 1.267E-05 | 2.082E-06 | 1.100E-05 | 1.500E-05 |
| 8440 | 422 | T422V | A | 9.4e-06 | 1.1e-05 | 1.1e-05 | 1.047E-05 | 9.238E-07 | 9.400E-06 | 1.100E-05 |
| 8441 | 423 | L423A | A | 1.6e-05 | 1.6e-05 | 1.4e-05 | 1.533E-05 | 1.155E-06 | 1.400E-05 | 1.600E-05 |
| 8442 | 423 | L423R | A | 8.3e-06 | 1.5e-05 | 1.5e-05 | 1.277E-05 | 3.868E-06 | 8.300E-06 | 1.500E-05 |
| 8443 | 423 | L423N | A | 1.4e-05 | 1.6e-05 | 1.6e-05 | 1.533E-05 | 1.155E-06 | 1.400E-05 | 1.600E-05 |
| 8444 | 423 | L423D | A | 1e-05 | 1.2e-05 | 1.7e-05 | 1.300E-05 | 3.606E-06 | 1.000E-05 | 1.700E-05 |
| 8445 | 423 | L423C | A | 1e-05 | 1.5e-05 | 1.7e-05 | 1.400E-05 | 3.606E-06 | 1.000E-05 | 1.700E-05 |
| 8446 | 423 | L423E | A | 1.7e-05 | 1.4e-05 | 1.4e-05 | 1.500E-05 | 1.732E-06 | 1.400E-05 | 1.700E-05 |
| 8447 | 423 | L423Q | A | 7.4e-06 | 1.5e-05 | 1.1e-05 | 1.113E-05 | 3.802E-06 | 7.400E-06 | 1.500E-05 |
| 8448 | 423 | L423G | A | 8.8e-06 | 1.5e-05 | 1.1e-05 | 1.160E-05 | 3.143E-06 | 8.800E-06 | 1.500E-05 |
| 8449 | 423 | L423H | A | 1.5e-05 | 1.7e-05 | 1e-05 | 1.400E-05 | 3.606E-06 | 1.000E-05 | 1.700E-05 |
| 8450 | 423 | L423I | A | 1.4e-05 | 1.4e-05 | 9.1e-06 | 1.237E-05 | 2.829E-06 | 9.100E-06 | 1.400E-05 |
| 8451 | 423 | L423L | A | 1.6e-05 | 1.5e-05 | 1.6e-05 | 1.567E-05 | 5.774E-07 | 1.500E-05 | 1.600E-05 |
| 8452 | 423 | L423K | A | 1.5e-05 | 1.2e-05 | 1.6e-05 | 1.433E-05 | 2.082E-06 | 1.200E-05 | 1.600E-05 |
| 8453 | 423 | L423M | A | 6.1e-06 | 7.5e-06 | 1e-05 | 7.867E-06 | 1.976E-06 | 6.100E-06 | 1.000E-05 |
| 8454 | 423 | L423F | A | 5.9e-06 | 9.2e-06 | 1.5e-05 | 1.003E-05 | 4.607E-06 | 5.900E-06 | 1.500E-05 |
| 8455 | 423 | L423P | A | 1.7e-05 | 1.6e-05 | 1.2e-05 | 1.500E-05 | 2.646E-06 | 1.200E-05 | 1.700E-05 |
| 8456 | 423 | L423S | A | 9.7e-06 | 1.3e-05 | 1.1e-05 | 1.123E-05 | 1.662E-06 | 9.700E-06 | 1.300E-05 |
| 8457 | 423 | L423T | A | 1.2e-05 | 1e-05 | 1.2e-05 | 1.133E-05 | 1.155E-06 | 1.000E-05 | 1.200E-05 |
| 8458 | 423 | L423W | A | 9.8e-06 | 1.5e-05 | 9.4e-06 | 1.140E-05 | 3.124E-06 | 9.400E-06 | 1.500E-05 |
| 8459 | 423 | L423Y | A | 1.5e-05 | 1.2e-05 | 1.4e-05 | 1.367E-05 | 1.528E-06 | 1.200E-05 | 1.500E-05 |
| 8460 | 423 | L423V | A | 1.3e-05 | 1.6e-05 | 1.1e-05 | 1.333E-05 | 2.517E-06 | 1.100E-05 | 1.600E-05 |
| 8461 | 424 | Q424A | A | 1.2e-05 | 9.3e-06 | 1.2e-05 | 1.110E-05 | 1.559E-06 | 9.300E-06 | 1.200E-05 |
| 8462 | 424 | Q424R | A | 1.4e-05 | 1.5e-05 | 7.8e-06 | 1.227E-05 | 3.900E-06 | 7.800E-06 | 1.500E-05 |
| 8463 | 424 | Q424N | A | 9.9e-06 | 1.3e-05 | 1e-05 | 1.097E-05 | 1.762E-06 | 9.900E-06 | 1.300E-05 |
| 8464 | 424 | Q424D | A | 1.3e-05 | 1e-05 | 1.3e-05 | 1.200E-05 | 1.732E-06 | 1.000E-05 | 1.300E-05 |
| 8465 | 424 | Q424C | A | 1e-05 | 9.2e-06 | 1.4e-05 | 1.107E-05 | 2.572E-06 | 9.200E-06 | 1.400E-05 |
| 8466 | 424 | Q424E | A | 1.5e-05 | 1.6e-05 | 1.1e-05 | 1.400E-05 | 2.646E-06 | 1.100E-05 | 1.600E-05 |
| 8467 | 424 | Q424Q | A | 1e-05 | 9.7e-06 | 1.6e-05 | 1.190E-05 | 3.554E-06 | 9.700E-06 | 1.600E-05 |
| 8468 | 424 | Q424G | A | 1.5e-05 | 1.1e-05 | 1.6e-05 | 1.400E-05 | 2.646E-06 | 1.100E-05 | 1.600E-05 |
| 8469 | 424 | Q424H | A | 8.9e-06 | 1.2e-05 | 1.4e-05 | 1.163E-05 | 2.570E-06 | 8.900E-06 | 1.400E-05 |
| 8470 | 424 | Q424I | A | 1.3e-05 | 7.6e-06 | 9.7e-06 | 1.010E-05 | 2.722E-06 | 7.600E-06 | 1.300E-05 |
| 8471 | 424 | Q424L | A | 1.1e-05 | 1.3e-05 | 1.2e-05 | 1.200E-05 | 1.000E-06 | 1.100E-05 | 1.300E-05 |
| 8472 | 424 | Q424K | A | 9.6e-06 | 1.4e-05 | 1.6e-05 | 1.320E-05 | 3.274E-06 | 9.600E-06 | 1.600E-05 |
| 8473 | 424 | Q424M | A | 1.3e-05 | 1.5e-05 | 1e-05 | 1.267E-05 | 2.517E-06 | 1.000E-05 | 1.500E-05 |
| 8474 | 424 | Q424F | A | 1e-05 | 9.7e-06 | 8.3e-06 | 9.333E-06 | 9.074E-07 | 8.300E-06 | 1.000E-05 |
| 8475 | 424 | Q424P | A | 1.5e-05 | 7.4e-06 | 6.7e-06 | 9.700E-06 | 4.603E-06 | 6.700E-06 | 1.500E-05 |
| 8476 | 424 | Q424S | A | 1.4e-05 | 1.1e-05 | 1.2e-05 | 1.233E-05 | 1.528E-06 | 1.100E-05 | 1.400E-05 |
| 8477 | 424 | Q424T | A | 1.4e-05 | 1.5e-05 | 1.6e-05 | 1.500E-05 | 1.000E-06 | 1.400E-05 | 1.600E-05 |
| 8478 | 424 | Q424W | A | 7.6e-06 | 1.4e-05 | 1e-05 | 1.053E-05 | 3.233E-06 | 7.600E-06 | 1.400E-05 |
| 8479 | 424 | Q424Y | A | 7.5e-06 | 1.6e-05 | 9.4e-06 | 1.097E-05 | 4.461E-06 | 7.500E-06 | 1.600E-05 |
| 8480 | 424 | Q424V | A | 1.1e-05 | 1.6e-05 | 1.3e-05 | 1.333E-05 | 2.517E-06 | 1.100E-05 | 1.600E-05 |
| 8481 | 425 | G425A | A | 1.2e-05 | 1.4e-05 | 9.7e-06 | 1.190E-05 | 2.152E-06 | 9.700E-06 | 1.400E-05 |
| 8482 | 425 | G425R | A | 1.4e-05 | 1.6e-05 | 1.6e-05 | 1.533E-05 | 1.155E-06 | 1.400E-05 | 1.600E-05 |
| 8483 | 425 | G425N | A | 1.4e-05 | 8.2e-06 | 1.1e-05 | 1.107E-05 | 2.901E-06 | 8.200E-06 | 1.400E-05 |
| 8484 | 425 | G425D | A | 1e-05 | 1.5e-05 | 9.3e-06 | 1.143E-05 | 3.109E-06 | 9.300E-06 | 1.500E-05 |
| 8485 | 425 | G425C | A | 1.5e-05 | 1e-05 | 1.4e-05 | 1.300E-05 | 2.646E-06 | 1.000E-05 | 1.500E-05 |
| 8486 | 425 | G425E | A | 1.6e-05 | 1.5e-05 | 1e-05 | 1.367E-05 | 3.215E-06 | 1.000E-05 | 1.600E-05 |
| 8487 | 425 | G425Q | A | 9.2e-06 | 1e-05 | 9.2e-06 | 9.467E-06 | 4.619E-07 | 9.200E-06 | 1.000E-05 |
| 8488 | 425 | G425G | A | 1.1e-05 | 1.1e-05 | 9.5e-06 | 1.050E-05 | 8.660E-07 | 9.500E-06 | 1.100E-05 |
| 8489 | 425 | G425H | A | 1.4e-05 | 1.1e-05 | 8.8e-06 | 1.127E-05 | 2.610E-06 | 8.800E-06 | 1.400E-05 |
| 8490 | 425 | G425I | A | 1.5e-05 | 1.6e-05 | 1e-05 | 1.367E-05 | 3.215E-06 | 1.000E-05 | 1.600E-05 |
| 8491 | 425 | G425L | A | 1.1e-05 | 1.6e-05 | 9.6e-06 | 1.220E-05 | 3.365E-06 | 9.600E-06 | 1.600E-05 |
| 8492 | 425 | G425K | A | 1.3e-05 | 9e-06 | 9.5e-06 | 1.050E-05 | 2.179E-06 | 9.000E-06 | 1.300E-05 |
| 8493 | 425 | G425M | A | 1.4e-05 | 1.4e-05 | 9.2e-06 | 1.240E-05 | 2.771E-06 | 9.200E-06 | 1.400E-05 |
| 8494 | 425 | G425F | A | 1.6e-05 | 1.7e-05 | 1.7e-05 | 1.667E-05 | 5.774E-07 | 1.600E-05 | 1.700E-05 |
| 8495 | 425 | G425P | A | 1.6e-05 | 1.2e-05 | 1.2e-05 | 1.333E-05 | 2.309E-06 | 1.200E-05 | 1.600E-05 |
| 8496 | 425 | G425S | A | 1.4e-05 | 1.4e-05 | 1.6e-05 | 1.467E-05 | 1.155E-06 | 1.400E-05 | 1.600E-05 |
| 8497 | 425 | G425T | A | 1.4e-05 | 1.4e-05 | 1.2e-05 | 1.333E-05 | 1.155E-06 | 1.200E-05 | 1.400E-05 |
| 8498 | 425 | G425W | A | 1.3e-05 | 1.5e-05 | 1.5e-05 | 1.433E-05 | 1.155E-06 | 1.300E-05 | 1.500E-05 |
| 8499 | 425 | G425Y | A | 1.6e-05 | 1.1e-05 | 1.2e-05 | 1.300E-05 | 2.646E-06 | 1.100E-05 | 1.600E-05 |

| | | | | | | | | | | |
|---|---|---|---|---|---|---|---|---|---|---|
| 8500 | 425 | G425V | A | 1e-05 | 1.8e-05 | 1.4e-05 | 1.400E-05 | 4.000E-06 | 1.000E-05 | 1.800E-05 |
| 8501 | 426 | L426A | A | 9.7e-06 | 1.1e-05 | 1.5e-05 | 1.190E-05 | 2.762E-06 | 9.700E-06 | 1.500E-05 |
| 8502 | 426 | L426R | A | 8.5e-06 | 1.4e-05 | 8.6e-06 | 1.037E-05 | 3.147E-06 | 8.500E-06 | 1.400E-05 |
| 8503 | 426 | L426N | A | 1.3e-05 | 1.3e-05 | 1e-05 | 1.200E-05 | 1.732E-06 | 1.000E-05 | 1.300E-05 |
| 8504 | 426 | L426D | A | 8.8e-06 | 1.1e-05 | 1.4e-05 | 1.127E-05 | 2.610E-06 | 8.800E-06 | 1.400E-05 |
| 8505 | 426 | L426C | A | 1.3e-05 | 1e-05 | 1.6e-05 | 1.300E-05 | 3.000E-06 | 1.000E-05 | 1.600E-05 |
| 8506 | 426 | L426E | A | 1.1e-05 | 1e-05 | 1.3e-05 | 1.133E-05 | 1.528E-06 | 1.000E-05 | 1.300E-05 |
| 8507 | 426 | L426Q | A | 1.4e-05 | 1e-05 | 1e-05 | 1.133E-05 | 2.309E-06 | 1.000E-05 | 1.400E-05 |
| 8508 | 426 | L426G | A | 8.8e-06 | 1.6e-05 | 1e-05 | 1.160E-05 | 3.857E-06 | 8.800E-06 | 1.600E-05 |
| 8509 | 426 | L426H | A | 1.4e-05 | 9.8e-06 | 9.2e-06 | 1.100E-05 | 2.615E-06 | 9.200E-06 | 1.400E-05 |
| 8510 | 426 | L426I | A | 1.6e-05 | 1.1e-05 | 1.6e-05 | 1.433E-05 | 2.887E-06 | 1.100E-05 | 1.600E-05 |
| 8511 | 426 | L426L | A | 1.4e-05 | 1.4e-05 | 1.1e-05 | 1.300E-05 | 1.732E-06 | 1.100E-05 | 1.400E-05 |
| 8512 | 426 | L426K | A | 1.3e-05 | 1.4e-05 | 1.5e-05 | 1.400E-05 | 1.000E-06 | 1.300E-05 | 1.500E-05 |
| 8513 | 426 | L426M | A | 1.1e-05 | 8e-06 | 9.9e-06 | 9.633E-06 | 1.518E-06 | 8.000E-06 | 1.100E-05 |
| 8514 | 426 | L426F | A | 1.4e-05 | 1.5e-05 | 1e-05 | 1.300E-05 | 2.646E-06 | 1.000E-05 | 1.500E-05 |
| 8515 | 426 | L426P | A | 1.3e-05 | 1e-05 | 1.5e-05 | 1.267E-05 | 2.517E-06 | 1.000E-05 | 1.500E-05 |
| 8516 | 426 | L426S | A | 1.6e-05 | 1.4e-05 | 9.3e-06 | 1.310E-05 | 3.439E-06 | 9.300E-06 | 1.600E-05 |
| 8517 | 426 | L426T | A | 1.1e-05 | 1.2e-05 | 1.7e-05 | 1.333E-05 | 3.215E-06 | 1.100E-05 | 1.700E-05 |
| 8518 | 426 | L426W | A | 1.5e-05 | 8e-06 | 1.5e-05 | 1.267E-05 | 4.041E-06 | 8.000E-06 | 1.500E-05 |
| 8519 | 426 | L426Y | A | 1.2e-05 | 1.5e-05 | 1.7e-05 | 1.467E-05 | 2.517E-06 | 1.200E-05 | 1.700E-05 |
| 8520 | 426 | L426V | A | 1.4e-05 | 1.4e-05 | 1.5e-05 | 1.433E-05 | 5.774E-07 | 1.400E-05 | 1.500E-05 |
| 8521 | 427 | G427A | A | 1.2e-05 | 1.3e-05 | 1.6e-05 | 1.367E-05 | 2.082E-06 | 1.200E-05 | 1.600E-05 |
| 8522 | 427 | G427R | A | 1.3e-05 | 1.5e-05 | 1.4e-05 | 1.400E-05 | 1.000E-06 | 1.300E-05 | 1.500E-05 |
| 8523 | 427 | G427N | A | 9.9e-06 | 1.1e-05 | 1.1e-05 | 1.063E-05 | 6.351E-07 | 9.900E-06 | 1.100E-05 |
| 8524 | 427 | G427D | A | 1.5e-05 | 1.5e-05 | 1.5e-05 | 1.500E-05 | 0.000E+00 | 1.500E-05 | 1.500E-05 |
| 8525 | 427 | G427C | A | 1.5e-05 | 1.5e-05 | 1.4e-05 | 1.467E-05 | 5.774E-07 | 1.400E-05 | 1.500E-05 |
| 8526 | 427 | G427E | A | 1.2e-05 | 1.2e-05 | 1.1e-05 | 1.167E-05 | 5.774E-07 | 1.100E-05 | 1.200E-05 |
| 8527 | 427 | G427Q | A | 1.3e-05 | 9.6e-06 | 1.3e-05 | 1.187E-05 | 1.963E-06 | 9.600E-06 | 1.300E-05 |
| 8528 | 427 | G427G | A | 1.3e-05 | 9.6e-06 | 1.5e-05 | 1.253E-05 | 2.730E-06 | 9.600E-06 | 1.500E-05 |
| 8529 | 427 | G427H | A | 1.4e-05 | 1.2e-05 | 1.5e-05 | 1.367E-05 | 1.528E-06 | 1.200E-05 | 1.500E-05 |
| 8530 | 427 | G427I | A | 1.6e-05 | 1.3e-05 | 1.2e-05 | 1.367E-05 | 2.082E-06 | 1.200E-05 | 1.600E-05 |
| 8531 | 427 | G427L | A | 1.5e-05 | 1.1e-05 | 1.5e-05 | 1.367E-05 | 2.309E-06 | 1.100E-05 | 1.500E-05 |
| 8532 | 427 | G427K | A | 1e-05 | 2.6e-05 | 1.5e-05 | 1.700E-05 | 8.185E-06 | 1.000E-05 | 2.600E-05 |
| 8533 | 427 | G427M | A | 1.3e-05 | 1.4e-05 | 1.6e-05 | 1.433E-05 | 1.528E-06 | 1.300E-05 | 1.600E-05 |
| 8534 | 427 | G427F | A | 8.2e-06 | 1.6e-05 | 1.3e-05 | 1.240E-05 | 3.934E-06 | 8.200E-06 | 1.600E-05 |
| 8535 | 427 | G427P | A | 1e-05 | 1.3e-05 | 9.6e-06 | 1.087E-05 | 1.858E-06 | 9.600E-06 | 1.300E-05 |
| 8536 | 427 | G427S | A | 9.9e-06 | 8.9e-06 | 9.8e-06 | 9.533E-06 | 5.508E-07 | 8.900E-06 | 9.900E-06 |
| 8537 | 427 | G427T | A | 1.4e-05 | 1.4e-05 | 1e-05 | 1.267E-05 | 2.309E-06 | 1.000E-05 | 1.400E-05 |
| 8538 | 427 | G427W | A | 8.9e-06 | 1e-05 | 1.5e-05 | 1.130E-05 | 3.251E-06 | 8.900E-06 | 1.500E-05 |
| 8539 | 427 | G427Y | A | 1.1e-05 | 1.2e-05 | 1.1e-05 | 1.133E-05 | 5.774E-07 | 1.100E-05 | 1.200E-05 |
| 8540 | 427 | G427V | A | 1e-05 | 1.3e-05 | 9.2e-06 | 1.073E-05 | 2.003E-06 | 9.200E-06 | 1.300E-05 |
| 8541 | 428 | I428A | A | 1.1e-05 | 1.5e-05 | 1.5e-05 | 1.367E-05 | 2.309E-06 | 1.100E-05 | 1.500E-05 |
| 8542 | 428 | I428R | A | 1.4e-05 | 9.5e-06 | 1.2e-05 | 1.183E-05 | 2.255E-06 | 9.500E-06 | 1.400E-05 |
| 8543 | 428 | I428N | A | 1.1e-05 | 1.6e-05 | 1.5e-05 | 1.400E-05 | 2.646E-06 | 1.100E-05 | 1.600E-05 |
| 8544 | 428 | I428D | A | 1.1e-05 | 1.6e-05 | 1.5e-05 | 1.400E-05 | 2.646E-06 | 1.100E-05 | 1.600E-05 |
| 8545 | 428 | I428C | A | 1.4e-05 | 1.5e-05 | 1.5e-05 | 1.467E-05 | 5.774E-07 | 1.400E-05 | 1.500E-05 |
| 8546 | 428 | I428E | A | 1.4e-05 | 9.6e-06 | 1.5e-05 | 1.287E-05 | 2.873E-06 | 9.600E-06 | 1.500E-05 |
| 8547 | 428 | I428Q | A | 1.5e-05 | 1.5e-05 | 1.3e-05 | 1.433E-05 | 1.155E-06 | 1.300E-05 | 1.500E-05 |
| 8548 | 428 | I428G | A | 1.6e-05 | 1.6e-05 | 1.3e-05 | 1.500E-05 | 1.732E-06 | 1.300E-05 | 1.600E-05 |
| 8549 | 428 | I428H | A | 1.4e-05 | 1.1e-05 | 1.3e-05 | 1.267E-05 | 1.528E-06 | 1.100E-05 | 1.400E-05 |
| 8550 | 428 | I428I | A | 1.3e-05 | 1.1e-05 | 1.6e-05 | 1.333E-05 | 2.517E-06 | 1.100E-05 | 1.600E-05 |
| 8551 | 428 | I428L | A | 1.1e-05 | 1e-05 | 1.1e-05 | 1.067E-05 | 5.774E-07 | 1.000E-05 | 1.100E-05 |
| 8552 | 428 | I428K | A | 1.1e-05 | 1.5e-05 | 1.4e-05 | 1.333E-05 | 2.082E-06 | 1.100E-05 | 1.500E-05 |
| 8553 | 428 | I428M | A | 1.7e-05 | 9.9e-06 | 9.4e-06 | 1.210E-05 | 4.251E-06 | 9.400E-06 | 1.700E-05 |
| 8554 | 428 | I428F | A | 1.7e-05 | 8e-06 | 1.1e-05 | 1.200E-05 | 4.583E-06 | 8.000E-06 | 1.700E-05 |
| 8555 | 428 | I428P | A | 1.5e-05 | 1.5e-05 | 8e-06 | 1.267E-05 | 4.041E-06 | 8.000E-06 | 1.500E-05 |
| 8556 | 428 | I428S | A | 1.6e-05 | 1.1e-05 | 9.9e-06 | 1.230E-05 | 3.251E-06 | 9.900E-06 | 1.600E-05 |
| 8557 | 428 | I428T | A | 1.4e-05 | 1.6e-05 | 7.7e-06 | 1.257E-05 | 4.332E-06 | 7.700E-06 | 1.600E-05 |
| 8558 | 428 | I428W | A | 9.8e-06 | 1.4e-05 | 1.1e-05 | 1.160E-05 | 2.163E-06 | 9.800E-06 | 1.400E-05 |
| 8559 | 428 | I428Y | A | 9.4e-06 | 1.2e-05 | 1.6e-05 | 1.247E-05 | 3.325E-06 | 9.400E-06 | 1.600E-05 |
| 8560 | 428 | I428V | A | 9.6e-06 | 1.3e-05 | 1.5e-05 | 1.253E-05 | 2.730E-06 | 9.600E-06 | 1.500E-05 |
| 8561 | 429 | S429A | A | 1.1e-05 | 2.8e-05 | 1.4e-05 | 1.767E-05 | 9.074E-06 | 1.100E-05 | 2.800E-05 |
| 8562 | 429 | S429R | A | 1.1e-05 | 8.4e-06 | 1.5e-05 | 1.147E-05 | 3.325E-06 | 8.400E-06 | 1.500E-05 |
| 8563 | 429 | S429N | A | 1.4e-05 | 1.5e-05 | 1e-05 | 1.300E-05 | 2.646E-06 | 1.000E-05 | 1.500E-05 |
| 8564 | 429 | S429D | A | 1.6e-05 | 1.5e-05 | 1.6e-05 | 1.567E-05 | 5.774E-07 | 1.500E-05 | 1.600E-05 |
| 8565 | 429 | S429C | A | 1.5e-05 | 1.2e-05 | 8.9e-06 | 1.197E-05 | 3.050E-06 | 8.900E-06 | 1.500E-05 |
| 8566 | 429 | S429E | A | 1.6e-05 | 9.7e-06 | 1.6e-05 | 1.390E-05 | 3.637E-06 | 9.700E-06 | 1.600E-05 |
| 8567 | 429 | S429Q | A | 9.7e-06 | 1.1e-05 | 1e-05 | 1.023E-05 | 6.807E-07 | 9.700E-06 | 1.100E-05 |
| 8568 | 429 | S429G | A | 1.5e-05 | 1.2e-05 | 1.2e-05 | 1.300E-05 | 1.732E-06 | 1.200E-05 | 1.500E-05 |
| 8569 | 429 | S429H | A | 1.6e-05 | 1.1e-05 | 1.7e-05 | 1.467E-05 | 3.215E-06 | 1.100E-05 | 1.700E-05 |
| 8570 | 429 | S429I | A | 1.8e-05 | 1.6e-05 | 1.8e-05 | 1.733E-05 | 1.155E-06 | 1.600E-05 | 1.800E-05 |
| 8571 | 429 | S429L | A | 1.6e-05 | 1.3e-05 | 1e-05 | 1.300E-05 | 3.000E-06 | 1.000E-05 | 1.600E-05 |
| 8572 | 429 | S429K | A | 1.6e-05 | 1e-05 | 1.4e-05 | 1.333E-05 | 3.055E-06 | 1.000E-05 | 1.600E-05 |
| 8573 | 429 | S429M | A | 1.3e-05 | 1e-05 | 9.9e-06 | 1.097E-05 | 1.762E-06 | 9.900E-06 | 1.300E-05 |
| 8574 | 429 | S429F | A | 1.8e-05 | 9.7e-06 | 1.1e-05 | 1.290E-05 | 4.464E-06 | 9.700E-06 | 1.800E-05 |
| 8575 | 429 | S429P | A | 1.2e-05 | 1.5e-05 | 1.1e-05 | 1.267E-05 | 2.082E-06 | 1.100E-05 | 1.500E-05 |
| 8576 | 429 | S429S | A | 9.6e-06 | 1.6e-05 | 1.4e-05 | 1.320E-05 | 3.274E-06 | 9.600E-06 | 1.600E-05 |
| 8577 | 429 | S429T | A | 1.4e-05 | 1.4e-05 | 1.6e-05 | 1.467E-05 | 1.155E-06 | 1.400E-05 | 1.600E-05 |
| 8578 | 429 | S429W | A | 1.1e-05 | 1.1e-05 | 9.2e-06 | 1.040E-05 | 1.039E-06 | 9.200E-06 | 1.100E-05 |
| 8579 | 429 | S429Y | A | 9.9e-06 | 1.6e-05 | 9.1e-06 | 1.167E-05 | 3.774E-06 | 9.100E-06 | 1.600E-05 |
| 8580 | 429 | S429V | A | 1.1e-05 | 1.3e-05 | 1.1e-05 | 1.167E-05 | 1.155E-06 | 1.100E-05 | 1.300E-05 |
| 8581 | 430 | W430A | A | 1.5e-05 | 1.6e-05 | 1.1e-05 | 1.400E-05 | 2.646E-06 | 1.100E-05 | 1.600E-05 |
| 8582 | 430 | W430R | A | 2.2e-05 | 1.1e-05 | 1.1e-05 | 1.467E-05 | 6.351E-06 | 1.100E-05 | 2.200E-05 |
| 8583 | 430 | W430N | A | 1.3e-05 | 1.4e-05 | 1.6e-05 | 1.433E-05 | 1.528E-06 | 1.300E-05 | 1.600E-05 |
| 8584 | 430 | W430D | A | 9.5e-06 | 1.5e-05 | 1.1e-05 | 1.183E-05 | 2.843E-06 | 9.500E-06 | 1.500E-05 |

| | | | | | | | | | | |
|---|---|---|---|---|---|---|---|---|---|---|
| 8585 | 430 | W430C | A | 1.3e-05 | 1e-05 | 1.2e-05 | 1.167E-05 | 1.528E-06 | 1.000E-05 | 1.300E-05 |
| 8586 | 430 | W430E | A | 1.2e-05 | 1.1e-05 | 1.4e-05 | 1.233E-05 | 1.528E-06 | 1.100E-05 | 1.400E-05 |
| 8587 | 430 | W430Q | A | 1.4e-05 | 1.5e-05 | 6.9e-06 | 1.197E-05 | 4.416E-06 | 6.900E-06 | 1.500E-05 |
| 8588 | 430 | W430G | A | 1.1e-05 | 1.8e-05 | 1e-05 | 1.300E-05 | 4.359E-06 | 1.000E-05 | 1.800E-05 |
| 8589 | 430 | W430H | A | 1.5e-05 | 1.6e-05 | 1.5e-05 | 1.533E-05 | 5.774E-07 | 1.500E-05 | 1.600E-05 |
| 8590 | 430 | W430I | A | 2.2e-05 | 7.2e-06 | 1.1e-05 | 1.340E-05 | 7.686E-06 | 7.200E-06 | 2.200E-05 |
| 8591 | 430 | W430L | A | 1.3e-05 | 1.6e-05 | 1.5e-05 | 1.467E-05 | 1.528E-06 | 1.300E-05 | 1.600E-05 |
| 8592 | 430 | W430K | A | 1.1e-05 | 1.6e-05 | 1.3e-05 | 1.333E-05 | 2.517E-06 | 1.100E-05 | 1.600E-05 |
| 8593 | 430 | W430M | A | 8.2e-06 | 1.3e-05 | 1.5e-05 | 1.207E-05 | 3.495E-06 | 8.200E-06 | 1.500E-05 |
| 8594 | 430 | W430F | A | 1e-05 | 8e-06 | 1.2e-05 | 1.000E-05 | 2.000E-06 | 8.000E-06 | 1.200E-05 |
| 8595 | 430 | W430P | A | 1.5e-05 | 1.5e-05 | 1.6e-05 | 1.533E-05 | 5.774E-07 | 1.500E-05 | 1.600E-05 |
| 8596 | 430 | W430S | A | 1.3e-05 | 9.8e-06 | 1.5e-05 | 1.260E-05 | 2.623E-06 | 9.800E-06 | 1.500E-05 |
| 8597 | 430 | W430T | A | 8.9e-06 | 1.2e-05 | 1.2e-05 | 1.097E-05 | 1.790E-06 | 8.900E-06 | 1.200E-05 |
| 8598 | 430 | W430W | A | 1.6e-05 | 9.8e-06 | 1e-05 | 1.193E-05 | 3.523E-06 | 9.800E-06 | 1.600E-05 |
| 8599 | 430 | W430Y | A | 1.4e-05 | 8.3e-06 | 1.4e-05 | 1.210E-05 | 3.291E-06 | 8.300E-06 | 1.400E-05 |
| 8600 | 430 | W430V | A | 1.4e-05 | 9.4e-06 | 1.6e-05 | 1.313E-05 | 3.384E-06 | 9.400E-06 | 1.600E-05 |
| 8601 | 431 | L431A | A | 1.6e-05 | 1.2e-05 | 1.6e-05 | 1.467E-05 | 2.309E-06 | 1.200E-05 | 1.600E-05 |
| 8602 | 431 | L431R | A | 1e-05 | 1.5e-05 | 1.4e-05 | 1.300E-05 | 2.646E-06 | 1.000E-05 | 1.500E-05 |
| 8603 | 431 | L431N | A | 1.7e-05 | 1.3e-05 | 1.1e-05 | 1.367E-05 | 3.055E-06 | 1.100E-05 | 1.700E-05 |
| 8604 | 431 | L431D | A | 1.4e-05 | 9.7e-06 | 7.4e-06 | 1.037E-05 | 3.350E-06 | 7.400E-06 | 1.400E-05 |
| 8605 | 431 | L431C | A | 9.8e-06 | 1e-05 | 1.4e-05 | 1.127E-05 | 2.369E-06 | 9.800E-06 | 1.400E-05 |
| 8606 | 431 | L431E | A | 1.2e-05 | 1.1e-05 | 1.2e-05 | 1.167E-05 | 5.774E-07 | 1.100E-05 | 1.200E-05 |
| 8607 | 431 | L431Q | A | 1.5e-05 | 1.1e-05 | 1.6e-05 | 1.400E-05 | 2.646E-06 | 1.100E-05 | 1.600E-05 |
| 8608 | 431 | L431G | A | 1.5e-05 | 1.1e-05 | 1.6e-05 | 1.400E-05 | 2.646E-06 | 1.100E-05 | 1.600E-05 |
| 8609 | 431 | L431H | A | 1.4e-05 | 1.2e-05 | 1.7e-05 | 1.433E-05 | 2.517E-06 | 1.200E-05 | 1.700E-05 |
| 8610 | 431 | L431I | A | 7.7e-06 | 1.6e-05 | 9e-06 | 1.090E-05 | 4.464E-06 | 7.700E-06 | 1.600E-05 |
| 8611 | 431 | L431L | A | 1.5e-05 | 1.5e-05 | 9.3e-06 | 1.310E-05 | 3.291E-06 | 9.300E-06 | 1.500E-05 |
| 8612 | 431 | L431K | A | 1.6e-05 | 1.5e-05 | 1.5e-05 | 1.533E-05 | 5.774E-07 | 1.500E-05 | 1.600E-05 |
| 8613 | 431 | L431M | A | 1e-05 | 8.8e-06 | 1.7e-05 | 1.193E-05 | 4.429E-06 | 8.800E-06 | 1.700E-05 |
| 8614 | 431 | L431F | A | 1.4e-05 | 1.5e-05 | 1.7e-05 | 1.533E-05 | 1.528E-06 | 1.400E-05 | 1.700E-05 |
| 8615 | 431 | L431P | A | 1.7e-05 | 1.1e-05 | 1.3e-05 | 1.367E-05 | 3.055E-06 | 1.100E-05 | 1.700E-05 |
| 8616 | 431 | L431S | A | 8.9e-06 | 1.5e-05 | 1.2e-05 | 1.197E-05 | 3.050E-06 | 8.900E-06 | 1.500E-05 |
| 8617 | 431 | L431T | A | 1.2e-05 | 1.6e-05 | 1.5e-05 | 1.433E-05 | 2.082E-06 | 1.200E-05 | 1.600E-05 |
| 8618 | 431 | L431W | A | 1e-05 | 1e-05 | 1.1e-05 | 1.033E-05 | 5.774E-07 | 1.000E-05 | 1.100E-05 |
| 8619 | 431 | L431Y | A | 1.5e-05 | 9.9e-06 | 1.6e-05 | 1.363E-05 | 3.272E-06 | 9.900E-06 | 1.600E-05 |
| 8620 | 431 | L431V | A | 1.7e-05 | 1.3e-05 | 9.6e-06 | 1.320E-05 | 3.704E-06 | 9.600E-06 | 1.700E-05 |
| 8621 | 432 | G432A | A | 1.7e-05 | 1.5e-05 | 2.1e-05 | 1.767E-05 | 3.055E-06 | 1.500E-05 | 2.100E-05 |
| 8622 | 432 | G432R | A | 1.4e-05 | 1e-05 | 1.4e-05 | 1.267E-05 | 2.309E-06 | 1.000E-05 | 1.400E-05 |
| 8623 | 432 | G432N | A | 1.6e-05 | 1.4e-05 | 1.2e-05 | 1.400E-05 | 2.000E-06 | 1.200E-05 | 1.600E-05 |
| 8624 | 432 | G432D | A | 1.4e-05 | 9.1e-06 | 1.4e-05 | 1.237E-05 | 2.829E-06 | 9.100E-06 | 1.400E-05 |
| 8625 | 432 | G432C | A | 6.5e-06 | 1.4e-05 | 1.4e-05 | 1.150E-05 | 4.330E-06 | 6.500E-06 | 1.400E-05 |
| 8626 | 432 | G432E | A | 9.8e-06 | 1.6e-05 | 1.1e-05 | 1.227E-05 | 3.288E-06 | 9.800E-06 | 1.600E-05 |
| 8627 | 432 | G432Q | A | 1.6e-05 | 1.4e-05 | 1.5e-05 | 1.500E-05 | 1.000E-06 | 1.400E-05 | 1.600E-05 |
| 8628 | 432 | G432G | A | 1.6e-05 | 1.5e-05 | 8.9e-06 | 1.330E-05 | 3.843E-06 | 8.900E-06 | 1.600E-05 |
| 8629 | 432 | G432H | A | 1.6e-05 | 1.3e-05 | 9e-06 | 1.267E-05 | 3.512E-06 | 9.000E-06 | 1.600E-05 |
| 8630 | 432 | G432I | A | 1e-05 | 1.6e-05 | 1.3e-05 | 1.300E-05 | 3.000E-06 | 1.000E-05 | 1.600E-05 |
| 8631 | 432 | G432L | A | 1e-05 | 1.1e-05 | 1e-05 | 1.033E-05 | 5.774E-07 | 1.000E-05 | 1.100E-05 |
| 8632 | 432 | G432K | A | 1.5e-05 | 8.9e-06 | 1.8e-05 | 1.397E-05 | 4.637E-06 | 8.900E-06 | 1.800E-05 |
| 8633 | 432 | G432M | A | 1.2e-05 | 1.1e-05 | 7.1e-06 | 1.003E-05 | 2.589E-06 | 7.100E-06 | 1.200E-05 |
| 8634 | 432 | G432F | A | 1e-05 | 1.2e-05 | 1.1e-05 | 1.100E-05 | 1.000E-06 | 1.000E-05 | 1.200E-05 |
| 8635 | 432 | G432P | A | 1.3e-05 | 8.2e-06 | 1.5e-05 | 1.207E-05 | 3.495E-06 | 8.200E-06 | 1.500E-05 |
| 8636 | 432 | G432S | A | 1.4e-05 | 1.4e-05 | 1.5e-05 | 1.433E-05 | 5.774E-07 | 1.400E-05 | 1.500E-05 |
| 8637 | 432 | G432T | A | 1.3e-05 | 1.6e-05 | 1.5e-05 | 1.467E-05 | 1.528E-06 | 1.300E-05 | 1.600E-05 |
| 8638 | 432 | G432W | A | 9.9e-06 | 1.4e-05 | 1e-05 | 1.130E-05 | 2.339E-06 | 9.900E-06 | 1.400E-05 |
| 8639 | 432 | G432Y | A | 1.2e-05 | 1.4e-05 | 8.6e-06 | 1.153E-05 | 2.730E-06 | 8.600E-06 | 1.400E-05 |
| 8640 | 432 | G432V | A | 1.6e-05 | 1.1e-05 | 1e-05 | 1.233E-05 | 3.215E-06 | 1.000E-05 | 1.600E-05 |
| 8641 | 433 | L433A | A | 1.7e-05 | 1.5e-05 | 9.5e-06 | 1.383E-05 | 3.884E-06 | 9.500E-06 | 1.700E-05 |
| 8642 | 433 | L433R | A | 9.6e-06 | 1.5e-05 | 1.3e-05 | 1.253E-05 | 2.730E-06 | 9.600E-06 | 1.500E-05 |
| 8643 | 433 | L433N | A | 1.4e-05 | 1.4e-05 | 1.7e-05 | 1.500E-05 | 1.732E-06 | 1.400E-05 | 1.700E-05 |
| 8644 | 433 | L433D | A | 1.4e-05 | 8.5e-06 | 1.6e-05 | 1.283E-05 | 3.884E-06 | 8.500E-06 | 1.600E-05 |
| 8645 | 433 | L433C | A | 1.4e-05 | 1e-05 | 1.3e-05 | 1.233E-05 | 2.082E-06 | 1.000E-05 | 1.400E-05 |
| 8646 | 433 | L433E | A | 1.3e-05 | 8.5e-06 | 1.4e-05 | 1.183E-05 | 2.930E-06 | 8.500E-06 | 1.400E-05 |
| 8647 | 433 | L433Q | A | 1.4e-05 | 1.1e-05 | 1.1e-05 | 1.200E-05 | 1.732E-06 | 1.100E-05 | 1.400E-05 |
| 8648 | 433 | L433G | A | 1.1e-05 | 1e-05 | 1.2e-05 | 1.100E-05 | 1.000E-06 | 1.000E-05 | 1.200E-05 |
| 8649 | 433 | L433H | A | 1.5e-05 | 1.6e-05 | 1.1e-05 | 1.400E-05 | 2.646E-06 | 1.100E-05 | 1.600E-05 |
| 8650 | 433 | L433I | A | 1.1e-05 | 1.5e-05 | 1e-05 | 1.200E-05 | 2.646E-06 | 1.000E-05 | 1.500E-05 |
| 8651 | 433 | L433L | A | 8.5e-06 | 1e-05 | 1.6e-05 | 1.150E-05 | 3.969E-06 | 8.500E-06 | 1.600E-05 |
| 8652 | 433 | L433K | A | 8e-06 | 1.5e-05 | 2.4e-05 | 1.567E-05 | 8.021E-06 | 8.000E-06 | 2.400E-05 |
| 8653 | 433 | L433M | A | 1.6e-05 | 9.6e-06 | 1.6e-05 | 1.387E-05 | 3.695E-06 | 9.600E-06 | 1.600E-05 |
| 8654 | 433 | L433F | A | 1.5e-05 | 2.8e-05 | 1.1e-05 | 1.800E-05 | 8.888E-06 | 1.100E-05 | 2.800E-05 |
| 8655 | 433 | L433P | A | 1e-05 | 1.5e-05 | 1.5e-05 | 1.333E-05 | 2.887E-06 | 1.000E-05 | 1.500E-05 |
| 8656 | 433 | L433S | A | 1.5e-05 | 1.6e-05 | 6.9e-06 | 1.263E-05 | 4.990E-06 | 6.900E-06 | 1.600E-05 |
| 8657 | 433 | L433T | A | 1.2e-05 | 1.5e-05 | 1e-05 | 1.233E-05 | 2.517E-06 | 1.000E-05 | 1.500E-05 |
| 8658 | 433 | L433W | A | 1.2e-05 | 1e-05 | 1e-05 | 1.067E-05 | 1.155E-06 | 1.000E-05 | 1.200E-05 |
| 8659 | 433 | L433Y | A | 1.5e-05 | 1.5e-05 | 9.5e-06 | 1.317E-05 | 3.175E-06 | 9.500E-06 | 1.500E-05 |
| 8660 | 433 | L433V | A | 1.4e-05 | 1.2e-05 | 1.5e-05 | 1.367E-05 | 1.528E-06 | 1.200E-05 | 1.500E-05 |
| 8661 | 434 | R434A | A | 1.4e-05 | 1.6e-05 | 1.6e-05 | 1.533E-05 | 1.155E-06 | 1.400E-05 | 1.600E-05 |
| 8662 | 434 | R434R | A | 1.4e-05 | 1.2e-05 | 1.3e-05 | 1.300E-05 | 1.000E-06 | 1.200E-05 | 1.400E-05 |
| 8663 | 434 | R434N | A | 1e-05 | 9.4e-06 | 1.1e-05 | 1.013E-05 | 8.083E-07 | 9.400E-06 | 1.100E-05 |
| 8664 | 434 | R434D | A | 1.5e-05 | 8.6e-06 | 1.6e-05 | 1.320E-05 | 4.015E-06 | 8.600E-06 | 1.600E-05 |
| 8665 | 434 | R434C | A | 1.7e-05 | 8.3e-06 | 1.6e-05 | 1.377E-05 | 4.761E-06 | 8.300E-06 | 1.700E-05 |
| 8666 | 434 | R434E | A | 1.7e-05 | 1.1e-05 | 9.1e-06 | 1.237E-05 | 4.124E-06 | 9.100E-06 | 1.700E-05 |
| 8667 | 434 | R434Q | A | 1.5e-05 | 1.6e-05 | 1.2e-05 | 1.433E-05 | 2.082E-06 | 1.200E-05 | 1.600E-05 |
| 8668 | 434 | R434G | A | 1.1e-05 | 1.6e-05 | 1.1e-05 | 1.267E-05 | 2.887E-06 | 1.100E-05 | 1.600E-05 |
| 8669 | 434 | R434H | A | 1.6e-05 | 1.7e-05 | 1.4e-05 | 1.567E-05 | 1.528E-06 | 1.400E-05 | 1.700E-05 |

| | | | | | | | | | | |
|---|---|---|---|---|---|---|---|---|---|---|
| 8670 | 434 | R434I | A | 1.3e-05 | 9.5e-06 | 1e-05 | 1.083E-05 | 1.893E-06 | 9.500E-06 | 1.300E-05 |
| 8671 | 434 | R434L | A | 1.4e-05 | 1.5e-05 | 1.2e-05 | 1.367E-05 | 1.528E-06 | 1.200E-05 | 1.500E-05 |
| 8672 | 434 | R434K | A | 1.6e-05 | 1.4e-05 | 1.5e-05 | 1.500E-05 | 1.000E-06 | 1.400E-05 | 1.600E-05 |
| 8673 | 434 | R434M | A | 1.5e-05 | 1e-05 | 1.5e-05 | 1.333E-05 | 2.887E-06 | 1.000E-05 | 1.500E-05 |
| 8674 | 434 | R434F | A | 9.9e-06 | 1.4e-05 | 1.7e-05 | 1.363E-05 | 3.564E-06 | 9.900E-06 | 1.700E-05 |
| 8675 | 434 | R434P | A | 1.6e-05 | 9.6e-06 | 9.7e-06 | 1.177E-05 | 3.667E-06 | 9.600E-06 | 1.600E-05 |
| 8676 | 434 | R434S | A | 1.5e-05 | 9.6e-06 | 1.4e-05 | 1.287E-05 | 2.873E-06 | 9.600E-06 | 1.500E-05 |
| 8677 | 434 | R434T | A | 8.8e-06 | 7.2e-06 | 1.3e-05 | 9.667E-06 | 2.996E-06 | 7.200E-06 | 1.300E-05 |
| 8678 | 434 | R434W | A | 1.3e-05 | 1.6e-05 | 1e-05 | 1.300E-05 | 3.000E-06 | 1.000E-05 | 1.600E-05 |
| 8679 | 434 | R434Y | A | 1.1e-05 | 9.1e-06 | 9.1e-06 | 9.733E-06 | 1.097E-06 | 9.100E-06 | 1.100E-05 |
| 8680 | 434 | R434V | A | 1.2e-05 | 1.6e-05 | 7.2e-06 | 1.173E-05 | 4.406E-06 | 7.200E-06 | 1.600E-05 |
| 8681 | 435 | S435A | A | 1.6e-05 | 1.3e-05 | 9.6e-06 | 1.287E-05 | 3.202E-06 | 9.600E-06 | 1.600E-05 |
| 8682 | 435 | S435R | A | 9.2e-06 | 1.2e-05 | 1.1e-05 | 1.073E-05 | 1.419E-06 | 9.200E-06 | 1.200E-05 |
| 8683 | 435 | S435N | A | 1.4e-05 | 1.5e-05 | 8.9e-06 | 1.263E-05 | 3.272E-06 | 8.900E-06 | 1.500E-05 |
| 8684 | 435 | S435D | A | 1e-05 | 9.9e-06 | 1.6e-05 | 1.197E-05 | 3.493E-06 | 9.900E-06 | 1.600E-05 |
| 8685 | 435 | S435C | A | 1.5e-05 | 2.3e-05 | 1.7e-05 | 1.833E-05 | 4.163E-06 | 1.500E-05 | 2.300E-05 |
| 8686 | 435 | S435E | A | 1.6e-05 | 1.6e-05 | 9.6e-06 | 1.387E-05 | 3.695E-06 | 9.600E-06 | 1.600E-05 |
| 8687 | 435 | S435Q | A | 1.5e-05 | 1e-05 | 8e-06 | 1.100E-05 | 3.606E-06 | 8.000E-06 | 1.500E-05 |
| 8688 | 435 | S435G | A | 8.3e-06 | 1.7e-05 | 1.1e-05 | 1.210E-05 | 4.453E-06 | 8.300E-06 | 1.700E-05 |
| 8689 | 435 | S435H | A | 8.3e-06 | 9.9e-06 | 1e-05 | 9.400E-06 | 9.539E-07 | 8.300E-06 | 1.000E-05 |
| 8690 | 435 | S435I | A | 1.4e-05 | 1.6e-05 | 1.3e-05 | 1.433E-05 | 1.528E-06 | 1.300E-05 | 1.600E-05 |
| 8691 | 435 | S435L | A | 1.7e-05 | 1.6e-05 | 1.6e-05 | 1.633E-05 | 5.774E-07 | 1.600E-05 | 1.700E-05 |
| 8692 | 435 | S435K | A | 1.4e-05 | 1.6e-05 | 1.1e-05 | 1.367E-05 | 2.517E-06 | 1.100E-05 | 1.600E-05 |
| 8693 | 435 | S435M | A | 1.7e-05 | 1.2e-05 | 1.5e-05 | 1.467E-05 | 2.517E-06 | 1.200E-05 | 1.700E-05 |
| 8694 | 435 | S435F | A | 1.1e-05 | 1.6e-05 | 1.6e-05 | 1.433E-05 | 2.887E-06 | 1.100E-05 | 1.600E-05 |
| 8695 | 435 | S435P | A | 1.5e-05 | 1e-05 | 1e-05 | 1.167E-05 | 2.887E-06 | 1.000E-05 | 1.500E-05 |
| 8696 | 435 | S435S | A | 1e-05 | 1e-05 | 1.5e-05 | 1.167E-05 | 2.887E-06 | 1.000E-05 | 1.500E-05 |
| 8697 | 435 | S435T | A | 1.1e-05 | 1.1e-05 | 8.3e-06 | 1.010E-05 | 1.559E-06 | 8.300E-06 | 1.100E-05 |
| 8698 | 435 | S435W | A | 1.7e-05 | 1.5e-05 | 1.3e-05 | 1.500E-05 | 2.000E-06 | 1.300E-05 | 1.700E-05 |
| 8699 | 435 | S435Y | A | 1.4e-05 | 1.6e-05 | 1.4e-05 | 1.467E-05 | 1.155E-06 | 1.400E-05 | 1.600E-05 |
| 8700 | 435 | S435V | A | 8.8e-06 | 1.1e-05 | 2.2e-05 | 1.393E-05 | 7.072E-06 | 8.800E-06 | 2.200E-05 |
| 8701 | 436 | L436A | A | 1.6e-05 | 1.1e-05 | 1.5e-05 | 1.400E-05 | 2.646E-06 | 1.100E-05 | 1.600E-05 |
| 8702 | 436 | L436R | A | 1.6e-05 | 1e-05 | 1e-05 | 1.200E-05 | 3.464E-06 | 1.000E-05 | 1.600E-05 |
| 8703 | 436 | L436N | A | 1.2e-05 | 8.7e-06 | 9.7e-06 | 1.013E-05 | 1.692E-06 | 8.700E-06 | 1.200E-05 |
| 8704 | 436 | L436D | A | 1e-05 | 1.8e-05 | 1.5e-05 | 1.433E-05 | 4.041E-06 | 1.000E-05 | 1.800E-05 |
| 8705 | 436 | L436C | A | 1.2e-05 | 1.6e-05 | 1.5e-05 | 1.433E-05 | 2.082E-06 | 1.200E-05 | 1.600E-05 |
| 8706 | 436 | L436E | A | 1.4e-05 | 1e-05 | 1.4e-05 | 1.267E-05 | 2.309E-06 | 1.000E-05 | 1.400E-05 |
| 8707 | 436 | L436Q | A | 1e-05 | 1.8e-05 | 1.3e-05 | 1.367E-05 | 4.041E-06 | 1.000E-05 | 1.800E-05 |
| 8708 | 436 | L436G | A | 9.8e-06 | 1.5e-05 | 1.6e-05 | 1.360E-05 | 3.329E-06 | 9.800E-06 | 1.600E-05 |
| 8709 | 436 | L436H | A | 1.4e-05 | 9.9e-06 | 1.4e-05 | 1.263E-05 | 2.367E-06 | 9.900E-06 | 1.400E-05 |
| 8710 | 436 | L436I | A | 1.1e-05 | 1.3e-05 | 1e-05 | 1.133E-05 | 1.528E-06 | 1.000E-05 | 1.300E-05 |
| 8711 | 436 | L436L | A | 1e-05 | 1.3e-05 | 8e-06 | 1.033E-05 | 2.517E-06 | 8.000E-06 | 1.300E-05 |
| 8712 | 436 | L436K | A | 1.5e-05 | 1.4e-05 | 1e-05 | 1.300E-05 | 2.646E-06 | 1.000E-05 | 1.500E-05 |
| 8713 | 436 | L436M | A | 1.2e-05 | 8.8e-06 | 1.7e-05 | 1.260E-05 | 4.133E-06 | 8.800E-06 | 1.700E-05 |
| 8714 | 436 | L436F | A | 8.6e-06 | 1e-05 | 1e-05 | 9.533E-06 | 8.083E-07 | 8.600E-06 | 1.000E-05 |
| 8715 | 436 | L436P | A | 2.6e-05 | 7.8e-06 | 1.5e-05 | 1.627E-05 | 9.166E-06 | 7.800E-06 | 2.600E-05 |
| 8716 | 436 | L436S | A | 1.8e-05 | 1.6e-05 | 1.2e-05 | 1.533E-05 | 3.055E-06 | 1.200E-05 | 1.800E-05 |
| 8717 | 436 | L436T | A | 1.5e-05 | 1.1e-05 | 2.3e-05 | 1.633E-05 | 6.110E-06 | 1.100E-05 | 2.300E-05 |
| 8718 | 436 | L436W | A | 1.6e-05 | 1.3e-05 | 1.4e-05 | 1.433E-05 | 1.528E-06 | 1.300E-05 | 1.600E-05 |
| 8719 | 436 | L436Y | A | 1.4e-05 | 9e-06 | 1.5e-05 | 1.267E-05 | 3.215E-06 | 9.000E-06 | 1.500E-05 |
| 8720 | 436 | L436V | A | 1.1e-05 | 8.9e-06 | 1.3e-05 | 1.097E-05 | 2.050E-06 | 8.900E-06 | 1.300E-05 |
| 8721 | 437 | R437A | A | 1.4e-05 | 7.3e-06 | 1.4e-05 | 1.177E-05 | 3.868E-06 | 7.300E-06 | 1.400E-05 |
| 8722 | 437 | R437R | A | 1.1e-05 | 1.5e-05 | 1.1e-05 | 1.233E-05 | 2.309E-06 | 1.100E-05 | 1.500E-05 |
| 8723 | 437 | R437N | A | 1.1e-05 | 1.3e-05 | 1.4e-05 | 1.267E-05 | 1.528E-06 | 1.100E-05 | 1.400E-05 |
| 8724 | 437 | R437D | A | 1.3e-05 | 1.7e-05 | 9.3e-06 | 1.310E-05 | 3.851E-06 | 9.300E-06 | 1.700E-05 |
| 8725 | 437 | R437C | A | 1.1e-05 | 1.4e-05 | 1.4e-05 | 1.300E-05 | 1.732E-06 | 1.100E-05 | 1.400E-05 |
| 8726 | 437 | R437E | A | 1.1e-05 | 1.7e-05 | 1.6e-05 | 1.467E-05 | 3.215E-06 | 1.100E-05 | 1.700E-05 |
| 8727 | 437 | R437Q | A | 9.2e-06 | 1.2e-05 | 1e-05 | 1.040E-05 | 1.442E-06 | 9.200E-06 | 1.200E-05 |
| 8728 | 437 | R437G | A | 9.3e-06 | 1.6e-05 | 8.7e-06 | 1.133E-05 | 4.053E-06 | 8.700E-06 | 1.600E-05 |
| 8729 | 437 | R437H | A | 1.5e-05 | 8.6e-06 | 1e-05 | 1.120E-05 | 3.365E-06 | 8.600E-06 | 1.500E-05 |
| 8730 | 437 | R437I | A | 1.6e-05 | 1.3e-05 | 1.4e-05 | 1.433E-05 | 1.528E-06 | 1.300E-05 | 1.600E-05 |
| 8731 | 437 | R437L | A | 9e-06 | 8.8e-06 | 1.5e-05 | 1.093E-05 | 3.523E-06 | 8.800E-06 | 1.500E-05 |
| 8732 | 437 | R437K | A | 1.4e-05 | 1e-05 | 1.5e-05 | 1.300E-05 | 2.646E-06 | 1.000E-05 | 1.500E-05 |
| 8733 | 437 | R437M | A | 1.6e-05 | 1.5e-05 | 1.1e-05 | 1.400E-05 | 2.646E-06 | 1.100E-05 | 1.600E-05 |
| 8734 | 437 | R437F | A | 1.2e-05 | 1e-05 | 1e-05 | 1.067E-05 | 1.155E-06 | 1.000E-05 | 1.200E-05 |
| 8735 | 437 | R437P | A | 1.4e-05 | 9.8e-06 | 1.5e-05 | 1.293E-05 | 2.759E-06 | 9.800E-06 | 1.500E-05 |
| 8736 | 437 | R437S | A | 1.5e-05 | 1.3e-05 | 1e-05 | 1.267E-05 | 2.517E-06 | 1.000E-05 | 1.500E-05 |
| 8737 | 437 | R437T | A | 1.5e-05 | 1.3e-05 | 9.6e-06 | 1.253E-05 | 2.730E-06 | 9.600E-06 | 1.500E-05 |
| 8738 | 437 | R437W | A | 1.4e-05 | 1.6e-05 | 1.3e-05 | 1.433E-05 | 1.528E-06 | 1.300E-05 | 1.600E-05 |
| 8739 | 437 | R437Y | A | 1.2e-05 | 1.1e-05 | 1.5e-05 | 1.267E-05 | 2.082E-06 | 1.100E-05 | 1.500E-05 |
| 8740 | 437 | R437V | A | 9.7e-06 | 1.1e-05 | 1e-05 | 1.023E-05 | 6.807E-07 | 9.700E-06 | 1.100E-05 |
| 8741 | 438 | E438A | A | 1.5e-05 | 1e-05 | 1.5e-05 | 1.333E-05 | 2.887E-06 | 1.000E-05 | 1.500E-05 |
| 8742 | 438 | E438R | A | 1.3e-05 | 1.3e-05 | 1.5e-05 | 1.367E-05 | 1.155E-06 | 1.300E-05 | 1.500E-05 |
| 8743 | 438 | E438N | A | 9.4e-06 | 9.8e-06 | 1.5e-05 | 1.140E-05 | 3.124E-06 | 9.400E-06 | 1.500E-05 |
| 8744 | 438 | E438D | A | 8.6e-06 | 1.1e-05 | 1.3e-05 | 1.087E-05 | 2.203E-06 | 8.600E-06 | 1.300E-05 |
| 8745 | 438 | E438C | A | 1.1e-05 | 1.2e-05 | 1.6e-05 | 1.300E-05 | 2.646E-06 | 1.100E-05 | 1.600E-05 |
| 8746 | 438 | E438E | A | 9.8e-06 | 1.1e-05 | 9.1e-06 | 9.967E-06 | 9.609E-07 | 9.100E-06 | 1.100E-05 |
| 8747 | 438 | E438Q | A | 1e-05 | 1.1e-05 | 1e-05 | 1.033E-05 | 5.774E-07 | 1.000E-05 | 1.100E-05 |
| 8748 | 438 | E438G | A | 1.4e-05 | 1.6e-05 | 1.7e-05 | 1.567E-05 | 1.528E-06 | 1.400E-05 | 1.700E-05 |
| 8749 | 438 | E438H | A | 9.4e-06 | 1.5e-05 | 1.1e-05 | 1.180E-05 | 2.884E-06 | 9.400E-06 | 1.500E-05 |
| 8750 | 438 | E438I | A | 1.3e-05 | 1e-05 | 1.1e-05 | 1.133E-05 | 1.528E-06 | 1.000E-05 | 1.300E-05 |
| 8751 | 438 | E438L | A | 1.5e-05 | 1e-05 | 9e-06 | 1.133E-05 | 3.215E-06 | 9.000E-06 | 1.500E-05 |
| 8752 | 438 | E438K | A | 1.1e-05 | 1.5e-05 | 1.8e-05 | 1.467E-05 | 3.512E-06 | 1.100E-05 | 1.800E-05 |
| 8753 | 438 | E438M | A | 1.4e-05 | 1.3e-05 | 1.6e-05 | 1.433E-05 | 1.528E-06 | 1.300E-05 | 1.600E-05 |
| 8754 | 438 | E438F | A | 1.6e-05 | 1.1e-05 | 1.6e-05 | 1.433E-05 | 2.887E-06 | 1.100E-05 | 1.600E-05 |

| | | | | | | | | | | |
|---|---|---|---|---|---|---|---|---|---|---|
| 8755 | 438 | E438P | A | 1e-05 | 1.6e-05 | 8.7e-06 | 1.157E-05 | 3.894E-06 | 8.700E-06 | 1.600E-05 |
| 8756 | 438 | E438S | A | 1e-05 | 1.5e-05 | 1e-05 | 1.167E-05 | 2.887E-06 | 1.000E-05 | 1.500E-05 |
| 8757 | 438 | E438T | A | 1.2e-05 | 9.5e-06 | 1.1e-05 | 1.083E-05 | 1.258E-06 | 9.500E-06 | 1.200E-05 |
| 8758 | 438 | E438W | A | 9.8e-06 | 1.5e-05 | 1.9e-05 | 1.460E-05 | 4.613E-06 | 9.800E-06 | 1.900E-05 |
| 8759 | 438 | E438Y | A | 1e-05 | 1.3e-05 | 1.1e-05 | 1.133E-05 | 1.528E-06 | 1.000E-05 | 1.300E-05 |
| 8760 | 438 | E438V | A | 9.4e-06 | 7.8e-06 | 1.6e-05 | 1.107E-05 | 4.347E-06 | 7.800E-06 | 1.600E-05 |
| 8761 | 439 | L439A | A | 1.6e-05 | 1.4e-05 | 1.5e-05 | 1.500E-05 | 1.000E-06 | 1.400E-05 | 1.600E-05 |
| 8762 | 439 | L439R | A | 1.4e-05 | 1.4e-05 | 1.4e-05 | 1.400E-05 | 0.000E+00 | 1.400E-05 | 1.400E-05 |
| 8763 | 439 | L439N | A | 1.3e-05 | 1.1e-05 | 1.4e-05 | 1.267E-05 | 1.528E-06 | 1.100E-05 | 1.400E-05 |
| 8764 | 439 | L439D | A | 1.8e-05 | 1.6e-05 | 9e-06 | 1.433E-05 | 4.726E-06 | 9.000E-06 | 1.800E-05 |
| 8765 | 439 | L439C | A | 1.4e-05 | 1e-05 | 1.2e-05 | 1.200E-05 | 2.000E-06 | 1.000E-05 | 1.400E-05 |
| 8766 | 439 | L439E | A | 1.6e-05 | 1e-05 | 1.1e-05 | 1.233E-05 | 3.215E-06 | 1.000E-05 | 1.600E-05 |
| 8767 | 439 | L439Q | A | 1.6e-05 | 1.2e-05 | 1.5e-05 | 1.433E-05 | 2.082E-06 | 1.200E-05 | 1.600E-05 |
| 8768 | 439 | L439G | A | 1.1e-05 | 9.9e-06 | 1.2e-05 | 1.097E-05 | 1.050E-06 | 9.900E-06 | 1.200E-05 |
| 8769 | 439 | L439H | A | 1.6e-05 | 1.4e-05 | 1.1e-05 | 1.367E-05 | 2.517E-06 | 1.100E-05 | 1.600E-05 |
| 8770 | 439 | L439I | A | 1.7e-05 | 1.5e-05 | 8.8e-06 | 1.360E-05 | 4.276E-06 | 8.800E-06 | 1.700E-05 |
| 8771 | 439 | L439L | A | 1.6e-05 | 1.5e-05 | 1.3e-05 | 1.467E-05 | 1.528E-06 | 1.300E-05 | 1.600E-05 |
| 8772 | 439 | L439K | A | 1.1e-05 | 1.8e-05 | 1.4e-05 | 1.433E-05 | 3.512E-06 | 1.100E-05 | 1.800E-05 |
| 8773 | 439 | L439M | A | 1.6e-05 | 9.6e-06 | 1e-05 | 1.187E-05 | 3.585E-06 | 9.600E-06 | 1.600E-05 |
| 8774 | 439 | L439F | A | 1.4e-05 | 1.4e-05 | 1.6e-05 | 1.467E-05 | 1.155E-06 | 1.400E-05 | 1.600E-05 |
| 8775 | 439 | L439P | A | 5.8e-06 | 1.7e-05 | 1e-05 | 1.093E-05 | 5.658E-06 | 5.800E-06 | 1.700E-05 |
| 8776 | 439 | L439S | A | 1.4e-05 | 1.6e-05 | 2.6e-05 | 1.867E-05 | 6.429E-06 | 1.400E-05 | 2.600E-05 |
| 8777 | 439 | L439T | A | 8.9e-06 | 9.7e-06 | 1.7e-05 | 1.187E-05 | 4.464E-06 | 8.900E-06 | 1.700E-05 |
| 8778 | 439 | L439W | A | 1e-05 | 1.6e-05 | 1.1e-05 | 1.233E-05 | 3.215E-06 | 1.000E-05 | 1.600E-05 |
| 8779 | 439 | L439Y | A | 1.2e-05 | 1.5e-05 | 1.5e-05 | 1.400E-05 | 1.732E-06 | 1.200E-05 | 1.500E-05 |
| 8780 | 439 | L439V | A | 1.1e-05 | 1.5e-05 | 1.1e-05 | 1.233E-05 | 2.309E-06 | 1.100E-05 | 1.500E-05 |
| 8781 | 440 | G440A | A | 1.6e-05 | 9.5e-06 | 1.1e-05 | 1.217E-05 | 3.403E-06 | 9.500E-06 | 1.600E-05 |
| 8782 | 440 | G440R | A | 1.1e-05 | 1e-05 | 1.5e-05 | 1.200E-05 | 2.646E-06 | 1.000E-05 | 1.500E-05 |
| 8783 | 440 | G440N | A | 9.3e-06 | 1.2e-05 | 1.6e-05 | 1.243E-05 | 3.371E-06 | 9.300E-06 | 1.600E-05 |
| 8784 | 440 | G440D | A | 1.1e-05 | 1.2e-05 | 1.6e-05 | 1.300E-05 | 2.646E-06 | 1.100E-05 | 1.600E-05 |
| 8785 | 440 | G440C | A | 1.7e-05 | 1.2e-05 | 1.5e-05 | 1.467E-05 | 2.517E-06 | 1.200E-05 | 1.700E-05 |
| 8786 | 440 | G440E | A | 1.5e-05 | 1.6e-05 | 1.4e-05 | 1.500E-05 | 1.000E-06 | 1.400E-05 | 1.600E-05 |
| 8787 | 440 | G440Q | A | 1e-05 | 1.1e-05 | 1e-05 | 1.033E-05 | 5.774E-07 | 1.000E-05 | 1.100E-05 |
| 8788 | 440 | G440G | A | 1.6e-05 | 1.6e-05 | 1.1e-05 | 1.433E-05 | 2.887E-06 | 1.100E-05 | 1.600E-05 |
| 8789 | 440 | G440H | A | 9.8e-06 | 1.6e-05 | 1.5e-05 | 1.360E-05 | 3.329E-06 | 9.800E-06 | 1.600E-05 |
| 8790 | 440 | G440I | A | 1.7e-05 | 1.2e-05 | 1.6e-05 | 1.500E-05 | 2.646E-06 | 1.200E-05 | 1.700E-05 |
| 8791 | 440 | G440L | A | 1.3e-05 | 9e-06 | 1.1e-05 | 1.100E-05 | 2.000E-06 | 9.000E-06 | 1.300E-05 |
| 8792 | 440 | G440K | A | 1.3e-05 | 1.6e-05 | 1e-05 | 1.300E-05 | 3.000E-06 | 1.000E-05 | 1.600E-05 |
| 8793 | 440 | G440M | A | 1.5e-05 | 1.8e-05 | 1.5e-05 | 1.600E-05 | 1.732E-06 | 1.500E-05 | 1.800E-05 |
| 8794 | 440 | G440F | A | 1.2e-05 | 1e-05 | 1.4e-05 | 1.200E-05 | 2.000E-06 | 1.000E-05 | 1.400E-05 |
| 8795 | 440 | G440P | A | 1.5e-05 | 1e-05 | 8.6e-06 | 1.120E-05 | 3.365E-06 | 8.600E-06 | 1.500E-05 |
| 8796 | 440 | G440S | A | 1.1e-05 | 1.1e-05 | 1.6e-05 | 1.267E-05 | 2.887E-06 | 1.100E-05 | 1.600E-05 |
| 8797 | 440 | G440T | A | 1.5e-05 | 1.5e-05 | 1.5e-05 | 1.500E-05 | 0.000E+00 | 1.500E-05 | 1.500E-05 |
| 8798 | 440 | G440W | A | 1.1e-05 | 9.3e-06 | 1.3e-05 | 1.110E-05 | 1.852E-06 | 9.300E-06 | 1.300E-05 |
| 8799 | 440 | G440Y | A | 1.3e-05 | 1e-05 | 1.5e-05 | 1.267E-05 | 2.517E-06 | 1.000E-05 | 1.500E-05 |
| 8800 | 440 | G440V | A | 1.4e-05 | 1.6e-05 | 1.2e-05 | 1.400E-05 | 2.000E-06 | 1.200E-05 | 1.600E-05 |
| 8801 | 441 | S441A | A | 1.5e-05 | 1.5e-05 | 1.6e-05 | 1.533E-05 | 5.774E-07 | 1.500E-05 | 1.600E-05 |
| 8802 | 441 | S441R | A | 1.5e-05 | 9.3e-06 | 1.4e-05 | 1.277E-05 | 3.044E-06 | 9.300E-06 | 1.500E-05 |
| 8803 | 441 | S441N | A | 1.5e-05 | 1.1e-05 | 1.4e-05 | 1.333E-05 | 2.082E-06 | 1.100E-05 | 1.500E-05 |
| 8804 | 441 | S441D | A | 1.5e-05 | 1.6e-05 | 1.1e-05 | 1.400E-05 | 2.646E-06 | 1.100E-05 | 1.600E-05 |
| 8805 | 441 | S441C | A | 1.4e-05 | 8.7e-06 | 1.5e-05 | 1.257E-05 | 3.386E-06 | 8.700E-06 | 1.500E-05 |
| 8806 | 441 | S441E | A | 1.5e-05 | 9.2e-06 | 1.1e-05 | 1.173E-05 | 2.969E-06 | 9.200E-06 | 1.500E-05 |
| 8807 | 441 | S441Q | A | 1.1e-05 | 1.3e-05 | 8.6e-06 | 1.087E-05 | 2.203E-06 | 8.600E-06 | 1.300E-05 |
| 8808 | 441 | S441G | A | 1.5e-05 | 8.7e-06 | 1.3e-05 | 1.223E-05 | 3.219E-06 | 8.700E-06 | 1.500E-05 |
| 8809 | 441 | S441H | A | 9.5e-06 | 1.4e-05 | 1.5e-05 | 1.283E-05 | 2.930E-06 | 9.500E-06 | 1.500E-05 |
| 8810 | 441 | S441I | A | 1.6e-05 | 1.2e-05 | 1.4e-05 | 1.400E-05 | 2.000E-06 | 1.200E-05 | 1.600E-05 |
| 8811 | 441 | S441L | A | 1.5e-05 | 1.1e-05 | 1.5e-05 | 1.367E-05 | 2.309E-06 | 1.100E-05 | 1.500E-05 |
| 8812 | 441 | S441K | A | 1.6e-05 | 1.6e-05 | 1.4e-05 | 1.533E-05 | 1.155E-06 | 1.400E-05 | 1.600E-05 |
| 8813 | 441 | S441M | A | 8.9e-06 | 1.4e-05 | 1e-05 | 1.097E-05 | 2.684E-06 | 8.900E-06 | 1.400E-05 |
| 8814 | 441 | S441F | A | 8.6e-06 | 1.1e-05 | 1.1e-05 | 1.020E-05 | 1.386E-06 | 8.600E-06 | 1.100E-05 |
| 8815 | 441 | S441P | A | 9.8e-06 | 7.1e-06 | 1.3e-05 | 9.967E-06 | 2.954E-06 | 7.100E-06 | 1.300E-05 |
| 8816 | 441 | S441S | A | 1.1e-05 | 1.5e-05 | 1.8e-05 | 1.467E-05 | 3.512E-06 | 1.100E-05 | 1.800E-05 |
| 8817 | 441 | S441T | A | 9.2e-06 | 1.1e-05 | 1e-05 | 1.007E-05 | 9.018E-07 | 9.200E-06 | 1.100E-05 |
| 8818 | 441 | S441W | A | 1.3e-05 | 1.2e-05 | 1.5e-05 | 1.333E-05 | 1.528E-06 | 1.200E-05 | 1.500E-05 |
| 8819 | 441 | S441Y | A | 1.5e-05 | 1.6e-05 | 1e-05 | 1.367E-05 | 3.215E-06 | 1.000E-05 | 1.600E-05 |
| 8820 | 441 | S441V | A | 1.6e-05 | 9.8e-06 | 1.5e-05 | 1.360E-05 | 3.329E-06 | 9.800E-06 | 1.600E-05 |
| 8821 | 442 | G442A | A | 8.7e-06 | 1e-05 | 7.1e-06 | 8.600E-06 | 1.453E-06 | 7.100E-06 | 1.000E-05 |
| 8822 | 442 | G442R | A | 1.8e-05 | 1.4e-05 | 1.4e-05 | 1.533E-05 | 2.309E-06 | 1.400E-05 | 1.800E-05 |
| 8823 | 442 | G442N | A | 1.4e-05 | 1e-05 | 1.4e-05 | 1.267E-05 | 2.309E-06 | 1.000E-05 | 1.400E-05 |
| 8824 | 442 | G442D | A | 9.6e-06 | 1e-05 | 9.4e-06 | 9.667E-06 | 3.055E-07 | 9.400E-06 | 1.000E-05 |
| 8825 | 442 | G442C | A | 1.1e-05 | 1.1e-05 | 1.3e-05 | 1.167E-05 | 1.155E-06 | 1.100E-05 | 1.300E-05 |
| 8826 | 442 | G442E | A | 1.5e-05 | 1.3e-05 | 9.2e-06 | 1.240E-05 | 2.946E-06 | 9.200E-06 | 1.500E-05 |
| 8827 | 442 | G442Q | A | 9.7e-06 | 1.4e-05 | 1.2e-05 | 1.190E-05 | 2.152E-06 | 9.700E-06 | 1.400E-05 |
| 8828 | 442 | G442G | A | 1.5e-05 | 1.6e-05 | 1.4e-05 | 1.500E-05 | 1.000E-06 | 1.400E-05 | 1.600E-05 |
| 8829 | 442 | G442H | A | 1.4e-05 | 8.2e-06 | 9.9e-06 | 1.070E-05 | 2.982E-06 | 8.200E-06 | 1.400E-05 |
| 8830 | 442 | G442I | A | 1.4e-05 | 8.7e-06 | 1.5e-05 | 1.257E-05 | 3.386E-06 | 8.700E-06 | 1.500E-05 |
| 8831 | 442 | G442L | A | 1.1e-05 | 1e-05 | 1.1e-05 | 1.067E-05 | 5.774E-07 | 1.000E-05 | 1.100E-05 |
| 8832 | 442 | G442K | A | 1.1e-05 | 1.4e-05 | 1.6e-05 | 1.367E-05 | 2.517E-06 | 1.100E-05 | 1.600E-05 |
| 8833 | 442 | G442M | A | 1.4e-05 | 9.9e-06 | 2.2e-05 | 1.530E-05 | 6.154E-06 | 9.900E-06 | 2.200E-05 |
| 8834 | 442 | G442F | A | 1.5e-05 | 9.2e-06 | 1.5e-05 | 1.307E-05 | 3.349E-06 | 9.200E-06 | 1.500E-05 |
| 8835 | 442 | G442P | A | 1.4e-05 | 1.5e-05 | 1e-05 | 1.300E-05 | 2.646E-06 | 1.000E-05 | 1.500E-05 |
| 8836 | 442 | G442S | A | 1e-05 | 1.5e-05 | 1.1e-05 | 1.200E-05 | 2.646E-06 | 1.000E-05 | 1.500E-05 |
| 8837 | 442 | G442T | A | 1.4e-05 | 9.8e-06 | 9.9e-06 | 1.123E-05 | 2.397E-06 | 9.800E-06 | 1.400E-05 |
| 8838 | 442 | G442W | A | 1e-05 | 1.5e-05 | 1.5e-05 | 1.333E-05 | 2.887E-06 | 1.000E-05 | 1.500E-05 |
| 8839 | 442 | G442Y | A | 1.5e-05 | 1.6e-05 | 1e-05 | 1.367E-05 | 3.215E-06 | 1.000E-05 | 1.600E-05 |

| | | | | | | | | | | |
|---|---|---|---|---|---|---|---|---|---|---|
| 8840 | 442 | G442V | A | 1.5e-05 | 1e-05 | 1.4e-05 | 1.300E-05 | 2.646E-06 | 1.000E-05 | 1.500E-05 |
| 8841 | 443 | L443A | A | 1e-05 | 7.3e-06 | 1.6e-05 | 1.110E-05 | 4.453E-06 | 7.300E-06 | 1.600E-05 |
| 8842 | 443 | L443R | A | 1.2e-05 | 1.2e-05 | 1.6e-05 | 1.333E-05 | 2.309E-06 | 1.200E-05 | 1.600E-05 |
| 8843 | 443 | L443N | A | 1.6e-05 | 1.3e-05 | 1.4e-05 | 1.433E-05 | 1.528E-06 | 1.300E-05 | 1.600E-05 |
| 8844 | 443 | L443D | A | 1.7e-05 | 1e-05 | 1.1e-05 | 1.267E-05 | 3.786E-06 | 1.000E-05 | 1.700E-05 |
| 8845 | 443 | L443C | A | 1.4e-05 | 1.4e-05 | 1e-05 | 1.267E-05 | 2.309E-06 | 1.000E-05 | 1.400E-05 |
| 8846 | 443 | L443E | A | 1.1e-05 | 1.5e-05 | 1.7e-05 | 1.433E-05 | 3.055E-06 | 1.100E-05 | 1.700E-05 |
| 8847 | 443 | L443Q | A | 1.5e-05 | 1.6e-05 | 1.3e-05 | 1.467E-05 | 1.528E-06 | 1.300E-05 | 1.600E-05 |
| 8848 | 443 | L443G | A | 1.1e-05 | 1.5e-05 | 1.5e-05 | 1.367E-05 | 2.309E-06 | 1.100E-05 | 1.500E-05 |
| 8849 | 443 | L443H | A | 1e-05 | 9.4e-06 | 9.1e-06 | 9.500E-06 | 4.583E-07 | 9.100E-06 | 1.000E-05 |
| 8850 | 443 | L443I | A | 9.2e-06 | 1.2e-05 | 1.1e-05 | 1.073E-05 | 1.419E-06 | 9.200E-06 | 1.200E-05 |
| 8851 | 443 | L443L | A | 1.1e-05 | 1.5e-05 | 1.3e-05 | 1.300E-05 | 2.000E-06 | 1.100E-05 | 1.500E-05 |
| 8852 | 443 | L443K | A | 1.5e-05 | 1.5e-05 | 1.3e-05 | 1.433E-05 | 1.155E-06 | 1.300E-05 | 1.500E-05 |
| 8853 | 443 | L443M | A | 1.3e-05 | 1.4e-05 | 1.4e-05 | 1.367E-05 | 5.774E-07 | 1.300E-05 | 1.400E-05 |
| 8854 | 443 | L443F | A | 1.4e-05 | 9.3e-06 | 1.7e-05 | 1.343E-05 | 3.881E-06 | 9.300E-06 | 1.700E-05 |
| 8855 | 443 | L443P | A | 9.3e-06 | 1.2e-05 | 9.9e-06 | 1.040E-05 | 1.418E-06 | 9.300E-06 | 1.200E-05 |
| 8856 | 443 | L443S | A | 1.1e-05 | 1.1e-05 | 1e-05 | 1.067E-05 | 5.774E-07 | 1.000E-05 | 1.100E-05 |
| 8857 | 443 | L443T | A | 1.4e-05 | 1.5e-05 | 1.6e-05 | 1.500E-05 | 1.000E-06 | 1.400E-05 | 1.600E-05 |
| 8858 | 443 | L443W | A | 1.1e-05 | 1.4e-05 | 1e-05 | 1.167E-05 | 2.082E-06 | 1.000E-05 | 1.400E-05 |
| 8859 | 443 | L443Y | A | 1.5e-05 | 4e-05 | 8.5e-06 | 2.117E-05 | 1.663E-05 | 8.500E-06 | 4.000E-05 |
| 8860 | 443 | L443V | A | 1.6e-05 | 1.1e-05 | 8.3e-06 | 1.177E-05 | 3.907E-06 | 8.300E-06 | 1.600E-05 |
| 8861 | 444 | A444A | A | 1.2e-05 | 9.8e-06 | 1.3e-05 | 1.160E-05 | 1.637E-06 | 9.800E-06 | 1.300E-05 |
| 8862 | 444 | A444R | A | 1.6e-05 | 1.1e-05 | 1.5e-05 | 1.400E-05 | 2.646E-06 | 1.100E-05 | 1.600E-05 |
| 8863 | 444 | A444N | A | 7e-06 | 1.7e-05 | 1.6e-05 | 1.333E-05 | 5.508E-06 | 7.000E-06 | 1.700E-05 |
| 8864 | 444 | A444D | A | 1.3e-05 | 1e-05 | 1e-05 | 1.100E-05 | 1.732E-06 | 1.000E-05 | 1.300E-05 |
| 8865 | 444 | A444C | A | 1.4e-05 | 1.3e-05 | 8.1e-06 | 1.170E-05 | 3.158E-06 | 8.100E-06 | 1.400E-05 |
| 8866 | 444 | A444E | A | 1.3e-05 | 1.5e-05 | 8.3e-06 | 1.210E-05 | 3.439E-06 | 8.300E-06 | 1.500E-05 |
| 8867 | 444 | A444Q | A | 1.6e-05 | 1.1e-05 | 1e-05 | 1.233E-05 | 3.215E-06 | 1.000E-05 | 1.600E-05 |
| 8868 | 444 | A444G | A | 9.6e-06 | 9.9e-06 | 9.9e-06 | 9.800E-06 | 1.732E-07 | 9.600E-06 | 9.900E-06 |
| 8869 | 444 | A444H | A | 1.1e-05 | 1.1e-05 | 1.4e-05 | 1.200E-05 | 1.732E-06 | 1.100E-05 | 1.400E-05 |
| 8870 | 444 | A444I | A | 1.5e-05 | 1.1e-05 | 1.6e-05 | 1.400E-05 | 2.646E-06 | 1.100E-05 | 1.600E-05 |
| 8871 | 444 | A444L | A | 1.6e-05 | 1e-05 | 1.6e-05 | 1.400E-05 | 3.464E-06 | 1.000E-05 | 1.600E-05 |
| 8872 | 444 | A444K | A | 1.7e-05 | 1.5e-05 | 1e-05 | 1.400E-05 | 3.606E-06 | 1.000E-05 | 1.700E-05 |
| 8873 | 444 | A444M | A | 1.6e-05 | 6.9e-06 | 1.5e-05 | 1.263E-05 | 4.990E-06 | 6.900E-06 | 1.600E-05 |
| 8874 | 444 | A444F | A | 1.4e-05 | 8.9e-06 | 1.4e-05 | 1.230E-05 | 2.944E-06 | 8.900E-06 | 1.400E-05 |
| 8875 | 444 | A444P | A | 1.1e-05 | 1.5e-05 | 1.3e-05 | 1.300E-05 | 2.000E-06 | 1.100E-05 | 1.500E-05 |
| 8876 | 444 | A444S | A | 1.3e-05 | 1.5e-05 | 1.1e-05 | 1.300E-05 | 2.000E-06 | 1.100E-05 | 1.500E-05 |
| 8877 | 444 | A444T | A | 1e-05 | 1.5e-05 | 1.4e-05 | 1.300E-05 | 2.646E-06 | 1.000E-05 | 1.500E-05 |
| 8878 | 444 | A444W | A | 1.1e-05 | 1.5e-05 | 9e-06 | 1.167E-05 | 3.055E-06 | 9.000E-06 | 1.500E-05 |
| 8879 | 444 | A444Y | A | 9.1e-06 | 1.3e-05 | 1.5e-05 | 1.237E-05 | 3.001E-06 | 9.100E-06 | 1.500E-05 |
| 8880 | 444 | A444V | A | 1.2e-05 | 1.5e-05 | 8.3e-06 | 1.177E-05 | 3.356E-06 | 8.300E-06 | 1.500E-05 |
| 8881 | 445 | L445A | A | 1.5e-05 | 1.5e-05 | 1.4e-05 | 1.467E-05 | 5.774E-07 | 1.400E-05 | 1.500E-05 |
| 8882 | 445 | L445R | A | 1.6e-05 | 1.1e-05 | 1.5e-05 | 1.400E-05 | 2.646E-06 | 1.100E-05 | 1.600E-05 |
| 8883 | 445 | L445N | A | 1.2e-05 | 1.3e-05 | 1.5e-05 | 1.333E-05 | 1.528E-06 | 1.200E-05 | 1.500E-05 |
| 8884 | 445 | L445D | A | 1.5e-05 | 1.4e-05 | 9.7e-06 | 1.290E-05 | 2.816E-06 | 9.700E-06 | 1.500E-05 |
| 8885 | 445 | L445C | A | 1.6e-05 | 1.6e-05 | 7.8e-06 | 1.327E-05 | 4.734E-06 | 7.800E-06 | 1.600E-05 |
| 8886 | 445 | L445E | A | 1e-05 | 1.4e-05 | 9e-06 | 1.100E-05 | 2.646E-06 | 9.000E-06 | 1.400E-05 |
| 8887 | 445 | L445Q | A | 1e-05 | 1.7e-05 | 1.1e-05 | 1.267E-05 | 3.786E-06 | 1.000E-05 | 1.700E-05 |
| 8888 | 445 | L445G | A | 1e-05 | 1.4e-05 | 1.5e-05 | 1.300E-05 | 2.646E-06 | 1.000E-05 | 1.500E-05 |
| 8889 | 445 | L445H | A | 1e-05 | 2.1e-05 | 1e-05 | 1.367E-05 | 6.351E-06 | 1.000E-05 | 2.100E-05 |
| 8890 | 445 | L445I | A | 1.1e-05 | 1.3e-05 | 1.4e-05 | 1.267E-05 | 1.528E-06 | 1.100E-05 | 1.400E-05 |
| 8891 | 445 | L445L | A | 1.6e-05 | 1.5e-05 | 1.3e-05 | 1.467E-05 | 1.528E-06 | 1.300E-05 | 1.600E-05 |
| 8892 | 445 | L445K | A | 8.8e-06 | 1.5e-05 | 1.5e-05 | 1.293E-05 | 3.580E-06 | 8.800E-06 | 1.500E-05 |
| 8893 | 445 | L445M | A | 1.1e-05 | 1.4e-05 | 1.7e-05 | 1.400E-05 | 3.000E-06 | 1.100E-05 | 1.700E-05 |
| 8894 | 445 | L445F | A | 1.6e-05 | 1.7e-05 | 1.1e-05 | 1.467E-05 | 3.215E-06 | 1.100E-05 | 1.700E-05 |
| 8895 | 445 | L445P | A | 7.3e-06 | 1.5e-05 | 1.4e-05 | 1.210E-05 | 4.187E-06 | 7.300E-06 | 1.500E-05 |
| 8896 | 445 | L445S | A | 1.6e-05 | 9.7e-06 | 1.5e-05 | 1.357E-05 | 3.386E-06 | 9.700E-06 | 1.600E-05 |
| 8897 | 445 | L445T | A | 1.5e-05 | 9.8e-06 | 1.5e-05 | 1.327E-05 | 3.002E-06 | 9.800E-06 | 1.500E-05 |
| 8898 | 445 | L445W | A | 1.6e-05 | 1.6e-05 | 1.3e-05 | 1.500E-05 | 1.732E-06 | 1.300E-05 | 1.600E-05 |
| 8899 | 445 | L445Y | A | 8.5e-06 | 1.4e-05 | 1.5e-05 | 1.250E-05 | 3.500E-06 | 8.500E-06 | 1.500E-05 |
| 8900 | 445 | L445V | A | 1.2e-05 | 1.4e-05 | 1.4e-05 | 1.333E-05 | 1.155E-06 | 1.200E-05 | 1.400E-05 |
| 8901 | 446 | I446A | A | 1.7e-05 | 1.5e-05 | 1.6e-05 | 1.600E-05 | 1.000E-06 | 1.500E-05 | 1.700E-05 |
| 8902 | 446 | I446R | A | 1.3e-05 | 1.4e-05 | 1.6e-05 | 1.433E-05 | 1.528E-06 | 1.300E-05 | 1.600E-05 |
| 8903 | 446 | I446N | A | 1.5e-05 | 1.2e-05 | 1.1e-05 | 1.267E-05 | 2.082E-06 | 1.100E-05 | 1.500E-05 |
| 8904 | 446 | I446D | A | 1.3e-05 | 1e-05 | 1.6e-05 | 1.300E-05 | 3.000E-06 | 1.000E-05 | 1.600E-05 |
| 8905 | 446 | I446C | A | 1.5e-05 | 1.1e-05 | 9.8e-06 | 1.193E-05 | 2.723E-06 | 9.800E-06 | 1.500E-05 |
| 8906 | 446 | I446E | A | 1.1e-05 | 1.3e-05 | 1.5e-05 | 1.300E-05 | 2.000E-06 | 1.100E-05 | 1.500E-05 |
| 8907 | 446 | I446Q | A | 1.2e-05 | 1.3e-05 | 1e-05 | 1.167E-05 | 1.528E-06 | 1.000E-05 | 1.300E-05 |
| 8908 | 446 | I446G | A | 1.4e-05 | 1.5e-05 | 1.1e-05 | 1.333E-05 | 2.082E-06 | 1.100E-05 | 1.500E-05 |
| 8909 | 446 | I446H | A | 1e-05 | 1.8e-05 | 1.3e-05 | 1.367E-05 | 4.041E-06 | 1.000E-05 | 1.800E-05 |
| 8910 | 446 | I446I | A | 1.6e-05 | 1.1e-05 | 1e-05 | 1.233E-05 | 3.215E-06 | 1.000E-05 | 1.600E-05 |
| 8911 | 446 | I446L | A | 1e-05 | 1e-05 | 1.4e-05 | 1.133E-05 | 2.309E-06 | 1.000E-05 | 1.400E-05 |
| 8912 | 446 | I446K | A | 9.6e-06 | 2.5e-05 | 1.6e-05 | 1.687E-05 | 7.736E-06 | 9.600E-06 | 2.500E-05 |
| 8913 | 446 | I446M | A | 8.6e-06 | 1.1e-05 | 9.7e-06 | 9.767E-06 | 1.201E-06 | 8.600E-06 | 1.100E-05 |
| 8914 | 446 | I446F | A | 6.4e-06 | 1.5e-05 | 1.3e-05 | 1.147E-05 | 4.500E-06 | 6.400E-06 | 1.500E-05 |
| 8915 | 446 | I446P | A | 1.6e-05 | 9.7e-06 | 1.3e-05 | 1.290E-05 | 3.151E-06 | 9.700E-06 | 1.600E-05 |
| 8916 | 446 | I446S | A | 9.2e-06 | 1.5e-05 | 1.7e-05 | 1.373E-05 | 4.051E-06 | 9.200E-06 | 1.700E-05 |
| 8917 | 446 | I446T | A | 1.5e-05 | 1.2e-05 | 1.6e-05 | 1.433E-05 | 2.082E-06 | 1.200E-05 | 1.600E-05 |
| 8918 | 446 | I446W | A | 1.5e-05 | 1.1e-05 | 8.3e-06 | 1.143E-05 | 3.371E-06 | 8.300E-06 | 1.500E-05 |
| 8919 | 446 | I446Y | A | 9.3e-06 | 1.7e-05 | 1.3e-05 | 1.310E-05 | 3.851E-06 | 9.300E-06 | 1.700E-05 |
| 8920 | 446 | I446V | A | 1.1e-05 | 1.6e-05 | 1.5e-05 | 1.400E-05 | 2.646E-06 | 1.100E-05 | 1.600E-05 |
| 8921 | 447 | H447A | A | 1.2e-05 | 1.6e-05 | 1.6e-05 | 1.467E-05 | 2.309E-06 | 1.200E-05 | 1.600E-05 |
| 8922 | 447 | H447R | A | 1e-05 | 1.6e-05 | 1.4e-05 | 1.333E-05 | 3.055E-06 | 1.000E-05 | 1.600E-05 |
| 8923 | 447 | H447N | A | 1e-05 | 1.5e-05 | 1.5e-05 | 1.333E-05 | 2.887E-06 | 1.000E-05 | 1.500E-05 |
| 8924 | 447 | H447D | A | 1.4e-05 | 1.1e-05 | 1.4e-05 | 1.300E-05 | 1.732E-06 | 1.100E-05 | 1.400E-05 |

| | | | | | | | | | | |
|---|---|---|---|---|---|---|---|---|---|---|
| 8925 | 447 | H447C | A | 1.4e-05 | 1.2e-05 | 1.1e-05 | 1.233E-05 | 1.528E-06 | 1.100E-05 | 1.400E-05 |
| 8926 | 447 | H447E | A | 1.5e-05 | 1.5e-05 | 1e-05 | 1.333E-05 | 2.887E-06 | 1.000E-05 | 1.500E-05 |
| 8927 | 447 | H447Q | A | 1e-05 | 1.4e-05 | 1.6e-05 | 1.333E-05 | 3.055E-06 | 1.000E-05 | 1.600E-05 |
| 8928 | 447 | H447G | A | 1.5e-05 | 1.3e-05 | 1.5e-05 | 1.433E-05 | 1.155E-06 | 1.300E-05 | 1.500E-05 |
| 8929 | 447 | H447H | A | 1.4e-05 | 1.4e-05 | 1.5e-05 | 1.433E-05 | 5.774E-07 | 1.400E-05 | 1.500E-05 |
| 8930 | 447 | H447I | A | 8.2e-06 | 1.3e-05 | 1.2e-05 | 1.107E-05 | 2.532E-06 | 8.200E-06 | 1.300E-05 |
| 8931 | 447 | H447L | A | 9.9e-06 | 1.2e-05 | 1.6e-05 | 1.263E-05 | 3.099E-06 | 9.900E-06 | 1.600E-05 |
| 8932 | 447 | H447K | A | 1.6e-05 | 1.1e-05 | 1.5e-05 | 1.400E-05 | 2.646E-06 | 1.100E-05 | 1.600E-05 |
| 8933 | 447 | H447M | A | 1e-05 | 1.4e-05 | 9.3e-06 | 1.110E-05 | 2.536E-06 | 9.300E-06 | 1.400E-05 |
| 8934 | 447 | H447F | A | 1.5e-05 | 1e-05 | 2.2e-05 | 1.567E-05 | 6.028E-06 | 1.000E-05 | 2.200E-05 |
| 8935 | 447 | H447P | A | 1.4e-05 | 1.1e-05 | 1.1e-05 | 1.200E-05 | 1.732E-06 | 1.100E-05 | 1.400E-05 |
| 8936 | 447 | H447S | A | 1.3e-05 | 1.1e-05 | 1.7e-05 | 1.367E-05 | 3.055E-06 | 1.100E-05 | 1.700E-05 |
| 8937 | 447 | H447T | A | 9.2e-06 | 1.1e-05 | 1e-05 | 1.007E-05 | 9.018E-07 | 9.200E-06 | 1.100E-05 |
| 8938 | 447 | H447W | A | 1.1e-05 | 1.4e-05 | 1.5e-05 | 1.333E-05 | 2.082E-06 | 1.100E-05 | 1.500E-05 |
| 8939 | 447 | H447Y | A | 8.9e-06 | 1e-05 | 1.6e-05 | 1.163E-05 | 3.821E-06 | 8.900E-06 | 1.600E-05 |
| 8940 | 447 | H447V | A | 1.7e-05 | 1e-05 | 8.2e-06 | 1.173E-05 | 4.649E-06 | 8.200E-06 | 1.700E-05 |
| 8941 | 448 | H448A | A | 1.4e-05 | 1.3e-05 | 9.6e-06 | 1.220E-05 | 2.307E-06 | 9.600E-06 | 1.400E-05 |
| 8942 | 448 | H448R | A | 7.9e-06 | 1.5e-05 | 1.6e-05 | 1.297E-05 | 4.416E-06 | 7.900E-06 | 1.600E-05 |
| 8943 | 448 | H448N | A | 8e-06 | 1.4e-05 | 1.7e-05 | 1.300E-05 | 4.583E-06 | 8.000E-06 | 1.700E-05 |
| 8944 | 448 | H448D | A | 1.5e-05 | 8.2e-06 | 1.1e-05 | 1.140E-05 | 3.418E-06 | 8.200E-06 | 1.500E-05 |
| 8945 | 448 | H448C | A | 1.1e-05 | 9.6e-06 | 1.4e-05 | 1.153E-05 | 2.248E-06 | 9.600E-06 | 1.400E-05 |
| 8946 | 448 | H448E | A | 9.9e-06 | 1.5e-05 | 1.3e-05 | 1.263E-05 | 2.570E-06 | 9.900E-06 | 1.500E-05 |
| 8947 | 448 | H448Q | A | 1.5e-05 | 1.5e-05 | 1.5e-05 | 1.500E-05 | 0.000E+00 | 1.500E-05 | 1.500E-05 |
| 8948 | 448 | H448G | A | 1.6e-05 | 1e-05 | 1.7e-05 | 1.433E-05 | 3.786E-06 | 1.000E-05 | 1.700E-05 |
| 8949 | 448 | H448H | A | 1.1e-05 | 9.3e-06 | 9.4e-06 | 9.900E-06 | 9.539E-07 | 9.300E-06 | 1.100E-05 |
| 8950 | 448 | H448I | A | 1.4e-05 | 1.8e-05 | 1.7e-05 | 1.633E-05 | 2.082E-06 | 1.400E-05 | 1.800E-05 |
| 8951 | 448 | H448L | A | 1.1e-05 | 1.5e-05 | 1.6e-05 | 1.400E-05 | 2.646E-06 | 1.100E-05 | 1.600E-05 |
| 8952 | 448 | H448K | A | 1.5e-05 | 9.6e-06 | 1.3e-05 | 1.253E-05 | 2.730E-06 | 9.600E-06 | 1.500E-05 |
| 8953 | 448 | H448M | A | 1.4e-05 | 1.6e-05 | 1e-05 | 1.333E-05 | 3.055E-06 | 1.000E-05 | 1.600E-05 |
| 8954 | 448 | H448F | A | 1.1e-05 | 1.2e-05 | 1.5e-05 | 1.267E-05 | 2.082E-06 | 1.100E-05 | 1.500E-05 |
| 8955 | 448 | H448P | A | 1.5e-05 | 1.4e-05 | 1.4e-05 | 1.433E-05 | 5.774E-07 | 1.400E-05 | 1.500E-05 |
| 8956 | 448 | H448S | A | 9.6e-06 | 1.5e-05 | 1.4e-05 | 1.287E-05 | 2.873E-06 | 9.600E-06 | 1.500E-05 |
| 8957 | 448 | H448T | A | 1.2e-05 | 1.2e-05 | 1.1e-05 | 1.167E-05 | 5.774E-07 | 1.100E-05 | 1.200E-05 |
| 8958 | 448 | H448W | A | 1.3e-05 | 1.1e-05 | 1.5e-05 | 1.300E-05 | 2.000E-06 | 1.100E-05 | 1.500E-05 |
| 8959 | 448 | H448Y | A | 1.5e-05 | 1.6e-05 | 1.4e-05 | 1.500E-05 | 1.000E-06 | 1.400E-05 | 1.600E-05 |
| 8960 | 448 | H448V | A | 1.5e-05 | 8.5e-06 | 1.3e-05 | 1.217E-05 | 3.329E-06 | 8.500E-06 | 1.500E-05 |
| 8961 | 449 | N449A | A | 9.6e-06 | 1.5e-05 | 1.2e-05 | 1.220E-05 | 2.706E-06 | 9.600E-06 | 1.500E-05 |
| 8962 | 449 | N449R | A | 1e-05 | 1.5e-05 | 1.3e-05 | 1.267E-05 | 2.517E-06 | 1.000E-05 | 1.500E-05 |
| 8963 | 449 | N449N | A | 1.2e-05 | 1e-05 | 1.4e-05 | 1.200E-05 | 2.000E-06 | 1.000E-05 | 1.400E-05 |
| 8964 | 449 | N449D | A | 1e-05 | 1.2e-05 | 1.5e-05 | 1.233E-05 | 2.517E-06 | 1.000E-05 | 1.500E-05 |
| 8965 | 449 | N449C | A | 1.5e-05 | 1.6e-05 | 1e-05 | 1.367E-05 | 3.215E-06 | 1.000E-05 | 1.600E-05 |
| 8966 | 449 | N449E | A | 9.1e-06 | 1.1e-05 | 1.4e-05 | 1.137E-05 | 2.470E-06 | 9.100E-06 | 1.400E-05 |
| 8967 | 449 | N449Q | A | 1.1e-05 | 1.1e-05 | 1.6e-05 | 1.267E-05 | 2.887E-06 | 1.100E-05 | 1.600E-05 |
| 8968 | 449 | N449G | A | 1.2e-05 | 1.5e-05 | 1.6e-05 | 1.433E-05 | 2.082E-06 | 1.200E-05 | 1.600E-05 |
| 8969 | 449 | N449H | A | 1.3e-05 | 1.5e-05 | 1.5e-05 | 1.433E-05 | 1.155E-06 | 1.300E-05 | 1.500E-05 |
| 8970 | 449 | N449I | A | 1.7e-05 | 1.1e-05 | 1e-05 | 1.267E-05 | 3.786E-06 | 1.000E-05 | 1.700E-05 |
| 8971 | 449 | N449L | A | 1.3e-05 | 1e-05 | 1.5e-05 | 1.267E-05 | 2.517E-06 | 1.000E-05 | 1.500E-05 |
| 8972 | 449 | N449K | A | 1.2e-05 | 1e-05 | 8.2e-06 | 1.007E-05 | 1.901E-06 | 8.200E-06 | 1.200E-05 |
| 8973 | 449 | N449M | A | 1.1e-05 | 1e-05 | 1.8e-05 | 1.300E-05 | 4.359E-06 | 1.000E-05 | 1.800E-05 |
| 8974 | 449 | N449F | A | 1.1e-05 | 1.7e-05 | 1.4e-05 | 1.400E-05 | 3.000E-06 | 1.100E-05 | 1.700E-05 |
| 8975 | 449 | N449P | A | 9.5e-06 | 1.5e-05 | 1.3e-05 | 1.250E-05 | 2.784E-06 | 9.500E-06 | 1.500E-05 |
| 8976 | 449 | N449S | A | 1.2e-05 | 1.5e-05 | 8.3e-06 | 1.177E-05 | 3.356E-06 | 8.300E-06 | 1.500E-05 |
| 8977 | 449 | N449T | A | 1.1e-05 | 1.6e-05 | 1.1e-05 | 1.267E-05 | 2.887E-06 | 1.100E-05 | 1.600E-05 |
| 8978 | 449 | N449W | A | 9.8e-06 | 1.5e-05 | 1.4e-05 | 1.293E-05 | 2.759E-06 | 9.800E-06 | 1.500E-05 |
| 8979 | 449 | N449Y | A | 1.5e-05 | 1.6e-05 | 1.1e-05 | 1.400E-05 | 2.646E-06 | 1.100E-05 | 1.600E-05 |
| 8980 | 449 | N449V | A | 9.4e-06 | 1.4e-05 | 1.4e-05 | 1.247E-05 | 2.656E-06 | 9.400E-06 | 1.400E-05 |
| 8981 | 450 | T450A | A | 1.3e-05 | 1.1e-05 | 1.4e-05 | 1.267E-05 | 1.528E-06 | 1.100E-05 | 1.400E-05 |
| 8982 | 450 | T450R | A | 1e-05 | 9.7e-06 | 1.4e-05 | 1.123E-05 | 2.401E-06 | 9.700E-06 | 1.400E-05 |
| 8983 | 450 | T450N | A | 8.5e-06 | 1.1e-05 | 1e-05 | 9.833E-06 | 1.258E-06 | 8.500E-06 | 1.100E-05 |
| 8984 | 450 | T450D | A | 1.5e-05 | 1.1e-05 | 1e-05 | 1.200E-05 | 2.646E-06 | 1.000E-05 | 1.500E-05 |
| 8985 | 450 | T450C | A | 1.8e-05 | 1.5e-05 | 9.5e-06 | 1.417E-05 | 4.311E-06 | 9.500E-06 | 1.800E-05 |
| 8986 | 450 | T450E | A | 1e-05 | 1.1e-05 | 1.6e-05 | 1.233E-05 | 3.215E-06 | 1.000E-05 | 1.600E-05 |
| 8987 | 450 | T450Q | A | 1.5e-05 | 1e-05 | 1.1e-05 | 1.200E-05 | 2.646E-06 | 1.000E-05 | 1.500E-05 |
| 8988 | 450 | T450G | A | 1.4e-05 | 1.1e-05 | 1.4e-05 | 1.300E-05 | 1.732E-06 | 1.100E-05 | 1.400E-05 |
| 8989 | 450 | T450H | A | 9.2e-06 | 1.5e-05 | 1.7e-05 | 1.373E-05 | 4.051E-06 | 9.200E-06 | 1.700E-05 |
| 8990 | 450 | T450I | A | 1.5e-05 | 1.6e-05 | 1.7e-05 | 1.600E-05 | 1.000E-06 | 1.500E-05 | 1.700E-05 |
| 8991 | 450 | T450L | A | 1.4e-05 | 8.6e-06 | 1.4e-05 | 1.220E-05 | 3.118E-06 | 8.600E-06 | 1.400E-05 |
| 8992 | 450 | T450K | A | 1.1e-05 | 9.3e-06 | 1e-05 | 1.010E-05 | 8.544E-07 | 9.300E-06 | 1.100E-05 |
| 8993 | 450 | T450M | A | 1.1e-05 | 1.6e-05 | 1.6e-05 | 1.433E-05 | 2.887E-06 | 1.100E-05 | 1.600E-05 |
| 8994 | 450 | T450F | A | 1.8e-05 | 1.1e-05 | 1.4e-05 | 1.433E-05 | 3.512E-06 | 1.100E-05 | 1.800E-05 |
| 8995 | 450 | T450P | A | 1.7e-05 | 1.6e-05 | 8.8e-06 | 1.393E-05 | 4.474E-06 | 8.800E-06 | 1.700E-05 |
| 8996 | 450 | T450S | A | 1.5e-05 | 1.1e-05 | 1.1e-05 | 1.233E-05 | 2.309E-06 | 1.100E-05 | 1.500E-05 |
| 8997 | 450 | T450T | A | 9.1e-06 | 1.5e-05 | 1.1e-05 | 1.170E-05 | 3.012E-06 | 9.100E-06 | 1.500E-05 |
| 8998 | 450 | T450W | A | 1.7e-05 | 1.6e-05 | 1.3e-05 | 1.533E-05 | 2.082E-06 | 1.300E-05 | 1.700E-05 |
| 8999 | 450 | T450Y | A | 1.1e-05 | 1.1e-05 | 1.1e-05 | 1.100E-05 | 2.075E-21 | 1.100E-05 | 1.100E-05 |
| 9000 | 450 | T450V | A | 1.6e-05 | 1.5e-05 | 1.3e-05 | 1.467E-05 | 1.528E-06 | 1.300E-05 | 1.600E-05 |
| 9001 | 451 | H451A | A | 1.8e-05 | 7.6e-06 | 1.4e-05 | 1.320E-05 | 5.246E-06 | 7.600E-06 | 1.800E-05 |
| 9002 | 451 | H451R | A | 1.1e-05 | 1.6e-05 | 1.6e-05 | 1.433E-05 | 2.887E-06 | 1.100E-05 | 1.600E-05 |
| 9003 | 451 | H451N | A | 1.1e-05 | 1.5e-05 | 1.1e-05 | 1.233E-05 | 2.309E-06 | 1.100E-05 | 1.500E-05 |
| 9004 | 451 | H451D | A | 1.3e-05 | 8.8e-06 | 9.9e-06 | 1.057E-05 | 2.178E-06 | 8.800E-06 | 1.300E-05 |
| 9005 | 451 | H451C | A | 1.1e-05 | 1.2e-05 | 1.5e-05 | 1.267E-05 | 2.082E-06 | 1.100E-05 | 1.500E-05 |
| 9006 | 451 | H451E | A | 1.5e-05 | 1.2e-05 | 1.4e-05 | 1.367E-05 | 1.528E-06 | 1.200E-05 | 1.500E-05 |
| 9007 | 451 | H451Q | A | 1.4e-05 | 1.6e-05 | 1.4e-05 | 1.467E-05 | 1.155E-06 | 1.400E-05 | 1.600E-05 |
| 9008 | 451 | H451G | A | 1.6e-05 | 1.2e-05 | 9.5e-06 | 1.250E-05 | 3.279E-06 | 9.500E-06 | 1.600E-05 |
| 9009 | 451 | H451H | A | 1e-05 | 1.4e-05 | 1.3e-05 | 1.233E-05 | 2.082E-06 | 1.000E-05 | 1.400E-05 |

| | | | | | | | | | | |
|---|---|---|---|---|---|---|---|---|---|---|
| 9010 | 451 | H451I | A | 1.6e-05 | 1.4e-05 | 1.5e-05 | 1.500E-05 | 1.000E-06 | 1.400E-05 | 1.600E-05 |
| 9011 | 451 | H451L | A | 1.4e-05 | 1.1e-05 | 1e-05 | 1.167E-05 | 2.082E-06 | 1.000E-05 | 1.400E-05 |
| 9012 | 451 | H451K | A | 1.2e-05 | 1.5e-05 | 1.6e-05 | 1.433E-05 | 2.082E-06 | 1.200E-05 | 1.600E-05 |
| 9013 | 451 | H451M | A | 1.4e-05 | 1.3e-05 | 9.9e-06 | 1.230E-05 | 2.138E-06 | 9.900E-06 | 1.400E-05 |
| 9014 | 451 | H451F | A | 1e-05 | 1.5e-05 | 1.5e-05 | 1.333E-05 | 2.887E-06 | 1.000E-05 | 1.500E-05 |
| 9015 | 451 | H451P | A | 1.5e-05 | 1.6e-05 | 1.4e-05 | 1.500E-05 | 1.000E-06 | 1.400E-05 | 1.600E-05 |
| 9016 | 451 | H451S | A | 1.6e-05 | 1.1e-05 | 9.4e-06 | 1.213E-05 | 3.443E-06 | 9.400E-06 | 1.600E-05 |
| 9017 | 451 | H451T | A | 1.4e-05 | 9.9e-06 | 1.5e-05 | 1.297E-05 | 2.702E-06 | 9.900E-06 | 1.500E-05 |
| 9018 | 451 | H451W | A | 1.3e-05 | 7.9e-06 | 8.4e-06 | 9.767E-06 | 2.811E-06 | 7.900E-06 | 1.300E-05 |
| 9019 | 451 | H451Y | A | 1.6e-05 | 1.5e-05 | 9.4e-06 | 1.347E-05 | 3.557E-06 | 9.400E-06 | 1.600E-05 |
| 9020 | 451 | H451V | A | 1.5e-05 | 1.3e-05 | 1.6e-05 | 1.467E-05 | 1.528E-06 | 1.300E-05 | 1.600E-05 |
| 9021 | 452 | L452A | A | 1.4e-05 | 1.1e-05 | 1e-05 | 1.167E-05 | 2.082E-06 | 1.000E-05 | 1.400E-05 |
| 9022 | 452 | L452R | A | 9.9e-06 | 1.1e-05 | 1.3e-05 | 1.130E-05 | 1.572E-06 | 9.900E-06 | 1.300E-05 |
| 9023 | 452 | L452N | A | 9.6e-06 | 1e-05 | 8.6e-06 | 9.400E-06 | 7.211E-07 | 8.600E-06 | 1.000E-05 |
| 9024 | 452 | L452D | A | 1.4e-05 | 1e-05 | 1.6e-05 | 1.333E-05 | 3.055E-06 | 1.000E-05 | 1.600E-05 |
| 9025 | 452 | L452C | A | 1e-05 | 1.4e-05 | 1.1e-05 | 1.167E-05 | 2.082E-06 | 1.000E-05 | 1.400E-05 |
| 9026 | 452 | L452E | A | 1e-05 | 1.7e-05 | 1.6e-05 | 1.433E-05 | 3.786E-06 | 1.000E-05 | 1.700E-05 |
| 9027 | 452 | L452Q | A | 1.6e-05 | 1.5e-05 | 1.6e-05 | 1.567E-05 | 5.774E-07 | 1.500E-05 | 1.600E-05 |
| 9028 | 452 | L452G | A | 1.4e-05 | 1.1e-05 | 1e-05 | 1.167E-05 | 2.082E-06 | 1.000E-05 | 1.400E-05 |
| 9029 | 452 | L452H | A | 1.7e-05 | 9.8e-06 | 8.6e-06 | 1.180E-05 | 4.543E-06 | 8.600E-06 | 1.700E-05 |
| 9030 | 452 | L452I | A | 1.2e-05 | 1.4e-05 | 1e-05 | 1.200E-05 | 2.000E-06 | 1.000E-05 | 1.400E-05 |
| 9031 | 452 | L452L | A | 9.9e-06 | 1.5e-05 | 8.5e-06 | 1.113E-05 | 3.421E-06 | 8.500E-06 | 1.500E-05 |
| 9032 | 452 | L452K | A | 1e-05 | 1.2e-05 | 9.4e-06 | 1.047E-05 | 1.361E-06 | 9.400E-06 | 1.200E-05 |
| 9033 | 452 | L452M | A | 1.1e-05 | 1e-05 | 1.5e-05 | 1.200E-05 | 2.646E-06 | 1.000E-05 | 1.500E-05 |
| 9034 | 452 | L452F | A | 1.6e-05 | 1.5e-05 | 8.8e-06 | 1.327E-05 | 3.900E-06 | 8.800E-06 | 1.600E-05 |
| 9035 | 452 | L452P | A | 1e-05 | 1.4e-05 | 1.5e-05 | 1.300E-05 | 2.646E-06 | 1.000E-05 | 1.500E-05 |
| 9036 | 452 | L452S | A | 1.5e-05 | 9.9e-06 | 1.2e-05 | 1.230E-05 | 2.563E-06 | 9.900E-06 | 1.500E-05 |
| 9037 | 452 | L452T | A | 1.1e-05 | 1.6e-05 | 1.2e-05 | 1.300E-05 | 2.646E-06 | 1.100E-05 | 1.600E-05 |
| 9038 | 452 | L452W | A | 1.5e-05 | 8.9e-06 | 1e-05 | 1.130E-05 | 3.251E-06 | 8.900E-06 | 1.500E-05 |
| 9039 | 452 | L452Y | A | 4.6e-05 | 1.5e-05 | 1.1e-05 | 2.400E-05 | 1.916E-05 | 1.100E-05 | 4.600E-05 |
| 9040 | 452 | L452V | A | 9.6e-06 | 1e-05 | 1.1e-05 | 1.020E-05 | 7.211E-07 | 9.600E-06 | 1.100E-05 |
| 9041 | 453 | C453A | A | 1.2e-05 | 1.6e-05 | 8.7e-06 | 1.223E-05 | 3.656E-06 | 8.700E-06 | 1.600E-05 |
| 9042 | 453 | C453R | A | 1.7e-05 | 1.5e-05 | 1e-05 | 1.400E-05 | 3.606E-06 | 1.000E-05 | 1.700E-05 |
| 9043 | 453 | C453N | A | 1.3e-05 | 1.1e-05 | 9.3e-06 | 1.110E-05 | 1.852E-06 | 9.300E-06 | 1.300E-05 |
| 9044 | 453 | C453D | A | 1.6e-05 | 1.6e-05 | 9.8e-06 | 1.393E-05 | 3.580E-06 | 9.800E-06 | 1.600E-05 |
| 9045 | 453 | C453C | A | 1.5e-05 | 1.4e-05 | 1.4e-05 | 1.433E-05 | 5.774E-07 | 1.400E-05 | 1.500E-05 |
| 9046 | 453 | C453E | A | 9.4e-06 | 9.1e-06 | 1.5e-05 | 1.117E-05 | 3.323E-06 | 9.100E-06 | 1.500E-05 |
| 9047 | 453 | C453Q | A | 1.6e-05 | 1.1e-05 | 1.6e-05 | 1.433E-05 | 2.887E-06 | 1.100E-05 | 1.600E-05 |
| 9048 | 453 | C453G | A | 1.5e-05 | 1.6e-05 | 1.3e-05 | 1.467E-05 | 1.528E-06 | 1.300E-05 | 1.600E-05 |
| 9049 | 453 | C453H | A | 1.5e-05 | 1.3e-05 | 1.5e-05 | 1.433E-05 | 1.155E-06 | 1.300E-05 | 1.500E-05 |
| 9050 | 453 | C453I | A | 1.5e-05 | 1.4e-05 | 1.6e-05 | 1.500E-05 | 1.000E-06 | 1.400E-05 | 1.600E-05 |
| 9051 | 453 | C453L | A | 1.4e-05 | 1.6e-05 | 1.1e-05 | 1.367E-05 | 2.517E-06 | 1.100E-05 | 1.600E-05 |
| 9052 | 453 | C453K | A | 1.5e-05 | 1.7e-05 | 1e-05 | 1.400E-05 | 3.606E-06 | 1.000E-05 | 1.700E-05 |
| 9053 | 453 | C453M | A | 1.3e-05 | 1.3e-05 | 1.4e-05 | 1.333E-05 | 5.774E-07 | 1.300E-05 | 1.400E-05 |
| 9054 | 453 | C453F | A | 1.3e-05 | 1.2e-05 | 1.1e-05 | 1.200E-05 | 1.000E-06 | 1.100E-05 | 1.300E-05 |
| 9055 | 453 | C453P | A | 1.5e-05 | 1.3e-05 | 1.1e-05 | 1.300E-05 | 2.000E-06 | 1.100E-05 | 1.500E-05 |
| 9056 | 453 | C453S | A | 1.4e-05 | 1.4e-05 | 1e-05 | 1.267E-05 | 2.309E-06 | 1.000E-05 | 1.400E-05 |
| 9057 | 453 | C453T | A | 1.3e-05 | 1.2e-05 | 9.4e-06 | 1.147E-05 | 1.858E-06 | 9.400E-06 | 1.300E-05 |
| 9058 | 453 | C453W | A | 1.3e-05 | 1.5e-05 | 1.6e-05 | 1.467E-05 | 1.528E-06 | 1.300E-05 | 1.600E-05 |
| 9059 | 453 | C453Y | A | 1e-05 | 1.4e-05 | 1.4e-05 | 1.267E-05 | 2.309E-06 | 1.000E-05 | 1.400E-05 |
| 9060 | 453 | C453V | A | 1.3e-05 | 1.7e-05 | 1.3e-05 | 1.433E-05 | 2.309E-06 | 1.300E-05 | 1.700E-05 |
| 9061 | 454 | F454A | A | 1.7e-05 | 1.7e-05 | 1.3e-05 | 1.567E-05 | 2.309E-06 | 1.300E-05 | 1.700E-05 |
| 9062 | 454 | F454R | A | 1.5e-05 | 1.5e-05 | 1.1e-05 | 1.367E-05 | 2.309E-06 | 1.100E-05 | 1.500E-05 |
| 9063 | 454 | F454N | A | 1.2e-05 | 9.9e-06 | 1.6e-05 | 1.263E-05 | 3.099E-06 | 9.900E-06 | 1.600E-05 |
| 9064 | 454 | F454D | A | 1.4e-05 | 1.5e-05 | 9e-06 | 1.267E-05 | 3.215E-06 | 9.000E-06 | 1.500E-05 |
| 9065 | 454 | F454C | A | 1.3e-05 | 1.3e-05 | 8.5e-06 | 1.150E-05 | 2.598E-06 | 8.500E-06 | 1.300E-05 |
| 9066 | 454 | F454E | A | 1.1e-05 | 8.9e-06 | 1.1e-05 | 1.030E-05 | 1.212E-06 | 8.900E-06 | 1.100E-05 |
| 9067 | 454 | F454Q | A | 1e-05 | 1.5e-05 | 9.7e-06 | 1.157E-05 | 2.977E-06 | 9.700E-06 | 1.500E-05 |
| 9068 | 454 | F454G | A | 1.5e-05 | 1.5e-05 | 1.4e-05 | 1.467E-05 | 5.774E-07 | 1.400E-05 | 1.500E-05 |
| 9069 | 454 | F454H | A | 1.9e-05 | 1.6e-05 | 1.1e-05 | 1.533E-05 | 4.041E-06 | 1.100E-05 | 1.900E-05 |
| 9070 | 454 | F454I | A | 1.2e-05 | 1.4e-05 | 1.6e-05 | 1.400E-05 | 2.000E-06 | 1.200E-05 | 1.600E-05 |
| 9071 | 454 | F454L | A | 1.3e-05 | 1.6e-05 | 1.3e-05 | 1.400E-05 | 1.732E-06 | 1.300E-05 | 1.600E-05 |
| 9072 | 454 | F454K | A | 1.6e-05 | 1e-05 | 1.3e-05 | 1.300E-05 | 3.000E-06 | 1.000E-05 | 1.600E-05 |
| 9073 | 454 | F454M | A | 1.7e-05 | 1.1e-05 | 1.5e-05 | 1.433E-05 | 3.055E-06 | 1.100E-05 | 1.700E-05 |
| 9074 | 454 | F454F | A | 1.1e-05 | 1.3e-05 | 1.1e-05 | 1.167E-05 | 1.155E-06 | 1.100E-05 | 1.300E-05 |
| 9075 | 454 | F454P | A | 8.1e-06 | 1.4e-05 | 1.5e-05 | 1.237E-05 | 3.729E-06 | 8.100E-06 | 1.500E-05 |
| 9076 | 454 | F454S | A | 1.5e-05 | 1.2e-05 | 1.5e-05 | 1.400E-05 | 1.732E-06 | 1.200E-05 | 1.500E-05 |
| 9077 | 454 | F454T | A | 6.8e-06 | 1.5e-05 | 1.6e-05 | 1.260E-05 | 5.048E-06 | 6.800E-06 | 1.600E-05 |
| 9078 | 454 | F454W | A | 1.4e-05 | 8.3e-06 | 9.2e-06 | 1.050E-05 | 3.064E-06 | 8.300E-06 | 1.400E-05 |
| 9079 | 454 | F454Y | A | 1.1e-05 | 1.2e-05 | 1.4e-05 | 1.233E-05 | 1.528E-06 | 1.100E-05 | 1.400E-05 |
| 9080 | 454 | F454V | A | 9.5e-06 | 1.6e-05 | 1.4e-05 | 1.317E-05 | 3.329E-06 | 9.500E-06 | 1.600E-05 |
| 9081 | 455 | V455A | A | 1.4e-05 | 1.5e-05 | 1.4e-05 | 1.433E-05 | 5.774E-07 | 1.400E-05 | 1.500E-05 |
| 9082 | 455 | V455R | A | 1.3e-05 | 1.4e-05 | 1.4e-05 | 1.367E-05 | 5.774E-07 | 1.300E-05 | 1.400E-05 |
| 9083 | 455 | V455N | A | 9.5e-06 | 1.5e-05 | 2.1e-05 | 1.517E-05 | 5.752E-06 | 9.500E-06 | 2.100E-05 |
| 9084 | 455 | V455D | A | 1.5e-05 | 9.4e-06 | 1.5e-05 | 1.313E-05 | 3.233E-06 | 9.400E-06 | 1.500E-05 |
| 9085 | 455 | V455C | A | 1.1e-05 | 1.1e-05 | 1.5e-05 | 1.233E-05 | 2.309E-06 | 1.100E-05 | 1.500E-05 |
| 9086 | 455 | V455E | A | 1.1e-05 | 8.4e-06 | 1.1e-05 | 1.013E-05 | 1.501E-06 | 8.400E-06 | 1.100E-05 |
| 9087 | 455 | V455Q | A | 1.6e-05 | 1.4e-05 | 1.1e-05 | 1.367E-05 | 2.517E-06 | 1.100E-05 | 1.600E-05 |
| 9088 | 455 | V455G | A | 1.5e-05 | 1.1e-05 | 1.5e-05 | 1.367E-05 | 2.309E-06 | 1.100E-05 | 1.500E-05 |
| 9089 | 455 | V455H | A | 1.3e-05 | 1.1e-05 | 1.6e-05 | 1.333E-05 | 2.517E-06 | 1.100E-05 | 1.600E-05 |
| 9090 | 455 | V455I | A | 1.6e-05 | 1e-05 | 1.6e-05 | 1.400E-05 | 3.464E-06 | 1.000E-05 | 1.600E-05 |
| 9091 | 455 | V455L | A | 1e-05 | 7.1e-06 | 1.5e-05 | 1.070E-05 | 3.996E-06 | 7.100E-06 | 1.500E-05 |
| 9092 | 455 | V455K | A | 1.3e-05 | 1.1e-05 | 1.7e-05 | 1.367E-05 | 3.055E-06 | 1.100E-05 | 1.700E-05 |
| 9093 | 455 | V455M | A | 1.5e-05 | 8e-06 | 1.2e-05 | 1.167E-05 | 3.512E-06 | 8.000E-06 | 1.500E-05 |
| 9094 | 455 | V455F | A | 1.5e-05 | 1e-05 | 1.1e-05 | 1.200E-05 | 2.646E-06 | 1.000E-05 | 1.500E-05 |

| | | | | | | | | | | |
|---|---|---|---|---|---|---|---|---|---|---|
| 9095 | 455 | V455P | A | 1.1e-05 | 1.5e-05 | 1.5e-05 | 1.367E-05 | 2.309E-06 | 1.100E-05 | 1.500E-05 |
| 9096 | 455 | V455S | A | 1.2e-05 | 1e-05 | 8.9e-06 | 1.030E-05 | 1.572E-06 | 8.900E-06 | 1.200E-05 |
| 9097 | 455 | V455T | A | 1.5e-05 | 1.3e-05 | 1.4e-05 | 1.400E-05 | 1.000E-06 | 1.300E-05 | 1.500E-05 |
| 9098 | 455 | V455W | A | 8.6e-06 | 1.4e-05 | 1e-05 | 1.087E-05 | 2.802E-06 | 8.600E-06 | 1.400E-05 |
| 9099 | 455 | V455Y | A | 9.2e-06 | 9.7e-06 | 1.5e-05 | 1.130E-05 | 3.214E-06 | 9.200E-06 | 1.500E-05 |
| 9100 | 455 | V455V | A | 1.4e-05 | 1.7e-05 | 1.4e-05 | 1.500E-05 | 1.732E-06 | 1.400E-05 | 1.700E-05 |
| 9101 | 456 | H456A | A | 1.1e-05 | 9e-06 | 1.2e-05 | 1.067E-05 | 1.528E-06 | 9.000E-06 | 1.200E-05 |
| 9102 | 456 | H456R | A | 1.4e-05 | 1.1e-05 | 9.7e-06 | 1.157E-05 | 2.205E-06 | 9.700E-06 | 1.400E-05 |
| 9103 | 456 | H456N | A | 1.5e-05 | 1.5e-05 | 1.5e-05 | 1.500E-05 | 0.000E+00 | 1.500E-05 | 1.500E-05 |
| 9104 | 456 | H456D | A | 1.6e-05 | 1e-05 | 8.7e-06 | 1.157E-05 | 3.894E-06 | 8.700E-06 | 1.600E-05 |
| 9105 | 456 | H456C | A | 8.5e-06 | 1e-05 | 1.4e-05 | 1.083E-05 | 2.843E-06 | 8.500E-06 | 1.400E-05 |
| 9106 | 456 | H456E | A | 1e-05 | 1.4e-05 | 9.4e-06 | 1.113E-05 | 2.501E-06 | 9.400E-06 | 1.400E-05 |
| 9107 | 456 | H456Q | A | 1e-05 | 1.5e-05 | 1.5e-05 | 1.333E-05 | 2.887E-06 | 1.000E-05 | 1.500E-05 |
| 9108 | 456 | H456G | A | 1e-05 | 1.7e-05 | 1e-05 | 1.233E-05 | 4.041E-06 | 1.000E-05 | 1.700E-05 |
| 9109 | 456 | H456H | A | 1.3e-05 | 1.5e-05 | 1.4e-05 | 1.400E-05 | 1.000E-06 | 1.300E-05 | 1.500E-05 |
| 9110 | 456 | H456I | A | 1.3e-05 | 1.3e-05 | 1.6e-05 | 1.400E-05 | 1.732E-06 | 1.300E-05 | 1.600E-05 |
| 9111 | 456 | H456L | A | 1.6e-05 | 1.6e-05 | 7.1e-06 | 1.303E-05 | 5.138E-06 | 7.100E-06 | 1.600E-05 |
| 9112 | 456 | H456K | A | 1.5e-05 | 1.6e-05 | 9.7e-06 | 1.357E-05 | 3.386E-06 | 9.700E-06 | 1.600E-05 |
| 9113 | 456 | H456M | A | 1.4e-05 | 1.5e-05 | 9.8e-06 | 1.293E-05 | 2.759E-06 | 9.800E-06 | 1.500E-05 |
| 9114 | 456 | H456F | A | 9.7e-06 | 1.5e-05 | 9.9e-06 | 1.153E-05 | 3.004E-06 | 9.700E-06 | 1.500E-05 |
| 9115 | 456 | H456P | A | 9.8e-06 | 1.1e-05 | 1e-05 | 1.027E-05 | 6.429E-07 | 9.800E-06 | 1.100E-05 |
| 9116 | 456 | H456S | A | 1.5e-05 | 1.6e-05 | 8.4e-06 | 1.313E-05 | 4.130E-06 | 8.400E-06 | 1.600E-05 |
| 9117 | 456 | H456T | A | 1.3e-05 | 8.5e-06 | 1.1e-05 | 1.083E-05 | 2.255E-06 | 8.500E-06 | 1.300E-05 |
| 9118 | 456 | H456W | A | 1.3e-05 | 1.4e-05 | 1.6e-05 | 1.433E-05 | 1.528E-06 | 1.300E-05 | 1.600E-05 |
| 9119 | 456 | H456Y | A | 1.2e-05 | 1.4e-05 | 8.7e-06 | 1.157E-05 | 2.676E-06 | 8.700E-06 | 1.400E-05 |
| 9120 | 456 | H456V | A | 1.7e-05 | 1.3e-05 | 1.6e-05 | 1.533E-05 | 2.082E-06 | 1.300E-05 | 1.700E-05 |
| 9121 | 457 | T457A | A | 1.1e-05 | 8.8e-06 | 1.7e-05 | 1.227E-05 | 4.244E-06 | 8.800E-06 | 1.700E-05 |
| 9122 | 457 | T457R | A | 1.3e-05 | 1.6e-05 | 1.3e-05 | 1.400E-05 | 1.732E-06 | 1.300E-05 | 1.600E-05 |
| 9123 | 457 | T457N | A | 9.2e-06 | 1e-05 | 1.6e-05 | 1.173E-05 | 3.717E-06 | 9.200E-06 | 1.600E-05 |
| 9124 | 457 | T457D | A | 1.6e-05 | 1.3e-05 | 1.4e-05 | 1.433E-05 | 1.528E-06 | 1.300E-05 | 1.600E-05 |
| 9125 | 457 | T457C | A | 1.4e-05 | 1.1e-05 | 1.2e-05 | 1.233E-05 | 1.528E-06 | 1.100E-05 | 1.400E-05 |
| 9126 | 457 | T457E | A | 9.1e-06 | 1.5e-05 | 1.3e-05 | 1.237E-05 | 3.001E-06 | 9.100E-06 | 1.500E-05 |
| 9127 | 457 | T457Q | A | 9.9e-06 | 1.5e-05 | 1.5e-05 | 1.330E-05 | 2.944E-06 | 9.900E-06 | 1.500E-05 |
| 9128 | 457 | T457G | A | 9e-06 | 1.3e-05 | 1.6e-05 | 1.267E-05 | 3.512E-06 | 9.000E-06 | 1.600E-05 |
| 9129 | 457 | T457H | A | 1.1e-05 | 1.1e-05 | 1.1e-05 | 1.100E-05 | 2.075E-21 | 1.100E-05 | 1.100E-05 |
| 9130 | 457 | T457I | A | 1.7e-05 | 1.6e-05 | 1.2e-05 | 1.500E-05 | 2.646E-06 | 1.200E-05 | 1.700E-05 |
| 9131 | 457 | T457L | A | 1.5e-05 | 1.5e-05 | 8.4e-06 | 1.280E-05 | 3.811E-06 | 8.400E-06 | 1.500E-05 |
| 9132 | 457 | T457K | A | 9.6e-06 | 1.2e-05 | 1.3e-05 | 1.153E-05 | 1.747E-06 | 9.600E-06 | 1.300E-05 |
| 9133 | 457 | T457M | A | 1.2e-05 | 1.1e-05 | 8.7e-06 | 1.057E-05 | 1.692E-06 | 8.700E-06 | 1.200E-05 |
| 9134 | 457 | T457F | A | 1e-05 | 2e-05 | 1.1e-05 | 1.367E-05 | 5.508E-06 | 1.000E-05 | 2.000E-05 |
| 9135 | 457 | T457P | A | 1.3e-05 | 1.5e-05 | 1.5e-05 | 1.433E-05 | 1.155E-06 | 1.300E-05 | 1.500E-05 |
| 9136 | 457 | T457S | A | 1e-05 | 1.6e-05 | 1.1e-05 | 1.233E-05 | 3.215E-06 | 1.000E-05 | 1.600E-05 |
| 9137 | 457 | T457T | A | 1.4e-05 | 1.3e-05 | 1.7e-05 | 1.467E-05 | 2.082E-06 | 1.300E-05 | 1.700E-05 |
| 9138 | 457 | T457W | A | 1.4e-05 | 1.7e-05 | 1.1e-05 | 1.400E-05 | 3.000E-06 | 1.100E-05 | 1.700E-05 |
| 9139 | 457 | T457Y | A | 1.3e-05 | 1.5e-05 | 1.2e-05 | 1.333E-05 | 1.528E-06 | 1.200E-05 | 1.500E-05 |
| 9140 | 457 | T457V | A | 1.3e-05 | 1.1e-05 | 1.6e-05 | 1.333E-05 | 2.517E-06 | 1.100E-05 | 1.600E-05 |
| 9141 | 458 | V458A | A | 9.6e-06 | 1.3e-05 | 1.7e-05 | 1.320E-05 | 3.704E-06 | 9.600E-06 | 1.700E-05 |
| 9142 | 458 | V458R | A | 1.5e-05 | 1.4e-05 | 1.4e-05 | 1.433E-05 | 5.774E-07 | 1.400E-05 | 1.500E-05 |
| 9143 | 458 | V458N | A | 1.7e-05 | 1.1e-05 | 1.4e-05 | 1.400E-05 | 3.000E-06 | 1.100E-05 | 1.700E-05 |
| 9144 | 458 | V458D | A | 8.7e-06 | 1e-05 | 1.5e-05 | 1.123E-05 | 3.326E-06 | 8.700E-06 | 1.500E-05 |
| 9145 | 458 | V458C | A | 1.4e-05 | 1.5e-05 | 1.4e-05 | 1.433E-05 | 5.774E-07 | 1.400E-05 | 1.500E-05 |
| 9146 | 458 | V458E | A | 1.1e-05 | 1.5e-05 | 1.4e-05 | 1.333E-05 | 2.082E-06 | 1.100E-05 | 1.500E-05 |
| 9147 | 458 | V458Q | A | 1.5e-05 | 1.5e-05 | 1.6e-05 | 1.533E-05 | 5.774E-07 | 1.500E-05 | 1.600E-05 |
| 9148 | 458 | V458G | A | 1e-05 | 1.4e-05 | 1e-05 | 1.133E-05 | 2.309E-06 | 1.000E-05 | 1.400E-05 |
| 9149 | 458 | V458H | A | 1.7e-05 | 1e-05 | 1.2e-05 | 1.300E-05 | 3.606E-06 | 1.000E-05 | 1.700E-05 |
| 9150 | 458 | V458I | A | 8.8e-06 | 1.1e-05 | 1.1e-05 | 1.027E-05 | 1.270E-06 | 8.800E-06 | 1.100E-05 |
| 9151 | 458 | V458L | A | 1.5e-05 | 1.1e-05 | 1.1e-05 | 1.233E-05 | 2.309E-06 | 1.100E-05 | 1.500E-05 |
| 9152 | 458 | V458K | A | 2.3e-05 | 1.5e-05 | 1.3e-05 | 1.700E-05 | 5.292E-06 | 1.300E-05 | 2.300E-05 |
| 9153 | 458 | V458M | A | 7.7e-06 | 1.7e-05 | 1.6e-05 | 1.357E-05 | 5.105E-06 | 7.700E-06 | 1.700E-05 |
| 9154 | 458 | V458F | A | 1.6e-05 | 1.5e-05 | 1.7e-05 | 1.600E-05 | 1.000E-06 | 1.500E-05 | 1.700E-05 |
| 9155 | 458 | V458P | A | 9.9e-06 | 1.6e-05 | 1.4e-05 | 1.330E-05 | 3.110E-06 | 9.900E-06 | 1.600E-05 |
| 9156 | 458 | V458S | A | 1.3e-05 | 1.3e-05 | 1.3e-05 | 1.300E-05 | 0.000E+00 | 1.300E-05 | 1.300E-05 |
| 9157 | 458 | V458T | A | 1.6e-05 | 1e-05 | 1.2e-05 | 1.267E-05 | 3.055E-06 | 1.000E-05 | 1.600E-05 |
| 9158 | 458 | V458W | A | 1.4e-05 | 1.5e-05 | 1.5e-05 | 1.467E-05 | 5.774E-07 | 1.400E-05 | 1.500E-05 |
| 9159 | 458 | V458Y | A | 1.6e-05 | 1.6e-05 | 1.5e-05 | 1.567E-05 | 5.774E-07 | 1.500E-05 | 1.600E-05 |
| 9160 | 458 | V458V | A | 1.3e-05 | 1.5e-05 | 1.7e-05 | 1.500E-05 | 2.000E-06 | 1.300E-05 | 1.700E-05 |
| 9161 | 459 | P459A | A | 1.1e-05 | 1.5e-05 | 1.8e-05 | 1.467E-05 | 3.512E-06 | 1.100E-05 | 1.800E-05 |
| 9162 | 459 | P459R | A | 1.6e-05 | 1.1e-05 | 1.2e-05 | 1.300E-05 | 2.646E-06 | 1.100E-05 | 1.600E-05 |
| 9163 | 459 | P459N | A | 1.3e-05 | 1.5e-05 | 1e-05 | 1.267E-05 | 2.517E-06 | 1.000E-05 | 1.500E-05 |
| 9164 | 459 | P459D | A | 1.5e-05 | 1.2e-05 | 9.8e-06 | 1.227E-05 | 2.610E-06 | 9.800E-06 | 1.500E-05 |
| 9165 | 459 | P459C | A | 1.5e-05 | 9.8e-06 | 7.8e-06 | 1.087E-05 | 3.717E-06 | 7.800E-06 | 1.500E-05 |
| 9166 | 459 | P459E | A | 1e-05 | 1.2e-05 | 1.5e-05 | 1.233E-05 | 2.517E-06 | 1.000E-05 | 1.500E-05 |
| 9167 | 459 | P459Q | A | 1.4e-05 | 1.2e-05 | 1e-05 | 1.200E-05 | 2.000E-06 | 1.000E-05 | 1.400E-05 |
| 9168 | 459 | P459G | A | 1.1e-05 | 1.1e-05 | 1.4e-05 | 1.200E-05 | 1.732E-06 | 1.100E-05 | 1.400E-05 |
| 9169 | 459 | P459H | A | 1.2e-05 | 8.1e-06 | 9.9e-06 | 1.000E-05 | 1.952E-06 | 8.100E-06 | 1.200E-05 |
| 9170 | 459 | P459I | A | 1.1e-05 | 9.5e-06 | 1.5e-05 | 1.183E-05 | 2.843E-06 | 9.500E-06 | 1.500E-05 |
| 9171 | 459 | P459L | A | 1.5e-05 | 1e-05 | 1.6e-05 | 1.367E-05 | 3.215E-06 | 1.000E-05 | 1.600E-05 |
| 9172 | 459 | P459K | A | 1.1e-05 | 8.5e-06 | 1.5e-05 | 1.150E-05 | 3.279E-06 | 8.500E-06 | 1.500E-05 |
| 9173 | 459 | P459M | A | 1.4e-05 | 1.5e-05 | 9.7e-06 | 1.290E-05 | 2.816E-06 | 9.700E-06 | 1.500E-05 |
| 9174 | 459 | P459F | A | 1.7e-05 | 1.1e-05 | 9.8e-06 | 1.260E-05 | 3.857E-06 | 9.800E-06 | 1.700E-05 |
| 9175 | 459 | P459P | A | 1.3e-05 | 1.5e-05 | 1.1e-05 | 1.300E-05 | 2.000E-06 | 1.100E-05 | 1.500E-05 |
| 9176 | 459 | P459S | A | 1.5e-05 | 1.9e-05 | 9.9e-06 | 1.463E-05 | 4.561E-06 | 9.900E-06 | 1.900E-05 |
| 9177 | 459 | P459T | A | 1.8e-05 | 9.2e-06 | 1.1e-05 | 1.273E-05 | 4.649E-06 | 9.200E-06 | 1.800E-05 |
| 9178 | 459 | P459W | A | 1.3e-05 | 1e-05 | 9.1e-06 | 1.070E-05 | 2.042E-06 | 9.100E-06 | 1.300E-05 |
| 9179 | 459 | P459Y | A | 9.2e-06 | 1e-05 | 1.4e-05 | 1.107E-05 | 2.572E-06 | 9.200E-06 | 1.400E-05 |

| | | | | | | | | | | |
|---|---|---|---|---|---|---|---|---|---|---|
| 9180 | 459 | P459V | A | 8.4e-06 | 8.4e-06 | 9.9e-06 | 8.900E-06 | 8.660E-07 | 8.400E-06 | 9.900E-06 |
| 9181 | 460 | W460A | A | 1e-05 | 1.5e-05 | 1.6e-05 | 1.367E-05 | 3.215E-06 | 1.000E-05 | 1.600E-05 |
| 9182 | 460 | W460R | A | 1.2e-05 | 1.6e-05 | 1e-05 | 1.267E-05 | 3.055E-06 | 1.000E-05 | 1.600E-05 |
| 9183 | 460 | W460N | A | 1.5e-05 | 8.7e-06 | 1.6e-05 | 1.323E-05 | 3.958E-06 | 8.700E-06 | 1.600E-05 |
| 9184 | 460 | W460D | A | 1.3e-05 | 1.7e-05 | 8.9e-06 | 1.297E-05 | 4.050E-06 | 8.900E-06 | 1.700E-05 |
| 9185 | 460 | W460C | A | 9e-06 | 1.4e-05 | 1.1e-05 | 1.133E-05 | 2.517E-06 | 9.000E-06 | 1.400E-05 |
| 9186 | 460 | W460E | A | 9.7e-06 | 1.2e-05 | 1.6e-05 | 1.257E-05 | 3.188E-06 | 9.700E-06 | 1.600E-05 |
| 9187 | 460 | W460Q | A | 1.2e-05 | 1.5e-05 | 1.6e-05 | 1.433E-05 | 2.082E-06 | 1.200E-05 | 1.600E-05 |
| 9188 | 460 | W460G | A | 1.6e-05 | 9.2e-06 | 1.2e-05 | 1.240E-05 | 3.418E-06 | 9.200E-06 | 1.600E-05 |
| 9189 | 460 | W460H | A | 1.5e-05 | 1.3e-05 | 9.9e-06 | 1.263E-05 | 2.570E-06 | 9.900E-06 | 1.500E-05 |
| 9190 | 460 | W460I | A | 9.7e-06 | 1.7e-05 | 1.1e-05 | 1.257E-05 | 3.894E-06 | 9.700E-06 | 1.700E-05 |
| 9191 | 460 | W460L | A | 1.7e-05 | 1.6e-05 | 1.6e-05 | 1.633E-05 | 5.774E-07 | 1.600E-05 | 1.700E-05 |
| 9192 | 460 | W460K | A | 1.8e-05 | 8.4e-06 | 1.3e-05 | 1.313E-05 | 4.801E-06 | 8.400E-06 | 1.800E-05 |
| 9193 | 460 | W460M | A | 1.6e-05 | 1.4e-05 | 1.8e-05 | 1.600E-05 | 2.000E-06 | 1.400E-05 | 1.800E-05 |
| 9194 | 460 | W460F | A | 1.1e-05 | 1.2e-05 | 1.5e-05 | 1.267E-05 | 2.082E-06 | 1.100E-05 | 1.500E-05 |
| 9195 | 460 | W460P | A | 1.7e-05 | 1.4e-05 | 1.5e-05 | 1.533E-05 | 1.528E-06 | 1.400E-05 | 1.700E-05 |
| 9196 | 460 | W460S | A | 1.7e-05 | 1.7e-05 | 1.7e-05 | 1.700E-05 | 0.000E+00 | 1.700E-05 | 1.700E-05 |
| 9197 | 460 | W460T | A | 9.6e-06 | 1.2e-05 | 1.6e-05 | 1.253E-05 | 3.233E-06 | 9.600E-06 | 1.600E-05 |
| 9198 | 460 | W460W | A | 1.5e-05 | 1.3e-05 | 1.1e-05 | 1.300E-05 | 2.000E-06 | 1.100E-05 | 1.500E-05 |
| 9199 | 460 | W460Y | A | 1.7e-05 | 1.4e-05 | 1.5e-05 | 1.533E-05 | 1.528E-06 | 1.400E-05 | 1.700E-05 |
| 9200 | 460 | W460V | A | 1.7e-05 | 1.7e-05 | 1.4e-05 | 1.600E-05 | 1.732E-06 | 1.400E-05 | 1.700E-05 |
| 9201 | 461 | D461A | A | 9.8e-06 | 1e-05 | 1.1e-05 | 1.027E-05 | 6.429E-07 | 9.800E-06 | 1.100E-05 |
| 9202 | 461 | D461R | A | 1.3e-05 | 1.6e-05 | 1.4e-05 | 1.433E-05 | 1.528E-06 | 1.300E-05 | 1.600E-05 |
| 9203 | 461 | D461N | A | 1.5e-05 | 1.6e-05 | 1.1e-05 | 1.400E-05 | 2.646E-06 | 1.100E-05 | 1.600E-05 |
| 9204 | 461 | D461D | A | 9.1e-06 | 9.9e-06 | 1.6e-05 | 1.167E-05 | 3.774E-06 | 9.100E-06 | 1.600E-05 |
| 9205 | 461 | D461C | A | 8e-06 | 1.3e-05 | 1.1e-05 | 1.067E-05 | 2.517E-06 | 8.000E-06 | 1.300E-05 |
| 9206 | 461 | D461E | A | 9.9e-06 | 9.5e-06 | 1.5e-05 | 1.147E-05 | 3.066E-06 | 9.500E-06 | 1.500E-05 |
| 9207 | 461 | D461Q | A | 1.7e-05 | 1.4e-05 | 1.7e-05 | 1.600E-05 | 1.732E-06 | 1.400E-05 | 1.700E-05 |
| 9208 | 461 | D461G | A | 9.8e-06 | 1.6e-05 | 9.8e-06 | 1.187E-05 | 3.580E-06 | 9.800E-06 | 1.600E-05 |
| 9209 | 461 | D461H | A | 1.4e-05 | 1e-05 | 1.3e-05 | 1.233E-05 | 2.082E-06 | 1.000E-05 | 1.400E-05 |
| 9210 | 461 | D461I | A | 1.1e-05 | 1.3e-05 | 1.3e-05 | 1.233E-05 | 1.155E-06 | 1.100E-05 | 1.300E-05 |
| 9211 | 461 | D461L | A | 1.5e-05 | 1.3e-05 | 1.2e-05 | 1.333E-05 | 1.528E-06 | 1.200E-05 | 1.500E-05 |
| 9212 | 461 | D461K | A | 1.4e-05 | 1.2e-05 | 5.8e-06 | 1.060E-05 | 4.276E-06 | 5.800E-06 | 1.400E-05 |
| 9213 | 461 | D461M | A | 1.5e-05 | 1.6e-05 | 1.2e-05 | 1.433E-05 | 2.082E-06 | 1.200E-05 | 1.600E-05 |
| 9214 | 461 | D461F | A | 1.6e-05 | 1.7e-05 | 1.3e-05 | 1.533E-05 | 2.082E-06 | 1.300E-05 | 1.700E-05 |
| 9215 | 461 | D461P | A | 1.1e-05 | 1.1e-05 | 9.5e-06 | 1.050E-05 | 8.660E-07 | 9.500E-06 | 1.100E-05 |
| 9216 | 461 | D461S | A | 1.1e-05 | 1.1e-05 | 1.6e-05 | 1.267E-05 | 2.887E-06 | 1.100E-05 | 1.600E-05 |
| 9217 | 461 | D461T | A | 1.4e-05 | 1.4e-05 | 9.2e-06 | 1.240E-05 | 2.771E-06 | 9.200E-06 | 1.400E-05 |
| 9218 | 461 | D461W | A | 1.3e-05 | 1.2e-05 | 1.4e-05 | 1.300E-05 | 1.000E-06 | 1.200E-05 | 1.400E-05 |
| 9219 | 461 | D461Y | A | 1.5e-05 | 1.2e-05 | 8.4e-06 | 1.180E-05 | 3.305E-06 | 8.400E-06 | 1.500E-05 |
| 9220 | 461 | D461V | A | 1.1e-05 | 9.7e-06 | 1.5e-05 | 1.190E-05 | 2.762E-06 | 9.700E-06 | 1.500E-05 |
| 9221 | 462 | Q462A | A | 1.4e-05 | 1.2e-05 | 1e-05 | 1.200E-05 | 2.000E-06 | 1.000E-05 | 1.400E-05 |
| 9222 | 462 | Q462R | A | 9.7e-06 | 1.4e-05 | 8.5e-06 | 1.073E-05 | 2.892E-06 | 8.500E-06 | 1.400E-05 |
| 9223 | 462 | Q462N | A | 1.8e-05 | 1.5e-05 | 1.7e-05 | 1.667E-05 | 1.528E-06 | 1.500E-05 | 1.800E-05 |
| 9224 | 462 | Q462D | A | 9.8e-06 | 1.1e-05 | 1.5e-05 | 1.193E-05 | 2.723E-06 | 9.800E-06 | 1.500E-05 |
| 9225 | 462 | Q462C | A | 1.4e-05 | 1.5e-05 | 1.1e-05 | 1.333E-05 | 2.082E-06 | 1.100E-05 | 1.500E-05 |
| 9226 | 462 | Q462E | A | 1.6e-05 | 9.9e-06 | 1.6e-05 | 1.397E-05 | 3.522E-06 | 9.900E-06 | 1.600E-05 |
| 9227 | 462 | Q462Q | A | 1.4e-05 | 6.3e-06 | 1.6e-05 | 1.210E-05 | 5.122E-06 | 6.300E-06 | 1.600E-05 |
| 9228 | 462 | Q462G | A | 1.5e-05 | 1e-05 | 1.1e-05 | 1.200E-05 | 2.646E-06 | 1.000E-05 | 1.500E-05 |
| 9229 | 462 | Q462H | A | 1.5e-05 | 1.5e-05 | 1.1e-05 | 1.367E-05 | 2.309E-06 | 1.100E-05 | 1.500E-05 |
| 9230 | 462 | Q462I | A | 1.5e-05 | 1.3e-05 | 1.5e-05 | 1.433E-05 | 1.155E-06 | 1.300E-05 | 1.500E-05 |
| 9231 | 462 | Q462L | A | 1.5e-05 | 9.1e-06 | 1.3e-05 | 1.237E-05 | 3.001E-06 | 9.100E-06 | 1.500E-05 |
| 9232 | 462 | Q462K | A | 1.2e-05 | 1e-05 | 1.5e-05 | 1.233E-05 | 2.517E-06 | 1.000E-05 | 1.500E-05 |
| 9233 | 462 | Q462M | A | 9e-06 | 9.3e-06 | 1.1e-05 | 9.767E-06 | 1.079E-06 | 9.000E-06 | 1.100E-05 |
| 9234 | 462 | Q462F | A | 7.2e-06 | 1.6e-05 | 1.4e-05 | 1.240E-05 | 4.613E-06 | 7.200E-06 | 1.600E-05 |
| 9235 | 462 | Q462P | A | 1e-05 | 6.9e-06 | 1.4e-05 | 1.030E-05 | 3.559E-06 | 6.900E-06 | 1.400E-05 |
| 9236 | 462 | Q462S | A | 1.1e-05 | 1.7e-05 | 1.5e-05 | 1.433E-05 | 3.055E-06 | 1.100E-05 | 1.700E-05 |
| 9237 | 462 | Q462T | A | 1.6e-05 | 1.1e-05 | 1e-05 | 1.233E-05 | 3.215E-06 | 1.000E-05 | 1.600E-05 |
| 9238 | 462 | Q462W | A | 1.1e-05 | 9.9e-06 | 1.3e-05 | 1.130E-05 | 1.572E-06 | 9.900E-06 | 1.300E-05 |
| 9239 | 462 | Q462Y | A | 1.7e-05 | 1.6e-05 | 1.2e-05 | 1.500E-05 | 2.646E-06 | 1.200E-05 | 1.700E-05 |
| 9240 | 462 | Q462V | A | 1.4e-05 | 1.5e-05 | 1.7e-05 | 1.533E-05 | 1.528E-06 | 1.400E-05 | 1.700E-05 |
| 9241 | 463 | L463A | A | 9.3e-06 | 1.1e-05 | 1e-05 | 1.010E-05 | 8.544E-07 | 9.300E-06 | 1.100E-05 |
| 9242 | 463 | L463R | A | 1e-05 | 1.1e-05 | 1.6e-05 | 1.233E-05 | 3.215E-06 | 1.000E-05 | 1.600E-05 |
| 9243 | 463 | L463N | A | 1.1e-05 | 1.4e-05 | 1e-05 | 1.167E-05 | 2.082E-06 | 1.000E-05 | 1.400E-05 |
| 9244 | 463 | L463D | A | 1.1e-05 | 1.3e-05 | 1.5e-05 | 1.300E-05 | 2.000E-06 | 1.100E-05 | 1.500E-05 |
| 9245 | 463 | L463C | A | 1.5e-05 | 9.7e-06 | 1.5e-05 | 1.323E-05 | 3.060E-06 | 9.700E-06 | 1.500E-05 |
| 9246 | 463 | L463E | A | 1.3e-05 | 1.5e-05 | 8.9e-06 | 1.230E-05 | 3.110E-06 | 8.900E-06 | 1.500E-05 |
| 9247 | 463 | L463Q | A | 1.9e-05 | 1.4e-05 | 1.5e-05 | 1.600E-05 | 2.646E-06 | 1.400E-05 | 1.900E-05 |
| 9248 | 463 | L463G | A | 1e-05 | 1.4e-05 | 1.1e-05 | 1.167E-05 | 2.082E-06 | 1.000E-05 | 1.400E-05 |
| 9249 | 463 | L463H | A | 1.6e-05 | 1.4e-05 | 1.3e-05 | 1.433E-05 | 1.528E-06 | 1.300E-05 | 1.600E-05 |
| 9250 | 463 | L463I | A | 1.5e-05 | 9.6e-06 | 1.7e-05 | 1.387E-05 | 3.828E-06 | 9.600E-06 | 1.700E-05 |
| 9251 | 463 | L463L | A | 1.1e-05 | 1.3e-05 | 1.2e-05 | 1.200E-05 | 1.000E-06 | 1.100E-05 | 1.300E-05 |
| 9252 | 463 | L463K | A | 1e-05 | 9.9e-06 | 1.6e-05 | 1.197E-05 | 3.493E-06 | 9.900E-06 | 1.600E-05 |
| 9253 | 463 | L463M | A | 1.3e-05 | 1.1e-05 | 1.5e-05 | 1.300E-05 | 2.000E-06 | 1.100E-05 | 1.500E-05 |
| 9254 | 463 | L463F | A | 1e-05 | 1.4e-05 | 1.4e-05 | 1.267E-05 | 2.309E-06 | 1.000E-05 | 1.400E-05 |
| 9255 | 463 | L463P | A | 1.4e-05 | 9.2e-06 | 8.8e-06 | 1.067E-05 | 2.894E-06 | 8.800E-06 | 1.400E-05 |
| 9256 | 463 | L463S | A | 1.4e-05 | 1.6e-05 | 1.3e-05 | 1.433E-05 | 1.528E-06 | 1.300E-05 | 1.600E-05 |
| 9257 | 463 | L463T | A | 1.1e-05 | 8.8e-06 | 1.4e-05 | 1.127E-05 | 2.610E-06 | 8.800E-06 | 1.400E-05 |
| 9258 | 463 | L463W | A | 1.6e-05 | 9.3e-06 | 1.2e-05 | 1.243E-05 | 3.371E-06 | 9.300E-06 | 1.600E-05 |
| 9259 | 463 | L463Y | A | 1.6e-05 | 1.6e-05 | 1e-05 | 1.400E-05 | 3.464E-06 | 1.000E-05 | 1.600E-05 |
| 9260 | 463 | L463V | A | 1.4e-05 | 1.3e-05 | 7.5e-06 | 1.150E-05 | 3.500E-06 | 7.500E-06 | 1.400E-05 |
| 9261 | 464 | F464A | A | 1.6e-05 | 1.6e-05 | 1.6e-05 | 1.600E-05 | 0.000E+00 | 1.600E-05 | 1.600E-05 |
| 9262 | 464 | F464R | A | 1.4e-05 | 1.1e-05 | 1e-05 | 1.167E-05 | 2.082E-06 | 1.000E-05 | 1.400E-05 |
| 9263 | 464 | F464N | A | 1.5e-05 | 1.1e-05 | 1.6e-05 | 1.400E-05 | 2.646E-06 | 1.100E-05 | 1.600E-05 |
| 9264 | 464 | F464D | A | 1.3e-05 | 1.6e-05 | 1e-05 | 1.300E-05 | 3.000E-06 | 1.000E-05 | 1.600E-05 |

| | | | | | | | | | | |
|---|---|---|---|---|---|---|---|---|---|---|
| 9265 | 464 | F464C | A | 9.5e-06 | 9.8e-06 | 1.6e-05 | 1.177E-05 | 3.669E-06 | 9.500E-06 | 1.600E-05 |
| 9266 | 464 | F464E | A | 1.4e-05 | 1.6e-05 | 1.1e-05 | 1.367E-05 | 2.517E-06 | 1.100E-05 | 1.600E-05 |
| 9267 | 464 | F464Q | A | 1.7e-05 | 9.5e-06 | 1.6e-05 | 1.417E-05 | 4.072E-06 | 9.500E-06 | 1.700E-05 |
| 9268 | 464 | F464G | A | 1.6e-05 | 7e-06 | 1.1e-05 | 1.133E-05 | 4.509E-06 | 7.000E-06 | 1.600E-05 |
| 9269 | 464 | F464H | A | 7.9e-06 | 8.2e-06 | 1.3e-05 | 9.700E-06 | 2.862E-06 | 7.900E-06 | 1.300E-05 |
| 9270 | 464 | F464I | A | 1.1e-05 | 1.1e-05 | 1.6e-05 | 1.267E-05 | 2.887E-06 | 1.100E-05 | 1.600E-05 |
| 9271 | 464 | F464L | A | 8.5e-06 | 1.3e-05 | 1.1e-05 | 1.083E-05 | 2.255E-06 | 8.500E-06 | 1.300E-05 |
| 9272 | 464 | F464K | A | 1.5e-05 | 1e-05 | 1.1e-05 | 1.200E-05 | 2.646E-06 | 1.000E-05 | 1.500E-05 |
| 9273 | 464 | F464M | A | 1.5e-05 | 1.6e-05 | 1e-05 | 1.367E-05 | 3.215E-06 | 1.000E-05 | 1.600E-05 |
| 9274 | 464 | F464F | A | 7.8e-06 | 1.4e-05 | 1.5e-05 | 1.227E-05 | 3.900E-06 | 7.800E-06 | 1.500E-05 |
| 9275 | 464 | F464P | A | 1.3e-05 | 1.7e-05 | 1.6e-05 | 1.533E-05 | 2.082E-06 | 1.300E-05 | 1.700E-05 |
| 9276 | 464 | F464S | A | 1.3e-05 | 1.5e-05 | 1.1e-05 | 1.300E-05 | 2.000E-06 | 1.100E-05 | 1.500E-05 |
| 9277 | 464 | F464T | A | 1.6e-05 | 1.6e-05 | 1.8e-05 | 1.667E-05 | 1.155E-06 | 1.600E-05 | 1.800E-05 |
| 9278 | 464 | F464W | A | 1.6e-05 | 1.1e-05 | 9.7e-06 | 1.223E-05 | 3.326E-06 | 9.700E-06 | 1.600E-05 |
| 9279 | 464 | F464Y | A | 1.2e-05 | 1.2e-05 | 1.5e-05 | 1.300E-05 | 1.732E-06 | 1.200E-05 | 1.500E-05 |
| 9280 | 464 | F464V | A | 1.1e-05 | 1e-05 | 1.5e-05 | 1.200E-05 | 2.646E-06 | 1.000E-05 | 1.500E-05 |
| 9281 | 465 | R465A | A | 1e-05 | 1.1e-05 | 9.7e-06 | 1.023E-05 | 6.807E-07 | 9.700E-06 | 1.100E-05 |
| 9282 | 465 | R465R | A | 9.7e-06 | 1.3e-05 | 1.6e-05 | 1.290E-05 | 3.151E-06 | 9.700E-06 | 1.600E-05 |
| 9283 | 465 | R465N | A | 9e-06 | 1.5e-05 | 1.7e-05 | 1.367E-05 | 4.163E-06 | 9.000E-06 | 1.700E-05 |
| 9284 | 465 | R465D | A | 9.8e-06 | 1.5e-05 | 1.4e-05 | 1.293E-05 | 2.759E-06 | 9.800E-06 | 1.500E-05 |
| 9285 | 465 | R465C | A | 1.7e-05 | 1.3e-05 | 1.5e-05 | 1.500E-05 | 2.000E-06 | 1.300E-05 | 1.700E-05 |
| 9286 | 465 | R465E | A | 8.9e-06 | 1.1e-05 | 1e-05 | 9.967E-06 | 1.050E-06 | 8.900E-06 | 1.100E-05 |
| 9287 | 465 | R465Q | A | 1.6e-05 | 1.6e-05 | 8.3e-06 | 1.343E-05 | 4.446E-06 | 8.300E-06 | 1.600E-05 |
| 9288 | 465 | R465G | A | 1.6e-05 | 1e-05 | 1.2e-05 | 1.267E-05 | 3.055E-06 | 1.000E-05 | 1.600E-05 |
| 9289 | 465 | R465H | A | 1.5e-05 | 1.6e-05 | 1.4e-05 | 1.500E-05 | 1.000E-06 | 1.400E-05 | 1.600E-05 |
| 9290 | 465 | R465I | A | 1.3e-05 | 1.5e-05 | 1.3e-05 | 1.367E-05 | 1.155E-06 | 1.300E-05 | 1.500E-05 |
| 9291 | 465 | R465L | A | 1e-05 | 1e-05 | 1e-05 | 1.000E-05 | 0.000E+00 | 1.000E-05 | 1.000E-05 |
| 9292 | 465 | R465K | A | 9.8e-06 | 1.3e-05 | 1.3e-05 | 1.193E-05 | 1.848E-06 | 9.800E-06 | 1.300E-05 |
| 9293 | 465 | R465M | A | 1.1e-05 | 1.1e-05 | 7.4e-06 | 9.800E-06 | 2.078E-06 | 7.400E-06 | 1.100E-05 |
| 9294 | 465 | R465F | A | 1.1e-05 | 1.4e-05 | 1.7e-05 | 1.400E-05 | 3.000E-06 | 1.100E-05 | 1.700E-05 |
| 9295 | 465 | R465P | A | 1.8e-05 | 1.6e-05 | 1.2e-05 | 1.533E-05 | 3.055E-06 | 1.200E-05 | 1.800E-05 |
| 9296 | 465 | R465S | A | 1.2e-05 | 1.5e-05 | 1e-05 | 1.233E-05 | 2.517E-06 | 1.000E-05 | 1.500E-05 |
| 9297 | 465 | R465T | A | 9.9e-06 | 1.2e-05 | 1.1e-05 | 1.097E-05 | 1.050E-06 | 9.900E-06 | 1.200E-05 |
| 9298 | 465 | R465W | A | 1.1e-05 | 9.6e-06 | 1.5e-05 | 1.187E-05 | 2.802E-06 | 9.600E-06 | 1.500E-05 |
| 9299 | 465 | R465Y | A | 1.1e-05 | 1.6e-05 | 1.3e-05 | 1.333E-05 | 2.517E-06 | 1.100E-05 | 1.600E-05 |
| 9300 | 465 | R465V | A | 1.2e-05 | 1.1e-05 | 1.6e-05 | 1.300E-05 | 2.646E-06 | 1.100E-05 | 1.600E-05 |
| 9301 | 466 | N466A | A | 8.6e-06 | 9.7e-06 | 1.6e-05 | 1.143E-05 | 3.993E-06 | 8.600E-06 | 1.600E-05 |
| 9302 | 466 | N466R | A | 1.1e-05 | 1.3e-05 | 1.4e-05 | 1.267E-05 | 1.528E-06 | 1.100E-05 | 1.400E-05 |
| 9303 | 466 | N466N | A | 1.4e-05 | 1.6e-05 | 1.4e-05 | 1.467E-05 | 1.155E-06 | 1.400E-05 | 1.600E-05 |
| 9304 | 466 | N466D | A | 1.1e-05 | 9.4e-06 | 1.6e-05 | 1.213E-05 | 3.443E-06 | 9.400E-06 | 1.600E-05 |
| 9305 | 466 | N466C | A | 9.6e-06 | 1.5e-05 | 1.1e-05 | 1.187E-05 | 2.802E-06 | 9.600E-06 | 1.500E-05 |
| 9306 | 466 | N466E | A | 1e-05 | 9.4e-06 | 1e-05 | 9.800E-06 | 3.464E-07 | 9.400E-06 | 1.000E-05 |
| 9307 | 466 | N466Q | A | 1.2e-05 | 1e-05 | 1.5e-05 | 1.233E-05 | 2.517E-06 | 1.000E-05 | 1.500E-05 |
| 9308 | 466 | N466G | A | 1.2e-05 | 1.6e-05 | 1.3e-05 | 1.367E-05 | 2.082E-06 | 1.200E-05 | 1.600E-05 |
| 9309 | 466 | N466H | A | 1.5e-05 | 1.7e-05 | 1e-05 | 1.400E-05 | 3.606E-06 | 1.000E-05 | 1.700E-05 |
| 9310 | 466 | N466I | A | 9.3e-06 | 1e-05 | 1e-05 | 9.767E-06 | 4.041E-07 | 9.300E-06 | 1.000E-05 |
| 9311 | 466 | N466L | A | 1.5e-05 | 1.2e-05 | 1.5e-05 | 1.400E-05 | 1.732E-06 | 1.200E-05 | 1.500E-05 |
| 9312 | 466 | N466K | A | 1.2e-05 | 1.7e-05 | 1.5e-05 | 1.467E-05 | 2.517E-06 | 1.200E-05 | 1.700E-05 |
| 9313 | 466 | N466M | A | 1.6e-05 | 1.5e-05 | 8.9e-06 | 1.330E-05 | 3.843E-06 | 8.900E-06 | 1.600E-05 |
| 9314 | 466 | N466F | A | 1.2e-05 | 1.1e-05 | 1.5e-05 | 1.267E-05 | 2.082E-06 | 1.100E-05 | 1.500E-05 |
| 9315 | 466 | N466P | A | 1.3e-05 | 1.3e-05 | 1.6e-05 | 1.400E-05 | 1.732E-06 | 1.300E-05 | 1.600E-05 |
| 9316 | 466 | N466S | A | 1.2e-05 | 1.5e-05 | 1.6e-05 | 1.433E-05 | 2.082E-06 | 1.200E-05 | 1.600E-05 |
| 9317 | 466 | N466T | A | 1e-05 | 8.3e-06 | 1.8e-05 | 1.210E-05 | 5.180E-06 | 8.300E-06 | 1.800E-05 |
| 9318 | 466 | N466W | A | 1.5e-05 | 1.6e-05 | 1.4e-05 | 1.500E-05 | 1.000E-06 | 1.400E-05 | 1.600E-05 |
| 9319 | 466 | N466Y | A | 1.5e-05 | 1.7e-05 | 9.6e-06 | 1.387E-05 | 3.828E-06 | 9.600E-06 | 1.700E-05 |
| 9320 | 466 | N466V | A | 1.6e-05 | 1.3e-05 | 9.9e-06 | 1.297E-05 | 3.050E-06 | 9.900E-06 | 1.600E-05 |
| 9321 | 467 | P467A | A | 1e-05 | 1.4e-05 | 1.6e-05 | 1.333E-05 | 3.055E-06 | 1.000E-05 | 1.600E-05 |
| 9322 | 467 | P467R | A | 8.2e-06 | 1.3e-05 | 7.6e-06 | 9.600E-06 | 2.960E-06 | 7.600E-06 | 1.300E-05 |
| 9323 | 467 | P467N | A | 1.5e-05 | 1.5e-05 | 1e-05 | 1.333E-05 | 2.887E-06 | 1.000E-05 | 1.500E-05 |
| 9324 | 467 | P467D | A | 1.4e-05 | 1.5e-05 | 9.3e-06 | 1.277E-05 | 3.044E-06 | 9.300E-06 | 1.500E-05 |
| 9325 | 467 | P467C | A | 1.5e-05 | 1.1e-05 | 1.1e-05 | 1.233E-05 | 2.309E-06 | 1.100E-05 | 1.500E-05 |
| 9326 | 467 | P467E | A | 1.7e-05 | 9.9e-06 | 9.9e-06 | 1.227E-05 | 4.099E-06 | 9.900E-06 | 1.700E-05 |
| 9327 | 467 | P467Q | A | 9.3e-06 | 1.6e-05 | 1.3e-05 | 1.277E-05 | 3.356E-06 | 9.300E-06 | 1.600E-05 |
| 9328 | 467 | P467G | A | 1e-05 | 1.3e-05 | 1.5e-05 | 1.267E-05 | 2.517E-06 | 1.000E-05 | 1.500E-05 |
| 9329 | 467 | P467H | A | 1e-05 | 9.6e-06 | 8.4e-06 | 9.333E-06 | 8.327E-07 | 8.400E-06 | 1.000E-05 |
| 9330 | 467 | P467I | A | 1.4e-05 | 1.5e-05 | 1.5e-05 | 1.467E-05 | 5.774E-07 | 1.400E-05 | 1.500E-05 |
| 9331 | 467 | P467L | A | 8.9e-06 | 1.1e-05 | 9.9e-06 | 9.933E-06 | 1.050E-06 | 8.900E-06 | 1.100E-05 |
| 9332 | 467 | P467K | A | 1.1e-05 | 1.4e-05 | 9.8e-06 | 1.160E-05 | 2.163E-06 | 9.800E-06 | 1.400E-05 |
| 9333 | 467 | P467M | A | 1.4e-05 | 1.1e-05 | 8.9e-06 | 1.130E-05 | 2.563E-06 | 8.900E-06 | 1.400E-05 |
| 9334 | 467 | P467F | A | 1.4e-05 | 1.6e-05 | 1e-05 | 1.333E-05 | 3.055E-06 | 1.000E-05 | 1.600E-05 |
| 9335 | 467 | P467P | A | 9.1e-06 | 1.6e-05 | 9.5e-06 | 1.153E-05 | 3.873E-06 | 9.100E-06 | 1.600E-05 |
| 9336 | 467 | P467S | A | 1e-05 | 1.3e-05 | 9.9e-06 | 1.097E-05 | 1.762E-06 | 9.900E-06 | 1.300E-05 |
| 9337 | 467 | P467T | A | 1.4e-05 | 9.1e-06 | 1.3e-05 | 1.203E-05 | 2.589E-06 | 9.100E-06 | 1.400E-05 |
| 9338 | 467 | P467W | A | 1.4e-05 | 1.6e-05 | 1e-05 | 1.333E-05 | 3.055E-06 | 1.000E-05 | 1.600E-05 |
| 9339 | 467 | P467Y | A | 1.3e-05 | 1.5e-05 | 1.3e-05 | 1.367E-05 | 1.155E-06 | 1.300E-05 | 1.500E-05 |
| 9340 | 467 | P467V | A | 9.5e-06 | 8.9e-06 | 1.5e-05 | 1.113E-05 | 3.362E-06 | 8.900E-06 | 1.500E-05 |
| 9341 | 468 | H468A | A | 1.6e-05 | 8.9e-06 | 1.5e-05 | 1.330E-05 | 3.843E-06 | 8.900E-06 | 1.600E-05 |
| 9342 | 468 | H468R | A | 1e-05 | 1.6e-05 | 1.2e-05 | 1.267E-05 | 3.055E-06 | 1.000E-05 | 1.600E-05 |
| 9343 | 468 | H468N | A | 1.3e-05 | 1e-05 | 1.4e-05 | 1.233E-05 | 2.082E-06 | 1.000E-05 | 1.400E-05 |
| 9344 | 468 | H468D | A | 1.1e-05 | 1.1e-05 | 1.3e-05 | 1.167E-05 | 1.155E-06 | 1.100E-05 | 1.300E-05 |
| 9345 | 468 | H468C | A | 1.5e-05 | 1.7e-05 | 9.9e-06 | 1.397E-05 | 3.661E-06 | 9.900E-06 | 1.700E-05 |
| 9346 | 468 | H468E | A | 1.6e-05 | 1.6e-05 | 1.2e-05 | 1.467E-05 | 2.309E-06 | 1.200E-05 | 1.600E-05 |
| 9347 | 468 | H468Q | A | 9.1e-06 | 1.2e-05 | 1.5e-05 | 1.203E-05 | 2.950E-06 | 9.100E-06 | 1.500E-05 |
| 9348 | 468 | H468G | A | 1.3e-05 | 1e-05 | 1.2e-05 | 1.167E-05 | 1.528E-06 | 1.000E-05 | 1.300E-05 |
| 9349 | 468 | H468H | A | 1.1e-05 | 1.6e-05 | 1.1e-05 | 1.267E-05 | 2.887E-06 | 1.100E-05 | 1.600E-05 |

| | | | | | | | | | | |
|---|---|---|---|---|---|---|---|---|---|---|
| 9350 | 468 | H468I | A | 1.1e-05 | 2e-05 | 8.7e-06 | 1.323E-05 | 5.972E-06 | 8.700E-06 | 2.000E-05 |
| 9351 | 468 | H468L | A | 1.3e-05 | 1.2e-05 | 1.6e-05 | 1.367E-05 | 2.082E-06 | 1.200E-05 | 1.600E-05 |
| 9352 | 468 | H468K | A | 1.2e-05 | 1.3e-05 | 1.1e-05 | 1.200E-05 | 1.000E-06 | 1.100E-05 | 1.300E-05 |
| 9353 | 468 | H468M | A | 1.9e-05 | 1.4e-05 | 1.3e-05 | 1.533E-05 | 3.215E-06 | 1.300E-05 | 1.900E-05 |
| 9354 | 468 | H468F | A | 1.4e-05 | 9.5e-06 | 1.7e-05 | 1.350E-05 | 3.775E-06 | 9.500E-06 | 1.700E-05 |
| 9355 | 468 | H468P | A | 1.5e-05 | 1.3e-05 | 1e-05 | 1.267E-05 | 2.517E-06 | 1.000E-05 | 1.500E-05 |
| 9356 | 468 | H468S | A | 1.6e-05 | 1.5e-05 | 9.1e-06 | 1.337E-05 | 3.729E-06 | 9.100E-06 | 1.600E-05 |
| 9357 | 468 | H468T | A | 8.9e-06 | 1e-05 | 1e-05 | 9.633E-06 | 6.351E-07 | 8.900E-06 | 1.000E-05 |
| 9358 | 468 | H468W | A | 1.3e-05 | 1.5e-05 | 1.1e-05 | 1.300E-05 | 2.000E-06 | 1.100E-05 | 1.500E-05 |
| 9359 | 468 | H468Y | A | 1.1e-05 | 1.6e-05 | 1e-05 | 1.233E-05 | 3.215E-06 | 1.000E-05 | 1.600E-05 |
| 9360 | 468 | H468V | A | 1.1e-05 | 1.1e-05 | 1.6e-05 | 1.267E-05 | 2.887E-06 | 1.100E-05 | 1.600E-05 |
| 9361 | 469 | Q469A | A | 1.4e-05 | 9.7e-06 | 1.5e-05 | 1.290E-05 | 2.816E-06 | 9.700E-06 | 1.500E-05 |
| 9362 | 469 | Q469R | A | 1.6e-05 | 1.7e-05 | 1.5e-05 | 1.600E-05 | 1.000E-06 | 1.500E-05 | 1.700E-05 |
| 9363 | 469 | Q469N | A | 1.1e-05 | 1e-05 | 1.4e-05 | 1.167E-05 | 2.082E-06 | 1.000E-05 | 1.400E-05 |
| 9364 | 469 | Q469D | A | 1.7e-05 | 1.1e-05 | 1.3e-05 | 1.367E-05 | 3.055E-06 | 1.100E-05 | 1.700E-05 |
| 9365 | 469 | Q469C | A | 8.9e-06 | 1.2e-05 | 1.6e-05 | 1.230E-05 | 3.559E-06 | 8.900E-06 | 1.600E-05 |
| 9366 | 469 | Q469E | A | 9.7e-06 | 1e-05 | 1.5e-05 | 1.157E-05 | 2.977E-06 | 9.700E-06 | 1.500E-05 |
| 9367 | 469 | Q469Q | A | 1.9e-05 | 1.1e-05 | 1.4e-05 | 1.467E-05 | 4.041E-06 | 1.100E-05 | 1.900E-05 |
| 9368 | 469 | Q469G | A | 1.5e-05 | 1e-05 | 1.5e-05 | 1.333E-05 | 2.887E-06 | 1.000E-05 | 1.500E-05 |
| 9369 | 469 | Q469H | A | 1.4e-05 | 1.8e-05 | 1.2e-05 | 1.467E-05 | 3.055E-06 | 1.200E-05 | 1.800E-05 |
| 9370 | 469 | Q469I | A | 1.8e-05 | 1.4e-05 | 1.1e-05 | 1.433E-05 | 3.512E-06 | 1.100E-05 | 1.800E-05 |
| 9371 | 469 | Q469L | A | 1.4e-05 | 1.8e-05 | 1.5e-05 | 1.567E-05 | 2.082E-06 | 1.400E-05 | 1.800E-05 |
| 9372 | 469 | Q469K | A | 1.6e-05 | 1.4e-05 | 1.4e-05 | 1.467E-05 | 1.155E-06 | 1.400E-05 | 1.600E-05 |
| 9373 | 469 | Q469M | A | 1.2e-05 | 1.4e-05 | 1.5e-05 | 1.367E-05 | 1.528E-06 | 1.200E-05 | 1.500E-05 |
| 9374 | 469 | Q469F | A | 1.5e-05 | 1.7e-05 | 1.4e-05 | 1.533E-05 | 1.528E-06 | 1.400E-05 | 1.700E-05 |
| 9375 | 469 | Q469P | A | 9.6e-06 | 1.7e-05 | 1.5e-05 | 1.387E-05 | 3.828E-06 | 9.600E-06 | 1.700E-05 |
| 9376 | 469 | Q469S | A | 1.1e-05 | 1.6e-05 | 1.6e-05 | 1.433E-05 | 2.887E-06 | 1.100E-05 | 1.600E-05 |
| 9377 | 469 | Q469T | A | 1.4e-05 | 1.7e-05 | 1.1e-05 | 1.400E-05 | 3.000E-06 | 1.100E-05 | 1.700E-05 |
| 9378 | 469 | Q469W | A | 9.9e-06 | 1.6e-05 | 1.3e-05 | 1.297E-05 | 3.050E-06 | 9.900E-06 | 1.600E-05 |
| 9379 | 469 | Q469Y | A | 1.7e-05 | 1.2e-05 | 1.5e-05 | 1.467E-05 | 2.517E-06 | 1.200E-05 | 1.700E-05 |
| 9380 | 469 | Q469V | A | 1.6e-05 | 1.7e-05 | 1.7e-05 | 1.667E-05 | 5.774E-07 | 1.600E-05 | 1.700E-05 |
| 9381 | 470 | A470A | A | 1.4e-05 | 1.2e-05 | 1.5e-05 | 1.367E-05 | 1.528E-06 | 1.200E-05 | 1.500E-05 |
| 9382 | 470 | A470R | A | 1.1e-05 | 1.6e-05 | 9.1e-06 | 1.203E-05 | 3.564E-06 | 9.100E-06 | 1.600E-05 |
| 9383 | 470 | A470N | A | 1.1e-05 | 1.7e-05 | 1.6e-05 | 1.467E-05 | 3.215E-06 | 1.100E-05 | 1.700E-05 |
| 9384 | 470 | A470D | A | 1.4e-05 | 1.6e-05 | 1.1e-05 | 1.367E-05 | 2.517E-06 | 1.100E-05 | 1.600E-05 |
| 9385 | 470 | A470C | A | 1.2e-05 | 1.7e-05 | 1e-05 | 1.300E-05 | 3.606E-06 | 1.000E-05 | 1.700E-05 |
| 9386 | 470 | A470E | A | 1.6e-05 | 1.3e-05 | 1.1e-05 | 1.333E-05 | 2.517E-06 | 1.100E-05 | 1.600E-05 |
| 9387 | 470 | A470Q | A | 9.8e-06 | 9.2e-06 | 9.4e-06 | 9.467E-06 | 3.055E-07 | 9.200E-06 | 9.800E-06 |
| 9388 | 470 | A470G | A | 1e-05 | 8.7e-06 | 1.6e-05 | 1.157E-05 | 3.894E-06 | 8.700E-06 | 1.600E-05 |
| 9389 | 470 | A470H | A | 1.1e-05 | 1.5e-05 | 1.2e-05 | 1.267E-05 | 2.082E-06 | 1.100E-05 | 1.500E-05 |
| 9390 | 470 | A470I | A | 1.5e-05 | 1.5e-05 | 1.8e-05 | 1.600E-05 | 1.732E-06 | 1.500E-05 | 1.800E-05 |
| 9391 | 470 | A470L | A | 1.3e-05 | 7.2e-06 | 1.6e-05 | 1.207E-05 | 4.474E-06 | 7.200E-06 | 1.600E-05 |
| 9392 | 470 | A470K | A | 1.1e-05 | 9e-06 | 1.5e-05 | 1.167E-05 | 3.055E-06 | 9.000E-06 | 1.500E-05 |
| 9393 | 470 | A470M | A | 1.3e-05 | 1.6e-05 | 1.1e-05 | 1.333E-05 | 2.517E-06 | 1.100E-05 | 1.600E-05 |
| 9394 | 470 | A470F | A | 1e-05 | 1.5e-05 | 1.2e-05 | 1.233E-05 | 2.517E-06 | 1.000E-05 | 1.500E-05 |
| 9395 | 470 | A470P | A | 1.1e-05 | 9.8e-06 | 1.6e-05 | 1.227E-05 | 3.288E-06 | 9.800E-06 | 1.600E-05 |
| 9396 | 470 | A470S | A | 1.6e-05 | 1.3e-05 | 1.2e-05 | 1.367E-05 | 2.082E-06 | 1.200E-05 | 1.600E-05 |
| 9397 | 470 | A470T | A | 1.5e-05 | 9e-06 | 9.9e-06 | 1.130E-05 | 3.236E-06 | 9.000E-06 | 1.500E-05 |
| 9398 | 470 | A470W | A | 1.6e-05 | 1.8e-05 | 1.4e-05 | 1.600E-05 | 2.000E-06 | 1.400E-05 | 1.800E-05 |
| 9399 | 470 | A470Y | A | 1.3e-05 | 1.6e-05 | 1.2e-05 | 1.367E-05 | 2.082E-06 | 1.200E-05 | 1.600E-05 |
| 9400 | 470 | A470V | A | 2.2e-05 | 9.7e-06 | 1.5e-05 | 1.557E-05 | 6.170E-06 | 9.700E-06 | 2.200E-05 |
| 9401 | 471 | L471A | A | 9.7e-06 | 1.3e-05 | 1.3e-05 | 1.190E-05 | 1.905E-06 | 9.700E-06 | 1.300E-05 |
| 9402 | 471 | L471R | A | 1.4e-05 | 8e-06 | 1.5e-05 | 1.233E-05 | 3.786E-06 | 8.000E-06 | 1.500E-05 |
| 9403 | 471 | L471N | A | 9.7e-06 | 1e-05 | 9.9e-06 | 9.867E-06 | 1.528E-07 | 9.700E-06 | 1.000E-05 |
| 9404 | 471 | L471D | A | 1.5e-05 | 1.5e-05 | 1e-05 | 1.333E-05 | 2.887E-06 | 1.000E-05 | 1.500E-05 |
| 9405 | 471 | L471C | A | 9.1e-06 | 1.3e-05 | 1e-05 | 1.070E-05 | 2.042E-06 | 9.100E-06 | 1.300E-05 |
| 9406 | 471 | L471E | A | 1.4e-05 | 1.2e-05 | 1.5e-05 | 1.367E-05 | 1.528E-06 | 1.200E-05 | 1.500E-05 |
| 9407 | 471 | L471Q | A | 1.2e-05 | 1.4e-05 | 1.2e-05 | 1.267E-05 | 1.155E-06 | 1.200E-05 | 1.400E-05 |
| 9408 | 471 | L471G | A | 1.1e-05 | 1.3e-05 | 2.2e-05 | 1.533E-05 | 5.859E-06 | 1.100E-05 | 2.200E-05 |
| 9409 | 471 | L471H | A | 1.4e-05 | 1.5e-05 | 1.1e-05 | 1.333E-05 | 2.082E-06 | 1.100E-05 | 1.500E-05 |
| 9410 | 471 | L471I | A | 1.5e-05 | 1.4e-05 | 1.5e-05 | 1.467E-05 | 5.774E-07 | 1.400E-05 | 1.500E-05 |
| 9411 | 471 | L471L | A | 1.4e-05 | 1.3e-05 | 1.6e-05 | 1.433E-05 | 1.528E-06 | 1.300E-05 | 1.600E-05 |
| 9412 | 471 | L471K | A | 8.6e-06 | 1.5e-05 | 1.1e-05 | 1.153E-05 | 3.233E-06 | 8.600E-06 | 1.500E-05 |
| 9413 | 471 | L471M | A | 1.6e-05 | 8.8e-06 | 1.6e-05 | 1.360E-05 | 4.157E-06 | 8.800E-06 | 1.600E-05 |
| 9414 | 471 | L471F | A | 1.1e-05 | 1.3e-05 | 9.6e-06 | 1.120E-05 | 1.709E-06 | 9.600E-06 | 1.300E-05 |
| 9415 | 471 | L471P | A | 1.1e-05 | 7.9e-06 | 1.6e-05 | 1.163E-05 | 4.087E-06 | 7.900E-06 | 1.600E-05 |
| 9416 | 471 | L471S | A | 1.6e-05 | 1e-05 | 9.1e-06 | 1.170E-05 | 3.751E-06 | 9.100E-06 | 1.600E-05 |
| 9417 | 471 | L471T | A | 1.4e-05 | 8.4e-06 | 1.1e-05 | 1.113E-05 | 2.802E-06 | 8.400E-06 | 1.400E-05 |
| 9418 | 471 | L471W | A | 1.4e-05 | 1.5e-05 | 1.6e-05 | 1.500E-05 | 1.000E-06 | 1.400E-05 | 1.600E-05 |
| 9419 | 471 | L471Y | A | 9.9e-06 | 1e-05 | 1.3e-05 | 1.097E-05 | 1.762E-06 | 9.900E-06 | 1.300E-05 |
| 9420 | 471 | L471V | A | 1.7e-05 | 1.4e-05 | 1.3e-05 | 1.467E-05 | 2.082E-06 | 1.300E-05 | 1.700E-05 |
| 9421 | 472 | L472A | A | 1.4e-05 | 1e-05 | 1.1e-05 | 1.167E-05 | 2.082E-06 | 1.000E-05 | 1.400E-05 |
| 9422 | 472 | L472R | A | 7.3e-06 | 1.1e-05 | 1.5e-05 | 1.110E-05 | 3.851E-06 | 7.300E-06 | 1.500E-05 |
| 9423 | 472 | L472N | A | 1e-05 | 1.3e-05 | 1.3e-05 | 1.200E-05 | 1.732E-06 | 1.000E-05 | 1.300E-05 |
| 9424 | 472 | L472D | A | 1.4e-05 | 1.5e-05 | 9.4e-06 | 1.280E-05 | 2.987E-06 | 9.400E-06 | 1.500E-05 |
| 9425 | 472 | L472C | A | 1.3e-05 | 1.5e-05 | 8.6e-06 | 1.220E-05 | 3.274E-06 | 8.600E-06 | 1.500E-05 |
| 9426 | 472 | L472E | A | 1.5e-05 | 1.6e-05 | 1.6e-05 | 1.567E-05 | 5.774E-07 | 1.500E-05 | 1.600E-05 |
| 9427 | 472 | L472Q | A | 1.1e-05 | 1.5e-05 | 1.3e-05 | 1.300E-05 | 2.000E-06 | 1.100E-05 | 1.500E-05 |
| 9428 | 472 | L472G | A | 8.4e-06 | 1.2e-05 | 7.9e-06 | 9.433E-06 | 2.237E-06 | 7.900E-06 | 1.200E-05 |
| 9429 | 472 | L472H | A | 1.5e-05 | 9.9e-06 | 7.5e-06 | 1.080E-05 | 3.830E-06 | 7.500E-06 | 1.500E-05 |
| 9430 | 472 | L472I | A | 1.3e-05 | 1.2e-05 | 1.3e-05 | 1.267E-05 | 5.774E-07 | 1.200E-05 | 1.300E-05 |
| 9431 | 472 | L472L | A | 1e-05 | 1.1e-05 | 1.4e-05 | 1.167E-05 | 2.082E-06 | 1.000E-05 | 1.400E-05 |
| 9432 | 472 | L472K | A | 1.6e-05 | 1.2e-05 | 1e-05 | 1.267E-05 | 3.055E-06 | 1.000E-05 | 1.600E-05 |
| 9433 | 472 | L472M | A | 1.1e-05 | 1.6e-05 | 8.7e-06 | 1.190E-05 | 3.732E-06 | 8.700E-06 | 1.600E-05 |
| 9434 | 472 | L472F | A | 9.7e-06 | 1.1e-05 | 9.9e-06 | 1.020E-05 | 7.000E-07 | 9.700E-06 | 1.100E-05 |

| | | | | | | | | | | |
|---|---|---|---|---|---|---|---|---|---|---|
| 9435 | 472 | L472P | A | 1e-05 | 1.1e-05 | 1.7e-05 | 1.267E-05 | 3.786E-06 | 1.000E-05 | 1.700E-05 |
| 9436 | 472 | L472S | A | 1.4e-05 | 1.1e-05 | 1.6e-05 | 1.367E-05 | 2.517E-06 | 1.100E-05 | 1.600E-05 |
| 9437 | 472 | L472T | A | 1.5e-05 | 9.5e-06 | 1.6e-05 | 1.350E-05 | 3.500E-06 | 9.500E-06 | 1.600E-05 |
| 9438 | 472 | L472W | A | 1.1e-05 | 1.6e-05 | 1.5e-05 | 1.400E-05 | 2.646E-06 | 1.100E-05 | 1.600E-05 |
| 9439 | 472 | L472Y | A | 1.4e-05 | 1.4e-05 | 9.8e-06 | 1.260E-05 | 2.425E-06 | 9.800E-06 | 1.400E-05 |
| 9440 | 472 | L472V | A | 8.2e-06 | 1.3e-05 | 1.6e-05 | 1.240E-05 | 3.934E-06 | 8.200E-06 | 1.600E-05 |
| 9441 | 473 | H473A | A | 1.3e-05 | 1.5e-05 | 1e-05 | 1.267E-05 | 2.517E-06 | 1.000E-05 | 1.500E-05 |
| 9442 | 473 | H473R | A | 8.5e-06 | 9.5e-06 | 8.6e-06 | 8.867E-06 | 5.508E-07 | 8.500E-06 | 9.500E-06 |
| 9443 | 473 | H473N | A | 1.6e-05 | 1.6e-05 | 1.6e-05 | 1.600E-05 | 0.000E+00 | 1.600E-05 | 1.600E-05 |
| 9444 | 473 | H473D | A | 1e-05 | 1.1e-05 | 1.4e-05 | 1.167E-05 | 2.082E-06 | 1.000E-05 | 1.400E-05 |
| 9445 | 473 | H473C | A | 1e-05 | 1e-05 | 8.8e-06 | 9.600E-06 | 6.928E-07 | 8.800E-06 | 1.000E-05 |
| 9446 | 473 | H473E | A | 8.9e-06 | 1e-05 | 1.1e-05 | 9.967E-06 | 1.050E-06 | 8.900E-06 | 1.100E-05 |
| 9447 | 473 | H473Q | A | 1.5e-05 | 1.2e-05 | 1.2e-05 | 1.300E-05 | 1.732E-06 | 1.200E-05 | 1.500E-05 |
| 9448 | 473 | H473G | A | 1.1e-05 | 1.4e-05 | 1.1e-05 | 1.200E-05 | 1.732E-06 | 1.100E-05 | 1.400E-05 |
| 9449 | 473 | H473H | A | 9.5e-06 | 1.1e-05 | 1.1e-05 | 1.050E-05 | 8.660E-07 | 9.500E-06 | 1.100E-05 |
| 9450 | 473 | H473I | A | 9.3e-06 | 9.3e-06 | 9.5e-06 | 9.367E-06 | 1.155E-07 | 9.300E-06 | 9.500E-06 |
| 9451 | 473 | H473L | A | 1.4e-05 | 1.5e-05 | 1.5e-05 | 1.467E-05 | 5.774E-07 | 1.400E-05 | 1.500E-05 |
| 9452 | 473 | H473K | A | 9.3e-06 | 1e-05 | 1.1e-05 | 1.010E-05 | 8.544E-07 | 9.300E-06 | 1.100E-05 |
| 9453 | 473 | H473M | A | 1.6e-05 | 1.6e-05 | 1.1e-05 | 1.433E-05 | 2.887E-06 | 1.100E-05 | 1.600E-05 |
| 9454 | 473 | H473F | A | 9.8e-06 | 9.8e-06 | 8.1e-06 | 9.233E-06 | 9.815E-07 | 8.100E-06 | 9.800E-06 |
| 9455 | 473 | H473P | A | 1.3e-05 | 1.5e-05 | 1.4e-05 | 1.400E-05 | 1.000E-06 | 1.300E-05 | 1.500E-05 |
| 9456 | 473 | H473S | A | 1.6e-05 | 9.6e-06 | 1.3e-05 | 1.287E-05 | 3.202E-06 | 9.600E-06 | 1.600E-05 |
| 9457 | 473 | H473T | A | 1.4e-05 | 1.5e-05 | 1.4e-05 | 1.433E-05 | 5.774E-07 | 1.400E-05 | 1.500E-05 |
| 9458 | 473 | H473W | A | 9.4e-06 | 1e-05 | 1.2e-05 | 1.047E-05 | 1.361E-06 | 9.400E-06 | 1.200E-05 |
| 9459 | 473 | H473Y | A | 1.4e-05 | 3.5e-05 | 9.5e-06 | 1.950E-05 | 1.361E-05 | 9.500E-06 | 3.500E-05 |
| 9460 | 473 | H473V | A | 1.3e-05 | 1.3e-05 | 1.5e-05 | 1.367E-05 | 1.155E-06 | 1.300E-05 | 1.500E-05 |
| 9461 | 474 | T474A | A | 1.5e-05 | 1.1e-05 | 9.5e-06 | 1.183E-05 | 2.843E-06 | 9.500E-06 | 1.500E-05 |
| 9462 | 474 | T474R | A | 1.5e-05 | 1.6e-05 | 1.4e-05 | 1.500E-05 | 1.000E-06 | 1.400E-05 | 1.600E-05 |
| 9463 | 474 | T474N | A | 1.5e-05 | 8.6e-06 | 1.8e-05 | 1.387E-05 | 4.801E-06 | 8.600E-06 | 1.800E-05 |
| 9464 | 474 | T474D | A | 1e-05 | 1.6e-05 | 1.5e-05 | 1.367E-05 | 3.215E-06 | 1.000E-05 | 1.600E-05 |
| 9465 | 474 | T474C | A | 1.1e-05 | 8.6e-06 | 7.3e-06 | 8.967E-06 | 1.877E-06 | 7.300E-06 | 1.100E-05 |
| 9466 | 474 | T474E | A | 1.4e-05 | 1.6e-05 | 1.2e-05 | 1.400E-05 | 2.000E-06 | 1.200E-05 | 1.600E-05 |
| 9467 | 474 | T474Q | A | 1.4e-05 | 1.4e-05 | 1.1e-05 | 1.300E-05 | 1.732E-06 | 1.100E-05 | 1.400E-05 |
| 9468 | 474 | T474G | A | 1.8e-05 | 1.4e-05 | 1.5e-05 | 1.567E-05 | 2.082E-06 | 1.400E-05 | 1.800E-05 |
| 9469 | 474 | T474H | A | 1e-05 | 1.6e-05 | 1.1e-05 | 1.233E-05 | 3.215E-06 | 1.000E-05 | 1.600E-05 |
| 9470 | 474 | T474I | A | 1.4e-05 | 1.1e-05 | 1.3e-05 | 1.267E-05 | 1.528E-06 | 1.100E-05 | 1.400E-05 |
| 9471 | 474 | T474L | A | 1.5e-05 | 1.5e-05 | 1.5e-05 | 1.500E-05 | 0.000E+00 | 1.500E-05 | 1.500E-05 |
| 9472 | 474 | T474K | A | 1.6e-05 | 8.2e-06 | 1.4e-05 | 1.273E-05 | 4.051E-06 | 8.200E-06 | 1.600E-05 |
| 9473 | 474 | T474M | A | 9.5e-06 | 1.1e-05 | 8.1e-06 | 9.533E-06 | 1.450E-06 | 8.100E-06 | 1.100E-05 |
| 9474 | 474 | T474F | A | 1.4e-05 | 1.7e-05 | 1.1e-05 | 1.400E-05 | 3.000E-06 | 1.100E-05 | 1.700E-05 |
| 9475 | 474 | T474P | A | 1e-05 | 1.4e-05 | 1.5e-05 | 1.300E-05 | 2.646E-06 | 1.000E-05 | 1.500E-05 |
| 9476 | 474 | T474S | A | 9.8e-06 | 1.3e-05 | 1.2e-05 | 1.160E-05 | 1.637E-06 | 9.800E-06 | 1.300E-05 |
| 9477 | 474 | T474T | A | 1.1e-05 | 1.7e-05 | 1.3e-05 | 1.367E-05 | 3.055E-06 | 1.100E-05 | 1.700E-05 |
| 9478 | 474 | T474W | A | 1.3e-05 | 1.3e-05 | 1e-05 | 1.200E-05 | 1.732E-06 | 1.000E-05 | 1.300E-05 |
| 9479 | 474 | T474Y | A | 6.8e-06 | 1e-05 | 1.5e-05 | 1.060E-05 | 4.133E-06 | 6.800E-06 | 1.500E-05 |
| 9480 | 474 | T474V | A | 1.5e-05 | 1.1e-05 | 1.5e-05 | 1.367E-05 | 2.309E-06 | 1.100E-05 | 1.500E-05 |
| 9481 | 475 | A475A | A | 1.5e-05 | 1.1e-05 | 1.7e-05 | 1.433E-05 | 3.055E-06 | 1.100E-05 | 1.700E-05 |
| 9482 | 475 | A475R | A | 1.1e-05 | 1.4e-05 | 1.4e-05 | 1.300E-05 | 1.732E-06 | 1.100E-05 | 1.400E-05 |
| 9483 | 475 | A475N | A | 1.5e-05 | 1e-05 | 8.9e-06 | 1.130E-05 | 3.251E-06 | 8.900E-06 | 1.500E-05 |
| 9484 | 475 | A475D | A | 2.1e-05 | 1.4e-05 | 1.6e-05 | 1.700E-05 | 3.606E-06 | 1.400E-05 | 2.100E-05 |
| 9485 | 475 | A475C | A | 1e-05 | 1e-05 | 1.1e-05 | 1.033E-05 | 5.774E-07 | 1.000E-05 | 1.100E-05 |
| 9486 | 475 | A475E | A | 9.9e-06 | 1.5e-05 | 1.5e-05 | 1.330E-05 | 2.944E-06 | 9.900E-06 | 1.500E-05 |
| 9487 | 475 | A475Q | A | 9.8e-06 | 1.1e-05 | 1.2e-05 | 1.093E-05 | 1.102E-06 | 9.800E-06 | 1.200E-05 |
| 9488 | 475 | A475G | A | 1.1e-05 | 1.6e-05 | 9.2e-06 | 1.207E-05 | 3.523E-06 | 9.200E-06 | 1.600E-05 |
| 9489 | 475 | A475H | A | 1.3e-05 | 1.1e-05 | 1.1e-05 | 1.167E-05 | 1.155E-06 | 1.100E-05 | 1.300E-05 |
| 9490 | 475 | A475I | A | 1.6e-05 | 1.3e-05 | 1.6e-05 | 1.500E-05 | 1.732E-06 | 1.300E-05 | 1.600E-05 |
| 9491 | 475 | A475L | A | 1.6e-05 | 1.4e-05 | 1.4e-05 | 1.467E-05 | 1.155E-06 | 1.400E-05 | 1.600E-05 |
| 9492 | 475 | A475K | A | 1.6e-05 | 1.5e-05 | 9.8e-06 | 1.360E-05 | 3.329E-06 | 9.800E-06 | 1.600E-05 |
| 9493 | 475 | A475M | A | 1.4e-05 | 1.1e-05 | 9.6e-06 | 1.153E-05 | 2.248E-06 | 9.600E-06 | 1.400E-05 |
| 9494 | 475 | A475F | A | 8.7e-06 | 1.5e-05 | 1.6e-05 | 1.323E-05 | 3.958E-06 | 8.700E-06 | 1.600E-05 |
| 9495 | 475 | A475P | A | 1.3e-05 | 1.4e-05 | 1.7e-05 | 1.467E-05 | 2.082E-06 | 1.300E-05 | 1.700E-05 |
| 9496 | 475 | A475S | A | 9e-06 | 1.5e-05 | 1.1e-05 | 1.167E-05 | 3.055E-06 | 9.000E-06 | 1.500E-05 |
| 9497 | 475 | A475T | A | 1.5e-05 | 1.5e-05 | 1.5e-05 | 1.500E-05 | 0.000E+00 | 1.500E-05 | 1.500E-05 |
| 9498 | 475 | A475W | A | 1e-05 | 1.5e-05 | 1.4e-05 | 1.300E-05 | 2.646E-06 | 1.000E-05 | 1.500E-05 |
| 9499 | 475 | A475Y | A | 1.6e-05 | 1.1e-05 | 1e-05 | 1.233E-05 | 3.215E-06 | 1.000E-05 | 1.600E-05 |
| 9500 | 475 | A475V | A | 1e-05 | 1.5e-05 | 1.7e-05 | 1.400E-05 | 3.606E-06 | 1.000E-05 | 1.700E-05 |
| 9501 | 476 | N476A | A | 1.4e-05 | 1.8e-05 | 1e-05 | 1.400E-05 | 4.000E-06 | 1.000E-05 | 1.800E-05 |
| 9502 | 476 | N476R | A | 1.5e-05 | 1.1e-05 | 9.5e-06 | 1.183E-05 | 2.843E-06 | 9.500E-06 | 1.500E-05 |
| 9503 | 476 | N476N | A | 1.5e-05 | 1.4e-05 | 1.4e-05 | 1.433E-05 | 5.774E-07 | 1.400E-05 | 1.500E-05 |
| 9504 | 476 | N476D | A | 1.7e-05 | 1.8e-05 | 1.3e-05 | 1.600E-05 | 2.646E-06 | 1.300E-05 | 1.800E-05 |
| 9505 | 476 | N476C | A | 8.8e-06 | 1.4e-05 | 1.6e-05 | 1.293E-05 | 3.717E-06 | 8.800E-06 | 1.600E-05 |
| 9506 | 476 | N476E | A | 1.2e-05 | 9.1e-06 | 1.1e-05 | 1.070E-05 | 1.473E-06 | 9.100E-06 | 1.200E-05 |
| 9507 | 476 | N476Q | A | 1.7e-05 | 1.4e-05 | 1.1e-05 | 1.400E-05 | 3.000E-06 | 1.100E-05 | 1.700E-05 |
| 9508 | 476 | N476G | A | 1.1e-05 | 1.1e-05 | 1e-05 | 1.067E-05 | 5.774E-07 | 1.000E-05 | 1.100E-05 |
| 9509 | 476 | N476H | A | 1.4e-05 | 1.3e-05 | 6.7e-06 | 1.123E-05 | 3.958E-06 | 6.700E-06 | 1.400E-05 |
| 9510 | 476 | N476I | A | 1.4e-05 | 1.1e-05 | 1.3e-05 | 1.267E-05 | 1.528E-06 | 1.100E-05 | 1.400E-05 |
| 9511 | 476 | N476L | A | 1.6e-05 | 8.8e-06 | 1.1e-05 | 1.193E-05 | 3.690E-06 | 8.800E-06 | 1.600E-05 |
| 9512 | 476 | N476K | A | 1.1e-05 | 1.6e-05 | 1.7e-05 | 1.467E-05 | 3.215E-06 | 1.100E-05 | 1.700E-05 |
| 9513 | 476 | N476M | A | 1.5e-05 | 1.3e-05 | 1.7e-05 | 1.500E-05 | 2.000E-06 | 1.300E-05 | 1.700E-05 |
| 9514 | 476 | N476F | A | 1.8e-05 | 1.2e-05 | 1.6e-05 | 1.533E-05 | 3.055E-06 | 1.200E-05 | 1.800E-05 |
| 9515 | 476 | N476P | A | 9.7e-06 | 1.1e-05 | 1.5e-05 | 1.190E-05 | 2.762E-06 | 9.700E-06 | 1.500E-05 |
| 9516 | 476 | N476S | A | 1.5e-05 | 1e-05 | 1.3e-05 | 1.267E-05 | 2.517E-06 | 1.000E-05 | 1.500E-05 |
| 9517 | 476 | N476T | A | 1.6e-05 | 1.2e-05 | 1.4e-05 | 1.400E-05 | 2.000E-06 | 1.200E-05 | 1.600E-05 |
| 9518 | 476 | N476W | A | 1.3e-05 | 1.4e-05 | 1.7e-05 | 1.467E-05 | 2.082E-06 | 1.300E-05 | 1.700E-05 |
| 9519 | 476 | N476Y | A | 1.2e-05 | 1.1e-05 | 9.9e-06 | 1.097E-05 | 1.050E-06 | 9.900E-06 | 1.200E-05 |

| | | | | | | | | | | |
|---|---|---|---|---|---|---|---|---|---|---|
| 9520 | 476 | N476V | A | 1.5e-05 | 1.7e-05 | 3e-05 | 2.067E-05 | 8.145E-06 | 1.500E-05 | 3.000E-05 |
| 9521 | 477 | R477A | A | 1.5e-05 | 1.3e-05 | 1.5e-05 | 1.433E-05 | 1.155E-06 | 1.300E-05 | 1.500E-05 |
| 9522 | 477 | R477R | A | 1.3e-05 | 9e-06 | 1.7e-05 | 1.300E-05 | 4.000E-06 | 9.000E-06 | 1.700E-05 |
| 9523 | 477 | R477N | A | 1e-05 | 1.5e-05 | 1.3e-05 | 1.267E-05 | 2.517E-06 | 1.000E-05 | 1.500E-05 |
| 9524 | 477 | R477D | A | 1.5e-05 | 1.4e-05 | 1.1e-05 | 1.333E-05 | 2.082E-06 | 1.100E-05 | 1.500E-05 |
| 9525 | 477 | R477C | A | 1.4e-05 | 1.6e-05 | 1.5e-05 | 1.500E-05 | 1.000E-06 | 1.400E-05 | 1.600E-05 |
| 9526 | 477 | R477E | A | 1.5e-05 | 1.6e-05 | 9.4e-06 | 1.347E-05 | 3.557E-06 | 9.400E-06 | 1.600E-05 |
| 9527 | 477 | R477Q | A | 1.5e-05 | 1.1e-05 | 8.2e-06 | 1.140E-05 | 3.418E-06 | 8.200E-06 | 1.500E-05 |
| 9528 | 477 | R477G | A | 1.5e-05 | 1.7e-05 | 1.5e-05 | 1.567E-05 | 1.155E-06 | 1.500E-05 | 1.700E-05 |
| 9529 | 477 | R477H | A | 9.8e-06 | 1.2e-05 | 9.5e-06 | 1.043E-05 | 1.365E-06 | 9.500E-06 | 1.200E-05 |
| 9530 | 477 | R477I | A | 1.3e-05 | 1.7e-05 | 1.6e-05 | 1.533E-05 | 2.082E-06 | 1.300E-05 | 1.700E-05 |
| 9531 | 477 | R477L | A | 1.5e-05 | 1.6e-05 | 1.3e-05 | 1.467E-05 | 1.528E-06 | 1.300E-05 | 1.600E-05 |
| 9532 | 477 | R477K | A | 1.4e-05 | 1.5e-05 | 1.3e-05 | 1.400E-05 | 1.000E-06 | 1.300E-05 | 1.500E-05 |
| 9533 | 477 | R477M | A | 1.5e-05 | 1.5e-05 | 1.3e-05 | 1.433E-05 | 1.155E-06 | 1.300E-05 | 1.500E-05 |
| 9534 | 477 | R477F | A | 9.4e-06 | 9e-06 | 1.2e-05 | 1.013E-05 | 1.629E-06 | 9.000E-06 | 1.200E-05 |
| 9535 | 477 | R477P | A | 1.5e-05 | 9.1e-06 | 1.1e-05 | 1.170E-05 | 3.012E-06 | 9.100E-06 | 1.500E-05 |
| 9536 | 477 | R477S | A | 1.5e-05 | 9.3e-06 | 9.9e-06 | 1.140E-05 | 3.132E-06 | 9.300E-06 | 1.500E-05 |
| 9537 | 477 | R477T | A | 1.7e-05 | 1.5e-05 | 1.4e-05 | 1.533E-05 | 1.528E-06 | 1.400E-05 | 1.700E-05 |
| 9538 | 477 | R477W | A | 9.4e-06 | 1.3e-05 | 9.4e-06 | 1.060E-05 | 2.078E-06 | 9.400E-06 | 1.300E-05 |
| 9539 | 477 | R477Y | A | 1.6e-05 | 1.4e-05 | 9.1e-06 | 1.303E-05 | 3.550E-06 | 9.100E-06 | 1.600E-05 |
| 9540 | 477 | R477V | A | 1.5e-05 | 1.6e-05 | 1.5e-05 | 1.533E-05 | 5.774E-07 | 1.500E-05 | 1.600E-05 |
| 9541 | 478 | P478A | A | 1.4e-05 | 1.5e-05 | 1e-05 | 1.300E-05 | 2.646E-06 | 1.000E-05 | 1.500E-05 |
| 9542 | 478 | P478R | A | 1.5e-05 | 1.3e-05 | 1.1e-05 | 1.300E-05 | 2.000E-06 | 1.100E-05 | 1.500E-05 |
| 9543 | 478 | P478N | A | 9e-06 | 9.7e-06 | 1.6e-05 | 1.157E-05 | 3.855E-06 | 9.000E-06 | 1.600E-05 |
| 9544 | 478 | P478D | A | 1.5e-05 | 1.4e-05 | 1.4e-05 | 1.433E-05 | 5.774E-07 | 1.400E-05 | 1.500E-05 |
| 9545 | 478 | P478C | A | 8.3e-06 | 1.4e-05 | 9e-06 | 1.043E-05 | 3.109E-06 | 8.300E-06 | 1.400E-05 |
| 9546 | 478 | P478E | A | 1.6e-05 | 9.1e-06 | 1.6e-05 | 1.370E-05 | 3.984E-06 | 9.100E-06 | 1.600E-05 |
| 9547 | 478 | P478Q | A | 1.1e-05 | 8e-06 | 9.5e-06 | 9.500E-06 | 1.500E-06 | 8.000E-06 | 1.100E-05 |
| 9548 | 478 | P478G | A | 1.2e-05 | 1.4e-05 | 1.9e-05 | 1.500E-05 | 3.606E-06 | 1.200E-05 | 1.900E-05 |
| 9549 | 478 | P478H | A | 1e-05 | 1.3e-05 | 8.3e-06 | 1.043E-05 | 2.380E-06 | 8.300E-06 | 1.300E-05 |
| 9550 | 478 | P478I | A | 1e-05 | 1.3e-05 | 1.4e-05 | 1.233E-05 | 2.082E-06 | 1.000E-05 | 1.400E-05 |
| 9551 | 478 | P478L | A | 1.2e-05 | 1.1e-05 | 1.4e-05 | 1.233E-05 | 1.528E-06 | 1.100E-05 | 1.400E-05 |
| 9552 | 478 | P478K | A | 1.3e-05 | 9.7e-06 | 9.5e-06 | 1.073E-05 | 1.966E-06 | 9.500E-06 | 1.300E-05 |
| 9553 | 478 | P478M | A | 1.6e-05 | 1.4e-05 | 8.7e-06 | 1.290E-05 | 3.772E-06 | 8.700E-06 | 1.600E-05 |
| 9554 | 478 | P478F | A | 1.1e-05 | 1.5e-05 | 1.5e-05 | 1.367E-05 | 2.309E-06 | 1.100E-05 | 1.500E-05 |
| 9555 | 478 | P478P | A | 1.2e-05 | 1.2e-05 | 1.4e-05 | 1.267E-05 | 1.155E-06 | 1.200E-05 | 1.400E-05 |
| 9556 | 478 | P478S | A | 1.4e-05 | 1.4e-05 | 1e-05 | 1.267E-05 | 2.309E-06 | 1.000E-05 | 1.400E-05 |
| 9557 | 478 | P478T | A | 1.5e-05 | 1.1e-05 | 1.2e-05 | 1.267E-05 | 2.082E-06 | 1.100E-05 | 1.500E-05 |
| 9558 | 478 | P478W | A | 1.2e-05 | 1.1e-05 | 1.6e-05 | 1.300E-05 | 2.646E-06 | 1.100E-05 | 1.600E-05 |
| 9559 | 478 | P478Y | A | 1e-05 | 1.5e-05 | 1e-05 | 1.167E-05 | 2.887E-06 | 1.000E-05 | 1.500E-05 |
| 9560 | 478 | P478V | A | 1.5e-05 | 9.2e-06 | 1.6e-05 | 1.340E-05 | 3.672E-06 | 9.200E-06 | 1.600E-05 |
| 9561 | 479 | E479A | A | 1.6e-05 | 9.4e-06 | 9.1e-06 | 1.150E-05 | 3.900E-06 | 9.100E-06 | 1.600E-05 |
| 9562 | 479 | E479R | A | 1.4e-05 | 1.4e-05 | 1e-05 | 1.267E-05 | 2.309E-06 | 1.000E-05 | 1.400E-05 |
| 9563 | 479 | E479N | A | 1.3e-05 | 1e-05 | 9.3e-06 | 1.077E-05 | 1.966E-06 | 9.300E-06 | 1.300E-05 |
| 9564 | 479 | E479D | A | 9.6e-06 | 9.4e-06 | 1.1e-05 | 1.000E-05 | 8.718E-07 | 9.400E-06 | 1.100E-05 |
| 9565 | 479 | E479C | A | 1.4e-05 | 9.1e-06 | 1.2e-05 | 1.170E-05 | 2.464E-06 | 9.100E-06 | 1.400E-05 |
| 9566 | 479 | E479E | A | 1e-05 | 1.5e-05 | 1.3e-05 | 1.267E-05 | 2.517E-06 | 1.000E-05 | 1.500E-05 |
| 9567 | 479 | E479Q | A | 1.2e-05 | 1e-05 | 1.5e-05 | 1.233E-05 | 2.517E-06 | 1.000E-05 | 1.500E-05 |
| 9568 | 479 | E479G | A | 1.6e-05 | 1.6e-05 | 1.6e-05 | 1.600E-05 | 0.000E+00 | 1.600E-05 | 1.600E-05 |
| 9569 | 479 | E479H | A | 1.6e-05 | 1.1e-05 | 9e-06 | 1.200E-05 | 3.606E-06 | 9.000E-06 | 1.600E-05 |
| 9570 | 479 | E479I | A | 1.2e-05 | 1.1e-05 | 6.1e-06 | 9.700E-06 | 3.158E-06 | 6.100E-06 | 1.200E-05 |
| 9571 | 479 | E479L | A | 1.1e-05 | 9.9e-06 | 1.5e-05 | 1.197E-05 | 2.684E-06 | 9.900E-06 | 1.500E-05 |
| 9572 | 479 | E479K | A | 1.3e-05 | 1.1e-05 | 1.6e-05 | 1.333E-05 | 2.517E-06 | 1.100E-05 | 1.600E-05 |
| 9573 | 479 | E479M | A | 1.3e-05 | 1e-05 | 1.3e-05 | 1.200E-05 | 1.732E-06 | 1.000E-05 | 1.300E-05 |
| 9574 | 479 | E479F | A | 7.2e-06 | 1.3e-05 | 1e-05 | 1.007E-05 | 2.901E-06 | 7.200E-06 | 1.300E-05 |
| 9575 | 479 | E479P | A | 1.6e-05 | 8.5e-06 | 1.1e-05 | 1.183E-05 | 3.819E-06 | 8.500E-06 | 1.600E-05 |
| 9576 | 479 | E479S | A | 1.4e-05 | 1e-05 | 1.4e-05 | 1.267E-05 | 2.309E-06 | 1.000E-05 | 1.400E-05 |
| 9577 | 479 | E479T | A | 1e-05 | 1.4e-05 | 1e-05 | 1.133E-05 | 2.309E-06 | 1.000E-05 | 1.400E-05 |
| 9578 | 479 | E479W | A | 1.4e-05 | 1.5e-05 | 1.5e-05 | 1.467E-05 | 5.774E-07 | 1.400E-05 | 1.500E-05 |
| 9579 | 479 | E479Y | A | 1.1e-05 | 1.4e-05 | 1.5e-05 | 1.333E-05 | 2.082E-06 | 1.100E-05 | 1.500E-05 |
| 9580 | 479 | E479V | A | 1.1e-05 | 1.5e-05 | 1.6e-05 | 1.400E-05 | 2.646E-06 | 1.100E-05 | 1.600E-05 |
| 9581 | 480 | D480A | A | 1.1e-05 | 1e-05 | 1.4e-05 | 1.167E-05 | 2.082E-06 | 1.000E-05 | 1.400E-05 |
| 9582 | 480 | D480R | A | 9.1e-06 | 1.2e-05 | 1.2e-05 | 1.103E-05 | 1.674E-06 | 9.100E-06 | 1.200E-05 |
| 9583 | 480 | D480N | A | 1.6e-05 | 1.3e-05 | 1.6e-05 | 1.500E-05 | 1.732E-06 | 1.300E-05 | 1.600E-05 |
| 9584 | 480 | D480D | A | 1.1e-05 | 1.3e-05 | 1.6e-05 | 1.333E-05 | 2.517E-06 | 1.100E-05 | 1.600E-05 |
| 9585 | 480 | D480C | A | 1.4e-05 | 9.7e-06 | 1.1e-05 | 1.157E-05 | 2.205E-06 | 9.700E-06 | 1.400E-05 |
| 9586 | 480 | D480E | A | 1.6e-05 | 1e-05 | 1.1e-05 | 1.233E-05 | 3.215E-06 | 1.000E-05 | 1.600E-05 |
| 9587 | 480 | D480Q | A | 9.3e-06 | 1.5e-05 | 1.3e-05 | 1.243E-05 | 2.892E-06 | 9.300E-06 | 1.500E-05 |
| 9588 | 480 | D480G | A | 8.9e-06 | 1.3e-05 | 1.1e-05 | 1.097E-05 | 2.050E-06 | 8.900E-06 | 1.300E-05 |
| 9589 | 480 | D480H | A | 1.5e-05 | 1.5e-05 | 1.3e-05 | 1.433E-05 | 1.155E-06 | 1.300E-05 | 1.500E-05 |
| 9590 | 480 | D480I | A | 1.3e-05 | 1.1e-05 | 1.4e-05 | 1.267E-05 | 1.528E-06 | 1.100E-05 | 1.400E-05 |
| 9591 | 480 | D480L | A | 9.1e-06 | 1.5e-05 | 1e-05 | 1.137E-05 | 3.179E-06 | 9.100E-06 | 1.500E-05 |
| 9592 | 480 | D480K | A | 9.9e-06 | 1.2e-05 | 1.4e-05 | 1.197E-05 | 2.050E-06 | 9.900E-06 | 1.400E-05 |
| 9593 | 480 | D480M | A | 1.6e-05 | 1.4e-05 | 1.4e-05 | 1.467E-05 | 1.155E-06 | 1.400E-05 | 1.600E-05 |
| 9594 | 480 | D480F | A | 1.5e-05 | 1e-05 | 1.3e-05 | 1.267E-05 | 2.517E-06 | 1.000E-05 | 1.500E-05 |
| 9595 | 480 | D480P | A | 1.5e-05 | 1.5e-05 | 1.5e-05 | 1.500E-05 | 0.000E+00 | 1.500E-05 | 1.500E-05 |
| 9596 | 480 | D480S | A | 8.1e-06 | 1.4e-05 | 1.5e-05 | 1.237E-05 | 3.729E-06 | 8.100E-06 | 1.500E-05 |
| 9597 | 480 | D480T | A | 1e-05 | 1.2e-05 | 1.5e-05 | 1.233E-05 | 2.517E-06 | 1.000E-05 | 1.500E-05 |
| 9598 | 480 | D480W | A | 1e-05 | 1e-05 | 1.5e-05 | 1.167E-05 | 2.887E-06 | 1.000E-05 | 1.500E-05 |
| 9599 | 480 | D480Y | A | 1.5e-05 | 1.6e-05 | 1.6e-05 | 1.567E-05 | 5.774E-07 | 1.500E-05 | 1.600E-05 |
| 9600 | 480 | D480V | A | 6.8e-06 | 1.6e-05 | 1.4e-05 | 1.227E-05 | 4.839E-06 | 6.800E-06 | 1.600E-05 |
| 9601 | 481 | E481A | A | 1e-05 | 1.2e-05 | 1.7e-05 | 1.300E-05 | 3.606E-06 | 1.000E-05 | 1.700E-05 |
| 9602 | 481 | E481R | A | 1.3e-05 | 1.6e-05 | 1.3e-05 | 1.400E-05 | 1.732E-06 | 1.300E-05 | 1.600E-05 |
| 9603 | 481 | E481N | A | 1.7e-05 | 1.6e-05 | 1.6e-05 | 1.633E-05 | 5.774E-07 | 1.600E-05 | 1.700E-05 |
| 9604 | 481 | E481D | A | 1.1e-05 | 1.4e-05 | 9.9e-06 | 1.163E-05 | 2.122E-06 | 9.900E-06 | 1.400E-05 |

| | | | | | | | | | | |
|---|---|---|---|---|---|---|---|---|---|---|
| 9605 | 481 | E481C | A | 1.5e-05 | 1.6e-05 | 1.1e-05 | 1.400E-05 | 2.646E-06 | 1.100E-05 | 1.600E-05 |
| 9606 | 481 | E481E | A | 1.5e-05 | 1.4e-05 | 1.4e-05 | 1.433E-05 | 5.774E-07 | 1.400E-05 | 1.500E-05 |
| 9607 | 481 | E481Q | A | 1.1e-05 | 1.5e-05 | 1.1e-05 | 1.233E-05 | 2.309E-06 | 1.100E-05 | 1.500E-05 |
| 9608 | 481 | E481G | A | 1.1e-05 | 1.6e-05 | 1.2e-05 | 1.300E-05 | 2.646E-06 | 1.100E-05 | 1.600E-05 |
| 9609 | 481 | E481H | A | 1.2e-05 | 9.7e-06 | 1.4e-05 | 1.190E-05 | 2.152E-06 | 9.700E-06 | 1.400E-05 |
| 9610 | 481 | E481I | A | 9.7e-06 | 1.5e-05 | 1.5e-05 | 1.323E-05 | 3.060E-06 | 9.700E-06 | 1.500E-05 |
| 9611 | 481 | E481L | A | 9.4e-06 | 1.1e-05 | 1.3e-05 | 1.113E-05 | 1.804E-06 | 9.400E-06 | 1.300E-05 |
| 9612 | 481 | E481K | A | 9.4e-06 | 9.8e-06 | 1.5e-05 | 1.140E-05 | 3.124E-06 | 9.400E-06 | 1.500E-05 |
| 9613 | 481 | E481M | A | 1.1e-05 | 1.3e-05 | 1.2e-05 | 1.200E-05 | 1.000E-06 | 1.100E-05 | 1.300E-05 |
| 9614 | 481 | E481F | A | 1.2e-05 | 1.5e-05 | 1e-05 | 1.233E-05 | 2.517E-06 | 1.000E-05 | 1.500E-05 |
| 9615 | 481 | E481P | A | 1.5e-05 | 1.8e-05 | 1.3e-05 | 1.533E-05 | 2.517E-06 | 1.300E-05 | 1.800E-05 |
| 9616 | 481 | E481S | A | 8.4e-06 | 1.5e-05 | 1.1e-05 | 1.147E-05 | 3.325E-06 | 8.400E-06 | 1.500E-05 |
| 9617 | 481 | E481T | A | 1.4e-05 | 1.1e-05 | 1.6e-05 | 1.367E-05 | 2.517E-06 | 1.100E-05 | 1.600E-05 |
| 9618 | 481 | E481W | A | 1.6e-05 | 1.6e-05 | 1.1e-05 | 1.433E-05 | 2.887E-06 | 1.100E-05 | 1.600E-05 |
| 9619 | 481 | E481Y | A | 1.6e-05 | 1.3e-05 | 1e-05 | 1.300E-05 | 3.000E-06 | 1.000E-05 | 1.600E-05 |
| 9620 | 481 | E481V | A | 1.4e-05 | 1.5e-05 | 1.2e-05 | 1.367E-05 | 1.528E-06 | 1.200E-05 | 1.500E-05 |
| 9621 | 482 | C482A | A | 1.5e-05 | 1.3e-05 | 1.5e-05 | 1.433E-05 | 1.155E-06 | 1.300E-05 | 1.500E-05 |
| 9622 | 482 | C482R | A | 1.4e-05 | 1.4e-05 | 1.6e-05 | 1.467E-05 | 1.155E-06 | 1.400E-05 | 1.600E-05 |
| 9623 | 482 | C482N | A | 1.4e-05 | 1.6e-05 | 7e-06 | 1.233E-05 | 4.726E-06 | 7.000E-06 | 1.600E-05 |
| 9624 | 482 | C482D | A | 1.3e-05 | 9.9e-06 | 1.3e-05 | 1.197E-05 | 1.790E-06 | 9.900E-06 | 1.300E-05 |
| 9625 | 482 | C482C | A | 1.3e-05 | 1.6e-05 | 9.5e-06 | 1.283E-05 | 3.253E-06 | 9.500E-06 | 1.600E-05 |
| 9626 | 482 | C482E | A | 1.1e-05 | 1.4e-05 | 9.6e-06 | 1.153E-05 | 2.248E-06 | 9.600E-06 | 1.400E-05 |
| 9627 | 482 | C482Q | A | 1.1e-05 | 9.4e-06 | 1.4e-05 | 1.147E-05 | 2.335E-06 | 9.400E-06 | 1.400E-05 |
| 9628 | 482 | C482G | A | 1.5e-05 | 1.3e-05 | 1.3e-05 | 1.367E-05 | 1.155E-06 | 1.300E-05 | 1.500E-05 |
| 9629 | 482 | C482H | A | 1.1e-05 | 1.5e-05 | 1.1e-05 | 1.233E-05 | 2.309E-06 | 1.100E-05 | 1.500E-05 |
| 9630 | 482 | C482I | A | 1.5e-05 | 1e-05 | 1.7e-05 | 1.400E-05 | 3.606E-06 | 1.000E-05 | 1.700E-05 |
| 9631 | 482 | C482L | A | 1.5e-05 | 1.5e-05 | 9.5e-06 | 1.317E-05 | 3.175E-06 | 9.500E-06 | 1.500E-05 |
| 9632 | 482 | C482K | A | 1.8e-05 | 1.7e-05 | 1.3e-05 | 1.600E-05 | 2.646E-06 | 1.300E-05 | 1.800E-05 |
| 9633 | 482 | C482M | A | 2e-05 | 1.3e-05 | 8.8e-06 | 1.393E-05 | 5.658E-06 | 8.800E-06 | 2.000E-05 |
| 9634 | 482 | C482F | A | 1.1e-05 | 8.9e-06 | 1.9e-05 | 1.297E-05 | 5.329E-06 | 8.900E-06 | 1.900E-05 |
| 9635 | 482 | C482P | A | 1.6e-05 | 1.4e-05 | 1.5e-05 | 1.500E-05 | 1.000E-06 | 1.400E-05 | 1.600E-05 |
| 9636 | 482 | C482S | A | 1.2e-05 | 1.3e-05 | 1.5e-05 | 1.333E-05 | 1.528E-06 | 1.200E-05 | 1.500E-05 |
| 9637 | 482 | C482T | A | 1.1e-05 | 1.6e-05 | 1e-05 | 1.233E-05 | 3.215E-06 | 1.000E-05 | 1.600E-05 |
| 9638 | 482 | C482W | A | 1.4e-05 | 1.7e-05 | 1.5e-05 | 1.533E-05 | 1.528E-06 | 1.400E-05 | 1.700E-05 |
| 9639 | 482 | C482Y | A | 1.1e-05 | 9.1e-06 | 1.4e-05 | 1.137E-05 | 2.470E-06 | 9.100E-06 | 1.400E-05 |
| 9640 | 482 | C482V | A | 8.4e-06 | 1.1e-05 | 1.2e-05 | 1.047E-05 | 1.858E-06 | 8.400E-06 | 1.200E-05 |
| 9641 | 483 | V483A | A | 9.6e-06 | 1.7e-05 | 1.5e-05 | 1.387E-05 | 3.828E-06 | 9.600E-06 | 1.700E-05 |
| 9642 | 483 | V483R | A | 1.6e-05 | 1.5e-05 | 1.6e-05 | 1.567E-05 | 5.774E-07 | 1.500E-05 | 1.600E-05 |
| 9643 | 483 | V483N | A | 1.4e-05 | 1.1e-05 | 9.8e-06 | 1.160E-05 | 2.163E-06 | 9.800E-06 | 1.400E-05 |
| 9644 | 483 | V483D | A | 1e-05 | 1.3e-05 | 1.2e-05 | 1.167E-05 | 1.528E-06 | 1.000E-05 | 1.300E-05 |
| 9645 | 483 | V483C | A | 1.1e-05 | 9.1e-06 | 1e-05 | 1.003E-05 | 9.504E-07 | 9.100E-06 | 1.100E-05 |
| 9646 | 483 | V483E | A | 1.5e-05 | 1.5e-05 | 1.7e-05 | 1.567E-05 | 1.155E-06 | 1.500E-05 | 1.700E-05 |
| 9647 | 483 | V483Q | A | 1.5e-05 | 1.6e-05 | 1e-05 | 1.367E-05 | 3.215E-06 | 1.000E-05 | 1.600E-05 |
| 9648 | 483 | V483G | A | 1.5e-05 | 1.6e-05 | 1.7e-05 | 1.600E-05 | 1.000E-06 | 1.500E-05 | 1.700E-05 |
| 9649 | 483 | V483H | A | 1.1e-05 | 1.6e-05 | 7.8e-06 | 1.160E-05 | 4.133E-06 | 7.800E-06 | 1.600E-05 |
| 9650 | 483 | V483I | A | 1.1e-05 | 9.2e-06 | 1.2e-05 | 1.073E-05 | 1.419E-06 | 9.200E-06 | 1.200E-05 |
| 9651 | 483 | V483L | A | 1.5e-05 | 1.1e-05 | 1.2e-05 | 1.267E-05 | 2.082E-06 | 1.100E-05 | 1.500E-05 |
| 9652 | 483 | V483K | A | 1.1e-05 | 9.9e-06 | 1.6e-05 | 1.230E-05 | 3.251E-06 | 9.900E-06 | 1.600E-05 |
| 9653 | 483 | V483M | A | 1.1e-05 | 1.5e-05 | 1.6e-05 | 1.400E-05 | 2.646E-06 | 1.100E-05 | 1.600E-05 |
| 9654 | 483 | V483F | A | 1.6e-05 | 1.7e-05 | 1.4e-05 | 1.567E-05 | 1.528E-06 | 1.400E-05 | 1.700E-05 |
| 9655 | 483 | V483P | A | 1.5e-05 | 1e-05 | 2.4e-05 | 1.633E-05 | 7.095E-06 | 1.000E-05 | 2.400E-05 |
| 9656 | 483 | V483S | A | 1e-05 | 1.5e-05 | 1.7e-05 | 1.400E-05 | 3.606E-06 | 1.000E-05 | 1.700E-05 |
| 9657 | 483 | V483T | A | 1.6e-05 | 1.6e-05 | 1.2e-05 | 1.467E-05 | 2.309E-06 | 1.200E-05 | 1.600E-05 |
| 9658 | 483 | V483W | A | 1.5e-05 | 1e-05 | 1.6e-05 | 1.367E-05 | 3.215E-06 | 1.000E-05 | 1.600E-05 |
| 9659 | 483 | V483Y | A | 1.3e-05 | 1.3e-05 | 1.5e-05 | 1.367E-05 | 1.155E-06 | 1.300E-05 | 1.500E-05 |
| 9660 | 483 | V483V | A | 1.4e-05 | 1.5e-05 | 1.1e-05 | 1.333E-05 | 2.082E-06 | 1.100E-05 | 1.500E-05 |
| 9661 | 484 | G484A | A | 1.4e-05 | 1.1e-05 | 1.4e-05 | 1.300E-05 | 1.732E-06 | 1.100E-05 | 1.400E-05 |
| 9662 | 484 | G484R | A | 1.1e-05 | 1.2e-05 | 1.7e-05 | 1.333E-05 | 3.215E-06 | 1.100E-05 | 1.700E-05 |
| 9663 | 484 | G484N | A | 9.6e-06 | 1e-05 | 9.9e-06 | 9.833E-06 | 2.082E-07 | 9.600E-06 | 1.000E-05 |
| 9664 | 484 | G484D | A | 1.3e-05 | 9.5e-06 | 1e-05 | 1.083E-05 | 1.893E-06 | 9.500E-06 | 1.300E-05 |
| 9665 | 484 | G484C | A | 1e-05 | 7.5e-06 | 1.6e-05 | 1.117E-05 | 4.368E-06 | 7.500E-06 | 1.600E-05 |
| 9666 | 484 | G484E | A | 1.1e-05 | 9.5e-06 | 9.3e-06 | 9.933E-06 | 9.292E-07 | 9.300E-06 | 1.100E-05 |
| 9667 | 484 | G484Q | A | 1.4e-05 | 9.2e-06 | 1.6e-05 | 1.307E-05 | 3.495E-06 | 9.200E-06 | 1.600E-05 |
| 9668 | 484 | G484G | A | 1.5e-05 | 1.5e-05 | 1.5e-05 | 1.500E-05 | 0.000E+00 | 1.500E-05 | 1.500E-05 |
| 9669 | 484 | G484H | A | 1.7e-05 | 1e-05 | 1e-05 | 1.233E-05 | 4.041E-06 | 1.000E-05 | 1.700E-05 |
| 9670 | 484 | G484I | A | 1.6e-05 | 1.1e-05 | 1.7e-05 | 1.467E-05 | 3.215E-06 | 1.100E-05 | 1.700E-05 |
| 9671 | 484 | G484L | A | 1.3e-05 | 5.9e-06 | 1.6e-05 | 1.163E-05 | 5.187E-06 | 5.900E-06 | 1.600E-05 |
| 9672 | 484 | G484K | A | 1.6e-05 | 1.2e-05 | 1.1e-05 | 1.300E-05 | 2.646E-06 | 1.100E-05 | 1.600E-05 |
| 9673 | 484 | G484M | A | 1.5e-05 | 1.3e-05 | 1.6e-05 | 1.467E-05 | 1.528E-06 | 1.300E-05 | 1.600E-05 |
| 9674 | 484 | G484F | A | 1.1e-05 | 1.6e-05 | 1.6e-05 | 1.433E-05 | 2.887E-06 | 1.100E-05 | 1.600E-05 |
| 9675 | 484 | G484P | A | 1.5e-05 | 9.5e-06 | 1.4e-05 | 1.283E-05 | 2.930E-06 | 9.500E-06 | 1.500E-05 |
| 9676 | 484 | G484S | A | 9e-06 | 1.4e-05 | 1.3e-05 | 1.200E-05 | 2.646E-06 | 9.000E-06 | 1.400E-05 |
| 9677 | 484 | G484T | A | 8.8e-06 | 1e-05 | 1.3e-05 | 1.060E-05 | 2.163E-06 | 8.800E-06 | 1.300E-05 |
| 9678 | 484 | G484W | A | 1.1e-05 | 8e-06 | 9.4e-06 | 9.467E-06 | 1.501E-06 | 8.000E-06 | 1.100E-05 |
| 9679 | 484 | G484Y | A | 1.1e-05 | 1.7e-05 | 1.3e-05 | 1.367E-05 | 3.055E-06 | 1.100E-05 | 1.700E-05 |
| 9680 | 484 | G484V | A | 1.3e-05 | 1e-05 | 1.5e-05 | 1.267E-05 | 2.517E-06 | 1.000E-05 | 1.500E-05 |
| 9681 | 485 | E485A | A | 1.5e-05 | 9.5e-06 | 1.5e-05 | 1.317E-05 | 3.175E-06 | 9.500E-06 | 1.500E-05 |
| 9682 | 485 | E485R | A | 1.4e-05 | 1.1e-05 | 1.2e-05 | 1.233E-05 | 1.528E-06 | 1.100E-05 | 1.400E-05 |
| 9683 | 485 | E485N | A | 1e-05 | 1.2e-05 | 1e-05 | 1.067E-05 | 1.155E-06 | 1.000E-05 | 1.200E-05 |
| 9684 | 485 | E485D | A | 1.3e-05 | 1.7e-05 | 9e-06 | 1.300E-05 | 4.000E-06 | 9.000E-06 | 1.700E-05 |
| 9685 | 485 | E485C | A | 1.6e-05 | 1.7e-05 | 1.3e-05 | 1.533E-05 | 2.082E-06 | 1.300E-05 | 1.700E-05 |
| 9686 | 485 | E485E | A | 8.4e-06 | 1.3e-05 | 1.5e-05 | 1.213E-05 | 3.384E-06 | 8.400E-06 | 1.500E-05 |
| 9687 | 485 | E485Q | A | 1.4e-05 | 1e-05 | 1.3e-05 | 1.233E-05 | 2.082E-06 | 1.000E-05 | 1.400E-05 |
| 9688 | 485 | E485G | A | 1.3e-05 | 1.3e-05 | 1.6e-05 | 1.400E-05 | 1.732E-06 | 1.300E-05 | 1.600E-05 |
| 9689 | 485 | E485H | A | 1e-05 | 1.6e-05 | 9.8e-06 | 1.193E-05 | 3.523E-06 | 9.800E-06 | 1.600E-05 |

| | | | | | | | | | | |
|---|---|---|---|---|---|---|---|---|---|---|
| 9690 | 485 | E485I | A | 1.1e-05 | 9.3e-06 | 1.5e-05 | 1.177E-05 | 2.926E-06 | 9.300E-06 | 1.500E-05 |
| 9691 | 485 | E485L | A | 8.4e-06 | 1.5e-05 | 1e-05 | 1.113E-05 | 3.443E-06 | 8.400E-06 | 1.500E-05 |
| 9692 | 485 | E485K | A | 1.1e-05 | 1.4e-05 | 1.1e-05 | 1.200E-05 | 1.732E-06 | 1.100E-05 | 1.400E-05 |
| 9693 | 485 | E485M | A | 1.4e-05 | 9.5e-06 | 1.4e-05 | 1.250E-05 | 2.598E-06 | 9.500E-06 | 1.400E-05 |
| 9694 | 485 | E485F | A | 1.3e-05 | 1.4e-05 | 6.4e-06 | 1.113E-05 | 4.130E-06 | 6.400E-06 | 1.400E-05 |
| 9695 | 485 | E485P | A | 1.1e-05 | 1e-05 | 1.1e-05 | 1.067E-05 | 5.774E-07 | 1.000E-05 | 1.100E-05 |
| 9696 | 485 | E485S | A | 9.2e-06 | 1.5e-05 | 1.3e-05 | 1.240E-05 | 2.946E-06 | 9.200E-06 | 1.500E-05 |
| 9697 | 485 | E485T | A | 1e-05 | 1.3e-05 | 1e-05 | 1.100E-05 | 1.732E-06 | 1.000E-05 | 1.300E-05 |
| 9698 | 485 | E485W | A | 1e-05 | 1.5e-05 | 1.1e-05 | 1.200E-05 | 2.646E-06 | 1.000E-05 | 1.500E-05 |
| 9699 | 485 | E485Y | A | 1.2e-05 | 1.4e-05 | 1.2e-05 | 1.267E-05 | 1.155E-06 | 1.200E-05 | 1.400E-05 |
| 9700 | 485 | E485V | A | 1.5e-05 | 1.2e-05 | 1e-05 | 1.233E-05 | 2.517E-06 | 1.000E-05 | 1.500E-05 |
| 9701 | 486 | G486A | A | 9.3e-06 | 1.6e-05 | 1.5e-05 | 1.343E-05 | 3.614E-06 | 9.300E-06 | 1.600E-05 |
| 9702 | 486 | G486R | A | 1.2e-05 | 1.7e-05 | 1e-05 | 1.300E-05 | 3.606E-06 | 1.000E-05 | 1.700E-05 |
| 9703 | 486 | G486N | A | 1.4e-05 | 1.5e-05 | 9.9e-06 | 1.297E-05 | 2.702E-06 | 9.900E-06 | 1.500E-05 |
| 9704 | 486 | G486D | A | 1.6e-05 | 1.5e-05 | 8.7e-06 | 1.323E-05 | 3.958E-06 | 8.700E-06 | 1.600E-05 |
| 9705 | 486 | G486C | A | 8.7e-06 | 1.4e-05 | 1.4e-05 | 1.223E-05 | 3.060E-06 | 8.700E-06 | 1.400E-05 |
| 9706 | 486 | G486E | A | 1.7e-05 | 1.4e-05 | 1.7e-05 | 1.600E-05 | 1.732E-06 | 1.400E-05 | 1.700E-05 |
| 9707 | 486 | G486Q | A | 8.4e-06 | 1.4e-05 | 1.4e-05 | 1.213E-05 | 3.233E-06 | 8.400E-06 | 1.400E-05 |
| 9708 | 486 | G486G | A | 9.1e-06 | 1.4e-05 | 1e-05 | 1.103E-05 | 2.608E-06 | 9.100E-06 | 1.400E-05 |
| 9709 | 486 | G486H | A | 1.4e-05 | 1.3e-05 | 9.6e-06 | 1.220E-05 | 2.307E-06 | 9.600E-06 | 1.400E-05 |
| 9710 | 486 | G486I | A | 1.3e-05 | 1.2e-05 | 1.6e-05 | 1.367E-05 | 2.082E-06 | 1.200E-05 | 1.600E-05 |
| 9711 | 486 | G486L | A | 1.5e-05 | 1.2e-05 | 1.4e-05 | 1.367E-05 | 1.528E-06 | 1.200E-05 | 1.500E-05 |
| 9712 | 486 | G486K | A | 7.8e-06 | 1e-05 | 8.7e-06 | 8.833E-06 | 1.106E-06 | 7.800E-06 | 1.000E-05 |
| 9713 | 486 | G486M | A | 8.7e-06 | 9e-06 | 1.3e-05 | 1.023E-05 | 2.401E-06 | 8.700E-06 | 1.300E-05 |
| 9714 | 486 | G486F | A | 1.7e-05 | 9.4e-06 | 1.5e-05 | 1.380E-05 | 3.940E-06 | 9.400E-06 | 1.700E-05 |
| 9715 | 486 | G486P | A | 1e-05 | 1.5e-05 | 9e-06 | 1.133E-05 | 3.215E-06 | 9.000E-06 | 1.500E-05 |
| 9716 | 486 | G486S | A | 1e-05 | 1.6e-05 | 9.8e-06 | 1.193E-05 | 3.523E-06 | 9.800E-06 | 1.600E-05 |
| 9717 | 486 | G486T | A | 1.4e-05 | 1.5e-05 | 1.2e-05 | 1.367E-05 | 1.528E-06 | 1.200E-05 | 1.500E-05 |
| 9718 | 486 | G486W | A | 1.3e-05 | 1.5e-05 | 1e-05 | 1.267E-05 | 2.517E-06 | 1.000E-05 | 1.500E-05 |
| 9719 | 486 | G486Y | A | 1.1e-05 | 9.2e-06 | 9.9e-06 | 1.003E-05 | 9.074E-07 | 9.200E-06 | 1.100E-05 |
| 9720 | 486 | G486V | A | 1.5e-05 | 1.6e-05 | 1.6e-05 | 1.567E-05 | 5.774E-07 | 1.500E-05 | 1.600E-05 |
| 9721 | 487 | L487A | A | 1.7e-05 | 1.7e-05 | 1.4e-05 | 1.600E-05 | 1.732E-06 | 1.400E-05 | 1.700E-05 |
| 9722 | 487 | L487R | A | 1.1e-05 | 1.6e-05 | 1.4e-05 | 1.367E-05 | 2.517E-06 | 1.100E-05 | 1.600E-05 |
| 9723 | 487 | L487N | A | 1.3e-05 | 9.4e-06 | 1.5e-05 | 1.247E-05 | 2.838E-06 | 9.400E-06 | 1.500E-05 |
| 9724 | 487 | L487D | A | 9.4e-06 | 1.4e-05 | 1e-05 | 1.113E-05 | 2.501E-06 | 9.400E-06 | 1.400E-05 |
| 9725 | 487 | L487C | A | 7.6e-06 | 1.2e-05 | 1.5e-05 | 1.153E-05 | 3.722E-06 | 7.600E-06 | 1.500E-05 |
| 9726 | 487 | L487E | A | 1.5e-05 | 1.2e-05 | 1.3e-05 | 1.333E-05 | 1.528E-06 | 1.200E-05 | 1.500E-05 |
| 9727 | 487 | L487Q | A | 1.3e-05 | 1.6e-05 | 1.5e-05 | 1.467E-05 | 1.528E-06 | 1.300E-05 | 1.600E-05 |
| 9728 | 487 | L487G | A | 1.1e-05 | 1.4e-05 | 9.7e-06 | 1.157E-05 | 2.205E-06 | 9.700E-06 | 1.400E-05 |
| 9729 | 487 | L487H | A | 1.6e-05 | 1.5e-05 | 1.5e-05 | 1.533E-05 | 5.774E-07 | 1.500E-05 | 1.600E-05 |
| 9730 | 487 | L487I | A | 1.6e-05 | 1.1e-05 | 1.5e-05 | 1.400E-05 | 2.646E-06 | 1.100E-05 | 1.600E-05 |
| 9731 | 487 | L487L | A | 1e-05 | 1.3e-05 | 1.4e-05 | 1.233E-05 | 2.082E-06 | 1.000E-05 | 1.400E-05 |
| 9732 | 487 | L487K | A | 1.4e-05 | 1.4e-05 | 1.4e-05 | 1.400E-05 | 0.000E+00 | 1.400E-05 | 1.400E-05 |
| 9733 | 487 | L487M | A | 8.6e-06 | 1.5e-05 | 1.6e-05 | 1.320E-05 | 4.015E-06 | 8.600E-06 | 1.600E-05 |
| 9734 | 487 | L487F | A | 1.4e-05 | 1.2e-05 | 1.4e-05 | 1.333E-05 | 1.155E-06 | 1.200E-05 | 1.400E-05 |
| 9735 | 487 | L487P | A | 9.9e-06 | 2.6e-05 | 1e-05 | 1.530E-05 | 9.267E-06 | 9.900E-06 | 2.600E-05 |
| 9736 | 487 | L487S | A | 1.5e-05 | 1.7e-05 | 1.5e-05 | 1.567E-05 | 1.155E-06 | 1.500E-05 | 1.700E-05 |
| 9737 | 487 | L487T | A | 1.4e-05 | 1.1e-05 | 1.5e-05 | 1.333E-05 | 2.082E-06 | 1.100E-05 | 1.500E-05 |
| 9738 | 487 | L487W | A | 1e-05 | 8e-06 | 1.1e-05 | 9.667E-06 | 1.528E-06 | 8.000E-06 | 1.100E-05 |
| 9739 | 487 | L487Y | A | 1.6e-05 | 1.3e-05 | 9.4e-06 | 1.280E-05 | 3.305E-06 | 9.400E-06 | 1.600E-05 |
| 9740 | 487 | L487V | A | 1e-05 | 1.7e-05 | 1.6e-05 | 1.433E-05 | 3.786E-06 | 1.000E-05 | 1.700E-05 |
| 9741 | 488 | A488A | A | 1.3e-05 | 1.6e-05 | 9.9e-06 | 1.297E-05 | 3.050E-06 | 9.900E-06 | 1.600E-05 |
| 9742 | 488 | A488R | A | 9.3e-06 | 1.6e-05 | 7.7e-06 | 1.100E-05 | 4.403E-06 | 7.700E-06 | 1.600E-05 |
| 9743 | 488 | A488N | A | 1.2e-05 | 1e-05 | 8.4e-06 | 1.013E-05 | 1.804E-06 | 8.400E-06 | 1.200E-05 |
| 9744 | 488 | A488D | A | 1.5e-05 | 9.6e-06 | 1.3e-05 | 1.253E-05 | 2.730E-06 | 9.600E-06 | 1.500E-05 |
| 9745 | 488 | A488C | A | 1e-05 | 1.2e-05 | 1.5e-05 | 1.233E-05 | 2.517E-06 | 1.000E-05 | 1.500E-05 |
| 9746 | 488 | A488E | A | 1.3e-05 | 1.1e-05 | 1.7e-05 | 1.367E-05 | 3.055E-06 | 1.100E-05 | 1.700E-05 |
| 9747 | 488 | A488Q | A | 1e-05 | 1.2e-05 | 1.1e-05 | 1.100E-05 | 1.000E-06 | 1.000E-05 | 1.200E-05 |
| 9748 | 488 | A488G | A | 1.5e-05 | 1.5e-05 | 1.4e-05 | 1.467E-05 | 5.774E-07 | 1.400E-05 | 1.500E-05 |
| 9749 | 488 | A488H | A | 1.7e-05 | 1.3e-05 | 1.1e-05 | 1.367E-05 | 3.055E-06 | 1.100E-05 | 1.700E-05 |
| 9750 | 488 | A488I | A | 9.9e-06 | 1e-05 | 1.4e-05 | 1.130E-05 | 2.339E-06 | 9.900E-06 | 1.400E-05 |
| 9751 | 488 | A488L | A | 1.5e-05 | 1.3e-05 | 9.4e-06 | 1.247E-05 | 2.838E-06 | 9.400E-06 | 1.500E-05 |
| 9752 | 488 | A488K | A | 8.9e-06 | 9.9e-06 | 1.3e-05 | 1.060E-05 | 2.138E-06 | 8.900E-06 | 1.300E-05 |
| 9753 | 488 | A488M | A | 1.5e-05 | 8.8e-06 | 1.1e-05 | 1.160E-05 | 3.143E-06 | 8.800E-06 | 1.500E-05 |
| 9754 | 488 | A488F | A | 1.1e-05 | 1.6e-05 | 9.7e-06 | 1.223E-05 | 3.326E-06 | 9.700E-06 | 1.600E-05 |
| 9755 | 488 | A488P | A | 1.6e-05 | 1.7e-05 | 1.4e-05 | 1.567E-05 | 1.528E-06 | 1.400E-05 | 1.700E-05 |
| 9756 | 488 | A488S | A | 9.7e-06 | 1.6e-05 | 1.1e-05 | 1.223E-05 | 3.326E-06 | 9.700E-06 | 1.600E-05 |
| 9757 | 488 | A488T | A | 1.1e-05 | 1.6e-05 | 1.4e-05 | 1.367E-05 | 2.517E-06 | 1.100E-05 | 1.600E-05 |
| 9758 | 488 | A488W | A | 1.2e-05 | 1e-05 | 1.4e-05 | 1.200E-05 | 2.000E-06 | 1.000E-05 | 1.400E-05 |
| 9759 | 488 | A488Y | A | 9.4e-06 | 1.7e-05 | 1.3e-05 | 1.313E-05 | 3.802E-06 | 9.400E-06 | 1.700E-05 |
| 9760 | 488 | A488V | A | 1.5e-05 | 9.7e-06 | 8e-06 | 1.090E-05 | 3.651E-06 | 8.000E-06 | 1.500E-05 |
| 9761 | 489 | C489A | A | 1.5e-05 | 1.4e-05 | 8.7e-06 | 1.257E-05 | 3.386E-06 | 8.700E-06 | 1.500E-05 |
| 9762 | 489 | C489R | A | 9.9e-06 | 1.4e-05 | 7e-06 | 1.030E-05 | 3.517E-06 | 7.000E-06 | 1.400E-05 |
| 9763 | 489 | C489N | A | 1.1e-05 | 1e-05 | 1.4e-05 | 1.167E-05 | 2.082E-06 | 1.000E-05 | 1.400E-05 |
| 9764 | 489 | C489D | A | 7.2e-06 | 1.5e-05 | 1.2e-05 | 1.140E-05 | 3.934E-06 | 7.200E-06 | 1.500E-05 |
| 9765 | 489 | C489C | A | 1.1e-05 | 1.6e-05 | 1.5e-05 | 1.400E-05 | 2.646E-06 | 1.100E-05 | 1.600E-05 |
| 9766 | 489 | C489E | A | 1.6e-05 | 1.4e-05 | 8.9e-06 | 1.297E-05 | 3.661E-06 | 8.900E-06 | 1.600E-05 |
| 9767 | 489 | C489Q | A | 1.5e-05 | 8e-06 | 1.1e-05 | 1.133E-05 | 3.512E-06 | 8.000E-06 | 1.500E-05 |
| 9768 | 489 | C489G | A | 1.1e-05 | 1.2e-05 | 1.1e-05 | 1.133E-05 | 5.774E-07 | 1.100E-05 | 1.200E-05 |
| 9769 | 489 | C489H | A | 1.4e-05 | 9.1e-06 | 9.9e-06 | 1.100E-05 | 2.629E-06 | 9.100E-06 | 1.400E-05 |
| 9770 | 489 | C489I | A | 1.1e-05 | 1.3e-05 | 9.6e-06 | 1.120E-05 | 1.709E-06 | 9.600E-06 | 1.300E-05 |
| 9771 | 489 | C489L | A | 1.3e-05 | 1.1e-05 | 9.8e-06 | 1.127E-05 | 1.617E-06 | 9.800E-06 | 1.300E-05 |
| 9772 | 489 | C489K | A | 1.3e-05 | 1.7e-05 | 1.1e-05 | 1.367E-05 | 3.055E-06 | 1.100E-05 | 1.700E-05 |
| 9773 | 489 | C489M | A | 1.7e-05 | 1.6e-05 | 1.1e-05 | 1.467E-05 | 3.215E-06 | 1.100E-05 | 1.700E-05 |
| 9774 | 489 | C489F | A | 1.1e-05 | 1.2e-05 | 1.4e-05 | 1.233E-05 | 1.528E-06 | 1.100E-05 | 1.400E-05 |

| | | | | | | | | | | |
|---|---|---|---|---|---|---|---|---|---|---|
| 9775 | 489 | C489P | A | 1.5e-05 | 9.9e-06 | 1.5e-05 | 1.330E-05 | 2.944E-06 | 9.900E-06 | 1.500E-05 |
| 9776 | 489 | C489S | A | 1.1e-05 | 1.2e-05 | 7.5e-06 | 1.017E-05 | 2.363E-06 | 7.500E-06 | 1.200E-05 |
| 9777 | 489 | C489T | A | 1.6e-05 | 8.5e-06 | 1.1e-05 | 1.183E-05 | 3.819E-06 | 8.500E-06 | 1.600E-05 |
| 9778 | 489 | C489W | A | 9.3e-06 | 1.4e-05 | 1.8e-05 | 1.377E-05 | 4.355E-06 | 9.300E-06 | 1.800E-05 |
| 9779 | 489 | C489Y | A | 1.6e-05 | 1.3e-05 | 1.3e-05 | 1.400E-05 | 1.732E-06 | 1.300E-05 | 1.600E-05 |
| 9780 | 489 | C489V | A | 1.5e-05 | 1.4e-05 | 1.5e-05 | 1.467E-05 | 5.774E-07 | 1.400E-05 | 1.500E-05 |
| 9781 | 490 | H490A | A | 1.9e-05 | 1.7e-05 | 1e-05 | 1.533E-05 | 4.726E-06 | 1.000E-05 | 1.900E-05 |
| 9782 | 490 | H490R | A | 8.1e-06 | 1.7e-05 | 1.5e-05 | 1.337E-05 | 4.669E-06 | 8.100E-06 | 1.700E-05 |
| 9783 | 490 | H490N | A | 1.5e-05 | 1.2e-05 | 1.4e-05 | 1.367E-05 | 1.528E-06 | 1.200E-05 | 1.500E-05 |
| 9784 | 490 | H490D | A | 9.3e-06 | 1.4e-05 | 1.6e-05 | 1.310E-05 | 3.439E-06 | 9.300E-06 | 1.600E-05 |
| 9785 | 490 | H490C | A | 3.5e-05 | 1.6e-05 | 1.5e-05 | 2.200E-05 | 1.127E-05 | 1.500E-05 | 3.500E-05 |
| 9786 | 490 | H490E | A | 6.9e-06 | 1.6e-05 | 1.3e-05 | 1.197E-05 | 4.637E-06 | 6.900E-06 | 1.600E-05 |
| 9787 | 490 | H490Q | A | 1e-05 | 9.9e-06 | 1.1e-05 | 1.030E-05 | 6.083E-07 | 9.900E-06 | 1.100E-05 |
| 9788 | 490 | H490G | A | 1.5e-05 | 1.4e-05 | 1.4e-05 | 1.433E-05 | 5.774E-07 | 1.400E-05 | 1.500E-05 |
| 9789 | 490 | H490H | A | 1.1e-05 | 1.4e-05 | 1.5e-05 | 1.333E-05 | 2.082E-06 | 1.100E-05 | 1.500E-05 |
| 9790 | 490 | H490I | A | 8.9e-06 | 1.6e-05 | 1.6e-05 | 1.363E-05 | 4.099E-06 | 8.900E-06 | 1.600E-05 |
| 9791 | 490 | H490L | A | 1.5e-05 | 1.8e-05 | 1.1e-05 | 1.467E-05 | 3.512E-06 | 1.100E-05 | 1.800E-05 |
| 9792 | 490 | H490K | A | 1.2e-05 | 1.6e-05 | 1.5e-05 | 1.433E-05 | 2.082E-06 | 1.200E-05 | 1.600E-05 |
| 9793 | 490 | H490M | A | 1.6e-05 | 9.8e-06 | 1e-05 | 1.193E-05 | 3.523E-06 | 9.800E-06 | 1.600E-05 |
| 9794 | 490 | H490F | A | 1.2e-05 | 1.2e-05 | 1.1e-05 | 1.167E-05 | 5.774E-07 | 1.100E-05 | 1.200E-05 |
| 9795 | 490 | H490P | A | 1.1e-05 | 9.4e-06 | 1.7e-05 | 1.247E-05 | 4.007E-06 | 9.400E-06 | 1.700E-05 |
| 9796 | 490 | H490S | A | 9.9e-06 | 1e-05 | 1.1e-05 | 1.030E-05 | 6.083E-07 | 9.900E-06 | 1.100E-05 |
| 9797 | 490 | H490T | A | 1.5e-05 | 1.7e-05 | 1.1e-05 | 1.433E-05 | 3.055E-06 | 1.100E-05 | 1.700E-05 |
| 9798 | 490 | H490W | A | 1.6e-05 | 1.6e-05 | 1.3e-05 | 1.500E-05 | 1.732E-06 | 1.300E-05 | 1.600E-05 |
| 9799 | 490 | H490Y | A | 1.3e-05 | 1.2e-05 | 1.4e-05 | 1.300E-05 | 1.000E-06 | 1.200E-05 | 1.400E-05 |
| 9800 | 490 | H490V | A | 1e-05 | 1.5e-05 | 1.1e-05 | 1.200E-05 | 2.646E-06 | 1.000E-05 | 1.500E-05 |
| 9801 | 491 | Q491A | A | 1.5e-05 | 1.5e-05 | 1e-05 | 1.333E-05 | 2.887E-06 | 1.000E-05 | 1.500E-05 |
| 9802 | 491 | Q491R | A | 1.6e-05 | 1.3e-05 | 1.7e-05 | 1.533E-05 | 2.082E-06 | 1.300E-05 | 1.700E-05 |
| 9803 | 491 | Q491N | A | 1.5e-05 | 1.4e-05 | 8.6e-06 | 1.253E-05 | 3.443E-06 | 8.600E-06 | 1.500E-05 |
| 9804 | 491 | Q491D | A | 1.1e-05 | 1.6e-05 | 1.4e-05 | 1.367E-05 | 2.517E-06 | 1.100E-05 | 1.600E-05 |
| 9805 | 491 | Q491C | A | 1.7e-05 | 1.4e-05 | 1e-05 | 1.367E-05 | 3.512E-06 | 1.000E-05 | 1.700E-05 |
| 9806 | 491 | Q491E | A | 1.6e-05 | 1e-05 | 1.1e-05 | 1.233E-05 | 3.215E-06 | 1.000E-05 | 1.600E-05 |
| 9807 | 491 | Q491Q | A | 1.6e-05 | 9.1e-06 | 1.2e-05 | 1.237E-05 | 3.465E-06 | 9.100E-06 | 1.600E-05 |
| 9808 | 491 | Q491G | A | 1.6e-05 | 1.1e-05 | 1.6e-05 | 1.433E-05 | 2.887E-06 | 1.100E-05 | 1.600E-05 |
| 9809 | 491 | Q491H | A | 1.5e-05 | 1.5e-05 | 1.4e-05 | 1.467E-05 | 5.774E-07 | 1.400E-05 | 1.500E-05 |
| 9810 | 491 | Q491I | A | 1.3e-05 | 9.6e-06 | 1.3e-05 | 1.187E-05 | 1.963E-06 | 9.600E-06 | 1.300E-05 |
| 9811 | 491 | Q491L | A | 1.4e-05 | 1.6e-05 | 1.3e-05 | 1.433E-05 | 1.528E-06 | 1.300E-05 | 1.600E-05 |
| 9812 | 491 | Q491K | A | 1.3e-05 | 9e-06 | 1.8e-05 | 1.333E-05 | 4.509E-06 | 9.000E-06 | 1.800E-05 |
| 9813 | 491 | Q491M | A | 1.6e-05 | 1.3e-05 | 1.5e-05 | 1.467E-05 | 1.528E-06 | 1.300E-05 | 1.600E-05 |
| 9814 | 491 | Q491F | A | 8.7e-06 | 1e-05 | 1.4e-05 | 1.090E-05 | 2.762E-06 | 8.700E-06 | 1.400E-05 |
| 9815 | 491 | Q491P | A | 1.2e-05 | 1e-05 | 1.6e-05 | 1.267E-05 | 3.055E-06 | 1.000E-05 | 1.600E-05 |
| 9816 | 491 | Q491S | A | 1.6e-05 | 1.2e-05 | 1.6e-05 | 1.467E-05 | 2.309E-06 | 1.200E-05 | 1.600E-05 |
| 9817 | 491 | Q491T | A | 1.6e-05 | 2e-05 | 1.5e-05 | 1.700E-05 | 2.646E-06 | 1.500E-05 | 2.000E-05 |
| 9818 | 491 | Q491W | A | 1.5e-05 | 1.4e-05 | 8.4e-06 | 1.247E-05 | 3.557E-06 | 8.400E-06 | 1.500E-05 |
| 9819 | 491 | Q491Y | A | 1.5e-05 | 1.2e-05 | 1.3e-05 | 1.333E-05 | 1.528E-06 | 1.200E-05 | 1.500E-05 |
| 9820 | 491 | Q491V | A | 1.5e-05 | 1.4e-05 | 8.4e-06 | 1.247E-05 | 3.557E-06 | 8.400E-06 | 1.500E-05 |
| 9821 | 492 | L492A | A | 1.1e-05 | 9.7e-06 | 1.4e-05 | 1.157E-05 | 2.205E-06 | 9.700E-06 | 1.400E-05 |
| 9822 | 492 | L492R | A | 1.5e-05 | 1.6e-05 | 1.5e-05 | 1.533E-05 | 5.774E-07 | 1.500E-05 | 1.600E-05 |
| 9823 | 492 | L492N | A | 1.1e-05 | 9.4e-06 | 1.1e-05 | 1.047E-05 | 9.238E-07 | 9.400E-06 | 1.100E-05 |
| 9824 | 492 | L492D | A | 1.2e-05 | 1.5e-05 | 1.4e-05 | 1.367E-05 | 1.528E-06 | 1.200E-05 | 1.500E-05 |
| 9825 | 492 | L492C | A | 6.8e-06 | 1.1e-05 | 1.4e-05 | 1.060E-05 | 3.617E-06 | 6.800E-06 | 1.400E-05 |
| 9826 | 492 | L492E | A | 8.8e-06 | 1.3e-05 | 1.3e-05 | 1.160E-05 | 2.425E-06 | 8.800E-06 | 1.300E-05 |
| 9827 | 492 | L492Q | A | 1.4e-05 | 9e-06 | 1.5e-05 | 1.267E-05 | 3.215E-06 | 9.000E-06 | 1.500E-05 |
| 9828 | 492 | L492G | A | 8.6e-06 | 1.2e-05 | 1.8e-05 | 1.287E-05 | 4.760E-06 | 8.600E-06 | 1.800E-05 |
| 9829 | 492 | L492H | A | 1.1e-05 | 1.4e-05 | 1.4e-05 | 1.300E-05 | 1.732E-06 | 1.100E-05 | 1.400E-05 |
| 9830 | 492 | L492I | A | 1.4e-05 | 1.1e-05 | 1.6e-05 | 1.367E-05 | 2.517E-06 | 1.100E-05 | 1.600E-05 |
| 9831 | 492 | L492L | A | 1e-05 | 1.1e-05 | 1.5e-05 | 1.200E-05 | 2.646E-06 | 1.000E-05 | 1.500E-05 |
| 9832 | 492 | L492K | A | 1e-05 | 1.4e-05 | 1.7e-05 | 1.367E-05 | 3.512E-06 | 1.000E-05 | 1.700E-05 |
| 9833 | 492 | L492M | A | 1.3e-05 | 1.6e-05 | 1.5e-05 | 1.467E-05 | 1.528E-06 | 1.300E-05 | 1.600E-05 |
| 9834 | 492 | L492F | A | 1.5e-05 | 1e-05 | 1.5e-05 | 1.333E-05 | 2.887E-06 | 1.000E-05 | 1.500E-05 |
| 9835 | 492 | L492P | A | 1.4e-05 | 9.5e-06 | 1.4e-05 | 1.250E-05 | 2.598E-06 | 9.500E-06 | 1.400E-05 |
| 9836 | 492 | L492S | A | 1.2e-05 | 9.5e-06 | 9.7e-06 | 1.040E-05 | 1.389E-06 | 9.500E-06 | 1.200E-05 |
| 9837 | 492 | L492T | A | 1.5e-05 | 1.4e-05 | 1.6e-05 | 1.500E-05 | 1.000E-06 | 1.400E-05 | 1.600E-05 |
| 9838 | 492 | L492W | A | 1.3e-05 | 1.4e-05 | 1.7e-05 | 1.467E-05 | 2.082E-06 | 1.300E-05 | 1.700E-05 |
| 9839 | 492 | L492Y | A | 1.4e-05 | 1.7e-05 | 1.3e-05 | 1.467E-05 | 2.082E-06 | 1.300E-05 | 1.700E-05 |
| 9840 | 492 | L492V | A | 1.3e-05 | 8.7e-06 | 1.6e-05 | 1.257E-05 | 3.669E-06 | 8.700E-06 | 1.600E-05 |
| 9841 | 493 | C493A | A | 1.1e-05 | 1e-05 | 1.5e-05 | 1.200E-05 | 2.646E-06 | 1.000E-05 | 1.500E-05 |
| 9842 | 493 | C493R | A | 1.5e-05 | 1.5e-05 | 1.5e-05 | 1.500E-05 | 0.000E+00 | 1.500E-05 | 1.500E-05 |
| 9843 | 493 | C493N | A | 7e-06 | 1e-05 | 1.1e-05 | 9.333E-06 | 2.082E-06 | 7.000E-06 | 1.100E-05 |
| 9844 | 493 | C493D | A | 1.1e-05 | 1.1e-05 | 1.9e-05 | 1.367E-05 | 4.619E-06 | 1.100E-05 | 1.900E-05 |
| 9845 | 493 | C493C | A | 1.5e-05 | 1.6e-05 | 1.8e-05 | 1.633E-05 | 1.528E-06 | 1.500E-05 | 1.800E-05 |
| 9846 | 493 | C493E | A | 1.6e-05 | 1.5e-05 | 1e-05 | 1.367E-05 | 3.215E-06 | 1.000E-05 | 1.600E-05 |
| 9847 | 493 | C493Q | A | 1.1e-05 | 1.5e-05 | 1.5e-05 | 1.367E-05 | 2.309E-06 | 1.100E-05 | 1.500E-05 |
| 9848 | 493 | C493G | A | 1.4e-05 | 9.3e-06 | 1.1e-05 | 1.143E-05 | 2.380E-06 | 9.300E-06 | 1.400E-05 |
| 9849 | 493 | C493H | A | 1.5e-05 | 1.8e-05 | 8.7e-06 | 1.390E-05 | 4.747E-06 | 8.700E-06 | 1.800E-05 |
| 9850 | 493 | C493I | A | 1.5e-05 | 1.6e-05 | 2.2e-05 | 1.767E-05 | 3.786E-06 | 1.500E-05 | 2.200E-05 |
| 9851 | 493 | C493L | A | 1.1e-05 | 1.5e-05 | 1.6e-05 | 1.400E-05 | 2.646E-06 | 1.100E-05 | 1.600E-05 |
| 9852 | 493 | C493K | A | 8.7e-06 | 1.6e-05 | 1e-05 | 1.157E-05 | 3.894E-06 | 8.700E-06 | 1.600E-05 |
| 9853 | 493 | C493M | A | 1.3e-05 | 1.4e-05 | 1.3e-05 | 1.333E-05 | 5.774E-07 | 1.300E-05 | 1.400E-05 |
| 9854 | 493 | C493F | A | 1.5e-05 | 1.5e-05 | 6.9e-06 | 1.230E-05 | 4.677E-06 | 6.900E-06 | 1.500E-05 |
| 9855 | 493 | C493P | A | 1.7e-05 | 1.4e-05 | 1.1e-05 | 1.400E-05 | 3.000E-06 | 1.100E-05 | 1.700E-05 |
| 9856 | 493 | C493S | A | 1.1e-05 | 1.6e-05 | 1.3e-05 | 1.333E-05 | 2.517E-06 | 1.100E-05 | 1.600E-05 |
| 9857 | 493 | C493T | A | 1.6e-05 | 1.2e-05 | 1.5e-05 | 1.433E-05 | 2.082E-06 | 1.200E-05 | 1.600E-05 |
| 9858 | 493 | C493W | A | 1.2e-05 | 1.2e-05 | 1e-05 | 1.133E-05 | 1.155E-06 | 1.000E-05 | 1.200E-05 |
| 9859 | 493 | C493Y | A | 1.4e-05 | 1.2e-05 | 1.3e-05 | 1.300E-05 | 1.000E-06 | 1.200E-05 | 1.400E-05 |

| | | | | | | | | | | |
|---|---|---|---|---|---|---|---|---|---|---|
| 9860 | 493 | C493V | A | 1.1e-05 | 1.1e-05 | 1.4e-05 | 1.200E-05 | 1.732E-06 | 1.100E-05 | 1.400E-05 |
| 9861 | 494 | A494A | A | 1.5e-05 | 1e-05 | 1.5e-05 | 1.333E-05 | 2.887E-06 | 1.000E-05 | 1.500E-05 |
| 9862 | 494 | A494R | A | 1e-05 | 1.3e-05 | 1.6e-05 | 1.300E-05 | 3.000E-06 | 1.000E-05 | 1.600E-05 |
| 9863 | 494 | A494N | A | 7.7e-06 | 1.1e-05 | 1.6e-05 | 1.157E-05 | 4.179E-06 | 7.700E-06 | 1.600E-05 |
| 9864 | 494 | A494D | A | 1.6e-05 | 5.8e-06 | 1.4e-05 | 1.193E-05 | 5.405E-06 | 5.800E-06 | 1.600E-05 |
| 9865 | 494 | A494C | A | 9e-06 | 1.4e-05 | 1.2e-05 | 1.167E-05 | 2.517E-06 | 9.000E-06 | 1.400E-05 |
| 9866 | 494 | A494E | A | 1.6e-05 | 9.4e-06 | 1.1e-05 | 1.213E-05 | 3.443E-06 | 9.400E-06 | 1.600E-05 |
| 9867 | 494 | A494Q | A | 1.4e-05 | 1.5e-05 | 1.4e-05 | 1.433E-05 | 5.774E-07 | 1.400E-05 | 1.500E-05 |
| 9868 | 494 | A494G | A | 8.2e-06 | 1.6e-05 | 1.2e-05 | 1.207E-05 | 3.900E-06 | 8.200E-06 | 1.600E-05 |
| 9869 | 494 | A494H | A | 1.6e-05 | 1.3e-05 | 7.9e-06 | 1.230E-05 | 4.095E-06 | 7.900E-06 | 1.600E-05 |
| 9870 | 494 | A494I | A | 1.3e-05 | 1.5e-05 | 1.1e-05 | 1.300E-05 | 2.000E-06 | 1.100E-05 | 1.500E-05 |
| 9871 | 494 | A494L | A | 1.3e-05 | 1.1e-05 | 7.6e-06 | 1.053E-05 | 2.730E-06 | 7.600E-06 | 1.300E-05 |
| 9872 | 494 | A494K | A | 1.5e-05 | 1.1e-05 | 1.2e-05 | 1.267E-05 | 2.082E-06 | 1.100E-05 | 1.500E-05 |
| 9873 | 494 | A494M | A | 1e-05 | 1.1e-05 | 1.1e-05 | 1.067E-05 | 5.774E-07 | 1.000E-05 | 1.100E-05 |
| 9874 | 494 | A494F | A | 1.5e-05 | 1.5e-05 | 1.5e-05 | 1.500E-05 | 0.000E+00 | 1.500E-05 | 1.500E-05 |
| 9875 | 494 | A494P | A | 1.2e-05 | 1.4e-05 | 1.2e-05 | 1.267E-05 | 1.155E-06 | 1.200E-05 | 1.400E-05 |
| 9876 | 494 | A494S | A | 1.5e-05 | 1.1e-05 | 9.4e-06 | 1.180E-05 | 2.884E-06 | 9.400E-06 | 1.500E-05 |
| 9877 | 494 | A494T | A | 1.4e-05 | 1.6e-05 | 1.3e-05 | 1.433E-05 | 1.528E-06 | 1.300E-05 | 1.600E-05 |
| 9878 | 494 | A494W | A | 1.3e-05 | 9.2e-06 | 9.9e-06 | 1.070E-05 | 2.022E-06 | 9.200E-06 | 1.300E-05 |
| 9879 | 494 | A494Y | A | 8.3e-06 | 8.7e-06 | 1.4e-05 | 1.033E-05 | 3.182E-06 | 8.300E-06 | 1.400E-05 |
| 9880 | 494 | A494V | A | 1.6e-05 | 1.4e-05 | 1.3e-05 | 1.433E-05 | 1.528E-06 | 1.300E-05 | 1.600E-05 |
| 9881 | 495 | R495A | A | 2.7e-05 | 1.1e-05 | 1.5e-05 | 1.767E-05 | 8.327E-06 | 1.100E-05 | 2.700E-05 |
| 9882 | 495 | R495R | A | 7e-06 | 1.5e-05 | 1.6e-05 | 1.267E-05 | 4.933E-06 | 7.000E-06 | 1.600E-05 |
| 9883 | 495 | R495N | A | 1.6e-05 | 1.3e-05 | 1.5e-05 | 1.467E-05 | 1.528E-06 | 1.300E-05 | 1.600E-05 |
| 9884 | 495 | R495D | A | 1.5e-05 | 9.2e-06 | 1.1e-05 | 1.173E-05 | 2.969E-06 | 9.200E-06 | 1.500E-05 |
| 9885 | 495 | R495C | A | 1.4e-05 | 9.3e-06 | 1.5e-05 | 1.277E-05 | 3.044E-06 | 9.300E-06 | 1.500E-05 |
| 9886 | 495 | R495E | A | 1.5e-05 | 1.3e-05 | 1e-05 | 1.267E-05 | 2.517E-06 | 1.000E-05 | 1.500E-05 |
| 9887 | 495 | R495Q | A | 1.6e-05 | 1.4e-05 | 9.3e-06 | 1.310E-05 | 3.439E-06 | 9.300E-06 | 1.600E-05 |
| 9888 | 495 | R495G | A | 1.5e-05 | 1.3e-05 | 1.2e-05 | 1.333E-05 | 1.528E-06 | 1.200E-05 | 1.500E-05 |
| 9889 | 495 | R495H | A | 1.6e-05 | 1.5e-05 | 1.2e-05 | 1.433E-05 | 2.082E-06 | 1.200E-05 | 1.600E-05 |
| 9890 | 495 | R495I | A | 1.2e-05 | 1.6e-05 | 1.5e-05 | 1.433E-05 | 2.082E-06 | 1.200E-05 | 1.600E-05 |
| 9891 | 495 | R495L | A | 1.5e-05 | 1.5e-05 | 1.4e-05 | 1.467E-05 | 5.774E-07 | 1.400E-05 | 1.500E-05 |
| 9892 | 495 | R495K | A | 1.4e-05 | 1.3e-05 | 9.8e-06 | 1.227E-05 | 2.194E-06 | 9.800E-06 | 1.400E-05 |
| 9893 | 495 | R495M | A | 1.4e-05 | 1.4e-05 | 1e-05 | 1.267E-05 | 2.309E-06 | 1.000E-05 | 1.400E-05 |
| 9894 | 495 | R495F | A | 1.4e-05 | 9.3e-06 | 9.8e-06 | 1.103E-05 | 2.581E-06 | 9.300E-06 | 1.400E-05 |
| 9895 | 495 | R495P | A | 9.4e-06 | 1.3e-05 | 1.3e-05 | 1.180E-05 | 2.078E-06 | 9.400E-06 | 1.300E-05 |
| 9896 | 495 | R495S | A | 1.6e-05 | 1.4e-05 | 1.4e-05 | 1.467E-05 | 1.155E-06 | 1.400E-05 | 1.600E-05 |
| 9897 | 495 | R495T | A | 1.5e-05 | 1.4e-05 | 1.5e-05 | 1.467E-05 | 5.774E-07 | 1.400E-05 | 1.500E-05 |
| 9898 | 495 | R495W | A | 9.3e-06 | 9.3e-06 | 1.1e-05 | 9.867E-06 | 9.815E-07 | 9.300E-06 | 1.100E-05 |
| 9899 | 495 | R495Y | A | 1.3e-05 | 1.5e-05 | 1.5e-05 | 1.433E-05 | 1.155E-06 | 1.300E-05 | 1.500E-05 |
| 9900 | 495 | R495V | A | 1.4e-05 | 1.3e-05 | 9.5e-06 | 1.217E-05 | 2.363E-06 | 9.500E-06 | 1.400E-05 |
| 9901 | 496 | G496A | A | 1e-05 | 1.5e-05 | 1.5e-05 | 1.333E-05 | 2.887E-06 | 1.000E-05 | 1.500E-05 |
| 9902 | 496 | G496R | A | 1.5e-05 | 1.6e-05 | 1.5e-05 | 1.533E-05 | 5.774E-07 | 1.500E-05 | 1.600E-05 |
| 9903 | 496 | G496N | A | 1.1e-05 | 1.4e-05 | 1.6e-05 | 1.367E-05 | 2.517E-06 | 1.100E-05 | 1.600E-05 |
| 9904 | 496 | G496D | A | 1e-05 | 9.2e-06 | 9.8e-06 | 9.667E-06 | 4.163E-07 | 9.200E-06 | 1.000E-05 |
| 9905 | 496 | G496C | A | 1.3e-05 | 1.2e-05 | 8e-06 | 1.100E-05 | 2.646E-06 | 8.000E-06 | 1.300E-05 |
| 9906 | 496 | G496E | A | 1e-05 | 8.3e-06 | 9.3e-06 | 9.200E-06 | 8.544E-07 | 8.300E-06 | 1.000E-05 |
| 9907 | 496 | G496Q | A | 1.3e-05 | 8.5e-06 | 9.8e-06 | 1.043E-05 | 2.316E-06 | 8.500E-06 | 1.300E-05 |
| 9908 | 496 | G496G | A | 1e-05 | 1.1e-05 | 6.4e-06 | 9.133E-06 | 2.419E-06 | 6.400E-06 | 1.100E-05 |
| 9909 | 496 | G496H | A | 8.7e-06 | 1.1e-05 | 1.6e-05 | 1.190E-05 | 3.732E-06 | 8.700E-06 | 1.600E-05 |
| 9910 | 496 | G496I | A | 1.3e-05 | 1.6e-05 | 9.9e-06 | 1.297E-05 | 3.050E-06 | 9.900E-06 | 1.600E-05 |
| 9911 | 496 | G496L | A | 8.8e-06 | 5.8e-06 | 1.4e-05 | 9.533E-06 | 4.149E-06 | 5.800E-06 | 1.400E-05 |
| 9912 | 496 | G496K | A | 1e-05 | 1.6e-05 | 8.6e-06 | 1.153E-05 | 3.931E-06 | 8.600E-06 | 1.600E-05 |
| 9913 | 496 | G496M | A | 1.3e-05 | 1.1e-05 | 1.3e-05 | 1.233E-05 | 1.155E-06 | 1.100E-05 | 1.300E-05 |
| 9914 | 496 | G496F | A | 6.4e-06 | 1.6e-05 | 1.6e-05 | 1.280E-05 | 5.543E-06 | 6.400E-06 | 1.600E-05 |
| 9915 | 496 | G496P | A | 1.3e-05 | 8.4e-06 | 1.6e-05 | 1.247E-05 | 3.828E-06 | 8.400E-06 | 1.600E-05 |
| 9916 | 496 | G496S | A | 1.5e-05 | 9.7e-06 | 8e-06 | 1.090E-05 | 3.651E-06 | 8.000E-06 | 1.500E-05 |
| 9917 | 496 | G496T | A | 1.4e-05 | 1.4e-05 | 1.3e-05 | 1.367E-05 | 5.774E-07 | 1.300E-05 | 1.400E-05 |
| 9918 | 496 | G496W | A | 9.7e-06 | 1.4e-05 | 1.7e-05 | 1.357E-05 | 3.669E-06 | 9.700E-06 | 1.700E-05 |
| 9919 | 496 | G496Y | A | 1.6e-05 | 8.6e-06 | 8.7e-06 | 1.110E-05 | 4.244E-06 | 8.600E-06 | 1.600E-05 |
| 9920 | 496 | G496V | A | 8.9e-06 | 1.5e-05 | 9.4e-06 | 1.110E-05 | 3.387E-06 | 8.900E-06 | 1.500E-05 |
| 9921 | 497 | H497A | A | 1.7e-05 | 1.1e-05 | 1.4e-05 | 1.400E-05 | 3.000E-06 | 1.100E-05 | 1.700E-05 |
| 9922 | 497 | H497R | A | 1.4e-05 | 1.4e-05 | 1.6e-05 | 1.467E-05 | 1.155E-06 | 1.400E-05 | 1.600E-05 |
| 9923 | 497 | H497N | A | 1.3e-05 | 1e-05 | 1.1e-05 | 1.133E-05 | 1.528E-06 | 1.000E-05 | 1.300E-05 |
| 9924 | 497 | H497D | A | 1.8e-05 | 1.6e-05 | 1.7e-05 | 1.700E-05 | 1.000E-06 | 1.600E-05 | 1.800E-05 |
| 9925 | 497 | H497C | A | 1.8e-05 | 1.7e-05 | 1.5e-05 | 1.667E-05 | 1.528E-06 | 1.500E-05 | 1.800E-05 |
| 9926 | 497 | H497E | A | 1.5e-05 | 1.5e-05 | 1.6e-05 | 1.533E-05 | 5.774E-07 | 1.500E-05 | 1.600E-05 |
| 9927 | 497 | H497Q | A | 8.9e-06 | 1e-05 | 1.2e-05 | 1.030E-05 | 1.572E-06 | 8.900E-06 | 1.200E-05 |
| 9928 | 497 | H497G | A | 1.5e-05 | 1.1e-05 | 1e-05 | 1.200E-05 | 2.646E-06 | 1.000E-05 | 1.500E-05 |
| 9929 | 497 | H497H | A | 1.4e-05 | 1e-05 | 1.1e-05 | 1.167E-05 | 2.082E-06 | 1.000E-05 | 1.400E-05 |
| 9930 | 497 | H497I | A | 1.2e-05 | 9.7e-06 | 1.7e-05 | 1.290E-05 | 3.732E-06 | 9.700E-06 | 1.700E-05 |
| 9931 | 497 | H497L | A | 7.9e-06 | 1.5e-05 | 1.1e-05 | 1.130E-05 | 3.559E-06 | 7.900E-06 | 1.500E-05 |
| 9932 | 497 | H497K | A | 1.7e-05 | 1.5e-05 | 1.1e-05 | 1.433E-05 | 3.055E-06 | 1.100E-05 | 1.700E-05 |
| 9933 | 497 | H497M | A | 1.5e-05 | 1.7e-05 | 1e-05 | 1.400E-05 | 3.606E-06 | 1.000E-05 | 1.700E-05 |
| 9934 | 497 | H497F | A | 2.2e-05 | 1e-05 | 1.1e-05 | 1.433E-05 | 6.658E-06 | 1.000E-05 | 2.200E-05 |
| 9935 | 497 | H497P | A | 1.3e-05 | 1.1e-05 | 1.3e-05 | 1.233E-05 | 1.155E-06 | 1.100E-05 | 1.300E-05 |
| 9936 | 497 | H497S | A | 7.2e-06 | 1.2e-05 | 9.4e-06 | 9.533E-06 | 2.403E-06 | 7.200E-06 | 1.200E-05 |
| 9937 | 497 | H497T | A | 1e-05 | 1.2e-05 | 1.7e-05 | 1.300E-05 | 3.606E-06 | 1.000E-05 | 1.700E-05 |
| 9938 | 497 | H497W | A | 1e-05 | 1e-05 | 1.1e-05 | 1.033E-05 | 5.774E-07 | 1.000E-05 | 1.100E-05 |
| 9939 | 497 | H497Y | A | 1.4e-05 | 1.5e-05 | 1.5e-05 | 1.467E-05 | 5.774E-07 | 1.400E-05 | 1.500E-05 |
| 9940 | 497 | H497V | A | 1.2e-05 | 1.6e-05 | 1.5e-05 | 1.433E-05 | 2.082E-06 | 1.200E-05 | 1.600E-05 |
| 9941 | 498 | C498A | A | 1.5e-05 | 1e-05 | 1.6e-05 | 1.367E-05 | 3.215E-06 | 1.000E-05 | 1.600E-05 |
| 9942 | 498 | C498R | A | 1.1e-05 | 8.4e-06 | 8e-06 | 9.133E-06 | 1.629E-06 | 8.000E-06 | 1.100E-05 |
| 9943 | 498 | C498N | A | 1.6e-05 | 2.4e-05 | 1.5e-05 | 1.833E-05 | 4.933E-06 | 1.500E-05 | 2.400E-05 |
| 9944 | 498 | C498D | A | 1.3e-05 | 8.6e-06 | 1e-05 | 1.053E-05 | 2.248E-06 | 8.600E-06 | 1.300E-05 |

| | | | | | | | | | | |
|---|---|---|---|---|---|---|---|---|---|---|
| 9945 | 498 | C498C | A | 1.1e-05 | 1.6e-05 | 1.3e-05 | 1.333E-05 | 2.517E-06 | 1.100E-05 | 1.600E-05 |
| 9946 | 498 | C498E | A | 9.2e-06 | 1e-05 | 1.5e-05 | 1.140E-05 | 3.143E-06 | 9.200E-06 | 1.500E-05 |
| 9947 | 498 | C498Q | A | 1.1e-05 | 1.4e-05 | 1.1e-05 | 1.200E-05 | 1.732E-06 | 1.100E-05 | 1.400E-05 |
| 9948 | 498 | C498G | A | 8.7e-06 | 1e-05 | 1.2e-05 | 1.023E-05 | 1.662E-06 | 8.700E-06 | 1.200E-05 |
| 9949 | 498 | C498H | A | 9.9e-06 | 9.5e-06 | 1.6e-05 | 1.180E-05 | 3.643E-06 | 9.500E-06 | 1.600E-05 |
| 9950 | 498 | C498I | A | 1e-05 | 1e-05 | 1e-05 | 1.000E-05 | 0.000E+00 | 1.000E-05 | 1.000E-05 |
| 9951 | 498 | C498L | A | 1.5e-05 | 1.4e-05 | 1.3e-05 | 1.400E-05 | 1.000E-06 | 1.300E-05 | 1.500E-05 |
| 9952 | 498 | C498K | A | 1.8e-05 | 1.7e-05 | 1.2e-05 | 1.567E-05 | 3.215E-06 | 1.200E-05 | 1.800E-05 |
| 9953 | 498 | C498M | A | 1e-05 | 1.2e-05 | 9.6e-06 | 1.053E-05 | 1.286E-06 | 9.600E-06 | 1.200E-05 |
| 9954 | 498 | C498F | A | 1.1e-05 | 9.4e-06 | 1.4e-05 | 1.147E-05 | 2.335E-06 | 9.400E-06 | 1.400E-05 |
| 9955 | 498 | C498P | A | 1.5e-05 | 1.3e-05 | 1.5e-05 | 1.433E-05 | 1.155E-06 | 1.300E-05 | 1.500E-05 |
| 9956 | 498 | C498S | A | 1.5e-05 | 9.9e-06 | 1.5e-05 | 1.330E-05 | 2.944E-06 | 9.900E-06 | 1.500E-05 |
| 9957 | 498 | C498T | A | 1.7e-05 | 1e-05 | 1.1e-05 | 1.267E-05 | 3.786E-06 | 1.000E-05 | 1.700E-05 |
| 9958 | 498 | C498W | A | 9.1e-06 | 9.3e-06 | 1e-05 | 9.467E-06 | 4.726E-07 | 9.100E-06 | 1.000E-05 |
| 9959 | 498 | C498Y | A | 1.5e-05 | 8.7e-06 | 9.1e-06 | 1.093E-05 | 3.528E-06 | 8.700E-06 | 1.500E-05 |
| 9960 | 498 | C498V | A | 1.3e-05 | 1.6e-05 | 1.1e-05 | 1.333E-05 | 2.517E-06 | 1.100E-05 | 1.600E-05 |
| 9961 | 499 | W499A | A | 8.9e-06 | 1.1e-05 | 1.1e-05 | 1.030E-05 | 1.212E-06 | 8.900E-06 | 1.100E-05 |
| 9962 | 499 | W499R | A | 1e-05 | 1.3e-05 | 1.6e-05 | 1.300E-05 | 3.000E-06 | 1.000E-05 | 1.600E-05 |
| 9963 | 499 | W499N | A | 1.5e-05 | 1.1e-05 | 1.6e-05 | 1.400E-05 | 2.646E-06 | 1.100E-05 | 1.600E-05 |
| 9964 | 499 | W499D | A | 1.5e-05 | 1.1e-05 | 1.6e-05 | 1.400E-05 | 2.646E-06 | 1.100E-05 | 1.600E-05 |
| 9965 | 499 | W499C | A | 9e-06 | 1.5e-05 | 1.6e-05 | 1.333E-05 | 3.786E-06 | 9.000E-06 | 1.600E-05 |
| 9966 | 499 | W499E | A | 1.2e-05 | 1.1e-05 | 9e-06 | 1.067E-05 | 1.528E-06 | 9.000E-06 | 1.200E-05 |
| 9967 | 499 | W499Q | A | 1.5e-05 | 1.6e-05 | 1.1e-05 | 1.400E-05 | 2.646E-06 | 1.100E-05 | 1.600E-05 |
| 9968 | 499 | W499G | A | 1.5e-05 | 1.6e-05 | 1.6e-05 | 1.567E-05 | 5.774E-07 | 1.500E-05 | 1.600E-05 |
| 9969 | 499 | W499H | A | 7.3e-06 | 1.8e-05 | 1.2e-05 | 1.243E-05 | 5.363E-06 | 7.300E-06 | 1.800E-05 |
| 9970 | 499 | W499I | A | 1.4e-05 | 1.8e-05 | 9.3e-06 | 1.377E-05 | 4.355E-06 | 9.300E-06 | 1.800E-05 |
| 9971 | 499 | W499L | A | 1.7e-05 | 1.4e-05 | 1.1e-05 | 1.400E-05 | 3.000E-06 | 1.100E-05 | 1.700E-05 |
| 9972 | 499 | W499K | A | 1.5e-05 | 1.1e-05 | 1.6e-05 | 1.400E-05 | 2.646E-06 | 1.100E-05 | 1.600E-05 |
| 9973 | 499 | W499M | A | 1.5e-05 | 1.4e-05 | 1.4e-05 | 1.433E-05 | 5.774E-07 | 1.400E-05 | 1.500E-05 |
| 9974 | 499 | W499F | A | 1.6e-05 | 1.8e-05 | 1.4e-05 | 1.600E-05 | 2.000E-06 | 1.400E-05 | 1.800E-05 |
| 9975 | 499 | W499P | A | 1.6e-05 | 1.6e-05 | 1.6e-05 | 1.600E-05 | 0.000E+00 | 1.600E-05 | 1.600E-05 |
| 9976 | 499 | W499S | A | 1.6e-05 | 1.1e-05 | 1.5e-05 | 1.400E-05 | 2.646E-06 | 1.100E-05 | 1.600E-05 |
| 9977 | 499 | W499T | A | 7.3e-06 | 1.5e-05 | 1e-05 | 1.077E-05 | 3.907E-06 | 7.300E-06 | 1.500E-05 |
| 9978 | 499 | W499W | A | 1.5e-05 | 1.5e-05 | 1.5e-05 | 1.500E-05 | 0.000E+00 | 1.500E-05 | 1.500E-05 |
| 9979 | 499 | W499Y | A | 1.4e-05 | 1.4e-05 | 1.1e-05 | 1.300E-05 | 1.732E-06 | 1.100E-05 | 1.400E-05 |
| 9980 | 499 | W499V | A | 1.5e-05 | 8.4e-06 | 1.2e-05 | 1.180E-05 | 3.305E-06 | 8.400E-06 | 1.500E-05 |
| 9981 | 500 | G500A | A | 1.4e-05 | 1.2e-05 | 8.4e-06 | 1.147E-05 | 2.838E-06 | 8.400E-06 | 1.400E-05 |
| 9982 | 500 | G500R | A | 1.3e-05 | 8.5e-06 | 1.5e-05 | 1.217E-05 | 3.329E-06 | 8.500E-06 | 1.500E-05 |
| 9983 | 500 | G500N | A | 1e-05 | 9.3e-06 | 8.6e-06 | 9.300E-06 | 7.000E-07 | 8.600E-06 | 1.000E-05 |
| 9984 | 500 | G500D | A | 9.7e-06 | 9.8e-06 | 6.2e-06 | 8.567E-06 | 2.050E-06 | 6.200E-06 | 9.800E-06 |
| 9985 | 500 | G500C | A | 1.1e-05 | 9.5e-06 | 9e-06 | 9.833E-06 | 1.041E-06 | 9.000E-06 | 1.100E-05 |
| 9986 | 500 | G500E | A | 1.1e-05 | 1.2e-05 | 9.8e-06 | 1.093E-05 | 1.102E-06 | 9.800E-06 | 1.200E-05 |
| 9987 | 500 | G500Q | A | 1.6e-05 | 1.2e-05 | 1e-05 | 1.267E-05 | 3.055E-06 | 1.000E-05 | 1.600E-05 |
| 9988 | 500 | G500G | A | 1.5e-05 | 9.6e-06 | 9.1e-06 | 1.123E-05 | 3.272E-06 | 9.100E-06 | 1.500E-05 |
| 9989 | 500 | G500H | A | 1.4e-05 | 1.5e-05 | 1.2e-05 | 1.367E-05 | 1.528E-06 | 1.200E-05 | 1.500E-05 |
| 9990 | 500 | G500I | A | 1.7e-05 | 1.4e-05 | 1.6e-05 | 1.567E-05 | 1.528E-06 | 1.400E-05 | 1.700E-05 |
| 9991 | 500 | G500L | A | 9.7e-06 | 9.9e-06 | 1.2e-05 | 1.053E-05 | 1.274E-06 | 9.700E-06 | 1.200E-05 |
| 9992 | 500 | G500K | A | 1.5e-05 | 1.3e-05 | 1.6e-05 | 1.467E-05 | 1.528E-06 | 1.300E-05 | 1.600E-05 |
| 9993 | 500 | G500M | A | 1.4e-05 | 9.7e-06 | 1.3e-05 | 1.223E-05 | 2.250E-06 | 9.700E-06 | 1.400E-05 |
| 9994 | 500 | G500F | A | 9.2e-06 | 1.5e-05 | 1e-05 | 1.140E-05 | 3.143E-06 | 9.200E-06 | 1.500E-05 |
| 9995 | 500 | G500P | A | 1.3e-05 | 1.1e-05 | 1.1e-05 | 1.167E-05 | 1.155E-06 | 1.100E-05 | 1.300E-05 |
| 9996 | 500 | G500S | A | 9.8e-06 | 1.6e-05 | 1.3e-05 | 1.293E-05 | 3.101E-06 | 9.800E-06 | 1.600E-05 |
| 9997 | 500 | G500T | A | 9.2e-06 | 1.6e-05 | 1.1e-05 | 1.207E-05 | 3.523E-06 | 9.200E-06 | 1.600E-05 |
| 9998 | 500 | G500W | A | 1e-05 | 1.6e-05 | 1.3e-05 | 1.300E-05 | 3.000E-06 | 1.000E-05 | 1.600E-05 |
| 9999 | 500 | G500Y | A | 1.4e-05 | 1.1e-05 | 9.6e-06 | 1.153E-05 | 2.248E-06 | 9.600E-06 | 1.400E-05 |
| 10000 | 500 | G500V | A | 1.6e-05 | 1.2e-05 | 1.1e-05 | 1.300E-05 | 2.646E-06 | 1.100E-05 | 1.600E-05 |
| 10001 | 501 | P501A | A | 1.6e-05 | 1.4e-05 | 1e-05 | 1.333E-05 | 3.055E-06 | 1.000E-05 | 1.600E-05 |
| 10002 | 501 | P501R | A | 1.1e-05 | 1.3e-05 | 9.8e-06 | 1.127E-05 | 1.617E-06 | 9.800E-06 | 1.300E-05 |
| 10003 | 501 | P501N | A | 1e-05 | 1.4e-05 | 1.6e-05 | 1.333E-05 | 3.055E-06 | 1.000E-05 | 1.600E-05 |
| 10004 | 501 | P501D | A | 1.6e-05 | 1.5e-05 | 1.7e-05 | 1.600E-05 | 1.000E-06 | 1.500E-05 | 1.700E-05 |
| 10005 | 501 | P501C | A | 1e-05 | 1.2e-05 | 8.5e-06 | 1.017E-05 | 1.756E-06 | 8.500E-06 | 1.200E-05 |
| 10006 | 501 | P501E | A | 1.6e-05 | 6.7e-06 | 1.5e-05 | 1.257E-05 | 5.105E-06 | 6.700E-06 | 1.600E-05 |
| 10007 | 501 | P501Q | A | 1.6e-05 | 1e-05 | 1.6e-05 | 1.400E-05 | 3.464E-06 | 1.000E-05 | 1.600E-05 |
| 10008 | 501 | P501G | A | 1.2e-05 | 1.5e-05 | 1.6e-05 | 1.433E-05 | 2.082E-06 | 1.200E-05 | 1.600E-05 |
| 10009 | 501 | P501H | A | 1.1e-05 | 1.5e-05 | 1.5e-05 | 1.367E-05 | 2.309E-06 | 1.100E-05 | 1.500E-05 |
| 10010 | 501 | P501I | A | 1e-05 | 9.6e-06 | 1.1e-05 | 1.020E-05 | 7.211E-07 | 9.600E-06 | 1.100E-05 |
| 10011 | 501 | P501L | A | 1.5e-05 | 9.7e-06 | 9e-06 | 1.123E-05 | 3.281E-06 | 9.000E-06 | 1.500E-05 |
| 10012 | 501 | P501K | A | 1.5e-05 | 1.1e-05 | 1.2e-05 | 1.267E-05 | 2.082E-06 | 1.100E-05 | 1.500E-05 |
| 10013 | 501 | P501M | A | 1.4e-05 | 1.3e-05 | 1.1e-05 | 1.267E-05 | 1.528E-06 | 1.100E-05 | 1.400E-05 |
| 10014 | 501 | P501F | A | 1.6e-05 | 1.2e-05 | 9.7e-06 | 1.257E-05 | 3.188E-06 | 9.700E-06 | 1.600E-05 |
| 10015 | 501 | P501P | A | 1e-05 | 1e-05 | 1e-05 | 1.000E-05 | 0.000E+00 | 1.000E-05 | 1.000E-05 |
| 10016 | 501 | P501S | A | 1.5e-05 | 9.7e-06 | 1.5e-05 | 1.323E-05 | 3.060E-06 | 9.700E-06 | 1.500E-05 |
| 10017 | 501 | P501T | A | 9.8e-06 | 1.6e-05 | 1.4e-05 | 1.327E-05 | 3.164E-06 | 9.800E-06 | 1.600E-05 |
| 10018 | 501 | P501W | A | 1.5e-05 | 1.5e-05 | 1.4e-05 | 1.467E-05 | 5.774E-07 | 1.400E-05 | 1.500E-05 |
| 10019 | 501 | P501Y | A | 1e-05 | 1.5e-05 | 1.5e-05 | 1.333E-05 | 2.887E-06 | 1.000E-05 | 1.500E-05 |
| 10020 | 501 | P501V | A | 7.6e-06 | 9.1e-06 | 1.6e-05 | 1.090E-05 | 4.480E-06 | 7.600E-06 | 1.600E-05 |
| 10021 | 502 | G502A | A | 1e-05 | 1e-05 | 1.1e-05 | 1.033E-05 | 5.774E-07 | 1.000E-05 | 1.100E-05 |
| 10022 | 502 | G502R | A | 1.8e-05 | 9.7e-06 | 1.1e-05 | 1.290E-05 | 4.464E-06 | 9.700E-06 | 1.800E-05 |
| 10023 | 502 | G502N | A | 1.4e-05 | 8.7e-06 | 9.2e-06 | 1.063E-05 | 2.926E-06 | 8.700E-06 | 1.400E-05 |
| 10024 | 502 | G502D | A | 9e-06 | 9e-06 | 1.5e-05 | 1.100E-05 | 3.464E-06 | 9.000E-06 | 1.500E-05 |
| 10025 | 502 | G502C | A | 1.4e-05 | 1.4e-05 | 1.5e-05 | 1.433E-05 | 5.774E-07 | 1.400E-05 | 1.500E-05 |
| 10026 | 502 | G502E | A | 1e-05 | 1e-05 | 1e-05 | 1.000E-05 | 0.000E+00 | 1.000E-05 | 1.000E-05 |
| 10027 | 502 | G502Q | A | 1.6e-05 | 9.9e-06 | 9.7e-06 | 1.187E-05 | 3.581E-06 | 9.700E-06 | 1.600E-05 |
| 10028 | 502 | G502G | A | 1.2e-05 | 1.3e-05 | 1.1e-05 | 1.200E-05 | 1.000E-06 | 1.100E-05 | 1.300E-05 |
| 10029 | 502 | G502H | A | 1.1e-05 | 1.3e-05 | 9.8e-06 | 1.127E-05 | 1.617E-06 | 9.800E-06 | 1.300E-05 |

| | | | | | | | | | | |
|---|---|---|---|---|---|---|---|---|---|---|
| 10030 | 502 | G502I | A | 1.4e-05 | 9.8e-06 | 1.4e-05 | 1.260E-05 | 2.425E-06 | 9.800E-06 | 1.400E-05 |
| 10031 | 502 | G502L | A | 1.7e-05 | 1.1e-05 | 1e-05 | 1.267E-05 | 3.786E-06 | 1.000E-05 | 1.700E-05 |
| 10032 | 502 | G502K | A | 1.5e-05 | 1.1e-05 | 1.4e-05 | 1.333E-05 | 2.082E-06 | 1.100E-05 | 1.500E-05 |
| 10033 | 502 | G502M | A | 1.4e-05 | 9.2e-06 | 1.5e-05 | 1.273E-05 | 3.101E-06 | 9.200E-06 | 1.500E-05 |
| 10034 | 502 | G502F | A | 1.5e-05 | 7.8e-06 | 1.2e-05 | 1.160E-05 | 3.617E-06 | 7.800E-06 | 1.500E-05 |
| 10035 | 502 | G502P | A | 1.4e-05 | 1.5e-05 | 1.4e-05 | 1.433E-05 | 5.774E-07 | 1.400E-05 | 1.500E-05 |
| 10036 | 502 | G502S | A | 9.1e-06 | 7.1e-06 | 1.6e-05 | 1.073E-05 | 4.669E-06 | 7.100E-06 | 1.600E-05 |
| 10037 | 502 | G502T | A | 1.1e-05 | 1.1e-05 | 1e-05 | 1.067E-05 | 5.774E-07 | 1.000E-05 | 1.100E-05 |
| 10038 | 502 | G502W | A | 1.1e-05 | 1.1e-05 | 1.4e-05 | 1.200E-05 | 1.732E-06 | 1.100E-05 | 1.400E-05 |
| 10039 | 502 | G502Y | A | 1.1e-05 | 1.3e-05 | 1.1e-05 | 1.167E-05 | 1.155E-06 | 1.100E-05 | 1.300E-05 |
| 10040 | 502 | G502V | A | 1.1e-05 | 1.1e-05 | 1.1e-05 | 1.100E-05 | 2.075E-21 | 1.100E-05 | 1.100E-05 |
| 10041 | 503 | P503A | A | 1.6e-05 | 1.6e-05 | 1.6e-05 | 1.600E-05 | 0.000E+00 | 1.600E-05 | 1.600E-05 |
| 10042 | 503 | P503R | A | 1e-05 | 1.2e-05 | 1.1e-05 | 1.100E-05 | 1.000E-06 | 1.000E-05 | 1.200E-05 |
| 10043 | 503 | P503N | A | 1e-05 | 1e-05 | 1.3e-05 | 1.100E-05 | 1.732E-06 | 1.000E-05 | 1.300E-05 |
| 10044 | 503 | P503D | A | 1.1e-05 | 1.5e-05 | 9e-06 | 1.167E-05 | 3.055E-06 | 9.000E-06 | 1.500E-05 |
| 10045 | 503 | P503C | A | 1e-05 | 1.4e-05 | 1.1e-05 | 1.167E-05 | 2.082E-06 | 1.000E-05 | 1.400E-05 |
| 10046 | 503 | P503E | A | 1.6e-05 | 7.4e-06 | 1.5e-05 | 1.280E-05 | 4.703E-06 | 7.400E-06 | 1.600E-05 |
| 10047 | 503 | P503Q | A | 1e-05 | 1.3e-05 | 1.4e-05 | 1.233E-05 | 2.082E-06 | 1.000E-05 | 1.400E-05 |
| 10048 | 503 | P503G | A | 1.1e-05 | 1.6e-05 | 1.5e-05 | 1.400E-05 | 2.646E-06 | 1.100E-05 | 1.600E-05 |
| 10049 | 503 | P503H | A | 1.3e-05 | 1.2e-05 | 1.5e-05 | 1.333E-05 | 1.528E-06 | 1.200E-05 | 1.500E-05 |
| 10050 | 503 | P503I | A | 1.3e-05 | 1.6e-05 | 1.6e-05 | 1.500E-05 | 1.732E-06 | 1.300E-05 | 1.600E-05 |
| 10051 | 503 | P503L | A | 1e-05 | 1.5e-05 | 1.1e-05 | 1.200E-05 | 2.646E-06 | 1.000E-05 | 1.500E-05 |
| 10052 | 503 | P503K | A | 1.3e-05 | 1.1e-05 | 1e-05 | 1.133E-05 | 1.528E-06 | 1.000E-05 | 1.300E-05 |
| 10053 | 503 | P503M | A | 1.1e-05 | 1.4e-05 | 1.1e-05 | 1.200E-05 | 1.732E-06 | 1.100E-05 | 1.400E-05 |
| 10054 | 503 | P503F | A | 1.1e-05 | 9.7e-06 | 1.6e-05 | 1.223E-05 | 3.326E-06 | 9.700E-06 | 1.600E-05 |
| 10055 | 503 | P503P | A | 1.1e-05 | 1.1e-05 | 1e-05 | 1.067E-05 | 5.774E-07 | 1.000E-05 | 1.100E-05 |
| 10056 | 503 | P503S | A | 9.1e-06 | 1.6e-05 | 1e-05 | 1.170E-05 | 3.751E-06 | 9.100E-06 | 1.600E-05 |
| 10057 | 503 | P503T | A | 1.4e-05 | 1.5e-05 | 1.4e-05 | 1.433E-05 | 5.774E-07 | 1.400E-05 | 1.500E-05 |
| 10058 | 503 | P503W | A | 1e-05 | 9.1e-06 | 1.7e-05 | 1.203E-05 | 4.325E-06 | 9.100E-06 | 1.700E-05 |
| 10059 | 503 | P503Y | A | 1.2e-05 | 1.5e-05 | 1.2e-05 | 1.300E-05 | 1.732E-06 | 1.200E-05 | 1.500E-05 |
| 10060 | 503 | P503V | A | 1.5e-05 | 1.1e-05 | 1.5e-05 | 1.367E-05 | 2.309E-06 | 1.100E-05 | 1.500E-05 |
| 10061 | 504 | T504A | A | 1e-05 | 1.7e-05 | 1.1e-05 | 1.267E-05 | 3.786E-06 | 1.000E-05 | 1.700E-05 |
| 10062 | 504 | T504R | A | 6.7e-06 | 1.7e-05 | 1.6e-05 | 1.323E-05 | 5.680E-06 | 6.700E-06 | 1.700E-05 |
| 10063 | 504 | T504N | A | 8.1e-06 | 1.5e-05 | 9.9e-06 | 1.100E-05 | 3.579E-06 | 8.100E-06 | 1.500E-05 |
| 10064 | 504 | T504D | A | 1.5e-05 | 1.5e-05 | 1.4e-05 | 1.467E-05 | 5.774E-07 | 1.400E-05 | 1.500E-05 |
| 10065 | 504 | T504C | A | 8e-06 | 1e-05 | 8.8e-06 | 8.933E-06 | 1.007E-06 | 8.000E-06 | 1.000E-05 |
| 10066 | 504 | T504E | A | 1e-05 | 1.1e-05 | 1.6e-05 | 1.233E-05 | 3.215E-06 | 1.000E-05 | 1.600E-05 |
| 10067 | 504 | T504Q | A | 1.6e-05 | 9.3e-06 | 7.4e-06 | 1.090E-05 | 4.518E-06 | 7.400E-06 | 1.600E-05 |
| 10068 | 504 | T504G | A | 1e-05 | 1.6e-05 | 1.6e-05 | 1.400E-05 | 3.464E-06 | 1.000E-05 | 1.600E-05 |
| 10069 | 504 | T504H | A | 1.5e-05 | 1.2e-05 | 9.7e-06 | 1.223E-05 | 2.658E-06 | 9.700E-06 | 1.500E-05 |
| 10070 | 504 | T504I | A | 1.5e-05 | 1.6e-05 | 9.6e-06 | 1.353E-05 | 3.443E-06 | 9.600E-06 | 1.600E-05 |
| 10071 | 504 | T504L | A | 1.1e-05 | 1.3e-05 | 1.6e-05 | 1.333E-05 | 2.517E-06 | 1.100E-05 | 1.600E-05 |
| 10072 | 504 | T504K | A | 1.6e-05 | 1e-05 | 8.2e-06 | 1.140E-05 | 4.084E-06 | 8.200E-06 | 1.600E-05 |
| 10073 | 504 | T504M | A | 1.4e-05 | 1.3e-05 | 1.8e-05 | 1.500E-05 | 2.646E-06 | 1.300E-05 | 1.800E-05 |
| 10074 | 504 | T504F | A | 1.1e-05 | 1.1e-05 | 1.1e-05 | 1.100E-05 | 2.075E-21 | 1.100E-05 | 1.100E-05 |
| 10075 | 504 | T504P | A | 1.1e-05 | 1.4e-05 | 1.1e-05 | 1.200E-05 | 1.732E-06 | 1.100E-05 | 1.400E-05 |
| 10076 | 504 | T504S | A | 1.4e-05 | 9.7e-06 | 1.1e-05 | 1.157E-05 | 2.205E-06 | 9.700E-06 | 1.400E-05 |
| 10077 | 504 | T504T | A | 1.5e-05 | 1.3e-05 | 1.7e-05 | 1.500E-05 | 2.000E-06 | 1.300E-05 | 1.700E-05 |
| 10078 | 504 | T504W | A | 1.2e-05 | 1.1e-05 | 1e-05 | 1.100E-05 | 1.000E-06 | 1.000E-05 | 1.200E-05 |
| 10079 | 504 | T504Y | A | 1e-05 | 1.7e-05 | 1.5e-05 | 1.400E-05 | 3.606E-06 | 1.000E-05 | 1.700E-05 |
| 10080 | 504 | T504V | A | 1.4e-05 | 1.5e-05 | 1.3e-05 | 1.400E-05 | 1.000E-06 | 1.300E-05 | 1.500E-05 |
| 10081 | 505 | Q505A | A | 1.7e-05 | 1.4e-05 | 1.4e-05 | 1.500E-05 | 1.732E-06 | 1.400E-05 | 1.700E-05 |
| 10082 | 505 | Q505R | A | 1.1e-05 | 1.1e-05 | 1.4e-05 | 1.200E-05 | 1.732E-06 | 1.100E-05 | 1.400E-05 |
| 10083 | 505 | Q505N | A | 1.1e-05 | 1.5e-05 | 1e-05 | 1.200E-05 | 2.646E-06 | 1.000E-05 | 1.500E-05 |
| 10084 | 505 | Q505D | A | 8.4e-06 | 1.6e-05 | 1.7e-05 | 1.380E-05 | 4.703E-06 | 8.400E-06 | 1.700E-05 |
| 10085 | 505 | Q505C | A | 8.3e-06 | 1.5e-05 | 1.5e-05 | 1.277E-05 | 3.868E-06 | 8.300E-06 | 1.500E-05 |
| 10086 | 505 | Q505E | A | 8.3e-06 | 1.2e-05 | 1.5e-05 | 1.177E-05 | 3.356E-06 | 8.300E-06 | 1.500E-05 |
| 10087 | 505 | Q505Q | A | 1.4e-05 | 1.5e-05 | 7.4e-06 | 1.213E-05 | 4.130E-06 | 7.400E-06 | 1.500E-05 |
| 10088 | 505 | Q505G | A | 1e-05 | 1.2e-05 | 1e-05 | 1.067E-05 | 1.155E-06 | 1.000E-05 | 1.200E-05 |
| 10089 | 505 | Q505H | A | 1.4e-05 | 1.6e-05 | 1.1e-05 | 1.367E-05 | 2.517E-06 | 1.100E-05 | 1.600E-05 |
| 10090 | 505 | Q505I | A | 1.6e-05 | 1.1e-05 | 1e-05 | 1.233E-05 | 3.215E-06 | 1.000E-05 | 1.600E-05 |
| 10091 | 505 | Q505L | A | 1.1e-05 | 1.5e-05 | 1.9e-05 | 1.500E-05 | 4.000E-06 | 1.100E-05 | 1.900E-05 |
| 10092 | 505 | Q505K | A | 1.5e-05 | 1.6e-05 | 7.9e-06 | 1.297E-05 | 4.416E-06 | 7.900E-06 | 1.600E-05 |
| 10093 | 505 | Q505M | A | 1.4e-05 | 1.5e-05 | 1.2e-05 | 1.367E-05 | 1.528E-06 | 1.200E-05 | 1.500E-05 |
| 10094 | 505 | Q505F | A | 1.4e-05 | 8.3e-06 | 1.6e-05 | 1.277E-05 | 3.995E-06 | 8.300E-06 | 1.600E-05 |
| 10095 | 505 | Q505P | A | 1.2e-05 | 9.8e-06 | 1.2e-05 | 1.127E-05 | 1.270E-06 | 9.800E-06 | 1.200E-05 |
| 10096 | 505 | Q505S | A | 1.6e-05 | 1.5e-05 | 1.8e-05 | 1.633E-05 | 1.528E-06 | 1.500E-05 | 1.800E-05 |
| 10097 | 505 | Q505T | A | 9.7e-06 | 1.7e-05 | 1e-05 | 1.223E-05 | 4.131E-06 | 9.700E-06 | 1.700E-05 |
| 10098 | 505 | Q505W | A | 9.3e-06 | 9.8e-06 | 1.2e-05 | 1.037E-05 | 1.436E-06 | 9.300E-06 | 1.200E-05 |
| 10099 | 505 | Q505Y | A | 1.6e-05 | 1.1e-05 | 1.4e-05 | 1.367E-05 | 2.517E-06 | 1.100E-05 | 1.600E-05 |
| 10100 | 505 | Q505V | A | 1.2e-05 | 1.5e-05 | 1.1e-05 | 1.267E-05 | 2.082E-06 | 1.100E-05 | 1.500E-05 |
| 10101 | 506 | C506A | A | 1.6e-05 | 1.7e-05 | 1.4e-05 | 1.567E-05 | 1.528E-06 | 1.400E-05 | 1.700E-05 |
| 10102 | 506 | C506R | A | 9.5e-06 | 1.2e-05 | 8.7e-06 | 1.007E-05 | 1.721E-06 | 8.700E-06 | 1.200E-05 |
| 10103 | 506 | C506N | A | 1.6e-05 | 1.5e-05 | 1.2e-05 | 1.433E-05 | 2.082E-06 | 1.200E-05 | 1.600E-05 |
| 10104 | 506 | C506D | A | 1e-05 | 1.4e-05 | 1.4e-05 | 1.267E-05 | 2.309E-06 | 1.000E-05 | 1.400E-05 |
| 10105 | 506 | C506C | A | 9.7e-06 | 1.4e-05 | 1.4e-05 | 1.257E-05 | 2.483E-06 | 9.700E-06 | 1.400E-05 |
| 10106 | 506 | C506E | A | 9.9e-06 | 1.5e-05 | 1.1e-05 | 1.197E-05 | 2.684E-06 | 9.900E-06 | 1.500E-05 |
| 10107 | 506 | C506Q | A | 1.2e-05 | 1.2e-05 | 1.1e-05 | 1.167E-05 | 5.774E-07 | 1.100E-05 | 1.200E-05 |
| 10108 | 506 | C506G | A | 1.5e-05 | 1.2e-05 | 1.7e-05 | 1.467E-05 | 2.517E-06 | 1.200E-05 | 1.700E-05 |
| 10109 | 506 | C506H | A | 1.6e-05 | 1.4e-05 | 1.1e-05 | 1.367E-05 | 2.517E-06 | 1.100E-05 | 1.600E-05 |
| 10110 | 506 | C506I | A | 9.2e-06 | 9.6e-06 | 1.5e-05 | 1.127E-05 | 3.239E-06 | 9.200E-06 | 1.500E-05 |
| 10111 | 506 | C506L | A | 8e-06 | 1e-05 | 1.5e-05 | 1.100E-05 | 3.606E-06 | 8.000E-06 | 1.500E-05 |
| 10112 | 506 | C506K | A | 1.7e-05 | 1.6e-05 | 9.4e-06 | 1.413E-05 | 4.130E-06 | 9.400E-06 | 1.700E-05 |
| 10113 | 506 | C506M | A | 1.3e-05 | 1.1e-05 | 1.3e-05 | 1.233E-05 | 1.155E-06 | 1.100E-05 | 1.300E-05 |
| 10114 | 506 | C506F | A | 1.4e-05 | 1.6e-05 | 1.5e-05 | 1.500E-05 | 1.000E-06 | 1.400E-05 | 1.600E-05 |

| | | | | | | | | | | |
|---|---|---|---|---|---|---|---|---|---|---|
| 10115 | 506 | C506P | A | 1.1e-05 | 1.3e-05 | 1.4e-05 | 1.267E-05 | 1.528E-06 | 1.100E-05 | 1.400E-05 |
| 10116 | 506 | C506S | A | 1.5e-05 | 1.6e-05 | 1.5e-05 | 1.533E-05 | 5.774E-07 | 1.500E-05 | 1.600E-05 |
| 10117 | 506 | C506T | A | 1.2e-05 | 1.2e-05 | 1e-05 | 1.133E-05 | 1.155E-06 | 1.000E-05 | 1.200E-05 |
| 10118 | 506 | C506W | A | 1.4e-05 | 8.5e-06 | 1.2e-05 | 1.150E-05 | 2.784E-06 | 8.500E-06 | 1.400E-05 |
| 10119 | 506 | C506Y | A | 1.1e-05 | 1.4e-05 | 1.4e-05 | 1.300E-05 | 1.732E-06 | 1.100E-05 | 1.400E-05 |
| 10120 | 506 | C506V | A | 1.5e-05 | 1.1e-05 | 1.7e-05 | 1.433E-05 | 3.055E-06 | 1.100E-05 | 1.700E-05 |
| 10121 | 507 | V507A | A | 1.3e-05 | 1.3e-05 | 1.5e-05 | 1.367E-05 | 1.155E-06 | 1.300E-05 | 1.500E-05 |
| 10122 | 507 | V507R | A | 1e-05 | 1.4e-05 | 1.4e-05 | 1.267E-05 | 2.309E-06 | 1.000E-05 | 1.400E-05 |
| 10123 | 507 | V507N | A | 1.1e-05 | 1e-05 | 9.3e-06 | 1.010E-05 | 8.544E-07 | 9.300E-06 | 1.100E-05 |
| 10124 | 507 | V507D | A | 1.5e-05 | 1.7e-05 | 1e-05 | 1.400E-05 | 3.606E-06 | 1.000E-05 | 1.700E-05 |
| 10125 | 507 | V507C | A | 1.4e-05 | 9.7e-06 | 9.3e-06 | 1.100E-05 | 2.606E-06 | 9.300E-06 | 1.400E-05 |
| 10126 | 507 | V507E | A | 1.6e-05 | 1.6e-05 | 1.5e-05 | 1.567E-05 | 5.774E-07 | 1.500E-05 | 1.600E-05 |
| 10127 | 507 | V507Q | A | 1.5e-05 | 9.1e-06 | 1.5e-05 | 1.303E-05 | 3.406E-06 | 9.100E-06 | 1.500E-05 |
| 10128 | 507 | V507G | A | 1e-05 | 9.7e-06 | 1.6e-05 | 1.190E-05 | 3.554E-06 | 9.700E-06 | 1.600E-05 |
| 10129 | 507 | V507H | A | 1e-05 | 8.5e-06 | 1.5e-05 | 1.117E-05 | 3.403E-06 | 8.500E-06 | 1.500E-05 |
| 10130 | 507 | V507I | A | 1.6e-05 | 1.5e-05 | 1.4e-05 | 1.500E-05 | 1.000E-06 | 1.400E-05 | 1.600E-05 |
| 10131 | 507 | V507L | A | 1.1e-05 | 1.6e-05 | 1.6e-05 | 1.433E-05 | 2.887E-06 | 1.100E-05 | 1.600E-05 |
| 10132 | 507 | V507K | A | 8.4e-06 | 1.5e-05 | 1.5e-05 | 1.280E-05 | 3.811E-06 | 8.400E-06 | 1.500E-05 |
| 10133 | 507 | V507M | A | 1.5e-05 | 1.1e-05 | 1.5e-05 | 1.367E-05 | 2.309E-06 | 1.100E-05 | 1.500E-05 |
| 10134 | 507 | V507F | A | 1.2e-05 | 1.5e-05 | 1.4e-05 | 1.367E-05 | 1.528E-06 | 1.200E-05 | 1.500E-05 |
| 10135 | 507 | V507P | A | 1e-05 | 1.3e-05 | 1.4e-05 | 1.233E-05 | 2.082E-06 | 1.000E-05 | 1.400E-05 |
| 10136 | 507 | V507S | A | 1.1e-05 | 1.6e-05 | 1.1e-05 | 1.267E-05 | 2.887E-06 | 1.100E-05 | 1.600E-05 |
| 10137 | 507 | V507T | A | 1.5e-05 | 8.1e-06 | 8.3e-06 | 1.047E-05 | 3.927E-06 | 8.100E-06 | 1.500E-05 |
| 10138 | 507 | V507W | A | 9.3e-06 | 1.1e-05 | 1e-05 | 1.010E-05 | 8.544E-07 | 9.300E-06 | 1.100E-05 |
| 10139 | 507 | V507Y | A | 8e-06 | 1.4e-05 | 1.3e-05 | 1.167E-05 | 3.215E-06 | 8.000E-06 | 1.400E-05 |
| 10140 | 507 | V507V | A | 1.5e-05 | 1.1e-05 | 1.1e-05 | 1.233E-05 | 2.309E-06 | 1.100E-05 | 1.500E-05 |
| 10141 | 508 | N508A | A | 1.6e-05 | 1.4e-05 | 9e-06 | 1.300E-05 | 3.606E-06 | 9.000E-06 | 1.600E-05 |
| 10142 | 508 | N508R | A | 1.5e-05 | 1.4e-05 | 1.5e-05 | 1.467E-05 | 5.774E-07 | 1.400E-05 | 1.500E-05 |
| 10143 | 508 | N508N | A | 1.4e-05 | 1.6e-05 | 1.1e-05 | 1.367E-05 | 2.517E-06 | 1.100E-05 | 1.600E-05 |
| 10144 | 508 | N508D | A | 1.4e-05 | 1e-05 | 1.1e-05 | 1.167E-05 | 2.082E-06 | 1.000E-05 | 1.400E-05 |
| 10145 | 508 | N508C | A | 1.5e-05 | 8.2e-06 | 1.4e-05 | 1.240E-05 | 3.672E-06 | 8.200E-06 | 1.500E-05 |
| 10146 | 508 | N508E | A | 1.3e-05 | 1.7e-05 | 1.4e-05 | 1.467E-05 | 2.082E-06 | 1.300E-05 | 1.700E-05 |
| 10147 | 508 | N508Q | A | 1.1e-05 | 1.7e-05 | 8.7e-06 | 1.223E-05 | 4.285E-06 | 8.700E-06 | 1.700E-05 |
| 10148 | 508 | N508G | A | 9.1e-06 | 9.8e-06 | 1.8e-05 | 1.230E-05 | 4.949E-06 | 9.100E-06 | 1.800E-05 |
| 10149 | 508 | N508H | A | 1.3e-05 | 1.3e-05 | 1.3e-05 | 1.300E-05 | 0.000E+00 | 1.300E-05 | 1.300E-05 |
| 10150 | 508 | N508I | A | 1.6e-05 | 1.6e-05 | 9.7e-06 | 1.390E-05 | 3.637E-06 | 9.700E-06 | 1.600E-05 |
| 10151 | 508 | N508L | A | 1e-05 | 1.6e-05 | 1.1e-05 | 1.233E-05 | 3.215E-06 | 1.000E-05 | 1.600E-05 |
| 10152 | 508 | N508K | A | 9.1e-06 | 1.4e-05 | 1.6e-05 | 1.303E-05 | 3.550E-06 | 9.100E-06 | 1.600E-05 |
| 10153 | 508 | N508M | A | 1.3e-05 | 8.8e-06 | 1e-05 | 1.060E-05 | 2.163E-06 | 8.800E-06 | 1.300E-05 |
| 10154 | 508 | N508F | A | 1e-05 | 1.2e-05 | 1.7e-05 | 1.300E-05 | 3.606E-06 | 1.000E-05 | 1.700E-05 |
| 10155 | 508 | N508P | A | 1.5e-05 | 1.5e-05 | 1.7e-05 | 1.567E-05 | 1.155E-06 | 1.500E-05 | 1.700E-05 |
| 10156 | 508 | N508S | A | 9.6e-06 | 8.7e-06 | 1.1e-05 | 9.767E-06 | 1.159E-06 | 8.700E-06 | 1.100E-05 |
| 10157 | 508 | N508T | A | 1.4e-05 | 1.5e-05 | 1.6e-05 | 1.500E-05 | 1.000E-06 | 1.400E-05 | 1.600E-05 |
| 10158 | 508 | N508W | A | 8.5e-06 | 6.3e-06 | 8.6e-06 | 7.800E-06 | 1.300E-06 | 6.300E-06 | 8.600E-06 |
| 10159 | 508 | N508Y | A | 1.4e-05 | 8.4e-06 | 9.4e-06 | 1.060E-05 | 2.987E-06 | 8.400E-06 | 1.400E-05 |
| 10160 | 508 | N508V | A | 1.6e-05 | 1.2e-05 | 1.3e-05 | 1.367E-05 | 2.082E-06 | 1.200E-05 | 1.600E-05 |
| 10161 | 509 | C509A | A | 1.6e-05 | 1.6e-05 | 2.3e-05 | 1.833E-05 | 4.041E-06 | 1.600E-05 | 2.300E-05 |
| 10162 | 509 | C509R | A | 1.6e-05 | 1e-05 | 1.2e-05 | 1.267E-05 | 3.055E-06 | 1.000E-05 | 1.600E-05 |
| 10163 | 509 | C509N | A | 1.2e-05 | 1.4e-05 | 1.5e-05 | 1.367E-05 | 1.528E-06 | 1.200E-05 | 1.500E-05 |
| 10164 | 509 | C509D | A | 1.5e-05 | 1.6e-05 | 1.6e-05 | 1.567E-05 | 5.774E-07 | 1.500E-05 | 1.600E-05 |
| 10165 | 509 | C509C | A | 1.2e-05 | 1.5e-05 | 1.4e-05 | 1.367E-05 | 1.528E-06 | 1.200E-05 | 1.500E-05 |
| 10166 | 509 | C509E | A | 1.4e-05 | 1.1e-05 | 1.4e-05 | 1.300E-05 | 1.732E-06 | 1.100E-05 | 1.400E-05 |
| 10167 | 509 | C509Q | A | 1.6e-05 | 1.5e-05 | 9.1e-06 | 1.337E-05 | 3.729E-06 | 9.100E-06 | 1.600E-05 |
| 10168 | 509 | C509G | A | 1.4e-05 | 1.2e-05 | 1.1e-05 | 1.233E-05 | 1.528E-06 | 1.100E-05 | 1.400E-05 |
| 10169 | 509 | C509H | A | 1.5e-05 | 1.5e-05 | 1.5e-05 | 1.500E-05 | 0.000E+00 | 1.500E-05 | 1.500E-05 |
| 10170 | 509 | C509I | A | 1.4e-05 | 1.5e-05 | 1.1e-05 | 1.333E-05 | 2.082E-06 | 1.100E-05 | 1.500E-05 |
| 10171 | 509 | C509L | A | 1.7e-05 | 1.6e-05 | 1e-05 | 1.433E-05 | 3.786E-06 | 1.000E-05 | 1.700E-05 |
| 10172 | 509 | C509K | A | 1.5e-05 | 1.6e-05 | 1.6e-05 | 1.567E-05 | 5.774E-07 | 1.500E-05 | 1.600E-05 |
| 10173 | 509 | C509M | A | 1.6e-05 | 1.5e-05 | 1.7e-05 | 1.600E-05 | 1.000E-06 | 1.500E-05 | 1.700E-05 |
| 10174 | 509 | C509F | A | 1.1e-05 | 9.4e-06 | 1e-05 | 1.013E-05 | 8.083E-07 | 9.400E-06 | 1.100E-05 |
| 10175 | 509 | C509P | A | 1.1e-05 | 1.5e-05 | 1.6e-05 | 1.400E-05 | 2.646E-06 | 1.100E-05 | 1.600E-05 |
| 10176 | 509 | C509S | A | 1.5e-05 | 1.1e-05 | 1.5e-05 | 1.367E-05 | 2.309E-06 | 1.100E-05 | 1.500E-05 |
| 10177 | 509 | C509T | A | 1.5e-05 | 1.2e-05 | 9.4e-06 | 1.213E-05 | 2.802E-06 | 9.400E-06 | 1.500E-05 |
| 10178 | 509 | C509W | A | 1.5e-05 | 9.1e-06 | 1.4e-05 | 1.270E-05 | 3.158E-06 | 9.100E-06 | 1.500E-05 |
| 10179 | 509 | C509Y | A | 1.6e-05 | 9.9e-06 | 1e-05 | 1.197E-05 | 3.493E-06 | 9.900E-06 | 1.600E-05 |
| 10180 | 509 | C509V | A | 1.6e-05 | 1.6e-05 | 1.7e-05 | 1.633E-05 | 5.774E-07 | 1.600E-05 | 1.700E-05 |
| 10181 | 510 | S510A | A | 9.3e-06 | 1.4e-05 | 1.3e-05 | 1.210E-05 | 2.476E-06 | 9.300E-06 | 1.400E-05 |
| 10182 | 510 | S510R | A | 1.1e-05 | 1.1e-05 | 1.8e-05 | 1.333E-05 | 4.041E-06 | 1.100E-05 | 1.800E-05 |
| 10183 | 510 | S510N | A | 1.5e-05 | 1.4e-05 | 1.1e-05 | 1.333E-05 | 2.082E-06 | 1.100E-05 | 1.500E-05 |
| 10184 | 510 | S510D | A | 9.8e-06 | 9.2e-06 | 1.4e-05 | 1.100E-05 | 2.615E-06 | 9.200E-06 | 1.400E-05 |
| 10185 | 510 | S510C | A | 1e-05 | 1.4e-05 | 1.2e-05 | 1.200E-05 | 2.000E-06 | 1.000E-05 | 1.400E-05 |
| 10186 | 510 | S510E | A | 1.1e-05 | 1.5e-05 | 1.5e-05 | 1.367E-05 | 2.309E-06 | 1.100E-05 | 1.500E-05 |
| 10187 | 510 | S510Q | A | 1.4e-05 | 1.1e-05 | 1.6e-05 | 1.367E-05 | 2.517E-06 | 1.100E-05 | 1.600E-05 |
| 10188 | 510 | S510G | A | 1.5e-05 | 1.1e-05 | 1.5e-05 | 1.367E-05 | 2.309E-06 | 1.100E-05 | 1.500E-05 |
| 10189 | 510 | S510H | A | 1e-05 | 1.4e-05 | 1.4e-05 | 1.267E-05 | 2.309E-06 | 1.000E-05 | 1.400E-05 |
| 10190 | 510 | S510I | A | 1.6e-05 | 1.5e-05 | 8e-06 | 1.300E-05 | 4.359E-06 | 8.000E-06 | 1.600E-05 |
| 10191 | 510 | S510L | A | 9.4e-06 | 1.1e-05 | 1.5e-05 | 1.180E-05 | 2.884E-06 | 9.400E-06 | 1.500E-05 |
| 10192 | 510 | S510K | A | 1.1e-05 | 1.5e-05 | 1.7e-05 | 1.433E-05 | 3.055E-06 | 1.100E-05 | 1.700E-05 |
| 10193 | 510 | S510M | A | 1.6e-05 | 1e-05 | 1.7e-05 | 1.433E-05 | 3.786E-06 | 1.000E-05 | 1.700E-05 |
| 10194 | 510 | S510F | A | 1.1e-05 | 1.3e-05 | 1.8e-05 | 1.400E-05 | 3.606E-06 | 1.100E-05 | 1.800E-05 |
| 10195 | 510 | S510P | A | 1.1e-05 | 1.7e-05 | 1.2e-05 | 1.333E-05 | 3.215E-06 | 1.100E-05 | 1.700E-05 |
| 10196 | 510 | S510S | A | 1.6e-05 | 2.3e-05 | 1.5e-05 | 1.800E-05 | 4.359E-06 | 1.500E-05 | 2.300E-05 |
| 10197 | 510 | S510T | A | 9.9e-06 | 1.5e-05 | 1.5e-05 | 1.330E-05 | 2.944E-06 | 9.900E-06 | 1.500E-05 |
| 10198 | 510 | S510W | A | 1e-05 | 1.5e-05 | 1.5e-05 | 1.333E-05 | 2.887E-06 | 1.000E-05 | 1.500E-05 |
| 10199 | 510 | S510Y | A | 1.2e-05 | 1.2e-05 | 1.3e-05 | 1.233E-05 | 5.774E-07 | 1.200E-05 | 1.300E-05 |

| | | | | | | | | | | |
|---|---|---|---|---|---|---|---|---|---|---|
| 10200 | 510 | S510V | A | 1.1e-05 | 1.5e-05 | 1.5e-05 | 1.367E-05 | 2.309E-06 | 1.100E-05 | 1.500E-05 |
| 10201 | 511 | Q511A | A | 1.5e-05 | 1.7e-05 | 8.7e-06 | 1.357E-05 | 4.332E-06 | 8.700E-06 | 1.700E-05 |
| 10202 | 511 | Q511R | A | 1.4e-05 | 1.3e-05 | 1.6e-05 | 1.433E-05 | 1.528E-06 | 1.300E-05 | 1.600E-05 |
| 10203 | 511 | Q511N | A | 9.4e-06 | 8.6e-06 | 1.2e-05 | 1.000E-05 | 1.778E-06 | 8.600E-06 | 1.200E-05 |
| 10204 | 511 | Q511D | A | 1.6e-05 | 1.6e-05 | 1.1e-05 | 1.433E-05 | 2.887E-06 | 1.100E-05 | 1.600E-05 |
| 10205 | 511 | Q511C | A | 1.6e-05 | 1.6e-05 | 8.5e-06 | 1.350E-05 | 4.330E-06 | 8.500E-06 | 1.600E-05 |
| 10206 | 511 | Q511E | A | 8.6e-06 | 1.5e-05 | 1.7e-05 | 1.353E-05 | 4.388E-06 | 8.600E-06 | 1.700E-05 |
| 10207 | 511 | Q511Q | A | 1.4e-05 | 1.6e-05 | 6e-06 | 1.200E-05 | 5.292E-06 | 6.000E-06 | 1.600E-05 |
| 10208 | 511 | Q511G | A | 1.6e-05 | 9.6e-06 | 1.2e-05 | 1.253E-05 | 3.233E-06 | 9.600E-06 | 1.600E-05 |
| 10209 | 511 | Q511H | A | 1.5e-05 | 1.5e-05 | 1e-05 | 1.333E-05 | 2.887E-06 | 1.000E-05 | 1.500E-05 |
| 10210 | 511 | Q511I | A | 1e-05 | 1.5e-05 | 1.4e-05 | 1.300E-05 | 2.646E-06 | 1.000E-05 | 1.500E-05 |
| 10211 | 511 | Q511L | A | 1.6e-05 | 1.6e-05 | 1.6e-05 | 1.600E-05 | 0.000E+00 | 1.600E-05 | 1.600E-05 |
| 10212 | 511 | Q511K | A | 1.6e-05 | 1.6e-05 | 1.2e-05 | 1.467E-05 | 2.309E-06 | 1.200E-05 | 1.600E-05 |
| 10213 | 511 | Q511M | A | 1.1e-05 | 1.4e-05 | 1.6e-05 | 1.367E-05 | 2.517E-06 | 1.100E-05 | 1.600E-05 |
| 10214 | 511 | Q511F | A | 1.6e-05 | 1.2e-05 | 1.3e-05 | 1.367E-05 | 2.082E-06 | 1.200E-05 | 1.600E-05 |
| 10215 | 511 | Q511P | A | 1.1e-05 | 1.6e-05 | 1e-05 | 1.233E-05 | 3.215E-06 | 1.000E-05 | 1.600E-05 |
| 10216 | 511 | Q511S | A | 8.6e-06 | 1.5e-05 | 1.7e-05 | 1.353E-05 | 4.388E-06 | 8.600E-06 | 1.700E-05 |
| 10217 | 511 | Q511T | A | 1.5e-05 | 1.3e-05 | 9.7e-06 | 1.257E-05 | 2.676E-06 | 9.700E-06 | 1.500E-05 |
| 10218 | 511 | Q511W | A | 1.3e-05 | 1e-05 | 1.6e-05 | 1.300E-05 | 3.000E-06 | 1.000E-05 | 1.600E-05 |
| 10219 | 511 | Q511Y | A | 1.6e-05 | 1.4e-05 | 9.8e-06 | 1.327E-05 | 3.164E-06 | 9.800E-06 | 1.600E-05 |
| 10220 | 511 | Q511V | A | 1.5e-05 | 1.2e-05 | 9e-06 | 1.200E-05 | 3.000E-06 | 9.000E-06 | 1.500E-05 |
| 10221 | 512 | F512A | A | 1.4e-05 | 1.4e-05 | 1.1e-05 | 1.300E-05 | 1.732E-06 | 1.100E-05 | 1.400E-05 |
| 10222 | 512 | F512R | A | 1.4e-05 | 9.9e-06 | 1.1e-05 | 1.163E-05 | 2.122E-06 | 9.900E-06 | 1.400E-05 |
| 10223 | 512 | F512N | A | 1.4e-05 | 1.1e-05 | 1.5e-05 | 1.333E-05 | 2.082E-06 | 1.100E-05 | 1.500E-05 |
| 10224 | 512 | F512D | A | 9.7e-06 | 9.3e-06 | 1.6e-05 | 1.167E-05 | 3.758E-06 | 9.300E-06 | 1.600E-05 |
| 10225 | 512 | F512C | A | 9.7e-06 | 1e-05 | 1e-05 | 9.900E-06 | 1.732E-07 | 9.700E-06 | 1.000E-05 |
| 10226 | 512 | F512E | A | 1.6e-05 | 1.4e-05 | 1e-05 | 1.333E-05 | 3.055E-06 | 1.000E-05 | 1.600E-05 |
| 10227 | 512 | F512Q | A | 9.7e-06 | 9.8e-06 | 1.4e-05 | 1.117E-05 | 2.454E-06 | 9.700E-06 | 1.400E-05 |
| 10228 | 512 | F512G | A | 1.1e-05 | 1.5e-05 | 1.3e-05 | 1.300E-05 | 2.000E-06 | 1.100E-05 | 1.500E-05 |
| 10229 | 512 | F512H | A | 1.4e-05 | 1.1e-05 | 1.8e-05 | 1.433E-05 | 3.512E-06 | 1.100E-05 | 1.800E-05 |
| 10230 | 512 | F512I | A | 1.4e-05 | 9.1e-06 | 8.7e-06 | 1.060E-05 | 2.951E-06 | 8.700E-06 | 1.400E-05 |
| 10231 | 512 | F512L | A | 1.3e-05 | 1.6e-05 | 1.3e-05 | 1.400E-05 | 1.732E-06 | 1.300E-05 | 1.600E-05 |
| 10232 | 512 | F512K | A | 1.1e-05 | 1.2e-05 | 9.3e-06 | 1.077E-05 | 1.365E-06 | 9.300E-06 | 1.200E-05 |
| 10233 | 512 | F512M | A | 1.1e-05 | 1.1e-05 | 1.6e-05 | 1.267E-05 | 2.887E-06 | 1.100E-05 | 1.600E-05 |
| 10234 | 512 | F512F | A | 2.8e-05 | 9.8e-06 | 1.2e-05 | 1.660E-05 | 9.934E-06 | 9.800E-06 | 2.800E-05 |
| 10235 | 512 | F512P | A | 1.6e-05 | 1.1e-05 | 9.7e-06 | 1.223E-05 | 3.326E-06 | 9.700E-06 | 1.600E-05 |
| 10236 | 512 | F512S | A | 1e-05 | 9.6e-06 | 8.5e-06 | 9.367E-06 | 7.767E-07 | 8.500E-06 | 1.000E-05 |
| 10237 | 512 | F512T | A | 1.4e-05 | 1.5e-05 | 1.4e-05 | 1.433E-05 | 5.774E-07 | 1.400E-05 | 1.500E-05 |
| 10238 | 512 | F512W | A | 1.5e-05 | 1.5e-05 | 1.6e-05 | 1.533E-05 | 5.774E-07 | 1.500E-05 | 1.600E-05 |
| 10239 | 512 | F512Y | A | 1.3e-05 | 1.2e-05 | 1.3e-05 | 1.267E-05 | 5.774E-07 | 1.200E-05 | 1.300E-05 |
| 10240 | 512 | F512V | A | 1.4e-05 | 1.2e-05 | 9.1e-06 | 1.170E-05 | 2.464E-06 | 9.100E-06 | 1.400E-05 |
| 10241 | 513 | L513A | A | 1.3e-05 | 1.5e-05 | 1.6e-05 | 1.467E-05 | 1.528E-06 | 1.300E-05 | 1.600E-05 |
| 10242 | 513 | L513R | A | 1.3e-05 | 1.5e-05 | 9.5e-06 | 1.250E-05 | 2.784E-06 | 9.500E-06 | 1.500E-05 |
| 10243 | 513 | L513N | A | 1.4e-05 | 1.5e-05 | 1.8e-05 | 1.567E-05 | 2.082E-06 | 1.400E-05 | 1.800E-05 |
| 10244 | 513 | L513D | A | 1.6e-05 | 1.3e-05 | 1.4e-05 | 1.433E-05 | 1.528E-06 | 1.300E-05 | 1.600E-05 |
| 10245 | 513 | L513C | A | 1.2e-05 | 9.3e-06 | 1.4e-05 | 1.177E-05 | 2.359E-06 | 9.300E-06 | 1.400E-05 |
| 10246 | 513 | L513E | A | 1.2e-05 | 9.5e-06 | 1.5e-05 | 1.217E-05 | 2.754E-06 | 9.500E-06 | 1.500E-05 |
| 10247 | 513 | L513Q | A | 1.2e-05 | 8.4e-06 | 1.4e-05 | 1.147E-05 | 2.838E-06 | 8.400E-06 | 1.400E-05 |
| 10248 | 513 | L513G | A | 1.4e-05 | 9.8e-06 | 9e-06 | 1.093E-05 | 2.686E-06 | 9.000E-06 | 1.400E-05 |
| 10249 | 513 | L513H | A | 1.2e-05 | 1.3e-05 | 1.2e-05 | 1.233E-05 | 5.774E-07 | 1.200E-05 | 1.300E-05 |
| 10250 | 513 | L513I | A | 9.5e-06 | 1.3e-05 | 1e-05 | 1.083E-05 | 1.893E-06 | 9.500E-06 | 1.300E-05 |
| 10251 | 513 | L513L | A | 9.8e-06 | 9.1e-06 | 8.7e-06 | 9.200E-06 | 5.568E-07 | 8.700E-06 | 9.800E-06 |
| 10252 | 513 | L513K | A | 1.3e-05 | 1.5e-05 | 1.1e-05 | 1.300E-05 | 2.000E-06 | 1.100E-05 | 1.500E-05 |
| 10253 | 513 | L513M | A | 1.3e-05 | 1.5e-05 | 8.2e-06 | 1.207E-05 | 3.495E-06 | 8.200E-06 | 1.500E-05 |
| 10254 | 513 | L513F | A | 1.5e-05 | 1e-05 | 1.4e-05 | 1.300E-05 | 2.646E-06 | 1.000E-05 | 1.500E-05 |
| 10255 | 513 | L513P | A | 1.5e-05 | 1e-05 | 1.1e-05 | 1.200E-05 | 2.646E-06 | 1.000E-05 | 1.500E-05 |
| 10256 | 513 | L513S | A | 1.7e-05 | 1.2e-05 | 1.4e-05 | 1.433E-05 | 2.517E-06 | 1.200E-05 | 1.700E-05 |
| 10257 | 513 | L513T | A | 1.6e-05 | 6.8e-06 | 1e-05 | 1.093E-05 | 4.670E-06 | 6.800E-06 | 1.600E-05 |
| 10258 | 513 | L513W | A | 9.3e-06 | 1.5e-05 | 1.7e-05 | 1.377E-05 | 3.995E-06 | 9.300E-06 | 1.700E-05 |
| 10259 | 513 | L513Y | A | 1.6e-05 | 1.5e-05 | 9.1e-06 | 1.337E-05 | 3.729E-06 | 9.100E-06 | 1.600E-05 |
| 10260 | 513 | L513V | A | 1.1e-05 | 9.2e-06 | 1.2e-05 | 1.073E-05 | 1.419E-06 | 9.200E-06 | 1.200E-05 |
| 10261 | 514 | R514A | A | 1.6e-05 | 8.8e-06 | 1.3e-05 | 1.260E-05 | 3.617E-06 | 8.800E-06 | 1.600E-05 |
| 10262 | 514 | R514R | A | 1.3e-05 | 1.5e-05 | 1.6e-05 | 1.467E-05 | 1.528E-06 | 1.300E-05 | 1.600E-05 |
| 10263 | 514 | R514N | A | 1.1e-05 | 1.1e-05 | 9.8e-06 | 1.060E-05 | 6.928E-07 | 9.800E-06 | 1.100E-05 |
| 10264 | 514 | R514D | A | 1.3e-05 | 1.4e-05 | 8.2e-06 | 1.173E-05 | 3.101E-06 | 8.200E-06 | 1.400E-05 |
| 10265 | 514 | R514C | A | 1.3e-05 | 1.5e-05 | 9.9e-06 | 1.263E-05 | 2.570E-06 | 9.900E-06 | 1.500E-05 |
| 10266 | 514 | R514E | A | 1.5e-05 | 1.5e-05 | 9.6e-06 | 1.320E-05 | 3.118E-06 | 9.600E-06 | 1.500E-05 |
| 10267 | 514 | R514Q | A | 1.1e-05 | 1.5e-05 | 1.3e-05 | 1.300E-05 | 2.000E-06 | 1.100E-05 | 1.500E-05 |
| 10268 | 514 | R514G | A | 1.5e-05 | 9.3e-06 | 1.5e-05 | 1.310E-05 | 3.291E-06 | 9.300E-06 | 1.500E-05 |
| 10269 | 514 | R514H | A | 1e-05 | 1.1e-05 | 1.1e-05 | 1.067E-05 | 5.774E-07 | 1.000E-05 | 1.100E-05 |
| 10270 | 514 | R514I | A | 9.1e-06 | 9e-06 | 1e-05 | 9.367E-06 | 5.508E-07 | 9.000E-06 | 1.000E-05 |
| 10271 | 514 | R514L | A | 1.5e-05 | 1.5e-05 | 1e-05 | 1.333E-05 | 2.887E-06 | 1.000E-05 | 1.500E-05 |
| 10272 | 514 | R514K | A | 8.9e-06 | 1e-05 | 1.2e-05 | 1.030E-05 | 1.572E-06 | 8.900E-06 | 1.200E-05 |
| 10273 | 514 | R514M | A | 1.5e-05 | 1.4e-05 | 1.4e-05 | 1.433E-05 | 5.774E-07 | 1.400E-05 | 1.500E-05 |
| 10274 | 514 | R514F | A | 7.7e-06 | 1.6e-05 | 1e-05 | 1.123E-05 | 4.285E-06 | 7.700E-06 | 1.600E-05 |
| 10275 | 514 | R514P | A | 1.5e-05 | 1.8e-05 | 1.1e-05 | 1.467E-05 | 3.512E-06 | 1.100E-05 | 1.800E-05 |
| 10276 | 514 | R514S | A | 1.3e-05 | 1.3e-05 | 1.1e-05 | 1.233E-05 | 1.155E-06 | 1.100E-05 | 1.300E-05 |
| 10277 | 514 | R514T | A | 1.4e-05 | 6.8e-06 | 1.5e-05 | 1.193E-05 | 4.474E-06 | 6.800E-06 | 1.500E-05 |
| 10278 | 514 | R514W | A | 1.3e-05 | 1.1e-05 | 1.4e-05 | 1.267E-05 | 1.528E-06 | 1.100E-05 | 1.400E-05 |
| 10279 | 514 | R514Y | A | 1.6e-05 | 1.3e-05 | 1.1e-05 | 1.333E-05 | 2.517E-06 | 1.100E-05 | 1.600E-05 |
| 10280 | 514 | R514V | A | 1.1e-05 | 1.1e-05 | 1.3e-05 | 1.167E-05 | 1.155E-06 | 1.100E-05 | 1.300E-05 |
| 10281 | 515 | G515A | A | 1.5e-05 | 1.1e-05 | 1.3e-05 | 1.300E-05 | 2.000E-06 | 1.100E-05 | 1.500E-05 |
| 10282 | 515 | G515R | A | 1.5e-05 | 1.5e-05 | 9.1e-06 | 1.303E-05 | 3.406E-06 | 9.100E-06 | 1.500E-05 |
| 10283 | 515 | G515N | A | 1e-05 | 1e-05 | 1.1e-05 | 1.033E-05 | 5.774E-07 | 1.000E-05 | 1.100E-05 |
| 10284 | 515 | G515D | A | 9.3e-06 | 1.3e-05 | 1.5e-05 | 1.243E-05 | 2.892E-06 | 9.300E-06 | 1.500E-05 |

| | | | | | | | | | | |
|---|---|---|---|---|---|---|---|---|---|---|
| 10285 | 515 | G515C | A | 1.4e-05 | 9.2e-06 | 1.3e-05 | 1.207E-05 | 2.532E-06 | 9.200E-06 | 1.400E-05 |
| 10286 | 515 | G515E | A | 9.7e-06 | 8.5e-06 | 1.6e-05 | 1.140E-05 | 4.029E-06 | 8.500E-06 | 1.600E-05 |
| 10287 | 515 | G515Q | A | 1.1e-05 | 1.7e-05 | 1e-05 | 1.267E-05 | 3.786E-06 | 1.000E-05 | 1.700E-05 |
| 10288 | 515 | G515G | A | 1.1e-05 | 1.1e-05 | 9.3e-06 | 1.043E-05 | 9.815E-07 | 9.300E-06 | 1.100E-05 |
| 10289 | 515 | G515H | A | 1.6e-05 | 1.1e-05 | 1.3e-05 | 1.333E-05 | 2.517E-06 | 1.100E-05 | 1.600E-05 |
| 10290 | 515 | G515I | A | 1.1e-05 | 1.4e-05 | 1.3e-05 | 1.267E-05 | 1.528E-06 | 1.100E-05 | 1.400E-05 |
| 10291 | 515 | G515L | A | 1.2e-05 | 1.1e-05 | 1.4e-05 | 1.233E-05 | 1.528E-06 | 1.100E-05 | 1.400E-05 |
| 10292 | 515 | G515K | A | 1.4e-05 | 1.1e-05 | 1.5e-05 | 1.333E-05 | 2.082E-06 | 1.100E-05 | 1.500E-05 |
| 10293 | 515 | G515M | A | 1.2e-05 | 1.5e-05 | 9.7e-06 | 1.223E-05 | 2.658E-06 | 9.700E-06 | 1.500E-05 |
| 10294 | 515 | G515F | A | 1.6e-05 | 1.4e-05 | 1.2e-05 | 1.400E-05 | 2.000E-06 | 1.200E-05 | 1.600E-05 |
| 10295 | 515 | G515P | A | 1.1e-05 | 1.6e-05 | 1.1e-05 | 1.267E-05 | 2.887E-06 | 1.100E-05 | 1.600E-05 |
| 10296 | 515 | G515S | A | 1.4e-05 | 1.1e-05 | 1.3e-05 | 1.267E-05 | 1.528E-06 | 1.100E-05 | 1.400E-05 |
| 10297 | 515 | G515T | A | 1.4e-05 | 1.5e-05 | 1e-05 | 1.300E-05 | 2.646E-06 | 1.000E-05 | 1.500E-05 |
| 10298 | 515 | G515W | A | 1.1e-05 | 9.1e-06 | 1.4e-05 | 1.137E-05 | 2.470E-06 | 9.100E-06 | 1.400E-05 |
| 10299 | 515 | G515Y | A | 1.1e-05 | 1.3e-05 | 1e-05 | 1.133E-05 | 1.528E-06 | 1.000E-05 | 1.300E-05 |
| 10300 | 515 | G515V | A | 1.4e-05 | 8.4e-06 | 1.1e-05 | 1.113E-05 | 2.802E-06 | 8.400E-06 | 1.400E-05 |
| 10301 | 516 | Q516A | A | 1.5e-05 | 1.5e-05 | 1.6e-05 | 1.533E-05 | 5.774E-07 | 1.500E-05 | 1.600E-05 |
| 10302 | 516 | Q516R | A | 1.2e-05 | 9.7e-06 | 8.1e-06 | 9.933E-06 | 1.960E-06 | 8.100E-06 | 1.200E-05 |
| 10303 | 516 | Q516N | A | 1.6e-05 | 1.3e-05 | 1.1e-05 | 1.333E-05 | 2.517E-06 | 1.100E-05 | 1.600E-05 |
| 10304 | 516 | Q516D | A | 1.2e-05 | 1.3e-05 | 1.1e-05 | 1.200E-05 | 1.000E-06 | 1.100E-05 | 1.300E-05 |
| 10305 | 516 | Q516C | A | 1.3e-05 | 1.1e-05 | 9e-06 | 1.100E-05 | 2.000E-06 | 9.000E-06 | 1.300E-05 |
| 10306 | 516 | Q516E | A | 1.6e-05 | 1.5e-05 | 1.6e-05 | 1.567E-05 | 5.774E-07 | 1.500E-05 | 1.600E-05 |
| 10307 | 516 | Q516Q | A | 1.5e-05 | 1.1e-05 | 1.4e-05 | 1.333E-05 | 2.082E-06 | 1.100E-05 | 1.500E-05 |
| 10308 | 516 | Q516G | A | 8.8e-06 | 1.1e-05 | 1.1e-05 | 1.027E-05 | 1.270E-06 | 8.800E-06 | 1.100E-05 |
| 10309 | 516 | Q516H | A | 8.1e-06 | 9.2e-06 | 1.7e-05 | 1.143E-05 | 4.852E-06 | 8.100E-06 | 1.700E-05 |
| 10310 | 516 | Q516I | A | 8.7e-06 | 1.2e-05 | 1.1e-05 | 1.057E-05 | 1.692E-06 | 8.700E-06 | 1.200E-05 |
| 10311 | 516 | Q516L | A | 1.6e-05 | 1.7e-05 | 1.6e-05 | 1.633E-05 | 5.774E-07 | 1.600E-05 | 1.700E-05 |
| 10312 | 516 | Q516K | A | 1.8e-05 | 1.5e-05 | 1.6e-05 | 1.633E-05 | 1.528E-06 | 1.500E-05 | 1.800E-05 |
| 10313 | 516 | Q516M | A | 9.5e-06 | 1.2e-05 | 1e-05 | 1.050E-05 | 1.323E-06 | 9.500E-06 | 1.200E-05 |
| 10314 | 516 | Q516F | A | 1.5e-05 | 9.4e-06 | 9.8e-06 | 1.140E-05 | 3.124E-06 | 9.400E-06 | 1.500E-05 |
| 10315 | 516 | Q516P | A | 1e-05 | 9.4e-06 | 1.4e-05 | 1.113E-05 | 2.501E-06 | 9.400E-06 | 1.400E-05 |
| 10316 | 516 | Q516S | A | 8.6e-06 | 1.8e-05 | 1e-05 | 1.220E-05 | 5.071E-06 | 8.600E-06 | 1.800E-05 |
| 10317 | 516 | Q516T | A | 1e-05 | 9.2e-06 | 9.9e-06 | 9.700E-06 | 4.359E-07 | 9.200E-06 | 1.000E-05 |
| 10318 | 516 | Q516W | A | 1.3e-05 | 1.6e-05 | 1.1e-05 | 1.333E-05 | 2.517E-06 | 1.100E-05 | 1.600E-05 |
| 10319 | 516 | Q516Y | A | 1.1e-05 | 1.6e-05 | 7.9e-06 | 1.163E-05 | 4.087E-06 | 7.900E-06 | 1.600E-05 |
| 10320 | 516 | Q516V | A | 1.6e-05 | 1.5e-05 | 9.1e-06 | 1.337E-05 | 3.729E-06 | 9.100E-06 | 1.600E-05 |
| 10321 | 517 | E517A | A | 1.4e-05 | 1.2e-05 | 1.1e-05 | 1.233E-05 | 1.528E-06 | 1.100E-05 | 1.400E-05 |
| 10322 | 517 | E517R | A | 9.2e-06 | 1.1e-05 | 1.1e-05 | 1.040E-05 | 1.039E-06 | 9.200E-06 | 1.100E-05 |
| 10323 | 517 | E517N | A | 1.4e-05 | 1.1e-05 | 1.5e-05 | 1.333E-05 | 2.082E-06 | 1.100E-05 | 1.500E-05 |
| 10324 | 517 | E517D | A | 1.1e-05 | 1.1e-05 | 1.4e-05 | 1.200E-05 | 1.732E-06 | 1.100E-05 | 1.400E-05 |
| 10325 | 517 | E517C | A | 1.4e-05 | 1.2e-05 | 9.8e-06 | 1.193E-05 | 2.101E-06 | 9.800E-06 | 1.400E-05 |
| 10326 | 517 | E517E | A | 1.6e-05 | 1.5e-05 | 1.6e-05 | 1.567E-05 | 5.774E-07 | 1.500E-05 | 1.600E-05 |
| 10327 | 517 | E517Q | A | 9e-06 | 1.7e-05 | 1.5e-05 | 1.367E-05 | 4.163E-06 | 9.000E-06 | 1.700E-05 |
| 10328 | 517 | E517G | A | 1.5e-05 | 1.4e-05 | 1.5e-05 | 1.467E-05 | 5.774E-07 | 1.400E-05 | 1.500E-05 |
| 10329 | 517 | E517H | A | 1.5e-05 | 1.5e-05 | 9.6e-06 | 1.320E-05 | 3.118E-06 | 9.600E-06 | 1.500E-05 |
| 10330 | 517 | E517I | A | 1.7e-05 | 1.5e-05 | 1.2e-05 | 1.467E-05 | 2.517E-06 | 1.200E-05 | 1.700E-05 |
| 10331 | 517 | E517L | A | 1.4e-05 | 1.3e-05 | 1.1e-05 | 1.267E-05 | 1.528E-06 | 1.100E-05 | 1.400E-05 |
| 10332 | 517 | E517K | A | 1.1e-05 | 1.2e-05 | 1.5e-05 | 1.267E-05 | 2.082E-06 | 1.100E-05 | 1.500E-05 |
| 10333 | 517 | E517M | A | 1.6e-05 | 1.4e-05 | 1.5e-05 | 1.500E-05 | 1.000E-06 | 1.400E-05 | 1.600E-05 |
| 10334 | 517 | E517F | A | 1.8e-05 | 9.7e-06 | 1.5e-05 | 1.423E-05 | 4.203E-06 | 9.700E-06 | 1.800E-05 |
| 10335 | 517 | E517P | A | 1.3e-05 | 9.2e-06 | 1.3e-05 | 1.173E-05 | 2.194E-06 | 9.200E-06 | 1.300E-05 |
| 10336 | 517 | E517S | A | 1.2e-05 | 1.5e-05 | 1.2e-05 | 1.300E-05 | 1.732E-06 | 1.200E-05 | 1.500E-05 |
| 10337 | 517 | E517T | A | 8.3e-06 | 1.4e-05 | 1.1e-05 | 1.110E-05 | 2.851E-06 | 8.300E-06 | 1.400E-05 |
| 10338 | 517 | E517W | A | 1.4e-05 | 1.1e-05 | 1.3e-05 | 1.267E-05 | 1.528E-06 | 1.100E-05 | 1.400E-05 |
| 10339 | 517 | E517Y | A | 1.4e-05 | 1.6e-05 | 1.3e-05 | 1.433E-05 | 1.528E-06 | 1.300E-05 | 1.600E-05 |
| 10340 | 517 | E517V | A | 1.3e-05 | 1.4e-05 | 1.9e-05 | 1.533E-05 | 3.215E-06 | 1.300E-05 | 1.900E-05 |
| 10341 | 518 | C518A | A | 1.1e-05 | 9.1e-06 | 8.8e-06 | 9.633E-06 | 1.193E-06 | 8.800E-06 | 1.100E-05 |
| 10342 | 518 | C518R | A | 1e-05 | 1.1e-05 | 1.7e-05 | 1.267E-05 | 3.786E-06 | 1.000E-05 | 1.700E-05 |
| 10343 | 518 | C518N | A | 1.2e-05 | 1.1e-05 | 1e-05 | 1.100E-05 | 1.000E-06 | 1.000E-05 | 1.200E-05 |
| 10344 | 518 | C518D | A | 1.5e-05 | 1.2e-05 | 8.8e-06 | 1.193E-05 | 3.101E-06 | 8.800E-06 | 1.500E-05 |
| 10345 | 518 | C518C | A | 1.5e-05 | 1.1e-05 | 1.6e-05 | 1.400E-05 | 2.646E-06 | 1.100E-05 | 1.600E-05 |
| 10346 | 518 | C518E | A | 1e-05 | 1.1e-05 | 1.3e-05 | 1.133E-05 | 1.528E-06 | 1.000E-05 | 1.300E-05 |
| 10347 | 518 | C518Q | A | 9.9e-06 | 1.5e-05 | 1.8e-05 | 1.430E-05 | 4.095E-06 | 9.900E-06 | 1.800E-05 |
| 10348 | 518 | C518G | A | 1.4e-05 | 9.6e-06 | 9.4e-06 | 1.100E-05 | 2.600E-06 | 9.400E-06 | 1.400E-05 |
| 10349 | 518 | C518H | A | 1.6e-05 | 1.1e-05 | 1.5e-05 | 1.400E-05 | 2.646E-06 | 1.100E-05 | 1.600E-05 |
| 10350 | 518 | C518I | A | 1.2e-05 | 1.7e-05 | 9.6e-06 | 1.287E-05 | 3.775E-06 | 9.600E-06 | 1.700E-05 |
| 10351 | 518 | C518L | A | 1.7e-05 | 1.5e-05 | 1e-05 | 1.400E-05 | 3.606E-06 | 1.000E-05 | 1.700E-05 |
| 10352 | 518 | C518K | A | 1.2e-05 | 1.1e-05 | 1.6e-05 | 1.300E-05 | 2.646E-06 | 1.100E-05 | 1.600E-05 |
| 10353 | 518 | C518M | A | 1.6e-05 | 1.1e-05 | 1.3e-05 | 1.333E-05 | 2.517E-06 | 1.100E-05 | 1.600E-05 |
| 10354 | 518 | C518F | A | 1.1e-05 | 1.6e-05 | 9.5e-06 | 1.217E-05 | 3.403E-06 | 9.500E-06 | 1.600E-05 |
| 10355 | 518 | C518P | A | 8.6e-06 | 2.5e-05 | 1.3e-05 | 1.553E-05 | 8.488E-06 | 8.600E-06 | 2.500E-05 |
| 10356 | 518 | C518S | A | 1.1e-05 | 8.5e-06 | 1.1e-05 | 1.017E-05 | 1.443E-06 | 8.500E-06 | 1.100E-05 |
| 10357 | 518 | C518T | A | 1.1e-05 | 7.3e-06 | 9.8e-06 | 9.367E-06 | 1.888E-06 | 7.300E-06 | 1.100E-05 |
| 10358 | 518 | C518W | A | 1.2e-05 | 1.5e-05 | 1.1e-05 | 1.267E-05 | 2.082E-06 | 1.100E-05 | 1.500E-05 |
| 10359 | 518 | C518Y | A | 1.1e-05 | 1.5e-05 | 1e-05 | 1.200E-05 | 2.646E-06 | 1.000E-05 | 1.500E-05 |
| 10360 | 518 | C518V | A | 1.2e-05 | 1.6e-05 | 1.6e-05 | 1.467E-05 | 2.309E-06 | 1.200E-05 | 1.600E-05 |
| 10361 | 519 | V519A | A | 9.6e-06 | 1.5e-05 | 9.3e-06 | 1.130E-05 | 3.208E-06 | 9.300E-06 | 1.500E-05 |
| 10362 | 519 | V519R | A | 1.1e-05 | 1.5e-05 | 1.1e-05 | 1.233E-05 | 2.309E-06 | 1.100E-05 | 1.500E-05 |
| 10363 | 519 | V519N | A | 1.5e-05 | 1.4e-05 | 1.3e-05 | 1.400E-05 | 1.000E-06 | 1.300E-05 | 1.500E-05 |
| 10364 | 519 | V519D | A | 1e-05 | 1e-05 | 1.6e-05 | 1.200E-05 | 3.464E-06 | 1.000E-05 | 1.600E-05 |
| 10365 | 519 | V519C | A | 1.1e-05 | 1.5e-05 | 1.6e-05 | 1.400E-05 | 2.646E-06 | 1.100E-05 | 1.600E-05 |
| 10366 | 519 | V519E | A | 1.5e-05 | 1.5e-05 | 8.9e-06 | 1.297E-05 | 3.522E-06 | 8.900E-06 | 1.500E-05 |
| 10367 | 519 | V519Q | A | 9.7e-06 | 9.7e-06 | 1e-05 | 9.800E-06 | 1.732E-07 | 9.700E-06 | 1.000E-05 |
| 10368 | 519 | V519G | A | 1.1e-05 | 1.1e-05 | 1.5e-05 | 1.233E-05 | 2.309E-06 | 1.100E-05 | 1.500E-05 |
| 10369 | 519 | V519H | A | 1.3e-05 | 1.3e-05 | 1.3e-05 | 1.300E-05 | 0.000E+00 | 1.300E-05 | 1.300E-05 |

| | | | | | | | | | | |
|---|---|---|---|---|---|---|---|---|---|---|
| 10370 | 519 | V519I | A | 1.2e-05 | 1.2e-05 | 1.6e-05 | 1.333E-05 | 2.309E-06 | 1.200E-05 | 1.600E-05 |
| 10371 | 519 | V519L | A | 1.5e-05 | 1.4e-05 | 1.7e-05 | 1.533E-05 | 1.528E-06 | 1.400E-05 | 1.700E-05 |
| 10372 | 519 | V519K | A | 7.7e-06 | 8.7e-06 | 8.7e-06 | 8.367E-06 | 5.774E-07 | 7.700E-06 | 8.700E-06 |
| 10373 | 519 | V519M | A | 1.6e-05 | 1e-05 | 1.4e-05 | 1.333E-05 | 3.055E-06 | 1.000E-05 | 1.600E-05 |
| 10374 | 519 | V519F | A | 1.5e-05 | 1.3e-05 | 9.8e-06 | 1.260E-05 | 2.623E-06 | 9.800E-06 | 1.500E-05 |
| 10375 | 519 | V519P | A | 1.1e-05 | 6.5e-06 | 1.4e-05 | 1.050E-05 | 3.775E-06 | 6.500E-06 | 1.400E-05 |
| 10376 | 519 | V519S | A | 9.9e-06 | 1.4e-05 | 1.3e-05 | 1.230E-05 | 2.138E-06 | 9.900E-06 | 1.400E-05 |
| 10377 | 519 | V519T | A | 8.6e-06 | 1.5e-05 | 1.6e-05 | 1.320E-05 | 4.015E-06 | 8.600E-06 | 1.600E-05 |
| 10378 | 519 | V519W | A | 1.6e-05 | 1.5e-05 | 1.4e-05 | 1.500E-05 | 1.000E-06 | 1.400E-05 | 1.600E-05 |
| 10379 | 519 | V519Y | A | 1.1e-05 | 1.4e-05 | 1.6e-05 | 1.367E-05 | 2.517E-06 | 1.100E-05 | 1.600E-05 |
| 10380 | 519 | V519V | A | 1.1e-05 | 8.9e-06 | 1.5e-05 | 1.163E-05 | 3.099E-06 | 8.900E-06 | 1.500E-05 |
| 10381 | 520 | E520A | A | 1.1e-05 | 1.6e-05 | 1.6e-05 | 1.433E-05 | 2.887E-06 | 1.100E-05 | 1.600E-05 |
| 10382 | 520 | E520R | A | 1.7e-05 | 1e-05 | 1.5e-05 | 1.400E-05 | 3.606E-06 | 1.000E-05 | 1.700E-05 |
| 10383 | 520 | E520N | A | 1.6e-05 | 1.6e-05 | 1.4e-05 | 1.533E-05 | 1.155E-06 | 1.400E-05 | 1.600E-05 |
| 10384 | 520 | E520D | A | 1.2e-05 | 1.4e-05 | 1.1e-05 | 1.233E-05 | 1.528E-06 | 1.100E-05 | 1.400E-05 |
| 10385 | 520 | E520C | A | 1.5e-05 | 1.4e-05 | 1.6e-05 | 1.500E-05 | 1.000E-06 | 1.400E-05 | 1.600E-05 |
| 10386 | 520 | E520E | A | 1e-05 | 6.8e-06 | 1.1e-05 | 9.267E-06 | 2.194E-06 | 6.800E-06 | 1.100E-05 |
| 10387 | 520 | E520Q | A | 8.9e-06 | 1.1e-05 | 1.4e-05 | 1.130E-05 | 2.563E-06 | 8.900E-06 | 1.400E-05 |
| 10388 | 520 | E520G | A | 8.6e-06 | 9.5e-06 | 1e-05 | 9.367E-06 | 7.095E-07 | 8.600E-06 | 1.000E-05 |
| 10389 | 520 | E520H | A | 9.2e-06 | 1e-05 | 1.5e-05 | 1.140E-05 | 3.143E-06 | 9.200E-06 | 1.500E-05 |
| 10390 | 520 | E520I | A | 1.1e-05 | 1.5e-05 | 1.1e-05 | 1.233E-05 | 2.309E-06 | 1.100E-05 | 1.500E-05 |
| 10391 | 520 | E520L | A | 9.5e-06 | 9.3e-06 | 1.7e-05 | 1.193E-05 | 4.389E-06 | 9.300E-06 | 1.700E-05 |
| 10392 | 520 | E520K | A | 1.3e-05 | 1.6e-05 | 1.2e-05 | 1.367E-05 | 2.082E-06 | 1.200E-05 | 1.600E-05 |
| 10393 | 520 | E520M | A | 1.8e-05 | 1e-05 | 1.4e-05 | 1.400E-05 | 4.000E-06 | 1.000E-05 | 1.800E-05 |
| 10394 | 520 | E520F | A | 1.1e-05 | 1.5e-05 | 8.3e-06 | 1.143E-05 | 3.371E-06 | 8.300E-06 | 1.500E-05 |
| 10395 | 520 | E520P | A | 1.5e-05 | 1.8e-05 | 1.1e-05 | 1.467E-05 | 3.512E-06 | 1.100E-05 | 1.800E-05 |
| 10396 | 520 | E520S | A | 8.7e-06 | 1.1e-05 | 8.5e-06 | 9.400E-06 | 1.389E-06 | 8.500E-06 | 1.100E-05 |
| 10397 | 520 | E520T | A | 1e-05 | 1e-05 | 7.4e-06 | 9.133E-06 | 1.501E-06 | 7.400E-06 | 1.000E-05 |
| 10398 | 520 | E520W | A | 1.2e-05 | 1.1e-05 | 1.1e-05 | 1.133E-05 | 5.774E-07 | 1.100E-05 | 1.200E-05 |
| 10399 | 520 | E520Y | A | 1.5e-05 | 1.1e-05 | 8.8e-06 | 1.160E-05 | 3.143E-06 | 8.800E-06 | 1.500E-05 |
| 10400 | 520 | E520V | A | 1.1e-05 | 1.4e-05 | 1.6e-05 | 1.367E-05 | 2.517E-06 | 1.100E-05 | 1.600E-05 |
| 10401 | 521 | E521A | A | 9.6e-06 | 1.6e-05 | 1.4e-05 | 1.320E-05 | 3.274E-06 | 9.600E-06 | 1.600E-05 |
| 10402 | 521 | E521R | A | 1.6e-05 | 1.5e-05 | 1.1e-05 | 1.400E-05 | 2.646E-06 | 1.100E-05 | 1.600E-05 |
| 10403 | 521 | E521N | A | 1.6e-05 | 1.6e-05 | 1.2e-05 | 1.467E-05 | 2.309E-06 | 1.200E-05 | 1.600E-05 |
| 10404 | 521 | E521D | A | 8.5e-06 | 1.6e-05 | 9.8e-06 | 1.143E-05 | 4.008E-06 | 8.500E-06 | 1.600E-05 |
| 10405 | 521 | E521C | A | 1.3e-05 | 1.8e-05 | 1e-05 | 1.367E-05 | 4.041E-06 | 1.000E-05 | 1.800E-05 |
| 10406 | 521 | E521E | A | 1.5e-05 | 7.3e-06 | 1.4e-05 | 1.210E-05 | 4.187E-06 | 7.300E-06 | 1.500E-05 |
| 10407 | 521 | E521Q | A | 1.6e-05 | 1e-05 | 1.1e-05 | 1.233E-05 | 3.215E-06 | 1.000E-05 | 1.600E-05 |
| 10408 | 521 | E521G | A | 1.4e-05 | 1.7e-05 | 1.2e-05 | 1.433E-05 | 2.517E-06 | 1.200E-05 | 1.700E-05 |
| 10409 | 521 | E521H | A | 1.5e-05 | 1.3e-05 | 1.6e-05 | 1.467E-05 | 1.528E-06 | 1.300E-05 | 1.600E-05 |
| 10410 | 521 | E521I | A | 1e-05 | 1.1e-05 | 8.3e-06 | 9.767E-06 | 1.365E-06 | 8.300E-06 | 1.100E-05 |
| 10411 | 521 | E521L | A | 1.1e-05 | 1.1e-05 | 1.5e-05 | 1.233E-05 | 2.309E-06 | 1.100E-05 | 1.500E-05 |
| 10412 | 521 | E521K | A | 1.5e-05 | 1.6e-05 | 1e-05 | 1.367E-05 | 3.215E-06 | 1.000E-05 | 1.600E-05 |
| 10413 | 521 | E521M | A | 1.7e-05 | 1.4e-05 | 2.2e-05 | 1.767E-05 | 4.041E-06 | 1.400E-05 | 2.200E-05 |
| 10414 | 521 | E521F | A | 8.7e-06 | 1.6e-05 | 1.4e-05 | 1.290E-05 | 3.772E-06 | 8.700E-06 | 1.600E-05 |
| 10415 | 521 | E521P | A | 1.5e-05 | 1.5e-05 | 1.1e-05 | 1.367E-05 | 2.309E-06 | 1.100E-05 | 1.500E-05 |
| 10416 | 521 | E521S | A | 1.5e-05 | 1.5e-05 | 1.1e-05 | 1.367E-05 | 2.309E-06 | 1.100E-05 | 1.500E-05 |
| 10417 | 521 | E521T | A | 1.7e-05 | 1.4e-05 | 8.3e-06 | 1.310E-05 | 4.419E-06 | 8.300E-06 | 1.700E-05 |
| 10418 | 521 | E521W | A | 8.9e-06 | 1.3e-05 | 9e-06 | 1.030E-05 | 2.339E-06 | 8.900E-06 | 1.300E-05 |
| 10419 | 521 | E521Y | A | 9.5e-06 | 1.7e-05 | 1.4e-05 | 1.350E-05 | 3.775E-06 | 9.500E-06 | 1.700E-05 |
| 10420 | 521 | E521V | A | 1.5e-05 | 1.5e-05 | 9.9e-06 | 1.330E-05 | 2.944E-06 | 9.900E-06 | 1.500E-05 |
| 10421 | 522 | C522A | A | 1.7e-05 | 1.6e-05 | 1.6e-05 | 1.633E-05 | 5.774E-07 | 1.600E-05 | 1.700E-05 |
| 10422 | 522 | C522R | A | 1.6e-05 | 1.2e-05 | 1.4e-05 | 1.400E-05 | 2.000E-06 | 1.200E-05 | 1.600E-05 |
| 10423 | 522 | C522N | A | 9.8e-06 | 1.1e-05 | 9.4e-06 | 1.007E-05 | 8.327E-07 | 9.400E-06 | 1.100E-05 |
| 10424 | 522 | C522D | A | 1.6e-05 | 1.4e-05 | 1.3e-05 | 1.433E-05 | 1.528E-06 | 1.300E-05 | 1.600E-05 |
| 10425 | 522 | C522C | A | 1.5e-05 | 1.4e-05 | 1.1e-05 | 1.333E-05 | 2.082E-06 | 1.100E-05 | 1.500E-05 |
| 10426 | 522 | C522E | A | 1.4e-05 | 1e-05 | 1.5e-05 | 1.300E-05 | 2.646E-06 | 1.000E-05 | 1.500E-05 |
| 10427 | 522 | C522Q | A | 1.3e-05 | 1.5e-05 | 1.1e-05 | 1.300E-05 | 2.000E-06 | 1.100E-05 | 1.500E-05 |
| 10428 | 522 | C522G | A | 1.6e-05 | 1.5e-05 | 1.5e-05 | 1.533E-05 | 5.774E-07 | 1.500E-05 | 1.600E-05 |
| 10429 | 522 | C522H | A | 1.4e-05 | 1.4e-05 | 1e-05 | 1.267E-05 | 2.309E-06 | 1.000E-05 | 1.400E-05 |
| 10430 | 522 | C522I | A | 9.7e-06 | 1e-05 | 1.3e-05 | 1.090E-05 | 1.825E-06 | 9.700E-06 | 1.300E-05 |
| 10431 | 522 | C522L | A | 1e-05 | 1.3e-05 | 6.9e-06 | 9.967E-06 | 3.050E-06 | 6.900E-06 | 1.300E-05 |
| 10432 | 522 | C522K | A | 9.2e-06 | 1.1e-05 | 1.4e-05 | 1.140E-05 | 2.425E-06 | 9.200E-06 | 1.400E-05 |
| 10433 | 522 | C522M | A | 1.1e-05 | 1e-05 | 1.6e-05 | 1.233E-05 | 3.215E-06 | 1.000E-05 | 1.600E-05 |
| 10434 | 522 | C522F | A | 9.3e-06 | 1.6e-05 | 9.3e-06 | 1.153E-05 | 3.868E-06 | 9.300E-06 | 1.600E-05 |
| 10435 | 522 | C522P | A | 1e-05 | 9.1e-06 | 1.6e-05 | 1.170E-05 | 3.751E-06 | 9.100E-06 | 1.600E-05 |
| 10436 | 522 | C522S | A | 1.1e-05 | 1.1e-05 | 1.4e-05 | 1.200E-05 | 1.732E-06 | 1.100E-05 | 1.400E-05 |
| 10437 | 522 | C522T | A | 1.5e-05 | 1e-05 | 1.5e-05 | 1.333E-05 | 2.887E-06 | 1.000E-05 | 1.500E-05 |
| 10438 | 522 | C522W | A | 1.4e-05 | 1.5e-05 | 1.4e-05 | 1.433E-05 | 5.774E-07 | 1.400E-05 | 1.500E-05 |
| 10439 | 522 | C522Y | A | 1.6e-05 | 8.5e-06 | 1.4e-05 | 1.283E-05 | 3.884E-06 | 8.500E-06 | 1.600E-05 |
| 10440 | 522 | C522V | A | 1.5e-05 | 1.4e-05 | 1e-05 | 1.300E-05 | 2.646E-06 | 1.000E-05 | 1.500E-05 |
| 10441 | 523 | R523A | A | 1.5e-05 | 1.5e-05 | 1.2e-05 | 1.400E-05 | 1.732E-06 | 1.200E-05 | 1.500E-05 |
| 10442 | 523 | R523R | A | 1.5e-05 | 1.3e-05 | 7.2e-06 | 1.173E-05 | 4.051E-06 | 7.200E-06 | 1.500E-05 |
| 10443 | 523 | R523N | A | 1.6e-05 | 1.3e-05 | 1e-05 | 1.300E-05 | 3.000E-06 | 1.000E-05 | 1.600E-05 |
| 10444 | 523 | R523D | A | 1.5e-05 | 7.1e-06 | 1.5e-05 | 1.237E-05 | 4.561E-06 | 7.100E-06 | 1.500E-05 |
| 10445 | 523 | R523C | A | 1.6e-05 | 1.6e-05 | 9e-06 | 1.367E-05 | 4.041E-06 | 9.000E-06 | 1.600E-05 |
| 10446 | 523 | R523E | A | 9e-06 | 9.9e-06 | 1.6e-05 | 1.163E-05 | 3.808E-06 | 9.000E-06 | 1.600E-05 |
| 10447 | 523 | R523Q | A | 1.5e-05 | 1.6e-05 | 1.5e-05 | 1.533E-05 | 5.774E-07 | 1.500E-05 | 1.600E-05 |
| 10448 | 523 | R523G | A | 1.1e-05 | 9.8e-06 | 1.2e-05 | 1.093E-05 | 1.102E-06 | 9.800E-06 | 1.200E-05 |
| 10449 | 523 | R523H | A | 7.1e-06 | 1.1e-05 | 9.2e-06 | 9.100E-06 | 1.952E-06 | 7.100E-06 | 1.100E-05 |
| 10450 | 523 | R523I | A | 1.6e-05 | 1.4e-05 | 1.6e-05 | 1.533E-05 | 1.155E-06 | 1.400E-05 | 1.600E-05 |
| 10451 | 523 | R523L | A | 9.1e-06 | 1.3e-05 | 1.1e-05 | 1.103E-05 | 1.950E-06 | 9.100E-06 | 1.300E-05 |
| 10452 | 523 | R523K | A | 1.5e-05 | 1.5e-05 | 1.2e-05 | 1.400E-05 | 1.732E-06 | 1.200E-05 | 1.500E-05 |
| 10453 | 523 | R523M | A | 1.5e-05 | 1.5e-05 | 1.3e-05 | 1.433E-05 | 1.155E-06 | 1.300E-05 | 1.500E-05 |
| 10454 | 523 | R523F | A | 1.5e-05 | 1e-05 | 1.7e-05 | 1.400E-05 | 3.606E-06 | 1.000E-05 | 1.700E-05 |

| | | | | | | | | | | |
|---|---|---|---|---|---|---|---|---|---|---|
| 10455 | 523 | R523P | A | 1.6e-05 | 8.2e-06 | 1.5e-05 | 1.307E-05 | 4.244E-06 | 8.200E-06 | 1.600E-05 |
| 10456 | 523 | R523S | A | 1.2e-05 | 1e-05 | 1.5e-05 | 1.233E-05 | 2.517E-06 | 1.000E-05 | 1.500E-05 |
| 10457 | 523 | R523T | A | 1.4e-05 | 1.4e-05 | 1.3e-05 | 1.367E-05 | 5.774E-07 | 1.300E-05 | 1.400E-05 |
| 10458 | 523 | R523W | A | 1.3e-05 | 1.3e-05 | 1e-05 | 1.200E-05 | 1.732E-06 | 1.000E-05 | 1.300E-05 |
| 10459 | 523 | R523Y | A | 1.3e-05 | 1.6e-05 | 1.6e-05 | 1.500E-05 | 1.732E-06 | 1.300E-05 | 1.600E-05 |
| 10460 | 523 | R523V | A | 1.6e-05 | 1.5e-05 | 1.6e-05 | 1.567E-05 | 5.774E-07 | 1.500E-05 | 1.600E-05 |
| 10461 | 524 | V524A | A | 1.3e-05 | 9.9e-06 | 1.2e-05 | 1.163E-05 | 1.582E-06 | 9.900E-06 | 1.300E-05 |
| 10462 | 524 | V524R | A | 1.5e-05 | 1.7e-05 | 1.3e-05 | 1.500E-05 | 2.000E-06 | 1.300E-05 | 1.700E-05 |
| 10463 | 524 | V524N | A | 8.4e-06 | 1.5e-05 | 1.5e-05 | 1.280E-05 | 3.811E-06 | 8.400E-06 | 1.500E-05 |
| 10464 | 524 | V524D | A | 8.8e-06 | 1.2e-05 | 1.2e-05 | 1.093E-05 | 1.848E-06 | 8.800E-06 | 1.200E-05 |
| 10465 | 524 | V524C | A | 1.5e-05 | 1.6e-05 | 9.3e-06 | 1.343E-05 | 3.614E-06 | 9.300E-06 | 1.600E-05 |
| 10466 | 524 | V524E | A | 7.8e-06 | 8.6e-06 | 1.4e-05 | 1.013E-05 | 3.372E-06 | 7.800E-06 | 1.400E-05 |
| 10467 | 524 | V524Q | A | 1.4e-05 | 1.5e-05 | 1.4e-05 | 1.433E-05 | 5.774E-07 | 1.400E-05 | 1.500E-05 |
| 10468 | 524 | V524G | A | 1e-05 | 1.1e-05 | 1.6e-05 | 1.233E-05 | 3.215E-06 | 1.000E-05 | 1.600E-05 |
| 10469 | 524 | V524H | A | 1.4e-05 | 1e-05 | 1.4e-05 | 1.267E-05 | 2.309E-06 | 1.000E-05 | 1.400E-05 |
| 10470 | 524 | V524I | A | 1e-05 | 1.6e-05 | 1.6e-05 | 1.400E-05 | 3.464E-06 | 1.000E-05 | 1.600E-05 |
| 10471 | 524 | V524L | A | 1.4e-05 | 1.5e-05 | 1.5e-05 | 1.467E-05 | 5.774E-07 | 1.400E-05 | 1.500E-05 |
| 10472 | 524 | V524K | A | 9.9e-06 | 9.8e-06 | 1.4e-05 | 1.123E-05 | 2.397E-06 | 9.800E-06 | 1.400E-05 |
| 10473 | 524 | V524M | A | 1e-05 | 9.2e-06 | 1.5e-05 | 1.140E-05 | 3.143E-06 | 9.200E-06 | 1.500E-05 |
| 10474 | 524 | V524F | A | 1e-05 | 1.1e-05 | 1.5e-05 | 1.200E-05 | 2.646E-06 | 1.000E-05 | 1.500E-05 |
| 10475 | 524 | V524P | A | 1.1e-05 | 1.6e-05 | 1.2e-05 | 1.300E-05 | 2.646E-06 | 1.100E-05 | 1.600E-05 |
| 10476 | 524 | V524S | A | 1.4e-05 | 1.4e-05 | 1.1e-05 | 1.300E-05 | 1.732E-06 | 1.100E-05 | 1.400E-05 |
| 10477 | 524 | V524T | A | 1.5e-05 | 1.6e-05 | 1.5e-05 | 1.533E-05 | 5.774E-07 | 1.500E-05 | 1.600E-05 |
| 10478 | 524 | V524W | A | 1.6e-05 | 1.5e-05 | 1.6e-05 | 1.567E-05 | 5.774E-07 | 1.500E-05 | 1.600E-05 |
| 10479 | 524 | V524Y | A | 1.5e-05 | 1.6e-05 | 8.3e-06 | 1.310E-05 | 4.187E-06 | 8.300E-06 | 1.600E-05 |
| 10480 | 524 | V524V | A | 1.2e-05 | 1.3e-05 | 1.4e-05 | 1.300E-05 | 1.000E-06 | 1.200E-05 | 1.400E-05 |
| 10481 | 525 | L525A | A | 1.1e-05 | 1.5e-05 | 1.5e-05 | 1.367E-05 | 2.309E-06 | 1.100E-05 | 1.500E-05 |
| 10482 | 525 | L525R | A | 1e-05 | 1.2e-05 | 1.4e-05 | 1.200E-05 | 2.000E-06 | 1.000E-05 | 1.400E-05 |
| 10483 | 525 | L525N | A | 1.2e-05 | 1.3e-05 | 1.1e-05 | 1.200E-05 | 1.000E-06 | 1.100E-05 | 1.300E-05 |
| 10484 | 525 | L525D | A | 8.5e-06 | 1.4e-05 | 1e-05 | 1.083E-05 | 2.843E-06 | 8.500E-06 | 1.400E-05 |
| 10485 | 525 | L525C | A | 1.3e-05 | 1.4e-05 | 1.2e-05 | 1.300E-05 | 1.000E-06 | 1.200E-05 | 1.400E-05 |
| 10486 | 525 | L525E | A | 1.6e-05 | 1.4e-05 | 1.1e-05 | 1.367E-05 | 2.517E-06 | 1.100E-05 | 1.600E-05 |
| 10487 | 525 | L525Q | A | 1.6e-05 | 1.6e-05 | 9e-06 | 1.367E-05 | 4.041E-06 | 9.000E-06 | 1.600E-05 |
| 10488 | 525 | L525G | A | 1.7e-05 | 7.8e-06 | 7.9e-06 | 1.090E-05 | 5.283E-06 | 7.800E-06 | 1.700E-05 |
| 10489 | 525 | L525H | A | 1.5e-05 | 1.2e-05 | 1.6e-05 | 1.433E-05 | 2.082E-06 | 1.200E-05 | 1.600E-05 |
| 10490 | 525 | L525I | A | 1e-05 | 1.6e-05 | 1.5e-05 | 1.367E-05 | 3.215E-06 | 1.000E-05 | 1.600E-05 |
| 10491 | 525 | L525L | A | 8.6e-06 | 1.6e-05 | 1.1e-05 | 1.187E-05 | 3.775E-06 | 8.600E-06 | 1.600E-05 |
| 10492 | 525 | L525K | A | 1.1e-05 | 1.6e-05 | 8.3e-06 | 1.177E-05 | 3.907E-06 | 8.300E-06 | 1.600E-05 |
| 10493 | 525 | L525M | A | 1e-05 | 1.1e-05 | 1e-05 | 1.033E-05 | 5.774E-07 | 1.000E-05 | 1.100E-05 |
| 10494 | 525 | L525F | A | 1.4e-05 | 1.5e-05 | 1e-05 | 1.300E-05 | 2.646E-06 | 1.000E-05 | 1.500E-05 |
| 10495 | 525 | L525P | A | 1.4e-05 | 1e-05 | 8.6e-06 | 1.087E-05 | 2.802E-06 | 8.600E-06 | 1.400E-05 |
| 10496 | 525 | L525S | A | 1.1e-05 | 1.5e-05 | 1.2e-05 | 1.267E-05 | 2.082E-06 | 1.100E-05 | 1.500E-05 |
| 10497 | 525 | L525T | A | 1.6e-05 | 1.6e-05 | 1.4e-05 | 1.533E-05 | 1.155E-06 | 1.400E-05 | 1.600E-05 |
| 10498 | 525 | L525W | A | 1e-05 | 1.2e-05 | 1.4e-05 | 1.200E-05 | 2.000E-06 | 1.000E-05 | 1.400E-05 |
| 10499 | 525 | L525Y | A | 1.1e-05 | 1.2e-05 | 1.5e-05 | 1.267E-05 | 2.082E-06 | 1.100E-05 | 1.500E-05 |
| 10500 | 525 | L525V | A | 1.4e-05 | 1.5e-05 | 8.4e-06 | 1.247E-05 | 3.557E-06 | 8.400E-06 | 1.500E-05 |
| 10501 | 526 | Q526A | A | 1.6e-05 | 1.4e-05 | 1.2e-05 | 1.400E-05 | 2.000E-06 | 1.200E-05 | 1.600E-05 |
| 10502 | 526 | Q526R | A | 1.5e-05 | 1.1e-05 | 1.7e-05 | 1.433E-05 | 3.055E-06 | 1.100E-05 | 1.700E-05 |
| 10503 | 526 | Q526N | A | 1e-05 | 1.7e-05 | 1.1e-05 | 1.267E-05 | 3.786E-06 | 1.000E-05 | 1.700E-05 |
| 10504 | 526 | Q526D | A | 1.5e-05 | 1.6e-05 | 1.6e-05 | 1.567E-05 | 5.774E-07 | 1.500E-05 | 1.600E-05 |
| 10505 | 526 | Q526C | A | 1.5e-05 | 9.7e-06 | 1.1e-05 | 1.190E-05 | 2.762E-06 | 9.700E-06 | 1.500E-05 |
| 10506 | 526 | Q526E | A | 9.5e-06 | 1.3e-05 | 1.4e-05 | 1.217E-05 | 2.363E-06 | 9.500E-06 | 1.400E-05 |
| 10507 | 526 | Q526Q | A | 1.5e-05 | 1.3e-05 | 1e-05 | 1.267E-05 | 2.517E-06 | 1.000E-05 | 1.500E-05 |
| 10508 | 526 | Q526G | A | 1.3e-05 | 9.2e-06 | 1.4e-05 | 1.207E-05 | 2.532E-06 | 9.200E-06 | 1.400E-05 |
| 10509 | 526 | Q526H | A | 1.6e-05 | 1.6e-05 | 1e-05 | 1.400E-05 | 3.464E-06 | 1.000E-05 | 1.600E-05 |
| 10510 | 526 | Q526I | A | 1.6e-05 | 1.2e-05 | 1.2e-05 | 1.333E-05 | 2.309E-06 | 1.200E-05 | 1.600E-05 |
| 10511 | 526 | Q526L | A | 1.3e-05 | 1e-05 | 1.2e-05 | 1.167E-05 | 1.528E-06 | 1.000E-05 | 1.300E-05 |
| 10512 | 526 | Q526K | A | 1.5e-05 | 1e-05 | 9.2e-06 | 1.140E-05 | 3.143E-06 | 9.200E-06 | 1.500E-05 |
| 10513 | 526 | Q526M | A | 1e-05 | 1.4e-05 | 8.6e-06 | 1.087E-05 | 2.802E-06 | 8.600E-06 | 1.400E-05 |
| 10514 | 526 | Q526F | A | 1e-05 | 1.5e-05 | 8.6e-06 | 1.120E-05 | 3.365E-06 | 8.600E-06 | 1.500E-05 |
| 10515 | 526 | Q526P | A | 1.5e-05 | 1.1e-05 | 9.6e-06 | 1.187E-05 | 2.802E-06 | 9.600E-06 | 1.500E-05 |
| 10516 | 526 | Q526S | A | 1.6e-05 | 1.3e-05 | 1.4e-05 | 1.433E-05 | 1.528E-06 | 1.300E-05 | 1.600E-05 |
| 10517 | 526 | Q526T | A | 1.5e-05 | 8.9e-06 | 1.4e-05 | 1.263E-05 | 3.272E-06 | 8.900E-06 | 1.500E-05 |
| 10518 | 526 | Q526W | A | 1.6e-05 | 1e-05 | 9.3e-06 | 1.177E-05 | 3.683E-06 | 9.300E-06 | 1.600E-05 |
| 10519 | 526 | Q526Y | A | 1e-05 | 1e-05 | 1.5e-05 | 1.167E-05 | 2.887E-06 | 1.000E-05 | 1.500E-05 |
| 10520 | 526 | Q526V | A | 1.6e-05 | 1.5e-05 | 1.7e-05 | 1.600E-05 | 1.000E-06 | 1.500E-05 | 1.700E-05 |
| 10521 | 527 | G527A | A | 9.8e-06 | 9.2e-06 | 1.7e-05 | 1.200E-05 | 4.341E-06 | 9.200E-06 | 1.700E-05 |
| 10522 | 527 | G527R | A | 1e-05 | 1.4e-05 | 1e-05 | 1.133E-05 | 2.309E-06 | 1.000E-05 | 1.400E-05 |
| 10523 | 527 | G527N | A | 1.3e-05 | 1.7e-05 | 8.5e-06 | 1.283E-05 | 4.252E-06 | 8.500E-06 | 1.700E-05 |
| 10524 | 527 | G527D | A | 1e-05 | 1.5e-05 | 1.5e-05 | 1.333E-05 | 2.887E-06 | 1.000E-05 | 1.500E-05 |
| 10525 | 527 | G527C | A | 9.5e-06 | 1.5e-05 | 1.5e-05 | 1.317E-05 | 3.175E-06 | 9.500E-06 | 1.500E-05 |
| 10526 | 527 | G527E | A | 1.1e-05 | 1.1e-05 | 1.8e-05 | 1.333E-05 | 4.041E-06 | 1.100E-05 | 1.800E-05 |
| 10527 | 527 | G527Q | A | 1e-05 | 1.3e-05 | 1.3e-05 | 1.200E-05 | 1.732E-06 | 1.000E-05 | 1.300E-05 |
| 10528 | 527 | G527G | A | 1e-05 | 1.3e-05 | 9.3e-06 | 1.077E-05 | 1.966E-06 | 9.300E-06 | 1.300E-05 |
| 10529 | 527 | G527H | A | 1e-05 | 9.1e-06 | 1.4e-05 | 1.103E-05 | 2.608E-06 | 9.100E-06 | 1.400E-05 |
| 10530 | 527 | G527I | A | 1e-05 | 1.1e-05 | 9.5e-06 | 1.017E-05 | 7.638E-07 | 9.500E-06 | 1.100E-05 |
| 10531 | 527 | G527L | A | 7.1e-06 | 1.7e-05 | 1.6e-05 | 1.337E-05 | 5.450E-06 | 7.100E-06 | 1.700E-05 |
| 10532 | 527 | G527K | A | 1.6e-05 | 1.2e-05 | 9.4e-06 | 1.247E-05 | 3.325E-06 | 9.400E-06 | 1.600E-05 |
| 10533 | 527 | G527M | A | 1.1e-05 | 1.4e-05 | 1e-05 | 1.167E-05 | 2.082E-06 | 1.000E-05 | 1.400E-05 |
| 10534 | 527 | G527F | A | 1.4e-05 | 9.6e-06 | 8.7e-06 | 1.077E-05 | 2.836E-06 | 8.700E-06 | 1.400E-05 |
| 10535 | 527 | G527P | A | 9e-06 | 1.5e-05 | 1.5e-05 | 1.300E-05 | 3.464E-06 | 9.000E-06 | 1.500E-05 |
| 10536 | 527 | G527S | A | 1.2e-05 | 1.7e-05 | 1.8e-05 | 1.567E-05 | 3.215E-06 | 1.200E-05 | 1.800E-05 |
| 10537 | 527 | G527T | A | 8.1e-06 | 1.4e-05 | 1e-05 | 1.070E-05 | 3.012E-06 | 8.100E-06 | 1.400E-05 |
| 10538 | 527 | G527W | A | 1.1e-05 | 9.2e-06 | 8.4e-06 | 9.533E-06 | 1.332E-06 | 8.400E-06 | 1.100E-05 |
| 10539 | 527 | G527Y | A | 1.2e-05 | 1e-05 | 8.3e-06 | 1.010E-05 | 1.852E-06 | 8.300E-06 | 1.200E-05 |

| | | | | | | | | | | |
|---|---|---|---|---|---|---|---|---|---|---|
| 10540 | 527 | G527V | A | 1.3e-05 | 1.5e-05 | 1.4e-05 | 1.400E-05 | 1.000E-06 | 1.300E-05 | 1.500E-05 |
| 10541 | 528 | L528A | A | 1.7e-05 | 1e-05 | 1.5e-05 | 1.400E-05 | 3.606E-06 | 1.000E-05 | 1.700E-05 |
| 10542 | 528 | L528R | A | 1.6e-05 | 1.5e-05 | 1.2e-05 | 1.433E-05 | 2.082E-06 | 1.200E-05 | 1.600E-05 |
| 10543 | 528 | L528N | A | 1.5e-05 | 1.3e-05 | 1.5e-05 | 1.433E-05 | 1.155E-06 | 1.300E-05 | 1.500E-05 |
| 10544 | 528 | L528D | A | 1.5e-05 | 1.4e-05 | 1.2e-05 | 1.367E-05 | 1.528E-06 | 1.200E-05 | 1.500E-05 |
| 10545 | 528 | L528C | A | 1.2e-05 | 1.5e-05 | 9.4e-06 | 1.213E-05 | 2.802E-06 | 9.400E-06 | 1.500E-05 |
| 10546 | 528 | L528E | A | 9.6e-06 | 1.5e-05 | 1.5e-05 | 1.320E-05 | 3.118E-06 | 9.600E-06 | 1.500E-05 |
| 10547 | 528 | L528Q | A | 1.1e-05 | 8.6e-06 | 1.1e-05 | 1.020E-05 | 1.386E-06 | 8.600E-06 | 1.100E-05 |
| 10548 | 528 | L528G | A | 1.6e-05 | 1.3e-05 | 1.2e-05 | 1.367E-05 | 2.082E-06 | 1.200E-05 | 1.600E-05 |
| 10549 | 528 | L528H | A | 1.3e-05 | 1e-05 | 1.2e-05 | 1.167E-05 | 1.528E-06 | 1.000E-05 | 1.300E-05 |
| 10550 | 528 | L528I | A | 1.4e-05 | 1.2e-05 | 8.4e-06 | 1.147E-05 | 2.838E-06 | 8.400E-06 | 1.400E-05 |
| 10551 | 528 | L528L | A | 1.6e-05 | 1.7e-05 | 1.5e-05 | 1.600E-05 | 1.000E-06 | 1.500E-05 | 1.700E-05 |
| 10552 | 528 | L528K | A | 8.8e-06 | 1.1e-05 | 1.4e-05 | 1.127E-05 | 2.610E-06 | 8.800E-06 | 1.400E-05 |
| 10553 | 528 | L528M | A | 1e-05 | 1.2e-05 | 9.6e-06 | 1.053E-05 | 1.286E-06 | 9.600E-06 | 1.200E-05 |
| 10554 | 528 | L528F | A | 1.4e-05 | 8.4e-06 | 1.1e-05 | 1.113E-05 | 2.802E-06 | 8.400E-06 | 1.400E-05 |
| 10555 | 528 | L528P | A | 1.6e-05 | 1.4e-05 | 1.4e-05 | 1.467E-05 | 1.155E-06 | 1.400E-05 | 1.600E-05 |
| 10556 | 528 | L528S | A | 1.2e-05 | 1.3e-05 | 1.2e-05 | 1.233E-05 | 5.774E-07 | 1.200E-05 | 1.300E-05 |
| 10557 | 528 | L528T | A | 9.3e-06 | 1.4e-05 | 1.6e-05 | 1.310E-05 | 3.439E-06 | 9.300E-06 | 1.600E-05 |
| 10558 | 528 | L528W | A | 1.2e-05 | 1.4e-05 | 1.5e-05 | 1.367E-05 | 1.528E-06 | 1.200E-05 | 1.500E-05 |
| 10559 | 528 | L528Y | A | 1.4e-05 | 1.2e-05 | 1e-05 | 1.200E-05 | 2.000E-06 | 1.000E-05 | 1.400E-05 |
| 10560 | 528 | L528V | A | 1.2e-05 | 1.5e-05 | 1e-05 | 1.233E-05 | 2.517E-06 | 1.000E-05 | 1.500E-05 |
| 10561 | 529 | P529A | A | 1.4e-05 | 1.6e-05 | 1.4e-05 | 1.467E-05 | 1.155E-06 | 1.400E-05 | 1.600E-05 |
| 10562 | 529 | P529R | A | 1.1e-05 | 1.5e-05 | 8.6e-06 | 1.153E-05 | 3.233E-06 | 8.600E-06 | 1.500E-05 |
| 10563 | 529 | P529N | A | 9.5e-06 | 8.2e-06 | 9.7e-06 | 9.133E-06 | 8.145E-07 | 8.200E-06 | 9.700E-06 |
| 10564 | 529 | P529D | A | 1.6e-05 | 1.1e-05 | 1.2e-05 | 1.300E-05 | 2.646E-06 | 1.100E-05 | 1.600E-05 |
| 10565 | 529 | P529C | A | 1.4e-05 | 1.5e-05 | 1.4e-05 | 1.433E-05 | 5.774E-07 | 1.400E-05 | 1.500E-05 |
| 10566 | 529 | P529E | A | 1.5e-05 | 1.4e-05 | 1.6e-05 | 1.500E-05 | 1.000E-06 | 1.400E-05 | 1.600E-05 |
| 10567 | 529 | P529Q | A | 9.1e-06 | 1.6e-05 | 2.1e-05 | 1.537E-05 | 5.975E-06 | 9.100E-06 | 2.100E-05 |
| 10568 | 529 | P529G | A | 1.8e-05 | 9.3e-06 | 1.5e-05 | 1.410E-05 | 4.419E-06 | 9.300E-06 | 1.800E-05 |
| 10569 | 529 | P529H | A | 1.1e-05 | 1.1e-05 | 1.5e-05 | 1.233E-05 | 2.309E-06 | 1.100E-05 | 1.500E-05 |
| 10570 | 529 | P529I | A | 9.9e-06 | 1e-05 | 1.7e-05 | 1.230E-05 | 4.071E-06 | 9.900E-06 | 1.700E-05 |
| 10571 | 529 | P529L | A | 1.1e-05 | 1.5e-05 | 1.4e-05 | 1.333E-05 | 2.082E-06 | 1.100E-05 | 1.500E-05 |
| 10572 | 529 | P529K | A | 1.5e-05 | 1.5e-05 | 1.4e-05 | 1.467E-05 | 5.774E-07 | 1.400E-05 | 1.500E-05 |
| 10573 | 529 | P529M | A | 1.1e-05 | 1.6e-05 | 1e-05 | 1.233E-05 | 3.215E-06 | 1.000E-05 | 1.600E-05 |
| 10574 | 529 | P529F | A | 7.4e-06 | 1.5e-05 | 1.3e-05 | 1.180E-05 | 3.940E-06 | 7.400E-06 | 1.500E-05 |
| 10575 | 529 | P529P | A | 1.5e-05 | 1.1e-05 | 1.4e-05 | 1.333E-05 | 2.082E-06 | 1.100E-05 | 1.500E-05 |
| 10576 | 529 | P529S | A | 1e-05 | 9.6e-06 | 1.2e-05 | 1.053E-05 | 1.286E-06 | 9.600E-06 | 1.200E-05 |
| 10577 | 529 | P529T | A | 1e-05 | 1.5e-05 | 8.1e-06 | 1.103E-05 | 3.564E-06 | 8.100E-06 | 1.500E-05 |
| 10578 | 529 | P529W | A | 1.4e-05 | 1.1e-05 | 1.4e-05 | 1.300E-05 | 1.732E-06 | 1.100E-05 | 1.400E-05 |
| 10579 | 529 | P529Y | A | 1.4e-05 | 1.3e-05 | 1.5e-05 | 1.400E-05 | 1.000E-06 | 1.300E-05 | 1.500E-05 |
| 10580 | 529 | P529V | A | 1e-05 | 1e-05 | 1.5e-05 | 1.167E-05 | 2.887E-06 | 1.000E-05 | 1.500E-05 |
| 10581 | 530 | R530A | A | 1.1e-05 | 8.9e-06 | 1.5e-05 | 1.163E-05 | 3.099E-06 | 8.900E-06 | 1.500E-05 |
| 10582 | 530 | R530R | A | 1.3e-05 | 8.5e-06 | 1.4e-05 | 1.183E-05 | 2.930E-06 | 8.500E-06 | 1.400E-05 |
| 10583 | 530 | R530N | A | 8e-06 | 9.3e-06 | 1.4e-05 | 1.043E-05 | 3.156E-06 | 8.000E-06 | 1.400E-05 |
| 10584 | 530 | R530D | A | 9.6e-06 | 1.4e-05 | 1.6e-05 | 1.320E-05 | 3.274E-06 | 9.600E-06 | 1.600E-05 |
| 10585 | 530 | R530C | A | 1.1e-05 | 1.2e-05 | 1.7e-05 | 1.333E-05 | 3.215E-06 | 1.100E-05 | 1.700E-05 |
| 10586 | 530 | R530E | A | 1.1e-05 | 1e-05 | 1.1e-05 | 1.067E-05 | 5.774E-07 | 1.000E-05 | 1.100E-05 |
| 10587 | 530 | R530Q | A | 9.4e-06 | 9.2e-06 | 1.6e-05 | 1.153E-05 | 3.870E-06 | 9.200E-06 | 1.600E-05 |
| 10588 | 530 | R530G | A | 9.4e-06 | 1.8e-05 | 9.3e-06 | 1.223E-05 | 4.994E-06 | 9.300E-06 | 1.800E-05 |
| 10589 | 530 | R530H | A | 1.1e-05 | 1.5e-05 | 1.2e-05 | 1.267E-05 | 2.082E-06 | 1.100E-05 | 1.500E-05 |
| 10590 | 530 | R530I | A | 1.4e-05 | 1.5e-05 | 1.1e-05 | 1.333E-05 | 2.082E-06 | 1.100E-05 | 1.500E-05 |
| 10591 | 530 | R530L | A | 1.3e-05 | 1.3e-05 | 1.5e-05 | 1.367E-05 | 1.155E-06 | 1.300E-05 | 1.500E-05 |
| 10592 | 530 | R530K | A | 1e-05 | 1e-05 | 9.9e-06 | 9.967E-06 | 5.774E-08 | 9.900E-06 | 1.000E-05 |
| 10593 | 530 | R530M | A | 1.1e-05 | 1.3e-05 | 8.4e-06 | 1.080E-05 | 2.307E-06 | 8.400E-06 | 1.300E-05 |
| 10594 | 530 | R530F | A | 1.1e-05 | 1.1e-05 | 9.2e-06 | 1.040E-05 | 1.039E-06 | 9.200E-06 | 1.100E-05 |
| 10595 | 530 | R530P | A | 1.5e-05 | 1.1e-05 | 1.6e-05 | 1.400E-05 | 2.646E-06 | 1.100E-05 | 1.600E-05 |
| 10596 | 530 | R530S | A | 8.9e-06 | 9.8e-06 | 1.1e-05 | 9.900E-06 | 1.054E-06 | 8.900E-06 | 1.100E-05 |
| 10597 | 530 | R530T | A | 1.5e-05 | 1.2e-05 | 1.6e-05 | 1.433E-05 | 2.082E-06 | 1.200E-05 | 1.600E-05 |
| 10598 | 530 | R530W | A | 9.4e-06 | 1e-05 | 1.4e-05 | 1.113E-05 | 2.501E-06 | 9.400E-06 | 1.400E-05 |
| 10599 | 530 | R530Y | A | 9.8e-06 | 1.6e-05 | 1e-05 | 1.193E-05 | 3.523E-06 | 9.800E-06 | 1.600E-05 |
| 10600 | 530 | R530V | A | 1.1e-05 | 9.6e-06 | 1.1e-05 | 1.053E-05 | 8.083E-07 | 9.600E-06 | 1.100E-05 |
| 10601 | 531 | E531A | A | 1.2e-05 | 1.3e-05 | 1.5e-05 | 1.333E-05 | 1.528E-06 | 1.200E-05 | 1.500E-05 |
| 10602 | 531 | E531R | A | 1.2e-05 | 1.5e-05 | 1.7e-05 | 1.467E-05 | 2.517E-06 | 1.200E-05 | 1.700E-05 |
| 10603 | 531 | E531N | A | 1.1e-05 | 6e-06 | 1e-05 | 9.000E-06 | 2.646E-06 | 6.000E-06 | 1.100E-05 |
| 10604 | 531 | E531D | A | 1.4e-05 | 1.1e-05 | 1.4e-05 | 1.300E-05 | 1.732E-06 | 1.100E-05 | 1.400E-05 |
| 10605 | 531 | E531C | A | 1.5e-05 | 1.1e-05 | 1.5e-05 | 1.367E-05 | 2.309E-06 | 1.100E-05 | 1.500E-05 |
| 10606 | 531 | E531E | A | 9.9e-06 | 1.3e-05 | 1e-05 | 1.097E-05 | 1.762E-06 | 9.900E-06 | 1.300E-05 |
| 10607 | 531 | E531Q | A | 1.5e-05 | 1e-05 | 1.1e-05 | 1.200E-05 | 2.646E-06 | 1.000E-05 | 1.500E-05 |
| 10608 | 531 | E531G | A | 1.1e-05 | 1.6e-05 | 1.5e-05 | 1.400E-05 | 2.646E-06 | 1.100E-05 | 1.600E-05 |
| 10609 | 531 | E531H | A | 1.5e-05 | 1.5e-05 | 1.4e-05 | 1.467E-05 | 5.774E-07 | 1.400E-05 | 1.500E-05 |
| 10610 | 531 | E531I | A | 1.1e-05 | 1.2e-05 | 1.3e-05 | 1.200E-05 | 1.000E-06 | 1.100E-05 | 1.300E-05 |
| 10611 | 531 | E531L | A | 1.7e-05 | 1.7e-05 | 1.6e-05 | 1.667E-05 | 5.774E-07 | 1.600E-05 | 1.700E-05 |
| 10612 | 531 | E531K | A | 1.6e-05 | 1.5e-05 | 9.8e-06 | 1.360E-05 | 3.329E-06 | 9.800E-06 | 1.600E-05 |
| 10613 | 531 | E531M | A | 1.7e-05 | 1e-05 | 9.5e-06 | 1.217E-05 | 4.193E-06 | 9.500E-06 | 1.700E-05 |
| 10614 | 531 | E531F | A | 1.4e-05 | 1.2e-05 | 1.1e-05 | 1.233E-05 | 1.528E-06 | 1.100E-05 | 1.400E-05 |
| 10615 | 531 | E531P | A | 1.5e-05 | 1.3e-05 | 1.3e-05 | 1.367E-05 | 1.155E-06 | 1.300E-05 | 1.500E-05 |
| 10616 | 531 | E531S | A | 1.4e-05 | 1.5e-05 | 1.1e-05 | 1.333E-05 | 2.082E-06 | 1.100E-05 | 1.500E-05 |
| 10617 | 531 | E531T | A | 7.7e-06 | 1.6e-05 | 1.5e-05 | 1.290E-05 | 4.531E-06 | 7.700E-06 | 1.600E-05 |
| 10618 | 531 | E531W | A | 1.4e-05 | 1.2e-05 | 9.4e-06 | 1.180E-05 | 2.307E-06 | 9.400E-06 | 1.400E-05 |
| 10619 | 531 | E531Y | A | 1.6e-05 | 1.1e-05 | 1.1e-05 | 1.267E-05 | 2.887E-06 | 1.100E-05 | 1.600E-05 |
| 10620 | 531 | E531V | A | 1.5e-05 | 1.4e-05 | 9.7e-06 | 1.290E-05 | 2.816E-06 | 9.700E-06 | 1.500E-05 |
| 10621 | 532 | Y532A | A | 1.5e-05 | 1.8e-05 | 1.5e-05 | 1.600E-05 | 1.732E-06 | 1.500E-05 | 1.800E-05 |
| 10622 | 532 | Y532R | A | 1.6e-05 | 1e-05 | 1e-05 | 1.200E-05 | 3.464E-06 | 1.000E-05 | 1.600E-05 |
| 10623 | 532 | Y532N | A | 1.1e-05 | 1.1e-05 | 1.1e-05 | 1.100E-05 | 2.075E-21 | 1.100E-05 | 1.100E-05 |
| 10624 | 532 | Y532D | A | 1.1e-05 | 1.6e-05 | 1.2e-05 | 1.300E-05 | 2.646E-06 | 1.100E-05 | 1.600E-05 |

| | | | | | | | | | | |
|---|---|---|---|---|---|---|---|---|---|---|
| 10625 | 532 | Y532C | A | 1.4e-05 | 1.5e-05 | 1e-05 | 1.300E-05 | 2.646E-06 | 1.000E-05 | 1.500E-05 |
| 10626 | 532 | Y532E | A | 9.3e-06 | 1.2e-05 | 1.2e-05 | 1.110E-05 | 1.559E-06 | 9.300E-06 | 1.200E-05 |
| 10627 | 532 | Y532Q | A | 1.1e-05 | 1.1e-05 | 1.5e-05 | 1.233E-05 | 2.309E-06 | 1.100E-05 | 1.500E-05 |
| 10628 | 532 | Y532G | A | 1.3e-05 | 1.1e-05 | 1.6e-05 | 1.333E-05 | 2.517E-06 | 1.100E-05 | 1.600E-05 |
| 10629 | 532 | Y532H | A | 7.3e-06 | 1.5e-05 | 9.6e-06 | 1.063E-05 | 3.953E-06 | 7.300E-06 | 1.500E-05 |
| 10630 | 532 | Y532I | A | 1.1e-05 | 1.1e-05 | 1e-05 | 1.067E-05 | 5.774E-07 | 1.000E-05 | 1.100E-05 |
| 10631 | 532 | Y532L | A | 1.2e-05 | 9.7e-06 | 1.7e-05 | 1.290E-05 | 3.732E-06 | 9.700E-06 | 1.700E-05 |
| 10632 | 532 | Y532K | A | 9.4e-06 | 1e-05 | 1.7e-05 | 1.213E-05 | 4.225E-06 | 9.400E-06 | 1.700E-05 |
| 10633 | 532 | Y532M | A | 7.8e-06 | 1.5e-05 | 7e-06 | 9.933E-06 | 4.406E-06 | 7.000E-06 | 1.500E-05 |
| 10634 | 532 | Y532F | A | 1.6e-05 | 1.4e-05 | 1.6e-05 | 1.533E-05 | 1.155E-06 | 1.400E-05 | 1.600E-05 |
| 10635 | 532 | Y532P | A | 1.6e-05 | 2e-05 | 1.2e-05 | 1.600E-05 | 4.000E-06 | 1.200E-05 | 2.000E-05 |
| 10636 | 532 | Y532S | A | 1.1e-05 | 9.7e-06 | 1.7e-05 | 1.257E-05 | 3.894E-06 | 9.700E-06 | 1.700E-05 |
| 10637 | 532 | Y532T | A | 1.4e-05 | 6.6e-06 | 1.5e-05 | 1.187E-05 | 4.588E-06 | 6.600E-06 | 1.500E-05 |
| 10638 | 532 | Y532W | A | 9.1e-06 | 9e-06 | 1.1e-05 | 9.700E-06 | 1.127E-06 | 9.000E-06 | 1.100E-05 |
| 10639 | 532 | Y532Y | A | 9.9e-06 | 1.4e-05 | 1.1e-05 | 1.163E-05 | 2.122E-06 | 9.900E-06 | 1.400E-05 |
| 10640 | 532 | Y532V | A | 8.4e-06 | 1.6e-05 | 1.3e-05 | 1.247E-05 | 3.828E-06 | 8.400E-06 | 1.600E-05 |
| 10641 | 533 | V533A | A | 1.6e-05 | 8.9e-06 | 1.5e-05 | 1.330E-05 | 3.843E-06 | 8.900E-06 | 1.600E-05 |
| 10642 | 533 | V533R | A | 1.6e-05 | 1.2e-05 | 1.3e-05 | 1.367E-05 | 2.082E-06 | 1.200E-05 | 1.600E-05 |
| 10643 | 533 | V533N | A | 1e-05 | 8.9e-06 | 9.4e-06 | 9.433E-06 | 5.508E-07 | 8.900E-06 | 1.000E-05 |
| 10644 | 533 | V533D | A | 1.2e-05 | 1e-05 | 1e-05 | 1.067E-05 | 1.155E-06 | 1.000E-05 | 1.200E-05 |
| 10645 | 533 | V533C | A | 8.9e-06 | 1.1e-05 | 1.3e-05 | 1.097E-05 | 2.050E-06 | 8.900E-06 | 1.300E-05 |
| 10646 | 533 | V533E | A | 1.5e-05 | 9e-06 | 9.5e-06 | 1.117E-05 | 3.329E-06 | 9.000E-06 | 1.500E-05 |
| 10647 | 533 | V533Q | A | 1e-05 | 9.1e-06 | 1.1e-05 | 1.003E-05 | 9.504E-07 | 9.100E-06 | 1.100E-05 |
| 10648 | 533 | V533G | A | 1.1e-05 | 1.1e-05 | 1.2e-05 | 1.133E-05 | 5.774E-07 | 1.100E-05 | 1.200E-05 |
| 10649 | 533 | V533H | A | 1.3e-05 | 1.3e-05 | 1.5e-05 | 1.367E-05 | 1.155E-06 | 1.300E-05 | 1.500E-05 |
| 10650 | 533 | V533I | A | 1.6e-05 | 1.1e-05 | 1.4e-05 | 1.367E-05 | 2.517E-06 | 1.100E-05 | 1.600E-05 |
| 10651 | 533 | V533L | A | 1.4e-05 | 8.6e-06 | 1.3e-05 | 1.187E-05 | 2.873E-06 | 8.600E-06 | 1.400E-05 |
| 10652 | 533 | V533K | A | 9.8e-06 | 9.8e-06 | 1.2e-05 | 1.053E-05 | 1.270E-06 | 9.800E-06 | 1.200E-05 |
| 10653 | 533 | V533M | A | 1.5e-05 | 8.8e-06 | 1.5e-05 | 1.293E-05 | 3.580E-06 | 8.800E-06 | 1.500E-05 |
| 10654 | 533 | V533F | A | 1.5e-05 | 1.4e-05 | 1e-05 | 1.300E-05 | 2.646E-06 | 1.000E-05 | 1.500E-05 |
| 10655 | 533 | V533P | A | 1.1e-05 | 1.1e-05 | 1.5e-05 | 1.233E-05 | 2.309E-06 | 1.100E-05 | 1.500E-05 |
| 10656 | 533 | V533S | A | 9.2e-06 | 1.6e-05 | 1.6e-05 | 1.373E-05 | 3.926E-06 | 9.200E-06 | 1.600E-05 |
| 10657 | 533 | V533T | A | 1.5e-05 | 1.4e-05 | 1.5e-05 | 1.467E-05 | 5.774E-07 | 1.400E-05 | 1.500E-05 |
| 10658 | 533 | V533W | A | 1e-05 | 1.1e-05 | 1.3e-05 | 1.133E-05 | 1.528E-06 | 1.000E-05 | 1.300E-05 |
| 10659 | 533 | V533Y | A | 1.5e-05 | 1.4e-05 | 1.1e-05 | 1.333E-05 | 2.082E-06 | 1.100E-05 | 1.500E-05 |
| 10660 | 533 | V533V | A | 1.8e-05 | 1.1e-05 | 1.1e-05 | 1.333E-05 | 4.041E-06 | 1.100E-05 | 1.800E-05 |
| 10661 | 534 | N534A | A | 1.3e-05 | 1.2e-05 | 1.4e-05 | 1.300E-05 | 1.000E-06 | 1.200E-05 | 1.400E-05 |
| 10662 | 534 | N534R | A | 1.4e-05 | 1.6e-05 | 1.1e-05 | 1.367E-05 | 2.517E-06 | 1.100E-05 | 1.600E-05 |
| 10663 | 534 | N534N | A | 1.5e-05 | 1e-05 | 8.2e-06 | 1.107E-05 | 3.523E-06 | 8.200E-06 | 1.500E-05 |
| 10664 | 534 | N534D | A | 1.7e-05 | 1.3e-05 | 1.1e-05 | 1.367E-05 | 3.055E-06 | 1.100E-05 | 1.700E-05 |
| 10665 | 534 | N534C | A | 9.2e-06 | 1.3e-05 | 1.3e-05 | 1.173E-05 | 2.194E-06 | 9.200E-06 | 1.300E-05 |
| 10666 | 534 | N534E | A | 1.6e-05 | 8.9e-06 | 1.6e-05 | 1.363E-05 | 4.099E-06 | 8.900E-06 | 1.600E-05 |
| 10667 | 534 | N534Q | A | 9.8e-06 | 8.8e-06 | 1.5e-05 | 1.120E-05 | 3.329E-06 | 8.800E-06 | 1.500E-05 |
| 10668 | 534 | N534G | A | 8.9e-06 | 1.6e-05 | 1.2e-05 | 1.230E-05 | 3.559E-06 | 8.900E-06 | 1.600E-05 |
| 10669 | 534 | N534H | A | 1.5e-05 | 1.4e-05 | 1.7e-05 | 1.533E-05 | 1.528E-06 | 1.400E-05 | 1.700E-05 |
| 10670 | 534 | N534I | A | 1.5e-05 | 1.5e-05 | 1.6e-05 | 1.533E-05 | 5.774E-07 | 1.500E-05 | 1.600E-05 |
| 10671 | 534 | N534L | A | 1e-05 | 1.7e-05 | 8.6e-06 | 1.187E-05 | 4.500E-06 | 8.600E-06 | 1.700E-05 |
| 10672 | 534 | N534K | A | 1.8e-05 | 1.1e-05 | 9.9e-06 | 1.297E-05 | 4.394E-06 | 9.900E-06 | 1.800E-05 |
| 10673 | 534 | N534M | A | 9.4e-06 | 1e-05 | 1.3e-05 | 1.080E-05 | 1.929E-06 | 9.400E-06 | 1.300E-05 |
| 10674 | 534 | N534F | A | 1e-05 | 1.3e-05 | 9.9e-06 | 1.097E-05 | 1.762E-06 | 9.900E-06 | 1.300E-05 |
| 10675 | 534 | N534P | A | 1.5e-05 | 1.1e-05 | 1.4e-05 | 1.333E-05 | 2.082E-06 | 1.100E-05 | 1.500E-05 |
| 10676 | 534 | N534S | A | 1.8e-05 | 9.6e-06 | 1e-05 | 1.253E-05 | 4.738E-06 | 9.600E-06 | 1.800E-05 |
| 10677 | 534 | N534T | A | 3.3e-05 | 1.6e-05 | 1.2e-05 | 2.033E-05 | 1.115E-05 | 1.200E-05 | 3.300E-05 |
| 10678 | 534 | N534W | A | 1.1e-05 | 1.6e-05 | 1.1e-05 | 1.267E-05 | 2.887E-06 | 1.100E-05 | 1.600E-05 |
| 10679 | 534 | N534Y | A | 1.4e-05 | 1.1e-05 | 9.7e-06 | 1.157E-05 | 2.205E-06 | 9.700E-06 | 1.400E-05 |
| 10680 | 534 | N534V | A | 1.5e-05 | 1.4e-05 | 1.6e-05 | 1.500E-05 | 1.000E-06 | 1.400E-05 | 1.600E-05 |
| 10681 | 535 | A535A | A | 1.4e-05 | 1e-05 | 1.5e-05 | 1.300E-05 | 2.646E-06 | 1.000E-05 | 1.500E-05 |
| 10682 | 535 | A535R | A | 1.1e-05 | 1.5e-05 | 7.9e-06 | 1.130E-05 | 3.559E-06 | 7.900E-06 | 1.500E-05 |
| 10683 | 535 | A535N | A | 1.1e-05 | 1.6e-05 | 1.4e-05 | 1.367E-05 | 2.517E-06 | 1.100E-05 | 1.600E-05 |
| 10684 | 535 | A535D | A | 1.5e-05 | 1.5e-05 | 2e-05 | 1.667E-05 | 2.887E-06 | 1.500E-05 | 2.000E-05 |
| 10685 | 535 | A535C | A | 1.3e-05 | 1.7e-05 | 1.4e-05 | 1.467E-05 | 2.082E-06 | 1.300E-05 | 1.700E-05 |
| 10686 | 535 | A535E | A | 1.3e-05 | 1.4e-05 | 1.1e-05 | 1.267E-05 | 1.528E-06 | 1.100E-05 | 1.400E-05 |
| 10687 | 535 | A535Q | A | 1.4e-05 | 8.2e-06 | 1.3e-05 | 1.173E-05 | 3.101E-06 | 8.200E-06 | 1.400E-05 |
| 10688 | 535 | A535G | A | 1e-05 | 1.4e-05 | 9.9e-06 | 1.130E-05 | 2.339E-06 | 9.900E-06 | 1.400E-05 |
| 10689 | 535 | A535H | A | 1.7e-05 | 1.1e-05 | 1e-05 | 1.267E-05 | 3.786E-06 | 1.000E-05 | 1.700E-05 |
| 10690 | 535 | A535I | A | 1.7e-05 | 1.6e-05 | 1e-05 | 1.433E-05 | 3.786E-06 | 1.000E-05 | 1.700E-05 |
| 10691 | 535 | A535L | A | 1.6e-05 | 1.5e-05 | 1.7e-05 | 1.600E-05 | 1.000E-06 | 1.500E-05 | 1.700E-05 |
| 10692 | 535 | A535K | A | 9.8e-06 | 1.2e-05 | 8.7e-06 | 1.017E-05 | 1.680E-06 | 8.700E-06 | 1.200E-05 |
| 10693 | 535 | A535M | A | 1.4e-05 | 1.3e-05 | 1.5e-05 | 1.400E-05 | 1.000E-06 | 1.300E-05 | 1.500E-05 |
| 10694 | 535 | A535F | A | 1.3e-05 | 1.1e-05 | 9.1e-06 | 1.103E-05 | 1.950E-06 | 9.100E-06 | 1.300E-05 |
| 10695 | 535 | A535P | A | 9.4e-06 | 1e-05 | 1.6e-05 | 1.180E-05 | 3.650E-06 | 9.400E-06 | 1.600E-05 |
| 10696 | 535 | A535S | A | 1.5e-05 | 1.4e-05 | 1.6e-05 | 1.500E-05 | 1.000E-06 | 1.400E-05 | 1.600E-05 |
| 10697 | 535 | A535T | A | 1.5e-05 | 1.1e-05 | 8e-06 | 1.133E-05 | 3.512E-06 | 8.000E-06 | 1.500E-05 |
| 10698 | 535 | A535W | A | 1.5e-05 | 9e-06 | 1.5e-05 | 1.300E-05 | 3.464E-06 | 9.000E-06 | 1.500E-05 |
| 10699 | 535 | A535Y | A | 1.1e-05 | 1.1e-05 | 1.4e-05 | 1.200E-05 | 1.732E-06 | 1.100E-05 | 1.400E-05 |
| 10700 | 535 | A535V | A | 1.6e-05 | 1.5e-05 | 1.4e-05 | 1.500E-05 | 1.000E-06 | 1.400E-05 | 1.600E-05 |
| 10701 | 536 | R536A | A | 1.1e-05 | 1.5e-05 | 1.2e-05 | 1.267E-05 | 2.082E-06 | 1.100E-05 | 1.500E-05 |
| 10702 | 536 | R536R | A | 1.5e-05 | 1.4e-05 | 1.1e-05 | 1.333E-05 | 2.082E-06 | 1.100E-05 | 1.500E-05 |
| 10703 | 536 | R536N | A | 1.6e-05 | 1e-05 | 9.9e-06 | 1.197E-05 | 3.493E-06 | 9.900E-06 | 1.600E-05 |
| 10704 | 536 | R536D | A | 1.1e-05 | 9e-06 | 9.9e-06 | 9.967E-06 | 1.002E-06 | 9.000E-06 | 1.100E-05 |
| 10705 | 536 | R536C | A | 1.5e-05 | 1.6e-05 | 1.1e-05 | 1.400E-05 | 2.646E-06 | 1.100E-05 | 1.600E-05 |
| 10706 | 536 | R536E | A | 1.4e-05 | 9.8e-06 | 1.5e-05 | 1.293E-05 | 2.759E-06 | 9.800E-06 | 1.500E-05 |
| 10707 | 536 | R536Q | A | 1.3e-05 | 1.1e-05 | 1.6e-05 | 1.333E-05 | 2.517E-06 | 1.100E-05 | 1.600E-05 |
| 10708 | 536 | R536G | A | 1.2e-05 | 1.3e-05 | 1.1e-05 | 1.200E-05 | 1.000E-06 | 1.100E-05 | 1.300E-05 |
| 10709 | 536 | R536H | A | 1.5e-05 | 1.1e-05 | 1.2e-05 | 1.267E-05 | 2.082E-06 | 1.100E-05 | 1.500E-05 |

| | | | | | | | | | | |
|---|---|---|---|---|---|---|---|---|---|---|
| 10710 | 536 | R536I | A | 8.5e-06 | 1.5e-05 | 1.4e-05 | 1.250E-05 | 3.500E-06 | 8.500E-06 | 1.500E-05 |
| 10711 | 536 | R536L | A | 9.8e-06 | 1.7e-05 | 1.1e-05 | 1.260E-05 | 3.857E-06 | 9.800E-06 | 1.700E-05 |
| 10712 | 536 | R536K | A | 1.1e-05 | 1.5e-05 | 8.5e-06 | 1.150E-05 | 3.279E-06 | 8.500E-06 | 1.500E-05 |
| 10713 | 536 | R536M | A | 1.1e-05 | 1.4e-05 | 1.5e-05 | 1.333E-05 | 2.082E-06 | 1.100E-05 | 1.500E-05 |
| 10714 | 536 | R536F | A | 1.1e-05 | 1.5e-05 | 1.1e-05 | 1.233E-05 | 2.309E-06 | 1.100E-05 | 1.500E-05 |
| 10715 | 536 | R536P | A | 1.2e-05 | 1.6e-05 | 9.7e-06 | 1.257E-05 | 3.188E-06 | 9.700E-06 | 1.600E-05 |
| 10716 | 536 | R536S | A | 1.1e-05 | 1.3e-05 | 9.6e-06 | 1.120E-05 | 1.709E-06 | 9.600E-06 | 1.300E-05 |
| 10717 | 536 | R536T | A | 1.4e-05 | 1e-05 | 7.2e-06 | 1.040E-05 | 3.418E-06 | 7.200E-06 | 1.400E-05 |
| 10718 | 536 | R536W | A | 9.2e-06 | 1.2e-05 | 1e-05 | 1.040E-05 | 1.442E-06 | 9.200E-06 | 1.200E-05 |
| 10719 | 536 | R536Y | A | 1.1e-05 | 8.9e-06 | 1.3e-05 | 1.097E-05 | 2.050E-06 | 8.900E-06 | 1.300E-05 |
| 10720 | 536 | R536V | A | 1.1e-05 | 1.1e-05 | 1.1e-05 | 1.100E-05 | 2.075E-21 | 1.100E-05 | 1.100E-05 |
| 10721 | 537 | H537A | A | 8.6e-06 | 9.9e-06 | 1.1e-05 | 9.833E-06 | 1.201E-06 | 8.600E-06 | 1.100E-05 |
| 10722 | 537 | H537R | A | 9.4e-06 | 1.9e-05 | 1.1e-05 | 1.313E-05 | 5.143E-06 | 9.400E-06 | 1.900E-05 |
| 10723 | 537 | H537N | A | 1.5e-05 | 1e-05 | 1.4e-05 | 1.300E-05 | 2.646E-06 | 1.000E-05 | 1.500E-05 |
| 10724 | 537 | H537D | A | 1.1e-05 | 1.7e-05 | 1e-05 | 1.267E-05 | 3.786E-06 | 1.000E-05 | 1.700E-05 |
| 10725 | 537 | H537C | A | 6.1e-06 | 1.4e-05 | 1.1e-05 | 1.037E-05 | 3.988E-06 | 6.100E-06 | 1.400E-05 |
| 10726 | 537 | H537E | A | 1.3e-05 | 1.1e-05 | 9.8e-06 | 1.127E-05 | 1.617E-06 | 9.800E-06 | 1.300E-05 |
| 10727 | 537 | H537Q | A | 1.1e-05 | 1e-05 | 1.2e-05 | 1.100E-05 | 1.000E-06 | 1.000E-05 | 1.200E-05 |
| 10728 | 537 | H537G | A | 1.1e-05 | 8.1e-06 | 1e-05 | 9.700E-06 | 1.473E-06 | 8.100E-06 | 1.100E-05 |
| 10729 | 537 | H537H | A | 9.9e-06 | 8.9e-06 | 9.3e-06 | 9.367E-06 | 5.033E-07 | 8.900E-06 | 9.900E-06 |
| 10730 | 537 | H537I | A | 1.6e-05 | 1.5e-05 | 1.1e-05 | 1.400E-05 | 2.646E-06 | 1.100E-05 | 1.600E-05 |
| 10731 | 537 | H537L | A | 1.3e-05 | 1.6e-05 | 1.6e-05 | 1.500E-05 | 1.732E-06 | 1.300E-05 | 1.600E-05 |
| 10732 | 537 | H537K | A | 1.5e-05 | 1.3e-05 | 1.5e-05 | 1.433E-05 | 1.155E-06 | 1.300E-05 | 1.500E-05 |
| 10733 | 537 | H537M | A | 1e-05 | 1e-05 | 1.2e-05 | 1.067E-05 | 1.155E-06 | 1.000E-05 | 1.200E-05 |
| 10734 | 537 | H537F | A | 1.6e-05 | 1.1e-05 | 1.2e-05 | 1.300E-05 | 2.646E-06 | 1.100E-05 | 1.600E-05 |
| 10735 | 537 | H537P | A | 1.6e-05 | 1.4e-05 | 1.8e-05 | 1.600E-05 | 2.000E-06 | 1.400E-05 | 1.800E-05 |
| 10736 | 537 | H537S | A | 1.5e-05 | 1.6e-05 | 9.7e-06 | 1.357E-05 | 3.386E-06 | 9.700E-06 | 1.600E-05 |
| 10737 | 537 | H537T | A | 9.9e-06 | 1.5e-05 | 1.4e-05 | 1.297E-05 | 2.702E-06 | 9.900E-06 | 1.500E-05 |
| 10738 | 537 | H537W | A | 1.1e-05 | 1.6e-05 | 1.4e-05 | 1.367E-05 | 2.517E-06 | 1.100E-05 | 1.600E-05 |
| 10739 | 537 | H537Y | A | 1.3e-05 | 1.1e-05 | 1e-05 | 1.133E-05 | 1.528E-06 | 1.000E-05 | 1.300E-05 |
| 10740 | 537 | H537V | A | 2e-05 | 1.6e-05 | 1.1e-05 | 1.567E-05 | 4.509E-06 | 1.100E-05 | 2.000E-05 |
| 10741 | 538 | C538A | A | 1.4e-05 | 1e-05 | 1.2e-05 | 1.200E-05 | 2.000E-06 | 1.000E-05 | 1.400E-05 |
| 10742 | 538 | C538R | A | 1.5e-05 | 1.2e-05 | 1.5e-05 | 1.400E-05 | 1.732E-06 | 1.200E-05 | 1.500E-05 |
| 10743 | 538 | C538N | A | 1e-05 | 9.8e-06 | 1e-05 | 9.933E-06 | 1.155E-07 | 9.800E-06 | 1.000E-05 |
| 10744 | 538 | C538D | A | 1.3e-05 | 1.4e-05 | 1.5e-05 | 1.400E-05 | 1.000E-06 | 1.300E-05 | 1.500E-05 |
| 10745 | 538 | C538C | A | 1.2e-05 | 1.1e-05 | 1.2e-05 | 1.167E-05 | 5.774E-07 | 1.100E-05 | 1.200E-05 |
| 10746 | 538 | C538E | A | 1.5e-05 | 1.4e-05 | 1.5e-05 | 1.467E-05 | 5.774E-07 | 1.400E-05 | 1.500E-05 |
| 10747 | 538 | C538Q | A | 1.1e-05 | 9.9e-06 | 1e-05 | 1.030E-05 | 6.083E-07 | 9.900E-06 | 1.100E-05 |
| 10748 | 538 | C538G | A | 1.6e-05 | 1.5e-05 | 1.5e-05 | 1.533E-05 | 5.774E-07 | 1.500E-05 | 1.600E-05 |
| 10749 | 538 | C538H | A | 9.2e-06 | 1.3e-05 | 1.5e-05 | 1.240E-05 | 2.946E-06 | 9.200E-06 | 1.500E-05 |
| 10750 | 538 | C538I | A | 9.3e-06 | 1e-05 | 1.3e-05 | 1.077E-05 | 1.966E-06 | 9.300E-06 | 1.300E-05 |
| 10751 | 538 | C538L | A | 1.1e-05 | 8.7e-06 | 1.5e-05 | 1.157E-05 | 3.188E-06 | 8.700E-06 | 1.500E-05 |
| 10752 | 538 | C538K | A | 1.2e-05 | 1.3e-05 | 1.9e-05 | 1.467E-05 | 3.786E-06 | 1.200E-05 | 1.900E-05 |
| 10753 | 538 | C538M | A | 1.5e-05 | 1.6e-05 | 1.5e-05 | 1.533E-05 | 5.774E-07 | 1.500E-05 | 1.600E-05 |
| 10754 | 538 | C538F | A | 1.7e-05 | 1.6e-05 | 1.5e-05 | 1.600E-05 | 1.000E-06 | 1.500E-05 | 1.700E-05 |
| 10755 | 538 | C538P | A | 1.1e-05 | 1.5e-05 | 1.4e-05 | 1.333E-05 | 2.082E-06 | 1.100E-05 | 1.500E-05 |
| 10756 | 538 | C538S | A | 1.5e-05 | 1.3e-05 | 1.3e-05 | 1.367E-05 | 1.155E-06 | 1.300E-05 | 1.500E-05 |
| 10757 | 538 | C538T | A | 1.7e-05 | 1.4e-05 | 1.6e-05 | 1.567E-05 | 1.528E-06 | 1.400E-05 | 1.700E-05 |
| 10758 | 538 | C538W | A | 1.5e-05 | 2.1e-05 | 1.9e-05 | 1.833E-05 | 3.055E-06 | 1.500E-05 | 2.100E-05 |
| 10759 | 538 | C538Y | A | 1.2e-05 | 1.4e-05 | 1.4e-05 | 1.333E-05 | 1.155E-06 | 1.200E-05 | 1.400E-05 |
| 10760 | 538 | C538V | A | 9.2e-06 | 8.5e-06 | 9.7e-06 | 9.133E-06 | 6.028E-07 | 8.500E-06 | 9.700E-06 |
| 10761 | 539 | L539A | A | 1.5e-05 | 1.3e-05 | 1.3e-05 | 1.367E-05 | 1.155E-06 | 1.300E-05 | 1.500E-05 |
| 10762 | 539 | L539R | A | 1.2e-05 | 8.9e-06 | 9.7e-06 | 1.020E-05 | 1.609E-06 | 8.900E-06 | 1.200E-05 |
| 10763 | 539 | L539N | A | 1.4e-05 | 1.1e-05 | 1.6e-05 | 1.367E-05 | 2.517E-06 | 1.100E-05 | 1.600E-05 |
| 10764 | 539 | L539D | A | 1.4e-05 | 1.5e-05 | 9.3e-06 | 1.277E-05 | 3.044E-06 | 9.300E-06 | 1.500E-05 |
| 10765 | 539 | L539C | A | 9.8e-06 | 1.5e-05 | 1e-05 | 1.160E-05 | 2.946E-06 | 9.800E-06 | 1.500E-05 |
| 10766 | 539 | L539E | A | 1.1e-05 | 1.4e-05 | 1.3e-05 | 1.267E-05 | 1.528E-06 | 1.100E-05 | 1.400E-05 |
| 10767 | 539 | L539Q | A | 9.7e-06 | 1.6e-05 | 1.6e-05 | 1.390E-05 | 3.637E-06 | 9.700E-06 | 1.600E-05 |
| 10768 | 539 | L539G | A | 1.3e-05 | 8.7e-06 | 9.1e-06 | 1.027E-05 | 2.376E-06 | 8.700E-06 | 1.300E-05 |
| 10769 | 539 | L539H | A | 1.2e-05 | 1.5e-05 | 1.4e-05 | 1.367E-05 | 1.528E-06 | 1.200E-05 | 1.500E-05 |
| 10770 | 539 | L539I | A | 1.6e-05 | 1.3e-05 | 1.8e-05 | 1.567E-05 | 2.517E-06 | 1.300E-05 | 1.800E-05 |
| 10771 | 539 | L539L | A | 9.8e-06 | 1.4e-05 | 1.1e-05 | 1.160E-05 | 2.163E-06 | 9.800E-06 | 1.400E-05 |
| 10772 | 539 | L539K | A | 1.6e-05 | 1.6e-05 | 9.6e-06 | 1.387E-05 | 3.695E-06 | 9.600E-06 | 1.600E-05 |
| 10773 | 539 | L539M | A | 2.4e-05 | 1.4e-05 | 9.9e-06 | 1.597E-05 | 7.253E-06 | 9.900E-06 | 2.400E-05 |
| 10774 | 539 | L539F | A | 1e-05 | 1.2e-05 | 1.5e-05 | 1.233E-05 | 2.517E-06 | 1.000E-05 | 1.500E-05 |
| 10775 | 539 | L539P | A | 9.9e-06 | 1.5e-05 | 1.4e-05 | 1.297E-05 | 2.702E-06 | 9.900E-06 | 1.500E-05 |
| 10776 | 539 | L539S | A | 1.4e-05 | 1.5e-05 | 1.4e-05 | 1.433E-05 | 5.774E-07 | 1.400E-05 | 1.500E-05 |
| 10777 | 539 | L539T | A | 1.6e-05 | 1.1e-05 | 1.1e-05 | 1.267E-05 | 2.887E-06 | 1.100E-05 | 1.600E-05 |
| 10778 | 539 | L539W | A | 1.1e-05 | 1.6e-05 | 8.4e-06 | 1.180E-05 | 3.863E-06 | 8.400E-06 | 1.600E-05 |
| 10779 | 539 | L539Y | A | 1e-05 | 9.7e-06 | 1.3e-05 | 1.090E-05 | 1.825E-06 | 9.700E-06 | 1.300E-05 |
| 10780 | 539 | L539V | A | 1.5e-05 | 1.7e-05 | 1.2e-05 | 1.467E-05 | 2.517E-06 | 1.200E-05 | 1.700E-05 |
| 10781 | 540 | P540A | A | 1.2e-05 | 1.1e-05 | 1.4e-05 | 1.233E-05 | 1.528E-06 | 1.100E-05 | 1.400E-05 |
| 10782 | 540 | P540R | A | 1.6e-05 | 1.5e-05 | 9.9e-06 | 1.363E-05 | 3.272E-06 | 9.900E-06 | 1.600E-05 |
| 10783 | 540 | P540N | A | 9.6e-06 | 9.7e-06 | 1e-05 | 9.767E-06 | 2.082E-07 | 9.600E-06 | 1.000E-05 |
| 10784 | 540 | P540D | A | 1.5e-05 | 1.6e-05 | 1.4e-05 | 1.500E-05 | 1.000E-06 | 1.400E-05 | 1.600E-05 |
| 10785 | 540 | P540C | A | 1.3e-05 | 1.3e-05 | 2e-05 | 1.533E-05 | 4.041E-06 | 1.300E-05 | 2.000E-05 |
| 10786 | 540 | P540E | A | 1.4e-05 | 1.6e-05 | 9.1e-06 | 1.303E-05 | 3.550E-06 | 9.100E-06 | 1.600E-05 |
| 10787 | 540 | P540Q | A | 1.6e-05 | 1.4e-05 | 1.8e-05 | 1.600E-05 | 2.000E-06 | 1.400E-05 | 1.800E-05 |
| 10788 | 540 | P540G | A | 1.1e-05 | 1.8e-05 | 1.7e-05 | 1.533E-05 | 3.786E-06 | 1.100E-05 | 1.800E-05 |
| 10789 | 540 | P540H | A | 1.3e-05 | 1.6e-05 | 1e-05 | 1.300E-05 | 3.000E-06 | 1.000E-05 | 1.600E-05 |
| 10790 | 540 | P540I | A | 9.3e-06 | 1.6e-05 | 1.2e-05 | 1.243E-05 | 3.371E-06 | 9.300E-06 | 1.600E-05 |
| 10791 | 540 | P540L | A | 1.6e-05 | 1.5e-05 | 1.5e-05 | 1.533E-05 | 5.774E-07 | 1.500E-05 | 1.600E-05 |
| 10792 | 540 | P540K | A | 9e-06 | 1.4e-05 | 1.3e-05 | 1.200E-05 | 2.646E-06 | 9.000E-06 | 1.400E-05 |
| 10793 | 540 | P540M | A | 1.5e-05 | 1.6e-05 | 1.5e-05 | 1.533E-05 | 5.774E-07 | 1.500E-05 | 1.600E-05 |
| 10794 | 540 | P540F | A | 1.2e-05 | 1.5e-05 | 1.6e-05 | 1.433E-05 | 2.082E-06 | 1.200E-05 | 1.600E-05 |

| | | | | | | | | | | |
|---|---|---|---|---|---|---|---|---|---|---|
| 10795 | 540 | P540P | A | 1.5e-05 | 1.6e-05 | 1.6e-05 | 1.567E-05 | 5.774E-07 | 1.500E-05 | 1.600E-05 |
| 10796 | 540 | P540S | A | 8.8e-06 | 1.4e-05 | 9.5e-06 | 1.077E-05 | 2.822E-06 | 8.800E-06 | 1.400E-05 |
| 10797 | 540 | P540T | A | 9.6e-06 | 1.7e-05 | 9.2e-06 | 1.193E-05 | 4.392E-06 | 9.200E-06 | 1.700E-05 |
| 10798 | 540 | P540W | A | 1.6e-05 | 1.7e-05 | 8.7e-06 | 1.390E-05 | 4.531E-06 | 8.700E-06 | 1.700E-05 |
| 10799 | 540 | P540Y | A | 1.5e-05 | 1.3e-05 | 9.1e-06 | 1.237E-05 | 3.001E-06 | 9.100E-06 | 1.500E-05 |
| 10800 | 540 | P540V | A | 9.3e-06 | 9.8e-06 | 1.4e-05 | 1.103E-05 | 2.581E-06 | 9.300E-06 | 1.400E-05 |
| 10801 | 541 | C541A | A | 1.1e-05 | 1.5e-05 | 8.5e-06 | 1.150E-05 | 3.279E-06 | 8.500E-06 | 1.500E-05 |
| 10802 | 541 | C541R | A | 1.5e-05 | 1.1e-05 | 1.6e-05 | 1.400E-05 | 2.646E-06 | 1.100E-05 | 1.600E-05 |
| 10803 | 541 | C541N | A | 1.3e-05 | 1.1e-05 | 1.5e-05 | 1.300E-05 | 2.000E-06 | 1.100E-05 | 1.500E-05 |
| 10804 | 541 | C541D | A | 1.7e-05 | 1e-05 | 1.2e-05 | 1.300E-05 | 3.606E-06 | 1.000E-05 | 1.700E-05 |
| 10805 | 541 | C541C | A | 1.4e-05 | 1.5e-05 | 1.5e-05 | 1.467E-05 | 5.774E-07 | 1.400E-05 | 1.500E-05 |
| 10806 | 541 | C541E | A | 1.5e-05 | 1.5e-05 | 1e-05 | 1.333E-05 | 2.887E-06 | 1.000E-05 | 1.500E-05 |
| 10807 | 541 | C541Q | A | 1.1e-05 | 1.1e-05 | 1.5e-05 | 1.233E-05 | 2.309E-06 | 1.100E-05 | 1.500E-05 |
| 10808 | 541 | C541G | A | 9.6e-06 | 1.5e-05 | 1.1e-05 | 1.187E-05 | 2.802E-06 | 9.600E-06 | 1.500E-05 |
| 10809 | 541 | C541H | A | 1.1e-05 | 1.7e-05 | 9.9e-06 | 1.263E-05 | 3.821E-06 | 9.900E-06 | 1.700E-05 |
| 10810 | 541 | C541I | A | 1.1e-05 | 1.6e-05 | 1.1e-05 | 1.267E-05 | 2.887E-06 | 1.100E-05 | 1.600E-05 |
| 10811 | 541 | C541L | A | 1.6e-05 | 1.2e-05 | 1.4e-05 | 1.400E-05 | 2.000E-06 | 1.200E-05 | 1.600E-05 |
| 10812 | 541 | C541K | A | 1.4e-05 | 8.8e-06 | 1e-05 | 1.093E-05 | 2.723E-06 | 8.800E-06 | 1.400E-05 |
| 10813 | 541 | C541M | A | 1e-05 | 1.5e-05 | 9.5e-06 | 1.150E-05 | 3.041E-06 | 9.500E-06 | 1.500E-05 |
| 10814 | 541 | C541F | A | 1.5e-05 | 1.7e-05 | 1.5e-05 | 1.567E-05 | 1.155E-06 | 1.500E-05 | 1.700E-05 |
| 10815 | 541 | C541P | A | 9.9e-06 | 1e-05 | 1e-05 | 9.967E-06 | 5.774E-08 | 9.900E-06 | 1.000E-05 |
| 10816 | 541 | C541S | A | 1.4e-05 | 1e-05 | 1.1e-05 | 1.167E-05 | 2.082E-06 | 1.000E-05 | 1.400E-05 |
| 10817 | 541 | C541T | A | 1e-05 | 9.5e-06 | 9e-06 | 9.500E-06 | 5.000E-07 | 9.000E-06 | 1.000E-05 |
| 10818 | 541 | C541W | A | 9.2e-06 | 1e-05 | 1.4e-05 | 1.107E-05 | 2.572E-06 | 9.200E-06 | 1.400E-05 |
| 10819 | 541 | C541Y | A | 1.6e-05 | 9.2e-06 | 1.4e-05 | 1.307E-05 | 3.495E-06 | 9.200E-06 | 1.600E-05 |
| 10820 | 541 | C541V | A | 1.1e-05 | 1.6e-05 | 1.1e-05 | 1.267E-05 | 2.887E-06 | 1.100E-05 | 1.600E-05 |
| 10821 | 542 | H542A | A | 1.1e-05 | 1.6e-05 | 1.4e-05 | 1.367E-05 | 2.517E-06 | 1.100E-05 | 1.600E-05 |
| 10822 | 542 | H542R | A | 1.1e-05 | 1.2e-05 | 1.4e-05 | 1.233E-05 | 1.528E-06 | 1.100E-05 | 1.400E-05 |
| 10823 | 542 | H542N | A | 7.8e-06 | 1.6e-05 | 1.5e-05 | 1.293E-05 | 4.474E-06 | 7.800E-06 | 1.600E-05 |
| 10824 | 542 | H542D | A | 1.6e-05 | 1.1e-05 | 1.1e-05 | 1.267E-05 | 2.887E-06 | 1.100E-05 | 1.600E-05 |
| 10825 | 542 | H542C | A | 9.6e-06 | 1.1e-05 | 1.1e-05 | 1.053E-05 | 8.083E-07 | 9.600E-06 | 1.100E-05 |
| 10826 | 542 | H542E | A | 1.2e-05 | 1.5e-05 | 1.1e-05 | 1.267E-05 | 2.082E-06 | 1.100E-05 | 1.500E-05 |
| 10827 | 542 | H542Q | A | 1.1e-05 | 1.3e-05 | 1.6e-05 | 1.333E-05 | 2.517E-06 | 1.100E-05 | 1.600E-05 |
| 10828 | 542 | H542G | A | 1.7e-05 | 9.9e-06 | 1.2e-05 | 1.297E-05 | 3.647E-06 | 9.900E-06 | 1.700E-05 |
| 10829 | 542 | H542H | A | 1.2e-05 | 1.3e-05 | 1.1e-05 | 1.200E-05 | 1.000E-06 | 1.100E-05 | 1.300E-05 |
| 10830 | 542 | H542I | A | 1.5e-05 | 1.6e-05 | 9.7e-06 | 1.357E-05 | 3.386E-06 | 9.700E-06 | 1.600E-05 |
| 10831 | 542 | H542L | A | 1.1e-05 | 1.5e-05 | 1.5e-05 | 1.367E-05 | 2.309E-06 | 1.100E-05 | 1.500E-05 |
| 10832 | 542 | H542K | A | 1e-05 | 1.1e-05 | 1e-05 | 1.033E-05 | 5.774E-07 | 1.000E-05 | 1.100E-05 |
| 10833 | 542 | H542M | A | 1.6e-05 | 1.4e-05 | 1.2e-05 | 1.400E-05 | 2.000E-06 | 1.200E-05 | 1.600E-05 |
| 10834 | 542 | H542F | A | 1.1e-05 | 1.9e-05 | 6.9e-06 | 1.230E-05 | 6.154E-06 | 6.900E-06 | 1.900E-05 |
| 10835 | 542 | H542P | A | 1.3e-05 | 1.4e-05 | 1.3e-05 | 1.333E-05 | 5.774E-07 | 1.300E-05 | 1.400E-05 |
| 10836 | 542 | H542S | A | 1.1e-05 | 1.1e-05 | 1.3e-05 | 1.167E-05 | 1.155E-06 | 1.100E-05 | 1.300E-05 |
| 10837 | 542 | H542T | A | 9.1e-06 | 1.6e-05 | 1.1e-05 | 1.203E-05 | 3.564E-06 | 9.100E-06 | 1.600E-05 |
| 10838 | 542 | H542W | A | 1.3e-05 | 1.6e-05 | 1e-05 | 1.300E-05 | 3.000E-06 | 1.000E-05 | 1.600E-05 |
| 10839 | 542 | H542Y | A | 9.2e-06 | 1e-05 | 1.5e-05 | 1.140E-05 | 3.143E-06 | 9.200E-06 | 1.500E-05 |
| 10840 | 542 | H542V | A | 1.4e-05 | 1.4e-05 | 1.6e-05 | 1.467E-05 | 1.155E-06 | 1.400E-05 | 1.600E-05 |
| 10841 | 543 | P543A | A | 1.4e-05 | 1.6e-05 | 1e-05 | 1.333E-05 | 3.055E-06 | 1.000E-05 | 1.600E-05 |
| 10842 | 543 | P543R | A | 1.2e-05 | 1.5e-05 | 1.2e-05 | 1.300E-05 | 1.732E-06 | 1.200E-05 | 1.500E-05 |
| 10843 | 543 | P543N | A | 1.5e-05 | 9.9e-06 | 8e-06 | 1.097E-05 | 3.620E-06 | 8.000E-06 | 1.500E-05 |
| 10844 | 543 | P543D | A | 1.7e-05 | 1.4e-05 | 1.5e-05 | 1.533E-05 | 1.528E-06 | 1.400E-05 | 1.700E-05 |
| 10845 | 543 | P543C | A | 1.4e-05 | 1.1e-05 | 1.3e-05 | 1.267E-05 | 1.528E-06 | 1.100E-05 | 1.400E-05 |
| 10846 | 543 | P543E | A | 9.7e-06 | 8.5e-06 | 1.4e-05 | 1.073E-05 | 2.892E-06 | 8.500E-06 | 1.400E-05 |
| 10847 | 543 | P543Q | A | 1.7e-05 | 1e-05 | 1e-05 | 1.233E-05 | 4.041E-06 | 1.000E-05 | 1.700E-05 |
| 10848 | 543 | P543G | A | 1.1e-05 | 1.7e-05 | 1.1e-05 | 1.300E-05 | 3.464E-06 | 1.100E-05 | 1.700E-05 |
| 10849 | 543 | P543H | A | 1.3e-05 | 1.5e-05 | 1.1e-05 | 1.300E-05 | 2.000E-06 | 1.100E-05 | 1.500E-05 |
| 10850 | 543 | P543I | A | 1.6e-05 | 1.1e-05 | 2.3e-05 | 1.667E-05 | 6.028E-06 | 1.100E-05 | 2.300E-05 |
| 10851 | 543 | P543L | A | 9e-06 | 1e-05 | 1e-05 | 9.667E-06 | 5.774E-07 | 9.000E-06 | 1.000E-05 |
| 10852 | 543 | P543K | A | 1.7e-05 | 1.5e-05 | 1.5e-05 | 1.567E-05 | 1.155E-06 | 1.500E-05 | 1.700E-05 |
| 10853 | 543 | P543M | A | 1.1e-05 | 1.3e-05 | 9.5e-06 | 1.117E-05 | 1.756E-06 | 9.500E-06 | 1.300E-05 |
| 10854 | 543 | P543F | A | 1.6e-05 | 1.5e-05 | 1.5e-05 | 1.533E-05 | 5.774E-07 | 1.500E-05 | 1.600E-05 |
| 10855 | 543 | P543P | A | 1.6e-05 | 1.6e-05 | 1.6e-05 | 1.600E-05 | 0.000E+00 | 1.600E-05 | 1.600E-05 |
| 10856 | 543 | P543S | A | 1.4e-05 | 1.3e-05 | 9.2e-06 | 1.207E-05 | 2.532E-06 | 9.200E-06 | 1.400E-05 |
| 10857 | 543 | P543T | A | 1.1e-05 | 1.4e-05 | 1.6e-05 | 1.367E-05 | 2.517E-06 | 1.100E-05 | 1.600E-05 |
| 10858 | 543 | P543W | A | 1.5e-05 | 1e-05 | 1.4e-05 | 1.300E-05 | 2.646E-06 | 1.000E-05 | 1.500E-05 |
| 10859 | 543 | P543Y | A | 1.2e-05 | 1.5e-05 | 1.5e-05 | 1.400E-05 | 1.732E-06 | 1.200E-05 | 1.500E-05 |
| 10860 | 543 | P543V | A | 1.7e-05 | 9.3e-06 | 1.3e-05 | 1.310E-05 | 3.851E-06 | 9.300E-06 | 1.700E-05 |
| 10861 | 544 | E544A | A | 1.5e-05 | 1.5e-05 | 1.4e-05 | 1.467E-05 | 5.774E-07 | 1.400E-05 | 1.500E-05 |
| 10862 | 544 | E544R | A | 1.5e-05 | 1.4e-05 | 6.9e-06 | 1.197E-05 | 4.416E-06 | 6.900E-06 | 1.500E-05 |
| 10863 | 544 | E544N | A | 1.5e-05 | 1.6e-05 | 1.1e-05 | 1.400E-05 | 2.646E-06 | 1.100E-05 | 1.600E-05 |
| 10864 | 544 | E544D | A | 1.1e-05 | 1.6e-05 | 9.9e-06 | 1.230E-05 | 3.251E-06 | 9.900E-06 | 1.600E-05 |
| 10865 | 544 | E544C | A | 1.6e-05 | 1.7e-05 | 1.6e-05 | 1.633E-05 | 5.774E-07 | 1.600E-05 | 1.700E-05 |
| 10866 | 544 | E544E | A | 1.1e-05 | 1.6e-05 | 7.2e-06 | 1.140E-05 | 4.414E-06 | 7.200E-06 | 1.600E-05 |
| 10867 | 544 | E544Q | A | 1e-05 | 1.3e-05 | 1.6e-05 | 1.300E-05 | 3.000E-06 | 1.000E-05 | 1.600E-05 |
| 10868 | 544 | E544G | A | 9.9e-06 | 1.6e-05 | 1.5e-05 | 1.363E-05 | 3.272E-06 | 9.900E-06 | 1.600E-05 |
| 10869 | 544 | E544H | A | 7e-06 | 9e-06 | 6.8e-06 | 7.600E-06 | 1.217E-06 | 6.800E-06 | 9.000E-06 |
| 10870 | 544 | E544I | A | 9.8e-06 | 1.3e-05 | 1e-05 | 1.093E-05 | 1.793E-06 | 9.800E-06 | 1.300E-05 |
| 10871 | 544 | E544L | A | 1.5e-05 | 1.3e-05 | 1.7e-05 | 1.500E-05 | 2.000E-06 | 1.300E-05 | 1.700E-05 |
| 10872 | 544 | E544K | A | 8.9e-06 | 1.4e-05 | 1.6e-05 | 1.297E-05 | 3.661E-06 | 8.900E-06 | 1.600E-05 |
| 10873 | 544 | E544M | A | 1.7e-05 | 1.1e-05 | 1.1e-05 | 1.300E-05 | 3.464E-06 | 1.100E-05 | 1.700E-05 |
| 10874 | 544 | E544F | A | 1.6e-05 | 9.4e-06 | 1.5e-05 | 1.347E-05 | 3.557E-06 | 9.400E-06 | 1.600E-05 |
| 10875 | 544 | E544P | A | 1.5e-05 | 1.2e-05 | 1.5e-05 | 1.400E-05 | 1.732E-06 | 1.200E-05 | 1.500E-05 |
| 10876 | 544 | E544S | A | 9.6e-06 | 9.9e-06 | 1.1e-05 | 1.017E-05 | 7.371E-07 | 9.600E-06 | 1.100E-05 |
| 10877 | 544 | E544T | A | 1.7e-05 | 1.4e-05 | 1.6e-05 | 1.567E-05 | 1.528E-06 | 1.400E-05 | 1.700E-05 |
| 10878 | 544 | E544W | A | 9.3e-06 | 6.8e-06 | 1e-05 | 8.700E-06 | 1.682E-06 | 6.800E-06 | 1.000E-05 |
| 10879 | 544 | E544Y | A | 1.1e-05 | 1e-05 | 1e-05 | 1.033E-05 | 5.774E-07 | 1.000E-05 | 1.100E-05 |

| | | | | | | | | | | |
|---|---|---|---|---|---|---|---|---|---|---|
| 10880 | 544 | E544V | A | 1.4e-05 | 1.6e-05 | 1e-05 | 1.333E-05 | 3.055E-06 | 1.000E-05 | 1.600E-05 |
| 10881 | 545 | C545A | A | 1.1e-05 | 9e-06 | 1e-05 | 1.000E-05 | 1.000E-06 | 9.000E-06 | 1.100E-05 |
| 10882 | 545 | C545R | A | 1.3e-05 | 1.5e-05 | 1.4e-05 | 1.400E-05 | 1.000E-06 | 1.300E-05 | 1.500E-05 |
| 10883 | 545 | C545N | A | 6.6e-06 | 1.5e-05 | 1.2e-05 | 1.120E-05 | 4.257E-06 | 6.600E-06 | 1.500E-05 |
| 10884 | 545 | C545D | A | 1.9e-05 | 1.1e-05 | 1.6e-05 | 1.533E-05 | 4.041E-06 | 1.100E-05 | 1.900E-05 |
| 10885 | 545 | C545C | A | 8.6e-06 | 1.3e-05 | 1.2e-05 | 1.120E-05 | 2.307E-06 | 8.600E-06 | 1.300E-05 |
| 10886 | 545 | C545E | A | 1e-05 | 1.6e-05 | 1.1e-05 | 1.233E-05 | 3.215E-06 | 1.000E-05 | 1.600E-05 |
| 10887 | 545 | C545Q | A | 1.7e-05 | 1.1e-05 | 1.3e-05 | 1.367E-05 | 3.055E-06 | 1.100E-05 | 1.700E-05 |
| 10888 | 545 | C545G | A | 1.3e-05 | 1.1e-05 | 8.7e-06 | 1.090E-05 | 2.152E-06 | 8.700E-06 | 1.300E-05 |
| 10889 | 545 | C545H | A | 1.2e-05 | 8.6e-06 | 1.2e-05 | 1.087E-05 | 1.963E-06 | 8.600E-06 | 1.200E-05 |
| 10890 | 545 | C545I | A | 1e-05 | 1.4e-05 | 1.5e-05 | 1.300E-05 | 2.646E-06 | 1.000E-05 | 1.500E-05 |
| 10891 | 545 | C545L | A | 1.1e-05 | 1.1e-05 | 1.5e-05 | 1.233E-05 | 2.309E-06 | 1.100E-05 | 1.500E-05 |
| 10892 | 545 | C545K | A | 1.4e-05 | 8.9e-06 | 1e-05 | 1.097E-05 | 2.684E-06 | 8.900E-06 | 1.400E-05 |
| 10893 | 545 | C545M | A | 9e-06 | 1.4e-05 | 1.8e-05 | 1.367E-05 | 4.509E-06 | 9.000E-06 | 1.800E-05 |
| 10894 | 545 | C545F | A | 1.6e-05 | 1.2e-05 | 1e-05 | 1.267E-05 | 3.055E-06 | 1.000E-05 | 1.600E-05 |
| 10895 | 545 | C545P | A | 1.7e-05 | 9.7e-06 | 1.5e-05 | 1.390E-05 | 3.772E-06 | 9.700E-06 | 1.700E-05 |
| 10896 | 545 | C545S | A | 1e-05 | 1.5e-05 | 1.6e-05 | 1.367E-05 | 3.215E-06 | 1.000E-05 | 1.600E-05 |
| 10897 | 545 | C545T | A | 1.4e-05 | 1.7e-05 | 1.3e-05 | 1.467E-05 | 2.082E-06 | 1.300E-05 | 1.700E-05 |
| 10898 | 545 | C545W | A | 1.5e-05 | 8.6e-06 | 1.5e-05 | 1.287E-05 | 3.695E-06 | 8.600E-06 | 1.500E-05 |
| 10899 | 545 | C545Y | A | 9.2e-06 | 1.5e-05 | 1.2e-05 | 1.207E-05 | 2.901E-06 | 9.200E-06 | 1.500E-05 |
| 10900 | 545 | C545V | A | 1.3e-05 | 1.4e-05 | 8.5e-06 | 1.183E-05 | 2.930E-06 | 8.500E-06 | 1.400E-05 |
| 10901 | 546 | Q546A | A | 1.6e-05 | 9e-06 | 6.9e-06 | 1.063E-05 | 4.765E-06 | 6.900E-06 | 1.600E-05 |
| 10902 | 546 | Q546R | A | 1.1e-05 | 8.3e-06 | 9.8e-06 | 9.700E-06 | 1.353E-06 | 8.300E-06 | 1.100E-05 |
| 10903 | 546 | Q546N | A | 1.5e-05 | 9.5e-06 | 1.1e-05 | 1.183E-05 | 2.843E-06 | 9.500E-06 | 1.500E-05 |
| 10904 | 546 | Q546D | A | 1.2e-05 | 1.6e-05 | 1.4e-05 | 1.400E-05 | 2.000E-06 | 1.200E-05 | 1.600E-05 |
| 10905 | 546 | Q546C | A | 8.8e-06 | 1.1e-05 | 7.9e-06 | 9.233E-06 | 1.595E-06 | 7.900E-06 | 1.100E-05 |
| 10906 | 546 | Q546E | A | 1.5e-05 | 1.3e-05 | 1.4e-05 | 1.400E-05 | 1.000E-06 | 1.300E-05 | 1.500E-05 |
| 10907 | 546 | Q546Q | A | 1e-05 | 1.1e-05 | 8.8e-06 | 9.933E-06 | 1.102E-06 | 8.800E-06 | 1.100E-05 |
| 10908 | 546 | Q546G | A | 1.6e-05 | 1.4e-05 | 9.5e-06 | 1.317E-05 | 3.329E-06 | 9.500E-06 | 1.600E-05 |
| 10909 | 546 | Q546H | A | 1.6e-05 | 1.6e-05 | 1.5e-05 | 1.567E-05 | 5.774E-07 | 1.500E-05 | 1.600E-05 |
| 10910 | 546 | Q546I | A | 1.1e-05 | 1.3e-05 | 1.2e-05 | 1.200E-05 | 1.000E-06 | 1.100E-05 | 1.300E-05 |
| 10911 | 546 | Q546L | A | 1.6e-05 | 1.6e-05 | 1.4e-05 | 1.533E-05 | 1.155E-06 | 1.400E-05 | 1.600E-05 |
| 10912 | 546 | Q546K | A | 1.4e-05 | 1.5e-05 | 9.3e-06 | 1.277E-05 | 3.044E-06 | 9.300E-06 | 1.500E-05 |
| 10913 | 546 | Q546M | A | 1.1e-05 | 1.1e-05 | 1.5e-05 | 1.233E-05 | 2.309E-06 | 1.100E-05 | 1.500E-05 |
| 10914 | 546 | Q546F | A | 1e-05 | 1.2e-05 | 1.7e-05 | 1.300E-05 | 3.606E-06 | 1.000E-05 | 1.700E-05 |
| 10915 | 546 | Q546P | A | 9e-06 | 9.2e-06 | 1.5e-05 | 1.107E-05 | 3.408E-06 | 9.000E-06 | 1.500E-05 |
| 10916 | 546 | Q546S | A | 1.4e-05 | 1.5e-05 | 1.5e-05 | 1.467E-05 | 5.774E-07 | 1.400E-05 | 1.500E-05 |
| 10917 | 546 | Q546T | A | 1.7e-05 | 1.4e-05 | 1.1e-05 | 1.400E-05 | 3.000E-06 | 1.100E-05 | 1.700E-05 |
| 10918 | 546 | Q546W | A | 9.7e-06 | 1e-05 | 1.7e-05 | 1.223E-05 | 4.131E-06 | 9.700E-06 | 1.700E-05 |
| 10919 | 546 | Q546Y | A | 1.5e-05 | 1e-05 | 9.3e-06 | 1.143E-05 | 3.109E-06 | 9.300E-06 | 1.500E-05 |
| 10920 | 546 | Q546V | A | 1.4e-05 | 1.1e-05 | 1.9e-05 | 1.467E-05 | 4.041E-06 | 1.100E-05 | 1.900E-05 |
| 10921 | 547 | P547A | A | 1e-05 | 1.5e-05 | 1.1e-05 | 1.200E-05 | 2.646E-06 | 1.000E-05 | 1.500E-05 |
| 10922 | 547 | P547R | A | 1.6e-05 | 8.4e-06 | 1.5e-05 | 1.313E-05 | 4.130E-06 | 8.400E-06 | 1.600E-05 |
| 10923 | 547 | P547N | A | 7.7e-06 | 1.2e-05 | 1.5e-05 | 1.157E-05 | 3.669E-06 | 7.700E-06 | 1.500E-05 |
| 10924 | 547 | P547D | A | 1e-05 | 1.5e-05 | 1.5e-05 | 1.333E-05 | 2.887E-06 | 1.000E-05 | 1.500E-05 |
| 10925 | 547 | P547C | A | 1.1e-05 | 1e-05 | 1.4e-05 | 1.167E-05 | 2.082E-06 | 1.000E-05 | 1.400E-05 |
| 10926 | 547 | P547E | A | 1.5e-05 | 1.2e-05 | 1.6e-05 | 1.433E-05 | 2.082E-06 | 1.200E-05 | 1.600E-05 |
| 10927 | 547 | P547Q | A | 1.5e-05 | 1.5e-05 | 1.5e-05 | 1.500E-05 | 0.000E+00 | 1.500E-05 | 1.500E-05 |
| 10928 | 547 | P547G | A | 1.1e-05 | 1.1e-05 | 1.6e-05 | 1.267E-05 | 2.887E-06 | 1.100E-05 | 1.600E-05 |
| 10929 | 547 | P547H | A | 1.8e-05 | 1.4e-05 | 1.6e-05 | 1.600E-05 | 2.000E-06 | 1.400E-05 | 1.800E-05 |
| 10930 | 547 | P547I | A | 1.4e-05 | 1.6e-05 | 1.1e-05 | 1.367E-05 | 2.517E-06 | 1.100E-05 | 1.600E-05 |
| 10931 | 547 | P547L | A | 8.2e-06 | 9.9e-06 | 1.1e-05 | 9.700E-06 | 1.411E-06 | 8.200E-06 | 1.100E-05 |
| 10932 | 547 | P547K | A | 1.3e-05 | 1.2e-05 | 8.5e-06 | 1.117E-05 | 2.363E-06 | 8.500E-06 | 1.300E-05 |
| 10933 | 547 | P547M | A | 1.3e-05 | 1.5e-05 | 1.6e-05 | 1.467E-05 | 1.528E-06 | 1.300E-05 | 1.600E-05 |
| 10934 | 547 | P547F | A | 8.1e-06 | 1.5e-05 | 2.3e-05 | 1.537E-05 | 7.457E-06 | 8.100E-06 | 2.300E-05 |
| 10935 | 547 | P547P | A | 1.7e-05 | 1.4e-05 | 1.7e-05 | 1.600E-05 | 1.732E-06 | 1.400E-05 | 1.700E-05 |
| 10936 | 547 | P547S | A | 1e-05 | 7.6e-06 | 1.1e-05 | 9.533E-06 | 1.747E-06 | 7.600E-06 | 1.100E-05 |
| 10937 | 547 | P547T | A | 1.4e-05 | 1.4e-05 | 1.1e-05 | 1.300E-05 | 1.732E-06 | 1.100E-05 | 1.400E-05 |
| 10938 | 547 | P547W | A | 1.1e-05 | 1.5e-05 | 1.2e-05 | 1.267E-05 | 2.082E-06 | 1.100E-05 | 1.500E-05 |
| 10939 | 547 | P547Y | A | 1.4e-05 | 1.6e-05 | 1.4e-05 | 1.467E-05 | 1.155E-06 | 1.400E-05 | 1.600E-05 |
| 10940 | 547 | P547V | A | 1.7e-05 | 1.5e-05 | 1.5e-05 | 1.567E-05 | 1.155E-06 | 1.500E-05 | 1.700E-05 |
| 10941 | 548 | Q548A | A | 1.1e-05 | 1.5e-05 | 1.3e-05 | 1.300E-05 | 2.000E-06 | 1.100E-05 | 1.500E-05 |
| 10942 | 548 | Q548R | A | 8.3e-06 | 1.4e-05 | 1.3e-05 | 1.177E-05 | 3.044E-06 | 8.300E-06 | 1.400E-05 |
| 10943 | 548 | Q548N | A | 1.1e-05 | 1.2e-05 | 1.5e-05 | 1.267E-05 | 2.082E-06 | 1.100E-05 | 1.500E-05 |
| 10944 | 548 | Q548D | A | 1.5e-05 | 1.5e-05 | 1.6e-05 | 1.533E-05 | 5.774E-07 | 1.500E-05 | 1.600E-05 |
| 10945 | 548 | Q548C | A | 1.7e-05 | 1.7e-05 | 1e-05 | 1.467E-05 | 4.041E-06 | 1.000E-05 | 1.700E-05 |
| 10946 | 548 | Q548E | A | 1.6e-05 | 1e-05 | 1.5e-05 | 1.367E-05 | 3.215E-06 | 1.000E-05 | 1.600E-05 |
| 10947 | 548 | Q548Q | A | 6.1e-06 | 1.1e-05 | 1.7e-05 | 1.137E-05 | 5.459E-06 | 6.100E-06 | 1.700E-05 |
| 10948 | 548 | Q548G | A | 1.1e-05 | 1.1e-05 | 1.7e-05 | 1.300E-05 | 3.464E-06 | 1.100E-05 | 1.700E-05 |
| 10949 | 548 | Q548H | A | 9.4e-06 | 1.5e-05 | 1.4e-05 | 1.280E-05 | 2.987E-06 | 9.400E-06 | 1.500E-05 |
| 10950 | 548 | Q548I | A | 1.7e-05 | 1.5e-05 | 1.6e-05 | 1.600E-05 | 1.000E-06 | 1.500E-05 | 1.700E-05 |
| 10951 | 548 | Q548L | A | 1.1e-05 | 1.5e-05 | 1.5e-05 | 1.367E-05 | 2.309E-06 | 1.100E-05 | 1.500E-05 |
| 10952 | 548 | Q548K | A | 1.1e-05 | 1.2e-05 | 1.5e-05 | 1.267E-05 | 2.082E-06 | 1.100E-05 | 1.500E-05 |
| 10953 | 548 | Q548M | A | 1.4e-05 | 1.6e-05 | 1.4e-05 | 1.467E-05 | 1.155E-06 | 1.400E-05 | 1.600E-05 |
| 10954 | 548 | Q548F | A | 9.9e-06 | 1.5e-05 | 1.2e-05 | 1.230E-05 | 2.563E-06 | 9.900E-06 | 1.500E-05 |
| 10955 | 548 | Q548P | A | 1.1e-05 | 9.1e-06 | 1e-05 | 1.003E-05 | 9.504E-07 | 9.100E-06 | 1.100E-05 |
| 10956 | 548 | Q548S | A | 1.6e-05 | 1.4e-05 | 1.1e-05 | 1.367E-05 | 2.517E-06 | 1.100E-05 | 1.600E-05 |
| 10957 | 548 | Q548T | A | 1.2e-05 | 1.7e-05 | 1.4e-05 | 1.433E-05 | 2.517E-06 | 1.200E-05 | 1.700E-05 |
| 10958 | 548 | Q548W | A | 9.4e-06 | 1.5e-05 | 1.4e-05 | 1.280E-05 | 2.987E-06 | 9.400E-06 | 1.500E-05 |
| 10959 | 548 | Q548Y | A | 9.7e-06 | 8.9e-06 | 1.5e-05 | 1.120E-05 | 3.315E-06 | 8.900E-06 | 1.500E-05 |
| 10960 | 548 | Q548V | A | 1.3e-05 | 8.9e-06 | 1.1e-05 | 1.097E-05 | 2.050E-06 | 8.900E-06 | 1.300E-05 |
| 10961 | 549 | N549A | A | 9.2e-06 | 8.5e-06 | 9.5e-06 | 9.067E-06 | 5.132E-07 | 8.500E-06 | 9.500E-06 |
| 10962 | 549 | N549R | A | 1.2e-05 | 1.3e-05 | 1.6e-05 | 1.367E-05 | 2.082E-06 | 1.200E-05 | 1.600E-05 |
| 10963 | 549 | N549N | A | 1.1e-05 | 7.7e-06 | 1.3e-05 | 1.057E-05 | 2.676E-06 | 7.700E-06 | 1.300E-05 |
| 10964 | 549 | N549D | A | 1.8e-05 | 1.1e-05 | 1.4e-05 | 1.433E-05 | 3.512E-06 | 1.100E-05 | 1.800E-05 |

| | | | | | | | | | | |
|---|---|---|---|---|---|---|---|---|---|---|
| 10965 | 549 | N549C | A | 7.1e-06 | 1.3e-05 | 9.5e-06 | 9.867E-06 | 2.967E-06 | 7.100E-06 | 1.300E-05 |
| 10966 | 549 | N549E | A | 1.3e-05 | 1e-05 | 1.1e-05 | 1.133E-05 | 1.528E-06 | 1.000E-05 | 1.300E-05 |
| 10967 | 549 | N549Q | A | 1.5e-05 | 1.6e-05 | 1.6e-05 | 1.567E-05 | 5.774E-07 | 1.500E-05 | 1.600E-05 |
| 10968 | 549 | N549G | A | 1.4e-05 | 1.6e-05 | 1e-05 | 1.333E-05 | 3.055E-06 | 1.000E-05 | 1.600E-05 |
| 10969 | 549 | N549H | A | 7.6e-06 | 1.5e-05 | 1.2e-05 | 1.153E-05 | 3.722E-06 | 7.600E-06 | 1.500E-05 |
| 10970 | 549 | N549I | A | 1.7e-05 | 1.5e-05 | 7.8e-06 | 1.327E-05 | 4.839E-06 | 7.800E-06 | 1.700E-05 |
| 10971 | 549 | N549L | A | 1.8e-05 | 1e-05 | 1.3e-05 | 1.367E-05 | 4.041E-06 | 1.000E-05 | 1.800E-05 |
| 10972 | 549 | N549K | A | 8.6e-06 | 1.5e-05 | 1e-05 | 1.120E-05 | 3.365E-06 | 8.600E-06 | 1.500E-05 |
| 10973 | 549 | N549M | A | 1.5e-05 | 1.1e-05 | 1.1e-05 | 1.233E-05 | 2.309E-06 | 1.100E-05 | 1.500E-05 |
| 10974 | 549 | N549F | A | 1.1e-05 | 1.3e-05 | 1e-05 | 1.133E-05 | 1.528E-06 | 1.000E-05 | 1.300E-05 |
| 10975 | 549 | N549P | A | 1.4e-05 | 1.5e-05 | 1e-05 | 1.300E-05 | 2.646E-06 | 1.000E-05 | 1.500E-05 |
| 10976 | 549 | N549S | A | 1.4e-05 | 1.5e-05 | 1.6e-05 | 1.500E-05 | 1.000E-06 | 1.400E-05 | 1.600E-05 |
| 10977 | 549 | N549T | A | 9.6e-06 | 1.5e-05 | 1.5e-05 | 1.320E-05 | 3.118E-06 | 9.600E-06 | 1.500E-05 |
| 10978 | 549 | N549W | A | 1.5e-05 | 1.2e-05 | 9.9e-06 | 1.230E-05 | 2.563E-06 | 9.900E-06 | 1.500E-05 |
| 10979 | 549 | N549Y | A | 9.7e-06 | 1e-05 | 1.4e-05 | 1.123E-05 | 2.401E-06 | 9.700E-06 | 1.400E-05 |
| 10980 | 549 | N549V | A | 1.5e-05 | 1.1e-05 | 1.6e-05 | 1.400E-05 | 2.646E-06 | 1.100E-05 | 1.600E-05 |
| 10981 | 550 | G550A | A | 1.4e-05 | 8.4e-06 | 1.7e-05 | 1.313E-05 | 4.365E-06 | 8.400E-06 | 1.700E-05 |
| 10982 | 550 | G550R | A | 1.1e-05 | 1.5e-05 | 1.2e-05 | 1.267E-05 | 2.082E-06 | 1.100E-05 | 1.500E-05 |
| 10983 | 550 | G550N | A | 1.5e-05 | 1.1e-05 | 9.8e-06 | 1.193E-05 | 2.723E-06 | 9.800E-06 | 1.500E-05 |
| 10984 | 550 | G550D | A | 1.3e-05 | 1.4e-05 | 8.8e-06 | 1.193E-05 | 2.759E-06 | 8.800E-06 | 1.400E-05 |
| 10985 | 550 | G550C | A | 1.9e-05 | 1.1e-05 | 1.4e-05 | 1.467E-05 | 4.041E-06 | 1.100E-05 | 1.900E-05 |
| 10986 | 550 | G550E | A | 1e-05 | 1.4e-05 | 1.6e-05 | 1.333E-05 | 3.055E-06 | 1.000E-05 | 1.600E-05 |
| 10987 | 550 | G550Q | A | 1.5e-05 | 1e-05 | 1.3e-05 | 1.267E-05 | 2.517E-06 | 1.000E-05 | 1.500E-05 |
| 10988 | 550 | G550G | A | 1.1e-05 | 9e-06 | 8.1e-06 | 9.367E-06 | 1.484E-06 | 8.100E-06 | 1.100E-05 |
| 10989 | 550 | G550H | A | 1.4e-05 | 1.4e-05 | 1.6e-05 | 1.467E-05 | 1.155E-06 | 1.400E-05 | 1.600E-05 |
| 10990 | 550 | G550I | A | 1.5e-05 | 1.7e-05 | 1e-05 | 1.400E-05 | 3.606E-06 | 1.000E-05 | 1.700E-05 |
| 10991 | 550 | G550L | A | 1.5e-05 | 1e-05 | 1.3e-05 | 1.267E-05 | 2.517E-06 | 1.000E-05 | 1.500E-05 |
| 10992 | 550 | G550K | A | 1.4e-05 | 1.4e-05 | 1.5e-05 | 1.433E-05 | 5.774E-07 | 1.400E-05 | 1.500E-05 |
| 10993 | 550 | G550M | A | 1.5e-05 | 9.5e-06 | 1.7e-05 | 1.383E-05 | 3.884E-06 | 9.500E-06 | 1.700E-05 |
| 10994 | 550 | G550F | A | 1.1e-05 | 9.6e-06 | 1.2e-05 | 1.087E-05 | 1.206E-06 | 9.600E-06 | 1.200E-05 |
| 10995 | 550 | G550P | A | 8.5e-06 | 1.3e-05 | 1e-05 | 1.050E-05 | 2.291E-06 | 8.500E-06 | 1.300E-05 |
| 10996 | 550 | G550S | A | 1.6e-05 | 1e-05 | 1.5e-05 | 1.367E-05 | 3.215E-06 | 1.000E-05 | 1.600E-05 |
| 10997 | 550 | G550T | A | 1e-05 | 1.3e-05 | 1.4e-05 | 1.233E-05 | 2.082E-06 | 1.000E-05 | 1.400E-05 |
| 10998 | 550 | G550W | A | 1.3e-05 | 2.6e-05 | 1.3e-05 | 1.733E-05 | 7.506E-06 | 1.300E-05 | 2.600E-05 |
| 10999 | 550 | G550Y | A | 1e-05 | 1.3e-05 | 1.5e-05 | 1.267E-05 | 2.517E-06 | 1.000E-05 | 1.500E-05 |
| 11000 | 550 | G550V | A | 8.2e-06 | 1.1e-05 | 1.4e-05 | 1.107E-05 | 2.901E-06 | 8.200E-06 | 1.400E-05 |
| 11001 | 551 | S551A | A | 1.6e-05 | 1.2e-05 | 1.2e-05 | 1.333E-05 | 2.309E-06 | 1.200E-05 | 1.600E-05 |
| 11002 | 551 | S551R | A | 2.2e-05 | 2.4e-05 | 1.5e-05 | 2.033E-05 | 4.726E-06 | 1.500E-05 | 2.400E-05 |
| 11003 | 551 | S551N | A | 1.5e-05 | 1.2e-05 | 1.1e-05 | 1.267E-05 | 2.082E-06 | 1.100E-05 | 1.500E-05 |
| 11004 | 551 | S551D | A | 1e-05 | 1.5e-05 | 1.3e-05 | 1.267E-05 | 2.517E-06 | 1.000E-05 | 1.500E-05 |
| 11005 | 551 | S551C | A | 1.5e-05 | 1.5e-05 | 1.4e-05 | 1.467E-05 | 5.774E-07 | 1.400E-05 | 1.500E-05 |
| 11006 | 551 | S551E | A | 1.3e-05 | 1.6e-05 | 1.7e-05 | 1.533E-05 | 2.082E-06 | 1.300E-05 | 1.700E-05 |
| 11007 | 551 | S551Q | A | 1.5e-05 | 9.3e-06 | 1.3e-05 | 1.243E-05 | 2.892E-06 | 9.300E-06 | 1.500E-05 |
| 11008 | 551 | S551G | A | 1.7e-05 | 1.4e-05 | 1.5e-05 | 1.533E-05 | 1.528E-06 | 1.400E-05 | 1.700E-05 |
| 11009 | 551 | S551H | A | 8.2e-06 | 1e-05 | 1.6e-05 | 1.140E-05 | 4.084E-06 | 8.200E-06 | 1.600E-05 |
| 11010 | 551 | S551I | A | 1.5e-05 | 1.4e-05 | 1.6e-05 | 1.500E-05 | 1.000E-06 | 1.400E-05 | 1.600E-05 |
| 11011 | 551 | S551L | A | 9.5e-06 | 1.7e-05 | 1.4e-05 | 1.350E-05 | 3.775E-06 | 9.500E-06 | 1.700E-05 |
| 11012 | 551 | S551K | A | 1.1e-05 | 1.5e-05 | 1.4e-05 | 1.333E-05 | 2.082E-06 | 1.100E-05 | 1.500E-05 |
| 11013 | 551 | S551M | A | 1.1e-05 | 1.7e-05 | 1.1e-05 | 1.300E-05 | 3.464E-06 | 1.100E-05 | 1.700E-05 |
| 11014 | 551 | S551F | A | 9e-06 | 1.7e-05 | 1.2e-05 | 1.267E-05 | 4.041E-06 | 9.000E-06 | 1.700E-05 |
| 11015 | 551 | S551P | A | 1.5e-05 | 1e-05 | 1.6e-05 | 1.367E-05 | 3.215E-06 | 1.000E-05 | 1.600E-05 |
| 11016 | 551 | S551S | A | 9.6e-06 | 1.5e-05 | 1.2e-05 | 1.220E-05 | 2.706E-06 | 9.600E-06 | 1.500E-05 |
| 11017 | 551 | S551T | A | 1.2e-05 | 1e-05 | 1.1e-05 | 1.100E-05 | 1.000E-06 | 1.000E-05 | 1.200E-05 |
| 11018 | 551 | S551W | A | 1.4e-05 | 1.6e-05 | 9.4e-06 | 1.313E-05 | 3.384E-06 | 9.400E-06 | 1.600E-05 |
| 11019 | 551 | S551Y | A | 1.4e-05 | 1.3e-05 | 1.7e-05 | 1.467E-05 | 2.082E-06 | 1.300E-05 | 1.700E-05 |
| 11020 | 551 | S551V | A | 9.3e-06 | 1.2e-05 | 1.5e-05 | 1.210E-05 | 2.851E-06 | 9.300E-06 | 1.500E-05 |
| 11021 | 552 | V552A | A | 1.4e-05 | 1.5e-05 | 1e-05 | 1.300E-05 | 2.646E-06 | 1.000E-05 | 1.500E-05 |
| 11022 | 552 | V552R | A | 1e-05 | 1.1e-05 | 1.3e-05 | 1.133E-05 | 1.528E-06 | 1.000E-05 | 1.300E-05 |
| 11023 | 552 | V552N | A | 1.5e-05 | 1.5e-05 | 1.5e-05 | 1.500E-05 | 0.000E+00 | 1.500E-05 | 1.500E-05 |
| 11024 | 552 | V552D | A | 1.1e-05 | 1.3e-05 | 1.3e-05 | 1.233E-05 | 1.155E-06 | 1.100E-05 | 1.300E-05 |
| 11025 | 552 | V552C | A | 1.1e-05 | 1.4e-05 | 1.2e-05 | 1.233E-05 | 1.528E-06 | 1.100E-05 | 1.400E-05 |
| 11026 | 552 | V552E | A | 1.5e-05 | 8.8e-06 | 9e-06 | 1.093E-05 | 3.523E-06 | 8.800E-06 | 1.500E-05 |
| 11027 | 552 | V552Q | A | 1.6e-05 | 1.6e-05 | 1e-05 | 1.400E-05 | 3.464E-06 | 1.000E-05 | 1.600E-05 |
| 11028 | 552 | V552G | A | 1.5e-05 | 9.5e-06 | 1.4e-05 | 1.283E-05 | 2.930E-06 | 9.500E-06 | 1.500E-05 |
| 11029 | 552 | V552H | A | 1e-05 | 9e-06 | 1.5e-05 | 1.133E-05 | 3.215E-06 | 9.000E-06 | 1.500E-05 |
| 11030 | 552 | V552I | A | 8.9e-06 | 1.4e-05 | 1.6e-05 | 1.297E-05 | 3.661E-06 | 8.900E-06 | 1.600E-05 |
| 11031 | 552 | V552L | A | 1.5e-05 | 1.6e-05 | 1.5e-05 | 1.533E-05 | 5.774E-07 | 1.500E-05 | 1.600E-05 |
| 11032 | 552 | V552K | A | 1.1e-05 | 9e-06 | 1e-05 | 1.000E-05 | 1.000E-06 | 9.000E-06 | 1.100E-05 |
| 11033 | 552 | V552M | A | 9.8e-06 | 1.7e-05 | 1.1e-05 | 1.260E-05 | 3.857E-06 | 9.800E-06 | 1.700E-05 |
| 11034 | 552 | V552F | A | 1.6e-05 | 1.7e-05 | 1.1e-05 | 1.467E-05 | 3.215E-06 | 1.100E-05 | 1.700E-05 |
| 11035 | 552 | V552P | A | 1.1e-05 | 1.5e-05 | 1.2e-05 | 1.267E-05 | 2.082E-06 | 1.100E-05 | 1.500E-05 |
| 11036 | 552 | V552S | A | 9.9e-06 | 1.6e-05 | 9e-06 | 1.163E-05 | 3.808E-06 | 9.000E-06 | 1.600E-05 |
| 11037 | 552 | V552T | A | 9.3e-06 | 1.2e-05 | 8e-06 | 9.767E-06 | 2.040E-06 | 8.000E-06 | 1.200E-05 |
| 11038 | 552 | V552W | A | 1.5e-05 | 1.5e-05 | 1e-05 | 1.333E-05 | 2.887E-06 | 1.000E-05 | 1.500E-05 |
| 11039 | 552 | V552Y | A | 1.4e-05 | 1.1e-05 | 9.6e-06 | 1.153E-05 | 2.248E-06 | 9.600E-06 | 1.400E-05 |
| 11040 | 552 | V552V | A | 1.6e-05 | 1.1e-05 | 8.2e-06 | 1.173E-05 | 3.951E-06 | 8.200E-06 | 1.600E-05 |
| 11041 | 553 | T553A | A | 1.2e-05 | 9.4e-06 | 1.2e-05 | 1.113E-05 | 1.501E-06 | 9.400E-06 | 1.200E-05 |
| 11042 | 553 | T553R | A | 1.2e-05 | 1.1e-05 | 9.5e-06 | 1.083E-05 | 1.258E-06 | 9.500E-06 | 1.200E-05 |
| 11043 | 553 | T553N | A | 1.1e-05 | 1.1e-05 | 1.1e-05 | 1.100E-05 | 2.075E-21 | 1.100E-05 | 1.100E-05 |
| 11044 | 553 | T553D | A | 1.3e-05 | 1.6e-05 | 1.3e-05 | 1.400E-05 | 1.732E-06 | 1.300E-05 | 1.600E-05 |
| 11045 | 553 | T553C | A | 1.1e-05 | 1.3e-05 | 1.4e-05 | 1.267E-05 | 1.528E-06 | 1.100E-05 | 1.400E-05 |
| 11046 | 553 | T553E | A | 1.5e-05 | 1.5e-05 | 1.4e-05 | 1.467E-05 | 5.774E-07 | 1.400E-05 | 1.500E-05 |
| 11047 | 553 | T553Q | A | 1.2e-05 | 9.9e-06 | 1e-05 | 1.063E-05 | 1.185E-06 | 9.900E-06 | 1.200E-05 |
| 11048 | 553 | T553G | A | 1.5e-05 | 1.4e-05 | 1.3e-05 | 1.400E-05 | 1.000E-06 | 1.300E-05 | 1.500E-05 |
| 11049 | 553 | T553H | A | 1.1e-05 | 1.5e-05 | 1.1e-05 | 1.233E-05 | 2.309E-06 | 1.100E-05 | 1.500E-05 |

| | | | | | | | | | | |
|---|---|---|---|---|---|---|---|---|---|---|
| 11050 | 553 | T553I | A | 1.1e-05 | 1.2e-05 | 1.5e-05 | 1.267E-05 | 2.082E-06 | 1.100E-05 | 1.500E-05 |
| 11051 | 553 | T553L | A | 1.5e-05 | 1.2e-05 | 1.2e-05 | 1.300E-05 | 1.732E-06 | 1.200E-05 | 1.500E-05 |
| 11052 | 553 | T553K | A | 1.4e-05 | 1.4e-05 | 1.4e-05 | 1.400E-05 | 0.000E+00 | 1.400E-05 | 1.400E-05 |
| 11053 | 553 | T553M | A | 1.8e-05 | 9.8e-06 | 1.1e-05 | 1.293E-05 | 4.429E-06 | 9.800E-06 | 1.800E-05 |
| 11054 | 553 | T553F | A | 1.5e-05 | 1.4e-05 | 1.6e-05 | 1.500E-05 | 1.000E-06 | 1.400E-05 | 1.600E-05 |
| 11055 | 553 | T553P | A | 8.6e-06 | 1.5e-05 | 8.3e-06 | 1.063E-05 | 3.785E-06 | 8.300E-06 | 1.500E-05 |
| 11056 | 553 | T553S | A | 1.1e-05 | 9.5e-06 | 1e-05 | 1.017E-05 | 7.638E-07 | 9.500E-06 | 1.100E-05 |
| 11057 | 553 | T553T | A | 1.5e-05 | 1.1e-05 | 1.1e-05 | 1.233E-05 | 2.309E-06 | 1.100E-05 | 1.500E-05 |
| 11058 | 553 | T553W | A | 1.5e-05 | 1.3e-05 | 1.4e-05 | 1.400E-05 | 1.000E-06 | 1.300E-05 | 1.500E-05 |
| 11059 | 553 | T553Y | A | 9.5e-06 | 9.9e-06 | 8.6e-06 | 9.333E-06 | 6.658E-07 | 8.600E-06 | 9.900E-06 |
| 11060 | 553 | T553V | A | 1.6e-05 | 1e-05 | 9.8e-06 | 1.193E-05 | 3.523E-06 | 9.800E-06 | 1.600E-05 |
| 11061 | 554 | C554A | A | 1e-05 | 1.5e-05 | 1.4e-05 | 1.300E-05 | 2.646E-06 | 1.000E-05 | 1.500E-05 |
| 11062 | 554 | C554R | A | 1.6e-05 | 9.5e-06 | 1.5e-05 | 1.350E-05 | 3.500E-06 | 9.500E-06 | 1.600E-05 |
| 11063 | 554 | C554N | A | 1.5e-05 | 1.2e-05 | 1.5e-05 | 1.400E-05 | 1.732E-06 | 1.200E-05 | 1.500E-05 |
| 11064 | 554 | C554D | A | 1.2e-05 | 1.1e-05 | 1.4e-05 | 1.233E-05 | 1.528E-06 | 1.100E-05 | 1.400E-05 |
| 11065 | 554 | C554C | A | 1.2e-05 | 1.6e-05 | 1.5e-05 | 1.433E-05 | 2.082E-06 | 1.200E-05 | 1.600E-05 |
| 11066 | 554 | C554E | A | 1.5e-05 | 9.2e-06 | 1.1e-05 | 1.173E-05 | 2.969E-06 | 9.200E-06 | 1.500E-05 |
| 11067 | 554 | C554Q | A | 9.1e-06 | 1e-05 | 1.4e-05 | 1.103E-05 | 2.608E-06 | 9.100E-06 | 1.400E-05 |
| 11068 | 554 | C554G | A | 1.6e-05 | 1.6e-05 | 1.2e-05 | 1.467E-05 | 2.309E-06 | 1.200E-05 | 1.600E-05 |
| 11069 | 554 | C554H | A | 1.3e-05 | 1.3e-05 | 1.6e-05 | 1.400E-05 | 1.732E-06 | 1.300E-05 | 1.600E-05 |
| 11070 | 554 | C554I | A | 9.6e-06 | 1.4e-05 | 1.1e-05 | 1.153E-05 | 2.248E-06 | 9.600E-06 | 1.400E-05 |
| 11071 | 554 | C554L | A | 1.4e-05 | 1.4e-05 | 1e-05 | 1.267E-05 | 2.309E-06 | 1.000E-05 | 1.400E-05 |
| 11072 | 554 | C554K | A | 1.2e-05 | 1.6e-05 | 9.6e-06 | 1.253E-05 | 3.233E-06 | 9.600E-06 | 1.600E-05 |
| 11073 | 554 | C554M | A | 1.5e-05 | 1.2e-05 | 1.7e-05 | 1.467E-05 | 2.517E-06 | 1.200E-05 | 1.700E-05 |
| 11074 | 554 | C554F | A | 1.6e-05 | 1.2e-05 | 1.5e-05 | 1.433E-05 | 2.082E-06 | 1.200E-05 | 1.600E-05 |
| 11075 | 554 | C554P | A | 1.6e-05 | 1.1e-05 | 1e-05 | 1.233E-05 | 3.215E-06 | 1.000E-05 | 1.600E-05 |
| 11076 | 554 | C554S | A | 1.4e-05 | 1.5e-05 | 1.5e-05 | 1.467E-05 | 5.774E-07 | 1.400E-05 | 1.500E-05 |
| 11077 | 554 | C554T | A | 1.4e-05 | 1.2e-05 | 1.3e-05 | 1.300E-05 | 1.000E-06 | 1.200E-05 | 1.400E-05 |
| 11078 | 554 | C554W | A | 1.1e-05 | 1e-05 | 8.9e-06 | 9.967E-06 | 1.050E-06 | 8.900E-06 | 1.100E-05 |
| 11079 | 554 | C554Y | A | 1.4e-05 | 1.6e-05 | 1.1e-05 | 1.367E-05 | 2.517E-06 | 1.100E-05 | 1.600E-05 |
| 11080 | 554 | C554V | A | 1e-05 | 1e-05 | 1.5e-05 | 1.167E-05 | 2.887E-06 | 1.000E-05 | 1.500E-05 |
| 11081 | 555 | F555A | A | 1.2e-05 | 1.1e-05 | 1e-05 | 1.100E-05 | 1.000E-06 | 1.000E-05 | 1.200E-05 |
| 11082 | 555 | F555R | A | 1.6e-05 | 1.1e-05 | 1.5e-05 | 1.400E-05 | 2.646E-06 | 1.100E-05 | 1.600E-05 |
| 11083 | 555 | F555N | A | 1.4e-05 | 7e-06 | 1.3e-05 | 1.133E-05 | 3.786E-06 | 7.000E-06 | 1.400E-05 |
| 11084 | 555 | F555D | A | 1e-05 | 1.8e-05 | 1.3e-05 | 1.367E-05 | 4.041E-06 | 1.000E-05 | 1.800E-05 |
| 11085 | 555 | F555C | A | 1e-05 | 1.4e-05 | 1.5e-05 | 1.300E-05 | 2.646E-06 | 1.000E-05 | 1.500E-05 |
| 11086 | 555 | F555E | A | 1.1e-05 | 1.6e-05 | 1.4e-05 | 1.367E-05 | 2.517E-06 | 1.100E-05 | 1.600E-05 |
| 11087 | 555 | F555Q | A | 1.3e-05 | 1.3e-05 | 1e-05 | 1.200E-05 | 1.732E-06 | 1.000E-05 | 1.300E-05 |
| 11088 | 555 | F555G | A | 5.8e-06 | 1.3e-05 | 1.6e-05 | 1.160E-05 | 5.242E-06 | 5.800E-06 | 1.600E-05 |
| 11089 | 555 | F555H | A | 1.3e-05 | 1.5e-05 | 7e-06 | 1.167E-05 | 4.163E-06 | 7.000E-06 | 1.500E-05 |
| 11090 | 555 | F555I | A | 9.7e-06 | 1.7e-05 | 1e-05 | 1.223E-05 | 4.131E-06 | 9.700E-06 | 1.700E-05 |
| 11091 | 555 | F555L | A | 1.4e-05 | 1.3e-05 | 9.4e-06 | 1.213E-05 | 2.419E-06 | 9.400E-06 | 1.400E-05 |
| 11092 | 555 | F555K | A | 1.3e-05 | 1e-05 | 1.6e-05 | 1.300E-05 | 3.000E-06 | 1.000E-05 | 1.600E-05 |
| 11093 | 555 | F555M | A | 1.5e-05 | 1e-05 | 1.2e-05 | 1.233E-05 | 2.517E-06 | 1.000E-05 | 1.500E-05 |
| 11094 | 555 | F555F | A | 9e-06 | 1.6e-05 | 1.7e-05 | 1.400E-05 | 4.359E-06 | 9.000E-06 | 1.700E-05 |
| 11095 | 555 | F555P | A | 1.7e-05 | 1.2e-05 | 1.5e-05 | 1.467E-05 | 2.517E-06 | 1.200E-05 | 1.700E-05 |
| 11096 | 555 | F555S | A | 1.4e-05 | 1e-05 | 1.4e-05 | 1.267E-05 | 2.309E-06 | 1.000E-05 | 1.400E-05 |
| 11097 | 555 | F555T | A | 1.5e-05 | 9.3e-06 | 1e-05 | 1.143E-05 | 3.109E-06 | 9.300E-06 | 1.500E-05 |
| 11098 | 555 | F555W | A | 1.6e-05 | 1.4e-05 | 2e-05 | 1.667E-05 | 3.055E-06 | 1.400E-05 | 2.000E-05 |
| 11099 | 555 | F555Y | A | 1.4e-05 | 8.1e-06 | 1e-05 | 1.070E-05 | 3.012E-06 | 8.100E-06 | 1.400E-05 |
| 11100 | 555 | F555V | A | 1.5e-05 | 1.1e-05 | 1.2e-05 | 1.267E-05 | 2.082E-06 | 1.100E-05 | 1.500E-05 |
| 11101 | 556 | G556A | A | 1.1e-05 | 1.3e-05 | 1e-05 | 1.133E-05 | 1.528E-06 | 1.000E-05 | 1.300E-05 |
| 11102 | 556 | G556R | A | 1.6e-05 | 1.5e-05 | 1.6e-05 | 1.567E-05 | 5.774E-07 | 1.500E-05 | 1.600E-05 |
| 11103 | 556 | G556N | A | 1.5e-05 | 7.9e-06 | 1.4e-05 | 1.230E-05 | 3.843E-06 | 7.900E-06 | 1.500E-05 |
| 11104 | 556 | G556D | A | 7.9e-06 | 1.5e-05 | 1.5e-05 | 1.263E-05 | 4.099E-06 | 7.900E-06 | 1.500E-05 |
| 11105 | 556 | G556C | A | 1.4e-05 | 1.3e-05 | 1.4e-05 | 1.367E-05 | 5.774E-07 | 1.300E-05 | 1.400E-05 |
| 11106 | 556 | G556E | A | 7.4e-06 | 1.1e-05 | 8e-06 | 8.800E-06 | 1.929E-06 | 7.400E-06 | 1.100E-05 |
| 11107 | 556 | G556Q | A | 1.4e-05 | 1e-05 | 1.4e-05 | 1.267E-05 | 2.309E-06 | 1.000E-05 | 1.400E-05 |
| 11108 | 556 | G556G | A | 1.1e-05 | 9.8e-06 | 1e-05 | 1.027E-05 | 6.429E-07 | 9.800E-06 | 1.100E-05 |
| 11109 | 556 | G556H | A | 1.5e-05 | 1.3e-05 | 1e-05 | 1.267E-05 | 2.517E-06 | 1.000E-05 | 1.500E-05 |
| 11110 | 556 | G556I | A | 1.4e-05 | 8.8e-06 | 1.3e-05 | 1.193E-05 | 2.759E-06 | 8.800E-06 | 1.400E-05 |
| 11111 | 556 | G556L | A | 1.4e-05 | 1.2e-05 | 1.5e-05 | 1.367E-05 | 1.528E-06 | 1.200E-05 | 1.500E-05 |
| 11112 | 556 | G556K | A | 7.9e-06 | 8.9e-06 | 1.5e-05 | 1.060E-05 | 3.843E-06 | 7.900E-06 | 1.500E-05 |
| 11113 | 556 | G556M | A | 1.7e-05 | 9.8e-06 | 1.6e-05 | 1.427E-05 | 3.900E-06 | 9.800E-06 | 1.700E-05 |
| 11114 | 556 | G556F | A | 1.5e-05 | 1.3e-05 | 1.5e-05 | 1.433E-05 | 1.155E-06 | 1.300E-05 | 1.500E-05 |
| 11115 | 556 | G556P | A | 1.2e-05 | 1.6e-05 | 1.5e-05 | 1.433E-05 | 2.082E-06 | 1.200E-05 | 1.600E-05 |
| 11116 | 556 | G556S | A | 1.2e-05 | 8.8e-06 | 1.8e-05 | 1.293E-05 | 4.670E-06 | 8.800E-06 | 1.800E-05 |
| 11117 | 556 | G556T | A | 1.2e-05 | 1.2e-05 | 1e-05 | 1.133E-05 | 1.155E-06 | 1.000E-05 | 1.200E-05 |
| 11118 | 556 | G556W | A | 1.1e-05 | 1.3e-05 | 1.4e-05 | 1.267E-05 | 1.528E-06 | 1.100E-05 | 1.400E-05 |
| 11119 | 556 | G556Y | A | 1e-05 | 9.4e-06 | 1.4e-05 | 1.113E-05 | 2.501E-06 | 9.400E-06 | 1.400E-05 |
| 11120 | 556 | G556V | A | 1.6e-05 | 1.3e-05 | 9e-06 | 1.267E-05 | 3.512E-06 | 9.000E-06 | 1.600E-05 |
| 11121 | 557 | P557A | A | 9.8e-06 | 1.5e-05 | 9.6e-06 | 1.147E-05 | 3.062E-06 | 9.600E-06 | 1.500E-05 |
| 11122 | 557 | P557R | A | 1.6e-05 | 1.1e-05 | 5.9e-06 | 1.097E-05 | 5.050E-06 | 5.900E-06 | 1.600E-05 |
| 11123 | 557 | P557N | A | 1.5e-05 | 9.9e-06 | 1.4e-05 | 1.297E-05 | 2.702E-06 | 9.900E-06 | 1.500E-05 |
| 11124 | 557 | P557D | A | 8.6e-06 | 1.3e-05 | 1.4e-05 | 1.187E-05 | 2.873E-06 | 8.600E-06 | 1.400E-05 |
| 11125 | 557 | P557C | A | 1.4e-05 | 1.2e-05 | 1.4e-05 | 1.333E-05 | 1.155E-06 | 1.200E-05 | 1.400E-05 |
| 11126 | 557 | P557E | A | 1.1e-05 | 1.4e-05 | 7.7e-06 | 1.090E-05 | 3.151E-06 | 7.700E-06 | 1.400E-05 |
| 11127 | 557 | P557Q | A | 1.1e-05 | 1.5e-05 | 1.4e-05 | 1.333E-05 | 2.082E-06 | 1.100E-05 | 1.500E-05 |
| 11128 | 557 | P557G | A | 1.2e-05 | 1.1e-05 | 9.7e-06 | 1.090E-05 | 1.153E-06 | 9.700E-06 | 1.200E-05 |
| 11129 | 557 | P557H | A | 8.8e-06 | 1e-05 | 1.5e-05 | 1.127E-05 | 3.288E-06 | 8.800E-06 | 1.500E-05 |
| 11130 | 557 | P557I | A | 1.1e-05 | 1.3e-05 | 1.6e-05 | 1.333E-05 | 2.517E-06 | 1.100E-05 | 1.600E-05 |
| 11131 | 557 | P557L | A | 9.6e-06 | 6.6e-06 | 1.5e-05 | 1.040E-05 | 4.257E-06 | 6.600E-06 | 1.500E-05 |
| 11132 | 557 | P557K | A | 1.6e-05 | 1.5e-05 | 1.6e-05 | 1.567E-05 | 5.774E-07 | 1.500E-05 | 1.600E-05 |
| 11133 | 557 | P557M | A | 1.5e-05 | 8.3e-06 | 1.1e-05 | 1.143E-05 | 3.371E-06 | 8.300E-06 | 1.500E-05 |
| 11134 | 557 | P557F | A | 8.8e-06 | 1e-05 | 9.6e-06 | 9.467E-06 | 6.110E-07 | 8.800E-06 | 1.000E-05 |

| | | | | | | | | | | |
|---|---|---|---|---|---|---|---|---|---|---|
| 11135 | 557 | P557P | A | 1.1e-05 | 1.5e-05 | 1.5e-05 | 1.367E-05 | 2.309E-06 | 1.100E-05 | 1.500E-05 |
| 11136 | 557 | P557S | A | 1.4e-05 | 1.6e-05 | 1.4e-05 | 1.467E-05 | 1.155E-06 | 1.400E-05 | 1.600E-05 |
| 11137 | 557 | P557T | A | 1.4e-05 | 1.5e-05 | 1.4e-05 | 1.433E-05 | 5.774E-07 | 1.400E-05 | 1.500E-05 |
| 11138 | 557 | P557W | A | 1.3e-05 | 1.2e-05 | 8e-06 | 1.100E-05 | 2.646E-06 | 8.000E-06 | 1.300E-05 |
| 11139 | 557 | P557Y | A | 9.4e-06 | 8.5e-06 | 9.6e-06 | 9.167E-06 | 5.859E-07 | 8.500E-06 | 9.600E-06 |
| 11140 | 557 | P557V | A | 1.5e-05 | 1.6e-05 | 1.3e-05 | 1.467E-05 | 1.528E-06 | 1.300E-05 | 1.600E-05 |
| 11141 | 558 | E558A | A | 9.9e-06 | 1.6e-05 | 8.5e-06 | 1.147E-05 | 3.988E-06 | 8.500E-06 | 1.600E-05 |
| 11142 | 558 | E558R | A | 1.5e-05 | 7.5e-06 | 1.6e-05 | 1.283E-05 | 4.646E-06 | 7.500E-06 | 1.600E-05 |
| 11143 | 558 | E558N | A | 1.6e-05 | 1.1e-05 | 1.5e-05 | 1.400E-05 | 2.646E-06 | 1.100E-05 | 1.600E-05 |
| 11144 | 558 | E558D | A | 1.6e-05 | 1.6e-05 | 1.6e-05 | 1.600E-05 | 0.000E+00 | 1.600E-05 | 1.600E-05 |
| 11145 | 558 | E558C | A | 8.4e-06 | 1.4e-05 | 1.5e-05 | 1.247E-05 | 3.557E-06 | 8.400E-06 | 1.500E-05 |
| 11146 | 558 | E558E | A | 1.3e-05 | 1.6e-05 | 1.4e-05 | 1.433E-05 | 1.528E-06 | 1.300E-05 | 1.600E-05 |
| 11147 | 558 | E558Q | A | 1.5e-05 | 1.2e-05 | 9.3e-06 | 1.210E-05 | 2.851E-06 | 9.300E-06 | 1.500E-05 |
| 11148 | 558 | E558G | A | 1.5e-05 | 8.8e-06 | 1.5e-05 | 1.293E-05 | 3.580E-06 | 8.800E-06 | 1.500E-05 |
| 11149 | 558 | E558H | A | 9.5e-06 | 8.3e-06 | 1.2e-05 | 9.933E-06 | 1.888E-06 | 8.300E-06 | 1.200E-05 |
| 11150 | 558 | E558I | A | 1.5e-05 | 1.6e-05 | 1.6e-05 | 1.567E-05 | 5.774E-07 | 1.500E-05 | 1.600E-05 |
| 11151 | 558 | E558L | A | 1.5e-05 | 1.4e-05 | 1e-05 | 1.300E-05 | 2.646E-06 | 1.000E-05 | 1.500E-05 |
| 11152 | 558 | E558K | A | 1.5e-05 | 1.1e-05 | 8.6e-06 | 1.153E-05 | 3.233E-06 | 8.600E-06 | 1.500E-05 |
| 11153 | 558 | E558M | A | 1.4e-05 | 1.1e-05 | 9.1e-06 | 1.137E-05 | 2.470E-06 | 9.100E-06 | 1.400E-05 |
| 11154 | 558 | E558F | A | 1.6e-05 | 1.1e-05 | 9.6e-06 | 1.220E-05 | 3.365E-06 | 9.600E-06 | 1.600E-05 |
| 11155 | 558 | E558P | A | 1.1e-05 | 1.6e-05 | 1.1e-05 | 1.267E-05 | 2.887E-06 | 1.100E-05 | 1.600E-05 |
| 11156 | 558 | E558S | A | 1.1e-05 | 1.1e-05 | 1.4e-05 | 1.200E-05 | 1.732E-06 | 1.100E-05 | 1.400E-05 |
| 11157 | 558 | E558T | A | 1.5e-05 | 1.6e-05 | 1.1e-05 | 1.400E-05 | 2.646E-06 | 1.100E-05 | 1.600E-05 |
| 11158 | 558 | E558W | A | 1.1e-05 | 1.2e-05 | 1.4e-05 | 1.233E-05 | 1.528E-06 | 1.100E-05 | 1.400E-05 |
| 11159 | 558 | E558Y | A | 8.8e-06 | 1.5e-05 | 9.7e-06 | 1.117E-05 | 3.350E-06 | 8.800E-06 | 1.500E-05 |
| 11160 | 558 | E558V | A | 1.2e-05 | 1e-05 | 1.3e-05 | 1.167E-05 | 1.528E-06 | 1.000E-05 | 1.300E-05 |
| 11161 | 559 | A559A | A | 1e-05 | 8.8e-06 | 1.1e-05 | 9.933E-06 | 1.102E-06 | 8.800E-06 | 1.100E-05 |
| 11162 | 559 | A559R | A | 8.5e-06 | 8.7e-06 | 9.5e-06 | 8.900E-06 | 5.292E-07 | 8.500E-06 | 9.500E-06 |
| 11163 | 559 | A559N | A | 8.9e-06 | 1.5e-05 | 1.4e-05 | 1.263E-05 | 3.272E-06 | 8.900E-06 | 1.500E-05 |
| 11164 | 559 | A559D | A | 1.5e-05 | 1.1e-05 | 1.1e-05 | 1.233E-05 | 2.309E-06 | 1.100E-05 | 1.500E-05 |
| 11165 | 559 | A559C | A | 2.6e-05 | 1.5e-05 | 1.5e-05 | 1.867E-05 | 6.351E-06 | 1.500E-05 | 2.600E-05 |
| 11166 | 559 | A559E | A | 1.4e-05 | 1.6e-05 | 9.4e-06 | 1.313E-05 | 3.384E-06 | 9.400E-06 | 1.600E-05 |
| 11167 | 559 | A559Q | A | 1e-05 | 1.5e-05 | 1.6e-05 | 1.367E-05 | 3.215E-06 | 1.000E-05 | 1.600E-05 |
| 11168 | 559 | A559G | A | 1.3e-05 | 1.4e-05 | 1.3e-05 | 1.333E-05 | 5.774E-07 | 1.300E-05 | 1.400E-05 |
| 11169 | 559 | A559H | A | 1.4e-05 | 1.3e-05 | 1e-05 | 1.233E-05 | 2.082E-06 | 1.000E-05 | 1.400E-05 |
| 11170 | 559 | A559I | A | 1.5e-05 | 1.1e-05 | 1.7e-05 | 1.433E-05 | 3.055E-06 | 1.100E-05 | 1.700E-05 |
| 11171 | 559 | A559L | A | 1.1e-05 | 1e-05 | 1.3e-05 | 1.133E-05 | 1.528E-06 | 1.000E-05 | 1.300E-05 |
| 11172 | 559 | A559K | A | 1.3e-05 | 1.1e-05 | 1.6e-05 | 1.333E-05 | 2.517E-06 | 1.100E-05 | 1.600E-05 |
| 11173 | 559 | A559M | A | 1.5e-05 | 9.7e-06 | 1.5e-05 | 1.323E-05 | 3.060E-06 | 9.700E-06 | 1.500E-05 |
| 11174 | 559 | A559F | A | 1.4e-05 | 1e-05 | 1e-05 | 1.133E-05 | 2.309E-06 | 1.000E-05 | 1.400E-05 |
| 11175 | 559 | A559P | A | 1.3e-05 | 1.5e-05 | 1.1e-05 | 1.300E-05 | 2.000E-06 | 1.100E-05 | 1.500E-05 |
| 11176 | 559 | A559S | A | 1.6e-05 | 1.6e-05 | 1.4e-05 | 1.533E-05 | 1.155E-06 | 1.400E-05 | 1.600E-05 |
| 11177 | 559 | A559T | A | 1.7e-05 | 1e-05 | 1.5e-05 | 1.400E-05 | 3.606E-06 | 1.000E-05 | 1.700E-05 |
| 11178 | 559 | A559W | A | 1.1e-05 | 1.3e-05 | 9.9e-06 | 1.130E-05 | 1.572E-06 | 9.900E-06 | 1.300E-05 |
| 11179 | 559 | A559Y | A | 1.2e-05 | 1.5e-05 | 1.6e-05 | 1.433E-05 | 2.082E-06 | 1.200E-05 | 1.600E-05 |
| 11180 | 559 | A559V | A | 1.3e-05 | 1.1e-05 | 1.2e-05 | 1.200E-05 | 1.000E-06 | 1.100E-05 | 1.300E-05 |
| 11181 | 560 | D560A | A | 1.2e-05 | 1.6e-05 | 1.7e-05 | 1.500E-05 | 2.646E-06 | 1.200E-05 | 1.700E-05 |
| 11182 | 560 | D560R | A | 8.9e-06 | 1.7e-05 | 8.8e-06 | 1.157E-05 | 4.706E-06 | 8.800E-06 | 1.700E-05 |
| 11183 | 560 | D560N | A | 1.4e-05 | 1.1e-05 | 1.4e-05 | 1.300E-05 | 1.732E-06 | 1.100E-05 | 1.400E-05 |
| 11184 | 560 | D560D | A | 8.2e-06 | 1.5e-05 | 1.5e-05 | 1.273E-05 | 3.926E-06 | 8.200E-06 | 1.500E-05 |
| 11185 | 560 | D560C | A | 1.1e-05 | 9.6e-06 | 1.3e-05 | 1.120E-05 | 1.709E-06 | 9.600E-06 | 1.300E-05 |
| 11186 | 560 | D560E | A | 1.5e-05 | 1.5e-05 | 1.6e-05 | 1.533E-05 | 5.774E-07 | 1.500E-05 | 1.600E-05 |
| 11187 | 560 | D560Q | A | 1e-05 | 9.8e-06 | 8.8e-06 | 9.533E-06 | 6.429E-07 | 8.800E-06 | 1.000E-05 |
| 11188 | 560 | D560G | A | 1.2e-05 | 1.3e-05 | 1.1e-05 | 1.200E-05 | 1.000E-06 | 1.100E-05 | 1.300E-05 |
| 11189 | 560 | D560H | A | 1.1e-05 | 1.6e-05 | 1.5e-05 | 1.400E-05 | 2.646E-06 | 1.100E-05 | 1.600E-05 |
| 11190 | 560 | D560I | A | 1.2e-05 | 9e-06 | 1.6e-05 | 1.233E-05 | 3.512E-06 | 9.000E-06 | 1.600E-05 |
| 11191 | 560 | D560L | A | 9.4e-06 | 1.1e-05 | 1.6e-05 | 1.213E-05 | 3.443E-06 | 9.400E-06 | 1.600E-05 |
| 11192 | 560 | D560K | A | 9.3e-06 | 1.5e-05 | 1.4e-05 | 1.277E-05 | 3.044E-06 | 9.300E-06 | 1.500E-05 |
| 11193 | 560 | D560M | A | 1.2e-05 | 1.3e-05 | 1e-05 | 1.167E-05 | 1.528E-06 | 1.000E-05 | 1.300E-05 |
| 11194 | 560 | D560F | A | 1.3e-05 | 1.6e-05 | 1.6e-05 | 1.500E-05 | 1.732E-06 | 1.300E-05 | 1.600E-05 |
| 11195 | 560 | D560P | A | 1.5e-05 | 1.1e-05 | 9.8e-06 | 1.193E-05 | 2.723E-06 | 9.800E-06 | 1.500E-05 |
| 11196 | 560 | D560S | A | 1.5e-05 | 8.8e-06 | 1.5e-05 | 1.293E-05 | 3.580E-06 | 8.800E-06 | 1.500E-05 |
| 11197 | 560 | D560T | A | 9.9e-06 | 1.3e-05 | 1e-05 | 1.097E-05 | 1.762E-06 | 9.900E-06 | 1.300E-05 |
| 11198 | 560 | D560W | A | 9.6e-06 | 8.6e-06 | 1.5e-05 | 1.107E-05 | 3.443E-06 | 8.600E-06 | 1.500E-05 |
| 11199 | 560 | D560Y | A | 1e-05 | 1.6e-05 | 1.5e-05 | 1.367E-05 | 3.215E-06 | 1.000E-05 | 1.600E-05 |
| 11200 | 560 | D560V | A | 1.5e-05 | 1.2e-05 | 1e-05 | 1.233E-05 | 2.517E-06 | 1.000E-05 | 1.500E-05 |
| 11201 | 561 | Q561A | A | 1.2e-05 | 9.3e-06 | 1.4e-05 | 1.177E-05 | 2.359E-06 | 9.300E-06 | 1.400E-05 |
| 11202 | 561 | Q561R | A | 1.7e-05 | 1.6e-05 | 1.6e-05 | 1.633E-05 | 5.774E-07 | 1.600E-05 | 1.700E-05 |
| 11203 | 561 | Q561N | A | 1.1e-05 | 9.2e-06 | 1.7e-05 | 1.240E-05 | 4.084E-06 | 9.200E-06 | 1.700E-05 |
| 11204 | 561 | Q561D | A | 1.6e-05 | 1.4e-05 | 1.5e-05 | 1.500E-05 | 1.000E-06 | 1.400E-05 | 1.600E-05 |
| 11205 | 561 | Q561C | A | 1.3e-05 | 1.6e-05 | 1e-05 | 1.300E-05 | 3.000E-06 | 1.000E-05 | 1.600E-05 |
| 11206 | 561 | Q561E | A | 1e-05 | 1.3e-05 | 1.5e-05 | 1.267E-05 | 2.517E-06 | 1.000E-05 | 1.500E-05 |
| 11207 | 561 | Q561Q | A | 1.7e-05 | 2.6e-05 | 1.1e-05 | 1.800E-05 | 7.550E-06 | 1.100E-05 | 2.600E-05 |
| 11208 | 561 | Q561G | A | 1e-05 | 1.7e-05 | 1.7e-05 | 1.467E-05 | 4.041E-06 | 1.000E-05 | 1.700E-05 |
| 11209 | 561 | Q561H | A | 9.2e-06 | 9.2e-06 | 1.6e-05 | 1.147E-05 | 3.926E-06 | 9.200E-06 | 1.600E-05 |
| 11210 | 561 | Q561I | A | 1.2e-05 | 1e-05 | 1.5e-05 | 1.233E-05 | 2.517E-06 | 1.000E-05 | 1.500E-05 |
| 11211 | 561 | Q561L | A | 1.1e-05 | 1.7e-05 | 1.5e-05 | 1.433E-05 | 3.055E-06 | 1.100E-05 | 1.700E-05 |
| 11212 | 561 | Q561K | A | 1.5e-05 | 1.6e-05 | 1.3e-05 | 1.467E-05 | 1.528E-06 | 1.300E-05 | 1.600E-05 |
| 11213 | 561 | Q561M | A | 1.2e-05 | 1.1e-05 | 1e-05 | 1.100E-05 | 1.000E-06 | 1.000E-05 | 1.200E-05 |
| 11214 | 561 | Q561F | A | 1.5e-05 | 1.5e-05 | 1.1e-05 | 1.367E-05 | 2.309E-06 | 1.100E-05 | 1.500E-05 |
| 11215 | 561 | Q561P | A | 1.7e-05 | 1.2e-05 | 1.1e-05 | 1.333E-05 | 3.215E-06 | 1.100E-05 | 1.700E-05 |
| 11216 | 561 | Q561S | A | 1.7e-05 | 1.5e-05 | 1.4e-05 | 1.533E-05 | 1.528E-06 | 1.400E-05 | 1.700E-05 |
| 11217 | 561 | Q561T | A | 1.4e-05 | 1.1e-05 | 1.6e-05 | 1.367E-05 | 2.517E-06 | 1.100E-05 | 1.600E-05 |
| 11218 | 561 | Q561W | A | 1.3e-05 | 1.1e-05 | 1e-05 | 1.133E-05 | 1.528E-06 | 1.000E-05 | 1.300E-05 |
| 11219 | 561 | Q561Y | A | 1.5e-05 | 8.9e-06 | 1.5e-05 | 1.297E-05 | 3.522E-06 | 8.900E-06 | 1.500E-05 |

| | | | | | | | | | | |
|---|---|---|---|---|---|---|---|---|---|---|
| 11220 | 561 | Q561V | A | 2.6e-05 | 9.3e-06 | 1.1e-05 | 1.543E-05 | 9.190E-06 | 9.300E-06 | 2.600E-05 |
| 11221 | 562 | C562A | A | 8.6e-06 | 1.1e-05 | 1.4e-05 | 1.120E-05 | 2.706E-06 | 8.600E-06 | 1.400E-05 |
| 11222 | 562 | C562R | A | 1.1e-05 | 1.4e-05 | 1.6e-05 | 1.367E-05 | 2.517E-06 | 1.100E-05 | 1.600E-05 |
| 11223 | 562 | C562N | A | 1.6e-05 | 1.1e-05 | 1.4e-05 | 1.367E-05 | 2.517E-06 | 1.100E-05 | 1.600E-05 |
| 11224 | 562 | C562D | A | 1.4e-05 | 1.2e-05 | 1.6e-05 | 1.400E-05 | 2.000E-06 | 1.200E-05 | 1.600E-05 |
| 11225 | 562 | C562C | A | 1.5e-05 | 9.8e-06 | 1.1e-05 | 1.193E-05 | 2.723E-06 | 9.800E-06 | 1.500E-05 |
| 11226 | 562 | C562E | A | 8e-06 | 1.5e-05 | 1.1e-05 | 1.133E-05 | 3.512E-06 | 8.000E-06 | 1.500E-05 |
| 11227 | 562 | C562Q | A | 8.3e-06 | 7.9e-06 | 1.6e-05 | 1.073E-05 | 4.565E-06 | 7.900E-06 | 1.600E-05 |
| 11228 | 562 | C562G | A | 1.4e-05 | 2e-05 | 1.2e-05 | 1.533E-05 | 4.163E-06 | 1.200E-05 | 2.000E-05 |
| 11229 | 562 | C562H | A | 9.8e-06 | 1.2e-05 | 1.6e-05 | 1.260E-05 | 3.143E-06 | 9.800E-06 | 1.600E-05 |
| 11230 | 562 | C562I | A | 1.4e-05 | 1.6e-05 | 1.5e-05 | 1.500E-05 | 1.000E-06 | 1.400E-05 | 1.600E-05 |
| 11231 | 562 | C562L | A | 8.9e-06 | 1.1e-05 | 1.3e-05 | 1.097E-05 | 2.050E-06 | 8.900E-06 | 1.300E-05 |
| 11232 | 562 | C562K | A | 1.6e-05 | 1.3e-05 | 1.4e-05 | 1.433E-05 | 1.528E-06 | 1.300E-05 | 1.600E-05 |
| 11233 | 562 | C562M | A | 1.4e-05 | 8.5e-06 | 1.3e-05 | 1.183E-05 | 2.930E-06 | 8.500E-06 | 1.400E-05 |
| 11234 | 562 | C562F | A | 1.6e-05 | 1.6e-05 | 1.3e-05 | 1.500E-05 | 1.732E-06 | 1.300E-05 | 1.600E-05 |
| 11235 | 562 | C562P | A | 1.6e-05 | 1.1e-05 | 9.4e-06 | 1.213E-05 | 3.443E-06 | 9.400E-06 | 1.600E-05 |
| 11236 | 562 | C562S | A | 1.2e-05 | 1.1e-05 | 1.1e-05 | 1.133E-05 | 5.774E-07 | 1.100E-05 | 1.200E-05 |
| 11237 | 562 | C562T | A | 9.9e-06 | 1.5e-05 | 1.6e-05 | 1.363E-05 | 3.272E-06 | 9.900E-06 | 1.600E-05 |
| 11238 | 562 | C562W | A | 9.6e-06 | 1.2e-05 | 1.3e-05 | 1.153E-05 | 1.747E-06 | 9.600E-06 | 1.300E-05 |
| 11239 | 562 | C562Y | A | 1.3e-05 | 1.6e-05 | 9.1e-06 | 1.270E-05 | 3.460E-06 | 9.100E-06 | 1.600E-05 |
| 11240 | 562 | C562V | A | 1.4e-05 | 9.6e-06 | 1e-05 | 1.120E-05 | 2.433E-06 | 9.600E-06 | 1.400E-05 |
| 11241 | 563 | V563A | A | 1.6e-05 | 1.3e-05 | 1.7e-05 | 1.533E-05 | 2.082E-06 | 1.300E-05 | 1.700E-05 |
| 11242 | 563 | V563R | A | 1e-05 | 1.5e-05 | 1.3e-05 | 1.267E-05 | 2.517E-06 | 1.000E-05 | 1.500E-05 |
| 11243 | 563 | V563N | A | 1.1e-05 | 1.5e-05 | 7.9e-06 | 1.130E-05 | 3.559E-06 | 7.900E-06 | 1.500E-05 |
| 11244 | 563 | V563D | A | 9.4e-06 | 1.5e-05 | 1e-05 | 1.147E-05 | 3.075E-06 | 9.400E-06 | 1.500E-05 |
| 11245 | 563 | V563C | A | 1.1e-05 | 1e-05 | 1.3e-05 | 1.133E-05 | 1.528E-06 | 1.000E-05 | 1.300E-05 |
| 11246 | 563 | V563E | A | 1.5e-05 | 1.5e-05 | 1.2e-05 | 1.400E-05 | 1.732E-06 | 1.200E-05 | 1.500E-05 |
| 11247 | 563 | V563Q | A | 1.6e-05 | 1.3e-05 | 1.1e-05 | 1.333E-05 | 2.517E-06 | 1.100E-05 | 1.600E-05 |
| 11248 | 563 | V563G | A | 1.7e-05 | 1.1e-05 | 9.8e-06 | 1.260E-05 | 3.857E-06 | 9.800E-06 | 1.700E-05 |
| 11249 | 563 | V563H | A | 1.4e-05 | 1.3e-05 | 8.4e-06 | 1.180E-05 | 2.987E-06 | 8.400E-06 | 1.400E-05 |
| 11250 | 563 | V563I | A | 1.3e-05 | 1.3e-05 | 1.4e-05 | 1.333E-05 | 5.774E-07 | 1.300E-05 | 1.400E-05 |
| 11251 | 563 | V563L | A | 1e-05 | 1.6e-05 | 1.6e-05 | 1.400E-05 | 3.464E-06 | 1.000E-05 | 1.600E-05 |
| 11252 | 563 | V563K | A | 1.6e-05 | 8.4e-06 | 1.3e-05 | 1.247E-05 | 3.828E-06 | 8.400E-06 | 1.600E-05 |
| 11253 | 563 | V563M | A | 9.9e-06 | 1.2e-05 | 1e-05 | 1.063E-05 | 1.185E-06 | 9.900E-06 | 1.200E-05 |
| 11254 | 563 | V563F | A | 1e-05 | 1.1e-05 | 1e-05 | 1.033E-05 | 5.774E-07 | 1.000E-05 | 1.100E-05 |
| 11255 | 563 | V563P | A | 1.2e-05 | 1e-05 | 1.7e-05 | 1.300E-05 | 3.606E-06 | 1.000E-05 | 1.700E-05 |
| 11256 | 563 | V563S | A | 1.6e-05 | 1.3e-05 | 1.6e-05 | 1.500E-05 | 1.732E-06 | 1.300E-05 | 1.600E-05 |
| 11257 | 563 | V563T | A | 1.5e-05 | 1.2e-05 | 1.1e-05 | 1.267E-05 | 2.082E-06 | 1.100E-05 | 1.500E-05 |
| 11258 | 563 | V563W | A | 1e-05 | 9.1e-06 | 1.4e-05 | 1.103E-05 | 2.608E-06 | 9.100E-06 | 1.400E-05 |
| 11259 | 563 | V563Y | A | 1.5e-05 | 1.4e-05 | 8.6e-06 | 1.253E-05 | 3.443E-06 | 8.600E-06 | 1.500E-05 |
| 11260 | 563 | V563V | A | 1.1e-05 | 1.4e-05 | 2.5e-05 | 1.667E-05 | 7.371E-06 | 1.100E-05 | 2.500E-05 |
| 11261 | 564 | A564A | A | 1e-05 | 1.6e-05 | 1.4e-05 | 1.333E-05 | 3.055E-06 | 1.000E-05 | 1.600E-05 |
| 11262 | 564 | A564R | A | 1.5e-05 | 1.5e-05 | 1.1e-05 | 1.367E-05 | 2.309E-06 | 1.100E-05 | 1.500E-05 |
| 11263 | 564 | A564N | A | 1.4e-05 | 1.4e-05 | 1.1e-05 | 1.300E-05 | 1.732E-06 | 1.100E-05 | 1.400E-05 |
| 11264 | 564 | A564D | A | 1.5e-05 | 1e-05 | 1e-05 | 1.167E-05 | 2.887E-06 | 1.000E-05 | 1.500E-05 |
| 11265 | 564 | A564C | A | 1.5e-05 | 1.1e-05 | 1.5e-05 | 1.367E-05 | 2.309E-06 | 1.100E-05 | 1.500E-05 |
| 11266 | 564 | A564E | A | 1.1e-05 | 1.6e-05 | 1.6e-05 | 1.433E-05 | 2.887E-06 | 1.100E-05 | 1.600E-05 |
| 11267 | 564 | A564Q | A | 1.3e-05 | 8.6e-06 | 1.6e-05 | 1.253E-05 | 3.722E-06 | 8.600E-06 | 1.600E-05 |
| 11268 | 564 | A564G | A | 1.6e-05 | 1.4e-05 | 1.4e-05 | 1.467E-05 | 1.155E-06 | 1.400E-05 | 1.600E-05 |
| 11269 | 564 | A564H | A | 1.6e-05 | 1.5e-05 | 1.5e-05 | 1.533E-05 | 5.774E-07 | 1.500E-05 | 1.600E-05 |
| 11270 | 564 | A564I | A | 9.1e-06 | 1.3e-05 | 1.3e-05 | 1.170E-05 | 2.252E-06 | 9.100E-06 | 1.300E-05 |
| 11271 | 564 | A564L | A | 7.9e-06 | 1e-05 | 1.4e-05 | 1.063E-05 | 3.099E-06 | 7.900E-06 | 1.400E-05 |
| 11272 | 564 | A564K | A | 9.1e-06 | 1.5e-05 | 1.8e-05 | 1.403E-05 | 4.528E-06 | 9.100E-06 | 1.800E-05 |
| 11273 | 564 | A564M | A | 1.5e-05 | 1.5e-05 | 1.1e-05 | 1.367E-05 | 2.309E-06 | 1.100E-05 | 1.500E-05 |
| 11274 | 564 | A564F | A | 1.5e-05 | 9.4e-06 | 1e-05 | 1.147E-05 | 3.075E-06 | 9.400E-06 | 1.500E-05 |
| 11275 | 564 | A564P | A | 1e-05 | 9.9e-06 | 9.6e-06 | 9.833E-06 | 2.082E-07 | 9.600E-06 | 1.000E-05 |
| 11276 | 564 | A564S | A | 1.1e-05 | 8.4e-06 | 1.6e-05 | 1.180E-05 | 3.863E-06 | 8.400E-06 | 1.600E-05 |
| 11277 | 564 | A564T | A | 1.1e-05 | 1.5e-05 | 1.4e-05 | 1.333E-05 | 2.082E-06 | 1.100E-05 | 1.500E-05 |
| 11278 | 564 | A564W | A | 1.4e-05 | 1.3e-05 | 1.2e-05 | 1.300E-05 | 1.000E-06 | 1.200E-05 | 1.400E-05 |
| 11279 | 564 | A564Y | A | 1e-05 | 1.5e-05 | 1.5e-05 | 1.333E-05 | 2.887E-06 | 1.000E-05 | 1.500E-05 |
| 11280 | 564 | A564V | A | 1e-05 | 1.5e-05 | 1.1e-05 | 1.200E-05 | 2.646E-06 | 1.000E-05 | 1.500E-05 |
| 11281 | 565 | C565A | A | 1.6e-05 | 1.2e-05 | 1.1e-05 | 1.300E-05 | 2.646E-06 | 1.100E-05 | 1.600E-05 |
| 11282 | 565 | C565R | A | 9.8e-06 | 8.8e-06 | 1.7e-05 | 1.187E-05 | 4.474E-06 | 8.800E-06 | 1.700E-05 |
| 11283 | 565 | C565N | A | 1.5e-05 | 9.8e-06 | 8.1e-06 | 1.097E-05 | 3.595E-06 | 8.100E-06 | 1.500E-05 |
| 11284 | 565 | C565D | A | 1.4e-05 | 1.4e-05 | 1.2e-05 | 1.333E-05 | 1.155E-06 | 1.200E-05 | 1.400E-05 |
| 11285 | 565 | C565C | A | 1.5e-05 | 1.5e-05 | 1.2e-05 | 1.400E-05 | 1.732E-06 | 1.200E-05 | 1.500E-05 |
| 11286 | 565 | C565E | A | 7.1e-06 | 1.4e-05 | 1.4e-05 | 1.170E-05 | 3.984E-06 | 7.100E-06 | 1.400E-05 |
| 11287 | 565 | C565Q | A | 2.3e-05 | 1.7e-05 | 1.1e-05 | 1.700E-05 | 6.000E-06 | 1.100E-05 | 2.300E-05 |
| 11288 | 565 | C565G | A | 1.4e-05 | 1e-05 | 1.4e-05 | 1.267E-05 | 2.309E-06 | 1.000E-05 | 1.400E-05 |
| 11289 | 565 | C565H | A | 1.1e-05 | 1.5e-05 | 1.5e-05 | 1.367E-05 | 2.309E-06 | 1.100E-05 | 1.500E-05 |
| 11290 | 565 | C565I | A | 1.6e-05 | 1.5e-05 | 1.5e-05 | 1.533E-05 | 5.774E-07 | 1.500E-05 | 1.600E-05 |
| 11291 | 565 | C565L | A | 1.6e-05 | 1e-05 | 9.4e-06 | 1.180E-05 | 3.650E-06 | 9.400E-06 | 1.600E-05 |
| 11292 | 565 | C565K | A | 1.5e-05 | 1.3e-05 | 9.9e-06 | 1.263E-05 | 2.570E-06 | 9.900E-06 | 1.500E-05 |
| 11293 | 565 | C565M | A | 1.2e-05 | 1e-05 | 1e-05 | 1.067E-05 | 1.155E-06 | 1.000E-05 | 1.200E-05 |
| 11294 | 565 | C565F | A | 9.2e-06 | 1.1e-05 | 1.5e-05 | 1.173E-05 | 2.969E-06 | 9.200E-06 | 1.500E-05 |
| 11295 | 565 | C565P | A | 1.1e-05 | 1.7e-05 | 1.3e-05 | 1.367E-05 | 3.055E-06 | 1.100E-05 | 1.700E-05 |
| 11296 | 565 | C565S | A | 9.5e-06 | 6.9e-06 | 1.3e-05 | 9.800E-06 | 3.061E-06 | 6.900E-06 | 1.300E-05 |
| 11297 | 565 | C565T | A | 1.5e-05 | 1.3e-05 | 9.7e-06 | 1.257E-05 | 2.676E-06 | 9.700E-06 | 1.500E-05 |
| 11298 | 565 | C565W | A | 9.5e-06 | 1.1e-05 | 2.4e-05 | 1.483E-05 | 7.974E-06 | 9.500E-06 | 2.400E-05 |
| 11299 | 565 | C565Y | A | 9.8e-06 | 1e-05 | 9.5e-06 | 9.767E-06 | 2.517E-07 | 9.500E-06 | 1.000E-05 |
| 11300 | 565 | C565V | A | 1.2e-05 | 1.1e-05 | 1.5e-05 | 1.267E-05 | 2.082E-06 | 1.100E-05 | 1.500E-05 |
| 11301 | 566 | A566A | A | 1.6e-05 | 9.1e-06 | 1.4e-05 | 1.303E-05 | 3.550E-06 | 9.100E-06 | 1.600E-05 |
| 11302 | 566 | A566R | A | 1e-05 | 1e-05 | 1e-05 | 1.000E-05 | 0.000E+00 | 1.000E-05 | 1.000E-05 |
| 11303 | 566 | A566N | A | 1e-05 | 1.2e-05 | 1.5e-05 | 1.233E-05 | 2.517E-06 | 1.000E-05 | 1.500E-05 |
| 11304 | 566 | A566D | A | 1.5e-05 | 7.9e-06 | 1.2e-05 | 1.163E-05 | 3.564E-06 | 7.900E-06 | 1.500E-05 |

| | | | | | | | | | | |
|---|---|---|---|---|---|---|---|---|---|---|
| 11305 | 566 | A566C | A | 9.4e-06 | 1.1e-05 | 1.5e-05 | 1.180E-05 | 2.884E-06 | 9.400E-06 | 1.500E-05 |
| 11306 | 566 | A566E | A | 1.4e-05 | 1.4e-05 | 1.5e-05 | 1.433E-05 | 5.774E-07 | 1.400E-05 | 1.500E-05 |
| 11307 | 566 | A566Q | A | 1e-05 | 1e-05 | 1.6e-05 | 1.200E-05 | 3.464E-06 | 1.000E-05 | 1.600E-05 |
| 11308 | 566 | A566G | A | 1.2e-05 | 1.6e-05 | 1.4e-05 | 1.400E-05 | 2.000E-06 | 1.200E-05 | 1.600E-05 |
| 11309 | 566 | A566H | A | 1.5e-05 | 1.2e-05 | 1.6e-05 | 1.433E-05 | 2.082E-06 | 1.200E-05 | 1.600E-05 |
| 11310 | 566 | A566I | A | 1.6e-05 | 1.6e-05 | 1.4e-05 | 1.533E-05 | 1.155E-06 | 1.400E-05 | 1.600E-05 |
| 11311 | 566 | A566L | A | 1.5e-05 | 1.5e-05 | 1.1e-05 | 1.367E-05 | 2.309E-06 | 1.100E-05 | 1.500E-05 |
| 11312 | 566 | A566K | A | 7.8e-06 | 1.2e-05 | 1e-05 | 9.933E-06 | 2.101E-06 | 7.800E-06 | 1.200E-05 |
| 11313 | 566 | A566M | A | 9.3e-06 | 1.6e-05 | 1e-05 | 1.177E-05 | 3.683E-06 | 9.300E-06 | 1.600E-05 |
| 11314 | 566 | A566F | A | 1.4e-05 | 1.1e-05 | 1.1e-05 | 1.200E-05 | 1.732E-06 | 1.100E-05 | 1.400E-05 |
| 11315 | 566 | A566P | A | 1.1e-05 | 1.3e-05 | 1.7e-05 | 1.367E-05 | 3.055E-06 | 1.100E-05 | 1.700E-05 |
| 11316 | 566 | A566S | A | 1.6e-05 | 1.5e-05 | 1.4e-05 | 1.500E-05 | 1.000E-06 | 1.400E-05 | 1.600E-05 |
| 11317 | 566 | A566T | A | 8.2e-06 | 5.5e-06 | 1.3e-05 | 8.900E-06 | 3.799E-06 | 5.500E-06 | 1.300E-05 |
| 11318 | 566 | A566W | A | 1.3e-05 | 9.5e-06 | 1.3e-05 | 1.183E-05 | 2.021E-06 | 9.500E-06 | 1.300E-05 |
| 11319 | 566 | A566Y | A | 9.3e-06 | 9.2e-06 | 1.5e-05 | 1.117E-05 | 3.320E-06 | 9.200E-06 | 1.500E-05 |
| 11320 | 566 | A566V | A | 9e-06 | 1.5e-05 | 1.5e-05 | 1.300E-05 | 3.464E-06 | 9.000E-06 | 1.500E-05 |
| 11321 | 567 | H567A | A | 1.2e-05 | 2.6e-05 | 1.5e-05 | 1.767E-05 | 7.371E-06 | 1.200E-05 | 2.600E-05 |
| 11322 | 567 | H567R | A | 1.6e-05 | 1.1e-05 | 9.4e-06 | 1.213E-05 | 3.443E-06 | 9.400E-06 | 1.600E-05 |
| 11323 | 567 | H567N | A | 1.3e-05 | 1.4e-05 | 1.1e-05 | 1.267E-05 | 1.528E-06 | 1.100E-05 | 1.400E-05 |
| 11324 | 567 | H567D | A | 1.1e-05 | 1.6e-05 | 1.6e-05 | 1.433E-05 | 2.887E-06 | 1.100E-05 | 1.600E-05 |
| 11325 | 567 | H567C | A | 1.1e-05 | 1e-05 | 1.2e-05 | 1.100E-05 | 1.000E-06 | 1.000E-05 | 1.200E-05 |
| 11326 | 567 | H567E | A | 1.2e-05 | 1.2e-05 | 1.5e-05 | 1.300E-05 | 1.732E-06 | 1.200E-05 | 1.500E-05 |
| 11327 | 567 | H567Q | A | 1.5e-05 | 1.7e-05 | 1.6e-05 | 1.600E-05 | 1.000E-06 | 1.500E-05 | 1.700E-05 |
| 11328 | 567 | H567G | A | 1.5e-05 | 9.5e-06 | 1.2e-05 | 1.217E-05 | 2.754E-06 | 9.500E-06 | 1.500E-05 |
| 11329 | 567 | H567H | A | 8.3e-06 | 1.1e-05 | 8.8e-06 | 9.367E-06 | 1.436E-06 | 8.300E-06 | 1.100E-05 |
| 11330 | 567 | H567I | A | 1.3e-05 | 1.3e-05 | 1e-05 | 1.200E-05 | 1.732E-06 | 1.000E-05 | 1.300E-05 |
| 11331 | 567 | H567L | A | 1.1e-05 | 1.1e-05 | 1.7e-05 | 1.300E-05 | 3.464E-06 | 1.100E-05 | 1.700E-05 |
| 11332 | 567 | H567K | A | 1.8e-05 | 1.7e-05 | 1.1e-05 | 1.533E-05 | 3.786E-06 | 1.100E-05 | 1.800E-05 |
| 11333 | 567 | H567M | A | 1.8e-05 | 1.6e-05 | 6.8e-06 | 1.360E-05 | 5.973E-06 | 6.800E-06 | 1.800E-05 |
| 11334 | 567 | H567F | A | 1e-05 | 1.5e-05 | 9.4e-06 | 1.147E-05 | 3.075E-06 | 9.400E-06 | 1.500E-05 |
| 11335 | 567 | H567P | A | 5.7e-06 | 1.2e-05 | 9.6e-06 | 9.100E-06 | 3.180E-06 | 5.700E-06 | 1.200E-05 |
| 11336 | 567 | H567S | A | 1.4e-05 | 1.3e-05 | 1e-05 | 1.233E-05 | 2.082E-06 | 1.000E-05 | 1.400E-05 |
| 11337 | 567 | H567T | A | 1.4e-05 | 9.7e-06 | 1.5e-05 | 1.290E-05 | 2.816E-06 | 9.700E-06 | 1.500E-05 |
| 11338 | 567 | H567W | A | 1.4e-05 | 1e-05 | 1.1e-05 | 1.167E-05 | 2.082E-06 | 1.000E-05 | 1.400E-05 |
| 11339 | 567 | H567Y | A | 1.4e-05 | 1.1e-05 | 1.1e-05 | 1.200E-05 | 1.732E-06 | 1.100E-05 | 1.400E-05 |
| 11340 | 567 | H567V | A | 1.5e-05 | 1.1e-05 | 1.4e-05 | 1.333E-05 | 2.082E-06 | 1.100E-05 | 1.500E-05 |
| 11341 | 568 | Y568A | A | 1.7e-05 | 1.8e-05 | 9.8e-06 | 1.493E-05 | 4.474E-06 | 9.800E-06 | 1.800E-05 |
| 11342 | 568 | Y568R | A | 9.9e-06 | 1.3e-05 | 1e-05 | 1.097E-05 | 1.762E-06 | 9.900E-06 | 1.300E-05 |
| 11343 | 568 | Y568N | A | 1.5e-05 | 1.3e-05 | 8.9e-06 | 1.230E-05 | 3.110E-06 | 8.900E-06 | 1.500E-05 |
| 11344 | 568 | Y568D | A | 1.2e-05 | 1.1e-05 | 1.5e-05 | 1.267E-05 | 2.082E-06 | 1.100E-05 | 1.500E-05 |
| 11345 | 568 | Y568C | A | 1.7e-05 | 9.8e-06 | 1.2e-05 | 1.293E-05 | 3.690E-06 | 9.800E-06 | 1.700E-05 |
| 11346 | 568 | Y568E | A | 1.5e-05 | 1.3e-05 | 1.7e-05 | 1.500E-05 | 2.000E-06 | 1.300E-05 | 1.700E-05 |
| 11347 | 568 | Y568Q | A | 1e-05 | 1.5e-05 | 9.8e-06 | 1.160E-05 | 2.946E-06 | 9.800E-06 | 1.500E-05 |
| 11348 | 568 | Y568G | A | 9.6e-06 | 1.2e-05 | 1.6e-05 | 1.253E-05 | 3.233E-06 | 9.600E-06 | 1.600E-05 |
| 11349 | 568 | Y568H | A | 1.4e-05 | 1.1e-05 | 1.7e-05 | 1.400E-05 | 3.000E-06 | 1.100E-05 | 1.700E-05 |
| 11350 | 568 | Y568I | A | 1.6e-05 | 1.5e-05 | 1.6e-05 | 1.567E-05 | 5.774E-07 | 1.500E-05 | 1.600E-05 |
| 11351 | 568 | Y568L | A | 1.2e-05 | 1e-05 | 1.4e-05 | 1.200E-05 | 2.000E-06 | 1.000E-05 | 1.400E-05 |
| 11352 | 568 | Y568K | A | 1.5e-05 | 1.5e-05 | 1.1e-05 | 1.367E-05 | 2.309E-06 | 1.100E-05 | 1.500E-05 |
| 11353 | 568 | Y568M | A | 9e-06 | 1.5e-05 | 1.5e-05 | 1.300E-05 | 3.464E-06 | 9.000E-06 | 1.500E-05 |
| 11354 | 568 | Y568F | A | 1.5e-05 | 1e-05 | 1.7e-05 | 1.400E-05 | 3.606E-06 | 1.000E-05 | 1.700E-05 |
| 11355 | 568 | Y568P | A | 1.1e-05 | 9.8e-06 | 1e-05 | 1.027E-05 | 6.429E-07 | 9.800E-06 | 1.100E-05 |
| 11356 | 568 | Y568S | A | 1.5e-05 | 1.1e-05 | 1e-05 | 1.200E-05 | 2.646E-06 | 1.000E-05 | 1.500E-05 |
| 11357 | 568 | Y568T | A | 1.4e-05 | 1.1e-05 | 1.2e-05 | 1.233E-05 | 1.528E-06 | 1.100E-05 | 1.400E-05 |
| 11358 | 568 | Y568W | A | 1e-05 | 9.9e-06 | 1.3e-05 | 1.097E-05 | 1.762E-06 | 9.900E-06 | 1.300E-05 |
| 11359 | 568 | Y568Y | A | 1.6e-05 | 1.1e-05 | 1.6e-05 | 1.433E-05 | 2.887E-06 | 1.100E-05 | 1.600E-05 |
| 11360 | 568 | Y568V | A | 1.6e-05 | 1.3e-05 | 1.7e-05 | 1.533E-05 | 2.082E-06 | 1.300E-05 | 1.700E-05 |
| 11361 | 569 | K569A | A | 1.5e-05 | 1.4e-05 | 1.6e-05 | 1.500E-05 | 1.000E-06 | 1.400E-05 | 1.600E-05 |
| 11362 | 569 | K569R | A | 1.5e-05 | 1.3e-05 | 1.3e-05 | 1.367E-05 | 1.155E-06 | 1.300E-05 | 1.500E-05 |
| 11363 | 569 | K569N | A | 1.5e-05 | 1.4e-05 | 8.9e-06 | 1.263E-05 | 3.272E-06 | 8.900E-06 | 1.500E-05 |
| 11364 | 569 | K569D | A | 1.1e-05 | 1.7e-05 | 1.5e-05 | 1.433E-05 | 3.055E-06 | 1.100E-05 | 1.700E-05 |
| 11365 | 569 | K569C | A | 8.7e-06 | 1e-05 | 1.2e-05 | 1.023E-05 | 1.662E-06 | 8.700E-06 | 1.200E-05 |
| 11366 | 569 | K569E | A | 9.5e-06 | 1.5e-05 | 1.5e-05 | 1.317E-05 | 3.175E-06 | 9.500E-06 | 1.500E-05 |
| 11367 | 569 | K569Q | A | 1.4e-05 | 1.2e-05 | 9.9e-06 | 1.197E-05 | 2.050E-06 | 9.900E-06 | 1.400E-05 |
| 11368 | 569 | K569G | A | 1.5e-05 | 1.6e-05 | 8.3e-06 | 1.310E-05 | 4.187E-06 | 8.300E-06 | 1.600E-05 |
| 11369 | 569 | K569H | A | 1e-05 | 1.6e-05 | 1.1e-05 | 1.233E-05 | 3.215E-06 | 1.000E-05 | 1.600E-05 |
| 11370 | 569 | K569I | A | 9.3e-06 | 1.5e-05 | 1.4e-05 | 1.277E-05 | 3.044E-06 | 9.300E-06 | 1.500E-05 |
| 11371 | 569 | K569L | A | 9.5e-06 | 1.5e-05 | 1.1e-05 | 1.183E-05 | 2.843E-06 | 9.500E-06 | 1.500E-05 |
| 11372 | 569 | K569K | A | 1.5e-05 | 1e-05 | 1.5e-05 | 1.333E-05 | 2.887E-06 | 1.000E-05 | 1.500E-05 |
| 11373 | 569 | K569M | A | 1.1e-05 | 9.7e-06 | 1e-05 | 1.023E-05 | 6.807E-07 | 9.700E-06 | 1.100E-05 |
| 11374 | 569 | K569F | A | 1.1e-05 | 1.5e-05 | 1.5e-05 | 1.367E-05 | 2.309E-06 | 1.100E-05 | 1.500E-05 |
| 11375 | 569 | K569P | A | 1.4e-05 | 1.6e-05 | 1.5e-05 | 1.500E-05 | 1.000E-06 | 1.400E-05 | 1.600E-05 |
| 11376 | 569 | K569S | A | 9.9e-06 | 1.2e-05 | 9.7e-06 | 1.053E-05 | 1.274E-06 | 9.700E-06 | 1.200E-05 |
| 11377 | 569 | K569T | A | 9.6e-06 | 1.1e-05 | 1.5e-05 | 1.187E-05 | 2.802E-06 | 9.600E-06 | 1.500E-05 |
| 11378 | 569 | K569W | A | 1.6e-05 | 1.3e-05 | 1.5e-05 | 1.467E-05 | 1.528E-06 | 1.300E-05 | 1.600E-05 |
| 11379 | 569 | K569Y | A | 1.1e-05 | 8.5e-06 | 9.1e-06 | 9.533E-06 | 1.305E-06 | 8.500E-06 | 1.100E-05 |
| 11380 | 569 | K569V | A | 1e-05 | 1e-05 | 1e-05 | 1.000E-05 | 0.000E+00 | 1.000E-05 | 1.000E-05 |
| 11381 | 570 | D570A | A | 1e-05 | 1.5e-05 | 1.7e-05 | 1.400E-05 | 3.606E-06 | 1.000E-05 | 1.700E-05 |
| 11382 | 570 | D570R | A | 1.5e-05 | 1.1e-05 | 2.1e-05 | 1.567E-05 | 5.033E-06 | 1.100E-05 | 2.100E-05 |
| 11383 | 570 | D570N | A | 1.6e-05 | 1.5e-05 | 1e-05 | 1.367E-05 | 3.215E-06 | 1.000E-05 | 1.600E-05 |
| 11384 | 570 | D570D | A | 9e-06 | 1e-05 | 7.2e-06 | 8.733E-06 | 1.419E-06 | 7.200E-06 | 1.000E-05 |
| 11385 | 570 | D570C | A | 9.2e-06 | 1.2e-05 | 1.3e-05 | 1.140E-05 | 1.970E-06 | 9.200E-06 | 1.300E-05 |
| 11386 | 570 | D570E | A | 1e-05 | 1.4e-05 | 1.4e-05 | 1.267E-05 | 2.309E-06 | 1.000E-05 | 1.400E-05 |
| 11387 | 570 | D570Q | A | 1.4e-05 | 1.5e-05 | 1.5e-05 | 1.467E-05 | 5.774E-07 | 1.400E-05 | 1.500E-05 |
| 11388 | 570 | D570G | A | 1.3e-05 | 9.1e-06 | 1.6e-05 | 1.270E-05 | 3.460E-06 | 9.100E-06 | 1.600E-05 |
| 11389 | 570 | D570H | A | 6.1e-06 | 1.4e-05 | 1e-05 | 1.003E-05 | 3.950E-06 | 6.100E-06 | 1.400E-05 |

| | | | | | | | | | | |
|---|---|---|---|---|---|---|---|---|---|---|
| 11390 | 570 | D570I | A | 1.2e-05 | 1.2e-05 | 1.6e-05 | 1.333E-05 | 2.309E-06 | 1.200E-05 | 1.600E-05 |
| 11391 | 570 | D570L | A | 1e-05 | 1e-05 | 1e-05 | 1.000E-05 | 0.000E+00 | 1.000E-05 | 1.000E-05 |
| 11392 | 570 | D570K | A | 1.5e-05 | 1.4e-05 | 8.5e-06 | 1.250E-05 | 3.500E-06 | 8.500E-06 | 1.500E-05 |
| 11393 | 570 | D570M | A | 6.3e-06 | 1e-05 | 1.7e-05 | 1.110E-05 | 5.434E-06 | 6.300E-06 | 1.700E-05 |
| 11394 | 570 | D570F | A | 8.5e-06 | 8.7e-06 | 9.2e-06 | 8.800E-06 | 3.606E-07 | 8.500E-06 | 9.200E-06 |
| 11395 | 570 | D570P | A | 1.1e-05 | 1.1e-05 | 1e-05 | 1.067E-05 | 5.774E-07 | 1.000E-05 | 1.100E-05 |
| 11396 | 570 | D570S | A | 1.2e-05 | 1e-05 | 1.7e-05 | 1.300E-05 | 3.606E-06 | 1.000E-05 | 1.700E-05 |
| 11397 | 570 | D570T | A | 1.3e-05 | 1.6e-05 | 8e-06 | 1.233E-05 | 4.041E-06 | 8.000E-06 | 1.600E-05 |
| 11398 | 570 | D570W | A | 8.5e-06 | 1.2e-05 | 1.3e-05 | 1.117E-05 | 2.363E-06 | 8.500E-06 | 1.300E-05 |
| 11399 | 570 | D570Y | A | 1.3e-05 | 7.7e-06 | 1.6e-05 | 1.223E-05 | 4.203E-06 | 7.700E-06 | 1.600E-05 |
| 11400 | 570 | D570V | A | 1.6e-05 | 9.8e-06 | 1.5e-05 | 1.360E-05 | 3.329E-06 | 9.800E-06 | 1.600E-05 |
| 11401 | 571 | P571A | A | 6.1e-06 | 1.7e-05 | 1.5e-05 | 1.270E-05 | 5.803E-06 | 6.100E-06 | 1.700E-05 |
| 11402 | 571 | P571R | A | 1.4e-05 | 1.3e-05 | 5e-06 | 1.067E-05 | 4.933E-06 | 5.000E-06 | 1.400E-05 |
| 11403 | 571 | P571N | A | 1.4e-05 | 9.9e-06 | 1e-05 | 1.130E-05 | 2.339E-06 | 9.900E-06 | 1.400E-05 |
| 11404 | 571 | P571D | A | 1.4e-05 | 1e-05 | 1.6e-05 | 1.333E-05 | 3.055E-06 | 1.000E-05 | 1.600E-05 |
| 11405 | 571 | P571C | A | 1.2e-05 | 8.9e-06 | 9.7e-06 | 1.020E-05 | 1.609E-06 | 8.900E-06 | 1.200E-05 |
| 11406 | 571 | P571E | A | 1.2e-05 | 1.2e-05 | 1.6e-05 | 1.333E-05 | 2.309E-06 | 1.200E-05 | 1.600E-05 |
| 11407 | 571 | P571Q | A | 1.2e-05 | 8.9e-06 | 1.9e-05 | 1.330E-05 | 5.174E-06 | 8.900E-06 | 1.900E-05 |
| 11408 | 571 | P571G | A | 1.5e-05 | 1.4e-05 | 1.5e-05 | 1.467E-05 | 5.774E-07 | 1.400E-05 | 1.500E-05 |
| 11409 | 571 | P571H | A | 1.1e-05 | 1e-05 | 2.1e-05 | 1.400E-05 | 6.083E-06 | 1.000E-05 | 2.100E-05 |
| 11410 | 571 | P571I | A | 1.6e-05 | 1.5e-05 | 1.4e-05 | 1.500E-05 | 1.000E-06 | 1.400E-05 | 1.600E-05 |
| 11411 | 571 | P571L | A | 1.6e-05 | 1.7e-05 | 1.2e-05 | 1.500E-05 | 2.646E-06 | 1.200E-05 | 1.700E-05 |
| 11412 | 571 | P571K | A | 1.3e-05 | 1e-05 | 1.1e-05 | 1.133E-05 | 1.528E-06 | 1.000E-05 | 1.300E-05 |
| 11413 | 571 | P571M | A | 1.2e-05 | 1e-05 | 1.6e-05 | 1.267E-05 | 3.055E-06 | 1.000E-05 | 1.600E-05 |
| 11414 | 571 | P571F | A | 1.3e-05 | 1.7e-05 | 9.9e-06 | 1.330E-05 | 3.559E-06 | 9.900E-06 | 1.700E-05 |
| 11415 | 571 | P571P | A | 1.1e-05 | 1.4e-05 | 1.5e-05 | 1.333E-05 | 2.082E-06 | 1.100E-05 | 1.500E-05 |
| 11416 | 571 | P571S | A | 1e-05 | 1.4e-05 | 1.4e-05 | 1.267E-05 | 2.309E-06 | 1.000E-05 | 1.400E-05 |
| 11417 | 571 | P571T | A | 1.6e-05 | 1.4e-05 | 1.6e-05 | 1.533E-05 | 1.155E-06 | 1.400E-05 | 1.600E-05 |
| 11418 | 571 | P571W | A | 1.2e-05 | 9.3e-06 | 1.5e-05 | 1.210E-05 | 2.851E-06 | 9.300E-06 | 1.500E-05 |
| 11419 | 571 | P571Y | A | 1.3e-05 | 1.5e-05 | 1.5e-05 | 1.433E-05 | 1.155E-06 | 1.300E-05 | 1.500E-05 |
| 11420 | 571 | P571V | A | 1e-05 | 1.4e-05 | 1e-05 | 1.133E-05 | 2.309E-06 | 1.000E-05 | 1.400E-05 |
| 11421 | 572 | P572A | A | 1.2e-05 | 1.5e-05 | 1e-05 | 1.233E-05 | 2.517E-06 | 1.000E-05 | 1.500E-05 |
| 11422 | 572 | P572R | A | 1e-05 | 1.7e-05 | 9.3e-06 | 1.210E-05 | 4.258E-06 | 9.300E-06 | 1.700E-05 |
| 11423 | 572 | P572N | A | 1.5e-05 | 8.4e-06 | 1.4e-05 | 1.247E-05 | 3.557E-06 | 8.400E-06 | 1.500E-05 |
| 11424 | 572 | P572D | A | 1e-05 | 1.5e-05 | 8.4e-06 | 1.113E-05 | 3.443E-06 | 8.400E-06 | 1.500E-05 |
| 11425 | 572 | P572C | A | 8.8e-06 | 1.4e-05 | 1.1e-05 | 1.127E-05 | 2.610E-06 | 8.800E-06 | 1.400E-05 |
| 11426 | 572 | P572E | A | 1.5e-05 | 1.1e-05 | 1.2e-05 | 1.267E-05 | 2.082E-06 | 1.100E-05 | 1.500E-05 |
| 11427 | 572 | P572Q | A | 9.8e-06 | 1.2e-05 | 1.4e-05 | 1.193E-05 | 2.101E-06 | 9.800E-06 | 1.400E-05 |
| 11428 | 572 | P572G | A | 1.5e-05 | 1.4e-05 | 2.4e-05 | 1.767E-05 | 5.508E-06 | 1.400E-05 | 2.400E-05 |
| 11429 | 572 | P572H | A | 1.5e-05 | 1.5e-05 | 1.6e-05 | 1.533E-05 | 5.774E-07 | 1.500E-05 | 1.600E-05 |
| 11430 | 572 | P572I | A | 1.1e-05 | 1e-05 | 1.7e-05 | 1.267E-05 | 3.786E-06 | 1.000E-05 | 1.700E-05 |
| 11431 | 572 | P572L | A | 9.8e-06 | 1.1e-05 | 1.5e-05 | 1.193E-05 | 2.723E-06 | 9.800E-06 | 1.500E-05 |
| 11432 | 572 | P572K | A | 1.1e-05 | 1e-05 | 1.6e-05 | 1.233E-05 | 3.215E-06 | 1.000E-05 | 1.600E-05 |
| 11433 | 572 | P572M | A | 1.6e-05 | 1e-05 | 1.5e-05 | 1.367E-05 | 3.215E-06 | 1.000E-05 | 1.600E-05 |
| 11434 | 572 | P572F | A | 1.5e-05 | 1.4e-05 | 9.8e-06 | 1.293E-05 | 2.759E-06 | 9.800E-06 | 1.500E-05 |
| 11435 | 572 | P572P | A | 1.3e-05 | 1e-05 | 1.5e-05 | 1.267E-05 | 2.517E-06 | 1.000E-05 | 1.500E-05 |
| 11436 | 572 | P572S | A | 1.5e-05 | 1.4e-05 | 1.4e-05 | 1.433E-05 | 5.774E-07 | 1.400E-05 | 1.500E-05 |
| 11437 | 572 | P572T | A | 1.6e-05 | 1.8e-05 | 1.6e-05 | 1.667E-05 | 1.155E-06 | 1.600E-05 | 1.800E-05 |
| 11438 | 572 | P572W | A | 1.6e-05 | 1.2e-05 | 1.5e-05 | 1.433E-05 | 2.082E-06 | 1.200E-05 | 1.600E-05 |
| 11439 | 572 | P572Y | A | 1.4e-05 | 1.4e-05 | 1e-05 | 1.267E-05 | 2.309E-06 | 1.000E-05 | 1.400E-05 |
| 11440 | 572 | P572V | A | 1.3e-05 | 1.5e-05 | 1.1e-05 | 1.300E-05 | 2.000E-06 | 1.100E-05 | 1.500E-05 |
| 11441 | 573 | F573A | A | 9.8e-06 | 1.5e-05 | 1.1e-05 | 1.193E-05 | 2.723E-06 | 9.800E-06 | 1.500E-05 |
| 11442 | 573 | F573R | A | 1.1e-05 | 1.5e-05 | 1.5e-05 | 1.367E-05 | 2.309E-06 | 1.100E-05 | 1.500E-05 |
| 11443 | 573 | F573N | A | 1.6e-05 | 1.1e-05 | 1.5e-05 | 1.400E-05 | 2.646E-06 | 1.100E-05 | 1.600E-05 |
| 11444 | 573 | F573D | A | 1.1e-05 | 1.5e-05 | 1.1e-05 | 1.233E-05 | 2.309E-06 | 1.100E-05 | 1.500E-05 |
| 11445 | 573 | F573C | A | 1.1e-05 | 1.4e-05 | 1e-05 | 1.167E-05 | 2.082E-06 | 1.000E-05 | 1.400E-05 |
| 11446 | 573 | F573E | A | 1e-05 | 2.5e-05 | 1.1e-05 | 1.533E-05 | 8.386E-06 | 1.000E-05 | 2.500E-05 |
| 11447 | 573 | F573Q | A | 1.5e-05 | 1.4e-05 | 1.5e-05 | 1.467E-05 | 5.774E-07 | 1.400E-05 | 1.500E-05 |
| 11448 | 573 | F573G | A | 1.5e-05 | 1.7e-05 | 1.6e-05 | 1.600E-05 | 1.000E-06 | 1.500E-05 | 1.700E-05 |
| 11449 | 573 | F573H | A | 1e-05 | 1.5e-05 | 8.7e-06 | 1.123E-05 | 3.326E-06 | 8.700E-06 | 1.500E-05 |
| 11450 | 573 | F573I | A | 8.3e-06 | 1.8e-05 | 8.8e-06 | 1.170E-05 | 5.462E-06 | 8.300E-06 | 1.800E-05 |
| 11451 | 573 | F573L | A | 1.4e-05 | 1.5e-05 | 1.5e-05 | 1.467E-05 | 5.774E-07 | 1.400E-05 | 1.500E-05 |
| 11452 | 573 | F573K | A | 1.4e-05 | 1.1e-05 | 1.4e-05 | 1.300E-05 | 1.732E-06 | 1.100E-05 | 1.400E-05 |
| 11453 | 573 | F573M | A | 1.3e-05 | 1.4e-05 | 1.5e-05 | 1.400E-05 | 1.000E-06 | 1.300E-05 | 1.500E-05 |
| 11454 | 573 | F573F | A | 8.3e-06 | 7e-06 | 9.3e-06 | 8.200E-06 | 1.153E-06 | 7.000E-06 | 9.300E-06 |
| 11455 | 573 | F573P | A | 9.5e-06 | 1e-05 | 1.4e-05 | 1.117E-05 | 2.466E-06 | 9.500E-06 | 1.400E-05 |
| 11456 | 573 | F573S | A | 1.4e-05 | 1.1e-05 | 9.9e-06 | 1.163E-05 | 2.122E-06 | 9.900E-06 | 1.400E-05 |
| 11457 | 573 | F573T | A | 1e-05 | 1.4e-05 | 1.5e-05 | 1.300E-05 | 2.646E-06 | 1.000E-05 | 1.500E-05 |
| 11458 | 573 | F573W | A | 1.5e-05 | 9.4e-06 | 1.2e-05 | 1.213E-05 | 2.802E-06 | 9.400E-06 | 1.500E-05 |
| 11459 | 573 | F573Y | A | 1.5e-05 | 1.4e-05 | 1.1e-05 | 1.333E-05 | 2.082E-06 | 1.100E-05 | 1.500E-05 |
| 11460 | 573 | F573V | A | 1.6e-05 | 1.4e-05 | 9.2e-06 | 1.307E-05 | 3.495E-06 | 9.200E-06 | 1.600E-05 |
| 11461 | 574 | C574A | A | 1.6e-05 | 1.2e-05 | 1.6e-05 | 1.467E-05 | 2.309E-06 | 1.200E-05 | 1.600E-05 |
| 11462 | 574 | C574R | A | 1.1e-05 | 1e-05 | 8.8e-06 | 9.933E-06 | 1.102E-06 | 8.800E-06 | 1.100E-05 |
| 11463 | 574 | C574N | A | 1.5e-05 | 8.1e-06 | 1.5e-05 | 1.270E-05 | 3.984E-06 | 8.100E-06 | 1.500E-05 |
| 11464 | 574 | C574D | A | 1.6e-05 | 1.4e-05 | 1.5e-05 | 1.500E-05 | 1.000E-06 | 1.400E-05 | 1.600E-05 |
| 11465 | 574 | C574C | A | 1.6e-05 | 1.4e-05 | 8.4e-06 | 1.280E-05 | 3.940E-06 | 8.400E-06 | 1.600E-05 |
| 11466 | 574 | C574E | A | 1.4e-05 | 1.1e-05 | 1.5e-05 | 1.333E-05 | 2.082E-06 | 1.100E-05 | 1.500E-05 |
| 11467 | 574 | C574Q | A | 1.8e-05 | 1.1e-05 | 1.5e-05 | 1.467E-05 | 3.512E-06 | 1.100E-05 | 1.800E-05 |
| 11468 | 574 | C574G | A | 1.3e-05 | 1.5e-05 | 1.3e-05 | 1.367E-05 | 1.155E-06 | 1.300E-05 | 1.500E-05 |
| 11469 | 574 | C574H | A | 1.4e-05 | 1.3e-05 | 1.2e-05 | 1.300E-05 | 1.000E-06 | 1.200E-05 | 1.400E-05 |
| 11470 | 574 | C574I | A | 1.5e-05 | 1.5e-05 | 1e-05 | 1.333E-05 | 2.887E-06 | 1.000E-05 | 1.500E-05 |
| 11471 | 574 | C574L | A | 1.5e-05 | 1e-05 | 8.8e-06 | 1.127E-05 | 3.288E-06 | 8.800E-06 | 1.500E-05 |
| 11472 | 574 | C574K | A | 1e-05 | 2e-05 | 1.6e-05 | 1.533E-05 | 5.033E-06 | 1.000E-05 | 2.000E-05 |
| 11473 | 574 | C574M | A | 1.5e-05 | 1.1e-05 | 1.6e-05 | 1.400E-05 | 2.646E-06 | 1.100E-05 | 1.600E-05 |
| 11474 | 574 | C574F | A | 1.2e-05 | 8e-06 | 1.1e-05 | 1.033E-05 | 2.082E-06 | 8.000E-06 | 1.200E-05 |

| | | | | | | | | | | |
|---|---|---|---|---|---|---|---|---|---|---|
| 11475 | 574 | C574P | A | 8.1e-06 | 9.2e-06 | 8.2e-06 | 8.500E-06 | 6.083E-07 | 8.100E-06 | 9.200E-06 |
| 11476 | 574 | C574S | A | 1.5e-05 | 1.6e-05 | 1.5e-05 | 1.533E-05 | 5.774E-07 | 1.500E-05 | 1.600E-05 |
| 11477 | 574 | C574T | A | 1.5e-05 | 1.4e-05 | 1.1e-05 | 1.333E-05 | 2.082E-06 | 1.100E-05 | 1.500E-05 |
| 11478 | 574 | C574W | A | 9.5e-06 | 1e-05 | 1.5e-05 | 1.150E-05 | 3.041E-06 | 9.500E-06 | 1.500E-05 |
| 11479 | 574 | C574Y | A | 1.1e-05 | 1.4e-05 | 1.7e-05 | 1.400E-05 | 3.000E-06 | 1.100E-05 | 1.700E-05 |
| 11480 | 574 | C574V | A | 1.5e-05 | 1.8e-05 | 1.3e-05 | 1.533E-05 | 2.517E-06 | 1.300E-05 | 1.800E-05 |
| 11481 | 575 | V575A | A | 1.1e-05 | 1.3e-05 | 1.4e-05 | 1.267E-05 | 1.528E-06 | 1.100E-05 | 1.400E-05 |
| 11482 | 575 | V575R | A | 8.5e-06 | 1.1e-05 | 1.2e-05 | 1.050E-05 | 1.803E-06 | 8.500E-06 | 1.200E-05 |
| 11483 | 575 | V575N | A | 1.4e-05 | 1.3e-05 | 1.5e-05 | 1.400E-05 | 1.000E-06 | 1.300E-05 | 1.500E-05 |
| 11484 | 575 | V575D | A | 1.5e-05 | 1.6e-05 | 1.2e-05 | 1.433E-05 | 2.082E-06 | 1.200E-05 | 1.600E-05 |
| 11485 | 575 | V575C | A | 1.3e-05 | 1.3e-05 | 1.3e-05 | 1.300E-05 | 0.000E+00 | 1.300E-05 | 1.300E-05 |
| 11486 | 575 | V575E | A | 1.4e-05 | 1.5e-05 | 1.5e-05 | 1.467E-05 | 5.774E-07 | 1.400E-05 | 1.500E-05 |
| 11487 | 575 | V575Q | A | 1.8e-05 | 1e-05 | 8.5e-06 | 1.217E-05 | 5.107E-06 | 8.500E-06 | 1.800E-05 |
| 11488 | 575 | V575G | A | 1.3e-05 | 8.6e-06 | 9.7e-06 | 1.043E-05 | 2.290E-06 | 8.600E-06 | 1.300E-05 |
| 11489 | 575 | V575H | A | 1.5e-05 | 1.4e-05 | 1.5e-05 | 1.467E-05 | 5.774E-07 | 1.400E-05 | 1.500E-05 |
| 11490 | 575 | V575I | A | 1.1e-05 | 1.1e-05 | 6.9e-06 | 9.633E-06 | 2.367E-06 | 6.900E-06 | 1.100E-05 |
| 11491 | 575 | V575L | A | 1.1e-05 | 1.7e-05 | 1.4e-05 | 1.400E-05 | 3.000E-06 | 1.100E-05 | 1.700E-05 |
| 11492 | 575 | V575K | A | 9.2e-06 | 1.4e-05 | 1.5e-05 | 1.273E-05 | 3.101E-06 | 9.200E-06 | 1.500E-05 |
| 11493 | 575 | V575M | A | 1.6e-05 | 1.7e-05 | 1.5e-05 | 1.600E-05 | 1.000E-06 | 1.500E-05 | 1.700E-05 |
| 11494 | 575 | V575F | A | 1.6e-05 | 1.5e-05 | 1.3e-05 | 1.467E-05 | 1.528E-06 | 1.300E-05 | 1.600E-05 |
| 11495 | 575 | V575P | A | 1.3e-05 | 1.7e-05 | 9.3e-06 | 1.310E-05 | 3.851E-06 | 9.300E-06 | 1.700E-05 |
| 11496 | 575 | V575S | A | 9.2e-06 | 1.3e-05 | 1.1e-05 | 1.107E-05 | 1.901E-06 | 9.200E-06 | 1.300E-05 |
| 11497 | 575 | V575T | A | 8.7e-06 | 1.5e-05 | 7.3e-06 | 1.033E-05 | 4.102E-06 | 7.300E-06 | 1.500E-05 |
| 11498 | 575 | V575W | A | 9.6e-06 | 9.9e-06 | 1.3e-05 | 1.083E-05 | 1.882E-06 | 9.600E-06 | 1.300E-05 |
| 11499 | 575 | V575Y | A | 1.4e-05 | 1.2e-05 | 1.3e-05 | 1.300E-05 | 1.000E-06 | 1.200E-05 | 1.400E-05 |
| 11500 | 575 | V575V | A | 1.2e-05 | 1.2e-05 | 1.4e-05 | 1.267E-05 | 1.155E-06 | 1.200E-05 | 1.400E-05 |
| 11501 | 576 | A576A | A | 9.8e-06 | 1.5e-05 | 1.1e-05 | 1.193E-05 | 2.723E-06 | 9.800E-06 | 1.500E-05 |
| 11502 | 576 | A576R | A | 1.5e-05 | 1.4e-05 | 1.5e-05 | 1.467E-05 | 5.774E-07 | 1.400E-05 | 1.500E-05 |
| 11503 | 576 | A576N | A | 1.6e-05 | 1.5e-05 | 9.9e-06 | 1.363E-05 | 3.272E-06 | 9.900E-06 | 1.600E-05 |
| 11504 | 576 | A576D | A | 1.4e-05 | 1.4e-05 | 1.4e-05 | 1.400E-05 | 0.000E+00 | 1.400E-05 | 1.400E-05 |
| 11505 | 576 | A576C | A | 8.9e-06 | 1.7e-05 | 9.4e-06 | 1.177E-05 | 4.539E-06 | 8.900E-06 | 1.700E-05 |
| 11506 | 576 | A576E | A | 1.2e-05 | 1.3e-05 | 1.3e-05 | 1.267E-05 | 5.774E-07 | 1.200E-05 | 1.300E-05 |
| 11507 | 576 | A576Q | A | 1.5e-05 | 1.1e-05 | 1.8e-05 | 1.467E-05 | 3.512E-06 | 1.100E-05 | 1.800E-05 |
| 11508 | 576 | A576G | A | 1.5e-05 | 1e-05 | 1.4e-05 | 1.300E-05 | 2.646E-06 | 1.000E-05 | 1.500E-05 |
| 11509 | 576 | A576H | A | 9.8e-06 | 1.3e-05 | 1e-05 | 1.093E-05 | 1.793E-06 | 9.800E-06 | 1.300E-05 |
| 11510 | 576 | A576I | A | 1.7e-05 | 1e-05 | 1.1e-05 | 1.267E-05 | 3.786E-06 | 1.000E-05 | 1.700E-05 |
| 11511 | 576 | A576L | A | 1.1e-05 | 1.4e-05 | 1.6e-05 | 1.367E-05 | 2.517E-06 | 1.100E-05 | 1.600E-05 |
| 11512 | 576 | A576K | A | 1.2e-05 | 1.5e-05 | 8.9e-06 | 1.197E-05 | 3.050E-06 | 8.900E-06 | 1.500E-05 |
| 11513 | 576 | A576M | A | 1.2e-05 | 1.3e-05 | 1.5e-05 | 1.333E-05 | 1.528E-06 | 1.200E-05 | 1.500E-05 |
| 11514 | 576 | A576F | A | 1e-05 | 9.8e-06 | 1.6e-05 | 1.193E-05 | 3.523E-06 | 9.800E-06 | 1.600E-05 |
| 11515 | 576 | A576P | A | 1.6e-05 | 9.4e-06 | 1.1e-05 | 1.213E-05 | 3.443E-06 | 9.400E-06 | 1.600E-05 |
| 11516 | 576 | A576S | A | 1.5e-05 | 8.3e-06 | 1.4e-05 | 1.243E-05 | 3.614E-06 | 8.300E-06 | 1.500E-05 |
| 11517 | 576 | A576T | A | 8.9e-06 | 1.1e-05 | 1e-05 | 9.967E-06 | 1.050E-06 | 8.900E-06 | 1.100E-05 |
| 11518 | 576 | A576W | A | 1.5e-05 | 9.7e-06 | 1e-05 | 1.157E-05 | 2.977E-06 | 9.700E-06 | 1.500E-05 |
| 11519 | 576 | A576Y | A | 1.5e-05 | 9.3e-06 | 9e-06 | 1.110E-05 | 3.381E-06 | 9.000E-06 | 1.500E-05 |
| 11520 | 576 | A576V | A | 8.7e-06 | 1.6e-05 | 8.6e-06 | 1.110E-05 | 4.244E-06 | 8.600E-06 | 1.600E-05 |
| 11521 | 577 | R577A | A | 9.2e-06 | 1.7e-05 | 1.1e-05 | 1.240E-05 | 4.084E-06 | 9.200E-06 | 1.700E-05 |
| 11522 | 577 | R577R | A | 1.2e-05 | 1.1e-05 | 1.4e-05 | 1.233E-05 | 1.528E-06 | 1.100E-05 | 1.400E-05 |
| 11523 | 577 | R577N | A | 1.2e-05 | 1.2e-05 | 9.9e-06 | 1.130E-05 | 1.212E-06 | 9.900E-06 | 1.200E-05 |
| 11524 | 577 | R577D | A | 1.3e-05 | 1.1e-05 | 1.6e-05 | 1.333E-05 | 2.517E-06 | 1.100E-05 | 1.600E-05 |
| 11525 | 577 | R577C | A | 9.6e-06 | 1.1e-05 | 1.2e-05 | 1.087E-05 | 1.206E-06 | 9.600E-06 | 1.200E-05 |
| 11526 | 577 | R577E | A | 9.2e-06 | 1.7e-05 | 1.4e-05 | 1.340E-05 | 3.934E-06 | 9.200E-06 | 1.700E-05 |
| 11527 | 577 | R577Q | A | 1.4e-05 | 8.2e-06 | 1.5e-05 | 1.240E-05 | 3.672E-06 | 8.200E-06 | 1.500E-05 |
| 11528 | 577 | R577G | A | 1.1e-05 | 1.5e-05 | 9.4e-06 | 1.180E-05 | 2.884E-06 | 9.400E-06 | 1.500E-05 |
| 11529 | 577 | R577H | A | 1.1e-05 | 9.3e-06 | 1.6e-05 | 1.210E-05 | 3.483E-06 | 9.300E-06 | 1.600E-05 |
| 11530 | 577 | R577I | A | 1.4e-05 | 1.4e-05 | 1.2e-05 | 1.333E-05 | 1.155E-06 | 1.200E-05 | 1.400E-05 |
| 11531 | 577 | R577L | A | 1.1e-05 | 9.5e-06 | 1.6e-05 | 1.217E-05 | 3.403E-06 | 9.500E-06 | 1.600E-05 |
| 11532 | 577 | R577K | A | 1.6e-05 | 1.4e-05 | 9.9e-06 | 1.330E-05 | 3.110E-06 | 9.900E-06 | 1.600E-05 |
| 11533 | 577 | R577M | A | 1.7e-05 | 8.1e-06 | 1.3e-05 | 1.270E-05 | 4.458E-06 | 8.100E-06 | 1.700E-05 |
| 11534 | 577 | R577F | A | 1.2e-05 | 1.5e-05 | 1.2e-05 | 1.300E-05 | 1.732E-06 | 1.200E-05 | 1.500E-05 |
| 11535 | 577 | R577P | A | 1.1e-05 | 1.5e-05 | 9.5e-06 | 1.183E-05 | 2.843E-06 | 9.500E-06 | 1.500E-05 |
| 11536 | 577 | R577S | A | 1.4e-05 | 1.5e-05 | 1.5e-05 | 1.467E-05 | 5.774E-07 | 1.400E-05 | 1.500E-05 |
| 11537 | 577 | R577T | A | 1e-05 | 1.3e-05 | 9.6e-06 | 1.087E-05 | 1.858E-06 | 9.600E-06 | 1.300E-05 |
| 11538 | 577 | R577W | A | 9.8e-06 | 9.8e-06 | 1.6e-05 | 1.187E-05 | 3.580E-06 | 9.800E-06 | 1.600E-05 |
| 11539 | 577 | R577Y | A | 1e-05 | 9.8e-06 | 9.5e-06 | 9.767E-06 | 2.517E-07 | 9.500E-06 | 1.000E-05 |
| 11540 | 577 | R577V | A | 9.7e-06 | 1.2e-05 | 1.6e-05 | 1.257E-05 | 3.188E-06 | 9.700E-06 | 1.600E-05 |
| 11541 | 578 | C578A | A | 1.6e-05 | 1.7e-05 | 1.2e-05 | 1.500E-05 | 2.646E-06 | 1.200E-05 | 1.700E-05 |
| 11542 | 578 | C578R | A | 1.1e-05 | 8.6e-06 | 1.4e-05 | 1.120E-05 | 2.706E-06 | 8.600E-06 | 1.400E-05 |
| 11543 | 578 | C578N | A | 1.4e-05 | 1.1e-05 | 1.5e-05 | 1.333E-05 | 2.082E-06 | 1.100E-05 | 1.500E-05 |
| 11544 | 578 | C578D | A | 1.3e-05 | 7.3e-06 | 1.5e-05 | 1.177E-05 | 3.995E-06 | 7.300E-06 | 1.500E-05 |
| 11545 | 578 | C578C | A | 1.1e-05 | 1.4e-05 | 1.4e-05 | 1.300E-05 | 1.732E-06 | 1.100E-05 | 1.400E-05 |
| 11546 | 578 | C578E | A | 1.3e-05 | 1.5e-05 | 1.6e-05 | 1.467E-05 | 1.528E-06 | 1.300E-05 | 1.600E-05 |
| 11547 | 578 | C578Q | A | 1.1e-05 | 1.6e-05 | 1e-05 | 1.233E-05 | 3.215E-06 | 1.000E-05 | 1.600E-05 |
| 11548 | 578 | C578G | A | 1.4e-05 | 1.1e-05 | 1.2e-05 | 1.233E-05 | 1.528E-06 | 1.100E-05 | 1.400E-05 |
| 11549 | 578 | C578H | A | 1.3e-05 | 1.9e-05 | 1.3e-05 | 1.500E-05 | 3.464E-06 | 1.300E-05 | 1.900E-05 |
| 11550 | 578 | C578I | A | 1.8e-05 | 9.6e-06 | 1.1e-05 | 1.287E-05 | 4.500E-06 | 9.600E-06 | 1.800E-05 |
| 11551 | 578 | C578L | A | 7.7e-06 | 1.3e-05 | 1.3e-05 | 1.123E-05 | 3.060E-06 | 7.700E-06 | 1.300E-05 |
| 11552 | 578 | C578K | A | 1e-05 | 1.3e-05 | 9.7e-06 | 1.090E-05 | 1.825E-06 | 9.700E-06 | 1.300E-05 |
| 11553 | 578 | C578M | A | 1.7e-05 | 1.1e-05 | 1.4e-05 | 1.400E-05 | 3.000E-06 | 1.100E-05 | 1.700E-05 |
| 11554 | 578 | C578F | A | 1.4e-05 | 1e-05 | 1.1e-05 | 1.167E-05 | 2.082E-06 | 1.000E-05 | 1.400E-05 |
| 11555 | 578 | C578P | A | 9.1e-06 | 1.4e-05 | 1.2e-05 | 1.170E-05 | 2.464E-06 | 9.100E-06 | 1.400E-05 |
| 11556 | 578 | C578S | A | 9.7e-06 | 1.5e-05 | 1.5e-05 | 1.323E-05 | 3.060E-06 | 9.700E-06 | 1.500E-05 |
| 11557 | 578 | C578T | A | 1.6e-05 | 1.6e-05 | 1.3e-05 | 1.500E-05 | 1.732E-06 | 1.300E-05 | 1.600E-05 |
| 11558 | 578 | C578W | A | 1.3e-05 | 1.4e-05 | 1.6e-05 | 1.433E-05 | 1.528E-06 | 1.300E-05 | 1.600E-05 |
| 11559 | 578 | C578Y | A | 1.5e-05 | 1.3e-05 | 1.6e-05 | 1.467E-05 | 1.528E-06 | 1.300E-05 | 1.600E-05 |

| | | | | | | | | | | |
|---|---|---|---|---|---|---|---|---|---|---|
| 11560 | 578 | C578V | A | 9.8e-06 | 1e-05 | 9.4e-06 | 9.733E-06 | 3.055E-07 | 9.400E-06 | 1.000E-05 |
| 11561 | 579 | P579A | A | 1.5e-05 | 1e-05 | 1.5e-05 | 1.333E-05 | 2.887E-06 | 1.000E-05 | 1.500E-05 |
| 11562 | 579 | P579R | A | 1.5e-05 | 8.9e-06 | 9.8e-06 | 1.123E-05 | 3.293E-06 | 8.900E-06 | 1.500E-05 |
| 11563 | 579 | P579N | A | 1.8e-05 | 1.4e-05 | 1e-05 | 1.400E-05 | 4.000E-06 | 1.000E-05 | 1.800E-05 |
| 11564 | 579 | P579D | A | 1.6e-05 | 1.1e-05 | 9.7e-06 | 1.223E-05 | 3.326E-06 | 9.700E-06 | 1.600E-05 |
| 11565 | 579 | P579C | A | 1.6e-05 | 1.4e-05 | 1.5e-05 | 1.500E-05 | 1.000E-06 | 1.400E-05 | 1.600E-05 |
| 11566 | 579 | P579E | A | 1e-05 | 1.4e-05 | 1.4e-05 | 1.267E-05 | 2.309E-06 | 1.000E-05 | 1.400E-05 |
| 11567 | 579 | P579Q | A | 1.4e-05 | 1.6e-05 | 9.6e-06 | 1.320E-05 | 3.274E-06 | 9.600E-06 | 1.600E-05 |
| 11568 | 579 | P579G | A | 1.1e-05 | 1.2e-05 | 1.6e-05 | 1.300E-05 | 2.646E-06 | 1.100E-05 | 1.600E-05 |
| 11569 | 579 | P579H | A | 1e-05 | 1.2e-05 | 8.3e-06 | 1.010E-05 | 1.852E-06 | 8.300E-06 | 1.200E-05 |
| 11570 | 579 | P579I | A | 1.5e-05 | 1.7e-05 | 8e-06 | 1.333E-05 | 4.726E-06 | 8.000E-06 | 1.700E-05 |
| 11571 | 579 | P579L | A | 8e-06 | 1.5e-05 | 1.6e-05 | 1.300E-05 | 4.359E-06 | 8.000E-06 | 1.600E-05 |
| 11572 | 579 | P579K | A | 2e-05 | 1.5e-05 | 1.3e-05 | 1.600E-05 | 3.606E-06 | 1.300E-05 | 2.000E-05 |
| 11573 | 579 | P579M | A | 1.1e-05 | 1.2e-05 | 1.4e-05 | 1.233E-05 | 1.528E-06 | 1.100E-05 | 1.400E-05 |
| 11574 | 579 | P579F | A | 9.9e-06 | 1e-05 | 1e-05 | 9.967E-06 | 5.774E-08 | 9.900E-06 | 1.000E-05 |
| 11575 | 579 | P579P | A | 1.1e-05 | 1.2e-05 | 1.6e-05 | 1.300E-05 | 2.646E-06 | 1.100E-05 | 1.600E-05 |
| 11576 | 579 | P579S | A | 1.2e-05 | 1.1e-05 | 1.4e-05 | 1.233E-05 | 1.528E-06 | 1.100E-05 | 1.400E-05 |
| 11577 | 579 | P579T | A | 1.4e-05 | 1.1e-05 | 9.9e-06 | 1.163E-05 | 2.122E-06 | 9.900E-06 | 1.400E-05 |
| 11578 | 579 | P579W | A | 8.8e-06 | 1.3e-05 | 9.8e-06 | 1.053E-05 | 2.194E-06 | 8.800E-06 | 1.300E-05 |
| 11579 | 579 | P579Y | A | 1.6e-05 | 9.8e-06 | 8.2e-06 | 1.133E-05 | 4.120E-06 | 8.200E-06 | 1.600E-05 |
| 11580 | 579 | P579V | A | 1.6e-05 | 9.7e-06 | 9.1e-06 | 1.160E-05 | 3.822E-06 | 9.100E-06 | 1.600E-05 |
| 11581 | 580 | S580A | A | 1e-05 | 1.4e-05 | 1.1e-05 | 1.167E-05 | 2.082E-06 | 1.000E-05 | 1.400E-05 |
| 11582 | 580 | S580R | A | 1.1e-05 | 1.5e-05 | 1.6e-05 | 1.400E-05 | 2.646E-06 | 1.100E-05 | 1.600E-05 |
| 11583 | 580 | S580N | A | 8.7e-06 | 7.8e-06 | 1.6e-05 | 1.083E-05 | 4.497E-06 | 7.800E-06 | 1.600E-05 |
| 11584 | 580 | S580D | A | 1.5e-05 | 1.3e-05 | 1.6e-05 | 1.467E-05 | 1.528E-06 | 1.300E-05 | 1.600E-05 |
| 11585 | 580 | S580C | A | 1.4e-05 | 1.3e-05 | 1.3e-05 | 1.333E-05 | 5.774E-07 | 1.300E-05 | 1.400E-05 |
| 11586 | 580 | S580E | A | 1e-05 | 9.9e-06 | 1.6e-05 | 1.197E-05 | 3.493E-06 | 9.900E-06 | 1.600E-05 |
| 11587 | 580 | S580Q | A | 1.5e-05 | 1e-05 | 1.1e-05 | 1.200E-05 | 2.646E-06 | 1.000E-05 | 1.500E-05 |
| 11588 | 580 | S580G | A | 1e-05 | 1.8e-05 | 1.4e-05 | 1.400E-05 | 4.000E-06 | 1.000E-05 | 1.800E-05 |
| 11589 | 580 | S580H | A | 1.1e-05 | 1.5e-05 | 1e-05 | 1.200E-05 | 2.646E-06 | 1.000E-05 | 1.500E-05 |
| 11590 | 580 | S580I | A | 1.5e-05 | 1.6e-05 | 9.9e-06 | 1.363E-05 | 3.272E-06 | 9.900E-06 | 1.600E-05 |
| 11591 | 580 | S580L | A | 1.4e-05 | 1.1e-05 | 1.3e-05 | 1.267E-05 | 1.528E-06 | 1.100E-05 | 1.400E-05 |
| 11592 | 580 | S580K | A | 1.5e-05 | 1.2e-05 | 7.7e-06 | 1.157E-05 | 3.669E-06 | 7.700E-06 | 1.500E-05 |
| 11593 | 580 | S580M | A | 1.7e-05 | 1e-05 | 9.9e-06 | 1.230E-05 | 4.071E-06 | 9.900E-06 | 1.700E-05 |
| 11594 | 580 | S580F | A | 1.3e-05 | 9.8e-06 | 1e-05 | 1.093E-05 | 1.793E-06 | 9.800E-06 | 1.300E-05 |
| 11595 | 580 | S580P | A | 1.1e-05 | 1.7e-05 | 1.5e-05 | 1.433E-05 | 3.055E-06 | 1.100E-05 | 1.700E-05 |
| 11596 | 580 | S580S | A | 1e-05 | 1.6e-05 | 1.4e-05 | 1.333E-05 | 3.055E-06 | 1.000E-05 | 1.600E-05 |
| 11597 | 580 | S580T | A | 1.3e-05 | 1.3e-05 | 1.1e-05 | 1.233E-05 | 1.155E-06 | 1.100E-05 | 1.300E-05 |
| 11598 | 580 | S580W | A | 1e-05 | 1.1e-05 | 1e-05 | 1.033E-05 | 5.774E-07 | 1.000E-05 | 1.100E-05 |
| 11599 | 580 | S580Y | A | 1.4e-05 | 1.4e-05 | 1.6e-05 | 1.467E-05 | 1.155E-06 | 1.400E-05 | 1.600E-05 |
| 11600 | 580 | S580V | A | 1.5e-05 | 1.4e-05 | 1.6e-05 | 1.500E-05 | 1.000E-06 | 1.400E-05 | 1.600E-05 |
| 11601 | 581 | G581A | A | 1.1e-05 | 1.4e-05 | 9.3e-06 | 1.143E-05 | 2.380E-06 | 9.300E-06 | 1.400E-05 |
| 11602 | 581 | G581R | A | 1.8e-05 | 1.5e-05 | 1.5e-05 | 1.600E-05 | 1.732E-06 | 1.500E-05 | 1.800E-05 |
| 11603 | 581 | G581N | A | 8.6e-06 | 1.3e-05 | 1.3e-05 | 1.153E-05 | 2.540E-06 | 8.600E-06 | 1.300E-05 |
| 11604 | 581 | G581D | A | 1.1e-05 | 1.4e-05 | 1.3e-05 | 1.267E-05 | 1.528E-06 | 1.100E-05 | 1.400E-05 |
| 11605 | 581 | G581C | A | 1.6e-05 | 1.2e-05 | 1.8e-05 | 1.533E-05 | 3.055E-06 | 1.200E-05 | 1.800E-05 |
| 11606 | 581 | G581E | A | 8.6e-06 | 1.6e-05 | 1.2e-05 | 1.220E-05 | 3.704E-06 | 8.600E-06 | 1.600E-05 |
| 11607 | 581 | G581Q | A | 1.4e-05 | 9.5e-06 | 1.1e-05 | 1.150E-05 | 2.291E-06 | 9.500E-06 | 1.400E-05 |
| 11608 | 581 | G581G | A | 9e-06 | 1.4e-05 | 1.6e-05 | 1.300E-05 | 3.606E-06 | 9.000E-06 | 1.600E-05 |
| 11609 | 581 | G581H | A | 6.8e-06 | 1e-05 | 1.4e-05 | 1.027E-05 | 3.607E-06 | 6.800E-06 | 1.400E-05 |
| 11610 | 581 | G581I | A | 7.9e-06 | 1.5e-05 | 1.5e-05 | 1.263E-05 | 4.099E-06 | 7.900E-06 | 1.500E-05 |
| 11611 | 581 | G581L | A | 9e-06 | 1.3e-05 | 1.4e-05 | 1.200E-05 | 2.646E-06 | 9.000E-06 | 1.400E-05 |
| 11612 | 581 | G581K | A | 8.9e-06 | 1.4e-05 | 1.6e-05 | 1.297E-05 | 3.661E-06 | 8.900E-06 | 1.600E-05 |
| 11613 | 581 | G581M | A | 1.4e-05 | 1.1e-05 | 1.3e-05 | 1.267E-05 | 1.528E-06 | 1.100E-05 | 1.400E-05 |
| 11614 | 581 | G581F | A | 9e-06 | 1.5e-05 | 1.5e-05 | 1.300E-05 | 3.464E-06 | 9.000E-06 | 1.500E-05 |
| 11615 | 581 | G581P | A | 8.8e-06 | 1.5e-05 | 7.9e-06 | 1.057E-05 | 3.866E-06 | 7.900E-06 | 1.500E-05 |
| 11616 | 581 | G581S | A | 1.4e-05 | 1.5e-05 | 9.1e-06 | 1.270E-05 | 3.158E-06 | 9.100E-06 | 1.500E-05 |
| 11617 | 581 | G581T | A | 1.7e-05 | 1.6e-05 | 1.5e-05 | 1.600E-05 | 1.000E-06 | 1.500E-05 | 1.700E-05 |
| 11618 | 581 | G581W | A | 1.6e-05 | 1.5e-05 | 9.9e-06 | 1.363E-05 | 3.272E-06 | 9.900E-06 | 1.600E-05 |
| 11619 | 581 | G581Y | A | 1.4e-05 | 7.7e-06 | 1.1e-05 | 1.090E-05 | 3.151E-06 | 7.700E-06 | 1.400E-05 |
| 11620 | 581 | G581V | A | 1e-05 | 1.1e-05 | 9.5e-06 | 1.017E-05 | 7.638E-07 | 9.500E-06 | 1.100E-05 |
| 11621 | 582 | V582A | A | 1e-05 | 1.2e-05 | 1.6e-05 | 1.267E-05 | 3.055E-06 | 1.000E-05 | 1.600E-05 |
| 11622 | 582 | V582R | A | 1.5e-05 | 1e-05 | 1.5e-05 | 1.333E-05 | 2.887E-06 | 1.000E-05 | 1.500E-05 |
| 11623 | 582 | V582N | A | 1.7e-05 | 1e-05 | 1.4e-05 | 1.367E-05 | 3.512E-06 | 1.000E-05 | 1.700E-05 |
| 11624 | 582 | V582D | A | 1.5e-05 | 1.1e-05 | 1.2e-05 | 1.267E-05 | 2.082E-06 | 1.100E-05 | 1.500E-05 |
| 11625 | 582 | V582C | A | 9.4e-06 | 1.5e-05 | 1.5e-05 | 1.313E-05 | 3.233E-06 | 9.400E-06 | 1.500E-05 |
| 11626 | 582 | V582E | A | 9.3e-06 | 9.6e-06 | 1.5e-05 | 1.130E-05 | 3.208E-06 | 9.300E-06 | 1.500E-05 |
| 11627 | 582 | V582Q | A | 1.5e-05 | 1.4e-05 | 1.3e-05 | 1.400E-05 | 1.000E-06 | 1.300E-05 | 1.500E-05 |
| 11628 | 582 | V582G | A | 1.4e-05 | 1.1e-05 | 1.3e-05 | 1.267E-05 | 1.528E-06 | 1.100E-05 | 1.400E-05 |
| 11629 | 582 | V582H | A | 1.1e-05 | 9.1e-06 | 8.2e-06 | 9.433E-06 | 1.429E-06 | 8.200E-06 | 1.100E-05 |
| 11630 | 582 | V582I | A | 1e-05 | 1.2e-05 | 9.1e-06 | 1.037E-05 | 1.484E-06 | 9.100E-06 | 1.200E-05 |
| 11631 | 582 | V582L | A | 1.2e-05 | 8.7e-06 | 1.2e-05 | 1.090E-05 | 1.905E-06 | 8.700E-06 | 1.200E-05 |
| 11632 | 582 | V582K | A | 1.7e-05 | 1.2e-05 | 1.1e-05 | 1.333E-05 | 3.215E-06 | 1.100E-05 | 1.700E-05 |
| 11633 | 582 | V582M | A | 1.3e-05 | 1.1e-05 | 1.6e-05 | 1.333E-05 | 2.517E-06 | 1.100E-05 | 1.600E-05 |
| 11634 | 582 | V582F | A | 9e-06 | 1e-05 | 1.4e-05 | 1.100E-05 | 2.646E-06 | 9.000E-06 | 1.400E-05 |
| 11635 | 582 | V582P | A | 1.2e-05 | 1.3e-05 | 1.1e-05 | 1.200E-05 | 1.000E-06 | 1.100E-05 | 1.300E-05 |
| 11636 | 582 | V582S | A | 1e-05 | 8.4e-06 | 1.3e-05 | 1.047E-05 | 2.335E-06 | 8.400E-06 | 1.300E-05 |
| 11637 | 582 | V582T | A | 1.5e-05 | 1.5e-05 | 8.6e-06 | 1.287E-05 | 3.695E-06 | 8.600E-06 | 1.500E-05 |
| 11638 | 582 | V582W | A | 8.9e-06 | 1.7e-05 | 1.2e-05 | 1.263E-05 | 4.087E-06 | 8.900E-06 | 1.700E-05 |
| 11639 | 582 | V582Y | A | 1.6e-05 | 1.2e-05 | 9.3e-06 | 1.243E-05 | 3.371E-06 | 9.300E-06 | 1.600E-05 |
| 11640 | 582 | V582V | A | 9.8e-06 | 7.7e-06 | 9.1e-06 | 8.867E-06 | 1.069E-06 | 7.700E-06 | 9.800E-06 |
| 11641 | 583 | K583A | A | 1.3e-05 | 1.5e-05 | 1.3e-05 | 1.367E-05 | 1.155E-06 | 1.300E-05 | 1.500E-05 |
| 11642 | 583 | K583R | A | 1.3e-05 | 1e-05 | 2.7e-05 | 1.667E-05 | 9.074E-06 | 1.000E-05 | 2.700E-05 |
| 11643 | 583 | K583N | A | 9.9e-06 | 1.8e-05 | 8.7e-06 | 1.220E-05 | 5.059E-06 | 8.700E-06 | 1.800E-05 |
| 11644 | 583 | K583D | A | 1.7e-05 | 1.5e-05 | 1.5e-05 | 1.567E-05 | 1.155E-06 | 1.500E-05 | 1.700E-05 |

| | | | | | | | | | | |
|---|---|---|---|---|---|---|---|---|---|---|
| 11645 | 583 | K583C | A | 1.3e-05 | 1.4e-05 | 1.1e-05 | 1.267E-05 | 1.528E-06 | 1.100E-05 | 1.400E-05 |
| 11646 | 583 | K583E | A | 1.6e-05 | 1.6e-05 | 1.5e-05 | 1.567E-05 | 5.774E-07 | 1.500E-05 | 1.600E-05 |
| 11647 | 583 | K583Q | A | 6.9e-06 | 1.2e-05 | 9.2e-06 | 9.367E-06 | 2.554E-06 | 6.900E-06 | 1.200E-05 |
| 11648 | 583 | K583G | A | 1.2e-05 | 1.5e-05 | 1.5e-05 | 1.400E-05 | 1.732E-06 | 1.200E-05 | 1.500E-05 |
| 11649 | 583 | K583H | A | 1.6e-05 | 9.6e-06 | 1.1e-05 | 1.220E-05 | 3.365E-06 | 9.600E-06 | 1.600E-05 |
| 11650 | 583 | K583I | A | 8.4e-06 | 1.5e-05 | 1.7e-05 | 1.347E-05 | 4.500E-06 | 8.400E-06 | 1.700E-05 |
| 11651 | 583 | K583L | A | 9.8e-06 | 1.1e-05 | 1.5e-05 | 1.193E-05 | 2.723E-06 | 9.800E-06 | 1.500E-05 |
| 11652 | 583 | K583K | A | 1.2e-05 | 1.5e-05 | 1.4e-05 | 1.367E-05 | 1.528E-06 | 1.200E-05 | 1.500E-05 |
| 11653 | 583 | K583M | A | 1.6e-05 | 7.5e-06 | 1.5e-05 | 1.283E-05 | 4.646E-06 | 7.500E-06 | 1.600E-05 |
| 11654 | 583 | K583F | A | 1.6e-05 | 1e-05 | 1.1e-05 | 1.233E-05 | 3.215E-06 | 1.000E-05 | 1.600E-05 |
| 11655 | 583 | K583P | A | 1.3e-05 | 1.5e-05 | 1.2e-05 | 1.333E-05 | 1.528E-06 | 1.200E-05 | 1.500E-05 |
| 11656 | 583 | K583S | A | 9.7e-06 | 9.8e-06 | 9.9e-06 | 9.800E-06 | 1.000E-07 | 9.700E-06 | 9.900E-06 |
| 11657 | 583 | K583T | A | 9.7e-06 | 1.5e-05 | 1.4e-05 | 1.290E-05 | 2.816E-06 | 9.700E-06 | 1.500E-05 |
| 11658 | 583 | K583W | A | 1.3e-05 | 1.4e-05 | 1.1e-05 | 1.267E-05 | 1.528E-06 | 1.100E-05 | 1.400E-05 |
| 11659 | 583 | K583Y | A | 1.1e-05 | 1.3e-05 | 1.3e-05 | 1.233E-05 | 1.155E-06 | 1.100E-05 | 1.300E-05 |
| 11660 | 583 | K583V | A | 1.5e-05 | 9.4e-06 | 9.3e-06 | 1.123E-05 | 3.262E-06 | 9.300E-06 | 1.500E-05 |
| 11661 | 584 | P584A | A | 1.4e-05 | 1.5e-05 | 1.4e-05 | 1.433E-05 | 5.774E-07 | 1.400E-05 | 1.500E-05 |
| 11662 | 584 | P584R | A | 1.5e-05 | 9.5e-06 | 1.7e-05 | 1.383E-05 | 3.884E-06 | 9.500E-06 | 1.700E-05 |
| 11663 | 584 | P584N | A | 1.4e-05 | 1.5e-05 | 1.3e-05 | 1.400E-05 | 1.000E-06 | 1.300E-05 | 1.500E-05 |
| 11664 | 584 | P584D | A | 1e-05 | 1e-05 | 1.2e-05 | 1.067E-05 | 1.155E-06 | 1.000E-05 | 1.200E-05 |
| 11665 | 584 | P584C | A | 6.2e-06 | 1.6e-05 | 1.2e-05 | 1.140E-05 | 4.927E-06 | 6.200E-06 | 1.600E-05 |
| 11666 | 584 | P584E | A | 1.5e-05 | 1.1e-05 | 1.6e-05 | 1.400E-05 | 2.646E-06 | 1.100E-05 | 1.600E-05 |
| 11667 | 584 | P584Q | A | 9.9e-06 | 1.4e-05 | 1.1e-05 | 1.163E-05 | 2.122E-06 | 9.900E-06 | 1.400E-05 |
| 11668 | 584 | P584G | A | 8.5e-06 | 1.5e-05 | 1.5e-05 | 1.283E-05 | 3.753E-06 | 8.500E-06 | 1.500E-05 |
| 11669 | 584 | P584H | A | 1.5e-05 | 1.5e-05 | 9.3e-06 | 1.310E-05 | 3.291E-06 | 9.300E-06 | 1.500E-05 |
| 11670 | 584 | P584I | A | 1.5e-05 | 1.7e-05 | 1.3e-05 | 1.500E-05 | 2.000E-06 | 1.300E-05 | 1.700E-05 |
| 11671 | 584 | P584L | A | 1.4e-05 | 1.4e-05 | 8.4e-06 | 1.213E-05 | 3.233E-06 | 8.400E-06 | 1.400E-05 |
| 11672 | 584 | P584K | A | 1e-05 | 1.6e-05 | 1.4e-05 | 1.333E-05 | 3.055E-06 | 1.000E-05 | 1.600E-05 |
| 11673 | 584 | P584M | A | 1.2e-05 | 9.7e-06 | 1.3e-05 | 1.157E-05 | 1.692E-06 | 9.700E-06 | 1.300E-05 |
| 11674 | 584 | P584F | A | 1.2e-05 | 1e-05 | 1.3e-05 | 1.167E-05 | 1.528E-06 | 1.000E-05 | 1.300E-05 |
| 11675 | 584 | P584P | A | 1.1e-05 | 1e-05 | 1.1e-05 | 1.067E-05 | 5.774E-07 | 1.000E-05 | 1.100E-05 |
| 11676 | 584 | P584S | A | 1.6e-05 | 1e-05 | 9.5e-06 | 1.183E-05 | 3.617E-06 | 9.500E-06 | 1.600E-05 |
| 11677 | 584 | P584T | A | 1e-05 | 1.3e-05 | 9.8e-06 | 1.093E-05 | 1.793E-06 | 9.800E-06 | 1.300E-05 |
| 11678 | 584 | P584W | A | 1.5e-05 | 9.2e-06 | 8.6e-06 | 1.093E-05 | 3.535E-06 | 8.600E-06 | 1.500E-05 |
| 11679 | 584 | P584Y | A | 1.4e-05 | 1.5e-05 | 1.4e-05 | 1.433E-05 | 5.774E-07 | 1.400E-05 | 1.500E-05 |
| 11680 | 584 | P584V | A | 9.5e-06 | 1.2e-05 | 1.2e-05 | 1.117E-05 | 1.443E-06 | 9.500E-06 | 1.200E-05 |
| 11681 | 585 | D585A | A | 9.9e-06 | 1.4e-05 | 1.1e-05 | 1.163E-05 | 2.122E-06 | 9.900E-06 | 1.400E-05 |
| 11682 | 585 | D585R | A | 2e-05 | 1.4e-05 | 9.4e-06 | 1.447E-05 | 5.315E-06 | 9.400E-06 | 2.000E-05 |
| 11683 | 585 | D585N | A | 1.6e-05 | 1e-05 | 1.6e-05 | 1.400E-05 | 3.464E-06 | 1.000E-05 | 1.600E-05 |
| 11684 | 585 | D585D | A | 9.1e-06 | 9.9e-06 | 1.1e-05 | 1.000E-05 | 9.539E-07 | 9.100E-06 | 1.100E-05 |
| 11685 | 585 | D585C | A | 1.5e-05 | 1.2e-05 | 1e-05 | 1.233E-05 | 2.517E-06 | 1.000E-05 | 1.500E-05 |
| 11686 | 585 | D585E | A | 1e-05 | 1.5e-05 | 1.5e-05 | 1.333E-05 | 2.887E-06 | 1.000E-05 | 1.500E-05 |
| 11687 | 585 | D585Q | A | 1.7e-05 | 1.4e-05 | 9.6e-06 | 1.353E-05 | 3.722E-06 | 9.600E-06 | 1.700E-05 |
| 11688 | 585 | D585G | A | 1.3e-05 | 1.4e-05 | 1.5e-05 | 1.400E-05 | 1.000E-06 | 1.300E-05 | 1.500E-05 |
| 11689 | 585 | D585H | A | 1.6e-05 | 1.3e-05 | 1.5e-05 | 1.467E-05 | 1.528E-06 | 1.300E-05 | 1.600E-05 |
| 11690 | 585 | D585I | A | 1.4e-05 | 1.3e-05 | 1.4e-05 | 1.367E-05 | 5.774E-07 | 1.300E-05 | 1.400E-05 |
| 11691 | 585 | D585L | A | 1.3e-05 | 1.5e-05 | 1.6e-05 | 1.467E-05 | 1.528E-06 | 1.300E-05 | 1.600E-05 |
| 11692 | 585 | D585K | A | 1e-05 | 1.7e-05 | 1.3e-05 | 1.333E-05 | 3.512E-06 | 1.000E-05 | 1.700E-05 |
| 11693 | 585 | D585M | A | 1.4e-05 | 1.3e-05 | 8.6e-06 | 1.187E-05 | 2.873E-06 | 8.600E-06 | 1.400E-05 |
| 11694 | 585 | D585F | A | 1e-05 | 1e-05 | 1.6e-05 | 1.200E-05 | 3.464E-06 | 1.000E-05 | 1.600E-05 |
| 11695 | 585 | D585P | A | 1e-05 | 1.1e-05 | 1.4e-05 | 1.167E-05 | 2.082E-06 | 1.000E-05 | 1.400E-05 |
| 11696 | 585 | D585S | A | 1.3e-05 | 9.7e-06 | 1.4e-05 | 1.223E-05 | 2.250E-06 | 9.700E-06 | 1.400E-05 |
| 11697 | 585 | D585T | A | 1.3e-05 | 1.4e-05 | 1.1e-05 | 1.267E-05 | 1.528E-06 | 1.100E-05 | 1.400E-05 |
| 11698 | 585 | D585W | A | 1e-05 | 1.3e-05 | 1.1e-05 | 1.133E-05 | 1.528E-06 | 1.000E-05 | 1.300E-05 |
| 11699 | 585 | D585Y | A | 1.3e-05 | 1.6e-05 | 9.9e-06 | 1.297E-05 | 3.050E-06 | 9.900E-06 | 1.600E-05 |
| 11700 | 585 | D585V | A | 1.6e-05 | 1.8e-05 | 1.4e-05 | 1.600E-05 | 2.000E-06 | 1.400E-05 | 1.800E-05 |
| 11701 | 586 | L586A | A | 1.7e-05 | 1e-05 | 9.4e-06 | 1.213E-05 | 4.225E-06 | 9.400E-06 | 1.700E-05 |
| 11702 | 586 | L586R | A | 1.1e-05 | 1.2e-05 | 1.6e-05 | 1.300E-05 | 2.646E-06 | 1.100E-05 | 1.600E-05 |
| 11703 | 586 | L586N | A | 1.6e-05 | 1.1e-05 | 9.5e-06 | 1.217E-05 | 3.403E-06 | 9.500E-06 | 1.600E-05 |
| 11704 | 586 | L586D | A | 7.6e-06 | 9.1e-06 | 1.4e-05 | 1.023E-05 | 3.347E-06 | 7.600E-06 | 1.400E-05 |
| 11705 | 586 | L586C | A | 9.8e-06 | 1.4e-05 | 1.4e-05 | 1.260E-05 | 2.425E-06 | 9.800E-06 | 1.400E-05 |
| 11706 | 586 | L586E | A | 1.1e-05 | 9.7e-06 | 8.8e-06 | 9.833E-06 | 1.106E-06 | 8.800E-06 | 1.100E-05 |
| 11707 | 586 | L586Q | A | 1.1e-05 | 1e-05 | 1.7e-05 | 1.267E-05 | 3.786E-06 | 1.000E-05 | 1.700E-05 |
| 11708 | 586 | L586G | A | 1.5e-05 | 9.6e-06 | 1.4e-05 | 1.287E-05 | 2.873E-06 | 9.600E-06 | 1.500E-05 |
| 11709 | 586 | L586H | A | 1.2e-05 | 1.5e-05 | 8.8e-06 | 1.193E-05 | 3.101E-06 | 8.800E-06 | 1.500E-05 |
| 11710 | 586 | L586I | A | 1.4e-05 | 1e-05 | 9.3e-06 | 1.110E-05 | 2.536E-06 | 9.300E-06 | 1.400E-05 |
| 11711 | 586 | L586L | A | 1.1e-05 | 1.1e-05 | 1.5e-05 | 1.233E-05 | 2.309E-06 | 1.100E-05 | 1.500E-05 |
| 11712 | 586 | L586K | A | 6.9e-06 | 1.3e-05 | 1.4e-05 | 1.130E-05 | 3.843E-06 | 6.900E-06 | 1.400E-05 |
| 11713 | 586 | L586M | A | 1.5e-05 | 1.6e-05 | 1.3e-05 | 1.467E-05 | 1.528E-06 | 1.300E-05 | 1.600E-05 |
| 11714 | 586 | L586F | A | 1e-05 | 1.4e-05 | 1.6e-05 | 1.333E-05 | 3.055E-06 | 1.000E-05 | 1.600E-05 |
| 11715 | 586 | L586P | A | 6.2e-06 | 1.3e-05 | 1.6e-05 | 1.173E-05 | 5.021E-06 | 6.200E-06 | 1.600E-05 |
| 11716 | 586 | L586S | A | 1.4e-05 | 1.5e-05 | 9.6e-06 | 1.287E-05 | 2.873E-06 | 9.600E-06 | 1.500E-05 |
| 11717 | 586 | L586T | A | 1.4e-05 | 1.4e-05 | 1.5e-05 | 1.433E-05 | 5.774E-07 | 1.400E-05 | 1.500E-05 |
| 11718 | 586 | L586W | A | 9.2e-06 | 1.4e-05 | 9.1e-06 | 1.077E-05 | 2.801E-06 | 9.100E-06 | 1.400E-05 |
| 11719 | 586 | L586Y | A | 1.5e-05 | 1.4e-05 | 1.3e-05 | 1.400E-05 | 1.000E-06 | 1.300E-05 | 1.500E-05 |
| 11720 | 586 | L586V | A | 1.5e-05 | 1e-05 | 1.1e-05 | 1.200E-05 | 2.646E-06 | 1.000E-05 | 1.500E-05 |
| 11721 | 587 | S587A | A | 1.2e-05 | 8.4e-06 | 1.5e-05 | 1.180E-05 | 3.305E-06 | 8.400E-06 | 1.500E-05 |
| 11722 | 587 | S587R | A | 1.1e-05 | 1.3e-05 | 8.5e-06 | 1.083E-05 | 2.255E-06 | 8.500E-06 | 1.300E-05 |
| 11723 | 587 | S587N | A | 1.3e-05 | 1.1e-05 | 9.1e-06 | 1.103E-05 | 1.950E-06 | 9.100E-06 | 1.300E-05 |
| 11724 | 587 | S587D | A | 1.8e-05 | 1.6e-05 | 1.4e-05 | 1.600E-05 | 2.000E-06 | 1.400E-05 | 1.800E-05 |
| 11725 | 587 | S587C | A | 1.1e-05 | 1.1e-05 | 1.6e-05 | 1.267E-05 | 2.887E-06 | 1.100E-05 | 1.600E-05 |
| 11726 | 587 | S587E | A | 8.9e-06 | 1.7e-05 | 1.6e-05 | 1.397E-05 | 4.416E-06 | 8.900E-06 | 1.700E-05 |
| 11727 | 587 | S587Q | A | 1.3e-05 | 1.1e-05 | 1.4e-05 | 1.267E-05 | 1.528E-06 | 1.100E-05 | 1.400E-05 |
| 11728 | 587 | S587G | A | 1.1e-05 | 1.4e-05 | 1.1e-05 | 1.200E-05 | 1.732E-06 | 1.100E-05 | 1.400E-05 |
| 11729 | 587 | S587H | A | 1.3e-05 | 1.5e-05 | 1.7e-05 | 1.500E-05 | 2.000E-06 | 1.300E-05 | 1.700E-05 |

| | | | | | | | | | | |
|---|---|---|---|---|---|---|---|---|---|---|
| 11730 | 587 | S587I | A | 9.8e-06 | 1.1e-05 | 1.2e-05 | 1.093E-05 | 1.102E-06 | 9.800E-06 | 1.200E-05 |
| 11731 | 587 | S587L | A | 1.5e-05 | 1.1e-05 | 1.6e-05 | 1.400E-05 | 2.646E-06 | 1.100E-05 | 1.600E-05 |
| 11732 | 587 | S587K | A | 1.6e-05 | 9.3e-06 | 1.4e-05 | 1.310E-05 | 3.439E-06 | 9.300E-06 | 1.600E-05 |
| 11733 | 587 | S587M | A | 1.8e-05 | 1.1e-05 | 1.4e-05 | 1.433E-05 | 3.512E-06 | 1.100E-05 | 1.800E-05 |
| 11734 | 587 | S587F | A | 1.1e-05 | 1.5e-05 | 1.8e-05 | 1.467E-05 | 3.512E-06 | 1.100E-05 | 1.800E-05 |
| 11735 | 587 | S587P | A | 1e-05 | 1.3e-05 | 1.6e-05 | 1.300E-05 | 3.000E-06 | 1.000E-05 | 1.600E-05 |
| 11736 | 587 | S587S | A | 8.7e-06 | 1.4e-05 | 1.4e-05 | 1.223E-05 | 3.060E-06 | 8.700E-06 | 1.400E-05 |
| 11737 | 587 | S587T | A | 1e-05 | 1.2e-05 | 9e-06 | 1.033E-05 | 1.528E-06 | 9.000E-06 | 1.200E-05 |
| 11738 | 587 | S587W | A | 1.4e-05 | 1.1e-05 | 1.6e-05 | 1.367E-05 | 2.517E-06 | 1.100E-05 | 1.600E-05 |
| 11739 | 587 | S587Y | A | 1.5e-05 | 1.5e-05 | 1.8e-05 | 1.600E-05 | 1.732E-06 | 1.500E-05 | 1.800E-05 |
| 11740 | 587 | S587V | A | 1.5e-05 | 1.1e-05 | 1.3e-05 | 1.300E-05 | 2.000E-06 | 1.100E-05 | 1.500E-05 |
| 11741 | 588 | Y588A | A | 1.5e-05 | 1.2e-05 | 1.1e-05 | 1.267E-05 | 2.082E-06 | 1.100E-05 | 1.500E-05 |
| 11742 | 588 | Y588R | A | 9.4e-06 | 1.1e-05 | 1.2e-05 | 1.080E-05 | 1.311E-06 | 9.400E-06 | 1.200E-05 |
| 11743 | 588 | Y588N | A | 1.1e-05 | 9.5e-06 | 1.1e-05 | 1.050E-05 | 8.660E-07 | 9.500E-06 | 1.100E-05 |
| 11744 | 588 | Y588D | A | 1e-05 | 1.5e-05 | 1.2e-05 | 1.233E-05 | 2.517E-06 | 1.000E-05 | 1.500E-05 |
| 11745 | 588 | Y588C | A | 1e-05 | 1.1e-05 | 1.2e-05 | 1.100E-05 | 1.000E-06 | 1.000E-05 | 1.200E-05 |
| 11746 | 588 | Y588E | A | 1.5e-05 | 1.3e-05 | 1.1e-05 | 1.300E-05 | 2.000E-06 | 1.100E-05 | 1.500E-05 |
| 11747 | 588 | Y588Q | A | 1.1e-05 | 1.4e-05 | 1.7e-05 | 1.400E-05 | 3.000E-06 | 1.100E-05 | 1.700E-05 |
| 11748 | 588 | Y588G | A | 1.1e-05 | 9.2e-06 | 1.5e-05 | 1.173E-05 | 2.969E-06 | 9.200E-06 | 1.500E-05 |
| 11749 | 588 | Y588H | A | 7.7e-06 | 1.5e-05 | 1.5e-05 | 1.257E-05 | 4.215E-06 | 7.700E-06 | 1.500E-05 |
| 11750 | 588 | Y588I | A | 7.9e-06 | 1.5e-05 | 1.5e-05 | 1.263E-05 | 4.099E-06 | 7.900E-06 | 1.500E-05 |
| 11751 | 588 | Y588L | A | 1.2e-05 | 1.1e-05 | 1e-05 | 1.100E-05 | 1.000E-06 | 1.000E-05 | 1.200E-05 |
| 11752 | 588 | Y588K | A | 1.2e-05 | 1.5e-05 | 1.4e-05 | 1.367E-05 | 1.528E-06 | 1.200E-05 | 1.500E-05 |
| 11753 | 588 | Y588M | A | 1.2e-05 | 1.4e-05 | 1e-05 | 1.200E-05 | 2.000E-06 | 1.000E-05 | 1.400E-05 |
| 11754 | 588 | Y588F | A | 1.5e-05 | 9.3e-06 | 1.4e-05 | 1.277E-05 | 3.044E-06 | 9.300E-06 | 1.500E-05 |
| 11755 | 588 | Y588P | A | 1.3e-05 | 1.5e-05 | 1.1e-05 | 1.300E-05 | 2.000E-06 | 1.100E-05 | 1.500E-05 |
| 11756 | 588 | Y588S | A | 1.5e-05 | 1.5e-05 | 1.6e-05 | 1.533E-05 | 5.774E-07 | 1.500E-05 | 1.600E-05 |
| 11757 | 588 | Y588T | A | 8.4e-06 | 1e-05 | 1.6e-05 | 1.147E-05 | 4.007E-06 | 8.400E-06 | 1.600E-05 |
| 11758 | 588 | Y588W | A | 9.6e-06 | 1.6e-05 | 9.2e-06 | 1.160E-05 | 3.816E-06 | 9.200E-06 | 1.600E-05 |
| 11759 | 588 | Y588Y | A | 1.7e-05 | 1.5e-05 | 1e-05 | 1.400E-05 | 3.606E-06 | 1.000E-05 | 1.700E-05 |
| 11760 | 588 | Y588V | A | 1.1e-05 | 1.1e-05 | 1e-05 | 1.067E-05 | 5.774E-07 | 1.000E-05 | 1.100E-05 |
| 11761 | 589 | M589A | A | 1.1e-05 | 1.7e-05 | 1.2e-05 | 1.333E-05 | 3.215E-06 | 1.100E-05 | 1.700E-05 |
| 11762 | 589 | M589R | A | 1.5e-05 | 1.1e-05 | 7.4e-06 | 1.113E-05 | 3.802E-06 | 7.400E-06 | 1.500E-05 |
| 11763 | 589 | M589N | A | 9.6e-06 | 9.4e-06 | 9.2e-06 | 9.400E-06 | 2.000E-07 | 9.200E-06 | 9.600E-06 |
| 11764 | 589 | M589D | A | 1.2e-05 | 1e-05 | 1.3e-05 | 1.167E-05 | 1.528E-06 | 1.000E-05 | 1.300E-05 |
| 11765 | 589 | M589C | A | 1.5e-05 | 9.3e-06 | 1.6e-05 | 1.343E-05 | 3.614E-06 | 9.300E-06 | 1.600E-05 |
| 11766 | 589 | M589E | A | 1e-05 | 1e-05 | 1.5e-05 | 1.167E-05 | 2.887E-06 | 1.000E-05 | 1.500E-05 |
| 11767 | 589 | M589Q | A | 1.3e-05 | 1e-05 | 8.4e-06 | 1.047E-05 | 2.335E-06 | 8.400E-06 | 1.300E-05 |
| 11768 | 589 | M589G | A | 1.6e-05 | 1.7e-05 | 1.6e-05 | 1.633E-05 | 5.774E-07 | 1.600E-05 | 1.700E-05 |
| 11769 | 589 | M589H | A | 6.5e-06 | 8.5e-06 | 9.6e-06 | 8.200E-06 | 1.572E-06 | 6.500E-06 | 9.600E-06 |
| 11770 | 589 | M589I | A | 1.6e-05 | 9.9e-06 | 1e-05 | 1.197E-05 | 3.493E-06 | 9.900E-06 | 1.600E-05 |
| 11771 | 589 | M589L | A | 9.9e-06 | 9.5e-06 | 1.2e-05 | 1.047E-05 | 1.343E-06 | 9.500E-06 | 1.200E-05 |
| 11772 | 589 | M589K | A | 9.8e-06 | 1.4e-05 | 1.1e-05 | 1.160E-05 | 2.163E-06 | 9.800E-06 | 1.400E-05 |
| 11773 | 589 | M589M | A | 9.7e-06 | 8.7e-06 | 1.1e-05 | 9.800E-06 | 1.153E-06 | 8.700E-06 | 1.100E-05 |
| 11774 | 589 | M589F | A | 1.3e-05 | 8.9e-06 | 1.4e-05 | 1.197E-05 | 2.702E-06 | 8.900E-06 | 1.400E-05 |
| 11775 | 589 | M589P | A | 1.5e-05 | 1e-05 | 1.5e-05 | 1.333E-05 | 2.887E-06 | 1.000E-05 | 1.500E-05 |
| 11776 | 589 | M589S | A | 8.8e-06 | 1e-05 | 1.2e-05 | 1.027E-05 | 1.617E-06 | 8.800E-06 | 1.200E-05 |
| 11777 | 589 | M589T | A | 1e-05 | 1.4e-05 | 1.4e-05 | 1.267E-05 | 2.309E-06 | 1.000E-05 | 1.400E-05 |
| 11778 | 589 | M589W | A | 1.3e-05 | 1.6e-05 | 2.7e-05 | 1.867E-05 | 7.371E-06 | 1.300E-05 | 2.700E-05 |
| 11779 | 589 | M589Y | A | 1.5e-05 | 1.5e-05 | 8.7e-06 | 1.290E-05 | 3.637E-06 | 8.700E-06 | 1.500E-05 |
| 11780 | 589 | M589V | A | 1.4e-05 | 8.3e-06 | 1.5e-05 | 1.243E-05 | 3.614E-06 | 8.300E-06 | 1.500E-05 |
| 11781 | 590 | P590A | A | 1.5e-05 | 1.7e-05 | 1.3e-05 | 1.500E-05 | 2.000E-06 | 1.300E-05 | 1.700E-05 |
| 11782 | 590 | P590R | A | 1.1e-05 | 1.4e-05 | 9.9e-06 | 1.163E-05 | 2.122E-06 | 9.900E-06 | 1.400E-05 |
| 11783 | 590 | P590N | A | 1.5e-05 | 1.4e-05 | 1.5e-05 | 1.467E-05 | 5.774E-07 | 1.400E-05 | 1.500E-05 |
| 11784 | 590 | P590D | A | 1e-05 | 1.6e-05 | 1.4e-05 | 1.333E-05 | 3.055E-06 | 1.000E-05 | 1.600E-05 |
| 11785 | 590 | P590C | A | 7.8e-06 | 1.5e-05 | 1.3e-05 | 1.193E-05 | 3.717E-06 | 7.800E-06 | 1.500E-05 |
| 11786 | 590 | P590E | A | 9.2e-06 | 1.3e-05 | 1.4e-05 | 1.207E-05 | 2.532E-06 | 9.200E-06 | 1.400E-05 |
| 11787 | 590 | P590Q | A | 1.7e-05 | 1.5e-05 | 1.3e-05 | 1.500E-05 | 2.000E-06 | 1.300E-05 | 1.700E-05 |
| 11788 | 590 | P590G | A | 8.5e-06 | 8.7e-06 | 1.2e-05 | 9.733E-06 | 1.966E-06 | 8.500E-06 | 1.200E-05 |
| 11789 | 590 | P590H | A | 1e-05 | 1.5e-05 | 1.6e-05 | 1.367E-05 | 3.215E-06 | 1.000E-05 | 1.600E-05 |
| 11790 | 590 | P590I | A | 9.7e-06 | 1.3e-05 | 9.3e-06 | 1.067E-05 | 2.031E-06 | 9.300E-06 | 1.300E-05 |
| 11791 | 590 | P590L | A | 1.4e-05 | 1.4e-05 | 9.5e-06 | 1.250E-05 | 2.598E-06 | 9.500E-06 | 1.400E-05 |
| 11792 | 590 | P590K | A | 1.2e-05 | 1.6e-05 | 1.6e-05 | 1.467E-05 | 2.309E-06 | 1.200E-05 | 1.600E-05 |
| 11793 | 590 | P590M | A | 6.2e-06 | 1.3e-05 | 1.3e-05 | 1.073E-05 | 3.926E-06 | 6.200E-06 | 1.300E-05 |
| 11794 | 590 | P590F | A | 1.7e-05 | 1.6e-05 | 9.6e-06 | 1.420E-05 | 4.015E-06 | 9.600E-06 | 1.700E-05 |
| 11795 | 590 | P590P | A | 1.6e-05 | 1.4e-05 | 1.2e-05 | 1.400E-05 | 2.000E-06 | 1.200E-05 | 1.600E-05 |
| 11796 | 590 | P590S | A | 2.1e-05 | 1.3e-05 | 1.4e-05 | 1.600E-05 | 4.359E-06 | 1.300E-05 | 2.100E-05 |
| 11797 | 590 | P590T | A | 1.3e-05 | 1.5e-05 | 1.3e-05 | 1.367E-05 | 1.155E-06 | 1.300E-05 | 1.500E-05 |
| 11798 | 590 | P590W | A | 9.9e-06 | 1.4e-05 | 1.3e-05 | 1.230E-05 | 2.138E-06 | 9.900E-06 | 1.400E-05 |
| 11799 | 590 | P590Y | A | 1.5e-05 | 1.2e-05 | 1.5e-05 | 1.400E-05 | 1.732E-06 | 1.200E-05 | 1.500E-05 |
| 11800 | 590 | P590V | A | 1.4e-05 | 1.6e-05 | 1.6e-05 | 1.533E-05 | 1.155E-06 | 1.400E-05 | 1.600E-05 |
| 11801 | 591 | I591A | A | 1.7e-05 | 9.6e-06 | 1.8e-05 | 1.487E-05 | 4.588E-06 | 9.600E-06 | 1.800E-05 |
| 11802 | 591 | I591R | A | 1.8e-05 | 1.9e-05 | 8.7e-06 | 1.523E-05 | 5.680E-06 | 8.700E-06 | 1.900E-05 |
| 11803 | 591 | I591N | A | 1.5e-05 | 1.4e-05 | 1.1e-05 | 1.333E-05 | 2.082E-06 | 1.100E-05 | 1.500E-05 |
| 11804 | 591 | I591D | A | 1.4e-05 | 1.5e-05 | 1.5e-05 | 1.467E-05 | 5.774E-07 | 1.400E-05 | 1.500E-05 |
| 11805 | 591 | I591C | A | 1.5e-05 | 1.5e-05 | 1.4e-05 | 1.467E-05 | 5.774E-07 | 1.400E-05 | 1.500E-05 |
| 11806 | 591 | I591E | A | 7.4e-06 | 1e-05 | 9.2e-06 | 8.867E-06 | 1.332E-06 | 7.400E-06 | 1.000E-05 |
| 11807 | 591 | I591Q | A | 1e-05 | 9.9e-06 | 1e-05 | 9.967E-06 | 5.774E-08 | 9.900E-06 | 1.000E-05 |
| 11808 | 591 | I591G | A | 1.1e-05 | 1.6e-05 | 1.6e-05 | 1.433E-05 | 2.887E-06 | 1.100E-05 | 1.600E-05 |
| 11809 | 591 | I591H | A | 1.5e-05 | 1.1e-05 | 1.5e-05 | 1.367E-05 | 2.309E-06 | 1.100E-05 | 1.500E-05 |
| 11810 | 591 | I591I | A | 1.1e-05 | 1.2e-05 | 1.4e-05 | 1.233E-05 | 1.528E-06 | 1.100E-05 | 1.400E-05 |
| 11811 | 591 | I591L | A | 1.5e-05 | 1.4e-05 | 1.5e-05 | 1.467E-05 | 5.774E-07 | 1.400E-05 | 1.500E-05 |
| 11812 | 591 | I591K | A | 9.3e-06 | 8.5e-06 | 9.9e-06 | 9.233E-06 | 7.024E-07 | 8.500E-06 | 9.900E-06 |
| 11813 | 591 | I591M | A | 1.1e-05 | 1.5e-05 | 1.5e-05 | 1.367E-05 | 2.309E-06 | 1.100E-05 | 1.500E-05 |
| 11814 | 591 | I591F | A | 1.7e-05 | 1.5e-05 | 9.4e-06 | 1.380E-05 | 3.940E-06 | 9.400E-06 | 1.700E-05 |

| | | | | | | | | | | |
|---|---|---|---|---|---|---|---|---|---|---|
| 11815 | 591 | I591P | A | 1.6e-05 | 1.6e-05 | 1.3e-05 | 1.500E-05 | 1.732E-06 | 1.300E-05 | 1.600E-05 |
| 11816 | 591 | I591S | A | 1.4e-05 | 9.9e-06 | 1.1e-05 | 1.163E-05 | 2.122E-06 | 9.900E-06 | 1.400E-05 |
| 11817 | 591 | I591T | A | 1.1e-05 | 9.9e-06 | 8.1e-06 | 9.667E-06 | 1.464E-06 | 8.100E-06 | 1.100E-05 |
| 11818 | 591 | I591W | A | 1.1e-05 | 8.8e-06 | 1.5e-05 | 1.160E-05 | 3.143E-06 | 8.800E-06 | 1.500E-05 |
| 11819 | 591 | I591Y | A | 1.1e-05 | 1.4e-05 | 1.5e-05 | 1.333E-05 | 2.082E-06 | 1.100E-05 | 1.500E-05 |
| 11820 | 591 | I591V | A | 1.7e-05 | 8.9e-06 | 1.5e-05 | 1.363E-05 | 4.219E-06 | 8.900E-06 | 1.700E-05 |
| 11821 | 592 | W592A | A | 9e-06 | 1.1e-05 | 9.5e-06 | 9.833E-06 | 1.041E-06 | 9.000E-06 | 1.100E-05 |
| 11822 | 592 | W592R | A | 1.1e-05 | 1.5e-05 | 1.6e-05 | 1.400E-05 | 2.646E-06 | 1.100E-05 | 1.600E-05 |
| 11823 | 592 | W592N | A | 9.5e-06 | 1.5e-05 | 1.1e-05 | 1.183E-05 | 2.843E-06 | 9.500E-06 | 1.500E-05 |
| 11824 | 592 | W592D | A | 1.1e-05 | 1.5e-05 | 1e-05 | 1.200E-05 | 2.646E-06 | 1.000E-05 | 1.500E-05 |
| 11825 | 592 | W592C | A | 1.8e-05 | 1.4e-05 | 1.3e-05 | 1.500E-05 | 2.646E-06 | 1.300E-05 | 1.800E-05 |
| 11826 | 592 | W592E | A | 1.5e-05 | 1.7e-05 | 1e-05 | 1.400E-05 | 3.606E-06 | 1.000E-05 | 1.700E-05 |
| 11827 | 592 | W592Q | A | 9.4e-06 | 1.5e-05 | 9.6e-06 | 1.133E-05 | 3.177E-06 | 9.400E-06 | 1.500E-05 |
| 11828 | 592 | W592G | A | 1.1e-05 | 1e-05 | 9.5e-06 | 1.017E-05 | 7.638E-07 | 9.500E-06 | 1.100E-05 |
| 11829 | 592 | W592H | A | 1.5e-05 | 9.3e-06 | 1.7e-05 | 1.377E-05 | 3.995E-06 | 9.300E-06 | 1.700E-05 |
| 11830 | 592 | W592I | A | 1.2e-05 | 1.5e-05 | 1.1e-05 | 1.267E-05 | 2.082E-06 | 1.100E-05 | 1.500E-05 |
| 11831 | 592 | W592L | A | 1.6e-05 | 1.3e-05 | 1e-05 | 1.300E-05 | 3.000E-06 | 1.000E-05 | 1.600E-05 |
| 11832 | 592 | W592K | A | 1.5e-05 | 1.4e-05 | 8.4e-06 | 1.247E-05 | 3.557E-06 | 8.400E-06 | 1.500E-05 |
| 11833 | 592 | W592M | A | 1.6e-05 | 1.4e-05 | 1.2e-05 | 1.400E-05 | 2.000E-06 | 1.200E-05 | 1.600E-05 |
| 11834 | 592 | W592F | A | 8.3e-06 | 1.1e-05 | 1.1e-05 | 1.010E-05 | 1.559E-06 | 8.300E-06 | 1.100E-05 |
| 11835 | 592 | W592P | A | 8.6e-06 | 1.6e-05 | 1.2e-05 | 1.220E-05 | 3.704E-06 | 8.600E-06 | 1.600E-05 |
| 11836 | 592 | W592S | A | 7.5e-06 | 1.6e-05 | 1.2e-05 | 1.183E-05 | 4.252E-06 | 7.500E-06 | 1.600E-05 |
| 11837 | 592 | W592T | A | 8.6e-06 | 9.6e-06 | 9e-06 | 9.067E-06 | 5.033E-07 | 8.600E-06 | 9.600E-06 |
| 11838 | 592 | W592W | A | 1.5e-05 | 7.7e-06 | 1.6e-05 | 1.290E-05 | 4.531E-06 | 7.700E-06 | 1.600E-05 |
| 11839 | 592 | W592Y | A | 1.6e-05 | 1.1e-05 | 1.5e-05 | 1.400E-05 | 2.646E-06 | 1.100E-05 | 1.600E-05 |
| 11840 | 592 | W592V | A | 9.9e-06 | 1e-05 | 1.4e-05 | 1.130E-05 | 2.339E-06 | 9.900E-06 | 1.400E-05 |
| 11841 | 593 | K593A | A | 1.4e-05 | 1.5e-05 | 1e-05 | 1.300E-05 | 2.646E-06 | 1.000E-05 | 1.500E-05 |
| 11842 | 593 | K593R | A | 1.1e-05 | 1.4e-05 | 1.5e-05 | 1.333E-05 | 2.082E-06 | 1.100E-05 | 1.500E-05 |
| 11843 | 593 | K593N | A | 1.5e-05 | 1.6e-05 | 1.6e-05 | 1.567E-05 | 5.774E-07 | 1.500E-05 | 1.600E-05 |
| 11844 | 593 | K593D | A | 9.4e-06 | 1.6e-05 | 1.5e-05 | 1.347E-05 | 3.557E-06 | 9.400E-06 | 1.600E-05 |
| 11845 | 593 | K593C | A | 8.8e-06 | 1.1e-05 | 9.4e-06 | 9.733E-06 | 1.137E-06 | 8.800E-06 | 1.100E-05 |
| 11846 | 593 | K593E | A | 1.6e-05 | 1e-05 | 9.4e-06 | 1.180E-05 | 3.650E-06 | 9.400E-06 | 1.600E-05 |
| 11847 | 593 | K593Q | A | 1.4e-05 | 1.1e-05 | 1.2e-05 | 1.233E-05 | 1.528E-06 | 1.100E-05 | 1.400E-05 |
| 11848 | 593 | K593G | A | 1.6e-05 | 1.4e-05 | 9.1e-06 | 1.303E-05 | 3.550E-06 | 9.100E-06 | 1.600E-05 |
| 11849 | 593 | K593H | A | 1.6e-05 | 9.6e-06 | 7e-06 | 1.087E-05 | 4.632E-06 | 7.000E-06 | 1.600E-05 |
| 11850 | 593 | K593I | A | 8.2e-06 | 9.5e-06 | 1.6e-05 | 1.123E-05 | 4.179E-06 | 8.200E-06 | 1.600E-05 |
| 11851 | 593 | K593L | A | 1e-05 | 1.3e-05 | 1.5e-05 | 1.267E-05 | 2.517E-06 | 1.000E-05 | 1.500E-05 |
| 11852 | 593 | K593K | A | 1.4e-05 | 9.2e-06 | 1.2e-05 | 1.173E-05 | 2.411E-06 | 9.200E-06 | 1.400E-05 |
| 11853 | 593 | K593M | A | 9.7e-06 | 1.5e-05 | 1.5e-05 | 1.323E-05 | 3.060E-06 | 9.700E-06 | 1.500E-05 |
| 11854 | 593 | K593F | A | 1.2e-05 | 1.1e-05 | 1.5e-05 | 1.267E-05 | 2.082E-06 | 1.100E-05 | 1.500E-05 |
| 11855 | 593 | K593P | A | 1.5e-05 | 1.7e-05 | 1.5e-05 | 1.567E-05 | 1.155E-06 | 1.500E-05 | 1.700E-05 |
| 11856 | 593 | K593S | A | 1e-05 | 1.4e-05 | 1.5e-05 | 1.300E-05 | 2.646E-06 | 1.000E-05 | 1.500E-05 |
| 11857 | 593 | K593T | A | 1.1e-05 | 1.6e-05 | 1.3e-05 | 1.333E-05 | 2.517E-06 | 1.100E-05 | 1.600E-05 |
| 11858 | 593 | K593W | A | 1.4e-05 | 1e-05 | 1.4e-05 | 1.267E-05 | 2.309E-06 | 1.000E-05 | 1.400E-05 |
| 11859 | 593 | K593Y | A | 1.5e-05 | 9.7e-06 | 1.3e-05 | 1.257E-05 | 2.676E-06 | 9.700E-06 | 1.500E-05 |
| 11860 | 593 | K593V | A | 1.5e-05 | 1e-05 | 1.4e-05 | 1.300E-05 | 2.646E-06 | 1.000E-05 | 1.500E-05 |
| 11861 | 594 | F594A | A | 1.5e-05 | 1.1e-05 | 1.3e-05 | 1.300E-05 | 2.000E-06 | 1.100E-05 | 1.500E-05 |
| 11862 | 594 | F594R | A | 1.1e-05 | 1.6e-05 | 1.4e-05 | 1.367E-05 | 2.517E-06 | 1.100E-05 | 1.600E-05 |
| 11863 | 594 | F594N | A | 9.1e-06 | 1.1e-05 | 1.6e-05 | 1.203E-05 | 3.564E-06 | 9.100E-06 | 1.600E-05 |
| 11864 | 594 | F594D | A | 1.4e-05 | 6.4e-06 | 1.2e-05 | 1.080E-05 | 3.940E-06 | 6.400E-06 | 1.400E-05 |
| 11865 | 594 | F594C | A | 1.1e-05 | 9.1e-06 | 9.7e-06 | 9.933E-06 | 9.713E-07 | 9.100E-06 | 1.100E-05 |
| 11866 | 594 | F594E | A | 1e-05 | 1.6e-05 | 1.4e-05 | 1.333E-05 | 3.055E-06 | 1.000E-05 | 1.600E-05 |
| 11867 | 594 | F594Q | A | 1.1e-05 | 9.9e-06 | 1.2e-05 | 1.097E-05 | 1.050E-06 | 9.900E-06 | 1.200E-05 |
| 11868 | 594 | F594G | A | 9.1e-06 | 1.7e-05 | 1.6e-05 | 1.403E-05 | 4.302E-06 | 9.100E-06 | 1.700E-05 |
| 11869 | 594 | F594H | A | 1e-05 | 1.5e-05 | 1.2e-05 | 1.233E-05 | 2.517E-06 | 1.000E-05 | 1.500E-05 |
| 11870 | 594 | F594I | A | 1.5e-05 | 1.5e-05 | 9.7e-06 | 1.323E-05 | 3.060E-06 | 9.700E-06 | 1.500E-05 |
| 11871 | 594 | F594L | A | 1.6e-05 | 1.6e-05 | 9.2e-06 | 1.373E-05 | 3.926E-06 | 9.200E-06 | 1.600E-05 |
| 11872 | 594 | F594K | A | 1.5e-05 | 1.4e-05 | 1.6e-05 | 1.500E-05 | 1.000E-06 | 1.400E-05 | 1.600E-05 |
| 11873 | 594 | F594M | A | 1.1e-05 | 1.6e-05 | 1.5e-05 | 1.400E-05 | 2.646E-06 | 1.100E-05 | 1.600E-05 |
| 11874 | 594 | F594F | A | 1.3e-05 | 9.2e-06 | 1e-05 | 1.073E-05 | 2.003E-06 | 9.200E-06 | 1.300E-05 |
| 11875 | 594 | F594P | A | 1.7e-05 | 1.6e-05 | 1e-05 | 1.433E-05 | 3.786E-06 | 1.000E-05 | 1.700E-05 |
| 11876 | 594 | F594S | A | 1.2e-05 | 9.9e-06 | 1.4e-05 | 1.197E-05 | 2.050E-06 | 9.900E-06 | 1.400E-05 |
| 11877 | 594 | F594T | A | 1.5e-05 | 1e-05 | 1.5e-05 | 1.333E-05 | 2.887E-06 | 1.000E-05 | 1.500E-05 |
| 11878 | 594 | F594W | A | 1.3e-05 | 9.5e-06 | 1.3e-05 | 1.183E-05 | 2.021E-06 | 9.500E-06 | 1.300E-05 |
| 11879 | 594 | F594Y | A | 1.6e-05 | 1.7e-05 | 1.6e-05 | 1.633E-05 | 5.774E-07 | 1.600E-05 | 1.700E-05 |
| 11880 | 594 | F594V | A | 1e-05 | 1.4e-05 | 1.2e-05 | 1.200E-05 | 2.000E-06 | 1.000E-05 | 1.400E-05 |
| 11881 | 595 | P595A | A | 1.4e-05 | 1e-05 | 1.1e-05 | 1.167E-05 | 2.082E-06 | 1.000E-05 | 1.400E-05 |
| 11882 | 595 | P595R | A | 1.4e-05 | 1.6e-05 | 1.1e-05 | 1.367E-05 | 2.517E-06 | 1.100E-05 | 1.600E-05 |
| 11883 | 595 | P595N | A | 1.1e-05 | 9.5e-06 | 9.1e-06 | 9.867E-06 | 1.002E-06 | 9.100E-06 | 1.100E-05 |
| 11884 | 595 | P595D | A | 1.1e-05 | 1.6e-05 | 1.3e-05 | 1.333E-05 | 2.517E-06 | 1.100E-05 | 1.600E-05 |
| 11885 | 595 | P595C | A | 1.6e-05 | 1.2e-05 | 8.5e-06 | 1.217E-05 | 3.753E-06 | 8.500E-06 | 1.600E-05 |
| 11886 | 595 | P595E | A | 1.4e-05 | 1.3e-05 | 1.2e-05 | 1.300E-05 | 1.000E-06 | 1.200E-05 | 1.400E-05 |
| 11887 | 595 | P595Q | A | 1.1e-05 | 1.1e-05 | 9.5e-06 | 1.050E-05 | 8.660E-07 | 9.500E-06 | 1.100E-05 |
| 11888 | 595 | P595G | A | 8.8e-06 | 1.5e-05 | 1.3e-05 | 1.227E-05 | 3.164E-06 | 8.800E-06 | 1.500E-05 |
| 11889 | 595 | P595H | A | 1.5e-05 | 1.3e-05 | 1.4e-05 | 1.400E-05 | 1.000E-06 | 1.300E-05 | 1.500E-05 |
| 11890 | 595 | P595I | A | 1.6e-05 | 1.7e-05 | 8.5e-06 | 1.383E-05 | 4.646E-06 | 8.500E-06 | 1.700E-05 |
| 11891 | 595 | P595L | A | 1.3e-05 | 1.7e-05 | 1.1e-05 | 1.367E-05 | 3.055E-06 | 1.100E-05 | 1.700E-05 |
| 11892 | 595 | P595K | A | 1.5e-05 | 8.5e-06 | 1e-05 | 1.117E-05 | 3.403E-06 | 8.500E-06 | 1.500E-05 |
| 11893 | 595 | P595M | A | 9.8e-06 | 1.5e-05 | 1e-05 | 1.160E-05 | 2.946E-06 | 9.800E-06 | 1.500E-05 |
| 11894 | 595 | P595F | A | 1.4e-05 | 1.3e-05 | 1.3e-05 | 1.333E-05 | 5.774E-07 | 1.300E-05 | 1.400E-05 |
| 11895 | 595 | P595P | A | 1.5e-05 | 1.2e-05 | 9.8e-06 | 1.227E-05 | 2.610E-06 | 9.800E-06 | 1.500E-05 |
| 11896 | 595 | P595S | A | 1.4e-05 | 8.7e-06 | 1.6e-05 | 1.290E-05 | 3.772E-06 | 8.700E-06 | 1.600E-05 |
| 11897 | 595 | P595T | A | 1.4e-05 | 9.6e-06 | 1.5e-05 | 1.287E-05 | 2.873E-06 | 9.600E-06 | 1.500E-05 |
| 11898 | 595 | P595W | A | 1.1e-05 | 1.5e-05 | 1.5e-05 | 1.367E-05 | 2.309E-06 | 1.100E-05 | 1.500E-05 |
| 11899 | 595 | P595Y | A | 1.4e-05 | 9.7e-06 | 1.5e-05 | 1.290E-05 | 2.816E-06 | 9.700E-06 | 1.500E-05 |

| | | | | | | | | | | |
|---|---|---|---|---|---|---|---|---|---|---|
| 11900 | 595 | P595V | A | 1.6e-05 | 1.4e-05 | 1.4e-05 | 1.467E-05 | 1.155E-06 | 1.400E-05 | 1.600E-05 |
| 11901 | 596 | D596A | A | 1e-05 | 1.3e-05 | 1.1e-05 | 1.133E-05 | 1.528E-06 | 1.000E-05 | 1.300E-05 |
| 11902 | 596 | D596R | A | 7.9e-06 | 1e-05 | 1.5e-05 | 1.097E-05 | 3.647E-06 | 7.900E-06 | 1.500E-05 |
| 11903 | 596 | D596N | A | 7.6e-06 | 1.5e-05 | 1.1e-05 | 1.120E-05 | 3.704E-06 | 7.600E-06 | 1.500E-05 |
| 11904 | 596 | D596D | A | 9.8e-06 | 1.6e-05 | 1e-05 | 1.193E-05 | 3.523E-06 | 9.800E-06 | 1.600E-05 |
| 11905 | 596 | D596C | A | 1.4e-05 | 1.6e-05 | 9.7e-06 | 1.323E-05 | 3.219E-06 | 9.700E-06 | 1.600E-05 |
| 11906 | 596 | D596E | A | 1.4e-05 | 1.4e-05 | 9.3e-06 | 1.243E-05 | 2.714E-06 | 9.300E-06 | 1.400E-05 |
| 11907 | 596 | D596Q | A | 9.3e-06 | 1.6e-05 | 1.1e-05 | 1.210E-05 | 3.483E-06 | 9.300E-06 | 1.600E-05 |
| 11908 | 596 | D596G | A | 1.1e-05 | 1.2e-05 | 1.4e-05 | 1.233E-05 | 1.528E-06 | 1.100E-05 | 1.400E-05 |
| 11909 | 596 | D596H | A | 1.5e-05 | 1.1e-05 | 9.9e-06 | 1.197E-05 | 2.684E-06 | 9.900E-06 | 1.500E-05 |
| 11910 | 596 | D596I | A | 8.8e-06 | 9.5e-06 | 1.1e-05 | 9.767E-06 | 1.124E-06 | 8.800E-06 | 1.100E-05 |
| 11911 | 596 | D596L | A | 1.6e-05 | 1.7e-05 | 1e-05 | 1.433E-05 | 3.786E-06 | 1.000E-05 | 1.700E-05 |
| 11912 | 596 | D596K | A | 1.4e-05 | 1.1e-05 | 1.3e-05 | 1.267E-05 | 1.528E-06 | 1.100E-05 | 1.400E-05 |
| 11913 | 596 | D596M | A | 1.1e-05 | 8.5e-06 | 9e-06 | 9.500E-06 | 1.323E-06 | 8.500E-06 | 1.100E-05 |
| 11914 | 596 | D596F | A | 8.3e-06 | 1.2e-05 | 1.2e-05 | 1.077E-05 | 2.136E-06 | 8.300E-06 | 1.200E-05 |
| 11915 | 596 | D596P | A | 1e-05 | 1.8e-05 | 1.1e-05 | 1.300E-05 | 4.359E-06 | 1.000E-05 | 1.800E-05 |
| 11916 | 596 | D596S | A | 1.5e-05 | 1.1e-05 | 1.3e-05 | 1.300E-05 | 2.000E-06 | 1.100E-05 | 1.500E-05 |
| 11917 | 596 | D596T | A | 1.5e-05 | 1e-05 | 8.2e-06 | 1.107E-05 | 3.523E-06 | 8.200E-06 | 1.500E-05 |
| 11918 | 596 | D596W | A | 6.4e-06 | 1.7e-05 | 1.6e-05 | 1.313E-05 | 5.853E-06 | 6.400E-06 | 1.700E-05 |
| 11919 | 596 | D596Y | A | 1.4e-05 | 1e-05 | 1.6e-05 | 1.333E-05 | 3.055E-06 | 1.000E-05 | 1.600E-05 |
| 11920 | 596 | D596V | A | 1.5e-05 | 9.7e-06 | 1.5e-05 | 1.323E-05 | 3.060E-06 | 9.700E-06 | 1.500E-05 |
| 11921 | 597 | E597A | A | 1.6e-05 | 1.4e-05 | 1.6e-05 | 1.533E-05 | 1.155E-06 | 1.400E-05 | 1.600E-05 |
| 11922 | 597 | E597R | A | 1.5e-05 | 1.5e-05 | 1.1e-05 | 1.367E-05 | 2.309E-06 | 1.100E-05 | 1.500E-05 |
| 11923 | 597 | E597N | A | 1.4e-05 | 1e-05 | 1.7e-05 | 1.367E-05 | 3.512E-06 | 1.000E-05 | 1.700E-05 |
| 11924 | 597 | E597D | A | 1.3e-05 | 1e-05 | 1.6e-05 | 1.300E-05 | 3.000E-06 | 1.000E-05 | 1.600E-05 |
| 11925 | 597 | E597C | A | 5.5e-06 | 9.4e-06 | 9.1e-06 | 8.000E-06 | 2.170E-06 | 5.500E-06 | 9.400E-06 |
| 11926 | 597 | E597E | A | 1.4e-05 | 1.5e-05 | 7.7e-06 | 1.223E-05 | 3.958E-06 | 7.700E-06 | 1.500E-05 |
| 11927 | 597 | E597Q | A | 1.5e-05 | 9.3e-06 | 8.3e-06 | 1.087E-05 | 3.614E-06 | 8.300E-06 | 1.500E-05 |
| 11928 | 597 | E597G | A | 8.2e-06 | 1.4e-05 | 1.1e-05 | 1.107E-05 | 2.901E-06 | 8.200E-06 | 1.400E-05 |
| 11929 | 597 | E597H | A | 1.5e-05 | 8.9e-06 | 1.5e-05 | 1.297E-05 | 3.522E-06 | 8.900E-06 | 1.500E-05 |
| 11930 | 597 | E597I | A | 1.3e-05 | 1.2e-05 | 1.1e-05 | 1.200E-05 | 1.000E-06 | 1.100E-05 | 1.300E-05 |
| 11931 | 597 | E597L | A | 8.6e-06 | 1.4e-05 | 8.8e-06 | 1.047E-05 | 3.062E-06 | 8.600E-06 | 1.400E-05 |
| 11932 | 597 | E597K | A | 1e-05 | 1e-05 | 1.5e-05 | 1.167E-05 | 2.887E-06 | 1.000E-05 | 1.500E-05 |
| 11933 | 597 | E597M | A | 7.9e-06 | 1.5e-05 | 8e-06 | 1.030E-05 | 4.071E-06 | 7.900E-06 | 1.500E-05 |
| 11934 | 597 | E597F | A | 1.4e-05 | 1.2e-05 | 1.1e-05 | 1.233E-05 | 1.528E-06 | 1.100E-05 | 1.400E-05 |
| 11935 | 597 | E597P | A | 1.6e-05 | 1.2e-05 | 1.4e-05 | 1.400E-05 | 2.000E-06 | 1.200E-05 | 1.600E-05 |
| 11936 | 597 | E597S | A | 1.5e-05 | 9.9e-06 | 1.6e-05 | 1.363E-05 | 3.272E-06 | 9.900E-06 | 1.600E-05 |
| 11937 | 597 | E597T | A | 1.6e-05 | 1.6e-05 | 1.1e-05 | 1.433E-05 | 2.887E-06 | 1.100E-05 | 1.600E-05 |
| 11938 | 597 | E597W | A | 1.1e-05 | 1.4e-05 | 1e-05 | 1.167E-05 | 2.082E-06 | 1.000E-05 | 1.400E-05 |
| 11939 | 597 | E597Y | A | 1.4e-05 | 1.3e-05 | 1.4e-05 | 1.367E-05 | 5.774E-07 | 1.300E-05 | 1.400E-05 |
| 11940 | 597 | E597V | A | 1.6e-05 | 1.6e-05 | 1.7e-05 | 1.633E-05 | 5.774E-07 | 1.600E-05 | 1.700E-05 |
| 11941 | 598 | E598A | A | 1.1e-05 | 1.6e-05 | 1.5e-05 | 1.400E-05 | 2.646E-06 | 1.100E-05 | 1.600E-05 |
| 11942 | 598 | E598R | A | 1.3e-05 | 1.4e-05 | 1.5e-05 | 1.400E-05 | 1.000E-06 | 1.300E-05 | 1.500E-05 |
| 11943 | 598 | E598N | A | 1.5e-05 | 1.4e-05 | 1.4e-05 | 1.433E-05 | 5.774E-07 | 1.400E-05 | 1.500E-05 |
| 11944 | 598 | E598D | A | 1.4e-05 | 8.4e-06 | 1.6e-05 | 1.280E-05 | 3.940E-06 | 8.400E-06 | 1.600E-05 |
| 11945 | 598 | E598C | A | 9.7e-06 | 8.4e-06 | 1.1e-05 | 9.700E-06 | 1.300E-06 | 8.400E-06 | 1.100E-05 |
| 11946 | 598 | E598E | A | 1.5e-05 | 1.1e-05 | 1.1e-05 | 1.233E-05 | 2.309E-06 | 1.100E-05 | 1.500E-05 |
| 11947 | 598 | E598Q | A | 1.5e-05 | 7.9e-06 | 9.9e-06 | 1.093E-05 | 3.661E-06 | 7.900E-06 | 1.500E-05 |
| 11948 | 598 | E598G | A | 1.2e-05 | 1.2e-05 | 1.1e-05 | 1.167E-05 | 5.774E-07 | 1.100E-05 | 1.200E-05 |
| 11949 | 598 | E598H | A | 1.6e-05 | 1.1e-05 | 1e-05 | 1.233E-05 | 3.215E-06 | 1.000E-05 | 1.600E-05 |
| 11950 | 598 | E598I | A | 9e-06 | 8.8e-06 | 1.4e-05 | 1.060E-05 | 2.946E-06 | 8.800E-06 | 1.400E-05 |
| 11951 | 598 | E598L | A | 1e-05 | 1.5e-05 | 1.8e-05 | 1.433E-05 | 4.041E-06 | 1.000E-05 | 1.800E-05 |
| 11952 | 598 | E598K | A | 1e-05 | 1.6e-05 | 1.3e-05 | 1.300E-05 | 3.000E-06 | 1.000E-05 | 1.600E-05 |
| 11953 | 598 | E598M | A | 1.5e-05 | 1.6e-05 | 1.1e-05 | 1.400E-05 | 2.646E-06 | 1.100E-05 | 1.600E-05 |
| 11954 | 598 | E598F | A | 9.7e-06 | 1.3e-05 | 1.5e-05 | 1.257E-05 | 2.676E-06 | 9.700E-06 | 1.500E-05 |
| 11955 | 598 | E598P | A | 1.6e-05 | 1.5e-05 | 8.1e-06 | 1.303E-05 | 4.302E-06 | 8.100E-06 | 1.600E-05 |
| 11956 | 598 | E598S | A | 9.1e-06 | 1.4e-05 | 1.4e-05 | 1.237E-05 | 2.829E-06 | 9.100E-06 | 1.400E-05 |
| 11957 | 598 | E598T | A | 1.4e-05 | 1.4e-05 | 1.3e-05 | 1.367E-05 | 5.774E-07 | 1.300E-05 | 1.400E-05 |
| 11958 | 598 | E598W | A | 1.1e-05 | 1.5e-05 | 9.5e-06 | 1.183E-05 | 2.843E-06 | 9.500E-06 | 1.500E-05 |
| 11959 | 598 | E598Y | A | 1.5e-05 | 1.5e-05 | 9.9e-06 | 1.330E-05 | 2.944E-06 | 9.900E-06 | 1.500E-05 |
| 11960 | 598 | E598V | A | 1.8e-05 | 1.7e-05 | 1.5e-05 | 1.667E-05 | 1.528E-06 | 1.500E-05 | 1.800E-05 |
| 11961 | 599 | G599A | A | 1.6e-05 | 1.4e-05 | 1e-05 | 1.333E-05 | 3.055E-06 | 1.000E-05 | 1.600E-05 |
| 11962 | 599 | G599R | A | 1.1e-05 | 1e-05 | 1.5e-05 | 1.200E-05 | 2.646E-06 | 1.000E-05 | 1.500E-05 |
| 11963 | 599 | G599N | A | 1.5e-05 | 9.4e-06 | 1.2e-05 | 1.213E-05 | 2.802E-06 | 9.400E-06 | 1.500E-05 |
| 11964 | 599 | G599D | A | 1.3e-05 | 1.2e-05 | 1.1e-05 | 1.200E-05 | 1.000E-06 | 1.100E-05 | 1.300E-05 |
| 11965 | 599 | G599C | A | 1.4e-05 | 9.6e-06 | 1.5e-05 | 1.287E-05 | 2.873E-06 | 9.600E-06 | 1.500E-05 |
| 11966 | 599 | G599E | A | 1.5e-05 | 1.2e-05 | 1e-05 | 1.233E-05 | 2.517E-06 | 1.000E-05 | 1.500E-05 |
| 11967 | 599 | G599Q | A | 1.7e-05 | 1.1e-05 | 1.7e-05 | 1.500E-05 | 3.464E-06 | 1.100E-05 | 1.700E-05 |
| 11968 | 599 | G599G | A | 1.6e-05 | 1.5e-05 | 1e-05 | 1.367E-05 | 3.215E-06 | 1.000E-05 | 1.600E-05 |
| 11969 | 599 | G599H | A | 8.2e-06 | 1.1e-05 | 1.3e-05 | 1.073E-05 | 2.411E-06 | 8.200E-06 | 1.300E-05 |
| 11970 | 599 | G599I | A | 9.4e-06 | 1e-05 | 8.1e-06 | 9.167E-06 | 9.713E-07 | 8.100E-06 | 1.000E-05 |
| 11971 | 599 | G599L | A | 1.4e-05 | 1.3e-05 | 1.2e-05 | 1.300E-05 | 1.000E-06 | 1.200E-05 | 1.400E-05 |
| 11972 | 599 | G599K | A | 1.1e-05 | 1e-05 | 9.4e-06 | 1.013E-05 | 8.083E-07 | 9.400E-06 | 1.100E-05 |
| 11973 | 599 | G599M | A | 1e-05 | 1.6e-05 | 1.7e-05 | 1.433E-05 | 3.786E-06 | 1.000E-05 | 1.700E-05 |
| 11974 | 599 | G599F | A | 9.5e-06 | 1.1e-05 | 1.2e-05 | 1.083E-05 | 1.258E-06 | 9.500E-06 | 1.200E-05 |
| 11975 | 599 | G599P | A | 1.4e-05 | 8.8e-06 | 1.5e-05 | 1.260E-05 | 3.329E-06 | 8.800E-06 | 1.500E-05 |
| 11976 | 599 | G599S | A | 9.5e-06 | 8.4e-06 | 1.5e-05 | 1.097E-05 | 3.536E-06 | 8.400E-06 | 1.500E-05 |
| 11977 | 599 | G599T | A | 1.3e-05 | 1.6e-05 | 1.3e-05 | 1.400E-05 | 1.732E-06 | 1.300E-05 | 1.600E-05 |
| 11978 | 599 | G599W | A | 1.1e-05 | 1.6e-05 | 9.3e-06 | 1.210E-05 | 3.483E-06 | 9.300E-06 | 1.600E-05 |
| 11979 | 599 | G599Y | A | 1.1e-05 | 1.3e-05 | 1.2e-05 | 1.200E-05 | 1.000E-06 | 1.100E-05 | 1.300E-05 |
| 11980 | 599 | G599V | A | 1.6e-05 | 1.3e-05 | 1.2e-05 | 1.367E-05 | 2.082E-06 | 1.200E-05 | 1.600E-05 |
| 11981 | 600 | A600A | A | 1.5e-05 | 1.7e-05 | 9.1e-06 | 1.370E-05 | 4.107E-06 | 9.100E-06 | 1.700E-05 |
| 11982 | 600 | A600R | A | 7.7e-06 | 9.6e-06 | 1.4e-05 | 1.043E-05 | 3.232E-06 | 7.700E-06 | 1.400E-05 |
| 11983 | 600 | A600N | A | 1.6e-05 | 1.1e-05 | 9.1e-06 | 1.203E-05 | 3.564E-06 | 9.100E-06 | 1.600E-05 |
| 11984 | 600 | A600D | A | 1.7e-05 | 9e-06 | 1e-05 | 1.200E-05 | 4.359E-06 | 9.000E-06 | 1.700E-05 |

| | | | | | | | | | | |
|---|---|---|---|---|---|---|---|---|---|---|
| 11985 | 600 | A600C | A | 1.3e-05 | 9.6e-06 | 8.5e-06 | 1.037E-05 | 2.346E-06 | 8.500E-06 | 1.300E-05 |
| 11986 | 600 | A600E | A | 1.3e-05 | 1.6e-05 | 9.2e-06 | 1.273E-05 | 3.408E-06 | 9.200E-06 | 1.600E-05 |
| 11987 | 600 | A600Q | A | 1.6e-05 | 1.5e-05 | 1.4e-05 | 1.500E-05 | 1.000E-06 | 1.400E-05 | 1.600E-05 |
| 11988 | 600 | A600G | A | 1.1e-05 | 1.1e-05 | 1.1e-05 | 1.100E-05 | 2.075E-21 | 1.100E-05 | 1.100E-05 |
| 11989 | 600 | A600H | A | 1.2e-05 | 1.3e-05 | 1.2e-05 | 1.233E-05 | 5.774E-07 | 1.200E-05 | 1.300E-05 |
| 11990 | 600 | A600I | A | 1.3e-05 | 1e-05 | 1.1e-05 | 1.133E-05 | 1.528E-06 | 1.000E-05 | 1.300E-05 |
| 11991 | 600 | A600L | A | 9.1e-06 | 1e-05 | 1.4e-05 | 1.103E-05 | 2.608E-06 | 9.100E-06 | 1.400E-05 |
| 11992 | 600 | A600K | A | 1.5e-05 | 1.1e-05 | 1e-05 | 1.200E-05 | 2.646E-06 | 1.000E-05 | 1.500E-05 |
| 11993 | 600 | A600M | A | 1.1e-05 | 1e-05 | 1.6e-05 | 1.233E-05 | 3.215E-06 | 1.000E-05 | 1.600E-05 |
| 11994 | 600 | A600F | A | 1.4e-05 | 1.7e-05 | 1.3e-05 | 1.467E-05 | 2.082E-06 | 1.300E-05 | 1.700E-05 |
| 11995 | 600 | A600P | A | 1.1e-05 | 1.4e-05 | 1.4e-05 | 1.300E-05 | 1.732E-06 | 1.100E-05 | 1.400E-05 |
| 11996 | 600 | A600S | A | 9.5e-06 | 1.5e-05 | 1e-05 | 1.150E-05 | 3.041E-06 | 9.500E-06 | 1.500E-05 |
| 11997 | 600 | A600T | A | 9.1e-06 | 8.8e-06 | 1.5e-05 | 1.097E-05 | 3.496E-06 | 8.800E-06 | 1.500E-05 |
| 11998 | 600 | A600W | A | 1.5e-05 | 1.5e-05 | 1.2e-05 | 1.400E-05 | 1.732E-06 | 1.200E-05 | 1.500E-05 |
| 11999 | 600 | A600Y | A | 1e-05 | 1.5e-05 | 1.1e-05 | 1.200E-05 | 2.646E-06 | 1.000E-05 | 1.500E-05 |
| 12000 | 600 | A600V | A | 9.2e-06 | 1.5e-05 | 1.5e-05 | 1.307E-05 | 3.349E-06 | 9.200E-06 | 1.500E-05 |
| 12001 | 601 | C601A | A | 1e-05 | 1.5e-05 | 9.5e-06 | 1.150E-05 | 3.041E-06 | 9.500E-06 | 1.500E-05 |
| 12002 | 601 | C601R | A | 9e-06 | 1e-05 | 9.6e-06 | 9.533E-06 | 5.033E-07 | 9.000E-06 | 1.000E-05 |
| 12003 | 601 | C601N | A | 1.2e-05 | 9.7e-06 | 1.6e-05 | 1.257E-05 | 3.188E-06 | 9.700E-06 | 1.600E-05 |
| 12004 | 601 | C601D | A | 1.2e-05 | 1.5e-05 | 1.3e-05 | 1.333E-05 | 1.528E-06 | 1.200E-05 | 1.500E-05 |
| 12005 | 601 | C601C | A | 1.1e-05 | 1e-05 | 8.7e-06 | 9.900E-06 | 1.153E-06 | 8.700E-06 | 1.100E-05 |
| 12006 | 601 | C601E | A | 1.7e-05 | 1.5e-05 | 9.7e-06 | 1.390E-05 | 3.772E-06 | 9.700E-06 | 1.700E-05 |
| 12007 | 601 | C601Q | A | 1.7e-05 | 1.1e-05 | 9.1e-06 | 1.237E-05 | 4.124E-06 | 9.100E-06 | 1.700E-05 |
| 12008 | 601 | C601G | A | 1.4e-05 | 1.1e-05 | 1.1e-05 | 1.200E-05 | 1.732E-06 | 1.100E-05 | 1.400E-05 |
| 12009 | 601 | C601H | A | 1.1e-05 | 1e-05 | 1.5e-05 | 1.200E-05 | 2.646E-06 | 1.000E-05 | 1.500E-05 |
| 12010 | 601 | C601I | A | 1.4e-05 | 1.1e-05 | 1.5e-05 | 1.333E-05 | 2.082E-06 | 1.100E-05 | 1.500E-05 |
| 12011 | 601 | C601L | A | 1.1e-05 | 1.5e-05 | 1.5e-05 | 1.367E-05 | 2.309E-06 | 1.100E-05 | 1.500E-05 |
| 12012 | 601 | C601K | A | 9.5e-06 | 1e-05 | 9.3e-06 | 9.600E-06 | 3.606E-07 | 9.300E-06 | 1.000E-05 |
| 12013 | 601 | C601M | A | 1.5e-05 | 1.5e-05 | 1.4e-05 | 1.467E-05 | 5.774E-07 | 1.400E-05 | 1.500E-05 |
| 12014 | 601 | C601F | A | 1.5e-05 | 1.4e-05 | 1.5e-05 | 1.467E-05 | 5.774E-07 | 1.400E-05 | 1.500E-05 |
| 12015 | 601 | C601P | A | 9.2e-06 | 1.5e-05 | 1.3e-05 | 1.240E-05 | 2.946E-06 | 9.200E-06 | 1.500E-05 |
| 12016 | 601 | C601S | A | 1.2e-05 | 1.5e-05 | 1.6e-05 | 1.433E-05 | 2.082E-06 | 1.200E-05 | 1.600E-05 |
| 12017 | 601 | C601T | A | 1.2e-05 | 1.1e-05 | 1.2e-05 | 1.167E-05 | 5.774E-07 | 1.100E-05 | 1.200E-05 |
| 12018 | 601 | C601W | A | 1.5e-05 | 1.5e-05 | 1e-05 | 1.333E-05 | 2.887E-06 | 1.000E-05 | 1.500E-05 |
| 12019 | 601 | C601Y | A | 1.1e-05 | 1.5e-05 | 1.5e-05 | 1.367E-05 | 2.309E-06 | 1.100E-05 | 1.500E-05 |
| 12020 | 601 | C601V | A | 1.5e-05 | 1e-05 | 8.7e-06 | 1.123E-05 | 3.326E-06 | 8.700E-06 | 1.500E-05 |
| 12021 | 602 | Q602A | A | 1.6e-05 | 1.1e-05 | 8.4e-06 | 1.180E-05 | 3.863E-06 | 8.400E-06 | 1.600E-05 |
| 12022 | 602 | Q602R | A | 1.6e-05 | 1.8e-05 | 1.1e-05 | 1.500E-05 | 3.606E-06 | 1.100E-05 | 1.800E-05 |
| 12023 | 602 | Q602N | A | 1.5e-05 | 9.6e-06 | 1.5e-05 | 1.320E-05 | 3.118E-06 | 9.600E-06 | 1.500E-05 |
| 12024 | 602 | Q602D | A | 1.5e-05 | 9.8e-06 | 1.3e-05 | 1.260E-05 | 2.623E-06 | 9.800E-06 | 1.500E-05 |
| 12025 | 602 | Q602C | A | 1.1e-05 | 1.1e-05 | 1.7e-05 | 1.300E-05 | 3.464E-06 | 1.100E-05 | 1.700E-05 |
| 12026 | 602 | Q602E | A | 7.6e-06 | 1.6e-05 | 1.1e-05 | 1.153E-05 | 4.225E-06 | 7.600E-06 | 1.600E-05 |
| 12027 | 602 | Q602Q | A | 9.3e-06 | 1.4e-05 | 1.7e-05 | 1.343E-05 | 3.881E-06 | 9.300E-06 | 1.700E-05 |
| 12028 | 602 | Q602G | A | 1.8e-05 | 1.5e-05 | 1.2e-05 | 1.500E-05 | 3.000E-06 | 1.200E-05 | 1.800E-05 |
| 12029 | 602 | Q602H | A | 1.4e-05 | 1.6e-05 | 1.1e-05 | 1.367E-05 | 2.517E-06 | 1.100E-05 | 1.600E-05 |
| 12030 | 602 | Q602I | A | 1.7e-05 | 8.9e-06 | 2.8e-05 | 1.797E-05 | 9.587E-06 | 8.900E-06 | 2.800E-05 |
| 12031 | 602 | Q602L | A | 1.5e-05 | 1e-05 | 1.3e-05 | 1.267E-05 | 2.517E-06 | 1.000E-05 | 1.500E-05 |
| 12032 | 602 | Q602K | A | 9.5e-06 | 1.7e-05 | 1.6e-05 | 1.417E-05 | 4.072E-06 | 9.500E-06 | 1.700E-05 |
| 12033 | 602 | Q602M | A | 9.1e-06 | 1.2e-05 | 8.6e-06 | 9.900E-06 | 1.836E-06 | 8.600E-06 | 1.200E-05 |
| 12034 | 602 | Q602F | A | 1.5e-05 | 1.5e-05 | 1.2e-05 | 1.400E-05 | 1.732E-06 | 1.200E-05 | 1.500E-05 |
| 12035 | 602 | Q602P | A | 1.3e-05 | 1.5e-05 | 8.2e-06 | 1.207E-05 | 3.495E-06 | 8.200E-06 | 1.500E-05 |
| 12036 | 602 | Q602S | A | 1.5e-05 | 9.8e-06 | 1.1e-05 | 1.193E-05 | 2.723E-06 | 9.800E-06 | 1.500E-05 |
| 12037 | 602 | Q602T | A | 1.4e-05 | 1.1e-05 | 1.1e-05 | 1.200E-05 | 1.732E-06 | 1.100E-05 | 1.400E-05 |
| 12038 | 602 | Q602W | A | 8.4e-06 | 1.6e-05 | 1.5e-05 | 1.313E-05 | 4.130E-06 | 8.400E-06 | 1.600E-05 |
| 12039 | 602 | Q602Y | A | 9.4e-06 | 9.9e-06 | 1.5e-05 | 1.143E-05 | 3.099E-06 | 9.400E-06 | 1.500E-05 |
| 12040 | 602 | Q602V | A | 1.5e-05 | 1.5e-05 | 1e-05 | 1.333E-05 | 2.887E-06 | 1.000E-05 | 1.500E-05 |
| 12041 | 603 | P603A | A | 1.2e-05 | 1.5e-05 | 1.1e-05 | 1.267E-05 | 2.082E-06 | 1.100E-05 | 1.500E-05 |
| 12042 | 603 | P603R | A | 8e-06 | 1e-05 | 1.4e-05 | 1.067E-05 | 3.055E-06 | 8.000E-06 | 1.400E-05 |
| 12043 | 603 | P603N | A | 1.6e-05 | 1.3e-05 | 1.4e-05 | 1.433E-05 | 1.528E-06 | 1.300E-05 | 1.600E-05 |
| 12044 | 603 | P603D | A | 9.5e-06 | 1.1e-05 | 1.4e-05 | 1.150E-05 | 2.291E-06 | 9.500E-06 | 1.400E-05 |
| 12045 | 603 | P603C | A | 1.5e-05 | 1.3e-05 | 1.5e-05 | 1.433E-05 | 1.155E-06 | 1.300E-05 | 1.500E-05 |
| 12046 | 603 | P603E | A | 8.5e-06 | 1.4e-05 | 1.3e-05 | 1.183E-05 | 2.930E-06 | 8.500E-06 | 1.400E-05 |
| 12047 | 603 | P603Q | A | 1.4e-05 | 1.1e-05 | 1.5e-05 | 1.333E-05 | 2.082E-06 | 1.100E-05 | 1.500E-05 |
| 12048 | 603 | P603G | A | 6.7e-06 | 2.5e-05 | 1.6e-05 | 1.590E-05 | 9.150E-06 | 6.700E-06 | 2.500E-05 |
| 12049 | 603 | P603H | A | 1.5e-05 | 1.2e-05 | 9.1e-06 | 1.203E-05 | 2.950E-06 | 9.100E-06 | 1.500E-05 |
| 12050 | 603 | P603I | A | 1e-05 | 1.6e-05 | 1.6e-05 | 1.400E-05 | 3.464E-06 | 1.000E-05 | 1.600E-05 |
| 12051 | 603 | P603L | A | 1.1e-05 | 1.5e-05 | 1.5e-05 | 1.367E-05 | 2.309E-06 | 1.100E-05 | 1.500E-05 |
| 12052 | 603 | P603K | A | 1.1e-05 | 1.4e-05 | 1.1e-05 | 1.200E-05 | 1.732E-06 | 1.100E-05 | 1.400E-05 |
| 12053 | 603 | P603M | A | 1.3e-05 | 1.5e-05 | 1e-05 | 1.267E-05 | 2.517E-06 | 1.000E-05 | 1.500E-05 |
| 12054 | 603 | P603F | A | 1.2e-05 | 1.6e-05 | 2.3e-05 | 1.700E-05 | 5.568E-06 | 1.200E-05 | 2.300E-05 |
| 12055 | 603 | P603P | A | 1.4e-05 | 1.1e-05 | 1.4e-05 | 1.300E-05 | 1.732E-06 | 1.100E-05 | 1.400E-05 |
| 12056 | 603 | P603S | A | 1e-05 | 1.3e-05 | 8e-06 | 1.033E-05 | 2.517E-06 | 8.000E-06 | 1.300E-05 |
| 12057 | 603 | P603T | A | 1e-05 | 1.1e-05 | 1.5e-05 | 1.200E-05 | 2.646E-06 | 1.000E-05 | 1.500E-05 |
| 12058 | 603 | P603W | A | 1.4e-05 | 9.8e-06 | 9.4e-06 | 1.107E-05 | 2.548E-06 | 9.400E-06 | 1.400E-05 |
| 12059 | 603 | P603Y | A | 9.3e-06 | 8.2e-06 | 1.6e-05 | 1.117E-05 | 4.222E-06 | 8.200E-06 | 1.600E-05 |
| 12060 | 603 | P603V | A | 1.2e-05 | 1.5e-05 | 1.5e-05 | 1.400E-05 | 1.732E-06 | 1.200E-05 | 1.500E-05 |
| 12061 | 604 | C604A | A | 1.6e-05 | 1.4e-05 | 1.4e-05 | 1.467E-05 | 1.155E-06 | 1.400E-05 | 1.600E-05 |
| 12062 | 604 | C604R | A | 8.4e-06 | 1.6e-05 | 1e-05 | 1.147E-05 | 4.007E-06 | 8.400E-06 | 1.600E-05 |
| 12063 | 604 | C604N | A | 1.1e-05 | 1.5e-05 | 1e-05 | 1.200E-05 | 2.646E-06 | 1.000E-05 | 1.500E-05 |
| 12064 | 604 | C604D | A | 1.6e-05 | 1.7e-05 | 1.1e-05 | 1.467E-05 | 3.215E-06 | 1.100E-05 | 1.700E-05 |
| 12065 | 604 | C604C | A | 1.2e-05 | 1.1e-05 | 2.4e-05 | 1.567E-05 | 7.234E-06 | 1.100E-05 | 2.400E-05 |
| 12066 | 604 | C604E | A | 1.1e-05 | 1.1e-05 | 1.4e-05 | 1.200E-05 | 1.732E-06 | 1.100E-05 | 1.400E-05 |
| 12067 | 604 | C604Q | A | 9.9e-06 | 9.6e-06 | 1.1e-05 | 1.017E-05 | 7.371E-07 | 9.600E-06 | 1.100E-05 |
| 12068 | 604 | C604G | A | 1.1e-05 | 1e-05 | 1.6e-05 | 1.233E-05 | 3.215E-06 | 1.000E-05 | 1.600E-05 |
| 12069 | 604 | C604H | A | 1.5e-05 | 1.5e-05 | 9.5e-06 | 1.317E-05 | 3.175E-06 | 9.500E-06 | 1.500E-05 |

| | | | | | | | | | | |
|---|---|---|---|---|---|---|---|---|---|---|
| 12070 | 604 | C604I | A | 1.2e-05 | 1.6e-05 | 1.5e-05 | 1.433E-05 | 2.082E-06 | 1.200E-05 | 1.600E-05 |
| 12071 | 604 | C604L | A | 1.1e-05 | 1e-05 | 1.1e-05 | 1.067E-05 | 5.774E-07 | 1.000E-05 | 1.100E-05 |
| 12072 | 604 | C604K | A | 1.3e-05 | 1.6e-05 | 9.9e-06 | 1.297E-05 | 3.050E-06 | 9.900E-06 | 1.600E-05 |
| 12073 | 604 | C604M | A | 1.5e-05 | 1.7e-05 | 1.5e-05 | 1.567E-05 | 1.155E-06 | 1.500E-05 | 1.700E-05 |
| 12074 | 604 | C604F | A | 1.3e-05 | 1.2e-05 | 9.7e-06 | 1.157E-05 | 1.692E-06 | 9.700E-06 | 1.300E-05 |
| 12075 | 604 | C604P | A | 1.6e-05 | 1.1e-05 | 1.1e-05 | 1.267E-05 | 2.887E-06 | 1.100E-05 | 1.600E-05 |
| 12076 | 604 | C604S | A | 1e-05 | 1.5e-05 | 1.4e-05 | 1.300E-05 | 2.646E-06 | 1.000E-05 | 1.500E-05 |
| 12077 | 604 | C604T | A | 1.1e-05 | 1.2e-05 | 1.5e-05 | 1.267E-05 | 2.082E-06 | 1.100E-05 | 1.500E-05 |
| 12078 | 604 | C604W | A | 1.4e-05 | 1.6e-05 | 1.7e-05 | 1.567E-05 | 1.528E-06 | 1.400E-05 | 1.700E-05 |
| 12079 | 604 | C604Y | A | 9.6e-06 | 1.4e-05 | 1.6e-05 | 1.320E-05 | 3.274E-06 | 9.600E-06 | 1.600E-05 |
| 12080 | 604 | C604V | A | 9.3e-06 | 1.5e-05 | 1.4e-05 | 1.277E-05 | 3.044E-06 | 9.300E-06 | 1.500E-05 |
| 12081 | 605 | P605A | A | 1.6e-05 | 1.6e-05 | 1.1e-05 | 1.433E-05 | 2.887E-06 | 1.100E-05 | 1.600E-05 |
| 12082 | 605 | P605R | A | 1.1e-05 | 9.9e-06 | 1.4e-05 | 1.163E-05 | 2.122E-06 | 9.900E-06 | 1.400E-05 |
| 12083 | 605 | P605N | A | 1.1e-05 | 1.7e-05 | 1.3e-05 | 1.367E-05 | 3.055E-06 | 1.100E-05 | 1.700E-05 |
| 12084 | 605 | P605D | A | 8.8e-06 | 1.4e-05 | 1.4e-05 | 1.227E-05 | 3.002E-06 | 8.800E-06 | 1.400E-05 |
| 12085 | 605 | P605C | A | 1.1e-05 | 1e-05 | 1.4e-05 | 1.167E-05 | 2.082E-06 | 1.000E-05 | 1.400E-05 |
| 12086 | 605 | P605E | A | 1e-05 | 1.4e-05 | 1.7e-05 | 1.367E-05 | 3.512E-06 | 1.000E-05 | 1.700E-05 |
| 12087 | 605 | P605Q | A | 1.3e-05 | 1.4e-05 | 1.1e-05 | 1.267E-05 | 1.528E-06 | 1.100E-05 | 1.400E-05 |
| 12088 | 605 | P605G | A | 1.6e-05 | 1.5e-05 | 1.3e-05 | 1.467E-05 | 1.528E-06 | 1.300E-05 | 1.600E-05 |
| 12089 | 605 | P605H | A | 2.1e-05 | 1e-05 | 1e-05 | 1.367E-05 | 6.351E-06 | 1.000E-05 | 2.100E-05 |
| 12090 | 605 | P605I | A | 1.1e-05 | 1.4e-05 | 9e-06 | 1.133E-05 | 2.517E-06 | 9.000E-06 | 1.400E-05 |
| 12091 | 605 | P605L | A | 1.6e-05 | 1.4e-05 | 1.5e-05 | 1.500E-05 | 1.000E-06 | 1.400E-05 | 1.600E-05 |
| 12092 | 605 | P605K | A | 1.6e-05 | 1.1e-05 | 1.1e-05 | 1.267E-05 | 2.887E-06 | 1.100E-05 | 1.600E-05 |
| 12093 | 605 | P605M | A | 1.1e-05 | 1.5e-05 | 1.5e-05 | 1.367E-05 | 2.309E-06 | 1.100E-05 | 1.500E-05 |
| 12094 | 605 | P605F | A | 9.6e-06 | 9e-06 | 8.3e-06 | 8.967E-06 | 6.506E-07 | 8.300E-06 | 9.600E-06 |
| 12095 | 605 | P605P | A | 1e-05 | 1e-05 | 7.5e-06 | 9.167E-06 | 1.443E-06 | 7.500E-06 | 1.000E-05 |
| 12096 | 605 | P605S | A | 1.5e-05 | 1e-05 | 1e-05 | 1.167E-05 | 2.887E-06 | 1.000E-05 | 1.500E-05 |
| 12097 | 605 | P605T | A | 8.3e-06 | 1.5e-05 | 1.1e-05 | 1.143E-05 | 3.371E-06 | 8.300E-06 | 1.500E-05 |
| 12098 | 605 | P605W | A | 1.4e-05 | 9.7e-06 | 6.9e-06 | 1.020E-05 | 3.576E-06 | 6.900E-06 | 1.400E-05 |
| 12099 | 605 | P605Y | A | 9.9e-06 | 7.1e-06 | 1.4e-05 | 1.033E-05 | 3.470E-06 | 7.100E-06 | 1.400E-05 |
| 12100 | 605 | P605V | A | 1.4e-05 | 1e-05 | 1.5e-05 | 1.300E-05 | 2.646E-06 | 1.000E-05 | 1.500E-05 |
| 12101 | 606 | I606A | A | 1.4e-05 | 1.6e-05 | 1.5e-05 | 1.500E-05 | 1.000E-06 | 1.400E-05 | 1.600E-05 |
| 12102 | 606 | I606R | A | 1.1e-05 | 1.1e-05 | 1.3e-05 | 1.167E-05 | 1.155E-06 | 1.100E-05 | 1.300E-05 |
| 12103 | 606 | I606N | A | 8.6e-06 | 1.1e-05 | 1e-05 | 9.867E-06 | 1.206E-06 | 8.600E-06 | 1.100E-05 |
| 12104 | 606 | I606D | A | 1.6e-05 | 1.7e-05 | 1.9e-05 | 1.733E-05 | 1.528E-06 | 1.600E-05 | 1.900E-05 |
| 12105 | 606 | I606C | A | 8.9e-06 | 1.5e-05 | 1e-05 | 1.130E-05 | 3.251E-06 | 8.900E-06 | 1.500E-05 |
| 12106 | 606 | I606E | A | 1.3e-05 | 1.2e-05 | 1.6e-05 | 1.367E-05 | 2.082E-06 | 1.200E-05 | 1.600E-05 |
| 12107 | 606 | I606Q | A | 1.3e-05 | 1.5e-05 | 1.6e-05 | 1.467E-05 | 1.528E-06 | 1.300E-05 | 1.600E-05 |
| 12108 | 606 | I606G | A | 1.2e-05 | 1.6e-05 | 1.1e-05 | 1.300E-05 | 2.646E-06 | 1.100E-05 | 1.600E-05 |
| 12109 | 606 | I606H | A | 1.4e-05 | 1e-05 | 9.3e-06 | 1.110E-05 | 2.536E-06 | 9.300E-06 | 1.400E-05 |
| 12110 | 606 | I606I | A | 1.6e-05 | 1.1e-05 | 1.1e-05 | 1.267E-05 | 2.887E-06 | 1.100E-05 | 1.600E-05 |
| 12111 | 606 | I606L | A | 1.4e-05 | 1.5e-05 | 1.2e-05 | 1.367E-05 | 1.528E-06 | 1.200E-05 | 1.500E-05 |
| 12112 | 606 | I606K | A | 1.1e-05 | 1.5e-05 | 1.4e-05 | 1.333E-05 | 2.082E-06 | 1.100E-05 | 1.500E-05 |
| 12113 | 606 | I606M | A | 1.2e-05 | 9.5e-06 | 1.6e-05 | 1.250E-05 | 3.279E-06 | 9.500E-06 | 1.600E-05 |
| 12114 | 606 | I606F | A | 9.4e-06 | 1.1e-05 | 1.1e-05 | 1.047E-05 | 9.238E-07 | 9.400E-06 | 1.100E-05 |
| 12115 | 606 | I606P | A | 1.1e-05 | 8.6e-06 | 1.5e-05 | 1.153E-05 | 3.233E-06 | 8.600E-06 | 1.500E-05 |
| 12116 | 606 | I606S | A | 1.4e-05 | 1.5e-05 | 1.1e-05 | 1.333E-05 | 2.082E-06 | 1.100E-05 | 1.500E-05 |
| 12117 | 606 | I606T | A | 1.4e-05 | 8.9e-06 | 1.1e-05 | 1.130E-05 | 2.563E-06 | 8.900E-06 | 1.400E-05 |
| 12118 | 606 | I606W | A | 1.1e-05 | 1.6e-05 | 1.5e-05 | 1.400E-05 | 2.646E-06 | 1.100E-05 | 1.600E-05 |
| 12119 | 606 | I606Y | A | 1.3e-05 | 1.1e-05 | 1e-05 | 1.133E-05 | 1.528E-06 | 1.000E-05 | 1.300E-05 |
| 12120 | 606 | I606V | A | 1.5e-05 | 9.8e-06 | 1.5e-05 | 1.327E-05 | 3.002E-06 | 9.800E-06 | 1.500E-05 |
| 12121 | 607 | N607A | A | 1e-05 | 1e-05 | 9.8e-06 | 9.933E-06 | 1.155E-07 | 9.800E-06 | 1.000E-05 |
| 12122 | 607 | N607R | A | 1.6e-05 | 1.6e-05 | 9.3e-06 | 1.377E-05 | 3.868E-06 | 9.300E-06 | 1.600E-05 |
| 12123 | 607 | N607N | A | 1.5e-05 | 1.3e-05 | 1.6e-05 | 1.467E-05 | 1.528E-06 | 1.300E-05 | 1.600E-05 |
| 12124 | 607 | N607D | A | 1.6e-05 | 1.6e-05 | 1.5e-05 | 1.567E-05 | 5.774E-07 | 1.500E-05 | 1.600E-05 |
| 12125 | 607 | N607C | A | 1.1e-05 | 8.7e-06 | 1e-05 | 9.900E-06 | 1.153E-06 | 8.700E-06 | 1.100E-05 |
| 12126 | 607 | N607E | A | 1.5e-05 | 1e-05 | 1.1e-05 | 1.200E-05 | 2.646E-06 | 1.000E-05 | 1.500E-05 |
| 12127 | 607 | N607Q | A | 1e-05 | 1.3e-05 | 1.1e-05 | 1.133E-05 | 1.528E-06 | 1.000E-05 | 1.300E-05 |
| 12128 | 607 | N607G | A | 1.1e-05 | 8.2e-06 | 1.2e-05 | 1.040E-05 | 1.970E-06 | 8.200E-06 | 1.200E-05 |
| 12129 | 607 | N607H | A | 1.1e-05 | 1.7e-05 | 1e-05 | 1.267E-05 | 3.786E-06 | 1.000E-05 | 1.700E-05 |
| 12130 | 607 | N607I | A | 1.2e-05 | 1.5e-05 | 1.1e-05 | 1.267E-05 | 2.082E-06 | 1.100E-05 | 1.500E-05 |
| 12131 | 607 | N607L | A | 1.5e-05 | 1e-05 | 1.6e-05 | 1.367E-05 | 3.215E-06 | 1.000E-05 | 1.600E-05 |
| 12132 | 607 | N607K | A | 1.1e-05 | 1.5e-05 | 1.8e-05 | 1.467E-05 | 3.512E-06 | 1.100E-05 | 1.800E-05 |
| 12133 | 607 | N607M | A | 1e-05 | 1.2e-05 | 1.2e-05 | 1.133E-05 | 1.155E-06 | 1.000E-05 | 1.200E-05 |
| 12134 | 607 | N607F | A | 1.3e-05 | 1.1e-05 | 1.4e-05 | 1.267E-05 | 1.528E-06 | 1.100E-05 | 1.400E-05 |
| 12135 | 607 | N607P | A | 1.6e-05 | 1.4e-05 | 1.7e-05 | 1.567E-05 | 1.528E-06 | 1.400E-05 | 1.700E-05 |
| 12136 | 607 | N607S | A | 1.1e-05 | 9.1e-06 | 7.8e-06 | 9.300E-06 | 1.609E-06 | 7.800E-06 | 1.100E-05 |
| 12137 | 607 | N607T | A | 1.1e-05 | 1.1e-05 | 1.5e-05 | 1.233E-05 | 2.309E-06 | 1.100E-05 | 1.500E-05 |
| 12138 | 607 | N607W | A | 1e-05 | 1.5e-05 | 1.1e-05 | 1.200E-05 | 2.646E-06 | 1.000E-05 | 1.500E-05 |
| 12139 | 607 | N607Y | A | 1.5e-05 | 1.1e-05 | 1.1e-05 | 1.233E-05 | 2.309E-06 | 1.100E-05 | 1.500E-05 |
| 12140 | 607 | N607V | A | 1.9e-05 | 1.6e-05 | 7e-06 | 1.400E-05 | 6.245E-06 | 7.000E-06 | 1.900E-05 |
| 12141 | 608 | C608A | A | 1.5e-05 | 1.1e-05 | 1.1e-05 | 1.233E-05 | 2.309E-06 | 1.100E-05 | 1.500E-05 |
| 12142 | 608 | C608R | A | 1e-05 | 1.1e-05 | 1.8e-05 | 1.300E-05 | 4.359E-06 | 1.000E-05 | 1.800E-05 |
| 12143 | 608 | C608N | A | 1e-05 | 1.6e-05 | 1.7e-05 | 1.433E-05 | 3.786E-06 | 1.000E-05 | 1.700E-05 |
| 12144 | 608 | C608D | A | 1.5e-05 | 1.2e-05 | 1.4e-05 | 1.367E-05 | 1.528E-06 | 1.200E-05 | 1.500E-05 |
| 12145 | 608 | C608C | A | 1.1e-05 | 8.8e-06 | 1.6e-05 | 1.193E-05 | 3.690E-06 | 8.800E-06 | 1.600E-05 |
| 12146 | 608 | C608E | A | 1.4e-05 | 9.9e-06 | 1.3e-05 | 1.230E-05 | 2.138E-06 | 9.900E-06 | 1.400E-05 |
| 12147 | 608 | C608Q | A | 2e-05 | 1.6e-05 | 1.5e-05 | 1.700E-05 | 2.646E-06 | 1.500E-05 | 2.000E-05 |
| 12148 | 608 | C608G | A | 1.1e-05 | 1.6e-05 | 1.1e-05 | 1.267E-05 | 2.887E-06 | 1.100E-05 | 1.600E-05 |
| 12149 | 608 | C608H | A | 1.5e-05 | 1.4e-05 | 1.3e-05 | 1.400E-05 | 1.000E-06 | 1.300E-05 | 1.500E-05 |
| 12150 | 608 | C608I | A | 1.5e-05 | 1.7e-05 | 1.4e-05 | 1.533E-05 | 1.528E-06 | 1.400E-05 | 1.700E-05 |
| 12151 | 608 | C608L | A | 1.6e-05 | 1e-05 | 1.1e-05 | 1.233E-05 | 3.215E-06 | 1.000E-05 | 1.600E-05 |
| 12152 | 608 | C608K | A | 1.8e-05 | 1.6e-05 | 1e-05 | 1.467E-05 | 4.163E-06 | 1.000E-05 | 1.800E-05 |
| 12153 | 608 | C608M | A | 1.6e-05 | 1.4e-05 | 1.3e-05 | 1.433E-05 | 1.528E-06 | 1.300E-05 | 1.600E-05 |
| 12154 | 608 | C608F | A | 1.3e-05 | 1.7e-05 | 1.1e-05 | 1.367E-05 | 3.055E-06 | 1.100E-05 | 1.700E-05 |

| | | | | | | | | | | |
|---|---|---|---|---|---|---|---|---|---|---|
| 12155 | 608 | C608P | A | 1.2e-05 | 1e-05 | 1.6e-05 | 1.267E-05 | 3.055E-06 | 1.000E-05 | 1.600E-05 |
| 12156 | 608 | C608S | A | 1.5e-05 | 9e-06 | 9.3e-06 | 1.110E-05 | 3.381E-06 | 9.000E-06 | 1.500E-05 |
| 12157 | 608 | C608T | A | 9.2e-06 | 1e-05 | 8.9e-06 | 9.367E-06 | 5.686E-07 | 8.900E-06 | 1.000E-05 |
| 12158 | 608 | C608W | A | 1e-05 | 1.6e-05 | 9.7e-06 | 1.190E-05 | 3.554E-06 | 9.700E-06 | 1.600E-05 |
| 12159 | 608 | C608Y | A | 1.3e-05 | 1.3e-05 | 1.4e-05 | 1.333E-05 | 5.774E-07 | 1.300E-05 | 1.400E-05 |
| 12160 | 608 | C608V | A | 1.1e-05 | 1.6e-05 | 9e-06 | 1.200E-05 | 3.606E-06 | 9.000E-06 | 1.600E-05 |
| 12161 | 609 | T609A | A | 1.3e-05 | 2e-05 | 1.3e-05 | 1.533E-05 | 4.041E-06 | 1.300E-05 | 2.000E-05 |
| 12162 | 609 | T609R | A | 1e-05 | 1e-05 | 1.4e-05 | 1.133E-05 | 2.309E-06 | 1.000E-05 | 1.400E-05 |
| 12163 | 609 | T609N | A | 8.8e-06 | 1.5e-05 | 1.6e-05 | 1.327E-05 | 3.900E-06 | 8.800E-06 | 1.600E-05 |
| 12164 | 609 | T609D | A | 9.7e-06 | 9.8e-06 | 7.2e-06 | 8.900E-06 | 1.473E-06 | 7.200E-06 | 9.800E-06 |
| 12165 | 609 | T609C | A | 1.5e-05 | 1.1e-05 | 1.4e-05 | 1.333E-05 | 2.082E-06 | 1.100E-05 | 1.500E-05 |
| 12166 | 609 | T609E | A | 1.1e-05 | 1.6e-05 | 1.7e-05 | 1.467E-05 | 3.215E-06 | 1.100E-05 | 1.700E-05 |
| 12167 | 609 | T609Q | A | 1.1e-05 | 8.4e-06 | 9.8e-06 | 9.733E-06 | 1.301E-06 | 8.400E-06 | 1.100E-05 |
| 12168 | 609 | T609G | A | 1.6e-05 | 1.6e-05 | 1.6e-05 | 1.600E-05 | 0.000E+00 | 1.600E-05 | 1.600E-05 |
| 12169 | 609 | T609H | A | 1.6e-05 | 5.8e-06 | 1.7e-05 | 1.293E-05 | 6.198E-06 | 5.800E-06 | 1.700E-05 |
| 12170 | 609 | T609I | A | 1.5e-05 | 1.6e-05 | 9.2e-06 | 1.340E-05 | 3.672E-06 | 9.200E-06 | 1.600E-05 |
| 12171 | 609 | T609L | A | 1.6e-05 | 1.6e-05 | 9e-06 | 1.367E-05 | 4.041E-06 | 9.000E-06 | 1.600E-05 |
| 12172 | 609 | T609K | A | 9.9e-06 | 1.6e-05 | 1.1e-05 | 1.230E-05 | 3.251E-06 | 9.900E-06 | 1.600E-05 |
| 12173 | 609 | T609M | A | 1.4e-05 | 6.1e-06 | 9.6e-06 | 9.900E-06 | 3.959E-06 | 6.100E-06 | 1.400E-05 |
| 12174 | 609 | T609F | A | 1.7e-05 | 1.4e-05 | 1.5e-05 | 1.533E-05 | 1.528E-06 | 1.400E-05 | 1.700E-05 |
| 12175 | 609 | T609P | A | 1.5e-05 | 1.3e-05 | 1.2e-05 | 1.333E-05 | 1.528E-06 | 1.200E-05 | 1.500E-05 |
| 12176 | 609 | T609S | A | 1.5e-05 | 1.5e-05 | 9.7e-06 | 1.323E-05 | 3.060E-06 | 9.700E-06 | 1.500E-05 |
| 12177 | 609 | T609T | A | 1e-05 | 1.3e-05 | 2.4e-05 | 1.567E-05 | 7.371E-06 | 1.000E-05 | 2.400E-05 |
| 12178 | 609 | T609W | A | 1.6e-05 | 1.6e-05 | 1.6e-05 | 1.600E-05 | 0.000E+00 | 1.600E-05 | 1.600E-05 |
| 12179 | 609 | T609Y | A | 8.1e-06 | 1.5e-05 | 1.2e-05 | 1.170E-05 | 3.460E-06 | 8.100E-06 | 1.500E-05 |
| 12180 | 609 | T609V | A | 1.3e-05 | 1.4e-05 | 1.5e-05 | 1.400E-05 | 1.000E-06 | 1.300E-05 | 1.500E-05 |
| 12181 | 610 | H610A | A | 1.6e-05 | 1.4e-05 | 1.2e-05 | 1.400E-05 | 2.000E-06 | 1.200E-05 | 1.600E-05 |
| 12182 | 610 | H610R | A | 9.9e-06 | 1.4e-05 | 1.5e-05 | 1.297E-05 | 2.702E-06 | 9.900E-06 | 1.500E-05 |
| 12183 | 610 | H610N | A | 8.7e-06 | 1.4e-05 | 1.2e-05 | 1.157E-05 | 2.676E-06 | 8.700E-06 | 1.400E-05 |
| 12184 | 610 | H610D | A | 1.5e-05 | 1e-05 | 1.6e-05 | 1.367E-05 | 3.215E-06 | 1.000E-05 | 1.600E-05 |
| 12185 | 610 | H610C | A | 1.1e-05 | 1.6e-05 | 1.6e-05 | 1.433E-05 | 2.887E-06 | 1.100E-05 | 1.600E-05 |
| 12186 | 610 | H610E | A | 1.4e-05 | 1.7e-05 | 1.5e-05 | 1.533E-05 | 1.528E-06 | 1.400E-05 | 1.700E-05 |
| 12187 | 610 | H610Q | A | 1.4e-05 | 1.3e-05 | 8.8e-06 | 1.193E-05 | 2.759E-06 | 8.800E-06 | 1.400E-05 |
| 12188 | 610 | H610G | A | 1.4e-05 | 9.7e-06 | 1.1e-05 | 1.157E-05 | 2.205E-06 | 9.700E-06 | 1.400E-05 |
| 12189 | 610 | H610H | A | 1.6e-05 | 1e-05 | 1e-05 | 1.200E-05 | 3.464E-06 | 1.000E-05 | 1.600E-05 |
| 12190 | 610 | H610I | A | 1e-05 | 1.5e-05 | 1.2e-05 | 1.233E-05 | 2.517E-06 | 1.000E-05 | 1.500E-05 |
| 12191 | 610 | H610L | A | 1.2e-05 | 1.5e-05 | 8.6e-06 | 1.187E-05 | 3.202E-06 | 8.600E-06 | 1.500E-05 |
| 12192 | 610 | H610K | A | 1.6e-05 | 1.7e-05 | 6.4e-06 | 1.313E-05 | 5.853E-06 | 6.400E-06 | 1.700E-05 |
| 12193 | 610 | H610M | A | 1.1e-05 | 1.6e-05 | 1e-05 | 1.233E-05 | 3.215E-06 | 1.000E-05 | 1.600E-05 |
| 12194 | 610 | H610F | A | 1.1e-05 | 1.1e-05 | 1.7e-05 | 1.300E-05 | 3.464E-06 | 1.100E-05 | 1.700E-05 |
| 12195 | 610 | H610P | A | 9.4e-06 | 1.4e-05 | 1.6e-05 | 1.313E-05 | 3.384E-06 | 9.400E-06 | 1.600E-05 |
| 12196 | 610 | H610S | A | 1.4e-05 | 1.3e-05 | 1.1e-05 | 1.267E-05 | 1.528E-06 | 1.100E-05 | 1.400E-05 |
| 12197 | 610 | H610T | A | 1.7e-05 | 1.1e-05 | 1.1e-05 | 1.300E-05 | 3.464E-06 | 1.100E-05 | 1.700E-05 |
| 12198 | 610 | H610W | A | 1e-05 | 6.7e-06 | 9.6e-06 | 8.767E-06 | 1.801E-06 | 6.700E-06 | 1.000E-05 |
| 12199 | 610 | H610Y | A | 1.6e-05 | 9.8e-06 | 1e-05 | 1.193E-05 | 3.523E-06 | 9.800E-06 | 1.600E-05 |
| 12200 | 610 | H610V | A | 8.8e-06 | 1.4e-05 | 1.5e-05 | 1.260E-05 | 3.329E-06 | 8.800E-06 | 1.500E-05 |
| 12201 | 611 | S611A | A | 1.1e-05 | 1.1e-05 | 1.6e-05 | 1.267E-05 | 2.887E-06 | 1.100E-05 | 1.600E-05 |
| 12202 | 611 | S611R | A | 1e-05 | 9.1e-06 | 1.3e-05 | 1.070E-05 | 2.042E-06 | 9.100E-06 | 1.300E-05 |
| 12203 | 611 | S611N | A | 1e-05 | 1e-05 | 1e-05 | 1.000E-05 | 0.000E+00 | 1.000E-05 | 1.000E-05 |
| 12204 | 611 | S611D | A | 1.3e-05 | 1e-05 | 1.3e-05 | 1.200E-05 | 1.732E-06 | 1.000E-05 | 1.300E-05 |
| 12205 | 611 | S611C | A | 1.7e-05 | 1.6e-05 | 1.4e-05 | 1.567E-05 | 1.528E-06 | 1.400E-05 | 1.700E-05 |
| 12206 | 611 | S611E | A | 9.7e-06 | 9.6e-06 | 1.5e-05 | 1.143E-05 | 3.089E-06 | 9.600E-06 | 1.500E-05 |
| 12207 | 611 | S611Q | A | 1.5e-05 | 1e-05 | 1.4e-05 | 1.300E-05 | 2.646E-06 | 1.000E-05 | 1.500E-05 |
| 12208 | 611 | S611G | A | 1e-05 | 1e-05 | 1.7e-05 | 1.233E-05 | 4.041E-06 | 1.000E-05 | 1.700E-05 |
| 12209 | 611 | S611H | A | 1.5e-05 | 1.6e-05 | 1.6e-05 | 1.567E-05 | 5.774E-07 | 1.500E-05 | 1.600E-05 |
| 12210 | 611 | S611I | A | 1.2e-05 | 1.6e-05 | 1.1e-05 | 1.300E-05 | 2.646E-06 | 1.100E-05 | 1.600E-05 |
| 12211 | 611 | S611L | A | 1.5e-05 | 8.3e-06 | 1.4e-05 | 1.243E-05 | 3.614E-06 | 8.300E-06 | 1.500E-05 |
| 12212 | 611 | S611K | A | 1.5e-05 | 1.1e-05 | 1.1e-05 | 1.233E-05 | 2.309E-06 | 1.100E-05 | 1.500E-05 |
| 12213 | 611 | S611M | A | 1.2e-05 | 7.8e-06 | 1.4e-05 | 1.127E-05 | 3.164E-06 | 7.800E-06 | 1.400E-05 |
| 12214 | 611 | S611F | A | 1.1e-05 | 1.4e-05 | 1.6e-05 | 1.367E-05 | 2.517E-06 | 1.100E-05 | 1.600E-05 |
| 12215 | 611 | S611P | A | 1.7e-05 | 1e-05 | 1.4e-05 | 1.367E-05 | 3.512E-06 | 1.000E-05 | 1.700E-05 |
| 12216 | 611 | S611S | A | 1.6e-05 | 1.1e-05 | 1.5e-05 | 1.400E-05 | 2.646E-06 | 1.100E-05 | 1.600E-05 |
| 12217 | 611 | S611T | A | 1.1e-05 | 1.2e-05 | 9.3e-06 | 1.077E-05 | 1.365E-06 | 9.300E-06 | 1.200E-05 |
| 12218 | 611 | S611W | A | 1.5e-05 | 1.6e-05 | 7.7e-06 | 1.290E-05 | 4.531E-06 | 7.700E-06 | 1.600E-05 |
| 12219 | 611 | S611Y | A | 1.6e-05 | 1.1e-05 | 8.5e-06 | 1.183E-05 | 3.819E-06 | 8.500E-06 | 1.600E-05 |
| 12220 | 611 | S611V | A | 1.4e-05 | 1.1e-05 | 1.6e-05 | 1.367E-05 | 2.517E-06 | 1.100E-05 | 1.600E-05 |
| 12221 | 612 | C612A | A | 1.6e-05 | 1.6e-05 | 1.6e-05 | 1.600E-05 | 0.000E+00 | 1.600E-05 | 1.600E-05 |
| 12222 | 612 | C612R | A | 1.7e-05 | 1e-05 | 1e-05 | 1.233E-05 | 4.041E-06 | 1.000E-05 | 1.700E-05 |
| 12223 | 612 | C612N | A | 1.3e-05 | 1.1e-05 | 1.5e-05 | 1.300E-05 | 2.000E-06 | 1.100E-05 | 1.500E-05 |
| 12224 | 612 | C612D | A | 1.5e-05 | 1e-05 | 1.5e-05 | 1.333E-05 | 2.887E-06 | 1.000E-05 | 1.500E-05 |
| 12225 | 612 | C612C | A | 1.2e-05 | 1.8e-05 | 1.6e-05 | 1.533E-05 | 3.055E-06 | 1.200E-05 | 1.800E-05 |
| 12226 | 612 | C612E | A | 1.4e-05 | 8.4e-06 | 9.9e-06 | 1.077E-05 | 2.899E-06 | 8.400E-06 | 1.400E-05 |
| 12227 | 612 | C612Q | A | 1.6e-05 | 1.5e-05 | 1e-05 | 1.367E-05 | 3.215E-06 | 1.000E-05 | 1.600E-05 |
| 12228 | 612 | C612G | A | 9.3e-06 | 1.5e-05 | 1.1e-05 | 1.177E-05 | 2.926E-06 | 9.300E-06 | 1.500E-05 |
| 12229 | 612 | C612H | A | 1.4e-05 | 1.4e-05 | 1.6e-05 | 1.467E-05 | 1.155E-06 | 1.400E-05 | 1.600E-05 |
| 12230 | 612 | C612I | A | 1.1e-05 | 1.1e-05 | 1.4e-05 | 1.200E-05 | 1.732E-06 | 1.100E-05 | 1.400E-05 |
| 12231 | 612 | C612L | A | 1.7e-05 | 1.5e-05 | 1.1e-05 | 1.433E-05 | 3.055E-06 | 1.100E-05 | 1.700E-05 |
| 12232 | 612 | C612K | A | 1.1e-05 | 1.3e-05 | 1.7e-05 | 1.367E-05 | 3.055E-06 | 1.100E-05 | 1.700E-05 |
| 12233 | 612 | C612M | A | 1.2e-05 | 1.7e-05 | 9.4e-06 | 1.280E-05 | 3.863E-06 | 9.400E-06 | 1.700E-05 |
| 12234 | 612 | C612F | A | 7.8e-06 | 1.7e-05 | 1.4e-05 | 1.293E-05 | 4.692E-06 | 7.800E-06 | 1.700E-05 |
| 12235 | 612 | C612P | A | 9.8e-06 | 1.5e-05 | 9.7e-06 | 1.150E-05 | 3.032E-06 | 9.700E-06 | 1.500E-05 |
| 12236 | 612 | C612S | A | 1.1e-05 | 1.3e-05 | 1.6e-05 | 1.333E-05 | 2.517E-06 | 1.100E-05 | 1.600E-05 |
| 12237 | 612 | C612T | A | 9.4e-06 | 1e-05 | 1.5e-05 | 1.147E-05 | 3.075E-06 | 9.400E-06 | 1.500E-05 |
| 12238 | 612 | C612W | A | 1.7e-05 | 1.2e-05 | 1.4e-05 | 1.433E-05 | 2.517E-06 | 1.200E-05 | 1.700E-05 |
| 12239 | 612 | C612Y | A | 1.5e-05 | 1.4e-05 | 1.5e-05 | 1.467E-05 | 5.774E-07 | 1.400E-05 | 1.500E-05 |

| | | | | | | | | | | |
|---|---|---|---|---|---|---|---|---|---|---|
| 12240 | 612 | C612V | A | 1.6e-05 | 1.5e-05 | 1e-05 | 1.367E-05 | 3.215E-06 | 1.000E-05 | 1.600E-05 |
| 12241 | 613 | V613A | A | 9.6e-06 | 1.6e-05 | 9.7e-06 | 1.177E-05 | 3.667E-06 | 9.600E-06 | 1.600E-05 |
| 12242 | 613 | V613R | A | 1e-05 | 1.5e-05 | 9.1e-06 | 1.137E-05 | 3.179E-06 | 9.100E-06 | 1.500E-05 |
| 12243 | 613 | V613N | A | 1.4e-05 | 7.1e-06 | 1.4e-05 | 1.170E-05 | 3.984E-06 | 7.100E-06 | 1.400E-05 |
| 12244 | 613 | V613D | A | 1.3e-05 | 1.6e-05 | 1.3e-05 | 1.400E-05 | 1.732E-06 | 1.300E-05 | 1.600E-05 |
| 12245 | 613 | V613C | A | 1e-05 | 2.1e-05 | 1.4e-05 | 1.500E-05 | 5.568E-06 | 1.000E-05 | 2.100E-05 |
| 12246 | 613 | V613E | A | 1.4e-05 | 1e-05 | 1.4e-05 | 1.267E-05 | 2.309E-06 | 1.000E-05 | 1.400E-05 |
| 12247 | 613 | V613Q | A | 9.8e-06 | 1.6e-05 | 1.1e-05 | 1.227E-05 | 3.288E-06 | 9.800E-06 | 1.600E-05 |
| 12248 | 613 | V613G | A | 1.5e-05 | 1.1e-05 | 1e-05 | 1.200E-05 | 2.646E-06 | 1.000E-05 | 1.500E-05 |
| 12249 | 613 | V613H | A | 1.1e-05 | 1.6e-05 | 1.5e-05 | 1.400E-05 | 2.646E-06 | 1.100E-05 | 1.600E-05 |
| 12250 | 613 | V613I | A | 9.9e-06 | 1.5e-05 | 1.5e-05 | 1.330E-05 | 2.944E-06 | 9.900E-06 | 1.500E-05 |
| 12251 | 613 | V613L | A | 1.5e-05 | 1.7e-05 | 1.5e-05 | 1.567E-05 | 1.155E-06 | 1.500E-05 | 1.700E-05 |
| 12252 | 613 | V613K | A | 1e-05 | 1.2e-05 | 1.4e-05 | 1.200E-05 | 2.000E-06 | 1.000E-05 | 1.400E-05 |
| 12253 | 613 | V613M | A | 1.5e-05 | 1.5e-05 | 1.7e-05 | 1.567E-05 | 1.155E-06 | 1.500E-05 | 1.700E-05 |
| 12254 | 613 | V613F | A | 8.7e-06 | 1.3e-05 | 1.1e-05 | 1.090E-05 | 2.152E-06 | 8.700E-06 | 1.300E-05 |
| 12255 | 613 | V613P | A | 1.6e-05 | 1.5e-05 | 1.3e-05 | 1.467E-05 | 1.528E-06 | 1.300E-05 | 1.600E-05 |
| 12256 | 613 | V613S | A | 1.2e-05 | 1.3e-05 | 1.1e-05 | 1.200E-05 | 1.000E-06 | 1.100E-05 | 1.300E-05 |
| 12257 | 613 | V613T | A | 1.1e-05 | 1.5e-05 | 1.5e-05 | 1.367E-05 | 2.309E-06 | 1.100E-05 | 1.500E-05 |
| 12258 | 613 | V613W | A | 1e-05 | 8.5e-06 | 1.6e-05 | 1.150E-05 | 3.969E-06 | 8.500E-06 | 1.600E-05 |
| 12259 | 613 | V613Y | A | 1.4e-05 | 1.6e-05 | 9.5e-06 | 1.317E-05 | 3.329E-06 | 9.500E-06 | 1.600E-05 |
| 12260 | 613 | V613V | A | 1.6e-05 | 1.4e-05 | 1.5e-05 | 1.500E-05 | 1.000E-06 | 1.400E-05 | 1.600E-05 |
| 12261 | 614 | D614A | A | 1.6e-05 | 1.5e-05 | 1.6e-05 | 1.567E-05 | 5.774E-07 | 1.500E-05 | 1.600E-05 |
| 12262 | 614 | D614R | A | 1.6e-05 | 1.6e-05 | 1.4e-05 | 1.533E-05 | 1.155E-06 | 1.400E-05 | 1.600E-05 |
| 12263 | 614 | D614N | A | 1.5e-05 | 1.4e-05 | 1.1e-05 | 1.333E-05 | 2.082E-06 | 1.100E-05 | 1.500E-05 |
| 12264 | 614 | D614D | A | 1.5e-05 | 1.2e-05 | 9.7e-06 | 1.223E-05 | 2.658E-06 | 9.700E-06 | 1.500E-05 |
| 12265 | 614 | D614C | A | 1e-05 | 1.3e-05 | 9.1e-06 | 1.070E-05 | 2.042E-06 | 9.100E-06 | 1.300E-05 |
| 12266 | 614 | D614E | A | 1.6e-05 | 1.6e-05 | 9.5e-06 | 1.383E-05 | 3.753E-06 | 9.500E-06 | 1.600E-05 |
| 12267 | 614 | D614Q | A | 1.1e-05 | 1.5e-05 | 1.5e-05 | 1.367E-05 | 2.309E-06 | 1.100E-05 | 1.500E-05 |
| 12268 | 614 | D614G | A | 1.4e-05 | 1.1e-05 | 1.6e-05 | 1.367E-05 | 2.517E-06 | 1.100E-05 | 1.600E-05 |
| 12269 | 614 | D614H | A | 9.3e-06 | 1.4e-05 | 1.4e-05 | 1.243E-05 | 2.714E-06 | 9.300E-06 | 1.400E-05 |
| 12270 | 614 | D614I | A | 1.5e-05 | 1e-05 | 1.4e-05 | 1.300E-05 | 2.646E-06 | 1.000E-05 | 1.500E-05 |
| 12271 | 614 | D614L | A | 1.3e-05 | 1.1e-05 | 9.6e-06 | 1.120E-05 | 1.709E-06 | 9.600E-06 | 1.300E-05 |
| 12272 | 614 | D614K | A | 1.1e-05 | 1.5e-05 | 9.9e-06 | 1.197E-05 | 2.684E-06 | 9.900E-06 | 1.500E-05 |
| 12273 | 614 | D614M | A | 1.5e-05 | 9.9e-06 | 1e-05 | 1.163E-05 | 2.916E-06 | 9.900E-06 | 1.500E-05 |
| 12274 | 614 | D614F | A | 1.6e-05 | 1.2e-05 | 1e-05 | 1.267E-05 | 3.055E-06 | 1.000E-05 | 1.600E-05 |
| 12275 | 614 | D614P | A | 1e-05 | 1.2e-05 | 7.7e-06 | 9.900E-06 | 2.152E-06 | 7.700E-06 | 1.200E-05 |
| 12276 | 614 | D614S | A | 1.5e-05 | 1.5e-05 | 9.2e-06 | 1.307E-05 | 3.349E-06 | 9.200E-06 | 1.500E-05 |
| 12277 | 614 | D614T | A | 1.6e-05 | 1.6e-05 | 1.6e-05 | 1.600E-05 | 0.000E+00 | 1.600E-05 | 1.600E-05 |
| 12278 | 614 | D614W | A | 9.2e-06 | 1.5e-05 | 1.6e-05 | 1.340E-05 | 3.672E-06 | 9.200E-06 | 1.600E-05 |
| 12279 | 614 | D614Y | A | 1.6e-05 | 1.5e-05 | 1e-05 | 1.367E-05 | 3.215E-06 | 1.000E-05 | 1.600E-05 |
| 12280 | 614 | D614V | A | 1.3e-05 | 9.8e-06 | 9.7e-06 | 1.083E-05 | 1.877E-06 | 9.700E-06 | 1.300E-05 |
| 12281 | 615 | L615A | A | 1.1e-05 | 1.3e-05 | 1.5e-05 | 1.300E-05 | 2.000E-06 | 1.100E-05 | 1.500E-05 |
| 12282 | 615 | L615R | A | 8.6e-06 | 9.6e-06 | 1.7e-05 | 1.173E-05 | 4.588E-06 | 8.600E-06 | 1.700E-05 |
| 12283 | 615 | L615N | A | 1.5e-05 | 9.8e-06 | 1.1e-05 | 1.193E-05 | 2.723E-06 | 9.800E-06 | 1.500E-05 |
| 12284 | 615 | L615D | A | 1e-05 | 1.1e-05 | 9.3e-06 | 1.010E-05 | 8.544E-07 | 9.300E-06 | 1.100E-05 |
| 12285 | 615 | L615C | A | 9.2e-06 | 1e-05 | 1.1e-05 | 1.007E-05 | 9.018E-07 | 9.200E-06 | 1.100E-05 |
| 12286 | 615 | L615E | A | 9.6e-06 | 1.4e-05 | 1e-05 | 1.120E-05 | 2.433E-06 | 9.600E-06 | 1.400E-05 |
| 12287 | 615 | L615Q | A | 1.1e-05 | 1.4e-05 | 1.4e-05 | 1.300E-05 | 1.732E-06 | 1.100E-05 | 1.400E-05 |
| 12288 | 615 | L615G | A | 1.6e-05 | 1.5e-05 | 9.8e-06 | 1.360E-05 | 3.329E-06 | 9.800E-06 | 1.600E-05 |
| 12289 | 615 | L615H | A | 9.2e-06 | 9.9e-06 | 1.3e-05 | 1.070E-05 | 2.022E-06 | 9.200E-06 | 1.300E-05 |
| 12290 | 615 | L615I | A | 1.6e-05 | 7.8e-06 | 8.6e-06 | 1.080E-05 | 4.521E-06 | 7.800E-06 | 1.600E-05 |
| 12291 | 615 | L615L | A | 9.2e-06 | 1.7e-05 | 1.4e-05 | 1.340E-05 | 3.934E-06 | 9.200E-06 | 1.700E-05 |
| 12292 | 615 | L615K | A | 1.2e-05 | 1.6e-05 | 9e-06 | 1.233E-05 | 3.512E-06 | 9.000E-06 | 1.600E-05 |
| 12293 | 615 | L615M | A | 1.8e-05 | 9.1e-06 | 1.1e-05 | 1.270E-05 | 4.687E-06 | 9.100E-06 | 1.800E-05 |
| 12294 | 615 | L615F | A | 1.2e-05 | 1.2e-05 | 1.2e-05 | 1.200E-05 | 0.000E+00 | 1.200E-05 | 1.200E-05 |
| 12295 | 615 | L615P | A | 1.3e-05 | 1.1e-05 | 8.7e-06 | 1.090E-05 | 2.152E-06 | 8.700E-06 | 1.300E-05 |
| 12296 | 615 | L615S | A | 1.5e-05 | 1.4e-05 | 1.5e-05 | 1.467E-05 | 5.774E-07 | 1.400E-05 | 1.500E-05 |
| 12297 | 615 | L615T | A | 1.5e-05 | 1e-05 | 1.4e-05 | 1.300E-05 | 2.646E-06 | 1.000E-05 | 1.500E-05 |
| 12298 | 615 | L615W | A | 1.4e-05 | 1.5e-05 | 1.2e-05 | 1.367E-05 | 1.528E-06 | 1.200E-05 | 1.500E-05 |
| 12299 | 615 | L615Y | A | 1.1e-05 | 8.2e-06 | 9.7e-06 | 9.633E-06 | 1.401E-06 | 8.200E-06 | 1.100E-05 |
| 12300 | 615 | L615V | A | 1.4e-05 | 1.2e-05 | 1.7e-05 | 1.433E-05 | 2.517E-06 | 1.200E-05 | 1.700E-05 |
| 12301 | 616 | D616A | A | 1.1e-05 | 1.3e-05 | 1.5e-05 | 1.300E-05 | 2.000E-06 | 1.100E-05 | 1.500E-05 |
| 12302 | 616 | D616R | A | 1.6e-05 | 8.2e-06 | 1.1e-05 | 1.173E-05 | 3.951E-06 | 8.200E-06 | 1.600E-05 |
| 12303 | 616 | D616N | A | 1.6e-05 | 1e-05 | 1.1e-05 | 1.233E-05 | 3.215E-06 | 1.000E-05 | 1.600E-05 |
| 12304 | 616 | D616D | A | 1.5e-05 | 9.8e-06 | 1.1e-05 | 1.193E-05 | 2.723E-06 | 9.800E-06 | 1.500E-05 |
| 12305 | 616 | D616C | A | 1.4e-05 | 1.1e-05 | 8e-06 | 1.100E-05 | 3.000E-06 | 8.000E-06 | 1.400E-05 |
| 12306 | 616 | D616E | A | 1.2e-05 | 1.7e-05 | 1.2e-05 | 1.367E-05 | 2.887E-06 | 1.200E-05 | 1.700E-05 |
| 12307 | 616 | D616Q | A | 1.3e-05 | 1e-05 | 1.5e-05 | 1.267E-05 | 2.517E-06 | 1.000E-05 | 1.500E-05 |
| 12308 | 616 | D616G | A | 9.1e-06 | 1.5e-05 | 1e-05 | 1.137E-05 | 3.179E-06 | 9.100E-06 | 1.500E-05 |
| 12309 | 616 | D616H | A | 1.4e-05 | 8.7e-06 | 9.2e-06 | 1.063E-05 | 2.926E-06 | 8.700E-06 | 1.400E-05 |
| 12310 | 616 | D616I | A | 1.3e-05 | 1.1e-05 | 9.4e-06 | 1.113E-05 | 1.804E-06 | 9.400E-06 | 1.300E-05 |
| 12311 | 616 | D616L | A | 1.5e-05 | 1.6e-05 | 8e-06 | 1.300E-05 | 4.359E-06 | 8.000E-06 | 1.600E-05 |
| 12312 | 616 | D616K | A | 1.5e-05 | 1.4e-05 | 1.2e-05 | 1.367E-05 | 1.528E-06 | 1.200E-05 | 1.500E-05 |
| 12313 | 616 | D616M | A | 1.8e-05 | 1.1e-05 | 1.1e-05 | 1.333E-05 | 4.041E-06 | 1.100E-05 | 1.800E-05 |
| 12314 | 616 | D616F | A | 1.5e-05 | 1.6e-05 | 1.5e-05 | 1.533E-05 | 5.774E-07 | 1.500E-05 | 1.600E-05 |
| 12315 | 616 | D616P | A | 1.6e-05 | 1e-05 | 1.3e-05 | 1.300E-05 | 3.000E-06 | 1.000E-05 | 1.600E-05 |
| 12316 | 616 | D616S | A | 1.5e-05 | 1e-05 | 1.1e-05 | 1.200E-05 | 2.646E-06 | 1.000E-05 | 1.500E-05 |
| 12317 | 616 | D616T | A | 9.8e-06 | 1e-05 | 1.1e-05 | 1.027E-05 | 6.429E-07 | 9.800E-06 | 1.100E-05 |
| 12318 | 616 | D616W | A | 8e-06 | 1e-05 | 8.6e-06 | 8.867E-06 | 1.026E-06 | 8.000E-06 | 1.000E-05 |
| 12319 | 616 | D616Y | A | 1.5e-05 | 1.3e-05 | 1.4e-05 | 1.400E-05 | 1.000E-06 | 1.300E-05 | 1.500E-05 |
| 12320 | 616 | D616V | A | 1.6e-05 | 1.4e-05 | 1.6e-05 | 1.533E-05 | 1.155E-06 | 1.400E-05 | 1.600E-05 |
| 12321 | 617 | D617A | A | 1.2e-05 | 1e-05 | 1.2e-05 | 1.133E-05 | 1.155E-06 | 1.000E-05 | 1.200E-05 |
| 12322 | 617 | D617R | A | 1.1e-05 | 1.5e-05 | 1.8e-05 | 1.467E-05 | 3.512E-06 | 1.100E-05 | 1.800E-05 |
| 12323 | 617 | D617N | A | 1.6e-05 | 1.7e-05 | 1.5e-05 | 1.600E-05 | 1.000E-06 | 1.500E-05 | 1.700E-05 |
| 12324 | 617 | D617D | A | 1.5e-05 | 8.9e-06 | 8.4e-06 | 1.077E-05 | 3.675E-06 | 8.400E-06 | 1.500E-05 |

| | | | | | | | | | | |
|---|---|---|---|---|---|---|---|---|---|---|
| 12325 | 617 | D617C | A | 1.2e-05 | 1.2e-05 | 1.5e-05 | 1.300E-05 | 1.732E-06 | 1.200E-05 | 1.500E-05 |
| 12326 | 617 | D617E | A | 9.4e-06 | 1.4e-05 | 1.3e-05 | 1.213E-05 | 2.419E-06 | 9.400E-06 | 1.400E-05 |
| 12327 | 617 | D617Q | A | 1.1e-05 | 1.4e-05 | 1.5e-05 | 1.333E-05 | 2.082E-06 | 1.100E-05 | 1.500E-05 |
| 12328 | 617 | D617G | A | 1.6e-05 | 1.5e-05 | 1.7e-05 | 1.600E-05 | 1.000E-06 | 1.500E-05 | 1.700E-05 |
| 12329 | 617 | D617H | A | 1e-05 | 1.5e-05 | 1.6e-05 | 1.367E-05 | 3.215E-06 | 1.000E-05 | 1.600E-05 |
| 12330 | 617 | D617I | A | 1.4e-05 | 1.1e-05 | 8e-06 | 1.100E-05 | 3.000E-06 | 8.000E-06 | 1.400E-05 |
| 12331 | 617 | D617L | A | 1.1e-05 | 1e-05 | 1.5e-05 | 1.200E-05 | 2.646E-06 | 1.000E-05 | 1.500E-05 |
| 12332 | 617 | D617K | A | 1.4e-05 | 1.3e-05 | 1.4e-05 | 1.367E-05 | 5.774E-07 | 1.300E-05 | 1.400E-05 |
| 12333 | 617 | D617M | A | 1.7e-05 | 1.1e-05 | 1.3e-05 | 1.367E-05 | 3.055E-06 | 1.100E-05 | 1.700E-05 |
| 12334 | 617 | D617F | A | 1.5e-05 | 1.7e-05 | 1e-05 | 1.400E-05 | 3.606E-06 | 1.000E-05 | 1.700E-05 |
| 12335 | 617 | D617P | A | 1.7e-05 | 9.7e-06 | 1.5e-05 | 1.390E-05 | 3.772E-06 | 9.700E-06 | 1.700E-05 |
| 12336 | 617 | D617S | A | 1.5e-05 | 1.1e-05 | 1e-05 | 1.200E-05 | 2.646E-06 | 1.000E-05 | 1.500E-05 |
| 12337 | 617 | D617T | A | 9.9e-06 | 9.7e-06 | 1.5e-05 | 1.153E-05 | 3.004E-06 | 9.700E-06 | 1.500E-05 |
| 12338 | 617 | D617W | A | 1.4e-05 | 1e-05 | 9.4e-06 | 1.113E-05 | 2.501E-06 | 9.400E-06 | 1.400E-05 |
| 12339 | 617 | D617Y | A | 1.3e-05 | 8.8e-06 | 9.2e-06 | 1.033E-05 | 2.318E-06 | 8.800E-06 | 1.300E-05 |
| 12340 | 617 | D617V | A | 1.1e-05 | 1.2e-05 | 1.4e-05 | 1.233E-05 | 1.528E-06 | 1.100E-05 | 1.400E-05 |
| 12341 | 618 | K618A | A | 1e-05 | 2.5e-05 | 9.1e-06 | 1.470E-05 | 8.931E-06 | 9.100E-06 | 2.500E-05 |
| 12342 | 618 | K618R | A | 1.2e-05 | 1.3e-05 | 1.4e-05 | 1.300E-05 | 1.000E-06 | 1.200E-05 | 1.400E-05 |
| 12343 | 618 | K618N | A | 1.2e-05 | 1.3e-05 | 1e-05 | 1.167E-05 | 1.528E-06 | 1.000E-05 | 1.300E-05 |
| 12344 | 618 | K618D | A | 1.5e-05 | 1.4e-05 | 1.6e-05 | 1.500E-05 | 1.000E-06 | 1.400E-05 | 1.600E-05 |
| 12345 | 618 | K618C | A | 1.5e-05 | 1e-05 | 1.3e-05 | 1.267E-05 | 2.517E-06 | 1.000E-05 | 1.500E-05 |
| 12346 | 618 | K618E | A | 9.6e-06 | 1.2e-05 | 1.4e-05 | 1.187E-05 | 2.203E-06 | 9.600E-06 | 1.400E-05 |
| 12347 | 618 | K618Q | A | 1.6e-05 | 1.6e-05 | 9.7e-06 | 1.390E-05 | 3.637E-06 | 9.700E-06 | 1.600E-05 |
| 12348 | 618 | K618G | A | 1.1e-05 | 1.4e-05 | 1.6e-05 | 1.367E-05 | 2.517E-06 | 1.100E-05 | 1.600E-05 |
| 12349 | 618 | K618H | A | 1.5e-05 | 9e-06 | 8.7e-06 | 1.090E-05 | 3.554E-06 | 8.700E-06 | 1.500E-05 |
| 12350 | 618 | K618I | A | 1.3e-05 | 2.4e-05 | 1.3e-05 | 1.667E-05 | 6.351E-06 | 1.300E-05 | 2.400E-05 |
| 12351 | 618 | K618L | A | 1.3e-05 | 1.2e-05 | 1.3e-05 | 1.267E-05 | 5.774E-07 | 1.200E-05 | 1.300E-05 |
| 12352 | 618 | K618K | A | 2.6e-05 | 1e-05 | 1.6e-05 | 1.733E-05 | 8.083E-06 | 1.000E-05 | 2.600E-05 |
| 12353 | 618 | K618M | A | 1.7e-05 | 1.4e-05 | 1.5e-05 | 1.533E-05 | 1.528E-06 | 1.400E-05 | 1.700E-05 |
| 12354 | 618 | K618F | A | 1.4e-05 | 1.5e-05 | 1e-05 | 1.300E-05 | 2.646E-06 | 1.000E-05 | 1.500E-05 |
| 12355 | 618 | K618P | A | 1e-05 | 1.4e-05 | 1.2e-05 | 1.200E-05 | 2.000E-06 | 1.000E-05 | 1.400E-05 |
| 12356 | 618 | K618S | A | 1.4e-05 | 9.3e-06 | 1.3e-05 | 1.210E-05 | 2.476E-06 | 9.300E-06 | 1.400E-05 |
| 12357 | 618 | K618T | A | 1.2e-05 | 1.6e-05 | 9.8e-06 | 1.260E-05 | 3.143E-06 | 9.800E-06 | 1.600E-05 |
| 12358 | 618 | K618W | A | 1.4e-05 | 1.5e-05 | 1.5e-05 | 1.467E-05 | 5.774E-07 | 1.400E-05 | 1.500E-05 |
| 12359 | 618 | K618Y | A | 1.4e-05 | 1.4e-05 | 1.3e-05 | 1.367E-05 | 5.774E-07 | 1.300E-05 | 1.400E-05 |
| 12360 | 618 | K618V | A | 1e-05 | 1.5e-05 | 1.6e-05 | 1.367E-05 | 3.215E-06 | 1.000E-05 | 1.600E-05 |
| 12361 | 619 | G619A | A | 1.5e-05 | 1.3e-05 | 1e-05 | 1.267E-05 | 2.517E-06 | 1.000E-05 | 1.500E-05 |
| 12362 | 619 | G619R | A | 9.3e-06 | 1.6e-05 | 1.3e-05 | 1.277E-05 | 3.356E-06 | 9.300E-06 | 1.600E-05 |
| 12363 | 619 | G619N | A | 1e-05 | 1.5e-05 | 1.5e-05 | 1.333E-05 | 2.887E-06 | 1.000E-05 | 1.500E-05 |
| 12364 | 619 | G619D | A | 1.8e-05 | 1e-05 | 1.5e-05 | 1.433E-05 | 4.041E-06 | 1.000E-05 | 1.800E-05 |
| 12365 | 619 | G619C | A | 6.5e-06 | 9.1e-06 | 1.4e-05 | 9.867E-06 | 3.808E-06 | 6.500E-06 | 1.400E-05 |
| 12366 | 619 | G619E | A | 1.3e-05 | 1.1e-05 | 9.8e-06 | 1.127E-05 | 1.617E-06 | 9.800E-06 | 1.300E-05 |
| 12367 | 619 | G619Q | A | 1.2e-05 | 1.5e-05 | 9.7e-06 | 1.223E-05 | 2.658E-06 | 9.700E-06 | 1.500E-05 |
| 12368 | 619 | G619G | A | 1.1e-05 | 1.2e-05 | 1.1e-05 | 1.133E-05 | 5.774E-07 | 1.100E-05 | 1.200E-05 |
| 12369 | 619 | G619H | A | 1.4e-05 | 1.4e-05 | 1e-05 | 1.267E-05 | 2.309E-06 | 1.000E-05 | 1.400E-05 |
| 12370 | 619 | G619I | A | 1.4e-05 | 1.5e-05 | 1.2e-05 | 1.367E-05 | 1.528E-06 | 1.200E-05 | 1.500E-05 |
| 12371 | 619 | G619L | A | 1.5e-05 | 1.4e-05 | 6.9e-06 | 1.197E-05 | 4.416E-06 | 6.900E-06 | 1.500E-05 |
| 12372 | 619 | G619K | A | 1.6e-05 | 1.4e-05 | 1.5e-05 | 1.500E-05 | 1.000E-06 | 1.400E-05 | 1.600E-05 |
| 12373 | 619 | G619M | A | 9.6e-06 | 1.5e-05 | 9.2e-06 | 1.127E-05 | 3.239E-06 | 9.200E-06 | 1.500E-05 |
| 12374 | 619 | G619F | A | 1.2e-05 | 1.3e-05 | 1.6e-05 | 1.367E-05 | 2.082E-06 | 1.200E-05 | 1.600E-05 |
| 12375 | 619 | G619P | A | 7.9e-06 | 1e-05 | 1.1e-05 | 9.633E-06 | 1.582E-06 | 7.900E-06 | 1.100E-05 |
| 12376 | 619 | G619S | A | 1.5e-05 | 1.5e-05 | 1.4e-05 | 1.467E-05 | 5.774E-07 | 1.400E-05 | 1.500E-05 |
| 12377 | 619 | G619T | A | 9.1e-06 | 1.8e-05 | 1.4e-05 | 1.370E-05 | 4.458E-06 | 9.100E-06 | 1.800E-05 |
| 12378 | 619 | G619W | A | 1e-05 | 1.1e-05 | 1.1e-05 | 1.067E-05 | 5.774E-07 | 1.000E-05 | 1.100E-05 |
| 12379 | 619 | G619Y | A | 1.5e-05 | 1.4e-05 | 9.8e-06 | 1.293E-05 | 2.759E-06 | 9.800E-06 | 1.500E-05 |
| 12380 | 619 | G619V | A | 1.4e-05 | 1.6e-05 | 1.6e-05 | 1.533E-05 | 1.155E-06 | 1.400E-05 | 1.600E-05 |
| 12381 | 620 | C620A | A | 1.4e-05 | 1.1e-05 | 1e-05 | 1.167E-05 | 2.082E-06 | 1.000E-05 | 1.400E-05 |
| 12382 | 620 | C620R | A | 1.1e-05 | 1.3e-05 | 9.6e-06 | 1.120E-05 | 1.709E-06 | 9.600E-06 | 1.300E-05 |
| 12383 | 620 | C620N | A | 1.1e-05 | 8.1e-06 | 1.3e-05 | 1.070E-05 | 2.464E-06 | 8.100E-06 | 1.300E-05 |
| 12384 | 620 | C620D | A | 1.1e-05 | 1.5e-05 | 1.5e-05 | 1.367E-05 | 2.309E-06 | 1.100E-05 | 1.500E-05 |
| 12385 | 620 | C620C | A | 1.1e-05 | 1.5e-05 | 1.1e-05 | 1.233E-05 | 2.309E-06 | 1.100E-05 | 1.500E-05 |
| 12386 | 620 | C620E | A | 1e-05 | 1.3e-05 | 1.5e-05 | 1.267E-05 | 2.517E-06 | 1.000E-05 | 1.500E-05 |
| 12387 | 620 | C620Q | A | 1e-05 | 7.2e-06 | 9.6e-06 | 8.933E-06 | 1.514E-06 | 7.200E-06 | 1.000E-05 |
| 12388 | 620 | C620G | A | 1.1e-05 | 1.7e-05 | 1.3e-05 | 1.367E-05 | 3.055E-06 | 1.100E-05 | 1.700E-05 |
| 12389 | 620 | C620H | A | 9.1e-06 | 1e-05 | 1.5e-05 | 1.137E-05 | 3.179E-06 | 9.100E-06 | 1.500E-05 |
| 12390 | 620 | C620I | A | 1.7e-05 | 1.1e-05 | 1.5e-05 | 1.433E-05 | 3.055E-06 | 1.100E-05 | 1.700E-05 |
| 12391 | 620 | C620L | A | 1.6e-05 | 1.5e-05 | 1.1e-05 | 1.400E-05 | 2.646E-06 | 1.100E-05 | 1.600E-05 |
| 12392 | 620 | C620K | A | 1.3e-05 | 1.5e-05 | 1.5e-05 | 1.433E-05 | 1.155E-06 | 1.300E-05 | 1.500E-05 |
| 12393 | 620 | C620M | A | 1.6e-05 | 1.6e-05 | 1e-05 | 1.400E-05 | 3.464E-06 | 1.000E-05 | 1.600E-05 |
| 12394 | 620 | C620F | A | 1.2e-05 | 1.2e-05 | 1e-05 | 1.133E-05 | 1.155E-06 | 1.000E-05 | 1.200E-05 |
| 12395 | 620 | C620P | A | 1.4e-05 | 9.3e-06 | 1.5e-05 | 1.277E-05 | 3.044E-06 | 9.300E-06 | 1.500E-05 |
| 12396 | 620 | C620S | A | 8.7e-06 | 1.4e-05 | 1.6e-05 | 1.290E-05 | 3.772E-06 | 8.700E-06 | 1.600E-05 |
| 12397 | 620 | C620T | A | 8.6e-06 | 1.4e-05 | 1.1e-05 | 1.120E-05 | 2.706E-06 | 8.600E-06 | 1.400E-05 |
| 12398 | 620 | C620W | A | 1e-05 | 1.4e-05 | 1.7e-05 | 1.367E-05 | 3.512E-06 | 1.000E-05 | 1.700E-05 |
| 12399 | 620 | C620Y | A | 1.3e-05 | 1.5e-05 | 1.3e-05 | 1.367E-05 | 1.155E-06 | 1.300E-05 | 1.500E-05 |
| 12400 | 620 | C620V | A | 1.1e-05 | 1.2e-05 | 1.3e-05 | 1.200E-05 | 1.000E-06 | 1.100E-05 | 1.300E-05 |
| 12401 | 621 | P621A | A | 1.5e-05 | 1.3e-05 | 1.6e-05 | 1.467E-05 | 1.528E-06 | 1.300E-05 | 1.600E-05 |
| 12402 | 621 | P621R | A | 1.6e-05 | 1e-05 | 9.1e-06 | 1.170E-05 | 3.751E-06 | 9.100E-06 | 1.600E-05 |
| 12403 | 621 | P621N | A | 1e-05 | 1e-05 | 1.5e-05 | 1.167E-05 | 2.887E-06 | 1.000E-05 | 1.500E-05 |
| 12404 | 621 | P621D | A | 1.1e-05 | 1e-05 | 1.4e-05 | 1.167E-05 | 2.082E-06 | 1.000E-05 | 1.400E-05 |
| 12405 | 621 | P621C | A | 8.4e-06 | 1.2e-05 | 1.2e-05 | 1.080E-05 | 2.078E-06 | 8.400E-06 | 1.200E-05 |
| 12406 | 621 | P621E | A | 8.8e-06 | 1.7e-05 | 1e-05 | 1.193E-05 | 4.429E-06 | 8.800E-06 | 1.700E-05 |
| 12407 | 621 | P621Q | A | 1.4e-05 | 1.6e-05 | 1.1e-05 | 1.367E-05 | 2.517E-06 | 1.100E-05 | 1.600E-05 |
| 12408 | 621 | P621G | A | 1.1e-05 | 1.6e-05 | 1.6e-05 | 1.433E-05 | 2.887E-06 | 1.100E-05 | 1.600E-05 |
| 12409 | 621 | P621H | A | 9.1e-06 | 1.6e-05 | 1.5e-05 | 1.337E-05 | 3.729E-06 | 9.100E-06 | 1.600E-05 |

| | | | | | | | | | | |
|---|---|---|---|---|---|---|---|---|---|---|
| 12410 | 621 | P621I | A | 1.1e-05 | 9.9e-06 | 1.5e-05 | 1.197E-05 | 2.684E-06 | 9.900E-06 | 1.500E-05 |
| 12411 | 621 | P621L | A | 1.3e-05 | 1.7e-05 | 9.3e-06 | 1.310E-05 | 3.851E-06 | 9.300E-06 | 1.700E-05 |
| 12412 | 621 | P621K | A | 1.1e-05 | 9.9e-06 | 1.5e-05 | 1.197E-05 | 2.684E-06 | 9.900E-06 | 1.500E-05 |
| 12413 | 621 | P621M | A | 1e-05 | 1.6e-05 | 1.4e-05 | 1.333E-05 | 3.055E-06 | 1.000E-05 | 1.600E-05 |
| 12414 | 621 | P621F | A | 1.5e-05 | 1e-05 | 1.7e-05 | 1.400E-05 | 3.606E-06 | 1.000E-05 | 1.700E-05 |
| 12415 | 621 | P621P | A | 1.4e-05 | 1.2e-05 | 1.2e-05 | 1.267E-05 | 1.155E-06 | 1.200E-05 | 1.400E-05 |
| 12416 | 621 | P621S | A | 1.5e-05 | 1.4e-05 | 2.4e-05 | 1.767E-05 | 5.508E-06 | 1.400E-05 | 2.400E-05 |
| 12417 | 621 | P621T | A | 1.1e-05 | 1.1e-05 | 1.6e-05 | 1.267E-05 | 2.887E-06 | 1.100E-05 | 1.600E-05 |
| 12418 | 621 | P621W | A | 1.4e-05 | 1.5e-05 | 9.3e-06 | 1.277E-05 | 3.044E-06 | 9.300E-06 | 1.500E-05 |
| 12419 | 621 | P621Y | A | 1.3e-05 | 1.5e-05 | 9.8e-06 | 1.260E-05 | 2.623E-06 | 9.800E-06 | 1.500E-05 |
| 12420 | 621 | P621V | A | 1.1e-05 | 1.6e-05 | 1.2e-05 | 1.300E-05 | 2.646E-06 | 1.100E-05 | 1.600E-05 |
| 12421 | 622 | A622A | A | 1.1e-05 | 8.8e-06 | 9.4e-06 | 9.733E-06 | 1.137E-06 | 8.800E-06 | 1.100E-05 |
| 12422 | 622 | A622R | A | 1.6e-05 | 1.4e-05 | 1.4e-05 | 1.467E-05 | 1.155E-06 | 1.400E-05 | 1.600E-05 |
| 12423 | 622 | A622N | A | 1.4e-05 | 1.4e-05 | 1e-05 | 1.267E-05 | 2.309E-06 | 1.000E-05 | 1.400E-05 |
| 12424 | 622 | A622D | A | 1.5e-05 | 1.4e-05 | 9.5e-06 | 1.283E-05 | 2.930E-06 | 9.500E-06 | 1.500E-05 |
| 12425 | 622 | A622C | A | 1e-05 | 1.4e-05 | 1.5e-05 | 1.300E-05 | 2.646E-06 | 1.000E-05 | 1.500E-05 |
| 12426 | 622 | A622E | A | 1.4e-05 | 1.1e-05 | 1.5e-05 | 1.333E-05 | 2.082E-06 | 1.100E-05 | 1.500E-05 |
| 12427 | 622 | A622Q | A | 1.4e-05 | 8.5e-06 | 9.2e-06 | 1.057E-05 | 2.994E-06 | 8.500E-06 | 1.400E-05 |
| 12428 | 622 | A622G | A | 1.6e-05 | 1.4e-05 | 1.5e-05 | 1.500E-05 | 1.000E-06 | 1.400E-05 | 1.600E-05 |
| 12429 | 622 | A622H | A | 1.5e-05 | 1.7e-05 | 1e-05 | 1.400E-05 | 3.606E-06 | 1.000E-05 | 1.700E-05 |
| 12430 | 622 | A622I | A | 1.5e-05 | 1e-05 | 1.1e-05 | 1.200E-05 | 2.646E-06 | 1.000E-05 | 1.500E-05 |
| 12431 | 622 | A622L | A | 9.7e-06 | 1.6e-05 | 9.1e-06 | 1.160E-05 | 3.822E-06 | 9.100E-06 | 1.600E-05 |
| 12432 | 622 | A622K | A | 1e-05 | 1.3e-05 | 8.6e-06 | 1.053E-05 | 2.248E-06 | 8.600E-06 | 1.300E-05 |
| 12433 | 622 | A622M | A | 1.5e-05 | 1.4e-05 | 1.6e-05 | 1.500E-05 | 1.000E-06 | 1.400E-05 | 1.600E-05 |
| 12434 | 622 | A622F | A | 9.9e-06 | 1e-05 | 1.5e-05 | 1.163E-05 | 2.916E-06 | 9.900E-06 | 1.500E-05 |
| 12435 | 622 | A622P | A | 1.1e-05 | 1.5e-05 | 1.5e-05 | 1.367E-05 | 2.309E-06 | 1.100E-05 | 1.500E-05 |
| 12436 | 622 | A622S | A | 1.6e-05 | 2.7e-05 | 1.5e-05 | 1.933E-05 | 6.658E-06 | 1.500E-05 | 2.700E-05 |
| 12437 | 622 | A622T | A | 1.6e-05 | 1.3e-05 | 1.4e-05 | 1.433E-05 | 1.528E-06 | 1.300E-05 | 1.600E-05 |
| 12438 | 622 | A622W | A | 1.6e-05 | 9.8e-06 | 1.3e-05 | 1.293E-05 | 3.101E-06 | 9.800E-06 | 1.600E-05 |
| 12439 | 622 | A622Y | A | 8.7e-06 | 1.1e-05 | 1.4e-05 | 1.123E-05 | 2.658E-06 | 8.700E-06 | 1.400E-05 |
| 12440 | 622 | A622V | A | 9e-06 | 1.5e-05 | 1.5e-05 | 1.300E-05 | 3.464E-06 | 9.000E-06 | 1.500E-05 |
| 12441 | 623 | D623A | B | 1.1e-05 | 1.2e-05 | 1.7e-05 | 1.333E-05 | 3.215E-06 | 1.100E-05 | 1.700E-05 |
| 12442 | 623 | D623R | B | 1.1e-05 | 1.2e-05 | 1e-05 | 1.100E-05 | 1.000E-06 | 1.000E-05 | 1.200E-05 |
| 12443 | 623 | D623N | B | 7.6e-06 | 1.5e-05 | 1.5e-05 | 1.253E-05 | 4.272E-06 | 7.600E-06 | 1.500E-05 |
| 12444 | 623 | D623D | B | 1.6e-05 | 1.5e-05 | 1.7e-05 | 1.600E-05 | 1.000E-06 | 1.500E-05 | 1.700E-05 |
| 12445 | 623 | D623C | B | 1.1e-05 | 1.5e-05 | 1.1e-05 | 1.233E-05 | 2.309E-06 | 1.100E-05 | 1.500E-05 |
| 12446 | 623 | D623E | B | 1e-05 | 1.2e-05 | 1.5e-05 | 1.233E-05 | 2.517E-06 | 1.000E-05 | 1.500E-05 |
| 12447 | 623 | D623Q | B | 9.6e-06 | 1e-05 | 1e-05 | 9.867E-06 | 2.309E-07 | 9.600E-06 | 1.000E-05 |
| 12448 | 623 | D623G | B | 1.5e-05 | 1.1e-05 | 1.4e-05 | 1.333E-05 | 2.082E-06 | 1.100E-05 | 1.500E-05 |
| 12449 | 623 | D623H | B | 1e-05 | 1.7e-05 | 8.5e-06 | 1.183E-05 | 4.537E-06 | 8.500E-06 | 1.700E-05 |
| 12450 | 623 | D623I | B | 1.5e-05 | 1.1e-05 | 1.5e-05 | 1.367E-05 | 2.309E-06 | 1.100E-05 | 1.500E-05 |
| 12451 | 623 | D623L | B | 1.4e-05 | 1.5e-05 | 1.5e-05 | 1.467E-05 | 5.774E-07 | 1.400E-05 | 1.500E-05 |
| 12452 | 623 | D623K | B | 9.9e-06 | 1.5e-05 | 1.5e-05 | 1.330E-05 | 2.944E-06 | 9.900E-06 | 1.500E-05 |
| 12453 | 623 | D623M | B | 6.1e-06 | 1.5e-05 | 1.8e-05 | 1.303E-05 | 6.189E-06 | 6.100E-06 | 1.800E-05 |
| 12454 | 623 | D623F | B | 1.1e-05 | 1.3e-05 | 1.5e-05 | 1.300E-05 | 2.000E-06 | 1.100E-05 | 1.500E-05 |
| 12455 | 623 | D623P | B | 1e-05 | 1.3e-05 | 1.2e-05 | 1.167E-05 | 1.528E-06 | 1.000E-05 | 1.300E-05 |
| 12456 | 623 | D623S | B | 1e-05 | 1.5e-05 | 1e-05 | 1.167E-05 | 2.887E-06 | 1.000E-05 | 1.500E-05 |
| 12457 | 623 | D623T | B | 1.6e-05 | 9.7e-06 | 1.1e-05 | 1.223E-05 | 3.326E-06 | 9.700E-06 | 1.600E-05 |
| 12458 | 623 | D623W | B | 1.4e-05 | 1e-05 | 9.2e-06 | 1.107E-05 | 2.572E-06 | 9.200E-06 | 1.400E-05 |
| 12459 | 623 | D623Y | B | 1e-05 | 8.8e-06 | 1.1e-05 | 9.933E-06 | 1.102E-06 | 8.800E-06 | 1.100E-05 |
| 12460 | 623 | D623V | B | 1.1e-05 | 1.4e-05 | 1.1e-05 | 1.200E-05 | 1.732E-06 | 1.100E-05 | 1.400E-05 |
| 12461 | 624 | I624A | B | 1.2e-05 | 1.6e-05 | 1.6e-05 | 1.467E-05 | 2.309E-06 | 1.200E-05 | 1.600E-05 |
| 12462 | 624 | I624R | B | 1.6e-05 | 9.5e-06 | 1.1e-05 | 1.217E-05 | 3.403E-06 | 9.500E-06 | 1.600E-05 |
| 12463 | 624 | I624N | B | 1.1e-05 | 1.4e-05 | 1.4e-05 | 1.300E-05 | 1.732E-06 | 1.100E-05 | 1.400E-05 |
| 12464 | 624 | I624D | B | 1.6e-05 | 1.1e-05 | 1.4e-05 | 1.367E-05 | 2.517E-06 | 1.100E-05 | 1.600E-05 |
| 12465 | 624 | I624C | B | 1.5e-05 | 1.7e-05 | 1.5e-05 | 1.567E-05 | 1.155E-06 | 1.500E-05 | 1.700E-05 |
| 12466 | 624 | I624E | B | 1.1e-05 | 1.5e-05 | 1.7e-05 | 1.433E-05 | 3.055E-06 | 1.100E-05 | 1.700E-05 |
| 12467 | 624 | I624Q | B | 1.1e-05 | 1e-05 | 1.6e-05 | 1.233E-05 | 3.215E-06 | 1.000E-05 | 1.600E-05 |
| 12468 | 624 | I624G | B | 1.1e-05 | 1.1e-05 | 9.7e-06 | 1.057E-05 | 7.506E-07 | 9.700E-06 | 1.100E-05 |
| 12469 | 624 | I624H | B | 1.6e-05 | 8.5e-06 | 1.5e-05 | 1.317E-05 | 4.072E-06 | 8.500E-06 | 1.600E-05 |
| 12470 | 624 | I624I | B | 1.6e-05 | 1.2e-05 | 1.5e-05 | 1.433E-05 | 2.082E-06 | 1.200E-05 | 1.600E-05 |
| 12471 | 624 | I624L | B | 1.4e-05 | 1.1e-05 | 1e-05 | 1.167E-05 | 2.082E-06 | 1.000E-05 | 1.400E-05 |
| 12472 | 624 | I624K | B | 1.4e-05 | 1.4e-05 | 1.2e-05 | 1.333E-05 | 1.155E-06 | 1.200E-05 | 1.400E-05 |
| 12473 | 624 | I624M | B | 1.7e-05 | 1.5e-05 | 1.5e-05 | 1.567E-05 | 1.155E-06 | 1.500E-05 | 1.700E-05 |
| 12474 | 624 | I624F | B | 1.5e-05 | 1.6e-05 | 1.6e-05 | 1.567E-05 | 5.774E-07 | 1.500E-05 | 1.600E-05 |
| 12475 | 624 | I624P | B | 1e-05 | 1.5e-05 | 1.5e-05 | 1.333E-05 | 2.887E-06 | 1.000E-05 | 1.500E-05 |
| 12476 | 624 | I624S | B | 1.1e-05 | 1.4e-05 | 1.5e-05 | 1.333E-05 | 2.082E-06 | 1.100E-05 | 1.500E-05 |
| 12477 | 624 | I624T | B | 1.8e-05 | 9e-06 | 1.5e-05 | 1.400E-05 | 4.583E-06 | 9.000E-06 | 1.800E-05 |
| 12478 | 624 | I624W | B | 1e-05 | 1.4e-05 | 9.2e-06 | 1.107E-05 | 2.572E-06 | 9.200E-06 | 1.400E-05 |
| 12479 | 624 | I624Y | B | 1.6e-05 | 1.5e-05 | 1.3e-05 | 1.467E-05 | 1.528E-06 | 1.300E-05 | 1.600E-05 |
| 12480 | 624 | I624V | B | 1.6e-05 | 1.6e-05 | 8.8e-06 | 1.360E-05 | 4.157E-06 | 8.800E-06 | 1.600E-05 |
| 12481 | 625 | Q625A | B | 1.1e-05 | 1.4e-05 | 1.5e-05 | 1.333E-05 | 2.082E-06 | 1.100E-05 | 1.500E-05 |
| 12482 | 625 | Q625R | B | 1.6e-05 | 1.6e-05 | 1e-05 | 1.400E-05 | 3.464E-06 | 1.000E-05 | 1.600E-05 |
| 12483 | 625 | Q625N | B | 1.1e-05 | 1.1e-05 | 8.4e-06 | 1.013E-05 | 1.501E-06 | 8.400E-06 | 1.100E-05 |
| 12484 | 625 | Q625D | B | 1.1e-05 | 1.4e-05 | 1.2e-05 | 1.233E-05 | 1.528E-06 | 1.100E-05 | 1.400E-05 |
| 12485 | 625 | Q625C | B | 1.6e-05 | 1e-05 | 1.4e-05 | 1.333E-05 | 3.055E-06 | 1.000E-05 | 1.600E-05 |
| 12486 | 625 | Q625E | B | 1.6e-05 | 1e-05 | 1e-05 | 1.200E-05 | 3.464E-06 | 1.000E-05 | 1.600E-05 |
| 12487 | 625 | Q625Q | B | 1.2e-05 | 1e-05 | 9.5e-06 | 1.050E-05 | 1.323E-06 | 9.500E-06 | 1.200E-05 |
| 12488 | 625 | Q625G | B | 1.7e-05 | 1.1e-05 | 1.3e-05 | 1.367E-05 | 3.055E-06 | 1.100E-05 | 1.700E-05 |
| 12489 | 625 | Q625H | B | 1.5e-05 | 1.5e-05 | 1.6e-05 | 1.533E-05 | 5.774E-07 | 1.500E-05 | 1.600E-05 |
| 12490 | 625 | Q625I | B | 8.7e-06 | 1.5e-05 | 1.5e-05 | 1.290E-05 | 3.637E-06 | 8.700E-06 | 1.500E-05 |
| 12491 | 625 | Q625L | B | 1.5e-05 | 1.7e-05 | 1.7e-05 | 1.633E-05 | 1.155E-06 | 1.500E-05 | 1.700E-05 |
| 12492 | 625 | Q625K | B | 1.6e-05 | 1.2e-05 | 9.7e-06 | 1.257E-05 | 3.188E-06 | 9.700E-06 | 1.600E-05 |
| 12493 | 625 | Q625M | B | 8.6e-06 | 1.4e-05 | 1.4e-05 | 1.220E-05 | 3.118E-06 | 8.600E-06 | 1.400E-05 |
| 12494 | 625 | Q625F | B | 8.8e-06 | 1.6e-05 | 1.2e-05 | 1.227E-05 | 3.607E-06 | 8.800E-06 | 1.600E-05 |

| | | | | | | | | | | |
|---|---|---|---|---|---|---|---|---|---|---|
| 12495 | 625 | Q625P | B | 1.2e-05 | 1.1e-05 | 1e-05 | 1.100E-05 | 1.000E-06 | 1.000E-05 | 1.200E-05 |
| 12496 | 625 | Q625S | B | 9.3e-06 | 1.1e-05 | 9.7e-06 | 1.000E-05 | 8.888E-07 | 9.300E-06 | 1.100E-05 |
| 12497 | 625 | Q625T | B | 1.5e-05 | 1.4e-05 | 1.6e-05 | 1.500E-05 | 1.000E-06 | 1.400E-05 | 1.600E-05 |
| 12498 | 625 | Q625W | B | 6.7e-06 | 1.5e-05 | 2.1e-05 | 1.423E-05 | 7.181E-06 | 6.700E-06 | 2.100E-05 |
| 12499 | 625 | Q625Y | B | 1.1e-05 | 1e-05 | 9.6e-06 | 1.020E-05 | 7.211E-07 | 9.600E-06 | 1.100E-05 |
| 12500 | 625 | Q625V | B | 1.5e-05 | 1e-05 | 1.7e-05 | 1.400E-05 | 3.606E-06 | 1.000E-05 | 1.700E-05 |
| 12501 | 626 | M626A | B | 1.6e-05 | 8.9e-06 | 1.4e-05 | 1.297E-05 | 3.661E-06 | 8.900E-06 | 1.600E-05 |
| 12502 | 626 | M626R | B | 8.3e-06 | 1.5e-05 | 1.4e-05 | 1.243E-05 | 3.614E-06 | 8.300E-06 | 1.500E-05 |
| 12503 | 626 | M626N | B | 8.2e-06 | 1.1e-05 | 1.4e-05 | 1.107E-05 | 2.901E-06 | 8.200E-06 | 1.400E-05 |
| 12504 | 626 | M626D | B | 1.6e-05 | 1.5e-05 | 1.5e-05 | 1.533E-05 | 5.774E-07 | 1.500E-05 | 1.600E-05 |
| 12505 | 626 | M626C | B | 1.3e-05 | 1.5e-05 | 9.8e-06 | 1.260E-05 | 2.623E-06 | 9.800E-06 | 1.500E-05 |
| 12506 | 626 | M626E | B | 1.6e-05 | 9.2e-06 | 1.4e-05 | 1.307E-05 | 3.495E-06 | 9.200E-06 | 1.600E-05 |
| 12507 | 626 | M626Q | B | 1.1e-05 | 1.4e-05 | 1.5e-05 | 1.333E-05 | 2.082E-06 | 1.100E-05 | 1.500E-05 |
| 12508 | 626 | M626G | B | 1e-05 | 1.5e-05 | 1e-05 | 1.167E-05 | 2.887E-06 | 1.000E-05 | 1.500E-05 |
| 12509 | 626 | M626H | B | 1.1e-05 | 9.9e-06 | 1.1e-05 | 1.063E-05 | 6.351E-07 | 9.900E-06 | 1.100E-05 |
| 12510 | 626 | M626I | B | 1.9e-05 | 1.3e-05 | 1e-05 | 1.400E-05 | 4.583E-06 | 1.000E-05 | 1.900E-05 |
| 12511 | 626 | M626L | B | 1e-05 | 1.7e-05 | 1.3e-05 | 1.333E-05 | 3.512E-06 | 1.000E-05 | 1.700E-05 |
| 12512 | 626 | M626K | B | 1.5e-05 | 1.1e-05 | 1.6e-05 | 1.400E-05 | 2.646E-06 | 1.100E-05 | 1.600E-05 |
| 12513 | 626 | M626M | B | 8.8e-06 | 1.2e-05 | 8.3e-06 | 9.700E-06 | 2.007E-06 | 8.300E-06 | 1.200E-05 |
| 12514 | 626 | M626F | B | 1.6e-05 | 1.2e-05 | 6.2e-06 | 1.140E-05 | 4.927E-06 | 6.200E-06 | 1.600E-05 |
| 12515 | 626 | M626P | B | 1.6e-05 | 1.5e-05 | 1.6e-05 | 1.567E-05 | 5.774E-07 | 1.500E-05 | 1.600E-05 |
| 12516 | 626 | M626S | B | 1.4e-05 | 8.5e-06 | 1e-05 | 1.083E-05 | 2.843E-06 | 8.500E-06 | 1.400E-05 |
| 12517 | 626 | M626T | B | 7.7e-06 | 1e-05 | 1e-05 | 9.233E-06 | 1.328E-06 | 7.700E-06 | 1.000E-05 |
| 12518 | 626 | M626W | B | 9e-06 | 1.1e-05 | 9.4e-06 | 9.800E-06 | 1.058E-06 | 9.000E-06 | 1.100E-05 |
| 12519 | 626 | M626Y | B | 1.3e-05 | 1e-05 | 9.9e-06 | 1.097E-05 | 1.762E-06 | 9.900E-06 | 1.300E-05 |
| 12520 | 626 | M626V | B | 1e-05 | 9.8e-06 | 1e-05 | 9.933E-06 | 1.155E-07 | 9.800E-06 | 1.000E-05 |
| 12521 | 627 | T627A | B | 1.4e-05 | 1.1e-05 | 1.5e-05 | 1.333E-05 | 2.082E-06 | 1.100E-05 | 1.500E-05 |
| 12522 | 627 | T627R | B | 1.8e-05 | 9.2e-06 | 1.5e-05 | 1.407E-05 | 4.474E-06 | 9.200E-06 | 1.800E-05 |
| 12523 | 627 | T627N | B | 1.6e-05 | 9.3e-06 | 1.2e-05 | 1.243E-05 | 3.371E-06 | 9.300E-06 | 1.600E-05 |
| 12524 | 627 | T627D | B | 1.5e-05 | 1.2e-05 | 1.6e-05 | 1.433E-05 | 2.082E-06 | 1.200E-05 | 1.600E-05 |
| 12525 | 627 | T627C | B | 1.1e-05 | 1.1e-05 | 1.2e-05 | 1.133E-05 | 5.774E-07 | 1.100E-05 | 1.200E-05 |
| 12526 | 627 | T627E | B | 1.6e-05 | 9.5e-06 | 1.5e-05 | 1.350E-05 | 3.500E-06 | 9.500E-06 | 1.600E-05 |
| 12527 | 627 | T627Q | B | 1.6e-05 | 1e-05 | 1.4e-05 | 1.333E-05 | 3.055E-06 | 1.000E-05 | 1.600E-05 |
| 12528 | 627 | T627G | B | 1.2e-05 | 1.6e-05 | 1.4e-05 | 1.400E-05 | 2.000E-06 | 1.200E-05 | 1.600E-05 |
| 12529 | 627 | T627H | B | 1.5e-05 | 1.2e-05 | 9.1e-06 | 1.203E-05 | 2.950E-06 | 9.100E-06 | 1.500E-05 |
| 12530 | 627 | T627I | B | 1.5e-05 | 1.2e-05 | 1.2e-05 | 1.300E-05 | 1.732E-06 | 1.200E-05 | 1.500E-05 |
| 12531 | 627 | T627L | B | 1.2e-05 | 9.8e-06 | 1e-05 | 1.060E-05 | 1.217E-06 | 9.800E-06 | 1.200E-05 |
| 12532 | 627 | T627K | B | 1.2e-05 | 9.8e-06 | 8.9e-06 | 1.023E-05 | 1.595E-06 | 8.900E-06 | 1.200E-05 |
| 12533 | 627 | T627M | B | 1.1e-05 | 5.6e-06 | 1e-05 | 8.867E-06 | 2.873E-06 | 5.600E-06 | 1.100E-05 |
| 12534 | 627 | T627F | B | 1e-05 | 1.6e-05 | 1.1e-05 | 1.233E-05 | 3.215E-06 | 1.000E-05 | 1.600E-05 |
| 12535 | 627 | T627P | B | 1.1e-05 | 9.6e-06 | 8.1e-06 | 9.567E-06 | 1.450E-06 | 8.100E-06 | 1.100E-05 |
| 12536 | 627 | T627S | B | 1.4e-05 | 1.2e-05 | 1.4e-05 | 1.333E-05 | 1.155E-06 | 1.200E-05 | 1.400E-05 |
| 12537 | 627 | T627T | B | 1.6e-05 | 1.1e-05 | 8.9e-06 | 1.197E-05 | 3.647E-06 | 8.900E-06 | 1.600E-05 |
| 12538 | 627 | T627W | B | 1.5e-05 | 1.6e-05 | 1.7e-05 | 1.600E-05 | 1.000E-06 | 1.500E-05 | 1.700E-05 |
| 12539 | 627 | T627Y | B | 1.6e-05 | 1.6e-05 | 1.4e-05 | 1.533E-05 | 1.155E-06 | 1.400E-05 | 1.600E-05 |
| 12540 | 627 | T627V | B | 1.4e-05 | 1.1e-05 | 1.6e-05 | 1.367E-05 | 2.517E-06 | 1.100E-05 | 1.600E-05 |
| 12541 | 628 | Q628A | B | 1.5e-05 | 9.9e-06 | 1.1e-05 | 1.197E-05 | 2.684E-06 | 9.900E-06 | 1.500E-05 |
| 12542 | 628 | Q628R | B | 9e-06 | 1.5e-05 | 1.4e-05 | 1.267E-05 | 3.215E-06 | 9.000E-06 | 1.500E-05 |
| 12543 | 628 | Q628N | B | 1.5e-05 | 1.1e-05 | 1.5e-05 | 1.367E-05 | 2.309E-06 | 1.100E-05 | 1.500E-05 |
| 12544 | 628 | Q628D | B | 1.3e-05 | 1.5e-05 | 1.4e-05 | 1.400E-05 | 1.000E-06 | 1.300E-05 | 1.500E-05 |
| 12545 | 628 | Q628C | B | 1.5e-05 | 1.1e-05 | 1.7e-05 | 1.433E-05 | 3.055E-06 | 1.100E-05 | 1.700E-05 |
| 12546 | 628 | Q628E | B | 9.5e-06 | 8.4e-06 | 1.6e-05 | 1.130E-05 | 4.107E-06 | 8.400E-06 | 1.600E-05 |
| 12547 | 628 | Q628Q | B | 9.2e-06 | 1.6e-05 | 1.4e-05 | 1.307E-05 | 3.495E-06 | 9.200E-06 | 1.600E-05 |
| 12548 | 628 | Q628G | B | 1.5e-05 | 9.8e-06 | 1.5e-05 | 1.327E-05 | 3.002E-06 | 9.800E-06 | 1.500E-05 |
| 12549 | 628 | Q628H | B | 7.9e-06 | 1.3e-05 | 1.5e-05 | 1.197E-05 | 3.661E-06 | 7.900E-06 | 1.500E-05 |
| 12550 | 628 | Q628I | B | 6.8e-06 | 1.1e-05 | 1e-05 | 9.267E-06 | 2.194E-06 | 6.800E-06 | 1.100E-05 |
| 12551 | 628 | Q628L | B | 1.5e-05 | 1.1e-05 | 1.6e-05 | 1.400E-05 | 2.646E-06 | 1.100E-05 | 1.600E-05 |
| 12552 | 628 | Q628K | B | 1e-05 | 1.1e-05 | 2e-05 | 1.367E-05 | 5.508E-06 | 1.000E-05 | 2.000E-05 |
| 12553 | 628 | Q628M | B | 1e-05 | 1.7e-05 | 1.5e-05 | 1.400E-05 | 3.606E-06 | 1.000E-05 | 1.700E-05 |
| 12554 | 628 | Q628F | B | 9.2e-06 | 1.2e-05 | 1.7e-05 | 1.273E-05 | 3.951E-06 | 9.200E-06 | 1.700E-05 |
| 12555 | 628 | Q628P | B | 1.5e-05 | 1.5e-05 | 9.8e-06 | 1.327E-05 | 3.002E-06 | 9.800E-06 | 1.500E-05 |
| 12556 | 628 | Q628S | B | 1.4e-05 | 1.3e-05 | 1.4e-05 | 1.367E-05 | 5.774E-07 | 1.300E-05 | 1.400E-05 |
| 12557 | 628 | Q628T | B | 1.5e-05 | 1.1e-05 | 1.3e-05 | 1.300E-05 | 2.000E-06 | 1.100E-05 | 1.500E-05 |
| 12558 | 628 | Q628W | B | 1.4e-05 | 1.4e-05 | 9.1e-06 | 1.237E-05 | 2.829E-06 | 9.100E-06 | 1.400E-05 |
| 12559 | 628 | Q628Y | B | 1.3e-05 | 1.7e-05 | 1.5e-05 | 1.500E-05 | 2.000E-06 | 1.300E-05 | 1.700E-05 |
| 12560 | 628 | Q628V | B | 1e-05 | 1.3e-05 | 1.1e-05 | 1.133E-05 | 1.528E-06 | 1.000E-05 | 1.300E-05 |
| 12561 | 629 | S629A | B | 1.1e-05 | 1.6e-05 | 1e-05 | 1.233E-05 | 3.215E-06 | 1.000E-05 | 1.600E-05 |
| 12562 | 629 | S629R | B | 1.6e-05 | 1.7e-05 | 1e-05 | 1.433E-05 | 3.786E-06 | 1.000E-05 | 1.700E-05 |
| 12563 | 629 | S629N | B | 1.1e-05 | 3.7e-05 | 2.1e-05 | 2.300E-05 | 1.311E-05 | 1.100E-05 | 3.700E-05 |
| 12564 | 629 | S629D | B | 8.5e-06 | 1.4e-05 | 1.7e-05 | 1.317E-05 | 4.311E-06 | 8.500E-06 | 1.700E-05 |
| 12565 | 629 | S629C | B | 1.5e-05 | 8.6e-06 | 1.6e-05 | 1.320E-05 | 4.015E-06 | 8.600E-06 | 1.600E-05 |
| 12566 | 629 | S629E | B | 1.5e-05 | 1.5e-05 | 1.4e-05 | 1.467E-05 | 5.774E-07 | 1.400E-05 | 1.500E-05 |
| 12567 | 629 | S629Q | B | 1.7e-05 | 1.7e-05 | 9.9e-06 | 1.463E-05 | 4.099E-06 | 9.900E-06 | 1.700E-05 |
| 12568 | 629 | S629G | B | 1.4e-05 | 1.5e-05 | 1.8e-05 | 1.567E-05 | 2.082E-06 | 1.400E-05 | 1.800E-05 |
| 12569 | 629 | S629H | B | 1.1e-05 | 1.3e-05 | 1.7e-05 | 1.367E-05 | 3.055E-06 | 1.100E-05 | 1.700E-05 |
| 12570 | 629 | S629I | B | 8.8e-06 | 1.1e-05 | 1.6e-05 | 1.193E-05 | 3.690E-06 | 8.800E-06 | 1.600E-05 |
| 12571 | 629 | S629L | B | 1.6e-05 | 1.2e-05 | 1.4e-05 | 1.400E-05 | 2.000E-06 | 1.200E-05 | 1.600E-05 |
| 12572 | 629 | S629K | B | 1.5e-05 | 1.2e-05 | 1.5e-05 | 1.400E-05 | 1.732E-06 | 1.200E-05 | 1.500E-05 |
| 12573 | 629 | S629M | B | 9.5e-06 | 1.4e-05 | 1.3e-05 | 1.217E-05 | 2.363E-06 | 9.500E-06 | 1.400E-05 |
| 12574 | 629 | S629F | B | 7.5e-06 | 9.1e-06 | 9.6e-06 | 8.733E-06 | 1.097E-06 | 7.500E-06 | 9.600E-06 |
| 12575 | 629 | S629P | B | 1.1e-05 | 1.5e-05 | 1.4e-05 | 1.333E-05 | 2.082E-06 | 1.100E-05 | 1.500E-05 |
| 12576 | 629 | S629S | B | 1.6e-05 | 1.1e-05 | 1e-05 | 1.233E-05 | 3.215E-06 | 1.000E-05 | 1.600E-05 |
| 12577 | 629 | S629T | B | 9.5e-06 | 1.4e-05 | 1.5e-05 | 1.283E-05 | 2.930E-06 | 9.500E-06 | 1.500E-05 |
| 12578 | 629 | S629W | B | 1.4e-05 | 1.3e-05 | 1.4e-05 | 1.367E-05 | 5.774E-07 | 1.300E-05 | 1.400E-05 |
| 12579 | 629 | S629Y | B | 1.3e-05 | 1.4e-05 | 1.6e-05 | 1.433E-05 | 1.528E-06 | 1.300E-05 | 1.600E-05 |

| | | | | | | | | | | |
|---|---|---|---|---|---|---|---|---|---|---|
| 12580 | 629 | S629V | B | 1.7e-05 | 1.5e-05 | 1.5e-05 | 1.567E-05 | 1.155E-06 | 1.500E-05 | 1.700E-05 |
| 12581 | 630 | P630A | B | 1.5e-05 | 1.3e-05 | 1.5e-05 | 1.433E-05 | 1.155E-06 | 1.300E-05 | 1.500E-05 |
| 12582 | 630 | P630R | B | 1.1e-05 | 1.8e-05 | 1.2e-05 | 1.367E-05 | 3.786E-06 | 1.100E-05 | 1.800E-05 |
| 12583 | 630 | P630N | B | 1.5e-05 | 9.7e-06 | 1e-05 | 1.157E-05 | 2.977E-06 | 9.700E-06 | 1.500E-05 |
| 12584 | 630 | P630D | B | 1.5e-05 | 8.8e-06 | 1.5e-05 | 1.293E-05 | 3.580E-06 | 8.800E-06 | 1.500E-05 |
| 12585 | 630 | P630C | B | 1.6e-05 | 9.8e-06 | 1.3e-05 | 1.293E-05 | 3.101E-06 | 9.800E-06 | 1.600E-05 |
| 12586 | 630 | P630E | B | 1.1e-05 | 8.1e-06 | 1.6e-05 | 1.170E-05 | 3.996E-06 | 8.100E-06 | 1.600E-05 |
| 12587 | 630 | P630Q | B | 1e-05 | 8.8e-06 | 1.6e-05 | 1.160E-05 | 3.857E-06 | 8.800E-06 | 1.600E-05 |
| 12588 | 630 | P630G | B | 9.5e-06 | 1.2e-05 | 1.2e-05 | 1.117E-05 | 1.443E-06 | 9.500E-06 | 1.200E-05 |
| 12589 | 630 | P630H | B | 8.7e-06 | 8.6e-06 | 1.6e-05 | 1.110E-05 | 4.244E-06 | 8.600E-06 | 1.600E-05 |
| 12590 | 630 | P630I | B | 1.6e-05 | 1.1e-05 | 9.2e-06 | 1.207E-05 | 3.523E-06 | 9.200E-06 | 1.600E-05 |
| 12591 | 630 | P630L | B | 9.1e-06 | 1.6e-05 | 1.2e-05 | 1.237E-05 | 3.465E-06 | 9.100E-06 | 1.600E-05 |
| 12592 | 630 | P630K | B | 1e-05 | 9.1e-06 | 1.6e-05 | 1.170E-05 | 3.751E-06 | 9.100E-06 | 1.600E-05 |
| 12593 | 630 | P630M | B | 1.1e-05 | 1e-05 | 1.4e-05 | 1.167E-05 | 2.082E-06 | 1.000E-05 | 1.400E-05 |
| 12594 | 630 | P630F | B | 1.6e-05 | 1.5e-05 | 1.6e-05 | 1.567E-05 | 5.774E-07 | 1.500E-05 | 1.600E-05 |
| 12595 | 630 | P630P | B | 1.1e-05 | 9.3e-06 | 9e-06 | 9.767E-06 | 1.079E-06 | 9.000E-06 | 1.100E-05 |
| 12596 | 630 | P630S | B | 1.1e-05 | 7.1e-06 | 9.8e-06 | 9.300E-06 | 1.997E-06 | 7.100E-06 | 1.100E-05 |
| 12597 | 630 | P630T | B | 1.6e-05 | 1.4e-05 | 7.5e-06 | 1.250E-05 | 4.444E-06 | 7.500E-06 | 1.600E-05 |
| 12598 | 630 | P630W | B | 1e-05 | 8.4e-06 | 9.5e-06 | 9.300E-06 | 8.185E-07 | 8.400E-06 | 1.000E-05 |
| 12599 | 630 | P630Y | B | 1.4e-05 | 1.6e-05 | 1.5e-05 | 1.500E-05 | 1.000E-06 | 1.400E-05 | 1.600E-05 |
| 12600 | 630 | P630V | B | 1e-05 | 1.7e-05 | 1.5e-05 | 1.400E-05 | 3.606E-06 | 1.000E-05 | 1.700E-05 |
| 12601 | 631 | S631A | B | 1.4e-05 | 1.1e-05 | 1.6e-05 | 1.367E-05 | 2.517E-06 | 1.100E-05 | 1.600E-05 |
| 12602 | 631 | S631R | B | 1.5e-05 | 1.4e-05 | 1.1e-05 | 1.333E-05 | 2.082E-06 | 1.100E-05 | 1.500E-05 |
| 12603 | 631 | S631N | B | 1.1e-05 | 1.3e-05 | 1.6e-05 | 1.333E-05 | 2.517E-06 | 1.100E-05 | 1.600E-05 |
| 12604 | 631 | S631D | B | 1e-05 | 1.5e-05 | 1.5e-05 | 1.333E-05 | 2.887E-06 | 1.000E-05 | 1.500E-05 |
| 12605 | 631 | S631C | B | 1.3e-05 | 1e-05 | 8.2e-06 | 1.040E-05 | 2.425E-06 | 8.200E-06 | 1.300E-05 |
| 12606 | 631 | S631E | B | 1.1e-05 | 7.6e-06 | 1.3e-05 | 1.053E-05 | 2.730E-06 | 7.600E-06 | 1.300E-05 |
| 12607 | 631 | S631Q | B | 1e-05 | 1.1e-05 | 1.2e-05 | 1.100E-05 | 1.000E-06 | 1.000E-05 | 1.200E-05 |
| 12608 | 631 | S631G | B | 1.6e-05 | 1.1e-05 | 1.6e-05 | 1.433E-05 | 2.887E-06 | 1.100E-05 | 1.600E-05 |
| 12609 | 631 | S631H | B | 1e-05 | 1.4e-05 | 1.4e-05 | 1.267E-05 | 2.309E-06 | 1.000E-05 | 1.400E-05 |
| 12610 | 631 | S631I | B | 1.1e-05 | 1e-05 | 1e-05 | 1.033E-05 | 5.774E-07 | 1.000E-05 | 1.100E-05 |
| 12611 | 631 | S631L | B | 1e-05 | 1.5e-05 | 1.5e-05 | 1.333E-05 | 2.887E-06 | 1.000E-05 | 1.500E-05 |
| 12612 | 631 | S631K | B | 1.6e-05 | 2.2e-05 | 1.7e-05 | 1.833E-05 | 3.215E-06 | 1.600E-05 | 2.200E-05 |
| 12613 | 631 | S631M | B | 1.1e-05 | 1.6e-05 | 1.5e-05 | 1.400E-05 | 2.646E-06 | 1.100E-05 | 1.600E-05 |
| 12614 | 631 | S631F | B | 1.6e-05 | 1.5e-05 | 1.1e-05 | 1.400E-05 | 2.646E-06 | 1.100E-05 | 1.600E-05 |
| 12615 | 631 | S631P | B | 1.8e-05 | 1.4e-05 | 1.3e-05 | 1.500E-05 | 2.646E-06 | 1.300E-05 | 1.800E-05 |
| 12616 | 631 | S631S | B | 1.1e-05 | 7.3e-06 | 1.4e-05 | 1.077E-05 | 3.356E-06 | 7.300E-06 | 1.400E-05 |
| 12617 | 631 | S631T | B | 1.4e-05 | 1.7e-05 | 1.2e-05 | 1.433E-05 | 2.517E-06 | 1.200E-05 | 1.700E-05 |
| 12618 | 631 | S631W | B | 1.3e-05 | 1.5e-05 | 1.5e-05 | 1.433E-05 | 1.155E-06 | 1.300E-05 | 1.500E-05 |
| 12619 | 631 | S631Y | B | 1.2e-05 | 1.4e-05 | 8.8e-06 | 1.160E-05 | 2.623E-06 | 8.800E-06 | 1.400E-05 |
| 12620 | 631 | S631V | B | 1.5e-05 | 1.1e-05 | 1.5e-05 | 1.367E-05 | 2.309E-06 | 1.100E-05 | 1.500E-05 |
| 12621 | 632 | S632A | B | 1.6e-05 | 1e-05 | 1.1e-05 | 1.233E-05 | 3.215E-06 | 1.000E-05 | 1.600E-05 |
| 12622 | 632 | S632R | B | 1.4e-05 | 1.4e-05 | 8.4e-06 | 1.213E-05 | 3.233E-06 | 8.400E-06 | 1.400E-05 |
| 12623 | 632 | S632N | B | 1.7e-05 | 1.6e-05 | 1.6e-05 | 1.633E-05 | 5.774E-07 | 1.600E-05 | 1.700E-05 |
| 12624 | 632 | S632D | B | 1.2e-05 | 9.3e-06 | 1.5e-05 | 1.210E-05 | 2.851E-06 | 9.300E-06 | 1.500E-05 |
| 12625 | 632 | S632C | B | 8.8e-06 | 1.6e-05 | 1.1e-05 | 1.193E-05 | 3.690E-06 | 8.800E-06 | 1.600E-05 |
| 12626 | 632 | S632E | B | 1.5e-05 | 1.6e-05 | 1.6e-05 | 1.567E-05 | 5.774E-07 | 1.500E-05 | 1.600E-05 |
| 12627 | 632 | S632Q | B | 1.4e-05 | 1.1e-05 | 8.6e-06 | 1.120E-05 | 2.706E-06 | 8.600E-06 | 1.400E-05 |
| 12628 | 632 | S632G | B | 1.6e-05 | 8.8e-06 | 1.2e-05 | 1.227E-05 | 3.607E-06 | 8.800E-06 | 1.600E-05 |
| 12629 | 632 | S632H | B | 1.3e-05 | 1.3e-05 | 1.1e-05 | 1.233E-05 | 1.155E-06 | 1.100E-05 | 1.300E-05 |
| 12630 | 632 | S632I | B | 1.6e-05 | 1.2e-05 | 1.5e-05 | 1.433E-05 | 2.082E-06 | 1.200E-05 | 1.600E-05 |
| 12631 | 632 | S632L | B | 1.6e-05 | 1.3e-05 | 1e-05 | 1.300E-05 | 3.000E-06 | 1.000E-05 | 1.600E-05 |
| 12632 | 632 | S632K | B | 1.6e-05 | 1.7e-05 | 1.5e-05 | 1.600E-05 | 1.000E-06 | 1.500E-05 | 1.700E-05 |
| 12633 | 632 | S632M | B | 9.5e-06 | 1.1e-05 | 9.4e-06 | 9.967E-06 | 8.963E-07 | 9.400E-06 | 1.100E-05 |
| 12634 | 632 | S632F | B | 1.2e-05 | 1.5e-05 | 9.3e-06 | 1.210E-05 | 2.851E-06 | 9.300E-06 | 1.500E-05 |
| 12635 | 632 | S632P | B | 1.5e-05 | 1.1e-05 | 1.3e-05 | 1.300E-05 | 2.000E-06 | 1.100E-05 | 1.500E-05 |
| 12636 | 632 | S632S | B | 1.3e-05 | 5.9e-06 | 1.1e-05 | 9.967E-06 | 3.661E-06 | 5.900E-06 | 1.300E-05 |
| 12637 | 632 | S632T | B | 1.1e-05 | 1.5e-05 | 1.1e-05 | 1.233E-05 | 2.309E-06 | 1.100E-05 | 1.500E-05 |
| 12638 | 632 | S632W | B | 1.1e-05 | 1e-05 | 1.5e-05 | 1.200E-05 | 2.646E-06 | 1.000E-05 | 1.500E-05 |
| 12639 | 632 | S632Y | B | 1.3e-05 | 8.8e-06 | 1.1e-05 | 1.093E-05 | 2.101E-06 | 8.800E-06 | 1.300E-05 |
| 12640 | 632 | S632V | B | 1.7e-05 | 1.6e-05 | 1.5e-05 | 1.600E-05 | 1.000E-06 | 1.500E-05 | 1.700E-05 |
| 12641 | 633 | L633A | B | 9.9e-06 | 1.3e-05 | 1.3e-05 | 1.197E-05 | 1.790E-06 | 9.900E-06 | 1.300E-05 |
| 12642 | 633 | L633R | B | 1.6e-05 | 9.3e-06 | 1.3e-05 | 1.277E-05 | 3.356E-06 | 9.300E-06 | 1.600E-05 |
| 12643 | 633 | L633N | B | 9.2e-06 | 1.5e-05 | 1.2e-05 | 1.207E-05 | 2.901E-06 | 9.200E-06 | 1.500E-05 |
| 12644 | 633 | L633D | B | 9.7e-06 | 1e-05 | 8.2e-06 | 9.300E-06 | 9.644E-07 | 8.200E-06 | 1.000E-05 |
| 12645 | 633 | L633C | B | 1.7e-05 | 1.5e-05 | 1.6e-05 | 1.600E-05 | 1.000E-06 | 1.500E-05 | 1.700E-05 |
| 12646 | 633 | L633E | B | 1e-05 | 1.5e-05 | 1.5e-05 | 1.333E-05 | 2.887E-06 | 1.000E-05 | 1.500E-05 |
| 12647 | 633 | L633Q | B | 9.7e-06 | 1.5e-05 | 1e-05 | 1.157E-05 | 2.977E-06 | 9.700E-06 | 1.500E-05 |
| 12648 | 633 | L633G | B | 9.9e-06 | 1.7e-05 | 1.5e-05 | 1.397E-05 | 3.661E-06 | 9.900E-06 | 1.700E-05 |
| 12649 | 633 | L633H | B | 1.5e-05 | 1.6e-05 | 1.4e-05 | 1.500E-05 | 1.000E-06 | 1.400E-05 | 1.600E-05 |
| 12650 | 633 | L633I | B | 1.2e-05 | 1.5e-05 | 9.3e-06 | 1.210E-05 | 2.851E-06 | 9.300E-06 | 1.500E-05 |
| 12651 | 633 | L633L | B | 1.2e-05 | 9.9e-06 | 1.1e-05 | 1.097E-05 | 1.050E-06 | 9.900E-06 | 1.200E-05 |
| 12652 | 633 | L633K | B | 1e-05 | 1.2e-05 | 1.2e-05 | 1.133E-05 | 1.155E-06 | 1.000E-05 | 1.200E-05 |
| 12653 | 633 | L633M | B | 1e-05 | 1.5e-05 | 1e-05 | 1.167E-05 | 2.887E-06 | 1.000E-05 | 1.500E-05 |
| 12654 | 633 | L633F | B | 1.6e-05 | 1.2e-05 | 1.1e-05 | 1.300E-05 | 2.646E-06 | 1.100E-05 | 1.600E-05 |
| 12655 | 633 | L633P | B | 1e-05 | 1.1e-05 | 1.4e-05 | 1.167E-05 | 2.082E-06 | 1.000E-05 | 1.400E-05 |
| 12656 | 633 | L633S | B | 1.5e-05 | 1.5e-05 | 1e-05 | 1.333E-05 | 2.887E-06 | 1.000E-05 | 1.500E-05 |
| 12657 | 633 | L633T | B | 1e-05 | 9.4e-06 | 1.4e-05 | 1.113E-05 | 2.501E-06 | 9.400E-06 | 1.400E-05 |
| 12658 | 633 | L633W | B | 1e-05 | 1.6e-05 | 1.5e-05 | 1.367E-05 | 3.215E-06 | 1.000E-05 | 1.600E-05 |
| 12659 | 633 | L633Y | B | 1.5e-05 | 1.6e-05 | 1.5e-05 | 1.533E-05 | 5.774E-07 | 1.500E-05 | 1.600E-05 |
| 12660 | 633 | L633V | B | 1.3e-05 | 1.4e-05 | 1.2e-05 | 1.300E-05 | 1.000E-06 | 1.200E-05 | 1.400E-05 |
| 12661 | 634 | S634A | B | 1.6e-05 | 1.3e-05 | 1.7e-05 | 1.533E-05 | 2.082E-06 | 1.300E-05 | 1.700E-05 |
| 12662 | 634 | S634R | B | 1.2e-05 | 1.5e-05 | 1.5e-05 | 1.400E-05 | 1.732E-06 | 1.200E-05 | 1.500E-05 |
| 12663 | 634 | S634N | B | 1.3e-05 | 1.7e-05 | 1.5e-05 | 1.500E-05 | 2.000E-06 | 1.300E-05 | 1.700E-05 |
| 12664 | 634 | S634D | B | 1.7e-05 | 1.5e-05 | 1.4e-05 | 1.533E-05 | 1.528E-06 | 1.400E-05 | 1.700E-05 |

| | | | | | | | | | | |
|---|---|---|---|---|---|---|---|---|---|---|
| 12665 | 634 | S634C | B | 1.5e-05 | 1.5e-05 | 1.3e-05 | 1.433E-05 | 1.155E-06 | 1.300E-05 | 1.500E-05 |
| 12666 | 634 | S634E | B | 1.3e-05 | 1.5e-05 | 1.1e-05 | 1.300E-05 | 2.000E-06 | 1.100E-05 | 1.500E-05 |
| 12667 | 634 | S634Q | B | 1.4e-05 | 1.2e-05 | 1.2e-05 | 1.267E-05 | 1.155E-06 | 1.200E-05 | 1.400E-05 |
| 12668 | 634 | S634G | B | 1.6e-05 | 1.1e-05 | 1.6e-05 | 1.433E-05 | 2.887E-06 | 1.100E-05 | 1.600E-05 |
| 12669 | 634 | S634H | B | 1.1e-05 | 1.4e-05 | 6.5e-06 | 1.050E-05 | 3.775E-06 | 6.500E-06 | 1.400E-05 |
| 12670 | 634 | S634I | B | 1e-05 | 1.2e-05 | 1.6e-05 | 1.267E-05 | 3.055E-06 | 1.000E-05 | 1.600E-05 |
| 12671 | 634 | S634L | B | 1.7e-05 | 8.5e-06 | 1.6e-05 | 1.383E-05 | 4.646E-06 | 8.500E-06 | 1.700E-05 |
| 12672 | 634 | S634K | B | 1.5e-05 | 1.7e-05 | 1.5e-05 | 1.567E-05 | 1.155E-06 | 1.500E-05 | 1.700E-05 |
| 12673 | 634 | S634M | B | 1.2e-05 | 1.6e-05 | 1.4e-05 | 1.400E-05 | 2.000E-06 | 1.200E-05 | 1.600E-05 |
| 12674 | 634 | S634F | B | 1.6e-05 | 1.7e-05 | 1.5e-05 | 1.600E-05 | 1.000E-06 | 1.500E-05 | 1.700E-05 |
| 12675 | 634 | S634P | B | 1.7e-05 | 1.6e-05 | 1.1e-05 | 1.467E-05 | 3.215E-06 | 1.100E-05 | 1.700E-05 |
| 12676 | 634 | S634S | B | 8.1e-06 | 1.1e-05 | 1.4e-05 | 1.103E-05 | 2.950E-06 | 8.100E-06 | 1.400E-05 |
| 12677 | 634 | S634T | B | 1.4e-05 | 1e-05 | 1.5e-05 | 1.300E-05 | 2.646E-06 | 1.000E-05 | 1.500E-05 |
| 12678 | 634 | S634W | B | 8.7e-06 | 1.1e-05 | 1.5e-05 | 1.157E-05 | 3.188E-06 | 8.700E-06 | 1.500E-05 |
| 12679 | 634 | S634Y | B | 8.9e-06 | 1.6e-05 | 1.6e-05 | 1.363E-05 | 4.099E-06 | 8.900E-06 | 1.600E-05 |
| 12680 | 634 | S634V | B | 1.6e-05 | 1.4e-05 | 1.4e-05 | 1.467E-05 | 1.155E-06 | 1.400E-05 | 1.600E-05 |
| 12681 | 635 | A635A | B | 1.6e-05 | 1.4e-05 | 1.4e-05 | 1.467E-05 | 1.155E-06 | 1.400E-05 | 1.600E-05 |
| 12682 | 635 | A635R | B | 1.6e-05 | 1.6e-05 | 9.8e-06 | 1.393E-05 | 3.580E-06 | 9.800E-06 | 1.600E-05 |
| 12683 | 635 | A635N | B | 1.3e-05 | 1.5e-05 | 8.5e-06 | 1.217E-05 | 3.329E-06 | 8.500E-06 | 1.500E-05 |
| 12684 | 635 | A635D | B | 1.3e-05 | 1.1e-05 | 1.7e-05 | 1.367E-05 | 3.055E-06 | 1.100E-05 | 1.700E-05 |
| 12685 | 635 | A635C | B | 1.5e-05 | 8.6e-06 | 9.6e-06 | 1.107E-05 | 3.443E-06 | 8.600E-06 | 1.500E-05 |
| 12686 | 635 | A635E | B | 9.1e-06 | 9.7e-06 | 1.5e-05 | 1.127E-05 | 3.247E-06 | 9.100E-06 | 1.500E-05 |
| 12687 | 635 | A635Q | B | 1e-05 | 1.5e-05 | 1.2e-05 | 1.233E-05 | 2.517E-06 | 1.000E-05 | 1.500E-05 |
| 12688 | 635 | A635G | B | 9.6e-06 | 1.1e-05 | 9.3e-06 | 9.967E-06 | 9.074E-07 | 9.300E-06 | 1.100E-05 |
| 12689 | 635 | A635H | B | 1.4e-05 | 1.6e-05 | 1.1e-05 | 1.367E-05 | 2.517E-06 | 1.100E-05 | 1.600E-05 |
| 12690 | 635 | A635I | B | 1.5e-05 | 8.8e-06 | 1.3e-05 | 1.227E-05 | 3.164E-06 | 8.800E-06 | 1.500E-05 |
| 12691 | 635 | A635L | B | 9.8e-06 | 1e-05 | 1.5e-05 | 1.160E-05 | 2.946E-06 | 9.800E-06 | 1.500E-05 |
| 12692 | 635 | A635K | B | 8.7e-06 | 1.4e-05 | 1.5e-05 | 1.257E-05 | 3.386E-06 | 8.700E-06 | 1.500E-05 |
| 12693 | 635 | A635M | B | 1.4e-05 | 1.4e-05 | 1.1e-05 | 1.300E-05 | 1.732E-06 | 1.100E-05 | 1.400E-05 |
| 12694 | 635 | A635F | B | 1.4e-05 | 1.3e-05 | 9.5e-06 | 1.217E-05 | 2.363E-06 | 9.500E-06 | 1.400E-05 |
| 12695 | 635 | A635P | B | 1.3e-05 | 1.4e-05 | 9.9e-06 | 1.230E-05 | 2.138E-06 | 9.900E-06 | 1.400E-05 |
| 12696 | 635 | A635S | B | 1.1e-05 | 8.9e-06 | 1.4e-05 | 1.130E-05 | 2.563E-06 | 8.900E-06 | 1.400E-05 |
| 12697 | 635 | A635T | B | 1.3e-05 | 1.5e-05 | 1.3e-05 | 1.367E-05 | 1.155E-06 | 1.300E-05 | 1.500E-05 |
| 12698 | 635 | A635W | B | 1.6e-05 | 1.4e-05 | 1.5e-05 | 1.500E-05 | 1.000E-06 | 1.400E-05 | 1.600E-05 |
| 12699 | 635 | A635Y | B | 1.2e-05 | 1.7e-05 | 1.5e-05 | 1.467E-05 | 2.517E-06 | 1.200E-05 | 1.700E-05 |
| 12700 | 635 | A635V | B | 1e-05 | 1.5e-05 | 9e-06 | 1.133E-05 | 3.215E-06 | 9.000E-06 | 1.500E-05 |
| 12701 | 636 | S636A | B | 1.5e-05 | 1.6e-05 | 1.4e-05 | 1.500E-05 | 1.000E-06 | 1.400E-05 | 1.600E-05 |
| 12702 | 636 | S636R | B | 1.5e-05 | 1e-05 | 1e-05 | 1.167E-05 | 2.887E-06 | 1.000E-05 | 1.500E-05 |
| 12703 | 636 | S636N | B | 1.7e-05 | 1.3e-05 | 1e-05 | 1.333E-05 | 3.512E-06 | 1.000E-05 | 1.700E-05 |
| 12704 | 636 | S636D | B | 1.4e-05 | 1.4e-05 | 1.1e-05 | 1.300E-05 | 1.732E-06 | 1.100E-05 | 1.400E-05 |
| 12705 | 636 | S636C | B | 1.5e-05 | 1.1e-05 | 1.7e-05 | 1.433E-05 | 3.055E-06 | 1.100E-05 | 1.700E-05 |
| 12706 | 636 | S636E | B | 1.4e-05 | 1.7e-05 | 8.4e-06 | 1.313E-05 | 4.365E-06 | 8.400E-06 | 1.700E-05 |
| 12707 | 636 | S636Q | B | 9.6e-06 | 1.6e-05 | 1.7e-05 | 1.420E-05 | 4.015E-06 | 9.600E-06 | 1.700E-05 |
| 12708 | 636 | S636G | B | 1e-05 | 9.9e-06 | 1.5e-05 | 1.163E-05 | 2.916E-06 | 9.900E-06 | 1.500E-05 |
| 12709 | 636 | S636H | B | 1.9e-05 | 1.4e-05 | 1.7e-05 | 1.667E-05 | 2.517E-06 | 1.400E-05 | 1.900E-05 |
| 12710 | 636 | S636I | B | 1.6e-05 | 1.1e-05 | 1.7e-05 | 1.467E-05 | 3.215E-06 | 1.100E-05 | 1.700E-05 |
| 12711 | 636 | S636L | B | 1.6e-05 | 1.3e-05 | 1.2e-05 | 1.367E-05 | 2.082E-06 | 1.200E-05 | 1.600E-05 |
| 12712 | 636 | S636K | B | 1.4e-05 | 1.4e-05 | 1.1e-05 | 1.300E-05 | 1.732E-06 | 1.100E-05 | 1.400E-05 |
| 12713 | 636 | S636M | B | 1.5e-05 | 1.6e-05 | 1.5e-05 | 1.533E-05 | 5.774E-07 | 1.500E-05 | 1.600E-05 |
| 12714 | 636 | S636F | B | 1.1e-05 | 1.3e-05 | 1e-05 | 1.133E-05 | 1.528E-06 | 1.000E-05 | 1.300E-05 |
| 12715 | 636 | S636P | B | 1.5e-05 | 1.6e-05 | 1.4e-05 | 1.500E-05 | 1.000E-06 | 1.400E-05 | 1.600E-05 |
| 12716 | 636 | S636S | B | 1.5e-05 | 1e-05 | 9.7e-06 | 1.157E-05 | 2.977E-06 | 9.700E-06 | 1.500E-05 |
| 12717 | 636 | S636T | B | 1.1e-05 | 1.3e-05 | 1.3e-05 | 1.233E-05 | 1.155E-06 | 1.100E-05 | 1.300E-05 |
| 12718 | 636 | S636W | B | 1.1e-05 | 1.1e-05 | 1.1e-05 | 1.100E-05 | 2.075E-21 | 1.100E-05 | 1.100E-05 |
| 12719 | 636 | S636Y | B | 1e-05 | 1.5e-05 | 1.1e-05 | 1.200E-05 | 2.646E-06 | 1.000E-05 | 1.500E-05 |
| 12720 | 636 | S636V | B | 1.7e-05 | 1.5e-05 | 1.5e-05 | 1.567E-05 | 1.155E-06 | 1.500E-05 | 1.700E-05 |
| 12721 | 637 | V637A | B | 1.4e-05 | 1.4e-05 | 8.1e-06 | 1.203E-05 | 3.406E-06 | 8.100E-06 | 1.400E-05 |
| 12722 | 637 | V637R | B | 1.7e-05 | 9.1e-06 | 1.6e-05 | 1.403E-05 | 4.302E-06 | 9.100E-06 | 1.700E-05 |
| 12723 | 637 | V637N | B | 1.5e-05 | 1.3e-05 | 1e-05 | 1.267E-05 | 2.517E-06 | 1.000E-05 | 1.500E-05 |
| 12724 | 637 | V637D | B | 1.1e-05 | 9.9e-06 | 1.3e-05 | 1.130E-05 | 1.572E-06 | 9.900E-06 | 1.300E-05 |
| 12725 | 637 | V637C | B | 1.2e-05 | 1.5e-05 | 1.4e-05 | 1.367E-05 | 1.528E-06 | 1.200E-05 | 1.500E-05 |
| 12726 | 637 | V637E | B | 1.5e-05 | 1.6e-05 | 9.1e-06 | 1.337E-05 | 3.729E-06 | 9.100E-06 | 1.600E-05 |
| 12727 | 637 | V637Q | B | 1.2e-05 | 9.6e-06 | 9.3e-06 | 1.030E-05 | 1.480E-06 | 9.300E-06 | 1.200E-05 |
| 12728 | 637 | V637G | B | 7.5e-06 | 1.1e-05 | 1.5e-05 | 1.117E-05 | 3.753E-06 | 7.500E-06 | 1.500E-05 |
| 12729 | 637 | V637H | B | 1.3e-05 | 9.2e-06 | 1.6e-05 | 1.273E-05 | 3.408E-06 | 9.200E-06 | 1.600E-05 |
| 12730 | 637 | V637I | B | 1e-05 | 1.4e-05 | 1.6e-05 | 1.333E-05 | 3.055E-06 | 1.000E-05 | 1.600E-05 |
| 12731 | 637 | V637L | B | 1.6e-05 | 1.1e-05 | 1.1e-05 | 1.267E-05 | 2.887E-06 | 1.100E-05 | 1.600E-05 |
| 12732 | 637 | V637K | B | 1.1e-05 | 1.4e-05 | 1.4e-05 | 1.300E-05 | 1.732E-06 | 1.100E-05 | 1.400E-05 |
| 12733 | 637 | V637M | B | 1.4e-05 | 1.3e-05 | 1.6e-05 | 1.433E-05 | 1.528E-06 | 1.300E-05 | 1.600E-05 |
| 12734 | 637 | V637F | B | 1.1e-05 | 1.5e-05 | 1.2e-05 | 1.267E-05 | 2.082E-06 | 1.100E-05 | 1.500E-05 |
| 12735 | 637 | V637P | B | 1.7e-05 | 1.6e-05 | 1.6e-05 | 1.633E-05 | 5.774E-07 | 1.600E-05 | 1.700E-05 |
| 12736 | 637 | V637S | B | 1e-05 | 9.6e-06 | 1.4e-05 | 1.120E-05 | 2.433E-06 | 9.600E-06 | 1.400E-05 |
| 12737 | 637 | V637T | B | 1.5e-05 | 1.2e-05 | 9.9e-06 | 1.230E-05 | 2.563E-06 | 9.900E-06 | 1.500E-05 |
| 12738 | 637 | V637W | B | 1.2e-05 | 7.9e-06 | 1.5e-05 | 1.163E-05 | 3.564E-06 | 7.900E-06 | 1.500E-05 |
| 12739 | 637 | V637Y | B | 9.9e-06 | 1.3e-05 | 1e-05 | 1.097E-05 | 1.762E-06 | 9.900E-06 | 1.300E-05 |
| 12740 | 637 | V637V | B | 1.7e-05 | 1.1e-05 | 8.4e-06 | 1.213E-05 | 4.411E-06 | 8.400E-06 | 1.700E-05 |
| 12741 | 638 | G638A | B | 9.3e-06 | 1.3e-05 | 1.1e-05 | 1.110E-05 | 1.852E-06 | 9.300E-06 | 1.300E-05 |
| 12742 | 638 | G638R | B | 1.7e-05 | 1.2e-05 | 1.5e-05 | 1.467E-05 | 2.517E-06 | 1.200E-05 | 1.700E-05 |
| 12743 | 638 | G638N | B | 1.3e-05 | 1.5e-05 | 1.5e-05 | 1.433E-05 | 1.155E-06 | 1.300E-05 | 1.500E-05 |
| 12744 | 638 | G638D | B | 9.3e-06 | 1e-05 | 1.7e-05 | 1.210E-05 | 4.258E-06 | 9.300E-06 | 1.700E-05 |
| 12745 | 638 | G638C | B | 1.6e-05 | 1.4e-05 | 1e-05 | 1.333E-05 | 3.055E-06 | 1.000E-05 | 1.600E-05 |
| 12746 | 638 | G638E | B | 9.8e-06 | 1.4e-05 | 1.5e-05 | 1.293E-05 | 2.759E-06 | 9.800E-06 | 1.500E-05 |
| 12747 | 638 | G638Q | B | 1.3e-05 | 9.7e-06 | 1.4e-05 | 1.223E-05 | 2.250E-06 | 9.700E-06 | 1.400E-05 |
| 12748 | 638 | G638G | B | 1.7e-05 | 7.5e-06 | 9.6e-06 | 1.137E-05 | 4.990E-06 | 7.500E-06 | 1.700E-05 |
| 12749 | 638 | G638H | B | 9.6e-06 | 1.4e-05 | 1e-05 | 1.120E-05 | 2.433E-06 | 9.600E-06 | 1.400E-05 |

| | | | | | | | | | | |
|---|---|---|---|---|---|---|---|---|---|---|
| 12750 | 638 | G638I | B | 1e-05 | 1.4e-05 | 1.2e-05 | 1.200E-05 | 2.000E-06 | 1.000E-05 | 1.400E-05 |
| 12751 | 638 | G638L | B | 1.1e-05 | 1e-05 | 1.3e-05 | 1.133E-05 | 1.528E-06 | 1.000E-05 | 1.300E-05 |
| 12752 | 638 | G638K | B | 1e-05 | 9.4e-06 | 1.5e-05 | 1.147E-05 | 3.075E-06 | 9.400E-06 | 1.500E-05 |
| 12753 | 638 | G638M | B | 1e-05 | 1.7e-05 | 9.9e-06 | 1.230E-05 | 4.071E-06 | 9.900E-06 | 1.700E-05 |
| 12754 | 638 | G638F | B | 1e-05 | 1e-05 | 1.7e-05 | 1.233E-05 | 4.041E-06 | 1.000E-05 | 1.700E-05 |
| 12755 | 638 | G638P | B | 1.7e-05 | 1.1e-05 | 1.5e-05 | 1.433E-05 | 3.055E-06 | 1.100E-05 | 1.700E-05 |
| 12756 | 638 | G638S | B | 8.1e-06 | 8.2e-06 | 9.4e-06 | 8.567E-06 | 7.234E-07 | 8.100E-06 | 9.400E-06 |
| 12757 | 638 | G638T | B | 1.1e-05 | 1.3e-05 | 1.1e-05 | 1.167E-05 | 1.155E-06 | 1.100E-05 | 1.300E-05 |
| 12758 | 638 | G638W | B | 1.4e-05 | 1.1e-05 | 9.4e-06 | 1.147E-05 | 2.335E-06 | 9.400E-06 | 1.400E-05 |
| 12759 | 638 | G638Y | B | 1.6e-05 | 1.1e-05 | 1e-05 | 1.233E-05 | 3.215E-06 | 1.000E-05 | 1.600E-05 |
| 12760 | 638 | G638V | B | 9.9e-06 | 8.8e-06 | 1e-05 | 9.567E-06 | 6.658E-07 | 8.800E-06 | 1.000E-05 |
| 12761 | 639 | D639A | B | 1.1e-05 | 1.4e-05 | 1.3e-05 | 1.267E-05 | 1.528E-06 | 1.100E-05 | 1.400E-05 |
| 12762 | 639 | D639R | B | 1.1e-05 | 1.3e-05 | 1e-05 | 1.133E-05 | 1.528E-06 | 1.000E-05 | 1.300E-05 |
| 12763 | 639 | D639N | B | 1e-05 | 1.4e-05 | 1.5e-05 | 1.300E-05 | 2.646E-06 | 1.000E-05 | 1.500E-05 |
| 12764 | 639 | D639D | B | 1.1e-05 | 9.4e-06 | 1.5e-05 | 1.180E-05 | 2.884E-06 | 9.400E-06 | 1.500E-05 |
| 12765 | 639 | D639C | B | 8.9e-06 | 1.1e-05 | 1e-05 | 9.967E-06 | 1.050E-06 | 8.900E-06 | 1.100E-05 |
| 12766 | 639 | D639E | B | 9.5e-06 | 1.4e-05 | 1.5e-05 | 1.283E-05 | 2.930E-06 | 9.500E-06 | 1.500E-05 |
| 12767 | 639 | D639Q | B | 1.5e-05 | 9.6e-06 | 9.1e-06 | 1.123E-05 | 3.272E-06 | 9.100E-06 | 1.500E-05 |
| 12768 | 639 | D639G | B | 1.1e-05 | 1.1e-05 | 1.5e-05 | 1.233E-05 | 2.309E-06 | 1.100E-05 | 1.500E-05 |
| 12769 | 639 | D639H | B | 1.4e-05 | 1.5e-05 | 1.3e-05 | 1.400E-05 | 1.000E-06 | 1.300E-05 | 1.500E-05 |
| 12770 | 639 | D639I | B | 1.7e-05 | 1.1e-05 | 1.6e-05 | 1.467E-05 | 3.215E-06 | 1.100E-05 | 1.700E-05 |
| 12771 | 639 | D639L | B | 1e-05 | 1.4e-05 | 1.2e-05 | 1.200E-05 | 2.000E-06 | 1.000E-05 | 1.400E-05 |
| 12772 | 639 | D639K | B | 1.5e-05 | 8.6e-06 | 1e-05 | 1.120E-05 | 3.365E-06 | 8.600E-06 | 1.500E-05 |
| 12773 | 639 | D639M | B | 8.4e-06 | 1.5e-05 | 9.5e-06 | 1.097E-05 | 3.536E-06 | 8.400E-06 | 1.500E-05 |
| 12774 | 639 | D639F | B | 1.6e-05 | 8.1e-06 | 9.7e-06 | 1.127E-05 | 4.177E-06 | 8.100E-06 | 1.600E-05 |
| 12775 | 639 | D639P | B | 1.5e-05 | 1.4e-05 | 1.7e-05 | 1.533E-05 | 1.528E-06 | 1.400E-05 | 1.700E-05 |
| 12776 | 639 | D639S | B | 9.3e-06 | 1.2e-05 | 1.1e-05 | 1.077E-05 | 1.365E-06 | 9.300E-06 | 1.200E-05 |
| 12777 | 639 | D639T | B | 9.7e-06 | 1.5e-05 | 1.8e-05 | 1.423E-05 | 4.203E-06 | 9.700E-06 | 1.800E-05 |
| 12778 | 639 | D639W | B | 1.5e-05 | 1.3e-05 | 1.6e-05 | 1.467E-05 | 1.528E-06 | 1.300E-05 | 1.600E-05 |
| 12779 | 639 | D639Y | B | 9e-06 | 1.4e-05 | 1e-05 | 1.100E-05 | 2.646E-06 | 9.000E-06 | 1.400E-05 |
| 12780 | 639 | D639V | B | 7.3e-06 | 1e-05 | 1.1e-05 | 9.433E-06 | 1.914E-06 | 7.300E-06 | 1.100E-05 |
| 12781 | 640 | R640A | B | 1.6e-05 | 1.5e-05 | 1.7e-05 | 1.600E-05 | 1.000E-06 | 1.500E-05 | 1.700E-05 |
| 12782 | 640 | R640R | B | 1.8e-05 | 1.3e-05 | 8.3e-06 | 1.310E-05 | 4.851E-06 | 8.300E-06 | 1.800E-05 |
| 12783 | 640 | R640N | B | 9.8e-06 | 1.5e-05 | 1.5e-05 | 1.327E-05 | 3.002E-06 | 9.800E-06 | 1.500E-05 |
| 12784 | 640 | R640D | B | 1.8e-05 | 1.4e-05 | 1.6e-05 | 1.600E-05 | 2.000E-06 | 1.400E-05 | 1.800E-05 |
| 12785 | 640 | R640C | B | 1.4e-05 | 1.3e-05 | 1.1e-05 | 1.267E-05 | 1.528E-06 | 1.100E-05 | 1.400E-05 |
| 12786 | 640 | R640E | B | 1.6e-05 | 1.4e-05 | 1.6e-05 | 1.533E-05 | 1.155E-06 | 1.400E-05 | 1.600E-05 |
| 12787 | 640 | R640Q | B | 1.7e-05 | 1.5e-05 | 1.5e-05 | 1.567E-05 | 1.155E-06 | 1.500E-05 | 1.700E-05 |
| 12788 | 640 | R640G | B | 1.4e-05 | 9.9e-06 | 9.3e-06 | 1.107E-05 | 2.558E-06 | 9.300E-06 | 1.400E-05 |
| 12789 | 640 | R640H | B | 1.8e-05 | 9.1e-06 | 1.3e-05 | 1.337E-05 | 4.461E-06 | 9.100E-06 | 1.800E-05 |
| 12790 | 640 | R640I | B | 9.2e-06 | 1.1e-05 | 1e-05 | 1.007E-05 | 9.018E-07 | 9.200E-06 | 1.100E-05 |
| 12791 | 640 | R640L | B | 1.6e-05 | 1e-05 | 1.5e-05 | 1.367E-05 | 3.215E-06 | 1.000E-05 | 1.600E-05 |
| 12792 | 640 | R640K | B | 1e-05 | 9.9e-06 | 1.4e-05 | 1.130E-05 | 2.339E-06 | 9.900E-06 | 1.400E-05 |
| 12793 | 640 | R640M | B | 8.4e-06 | 1.3e-05 | 1e-05 | 1.047E-05 | 2.335E-06 | 8.400E-06 | 1.300E-05 |
| 12794 | 640 | R640F | B | 1.1e-05 | 1.5e-05 | 1.1e-05 | 1.233E-05 | 2.309E-06 | 1.100E-05 | 1.500E-05 |
| 12795 | 640 | R640P | B | 9.7e-06 | 7.7e-06 | 1e-05 | 9.133E-06 | 1.250E-06 | 7.700E-06 | 1.000E-05 |
| 12796 | 640 | R640S | B | 9.8e-06 | 1.2e-05 | 1.5e-05 | 1.227E-05 | 2.610E-06 | 9.800E-06 | 1.500E-05 |
| 12797 | 640 | R640T | B | 1.1e-05 | 1.3e-05 | 1e-05 | 1.133E-05 | 1.528E-06 | 1.000E-05 | 1.300E-05 |
| 12798 | 640 | R640W | B | 1.1e-05 | 1.4e-05 | 1e-05 | 1.167E-05 | 2.082E-06 | 1.000E-05 | 1.400E-05 |
| 12799 | 640 | R640Y | B | 1.5e-05 | 1.1e-05 | 1.1e-05 | 1.233E-05 | 2.309E-06 | 1.100E-05 | 1.500E-05 |
| 12800 | 640 | R640V | B | 1.5e-05 | 8.9e-06 | 1.4e-05 | 1.263E-05 | 3.272E-06 | 8.900E-06 | 1.500E-05 |
| 12801 | 641 | V641A | B | 1.5e-05 | 1.6e-05 | 1.1e-05 | 1.400E-05 | 2.646E-06 | 1.100E-05 | 1.600E-05 |
| 12802 | 641 | V641R | B | 1.6e-05 | 1.5e-05 | 1.6e-05 | 1.567E-05 | 5.774E-07 | 1.500E-05 | 1.600E-05 |
| 12803 | 641 | V641N | B | 9.8e-06 | 1.4e-05 | 1.5e-05 | 1.293E-05 | 2.759E-06 | 9.800E-06 | 1.500E-05 |
| 12804 | 641 | V641D | B | 1.5e-05 | 1.5e-05 | 1.3e-05 | 1.433E-05 | 1.155E-06 | 1.300E-05 | 1.500E-05 |
| 12805 | 641 | V641C | B | 1.1e-05 | 8.7e-06 | 9.9e-06 | 9.867E-06 | 1.150E-06 | 8.700E-06 | 1.100E-05 |
| 12806 | 641 | V641E | B | 1.3e-05 | 1.3e-05 | 1.3e-05 | 1.300E-05 | 0.000E+00 | 1.300E-05 | 1.300E-05 |
| 12807 | 641 | V641Q | B | 1e-05 | 1.3e-05 | 1e-05 | 1.100E-05 | 1.732E-06 | 1.000E-05 | 1.300E-05 |
| 12808 | 641 | V641G | B | 1.5e-05 | 1.2e-05 | 1.6e-05 | 1.433E-05 | 2.082E-06 | 1.200E-05 | 1.600E-05 |
| 12809 | 641 | V641H | B | 8.5e-06 | 1.1e-05 | 1.6e-05 | 1.183E-05 | 3.819E-06 | 8.500E-06 | 1.600E-05 |
| 12810 | 641 | V641I | B | 1e-05 | 1.1e-05 | 1.4e-05 | 1.167E-05 | 2.082E-06 | 1.000E-05 | 1.400E-05 |
| 12811 | 641 | V641L | B | 1.5e-05 | 9.7e-06 | 1.3e-05 | 1.257E-05 | 2.676E-06 | 9.700E-06 | 1.500E-05 |
| 12812 | 641 | V641K | B | 1.4e-05 | 1e-05 | 1.4e-05 | 1.267E-05 | 2.309E-06 | 1.000E-05 | 1.400E-05 |
| 12813 | 641 | V641M | B | 1.4e-05 | 9.4e-06 | 1.1e-05 | 1.147E-05 | 2.335E-06 | 9.400E-06 | 1.400E-05 |
| 12814 | 641 | V641F | B | 9e-06 | 1.5e-05 | 1.1e-05 | 1.167E-05 | 3.055E-06 | 9.000E-06 | 1.500E-05 |
| 12815 | 641 | V641P | B | 1e-05 | 1.3e-05 | 1.6e-05 | 1.300E-05 | 3.000E-06 | 1.000E-05 | 1.600E-05 |
| 12816 | 641 | V641S | B | 1.4e-05 | 1.4e-05 | 1.3e-05 | 1.367E-05 | 5.774E-07 | 1.300E-05 | 1.400E-05 |
| 12817 | 641 | V641T | B | 1.4e-05 | 1.5e-05 | 1.6e-05 | 1.500E-05 | 1.000E-06 | 1.400E-05 | 1.600E-05 |
| 12818 | 641 | V641W | B | 1.6e-05 | 7.5e-06 | 1.5e-05 | 1.283E-05 | 4.646E-06 | 7.500E-06 | 1.600E-05 |
| 12819 | 641 | V641Y | B | 1.4e-05 | 1e-05 | 1.5e-05 | 1.300E-05 | 2.646E-06 | 1.000E-05 | 1.500E-05 |
| 12820 | 641 | V641V | B | 7.4e-06 | 1.5e-05 | 1.6e-05 | 1.280E-05 | 4.703E-06 | 7.400E-06 | 1.600E-05 |
| 12821 | 642 | T642A | B | 1.2e-05 | 1.6e-05 | 1.2e-05 | 1.333E-05 | 2.309E-06 | 1.200E-05 | 1.600E-05 |
| 12822 | 642 | T642R | B | 1.5e-05 | 1e-05 | 1.3e-05 | 1.267E-05 | 2.517E-06 | 1.000E-05 | 1.500E-05 |
| 12823 | 642 | T642N | B | 9.4e-06 | 1.5e-05 | 1.3e-05 | 1.247E-05 | 2.838E-06 | 9.400E-06 | 1.500E-05 |
| 12824 | 642 | T642D | B | 1.5e-05 | 8.2e-06 | 1.1e-05 | 1.140E-05 | 3.418E-06 | 8.200E-06 | 1.500E-05 |
| 12825 | 642 | T642C | B | 1.5e-05 | 9.3e-06 | 1.8e-05 | 1.410E-05 | 4.419E-06 | 9.300E-06 | 1.800E-05 |
| 12826 | 642 | T642E | B | 1.6e-05 | 1.7e-05 | 1.6e-05 | 1.633E-05 | 5.774E-07 | 1.600E-05 | 1.700E-05 |
| 12827 | 642 | T642Q | B | 1.5e-05 | 1.4e-05 | 1.5e-05 | 1.467E-05 | 5.774E-07 | 1.400E-05 | 1.500E-05 |
| 12828 | 642 | T642G | B | 9.2e-06 | 1.6e-05 | 1e-05 | 1.173E-05 | 3.717E-06 | 9.200E-06 | 1.600E-05 |
| 12829 | 642 | T642H | B | 8.7e-06 | 1.5e-05 | 1.5e-05 | 1.290E-05 | 3.637E-06 | 8.700E-06 | 1.500E-05 |
| 12830 | 642 | T642I | B | 1.8e-05 | 9.4e-06 | 1.4e-05 | 1.380E-05 | 4.303E-06 | 9.400E-06 | 1.800E-05 |
| 12831 | 642 | T642L | B | 1.1e-05 | 1.5e-05 | 1.5e-05 | 1.367E-05 | 2.309E-06 | 1.100E-05 | 1.500E-05 |
| 12832 | 642 | T642K | B | 1e-05 | 1.6e-05 | 1.1e-05 | 1.233E-05 | 3.215E-06 | 1.000E-05 | 1.600E-05 |
| 12833 | 642 | T642M | B | 1.1e-05 | 1.1e-05 | 1.1e-05 | 1.100E-05 | 2.075E-21 | 1.100E-05 | 1.100E-05 |
| 12834 | 642 | T642F | B | 8.7e-06 | 1.1e-05 | 6.9e-06 | 8.867E-06 | 2.055E-06 | 6.900E-06 | 1.100E-05 |

| | | | | | | | | | | |
|---|---|---|---|---|---|---|---|---|---|---|
| 12835 | 642 | T642P | B | 1.6e-05 | 1.2e-05 | 9.9e-06 | 1.263E-05 | 3.099E-06 | 9.900E-06 | 1.600E-05 |
| 12836 | 642 | T642S | B | 1e-05 | 1.5e-05 | 1e-05 | 1.167E-05 | 2.887E-06 | 1.000E-05 | 1.500E-05 |
| 12837 | 642 | T642T | B | 1.5e-05 | 1e-05 | 1.7e-05 | 1.400E-05 | 3.606E-06 | 1.000E-05 | 1.700E-05 |
| 12838 | 642 | T642W | B | 1.2e-05 | 1.2e-05 | 1e-05 | 1.133E-05 | 1.155E-06 | 1.000E-05 | 1.200E-05 |
| 12839 | 642 | T642Y | B | 9.8e-06 | 8.8e-06 | 1e-05 | 9.533E-06 | 6.429E-07 | 8.800E-06 | 1.000E-05 |
| 12840 | 642 | T642V | B | 1.6e-05 | 1.1e-05 | 1.4e-05 | 1.367E-05 | 2.517E-06 | 1.100E-05 | 1.600E-05 |
| 12841 | 643 | I643A | B | 8.6e-06 | 1.2e-05 | 9.7e-06 | 1.010E-05 | 1.735E-06 | 8.600E-06 | 1.200E-05 |
| 12842 | 643 | I643R | B | 8.4e-06 | 1.1e-05 | 1.2e-05 | 1.047E-05 | 1.858E-06 | 8.400E-06 | 1.200E-05 |
| 12843 | 643 | I643N | B | 1.5e-05 | 8.7e-06 | 1.4e-05 | 1.257E-05 | 3.386E-06 | 8.700E-06 | 1.500E-05 |
| 12844 | 643 | I643D | B | 1.5e-05 | 1.6e-05 | 1.2e-05 | 1.433E-05 | 2.082E-06 | 1.200E-05 | 1.600E-05 |
| 12845 | 643 | I643C | B | 9.7e-06 | 1.4e-05 | 8.4e-06 | 1.070E-05 | 2.931E-06 | 8.400E-06 | 1.400E-05 |
| 12846 | 643 | I643E | B | 1.7e-05 | 1e-05 | 1.3e-05 | 1.333E-05 | 3.512E-06 | 1.000E-05 | 1.700E-05 |
| 12847 | 643 | I643Q | B | 6.8e-06 | 1e-05 | 9.9e-06 | 8.900E-06 | 1.819E-06 | 6.800E-06 | 1.000E-05 |
| 12848 | 643 | I643G | B | 1e-05 | 1.4e-05 | 1.4e-05 | 1.267E-05 | 2.309E-06 | 1.000E-05 | 1.400E-05 |
| 12849 | 643 | I643H | B | 1.2e-05 | 1.1e-05 | 9.8e-06 | 1.093E-05 | 1.102E-06 | 9.800E-06 | 1.200E-05 |
| 12850 | 643 | I643I | B | 1e-05 | 1.5e-05 | 8.5e-06 | 1.117E-05 | 3.403E-06 | 8.500E-06 | 1.500E-05 |
| 12851 | 643 | I643L | B | 1.5e-05 | 1.4e-05 | 1.4e-05 | 1.433E-05 | 5.774E-07 | 1.400E-05 | 1.500E-05 |
| 12852 | 643 | I643K | B | 1e-05 | 1.5e-05 | 1.4e-05 | 1.300E-05 | 2.646E-06 | 1.000E-05 | 1.500E-05 |
| 12853 | 643 | I643M | B | 1.6e-05 | 1.1e-05 | 1.2e-05 | 1.300E-05 | 2.646E-06 | 1.100E-05 | 1.600E-05 |
| 12854 | 643 | I643F | B | 9.1e-06 | 1.2e-05 | 1e-05 | 1.037E-05 | 1.484E-06 | 9.100E-06 | 1.200E-05 |
| 12855 | 643 | I643P | B | 9.1e-06 | 1.4e-05 | 1.7e-05 | 1.337E-05 | 3.988E-06 | 9.100E-06 | 1.700E-05 |
| 12856 | 643 | I643S | B | 1.5e-05 | 1.5e-05 | 1.7e-05 | 1.567E-05 | 1.155E-06 | 1.500E-05 | 1.700E-05 |
| 12857 | 643 | I643T | B | 1e-05 | 1.5e-05 | 1.5e-05 | 1.333E-05 | 2.887E-06 | 1.000E-05 | 1.500E-05 |
| 12858 | 643 | I643W | B | 9.8e-06 | 1.3e-05 | 1.5e-05 | 1.260E-05 | 2.623E-06 | 9.800E-06 | 1.500E-05 |
| 12859 | 643 | I643Y | B | 1.5e-05 | 1.5e-05 | 1.5e-05 | 1.500E-05 | 0.000E+00 | 1.500E-05 | 1.500E-05 |
| 12860 | 643 | I643V | B | 1.4e-05 | 1.8e-05 | 1.2e-05 | 1.467E-05 | 3.055E-06 | 1.200E-05 | 1.800E-05 |
| 12861 | 644 | T644A | B | 8.3e-06 | 1.5e-05 | 1.2e-05 | 1.177E-05 | 3.356E-06 | 8.300E-06 | 1.500E-05 |
| 12862 | 644 | T644R | B | 1.5e-05 | 1.5e-05 | 1.2e-05 | 1.400E-05 | 1.732E-06 | 1.200E-05 | 1.500E-05 |
| 12863 | 644 | T644N | B | 1.5e-05 | 1.6e-05 | 1.3e-05 | 1.467E-05 | 1.528E-06 | 1.300E-05 | 1.600E-05 |
| 12864 | 644 | T644D | B | 9.2e-06 | 1.5e-05 | 1.1e-05 | 1.173E-05 | 2.969E-06 | 9.200E-06 | 1.500E-05 |
| 12865 | 644 | T644C | B | 8e-06 | 1.5e-05 | 1.5e-05 | 1.267E-05 | 4.041E-06 | 8.000E-06 | 1.500E-05 |
| 12866 | 644 | T644E | B | 1.5e-05 | 1.1e-05 | 1.8e-05 | 1.467E-05 | 3.512E-06 | 1.100E-05 | 1.800E-05 |
| 12867 | 644 | T644Q | B | 1.5e-05 | 1.5e-05 | 1.1e-05 | 1.367E-05 | 2.309E-06 | 1.100E-05 | 1.500E-05 |
| 12868 | 644 | T644G | B | 5.9e-06 | 9e-06 | 9.3e-06 | 8.067E-06 | 1.882E-06 | 5.900E-06 | 9.300E-06 |
| 12869 | 644 | T644H | B | 1e-05 | 1.1e-05 | 1.1e-05 | 1.067E-05 | 5.774E-07 | 1.000E-05 | 1.100E-05 |
| 12870 | 644 | T644I | B | 1.5e-05 | 1.3e-05 | 8.2e-06 | 1.207E-05 | 3.495E-06 | 8.200E-06 | 1.500E-05 |
| 12871 | 644 | T644L | B | 1e-05 | 7.5e-06 | 1.5e-05 | 1.083E-05 | 3.819E-06 | 7.500E-06 | 1.500E-05 |
| 12872 | 644 | T644K | B | 1.5e-05 | 1.4e-05 | 9e-06 | 1.267E-05 | 3.215E-06 | 9.000E-06 | 1.500E-05 |
| 12873 | 644 | T644M | B | 1e-05 | 1.4e-05 | 1.8e-05 | 1.400E-05 | 4.000E-06 | 1.000E-05 | 1.800E-05 |
| 12874 | 644 | T644F | B | 1.3e-05 | 1.6e-05 | 1.2e-05 | 1.367E-05 | 2.082E-06 | 1.200E-05 | 1.600E-05 |
| 12875 | 644 | T644P | B | 1.6e-05 | 1.6e-05 | 1.1e-05 | 1.433E-05 | 2.887E-06 | 1.100E-05 | 1.600E-05 |
| 12876 | 644 | T644S | B | 1.2e-05 | 9.1e-06 | 1.1e-05 | 1.070E-05 | 1.473E-06 | 9.100E-06 | 1.200E-05 |
| 12877 | 644 | T644T | B | 1.6e-05 | 1.5e-05 | 1.5e-05 | 1.533E-05 | 5.774E-07 | 1.500E-05 | 1.600E-05 |
| 12878 | 644 | T644W | B | 1.3e-05 | 1.6e-05 | 1.3e-05 | 1.400E-05 | 1.732E-06 | 1.300E-05 | 1.600E-05 |
| 12879 | 644 | T644Y | B | 1.1e-05 | 1.7e-05 | 1.7e-05 | 1.500E-05 | 3.464E-06 | 1.100E-05 | 1.700E-05 |
| 12880 | 644 | T644V | B | 1.7e-05 | 1.4e-05 | 1.5e-05 | 1.533E-05 | 1.528E-06 | 1.400E-05 | 1.700E-05 |
| 12881 | 645 | C645A | B | 1.2e-05 | 9.3e-06 | 1e-05 | 1.043E-05 | 1.401E-06 | 9.300E-06 | 1.200E-05 |
| 12882 | 645 | C645R | B | 1.5e-05 | 1.1e-05 | 1.7e-05 | 1.433E-05 | 3.055E-06 | 1.100E-05 | 1.700E-05 |
| 12883 | 645 | C645N | B | 1.8e-05 | 1.4e-05 | 1.5e-05 | 1.567E-05 | 2.082E-06 | 1.400E-05 | 1.800E-05 |
| 12884 | 645 | C645D | B | 1.4e-05 | 9.8e-06 | 1.7e-05 | 1.360E-05 | 3.617E-06 | 9.800E-06 | 1.700E-05 |
| 12885 | 645 | C645C | B | 1.4e-05 | 8.9e-06 | 1.6e-05 | 1.297E-05 | 3.661E-06 | 8.900E-06 | 1.600E-05 |
| 12886 | 645 | C645E | B | 1.2e-05 | 1.6e-05 | 1.4e-05 | 1.400E-05 | 2.000E-06 | 1.200E-05 | 1.600E-05 |
| 12887 | 645 | C645Q | B | 1e-05 | 1.1e-05 | 1e-05 | 1.033E-05 | 5.774E-07 | 1.000E-05 | 1.100E-05 |
| 12888 | 645 | C645G | B | 1.2e-05 | 1.1e-05 | 1.6e-05 | 1.300E-05 | 2.646E-06 | 1.100E-05 | 1.600E-05 |
| 12889 | 645 | C645H | B | 1.5e-05 | 8.4e-06 | 1.2e-05 | 1.180E-05 | 3.305E-06 | 8.400E-06 | 1.500E-05 |
| 12890 | 645 | C645I | B | 1.5e-05 | 8.6e-06 | 1.5e-05 | 1.287E-05 | 3.695E-06 | 8.600E-06 | 1.500E-05 |
| 12891 | 645 | C645L | B | 9.8e-06 | 9.7e-06 | 9.2e-06 | 9.567E-06 | 3.215E-07 | 9.200E-06 | 9.800E-06 |
| 12892 | 645 | C645K | B | 1.1e-05 | 1.5e-05 | 1.5e-05 | 1.367E-05 | 2.309E-06 | 1.100E-05 | 1.500E-05 |
| 12893 | 645 | C645M | B | 7.2e-06 | 1.1e-05 | 5.9e-06 | 8.033E-06 | 2.650E-06 | 5.900E-06 | 1.100E-05 |
| 12894 | 645 | C645F | B | 1.4e-05 | 1.6e-05 | 1.7e-05 | 1.567E-05 | 1.528E-06 | 1.400E-05 | 1.700E-05 |
| 12895 | 645 | C645P | B | 1.5e-05 | 1.2e-05 | 1.5e-05 | 1.400E-05 | 1.732E-06 | 1.200E-05 | 1.500E-05 |
| 12896 | 645 | C645S | B | 1.6e-05 | 1.6e-05 | 1e-05 | 1.400E-05 | 3.464E-06 | 1.000E-05 | 1.600E-05 |
| 12897 | 645 | C645T | B | 1.1e-05 | 1.5e-05 | 1.3e-05 | 1.300E-05 | 2.000E-06 | 1.100E-05 | 1.500E-05 |
| 12898 | 645 | C645W | B | 1e-05 | 9.6e-06 | 9.6e-06 | 9.733E-06 | 2.309E-07 | 9.600E-06 | 1.000E-05 |
| 12899 | 645 | C645Y | B | 1.5e-05 | 1.5e-05 | 8.2e-06 | 1.273E-05 | 3.926E-06 | 8.200E-06 | 1.500E-05 |
| 12900 | 645 | C645V | B | 1.4e-05 | 1.5e-05 | 1.7e-05 | 1.533E-05 | 1.528E-06 | 1.400E-05 | 1.700E-05 |
| 12901 | 646 | K646A | B | 1.1e-05 | 1.8e-05 | 9.2e-06 | 1.273E-05 | 4.649E-06 | 9.200E-06 | 1.800E-05 |
| 12902 | 646 | K646R | B | 1.8e-05 | 1.3e-05 | 1.5e-05 | 1.533E-05 | 2.517E-06 | 1.300E-05 | 1.800E-05 |
| 12903 | 646 | K646N | B | 1e-05 | 1e-05 | 1.2e-05 | 1.067E-05 | 1.155E-06 | 1.000E-05 | 1.200E-05 |
| 12904 | 646 | K646D | B | 1.3e-05 | 1.1e-05 | 1.4e-05 | 1.267E-05 | 1.528E-06 | 1.100E-05 | 1.400E-05 |
| 12905 | 646 | K646C | B | 1.4e-05 | 1.5e-05 | 9.8e-06 | 1.293E-05 | 2.759E-06 | 9.800E-06 | 1.500E-05 |
| 12906 | 646 | K646E | B | 1.5e-05 | 1.1e-05 | 8.4e-06 | 1.147E-05 | 3.325E-06 | 8.400E-06 | 1.500E-05 |
| 12907 | 646 | K646Q | B | 1e-05 | 1.1e-05 | 1.3e-05 | 1.133E-05 | 1.528E-06 | 1.000E-05 | 1.300E-05 |
| 12908 | 646 | K646G | B | 1.4e-05 | 1.5e-05 | 1.3e-05 | 1.400E-05 | 1.000E-06 | 1.300E-05 | 1.500E-05 |
| 12909 | 646 | K646H | B | 1.3e-05 | 1.2e-05 | 1.3e-05 | 1.267E-05 | 5.774E-07 | 1.200E-05 | 1.300E-05 |
| 12910 | 646 | K646I | B | 1.6e-05 | 1e-05 | 1.2e-05 | 1.267E-05 | 3.055E-06 | 1.000E-05 | 1.600E-05 |
| 12911 | 646 | K646L | B | 1.1e-05 | 1.4e-05 | 8.4e-06 | 1.113E-05 | 2.802E-06 | 8.400E-06 | 1.400E-05 |
| 12912 | 646 | K646K | B | 1.5e-05 | 1.6e-05 | 1.1e-05 | 1.400E-05 | 2.646E-06 | 1.100E-05 | 1.600E-05 |
| 12913 | 646 | K646M | B | 1.2e-05 | 1.4e-05 | 1.6e-05 | 1.400E-05 | 2.000E-06 | 1.200E-05 | 1.600E-05 |
| 12914 | 646 | K646F | B | 1.5e-05 | 1.4e-05 | 1.5e-05 | 1.467E-05 | 5.774E-07 | 1.400E-05 | 1.500E-05 |
| 12915 | 646 | K646P | B | 1.2e-05 | 1.5e-05 | 1.5e-05 | 1.400E-05 | 1.732E-06 | 1.200E-05 | 1.500E-05 |
| 12916 | 646 | K646S | B | 8.4e-06 | 6.3e-06 | 1.6e-05 | 1.023E-05 | 5.103E-06 | 6.300E-06 | 1.600E-05 |
| 12917 | 646 | K646T | B | 1.1e-05 | 1.1e-05 | 1.1e-05 | 1.100E-05 | 2.075E-21 | 1.100E-05 | 1.100E-05 |
| 12918 | 646 | K646W | B | 1e-05 | 1.6e-05 | 1.6e-05 | 1.400E-05 | 3.464E-06 | 1.000E-05 | 1.600E-05 |
| 12919 | 646 | K646Y | B | 9.8e-06 | 1.6e-05 | 1.5e-05 | 1.360E-05 | 3.329E-06 | 9.800E-06 | 1.600E-05 |

| | | | | | | | | | | |
|---|---|---|---|---|---|---|---|---|---|---|
| 12920 | 646 | K646V | B | 1.2e-05 | 1e-05 | 1.5e-05 | 1.233E-05 | 2.517E-06 | 1.000E-05 | 1.500E-05 |
| 12921 | 647 | A647A | B | 1e-05 | 8.9e-06 | 1.7e-05 | 1.197E-05 | 4.394E-06 | 8.900E-06 | 1.700E-05 |
| 12922 | 647 | A647R | B | 1.7e-05 | 1.6e-05 | 1.3e-05 | 1.533E-05 | 2.082E-06 | 1.300E-05 | 1.700E-05 |
| 12923 | 647 | A647N | B | 1.6e-05 | 1.3e-05 | 1.1e-05 | 1.333E-05 | 2.517E-06 | 1.100E-05 | 1.600E-05 |
| 12924 | 647 | A647D | B | 1.5e-05 | 1.4e-05 | 1e-05 | 1.300E-05 | 2.646E-06 | 1.000E-05 | 1.500E-05 |
| 12925 | 647 | A647C | B | 1e-05 | 1.1e-05 | 1.6e-05 | 1.233E-05 | 3.215E-06 | 1.000E-05 | 1.600E-05 |
| 12926 | 647 | A647E | B | 1.5e-05 | 7.6e-06 | 1.3e-05 | 1.187E-05 | 3.828E-06 | 7.600E-06 | 1.500E-05 |
| 12927 | 647 | A647Q | B | 1.5e-05 | 1.2e-05 | 1.5e-05 | 1.400E-05 | 1.732E-06 | 1.200E-05 | 1.500E-05 |
| 12928 | 647 | A647G | B | 1.6e-05 | 1.1e-05 | 1.4e-05 | 1.367E-05 | 2.517E-06 | 1.100E-05 | 1.600E-05 |
| 12929 | 647 | A647H | B | 1e-05 | 1.4e-05 | 1e-05 | 1.133E-05 | 2.309E-06 | 1.000E-05 | 1.400E-05 |
| 12930 | 647 | A647I | B | 1.7e-05 | 1.6e-05 | 1.4e-05 | 1.567E-05 | 1.528E-06 | 1.400E-05 | 1.700E-05 |
| 12931 | 647 | A647L | B | 1.5e-05 | 1.5e-05 | 1.1e-05 | 1.367E-05 | 2.309E-06 | 1.100E-05 | 1.500E-05 |
| 12932 | 647 | A647K | B | 1.3e-05 | 1.5e-05 | 1.6e-05 | 1.467E-05 | 1.528E-06 | 1.300E-05 | 1.600E-05 |
| 12933 | 647 | A647M | B | 9.8e-06 | 1e-05 | 1.3e-05 | 1.093E-05 | 1.793E-06 | 9.800E-06 | 1.300E-05 |
| 12934 | 647 | A647F | B | 1.6e-05 | 1.1e-05 | 1.5e-05 | 1.400E-05 | 2.646E-06 | 1.100E-05 | 1.600E-05 |
| 12935 | 647 | A647P | B | 1.5e-05 | 1.5e-05 | 1.1e-05 | 1.367E-05 | 2.309E-06 | 1.100E-05 | 1.500E-05 |
| 12936 | 647 | A647S | B | 1.7e-05 | 1.5e-05 | 1.5e-05 | 1.567E-05 | 1.155E-06 | 1.500E-05 | 1.700E-05 |
| 12937 | 647 | A647T | B | 1.3e-05 | 1.5e-05 | 1.4e-05 | 1.400E-05 | 1.000E-06 | 1.300E-05 | 1.500E-05 |
| 12938 | 647 | A647W | B | 1.5e-05 | 1.1e-05 | 1.1e-05 | 1.233E-05 | 2.309E-06 | 1.100E-05 | 1.500E-05 |
| 12939 | 647 | A647Y | B | 8.4e-06 | 1.1e-05 | 1.4e-05 | 1.113E-05 | 2.802E-06 | 8.400E-06 | 1.400E-05 |
| 12940 | 647 | A647V | B | 1.6e-05 | 1.1e-05 | 1.6e-05 | 1.433E-05 | 2.887E-06 | 1.100E-05 | 1.600E-05 |
| 12941 | 648 | S648A | B | 1.6e-05 | 1.6e-05 | 1e-05 | 1.400E-05 | 3.464E-06 | 1.000E-05 | 1.600E-05 |
| 12942 | 648 | S648R | B | 1.6e-05 | 1.3e-05 | 1.4e-05 | 1.433E-05 | 1.528E-06 | 1.300E-05 | 1.600E-05 |
| 12943 | 648 | S648N | B | 1.6e-05 | 1.6e-05 | 1.6e-05 | 1.600E-05 | 0.000E+00 | 1.600E-05 | 1.600E-05 |
| 12944 | 648 | S648D | B | 1e-05 | 1.4e-05 | 1.5e-05 | 1.300E-05 | 2.646E-06 | 1.000E-05 | 1.500E-05 |
| 12945 | 648 | S648C | B | 1.9e-05 | 1e-05 | 1.6e-05 | 1.500E-05 | 4.583E-06 | 1.000E-05 | 1.900E-05 |
| 12946 | 648 | S648E | B | 1.4e-05 | 1.2e-05 | 1.1e-05 | 1.233E-05 | 1.528E-06 | 1.100E-05 | 1.400E-05 |
| 12947 | 648 | S648Q | B | 8.6e-06 | 1.1e-05 | 1e-05 | 9.867E-06 | 1.206E-06 | 8.600E-06 | 1.100E-05 |
| 12948 | 648 | S648G | B | 9.9e-06 | 1.8e-05 | 1.5e-05 | 1.430E-05 | 4.095E-06 | 9.900E-06 | 1.800E-05 |
| 12949 | 648 | S648H | B | 9.5e-06 | 1.6e-05 | 1.8e-05 | 1.450E-05 | 4.444E-06 | 9.500E-06 | 1.800E-05 |
| 12950 | 648 | S648I | B | 8.7e-06 | 1.6e-05 | 8.4e-06 | 1.103E-05 | 4.304E-06 | 8.400E-06 | 1.600E-05 |
| 12951 | 648 | S648L | B | 1.6e-05 | 8.4e-06 | 1.6e-05 | 1.347E-05 | 4.388E-06 | 8.400E-06 | 1.600E-05 |
| 12952 | 648 | S648K | B | 1.3e-05 | 1.6e-05 | 1.4e-05 | 1.433E-05 | 1.528E-06 | 1.300E-05 | 1.600E-05 |
| 12953 | 648 | S648M | B | 8.1e-06 | 1.1e-05 | 2.4e-05 | 1.437E-05 | 8.468E-06 | 8.100E-06 | 2.400E-05 |
| 12954 | 648 | S648F | B | 1.2e-05 | 1.6e-05 | 1.1e-05 | 1.300E-05 | 2.646E-06 | 1.100E-05 | 1.600E-05 |
| 12955 | 648 | S648P | B | 1.5e-05 | 1.3e-05 | 1e-05 | 1.267E-05 | 2.517E-06 | 1.000E-05 | 1.500E-05 |
| 12956 | 648 | S648S | B | 1.2e-05 | 1.1e-05 | 1.6e-05 | 1.300E-05 | 2.646E-06 | 1.100E-05 | 1.600E-05 |
| 12957 | 648 | S648T | B | 1.3e-05 | 1.1e-05 | 1.6e-05 | 1.333E-05 | 2.517E-06 | 1.100E-05 | 1.600E-05 |
| 12958 | 648 | S648W | B | 1.5e-05 | 1.1e-05 | 9.3e-06 | 1.177E-05 | 2.926E-06 | 9.300E-06 | 1.500E-05 |
| 12959 | 648 | S648Y | B | 1.3e-05 | 9.8e-06 | 9.3e-06 | 1.070E-05 | 2.007E-06 | 9.300E-06 | 1.300E-05 |
| 12960 | 648 | S648V | B | 1.6e-05 | 1e-05 | 1.6e-05 | 1.400E-05 | 3.464E-06 | 1.000E-05 | 1.600E-05 |
| 12961 | 649 | Q649A | B | 1.1e-05 | 1.2e-05 | 1.3e-05 | 1.200E-05 | 1.000E-06 | 1.100E-05 | 1.300E-05 |
| 12962 | 649 | Q649R | B | 1.4e-05 | 1.1e-05 | 1.4e-05 | 1.300E-05 | 1.732E-06 | 1.100E-05 | 1.400E-05 |
| 12963 | 649 | Q649N | B | 1.3e-05 | 1e-05 | 8.7e-06 | 1.057E-05 | 2.205E-06 | 8.700E-06 | 1.300E-05 |
| 12964 | 649 | Q649D | B | 1.6e-05 | 1.5e-05 | 1.6e-05 | 1.567E-05 | 5.774E-07 | 1.500E-05 | 1.600E-05 |
| 12965 | 649 | Q649C | B | 9.2e-06 | 1e-05 | 1e-05 | 9.733E-06 | 4.619E-07 | 9.200E-06 | 1.000E-05 |
| 12966 | 649 | Q649E | B | 1.4e-05 | 1.5e-05 | 1.6e-05 | 1.500E-05 | 1.000E-06 | 1.400E-05 | 1.600E-05 |
| 12967 | 649 | Q649Q | B | 1.1e-05 | 1.5e-05 | 1.5e-05 | 1.367E-05 | 2.309E-06 | 1.100E-05 | 1.500E-05 |
| 12968 | 649 | Q649G | B | 9.3e-06 | 1.6e-05 | 1.5e-05 | 1.343E-05 | 3.614E-06 | 9.300E-06 | 1.600E-05 |
| 12969 | 649 | Q649H | B | 1.3e-05 | 9.6e-06 | 1.1e-05 | 1.120E-05 | 1.709E-06 | 9.600E-06 | 1.300E-05 |
| 12970 | 649 | Q649I | B | 1.3e-05 | 1.5e-05 | 1.5e-05 | 1.433E-05 | 1.155E-06 | 1.300E-05 | 1.500E-05 |
| 12971 | 649 | Q649L | B | 1.3e-05 | 1.5e-05 | 1.6e-05 | 1.467E-05 | 1.528E-06 | 1.300E-05 | 1.600E-05 |
| 12972 | 649 | Q649K | B | 9.8e-06 | 1.6e-05 | 1.1e-05 | 1.227E-05 | 3.288E-06 | 9.800E-06 | 1.600E-05 |
| 12973 | 649 | Q649M | B | 1.1e-05 | 1.5e-05 | 9.9e-06 | 1.197E-05 | 2.684E-06 | 9.900E-06 | 1.500E-05 |
| 12974 | 649 | Q649F | B | 1.2e-05 | 1.5e-05 | 1.6e-05 | 1.433E-05 | 2.082E-06 | 1.200E-05 | 1.600E-05 |
| 12975 | 649 | Q649P | B | 9.1e-06 | 1.5e-05 | 1.3e-05 | 1.237E-05 | 3.001E-06 | 9.100E-06 | 1.500E-05 |
| 12976 | 649 | Q649S | B | 1.5e-05 | 1.1e-05 | 1.6e-05 | 1.400E-05 | 2.646E-06 | 1.100E-05 | 1.600E-05 |
| 12977 | 649 | Q649T | B | 1.7e-05 | 1.5e-05 | 1.2e-05 | 1.467E-05 | 2.517E-06 | 1.200E-05 | 1.700E-05 |
| 12978 | 649 | Q649W | B | 1.4e-05 | 1e-05 | 1.5e-05 | 1.300E-05 | 2.646E-06 | 1.000E-05 | 1.500E-05 |
| 12979 | 649 | Q649Y | B | 1.5e-05 | 1.6e-05 | 1.2e-05 | 1.433E-05 | 2.082E-06 | 1.200E-05 | 1.600E-05 |
| 12980 | 649 | Q649V | B | 1.1e-05 | 7.6e-06 | 1e-05 | 9.533E-06 | 1.747E-06 | 7.600E-06 | 1.100E-05 |
| 12981 | 650 | D650A | B | 1e-05 | 6.6e-06 | 1.1e-05 | 9.200E-06 | 2.307E-06 | 6.600E-06 | 1.100E-05 |
| 12982 | 650 | D650R | B | 1e-05 | 1.1e-05 | 1.1e-05 | 1.067E-05 | 5.774E-07 | 1.000E-05 | 1.100E-05 |
| 12983 | 650 | D650N | B | 1.6e-05 | 1.7e-05 | 9.9e-06 | 1.430E-05 | 3.843E-06 | 9.900E-06 | 1.700E-05 |
| 12984 | 650 | D650D | B | 1.4e-05 | 2.4e-05 | 1e-05 | 1.600E-05 | 7.211E-06 | 1.000E-05 | 2.400E-05 |
| 12985 | 650 | D650C | B | 1.3e-05 | 1.4e-05 | 1.4e-05 | 1.367E-05 | 5.774E-07 | 1.300E-05 | 1.400E-05 |
| 12986 | 650 | D650E | B | 1.5e-05 | 1.7e-05 | 1.2e-05 | 1.467E-05 | 2.517E-06 | 1.200E-05 | 1.700E-05 |
| 12987 | 650 | D650Q | B | 9.7e-06 | 8.8e-06 | 2.1e-05 | 1.317E-05 | 6.799E-06 | 8.800E-06 | 2.100E-05 |
| 12988 | 650 | D650G | B | 1.5e-05 | 1.6e-05 | 1.1e-05 | 1.400E-05 | 2.646E-06 | 1.100E-05 | 1.600E-05 |
| 12989 | 650 | D650H | B | 6.9e-06 | 1.5e-05 | 1.5e-05 | 1.230E-05 | 4.677E-06 | 6.900E-06 | 1.500E-05 |
| 12990 | 650 | D650I | B | 1.6e-05 | 1.5e-05 | 1.1e-05 | 1.400E-05 | 2.646E-06 | 1.100E-05 | 1.600E-05 |
| 12991 | 650 | D650L | B | 1.7e-05 | 9.9e-06 | 1.4e-05 | 1.363E-05 | 3.564E-06 | 9.900E-06 | 1.700E-05 |
| 12992 | 650 | D650K | B | 1.6e-05 | 1.5e-05 | 7.3e-06 | 1.277E-05 | 4.761E-06 | 7.300E-06 | 1.600E-05 |
| 12993 | 650 | D650M | B | 1.4e-05 | 1.3e-05 | 9.9e-06 | 1.230E-05 | 2.138E-06 | 9.900E-06 | 1.400E-05 |
| 12994 | 650 | D650F | B | 1e-05 | 1.5e-05 | 9.7e-06 | 1.157E-05 | 2.977E-06 | 9.700E-06 | 1.500E-05 |
| 12995 | 650 | D650P | B | 9.3e-06 | 1.1e-05 | 1.6e-05 | 1.210E-05 | 3.483E-06 | 9.300E-06 | 1.600E-05 |
| 12996 | 650 | D650S | B | 1.4e-05 | 1.1e-05 | 1.1e-05 | 1.200E-05 | 1.732E-06 | 1.100E-05 | 1.400E-05 |
| 12997 | 650 | D650T | B | 8e-06 | 1.5e-05 | 1.6e-05 | 1.300E-05 | 4.359E-06 | 8.000E-06 | 1.600E-05 |
| 12998 | 650 | D650W | B | 1.5e-05 | 1.5e-05 | 1.1e-05 | 1.367E-05 | 2.309E-06 | 1.100E-05 | 1.500E-05 |
| 12999 | 650 | D650Y | B | 1.6e-05 | 1.1e-05 | 1.5e-05 | 1.400E-05 | 2.646E-06 | 1.100E-05 | 1.600E-05 |
| 13000 | 650 | D650V | B | 1.7e-05 | 1.5e-05 | 1.6e-05 | 1.600E-05 | 1.000E-06 | 1.500E-05 | 1.700E-05 |
| 13001 | 651 | V651A | B | 1.5e-05 | 9.4e-06 | 1.1e-05 | 1.180E-05 | 2.884E-06 | 9.400E-06 | 1.500E-05 |
| 13002 | 651 | V651R | B | 1.1e-05 | 1.5e-05 | 1.3e-05 | 1.300E-05 | 2.000E-06 | 1.100E-05 | 1.500E-05 |
| 13003 | 651 | V651N | B | 9.2e-06 | 1.5e-05 | 1.4e-05 | 1.273E-05 | 3.101E-06 | 9.200E-06 | 1.500E-05 |
| 13004 | 651 | V651D | B | 1.5e-05 | 1.6e-05 | 1.5e-05 | 1.533E-05 | 5.774E-07 | 1.500E-05 | 1.600E-05 |

| | | | | | | | | | | |
|---|---|---|---|---|---|---|---|---|---|---|
| 13005 | 651 | V651C | B | 9.7e-06 | 1.6e-05 | 1.4e-05 | 1.323E-05 | 3.219E-06 | 9.700E-06 | 1.600E-05 |
| 13006 | 651 | V651E | B | 1.5e-05 | 1.5e-05 | 1.6e-05 | 1.533E-05 | 5.774E-07 | 1.500E-05 | 1.600E-05 |
| 13007 | 651 | V651Q | B | 1.5e-05 | 1e-05 | 1.5e-05 | 1.333E-05 | 2.887E-06 | 1.000E-05 | 1.500E-05 |
| 13008 | 651 | V651G | B | 1.7e-05 | 1.6e-05 | 1.2e-05 | 1.500E-05 | 2.646E-06 | 1.200E-05 | 1.700E-05 |
| 13009 | 651 | V651H | B | 1.5e-05 | 8.8e-06 | 9.9e-06 | 1.123E-05 | 3.308E-06 | 8.800E-06 | 1.500E-05 |
| 13010 | 651 | V651I | B | 1.5e-05 | 1.6e-05 | 9.7e-06 | 1.357E-05 | 3.386E-06 | 9.700E-06 | 1.600E-05 |
| 13011 | 651 | V651L | B | 1.7e-05 | 1.5e-05 | 1.6e-05 | 1.600E-05 | 1.000E-06 | 1.500E-05 | 1.700E-05 |
| 13012 | 651 | V651K | B | 1.5e-05 | 1.5e-05 | 1.1e-05 | 1.367E-05 | 2.309E-06 | 1.100E-05 | 1.500E-05 |
| 13013 | 651 | V651M | B | 1.5e-05 | 1.4e-05 | 1.7e-05 | 1.533E-05 | 1.528E-06 | 1.400E-05 | 1.700E-05 |
| 13014 | 651 | V651F | B | 1.6e-05 | 9.3e-06 | 1.2e-05 | 1.243E-05 | 3.371E-06 | 9.300E-06 | 1.600E-05 |
| 13015 | 651 | V651P | B | 1.7e-05 | 1.5e-05 | 1.5e-05 | 1.567E-05 | 1.155E-06 | 1.500E-05 | 1.700E-05 |
| 13016 | 651 | V651S | B | 1.2e-05 | 1.5e-05 | 2.5e-05 | 1.733E-05 | 6.807E-06 | 1.200E-05 | 2.500E-05 |
| 13017 | 651 | V651T | B | 1e-05 | 1.5e-05 | 1e-05 | 1.167E-05 | 2.887E-06 | 1.000E-05 | 1.500E-05 |
| 13018 | 651 | V651W | B | 1.5e-05 | 1e-05 | 1.6e-05 | 1.367E-05 | 3.215E-06 | 1.000E-05 | 1.600E-05 |
| 13019 | 651 | V651Y | B | 1.4e-05 | 1.1e-05 | 1.5e-05 | 1.333E-05 | 2.082E-06 | 1.100E-05 | 1.500E-05 |
| 13020 | 651 | V651V | B | 1.5e-05 | 1.1e-05 | 2.3e-05 | 1.633E-05 | 6.110E-06 | 1.100E-05 | 2.300E-05 |
| 13021 | 652 | S652A | B | 1e-05 | 9.6e-06 | 1.6e-05 | 1.187E-05 | 3.585E-06 | 9.600E-06 | 1.600E-05 |
| 13022 | 652 | S652R | B | 9.4e-06 | 1.6e-05 | 1.5e-05 | 1.347E-05 | 3.557E-06 | 9.400E-06 | 1.600E-05 |
| 13023 | 652 | S652N | B | 1.3e-05 | 1.5e-05 | 1.1e-05 | 1.300E-05 | 2.000E-06 | 1.100E-05 | 1.500E-05 |
| 13024 | 652 | S652D | B | 1.5e-05 | 1.5e-05 | 1.2e-05 | 1.400E-05 | 1.732E-06 | 1.200E-05 | 1.500E-05 |
| 13025 | 652 | S652C | B | 7.8e-06 | 1.5e-05 | 1.4e-05 | 1.227E-05 | 3.900E-06 | 7.800E-06 | 1.500E-05 |
| 13026 | 652 | S652E | B | 1.5e-05 | 9.3e-06 | 9.8e-06 | 1.137E-05 | 3.156E-06 | 9.300E-06 | 1.500E-05 |
| 13027 | 652 | S652Q | B | 1.6e-05 | 9.2e-06 | 1.6e-05 | 1.373E-05 | 3.926E-06 | 9.200E-06 | 1.600E-05 |
| 13028 | 652 | S652G | B | 1e-05 | 9.8e-06 | 2.2e-05 | 1.393E-05 | 6.987E-06 | 9.800E-06 | 2.200E-05 |
| 13029 | 652 | S652H | B | 1e-05 | 1.2e-05 | 1.1e-05 | 1.100E-05 | 1.000E-06 | 1.000E-05 | 1.200E-05 |
| 13030 | 652 | S652I | B | 6.3e-06 | 1.3e-05 | 1e-05 | 9.767E-06 | 3.356E-06 | 6.300E-06 | 1.300E-05 |
| 13031 | 652 | S652L | B | 9.1e-06 | 1.2e-05 | 1.5e-05 | 1.203E-05 | 2.950E-06 | 9.100E-06 | 1.500E-05 |
| 13032 | 652 | S652K | B | 1.3e-05 | 1.1e-05 | 1.5e-05 | 1.300E-05 | 2.000E-06 | 1.100E-05 | 1.500E-05 |
| 13033 | 652 | S652M | B | 1.5e-05 | 1.2e-05 | 1.8e-05 | 1.500E-05 | 3.000E-06 | 1.200E-05 | 1.800E-05 |
| 13034 | 652 | S652F | B | 1.5e-05 | 1.2e-05 | 1.1e-05 | 1.267E-05 | 2.082E-06 | 1.100E-05 | 1.500E-05 |
| 13035 | 652 | S652P | B | 1.5e-05 | 1.8e-05 | 1.7e-05 | 1.667E-05 | 1.528E-06 | 1.500E-05 | 1.800E-05 |
| 13036 | 652 | S652S | B | 8e-06 | 1.5e-05 | 1.1e-05 | 1.133E-05 | 3.512E-06 | 8.000E-06 | 1.500E-05 |
| 13037 | 652 | S652T | B | 9.7e-06 | 9.9e-06 | 1.4e-05 | 1.120E-05 | 2.427E-06 | 9.700E-06 | 1.400E-05 |
| 13038 | 652 | S652W | B | 1.1e-05 | 1.3e-05 | 1.6e-05 | 1.333E-05 | 2.517E-06 | 1.100E-05 | 1.600E-05 |
| 13039 | 652 | S652Y | B | 1.5e-05 | 1.4e-05 | 1e-05 | 1.300E-05 | 2.646E-06 | 1.000E-05 | 1.500E-05 |
| 13040 | 652 | S652V | B | 1.2e-05 | 1.1e-05 | 9.9e-06 | 1.097E-05 | 1.050E-06 | 9.900E-06 | 1.200E-05 |
| 13041 | 653 | I653A | B | 1.2e-05 | 9.9e-06 | 9.6e-06 | 1.050E-05 | 1.308E-06 | 9.600E-06 | 1.200E-05 |
| 13042 | 653 | I653R | B | 1.2e-05 | 1.4e-05 | 1.3e-05 | 1.300E-05 | 1.000E-06 | 1.200E-05 | 1.400E-05 |
| 13043 | 653 | I653N | B | 9.5e-06 | 1.5e-05 | 1.2e-05 | 1.217E-05 | 2.754E-06 | 9.500E-06 | 1.500E-05 |
| 13044 | 653 | I653D | B | 1.5e-05 | 1.5e-05 | 1.2e-05 | 1.400E-05 | 1.732E-06 | 1.200E-05 | 1.500E-05 |
| 13045 | 653 | I653C | B | 1.3e-05 | 1.5e-05 | 1e-05 | 1.267E-05 | 2.517E-06 | 1.000E-05 | 1.500E-05 |
| 13046 | 653 | I653E | B | 1.1e-05 | 1.5e-05 | 7.3e-06 | 1.110E-05 | 3.851E-06 | 7.300E-06 | 1.500E-05 |
| 13047 | 653 | I653Q | B | 9.5e-06 | 1.2e-05 | 1.1e-05 | 1.083E-05 | 1.258E-06 | 9.500E-06 | 1.200E-05 |
| 13048 | 653 | I653G | B | 1.5e-05 | 1.2e-05 | 1.4e-05 | 1.367E-05 | 1.528E-06 | 1.200E-05 | 1.500E-05 |
| 13049 | 653 | I653H | B | 8.6e-06 | 1.4e-05 | 1.1e-05 | 1.120E-05 | 2.706E-06 | 8.600E-06 | 1.400E-05 |
| 13050 | 653 | I653I | B | 1.5e-05 | 1.1e-05 | 9.3e-06 | 1.177E-05 | 2.926E-06 | 9.300E-06 | 1.500E-05 |
| 13051 | 653 | I653L | B | 1.1e-05 | 1.4e-05 | 8.3e-06 | 1.110E-05 | 2.851E-06 | 8.300E-06 | 1.400E-05 |
| 13052 | 653 | I653K | B | 1.3e-05 | 8.8e-06 | 1.3e-05 | 1.160E-05 | 2.425E-06 | 8.800E-06 | 1.300E-05 |
| 13053 | 653 | I653M | B | 1.1e-05 | 1.7e-05 | 6e-06 | 1.133E-05 | 5.508E-06 | 6.000E-06 | 1.700E-05 |
| 13054 | 653 | I653F | B | 1e-05 | 1e-05 | 9.9e-06 | 9.967E-06 | 5.774E-08 | 9.900E-06 | 1.000E-05 |
| 13055 | 653 | I653P | B | 1.7e-05 | 8.8e-06 | 1.4e-05 | 1.327E-05 | 4.149E-06 | 8.800E-06 | 1.700E-05 |
| 13056 | 653 | I653S | B | 1.3e-05 | 1.9e-05 | 1e-05 | 1.400E-05 | 4.583E-06 | 1.000E-05 | 1.900E-05 |
| 13057 | 653 | I653T | B | 1.4e-05 | 9e-06 | 1.6e-05 | 1.300E-05 | 3.606E-06 | 9.000E-06 | 1.600E-05 |
| 13058 | 653 | I653W | B | 7.2e-06 | 9.7e-06 | 1.5e-05 | 1.063E-05 | 3.983E-06 | 7.200E-06 | 1.500E-05 |
| 13059 | 653 | I653Y | B | 1e-05 | 8.2e-06 | 2.1e-05 | 1.307E-05 | 6.929E-06 | 8.200E-06 | 2.100E-05 |
| 13060 | 653 | I653V | B | 1.1e-05 | 1.1e-05 | 1.3e-05 | 1.167E-05 | 1.155E-06 | 1.100E-05 | 1.300E-05 |
| 13061 | 654 | G654A | B | 1e-05 | 8.9e-06 | 1.2e-05 | 1.030E-05 | 1.572E-06 | 8.900E-06 | 1.200E-05 |
| 13062 | 654 | G654R | B | 9.3e-06 | 8.7e-06 | 1.2e-05 | 1.000E-05 | 1.758E-06 | 8.700E-06 | 1.200E-05 |
| 13063 | 654 | G654N | B | 1e-05 | 9.3e-06 | 1.1e-05 | 1.010E-05 | 8.544E-07 | 9.300E-06 | 1.100E-05 |
| 13064 | 654 | G654D | B | 9.1e-06 | 1.2e-05 | 1.1e-05 | 1.070E-05 | 1.473E-06 | 9.100E-06 | 1.200E-05 |
| 13065 | 654 | G654C | B | 8.5e-06 | 1e-05 | 1.7e-05 | 1.183E-05 | 4.537E-06 | 8.500E-06 | 1.700E-05 |
| 13066 | 654 | G654E | B | 1.5e-05 | 9.4e-06 | 1.5e-05 | 1.313E-05 | 3.233E-06 | 9.400E-06 | 1.500E-05 |
| 13067 | 654 | G654Q | B | 1.3e-05 | 1.5e-05 | 1.5e-05 | 1.433E-05 | 1.155E-06 | 1.300E-05 | 1.500E-05 |
| 13068 | 654 | G654G | B | 5.3e-06 | 1.2e-05 | 1e-05 | 9.100E-06 | 3.439E-06 | 5.300E-06 | 1.200E-05 |
| 13069 | 654 | G654H | B | 1.5e-05 | 9.9e-06 | 1.9e-05 | 1.463E-05 | 4.561E-06 | 9.900E-06 | 1.900E-05 |
| 13070 | 654 | G654I | B | 1.1e-05 | 1.4e-05 | 1.4e-05 | 1.300E-05 | 1.732E-06 | 1.100E-05 | 1.400E-05 |
| 13071 | 654 | G654L | B | 1.1e-05 | 1.4e-05 | 1.5e-05 | 1.333E-05 | 2.082E-06 | 1.100E-05 | 1.500E-05 |
| 13072 | 654 | G654K | B | 1.1e-05 | 1.6e-05 | 6.9e-06 | 1.130E-05 | 4.557E-06 | 6.900E-06 | 1.600E-05 |
| 13073 | 654 | G654M | B | 1.6e-05 | 1.6e-05 | 1.5e-05 | 1.567E-05 | 5.774E-07 | 1.500E-05 | 1.600E-05 |
| 13074 | 654 | G654F | B | 1.1e-05 | 1.6e-05 | 1.6e-05 | 1.433E-05 | 2.887E-06 | 1.100E-05 | 1.600E-05 |
| 13075 | 654 | G654P | B | 1.1e-05 | 1.7e-05 | 1.1e-05 | 1.300E-05 | 3.464E-06 | 1.100E-05 | 1.700E-05 |
| 13076 | 654 | G654S | B | 8.8e-06 | 1.4e-05 | 1.1e-05 | 1.127E-05 | 2.610E-06 | 8.800E-06 | 1.400E-05 |
| 13077 | 654 | G654T | B | 1.5e-05 | 1.6e-05 | 1.5e-05 | 1.533E-05 | 5.774E-07 | 1.500E-05 | 1.600E-05 |
| 13078 | 654 | G654W | B | 1.5e-05 | 1.6e-05 | 7.2e-06 | 1.273E-05 | 4.818E-06 | 7.200E-06 | 1.600E-05 |
| 13079 | 654 | G654Y | B | 9.7e-06 | 9.3e-06 | 1.4e-05 | 1.100E-05 | 2.606E-06 | 9.300E-06 | 1.400E-05 |
| 13080 | 654 | G654V | B | 1.5e-05 | 1.4e-05 | 1.2e-05 | 1.367E-05 | 1.528E-06 | 1.200E-05 | 1.500E-05 |
| 13081 | 655 | V655A | B | 1.5e-05 | 1.2e-05 | 1.4e-05 | 1.367E-05 | 1.528E-06 | 1.200E-05 | 1.500E-05 |
| 13082 | 655 | V655R | B | 1.4e-05 | 1.2e-05 | 1.5e-05 | 1.367E-05 | 1.528E-06 | 1.200E-05 | 1.500E-05 |
| 13083 | 655 | V655N | B | 1.3e-05 | 1.1e-05 | 1.6e-05 | 1.333E-05 | 2.517E-06 | 1.100E-05 | 1.600E-05 |
| 13084 | 655 | V655D | B | 1.5e-05 | 1.6e-05 | 9.9e-06 | 1.363E-05 | 3.272E-06 | 9.900E-06 | 1.600E-05 |
| 13085 | 655 | V655C | B | 1.5e-05 | 8.1e-06 | 1.6e-05 | 1.303E-05 | 4.302E-06 | 8.100E-06 | 1.600E-05 |
| 13086 | 655 | V655E | B | 9.8e-06 | 1.4e-05 | 1.4e-05 | 1.260E-05 | 2.425E-06 | 9.800E-06 | 1.400E-05 |
| 13087 | 655 | V655Q | B | 1.3e-05 | 1.7e-05 | 1.6e-05 | 1.533E-05 | 2.082E-06 | 1.300E-05 | 1.700E-05 |
| 13088 | 655 | V655G | B | 1.7e-05 | 1.5e-05 | 1.5e-05 | 1.567E-05 | 1.155E-06 | 1.500E-05 | 1.700E-05 |
| 13089 | 655 | V655H | B | 7.6e-06 | 9.2e-06 | 1.7e-05 | 1.127E-05 | 5.029E-06 | 7.600E-06 | 1.700E-05 |

| | | | | | | | | | | |
|---|---|---|---|---|---|---|---|---|---|---|
| 13090 | 655 | V655I | B | 1.2e-05 | 1.5e-05 | 1e-05 | 1.233E-05 | 2.517E-06 | 1.000E-05 | 1.500E-05 |
| 13091 | 655 | V655L | B | 1.6e-05 | 1.1e-05 | 1.6e-05 | 1.433E-05 | 2.887E-06 | 1.100E-05 | 1.600E-05 |
| 13092 | 655 | V655K | B | 1.6e-05 | 1.1e-05 | 1.2e-05 | 1.300E-05 | 2.646E-06 | 1.100E-05 | 1.600E-05 |
| 13093 | 655 | V655M | B | 8.9e-06 | 1.7e-05 | 1.4e-05 | 1.330E-05 | 4.095E-06 | 8.900E-06 | 1.700E-05 |
| 13094 | 655 | V655F | B | 1.2e-05 | 1e-05 | 1.5e-05 | 1.233E-05 | 2.517E-06 | 1.000E-05 | 1.500E-05 |
| 13095 | 655 | V655P | B | 1.2e-05 | 1e-05 | 1.5e-05 | 1.233E-05 | 2.517E-06 | 1.000E-05 | 1.500E-05 |
| 13096 | 655 | V655S | B | 1.5e-05 | 1.1e-05 | 1.5e-05 | 1.367E-05 | 2.309E-06 | 1.100E-05 | 1.500E-05 |
| 13097 | 655 | V655T | B | 1.5e-05 | 1e-05 | 1.6e-05 | 1.367E-05 | 3.215E-06 | 1.000E-05 | 1.600E-05 |
| 13098 | 655 | V655W | B | 1e-05 | 1.1e-05 | 1.8e-05 | 1.300E-05 | 4.359E-06 | 1.000E-05 | 1.800E-05 |
| 13099 | 655 | V655Y | B | 1.4e-05 | 1.6e-05 | 1.5e-05 | 1.500E-05 | 1.000E-06 | 1.400E-05 | 1.600E-05 |
| 13100 | 655 | V655V | B | 1.4e-05 | 1.4e-05 | 1.5e-05 | 1.433E-05 | 5.774E-07 | 1.400E-05 | 1.500E-05 |
| 13101 | 656 | A656A | B | 1.1e-05 | 9.5e-06 | 1.7e-05 | 1.250E-05 | 3.969E-06 | 9.500E-06 | 1.700E-05 |
| 13102 | 656 | A656R | B | 9.2e-06 | 1.3e-05 | 8.9e-06 | 1.037E-05 | 2.285E-06 | 8.900E-06 | 1.300E-05 |
| 13103 | 656 | A656N | B | 1.4e-05 | 1.4e-05 | 1.6e-05 | 1.467E-05 | 1.155E-06 | 1.400E-05 | 1.600E-05 |
| 13104 | 656 | A656D | B | 1.3e-05 | 8.8e-06 | 1.4e-05 | 1.193E-05 | 2.759E-06 | 8.800E-06 | 1.400E-05 |
| 13105 | 656 | A656C | B | 1.5e-05 | 1e-05 | 1.1e-05 | 1.200E-05 | 2.646E-06 | 1.000E-05 | 1.500E-05 |
| 13106 | 656 | A656E | B | 2.1e-05 | 1.5e-05 | 1.4e-05 | 1.667E-05 | 3.786E-06 | 1.400E-05 | 2.100E-05 |
| 13107 | 656 | A656Q | B | 1.4e-05 | 1.1e-05 | 1.4e-05 | 1.300E-05 | 1.732E-06 | 1.100E-05 | 1.400E-05 |
| 13108 | 656 | A656G | B | 9.1e-06 | 1.5e-05 | 1.5e-05 | 1.303E-05 | 3.406E-06 | 9.100E-06 | 1.500E-05 |
| 13109 | 656 | A656H | B | 1.3e-05 | 7.9e-06 | 1.2e-05 | 1.097E-05 | 2.702E-06 | 7.900E-06 | 1.300E-05 |
| 13110 | 656 | A656I | B | 1.2e-05 | 9.6e-06 | 1.5e-05 | 1.220E-05 | 2.706E-06 | 9.600E-06 | 1.500E-05 |
| 13111 | 656 | A656L | B | 1.5e-05 | 1.6e-05 | 1.6e-05 | 1.567E-05 | 5.774E-07 | 1.500E-05 | 1.600E-05 |
| 13112 | 656 | A656K | B | 1.2e-05 | 1.7e-05 | 1.3e-05 | 1.400E-05 | 2.646E-06 | 1.200E-05 | 1.700E-05 |
| 13113 | 656 | A656M | B | 9.5e-06 | 1e-05 | 1.4e-05 | 1.117E-05 | 2.466E-06 | 9.500E-06 | 1.400E-05 |
| 13114 | 656 | A656F | B | 1.7e-05 | 1.6e-05 | 1e-05 | 1.433E-05 | 3.786E-06 | 1.000E-05 | 1.700E-05 |
| 13115 | 656 | A656P | B | 1e-05 | 6.5e-06 | 1.5e-05 | 1.050E-05 | 4.272E-06 | 6.500E-06 | 1.500E-05 |
| 13116 | 656 | A656S | B | 1.5e-05 | 1.1e-05 | 1.6e-05 | 1.400E-05 | 2.646E-06 | 1.100E-05 | 1.600E-05 |
| 13117 | 656 | A656T | B | 1.2e-05 | 1.4e-05 | 7.4e-06 | 1.113E-05 | 3.384E-06 | 7.400E-06 | 1.400E-05 |
| 13118 | 656 | A656W | B | 9.2e-06 | 1.3e-05 | 1.3e-05 | 1.173E-05 | 2.194E-06 | 9.200E-06 | 1.300E-05 |
| 13119 | 656 | A656Y | B | 1.1e-05 | 8.6e-06 | 1.4e-05 | 1.120E-05 | 2.706E-06 | 8.600E-06 | 1.400E-05 |
| 13120 | 656 | A656V | B | 1.5e-05 | 1e-05 | 1.4e-05 | 1.300E-05 | 2.646E-06 | 1.000E-05 | 1.500E-05 |
| 13121 | 657 | W657A | B | 1.6e-05 | 1.2e-05 | 9.6e-06 | 1.253E-05 | 3.233E-06 | 9.600E-06 | 1.600E-05 |
| 13122 | 657 | W657R | B | 8.8e-06 | 1.1e-05 | 1e-05 | 9.933E-06 | 1.102E-06 | 8.800E-06 | 1.100E-05 |
| 13123 | 657 | W657N | B | 1.4e-05 | 1.6e-05 | 1.3e-05 | 1.433E-05 | 1.528E-06 | 1.300E-05 | 1.600E-05 |
| 13124 | 657 | W657D | B | 1.3e-05 | 1e-05 | 1.2e-05 | 1.167E-05 | 1.528E-06 | 1.000E-05 | 1.300E-05 |
| 13125 | 657 | W657C | B | 1.1e-05 | 1.1e-05 | 1.6e-05 | 1.267E-05 | 2.887E-06 | 1.100E-05 | 1.600E-05 |
| 13126 | 657 | W657E | B | 1.2e-05 | 1.3e-05 | 1.5e-05 | 1.333E-05 | 1.528E-06 | 1.200E-05 | 1.500E-05 |
| 13127 | 657 | W657Q | B | 1.7e-05 | 1.1e-05 | 1.2e-05 | 1.333E-05 | 3.215E-06 | 1.100E-05 | 1.700E-05 |
| 13128 | 657 | W657G | B | 1.1e-05 | 1e-05 | 1.8e-05 | 1.300E-05 | 4.359E-06 | 1.000E-05 | 1.800E-05 |
| 13129 | 657 | W657H | B | 1e-05 | 1.3e-05 | 9.4e-06 | 1.080E-05 | 1.929E-06 | 9.400E-06 | 1.300E-05 |
| 13130 | 657 | W657I | B | 7.6e-06 | 9.5e-06 | 9.5e-06 | 8.867E-06 | 1.097E-06 | 7.600E-06 | 9.500E-06 |
| 13131 | 657 | W657L | B | 1.1e-05 | 1.6e-05 | 1.9e-05 | 1.533E-05 | 4.041E-06 | 1.100E-05 | 1.900E-05 |
| 13132 | 657 | W657K | B | 1.1e-05 | 8.2e-06 | 1.3e-05 | 1.073E-05 | 2.411E-06 | 8.200E-06 | 1.300E-05 |
| 13133 | 657 | W657M | B | 1e-05 | 1.6e-05 | 1.6e-05 | 1.400E-05 | 3.464E-06 | 1.000E-05 | 1.600E-05 |
| 13134 | 657 | W657F | B | 1.3e-05 | 1.1e-05 | 1e-05 | 1.133E-05 | 1.528E-06 | 1.000E-05 | 1.300E-05 |
| 13135 | 657 | W657P | B | 8.6e-06 | 1.1e-05 | 1.7e-05 | 1.220E-05 | 4.327E-06 | 8.600E-06 | 1.700E-05 |
| 13136 | 657 | W657S | B | 9.8e-06 | 9.9e-06 | 1.5e-05 | 1.157E-05 | 2.974E-06 | 9.800E-06 | 1.500E-05 |
| 13137 | 657 | W657T | B | 1.5e-05 | 1e-05 | 8.9e-06 | 1.130E-05 | 3.251E-06 | 8.900E-06 | 1.500E-05 |
| 13138 | 657 | W657W | B | 1.4e-05 | 1e-05 | 1.5e-05 | 1.300E-05 | 2.646E-06 | 1.000E-05 | 1.500E-05 |
| 13139 | 657 | W657Y | B | 1.1e-05 | 1.3e-05 | 1.5e-05 | 1.300E-05 | 2.000E-06 | 1.100E-05 | 1.500E-05 |
| 13140 | 657 | W657V | B | 1.6e-05 | 1.3e-05 | 1.1e-05 | 1.333E-05 | 2.517E-06 | 1.100E-05 | 1.600E-05 |
| 13141 | 658 | Y658A | B | 1.7e-05 | 1e-05 | 1.1e-05 | 1.267E-05 | 3.786E-06 | 1.000E-05 | 1.700E-05 |
| 13142 | 658 | Y658R | B | 1.7e-05 | 1.5e-05 | 1.1e-05 | 1.433E-05 | 3.055E-06 | 1.100E-05 | 1.700E-05 |
| 13143 | 658 | Y658N | B | 1.2e-05 | 1.5e-05 | 1.3e-05 | 1.333E-05 | 1.528E-06 | 1.200E-05 | 1.500E-05 |
| 13144 | 658 | Y658D | B | 1.4e-05 | 1e-05 | 1.5e-05 | 1.300E-05 | 2.646E-06 | 1.000E-05 | 1.500E-05 |
| 13145 | 658 | Y658C | B | 9.5e-06 | 1.4e-05 | 1.8e-05 | 1.383E-05 | 4.252E-06 | 9.500E-06 | 1.800E-05 |
| 13146 | 658 | Y658E | B | 1.5e-05 | 8.6e-06 | 8.1e-06 | 1.057E-05 | 3.848E-06 | 8.100E-06 | 1.500E-05 |
| 13147 | 658 | Y658Q | B | 1.4e-05 | 9.7e-06 | 9e-06 | 1.090E-05 | 2.707E-06 | 9.000E-06 | 1.400E-05 |
| 13148 | 658 | Y658G | B | 1.5e-05 | 9.7e-06 | 1.5e-05 | 1.323E-05 | 3.060E-06 | 9.700E-06 | 1.500E-05 |
| 13149 | 658 | Y658H | B | 1.1e-05 | 1.5e-05 | 8.2e-06 | 1.140E-05 | 3.418E-06 | 8.200E-06 | 1.500E-05 |
| 13150 | 658 | Y658I | B | 1.3e-05 | 1.6e-05 | 1.4e-05 | 1.433E-05 | 1.528E-06 | 1.300E-05 | 1.600E-05 |
| 13151 | 658 | Y658L | B | 1.7e-05 | 1e-05 | 1.6e-05 | 1.433E-05 | 3.786E-06 | 1.000E-05 | 1.700E-05 |
| 13152 | 658 | Y658K | B | 1.5e-05 | 9.4e-06 | 1.3e-05 | 1.247E-05 | 2.838E-06 | 9.400E-06 | 1.500E-05 |
| 13153 | 658 | Y658M | B | 1.6e-05 | 1.6e-05 | 1.6e-05 | 1.600E-05 | 0.000E+00 | 1.600E-05 | 1.600E-05 |
| 13154 | 658 | Y658F | B | 1.6e-05 | 1.2e-05 | 1.2e-05 | 1.333E-05 | 2.309E-06 | 1.200E-05 | 1.600E-05 |
| 13155 | 658 | Y658P | B | 1.6e-05 | 1.2e-05 | 1.5e-05 | 1.433E-05 | 2.082E-06 | 1.200E-05 | 1.600E-05 |
| 13156 | 658 | Y658S | B | 1.6e-05 | 1.1e-05 | 1.7e-05 | 1.467E-05 | 3.215E-06 | 1.100E-05 | 1.700E-05 |
| 13157 | 658 | Y658T | B | 1.6e-05 | 1.4e-05 | 1e-05 | 1.333E-05 | 3.055E-06 | 1.000E-05 | 1.600E-05 |
| 13158 | 658 | Y658W | B | 1.1e-05 | 1.1e-05 | 1.4e-05 | 1.200E-05 | 1.732E-06 | 1.100E-05 | 1.400E-05 |
| 13159 | 658 | Y658Y | B | 1e-05 | 1.4e-05 | 9.5e-06 | 1.117E-05 | 2.466E-06 | 9.500E-06 | 1.400E-05 |
| 13160 | 658 | Y658V | B | 1.5e-05 | 1.6e-05 | 1.4e-05 | 1.500E-05 | 1.000E-06 | 1.400E-05 | 1.600E-05 |
| 13161 | 659 | Q659A | B | 1.3e-05 | 1.4e-05 | 1e-05 | 1.233E-05 | 2.082E-06 | 1.000E-05 | 1.400E-05 |
| 13162 | 659 | Q659R | B | 8.9e-06 | 1.5e-05 | 8.5e-06 | 1.080E-05 | 3.643E-06 | 8.500E-06 | 1.500E-05 |
| 13163 | 659 | Q659N | B | 1.4e-05 | 8.5e-06 | 1.5e-05 | 1.250E-05 | 3.500E-06 | 8.500E-06 | 1.500E-05 |
| 13164 | 659 | Q659D | B | 1.7e-05 | 1.7e-05 | 1.1e-05 | 1.500E-05 | 3.464E-06 | 1.100E-05 | 1.700E-05 |
| 13165 | 659 | Q659C | B | 1.8e-05 | 1.4e-05 | 1.4e-05 | 1.533E-05 | 2.309E-06 | 1.400E-05 | 1.800E-05 |
| 13166 | 659 | Q659E | B | 1e-05 | 1.5e-05 | 1.5e-05 | 1.333E-05 | 2.887E-06 | 1.000E-05 | 1.500E-05 |
| 13167 | 659 | Q659Q | B | 1.7e-05 | 1.3e-05 | 1.4e-05 | 1.467E-05 | 2.082E-06 | 1.300E-05 | 1.700E-05 |
| 13168 | 659 | Q659G | B | 1.4e-05 | 1e-05 | 1.6e-05 | 1.333E-05 | 3.055E-06 | 1.000E-05 | 1.600E-05 |
| 13169 | 659 | Q659H | B | 8.1e-06 | 1.7e-05 | 1.1e-05 | 1.203E-05 | 4.539E-06 | 8.100E-06 | 1.700E-05 |
| 13170 | 659 | Q659I | B | 8.9e-06 | 1.6e-05 | 9.6e-06 | 1.150E-05 | 3.913E-06 | 8.900E-06 | 1.600E-05 |
| 13171 | 659 | Q659L | B | 1.5e-05 | 1.5e-05 | 1.3e-05 | 1.433E-05 | 1.155E-06 | 1.300E-05 | 1.500E-05 |
| 13172 | 659 | Q659K | B | 9.3e-06 | 1.4e-05 | 1.6e-05 | 1.310E-05 | 3.439E-06 | 9.300E-06 | 1.600E-05 |
| 13173 | 659 | Q659M | B | 1.2e-05 | 1e-05 | 9.1e-06 | 1.037E-05 | 1.484E-06 | 9.100E-06 | 1.200E-05 |
| 13174 | 659 | Q659F | B | 1.6e-05 | 9.7e-06 | 1.7e-05 | 1.423E-05 | 3.958E-06 | 9.700E-06 | 1.700E-05 |

| | | | | | | | | | | |
|---|---|---|---|---|---|---|---|---|---|---|
| 13175 | 659 | Q659P | B | 1.2e-05 | 1.5e-05 | 9.4e-06 | 1.213E-05 | 2.802E-06 | 9.400E-06 | 1.500E-05 |
| 13176 | 659 | Q659S | B | 1.1e-05 | 1e-05 | 9.8e-06 | 1.027E-05 | 6.429E-07 | 9.800E-06 | 1.100E-05 |
| 13177 | 659 | Q659T | B | 9.9e-06 | 1.5e-05 | 1.6e-05 | 1.363E-05 | 3.272E-06 | 9.900E-06 | 1.600E-05 |
| 13178 | 659 | Q659W | B | 1e-05 | 1e-05 | 1.6e-05 | 1.200E-05 | 3.464E-06 | 1.000E-05 | 1.600E-05 |
| 13179 | 659 | Q659Y | B | 1.3e-05 | 1.2e-05 | 1.7e-05 | 1.400E-05 | 2.646E-06 | 1.200E-05 | 1.700E-05 |
| 13180 | 659 | Q659V | B | 1.4e-05 | 1.4e-05 | 1.5e-05 | 1.433E-05 | 5.774E-07 | 1.400E-05 | 1.500E-05 |
| 13181 | 660 | Q660A | B | 1.5e-05 | 1.4e-05 | 1.4e-05 | 1.433E-05 | 5.774E-07 | 1.400E-05 | 1.500E-05 |
| 13182 | 660 | Q660R | B | 1.7e-05 | 1.7e-05 | 1.1e-05 | 1.500E-05 | 3.464E-06 | 1.100E-05 | 1.700E-05 |
| 13183 | 660 | Q660N | B | 1.6e-05 | 9.2e-06 | 9.4e-06 | 1.153E-05 | 3.870E-06 | 9.200E-06 | 1.600E-05 |
| 13184 | 660 | Q660D | B | 1.5e-05 | 1.5e-05 | 9.9e-06 | 1.330E-05 | 2.944E-06 | 9.900E-06 | 1.500E-05 |
| 13185 | 660 | Q660C | B | 1.1e-05 | 8e-06 | 9.9e-06 | 9.633E-06 | 1.518E-06 | 8.000E-06 | 1.100E-05 |
| 13186 | 660 | Q660E | B | 1.1e-05 | 8.8e-06 | 8.9e-06 | 9.567E-06 | 1.242E-06 | 8.800E-06 | 1.100E-05 |
| 13187 | 660 | Q660Q | B | 9.9e-06 | 1.4e-05 | 1e-05 | 1.130E-05 | 2.339E-06 | 9.900E-06 | 1.400E-05 |
| 13188 | 660 | Q660G | B | 1.6e-05 | 1.3e-05 | 1.1e-05 | 1.333E-05 | 2.517E-06 | 1.100E-05 | 1.600E-05 |
| 13189 | 660 | Q660H | B | 1.5e-05 | 1.5e-05 | 1e-05 | 1.333E-05 | 2.887E-06 | 1.000E-05 | 1.500E-05 |
| 13190 | 660 | Q660I | B | 1.4e-05 | 1.7e-05 | 1.5e-05 | 1.533E-05 | 1.528E-06 | 1.400E-05 | 1.700E-05 |
| 13191 | 660 | Q660L | B | 1.6e-05 | 1.2e-05 | 1.6e-05 | 1.467E-05 | 2.309E-06 | 1.200E-05 | 1.600E-05 |
| 13192 | 660 | Q660K | B | 1.5e-05 | 1.1e-05 | 1.2e-05 | 1.267E-05 | 2.082E-06 | 1.100E-05 | 1.500E-05 |
| 13193 | 660 | Q660M | B | 1.2e-05 | 8.8e-06 | 1.6e-05 | 1.227E-05 | 3.607E-06 | 8.800E-06 | 1.600E-05 |
| 13194 | 660 | Q660F | B | 1.3e-05 | 1.1e-05 | 1.5e-05 | 1.300E-05 | 2.000E-06 | 1.100E-05 | 1.500E-05 |
| 13195 | 660 | Q660P | B | 1.1e-05 | 1.5e-05 | 1.6e-05 | 1.400E-05 | 2.646E-06 | 1.100E-05 | 1.600E-05 |
| 13196 | 660 | Q660S | B | 1e-05 | 1.4e-05 | 1.4e-05 | 1.267E-05 | 2.309E-06 | 1.000E-05 | 1.400E-05 |
| 13197 | 660 | Q660T | B | 1.7e-05 | 8.2e-06 | 1.4e-05 | 1.307E-05 | 4.474E-06 | 8.200E-06 | 1.700E-05 |
| 13198 | 660 | Q660W | B | 1.1e-05 | 1e-05 | 1.3e-05 | 1.133E-05 | 1.528E-06 | 1.000E-05 | 1.300E-05 |
| 13199 | 660 | Q660Y | B | 1.4e-05 | 1.3e-05 | 1.4e-05 | 1.367E-05 | 5.774E-07 | 1.300E-05 | 1.400E-05 |
| 13200 | 660 | Q660V | B | 1.5e-05 | 1.6e-05 | 1.7e-05 | 1.600E-05 | 1.000E-06 | 1.500E-05 | 1.700E-05 |
| 13201 | 661 | K661A | B | 1.5e-05 | 1e-05 | 1.4e-05 | 1.300E-05 | 2.646E-06 | 1.000E-05 | 1.500E-05 |
| 13202 | 661 | K661R | B | 1.1e-05 | 1.5e-05 | 1.5e-05 | 1.367E-05 | 2.309E-06 | 1.100E-05 | 1.500E-05 |
| 13203 | 661 | K661N | B | 1e-05 | 1.6e-05 | 1.5e-05 | 1.367E-05 | 3.215E-06 | 1.000E-05 | 1.600E-05 |
| 13204 | 661 | K661D | B | 1.5e-05 | 1.3e-05 | 1.1e-05 | 1.300E-05 | 2.000E-06 | 1.100E-05 | 1.500E-05 |
| 13205 | 661 | K661C | B | 1.6e-05 | 1.5e-05 | 1.3e-05 | 1.467E-05 | 1.528E-06 | 1.300E-05 | 1.600E-05 |
| 13206 | 661 | K661E | B | 1.6e-05 | 1.3e-05 | 1.5e-05 | 1.467E-05 | 1.528E-06 | 1.300E-05 | 1.600E-05 |
| 13207 | 661 | K661Q | B | 1.6e-05 | 1.2e-05 | 1.1e-05 | 1.300E-05 | 2.646E-06 | 1.100E-05 | 1.600E-05 |
| 13208 | 661 | K661G | B | 1e-05 | 1.5e-05 | 1.6e-05 | 1.367E-05 | 3.215E-06 | 1.000E-05 | 1.600E-05 |
| 13209 | 661 | K661H | B | 1.2e-05 | 1.1e-05 | 1.4e-05 | 1.233E-05 | 1.528E-06 | 1.100E-05 | 1.400E-05 |
| 13210 | 661 | K661I | B | 1.5e-05 | 1e-05 | 1.5e-05 | 1.333E-05 | 2.887E-06 | 1.000E-05 | 1.500E-05 |
| 13211 | 661 | K661L | B | 9.3e-06 | 9.9e-06 | 9.2e-06 | 9.467E-06 | 3.786E-07 | 9.200E-06 | 9.900E-06 |
| 13212 | 661 | K661K | B | 7.4e-06 | 1.5e-05 | 1.6e-05 | 1.280E-05 | 4.703E-06 | 7.400E-06 | 1.600E-05 |
| 13213 | 661 | K661M | B | 1e-05 | 1.1e-05 | 1.1e-05 | 1.067E-05 | 5.774E-07 | 1.000E-05 | 1.100E-05 |
| 13214 | 661 | K661F | B | 8.2e-06 | 9.3e-06 | 1.8e-05 | 1.183E-05 | 5.369E-06 | 8.200E-06 | 1.800E-05 |
| 13215 | 661 | K661P | B | 1.5e-05 | 1.4e-05 | 1.6e-05 | 1.500E-05 | 1.000E-06 | 1.400E-05 | 1.600E-05 |
| 13216 | 661 | K661S | B | 8.9e-06 | 8.3e-06 | 1.2e-05 | 9.733E-06 | 1.986E-06 | 8.300E-06 | 1.200E-05 |
| 13217 | 661 | K661T | B | 8.8e-06 | 1.6e-05 | 1.4e-05 | 1.293E-05 | 3.717E-06 | 8.800E-06 | 1.600E-05 |
| 13218 | 661 | K661W | B | 1e-05 | 1.2e-05 | 1.2e-05 | 1.133E-05 | 1.155E-06 | 1.000E-05 | 1.200E-05 |
| 13219 | 661 | K661Y | B | 1.4e-05 | 1.4e-05 | 1e-05 | 1.267E-05 | 2.309E-06 | 1.000E-05 | 1.400E-05 |
| 13220 | 661 | K661V | B | 1.5e-05 | 1.3e-05 | 1.4e-05 | 1.400E-05 | 1.000E-06 | 1.300E-05 | 1.500E-05 |
| 13221 | 662 | P662A | B | 1.4e-05 | 1.2e-05 | 1.4e-05 | 1.333E-05 | 1.155E-06 | 1.200E-05 | 1.400E-05 |
| 13222 | 662 | P662R | B | 1.4e-05 | 1.5e-05 | 1.3e-05 | 1.400E-05 | 1.000E-06 | 1.300E-05 | 1.500E-05 |
| 13223 | 662 | P662N | B | 1.1e-05 | 8.6e-06 | 1.3e-05 | 1.087E-05 | 2.203E-06 | 8.600E-06 | 1.300E-05 |
| 13224 | 662 | P662D | B | 1.6e-05 | 1.6e-05 | 1.5e-05 | 1.567E-05 | 5.774E-07 | 1.500E-05 | 1.600E-05 |
| 13225 | 662 | P662C | B | 9.2e-06 | 8.4e-06 | 9.9e-06 | 9.167E-06 | 7.506E-07 | 8.400E-06 | 9.900E-06 |
| 13226 | 662 | P662E | B | 1e-05 | 1.7e-05 | 1.5e-05 | 1.400E-05 | 3.606E-06 | 1.000E-05 | 1.700E-05 |
| 13227 | 662 | P662Q | B | 1.2e-05 | 1.1e-05 | 1e-05 | 1.100E-05 | 1.000E-06 | 1.000E-05 | 1.200E-05 |
| 13228 | 662 | P662G | B | 1.5e-05 | 1.2e-05 | 1.3e-05 | 1.333E-05 | 1.528E-06 | 1.200E-05 | 1.500E-05 |
| 13229 | 662 | P662H | B | 1.6e-05 | 1e-05 | 9.7e-06 | 1.190E-05 | 3.554E-06 | 9.700E-06 | 1.600E-05 |
| 13230 | 662 | P662I | B | 1.6e-05 | 1.4e-05 | 9.9e-06 | 1.330E-05 | 3.110E-06 | 9.900E-06 | 1.600E-05 |
| 13231 | 662 | P662L | B | 1.5e-05 | 1.6e-05 | 1.6e-05 | 1.567E-05 | 5.774E-07 | 1.500E-05 | 1.600E-05 |
| 13232 | 662 | P662K | B | 1.5e-05 | 9.3e-06 | 1.4e-05 | 1.277E-05 | 3.044E-06 | 9.300E-06 | 1.500E-05 |
| 13233 | 662 | P662M | B | 1.3e-05 | 1.6e-05 | 1.1e-05 | 1.333E-05 | 2.517E-06 | 1.100E-05 | 1.600E-05 |
| 13234 | 662 | P662F | B | 1.3e-05 | 1.2e-05 | 1.5e-05 | 1.333E-05 | 1.528E-06 | 1.200E-05 | 1.500E-05 |
| 13235 | 662 | P662P | B | 1.4e-05 | 1.6e-05 | 1.6e-05 | 1.533E-05 | 1.155E-06 | 1.400E-05 | 1.600E-05 |
| 13236 | 662 | P662S | B | 1.5e-05 | 1e-05 | 1.2e-05 | 1.233E-05 | 2.517E-06 | 1.000E-05 | 1.500E-05 |
| 13237 | 662 | P662T | B | 1e-05 | 9.9e-06 | 1.5e-05 | 1.163E-05 | 2.916E-06 | 9.900E-06 | 1.500E-05 |
| 13238 | 662 | P662W | B | 9.7e-06 | 1.5e-05 | 1.4e-05 | 1.290E-05 | 2.816E-06 | 9.700E-06 | 1.500E-05 |
| 13239 | 662 | P662Y | B | 1.5e-05 | 1.4e-05 | 1.1e-05 | 1.333E-05 | 2.082E-06 | 1.100E-05 | 1.500E-05 |
| 13240 | 662 | P662V | B | 1.2e-05 | 9.3e-06 | 1e-05 | 1.043E-05 | 1.401E-06 | 9.300E-06 | 1.200E-05 |
| 13241 | 663 | G663A | B | 1.1e-05 | 1.5e-05 | 1.7e-05 | 1.433E-05 | 3.055E-06 | 1.100E-05 | 1.700E-05 |
| 13242 | 663 | G663R | B | 1.6e-05 | 1.2e-05 | 1.2e-05 | 1.333E-05 | 2.309E-06 | 1.200E-05 | 1.600E-05 |
| 13243 | 663 | G663N | B | 1.4e-05 | 9.9e-06 | 1.5e-05 | 1.297E-05 | 2.702E-06 | 9.900E-06 | 1.500E-05 |
| 13244 | 663 | G663D | B | 6.7e-06 | 1.1e-05 | 1.1e-05 | 9.567E-06 | 2.483E-06 | 6.700E-06 | 1.100E-05 |
| 13245 | 663 | G663C | B | 8.2e-06 | 9.4e-06 | 1.5e-05 | 1.087E-05 | 3.630E-06 | 8.200E-06 | 1.500E-05 |
| 13246 | 663 | G663E | B | 1.4e-05 | 1e-05 | 1.1e-05 | 1.167E-05 | 2.082E-06 | 1.000E-05 | 1.400E-05 |
| 13247 | 663 | G663Q | B | 1.4e-05 | 1.1e-05 | 1.4e-05 | 1.300E-05 | 1.732E-06 | 1.100E-05 | 1.400E-05 |
| 13248 | 663 | G663G | B | 1.1e-05 | 1.1e-05 | 1.1e-05 | 1.100E-05 | 2.075E-21 | 1.100E-05 | 1.100E-05 |
| 13249 | 663 | G663H | B | 8.5e-06 | 1e-05 | 1.4e-05 | 1.083E-05 | 2.843E-06 | 8.500E-06 | 1.400E-05 |
| 13250 | 663 | G663I | B | 9.5e-06 | 1.8e-05 | 1.4e-05 | 1.383E-05 | 4.252E-06 | 9.500E-06 | 1.800E-05 |
| 13251 | 663 | G663L | B | 1.4e-05 | 8.6e-06 | 1.1e-05 | 1.120E-05 | 2.706E-06 | 8.600E-06 | 1.400E-05 |
| 13252 | 663 | G663K | B | 1.6e-05 | 1.5e-05 | 1.2e-05 | 1.433E-05 | 2.082E-06 | 1.200E-05 | 1.600E-05 |
| 13253 | 663 | G663M | B | 1.7e-05 | 1.2e-05 | 1.5e-05 | 1.467E-05 | 2.517E-06 | 1.200E-05 | 1.700E-05 |
| 13254 | 663 | G663F | B | 1.5e-05 | 1.2e-05 | 1e-05 | 1.233E-05 | 2.517E-06 | 1.000E-05 | 1.500E-05 |
| 13255 | 663 | G663P | B | 1e-05 | 1.1e-05 | 8.5e-06 | 9.833E-06 | 1.258E-06 | 8.500E-06 | 1.100E-05 |
| 13256 | 663 | G663S | B | 1.4e-05 | 1.5e-05 | 1e-05 | 1.300E-05 | 2.646E-06 | 1.000E-05 | 1.500E-05 |
| 13257 | 663 | G663T | B | 1.4e-05 | 1.1e-05 | 1e-05 | 1.167E-05 | 2.082E-06 | 1.000E-05 | 1.400E-05 |
| 13258 | 663 | G663W | B | 8.5e-06 | 9.2e-06 | 1.2e-05 | 9.900E-06 | 1.852E-06 | 8.500E-06 | 1.200E-05 |
| 13259 | 663 | G663Y | B | 1.2e-05 | 1.2e-05 | 9.8e-06 | 1.127E-05 | 1.270E-06 | 9.800E-06 | 1.200E-05 |

| | | | | | | | | | | |
|---|---|---|---|---|---|---|---|---|---|---|
| 13260 | 663 | G663V | B | 1.1e-05 | 5.6e-06 | 1.6e-05 | 1.087E-05 | 5.201E-06 | 5.600E-06 | 1.600E-05 |
| 13261 | 664 | K664A | B | 1.6e-05 | 7.6e-06 | 9.4e-06 | 1.100E-05 | 4.423E-06 | 7.600E-06 | 1.600E-05 |
| 13262 | 664 | K664R | B | 1.1e-05 | 8.2e-06 | 1.2e-05 | 1.040E-05 | 1.970E-06 | 8.200E-06 | 1.200E-05 |
| 13263 | 664 | K664N | B | 1.3e-05 | 1.1e-05 | 8.5e-06 | 1.083E-05 | 2.255E-06 | 8.500E-06 | 1.300E-05 |
| 13264 | 664 | K664D | B | 9.8e-06 | 1.4e-05 | 1.6e-05 | 1.327E-05 | 3.164E-06 | 9.800E-06 | 1.600E-05 |
| 13265 | 664 | K664C | B | 1.6e-05 | 6.1e-06 | 1.5e-05 | 1.237E-05 | 5.450E-06 | 6.100E-06 | 1.600E-05 |
| 13266 | 664 | K664E | B | 1.4e-05 | 1.6e-05 | 1.5e-05 | 1.500E-05 | 1.000E-06 | 1.400E-05 | 1.600E-05 |
| 13267 | 664 | K664Q | B | 1.2e-05 | 1.5e-05 | 1e-05 | 1.233E-05 | 2.517E-06 | 1.000E-05 | 1.500E-05 |
| 13268 | 664 | K664G | B | 1.5e-05 | 8.7e-06 | 7.5e-06 | 1.040E-05 | 4.029E-06 | 7.500E-06 | 1.500E-05 |
| 13269 | 664 | K664H | B | 9.9e-06 | 1.3e-05 | 1.1e-05 | 1.130E-05 | 1.572E-06 | 9.900E-06 | 1.300E-05 |
| 13270 | 664 | K664I | B | 1.5e-05 | 1.1e-05 | 1.5e-05 | 1.367E-05 | 2.309E-06 | 1.100E-05 | 1.500E-05 |
| 13271 | 664 | K664L | B | 1.1e-05 | 1.2e-05 | 1.3e-05 | 1.200E-05 | 1.000E-06 | 1.100E-05 | 1.300E-05 |
| 13272 | 664 | K664K | B | 1.5e-05 | 1.2e-05 | 1.4e-05 | 1.367E-05 | 1.528E-06 | 1.200E-05 | 1.500E-05 |
| 13273 | 664 | K664M | B | 9.2e-06 | 1.1e-05 | 1.5e-05 | 1.173E-05 | 2.969E-06 | 9.200E-06 | 1.500E-05 |
| 13274 | 664 | K664F | B | 1.5e-05 | 1.6e-05 | 1.3e-05 | 1.467E-05 | 1.528E-06 | 1.300E-05 | 1.600E-05 |
| 13275 | 664 | K664P | B | 1e-05 | 1.1e-05 | 9.3e-06 | 1.010E-05 | 8.544E-07 | 9.300E-06 | 1.100E-05 |
| 13276 | 664 | K664S | B | 9.9e-06 | 1.3e-05 | 1.1e-05 | 1.130E-05 | 1.572E-06 | 9.900E-06 | 1.300E-05 |
| 13277 | 664 | K664T | B | 1.5e-05 | 1.5e-05 | 1.3e-05 | 1.433E-05 | 1.155E-06 | 1.300E-05 | 1.500E-05 |
| 13278 | 664 | K664W | B | 9.2e-06 | 9.8e-06 | 8.7e-06 | 9.233E-06 | 5.508E-07 | 8.700E-06 | 9.800E-06 |
| 13279 | 664 | K664Y | B | 1e-05 | 1.3e-05 | 1.4e-05 | 1.233E-05 | 2.082E-06 | 1.000E-05 | 1.400E-05 |
| 13280 | 664 | K664V | B | 1e-05 | 1.4e-05 | 1.5e-05 | 1.300E-05 | 2.646E-06 | 1.000E-05 | 1.500E-05 |
| 13281 | 665 | A665A | B | 1e-05 | 1.1e-05 | 1.4e-05 | 1.167E-05 | 2.082E-06 | 1.000E-05 | 1.400E-05 |
| 13282 | 665 | A665R | B | 1.2e-05 | 1.1e-05 | 1.5e-05 | 1.267E-05 | 2.082E-06 | 1.100E-05 | 1.500E-05 |
| 13283 | 665 | A665N | B | 1e-05 | 1.6e-05 | 1e-05 | 1.200E-05 | 3.464E-06 | 1.000E-05 | 1.600E-05 |
| 13284 | 665 | A665D | B | 1.5e-05 | 1.7e-05 | 1.5e-05 | 1.567E-05 | 1.155E-06 | 1.500E-05 | 1.700E-05 |
| 13285 | 665 | A665C | B | 9e-06 | 1.1e-05 | 1.7e-05 | 1.233E-05 | 4.163E-06 | 9.000E-06 | 1.700E-05 |
| 13286 | 665 | A665E | B | 1.5e-05 | 1.1e-05 | 1.4e-05 | 1.333E-05 | 2.082E-06 | 1.100E-05 | 1.500E-05 |
| 13287 | 665 | A665Q | B | 1.6e-05 | 9.9e-06 | 8.6e-06 | 1.150E-05 | 3.951E-06 | 8.600E-06 | 1.600E-05 |
| 13288 | 665 | A665G | B | 1.3e-05 | 1.3e-05 | 1.5e-05 | 1.367E-05 | 1.155E-06 | 1.300E-05 | 1.500E-05 |
| 13289 | 665 | A665H | B | 9.4e-06 | 1.4e-05 | 1.6e-05 | 1.313E-05 | 3.384E-06 | 9.400E-06 | 1.600E-05 |
| 13290 | 665 | A665I | B | 8.8e-06 | 1.4e-05 | 9.8e-06 | 1.087E-05 | 2.759E-06 | 8.800E-06 | 1.400E-05 |
| 13291 | 665 | A665L | B | 1e-05 | 7.4e-06 | 1.6e-05 | 1.113E-05 | 4.411E-06 | 7.400E-06 | 1.600E-05 |
| 13292 | 665 | A665K | B | 1.3e-05 | 9.5e-06 | 9.6e-06 | 1.070E-05 | 1.992E-06 | 9.500E-06 | 1.300E-05 |
| 13293 | 665 | A665M | B | 1.2e-05 | 1.6e-05 | 1e-05 | 1.267E-05 | 3.055E-06 | 1.000E-05 | 1.600E-05 |
| 13294 | 665 | A665F | B | 1.4e-05 | 1.1e-05 | 9.6e-06 | 1.153E-05 | 2.248E-06 | 9.600E-06 | 1.400E-05 |
| 13295 | 665 | A665P | B | 1.8e-05 | 9.7e-06 | 2.1e-05 | 1.623E-05 | 5.853E-06 | 9.700E-06 | 2.100E-05 |
| 13296 | 665 | A665S | B | 1.4e-05 | 1.6e-05 | 1.7e-05 | 1.567E-05 | 1.528E-06 | 1.400E-05 | 1.700E-05 |
| 13297 | 665 | A665T | B | 1.3e-05 | 1e-05 | 1e-05 | 1.100E-05 | 1.732E-06 | 1.000E-05 | 1.300E-05 |
| 13298 | 665 | A665W | B | 6.9e-06 | 1.4e-05 | 1.2e-05 | 1.097E-05 | 3.661E-06 | 6.900E-06 | 1.400E-05 |
| 13299 | 665 | A665Y | B | 1.5e-05 | 1.2e-05 | 1.5e-05 | 1.400E-05 | 1.732E-06 | 1.200E-05 | 1.500E-05 |
| 13300 | 665 | A665V | B | 1.2e-05 | 1.2e-05 | 1.4e-05 | 1.267E-05 | 1.155E-06 | 1.200E-05 | 1.400E-05 |
| 13301 | 666 | P666A | B | 1.1e-05 | 9.4e-06 | 1.3e-05 | 1.113E-05 | 1.804E-06 | 9.400E-06 | 1.300E-05 |
| 13302 | 666 | P666R | B | 1.1e-05 | 1.5e-05 | 9.5e-06 | 1.183E-05 | 2.843E-06 | 9.500E-06 | 1.500E-05 |
| 13303 | 666 | P666N | B | 1.4e-05 | 8.7e-06 | 1.7e-05 | 1.323E-05 | 4.203E-06 | 8.700E-06 | 1.700E-05 |
| 13304 | 666 | P666D | B | 1.6e-05 | 1.1e-05 | 1e-05 | 1.233E-05 | 3.215E-06 | 1.000E-05 | 1.600E-05 |
| 13305 | 666 | P666C | B | 9.2e-06 | 1.6e-05 | 1.1e-05 | 1.207E-05 | 3.523E-06 | 9.200E-06 | 1.600E-05 |
| 13306 | 666 | P666E | B | 1e-05 | 1.3e-05 | 1.1e-05 | 1.133E-05 | 1.528E-06 | 1.000E-05 | 1.300E-05 |
| 13307 | 666 | P666Q | B | 1e-05 | 1.1e-05 | 1.1e-05 | 1.067E-05 | 5.774E-07 | 1.000E-05 | 1.100E-05 |
| 13308 | 666 | P666G | B | 1.6e-05 | 1.5e-05 | 8.5e-06 | 1.317E-05 | 4.072E-06 | 8.500E-06 | 1.600E-05 |
| 13309 | 666 | P666H | B | 1e-05 | 1.6e-05 | 1.5e-05 | 1.367E-05 | 3.215E-06 | 1.000E-05 | 1.600E-05 |
| 13310 | 666 | P666I | B | 1.6e-05 | 1.2e-05 | 1e-05 | 1.267E-05 | 3.055E-06 | 1.000E-05 | 1.600E-05 |
| 13311 | 666 | P666L | B | 1.3e-05 | 9.5e-06 | 1.5e-05 | 1.250E-05 | 2.784E-06 | 9.500E-06 | 1.500E-05 |
| 13312 | 666 | P666K | B | 9.6e-06 | 1.4e-05 | 1.5e-05 | 1.287E-05 | 2.873E-06 | 9.600E-06 | 1.500E-05 |
| 13313 | 666 | P666M | B | 1.4e-05 | 9.4e-06 | 9.1e-06 | 1.083E-05 | 2.747E-06 | 9.100E-06 | 1.400E-05 |
| 13314 | 666 | P666F | B | 2e-05 | 1.2e-05 | 1.1e-05 | 1.433E-05 | 4.933E-06 | 1.100E-05 | 2.000E-05 |
| 13315 | 666 | P666P | B | 1.3e-05 | 1.1e-05 | 1.5e-05 | 1.300E-05 | 2.000E-06 | 1.100E-05 | 1.500E-05 |
| 13316 | 666 | P666S | B | 9.5e-06 | 1.5e-05 | 9.4e-06 | 1.130E-05 | 3.205E-06 | 9.400E-06 | 1.500E-05 |
| 13317 | 666 | P666T | B | 1.5e-05 | 1.5e-05 | 1.1e-05 | 1.367E-05 | 2.309E-06 | 1.100E-05 | 1.500E-05 |
| 13318 | 666 | P666W | B | 1.4e-05 | 1.5e-05 | 1.4e-05 | 1.433E-05 | 5.774E-07 | 1.400E-05 | 1.500E-05 |
| 13319 | 666 | P666Y | B | 1.5e-05 | 1.1e-05 | 1.6e-05 | 1.400E-05 | 2.646E-06 | 1.100E-05 | 1.600E-05 |
| 13320 | 666 | P666V | B | 1.3e-05 | 1.4e-05 | 1.4e-05 | 1.367E-05 | 5.774E-07 | 1.300E-05 | 1.400E-05 |
| 13321 | 667 | K667A | B | 1.6e-05 | 1.6e-05 | 1.5e-05 | 1.567E-05 | 5.774E-07 | 1.500E-05 | 1.600E-05 |
| 13322 | 667 | K667R | B | 1.1e-05 | 1.2e-05 | 1.5e-05 | 1.267E-05 | 2.082E-06 | 1.100E-05 | 1.500E-05 |
| 13323 | 667 | K667N | B | 6.7e-06 | 1.4e-05 | 1.7e-05 | 1.257E-05 | 5.297E-06 | 6.700E-06 | 1.700E-05 |
| 13324 | 667 | K667D | B | 9.9e-06 | 1.4e-05 | 1.5e-05 | 1.297E-05 | 2.702E-06 | 9.900E-06 | 1.500E-05 |
| 13325 | 667 | K667C | B | 1.5e-05 | 1.3e-05 | 1.2e-05 | 1.333E-05 | 1.528E-06 | 1.200E-05 | 1.500E-05 |
| 13326 | 667 | K667E | B | 1.2e-05 | 1.4e-05 | 1.4e-05 | 1.333E-05 | 1.155E-06 | 1.200E-05 | 1.400E-05 |
| 13327 | 667 | K667Q | B | 1.1e-05 | 1.4e-05 | 1.6e-05 | 1.367E-05 | 2.517E-06 | 1.100E-05 | 1.600E-05 |
| 13328 | 667 | K667G | B | 1.1e-05 | 1.4e-05 | 1.5e-05 | 1.333E-05 | 2.082E-06 | 1.100E-05 | 1.500E-05 |
| 13329 | 667 | K667H | B | 9.4e-06 | 1.5e-05 | 1.4e-05 | 1.280E-05 | 2.987E-06 | 9.400E-06 | 1.500E-05 |
| 13330 | 667 | K667I | B | 1.3e-05 | 1.6e-05 | 1.1e-05 | 1.333E-05 | 2.517E-06 | 1.100E-05 | 1.600E-05 |
| 13331 | 667 | K667L | B | 8.9e-06 | 1.3e-05 | 1.6e-05 | 1.263E-05 | 3.564E-06 | 8.900E-06 | 1.600E-05 |
| 13332 | 667 | K667K | B | 8.8e-06 | 8.3e-06 | 1.5e-05 | 1.070E-05 | 3.732E-06 | 8.300E-06 | 1.500E-05 |
| 13333 | 667 | K667M | B | 1.2e-05 | 1.2e-05 | 1.5e-05 | 1.300E-05 | 1.732E-06 | 1.200E-05 | 1.500E-05 |
| 13334 | 667 | K667F | B | 1.1e-05 | 1.4e-05 | 1.4e-05 | 1.300E-05 | 1.732E-06 | 1.100E-05 | 1.400E-05 |
| 13335 | 667 | K667P | B | 1.1e-05 | 1.1e-05 | 1.5e-05 | 1.233E-05 | 2.309E-06 | 1.100E-05 | 1.500E-05 |
| 13336 | 667 | K667S | B | 1.3e-05 | 1.1e-05 | 1.1e-05 | 1.167E-05 | 1.155E-06 | 1.100E-05 | 1.300E-05 |
| 13337 | 667 | K667T | B | 1.4e-05 | 1.4e-05 | 1.5e-05 | 1.433E-05 | 5.774E-07 | 1.400E-05 | 1.500E-05 |
| 13338 | 667 | K667W | B | 1.4e-05 | 1.1e-05 | 1.5e-05 | 1.333E-05 | 2.082E-06 | 1.100E-05 | 1.500E-05 |
| 13339 | 667 | K667Y | B | 1.4e-05 | 1.5e-05 | 8.3e-06 | 1.243E-05 | 3.614E-06 | 8.300E-06 | 1.500E-05 |
| 13340 | 667 | K667V | B | 1.1e-05 | 9.7e-06 | 8.9e-06 | 9.867E-06 | 1.060E-06 | 8.900E-06 | 1.100E-05 |
| 13341 | 668 | L668A | B | 1e-05 | 9.9e-06 | 1.1e-05 | 1.030E-05 | 6.083E-07 | 9.900E-06 | 1.100E-05 |
| 13342 | 668 | L668R | B | 1.1e-05 | 8.9e-06 | 1.5e-05 | 1.163E-05 | 3.099E-06 | 8.900E-06 | 1.500E-05 |
| 13343 | 668 | L668N | B | 1.1e-05 | 1.4e-05 | 1.4e-05 | 1.300E-05 | 1.732E-06 | 1.100E-05 | 1.400E-05 |
| 13344 | 668 | L668D | B | 1.4e-05 | 9.7e-06 | 1.6e-05 | 1.323E-05 | 3.219E-06 | 9.700E-06 | 1.600E-05 |

| | | | | | | | | | | |
|---|---|---|---|---|---|---|---|---|---|---|
| 13345 | 668 | L668C | B | 1.6e-05 | 1.3e-05 | 1.1e-05 | 1.333E-05 | 2.517E-06 | 1.100E-05 | 1.600E-05 |
| 13346 | 668 | L668E | B | 1.2e-05 | 1.3e-05 | 1.4e-05 | 1.300E-05 | 1.000E-06 | 1.200E-05 | 1.400E-05 |
| 13347 | 668 | L668Q | B | 1.4e-05 | 1.5e-05 | 1.3e-05 | 1.400E-05 | 1.000E-06 | 1.300E-05 | 1.500E-05 |
| 13348 | 668 | L668G | B | 1.5e-05 | 1.5e-05 | 1.6e-05 | 1.533E-05 | 5.774E-07 | 1.500E-05 | 1.600E-05 |
| 13349 | 668 | L668H | B | 1.4e-05 | 1.6e-05 | 1.5e-05 | 1.500E-05 | 1.000E-06 | 1.400E-05 | 1.600E-05 |
| 13350 | 668 | L668I | B | 1.5e-05 | 1e-05 | 1.1e-05 | 1.200E-05 | 2.646E-06 | 1.000E-05 | 1.500E-05 |
| 13351 | 668 | L668L | B | 1.6e-05 | 1.1e-05 | 1.2e-05 | 1.300E-05 | 2.646E-06 | 1.100E-05 | 1.600E-05 |
| 13352 | 668 | L668K | B | 1.2e-05 | 1.5e-05 | 1.4e-05 | 1.367E-05 | 1.528E-06 | 1.200E-05 | 1.500E-05 |
| 13353 | 668 | L668M | B | 1.3e-05 | 1.5e-05 | 1.3e-05 | 1.367E-05 | 1.155E-06 | 1.300E-05 | 1.500E-05 |
| 13354 | 668 | L668F | B | 1.6e-05 | 1e-05 | 1.5e-05 | 1.367E-05 | 3.215E-06 | 1.000E-05 | 1.600E-05 |
| 13355 | 668 | L668P | B | 1.1e-05 | 1.1e-05 | 1.3e-05 | 1.167E-05 | 1.155E-06 | 1.100E-05 | 1.300E-05 |
| 13356 | 668 | L668S | B | 1.7e-05 | 1e-05 | 1e-05 | 1.233E-05 | 4.041E-06 | 1.000E-05 | 1.700E-05 |
| 13357 | 668 | L668T | B | 1.5e-05 | 9.7e-06 | 1.1e-05 | 1.190E-05 | 2.762E-06 | 9.700E-06 | 1.500E-05 |
| 13358 | 668 | L668W | B | 8.5e-06 | 9.3e-06 | 1.5e-05 | 1.093E-05 | 3.544E-06 | 8.500E-06 | 1.500E-05 |
| 13359 | 668 | L668Y | B | 1.5e-05 | 1.4e-05 | 1.2e-05 | 1.367E-05 | 1.528E-06 | 1.200E-05 | 1.500E-05 |
| 13360 | 668 | L668V | B | 1.6e-05 | 1.5e-05 | 1.6e-05 | 1.567E-05 | 5.774E-07 | 1.500E-05 | 1.600E-05 |
| 13361 | 669 | L669A | B | 1.8e-05 | 1.1e-05 | 8.2e-06 | 1.240E-05 | 5.048E-06 | 8.200E-06 | 1.800E-05 |
| 13362 | 669 | L669R | B | 1.1e-05 | 1.4e-05 | 1.8e-05 | 1.433E-05 | 3.512E-06 | 1.100E-05 | 1.800E-05 |
| 13363 | 669 | L669N | B | 1.1e-05 | 9.1e-06 | 1.1e-05 | 1.037E-05 | 1.097E-06 | 9.100E-06 | 1.100E-05 |
| 13364 | 669 | L669D | B | 1.5e-05 | 1.6e-05 | 1.7e-05 | 1.600E-05 | 1.000E-06 | 1.500E-05 | 1.700E-05 |
| 13365 | 669 | L669C | B | 9.3e-06 | 1.4e-05 | 1.3e-05 | 1.210E-05 | 2.476E-06 | 9.300E-06 | 1.400E-05 |
| 13366 | 669 | L669E | B | 1.4e-05 | 1.3e-05 | 1.6e-05 | 1.433E-05 | 1.528E-06 | 1.300E-05 | 1.600E-05 |
| 13367 | 669 | L669Q | B | 1e-05 | 1.4e-05 | 1.1e-05 | 1.167E-05 | 2.082E-06 | 1.000E-05 | 1.400E-05 |
| 13368 | 669 | L669G | B | 1.9e-05 | 1e-05 | 1.1e-05 | 1.333E-05 | 4.933E-06 | 1.000E-05 | 1.900E-05 |
| 13369 | 669 | L669H | B | 1.1e-05 | 1e-05 | 1.6e-05 | 1.233E-05 | 3.215E-06 | 1.000E-05 | 1.600E-05 |
| 13370 | 669 | L669I | B | 1.1e-05 | 1.1e-05 | 1.6e-05 | 1.267E-05 | 2.887E-06 | 1.100E-05 | 1.600E-05 |
| 13371 | 669 | L669L | B | 1.6e-05 | 1e-05 | 1.6e-05 | 1.400E-05 | 3.464E-06 | 1.000E-05 | 1.600E-05 |
| 13372 | 669 | L669K | B | 1.5e-05 | 1.1e-05 | 1e-05 | 1.200E-05 | 2.646E-06 | 1.000E-05 | 1.500E-05 |
| 13373 | 669 | L669M | B | 1.5e-05 | 1.7e-05 | 1.1e-05 | 1.433E-05 | 3.055E-06 | 1.100E-05 | 1.700E-05 |
| 13374 | 669 | L669F | B | 9.4e-06 | 1.4e-05 | 1.7e-05 | 1.347E-05 | 3.828E-06 | 9.400E-06 | 1.700E-05 |
| 13375 | 669 | L669P | B | 1.6e-05 | 1.6e-05 | 1.6e-05 | 1.600E-05 | 0.000E+00 | 1.600E-05 | 1.600E-05 |
| 13376 | 669 | L669S | B | 1.5e-05 | 1.5e-05 | 1.1e-05 | 1.367E-05 | 2.309E-06 | 1.100E-05 | 1.500E-05 |
| 13377 | 669 | L669T | B | 1.6e-05 | 1.1e-05 | 1.6e-05 | 1.433E-05 | 2.887E-06 | 1.100E-05 | 1.600E-05 |
| 13378 | 669 | L669W | B | 1.6e-05 | 1.3e-05 | 1.5e-05 | 1.467E-05 | 1.528E-06 | 1.300E-05 | 1.600E-05 |
| 13379 | 669 | L669Y | B | 1.3e-05 | 2.2e-05 | 1.1e-05 | 1.533E-05 | 5.859E-06 | 1.100E-05 | 2.200E-05 |
| 13380 | 669 | L669V | B | 1.6e-05 | 1.4e-05 | 1.6e-05 | 1.533E-05 | 1.155E-06 | 1.400E-05 | 1.600E-05 |
| 13381 | 670 | I670A | B | 1.2e-05 | 1.5e-05 | 1e-05 | 1.233E-05 | 2.517E-06 | 1.000E-05 | 1.500E-05 |
| 13382 | 670 | I670R | B | 6.3e-06 | 1.4e-05 | 9.8e-06 | 1.003E-05 | 3.855E-06 | 6.300E-06 | 1.400E-05 |
| 13383 | 670 | I670N | B | 1.4e-05 | 1.5e-05 | 1e-05 | 1.300E-05 | 2.646E-06 | 1.000E-05 | 1.500E-05 |
| 13384 | 670 | I670D | B | 1.1e-05 | 8.7e-06 | 1.3e-05 | 1.090E-05 | 2.152E-06 | 8.700E-06 | 1.300E-05 |
| 13385 | 670 | I670C | B | 9.2e-06 | 8.2e-06 | 1.1e-05 | 9.467E-06 | 1.419E-06 | 8.200E-06 | 1.100E-05 |
| 13386 | 670 | I670E | B | 1.7e-05 | 1.4e-05 | 1e-05 | 1.367E-05 | 3.512E-06 | 1.000E-05 | 1.700E-05 |
| 13387 | 670 | I670Q | B | 1.4e-05 | 1.5e-05 | 1.5e-05 | 1.467E-05 | 5.774E-07 | 1.400E-05 | 1.500E-05 |
| 13388 | 670 | I670G | B | 1.1e-05 | 1.1e-05 | 1.3e-05 | 1.167E-05 | 1.155E-06 | 1.100E-05 | 1.300E-05 |
| 13389 | 670 | I670H | B | 1.1e-05 | 1.7e-05 | 1.6e-05 | 1.467E-05 | 3.215E-06 | 1.100E-05 | 1.700E-05 |
| 13390 | 670 | I670I | B | 1.5e-05 | 1.1e-05 | 1.5e-05 | 1.367E-05 | 2.309E-06 | 1.100E-05 | 1.500E-05 |
| 13391 | 670 | I670L | B | 1.4e-05 | 1.5e-05 | 9.7e-06 | 1.290E-05 | 2.816E-06 | 9.700E-06 | 1.500E-05 |
| 13392 | 670 | I670K | B | 7.1e-06 | 1.5e-05 | 6.9e-06 | 9.667E-06 | 4.620E-06 | 6.900E-06 | 1.500E-05 |
| 13393 | 670 | I670M | B | 1e-05 | 1.1e-05 | 8.7e-06 | 9.900E-06 | 1.153E-06 | 8.700E-06 | 1.100E-05 |
| 13394 | 670 | I670F | B | 1.5e-05 | 1.6e-05 | 1.5e-05 | 1.533E-05 | 5.774E-07 | 1.500E-05 | 1.600E-05 |
| 13395 | 670 | I670P | B | 1e-05 | 1.4e-05 | 1.4e-05 | 1.267E-05 | 2.309E-06 | 1.000E-05 | 1.400E-05 |
| 13396 | 670 | I670S | B | 1.5e-05 | 1.1e-05 | 1.5e-05 | 1.367E-05 | 2.309E-06 | 1.100E-05 | 1.500E-05 |
| 13397 | 670 | I670T | B | 1.5e-05 | 1.4e-05 | 1e-05 | 1.300E-05 | 2.646E-06 | 1.000E-05 | 1.500E-05 |
| 13398 | 670 | I670W | B | 1.5e-05 | 1.4e-05 | 9.8e-06 | 1.293E-05 | 2.759E-06 | 9.800E-06 | 1.500E-05 |
| 13399 | 670 | I670Y | B | 1.1e-05 | 9.7e-06 | 1.6e-05 | 1.223E-05 | 3.326E-06 | 9.700E-06 | 1.600E-05 |
| 13400 | 670 | I670V | B | 1.3e-05 | 1.2e-05 | 8.7e-06 | 1.123E-05 | 2.250E-06 | 8.700E-06 | 1.300E-05 |
| 13401 | 671 | Y671A | B | 2.2e-05 | 2.1e-05 | 2.6e-05 | 2.300E-05 | 2.646E-06 | 2.100E-05 | 2.600E-05 |
| 13402 | 671 | Y671R | B | 9.3e-06 | 1.6e-05 | 1.6e-05 | 1.377E-05 | 3.868E-06 | 9.300E-06 | 1.600E-05 |
| 13403 | 671 | Y671N | B | 2.6e-05 | 2e-05 | 2.9e-05 | 2.500E-05 | 4.583E-06 | 2.000E-05 | 2.900E-05 |
| 13404 | 671 | Y671D | B | 1.9e-05 | 1.8e-05 | 2e-05 | 1.900E-05 | 1.000E-06 | 1.800E-05 | 2.000E-05 |
| 13405 | 671 | Y671C | B | 2.3e-05 | 2.2e-05 | 2.4e-05 | 2.300E-05 | 1.000E-06 | 2.200E-05 | 2.400E-05 |
| 13406 | 671 | Y671E | B | 1.6e-05 | 2.5e-05 | 3e-05 | 2.367E-05 | 7.095E-06 | 1.600E-05 | 3.000E-05 |
| 13407 | 671 | Y671Q | B | 1.7e-05 | 2.8e-05 | 2.9e-05 | 2.467E-05 | 6.658E-06 | 1.700E-05 | 2.900E-05 |
| 13408 | 671 | Y671G | B | 2.1e-05 | 2.5e-05 | 2.3e-05 | 2.300E-05 | 2.000E-06 | 2.100E-05 | 2.500E-05 |
| 13409 | 671 | Y671H | B | 1.3e-05 | 1.4e-05 | 1e-05 | 1.233E-05 | 2.082E-06 | 1.000E-05 | 1.400E-05 |
| 13410 | 671 | Y671I | B | 2.7e-05 | 1.3e-05 | 2.9e-05 | 2.300E-05 | 8.718E-06 | 1.300E-05 | 2.900E-05 |
| 13411 | 671 | Y671L | B | 3e-05 | 1.5e-05 | 2.7e-05 | 2.400E-05 | 7.937E-06 | 1.500E-05 | 3.000E-05 |
| 13412 | 671 | Y671K | B | 2.3e-05 | 2.7e-05 | 1e-05 | 2.000E-05 | 8.888E-06 | 1.000E-05 | 2.700E-05 |
| 13413 | 671 | Y671M | B | 2.2e-05 | 2.4e-05 | 2.5e-05 | 2.367E-05 | 1.528E-06 | 2.200E-05 | 2.500E-05 |
| 13414 | 671 | Y671F | B | 1.3e-05 | 1.5e-05 | 1.4e-05 | 1.400E-05 | 1.000E-06 | 1.300E-05 | 1.500E-05 |
| 13415 | 671 | Y671P | B | 2.2e-05 | 2.3e-05 | 1.8e-05 | 2.100E-05 | 2.646E-06 | 1.800E-05 | 2.300E-05 |
| 13416 | 671 | Y671S | B | 1.8e-05 | 3.4e-05 | 1.8e-05 | 2.333E-05 | 9.238E-06 | 1.800E-05 | 3.400E-05 |
| 13417 | 671 | Y671T | B | 2.2e-05 | 2.4e-05 | 2.7e-05 | 2.433E-05 | 2.517E-06 | 2.200E-05 | 2.700E-05 |
| 13418 | 671 | Y671W | B | 1.4e-05 | 1.4e-05 | 1.1e-05 | 1.300E-05 | 1.732E-06 | 1.100E-05 | 1.400E-05 |
| 13419 | 671 | Y671Y | B | 1.6e-05 | 1.8e-05 | 1.2e-05 | 1.533E-05 | 3.055E-06 | 1.200E-05 | 1.800E-05 |
| 13420 | 671 | Y671V | B | 2.2e-05 | 1.8e-05 | 2.1e-05 | 2.033E-05 | 2.082E-06 | 1.800E-05 | 2.200E-05 |
| 13421 | 672 | S672A | B | 1.2e-05 | 4.4e-06 | 7.9e-06 | 8.100E-06 | 3.804E-06 | 4.400E-06 | 1.200E-05 |
| 13422 | 672 | S672R | B | 5.4e-06 | 7.9e-06 | 8.5e-06 | 7.267E-06 | 1.644E-06 | 5.400E-06 | 8.500E-06 |
| 13423 | 672 | S672N | B | 9.6e-06 | 1.6e-05 | 1.5e-05 | 1.353E-05 | 3.443E-06 | 9.600E-06 | 1.600E-05 |
| 13424 | 672 | S672D | B | 7.6e-06 | 1.1e-05 | 7e-06 | 8.533E-06 | 2.157E-06 | 7.000E-06 | 1.100E-05 |
| 13425 | 672 | S672C | B | 5.8e-06 | 7.1e-06 | 5.8e-06 | 6.233E-06 | 7.506E-07 | 5.800E-06 | 7.100E-06 |
| 13426 | 672 | S672E | B | 9.7e-06 | 7.8e-06 | 8.8e-06 | 8.767E-06 | 9.504E-07 | 7.800E-06 | 9.700E-06 |
| 13427 | 672 | S672Q | B | 1.2e-05 | 9.2e-06 | 4.7e-06 | 8.633E-06 | 3.683E-06 | 4.700E-06 | 1.200E-05 |
| 13428 | 672 | S672G | B | 1.1e-05 | 9.5e-06 | 8.3e-06 | 9.600E-06 | 1.353E-06 | 8.300E-06 | 1.100E-05 |
| 13429 | 672 | S672H | B | 9.5e-06 | 7.5e-06 | 8.9e-06 | 8.633E-06 | 1.026E-06 | 7.500E-06 | 9.500E-06 |

| | | | | | | | | | | |
|---|---|---|---|---|---|---|---|---|---|---|
| 13430 | 672 | S672I | B | 8.6e-06 | 4.5e-06 | 7.9e-06 | 7.000E-06 | 2.193E-06 | 4.500E-06 | 8.600E-06 |
| 13431 | 672 | S672L | B | 6.3e-06 | 7.5e-06 | 5e-06 | 6.267E-06 | 1.250E-06 | 5.000E-06 | 7.500E-06 |
| 13432 | 672 | S672K | B | 1e-05 | 7.4e-06 | 6.7e-06 | 8.033E-06 | 1.739E-06 | 6.700E-06 | 1.000E-05 |
| 13433 | 672 | S672M | B | 6.4e-06 | 3.5e-06 | 6.7e-06 | 5.533E-06 | 1.767E-06 | 3.500E-06 | 6.700E-06 |
| 13434 | 672 | S672F | B | 5.5e-06 | 5.8e-06 | 7.7e-06 | 6.333E-06 | 1.193E-06 | 5.500E-06 | 7.700E-06 |
| 13435 | 672 | S672P | B | 5.5e-06 | 5.2e-06 | 7.3e-06 | 6.000E-06 | 1.136E-06 | 5.200E-06 | 7.300E-06 |
| 13436 | 672 | S672S | B | 1e-05 | 1.1e-05 | 1.6e-05 | 1.233E-05 | 3.215E-06 | 1.000E-05 | 1.600E-05 |
| 13437 | 672 | S672T | B | 1.6e-05 | 9.2e-06 | 1.4e-05 | 1.307E-05 | 3.495E-06 | 9.200E-06 | 1.600E-05 |
| 13438 | 672 | S672W | B | 5.2e-06 | 7.8e-06 | 5e-06 | 6.000E-06 | 1.562E-06 | 5.000E-06 | 7.800E-06 |
| 13439 | 672 | S672Y | B | 8.4e-06 | 7.4e-06 | 5.5e-06 | 7.100E-06 | 1.473E-06 | 5.500E-06 | 8.400E-06 |
| 13440 | 672 | S672V | B | 4.3e-06 | 5.7e-06 | 6.2e-06 | 5.400E-06 | 9.849E-07 | 4.300E-06 | 6.200E-06 |
| 13441 | 673 | A673A | B | 1.1e-05 | 1.2e-05 | 1.7e-05 | 1.333E-05 | 3.215E-06 | 1.100E-05 | 1.700E-05 |
| 13442 | 673 | A673R | B | 1e-05 | 1.4e-05 | 1.7e-05 | 1.367E-05 | 3.512E-06 | 1.000E-05 | 1.700E-05 |
| 13443 | 673 | A673N | B | 1.3e-05 | 9.1e-06 | 2e-05 | 1.403E-05 | 5.523E-06 | 9.100E-06 | 2.000E-05 |
| 13444 | 673 | A673D | B | 1.5e-05 | 1.2e-05 | 1.7e-05 | 1.467E-05 | 2.517E-06 | 1.200E-05 | 1.700E-05 |
| 13445 | 673 | A673C | B | 1.4e-05 | 1.1e-05 | 1.5e-05 | 1.333E-05 | 2.082E-06 | 1.100E-05 | 1.500E-05 |
| 13446 | 673 | A673E | B | 8.9e-06 | 1e-05 | 9e-06 | 9.300E-06 | 6.083E-07 | 8.900E-06 | 1.000E-05 |
| 13447 | 673 | A673Q | B | 1.3e-05 | 1e-05 | 1.5e-05 | 1.267E-05 | 2.517E-06 | 1.000E-05 | 1.500E-05 |
| 13448 | 673 | A673G | B | 1.4e-05 | 1.1e-05 | 1.5e-05 | 1.333E-05 | 2.082E-06 | 1.100E-05 | 1.500E-05 |
| 13449 | 673 | A673H | B | 1.5e-05 | 1.6e-05 | 1.5e-05 | 1.533E-05 | 5.774E-07 | 1.500E-05 | 1.600E-05 |
| 13450 | 673 | A673I | B | 1.5e-05 | 1.5e-05 | 1e-05 | 1.333E-05 | 2.887E-06 | 1.000E-05 | 1.500E-05 |
| 13451 | 673 | A673L | B | 1.6e-05 | 1.3e-05 | 1e-05 | 1.300E-05 | 3.000E-06 | 1.000E-05 | 1.600E-05 |
| 13452 | 673 | A673K | B | 1.2e-05 | 1e-05 | 1.2e-05 | 1.133E-05 | 1.155E-06 | 1.000E-05 | 1.200E-05 |
| 13453 | 673 | A673M | B | 1.5e-05 | 9.3e-06 | 2.1e-05 | 1.510E-05 | 5.851E-06 | 9.300E-06 | 2.100E-05 |
| 13454 | 673 | A673F | B | 7.2e-06 | 9.9e-06 | 1.1e-05 | 9.367E-06 | 1.955E-06 | 7.200E-06 | 1.100E-05 |
| 13455 | 673 | A673P | B | 1.2e-05 | 1.4e-05 | 1e-05 | 1.200E-05 | 2.000E-06 | 1.000E-05 | 1.400E-05 |
| 13456 | 673 | A673S | B | 1.3e-05 | 1.6e-05 | 1.1e-05 | 1.333E-05 | 2.517E-06 | 1.100E-05 | 1.600E-05 |
| 13457 | 673 | A673T | B | 8.8e-06 | 7.6e-06 | 1.4e-05 | 1.013E-05 | 3.402E-06 | 7.600E-06 | 1.400E-05 |
| 13458 | 673 | A673W | B | 1.1e-05 | 1.5e-05 | 9.8e-06 | 1.193E-05 | 2.723E-06 | 9.800E-06 | 1.500E-05 |
| 13459 | 673 | A673Y | B | 1.3e-05 | 1.3e-05 | 1.7e-05 | 1.433E-05 | 2.309E-06 | 1.300E-05 | 1.700E-05 |
| 13460 | 673 | A673V | B | 9.9e-06 | 1.1e-05 | 9.1e-06 | 1.000E-05 | 9.539E-07 | 9.100E-06 | 1.100E-05 |
| 13461 | 674 | S674A | B | 1.5e-05 | 1.6e-05 | 1.4e-05 | 1.500E-05 | 1.000E-06 | 1.400E-05 | 1.600E-05 |
| 13462 | 674 | S674R | B | 9.7e-06 | 1e-05 | 1e-05 | 9.900E-06 | 1.732E-07 | 9.700E-06 | 1.000E-05 |
| 13463 | 674 | S674N | B | 1.6e-05 | 1.2e-05 | 8.7e-06 | 1.223E-05 | 3.656E-06 | 8.700E-06 | 1.600E-05 |
| 13464 | 674 | S674D | B | 9.7e-06 | 1.5e-05 | 6.9e-06 | 1.053E-05 | 4.114E-06 | 6.900E-06 | 1.500E-05 |
| 13465 | 674 | S674C | B | 1.5e-05 | 1.4e-05 | 1.5e-05 | 1.467E-05 | 5.774E-07 | 1.400E-05 | 1.500E-05 |
| 13466 | 674 | S674E | B | 1.4e-05 | 1.6e-05 | 1.7e-05 | 1.567E-05 | 1.528E-06 | 1.400E-05 | 1.700E-05 |
| 13467 | 674 | S674Q | B | 1.4e-05 | 1.2e-05 | 1.1e-05 | 1.233E-05 | 1.528E-06 | 1.100E-05 | 1.400E-05 |
| 13468 | 674 | S674G | B | 1e-05 | 8.7e-06 | 1.6e-05 | 1.157E-05 | 3.894E-06 | 8.700E-06 | 1.600E-05 |
| 13469 | 674 | S674H | B | 1.5e-05 | 1.5e-05 | 1.5e-05 | 1.500E-05 | 0.000E+00 | 1.500E-05 | 1.500E-05 |
| 13470 | 674 | S674I | B | 9.3e-06 | 1.1e-05 | 7.2e-06 | 9.167E-06 | 1.904E-06 | 7.200E-06 | 1.100E-05 |
| 13471 | 674 | S674L | B | 1.7e-05 | 1e-05 | 1.2e-05 | 1.300E-05 | 3.606E-06 | 1.000E-05 | 1.700E-05 |
| 13472 | 674 | S674K | B | 9.1e-06 | 1.6e-05 | 1.1e-05 | 1.203E-05 | 3.564E-06 | 9.100E-06 | 1.600E-05 |
| 13473 | 674 | S674M | B | 8.6e-06 | 1.1e-05 | 1.3e-05 | 1.087E-05 | 2.203E-06 | 8.600E-06 | 1.300E-05 |
| 13474 | 674 | S674F | B | 1.1e-05 | 1.5e-05 | 1e-05 | 1.200E-05 | 2.646E-06 | 1.000E-05 | 1.500E-05 |
| 13475 | 674 | S674P | B | 9.9e-06 | 1.2e-05 | 1.7e-05 | 1.297E-05 | 3.647E-06 | 9.900E-06 | 1.700E-05 |
| 13476 | 674 | S674S | B | 1.5e-05 | 9.8e-06 | 9.3e-06 | 1.137E-05 | 3.156E-06 | 9.300E-06 | 1.500E-05 |
| 13477 | 674 | S674T | B | 9.8e-06 | 1.1e-05 | 1.4e-05 | 1.160E-05 | 2.163E-06 | 9.800E-06 | 1.400E-05 |
| 13478 | 674 | S674W | B | 1.7e-05 | 1.3e-05 | 8.8e-06 | 1.293E-05 | 4.100E-06 | 8.800E-06 | 1.700E-05 |
| 13479 | 674 | S674Y | B | 1.2e-05 | 1.7e-05 | 1e-05 | 1.300E-05 | 3.606E-06 | 1.000E-05 | 1.700E-05 |
| 13480 | 674 | S674V | B | 1.1e-05 | 1.8e-05 | 1.1e-05 | 1.333E-05 | 4.041E-06 | 1.100E-05 | 1.800E-05 |
| 13481 | 675 | Y675A | B | 1.5e-05 | 1.4e-05 | 2.2e-05 | 1.700E-05 | 4.359E-06 | 1.400E-05 | 2.200E-05 |
| 13482 | 675 | Y675R | B | 2e-05 | 2e-05 | 1.8e-05 | 1.933E-05 | 1.155E-06 | 1.800E-05 | 2.000E-05 |
| 13483 | 675 | Y675N | B | 1.9e-05 | 2.4e-05 | 1.8e-05 | 2.033E-05 | 3.215E-06 | 1.800E-05 | 2.400E-05 |
| 13484 | 675 | Y675D | B | 1.9e-05 | 1.2e-05 | 1.6e-05 | 1.567E-05 | 3.512E-06 | 1.200E-05 | 1.900E-05 |
| 13485 | 675 | Y675C | B | 2e-05 | 9.1e-06 | 1.2e-05 | 1.370E-05 | 5.645E-06 | 9.100E-06 | 2.000E-05 |
| 13486 | 675 | Y675E | B | 1.7e-05 | 1.3e-05 | 1.8e-05 | 1.600E-05 | 2.646E-06 | 1.300E-05 | 1.800E-05 |
| 13487 | 675 | Y675Q | B | 2.8e-05 | 1.6e-05 | 1.7e-05 | 2.033E-05 | 6.658E-06 | 1.600E-05 | 2.800E-05 |
| 13488 | 675 | Y675G | B | 1.6e-05 | 1.2e-05 | 2.4e-05 | 1.733E-05 | 6.110E-06 | 1.200E-05 | 2.400E-05 |
| 13489 | 675 | Y675H | B | 1.6e-05 | 1.3e-05 | 1.3e-05 | 1.400E-05 | 1.732E-06 | 1.300E-05 | 1.600E-05 |
| 13490 | 675 | Y675I | B | 1.3e-05 | 1.1e-05 | 1.8e-05 | 1.400E-05 | 3.606E-06 | 1.100E-05 | 1.800E-05 |
| 13491 | 675 | Y675L | B | 1.2e-05 | 1.3e-05 | 1.4e-05 | 1.300E-05 | 1.000E-06 | 1.200E-05 | 1.400E-05 |
| 13492 | 675 | Y675K | B | 1.3e-05 | 2e-05 | 1.6e-05 | 1.633E-05 | 3.512E-06 | 1.300E-05 | 2.000E-05 |
| 13493 | 675 | Y675M | B | 1.5e-05 | 9.3e-06 | 1.4e-05 | 1.277E-05 | 3.044E-06 | 9.300E-06 | 1.500E-05 |
| 13494 | 675 | Y675F | B | 1.6e-05 | 8.5e-06 | 1.6e-05 | 1.350E-05 | 4.330E-06 | 8.500E-06 | 1.600E-05 |
| 13495 | 675 | Y675P | B | 1.7e-05 | 2.7e-05 | 2e-05 | 2.133E-05 | 5.132E-06 | 1.700E-05 | 2.700E-05 |
| 13496 | 675 | Y675S | B | 1.8e-05 | 1.2e-05 | 2.5e-05 | 1.833E-05 | 6.506E-06 | 1.200E-05 | 2.500E-05 |
| 13497 | 675 | Y675T | B | 1.2e-05 | 1.1e-05 | 1.5e-05 | 1.267E-05 | 2.082E-06 | 1.100E-05 | 1.500E-05 |
| 13498 | 675 | Y675W | B | 1.4e-05 | 1.6e-05 | 2.6e-05 | 1.867E-05 | 6.429E-06 | 1.400E-05 | 2.600E-05 |
| 13499 | 675 | Y675Y | B | 1.5e-05 | 1.5e-05 | 1.5e-05 | 1.500E-05 | 0.000E+00 | 1.500E-05 | 1.500E-05 |
| 13500 | 675 | Y675V | B | 2.1e-05 | 2.1e-05 | 2.1e-05 | 2.100E-05 | 0.000E+00 | 2.100E-05 | 2.100E-05 |
| 13501 | 676 | R676A | B | 9.1e-06 | 1.5e-05 | 1.4e-05 | 1.270E-05 | 3.158E-06 | 9.100E-06 | 1.500E-05 |
| 13502 | 676 | R676R | B | 6.5e-06 | 1e-05 | 1.2e-05 | 9.500E-06 | 2.784E-06 | 6.500E-06 | 1.200E-05 |
| 13503 | 676 | R676N | B | 1.5e-05 | 1.6e-05 | 8.7e-06 | 1.323E-05 | 3.958E-06 | 8.700E-06 | 1.600E-05 |
| 13504 | 676 | R676D | B | 1.6e-05 | 1.6e-05 | 1.3e-05 | 1.500E-05 | 1.732E-06 | 1.300E-05 | 1.600E-05 |
| 13505 | 676 | R676C | B | 1.5e-05 | 1.4e-05 | 1.2e-05 | 1.367E-05 | 1.528E-06 | 1.200E-05 | 1.500E-05 |
| 13506 | 676 | R676E | B | 1.5e-05 | 1.2e-05 | 9.2e-06 | 1.207E-05 | 2.901E-06 | 9.200E-06 | 1.500E-05 |
| 13507 | 676 | R676Q | B | 1e-05 | 1.5e-05 | 1.6e-05 | 1.367E-05 | 3.215E-06 | 1.000E-05 | 1.600E-05 |
| 13508 | 676 | R676G | B | 1.4e-05 | 1.2e-05 | 1.4e-05 | 1.333E-05 | 1.155E-06 | 1.200E-05 | 1.400E-05 |
| 13509 | 676 | R676H | B | 1.1e-05 | 1.6e-05 | 1e-05 | 1.233E-05 | 3.215E-06 | 1.000E-05 | 1.600E-05 |
| 13510 | 676 | R676I | B | 1.5e-05 | 1e-05 | 1.6e-05 | 1.367E-05 | 3.215E-06 | 1.000E-05 | 1.600E-05 |
| 13511 | 676 | R676L | B | 1.5e-05 | 8.8e-06 | 1.3e-05 | 1.227E-05 | 3.164E-06 | 8.800E-06 | 1.500E-05 |
| 13512 | 676 | R676K | B | 1.4e-05 | 1.7e-05 | 1.3e-05 | 1.467E-05 | 2.082E-06 | 1.300E-05 | 1.700E-05 |
| 13513 | 676 | R676M | B | 1.6e-05 | 1.4e-05 | 1.2e-05 | 1.400E-05 | 2.000E-06 | 1.200E-05 | 1.600E-05 |
| 13514 | 676 | R676F | B | 1.2e-05 | 9.9e-06 | 1.4e-05 | 1.197E-05 | 2.050E-06 | 9.900E-06 | 1.400E-05 |

| | | | | | | | | | | |
|---|---|---|---|---|---|---|---|---|---|---|
| 13515 | 676 | R676P | B | 1e-05 | 7.9e-06 | 1.2e-05 | 9.967E-06 | 2.050E-06 | 7.900E-06 | 1.200E-05 |
| 13516 | 676 | R676S | B | 1.5e-05 | 1e-05 | 1.5e-05 | 1.333E-05 | 2.887E-06 | 1.000E-05 | 1.500E-05 |
| 13517 | 676 | R676T | B | 1.5e-05 | 1.1e-05 | 1.5e-05 | 1.367E-05 | 2.309E-06 | 1.100E-05 | 1.500E-05 |
| 13518 | 676 | R676W | B | 9.7e-06 | 1.5e-05 | 1.3e-05 | 1.257E-05 | 2.676E-06 | 9.700E-06 | 1.500E-05 |
| 13519 | 676 | R676Y | B | 1.1e-05 | 1.4e-05 | 1.6e-05 | 1.367E-05 | 2.517E-06 | 1.100E-05 | 1.600E-05 |
| 13520 | 676 | R676V | B | 1.2e-05 | 1.6e-05 | 9.9e-06 | 1.263E-05 | 3.099E-06 | 9.900E-06 | 1.600E-05 |
| 13521 | 677 | Y677A | B | 1.3e-05 | 1.6e-05 | 1.4e-05 | 1.433E-05 | 1.528E-06 | 1.300E-05 | 1.600E-05 |
| 13522 | 677 | Y677R | B | 1.6e-05 | 1.6e-05 | 1.8e-05 | 1.667E-05 | 1.155E-06 | 1.600E-05 | 1.800E-05 |
| 13523 | 677 | Y677N | B | 1.1e-05 | 2e-05 | 1.8e-05 | 1.633E-05 | 4.726E-06 | 1.100E-05 | 2.000E-05 |
| 13524 | 677 | Y677D | B | 1.2e-05 | 1.5e-05 | 1.2e-05 | 1.300E-05 | 1.732E-06 | 1.200E-05 | 1.500E-05 |
| 13525 | 677 | Y677C | B | 2e-05 | 1.3e-05 | 1.1e-05 | 1.467E-05 | 4.726E-06 | 1.100E-05 | 2.000E-05 |
| 13526 | 677 | Y677E | B | 1.8e-05 | 1.4e-05 | 9e-06 | 1.367E-05 | 4.509E-06 | 9.000E-06 | 1.800E-05 |
| 13527 | 677 | Y677Q | B | 1.2e-05 | 7.5e-06 | 1.6e-05 | 1.183E-05 | 4.252E-06 | 7.500E-06 | 1.600E-05 |
| 13528 | 677 | Y677G | B | 1.9e-05 | 1.3e-05 | 1.9e-05 | 1.700E-05 | 3.464E-06 | 1.300E-05 | 1.900E-05 |
| 13529 | 677 | Y677H | B | 1.4e-05 | 9.3e-06 | 1.3e-05 | 1.210E-05 | 2.476E-06 | 9.300E-06 | 1.400E-05 |
| 13530 | 677 | Y677I | B | 1.6e-05 | 1.1e-05 | 1.6e-05 | 1.433E-05 | 2.887E-06 | 1.100E-05 | 1.600E-05 |
| 13531 | 677 | Y677L | B | 1.7e-05 | 1.6e-05 | 1.1e-05 | 1.467E-05 | 3.215E-06 | 1.100E-05 | 1.700E-05 |
| 13532 | 677 | Y677K | B | 9.3e-06 | 1.1e-05 | 1.3e-05 | 1.110E-05 | 1.852E-06 | 9.300E-06 | 1.300E-05 |
| 13533 | 677 | Y677M | B | 1.1e-05 | 2e-05 | 2e-05 | 1.700E-05 | 5.196E-06 | 1.100E-05 | 2.000E-05 |
| 13534 | 677 | Y677F | B | 1.3e-05 | 1.4e-05 | 1.1e-05 | 1.267E-05 | 1.528E-06 | 1.100E-05 | 1.400E-05 |
| 13535 | 677 | Y677P | B | 1.8e-05 | 1.2e-05 | 1.8e-05 | 1.600E-05 | 3.464E-06 | 1.200E-05 | 1.800E-05 |
| 13536 | 677 | Y677S | B | 1.5e-05 | 1.7e-05 | 1.7e-05 | 1.633E-05 | 1.155E-06 | 1.500E-05 | 1.700E-05 |
| 13537 | 677 | Y677T | B | 1.4e-05 | 1.8e-05 | 1.3e-05 | 1.500E-05 | 2.646E-06 | 1.300E-05 | 1.800E-05 |
| 13538 | 677 | Y677W | B | 1.5e-05 | 1.3e-05 | 1.7e-05 | 1.500E-05 | 2.000E-06 | 1.300E-05 | 1.700E-05 |
| 13539 | 677 | Y677Y | B | 1.1e-05 | 1e-05 | 1e-05 | 1.033E-05 | 5.774E-07 | 1.000E-05 | 1.100E-05 |
| 13540 | 677 | Y677V | B | 1.7e-05 | 1.7e-05 | 1.4e-05 | 1.600E-05 | 1.732E-06 | 1.400E-05 | 1.700E-05 |
| 13541 | 678 | T678A | B | 7.1e-06 | 1.5e-05 | 1.3e-05 | 1.170E-05 | 4.107E-06 | 7.100E-06 | 1.500E-05 |
| 13542 | 678 | T678R | B | 9.2e-06 | 8.7e-06 | 1.1e-05 | 9.633E-06 | 1.210E-06 | 8.700E-06 | 1.100E-05 |
| 13543 | 678 | T678N | B | 1.6e-05 | 1.4e-05 | 1.5e-05 | 1.500E-05 | 1.000E-06 | 1.400E-05 | 1.600E-05 |
| 13544 | 678 | T678D | B | 9e-06 | 1.3e-05 | 1.3e-05 | 1.167E-05 | 2.309E-06 | 9.000E-06 | 1.300E-05 |
| 13545 | 678 | T678C | B | 1.3e-05 | 1.2e-05 | 1.5e-05 | 1.333E-05 | 1.528E-06 | 1.200E-05 | 1.500E-05 |
| 13546 | 678 | T678E | B | 4.3e-06 | 1.5e-05 | 1.6e-05 | 1.177E-05 | 6.486E-06 | 4.300E-06 | 1.600E-05 |
| 13547 | 678 | T678Q | B | 7.6e-06 | 1.2e-05 | 5.5e-06 | 8.367E-06 | 3.317E-06 | 5.500E-06 | 1.200E-05 |
| 13548 | 678 | T678G | B | 7.7e-06 | 1.6e-05 | 1.2e-05 | 1.190E-05 | 4.151E-06 | 7.700E-06 | 1.600E-05 |
| 13549 | 678 | T678H | B | 1.5e-05 | 9e-06 | 1.6e-05 | 1.333E-05 | 3.786E-06 | 9.000E-06 | 1.600E-05 |
| 13550 | 678 | T678I | B | 1.1e-05 | 1.3e-05 | 1.3e-05 | 1.233E-05 | 1.155E-06 | 1.100E-05 | 1.300E-05 |
| 13551 | 678 | T678L | B | 1.4e-05 | 1.3e-05 | 1.7e-05 | 1.467E-05 | 2.082E-06 | 1.300E-05 | 1.700E-05 |
| 13552 | 678 | T678K | B | 1.1e-05 | 6.7e-06 | 1.3e-05 | 1.023E-05 | 3.219E-06 | 6.700E-06 | 1.300E-05 |
| 13553 | 678 | T678M | B | 8.2e-06 | 1e-05 | 1.6e-05 | 1.140E-05 | 4.084E-06 | 8.200E-06 | 1.600E-05 |
| 13554 | 678 | T678F | B | 1e-05 | 1.1e-05 | 1.2e-05 | 1.100E-05 | 1.000E-06 | 1.000E-05 | 1.200E-05 |
| 13555 | 678 | T678P | B | 7.8e-06 | 1.3e-05 | 1.3e-05 | 1.127E-05 | 3.002E-06 | 7.800E-06 | 1.300E-05 |
| 13556 | 678 | T678S | B | 9.6e-06 | 9.7e-06 | 1.1e-05 | 1.010E-05 | 7.810E-07 | 9.600E-06 | 1.100E-05 |
| 13557 | 678 | T678T | B | 1e-05 | 5.8e-06 | 1.4e-05 | 9.933E-06 | 4.100E-06 | 5.800E-06 | 1.400E-05 |
| 13558 | 678 | T678W | B | 1.4e-05 | 7e-06 | 1.2e-05 | 1.100E-05 | 3.606E-06 | 7.000E-06 | 1.400E-05 |
| 13559 | 678 | T678Y | B | 1.2e-05 | 1.3e-05 | 1.5e-05 | 1.333E-05 | 1.528E-06 | 1.200E-05 | 1.500E-05 |
| 13560 | 678 | T678V | B | 1.1e-05 | 1.6e-05 | 1e-05 | 1.233E-05 | 3.215E-06 | 1.000E-05 | 1.600E-05 |
| 13561 | 679 | G679A | B | 1.6e-05 | 1.4e-05 | 1.4e-05 | 1.467E-05 | 1.155E-06 | 1.400E-05 | 1.600E-05 |
| 13562 | 679 | G679R | B | 1.1e-05 | 1.5e-05 | 1.1e-05 | 1.233E-05 | 2.309E-06 | 1.100E-05 | 1.500E-05 |
| 13563 | 679 | G679N | B | 1.1e-05 | 8.9e-06 | 1.5e-05 | 1.163E-05 | 3.099E-06 | 8.900E-06 | 1.500E-05 |
| 13564 | 679 | G679D | B | 1.4e-05 | 1.4e-05 | 1.5e-05 | 1.433E-05 | 5.774E-07 | 1.400E-05 | 1.500E-05 |
| 13565 | 679 | G679C | B | 1.2e-05 | 1.1e-05 | 1.5e-05 | 1.267E-05 | 2.082E-06 | 1.100E-05 | 1.500E-05 |
| 13566 | 679 | G679E | B | 1.5e-05 | 1.4e-05 | 1e-05 | 1.300E-05 | 2.646E-06 | 1.000E-05 | 1.500E-05 |
| 13567 | 679 | G679Q | B | 7.8e-06 | 1.4e-05 | 1e-05 | 1.060E-05 | 3.143E-06 | 7.800E-06 | 1.400E-05 |
| 13568 | 679 | G679G | B | 1.6e-05 | 1e-05 | 1.6e-05 | 1.400E-05 | 3.464E-06 | 1.000E-05 | 1.600E-05 |
| 13569 | 679 | G679H | B | 1.2e-05 | 1.6e-05 | 1e-05 | 1.267E-05 | 3.055E-06 | 1.000E-05 | 1.600E-05 |
| 13570 | 679 | G679I | B | 1.1e-05 | 1e-05 | 1e-05 | 1.033E-05 | 5.774E-07 | 1.000E-05 | 1.100E-05 |
| 13571 | 679 | G679L | B | 1e-05 | 6.5e-06 | 1.5e-05 | 1.050E-05 | 4.272E-06 | 6.500E-06 | 1.500E-05 |
| 13572 | 679 | G679K | B | 1.3e-05 | 1.6e-05 | 1e-05 | 1.300E-05 | 3.000E-06 | 1.000E-05 | 1.600E-05 |
| 13573 | 679 | G679M | B | 1.2e-05 | 1.5e-05 | 1.5e-05 | 1.400E-05 | 1.732E-06 | 1.200E-05 | 1.500E-05 |
| 13574 | 679 | G679F | B | 1.5e-05 | 1.3e-05 | 8.1e-06 | 1.203E-05 | 3.550E-06 | 8.100E-06 | 1.500E-05 |
| 13575 | 679 | G679P | B | 8.6e-06 | 1e-05 | 8e-06 | 8.867E-06 | 1.026E-06 | 8.000E-06 | 1.000E-05 |
| 13576 | 679 | G679S | B | 1.6e-05 | 1.1e-05 | 1e-05 | 1.233E-05 | 3.215E-06 | 1.000E-05 | 1.600E-05 |
| 13577 | 679 | G679T | B | 1.1e-05 | 1.6e-05 | 9e-06 | 1.200E-05 | 3.606E-06 | 9.000E-06 | 1.600E-05 |
| 13578 | 679 | G679W | B | 1.4e-05 | 1.7e-05 | 1e-05 | 1.367E-05 | 3.512E-06 | 1.000E-05 | 1.700E-05 |
| 13579 | 679 | G679Y | B | 1.3e-05 | 1.6e-05 | 1.5e-05 | 1.467E-05 | 1.528E-06 | 1.300E-05 | 1.600E-05 |
| 13580 | 679 | G679V | B | 1.4e-05 | 1.2e-05 | 1.6e-05 | 1.400E-05 | 2.000E-06 | 1.200E-05 | 1.600E-05 |
| 13581 | 680 | V680A | B | 1.1e-05 | 1.6e-05 | 1e-05 | 1.233E-05 | 3.215E-06 | 1.000E-05 | 1.600E-05 |
| 13582 | 680 | V680R | B | 1.4e-05 | 1.7e-05 | 9e-06 | 1.333E-05 | 4.041E-06 | 9.000E-06 | 1.700E-05 |
| 13583 | 680 | V680N | B | 1.3e-05 | 1.8e-05 | 1.2e-05 | 1.433E-05 | 3.215E-06 | 1.200E-05 | 1.800E-05 |
| 13584 | 680 | V680D | B | 1.3e-05 | 1e-05 | 1.4e-05 | 1.233E-05 | 2.082E-06 | 1.000E-05 | 1.400E-05 |
| 13585 | 680 | V680C | B | 9.5e-06 | 1.1e-05 | 1.1e-05 | 1.050E-05 | 8.660E-07 | 9.500E-06 | 1.100E-05 |
| 13586 | 680 | V680E | B | 1.5e-05 | 1.6e-05 | 8.5e-06 | 1.317E-05 | 4.072E-06 | 8.500E-06 | 1.600E-05 |
| 13587 | 680 | V680Q | B | 1.1e-05 | 1.1e-05 | 7.9e-06 | 9.967E-06 | 1.790E-06 | 7.900E-06 | 1.100E-05 |
| 13588 | 680 | V680G | B | 1.5e-05 | 1.1e-05 | 1.6e-05 | 1.400E-05 | 2.646E-06 | 1.100E-05 | 1.600E-05 |
| 13589 | 680 | V680H | B | 1.5e-05 | 1.5e-05 | 1.2e-05 | 1.400E-05 | 1.732E-06 | 1.200E-05 | 1.500E-05 |
| 13590 | 680 | V680I | B | 1.5e-05 | 1.6e-05 | 9.9e-06 | 1.363E-05 | 3.272E-06 | 9.900E-06 | 1.600E-05 |
| 13591 | 680 | V680L | B | 1.4e-05 | 1.7e-05 | 1.1e-05 | 1.400E-05 | 3.000E-06 | 1.100E-05 | 1.700E-05 |
| 13592 | 680 | V680K | B | 1.2e-05 | 1.6e-05 | 5.8e-06 | 1.127E-05 | 5.139E-06 | 5.800E-06 | 1.600E-05 |
| 13593 | 680 | V680M | B | 1.5e-05 | 1.4e-05 | 1.6e-05 | 1.500E-05 | 1.000E-06 | 1.400E-05 | 1.600E-05 |
| 13594 | 680 | V680F | B | 1.4e-05 | 1.5e-05 | 1.7e-05 | 1.533E-05 | 1.528E-06 | 1.400E-05 | 1.700E-05 |
| 13595 | 680 | V680P | B | 1.2e-05 | 1.4e-05 | 1.1e-05 | 1.233E-05 | 1.528E-06 | 1.100E-05 | 1.400E-05 |
| 13596 | 680 | V680S | B | 1e-05 | 9.5e-06 | 1.6e-05 | 1.183E-05 | 3.617E-06 | 9.500E-06 | 1.600E-05 |
| 13597 | 680 | V680T | B | 1.5e-05 | 9.9e-06 | 1.4e-05 | 1.297E-05 | 2.702E-06 | 9.900E-06 | 1.500E-05 |
| 13598 | 680 | V680W | B | 1.3e-05 | 1.4e-05 | 1.3e-05 | 1.333E-05 | 5.774E-07 | 1.300E-05 | 1.400E-05 |
| 13599 | 680 | V680Y | B | 9.1e-06 | 1.4e-05 | 9.3e-06 | 1.080E-05 | 2.773E-06 | 9.100E-06 | 1.400E-05 |

| | | | | | | | | | | |
|---|---|---|---|---|---|---|---|---|---|---|
| 13600 | 680 | V680V | B | 1.5e-05 | 1e-05 | 1e-05 | 1.167E-05 | 2.887E-06 | 1.000E-05 | 1.500E-05 |
| 13601 | 681 | P681A | B | 1.1e-05 | 1.7e-05 | 8.3e-06 | 1.210E-05 | 4.453E-06 | 8.300E-06 | 1.700E-05 |
| 13602 | 681 | P681R | B | 1e-05 | 1.4e-05 | 9.5e-06 | 1.117E-05 | 2.466E-06 | 9.500E-06 | 1.400E-05 |
| 13603 | 681 | P681N | B | 1.2e-05 | 1.3e-05 | 1.2e-05 | 1.233E-05 | 5.774E-07 | 1.200E-05 | 1.300E-05 |
| 13604 | 681 | P681D | B | 1.7e-05 | 9.4e-06 | 1.5e-05 | 1.380E-05 | 3.940E-06 | 9.400E-06 | 1.700E-05 |
| 13605 | 681 | P681C | B | 1.5e-05 | 1.4e-05 | 8.5e-06 | 1.250E-05 | 3.500E-06 | 8.500E-06 | 1.500E-05 |
| 13606 | 681 | P681E | B | 1.3e-05 | 9.7e-06 | 1.2e-05 | 1.157E-05 | 1.692E-06 | 9.700E-06 | 1.300E-05 |
| 13607 | 681 | P681Q | B | 1.4e-05 | 1.1e-05 | 1.3e-05 | 1.267E-05 | 1.528E-06 | 1.100E-05 | 1.400E-05 |
| 13608 | 681 | P681G | B | 1.2e-05 | 1e-05 | 1e-05 | 1.067E-05 | 1.155E-06 | 1.000E-05 | 1.200E-05 |
| 13609 | 681 | P681H | B | 1e-05 | 1.6e-05 | 1.2e-05 | 1.267E-05 | 3.055E-06 | 1.000E-05 | 1.600E-05 |
| 13610 | 681 | P681I | B | 1.7e-05 | 1.4e-05 | 1.5e-05 | 1.533E-05 | 1.528E-06 | 1.400E-05 | 1.700E-05 |
| 13611 | 681 | P681L | B | 8.9e-06 | 1.1e-05 | 1.4e-05 | 1.130E-05 | 2.563E-06 | 8.900E-06 | 1.400E-05 |
| 13612 | 681 | P681K | B | 9.1e-06 | 1.7e-05 | 1e-05 | 1.203E-05 | 4.325E-06 | 9.100E-06 | 1.700E-05 |
| 13613 | 681 | P681M | B | 1e-05 | 1.4e-05 | 9.5e-06 | 1.117E-05 | 2.466E-06 | 9.500E-06 | 1.400E-05 |
| 13614 | 681 | P681F | B | 1.5e-05 | 1e-05 | 1.2e-05 | 1.233E-05 | 2.517E-06 | 1.000E-05 | 1.500E-05 |
| 13615 | 681 | P681P | B | 1.4e-05 | 1.5e-05 | 1.1e-05 | 1.333E-05 | 2.082E-06 | 1.100E-05 | 1.500E-05 |
| 13616 | 681 | P681S | B | 1.1e-05 | 1e-05 | 9.7e-06 | 1.023E-05 | 6.807E-07 | 9.700E-06 | 1.100E-05 |
| 13617 | 681 | P681T | B | 1e-05 | 1.5e-05 | 1.6e-05 | 1.367E-05 | 3.215E-06 | 1.000E-05 | 1.600E-05 |
| 13618 | 681 | P681W | B | 1.5e-05 | 1.4e-05 | 8.8e-06 | 1.260E-05 | 3.329E-06 | 8.800E-06 | 1.500E-05 |
| 13619 | 681 | P681Y | B | 1.5e-05 | 1e-05 | 1.5e-05 | 1.333E-05 | 2.887E-06 | 1.000E-05 | 1.500E-05 |
| 13620 | 681 | P681V | B | 1.1e-05 | 7.3e-06 | 1.1e-05 | 9.767E-06 | 2.136E-06 | 7.300E-06 | 1.100E-05 |
| 13621 | 682 | S682A | B | 1.4e-05 | 1.5e-05 | 9.2e-06 | 1.273E-05 | 3.101E-06 | 9.200E-06 | 1.500E-05 |
| 13622 | 682 | S682R | B | 1.5e-05 | 1.1e-05 | 1.7e-05 | 1.433E-05 | 3.055E-06 | 1.100E-05 | 1.700E-05 |
| 13623 | 682 | S682N | B | 1.1e-05 | 7.7e-06 | 1.6e-05 | 1.157E-05 | 4.179E-06 | 7.700E-06 | 1.600E-05 |
| 13624 | 682 | S682D | B | 1.3e-05 | 8e-06 | 1.3e-05 | 1.133E-05 | 2.887E-06 | 8.000E-06 | 1.300E-05 |
| 13625 | 682 | S682C | B | 1.2e-05 | 1.7e-05 | 1.2e-05 | 1.367E-05 | 2.887E-06 | 1.200E-05 | 1.700E-05 |
| 13626 | 682 | S682E | B | 9.6e-06 | 1.5e-05 | 1.5e-05 | 1.320E-05 | 3.118E-06 | 9.600E-06 | 1.500E-05 |
| 13627 | 682 | S682Q | B | 1.2e-05 | 1.5e-05 | 1.4e-05 | 1.367E-05 | 1.528E-06 | 1.200E-05 | 1.500E-05 |
| 13628 | 682 | S682G | B | 1.4e-05 | 9.2e-06 | 1.1e-05 | 1.140E-05 | 2.425E-06 | 9.200E-06 | 1.400E-05 |
| 13629 | 682 | S682H | B | 1.4e-05 | 1e-05 | 1.2e-05 | 1.200E-05 | 2.000E-06 | 1.000E-05 | 1.400E-05 |
| 13630 | 682 | S682I | B | 1.7e-05 | 8e-06 | 1.3e-05 | 1.267E-05 | 4.509E-06 | 8.000E-06 | 1.700E-05 |
| 13631 | 682 | S682L | B | 1.4e-05 | 1.7e-05 | 1.1e-05 | 1.400E-05 | 3.000E-06 | 1.100E-05 | 1.700E-05 |
| 13632 | 682 | S682K | B | 1.1e-05 | 1.7e-05 | 9.6e-06 | 1.253E-05 | 3.931E-06 | 9.600E-06 | 1.700E-05 |
| 13633 | 682 | S682M | B | 1.5e-05 | 9.9e-06 | 1.2e-05 | 1.230E-05 | 2.563E-06 | 9.900E-06 | 1.500E-05 |
| 13634 | 682 | S682F | B | 8.8e-06 | 1.5e-05 | 1.1e-05 | 1.160E-05 | 3.143E-06 | 8.800E-06 | 1.500E-05 |
| 13635 | 682 | S682P | B | 1.1e-05 | 1.1e-05 | 9.4e-06 | 1.047E-05 | 9.238E-07 | 9.400E-06 | 1.100E-05 |
| 13636 | 682 | S682S | B | 1.4e-05 | 1.5e-05 | 1.5e-05 | 1.467E-05 | 5.774E-07 | 1.400E-05 | 1.500E-05 |
| 13637 | 682 | S682T | B | 1.5e-05 | 1.5e-05 | 1e-05 | 1.333E-05 | 2.887E-06 | 1.000E-05 | 1.500E-05 |
| 13638 | 682 | S682W | B | 1e-05 | 7.9e-06 | 1.4e-05 | 1.063E-05 | 3.099E-06 | 7.900E-06 | 1.400E-05 |
| 13639 | 682 | S682Y | B | 1.5e-05 | 8.2e-06 | 1.5e-05 | 1.273E-05 | 3.926E-06 | 8.200E-06 | 1.500E-05 |
| 13640 | 682 | S682V | B | 1.2e-05 | 1.2e-05 | 1.5e-05 | 1.300E-05 | 1.732E-06 | 1.200E-05 | 1.500E-05 |
| 13641 | 683 | R683A | B | 1e-05 | 1.6e-05 | 1.6e-05 | 1.400E-05 | 3.464E-06 | 1.000E-05 | 1.600E-05 |
| 13642 | 683 | R683R | B | 1.6e-05 | 1.4e-05 | 1.8e-05 | 1.600E-05 | 2.000E-06 | 1.400E-05 | 1.800E-05 |
| 13643 | 683 | R683N | B | 1e-05 | 1.2e-05 | 9.2e-06 | 1.040E-05 | 1.442E-06 | 9.200E-06 | 1.200E-05 |
| 13644 | 683 | R683D | B | 1.5e-05 | 1.7e-05 | 1.5e-05 | 1.567E-05 | 1.155E-06 | 1.500E-05 | 1.700E-05 |
| 13645 | 683 | R683C | B | 1.1e-05 | 1.4e-05 | 1.2e-05 | 1.233E-05 | 1.528E-06 | 1.100E-05 | 1.400E-05 |
| 13646 | 683 | R683E | B | 1.6e-05 | 1.4e-05 | 1.1e-05 | 1.367E-05 | 2.517E-06 | 1.100E-05 | 1.600E-05 |
| 13647 | 683 | R683Q | B | 1.6e-05 | 1.5e-05 | 1e-05 | 1.367E-05 | 3.215E-06 | 1.000E-05 | 1.600E-05 |
| 13648 | 683 | R683G | B | 1.2e-05 | 1.1e-05 | 1.6e-05 | 1.300E-05 | 2.646E-06 | 1.100E-05 | 1.600E-05 |
| 13649 | 683 | R683H | B | 1.5e-05 | 9.3e-06 | 7.4e-06 | 1.057E-05 | 3.955E-06 | 7.400E-06 | 1.500E-05 |
| 13650 | 683 | R683I | B | 1.3e-05 | 1.1e-05 | 9.4e-06 | 1.113E-05 | 1.804E-06 | 9.400E-06 | 1.300E-05 |
| 13651 | 683 | R683L | B | 1.1e-05 | 1.1e-05 | 1.1e-05 | 1.100E-05 | 2.075E-21 | 1.100E-05 | 1.100E-05 |
| 13652 | 683 | R683K | B | 8.4e-06 | 1.3e-05 | 1.1e-05 | 1.080E-05 | 2.307E-06 | 8.400E-06 | 1.300E-05 |
| 13653 | 683 | R683M | B | 1.7e-05 | 1.2e-05 | 1.5e-05 | 1.467E-05 | 2.517E-06 | 1.200E-05 | 1.700E-05 |
| 13654 | 683 | R683F | B | 1.3e-05 | 1.1e-05 | 1.4e-05 | 1.267E-05 | 1.528E-06 | 1.100E-05 | 1.400E-05 |
| 13655 | 683 | R683P | B | 1.5e-05 | 1.4e-05 | 1.7e-05 | 1.533E-05 | 1.528E-06 | 1.400E-05 | 1.700E-05 |
| 13656 | 683 | R683S | B | 1.3e-05 | 1.4e-05 | 1.5e-05 | 1.400E-05 | 1.000E-06 | 1.300E-05 | 1.500E-05 |
| 13657 | 683 | R683T | B | 1.3e-05 | 9.7e-06 | 9.4e-06 | 1.070E-05 | 1.997E-06 | 9.400E-06 | 1.300E-05 |
| 13658 | 683 | R683W | B | 1.1e-05 | 1.6e-05 | 8.7e-06 | 1.190E-05 | 3.732E-06 | 8.700E-06 | 1.600E-05 |
| 13659 | 683 | R683Y | B | 1.5e-05 | 9.5e-06 | 1.5e-05 | 1.317E-05 | 3.175E-06 | 9.500E-06 | 1.500E-05 |
| 13660 | 683 | R683V | B | 1.4e-05 | 1e-05 | 1.4e-05 | 1.267E-05 | 2.309E-06 | 1.000E-05 | 1.400E-05 |
| 13661 | 684 | F684A | B | 1.4e-05 | 1.4e-05 | 1.4e-05 | 1.400E-05 | 0.000E+00 | 1.400E-05 | 1.400E-05 |
| 13662 | 684 | F684R | B | 1.4e-05 | 8.6e-06 | 1.6e-05 | 1.287E-05 | 3.828E-06 | 8.600E-06 | 1.600E-05 |
| 13663 | 684 | F684N | B | 9.9e-06 | 1.4e-05 | 1.1e-05 | 1.163E-05 | 2.122E-06 | 9.900E-06 | 1.400E-05 |
| 13664 | 684 | F684D | B | 1.1e-05 | 1.6e-05 | 8.1e-06 | 1.170E-05 | 3.996E-06 | 8.100E-06 | 1.600E-05 |
| 13665 | 684 | F684C | B | 1.6e-05 | 1.3e-05 | 1.1e-05 | 1.333E-05 | 2.517E-06 | 1.100E-05 | 1.600E-05 |
| 13666 | 684 | F684E | B | 1.5e-05 | 1.6e-05 | 1.2e-05 | 1.433E-05 | 2.082E-06 | 1.200E-05 | 1.600E-05 |
| 13667 | 684 | F684Q | B | 1.6e-05 | 1e-05 | 1.6e-05 | 1.400E-05 | 3.464E-06 | 1.000E-05 | 1.600E-05 |
| 13668 | 684 | F684G | B | 9.3e-06 | 1.6e-05 | 1.2e-05 | 1.243E-05 | 3.371E-06 | 9.300E-06 | 1.600E-05 |
| 13669 | 684 | F684H | B | 1.4e-05 | 1.2e-05 | 1e-05 | 1.200E-05 | 2.000E-06 | 1.000E-05 | 1.400E-05 |
| 13670 | 684 | F684I | B | 1.1e-05 | 9.6e-06 | 1.1e-05 | 1.053E-05 | 8.083E-07 | 9.600E-06 | 1.100E-05 |
| 13671 | 684 | F684L | B | 1.1e-05 | 1.7e-05 | 1.3e-05 | 1.367E-05 | 3.055E-06 | 1.100E-05 | 1.700E-05 |
| 13672 | 684 | F684K | B | 1.4e-05 | 1.5e-05 | 7.7e-06 | 1.223E-05 | 3.958E-06 | 7.700E-06 | 1.500E-05 |
| 13673 | 684 | F684M | B | 1.3e-05 | 9.6e-06 | 1.4e-05 | 1.220E-05 | 2.307E-06 | 9.600E-06 | 1.400E-05 |
| 13674 | 684 | F684F | B | 1.5e-05 | 1.5e-05 | 1.5e-05 | 1.500E-05 | 0.000E+00 | 1.500E-05 | 1.500E-05 |
| 13675 | 684 | F684P | B | 1.1e-05 | 1.4e-05 | 1.5e-05 | 1.333E-05 | 2.082E-06 | 1.100E-05 | 1.500E-05 |
| 13676 | 684 | F684S | B | 1.6e-05 | 1.5e-05 | 8.2e-06 | 1.307E-05 | 4.244E-06 | 8.200E-06 | 1.600E-05 |
| 13677 | 684 | F684T | B | 1.5e-05 | 1.1e-05 | 9.3e-06 | 1.177E-05 | 2.926E-06 | 9.300E-06 | 1.500E-05 |
| 13678 | 684 | F684W | B | 1.1e-05 | 1.2e-05 | 9.5e-06 | 1.083E-05 | 1.258E-06 | 9.500E-06 | 1.200E-05 |
| 13679 | 684 | F684Y | B | 2.5e-05 | 1.6e-05 | 1.4e-05 | 1.833E-05 | 5.859E-06 | 1.400E-05 | 2.500E-05 |
| 13680 | 684 | F684V | B | 1e-05 | 1.1e-05 | 8.3e-06 | 9.767E-06 | 1.365E-06 | 8.300E-06 | 1.100E-05 |
| 13681 | 685 | S685A | B | 1.6e-05 | 1.6e-05 | 1.5e-05 | 1.567E-05 | 5.774E-07 | 1.500E-05 | 1.600E-05 |
| 13682 | 685 | S685R | B | 1.4e-05 | 1.2e-05 | 1.5e-05 | 1.367E-05 | 1.528E-06 | 1.200E-05 | 1.500E-05 |
| 13683 | 685 | S685N | B | 1e-05 | 1.7e-05 | 1.5e-05 | 1.400E-05 | 3.606E-06 | 1.000E-05 | 1.700E-05 |
| 13684 | 685 | S685D | B | 1.1e-05 | 1.1e-05 | 1.7e-05 | 1.300E-05 | 3.464E-06 | 1.100E-05 | 1.700E-05 |

| | | | | | | | | | | |
|---|---|---|---|---|---|---|---|---|---|---|
| 13685 | 685 | S685C | B | 1.4e-05 | 1.3e-05 | 1.1e-05 | 1.267E-05 | 1.528E-06 | 1.100E-05 | 1.400E-05 |
| 13686 | 685 | S685E | B | 1.7e-05 | 2.2e-05 | 1.6e-05 | 1.833E-05 | 3.215E-06 | 1.600E-05 | 2.200E-05 |
| 13687 | 685 | S685Q | B | 1.5e-05 | 1.6e-05 | 9.7e-06 | 1.357E-05 | 3.386E-06 | 9.700E-06 | 1.600E-05 |
| 13688 | 685 | S685G | B | 1.7e-05 | 1.3e-05 | 1.2e-05 | 1.400E-05 | 2.646E-06 | 1.200E-05 | 1.700E-05 |
| 13689 | 685 | S685H | B | 9.7e-06 | 1.1e-05 | 1.6e-05 | 1.223E-05 | 3.326E-06 | 9.700E-06 | 1.600E-05 |
| 13690 | 685 | S685I | B | 1.5e-05 | 1.5e-05 | 1e-05 | 1.333E-05 | 2.887E-06 | 1.000E-05 | 1.500E-05 |
| 13691 | 685 | S685L | B | 1.6e-05 | 1.5e-05 | 1e-05 | 1.367E-05 | 3.215E-06 | 1.000E-05 | 1.600E-05 |
| 13692 | 685 | S685K | B | 1.5e-05 | 1.3e-05 | 9.1e-06 | 1.237E-05 | 3.001E-06 | 9.100E-06 | 1.500E-05 |
| 13693 | 685 | S685M | B | 1.5e-05 | 1.4e-05 | 1.3e-05 | 1.400E-05 | 1.000E-06 | 1.300E-05 | 1.500E-05 |
| 13694 | 685 | S685F | B | 1.6e-05 | 1.1e-05 | 1.5e-05 | 1.400E-05 | 2.646E-06 | 1.100E-05 | 1.600E-05 |
| 13695 | 685 | S685P | B | 8e-06 | 1.8e-05 | 1e-05 | 1.200E-05 | 5.292E-06 | 8.000E-06 | 1.800E-05 |
| 13696 | 685 | S685S | B | 1e-05 | 1.5e-05 | 1.7e-05 | 1.400E-05 | 3.606E-06 | 1.000E-05 | 1.700E-05 |
| 13697 | 685 | S685T | B | 9.1e-06 | 1.1e-05 | 1e-05 | 1.003E-05 | 9.504E-07 | 9.100E-06 | 1.100E-05 |
| 13698 | 685 | S685W | B | 9.8e-06 | 1e-05 | 1.4e-05 | 1.127E-05 | 2.369E-06 | 9.800E-06 | 1.400E-05 |
| 13699 | 685 | S685Y | B | 1e-05 | 1.1e-05 | 1.3e-05 | 1.133E-05 | 1.528E-06 | 1.000E-05 | 1.300E-05 |
| 13700 | 685 | S685V | B | 1.6e-05 | 1.1e-05 | 1e-05 | 1.233E-05 | 3.215E-06 | 1.000E-05 | 1.600E-05 |
| 13701 | 686 | G686A | B | 1.1e-05 | 9.7e-06 | 1.6e-05 | 1.223E-05 | 3.326E-06 | 9.700E-06 | 1.600E-05 |
| 13702 | 686 | G686R | B | 1.6e-05 | 1.4e-05 | 1.3e-05 | 1.433E-05 | 1.528E-06 | 1.300E-05 | 1.600E-05 |
| 13703 | 686 | G686N | B | 1.6e-05 | 1.5e-05 | 1.5e-05 | 1.533E-05 | 5.774E-07 | 1.500E-05 | 1.600E-05 |
| 13704 | 686 | G686D | B | 1.5e-05 | 1e-05 | 1.1e-05 | 1.200E-05 | 2.646E-06 | 1.000E-05 | 1.500E-05 |
| 13705 | 686 | G686C | B | 1.3e-05 | 1e-05 | 1.4e-05 | 1.233E-05 | 2.082E-06 | 1.000E-05 | 1.400E-05 |
| 13706 | 686 | G686E | B | 1.5e-05 | 9.2e-06 | 4e-05 | 2.140E-05 | 1.637E-05 | 9.200E-06 | 4.000E-05 |
| 13707 | 686 | G686Q | B | 9e-06 | 1.1e-05 | 1.1e-05 | 1.033E-05 | 1.155E-06 | 9.000E-06 | 1.100E-05 |
| 13708 | 686 | G686G | B | 1.1e-05 | 1.5e-05 | 1.1e-05 | 1.233E-05 | 2.309E-06 | 1.100E-05 | 1.500E-05 |
| 13709 | 686 | G686H | B | 1.6e-05 | 1.4e-05 | 1.5e-05 | 1.500E-05 | 1.000E-06 | 1.400E-05 | 1.600E-05 |
| 13710 | 686 | G686I | B | 1.6e-05 | 1.6e-05 | 8.1e-06 | 1.337E-05 | 4.561E-06 | 8.100E-06 | 1.600E-05 |
| 13711 | 686 | G686L | B | 1.4e-05 | 1e-05 | 1.1e-05 | 1.167E-05 | 2.082E-06 | 1.000E-05 | 1.400E-05 |
| 13712 | 686 | G686K | B | 1.5e-05 | 1.6e-05 | 1.7e-05 | 1.600E-05 | 1.000E-06 | 1.500E-05 | 1.700E-05 |
| 13713 | 686 | G686M | B | 1.7e-05 | 1e-05 | 1.2e-05 | 1.300E-05 | 3.606E-06 | 1.000E-05 | 1.700E-05 |
| 13714 | 686 | G686F | B | 1.5e-05 | 1.3e-05 | 1e-05 | 1.267E-05 | 2.517E-06 | 1.000E-05 | 1.500E-05 |
| 13715 | 686 | G686P | B | 1.4e-05 | 1.6e-05 | 1.1e-05 | 1.367E-05 | 2.517E-06 | 1.100E-05 | 1.600E-05 |
| 13716 | 686 | G686S | B | 9.8e-06 | 1.4e-05 | 1e-05 | 1.127E-05 | 2.369E-06 | 9.800E-06 | 1.400E-05 |
| 13717 | 686 | G686T | B | 6.8e-06 | 9.3e-06 | 9.9e-06 | 8.667E-06 | 1.644E-06 | 6.800E-06 | 9.900E-06 |
| 13718 | 686 | G686W | B | 1.5e-05 | 1.5e-05 | 1.4e-05 | 1.467E-05 | 5.774E-07 | 1.400E-05 | 1.500E-05 |
| 13719 | 686 | G686Y | B | 1.5e-05 | 1e-05 | 1.2e-05 | 1.233E-05 | 2.517E-06 | 1.000E-05 | 1.500E-05 |
| 13720 | 686 | G686V | B | 1.4e-05 | 1.6e-05 | 1.6e-05 | 1.533E-05 | 1.155E-06 | 1.400E-05 | 1.600E-05 |
| 13721 | 687 | S687A | B | 1.1e-05 | 9e-06 | 2e-05 | 1.333E-05 | 5.859E-06 | 9.000E-06 | 2.000E-05 |
| 13722 | 687 | S687R | B | 1.2e-05 | 1.2e-05 | 1.5e-05 | 1.300E-05 | 1.732E-06 | 1.200E-05 | 1.500E-05 |
| 13723 | 687 | S687N | B | 1.6e-05 | 9.6e-06 | 9.7e-06 | 1.177E-05 | 3.667E-06 | 9.600E-06 | 1.600E-05 |
| 13724 | 687 | S687D | B | 1.2e-05 | 1.4e-05 | 9.9e-06 | 1.197E-05 | 2.050E-06 | 9.900E-06 | 1.400E-05 |
| 13725 | 687 | S687C | B | 9.7e-06 | 1.4e-05 | 9.2e-06 | 1.097E-05 | 2.639E-06 | 9.200E-06 | 1.400E-05 |
| 13726 | 687 | S687E | B | 1.3e-05 | 1e-05 | 1.6e-05 | 1.300E-05 | 3.000E-06 | 1.000E-05 | 1.600E-05 |
| 13727 | 687 | S687Q | B | 6.2e-06 | 1.5e-05 | 6.7e-06 | 9.300E-06 | 4.943E-06 | 6.200E-06 | 1.500E-05 |
| 13728 | 687 | S687G | B | 1.8e-05 | 1.6e-05 | 1.2e-05 | 1.533E-05 | 3.055E-06 | 1.200E-05 | 1.800E-05 |
| 13729 | 687 | S687H | B | 1e-05 | 1.6e-05 | 9.5e-06 | 1.183E-05 | 3.617E-06 | 9.500E-06 | 1.600E-05 |
| 13730 | 687 | S687I | B | 1e-05 | 1.7e-05 | 2.4e-05 | 1.700E-05 | 7.000E-06 | 1.000E-05 | 2.400E-05 |
| 13731 | 687 | S687L | B | 9.3e-06 | 1e-05 | 1.1e-05 | 1.010E-05 | 8.544E-07 | 9.300E-06 | 1.100E-05 |
| 13732 | 687 | S687K | B | 1.5e-05 | 1.4e-05 | 9.9e-06 | 1.297E-05 | 2.702E-06 | 9.900E-06 | 1.500E-05 |
| 13733 | 687 | S687M | B | 1.7e-05 | 1e-05 | 1.6e-05 | 1.433E-05 | 3.786E-06 | 1.000E-05 | 1.700E-05 |
| 13734 | 687 | S687F | B | 1.5e-05 | 1.5e-05 | 1.6e-05 | 1.533E-05 | 5.774E-07 | 1.500E-05 | 1.600E-05 |
| 13735 | 687 | S687P | B | 1.2e-05 | 1e-05 | 1.4e-05 | 1.200E-05 | 2.000E-06 | 1.000E-05 | 1.400E-05 |
| 13736 | 687 | S687S | B | 1.8e-05 | 1.7e-05 | 1.5e-05 | 1.667E-05 | 1.528E-06 | 1.500E-05 | 1.800E-05 |
| 13737 | 687 | S687T | B | 1.7e-05 | 9.8e-06 | 9.3e-06 | 1.203E-05 | 4.309E-06 | 9.300E-06 | 1.700E-05 |
| 13738 | 687 | S687W | B | 8e-06 | 1.5e-05 | 1.4e-05 | 1.233E-05 | 3.786E-06 | 8.000E-06 | 1.500E-05 |
| 13739 | 687 | S687Y | B | 1.6e-05 | 1.4e-05 | 1.2e-05 | 1.400E-05 | 2.000E-06 | 1.200E-05 | 1.600E-05 |
| 13740 | 687 | S687V | B | 1.5e-05 | 1.1e-05 | 1e-05 | 1.200E-05 | 2.646E-06 | 1.000E-05 | 1.500E-05 |
| 13741 | 688 | G688A | B | 1.4e-05 | 1.3e-05 | 1.5e-05 | 1.400E-05 | 1.000E-06 | 1.300E-05 | 1.500E-05 |
| 13742 | 688 | G688R | B | 1.1e-05 | 8e-06 | 8.1e-06 | 9.033E-06 | 1.704E-06 | 8.000E-06 | 1.100E-05 |
| 13743 | 688 | G688N | B | 1.5e-05 | 9.6e-06 | 1.1e-05 | 1.187E-05 | 2.802E-06 | 9.600E-06 | 1.500E-05 |
| 13744 | 688 | G688D | B | 8.8e-06 | 1.4e-05 | 1e-05 | 1.093E-05 | 2.723E-06 | 8.800E-06 | 1.400E-05 |
| 13745 | 688 | G688C | B | 9.5e-06 | 1.4e-05 | 1e-05 | 1.117E-05 | 2.466E-06 | 9.500E-06 | 1.400E-05 |
| 13746 | 688 | G688E | B | 1.4e-05 | 1.5e-05 | 1.1e-05 | 1.333E-05 | 2.082E-06 | 1.100E-05 | 1.500E-05 |
| 13747 | 688 | G688Q | B | 1.2e-05 | 1.2e-05 | 9.9e-06 | 1.130E-05 | 1.212E-06 | 9.900E-06 | 1.200E-05 |
| 13748 | 688 | G688G | B | 1.5e-05 | 1e-05 | 9.1e-06 | 1.137E-05 | 3.179E-06 | 9.100E-06 | 1.500E-05 |
| 13749 | 688 | G688H | B | 1.2e-05 | 1.5e-05 | 1.1e-05 | 1.267E-05 | 2.082E-06 | 1.100E-05 | 1.500E-05 |
| 13750 | 688 | G688I | B | 6.2e-06 | 8.6e-06 | 1.7e-05 | 1.060E-05 | 5.671E-06 | 6.200E-06 | 1.700E-05 |
| 13751 | 688 | G688L | B | 8.6e-06 | 1.5e-05 | 1.1e-05 | 1.153E-05 | 3.233E-06 | 8.600E-06 | 1.500E-05 |
| 13752 | 688 | G688K | B | 1.1e-05 | 9.1e-06 | 1.4e-05 | 1.137E-05 | 2.470E-06 | 9.100E-06 | 1.400E-05 |
| 13753 | 688 | G688M | B | 8e-06 | 1.5e-05 | 1.3e-05 | 1.200E-05 | 3.606E-06 | 8.000E-06 | 1.500E-05 |
| 13754 | 688 | G688F | B | 1.1e-05 | 8e-06 | 1.3e-05 | 1.067E-05 | 2.517E-06 | 8.000E-06 | 1.300E-05 |
| 13755 | 688 | G688P | B | 1.3e-05 | 1.5e-05 | 1.5e-05 | 1.433E-05 | 1.155E-06 | 1.300E-05 | 1.500E-05 |
| 13756 | 688 | G688S | B | 1.4e-05 | 1.5e-05 | 1.4e-05 | 1.433E-05 | 5.774E-07 | 1.400E-05 | 1.500E-05 |
| 13757 | 688 | G688T | B | 1.3e-05 | 1.5e-05 | 1e-05 | 1.267E-05 | 2.517E-06 | 1.000E-05 | 1.500E-05 |
| 13758 | 688 | G688W | B | 1.4e-05 | 1.1e-05 | 1.6e-05 | 1.367E-05 | 2.517E-06 | 1.100E-05 | 1.600E-05 |
| 13759 | 688 | G688Y | B | 1e-05 | 1.4e-05 | 1.4e-05 | 1.267E-05 | 2.309E-06 | 1.000E-05 | 1.400E-05 |
| 13760 | 688 | G688V | B | 1.6e-05 | 1e-05 | 1.2e-05 | 1.267E-05 | 3.055E-06 | 1.000E-05 | 1.600E-05 |
| 13761 | 689 | S689A | B | 7.2e-06 | 1.5e-05 | 1.1e-05 | 1.107E-05 | 3.900E-06 | 7.200E-06 | 1.500E-05 |
| 13762 | 689 | S689R | B | 1.1e-05 | 1.4e-05 | 1.7e-05 | 1.400E-05 | 3.000E-06 | 1.100E-05 | 1.700E-05 |
| 13763 | 689 | S689N | B | 1.6e-05 | 8.5e-06 | 1.2e-05 | 1.217E-05 | 3.753E-06 | 8.500E-06 | 1.600E-05 |
| 13764 | 689 | S689D | B | 1e-05 | 1.1e-05 | 1.5e-05 | 1.200E-05 | 2.646E-06 | 1.000E-05 | 1.500E-05 |
| 13765 | 689 | S689C | B | 8.7e-06 | 1.6e-05 | 1.2e-05 | 1.223E-05 | 3.656E-06 | 8.700E-06 | 1.600E-05 |
| 13766 | 689 | S689E | B | 1.5e-05 | 1e-05 | 8e-06 | 1.100E-05 | 3.606E-06 | 8.000E-06 | 1.500E-05 |
| 13767 | 689 | S689Q | B | 1.3e-05 | 1.6e-05 | 1.1e-05 | 1.333E-05 | 2.517E-06 | 1.100E-05 | 1.600E-05 |
| 13768 | 689 | S689G | B | 1.2e-05 | 1.2e-05 | 1.3e-05 | 1.233E-05 | 5.774E-07 | 1.200E-05 | 1.300E-05 |
| 13769 | 689 | S689H | B | 1.6e-05 | 1.6e-05 | 1.4e-05 | 1.533E-05 | 1.155E-06 | 1.400E-05 | 1.600E-05 |

| | | | | | | | | | | |
|---|---|---|---|---|---|---|---|---|---|---|
| 13770 | 689 | S689I | B | 1.1e-05 | 1.7e-05 | 1.4e-05 | 1.400E-05 | 3.000E-06 | 1.100E-05 | 1.700E-05 |
| 13771 | 689 | S689L | B | 1.1e-05 | 1.1e-05 | 1.5e-05 | 1.233E-05 | 2.309E-06 | 1.100E-05 | 1.500E-05 |
| 13772 | 689 | S689K | B | 1.1e-05 | 1.4e-05 | 1e-05 | 1.167E-05 | 2.082E-06 | 1.000E-05 | 1.400E-05 |
| 13773 | 689 | S689M | B | 1.7e-05 | 1.4e-05 | 1.7e-05 | 1.600E-05 | 1.732E-06 | 1.400E-05 | 1.700E-05 |
| 13774 | 689 | S689F | B | 1.5e-05 | 1.3e-05 | 1.3e-05 | 1.367E-05 | 1.155E-06 | 1.300E-05 | 1.500E-05 |
| 13775 | 689 | S689P | B | 9.7e-06 | 1.4e-05 | 1.8e-05 | 1.390E-05 | 4.151E-06 | 9.700E-06 | 1.800E-05 |
| 13776 | 689 | S689S | B | 1.1e-05 | 9.2e-06 | 1.3e-05 | 1.107E-05 | 1.901E-06 | 9.200E-06 | 1.300E-05 |
| 13777 | 689 | S689T | B | 1.9e-05 | 1.6e-05 | 1.7e-05 | 1.733E-05 | 1.528E-06 | 1.600E-05 | 1.900E-05 |
| 13778 | 689 | S689W | B | 1.4e-05 | 1.6e-05 | 1.2e-05 | 1.400E-05 | 2.000E-06 | 1.200E-05 | 1.600E-05 |
| 13779 | 689 | S689Y | B | 1.6e-05 | 1.5e-05 | 1.4e-05 | 1.500E-05 | 1.000E-06 | 1.400E-05 | 1.600E-05 |
| 13780 | 689 | S689V | B | 1.8e-05 | 1e-05 | 1.5e-05 | 1.433E-05 | 4.041E-06 | 1.000E-05 | 1.800E-05 |
| 13781 | 690 | G690A | B | 1.7e-05 | 1.3e-05 | 1.5e-05 | 1.500E-05 | 2.000E-06 | 1.300E-05 | 1.700E-05 |
| 13782 | 690 | G690R | B | 1.4e-05 | 8.6e-06 | 1.5e-05 | 1.253E-05 | 3.443E-06 | 8.600E-06 | 1.500E-05 |
| 13783 | 690 | G690N | B | 9.9e-06 | 1.1e-05 | 1.4e-05 | 1.163E-05 | 2.122E-06 | 9.900E-06 | 1.400E-05 |
| 13784 | 690 | G690D | B | 1.6e-05 | 1.7e-05 | 1.5e-05 | 1.600E-05 | 1.000E-06 | 1.500E-05 | 1.700E-05 |
| 13785 | 690 | G690C | B | 1.6e-05 | 1.5e-05 | 9.8e-06 | 1.360E-05 | 3.329E-06 | 9.800E-06 | 1.600E-05 |
| 13786 | 690 | G690E | B | 1.5e-05 | 1.4e-05 | 1.5e-05 | 1.467E-05 | 5.774E-07 | 1.400E-05 | 1.500E-05 |
| 13787 | 690 | G690Q | B | 9.6e-06 | 1e-05 | 1.4e-05 | 1.120E-05 | 2.433E-06 | 9.600E-06 | 1.400E-05 |
| 13788 | 690 | G690G | B | 1.8e-05 | 1.4e-05 | 1.6e-05 | 1.600E-05 | 2.000E-06 | 1.400E-05 | 1.800E-05 |
| 13789 | 690 | G690H | B | 8.9e-06 | 1e-05 | 1.5e-05 | 1.130E-05 | 3.251E-06 | 8.900E-06 | 1.500E-05 |
| 13790 | 690 | G690I | B | 1.6e-05 | 1.6e-05 | 1.5e-05 | 1.567E-05 | 5.774E-07 | 1.500E-05 | 1.600E-05 |
| 13791 | 690 | G690L | B | 1.6e-05 | 1.3e-05 | 1.5e-05 | 1.467E-05 | 1.528E-06 | 1.300E-05 | 1.600E-05 |
| 13792 | 690 | G690K | B | 1.6e-05 | 1.4e-05 | 8.2e-06 | 1.273E-05 | 4.051E-06 | 8.200E-06 | 1.600E-05 |
| 13793 | 690 | G690M | B | 9.2e-06 | 1.2e-05 | 8.2e-06 | 9.800E-06 | 1.970E-06 | 8.200E-06 | 1.200E-05 |
| 13794 | 690 | G690F | B | 1.6e-05 | 1.1e-05 | 1.2e-05 | 1.300E-05 | 2.646E-06 | 1.100E-05 | 1.600E-05 |
| 13795 | 690 | G690P | B | 1.1e-05 | 1.6e-05 | 1.5e-05 | 1.400E-05 | 2.646E-06 | 1.100E-05 | 1.600E-05 |
| 13796 | 690 | G690S | B | 1.2e-05 | 1.4e-05 | 1.1e-05 | 1.233E-05 | 1.528E-06 | 1.100E-05 | 1.400E-05 |
| 13797 | 690 | G690T | B | 1.3e-05 | 9.8e-06 | 1.1e-05 | 1.127E-05 | 1.617E-06 | 9.800E-06 | 1.300E-05 |
| 13798 | 690 | G690W | B | 1.1e-05 | 1.4e-05 | 1.5e-05 | 1.333E-05 | 2.082E-06 | 1.100E-05 | 1.500E-05 |
| 13799 | 690 | G690Y | B | 1.6e-05 | 1.5e-05 | 1.5e-05 | 1.533E-05 | 5.774E-07 | 1.500E-05 | 1.600E-05 |
| 13800 | 690 | G690V | B | 9.1e-06 | 1.1e-05 | 1.4e-05 | 1.137E-05 | 2.470E-06 | 9.100E-06 | 1.400E-05 |
| 13801 | 691 | T691A | B | 1.5e-05 | 9.3e-06 | 1.2e-05 | 1.210E-05 | 2.851E-06 | 9.300E-06 | 1.500E-05 |
| 13802 | 691 | T691R | B | 1e-05 | 1.6e-05 | 1.2e-05 | 1.267E-05 | 3.055E-06 | 1.000E-05 | 1.600E-05 |
| 13803 | 691 | T691N | B | 1.5e-05 | 9.8e-06 | 1.3e-05 | 1.260E-05 | 2.623E-06 | 9.800E-06 | 1.500E-05 |
| 13804 | 691 | T691D | B | 1.5e-05 | 1.5e-05 | 1.1e-05 | 1.367E-05 | 2.309E-06 | 1.100E-05 | 1.500E-05 |
| 13805 | 691 | T691C | B | 1.4e-05 | 9.6e-06 | 1.1e-05 | 1.153E-05 | 2.248E-06 | 9.600E-06 | 1.400E-05 |
| 13806 | 691 | T691E | B | 1e-05 | 1.4e-05 | 1.1e-05 | 1.167E-05 | 2.082E-06 | 1.000E-05 | 1.400E-05 |
| 13807 | 691 | T691Q | B | 7e-06 | 1.6e-05 | 1.3e-05 | 1.200E-05 | 4.583E-06 | 7.000E-06 | 1.600E-05 |
| 13808 | 691 | T691G | B | 1.4e-05 | 1.3e-05 | 1.1e-05 | 1.267E-05 | 1.528E-06 | 1.100E-05 | 1.400E-05 |
| 13809 | 691 | T691H | B | 1.5e-05 | 1.5e-05 | 1.8e-05 | 1.600E-05 | 1.732E-06 | 1.500E-05 | 1.800E-05 |
| 13810 | 691 | T691I | B | 1.6e-05 | 1.5e-05 | 1.1e-05 | 1.400E-05 | 2.646E-06 | 1.100E-05 | 1.600E-05 |
| 13811 | 691 | T691L | B | 1e-05 | 9.2e-06 | 1.5e-05 | 1.140E-05 | 3.143E-06 | 9.200E-06 | 1.500E-05 |
| 13812 | 691 | T691K | B | 1.2e-05 | 9.1e-06 | 1.1e-05 | 1.070E-05 | 1.473E-06 | 9.100E-06 | 1.200E-05 |
| 13813 | 691 | T691M | B | 1.4e-05 | 1e-05 | 1.5e-05 | 1.300E-05 | 2.646E-06 | 1.000E-05 | 1.500E-05 |
| 13814 | 691 | T691F | B | 1.7e-05 | 7.7e-06 | 1e-05 | 1.157E-05 | 4.844E-06 | 7.700E-06 | 1.700E-05 |
| 13815 | 691 | T691P | B | 1.2e-05 | 1.1e-05 | 1.2e-05 | 1.167E-05 | 5.774E-07 | 1.100E-05 | 1.200E-05 |
| 13816 | 691 | T691S | B | 1.3e-05 | 1.6e-05 | 1e-05 | 1.300E-05 | 3.000E-06 | 1.000E-05 | 1.600E-05 |
| 13817 | 691 | T691T | B | 1.4e-05 | 8.4e-06 | 1.6e-05 | 1.280E-05 | 3.940E-06 | 8.400E-06 | 1.600E-05 |
| 13818 | 691 | T691W | B | 1.1e-05 | 1.1e-05 | 1.2e-05 | 1.133E-05 | 5.774E-07 | 1.100E-05 | 1.200E-05 |
| 13819 | 691 | T691Y | B | 8.8e-06 | 9.6e-06 | 1.7e-05 | 1.180E-05 | 4.521E-06 | 8.800E-06 | 1.700E-05 |
| 13820 | 691 | T691V | B | 1.7e-05 | 1.1e-05 | 1.5e-05 | 1.433E-05 | 3.055E-06 | 1.100E-05 | 1.700E-05 |
| 13821 | 692 | D692A | B | 1.2e-05 | 1.7e-05 | 1e-05 | 1.300E-05 | 3.606E-06 | 1.000E-05 | 1.700E-05 |
| 13822 | 692 | D692R | B | 1.6e-05 | 1.5e-05 | 1.5e-05 | 1.533E-05 | 5.774E-07 | 1.500E-05 | 1.600E-05 |
| 13823 | 692 | D692N | B | 1.2e-05 | 1.2e-05 | 1e-05 | 1.133E-05 | 1.155E-06 | 1.000E-05 | 1.200E-05 |
| 13824 | 692 | D692D | B | 1.5e-05 | 1.5e-05 | 1.7e-05 | 1.567E-05 | 1.155E-06 | 1.500E-05 | 1.700E-05 |
| 13825 | 692 | D692C | B | 9.6e-06 | 1.5e-05 | 1.2e-05 | 1.220E-05 | 2.706E-06 | 9.600E-06 | 1.500E-05 |
| 13826 | 692 | D692E | B | 1.5e-05 | 1.1e-05 | 9.6e-06 | 1.187E-05 | 2.802E-06 | 9.600E-06 | 1.500E-05 |
| 13827 | 692 | D692Q | B | 1.4e-05 | 1.3e-05 | 1e-05 | 1.233E-05 | 2.082E-06 | 1.000E-05 | 1.400E-05 |
| 13828 | 692 | D692G | B | 1e-05 | 1.8e-05 | 1.7e-05 | 1.500E-05 | 4.359E-06 | 1.000E-05 | 1.800E-05 |
| 13829 | 692 | D692H | B | 1.1e-05 | 1.1e-05 | 1.1e-05 | 1.100E-05 | 2.075E-21 | 1.100E-05 | 1.100E-05 |
| 13830 | 692 | D692I | B | 1e-05 | 1.4e-05 | 9.2e-06 | 1.107E-05 | 2.572E-06 | 9.200E-06 | 1.400E-05 |
| 13831 | 692 | D692L | B | 1.4e-05 | 9.7e-06 | 1.4e-05 | 1.257E-05 | 2.483E-06 | 9.700E-06 | 1.400E-05 |
| 13832 | 692 | D692K | B | 1.6e-05 | 1.4e-05 | 1.1e-05 | 1.367E-05 | 2.517E-06 | 1.100E-05 | 1.600E-05 |
| 13833 | 692 | D692M | B | 1.6e-05 | 1.7e-05 | 1.6e-05 | 1.633E-05 | 5.774E-07 | 1.600E-05 | 1.700E-05 |
| 13834 | 692 | D692F | B | 1.1e-05 | 1.5e-05 | 1.3e-05 | 1.300E-05 | 2.000E-06 | 1.100E-05 | 1.500E-05 |
| 13835 | 692 | D692P | B | 8.9e-06 | 1.2e-05 | 1.6e-05 | 1.230E-05 | 3.559E-06 | 8.900E-06 | 1.600E-05 |
| 13836 | 692 | D692S | B | 9.5e-06 | 9.4e-06 | 1.4e-05 | 1.097E-05 | 2.627E-06 | 9.400E-06 | 1.400E-05 |
| 13837 | 692 | D692T | B | 9.3e-06 | 9.1e-06 | 1.3e-05 | 1.047E-05 | 2.196E-06 | 9.100E-06 | 1.300E-05 |
| 13838 | 692 | D692W | B | 8.9e-06 | 1.5e-05 | 1.4e-05 | 1.263E-05 | 3.272E-06 | 8.900E-06 | 1.500E-05 |
| 13839 | 692 | D692Y | B | 1.5e-05 | 1.4e-05 | 8.6e-06 | 1.253E-05 | 3.443E-06 | 8.600E-06 | 1.500E-05 |
| 13840 | 692 | D692V | B | 1e-05 | 1.6e-05 | 9.6e-06 | 1.187E-05 | 3.585E-06 | 9.600E-06 | 1.600E-05 |
| 13841 | 693 | F693A | B | 1.7e-05 | 9.7e-06 | 1.1e-05 | 1.257E-05 | 3.894E-06 | 9.700E-06 | 1.700E-05 |
| 13842 | 693 | F693R | B | 9.5e-06 | 1.5e-05 | 1.5e-05 | 1.317E-05 | 3.175E-06 | 9.500E-06 | 1.500E-05 |
| 13843 | 693 | F693N | B | 1.4e-05 | 9.4e-06 | 1.1e-05 | 1.147E-05 | 2.335E-06 | 9.400E-06 | 1.400E-05 |
| 13844 | 693 | F693D | B | 1.1e-05 | 1e-05 | 1.5e-05 | 1.200E-05 | 2.646E-06 | 1.000E-05 | 1.500E-05 |
| 13845 | 693 | F693C | B | 1.2e-05 | 2.6e-05 | 1.5e-05 | 1.767E-05 | 7.371E-06 | 1.200E-05 | 2.600E-05 |
| 13846 | 693 | F693E | B | 1.5e-05 | 1.2e-05 | 1.5e-05 | 1.400E-05 | 1.732E-06 | 1.200E-05 | 1.500E-05 |
| 13847 | 693 | F693Q | B | 1.1e-05 | 9.1e-06 | 1.1e-05 | 1.037E-05 | 1.097E-06 | 9.100E-06 | 1.100E-05 |
| 13848 | 693 | F693G | B | 1.6e-05 | 1.6e-05 | 4.2e-05 | 2.467E-05 | 1.501E-05 | 1.600E-05 | 4.200E-05 |
| 13849 | 693 | F693H | B | 1.4e-05 | 1.4e-05 | 9.3e-06 | 1.243E-05 | 2.714E-06 | 9.300E-06 | 1.400E-05 |
| 13850 | 693 | F693I | B | 9.6e-06 | 1.6e-05 | 1.5e-05 | 1.353E-05 | 3.443E-06 | 9.600E-06 | 1.600E-05 |
| 13851 | 693 | F693L | B | 1.7e-05 | 8.6e-06 | 9.3e-06 | 1.163E-05 | 4.661E-06 | 8.600E-06 | 1.700E-05 |
| 13852 | 693 | F693K | B | 1.5e-05 | 1.6e-05 | 1.1e-05 | 1.400E-05 | 2.646E-06 | 1.100E-05 | 1.600E-05 |
| 13853 | 693 | F693M | B | 1e-05 | 1e-05 | 1.3e-05 | 1.100E-05 | 1.732E-06 | 1.000E-05 | 1.300E-05 |
| 13854 | 693 | F693F | B | 7e-06 | 1.6e-05 | 1.4e-05 | 1.233E-05 | 4.726E-06 | 7.000E-06 | 1.600E-05 |

| | | | | | | | | | | |
|---|---|---|---|---|---|---|---|---|---|---|
| 13855 | 693 | F693P | B | 8.1e-06 | 1.5e-05 | 9e-06 | 1.070E-05 | 3.751E-06 | 8.100E-06 | 1.500E-05 |
| 13856 | 693 | F693S | B | 1.6e-05 | 1.5e-05 | 1.1e-05 | 1.400E-05 | 2.646E-06 | 1.100E-05 | 1.600E-05 |
| 13857 | 693 | F693T | B | 1.6e-05 | 2.4e-05 | 1.1e-05 | 1.700E-05 | 6.557E-06 | 1.100E-05 | 2.400E-05 |
| 13858 | 693 | F693W | B | 1.4e-05 | 1.2e-05 | 1.1e-05 | 1.233E-05 | 1.528E-06 | 1.100E-05 | 1.400E-05 |
| 13859 | 693 | F693Y | B | 1.2e-05 | 1.2e-05 | 1.6e-05 | 1.333E-05 | 2.309E-06 | 1.200E-05 | 1.600E-05 |
| 13860 | 693 | F693V | B | 9.9e-06 | 1.6e-05 | 8.5e-06 | 1.147E-05 | 3.988E-06 | 8.500E-06 | 1.600E-05 |
| 13861 | 694 | T694A | B | 1.4e-05 | 9.6e-06 | 1.6e-05 | 1.320E-05 | 3.274E-06 | 9.600E-06 | 1.600E-05 |
| 13862 | 694 | T694R | B | 1.6e-05 | 1e-05 | 1.3e-05 | 1.300E-05 | 3.000E-06 | 1.000E-05 | 1.600E-05 |
| 13863 | 694 | T694N | B | 1.4e-05 | 1.1e-05 | 1.3e-05 | 1.267E-05 | 1.528E-06 | 1.100E-05 | 1.400E-05 |
| 13864 | 694 | T694D | B | 9.2e-06 | 8.8e-06 | 9.6e-06 | 9.200E-06 | 4.000E-07 | 8.800E-06 | 9.600E-06 |
| 13865 | 694 | T694C | B | 1.4e-05 | 1e-05 | 1e-05 | 1.133E-05 | 2.309E-06 | 1.000E-05 | 1.400E-05 |
| 13866 | 694 | T694E | B | 1.6e-05 | 1.2e-05 | 1.6e-05 | 1.467E-05 | 2.309E-06 | 1.200E-05 | 1.600E-05 |
| 13867 | 694 | T694Q | B | 1.2e-05 | 9.4e-06 | 1e-05 | 1.047E-05 | 1.361E-06 | 9.400E-06 | 1.200E-05 |
| 13868 | 694 | T694G | B | 1.6e-05 | 1.5e-05 | 1.8e-05 | 1.633E-05 | 1.528E-06 | 1.500E-05 | 1.800E-05 |
| 13869 | 694 | T694H | B | 1.5e-05 | 1.6e-05 | 1.1e-05 | 1.400E-05 | 2.646E-06 | 1.100E-05 | 1.600E-05 |
| 13870 | 694 | T694I | B | 1.5e-05 | 1.5e-05 | 1.2e-05 | 1.400E-05 | 1.732E-06 | 1.200E-05 | 1.500E-05 |
| 13871 | 694 | T694L | B | 1.6e-05 | 1.1e-05 | 1.2e-05 | 1.300E-05 | 2.646E-06 | 1.100E-05 | 1.600E-05 |
| 13872 | 694 | T694K | B | 1.7e-05 | 1.1e-05 | 1.4e-05 | 1.400E-05 | 3.000E-06 | 1.100E-05 | 1.700E-05 |
| 13873 | 694 | T694M | B | 1.5e-05 | 1.6e-05 | 1.7e-05 | 1.600E-05 | 1.000E-06 | 1.500E-05 | 1.700E-05 |
| 13874 | 694 | T694F | B | 1e-05 | 1.5e-05 | 9.7e-06 | 1.157E-05 | 2.977E-06 | 9.700E-06 | 1.500E-05 |
| 13875 | 694 | T694P | B | 1.1e-05 | 1.2e-05 | 1.7e-05 | 1.333E-05 | 3.215E-06 | 1.100E-05 | 1.700E-05 |
| 13876 | 694 | T694S | B | 1.5e-05 | 9.9e-06 | 1.5e-05 | 1.330E-05 | 2.944E-06 | 9.900E-06 | 1.500E-05 |
| 13877 | 694 | T694T | B | 1.7e-05 | 9.9e-06 | 1.5e-05 | 1.397E-05 | 3.661E-06 | 9.900E-06 | 1.700E-05 |
| 13878 | 694 | T694W | B | 9.2e-06 | 1.4e-05 | 1.1e-05 | 1.140E-05 | 2.425E-06 | 9.200E-06 | 1.400E-05 |
| 13879 | 694 | T694Y | B | 1.6e-05 | 2.2e-05 | 9.9e-06 | 1.597E-05 | 6.050E-06 | 9.900E-06 | 2.200E-05 |
| 13880 | 694 | T694V | B | 1e-05 | 7.7e-06 | 1.1e-05 | 9.567E-06 | 1.692E-06 | 7.700E-06 | 1.100E-05 |
| 13881 | 695 | L695A | B | 1.4e-05 | 1e-05 | 1.5e-05 | 1.300E-05 | 2.646E-06 | 1.000E-05 | 1.500E-05 |
| 13882 | 695 | L695R | B | 9.3e-06 | 1.2e-05 | 1e-05 | 1.043E-05 | 1.401E-06 | 9.300E-06 | 1.200E-05 |
| 13883 | 695 | L695N | B | 1.5e-05 | 1.5e-05 | 1.5e-05 | 1.500E-05 | 0.000E+00 | 1.500E-05 | 1.500E-05 |
| 13884 | 695 | L695D | B | 1.5e-05 | 1.1e-05 | 1.5e-05 | 1.367E-05 | 2.309E-06 | 1.100E-05 | 1.500E-05 |
| 13885 | 695 | L695C | B | 1.3e-05 | 1.5e-05 | 1.1e-05 | 1.300E-05 | 2.000E-06 | 1.100E-05 | 1.500E-05 |
| 13886 | 695 | L695E | B | 1.7e-05 | 8.9e-06 | 1.5e-05 | 1.363E-05 | 4.219E-06 | 8.900E-06 | 1.700E-05 |
| 13887 | 695 | L695Q | B | 9.4e-06 | 1.1e-05 | 9.8e-06 | 1.007E-05 | 8.327E-07 | 9.400E-06 | 1.100E-05 |
| 13888 | 695 | L695G | B | 1.6e-05 | 1.1e-05 | 1.4e-05 | 1.367E-05 | 2.517E-06 | 1.100E-05 | 1.600E-05 |
| 13889 | 695 | L695H | B | 1.4e-05 | 1.3e-05 | 1.1e-05 | 1.267E-05 | 1.528E-06 | 1.100E-05 | 1.400E-05 |
| 13890 | 695 | L695I | B | 1.5e-05 | 1e-05 | 1.2e-05 | 1.233E-05 | 2.517E-06 | 1.000E-05 | 1.500E-05 |
| 13891 | 695 | L695L | B | 1e-05 | 7.9e-06 | 1.6e-05 | 1.130E-05 | 4.204E-06 | 7.900E-06 | 1.600E-05 |
| 13892 | 695 | L695K | B | 1.6e-05 | 6.9e-06 | 1.6e-05 | 1.297E-05 | 5.254E-06 | 6.900E-06 | 1.600E-05 |
| 13893 | 695 | L695M | B | 1.3e-05 | 7.8e-06 | 1.5e-05 | 1.193E-05 | 3.717E-06 | 7.800E-06 | 1.500E-05 |
| 13894 | 695 | L695F | B | 1.4e-05 | 1.4e-05 | 1.8e-05 | 1.533E-05 | 2.309E-06 | 1.400E-05 | 1.800E-05 |
| 13895 | 695 | L695P | B | 1.4e-05 | 1e-05 | 1.6e-05 | 1.333E-05 | 3.055E-06 | 1.000E-05 | 1.600E-05 |
| 13896 | 695 | L695S | B | 1.6e-05 | 1.5e-05 | 1.4e-05 | 1.500E-05 | 1.000E-06 | 1.400E-05 | 1.600E-05 |
| 13897 | 695 | L695T | B | 1e-05 | 1.5e-05 | 1.5e-05 | 1.333E-05 | 2.887E-06 | 1.000E-05 | 1.500E-05 |
| 13898 | 695 | L695W | B | 9.8e-06 | 8.1e-06 | 1.1e-05 | 9.633E-06 | 1.457E-06 | 8.100E-06 | 1.100E-05 |
| 13899 | 695 | L695Y | B | 1.1e-05 | 1.5e-05 | 1.5e-05 | 1.367E-05 | 2.309E-06 | 1.100E-05 | 1.500E-05 |
| 13900 | 695 | L695V | B | 1.5e-05 | 1e-05 | 1.5e-05 | 1.333E-05 | 2.887E-06 | 1.000E-05 | 1.500E-05 |
| 13901 | 696 | T696A | B | 1.2e-05 | 1.5e-05 | 1.7e-05 | 1.467E-05 | 2.517E-06 | 1.200E-05 | 1.700E-05 |
| 13902 | 696 | T696R | B | 1.3e-05 | 1.5e-05 | 1.2e-05 | 1.333E-05 | 1.528E-06 | 1.200E-05 | 1.500E-05 |
| 13903 | 696 | T696N | B | 9.9e-06 | 1.3e-05 | 1.1e-05 | 1.130E-05 | 1.572E-06 | 9.900E-06 | 1.300E-05 |
| 13904 | 696 | T696D | B | 1.1e-05 | 9.1e-06 | 1.1e-05 | 1.037E-05 | 1.097E-06 | 9.100E-06 | 1.100E-05 |
| 13905 | 696 | T696C | B | 1.7e-05 | 1e-05 | 9.7e-06 | 1.223E-05 | 4.131E-06 | 9.700E-06 | 1.700E-05 |
| 13906 | 696 | T696E | B | 1.6e-05 | 1.7e-05 | 1.1e-05 | 1.467E-05 | 3.215E-06 | 1.100E-05 | 1.700E-05 |
| 13907 | 696 | T696Q | B | 1.5e-05 | 1e-05 | 1.6e-05 | 1.367E-05 | 3.215E-06 | 1.000E-05 | 1.600E-05 |
| 13908 | 696 | T696G | B | 1.1e-05 | 9.7e-06 | 1.3e-05 | 1.123E-05 | 1.662E-06 | 9.700E-06 | 1.300E-05 |
| 13909 | 696 | T696H | B | 1.3e-05 | 1e-05 | 1.4e-05 | 1.233E-05 | 2.082E-06 | 1.000E-05 | 1.400E-05 |
| 13910 | 696 | T696I | B | 1e-05 | 1.5e-05 | 1.5e-05 | 1.333E-05 | 2.887E-06 | 1.000E-05 | 1.500E-05 |
| 13911 | 696 | T696L | B | 1.4e-05 | 1.5e-05 | 8.9e-06 | 1.263E-05 | 3.272E-06 | 8.900E-06 | 1.500E-05 |
| 13912 | 696 | T696K | B | 1.6e-05 | 1.7e-05 | 1.5e-05 | 1.600E-05 | 1.000E-06 | 1.500E-05 | 1.700E-05 |
| 13913 | 696 | T696M | B | 1.4e-05 | 1.5e-05 | 1e-05 | 1.300E-05 | 2.646E-06 | 1.000E-05 | 1.500E-05 |
| 13914 | 696 | T696F | B | 1.5e-05 | 1.4e-05 | 1.5e-05 | 1.467E-05 | 5.774E-07 | 1.400E-05 | 1.500E-05 |
| 13915 | 696 | T696P | B | 1.1e-05 | 1.4e-05 | 8.3e-06 | 1.110E-05 | 2.851E-06 | 8.300E-06 | 1.400E-05 |
| 13916 | 696 | T696S | B | 1.7e-05 | 1.3e-05 | 1.2e-05 | 1.400E-05 | 2.646E-06 | 1.200E-05 | 1.700E-05 |
| 13917 | 696 | T696T | B | 1.3e-05 | 9.8e-06 | 1.1e-05 | 1.127E-05 | 1.617E-06 | 9.800E-06 | 1.300E-05 |
| 13918 | 696 | T696W | B | 8.7e-06 | 1.7e-05 | 1.4e-05 | 1.323E-05 | 4.203E-06 | 8.700E-06 | 1.700E-05 |
| 13919 | 696 | T696Y | B | 8.5e-06 | 9.3e-06 | 1.4e-05 | 1.060E-05 | 2.972E-06 | 8.500E-06 | 1.400E-05 |
| 13920 | 696 | T696V | B | 1.2e-05 | 1.6e-05 | 1.4e-05 | 1.400E-05 | 2.000E-06 | 1.200E-05 | 1.600E-05 |
| 13921 | 697 | I697A | B | 1.6e-05 | 9.8e-06 | 1.1e-05 | 1.227E-05 | 3.288E-06 | 9.800E-06 | 1.600E-05 |
| 13922 | 697 | I697R | B | 1.4e-05 | 1.5e-05 | 1.5e-05 | 1.467E-05 | 5.774E-07 | 1.400E-05 | 1.500E-05 |
| 13923 | 697 | I697N | B | 1.1e-05 | 1.5e-05 | 1e-05 | 1.200E-05 | 2.646E-06 | 1.000E-05 | 1.500E-05 |
| 13924 | 697 | I697D | B | 1.1e-05 | 1.7e-05 | 1.5e-05 | 1.433E-05 | 3.055E-06 | 1.100E-05 | 1.700E-05 |
| 13925 | 697 | I697C | B | 1.4e-05 | 1.6e-05 | 1e-05 | 1.333E-05 | 3.055E-06 | 1.000E-05 | 1.600E-05 |
| 13926 | 697 | I697E | B | 1.4e-05 | 1.5e-05 | 1e-05 | 1.300E-05 | 2.646E-06 | 1.000E-05 | 1.500E-05 |
| 13927 | 697 | I697Q | B | 1.5e-05 | 1.4e-05 | 1.4e-05 | 1.433E-05 | 5.774E-07 | 1.400E-05 | 1.500E-05 |
| 13928 | 697 | I697G | B | 9.9e-06 | 1.5e-05 | 1.1e-05 | 1.197E-05 | 2.684E-06 | 9.900E-06 | 1.500E-05 |
| 13929 | 697 | I697H | B | 8.8e-06 | 1.5e-05 | 1.3e-05 | 1.227E-05 | 3.164E-06 | 8.800E-06 | 1.500E-05 |
| 13930 | 697 | I697I | B | 1e-05 | 9.8e-06 | 1.1e-05 | 1.027E-05 | 6.429E-07 | 9.800E-06 | 1.100E-05 |
| 13931 | 697 | I697L | B | 2.4e-05 | 1.6e-05 | 1.1e-05 | 1.700E-05 | 6.557E-06 | 1.100E-05 | 2.400E-05 |
| 13932 | 697 | I697K | B | 1e-05 | 8.2e-06 | 1.4e-05 | 1.073E-05 | 2.969E-06 | 8.200E-06 | 1.400E-05 |
| 13933 | 697 | I697M | B | 1.9e-05 | 1.6e-05 | 1.5e-05 | 1.667E-05 | 2.082E-06 | 1.500E-05 | 1.900E-05 |
| 13934 | 697 | I697F | B | 1.1e-05 | 1.6e-05 | 1.5e-05 | 1.400E-05 | 2.646E-06 | 1.100E-05 | 1.600E-05 |
| 13935 | 697 | I697P | B | 1.1e-05 | 1.6e-05 | 1e-05 | 1.233E-05 | 3.215E-06 | 1.000E-05 | 1.600E-05 |
| 13936 | 697 | I697S | B | 1.4e-05 | 1e-05 | 1.4e-05 | 1.267E-05 | 2.309E-06 | 1.000E-05 | 1.400E-05 |
| 13937 | 697 | I697T | B | 1.1e-05 | 1.3e-05 | 1.3e-05 | 1.233E-05 | 1.155E-06 | 1.100E-05 | 1.300E-05 |
| 13938 | 697 | I697W | B | 9.1e-06 | 1.7e-05 | 1.5e-05 | 1.370E-05 | 4.107E-06 | 9.100E-06 | 1.700E-05 |
| 13939 | 697 | I697Y | B | 8.6e-06 | 1e-05 | 1.3e-05 | 1.053E-05 | 2.248E-06 | 8.600E-06 | 1.300E-05 |

| | | | | | | | | | | |
|---|---|---|---|---|---|---|---|---|---|---|
| 13940 | 697 | I697V | B | 1.4e-05 | 1.3e-05 | 1.5e-05 | 1.400E-05 | 1.000E-06 | 1.300E-05 | 1.500E-05 |
| 13941 | 698 | S698A | B | 1.2e-05 | 1.7e-05 | 1.4e-05 | 1.433E-05 | 2.517E-06 | 1.200E-05 | 1.700E-05 |
| 13942 | 698 | S698R | B | 1.5e-05 | 1.6e-05 | 1e-05 | 1.367E-05 | 3.215E-06 | 1.000E-05 | 1.600E-05 |
| 13943 | 698 | S698N | B | 1.1e-05 | 1.6e-05 | 1.3e-05 | 1.333E-05 | 2.517E-06 | 1.100E-05 | 1.600E-05 |
| 13944 | 698 | S698D | B | 1.6e-05 | 1.6e-05 | 1.2e-05 | 1.467E-05 | 2.309E-06 | 1.200E-05 | 1.600E-05 |
| 13945 | 698 | S698C | B | 1.6e-05 | 1e-05 | 1.2e-05 | 1.267E-05 | 3.055E-06 | 1.000E-05 | 1.600E-05 |
| 13946 | 698 | S698E | B | 1.5e-05 | 1.5e-05 | 1e-05 | 1.333E-05 | 2.887E-06 | 1.000E-05 | 1.500E-05 |
| 13947 | 698 | S698Q | B | 1.5e-05 | 8.4e-06 | 1.4e-05 | 1.247E-05 | 3.557E-06 | 8.400E-06 | 1.500E-05 |
| 13948 | 698 | S698G | B | 1.4e-05 | 1.4e-05 | 1.2e-05 | 1.333E-05 | 1.155E-06 | 1.200E-05 | 1.400E-05 |
| 13949 | 698 | S698H | B | 9.3e-06 | 1.2e-05 | 1.8e-05 | 1.310E-05 | 4.453E-06 | 9.300E-06 | 1.800E-05 |
| 13950 | 698 | S698I | B | 1.5e-05 | 1.1e-05 | 1.3e-05 | 1.300E-05 | 2.000E-06 | 1.100E-05 | 1.500E-05 |
| 13951 | 698 | S698L | B | 1.4e-05 | 1.5e-05 | 1.6e-05 | 1.500E-05 | 1.000E-06 | 1.400E-05 | 1.600E-05 |
| 13952 | 698 | S698K | B | 1.1e-05 | 1.1e-05 | 1.1e-05 | 1.100E-05 | 2.075E-21 | 1.100E-05 | 1.100E-05 |
| 13953 | 698 | S698M | B | 1.5e-05 | 9.1e-06 | 1e-05 | 1.137E-05 | 3.179E-06 | 9.100E-06 | 1.500E-05 |
| 13954 | 698 | S698F | B | 1.4e-05 | 1.1e-05 | 1.4e-05 | 1.300E-05 | 1.732E-06 | 1.100E-05 | 1.400E-05 |
| 13955 | 698 | S698P | B | 1.5e-05 | 1.1e-05 | 1e-05 | 1.200E-05 | 2.646E-06 | 1.000E-05 | 1.500E-05 |
| 13956 | 698 | S698S | B | 1.1e-05 | 1.4e-05 | 1.7e-05 | 1.400E-05 | 3.000E-06 | 1.100E-05 | 1.700E-05 |
| 13957 | 698 | S698T | B | 9.1e-06 | 1.3e-05 | 1.6e-05 | 1.270E-05 | 3.460E-06 | 9.100E-06 | 1.600E-05 |
| 13958 | 698 | S698W | B | 1.1e-05 | 1.6e-05 | 1.5e-05 | 1.400E-05 | 2.646E-06 | 1.100E-05 | 1.600E-05 |
| 13959 | 698 | S698Y | B | 1.5e-05 | 1e-05 | 1.8e-05 | 1.433E-05 | 4.041E-06 | 1.000E-05 | 1.800E-05 |
| 13960 | 698 | S698V | B | 1.5e-05 | 1e-05 | 1.2e-05 | 1.233E-05 | 2.517E-06 | 1.000E-05 | 1.500E-05 |
| 13961 | 699 | S699A | B | 1.2e-05 | 1.2e-05 | 1e-05 | 1.133E-05 | 1.155E-06 | 1.000E-05 | 1.200E-05 |
| 13962 | 699 | S699R | B | 1.3e-05 | 1e-05 | 1.6e-05 | 1.300E-05 | 3.000E-06 | 1.000E-05 | 1.600E-05 |
| 13963 | 699 | S699N | B | 1.2e-05 | 1.3e-05 | 1.1e-05 | 1.200E-05 | 1.000E-06 | 1.100E-05 | 1.300E-05 |
| 13964 | 699 | S699D | B | 1.5e-05 | 1.7e-05 | 1e-05 | 1.400E-05 | 3.606E-06 | 1.000E-05 | 1.700E-05 |
| 13965 | 699 | S699C | B | 1.2e-05 | 1.6e-05 | 7.2e-06 | 1.173E-05 | 4.406E-06 | 7.200E-06 | 1.600E-05 |
| 13966 | 699 | S699E | B | 1.2e-05 | 1.1e-05 | 1.4e-05 | 1.233E-05 | 1.528E-06 | 1.100E-05 | 1.400E-05 |
| 13967 | 699 | S699Q | B | 1.3e-05 | 1.7e-05 | 1e-05 | 1.333E-05 | 3.512E-06 | 1.000E-05 | 1.700E-05 |
| 13968 | 699 | S699G | B | 1.6e-05 | 1.4e-05 | 1.1e-05 | 1.367E-05 | 2.517E-06 | 1.100E-05 | 1.600E-05 |
| 13969 | 699 | S699H | B | 1.1e-05 | 1.6e-05 | 1.1e-05 | 1.267E-05 | 2.887E-06 | 1.100E-05 | 1.600E-05 |
| 13970 | 699 | S699I | B | 1.4e-05 | 1.4e-05 | 1e-05 | 1.267E-05 | 2.309E-06 | 1.000E-05 | 1.400E-05 |
| 13971 | 699 | S699L | B | 1.6e-05 | 1.5e-05 | 1.3e-05 | 1.467E-05 | 1.528E-06 | 1.300E-05 | 1.600E-05 |
| 13972 | 699 | S699K | B | 1e-05 | 1.6e-05 | 1.1e-05 | 1.233E-05 | 3.215E-06 | 1.000E-05 | 1.600E-05 |
| 13973 | 699 | S699M | B | 1.3e-05 | 1e-05 | 8.8e-06 | 1.060E-05 | 2.163E-06 | 8.800E-06 | 1.300E-05 |
| 13974 | 699 | S699F | B | 1.1e-05 | 9.1e-06 | 1.7e-05 | 1.237E-05 | 4.124E-06 | 9.100E-06 | 1.700E-05 |
| 13975 | 699 | S699P | B | 1.1e-05 | 1.8e-05 | 9.5e-06 | 1.283E-05 | 4.537E-06 | 9.500E-06 | 1.800E-05 |
| 13976 | 699 | S699S | B | 1.1e-05 | 1e-05 | 9.7e-06 | 1.023E-05 | 6.807E-07 | 9.700E-06 | 1.100E-05 |
| 13977 | 699 | S699T | B | 1.4e-05 | 1.4e-05 | 8.8e-06 | 1.227E-05 | 3.002E-06 | 8.800E-06 | 1.400E-05 |
| 13978 | 699 | S699W | B | 1.3e-05 | 1.6e-05 | 1.5e-05 | 1.467E-05 | 1.528E-06 | 1.300E-05 | 1.600E-05 |
| 13979 | 699 | S699Y | B | 6.7e-06 | 9.7e-06 | 1.2e-05 | 9.467E-06 | 2.658E-06 | 6.700E-06 | 1.200E-05 |
| 13980 | 699 | S699V | B | 1.6e-05 | 1.4e-05 | 1.5e-05 | 1.500E-05 | 1.000E-06 | 1.400E-05 | 1.600E-05 |
| 13981 | 700 | L700A | B | 1.7e-05 | 1.8e-05 | 1.1e-05 | 1.533E-05 | 3.786E-06 | 1.100E-05 | 1.800E-05 |
| 13982 | 700 | L700R | B | 1.1e-05 | 1.5e-05 | 1.1e-05 | 1.233E-05 | 2.309E-06 | 1.100E-05 | 1.500E-05 |
| 13983 | 700 | L700N | B | 1e-05 | 1.6e-05 | 8.7e-06 | 1.157E-05 | 3.894E-06 | 8.700E-06 | 1.600E-05 |
| 13984 | 700 | L700D | B | 1.5e-05 | 1.4e-05 | 1.5e-05 | 1.467E-05 | 5.774E-07 | 1.400E-05 | 1.500E-05 |
| 13985 | 700 | L700C | B | 1.5e-05 | 1e-05 | 8.8e-06 | 1.127E-05 | 3.288E-06 | 8.800E-06 | 1.500E-05 |
| 13986 | 700 | L700E | B | 9.8e-06 | 1.8e-05 | 1.5e-05 | 1.427E-05 | 4.149E-06 | 9.800E-06 | 1.800E-05 |
| 13987 | 700 | L700Q | B | 1.2e-05 | 1.6e-05 | 1.1e-05 | 1.300E-05 | 2.646E-06 | 1.100E-05 | 1.600E-05 |
| 13988 | 700 | L700G | B | 1.6e-05 | 1.6e-05 | 1.1e-05 | 1.433E-05 | 2.887E-06 | 1.100E-05 | 1.600E-05 |
| 13989 | 700 | L700H | B | 1.3e-05 | 1.1e-05 | 1.1e-05 | 1.167E-05 | 1.155E-06 | 1.100E-05 | 1.300E-05 |
| 13990 | 700 | L700I | B | 1.2e-05 | 1e-05 | 1.3e-05 | 1.167E-05 | 1.528E-06 | 1.000E-05 | 1.300E-05 |
| 13991 | 700 | L700L | B | 1.4e-05 | 9.7e-06 | 1.1e-05 | 1.157E-05 | 2.205E-06 | 9.700E-06 | 1.400E-05 |
| 13992 | 700 | L700K | B | 1.5e-05 | 9.7e-06 | 1.1e-05 | 1.190E-05 | 2.762E-06 | 9.700E-06 | 1.500E-05 |
| 13993 | 700 | L700M | B | 1.1e-05 | 1.5e-05 | 1.1e-05 | 1.233E-05 | 2.309E-06 | 1.100E-05 | 1.500E-05 |
| 13994 | 700 | L700F | B | 8.2e-06 | 1.3e-05 | 1.2e-05 | 1.107E-05 | 2.532E-06 | 8.200E-06 | 1.300E-05 |
| 13995 | 700 | L700P | B | 9.5e-06 | 9.8e-06 | 1.1e-05 | 1.010E-05 | 7.937E-07 | 9.500E-06 | 1.100E-05 |
| 13996 | 700 | L700S | B | 1.1e-05 | 9.7e-06 | 1e-05 | 1.023E-05 | 6.807E-07 | 9.700E-06 | 1.100E-05 |
| 13997 | 700 | L700T | B | 1.1e-05 | 1.6e-05 | 1.4e-05 | 1.367E-05 | 2.517E-06 | 1.100E-05 | 1.600E-05 |
| 13998 | 700 | L700W | B | 1.1e-05 | 1.5e-05 | 9.8e-06 | 1.193E-05 | 2.723E-06 | 9.800E-06 | 1.500E-05 |
| 13999 | 700 | L700Y | B | 1.5e-05 | 1.1e-05 | 1.2e-05 | 1.267E-05 | 2.082E-06 | 1.100E-05 | 1.500E-05 |
| 14000 | 700 | L700V | B | 1.3e-05 | 1.4e-05 | 9.8e-06 | 1.227E-05 | 2.194E-06 | 9.800E-06 | 1.400E-05 |
| 14001 | 701 | Q701A | B | 1.5e-05 | 1.6e-05 | 1.1e-05 | 1.400E-05 | 2.646E-06 | 1.100E-05 | 1.600E-05 |
| 14002 | 701 | Q701R | B | 1.3e-05 | 1.2e-05 | 1.2e-05 | 1.233E-05 | 5.774E-07 | 1.200E-05 | 1.300E-05 |
| 14003 | 701 | Q701N | B | 1.4e-05 | 9.9e-06 | 8.8e-06 | 1.090E-05 | 2.740E-06 | 8.800E-06 | 1.400E-05 |
| 14004 | 701 | Q701D | B | 1.7e-05 | 1.6e-05 | 1.5e-05 | 1.600E-05 | 1.000E-06 | 1.500E-05 | 1.700E-05 |
| 14005 | 701 | Q701C | B | 1.1e-05 | 1.5e-05 | 1.3e-05 | 1.300E-05 | 2.000E-06 | 1.100E-05 | 1.500E-05 |
| 14006 | 701 | Q701E | B | 1.1e-05 | 1.6e-05 | 1.2e-05 | 1.300E-05 | 2.646E-06 | 1.100E-05 | 1.600E-05 |
| 14007 | 701 | Q701Q | B | 8.2e-06 | 1.4e-05 | 2e-05 | 1.407E-05 | 5.900E-06 | 8.200E-06 | 2.000E-05 |
| 14008 | 701 | Q701G | B | 1.1e-05 | 1.8e-05 | 1.4e-05 | 1.433E-05 | 3.512E-06 | 1.100E-05 | 1.800E-05 |
| 14009 | 701 | Q701H | B | 9.4e-06 | 1.8e-05 | 1.4e-05 | 1.380E-05 | 4.303E-06 | 9.400E-06 | 1.800E-05 |
| 14010 | 701 | Q701I | B | 1e-05 | 1.4e-05 | 1.2e-05 | 1.200E-05 | 2.000E-06 | 1.000E-05 | 1.400E-05 |
| 14011 | 701 | Q701L | B | 1.6e-05 | 1.2e-05 | 1e-05 | 1.267E-05 | 3.055E-06 | 1.000E-05 | 1.600E-05 |
| 14012 | 701 | Q701K | B | 1.5e-05 | 1.7e-05 | 1.2e-05 | 1.467E-05 | 2.517E-06 | 1.200E-05 | 1.700E-05 |
| 14013 | 701 | Q701M | B | 7.4e-06 | 9.6e-06 | 1.4e-05 | 1.033E-05 | 3.361E-06 | 7.400E-06 | 1.400E-05 |
| 14014 | 701 | Q701F | B | 1.6e-05 | 7.5e-06 | 1e-05 | 1.117E-05 | 4.368E-06 | 7.500E-06 | 1.600E-05 |
| 14015 | 701 | Q701P | B | 1.2e-05 | 1.1e-05 | 1e-05 | 1.100E-05 | 1.000E-06 | 1.000E-05 | 1.200E-05 |
| 14016 | 701 | Q701S | B | 1.6e-05 | 1.5e-05 | 1.3e-05 | 1.467E-05 | 1.528E-06 | 1.300E-05 | 1.600E-05 |
| 14017 | 701 | Q701T | B | 1.6e-05 | 1e-05 | 1.2e-05 | 1.267E-05 | 3.055E-06 | 1.000E-05 | 1.600E-05 |
| 14018 | 701 | Q701W | B | 1.5e-05 | 1.5e-05 | 9.5e-06 | 1.317E-05 | 3.175E-06 | 9.500E-06 | 1.500E-05 |
| 14019 | 701 | Q701Y | B | 1.4e-05 | 1.6e-05 | 1.1e-05 | 1.367E-05 | 2.517E-06 | 1.100E-05 | 1.600E-05 |
| 14020 | 701 | Q701V | B | 1.1e-05 | 1.7e-05 | 1.2e-05 | 1.333E-05 | 3.215E-06 | 1.100E-05 | 1.700E-05 |
| 14021 | 702 | P702A | B | 1.4e-05 | 1.1e-05 | 1.4e-05 | 1.300E-05 | 1.732E-06 | 1.100E-05 | 1.400E-05 |
| 14022 | 702 | P702R | B | 1.5e-05 | 1.1e-05 | 1.4e-05 | 1.333E-05 | 2.082E-06 | 1.100E-05 | 1.500E-05 |
| 14023 | 702 | P702N | B | 1.1e-05 | 9.9e-06 | 1.6e-05 | 1.230E-05 | 3.251E-06 | 9.900E-06 | 1.600E-05 |
| 14024 | 702 | P702D | B | 1.6e-05 | 1.6e-05 | 1.3e-05 | 1.500E-05 | 1.732E-06 | 1.300E-05 | 1.600E-05 |

| | | | | | | | | | | |
|---|---|---|---|---|---|---|---|---|---|---|
| 14025 | 702 | P702C | B | 1.5e-05 | 1.5e-05 | 1.1e-05 | 1.367E-05 | 2.309E-06 | 1.100E-05 | 1.500E-05 |
| 14026 | 702 | P702E | B | 8.4e-06 | 1.5e-05 | 1.2e-05 | 1.180E-05 | 3.305E-06 | 8.400E-06 | 1.500E-05 |
| 14027 | 702 | P702Q | B | 1.5e-05 | 8.7e-06 | 1.5e-05 | 1.290E-05 | 3.637E-06 | 8.700E-06 | 1.500E-05 |
| 14028 | 702 | P702G | B | 1.8e-05 | 1.4e-05 | 1e-05 | 1.400E-05 | 4.000E-06 | 1.000E-05 | 1.800E-05 |
| 14029 | 702 | P702H | B | 1.7e-05 | 1.2e-05 | 1.7e-05 | 1.533E-05 | 2.887E-06 | 1.200E-05 | 1.700E-05 |
| 14030 | 702 | P702I | B | 1.6e-05 | 1.1e-05 | 1e-05 | 1.233E-05 | 3.215E-06 | 1.000E-05 | 1.600E-05 |
| 14031 | 702 | P702L | B | 1.6e-05 | 1.5e-05 | 1.5e-05 | 1.533E-05 | 5.774E-07 | 1.500E-05 | 1.600E-05 |
| 14032 | 702 | P702K | B | 1.1e-05 | 1.6e-05 | 1.3e-05 | 1.333E-05 | 2.517E-06 | 1.100E-05 | 1.600E-05 |
| 14033 | 702 | P702M | B | 1.2e-05 | 1.5e-05 | 1.6e-05 | 1.433E-05 | 2.082E-06 | 1.200E-05 | 1.600E-05 |
| 14034 | 702 | P702F | B | 1.5e-05 | 1.1e-05 | 9.2e-06 | 1.173E-05 | 2.969E-06 | 9.200E-06 | 1.500E-05 |
| 14035 | 702 | P702P | B | 1.1e-05 | 1e-05 | 8.3e-06 | 9.767E-06 | 1.365E-06 | 8.300E-06 | 1.100E-05 |
| 14036 | 702 | P702S | B | 1.1e-05 | 1.6e-05 | 1.1e-05 | 1.267E-05 | 2.887E-06 | 1.100E-05 | 1.600E-05 |
| 14037 | 702 | P702T | B | 1e-05 | 9.6e-06 | 1.5e-05 | 1.153E-05 | 3.009E-06 | 9.600E-06 | 1.500E-05 |
| 14038 | 702 | P702W | B | 1.6e-05 | 1.1e-05 | 1.6e-05 | 1.433E-05 | 2.887E-06 | 1.100E-05 | 1.600E-05 |
| 14039 | 702 | P702Y | B | 1.5e-05 | 1.4e-05 | 1e-05 | 1.300E-05 | 2.646E-06 | 1.000E-05 | 1.500E-05 |
| 14040 | 702 | P702V | B | 1.1e-05 | 1.4e-05 | 9.9e-06 | 1.163E-05 | 2.122E-06 | 9.900E-06 | 1.400E-05 |
| 14041 | 703 | E703A | B | 1e-05 | 1.1e-05 | 1.2e-05 | 1.100E-05 | 1.000E-06 | 1.000E-05 | 1.200E-05 |
| 14042 | 703 | E703R | B | 1.1e-05 | 1.1e-05 | 1.8e-05 | 1.333E-05 | 4.041E-06 | 1.100E-05 | 1.800E-05 |
| 14043 | 703 | E703N | B | 9.5e-06 | 1.4e-05 | 1.2e-05 | 1.183E-05 | 2.255E-06 | 9.500E-06 | 1.400E-05 |
| 14044 | 703 | E703D | B | 1.1e-05 | 1.1e-05 | 1.5e-05 | 1.233E-05 | 2.309E-06 | 1.100E-05 | 1.500E-05 |
| 14045 | 703 | E703C | B | 1.5e-05 | 1.4e-05 | 9e-06 | 1.267E-05 | 3.215E-06 | 9.000E-06 | 1.500E-05 |
| 14046 | 703 | E703E | B | 1.2e-05 | 7e-06 | 9.2e-06 | 9.400E-06 | 2.506E-06 | 7.000E-06 | 1.200E-05 |
| 14047 | 703 | E703Q | B | 8.6e-06 | 1.1e-05 | 1.5e-05 | 1.153E-05 | 3.233E-06 | 8.600E-06 | 1.500E-05 |
| 14048 | 703 | E703G | B | 1.5e-05 | 1.6e-05 | 1.5e-05 | 1.533E-05 | 5.774E-07 | 1.500E-05 | 1.600E-05 |
| 14049 | 703 | E703H | B | 1.4e-05 | 1.4e-05 | 1.5e-05 | 1.433E-05 | 5.774E-07 | 1.400E-05 | 1.500E-05 |
| 14050 | 703 | E703I | B | 1.2e-05 | 9.2e-06 | 8.6e-06 | 9.933E-06 | 1.815E-06 | 8.600E-06 | 1.200E-05 |
| 14051 | 703 | E703L | B | 1.5e-05 | 1e-05 | 1.4e-05 | 1.300E-05 | 2.646E-06 | 1.000E-05 | 1.500E-05 |
| 14052 | 703 | E703K | B | 1.5e-05 | 1.5e-05 | 1.1e-05 | 1.367E-05 | 2.309E-06 | 1.100E-05 | 1.500E-05 |
| 14053 | 703 | E703M | B | 1.6e-05 | 1.6e-05 | 1.3e-05 | 1.500E-05 | 1.732E-06 | 1.300E-05 | 1.600E-05 |
| 14054 | 703 | E703F | B | 1.7e-05 | 1.3e-05 | 1.5e-05 | 1.500E-05 | 2.000E-06 | 1.300E-05 | 1.700E-05 |
| 14055 | 703 | E703P | B | 9.3e-06 | 8.7e-06 | 1.1e-05 | 9.667E-06 | 1.193E-06 | 8.700E-06 | 1.100E-05 |
| 14056 | 703 | E703S | B | 1.6e-05 | 1.1e-05 | 1.3e-05 | 1.333E-05 | 2.517E-06 | 1.100E-05 | 1.600E-05 |
| 14057 | 703 | E703T | B | 1.5e-05 | 9.3e-06 | 7.7e-06 | 1.067E-05 | 3.837E-06 | 7.700E-06 | 1.500E-05 |
| 14058 | 703 | E703W | B | 1.4e-05 | 1.3e-05 | 1.5e-05 | 1.400E-05 | 1.000E-06 | 1.300E-05 | 1.500E-05 |
| 14059 | 703 | E703Y | B | 1.4e-05 | 1.6e-05 | 1.3e-05 | 1.433E-05 | 1.528E-06 | 1.300E-05 | 1.600E-05 |
| 14060 | 703 | E703V | B | 1.3e-05 | 1.4e-05 | 1e-05 | 1.233E-05 | 2.082E-06 | 1.000E-05 | 1.400E-05 |
| 14061 | 704 | D704A | B | 1.1e-05 | 1.3e-05 | 1.3e-05 | 1.233E-05 | 1.155E-06 | 1.100E-05 | 1.300E-05 |
| 14062 | 704 | D704R | B | 1.7e-05 | 8.4e-06 | 1.5e-05 | 1.347E-05 | 4.500E-06 | 8.400E-06 | 1.700E-05 |
| 14063 | 704 | D704N | B | 7.3e-06 | 1.4e-05 | 1e-05 | 1.043E-05 | 3.371E-06 | 7.300E-06 | 1.400E-05 |
| 14064 | 704 | D704D | B | 9.1e-06 | 1e-05 | 1.4e-05 | 1.103E-05 | 2.608E-06 | 9.100E-06 | 1.400E-05 |
| 14065 | 704 | D704C | B | 1.1e-05 | 1.4e-05 | 9.6e-06 | 1.153E-05 | 2.248E-06 | 9.600E-06 | 1.400E-05 |
| 14066 | 704 | D704E | B | 1.2e-05 | 1.2e-05 | 8.7e-06 | 1.090E-05 | 1.905E-06 | 8.700E-06 | 1.200E-05 |
| 14067 | 704 | D704Q | B | 1.1e-05 | 1.3e-05 | 1.1e-05 | 1.167E-05 | 1.155E-06 | 1.100E-05 | 1.300E-05 |
| 14068 | 704 | D704G | B | 1e-05 | 1.6e-05 | 1.1e-05 | 1.233E-05 | 3.215E-06 | 1.000E-05 | 1.600E-05 |
| 14069 | 704 | D704H | B | 1.5e-05 | 1.3e-05 | 1.3e-05 | 1.367E-05 | 1.155E-06 | 1.300E-05 | 1.500E-05 |
| 14070 | 704 | D704I | B | 1.4e-05 | 1.5e-05 | 1.1e-05 | 1.333E-05 | 2.082E-06 | 1.100E-05 | 1.500E-05 |
| 14071 | 704 | D704L | B | 1e-05 | 1.2e-05 | 1.5e-05 | 1.233E-05 | 2.517E-06 | 1.000E-05 | 1.500E-05 |
| 14072 | 704 | D704K | B | 1.1e-05 | 1.7e-05 | 1e-05 | 1.267E-05 | 3.786E-06 | 1.000E-05 | 1.700E-05 |
| 14073 | 704 | D704M | B | 1.6e-05 | 1.6e-05 | 1.4e-05 | 1.533E-05 | 1.155E-06 | 1.400E-05 | 1.600E-05 |
| 14074 | 704 | D704F | B | 8.6e-06 | 1.4e-05 | 1.5e-05 | 1.253E-05 | 3.443E-06 | 8.600E-06 | 1.500E-05 |
| 14075 | 704 | D704P | B | 1.1e-05 | 1.6e-05 | 6.7e-06 | 1.123E-05 | 4.654E-06 | 6.700E-06 | 1.600E-05 |
| 14076 | 704 | D704S | B | 1.3e-05 | 9.7e-06 | 1.2e-05 | 1.157E-05 | 1.692E-06 | 9.700E-06 | 1.300E-05 |
| 14077 | 704 | D704T | B | 1.5e-05 | 1.7e-05 | 1.4e-05 | 1.533E-05 | 1.528E-06 | 1.400E-05 | 1.700E-05 |
| 14078 | 704 | D704W | B | 1.7e-05 | 1.2e-05 | 9.8e-06 | 1.293E-05 | 3.690E-06 | 9.800E-06 | 1.700E-05 |
| 14079 | 704 | D704Y | B | 8.5e-06 | 1e-05 | 1.1e-05 | 9.833E-06 | 1.258E-06 | 8.500E-06 | 1.100E-05 |
| 14080 | 704 | D704V | B | 1.2e-05 | 1.2e-05 | 1.5e-05 | 1.300E-05 | 1.732E-06 | 1.200E-05 | 1.500E-05 |
| 14081 | 705 | F705A | B | 1.4e-05 | 1.6e-05 | 1.2e-05 | 1.400E-05 | 2.000E-06 | 1.200E-05 | 1.600E-05 |
| 14082 | 705 | F705R | B | 1.1e-05 | 1e-05 | 1.1e-05 | 1.067E-05 | 5.774E-07 | 1.000E-05 | 1.100E-05 |
| 14083 | 705 | F705N | B | 1.5e-05 | 1.5e-05 | 1.5e-05 | 1.500E-05 | 0.000E+00 | 1.500E-05 | 1.500E-05 |
| 14084 | 705 | F705D | B | 1.1e-05 | 1.1e-05 | 8.7e-06 | 1.023E-05 | 1.328E-06 | 8.700E-06 | 1.100E-05 |
| 14085 | 705 | F705C | B | 9.1e-06 | 1.3e-05 | 1.2e-05 | 1.137E-05 | 2.026E-06 | 9.100E-06 | 1.300E-05 |
| 14086 | 705 | F705E | B | 1.4e-05 | 1.1e-05 | 1.1e-05 | 1.200E-05 | 1.732E-06 | 1.100E-05 | 1.400E-05 |
| 14087 | 705 | F705Q | B | 1.4e-05 | 9.8e-06 | 1.1e-05 | 1.160E-05 | 2.163E-06 | 9.800E-06 | 1.400E-05 |
| 14088 | 705 | F705G | B | 1.2e-05 | 1e-05 | 1.1e-05 | 1.100E-05 | 1.000E-06 | 1.000E-05 | 1.200E-05 |
| 14089 | 705 | F705H | B | 1.5e-05 | 1.6e-05 | 1.5e-05 | 1.533E-05 | 5.774E-07 | 1.500E-05 | 1.600E-05 |
| 14090 | 705 | F705I | B | 8.9e-06 | 1.2e-05 | 1.4e-05 | 1.163E-05 | 2.570E-06 | 8.900E-06 | 1.400E-05 |
| 14091 | 705 | F705L | B | 9.2e-06 | 1.1e-05 | 9e-06 | 9.733E-06 | 1.102E-06 | 9.000E-06 | 1.100E-05 |
| 14092 | 705 | F705K | B | 9e-06 | 9.9e-06 | 1.2e-05 | 1.030E-05 | 1.539E-06 | 9.000E-06 | 1.200E-05 |
| 14093 | 705 | F705M | B | 1.4e-05 | 1e-05 | 1.8e-05 | 1.400E-05 | 4.000E-06 | 1.000E-05 | 1.800E-05 |
| 14094 | 705 | F705F | B | 1.5e-05 | 1.3e-05 | 1.4e-05 | 1.400E-05 | 1.000E-06 | 1.300E-05 | 1.500E-05 |
| 14095 | 705 | F705P | B | 1.4e-05 | 1.7e-05 | 1e-05 | 1.367E-05 | 3.512E-06 | 1.000E-05 | 1.700E-05 |
| 14096 | 705 | F705S | B | 6.2e-06 | 1.1e-05 | 1.5e-05 | 1.073E-05 | 4.406E-06 | 6.200E-06 | 1.500E-05 |
| 14097 | 705 | F705T | B | 1.5e-05 | 1.1e-05 | 1.7e-05 | 1.433E-05 | 3.055E-06 | 1.100E-05 | 1.700E-05 |
| 14098 | 705 | F705W | B | 9.5e-06 | 1.2e-05 | 1.6e-05 | 1.250E-05 | 3.279E-06 | 9.500E-06 | 1.600E-05 |
| 14099 | 705 | F705Y | B | 1.2e-05 | 1.5e-05 | 1.1e-05 | 1.267E-05 | 2.082E-06 | 1.100E-05 | 1.500E-05 |
| 14100 | 705 | F705V | B | 1.1e-05 | 1.6e-05 | 1.2e-05 | 1.300E-05 | 2.646E-06 | 1.100E-05 | 1.600E-05 |
| 14101 | 706 | A706A | B | 1e-05 | 1.4e-05 | 1.3e-05 | 1.233E-05 | 2.082E-06 | 1.000E-05 | 1.400E-05 |
| 14102 | 706 | A706R | B | 1.1e-05 | 1.6e-05 | 1.4e-05 | 1.367E-05 | 2.517E-06 | 1.100E-05 | 1.600E-05 |
| 14103 | 706 | A706N | B | 9.8e-06 | 1.4e-05 | 1.3e-05 | 1.227E-05 | 2.194E-06 | 9.800E-06 | 1.400E-05 |
| 14104 | 706 | A706D | B | 1.5e-05 | 1.6e-05 | 1.2e-05 | 1.433E-05 | 2.082E-06 | 1.200E-05 | 1.600E-05 |
| 14105 | 706 | A706C | B | 1.3e-05 | 1.5e-05 | 1.4e-05 | 1.400E-05 | 1.000E-06 | 1.300E-05 | 1.500E-05 |
| 14106 | 706 | A706E | B | 9.9e-06 | 1e-05 | 1.4e-05 | 1.130E-05 | 2.339E-06 | 9.900E-06 | 1.400E-05 |
| 14107 | 706 | A706Q | B | 1.8e-05 | 1.6e-05 | 1.6e-05 | 1.667E-05 | 1.155E-06 | 1.600E-05 | 1.800E-05 |
| 14108 | 706 | A706G | B | 1.5e-05 | 9.8e-06 | 1e-05 | 1.160E-05 | 2.946E-06 | 9.800E-06 | 1.500E-05 |
| 14109 | 706 | A706H | B | 8.7e-06 | 2.2e-05 | 1.2e-05 | 1.423E-05 | 6.926E-06 | 8.700E-06 | 2.200E-05 |

| | | | | | | | | | | |
|---|---|---|---|---|---|---|---|---|---|---|
| 14110 | 706 | A706I | B | 9.2e-06 | 9.8e-06 | 1e-05 | 9.667E-06 | 4.163E-07 | 9.200E-06 | 1.000E-05 |
| 14111 | 706 | A706L | B | 1.3e-05 | 1.4e-05 | 1.6e-05 | 1.433E-05 | 1.528E-06 | 1.300E-05 | 1.600E-05 |
| 14112 | 706 | A706K | B | 1.1e-05 | 1.4e-05 | 1.5e-05 | 1.333E-05 | 2.082E-06 | 1.100E-05 | 1.500E-05 |
| 14113 | 706 | A706M | B | 1.5e-05 | 1.5e-05 | 1e-05 | 1.333E-05 | 2.887E-06 | 1.000E-05 | 1.500E-05 |
| 14114 | 706 | A706F | B | 1.7e-05 | 1.1e-05 | 1.3e-05 | 1.367E-05 | 3.055E-06 | 1.100E-05 | 1.700E-05 |
| 14115 | 706 | A706P | B | 1.4e-05 | 1.5e-05 | 1.7e-05 | 1.533E-05 | 1.528E-06 | 1.400E-05 | 1.700E-05 |
| 14116 | 706 | A706S | B | 9.1e-06 | 1.3e-05 | 9.5e-06 | 1.053E-05 | 2.146E-06 | 9.100E-06 | 1.300E-05 |
| 14117 | 706 | A706T | B | 1.6e-05 | 1e-05 | 8.5e-06 | 1.150E-05 | 3.969E-06 | 8.500E-06 | 1.600E-05 |
| 14118 | 706 | A706W | B | 1.6e-05 | 2.4e-05 | 8.6e-06 | 1.620E-05 | 7.702E-06 | 8.600E-06 | 2.400E-05 |
| 14119 | 706 | A706Y | B | 1.8e-05 | 1.1e-05 | 1.6e-05 | 1.500E-05 | 3.606E-06 | 1.100E-05 | 1.800E-05 |
| 14120 | 706 | A706V | B | 1.1e-05 | 1.4e-05 | 1.6e-05 | 1.367E-05 | 2.517E-06 | 1.100E-05 | 1.600E-05 |
| 14121 | 707 | T707A | B | 1.1e-05 | 1e-05 | 1.7e-05 | 1.267E-05 | 3.786E-06 | 1.000E-05 | 1.700E-05 |
| 14122 | 707 | T707R | B | 1.5e-05 | 1.7e-05 | 9.3e-06 | 1.377E-05 | 3.995E-06 | 9.300E-06 | 1.700E-05 |
| 14123 | 707 | T707N | B | 1e-05 | 1.6e-05 | 1.6e-05 | 1.400E-05 | 3.464E-06 | 1.000E-05 | 1.600E-05 |
| 14124 | 707 | T707D | B | 1.1e-05 | 1.1e-05 | 1e-05 | 1.067E-05 | 5.774E-07 | 1.000E-05 | 1.100E-05 |
| 14125 | 707 | T707C | B | 1e-05 | 1e-05 | 1.4e-05 | 1.133E-05 | 2.309E-06 | 1.000E-05 | 1.400E-05 |
| 14126 | 707 | T707E | B | 1.2e-05 | 1.5e-05 | 8.9e-06 | 1.197E-05 | 3.050E-06 | 8.900E-06 | 1.500E-05 |
| 14127 | 707 | T707Q | B | 1e-05 | 1.4e-05 | 8.7e-06 | 1.090E-05 | 2.762E-06 | 8.700E-06 | 1.400E-05 |
| 14128 | 707 | T707G | B | 1e-05 | 1.6e-05 | 1.1e-05 | 1.233E-05 | 3.215E-06 | 1.000E-05 | 1.600E-05 |
| 14129 | 707 | T707H | B | 1.4e-05 | 9.1e-06 | 1.1e-05 | 1.137E-05 | 2.470E-06 | 9.100E-06 | 1.400E-05 |
| 14130 | 707 | T707I | B | 1.6e-05 | 1e-05 | 1.7e-05 | 1.433E-05 | 3.786E-06 | 1.000E-05 | 1.700E-05 |
| 14131 | 707 | T707L | B | 1.5e-05 | 1.5e-05 | 1.4e-05 | 1.467E-05 | 5.774E-07 | 1.400E-05 | 1.500E-05 |
| 14132 | 707 | T707K | B | 1.4e-05 | 1.5e-05 | 1.7e-05 | 1.533E-05 | 1.528E-06 | 1.400E-05 | 1.700E-05 |
| 14133 | 707 | T707M | B | 1.1e-05 | 1.7e-05 | 1.2e-05 | 1.333E-05 | 3.215E-06 | 1.100E-05 | 1.700E-05 |
| 14134 | 707 | T707F | B | 1.4e-05 | 1.5e-05 | 1.7e-05 | 1.533E-05 | 1.528E-06 | 1.400E-05 | 1.700E-05 |
| 14135 | 707 | T707P | B | 1.3e-05 | 1.5e-05 | 1.1e-05 | 1.300E-05 | 2.000E-06 | 1.100E-05 | 1.500E-05 |
| 14136 | 707 | T707S | B | 1.1e-05 | 1.5e-05 | 1.4e-05 | 1.333E-05 | 2.082E-06 | 1.100E-05 | 1.500E-05 |
| 14137 | 707 | T707T | B | 1.5e-05 | 1.5e-05 | 1.7e-05 | 1.567E-05 | 1.155E-06 | 1.500E-05 | 1.700E-05 |
| 14138 | 707 | T707W | B | 1.1e-05 | 1.1e-05 | 1.1e-05 | 1.100E-05 | 2.075E-21 | 1.100E-05 | 1.100E-05 |
| 14139 | 707 | T707Y | B | 9.8e-06 | 1.3e-05 | 1e-05 | 1.093E-05 | 1.793E-06 | 9.800E-06 | 1.300E-05 |
| 14140 | 707 | T707V | B | 1.6e-05 | 9.7e-06 | 1.1e-05 | 1.223E-05 | 3.326E-06 | 9.700E-06 | 1.600E-05 |
| 14141 | 708 | Y708A | B | 1.5e-05 | 1.7e-05 | 1.6e-05 | 1.600E-05 | 1.000E-06 | 1.500E-05 | 1.700E-05 |
| 14142 | 708 | Y708R | B | 8.7e-06 | 1.5e-05 | 1.1e-05 | 1.157E-05 | 3.188E-06 | 8.700E-06 | 1.500E-05 |
| 14143 | 708 | Y708N | B | 9.6e-06 | 1.4e-05 | 1.5e-05 | 1.287E-05 | 2.873E-06 | 9.600E-06 | 1.500E-05 |
| 14144 | 708 | Y708D | B | 1.4e-05 | 1.1e-05 | 1.7e-05 | 1.400E-05 | 3.000E-06 | 1.100E-05 | 1.700E-05 |
| 14145 | 708 | Y708C | B | 1.4e-05 | 1.4e-05 | 1.5e-05 | 1.433E-05 | 5.774E-07 | 1.400E-05 | 1.500E-05 |
| 14146 | 708 | Y708E | B | 1.2e-05 | 1.4e-05 | 1.6e-05 | 1.400E-05 | 2.000E-06 | 1.200E-05 | 1.600E-05 |
| 14147 | 708 | Y708Q | B | 1.3e-05 | 1.5e-05 | 1.5e-05 | 1.433E-05 | 1.155E-06 | 1.300E-05 | 1.500E-05 |
| 14148 | 708 | Y708G | B | 8.8e-06 | 1.5e-05 | 1.1e-05 | 1.160E-05 | 3.143E-06 | 8.800E-06 | 1.500E-05 |
| 14149 | 708 | Y708H | B | 1.5e-05 | 1.5e-05 | 1.1e-05 | 1.367E-05 | 2.309E-06 | 1.100E-05 | 1.500E-05 |
| 14150 | 708 | Y708I | B | 1.6e-05 | 9.5e-06 | 1.7e-05 | 1.417E-05 | 4.072E-06 | 9.500E-06 | 1.700E-05 |
| 14151 | 708 | Y708L | B | 1e-05 | 1.1e-05 | 9.5e-06 | 1.017E-05 | 7.638E-07 | 9.500E-06 | 1.100E-05 |
| 14152 | 708 | Y708K | B | 1.5e-05 | 1.3e-05 | 1.3e-05 | 1.367E-05 | 1.155E-06 | 1.300E-05 | 1.500E-05 |
| 14153 | 708 | Y708M | B | 1e-05 | 1.5e-05 | 1.5e-05 | 1.333E-05 | 2.887E-06 | 1.000E-05 | 1.500E-05 |
| 14154 | 708 | Y708F | B | 1.4e-05 | 1.4e-05 | 1.6e-05 | 1.467E-05 | 1.155E-06 | 1.400E-05 | 1.600E-05 |
| 14155 | 708 | Y708P | B | 9.2e-06 | 9.4e-06 | 1.3e-05 | 1.053E-05 | 2.139E-06 | 9.200E-06 | 1.300E-05 |
| 14156 | 708 | Y708S | B | 1.1e-05 | 1e-05 | 1.4e-05 | 1.167E-05 | 2.082E-06 | 1.000E-05 | 1.400E-05 |
| 14157 | 708 | Y708T | B | 8.4e-06 | 8.9e-06 | 1e-05 | 9.100E-06 | 8.185E-07 | 8.400E-06 | 1.000E-05 |
| 14158 | 708 | Y708W | B | 1.1e-05 | 1.5e-05 | 1.1e-05 | 1.233E-05 | 2.309E-06 | 1.100E-05 | 1.500E-05 |
| 14159 | 708 | Y708Y | B | 1.4e-05 | 1.5e-05 | 1.3e-05 | 1.400E-05 | 1.000E-06 | 1.300E-05 | 1.500E-05 |
| 14160 | 708 | Y708V | B | 1.2e-05 | 1.5e-05 | 1e-05 | 1.233E-05 | 2.517E-06 | 1.000E-05 | 1.500E-05 |
| 14161 | 709 | Y709A | B | 1.1e-05 | 1e-05 | 1.5e-05 | 1.200E-05 | 2.646E-06 | 1.000E-05 | 1.500E-05 |
| 14162 | 709 | Y709R | B | 1.2e-05 | 1.4e-05 | 1.1e-05 | 1.233E-05 | 1.528E-06 | 1.100E-05 | 1.400E-05 |
| 14163 | 709 | Y709N | B | 1.4e-05 | 1.1e-05 | 1.3e-05 | 1.267E-05 | 1.528E-06 | 1.100E-05 | 1.400E-05 |
| 14164 | 709 | Y709D | B | 1.1e-05 | 1.4e-05 | 9.8e-06 | 1.160E-05 | 2.163E-06 | 9.800E-06 | 1.400E-05 |
| 14165 | 709 | Y709C | B | 1.3e-05 | 9.3e-06 | 1.3e-05 | 1.177E-05 | 2.136E-06 | 9.300E-06 | 1.300E-05 |
| 14166 | 709 | Y709E | B | 1.6e-05 | 1.1e-05 | 1e-05 | 1.233E-05 | 3.215E-06 | 1.000E-05 | 1.600E-05 |
| 14167 | 709 | Y709Q | B | 1.5e-05 | 1.5e-05 | 1.1e-05 | 1.367E-05 | 2.309E-06 | 1.100E-05 | 1.500E-05 |
| 14168 | 709 | Y709G | B | 1.4e-05 | 1.6e-05 | 1.5e-05 | 1.500E-05 | 1.000E-06 | 1.400E-05 | 1.600E-05 |
| 14169 | 709 | Y709H | B | 1.2e-05 | 1.5e-05 | 1.4e-05 | 1.367E-05 | 1.528E-06 | 1.200E-05 | 1.500E-05 |
| 14170 | 709 | Y709I | B | 1e-05 | 9.3e-06 | 1.2e-05 | 1.043E-05 | 1.401E-06 | 9.300E-06 | 1.200E-05 |
| 14171 | 709 | Y709L | B | 1.6e-05 | 8.9e-06 | 1.5e-05 | 1.330E-05 | 3.843E-06 | 8.900E-06 | 1.600E-05 |
| 14172 | 709 | Y709K | B | 1.2e-05 | 1.1e-05 | 1.1e-05 | 1.133E-05 | 5.774E-07 | 1.100E-05 | 1.200E-05 |
| 14173 | 709 | Y709M | B | 1.6e-05 | 1.1e-05 | 1.1e-05 | 1.267E-05 | 2.887E-06 | 1.100E-05 | 1.600E-05 |
| 14174 | 709 | Y709F | B | 8.8e-06 | 9.3e-06 | 1.2e-05 | 1.003E-05 | 1.721E-06 | 8.800E-06 | 1.200E-05 |
| 14175 | 709 | Y709P | B | 1.1e-05 | 1.3e-05 | 1.6e-05 | 1.333E-05 | 2.517E-06 | 1.100E-05 | 1.600E-05 |
| 14176 | 709 | Y709S | B | 1.6e-05 | 1.6e-05 | 1.5e-05 | 1.567E-05 | 5.774E-07 | 1.500E-05 | 1.600E-05 |
| 14177 | 709 | Y709T | B | 1.3e-05 | 1.6e-05 | 1.2e-05 | 1.367E-05 | 2.082E-06 | 1.200E-05 | 1.600E-05 |
| 14178 | 709 | Y709W | B | 1.2e-05 | 1.1e-05 | 1.4e-05 | 1.233E-05 | 1.528E-06 | 1.100E-05 | 1.400E-05 |
| 14179 | 709 | Y709Y | B | 9.4e-06 | 9.9e-06 | 1.3e-05 | 1.077E-05 | 1.950E-06 | 9.400E-06 | 1.300E-05 |
| 14180 | 709 | Y709V | B | 1.6e-05 | 1.6e-05 | 1.5e-05 | 1.567E-05 | 5.774E-07 | 1.500E-05 | 1.600E-05 |
| 14181 | 710 | C710A | B | 1e-05 | 1.3e-05 | 1.5e-05 | 1.267E-05 | 2.517E-06 | 1.000E-05 | 1.500E-05 |
| 14182 | 710 | C710R | B | 1.3e-05 | 9.3e-06 | 1.1e-05 | 1.110E-05 | 1.852E-06 | 9.300E-06 | 1.300E-05 |
| 14183 | 710 | C710N | B | 1.7e-05 | 1.5e-05 | 1.1e-05 | 1.433E-05 | 3.055E-06 | 1.100E-05 | 1.700E-05 |
| 14184 | 710 | C710D | B | 1.5e-05 | 1.1e-05 | 1.5e-05 | 1.367E-05 | 2.309E-06 | 1.100E-05 | 1.500E-05 |
| 14185 | 710 | C710C | B | 1.6e-05 | 1e-05 | 1.5e-05 | 1.367E-05 | 3.215E-06 | 1.000E-05 | 1.600E-05 |
| 14186 | 710 | C710E | B | 1.3e-05 | 9.6e-06 | 1e-05 | 1.087E-05 | 1.858E-06 | 9.600E-06 | 1.300E-05 |
| 14187 | 710 | C710Q | B | 1.7e-05 | 1.3e-05 | 1.5e-05 | 1.500E-05 | 2.000E-06 | 1.300E-05 | 1.700E-05 |
| 14188 | 710 | C710G | B | 1.6e-05 | 1.7e-05 | 7.9e-06 | 1.363E-05 | 4.990E-06 | 7.900E-06 | 1.700E-05 |
| 14189 | 710 | C710H | B | 1.4e-05 | 9.2e-06 | 1.6e-05 | 1.307E-05 | 3.495E-06 | 9.200E-06 | 1.600E-05 |
| 14190 | 710 | C710I | B | 1e-05 | 1.3e-05 | 1.5e-05 | 1.267E-05 | 2.517E-06 | 1.000E-05 | 1.500E-05 |
| 14191 | 710 | C710L | B | 1.2e-05 | 1.4e-05 | 9.9e-06 | 1.197E-05 | 2.050E-06 | 9.900E-06 | 1.400E-05 |
| 14192 | 710 | C710K | B | 1.4e-05 | 1.6e-05 | 1.4e-05 | 1.467E-05 | 1.155E-06 | 1.400E-05 | 1.600E-05 |
| 14193 | 710 | C710M | B | 1.1e-05 | 1.5e-05 | 1.4e-05 | 1.333E-05 | 2.082E-06 | 1.100E-05 | 1.500E-05 |
| 14194 | 710 | C710F | B | 1.4e-05 | 1.4e-05 | 1.7e-05 | 1.500E-05 | 1.732E-06 | 1.400E-05 | 1.700E-05 |

| | | | | | | | | | | |
|---|---|---|---|---|---|---|---|---|---|---|
| 14195 | 710 | C710P | B | 9.1e-06 | 1.7e-05 | 1.1e-05 | 1.237E-05 | 4.124E-06 | 9.100E-06 | 1.700E-05 |
| 14196 | 710 | C710S | B | 1.5e-05 | 1.5e-05 | 1.5e-05 | 1.500E-05 | 0.000E+00 | 1.500E-05 | 1.500E-05 |
| 14197 | 710 | C710T | B | 1.4e-05 | 1.1e-05 | 8.8e-06 | 1.127E-05 | 2.610E-06 | 8.800E-06 | 1.400E-05 |
| 14198 | 710 | C710W | B | 1.5e-05 | 1.5e-05 | 1e-05 | 1.333E-05 | 2.887E-06 | 1.000E-05 | 1.500E-05 |
| 14199 | 710 | C710Y | B | 1.3e-05 | 8.4e-06 | 1e-05 | 1.047E-05 | 2.335E-06 | 8.400E-06 | 1.300E-05 |
| 14200 | 710 | C710V | B | 1.4e-05 | 1.1e-05 | 1.5e-05 | 1.333E-05 | 2.082E-06 | 1.100E-05 | 1.500E-05 |
| 14201 | 711 | Q711A | B | 8.4e-06 | 1.4e-05 | 9.6e-06 | 1.067E-05 | 2.948E-06 | 8.400E-06 | 1.400E-05 |
| 14202 | 711 | Q711R | B | 1.2e-05 | 1e-05 | 1.4e-05 | 1.200E-05 | 2.000E-06 | 1.000E-05 | 1.400E-05 |
| 14203 | 711 | Q711N | B | 9.2e-06 | 9.8e-06 | 1.1e-05 | 1.000E-05 | 9.165E-07 | 9.200E-06 | 1.100E-05 |
| 14204 | 711 | Q711D | B | 1.6e-05 | 1.4e-05 | 9.8e-06 | 1.327E-05 | 3.164E-06 | 9.800E-06 | 1.600E-05 |
| 14205 | 711 | Q711C | B | 1.6e-05 | 1.5e-05 | 9.4e-06 | 1.347E-05 | 3.557E-06 | 9.400E-06 | 1.600E-05 |
| 14206 | 711 | Q711E | B | 1.6e-05 | 1.4e-05 | 1.5e-05 | 1.500E-05 | 1.000E-06 | 1.400E-05 | 1.600E-05 |
| 14207 | 711 | Q711Q | B | 1.1e-05 | 9.3e-06 | 1.2e-05 | 1.077E-05 | 1.365E-06 | 9.300E-06 | 1.200E-05 |
| 14208 | 711 | Q711G | B | 1.2e-05 | 1.2e-05 | 6.6e-06 | 1.020E-05 | 3.118E-06 | 6.600E-06 | 1.200E-05 |
| 14209 | 711 | Q711H | B | 9.2e-06 | 1.3e-05 | 1e-05 | 1.073E-05 | 2.003E-06 | 9.200E-06 | 1.300E-05 |
| 14210 | 711 | Q711I | B | 1.6e-05 | 1.7e-05 | 6.3e-06 | 1.310E-05 | 5.910E-06 | 6.300E-06 | 1.700E-05 |
| 14211 | 711 | Q711L | B | 1.2e-05 | 1e-05 | 9.2e-06 | 1.040E-05 | 1.442E-06 | 9.200E-06 | 1.200E-05 |
| 14212 | 711 | Q711K | B | 1e-05 | 9.7e-06 | 1.5e-05 | 1.157E-05 | 2.977E-06 | 9.700E-06 | 1.500E-05 |
| 14213 | 711 | Q711M | B | 1.4e-05 | 1.6e-05 | 1.2e-05 | 1.400E-05 | 2.000E-06 | 1.200E-05 | 1.600E-05 |
| 14214 | 711 | Q711F | B | 8.4e-06 | 1.4e-05 | 8.9e-06 | 1.043E-05 | 3.099E-06 | 8.400E-06 | 1.400E-05 |
| 14215 | 711 | Q711P | B | 1.2e-05 | 1e-05 | 1e-05 | 1.067E-05 | 1.155E-06 | 1.000E-05 | 1.200E-05 |
| 14216 | 711 | Q711S | B | 1.1e-05 | 1.1e-05 | 1.6e-05 | 1.267E-05 | 2.887E-06 | 1.100E-05 | 1.600E-05 |
| 14217 | 711 | Q711T | B | 1.5e-05 | 1e-05 | 7.1e-06 | 1.070E-05 | 3.996E-06 | 7.100E-06 | 1.500E-05 |
| 14218 | 711 | Q711W | B | 1.1e-05 | 1.3e-05 | 1.1e-05 | 1.167E-05 | 1.155E-06 | 1.100E-05 | 1.300E-05 |
| 14219 | 711 | Q711Y | B | 1.3e-05 | 1.1e-05 | 1.5e-05 | 1.300E-05 | 2.000E-06 | 1.100E-05 | 1.500E-05 |
| 14220 | 711 | Q711V | B | 1.5e-05 | 1.3e-05 | 1.6e-05 | 1.467E-05 | 1.528E-06 | 1.300E-05 | 1.600E-05 |
| 14221 | 712 | Q712A | B | 1e-05 | 1.3e-05 | 1.8e-05 | 1.367E-05 | 4.041E-06 | 1.000E-05 | 1.800E-05 |
| 14222 | 712 | Q712R | B | 1.4e-05 | 1.8e-05 | 1.5e-05 | 1.567E-05 | 2.082E-06 | 1.400E-05 | 1.800E-05 |
| 14223 | 712 | Q712N | B | 1.5e-05 | 1.4e-05 | 1.6e-05 | 1.500E-05 | 1.000E-06 | 1.400E-05 | 1.600E-05 |
| 14224 | 712 | Q712D | B | 1.2e-05 | 1.4e-05 | 1.3e-05 | 1.300E-05 | 1.000E-06 | 1.200E-05 | 1.400E-05 |
| 14225 | 712 | Q712C | B | 1.2e-05 | 1.4e-05 | 1.5e-05 | 1.367E-05 | 1.528E-06 | 1.200E-05 | 1.500E-05 |
| 14226 | 712 | Q712E | B | 8.7e-06 | 1.4e-05 | 1e-05 | 1.090E-05 | 2.762E-06 | 8.700E-06 | 1.400E-05 |
| 14227 | 712 | Q712Q | B | 1.3e-05 | 1.5e-05 | 1e-05 | 1.267E-05 | 2.517E-06 | 1.000E-05 | 1.500E-05 |
| 14228 | 712 | Q712G | B | 1.8e-05 | 1.4e-05 | 1.7e-05 | 1.633E-05 | 2.082E-06 | 1.400E-05 | 1.800E-05 |
| 14229 | 712 | Q712H | B | 1.3e-05 | 1.1e-05 | 1.5e-05 | 1.300E-05 | 2.000E-06 | 1.100E-05 | 1.500E-05 |
| 14230 | 712 | Q712I | B | 1.6e-05 | 1.1e-05 | 7.5e-06 | 1.150E-05 | 4.272E-06 | 7.500E-06 | 1.600E-05 |
| 14231 | 712 | Q712L | B | 1.4e-05 | 1.1e-05 | 1.4e-05 | 1.300E-05 | 1.732E-06 | 1.100E-05 | 1.400E-05 |
| 14232 | 712 | Q712K | B | 5.9e-06 | 1.6e-05 | 1.1e-05 | 1.097E-05 | 5.050E-06 | 5.900E-06 | 1.600E-05 |
| 14233 | 712 | Q712M | B | 9e-06 | 1.5e-05 | 1.5e-05 | 1.300E-05 | 3.464E-06 | 9.000E-06 | 1.500E-05 |
| 14234 | 712 | Q712F | B | 1.2e-05 | 1.5e-05 | 1.6e-05 | 1.433E-05 | 2.082E-06 | 1.200E-05 | 1.600E-05 |
| 14235 | 712 | Q712P | B | 1.5e-05 | 1.4e-05 | 6.1e-06 | 1.170E-05 | 4.875E-06 | 6.100E-06 | 1.500E-05 |
| 14236 | 712 | Q712S | B | 1.1e-05 | 7.4e-06 | 8.1e-06 | 8.833E-06 | 1.909E-06 | 7.400E-06 | 1.100E-05 |
| 14237 | 712 | Q712T | B | 1.6e-05 | 1.6e-05 | 1.5e-05 | 1.567E-05 | 5.774E-07 | 1.500E-05 | 1.600E-05 |
| 14238 | 712 | Q712W | B | 1.2e-05 | 1e-05 | 9.4e-06 | 1.047E-05 | 1.361E-06 | 9.400E-06 | 1.200E-05 |
| 14239 | 712 | Q712Y | B | 1.1e-05 | 1.2e-05 | 9.7e-06 | 1.090E-05 | 1.153E-06 | 9.700E-06 | 1.200E-05 |
| 14240 | 712 | Q712V | B | 1.5e-05 | 9.5e-06 | 1.5e-05 | 1.317E-05 | 3.175E-06 | 9.500E-06 | 1.500E-05 |
| 14241 | 713 | Y713A | B | 1.5e-05 | 1.2e-05 | 1.6e-05 | 1.433E-05 | 2.082E-06 | 1.200E-05 | 1.600E-05 |
| 14242 | 713 | Y713R | B | 1.4e-05 | 1.4e-05 | 1.8e-05 | 1.533E-05 | 2.309E-06 | 1.400E-05 | 1.800E-05 |
| 14243 | 713 | Y713N | B | 8.9e-06 | 1.7e-05 | 1.4e-05 | 1.330E-05 | 4.095E-06 | 8.900E-06 | 1.700E-05 |
| 14244 | 713 | Y713D | B | 1.3e-05 | 1.4e-05 | 1.6e-05 | 1.433E-05 | 1.528E-06 | 1.300E-05 | 1.600E-05 |
| 14245 | 713 | Y713C | B | 1.6e-05 | 1.3e-05 | 1.5e-05 | 1.467E-05 | 1.528E-06 | 1.300E-05 | 1.600E-05 |
| 14246 | 713 | Y713E | B | 1.5e-05 | 1.4e-05 | 1.6e-05 | 1.500E-05 | 1.000E-06 | 1.400E-05 | 1.600E-05 |
| 14247 | 713 | Y713Q | B | 1e-05 | 1.5e-05 | 9.9e-06 | 1.163E-05 | 2.916E-06 | 9.900E-06 | 1.500E-05 |
| 14248 | 713 | Y713G | B | 1.6e-05 | 1.6e-05 | 1.2e-05 | 1.467E-05 | 2.309E-06 | 1.200E-05 | 1.600E-05 |
| 14249 | 713 | Y713H | B | 1.7e-05 | 1e-05 | 1.5e-05 | 1.400E-05 | 3.606E-06 | 1.000E-05 | 1.700E-05 |
| 14250 | 713 | Y713I | B | 1.6e-05 | 1.2e-05 | 1e-05 | 1.267E-05 | 3.055E-06 | 1.000E-05 | 1.600E-05 |
| 14251 | 713 | Y713L | B | 1.5e-05 | 1.5e-05 | 1.5e-05 | 1.500E-05 | 0.000E+00 | 1.500E-05 | 1.500E-05 |
| 14252 | 713 | Y713K | B | 1.1e-05 | 8.9e-06 | 6.8e-06 | 8.900E-06 | 2.100E-06 | 6.800E-06 | 1.100E-05 |
| 14253 | 713 | Y713M | B | 1.5e-05 | 1.5e-05 | 1.1e-05 | 1.367E-05 | 2.309E-06 | 1.100E-05 | 1.500E-05 |
| 14254 | 713 | Y713F | B | 9.5e-06 | 1.3e-05 | 1.3e-05 | 1.183E-05 | 2.021E-06 | 9.500E-06 | 1.300E-05 |
| 14255 | 713 | Y713P | B | 1.5e-05 | 9.9e-06 | 1.6e-05 | 1.363E-05 | 3.272E-06 | 9.900E-06 | 1.600E-05 |
| 14256 | 713 | Y713S | B | 1.8e-05 | 1e-05 | 1.5e-05 | 1.433E-05 | 4.041E-06 | 1.000E-05 | 1.800E-05 |
| 14257 | 713 | Y713T | B | 1.2e-05 | 1.6e-05 | 1.6e-05 | 1.467E-05 | 2.309E-06 | 1.200E-05 | 1.600E-05 |
| 14258 | 713 | Y713W | B | 1.6e-05 | 1.3e-05 | 9.3e-06 | 1.277E-05 | 3.356E-06 | 9.300E-06 | 1.600E-05 |
| 14259 | 713 | Y713Y | B | 1.6e-05 | 1e-05 | 7e-06 | 1.100E-05 | 4.583E-06 | 7.000E-06 | 1.600E-05 |
| 14260 | 713 | Y713V | B | 1.4e-05 | 1.1e-05 | 1.6e-05 | 1.367E-05 | 2.517E-06 | 1.100E-05 | 1.600E-05 |
| 14261 | 714 | Y714A | B | 1.7e-05 | 1.7e-05 | 2.3e-05 | 1.900E-05 | 3.464E-06 | 1.700E-05 | 2.300E-05 |
| 14262 | 714 | Y714R | B | 1.2e-05 | 1.1e-05 | 1.2e-05 | 1.167E-05 | 5.774E-07 | 1.100E-05 | 1.200E-05 |
| 14263 | 714 | Y714N | B | 9.9e-06 | 1.7e-05 | 1.5e-05 | 1.397E-05 | 3.661E-06 | 9.900E-06 | 1.700E-05 |
| 14264 | 714 | Y714D | B | 1.1e-05 | 1.5e-05 | 1.5e-05 | 1.367E-05 | 2.309E-06 | 1.100E-05 | 1.500E-05 |
| 14265 | 714 | Y714C | B | 1.7e-05 | 1.4e-05 | 1.6e-05 | 1.567E-05 | 1.528E-06 | 1.400E-05 | 1.700E-05 |
| 14266 | 714 | Y714E | B | 1.5e-05 | 1.3e-05 | 1.6e-05 | 1.467E-05 | 1.528E-06 | 1.300E-05 | 1.600E-05 |
| 14267 | 714 | Y714Q | B | 1e-05 | 1.4e-05 | 1.4e-05 | 1.267E-05 | 2.309E-06 | 1.000E-05 | 1.400E-05 |
| 14268 | 714 | Y714G | B | 1.4e-05 | 1.6e-05 | 1.2e-05 | 1.400E-05 | 2.000E-06 | 1.200E-05 | 1.600E-05 |
| 14269 | 714 | Y714H | B | 1.4e-05 | 9.9e-06 | 1.5e-05 | 1.297E-05 | 2.702E-06 | 9.900E-06 | 1.500E-05 |
| 14270 | 714 | Y714I | B | 1.6e-05 | 1e-05 | 1.1e-05 | 1.233E-05 | 3.215E-06 | 1.000E-05 | 1.600E-05 |
| 14271 | 714 | Y714L | B | 9.2e-06 | 1.3e-05 | 1.5e-05 | 1.240E-05 | 2.946E-06 | 9.200E-06 | 1.500E-05 |
| 14272 | 714 | Y714K | B | 1.5e-05 | 9.6e-06 | 1.5e-05 | 1.320E-05 | 3.118E-06 | 9.600E-06 | 1.500E-05 |
| 14273 | 714 | Y714M | B | 1.1e-05 | 8.7e-06 | 1e-05 | 9.900E-06 | 1.153E-06 | 8.700E-06 | 1.100E-05 |
| 14274 | 714 | Y714F | B | 1e-05 | 8.5e-06 | 1e-05 | 9.500E-06 | 8.660E-07 | 8.500E-06 | 1.000E-05 |
| 14275 | 714 | Y714P | B | 1.6e-05 | 1.2e-05 | 1.2e-05 | 1.333E-05 | 2.309E-06 | 1.200E-05 | 1.600E-05 |
| 14276 | 714 | Y714S | B | 9.6e-06 | 1.4e-05 | 1.3e-05 | 1.220E-05 | 2.307E-06 | 9.600E-06 | 1.400E-05 |
| 14277 | 714 | Y714T | B | 8.2e-06 | 1.1e-05 | 1e-05 | 9.733E-06 | 1.419E-06 | 8.200E-06 | 1.100E-05 |
| 14278 | 714 | Y714W | B | 1.5e-05 | 9.7e-06 | 1.2e-05 | 1.223E-05 | 2.658E-06 | 9.700E-06 | 1.500E-05 |
| 14279 | 714 | Y714Y | B | 1.1e-05 | 1.1e-05 | 1.5e-05 | 1.233E-05 | 2.309E-06 | 1.100E-05 | 1.500E-05 |

| | | | | | | | | | | |
|---|---|---|---|---|---|---|---|---|---|---|
| 14280 | 714 | Y714V | B | 1.4e-05 | 1.5e-05 | 1.1e-05 | 1.333E-05 | 2.082E-06 | 1.100E-05 | 1.500E-05 |
| 14281 | 715 | I715A | B | 1.5e-05 | 1.6e-05 | 1.3e-05 | 1.467E-05 | 1.528E-06 | 1.300E-05 | 1.600E-05 |
| 14282 | 715 | I715R | B | 9.9e-06 | 9.2e-06 | 1.1e-05 | 1.003E-05 | 9.074E-07 | 9.200E-06 | 1.100E-05 |
| 14283 | 715 | I715N | B | 2.3e-05 | 1e-05 | 1.1e-05 | 1.467E-05 | 7.234E-06 | 1.000E-05 | 2.300E-05 |
| 14284 | 715 | I715D | B | 1.5e-05 | 1.1e-05 | 1.5e-05 | 1.367E-05 | 2.309E-06 | 1.100E-05 | 1.500E-05 |
| 14285 | 715 | I715C | B | 9.8e-06 | 1.4e-05 | 1.3e-05 | 1.227E-05 | 2.194E-06 | 9.800E-06 | 1.400E-05 |
| 14286 | 715 | I715E | B | 1.3e-05 | 7.8e-06 | 1.4e-05 | 1.160E-05 | 3.329E-06 | 7.800E-06 | 1.400E-05 |
| 14287 | 715 | I715Q | B | 1e-05 | 1.5e-05 | 1.1e-05 | 1.200E-05 | 2.646E-06 | 1.000E-05 | 1.500E-05 |
| 14288 | 715 | I715G | B | 8.9e-06 | 1.2e-05 | 9.1e-06 | 1.000E-05 | 1.735E-06 | 8.900E-06 | 1.200E-05 |
| 14289 | 715 | I715H | B | 1e-05 | 9.3e-06 | 1.4e-05 | 1.110E-05 | 2.536E-06 | 9.300E-06 | 1.400E-05 |
| 14290 | 715 | I715I | B | 1e-05 | 1.2e-05 | 8.5e-06 | 1.017E-05 | 1.756E-06 | 8.500E-06 | 1.200E-05 |
| 14291 | 715 | I715L | B | 1.5e-05 | 1.6e-05 | 1.6e-05 | 1.567E-05 | 5.774E-07 | 1.500E-05 | 1.600E-05 |
| 14292 | 715 | I715K | B | 1.1e-05 | 1.4e-05 | 2.1e-05 | 1.533E-05 | 5.132E-06 | 1.100E-05 | 2.100E-05 |
| 14293 | 715 | I715M | B | 1.4e-05 | 1.4e-05 | 1.3e-05 | 1.367E-05 | 5.774E-07 | 1.300E-05 | 1.400E-05 |
| 14294 | 715 | I715F | B | 8.6e-06 | 1.5e-05 | 1.6e-05 | 1.320E-05 | 4.015E-06 | 8.600E-06 | 1.600E-05 |
| 14295 | 715 | I715P | B | 1.2e-05 | 1.4e-05 | 1.6e-05 | 1.400E-05 | 2.000E-06 | 1.200E-05 | 1.600E-05 |
| 14296 | 715 | I715S | B | 8.6e-06 | 9.6e-06 | 1.3e-05 | 1.040E-05 | 2.307E-06 | 8.600E-06 | 1.300E-05 |
| 14297 | 715 | I715T | B | 1e-05 | 1.4e-05 | 1.5e-05 | 1.300E-05 | 2.646E-06 | 1.000E-05 | 1.500E-05 |
| 14298 | 715 | I715W | B | 1.5e-05 | 1.2e-05 | 1.5e-05 | 1.400E-05 | 1.732E-06 | 1.200E-05 | 1.500E-05 |
| 14299 | 715 | I715Y | B | 8.7e-06 | 1.3e-05 | 8.6e-06 | 1.010E-05 | 2.512E-06 | 8.600E-06 | 1.300E-05 |
| 14300 | 715 | I715V | B | 1e-05 | 1.9e-05 | 9.3e-06 | 1.277E-05 | 5.410E-06 | 9.300E-06 | 1.900E-05 |
| 14301 | 716 | Y716A | B | 1.6e-05 | 1.7e-05 | 1.7e-05 | 1.667E-05 | 5.774E-07 | 1.600E-05 | 1.700E-05 |
| 14302 | 716 | Y716R | B | 1.9e-05 | 1.8e-05 | 1.8e-05 | 1.833E-05 | 5.774E-07 | 1.800E-05 | 1.900E-05 |
| 14303 | 716 | Y716N | B | 2.4e-05 | 2.2e-05 | 1.7e-05 | 2.100E-05 | 3.606E-06 | 1.700E-05 | 2.400E-05 |
| 14304 | 716 | Y716D | B | 1.7e-05 | 1.6e-05 | 1.5e-05 | 1.600E-05 | 1.000E-06 | 1.500E-05 | 1.700E-05 |
| 14305 | 716 | Y716C | B | 1.9e-05 | 1.8e-05 | 1.5e-05 | 1.733E-05 | 2.082E-06 | 1.500E-05 | 1.900E-05 |
| 14306 | 716 | Y716E | B | 1.9e-05 | 1.6e-05 | 1.6e-05 | 1.700E-05 | 1.732E-06 | 1.600E-05 | 1.900E-05 |
| 14307 | 716 | Y716Q | B | 2e-05 | 2.4e-05 | 2.4e-05 | 2.267E-05 | 2.309E-06 | 2.000E-05 | 2.400E-05 |
| 14308 | 716 | Y716G | B | 2.6e-05 | 2.5e-05 | 2.6e-05 | 2.567E-05 | 5.774E-07 | 2.500E-05 | 2.600E-05 |
| 14309 | 716 | Y716H | B | 2.2e-05 | 1.7e-05 | 1.8e-05 | 1.900E-05 | 2.646E-06 | 1.700E-05 | 2.200E-05 |
| 14310 | 716 | Y716I | B | 1.5e-05 | 1.6e-05 | 1.5e-05 | 1.533E-05 | 5.774E-07 | 1.500E-05 | 1.600E-05 |
| 14311 | 716 | Y716L | B | 1.5e-05 | 1.6e-05 | 1.7e-05 | 1.600E-05 | 1.000E-06 | 1.500E-05 | 1.700E-05 |
| 14312 | 716 | Y716K | B | 2e-05 | 2.1e-05 | 1.8e-05 | 1.967E-05 | 1.528E-06 | 1.800E-05 | 2.100E-05 |
| 14313 | 716 | Y716M | B | 1.6e-05 | 1e-05 | 1.3e-05 | 1.300E-05 | 3.000E-06 | 1.000E-05 | 1.600E-05 |
| 14314 | 716 | Y716F | B | 1.6e-05 | 1.4e-05 | 1.6e-05 | 1.533E-05 | 1.155E-06 | 1.400E-05 | 1.600E-05 |
| 14315 | 716 | Y716P | B | 1.6e-05 | 1.9e-05 | 1.8e-05 | 1.767E-05 | 1.528E-06 | 1.600E-05 | 1.900E-05 |
| 14316 | 716 | Y716S | B | 2.3e-05 | 2.4e-05 | 2.2e-05 | 2.300E-05 | 1.000E-06 | 2.200E-05 | 2.400E-05 |
| 14317 | 716 | Y716T | B | 2.5e-05 | 1.5e-05 | 2.3e-05 | 2.100E-05 | 5.292E-06 | 1.500E-05 | 2.500E-05 |
| 14318 | 716 | Y716W | B | 1.1e-05 | 9.3e-06 | 7.9e-06 | 9.400E-06 | 1.552E-06 | 7.900E-06 | 1.100E-05 |
| 14319 | 716 | Y716Y | B | 1.5e-05 | 1e-05 | 1.1e-05 | 1.200E-05 | 2.646E-06 | 1.000E-05 | 1.500E-05 |
| 14320 | 716 | Y716V | B | 1.5e-05 | 1.7e-05 | 1.8e-05 | 1.667E-05 | 1.528E-06 | 1.500E-05 | 1.800E-05 |
| 14321 | 717 | P717A | B | 1.5e-05 | 1.4e-05 | 1.2e-05 | 1.367E-05 | 1.528E-06 | 1.200E-05 | 1.500E-05 |
| 14322 | 717 | P717R | B | 9.7e-06 | 1.4e-05 | 1.2e-05 | 1.190E-05 | 2.152E-06 | 9.700E-06 | 1.400E-05 |
| 14323 | 717 | P717N | B | 1.5e-05 | 1.6e-05 | 9.6e-06 | 1.353E-05 | 3.443E-06 | 9.600E-06 | 1.600E-05 |
| 14324 | 717 | P717D | B | 1e-05 | 1.4e-05 | 1.1e-05 | 1.167E-05 | 2.082E-06 | 1.000E-05 | 1.400E-05 |
| 14325 | 717 | P717C | B | 1.4e-05 | 1.1e-05 | 1.4e-05 | 1.300E-05 | 1.732E-06 | 1.100E-05 | 1.400E-05 |
| 14326 | 717 | P717E | B | 1.5e-05 | 1e-05 | 1.3e-05 | 1.267E-05 | 2.517E-06 | 1.000E-05 | 1.500E-05 |
| 14327 | 717 | P717Q | B | 1.4e-05 | 1.6e-05 | 9.3e-06 | 1.310E-05 | 3.439E-06 | 9.300E-06 | 1.600E-05 |
| 14328 | 717 | P717G | B | 1.6e-05 | 1.4e-05 | 1.8e-05 | 1.600E-05 | 2.000E-06 | 1.400E-05 | 1.800E-05 |
| 14329 | 717 | P717H | B | 1.1e-05 | 1.1e-05 | 1.5e-05 | 1.233E-05 | 2.309E-06 | 1.100E-05 | 1.500E-05 |
| 14330 | 717 | P717I | B | 1e-05 | 8.8e-06 | 1.5e-05 | 1.127E-05 | 3.288E-06 | 8.800E-06 | 1.500E-05 |
| 14331 | 717 | P717L | B | 1e-05 | 1e-05 | 1e-05 | 1.000E-05 | 0.000E+00 | 1.000E-05 | 1.000E-05 |
| 14332 | 717 | P717K | B | 1.1e-05 | 7.5e-06 | 1.7e-05 | 1.183E-05 | 4.805E-06 | 7.500E-06 | 1.700E-05 |
| 14333 | 717 | P717M | B | 1.3e-05 | 1.1e-05 | 1.6e-05 | 1.333E-05 | 2.517E-06 | 1.100E-05 | 1.600E-05 |
| 14334 | 717 | P717F | B | 1e-05 | 1.4e-05 | 1.4e-05 | 1.267E-05 | 2.309E-06 | 1.000E-05 | 1.400E-05 |
| 14335 | 717 | P717P | B | 1e-05 | 1.1e-05 | 9.1e-06 | 1.003E-05 | 9.504E-07 | 9.100E-06 | 1.100E-05 |
| 14336 | 717 | P717S | B | 1e-05 | 9.1e-06 | 1e-05 | 9.700E-06 | 5.196E-07 | 9.100E-06 | 1.000E-05 |
| 14337 | 717 | P717T | B | 1.4e-05 | 1.1e-05 | 1e-05 | 1.167E-05 | 2.082E-06 | 1.000E-05 | 1.400E-05 |
| 14338 | 717 | P717W | B | 1.5e-05 | 1e-05 | 1.2e-05 | 1.233E-05 | 2.517E-06 | 1.000E-05 | 1.500E-05 |
| 14339 | 717 | P717Y | B | 2.5e-05 | 1.5e-05 | 1.1e-05 | 1.700E-05 | 7.211E-06 | 1.100E-05 | 2.500E-05 |
| 14340 | 717 | P717V | B | 1.1e-05 | 1.5e-05 | 1.5e-05 | 1.367E-05 | 2.309E-06 | 1.100E-05 | 1.500E-05 |
| 14341 | 718 | Y718A | B | 1.1e-05 | 1.6e-05 | 1.3e-05 | 1.333E-05 | 2.517E-06 | 1.100E-05 | 1.600E-05 |
| 14342 | 718 | Y718R | B | 8.1e-06 | 1.2e-05 | 1.7e-05 | 1.237E-05 | 4.461E-06 | 8.100E-06 | 1.700E-05 |
| 14343 | 718 | Y718N | B | 1.3e-05 | 1.4e-05 | 1.4e-05 | 1.367E-05 | 5.774E-07 | 1.300E-05 | 1.400E-05 |
| 14344 | 718 | Y718D | B | 1.7e-05 | 1.5e-05 | 1.6e-05 | 1.600E-05 | 1.000E-06 | 1.500E-05 | 1.700E-05 |
| 14345 | 718 | Y718C | B | 1e-05 | 9.8e-06 | 1.4e-05 | 1.127E-05 | 2.369E-06 | 9.800E-06 | 1.400E-05 |
| 14346 | 718 | Y718E | B | 1.4e-05 | 1.7e-05 | 1.1e-05 | 1.400E-05 | 3.000E-06 | 1.100E-05 | 1.700E-05 |
| 14347 | 718 | Y718Q | B | 9e-06 | 1.2e-05 | 1e-05 | 1.033E-05 | 1.528E-06 | 9.000E-06 | 1.200E-05 |
| 14348 | 718 | Y718G | B | 8.7e-06 | 1.5e-05 | 1.5e-05 | 1.290E-05 | 3.637E-06 | 8.700E-06 | 1.500E-05 |
| 14349 | 718 | Y718H | B | 1.5e-05 | 1.6e-05 | 1.4e-05 | 1.500E-05 | 1.000E-06 | 1.400E-05 | 1.600E-05 |
| 14350 | 718 | Y718I | B | 1.7e-05 | 8.9e-06 | 9.1e-06 | 1.167E-05 | 4.620E-06 | 8.900E-06 | 1.700E-05 |
| 14351 | 718 | Y718L | B | 1.7e-05 | 1.5e-05 | 1.6e-05 | 1.600E-05 | 1.000E-06 | 1.500E-05 | 1.700E-05 |
| 14352 | 718 | Y718K | B | 1.5e-05 | 1.4e-05 | 1.4e-05 | 1.433E-05 | 5.774E-07 | 1.400E-05 | 1.500E-05 |
| 14353 | 718 | Y718M | B | 1.3e-05 | 1.1e-05 | 1.5e-05 | 1.300E-05 | 2.000E-06 | 1.100E-05 | 1.500E-05 |
| 14354 | 718 | Y718F | B | 1.1e-05 | 1.4e-05 | 1.2e-05 | 1.233E-05 | 1.528E-06 | 1.100E-05 | 1.400E-05 |
| 14355 | 718 | Y718P | B | 7.6e-06 | 1.1e-05 | 1.2e-05 | 1.020E-05 | 2.307E-06 | 7.600E-06 | 1.200E-05 |
| 14356 | 718 | Y718S | B | 1.1e-05 | 1.6e-05 | 1.2e-05 | 1.300E-05 | 2.646E-06 | 1.100E-05 | 1.600E-05 |
| 14357 | 718 | Y718T | B | 1.3e-05 | 1e-05 | 1.6e-05 | 1.300E-05 | 3.000E-06 | 1.000E-05 | 1.600E-05 |
| 14358 | 718 | Y718W | B | 1.4e-05 | 9.5e-06 | 1.1e-05 | 1.150E-05 | 2.291E-06 | 9.500E-06 | 1.400E-05 |
| 14359 | 718 | Y718Y | B | 9.8e-06 | 1.4e-05 | 7.3e-06 | 1.037E-05 | 3.386E-06 | 7.300E-06 | 1.400E-05 |
| 14360 | 718 | Y718V | B | 1.1e-05 | 1.5e-05 | 1.4e-05 | 1.333E-05 | 2.082E-06 | 1.100E-05 | 1.500E-05 |
| 14361 | 719 | T719A | B | 1.5e-05 | 1.8e-05 | 9.8e-06 | 1.427E-05 | 4.149E-06 | 9.800E-06 | 1.800E-05 |
| 14362 | 719 | T719R | B | 8.1e-06 | 1e-05 | 1.1e-05 | 9.700E-06 | 1.473E-06 | 8.100E-06 | 1.100E-05 |
| 14363 | 719 | T719N | B | 1.2e-05 | 1.5e-05 | 9.3e-06 | 1.210E-05 | 2.851E-06 | 9.300E-06 | 1.500E-05 |
| 14364 | 719 | T719D | B | 1.5e-05 | 1.5e-05 | 1.4e-05 | 1.467E-05 | 5.774E-07 | 1.400E-05 | 1.500E-05 |

| | | | | | | | | | | |
|---|---|---|---|---|---|---|---|---|---|---|
| 14365 | 719 | T719C | B | 1.6e-05 | 1.1e-05 | 1e-05 | 1.233E-05 | 3.215E-06 | 1.000E-05 | 1.600E-05 |
| 14366 | 719 | T719E | B | 1.3e-05 | 1.4e-05 | 1.5e-05 | 1.400E-05 | 1.000E-06 | 1.300E-05 | 1.500E-05 |
| 14367 | 719 | T719Q | B | 1.1e-05 | 8e-06 | 1.3e-05 | 1.067E-05 | 2.517E-06 | 8.000E-06 | 1.300E-05 |
| 14368 | 719 | T719G | B | 9.5e-06 | 1.6e-05 | 1.1e-05 | 1.217E-05 | 3.403E-06 | 9.500E-06 | 1.600E-05 |
| 14369 | 719 | T719H | B | 1e-05 | 1.4e-05 | 6.2e-06 | 1.007E-05 | 3.900E-06 | 6.200E-06 | 1.400E-05 |
| 14370 | 719 | T719I | B | 1.5e-05 | 1.2e-05 | 1.6e-05 | 1.433E-05 | 2.082E-06 | 1.200E-05 | 1.600E-05 |
| 14371 | 719 | T719L | B | 1.7e-05 | 1.6e-05 | 1.1e-05 | 1.467E-05 | 3.215E-06 | 1.100E-05 | 1.700E-05 |
| 14372 | 719 | T719K | B | 1.4e-05 | 1.2e-05 | 1.7e-05 | 1.433E-05 | 2.517E-06 | 1.200E-05 | 1.700E-05 |
| 14373 | 719 | T719M | B | 9.4e-06 | 1.1e-05 | 1.2e-05 | 1.080E-05 | 1.311E-06 | 9.400E-06 | 1.200E-05 |
| 14374 | 719 | T719F | B | 1.1e-05 | 1.3e-05 | 9.8e-06 | 1.127E-05 | 1.617E-06 | 9.800E-06 | 1.300E-05 |
| 14375 | 719 | T719P | B | 1.4e-05 | 9.4e-06 | 1.1e-05 | 1.147E-05 | 2.335E-06 | 9.400E-06 | 1.400E-05 |
| 14376 | 719 | T719S | B | 1.5e-05 | 8.3e-06 | 1.7e-05 | 1.343E-05 | 4.557E-06 | 8.300E-06 | 1.700E-05 |
| 14377 | 719 | T719T | B | 7.6e-06 | 1.1e-05 | 1.5e-05 | 1.120E-05 | 3.704E-06 | 7.600E-06 | 1.500E-05 |
| 14378 | 719 | T719W | B | 1e-05 | 1e-05 | 9.9e-06 | 9.967E-06 | 5.774E-08 | 9.900E-06 | 1.000E-05 |
| 14379 | 719 | T719Y | B | 1.1e-05 | 8.6e-06 | 1.4e-05 | 1.120E-05 | 2.706E-06 | 8.600E-06 | 1.400E-05 |
| 14380 | 719 | T719V | B | 1.1e-05 | 1.1e-05 | 1.3e-05 | 1.167E-05 | 1.155E-06 | 1.100E-05 | 1.300E-05 |
| 14381 | 720 | F720A | B | 1.5e-05 | 1.5e-05 | 1.3e-05 | 1.433E-05 | 1.155E-06 | 1.300E-05 | 1.500E-05 |
| 14382 | 720 | F720R | B | 1.2e-05 | 1.5e-05 | 1e-05 | 1.233E-05 | 2.517E-06 | 1.000E-05 | 1.500E-05 |
| 14383 | 720 | F720N | B | 9.7e-06 | 1.5e-05 | 1e-05 | 1.157E-05 | 2.977E-06 | 9.700E-06 | 1.500E-05 |
| 14384 | 720 | F720D | B | 1.5e-05 | 1.5e-05 | 1.5e-05 | 1.500E-05 | 0.000E+00 | 1.500E-05 | 1.500E-05 |
| 14385 | 720 | F720C | B | 1.3e-05 | 1e-05 | 1.7e-05 | 1.333E-05 | 3.512E-06 | 1.000E-05 | 1.700E-05 |
| 14386 | 720 | F720E | B | 1.1e-05 | 1.6e-05 | 1e-05 | 1.233E-05 | 3.215E-06 | 1.000E-05 | 1.600E-05 |
| 14387 | 720 | F720Q | B | 1.5e-05 | 1.1e-05 | 1.1e-05 | 1.233E-05 | 2.309E-06 | 1.100E-05 | 1.500E-05 |
| 14388 | 720 | F720G | B | 1e-05 | 1e-05 | 1.6e-05 | 1.200E-05 | 3.464E-06 | 1.000E-05 | 1.600E-05 |
| 14389 | 720 | F720H | B | 9.5e-06 | 1e-05 | 1.4e-05 | 1.117E-05 | 2.466E-06 | 9.500E-06 | 1.400E-05 |
| 14390 | 720 | F720I | B | 1.3e-05 | 9.9e-06 | 1.2e-05 | 1.163E-05 | 1.582E-06 | 9.900E-06 | 1.300E-05 |
| 14391 | 720 | F720L | B | 1.3e-05 | 9.7e-06 | 9.8e-06 | 1.083E-05 | 1.877E-06 | 9.700E-06 | 1.300E-05 |
| 14392 | 720 | F720K | B | 1.6e-05 | 1.1e-05 | 9.5e-06 | 1.217E-05 | 3.403E-06 | 9.500E-06 | 1.600E-05 |
| 14393 | 720 | F720M | B | 9.4e-06 | 1.5e-05 | 1.5e-05 | 1.313E-05 | 3.233E-06 | 9.400E-06 | 1.500E-05 |
| 14394 | 720 | F720F | B | 1.1e-05 | 1e-05 | 1.1e-05 | 1.067E-05 | 5.774E-07 | 1.000E-05 | 1.100E-05 |
| 14395 | 720 | F720P | B | 1.2e-05 | 1.3e-05 | 1e-05 | 1.167E-05 | 1.528E-06 | 1.000E-05 | 1.300E-05 |
| 14396 | 720 | F720S | B | 1.4e-05 | 1.6e-05 | 8.2e-06 | 1.273E-05 | 4.051E-06 | 8.200E-06 | 1.600E-05 |
| 14397 | 720 | F720T | B | 1.3e-05 | 1e-05 | 8.4e-06 | 1.047E-05 | 2.335E-06 | 8.400E-06 | 1.300E-05 |
| 14398 | 720 | F720W | B | 1e-05 | 1.5e-05 | 1e-05 | 1.167E-05 | 2.887E-06 | 1.000E-05 | 1.500E-05 |
| 14399 | 720 | F720Y | B | 2.3e-05 | 1.5e-05 | 1.4e-05 | 1.733E-05 | 4.933E-06 | 1.400E-05 | 2.300E-05 |
| 14400 | 720 | F720V | B | 7.8e-06 | 7.8e-06 | 1.1e-05 | 8.867E-06 | 1.848E-06 | 7.800E-06 | 1.100E-05 |
| 14401 | 721 | G721A | B | 1.5e-05 | 1.5e-05 | 1.5e-05 | 1.500E-05 | 0.000E+00 | 1.500E-05 | 1.500E-05 |
| 14402 | 721 | G721R | B | 1.1e-05 | 9.3e-06 | 1.6e-05 | 1.210E-05 | 3.483E-06 | 9.300E-06 | 1.600E-05 |
| 14403 | 721 | G721N | B | 1.1e-05 | 1.4e-05 | 1.1e-05 | 1.200E-05 | 1.732E-06 | 1.100E-05 | 1.400E-05 |
| 14404 | 721 | G721D | B | 1.5e-05 | 1.4e-05 | 1.7e-05 | 1.533E-05 | 1.528E-06 | 1.400E-05 | 1.700E-05 |
| 14405 | 721 | G721C | B | 1.5e-05 | 1.5e-05 | 1.3e-05 | 1.433E-05 | 1.155E-06 | 1.300E-05 | 1.500E-05 |
| 14406 | 721 | G721E | B | 1.3e-05 | 1e-05 | 1.3e-05 | 1.200E-05 | 1.732E-06 | 1.000E-05 | 1.300E-05 |
| 14407 | 721 | G721Q | B | 1.1e-05 | 1.5e-05 | 1.4e-05 | 1.333E-05 | 2.082E-06 | 1.100E-05 | 1.500E-05 |
| 14408 | 721 | G721G | B | 1.1e-05 | 1.6e-05 | 1.4e-05 | 1.367E-05 | 2.517E-06 | 1.100E-05 | 1.600E-05 |
| 14409 | 721 | G721H | B | 1e-05 | 1.5e-05 | 1.4e-05 | 1.300E-05 | 2.646E-06 | 1.000E-05 | 1.500E-05 |
| 14410 | 721 | G721I | B | 1.6e-05 | 1e-05 | 8.9e-06 | 1.163E-05 | 3.821E-06 | 8.900E-06 | 1.600E-05 |
| 14411 | 721 | G721L | B | 1.6e-05 | 1.4e-05 | 1.1e-05 | 1.367E-05 | 2.517E-06 | 1.100E-05 | 1.600E-05 |
| 14412 | 721 | G721K | B | 9.1e-06 | 1e-05 | 1.5e-05 | 1.137E-05 | 3.179E-06 | 9.100E-06 | 1.500E-05 |
| 14413 | 721 | G721M | B | 1e-05 | 1e-05 | 8.6e-06 | 9.533E-06 | 8.083E-07 | 8.600E-06 | 1.000E-05 |
| 14414 | 721 | G721F | B | 9.6e-06 | 9.6e-06 | 8.7e-06 | 9.300E-06 | 5.196E-07 | 8.700E-06 | 9.600E-06 |
| 14415 | 721 | G721P | B | 1.6e-05 | 1.1e-05 | 1.5e-05 | 1.400E-05 | 2.646E-06 | 1.100E-05 | 1.600E-05 |
| 14416 | 721 | G721S | B | 9e-06 | 9.8e-06 | 1.2e-05 | 1.027E-05 | 1.553E-06 | 9.000E-06 | 1.200E-05 |
| 14417 | 721 | G721T | B | 1.3e-05 | 1.5e-05 | 1.4e-05 | 1.400E-05 | 1.000E-06 | 1.300E-05 | 1.500E-05 |
| 14418 | 721 | G721W | B | 1.2e-05 | 8.7e-06 | 1.5e-05 | 1.190E-05 | 3.151E-06 | 8.700E-06 | 1.500E-05 |
| 14419 | 721 | G721Y | B | 1.5e-05 | 1.2e-05 | 1e-05 | 1.233E-05 | 2.517E-06 | 1.000E-05 | 1.500E-05 |
| 14420 | 721 | G721V | B | 9.9e-06 | 9.7e-06 | 1.6e-05 | 1.187E-05 | 3.581E-06 | 9.700E-06 | 1.600E-05 |
| 14421 | 722 | Q722A | B | 1.1e-05 | 9.3e-06 | 1.7e-05 | 1.243E-05 | 4.045E-06 | 9.300E-06 | 1.700E-05 |
| 14422 | 722 | Q722R | B | 9.7e-06 | 1.1e-05 | 1.6e-05 | 1.223E-05 | 3.326E-06 | 9.700E-06 | 1.600E-05 |
| 14423 | 722 | Q722N | B | 1.7e-05 | 1.2e-05 | 1.6e-05 | 1.500E-05 | 2.646E-06 | 1.200E-05 | 1.700E-05 |
| 14424 | 722 | Q722D | B | 1.6e-05 | 1.7e-05 | 9.7e-06 | 1.423E-05 | 3.958E-06 | 9.700E-06 | 1.700E-05 |
| 14425 | 722 | Q722C | B | 1.3e-05 | 9.4e-06 | 1.5e-05 | 1.247E-05 | 2.838E-06 | 9.400E-06 | 1.500E-05 |
| 14426 | 722 | Q722E | B | 1.5e-05 | 9.7e-06 | 7.8e-06 | 1.083E-05 | 3.731E-06 | 7.800E-06 | 1.500E-05 |
| 14427 | 722 | Q722Q | B | 1.2e-05 | 1.1e-05 | 1.2e-05 | 1.167E-05 | 5.774E-07 | 1.100E-05 | 1.200E-05 |
| 14428 | 722 | Q722G | B | 1.5e-05 | 1.3e-05 | 1.6e-05 | 1.467E-05 | 1.528E-06 | 1.300E-05 | 1.600E-05 |
| 14429 | 722 | Q722H | B | 1.1e-05 | 1e-05 | 1.4e-05 | 1.167E-05 | 2.082E-06 | 1.000E-05 | 1.400E-05 |
| 14430 | 722 | Q722I | B | 1.4e-05 | 1.7e-05 | 1.1e-05 | 1.400E-05 | 3.000E-06 | 1.100E-05 | 1.700E-05 |
| 14431 | 722 | Q722L | B | 1.7e-05 | 1.5e-05 | 1e-05 | 1.400E-05 | 3.606E-06 | 1.000E-05 | 1.700E-05 |
| 14432 | 722 | Q722K | B | 1.3e-05 | 1.8e-05 | 1.1e-05 | 1.400E-05 | 3.606E-06 | 1.100E-05 | 1.800E-05 |
| 14433 | 722 | Q722M | B | 1e-05 | 1e-05 | 1.2e-05 | 1.067E-05 | 1.155E-06 | 1.000E-05 | 1.200E-05 |
| 14434 | 722 | Q722F | B | 9.4e-06 | 1.5e-05 | 8.1e-06 | 1.083E-05 | 3.667E-06 | 8.100E-06 | 1.500E-05 |
| 14435 | 722 | Q722P | B | 1.6e-05 | 1.6e-05 | 1.1e-05 | 1.433E-05 | 2.887E-06 | 1.100E-05 | 1.600E-05 |
| 14436 | 722 | Q722S | B | 1.4e-05 | 1.1e-05 | 1.4e-05 | 1.300E-05 | 1.732E-06 | 1.100E-05 | 1.400E-05 |
| 14437 | 722 | Q722T | B | 1.5e-05 | 1.3e-05 | 1.4e-05 | 1.400E-05 | 1.000E-06 | 1.300E-05 | 1.500E-05 |
| 14438 | 722 | Q722W | B | 1.3e-05 | 1.4e-05 | 1.7e-05 | 1.467E-05 | 2.082E-06 | 1.300E-05 | 1.700E-05 |
| 14439 | 722 | Q722Y | B | 9.8e-06 | 1.4e-05 | 1.5e-05 | 1.293E-05 | 2.759E-06 | 9.800E-06 | 1.500E-05 |
| 14440 | 722 | Q722V | B | 1.1e-05 | 9.9e-06 | 1.4e-05 | 1.163E-05 | 2.122E-06 | 9.900E-06 | 1.400E-05 |
| 14441 | 723 | G723A | B | 1.4e-05 | 1.1e-05 | 9.5e-06 | 1.150E-05 | 2.291E-06 | 9.500E-06 | 1.400E-05 |
| 14442 | 723 | G723R | B | 1.5e-05 | 1.2e-05 | 8.4e-06 | 1.180E-05 | 3.305E-06 | 8.400E-06 | 1.500E-05 |
| 14443 | 723 | G723N | B | 1.5e-05 | 1.6e-05 | 1.6e-05 | 1.567E-05 | 5.774E-07 | 1.500E-05 | 1.600E-05 |
| 14444 | 723 | G723D | B | 1.5e-05 | 1.1e-05 | 1.6e-05 | 1.400E-05 | 2.646E-06 | 1.100E-05 | 1.600E-05 |
| 14445 | 723 | G723C | B | 9.2e-06 | 1.4e-05 | 1.6e-05 | 1.307E-05 | 3.495E-06 | 9.200E-06 | 1.600E-05 |
| 14446 | 723 | G723E | B | 1.6e-05 | 1.8e-05 | 1e-05 | 1.467E-05 | 4.163E-06 | 1.000E-05 | 1.800E-05 |
| 14447 | 723 | G723Q | B | 1.7e-05 | 1.6e-05 | 1.5e-05 | 1.600E-05 | 1.000E-06 | 1.500E-05 | 1.700E-05 |
| 14448 | 723 | G723G | B | 7.5e-06 | 1.2e-05 | 1.6e-05 | 1.183E-05 | 4.252E-06 | 7.500E-06 | 1.600E-05 |
| 14449 | 723 | G723H | B | 1e-05 | 1.4e-05 | 1.5e-05 | 1.300E-05 | 2.646E-06 | 1.000E-05 | 1.500E-05 |

| | | | | | | | | | | |
|---|---|---|---|---|---|---|---|---|---|---|
| 14450 | 723 | G723I | B | 1.1e-05 | 1.5e-05 | 1.5e-05 | 1.367E-05 | 2.309E-06 | 1.100E-05 | 1.500E-05 |
| 14451 | 723 | G723L | B | 1.4e-05 | 8.6e-06 | 1.4e-05 | 1.220E-05 | 3.118E-06 | 8.600E-06 | 1.400E-05 |
| 14452 | 723 | G723K | B | 1.1e-05 | 1.1e-05 | 1.5e-05 | 1.233E-05 | 2.309E-06 | 1.100E-05 | 1.500E-05 |
| 14453 | 723 | G723M | B | 1.5e-05 | 1.3e-05 | 7.6e-06 | 1.187E-05 | 3.828E-06 | 7.600E-06 | 1.500E-05 |
| 14454 | 723 | G723F | B | 1.4e-05 | 9.6e-06 | 1.6e-05 | 1.320E-05 | 3.274E-06 | 9.600E-06 | 1.600E-05 |
| 14455 | 723 | G723P | B | 1.1e-05 | 1e-05 | 1.6e-05 | 1.233E-05 | 3.215E-06 | 1.000E-05 | 1.600E-05 |
| 14456 | 723 | G723S | B | 1.1e-05 | 1.6e-05 | 9.5e-06 | 1.217E-05 | 3.403E-06 | 9.500E-06 | 1.600E-05 |
| 14457 | 723 | G723T | B | 1.6e-05 | 8.7e-06 | 1.7e-05 | 1.390E-05 | 4.531E-06 | 8.700E-06 | 1.700E-05 |
| 14458 | 723 | G723W | B | 1.4e-05 | 1.6e-05 | 1.1e-05 | 1.367E-05 | 2.517E-06 | 1.100E-05 | 1.600E-05 |
| 14459 | 723 | G723Y | B | 1.4e-05 | 1.5e-05 | 1.6e-05 | 1.500E-05 | 1.000E-06 | 1.400E-05 | 1.600E-05 |
| 14460 | 723 | G723V | B | 1.1e-05 | 1.5e-05 | 1.6e-05 | 1.400E-05 | 2.646E-06 | 1.100E-05 | 1.600E-05 |
| 14461 | 724 | T724A | B | 1.9e-05 | 1.6e-05 | 1.5e-05 | 1.667E-05 | 2.082E-06 | 1.500E-05 | 1.900E-05 |
| 14462 | 724 | T724R | B | 1.9e-05 | 1.2e-05 | 9.7e-06 | 1.357E-05 | 4.844E-06 | 9.700E-06 | 1.900E-05 |
| 14463 | 724 | T724N | B | 1.5e-05 | 1.2e-05 | 8.7e-06 | 1.190E-05 | 3.151E-06 | 8.700E-06 | 1.500E-05 |
| 14464 | 724 | T724D | B | 1.5e-05 | 1.4e-05 | 1.7e-05 | 1.533E-05 | 1.528E-06 | 1.400E-05 | 1.700E-05 |
| 14465 | 724 | T724C | B | 1.5e-05 | 1.6e-05 | 1.5e-05 | 1.533E-05 | 5.774E-07 | 1.500E-05 | 1.600E-05 |
| 14466 | 724 | T724E | B | 1.6e-05 | 1.3e-05 | 1.5e-05 | 1.467E-05 | 1.528E-06 | 1.300E-05 | 1.600E-05 |
| 14467 | 724 | T724Q | B | 1.5e-05 | 9.5e-06 | 8.6e-06 | 1.103E-05 | 3.465E-06 | 8.600E-06 | 1.500E-05 |
| 14468 | 724 | T724G | B | 1.3e-05 | 1.6e-05 | 1.4e-05 | 1.433E-05 | 1.528E-06 | 1.300E-05 | 1.600E-05 |
| 14469 | 724 | T724H | B | 1.1e-05 | 9.7e-06 | 1.1e-05 | 1.057E-05 | 7.506E-07 | 9.700E-06 | 1.100E-05 |
| 14470 | 724 | T724I | B | 1.5e-05 | 1e-05 | 9.6e-06 | 1.153E-05 | 3.009E-06 | 9.600E-06 | 1.500E-05 |
| 14471 | 724 | T724L | B | 1.4e-05 | 1.1e-05 | 1.4e-05 | 1.300E-05 | 1.732E-06 | 1.100E-05 | 1.400E-05 |
| 14472 | 724 | T724K | B | 9.7e-06 | 8.2e-06 | 1.6e-05 | 1.130E-05 | 4.139E-06 | 8.200E-06 | 1.600E-05 |
| 14473 | 724 | T724M | B | 1.2e-05 | 1.1e-05 | 9e-06 | 1.067E-05 | 1.528E-06 | 9.000E-06 | 1.200E-05 |
| 14474 | 724 | T724F | B | 1e-05 | 1e-05 | 1.5e-05 | 1.167E-05 | 2.887E-06 | 1.000E-05 | 1.500E-05 |
| 14475 | 724 | T724P | B | 1e-05 | 1.4e-05 | 1.1e-05 | 1.167E-05 | 2.082E-06 | 1.000E-05 | 1.400E-05 |
| 14476 | 724 | T724S | B | 9.3e-06 | 1.6e-05 | 1.6e-05 | 1.377E-05 | 3.868E-06 | 9.300E-06 | 1.600E-05 |
| 14477 | 724 | T724T | B | 1.6e-05 | 1.1e-05 | 1e-05 | 1.233E-05 | 3.215E-06 | 1.000E-05 | 1.600E-05 |
| 14478 | 724 | T724W | B | 9.9e-06 | 1.5e-05 | 7.5e-06 | 1.080E-05 | 3.830E-06 | 7.500E-06 | 1.500E-05 |
| 14479 | 724 | T724Y | B | 9.1e-06 | 1.4e-05 | 8.1e-06 | 1.040E-05 | 3.158E-06 | 8.100E-06 | 1.400E-05 |
| 14480 | 724 | T724V | B | 2.5e-05 | 1.1e-05 | 1.5e-05 | 1.700E-05 | 7.211E-06 | 1.100E-05 | 2.500E-05 |
| 14481 | 725 | K725A | B | 1.6e-05 | 1e-05 | 1.1e-05 | 1.233E-05 | 3.215E-06 | 1.000E-05 | 1.600E-05 |
| 14482 | 725 | K725R | B | 1.3e-05 | 1.2e-05 | 1.5e-05 | 1.333E-05 | 1.528E-06 | 1.200E-05 | 1.500E-05 |
| 14483 | 725 | K725N | B | 1.5e-05 | 1.4e-05 | 1.4e-05 | 1.433E-05 | 5.774E-07 | 1.400E-05 | 1.500E-05 |
| 14484 | 725 | K725D | B | 1.6e-05 | 1.1e-05 | 1e-05 | 1.233E-05 | 3.215E-06 | 1.000E-05 | 1.600E-05 |
| 14485 | 725 | K725C | B | 1.5e-05 | 1.1e-05 | 1e-05 | 1.200E-05 | 2.646E-06 | 1.000E-05 | 1.500E-05 |
| 14486 | 725 | K725E | B | 1.4e-05 | 1.8e-05 | 1.5e-05 | 1.567E-05 | 2.082E-06 | 1.400E-05 | 1.800E-05 |
| 14487 | 725 | K725Q | B | 9.6e-06 | 1.4e-05 | 1.6e-05 | 1.320E-05 | 3.274E-06 | 9.600E-06 | 1.600E-05 |
| 14488 | 725 | K725G | B | 1e-05 | 1.1e-05 | 9.9e-06 | 1.030E-05 | 6.083E-07 | 9.900E-06 | 1.100E-05 |
| 14489 | 725 | K725H | B | 1.7e-05 | 1.4e-05 | 1.1e-05 | 1.400E-05 | 3.000E-06 | 1.100E-05 | 1.700E-05 |
| 14490 | 725 | K725I | B | 8.8e-06 | 1.4e-05 | 1.2e-05 | 1.160E-05 | 2.623E-06 | 8.800E-06 | 1.400E-05 |
| 14491 | 725 | K725L | B | 1.2e-05 | 1.1e-05 | 1.6e-05 | 1.300E-05 | 2.646E-06 | 1.100E-05 | 1.600E-05 |
| 14492 | 725 | K725K | B | 1.1e-05 | 1.5e-05 | 1e-05 | 1.200E-05 | 2.646E-06 | 1.000E-05 | 1.500E-05 |
| 14493 | 725 | K725M | B | 1.4e-05 | 1.2e-05 | 9.6e-06 | 1.187E-05 | 2.203E-06 | 9.600E-06 | 1.400E-05 |
| 14494 | 725 | K725F | B | 1e-05 | 1.1e-05 | 1.1e-05 | 1.067E-05 | 5.774E-07 | 1.000E-05 | 1.100E-05 |
| 14495 | 725 | K725P | B | 1.2e-05 | 1.3e-05 | 8.9e-06 | 1.130E-05 | 2.138E-06 | 8.900E-06 | 1.300E-05 |
| 14496 | 725 | K725S | B | 9.7e-06 | 9.7e-06 | 1.3e-05 | 1.080E-05 | 1.905E-06 | 9.700E-06 | 1.300E-05 |
| 14497 | 725 | K725T | B | 1.2e-05 | 8.4e-06 | 1e-05 | 1.013E-05 | 1.804E-06 | 8.400E-06 | 1.200E-05 |
| 14498 | 725 | K725W | B | 2.1e-05 | 1.3e-05 | 1.1e-05 | 1.500E-05 | 5.292E-06 | 1.100E-05 | 2.100E-05 |
| 14499 | 725 | K725Y | B | 1.6e-05 | 1.4e-05 | 1.1e-05 | 1.367E-05 | 2.517E-06 | 1.100E-05 | 1.600E-05 |
| 14500 | 725 | K725V | B | 1.4e-05 | 1.1e-05 | 1e-05 | 1.167E-05 | 2.082E-06 | 1.000E-05 | 1.400E-05 |
| 14501 | 726 | V726A | B | 8.2e-06 | 1e-05 | 1.4e-05 | 1.073E-05 | 2.969E-06 | 8.200E-06 | 1.400E-05 |
| 14502 | 726 | V726R | B | 1.2e-05 | 1.1e-05 | 1e-05 | 1.100E-05 | 1.000E-06 | 1.000E-05 | 1.200E-05 |
| 14503 | 726 | V726N | B | 8.1e-06 | 1.1e-05 | 1.5e-05 | 1.137E-05 | 3.465E-06 | 8.100E-06 | 1.500E-05 |
| 14504 | 726 | V726D | B | 1.1e-05 | 1.8e-05 | 1.3e-05 | 1.400E-05 | 3.606E-06 | 1.100E-05 | 1.800E-05 |
| 14505 | 726 | V726C | B | 1.3e-05 | 8.9e-06 | 9.5e-06 | 1.047E-05 | 2.214E-06 | 8.900E-06 | 1.300E-05 |
| 14506 | 726 | V726E | B | 1.7e-05 | 1.1e-05 | 8.3e-06 | 1.210E-05 | 4.453E-06 | 8.300E-06 | 1.700E-05 |
| 14507 | 726 | V726Q | B | 1.2e-05 | 1.2e-05 | 9.9e-06 | 1.130E-05 | 1.212E-06 | 9.900E-06 | 1.200E-05 |
| 14508 | 726 | V726G | B | 1.2e-05 | 1.5e-05 | 1.5e-05 | 1.400E-05 | 1.732E-06 | 1.200E-05 | 1.500E-05 |
| 14509 | 726 | V726H | B | 1.6e-05 | 1.4e-05 | 2.4e-05 | 1.800E-05 | 5.292E-06 | 1.400E-05 | 2.400E-05 |
| 14510 | 726 | V726I | B | 1.7e-05 | 6.3e-06 | 1.6e-05 | 1.310E-05 | 5.910E-06 | 6.300E-06 | 1.700E-05 |
| 14511 | 726 | V726L | B | 1.5e-05 | 1.4e-05 | 1.2e-05 | 1.367E-05 | 1.528E-06 | 1.200E-05 | 1.500E-05 |
| 14512 | 726 | V726K | B | 1e-05 | 1e-05 | 7.5e-06 | 9.167E-06 | 1.443E-06 | 7.500E-06 | 1.000E-05 |
| 14513 | 726 | V726M | B | 1.6e-05 | 9.7e-06 | 1.4e-05 | 1.323E-05 | 3.219E-06 | 9.700E-06 | 1.600E-05 |
| 14514 | 726 | V726F | B | 1.1e-05 | 9.3e-06 | 7.1e-06 | 9.133E-06 | 1.955E-06 | 7.100E-06 | 1.100E-05 |
| 14515 | 726 | V726P | B | 8.8e-06 | 1.5e-05 | 9e-06 | 1.093E-05 | 3.523E-06 | 8.800E-06 | 1.500E-05 |
| 14516 | 726 | V726S | B | 1.6e-05 | 9.6e-06 | 1.4e-05 | 1.320E-05 | 3.274E-06 | 9.600E-06 | 1.600E-05 |
| 14517 | 726 | V726T | B | 1e-05 | 9.9e-06 | 1.2e-05 | 1.063E-05 | 1.185E-06 | 9.900E-06 | 1.200E-05 |
| 14518 | 726 | V726W | B | 1.6e-05 | 1.3e-05 | 1.5e-05 | 1.467E-05 | 1.528E-06 | 1.300E-05 | 1.600E-05 |
| 14519 | 726 | V726Y | B | 8e-06 | 9.9e-06 | 8.4e-06 | 8.767E-06 | 1.002E-06 | 8.000E-06 | 9.900E-06 |
| 14520 | 726 | V726V | B | 9.4e-06 | 1e-05 | 1.4e-05 | 1.113E-05 | 2.501E-06 | 9.400E-06 | 1.400E-05 |
| 14521 | 727 | E727A | B | 1.5e-05 | 1.3e-05 | 1.5e-05 | 1.433E-05 | 1.155E-06 | 1.300E-05 | 1.500E-05 |
| 14522 | 727 | E727R | B | 2e-05 | 9.1e-06 | 1e-05 | 1.303E-05 | 6.050E-06 | 9.100E-06 | 2.000E-05 |
| 14523 | 727 | E727N | B | 9.5e-06 | 1.4e-05 | 1.5e-05 | 1.283E-05 | 2.930E-06 | 9.500E-06 | 1.500E-05 |
| 14524 | 727 | E727D | B | 8.8e-06 | 1.5e-05 | 1.2e-05 | 1.193E-05 | 3.101E-06 | 8.800E-06 | 1.500E-05 |
| 14525 | 727 | E727C | B | 1.5e-05 | 8.3e-06 | 1.5e-05 | 1.277E-05 | 3.868E-06 | 8.300E-06 | 1.500E-05 |
| 14526 | 727 | E727E | B | 1.4e-05 | 8.8e-06 | 1.5e-05 | 1.260E-05 | 3.329E-06 | 8.800E-06 | 1.500E-05 |
| 14527 | 727 | E727Q | B | 1.1e-05 | 7.7e-06 | 1.5e-05 | 1.123E-05 | 3.656E-06 | 7.700E-06 | 1.500E-05 |
| 14528 | 727 | E727G | B | 1.5e-05 | 1e-05 | 1.1e-05 | 1.200E-05 | 2.646E-06 | 1.000E-05 | 1.500E-05 |
| 14529 | 727 | E727H | B | 1.5e-05 | 1.1e-05 | 1.2e-05 | 1.267E-05 | 2.082E-06 | 1.100E-05 | 1.500E-05 |
| 14530 | 727 | E727I | B | 1.1e-05 | 1.6e-05 | 1.6e-05 | 1.433E-05 | 2.887E-06 | 1.100E-05 | 1.600E-05 |
| 14531 | 727 | E727L | B | 1e-05 | 1.5e-05 | 9.2e-06 | 1.140E-05 | 3.143E-06 | 9.200E-06 | 1.500E-05 |
| 14532 | 727 | E727K | B | 1.7e-05 | 9.7e-06 | 1.1e-05 | 1.257E-05 | 3.894E-06 | 9.700E-06 | 1.700E-05 |
| 14533 | 727 | E727M | B | 1.5e-05 | 1e-05 | 1.5e-05 | 1.333E-05 | 2.887E-06 | 1.000E-05 | 1.500E-05 |
| 14534 | 727 | E727F | B | 1.1e-05 | 1.2e-05 | 1.1e-05 | 1.133E-05 | 5.774E-07 | 1.100E-05 | 1.200E-05 |

| | | | | | | | | | | |
|---|---|---|---|---|---|---|---|---|---|---|
| 14535 | 727 | E727P | B | 7.9e-06 | 1.4e-05 | 1.5e-05 | 1.230E-05 | 3.843E-06 | 7.900E-06 | 1.500E-05 |
| 14536 | 727 | E727S | B | 1.4e-05 | 1.5e-05 | 1.3e-05 | 1.400E-05 | 1.000E-06 | 1.300E-05 | 1.500E-05 |
| 14537 | 727 | E727T | B | 1e-05 | 1.5e-05 | 1.4e-05 | 1.300E-05 | 2.646E-06 | 1.000E-05 | 1.500E-05 |
| 14538 | 727 | E727W | B | 6.1e-06 | 1e-05 | 1.4e-05 | 1.003E-05 | 3.950E-06 | 6.100E-06 | 1.400E-05 |
| 14539 | 727 | E727Y | B | 9.9e-06 | 1e-05 | 1.6e-05 | 1.197E-05 | 3.493E-06 | 9.900E-06 | 1.600E-05 |
| 14540 | 727 | E727V | B | 1.3e-05 | 1.4e-05 | 1e-05 | 1.233E-05 | 2.082E-06 | 1.000E-05 | 1.400E-05 |
| 14541 | 728 | I728A | B | 1.4e-05 | 1.1e-05 | 9.9e-06 | 1.163E-05 | 2.122E-06 | 9.900E-06 | 1.400E-05 |
| 14542 | 728 | I728R | B | 7.2e-06 | 1.7e-05 | 9.5e-06 | 1.123E-05 | 5.125E-06 | 7.200E-06 | 1.700E-05 |
| 14543 | 728 | I728N | B | 1.5e-05 | 1.1e-05 | 1.2e-05 | 1.267E-05 | 2.082E-06 | 1.100E-05 | 1.500E-05 |
| 14544 | 728 | I728D | B | 1.3e-05 | 1.7e-05 | 1.3e-05 | 1.433E-05 | 2.309E-06 | 1.300E-05 | 1.700E-05 |
| 14545 | 728 | I728C | B | 1.5e-05 | 1e-05 | 1.1e-05 | 1.200E-05 | 2.646E-06 | 1.000E-05 | 1.500E-05 |
| 14546 | 728 | I728E | B | 1.3e-05 | 1.3e-05 | 7.7e-06 | 1.123E-05 | 3.060E-06 | 7.700E-06 | 1.300E-05 |
| 14547 | 728 | I728Q | B | 1.1e-05 | 1.1e-05 | 8.2e-06 | 1.007E-05 | 1.617E-06 | 8.200E-06 | 1.100E-05 |
| 14548 | 728 | I728G | B | 1.5e-05 | 1.4e-05 | 1.5e-05 | 1.467E-05 | 5.774E-07 | 1.400E-05 | 1.500E-05 |
| 14549 | 728 | I728H | B | 1.5e-05 | 9.6e-06 | 9.5e-06 | 1.137E-05 | 3.147E-06 | 9.500E-06 | 1.500E-05 |
| 14550 | 728 | I728I | B | 9.2e-06 | 1.4e-05 | 1.6e-05 | 1.307E-05 | 3.495E-06 | 9.200E-06 | 1.600E-05 |
| 14551 | 728 | I728L | B | 1.6e-05 | 1.5e-05 | 9.4e-06 | 1.347E-05 | 3.557E-06 | 9.400E-06 | 1.600E-05 |
| 14552 | 728 | I728K | B | 1e-05 | 1.5e-05 | 1.2e-05 | 1.233E-05 | 2.517E-06 | 1.000E-05 | 1.500E-05 |
| 14553 | 728 | I728M | B | 9.8e-06 | 2.3e-05 | 1.5e-05 | 1.593E-05 | 6.649E-06 | 9.800E-06 | 2.300E-05 |
| 14554 | 728 | I728F | B | 1.8e-05 | 8.5e-06 | 1.5e-05 | 1.383E-05 | 4.856E-06 | 8.500E-06 | 1.800E-05 |
| 14555 | 728 | I728P | B | 1.2e-05 | 1.1e-05 | 1.6e-05 | 1.300E-05 | 2.646E-06 | 1.100E-05 | 1.600E-05 |
| 14556 | 728 | I728S | B | 1.2e-05 | 1.6e-05 | 1.5e-05 | 1.433E-05 | 2.082E-06 | 1.200E-05 | 1.600E-05 |
| 14557 | 728 | I728T | B | 9.4e-06 | 1.3e-05 | 1.7e-05 | 1.313E-05 | 3.802E-06 | 9.400E-06 | 1.700E-05 |
| 14558 | 728 | I728W | B | 1.4e-05 | 9.7e-06 | 1e-05 | 1.123E-05 | 2.401E-06 | 9.700E-06 | 1.400E-05 |
| 14559 | 728 | I728Y | B | 1.5e-05 | 1.4e-05 | 1.4e-05 | 1.433E-05 | 5.774E-07 | 1.400E-05 | 1.500E-05 |
| 14560 | 728 | I728V | B | 1e-05 | 1.7e-05 | 1.7e-05 | 1.467E-05 | 4.041E-06 | 1.000E-05 | 1.700E-05 |
| 14561 | 729 | K729A | B | 1e-05 | 1.3e-05 | 1.1e-05 | 1.133E-05 | 1.528E-06 | 1.000E-05 | 1.300E-05 |
| 14562 | 729 | K729R | B | 1.8e-05 | 9.2e-06 | 1.2e-05 | 1.307E-05 | 4.496E-06 | 9.200E-06 | 1.800E-05 |
| 14563 | 729 | K729N | B | 1.4e-05 | 1.2e-05 | 1.1e-05 | 1.233E-05 | 1.528E-06 | 1.100E-05 | 1.400E-05 |
| 14564 | 729 | K729D | B | 1.3e-05 | 9.6e-06 | 1e-05 | 1.087E-05 | 1.858E-06 | 9.600E-06 | 1.300E-05 |
| 14565 | 729 | K729C | B | 1e-05 | 1e-05 | 9.5e-06 | 9.833E-06 | 2.887E-07 | 9.500E-06 | 1.000E-05 |
| 14566 | 729 | K729E | B | 1e-05 | 8.2e-06 | 1.1e-05 | 9.733E-06 | 1.419E-06 | 8.200E-06 | 1.100E-05 |
| 14567 | 729 | K729Q | B | 1.3e-05 | 9.9e-06 | 1.2e-05 | 1.163E-05 | 1.582E-06 | 9.900E-06 | 1.300E-05 |
| 14568 | 729 | K729G | B | 9.8e-06 | 1.3e-05 | 1e-05 | 1.093E-05 | 1.793E-06 | 9.800E-06 | 1.300E-05 |
| 14569 | 729 | K729H | B | 8.3e-06 | 1.6e-05 | 1.2e-05 | 1.210E-05 | 3.851E-06 | 8.300E-06 | 1.600E-05 |
| 14570 | 729 | K729I | B | 1.5e-05 | 1.5e-05 | 1.6e-05 | 1.533E-05 | 5.774E-07 | 1.500E-05 | 1.600E-05 |
| 14571 | 729 | K729L | B | 8.6e-06 | 1.1e-05 | 1.5e-05 | 1.153E-05 | 3.233E-06 | 8.600E-06 | 1.500E-05 |
| 14572 | 729 | K729K | B | 9.4e-06 | 8.6e-06 | 1.4e-05 | 1.067E-05 | 2.914E-06 | 8.600E-06 | 1.400E-05 |
| 14573 | 729 | K729M | B | 1.5e-05 | 9.8e-06 | 1.6e-05 | 1.360E-05 | 3.329E-06 | 9.800E-06 | 1.600E-05 |
| 14574 | 729 | K729F | B | 1.3e-05 | 1.5e-05 | 9.7e-06 | 1.257E-05 | 2.676E-06 | 9.700E-06 | 1.500E-05 |
| 14575 | 729 | K729P | B | 6.8e-06 | 1.3e-05 | 9.7e-06 | 9.833E-06 | 3.102E-06 | 6.800E-06 | 1.300E-05 |
| 14576 | 729 | K729S | B | 1.1e-05 | 1.3e-05 | 1.1e-05 | 1.167E-05 | 1.155E-06 | 1.100E-05 | 1.300E-05 |
| 14577 | 729 | K729T | B | 1.4e-05 | 1.1e-05 | 1.5e-05 | 1.333E-05 | 2.082E-06 | 1.100E-05 | 1.500E-05 |
| 14578 | 729 | K729W | B | 1.4e-05 | 1.4e-05 | 1e-05 | 1.267E-05 | 2.309E-06 | 1.000E-05 | 1.400E-05 |
| 14579 | 729 | K729Y | B | 6e-06 | 1e-05 | 1.4e-05 | 1.000E-05 | 4.000E-06 | 6.000E-06 | 1.400E-05 |
| 14580 | 729 | K729V | B | 8.5e-06 | 9.8e-06 | 1.6e-05 | 1.143E-05 | 4.008E-06 | 8.500E-06 | 1.600E-05 |
| 14581 | 730 | R730A | B | 1.2e-05 | 9.2e-06 | 1.6e-05 | 1.240E-05 | 3.418E-06 | 9.200E-06 | 1.600E-05 |
| 14582 | 730 | R730R | B | 1.3e-05 | 1.6e-05 | 1.1e-05 | 1.333E-05 | 2.517E-06 | 1.100E-05 | 1.600E-05 |
| 14583 | 730 | R730N | B | 9e-06 | 1.4e-05 | 1.1e-05 | 1.133E-05 | 2.517E-06 | 9.000E-06 | 1.400E-05 |
| 14584 | 730 | R730D | B | 9.1e-06 | 7.9e-06 | 1.1e-05 | 9.333E-06 | 1.563E-06 | 7.900E-06 | 1.100E-05 |
| 14585 | 730 | R730C | B | 9.8e-06 | 1.4e-05 | 1.2e-05 | 1.193E-05 | 2.101E-06 | 9.800E-06 | 1.400E-05 |
| 14586 | 730 | R730E | B | 1.2e-05 | 1e-05 | 1.1e-05 | 1.100E-05 | 1.000E-06 | 1.000E-05 | 1.200E-05 |
| 14587 | 730 | R730Q | B | 9.3e-06 | 1.5e-05 | 1.5e-05 | 1.310E-05 | 3.291E-06 | 9.300E-06 | 1.500E-05 |
| 14588 | 730 | R730G | B | 9.8e-06 | 1.5e-05 | 1.2e-05 | 1.227E-05 | 2.610E-06 | 9.800E-06 | 1.500E-05 |
| 14589 | 730 | R730H | B | 1.5e-05 | 1.4e-05 | 1.5e-05 | 1.467E-05 | 5.774E-07 | 1.400E-05 | 1.500E-05 |
| 14590 | 730 | R730I | B | 1.1e-05 | 1.1e-05 | 7.6e-06 | 9.867E-06 | 1.963E-06 | 7.600E-06 | 1.100E-05 |
| 14591 | 730 | R730L | B | 8.8e-06 | 1.3e-05 | 8.8e-06 | 1.020E-05 | 2.425E-06 | 8.800E-06 | 1.300E-05 |
| 14592 | 730 | R730K | B | 1e-05 | 1.9e-05 | 1.4e-05 | 1.433E-05 | 4.509E-06 | 1.000E-05 | 1.900E-05 |
| 14593 | 730 | R730M | B | 1.6e-05 | 1.6e-05 | 1.4e-05 | 1.533E-05 | 1.155E-06 | 1.400E-05 | 1.600E-05 |
| 14594 | 730 | R730F | B | 1.1e-05 | 1e-05 | 1.5e-05 | 1.200E-05 | 2.646E-06 | 1.000E-05 | 1.500E-05 |
| 14595 | 730 | R730P | B | 1.6e-05 | 1.7e-05 | 1.4e-05 | 1.567E-05 | 1.528E-06 | 1.400E-05 | 1.700E-05 |
| 14596 | 730 | R730S | B | 9.8e-06 | 1.6e-05 | 1.3e-05 | 1.293E-05 | 3.101E-06 | 9.800E-06 | 1.600E-05 |
| 14597 | 730 | R730T | B | 1.5e-05 | 1.4e-05 | 1.2e-05 | 1.367E-05 | 1.528E-06 | 1.200E-05 | 1.500E-05 |
| 14598 | 730 | R730W | B | 1.1e-05 | 7.9e-06 | 1.6e-05 | 1.163E-05 | 4.087E-06 | 7.900E-06 | 1.600E-05 |
| 14599 | 730 | R730Y | B | 1.3e-05 | 1.1e-05 | 1.5e-05 | 1.300E-05 | 2.000E-06 | 1.100E-05 | 1.500E-05 |
| 14600 | 730 | R730V | B | 1.6e-05 | 1.4e-05 | 1.1e-05 | 1.367E-05 | 2.517E-06 | 1.100E-05 | 1.600E-05 |
| 14601 | 731 | T731A | B | 1.5e-05 | 8.2e-06 | 1.3e-05 | 1.207E-05 | 3.495E-06 | 8.200E-06 | 1.500E-05 |
| 14602 | 731 | T731R | B | 1.1e-05 | 1.7e-05 | 1.2e-05 | 1.333E-05 | 3.215E-06 | 1.100E-05 | 1.700E-05 |
| 14603 | 731 | T731N | B | 9.4e-06 | 8.9e-06 | 1.5e-05 | 1.110E-05 | 3.387E-06 | 8.900E-06 | 1.500E-05 |
| 14604 | 731 | T731D | B | 1e-05 | 9.3e-06 | 1.1e-05 | 1.010E-05 | 8.544E-07 | 9.300E-06 | 1.100E-05 |
| 14605 | 731 | T731C | B | 1e-05 | 1.1e-05 | 9.8e-06 | 1.027E-05 | 6.429E-07 | 9.800E-06 | 1.100E-05 |
| 14606 | 731 | T731E | B | 1.3e-05 | 1.1e-05 | 9.5e-06 | 1.117E-05 | 1.756E-06 | 9.500E-06 | 1.300E-05 |
| 14607 | 731 | T731Q | B | 1.5e-05 | 1.5e-05 | 1.5e-05 | 1.500E-05 | 0.000E+00 | 1.500E-05 | 1.500E-05 |
| 14608 | 731 | T731G | B | 1.2e-05 | 1e-05 | 1.1e-05 | 1.100E-05 | 1.000E-06 | 1.000E-05 | 1.200E-05 |
| 14609 | 731 | T731H | B | 1.3e-05 | 9.8e-06 | 9.9e-06 | 1.090E-05 | 1.819E-06 | 9.800E-06 | 1.300E-05 |
| 14610 | 731 | T731I | B | 1.6e-05 | 1.6e-05 | 1.7e-05 | 1.633E-05 | 5.774E-07 | 1.600E-05 | 1.700E-05 |
| 14611 | 731 | T731L | B | 1e-05 | 1.5e-05 | 1.4e-05 | 1.300E-05 | 2.646E-06 | 1.000E-05 | 1.500E-05 |
| 14612 | 731 | T731K | B | 1.4e-05 | 1.1e-05 | 1.1e-05 | 1.200E-05 | 1.732E-06 | 1.100E-05 | 1.400E-05 |
| 14613 | 731 | T731M | B | 1.2e-05 | 1.6e-05 | 1.3e-05 | 1.367E-05 | 2.082E-06 | 1.200E-05 | 1.600E-05 |
| 14614 | 731 | T731F | B | 1.6e-05 | 1.1e-05 | 1.6e-05 | 1.433E-05 | 2.887E-06 | 1.100E-05 | 1.600E-05 |
| 14615 | 731 | T731P | B | 1.5e-05 | 1.5e-05 | 9.9e-06 | 1.330E-05 | 2.944E-06 | 9.900E-06 | 1.500E-05 |
| 14616 | 731 | T731S | B | 1.6e-05 | 1e-05 | 1.5e-05 | 1.367E-05 | 3.215E-06 | 1.000E-05 | 1.600E-05 |
| 14617 | 731 | T731T | B | 1e-05 | 9.7e-06 | 1e-05 | 9.900E-06 | 1.732E-07 | 9.700E-06 | 1.000E-05 |
| 14618 | 731 | T731W | B | 1.6e-05 | 1.7e-05 | 1.2e-05 | 1.500E-05 | 2.646E-06 | 1.200E-05 | 1.700E-05 |
| 14619 | 731 | T731Y | B | 1.1e-05 | 1.6e-05 | 1.7e-05 | 1.467E-05 | 3.215E-06 | 1.100E-05 | 1.700E-05 |

| | | | | | | | | | | |
|---|---|---|---|---|---|---|---|---|---|---|
| 14620 | 731 | T731V | B | 1.5e-05 | 1.5e-05 | 1.3e-05 | 1.433E-05 | 1.155E-06 | 1.300E-05 | 1.500E-05 |
| 14621 | 732 | V732A | B | 8.5e-06 | 1.6e-05 | 9.2e-06 | 1.123E-05 | 4.143E-06 | 8.500E-06 | 1.600E-05 |
| 14622 | 732 | V732R | B | 1.4e-05 | 1.6e-05 | 1e-05 | 1.333E-05 | 3.055E-06 | 1.000E-05 | 1.600E-05 |
| 14623 | 732 | V732N | B | 1.2e-05 | 9.5e-06 | 1.6e-05 | 1.250E-05 | 3.279E-06 | 9.500E-06 | 1.600E-05 |
| 14624 | 732 | V732D | B | 1.4e-05 | 1.3e-05 | 1.4e-05 | 1.367E-05 | 5.774E-07 | 1.300E-05 | 1.400E-05 |
| 14625 | 732 | V732C | B | 1.3e-05 | 1.6e-05 | 1.5e-05 | 1.467E-05 | 1.528E-06 | 1.300E-05 | 1.600E-05 |
| 14626 | 732 | V732E | B | 1.7e-05 | 1.4e-05 | 4.9e-06 | 1.197E-05 | 6.301E-06 | 4.900E-06 | 1.700E-05 |
| 14627 | 732 | V732Q | B | 9.5e-06 | 1.4e-05 | 1.5e-05 | 1.283E-05 | 2.930E-06 | 9.500E-06 | 1.500E-05 |
| 14628 | 732 | V732G | B | 1.5e-05 | 1.4e-05 | 1.5e-05 | 1.467E-05 | 5.774E-07 | 1.400E-05 | 1.500E-05 |
| 14629 | 732 | V732H | B | 1.6e-05 | 1.5e-05 | 1.5e-05 | 1.533E-05 | 5.774E-07 | 1.500E-05 | 1.600E-05 |
| 14630 | 732 | V732I | B | 1.2e-05 | 9.3e-06 | 9.6e-06 | 1.030E-05 | 1.480E-06 | 9.300E-06 | 1.200E-05 |
| 14631 | 732 | V732L | B | 1.6e-05 | 1e-05 | 1e-05 | 1.200E-05 | 3.464E-06 | 1.000E-05 | 1.600E-05 |
| 14632 | 732 | V732K | B | 1.2e-05 | 1.4e-05 | 1.4e-05 | 1.333E-05 | 1.155E-06 | 1.200E-05 | 1.400E-05 |
| 14633 | 732 | V732M | B | 8.6e-06 | 1e-05 | 1.1e-05 | 9.867E-06 | 1.206E-06 | 8.600E-06 | 1.100E-05 |
| 14634 | 732 | V732F | B | 1.5e-05 | 1.1e-05 | 1.6e-05 | 1.400E-05 | 2.646E-06 | 1.100E-05 | 1.600E-05 |
| 14635 | 732 | V732P | B | 1.2e-05 | 9.9e-06 | 1.3e-05 | 1.163E-05 | 1.582E-06 | 9.900E-06 | 1.300E-05 |
| 14636 | 732 | V732S | B | 1.1e-05 | 1.6e-05 | 9.8e-06 | 1.227E-05 | 3.288E-06 | 9.800E-06 | 1.600E-05 |
| 14637 | 732 | V732T | B | 9.1e-06 | 1.5e-05 | 1.5e-05 | 1.303E-05 | 3.406E-06 | 9.100E-06 | 1.500E-05 |
| 14638 | 732 | V732W | B | 1.9e-05 | 1.5e-05 | 1.5e-05 | 1.633E-05 | 2.309E-06 | 1.500E-05 | 1.900E-05 |
| 14639 | 732 | V732Y | B | 1.7e-05 | 1.5e-05 | 8.6e-06 | 1.353E-05 | 4.388E-06 | 8.600E-06 | 1.700E-05 |
| 14640 | 732 | V732V | B | 1.5e-05 | 1.4e-05 | 9.6e-06 | 1.287E-05 | 2.873E-06 | 9.600E-06 | 1.500E-05 |
| 14641 | 733 | A733A | B | 1.1e-05 | 1.4e-05 | 1.2e-05 | 1.233E-05 | 1.528E-06 | 1.100E-05 | 1.400E-05 |
| 14642 | 733 | A733R | B | 1.4e-05 | 1.5e-05 | 1.6e-05 | 1.500E-05 | 1.000E-06 | 1.400E-05 | 1.600E-05 |
| 14643 | 733 | A733N | B | 1.5e-05 | 9.1e-06 | 1.7e-05 | 1.370E-05 | 4.107E-06 | 9.100E-06 | 1.700E-05 |
| 14644 | 733 | A733D | B | 1.3e-05 | 9.2e-06 | 1e-05 | 1.073E-05 | 2.003E-06 | 9.200E-06 | 1.300E-05 |
| 14645 | 733 | A733C | B | 1.5e-05 | 1e-05 | 1e-05 | 1.167E-05 | 2.887E-06 | 1.000E-05 | 1.500E-05 |
| 14646 | 733 | A733E | B | 2.2e-05 | 1.4e-05 | 8.3e-06 | 1.477E-05 | 6.882E-06 | 8.300E-06 | 2.200E-05 |
| 14647 | 733 | A733Q | B | 1.1e-05 | 1.3e-05 | 9.8e-06 | 1.127E-05 | 1.617E-06 | 9.800E-06 | 1.300E-05 |
| 14648 | 733 | A733G | B | 1.2e-05 | 1.7e-05 | 1.2e-05 | 1.367E-05 | 2.887E-06 | 1.200E-05 | 1.700E-05 |
| 14649 | 733 | A733H | B | 1e-05 | 8.8e-06 | 1.4e-05 | 1.093E-05 | 2.723E-06 | 8.800E-06 | 1.400E-05 |
| 14650 | 733 | A733I | B | 1.6e-05 | 1e-05 | 7.8e-06 | 1.127E-05 | 4.244E-06 | 7.800E-06 | 1.600E-05 |
| 14651 | 733 | A733L | B | 1.1e-05 | 1.7e-05 | 9.3e-06 | 1.243E-05 | 4.045E-06 | 9.300E-06 | 1.700E-05 |
| 14652 | 733 | A733K | B | 1.1e-05 | 9.9e-06 | 1.5e-05 | 1.197E-05 | 2.684E-06 | 9.900E-06 | 1.500E-05 |
| 14653 | 733 | A733M | B | 1.5e-05 | 1.1e-05 | 7.8e-06 | 1.127E-05 | 3.607E-06 | 7.800E-06 | 1.500E-05 |
| 14654 | 733 | A733F | B | 1.5e-05 | 1.2e-05 | 1.5e-05 | 1.400E-05 | 1.732E-06 | 1.200E-05 | 1.500E-05 |
| 14655 | 733 | A733P | B | 1.5e-05 | 1.2e-05 | 1.4e-05 | 1.367E-05 | 1.528E-06 | 1.200E-05 | 1.500E-05 |
| 14656 | 733 | A733S | B | 9.1e-06 | 1.5e-05 | 1.6e-05 | 1.337E-05 | 3.729E-06 | 9.100E-06 | 1.600E-05 |
| 14657 | 733 | A733T | B | 1e-05 | 1.6e-05 | 1e-05 | 1.200E-05 | 3.464E-06 | 1.000E-05 | 1.600E-05 |
| 14658 | 733 | A733W | B | 1.4e-05 | 1.6e-05 | 1e-05 | 1.333E-05 | 3.055E-06 | 1.000E-05 | 1.600E-05 |
| 14659 | 733 | A733Y | B | 9.3e-06 | 8.9e-06 | 1.6e-05 | 1.140E-05 | 3.989E-06 | 8.900E-06 | 1.600E-05 |
| 14660 | 733 | A733V | B | 1.1e-05 | 1.4e-05 | 1.5e-05 | 1.333E-05 | 2.082E-06 | 1.100E-05 | 1.500E-05 |
| 14661 | 734 | A734A | B | 1.8e-05 | 1.5e-05 | 1.6e-05 | 1.633E-05 | 1.528E-06 | 1.500E-05 | 1.800E-05 |
| 14662 | 734 | A734R | B | 1e-05 | 1.5e-05 | 1.6e-05 | 1.367E-05 | 3.215E-06 | 1.000E-05 | 1.600E-05 |
| 14663 | 734 | A734N | B | 8.6e-06 | 1.6e-05 | 1.3e-05 | 1.253E-05 | 3.722E-06 | 8.600E-06 | 1.600E-05 |
| 14664 | 734 | A734D | B | 1.5e-05 | 1.1e-05 | 1.5e-05 | 1.367E-05 | 2.309E-06 | 1.100E-05 | 1.500E-05 |
| 14665 | 734 | A734C | B | 9.5e-06 | 1.1e-05 | 1.5e-05 | 1.183E-05 | 2.843E-06 | 9.500E-06 | 1.500E-05 |
| 14666 | 734 | A734E | B | 1.6e-05 | 1.5e-05 | 1.1e-05 | 1.400E-05 | 2.646E-06 | 1.100E-05 | 1.600E-05 |
| 14667 | 734 | A734Q | B | 1.6e-05 | 1.3e-05 | 1.5e-05 | 1.467E-05 | 1.528E-06 | 1.300E-05 | 1.600E-05 |
| 14668 | 734 | A734G | B | 1.1e-05 | 1e-05 | 1.2e-05 | 1.100E-05 | 1.000E-06 | 1.000E-05 | 1.200E-05 |
| 14669 | 734 | A734H | B | 9e-06 | 9e-06 | 9.4e-06 | 9.133E-06 | 2.309E-07 | 9.000E-06 | 9.400E-06 |
| 14670 | 734 | A734I | B | 1.6e-05 | 9.4e-06 | 1.1e-05 | 1.213E-05 | 3.443E-06 | 9.400E-06 | 1.600E-05 |
| 14671 | 734 | A734L | B | 1.6e-05 | 9.9e-06 | 8.5e-06 | 1.147E-05 | 3.988E-06 | 8.500E-06 | 1.600E-05 |
| 14672 | 734 | A734K | B | 1.6e-05 | 1.1e-05 | 1e-05 | 1.233E-05 | 3.215E-06 | 1.000E-05 | 1.600E-05 |
| 14673 | 734 | A734M | B | 1.5e-05 | 8.3e-06 | 1.6e-05 | 1.310E-05 | 4.187E-06 | 8.300E-06 | 1.600E-05 |
| 14674 | 734 | A734F | B | 1.3e-05 | 1.4e-05 | 1.1e-05 | 1.267E-05 | 1.528E-06 | 1.100E-05 | 1.400E-05 |
| 14675 | 734 | A734P | B | 1.2e-05 | 1.6e-05 | 1.5e-05 | 1.433E-05 | 2.082E-06 | 1.200E-05 | 1.600E-05 |
| 14676 | 734 | A734S | B | 1.7e-05 | 1e-05 | 1.2e-05 | 1.300E-05 | 3.606E-06 | 1.000E-05 | 1.700E-05 |
| 14677 | 734 | A734T | B | 1.4e-05 | 9.9e-06 | 9.4e-06 | 1.110E-05 | 2.524E-06 | 9.400E-06 | 1.400E-05 |
| 14678 | 734 | A734W | B | 1.2e-05 | 9.7e-06 | 1.3e-05 | 1.157E-05 | 1.692E-06 | 9.700E-06 | 1.300E-05 |
| 14679 | 734 | A734Y | B | 1.1e-05 | 1.4e-05 | 1.1e-05 | 1.200E-05 | 1.732E-06 | 1.100E-05 | 1.400E-05 |
| 14680 | 734 | A734V | B | 1.5e-05 | 9.8e-06 | 8e-06 | 1.093E-05 | 3.635E-06 | 8.000E-06 | 1.500E-05 |
| 14681 | 735 | P735A | B | 1.4e-05 | 1.3e-05 | 1.1e-05 | 1.267E-05 | 1.528E-06 | 1.100E-05 | 1.400E-05 |
| 14682 | 735 | P735R | B | 1.2e-05 | 1.1e-05 | 1.2e-05 | 1.167E-05 | 5.774E-07 | 1.100E-05 | 1.200E-05 |
| 14683 | 735 | P735N | B | 1e-05 | 1e-05 | 1.3e-05 | 1.100E-05 | 1.732E-06 | 1.000E-05 | 1.300E-05 |
| 14684 | 735 | P735D | B | 1e-05 | 1e-05 | 1.5e-05 | 1.167E-05 | 2.887E-06 | 1.000E-05 | 1.500E-05 |
| 14685 | 735 | P735C | B | 1.2e-05 | 1e-05 | 1.4e-05 | 1.200E-05 | 2.000E-06 | 1.000E-05 | 1.400E-05 |
| 14686 | 735 | P735E | B | 1.6e-05 | 1.4e-05 | 1.5e-05 | 1.500E-05 | 1.000E-06 | 1.400E-05 | 1.600E-05 |
| 14687 | 735 | P735Q | B | 1e-05 | 9.7e-06 | 1.1e-05 | 1.023E-05 | 6.807E-07 | 9.700E-06 | 1.100E-05 |
| 14688 | 735 | P735G | B | 1.6e-05 | 8.3e-06 | 9.7e-06 | 1.133E-05 | 4.102E-06 | 8.300E-06 | 1.600E-05 |
| 14689 | 735 | P735H | B | 1.2e-05 | 1.1e-05 | 1.3e-05 | 1.200E-05 | 1.000E-06 | 1.100E-05 | 1.300E-05 |
| 14690 | 735 | P735I | B | 1.6e-05 | 1.5e-05 | 1.5e-05 | 1.533E-05 | 5.774E-07 | 1.500E-05 | 1.600E-05 |
| 14691 | 735 | P735L | B | 1e-05 | 9.2e-06 | 1.4e-05 | 1.107E-05 | 2.572E-06 | 9.200E-06 | 1.400E-05 |
| 14692 | 735 | P735K | B | 1e-05 | 1e-05 | 1.4e-05 | 1.133E-05 | 2.309E-06 | 1.000E-05 | 1.400E-05 |
| 14693 | 735 | P735M | B | 1.6e-05 | 6.5e-06 | 1e-05 | 1.083E-05 | 4.805E-06 | 6.500E-06 | 1.600E-05 |
| 14694 | 735 | P735F | B | 1.1e-05 | 1.6e-05 | 1.1e-05 | 1.267E-05 | 2.887E-06 | 1.100E-05 | 1.600E-05 |
| 14695 | 735 | P735P | B | 6.2e-06 | 9.8e-06 | 1e-05 | 8.667E-06 | 2.139E-06 | 6.200E-06 | 1.000E-05 |
| 14696 | 735 | P735S | B | 8.4e-06 | 1.5e-05 | 1.7e-05 | 1.347E-05 | 4.500E-06 | 8.400E-06 | 1.700E-05 |
| 14697 | 735 | P735T | B | 9.8e-06 | 1.3e-05 | 1.6e-05 | 1.293E-05 | 3.101E-06 | 9.800E-06 | 1.600E-05 |
| 14698 | 735 | P735W | B | 9.5e-06 | 8.7e-06 | 1.5e-05 | 1.107E-05 | 3.430E-06 | 8.700E-06 | 1.500E-05 |
| 14699 | 735 | P735Y | B | 9.9e-06 | 1.2e-05 | 1.6e-05 | 1.263E-05 | 3.099E-06 | 9.900E-06 | 1.600E-05 |
| 14700 | 735 | P735V | B | 1.3e-05 | 1.8e-05 | 1.6e-05 | 1.567E-05 | 2.517E-06 | 1.300E-05 | 1.800E-05 |
| 14701 | 736 | S736A | B | 1.5e-05 | 1.1e-05 | 1.3e-05 | 1.300E-05 | 2.000E-06 | 1.100E-05 | 1.500E-05 |
| 14702 | 736 | S736R | B | 1.4e-05 | 1.5e-05 | 9.1e-06 | 1.270E-05 | 3.158E-06 | 9.100E-06 | 1.500E-05 |
| 14703 | 736 | S736N | B | 1.1e-05 | 1.5e-05 | 1.6e-05 | 1.400E-05 | 2.646E-06 | 1.100E-05 | 1.600E-05 |
| 14704 | 736 | S736D | B | 1.5e-05 | 8.2e-06 | 1.2e-05 | 1.173E-05 | 3.408E-06 | 8.200E-06 | 1.500E-05 |

| | | | | | | | | | | |
|---|---|---|---|---|---|---|---|---|---|---|
| 14705 | 736 | S736C | B | 1.5e-05 | 1.3e-05 | 1e-05 | 1.267E-05 | 2.517E-06 | 1.000E-05 | 1.500E-05 |
| 14706 | 736 | S736E | B | 9.5e-06 | 1.4e-05 | 9.5e-06 | 1.100E-05 | 2.598E-06 | 9.500E-06 | 1.400E-05 |
| 14707 | 736 | S736Q | B | 1.2e-05 | 1.1e-05 | 1.7e-05 | 1.333E-05 | 3.215E-06 | 1.100E-05 | 1.700E-05 |
| 14708 | 736 | S736G | B | 1.6e-05 | 1.5e-05 | 1.1e-05 | 1.400E-05 | 2.646E-06 | 1.100E-05 | 1.600E-05 |
| 14709 | 736 | S736H | B | 1.1e-05 | 1e-05 | 1.4e-05 | 1.167E-05 | 2.082E-06 | 1.000E-05 | 1.400E-05 |
| 14710 | 736 | S736I | B | 1e-05 | 1.4e-05 | 1.5e-05 | 1.300E-05 | 2.646E-06 | 1.000E-05 | 1.500E-05 |
| 14711 | 736 | S736L | B | 1e-05 | 1.6e-05 | 1.7e-05 | 1.433E-05 | 3.786E-06 | 1.000E-05 | 1.700E-05 |
| 14712 | 736 | S736K | B | 1.3e-05 | 1.1e-05 | 1e-05 | 1.133E-05 | 1.528E-06 | 1.000E-05 | 1.300E-05 |
| 14713 | 736 | S736M | B | 1.6e-05 | 1.1e-05 | 1.2e-05 | 1.300E-05 | 2.646E-06 | 1.100E-05 | 1.600E-05 |
| 14714 | 736 | S736F | B | 1.1e-05 | 1.7e-05 | 1.3e-05 | 1.367E-05 | 3.055E-06 | 1.100E-05 | 1.700E-05 |
| 14715 | 736 | S736P | B | 1.6e-05 | 1.2e-05 | 8.7e-06 | 1.223E-05 | 3.656E-06 | 8.700E-06 | 1.600E-05 |
| 14716 | 736 | S736S | B | 1.6e-05 | 1.1e-05 | 1.6e-05 | 1.433E-05 | 2.887E-06 | 1.100E-05 | 1.600E-05 |
| 14717 | 736 | S736T | B | 1.1e-05 | 1.6e-05 | 1.4e-05 | 1.367E-05 | 2.517E-06 | 1.100E-05 | 1.600E-05 |
| 14718 | 736 | S736W | B | 1.5e-05 | 1.1e-05 | 9.8e-06 | 1.193E-05 | 2.723E-06 | 9.800E-06 | 1.500E-05 |
| 14719 | 736 | S736Y | B | 1.3e-05 | 1e-05 | 1.5e-05 | 1.267E-05 | 2.517E-06 | 1.000E-05 | 1.500E-05 |
| 14720 | 736 | S736V | B | 6e-06 | 9.8e-06 | 1.1e-05 | 8.933E-06 | 2.610E-06 | 6.000E-06 | 1.100E-05 |
| 14721 | 737 | V737A | B | 1e-05 | 1.6e-05 | 1.5e-05 | 1.367E-05 | 3.215E-06 | 1.000E-05 | 1.600E-05 |
| 14722 | 737 | V737R | B | 1.4e-05 | 1.1e-05 | 1.1e-05 | 1.200E-05 | 1.732E-06 | 1.100E-05 | 1.400E-05 |
| 14723 | 737 | V737N | B | 1.5e-05 | 1.5e-05 | 1.5e-05 | 1.500E-05 | 0.000E+00 | 1.500E-05 | 1.500E-05 |
| 14724 | 737 | V737D | B | 9.7e-06 | 1.4e-05 | 1.2e-05 | 1.190E-05 | 2.152E-06 | 9.700E-06 | 1.400E-05 |
| 14725 | 737 | V737C | B | 1.6e-05 | 1.1e-05 | 1.5e-05 | 1.400E-05 | 2.646E-06 | 1.100E-05 | 1.600E-05 |
| 14726 | 737 | V737E | B | 1.2e-05 | 1.5e-05 | 1.4e-05 | 1.367E-05 | 1.528E-06 | 1.200E-05 | 1.500E-05 |
| 14727 | 737 | V737Q | B | 1.1e-05 | 1.3e-05 | 1.4e-05 | 1.267E-05 | 1.528E-06 | 1.100E-05 | 1.400E-05 |
| 14728 | 737 | V737G | B | 1e-05 | 1e-05 | 1.1e-05 | 1.033E-05 | 5.774E-07 | 1.000E-05 | 1.100E-05 |
| 14729 | 737 | V737H | B | 1.6e-05 | 1.4e-05 | 1.3e-05 | 1.433E-05 | 1.528E-06 | 1.300E-05 | 1.600E-05 |
| 14730 | 737 | V737I | B | 1e-05 | 1.5e-05 | 9.8e-06 | 1.160E-05 | 2.946E-06 | 9.800E-06 | 1.500E-05 |
| 14731 | 737 | V737L | B | 1.3e-05 | 1e-05 | 1.1e-05 | 1.133E-05 | 1.528E-06 | 1.000E-05 | 1.300E-05 |
| 14732 | 737 | V737K | B | 1.5e-05 | 8.7e-06 | 1e-05 | 1.123E-05 | 3.326E-06 | 8.700E-06 | 1.500E-05 |
| 14733 | 737 | V737M | B | 1.3e-05 | 1.2e-05 | 1e-05 | 1.167E-05 | 1.528E-06 | 1.000E-05 | 1.300E-05 |
| 14734 | 737 | V737F | B | 9.2e-06 | 1.3e-05 | 8.5e-06 | 1.023E-05 | 2.421E-06 | 8.500E-06 | 1.300E-05 |
| 14735 | 737 | V737P | B | 1.5e-05 | 1.1e-05 | 9.9e-06 | 1.197E-05 | 2.684E-06 | 9.900E-06 | 1.500E-05 |
| 14736 | 737 | V737S | B | 1e-05 | 1.1e-05 | 1e-05 | 1.033E-05 | 5.774E-07 | 1.000E-05 | 1.100E-05 |
| 14737 | 737 | V737T | B | 1.5e-05 | 1.5e-05 | 1.5e-05 | 1.500E-05 | 0.000E+00 | 1.500E-05 | 1.500E-05 |
| 14738 | 737 | V737W | B | 1e-05 | 9.6e-06 | 1.4e-05 | 1.120E-05 | 2.433E-06 | 9.600E-06 | 1.400E-05 |
| 14739 | 737 | V737Y | B | 1.2e-05 | 1.1e-05 | 9.8e-06 | 1.093E-05 | 1.102E-06 | 9.800E-06 | 1.200E-05 |
| 14740 | 737 | V737V | B | 1e-05 | 1.3e-05 | 1.3e-05 | 1.200E-05 | 1.732E-06 | 1.000E-05 | 1.300E-05 |
| 14741 | 738 | F738A | B | 1.5e-05 | 9.5e-06 | 1.3e-05 | 1.250E-05 | 2.784E-06 | 9.500E-06 | 1.500E-05 |
| 14742 | 738 | F738R | B | 1.3e-05 | 9.2e-06 | 1.4e-05 | 1.207E-05 | 2.532E-06 | 9.200E-06 | 1.400E-05 |
| 14743 | 738 | F738N | B | 1e-05 | 1.2e-05 | 1.5e-05 | 1.233E-05 | 2.517E-06 | 1.000E-05 | 1.500E-05 |
| 14744 | 738 | F738D | B | 1e-05 | 9.5e-06 | 1.1e-05 | 1.017E-05 | 7.638E-07 | 9.500E-06 | 1.100E-05 |
| 14745 | 738 | F738C | B | 9.5e-06 | 9e-06 | 1e-05 | 9.500E-06 | 5.000E-07 | 9.000E-06 | 1.000E-05 |
| 14746 | 738 | F738E | B | 1.1e-05 | 8e-06 | 9.1e-06 | 9.367E-06 | 1.518E-06 | 8.000E-06 | 1.100E-05 |
| 14747 | 738 | F738Q | B | 1e-05 | 1.3e-05 | 1.3e-05 | 1.200E-05 | 1.732E-06 | 1.000E-05 | 1.300E-05 |
| 14748 | 738 | F738G | B | 1e-05 | 1.7e-05 | 1.5e-05 | 1.400E-05 | 3.606E-06 | 1.000E-05 | 1.700E-05 |
| 14749 | 738 | F738H | B | 1.4e-05 | 9.3e-06 | 1e-05 | 1.110E-05 | 2.536E-06 | 9.300E-06 | 1.400E-05 |
| 14750 | 738 | F738I | B | 1.6e-05 | 1.5e-05 | 1.2e-05 | 1.433E-05 | 2.082E-06 | 1.200E-05 | 1.600E-05 |
| 14751 | 738 | F738L | B | 1.2e-05 | 9.9e-06 | 1.3e-05 | 1.163E-05 | 1.582E-06 | 9.900E-06 | 1.300E-05 |
| 14752 | 738 | F738K | B | 1.5e-05 | 7e-06 | 1.6e-05 | 1.267E-05 | 4.933E-06 | 7.000E-06 | 1.600E-05 |
| 14753 | 738 | F738M | B | 9.8e-06 | 8.8e-06 | 1.1e-05 | 9.867E-06 | 1.102E-06 | 8.800E-06 | 1.100E-05 |
| 14754 | 738 | F738F | B | 1.6e-05 | 1.6e-05 | 1.4e-05 | 1.533E-05 | 1.155E-06 | 1.400E-05 | 1.600E-05 |
| 14755 | 738 | F738P | B | 1.2e-05 | 1.5e-05 | 1e-05 | 1.233E-05 | 2.517E-06 | 1.000E-05 | 1.500E-05 |
| 14756 | 738 | F738S | B | 1.5e-05 | 8.3e-06 | 1.3e-05 | 1.210E-05 | 3.439E-06 | 8.300E-06 | 1.500E-05 |
| 14757 | 738 | F738T | B | 1.5e-05 | 1.5e-05 | 1.5e-05 | 1.500E-05 | 0.000E+00 | 1.500E-05 | 1.500E-05 |
| 14758 | 738 | F738W | B | 9.4e-06 | 1e-05 | 1e-05 | 9.800E-06 | 3.464E-07 | 9.400E-06 | 1.000E-05 |
| 14759 | 738 | F738Y | B | 1.3e-05 | 1e-05 | 1.7e-05 | 1.333E-05 | 3.512E-06 | 1.000E-05 | 1.700E-05 |
| 14760 | 738 | F738V | B | 1.3e-05 | 1.6e-05 | 9.2e-06 | 1.273E-05 | 3.408E-06 | 9.200E-06 | 1.600E-05 |
| 14761 | 739 | I739A | B | 1.5e-05 | 1.4e-05 | 1.5e-05 | 1.467E-05 | 5.774E-07 | 1.400E-05 | 1.500E-05 |
| 14762 | 739 | I739R | B | 1.6e-05 | 1.7e-05 | 7.8e-06 | 1.360E-05 | 5.048E-06 | 7.800E-06 | 1.700E-05 |
| 14763 | 739 | I739N | B | 1.3e-05 | 1.5e-05 | 1.6e-05 | 1.467E-05 | 1.528E-06 | 1.300E-05 | 1.600E-05 |
| 14764 | 739 | I739D | B | 1.1e-05 | 1.5e-05 | 1.6e-05 | 1.400E-05 | 2.646E-06 | 1.100E-05 | 1.600E-05 |
| 14765 | 739 | I739C | B | 1.5e-05 | 1.5e-05 | 1.4e-05 | 1.467E-05 | 5.774E-07 | 1.400E-05 | 1.500E-05 |
| 14766 | 739 | I739E | B | 1e-05 | 8.8e-06 | 9.3e-06 | 9.367E-06 | 6.028E-07 | 8.800E-06 | 1.000E-05 |
| 14767 | 739 | I739Q | B | 1.6e-05 | 1.4e-05 | 1.5e-05 | 1.500E-05 | 1.000E-06 | 1.400E-05 | 1.600E-05 |
| 14768 | 739 | I739G | B | 1.1e-05 | 1.1e-05 | 1.1e-05 | 1.100E-05 | 2.075E-21 | 1.100E-05 | 1.100E-05 |
| 14769 | 739 | I739H | B | 9.4e-06 | 1.6e-05 | 1.2e-05 | 1.247E-05 | 3.325E-06 | 9.400E-06 | 1.600E-05 |
| 14770 | 739 | I739I | B | 8.5e-06 | 1.4e-05 | 1.5e-05 | 1.250E-05 | 3.500E-06 | 8.500E-06 | 1.500E-05 |
| 14771 | 739 | I739L | B | 1.7e-05 | 1.4e-05 | 1.6e-05 | 1.567E-05 | 1.528E-06 | 1.400E-05 | 1.700E-05 |
| 14772 | 739 | I739K | B | 1.6e-05 | 1.5e-05 | 1.5e-05 | 1.533E-05 | 5.774E-07 | 1.500E-05 | 1.600E-05 |
| 14773 | 739 | I739M | B | 1.3e-05 | 1.5e-05 | 1.5e-05 | 1.433E-05 | 1.155E-06 | 1.300E-05 | 1.500E-05 |
| 14774 | 739 | I739F | B | 1.3e-05 | 9.8e-06 | 1e-05 | 1.093E-05 | 1.793E-06 | 9.800E-06 | 1.300E-05 |
| 14775 | 739 | I739P | B | 8.6e-06 | 9.8e-06 | 1e-05 | 9.467E-06 | 7.572E-07 | 8.600E-06 | 1.000E-05 |
| 14776 | 739 | I739S | B | 1.6e-05 | 8.9e-06 | 1.4e-05 | 1.297E-05 | 3.661E-06 | 8.900E-06 | 1.600E-05 |
| 14777 | 739 | I739T | B | 1.4e-05 | 1.6e-05 | 1.5e-05 | 1.500E-05 | 1.000E-06 | 1.400E-05 | 1.600E-05 |
| 14778 | 739 | I739W | B | 8.9e-06 | 1.6e-05 | 1.4e-05 | 1.297E-05 | 3.661E-06 | 8.900E-06 | 1.600E-05 |
| 14779 | 739 | I739Y | B | 8.2e-06 | 2.4e-05 | 9.6e-06 | 1.393E-05 | 8.746E-06 | 8.200E-06 | 2.400E-05 |
| 14780 | 739 | I739V | B | 1.5e-05 | 1.6e-05 | 1.6e-05 | 1.567E-05 | 5.774E-07 | 1.500E-05 | 1.600E-05 |
| 14781 | 740 | F740A | B | 1.1e-05 | 1.6e-05 | 2e-05 | 1.567E-05 | 4.509E-06 | 1.100E-05 | 2.000E-05 |
| 14782 | 740 | F740R | B | 1.4e-05 | 1e-05 | 1.3e-05 | 1.233E-05 | 2.082E-06 | 1.000E-05 | 1.400E-05 |
| 14783 | 740 | F740N | B | 1.5e-05 | 1.2e-05 | 9.4e-06 | 1.213E-05 | 2.802E-06 | 9.400E-06 | 1.500E-05 |
| 14784 | 740 | F740D | B | 1.4e-05 | 1.5e-05 | 9.7e-06 | 1.290E-05 | 2.816E-06 | 9.700E-06 | 1.500E-05 |
| 14785 | 740 | F740C | B | 9.6e-06 | 1.3e-05 | 1.4e-05 | 1.220E-05 | 2.307E-06 | 9.600E-06 | 1.400E-05 |
| 14786 | 740 | F740E | B | 1e-05 | 6.6e-06 | 1.7e-05 | 1.120E-05 | 5.303E-06 | 6.600E-06 | 1.700E-05 |
| 14787 | 740 | F740Q | B | 9.1e-06 | 1e-05 | 9.4e-06 | 9.500E-06 | 4.583E-07 | 9.100E-06 | 1.000E-05 |
| 14788 | 740 | F740G | B | 1.1e-05 | 1.5e-05 | 1.6e-05 | 1.400E-05 | 2.646E-06 | 1.100E-05 | 1.600E-05 |
| 14789 | 740 | F740H | B | 1.4e-05 | 9.6e-06 | 9.9e-06 | 1.117E-05 | 2.458E-06 | 9.600E-06 | 1.400E-05 |

| | | | | | | | | | | |
|---|---|---|---|---|---|---|---|---|---|---|
| 14790 | 740 | F740I | B | 1.6e-05 | 1.1e-05 | 1e-05 | 1.233E-05 | 3.215E-06 | 1.000E-05 | 1.600E-05 |
| 14791 | 740 | F740L | B | 1.1e-05 | 1.5e-05 | 1.7e-05 | 1.433E-05 | 3.055E-06 | 1.100E-05 | 1.700E-05 |
| 14792 | 740 | F740K | B | 5.7e-06 | 1.3e-05 | 1.3e-05 | 1.057E-05 | 4.215E-06 | 5.700E-06 | 1.300E-05 |
| 14793 | 740 | F740M | B | 1.3e-05 | 1.4e-05 | 1.4e-05 | 1.367E-05 | 5.774E-07 | 1.300E-05 | 1.400E-05 |
| 14794 | 740 | F740F | B | 8.9e-06 | 1.2e-05 | 1.5e-05 | 1.197E-05 | 3.050E-06 | 8.900E-06 | 1.500E-05 |
| 14795 | 740 | F740P | B | 1.5e-05 | 1e-05 | 1e-05 | 1.167E-05 | 2.887E-06 | 1.000E-05 | 1.500E-05 |
| 14796 | 740 | F740S | B | 9.7e-06 | 1.3e-05 | 8.1e-06 | 1.027E-05 | 2.499E-06 | 8.100E-06 | 1.300E-05 |
| 14797 | 740 | F740T | B | 1.4e-05 | 9.2e-06 | 1.6e-05 | 1.307E-05 | 3.495E-06 | 9.200E-06 | 1.600E-05 |
| 14798 | 740 | F740W | B | 1.2e-05 | 1.5e-05 | 1.5e-05 | 1.400E-05 | 1.732E-06 | 1.200E-05 | 1.500E-05 |
| 14799 | 740 | F740Y | B | 1.5e-05 | 1e-05 | 7.9e-06 | 1.097E-05 | 3.647E-06 | 7.900E-06 | 1.500E-05 |
| 14800 | 740 | F740V | B | 1.6e-05 | 1.1e-05 | 1.5e-05 | 1.400E-05 | 2.646E-06 | 1.100E-05 | 1.600E-05 |
| 14801 | 741 | P741A | B | 1.6e-05 | 1.3e-05 | 1.1e-05 | 1.333E-05 | 2.517E-06 | 1.100E-05 | 1.600E-05 |
| 14802 | 741 | P741R | B | 9.4e-06 | 1.1e-05 | 1.1e-05 | 1.047E-05 | 9.238E-07 | 9.400E-06 | 1.100E-05 |
| 14803 | 741 | P741N | B | 1e-05 | 7.6e-06 | 1.1e-05 | 9.533E-06 | 1.747E-06 | 7.600E-06 | 1.100E-05 |
| 14804 | 741 | P741D | B | 6.9e-06 | 1.1e-05 | 1.4e-05 | 1.063E-05 | 3.564E-06 | 6.900E-06 | 1.400E-05 |
| 14805 | 741 | P741C | B | 1.6e-05 | 1.1e-05 | 1.2e-05 | 1.300E-05 | 2.646E-06 | 1.100E-05 | 1.600E-05 |
| 14806 | 741 | P741E | B | 1.1e-05 | 9.8e-06 | 1e-05 | 1.027E-05 | 6.429E-07 | 9.800E-06 | 1.100E-05 |
| 14807 | 741 | P741Q | B | 1.6e-05 | 1.5e-05 | 1.3e-05 | 1.467E-05 | 1.528E-06 | 1.300E-05 | 1.600E-05 |
| 14808 | 741 | P741G | B | 1.1e-05 | 8e-06 | 1.5e-05 | 1.133E-05 | 3.512E-06 | 8.000E-06 | 1.500E-05 |
| 14809 | 741 | P741H | B | 1.4e-05 | 1.4e-05 | 1.4e-05 | 1.400E-05 | 0.000E+00 | 1.400E-05 | 1.400E-05 |
| 14810 | 741 | P741I | B | 1.6e-05 | 8.8e-06 | 1.2e-05 | 1.227E-05 | 3.607E-06 | 8.800E-06 | 1.600E-05 |
| 14811 | 741 | P741L | B | 1.2e-05 | 1.4e-05 | 1.6e-05 | 1.400E-05 | 2.000E-06 | 1.200E-05 | 1.600E-05 |
| 14812 | 741 | P741K | B | 1.4e-05 | 9.4e-06 | 7.5e-06 | 1.030E-05 | 3.342E-06 | 7.500E-06 | 1.400E-05 |
| 14813 | 741 | P741M | B | 1.4e-05 | 1.4e-05 | 9.8e-06 | 1.260E-05 | 2.425E-06 | 9.800E-06 | 1.400E-05 |
| 14814 | 741 | P741F | B | 1.6e-05 | 1.1e-05 | 1.1e-05 | 1.267E-05 | 2.887E-06 | 1.100E-05 | 1.600E-05 |
| 14815 | 741 | P741P | B | 1.4e-05 | 9.2e-06 | 9.3e-06 | 1.083E-05 | 2.743E-06 | 9.200E-06 | 1.400E-05 |
| 14816 | 741 | P741S | B | 1e-05 | 9.5e-06 | 1.5e-05 | 1.150E-05 | 3.041E-06 | 9.500E-06 | 1.500E-05 |
| 14817 | 741 | P741T | B | 1.6e-05 | 1.2e-05 | 1.2e-05 | 1.333E-05 | 2.309E-06 | 1.200E-05 | 1.600E-05 |
| 14818 | 741 | P741W | B | 1.1e-05 | 1.5e-05 | 1.5e-05 | 1.367E-05 | 2.309E-06 | 1.100E-05 | 1.500E-05 |
| 14819 | 741 | P741Y | B | 1e-05 | 1e-05 | 1.5e-05 | 1.167E-05 | 2.887E-06 | 1.000E-05 | 1.500E-05 |
| 14820 | 741 | P741V | B | 1.4e-05 | 9.8e-06 | 1e-05 | 1.127E-05 | 2.369E-06 | 9.800E-06 | 1.400E-05 |
| 14821 | 742 | P742A | B | 1e-05 | 1.6e-05 | 9.2e-06 | 1.173E-05 | 3.717E-06 | 9.200E-06 | 1.600E-05 |
| 14822 | 742 | P742R | B | 1.6e-05 | 9.5e-06 | 1.5e-05 | 1.350E-05 | 3.500E-06 | 9.500E-06 | 1.600E-05 |
| 14823 | 742 | P742N | B | 1.4e-05 | 1.3e-05 | 1.3e-05 | 1.333E-05 | 5.774E-07 | 1.300E-05 | 1.400E-05 |
| 14824 | 742 | P742D | B | 9.8e-06 | 1.1e-05 | 9.8e-06 | 1.020E-05 | 6.928E-07 | 9.800E-06 | 1.100E-05 |
| 14825 | 742 | P742C | B | 1.4e-05 | 8.9e-06 | 1.2e-05 | 1.163E-05 | 2.570E-06 | 8.900E-06 | 1.400E-05 |
| 14826 | 742 | P742E | B | 1e-05 | 1.5e-05 | 1.5e-05 | 1.333E-05 | 2.887E-06 | 1.000E-05 | 1.500E-05 |
| 14827 | 742 | P742Q | B | 1.3e-05 | 8.5e-06 | 8.4e-06 | 9.967E-06 | 2.627E-06 | 8.400E-06 | 1.300E-05 |
| 14828 | 742 | P742G | B | 8.6e-06 | 1.6e-05 | 1.1e-05 | 1.187E-05 | 3.775E-06 | 8.600E-06 | 1.600E-05 |
| 14829 | 742 | P742H | B | 1.5e-05 | 8.2e-06 | 1.4e-05 | 1.240E-05 | 3.672E-06 | 8.200E-06 | 1.500E-05 |
| 14830 | 742 | P742I | B | 1.6e-05 | 1.4e-05 | 1.6e-05 | 1.533E-05 | 1.155E-06 | 1.400E-05 | 1.600E-05 |
| 14831 | 742 | P742L | B | 1.4e-05 | 1.7e-05 | 9.7e-06 | 1.357E-05 | 3.669E-06 | 9.700E-06 | 1.700E-05 |
| 14832 | 742 | P742K | B | 1.3e-05 | 1.5e-05 | 1.5e-05 | 1.433E-05 | 1.155E-06 | 1.300E-05 | 1.500E-05 |
| 14833 | 742 | P742M | B | 1.5e-05 | 1.1e-05 | 1.5e-05 | 1.367E-05 | 2.309E-06 | 1.100E-05 | 1.500E-05 |
| 14834 | 742 | P742F | B | 1e-05 | 1.5e-05 | 1.2e-05 | 1.233E-05 | 2.517E-06 | 1.000E-05 | 1.500E-05 |
| 14835 | 742 | P742P | B | 1.1e-05 | 1.6e-05 | 1.8e-05 | 1.500E-05 | 3.606E-06 | 1.100E-05 | 1.800E-05 |
| 14836 | 742 | P742S | B | 1.4e-05 | 8.3e-06 | 1.4e-05 | 1.210E-05 | 3.291E-06 | 8.300E-06 | 1.400E-05 |
| 14837 | 742 | P742T | B | 1.4e-05 | 1.5e-05 | 1.1e-05 | 1.333E-05 | 2.082E-06 | 1.100E-05 | 1.500E-05 |
| 14838 | 742 | P742W | B | 1.1e-05 | 1.4e-05 | 9.1e-06 | 1.137E-05 | 2.470E-06 | 9.100E-06 | 1.400E-05 |
| 14839 | 742 | P742Y | B | 1.5e-05 | 1.1e-05 | 1.5e-05 | 1.367E-05 | 2.309E-06 | 1.100E-05 | 1.500E-05 |
| 14840 | 742 | P742V | B | 1.2e-05 | 8.4e-06 | 1e-05 | 1.013E-05 | 1.804E-06 | 8.400E-06 | 1.200E-05 |
| 14841 | 743 | S743A | B | 1.6e-05 | 1.5e-05 | 1.4e-05 | 1.500E-05 | 1.000E-06 | 1.400E-05 | 1.600E-05 |
| 14842 | 743 | S743R | B | 1.4e-05 | 1.1e-05 | 1.1e-05 | 1.200E-05 | 1.732E-06 | 1.100E-05 | 1.400E-05 |
| 14843 | 743 | S743N | B | 1.7e-05 | 1.1e-05 | 1.4e-05 | 1.400E-05 | 3.000E-06 | 1.100E-05 | 1.700E-05 |
| 14844 | 743 | S743D | B | 1.5e-05 | 1.4e-05 | 1.5e-05 | 1.467E-05 | 5.774E-07 | 1.400E-05 | 1.500E-05 |
| 14845 | 743 | S743C | B | 1.3e-05 | 1.5e-05 | 1.1e-05 | 1.300E-05 | 2.000E-06 | 1.100E-05 | 1.500E-05 |
| 14846 | 743 | S743E | B | 1.5e-05 | 1.8e-05 | 1.4e-05 | 1.567E-05 | 2.082E-06 | 1.400E-05 | 1.800E-05 |
| 14847 | 743 | S743Q | B | 1.5e-05 | 1e-05 | 1.6e-05 | 1.367E-05 | 3.215E-06 | 1.000E-05 | 1.600E-05 |
| 14848 | 743 | S743G | B | 1.6e-05 | 9.1e-06 | 1.6e-05 | 1.370E-05 | 3.984E-06 | 9.100E-06 | 1.600E-05 |
| 14849 | 743 | S743H | B | 9.8e-06 | 8.3e-06 | 1e-05 | 9.367E-06 | 9.292E-07 | 8.300E-06 | 1.000E-05 |
| 14850 | 743 | S743I | B | 8.9e-06 | 1.4e-05 | 1.4e-05 | 1.230E-05 | 2.944E-06 | 8.900E-06 | 1.400E-05 |
| 14851 | 743 | S743L | B | 1e-05 | 1.7e-05 | 1.3e-05 | 1.333E-05 | 3.512E-06 | 1.000E-05 | 1.700E-05 |
| 14852 | 743 | S743K | B | 1.2e-05 | 1.7e-05 | 9.6e-06 | 1.287E-05 | 3.775E-06 | 9.600E-06 | 1.700E-05 |
| 14853 | 743 | S743M | B | 9.4e-06 | 1.3e-05 | 1.1e-05 | 1.113E-05 | 1.804E-06 | 9.400E-06 | 1.300E-05 |
| 14854 | 743 | S743F | B | 8.4e-06 | 1.1e-05 | 1e-05 | 9.800E-06 | 1.311E-06 | 8.400E-06 | 1.100E-05 |
| 14855 | 743 | S743P | B | 1.6e-05 | 1.1e-05 | 1.1e-05 | 1.267E-05 | 2.887E-06 | 1.100E-05 | 1.600E-05 |
| 14856 | 743 | S743S | B | 1.1e-05 | 1.4e-05 | 1e-05 | 1.167E-05 | 2.082E-06 | 1.000E-05 | 1.400E-05 |
| 14857 | 743 | S743T | B | 1e-05 | 1.4e-05 | 1.4e-05 | 1.267E-05 | 2.309E-06 | 1.000E-05 | 1.400E-05 |
| 14858 | 743 | S743W | B | 9.2e-06 | 1.3e-05 | 1.5e-05 | 1.240E-05 | 2.946E-06 | 9.200E-06 | 1.500E-05 |
| 14859 | 743 | S743Y | B | 1e-05 | 1e-05 | 1e-05 | 1.000E-05 | 0.000E+00 | 1.000E-05 | 1.000E-05 |
| 14860 | 743 | S743V | B | 1.7e-05 | 9e-06 | 1.3e-05 | 1.300E-05 | 4.000E-06 | 9.000E-06 | 1.700E-05 |
| 14861 | 744 | D744A | B | 1.2e-05 | 1.5e-05 | 1.6e-05 | 1.433E-05 | 2.082E-06 | 1.200E-05 | 1.600E-05 |
| 14862 | 744 | D744R | B | 1.4e-05 | 1e-05 | 1.7e-05 | 1.367E-05 | 3.512E-06 | 1.000E-05 | 1.700E-05 |
| 14863 | 744 | D744N | B | 1.3e-05 | 1.5e-05 | 1.5e-05 | 1.433E-05 | 1.155E-06 | 1.300E-05 | 1.500E-05 |
| 14864 | 744 | D744D | B | 1.3e-05 | 1.4e-05 | 1e-05 | 1.233E-05 | 2.082E-06 | 1.000E-05 | 1.400E-05 |
| 14865 | 744 | D744C | B | 1e-05 | 1.6e-05 | 1.7e-05 | 1.433E-05 | 3.786E-06 | 1.000E-05 | 1.700E-05 |
| 14866 | 744 | D744E | B | 1.6e-05 | 1.6e-05 | 1.4e-05 | 1.533E-05 | 1.155E-06 | 1.400E-05 | 1.600E-05 |
| 14867 | 744 | D744Q | B | 1e-05 | 8.8e-06 | 1.3e-05 | 1.060E-05 | 2.163E-06 | 8.800E-06 | 1.300E-05 |
| 14868 | 744 | D744G | B | 8.5e-06 | 1.2e-05 | 1.6e-05 | 1.217E-05 | 3.753E-06 | 8.500E-06 | 1.600E-05 |
| 14869 | 744 | D744H | B | 1e-05 | 1e-05 | 1.5e-05 | 1.167E-05 | 2.887E-06 | 1.000E-05 | 1.500E-05 |
| 14870 | 744 | D744I | B | 1.5e-05 | 1.6e-05 | 1e-05 | 1.367E-05 | 3.215E-06 | 1.000E-05 | 1.600E-05 |
| 14871 | 744 | D744L | B | 1.5e-05 | 1.7e-05 | 1.7e-05 | 1.633E-05 | 1.155E-06 | 1.500E-05 | 1.700E-05 |
| 14872 | 744 | D744K | B | 9.9e-06 | 1.5e-05 | 1e-05 | 1.163E-05 | 2.916E-06 | 9.900E-06 | 1.500E-05 |
| 14873 | 744 | D744M | B | 1.4e-05 | 1.4e-05 | 1.5e-05 | 1.433E-05 | 5.774E-07 | 1.400E-05 | 1.500E-05 |
| 14874 | 744 | D744F | B | 1e-05 | 1e-05 | 1.4e-05 | 1.133E-05 | 2.309E-06 | 1.000E-05 | 1.400E-05 |

| | | | | | | | | | | |
|---|---|---|---|---|---|---|---|---|---|---|
| 14875 | 744 | D744P | B | 5.8e-06 | 1.3e-05 | 1.3e-05 | 1.060E-05 | 4.157E-06 | 5.800E-06 | 1.300E-05 |
| 14876 | 744 | D744S | B | 1.6e-05 | 1.1e-05 | 1e-05 | 1.233E-05 | 3.215E-06 | 1.000E-05 | 1.600E-05 |
| 14877 | 744 | D744T | B | 1.4e-05 | 1.2e-05 | 1.6e-05 | 1.400E-05 | 2.000E-06 | 1.200E-05 | 1.600E-05 |
| 14878 | 744 | D744W | B | 1.4e-05 | 1.1e-05 | 1.5e-05 | 1.333E-05 | 2.082E-06 | 1.100E-05 | 1.500E-05 |
| 14879 | 744 | D744Y | B | 1e-05 | 9e-06 | 1.4e-05 | 1.100E-05 | 2.646E-06 | 9.000E-06 | 1.400E-05 |
| 14880 | 744 | D744V | B | 1.1e-05 | 1.1e-05 | 1.1e-05 | 1.100E-05 | 2.075E-21 | 1.100E-05 | 1.100E-05 |
| 14881 | 745 | E745A | B | 1.4e-05 | 1.1e-05 | 9.9e-06 | 1.163E-05 | 2.122E-06 | 9.900E-06 | 1.400E-05 |
| 14882 | 745 | E745R | B | 9.8e-06 | 1.5e-05 | 1.1e-05 | 1.193E-05 | 2.723E-06 | 9.800E-06 | 1.500E-05 |
| 14883 | 745 | E745N | B | 1.5e-05 | 1.3e-05 | 1.6e-05 | 1.467E-05 | 1.528E-06 | 1.300E-05 | 1.600E-05 |
| 14884 | 745 | E745D | B | 1.5e-05 | 9.2e-06 | 1.1e-05 | 1.173E-05 | 2.969E-06 | 9.200E-06 | 1.500E-05 |
| 14885 | 745 | E745C | B | 1.1e-05 | 1.3e-05 | 1.6e-05 | 1.333E-05 | 2.517E-06 | 1.100E-05 | 1.600E-05 |
| 14886 | 745 | E745E | B | 1.4e-05 | 8.2e-06 | 1.4e-05 | 1.207E-05 | 3.349E-06 | 8.200E-06 | 1.400E-05 |
| 14887 | 745 | E745Q | B | 1.3e-05 | 8.7e-06 | 1.1e-05 | 1.090E-05 | 2.152E-06 | 8.700E-06 | 1.300E-05 |
| 14888 | 745 | E745G | B | 1.5e-05 | 1.6e-05 | 1e-05 | 1.367E-05 | 3.215E-06 | 1.000E-05 | 1.600E-05 |
| 14889 | 745 | E745H | B | 1.5e-05 | 9.9e-06 | 1.1e-05 | 1.197E-05 | 2.684E-06 | 9.900E-06 | 1.500E-05 |
| 14890 | 745 | E745I | B | 1.8e-05 | 1.5e-05 | 1.4e-05 | 1.567E-05 | 2.082E-06 | 1.400E-05 | 1.800E-05 |
| 14891 | 745 | E745L | B | 1.4e-05 | 1.6e-05 | 9e-06 | 1.300E-05 | 3.606E-06 | 9.000E-06 | 1.600E-05 |
| 14892 | 745 | E745K | B | 1e-05 | 1e-05 | 1.1e-05 | 1.033E-05 | 5.774E-07 | 1.000E-05 | 1.100E-05 |
| 14893 | 745 | E745M | B | 1.1e-05 | 9.8e-06 | 1.4e-05 | 1.160E-05 | 2.163E-06 | 9.800E-06 | 1.400E-05 |
| 14894 | 745 | E745F | B | 1.2e-05 | 6.5e-06 | 9.7e-06 | 9.400E-06 | 2.762E-06 | 6.500E-06 | 1.200E-05 |
| 14895 | 745 | E745P | B | 9.1e-06 | 1.3e-05 | 1.5e-05 | 1.237E-05 | 3.001E-06 | 9.100E-06 | 1.500E-05 |
| 14896 | 745 | E745S | B | 1.1e-05 | 1.5e-05 | 1.5e-05 | 1.367E-05 | 2.309E-06 | 1.100E-05 | 1.500E-05 |
| 14897 | 745 | E745T | B | 1.4e-05 | 1e-05 | 9.3e-06 | 1.110E-05 | 2.536E-06 | 9.300E-06 | 1.400E-05 |
| 14898 | 745 | E745W | B | 1e-05 | 9.5e-06 | 1.5e-05 | 1.150E-05 | 3.041E-06 | 9.500E-06 | 1.500E-05 |
| 14899 | 745 | E745Y | B | 1.5e-05 | 1e-05 | 1.5e-05 | 1.333E-05 | 2.887E-06 | 1.000E-05 | 1.500E-05 |
| 14900 | 745 | E745V | B | 1.2e-05 | 1.5e-05 | 1.1e-05 | 1.267E-05 | 2.082E-06 | 1.100E-05 | 1.500E-05 |
| 14901 | 746 | Q746A | B | 1.6e-05 | 1.5e-05 | 1e-05 | 1.367E-05 | 3.215E-06 | 1.000E-05 | 1.600E-05 |
| 14902 | 746 | Q746R | B | 1e-05 | 1.5e-05 | 1e-05 | 1.167E-05 | 2.887E-06 | 1.000E-05 | 1.500E-05 |
| 14903 | 746 | Q746N | B | 1.4e-05 | 1.5e-05 | 1.5e-05 | 1.467E-05 | 5.774E-07 | 1.400E-05 | 1.500E-05 |
| 14904 | 746 | Q746D | B | 1.4e-05 | 1.5e-05 | 1e-05 | 1.300E-05 | 2.646E-06 | 1.000E-05 | 1.500E-05 |
| 14905 | 746 | Q746C | B | 9.6e-06 | 1.5e-05 | 1.3e-05 | 1.253E-05 | 2.730E-06 | 9.600E-06 | 1.500E-05 |
| 14906 | 746 | Q746E | B | 9.5e-06 | 1.6e-05 | 1.3e-05 | 1.283E-05 | 3.253E-06 | 9.500E-06 | 1.600E-05 |
| 14907 | 746 | Q746Q | B | 1.8e-05 | 1.4e-05 | 1.5e-05 | 1.567E-05 | 2.082E-06 | 1.400E-05 | 1.800E-05 |
| 14908 | 746 | Q746G | B | 1.5e-05 | 1.4e-05 | 1.2e-05 | 1.367E-05 | 1.528E-06 | 1.200E-05 | 1.500E-05 |
| 14909 | 746 | Q746H | B | 1.3e-05 | 1.5e-05 | 9.1e-06 | 1.237E-05 | 3.001E-06 | 9.100E-06 | 1.500E-05 |
| 14910 | 746 | Q746I | B | 1.2e-05 | 1e-05 | 1e-05 | 1.067E-05 | 1.155E-06 | 1.000E-05 | 1.200E-05 |
| 14911 | 746 | Q746L | B | 1.1e-05 | 1.3e-05 | 1.2e-05 | 1.200E-05 | 1.000E-06 | 1.100E-05 | 1.300E-05 |
| 14912 | 746 | Q746K | B | 1e-05 | 1.5e-05 | 1.1e-05 | 1.200E-05 | 2.646E-06 | 1.000E-05 | 1.500E-05 |
| 14913 | 746 | Q746M | B | 1.5e-05 | 1.4e-05 | 1.3e-05 | 1.400E-05 | 1.000E-06 | 1.300E-05 | 1.500E-05 |
| 14914 | 746 | Q746F | B | 1e-05 | 1.6e-05 | 1.4e-05 | 1.333E-05 | 3.055E-06 | 1.000E-05 | 1.600E-05 |
| 14915 | 746 | Q746P | B | 1e-05 | 1.7e-05 | 1.1e-05 | 1.267E-05 | 3.786E-06 | 1.000E-05 | 1.700E-05 |
| 14916 | 746 | Q746S | B | 1e-05 | 1.6e-05 | 1.3e-05 | 1.300E-05 | 3.000E-06 | 1.000E-05 | 1.600E-05 |
| 14917 | 746 | Q746T | B | 2e-05 | 1.4e-05 | 1e-05 | 1.467E-05 | 5.033E-06 | 1.000E-05 | 2.000E-05 |
| 14918 | 746 | Q746W | B | 1.6e-05 | 9.6e-06 | 9.5e-06 | 1.170E-05 | 3.724E-06 | 9.500E-06 | 1.600E-05 |
| 14919 | 746 | Q746Y | B | 1.2e-05 | 8.9e-06 | 9.2e-06 | 1.003E-05 | 1.710E-06 | 8.900E-06 | 1.200E-05 |
| 14920 | 746 | Q746V | B | 2e-05 | 1.5e-05 | 1.7e-05 | 1.733E-05 | 2.517E-06 | 1.500E-05 | 2.000E-05 |
| 14921 | 747 | L747A | B | 9.7e-06 | 1.4e-05 | 9.9e-06 | 1.120E-05 | 2.427E-06 | 9.700E-06 | 1.400E-05 |
| 14922 | 747 | L747R | B | 1.1e-05 | 9.9e-06 | 1.1e-05 | 1.063E-05 | 6.351E-07 | 9.900E-06 | 1.100E-05 |
| 14923 | 747 | L747N | B | 1.1e-05 | 1.4e-05 | 1e-05 | 1.167E-05 | 2.082E-06 | 1.000E-05 | 1.400E-05 |
| 14924 | 747 | L747D | B | 1.5e-05 | 1.4e-05 | 1.2e-05 | 1.367E-05 | 1.528E-06 | 1.200E-05 | 1.500E-05 |
| 14925 | 747 | L747C | B | 1.3e-05 | 1.5e-05 | 1e-05 | 1.267E-05 | 2.517E-06 | 1.000E-05 | 1.500E-05 |
| 14926 | 747 | L747E | B | 9.7e-06 | 1.4e-05 | 9.4e-06 | 1.103E-05 | 2.574E-06 | 9.400E-06 | 1.400E-05 |
| 14927 | 747 | L747Q | B | 1.4e-05 | 9.5e-06 | 1.4e-05 | 1.250E-05 | 2.598E-06 | 9.500E-06 | 1.400E-05 |
| 14928 | 747 | L747G | B | 1.3e-05 | 1.2e-05 | 1.5e-05 | 1.333E-05 | 1.528E-06 | 1.200E-05 | 1.500E-05 |
| 14929 | 747 | L747H | B | 1.4e-05 | 8.6e-06 | 1.3e-05 | 1.187E-05 | 2.873E-06 | 8.600E-06 | 1.400E-05 |
| 14930 | 747 | L747I | B | 1.6e-05 | 1.3e-05 | 1.5e-05 | 1.467E-05 | 1.528E-06 | 1.300E-05 | 1.600E-05 |
| 14931 | 747 | L747L | B | 1.5e-05 | 1.5e-05 | 8.5e-06 | 1.283E-05 | 3.753E-06 | 8.500E-06 | 1.500E-05 |
| 14932 | 747 | L747K | B | 1.5e-05 | 6.2e-06 | 1.5e-05 | 1.207E-05 | 5.081E-06 | 6.200E-06 | 1.500E-05 |
| 14933 | 747 | L747M | B | 9.3e-06 | 1.1e-05 | 1.4e-05 | 1.143E-05 | 2.380E-06 | 9.300E-06 | 1.400E-05 |
| 14934 | 747 | L747F | B | 1e-05 | 1.4e-05 | 2.4e-05 | 1.600E-05 | 7.211E-06 | 1.000E-05 | 2.400E-05 |
| 14935 | 747 | L747P | B | 1.3e-05 | 1e-05 | 1e-05 | 1.100E-05 | 1.732E-06 | 1.000E-05 | 1.300E-05 |
| 14936 | 747 | L747S | B | 1e-05 | 9.4e-06 | 1e-05 | 9.800E-06 | 3.464E-07 | 9.400E-06 | 1.000E-05 |
| 14937 | 747 | L747T | B | 1.4e-05 | 1.5e-05 | 1.6e-05 | 1.500E-05 | 1.000E-06 | 1.400E-05 | 1.600E-05 |
| 14938 | 747 | L747W | B | 1e-05 | 1.5e-05 | 1e-05 | 1.167E-05 | 2.887E-06 | 1.000E-05 | 1.500E-05 |
| 14939 | 747 | L747Y | B | 1.1e-05 | 1.3e-05 | 1.1e-05 | 1.167E-05 | 1.155E-06 | 1.100E-05 | 1.300E-05 |
| 14940 | 747 | L747V | B | 1.3e-05 | 1.3e-05 | 1.4e-05 | 1.333E-05 | 5.774E-07 | 1.300E-05 | 1.400E-05 |
| 14941 | 748 | K748A | B | 1.6e-05 | 1.6e-05 | 1.4e-05 | 1.533E-05 | 1.155E-06 | 1.400E-05 | 1.600E-05 |
| 14942 | 748 | K748R | B | 8.9e-06 | 1e-05 | 1.7e-05 | 1.197E-05 | 4.394E-06 | 8.900E-06 | 1.700E-05 |
| 14943 | 748 | K748N | B | 1.5e-05 | 1.1e-05 | 9.6e-06 | 1.187E-05 | 2.802E-06 | 9.600E-06 | 1.500E-05 |
| 14944 | 748 | K748D | B | 1.5e-05 | 1.5e-05 | 1.7e-05 | 1.567E-05 | 1.155E-06 | 1.500E-05 | 1.700E-05 |
| 14945 | 748 | K748C | B | 1.4e-05 | 8.7e-06 | 1.3e-05 | 1.190E-05 | 2.816E-06 | 8.700E-06 | 1.400E-05 |
| 14946 | 748 | K748E | B | 1.4e-05 | 1.6e-05 | 1.5e-05 | 1.500E-05 | 1.000E-06 | 1.400E-05 | 1.600E-05 |
| 14947 | 748 | K748Q | B | 1.4e-05 | 9.6e-06 | 1.1e-05 | 1.153E-05 | 2.248E-06 | 9.600E-06 | 1.400E-05 |
| 14948 | 748 | K748G | B | 1e-05 | 1.1e-05 | 9.2e-06 | 1.007E-05 | 9.018E-07 | 9.200E-06 | 1.100E-05 |
| 14949 | 748 | K748H | B | 1.1e-05 | 1.5e-05 | 1.4e-05 | 1.333E-05 | 2.082E-06 | 1.100E-05 | 1.500E-05 |
| 14950 | 748 | K748I | B | 1.1e-05 | 1.6e-05 | 1e-05 | 1.233E-05 | 3.215E-06 | 1.000E-05 | 1.600E-05 |
| 14951 | 748 | K748L | B | 1.6e-05 | 1.7e-05 | 1.4e-05 | 1.567E-05 | 1.528E-06 | 1.400E-05 | 1.700E-05 |
| 14952 | 748 | K748K | B | 1.3e-05 | 1.3e-05 | 1e-05 | 1.200E-05 | 1.732E-06 | 1.000E-05 | 1.300E-05 |
| 14953 | 748 | K748M | B | 9.7e-06 | 1.7e-05 | 1.4e-05 | 1.357E-05 | 3.669E-06 | 9.700E-06 | 1.700E-05 |
| 14954 | 748 | K748F | B | 1.6e-05 | 1.4e-05 | 1e-05 | 1.333E-05 | 3.055E-06 | 1.000E-05 | 1.600E-05 |
| 14955 | 748 | K748P | B | 1e-05 | 1.6e-05 | 1.4e-05 | 1.333E-05 | 3.055E-06 | 1.000E-05 | 1.600E-05 |
| 14956 | 748 | K748S | B | 1e-05 | 9.4e-06 | 1.1e-05 | 1.013E-05 | 8.083E-07 | 9.400E-06 | 1.100E-05 |
| 14957 | 748 | K748T | B | 1.1e-05 | 1.2e-05 | 1.5e-05 | 1.267E-05 | 2.082E-06 | 1.100E-05 | 1.500E-05 |
| 14958 | 748 | K748W | B | 1.4e-05 | 9.7e-06 | 1.5e-05 | 1.290E-05 | 2.816E-06 | 9.700E-06 | 1.500E-05 |
| 14959 | 748 | K748Y | B | 9.9e-06 | 1.5e-05 | 9.2e-06 | 1.137E-05 | 3.166E-06 | 9.200E-06 | 1.500E-05 |

| | | | | | | | | | | |
|---|---|---|---|---|---|---|---|---|---|---|
| 14960 | 748 | K748V | B | 1.6e-05 | 1.5e-05 | 1.4e-05 | 1.500E-05 | 1.000E-06 | 1.400E-05 | 1.600E-05 |
| 14961 | 749 | S749A | B | 1.6e-05 | 1.5e-05 | 1.3e-05 | 1.467E-05 | 1.528E-06 | 1.300E-05 | 1.600E-05 |
| 14962 | 749 | S749R | B | 1.6e-05 | 1.6e-05 | 1.1e-05 | 1.433E-05 | 2.887E-06 | 1.100E-05 | 1.600E-05 |
| 14963 | 749 | S749N | B | 1.5e-05 | 8.7e-06 | 1.5e-05 | 1.290E-05 | 3.637E-06 | 8.700E-06 | 1.500E-05 |
| 14964 | 749 | S749D | B | 1.2e-05 | 1.1e-05 | 1.4e-05 | 1.233E-05 | 1.528E-06 | 1.100E-05 | 1.400E-05 |
| 14965 | 749 | S749C | B | 1.6e-05 | 1.7e-05 | 1.1e-05 | 1.467E-05 | 3.215E-06 | 1.100E-05 | 1.700E-05 |
| 14966 | 749 | S749E | B | 1e-05 | 1.4e-05 | 1.5e-05 | 1.300E-05 | 2.646E-06 | 1.000E-05 | 1.500E-05 |
| 14967 | 749 | S749Q | B | 1.4e-05 | 1.4e-05 | 1e-05 | 1.267E-05 | 2.309E-06 | 1.000E-05 | 1.400E-05 |
| 14968 | 749 | S749G | B | 1.5e-05 | 1.6e-05 | 1.5e-05 | 1.533E-05 | 5.774E-07 | 1.500E-05 | 1.600E-05 |
| 14969 | 749 | S749H | B | 8.1e-06 | 9.5e-06 | 1.1e-05 | 9.533E-06 | 1.450E-06 | 8.100E-06 | 1.100E-05 |
| 14970 | 749 | S749I | B | 1.2e-05 | 1.4e-05 | 1e-05 | 1.200E-05 | 2.000E-06 | 1.000E-05 | 1.400E-05 |
| 14971 | 749 | S749L | B | 1.1e-05 | 1.4e-05 | 1.6e-05 | 1.367E-05 | 2.517E-06 | 1.100E-05 | 1.600E-05 |
| 14972 | 749 | S749K | B | 1.6e-05 | 1e-05 | 1.5e-05 | 1.367E-05 | 3.215E-06 | 1.000E-05 | 1.600E-05 |
| 14973 | 749 | S749M | B | 1.5e-05 | 1e-05 | 1.5e-05 | 1.333E-05 | 2.887E-06 | 1.000E-05 | 1.500E-05 |
| 14974 | 749 | S749F | B | 8.9e-06 | 1.1e-05 | 1.1e-05 | 1.030E-05 | 1.212E-06 | 8.900E-06 | 1.100E-05 |
| 14975 | 749 | S749P | B | 1e-05 | 1.3e-05 | 1.1e-05 | 1.133E-05 | 1.528E-06 | 1.000E-05 | 1.300E-05 |
| 14976 | 749 | S749S | B | 1.5e-05 | 1.4e-05 | 1.4e-05 | 1.433E-05 | 5.774E-07 | 1.400E-05 | 1.500E-05 |
| 14977 | 749 | S749T | B | 9.6e-06 | 1.3e-05 | 9.1e-06 | 1.057E-05 | 2.122E-06 | 9.100E-06 | 1.300E-05 |
| 14978 | 749 | S749W | B | 1e-05 | 1.8e-05 | 1.3e-05 | 1.367E-05 | 4.041E-06 | 1.000E-05 | 1.800E-05 |
| 14979 | 749 | S749Y | B | 1.8e-05 | 1e-05 | 1.6e-05 | 1.467E-05 | 4.163E-06 | 1.000E-05 | 1.800E-05 |
| 14980 | 749 | S749V | B | 1.1e-05 | 1.5e-05 | 1.1e-05 | 1.233E-05 | 2.309E-06 | 1.100E-05 | 1.500E-05 |
| 14981 | 750 | G750A | B | 1.1e-05 | 1e-05 | 1.6e-05 | 1.233E-05 | 3.215E-06 | 1.000E-05 | 1.600E-05 |
| 14982 | 750 | G750R | B | 9.1e-06 | 1.4e-05 | 9.9e-06 | 1.100E-05 | 2.629E-06 | 9.100E-06 | 1.400E-05 |
| 14983 | 750 | G750N | B | 1e-05 | 1.6e-05 | 1.6e-05 | 1.400E-05 | 3.464E-06 | 1.000E-05 | 1.600E-05 |
| 14984 | 750 | G750D | B | 2.2e-05 | 1.4e-05 | 9.5e-06 | 1.517E-05 | 6.331E-06 | 9.500E-06 | 2.200E-05 |
| 14985 | 750 | G750C | B | 1.3e-05 | 1.5e-05 | 1.5e-05 | 1.433E-05 | 1.155E-06 | 1.300E-05 | 1.500E-05 |
| 14986 | 750 | G750E | B | 1.6e-05 | 1.5e-05 | 1.3e-05 | 1.467E-05 | 1.528E-06 | 1.300E-05 | 1.600E-05 |
| 14987 | 750 | G750Q | B | 1.6e-05 | 1.6e-05 | 1.3e-05 | 1.500E-05 | 1.732E-06 | 1.300E-05 | 1.600E-05 |
| 14988 | 750 | G750G | B | 1.5e-05 | 1.4e-05 | 1.7e-05 | 1.533E-05 | 1.528E-06 | 1.400E-05 | 1.700E-05 |
| 14989 | 750 | G750H | B | 1.1e-05 | 1.1e-05 | 1.6e-05 | 1.267E-05 | 2.887E-06 | 1.100E-05 | 1.600E-05 |
| 14990 | 750 | G750I | B | 1.4e-05 | 1.6e-05 | 1.9e-05 | 1.633E-05 | 2.517E-06 | 1.400E-05 | 1.900E-05 |
| 14991 | 750 | G750L | B | 9.8e-06 | 1.5e-05 | 1.5e-05 | 1.327E-05 | 3.002E-06 | 9.800E-06 | 1.500E-05 |
| 14992 | 750 | G750K | B | 1.7e-05 | 1.7e-05 | 8.5e-06 | 1.417E-05 | 4.907E-06 | 8.500E-06 | 1.700E-05 |
| 14993 | 750 | G750M | B | 2e-05 | 1.5e-05 | 9.4e-06 | 1.480E-05 | 5.303E-06 | 9.400E-06 | 2.000E-05 |
| 14994 | 750 | G750F | B | 1.2e-05 | 1.5e-05 | 1.1e-05 | 1.267E-05 | 2.082E-06 | 1.100E-05 | 1.500E-05 |
| 14995 | 750 | G750P | B | 1e-05 | 2.2e-05 | 1.6e-05 | 1.600E-05 | 6.000E-06 | 1.000E-05 | 2.200E-05 |
| 14996 | 750 | G750S | B | 5.8e-06 | 1.2e-05 | 1.4e-05 | 1.060E-05 | 4.276E-06 | 5.800E-06 | 1.400E-05 |
| 14997 | 750 | G750T | B | 1.5e-05 | 1e-05 | 1.4e-05 | 1.300E-05 | 2.646E-06 | 1.000E-05 | 1.500E-05 |
| 14998 | 750 | G750W | B | 1e-05 | 1e-05 | 9.3e-06 | 9.767E-06 | 4.041E-07 | 9.300E-06 | 1.000E-05 |
| 14999 | 750 | G750Y | B | 9.3e-06 | 1.4e-05 | 1.6e-05 | 1.310E-05 | 3.439E-06 | 9.300E-06 | 1.600E-05 |
| 15000 | 750 | G750V | B | 1.3e-05 | 1.2e-05 | 1.6e-05 | 1.367E-05 | 2.082E-06 | 1.200E-05 | 1.600E-05 |
| 15001 | 751 | T751A | B | 1.2e-05 | 1.2e-05 | 8.9e-06 | 1.097E-05 | 1.790E-06 | 8.900E-06 | 1.200E-05 |
| 15002 | 751 | T751R | B | 1.2e-05 | 1.1e-05 | 1.4e-05 | 1.233E-05 | 1.528E-06 | 1.100E-05 | 1.400E-05 |
| 15003 | 751 | T751N | B | 1.5e-05 | 9.8e-06 | 1.6e-05 | 1.360E-05 | 3.329E-06 | 9.800E-06 | 1.600E-05 |
| 15004 | 751 | T751D | B | 1.6e-05 | 8.9e-06 | 1.2e-05 | 1.230E-05 | 3.559E-06 | 8.900E-06 | 1.600E-05 |
| 15005 | 751 | T751C | B | 1.3e-05 | 1.5e-05 | 1.6e-05 | 1.467E-05 | 1.528E-06 | 1.300E-05 | 1.600E-05 |
| 15006 | 751 | T751E | B | 1.6e-05 | 1e-05 | 1.5e-05 | 1.367E-05 | 3.215E-06 | 1.000E-05 | 1.600E-05 |
| 15007 | 751 | T751Q | B | 8.6e-06 | 1.2e-05 | 1.7e-05 | 1.253E-05 | 4.225E-06 | 8.600E-06 | 1.700E-05 |
| 15008 | 751 | T751G | B | 1.7e-05 | 1.7e-05 | 1.4e-05 | 1.600E-05 | 1.732E-06 | 1.400E-05 | 1.700E-05 |
| 15009 | 751 | T751H | B | 1.1e-05 | 1.4e-05 | 1.5e-05 | 1.333E-05 | 2.082E-06 | 1.100E-05 | 1.500E-05 |
| 15010 | 751 | T751I | B | 1.6e-05 | 1.8e-05 | 1.1e-05 | 1.500E-05 | 3.606E-06 | 1.100E-05 | 1.800E-05 |
| 15011 | 751 | T751L | B | 1.1e-05 | 1.5e-05 | 1e-05 | 1.200E-05 | 2.646E-06 | 1.000E-05 | 1.500E-05 |
| 15012 | 751 | T751K | B | 9.8e-06 | 9.9e-06 | 1e-05 | 9.900E-06 | 1.000E-07 | 9.800E-06 | 1.000E-05 |
| 15013 | 751 | T751M | B | 1.3e-05 | 9.3e-06 | 1.6e-05 | 1.277E-05 | 3.356E-06 | 9.300E-06 | 1.600E-05 |
| 15014 | 751 | T751F | B | 1.5e-05 | 1.7e-05 | 8.9e-06 | 1.363E-05 | 4.219E-06 | 8.900E-06 | 1.700E-05 |
| 15015 | 751 | T751P | B | 1.5e-05 | 1.4e-05 | 1.5e-05 | 1.467E-05 | 5.774E-07 | 1.400E-05 | 1.500E-05 |
| 15016 | 751 | T751S | B | 1.7e-05 | 1.4e-05 | 1.1e-05 | 1.400E-05 | 3.000E-06 | 1.100E-05 | 1.700E-05 |
| 15017 | 751 | T751T | B | 1.5e-05 | 1.3e-05 | 7.1e-06 | 1.170E-05 | 4.107E-06 | 7.100E-06 | 1.500E-05 |
| 15018 | 751 | T751W | B | 9.2e-06 | 1.2e-05 | 1.3e-05 | 1.140E-05 | 1.970E-06 | 9.200E-06 | 1.300E-05 |
| 15019 | 751 | T751Y | B | 1.8e-05 | 1.6e-05 | 5.5e-06 | 1.317E-05 | 6.714E-06 | 5.500E-06 | 1.800E-05 |
| 15020 | 751 | T751V | B | 1.4e-05 | 1.2e-05 | 8.5e-06 | 1.150E-05 | 2.784E-06 | 8.500E-06 | 1.400E-05 |
| 15021 | 752 | A752A | B | 1.1e-05 | 1.1e-05 | 1.1e-05 | 1.100E-05 | 2.075E-21 | 1.100E-05 | 1.100E-05 |
| 15022 | 752 | A752R | B | 1.4e-05 | 1.6e-05 | 1.1e-05 | 1.367E-05 | 2.517E-06 | 1.100E-05 | 1.600E-05 |
| 15023 | 752 | A752N | B | 1e-05 | 1.7e-05 | 1.6e-05 | 1.433E-05 | 3.786E-06 | 1.000E-05 | 1.700E-05 |
| 15024 | 752 | A752D | B | 1.3e-05 | 1.6e-05 | 1.5e-05 | 1.467E-05 | 1.528E-06 | 1.300E-05 | 1.600E-05 |
| 15025 | 752 | A752C | B | 1.7e-05 | 1.1e-05 | 8.3e-06 | 1.210E-05 | 4.453E-06 | 8.300E-06 | 1.700E-05 |
| 15026 | 752 | A752E | B | 9.9e-06 | 1.5e-05 | 1.5e-05 | 1.330E-05 | 2.944E-06 | 9.900E-06 | 1.500E-05 |
| 15027 | 752 | A752Q | B | 1.7e-05 | 9.2e-06 | 1.2e-05 | 1.273E-05 | 3.951E-06 | 9.200E-06 | 1.700E-05 |
| 15028 | 752 | A752G | B | 1.1e-05 | 1.4e-05 | 1.4e-05 | 1.300E-05 | 1.732E-06 | 1.100E-05 | 1.400E-05 |
| 15029 | 752 | A752H | B | 8.8e-06 | 1.1e-05 | 1.1e-05 | 1.027E-05 | 1.270E-06 | 8.800E-06 | 1.100E-05 |
| 15030 | 752 | A752I | B | 1.6e-05 | 1e-05 | 9.2e-06 | 1.173E-05 | 3.717E-06 | 9.200E-06 | 1.600E-05 |
| 15031 | 752 | A752L | B | 3.2e-05 | 1.7e-05 | 1.1e-05 | 2.000E-05 | 1.082E-05 | 1.100E-05 | 3.200E-05 |
| 15032 | 752 | A752K | B | 9.5e-06 | 1.6e-05 | 1.2e-05 | 1.250E-05 | 3.279E-06 | 9.500E-06 | 1.600E-05 |
| 15033 | 752 | A752M | B | 1.5e-05 | 1.2e-05 | 1.4e-05 | 1.367E-05 | 1.528E-06 | 1.200E-05 | 1.500E-05 |
| 15034 | 752 | A752F | B | 1.1e-05 | 1.4e-05 | 9.7e-06 | 1.157E-05 | 2.205E-06 | 9.700E-06 | 1.400E-05 |
| 15035 | 752 | A752P | B | 2.6e-05 | 9.8e-06 | 1e-05 | 1.527E-05 | 9.296E-06 | 9.800E-06 | 2.600E-05 |
| 15036 | 752 | A752S | B | 1.3e-05 | 1.5e-05 | 1.3e-05 | 1.367E-05 | 1.155E-06 | 1.300E-05 | 1.500E-05 |
| 15037 | 752 | A752T | B | 1.4e-05 | 1.6e-05 | 1.7e-05 | 1.567E-05 | 1.528E-06 | 1.400E-05 | 1.700E-05 |
| 15038 | 752 | A752W | B | 1e-05 | 1.2e-05 | 1.3e-05 | 1.167E-05 | 1.528E-06 | 1.000E-05 | 1.300E-05 |
| 15039 | 752 | A752Y | B | 1.1e-05 | 1.1e-05 | 1e-05 | 1.067E-05 | 5.774E-07 | 1.000E-05 | 1.100E-05 |
| 15040 | 752 | A752V | B | 1.4e-05 | 1.3e-05 | 1.4e-05 | 1.367E-05 | 5.774E-07 | 1.300E-05 | 1.400E-05 |
| 15041 | 753 | S753A | B | 9.7e-06 | 1e-05 | 1.4e-05 | 1.123E-05 | 2.401E-06 | 9.700E-06 | 1.400E-05 |
| 15042 | 753 | S753R | B | 9.2e-06 | 1.2e-05 | 1.1e-05 | 1.073E-05 | 1.419E-06 | 9.200E-06 | 1.200E-05 |
| 15043 | 753 | S753N | B | 8.8e-06 | 1e-05 | 1e-05 | 9.600E-06 | 6.928E-07 | 8.800E-06 | 1.000E-05 |
| 15044 | 753 | S753D | B | 1.5e-05 | 9.6e-06 | 1.8e-05 | 1.420E-05 | 4.257E-06 | 9.600E-06 | 1.800E-05 |

| | | | | | | | | | | |
|---|---|---|---|---|---|---|---|---|---|---|
| 15045 | 753 | S753C | B | 1.1e-05 | 1.1e-05 | 1.1e-05 | 1.100E-05 | 2.075E-21 | 1.100E-05 | 1.100E-05 |
| 15046 | 753 | S753E | B | 1.5e-05 | 9.4e-06 | 1.6e-05 | 1.347E-05 | 3.557E-06 | 9.400E-06 | 1.600E-05 |
| 15047 | 753 | S753Q | B | 1.3e-05 | 1.6e-05 | 1.4e-05 | 1.433E-05 | 1.528E-06 | 1.300E-05 | 1.600E-05 |
| 15048 | 753 | S753G | B | 1.5e-05 | 8.8e-06 | 1.5e-05 | 1.293E-05 | 3.580E-06 | 8.800E-06 | 1.500E-05 |
| 15049 | 753 | S753H | B | 1.5e-05 | 1.2e-05 | 9.4e-06 | 1.213E-05 | 2.802E-06 | 9.400E-06 | 1.500E-05 |
| 15050 | 753 | S753I | B | 1.5e-05 | 1.1e-05 | 1.9e-05 | 1.500E-05 | 4.000E-06 | 1.100E-05 | 1.900E-05 |
| 15051 | 753 | S753L | B | 1.4e-05 | 1.6e-05 | 1e-05 | 1.333E-05 | 3.055E-06 | 1.000E-05 | 1.600E-05 |
| 15052 | 753 | S753K | B | 1.4e-05 | 1.6e-05 | 1.1e-05 | 1.367E-05 | 2.517E-06 | 1.100E-05 | 1.600E-05 |
| 15053 | 753 | S753M | B | 1e-05 | 1.5e-05 | 1.1e-05 | 1.200E-05 | 2.646E-06 | 1.000E-05 | 1.500E-05 |
| 15054 | 753 | S753F | B | 1.4e-05 | 1.5e-05 | 1.3e-05 | 1.400E-05 | 1.000E-06 | 1.300E-05 | 1.500E-05 |
| 15055 | 753 | S753P | B | 1.1e-05 | 9.9e-06 | 1.3e-05 | 1.130E-05 | 1.572E-06 | 9.900E-06 | 1.300E-05 |
| 15056 | 753 | S753S | B | 1.5e-05 | 1.5e-05 | 8.5e-06 | 1.283E-05 | 3.753E-06 | 8.500E-06 | 1.500E-05 |
| 15057 | 753 | S753T | B | 1.6e-05 | 1.1e-05 | 1.1e-05 | 1.267E-05 | 2.887E-06 | 1.100E-05 | 1.600E-05 |
| 15058 | 753 | S753W | B | 1.6e-05 | 1.3e-05 | 8.1e-06 | 1.237E-05 | 3.988E-06 | 8.100E-06 | 1.600E-05 |
| 15059 | 753 | S753Y | B | 1.5e-05 | 1e-05 | 1.5e-05 | 1.333E-05 | 2.887E-06 | 1.000E-05 | 1.500E-05 |
| 15060 | 753 | S753V | B | 1.4e-05 | 9.2e-06 | 1.1e-05 | 1.140E-05 | 2.425E-06 | 9.200E-06 | 1.400E-05 |
| 15061 | 754 | V754A | B | 1.2e-05 | 1.4e-05 | 1.6e-05 | 1.400E-05 | 2.000E-06 | 1.200E-05 | 1.600E-05 |
| 15062 | 754 | V754R | B | 1.5e-05 | 1.6e-05 | 1.3e-05 | 1.467E-05 | 1.528E-06 | 1.300E-05 | 1.600E-05 |
| 15063 | 754 | V754N | B | 8.3e-06 | 1.5e-05 | 1.3e-05 | 1.210E-05 | 3.439E-06 | 8.300E-06 | 1.500E-05 |
| 15064 | 754 | V754D | B | 1.7e-05 | 8.2e-06 | 1.5e-05 | 1.340E-05 | 4.613E-06 | 8.200E-06 | 1.700E-05 |
| 15065 | 754 | V754C | B | 1.4e-05 | 1.3e-05 | 8.8e-06 | 1.193E-05 | 2.759E-06 | 8.800E-06 | 1.400E-05 |
| 15066 | 754 | V754E | B | 9.5e-06 | 1.1e-05 | 9.6e-06 | 1.003E-05 | 8.386E-07 | 9.500E-06 | 1.100E-05 |
| 15067 | 754 | V754Q | B | 1.1e-05 | 1.7e-05 | 9.7e-06 | 1.257E-05 | 3.894E-06 | 9.700E-06 | 1.700E-05 |
| 15068 | 754 | V754G | B | 1.4e-05 | 1e-05 | 6.2e-06 | 1.007E-05 | 3.900E-06 | 6.200E-06 | 1.400E-05 |
| 15069 | 754 | V754H | B | 1.4e-05 | 8.9e-06 | 7.7e-06 | 1.020E-05 | 3.345E-06 | 7.700E-06 | 1.400E-05 |
| 15070 | 754 | V754I | B | 9.6e-06 | 1.6e-05 | 8.9e-06 | 1.150E-05 | 3.913E-06 | 8.900E-06 | 1.600E-05 |
| 15071 | 754 | V754L | B | 1.2e-05 | 1.4e-05 | 1e-05 | 1.200E-05 | 2.000E-06 | 1.000E-05 | 1.400E-05 |
| 15072 | 754 | V754K | B | 1.4e-05 | 8.2e-06 | 1.1e-05 | 1.107E-05 | 2.901E-06 | 8.200E-06 | 1.400E-05 |
| 15073 | 754 | V754M | B | 9.7e-06 | 1.6e-05 | 1.4e-05 | 1.323E-05 | 3.219E-06 | 9.700E-06 | 1.600E-05 |
| 15074 | 754 | V754F | B | 1.5e-05 | 9.5e-06 | 2.2e-05 | 1.550E-05 | 6.265E-06 | 9.500E-06 | 2.200E-05 |
| 15075 | 754 | V754P | B | 1.1e-05 | 1.1e-05 | 1.1e-05 | 1.100E-05 | 2.075E-21 | 1.100E-05 | 1.100E-05 |
| 15076 | 754 | V754S | B | 1.5e-05 | 1.4e-05 | 1.1e-05 | 1.333E-05 | 2.082E-06 | 1.100E-05 | 1.500E-05 |
| 15077 | 754 | V754T | B | 1.1e-05 | 1.3e-05 | 9.9e-06 | 1.130E-05 | 1.572E-06 | 9.900E-06 | 1.300E-05 |
| 15078 | 754 | V754W | B | 1.1e-05 | 1.1e-05 | 1.8e-05 | 1.333E-05 | 4.041E-06 | 1.100E-05 | 1.800E-05 |
| 15079 | 754 | V754Y | B | 1.2e-05 | 1.5e-05 | 1.5e-05 | 1.400E-05 | 1.732E-06 | 1.200E-05 | 1.500E-05 |
| 15080 | 754 | V754V | B | 1.1e-05 | 1.1e-05 | 1.4e-05 | 1.200E-05 | 1.732E-06 | 1.100E-05 | 1.400E-05 |
| 15081 | 755 | V755A | B | 1e-05 | 9.2e-06 | 1.3e-05 | 1.073E-05 | 2.003E-06 | 9.200E-06 | 1.300E-05 |
| 15082 | 755 | V755R | B | 8.2e-06 | 1e-05 | 1.7e-05 | 1.173E-05 | 4.649E-06 | 8.200E-06 | 1.700E-05 |
| 15083 | 755 | V755N | B | 1.2e-05 | 1.2e-05 | 1.3e-05 | 1.233E-05 | 5.774E-07 | 1.200E-05 | 1.300E-05 |
| 15084 | 755 | V755D | B | 1.6e-05 | 1.3e-05 | 1.6e-05 | 1.500E-05 | 1.732E-06 | 1.300E-05 | 1.600E-05 |
| 15085 | 755 | V755C | B | 1.5e-05 | 3.8e-05 | 1.1e-05 | 2.133E-05 | 1.457E-05 | 1.100E-05 | 3.800E-05 |
| 15086 | 755 | V755E | B | 1.2e-05 | 9.8e-06 | 1.3e-05 | 1.160E-05 | 1.637E-06 | 9.800E-06 | 1.300E-05 |
| 15087 | 755 | V755Q | B | 1.3e-05 | 1e-05 | 1.3e-05 | 1.200E-05 | 1.732E-06 | 1.000E-05 | 1.300E-05 |
| 15088 | 755 | V755G | B | 1.6e-05 | 1.6e-05 | 1.1e-05 | 1.433E-05 | 2.887E-06 | 1.100E-05 | 1.600E-05 |
| 15089 | 755 | V755H | B | 1.7e-05 | 1e-05 | 1e-05 | 1.233E-05 | 4.041E-06 | 1.000E-05 | 1.700E-05 |
| 15090 | 755 | V755I | B | 1e-05 | 8.9e-06 | 1.3e-05 | 1.063E-05 | 2.122E-06 | 8.900E-06 | 1.300E-05 |
| 15091 | 755 | V755L | B | 1.5e-05 | 1.5e-05 | 1.1e-05 | 1.367E-05 | 2.309E-06 | 1.100E-05 | 1.500E-05 |
| 15092 | 755 | V755K | B | 1.1e-05 | 1.1e-05 | 1.2e-05 | 1.133E-05 | 5.774E-07 | 1.100E-05 | 1.200E-05 |
| 15093 | 755 | V755M | B | 1.5e-05 | 1.2e-05 | 1.5e-05 | 1.400E-05 | 1.732E-06 | 1.200E-05 | 1.500E-05 |
| 15094 | 755 | V755F | B | 1.7e-05 | 1.6e-05 | 1.1e-05 | 1.467E-05 | 3.215E-06 | 1.100E-05 | 1.700E-05 |
| 15095 | 755 | V755P | B | 1.6e-05 | 1.5e-05 | 1.5e-05 | 1.533E-05 | 5.774E-07 | 1.500E-05 | 1.600E-05 |
| 15096 | 755 | V755S | B | 1.4e-05 | 1e-05 | 1.3e-05 | 1.233E-05 | 2.082E-06 | 1.000E-05 | 1.400E-05 |
| 15097 | 755 | V755T | B | 1e-05 | 1.4e-05 | 1.6e-05 | 1.333E-05 | 3.055E-06 | 1.000E-05 | 1.600E-05 |
| 15098 | 755 | V755W | B | 1.6e-05 | 1.6e-05 | 1e-05 | 1.400E-05 | 3.464E-06 | 1.000E-05 | 1.600E-05 |
| 15099 | 755 | V755Y | B | 2.3e-05 | 1.6e-05 | 1.4e-05 | 1.767E-05 | 4.726E-06 | 1.400E-05 | 2.300E-05 |
| 15100 | 755 | V755V | B | 1.5e-05 | 1.5e-05 | 1.1e-05 | 1.367E-05 | 2.309E-06 | 1.100E-05 | 1.500E-05 |
| 15101 | 756 | C756A | B | 1.3e-05 | 1.6e-05 | 1.1e-05 | 1.333E-05 | 2.517E-06 | 1.100E-05 | 1.600E-05 |
| 15102 | 756 | C756R | B | 1.4e-05 | 1.3e-05 | 7.5e-06 | 1.150E-05 | 3.500E-06 | 7.500E-06 | 1.400E-05 |
| 15103 | 756 | C756N | B | 8.5e-06 | 1.6e-05 | 1e-05 | 1.150E-05 | 3.969E-06 | 8.500E-06 | 1.600E-05 |
| 15104 | 756 | C756D | B | 1.1e-05 | 1.6e-05 | 1.5e-05 | 1.400E-05 | 2.646E-06 | 1.100E-05 | 1.600E-05 |
| 15105 | 756 | C756C | B | 1.5e-05 | 1.2e-05 | 1.4e-05 | 1.367E-05 | 1.528E-06 | 1.200E-05 | 1.500E-05 |
| 15106 | 756 | C756E | B | 1.1e-05 | 1.5e-05 | 1.2e-05 | 1.267E-05 | 2.082E-06 | 1.100E-05 | 1.500E-05 |
| 15107 | 756 | C756Q | B | 1.2e-05 | 7.4e-06 | 1.4e-05 | 1.113E-05 | 3.384E-06 | 7.400E-06 | 1.400E-05 |
| 15108 | 756 | C756G | B | 1.4e-05 | 1.4e-05 | 1.3e-05 | 1.367E-05 | 5.774E-07 | 1.300E-05 | 1.400E-05 |
| 15109 | 756 | C756H | B | 1.5e-05 | 8.9e-06 | 1.1e-05 | 1.163E-05 | 3.099E-06 | 8.900E-06 | 1.500E-05 |
| 15110 | 756 | C756I | B | 1.1e-05 | 1.5e-05 | 8.6e-06 | 1.153E-05 | 3.233E-06 | 8.600E-06 | 1.500E-05 |
| 15111 | 756 | C756L | B | 1.3e-05 | 1.4e-05 | 1.4e-05 | 1.367E-05 | 5.774E-07 | 1.300E-05 | 1.400E-05 |
| 15112 | 756 | C756K | B | 1.5e-05 | 1.5e-05 | 8.3e-06 | 1.277E-05 | 3.868E-06 | 8.300E-06 | 1.500E-05 |
| 15113 | 756 | C756M | B | 9.8e-06 | 1.4e-05 | 9.3e-06 | 1.103E-05 | 2.581E-06 | 9.300E-06 | 1.400E-05 |
| 15114 | 756 | C756F | B | 1.5e-05 | 1.2e-05 | 1.7e-05 | 1.467E-05 | 2.517E-06 | 1.200E-05 | 1.700E-05 |
| 15115 | 756 | C756P | B | 1.3e-05 | 1.3e-05 | 1.4e-05 | 1.333E-05 | 5.774E-07 | 1.300E-05 | 1.400E-05 |
| 15116 | 756 | C756S | B | 1.1e-05 | 9.8e-06 | 1.6e-05 | 1.227E-05 | 3.288E-06 | 9.800E-06 | 1.600E-05 |
| 15117 | 756 | C756T | B | 1.1e-05 | 1.1e-05 | 1.3e-05 | 1.167E-05 | 1.155E-06 | 1.100E-05 | 1.300E-05 |
| 15118 | 756 | C756W | B | 1.5e-05 | 7.6e-06 | 1e-05 | 1.087E-05 | 3.775E-06 | 7.600E-06 | 1.500E-05 |
| 15119 | 756 | C756Y | B | 1.4e-05 | 1.4e-05 | 1.5e-05 | 1.433E-05 | 5.774E-07 | 1.400E-05 | 1.500E-05 |
| 15120 | 756 | C756V | B | 1e-05 | 1e-05 | 1e-05 | 1.000E-05 | 0.000E+00 | 1.000E-05 | 1.000E-05 |
| 15121 | 757 | L757A | B | 1.2e-05 | 1.7e-05 | 1.3e-05 | 1.400E-05 | 2.646E-06 | 1.200E-05 | 1.700E-05 |
| 15122 | 757 | L757R | B | 1.6e-05 | 1.6e-05 | 1.4e-05 | 1.533E-05 | 1.155E-06 | 1.400E-05 | 1.600E-05 |
| 15123 | 757 | L757N | B | 1.3e-05 | 1.1e-05 | 1.5e-05 | 1.300E-05 | 2.000E-06 | 1.100E-05 | 1.500E-05 |
| 15124 | 757 | L757D | B | 1.5e-05 | 1.5e-05 | 1.1e-05 | 1.367E-05 | 2.309E-06 | 1.100E-05 | 1.500E-05 |
| 15125 | 757 | L757C | B | 1.3e-05 | 1.1e-05 | 9.4e-06 | 1.113E-05 | 1.804E-06 | 9.400E-06 | 1.300E-05 |
| 15126 | 757 | L757E | B | 1e-05 | 1.4e-05 | 1.1e-05 | 1.167E-05 | 2.082E-06 | 1.000E-05 | 1.400E-05 |
| 15127 | 757 | L757Q | B | 1.4e-05 | 1.1e-05 | 1.6e-05 | 1.367E-05 | 2.517E-06 | 1.100E-05 | 1.600E-05 |
| 15128 | 757 | L757G | B | 1.5e-05 | 1.4e-05 | 1.5e-05 | 1.467E-05 | 5.774E-07 | 1.400E-05 | 1.500E-05 |
| 15129 | 757 | L757H | B | 1.5e-05 | 9.6e-06 | 2e-05 | 1.487E-05 | 5.201E-06 | 9.600E-06 | 2.000E-05 |

| | | | | | | | | | | |
|---|---|---|---|---|---|---|---|---|---|---|
| 15130 | 757 | L757I | B | 1e-05 | 1e-05 | 1.4e-05 | 1.133E-05 | 2.309E-06 | 1.000E-05 | 1.400E-05 |
| 15131 | 757 | L757L | B | 1.4e-05 | 1.1e-05 | 1.1e-05 | 1.200E-05 | 1.732E-06 | 1.100E-05 | 1.400E-05 |
| 15132 | 757 | L757K | B | 1.4e-05 | 1.4e-05 | 9.9e-06 | 1.263E-05 | 2.367E-06 | 9.900E-06 | 1.400E-05 |
| 15133 | 757 | L757M | B | 1.7e-05 | 1.6e-05 | 9.2e-06 | 1.407E-05 | 4.244E-06 | 9.200E-06 | 1.700E-05 |
| 15134 | 757 | L757F | B | 1e-05 | 1.3e-05 | 9.6e-06 | 1.087E-05 | 1.858E-06 | 9.600E-06 | 1.300E-05 |
| 15135 | 757 | L757P | B | 1.6e-05 | 1.3e-05 | 1.5e-05 | 1.467E-05 | 1.528E-06 | 1.300E-05 | 1.600E-05 |
| 15136 | 757 | L757S | B | 1.2e-05 | 9e-06 | 1.1e-05 | 1.067E-05 | 1.528E-06 | 9.000E-06 | 1.200E-05 |
| 15137 | 757 | L757T | B | 1.3e-05 | 1.4e-05 | 1.4e-05 | 1.367E-05 | 5.774E-07 | 1.300E-05 | 1.400E-05 |
| 15138 | 757 | L757W | B | 1.5e-05 | 1.5e-05 | 9.7e-06 | 1.323E-05 | 3.060E-06 | 9.700E-06 | 1.500E-05 |
| 15139 | 757 | L757Y | B | 9.5e-06 | 7.7e-06 | 1.2e-05 | 9.733E-06 | 2.159E-06 | 7.700E-06 | 1.200E-05 |
| 15140 | 757 | L757V | B | 1.1e-05 | 1.4e-05 | 1.4e-05 | 1.300E-05 | 1.732E-06 | 1.100E-05 | 1.400E-05 |
| 15141 | 758 | L758A | B | 9.3e-06 | 1.4e-05 | 9.5e-06 | 1.093E-05 | 2.658E-06 | 9.300E-06 | 1.400E-05 |
| 15142 | 758 | L758R | B | 1.1e-05 | 7.1e-06 | 9.9e-06 | 9.333E-06 | 2.011E-06 | 7.100E-06 | 1.100E-05 |
| 15143 | 758 | L758N | B | 1.3e-05 | 1.4e-05 | 1.6e-05 | 1.433E-05 | 1.528E-06 | 1.300E-05 | 1.600E-05 |
| 15144 | 758 | L758D | B | 1.2e-05 | 1.7e-05 | 1.4e-05 | 1.433E-05 | 2.517E-06 | 1.200E-05 | 1.700E-05 |
| 15145 | 758 | L758C | B | 9.7e-06 | 1e-05 | 1.2e-05 | 1.057E-05 | 1.250E-06 | 9.700E-06 | 1.200E-05 |
| 15146 | 758 | L758E | B | 9.9e-06 | 9.4e-06 | 9.7e-06 | 9.667E-06 | 2.517E-07 | 9.400E-06 | 9.900E-06 |
| 15147 | 758 | L758Q | B | 9.9e-06 | 1.2e-05 | 1.5e-05 | 1.230E-05 | 2.563E-06 | 9.900E-06 | 1.500E-05 |
| 15148 | 758 | L758G | B | 1.4e-05 | 1e-05 | 1.4e-05 | 1.267E-05 | 2.309E-06 | 1.000E-05 | 1.400E-05 |
| 15149 | 758 | L758H | B | 8.5e-06 | 8.7e-06 | 1.4e-05 | 1.040E-05 | 3.119E-06 | 8.500E-06 | 1.400E-05 |
| 15150 | 758 | L758I | B | 9.4e-06 | 1.7e-05 | 1.3e-05 | 1.313E-05 | 3.802E-06 | 9.400E-06 | 1.700E-05 |
| 15151 | 758 | L758L | B | 1e-05 | 9.4e-06 | 1.4e-05 | 1.113E-05 | 2.501E-06 | 9.400E-06 | 1.400E-05 |
| 15152 | 758 | L758K | B | 1.5e-05 | 1e-05 | 1.4e-05 | 1.300E-05 | 2.646E-06 | 1.000E-05 | 1.500E-05 |
| 15153 | 758 | L758M | B | 9.9e-06 | 1.3e-05 | 8.9e-06 | 1.060E-05 | 2.138E-06 | 8.900E-06 | 1.300E-05 |
| 15154 | 758 | L758F | B | 1.2e-05 | 1.6e-05 | 2.2e-05 | 1.667E-05 | 5.033E-06 | 1.200E-05 | 2.200E-05 |
| 15155 | 758 | L758P | B | 1.4e-05 | 1.7e-05 | 1.6e-05 | 1.567E-05 | 1.528E-06 | 1.400E-05 | 1.700E-05 |
| 15156 | 758 | L758S | B | 9.8e-06 | 1.4e-05 | 1.6e-05 | 1.327E-05 | 3.164E-06 | 9.800E-06 | 1.600E-05 |
| 15157 | 758 | L758T | B | 9.7e-06 | 1.1e-05 | 1e-05 | 1.023E-05 | 6.807E-07 | 9.700E-06 | 1.100E-05 |
| 15158 | 758 | L758W | B | 1.1e-05 | 7.7e-06 | 9.9e-06 | 9.533E-06 | 1.680E-06 | 7.700E-06 | 1.100E-05 |
| 15159 | 758 | L758Y | B | 1.5e-05 | 1.3e-05 | 1.5e-05 | 1.433E-05 | 1.155E-06 | 1.300E-05 | 1.500E-05 |
| 15160 | 758 | L758V | B | 1.7e-05 | 1.6e-05 | 1.1e-05 | 1.467E-05 | 3.215E-06 | 1.100E-05 | 1.700E-05 |
| 15161 | 759 | N759A | B | 1.2e-05 | 9.5e-06 | 1.5e-05 | 1.217E-05 | 2.754E-06 | 9.500E-06 | 1.500E-05 |
| 15162 | 759 | N759R | B | 1.6e-05 | 1.1e-05 | 1e-05 | 1.233E-05 | 3.215E-06 | 1.000E-05 | 1.600E-05 |
| 15163 | 759 | N759N | B | 8.2e-06 | 1.5e-05 | 1.4e-05 | 1.240E-05 | 3.672E-06 | 8.200E-06 | 1.500E-05 |
| 15164 | 759 | N759D | B | 1.5e-05 | 8.9e-06 | 1.3e-05 | 1.230E-05 | 3.110E-06 | 8.900E-06 | 1.500E-05 |
| 15165 | 759 | N759C | B | 1.4e-05 | 1.7e-05 | 1.4e-05 | 1.500E-05 | 1.732E-06 | 1.400E-05 | 1.700E-05 |
| 15166 | 759 | N759E | B | 1.1e-05 | 1.5e-05 | 9.8e-06 | 1.193E-05 | 2.723E-06 | 9.800E-06 | 1.500E-05 |
| 15167 | 759 | N759Q | B | 1.1e-05 | 1e-05 | 9.4e-06 | 1.013E-05 | 8.083E-07 | 9.400E-06 | 1.100E-05 |
| 15168 | 759 | N759G | B | 1.5e-05 | 1.2e-05 | 1.1e-05 | 1.267E-05 | 2.082E-06 | 1.100E-05 | 1.500E-05 |
| 15169 | 759 | N759H | B | 1.2e-05 | 1.3e-05 | 1.5e-05 | 1.333E-05 | 1.528E-06 | 1.200E-05 | 1.500E-05 |
| 15170 | 759 | N759I | B | 1.5e-05 | 1.5e-05 | 7.7e-06 | 1.257E-05 | 4.215E-06 | 7.700E-06 | 1.500E-05 |
| 15171 | 759 | N759L | B | 9.9e-06 | 1.1e-05 | 1.1e-05 | 1.063E-05 | 6.351E-07 | 9.900E-06 | 1.100E-05 |
| 15172 | 759 | N759K | B | 1.5e-05 | 1e-05 | 1.6e-05 | 1.367E-05 | 3.215E-06 | 1.000E-05 | 1.600E-05 |
| 15173 | 759 | N759M | B | 1.5e-05 | 1.6e-05 | 1.6e-05 | 1.567E-05 | 5.774E-07 | 1.500E-05 | 1.600E-05 |
| 15174 | 759 | N759F | B | 1.3e-05 | 1.4e-05 | 1.2e-05 | 1.300E-05 | 1.000E-06 | 1.200E-05 | 1.400E-05 |
| 15175 | 759 | N759P | B | 1.6e-05 | 1.2e-05 | 1.2e-05 | 1.333E-05 | 2.309E-06 | 1.200E-05 | 1.600E-05 |
| 15176 | 759 | N759S | B | 1e-05 | 1.4e-05 | 1.5e-05 | 1.300E-05 | 2.646E-06 | 1.000E-05 | 1.500E-05 |
| 15177 | 759 | N759T | B | 1.5e-05 | 9.4e-06 | 1.5e-05 | 1.313E-05 | 3.233E-06 | 9.400E-06 | 1.500E-05 |
| 15178 | 759 | N759W | B | 1e-05 | 1.5e-05 | 1.6e-05 | 1.367E-05 | 3.215E-06 | 1.000E-05 | 1.600E-05 |
| 15179 | 759 | N759Y | B | 1.6e-05 | 1.3e-05 | 9.4e-06 | 1.280E-05 | 3.305E-06 | 9.400E-06 | 1.600E-05 |
| 15180 | 759 | N759V | B | 1.4e-05 | 1.4e-05 | 1.3e-05 | 1.367E-05 | 5.774E-07 | 1.300E-05 | 1.400E-05 |
| 15181 | 760 | N760A | B | 1.6e-05 | 1.3e-05 | 1.1e-05 | 1.333E-05 | 2.517E-06 | 1.100E-05 | 1.600E-05 |
| 15182 | 760 | N760R | B | 1.8e-05 | 1.8e-05 | 1.2e-05 | 1.600E-05 | 3.464E-06 | 1.200E-05 | 1.800E-05 |
| 15183 | 760 | N760N | B | 1.1e-05 | 9.5e-06 | 1.4e-05 | 1.150E-05 | 2.291E-06 | 9.500E-06 | 1.400E-05 |
| 15184 | 760 | N760D | B | 8.5e-06 | 1.3e-05 | 1.4e-05 | 1.183E-05 | 2.930E-06 | 8.500E-06 | 1.400E-05 |
| 15185 | 760 | N760C | B | 1.5e-05 | 1.2e-05 | 1e-05 | 1.233E-05 | 2.517E-06 | 1.000E-05 | 1.500E-05 |
| 15186 | 760 | N760E | B | 1.7e-05 | 1.1e-05 | 1.2e-05 | 1.333E-05 | 3.215E-06 | 1.100E-05 | 1.700E-05 |
| 15187 | 760 | N760Q | B | 1e-05 | 1.5e-05 | 1.5e-05 | 1.333E-05 | 2.887E-06 | 1.000E-05 | 1.500E-05 |
| 15188 | 760 | N760G | B | 1.1e-05 | 1.2e-05 | 1.3e-05 | 1.200E-05 | 1.000E-06 | 1.100E-05 | 1.300E-05 |
| 15189 | 760 | N760H | B | 9.3e-06 | 1e-05 | 1.5e-05 | 1.143E-05 | 3.109E-06 | 9.300E-06 | 1.500E-05 |
| 15190 | 760 | N760I | B | 1.1e-05 | 1.2e-05 | 1e-05 | 1.100E-05 | 1.000E-06 | 1.000E-05 | 1.200E-05 |
| 15191 | 760 | N760L | B | 1.1e-05 | 1.1e-05 | 1.3e-05 | 1.167E-05 | 1.155E-06 | 1.100E-05 | 1.300E-05 |
| 15192 | 760 | N760K | B | 9.9e-06 | 1.1e-05 | 1.4e-05 | 1.163E-05 | 2.122E-06 | 9.900E-06 | 1.400E-05 |
| 15193 | 760 | N760M | B | 1.3e-05 | 1.1e-05 | 1.7e-05 | 1.367E-05 | 3.055E-06 | 1.100E-05 | 1.700E-05 |
| 15194 | 760 | N760F | B | 9.9e-06 | 1.6e-05 | 1.1e-05 | 1.230E-05 | 3.251E-06 | 9.900E-06 | 1.600E-05 |
| 15195 | 760 | N760P | B | 1.3e-05 | 1.7e-05 | 1.1e-05 | 1.367E-05 | 3.055E-06 | 1.100E-05 | 1.700E-05 |
| 15196 | 760 | N760S | B | 1e-05 | 1.5e-05 | 1.6e-05 | 1.367E-05 | 3.215E-06 | 1.000E-05 | 1.600E-05 |
| 15197 | 760 | N760T | B | 1.5e-05 | 1.7e-05 | 9.7e-06 | 1.390E-05 | 3.772E-06 | 9.700E-06 | 1.700E-05 |
| 15198 | 760 | N760W | B | 1.5e-05 | 8.8e-06 | 1.5e-05 | 1.293E-05 | 3.580E-06 | 8.800E-06 | 1.500E-05 |
| 15199 | 760 | N760Y | B | 1.6e-05 | 1.2e-05 | 1.4e-05 | 1.400E-05 | 2.000E-06 | 1.200E-05 | 1.600E-05 |
| 15200 | 760 | N760V | B | 1.6e-05 | 1.5e-05 | 9.7e-06 | 1.357E-05 | 3.386E-06 | 9.700E-06 | 1.600E-05 |
| 15201 | 761 | F761A | B | 1.4e-05 | 9.4e-06 | 1.1e-05 | 1.147E-05 | 2.335E-06 | 9.400E-06 | 1.400E-05 |
| 15202 | 761 | F761R | B | 1.3e-05 | 8.3e-06 | 1.1e-05 | 1.077E-05 | 2.359E-06 | 8.300E-06 | 1.300E-05 |
| 15203 | 761 | F761N | B | 8.4e-06 | 1.7e-05 | 1.4e-05 | 1.313E-05 | 4.365E-06 | 8.400E-06 | 1.700E-05 |
| 15204 | 761 | F761D | B | 1.5e-05 | 1.5e-05 | 1.4e-05 | 1.467E-05 | 5.774E-07 | 1.400E-05 | 1.500E-05 |
| 15205 | 761 | F761C | B | 1e-05 | 1e-05 | 9.6e-06 | 9.867E-06 | 2.309E-07 | 9.600E-06 | 1.000E-05 |
| 15206 | 761 | F761E | B | 1.1e-05 | 1.7e-05 | 9.4e-06 | 1.247E-05 | 4.007E-06 | 9.400E-06 | 1.700E-05 |
| 15207 | 761 | F761Q | B | 1e-05 | 1.3e-05 | 9.5e-06 | 1.083E-05 | 1.893E-06 | 9.500E-06 | 1.300E-05 |
| 15208 | 761 | F761G | B | 1.5e-05 | 1.5e-05 | 9.6e-06 | 1.320E-05 | 3.118E-06 | 9.600E-06 | 1.500E-05 |
| 15209 | 761 | F761H | B | 1.1e-05 | 9.4e-06 | 1.4e-05 | 1.147E-05 | 2.335E-06 | 9.400E-06 | 1.400E-05 |
| 15210 | 761 | F761I | B | 1.6e-05 | 1.6e-05 | 1.8e-05 | 1.667E-05 | 1.155E-06 | 1.600E-05 | 1.800E-05 |
| 15211 | 761 | F761L | B | 8.2e-06 | 1.5e-05 | 9.9e-06 | 1.103E-05 | 3.539E-06 | 8.200E-06 | 1.500E-05 |
| 15212 | 761 | F761K | B | 9.2e-06 | 1.2e-05 | 1.1e-05 | 1.073E-05 | 1.419E-06 | 9.200E-06 | 1.200E-05 |
| 15213 | 761 | F761M | B | 1.7e-05 | 1.6e-05 | 1.5e-05 | 1.600E-05 | 1.000E-06 | 1.500E-05 | 1.700E-05 |
| 15214 | 761 | F761F | B | 1e-05 | 1.3e-05 | 1.4e-05 | 1.233E-05 | 2.082E-06 | 1.000E-05 | 1.400E-05 |

| | | | | | | | | | | |
|---|---|---|---|---|---|---|---|---|---|---|
| 15215 | 761 | F761P | B | 8.4e-06 | 1.4e-05 | 8.8e-06 | 1.040E-05 | 3.124E-06 | 8.400E-06 | 1.400E-05 |
| 15216 | 761 | F761S | B | 1.5e-05 | 1.5e-05 | 1.2e-05 | 1.400E-05 | 1.732E-06 | 1.200E-05 | 1.500E-05 |
| 15217 | 761 | F761T | B | 1.4e-05 | 1.4e-05 | 1.6e-05 | 1.467E-05 | 1.155E-06 | 1.400E-05 | 1.600E-05 |
| 15218 | 761 | F761W | B | 1.5e-05 | 8.7e-06 | 1.4e-05 | 1.257E-05 | 3.386E-06 | 8.700E-06 | 1.500E-05 |
| 15219 | 761 | F761Y | B | 1e-05 | 1.1e-05 | 1.7e-05 | 1.267E-05 | 3.786E-06 | 1.000E-05 | 1.700E-05 |
| 15220 | 761 | F761V | B | 1.5e-05 | 9.9e-06 | 1.6e-05 | 1.363E-05 | 3.272E-06 | 9.900E-06 | 1.600E-05 |
| 15221 | 762 | Y762A | B | 9.9e-06 | 1.1e-05 | 1.4e-05 | 1.163E-05 | 2.122E-06 | 9.900E-06 | 1.400E-05 |
| 15222 | 762 | Y762R | B | 1.1e-05 | 7.4e-06 | 8.8e-06 | 9.067E-06 | 1.815E-06 | 7.400E-06 | 1.100E-05 |
| 15223 | 762 | Y762N | B | 1e-05 | 1e-05 | 1.7e-05 | 1.233E-05 | 4.041E-06 | 1.000E-05 | 1.700E-05 |
| 15224 | 762 | Y762D | B | 1.1e-05 | 1.4e-05 | 1.5e-05 | 1.333E-05 | 2.082E-06 | 1.100E-05 | 1.500E-05 |
| 15225 | 762 | Y762C | B | 1.6e-05 | 9.3e-06 | 1.5e-05 | 1.343E-05 | 3.614E-06 | 9.300E-06 | 1.600E-05 |
| 15226 | 762 | Y762E | B | 1.6e-05 | 9.1e-06 | 1.4e-05 | 1.303E-05 | 3.550E-06 | 9.100E-06 | 1.600E-05 |
| 15227 | 762 | Y762Q | B | 1.3e-05 | 1.2e-05 | 1.4e-05 | 1.300E-05 | 1.000E-06 | 1.200E-05 | 1.400E-05 |
| 15228 | 762 | Y762G | B | 1e-05 | 1.1e-05 | 9.9e-06 | 1.030E-05 | 6.083E-07 | 9.900E-06 | 1.100E-05 |
| 15229 | 762 | Y762H | B | 6.2e-06 | 7.1e-06 | 1.5e-05 | 9.433E-06 | 4.842E-06 | 6.200E-06 | 1.500E-05 |
| 15230 | 762 | Y762I | B | 1.5e-05 | 1.6e-05 | 1.3e-05 | 1.467E-05 | 1.528E-06 | 1.300E-05 | 1.600E-05 |
| 15231 | 762 | Y762L | B | 1e-05 | 1.1e-05 | 1.6e-05 | 1.233E-05 | 3.215E-06 | 1.000E-05 | 1.600E-05 |
| 15232 | 762 | Y762K | B | 1.3e-05 | 1e-05 | 1.3e-05 | 1.200E-05 | 1.732E-06 | 1.000E-05 | 1.300E-05 |
| 15233 | 762 | Y762M | B | 1.5e-05 | 1.3e-05 | 1.1e-05 | 1.300E-05 | 2.000E-06 | 1.100E-05 | 1.500E-05 |
| 15234 | 762 | Y762F | B | 1.7e-05 | 1.3e-05 | 1e-05 | 1.333E-05 | 3.512E-06 | 1.000E-05 | 1.700E-05 |
| 15235 | 762 | Y762P | B | 9.1e-06 | 1.1e-05 | 1.6e-05 | 1.203E-05 | 3.564E-06 | 9.100E-06 | 1.600E-05 |
| 15236 | 762 | Y762S | B | 1.5e-05 | 1.4e-05 | 1e-05 | 1.300E-05 | 2.646E-06 | 1.000E-05 | 1.500E-05 |
| 15237 | 762 | Y762T | B | 1.4e-05 | 1.3e-05 | 1.5e-05 | 1.400E-05 | 1.000E-06 | 1.300E-05 | 1.500E-05 |
| 15238 | 762 | Y762W | B | 1.3e-05 | 1.5e-05 | 1.4e-05 | 1.400E-05 | 1.000E-06 | 1.300E-05 | 1.500E-05 |
| 15239 | 762 | Y762Y | B | 1.3e-05 | 9.4e-06 | 1.6e-05 | 1.280E-05 | 3.305E-06 | 9.400E-06 | 1.600E-05 |
| 15240 | 762 | Y762V | B | 1.1e-05 | 1e-05 | 1.1e-05 | 1.067E-05 | 5.774E-07 | 1.000E-05 | 1.100E-05 |
| 15241 | 763 | P763A | B | 1.5e-05 | 1.5e-05 | 1.1e-05 | 1.367E-05 | 2.309E-06 | 1.100E-05 | 1.500E-05 |
| 15242 | 763 | P763R | B | 1.5e-05 | 1.6e-05 | 1.6e-05 | 1.567E-05 | 5.774E-07 | 1.500E-05 | 1.600E-05 |
| 15243 | 763 | P763N | B | 1.5e-05 | 1.6e-05 | 9.2e-06 | 1.340E-05 | 3.672E-06 | 9.200E-06 | 1.600E-05 |
| 15244 | 763 | P763D | B | 9.1e-06 | 1.4e-05 | 1.2e-05 | 1.170E-05 | 2.464E-06 | 9.100E-06 | 1.400E-05 |
| 15245 | 763 | P763C | B | 9.8e-06 | 1.5e-05 | 1.5e-05 | 1.327E-05 | 3.002E-06 | 9.800E-06 | 1.500E-05 |
| 15246 | 763 | P763E | B | 1.5e-05 | 1.6e-05 | 1.4e-05 | 1.500E-05 | 1.000E-06 | 1.400E-05 | 1.600E-05 |
| 15247 | 763 | P763Q | B | 1.6e-05 | 1.2e-05 | 1.1e-05 | 1.300E-05 | 2.646E-06 | 1.100E-05 | 1.600E-05 |
| 15248 | 763 | P763G | B | 8.4e-06 | 7.8e-06 | 1.5e-05 | 1.040E-05 | 3.995E-06 | 7.800E-06 | 1.500E-05 |
| 15249 | 763 | P763H | B | 9.3e-06 | 8.7e-06 | 1.5e-05 | 1.100E-05 | 3.477E-06 | 8.700E-06 | 1.500E-05 |
| 15250 | 763 | P763I | B | 1.8e-05 | 1.4e-05 | 1.4e-05 | 1.533E-05 | 2.309E-06 | 1.400E-05 | 1.800E-05 |
| 15251 | 763 | P763L | B | 9.9e-06 | 1e-05 | 1.5e-05 | 1.163E-05 | 2.916E-06 | 9.900E-06 | 1.500E-05 |
| 15252 | 763 | P763K | B | 9.7e-06 | 9.9e-06 | 9.4e-06 | 9.667E-06 | 2.517E-07 | 9.400E-06 | 9.900E-06 |
| 15253 | 763 | P763M | B | 1.2e-05 | 9e-06 | 1.5e-05 | 1.200E-05 | 3.000E-06 | 9.000E-06 | 1.500E-05 |
| 15254 | 763 | P763F | B | 1.5e-05 | 1.3e-05 | 1.3e-05 | 1.367E-05 | 1.155E-06 | 1.300E-05 | 1.500E-05 |
| 15255 | 763 | P763P | B | 1.4e-05 | 1e-05 | 8.3e-06 | 1.077E-05 | 2.926E-06 | 8.300E-06 | 1.400E-05 |
| 15256 | 763 | P763S | B | 1.4e-05 | 1.1e-05 | 1.2e-05 | 1.233E-05 | 1.528E-06 | 1.100E-05 | 1.400E-05 |
| 15257 | 763 | P763T | B | 9.9e-06 | 1.5e-05 | 1.3e-05 | 1.263E-05 | 2.570E-06 | 9.900E-06 | 1.500E-05 |
| 15258 | 763 | P763W | B | 9.6e-06 | 1.5e-05 | 6.7e-06 | 1.043E-05 | 4.212E-06 | 6.700E-06 | 1.500E-05 |
| 15259 | 763 | P763Y | B | 1.5e-05 | 1.5e-05 | 1.4e-05 | 1.467E-05 | 5.774E-07 | 1.400E-05 | 1.500E-05 |
| 15260 | 763 | P763V | B | 1.1e-05 | 1.1e-05 | 1.1e-05 | 1.100E-05 | 2.075E-21 | 1.100E-05 | 1.100E-05 |
| 15261 | 764 | R764A | B | 1.4e-05 | 1.3e-05 | 1.3e-05 | 1.333E-05 | 5.774E-07 | 1.300E-05 | 1.400E-05 |
| 15262 | 764 | R764R | B | 1e-05 | 1.3e-05 | 1e-05 | 1.100E-05 | 1.732E-06 | 1.000E-05 | 1.300E-05 |
| 15263 | 764 | R764N | B | 1.5e-05 | 1.4e-05 | 1.4e-05 | 1.433E-05 | 5.774E-07 | 1.400E-05 | 1.500E-05 |
| 15264 | 764 | R764D | B | 1e-05 | 8.2e-06 | 7.7e-06 | 8.633E-06 | 1.210E-06 | 7.700E-06 | 1.000E-05 |
| 15265 | 764 | R764C | B | 1.5e-05 | 1.5e-05 | 1.5e-05 | 1.500E-05 | 0.000E+00 | 1.500E-05 | 1.500E-05 |
| 15266 | 764 | R764E | B | 1.4e-05 | 1.4e-05 | 1e-05 | 1.267E-05 | 2.309E-06 | 1.000E-05 | 1.400E-05 |
| 15267 | 764 | R764Q | B | 1e-05 | 1.5e-05 | 9.8e-06 | 1.160E-05 | 2.946E-06 | 9.800E-06 | 1.500E-05 |
| 15268 | 764 | R764G | B | 1.1e-05 | 1.9e-05 | 1.5e-05 | 1.500E-05 | 4.000E-06 | 1.100E-05 | 1.900E-05 |
| 15269 | 764 | R764H | B | 1.4e-05 | 1.2e-05 | 1.6e-05 | 1.400E-05 | 2.000E-06 | 1.200E-05 | 1.600E-05 |
| 15270 | 764 | R764I | B | 1.1e-05 | 1e-05 | 1.7e-05 | 1.267E-05 | 3.786E-06 | 1.000E-05 | 1.700E-05 |
| 15271 | 764 | R764L | B | 1.6e-05 | 8.9e-06 | 1.5e-05 | 1.330E-05 | 3.843E-06 | 8.900E-06 | 1.600E-05 |
| 15272 | 764 | R764K | B | 1.4e-05 | 1.4e-05 | 1.4e-05 | 1.400E-05 | 0.000E+00 | 1.400E-05 | 1.400E-05 |
| 15273 | 764 | R764M | B | 1.5e-05 | 6.5e-06 | 1.4e-05 | 1.183E-05 | 4.646E-06 | 6.500E-06 | 1.500E-05 |
| 15274 | 764 | R764F | B | 1.5e-05 | 1.3e-05 | 2.6e-05 | 1.800E-05 | 7.000E-06 | 1.300E-05 | 2.600E-05 |
| 15275 | 764 | R764P | B | 1.5e-05 | 1.1e-05 | 1.6e-05 | 1.400E-05 | 2.646E-06 | 1.100E-05 | 1.600E-05 |
| 15276 | 764 | R764S | B | 1.3e-05 | 1.5e-05 | 1.1e-05 | 1.300E-05 | 2.000E-06 | 1.100E-05 | 1.500E-05 |
| 15277 | 764 | R764T | B | 1.4e-05 | 1.6e-05 | 1.1e-05 | 1.367E-05 | 2.517E-06 | 1.100E-05 | 1.600E-05 |
| 15278 | 764 | R764W | B | 1e-05 | 1.5e-05 | 1.4e-05 | 1.300E-05 | 2.646E-06 | 1.000E-05 | 1.500E-05 |
| 15279 | 764 | R764Y | B | 1.3e-05 | 1.6e-05 | 1.4e-05 | 1.433E-05 | 1.528E-06 | 1.300E-05 | 1.600E-05 |
| 15280 | 764 | R764V | B | 1.1e-05 | 9.6e-06 | 1.1e-05 | 1.053E-05 | 8.083E-07 | 9.600E-06 | 1.100E-05 |
| 15281 | 765 | E765A | B | 1.2e-05 | 1.3e-05 | 8.2e-06 | 1.107E-05 | 2.532E-06 | 8.200E-06 | 1.300E-05 |
| 15282 | 765 | E765R | B | 1.5e-05 | 8.7e-06 | 9.2e-06 | 1.097E-05 | 3.502E-06 | 8.700E-06 | 1.500E-05 |
| 15283 | 765 | E765N | B | 1e-05 | 1.1e-05 | 1.2e-05 | 1.100E-05 | 1.000E-06 | 1.000E-05 | 1.200E-05 |
| 15284 | 765 | E765D | B | 1e-05 | 1.5e-05 | 1.3e-05 | 1.267E-05 | 2.517E-06 | 1.000E-05 | 1.500E-05 |
| 15285 | 765 | E765C | B | 9.2e-06 | 1.3e-05 | 8.5e-06 | 1.023E-05 | 2.421E-06 | 8.500E-06 | 1.300E-05 |
| 15286 | 765 | E765E | B | 9.1e-06 | 1.6e-05 | 1.3e-05 | 1.270E-05 | 3.460E-06 | 9.100E-06 | 1.600E-05 |
| 15287 | 765 | E765Q | B | 1.6e-05 | 9.1e-06 | 1.4e-05 | 1.303E-05 | 3.550E-06 | 9.100E-06 | 1.600E-05 |
| 15288 | 765 | E765G | B | 7.4e-06 | 1.4e-05 | 1.7e-05 | 1.280E-05 | 4.911E-06 | 7.400E-06 | 1.700E-05 |
| 15289 | 765 | E765H | B | 9.4e-06 | 1.1e-05 | 1.5e-05 | 1.180E-05 | 2.884E-06 | 9.400E-06 | 1.500E-05 |
| 15290 | 765 | E765I | B | 1.1e-05 | 1.5e-05 | 1.1e-05 | 1.233E-05 | 2.309E-06 | 1.100E-05 | 1.500E-05 |
| 15291 | 765 | E765L | B | 9.4e-06 | 1.5e-05 | 9.9e-06 | 1.143E-05 | 3.099E-06 | 9.400E-06 | 1.500E-05 |
| 15292 | 765 | E765K | B | 1.4e-05 | 1.6e-05 | 1.3e-05 | 1.433E-05 | 1.528E-06 | 1.300E-05 | 1.600E-05 |
| 15293 | 765 | E765M | B | 1.3e-05 | 1.2e-05 | 1.2e-05 | 1.233E-05 | 5.774E-07 | 1.200E-05 | 1.300E-05 |
| 15294 | 765 | E765F | B | 1.5e-05 | 1.2e-05 | 1.3e-05 | 1.333E-05 | 1.528E-06 | 1.200E-05 | 1.500E-05 |
| 15295 | 765 | E765P | B | 1.6e-05 | 1e-05 | 1e-05 | 1.200E-05 | 3.464E-06 | 1.000E-05 | 1.600E-05 |
| 15296 | 765 | E765S | B | 7.7e-06 | 1.6e-05 | 1.5e-05 | 1.290E-05 | 4.531E-06 | 7.700E-06 | 1.600E-05 |
| 15297 | 765 | E765T | B | 9.6e-06 | 1.6e-05 | 1e-05 | 1.187E-05 | 3.585E-06 | 9.600E-06 | 1.600E-05 |
| 15298 | 765 | E765W | B | 1.5e-05 | 1e-05 | 1.1e-05 | 1.200E-05 | 2.646E-06 | 1.000E-05 | 1.500E-05 |
| 15299 | 765 | E765Y | B | 1e-05 | 1.4e-05 | 1.5e-05 | 1.300E-05 | 2.646E-06 | 1.000E-05 | 1.500E-05 |

| | | | | | | | | | | |
|---|---|---|---|---|---|---|---|---|---|---|
| 15300 | 765 | E765V | B | 1e-05 | 8.8e-06 | 1e-05 | 9.600E-06 | 6.928E-07 | 8.800E-06 | 1.000E-05 |
| 15301 | 766 | A766A | B | 8.5e-06 | 1.3e-05 | 1.6e-05 | 1.250E-05 | 3.775E-06 | 8.500E-06 | 1.600E-05 |
| 15302 | 766 | A766R | B | 1.2e-05 | 9.6e-06 | 1.1e-05 | 1.087E-05 | 1.206E-06 | 9.600E-06 | 1.200E-05 |
| 15303 | 766 | A766N | B | 1.6e-05 | 1.5e-05 | 1.4e-05 | 1.500E-05 | 1.000E-06 | 1.400E-05 | 1.600E-05 |
| 15304 | 766 | A766D | B | 1e-05 | 1.6e-05 | 1.4e-05 | 1.333E-05 | 3.055E-06 | 1.000E-05 | 1.600E-05 |
| 15305 | 766 | A766C | B | 9.7e-06 | 9.9e-06 | 1.5e-05 | 1.153E-05 | 3.004E-06 | 9.700E-06 | 1.500E-05 |
| 15306 | 766 | A766E | B | 1.1e-05 | 1e-05 | 7.8e-06 | 9.600E-06 | 1.637E-06 | 7.800E-06 | 1.100E-05 |
| 15307 | 766 | A766Q | B | 1.6e-05 | 1e-05 | 1.1e-05 | 1.233E-05 | 3.215E-06 | 1.000E-05 | 1.600E-05 |
| 15308 | 766 | A766G | B | 1.5e-05 | 1e-05 | 1.1e-05 | 1.200E-05 | 2.646E-06 | 1.000E-05 | 1.500E-05 |
| 15309 | 766 | A766H | B | 1.2e-05 | 1.4e-05 | 9.1e-06 | 1.170E-05 | 2.464E-06 | 9.100E-06 | 1.400E-05 |
| 15310 | 766 | A766I | B | 1.4e-05 | 1.6e-05 | 1.1e-05 | 1.367E-05 | 2.517E-06 | 1.100E-05 | 1.600E-05 |
| 15311 | 766 | A766L | B | 1.1e-05 | 1.7e-05 | 8.9e-06 | 1.230E-05 | 4.204E-06 | 8.900E-06 | 1.700E-05 |
| 15312 | 766 | A766K | B | 1e-05 | 1.5e-05 | 1.6e-05 | 1.367E-05 | 3.215E-06 | 1.000E-05 | 1.600E-05 |
| 15313 | 766 | A766M | B | 7.6e-06 | 1.6e-05 | 8.8e-06 | 1.080E-05 | 4.543E-06 | 7.600E-06 | 1.600E-05 |
| 15314 | 766 | A766F | B | 1.6e-05 | 1.2e-05 | 1.2e-05 | 1.333E-05 | 2.309E-06 | 1.200E-05 | 1.600E-05 |
| 15315 | 766 | A766P | B | 1e-05 | 1.1e-05 | 1.5e-05 | 1.200E-05 | 2.646E-06 | 1.000E-05 | 1.500E-05 |
| 15316 | 766 | A766S | B | 2.3e-05 | 9.3e-06 | 1.2e-05 | 1.477E-05 | 7.257E-06 | 9.300E-06 | 2.300E-05 |
| 15317 | 766 | A766T | B | 1.7e-05 | 1.5e-05 | 9.9e-06 | 1.397E-05 | 3.661E-06 | 9.900E-06 | 1.700E-05 |
| 15318 | 766 | A766W | B | 8.2e-06 | 1.4e-05 | 1.4e-05 | 1.207E-05 | 3.349E-06 | 8.200E-06 | 1.400E-05 |
| 15319 | 766 | A766Y | B | 1.5e-05 | 1.5e-05 | 2.1e-05 | 1.700E-05 | 3.464E-06 | 1.500E-05 | 2.100E-05 |
| 15320 | 766 | A766V | B | 1.6e-05 | 1.2e-05 | 6.6e-06 | 1.153E-05 | 4.717E-06 | 6.600E-06 | 1.600E-05 |
| 15321 | 767 | K767A | B | 1.5e-05 | 1.1e-05 | 8.1e-06 | 1.137E-05 | 3.465E-06 | 8.100E-06 | 1.500E-05 |
| 15322 | 767 | K767R | B | 1.4e-05 | 1.6e-05 | 1e-05 | 1.333E-05 | 3.055E-06 | 1.000E-05 | 1.600E-05 |
| 15323 | 767 | K767N | B | 7.2e-06 | 9.2e-06 | 8.7e-06 | 8.367E-06 | 1.041E-06 | 7.200E-06 | 9.200E-06 |
| 15324 | 767 | K767D | B | 1.2e-05 | 1.1e-05 | 1.4e-05 | 1.233E-05 | 1.528E-06 | 1.100E-05 | 1.400E-05 |
| 15325 | 767 | K767C | B | 1.6e-05 | 1.3e-05 | 1.4e-05 | 1.433E-05 | 1.528E-06 | 1.300E-05 | 1.600E-05 |
| 15326 | 767 | K767E | B | 8.8e-06 | 1.6e-05 | 1.6e-05 | 1.360E-05 | 4.157E-06 | 8.800E-06 | 1.600E-05 |
| 15327 | 767 | K767Q | B | 9.2e-06 | 1.6e-05 | 1.2e-05 | 1.240E-05 | 3.418E-06 | 9.200E-06 | 1.600E-05 |
| 15328 | 767 | K767G | B | 9.4e-06 | 1e-05 | 1.4e-05 | 1.113E-05 | 2.501E-06 | 9.400E-06 | 1.400E-05 |
| 15329 | 767 | K767H | B | 1.4e-05 | 1.5e-05 | 1.4e-05 | 1.433E-05 | 5.774E-07 | 1.400E-05 | 1.500E-05 |
| 15330 | 767 | K767I | B | 1.4e-05 | 1.2e-05 | 1.1e-05 | 1.233E-05 | 1.528E-06 | 1.100E-05 | 1.400E-05 |
| 15331 | 767 | K767L | B | 1.5e-05 | 1.6e-05 | 1.6e-05 | 1.567E-05 | 5.774E-07 | 1.500E-05 | 1.600E-05 |
| 15332 | 767 | K767K | B | 1.7e-05 | 1.1e-05 | 1.1e-05 | 1.300E-05 | 3.464E-06 | 1.100E-05 | 1.700E-05 |
| 15333 | 767 | K767M | B | 8.8e-06 | 1.4e-05 | 1.6e-05 | 1.293E-05 | 3.717E-06 | 8.800E-06 | 1.600E-05 |
| 15334 | 767 | K767F | B | 8.6e-06 | 1.6e-05 | 1.6e-05 | 1.353E-05 | 4.272E-06 | 8.600E-06 | 1.600E-05 |
| 15335 | 767 | K767P | B | 1.8e-05 | 1.5e-05 | 9.6e-06 | 1.420E-05 | 4.257E-06 | 9.600E-06 | 1.800E-05 |
| 15336 | 767 | K767S | B | 9.3e-06 | 1.4e-05 | 1.4e-05 | 1.243E-05 | 2.714E-06 | 9.300E-06 | 1.400E-05 |
| 15337 | 767 | K767T | B | 1.6e-05 | 7e-06 | 9.3e-06 | 1.077E-05 | 4.676E-06 | 7.000E-06 | 1.600E-05 |
| 15338 | 767 | K767W | B | 1.3e-05 | 9.1e-06 | 1.3e-05 | 1.170E-05 | 2.252E-06 | 9.100E-06 | 1.300E-05 |
| 15339 | 767 | K767Y | B | 9.9e-06 | 1e-05 | 1e-05 | 9.967E-06 | 5.774E-08 | 9.900E-06 | 1.000E-05 |
| 15340 | 767 | K767V | B | 1.5e-05 | 1.1e-05 | 9e-06 | 1.167E-05 | 3.055E-06 | 9.000E-06 | 1.500E-05 |
| 15341 | 768 | V768A | B | 1.6e-05 | 1e-05 | 1.5e-05 | 1.367E-05 | 3.215E-06 | 1.000E-05 | 1.600E-05 |
| 15342 | 768 | V768R | B | 1.7e-05 | 1.6e-05 | 1.3e-05 | 1.533E-05 | 2.082E-06 | 1.300E-05 | 1.700E-05 |
| 15343 | 768 | V768N | B | 1.2e-05 | 1.5e-05 | 9.4e-06 | 1.213E-05 | 2.802E-06 | 9.400E-06 | 1.500E-05 |
| 15344 | 768 | V768D | B | 9.5e-06 | 1.3e-05 | 1e-05 | 1.083E-05 | 1.893E-06 | 9.500E-06 | 1.300E-05 |
| 15345 | 768 | V768C | B | 6e-06 | 1e-05 | 1.5e-05 | 1.033E-05 | 4.509E-06 | 6.000E-06 | 1.500E-05 |
| 15346 | 768 | V768E | B | 1.4e-05 | 1.2e-05 | 1.2e-05 | 1.267E-05 | 1.155E-06 | 1.200E-05 | 1.400E-05 |
| 15347 | 768 | V768Q | B | 1.3e-05 | 1.4e-05 | 1.4e-05 | 1.367E-05 | 5.774E-07 | 1.300E-05 | 1.400E-05 |
| 15348 | 768 | V768G | B | 1e-05 | 1.6e-05 | 1.1e-05 | 1.233E-05 | 3.215E-06 | 1.000E-05 | 1.600E-05 |
| 15349 | 768 | V768H | B | 1.3e-05 | 9.4e-06 | 1.9e-05 | 1.380E-05 | 4.850E-06 | 9.400E-06 | 1.900E-05 |
| 15350 | 768 | V768I | B | 1.5e-05 | 1.5e-05 | 1.4e-05 | 1.467E-05 | 5.774E-07 | 1.400E-05 | 1.500E-05 |
| 15351 | 768 | V768L | B | 1.2e-05 | 1.1e-05 | 1e-05 | 1.100E-05 | 1.000E-06 | 1.000E-05 | 1.200E-05 |
| 15352 | 768 | V768K | B | 9.6e-06 | 1e-05 | 1.6e-05 | 1.187E-05 | 3.585E-06 | 9.600E-06 | 1.600E-05 |
| 15353 | 768 | V768M | B | 9.7e-06 | 1.6e-05 | 8.8e-06 | 1.150E-05 | 3.923E-06 | 8.800E-06 | 1.600E-05 |
| 15354 | 768 | V768F | B | 1.6e-05 | 1.1e-05 | 1.7e-05 | 1.467E-05 | 3.215E-06 | 1.100E-05 | 1.700E-05 |
| 15355 | 768 | V768P | B | 6.5e-06 | 2.6e-05 | 1e-05 | 1.417E-05 | 1.040E-05 | 6.500E-06 | 2.600E-05 |
| 15356 | 768 | V768S | B | 1.4e-05 | 1.4e-05 | 1.6e-05 | 1.467E-05 | 1.155E-06 | 1.400E-05 | 1.600E-05 |
| 15357 | 768 | V768T | B | 8.2e-06 | 1.3e-05 | 1.1e-05 | 1.073E-05 | 2.411E-06 | 8.200E-06 | 1.300E-05 |
| 15358 | 768 | V768W | B | 1.4e-05 | 1.4e-05 | 1e-05 | 1.267E-05 | 2.309E-06 | 1.000E-05 | 1.400E-05 |
| 15359 | 768 | V768Y | B | 1.3e-05 | 1.1e-05 | 1e-05 | 1.133E-05 | 1.528E-06 | 1.000E-05 | 1.300E-05 |
| 15360 | 768 | V768V | B | 1e-05 | 1e-05 | 1.2e-05 | 1.067E-05 | 1.155E-06 | 1.000E-05 | 1.200E-05 |
| 15361 | 769 | Q769A | B | 1e-05 | 9.1e-06 | 1.6e-05 | 1.170E-05 | 3.751E-06 | 9.100E-06 | 1.600E-05 |
| 15362 | 769 | Q769R | B | 1.6e-05 | 1.2e-05 | 9.8e-06 | 1.260E-05 | 3.143E-06 | 9.800E-06 | 1.600E-05 |
| 15363 | 769 | Q769N | B | 1.4e-05 | 1.1e-05 | 1.7e-05 | 1.400E-05 | 3.000E-06 | 1.100E-05 | 1.700E-05 |
| 15364 | 769 | Q769D | B | 9.6e-06 | 1.5e-05 | 9.1e-06 | 1.123E-05 | 3.272E-06 | 9.100E-06 | 1.500E-05 |
| 15365 | 769 | Q769C | B | 1.4e-05 | 1.5e-05 | 1.1e-05 | 1.333E-05 | 2.082E-06 | 1.100E-05 | 1.500E-05 |
| 15366 | 769 | Q769E | B | 1e-05 | 1.6e-05 | 1.2e-05 | 1.267E-05 | 3.055E-06 | 1.000E-05 | 1.600E-05 |
| 15367 | 769 | Q769Q | B | 1.5e-05 | 6.5e-06 | 1e-05 | 1.050E-05 | 4.272E-06 | 6.500E-06 | 1.500E-05 |
| 15368 | 769 | Q769G | B | 1.5e-05 | 1.7e-05 | 1.6e-05 | 1.600E-05 | 1.000E-06 | 1.500E-05 | 1.700E-05 |
| 15369 | 769 | Q769H | B | 2.3e-05 | 1.4e-05 | 1.1e-05 | 1.600E-05 | 6.245E-06 | 1.100E-05 | 2.300E-05 |
| 15370 | 769 | Q769I | B | 1e-05 | 1.4e-05 | 1.6e-05 | 1.333E-05 | 3.055E-06 | 1.000E-05 | 1.600E-05 |
| 15371 | 769 | Q769L | B | 1.3e-05 | 6.1e-06 | 1.3e-05 | 1.070E-05 | 3.984E-06 | 6.100E-06 | 1.300E-05 |
| 15372 | 769 | Q769K | B | 1.1e-05 | 9.5e-06 | 1.4e-05 | 1.150E-05 | 2.291E-06 | 9.500E-06 | 1.400E-05 |
| 15373 | 769 | Q769M | B | 1.5e-05 | 1.5e-05 | 1.1e-05 | 1.367E-05 | 2.309E-06 | 1.100E-05 | 1.500E-05 |
| 15374 | 769 | Q769F | B | 1.1e-05 | 9e-06 | 9.6e-06 | 9.867E-06 | 1.026E-06 | 9.000E-06 | 1.100E-05 |
| 15375 | 769 | Q769P | B | 1.6e-05 | 1.6e-05 | 1.3e-05 | 1.500E-05 | 1.732E-06 | 1.300E-05 | 1.600E-05 |
| 15376 | 769 | Q769S | B | 1.5e-05 | 1.4e-05 | 9.2e-06 | 1.273E-05 | 3.101E-06 | 9.200E-06 | 1.500E-05 |
| 15377 | 769 | Q769T | B | 1e-05 | 1e-05 | 1.4e-05 | 1.133E-05 | 2.309E-06 | 1.000E-05 | 1.400E-05 |
| 15378 | 769 | Q769W | B | 1.3e-05 | 1.4e-05 | 9.3e-06 | 1.210E-05 | 2.476E-06 | 9.300E-06 | 1.400E-05 |
| 15379 | 769 | Q769Y | B | 8.3e-06 | 1.4e-05 | 2.4e-05 | 1.543E-05 | 7.948E-06 | 8.300E-06 | 2.400E-05 |
| 15380 | 769 | Q769V | B | 1.4e-05 | 1.5e-05 | 1.5e-05 | 1.467E-05 | 5.774E-07 | 1.400E-05 | 1.500E-05 |
| 15381 | 770 | W770A | B | 1e-05 | 1.5e-05 | 1.3e-05 | 1.267E-05 | 2.517E-06 | 1.000E-05 | 1.500E-05 |
| 15382 | 770 | W770R | B | 8.9e-06 | 1.5e-05 | 1.5e-05 | 1.297E-05 | 3.522E-06 | 8.900E-06 | 1.500E-05 |
| 15383 | 770 | W770N | B | 1.1e-05 | 1.4e-05 | 1.6e-05 | 1.367E-05 | 2.517E-06 | 1.100E-05 | 1.600E-05 |
| 15384 | 770 | W770D | B | 1.3e-05 | 1.4e-05 | 1.7e-05 | 1.467E-05 | 2.082E-06 | 1.300E-05 | 1.700E-05 |

| | | | | | | | | | | |
|---|---|---|---|---|---|---|---|---|---|---|
| 15385 | 770 | W770C | B | 1e-05 | 1.2e-05 | 1.5e-05 | 1.233E-05 | 2.517E-06 | 1.000E-05 | 1.500E-05 |
| 15386 | 770 | W770E | B | 1.4e-05 | 1.3e-05 | 1.2e-05 | 1.300E-05 | 1.000E-06 | 1.200E-05 | 1.400E-05 |
| 15387 | 770 | W770Q | B | 1.3e-05 | 8.8e-06 | 1.5e-05 | 1.227E-05 | 3.164E-06 | 8.800E-06 | 1.500E-05 |
| 15388 | 770 | W770G | B | 1e-05 | 1.2e-05 | 1.4e-05 | 1.200E-05 | 2.000E-06 | 1.000E-05 | 1.400E-05 |
| 15389 | 770 | W770H | B | 1.1e-05 | 1.1e-05 | 1.5e-05 | 1.233E-05 | 2.309E-06 | 1.100E-05 | 1.500E-05 |
| 15390 | 770 | W770I | B | 1.5e-05 | 1.1e-05 | 1.6e-05 | 1.400E-05 | 2.646E-06 | 1.100E-05 | 1.600E-05 |
| 15391 | 770 | W770L | B | 1.1e-05 | 1.6e-05 | 1.1e-05 | 1.267E-05 | 2.887E-06 | 1.100E-05 | 1.600E-05 |
| 15392 | 770 | W770K | B | 1.5e-05 | 1.7e-05 | 1.1e-05 | 1.433E-05 | 3.055E-06 | 1.100E-05 | 1.700E-05 |
| 15393 | 770 | W770M | B | 1.4e-05 | 9.8e-06 | 1.4e-05 | 1.260E-05 | 2.425E-06 | 9.800E-06 | 1.400E-05 |
| 15394 | 770 | W770F | B | 1.6e-05 | 1.7e-05 | 1.1e-05 | 1.467E-05 | 3.215E-06 | 1.100E-05 | 1.700E-05 |
| 15395 | 770 | W770P | B | 1.5e-05 | 1.5e-05 | 1.3e-05 | 1.433E-05 | 1.155E-06 | 1.300E-05 | 1.500E-05 |
| 15396 | 770 | W770S | B | 1.4e-05 | 9.7e-06 | 1.5e-05 | 1.290E-05 | 2.816E-06 | 9.700E-06 | 1.500E-05 |
| 15397 | 770 | W770T | B | 8.4e-06 | 1e-05 | 1.6e-05 | 1.147E-05 | 4.007E-06 | 8.400E-06 | 1.600E-05 |
| 15398 | 770 | W770W | B | 1e-05 | 1.5e-05 | 1e-05 | 1.167E-05 | 2.887E-06 | 1.000E-05 | 1.500E-05 |
| 15399 | 770 | W770Y | B | 8.5e-06 | 1.3e-05 | 9.6e-06 | 1.037E-05 | 2.346E-06 | 8.500E-06 | 1.300E-05 |
| 15400 | 770 | W770V | B | 1.4e-05 | 1.5e-05 | 1.4e-05 | 1.433E-05 | 5.774E-07 | 1.400E-05 | 1.500E-05 |
| 15401 | 771 | K771A | B | 1.1e-05 | 1.1e-05 | 1.4e-05 | 1.200E-05 | 1.732E-06 | 1.100E-05 | 1.400E-05 |
| 15402 | 771 | K771R | B | 1.4e-05 | 9e-06 | 1.3e-05 | 1.200E-05 | 2.646E-06 | 9.000E-06 | 1.400E-05 |
| 15403 | 771 | K771N | B | 1.6e-05 | 1.3e-05 | 9.4e-06 | 1.280E-05 | 3.305E-06 | 9.400E-06 | 1.600E-05 |
| 15404 | 771 | K771D | B | 1.6e-05 | 1.2e-05 | 9e-06 | 1.233E-05 | 3.512E-06 | 9.000E-06 | 1.600E-05 |
| 15405 | 771 | K771C | B | 8.2e-06 | 9.1e-06 | 1.2e-05 | 9.767E-06 | 1.986E-06 | 8.200E-06 | 1.200E-05 |
| 15406 | 771 | K771E | B | 1.5e-05 | 1.6e-05 | 1.2e-05 | 1.433E-05 | 2.082E-06 | 1.200E-05 | 1.600E-05 |
| 15407 | 771 | K771Q | B | 1.4e-05 | 1e-05 | 1e-05 | 1.133E-05 | 2.309E-06 | 1.000E-05 | 1.400E-05 |
| 15408 | 771 | K771G | B | 1.4e-05 | 1.6e-05 | 9.8e-06 | 1.327E-05 | 3.164E-06 | 9.800E-06 | 1.600E-05 |
| 15409 | 771 | K771H | B | 1.3e-05 | 1.5e-05 | 1.1e-05 | 1.300E-05 | 2.000E-06 | 1.100E-05 | 1.500E-05 |
| 15410 | 771 | K771I | B | 9.9e-06 | 1.6e-05 | 1.3e-05 | 1.297E-05 | 3.050E-06 | 9.900E-06 | 1.600E-05 |
| 15411 | 771 | K771L | B | 1e-05 | 9.5e-06 | 8.9e-06 | 9.467E-06 | 5.508E-07 | 8.900E-06 | 1.000E-05 |
| 15412 | 771 | K771K | B | 1.6e-05 | 1.3e-05 | 1.6e-05 | 1.500E-05 | 1.732E-06 | 1.300E-05 | 1.600E-05 |
| 15413 | 771 | K771M | B | 9.5e-06 | 1.5e-05 | 1.5e-05 | 1.317E-05 | 3.175E-06 | 9.500E-06 | 1.500E-05 |
| 15414 | 771 | K771F | B | 1.4e-05 | 1.5e-05 | 9e-06 | 1.267E-05 | 3.215E-06 | 9.000E-06 | 1.500E-05 |
| 15415 | 771 | K771P | B | 1.6e-05 | 1.6e-05 | 1.5e-05 | 1.567E-05 | 5.774E-07 | 1.500E-05 | 1.600E-05 |
| 15416 | 771 | K771S | B | 8e-06 | 8e-06 | 1.2e-05 | 9.333E-06 | 2.309E-06 | 8.000E-06 | 1.200E-05 |
| 15417 | 771 | K771T | B | 8.3e-06 | 1.5e-05 | 1.7e-05 | 1.343E-05 | 4.557E-06 | 8.300E-06 | 1.700E-05 |
| 15418 | 771 | K771W | B | 1.4e-05 | 1.5e-05 | 1.3e-05 | 1.400E-05 | 1.000E-06 | 1.300E-05 | 1.500E-05 |
| 15419 | 771 | K771Y | B | 1e-05 | 1e-05 | 1.2e-05 | 1.067E-05 | 1.155E-06 | 1.000E-05 | 1.200E-05 |
| 15420 | 771 | K771V | B | 1.5e-05 | 1.5e-05 | 1.5e-05 | 1.500E-05 | 0.000E+00 | 1.500E-05 | 1.500E-05 |
| 15421 | 772 | V772A | B | 1.3e-05 | 1.5e-05 | 1.5e-05 | 1.433E-05 | 1.155E-06 | 1.300E-05 | 1.500E-05 |
| 15422 | 772 | V772R | B | 8.7e-06 | 1.5e-05 | 1.5e-05 | 1.290E-05 | 3.637E-06 | 8.700E-06 | 1.500E-05 |
| 15423 | 772 | V772N | B | 1.6e-05 | 1.6e-05 | 1.4e-05 | 1.533E-05 | 1.155E-06 | 1.400E-05 | 1.600E-05 |
| 15424 | 772 | V772D | B | 1.8e-05 | 9.5e-06 | 8.6e-06 | 1.203E-05 | 5.187E-06 | 8.600E-06 | 1.800E-05 |
| 15425 | 772 | V772C | B | 1.1e-05 | 1.5e-05 | 1e-05 | 1.200E-05 | 2.646E-06 | 1.000E-05 | 1.500E-05 |
| 15426 | 772 | V772E | B | 1e-05 | 9.4e-06 | 9.3e-06 | 9.567E-06 | 3.786E-07 | 9.300E-06 | 1.000E-05 |
| 15427 | 772 | V772Q | B | 1.5e-05 | 9.8e-06 | 1.5e-05 | 1.327E-05 | 3.002E-06 | 9.800E-06 | 1.500E-05 |
| 15428 | 772 | V772G | B | 1.4e-05 | 1.7e-05 | 1.7e-05 | 1.600E-05 | 1.732E-06 | 1.400E-05 | 1.700E-05 |
| 15429 | 772 | V772H | B | 1.3e-05 | 1.6e-05 | 1e-05 | 1.300E-05 | 3.000E-06 | 1.000E-05 | 1.600E-05 |
| 15430 | 772 | V772I | B | 1.5e-05 | 1.2e-05 | 1e-05 | 1.233E-05 | 2.517E-06 | 1.000E-05 | 1.500E-05 |
| 15431 | 772 | V772L | B | 1e-05 | 9.6e-06 | 1.4e-05 | 1.120E-05 | 2.433E-06 | 9.600E-06 | 1.400E-05 |
| 15432 | 772 | V772K | B | 1.4e-05 | 1e-05 | 9e-06 | 1.100E-05 | 2.646E-06 | 9.000E-06 | 1.400E-05 |
| 15433 | 772 | V772M | B | 9.5e-06 | 1.4e-05 | 1.5e-05 | 1.283E-05 | 2.930E-06 | 9.500E-06 | 1.500E-05 |
| 15434 | 772 | V772F | B | 1.1e-05 | 9.3e-06 | 9.5e-06 | 9.933E-06 | 9.292E-07 | 9.300E-06 | 1.100E-05 |
| 15435 | 772 | V772P | B | 9e-06 | 1.1e-05 | 1.5e-05 | 1.167E-05 | 3.055E-06 | 9.000E-06 | 1.500E-05 |
| 15436 | 772 | V772S | B | 1.4e-05 | 1.1e-05 | 1.2e-05 | 1.233E-05 | 1.528E-06 | 1.100E-05 | 1.400E-05 |
| 15437 | 772 | V772T | B | 7.9e-06 | 1.4e-05 | 1e-05 | 1.063E-05 | 3.099E-06 | 7.900E-06 | 1.400E-05 |
| 15438 | 772 | V772W | B | 1.3e-05 | 1.6e-05 | 1.5e-05 | 1.467E-05 | 1.528E-06 | 1.300E-05 | 1.600E-05 |
| 15439 | 772 | V772Y | B | 1.7e-05 | 8.6e-06 | 9.5e-06 | 1.170E-05 | 4.612E-06 | 8.600E-06 | 1.700E-05 |
| 15440 | 772 | V772V | B | 1.3e-05 | 6.1e-06 | 1.6e-05 | 1.170E-05 | 5.076E-06 | 6.100E-06 | 1.600E-05 |
| 15441 | 773 | D773A | B | 1e-05 | 1.6e-05 | 1.6e-05 | 1.400E-05 | 3.464E-06 | 1.000E-05 | 1.600E-05 |
| 15442 | 773 | D773R | B | 8.4e-06 | 1.1e-05 | 1.1e-05 | 1.013E-05 | 1.501E-06 | 8.400E-06 | 1.100E-05 |
| 15443 | 773 | D773N | B | 1.5e-05 | 1.5e-05 | 1.1e-05 | 1.367E-05 | 2.309E-06 | 1.100E-05 | 1.500E-05 |
| 15444 | 773 | D773D | B | 1.1e-05 | 1e-05 | 1.1e-05 | 1.067E-05 | 5.774E-07 | 1.000E-05 | 1.100E-05 |
| 15445 | 773 | D773C | B | 1.4e-05 | 9.5e-06 | 1.2e-05 | 1.183E-05 | 2.255E-06 | 9.500E-06 | 1.400E-05 |
| 15446 | 773 | D773E | B | 1.7e-05 | 1.5e-05 | 1.4e-05 | 1.533E-05 | 1.528E-06 | 1.400E-05 | 1.700E-05 |
| 15447 | 773 | D773Q | B | 1.4e-05 | 1.1e-05 | 8.8e-06 | 1.127E-05 | 2.610E-06 | 8.800E-06 | 1.400E-05 |
| 15448 | 773 | D773G | B | 7.6e-06 | 9.6e-06 | 1.6e-05 | 1.107E-05 | 4.388E-06 | 7.600E-06 | 1.600E-05 |
| 15449 | 773 | D773H | B | 1.1e-05 | 1.5e-05 | 9.8e-06 | 1.193E-05 | 2.723E-06 | 9.800E-06 | 1.500E-05 |
| 15450 | 773 | D773I | B | 1.1e-05 | 7.5e-06 | 9.8e-06 | 9.433E-06 | 1.779E-06 | 7.500E-06 | 1.100E-05 |
| 15451 | 773 | D773L | B | 8.9e-06 | 9.6e-06 | 1.1e-05 | 9.833E-06 | 1.069E-06 | 8.900E-06 | 1.100E-05 |
| 15452 | 773 | D773K | B | 1e-05 | 1.3e-05 | 1.5e-05 | 1.267E-05 | 2.517E-06 | 1.000E-05 | 1.500E-05 |
| 15453 | 773 | D773M | B | 1.5e-05 | 1e-05 | 8.6e-06 | 1.120E-05 | 3.365E-06 | 8.600E-06 | 1.500E-05 |
| 15454 | 773 | D773F | B | 1.6e-05 | 1.3e-05 | 1.6e-05 | 1.500E-05 | 1.732E-06 | 1.300E-05 | 1.600E-05 |
| 15455 | 773 | D773P | B | 1.6e-05 | 1.2e-05 | 9.7e-06 | 1.257E-05 | 3.188E-06 | 9.700E-06 | 1.600E-05 |
| 15456 | 773 | D773S | B | 1.4e-05 | 1.1e-05 | 9.2e-06 | 1.140E-05 | 2.425E-06 | 9.200E-06 | 1.400E-05 |
| 15457 | 773 | D773T | B | 1.5e-05 | 1e-05 | 7.3e-06 | 1.077E-05 | 3.907E-06 | 7.300E-06 | 1.500E-05 |
| 15458 | 773 | D773W | B | 1.1e-05 | 1.2e-05 | 1.6e-05 | 1.300E-05 | 2.646E-06 | 1.100E-05 | 1.600E-05 |
| 15459 | 773 | D773Y | B | 8.1e-06 | 1.1e-05 | 1.4e-05 | 1.103E-05 | 2.950E-06 | 8.100E-06 | 1.400E-05 |
| 15460 | 773 | D773V | B | 1.1e-05 | 7.1e-06 | 1.1e-05 | 9.700E-06 | 2.252E-06 | 7.100E-06 | 1.100E-05 |
| 15461 | 774 | N774A | B | 9.2e-06 | 9.1e-06 | 1.2e-05 | 1.010E-05 | 1.646E-06 | 9.100E-06 | 1.200E-05 |
| 15462 | 774 | N774R | B | 1.1e-05 | 9.3e-06 | 1.5e-05 | 1.177E-05 | 2.926E-06 | 9.300E-06 | 1.500E-05 |
| 15463 | 774 | N774N | B | 1.1e-05 | 1.4e-05 | 1.3e-05 | 1.267E-05 | 1.528E-06 | 1.100E-05 | 1.400E-05 |
| 15464 | 774 | N774D | B | 1.5e-05 | 1.4e-05 | 1.6e-05 | 1.500E-05 | 1.000E-06 | 1.400E-05 | 1.600E-05 |
| 15465 | 774 | N774C | B | 1.5e-05 | 1.1e-05 | 1.6e-05 | 1.400E-05 | 2.646E-06 | 1.100E-05 | 1.600E-05 |
| 15466 | 774 | N774E | B | 1.5e-05 | 1e-05 | 1.6e-05 | 1.367E-05 | 3.215E-06 | 1.000E-05 | 1.600E-05 |
| 15467 | 774 | N774Q | B | 1e-05 | 1.1e-05 | 9.7e-06 | 1.023E-05 | 6.807E-07 | 9.700E-06 | 1.100E-05 |
| 15468 | 774 | N774G | B | 1.6e-05 | 1.4e-05 | 1.7e-05 | 1.567E-05 | 1.528E-06 | 1.400E-05 | 1.700E-05 |
| 15469 | 774 | N774H | B | 1e-05 | 9.4e-06 | 1.6e-05 | 1.180E-05 | 3.650E-06 | 9.400E-06 | 1.600E-05 |

| | | | | | | | | | | |
|---|---|---|---|---|---|---|---|---|---|---|
| 15470 | 774 | N774I | B | 1.2e-05 | 1.4e-05 | 1.5e-05 | 1.367E-05 | 1.528E-06 | 1.200E-05 | 1.500E-05 |
| 15471 | 774 | N774L | B | 1.1e-05 | 1.6e-05 | 1.3e-05 | 1.333E-05 | 2.517E-06 | 1.100E-05 | 1.600E-05 |
| 15472 | 774 | N774K | B | 1.2e-05 | 1.5e-05 | 1.2e-05 | 1.300E-05 | 1.732E-06 | 1.200E-05 | 1.500E-05 |
| 15473 | 774 | N774M | B | 1.7e-05 | 1.3e-05 | 9.7e-06 | 1.323E-05 | 3.656E-06 | 9.700E-06 | 1.700E-05 |
| 15474 | 774 | N774F | B | 1.7e-05 | 1.5e-05 | 9.1e-06 | 1.370E-05 | 4.107E-06 | 9.100E-06 | 1.700E-05 |
| 15475 | 774 | N774P | B | 1e-05 | 1.7e-05 | 1.5e-05 | 1.400E-05 | 3.606E-06 | 1.000E-05 | 1.700E-05 |
| 15476 | 774 | N774S | B | 1.1e-05 | 1.7e-05 | 1.1e-05 | 1.300E-05 | 3.464E-06 | 1.100E-05 | 1.700E-05 |
| 15477 | 774 | N774T | B | 1.3e-05 | 1.5e-05 | 1e-05 | 1.267E-05 | 2.517E-06 | 1.000E-05 | 1.500E-05 |
| 15478 | 774 | N774W | B | 1e-05 | 8.9e-06 | 1.1e-05 | 9.967E-06 | 1.050E-06 | 8.900E-06 | 1.100E-05 |
| 15479 | 774 | N774Y | B | 1.5e-05 | 8e-06 | 1.4e-05 | 1.233E-05 | 3.786E-06 | 8.000E-06 | 1.500E-05 |
| 15480 | 774 | N774V | B | 1e-05 | 6.6e-06 | 1.6e-05 | 1.087E-05 | 4.760E-06 | 6.600E-06 | 1.600E-05 |
| 15481 | 775 | A775A | B | 1.7e-05 | 1e-05 | 7.8e-06 | 1.160E-05 | 4.804E-06 | 7.800E-06 | 1.700E-05 |
| 15482 | 775 | A775R | B | 1.2e-05 | 1.5e-05 | 1.3e-05 | 1.333E-05 | 1.528E-06 | 1.200E-05 | 1.500E-05 |
| 15483 | 775 | A775N | B | 1.4e-05 | 1e-05 | 9.1e-06 | 1.103E-05 | 2.608E-06 | 9.100E-06 | 1.400E-05 |
| 15484 | 775 | A775D | B | 9.9e-06 | 1.4e-05 | 1.1e-05 | 1.163E-05 | 2.122E-06 | 9.900E-06 | 1.400E-05 |
| 15485 | 775 | A775C | B | 1.1e-05 | 1.1e-05 | 1.1e-05 | 1.100E-05 | 2.075E-21 | 1.100E-05 | 1.100E-05 |
| 15486 | 775 | A775E | B | 1.1e-05 | 9.7e-06 | 1e-05 | 1.023E-05 | 6.807E-07 | 9.700E-06 | 1.100E-05 |
| 15487 | 775 | A775Q | B | 1.3e-05 | 1.5e-05 | 1.2e-05 | 1.333E-05 | 1.528E-06 | 1.200E-05 | 1.500E-05 |
| 15488 | 775 | A775G | B | 1.2e-05 | 1.1e-05 | 1.1e-05 | 1.133E-05 | 5.774E-07 | 1.100E-05 | 1.200E-05 |
| 15489 | 775 | A775H | B | 7.8e-06 | 1.5e-05 | 1.4e-05 | 1.227E-05 | 3.900E-06 | 7.800E-06 | 1.500E-05 |
| 15490 | 775 | A775I | B | 6.7e-06 | 1.2e-05 | 9.5e-06 | 9.400E-06 | 2.651E-06 | 6.700E-06 | 1.200E-05 |
| 15491 | 775 | A775L | B | 1.1e-05 | 1.5e-05 | 1.6e-05 | 1.400E-05 | 2.646E-06 | 1.100E-05 | 1.600E-05 |
| 15492 | 775 | A775K | B | 9e-06 | 1.5e-05 | 1.5e-05 | 1.300E-05 | 3.464E-06 | 9.000E-06 | 1.500E-05 |
| 15493 | 775 | A775M | B | 1.4e-05 | 9.4e-06 | 1.3e-05 | 1.213E-05 | 2.419E-06 | 9.400E-06 | 1.400E-05 |
| 15494 | 775 | A775F | B | 1.5e-05 | 9.2e-06 | 9.8e-06 | 1.133E-05 | 3.190E-06 | 9.200E-06 | 1.500E-05 |
| 15495 | 775 | A775P | B | 1.1e-05 | 1.5e-05 | 8.7e-06 | 1.157E-05 | 3.188E-06 | 8.700E-06 | 1.500E-05 |
| 15496 | 775 | A775S | B | 1e-05 | 1.3e-05 | 1.5e-05 | 1.267E-05 | 2.517E-06 | 1.000E-05 | 1.500E-05 |
| 15497 | 775 | A775T | B | 1.5e-05 | 1.5e-05 | 1.4e-05 | 1.467E-05 | 5.774E-07 | 1.400E-05 | 1.500E-05 |
| 15498 | 775 | A775W | B | 1.6e-05 | 1.4e-05 | 9.8e-06 | 1.327E-05 | 3.164E-06 | 9.800E-06 | 1.600E-05 |
| 15499 | 775 | A775Y | B | 1.4e-05 | 1e-05 | 1.5e-05 | 1.300E-05 | 2.646E-06 | 1.000E-05 | 1.500E-05 |
| 15500 | 775 | A775V | B | 2.2e-05 | 1.6e-05 | 8.1e-06 | 1.537E-05 | 6.972E-06 | 8.100E-06 | 2.200E-05 |
| 15501 | 776 | L776A | B | 1.4e-05 | 1.2e-05 | 1.2e-05 | 1.267E-05 | 1.155E-06 | 1.200E-05 | 1.400E-05 |
| 15502 | 776 | L776R | B | 1.5e-05 | 1.5e-05 | 1e-05 | 1.333E-05 | 2.887E-06 | 1.000E-05 | 1.500E-05 |
| 15503 | 776 | L776N | B | 1.5e-05 | 8e-06 | 9.8e-06 | 1.093E-05 | 3.635E-06 | 8.000E-06 | 1.500E-05 |
| 15504 | 776 | L776D | B | 8.3e-06 | 1.7e-05 | 1.1e-05 | 1.210E-05 | 4.453E-06 | 8.300E-06 | 1.700E-05 |
| 15505 | 776 | L776C | B | 1.5e-05 | 9.2e-06 | 1.3e-05 | 1.240E-05 | 2.946E-06 | 9.200E-06 | 1.500E-05 |
| 15506 | 776 | L776E | B | 1.4e-05 | 1.4e-05 | 9e-06 | 1.233E-05 | 2.887E-06 | 9.000E-06 | 1.400E-05 |
| 15507 | 776 | L776Q | B | 9.1e-06 | 1.3e-05 | 1.5e-05 | 1.237E-05 | 3.001E-06 | 9.100E-06 | 1.500E-05 |
| 15508 | 776 | L776G | B | 1e-05 | 1.7e-05 | 1.4e-05 | 1.367E-05 | 3.512E-06 | 1.000E-05 | 1.700E-05 |
| 15509 | 776 | L776H | B | 1.1e-05 | 1.4e-05 | 1.4e-05 | 1.300E-05 | 1.732E-06 | 1.100E-05 | 1.400E-05 |
| 15510 | 776 | L776I | B | 1.5e-05 | 1.1e-05 | 9.2e-06 | 1.173E-05 | 2.969E-06 | 9.200E-06 | 1.500E-05 |
| 15511 | 776 | L776L | B | 1.5e-05 | 1.5e-05 | 1.1e-05 | 1.367E-05 | 2.309E-06 | 1.100E-05 | 1.500E-05 |
| 15512 | 776 | L776K | B | 9.6e-06 | 1.4e-05 | 1.4e-05 | 1.253E-05 | 2.540E-06 | 9.600E-06 | 1.400E-05 |
| 15513 | 776 | L776M | B | 1.3e-05 | 1.3e-05 | 1.5e-05 | 1.367E-05 | 1.155E-06 | 1.300E-05 | 1.500E-05 |
| 15514 | 776 | L776F | B | 1.1e-05 | 1.5e-05 | 1.1e-05 | 1.233E-05 | 2.309E-06 | 1.100E-05 | 1.500E-05 |
| 15515 | 776 | L776P | B | 1.1e-05 | 1.5e-05 | 1.6e-05 | 1.400E-05 | 2.646E-06 | 1.100E-05 | 1.600E-05 |
| 15516 | 776 | L776S | B | 1e-05 | 1.2e-05 | 1.1e-05 | 1.100E-05 | 1.000E-06 | 1.000E-05 | 1.200E-05 |
| 15517 | 776 | L776T | B | 9.5e-06 | 1.4e-05 | 8.6e-06 | 1.070E-05 | 2.893E-06 | 8.600E-06 | 1.400E-05 |
| 15518 | 776 | L776W | B | 1.4e-05 | 1.5e-05 | 1.1e-05 | 1.333E-05 | 2.082E-06 | 1.100E-05 | 1.500E-05 |
| 15519 | 776 | L776Y | B | 1.3e-05 | 8.5e-06 | 9.6e-06 | 1.037E-05 | 2.346E-06 | 8.500E-06 | 1.300E-05 |
| 15520 | 776 | L776V | B | 9.6e-06 | 1.6e-05 | 1e-05 | 1.187E-05 | 3.585E-06 | 9.600E-06 | 1.600E-05 |
| 15521 | 777 | Q777A | B | 1.1e-05 | 9.1e-06 | 1.1e-05 | 1.037E-05 | 1.097E-06 | 9.100E-06 | 1.100E-05 |
| 15522 | 777 | Q777R | B | 1.4e-05 | 1.1e-05 | 1.5e-05 | 1.333E-05 | 2.082E-06 | 1.100E-05 | 1.500E-05 |
| 15523 | 777 | Q777N | B | 1.4e-05 | 1.3e-05 | 1e-05 | 1.233E-05 | 2.082E-06 | 1.000E-05 | 1.400E-05 |
| 15524 | 777 | Q777D | B | 1.5e-05 | 1.2e-05 | 1e-05 | 1.233E-05 | 2.517E-06 | 1.000E-05 | 1.500E-05 |
| 15525 | 777 | Q777C | B | 1.7e-05 | 1.1e-05 | 1.5e-05 | 1.433E-05 | 3.055E-06 | 1.100E-05 | 1.700E-05 |
| 15526 | 777 | Q777E | B | 1e-05 | 1.5e-05 | 1.5e-05 | 1.333E-05 | 2.887E-06 | 1.000E-05 | 1.500E-05 |
| 15527 | 777 | Q777Q | B | 1.1e-05 | 1.1e-05 | 1.4e-05 | 1.200E-05 | 1.732E-06 | 1.100E-05 | 1.400E-05 |
| 15528 | 777 | Q777G | B | 1.5e-05 | 1.1e-05 | 1.8e-05 | 1.467E-05 | 3.512E-06 | 1.100E-05 | 1.800E-05 |
| 15529 | 777 | Q777H | B | 1.2e-05 | 1.6e-05 | 1.3e-05 | 1.367E-05 | 2.082E-06 | 1.200E-05 | 1.600E-05 |
| 15530 | 777 | Q777I | B | 1.4e-05 | 1.5e-05 | 1e-05 | 1.300E-05 | 2.646E-06 | 1.000E-05 | 1.500E-05 |
| 15531 | 777 | Q777L | B | 1.7e-05 | 1e-05 | 1.2e-05 | 1.300E-05 | 3.606E-06 | 1.000E-05 | 1.700E-05 |
| 15532 | 777 | Q777K | B | 8.8e-06 | 1.5e-05 | 1.2e-05 | 1.193E-05 | 3.101E-06 | 8.800E-06 | 1.500E-05 |
| 15533 | 777 | Q777M | B | 9e-06 | 1.4e-05 | 1.1e-05 | 1.133E-05 | 2.517E-06 | 9.000E-06 | 1.400E-05 |
| 15534 | 777 | Q777F | B | 1.2e-05 | 1.5e-05 | 7.3e-06 | 1.143E-05 | 3.881E-06 | 7.300E-06 | 1.500E-05 |
| 15535 | 777 | Q777P | B | 1.6e-05 | 1.7e-05 | 1.4e-05 | 1.567E-05 | 1.528E-06 | 1.400E-05 | 1.700E-05 |
| 15536 | 777 | Q777S | B | 1.5e-05 | 1.3e-05 | 1e-05 | 1.267E-05 | 2.517E-06 | 1.000E-05 | 1.500E-05 |
| 15537 | 777 | Q777T | B | 1.1e-05 | 1.5e-05 | 1.4e-05 | 1.333E-05 | 2.082E-06 | 1.100E-05 | 1.500E-05 |
| 15538 | 777 | Q777W | B | 1.4e-05 | 9.8e-06 | 1e-05 | 1.127E-05 | 2.369E-06 | 9.800E-06 | 1.400E-05 |
| 15539 | 777 | Q777Y | B | 1e-05 | 7.8e-06 | 9.2e-06 | 9.000E-06 | 1.114E-06 | 7.800E-06 | 1.000E-05 |
| 15540 | 777 | Q777V | B | 1.6e-05 | 1.6e-05 | 1.5e-05 | 1.567E-05 | 5.774E-07 | 1.500E-05 | 1.600E-05 |
| 15541 | 778 | S778A | B | 1.5e-05 | 1.7e-05 | 1.1e-05 | 1.433E-05 | 3.055E-06 | 1.100E-05 | 1.700E-05 |
| 15542 | 778 | S778R | B | 1.5e-05 | 1.6e-05 | 1.5e-05 | 1.533E-05 | 5.774E-07 | 1.500E-05 | 1.600E-05 |
| 15543 | 778 | S778N | B | 1.6e-05 | 1.4e-05 | 1.1e-05 | 1.367E-05 | 2.517E-06 | 1.100E-05 | 1.600E-05 |
| 15544 | 778 | S778D | B | 9.5e-06 | 6.9e-06 | 1.7e-05 | 1.113E-05 | 5.244E-06 | 6.900E-06 | 1.700E-05 |
| 15545 | 778 | S778C | B | 8.9e-06 | 8.7e-06 | 1.1e-05 | 9.533E-06 | 1.274E-06 | 8.700E-06 | 1.100E-05 |
| 15546 | 778 | S778E | B | 1.2e-05 | 1.2e-05 | 1.2e-05 | 1.200E-05 | 0.000E+00 | 1.200E-05 | 1.200E-05 |
| 15547 | 778 | S778Q | B | 6.4e-06 | 1.4e-05 | 1.5e-05 | 1.180E-05 | 4.703E-06 | 6.400E-06 | 1.500E-05 |
| 15548 | 778 | S778G | B | 1.3e-05 | 9.1e-06 | 1.1e-05 | 1.103E-05 | 1.950E-06 | 9.100E-06 | 1.300E-05 |
| 15549 | 778 | S778H | B | 1.5e-05 | 8.7e-06 | 1.6e-05 | 1.323E-05 | 3.958E-06 | 8.700E-06 | 1.600E-05 |
| 15550 | 778 | S778I | B | 1.3e-05 | 1.4e-05 | 1.6e-05 | 1.433E-05 | 1.528E-06 | 1.300E-05 | 1.600E-05 |
| 15551 | 778 | S778L | B | 1.7e-05 | 8.2e-06 | 1e-05 | 1.173E-05 | 4.649E-06 | 8.200E-06 | 1.700E-05 |
| 15552 | 778 | S778K | B | 1.6e-05 | 1.4e-05 | 1.5e-05 | 1.500E-05 | 1.000E-06 | 1.400E-05 | 1.600E-05 |
| 15553 | 778 | S778M | B | 1.1e-05 | 1.2e-05 | 1.5e-05 | 1.267E-05 | 2.082E-06 | 1.100E-05 | 1.500E-05 |
| 15554 | 778 | S778F | B | 1e-05 | 1.5e-05 | 1.7e-05 | 1.400E-05 | 3.606E-06 | 1.000E-05 | 1.700E-05 |

| | | | | | | | | | | |
|---|---|---|---|---|---|---|---|---|---|---|
| 15555 | 778 | S778P | B | 8.4e-06 | 1.7e-05 | 9.4e-06 | 1.160E-05 | 4.703E-06 | 8.400E-06 | 1.700E-05 |
| 15556 | 778 | S778S | B | 1.4e-05 | 1.1e-05 | 1.2e-05 | 1.233E-05 | 1.528E-06 | 1.100E-05 | 1.400E-05 |
| 15557 | 778 | S778T | B | 1.2e-05 | 1.4e-05 | 1.4e-05 | 1.333E-05 | 1.155E-06 | 1.200E-05 | 1.400E-05 |
| 15558 | 778 | S778W | B | 1e-05 | 1.6e-05 | 1.7e-05 | 1.433E-05 | 3.786E-06 | 1.000E-05 | 1.700E-05 |
| 15559 | 778 | S778Y | B | 1e-05 | 1.2e-05 | 1.2e-05 | 1.133E-05 | 1.155E-06 | 1.000E-05 | 1.200E-05 |
| 15560 | 778 | S778V | B | 1.7e-05 | 1e-05 | 1e-05 | 1.233E-05 | 4.041E-06 | 1.000E-05 | 1.700E-05 |
| 15561 | 779 | G779A | B | 1.5e-05 | 1.2e-05 | 1.1e-05 | 1.267E-05 | 2.082E-06 | 1.100E-05 | 1.500E-05 |
| 15562 | 779 | G779R | B | 1.8e-05 | 1.4e-05 | 1.1e-05 | 1.433E-05 | 3.512E-06 | 1.100E-05 | 1.800E-05 |
| 15563 | 779 | G779N | B | 1e-05 | 1.5e-05 | 9.6e-06 | 1.153E-05 | 3.009E-06 | 9.600E-06 | 1.500E-05 |
| 15564 | 779 | G779D | B | 1.5e-05 | 9.7e-06 | 1e-05 | 1.157E-05 | 2.977E-06 | 9.700E-06 | 1.500E-05 |
| 15565 | 779 | G779C | B | 1.1e-05 | 9.7e-06 | 1.4e-05 | 1.157E-05 | 2.205E-06 | 9.700E-06 | 1.400E-05 |
| 15566 | 779 | G779E | B | 1.3e-05 | 9.7e-06 | 7.8e-06 | 1.017E-05 | 2.631E-06 | 7.800E-06 | 1.300E-05 |
| 15567 | 779 | G779Q | B | 1.3e-05 | 1.4e-05 | 6.6e-06 | 1.120E-05 | 4.015E-06 | 6.600E-06 | 1.400E-05 |
| 15568 | 779 | G779G | B | 1e-05 | 1.6e-05 | 1.6e-05 | 1.400E-05 | 3.464E-06 | 1.000E-05 | 1.600E-05 |
| 15569 | 779 | G779H | B | 8.9e-06 | 9.6e-06 | 1e-05 | 9.500E-06 | 5.568E-07 | 8.900E-06 | 1.000E-05 |
| 15570 | 779 | G779I | B | 1.6e-05 | 1.1e-05 | 1.2e-05 | 1.300E-05 | 2.646E-06 | 1.100E-05 | 1.600E-05 |
| 15571 | 779 | G779L | B | 1.2e-05 | 1.4e-05 | 7.4e-06 | 1.113E-05 | 3.384E-06 | 7.400E-06 | 1.400E-05 |
| 15572 | 779 | G779K | B | 1.6e-05 | 8.7e-06 | 1e-05 | 1.157E-05 | 3.894E-06 | 8.700E-06 | 1.600E-05 |
| 15573 | 779 | G779M | B | 1.7e-05 | 1.4e-05 | 1.6e-05 | 1.567E-05 | 1.528E-06 | 1.400E-05 | 1.700E-05 |
| 15574 | 779 | G779F | B | 1.3e-05 | 1.5e-05 | 1.1e-05 | 1.300E-05 | 2.000E-06 | 1.100E-05 | 1.500E-05 |
| 15575 | 779 | G779P | B | 1.5e-05 | 9.8e-06 | 1.6e-05 | 1.360E-05 | 3.329E-06 | 9.800E-06 | 1.600E-05 |
| 15576 | 779 | G779S | B | 1.5e-05 | 1.5e-05 | 1.2e-05 | 1.400E-05 | 1.732E-06 | 1.200E-05 | 1.500E-05 |
| 15577 | 779 | G779T | B | 1.3e-05 | 1e-05 | 1.3e-05 | 1.200E-05 | 1.732E-06 | 1.000E-05 | 1.300E-05 |
| 15578 | 779 | G779W | B | 9.7e-06 | 1.3e-05 | 1.4e-05 | 1.223E-05 | 2.250E-06 | 9.700E-06 | 1.400E-05 |
| 15579 | 779 | G779Y | B | 1.2e-05 | 9.9e-06 | 1.4e-05 | 1.197E-05 | 2.050E-06 | 9.900E-06 | 1.400E-05 |
| 15580 | 779 | G779V | B | 1.5e-05 | 9e-06 | 9.4e-06 | 1.113E-05 | 3.355E-06 | 9.000E-06 | 1.500E-05 |
| 15581 | 780 | N780A | B | 9.3e-06 | 1.4e-05 | 1.5e-05 | 1.277E-05 | 3.044E-06 | 9.300E-06 | 1.500E-05 |
| 15582 | 780 | N780R | B | 1e-05 | 1.1e-05 | 9.9e-06 | 1.030E-05 | 6.083E-07 | 9.900E-06 | 1.100E-05 |
| 15583 | 780 | N780N | B | 1.2e-05 | 1.1e-05 | 1.5e-05 | 1.267E-05 | 2.082E-06 | 1.100E-05 | 1.500E-05 |
| 15584 | 780 | N780D | B | 1.6e-05 | 1.4e-05 | 7e-06 | 1.233E-05 | 4.726E-06 | 7.000E-06 | 1.600E-05 |
| 15585 | 780 | N780C | B | 9.7e-06 | 1.3e-05 | 1.4e-05 | 1.223E-05 | 2.250E-06 | 9.700E-06 | 1.400E-05 |
| 15586 | 780 | N780E | B | 1.5e-05 | 1.7e-05 | 1.6e-05 | 1.600E-05 | 1.000E-06 | 1.500E-05 | 1.700E-05 |
| 15587 | 780 | N780Q | B | 1.1e-05 | 9.6e-06 | 1.1e-05 | 1.053E-05 | 8.083E-07 | 9.600E-06 | 1.100E-05 |
| 15588 | 780 | N780G | B | 1.2e-05 | 1.4e-05 | 1.6e-05 | 1.400E-05 | 2.000E-06 | 1.200E-05 | 1.600E-05 |
| 15589 | 780 | N780H | B | 1.7e-05 | 1e-05 | 1.3e-05 | 1.333E-05 | 3.512E-06 | 1.000E-05 | 1.700E-05 |
| 15590 | 780 | N780I | B | 1.1e-05 | 1.6e-05 | 1.4e-05 | 1.367E-05 | 2.517E-06 | 1.100E-05 | 1.600E-05 |
| 15591 | 780 | N780L | B | 1.5e-05 | 1.7e-05 | 1e-05 | 1.400E-05 | 3.606E-06 | 1.000E-05 | 1.700E-05 |
| 15592 | 780 | N780K | B | 1.2e-05 | 1.4e-05 | 9.3e-06 | 1.177E-05 | 2.359E-06 | 9.300E-06 | 1.400E-05 |
| 15593 | 780 | N780M | B | 1.3e-05 | 1.7e-05 | 1.8e-05 | 1.600E-05 | 2.646E-06 | 1.300E-05 | 1.800E-05 |
| 15594 | 780 | N780F | B | 1.7e-05 | 1.5e-05 | 1.3e-05 | 1.500E-05 | 2.000E-06 | 1.300E-05 | 1.700E-05 |
| 15595 | 780 | N780P | B | 1.2e-05 | 8.8e-06 | 1.5e-05 | 1.193E-05 | 3.101E-06 | 8.800E-06 | 1.500E-05 |
| 15596 | 780 | N780S | B | 1.4e-05 | 1.5e-05 | 1.5e-05 | 1.467E-05 | 5.774E-07 | 1.400E-05 | 1.500E-05 |
| 15597 | 780 | N780T | B | 1.2e-05 | 1.2e-05 | 1.1e-05 | 1.167E-05 | 5.774E-07 | 1.100E-05 | 1.200E-05 |
| 15598 | 780 | N780W | B | 1.5e-05 | 1.5e-05 | 1.3e-05 | 1.433E-05 | 1.155E-06 | 1.300E-05 | 1.500E-05 |
| 15599 | 780 | N780Y | B | 1.6e-05 | 1.6e-05 | 1.1e-05 | 1.433E-05 | 2.887E-06 | 1.100E-05 | 1.600E-05 |
| 15600 | 780 | N780V | B | 1e-05 | 1.8e-05 | 1.2e-05 | 1.333E-05 | 4.163E-06 | 1.000E-05 | 1.800E-05 |
| 15601 | 781 | S781A | B | 1.5e-05 | 1.6e-05 | 1.1e-05 | 1.400E-05 | 2.646E-06 | 1.100E-05 | 1.600E-05 |
| 15602 | 781 | S781R | B | 1.6e-05 | 9.4e-06 | 1.2e-05 | 1.247E-05 | 3.325E-06 | 9.400E-06 | 1.600E-05 |
| 15603 | 781 | S781N | B | 1.4e-05 | 1.4e-05 | 1.6e-05 | 1.467E-05 | 1.155E-06 | 1.400E-05 | 1.600E-05 |
| 15604 | 781 | S781D | B | 1.2e-05 | 1.7e-05 | 1.1e-05 | 1.333E-05 | 3.215E-06 | 1.100E-05 | 1.700E-05 |
| 15605 | 781 | S781C | B | 1.4e-05 | 1.2e-05 | 1.4e-05 | 1.333E-05 | 1.155E-06 | 1.200E-05 | 1.400E-05 |
| 15606 | 781 | S781E | B | 1.3e-05 | 1.6e-05 | 1.1e-05 | 1.333E-05 | 2.517E-06 | 1.100E-05 | 1.600E-05 |
| 15607 | 781 | S781Q | B | 1.4e-05 | 1.6e-05 | 7.2e-06 | 1.240E-05 | 4.613E-06 | 7.200E-06 | 1.600E-05 |
| 15608 | 781 | S781G | B | 1.5e-05 | 1.5e-05 | 1.3e-05 | 1.433E-05 | 1.155E-06 | 1.300E-05 | 1.500E-05 |
| 15609 | 781 | S781H | B | 1.4e-05 | 1.6e-05 | 1.5e-05 | 1.500E-05 | 1.000E-06 | 1.400E-05 | 1.600E-05 |
| 15610 | 781 | S781I | B | 1.6e-05 | 1.5e-05 | 1.4e-05 | 1.500E-05 | 1.000E-06 | 1.400E-05 | 1.600E-05 |
| 15611 | 781 | S781L | B | 1.3e-05 | 1.6e-05 | 9.6e-06 | 1.287E-05 | 3.202E-06 | 9.600E-06 | 1.600E-05 |
| 15612 | 781 | S781K | B | 1e-05 | 1.5e-05 | 1e-05 | 1.167E-05 | 2.887E-06 | 1.000E-05 | 1.500E-05 |
| 15613 | 781 | S781M | B | 1.4e-05 | 1.7e-05 | 1e-05 | 1.367E-05 | 3.512E-06 | 1.000E-05 | 1.700E-05 |
| 15614 | 781 | S781F | B | 1.7e-05 | 9.5e-06 | 1e-05 | 1.217E-05 | 4.193E-06 | 9.500E-06 | 1.700E-05 |
| 15615 | 781 | S781P | B | 1.7e-05 | 1.1e-05 | 1e-05 | 1.267E-05 | 3.786E-06 | 1.000E-05 | 1.700E-05 |
| 15616 | 781 | S781S | B | 1.3e-05 | 8.3e-06 | 9.2e-06 | 1.017E-05 | 2.495E-06 | 8.300E-06 | 1.300E-05 |
| 15617 | 781 | S781T | B | 2e-05 | 1.4e-05 | 8.2e-06 | 1.407E-05 | 5.900E-06 | 8.200E-06 | 2.000E-05 |
| 15618 | 781 | S781W | B | 8.9e-06 | 1.6e-05 | 1.2e-05 | 1.230E-05 | 3.559E-06 | 8.900E-06 | 1.600E-05 |
| 15619 | 781 | S781Y | B | 1.1e-05 | 1.7e-05 | 1e-05 | 1.267E-05 | 3.786E-06 | 1.000E-05 | 1.700E-05 |
| 15620 | 781 | S781V | B | 1.1e-05 | 1.1e-05 | 1.4e-05 | 1.200E-05 | 1.732E-06 | 1.100E-05 | 1.400E-05 |
| 15621 | 782 | Q782A | B | 1.4e-05 | 1.5e-05 | 1.4e-05 | 1.433E-05 | 5.774E-07 | 1.400E-05 | 1.500E-05 |
| 15622 | 782 | Q782R | B | 1.5e-05 | 1.6e-05 | 1.1e-05 | 1.400E-05 | 2.646E-06 | 1.100E-05 | 1.600E-05 |
| 15623 | 782 | Q782N | B | 1e-05 | 9.5e-06 | 1.5e-05 | 1.150E-05 | 3.041E-06 | 9.500E-06 | 1.500E-05 |
| 15624 | 782 | Q782D | B | 1.3e-05 | 1.3e-05 | 1.4e-05 | 1.333E-05 | 5.774E-07 | 1.300E-05 | 1.400E-05 |
| 15625 | 782 | Q782C | B | 1.3e-05 | 1.1e-05 | 1.6e-05 | 1.333E-05 | 2.517E-06 | 1.100E-05 | 1.600E-05 |
| 15626 | 782 | Q782E | B | 1.1e-05 | 1e-05 | 1.4e-05 | 1.167E-05 | 2.082E-06 | 1.000E-05 | 1.400E-05 |
| 15627 | 782 | Q782Q | B | 9e-06 | 1.5e-05 | 1.1e-05 | 1.167E-05 | 3.055E-06 | 9.000E-06 | 1.500E-05 |
| 15628 | 782 | Q782G | B | 9.9e-06 | 1.1e-05 | 1.5e-05 | 1.197E-05 | 2.684E-06 | 9.900E-06 | 1.500E-05 |
| 15629 | 782 | Q782H | B | 1.1e-05 | 1e-05 | 1.6e-05 | 1.233E-05 | 3.215E-06 | 1.000E-05 | 1.600E-05 |
| 15630 | 782 | Q782I | B | 1.4e-05 | 1.3e-05 | 1.6e-05 | 1.433E-05 | 1.528E-06 | 1.300E-05 | 1.600E-05 |
| 15631 | 782 | Q782L | B | 1.1e-05 | 1.6e-05 | 1.1e-05 | 1.267E-05 | 2.887E-06 | 1.100E-05 | 1.600E-05 |
| 15632 | 782 | Q782K | B | 1.1e-05 | 9e-06 | 1e-05 | 1.000E-05 | 1.000E-06 | 9.000E-06 | 1.100E-05 |
| 15633 | 782 | Q782M | B | 8.8e-06 | 8.9e-06 | 1.4e-05 | 1.057E-05 | 2.974E-06 | 8.800E-06 | 1.400E-05 |
| 15634 | 782 | Q782F | B | 1.5e-05 | 1.4e-05 | 1e-05 | 1.300E-05 | 2.646E-06 | 1.000E-05 | 1.500E-05 |
| 15635 | 782 | Q782P | B | 1.3e-05 | 1.5e-05 | 1.5e-05 | 1.433E-05 | 1.155E-06 | 1.300E-05 | 1.500E-05 |
| 15636 | 782 | Q782S | B | 1.2e-05 | 1.4e-05 | 1.5e-05 | 1.367E-05 | 1.528E-06 | 1.200E-05 | 1.500E-05 |
| 15637 | 782 | Q782T | B | 9.6e-06 | 1.4e-05 | 1.6e-05 | 1.320E-05 | 3.274E-06 | 9.600E-06 | 1.600E-05 |
| 15638 | 782 | Q782W | B | 1.6e-05 | 1.6e-05 | 1.5e-05 | 1.567E-05 | 5.774E-07 | 1.500E-05 | 1.600E-05 |
| 15639 | 782 | Q782Y | B | 1.6e-05 | 1.5e-05 | 1.5e-05 | 1.533E-05 | 5.774E-07 | 1.500E-05 | 1.600E-05 |

| | | | | | | | | | | |
|---|---|---|---|---|---|---|---|---|---|---|
| 15640 | 782 | Q782V | B | 9.9e-06 | 1.4e-05 | 1.6e-05 | 1.330E-05 | 3.110E-06 | 9.900E-06 | 1.600E-05 |
| 15641 | 783 | E783A | B | 1.7e-05 | 1.1e-05 | 1.5e-05 | 1.433E-05 | 3.055E-06 | 1.100E-05 | 1.700E-05 |
| 15642 | 783 | E783R | B | 8.2e-06 | 1e-05 | 1.4e-05 | 1.073E-05 | 2.969E-06 | 8.200E-06 | 1.400E-05 |
| 15643 | 783 | E783N | B | 1.5e-05 | 1.4e-05 | 8.1e-06 | 1.237E-05 | 3.729E-06 | 8.100E-06 | 1.500E-05 |
| 15644 | 783 | E783D | B | 1.1e-05 | 1.5e-05 | 1.1e-05 | 1.233E-05 | 2.309E-06 | 1.100E-05 | 1.500E-05 |
| 15645 | 783 | E783C | B | 1.5e-05 | 7.1e-06 | 1.1e-05 | 1.103E-05 | 3.950E-06 | 7.100E-06 | 1.500E-05 |
| 15646 | 783 | E783E | B | 1.3e-05 | 1.1e-05 | 1.5e-05 | 1.300E-05 | 2.000E-06 | 1.100E-05 | 1.500E-05 |
| 15647 | 783 | E783Q | B | 1.5e-05 | 6.8e-06 | 1.3e-05 | 1.160E-05 | 4.276E-06 | 6.800E-06 | 1.500E-05 |
| 15648 | 783 | E783G | B | 1.4e-05 | 1.2e-05 | 1.6e-05 | 1.400E-05 | 2.000E-06 | 1.200E-05 | 1.600E-05 |
| 15649 | 783 | E783H | B | 1.5e-05 | 1.5e-05 | 1e-05 | 1.333E-05 | 2.887E-06 | 1.000E-05 | 1.500E-05 |
| 15650 | 783 | E783I | B | 8.9e-06 | 9.9e-06 | 1.1e-05 | 9.933E-06 | 1.050E-06 | 8.900E-06 | 1.100E-05 |
| 15651 | 783 | E783L | B | 8.9e-06 | 1.6e-05 | 1.6e-05 | 1.363E-05 | 4.099E-06 | 8.900E-06 | 1.600E-05 |
| 15652 | 783 | E783K | B | 1.1e-05 | 1.3e-05 | 1.5e-05 | 1.300E-05 | 2.000E-06 | 1.100E-05 | 1.500E-05 |
| 15653 | 783 | E783M | B | 1e-05 | 9e-06 | 1.5e-05 | 1.133E-05 | 3.215E-06 | 9.000E-06 | 1.500E-05 |
| 15654 | 783 | E783F | B | 9.4e-06 | 7.6e-06 | 9.7e-06 | 8.900E-06 | 1.136E-06 | 7.600E-06 | 9.700E-06 |
| 15655 | 783 | E783P | B | 9.4e-06 | 1.5e-05 | 1.1e-05 | 1.180E-05 | 2.884E-06 | 9.400E-06 | 1.500E-05 |
| 15656 | 783 | E783S | B | 1.3e-05 | 1e-05 | 1e-05 | 1.100E-05 | 1.732E-06 | 1.000E-05 | 1.300E-05 |
| 15657 | 783 | E783T | B | 1.2e-05 | 8.4e-06 | 1.6e-05 | 1.213E-05 | 3.802E-06 | 8.400E-06 | 1.600E-05 |
| 15658 | 783 | E783W | B | 1.4e-05 | 1.4e-05 | 1.3e-05 | 1.367E-05 | 5.774E-07 | 1.300E-05 | 1.400E-05 |
| 15659 | 783 | E783Y | B | 9.6e-06 | 1.2e-05 | 1.5e-05 | 1.220E-05 | 2.706E-06 | 9.600E-06 | 1.500E-05 |
| 15660 | 783 | E783V | B | 9.9e-06 | 9.1e-06 | 1e-05 | 9.667E-06 | 4.933E-07 | 9.100E-06 | 1.000E-05 |
| 15661 | 784 | S784A | B | 1.1e-05 | 1.1e-05 | 1.1e-05 | 1.100E-05 | 2.075E-21 | 1.100E-05 | 1.100E-05 |
| 15662 | 784 | S784R | B | 1.4e-05 | 1.6e-05 | 9.5e-06 | 1.317E-05 | 3.329E-06 | 9.500E-06 | 1.600E-05 |
| 15663 | 784 | S784N | B | 1.5e-05 | 1.5e-05 | 8.2e-06 | 1.273E-05 | 3.926E-06 | 8.200E-06 | 1.500E-05 |
| 15664 | 784 | S784D | B | 9.4e-06 | 1.4e-05 | 1.6e-05 | 1.313E-05 | 3.384E-06 | 9.400E-06 | 1.600E-05 |
| 15665 | 784 | S784C | B | 1.4e-05 | 1.4e-05 | 1.4e-05 | 1.400E-05 | 0.000E+00 | 1.400E-05 | 1.400E-05 |
| 15666 | 784 | S784E | B | 1.5e-05 | 1.1e-05 | 1.6e-05 | 1.400E-05 | 2.646E-06 | 1.100E-05 | 1.600E-05 |
| 15667 | 784 | S784Q | B | 1.7e-05 | 1e-05 | 1.1e-05 | 1.267E-05 | 3.786E-06 | 1.000E-05 | 1.700E-05 |
| 15668 | 784 | S784G | B | 9.8e-06 | 1e-05 | 1.1e-05 | 1.027E-05 | 6.429E-07 | 9.800E-06 | 1.100E-05 |
| 15669 | 784 | S784H | B | 9.7e-06 | 1.3e-05 | 1.5e-05 | 1.257E-05 | 2.676E-06 | 9.700E-06 | 1.500E-05 |
| 15670 | 784 | S784I | B | 1.5e-05 | 1.6e-05 | 1e-05 | 1.367E-05 | 3.215E-06 | 1.000E-05 | 1.600E-05 |
| 15671 | 784 | S784L | B | 1e-05 | 1.6e-05 | 1.2e-05 | 1.267E-05 | 3.055E-06 | 1.000E-05 | 1.600E-05 |
| 15672 | 784 | S784K | B | 1.1e-05 | 1.1e-05 | 1.1e-05 | 1.100E-05 | 2.075E-21 | 1.100E-05 | 1.100E-05 |
| 15673 | 784 | S784M | B | 1.4e-05 | 1.4e-05 | 1.6e-05 | 1.467E-05 | 1.155E-06 | 1.400E-05 | 1.600E-05 |
| 15674 | 784 | S784F | B | 9e-06 | 7.9e-06 | 1.6e-05 | 1.097E-05 | 4.394E-06 | 7.900E-06 | 1.600E-05 |
| 15675 | 784 | S784P | B | 1.5e-05 | 1.6e-05 | 1.5e-05 | 1.533E-05 | 5.774E-07 | 1.500E-05 | 1.600E-05 |
| 15676 | 784 | S784S | B | 1.7e-05 | 1.6e-05 | 1.2e-05 | 1.500E-05 | 2.646E-06 | 1.200E-05 | 1.700E-05 |
| 15677 | 784 | S784T | B | 1.2e-05 | 1.7e-05 | 1e-05 | 1.300E-05 | 3.606E-06 | 1.000E-05 | 1.700E-05 |
| 15678 | 784 | S784W | B | 1.6e-05 | 7.3e-06 | 1.5e-05 | 1.277E-05 | 4.761E-06 | 7.300E-06 | 1.600E-05 |
| 15679 | 784 | S784Y | B | 1.3e-05 | 1e-05 | 1.1e-05 | 1.133E-05 | 1.528E-06 | 1.000E-05 | 1.300E-05 |
| 15680 | 784 | S784V | B | 1e-05 | 1.5e-05 | 1.4e-05 | 1.300E-05 | 2.646E-06 | 1.000E-05 | 1.500E-05 |
| 15681 | 785 | V785A | B | 1.3e-05 | 1.3e-05 | 1e-05 | 1.200E-05 | 1.732E-06 | 1.000E-05 | 1.300E-05 |
| 15682 | 785 | V785R | B | 1.1e-05 | 1.4e-05 | 1.1e-05 | 1.200E-05 | 1.732E-06 | 1.100E-05 | 1.400E-05 |
| 15683 | 785 | V785N | B | 9.7e-06 | 8.8e-06 | 1.4e-05 | 1.083E-05 | 2.779E-06 | 8.800E-06 | 1.400E-05 |
| 15684 | 785 | V785D | B | 9.8e-06 | 1.7e-05 | 1.5e-05 | 1.393E-05 | 3.717E-06 | 9.800E-06 | 1.700E-05 |
| 15685 | 785 | V785C | B | 8.9e-06 | 1.3e-05 | 6.9e-06 | 9.600E-06 | 3.110E-06 | 6.900E-06 | 1.300E-05 |
| 15686 | 785 | V785E | B | 1.3e-05 | 1e-05 | 1.2e-05 | 1.167E-05 | 1.528E-06 | 1.000E-05 | 1.300E-05 |
| 15687 | 785 | V785Q | B | 1.1e-05 | 1.4e-05 | 1.4e-05 | 1.300E-05 | 1.732E-06 | 1.100E-05 | 1.400E-05 |
| 15688 | 785 | V785G | B | 1.5e-05 | 1.4e-05 | 1.6e-05 | 1.500E-05 | 1.000E-06 | 1.400E-05 | 1.600E-05 |
| 15689 | 785 | V785H | B | 1.4e-05 | 1.4e-05 | 1.6e-05 | 1.467E-05 | 1.155E-06 | 1.400E-05 | 1.600E-05 |
| 15690 | 785 | V785I | B | 1.5e-05 | 1.6e-05 | 1.5e-05 | 1.533E-05 | 5.774E-07 | 1.500E-05 | 1.600E-05 |
| 15691 | 785 | V785L | B | 1.5e-05 | 1.5e-05 | 1.7e-05 | 1.567E-05 | 1.155E-06 | 1.500E-05 | 1.700E-05 |
| 15692 | 785 | V785K | B | 1e-05 | 1.7e-05 | 1e-05 | 1.233E-05 | 4.041E-06 | 1.000E-05 | 1.700E-05 |
| 15693 | 785 | V785M | B | 1.6e-05 | 1e-05 | 7.4e-06 | 1.113E-05 | 4.411E-06 | 7.400E-06 | 1.600E-05 |
| 15694 | 785 | V785F | B | 1.5e-05 | 1.1e-05 | 1.1e-05 | 1.233E-05 | 2.309E-06 | 1.100E-05 | 1.500E-05 |
| 15695 | 785 | V785P | B | 1.6e-05 | 1.3e-05 | 1.5e-05 | 1.467E-05 | 1.528E-06 | 1.300E-05 | 1.600E-05 |
| 15696 | 785 | V785S | B | 1.4e-05 | 1.5e-05 | 8.1e-06 | 1.237E-05 | 3.729E-06 | 8.100E-06 | 1.500E-05 |
| 15697 | 785 | V785T | B | 9.8e-06 | 9.5e-06 | 1.4e-05 | 1.110E-05 | 2.516E-06 | 9.500E-06 | 1.400E-05 |
| 15698 | 785 | V785W | B | 1.4e-05 | 1.4e-05 | 1.1e-05 | 1.300E-05 | 1.732E-06 | 1.100E-05 | 1.400E-05 |
| 15699 | 785 | V785Y | B | 1.1e-05 | 2e-05 | 1.1e-05 | 1.400E-05 | 5.196E-06 | 1.100E-05 | 2.000E-05 |
| 15700 | 785 | V785V | B | 1.1e-05 | 1.4e-05 | 1e-05 | 1.167E-05 | 2.082E-06 | 1.000E-05 | 1.400E-05 |
| 15701 | 786 | T786A | B | 1.4e-05 | 1e-05 | 1.3e-05 | 1.233E-05 | 2.082E-06 | 1.000E-05 | 1.400E-05 |
| 15702 | 786 | T786R | B | 9.3e-06 | 1.7e-05 | 1.5e-05 | 1.377E-05 | 3.995E-06 | 9.300E-06 | 1.700E-05 |
| 15703 | 786 | T786N | B | 1.1e-05 | 1.5e-05 | 1.1e-05 | 1.233E-05 | 2.309E-06 | 1.100E-05 | 1.500E-05 |
| 15704 | 786 | T786D | B | 1.6e-05 | 1.2e-05 | 1e-05 | 1.267E-05 | 3.055E-06 | 1.000E-05 | 1.600E-05 |
| 15705 | 786 | T786C | B | 9.9e-06 | 1.3e-05 | 1.1e-05 | 1.130E-05 | 1.572E-06 | 9.900E-06 | 1.300E-05 |
| 15706 | 786 | T786E | B | 9.6e-06 | 1.2e-05 | 1e-05 | 1.053E-05 | 1.286E-06 | 9.600E-06 | 1.200E-05 |
| 15707 | 786 | T786Q | B | 1.1e-05 | 1e-05 | 1.5e-05 | 1.200E-05 | 2.646E-06 | 1.000E-05 | 1.500E-05 |
| 15708 | 786 | T786G | B | 1.4e-05 | 1.5e-05 | 1.1e-05 | 1.333E-05 | 2.082E-06 | 1.100E-05 | 1.500E-05 |
| 15709 | 786 | T786H | B | 1e-05 | 1.5e-05 | 1.4e-05 | 1.300E-05 | 2.646E-06 | 1.000E-05 | 1.500E-05 |
| 15710 | 786 | T786I | B | 1.5e-05 | 1.6e-05 | 1.2e-05 | 1.433E-05 | 2.082E-06 | 1.200E-05 | 1.600E-05 |
| 15711 | 786 | T786L | B | 1.6e-05 | 1.8e-05 | 1.1e-05 | 1.500E-05 | 3.606E-06 | 1.100E-05 | 1.800E-05 |
| 15712 | 786 | T786K | B | 1.5e-05 | 1.5e-05 | 1.1e-05 | 1.367E-05 | 2.309E-06 | 1.100E-05 | 1.500E-05 |
| 15713 | 786 | T786M | B | 8.1e-06 | 1.4e-05 | 1.2e-05 | 1.137E-05 | 3.001E-06 | 8.100E-06 | 1.400E-05 |
| 15714 | 786 | T786F | B | 1.1e-05 | 8.1e-06 | 8.8e-06 | 9.300E-06 | 1.513E-06 | 8.100E-06 | 1.100E-05 |
| 15715 | 786 | T786P | B | 1.1e-05 | 1e-05 | 1.1e-05 | 1.067E-05 | 5.774E-07 | 1.000E-05 | 1.100E-05 |
| 15716 | 786 | T786S | B | 2.6e-05 | 1.7e-05 | 7.5e-06 | 1.683E-05 | 9.251E-06 | 7.500E-06 | 2.600E-05 |
| 15717 | 786 | T786T | B | 1e-05 | 1.5e-05 | 1.1e-05 | 1.200E-05 | 2.646E-06 | 1.000E-05 | 1.500E-05 |
| 15718 | 786 | T786W | B | 1e-05 | 1.6e-05 | 7.2e-06 | 1.107E-05 | 4.496E-06 | 7.200E-06 | 1.600E-05 |
| 15719 | 786 | T786Y | B | 1.7e-05 | 8.5e-06 | 1.1e-05 | 1.217E-05 | 4.368E-06 | 8.500E-06 | 1.700E-05 |
| 15720 | 786 | T786V | B | 1.7e-05 | 8.9e-06 | 8.2e-06 | 1.137E-05 | 4.891E-06 | 8.200E-06 | 1.700E-05 |
| 15721 | 787 | E787A | B | 1.5e-05 | 1.2e-05 | 1.6e-05 | 1.433E-05 | 2.082E-06 | 1.200E-05 | 1.600E-05 |
| 15722 | 787 | E787R | B | 1.5e-05 | 1.2e-05 | 1.6e-05 | 1.433E-05 | 2.082E-06 | 1.200E-05 | 1.600E-05 |
| 15723 | 787 | E787N | B | 1.6e-05 | 1.5e-05 | 1.6e-05 | 1.567E-05 | 5.774E-07 | 1.500E-05 | 1.600E-05 |
| 15724 | 787 | E787D | B | 1.6e-05 | 9.3e-06 | 1.1e-05 | 1.210E-05 | 3.483E-06 | 9.300E-06 | 1.600E-05 |

| | | | | | | | | | | |
|---|---|---|---|---|---|---|---|---|---|---|
| 15725 | 787 | E787C | B | 1.7e-05 | 1e-05 | 1e-05 | 1.233E-05 | 4.041E-06 | 1.000E-05 | 1.700E-05 |
| 15726 | 787 | E787E | B | 6.8e-06 | 1.7e-05 | 1.4e-05 | 1.260E-05 | 5.242E-06 | 6.800E-06 | 1.700E-05 |
| 15727 | 787 | E787Q | B | 1.5e-05 | 9.8e-06 | 1.4e-05 | 1.293E-05 | 2.759E-06 | 9.800E-06 | 1.500E-05 |
| 15728 | 787 | E787G | B | 9.1e-06 | 1.1e-05 | 8.9e-06 | 9.667E-06 | 1.159E-06 | 8.900E-06 | 1.100E-05 |
| 15729 | 787 | E787H | B | 1.5e-05 | 9.4e-06 | 1.1e-05 | 1.180E-05 | 2.884E-06 | 9.400E-06 | 1.500E-05 |
| 15730 | 787 | E787I | B | 8.7e-06 | 8.8e-06 | 1.5e-05 | 1.083E-05 | 3.609E-06 | 8.700E-06 | 1.500E-05 |
| 15731 | 787 | E787L | B | 1e-05 | 1.5e-05 | 1.6e-05 | 1.367E-05 | 3.215E-06 | 1.000E-05 | 1.600E-05 |
| 15732 | 787 | E787K | B | 1e-05 | 8.9e-06 | 1.4e-05 | 1.097E-05 | 2.684E-06 | 8.900E-06 | 1.400E-05 |
| 15733 | 787 | E787M | B | 1.5e-05 | 1.4e-05 | 1.1e-05 | 1.333E-05 | 2.082E-06 | 1.100E-05 | 1.500E-05 |
| 15734 | 787 | E787F | B | 1.6e-05 | 1.3e-05 | 1.6e-05 | 1.500E-05 | 1.732E-06 | 1.300E-05 | 1.600E-05 |
| 15735 | 787 | E787P | B | 1.1e-05 | 1.6e-05 | 1.3e-05 | 1.333E-05 | 2.517E-06 | 1.100E-05 | 1.600E-05 |
| 15736 | 787 | E787S | B | 1.4e-05 | 1.4e-05 | 1.1e-05 | 1.300E-05 | 1.732E-06 | 1.100E-05 | 1.400E-05 |
| 15737 | 787 | E787T | B | 1.4e-05 | 1e-05 | 9.5e-06 | 1.117E-05 | 2.466E-06 | 9.500E-06 | 1.400E-05 |
| 15738 | 787 | E787W | B | 1.5e-05 | 1.5e-05 | 1.1e-05 | 1.367E-05 | 2.309E-06 | 1.100E-05 | 1.500E-05 |
| 15739 | 787 | E787Y | B | 1.2e-05 | 5.9e-06 | 8.7e-06 | 8.867E-06 | 3.053E-06 | 5.900E-06 | 1.200E-05 |
| 15740 | 787 | E787V | B | 1.1e-05 | 1.5e-05 | 8.7e-06 | 1.157E-05 | 3.188E-06 | 8.700E-06 | 1.500E-05 |
| 15741 | 788 | Q788A | B | 1.5e-05 | 1.6e-05 | 9.2e-06 | 1.340E-05 | 3.672E-06 | 9.200E-06 | 1.600E-05 |
| 15742 | 788 | Q788R | B | 1.1e-05 | 1.6e-05 | 1.2e-05 | 1.300E-05 | 2.646E-06 | 1.100E-05 | 1.600E-05 |
| 15743 | 788 | Q788N | B | 1.5e-05 | 1e-05 | 8.6e-06 | 1.120E-05 | 3.365E-06 | 8.600E-06 | 1.500E-05 |
| 15744 | 788 | Q788D | B | 1.5e-05 | 7.2e-06 | 1.2e-05 | 1.140E-05 | 3.934E-06 | 7.200E-06 | 1.500E-05 |
| 15745 | 788 | Q788C | B | 1.5e-05 | 1.5e-05 | 1e-05 | 1.333E-05 | 2.887E-06 | 1.000E-05 | 1.500E-05 |
| 15746 | 788 | Q788E | B | 1.4e-05 | 1.1e-05 | 1.6e-05 | 1.367E-05 | 2.517E-06 | 1.100E-05 | 1.600E-05 |
| 15747 | 788 | Q788Q | B | 1.1e-05 | 1.7e-05 | 1.3e-05 | 1.367E-05 | 3.055E-06 | 1.100E-05 | 1.700E-05 |
| 15748 | 788 | Q788G | B | 1.2e-05 | 9.9e-06 | 1.2e-05 | 1.130E-05 | 1.212E-06 | 9.900E-06 | 1.200E-05 |
| 15749 | 788 | Q788H | B | 9.6e-06 | 1.5e-05 | 1.6e-05 | 1.353E-05 | 3.443E-06 | 9.600E-06 | 1.600E-05 |
| 15750 | 788 | Q788I | B | 1.6e-05 | 1.6e-05 | 1.1e-05 | 1.433E-05 | 2.887E-06 | 1.100E-05 | 1.600E-05 |
| 15751 | 788 | Q788L | B | 1.6e-05 | 8.7e-06 | 1e-05 | 1.157E-05 | 3.894E-06 | 8.700E-06 | 1.600E-05 |
| 15752 | 788 | Q788K | B | 1.6e-05 | 1.1e-05 | 1.5e-05 | 1.400E-05 | 2.646E-06 | 1.100E-05 | 1.600E-05 |
| 15753 | 788 | Q788M | B | 1.6e-05 | 1.1e-05 | 1.5e-05 | 1.400E-05 | 2.646E-06 | 1.100E-05 | 1.600E-05 |
| 15754 | 788 | Q788F | B | 1.6e-05 | 1.6e-05 | 1.6e-05 | 1.600E-05 | 0.000E+00 | 1.600E-05 | 1.600E-05 |
| 15755 | 788 | Q788P | B | 1.2e-05 | 1.6e-05 | 1.1e-05 | 1.300E-05 | 2.646E-06 | 1.100E-05 | 1.600E-05 |
| 15756 | 788 | Q788S | B | 1.3e-05 | 1.4e-05 | 1.4e-05 | 1.367E-05 | 5.774E-07 | 1.300E-05 | 1.400E-05 |
| 15757 | 788 | Q788T | B | 1.3e-05 | 1.1e-05 | 1.4e-05 | 1.267E-05 | 1.528E-06 | 1.100E-05 | 1.400E-05 |
| 15758 | 788 | Q788W | B | 1.3e-05 | 1.2e-05 | 1.4e-05 | 1.300E-05 | 1.000E-06 | 1.200E-05 | 1.400E-05 |
| 15759 | 788 | Q788Y | B | 1.7e-05 | 1.7e-05 | 1.2e-05 | 1.533E-05 | 2.887E-06 | 1.200E-05 | 1.700E-05 |
| 15760 | 788 | Q788V | B | 9.8e-06 | 1.7e-05 | 1.1e-05 | 1.260E-05 | 3.857E-06 | 9.800E-06 | 1.700E-05 |
| 15761 | 789 | D789A | B | 1.1e-05 | 9.7e-06 | 1.4e-05 | 1.157E-05 | 2.205E-06 | 9.700E-06 | 1.400E-05 |
| 15762 | 789 | D789R | B | 1.4e-05 | 1.4e-05 | 1.4e-05 | 1.400E-05 | 0.000E+00 | 1.400E-05 | 1.400E-05 |
| 15763 | 789 | D789N | B | 9.8e-06 | 9e-06 | 1.6e-05 | 1.160E-05 | 3.831E-06 | 9.000E-06 | 1.600E-05 |
| 15764 | 789 | D789D | B | 1.4e-05 | 1.3e-05 | 1.4e-05 | 1.367E-05 | 5.774E-07 | 1.300E-05 | 1.400E-05 |
| 15765 | 789 | D789C | B | 1.6e-05 | 1.4e-05 | 8.8e-06 | 1.293E-05 | 3.717E-06 | 8.800E-06 | 1.600E-05 |
| 15766 | 789 | D789E | B | 1.1e-05 | 6.5e-06 | 9.1e-06 | 8.867E-06 | 2.259E-06 | 6.500E-06 | 1.100E-05 |
| 15767 | 789 | D789Q | B | 1.6e-05 | 1.4e-05 | 1.6e-05 | 1.533E-05 | 1.155E-06 | 1.400E-05 | 1.600E-05 |
| 15768 | 789 | D789G | B | 1.6e-05 | 1.7e-05 | 1.2e-05 | 1.500E-05 | 2.646E-06 | 1.200E-05 | 1.700E-05 |
| 15769 | 789 | D789H | B | 9.3e-06 | 1e-05 | 1.5e-05 | 1.143E-05 | 3.109E-06 | 9.300E-06 | 1.500E-05 |
| 15770 | 789 | D789I | B | 1.3e-05 | 8.4e-06 | 1.3e-05 | 1.147E-05 | 2.656E-06 | 8.400E-06 | 1.300E-05 |
| 15771 | 789 | D789L | B | 1.6e-05 | 1.5e-05 | 1.1e-05 | 1.400E-05 | 2.646E-06 | 1.100E-05 | 1.600E-05 |
| 15772 | 789 | D789K | B | 1.4e-05 | 1.4e-05 | 1.5e-05 | 1.433E-05 | 5.774E-07 | 1.400E-05 | 1.500E-05 |
| 15773 | 789 | D789M | B | 1.6e-05 | 1.2e-05 | 9.8e-06 | 1.260E-05 | 3.143E-06 | 9.800E-06 | 1.600E-05 |
| 15774 | 789 | D789F | B | 1.6e-05 | 1.5e-05 | 1.7e-05 | 1.600E-05 | 1.000E-06 | 1.500E-05 | 1.700E-05 |
| 15775 | 789 | D789P | B | 1.4e-05 | 1.2e-05 | 1.1e-05 | 1.233E-05 | 1.528E-06 | 1.100E-05 | 1.400E-05 |
| 15776 | 789 | D789S | B | 1.2e-05 | 1.4e-05 | 1.3e-05 | 1.300E-05 | 1.000E-06 | 1.200E-05 | 1.400E-05 |
| 15777 | 789 | D789T | B | 1.4e-05 | 1.6e-05 | 1.4e-05 | 1.467E-05 | 1.155E-06 | 1.400E-05 | 1.600E-05 |
| 15778 | 789 | D789W | B | 1.4e-05 | 1.1e-05 | 1e-05 | 1.167E-05 | 2.082E-06 | 1.000E-05 | 1.400E-05 |
| 15779 | 789 | D789Y | B | 1.1e-05 | 7.9e-06 | 1.2e-05 | 1.030E-05 | 2.138E-06 | 7.900E-06 | 1.200E-05 |
| 15780 | 789 | D789V | B | 1.1e-05 | 1.7e-05 | 1.4e-05 | 1.400E-05 | 3.000E-06 | 1.100E-05 | 1.700E-05 |
| 15781 | 790 | S790A | B | 1.6e-05 | 1.1e-05 | 1.4e-05 | 1.367E-05 | 2.517E-06 | 1.100E-05 | 1.600E-05 |
| 15782 | 790 | S790R | B | 1.3e-05 | 1.6e-05 | 1.1e-05 | 1.333E-05 | 2.517E-06 | 1.100E-05 | 1.600E-05 |
| 15783 | 790 | S790N | B | 1.5e-05 | 1.5e-05 | 1.5e-05 | 1.500E-05 | 0.000E+00 | 1.500E-05 | 1.500E-05 |
| 15784 | 790 | S790D | B | 1.5e-05 | 1.1e-05 | 1.4e-05 | 1.333E-05 | 2.082E-06 | 1.100E-05 | 1.500E-05 |
| 15785 | 790 | S790C | B | 1.1e-05 | 1.6e-05 | 9.9e-06 | 1.230E-05 | 3.251E-06 | 9.900E-06 | 1.600E-05 |
| 15786 | 790 | S790E | B | 1.4e-05 | 1.1e-05 | 1e-05 | 1.167E-05 | 2.082E-06 | 1.000E-05 | 1.400E-05 |
| 15787 | 790 | S790Q | B | 1.6e-05 | 1.7e-05 | 1.4e-05 | 1.567E-05 | 1.528E-06 | 1.400E-05 | 1.700E-05 |
| 15788 | 790 | S790G | B | 1e-05 | 1.8e-05 | 1.1e-05 | 1.300E-05 | 4.359E-06 | 1.000E-05 | 1.800E-05 |
| 15789 | 790 | S790H | B | 1.3e-05 | 1e-05 | 1.5e-05 | 1.267E-05 | 2.517E-06 | 1.000E-05 | 1.500E-05 |
| 15790 | 790 | S790I | B | 1e-05 | 1.6e-05 | 1.3e-05 | 1.300E-05 | 3.000E-06 | 1.000E-05 | 1.600E-05 |
| 15791 | 790 | S790L | B | 1.1e-05 | 1.1e-05 | 1.4e-05 | 1.200E-05 | 1.732E-06 | 1.100E-05 | 1.400E-05 |
| 15792 | 790 | S790K | B | 1.7e-05 | 1.4e-05 | 1.7e-05 | 1.600E-05 | 1.732E-06 | 1.400E-05 | 1.700E-05 |
| 15793 | 790 | S790M | B | 1.4e-05 | 1.5e-05 | 9.7e-06 | 1.290E-05 | 2.816E-06 | 9.700E-06 | 1.500E-05 |
| 15794 | 790 | S790F | B | 1.4e-05 | 1.6e-05 | 1e-05 | 1.333E-05 | 3.055E-06 | 1.000E-05 | 1.600E-05 |
| 15795 | 790 | S790P | B | 1.6e-05 | 1.7e-05 | 7.8e-06 | 1.360E-05 | 5.048E-06 | 7.800E-06 | 1.700E-05 |
| 15796 | 790 | S790S | B | 1.8e-05 | 1.1e-05 | 1.5e-05 | 1.467E-05 | 3.512E-06 | 1.100E-05 | 1.800E-05 |
| 15797 | 790 | S790T | B | 8.9e-06 | 1.6e-05 | 9.6e-06 | 1.150E-05 | 3.913E-06 | 8.900E-06 | 1.600E-05 |
| 15798 | 790 | S790W | B | 1.3e-05 | 8.8e-06 | 1.3e-05 | 1.160E-05 | 2.425E-06 | 8.800E-06 | 1.300E-05 |
| 15799 | 790 | S790Y | B | 9e-06 | 1.2e-05 | 1.4e-05 | 1.167E-05 | 2.517E-06 | 9.000E-06 | 1.400E-05 |
| 15800 | 790 | S790V | B | 1.5e-05 | 9.9e-06 | 1.2e-05 | 1.230E-05 | 2.563E-06 | 9.900E-06 | 1.500E-05 |
| 15801 | 791 | K791A | B | 1.7e-05 | 1e-05 | 1.3e-05 | 1.333E-05 | 3.512E-06 | 1.000E-05 | 1.700E-05 |
| 15802 | 791 | K791R | B | 9.1e-06 | 1.5e-05 | 1e-05 | 1.137E-05 | 3.179E-06 | 9.100E-06 | 1.500E-05 |
| 15803 | 791 | K791N | B | 1.2e-05 | 1.5e-05 | 9.8e-06 | 1.227E-05 | 2.610E-06 | 9.800E-06 | 1.500E-05 |
| 15804 | 791 | K791D | B | 9.2e-06 | 1e-05 | 1.3e-05 | 1.073E-05 | 2.003E-06 | 9.200E-06 | 1.300E-05 |
| 15805 | 791 | K791C | B | 1.3e-05 | 1.5e-05 | 1.4e-05 | 1.400E-05 | 1.000E-06 | 1.300E-05 | 1.500E-05 |
| 15806 | 791 | K791E | B | 1.5e-05 | 9.3e-06 | 1.5e-05 | 1.310E-05 | 3.291E-06 | 9.300E-06 | 1.500E-05 |
| 15807 | 791 | K791Q | B | 9.7e-06 | 1e-05 | 1.1e-05 | 1.023E-05 | 6.807E-07 | 9.700E-06 | 1.100E-05 |
| 15808 | 791 | K791G | B | 9.6e-06 | 7.1e-06 | 1.3e-05 | 9.900E-06 | 2.961E-06 | 7.100E-06 | 1.300E-05 |
| 15809 | 791 | K791H | B | 1.5e-05 | 1.2e-05 | 9.8e-06 | 1.227E-05 | 2.610E-06 | 9.800E-06 | 1.500E-05 |

| | | | | | | | | | | |
|---|---|---|---|---|---|---|---|---|---|---|
| 15810 | 791 | K791I | B | 1.4e-05 | 1.4e-05 | 1.2e-05 | 1.333E-05 | 1.155E-06 | 1.200E-05 | 1.400E-05 |
| 15811 | 791 | K791L | B | 8.2e-06 | 1e-05 | 1.3e-05 | 1.040E-05 | 2.425E-06 | 8.200E-06 | 1.300E-05 |
| 15812 | 791 | K791K | B | 9.7e-06 | 1.1e-05 | 1.1e-05 | 1.057E-05 | 7.506E-07 | 9.700E-06 | 1.100E-05 |
| 15813 | 791 | K791M | B | 9.7e-06 | 1e-05 | 1.2e-05 | 1.057E-05 | 1.250E-06 | 9.700E-06 | 1.200E-05 |
| 15814 | 791 | K791F | B | 8.4e-06 | 1e-05 | 2.6e-05 | 1.480E-05 | 9.732E-06 | 8.400E-06 | 2.600E-05 |
| 15815 | 791 | K791P | B | 1e-05 | 1.6e-05 | 7.7e-06 | 1.123E-05 | 4.285E-06 | 7.700E-06 | 1.600E-05 |
| 15816 | 791 | K791S | B | 9.5e-06 | 1.5e-05 | 8.4e-06 | 1.097E-05 | 3.536E-06 | 8.400E-06 | 1.500E-05 |
| 15817 | 791 | K791T | B | 1e-05 | 1.2e-05 | 1.6e-05 | 1.267E-05 | 3.055E-06 | 1.000E-05 | 1.600E-05 |
| 15818 | 791 | K791W | B | 1.8e-05 | 9.3e-06 | 1e-05 | 1.243E-05 | 4.834E-06 | 9.300E-06 | 1.800E-05 |
| 15819 | 791 | K791Y | B | 1.3e-05 | 1.5e-05 | 1.3e-05 | 1.367E-05 | 1.155E-06 | 1.300E-05 | 1.500E-05 |
| 15820 | 791 | K791V | B | 1e-05 | 1.4e-05 | 1e-05 | 1.133E-05 | 2.309E-06 | 1.000E-05 | 1.400E-05 |
| 15821 | 792 | D792A | B | 1.4e-05 | 1e-05 | 1e-05 | 1.133E-05 | 2.309E-06 | 1.000E-05 | 1.400E-05 |
| 15822 | 792 | D792R | B | 1.5e-05 | 1.7e-05 | 1.2e-05 | 1.467E-05 | 2.517E-06 | 1.200E-05 | 1.700E-05 |
| 15823 | 792 | D792N | B | 1.3e-05 | 1.3e-05 | 1e-05 | 1.200E-05 | 1.732E-06 | 1.000E-05 | 1.300E-05 |
| 15824 | 792 | D792D | B | 1.6e-05 | 9.3e-06 | 1.1e-05 | 1.210E-05 | 3.483E-06 | 9.300E-06 | 1.600E-05 |
| 15825 | 792 | D792C | B | 9.2e-06 | 1e-05 | 1.3e-05 | 1.073E-05 | 2.003E-06 | 9.200E-06 | 1.300E-05 |
| 15826 | 792 | D792E | B | 1.6e-05 | 1.6e-05 | 1.2e-05 | 1.467E-05 | 2.309E-06 | 1.200E-05 | 1.600E-05 |
| 15827 | 792 | D792Q | B | 1e-05 | 1.5e-05 | 1.2e-05 | 1.233E-05 | 2.517E-06 | 1.000E-05 | 1.500E-05 |
| 15828 | 792 | D792G | B | 1.5e-05 | 1.1e-05 | 1.4e-05 | 1.333E-05 | 2.082E-06 | 1.100E-05 | 1.500E-05 |
| 15829 | 792 | D792H | B | 8.5e-06 | 9e-06 | 1.2e-05 | 9.833E-06 | 1.893E-06 | 8.500E-06 | 1.200E-05 |
| 15830 | 792 | D792I | B | 1.6e-05 | 1.7e-05 | 1.4e-05 | 1.567E-05 | 1.528E-06 | 1.400E-05 | 1.700E-05 |
| 15831 | 792 | D792L | B | 1.5e-05 | 9.4e-06 | 1.2e-05 | 1.213E-05 | 2.802E-06 | 9.400E-06 | 1.500E-05 |
| 15832 | 792 | D792K | B | 1.5e-05 | 1.5e-05 | 1.1e-05 | 1.367E-05 | 2.309E-06 | 1.100E-05 | 1.500E-05 |
| 15833 | 792 | D792M | B | 1.5e-05 | 1.5e-05 | 1.4e-05 | 1.467E-05 | 5.774E-07 | 1.400E-05 | 1.500E-05 |
| 15834 | 792 | D792F | B | 1.2e-05 | 9.6e-06 | 1e-05 | 1.053E-05 | 1.286E-06 | 9.600E-06 | 1.200E-05 |
| 15835 | 792 | D792P | B | 1.4e-05 | 6.8e-06 | 1.7e-05 | 1.260E-05 | 5.242E-06 | 6.800E-06 | 1.700E-05 |
| 15836 | 792 | D792S | B | 1e-05 | 9.9e-06 | 1.1e-05 | 1.030E-05 | 6.083E-07 | 9.900E-06 | 1.100E-05 |
| 15837 | 792 | D792T | B | 1e-05 | 1.4e-05 | 1.2e-05 | 1.200E-05 | 2.000E-06 | 1.000E-05 | 1.400E-05 |
| 15838 | 792 | D792W | B | 1.1e-05 | 1e-05 | 1.2e-05 | 1.100E-05 | 1.000E-06 | 1.000E-05 | 1.200E-05 |
| 15839 | 792 | D792Y | B | 1.7e-05 | 1.4e-05 | 1e-05 | 1.367E-05 | 3.512E-06 | 1.000E-05 | 1.700E-05 |
| 15840 | 792 | D792V | B | 1.7e-05 | 1.6e-05 | 7.1e-06 | 1.337E-05 | 5.450E-06 | 7.100E-06 | 1.700E-05 |
| 15841 | 793 | S793A | B | 1.5e-05 | 1.1e-05 | 1.4e-05 | 1.333E-05 | 2.082E-06 | 1.100E-05 | 1.500E-05 |
| 15842 | 793 | S793R | B | 1e-05 | 1.1e-05 | 1e-05 | 1.033E-05 | 5.774E-07 | 1.000E-05 | 1.100E-05 |
| 15843 | 793 | S793N | B | 1.1e-05 | 1e-05 | 8e-06 | 9.667E-06 | 1.528E-06 | 8.000E-06 | 1.100E-05 |
| 15844 | 793 | S793D | B | 1.4e-05 | 1.5e-05 | 1.2e-05 | 1.367E-05 | 1.528E-06 | 1.200E-05 | 1.500E-05 |
| 15845 | 793 | S793C | B | 1.5e-05 | 1.2e-05 | 1.5e-05 | 1.400E-05 | 1.732E-06 | 1.200E-05 | 1.500E-05 |
| 15846 | 793 | S793E | B | 1.3e-05 | 1.4e-05 | 9.6e-06 | 1.220E-05 | 2.307E-06 | 9.600E-06 | 1.400E-05 |
| 15847 | 793 | S793Q | B | 1.3e-05 | 1.4e-05 | 1.3e-05 | 1.333E-05 | 5.774E-07 | 1.300E-05 | 1.400E-05 |
| 15848 | 793 | S793G | B | 1.6e-05 | 1.6e-05 | 1.4e-05 | 1.533E-05 | 1.155E-06 | 1.400E-05 | 1.600E-05 |
| 15849 | 793 | S793H | B | 1.7e-05 | 1.5e-05 | 8.3e-06 | 1.343E-05 | 4.557E-06 | 8.300E-06 | 1.700E-05 |
| 15850 | 793 | S793I | B | 1.8e-05 | 1.4e-05 | 1.5e-05 | 1.567E-05 | 2.082E-06 | 1.400E-05 | 1.800E-05 |
| 15851 | 793 | S793L | B | 1.4e-05 | 1.2e-05 | 1.6e-05 | 1.400E-05 | 2.000E-06 | 1.200E-05 | 1.600E-05 |
| 15852 | 793 | S793K | B | 1.6e-05 | 1.7e-05 | 1.5e-05 | 1.600E-05 | 1.000E-06 | 1.500E-05 | 1.700E-05 |
| 15853 | 793 | S793M | B | 1.1e-05 | 1.2e-05 | 1.5e-05 | 1.267E-05 | 2.082E-06 | 1.100E-05 | 1.500E-05 |
| 15854 | 793 | S793F | B | 1.1e-05 | 9.9e-06 | 1.6e-05 | 1.230E-05 | 3.251E-06 | 9.900E-06 | 1.600E-05 |
| 15855 | 793 | S793P | B | 9.6e-06 | 1.5e-05 | 1e-05 | 1.153E-05 | 3.009E-06 | 9.600E-06 | 1.500E-05 |
| 15856 | 793 | S793S | B | 1.5e-05 | 1.6e-05 | 1.6e-05 | 1.567E-05 | 5.774E-07 | 1.500E-05 | 1.600E-05 |
| 15857 | 793 | S793T | B | 1.6e-05 | 1.1e-05 | 1.6e-05 | 1.433E-05 | 2.887E-06 | 1.100E-05 | 1.600E-05 |
| 15858 | 793 | S793W | B | 1.4e-05 | 1e-05 | 1.5e-05 | 1.300E-05 | 2.646E-06 | 1.000E-05 | 1.500E-05 |
| 15859 | 793 | S793Y | B | 1e-05 | 1.4e-05 | 1.1e-05 | 1.167E-05 | 2.082E-06 | 1.000E-05 | 1.400E-05 |
| 15860 | 793 | S793V | B | 2.1e-05 | 1.5e-05 | 1e-05 | 1.533E-05 | 5.508E-06 | 1.000E-05 | 2.100E-05 |
| 15861 | 794 | T794A | B | 1.3e-05 | 8.2e-06 | 1.4e-05 | 1.173E-05 | 3.101E-06 | 8.200E-06 | 1.400E-05 |
| 15862 | 794 | T794R | B | 1.6e-05 | 9.5e-06 | 1.1e-05 | 1.217E-05 | 3.403E-06 | 9.500E-06 | 1.600E-05 |
| 15863 | 794 | T794N | B | 1.6e-05 | 1.5e-05 | 1.5e-05 | 1.533E-05 | 5.774E-07 | 1.500E-05 | 1.600E-05 |
| 15864 | 794 | T794D | B | 1.2e-05 | 1.5e-05 | 1.6e-05 | 1.433E-05 | 2.082E-06 | 1.200E-05 | 1.600E-05 |
| 15865 | 794 | T794C | B | 1.4e-05 | 1e-05 | 1.4e-05 | 1.267E-05 | 2.309E-06 | 1.000E-05 | 1.400E-05 |
| 15866 | 794 | T794E | B | 1.6e-05 | 1.4e-05 | 1.2e-05 | 1.400E-05 | 2.000E-06 | 1.200E-05 | 1.600E-05 |
| 15867 | 794 | T794Q | B | 1.5e-05 | 1.5e-05 | 1.2e-05 | 1.400E-05 | 1.732E-06 | 1.200E-05 | 1.500E-05 |
| 15868 | 794 | T794G | B | 1.5e-05 | 9.9e-06 | 9e-06 | 1.130E-05 | 3.236E-06 | 9.000E-06 | 1.500E-05 |
| 15869 | 794 | T794H | B | 1.6e-05 | 1e-05 | 8.5e-06 | 1.150E-05 | 3.969E-06 | 8.500E-06 | 1.600E-05 |
| 15870 | 794 | T794I | B | 1.2e-05 | 1e-05 | 1.4e-05 | 1.200E-05 | 2.000E-06 | 1.000E-05 | 1.400E-05 |
| 15871 | 794 | T794L | B | 6.7e-06 | 1.1e-05 | 1.6e-05 | 1.123E-05 | 4.654E-06 | 6.700E-06 | 1.600E-05 |
| 15872 | 794 | T794K | B | 1.3e-05 | 1.4e-05 | 1.1e-05 | 1.267E-05 | 1.528E-06 | 1.100E-05 | 1.400E-05 |
| 15873 | 794 | T794M | B | 1.6e-05 | 9.3e-06 | 1.3e-05 | 1.277E-05 | 3.356E-06 | 9.300E-06 | 1.600E-05 |
| 15874 | 794 | T794F | B | 1.3e-05 | 1.6e-05 | 1.6e-05 | 1.500E-05 | 1.732E-06 | 1.300E-05 | 1.600E-05 |
| 15875 | 794 | T794P | B | 1.5e-05 | 1e-05 | 1.3e-05 | 1.267E-05 | 2.517E-06 | 1.000E-05 | 1.500E-05 |
| 15876 | 794 | T794S | B | 1.6e-05 | 1.5e-05 | 1.6e-05 | 1.567E-05 | 5.774E-07 | 1.500E-05 | 1.600E-05 |
| 15877 | 794 | T794T | B | 1.6e-05 | 1e-05 | 1.1e-05 | 1.233E-05 | 3.215E-06 | 1.000E-05 | 1.600E-05 |
| 15878 | 794 | T794W | B | 1.6e-05 | 1.3e-05 | 1.5e-05 | 1.467E-05 | 1.528E-06 | 1.300E-05 | 1.600E-05 |
| 15879 | 794 | T794Y | B | 1.1e-05 | 1.6e-05 | 1.7e-05 | 1.467E-05 | 3.215E-06 | 1.100E-05 | 1.700E-05 |
| 15880 | 794 | T794V | B | 1e-05 | 1.2e-05 | 9.1e-06 | 1.037E-05 | 1.484E-06 | 9.100E-06 | 1.200E-05 |
| 15881 | 795 | Y795A | B | 8e-06 | 9.9e-06 | 1.1e-05 | 9.633E-06 | 1.518E-06 | 8.000E-06 | 1.100E-05 |
| 15882 | 795 | Y795R | B | 1.6e-05 | 1.6e-05 | 1.4e-05 | 1.533E-05 | 1.155E-06 | 1.400E-05 | 1.600E-05 |
| 15883 | 795 | Y795N | B | 1.6e-05 | 1.7e-05 | 1.1e-05 | 1.467E-05 | 3.215E-06 | 1.100E-05 | 1.700E-05 |
| 15884 | 795 | Y795D | B | 1.5e-05 | 1.1e-05 | 1.3e-05 | 1.300E-05 | 2.000E-06 | 1.100E-05 | 1.500E-05 |
| 15885 | 795 | Y795C | B | 1.5e-05 | 1e-05 | 1.2e-05 | 1.233E-05 | 2.517E-06 | 1.000E-05 | 1.500E-05 |
| 15886 | 795 | Y795E | B | 1.6e-05 | 1.1e-05 | 1.2e-05 | 1.300E-05 | 2.646E-06 | 1.100E-05 | 1.600E-05 |
| 15887 | 795 | Y795Q | B | 1.1e-05 | 1.4e-05 | 1.6e-05 | 1.367E-05 | 2.517E-06 | 1.100E-05 | 1.600E-05 |
| 15888 | 795 | Y795G | B | 1.2e-05 | 1.1e-05 | 9.7e-06 | 1.090E-05 | 1.153E-06 | 9.700E-06 | 1.200E-05 |
| 15889 | 795 | Y795H | B | 1e-05 | 1.1e-05 | 8.7e-06 | 9.900E-06 | 1.153E-06 | 8.700E-06 | 1.100E-05 |
| 15890 | 795 | Y795I | B | 1.6e-05 | 1.8e-05 | 1.6e-05 | 1.667E-05 | 1.155E-06 | 1.600E-05 | 1.800E-05 |
| 15891 | 795 | Y795L | B | 1.4e-05 | 1.4e-05 | 8.9e-06 | 1.230E-05 | 2.944E-06 | 8.900E-06 | 1.400E-05 |
| 15892 | 795 | Y795K | B | 1.3e-05 | 8.6e-06 | 9.5e-06 | 1.037E-05 | 2.325E-06 | 8.600E-06 | 1.300E-05 |
| 15893 | 795 | Y795M | B | 1.6e-05 | 1e-05 | 1.7e-05 | 1.433E-05 | 3.786E-06 | 1.000E-05 | 1.700E-05 |
| 15894 | 795 | Y795F | B | 1.2e-05 | 1.4e-05 | 8.8e-06 | 1.160E-05 | 2.623E-06 | 8.800E-06 | 1.400E-05 |

| | | | | | | | | | | |
|---|---|---|---|---|---|---|---|---|---|---|
| 15895 | 795 | Y795P | B | 1.6e-05 | 1.2e-05 | 1.4e-05 | 1.400E-05 | 2.000E-06 | 1.200E-05 | 1.600E-05 |
| 15896 | 795 | Y795S | B | 1.5e-05 | 1.3e-05 | 1e-05 | 1.267E-05 | 2.517E-06 | 1.000E-05 | 1.500E-05 |
| 15897 | 795 | Y795T | B | 1.4e-05 | 9.6e-06 | 9.1e-06 | 1.090E-05 | 2.696E-06 | 9.100E-06 | 1.400E-05 |
| 15898 | 795 | Y795W | B | 9.6e-06 | 1.4e-05 | 1.5e-05 | 1.287E-05 | 2.873E-06 | 9.600E-06 | 1.500E-05 |
| 15899 | 795 | Y795Y | B | 8.7e-06 | 1.6e-05 | 1.1e-05 | 1.190E-05 | 3.732E-06 | 8.700E-06 | 1.600E-05 |
| 15900 | 795 | Y795V | B | 1.3e-05 | 1.1e-05 | 9.6e-06 | 1.120E-05 | 1.709E-06 | 9.600E-06 | 1.300E-05 |
| 15901 | 796 | S796A | B | 1.5e-05 | 9.8e-06 | 1.6e-05 | 1.360E-05 | 3.329E-06 | 9.800E-06 | 1.600E-05 |
| 15902 | 796 | S796R | B | 1.4e-05 | 1e-05 | 1e-05 | 1.133E-05 | 2.309E-06 | 1.000E-05 | 1.400E-05 |
| 15903 | 796 | S796N | B | 9.8e-06 | 1.5e-05 | 9.2e-06 | 1.133E-05 | 3.190E-06 | 9.200E-06 | 1.500E-05 |
| 15904 | 796 | S796D | B | 1.8e-05 | 1e-05 | 2.3e-05 | 1.700E-05 | 6.557E-06 | 1.000E-05 | 2.300E-05 |
| 15905 | 796 | S796C | B | 1.4e-05 | 9.3e-06 | 8.5e-06 | 1.060E-05 | 2.972E-06 | 8.500E-06 | 1.400E-05 |
| 15906 | 796 | S796E | B | 1.1e-05 | 1.5e-05 | 1.5e-05 | 1.367E-05 | 2.309E-06 | 1.100E-05 | 1.500E-05 |
| 15907 | 796 | S796Q | B | 8.8e-06 | 1.4e-05 | 1.6e-05 | 1.293E-05 | 3.717E-06 | 8.800E-06 | 1.600E-05 |
| 15908 | 796 | S796G | B | 1.5e-05 | 1.3e-05 | 1.4e-05 | 1.400E-05 | 1.000E-06 | 1.300E-05 | 1.500E-05 |
| 15909 | 796 | S796H | B | 1e-05 | 1.6e-05 | 1.1e-05 | 1.233E-05 | 3.215E-06 | 1.000E-05 | 1.600E-05 |
| 15910 | 796 | S796I | B | 1.3e-05 | 1.5e-05 | 1.7e-05 | 1.500E-05 | 2.000E-06 | 1.300E-05 | 1.700E-05 |
| 15911 | 796 | S796L | B | 1.1e-05 | 9.9e-06 | 1e-05 | 1.030E-05 | 6.083E-07 | 9.900E-06 | 1.100E-05 |
| 15912 | 796 | S796K | B | 9.8e-06 | 1e-05 | 1.3e-05 | 1.093E-05 | 1.793E-06 | 9.800E-06 | 1.300E-05 |
| 15913 | 796 | S796M | B | 9.3e-06 | 1.3e-05 | 1.5e-05 | 1.243E-05 | 2.892E-06 | 9.300E-06 | 1.500E-05 |
| 15914 | 796 | S796F | B | 9.8e-06 | 1.6e-05 | 1.4e-05 | 1.327E-05 | 3.164E-06 | 9.800E-06 | 1.600E-05 |
| 15915 | 796 | S796P | B | 1.4e-05 | 9.7e-06 | 1.5e-05 | 1.290E-05 | 2.816E-06 | 9.700E-06 | 1.500E-05 |
| 15916 | 796 | S796S | B | 1e-05 | 1.6e-05 | 1.6e-05 | 1.400E-05 | 3.464E-06 | 1.000E-05 | 1.600E-05 |
| 15917 | 796 | S796T | B | 1.5e-05 | 1.3e-05 | 1.5e-05 | 1.433E-05 | 1.155E-06 | 1.300E-05 | 1.500E-05 |
| 15918 | 796 | S796W | B | 1.3e-05 | 1.6e-05 | 1.3e-05 | 1.400E-05 | 1.732E-06 | 1.300E-05 | 1.600E-05 |
| 15919 | 796 | S796Y | B | 1.6e-05 | 1.3e-05 | 1.5e-05 | 1.467E-05 | 1.528E-06 | 1.300E-05 | 1.600E-05 |
| 15920 | 796 | S796V | B | 1.4e-05 | 1.2e-05 | 1.4e-05 | 1.333E-05 | 1.155E-06 | 1.200E-05 | 1.400E-05 |
| 15921 | 797 | L797A | B | 1.1e-05 | 1.4e-05 | 8.5e-06 | 1.117E-05 | 2.754E-06 | 8.500E-06 | 1.400E-05 |
| 15922 | 797 | L797R | B | 1.5e-05 | 1.2e-05 | 1.2e-05 | 1.300E-05 | 1.732E-06 | 1.200E-05 | 1.500E-05 |
| 15923 | 797 | L797N | B | 1.5e-05 | 1.4e-05 | 1.5e-05 | 1.467E-05 | 5.774E-07 | 1.400E-05 | 1.500E-05 |
| 15924 | 797 | L797D | B | 8.8e-06 | 1e-05 | 9.3e-06 | 9.367E-06 | 6.028E-07 | 8.800E-06 | 1.000E-05 |
| 15925 | 797 | L797C | B | 1.4e-05 | 1.3e-05 | 1.4e-05 | 1.367E-05 | 5.774E-07 | 1.300E-05 | 1.400E-05 |
| 15926 | 797 | L797E | B | 1.1e-05 | 1.5e-05 | 5.6e-06 | 1.053E-05 | 4.717E-06 | 5.600E-06 | 1.500E-05 |
| 15927 | 797 | L797Q | B | 1.4e-05 | 9.6e-06 | 1.4e-05 | 1.253E-05 | 2.540E-06 | 9.600E-06 | 1.400E-05 |
| 15928 | 797 | L797G | B | 1e-05 | 1.7e-05 | 9.7e-06 | 1.223E-05 | 4.131E-06 | 9.700E-06 | 1.700E-05 |
| 15929 | 797 | L797H | B | 8.8e-06 | 9e-06 | 1.2e-05 | 9.933E-06 | 1.793E-06 | 8.800E-06 | 1.200E-05 |
| 15930 | 797 | L797I | B | 1.5e-05 | 1.1e-05 | 1e-05 | 1.200E-05 | 2.646E-06 | 1.000E-05 | 1.500E-05 |
| 15931 | 797 | L797L | B | 1.6e-05 | 1.5e-05 | 1.1e-05 | 1.400E-05 | 2.646E-06 | 1.100E-05 | 1.600E-05 |
| 15932 | 797 | L797K | B | 9.8e-06 | 1.5e-05 | 1.1e-05 | 1.193E-05 | 2.723E-06 | 9.800E-06 | 1.500E-05 |
| 15933 | 797 | L797M | B | 1.6e-05 | 9.5e-06 | 1.1e-05 | 1.217E-05 | 3.403E-06 | 9.500E-06 | 1.600E-05 |
| 15934 | 797 | L797F | B | 9.3e-06 | 1.6e-05 | 1e-05 | 1.177E-05 | 3.683E-06 | 9.300E-06 | 1.600E-05 |
| 15935 | 797 | L797P | B | 9.7e-06 | 9.3e-06 | 1.4e-05 | 1.100E-05 | 2.606E-06 | 9.300E-06 | 1.400E-05 |
| 15936 | 797 | L797S | B | 1.4e-05 | 1.6e-05 | 9.6e-06 | 1.320E-05 | 3.274E-06 | 9.600E-06 | 1.600E-05 |
| 15937 | 797 | L797T | B | 1e-05 | 1.4e-05 | 1.1e-05 | 1.167E-05 | 2.082E-06 | 1.000E-05 | 1.400E-05 |
| 15938 | 797 | L797W | B | 1.5e-05 | 1.5e-05 | 1.4e-05 | 1.467E-05 | 5.774E-07 | 1.400E-05 | 1.500E-05 |
| 15939 | 797 | L797Y | B | 1.5e-05 | 1.5e-05 | 1.5e-05 | 1.500E-05 | 0.000E+00 | 1.500E-05 | 1.500E-05 |
| 15940 | 797 | L797V | B | 2.6e-05 | 1.7e-05 | 1.4e-05 | 1.900E-05 | 6.245E-06 | 1.400E-05 | 2.600E-05 |
| 15941 | 798 | S798A | B | 9.7e-06 | 7.9e-06 | 1.5e-05 | 1.087E-05 | 3.691E-06 | 7.900E-06 | 1.500E-05 |
| 15942 | 798 | S798R | B | 1.5e-05 | 1.6e-05 | 1.5e-05 | 1.533E-05 | 5.774E-07 | 1.500E-05 | 1.600E-05 |
| 15943 | 798 | S798N | B | 1e-05 | 1.5e-05 | 1.6e-05 | 1.367E-05 | 3.215E-06 | 1.000E-05 | 1.600E-05 |
| 15944 | 798 | S798D | B | 1.4e-05 | 1.2e-05 | 1.1e-05 | 1.233E-05 | 1.528E-06 | 1.100E-05 | 1.400E-05 |
| 15945 | 798 | S798C | B | 1e-05 | 1.5e-05 | 9.6e-06 | 1.153E-05 | 3.009E-06 | 9.600E-06 | 1.500E-05 |
| 15946 | 798 | S798E | B | 1.5e-05 | 1.3e-05 | 1.5e-05 | 1.433E-05 | 1.155E-06 | 1.300E-05 | 1.500E-05 |
| 15947 | 798 | S798Q | B | 1.1e-05 | 9e-06 | 9.3e-06 | 9.767E-06 | 1.079E-06 | 9.000E-06 | 1.100E-05 |
| 15948 | 798 | S798G | B | 1.4e-05 | 1.1e-05 | 1.5e-05 | 1.333E-05 | 2.082E-06 | 1.100E-05 | 1.500E-05 |
| 15949 | 798 | S798H | B | 1e-05 | 1.1e-05 | 1.4e-05 | 1.167E-05 | 2.082E-06 | 1.000E-05 | 1.400E-05 |
| 15950 | 798 | S798I | B | 1.5e-05 | 8.6e-06 | 1.6e-05 | 1.320E-05 | 4.015E-06 | 8.600E-06 | 1.600E-05 |
| 15951 | 798 | S798L | B | 6.5e-06 | 1.7e-05 | 1.5e-05 | 1.283E-05 | 5.575E-06 | 6.500E-06 | 1.700E-05 |
| 15952 | 798 | S798K | B | 1.1e-05 | 1.5e-05 | 9.6e-06 | 1.187E-05 | 2.802E-06 | 9.600E-06 | 1.500E-05 |
| 15953 | 798 | S798M | B | 8.5e-06 | 1.1e-05 | 1e-05 | 9.833E-06 | 1.258E-06 | 8.500E-06 | 1.100E-05 |
| 15954 | 798 | S798F | B | 1.6e-05 | 9.4e-06 | 2.2e-05 | 1.580E-05 | 6.302E-06 | 9.400E-06 | 2.200E-05 |
| 15955 | 798 | S798P | B | 1.2e-05 | 1e-05 | 1.7e-05 | 1.300E-05 | 3.606E-06 | 1.000E-05 | 1.700E-05 |
| 15956 | 798 | S798S | B | 1.1e-05 | 1.1e-05 | 8.8e-06 | 1.027E-05 | 1.270E-06 | 8.800E-06 | 1.100E-05 |
| 15957 | 798 | S798T | B | 1.4e-05 | 1.7e-05 | 1.6e-05 | 1.567E-05 | 1.528E-06 | 1.400E-05 | 1.700E-05 |
| 15958 | 798 | S798W | B | 1.3e-05 | 1.4e-05 | 1e-05 | 1.233E-05 | 2.082E-06 | 1.000E-05 | 1.400E-05 |
| 15959 | 798 | S798Y | B | 9.8e-06 | 1.1e-05 | 1.7e-05 | 1.260E-05 | 3.857E-06 | 9.800E-06 | 1.700E-05 |
| 15960 | 798 | S798V | B | 6.2e-06 | 1.6e-05 | 1.6e-05 | 1.273E-05 | 5.658E-06 | 6.200E-06 | 1.600E-05 |
| 15961 | 799 | S799A | B | 1.6e-05 | 1.3e-05 | 1e-05 | 1.300E-05 | 3.000E-06 | 1.000E-05 | 1.600E-05 |
| 15962 | 799 | S799R | B | 1.1e-05 | 1e-05 | 1.4e-05 | 1.167E-05 | 2.082E-06 | 1.000E-05 | 1.400E-05 |
| 15963 | 799 | S799N | B | 1.5e-05 | 1.3e-05 | 1.6e-05 | 1.467E-05 | 1.528E-06 | 1.300E-05 | 1.600E-05 |
| 15964 | 799 | S799D | B | 1.2e-05 | 1.1e-05 | 1e-05 | 1.100E-05 | 1.000E-06 | 1.000E-05 | 1.200E-05 |
| 15965 | 799 | S799C | B | 1e-05 | 1e-05 | 1.4e-05 | 1.133E-05 | 2.309E-06 | 1.000E-05 | 1.400E-05 |
| 15966 | 799 | S799E | B | 1.5e-05 | 1.6e-05 | 1.7e-05 | 1.600E-05 | 1.000E-06 | 1.500E-05 | 1.700E-05 |
| 15967 | 799 | S799Q | B | 1.7e-05 | 9.6e-06 | 9.8e-06 | 1.213E-05 | 4.216E-06 | 9.600E-06 | 1.700E-05 |
| 15968 | 799 | S799G | B | 1.6e-05 | 1.5e-05 | 9.1e-06 | 1.337E-05 | 3.729E-06 | 9.100E-06 | 1.600E-05 |
| 15969 | 799 | S799H | B | 1e-05 | 1.6e-05 | 9.1e-06 | 1.170E-05 | 3.751E-06 | 9.100E-06 | 1.600E-05 |
| 15970 | 799 | S799I | B | 9.9e-06 | 1.5e-05 | 1.2e-05 | 1.230E-05 | 2.563E-06 | 9.900E-06 | 1.500E-05 |
| 15971 | 799 | S799L | B | 1.1e-05 | 9.6e-06 | 9.3e-06 | 9.967E-06 | 9.074E-07 | 9.300E-06 | 1.100E-05 |
| 15972 | 799 | S799K | B | 1.1e-05 | 1.1e-05 | 1.1e-05 | 1.100E-05 | 2.075E-21 | 1.100E-05 | 1.100E-05 |
| 15973 | 799 | S799M | B | 1.1e-05 | 9.7e-06 | 1.1e-05 | 1.057E-05 | 7.506E-07 | 9.700E-06 | 1.100E-05 |
| 15974 | 799 | S799F | B | 1.5e-05 | 9.4e-06 | 1.1e-05 | 1.180E-05 | 2.884E-06 | 9.400E-06 | 1.500E-05 |
| 15975 | 799 | S799P | B | 1.8e-05 | 9.8e-06 | 8.8e-06 | 1.220E-05 | 5.048E-06 | 8.800E-06 | 1.800E-05 |
| 15976 | 799 | S799S | B | 1.7e-05 | 1.6e-05 | 1.3e-05 | 1.533E-05 | 2.082E-06 | 1.300E-05 | 1.700E-05 |
| 15977 | 799 | S799T | B | 1.2e-05 | 1.3e-05 | 1.5e-05 | 1.333E-05 | 1.528E-06 | 1.200E-05 | 1.500E-05 |
| 15978 | 799 | S799W | B | 1.2e-05 | 1.6e-05 | 1.1e-05 | 1.300E-05 | 2.646E-06 | 1.100E-05 | 1.600E-05 |
| 15979 | 799 | S799Y | B | 1.4e-05 | 1.1e-05 | 1.2e-05 | 1.233E-05 | 1.528E-06 | 1.100E-05 | 1.400E-05 |

| | | | | | | | | | | |
|---|---|---|---|---|---|---|---|---|---|---|
| 15980 | 799 | S799V | B | 1.7e-05 | 1.5e-05 | 1e-05 | 1.400E-05 | 3.606E-06 | 1.000E-05 | 1.700E-05 |
| 15981 | 800 | T800A | B | 1.1e-05 | 1.5e-05 | 9.9e-06 | 1.197E-05 | 2.684E-06 | 9.900E-06 | 1.500E-05 |
| 15982 | 800 | T800R | B | 1e-05 | 1.3e-05 | 1e-05 | 1.100E-05 | 1.732E-06 | 1.000E-05 | 1.300E-05 |
| 15983 | 800 | T800N | B | 1e-05 | 1.6e-05 | 8.6e-06 | 1.153E-05 | 3.931E-06 | 8.600E-06 | 1.600E-05 |
| 15984 | 800 | T800D | B | 1e-05 | 1.5e-05 | 9e-06 | 1.133E-05 | 3.215E-06 | 9.000E-06 | 1.500E-05 |
| 15985 | 800 | T800C | B | 1.8e-05 | 1.1e-05 | 1.2e-05 | 1.367E-05 | 3.786E-06 | 1.100E-05 | 1.800E-05 |
| 15986 | 800 | T800E | B | 1.5e-05 | 1.1e-05 | 1.7e-05 | 1.433E-05 | 3.055E-06 | 1.100E-05 | 1.700E-05 |
| 15987 | 800 | T800Q | B | 1.5e-05 | 1.2e-05 | 1.4e-05 | 1.367E-05 | 1.528E-06 | 1.200E-05 | 1.500E-05 |
| 15988 | 800 | T800G | B | 7.8e-06 | 1.1e-05 | 9.9e-06 | 9.567E-06 | 1.626E-06 | 7.800E-06 | 1.100E-05 |
| 15989 | 800 | T800H | B | 1e-05 | 1.1e-05 | 1.1e-05 | 1.067E-05 | 5.774E-07 | 1.000E-05 | 1.100E-05 |
| 15990 | 800 | T800I | B | 9.3e-06 | 8.4e-06 | 1.2e-05 | 9.900E-06 | 1.873E-06 | 8.400E-06 | 1.200E-05 |
| 15991 | 800 | T800L | B | 1.4e-05 | 1.3e-05 | 1.1e-05 | 1.267E-05 | 1.528E-06 | 1.100E-05 | 1.400E-05 |
| 15992 | 800 | T800K | B | 2.1e-05 | 1.3e-05 | 1.3e-05 | 1.567E-05 | 4.619E-06 | 1.300E-05 | 2.100E-05 |
| 15993 | 800 | T800M | B | 1.4e-05 | 1.1e-05 | 1e-05 | 1.167E-05 | 2.082E-06 | 1.000E-05 | 1.400E-05 |
| 15994 | 800 | T800F | B | 1.5e-05 | 1.2e-05 | 1.1e-05 | 1.267E-05 | 2.082E-06 | 1.100E-05 | 1.500E-05 |
| 15995 | 800 | T800P | B | 1.1e-05 | 1.5e-05 | 1.7e-05 | 1.433E-05 | 3.055E-06 | 1.100E-05 | 1.700E-05 |
| 15996 | 800 | T800S | B | 1e-05 | 1.1e-05 | 1.1e-05 | 1.067E-05 | 5.774E-07 | 1.000E-05 | 1.100E-05 |
| 15997 | 800 | T800T | B | 1.3e-05 | 1.2e-05 | 1.2e-05 | 1.233E-05 | 5.774E-07 | 1.200E-05 | 1.300E-05 |
| 15998 | 800 | T800W | B | 1.1e-05 | 1.2e-05 | 9.8e-06 | 1.093E-05 | 1.102E-06 | 9.800E-06 | 1.200E-05 |
| 15999 | 800 | T800Y | B | 6.7e-06 | 1.5e-05 | 9.6e-06 | 1.043E-05 | 4.212E-06 | 6.700E-06 | 1.500E-05 |
| 16000 | 800 | T800V | B | 1.6e-05 | 1.4e-05 | 1.4e-05 | 1.467E-05 | 1.155E-06 | 1.400E-05 | 1.600E-05 |
| 16001 | 801 | L801A | B | 1.4e-05 | 9.2e-06 | 9.5e-06 | 1.090E-05 | 2.689E-06 | 9.200E-06 | 1.400E-05 |
| 16002 | 801 | L801R | B | 1.3e-05 | 1.2e-05 | 8.9e-06 | 1.130E-05 | 2.138E-06 | 8.900E-06 | 1.300E-05 |
| 16003 | 801 | L801N | B | 1.4e-05 | 1.6e-05 | 1e-05 | 1.333E-05 | 3.055E-06 | 1.000E-05 | 1.600E-05 |
| 16004 | 801 | L801D | B | 1.2e-05 | 1.3e-05 | 1.6e-05 | 1.367E-05 | 2.082E-06 | 1.200E-05 | 1.600E-05 |
| 16005 | 801 | L801C | B | 1.1e-05 | 1.2e-05 | 1.5e-05 | 1.267E-05 | 2.082E-06 | 1.100E-05 | 1.500E-05 |
| 16006 | 801 | L801E | B | 1.7e-05 | 1.6e-05 | 1.2e-05 | 1.500E-05 | 2.646E-06 | 1.200E-05 | 1.700E-05 |
| 16007 | 801 | L801Q | B | 8.9e-06 | 1.5e-05 | 9.7e-06 | 1.120E-05 | 3.315E-06 | 8.900E-06 | 1.500E-05 |
| 16008 | 801 | L801G | B | 1.1e-05 | 1.6e-05 | 1.5e-05 | 1.400E-05 | 2.646E-06 | 1.100E-05 | 1.600E-05 |
| 16009 | 801 | L801H | B | 1.6e-05 | 1.4e-05 | 1.4e-05 | 1.467E-05 | 1.155E-06 | 1.400E-05 | 1.600E-05 |
| 16010 | 801 | L801I | B | 1.5e-05 | 1.3e-05 | 1.6e-05 | 1.467E-05 | 1.528E-06 | 1.300E-05 | 1.600E-05 |
| 16011 | 801 | L801L | B | 1.8e-05 | 1.1e-05 | 1.4e-05 | 1.433E-05 | 3.512E-06 | 1.100E-05 | 1.800E-05 |
| 16012 | 801 | L801K | B | 1.3e-05 | 1e-05 | 1.3e-05 | 1.200E-05 | 1.732E-06 | 1.000E-05 | 1.300E-05 |
| 16013 | 801 | L801M | B | 9.7e-06 | 9.2e-06 | 1.6e-05 | 1.163E-05 | 3.790E-06 | 9.200E-06 | 1.600E-05 |
| 16014 | 801 | L801F | B | 2.6e-05 | 1.3e-05 | 1.4e-05 | 1.767E-05 | 7.234E-06 | 1.300E-05 | 2.600E-05 |
| 16015 | 801 | L801P | B | 1.3e-05 | 8.6e-06 | 1e-05 | 1.053E-05 | 2.248E-06 | 8.600E-06 | 1.300E-05 |
| 16016 | 801 | L801S | B | 1.6e-05 | 1.1e-05 | 1e-05 | 1.233E-05 | 3.215E-06 | 1.000E-05 | 1.600E-05 |
| 16017 | 801 | L801T | B | 1.1e-05 | 1.4e-05 | 1.5e-05 | 1.333E-05 | 2.082E-06 | 1.100E-05 | 1.500E-05 |
| 16018 | 801 | L801W | B | 1.5e-05 | 1.1e-05 | 1.5e-05 | 1.367E-05 | 2.309E-06 | 1.100E-05 | 1.500E-05 |
| 16019 | 801 | L801Y | B | 9.1e-06 | 1e-05 | 2.2e-05 | 1.370E-05 | 7.202E-06 | 9.100E-06 | 2.200E-05 |
| 16020 | 801 | L801V | B | 1.4e-05 | 1.6e-05 | 7.2e-06 | 1.240E-05 | 4.613E-06 | 7.200E-06 | 1.600E-05 |
| 16021 | 802 | T802A | B | 9.2e-06 | 7.5e-06 | 9e-06 | 8.567E-06 | 9.292E-07 | 7.500E-06 | 9.200E-06 |
| 16022 | 802 | T802R | B | 1.5e-05 | 1.1e-05 | 1.4e-05 | 1.333E-05 | 2.082E-06 | 1.100E-05 | 1.500E-05 |
| 16023 | 802 | T802N | B | 1.4e-05 | 1.3e-05 | 9.4e-06 | 1.213E-05 | 2.419E-06 | 9.400E-06 | 1.400E-05 |
| 16024 | 802 | T802D | B | 9.3e-06 | 1.1e-05 | 1.5e-05 | 1.177E-05 | 2.926E-06 | 9.300E-06 | 1.500E-05 |
| 16025 | 802 | T802C | B | 9e-06 | 1.4e-05 | 1.1e-05 | 1.133E-05 | 2.517E-06 | 9.000E-06 | 1.400E-05 |
| 16026 | 802 | T802E | B | 1.4e-05 | 1.5e-05 | 1.4e-05 | 1.433E-05 | 5.774E-07 | 1.400E-05 | 1.500E-05 |
| 16027 | 802 | T802Q | B | 9.8e-06 | 1.5e-05 | 1.5e-05 | 1.327E-05 | 3.002E-06 | 9.800E-06 | 1.500E-05 |
| 16028 | 802 | T802G | B | 1.7e-05 | 2e-05 | 1.2e-05 | 1.633E-05 | 4.041E-06 | 1.200E-05 | 2.000E-05 |
| 16029 | 802 | T802H | B | 1.6e-05 | 1.4e-05 | 1.5e-05 | 1.500E-05 | 1.000E-06 | 1.400E-05 | 1.600E-05 |
| 16030 | 802 | T802I | B | 1.6e-05 | 1.2e-05 | 1.3e-05 | 1.367E-05 | 2.082E-06 | 1.200E-05 | 1.600E-05 |
| 16031 | 802 | T802L | B | 1.9e-05 | 1.6e-05 | 1.6e-05 | 1.700E-05 | 1.732E-06 | 1.600E-05 | 1.900E-05 |
| 16032 | 802 | T802K | B | 1.6e-05 | 1e-05 | 1.5e-05 | 1.367E-05 | 3.215E-06 | 1.000E-05 | 1.600E-05 |
| 16033 | 802 | T802M | B | 1.7e-05 | 1.7e-05 | 1.6e-05 | 1.667E-05 | 5.774E-07 | 1.600E-05 | 1.700E-05 |
| 16034 | 802 | T802F | B | 1.1e-05 | 1.1e-05 | 1.5e-05 | 1.233E-05 | 2.309E-06 | 1.100E-05 | 1.500E-05 |
| 16035 | 802 | T802P | B | 1.1e-05 | 1.1e-05 | 1.6e-05 | 1.267E-05 | 2.887E-06 | 1.100E-05 | 1.600E-05 |
| 16036 | 802 | T802S | B | 1.4e-05 | 1.5e-05 | 1.4e-05 | 1.433E-05 | 5.774E-07 | 1.400E-05 | 1.500E-05 |
| 16037 | 802 | T802T | B | 1.7e-05 | 1.3e-05 | 8e-06 | 1.267E-05 | 4.509E-06 | 8.000E-06 | 1.700E-05 |
| 16038 | 802 | T802W | B | 1.1e-05 | 1e-05 | 1.4e-05 | 1.167E-05 | 2.082E-06 | 1.000E-05 | 1.400E-05 |
| 16039 | 802 | T802Y | B | 1.5e-05 | 1e-05 | 1.6e-05 | 1.367E-05 | 3.215E-06 | 1.000E-05 | 1.600E-05 |
| 16040 | 802 | T802V | B | 1.2e-05 | 9.8e-06 | 1.4e-05 | 1.193E-05 | 2.101E-06 | 9.800E-06 | 1.400E-05 |
| 16041 | 803 | L803A | B | 1.1e-05 | 1.4e-05 | 1e-05 | 1.167E-05 | 2.082E-06 | 1.000E-05 | 1.400E-05 |
| 16042 | 803 | L803R | B | 1.1e-05 | 1.3e-05 | 1.3e-05 | 1.233E-05 | 1.155E-06 | 1.100E-05 | 1.300E-05 |
| 16043 | 803 | L803N | B | 1e-05 | 9.2e-06 | 1.4e-05 | 1.107E-05 | 2.572E-06 | 9.200E-06 | 1.400E-05 |
| 16044 | 803 | L803D | B | 1.6e-05 | 9.9e-06 | 1.4e-05 | 1.330E-05 | 3.110E-06 | 9.900E-06 | 1.600E-05 |
| 16045 | 803 | L803C | B | 1.5e-05 | 1e-05 | 9.3e-06 | 1.143E-05 | 3.109E-06 | 9.300E-06 | 1.500E-05 |
| 16046 | 803 | L803E | B | 1.1e-05 | 1.5e-05 | 1.6e-05 | 1.400E-05 | 2.646E-06 | 1.100E-05 | 1.600E-05 |
| 16047 | 803 | L803Q | B | 1.4e-05 | 1e-05 | 1.5e-05 | 1.300E-05 | 2.646E-06 | 1.000E-05 | 1.500E-05 |
| 16048 | 803 | L803G | B | 1.2e-05 | 1.5e-05 | 1.6e-05 | 1.433E-05 | 2.082E-06 | 1.200E-05 | 1.600E-05 |
| 16049 | 803 | L803H | B | 9.6e-06 | 1.3e-05 | 1e-05 | 1.087E-05 | 1.858E-06 | 9.600E-06 | 1.300E-05 |
| 16050 | 803 | L803I | B | 1.1e-05 | 9.7e-06 | 1.4e-05 | 1.157E-05 | 2.205E-06 | 9.700E-06 | 1.400E-05 |
| 16051 | 803 | L803L | B | 1e-05 | 1.4e-05 | 1.6e-05 | 1.333E-05 | 3.055E-06 | 1.000E-05 | 1.600E-05 |
| 16052 | 803 | L803K | B | 1.1e-05 | 9.4e-06 | 8.9e-06 | 9.767E-06 | 1.097E-06 | 8.900E-06 | 1.100E-05 |
| 16053 | 803 | L803M | B | 1.3e-05 | 1.1e-05 | 1.5e-05 | 1.300E-05 | 2.000E-06 | 1.100E-05 | 1.500E-05 |
| 16054 | 803 | L803F | B | 1.4e-05 | 1.1e-05 | 1.6e-05 | 1.367E-05 | 2.517E-06 | 1.100E-05 | 1.600E-05 |
| 16055 | 803 | L803P | B | 8.6e-06 | 1.3e-05 | 1.6e-05 | 1.253E-05 | 3.722E-06 | 8.600E-06 | 1.600E-05 |
| 16056 | 803 | L803S | B | 1.5e-05 | 1.2e-05 | 1.5e-05 | 1.400E-05 | 1.732E-06 | 1.200E-05 | 1.500E-05 |
| 16057 | 803 | L803T | B | 1.3e-05 | 9.5e-06 | 1.4e-05 | 1.217E-05 | 2.363E-06 | 9.500E-06 | 1.400E-05 |
| 16058 | 803 | L803W | B | 1e-05 | 1.6e-05 | 9.1e-06 | 1.170E-05 | 3.751E-06 | 9.100E-06 | 1.600E-05 |
| 16059 | 803 | L803Y | B | 1.3e-05 | 1.5e-05 | 7.7e-06 | 1.190E-05 | 3.772E-06 | 7.700E-06 | 1.500E-05 |
| 16060 | 803 | L803V | B | 1.6e-05 | 7.8e-06 | 1.3e-05 | 1.227E-05 | 4.149E-06 | 7.800E-06 | 1.600E-05 |
| 16061 | 804 | S804A | B | 1.1e-05 | 1.5e-05 | 9.6e-06 | 1.187E-05 | 2.802E-06 | 9.600E-06 | 1.500E-05 |
| 16062 | 804 | S804R | B | 9e-06 | 1.5e-05 | 1.4e-05 | 1.267E-05 | 3.215E-06 | 9.000E-06 | 1.500E-05 |
| 16063 | 804 | S804N | B | 1.3e-05 | 1.1e-05 | 1.3e-05 | 1.233E-05 | 1.155E-06 | 1.100E-05 | 1.300E-05 |
| 16064 | 804 | S804D | B | 1.7e-05 | 1.7e-05 | 1.1e-05 | 1.500E-05 | 3.464E-06 | 1.100E-05 | 1.700E-05 |

| | | | | | | | | | | |
|---|---|---|---|---|---|---|---|---|---|---|
| 16065 | 804 | S804C | B | 6.7e-06 | 1.6e-05 | 1.5e-05 | 1.257E-05 | 5.105E-06 | 6.700E-06 | 1.600E-05 |
| 16066 | 804 | S804E | B | 1.1e-05 | 1.1e-05 | 1.1e-05 | 1.100E-05 | 2.075E-21 | 1.100E-05 | 1.100E-05 |
| 16067 | 804 | S804Q | B | 6.3e-06 | 9.8e-06 | 1.6e-05 | 1.070E-05 | 4.912E-06 | 6.300E-06 | 1.600E-05 |
| 16068 | 804 | S804G | B | 1.6e-05 | 9.5e-06 | 1.5e-05 | 1.350E-05 | 3.500E-06 | 9.500E-06 | 1.600E-05 |
| 16069 | 804 | S804H | B | 1.1e-05 | 1.4e-05 | 1.5e-05 | 1.333E-05 | 2.082E-06 | 1.100E-05 | 1.500E-05 |
| 16070 | 804 | S804I | B | 1.6e-05 | 1.5e-05 | 9.6e-06 | 1.353E-05 | 3.443E-06 | 9.600E-06 | 1.600E-05 |
| 16071 | 804 | S804L | B | 9.8e-06 | 1.1e-05 | 1.2e-05 | 1.093E-05 | 1.102E-06 | 9.800E-06 | 1.200E-05 |
| 16072 | 804 | S804K | B | 1.3e-05 | 1.1e-05 | 1.1e-05 | 1.167E-05 | 1.155E-06 | 1.100E-05 | 1.300E-05 |
| 16073 | 804 | S804M | B | 1.5e-05 | 1.4e-05 | 8.3e-06 | 1.243E-05 | 3.614E-06 | 8.300E-06 | 1.500E-05 |
| 16074 | 804 | S804F | B | 1.1e-05 | 1.4e-05 | 1.1e-05 | 1.200E-05 | 1.732E-06 | 1.100E-05 | 1.400E-05 |
| 16075 | 804 | S804P | B | 1.5e-05 | 1.5e-05 | 1.5e-05 | 1.500E-05 | 0.000E+00 | 1.500E-05 | 1.500E-05 |
| 16076 | 804 | S804S | B | 8.7e-06 | 1.6e-05 | 1.5e-05 | 1.323E-05 | 3.958E-06 | 8.700E-06 | 1.600E-05 |
| 16077 | 804 | S804T | B | 1.1e-05 | 1.1e-05 | 1.6e-05 | 1.267E-05 | 2.887E-06 | 1.100E-05 | 1.600E-05 |
| 16078 | 804 | S804W | B | 1.5e-05 | 1.2e-05 | 9.6e-06 | 1.220E-05 | 2.706E-06 | 9.600E-06 | 1.500E-05 |
| 16079 | 804 | S804Y | B | 1.4e-05 | 1.1e-05 | 1.6e-05 | 1.367E-05 | 2.517E-06 | 1.100E-05 | 1.600E-05 |
| 16080 | 804 | S804V | B | 1.5e-05 | 1.2e-05 | 1.1e-05 | 1.267E-05 | 2.082E-06 | 1.100E-05 | 1.500E-05 |
| 16081 | 805 | K805A | B | 1.5e-05 | 1.2e-05 | 1.1e-05 | 1.267E-05 | 2.082E-06 | 1.100E-05 | 1.500E-05 |
| 16082 | 805 | K805R | B | 2.6e-05 | 1.5e-05 | 1.5e-05 | 1.867E-05 | 6.351E-06 | 1.500E-05 | 2.600E-05 |
| 16083 | 805 | K805N | B | 9.5e-06 | 1.4e-05 | 1.5e-05 | 1.283E-05 | 2.930E-06 | 9.500E-06 | 1.500E-05 |
| 16084 | 805 | K805D | B | 1.1e-05 | 1.1e-05 | 1.5e-05 | 1.233E-05 | 2.309E-06 | 1.100E-05 | 1.500E-05 |
| 16085 | 805 | K805C | B | 8.7e-06 | 1.5e-05 | 9.9e-06 | 1.120E-05 | 3.345E-06 | 8.700E-06 | 1.500E-05 |
| 16086 | 805 | K805E | B | 1.5e-05 | 1e-05 | 8.8e-06 | 1.127E-05 | 3.288E-06 | 8.800E-06 | 1.500E-05 |
| 16087 | 805 | K805Q | B | 8.6e-06 | 1.5e-05 | 1.2e-05 | 1.187E-05 | 3.202E-06 | 8.600E-06 | 1.500E-05 |
| 16088 | 805 | K805G | B | 1.5e-05 | 8e-06 | 9.4e-06 | 1.080E-05 | 3.704E-06 | 8.000E-06 | 1.500E-05 |
| 16089 | 805 | K805H | B | 1e-05 | 1.5e-05 | 9.8e-06 | 1.160E-05 | 2.946E-06 | 9.800E-06 | 1.500E-05 |
| 16090 | 805 | K805I | B | 9e-06 | 1e-05 | 1.6e-05 | 1.167E-05 | 3.786E-06 | 9.000E-06 | 1.600E-05 |
| 16091 | 805 | K805L | B | 1.5e-05 | 1.2e-05 | 1.1e-05 | 1.267E-05 | 2.082E-06 | 1.100E-05 | 1.500E-05 |
| 16092 | 805 | K805K | B | 1.4e-05 | 1.5e-05 | 1.7e-05 | 1.533E-05 | 1.528E-06 | 1.400E-05 | 1.700E-05 |
| 16093 | 805 | K805M | B | 9.4e-06 | 1.1e-05 | 1.8e-05 | 1.280E-05 | 4.574E-06 | 9.400E-06 | 1.800E-05 |
| 16094 | 805 | K805F | B | 1e-05 | 1.5e-05 | 1.4e-05 | 1.300E-05 | 2.646E-06 | 1.000E-05 | 1.500E-05 |
| 16095 | 805 | K805P | B | 1e-05 | 1.5e-05 | 1.5e-05 | 1.333E-05 | 2.887E-06 | 1.000E-05 | 1.500E-05 |
| 16096 | 805 | K805S | B | 1.4e-05 | 1.5e-05 | 1.3e-05 | 1.400E-05 | 1.000E-06 | 1.300E-05 | 1.500E-05 |
| 16097 | 805 | K805T | B | 1.2e-05 | 1.4e-05 | 1.5e-05 | 1.367E-05 | 1.528E-06 | 1.200E-05 | 1.500E-05 |
| 16098 | 805 | K805W | B | 1.2e-05 | 1e-05 | 1.6e-05 | 1.267E-05 | 3.055E-06 | 1.000E-05 | 1.600E-05 |
| 16099 | 805 | K805Y | B | 1.1e-05 | 1e-05 | 1.3e-05 | 1.133E-05 | 1.528E-06 | 1.000E-05 | 1.300E-05 |
| 16100 | 805 | K805V | B | 1.4e-05 | 1.6e-05 | 9.9e-06 | 1.330E-05 | 3.110E-06 | 9.900E-06 | 1.600E-05 |
| 16101 | 806 | A806A | B | 6.8e-06 | 1.7e-05 | 9.6e-06 | 1.113E-05 | 5.270E-06 | 6.800E-06 | 1.700E-05 |
| 16102 | 806 | A806R | B | 1.5e-05 | 1.5e-05 | 9.7e-06 | 1.323E-05 | 3.060E-06 | 9.700E-06 | 1.500E-05 |
| 16103 | 806 | A806N | B | 1e-05 | 1.5e-05 | 1.4e-05 | 1.300E-05 | 2.646E-06 | 1.000E-05 | 1.500E-05 |
| 16104 | 806 | A806D | B | 1.4e-05 | 1.5e-05 | 1.1e-05 | 1.333E-05 | 2.082E-06 | 1.100E-05 | 1.500E-05 |
| 16105 | 806 | A806C | B | 1.5e-05 | 1.2e-05 | 1.6e-05 | 1.433E-05 | 2.082E-06 | 1.200E-05 | 1.600E-05 |
| 16106 | 806 | A806E | B | 9.1e-06 | 1.5e-05 | 8.4e-06 | 1.083E-05 | 3.625E-06 | 8.400E-06 | 1.500E-05 |
| 16107 | 806 | A806Q | B | 9.5e-06 | 1.4e-05 | 9.3e-06 | 1.093E-05 | 2.658E-06 | 9.300E-06 | 1.400E-05 |
| 16108 | 806 | A806G | B | 1.1e-05 | 1.8e-05 | 9.8e-06 | 1.293E-05 | 4.429E-06 | 9.800E-06 | 1.800E-05 |
| 16109 | 806 | A806H | B | 1.6e-05 | 1.4e-05 | 1e-05 | 1.333E-05 | 3.055E-06 | 1.000E-05 | 1.600E-05 |
| 16110 | 806 | A806I | B | 1.3e-05 | 1.2e-05 | 1e-05 | 1.167E-05 | 1.528E-06 | 1.000E-05 | 1.300E-05 |
| 16111 | 806 | A806L | B | 1.4e-05 | 1.5e-05 | 1.5e-05 | 1.467E-05 | 5.774E-07 | 1.400E-05 | 1.500E-05 |
| 16112 | 806 | A806K | B | 1.3e-05 | 1.1e-05 | 1.2e-05 | 1.200E-05 | 1.000E-06 | 1.100E-05 | 1.300E-05 |
| 16113 | 806 | A806M | B | 9e-06 | 1.1e-05 | 1.5e-05 | 1.167E-05 | 3.055E-06 | 9.000E-06 | 1.500E-05 |
| 16114 | 806 | A806F | B | 9.5e-06 | 1.2e-05 | 1.2e-05 | 1.117E-05 | 1.443E-06 | 9.500E-06 | 1.200E-05 |
| 16115 | 806 | A806P | B | 1.5e-05 | 1.5e-05 | 1.6e-05 | 1.533E-05 | 5.774E-07 | 1.500E-05 | 1.600E-05 |
| 16116 | 806 | A806S | B | 1.5e-05 | 1.2e-05 | 1.3e-05 | 1.333E-05 | 1.528E-06 | 1.200E-05 | 1.500E-05 |
| 16117 | 806 | A806T | B | 1.4e-05 | 1.1e-05 | 9.8e-06 | 1.160E-05 | 2.163E-06 | 9.800E-06 | 1.400E-05 |
| 16118 | 806 | A806W | B | 1e-05 | 8.2e-06 | 1.4e-05 | 1.073E-05 | 2.969E-06 | 8.200E-06 | 1.400E-05 |
| 16119 | 806 | A806Y | B | 1.4e-05 | 1.4e-05 | 1.4e-05 | 1.400E-05 | 0.000E+00 | 1.400E-05 | 1.400E-05 |
| 16120 | 806 | A806V | B | 8.8e-06 | 8.8e-06 | 1.4e-05 | 1.053E-05 | 3.002E-06 | 8.800E-06 | 1.400E-05 |
| 16121 | 807 | D807A | B | 9.7e-06 | 1.2e-05 | 1.6e-05 | 1.257E-05 | 3.188E-06 | 9.700E-06 | 1.600E-05 |
| 16122 | 807 | D807R | B | 1.6e-05 | 1.6e-05 | 8.2e-06 | 1.340E-05 | 4.503E-06 | 8.200E-06 | 1.600E-05 |
| 16123 | 807 | D807N | B | 1.5e-05 | 1.6e-05 | 1.6e-05 | 1.567E-05 | 5.774E-07 | 1.500E-05 | 1.600E-05 |
| 16124 | 807 | D807D | B | 1.7e-05 | 1.2e-05 | 1.5e-05 | 1.467E-05 | 2.517E-06 | 1.200E-05 | 1.700E-05 |
| 16125 | 807 | D807C | B | 1.6e-05 | 1.3e-05 | 1.3e-05 | 1.400E-05 | 1.732E-06 | 1.300E-05 | 1.600E-05 |
| 16126 | 807 | D807E | B | 1.4e-05 | 1.1e-05 | 1.6e-05 | 1.367E-05 | 2.517E-06 | 1.100E-05 | 1.600E-05 |
| 16127 | 807 | D807Q | B | 1.1e-05 | 1.6e-05 | 1.1e-05 | 1.267E-05 | 2.887E-06 | 1.100E-05 | 1.600E-05 |
| 16128 | 807 | D807G | B | 1.3e-05 | 1.4e-05 | 1.3e-05 | 1.333E-05 | 5.774E-07 | 1.300E-05 | 1.400E-05 |
| 16129 | 807 | D807H | B | 1.5e-05 | 1e-05 | 1.5e-05 | 1.333E-05 | 2.887E-06 | 1.000E-05 | 1.500E-05 |
| 16130 | 807 | D807I | B | 1.7e-05 | 1.1e-05 | 1.2e-05 | 1.333E-05 | 3.215E-06 | 1.100E-05 | 1.700E-05 |
| 16131 | 807 | D807L | B | 1.4e-05 | 1.2e-05 | 1.5e-05 | 1.367E-05 | 1.528E-06 | 1.200E-05 | 1.500E-05 |
| 16132 | 807 | D807K | B | 1.5e-05 | 9.5e-06 | 1.5e-05 | 1.317E-05 | 3.175E-06 | 9.500E-06 | 1.500E-05 |
| 16133 | 807 | D807M | B | 1.1e-05 | 1.7e-05 | 9.2e-06 | 1.240E-05 | 4.084E-06 | 9.200E-06 | 1.700E-05 |
| 16134 | 807 | D807F | B | 1.1e-05 | 9.9e-06 | 1.6e-05 | 1.230E-05 | 3.251E-06 | 9.900E-06 | 1.600E-05 |
| 16135 | 807 | D807P | B | 9e-06 | 1.7e-05 | 1.3e-05 | 1.300E-05 | 4.000E-06 | 9.000E-06 | 1.700E-05 |
| 16136 | 807 | D807S | B | 8.3e-06 | 1.5e-05 | 1.1e-05 | 1.143E-05 | 3.371E-06 | 8.300E-06 | 1.500E-05 |
| 16137 | 807 | D807T | B | 1.2e-05 | 1.6e-05 | 1.5e-05 | 1.433E-05 | 2.082E-06 | 1.200E-05 | 1.600E-05 |
| 16138 | 807 | D807W | B | 1.4e-05 | 1.6e-05 | 1.4e-05 | 1.467E-05 | 1.155E-06 | 1.400E-05 | 1.600E-05 |
| 16139 | 807 | D807Y | B | 8.8e-06 | 1.4e-05 | 1.4e-05 | 1.227E-05 | 3.002E-06 | 8.800E-06 | 1.400E-05 |
| 16140 | 807 | D807V | B | 1.1e-05 | 1.6e-05 | 1.6e-05 | 1.433E-05 | 2.887E-06 | 1.100E-05 | 1.600E-05 |
| 16141 | 808 | Y808A | B | 1.6e-05 | 9.4e-06 | 1.4e-05 | 1.313E-05 | 3.384E-06 | 9.400E-06 | 1.600E-05 |
| 16142 | 808 | Y808R | B | 1.6e-05 | 1.6e-05 | 1.6e-05 | 1.600E-05 | 0.000E+00 | 1.600E-05 | 1.600E-05 |
| 16143 | 808 | Y808N | B | 7.9e-06 | 1.3e-05 | 8.1e-06 | 9.667E-06 | 2.888E-06 | 7.900E-06 | 1.300E-05 |
| 16144 | 808 | Y808D | B | 8.8e-06 | 1.1e-05 | 1.1e-05 | 1.027E-05 | 1.270E-06 | 8.800E-06 | 1.100E-05 |
| 16145 | 808 | Y808C | B | 1.6e-05 | 9.7e-06 | 5.8e-06 | 1.050E-05 | 5.147E-06 | 5.800E-06 | 1.600E-05 |
| 16146 | 808 | Y808E | B | 1e-05 | 1.4e-05 | 1.2e-05 | 1.200E-05 | 2.000E-06 | 1.000E-05 | 1.400E-05 |
| 16147 | 808 | Y808Q | B | 1.7e-05 | 1.2e-05 | 1.1e-05 | 1.333E-05 | 3.215E-06 | 1.100E-05 | 1.700E-05 |
| 16148 | 808 | Y808G | B | 1.6e-05 | 6.1e-06 | 1.1e-05 | 1.103E-05 | 4.950E-06 | 6.100E-06 | 1.600E-05 |
| 16149 | 808 | Y808H | B | 1.3e-05 | 1.6e-05 | 1.5e-05 | 1.467E-05 | 1.528E-06 | 1.300E-05 | 1.600E-05 |

| | | | | | | | | | | |
|---|---|---|---|---|---|---|---|---|---|---|
| 16150 | 808 | Y808I | B | 1.2e-05 | 1.6e-05 | 1.7e-05 | 1.500E-05 | 2.646E-06 | 1.200E-05 | 1.700E-05 |
| 16151 | 808 | Y808L | B | 1.7e-05 | 1.3e-05 | 1.1e-05 | 1.367E-05 | 3.055E-06 | 1.100E-05 | 1.700E-05 |
| 16152 | 808 | Y808K | B | 1e-05 | 1.4e-05 | 1.3e-05 | 1.233E-05 | 2.082E-06 | 1.000E-05 | 1.400E-05 |
| 16153 | 808 | Y808M | B | 1.6e-05 | 1.6e-05 | 1e-05 | 1.400E-05 | 3.464E-06 | 1.000E-05 | 1.600E-05 |
| 16154 | 808 | Y808F | B | 1.5e-05 | 1.5e-05 | 1.1e-05 | 1.367E-05 | 2.309E-06 | 1.100E-05 | 1.500E-05 |
| 16155 | 808 | Y808P | B | 1.6e-05 | 1.3e-05 | 1.3e-05 | 1.400E-05 | 1.732E-06 | 1.300E-05 | 1.600E-05 |
| 16156 | 808 | Y808S | B | 1.4e-05 | 7.9e-06 | 1.3e-05 | 1.163E-05 | 3.272E-06 | 7.900E-06 | 1.400E-05 |
| 16157 | 808 | Y808T | B | 1.1e-05 | 1e-05 | 1e-05 | 1.033E-05 | 5.774E-07 | 1.000E-05 | 1.100E-05 |
| 16158 | 808 | Y808W | B | 1e-05 | 1.7e-05 | 1.4e-05 | 1.367E-05 | 3.512E-06 | 1.000E-05 | 1.700E-05 |
| 16159 | 808 | Y808Y | B | 1.6e-05 | 1.5e-05 | 1.3e-05 | 1.467E-05 | 1.528E-06 | 1.300E-05 | 1.600E-05 |
| 16160 | 808 | Y808V | B | 1.2e-05 | 1e-05 | 1e-05 | 1.067E-05 | 1.155E-06 | 1.000E-05 | 1.200E-05 |
| 16161 | 809 | E809A | B | 1.5e-05 | 1.6e-05 | 8.7e-06 | 1.323E-05 | 3.958E-06 | 8.700E-06 | 1.600E-05 |
| 16162 | 809 | E809R | B | 1e-05 | 1.5e-05 | 9.5e-06 | 1.150E-05 | 3.041E-06 | 9.500E-06 | 1.500E-05 |
| 16163 | 809 | E809N | B | 1.8e-05 | 1.5e-05 | 1.1e-05 | 1.467E-05 | 3.512E-06 | 1.100E-05 | 1.800E-05 |
| 16164 | 809 | E809D | B | 1.5e-05 | 1.3e-05 | 1.1e-05 | 1.300E-05 | 2.000E-06 | 1.100E-05 | 1.500E-05 |
| 16165 | 809 | E809C | B | 1.5e-05 | 1.3e-05 | 1.6e-05 | 1.467E-05 | 1.528E-06 | 1.300E-05 | 1.600E-05 |
| 16166 | 809 | E809E | B | 1.5e-05 | 1.6e-05 | 9.2e-06 | 1.340E-05 | 3.672E-06 | 9.200E-06 | 1.600E-05 |
| 16167 | 809 | E809Q | B | 1.5e-05 | 8.4e-06 | 1.1e-05 | 1.147E-05 | 3.325E-06 | 8.400E-06 | 1.500E-05 |
| 16168 | 809 | E809G | B | 1.4e-05 | 1.6e-05 | 1.1e-05 | 1.367E-05 | 2.517E-06 | 1.100E-05 | 1.600E-05 |
| 16169 | 809 | E809H | B | 1.4e-05 | 9.1e-06 | 1.1e-05 | 1.137E-05 | 2.470E-06 | 9.100E-06 | 1.400E-05 |
| 16170 | 809 | E809I | B | 1.5e-05 | 1.5e-05 | 1.8e-05 | 1.600E-05 | 1.732E-06 | 1.500E-05 | 1.800E-05 |
| 16171 | 809 | E809L | B | 1e-05 | 1.6e-05 | 1.6e-05 | 1.400E-05 | 3.464E-06 | 1.000E-05 | 1.600E-05 |
| 16172 | 809 | E809K | B | 1.1e-05 | 1.1e-05 | 9.6e-06 | 1.053E-05 | 8.083E-07 | 9.600E-06 | 1.100E-05 |
| 16173 | 809 | E809M | B | 9.9e-06 | 1.1e-05 | 1.3e-05 | 1.130E-05 | 1.572E-06 | 9.900E-06 | 1.300E-05 |
| 16174 | 809 | E809F | B | 1.2e-05 | 1.6e-05 | 1.6e-05 | 1.467E-05 | 2.309E-06 | 1.200E-05 | 1.600E-05 |
| 16175 | 809 | E809P | B | 9.7e-06 | 1.1e-05 | 1.1e-05 | 1.057E-05 | 7.506E-07 | 9.700E-06 | 1.100E-05 |
| 16176 | 809 | E809S | B | 1.8e-05 | 1.5e-05 | 1.5e-05 | 1.600E-05 | 1.732E-06 | 1.500E-05 | 1.800E-05 |
| 16177 | 809 | E809T | B | 9.7e-06 | 1.4e-05 | 1e-05 | 1.123E-05 | 2.401E-06 | 9.700E-06 | 1.400E-05 |
| 16178 | 809 | E809W | B | 1.2e-05 | 1.6e-05 | 1.2e-05 | 1.333E-05 | 2.309E-06 | 1.200E-05 | 1.600E-05 |
| 16179 | 809 | E809Y | B | 1.5e-05 | 1.7e-05 | 1.5e-05 | 1.567E-05 | 1.155E-06 | 1.500E-05 | 1.700E-05 |
| 16180 | 809 | E809V | B | 1e-05 | 7.9e-06 | 8.9e-06 | 8.933E-06 | 1.050E-06 | 7.900E-06 | 1.000E-05 |
| 16181 | 810 | K810A | B | 1.2e-05 | 1e-05 | 1.4e-05 | 1.200E-05 | 2.000E-06 | 1.000E-05 | 1.400E-05 |
| 16182 | 810 | K810R | B | 1.4e-05 | 9.6e-06 | 1.8e-05 | 1.387E-05 | 4.202E-06 | 9.600E-06 | 1.800E-05 |
| 16183 | 810 | K810N | B | 1.2e-05 | 1.5e-05 | 1.6e-05 | 1.433E-05 | 2.082E-06 | 1.200E-05 | 1.600E-05 |
| 16184 | 810 | K810D | B | 1.6e-05 | 1.5e-05 | 1e-05 | 1.367E-05 | 3.215E-06 | 1.000E-05 | 1.600E-05 |
| 16185 | 810 | K810C | B | 1.1e-05 | 1.6e-05 | 8.4e-06 | 1.180E-05 | 3.863E-06 | 8.400E-06 | 1.600E-05 |
| 16186 | 810 | K810E | B | 9.9e-06 | 1.2e-05 | 1.5e-05 | 1.230E-05 | 2.563E-06 | 9.900E-06 | 1.500E-05 |
| 16187 | 810 | K810Q | B | 8.7e-06 | 1.5e-05 | 1.2e-05 | 1.190E-05 | 3.151E-06 | 8.700E-06 | 1.500E-05 |
| 16188 | 810 | K810G | B | 1e-05 | 1.5e-05 | 1.5e-05 | 1.333E-05 | 2.887E-06 | 1.000E-05 | 1.500E-05 |
| 16189 | 810 | K810H | B | 1.5e-05 | 1.9e-05 | 1e-05 | 1.467E-05 | 4.509E-06 | 1.000E-05 | 1.900E-05 |
| 16190 | 810 | K810I | B | 9e-06 | 1.5e-05 | 1.6e-05 | 1.333E-05 | 3.786E-06 | 9.000E-06 | 1.600E-05 |
| 16191 | 810 | K810L | B | 1.1e-05 | 1e-05 | 1.1e-05 | 1.067E-05 | 5.774E-07 | 1.000E-05 | 1.100E-05 |
| 16192 | 810 | K810K | B | 6.2e-06 | 1.2e-05 | 1.9e-05 | 1.240E-05 | 6.409E-06 | 6.200E-06 | 1.900E-05 |
| 16193 | 810 | K810M | B | 1e-05 | 1.3e-05 | 8.9e-06 | 1.063E-05 | 2.122E-06 | 8.900E-06 | 1.300E-05 |
| 16194 | 810 | K810F | B | 1.6e-05 | 1.2e-05 | 1.2e-05 | 1.333E-05 | 2.309E-06 | 1.200E-05 | 1.600E-05 |
| 16195 | 810 | K810P | B | 1.4e-05 | 1.1e-05 | 9.1e-06 | 1.137E-05 | 2.470E-06 | 9.100E-06 | 1.400E-05 |
| 16196 | 810 | K810S | B | 1.4e-05 | 8.9e-06 | 1.4e-05 | 1.230E-05 | 2.944E-06 | 8.900E-06 | 1.400E-05 |
| 16197 | 810 | K810T | B | 1.6e-05 | 1.3e-05 | 6.8e-06 | 1.193E-05 | 4.692E-06 | 6.800E-06 | 1.600E-05 |
| 16198 | 810 | K810W | B | 1.6e-05 | 1.4e-05 | 9.9e-06 | 1.330E-05 | 3.110E-06 | 9.900E-06 | 1.600E-05 |
| 16199 | 810 | K810Y | B | 1.6e-05 | 3.8e-05 | 9e-06 | 2.100E-05 | 1.513E-05 | 9.000E-06 | 3.800E-05 |
| 16200 | 810 | K810V | B | 1.5e-05 | 9.8e-06 | 1.4e-05 | 1.293E-05 | 2.759E-06 | 9.800E-06 | 1.500E-05 |
| 16201 | 811 | H811A | B | 1e-05 | 1.5e-05 | 1.5e-05 | 1.333E-05 | 2.887E-06 | 1.000E-05 | 1.500E-05 |
| 16202 | 811 | H811R | B | 8.3e-06 | 1.6e-05 | 1.3e-05 | 1.243E-05 | 3.881E-06 | 8.300E-06 | 1.600E-05 |
| 16203 | 811 | H811N | B | 1.1e-05 | 1.4e-05 | 1.1e-05 | 1.200E-05 | 1.732E-06 | 1.100E-05 | 1.400E-05 |
| 16204 | 811 | H811D | B | 1.5e-05 | 1e-05 | 1.1e-05 | 1.200E-05 | 2.646E-06 | 1.000E-05 | 1.500E-05 |
| 16205 | 811 | H811C | B | 8.8e-06 | 1.6e-05 | 1.4e-05 | 1.293E-05 | 3.717E-06 | 8.800E-06 | 1.600E-05 |
| 16206 | 811 | H811E | B | 9.9e-06 | 1.6e-05 | 1.7e-05 | 1.430E-05 | 3.843E-06 | 9.900E-06 | 1.700E-05 |
| 16207 | 811 | H811Q | B | 1.6e-05 | 1.3e-05 | 8.5e-06 | 1.250E-05 | 3.775E-06 | 8.500E-06 | 1.600E-05 |
| 16208 | 811 | H811G | B | 9.6e-06 | 9.8e-06 | 1.6e-05 | 1.180E-05 | 3.639E-06 | 9.600E-06 | 1.600E-05 |
| 16209 | 811 | H811H | B | 8.6e-06 | 1.5e-05 | 1.4e-05 | 1.253E-05 | 3.443E-06 | 8.600E-06 | 1.500E-05 |
| 16210 | 811 | H811I | B | 1.5e-05 | 1.5e-05 | 1.2e-05 | 1.400E-05 | 1.732E-06 | 1.200E-05 | 1.500E-05 |
| 16211 | 811 | H811L | B | 1.6e-05 | 1.5e-05 | 1.4e-05 | 1.500E-05 | 1.000E-06 | 1.400E-05 | 1.600E-05 |
| 16212 | 811 | H811K | B | 2e-05 | 1.1e-05 | 1.4e-05 | 1.500E-05 | 4.583E-06 | 1.100E-05 | 2.000E-05 |
| 16213 | 811 | H811M | B | 8.9e-06 | 1.5e-05 | 9.1e-06 | 1.100E-05 | 3.466E-06 | 8.900E-06 | 1.500E-05 |
| 16214 | 811 | H811F | B | 1.2e-05 | 1.4e-05 | 1.6e-05 | 1.400E-05 | 2.000E-06 | 1.200E-05 | 1.600E-05 |
| 16215 | 811 | H811P | B | 1.2e-05 | 1.5e-05 | 9.4e-06 | 1.213E-05 | 2.802E-06 | 9.400E-06 | 1.500E-05 |
| 16216 | 811 | H811S | B | 1.1e-05 | 1.4e-05 | 1.1e-05 | 1.200E-05 | 1.732E-06 | 1.100E-05 | 1.400E-05 |
| 16217 | 811 | H811T | B | 1.6e-05 | 1e-05 | 1.1e-05 | 1.233E-05 | 3.215E-06 | 1.000E-05 | 1.600E-05 |
| 16218 | 811 | H811W | B | 1.1e-05 | 1.1e-05 | 2.2e-05 | 1.467E-05 | 6.351E-06 | 1.100E-05 | 2.200E-05 |
| 16219 | 811 | H811Y | B | 1.6e-05 | 9.8e-06 | 1.5e-05 | 1.360E-05 | 3.329E-06 | 9.800E-06 | 1.600E-05 |
| 16220 | 811 | H811V | B | 1.6e-05 | 1.3e-05 | 1.7e-05 | 1.533E-05 | 2.082E-06 | 1.300E-05 | 1.700E-05 |
| 16221 | 812 | K812A | B | 1.4e-05 | 1.6e-05 | 1.6e-05 | 1.533E-05 | 1.155E-06 | 1.400E-05 | 1.600E-05 |
| 16222 | 812 | K812R | B | 1e-05 | 1.6e-05 | 1.6e-05 | 1.400E-05 | 3.464E-06 | 1.000E-05 | 1.600E-05 |
| 16223 | 812 | K812N | B | 8.9e-06 | 6.8e-06 | 1.4e-05 | 9.900E-06 | 3.703E-06 | 6.800E-06 | 1.400E-05 |
| 16224 | 812 | K812D | B | 1e-05 | 1.1e-05 | 1.4e-05 | 1.167E-05 | 2.082E-06 | 1.000E-05 | 1.400E-05 |
| 16225 | 812 | K812C | B | 1.2e-05 | 1.5e-05 | 1.7e-05 | 1.467E-05 | 2.517E-06 | 1.200E-05 | 1.700E-05 |
| 16226 | 812 | K812E | B | 1.5e-05 | 9.9e-06 | 1.6e-05 | 1.363E-05 | 3.272E-06 | 9.900E-06 | 1.600E-05 |
| 16227 | 812 | K812Q | B | 1.4e-05 | 8.6e-06 | 1.3e-05 | 1.187E-05 | 2.873E-06 | 8.600E-06 | 1.400E-05 |
| 16228 | 812 | K812G | B | 1.1e-05 | 1.5e-05 | 1.5e-05 | 1.367E-05 | 2.309E-06 | 1.100E-05 | 1.500E-05 |
| 16229 | 812 | K812H | B | 1.1e-05 | 1.2e-05 | 1.5e-05 | 1.267E-05 | 2.082E-06 | 1.100E-05 | 1.500E-05 |
| 16230 | 812 | K812I | B | 1.6e-05 | 8.5e-06 | 1.2e-05 | 1.217E-05 | 3.753E-06 | 8.500E-06 | 1.600E-05 |
| 16231 | 812 | K812L | B | 9e-06 | 1.3e-05 | 1e-05 | 1.067E-05 | 2.082E-06 | 9.000E-06 | 1.300E-05 |
| 16232 | 812 | K812K | B | 1.5e-05 | 1.4e-05 | 1.7e-05 | 1.533E-05 | 1.528E-06 | 1.400E-05 | 1.700E-05 |
| 16233 | 812 | K812M | B | 1.6e-05 | 1e-05 | 1.5e-05 | 1.367E-05 | 3.215E-06 | 1.000E-05 | 1.600E-05 |
| 16234 | 812 | K812F | B | 1e-05 | 1.3e-05 | 1.4e-05 | 1.233E-05 | 2.082E-06 | 1.000E-05 | 1.400E-05 |

| | | | | | | | | | | |
|---|---|---|---|---|---|---|---|---|---|---|
| 16235 | 812 | K812P | B | 1.7e-05 | 1.3e-05 | 1.5e-05 | 1.500E-05 | 2.000E-06 | 1.300E-05 | 1.700E-05 |
| 16236 | 812 | K812S | B | 1.3e-05 | 1.5e-05 | 8.4e-06 | 1.213E-05 | 3.384E-06 | 8.400E-06 | 1.500E-05 |
| 16237 | 812 | K812T | B | 1.6e-05 | 9.6e-06 | 1.2e-05 | 1.253E-05 | 3.233E-06 | 9.600E-06 | 1.600E-05 |
| 16238 | 812 | K812W | B | 1.3e-05 | 1.5e-05 | 1.4e-05 | 1.400E-05 | 1.000E-06 | 1.300E-05 | 1.500E-05 |
| 16239 | 812 | K812Y | B | 1.2e-05 | 1.6e-05 | 1.3e-05 | 1.367E-05 | 2.082E-06 | 1.200E-05 | 1.600E-05 |
| 16240 | 812 | K812V | B | 9.2e-06 | 1.6e-05 | 1.3e-05 | 1.273E-05 | 3.408E-06 | 9.200E-06 | 1.600E-05 |
| 16241 | 813 | V813A | B | 1.4e-05 | 1.6e-05 | 1.3e-05 | 1.433E-05 | 1.528E-06 | 1.300E-05 | 1.600E-05 |
| 16242 | 813 | V813R | B | 1.4e-05 | 1.2e-05 | 1.5e-05 | 1.367E-05 | 1.528E-06 | 1.200E-05 | 1.500E-05 |
| 16243 | 813 | V813N | B | 1.3e-05 | 8.6e-06 | 1.2e-05 | 1.120E-05 | 2.307E-06 | 8.600E-06 | 1.300E-05 |
| 16244 | 813 | V813D | B | 1.1e-05 | 1.5e-05 | 1.5e-05 | 1.367E-05 | 2.309E-06 | 1.100E-05 | 1.500E-05 |
| 16245 | 813 | V813C | B | 1.5e-05 | 1.5e-05 | 1.6e-05 | 1.533E-05 | 5.774E-07 | 1.500E-05 | 1.600E-05 |
| 16246 | 813 | V813E | B | 1.6e-05 | 1.2e-05 | 1.2e-05 | 1.333E-05 | 2.309E-06 | 1.200E-05 | 1.600E-05 |
| 16247 | 813 | V813Q | B | 1.4e-05 | 1e-05 | 1.6e-05 | 1.333E-05 | 3.055E-06 | 1.000E-05 | 1.600E-05 |
| 16248 | 813 | V813G | B | 1.4e-05 | 1.3e-05 | 1.4e-05 | 1.367E-05 | 5.774E-07 | 1.300E-05 | 1.400E-05 |
| 16249 | 813 | V813H | B | 1.6e-05 | 1.3e-05 | 8.8e-06 | 1.260E-05 | 3.617E-06 | 8.800E-06 | 1.600E-05 |
| 16250 | 813 | V813I | B | 1.6e-05 | 1.6e-05 | 1e-05 | 1.400E-05 | 3.464E-06 | 1.000E-05 | 1.600E-05 |
| 16251 | 813 | V813L | B | 1.3e-05 | 8.2e-06 | 1.6e-05 | 1.240E-05 | 3.934E-06 | 8.200E-06 | 1.600E-05 |
| 16252 | 813 | V813K | B | 1.2e-05 | 1.4e-05 | 1.5e-05 | 1.367E-05 | 1.528E-06 | 1.200E-05 | 1.500E-05 |
| 16253 | 813 | V813M | B | 1.1e-05 | 1.7e-05 | 8e-06 | 1.200E-05 | 4.583E-06 | 8.000E-06 | 1.700E-05 |
| 16254 | 813 | V813F | B | 1.6e-05 | 1e-05 | 9.1e-06 | 1.170E-05 | 3.751E-06 | 9.100E-06 | 1.600E-05 |
| 16255 | 813 | V813P | B | 1.7e-05 | 1.4e-05 | 1.5e-05 | 1.533E-05 | 1.528E-06 | 1.400E-05 | 1.700E-05 |
| 16256 | 813 | V813S | B | 1e-05 | 6e-06 | 1e-05 | 8.667E-06 | 2.309E-06 | 6.000E-06 | 1.000E-05 |
| 16257 | 813 | V813T | B | 1.1e-05 | 1.7e-05 | 1e-05 | 1.267E-05 | 3.786E-06 | 1.000E-05 | 1.700E-05 |
| 16258 | 813 | V813W | B | 2e-05 | 1.4e-05 | 8.1e-06 | 1.403E-05 | 5.950E-06 | 8.100E-06 | 2.000E-05 |
| 16259 | 813 | V813Y | B | 1e-05 | 1.4e-05 | 1.6e-05 | 1.333E-05 | 3.055E-06 | 1.000E-05 | 1.600E-05 |
| 16260 | 813 | V813V | B | 8.9e-06 | 1.3e-05 | 1.5e-05 | 1.230E-05 | 3.110E-06 | 8.900E-06 | 1.500E-05 |
| 16261 | 814 | Y814A | B | 1.2e-05 | 1.4e-05 | 1.2e-05 | 1.267E-05 | 1.155E-06 | 1.200E-05 | 1.400E-05 |
| 16262 | 814 | Y814R | B | 1.7e-05 | 1.1e-05 | 1.5e-05 | 1.433E-05 | 3.055E-06 | 1.100E-05 | 1.700E-05 |
| 16263 | 814 | Y814N | B | 1.1e-05 | 1.2e-05 | 1.1e-05 | 1.133E-05 | 5.774E-07 | 1.100E-05 | 1.200E-05 |
| 16264 | 814 | Y814D | B | 1.5e-05 | 1.5e-05 | 1.1e-05 | 1.367E-05 | 2.309E-06 | 1.100E-05 | 1.500E-05 |
| 16265 | 814 | Y814C | B | 8.9e-06 | 1.4e-05 | 1.6e-05 | 1.297E-05 | 3.661E-06 | 8.900E-06 | 1.600E-05 |
| 16266 | 814 | Y814E | B | 1.5e-05 | 1e-05 | 1.5e-05 | 1.333E-05 | 2.887E-06 | 1.000E-05 | 1.500E-05 |
| 16267 | 814 | Y814Q | B | 8.3e-06 | 8.2e-06 | 1.6e-05 | 1.083E-05 | 4.475E-06 | 8.200E-06 | 1.600E-05 |
| 16268 | 814 | Y814G | B | 1.9e-05 | 1.5e-05 | 1e-05 | 1.467E-05 | 4.509E-06 | 1.000E-05 | 1.900E-05 |
| 16269 | 814 | Y814H | B | 1.3e-05 | 1e-05 | 6.4e-06 | 9.800E-06 | 3.305E-06 | 6.400E-06 | 1.300E-05 |
| 16270 | 814 | Y814I | B | 1.1e-05 | 1.2e-05 | 1.5e-05 | 1.267E-05 | 2.082E-06 | 1.100E-05 | 1.500E-05 |
| 16271 | 814 | Y814L | B | 8.9e-06 | 1.3e-05 | 1.1e-05 | 1.097E-05 | 2.050E-06 | 8.900E-06 | 1.300E-05 |
| 16272 | 814 | Y814K | B | 1.5e-05 | 1e-05 | 9.9e-06 | 1.163E-05 | 2.916E-06 | 9.900E-06 | 1.500E-05 |
| 16273 | 814 | Y814M | B | 1.6e-05 | 9.5e-06 | 8.9e-06 | 1.147E-05 | 3.937E-06 | 8.900E-06 | 1.600E-05 |
| 16274 | 814 | Y814F | B | 9.9e-06 | 1.6e-05 | 1e-05 | 1.197E-05 | 3.493E-06 | 9.900E-06 | 1.600E-05 |
| 16275 | 814 | Y814P | B | 8.1e-06 | 1.6e-05 | 1.6e-05 | 1.337E-05 | 4.561E-06 | 8.100E-06 | 1.600E-05 |
| 16276 | 814 | Y814S | B | 1.8e-05 | 7.3e-06 | 9.6e-06 | 1.163E-05 | 5.632E-06 | 7.300E-06 | 1.800E-05 |
| 16277 | 814 | Y814T | B | 9e-06 | 1.1e-05 | 1.6e-05 | 1.200E-05 | 3.606E-06 | 9.000E-06 | 1.600E-05 |
| 16278 | 814 | Y814W | B | 9.2e-06 | 1e-05 | 1.1e-05 | 1.007E-05 | 9.018E-07 | 9.200E-06 | 1.100E-05 |
| 16279 | 814 | Y814Y | B | 8.8e-06 | 1.4e-05 | 1.1e-05 | 1.127E-05 | 2.610E-06 | 8.800E-06 | 1.400E-05 |
| 16280 | 814 | Y814V | B | 1.1e-05 | 1.5e-05 | 1.5e-05 | 1.367E-05 | 2.309E-06 | 1.100E-05 | 1.500E-05 |
| 16281 | 815 | A815A | B | 1.2e-05 | 1.5e-05 | 1.6e-05 | 1.433E-05 | 2.082E-06 | 1.200E-05 | 1.600E-05 |
| 16282 | 815 | A815R | B | 1.7e-05 | 1.4e-05 | 9.7e-06 | 1.357E-05 | 3.669E-06 | 9.700E-06 | 1.700E-05 |
| 16283 | 815 | A815N | B | 9.9e-06 | 1.3e-05 | 1.3e-05 | 1.197E-05 | 1.790E-06 | 9.900E-06 | 1.300E-05 |
| 16284 | 815 | A815D | B | 1.1e-05 | 1.5e-05 | 1.7e-05 | 1.433E-05 | 3.055E-06 | 1.100E-05 | 1.700E-05 |
| 16285 | 815 | A815C | B | 1.4e-05 | 1.4e-05 | 1.6e-05 | 1.467E-05 | 1.155E-06 | 1.400E-05 | 1.600E-05 |
| 16286 | 815 | A815E | B | 1.5e-05 | 1.5e-05 | 1.5e-05 | 1.500E-05 | 0.000E+00 | 1.500E-05 | 1.500E-05 |
| 16287 | 815 | A815Q | B | 1.3e-05 | 1.2e-05 | 1.1e-05 | 1.200E-05 | 1.000E-06 | 1.100E-05 | 1.300E-05 |
| 16288 | 815 | A815G | B | 1.5e-05 | 1.1e-05 | 1.1e-05 | 1.233E-05 | 2.309E-06 | 1.100E-05 | 1.500E-05 |
| 16289 | 815 | A815H | B | 1.5e-05 | 1.3e-05 | 9.9e-06 | 1.263E-05 | 2.570E-06 | 9.900E-06 | 1.500E-05 |
| 16290 | 815 | A815I | B | 1.3e-05 | 1.1e-05 | 1.6e-05 | 1.333E-05 | 2.517E-06 | 1.100E-05 | 1.600E-05 |
| 16291 | 815 | A815L | B | 1.5e-05 | 1.4e-05 | 6e-06 | 1.167E-05 | 4.933E-06 | 6.000E-06 | 1.500E-05 |
| 16292 | 815 | A815K | B | 1.5e-05 | 1.7e-05 | 1.7e-05 | 1.633E-05 | 1.155E-06 | 1.500E-05 | 1.700E-05 |
| 16293 | 815 | A815M | B | 1.5e-05 | 1.5e-05 | 1.5e-05 | 1.500E-05 | 0.000E+00 | 1.500E-05 | 1.500E-05 |
| 16294 | 815 | A815F | B | 1.5e-05 | 1.5e-05 | 1.1e-05 | 1.367E-05 | 2.309E-06 | 1.100E-05 | 1.500E-05 |
| 16295 | 815 | A815P | B | 1.5e-05 | 1.6e-05 | 1e-05 | 1.367E-05 | 3.215E-06 | 1.000E-05 | 1.600E-05 |
| 16296 | 815 | A815S | B | 1.1e-05 | 1.5e-05 | 1.2e-05 | 1.267E-05 | 2.082E-06 | 1.100E-05 | 1.500E-05 |
| 16297 | 815 | A815T | B | 1.1e-05 | 1.3e-05 | 7.8e-06 | 1.060E-05 | 2.623E-06 | 7.800E-06 | 1.300E-05 |
| 16298 | 815 | A815W | B | 1.4e-05 | 1.1e-05 | 1e-05 | 1.167E-05 | 2.082E-06 | 1.000E-05 | 1.400E-05 |
| 16299 | 815 | A815Y | B | 1.4e-05 | 1e-05 | 1.3e-05 | 1.233E-05 | 2.082E-06 | 1.000E-05 | 1.400E-05 |
| 16300 | 815 | A815V | B | 1.1e-05 | 8.8e-06 | 1.6e-05 | 1.193E-05 | 3.690E-06 | 8.800E-06 | 1.600E-05 |
| 16301 | 816 | C816A | B | 1.4e-05 | 1.3e-05 | 1.5e-05 | 1.400E-05 | 1.000E-06 | 1.300E-05 | 1.500E-05 |
| 16302 | 816 | C816R | B | 1.5e-05 | 1.3e-05 | 9.7e-06 | 1.257E-05 | 2.676E-06 | 9.700E-06 | 1.500E-05 |
| 16303 | 816 | C816N | B | 1.2e-05 | 1.5e-05 | 1.5e-05 | 1.400E-05 | 1.732E-06 | 1.200E-05 | 1.500E-05 |
| 16304 | 816 | C816D | B | 8.2e-06 | 9.8e-06 | 8.5e-06 | 8.833E-06 | 8.505E-07 | 8.200E-06 | 9.800E-06 |
| 16305 | 816 | C816C | B | 2.1e-05 | 1.1e-05 | 1.6e-05 | 1.600E-05 | 5.000E-06 | 1.100E-05 | 2.100E-05 |
| 16306 | 816 | C816E | B | 1.5e-05 | 1.4e-05 | 1.5e-05 | 1.467E-05 | 5.774E-07 | 1.400E-05 | 1.500E-05 |
| 16307 | 816 | C816Q | B | 1e-05 | 8.7e-06 | 1.7e-05 | 1.190E-05 | 4.464E-06 | 8.700E-06 | 1.700E-05 |
| 16308 | 816 | C816G | B | 8.9e-06 | 1.5e-05 | 1.6e-05 | 1.330E-05 | 3.843E-06 | 8.900E-06 | 1.600E-05 |
| 16309 | 816 | C816H | B | 1.3e-05 | 1.7e-05 | 9.9e-06 | 1.330E-05 | 3.559E-06 | 9.900E-06 | 1.700E-05 |
| 16310 | 816 | C816I | B | 1.3e-05 | 1.4e-05 | 1.6e-05 | 1.433E-05 | 1.528E-06 | 1.300E-05 | 1.600E-05 |
| 16311 | 816 | C816L | B | 1e-05 | 1.4e-05 | 1.1e-05 | 1.167E-05 | 2.082E-06 | 1.000E-05 | 1.400E-05 |
| 16312 | 816 | C816K | B | 1.5e-05 | 1.9e-05 | 1.5e-05 | 1.633E-05 | 2.309E-06 | 1.500E-05 | 1.900E-05 |
| 16313 | 816 | C816M | B | 1.5e-05 | 1.6e-05 | 1.2e-05 | 1.433E-05 | 2.082E-06 | 1.200E-05 | 1.600E-05 |
| 16314 | 816 | C816F | B | 9.6e-06 | 1.5e-05 | 1.5e-05 | 1.320E-05 | 3.118E-06 | 9.600E-06 | 1.500E-05 |
| 16315 | 816 | C816P | B | 9.8e-06 | 1.4e-05 | 1.4e-05 | 1.260E-05 | 2.425E-06 | 9.800E-06 | 1.400E-05 |
| 16316 | 816 | C816S | B | 1.4e-05 | 1.1e-05 | 1.2e-05 | 1.233E-05 | 1.528E-06 | 1.100E-05 | 1.400E-05 |
| 16317 | 816 | C816T | B | 1.4e-05 | 1e-05 | 7.4e-06 | 1.047E-05 | 3.325E-06 | 7.400E-06 | 1.400E-05 |
| 16318 | 816 | C816W | B | 1.2e-05 | 1.3e-05 | 1.7e-05 | 1.400E-05 | 2.646E-06 | 1.200E-05 | 1.700E-05 |
| 16319 | 816 | C816Y | B | 1.3e-05 | 1.5e-05 | 1e-05 | 1.267E-05 | 2.517E-06 | 1.000E-05 | 1.500E-05 |

| | | | | | | | | | | |
|---|---|---|---|---|---|---|---|---|---|---|
| 16320 | 816 | C816V | B | 1.6e-05 | 9.9e-06 | 9.2e-06 | 1.170E-05 | 3.740E-06 | 9.200E-06 | 1.600E-05 |
| 16321 | 817 | E817A | B | 1.4e-05 | 1.6e-05 | 1.4e-05 | 1.467E-05 | 1.155E-06 | 1.400E-05 | 1.600E-05 |
| 16322 | 817 | E817R | B | 1.5e-05 | 1.5e-05 | 1.5e-05 | 1.500E-05 | 0.000E+00 | 1.500E-05 | 1.500E-05 |
| 16323 | 817 | E817N | B | 1.4e-05 | 1.1e-05 | 1e-05 | 1.167E-05 | 2.082E-06 | 1.000E-05 | 1.400E-05 |
| 16324 | 817 | E817D | B | 1.3e-05 | 1.1e-05 | 1.3e-05 | 1.233E-05 | 1.155E-06 | 1.100E-05 | 1.300E-05 |
| 16325 | 817 | E817C | B | 1.4e-05 | 1.6e-05 | 1.1e-05 | 1.367E-05 | 2.517E-06 | 1.100E-05 | 1.600E-05 |
| 16326 | 817 | E817E | B | 8.3e-06 | 1.3e-05 | 1.2e-05 | 1.110E-05 | 2.476E-06 | 8.300E-06 | 1.300E-05 |
| 16327 | 817 | E817Q | B | 8.9e-06 | 1e-05 | 1.7e-05 | 1.197E-05 | 4.394E-06 | 8.900E-06 | 1.700E-05 |
| 16328 | 817 | E817G | B | 9.9e-06 | 1.3e-05 | 1.1e-05 | 1.130E-05 | 1.572E-06 | 9.900E-06 | 1.300E-05 |
| 16329 | 817 | E817H | B | 1.1e-05 | 1.4e-05 | 1.4e-05 | 1.300E-05 | 1.732E-06 | 1.100E-05 | 1.400E-05 |
| 16330 | 817 | E817I | B | 1.2e-05 | 7.9e-06 | 1.4e-05 | 1.130E-05 | 3.110E-06 | 7.900E-06 | 1.400E-05 |
| 16331 | 817 | E817L | B | 1.6e-05 | 1.6e-05 | 1.5e-05 | 1.567E-05 | 5.774E-07 | 1.500E-05 | 1.600E-05 |
| 16332 | 817 | E817K | B | 1.4e-05 | 8.6e-06 | 1.6e-05 | 1.287E-05 | 3.828E-06 | 8.600E-06 | 1.600E-05 |
| 16333 | 817 | E817M | B | 1.6e-05 | 1.5e-05 | 9.1e-06 | 1.337E-05 | 3.729E-06 | 9.100E-06 | 1.600E-05 |
| 16334 | 817 | E817F | B | 1.6e-05 | 1.5e-05 | 9.7e-06 | 1.357E-05 | 3.386E-06 | 9.700E-06 | 1.600E-05 |
| 16335 | 817 | E817P | B | 9.5e-06 | 1.6e-05 | 8.6e-06 | 1.137E-05 | 4.038E-06 | 8.600E-06 | 1.600E-05 |
| 16336 | 817 | E817S | B | 1.5e-05 | 9.7e-06 | 1.7e-05 | 1.390E-05 | 3.772E-06 | 9.700E-06 | 1.700E-05 |
| 16337 | 817 | E817T | B | 1.4e-05 | 1.3e-05 | 1.1e-05 | 1.267E-05 | 1.528E-06 | 1.100E-05 | 1.400E-05 |
| 16338 | 817 | E817W | B | 9.7e-06 | 1.4e-05 | 8.9e-06 | 1.087E-05 | 2.743E-06 | 8.900E-06 | 1.400E-05 |
| 16339 | 817 | E817Y | B | 1.7e-05 | 9.8e-06 | 1e-05 | 1.227E-05 | 4.100E-06 | 9.800E-06 | 1.700E-05 |
| 16340 | 817 | E817V | B | 1.1e-05 | 1.4e-05 | 1.3e-05 | 1.267E-05 | 1.528E-06 | 1.100E-05 | 1.400E-05 |
| 16341 | 818 | V818A | B | 8.1e-06 | 7e-06 | 1.5e-05 | 1.003E-05 | 4.336E-06 | 7.000E-06 | 1.500E-05 |
| 16342 | 818 | V818R | B | 9.7e-06 | 1.3e-05 | 7.7e-06 | 1.013E-05 | 2.676E-06 | 7.700E-06 | 1.300E-05 |
| 16343 | 818 | V818N | B | 1.6e-05 | 1.4e-05 | 1.3e-05 | 1.433E-05 | 1.528E-06 | 1.300E-05 | 1.600E-05 |
| 16344 | 818 | V818D | B | 1e-05 | 1.7e-05 | 2.3e-05 | 1.667E-05 | 6.506E-06 | 1.000E-05 | 2.300E-05 |
| 16345 | 818 | V818C | B | 1.3e-05 | 1.7e-05 | 2.6e-05 | 1.867E-05 | 6.658E-06 | 1.300E-05 | 2.600E-05 |
| 16346 | 818 | V818E | B | 1.1e-05 | 6.5e-06 | 1.8e-05 | 1.183E-05 | 5.795E-06 | 6.500E-06 | 1.800E-05 |
| 16347 | 818 | V818Q | B | 1.4e-05 | 1.1e-05 | 1.5e-05 | 1.333E-05 | 2.082E-06 | 1.100E-05 | 1.500E-05 |
| 16348 | 818 | V818G | B | 1e-05 | 1.4e-05 | 1.6e-05 | 1.333E-05 | 3.055E-06 | 1.000E-05 | 1.600E-05 |
| 16349 | 818 | V818H | B | 1.4e-05 | 1.5e-05 | 1.5e-05 | 1.467E-05 | 5.774E-07 | 1.400E-05 | 1.500E-05 |
| 16350 | 818 | V818I | B | 1.6e-05 | 1.3e-05 | 1.6e-05 | 1.500E-05 | 1.732E-06 | 1.300E-05 | 1.600E-05 |
| 16351 | 818 | V818L | B | 9.9e-06 | 1.6e-05 | 8.4e-06 | 1.143E-05 | 4.025E-06 | 8.400E-06 | 1.600E-05 |
| 16352 | 818 | V818K | B | 1.5e-05 | 1.1e-05 | 6.9e-06 | 1.097E-05 | 4.050E-06 | 6.900E-06 | 1.500E-05 |
| 16353 | 818 | V818M | B | 1.7e-05 | 1.6e-05 | 9.8e-06 | 1.427E-05 | 3.900E-06 | 9.800E-06 | 1.700E-05 |
| 16354 | 818 | V818F | B | 1.5e-05 | 1.1e-05 | 1.3e-05 | 1.300E-05 | 2.000E-06 | 1.100E-05 | 1.500E-05 |
| 16355 | 818 | V818P | B | 1.5e-05 | 1.4e-05 | 1.5e-05 | 1.467E-05 | 5.774E-07 | 1.400E-05 | 1.500E-05 |
| 16356 | 818 | V818S | B | 8.7e-06 | 1.1e-05 | 1.1e-05 | 1.023E-05 | 1.328E-06 | 8.700E-06 | 1.100E-05 |
| 16357 | 818 | V818T | B | 1.4e-05 | 1.6e-05 | 1.1e-05 | 1.367E-05 | 2.517E-06 | 1.100E-05 | 1.600E-05 |
| 16358 | 818 | V818W | B | 1.5e-05 | 1.4e-05 | 1.5e-05 | 1.467E-05 | 5.774E-07 | 1.400E-05 | 1.500E-05 |
| 16359 | 818 | V818Y | B | 9.7e-06 | 1.1e-05 | 1.5e-05 | 1.190E-05 | 2.762E-06 | 9.700E-06 | 1.500E-05 |
| 16360 | 818 | V818V | B | 1.5e-05 | 1.5e-05 | 1.5e-05 | 1.500E-05 | 0.000E+00 | 1.500E-05 | 1.500E-05 |
| 16361 | 819 | T819A | B | 1.6e-05 | 1.4e-05 | 1.6e-05 | 1.533E-05 | 1.155E-06 | 1.400E-05 | 1.600E-05 |
| 16362 | 819 | T819R | B | 1.4e-05 | 1.5e-05 | 1e-05 | 1.300E-05 | 2.646E-06 | 1.000E-05 | 1.500E-05 |
| 16363 | 819 | T819N | B | 1.5e-05 | 8.6e-06 | 1.5e-05 | 1.287E-05 | 3.695E-06 | 8.600E-06 | 1.500E-05 |
| 16364 | 819 | T819D | B | 1.1e-05 | 1.4e-05 | 1.5e-05 | 1.333E-05 | 2.082E-06 | 1.100E-05 | 1.500E-05 |
| 16365 | 819 | T819C | B | 1.4e-05 | 1.1e-05 | 1e-05 | 1.167E-05 | 2.082E-06 | 1.000E-05 | 1.400E-05 |
| 16366 | 819 | T819E | B | 9.9e-06 | 1.6e-05 | 1.6e-05 | 1.397E-05 | 3.522E-06 | 9.900E-06 | 1.600E-05 |
| 16367 | 819 | T819Q | B | 1e-05 | 1.6e-05 | 1.7e-05 | 1.433E-05 | 3.786E-06 | 1.000E-05 | 1.700E-05 |
| 16368 | 819 | T819G | B | 1.3e-05 | 1.1e-05 | 1.9e-05 | 1.433E-05 | 4.163E-06 | 1.100E-05 | 1.900E-05 |
| 16369 | 819 | T819H | B | 1.6e-05 | 9.9e-06 | 1.1e-05 | 1.230E-05 | 3.251E-06 | 9.900E-06 | 1.600E-05 |
| 16370 | 819 | T819I | B | 1.1e-05 | 1.7e-05 | 8.9e-06 | 1.230E-05 | 4.204E-06 | 8.900E-06 | 1.700E-05 |
| 16371 | 819 | T819L | B | 1.5e-05 | 1.1e-05 | 1.1e-05 | 1.233E-05 | 2.309E-06 | 1.100E-05 | 1.500E-05 |
| 16372 | 819 | T819K | B | 7.6e-06 | 1e-05 | 1.2e-05 | 9.867E-06 | 2.203E-06 | 7.600E-06 | 1.200E-05 |
| 16373 | 819 | T819M | B | 1.5e-05 | 1.5e-05 | 1e-05 | 1.333E-05 | 2.887E-06 | 1.000E-05 | 1.500E-05 |
| 16374 | 819 | T819F | B | 1.6e-05 | 1.4e-05 | 9.5e-06 | 1.317E-05 | 3.329E-06 | 9.500E-06 | 1.600E-05 |
| 16375 | 819 | T819P | B | 8.1e-06 | 8.8e-06 | 1.2e-05 | 9.633E-06 | 2.079E-06 | 8.100E-06 | 1.200E-05 |
| 16376 | 819 | T819S | B | 1.2e-05 | 9.6e-06 | 1.5e-05 | 1.220E-05 | 2.706E-06 | 9.600E-06 | 1.500E-05 |
| 16377 | 819 | T819T | B | 1.2e-05 | 9e-06 | 9.4e-06 | 1.013E-05 | 1.629E-06 | 9.000E-06 | 1.200E-05 |
| 16378 | 819 | T819W | B | 1e-05 | 1.5e-05 | 1.4e-05 | 1.300E-05 | 2.646E-06 | 1.000E-05 | 1.500E-05 |
| 16379 | 819 | T819Y | B | 1e-05 | 9.6e-06 | 1.1e-05 | 1.020E-05 | 7.211E-07 | 9.600E-06 | 1.100E-05 |
| 16380 | 819 | T819V | B | 1.6e-05 | 1.1e-05 | 1.7e-05 | 1.467E-05 | 3.215E-06 | 1.100E-05 | 1.700E-05 |
| 16381 | 820 | H820A | B | 9.3e-06 | 1e-05 | 1.1e-05 | 1.010E-05 | 8.544E-07 | 9.300E-06 | 1.100E-05 |
| 16382 | 820 | H820R | B | 1e-05 | 1.2e-05 | 1.6e-05 | 1.267E-05 | 3.055E-06 | 1.000E-05 | 1.600E-05 |
| 16383 | 820 | H820N | B | 1.3e-05 | 1.5e-05 | 1.7e-05 | 1.500E-05 | 2.000E-06 | 1.300E-05 | 1.700E-05 |
| 16384 | 820 | H820D | B | 1.7e-05 | 1.4e-05 | 1.3e-05 | 1.467E-05 | 2.082E-06 | 1.300E-05 | 1.700E-05 |
| 16385 | 820 | H820C | B | 1.5e-05 | 2.8e-05 | 1.3e-05 | 1.867E-05 | 8.145E-06 | 1.300E-05 | 2.800E-05 |
| 16386 | 820 | H820E | B | 9.9e-06 | 1.1e-05 | 1.3e-05 | 1.130E-05 | 1.572E-06 | 9.900E-06 | 1.300E-05 |
| 16387 | 820 | H820Q | B | 9.2e-06 | 1e-05 | 1.1e-05 | 1.007E-05 | 9.018E-07 | 9.200E-06 | 1.100E-05 |
| 16388 | 820 | H820G | B | 1.4e-05 | 1.2e-05 | 1.1e-05 | 1.233E-05 | 1.528E-06 | 1.100E-05 | 1.400E-05 |
| 16389 | 820 | H820H | B | 1.2e-05 | 1.6e-05 | 1.4e-05 | 1.400E-05 | 2.000E-06 | 1.200E-05 | 1.600E-05 |
| 16390 | 820 | H820I | B | 1.5e-05 | 1.2e-05 | 1.7e-05 | 1.467E-05 | 2.517E-06 | 1.200E-05 | 1.700E-05 |
| 16391 | 820 | H820L | B | 9.6e-06 | 1e-05 | 1e-05 | 9.867E-06 | 2.309E-07 | 9.600E-06 | 1.000E-05 |
| 16392 | 820 | H820K | B | 1.6e-05 | 1.5e-05 | 1.8e-05 | 1.633E-05 | 1.528E-06 | 1.500E-05 | 1.800E-05 |
| 16393 | 820 | H820M | B | 1.1e-05 | 1.4e-05 | 9.3e-06 | 1.143E-05 | 2.380E-06 | 9.300E-06 | 1.400E-05 |
| 16394 | 820 | H820F | B | 1.5e-05 | 9.2e-06 | 1.8e-05 | 1.407E-05 | 4.474E-06 | 9.200E-06 | 1.800E-05 |
| 16395 | 820 | H820P | B | 1.3e-05 | 1.1e-05 | 1.2e-05 | 1.200E-05 | 1.000E-06 | 1.100E-05 | 1.300E-05 |
| 16396 | 820 | H820S | B | 1e-05 | 1.5e-05 | 1.1e-05 | 1.200E-05 | 2.646E-06 | 1.000E-05 | 1.500E-05 |
| 16397 | 820 | H820T | B | 1.7e-05 | 1.3e-05 | 1.2e-05 | 1.400E-05 | 2.646E-06 | 1.200E-05 | 1.700E-05 |
| 16398 | 820 | H820W | B | 7.4e-06 | 1.6e-05 | 1.5e-05 | 1.280E-05 | 4.703E-06 | 7.400E-06 | 1.600E-05 |
| 16399 | 820 | H820Y | B | 1.3e-05 | 1.6e-05 | 9.3e-06 | 1.277E-05 | 3.356E-06 | 9.300E-06 | 1.600E-05 |
| 16400 | 820 | H820V | B | 1.5e-05 | 1.3e-05 | 1.5e-05 | 1.433E-05 | 1.155E-06 | 1.300E-05 | 1.500E-05 |
| 16401 | 821 | Q821A | B | 1.7e-05 | 1e-05 | 1.6e-05 | 1.433E-05 | 3.786E-06 | 1.000E-05 | 1.700E-05 |
| 16402 | 821 | Q821R | B | 9.7e-06 | 1.6e-05 | 1.5e-05 | 1.357E-05 | 3.386E-06 | 9.700E-06 | 1.600E-05 |
| 16403 | 821 | Q821N | B | 1.1e-05 | 7.5e-06 | 1.6e-05 | 1.150E-05 | 4.272E-06 | 7.500E-06 | 1.600E-05 |
| 16404 | 821 | Q821D | B | 1.2e-05 | 9.9e-06 | 1.1e-05 | 1.097E-05 | 1.050E-06 | 9.900E-06 | 1.200E-05 |

| | | | | | | | | | | |
|---|---|---|---|---|---|---|---|---|---|---|
| 16405 | 821 | Q821C | B | 1.5e-05 | 9.3e-06 | 1.6e-05 | 1.343E-05 | 3.614E-06 | 9.300E-06 | 1.600E-05 |
| 16406 | 821 | Q821E | B | 9.7e-06 | 1.5e-05 | 1.5e-05 | 1.323E-05 | 3.060E-06 | 9.700E-06 | 1.500E-05 |
| 16407 | 821 | Q821Q | B | 1.7e-05 | 1.5e-05 | 1.1e-05 | 1.433E-05 | 3.055E-06 | 1.100E-05 | 1.700E-05 |
| 16408 | 821 | Q821G | B | 1.3e-05 | 1e-05 | 1.1e-05 | 1.133E-05 | 1.528E-06 | 1.000E-05 | 1.300E-05 |
| 16409 | 821 | Q821H | B | 1.3e-05 | 1.9e-05 | 1.5e-05 | 1.567E-05 | 3.055E-06 | 1.300E-05 | 1.900E-05 |
| 16410 | 821 | Q821I | B | 1.6e-05 | 1.3e-05 | 1.3e-05 | 1.400E-05 | 1.732E-06 | 1.300E-05 | 1.600E-05 |
| 16411 | 821 | Q821L | B | 1.7e-05 | 1.6e-05 | 1.1e-05 | 1.467E-05 | 3.215E-06 | 1.100E-05 | 1.700E-05 |
| 16412 | 821 | Q821K | B | 1e-05 | 9.3e-06 | 8.9e-06 | 9.400E-06 | 5.568E-07 | 8.900E-06 | 1.000E-05 |
| 16413 | 821 | Q821M | B | 1.6e-05 | 1.9e-05 | 1.4e-05 | 1.633E-05 | 2.517E-06 | 1.400E-05 | 1.900E-05 |
| 16414 | 821 | Q821F | B | 1.1e-05 | 8.4e-06 | 1e-05 | 9.800E-06 | 1.311E-06 | 8.400E-06 | 1.100E-05 |
| 16415 | 821 | Q821P | B | 1.1e-05 | 1.4e-05 | 1.6e-05 | 1.367E-05 | 2.517E-06 | 1.100E-05 | 1.600E-05 |
| 16416 | 821 | Q821S | B | 1.5e-05 | 1.2e-05 | 1.2e-05 | 1.300E-05 | 1.732E-06 | 1.200E-05 | 1.500E-05 |
| 16417 | 821 | Q821T | B | 1.1e-05 | 1.5e-05 | 1.4e-05 | 1.333E-05 | 2.082E-06 | 1.100E-05 | 1.500E-05 |
| 16418 | 821 | Q821W | B | 9.8e-06 | 1.2e-05 | 1.1e-05 | 1.093E-05 | 1.102E-06 | 9.800E-06 | 1.200E-05 |
| 16419 | 821 | Q821Y | B | 1.1e-05 | 1.2e-05 | 7.1e-06 | 1.003E-05 | 2.589E-06 | 7.100E-06 | 1.200E-05 |
| 16420 | 821 | Q821V | B | 1e-05 | 1.7e-05 | 1.6e-05 | 1.433E-05 | 3.786E-06 | 1.000E-05 | 1.700E-05 |
| 16421 | 822 | G822A | B | 1.5e-05 | 1.6e-05 | 1e-05 | 1.367E-05 | 3.215E-06 | 1.000E-05 | 1.600E-05 |
| 16422 | 822 | G822R | B | 1.4e-05 | 9.7e-06 | 1.4e-05 | 1.257E-05 | 2.483E-06 | 9.700E-06 | 1.400E-05 |
| 16423 | 822 | G822N | B | 1.2e-05 | 1e-05 | 8e-06 | 1.000E-05 | 2.000E-06 | 8.000E-06 | 1.200E-05 |
| 16424 | 822 | G822D | B | 1.2e-05 | 1.4e-05 | 9.3e-06 | 1.177E-05 | 2.359E-06 | 9.300E-06 | 1.400E-05 |
| 16425 | 822 | G822C | B | 1.2e-05 | 1.5e-05 | 1.6e-05 | 1.433E-05 | 2.082E-06 | 1.200E-05 | 1.600E-05 |
| 16426 | 822 | G822E | B | 1e-05 | 1e-05 | 1.6e-05 | 1.200E-05 | 3.464E-06 | 1.000E-05 | 1.600E-05 |
| 16427 | 822 | G822Q | B | 1.2e-05 | 1.5e-05 | 9.2e-06 | 1.207E-05 | 2.901E-06 | 9.200E-06 | 1.500E-05 |
| 16428 | 822 | G822G | B | 1e-05 | 1.1e-05 | 1.4e-05 | 1.167E-05 | 2.082E-06 | 1.000E-05 | 1.400E-05 |
| 16429 | 822 | G822H | B | 8.9e-06 | 1.4e-05 | 1.5e-05 | 1.263E-05 | 3.272E-06 | 8.900E-06 | 1.500E-05 |
| 16430 | 822 | G822I | B | 1.1e-05 | 1.6e-05 | 1.3e-05 | 1.333E-05 | 2.517E-06 | 1.100E-05 | 1.600E-05 |
| 16431 | 822 | G822L | B | 1.2e-05 | 1.1e-05 | 1.4e-05 | 1.233E-05 | 1.528E-06 | 1.100E-05 | 1.400E-05 |
| 16432 | 822 | G822K | B | 1.1e-05 | 1.1e-05 | 1.4e-05 | 1.200E-05 | 1.732E-06 | 1.100E-05 | 1.400E-05 |
| 16433 | 822 | G822M | B | 9.8e-06 | 1e-05 | 7.4e-06 | 9.067E-06 | 1.447E-06 | 7.400E-06 | 1.000E-05 |
| 16434 | 822 | G822F | B | 1.3e-05 | 9.8e-06 | 1.1e-05 | 1.127E-05 | 1.617E-06 | 9.800E-06 | 1.300E-05 |
| 16435 | 822 | G822P | B | 1e-05 | 1.6e-05 | 1.4e-05 | 1.333E-05 | 3.055E-06 | 1.000E-05 | 1.600E-05 |
| 16436 | 822 | G822S | B | 1.2e-05 | 1.1e-05 | 1e-05 | 1.100E-05 | 1.000E-06 | 1.000E-05 | 1.200E-05 |
| 16437 | 822 | G822T | B | 1.6e-05 | 1.4e-05 | 1.3e-05 | 1.433E-05 | 1.528E-06 | 1.300E-05 | 1.600E-05 |
| 16438 | 822 | G822W | B | 1.1e-05 | 1e-05 | 1.1e-05 | 1.067E-05 | 5.774E-07 | 1.000E-05 | 1.100E-05 |
| 16439 | 822 | G822Y | B | 1.5e-05 | 9.1e-06 | 1e-05 | 1.137E-05 | 3.179E-06 | 9.100E-06 | 1.500E-05 |
| 16440 | 822 | G822V | B | 8.3e-06 | 7.7e-06 | 1e-05 | 8.667E-06 | 1.193E-06 | 7.700E-06 | 1.000E-05 |
| 16441 | 823 | L823A | B | 1.1e-05 | 1.6e-05 | 8.9e-06 | 1.197E-05 | 3.647E-06 | 8.900E-06 | 1.600E-05 |
| 16442 | 823 | L823R | B | 1.6e-05 | 1.1e-05 | 1.5e-05 | 1.400E-05 | 2.646E-06 | 1.100E-05 | 1.600E-05 |
| 16443 | 823 | L823N | B | 9.7e-06 | 1.4e-05 | 1e-05 | 1.123E-05 | 2.401E-06 | 9.700E-06 | 1.400E-05 |
| 16444 | 823 | L823D | B | 6.4e-06 | 1.4e-05 | 1e-05 | 1.013E-05 | 3.802E-06 | 6.400E-06 | 1.400E-05 |
| 16445 | 823 | L823C | B | 1.2e-05 | 1.1e-05 | 1.5e-05 | 1.267E-05 | 2.082E-06 | 1.100E-05 | 1.500E-05 |
| 16446 | 823 | L823E | B | 1.2e-05 | 1.6e-05 | 1.5e-05 | 1.433E-05 | 2.082E-06 | 1.200E-05 | 1.600E-05 |
| 16447 | 823 | L823Q | B | 1.1e-05 | 1.3e-05 | 1.7e-05 | 1.367E-05 | 3.055E-06 | 1.100E-05 | 1.700E-05 |
| 16448 | 823 | L823G | B | 9.6e-06 | 1.3e-05 | 1.5e-05 | 1.253E-05 | 2.730E-06 | 9.600E-06 | 1.500E-05 |
| 16449 | 823 | L823H | B | 1.5e-05 | 1e-05 | 8.3e-06 | 1.110E-05 | 3.483E-06 | 8.300E-06 | 1.500E-05 |
| 16450 | 823 | L823I | B | 1e-05 | 1.6e-05 | 1.3e-05 | 1.300E-05 | 3.000E-06 | 1.000E-05 | 1.600E-05 |
| 16451 | 823 | L823L | B | 1.3e-05 | 1e-05 | 1.3e-05 | 1.200E-05 | 1.732E-06 | 1.000E-05 | 1.300E-05 |
| 16452 | 823 | L823K | B | 1.1e-05 | 1.1e-05 | 1.5e-05 | 1.233E-05 | 2.309E-06 | 1.100E-05 | 1.500E-05 |
| 16453 | 823 | L823M | B | 6.5e-06 | 1.6e-05 | 1.4e-05 | 1.217E-05 | 5.008E-06 | 6.500E-06 | 1.600E-05 |
| 16454 | 823 | L823F | B | 1.7e-05 | 1.4e-05 | 1.5e-05 | 1.533E-05 | 1.528E-06 | 1.400E-05 | 1.700E-05 |
| 16455 | 823 | L823P | B | 1.4e-05 | 1.4e-05 | 1.2e-05 | 1.333E-05 | 1.155E-06 | 1.200E-05 | 1.400E-05 |
| 16456 | 823 | L823S | B | 1e-05 | 1.4e-05 | 2.1e-05 | 1.500E-05 | 5.568E-06 | 1.000E-05 | 2.100E-05 |
| 16457 | 823 | L823T | B | 1.3e-05 | 1.6e-05 | 1.3e-05 | 1.400E-05 | 1.732E-06 | 1.300E-05 | 1.600E-05 |
| 16458 | 823 | L823W | B | 1.4e-05 | 1.7e-05 | 9.6e-06 | 1.353E-05 | 3.722E-06 | 9.600E-06 | 1.700E-05 |
| 16459 | 823 | L823Y | B | 1.6e-05 | 1.1e-05 | 9.5e-06 | 1.217E-05 | 3.403E-06 | 9.500E-06 | 1.600E-05 |
| 16460 | 823 | L823V | B | 1e-05 | 1.3e-05 | 1.5e-05 | 1.267E-05 | 2.517E-06 | 1.000E-05 | 1.500E-05 |
| 16461 | 824 | S824A | B | 9.6e-06 | 1.1e-05 | 1.6e-05 | 1.220E-05 | 3.365E-06 | 9.600E-06 | 1.600E-05 |
| 16462 | 824 | S824R | B | 1.1e-05 | 1.1e-05 | 9.4e-06 | 1.047E-05 | 9.238E-07 | 9.400E-06 | 1.100E-05 |
| 16463 | 824 | S824N | B | 1.1e-05 | 9.2e-06 | 1.5e-05 | 1.173E-05 | 2.969E-06 | 9.200E-06 | 1.500E-05 |
| 16464 | 824 | S824D | B | 1.4e-05 | 9.4e-06 | 8.6e-06 | 1.067E-05 | 2.914E-06 | 8.600E-06 | 1.400E-05 |
| 16465 | 824 | S824C | B | 1.5e-05 | 1.6e-05 | 2.8e-05 | 1.967E-05 | 7.234E-06 | 1.500E-05 | 2.800E-05 |
| 16466 | 824 | S824E | B | 9.8e-06 | 1.2e-05 | 1.5e-05 | 1.227E-05 | 2.610E-06 | 9.800E-06 | 1.500E-05 |
| 16467 | 824 | S824Q | B | 6.7e-06 | 1.4e-05 | 1.1e-05 | 1.057E-05 | 3.669E-06 | 6.700E-06 | 1.400E-05 |
| 16468 | 824 | S824G | B | 1.4e-05 | 1.7e-05 | 1.6e-05 | 1.567E-05 | 1.528E-06 | 1.400E-05 | 1.700E-05 |
| 16469 | 824 | S824H | B | 1.6e-05 | 9.9e-06 | 1.5e-05 | 1.363E-05 | 3.272E-06 | 9.900E-06 | 1.600E-05 |
| 16470 | 824 | S824I | B | 1.6e-05 | 9.9e-06 | 1.1e-05 | 1.230E-05 | 3.251E-06 | 9.900E-06 | 1.600E-05 |
| 16471 | 824 | S824L | B | 1.6e-05 | 1.1e-05 | 1.7e-05 | 1.467E-05 | 3.215E-06 | 1.100E-05 | 1.700E-05 |
| 16472 | 824 | S824K | B | 1.2e-05 | 1e-05 | 1.6e-05 | 1.267E-05 | 3.055E-06 | 1.000E-05 | 1.600E-05 |
| 16473 | 824 | S824M | B | 1e-05 | 1e-05 | 1.5e-05 | 1.167E-05 | 2.887E-06 | 1.000E-05 | 1.500E-05 |
| 16474 | 824 | S824F | B | 9.3e-06 | 1.5e-05 | 1e-05 | 1.143E-05 | 3.109E-06 | 9.300E-06 | 1.500E-05 |
| 16475 | 824 | S824P | B | 1.5e-05 | 1.6e-05 | 8.5e-06 | 1.317E-05 | 4.072E-06 | 8.500E-06 | 1.600E-05 |
| 16476 | 824 | S824S | B | 1e-05 | 1.4e-05 | 1.1e-05 | 1.167E-05 | 2.082E-06 | 1.000E-05 | 1.400E-05 |
| 16477 | 824 | S824T | B | 1.3e-05 | 1.4e-05 | 8.9e-06 | 1.197E-05 | 2.702E-06 | 8.900E-06 | 1.400E-05 |
| 16478 | 824 | S824W | B | 9.5e-06 | 9.7e-06 | 1.1e-05 | 1.007E-05 | 8.145E-07 | 9.500E-06 | 1.100E-05 |
| 16479 | 824 | S824Y | B | 1.6e-05 | 1e-05 | 1.4e-05 | 1.333E-05 | 3.055E-06 | 1.000E-05 | 1.600E-05 |
| 16480 | 824 | S824V | B | 6.3e-06 | 1.2e-05 | 1.1e-05 | 9.767E-06 | 3.044E-06 | 6.300E-06 | 1.200E-05 |
| 16481 | 825 | S825A | B | 1.3e-05 | 1.2e-05 | 1.2e-05 | 1.233E-05 | 5.774E-07 | 1.200E-05 | 1.300E-05 |
| 16482 | 825 | S825R | B | 1.3e-05 | 1.4e-05 | 1.5e-05 | 1.400E-05 | 1.000E-06 | 1.300E-05 | 1.500E-05 |
| 16483 | 825 | S825N | B | 9.9e-06 | 1e-05 | 1.2e-05 | 1.063E-05 | 1.185E-06 | 9.900E-06 | 1.200E-05 |
| 16484 | 825 | S825D | B | 1.4e-05 | 1.1e-05 | 1.5e-05 | 1.333E-05 | 2.082E-06 | 1.100E-05 | 1.500E-05 |
| 16485 | 825 | S825C | B | 1.5e-05 | 1.1e-05 | 1.7e-05 | 1.433E-05 | 3.055E-06 | 1.100E-05 | 1.700E-05 |
| 16486 | 825 | S825E | B | 1.4e-05 | 1.5e-05 | 1.5e-05 | 1.467E-05 | 5.774E-07 | 1.400E-05 | 1.500E-05 |
| 16487 | 825 | S825Q | B | 9e-06 | 1.4e-05 | 1.4e-05 | 1.233E-05 | 2.887E-06 | 9.000E-06 | 1.400E-05 |
| 16488 | 825 | S825G | B | 1e-05 | 1.1e-05 | 1.6e-05 | 1.233E-05 | 3.215E-06 | 1.000E-05 | 1.600E-05 |
| 16489 | 825 | S825H | B | 1e-05 | 1.6e-05 | 1.4e-05 | 1.333E-05 | 3.055E-06 | 1.000E-05 | 1.600E-05 |

| | | | | | | | | | | |
|---|---|---|---|---|---|---|---|---|---|---|
| 16490 | 825 | S825I | B | 1e-05 | 1.4e-05 | 6.8e-06 | 1.027E-05 | 3.607E-06 | 6.800E-06 | 1.400E-05 |
| 16491 | 825 | S825L | B | 1.3e-05 | 9.5e-06 | 1.5e-05 | 1.250E-05 | 2.784E-06 | 9.500E-06 | 1.500E-05 |
| 16492 | 825 | S825K | B | 1.1e-05 | 1.1e-05 | 1e-05 | 1.067E-05 | 5.774E-07 | 1.000E-05 | 1.100E-05 |
| 16493 | 825 | S825M | B | 1.5e-05 | 1.4e-05 | 1.6e-05 | 1.500E-05 | 1.000E-06 | 1.400E-05 | 1.600E-05 |
| 16494 | 825 | S825F | B | 1e-05 | 1.3e-05 | 1.6e-05 | 1.300E-05 | 3.000E-06 | 1.000E-05 | 1.600E-05 |
| 16495 | 825 | S825P | B | 1.5e-05 | 6.3e-06 | 8.7e-06 | 1.000E-05 | 4.493E-06 | 6.300E-06 | 1.500E-05 |
| 16496 | 825 | S825S | B | 1.6e-05 | 9.1e-06 | 1.1e-05 | 1.203E-05 | 3.564E-06 | 9.100E-06 | 1.600E-05 |
| 16497 | 825 | S825T | B | 1.1e-05 | 1.5e-05 | 1.5e-05 | 1.367E-05 | 2.309E-06 | 1.100E-05 | 1.500E-05 |
| 16498 | 825 | S825W | B | 1.4e-05 | 1.4e-05 | 9.6e-06 | 1.253E-05 | 2.540E-06 | 9.600E-06 | 1.400E-05 |
| 16499 | 825 | S825Y | B | 1.2e-05 | 1.3e-05 | 9.5e-06 | 1.150E-05 | 1.803E-06 | 9.500E-06 | 1.300E-05 |
| 16500 | 825 | S825V | B | 1.7e-05 | 1.5e-05 | 1.6e-05 | 1.600E-05 | 1.000E-06 | 1.500E-05 | 1.700E-05 |
| 16501 | 826 | P826A | B | 1.7e-05 | 1.5e-05 | 1e-05 | 1.400E-05 | 3.606E-06 | 1.000E-05 | 1.700E-05 |
| 16502 | 826 | P826R | B | 8.4e-06 | 1e-05 | 1.1e-05 | 9.800E-06 | 1.311E-06 | 8.400E-06 | 1.100E-05 |
| 16503 | 826 | P826N | B | 9.1e-06 | 1.6e-05 | 1.4e-05 | 1.303E-05 | 3.550E-06 | 9.100E-06 | 1.600E-05 |
| 16504 | 826 | P826D | B | 1.6e-05 | 1.5e-05 | 1.5e-05 | 1.533E-05 | 5.774E-07 | 1.500E-05 | 1.600E-05 |
| 16505 | 826 | P826C | B | 1.1e-05 | 1.1e-05 | 1.7e-05 | 1.300E-05 | 3.464E-06 | 1.100E-05 | 1.700E-05 |
| 16506 | 826 | P826E | B | 1e-05 | 1.4e-05 | 1.4e-05 | 1.267E-05 | 2.309E-06 | 1.000E-05 | 1.400E-05 |
| 16507 | 826 | P826Q | B | 1.2e-05 | 1.3e-05 | 1.6e-05 | 1.367E-05 | 2.082E-06 | 1.200E-05 | 1.600E-05 |
| 16508 | 826 | P826G | B | 1.3e-05 | 1.5e-05 | 1.7e-05 | 1.500E-05 | 2.000E-06 | 1.300E-05 | 1.700E-05 |
| 16509 | 826 | P826H | B | 1.4e-05 | 9.2e-06 | 1.4e-05 | 1.240E-05 | 2.771E-06 | 9.200E-06 | 1.400E-05 |
| 16510 | 826 | P826I | B | 1.5e-05 | 1.4e-05 | 1.6e-05 | 1.500E-05 | 1.000E-06 | 1.400E-05 | 1.600E-05 |
| 16511 | 826 | P826L | B | 1.5e-05 | 1.3e-05 | 9.2e-06 | 1.240E-05 | 2.946E-06 | 9.200E-06 | 1.500E-05 |
| 16512 | 826 | P826K | B | 1.4e-05 | 9.3e-06 | 1.4e-05 | 1.243E-05 | 2.714E-06 | 9.300E-06 | 1.400E-05 |
| 16513 | 826 | P826M | B | 1.4e-05 | 9.5e-06 | 1.3e-05 | 1.217E-05 | 2.363E-06 | 9.500E-06 | 1.400E-05 |
| 16514 | 826 | P826F | B | 8.1e-06 | 1.7e-05 | 9.5e-06 | 1.153E-05 | 4.786E-06 | 8.100E-06 | 1.700E-05 |
| 16515 | 826 | P826P | B | 2.1e-05 | 1.4e-05 | 1.1e-05 | 1.533E-05 | 5.132E-06 | 1.100E-05 | 2.100E-05 |
| 16516 | 826 | P826S | B | 1.1e-05 | 1.5e-05 | 1.4e-05 | 1.333E-05 | 2.082E-06 | 1.100E-05 | 1.500E-05 |
| 16517 | 826 | P826T | B | 1.5e-05 | 1.2e-05 | 9.7e-06 | 1.223E-05 | 2.658E-06 | 9.700E-06 | 1.500E-05 |
| 16518 | 826 | P826W | B | 1.4e-05 | 9e-06 | 1.6e-05 | 1.300E-05 | 3.606E-06 | 9.000E-06 | 1.600E-05 |
| 16519 | 826 | P826Y | B | 1e-05 | 1.2e-05 | 1.5e-05 | 1.233E-05 | 2.517E-06 | 1.000E-05 | 1.500E-05 |
| 16520 | 826 | P826V | B | 1.5e-05 | 1e-05 | 1.6e-05 | 1.367E-05 | 3.215E-06 | 1.000E-05 | 1.600E-05 |
| 16521 | 827 | V827A | B | 1.1e-05 | 1.6e-05 | 1.3e-05 | 1.333E-05 | 2.517E-06 | 1.100E-05 | 1.600E-05 |
| 16522 | 827 | V827R | B | 1e-05 | 9.8e-06 | 9.5e-06 | 9.767E-06 | 2.517E-07 | 9.500E-06 | 1.000E-05 |
| 16523 | 827 | V827N | B | 1.4e-05 | 9.4e-06 | 1.3e-05 | 1.213E-05 | 2.419E-06 | 9.400E-06 | 1.400E-05 |
| 16524 | 827 | V827D | B | 8.4e-06 | 1.5e-05 | 1.6e-05 | 1.313E-05 | 4.130E-06 | 8.400E-06 | 1.600E-05 |
| 16525 | 827 | V827C | B | 1.6e-05 | 9.7e-06 | 1.4e-05 | 1.323E-05 | 3.219E-06 | 9.700E-06 | 1.600E-05 |
| 16526 | 827 | V827E | B | 1.3e-05 | 9.5e-06 | 1.4e-05 | 1.217E-05 | 2.363E-06 | 9.500E-06 | 1.400E-05 |
| 16527 | 827 | V827Q | B | 1.6e-05 | 1.1e-05 | 8.5e-06 | 1.183E-05 | 3.819E-06 | 8.500E-06 | 1.600E-05 |
| 16528 | 827 | V827G | B | 1.4e-05 | 1.6e-05 | 1.5e-05 | 1.500E-05 | 1.000E-06 | 1.400E-05 | 1.600E-05 |
| 16529 | 827 | V827H | B | 1.5e-05 | 6.7e-06 | 1e-05 | 1.057E-05 | 4.179E-06 | 6.700E-06 | 1.500E-05 |
| 16530 | 827 | V827I | B | 1.5e-05 | 1.4e-05 | 1.3e-05 | 1.400E-05 | 1.000E-06 | 1.300E-05 | 1.500E-05 |
| 16531 | 827 | V827L | B | 1.5e-05 | 1.1e-05 | 9.5e-06 | 1.183E-05 | 2.843E-06 | 9.500E-06 | 1.500E-05 |
| 16532 | 827 | V827K | B | 8.1e-06 | 9e-06 | 1.3e-05 | 1.003E-05 | 2.608E-06 | 8.100E-06 | 1.300E-05 |
| 16533 | 827 | V827M | B | 3.6e-05 | 1.3e-05 | 1.6e-05 | 2.167E-05 | 1.250E-05 | 1.300E-05 | 3.600E-05 |
| 16534 | 827 | V827F | B | 1.3e-05 | 1e-05 | 1.6e-05 | 1.300E-05 | 3.000E-06 | 1.000E-05 | 1.600E-05 |
| 16535 | 827 | V827P | B | 1.6e-05 | 1.5e-05 | 1.4e-05 | 1.500E-05 | 1.000E-06 | 1.400E-05 | 1.600E-05 |
| 16536 | 827 | V827S | B | 1.4e-05 | 1.5e-05 | 1.6e-05 | 1.500E-05 | 1.000E-06 | 1.400E-05 | 1.600E-05 |
| 16537 | 827 | V827T | B | 8e-06 | 1.4e-05 | 1.2e-05 | 1.133E-05 | 3.055E-06 | 8.000E-06 | 1.400E-05 |
| 16538 | 827 | V827W | B | 1.4e-05 | 8.7e-06 | 1.5e-05 | 1.257E-05 | 3.386E-06 | 8.700E-06 | 1.500E-05 |
| 16539 | 827 | V827Y | B | 1.5e-05 | 1.4e-05 | 1e-05 | 1.300E-05 | 2.646E-06 | 1.000E-05 | 1.500E-05 |
| 16540 | 827 | V827V | B | 1.4e-05 | 1.6e-05 | 1.4e-05 | 1.467E-05 | 1.155E-06 | 1.400E-05 | 1.600E-05 |
| 16541 | 828 | T828A | B | 9.7e-06 | 1.6e-05 | 1e-05 | 1.190E-05 | 3.554E-06 | 9.700E-06 | 1.600E-05 |
| 16542 | 828 | T828R | B | 1.5e-05 | 1.2e-05 | 1.5e-05 | 1.400E-05 | 1.732E-06 | 1.200E-05 | 1.500E-05 |
| 16543 | 828 | T828N | B | 9.9e-06 | 9.6e-06 | 1e-05 | 9.833E-06 | 2.082E-07 | 9.600E-06 | 1.000E-05 |
| 16544 | 828 | T828D | B | 9.9e-06 | 1e-05 | 7.6e-06 | 9.167E-06 | 1.358E-06 | 7.600E-06 | 1.000E-05 |
| 16545 | 828 | T828C | B | 1.3e-05 | 1.2e-05 | 1.5e-05 | 1.333E-05 | 1.528E-06 | 1.200E-05 | 1.500E-05 |
| 16546 | 828 | T828E | B | 1.5e-05 | 1.6e-05 | 1.1e-05 | 1.400E-05 | 2.646E-06 | 1.100E-05 | 1.600E-05 |
| 16547 | 828 | T828Q | B | 1.5e-05 | 9.3e-06 | 1.6e-05 | 1.343E-05 | 3.614E-06 | 9.300E-06 | 1.600E-05 |
| 16548 | 828 | T828G | B | 1e-05 | 1.5e-05 | 1.3e-05 | 1.267E-05 | 2.517E-06 | 1.000E-05 | 1.500E-05 |
| 16549 | 828 | T828H | B | 1.1e-05 | 1.2e-05 | 1.4e-05 | 1.233E-05 | 1.528E-06 | 1.100E-05 | 1.400E-05 |
| 16550 | 828 | T828I | B | 1.3e-05 | 1.4e-05 | 1.6e-05 | 1.433E-05 | 1.528E-06 | 1.300E-05 | 1.600E-05 |
| 16551 | 828 | T828L | B | 1e-05 | 1.6e-05 | 1.5e-05 | 1.367E-05 | 3.215E-06 | 1.000E-05 | 1.600E-05 |
| 16552 | 828 | T828K | B | 1e-05 | 1e-05 | 1.4e-05 | 1.133E-05 | 2.309E-06 | 1.000E-05 | 1.400E-05 |
| 16553 | 828 | T828M | B | 1.5e-05 | 1e-05 | 9.8e-06 | 1.160E-05 | 2.946E-06 | 9.800E-06 | 1.500E-05 |
| 16554 | 828 | T828F | B | 1e-05 | 1.1e-05 | 1.4e-05 | 1.167E-05 | 2.082E-06 | 1.000E-05 | 1.400E-05 |
| 16555 | 828 | T828P | B | 1.2e-05 | 1.1e-05 | 1.2e-05 | 1.167E-05 | 5.774E-07 | 1.100E-05 | 1.200E-05 |
| 16556 | 828 | T828S | B | 7.7e-06 | 1.5e-05 | 9.4e-06 | 1.070E-05 | 3.820E-06 | 7.700E-06 | 1.500E-05 |
| 16557 | 828 | T828T | B | 1.6e-05 | 1.6e-05 | 1.5e-05 | 1.567E-05 | 5.774E-07 | 1.500E-05 | 1.600E-05 |
| 16558 | 828 | T828W | B | 1.7e-05 | 1.3e-05 | 1.5e-05 | 1.500E-05 | 2.000E-06 | 1.300E-05 | 1.700E-05 |
| 16559 | 828 | T828Y | B | 8.3e-06 | 1e-05 | 8.4e-06 | 8.900E-06 | 9.539E-07 | 8.300E-06 | 1.000E-05 |
| 16560 | 828 | T828V | B | 1e-05 | 1.5e-05 | 1.5e-05 | 1.333E-05 | 2.887E-06 | 1.000E-05 | 1.500E-05 |
| 16561 | 829 | K829A | B | 1.2e-05 | 1.6e-05 | 1.5e-05 | 1.433E-05 | 2.082E-06 | 1.200E-05 | 1.600E-05 |
| 16562 | 829 | K829R | B | 1e-05 | 8.6e-06 | 1.3e-05 | 1.053E-05 | 2.248E-06 | 8.600E-06 | 1.300E-05 |
| 16563 | 829 | K829N | B | 8.9e-06 | 1.2e-05 | 1e-05 | 1.030E-05 | 1.572E-06 | 8.900E-06 | 1.200E-05 |
| 16564 | 829 | K829D | B | 9.9e-06 | 1e-05 | 8.8e-06 | 9.567E-06 | 6.658E-07 | 8.800E-06 | 1.000E-05 |
| 16565 | 829 | K829C | B | 1e-05 | 1e-05 | 1.6e-05 | 1.200E-05 | 3.464E-06 | 1.000E-05 | 1.600E-05 |
| 16566 | 829 | K829E | B | 9.6e-06 | 1.1e-05 | 1.6e-05 | 1.220E-05 | 3.365E-06 | 9.600E-06 | 1.600E-05 |
| 16567 | 829 | K829Q | B | 1.7e-05 | 1.2e-05 | 7e-06 | 1.200E-05 | 5.000E-06 | 7.000E-06 | 1.700E-05 |
| 16568 | 829 | K829G | B | 1.2e-05 | 1.2e-05 | 1.2e-05 | 1.200E-05 | 0.000E+00 | 1.200E-05 | 1.200E-05 |
| 16569 | 829 | K829H | B | 1.2e-05 | 1.4e-05 | 1.2e-05 | 1.267E-05 | 1.155E-06 | 1.200E-05 | 1.400E-05 |
| 16570 | 829 | K829I | B | 1.7e-05 | 1.5e-05 | 1.5e-05 | 1.567E-05 | 1.155E-06 | 1.500E-05 | 1.700E-05 |
| 16571 | 829 | K829L | B | 1e-05 | 1.5e-05 | 1.5e-05 | 1.333E-05 | 2.887E-06 | 1.000E-05 | 1.500E-05 |
| 16572 | 829 | K829K | B | 9.9e-06 | 1.5e-05 | 6.8e-06 | 1.057E-05 | 4.140E-06 | 6.800E-06 | 1.500E-05 |
| 16573 | 829 | K829M | B | 1.1e-05 | 1.3e-05 | 9.8e-06 | 1.127E-05 | 1.617E-06 | 9.800E-06 | 1.300E-05 |
| 16574 | 829 | K829F | B | 1.7e-05 | 1.1e-05 | 9.9e-06 | 1.263E-05 | 3.821E-06 | 9.900E-06 | 1.700E-05 |

| | | | | | | | | | | |
|---|---|---|---|---|---|---|---|---|---|---|
| 16575 | 829 | K829P | B | 1.6e-05 | 1.4e-05 | 1.5e-05 | 1.500E-05 | 1.000E-06 | 1.400E-05 | 1.600E-05 |
| 16576 | 829 | K829S | B | 1.3e-05 | 1.4e-05 | 1.4e-05 | 1.367E-05 | 5.774E-07 | 1.300E-05 | 1.400E-05 |
| 16577 | 829 | K829T | B | 9.3e-06 | 1.4e-05 | 8.6e-06 | 1.063E-05 | 2.937E-06 | 8.600E-06 | 1.400E-05 |
| 16578 | 829 | K829W | B | 1.1e-05 | 1.4e-05 | 1.2e-05 | 1.233E-05 | 1.528E-06 | 1.100E-05 | 1.400E-05 |
| 16579 | 829 | K829Y | B | 1.4e-05 | 1.6e-05 | 1.3e-05 | 1.433E-05 | 1.528E-06 | 1.300E-05 | 1.600E-05 |
| 16580 | 829 | K829V | B | 1.4e-05 | 1.4e-05 | 1.5e-05 | 1.433E-05 | 5.774E-07 | 1.400E-05 | 1.500E-05 |
| 16581 | 830 | S830A | B | 1.5e-05 | 1.3e-05 | 1.2e-05 | 1.333E-05 | 1.528E-06 | 1.200E-05 | 1.500E-05 |
| 16582 | 830 | S830R | B | 1.1e-05 | 1.5e-05 | 9.8e-06 | 1.193E-05 | 2.723E-06 | 9.800E-06 | 1.500E-05 |
| 16583 | 830 | S830N | B | 9.9e-06 | 1.1e-05 | 1.6e-05 | 1.230E-05 | 3.251E-06 | 9.900E-06 | 1.600E-05 |
| 16584 | 830 | S830D | B | 1.5e-05 | 1.4e-05 | 1.5e-05 | 1.467E-05 | 5.774E-07 | 1.400E-05 | 1.500E-05 |
| 16585 | 830 | S830C | B | 1.6e-05 | 1.4e-05 | 1.6e-05 | 1.533E-05 | 1.155E-06 | 1.400E-05 | 1.600E-05 |
| 16586 | 830 | S830E | B | 1.4e-05 | 9.6e-06 | 9.7e-06 | 1.110E-05 | 2.512E-06 | 9.600E-06 | 1.400E-05 |
| 16587 | 830 | S830Q | B | 9.9e-06 | 1.2e-05 | 1.1e-05 | 1.097E-05 | 1.050E-06 | 9.900E-06 | 1.200E-05 |
| 16588 | 830 | S830G | B | 1.6e-05 | 9.3e-06 | 1.7e-05 | 1.410E-05 | 4.187E-06 | 9.300E-06 | 1.700E-05 |
| 16589 | 830 | S830H | B | 1.3e-05 | 1.5e-05 | 1.6e-05 | 1.467E-05 | 1.528E-06 | 1.300E-05 | 1.600E-05 |
| 16590 | 830 | S830I | B | 1.2e-05 | 1.1e-05 | 1.8e-05 | 1.367E-05 | 3.786E-06 | 1.100E-05 | 1.800E-05 |
| 16591 | 830 | S830L | B | 1.5e-05 | 8.7e-06 | 9.7e-06 | 1.113E-05 | 3.386E-06 | 8.700E-06 | 1.500E-05 |
| 16592 | 830 | S830K | B | 1.4e-05 | 1.1e-05 | 1.6e-05 | 1.367E-05 | 2.517E-06 | 1.100E-05 | 1.600E-05 |
| 16593 | 830 | S830M | B | 1.6e-05 | 1.5e-05 | 1.6e-05 | 1.567E-05 | 5.774E-07 | 1.500E-05 | 1.600E-05 |
| 16594 | 830 | S830F | B | 1.4e-05 | 1.6e-05 | 1e-05 | 1.333E-05 | 3.055E-06 | 1.000E-05 | 1.600E-05 |
| 16595 | 830 | S830P | B | 1.2e-05 | 1.1e-05 | 1.2e-05 | 1.167E-05 | 5.774E-07 | 1.100E-05 | 1.200E-05 |
| 16596 | 830 | S830S | B | 1e-05 | 1.4e-05 | 1.5e-05 | 1.300E-05 | 2.646E-06 | 1.000E-05 | 1.500E-05 |
| 16597 | 830 | S830T | B | 1.6e-05 | 1.7e-05 | 1.6e-05 | 1.633E-05 | 5.774E-07 | 1.600E-05 | 1.700E-05 |
| 16598 | 830 | S830W | B | 1.5e-05 | 7.9e-06 | 1.4e-05 | 1.230E-05 | 3.843E-06 | 7.900E-06 | 1.500E-05 |
| 16599 | 830 | S830Y | B | 1e-05 | 1.1e-05 | 1.5e-05 | 1.200E-05 | 2.646E-06 | 1.000E-05 | 1.500E-05 |
| 16600 | 830 | S830V | B | 9.9e-06 | 1.7e-05 | 1.1e-05 | 1.263E-05 | 3.821E-06 | 9.900E-06 | 1.700E-05 |
| 16601 | 831 | F831A | B | 1.5e-05 | 1.5e-05 | 1.7e-05 | 1.567E-05 | 1.155E-06 | 1.500E-05 | 1.700E-05 |
| 16602 | 831 | F831R | B | 1.5e-05 | 1.8e-05 | 8.8e-06 | 1.393E-05 | 4.692E-06 | 8.800E-06 | 1.800E-05 |
| 16603 | 831 | F831N | B | 1e-05 | 1e-05 | 1.3e-05 | 1.100E-05 | 1.732E-06 | 1.000E-05 | 1.300E-05 |
| 16604 | 831 | F831D | B | 1.6e-05 | 1.1e-05 | 1.7e-05 | 1.467E-05 | 3.215E-06 | 1.100E-05 | 1.700E-05 |
| 16605 | 831 | F831C | B | 1.5e-05 | 1.5e-05 | 1.4e-05 | 1.467E-05 | 5.774E-07 | 1.400E-05 | 1.500E-05 |
| 16606 | 831 | F831E | B | 1.4e-05 | 1.5e-05 | 1.1e-05 | 1.333E-05 | 2.082E-06 | 1.100E-05 | 1.500E-05 |
| 16607 | 831 | F831Q | B | 1.2e-05 | 1e-05 | 9.8e-06 | 1.060E-05 | 1.217E-06 | 9.800E-06 | 1.200E-05 |
| 16608 | 831 | F831G | B | 1.4e-05 | 1.1e-05 | 1.7e-05 | 1.400E-05 | 3.000E-06 | 1.100E-05 | 1.700E-05 |
| 16609 | 831 | F831H | B | 1.3e-05 | 9.4e-06 | 1e-05 | 1.080E-05 | 1.929E-06 | 9.400E-06 | 1.300E-05 |
| 16610 | 831 | F831I | B | 1.3e-05 | 1.5e-05 | 9.4e-06 | 1.247E-05 | 2.838E-06 | 9.400E-06 | 1.500E-05 |
| 16611 | 831 | F831L | B | 9.7e-06 | 1.5e-05 | 1.5e-05 | 1.323E-05 | 3.060E-06 | 9.700E-06 | 1.500E-05 |
| 16612 | 831 | F831K | B | 1.1e-05 | 1.5e-05 | 1.5e-05 | 1.367E-05 | 2.309E-06 | 1.100E-05 | 1.500E-05 |
| 16613 | 831 | F831M | B | 1.1e-05 | 1.6e-05 | 1.6e-05 | 1.433E-05 | 2.887E-06 | 1.100E-05 | 1.600E-05 |
| 16614 | 831 | F831F | B | 1.4e-05 | 1.5e-05 | 9.4e-06 | 1.280E-05 | 2.987E-06 | 9.400E-06 | 1.500E-05 |
| 16615 | 831 | F831P | B | 9.2e-06 | 1.6e-05 | 9.5e-06 | 1.157E-05 | 3.842E-06 | 9.200E-06 | 1.600E-05 |
| 16616 | 831 | F831S | B | 1.3e-05 | 1e-05 | 1.2e-05 | 1.167E-05 | 1.528E-06 | 1.000E-05 | 1.300E-05 |
| 16617 | 831 | F831T | B | 1.3e-05 | 1.1e-05 | 1e-05 | 1.133E-05 | 1.528E-06 | 1.000E-05 | 1.300E-05 |
| 16618 | 831 | F831W | B | 1.3e-05 | 9.9e-06 | 9.4e-06 | 1.077E-05 | 1.950E-06 | 9.400E-06 | 1.300E-05 |
| 16619 | 831 | F831Y | B | 1.4e-05 | 1.4e-05 | 1.1e-05 | 1.300E-05 | 1.732E-06 | 1.100E-05 | 1.400E-05 |
| 16620 | 831 | F831V | B | 1.5e-05 | 1.7e-05 | 1.6e-05 | 1.600E-05 | 1.000E-06 | 1.500E-05 | 1.700E-05 |
| 16621 | 832 | N832A | B | 9.6e-06 | 1.5e-05 | 1.4e-05 | 1.287E-05 | 2.873E-06 | 9.600E-06 | 1.500E-05 |
| 16622 | 832 | N832R | B | 1.1e-05 | 1.6e-05 | 1.4e-05 | 1.367E-05 | 2.517E-06 | 1.100E-05 | 1.600E-05 |
| 16623 | 832 | N832N | B | 2.2e-05 | 1.5e-05 | 1.5e-05 | 1.733E-05 | 4.041E-06 | 1.500E-05 | 2.200E-05 |
| 16624 | 832 | N832D | B | 1.1e-05 | 9.2e-06 | 1.6e-05 | 1.207E-05 | 3.523E-06 | 9.200E-06 | 1.600E-05 |
| 16625 | 832 | N832C | B | 9.9e-06 | 1.1e-05 | 2.4e-05 | 1.497E-05 | 7.842E-06 | 9.900E-06 | 2.400E-05 |
| 16626 | 832 | N832E | B | 1e-05 | 9.5e-06 | 9.6e-06 | 9.700E-06 | 2.646E-07 | 9.500E-06 | 1.000E-05 |
| 16627 | 832 | N832Q | B | 1.4e-05 | 1.2e-05 | 1.5e-05 | 1.367E-05 | 1.528E-06 | 1.200E-05 | 1.500E-05 |
| 16628 | 832 | N832G | B | 1.2e-05 | 1.2e-05 | 1.6e-05 | 1.333E-05 | 2.309E-06 | 1.200E-05 | 1.600E-05 |
| 16629 | 832 | N832H | B | 9.6e-06 | 1.5e-05 | 1.1e-05 | 1.187E-05 | 2.802E-06 | 9.600E-06 | 1.500E-05 |
| 16630 | 832 | N832I | B | 8.7e-06 | 1.4e-05 | 1e-05 | 1.090E-05 | 2.762E-06 | 8.700E-06 | 1.400E-05 |
| 16631 | 832 | N832L | B | 9.2e-06 | 1.2e-05 | 1.6e-05 | 1.240E-05 | 3.418E-06 | 9.200E-06 | 1.600E-05 |
| 16632 | 832 | N832K | B | 1.2e-05 | 9.5e-06 | 1.2e-05 | 1.117E-05 | 1.443E-06 | 9.500E-06 | 1.200E-05 |
| 16633 | 832 | N832M | B | 1.4e-05 | 1.1e-05 | 1.5e-05 | 1.333E-05 | 2.082E-06 | 1.100E-05 | 1.500E-05 |
| 16634 | 832 | N832F | B | 1.4e-05 | 1.1e-05 | 1e-05 | 1.167E-05 | 2.082E-06 | 1.000E-05 | 1.400E-05 |
| 16635 | 832 | N832P | B | 1.8e-05 | 1.5e-05 | 1e-05 | 1.433E-05 | 4.041E-06 | 1.000E-05 | 1.800E-05 |
| 16636 | 832 | N832S | B | 1.5e-05 | 1.4e-05 | 1.7e-05 | 1.533E-05 | 1.528E-06 | 1.400E-05 | 1.700E-05 |
| 16637 | 832 | N832T | B | 1.4e-05 | 1.2e-05 | 1.2e-05 | 1.267E-05 | 1.155E-06 | 1.200E-05 | 1.400E-05 |
| 16638 | 832 | N832W | B | 1.5e-05 | 1e-05 | 1e-05 | 1.167E-05 | 2.887E-06 | 1.000E-05 | 1.500E-05 |
| 16639 | 832 | N832Y | B | 1.6e-05 | 1.5e-05 | 1e-05 | 1.367E-05 | 3.215E-06 | 1.000E-05 | 1.600E-05 |
| 16640 | 832 | N832V | B | 1.6e-05 | 1.5e-05 | 1e-05 | 1.367E-05 | 3.215E-06 | 1.000E-05 | 1.600E-05 |
| 16641 | 833 | R833A | B | 1.1e-05 | 1e-05 | 1.3e-05 | 1.133E-05 | 1.528E-06 | 1.000E-05 | 1.300E-05 |
| 16642 | 833 | R833R | B | 1.4e-05 | 1.4e-05 | 1.1e-05 | 1.300E-05 | 1.732E-06 | 1.100E-05 | 1.400E-05 |
| 16643 | 833 | R833N | B | 1.1e-05 | 1.1e-05 | 1.3e-05 | 1.167E-05 | 1.155E-06 | 1.100E-05 | 1.300E-05 |
| 16644 | 833 | R833D | B | 1.4e-05 | 9.2e-06 | 1.5e-05 | 1.273E-05 | 3.101E-06 | 9.200E-06 | 1.500E-05 |
| 16645 | 833 | R833C | B | 1.4e-05 | 8.2e-06 | 7.5e-06 | 9.900E-06 | 3.568E-06 | 7.500E-06 | 1.400E-05 |
| 16646 | 833 | R833E | B | 1.3e-05 | 9.6e-06 | 7.3e-06 | 9.967E-06 | 2.868E-06 | 7.300E-06 | 1.300E-05 |
| 16647 | 833 | R833Q | B | 9.4e-06 | 1.2e-05 | 8.5e-06 | 9.967E-06 | 1.818E-06 | 8.500E-06 | 1.200E-05 |
| 16648 | 833 | R833G | B | 8.7e-06 | 1.3e-05 | 1.5e-05 | 1.223E-05 | 3.219E-06 | 8.700E-06 | 1.500E-05 |
| 16649 | 833 | R833H | B | 1.1e-05 | 1.5e-05 | 1.5e-05 | 1.367E-05 | 2.309E-06 | 1.100E-05 | 1.500E-05 |
| 16650 | 833 | R833I | B | 8.5e-06 | 1.7e-05 | 1.1e-05 | 1.217E-05 | 4.368E-06 | 8.500E-06 | 1.700E-05 |
| 16651 | 833 | R833L | B | 1.6e-05 | 1.5e-05 | 9.7e-06 | 1.357E-05 | 3.386E-06 | 9.700E-06 | 1.600E-05 |
| 16652 | 833 | R833K | B | 1.3e-05 | 1.1e-05 | 9.8e-06 | 1.127E-05 | 1.617E-06 | 9.800E-06 | 1.300E-05 |
| 16653 | 833 | R833M | B | 1.1e-05 | 1e-05 | 1.1e-05 | 1.067E-05 | 5.774E-07 | 1.000E-05 | 1.100E-05 |
| 16654 | 833 | R833F | B | 9.9e-06 | 1.5e-05 | 8.4e-06 | 1.110E-05 | 3.460E-06 | 8.400E-06 | 1.500E-05 |
| 16655 | 833 | R833P | B | 1.4e-05 | 1e-05 | 1.3e-05 | 1.233E-05 | 2.082E-06 | 1.000E-05 | 1.400E-05 |
| 16656 | 833 | R833S | B | 1.4e-05 | 1.5e-05 | 1.1e-05 | 1.333E-05 | 2.082E-06 | 1.100E-05 | 1.500E-05 |
| 16657 | 833 | R833T | B | 9.1e-06 | 1.3e-05 | 9.6e-06 | 1.057E-05 | 2.122E-06 | 9.100E-06 | 1.300E-05 |
| 16658 | 833 | R833W | B | 1.5e-05 | 1.4e-05 | 1.5e-05 | 1.467E-05 | 5.774E-07 | 1.400E-05 | 1.500E-05 |
| 16659 | 833 | R833Y | B | 1.1e-05 | 1e-05 | 1.5e-05 | 1.200E-05 | 2.646E-06 | 1.000E-05 | 1.500E-05 |

| | | | | | | | | | | |
|---|---|---|---|---|---|---|---|---|---|---|
| 16660 | 833 | R833V | B | 1.1e-05 | 1.6e-05 | 8.8e-06 | 1.193E-05 | 3.690E-06 | 8.800E-06 | 1.600E-05 |
| 16661 | 834 | G834A | B | 1e-05 | 1.1e-05 | 1.1e-05 | 1.067E-05 | 5.774E-07 | 1.000E-05 | 1.100E-05 |
| 16662 | 834 | G834R | B | 8.4e-06 | 1.5e-05 | 1.5e-05 | 1.280E-05 | 3.811E-06 | 8.400E-06 | 1.500E-05 |
| 16663 | 834 | G834N | B | 1.1e-05 | 1.3e-05 | 1.5e-05 | 1.300E-05 | 2.000E-06 | 1.100E-05 | 1.500E-05 |
| 16664 | 834 | G834D | B | 1.4e-05 | 1e-05 | 7.8e-06 | 1.060E-05 | 3.143E-06 | 7.800E-06 | 1.400E-05 |
| 16665 | 834 | G834C | B | 1.6e-05 | 1.4e-05 | 1.5e-05 | 1.500E-05 | 1.000E-06 | 1.400E-05 | 1.600E-05 |
| 16666 | 834 | G834E | B | 1.3e-05 | 1.6e-05 | 9.2e-06 | 1.273E-05 | 3.408E-06 | 9.200E-06 | 1.600E-05 |
| 16667 | 834 | G834Q | B | 1.3e-05 | 1.4e-05 | 9.9e-06 | 1.230E-05 | 2.138E-06 | 9.900E-06 | 1.400E-05 |
| 16668 | 834 | G834G | B | 1.1e-05 | 1.1e-05 | 1.5e-05 | 1.233E-05 | 2.309E-06 | 1.100E-05 | 1.500E-05 |
| 16669 | 834 | G834H | B | 7.2e-06 | 1e-05 | 1.5e-05 | 1.073E-05 | 3.951E-06 | 7.200E-06 | 1.500E-05 |
| 16670 | 834 | G834I | B | 1.4e-05 | 1.6e-05 | 1.5e-05 | 1.500E-05 | 1.000E-06 | 1.400E-05 | 1.600E-05 |
| 16671 | 834 | G834L | B | 1.5e-05 | 9.6e-06 | 8.9e-06 | 1.117E-05 | 3.338E-06 | 8.900E-06 | 1.500E-05 |
| 16672 | 834 | G834K | B | 9.9e-06 | 1.1e-05 | 1.4e-05 | 1.163E-05 | 2.122E-06 | 9.900E-06 | 1.400E-05 |
| 16673 | 834 | G834M | B | 1.1e-05 | 1.1e-05 | 1.2e-05 | 1.133E-05 | 5.774E-07 | 1.100E-05 | 1.200E-05 |
| 16674 | 834 | G834F | B | 1.6e-05 | 1.1e-05 | 1.6e-05 | 1.433E-05 | 2.887E-06 | 1.100E-05 | 1.600E-05 |
| 16675 | 834 | G834P | B | 1.1e-05 | 1.3e-05 | 1.3e-05 | 1.233E-05 | 1.155E-06 | 1.100E-05 | 1.300E-05 |
| 16676 | 834 | G834S | B | 1.6e-05 | 1.3e-05 | 1e-05 | 1.300E-05 | 3.000E-06 | 1.000E-05 | 1.600E-05 |
| 16677 | 834 | G834T | B | 1.7e-05 | 9.3e-06 | 7.5e-06 | 1.127E-05 | 5.046E-06 | 7.500E-06 | 1.700E-05 |
| 16678 | 834 | G834W | B | 1.1e-05 | 1.1e-05 | 1.5e-05 | 1.233E-05 | 2.309E-06 | 1.100E-05 | 1.500E-05 |
| 16679 | 834 | G834Y | B | 1.5e-05 | 1.6e-05 | 1.1e-05 | 1.400E-05 | 2.646E-06 | 1.100E-05 | 1.600E-05 |
| 16680 | 834 | G834V | B | 1.3e-05 | 1.6e-05 | 1.3e-05 | 1.400E-05 | 1.732E-06 | 1.300E-05 | 1.600E-05 |
| 16681 | 835 | E835A | B | 1.4e-05 | 1.3e-05 | 1e-05 | 1.233E-05 | 2.082E-06 | 1.000E-05 | 1.400E-05 |
| 16682 | 835 | E835R | B | 2.8e-05 | 1.5e-05 | 1.2e-05 | 1.833E-05 | 8.505E-06 | 1.200E-05 | 2.800E-05 |
| 16683 | 835 | E835N | B | 1.6e-05 | 1.5e-05 | 1.2e-05 | 1.433E-05 | 2.082E-06 | 1.200E-05 | 1.600E-05 |
| 16684 | 835 | E835D | B | 1.2e-05 | 1.7e-05 | 1.4e-05 | 1.433E-05 | 2.517E-06 | 1.200E-05 | 1.700E-05 |
| 16685 | 835 | E835C | B | 1.4e-05 | 1.4e-05 | 1.3e-05 | 1.367E-05 | 5.774E-07 | 1.300E-05 | 1.400E-05 |
| 16686 | 835 | E835E | B | 1.5e-05 | 1.1e-05 | 1.6e-05 | 1.400E-05 | 2.646E-06 | 1.100E-05 | 1.600E-05 |
| 16687 | 835 | E835Q | B | 6.2e-06 | 1.1e-05 | 9.6e-06 | 8.933E-06 | 2.468E-06 | 6.200E-06 | 1.100E-05 |
| 16688 | 835 | E835G | B | 1.6e-05 | 1.3e-05 | 1.1e-05 | 1.333E-05 | 2.517E-06 | 1.100E-05 | 1.600E-05 |
| 16689 | 835 | E835H | B | 1.3e-05 | 1.1e-05 | 1e-05 | 1.133E-05 | 1.528E-06 | 1.000E-05 | 1.300E-05 |
| 16690 | 835 | E835I | B | 1.6e-05 | 1.3e-05 | 1.1e-05 | 1.333E-05 | 2.517E-06 | 1.100E-05 | 1.600E-05 |
| 16691 | 835 | E835L | B | 1.1e-05 | 1.7e-05 | 1.5e-05 | 1.433E-05 | 3.055E-06 | 1.100E-05 | 1.700E-05 |
| 16692 | 835 | E835K | B | 1.6e-05 | 1e-05 | 1.1e-05 | 1.233E-05 | 3.215E-06 | 1.000E-05 | 1.600E-05 |
| 16693 | 835 | E835M | B | 1.5e-05 | 1.4e-05 | 1e-05 | 1.300E-05 | 2.646E-06 | 1.000E-05 | 1.500E-05 |
| 16694 | 835 | E835F | B | 1e-05 | 1.1e-05 | 9e-06 | 1.000E-05 | 1.000E-06 | 9.000E-06 | 1.100E-05 |
| 16695 | 835 | E835P | B | 1.1e-05 | 1.5e-05 | 1.6e-05 | 1.400E-05 | 2.646E-06 | 1.100E-05 | 1.600E-05 |
| 16696 | 835 | E835S | B | 1.4e-05 | 8.8e-06 | 1e-05 | 1.093E-05 | 2.723E-06 | 8.800E-06 | 1.400E-05 |
| 16697 | 835 | E835T | B | 1.4e-05 | 1.1e-05 | 1.1e-05 | 1.200E-05 | 1.732E-06 | 1.100E-05 | 1.400E-05 |
| 16698 | 835 | E835W | B | 1.5e-05 | 1.7e-05 | 1.4e-05 | 1.533E-05 | 1.528E-06 | 1.400E-05 | 1.700E-05 |
| 16699 | 835 | E835Y | B | 9.8e-06 | 1.1e-05 | 1.4e-05 | 1.160E-05 | 2.163E-06 | 9.800E-06 | 1.400E-05 |
| 16700 | 835 | E835V | B | 1.4e-05 | 1.6e-05 | 1.5e-05 | 1.500E-05 | 1.000E-06 | 1.400E-05 | 1.600E-05 |
| 16701 | 836 | C836A | B | 1.3e-05 | 9.3e-06 | 1.4e-05 | 1.210E-05 | 2.476E-06 | 9.300E-06 | 1.400E-05 |
| 16702 | 836 | C836R | B | 1.1e-05 | 1.5e-05 | 9.6e-06 | 1.187E-05 | 2.802E-06 | 9.600E-06 | 1.500E-05 |
| 16703 | 836 | C836N | B | 1.5e-05 | 9.8e-06 | 1.7e-05 | 1.393E-05 | 3.717E-06 | 9.800E-06 | 1.700E-05 |
| 16704 | 836 | C836D | B | 7.8e-06 | 1.4e-05 | 8.7e-06 | 1.017E-05 | 3.350E-06 | 7.800E-06 | 1.400E-05 |
| 16705 | 836 | C836C | B | 1.4e-05 | 1e-05 | 1.2e-05 | 1.200E-05 | 2.000E-06 | 1.000E-05 | 1.400E-05 |
| 16706 | 836 | C836E | B | 1.5e-05 | 7.4e-06 | 1.5e-05 | 1.247E-05 | 4.388E-06 | 7.400E-06 | 1.500E-05 |
| 16707 | 836 | C836Q | B | 1.3e-05 | 1e-05 | 1.5e-05 | 1.267E-05 | 2.517E-06 | 1.000E-05 | 1.500E-05 |
| 16708 | 836 | C836G | B | 1.1e-05 | 7.2e-06 | 1.2e-05 | 1.007E-05 | 2.532E-06 | 7.200E-06 | 1.200E-05 |
| 16709 | 836 | C836H | B | 1.3e-05 | 9.5e-06 | 1e-05 | 1.083E-05 | 1.893E-06 | 9.500E-06 | 1.300E-05 |
| 16710 | 836 | C836I | B | 7.9e-06 | 9e-06 | 1.1e-05 | 9.300E-06 | 1.572E-06 | 7.900E-06 | 1.100E-05 |
| 16711 | 836 | C836L | B | 1.6e-05 | 1.7e-05 | 1.3e-05 | 1.533E-05 | 2.082E-06 | 1.300E-05 | 1.700E-05 |
| 16712 | 836 | C836K | B | 1.5e-05 | 9.2e-06 | 1.2e-05 | 1.207E-05 | 2.901E-06 | 9.200E-06 | 1.500E-05 |
| 16713 | 836 | C836M | B | 1.1e-05 | 8.1e-06 | 1.3e-05 | 1.070E-05 | 2.464E-06 | 8.100E-06 | 1.300E-05 |
| 16714 | 836 | C836F | B | 1.6e-05 | 1.6e-05 | 1.8e-05 | 1.667E-05 | 1.155E-06 | 1.600E-05 | 1.800E-05 |
| 16715 | 836 | C836P | B | 1.4e-05 | 1.6e-05 | 1.5e-05 | 1.500E-05 | 1.000E-06 | 1.400E-05 | 1.600E-05 |
| 16716 | 836 | C836S | B | 1.6e-05 | 1.6e-05 | 1.6e-05 | 1.600E-05 | 0.000E+00 | 1.600E-05 | 1.600E-05 |
| 16717 | 836 | C836T | B | 1.2e-05 | 1.5e-05 | 1.1e-05 | 1.267E-05 | 2.082E-06 | 1.100E-05 | 1.500E-05 |
| 16718 | 836 | C836W | B | 1.3e-05 | 1.5e-05 | 1.1e-05 | 1.300E-05 | 2.000E-06 | 1.100E-05 | 1.500E-05 |
| 16719 | 836 | C836Y | B | 1.4e-05 | 1.4e-05 | 1.4e-05 | 1.400E-05 | 0.000E+00 | 1.400E-05 | 1.400E-05 |
| 16720 | 836 | C836V | B | 1.3e-05 | 1.7e-05 | 1.3e-05 | 1.433E-05 | 2.309E-06 | 1.300E-05 | 1.700E-05 |
| 16721 | 837 | E837A | C | 1.6e-05 | 1.6e-05 | 1.4e-05 | 1.533E-05 | 1.155E-06 | 1.400E-05 | 1.600E-05 |
| 16722 | 837 | E837R | C | 1.5e-05 | 1.5e-05 | 7.7e-06 | 1.257E-05 | 4.215E-06 | 7.700E-06 | 1.500E-05 |
| 16723 | 837 | E837N | C | 1.5e-05 | 9.6e-06 | 9.4e-06 | 1.133E-05 | 3.177E-06 | 9.400E-06 | 1.500E-05 |
| 16724 | 837 | E837D | C | 8.3e-06 | 1.6e-05 | 1.1e-05 | 1.177E-05 | 3.907E-06 | 8.300E-06 | 1.600E-05 |
| 16725 | 837 | E837C | C | 1.2e-05 | 1.4e-05 | 1.3e-05 | 1.300E-05 | 1.000E-06 | 1.200E-05 | 1.400E-05 |
| 16726 | 837 | E837E | C | 1.3e-05 | 1.6e-05 | 9.1e-06 | 1.270E-05 | 3.460E-06 | 9.100E-06 | 1.600E-05 |
| 16727 | 837 | E837Q | C | 9.9e-06 | 7.4e-06 | 9e-06 | 8.767E-06 | 1.266E-06 | 7.400E-06 | 9.900E-06 |
| 16728 | 837 | E837G | C | 1.7e-05 | 1.4e-05 | 1.3e-05 | 1.467E-05 | 2.082E-06 | 1.300E-05 | 1.700E-05 |
| 16729 | 837 | E837H | C | 8.4e-06 | 1.4e-05 | 1.1e-05 | 1.113E-05 | 2.802E-06 | 8.400E-06 | 1.400E-05 |
| 16730 | 837 | E837I | C | 9e-06 | 1.1e-05 | 8.7e-06 | 9.567E-06 | 1.250E-06 | 8.700E-06 | 1.100E-05 |
| 16731 | 837 | E837L | C | 2.8e-05 | 1.6e-05 | 1.4e-05 | 1.933E-05 | 7.572E-06 | 1.400E-05 | 2.800E-05 |
| 16732 | 837 | E837K | C | 6e-06 | 1.4e-05 | 9.4e-06 | 9.800E-06 | 4.015E-06 | 6.000E-06 | 1.400E-05 |
| 16733 | 837 | E837M | C | 1.6e-05 | 1.4e-05 | 1.6e-05 | 1.533E-05 | 1.155E-06 | 1.400E-05 | 1.600E-05 |
| 16734 | 837 | E837F | C | 1.1e-05 | 1.4e-05 | 1.6e-05 | 1.367E-05 | 2.517E-06 | 1.100E-05 | 1.600E-05 |
| 16735 | 837 | E837P | C | 1.1e-05 | 1.2e-05 | 1.4e-05 | 1.233E-05 | 1.528E-06 | 1.100E-05 | 1.400E-05 |
| 16736 | 837 | E837S | C | 1e-05 | 1.6e-05 | 1.5e-05 | 1.367E-05 | 3.215E-06 | 1.000E-05 | 1.600E-05 |
| 16737 | 837 | E837T | C | 1e-05 | 1.3e-05 | 1.3e-05 | 1.200E-05 | 1.732E-06 | 1.000E-05 | 1.300E-05 |
| 16738 | 837 | E837W | C | 1.1e-05 | 1.2e-05 | 1e-05 | 1.100E-05 | 1.000E-06 | 1.000E-05 | 1.200E-05 |
| 16739 | 837 | E837Y | C | 9.9e-06 | 1.5e-05 | 9.9e-06 | 1.160E-05 | 2.944E-06 | 9.900E-06 | 1.500E-05 |
| 16740 | 837 | E837V | C | 1.6e-05 | 1e-05 | 1.2e-05 | 1.267E-05 | 3.055E-06 | 1.000E-05 | 1.600E-05 |
| 16741 | 838 | V838A | C | 1.1e-05 | 1.4e-05 | 1.1e-05 | 1.200E-05 | 1.732E-06 | 1.100E-05 | 1.400E-05 |
| 16742 | 838 | V838R | C | 1.4e-05 | 8.4e-06 | 1.7e-05 | 1.313E-05 | 4.365E-06 | 8.400E-06 | 1.700E-05 |
| 16743 | 838 | V838N | C | 1.6e-05 | 1.5e-05 | 1e-05 | 1.367E-05 | 3.215E-06 | 1.000E-05 | 1.600E-05 |
| 16744 | 838 | V838D | C | 1.1e-05 | 1.4e-05 | 1.5e-05 | 1.333E-05 | 2.082E-06 | 1.100E-05 | 1.500E-05 |

| | | | | | | | | | | |
|---|---|---|---|---|---|---|---|---|---|---|
| 16745 | 838 | V838C | C | 1.3e-05 | 1.1e-05 | 1e-05 | 1.133E-05 | 1.528E-06 | 1.000E-05 | 1.300E-05 |
| 16746 | 838 | V838E | C | 8.8e-06 | 1.6e-05 | 1.6e-05 | 1.360E-05 | 4.157E-06 | 8.800E-06 | 1.600E-05 |
| 16747 | 838 | V838Q | C | 1.1e-05 | 1e-05 | 1.5e-05 | 1.200E-05 | 2.646E-06 | 1.000E-05 | 1.500E-05 |
| 16748 | 838 | V838G | C | 1e-05 | 1.1e-05 | 1.4e-05 | 1.167E-05 | 2.082E-06 | 1.000E-05 | 1.400E-05 |
| 16749 | 838 | V838H | C | 8.5e-06 | 8.7e-06 | 1.4e-05 | 1.040E-05 | 3.119E-06 | 8.500E-06 | 1.400E-05 |
| 16750 | 838 | V838I | C | 1.6e-05 | 9.3e-06 | 1.5e-05 | 1.343E-05 | 3.614E-06 | 9.300E-06 | 1.600E-05 |
| 16751 | 838 | V838L | C | 1.6e-05 | 9.7e-06 | 1.3e-05 | 1.290E-05 | 3.151E-06 | 9.700E-06 | 1.600E-05 |
| 16752 | 838 | V838K | C | 1.6e-05 | 1.6e-05 | 1.5e-05 | 1.567E-05 | 5.774E-07 | 1.500E-05 | 1.600E-05 |
| 16753 | 838 | V838M | C | 1.1e-05 | 1.3e-05 | 1.3e-05 | 1.233E-05 | 1.155E-06 | 1.100E-05 | 1.300E-05 |
| 16754 | 838 | V838F | C | 9.5e-06 | 1.5e-05 | 1.2e-05 | 1.217E-05 | 2.754E-06 | 9.500E-06 | 1.500E-05 |
| 16755 | 838 | V838P | C | 9.8e-06 | 1.5e-05 | 1e-05 | 1.160E-05 | 2.946E-06 | 9.800E-06 | 1.500E-05 |
| 16756 | 838 | V838S | C | 1.1e-05 | 1e-05 | 9.9e-06 | 1.030E-05 | 6.083E-07 | 9.900E-06 | 1.100E-05 |
| 16757 | 838 | V838T | C | 1.6e-05 | 1.1e-05 | 9.5e-06 | 1.217E-05 | 3.403E-06 | 9.500E-06 | 1.600E-05 |
| 16758 | 838 | V838W | C | 1.6e-05 | 9.3e-06 | 1.1e-05 | 1.210E-05 | 3.483E-06 | 9.300E-06 | 1.600E-05 |
| 16759 | 838 | V838Y | C | 1e-05 | 1.4e-05 | 6.9e-06 | 1.030E-05 | 3.559E-06 | 6.900E-06 | 1.400E-05 |
| 16760 | 838 | V838V | C | 1.3e-05 | 1.6e-05 | 1.5e-05 | 1.467E-05 | 1.528E-06 | 1.300E-05 | 1.600E-05 |
| 16761 | 839 | Q839A | C | 9e-06 | 1.1e-05 | 1.5e-05 | 1.167E-05 | 3.055E-06 | 9.000E-06 | 1.500E-05 |
| 16762 | 839 | Q839R | C | 1.6e-05 | 1.4e-05 | 1.1e-05 | 1.367E-05 | 2.517E-06 | 1.100E-05 | 1.600E-05 |
| 16763 | 839 | Q839N | C | 1.1e-05 | 1.2e-05 | 1.5e-05 | 1.267E-05 | 2.082E-06 | 1.100E-05 | 1.500E-05 |
| 16764 | 839 | Q839D | C | 1.1e-05 | 1.5e-05 | 9.6e-06 | 1.187E-05 | 2.802E-06 | 9.600E-06 | 1.500E-05 |
| 16765 | 839 | Q839C | C | 1.4e-05 | 1.5e-05 | 9.7e-06 | 1.290E-05 | 2.816E-06 | 9.700E-06 | 1.500E-05 |
| 16766 | 839 | Q839E | C | 1e-05 | 9.7e-06 | 1e-05 | 9.900E-06 | 1.732E-07 | 9.700E-06 | 1.000E-05 |
| 16767 | 839 | Q839Q | C | 1e-05 | 1.1e-05 | 1.1e-05 | 1.067E-05 | 5.774E-07 | 1.000E-05 | 1.100E-05 |
| 16768 | 839 | Q839G | C | 1.1e-05 | 1.2e-05 | 1.4e-05 | 1.233E-05 | 1.528E-06 | 1.100E-05 | 1.400E-05 |
| 16769 | 839 | Q839H | C | 1e-05 | 1.6e-05 | 1.2e-05 | 1.267E-05 | 3.055E-06 | 1.000E-05 | 1.600E-05 |
| 16770 | 839 | Q839I | C | 1.1e-05 | 1.5e-05 | 1.6e-05 | 1.400E-05 | 2.646E-06 | 1.100E-05 | 1.600E-05 |
| 16771 | 839 | Q839L | C | 1.7e-05 | 1.3e-05 | 1.1e-05 | 1.367E-05 | 3.055E-06 | 1.100E-05 | 1.700E-05 |
| 16772 | 839 | Q839K | C | 1.6e-05 | 1.2e-05 | 1.1e-05 | 1.300E-05 | 2.646E-06 | 1.100E-05 | 1.600E-05 |
| 16773 | 839 | Q839M | C | 1.4e-05 | 9.6e-06 | 9e-06 | 1.087E-05 | 2.730E-06 | 9.000E-06 | 1.400E-05 |
| 16774 | 839 | Q839F | C | 9.7e-06 | 1.4e-05 | 1.3e-05 | 1.223E-05 | 2.250E-06 | 9.700E-06 | 1.400E-05 |
| 16775 | 839 | Q839P | C | 1.6e-05 | 1.5e-05 | 1.5e-05 | 1.533E-05 | 5.774E-07 | 1.500E-05 | 1.600E-05 |
| 16776 | 839 | Q839S | C | 1.2e-05 | 1.3e-05 | 1.3e-05 | 1.267E-05 | 5.774E-07 | 1.200E-05 | 1.300E-05 |
| 16777 | 839 | Q839T | C | 1.7e-05 | 9.5e-06 | 1.5e-05 | 1.383E-05 | 3.884E-06 | 9.500E-06 | 1.700E-05 |
| 16778 | 839 | Q839W | C | 1.1e-05 | 1.2e-05 | 8.5e-06 | 1.050E-05 | 1.803E-06 | 8.500E-06 | 1.200E-05 |
| 16779 | 839 | Q839Y | C | 1.4e-05 | 1.7e-05 | 9.7e-06 | 1.357E-05 | 3.669E-06 | 9.700E-06 | 1.700E-05 |
| 16780 | 839 | Q839V | C | 1.2e-05 | 1.7e-05 | 1.4e-05 | 1.433E-05 | 2.517E-06 | 1.200E-05 | 1.700E-05 |
| 16781 | 840 | L840A | C | 9.8e-06 | 9.9e-06 | 1.2e-05 | 1.057E-05 | 1.242E-06 | 9.800E-06 | 1.200E-05 |
| 16782 | 840 | L840R | C | 8.1e-06 | 1.7e-05 | 9.4e-06 | 1.150E-05 | 4.807E-06 | 8.100E-06 | 1.700E-05 |
| 16783 | 840 | L840N | C | 1e-05 | 1e-05 | 1.3e-05 | 1.100E-05 | 1.732E-06 | 1.000E-05 | 1.300E-05 |
| 16784 | 840 | L840D | C | 1.4e-05 | 1.3e-05 | 1.6e-05 | 1.433E-05 | 1.528E-06 | 1.300E-05 | 1.600E-05 |
| 16785 | 840 | L840C | C | 1.5e-05 | 1.4e-05 | 1.2e-05 | 1.367E-05 | 1.528E-06 | 1.200E-05 | 1.500E-05 |
| 16786 | 840 | L840E | C | 1.5e-05 | 1.4e-05 | 1.5e-05 | 1.467E-05 | 5.774E-07 | 1.400E-05 | 1.500E-05 |
| 16787 | 840 | L840Q | C | 1.4e-05 | 1.5e-05 | 1.4e-05 | 1.433E-05 | 5.774E-07 | 1.400E-05 | 1.500E-05 |
| 16788 | 840 | L840G | C | 1.1e-05 | 1.5e-05 | 1.5e-05 | 1.367E-05 | 2.309E-06 | 1.100E-05 | 1.500E-05 |
| 16789 | 840 | L840H | C | 9.4e-06 | 9.8e-06 | 1e-05 | 9.733E-06 | 3.055E-07 | 9.400E-06 | 1.000E-05 |
| 16790 | 840 | L840I | C | 1.5e-05 | 9.1e-06 | 1.5e-05 | 1.303E-05 | 3.406E-06 | 9.100E-06 | 1.500E-05 |
| 16791 | 840 | L840L | C | 1.1e-05 | 1.6e-05 | 1.4e-05 | 1.367E-05 | 2.517E-06 | 1.100E-05 | 1.600E-05 |
| 16792 | 840 | L840K | C | 5.8e-06 | 1e-05 | 1.1e-05 | 8.933E-06 | 2.759E-06 | 5.800E-06 | 1.100E-05 |
| 16793 | 840 | L840M | C | 1.6e-05 | 1.5e-05 | 1.4e-05 | 1.500E-05 | 1.000E-06 | 1.400E-05 | 1.600E-05 |
| 16794 | 840 | L840F | C | 1.1e-05 | 1.2e-05 | 1.2e-05 | 1.167E-05 | 5.774E-07 | 1.100E-05 | 1.200E-05 |
| 16795 | 840 | L840P | C | 9.1e-06 | 1.5e-05 | 8.9e-06 | 1.100E-05 | 3.466E-06 | 8.900E-06 | 1.500E-05 |
| 16796 | 840 | L840S | C | 1.9e-05 | 1.1e-05 | 1.4e-05 | 1.467E-05 | 4.041E-06 | 1.100E-05 | 1.900E-05 |
| 16797 | 840 | L840T | C | 9.9e-06 | 1.4e-05 | 9e-06 | 1.097E-05 | 2.665E-06 | 9.000E-06 | 1.400E-05 |
| 16798 | 840 | L840W | C | 9.6e-06 | 9.7e-06 | 1.5e-05 | 1.143E-05 | 3.089E-06 | 9.600E-06 | 1.500E-05 |
| 16799 | 840 | L840Y | C | 1e-05 | 1.6e-05 | 9.6e-06 | 1.187E-05 | 3.585E-06 | 9.600E-06 | 1.600E-05 |
| 16800 | 840 | L840V | C | 8.1e-06 | 1.5e-05 | 1.1e-05 | 1.137E-05 | 3.465E-06 | 8.100E-06 | 1.500E-05 |
| 16801 | 841 | V841A | C | 1.1e-05 | 1.6e-05 | 1.6e-05 | 1.433E-05 | 2.887E-06 | 1.100E-05 | 1.600E-05 |
| 16802 | 841 | V841R | C | 9e-06 | 1.5e-05 | 1.6e-05 | 1.333E-05 | 3.786E-06 | 9.000E-06 | 1.600E-05 |
| 16803 | 841 | V841N | C | 1.4e-05 | 1.4e-05 | 1.5e-05 | 1.433E-05 | 5.774E-07 | 1.400E-05 | 1.500E-05 |
| 16804 | 841 | V841D | C | 9.7e-06 | 1.5e-05 | 9.3e-06 | 1.133E-05 | 3.182E-06 | 9.300E-06 | 1.500E-05 |
| 16805 | 841 | V841C | C | 1.1e-05 | 1.3e-05 | 1.6e-05 | 1.333E-05 | 2.517E-06 | 1.100E-05 | 1.600E-05 |
| 16806 | 841 | V841E | C | 1.3e-05 | 1.4e-05 | 1.3e-05 | 1.333E-05 | 5.774E-07 | 1.300E-05 | 1.400E-05 |
| 16807 | 841 | V841Q | C | 1.5e-05 | 1e-05 | 1.1e-05 | 1.200E-05 | 2.646E-06 | 1.000E-05 | 1.500E-05 |
| 16808 | 841 | V841G | C | 9.4e-06 | 1.1e-05 | 1.5e-05 | 1.180E-05 | 2.884E-06 | 9.400E-06 | 1.500E-05 |
| 16809 | 841 | V841H | C | 9.5e-06 | 6.1e-06 | 1.5e-05 | 1.020E-05 | 4.491E-06 | 6.100E-06 | 1.500E-05 |
| 16810 | 841 | V841I | C | 1.5e-05 | 1.4e-05 | 8.8e-06 | 1.260E-05 | 3.329E-06 | 8.800E-06 | 1.500E-05 |
| 16811 | 841 | V841L | C | 8.8e-06 | 1.1e-05 | 1.1e-05 | 1.027E-05 | 1.270E-06 | 8.800E-06 | 1.100E-05 |
| 16812 | 841 | V841K | C | 1.4e-05 | 1.1e-05 | 1e-05 | 1.167E-05 | 2.082E-06 | 1.000E-05 | 1.400E-05 |
| 16813 | 841 | V841M | C | 8.6e-06 | 1.3e-05 | 9.6e-06 | 1.040E-05 | 2.307E-06 | 8.600E-06 | 1.300E-05 |
| 16814 | 841 | V841F | C | 1.5e-05 | 1.5e-05 | 1.5e-05 | 1.500E-05 | 0.000E+00 | 1.500E-05 | 1.500E-05 |
| 16815 | 841 | V841P | C | 1.2e-05 | 9.8e-06 | 1.5e-05 | 1.227E-05 | 2.610E-06 | 9.800E-06 | 1.500E-05 |
| 16816 | 841 | V841S | C | 1.2e-05 | 7.8e-06 | 9.1e-06 | 9.633E-06 | 2.150E-06 | 7.800E-06 | 1.200E-05 |
| 16817 | 841 | V841T | C | 2.1e-05 | 1.4e-05 | 1.5e-05 | 1.667E-05 | 3.786E-06 | 1.400E-05 | 2.100E-05 |
| 16818 | 841 | V841W | C | 1.5e-05 | 1e-05 | 1.7e-05 | 1.400E-05 | 3.606E-06 | 1.000E-05 | 1.700E-05 |
| 16819 | 841 | V841Y | C | 1.1e-05 | 1.1e-05 | 1e-05 | 1.067E-05 | 5.774E-07 | 1.000E-05 | 1.100E-05 |
| 16820 | 841 | V841V | C | 9.2e-06 | 1.3e-05 | 8e-06 | 1.007E-05 | 2.610E-06 | 8.000E-06 | 1.300E-05 |
| 16821 | 842 | E842A | C | 1.1e-05 | 1.3e-05 | 1.4e-05 | 1.267E-05 | 1.528E-06 | 1.100E-05 | 1.400E-05 |
| 16822 | 842 | E842R | C | 1.3e-05 | 1.5e-05 | 1e-05 | 1.267E-05 | 2.517E-06 | 1.000E-05 | 1.500E-05 |
| 16823 | 842 | E842N | C | 1.4e-05 | 1.5e-05 | 1.2e-05 | 1.367E-05 | 1.528E-06 | 1.200E-05 | 1.500E-05 |
| 16824 | 842 | E842D | C | 1.7e-05 | 1.4e-05 | 1.4e-05 | 1.500E-05 | 1.732E-06 | 1.400E-05 | 1.700E-05 |
| 16825 | 842 | E842C | C | 9.4e-06 | 1.1e-05 | 1.3e-05 | 1.113E-05 | 1.804E-06 | 9.400E-06 | 1.300E-05 |
| 16826 | 842 | E842E | C | 1.1e-05 | 6.3e-06 | 1.2e-05 | 9.767E-06 | 3.044E-06 | 6.300E-06 | 1.200E-05 |
| 16827 | 842 | E842Q | C | 1.6e-05 | 1.6e-05 | 1.5e-05 | 1.567E-05 | 5.774E-07 | 1.500E-05 | 1.600E-05 |
| 16828 | 842 | E842G | C | 1e-05 | 2.6e-05 | 1.4e-05 | 1.667E-05 | 8.327E-06 | 1.000E-05 | 2.600E-05 |
| 16829 | 842 | E842H | C | 8.4e-06 | 1.1e-05 | 9.2e-06 | 9.533E-06 | 1.332E-06 | 8.400E-06 | 1.100E-05 |

| | | | | | | | | | | |
|---|---|---|---|---|---|---|---|---|---|---|
| 16830 | 842 | E842I | C | 7.4e-06 | 1.1e-05 | 1.5e-05 | 1.113E-05 | 3.802E-06 | 7.400E-06 | 1.500E-05 |
| 16831 | 842 | E842L | C | 1e-05 | 1.6e-05 | 1.2e-05 | 1.267E-05 | 3.055E-06 | 1.000E-05 | 1.600E-05 |
| 16832 | 842 | E842K | C | 1.1e-05 | 9e-06 | 1.6e-05 | 1.200E-05 | 3.606E-06 | 9.000E-06 | 1.600E-05 |
| 16833 | 842 | E842M | C | 1e-05 | 1.6e-05 | 1.5e-05 | 1.367E-05 | 3.215E-06 | 1.000E-05 | 1.600E-05 |
| 16834 | 842 | E842F | C | 1.1e-05 | 1.3e-05 | 1.3e-05 | 1.233E-05 | 1.155E-06 | 1.100E-05 | 1.300E-05 |
| 16835 | 842 | E842P | C | 8.3e-06 | 1.4e-05 | 1.7e-05 | 1.310E-05 | 4.419E-06 | 8.300E-06 | 1.700E-05 |
| 16836 | 842 | E842S | C | 1.3e-05 | 1e-05 | 1.6e-05 | 1.300E-05 | 3.000E-06 | 1.000E-05 | 1.600E-05 |
| 16837 | 842 | E842T | C | 1.2e-05 | 1.1e-05 | 1.3e-05 | 1.200E-05 | 1.000E-06 | 1.100E-05 | 1.300E-05 |
| 16838 | 842 | E842W | C | 1.3e-05 | 7e-06 | 1.6e-05 | 1.200E-05 | 4.583E-06 | 7.000E-06 | 1.600E-05 |
| 16839 | 842 | E842Y | C | 1.5e-05 | 1.1e-05 | 1.3e-05 | 1.300E-05 | 2.000E-06 | 1.100E-05 | 1.500E-05 |
| 16840 | 842 | E842V | C | 1.7e-05 | 1.1e-05 | 1e-05 | 1.267E-05 | 3.786E-06 | 1.000E-05 | 1.700E-05 |
| 16841 | 843 | S843A | C | 7.2e-06 | 1.1e-05 | 1.6e-05 | 1.140E-05 | 4.414E-06 | 7.200E-06 | 1.600E-05 |
| 16842 | 843 | S843R | C | 1.3e-05 | 1.7e-05 | 9.8e-06 | 1.327E-05 | 3.607E-06 | 9.800E-06 | 1.700E-05 |
| 16843 | 843 | S843N | C | 8.2e-06 | 9.9e-06 | 1.5e-05 | 1.103E-05 | 3.539E-06 | 8.200E-06 | 1.500E-05 |
| 16844 | 843 | S843D | C | 1.2e-05 | 9.5e-06 | 1.4e-05 | 1.183E-05 | 2.255E-06 | 9.500E-06 | 1.400E-05 |
| 16845 | 843 | S843C | C | 1.5e-05 | 9.2e-06 | 1.1e-05 | 1.173E-05 | 2.969E-06 | 9.200E-06 | 1.500E-05 |
| 16846 | 843 | S843E | C | 9.6e-06 | 1.6e-05 | 1.3e-05 | 1.287E-05 | 3.202E-06 | 9.600E-06 | 1.600E-05 |
| 16847 | 843 | S843Q | C | 1.6e-05 | 1.5e-05 | 1.1e-05 | 1.400E-05 | 2.646E-06 | 1.100E-05 | 1.600E-05 |
| 16848 | 843 | S843G | C | 1.7e-05 | 1e-05 | 1.1e-05 | 1.267E-05 | 3.786E-06 | 1.000E-05 | 1.700E-05 |
| 16849 | 843 | S843H | C | 1.5e-05 | 9.7e-06 | 1.2e-05 | 1.223E-05 | 2.658E-06 | 9.700E-06 | 1.500E-05 |
| 16850 | 843 | S843I | C | 1.1e-05 | 1.6e-05 | 1.5e-05 | 1.400E-05 | 2.646E-06 | 1.100E-05 | 1.600E-05 |
| 16851 | 843 | S843L | C | 1.6e-05 | 1.1e-05 | 1.9e-05 | 1.533E-05 | 4.041E-06 | 1.100E-05 | 1.900E-05 |
| 16852 | 843 | S843K | C | 1.3e-05 | 1e-05 | 1.8e-05 | 1.367E-05 | 4.041E-06 | 1.000E-05 | 1.800E-05 |
| 16853 | 843 | S843M | C | 9.2e-06 | 1.2e-05 | 1.3e-05 | 1.140E-05 | 1.970E-06 | 9.200E-06 | 1.300E-05 |
| 16854 | 843 | S843F | C | 1.6e-05 | 1e-05 | 1e-05 | 1.200E-05 | 3.464E-06 | 1.000E-05 | 1.600E-05 |
| 16855 | 843 | S843P | C | 1.1e-05 | 1.5e-05 | 1.5e-05 | 1.367E-05 | 2.309E-06 | 1.100E-05 | 1.500E-05 |
| 16856 | 843 | S843S | C | 1.4e-05 | 1.5e-05 | 1.4e-05 | 1.433E-05 | 5.774E-07 | 1.400E-05 | 1.500E-05 |
| 16857 | 843 | S843T | C | 1.3e-05 | 1.3e-05 | 1.1e-05 | 1.233E-05 | 1.155E-06 | 1.100E-05 | 1.300E-05 |
| 16858 | 843 | S843W | C | 1e-05 | 1.4e-05 | 1.1e-05 | 1.167E-05 | 2.082E-06 | 1.000E-05 | 1.400E-05 |
| 16859 | 843 | S843Y | C | 1.5e-05 | 8.7e-06 | 1.5e-05 | 1.290E-05 | 3.637E-06 | 8.700E-06 | 1.500E-05 |
| 16860 | 843 | S843V | C | 1.6e-05 | 1.5e-05 | 1.5e-05 | 1.533E-05 | 5.774E-07 | 1.500E-05 | 1.600E-05 |
| 16861 | 844 | G844A | C | 1.5e-05 | 6.6e-06 | 9.4e-06 | 1.033E-05 | 4.277E-06 | 6.600E-06 | 1.500E-05 |
| 16862 | 844 | G844R | C | 9.8e-06 | 1.4e-05 | 1.1e-05 | 1.160E-05 | 2.163E-06 | 9.800E-06 | 1.400E-05 |
| 16863 | 844 | G844N | C | 1.3e-05 | 8e-06 | 1.4e-05 | 1.167E-05 | 3.215E-06 | 8.000E-06 | 1.400E-05 |
| 16864 | 844 | G844D | C | 1.4e-05 | 9.5e-06 | 1.5e-05 | 1.283E-05 | 2.930E-06 | 9.500E-06 | 1.500E-05 |
| 16865 | 844 | G844C | C | 1.4e-05 | 1.4e-05 | 1.6e-05 | 1.467E-05 | 1.155E-06 | 1.400E-05 | 1.600E-05 |
| 16866 | 844 | G844E | C | 1.1e-05 | 1.2e-05 | 1.3e-05 | 1.200E-05 | 1.000E-06 | 1.100E-05 | 1.300E-05 |
| 16867 | 844 | G844Q | C | 1.5e-05 | 1.4e-05 | 1.4e-05 | 1.433E-05 | 5.774E-07 | 1.400E-05 | 1.500E-05 |
| 16868 | 844 | G844G | C | 1.5e-05 | 1.1e-05 | 1e-05 | 1.200E-05 | 2.646E-06 | 1.000E-05 | 1.500E-05 |
| 16869 | 844 | G844H | C | 8.5e-06 | 1.6e-05 | 1.5e-05 | 1.317E-05 | 4.072E-06 | 8.500E-06 | 1.600E-05 |
| 16870 | 844 | G844I | C | 9.5e-06 | 1.5e-05 | 1.1e-05 | 1.183E-05 | 2.843E-06 | 9.500E-06 | 1.500E-05 |
| 16871 | 844 | G844L | C | 1.2e-05 | 1e-05 | 1.4e-05 | 1.200E-05 | 2.000E-06 | 1.000E-05 | 1.400E-05 |
| 16872 | 844 | G844K | C | 1.6e-05 | 1e-05 | 1.4e-05 | 1.333E-05 | 3.055E-06 | 1.000E-05 | 1.600E-05 |
| 16873 | 844 | G844M | C | 1.1e-05 | 1.6e-05 | 1.1e-05 | 1.267E-05 | 2.887E-06 | 1.100E-05 | 1.600E-05 |
| 16874 | 844 | G844F | C | 1.6e-05 | 9.6e-06 | 1.5e-05 | 1.353E-05 | 3.443E-06 | 9.600E-06 | 1.600E-05 |
| 16875 | 844 | G844P | C | 1.8e-05 | 9.6e-06 | 1.5e-05 | 1.420E-05 | 4.257E-06 | 9.600E-06 | 1.800E-05 |
| 16876 | 844 | G844S | C | 1.7e-05 | 1.1e-05 | 1.3e-05 | 1.367E-05 | 3.055E-06 | 1.100E-05 | 1.700E-05 |
| 16877 | 844 | G844T | C | 9.5e-06 | 5.7e-06 | 1e-05 | 8.400E-06 | 2.352E-06 | 5.700E-06 | 1.000E-05 |
| 16878 | 844 | G844W | C | 1.1e-05 | 1.4e-05 | 1.4e-05 | 1.300E-05 | 1.732E-06 | 1.100E-05 | 1.400E-05 |
| 16879 | 844 | G844Y | C | 1.3e-05 | 1.1e-05 | 1.2e-05 | 1.200E-05 | 1.000E-06 | 1.100E-05 | 1.300E-05 |
| 16880 | 844 | G844V | C | 6.6e-06 | 1.5e-05 | 9.9e-06 | 1.050E-05 | 4.232E-06 | 6.600E-06 | 1.500E-05 |
| 16881 | 845 | G845A | C | 9.5e-06 | 1.1e-05 | 1.2e-05 | 1.083E-05 | 1.258E-06 | 9.500E-06 | 1.200E-05 |
| 16882 | 845 | G845R | C | 9.5e-06 | 9.2e-06 | 1e-05 | 9.567E-06 | 4.041E-07 | 9.200E-06 | 1.000E-05 |
| 16883 | 845 | G845N | C | 9.7e-06 | 1.6e-05 | 1e-05 | 1.190E-05 | 3.554E-06 | 9.700E-06 | 1.600E-05 |
| 16884 | 845 | G845D | C | 1.4e-05 | 1.4e-05 | 1.4e-05 | 1.400E-05 | 0.000E+00 | 1.400E-05 | 1.400E-05 |
| 16885 | 845 | G845C | C | 1.3e-05 | 1.4e-05 | 8.5e-06 | 1.183E-05 | 2.930E-06 | 8.500E-06 | 1.400E-05 |
| 16886 | 845 | G845E | C | 1.3e-05 | 9.3e-06 | 9.3e-06 | 1.053E-05 | 2.136E-06 | 9.300E-06 | 1.300E-05 |
| 16887 | 845 | G845Q | C | 1.5e-05 | 1.5e-05 | 1e-05 | 1.333E-05 | 2.887E-06 | 1.000E-05 | 1.500E-05 |
| 16888 | 845 | G845G | C | 1.2e-05 | 2.4e-05 | 1.1e-05 | 1.567E-05 | 7.234E-06 | 1.100E-05 | 2.400E-05 |
| 16889 | 845 | G845H | C | 7.6e-06 | 1e-05 | 1.3e-05 | 1.020E-05 | 2.706E-06 | 7.600E-06 | 1.300E-05 |
| 16890 | 845 | G845I | C | 9.7e-06 | 1e-05 | 1.4e-05 | 1.123E-05 | 2.401E-06 | 9.700E-06 | 1.400E-05 |
| 16891 | 845 | G845L | C | 1.4e-05 | 1.3e-05 | 1.7e-05 | 1.467E-05 | 2.082E-06 | 1.300E-05 | 1.700E-05 |
| 16892 | 845 | G845K | C | 1.2e-05 | 1.4e-05 | 1.1e-05 | 1.233E-05 | 1.528E-06 | 1.100E-05 | 1.400E-05 |
| 16893 | 845 | G845M | C | 1.4e-05 | 1e-05 | 6.1e-06 | 1.003E-05 | 3.950E-06 | 6.100E-06 | 1.400E-05 |
| 16894 | 845 | G845F | C | 1.8e-05 | 1e-05 | 1.7e-05 | 1.500E-05 | 4.359E-06 | 1.000E-05 | 1.800E-05 |
| 16895 | 845 | G845P | C | 1e-05 | 1.2e-05 | 1.5e-05 | 1.233E-05 | 2.517E-06 | 1.000E-05 | 1.500E-05 |
| 16896 | 845 | G845S | C | 8.1e-06 | 1.4e-05 | 1.5e-05 | 1.237E-05 | 3.729E-06 | 8.100E-06 | 1.500E-05 |
| 16897 | 845 | G845T | C | 1e-05 | 1.2e-05 | 1.6e-05 | 1.267E-05 | 3.055E-06 | 1.000E-05 | 1.600E-05 |
| 16898 | 845 | G845W | C | 1e-05 | 9.5e-06 | 9.2e-06 | 9.567E-06 | 4.041E-07 | 9.200E-06 | 1.000E-05 |
| 16899 | 845 | G845Y | C | 1.1e-05 | 1.3e-05 | 1e-05 | 1.133E-05 | 1.528E-06 | 1.000E-05 | 1.300E-05 |
| 16900 | 845 | G845V | C | 8.6e-06 | 9.3e-06 | 1.7e-05 | 1.163E-05 | 4.661E-06 | 8.600E-06 | 1.700E-05 |
| 16901 | 846 | G846A | C | 1.3e-05 | 1.5e-05 | 1.6e-05 | 1.467E-05 | 1.528E-06 | 1.300E-05 | 1.600E-05 |
| 16902 | 846 | G846R | C | 1.5e-05 | 1.4e-05 | 1.5e-05 | 1.467E-05 | 5.774E-07 | 1.400E-05 | 1.500E-05 |
| 16903 | 846 | G846N | C | 1.1e-05 | 8.6e-06 | 1.1e-05 | 1.020E-05 | 1.386E-06 | 8.600E-06 | 1.100E-05 |
| 16904 | 846 | G846D | C | 1.6e-05 | 6e-06 | 8.4e-06 | 1.013E-05 | 5.220E-06 | 6.000E-06 | 1.600E-05 |
| 16905 | 846 | G846C | C | 9.8e-06 | 1.5e-05 | 7.2e-06 | 1.067E-05 | 3.972E-06 | 7.200E-06 | 1.500E-05 |
| 16906 | 846 | G846E | C | 8.6e-06 | 1.3e-05 | 1.6e-05 | 1.253E-05 | 3.722E-06 | 8.600E-06 | 1.600E-05 |
| 16907 | 846 | G846Q | C | 1.1e-05 | 1.4e-05 | 1.5e-05 | 1.333E-05 | 2.082E-06 | 1.100E-05 | 1.500E-05 |
| 16908 | 846 | G846G | C | 9.8e-06 | 1.1e-05 | 1.5e-05 | 1.193E-05 | 2.723E-06 | 9.800E-06 | 1.500E-05 |
| 16909 | 846 | G846H | C | 6.9e-06 | 9.9e-06 | 9.4e-06 | 8.733E-06 | 1.607E-06 | 6.900E-06 | 9.900E-06 |
| 16910 | 846 | G846I | C | 6.8e-06 | 1.4e-05 | 9.7e-06 | 1.017E-05 | 3.623E-06 | 6.800E-06 | 1.400E-05 |
| 16911 | 846 | G846L | C | 1.1e-05 | 1.7e-05 | 1.1e-05 | 1.300E-05 | 3.464E-06 | 1.100E-05 | 1.700E-05 |
| 16912 | 846 | G846K | C | 1e-05 | 1.3e-05 | 9.3e-06 | 1.077E-05 | 1.966E-06 | 9.300E-06 | 1.300E-05 |
| 16913 | 846 | G846M | C | 1e-05 | 1e-05 | 1.1e-05 | 1.033E-05 | 5.774E-07 | 1.000E-05 | 1.100E-05 |
| 16914 | 846 | G846F | C | 1.1e-05 | 1.2e-05 | 1.7e-05 | 1.333E-05 | 3.215E-06 | 1.100E-05 | 1.700E-05 |

| | | | | | | | | | | |
|---|---|---|---|---|---|---|---|---|---|---|
| 16915 | 846 | G846P | C | 1.4e-05 | 1.5e-05 | 1.4e-05 | 1.433E-05 | 5.774E-07 | 1.400E-05 | 1.500E-05 |
| 16916 | 846 | G846S | C | 1.5e-05 | 1.2e-05 | 1.5e-05 | 1.400E-05 | 1.732E-06 | 1.200E-05 | 1.500E-05 |
| 16917 | 846 | G846T | C | 1.3e-05 | 1.1e-05 | 1.3e-05 | 1.233E-05 | 1.155E-06 | 1.100E-05 | 1.300E-05 |
| 16918 | 846 | G846W | C | 1.5e-05 | 1.5e-05 | 1.4e-05 | 1.467E-05 | 5.774E-07 | 1.400E-05 | 1.500E-05 |
| 16919 | 846 | G846Y | C | 1.1e-05 | 1.6e-05 | 1.1e-05 | 1.267E-05 | 2.887E-06 | 1.100E-05 | 1.600E-05 |
| 16920 | 846 | G846V | C | 1.1e-05 | 1.4e-05 | 1.3e-05 | 1.267E-05 | 1.528E-06 | 1.100E-05 | 1.400E-05 |
| 16921 | 847 | L847A | C | 1.6e-05 | 1.2e-05 | 1.4e-05 | 1.400E-05 | 2.000E-06 | 1.200E-05 | 1.600E-05 |
| 16922 | 847 | L847R | C | 2e-05 | 1.5e-05 | 1.7e-05 | 1.733E-05 | 2.517E-06 | 1.500E-05 | 2.000E-05 |
| 16923 | 847 | L847N | C | 1.3e-05 | 7.3e-06 | 1.4e-05 | 1.143E-05 | 3.614E-06 | 7.300E-06 | 1.400E-05 |
| 16924 | 847 | L847D | C | 1.5e-05 | 1e-05 | 1e-05 | 1.167E-05 | 2.887E-06 | 1.000E-05 | 1.500E-05 |
| 16925 | 847 | L847C | C | 1.5e-05 | 8.4e-06 | 1.4e-05 | 1.247E-05 | 3.557E-06 | 8.400E-06 | 1.500E-05 |
| 16926 | 847 | L847E | C | 1e-05 | 1.5e-05 | 9.8e-06 | 1.160E-05 | 2.946E-06 | 9.800E-06 | 1.500E-05 |
| 16927 | 847 | L847Q | C | 1.4e-05 | 9.3e-06 | 1.7e-05 | 1.343E-05 | 3.881E-06 | 9.300E-06 | 1.700E-05 |
| 16928 | 847 | L847G | C | 9.7e-06 | 1.4e-05 | 1.2e-05 | 1.190E-05 | 2.152E-06 | 9.700E-06 | 1.400E-05 |
| 16929 | 847 | L847H | C | 1.1e-05 | 9.7e-06 | 9.5e-06 | 1.007E-05 | 8.145E-07 | 9.500E-06 | 1.100E-05 |
| 16930 | 847 | L847I | C | 1.6e-05 | 7.9e-06 | 1.4e-05 | 1.263E-05 | 4.219E-06 | 7.900E-06 | 1.600E-05 |
| 16931 | 847 | L847L | C | 1.1e-05 | 1.6e-05 | 1.4e-05 | 1.367E-05 | 2.517E-06 | 1.100E-05 | 1.600E-05 |
| 16932 | 847 | L847K | C | 1.5e-05 | 1.6e-05 | 1.2e-05 | 1.433E-05 | 2.082E-06 | 1.200E-05 | 1.600E-05 |
| 16933 | 847 | L847M | C | 8.2e-06 | 1.5e-05 | 9e-06 | 1.073E-05 | 3.717E-06 | 8.200E-06 | 1.500E-05 |
| 16934 | 847 | L847F | C | 1.7e-05 | 1e-05 | 1.1e-05 | 1.267E-05 | 3.786E-06 | 1.000E-05 | 1.700E-05 |
| 16935 | 847 | L847P | C | 1.4e-05 | 1e-05 | 1.4e-05 | 1.267E-05 | 2.309E-06 | 1.000E-05 | 1.400E-05 |
| 16936 | 847 | L847S | C | 8.4e-06 | 1.5e-05 | 1.1e-05 | 1.147E-05 | 3.325E-06 | 8.400E-06 | 1.500E-05 |
| 16937 | 847 | L847T | C | 1.5e-05 | 1.5e-05 | 1.1e-05 | 1.367E-05 | 2.309E-06 | 1.100E-05 | 1.500E-05 |
| 16938 | 847 | L847W | C | 1e-05 | 1.7e-05 | 1.4e-05 | 1.367E-05 | 3.512E-06 | 1.000E-05 | 1.700E-05 |
| 16939 | 847 | L847Y | C | 1.1e-05 | 1.5e-05 | 1.1e-05 | 1.233E-05 | 2.309E-06 | 1.100E-05 | 1.500E-05 |
| 16940 | 847 | L847V | C | 1e-05 | 1.5e-05 | 1.6e-05 | 1.367E-05 | 3.215E-06 | 1.000E-05 | 1.600E-05 |
| 16941 | 848 | V848A | C | 1.2e-05 | 1.4e-05 | 8.2e-06 | 1.140E-05 | 2.946E-06 | 8.200E-06 | 1.400E-05 |
| 16942 | 848 | V848R | C | 1.4e-05 | 1.4e-05 | 1.6e-05 | 1.467E-05 | 1.155E-06 | 1.400E-05 | 1.600E-05 |
| 16943 | 848 | V848N | C | 1.7e-05 | 1.5e-05 | 9.9e-06 | 1.397E-05 | 3.661E-06 | 9.900E-06 | 1.700E-05 |
| 16944 | 848 | V848D | C | 1.1e-05 | 1.5e-05 | 1.2e-05 | 1.267E-05 | 2.082E-06 | 1.100E-05 | 1.500E-05 |
| 16945 | 848 | V848C | C | 1e-05 | 1.6e-05 | 1.5e-05 | 1.367E-05 | 3.215E-06 | 1.000E-05 | 1.600E-05 |
| 16946 | 848 | V848E | C | 1.1e-05 | 8.7e-06 | 9.7e-06 | 9.800E-06 | 1.153E-06 | 8.700E-06 | 1.100E-05 |
| 16947 | 848 | V848Q | C | 9.1e-06 | 1.6e-05 | 9.7e-06 | 1.160E-05 | 3.822E-06 | 9.100E-06 | 1.600E-05 |
| 16948 | 848 | V848G | C | 1.3e-05 | 1.7e-05 | 1.8e-05 | 1.600E-05 | 2.646E-06 | 1.300E-05 | 1.800E-05 |
| 16949 | 848 | V848H | C | 8.6e-06 | 1.6e-05 | 9.8e-06 | 1.147E-05 | 3.972E-06 | 8.600E-06 | 1.600E-05 |
| 16950 | 848 | V848I | C | 1.1e-05 | 1.1e-05 | 1.2e-05 | 1.133E-05 | 5.774E-07 | 1.100E-05 | 1.200E-05 |
| 16951 | 848 | V848L | C | 1e-05 | 1e-05 | 1.3e-05 | 1.100E-05 | 1.732E-06 | 1.000E-05 | 1.300E-05 |
| 16952 | 848 | V848K | C | 1.3e-05 | 1.5e-05 | 1.7e-05 | 1.500E-05 | 2.000E-06 | 1.300E-05 | 1.700E-05 |
| 16953 | 848 | V848M | C | 1.5e-05 | 1e-05 | 1.1e-05 | 1.200E-05 | 2.646E-06 | 1.000E-05 | 1.500E-05 |
| 16954 | 848 | V848F | C | 1.3e-05 | 1.4e-05 | 1.6e-05 | 1.433E-05 | 1.528E-06 | 1.300E-05 | 1.600E-05 |
| 16955 | 848 | V848P | C | 1.1e-05 | 9.4e-06 | 1.1e-05 | 1.047E-05 | 9.238E-07 | 9.400E-06 | 1.100E-05 |
| 16956 | 848 | V848S | C | 9.6e-06 | 1.2e-05 | 9.5e-06 | 1.037E-05 | 1.415E-06 | 9.500E-06 | 1.200E-05 |
| 16957 | 848 | V848T | C | 1.5e-05 | 6.8e-06 | 6.8e-06 | 9.533E-06 | 4.734E-06 | 6.800E-06 | 1.500E-05 |
| 16958 | 848 | V848W | C | 8.9e-06 | 1.1e-05 | 1.5e-05 | 1.163E-05 | 3.099E-06 | 8.900E-06 | 1.500E-05 |
| 16959 | 848 | V848Y | C | 1.3e-05 | 1e-05 | 1.4e-05 | 1.233E-05 | 2.082E-06 | 1.000E-05 | 1.400E-05 |
| 16960 | 848 | V848V | C | 1.1e-05 | 9.9e-06 | 1e-05 | 1.030E-05 | 6.083E-07 | 9.900E-06 | 1.100E-05 |
| 16961 | 849 | Q849A | C | 1.5e-05 | 1.1e-05 | 1.1e-05 | 1.233E-05 | 2.309E-06 | 1.100E-05 | 1.500E-05 |
| 16962 | 849 | Q849R | C | 1.5e-05 | 1.9e-05 | 1.2e-05 | 1.533E-05 | 3.512E-06 | 1.200E-05 | 1.900E-05 |
| 16963 | 849 | Q849N | C | 8.8e-06 | 1e-05 | 7.9e-06 | 8.900E-06 | 1.054E-06 | 7.900E-06 | 1.000E-05 |
| 16964 | 849 | Q849D | C | 1.1e-05 | 1.1e-05 | 8e-06 | 1.000E-05 | 1.732E-06 | 8.000E-06 | 1.100E-05 |
| 16965 | 849 | Q849C | C | 1.4e-05 | 1.1e-05 | 9.8e-06 | 1.160E-05 | 2.163E-06 | 9.800E-06 | 1.400E-05 |
| 16966 | 849 | Q849E | C | 1.4e-05 | 8.9e-06 | 1.7e-05 | 1.330E-05 | 4.095E-06 | 8.900E-06 | 1.700E-05 |
| 16967 | 849 | Q849Q | C | 1e-05 | 1e-05 | 9.7e-06 | 9.900E-06 | 1.732E-07 | 9.700E-06 | 1.000E-05 |
| 16968 | 849 | Q849G | C | 1.4e-05 | 1e-05 | 1.2e-05 | 1.200E-05 | 2.000E-06 | 1.000E-05 | 1.400E-05 |
| 16969 | 849 | Q849H | C | 1.5e-05 | 1.6e-05 | 1.6e-05 | 1.567E-05 | 5.774E-07 | 1.500E-05 | 1.600E-05 |
| 16970 | 849 | Q849I | C | 1.3e-05 | 1.1e-05 | 1.1e-05 | 1.167E-05 | 1.155E-06 | 1.100E-05 | 1.300E-05 |
| 16971 | 849 | Q849L | C | 1.2e-05 | 1.1e-05 | 1.3e-05 | 1.200E-05 | 1.000E-06 | 1.100E-05 | 1.300E-05 |
| 16972 | 849 | Q849K | C | 1.7e-05 | 1.5e-05 | 1.6e-05 | 1.600E-05 | 1.000E-06 | 1.500E-05 | 1.700E-05 |
| 16973 | 849 | Q849M | C | 1.1e-05 | 8.5e-06 | 1.1e-05 | 1.017E-05 | 1.443E-06 | 8.500E-06 | 1.100E-05 |
| 16974 | 849 | Q849F | C | 9.4e-06 | 1.7e-05 | 9.4e-06 | 1.193E-05 | 4.388E-06 | 9.400E-06 | 1.700E-05 |
| 16975 | 849 | Q849P | C | 1.1e-05 | 9.8e-06 | 1.6e-05 | 1.227E-05 | 3.288E-06 | 9.800E-06 | 1.600E-05 |
| 16976 | 849 | Q849S | C | 1.5e-05 | 8.7e-06 | 1.4e-05 | 1.257E-05 | 3.386E-06 | 8.700E-06 | 1.500E-05 |
| 16977 | 849 | Q849T | C | 1.4e-05 | 1.2e-05 | 1.1e-05 | 1.233E-05 | 1.528E-06 | 1.100E-05 | 1.400E-05 |
| 16978 | 849 | Q849W | C | 9.9e-06 | 1.6e-05 | 1.4e-05 | 1.330E-05 | 3.110E-06 | 9.900E-06 | 1.600E-05 |
| 16979 | 849 | Q849Y | C | 1.1e-05 | 1e-05 | 1e-05 | 1.033E-05 | 5.774E-07 | 1.000E-05 | 1.100E-05 |
| 16980 | 849 | Q849V | C | 1.5e-05 | 1.6e-05 | 1.4e-05 | 1.500E-05 | 1.000E-06 | 1.400E-05 | 1.600E-05 |
| 16981 | 850 | P850A | C | 9.4e-06 | 1.2e-05 | 1.5e-05 | 1.213E-05 | 2.802E-06 | 9.400E-06 | 1.500E-05 |
| 16982 | 850 | P850R | C | 5.9e-06 | 8.8e-06 | 1.4e-05 | 9.567E-06 | 4.104E-06 | 5.900E-06 | 1.400E-05 |
| 16983 | 850 | P850N | C | 1.1e-05 | 1.6e-05 | 1.3e-05 | 1.333E-05 | 2.517E-06 | 1.100E-05 | 1.600E-05 |
| 16984 | 850 | P850D | C | 1.2e-05 | 1.1e-05 | 2e-05 | 1.433E-05 | 4.933E-06 | 1.100E-05 | 2.000E-05 |
| 16985 | 850 | P850C | C | 1.6e-05 | 8.6e-06 | 1.5e-05 | 1.320E-05 | 4.015E-06 | 8.600E-06 | 1.600E-05 |
| 16986 | 850 | P850E | C | 1.2e-05 | 1.6e-05 | 1.7e-05 | 1.500E-05 | 2.646E-06 | 1.200E-05 | 1.700E-05 |
| 16987 | 850 | P850Q | C | 1.5e-05 | 1.3e-05 | 1.5e-05 | 1.433E-05 | 1.155E-06 | 1.300E-05 | 1.500E-05 |
| 16988 | 850 | P850G | C | 1.5e-05 | 6.4e-06 | 1.1e-05 | 1.080E-05 | 4.303E-06 | 6.400E-06 | 1.500E-05 |
| 16989 | 850 | P850H | C | 1.5e-05 | 2e-05 | 1.4e-05 | 1.633E-05 | 3.215E-06 | 1.400E-05 | 2.000E-05 |
| 16990 | 850 | P850I | C | 1.7e-05 | 1.6e-05 | 6.5e-06 | 1.317E-05 | 5.795E-06 | 6.500E-06 | 1.700E-05 |
| 16991 | 850 | P850L | C | 8.9e-06 | 1.1e-05 | 1.6e-05 | 1.197E-05 | 3.647E-06 | 8.900E-06 | 1.600E-05 |
| 16992 | 850 | P850K | C | 1.7e-05 | 1.6e-05 | 1.5e-05 | 1.600E-05 | 1.000E-06 | 1.500E-05 | 1.700E-05 |
| 16993 | 850 | P850M | C | 1.1e-05 | 1.1e-05 | 1e-05 | 1.067E-05 | 5.774E-07 | 1.000E-05 | 1.100E-05 |
| 16994 | 850 | P850F | C | 1.4e-05 | 1.6e-05 | 8.9e-06 | 1.297E-05 | 3.661E-06 | 8.900E-06 | 1.600E-05 |
| 16995 | 850 | P850P | C | 1e-05 | 8.7e-06 | 1.4e-05 | 1.090E-05 | 2.762E-06 | 8.700E-06 | 1.400E-05 |
| 16996 | 850 | P850S | C | 1.3e-05 | 1.2e-05 | 1.6e-05 | 1.367E-05 | 2.082E-06 | 1.200E-05 | 1.600E-05 |
| 16997 | 850 | P850T | C | 1.3e-05 | 1.4e-05 | 1.3e-05 | 1.333E-05 | 5.774E-07 | 1.300E-05 | 1.400E-05 |
| 16998 | 850 | P850W | C | 1.3e-05 | 1.6e-05 | 1.3e-05 | 1.400E-05 | 1.732E-06 | 1.300E-05 | 1.600E-05 |
| 16999 | 850 | P850Y | C | 1e-05 | 1.6e-05 | 1.4e-05 | 1.333E-05 | 3.055E-06 | 1.000E-05 | 1.600E-05 |

| | | | | | | | | | | |
|---|---|---|---|---|---|---|---|---|---|---|
| 17000 | 850 | P850V | C | 1.4e-05 | 8.7e-06 | 1.2e-05 | 1.157E-05 | 2.676E-06 | 8.700E-06 | 1.400E-05 |
| 17001 | 851 | G851A | C | 1.1e-05 | 1.4e-05 | 1.1e-05 | 1.200E-05 | 1.732E-06 | 1.100E-05 | 1.400E-05 |
| 17002 | 851 | G851R | C | 1.6e-05 | 1.6e-05 | 8.3e-06 | 1.343E-05 | 4.446E-06 | 8.300E-06 | 1.600E-05 |
| 17003 | 851 | G851N | C | 1.5e-05 | 1.1e-05 | 8.2e-06 | 1.140E-05 | 3.418E-06 | 8.200E-06 | 1.500E-05 |
| 17004 | 851 | G851D | C | 9.6e-06 | 1.3e-05 | 1.5e-05 | 1.253E-05 | 2.730E-06 | 9.600E-06 | 1.500E-05 |
| 17005 | 851 | G851C | C | 1.1e-05 | 1e-05 | 8.7e-06 | 9.900E-06 | 1.153E-06 | 8.700E-06 | 1.100E-05 |
| 17006 | 851 | G851E | C | 9.2e-06 | 9.1e-06 | 1.6e-05 | 1.143E-05 | 3.955E-06 | 9.100E-06 | 1.600E-05 |
| 17007 | 851 | G851Q | C | 8.6e-06 | 1.1e-05 | 1.7e-05 | 1.220E-05 | 4.327E-06 | 8.600E-06 | 1.700E-05 |
| 17008 | 851 | G851G | C | 1.1e-05 | 9.9e-06 | 1.5e-05 | 1.197E-05 | 2.684E-06 | 9.900E-06 | 1.500E-05 |
| 17009 | 851 | G851H | C | 1.5e-05 | 1.3e-05 | 1.4e-05 | 1.400E-05 | 1.000E-06 | 1.300E-05 | 1.500E-05 |
| 17010 | 851 | G851I | C | 1.6e-05 | 1.3e-05 | 1e-05 | 1.300E-05 | 3.000E-06 | 1.000E-05 | 1.600E-05 |
| 17011 | 851 | G851L | C | 9.8e-06 | 1.6e-05 | 1.5e-05 | 1.360E-05 | 3.329E-06 | 9.800E-06 | 1.600E-05 |
| 17012 | 851 | G851K | C | 8.8e-06 | 7.7e-06 | 1.7e-05 | 1.117E-05 | 5.082E-06 | 7.700E-06 | 1.700E-05 |
| 17013 | 851 | G851M | C | 9.3e-06 | 1.5e-05 | 1.6e-05 | 1.343E-05 | 3.614E-06 | 9.300E-06 | 1.600E-05 |
| 17014 | 851 | G851F | C | 1.1e-05 | 9.7e-06 | 1.5e-05 | 1.190E-05 | 2.762E-06 | 9.700E-06 | 1.500E-05 |
| 17015 | 851 | G851P | C | 1.2e-05 | 1.6e-05 | 1.4e-05 | 1.400E-05 | 2.000E-06 | 1.200E-05 | 1.600E-05 |
| 17016 | 851 | G851S | C | 1.6e-05 | 1.3e-05 | 9.4e-06 | 1.280E-05 | 3.305E-06 | 9.400E-06 | 1.600E-05 |
| 17017 | 851 | G851T | C | 1.5e-05 | 1.3e-05 | 9.7e-06 | 1.257E-05 | 2.676E-06 | 9.700E-06 | 1.500E-05 |
| 17018 | 851 | G851W | C | 1.3e-05 | 1.1e-05 | 1.1e-05 | 1.167E-05 | 1.155E-06 | 1.100E-05 | 1.300E-05 |
| 17019 | 851 | G851Y | C | 1.5e-05 | 1.1e-05 | 9.4e-06 | 1.180E-05 | 2.884E-06 | 9.400E-06 | 1.500E-05 |
| 17020 | 851 | G851V | C | 8.3e-06 | 1.1e-05 | 9.3e-06 | 9.533E-06 | 1.365E-06 | 8.300E-06 | 1.100E-05 |
| 17021 | 852 | G852A | C | 1.2e-05 | 1e-05 | 1e-05 | 1.067E-05 | 1.155E-06 | 1.000E-05 | 1.200E-05 |
| 17022 | 852 | G852R | C | 1.6e-05 | 1.5e-05 | 1.4e-05 | 1.500E-05 | 1.000E-06 | 1.400E-05 | 1.600E-05 |
| 17023 | 852 | G852N | C | 8.6e-06 | 1.1e-05 | 1.5e-05 | 1.153E-05 | 3.233E-06 | 8.600E-06 | 1.500E-05 |
| 17024 | 852 | G852D | C | 1.1e-05 | 7.5e-06 | 1.5e-05 | 1.117E-05 | 3.753E-06 | 7.500E-06 | 1.500E-05 |
| 17025 | 852 | G852C | C | 1.7e-05 | 1.1e-05 | 1.4e-05 | 1.400E-05 | 3.000E-06 | 1.100E-05 | 1.700E-05 |
| 17026 | 852 | G852E | C | 1.5e-05 | 1.5e-05 | 1.2e-05 | 1.400E-05 | 1.732E-06 | 1.200E-05 | 1.500E-05 |
| 17027 | 852 | G852Q | C | 1.4e-05 | 1.2e-05 | 1.5e-05 | 1.367E-05 | 1.528E-06 | 1.200E-05 | 1.500E-05 |
| 17028 | 852 | G852G | C | 1.4e-05 | 1.3e-05 | 1.5e-05 | 1.400E-05 | 1.000E-06 | 1.300E-05 | 1.500E-05 |
| 17029 | 852 | G852H | C | 9.8e-06 | 1.4e-05 | 1.3e-05 | 1.227E-05 | 2.194E-06 | 9.800E-06 | 1.400E-05 |
| 17030 | 852 | G852I | C | 8.5e-06 | 9.8e-06 | 1.3e-05 | 1.043E-05 | 2.316E-06 | 8.500E-06 | 1.300E-05 |
| 17031 | 852 | G852L | C | 1.1e-05 | 1.6e-05 | 1.1e-05 | 1.267E-05 | 2.887E-06 | 1.100E-05 | 1.600E-05 |
| 17032 | 852 | G852K | C | 1.3e-05 | 1.4e-05 | 1.6e-05 | 1.433E-05 | 1.528E-06 | 1.300E-05 | 1.600E-05 |
| 17033 | 852 | G852M | C | 1.1e-05 | 1.5e-05 | 1.4e-05 | 1.333E-05 | 2.082E-06 | 1.100E-05 | 1.500E-05 |
| 17034 | 852 | G852F | C | 1.6e-05 | 1e-05 | 1.3e-05 | 1.300E-05 | 3.000E-06 | 1.000E-05 | 1.600E-05 |
| 17035 | 852 | G852P | C | 1e-05 | 1.1e-05 | 1.2e-05 | 1.100E-05 | 1.000E-06 | 1.000E-05 | 1.200E-05 |
| 17036 | 852 | G852S | C | 1.5e-05 | 1.4e-05 | 1.1e-05 | 1.333E-05 | 2.082E-06 | 1.100E-05 | 1.500E-05 |
| 17037 | 852 | G852T | C | 1.1e-05 | 1.5e-05 | 1.5e-05 | 1.367E-05 | 2.309E-06 | 1.100E-05 | 1.500E-05 |
| 17038 | 852 | G852W | C | 1.2e-05 | 9.5e-06 | 1.4e-05 | 1.183E-05 | 2.255E-06 | 9.500E-06 | 1.400E-05 |
| 17039 | 852 | G852Y | C | 9.4e-06 | 1.3e-05 | 1.1e-05 | 1.113E-05 | 1.804E-06 | 9.400E-06 | 1.300E-05 |
| 17040 | 852 | G852V | C | 1.5e-05 | 1.6e-05 | 1.2e-05 | 1.433E-05 | 2.082E-06 | 1.200E-05 | 1.600E-05 |
| 17041 | 853 | S853A | C | 9.7e-06 | 1.1e-05 | 1.2e-05 | 1.090E-05 | 1.153E-06 | 9.700E-06 | 1.200E-05 |
| 17042 | 853 | S853R | C | 1.3e-05 | 1.6e-05 | 1e-05 | 1.300E-05 | 3.000E-06 | 1.000E-05 | 1.600E-05 |
| 17043 | 853 | S853N | C | 1.6e-05 | 8.4e-06 | 1.4e-05 | 1.280E-05 | 3.940E-06 | 8.400E-06 | 1.600E-05 |
| 17044 | 853 | S853D | C | 1.1e-05 | 1.4e-05 | 1.1e-05 | 1.200E-05 | 1.732E-06 | 1.100E-05 | 1.400E-05 |
| 17045 | 853 | S853C | C | 1.7e-05 | 1.5e-05 | 1.1e-05 | 1.433E-05 | 3.055E-06 | 1.100E-05 | 1.700E-05 |
| 17046 | 853 | S853E | C | 8.2e-06 | 1e-05 | 1.1e-05 | 9.733E-06 | 1.419E-06 | 8.200E-06 | 1.100E-05 |
| 17047 | 853 | S853Q | C | 1.4e-05 | 1e-05 | 1.2e-05 | 1.200E-05 | 2.000E-06 | 1.000E-05 | 1.400E-05 |
| 17048 | 853 | S853G | C | 1.2e-05 | 1.3e-05 | 1.2e-05 | 1.233E-05 | 5.774E-07 | 1.200E-05 | 1.300E-05 |
| 17049 | 853 | S853H | C | 1.5e-05 | 9.9e-06 | 1.5e-05 | 1.330E-05 | 2.944E-06 | 9.900E-06 | 1.500E-05 |
| 17050 | 853 | S853I | C | 1.5e-05 | 1e-05 | 1.4e-05 | 1.300E-05 | 2.646E-06 | 1.000E-05 | 1.500E-05 |
| 17051 | 853 | S853L | C | 1.6e-05 | 1e-05 | 8.7e-06 | 1.157E-05 | 3.894E-06 | 8.700E-06 | 1.600E-05 |
| 17052 | 853 | S853K | C | 1.2e-05 | 1.6e-05 | 1.1e-05 | 1.300E-05 | 2.646E-06 | 1.100E-05 | 1.600E-05 |
| 17053 | 853 | S853M | C | 1.3e-05 | 1.6e-05 | 1.7e-05 | 1.533E-05 | 2.082E-06 | 1.300E-05 | 1.700E-05 |
| 17054 | 853 | S853F | C | 1.5e-05 | 1.2e-05 | 1e-05 | 1.233E-05 | 2.517E-06 | 1.000E-05 | 1.500E-05 |
| 17055 | 853 | S853P | C | 1.1e-05 | 1.5e-05 | 1.1e-05 | 1.233E-05 | 2.309E-06 | 1.100E-05 | 1.500E-05 |
| 17056 | 853 | S853S | C | 1e-05 | 1e-05 | 1.5e-05 | 1.167E-05 | 2.887E-06 | 1.000E-05 | 1.500E-05 |
| 17057 | 853 | S853T | C | 1.5e-05 | 6.5e-06 | 1.5e-05 | 1.217E-05 | 4.907E-06 | 6.500E-06 | 1.500E-05 |
| 17058 | 853 | S853W | C | 1.4e-05 | 1.5e-05 | 1.5e-05 | 1.467E-05 | 5.774E-07 | 1.400E-05 | 1.500E-05 |
| 17059 | 853 | S853Y | C | 1.1e-05 | 1.5e-05 | 1.5e-05 | 1.367E-05 | 2.309E-06 | 1.100E-05 | 1.500E-05 |
| 17060 | 853 | S853V | C | 5.9e-06 | 9.6e-06 | 9.4e-06 | 8.300E-06 | 2.081E-06 | 5.900E-06 | 9.600E-06 |
| 17061 | 854 | L854A | C | 1.4e-05 | 1.1e-05 | 1.6e-05 | 1.367E-05 | 2.517E-06 | 1.100E-05 | 1.600E-05 |
| 17062 | 854 | L854R | C | 1.5e-05 | 9.7e-06 | 1.1e-05 | 1.190E-05 | 2.762E-06 | 9.700E-06 | 1.500E-05 |
| 17063 | 854 | L854N | C | 1.4e-05 | 1.2e-05 | 1e-05 | 1.200E-05 | 2.000E-06 | 1.000E-05 | 1.400E-05 |
| 17064 | 854 | L854D | C | 1.1e-05 | 1.5e-05 | 8.7e-06 | 1.157E-05 | 3.188E-06 | 8.700E-06 | 1.500E-05 |
| 17065 | 854 | L854C | C | 8.4e-06 | 1.1e-05 | 9.9e-06 | 9.767E-06 | 1.305E-06 | 8.400E-06 | 1.100E-05 |
| 17066 | 854 | L854E | C | 1.4e-05 | 1.1e-05 | 1.6e-05 | 1.367E-05 | 2.517E-06 | 1.100E-05 | 1.600E-05 |
| 17067 | 854 | L854Q | C | 1.4e-05 | 8.6e-06 | 8.9e-06 | 1.050E-05 | 3.035E-06 | 8.600E-06 | 1.400E-05 |
| 17068 | 854 | L854G | C | 1.1e-05 | 1.6e-05 | 1.5e-05 | 1.400E-05 | 2.646E-06 | 1.100E-05 | 1.600E-05 |
| 17069 | 854 | L854H | C | 1.1e-05 | 1.4e-05 | 1.1e-05 | 1.200E-05 | 1.732E-06 | 1.100E-05 | 1.400E-05 |
| 17070 | 854 | L854I | C | 8.3e-06 | 9.2e-06 | 1.3e-05 | 1.017E-05 | 2.495E-06 | 8.300E-06 | 1.300E-05 |
| 17071 | 854 | L854L | C | 9.6e-06 | 9.4e-06 | 1.5e-05 | 1.133E-05 | 3.177E-06 | 9.400E-06 | 1.500E-05 |
| 17072 | 854 | L854K | C | 9.9e-06 | 1.3e-05 | 9.2e-06 | 1.070E-05 | 2.022E-06 | 9.200E-06 | 1.300E-05 |
| 17073 | 854 | L854M | C | 8.7e-06 | 1.6e-05 | 1.2e-05 | 1.223E-05 | 3.656E-06 | 8.700E-06 | 1.600E-05 |
| 17074 | 854 | L854F | C | 1.6e-05 | 1.2e-05 | 1.6e-05 | 1.467E-05 | 2.309E-06 | 1.200E-05 | 1.600E-05 |
| 17075 | 854 | L854P | C | 1e-05 | 1.5e-05 | 1.1e-05 | 1.200E-05 | 2.646E-06 | 1.000E-05 | 1.500E-05 |
| 17076 | 854 | L854S | C | 1.4e-05 | 1.1e-05 | 1.1e-05 | 1.200E-05 | 1.732E-06 | 1.100E-05 | 1.400E-05 |
| 17077 | 854 | L854T | C | 1.5e-05 | 1.1e-05 | 8.5e-06 | 1.150E-05 | 3.279E-06 | 8.500E-06 | 1.500E-05 |
| 17078 | 854 | L854W | C | 1.4e-05 | 1.4e-05 | 1.6e-05 | 1.467E-05 | 1.155E-06 | 1.400E-05 | 1.600E-05 |
| 17079 | 854 | L854Y | C | 1e-05 | 1.1e-05 | 1.4e-05 | 1.167E-05 | 2.082E-06 | 1.000E-05 | 1.400E-05 |
| 17080 | 854 | L854V | C | 1.1e-05 | 1.2e-05 | 1.3e-05 | 1.200E-05 | 1.000E-06 | 1.100E-05 | 1.300E-05 |
| 17081 | 855 | R855A | C | 1.6e-05 | 1.1e-05 | 1e-05 | 1.233E-05 | 3.215E-06 | 1.000E-05 | 1.600E-05 |
| 17082 | 855 | R855R | C | 1.4e-05 | 1.6e-05 | 1.5e-05 | 1.500E-05 | 1.000E-06 | 1.400E-05 | 1.600E-05 |
| 17083 | 855 | R855N | C | 1.1e-05 | 1.6e-05 | 1e-05 | 1.233E-05 | 3.215E-06 | 1.000E-05 | 1.600E-05 |
| 17084 | 855 | R855D | C | 7.2e-06 | 1.4e-05 | 1.6e-05 | 1.240E-05 | 4.613E-06 | 7.200E-06 | 1.600E-05 |

| | | | | | | | | | | |
|---|---|---|---|---|---|---|---|---|---|---|
| 17085 | 855 | R855C | C | 1.3e-05 | 9.2e-06 | 1.4e-05 | 1.207E-05 | 2.532E-06 | 9.200E-06 | 1.400E-05 |
| 17086 | 855 | R855E | C | 1.1e-05 | 1.5e-05 | 8.9e-06 | 1.163E-05 | 3.099E-06 | 8.900E-06 | 1.500E-05 |
| 17087 | 855 | R855Q | C | 1e-05 | 1.5e-05 | 1.6e-05 | 1.367E-05 | 3.215E-06 | 1.000E-05 | 1.600E-05 |
| 17088 | 855 | R855G | C | 1.7e-05 | 1.1e-05 | 1.4e-05 | 1.400E-05 | 3.000E-06 | 1.100E-05 | 1.700E-05 |
| 17089 | 855 | R855H | C | 9.5e-06 | 1.6e-05 | 1.6e-05 | 1.383E-05 | 3.753E-06 | 9.500E-06 | 1.600E-05 |
| 17090 | 855 | R855I | C | 9.1e-06 | 1.2e-05 | 9.2e-06 | 1.010E-05 | 1.646E-06 | 9.100E-06 | 1.200E-05 |
| 17091 | 855 | R855L | C | 9.4e-06 | 1.7e-05 | 1e-05 | 1.213E-05 | 4.225E-06 | 9.400E-06 | 1.700E-05 |
| 17092 | 855 | R855K | C | 1.4e-05 | 1.5e-05 | 1.1e-05 | 1.333E-05 | 2.082E-06 | 1.100E-05 | 1.500E-05 |
| 17093 | 855 | R855M | C | 1.5e-05 | 1.1e-05 | 1.5e-05 | 1.367E-05 | 2.309E-06 | 1.100E-05 | 1.500E-05 |
| 17094 | 855 | R855F | C | 1.6e-05 | 1.5e-05 | 1.1e-05 | 1.400E-05 | 2.646E-06 | 1.100E-05 | 1.600E-05 |
| 17095 | 855 | R855P | C | 1e-05 | 1.5e-05 | 1.3e-05 | 1.267E-05 | 2.517E-06 | 1.000E-05 | 1.500E-05 |
| 17096 | 855 | R855S | C | 1.5e-05 | 1.3e-05 | 1.4e-05 | 1.400E-05 | 1.000E-06 | 1.300E-05 | 1.500E-05 |
| 17097 | 855 | R855T | C | 9e-06 | 1.1e-05 | 1.2e-05 | 1.067E-05 | 1.528E-06 | 9.000E-06 | 1.200E-05 |
| 17098 | 855 | R855W | C | 1e-05 | 1e-05 | 1.1e-05 | 1.033E-05 | 5.774E-07 | 1.000E-05 | 1.100E-05 |
| 17099 | 855 | R855Y | C | 1.3e-05 | 1.1e-05 | 1e-05 | 1.133E-05 | 1.528E-06 | 1.000E-05 | 1.300E-05 |
| 17100 | 855 | R855V | C | 1.6e-05 | 1.3e-05 | 9.3e-06 | 1.277E-05 | 3.356E-06 | 9.300E-06 | 1.600E-05 |
| 17101 | 856 | L856A | C | 1.1e-05 | 1.3e-05 | 1.2e-05 | 1.200E-05 | 1.000E-06 | 1.100E-05 | 1.300E-05 |
| 17102 | 856 | L856R | C | 1e-05 | 7e-06 | 9.2e-06 | 8.733E-06 | 1.553E-06 | 7.000E-06 | 1.000E-05 |
| 17103 | 856 | L856N | C | 1.6e-05 | 1.4e-05 | 1.3e-05 | 1.433E-05 | 1.528E-06 | 1.300E-05 | 1.600E-05 |
| 17104 | 856 | L856D | C | 1.6e-05 | 1.5e-05 | 1.5e-05 | 1.533E-05 | 5.774E-07 | 1.500E-05 | 1.600E-05 |
| 17105 | 856 | L856C | C | 1.7e-05 | 9.4e-06 | 9e-06 | 1.180E-05 | 4.508E-06 | 9.000E-06 | 1.700E-05 |
| 17106 | 856 | L856E | C | 1.5e-05 | 1.7e-05 | 1.3e-05 | 1.500E-05 | 2.000E-06 | 1.300E-05 | 1.700E-05 |
| 17107 | 856 | L856Q | C | 1.7e-05 | 9.3e-06 | 1.4e-05 | 1.343E-05 | 3.881E-06 | 9.300E-06 | 1.700E-05 |
| 17108 | 856 | L856G | C | 1.5e-05 | 1.5e-05 | 1.2e-05 | 1.400E-05 | 1.732E-06 | 1.200E-05 | 1.500E-05 |
| 17109 | 856 | L856H | C | 1.4e-05 | 1.6e-05 | 9.3e-06 | 1.310E-05 | 3.439E-06 | 9.300E-06 | 1.600E-05 |
| 17110 | 856 | L856I | C | 1.3e-05 | 1.6e-05 | 1.6e-05 | 1.500E-05 | 1.732E-06 | 1.300E-05 | 1.600E-05 |
| 17111 | 856 | L856L | C | 1.5e-05 | 1.5e-05 | 1.5e-05 | 1.500E-05 | 0.000E+00 | 1.500E-05 | 1.500E-05 |
| 17112 | 856 | L856K | C | 8.7e-06 | 9e-06 | 1.5e-05 | 1.090E-05 | 3.554E-06 | 8.700E-06 | 1.500E-05 |
| 17113 | 856 | L856M | C | 1.4e-05 | 1.3e-05 | 1.4e-05 | 1.367E-05 | 5.774E-07 | 1.300E-05 | 1.400E-05 |
| 17114 | 856 | L856F | C | 1.4e-05 | 1.5e-05 | 1.5e-05 | 1.467E-05 | 5.774E-07 | 1.400E-05 | 1.500E-05 |
| 17115 | 856 | L856P | C | 1.1e-05 | 1.8e-05 | 1.5e-05 | 1.467E-05 | 3.512E-06 | 1.100E-05 | 1.800E-05 |
| 17116 | 856 | L856S | C | 9.1e-06 | 1.1e-05 | 1.5e-05 | 1.170E-05 | 3.012E-06 | 9.100E-06 | 1.500E-05 |
| 17117 | 856 | L856T | C | 1.1e-05 | 1.2e-05 | 1.3e-05 | 1.200E-05 | 1.000E-06 | 1.100E-05 | 1.300E-05 |
| 17118 | 856 | L856W | C | 1.3e-05 | 1e-05 | 1.1e-05 | 1.133E-05 | 1.528E-06 | 1.000E-05 | 1.300E-05 |
| 17119 | 856 | L856Y | C | 1e-05 | 1.6e-05 | 1.6e-05 | 1.400E-05 | 3.464E-06 | 1.000E-05 | 1.600E-05 |
| 17120 | 856 | L856V | C | 1.8e-05 | 1.1e-05 | 1e-05 | 1.300E-05 | 4.359E-06 | 1.000E-05 | 1.800E-05 |
| 17121 | 857 | S857A | C | 1.1e-05 | 1.5e-05 | 9.8e-06 | 1.193E-05 | 2.723E-06 | 9.800E-06 | 1.500E-05 |
| 17122 | 857 | S857R | C | 1.1e-05 | 1.6e-05 | 9e-06 | 1.200E-05 | 3.606E-06 | 9.000E-06 | 1.600E-05 |
| 17123 | 857 | S857N | C | 1e-05 | 9.1e-06 | 1.5e-05 | 1.137E-05 | 3.179E-06 | 9.100E-06 | 1.500E-05 |
| 17124 | 857 | S857D | C | 7.7e-06 | 1.4e-05 | 1.5e-05 | 1.223E-05 | 3.958E-06 | 7.700E-06 | 1.500E-05 |
| 17125 | 857 | S857C | C | 9e-06 | 1e-05 | 1e-05 | 9.667E-06 | 5.774E-07 | 9.000E-06 | 1.000E-05 |
| 17126 | 857 | S857E | C | 1.4e-05 | 1.3e-05 | 1.4e-05 | 1.367E-05 | 5.774E-07 | 1.300E-05 | 1.400E-05 |
| 17127 | 857 | S857Q | C | 1e-05 | 9.2e-06 | 1.4e-05 | 1.107E-05 | 2.572E-06 | 9.200E-06 | 1.400E-05 |
| 17128 | 857 | S857G | C | 1.2e-05 | 9.2e-06 | 1.5e-05 | 1.207E-05 | 2.901E-06 | 9.200E-06 | 1.500E-05 |
| 17129 | 857 | S857H | C | 1.2e-05 | 6.2e-06 | 1.5e-05 | 1.107E-05 | 4.474E-06 | 6.200E-06 | 1.500E-05 |
| 17130 | 857 | S857I | C | 1.3e-05 | 1.6e-05 | 1.2e-05 | 1.367E-05 | 2.082E-06 | 1.200E-05 | 1.600E-05 |
| 17131 | 857 | S857L | C | 1.5e-05 | 1.1e-05 | 1.1e-05 | 1.233E-05 | 2.309E-06 | 1.100E-05 | 1.500E-05 |
| 17132 | 857 | S857K | C | 1.6e-05 | 1.1e-05 | 1.8e-05 | 1.500E-05 | 3.606E-06 | 1.100E-05 | 1.800E-05 |
| 17133 | 857 | S857M | C | 1.5e-05 | 1e-05 | 1.1e-05 | 1.200E-05 | 2.646E-06 | 1.000E-05 | 1.500E-05 |
| 17134 | 857 | S857F | C | 1.7e-05 | 1.4e-05 | 1.6e-05 | 1.567E-05 | 1.528E-06 | 1.400E-05 | 1.700E-05 |
| 17135 | 857 | S857P | C | 1.1e-05 | 1.6e-05 | 1e-05 | 1.233E-05 | 3.215E-06 | 1.000E-05 | 1.600E-05 |
| 17136 | 857 | S857S | C | 1.6e-05 | 9.8e-06 | 1.2e-05 | 1.260E-05 | 3.143E-06 | 9.800E-06 | 1.600E-05 |
| 17137 | 857 | S857T | C | 1e-05 | 5.9e-06 | 1.5e-05 | 1.030E-05 | 4.557E-06 | 5.900E-06 | 1.500E-05 |
| 17138 | 857 | S857W | C | 9.5e-06 | 1.1e-05 | 1.4e-05 | 1.150E-05 | 2.291E-06 | 9.500E-06 | 1.400E-05 |
| 17139 | 857 | S857Y | C | 1e-05 | 9.6e-06 | 1.4e-05 | 1.120E-05 | 2.433E-06 | 9.600E-06 | 1.400E-05 |
| 17140 | 857 | S857V | C | 1e-05 | 1.1e-05 | 1.1e-05 | 1.067E-05 | 5.774E-07 | 1.000E-05 | 1.100E-05 |
| 17141 | 858 | C858A | C | 1.4e-05 | 1.4e-05 | 1e-05 | 1.267E-05 | 2.309E-06 | 1.000E-05 | 1.400E-05 |
| 17142 | 858 | C858R | C | 1.5e-05 | 1.4e-05 | 1.5e-05 | 1.467E-05 | 5.774E-07 | 1.400E-05 | 1.500E-05 |
| 17143 | 858 | C858N | C | 1e-05 | 1.5e-05 | 9e-06 | 1.133E-05 | 3.215E-06 | 9.000E-06 | 1.500E-05 |
| 17144 | 858 | C858D | C | 8.7e-06 | 1.4e-05 | 9.8e-06 | 1.083E-05 | 2.797E-06 | 8.700E-06 | 1.400E-05 |
| 17145 | 858 | C858C | C | 1.6e-05 | 1.4e-05 | 1.3e-05 | 1.433E-05 | 1.528E-06 | 1.300E-05 | 1.600E-05 |
| 17146 | 858 | C858E | C | 9.3e-06 | 1.4e-05 | 1.4e-05 | 1.243E-05 | 2.714E-06 | 9.300E-06 | 1.400E-05 |
| 17147 | 858 | C858Q | C | 1.5e-05 | 1.2e-05 | 1.3e-05 | 1.333E-05 | 1.528E-06 | 1.200E-05 | 1.500E-05 |
| 17148 | 858 | C858G | C | 9.3e-06 | 1.7e-05 | 1.1e-05 | 1.243E-05 | 4.045E-06 | 9.300E-06 | 1.700E-05 |
| 17149 | 858 | C858H | C | 1.8e-05 | 1e-05 | 1e-05 | 1.267E-05 | 4.619E-06 | 1.000E-05 | 1.800E-05 |
| 17150 | 858 | C858I | C | 1.6e-05 | 9.5e-06 | 8.9e-06 | 1.147E-05 | 3.937E-06 | 8.900E-06 | 1.600E-05 |
| 17151 | 858 | C858L | C | 1.4e-05 | 1.7e-05 | 1.4e-05 | 1.500E-05 | 1.732E-06 | 1.400E-05 | 1.700E-05 |
| 17152 | 858 | C858K | C | 1.6e-05 | 1.5e-05 | 9.5e-06 | 1.350E-05 | 3.500E-06 | 9.500E-06 | 1.600E-05 |
| 17153 | 858 | C858M | C | 1.3e-05 | 1.4e-05 | 1.1e-05 | 1.267E-05 | 1.528E-06 | 1.100E-05 | 1.400E-05 |
| 17154 | 858 | C858F | C | 1.7e-05 | 1.6e-05 | 1.1e-05 | 1.467E-05 | 3.215E-06 | 1.100E-05 | 1.700E-05 |
| 17155 | 858 | C858P | C | 1.3e-05 | 1.3e-05 | 1.5e-05 | 1.367E-05 | 1.155E-06 | 1.300E-05 | 1.500E-05 |
| 17156 | 858 | C858S | C | 1.9e-05 | 1.6e-05 | 9.1e-06 | 1.470E-05 | 5.076E-06 | 9.100E-06 | 1.900E-05 |
| 17157 | 858 | C858T | C | 1.5e-05 | 1.5e-05 | 1.5e-05 | 1.500E-05 | 0.000E+00 | 1.500E-05 | 1.500E-05 |
| 17158 | 858 | C858W | C | 1.6e-05 | 1.3e-05 | 9.3e-06 | 1.277E-05 | 3.356E-06 | 9.300E-06 | 1.600E-05 |
| 17159 | 858 | C858Y | C | 1.3e-05 | 1.6e-05 | 9.3e-06 | 1.277E-05 | 3.356E-06 | 9.300E-06 | 1.600E-05 |
| 17160 | 858 | C858V | C | 4e-05 | 9.9e-06 | 1.6e-05 | 2.197E-05 | 1.591E-05 | 9.900E-06 | 4.000E-05 |
| 17161 | 859 | A859A | C | 8.8e-06 | 1.2e-05 | 7.2e-06 | 9.333E-06 | 2.444E-06 | 7.200E-06 | 1.200E-05 |
| 17162 | 859 | A859R | C | 1.2e-05 | 7.8e-06 | 1.7e-05 | 1.227E-05 | 4.606E-06 | 7.800E-06 | 1.700E-05 |
| 17163 | 859 | A859N | C | 1.5e-05 | 1.1e-05 | 1e-05 | 1.200E-05 | 2.646E-06 | 1.000E-05 | 1.500E-05 |
| 17164 | 859 | A859D | C | 1.6e-05 | 1.1e-05 | 1.1e-05 | 1.267E-05 | 2.887E-06 | 1.100E-05 | 1.600E-05 |
| 17165 | 859 | A859C | C | 1.4e-05 | 1e-05 | 6.2e-06 | 1.007E-05 | 3.900E-06 | 6.200E-06 | 1.400E-05 |
| 17166 | 859 | A859E | C | 1e-05 | 1.5e-05 | 1.4e-05 | 1.300E-05 | 2.646E-06 | 1.000E-05 | 1.500E-05 |
| 17167 | 859 | A859Q | C | 9.9e-06 | 1.1e-05 | 1.3e-05 | 1.130E-05 | 1.572E-06 | 9.900E-06 | 1.300E-05 |
| 17168 | 859 | A859G | C | 1.5e-05 | 1.7e-05 | 9.7e-06 | 1.390E-05 | 3.772E-06 | 9.700E-06 | 1.700E-05 |
| 17169 | 859 | A859H | C | 1.2e-05 | 1.4e-05 | 1.5e-05 | 1.367E-05 | 1.528E-06 | 1.200E-05 | 1.500E-05 |

| | | | | | | | | | | |
|---|---|---|---|---|---|---|---|---|---|---|
| 17170 | 859 | A859I | C | 1.5e-05 | 8.8e-06 | 9.6e-06 | 1.113E-05 | 3.372E-06 | 8.800E-06 | 1.500E-05 |
| 17171 | 859 | A859L | C | 8.4e-06 | 9.8e-06 | 8.8e-06 | 9.000E-06 | 7.211E-07 | 8.400E-06 | 9.800E-06 |
| 17172 | 859 | A859K | C | 1.6e-05 | 1.1e-05 | 1.1e-05 | 1.267E-05 | 2.887E-06 | 1.100E-05 | 1.600E-05 |
| 17173 | 859 | A859M | C | 9.9e-06 | 1.1e-05 | 1.6e-05 | 1.230E-05 | 3.251E-06 | 9.900E-06 | 1.600E-05 |
| 17174 | 859 | A859F | C | 1.6e-05 | 1.1e-05 | 1.5e-05 | 1.400E-05 | 2.646E-06 | 1.100E-05 | 1.600E-05 |
| 17175 | 859 | A859P | C | 1.3e-05 | 1.7e-05 | 1.2e-05 | 1.400E-05 | 2.646E-06 | 1.200E-05 | 1.700E-05 |
| 17176 | 859 | A859S | C | 1.3e-05 | 1.4e-05 | 1.3e-05 | 1.333E-05 | 5.774E-07 | 1.300E-05 | 1.400E-05 |
| 17177 | 859 | A859T | C | 1.4e-05 | 1.5e-05 | 7.2e-06 | 1.207E-05 | 4.244E-06 | 7.200E-06 | 1.500E-05 |
| 17178 | 859 | A859W | C | 1.2e-05 | 1.2e-05 | 1.5e-05 | 1.300E-05 | 1.732E-06 | 1.200E-05 | 1.500E-05 |
| 17179 | 859 | A859Y | C | 9.7e-06 | 1.1e-05 | 1.6e-05 | 1.223E-05 | 3.326E-06 | 9.700E-06 | 1.600E-05 |
| 17180 | 859 | A859V | C | 1.1e-05 | 1.6e-05 | 1.5e-05 | 1.400E-05 | 2.646E-06 | 1.100E-05 | 1.600E-05 |
| 17181 | 860 | A860A | C | 1.5e-05 | 1e-05 | 1.4e-05 | 1.300E-05 | 2.646E-06 | 1.000E-05 | 1.500E-05 |
| 17182 | 860 | A860R | C | 8.8e-06 | 1.6e-05 | 1.7e-05 | 1.393E-05 | 4.474E-06 | 8.800E-06 | 1.700E-05 |
| 17183 | 860 | A860N | C | 1.5e-05 | 1e-05 | 1.3e-05 | 1.267E-05 | 2.517E-06 | 1.000E-05 | 1.500E-05 |
| 17184 | 860 | A860D | C | 1e-05 | 8.5e-06 | 1.4e-05 | 1.083E-05 | 2.843E-06 | 8.500E-06 | 1.400E-05 |
| 17185 | 860 | A860C | C | 1.1e-05 | 1e-05 | 1e-05 | 1.033E-05 | 5.774E-07 | 1.000E-05 | 1.100E-05 |
| 17186 | 860 | A860E | C | 7.5e-06 | 1.6e-05 | 1.5e-05 | 1.283E-05 | 4.646E-06 | 7.500E-06 | 1.600E-05 |
| 17187 | 860 | A860Q | C | 1e-05 | 9.4e-06 | 1.6e-05 | 1.180E-05 | 3.650E-06 | 9.400E-06 | 1.600E-05 |
| 17188 | 860 | A860G | C | 1.1e-05 | 1e-05 | 1.4e-05 | 1.167E-05 | 2.082E-06 | 1.000E-05 | 1.400E-05 |
| 17189 | 860 | A860H | C | 1.1e-05 | 1.5e-05 | 9.4e-06 | 1.180E-05 | 2.884E-06 | 9.400E-06 | 1.500E-05 |
| 17190 | 860 | A860I | C | 1.6e-05 | 1.4e-05 | 1.2e-05 | 1.400E-05 | 2.000E-06 | 1.200E-05 | 1.600E-05 |
| 17191 | 860 | A860L | C | 1.4e-05 | 1.6e-05 | 1e-05 | 1.333E-05 | 3.055E-06 | 1.000E-05 | 1.600E-05 |
| 17192 | 860 | A860K | C | 1.5e-05 | 1.1e-05 | 1.6e-05 | 1.400E-05 | 2.646E-06 | 1.100E-05 | 1.600E-05 |
| 17193 | 860 | A860M | C | 1.5e-05 | 1.1e-05 | 1.5e-05 | 1.367E-05 | 2.309E-06 | 1.100E-05 | 1.500E-05 |
| 17194 | 860 | A860F | C | 1.5e-05 | 1e-05 | 1.1e-05 | 1.200E-05 | 2.646E-06 | 1.000E-05 | 1.500E-05 |
| 17195 | 860 | A860P | C | 1.5e-05 | 1.6e-05 | 1e-05 | 1.367E-05 | 3.215E-06 | 1.000E-05 | 1.600E-05 |
| 17196 | 860 | A860S | C | 9.9e-06 | 1.1e-05 | 1.1e-05 | 1.063E-05 | 6.351E-07 | 9.900E-06 | 1.100E-05 |
| 17197 | 860 | A860T | C | 9.2e-06 | 9.5e-06 | 1.3e-05 | 1.057E-05 | 2.113E-06 | 9.200E-06 | 1.300E-05 |
| 17198 | 860 | A860W | C | 8.8e-06 | 1.4e-05 | 1.5e-05 | 1.260E-05 | 3.329E-06 | 8.800E-06 | 1.500E-05 |
| 17199 | 860 | A860Y | C | 1.6e-05 | 1.2e-05 | 1.3e-05 | 1.367E-05 | 2.082E-06 | 1.200E-05 | 1.600E-05 |
| 17200 | 860 | A860V | C | 1.1e-05 | 1.5e-05 | 1.3e-05 | 1.300E-05 | 2.000E-06 | 1.100E-05 | 1.500E-05 |
| 17201 | 861 | S861A | C | 1.3e-05 | 1.5e-05 | 1.4e-05 | 1.400E-05 | 1.000E-06 | 1.300E-05 | 1.500E-05 |
| 17202 | 861 | S861R | C | 1.5e-05 | 1.4e-05 | 9.8e-06 | 1.293E-05 | 2.759E-06 | 9.800E-06 | 1.500E-05 |
| 17203 | 861 | S861N | C | 1.5e-05 | 1.2e-05 | 1e-05 | 1.233E-05 | 2.517E-06 | 1.000E-05 | 1.500E-05 |
| 17204 | 861 | S861D | C | 9.5e-06 | 9.7e-06 | 1.7e-05 | 1.207E-05 | 4.274E-06 | 9.500E-06 | 1.700E-05 |
| 17205 | 861 | S861C | C | 1.3e-05 | 1e-05 | 1.6e-05 | 1.300E-05 | 3.000E-06 | 1.000E-05 | 1.600E-05 |
| 17206 | 861 | S861E | C | 9.7e-06 | 1.6e-05 | 9.9e-06 | 1.187E-05 | 3.581E-06 | 9.700E-06 | 1.600E-05 |
| 17207 | 861 | S861Q | C | 1.4e-05 | 1.5e-05 | 1.4e-05 | 1.433E-05 | 5.774E-07 | 1.400E-05 | 1.500E-05 |
| 17208 | 861 | S861G | C | 1.1e-05 | 8.9e-06 | 1.1e-05 | 1.030E-05 | 1.212E-06 | 8.900E-06 | 1.100E-05 |
| 17209 | 861 | S861H | C | 1.4e-05 | 1.6e-05 | 1.1e-05 | 1.367E-05 | 2.517E-06 | 1.100E-05 | 1.600E-05 |
| 17210 | 861 | S861I | C | 1.1e-05 | 1.5e-05 | 9.7e-06 | 1.190E-05 | 2.762E-06 | 9.700E-06 | 1.500E-05 |
| 17211 | 861 | S861L | C | 1e-05 | 1.8e-05 | 9.6e-06 | 1.253E-05 | 4.738E-06 | 9.600E-06 | 1.800E-05 |
| 17212 | 861 | S861K | C | 1.3e-05 | 1.1e-05 | 9.6e-06 | 1.120E-05 | 1.709E-06 | 9.600E-06 | 1.300E-05 |
| 17213 | 861 | S861M | C | 1.5e-05 | 1.4e-05 | 1.6e-05 | 1.500E-05 | 1.000E-06 | 1.400E-05 | 1.600E-05 |
| 17214 | 861 | S861F | C | 1.5e-05 | 1e-05 | 1.1e-05 | 1.200E-05 | 2.646E-06 | 1.000E-05 | 1.500E-05 |
| 17215 | 861 | S861P | C | 1.7e-05 | 1.5e-05 | 1.1e-05 | 1.433E-05 | 3.055E-06 | 1.100E-05 | 1.700E-05 |
| 17216 | 861 | S861S | C | 1.5e-05 | 1e-05 | 1.4e-05 | 1.300E-05 | 2.646E-06 | 1.000E-05 | 1.500E-05 |
| 17217 | 861 | S861T | C | 9.5e-06 | 1.1e-05 | 1.7e-05 | 1.250E-05 | 3.969E-06 | 9.500E-06 | 1.700E-05 |
| 17218 | 861 | S861W | C | 1.6e-05 | 9.5e-06 | 8.7e-06 | 1.140E-05 | 4.004E-06 | 8.700E-06 | 1.600E-05 |
| 17219 | 861 | S861Y | C | 1.2e-05 | 1.3e-05 | 9.5e-06 | 1.150E-05 | 1.803E-06 | 9.500E-06 | 1.300E-05 |
| 17220 | 861 | S861V | C | 1.2e-05 | 1.1e-05 | 1e-05 | 1.100E-05 | 1.000E-06 | 1.000E-05 | 1.200E-05 |
| 17221 | 862 | G862A | C | 1.5e-05 | 8.6e-06 | 1.1e-05 | 1.153E-05 | 3.233E-06 | 8.600E-06 | 1.500E-05 |
| 17222 | 862 | G862R | C | 1.4e-05 | 1.5e-05 | 1e-05 | 1.300E-05 | 2.646E-06 | 1.000E-05 | 1.500E-05 |
| 17223 | 862 | G862N | C | 1.4e-05 | 9.9e-06 | 1.5e-05 | 1.297E-05 | 2.702E-06 | 9.900E-06 | 1.500E-05 |
| 17224 | 862 | G862D | C | 1.5e-05 | 1.5e-05 | 1.4e-05 | 1.467E-05 | 5.774E-07 | 1.400E-05 | 1.500E-05 |
| 17225 | 862 | G862C | C | 1e-05 | 1.5e-05 | 1.4e-05 | 1.300E-05 | 2.646E-06 | 1.000E-05 | 1.500E-05 |
| 17226 | 862 | G862E | C | 1.7e-05 | 1.5e-05 | 1.3e-05 | 1.500E-05 | 2.000E-06 | 1.300E-05 | 1.700E-05 |
| 17227 | 862 | G862Q | C | 9.1e-06 | 1.7e-05 | 1.1e-05 | 1.237E-05 | 4.124E-06 | 9.100E-06 | 1.700E-05 |
| 17228 | 862 | G862G | C | 1.5e-05 | 7.9e-06 | 1.7e-05 | 1.330E-05 | 4.782E-06 | 7.900E-06 | 1.700E-05 |
| 17229 | 862 | G862H | C | 1.5e-05 | 1.2e-05 | 1e-05 | 1.233E-05 | 2.517E-06 | 1.000E-05 | 1.500E-05 |
| 17230 | 862 | G862I | C | 9.1e-06 | 1.1e-05 | 1.1e-05 | 1.037E-05 | 1.097E-06 | 9.100E-06 | 1.100E-05 |
| 17231 | 862 | G862L | C | 7.9e-06 | 1.1e-05 | 1.2e-05 | 1.030E-05 | 2.138E-06 | 7.900E-06 | 1.200E-05 |
| 17232 | 862 | G862K | C | 1.7e-05 | 1.4e-05 | 1.2e-05 | 1.433E-05 | 2.517E-06 | 1.200E-05 | 1.700E-05 |
| 17233 | 862 | G862M | C | 1.6e-05 | 1.2e-05 | 9.8e-06 | 1.260E-05 | 3.143E-06 | 9.800E-06 | 1.600E-05 |
| 17234 | 862 | G862F | C | 1.4e-05 | 1.7e-05 | 1.6e-05 | 1.567E-05 | 1.528E-06 | 1.400E-05 | 1.700E-05 |
| 17235 | 862 | G862P | C | 1.5e-05 | 1.1e-05 | 1.5e-05 | 1.367E-05 | 2.309E-06 | 1.100E-05 | 1.500E-05 |
| 17236 | 862 | G862S | C | 9.8e-06 | 1.3e-05 | 1.4e-05 | 1.227E-05 | 2.194E-06 | 9.800E-06 | 1.400E-05 |
| 17237 | 862 | G862T | C | 1.1e-05 | 1.6e-05 | 1e-05 | 1.233E-05 | 3.215E-06 | 1.000E-05 | 1.600E-05 |
| 17238 | 862 | G862W | C | 1.4e-05 | 1.4e-05 | 1.3e-05 | 1.367E-05 | 5.774E-07 | 1.300E-05 | 1.400E-05 |
| 17239 | 862 | G862Y | C | 9.8e-06 | 9.2e-06 | 8.9e-06 | 9.300E-06 | 4.583E-07 | 8.900E-06 | 9.800E-06 |
| 17240 | 862 | G862V | C | 1.5e-05 | 1.1e-05 | 1e-05 | 1.200E-05 | 2.646E-06 | 1.000E-05 | 1.500E-05 |
| 17241 | 863 | F863A | C | 9.3e-06 | 1e-05 | 9e-06 | 9.433E-06 | 5.132E-07 | 9.000E-06 | 1.000E-05 |
| 17242 | 863 | F863R | C | 1e-05 | 1.1e-05 | 9.8e-06 | 1.027E-05 | 6.429E-07 | 9.800E-06 | 1.100E-05 |
| 17243 | 863 | F863N | C | 1.5e-05 | 1.4e-05 | 1.1e-05 | 1.333E-05 | 2.082E-06 | 1.100E-05 | 1.500E-05 |
| 17244 | 863 | F863D | C | 1.6e-05 | 1.4e-05 | 9e-06 | 1.300E-05 | 3.606E-06 | 9.000E-06 | 1.600E-05 |
| 17245 | 863 | F863C | C | 1.5e-05 | 1.3e-05 | 1.5e-05 | 1.433E-05 | 1.155E-06 | 1.300E-05 | 1.500E-05 |
| 17246 | 863 | F863E | C | 1.5e-05 | 9.1e-06 | 1.1e-05 | 1.170E-05 | 3.012E-06 | 9.100E-06 | 1.500E-05 |
| 17247 | 863 | F863Q | C | 9.8e-06 | 1.5e-05 | 9.1e-06 | 1.130E-05 | 3.223E-06 | 9.100E-06 | 1.500E-05 |
| 17248 | 863 | F863G | C | 1.6e-05 | 1.5e-05 | 1.7e-05 | 1.600E-05 | 1.000E-06 | 1.500E-05 | 1.700E-05 |
| 17249 | 863 | F863H | C | 8.5e-06 | 1.1e-05 | 1.5e-05 | 1.150E-05 | 3.279E-06 | 8.500E-06 | 1.500E-05 |
| 17250 | 863 | F863I | C | 1e-05 | 1.1e-05 | 1.5e-05 | 1.200E-05 | 2.646E-06 | 1.000E-05 | 1.500E-05 |
| 17251 | 863 | F863L | C | 8.1e-06 | 1.4e-05 | 1.4e-05 | 1.203E-05 | 3.406E-06 | 8.100E-06 | 1.400E-05 |
| 17252 | 863 | F863K | C | 9.3e-06 | 1.5e-05 | 1.1e-05 | 1.177E-05 | 2.926E-06 | 9.300E-06 | 1.500E-05 |
| 17253 | 863 | F863M | C | 1.1e-05 | 1.5e-05 | 1.3e-05 | 1.300E-05 | 2.000E-06 | 1.100E-05 | 1.500E-05 |
| 17254 | 863 | F863F | C | 1.2e-05 | 1.1e-05 | 1.4e-05 | 1.233E-05 | 1.528E-06 | 1.100E-05 | 1.400E-05 |

| | | | | | | | | | | |
|---|---|---|---|---|---|---|---|---|---|---|
| 17255 | 863 | F863P | C | 7.8e-06 | 1.5e-05 | 1.6e-05 | 1.293E-05 | 4.474E-06 | 7.800E-06 | 1.600E-05 |
| 17256 | 863 | F863S | C | 1.5e-05 | 8.8e-06 | 1.2e-05 | 1.193E-05 | 3.101E-06 | 8.800E-06 | 1.500E-05 |
| 17257 | 863 | F863T | C | 1.7e-05 | 1.4e-05 | 9.5e-06 | 1.350E-05 | 3.775E-06 | 9.500E-06 | 1.700E-05 |
| 17258 | 863 | F863W | C | 1.4e-05 | 1.3e-05 | 8.4e-06 | 1.180E-05 | 2.987E-06 | 8.400E-06 | 1.400E-05 |
| 17259 | 863 | F863Y | C | 1.7e-05 | 1.6e-05 | 1.1e-05 | 1.467E-05 | 3.215E-06 | 1.100E-05 | 1.700E-05 |
| 17260 | 863 | F863V | C | 1.5e-05 | 1.5e-05 | 1.6e-05 | 1.533E-05 | 5.774E-07 | 1.500E-05 | 1.600E-05 |
| 17261 | 864 | T864A | C | 1.2e-05 | 1e-05 | 1.3e-05 | 1.167E-05 | 1.528E-06 | 1.000E-05 | 1.300E-05 |
| 17262 | 864 | T864R | C | 1.5e-05 | 1.2e-05 | 1.7e-05 | 1.467E-05 | 2.517E-06 | 1.200E-05 | 1.700E-05 |
| 17263 | 864 | T864N | C | 1e-05 | 1.5e-05 | 9.7e-06 | 1.157E-05 | 2.977E-06 | 9.700E-06 | 1.500E-05 |
| 17264 | 864 | T864D | C | 9.6e-06 | 1.6e-05 | 9.1e-06 | 1.157E-05 | 3.848E-06 | 9.100E-06 | 1.600E-05 |
| 17265 | 864 | T864C | C | 1.4e-05 | 1.7e-05 | 1.5e-05 | 1.533E-05 | 1.528E-06 | 1.400E-05 | 1.700E-05 |
| 17266 | 864 | T864E | C | 1.5e-05 | 1.1e-05 | 1.5e-05 | 1.367E-05 | 2.309E-06 | 1.100E-05 | 1.500E-05 |
| 17267 | 864 | T864Q | C | 1.6e-05 | 1.1e-05 | 1.1e-05 | 1.267E-05 | 2.887E-06 | 1.100E-05 | 1.600E-05 |
| 17268 | 864 | T864G | C | 1.3e-05 | 1.1e-05 | 1.1e-05 | 1.167E-05 | 1.155E-06 | 1.100E-05 | 1.300E-05 |
| 17269 | 864 | T864H | C | 8.3e-06 | 1.4e-05 | 1.1e-05 | 1.110E-05 | 2.851E-06 | 8.300E-06 | 1.400E-05 |
| 17270 | 864 | T864I | C | 1.1e-05 | 1.6e-05 | 1.6e-05 | 1.433E-05 | 2.887E-06 | 1.100E-05 | 1.600E-05 |
| 17271 | 864 | T864L | C | 1.5e-05 | 9.9e-06 | 1.1e-05 | 1.197E-05 | 2.684E-06 | 9.900E-06 | 1.500E-05 |
| 17272 | 864 | T864K | C | 1.1e-05 | 1.2e-05 | 1.1e-05 | 1.133E-05 | 5.774E-07 | 1.100E-05 | 1.200E-05 |
| 17273 | 864 | T864M | C | 1.5e-05 | 1.1e-05 | 1.5e-05 | 1.367E-05 | 2.309E-06 | 1.100E-05 | 1.500E-05 |
| 17274 | 864 | T864F | C | 1.1e-05 | 1.6e-05 | 1.6e-05 | 1.433E-05 | 2.887E-06 | 1.100E-05 | 1.600E-05 |
| 17275 | 864 | T864P | C | 1.2e-05 | 1.6e-05 | 1.3e-05 | 1.367E-05 | 2.082E-06 | 1.200E-05 | 1.600E-05 |
| 17276 | 864 | T864S | C | 1.5e-05 | 1.1e-05 | 9.5e-06 | 1.183E-05 | 2.843E-06 | 9.500E-06 | 1.500E-05 |
| 17277 | 864 | T864T | C | 1.2e-05 | 1.5e-05 | 1.6e-05 | 1.433E-05 | 2.082E-06 | 1.200E-05 | 1.600E-05 |
| 17278 | 864 | T864W | C | 1e-05 | 1.2e-05 | 1.4e-05 | 1.200E-05 | 2.000E-06 | 1.000E-05 | 1.400E-05 |
| 17279 | 864 | T864Y | C | 1.5e-05 | 1.7e-05 | 1.4e-05 | 1.533E-05 | 1.528E-06 | 1.400E-05 | 1.700E-05 |
| 17280 | 864 | T864V | C | 1.1e-05 | 1.2e-05 | 7e-06 | 1.000E-05 | 2.646E-06 | 7.000E-06 | 1.200E-05 |
| 17281 | 865 | F865A | C | 9.3e-06 | 1.4e-05 | 9.8e-06 | 1.103E-05 | 2.581E-06 | 9.300E-06 | 1.400E-05 |
| 17282 | 865 | F865R | C | 1.7e-05 | 9e-06 | 1.4e-05 | 1.333E-05 | 4.041E-06 | 9.000E-06 | 1.700E-05 |
| 17283 | 865 | F865N | C | 1.4e-05 | 1.2e-05 | 1.3e-05 | 1.300E-05 | 1.000E-06 | 1.200E-05 | 1.400E-05 |
| 17284 | 865 | F865D | C | 8e-06 | 1.6e-05 | 1.6e-05 | 1.333E-05 | 4.619E-06 | 8.000E-06 | 1.600E-05 |
| 17285 | 865 | F865C | C | 1.6e-05 | 1.7e-05 | 6.9e-06 | 1.330E-05 | 5.565E-06 | 6.900E-06 | 1.700E-05 |
| 17286 | 865 | F865E | C | 1.4e-05 | 1e-05 | 1.1e-05 | 1.167E-05 | 2.082E-06 | 1.000E-05 | 1.400E-05 |
| 17287 | 865 | F865Q | C | 8.9e-06 | 1.6e-05 | 1.1e-05 | 1.197E-05 | 3.647E-06 | 8.900E-06 | 1.600E-05 |
| 17288 | 865 | F865G | C | 1.5e-05 | 7.4e-06 | 1.5e-05 | 1.247E-05 | 4.388E-06 | 7.400E-06 | 1.500E-05 |
| 17289 | 865 | F865H | C | 1.6e-05 | 1e-05 | 1.6e-05 | 1.400E-05 | 3.464E-06 | 1.000E-05 | 1.600E-05 |
| 17290 | 865 | F865I | C | 1.1e-05 | 1.7e-05 | 1.5e-05 | 1.433E-05 | 3.055E-06 | 1.100E-05 | 1.700E-05 |
| 17291 | 865 | F865L | C | 1.7e-05 | 1.5e-05 | 1.1e-05 | 1.433E-05 | 3.055E-06 | 1.100E-05 | 1.700E-05 |
| 17292 | 865 | F865K | C | 1.6e-05 | 1.1e-05 | 1.1e-05 | 1.267E-05 | 2.887E-06 | 1.100E-05 | 1.600E-05 |
| 17293 | 865 | F865M | C | 1.6e-05 | 1.4e-05 | 1.1e-05 | 1.367E-05 | 2.517E-06 | 1.100E-05 | 1.600E-05 |
| 17294 | 865 | F865F | C | 1.5e-05 | 1.4e-05 | 9.8e-06 | 1.293E-05 | 2.759E-06 | 9.800E-06 | 1.500E-05 |
| 17295 | 865 | F865P | C | 1.5e-05 | 7.7e-06 | 9.5e-06 | 1.073E-05 | 3.803E-06 | 7.700E-06 | 1.500E-05 |
| 17296 | 865 | F865S | C | 1e-05 | 1.7e-05 | 1.5e-05 | 1.400E-05 | 3.606E-06 | 1.000E-05 | 1.700E-05 |
| 17297 | 865 | F865T | C | 1.2e-05 | 1.2e-05 | 1.5e-05 | 1.300E-05 | 1.732E-06 | 1.200E-05 | 1.500E-05 |
| 17298 | 865 | F865W | C | 1.4e-05 | 8.7e-06 | 1.1e-05 | 1.123E-05 | 2.658E-06 | 8.700E-06 | 1.400E-05 |
| 17299 | 865 | F865Y | C | 9.1e-06 | 1.3e-05 | 1.1e-05 | 1.103E-05 | 1.950E-06 | 9.100E-06 | 1.300E-05 |
| 17300 | 865 | F865V | C | 1.2e-05 | 9.8e-06 | 1.4e-05 | 1.193E-05 | 2.101E-06 | 9.800E-06 | 1.400E-05 |
| 17301 | 866 | T866A | C | 1.6e-05 | 1.2e-05 | 1.1e-05 | 1.300E-05 | 2.646E-06 | 1.100E-05 | 1.600E-05 |
| 17302 | 866 | T866R | C | 1.3e-05 | 1.6e-05 | 1.3e-05 | 1.400E-05 | 1.732E-06 | 1.300E-05 | 1.600E-05 |
| 17303 | 866 | T866N | C | 1.3e-05 | 1.6e-05 | 8.6e-06 | 1.253E-05 | 3.722E-06 | 8.600E-06 | 1.600E-05 |
| 17304 | 866 | T866D | C | 1.4e-05 | 9.7e-06 | 1.7e-05 | 1.357E-05 | 3.669E-06 | 9.700E-06 | 1.700E-05 |
| 17305 | 866 | T866C | C | 9.5e-06 | 1.3e-05 | 1.3e-05 | 1.183E-05 | 2.021E-06 | 9.500E-06 | 1.300E-05 |
| 17306 | 866 | T866E | C | 1.1e-05 | 1e-05 | 9.1e-06 | 1.003E-05 | 9.504E-07 | 9.100E-06 | 1.100E-05 |
| 17307 | 866 | T866Q | C | 1.7e-05 | 1.5e-05 | 1.3e-05 | 1.500E-05 | 2.000E-06 | 1.300E-05 | 1.700E-05 |
| 17308 | 866 | T866G | C | 1.6e-05 | 1.4e-05 | 8.8e-06 | 1.293E-05 | 3.717E-06 | 8.800E-06 | 1.600E-05 |
| 17309 | 866 | T866H | C | 1.4e-05 | 1.4e-05 | 1.1e-05 | 1.300E-05 | 1.732E-06 | 1.100E-05 | 1.400E-05 |
| 17310 | 866 | T866I | C | 8.6e-06 | 1.5e-05 | 1e-05 | 1.120E-05 | 3.365E-06 | 8.600E-06 | 1.500E-05 |
| 17311 | 866 | T866L | C | 1e-05 | 1.1e-05 | 1.4e-05 | 1.167E-05 | 2.082E-06 | 1.000E-05 | 1.400E-05 |
| 17312 | 866 | T866K | C | 1.5e-05 | 1e-05 | 1.7e-05 | 1.400E-05 | 3.606E-06 | 1.000E-05 | 1.700E-05 |
| 17313 | 866 | T866M | C | 1.4e-05 | 1.4e-05 | 1.2e-05 | 1.333E-05 | 1.155E-06 | 1.200E-05 | 1.400E-05 |
| 17314 | 866 | T866F | C | 1.5e-05 | 1.5e-05 | 1.1e-05 | 1.367E-05 | 2.309E-06 | 1.100E-05 | 1.500E-05 |
| 17315 | 866 | T866P | C | 1.6e-05 | 1.5e-05 | 1.7e-05 | 1.600E-05 | 1.000E-06 | 1.500E-05 | 1.700E-05 |
| 17316 | 866 | T866S | C | 1.5e-05 | 1.6e-05 | 1.2e-05 | 1.433E-05 | 2.082E-06 | 1.200E-05 | 1.600E-05 |
| 17317 | 866 | T866T | C | 1.7e-05 | 9.3e-06 | 1.5e-05 | 1.377E-05 | 3.995E-06 | 9.300E-06 | 1.700E-05 |
| 17318 | 866 | T866W | C | 1.1e-05 | 9.8e-06 | 1.5e-05 | 1.193E-05 | 2.723E-06 | 9.800E-06 | 1.500E-05 |
| 17319 | 866 | T866Y | C | 8.8e-06 | 1.7e-05 | 1.6e-05 | 1.393E-05 | 4.474E-06 | 8.800E-06 | 1.700E-05 |
| 17320 | 866 | T866V | C | 1.1e-05 | 1.5e-05 | 1e-05 | 1.200E-05 | 2.646E-06 | 1.000E-05 | 1.500E-05 |
| 17321 | 867 | D867A | C | 1.4e-05 | 1.5e-05 | 1e-05 | 1.300E-05 | 2.646E-06 | 1.000E-05 | 1.500E-05 |
| 17322 | 867 | D867R | C | 1.6e-05 | 8.9e-06 | 1.6e-05 | 1.363E-05 | 4.099E-06 | 8.900E-06 | 1.600E-05 |
| 17323 | 867 | D867N | C | 1.6e-05 | 9.8e-06 | 1.3e-05 | 1.293E-05 | 3.101E-06 | 9.800E-06 | 1.600E-05 |
| 17324 | 867 | D867D | C | 7.7e-06 | 1.4e-05 | 1.3e-05 | 1.157E-05 | 3.386E-06 | 7.700E-06 | 1.400E-05 |
| 17325 | 867 | D867C | C | 1.6e-05 | 1e-05 | 1.5e-05 | 1.367E-05 | 3.215E-06 | 1.000E-05 | 1.600E-05 |
| 17326 | 867 | D867E | C | 1.5e-05 | 1.5e-05 | 1.3e-05 | 1.433E-05 | 1.155E-06 | 1.300E-05 | 1.500E-05 |
| 17327 | 867 | D867Q | C | 1.4e-05 | 1.6e-05 | 1.3e-05 | 1.433E-05 | 1.528E-06 | 1.300E-05 | 1.600E-05 |
| 17328 | 867 | D867G | C | 1.1e-05 | 1.4e-05 | 8.6e-06 | 1.120E-05 | 2.706E-06 | 8.600E-06 | 1.400E-05 |
| 17329 | 867 | D867H | C | 8.5e-06 | 1.5e-05 | 9.8e-06 | 1.110E-05 | 3.439E-06 | 8.500E-06 | 1.500E-05 |
| 17330 | 867 | D867I | C | 1.1e-05 | 1.6e-05 | 9.9e-06 | 1.230E-05 | 3.251E-06 | 9.900E-06 | 1.600E-05 |
| 17331 | 867 | D867L | C | 1.1e-05 | 9.9e-06 | 1.5e-05 | 1.197E-05 | 2.684E-06 | 9.900E-06 | 1.500E-05 |
| 17332 | 867 | D867K | C | 1.1e-05 | 1.4e-05 | 1.1e-05 | 1.200E-05 | 1.732E-06 | 1.100E-05 | 1.400E-05 |
| 17333 | 867 | D867M | C | 1.4e-05 | 1.2e-05 | 1.1e-05 | 1.233E-05 | 1.528E-06 | 1.100E-05 | 1.400E-05 |
| 17334 | 867 | D867F | C | 1.6e-05 | 1.7e-05 | 1.6e-05 | 1.633E-05 | 5.774E-07 | 1.600E-05 | 1.700E-05 |
| 17335 | 867 | D867P | C | 1e-05 | 1.2e-05 | 1.4e-05 | 1.200E-05 | 2.000E-06 | 1.000E-05 | 1.400E-05 |
| 17336 | 867 | D867S | C | 1.4e-05 | 1e-05 | 1e-05 | 1.133E-05 | 2.309E-06 | 1.000E-05 | 1.400E-05 |
| 17337 | 867 | D867T | C | 1.4e-05 | 1.5e-05 | 1.1e-05 | 1.333E-05 | 2.082E-06 | 1.100E-05 | 1.500E-05 |
| 17338 | 867 | D867W | C | 1.6e-05 | 1.6e-05 | 1.2e-05 | 1.467E-05 | 2.309E-06 | 1.200E-05 | 1.600E-05 |
| 17339 | 867 | D867Y | C | 1e-05 | 9.6e-06 | 1.3e-05 | 1.087E-05 | 1.858E-06 | 9.600E-06 | 1.300E-05 |

| | | | | | | | | | | |
|---|---|---|---|---|---|---|---|---|---|---|
| 17340 | 867 | D867V | C | 1e-05 | 1.4e-05 | 1.5e-05 | 1.300E-05 | 2.646E-06 | 1.000E-05 | 1.500E-05 |
| 17341 | 868 | Y868A | C | 1.5e-05 | 9e-06 | 1.7e-05 | 1.367E-05 | 4.163E-06 | 9.000E-06 | 1.700E-05 |
| 17342 | 868 | Y868R | C | 1.7e-05 | 1e-05 | 1e-05 | 1.233E-05 | 4.041E-06 | 1.000E-05 | 1.700E-05 |
| 17343 | 868 | Y868N | C | 1.5e-05 | 1.3e-05 | 1.6e-05 | 1.467E-05 | 1.528E-06 | 1.300E-05 | 1.600E-05 |
| 17344 | 868 | Y868D | C | 1.6e-05 | 1.1e-05 | 1.3e-05 | 1.333E-05 | 2.517E-06 | 1.100E-05 | 1.600E-05 |
| 17345 | 868 | Y868C | C | 9.6e-06 | 1.1e-05 | 9.9e-06 | 1.017E-05 | 7.371E-07 | 9.600E-06 | 1.100E-05 |
| 17346 | 868 | Y868E | C | 1.4e-05 | 1.2e-05 | 1e-05 | 1.200E-05 | 2.000E-06 | 1.000E-05 | 1.400E-05 |
| 17347 | 868 | Y868Q | C | 1.3e-05 | 1.2e-05 | 1.4e-05 | 1.300E-05 | 1.000E-06 | 1.200E-05 | 1.400E-05 |
| 17348 | 868 | Y868G | C | 1.6e-05 | 1.5e-05 | 1.6e-05 | 1.567E-05 | 5.774E-07 | 1.500E-05 | 1.600E-05 |
| 17349 | 868 | Y868H | C | 9.8e-06 | 9.8e-06 | 1.6e-05 | 1.187E-05 | 3.580E-06 | 9.800E-06 | 1.600E-05 |
| 17350 | 868 | Y868I | C | 1.4e-05 | 1.5e-05 | 9.3e-06 | 1.277E-05 | 3.044E-06 | 9.300E-06 | 1.500E-05 |
| 17351 | 868 | Y868L | C | 1.6e-05 | 9.8e-06 | 9.7e-06 | 1.183E-05 | 3.609E-06 | 9.700E-06 | 1.600E-05 |
| 17352 | 868 | Y868K | C | 1.5e-05 | 1.6e-05 | 1.4e-05 | 1.500E-05 | 1.000E-06 | 1.400E-05 | 1.600E-05 |
| 17353 | 868 | Y868M | C | 1.7e-05 | 1.5e-05 | 1.4e-05 | 1.533E-05 | 1.528E-06 | 1.400E-05 | 1.700E-05 |
| 17354 | 868 | Y868F | C | 8.5e-06 | 1.5e-05 | 1.6e-05 | 1.317E-05 | 4.072E-06 | 8.500E-06 | 1.600E-05 |
| 17355 | 868 | Y868P | C | 1.7e-05 | 1e-05 | 1.2e-05 | 1.300E-05 | 3.606E-06 | 1.000E-05 | 1.700E-05 |
| 17356 | 868 | Y868S | C | 1.8e-05 | 1.5e-05 | 1.5e-05 | 1.600E-05 | 1.732E-06 | 1.500E-05 | 1.800E-05 |
| 17357 | 868 | Y868T | C | 7.6e-06 | 1e-05 | 1.6e-05 | 1.120E-05 | 4.327E-06 | 7.600E-06 | 1.600E-05 |
| 17358 | 868 | Y868W | C | 9.6e-06 | 1.3e-05 | 1.6e-05 | 1.287E-05 | 3.202E-06 | 9.600E-06 | 1.600E-05 |
| 17359 | 868 | Y868Y | C | 1.8e-05 | 9.2e-06 | 1.4e-05 | 1.373E-05 | 4.406E-06 | 9.200E-06 | 1.800E-05 |
| 17360 | 868 | Y868V | C | 1.6e-05 | 1.2e-05 | 1.5e-05 | 1.433E-05 | 2.082E-06 | 1.200E-05 | 1.600E-05 |
| 17361 | 869 | T869A | C | 1.5e-05 | 1.1e-05 | 1.7e-05 | 1.433E-05 | 3.055E-06 | 1.100E-05 | 1.700E-05 |
| 17362 | 869 | T869R | C | 1.5e-05 | 1.5e-05 | 1.6e-05 | 1.533E-05 | 5.774E-07 | 1.500E-05 | 1.600E-05 |
| 17363 | 869 | T869N | C | 1.5e-05 | 1.5e-05 | 1.4e-05 | 1.467E-05 | 5.774E-07 | 1.400E-05 | 1.500E-05 |
| 17364 | 869 | T869D | C | 1.6e-05 | 1.4e-05 | 1.6e-05 | 1.533E-05 | 1.155E-06 | 1.400E-05 | 1.600E-05 |
| 17365 | 869 | T869C | C | 1.7e-05 | 1.1e-05 | 1.1e-05 | 1.300E-05 | 3.464E-06 | 1.100E-05 | 1.700E-05 |
| 17366 | 869 | T869E | C | 1.8e-05 | 1.2e-05 | 1.4e-05 | 1.467E-05 | 3.055E-06 | 1.200E-05 | 1.800E-05 |
| 17367 | 869 | T869Q | C | 1.5e-05 | 1.1e-05 | 1.1e-05 | 1.233E-05 | 2.309E-06 | 1.100E-05 | 1.500E-05 |
| 17368 | 869 | T869G | C | 1.1e-05 | 1e-05 | 1.1e-05 | 1.067E-05 | 5.774E-07 | 1.000E-05 | 1.100E-05 |
| 17369 | 869 | T869H | C | 1.6e-05 | 1.6e-05 | 1.5e-05 | 1.567E-05 | 5.774E-07 | 1.500E-05 | 1.600E-05 |
| 17370 | 869 | T869I | C | 1.5e-05 | 1.7e-05 | 1.6e-05 | 1.600E-05 | 1.000E-06 | 1.500E-05 | 1.700E-05 |
| 17371 | 869 | T869L | C | 1.8e-05 | 1.4e-05 | 1.2e-05 | 1.467E-05 | 3.055E-06 | 1.200E-05 | 1.800E-05 |
| 17372 | 869 | T869K | C | 8.9e-06 | 1.3e-05 | 1.5e-05 | 1.230E-05 | 3.110E-06 | 8.900E-06 | 1.500E-05 |
| 17373 | 869 | T869M | C | 1.2e-05 | 1.2e-05 | 1.4e-05 | 1.267E-05 | 1.155E-06 | 1.200E-05 | 1.400E-05 |
| 17374 | 869 | T869F | C | 1.6e-05 | 1.3e-05 | 8.1e-06 | 1.237E-05 | 3.988E-06 | 8.100E-06 | 1.600E-05 |
| 17375 | 869 | T869P | C | 1.6e-05 | 1.5e-05 | 1.7e-05 | 1.600E-05 | 1.000E-06 | 1.500E-05 | 1.700E-05 |
| 17376 | 869 | T869S | C | 1.6e-05 | 1.6e-05 | 9.7e-06 | 1.390E-05 | 3.637E-06 | 9.700E-06 | 1.600E-05 |
| 17377 | 869 | T869T | C | 1.4e-05 | 1.5e-05 | 1.7e-05 | 1.533E-05 | 1.528E-06 | 1.400E-05 | 1.700E-05 |
| 17378 | 869 | T869W | C | 1.3e-05 | 9.1e-06 | 1.7e-05 | 1.303E-05 | 3.950E-06 | 9.100E-06 | 1.700E-05 |
| 17379 | 869 | T869Y | C | 1.4e-05 | 1.3e-05 | 1e-05 | 1.233E-05 | 2.082E-06 | 1.000E-05 | 1.400E-05 |
| 17380 | 869 | T869V | C | 7.9e-06 | 8.5e-06 | 1.6e-05 | 1.080E-05 | 4.513E-06 | 7.900E-06 | 1.600E-05 |
| 17381 | 870 | M870A | C | 1.1e-05 | 1.7e-05 | 1.7e-05 | 1.500E-05 | 3.464E-06 | 1.100E-05 | 1.700E-05 |
| 17382 | 870 | M870R | C | 1.4e-05 | 1.1e-05 | 1e-05 | 1.167E-05 | 2.082E-06 | 1.000E-05 | 1.400E-05 |
| 17383 | 870 | M870N | C | 1.3e-05 | 1.1e-05 | 9.6e-06 | 1.120E-05 | 1.709E-06 | 9.600E-06 | 1.300E-05 |
| 17384 | 870 | M870D | C | 1.6e-05 | 1.4e-05 | 1.3e-05 | 1.433E-05 | 1.528E-06 | 1.300E-05 | 1.600E-05 |
| 17385 | 870 | M870C | C | 1.6e-05 | 1.5e-05 | 1.2e-05 | 1.433E-05 | 2.082E-06 | 1.200E-05 | 1.600E-05 |
| 17386 | 870 | M870E | C | 1.5e-05 | 1.3e-05 | 1.7e-05 | 1.500E-05 | 2.000E-06 | 1.300E-05 | 1.700E-05 |
| 17387 | 870 | M870Q | C | 1.4e-05 | 1.5e-05 | 1.3e-05 | 1.400E-05 | 1.000E-06 | 1.300E-05 | 1.500E-05 |
| 17388 | 870 | M870G | C | 1.2e-05 | 1.6e-05 | 1.6e-05 | 1.467E-05 | 2.309E-06 | 1.200E-05 | 1.600E-05 |
| 17389 | 870 | M870H | C | 1.3e-05 | 1.4e-05 | 9.4e-06 | 1.213E-05 | 2.419E-06 | 9.400E-06 | 1.400E-05 |
| 17390 | 870 | M870I | C | 1.6e-05 | 9.8e-06 | 8.9e-06 | 1.157E-05 | 3.866E-06 | 8.900E-06 | 1.600E-05 |
| 17391 | 870 | M870L | C | 1.2e-05 | 6.7e-06 | 1.7e-05 | 1.190E-05 | 5.151E-06 | 6.700E-06 | 1.700E-05 |
| 17392 | 870 | M870K | C | 1e-05 | 1.5e-05 | 1.6e-05 | 1.367E-05 | 3.215E-06 | 1.000E-05 | 1.600E-05 |
| 17393 | 870 | M870M | C | 1.5e-05 | 9.4e-06 | 1e-05 | 1.147E-05 | 3.075E-06 | 9.400E-06 | 1.500E-05 |
| 17394 | 870 | M870F | C | 1.6e-05 | 1.7e-05 | 1.5e-05 | 1.600E-05 | 1.000E-06 | 1.500E-05 | 1.700E-05 |
| 17395 | 870 | M870P | C | 9.4e-06 | 1e-05 | 9.3e-06 | 9.567E-06 | 3.786E-07 | 9.300E-06 | 1.000E-05 |
| 17396 | 870 | M870S | C | 1.1e-05 | 1.5e-05 | 1.3e-05 | 1.300E-05 | 2.000E-06 | 1.100E-05 | 1.500E-05 |
| 17397 | 870 | M870T | C | 1e-05 | 2.2e-05 | 1.5e-05 | 1.567E-05 | 6.028E-06 | 1.000E-05 | 2.200E-05 |
| 17398 | 870 | M870W | C | 1e-05 | 1.5e-05 | 1.2e-05 | 1.233E-05 | 2.517E-06 | 1.000E-05 | 1.500E-05 |
| 17399 | 870 | M870Y | C | 1e-05 | 1.2e-05 | 9.7e-06 | 1.057E-05 | 1.250E-06 | 9.700E-06 | 1.200E-05 |
| 17400 | 870 | M870V | C | 1.4e-05 | 1.4e-05 | 1e-05 | 1.267E-05 | 2.309E-06 | 1.000E-05 | 1.400E-05 |
| 17401 | 871 | D871A | C | 1.4e-05 | 1.2e-05 | 1.5e-05 | 1.367E-05 | 1.528E-06 | 1.200E-05 | 1.500E-05 |
| 17402 | 871 | D871R | C | 1.1e-05 | 1.6e-05 | 1.1e-05 | 1.267E-05 | 2.887E-06 | 1.100E-05 | 1.600E-05 |
| 17403 | 871 | D871N | C | 1.2e-05 | 1e-05 | 1.5e-05 | 1.233E-05 | 2.517E-06 | 1.000E-05 | 1.500E-05 |
| 17404 | 871 | D871D | C | 1.8e-05 | 1e-05 | 2.1e-05 | 1.633E-05 | 5.686E-06 | 1.000E-05 | 2.100E-05 |
| 17405 | 871 | D871C | C | 1.7e-05 | 1.5e-05 | 1.5e-05 | 1.567E-05 | 1.155E-06 | 1.500E-05 | 1.700E-05 |
| 17406 | 871 | D871E | C | 1.2e-05 | 1.3e-05 | 1e-05 | 1.167E-05 | 1.528E-06 | 1.000E-05 | 1.300E-05 |
| 17407 | 871 | D871Q | C | 1.5e-05 | 1e-05 | 1e-05 | 1.167E-05 | 2.887E-06 | 1.000E-05 | 1.500E-05 |
| 17408 | 871 | D871G | C | 8.5e-06 | 1.5e-05 | 9.8e-06 | 1.110E-05 | 3.439E-06 | 8.500E-06 | 1.500E-05 |
| 17409 | 871 | D871H | C | 1.6e-05 | 1.3e-05 | 1.2e-05 | 1.367E-05 | 2.082E-06 | 1.200E-05 | 1.600E-05 |
| 17410 | 871 | D871I | C | 1e-05 | 5.8e-06 | 1.7e-05 | 1.093E-05 | 5.658E-06 | 5.800E-06 | 1.700E-05 |
| 17411 | 871 | D871L | C | 1.1e-05 | 1.5e-05 | 1e-05 | 1.200E-05 | 2.646E-06 | 1.000E-05 | 1.500E-05 |
| 17412 | 871 | D871K | C | 1.2e-05 | 8.6e-06 | 1.4e-05 | 1.153E-05 | 2.730E-06 | 8.600E-06 | 1.400E-05 |
| 17413 | 871 | D871M | C | 1.2e-05 | 1.1e-05 | 1.2e-05 | 1.167E-05 | 5.774E-07 | 1.100E-05 | 1.200E-05 |
| 17414 | 871 | D871F | C | 1.1e-05 | 1e-05 | 1.5e-05 | 1.200E-05 | 2.646E-06 | 1.000E-05 | 1.500E-05 |
| 17415 | 871 | D871P | C | 1.1e-05 | 1.1e-05 | 1.4e-05 | 1.200E-05 | 1.732E-06 | 1.100E-05 | 1.400E-05 |
| 17416 | 871 | D871S | C | 9.7e-06 | 1.3e-05 | 9.2e-06 | 1.063E-05 | 2.065E-06 | 9.200E-06 | 1.300E-05 |
| 17417 | 871 | D871T | C | 1.5e-05 | 1e-05 | 1.4e-05 | 1.300E-05 | 2.646E-06 | 1.000E-05 | 1.500E-05 |
| 17418 | 871 | D871W | C | 9.5e-06 | 1.4e-05 | 1.5e-05 | 1.283E-05 | 2.930E-06 | 9.500E-06 | 1.500E-05 |
| 17419 | 871 | D871Y | C | 1e-05 | 1.2e-05 | 1.1e-05 | 1.100E-05 | 1.000E-06 | 1.000E-05 | 1.200E-05 |
| 17420 | 871 | D871V | C | 1.5e-05 | 1.5e-05 | 1.6e-05 | 1.533E-05 | 5.774E-07 | 1.500E-05 | 1.600E-05 |
| 17421 | 872 | W872A | C | 1.2e-05 | 1.6e-05 | 1.5e-05 | 1.433E-05 | 2.082E-06 | 1.200E-05 | 1.600E-05 |
| 17422 | 872 | W872R | C | 1.3e-05 | 1.5e-05 | 1.4e-05 | 1.400E-05 | 1.000E-06 | 1.300E-05 | 1.500E-05 |
| 17423 | 872 | W872N | C | 1.3e-05 | 1.6e-05 | 1e-05 | 1.300E-05 | 3.000E-06 | 1.000E-05 | 1.600E-05 |
| 17424 | 872 | W872D | C | 1.4e-05 | 1e-05 | 1.1e-05 | 1.167E-05 | 2.082E-06 | 1.000E-05 | 1.400E-05 |

| | | | | | | | | | | |
|---|---|---|---|---|---|---|---|---|---|---|
| 17425 | 872 | W872C | C | 1.5e-05 | 1.4e-05 | 1.1e-05 | 1.333E-05 | 2.082E-06 | 1.100E-05 | 1.500E-05 |
| 17426 | 872 | W872E | C | 1.5e-05 | 1.1e-05 | 1.4e-05 | 1.333E-05 | 2.082E-06 | 1.100E-05 | 1.500E-05 |
| 17427 | 872 | W872Q | C | 1.1e-05 | 9.5e-06 | 1.2e-05 | 1.083E-05 | 1.258E-06 | 9.500E-06 | 1.200E-05 |
| 17428 | 872 | W872G | C | 1.5e-05 | 9.7e-06 | 7.5e-06 | 1.073E-05 | 3.855E-06 | 7.500E-06 | 1.500E-05 |
| 17429 | 872 | W872H | C | 1.1e-05 | 1.3e-05 | 8.7e-06 | 1.090E-05 | 2.152E-06 | 8.700E-06 | 1.300E-05 |
| 17430 | 872 | W872I | C | 6.3e-06 | 1.7e-05 | 1.1e-05 | 1.143E-05 | 5.363E-06 | 6.300E-06 | 1.700E-05 |
| 17431 | 872 | W872L | C | 9.1e-06 | 1.1e-05 | 1.6e-05 | 1.203E-05 | 3.564E-06 | 9.100E-06 | 1.600E-05 |
| 17432 | 872 | W872K | C | 9.6e-06 | 1.3e-05 | 1e-05 | 1.087E-05 | 1.858E-06 | 9.600E-06 | 1.300E-05 |
| 17433 | 872 | W872M | C | 8.3e-06 | 1.2e-05 | 1.4e-05 | 1.143E-05 | 2.892E-06 | 8.300E-06 | 1.400E-05 |
| 17434 | 872 | W872F | C | 1.1e-05 | 9.6e-06 | 1.4e-05 | 1.153E-05 | 2.248E-06 | 9.600E-06 | 1.400E-05 |
| 17435 | 872 | W872P | C | 1.5e-05 | 1.5e-05 | 1.6e-05 | 1.533E-05 | 5.774E-07 | 1.500E-05 | 1.600E-05 |
| 17436 | 872 | W872S | C | 1.7e-05 | 1.1e-05 | 1.5e-05 | 1.433E-05 | 3.055E-06 | 1.100E-05 | 1.700E-05 |
| 17437 | 872 | W872T | C | 1.4e-05 | 9.4e-06 | 1.5e-05 | 1.280E-05 | 2.987E-06 | 9.400E-06 | 1.500E-05 |
| 17438 | 872 | W872W | C | 1.1e-05 | 1.6e-05 | 1.5e-05 | 1.400E-05 | 2.646E-06 | 1.100E-05 | 1.600E-05 |
| 17439 | 872 | W872Y | C | 1.6e-05 | 1e-05 | 8.7e-06 | 1.157E-05 | 3.894E-06 | 8.700E-06 | 1.600E-05 |
| 17440 | 872 | W872V | C | 9.2e-06 | 1.2e-05 | 1.5e-05 | 1.207E-05 | 2.901E-06 | 9.200E-06 | 1.500E-05 |
| 17441 | 873 | V873A | C | 9.5e-06 | 7.1e-06 | 1.4e-05 | 1.020E-05 | 3.503E-06 | 7.100E-06 | 1.400E-05 |
| 17442 | 873 | V873R | C | 1e-05 | 1.1e-05 | 1.5e-05 | 1.200E-05 | 2.646E-06 | 1.000E-05 | 1.500E-05 |
| 17443 | 873 | V873N | C | 1.2e-05 | 1.6e-05 | 7.4e-06 | 1.180E-05 | 4.303E-06 | 7.400E-06 | 1.600E-05 |
| 17444 | 873 | V873D | C | 1e-05 | 1.1e-05 | 9.7e-06 | 1.023E-05 | 6.807E-07 | 9.700E-06 | 1.100E-05 |
| 17445 | 873 | V873C | C | 1.4e-05 | 1.6e-05 | 1.1e-05 | 1.367E-05 | 2.517E-06 | 1.100E-05 | 1.600E-05 |
| 17446 | 873 | V873E | C | 1.4e-05 | 1.4e-05 | 8.3e-06 | 1.210E-05 | 3.291E-06 | 8.300E-06 | 1.400E-05 |
| 17447 | 873 | V873Q | C | 1.2e-05 | 1.5e-05 | 9.6e-06 | 1.220E-05 | 2.706E-06 | 9.600E-06 | 1.500E-05 |
| 17448 | 873 | V873G | C | 1e-05 | 9.6e-06 | 1.5e-05 | 1.153E-05 | 3.009E-06 | 9.600E-06 | 1.500E-05 |
| 17449 | 873 | V873H | C | 1.5e-05 | 1.5e-05 | 1.2e-05 | 1.400E-05 | 1.732E-06 | 1.200E-05 | 1.500E-05 |
| 17450 | 873 | V873I | C | 1.1e-05 | 1.3e-05 | 1.6e-05 | 1.333E-05 | 2.517E-06 | 1.100E-05 | 1.600E-05 |
| 17451 | 873 | V873L | C | 1.6e-05 | 1.4e-05 | 1.4e-05 | 1.467E-05 | 1.155E-06 | 1.400E-05 | 1.600E-05 |
| 17452 | 873 | V873K | C | 9.7e-06 | 1.1e-05 | 1.1e-05 | 1.057E-05 | 7.506E-07 | 9.700E-06 | 1.100E-05 |
| 17453 | 873 | V873M | C | 1e-05 | 1e-05 | 9.4e-06 | 9.800E-06 | 3.464E-07 | 9.400E-06 | 1.000E-05 |
| 17454 | 873 | V873F | C | 7.9e-06 | 1.5e-05 | 1.5e-05 | 1.263E-05 | 4.099E-06 | 7.900E-06 | 1.500E-05 |
| 17455 | 873 | V873P | C | 1.1e-05 | 1.5e-05 | 1.6e-05 | 1.400E-05 | 2.646E-06 | 1.100E-05 | 1.600E-05 |
| 17456 | 873 | V873S | C | 1e-05 | 1.3e-05 | 1.6e-05 | 1.300E-05 | 3.000E-06 | 1.000E-05 | 1.600E-05 |
| 17457 | 873 | V873T | C | 1.1e-05 | 1.5e-05 | 1.4e-05 | 1.333E-05 | 2.082E-06 | 1.100E-05 | 1.500E-05 |
| 17458 | 873 | V873W | C | 9.9e-06 | 1.5e-05 | 1.5e-05 | 1.330E-05 | 2.944E-06 | 9.900E-06 | 1.500E-05 |
| 17459 | 873 | V873Y | C | 1.4e-05 | 1.6e-05 | 1.1e-05 | 1.367E-05 | 2.517E-06 | 1.100E-05 | 1.600E-05 |
| 17460 | 873 | V873V | C | 1.4e-05 | 1.7e-05 | 9.6e-06 | 1.353E-05 | 3.722E-06 | 9.600E-06 | 1.700E-05 |
| 17461 | 874 | R874A | C | 1.1e-05 | 1.5e-05 | 1.5e-05 | 1.367E-05 | 2.309E-06 | 1.100E-05 | 1.500E-05 |
| 17462 | 874 | R874R | C | 1.4e-05 | 1.6e-05 | 1.6e-05 | 1.533E-05 | 1.155E-06 | 1.400E-05 | 1.600E-05 |
| 17463 | 874 | R874N | C | 1.7e-05 | 1.7e-05 | 8.6e-06 | 1.420E-05 | 4.850E-06 | 8.600E-06 | 1.700E-05 |
| 17464 | 874 | R874D | C | 9.9e-06 | 1.4e-05 | 8.5e-06 | 1.080E-05 | 2.858E-06 | 8.500E-06 | 1.400E-05 |
| 17465 | 874 | R874C | C | 1.1e-05 | 1.6e-05 | 9.6e-06 | 1.220E-05 | 3.365E-06 | 9.600E-06 | 1.600E-05 |
| 17466 | 874 | R874E | C | 1.1e-05 | 9.3e-06 | 1.6e-05 | 1.210E-05 | 3.483E-06 | 9.300E-06 | 1.600E-05 |
| 17467 | 874 | R874Q | C | 9.8e-06 | 1.6e-05 | 1.4e-05 | 1.327E-05 | 3.164E-06 | 9.800E-06 | 1.600E-05 |
| 17468 | 874 | R874G | C | 1.1e-05 | 1.5e-05 | 2.5e-05 | 1.700E-05 | 7.211E-06 | 1.100E-05 | 2.500E-05 |
| 17469 | 874 | R874H | C | 1.5e-05 | 1.7e-05 | 1.2e-05 | 1.467E-05 | 2.517E-06 | 1.200E-05 | 1.700E-05 |
| 17470 | 874 | R874I | C | 1.4e-05 | 1.5e-05 | 8.9e-06 | 1.263E-05 | 3.272E-06 | 8.900E-06 | 1.500E-05 |
| 17471 | 874 | R874L | C | 1.6e-05 | 1.3e-05 | 1.4e-05 | 1.433E-05 | 1.528E-06 | 1.300E-05 | 1.600E-05 |
| 17472 | 874 | R874K | C | 9.5e-06 | 9.9e-06 | 1.4e-05 | 1.113E-05 | 2.491E-06 | 9.500E-06 | 1.400E-05 |
| 17473 | 874 | R874M | C | 1.3e-05 | 1e-05 | 1.5e-05 | 1.267E-05 | 2.517E-06 | 1.000E-05 | 1.500E-05 |
| 17474 | 874 | R874F | C | 1.1e-05 | 1.2e-05 | 1e-05 | 1.100E-05 | 1.000E-06 | 1.000E-05 | 1.200E-05 |
| 17475 | 874 | R874P | C | 1.6e-05 | 1.5e-05 | 1.7e-05 | 1.600E-05 | 1.000E-06 | 1.500E-05 | 1.700E-05 |
| 17476 | 874 | R874S | C | 1e-05 | 1.6e-05 | 1.5e-05 | 1.367E-05 | 3.215E-06 | 1.000E-05 | 1.600E-05 |
| 17477 | 874 | R874T | C | 1e-05 | 1.2e-05 | 1.6e-05 | 1.267E-05 | 3.055E-06 | 1.000E-05 | 1.600E-05 |
| 17478 | 874 | R874W | C | 8.3e-06 | 1.5e-05 | 1.4e-05 | 1.243E-05 | 3.614E-06 | 8.300E-06 | 1.500E-05 |
| 17479 | 874 | R874Y | C | 1.4e-05 | 9.8e-06 | 9.3e-06 | 1.103E-05 | 2.581E-06 | 9.300E-06 | 1.400E-05 |
| 17480 | 874 | R874V | C | 1.4e-05 | 1.1e-05 | 1.2e-05 | 1.233E-05 | 1.528E-06 | 1.100E-05 | 1.400E-05 |
| 17481 | 875 | Q875A | C | 1.6e-05 | 1.1e-05 | 1.5e-05 | 1.400E-05 | 2.646E-06 | 1.100E-05 | 1.600E-05 |
| 17482 | 875 | Q875R | C | 1.6e-05 | 1.5e-05 | 1.4e-05 | 1.500E-05 | 1.000E-06 | 1.400E-05 | 1.600E-05 |
| 17483 | 875 | Q875N | C | 1.5e-05 | 1.5e-05 | 8.9e-06 | 1.297E-05 | 3.522E-06 | 8.900E-06 | 1.500E-05 |
| 17484 | 875 | Q875D | C | 1.5e-05 | 7.5e-06 | 9.4e-06 | 1.063E-05 | 3.899E-06 | 7.500E-06 | 1.500E-05 |
| 17485 | 875 | Q875C | C | 1.3e-05 | 1.5e-05 | 1.4e-05 | 1.400E-05 | 1.000E-06 | 1.300E-05 | 1.500E-05 |
| 17486 | 875 | Q875E | C | 1.5e-05 | 1.3e-05 | 1.4e-05 | 1.400E-05 | 1.000E-06 | 1.300E-05 | 1.500E-05 |
| 17487 | 875 | Q875Q | C | 1.4e-05 | 1.5e-05 | 1e-05 | 1.300E-05 | 2.646E-06 | 1.000E-05 | 1.500E-05 |
| 17488 | 875 | Q875G | C | 1.2e-05 | 9.9e-06 | 1.1e-05 | 1.097E-05 | 1.050E-06 | 9.900E-06 | 1.200E-05 |
| 17489 | 875 | Q875H | C | 1.1e-05 | 1.6e-05 | 1.7e-05 | 1.467E-05 | 3.215E-06 | 1.100E-05 | 1.700E-05 |
| 17490 | 875 | Q875I | C | 1.6e-05 | 1.6e-05 | 8.8e-06 | 1.360E-05 | 4.157E-06 | 8.800E-06 | 1.600E-05 |
| 17491 | 875 | Q875L | C | 9e-06 | 1.4e-05 | 1.6e-05 | 1.300E-05 | 3.606E-06 | 9.000E-06 | 1.600E-05 |
| 17492 | 875 | Q875K | C | 1.5e-05 | 1.2e-05 | 7.9e-06 | 1.163E-05 | 3.564E-06 | 7.900E-06 | 1.500E-05 |
| 17493 | 875 | Q875M | C | 1.4e-05 | 1.1e-05 | 1.3e-05 | 1.267E-05 | 1.528E-06 | 1.100E-05 | 1.400E-05 |
| 17494 | 875 | Q875F | C | 1e-05 | 1.5e-05 | 1.4e-05 | 1.300E-05 | 2.646E-06 | 1.000E-05 | 1.500E-05 |
| 17495 | 875 | Q875P | C | 7.7e-06 | 1.5e-05 | 1.1e-05 | 1.123E-05 | 3.656E-06 | 7.700E-06 | 1.500E-05 |
| 17496 | 875 | Q875S | C | 1e-05 | 8.9e-06 | 1.5e-05 | 1.130E-05 | 3.251E-06 | 8.900E-06 | 1.500E-05 |
| 17497 | 875 | Q875T | C | 1.4e-05 | 1.4e-05 | 1.6e-05 | 1.467E-05 | 1.155E-06 | 1.400E-05 | 1.600E-05 |
| 17498 | 875 | Q875W | C | 1.5e-05 | 1.6e-05 | 1.4e-05 | 1.500E-05 | 1.000E-06 | 1.400E-05 | 1.600E-05 |
| 17499 | 875 | Q875Y | C | 1.5e-05 | 1.4e-05 | 1.5e-05 | 1.467E-05 | 5.774E-07 | 1.400E-05 | 1.500E-05 |
| 17500 | 875 | Q875V | C | 8.1e-06 | 1.7e-05 | 7.5e-06 | 1.087E-05 | 5.320E-06 | 7.500E-06 | 1.700E-05 |
| 17501 | 876 | A876A | C | 1e-05 | 1e-05 | 1.4e-05 | 1.133E-05 | 2.309E-06 | 1.000E-05 | 1.400E-05 |
| 17502 | 876 | A876R | C | 1e-05 | 1.4e-05 | 9.3e-06 | 1.110E-05 | 2.536E-06 | 9.300E-06 | 1.400E-05 |
| 17503 | 876 | A876N | C | 1e-05 | 1.2e-05 | 1.6e-05 | 1.267E-05 | 3.055E-06 | 1.000E-05 | 1.600E-05 |
| 17504 | 876 | A876D | C | 1.2e-05 | 1.1e-05 | 1.2e-05 | 1.167E-05 | 5.774E-07 | 1.100E-05 | 1.200E-05 |
| 17505 | 876 | A876C | C | 1.6e-05 | 1.5e-05 | 1.4e-05 | 1.500E-05 | 1.000E-06 | 1.400E-05 | 1.600E-05 |
| 17506 | 876 | A876E | C | 7.9e-06 | 1.2e-05 | 1e-05 | 9.967E-06 | 2.050E-06 | 7.900E-06 | 1.200E-05 |
| 17507 | 876 | A876Q | C | 1.4e-05 | 1.4e-05 | 1.1e-05 | 1.300E-05 | 1.732E-06 | 1.100E-05 | 1.400E-05 |
| 17508 | 876 | A876G | C | 9.7e-06 | 1.5e-05 | 1.4e-05 | 1.290E-05 | 2.816E-06 | 9.700E-06 | 1.500E-05 |
| 17509 | 876 | A876H | C | 1.7e-05 | 1e-05 | 1e-05 | 1.233E-05 | 4.041E-06 | 1.000E-05 | 1.700E-05 |

| | | | | | | | | | | |
|---|---|---|---|---|---|---|---|---|---|---|
| 17510 | 876 | A876I | C | 1.1e-05 | 1.1e-05 | 8.9e-06 | 1.030E-05 | 1.212E-06 | 8.900E-06 | 1.100E-05 |
| 17511 | 876 | A876L | C | 1e-05 | 1.2e-05 | 1.5e-05 | 1.233E-05 | 2.517E-06 | 1.000E-05 | 1.500E-05 |
| 17512 | 876 | A876K | C | 1.1e-05 | 1.4e-05 | 1.1e-05 | 1.200E-05 | 1.732E-06 | 1.100E-05 | 1.400E-05 |
| 17513 | 876 | A876M | C | 1.6e-05 | 1.6e-05 | 1e-05 | 1.400E-05 | 3.464E-06 | 1.000E-05 | 1.600E-05 |
| 17514 | 876 | A876F | C | 1.1e-05 | 1.4e-05 | 1.6e-05 | 1.367E-05 | 2.517E-06 | 1.100E-05 | 1.600E-05 |
| 17515 | 876 | A876P | C | 1.6e-05 | 1e-05 | 1.3e-05 | 1.300E-05 | 3.000E-06 | 1.000E-05 | 1.600E-05 |
| 17516 | 876 | A876S | C | 9.8e-06 | 1.6e-05 | 1.3e-05 | 1.293E-05 | 3.101E-06 | 9.800E-06 | 1.600E-05 |
| 17517 | 876 | A876T | C | 1.5e-05 | 1.4e-05 | 9.9e-06 | 1.297E-05 | 2.702E-06 | 9.900E-06 | 1.500E-05 |
| 17518 | 876 | A876W | C | 5.8e-06 | 6.7e-06 | 1e-05 | 7.500E-06 | 2.211E-06 | 5.800E-06 | 1.000E-05 |
| 17519 | 876 | A876Y | C | 8.9e-06 | 1.2e-05 | 9e-06 | 9.967E-06 | 1.762E-06 | 8.900E-06 | 1.200E-05 |
| 17520 | 876 | A876V | C | 1.5e-05 | 1e-05 | 1.5e-05 | 1.333E-05 | 2.887E-06 | 1.000E-05 | 1.500E-05 |
| 17521 | 877 | P877A | C | 1.6e-05 | 1.2e-05 | 1.6e-05 | 1.467E-05 | 2.309E-06 | 1.200E-05 | 1.600E-05 |
| 17522 | 877 | P877R | C | 1.6e-05 | 1.3e-05 | 1.5e-05 | 1.467E-05 | 1.528E-06 | 1.300E-05 | 1.600E-05 |
| 17523 | 877 | P877N | C | 1.3e-05 | 1e-05 | 1.5e-05 | 1.267E-05 | 2.517E-06 | 1.000E-05 | 1.500E-05 |
| 17524 | 877 | P877D | C | 1e-05 | 9e-06 | 1.6e-05 | 1.167E-05 | 3.786E-06 | 9.000E-06 | 1.600E-05 |
| 17525 | 877 | P877C | C | 1e-05 | 1.6e-05 | 1.7e-05 | 1.433E-05 | 3.786E-06 | 1.000E-05 | 1.700E-05 |
| 17526 | 877 | P877E | C | 1.6e-05 | 1.5e-05 | 1.1e-05 | 1.400E-05 | 2.646E-06 | 1.100E-05 | 1.600E-05 |
| 17527 | 877 | P877Q | C | 9.9e-06 | 1.6e-05 | 1.5e-05 | 1.363E-05 | 3.272E-06 | 9.900E-06 | 1.600E-05 |
| 17528 | 877 | P877G | C | 1.2e-05 | 1.6e-05 | 1.6e-05 | 1.467E-05 | 2.309E-06 | 1.200E-05 | 1.600E-05 |
| 17529 | 877 | P877H | C | 7.9e-06 | 1.2e-05 | 8.9e-06 | 9.600E-06 | 2.138E-06 | 7.900E-06 | 1.200E-05 |
| 17530 | 877 | P877I | C | 1.4e-05 | 1.4e-05 | 1e-05 | 1.267E-05 | 2.309E-06 | 1.000E-05 | 1.400E-05 |
| 17531 | 877 | P877L | C | 1.1e-05 | 1.2e-05 | 9.8e-06 | 1.093E-05 | 1.102E-06 | 9.800E-06 | 1.200E-05 |
| 17532 | 877 | P877K | C | 1.5e-05 | 8.2e-06 | 9.4e-06 | 1.087E-05 | 3.630E-06 | 8.200E-06 | 1.500E-05 |
| 17533 | 877 | P877M | C | 1.1e-05 | 1.1e-05 | 1.1e-05 | 1.100E-05 | 2.075E-21 | 1.100E-05 | 1.100E-05 |
| 17534 | 877 | P877F | C | 1.1e-05 | 1e-05 | 1.6e-05 | 1.233E-05 | 3.215E-06 | 1.000E-05 | 1.600E-05 |
| 17535 | 877 | P877P | C | 1.5e-05 | 1.5e-05 | 9.7e-06 | 1.323E-05 | 3.060E-06 | 9.700E-06 | 1.500E-05 |
| 17536 | 877 | P877S | C | 1e-05 | 8.9e-06 | 9.5e-06 | 9.467E-06 | 5.508E-07 | 8.900E-06 | 1.000E-05 |
| 17537 | 877 | P877T | C | 1.1e-05 | 1.5e-05 | 1.4e-05 | 1.333E-05 | 2.082E-06 | 1.100E-05 | 1.500E-05 |
| 17538 | 877 | P877W | C | 1.1e-05 | 1.3e-05 | 1.1e-05 | 1.167E-05 | 1.155E-06 | 1.100E-05 | 1.300E-05 |
| 17539 | 877 | P877Y | C | 1.7e-05 | 1.2e-05 | 9.1e-06 | 1.270E-05 | 3.996E-06 | 9.100E-06 | 1.700E-05 |
| 17540 | 877 | P877V | C | 1.5e-05 | 1.3e-05 | 1.4e-05 | 1.400E-05 | 1.000E-06 | 1.300E-05 | 1.500E-05 |
| 17541 | 878 | G878A | C | 1.1e-05 | 1.8e-05 | 1.8e-05 | 1.567E-05 | 4.041E-06 | 1.100E-05 | 1.800E-05 |
| 17542 | 878 | G878R | C | 1.1e-05 | 1.5e-05 | 1.6e-05 | 1.400E-05 | 2.646E-06 | 1.100E-05 | 1.600E-05 |
| 17543 | 878 | G878N | C | 1.6e-05 | 8.8e-06 | 8.7e-06 | 1.117E-05 | 4.186E-06 | 8.700E-06 | 1.600E-05 |
| 17544 | 878 | G878D | C | 1.1e-05 | 1.5e-05 | 8.3e-06 | 1.143E-05 | 3.371E-06 | 8.300E-06 | 1.500E-05 |
| 17545 | 878 | G878C | C | 1.5e-05 | 1.1e-05 | 8.6e-06 | 1.153E-05 | 3.233E-06 | 8.600E-06 | 1.500E-05 |
| 17546 | 878 | G878E | C | 1.4e-05 | 1.6e-05 | 1.1e-05 | 1.367E-05 | 2.517E-06 | 1.100E-05 | 1.600E-05 |
| 17547 | 878 | G878Q | C | 1.6e-05 | 1.3e-05 | 8.5e-06 | 1.250E-05 | 3.775E-06 | 8.500E-06 | 1.600E-05 |
| 17548 | 878 | G878G | C | 1.6e-05 | 1.2e-05 | 7.7e-06 | 1.190E-05 | 4.151E-06 | 7.700E-06 | 1.600E-05 |
| 17549 | 878 | G878H | C | 1.2e-05 | 1.4e-05 | 8.4e-06 | 1.147E-05 | 2.838E-06 | 8.400E-06 | 1.400E-05 |
| 17550 | 878 | G878I | C | 1.6e-05 | 1.6e-05 | 1.8e-05 | 1.667E-05 | 1.155E-06 | 1.600E-05 | 1.800E-05 |
| 17551 | 878 | G878L | C | 9.7e-06 | 1.5e-05 | 9.9e-06 | 1.153E-05 | 3.004E-06 | 9.700E-06 | 1.500E-05 |
| 17552 | 878 | G878K | C | 9.5e-06 | 1.1e-05 | 1.4e-05 | 1.150E-05 | 2.291E-06 | 9.500E-06 | 1.400E-05 |
| 17553 | 878 | G878M | C | 1.6e-05 | 1.2e-05 | 1.6e-05 | 1.467E-05 | 2.309E-06 | 1.200E-05 | 1.600E-05 |
| 17554 | 878 | G878F | C | 1.4e-05 | 1.1e-05 | 1.4e-05 | 1.300E-05 | 1.732E-06 | 1.100E-05 | 1.400E-05 |
| 17555 | 878 | G878P | C | 1.6e-05 | 1.4e-05 | 1.5e-05 | 1.500E-05 | 1.000E-06 | 1.400E-05 | 1.600E-05 |
| 17556 | 878 | G878S | C | 7.7e-06 | 1.3e-05 | 1.6e-05 | 1.223E-05 | 4.203E-06 | 7.700E-06 | 1.600E-05 |
| 17557 | 878 | G878T | C | 1.5e-05 | 1.7e-05 | 1e-05 | 1.400E-05 | 3.606E-06 | 1.000E-05 | 1.700E-05 |
| 17558 | 878 | G878W | C | 1.6e-05 | 1.5e-05 | 1.3e-05 | 1.467E-05 | 1.528E-06 | 1.300E-05 | 1.600E-05 |
| 17559 | 878 | G878Y | C | 1e-05 | 1.6e-05 | 1.4e-05 | 1.333E-05 | 3.055E-06 | 1.000E-05 | 1.600E-05 |
| 17560 | 878 | G878V | C | 1.7e-05 | 1.3e-05 | 1.3e-05 | 1.433E-05 | 2.309E-06 | 1.300E-05 | 1.700E-05 |
| 17561 | 879 | K879A | C | 1.6e-05 | 1.1e-05 | 1.5e-05 | 1.400E-05 | 2.646E-06 | 1.100E-05 | 1.600E-05 |
| 17562 | 879 | K879R | C | 1.4e-05 | 1.2e-05 | 9.6e-06 | 1.187E-05 | 2.203E-06 | 9.600E-06 | 1.400E-05 |
| 17563 | 879 | K879N | C | 1.3e-05 | 1e-05 | 1.5e-05 | 1.267E-05 | 2.517E-06 | 1.000E-05 | 1.500E-05 |
| 17564 | 879 | K879D | C | 1.5e-05 | 1.1e-05 | 1.4e-05 | 1.333E-05 | 2.082E-06 | 1.100E-05 | 1.500E-05 |
| 17565 | 879 | K879C | C | 1.5e-05 | 8.8e-06 | 1e-05 | 1.127E-05 | 3.288E-06 | 8.800E-06 | 1.500E-05 |
| 17566 | 879 | K879E | C | 1.2e-05 | 1.3e-05 | 1.3e-05 | 1.267E-05 | 5.774E-07 | 1.200E-05 | 1.300E-05 |
| 17567 | 879 | K879Q | C | 8e-06 | 9.4e-06 | 1e-05 | 9.133E-06 | 1.026E-06 | 8.000E-06 | 1.000E-05 |
| 17568 | 879 | K879G | C | 1.5e-05 | 1.4e-05 | 1.5e-05 | 1.467E-05 | 5.774E-07 | 1.400E-05 | 1.500E-05 |
| 17569 | 879 | K879H | C | 1.3e-05 | 1.5e-05 | 1.5e-05 | 1.433E-05 | 1.155E-06 | 1.300E-05 | 1.500E-05 |
| 17570 | 879 | K879I | C | 1.5e-05 | 1.1e-05 | 1.3e-05 | 1.300E-05 | 2.000E-06 | 1.100E-05 | 1.500E-05 |
| 17571 | 879 | K879L | C | 1.1e-05 | 8.8e-06 | 9.6e-06 | 9.800E-06 | 1.114E-06 | 8.800E-06 | 1.100E-05 |
| 17572 | 879 | K879K | C | 1.5e-05 | 1e-05 | 1.5e-05 | 1.333E-05 | 2.887E-06 | 1.000E-05 | 1.500E-05 |
| 17573 | 879 | K879M | C | 1e-05 | 1.5e-05 | 1.3e-05 | 1.267E-05 | 2.517E-06 | 1.000E-05 | 1.500E-05 |
| 17574 | 879 | K879F | C | 1.5e-05 | 1.4e-05 | 1.6e-05 | 1.500E-05 | 1.000E-06 | 1.400E-05 | 1.600E-05 |
| 17575 | 879 | K879P | C | 1.5e-05 | 1.1e-05 | 1.4e-05 | 1.333E-05 | 2.082E-06 | 1.100E-05 | 1.500E-05 |
| 17576 | 879 | K879S | C | 1.5e-05 | 8.9e-06 | 1.1e-05 | 1.163E-05 | 3.099E-06 | 8.900E-06 | 1.500E-05 |
| 17577 | 879 | K879T | C | 9.5e-06 | 1.3e-05 | 1.6e-05 | 1.283E-05 | 3.253E-06 | 9.500E-06 | 1.600E-05 |
| 17578 | 879 | K879W | C | 1.6e-05 | 1.2e-05 | 1.3e-05 | 1.367E-05 | 2.082E-06 | 1.200E-05 | 1.600E-05 |
| 17579 | 879 | K879Y | C | 1e-05 | 1.5e-05 | 1.2e-05 | 1.233E-05 | 2.517E-06 | 1.000E-05 | 1.500E-05 |
| 17580 | 879 | K879V | C | 1.4e-05 | 1.1e-05 | 8.3e-06 | 1.110E-05 | 2.851E-06 | 8.300E-06 | 1.400E-05 |
| 17581 | 880 | G880A | C | 1.5e-05 | 9.3e-06 | 1e-05 | 1.143E-05 | 3.109E-06 | 9.300E-06 | 1.500E-05 |
| 17582 | 880 | G880R | C | 1.5e-05 | 1.1e-05 | 1.4e-05 | 1.333E-05 | 2.082E-06 | 1.100E-05 | 1.500E-05 |
| 17583 | 880 | G880N | C | 1.2e-05 | 9.1e-06 | 1.2e-05 | 1.103E-05 | 1.674E-06 | 9.100E-06 | 1.200E-05 |
| 17584 | 880 | G880D | C | 1.2e-05 | 8.8e-06 | 1.4e-05 | 1.160E-05 | 2.623E-06 | 8.800E-06 | 1.400E-05 |
| 17585 | 880 | G880C | C | 1.1e-05 | 1.1e-05 | 1e-05 | 1.067E-05 | 5.774E-07 | 1.000E-05 | 1.100E-05 |
| 17586 | 880 | G880E | C | 1.2e-05 | 1.4e-05 | 9.2e-06 | 1.173E-05 | 2.411E-06 | 9.200E-06 | 1.400E-05 |
| 17587 | 880 | G880Q | C | 1e-05 | 8.3e-06 | 1e-05 | 9.433E-06 | 9.815E-07 | 8.300E-06 | 1.000E-05 |
| 17588 | 880 | G880G | C | 1.3e-05 | 1.6e-05 | 8.5e-06 | 1.250E-05 | 3.775E-06 | 8.500E-06 | 1.600E-05 |
| 17589 | 880 | G880H | C | 6.7e-06 | 1.2e-05 | 1.4e-05 | 1.090E-05 | 3.772E-06 | 6.700E-06 | 1.400E-05 |
| 17590 | 880 | G880I | C | 1.4e-05 | 1e-05 | 1.1e-05 | 1.167E-05 | 2.082E-06 | 1.000E-05 | 1.400E-05 |
| 17591 | 880 | G880L | C | 1.4e-05 | 1e-05 | 1.5e-05 | 1.300E-05 | 2.646E-06 | 1.000E-05 | 1.500E-05 |
| 17592 | 880 | G880K | C | 1.4e-05 | 9.9e-06 | 1.4e-05 | 1.263E-05 | 2.367E-06 | 9.900E-06 | 1.400E-05 |
| 17593 | 880 | G880M | C | 1.4e-05 | 1.2e-05 | 9e-06 | 1.167E-05 | 2.517E-06 | 9.000E-06 | 1.400E-05 |
| 17594 | 880 | G880F | C | 1.6e-05 | 1.4e-05 | 2.3e-05 | 1.767E-05 | 4.726E-06 | 1.400E-05 | 2.300E-05 |

| | | | | | | | | | | |
|---|---|---|---|---|---|---|---|---|---|---|
| 17595 | 880 | G880P | C | 1.2e-05 | 9e-06 | 1.1e-05 | 1.067E-05 | 1.528E-06 | 9.000E-06 | 1.200E-05 |
| 17596 | 880 | G880S | C | 1.5e-05 | 1.4e-05 | 1e-05 | 1.300E-05 | 2.646E-06 | 1.000E-05 | 1.500E-05 |
| 17597 | 880 | G880T | C | 1.7e-05 | 1e-05 | 1.1e-05 | 1.267E-05 | 3.786E-06 | 1.000E-05 | 1.700E-05 |
| 17598 | 880 | G880W | C | 1.4e-05 | 1.2e-05 | 1e-05 | 1.200E-05 | 2.000E-06 | 1.000E-05 | 1.400E-05 |
| 17599 | 880 | G880Y | C | 1.5e-05 | 1.5e-05 | 9.7e-06 | 1.323E-05 | 3.060E-06 | 9.700E-06 | 1.500E-05 |
| 17600 | 880 | G880V | C | 9.3e-06 | 1.7e-05 | 1.2e-05 | 1.277E-05 | 3.907E-06 | 9.300E-06 | 1.700E-05 |
| 17601 | 881 | L881A | C | 1.1e-05 | 1.6e-05 | 6.7e-06 | 1.123E-05 | 4.654E-06 | 6.700E-06 | 1.600E-05 |
| 17602 | 881 | L881R | C | 1.1e-05 | 1e-05 | 1e-05 | 1.033E-05 | 5.774E-07 | 1.000E-05 | 1.100E-05 |
| 17603 | 881 | L881N | C | 1.1e-05 | 1e-05 | 9.9e-06 | 1.030E-05 | 6.083E-07 | 9.900E-06 | 1.100E-05 |
| 17604 | 881 | L881D | C | 1.6e-05 | 1.3e-05 | 1.6e-05 | 1.500E-05 | 1.732E-06 | 1.300E-05 | 1.600E-05 |
| 17605 | 881 | L881C | C | 1.3e-05 | 1.3e-05 | 1e-05 | 1.200E-05 | 1.732E-06 | 1.000E-05 | 1.300E-05 |
| 17606 | 881 | L881E | C | 1.6e-05 | 9.8e-06 | 1e-05 | 1.193E-05 | 3.523E-06 | 9.800E-06 | 1.600E-05 |
| 17607 | 881 | L881Q | C | 1.5e-05 | 1e-05 | 9.6e-06 | 1.153E-05 | 3.009E-06 | 9.600E-06 | 1.500E-05 |
| 17608 | 881 | L881G | C | 1.5e-05 | 1e-05 | 1.7e-05 | 1.400E-05 | 3.606E-06 | 1.000E-05 | 1.700E-05 |
| 17609 | 881 | L881H | C | 1.4e-05 | 1.4e-05 | 1e-05 | 1.267E-05 | 2.309E-06 | 1.000E-05 | 1.400E-05 |
| 17610 | 881 | L881I | C | 1.2e-05 | 9.9e-06 | 8.2e-06 | 1.003E-05 | 1.904E-06 | 8.200E-06 | 1.200E-05 |
| 17611 | 881 | L881L | C | 1.1e-05 | 9.7e-06 | 1e-05 | 1.023E-05 | 6.807E-07 | 9.700E-06 | 1.100E-05 |
| 17612 | 881 | L881K | C | 1.3e-05 | 1.4e-05 | 1.6e-05 | 1.433E-05 | 1.528E-06 | 1.300E-05 | 1.600E-05 |
| 17613 | 881 | L881M | C | 1.4e-05 | 1.2e-05 | 1.6e-05 | 1.400E-05 | 2.000E-06 | 1.200E-05 | 1.600E-05 |
| 17614 | 881 | L881F | C | 1e-05 | 1.4e-05 | 1.5e-05 | 1.300E-05 | 2.646E-06 | 1.000E-05 | 1.500E-05 |
| 17615 | 881 | L881P | C | 1.6e-05 | 1.4e-05 | 1.1e-05 | 1.367E-05 | 2.517E-06 | 1.100E-05 | 1.600E-05 |
| 17616 | 881 | L881S | C | 8.7e-06 | 1.4e-05 | 1.4e-05 | 1.223E-05 | 3.060E-06 | 8.700E-06 | 1.400E-05 |
| 17617 | 881 | L881T | C | 1.5e-05 | 1.1e-05 | 1.5e-05 | 1.367E-05 | 2.309E-06 | 1.100E-05 | 1.500E-05 |
| 17618 | 881 | L881W | C | 1.6e-05 | 1.3e-05 | 1.5e-05 | 1.467E-05 | 1.528E-06 | 1.300E-05 | 1.600E-05 |
| 17619 | 881 | L881Y | C | 1.5e-05 | 1.5e-05 | 1.5e-05 | 1.500E-05 | 0.000E+00 | 1.500E-05 | 1.500E-05 |
| 17620 | 881 | L881V | C | 1.2e-05 | 1.5e-05 | 1.3e-05 | 1.333E-05 | 1.528E-06 | 1.200E-05 | 1.500E-05 |
| 17621 | 882 | E882A | C | 1.6e-05 | 9.6e-06 | 1e-05 | 1.187E-05 | 3.585E-06 | 9.600E-06 | 1.600E-05 |
| 17622 | 882 | E882R | C | 1e-05 | 1.6e-05 | 8.5e-06 | 1.150E-05 | 3.969E-06 | 8.500E-06 | 1.600E-05 |
| 17623 | 882 | E882N | C | 1.5e-05 | 1.5e-05 | 1e-05 | 1.333E-05 | 2.887E-06 | 1.000E-05 | 1.500E-05 |
| 17624 | 882 | E882D | C | 1e-05 | 1.4e-05 | 1e-05 | 1.133E-05 | 2.309E-06 | 1.000E-05 | 1.400E-05 |
| 17625 | 882 | E882C | C | 8.2e-06 | 1.5e-05 | 7.6e-06 | 1.027E-05 | 4.110E-06 | 7.600E-06 | 1.500E-05 |
| 17626 | 882 | E882E | C | 8.9e-06 | 1.2e-05 | 1.3e-05 | 1.130E-05 | 2.138E-06 | 8.900E-06 | 1.300E-05 |
| 17627 | 882 | E882Q | C | 1.4e-05 | 1.1e-05 | 1.4e-05 | 1.300E-05 | 1.732E-06 | 1.100E-05 | 1.400E-05 |
| 17628 | 882 | E882G | C | 9.1e-06 | 1.1e-05 | 9.5e-06 | 9.867E-06 | 1.002E-06 | 9.100E-06 | 1.100E-05 |
| 17629 | 882 | E882H | C | 1.4e-05 | 1.2e-05 | 1.5e-05 | 1.367E-05 | 1.528E-06 | 1.200E-05 | 1.500E-05 |
| 17630 | 882 | E882I | C | 1.6e-05 | 1.3e-05 | 1.7e-05 | 1.533E-05 | 2.082E-06 | 1.300E-05 | 1.700E-05 |
| 17631 | 882 | E882L | C | 1.1e-05 | 1.2e-05 | 1.1e-05 | 1.133E-05 | 5.774E-07 | 1.100E-05 | 1.200E-05 |
| 17632 | 882 | E882K | C | 1.7e-05 | 8e-06 | 1.8e-05 | 1.433E-05 | 5.508E-06 | 8.000E-06 | 1.800E-05 |
| 17633 | 882 | E882M | C | 8.7e-06 | 1.5e-05 | 1.1e-05 | 1.157E-05 | 3.188E-06 | 8.700E-06 | 1.500E-05 |
| 17634 | 882 | E882F | C | 1.3e-05 | 1.5e-05 | 5.9e-06 | 1.130E-05 | 4.782E-06 | 5.900E-06 | 1.500E-05 |
| 17635 | 882 | E882P | C | 9.3e-06 | 1.1e-05 | 1.7e-05 | 1.243E-05 | 4.045E-06 | 9.300E-06 | 1.700E-05 |
| 17636 | 882 | E882S | C | 1.2e-05 | 9.2e-06 | 8.6e-06 | 9.933E-06 | 1.815E-06 | 8.600E-06 | 1.200E-05 |
| 17637 | 882 | E882T | C | 9.4e-06 | 1.6e-05 | 1.5e-05 | 1.347E-05 | 3.557E-06 | 9.400E-06 | 1.600E-05 |
| 17638 | 882 | E882W | C | 1.5e-05 | 8.1e-06 | 1.6e-05 | 1.303E-05 | 4.302E-06 | 8.100E-06 | 1.600E-05 |
| 17639 | 882 | E882Y | C | 1.5e-05 | 1.4e-05 | 1e-05 | 1.300E-05 | 2.646E-06 | 1.000E-05 | 1.500E-05 |
| 17640 | 882 | E882V | C | 1.5e-05 | 1e-05 | 1.5e-05 | 1.333E-05 | 2.887E-06 | 1.000E-05 | 1.500E-05 |
| 17641 | 883 | W883A | C | 1.1e-05 | 8.8e-06 | 1.6e-05 | 1.193E-05 | 3.690E-06 | 8.800E-06 | 1.600E-05 |
| 17642 | 883 | W883R | C | 1e-05 | 9.2e-06 | 1.3e-05 | 1.073E-05 | 2.003E-06 | 9.200E-06 | 1.300E-05 |
| 17643 | 883 | W883N | C | 1.3e-05 | 1.5e-05 | 1.5e-05 | 1.433E-05 | 1.155E-06 | 1.300E-05 | 1.500E-05 |
| 17644 | 883 | W883D | C | 1.7e-05 | 1.1e-05 | 1.6e-05 | 1.467E-05 | 3.215E-06 | 1.100E-05 | 1.700E-05 |
| 17645 | 883 | W883C | C | 1e-05 | 1.5e-05 | 9.9e-06 | 1.163E-05 | 2.916E-06 | 9.900E-06 | 1.500E-05 |
| 17646 | 883 | W883E | C | 1e-05 | 1.6e-05 | 9.7e-06 | 1.190E-05 | 3.554E-06 | 9.700E-06 | 1.600E-05 |
| 17647 | 883 | W883Q | C | 1.1e-05 | 2.2e-05 | 1.5e-05 | 1.600E-05 | 5.568E-06 | 1.100E-05 | 2.200E-05 |
| 17648 | 883 | W883G | C | 1.2e-05 | 1.5e-05 | 1.6e-05 | 1.433E-05 | 2.082E-06 | 1.200E-05 | 1.600E-05 |
| 17649 | 883 | W883H | C | 8.4e-06 | 8.9e-06 | 8.6e-06 | 8.633E-06 | 2.517E-07 | 8.400E-06 | 8.900E-06 |
| 17650 | 883 | W883I | C | 9.4e-06 | 1e-05 | 1.1e-05 | 1.013E-05 | 8.083E-07 | 9.400E-06 | 1.100E-05 |
| 17651 | 883 | W883L | C | 1.2e-05 | 1.6e-05 | 1.5e-05 | 1.433E-05 | 2.082E-06 | 1.200E-05 | 1.600E-05 |
| 17652 | 883 | W883K | C | 1.6e-05 | 1.6e-05 | 1.7e-05 | 1.633E-05 | 5.774E-07 | 1.600E-05 | 1.700E-05 |
| 17653 | 883 | W883M | C | 1.5e-05 | 1.7e-05 | 1.6e-05 | 1.600E-05 | 1.000E-06 | 1.500E-05 | 1.700E-05 |
| 17654 | 883 | W883F | C | 8.4e-06 | 1.7e-05 | 9e-06 | 1.147E-05 | 4.801E-06 | 8.400E-06 | 1.700E-05 |
| 17655 | 883 | W883P | C | 1.4e-05 | 1.3e-05 | 2e-05 | 1.567E-05 | 3.786E-06 | 1.300E-05 | 2.000E-05 |
| 17656 | 883 | W883S | C | 1.4e-05 | 1.4e-05 | 1.1e-05 | 1.300E-05 | 1.732E-06 | 1.100E-05 | 1.400E-05 |
| 17657 | 883 | W883T | C | 1.4e-05 | 1.6e-05 | 1.1e-05 | 1.367E-05 | 2.517E-06 | 1.100E-05 | 1.600E-05 |
| 17658 | 883 | W883W | C | 1.4e-05 | 1.5e-05 | 1.2e-05 | 1.367E-05 | 1.528E-06 | 1.200E-05 | 1.500E-05 |
| 17659 | 883 | W883Y | C | 1.5e-05 | 1.1e-05 | 1.1e-05 | 1.233E-05 | 2.309E-06 | 1.100E-05 | 1.500E-05 |
| 17660 | 883 | W883V | C | 1.7e-05 | 1.6e-05 | 1.7e-05 | 1.667E-05 | 5.774E-07 | 1.600E-05 | 1.700E-05 |
| 17661 | 884 | V884A | C | 1.5e-05 | 1.4e-05 | 1.4e-05 | 1.433E-05 | 5.774E-07 | 1.400E-05 | 1.500E-05 |
| 17662 | 884 | V884R | C | 9.8e-06 | 1.1e-05 | 1e-05 | 1.027E-05 | 6.429E-07 | 9.800E-06 | 1.100E-05 |
| 17663 | 884 | V884N | C | 1.4e-05 | 1.2e-05 | 8.3e-06 | 1.143E-05 | 2.892E-06 | 8.300E-06 | 1.400E-05 |
| 17664 | 884 | V884D | C | 1.4e-05 | 1.2e-05 | 1.7e-05 | 1.433E-05 | 2.517E-06 | 1.200E-05 | 1.700E-05 |
| 17665 | 884 | V884C | C | 1.4e-05 | 1.4e-05 | 1.6e-05 | 1.467E-05 | 1.155E-06 | 1.400E-05 | 1.600E-05 |
| 17666 | 884 | V884E | C | 1.5e-05 | 1.2e-05 | 1.7e-05 | 1.467E-05 | 2.517E-06 | 1.200E-05 | 1.700E-05 |
| 17667 | 884 | V884Q | C | 1.4e-05 | 9.9e-06 | 1.4e-05 | 1.263E-05 | 2.367E-06 | 9.900E-06 | 1.400E-05 |
| 17668 | 884 | V884G | C | 1.5e-05 | 1.5e-05 | 1.6e-05 | 1.533E-05 | 5.774E-07 | 1.500E-05 | 1.600E-05 |
| 17669 | 884 | V884H | C | 1.5e-05 | 1.3e-05 | 8.3e-06 | 1.210E-05 | 3.439E-06 | 8.300E-06 | 1.500E-05 |
| 17670 | 884 | V884I | C | 1.3e-05 | 1.5e-05 | 1.1e-05 | 1.300E-05 | 2.000E-06 | 1.100E-05 | 1.500E-05 |
| 17671 | 884 | V884L | C | 7.1e-06 | 1.5e-05 | 1.3e-05 | 1.170E-05 | 4.107E-06 | 7.100E-06 | 1.500E-05 |
| 17672 | 884 | V884K | C | 1.1e-05 | 1.1e-05 | 1.1e-05 | 1.100E-05 | 2.075E-21 | 1.100E-05 | 1.100E-05 |
| 17673 | 884 | V884M | C | 1.5e-05 | 1.5e-05 | 1.2e-05 | 1.400E-05 | 1.732E-06 | 1.200E-05 | 1.500E-05 |
| 17674 | 884 | V884F | C | 1e-05 | 1.5e-05 | 7.7e-06 | 1.090E-05 | 3.732E-06 | 7.700E-06 | 1.500E-05 |
| 17675 | 884 | V884P | C | 9.8e-06 | 1.3e-05 | 2.1e-05 | 1.460E-05 | 5.769E-06 | 9.800E-06 | 2.100E-05 |
| 17676 | 884 | V884S | C | 1.5e-05 | 1.1e-05 | 1.4e-05 | 1.333E-05 | 2.082E-06 | 1.100E-05 | 1.500E-05 |
| 17677 | 884 | V884T | C | 1.4e-05 | 1e-05 | 1.4e-05 | 1.267E-05 | 2.309E-06 | 1.000E-05 | 1.400E-05 |
| 17678 | 884 | V884W | C | 1.4e-05 | 2.2e-05 | 1.1e-05 | 1.567E-05 | 5.686E-06 | 1.100E-05 | 2.200E-05 |
| 17679 | 884 | V884Y | C | 7.8e-06 | 1.6e-05 | 1.1e-05 | 1.160E-05 | 4.133E-06 | 7.800E-06 | 1.600E-05 |

| | | | | | | | | | | |
|---|---|---|---|---|---|---|---|---|---|---|
| 17680 | 884 | V884V | C | 9.4e-06 | 9.9e-06 | 8.2e-06 | 9.167E-06 | 8.737E-07 | 8.200E-06 | 9.900E-06 |
| 17681 | 885 | A885A | C | 6.7e-06 | 1.1e-05 | 1.7e-05 | 1.157E-05 | 5.173E-06 | 6.700E-06 | 1.700E-05 |
| 17682 | 885 | A885R | C | 1.3e-05 | 1.4e-05 | 1e-05 | 1.233E-05 | 2.082E-06 | 1.000E-05 | 1.400E-05 |
| 17683 | 885 | A885N | C | 8.6e-06 | 1.2e-05 | 1.3e-05 | 1.120E-05 | 2.307E-06 | 8.600E-06 | 1.300E-05 |
| 17684 | 885 | A885D | C | 9.1e-06 | 1e-05 | 1.1e-05 | 1.003E-05 | 9.504E-07 | 9.100E-06 | 1.100E-05 |
| 17685 | 885 | A885C | C | 1.5e-05 | 9.2e-06 | 1.1e-05 | 1.173E-05 | 2.969E-06 | 9.200E-06 | 1.500E-05 |
| 17686 | 885 | A885E | C | 1.5e-05 | 9.6e-06 | 1e-05 | 1.153E-05 | 3.009E-06 | 9.600E-06 | 1.500E-05 |
| 17687 | 885 | A885Q | C | 1.4e-05 | 1.4e-05 | 1.5e-05 | 1.433E-05 | 5.774E-07 | 1.400E-05 | 1.500E-05 |
| 17688 | 885 | A885G | C | 1.6e-05 | 1.1e-05 | 1.2e-05 | 1.300E-05 | 2.646E-06 | 1.100E-05 | 1.600E-05 |
| 17689 | 885 | A885H | C | 1.5e-05 | 9e-06 | 9.6e-06 | 1.120E-05 | 3.305E-06 | 9.000E-06 | 1.500E-05 |
| 17690 | 885 | A885I | C | 1.6e-05 | 1.2e-05 | 1.2e-05 | 1.333E-05 | 2.309E-06 | 1.200E-05 | 1.600E-05 |
| 17691 | 885 | A885L | C | 1.6e-05 | 1.1e-05 | 9.8e-06 | 1.227E-05 | 3.288E-06 | 9.800E-06 | 1.600E-05 |
| 17692 | 885 | A885K | C | 1e-05 | 1.6e-05 | 1.5e-05 | 1.367E-05 | 3.215E-06 | 1.000E-05 | 1.600E-05 |
| 17693 | 885 | A885M | C | 1.4e-05 | 1.1e-05 | 1.2e-05 | 1.233E-05 | 1.528E-06 | 1.100E-05 | 1.400E-05 |
| 17694 | 885 | A885F | C | 1e-05 | 3.1e-05 | 1.4e-05 | 1.833E-05 | 1.115E-05 | 1.000E-05 | 3.100E-05 |
| 17695 | 885 | A885P | C | 1.5e-05 | 9.3e-06 | 9.2e-06 | 1.117E-05 | 3.320E-06 | 9.200E-06 | 1.500E-05 |
| 17696 | 885 | A885S | C | 1.6e-05 | 1.1e-05 | 8.8e-06 | 1.193E-05 | 3.690E-06 | 8.800E-06 | 1.600E-05 |
| 17697 | 885 | A885T | C | 1.5e-05 | 1.5e-05 | 1.6e-05 | 1.533E-05 | 5.774E-07 | 1.500E-05 | 1.600E-05 |
| 17698 | 885 | A885W | C | 1.7e-05 | 1.3e-05 | 1.7e-05 | 1.567E-05 | 2.309E-06 | 1.300E-05 | 1.700E-05 |
| 17699 | 885 | A885Y | C | 9.2e-06 | 1e-05 | 1.5e-05 | 1.140E-05 | 3.143E-06 | 9.200E-06 | 1.500E-05 |
| 17700 | 885 | A885V | C | 1.6e-05 | 1.3e-05 | 7e-06 | 1.200E-05 | 4.583E-06 | 7.000E-06 | 1.600E-05 |
| 17701 | 886 | D886A | C | 9.8e-06 | 1.4e-05 | 9.7e-06 | 1.117E-05 | 2.454E-06 | 9.700E-06 | 1.400E-05 |
| 17702 | 886 | D886R | C | 1.3e-05 | 9.2e-06 | 1.4e-05 | 1.207E-05 | 2.532E-06 | 9.200E-06 | 1.400E-05 |
| 17703 | 886 | D886N | C | 1.6e-05 | 1e-05 | 1.4e-05 | 1.333E-05 | 3.055E-06 | 1.000E-05 | 1.600E-05 |
| 17704 | 886 | D886D | C | 1.5e-05 | 1.5e-05 | 1.4e-05 | 1.467E-05 | 5.774E-07 | 1.400E-05 | 1.500E-05 |
| 17705 | 886 | D886C | C | 1e-05 | 9.7e-06 | 1.1e-05 | 1.023E-05 | 6.807E-07 | 9.700E-06 | 1.100E-05 |
| 17706 | 886 | D886E | C | 1.3e-05 | 1.5e-05 | 1.5e-05 | 1.433E-05 | 1.155E-06 | 1.300E-05 | 1.500E-05 |
| 17707 | 886 | D886Q | C | 1.5e-05 | 1.4e-05 | 1e-05 | 1.300E-05 | 2.646E-06 | 1.000E-05 | 1.500E-05 |
| 17708 | 886 | D886G | C | 1.4e-05 | 1.3e-05 | 1.5e-05 | 1.400E-05 | 1.000E-06 | 1.300E-05 | 1.500E-05 |
| 17709 | 886 | D886H | C | 9.8e-06 | 1.3e-05 | 1.1e-05 | 1.127E-05 | 1.617E-06 | 9.800E-06 | 1.300E-05 |
| 17710 | 886 | D886I | C | 1e-05 | 1e-05 | 1.4e-05 | 1.133E-05 | 2.309E-06 | 1.000E-05 | 1.400E-05 |
| 17711 | 886 | D886L | C | 1.5e-05 | 9.8e-06 | 1.2e-05 | 1.227E-05 | 2.610E-06 | 9.800E-06 | 1.500E-05 |
| 17712 | 886 | D886K | C | 1.4e-05 | 1.1e-05 | 1.1e-05 | 1.200E-05 | 1.732E-06 | 1.100E-05 | 1.400E-05 |
| 17713 | 886 | D886M | C | 1.4e-05 | 1e-05 | 1.7e-05 | 1.367E-05 | 3.512E-06 | 1.000E-05 | 1.700E-05 |
| 17714 | 886 | D886F | C | 1.5e-05 | 2.1e-05 | 1.4e-05 | 1.667E-05 | 3.786E-06 | 1.400E-05 | 2.100E-05 |
| 17715 | 886 | D886P | C | 9e-06 | 1.1e-05 | 1.5e-05 | 1.167E-05 | 3.055E-06 | 9.000E-06 | 1.500E-05 |
| 17716 | 886 | D886S | C | 1.5e-05 | 1.3e-05 | 2.2e-05 | 1.667E-05 | 4.726E-06 | 1.300E-05 | 2.200E-05 |
| 17717 | 886 | D886T | C | 1.5e-05 | 1.4e-05 | 1.1e-05 | 1.333E-05 | 2.082E-06 | 1.100E-05 | 1.500E-05 |
| 17718 | 886 | D886W | C | 1.1e-05 | 1.6e-05 | 1.8e-05 | 1.500E-05 | 3.606E-06 | 1.100E-05 | 1.800E-05 |
| 17719 | 886 | D886Y | C | 1e-05 | 1.2e-05 | 1.4e-05 | 1.200E-05 | 2.000E-06 | 1.000E-05 | 1.400E-05 |
| 17720 | 886 | D886V | C | 1.6e-05 | 7.9e-06 | 1.6e-05 | 1.330E-05 | 4.677E-06 | 7.900E-06 | 1.600E-05 |
| 17721 | 887 | V887A | C | 9.2e-06 | 8.2e-06 | 1.2e-05 | 9.800E-06 | 1.970E-06 | 8.200E-06 | 1.200E-05 |
| 17722 | 887 | V887R | C | 9.5e-06 | 1e-05 | 1.3e-05 | 1.083E-05 | 1.893E-06 | 9.500E-06 | 1.300E-05 |
| 17723 | 887 | V887N | C | 1.4e-05 | 9.8e-06 | 1.7e-05 | 1.360E-05 | 3.617E-06 | 9.800E-06 | 1.700E-05 |
| 17724 | 887 | V887D | C | 9.7e-06 | 1.5e-05 | 1.6e-05 | 1.357E-05 | 3.386E-06 | 9.700E-06 | 1.600E-05 |
| 17725 | 887 | V887C | C | 9.5e-06 | 1.1e-05 | 1e-05 | 1.017E-05 | 7.638E-07 | 9.500E-06 | 1.100E-05 |
| 17726 | 887 | V887E | C | 1.5e-05 | 8.7e-06 | 1.1e-05 | 1.157E-05 | 3.188E-06 | 8.700E-06 | 1.500E-05 |
| 17727 | 887 | V887Q | C | 1.2e-05 | 9.7e-06 | 1.5e-05 | 1.223E-05 | 2.658E-06 | 9.700E-06 | 1.500E-05 |
| 17728 | 887 | V887G | C | 1.5e-05 | 1.1e-05 | 1.5e-05 | 1.367E-05 | 2.309E-06 | 1.100E-05 | 1.500E-05 |
| 17729 | 887 | V887H | C | 1e-05 | 1.6e-05 | 1.4e-05 | 1.333E-05 | 3.055E-06 | 1.000E-05 | 1.600E-05 |
| 17730 | 887 | V887I | C | 1e-05 | 1.1e-05 | 1.3e-05 | 1.133E-05 | 1.528E-06 | 1.000E-05 | 1.300E-05 |
| 17731 | 887 | V887L | C | 1.1e-05 | 1.4e-05 | 1.6e-05 | 1.367E-05 | 2.517E-06 | 1.100E-05 | 1.600E-05 |
| 17732 | 887 | V887K | C | 1.6e-05 | 1.6e-05 | 1e-05 | 1.400E-05 | 3.464E-06 | 1.000E-05 | 1.600E-05 |
| 17733 | 887 | V887M | C | 1.5e-05 | 1.5e-05 | 1.5e-05 | 1.500E-05 | 0.000E+00 | 1.500E-05 | 1.500E-05 |
| 17734 | 887 | V887F | C | 1e-05 | 1e-05 | 1.3e-05 | 1.100E-05 | 1.732E-06 | 1.000E-05 | 1.300E-05 |
| 17735 | 887 | V887P | C | 1.1e-05 | 1.1e-05 | 1.2e-05 | 1.133E-05 | 5.774E-07 | 1.100E-05 | 1.200E-05 |
| 17736 | 887 | V887S | C | 1.1e-05 | 1.1e-05 | 1.7e-05 | 1.300E-05 | 3.464E-06 | 1.100E-05 | 1.700E-05 |
| 17737 | 887 | V887T | C | 1.3e-05 | 1.1e-05 | 8.5e-06 | 1.083E-05 | 2.255E-06 | 8.500E-06 | 1.300E-05 |
| 17738 | 887 | V887W | C | 1e-05 | 1.4e-05 | 1.6e-05 | 1.333E-05 | 3.055E-06 | 1.000E-05 | 1.600E-05 |
| 17739 | 887 | V887Y | C | 1.1e-05 | 1.6e-05 | 1.3e-05 | 1.333E-05 | 2.517E-06 | 1.100E-05 | 1.600E-05 |
| 17740 | 887 | V887V | C | 1.4e-05 | 1e-05 | 1.5e-05 | 1.300E-05 | 2.646E-06 | 1.000E-05 | 1.500E-05 |
| 17741 | 888 | N888A | C | 1.1e-05 | 1.3e-05 | 1.5e-05 | 1.300E-05 | 2.000E-06 | 1.100E-05 | 1.500E-05 |
| 17742 | 888 | N888R | C | 1.5e-05 | 1.5e-05 | 1e-05 | 1.333E-05 | 2.887E-06 | 1.000E-05 | 1.500E-05 |
| 17743 | 888 | N888N | C | 1.2e-05 | 8.9e-06 | 1.1e-05 | 1.063E-05 | 1.582E-06 | 8.900E-06 | 1.200E-05 |
| 17744 | 888 | N888D | C | 1.4e-05 | 1.4e-05 | 6.9e-06 | 1.163E-05 | 4.099E-06 | 6.900E-06 | 1.400E-05 |
| 17745 | 888 | N888C | C | 1.5e-05 | 1.1e-05 | 1.6e-05 | 1.400E-05 | 2.646E-06 | 1.100E-05 | 1.600E-05 |
| 17746 | 888 | N888E | C | 1.3e-05 | 1.3e-05 | 1.6e-05 | 1.400E-05 | 1.732E-06 | 1.300E-05 | 1.600E-05 |
| 17747 | 888 | N888Q | C | 1.3e-05 | 1e-05 | 1.1e-05 | 1.133E-05 | 1.528E-06 | 1.000E-05 | 1.300E-05 |
| 17748 | 888 | N888G | C | 1.6e-05 | 1.1e-05 | 1e-05 | 1.233E-05 | 3.215E-06 | 1.000E-05 | 1.600E-05 |
| 17749 | 888 | N888H | C | 1.5e-05 | 1.4e-05 | 1e-05 | 1.300E-05 | 2.646E-06 | 1.000E-05 | 1.500E-05 |
| 17750 | 888 | N888I | C | 1.6e-05 | 1.1e-05 | 1.4e-05 | 1.367E-05 | 2.517E-06 | 1.100E-05 | 1.600E-05 |
| 17751 | 888 | N888L | C | 1.1e-05 | 1.4e-05 | 8.2e-06 | 1.107E-05 | 2.901E-06 | 8.200E-06 | 1.400E-05 |
| 17752 | 888 | N888K | C | 1.5e-05 | 1.6e-05 | 9.8e-06 | 1.360E-05 | 3.329E-06 | 9.800E-06 | 1.600E-05 |
| 17753 | 888 | N888M | C | 1.1e-05 | 1.2e-05 | 9.9e-06 | 1.097E-05 | 1.050E-06 | 9.900E-06 | 1.200E-05 |
| 17754 | 888 | N888F | C | 1.5e-05 | 1.9e-05 | 1.5e-05 | 1.633E-05 | 2.309E-06 | 1.500E-05 | 1.900E-05 |
| 17755 | 888 | N888P | C | 1.1e-05 | 6.6e-06 | 1.7e-05 | 1.153E-05 | 5.220E-06 | 6.600E-06 | 1.700E-05 |
| 17756 | 888 | N888S | C | 1.5e-05 | 9.3e-06 | 1.5e-05 | 1.310E-05 | 3.291E-06 | 9.300E-06 | 1.500E-05 |
| 17757 | 888 | N888T | C | 1.3e-05 | 1.1e-05 | 1.7e-05 | 1.367E-05 | 3.055E-06 | 1.100E-05 | 1.700E-05 |
| 17758 | 888 | N888W | C | 1.5e-05 | 2.9e-05 | 1.5e-05 | 1.967E-05 | 8.083E-06 | 1.500E-05 | 2.900E-05 |
| 17759 | 888 | N888Y | C | 1.5e-05 | 1.1e-05 | 1.5e-05 | 1.367E-05 | 2.309E-06 | 1.100E-05 | 1.500E-05 |
| 17760 | 888 | N888V | C | 1.1e-05 | 9.5e-06 | 1.4e-05 | 1.150E-05 | 2.291E-06 | 9.500E-06 | 1.400E-05 |
| 17761 | 889 | P889A | C | 1e-05 | 9.6e-06 | 1.8e-05 | 1.253E-05 | 4.738E-06 | 9.600E-06 | 1.800E-05 |
| 17762 | 889 | P889R | C | 1.6e-05 | 1.4e-05 | 1.1e-05 | 1.367E-05 | 2.517E-06 | 1.100E-05 | 1.600E-05 |
| 17763 | 889 | P889N | C | 9.8e-06 | 1.3e-05 | 1.5e-05 | 1.260E-05 | 2.623E-06 | 9.800E-06 | 1.500E-05 |
| 17764 | 889 | P889D | C | 1.7e-05 | 1.6e-05 | 1.6e-05 | 1.633E-05 | 5.774E-07 | 1.600E-05 | 1.700E-05 |

| | | | | | | | | | | |
|---|---|---|---|---|---|---|---|---|---|---|
| 17765 | 889 | P889C | C | 1.5e-05 | 1.5e-05 | 1.2e-05 | 1.400E-05 | 1.732E-06 | 1.200E-05 | 1.500E-05 |
| 17766 | 889 | P889E | C | 1.4e-05 | 1.4e-05 | 1e-05 | 1.267E-05 | 2.309E-06 | 1.000E-05 | 1.400E-05 |
| 17767 | 889 | P889Q | C | 1.6e-05 | 1.6e-05 | 9.9e-06 | 1.397E-05 | 3.522E-06 | 9.900E-06 | 1.600E-05 |
| 17768 | 889 | P889G | C | 1.1e-05 | 1.6e-05 | 1.4e-05 | 1.367E-05 | 2.517E-06 | 1.100E-05 | 1.600E-05 |
| 17769 | 889 | P889H | C | 1.2e-05 | 1.3e-05 | 1.1e-05 | 1.200E-05 | 1.000E-06 | 1.100E-05 | 1.300E-05 |
| 17770 | 889 | P889I | C | 1.5e-05 | 1.5e-05 | 1.1e-05 | 1.367E-05 | 2.309E-06 | 1.100E-05 | 1.500E-05 |
| 17771 | 889 | P889L | C | 1.5e-05 | 1e-05 | 1.3e-05 | 1.267E-05 | 2.517E-06 | 1.000E-05 | 1.500E-05 |
| 17772 | 889 | P889K | C | 1.1e-05 | 1.2e-05 | 1.2e-05 | 1.167E-05 | 5.774E-07 | 1.100E-05 | 1.200E-05 |
| 17773 | 889 | P889M | C | 1.2e-05 | 1.4e-05 | 1.5e-05 | 1.367E-05 | 1.528E-06 | 1.200E-05 | 1.500E-05 |
| 17774 | 889 | P889F | C | 1.1e-05 | 1.5e-05 | 1.1e-05 | 1.233E-05 | 2.309E-06 | 1.100E-05 | 1.500E-05 |
| 17775 | 889 | P889P | C | 1.3e-05 | 1e-05 | 1.5e-05 | 1.267E-05 | 2.517E-06 | 1.000E-05 | 1.500E-05 |
| 17776 | 889 | P889S | C | 1.5e-05 | 1.5e-05 | 1.4e-05 | 1.467E-05 | 5.774E-07 | 1.400E-05 | 1.500E-05 |
| 17777 | 889 | P889T | C | 1.1e-05 | 1.1e-05 | 1.1e-05 | 1.100E-05 | 2.075E-21 | 1.100E-05 | 1.100E-05 |
| 17778 | 889 | P889W | C | 1.3e-05 | 1.6e-05 | 1.1e-05 | 1.333E-05 | 2.517E-06 | 1.100E-05 | 1.600E-05 |
| 17779 | 889 | P889Y | C | 1.4e-05 | 1.5e-05 | 1.5e-05 | 1.467E-05 | 5.774E-07 | 1.400E-05 | 1.500E-05 |
| 17780 | 889 | P889V | C | 1e-05 | 1.2e-05 | 1.5e-05 | 1.233E-05 | 2.517E-06 | 1.000E-05 | 1.500E-05 |
| 17781 | 890 | N890A | C | 1.1e-05 | 1.1e-05 | 1.5e-05 | 1.233E-05 | 2.309E-06 | 1.100E-05 | 1.500E-05 |
| 17782 | 890 | N890R | C | 9.1e-06 | 1.6e-05 | 1.6e-05 | 1.370E-05 | 3.984E-06 | 9.100E-06 | 1.600E-05 |
| 17783 | 890 | N890N | C | 1.1e-05 | 1.2e-05 | 1.4e-05 | 1.233E-05 | 1.528E-06 | 1.100E-05 | 1.400E-05 |
| 17784 | 890 | N890D | C | 1.5e-05 | 1.9e-05 | 1.7e-05 | 1.700E-05 | 2.000E-06 | 1.500E-05 | 1.900E-05 |
| 17785 | 890 | N890C | C | 1.2e-05 | 1.3e-05 | 1.5e-05 | 1.333E-05 | 1.528E-06 | 1.200E-05 | 1.500E-05 |
| 17786 | 890 | N890E | C | 9.7e-06 | 1.2e-05 | 1e-05 | 1.057E-05 | 1.250E-06 | 9.700E-06 | 1.200E-05 |
| 17787 | 890 | N890Q | C | 1.1e-05 | 1.1e-05 | 9.2e-06 | 1.040E-05 | 1.039E-06 | 9.200E-06 | 1.100E-05 |
| 17788 | 890 | N890G | C | 9.5e-06 | 7.7e-06 | 1.4e-05 | 1.040E-05 | 3.245E-06 | 7.700E-06 | 1.400E-05 |
| 17789 | 890 | N890H | C | 1.1e-05 | 1.5e-05 | 1.5e-05 | 1.367E-05 | 2.309E-06 | 1.100E-05 | 1.500E-05 |
| 17790 | 890 | N890I | C | 8.1e-06 | 1.5e-05 | 9.9e-06 | 1.100E-05 | 3.579E-06 | 8.100E-06 | 1.500E-05 |
| 17791 | 890 | N890L | C | 1.4e-05 | 1.1e-05 | 1.3e-05 | 1.267E-05 | 1.528E-06 | 1.100E-05 | 1.400E-05 |
| 17792 | 890 | N890K | C | 1.2e-05 | 1.1e-05 | 1.5e-05 | 1.267E-05 | 2.082E-06 | 1.100E-05 | 1.500E-05 |
| 17793 | 890 | N890M | C | 1e-05 | 1.7e-05 | 1.6e-05 | 1.433E-05 | 3.786E-06 | 1.000E-05 | 1.700E-05 |
| 17794 | 890 | N890F | C | 1.5e-05 | 1.6e-05 | 1.7e-05 | 1.600E-05 | 1.000E-06 | 1.500E-05 | 1.700E-05 |
| 17795 | 890 | N890P | C | 1e-05 | 1.5e-05 | 1.4e-05 | 1.300E-05 | 2.646E-06 | 1.000E-05 | 1.500E-05 |
| 17796 | 890 | N890S | C | 1e-05 | 8.4e-06 | 9.9e-06 | 9.433E-06 | 8.963E-07 | 8.400E-06 | 1.000E-05 |
| 17797 | 890 | N890T | C | 1.5e-05 | 1.2e-05 | 1.6e-05 | 1.433E-05 | 2.082E-06 | 1.200E-05 | 1.600E-05 |
| 17798 | 890 | N890W | C | 1.3e-05 | 1.6e-05 | 1.3e-05 | 1.400E-05 | 1.732E-06 | 1.300E-05 | 1.600E-05 |
| 17799 | 890 | N890Y | C | 1.4e-05 | 9.6e-06 | 9.5e-06 | 1.103E-05 | 2.570E-06 | 9.500E-06 | 1.400E-05 |
| 17800 | 890 | N890V | C | 8.8e-06 | 1.3e-05 | 1.2e-05 | 1.127E-05 | 2.194E-06 | 8.800E-06 | 1.300E-05 |
| 17801 | 891 | S891A | C | 1.6e-05 | 1.7e-05 | 1.1e-05 | 1.467E-05 | 3.215E-06 | 1.100E-05 | 1.700E-05 |
| 17802 | 891 | S891R | C | 1.7e-05 | 1.1e-05 | 1.7e-05 | 1.500E-05 | 3.464E-06 | 1.100E-05 | 1.700E-05 |
| 17803 | 891 | S891N | C | 1.1e-05 | 1.6e-05 | 9.1e-06 | 1.203E-05 | 3.564E-06 | 9.100E-06 | 1.600E-05 |
| 17804 | 891 | S891D | C | 1.5e-05 | 1.9e-05 | 1e-05 | 1.467E-05 | 4.509E-06 | 1.000E-05 | 1.900E-05 |
| 17805 | 891 | S891C | C | 1.1e-05 | 1e-05 | 9.2e-06 | 1.007E-05 | 9.018E-07 | 9.200E-06 | 1.100E-05 |
| 17806 | 891 | S891E | C | 1.6e-05 | 1.4e-05 | 1.6e-05 | 1.533E-05 | 1.155E-06 | 1.400E-05 | 1.600E-05 |
| 17807 | 891 | S891Q | C | 8.5e-06 | 1e-05 | 1.1e-05 | 9.833E-06 | 1.258E-06 | 8.500E-06 | 1.100E-05 |
| 17808 | 891 | S891G | C | 1.7e-05 | 1.6e-05 | 1.2e-05 | 1.500E-05 | 2.646E-06 | 1.200E-05 | 1.700E-05 |
| 17809 | 891 | S891H | C | 1.1e-05 | 1.1e-05 | 1.5e-05 | 1.233E-05 | 2.309E-06 | 1.100E-05 | 1.500E-05 |
| 17810 | 891 | S891I | C | 1.7e-05 | 1e-05 | 9.9e-06 | 1.230E-05 | 4.071E-06 | 9.900E-06 | 1.700E-05 |
| 17811 | 891 | S891L | C | 1.5e-05 | 1.1e-05 | 1.6e-05 | 1.400E-05 | 2.646E-06 | 1.100E-05 | 1.600E-05 |
| 17812 | 891 | S891K | C | 1.5e-05 | 1.3e-05 | 9.4e-06 | 1.247E-05 | 2.838E-06 | 9.400E-06 | 1.500E-05 |
| 17813 | 891 | S891M | C | 1.6e-05 | 1.3e-05 | 1.7e-05 | 1.533E-05 | 2.082E-06 | 1.300E-05 | 1.700E-05 |
| 17814 | 891 | S891F | C | 1.5e-05 | 9.5e-06 | 1.3e-05 | 1.250E-05 | 2.784E-06 | 9.500E-06 | 1.500E-05 |
| 17815 | 891 | S891P | C | 1.7e-05 | 1.4e-05 | 1.5e-05 | 1.533E-05 | 1.528E-06 | 1.400E-05 | 1.700E-05 |
| 17816 | 891 | S891S | C | 1.3e-05 | 1.6e-05 | 1.5e-05 | 1.467E-05 | 1.528E-06 | 1.300E-05 | 1.600E-05 |
| 17817 | 891 | S891T | C | 1.5e-05 | 1.1e-05 | 1.1e-05 | 1.233E-05 | 2.309E-06 | 1.100E-05 | 1.500E-05 |
| 17818 | 891 | S891W | C | 1e-05 | 1.6e-05 | 1.5e-05 | 1.367E-05 | 3.215E-06 | 1.000E-05 | 1.600E-05 |
| 17819 | 891 | S891Y | C | 1.6e-05 | 1.2e-05 | 1.1e-05 | 1.300E-05 | 2.646E-06 | 1.100E-05 | 1.600E-05 |
| 17820 | 891 | S891V | C | 1.6e-05 | 1.4e-05 | 7.5e-06 | 1.250E-05 | 4.444E-06 | 7.500E-06 | 1.600E-05 |
| 17821 | 892 | G892A | C | 1.5e-05 | 1.5e-05 | 1.1e-05 | 1.367E-05 | 2.309E-06 | 1.100E-05 | 1.500E-05 |
| 17822 | 892 | G892R | C | 1.1e-05 | 1.1e-05 | 1.2e-05 | 1.133E-05 | 5.774E-07 | 1.100E-05 | 1.200E-05 |
| 17823 | 892 | G892N | C | 1e-05 | 1.1e-05 | 1.4e-05 | 1.167E-05 | 2.082E-06 | 1.000E-05 | 1.400E-05 |
| 17824 | 892 | G892D | C | 6.6e-06 | 1e-05 | 9.5e-06 | 8.700E-06 | 1.836E-06 | 6.600E-06 | 1.000E-05 |
| 17825 | 892 | G892C | C | 1.3e-05 | 9.1e-06 | 1.5e-05 | 1.237E-05 | 3.001E-06 | 9.100E-06 | 1.500E-05 |
| 17826 | 892 | G892E | C | 8.4e-06 | 1.6e-05 | 1.1e-05 | 1.180E-05 | 3.863E-06 | 8.400E-06 | 1.600E-05 |
| 17827 | 892 | G892Q | C | 1e-05 | 1.2e-05 | 1.1e-05 | 1.100E-05 | 1.000E-06 | 1.000E-05 | 1.200E-05 |
| 17828 | 892 | G892G | C | 9.6e-06 | 1.4e-05 | 1.6e-05 | 1.320E-05 | 3.274E-06 | 9.600E-06 | 1.600E-05 |
| 17829 | 892 | G892H | C | 1.1e-05 | 1.3e-05 | 9.1e-06 | 1.103E-05 | 1.950E-06 | 9.100E-06 | 1.300E-05 |
| 17830 | 892 | G892I | C | 1.4e-05 | 1.1e-05 | 1.6e-05 | 1.367E-05 | 2.517E-06 | 1.100E-05 | 1.600E-05 |
| 17831 | 892 | G892L | C | 1.6e-05 | 1e-05 | 1.4e-05 | 1.333E-05 | 3.055E-06 | 1.000E-05 | 1.600E-05 |
| 17832 | 892 | G892K | C | 1.5e-05 | 1.5e-05 | 7.5e-06 | 1.250E-05 | 4.330E-06 | 7.500E-06 | 1.500E-05 |
| 17833 | 892 | G892M | C | 9.8e-06 | 1e-05 | 1.8e-05 | 1.260E-05 | 4.678E-06 | 9.800E-06 | 1.800E-05 |
| 17834 | 892 | G892F | C | 1.4e-05 | 1.6e-05 | 1.3e-05 | 1.433E-05 | 1.528E-06 | 1.300E-05 | 1.600E-05 |
| 17835 | 892 | G892P | C | 1.1e-05 | 1e-05 | 1.4e-05 | 1.167E-05 | 2.082E-06 | 1.000E-05 | 1.400E-05 |
| 17836 | 892 | G892S | C | 1.4e-05 | 1.7e-05 | 1.1e-05 | 1.400E-05 | 3.000E-06 | 1.100E-05 | 1.700E-05 |
| 17837 | 892 | G892T | C | 9.6e-06 | 1e-05 | 9.2e-06 | 9.600E-06 | 4.000E-07 | 9.200E-06 | 1.000E-05 |
| 17838 | 892 | G892W | C | 1.6e-05 | 8.8e-06 | 1.5e-05 | 1.327E-05 | 3.900E-06 | 8.800E-06 | 1.600E-05 |
| 17839 | 892 | G892Y | C | 1.5e-05 | 1.2e-05 | 1e-05 | 1.233E-05 | 2.517E-06 | 1.000E-05 | 1.500E-05 |
| 17840 | 892 | G892V | C | 9.9e-06 | 1.5e-05 | 1.5e-05 | 1.330E-05 | 2.944E-06 | 9.900E-06 | 1.500E-05 |
| 17841 | 893 | G893A | C | 1.1e-05 | 1.6e-05 | 9.4e-06 | 1.213E-05 | 3.443E-06 | 9.400E-06 | 1.600E-05 |
| 17842 | 893 | G893R | C | 1.4e-05 | 1.1e-05 | 9.5e-06 | 1.150E-05 | 2.291E-06 | 9.500E-06 | 1.400E-05 |
| 17843 | 893 | G893N | C | 1.8e-05 | 9.7e-06 | 1.4e-05 | 1.390E-05 | 4.151E-06 | 9.700E-06 | 1.800E-05 |
| 17844 | 893 | G893D | C | 9.4e-06 | 1.4e-05 | 1.2e-05 | 1.180E-05 | 2.307E-06 | 9.400E-06 | 1.400E-05 |
| 17845 | 893 | G893C | C | 1.2e-05 | 1.4e-05 | 1.3e-05 | 1.300E-05 | 1.000E-06 | 1.200E-05 | 1.400E-05 |
| 17846 | 893 | G893E | C | 1.2e-05 | 1.3e-05 | 9.3e-06 | 1.143E-05 | 1.914E-06 | 9.300E-06 | 1.300E-05 |
| 17847 | 893 | G893Q | C | 1.4e-05 | 1.1e-05 | 9.3e-06 | 1.143E-05 | 2.380E-06 | 9.300E-06 | 1.400E-05 |
| 17848 | 893 | G893G | C | 1.3e-05 | 9.5e-06 | 1.1e-05 | 1.117E-05 | 1.756E-06 | 9.500E-06 | 1.300E-05 |
| 17849 | 893 | G893H | C | 9.8e-06 | 9.4e-06 | 1.1e-05 | 1.007E-05 | 8.327E-07 | 9.400E-06 | 1.100E-05 |

| | | | | | | | | | | |
|---|---|---|---|---|---|---|---|---|---|---|
| 17850 | 893 | G893I | C | 9.7e-06 | 9.2e-06 | 1.7e-05 | 1.197E-05 | 4.366E-06 | 9.200E-06 | 1.700E-05 |
| 17851 | 893 | G893L | C | 1e-05 | 1.1e-05 | 1.6e-05 | 1.233E-05 | 3.215E-06 | 1.000E-05 | 1.600E-05 |
| 17852 | 893 | G893K | C | 1.5e-05 | 1.3e-05 | 1e-05 | 1.267E-05 | 2.517E-06 | 1.000E-05 | 1.500E-05 |
| 17853 | 893 | G893M | C | 1.8e-05 | 1.1e-05 | 1e-05 | 1.300E-05 | 4.359E-06 | 1.000E-05 | 1.800E-05 |
| 17854 | 893 | G893F | C | 1.6e-05 | 1.6e-05 | 1.1e-05 | 1.433E-05 | 2.887E-06 | 1.100E-05 | 1.600E-05 |
| 17855 | 893 | G893P | C | 1.2e-05 | 5.8e-06 | 1e-05 | 9.267E-06 | 3.164E-06 | 5.800E-06 | 1.200E-05 |
| 17856 | 893 | G893S | C | 1e-05 | 1.6e-05 | 6.8e-06 | 1.093E-05 | 4.670E-06 | 6.800E-06 | 1.600E-05 |
| 17857 | 893 | G893T | C | 1.4e-05 | 1.1e-05 | 8.3e-06 | 1.110E-05 | 2.851E-06 | 8.300E-06 | 1.400E-05 |
| 17858 | 893 | G893W | C | 1.4e-05 | 1.5e-05 | 1.4e-05 | 1.433E-05 | 5.774E-07 | 1.400E-05 | 1.500E-05 |
| 17859 | 893 | G893Y | C | 1.4e-05 | 1.1e-05 | 1.5e-05 | 1.333E-05 | 2.082E-06 | 1.100E-05 | 1.500E-05 |
| 17860 | 893 | G893V | C | 1.5e-05 | 1.8e-05 | 1.4e-05 | 1.567E-05 | 2.082E-06 | 1.400E-05 | 1.800E-05 |
| 17861 | 894 | S894A | C | 1.7e-05 | 7.1e-06 | 1.1e-05 | 1.170E-05 | 4.987E-06 | 7.100E-06 | 1.700E-05 |
| 17862 | 894 | S894R | C | 1.6e-05 | 9.2e-06 | 1.4e-05 | 1.307E-05 | 3.495E-06 | 9.200E-06 | 1.600E-05 |
| 17863 | 894 | S894N | C | 1.1e-05 | 8.1e-06 | 1.2e-05 | 1.037E-05 | 2.026E-06 | 8.100E-06 | 1.200E-05 |
| 17864 | 894 | S894D | C | 1.5e-05 | 9.4e-06 | 1.4e-05 | 1.280E-05 | 2.987E-06 | 9.400E-06 | 1.500E-05 |
| 17865 | 894 | S894C | C | 9.8e-06 | 1.2e-05 | 1e-05 | 1.060E-05 | 1.217E-06 | 9.800E-06 | 1.200E-05 |
| 17866 | 894 | S894E | C | 1e-05 | 1e-05 | 1e-05 | 1.000E-05 | 0.000E+00 | 1.000E-05 | 1.000E-05 |
| 17867 | 894 | S894Q | C | 1.3e-05 | 1.3e-05 | 1.2e-05 | 1.267E-05 | 5.774E-07 | 1.200E-05 | 1.300E-05 |
| 17868 | 894 | S894G | C | 1.6e-05 | 1.2e-05 | 1.8e-05 | 1.533E-05 | 3.055E-06 | 1.200E-05 | 1.800E-05 |
| 17869 | 894 | S894H | C | 8.9e-06 | 1.1e-05 | 1.4e-05 | 1.130E-05 | 2.563E-06 | 8.900E-06 | 1.400E-05 |
| 17870 | 894 | S894I | C | 1.5e-05 | 9e-06 | 1.6e-05 | 1.333E-05 | 3.786E-06 | 9.000E-06 | 1.600E-05 |
| 17871 | 894 | S894L | C | 1.6e-05 | 1.1e-05 | 1e-05 | 1.233E-05 | 3.215E-06 | 1.000E-05 | 1.600E-05 |
| 17872 | 894 | S894K | C | 1.4e-05 | 1.5e-05 | 1e-05 | 1.300E-05 | 2.646E-06 | 1.000E-05 | 1.500E-05 |
| 17873 | 894 | S894M | C | 1.1e-05 | 1.6e-05 | 1.6e-05 | 1.433E-05 | 2.887E-06 | 1.100E-05 | 1.600E-05 |
| 17874 | 894 | S894F | C | 1.2e-05 | 1.2e-05 | 1.5e-05 | 1.300E-05 | 1.732E-06 | 1.200E-05 | 1.500E-05 |
| 17875 | 894 | S894P | C | 1.8e-05 | 1.4e-05 | 1.2e-05 | 1.467E-05 | 3.055E-06 | 1.200E-05 | 1.800E-05 |
| 17876 | 894 | S894S | C | 9.4e-06 | 1e-05 | 1.7e-05 | 1.213E-05 | 4.225E-06 | 9.400E-06 | 1.700E-05 |
| 17877 | 894 | S894T | C | 1.1e-05 | 1e-05 | 1.5e-05 | 1.200E-05 | 2.646E-06 | 1.000E-05 | 1.500E-05 |
| 17878 | 894 | S894W | C | 1.4e-05 | 1.5e-05 | 1.3e-05 | 1.400E-05 | 1.000E-06 | 1.300E-05 | 1.500E-05 |
| 17879 | 894 | S894Y | C | 8.9e-06 | 1.1e-05 | 8.8e-06 | 9.567E-06 | 1.242E-06 | 8.800E-06 | 1.100E-05 |
| 17880 | 894 | S894V | C | 1.1e-05 | 1.4e-05 | 1.3e-05 | 1.267E-05 | 1.528E-06 | 1.100E-05 | 1.400E-05 |
| 17881 | 895 | I895A | C | 1.1e-05 | 9e-06 | 1.2e-05 | 1.067E-05 | 1.528E-06 | 9.000E-06 | 1.200E-05 |
| 17882 | 895 | I895R | C | 1e-05 | 8.8e-06 | 6.7e-06 | 8.500E-06 | 1.670E-06 | 6.700E-06 | 1.000E-05 |
| 17883 | 895 | I895N | C | 1.1e-05 | 9.9e-06 | 1.1e-05 | 1.063E-05 | 6.351E-07 | 9.900E-06 | 1.100E-05 |
| 17884 | 895 | I895D | C | 1.4e-05 | 1.1e-05 | 1.3e-05 | 1.267E-05 | 1.528E-06 | 1.100E-05 | 1.400E-05 |
| 17885 | 895 | I895C | C | 1.3e-05 | 1.2e-05 | 7.3e-06 | 1.077E-05 | 3.044E-06 | 7.300E-06 | 1.300E-05 |
| 17886 | 895 | I895E | C | 9.5e-06 | 6.9e-06 | 1.2e-05 | 9.467E-06 | 2.550E-06 | 6.900E-06 | 1.200E-05 |
| 17887 | 895 | I895Q | C | 1.5e-05 | 9.8e-06 | 1.5e-05 | 1.327E-05 | 3.002E-06 | 9.800E-06 | 1.500E-05 |
| 17888 | 895 | I895G | C | 1.3e-05 | 1.1e-05 | 1.3e-05 | 1.233E-05 | 1.155E-06 | 1.100E-05 | 1.300E-05 |
| 17889 | 895 | I895H | C | 1.6e-05 | 1.3e-05 | 8.3e-06 | 1.243E-05 | 3.881E-06 | 8.300E-06 | 1.600E-05 |
| 17890 | 895 | I895I | C | 1.2e-05 | 1.7e-05 | 1.1e-05 | 1.333E-05 | 3.215E-06 | 1.100E-05 | 1.700E-05 |
| 17891 | 895 | I895L | C | 1.1e-05 | 6.6e-06 | 1.2e-05 | 9.867E-06 | 2.873E-06 | 6.600E-06 | 1.200E-05 |
| 17892 | 895 | I895K | C | 1.6e-05 | 1e-05 | 1.4e-05 | 1.333E-05 | 3.055E-06 | 1.000E-05 | 1.600E-05 |
| 17893 | 895 | I895M | C | 6.8e-06 | 1.7e-05 | 1.4e-05 | 1.260E-05 | 5.242E-06 | 6.800E-06 | 1.700E-05 |
| 17894 | 895 | I895F | C | 1.6e-05 | 1.2e-05 | 1.1e-05 | 1.300E-05 | 2.646E-06 | 1.100E-05 | 1.600E-05 |
| 17895 | 895 | I895P | C | 1e-05 | 7.5e-06 | 1.3e-05 | 1.017E-05 | 2.754E-06 | 7.500E-06 | 1.300E-05 |
| 17896 | 895 | I895S | C | 1.4e-05 | 9.5e-06 | 1.2e-05 | 1.183E-05 | 2.255E-06 | 9.500E-06 | 1.400E-05 |
| 17897 | 895 | I895T | C | 1.2e-05 | 1.3e-05 | 1.2e-05 | 1.233E-05 | 5.774E-07 | 1.200E-05 | 1.300E-05 |
| 17898 | 895 | I895W | C | 1.3e-05 | 1.4e-05 | 1.5e-05 | 1.400E-05 | 1.000E-06 | 1.300E-05 | 1.500E-05 |
| 17899 | 895 | I895Y | C | 1e-05 | 9.8e-06 | 9.4e-06 | 9.733E-06 | 3.055E-07 | 9.400E-06 | 1.000E-05 |
| 17900 | 895 | I895V | C | 1.5e-05 | 1.6e-05 | 8.6e-06 | 1.320E-05 | 4.015E-06 | 8.600E-06 | 1.600E-05 |
| 17901 | 896 | Y896A | C | 1.5e-05 | 1.1e-05 | 1.1e-05 | 1.233E-05 | 2.309E-06 | 1.100E-05 | 1.500E-05 |
| 17902 | 896 | Y896R | C | 9.9e-06 | 1.6e-05 | 1.4e-05 | 1.330E-05 | 3.110E-06 | 9.900E-06 | 1.600E-05 |
| 17903 | 896 | Y896N | C | 9e-06 | 1.3e-05 | 1e-05 | 1.067E-05 | 2.082E-06 | 9.000E-06 | 1.300E-05 |
| 17904 | 896 | Y896D | C | 1.7e-05 | 1.1e-05 | 1.5e-05 | 1.433E-05 | 3.055E-06 | 1.100E-05 | 1.700E-05 |
| 17905 | 896 | Y896C | C | 1.1e-05 | 1.6e-05 | 1.6e-05 | 1.433E-05 | 2.887E-06 | 1.100E-05 | 1.600E-05 |
| 17906 | 896 | Y896E | C | 1.3e-05 | 1.1e-05 | 1.4e-05 | 1.267E-05 | 1.528E-06 | 1.100E-05 | 1.400E-05 |
| 17907 | 896 | Y896Q | C | 1.1e-05 | 1.6e-05 | 1e-05 | 1.233E-05 | 3.215E-06 | 1.000E-05 | 1.600E-05 |
| 17908 | 896 | Y896G | C | 1.6e-05 | 1e-05 | 1.1e-05 | 1.233E-05 | 3.215E-06 | 1.000E-05 | 1.600E-05 |
| 17909 | 896 | Y896H | C | 7.9e-06 | 1.6e-05 | 1.1e-05 | 1.163E-05 | 4.087E-06 | 7.900E-06 | 1.600E-05 |
| 17910 | 896 | Y896I | C | 1.4e-05 | 1.7e-05 | 1.7e-05 | 1.600E-05 | 1.732E-06 | 1.400E-05 | 1.700E-05 |
| 17911 | 896 | Y896L | C | 1.7e-05 | 9.9e-06 | 1.7e-05 | 1.463E-05 | 4.099E-06 | 9.900E-06 | 1.700E-05 |
| 17912 | 896 | Y896K | C | 9.8e-06 | 1.6e-05 | 1.1e-05 | 1.227E-05 | 3.288E-06 | 9.800E-06 | 1.600E-05 |
| 17913 | 896 | Y896M | C | 1.6e-05 | 1e-05 | 8.5e-06 | 1.150E-05 | 3.969E-06 | 8.500E-06 | 1.600E-05 |
| 17914 | 896 | Y896F | C | 1.5e-05 | 1.1e-05 | 1.6e-05 | 1.400E-05 | 2.646E-06 | 1.100E-05 | 1.600E-05 |
| 17915 | 896 | Y896P | C | 1.1e-05 | 1.2e-05 | 1.4e-05 | 1.233E-05 | 1.528E-06 | 1.100E-05 | 1.400E-05 |
| 17916 | 896 | Y896S | C | 1.1e-05 | 1.8e-05 | 1.7e-05 | 1.533E-05 | 3.786E-06 | 1.100E-05 | 1.800E-05 |
| 17917 | 896 | Y896T | C | 9.8e-06 | 1.2e-05 | 1.3e-05 | 1.160E-05 | 1.637E-06 | 9.800E-06 | 1.300E-05 |
| 17918 | 896 | Y896W | C | 1.6e-05 | 1e-05 | 9.5e-06 | 1.183E-05 | 3.617E-06 | 9.500E-06 | 1.600E-05 |
| 17919 | 896 | Y896Y | C | 1.6e-05 | 1.1e-05 | 1.1e-05 | 1.267E-05 | 2.887E-06 | 1.100E-05 | 1.600E-05 |
| 17920 | 896 | Y896V | C | 1.3e-05 | 8.8e-06 | 1.5e-05 | 1.227E-05 | 3.164E-06 | 8.800E-06 | 1.500E-05 |
| 17921 | 897 | N897A | C | 1.1e-05 | 1e-05 | 1.5e-05 | 1.200E-05 | 2.646E-06 | 1.000E-05 | 1.500E-05 |
| 17922 | 897 | N897R | C | 1.3e-05 | 1.7e-05 | 1.7e-05 | 1.567E-05 | 2.309E-06 | 1.300E-05 | 1.700E-05 |
| 17923 | 897 | N897N | C | 1.4e-05 | 1.2e-05 | 1.1e-05 | 1.233E-05 | 1.528E-06 | 1.100E-05 | 1.400E-05 |
| 17924 | 897 | N897D | C | 1.5e-05 | 1.4e-05 | 1.5e-05 | 1.467E-05 | 5.774E-07 | 1.400E-05 | 1.500E-05 |
| 17925 | 897 | N897C | C | 1.1e-05 | 1.4e-05 | 1.4e-05 | 1.300E-05 | 1.732E-06 | 1.100E-05 | 1.400E-05 |
| 17926 | 897 | N897E | C | 9.6e-06 | 1e-05 | 1.1e-05 | 1.020E-05 | 7.211E-07 | 9.600E-06 | 1.100E-05 |
| 17927 | 897 | N897Q | C | 1.1e-05 | 1.5e-05 | 7.7e-06 | 1.123E-05 | 3.656E-06 | 7.700E-06 | 1.500E-05 |
| 17928 | 897 | N897G | C | 1.3e-05 | 1.5e-05 | 1.7e-05 | 1.500E-05 | 2.000E-06 | 1.300E-05 | 1.700E-05 |
| 17929 | 897 | N897H | C | 1.7e-05 | 1.3e-05 | 9.3e-06 | 1.310E-05 | 3.851E-06 | 9.300E-06 | 1.700E-05 |
| 17930 | 897 | N897I | C | 1.1e-05 | 1.1e-05 | 1.5e-05 | 1.233E-05 | 2.309E-06 | 1.100E-05 | 1.500E-05 |
| 17931 | 897 | N897L | C | 1.1e-05 | 1.7e-05 | 1.1e-05 | 1.300E-05 | 3.464E-06 | 1.100E-05 | 1.700E-05 |
| 17932 | 897 | N897K | C | 1.2e-05 | 9.6e-06 | 1.6e-05 | 1.253E-05 | 3.233E-06 | 9.600E-06 | 1.600E-05 |
| 17933 | 897 | N897M | C | 1.4e-05 | 1e-05 | 1.1e-05 | 1.167E-05 | 2.082E-06 | 1.000E-05 | 1.400E-05 |
| 17934 | 897 | N897F | C | 1.5e-05 | 1.3e-05 | 1.6e-05 | 1.467E-05 | 1.528E-06 | 1.300E-05 | 1.600E-05 |

| | | | | | | | | | | |
|---|---|---|---|---|---|---|---|---|---|---|
| 17935 | 897 | N897P | C | 1.7e-05 | 1.4e-05 | 9.8e-06 | 1.360E-05 | 3.617E-06 | 9.800E-06 | 1.700E-05 |
| 17936 | 897 | N897S | C | 9.6e-06 | 9.2e-06 | 1.6e-05 | 1.160E-05 | 3.816E-06 | 9.200E-06 | 1.600E-05 |
| 17937 | 897 | N897T | C | 1.4e-05 | 1.4e-05 | 1e-05 | 1.267E-05 | 2.309E-06 | 1.000E-05 | 1.400E-05 |
| 17938 | 897 | N897W | C | 1.6e-05 | 9.9e-06 | 1.5e-05 | 1.363E-05 | 3.272E-06 | 9.900E-06 | 1.600E-05 |
| 17939 | 897 | N897Y | C | 1.1e-05 | 9.3e-06 | 1.5e-05 | 1.177E-05 | 2.926E-06 | 9.300E-06 | 1.500E-05 |
| 17940 | 897 | N897V | C | 1.5e-05 | 1.4e-05 | 1.6e-05 | 1.500E-05 | 1.000E-06 | 1.400E-05 | 1.600E-05 |
| 17941 | 898 | Q898A | C | 1.7e-05 | 1.5e-05 | 1.2e-05 | 1.467E-05 | 2.517E-06 | 1.200E-05 | 1.700E-05 |
| 17942 | 898 | Q898R | C | 9.7e-06 | 1.5e-05 | 1e-05 | 1.157E-05 | 2.977E-06 | 9.700E-06 | 1.500E-05 |
| 17943 | 898 | Q898N | C | 1.1e-05 | 1.6e-05 | 1.1e-05 | 1.267E-05 | 2.887E-06 | 1.100E-05 | 1.600E-05 |
| 17944 | 898 | Q898D | C | 9.4e-06 | 1.1e-05 | 8.8e-06 | 9.733E-06 | 1.137E-06 | 8.800E-06 | 1.100E-05 |
| 17945 | 898 | Q898C | C | 1.7e-05 | 1.1e-05 | 1.1e-05 | 1.300E-05 | 3.464E-06 | 1.100E-05 | 1.700E-05 |
| 17946 | 898 | Q898E | C | 1.1e-05 | 1.5e-05 | 1.3e-05 | 1.300E-05 | 2.000E-06 | 1.100E-05 | 1.500E-05 |
| 17947 | 898 | Q898Q | C | 8.8e-06 | 1.5e-05 | 1.6e-05 | 1.327E-05 | 3.900E-06 | 8.800E-06 | 1.600E-05 |
| 17948 | 898 | Q898G | C | 7.5e-06 | 1e-05 | 1.5e-05 | 1.083E-05 | 3.819E-06 | 7.500E-06 | 1.500E-05 |
| 17949 | 898 | Q898H | C | 1.7e-05 | 1.4e-05 | 1.1e-05 | 1.400E-05 | 3.000E-06 | 1.100E-05 | 1.700E-05 |
| 17950 | 898 | Q898I | C | 2.2e-05 | 1.1e-05 | 1.2e-05 | 1.500E-05 | 6.083E-06 | 1.100E-05 | 2.200E-05 |
| 17951 | 898 | Q898L | C | 1.6e-05 | 1.5e-05 | 1.5e-05 | 1.533E-05 | 5.774E-07 | 1.500E-05 | 1.600E-05 |
| 17952 | 898 | Q898K | C | 1.6e-05 | 1e-05 | 1.1e-05 | 1.233E-05 | 3.215E-06 | 1.000E-05 | 1.600E-05 |
| 17953 | 898 | Q898M | C | 1.5e-05 | 1.2e-05 | 1.4e-05 | 1.367E-05 | 1.528E-06 | 1.200E-05 | 1.500E-05 |
| 17954 | 898 | Q898F | C | 1e-05 | 1.6e-05 | 1e-05 | 1.200E-05 | 3.464E-06 | 1.000E-05 | 1.600E-05 |
| 17955 | 898 | Q898P | C | 1.6e-05 | 1e-05 | 1.2e-05 | 1.267E-05 | 3.055E-06 | 1.000E-05 | 1.600E-05 |
| 17956 | 898 | Q898S | C | 1.5e-05 | 1e-05 | 1e-05 | 1.167E-05 | 2.887E-06 | 1.000E-05 | 1.500E-05 |
| 17957 | 898 | Q898T | C | 1.1e-05 | 1.1e-05 | 1.4e-05 | 1.200E-05 | 1.732E-06 | 1.100E-05 | 1.400E-05 |
| 17958 | 898 | Q898W | C | 9.7e-06 | 7.7e-06 | 1.3e-05 | 1.013E-05 | 2.676E-06 | 7.700E-06 | 1.300E-05 |
| 17959 | 898 | Q898Y | C | 1.7e-05 | 1.5e-05 | 1.5e-05 | 1.567E-05 | 1.155E-06 | 1.500E-05 | 1.700E-05 |
| 17960 | 898 | Q898V | C | 9.7e-06 | 9.1e-06 | 1.4e-05 | 1.093E-05 | 2.673E-06 | 9.100E-06 | 1.400E-05 |
| 17961 | 899 | R899A | C | 1.1e-05 | 8.3e-06 | 1.3e-05 | 1.077E-05 | 2.359E-06 | 8.300E-06 | 1.300E-05 |
| 17962 | 899 | R899R | C | 1e-05 | 1.5e-05 | 9.2e-06 | 1.140E-05 | 3.143E-06 | 9.200E-06 | 1.500E-05 |
| 17963 | 899 | R899N | C | 1.4e-05 | 1.5e-05 | 1.5e-05 | 1.467E-05 | 5.774E-07 | 1.400E-05 | 1.500E-05 |
| 17964 | 899 | R899D | C | 1.6e-05 | 1e-05 | 1e-05 | 1.200E-05 | 3.464E-06 | 1.000E-05 | 1.600E-05 |
| 17965 | 899 | R899C | C | 1.5e-05 | 1.7e-05 | 1.2e-05 | 1.467E-05 | 2.517E-06 | 1.200E-05 | 1.700E-05 |
| 17966 | 899 | R899E | C | 1.1e-05 | 1.7e-05 | 1e-05 | 1.267E-05 | 3.786E-06 | 1.000E-05 | 1.700E-05 |
| 17967 | 899 | R899Q | C | 1e-05 | 1.6e-05 | 1e-05 | 1.200E-05 | 3.464E-06 | 1.000E-05 | 1.600E-05 |
| 17968 | 899 | R899G | C | 1.5e-05 | 1.1e-05 | 1.6e-05 | 1.400E-05 | 2.646E-06 | 1.100E-05 | 1.600E-05 |
| 17969 | 899 | R899H | C | 1.3e-05 | 1.5e-05 | 1.5e-05 | 1.433E-05 | 1.155E-06 | 1.300E-05 | 1.500E-05 |
| 17970 | 899 | R899I | C | 1.8e-05 | 1.5e-05 | 1.4e-05 | 1.567E-05 | 2.082E-06 | 1.400E-05 | 1.800E-05 |
| 17971 | 899 | R899L | C | 9.4e-06 | 1.5e-05 | 1e-05 | 1.147E-05 | 3.075E-06 | 9.400E-06 | 1.500E-05 |
| 17972 | 899 | R899K | C | 1.4e-05 | 1.1e-05 | 1e-05 | 1.167E-05 | 2.082E-06 | 1.000E-05 | 1.400E-05 |
| 17973 | 899 | R899M | C | 1.5e-05 | 9.1e-06 | 1.1e-05 | 1.170E-05 | 3.012E-06 | 9.100E-06 | 1.500E-05 |
| 17974 | 899 | R899F | C | 1.1e-05 | 1.8e-05 | 9.8e-06 | 1.293E-05 | 4.429E-06 | 9.800E-06 | 1.800E-05 |
| 17975 | 899 | R899P | C | 1e-05 | 1e-05 | 1.6e-05 | 1.200E-05 | 3.464E-06 | 1.000E-05 | 1.600E-05 |
| 17976 | 899 | R899S | C | 1.2e-05 | 1.6e-05 | 1.4e-05 | 1.400E-05 | 2.000E-06 | 1.200E-05 | 1.600E-05 |
| 17977 | 899 | R899T | C | 1.1e-05 | 1.5e-05 | 1.4e-05 | 1.333E-05 | 2.082E-06 | 1.100E-05 | 1.500E-05 |
| 17978 | 899 | R899W | C | 8.2e-06 | 8.3e-06 | 1.4e-05 | 1.017E-05 | 3.320E-06 | 8.200E-06 | 1.400E-05 |
| 17979 | 899 | R899Y | C | 1.3e-05 | 9.4e-06 | 1e-05 | 1.080E-05 | 1.929E-06 | 9.400E-06 | 1.300E-05 |
| 17980 | 899 | R899V | C | 1.7e-05 | 1e-05 | 9.3e-06 | 1.210E-05 | 4.258E-06 | 9.300E-06 | 1.700E-05 |
| 17981 | 900 | F900A | C | 1.5e-05 | 1.4e-05 | 1.1e-05 | 1.333E-05 | 2.082E-06 | 1.100E-05 | 1.500E-05 |
| 17982 | 900 | F900R | C | 1.4e-05 | 1.5e-05 | 8.6e-06 | 1.253E-05 | 3.443E-06 | 8.600E-06 | 1.500E-05 |
| 17983 | 900 | F900N | C | 9.8e-06 | 1.5e-05 | 1.7e-05 | 1.393E-05 | 3.717E-06 | 9.800E-06 | 1.700E-05 |
| 17984 | 900 | F900D | C | 1.1e-05 | 1.1e-05 | 1.8e-05 | 1.333E-05 | 4.041E-06 | 1.100E-05 | 1.800E-05 |
| 17985 | 900 | F900C | C | 1.6e-05 | 1.1e-05 | 9.3e-06 | 1.210E-05 | 3.483E-06 | 9.300E-06 | 1.600E-05 |
| 17986 | 900 | F900E | C | 1.4e-05 | 1.6e-05 | 1.4e-05 | 1.467E-05 | 1.155E-06 | 1.400E-05 | 1.600E-05 |
| 17987 | 900 | F900Q | C | 1.5e-05 | 1.3e-05 | 1e-05 | 1.267E-05 | 2.517E-06 | 1.000E-05 | 1.500E-05 |
| 17988 | 900 | F900G | C | 1e-05 | 1.5e-05 | 1.5e-05 | 1.333E-05 | 2.887E-06 | 1.000E-05 | 1.500E-05 |
| 17989 | 900 | F900H | C | 1.1e-05 | 1.1e-05 | 1.7e-05 | 1.300E-05 | 3.464E-06 | 1.100E-05 | 1.700E-05 |
| 17990 | 900 | F900I | C | 1.3e-05 | 1.4e-05 | 1.1e-05 | 1.267E-05 | 1.528E-06 | 1.100E-05 | 1.400E-05 |
| 17991 | 900 | F900L | C | 1.6e-05 | 1.4e-05 | 1.4e-05 | 1.467E-05 | 1.155E-06 | 1.400E-05 | 1.600E-05 |
| 17992 | 900 | F900K | C | 1.5e-05 | 1.1e-05 | 1.5e-05 | 1.367E-05 | 2.309E-06 | 1.100E-05 | 1.500E-05 |
| 17993 | 900 | F900M | C | 1.7e-05 | 8.9e-06 | 1e-05 | 1.197E-05 | 4.394E-06 | 8.900E-06 | 1.700E-05 |
| 17994 | 900 | F900F | C | 9e-06 | 8.9e-06 | 9.3e-06 | 9.067E-06 | 2.082E-07 | 8.900E-06 | 9.300E-06 |
| 17995 | 900 | F900P | C | 1.6e-05 | 1.2e-05 | 1.6e-05 | 1.467E-05 | 2.309E-06 | 1.200E-05 | 1.600E-05 |
| 17996 | 900 | F900S | C | 1.1e-05 | 1.3e-05 | 9.7e-06 | 1.123E-05 | 1.662E-06 | 9.700E-06 | 1.300E-05 |
| 17997 | 900 | F900T | C | 8.5e-06 | 1.7e-05 | 1.7e-05 | 1.417E-05 | 4.907E-06 | 8.500E-06 | 1.700E-05 |
| 17998 | 900 | F900W | C | 1.3e-05 | 1.5e-05 | 1e-05 | 1.267E-05 | 2.517E-06 | 1.000E-05 | 1.500E-05 |
| 17999 | 900 | F900Y | C | 1.5e-05 | 3e-05 | 1.5e-05 | 2.000E-05 | 8.660E-06 | 1.500E-05 | 3.000E-05 |
| 18000 | 900 | F900V | C | 1.5e-05 | 9.6e-06 | 1.1e-05 | 1.187E-05 | 2.802E-06 | 9.600E-06 | 1.500E-05 |
| 18001 | 901 | K901A | C | 9.1e-06 | 1.1e-05 | 1.6e-05 | 1.203E-05 | 3.564E-06 | 9.100E-06 | 1.600E-05 |
| 18002 | 901 | K901R | C | 1.1e-05 | 1.6e-05 | 1.1e-05 | 1.267E-05 | 2.887E-06 | 1.100E-05 | 1.600E-05 |
| 18003 | 901 | K901N | C | 1.4e-05 | 7.4e-06 | 1.1e-05 | 1.080E-05 | 3.305E-06 | 7.400E-06 | 1.400E-05 |
| 18004 | 901 | K901D | C | 1.2e-05 | 9.8e-06 | 1.7e-05 | 1.293E-05 | 3.690E-06 | 9.800E-06 | 1.700E-05 |
| 18005 | 901 | K901C | C | 1.5e-05 | 1.1e-05 | 9.8e-06 | 1.193E-05 | 2.723E-06 | 9.800E-06 | 1.500E-05 |
| 18006 | 901 | K901E | C | 1.1e-05 | 1.4e-05 | 1.5e-05 | 1.333E-05 | 2.082E-06 | 1.100E-05 | 1.500E-05 |
| 18007 | 901 | K901Q | C | 1.4e-05 | 1.1e-05 | 1.4e-05 | 1.300E-05 | 1.732E-06 | 1.100E-05 | 1.400E-05 |
| 18008 | 901 | K901G | C | 1.5e-05 | 1.5e-05 | 1.4e-05 | 1.467E-05 | 5.774E-07 | 1.400E-05 | 1.500E-05 |
| 18009 | 901 | K901H | C | 1.5e-05 | 1.5e-05 | 9.9e-06 | 1.330E-05 | 2.944E-06 | 9.900E-06 | 1.500E-05 |
| 18010 | 901 | K901I | C | 1.4e-05 | 1.6e-05 | 1.1e-05 | 1.367E-05 | 2.517E-06 | 1.100E-05 | 1.600E-05 |
| 18011 | 901 | K901L | C | 1.4e-05 | 1.6e-05 | 9.9e-06 | 1.330E-05 | 3.110E-06 | 9.900E-06 | 1.600E-05 |
| 18012 | 901 | K901K | C | 1e-05 | 9.5e-06 | 1.7e-05 | 1.217E-05 | 4.193E-06 | 9.500E-06 | 1.700E-05 |
| 18013 | 901 | K901M | C | 1.6e-05 | 1.5e-05 | 2.4e-05 | 1.833E-05 | 4.933E-06 | 1.500E-05 | 2.400E-05 |
| 18014 | 901 | K901F | C | 1.4e-05 | 9.6e-06 | 9.8e-06 | 1.113E-05 | 2.485E-06 | 9.600E-06 | 1.400E-05 |
| 18015 | 901 | K901P | C | 9.5e-06 | 1.7e-05 | 1e-05 | 1.217E-05 | 4.193E-06 | 9.500E-06 | 1.700E-05 |
| 18016 | 901 | K901S | C | 1.4e-05 | 1.5e-05 | 1.5e-05 | 1.467E-05 | 5.774E-07 | 1.400E-05 | 1.500E-05 |
| 18017 | 901 | K901T | C | 1.4e-05 | 1.5e-05 | 1.7e-05 | 1.533E-05 | 1.528E-06 | 1.400E-05 | 1.700E-05 |
| 18018 | 901 | K901W | C | 1.4e-05 | 1.5e-05 | 1.5e-05 | 1.467E-05 | 5.774E-07 | 1.400E-05 | 1.500E-05 |
| 18019 | 901 | K901Y | C | 1.5e-05 | 1.7e-05 | 9.7e-06 | 1.390E-05 | 3.772E-06 | 9.700E-06 | 1.700E-05 |

| | | | | | | | | | | |
|---|---|---|---|---|---|---|---|---|---|---|
| 18020 | 901 | K901V | C | 9.9e-06 | 9.9e-06 | 1.5e-05 | 1.160E-05 | 2.944E-06 | 9.900E-06 | 1.500E-05 |
| 18021 | 902 | G902A | C | 1.5e-05 | 1.1e-05 | 1.6e-05 | 1.400E-05 | 2.646E-06 | 1.100E-05 | 1.600E-05 |
| 18022 | 902 | G902R | C | 1.8e-05 | 1.5e-05 | 1.4e-05 | 1.567E-05 | 2.082E-06 | 1.400E-05 | 1.800E-05 |
| 18023 | 902 | G902N | C | 9e-06 | 1.1e-05 | 1.4e-05 | 1.133E-05 | 2.517E-06 | 9.000E-06 | 1.400E-05 |
| 18024 | 902 | G902D | C | 7.6e-06 | 1.5e-05 | 1.1e-05 | 1.120E-05 | 3.704E-06 | 7.600E-06 | 1.500E-05 |
| 18025 | 902 | G902C | C | 1.5e-05 | 1.3e-05 | 1.4e-05 | 1.400E-05 | 1.000E-06 | 1.300E-05 | 1.500E-05 |
| 18026 | 902 | G902E | C | 1.4e-05 | 1.4e-05 | 1.3e-05 | 1.367E-05 | 5.774E-07 | 1.300E-05 | 1.400E-05 |
| 18027 | 902 | G902Q | C | 1.6e-05 | 1.1e-05 | 1.5e-05 | 1.400E-05 | 2.646E-06 | 1.100E-05 | 1.600E-05 |
| 18028 | 902 | G902G | C | 1.6e-05 | 1.4e-05 | 1.7e-05 | 1.567E-05 | 1.528E-06 | 1.400E-05 | 1.700E-05 |
| 18029 | 902 | G902H | C | 1.4e-05 | 1.4e-05 | 1.5e-05 | 1.433E-05 | 5.774E-07 | 1.400E-05 | 1.500E-05 |
| 18030 | 902 | G902I | C | 1.2e-05 | 1.5e-05 | 1.5e-05 | 1.400E-05 | 1.732E-06 | 1.200E-05 | 1.500E-05 |
| 18031 | 902 | G902L | C | 1.1e-05 | 1.1e-05 | 1.1e-05 | 1.100E-05 | 2.075E-21 | 1.100E-05 | 1.100E-05 |
| 18032 | 902 | G902K | C | 1.4e-05 | 8.6e-06 | 1.1e-05 | 1.120E-05 | 2.706E-06 | 8.600E-06 | 1.400E-05 |
| 18033 | 902 | G902M | C | 1.1e-05 | 1.1e-05 | 1.5e-05 | 1.233E-05 | 2.309E-06 | 1.100E-05 | 1.500E-05 |
| 18034 | 902 | G902F | C | 8.9e-06 | 1.7e-05 | 1.3e-05 | 1.297E-05 | 4.050E-06 | 8.900E-06 | 1.700E-05 |
| 18035 | 902 | G902P | C | 1.2e-05 | 9.4e-06 | 1.4e-05 | 1.180E-05 | 2.307E-06 | 9.400E-06 | 1.400E-05 |
| 18036 | 902 | G902S | C | 1.2e-05 | 8.9e-06 | 1.9e-05 | 1.330E-05 | 5.174E-06 | 8.900E-06 | 1.900E-05 |
| 18037 | 902 | G902T | C | 1.6e-05 | 1.4e-05 | 1.2e-05 | 1.400E-05 | 2.000E-06 | 1.200E-05 | 1.600E-05 |
| 18038 | 902 | G902W | C | 1.5e-05 | 1.3e-05 | 9.7e-06 | 1.257E-05 | 2.676E-06 | 9.700E-06 | 1.500E-05 |
| 18039 | 902 | G902Y | C | 1.6e-05 | 1.5e-05 | 1.2e-05 | 1.433E-05 | 2.082E-06 | 1.200E-05 | 1.600E-05 |
| 18040 | 902 | G902V | C | 1.2e-05 | 1.5e-05 | 1.5e-05 | 1.400E-05 | 1.732E-06 | 1.200E-05 | 1.500E-05 |
| 18041 | 903 | R903A | C | 1.6e-05 | 1.5e-05 | 8.3e-06 | 1.310E-05 | 4.187E-06 | 8.300E-06 | 1.600E-05 |
| 18042 | 903 | R903R | C | 1.5e-05 | 9.7e-06 | 1.1e-05 | 1.190E-05 | 2.762E-06 | 9.700E-06 | 1.500E-05 |
| 18043 | 903 | R903N | C | 1.4e-05 | 1.3e-05 | 1.4e-05 | 1.367E-05 | 5.774E-07 | 1.300E-05 | 1.400E-05 |
| 18044 | 903 | R903D | C | 1e-05 | 1.1e-05 | 1.1e-05 | 1.067E-05 | 5.774E-07 | 1.000E-05 | 1.100E-05 |
| 18045 | 903 | R903C | C | 1e-05 | 1.3e-05 | 1e-05 | 1.100E-05 | 1.732E-06 | 1.000E-05 | 1.300E-05 |
| 18046 | 903 | R903E | C | 1.5e-05 | 1.6e-05 | 1.4e-05 | 1.500E-05 | 1.000E-06 | 1.400E-05 | 1.600E-05 |
| 18047 | 903 | R903Q | C | 1.3e-05 | 9.7e-06 | 1.5e-05 | 1.257E-05 | 2.676E-06 | 9.700E-06 | 1.500E-05 |
| 18048 | 903 | R903G | C | 1.4e-05 | 1.5e-05 | 1e-05 | 1.300E-05 | 2.646E-06 | 1.000E-05 | 1.500E-05 |
| 18049 | 903 | R903H | C | 1.1e-05 | 1.1e-05 | 1.5e-05 | 1.233E-05 | 2.309E-06 | 1.100E-05 | 1.500E-05 |
| 18050 | 903 | R903I | C | 1.1e-05 | 9.6e-06 | 1.1e-05 | 1.053E-05 | 8.083E-07 | 9.600E-06 | 1.100E-05 |
| 18051 | 903 | R903L | C | 1.6e-05 | 9.7e-06 | 1.6e-05 | 1.390E-05 | 3.637E-06 | 9.700E-06 | 1.600E-05 |
| 18052 | 903 | R903K | C | 5.8e-06 | 9.5e-06 | 1.6e-05 | 1.043E-05 | 5.164E-06 | 5.800E-06 | 1.600E-05 |
| 18053 | 903 | R903M | C | 1.1e-05 | 1.4e-05 | 1.1e-05 | 1.200E-05 | 1.732E-06 | 1.100E-05 | 1.400E-05 |
| 18054 | 903 | R903F | C | 1.4e-05 | 1.5e-05 | 1.1e-05 | 1.333E-05 | 2.082E-06 | 1.100E-05 | 1.500E-05 |
| 18055 | 903 | R903P | C | 8.9e-06 | 1.5e-05 | 1.1e-05 | 1.163E-05 | 3.099E-06 | 8.900E-06 | 1.500E-05 |
| 18056 | 903 | R903S | C | 1.1e-05 | 1.3e-05 | 1.6e-05 | 1.333E-05 | 2.517E-06 | 1.100E-05 | 1.600E-05 |
| 18057 | 903 | R903T | C | 1e-05 | 1.2e-05 | 1.7e-05 | 1.300E-05 | 3.606E-06 | 1.000E-05 | 1.700E-05 |
| 18058 | 903 | R903W | C | 9.7e-06 | 1.6e-05 | 1.4e-05 | 1.323E-05 | 3.219E-06 | 9.700E-06 | 1.600E-05 |
| 18059 | 903 | R903Y | C | 1.5e-05 | 1e-05 | 7.2e-06 | 1.073E-05 | 3.951E-06 | 7.200E-06 | 1.500E-05 |
| 18060 | 903 | R903V | C | 1.4e-05 | 8.3e-06 | 1.4e-05 | 1.210E-05 | 3.291E-06 | 8.300E-06 | 1.400E-05 |
| 18061 | 904 | F904A | C | 1.4e-05 | 1e-05 | 1.1e-05 | 1.167E-05 | 2.082E-06 | 1.000E-05 | 1.400E-05 |
| 18062 | 904 | F904R | C | 7.6e-06 | 1.1e-05 | 1.1e-05 | 9.867E-06 | 1.963E-06 | 7.600E-06 | 1.100E-05 |
| 18063 | 904 | F904N | C | 1.4e-05 | 1.4e-05 | 1e-05 | 1.267E-05 | 2.309E-06 | 1.000E-05 | 1.400E-05 |
| 18064 | 904 | F904D | C | 1.4e-05 | 1.3e-05 | 1.1e-05 | 1.267E-05 | 1.528E-06 | 1.100E-05 | 1.400E-05 |
| 18065 | 904 | F904C | C | 1.7e-05 | 1.4e-05 | 1.5e-05 | 1.533E-05 | 1.528E-06 | 1.400E-05 | 1.700E-05 |
| 18066 | 904 | F904E | C | 9.7e-06 | 1.2e-05 | 1.6e-05 | 1.257E-05 | 3.188E-06 | 9.700E-06 | 1.600E-05 |
| 18067 | 904 | F904Q | C | 1.2e-05 | 1.6e-05 | 1.6e-05 | 1.467E-05 | 2.309E-06 | 1.200E-05 | 1.600E-05 |
| 18068 | 904 | F904G | C | 1.5e-05 | 1.6e-05 | 1.4e-05 | 1.500E-05 | 1.000E-06 | 1.400E-05 | 1.600E-05 |
| 18069 | 904 | F904H | C | 1.6e-05 | 1.4e-05 | 1.7e-05 | 1.567E-05 | 1.528E-06 | 1.400E-05 | 1.700E-05 |
| 18070 | 904 | F904I | C | 9.6e-06 | 1.5e-05 | 1.7e-05 | 1.387E-05 | 3.828E-06 | 9.600E-06 | 1.700E-05 |
| 18071 | 904 | F904L | C | 1.4e-05 | 1.6e-05 | 1.5e-05 | 1.500E-05 | 1.000E-06 | 1.400E-05 | 1.600E-05 |
| 18072 | 904 | F904K | C | 1e-05 | 9.2e-06 | 1e-05 | 9.733E-06 | 4.619E-07 | 9.200E-06 | 1.000E-05 |
| 18073 | 904 | F904M | C | 1.6e-05 | 9.2e-06 | 1.1e-05 | 1.207E-05 | 3.523E-06 | 9.200E-06 | 1.600E-05 |
| 18074 | 904 | F904F | C | 1.5e-05 | 8.5e-06 | 1.5e-05 | 1.283E-05 | 3.753E-06 | 8.500E-06 | 1.500E-05 |
| 18075 | 904 | F904P | C | 1.7e-05 | 1.4e-05 | 1.4e-05 | 1.500E-05 | 1.732E-06 | 1.400E-05 | 1.700E-05 |
| 18076 | 904 | F904S | C | 1e-05 | 1.1e-05 | 1.1e-05 | 1.067E-05 | 5.774E-07 | 1.000E-05 | 1.100E-05 |
| 18077 | 904 | F904T | C | 1.3e-05 | 1.5e-05 | 1.3e-05 | 1.367E-05 | 1.155E-06 | 1.300E-05 | 1.500E-05 |
| 18078 | 904 | F904W | C | 1.4e-05 | 1.5e-05 | 1.2e-05 | 1.367E-05 | 1.528E-06 | 1.200E-05 | 1.500E-05 |
| 18079 | 904 | F904Y | C | 1e-05 | 1.4e-05 | 1.3e-05 | 1.233E-05 | 2.082E-06 | 1.000E-05 | 1.400E-05 |
| 18080 | 904 | F904V | C | 1.5e-05 | 1.6e-05 | 1.4e-05 | 1.500E-05 | 1.000E-06 | 1.400E-05 | 1.600E-05 |
| 18081 | 905 | T905A | C | 1.4e-05 | 1.1e-05 | 1.3e-05 | 1.267E-05 | 1.528E-06 | 1.100E-05 | 1.400E-05 |
| 18082 | 905 | T905R | C | 1.5e-05 | 1.7e-05 | 1.6e-05 | 1.600E-05 | 1.000E-06 | 1.500E-05 | 1.700E-05 |
| 18083 | 905 | T905N | C | 1.4e-05 | 9.4e-06 | 1.1e-05 | 1.147E-05 | 2.335E-06 | 9.400E-06 | 1.400E-05 |
| 18084 | 905 | T905D | C | 1.1e-05 | 1.6e-05 | 1.6e-05 | 1.433E-05 | 2.887E-06 | 1.100E-05 | 1.600E-05 |
| 18085 | 905 | T905C | C | 1.5e-05 | 1.6e-05 | 9.8e-06 | 1.360E-05 | 3.329E-06 | 9.800E-06 | 1.600E-05 |
| 18086 | 905 | T905E | C | 1.3e-05 | 1.5e-05 | 1e-05 | 1.267E-05 | 2.517E-06 | 1.000E-05 | 1.500E-05 |
| 18087 | 905 | T905Q | C | 1.5e-05 | 1.4e-05 | 9.6e-06 | 1.287E-05 | 2.873E-06 | 9.600E-06 | 1.500E-05 |
| 18088 | 905 | T905G | C | 1e-05 | 1.3e-05 | 1.5e-05 | 1.267E-05 | 2.517E-06 | 1.000E-05 | 1.500E-05 |
| 18089 | 905 | T905H | C | 1.6e-05 | 1e-05 | 1.6e-05 | 1.400E-05 | 3.464E-06 | 1.000E-05 | 1.600E-05 |
| 18090 | 905 | T905I | C | 1.6e-05 | 1e-05 | 1.4e-05 | 1.333E-05 | 3.055E-06 | 1.000E-05 | 1.600E-05 |
| 18091 | 905 | T905L | C | 1.4e-05 | 1.1e-05 | 1.3e-05 | 1.267E-05 | 1.528E-06 | 1.100E-05 | 1.400E-05 |
| 18092 | 905 | T905K | C | 1.2e-05 | 1.1e-05 | 1.6e-05 | 1.300E-05 | 2.646E-06 | 1.100E-05 | 1.600E-05 |
| 18093 | 905 | T905M | C | 1.6e-05 | 1.1e-05 | 9.9e-06 | 1.230E-05 | 3.251E-06 | 9.900E-06 | 1.600E-05 |
| 18094 | 905 | T905F | C | 1.6e-05 | 1.3e-05 | 2.2e-05 | 1.700E-05 | 4.583E-06 | 1.300E-05 | 2.200E-05 |
| 18095 | 905 | T905P | C | 1.6e-05 | 1.5e-05 | 1e-05 | 1.367E-05 | 3.215E-06 | 1.000E-05 | 1.600E-05 |
| 18096 | 905 | T905S | C | 9.4e-06 | 6.4e-06 | 1.1e-05 | 8.933E-06 | 2.335E-06 | 6.400E-06 | 1.100E-05 |
| 18097 | 905 | T905T | C | 1e-05 | 1.6e-05 | 1.6e-05 | 1.400E-05 | 3.464E-06 | 1.000E-05 | 1.600E-05 |
| 18098 | 905 | T905W | C | 1.5e-05 | 1e-05 | 1.4e-05 | 1.300E-05 | 2.646E-06 | 1.000E-05 | 1.500E-05 |
| 18099 | 905 | T905Y | C | 1e-05 | 9.5e-06 | 9.8e-06 | 9.767E-06 | 2.517E-07 | 9.500E-06 | 1.000E-05 |
| 18100 | 905 | T905V | C | 2.3e-05 | 1.6e-05 | 1.1e-05 | 1.667E-05 | 6.028E-06 | 1.100E-05 | 2.300E-05 |
| 18101 | 906 | L906A | C | 1.1e-05 | 1.1e-05 | 1.6e-05 | 1.267E-05 | 2.887E-06 | 1.100E-05 | 1.600E-05 |
| 18102 | 906 | L906R | C | 1.1e-05 | 1e-05 | 9e-06 | 1.000E-05 | 1.000E-06 | 9.000E-06 | 1.100E-05 |
| 18103 | 906 | L906N | C | 1.1e-05 | 1.5e-05 | 1.2e-05 | 1.267E-05 | 2.082E-06 | 1.100E-05 | 1.500E-05 |
| 18104 | 906 | L906D | C | 1.6e-05 | 1.5e-05 | 8.9e-06 | 1.330E-05 | 3.843E-06 | 8.900E-06 | 1.600E-05 |

| | | | | | | | | | | |
|---|---|---|---|---|---|---|---|---|---|---|
| 18105 | 906 | L906C | C | 1.4e-05 | 1.1e-05 | 1.1e-05 | 1.200E-05 | 1.732E-06 | 1.100E-05 | 1.400E-05 |
| 18106 | 906 | L906E | C | 1.6e-05 | 1.2e-05 | 1.2e-05 | 1.333E-05 | 2.309E-06 | 1.200E-05 | 1.600E-05 |
| 18107 | 906 | L906Q | C | 1.3e-05 | 1.4e-05 | 8.3e-06 | 1.177E-05 | 3.044E-06 | 8.300E-06 | 1.400E-05 |
| 18108 | 906 | L906G | C | 1e-05 | 1.5e-05 | 1.1e-05 | 1.200E-05 | 2.646E-06 | 1.000E-05 | 1.500E-05 |
| 18109 | 906 | L906H | C | 1.6e-05 | 1.4e-05 | 1.4e-05 | 1.467E-05 | 1.155E-06 | 1.400E-05 | 1.600E-05 |
| 18110 | 906 | L906I | C | 1.6e-05 | 1.1e-05 | 1.5e-05 | 1.400E-05 | 2.646E-06 | 1.100E-05 | 1.600E-05 |
| 18111 | 906 | L906L | C | 1.4e-05 | 1.4e-05 | 1.5e-05 | 1.433E-05 | 5.774E-07 | 1.400E-05 | 1.500E-05 |
| 18112 | 906 | L906K | C | 1.3e-05 | 9.6e-06 | 1.4e-05 | 1.220E-05 | 2.307E-06 | 9.600E-06 | 1.400E-05 |
| 18113 | 906 | L906M | C | 9.2e-06 | 1.5e-05 | 1.2e-05 | 1.207E-05 | 2.901E-06 | 9.200E-06 | 1.500E-05 |
| 18114 | 906 | L906F | C | 1.6e-05 | 9.9e-06 | 1.2e-05 | 1.263E-05 | 3.099E-06 | 9.900E-06 | 1.600E-05 |
| 18115 | 906 | L906P | C | 1.6e-05 | 1.1e-05 | 1.2e-05 | 1.300E-05 | 2.646E-06 | 1.100E-05 | 1.600E-05 |
| 18116 | 906 | L906S | C | 9.2e-06 | 2e-05 | 1.5e-05 | 1.473E-05 | 5.405E-06 | 9.200E-06 | 2.000E-05 |
| 18117 | 906 | L906T | C | 1.4e-05 | 7.7e-06 | 1.6e-05 | 1.257E-05 | 4.332E-06 | 7.700E-06 | 1.600E-05 |
| 18118 | 906 | L906W | C | 1.5e-05 | 9.3e-06 | 1.4e-05 | 1.277E-05 | 3.044E-06 | 9.300E-06 | 1.500E-05 |
| 18119 | 906 | L906Y | C | 1.5e-05 | 1.4e-05 | 1.1e-05 | 1.333E-05 | 2.082E-06 | 1.100E-05 | 1.500E-05 |
| 18120 | 906 | L906V | C | 1e-05 | 1.1e-05 | 9.4e-06 | 1.013E-05 | 8.083E-07 | 9.400E-06 | 1.100E-05 |
| 18121 | 907 | S907A | C | 1.6e-05 | 1.1e-05 | 1.4e-05 | 1.367E-05 | 2.517E-06 | 1.100E-05 | 1.600E-05 |
| 18122 | 907 | S907R | C | 1e-05 | 9.6e-06 | 1.1e-05 | 1.020E-05 | 7.211E-07 | 9.600E-06 | 1.100E-05 |
| 18123 | 907 | S907N | C | 1.2e-05 | 1.4e-05 | 1.1e-05 | 1.233E-05 | 1.528E-06 | 1.100E-05 | 1.400E-05 |
| 18124 | 907 | S907D | C | 1e-05 | 1.1e-05 | 1e-05 | 1.033E-05 | 5.774E-07 | 1.000E-05 | 1.100E-05 |
| 18125 | 907 | S907C | C | 1.3e-05 | 1.2e-05 | 1.4e-05 | 1.300E-05 | 1.000E-06 | 1.200E-05 | 1.400E-05 |
| 18126 | 907 | S907E | C | 1.5e-05 | 7.8e-06 | 1.7e-05 | 1.327E-05 | 4.839E-06 | 7.800E-06 | 1.700E-05 |
| 18127 | 907 | S907Q | C | 9.2e-06 | 1.5e-05 | 2.5e-05 | 1.640E-05 | 7.992E-06 | 9.200E-06 | 2.500E-05 |
| 18128 | 907 | S907G | C | 1.4e-05 | 1.3e-05 | 1.1e-05 | 1.267E-05 | 1.528E-06 | 1.100E-05 | 1.400E-05 |
| 18129 | 907 | S907H | C | 1.5e-05 | 1.4e-05 | 1e-05 | 1.300E-05 | 2.646E-06 | 1.000E-05 | 1.500E-05 |
| 18130 | 907 | S907I | C | 1.8e-05 | 1.1e-05 | 1.2e-05 | 1.367E-05 | 3.786E-06 | 1.100E-05 | 1.800E-05 |
| 18131 | 907 | S907L | C | 1.8e-05 | 1.1e-05 | 1.8e-05 | 1.567E-05 | 4.041E-06 | 1.100E-05 | 1.800E-05 |
| 18132 | 907 | S907K | C | 1e-05 | 1.1e-05 | 7.6e-06 | 9.533E-06 | 1.747E-06 | 7.600E-06 | 1.100E-05 |
| 18133 | 907 | S907M | C | 1.6e-05 | 1.4e-05 | 1.4e-05 | 1.467E-05 | 1.155E-06 | 1.400E-05 | 1.600E-05 |
| 18134 | 907 | S907F | C | 1.7e-05 | 1.6e-05 | 1.4e-05 | 1.567E-05 | 1.528E-06 | 1.400E-05 | 1.700E-05 |
| 18135 | 907 | S907P | C | 1.5e-05 | 8.9e-06 | 1.7e-05 | 1.363E-05 | 4.219E-06 | 8.900E-06 | 1.700E-05 |
| 18136 | 907 | S907S | C | 9e-06 | 1.1e-05 | 9.2e-06 | 9.733E-06 | 1.102E-06 | 9.000E-06 | 1.100E-05 |
| 18137 | 907 | S907T | C | 1.5e-05 | 1e-05 | 1e-05 | 1.167E-05 | 2.887E-06 | 1.000E-05 | 1.500E-05 |
| 18138 | 907 | S907W | C | 1.7e-05 | 1.4e-05 | 1.1e-05 | 1.400E-05 | 3.000E-06 | 1.100E-05 | 1.700E-05 |
| 18139 | 907 | S907Y | C | 1.4e-05 | 8.4e-06 | 9.2e-06 | 1.053E-05 | 3.029E-06 | 8.400E-06 | 1.400E-05 |
| 18140 | 907 | S907V | C | 1.2e-05 | 1.1e-05 | 1.8e-05 | 1.367E-05 | 3.786E-06 | 1.100E-05 | 1.800E-05 |
| 18141 | 908 | V908A | C | 1.4e-05 | 1.5e-05 | 1.1e-05 | 1.333E-05 | 2.082E-06 | 1.100E-05 | 1.500E-05 |
| 18142 | 908 | V908R | C | 1.4e-05 | 1.1e-05 | 8.8e-06 | 1.127E-05 | 2.610E-06 | 8.800E-06 | 1.400E-05 |
| 18143 | 908 | V908N | C | 1.4e-05 | 1e-05 | 1e-05 | 1.133E-05 | 2.309E-06 | 1.000E-05 | 1.400E-05 |
| 18144 | 908 | V908D | C | 1.6e-05 | 1.4e-05 | 1.5e-05 | 1.500E-05 | 1.000E-06 | 1.400E-05 | 1.600E-05 |
| 18145 | 908 | V908C | C | 1.5e-05 | 1e-05 | 1.5e-05 | 1.333E-05 | 2.887E-06 | 1.000E-05 | 1.500E-05 |
| 18146 | 908 | V908E | C | 1.3e-05 | 1.6e-05 | 1.5e-05 | 1.467E-05 | 1.528E-06 | 1.300E-05 | 1.600E-05 |
| 18147 | 908 | V908Q | C | 1.5e-05 | 9.2e-06 | 1.4e-05 | 1.273E-05 | 3.101E-06 | 9.200E-06 | 1.500E-05 |
| 18148 | 908 | V908G | C | 9e-06 | 8.3e-06 | 1.1e-05 | 9.433E-06 | 1.401E-06 | 8.300E-06 | 1.100E-05 |
| 18149 | 908 | V908H | C | 8.3e-06 | 1.5e-05 | 1e-05 | 1.110E-05 | 3.483E-06 | 8.300E-06 | 1.500E-05 |
| 18150 | 908 | V908I | C | 1.5e-05 | 1.5e-05 | 9.6e-06 | 1.320E-05 | 3.118E-06 | 9.600E-06 | 1.500E-05 |
| 18151 | 908 | V908L | C | 1.1e-05 | 1.1e-05 | 9.9e-06 | 1.063E-05 | 6.351E-07 | 9.900E-06 | 1.100E-05 |
| 18152 | 908 | V908K | C | 1.4e-05 | 1.5e-05 | 1.4e-05 | 1.433E-05 | 5.774E-07 | 1.400E-05 | 1.500E-05 |
| 18153 | 908 | V908M | C | 1.4e-05 | 1.4e-05 | 1.4e-05 | 1.400E-05 | 0.000E+00 | 1.400E-05 | 1.400E-05 |
| 18154 | 908 | V908F | C | 1.4e-05 | 1e-05 | 1.8e-05 | 1.400E-05 | 4.000E-06 | 1.000E-05 | 1.800E-05 |
| 18155 | 908 | V908P | C | 1.5e-05 | 1.5e-05 | 1e-05 | 1.333E-05 | 2.887E-06 | 1.000E-05 | 1.500E-05 |
| 18156 | 908 | V908S | C | 1.5e-05 | 1.6e-05 | 1.5e-05 | 1.533E-05 | 5.774E-07 | 1.500E-05 | 1.600E-05 |
| 18157 | 908 | V908T | C | 1.3e-05 | 1.1e-05 | 1.3e-05 | 1.233E-05 | 1.155E-06 | 1.100E-05 | 1.300E-05 |
| 18158 | 908 | V908W | C | 1.5e-05 | 9.7e-06 | 9e-06 | 1.123E-05 | 3.281E-06 | 9.000E-06 | 1.500E-05 |
| 18159 | 908 | V908Y | C | 1.6e-05 | 8.2e-06 | 1.3e-05 | 1.240E-05 | 3.934E-06 | 8.200E-06 | 1.600E-05 |
| 18160 | 908 | V908V | C | 1.5e-05 | 1.4e-05 | 1.6e-05 | 1.500E-05 | 1.000E-06 | 1.400E-05 | 1.600E-05 |
| 18161 | 909 | D909A | C | 1.4e-05 | 1.1e-05 | 1e-05 | 1.167E-05 | 2.082E-06 | 1.000E-05 | 1.400E-05 |
| 18162 | 909 | D909R | C | 1.5e-05 | 1.2e-05 | 1.1e-05 | 1.267E-05 | 2.082E-06 | 1.100E-05 | 1.500E-05 |
| 18163 | 909 | D909N | C | 1.4e-05 | 1.4e-05 | 1.5e-05 | 1.433E-05 | 5.774E-07 | 1.400E-05 | 1.500E-05 |
| 18164 | 909 | D909D | C | 1.7e-05 | 1.5e-05 | 9.5e-06 | 1.383E-05 | 3.884E-06 | 9.500E-06 | 1.700E-05 |
| 18165 | 909 | D909C | C | 1.4e-05 | 9.6e-06 | 1.2e-05 | 1.187E-05 | 2.203E-06 | 9.600E-06 | 1.400E-05 |
| 18166 | 909 | D909E | C | 1e-05 | 1.5e-05 | 1.5e-05 | 1.333E-05 | 2.887E-06 | 1.000E-05 | 1.500E-05 |
| 18167 | 909 | D909Q | C | 9.9e-06 | 1.4e-05 | 8.4e-06 | 1.077E-05 | 2.899E-06 | 8.400E-06 | 1.400E-05 |
| 18168 | 909 | D909G | C | 1.7e-05 | 1e-05 | 1.3e-05 | 1.333E-05 | 3.512E-06 | 1.000E-05 | 1.700E-05 |
| 18169 | 909 | D909H | C | 1.5e-05 | 1.5e-05 | 1.4e-05 | 1.467E-05 | 5.774E-07 | 1.400E-05 | 1.500E-05 |
| 18170 | 909 | D909I | C | 1.6e-05 | 1.6e-05 | 1.1e-05 | 1.433E-05 | 2.887E-06 | 1.100E-05 | 1.600E-05 |
| 18171 | 909 | D909L | C | 9.5e-06 | 8.7e-06 | 1.6e-05 | 1.140E-05 | 4.004E-06 | 8.700E-06 | 1.600E-05 |
| 18172 | 909 | D909K | C | 8.1e-06 | 1.7e-05 | 1.7e-05 | 1.403E-05 | 5.138E-06 | 8.100E-06 | 1.700E-05 |
| 18173 | 909 | D909M | C | 1.6e-05 | 1.1e-05 | 1.1e-05 | 1.267E-05 | 2.887E-06 | 1.100E-05 | 1.600E-05 |
| 18174 | 909 | D909F | C | 1.1e-05 | 1.3e-05 | 1.4e-05 | 1.267E-05 | 1.528E-06 | 1.100E-05 | 1.400E-05 |
| 18175 | 909 | D909P | C | 1.1e-05 | 9.1e-06 | 1.3e-05 | 1.103E-05 | 1.950E-06 | 9.100E-06 | 1.300E-05 |
| 18176 | 909 | D909S | C | 2.1e-05 | 1e-05 | 1e-05 | 1.367E-05 | 6.351E-06 | 1.000E-05 | 2.100E-05 |
| 18177 | 909 | D909T | C | 1.3e-05 | 1.4e-05 | 1.6e-05 | 1.433E-05 | 1.528E-06 | 1.300E-05 | 1.600E-05 |
| 18178 | 909 | D909W | C | 1.6e-05 | 1.4e-05 | 1.4e-05 | 1.467E-05 | 1.155E-06 | 1.400E-05 | 1.600E-05 |
| 18179 | 909 | D909Y | C | 8.3e-06 | 1.1e-05 | 1.6e-05 | 1.177E-05 | 3.907E-06 | 8.300E-06 | 1.600E-05 |
| 18180 | 909 | D909V | C | 1.1e-05 | 1.5e-05 | 1.1e-05 | 1.233E-05 | 2.309E-06 | 1.100E-05 | 1.500E-05 |
| 18181 | 910 | R910A | C | 1.1e-05 | 8.9e-06 | 1.6e-05 | 1.197E-05 | 3.647E-06 | 8.900E-06 | 1.600E-05 |
| 18182 | 910 | R910R | C | 8.6e-06 | 1.4e-05 | 1.4e-05 | 1.220E-05 | 3.118E-06 | 8.600E-06 | 1.400E-05 |
| 18183 | 910 | R910N | C | 1.5e-05 | 5.2e-06 | 9.2e-06 | 9.800E-06 | 4.927E-06 | 5.200E-06 | 1.500E-05 |
| 18184 | 910 | R910D | C | 1.5e-05 | 1e-05 | 1.2e-05 | 1.233E-05 | 2.517E-06 | 1.000E-05 | 1.500E-05 |
| 18185 | 910 | R910C | C | 1.5e-05 | 1.2e-05 | 1.3e-05 | 1.333E-05 | 1.528E-06 | 1.200E-05 | 1.500E-05 |
| 18186 | 910 | R910E | C | 1.5e-05 | 1e-05 | 1.3e-05 | 1.267E-05 | 2.517E-06 | 1.000E-05 | 1.500E-05 |
| 18187 | 910 | R910Q | C | 1.2e-05 | 1.5e-05 | 1.5e-05 | 1.400E-05 | 1.732E-06 | 1.200E-05 | 1.500E-05 |
| 18188 | 910 | R910G | C | 1.5e-05 | 9.6e-06 | 1.5e-05 | 1.320E-05 | 3.118E-06 | 9.600E-06 | 1.500E-05 |
| 18189 | 910 | R910H | C | 1.5e-05 | 1.3e-05 | 8.3e-06 | 1.210E-05 | 3.439E-06 | 8.300E-06 | 1.500E-05 |

| | | | | | | | | | | |
|---|---|---|---|---|---|---|---|---|---|---|
| 18190 | 910 | R910I | C | 1.4e-05 | 6.3e-06 | 1.6e-05 | 1.210E-05 | 5.122E-06 | 6.300E-06 | 1.600E-05 |
| 18191 | 910 | R910L | C | 1.1e-05 | 1.6e-05 | 1.3e-05 | 1.333E-05 | 2.517E-06 | 1.100E-05 | 1.600E-05 |
| 18192 | 910 | R910K | C | 1.1e-05 | 1.4e-05 | 1.4e-05 | 1.300E-05 | 1.732E-06 | 1.100E-05 | 1.400E-05 |
| 18193 | 910 | R910M | C | 4.1e-05 | 1.1e-05 | 1e-05 | 2.067E-05 | 1.762E-05 | 1.000E-05 | 4.100E-05 |
| 18194 | 910 | R910F | C | 1.2e-05 | 1.6e-05 | 1.6e-05 | 1.467E-05 | 2.309E-06 | 1.200E-05 | 1.600E-05 |
| 18195 | 910 | R910P | C | 1.5e-05 | 7.3e-06 | 1.5e-05 | 1.243E-05 | 4.446E-06 | 7.300E-06 | 1.500E-05 |
| 18196 | 910 | R910S | C | 8.5e-06 | 1.5e-05 | 1.1e-05 | 1.150E-05 | 3.279E-06 | 8.500E-06 | 1.500E-05 |
| 18197 | 910 | R910T | C | 7.7e-06 | 1e-05 | 1.5e-05 | 1.090E-05 | 3.732E-06 | 7.700E-06 | 1.500E-05 |
| 18198 | 910 | R910W | C | 8.4e-06 | 1.1e-05 | 1.2e-05 | 1.047E-05 | 1.858E-06 | 8.400E-06 | 1.200E-05 |
| 18199 | 910 | R910Y | C | 1.4e-05 | 1.5e-05 | 1.7e-05 | 1.533E-05 | 1.528E-06 | 1.400E-05 | 1.700E-05 |
| 18200 | 910 | R910V | C | 1.5e-05 | 9.4e-06 | 1.2e-05 | 1.213E-05 | 2.802E-06 | 9.400E-06 | 1.500E-05 |
| 18201 | 911 | S911A | C | 1.5e-05 | 1.5e-05 | 9.8e-06 | 1.327E-05 | 3.002E-06 | 9.800E-06 | 1.500E-05 |
| 18202 | 911 | S911R | C | 1.4e-05 | 7.6e-06 | 9.8e-06 | 1.047E-05 | 3.252E-06 | 7.600E-06 | 1.400E-05 |
| 18203 | 911 | S911N | C | 1.5e-05 | 1.6e-05 | 1.5e-05 | 1.533E-05 | 5.774E-07 | 1.500E-05 | 1.600E-05 |
| 18204 | 911 | S911D | C | 1.4e-05 | 1.1e-05 | 1.3e-05 | 1.267E-05 | 1.528E-06 | 1.100E-05 | 1.400E-05 |
| 18205 | 911 | S911C | C | 1.5e-05 | 1.9e-05 | 1.6e-05 | 1.667E-05 | 2.082E-06 | 1.500E-05 | 1.900E-05 |
| 18206 | 911 | S911E | C | 1.1e-05 | 1.2e-05 | 1.5e-05 | 1.267E-05 | 2.082E-06 | 1.100E-05 | 1.500E-05 |
| 18207 | 911 | S911Q | C | 1.6e-05 | 9e-06 | 1.6e-05 | 1.367E-05 | 4.041E-06 | 9.000E-06 | 1.600E-05 |
| 18208 | 911 | S911G | C | 1.3e-05 | 1.6e-05 | 1e-05 | 1.300E-05 | 3.000E-06 | 1.000E-05 | 1.600E-05 |
| 18209 | 911 | S911H | C | 1.4e-05 | 1.5e-05 | 1.5e-05 | 1.467E-05 | 5.774E-07 | 1.400E-05 | 1.500E-05 |
| 18210 | 911 | S911I | C | 1.1e-05 | 1.1e-05 | 1.5e-05 | 1.233E-05 | 2.309E-06 | 1.100E-05 | 1.500E-05 |
| 18211 | 911 | S911L | C | 1.1e-05 | 1.1e-05 | 1.1e-05 | 1.100E-05 | 2.075E-21 | 1.100E-05 | 1.100E-05 |
| 18212 | 911 | S911K | C | 1.1e-05 | 1.1e-05 | 1.5e-05 | 1.233E-05 | 2.309E-06 | 1.100E-05 | 1.500E-05 |
| 18213 | 911 | S911M | C | 1.4e-05 | 1.3e-05 | 8.7e-06 | 1.190E-05 | 2.816E-06 | 8.700E-06 | 1.400E-05 |
| 18214 | 911 | S911F | C | 1.4e-05 | 1.2e-05 | 1.2e-05 | 1.267E-05 | 1.155E-06 | 1.200E-05 | 1.400E-05 |
| 18215 | 911 | S911P | C | 1.4e-05 | 1.6e-05 | 1.6e-05 | 1.533E-05 | 1.155E-06 | 1.400E-05 | 1.600E-05 |
| 18216 | 911 | S911S | C | 1.1e-05 | 1.5e-05 | 1.6e-05 | 1.400E-05 | 2.646E-06 | 1.100E-05 | 1.600E-05 |
| 18217 | 911 | S911T | C | 1.5e-05 | 1.6e-05 | 1.5e-05 | 1.533E-05 | 5.774E-07 | 1.500E-05 | 1.600E-05 |
| 18218 | 911 | S911W | C | 1.5e-05 | 1.3e-05 | 1.4e-05 | 1.400E-05 | 1.000E-06 | 1.300E-05 | 1.500E-05 |
| 18219 | 911 | S911Y | C | 9.9e-06 | 1.7e-05 | 1.2e-05 | 1.297E-05 | 3.647E-06 | 9.900E-06 | 1.700E-05 |
| 18220 | 911 | S911V | C | 1.6e-05 | 7.6e-06 | 1.6e-05 | 1.320E-05 | 4.850E-06 | 7.600E-06 | 1.600E-05 |
| 18221 | 912 | K912A | C | 2e-05 | 1e-05 | 1.5e-05 | 1.500E-05 | 5.000E-06 | 1.000E-05 | 2.000E-05 |
| 18222 | 912 | K912R | C | 8.6e-06 | 1.3e-05 | 1.5e-05 | 1.220E-05 | 3.274E-06 | 8.600E-06 | 1.500E-05 |
| 18223 | 912 | K912N | C | 9e-06 | 1.5e-05 | 1.3e-05 | 1.233E-05 | 3.055E-06 | 9.000E-06 | 1.500E-05 |
| 18224 | 912 | K912D | C | 9.9e-06 | 8.7e-06 | 1.5e-05 | 1.120E-05 | 3.345E-06 | 8.700E-06 | 1.500E-05 |
| 18225 | 912 | K912C | C | 9e-06 | 1.1e-05 | 9e-06 | 9.667E-06 | 1.155E-06 | 9.000E-06 | 1.100E-05 |
| 18226 | 912 | K912E | C | 1.3e-05 | 1.2e-05 | 1.1e-05 | 1.200E-05 | 1.000E-06 | 1.100E-05 | 1.300E-05 |
| 18227 | 912 | K912Q | C | 9e-06 | 1.8e-05 | 6.7e-06 | 1.123E-05 | 5.972E-06 | 6.700E-06 | 1.800E-05 |
| 18228 | 912 | K912G | C | 1.5e-05 | 1.4e-05 | 1e-05 | 1.300E-05 | 2.646E-06 | 1.000E-05 | 1.500E-05 |
| 18229 | 912 | K912H | C | 1.5e-05 | 1.1e-05 | 1.5e-05 | 1.367E-05 | 2.309E-06 | 1.100E-05 | 1.500E-05 |
| 18230 | 912 | K912I | C | 8.4e-06 | 1.3e-05 | 1e-05 | 1.047E-05 | 2.335E-06 | 8.400E-06 | 1.300E-05 |
| 18231 | 912 | K912L | C | 1.3e-05 | 1.1e-05 | 1e-05 | 1.133E-05 | 1.528E-06 | 1.000E-05 | 1.300E-05 |
| 18232 | 912 | K912K | C | 1.1e-05 | 1.4e-05 | 9.6e-06 | 1.153E-05 | 2.248E-06 | 9.600E-06 | 1.400E-05 |
| 18233 | 912 | K912M | C | 1.4e-05 | 1e-05 | 1e-05 | 1.133E-05 | 2.309E-06 | 1.000E-05 | 1.400E-05 |
| 18234 | 912 | K912F | C | 1e-05 | 9.3e-06 | 1.5e-05 | 1.143E-05 | 3.109E-06 | 9.300E-06 | 1.500E-05 |
| 18235 | 912 | K912P | C | 1.4e-05 | 1.5e-05 | 1.5e-05 | 1.467E-05 | 5.774E-07 | 1.400E-05 | 1.500E-05 |
| 18236 | 912 | K912S | C | 1.2e-05 | 8.7e-06 | 1.4e-05 | 1.157E-05 | 2.676E-06 | 8.700E-06 | 1.400E-05 |
| 18237 | 912 | K912T | C | 1.4e-05 | 1.2e-05 | 1.5e-05 | 1.367E-05 | 1.528E-06 | 1.200E-05 | 1.500E-05 |
| 18238 | 912 | K912W | C | 1e-05 | 9e-06 | 1.2e-05 | 1.033E-05 | 1.528E-06 | 9.000E-06 | 1.200E-05 |
| 18239 | 912 | K912Y | C | 1.1e-05 | 1e-05 | 1.1e-05 | 1.067E-05 | 5.774E-07 | 1.000E-05 | 1.100E-05 |
| 18240 | 912 | K912V | C | 9.5e-06 | 1e-05 | 9.4e-06 | 9.633E-06 | 3.215E-07 | 9.400E-06 | 1.000E-05 |
| 18241 | 913 | N913A | C | 7.2e-06 | 9.2e-06 | 1e-05 | 8.800E-06 | 1.442E-06 | 7.200E-06 | 1.000E-05 |
| 18242 | 913 | N913R | C | 9.2e-06 | 1.4e-05 | 1.6e-05 | 1.307E-05 | 3.495E-06 | 9.200E-06 | 1.600E-05 |
| 18243 | 913 | N913N | C | 1.2e-05 | 1.4e-05 | 1.2e-05 | 1.267E-05 | 1.155E-06 | 1.200E-05 | 1.400E-05 |
| 18244 | 913 | N913D | C | 1e-05 | 1.7e-05 | 1.4e-05 | 1.367E-05 | 3.512E-06 | 1.000E-05 | 1.700E-05 |
| 18245 | 913 | N913C | C | 1.1e-05 | 1.5e-05 | 8.9e-06 | 1.163E-05 | 3.099E-06 | 8.900E-06 | 1.500E-05 |
| 18246 | 913 | N913E | C | 1.1e-05 | 1.6e-05 | 1.3e-05 | 1.333E-05 | 2.517E-06 | 1.100E-05 | 1.600E-05 |
| 18247 | 913 | N913Q | C | 1.1e-05 | 9e-06 | 1.6e-05 | 1.200E-05 | 3.606E-06 | 9.000E-06 | 1.600E-05 |
| 18248 | 913 | N913G | C | 9.9e-06 | 1.1e-05 | 1.1e-05 | 1.063E-05 | 6.351E-07 | 9.900E-06 | 1.100E-05 |
| 18249 | 913 | N913H | C | 1.2e-05 | 1.4e-05 | 8.7e-06 | 1.157E-05 | 2.676E-06 | 8.700E-06 | 1.400E-05 |
| 18250 | 913 | N913I | C | 9.5e-06 | 1e-05 | 1.6e-05 | 1.183E-05 | 3.617E-06 | 9.500E-06 | 1.600E-05 |
| 18251 | 913 | N913L | C | 1.2e-05 | 1.1e-05 | 1.1e-05 | 1.133E-05 | 5.774E-07 | 1.100E-05 | 1.200E-05 |
| 18252 | 913 | N913K | C | 1.6e-05 | 1.1e-05 | 1.5e-05 | 1.400E-05 | 2.646E-06 | 1.100E-05 | 1.600E-05 |
| 18253 | 913 | N913M | C | 1.3e-05 | 1.5e-05 | 1.5e-05 | 1.433E-05 | 1.155E-06 | 1.300E-05 | 1.500E-05 |
| 18254 | 913 | N913F | C | 1.6e-05 | 1.4e-05 | 1.1e-05 | 1.367E-05 | 2.517E-06 | 1.100E-05 | 1.600E-05 |
| 18255 | 913 | N913P | C | 1e-05 | 8.6e-06 | 1.1e-05 | 9.867E-06 | 1.206E-06 | 8.600E-06 | 1.100E-05 |
| 18256 | 913 | N913S | C | 1.3e-05 | 1.1e-05 | 1.4e-05 | 1.267E-05 | 1.528E-06 | 1.100E-05 | 1.400E-05 |
| 18257 | 913 | N913T | C | 9.7e-06 | 9.1e-06 | 1.4e-05 | 1.093E-05 | 2.673E-06 | 9.100E-06 | 1.400E-05 |
| 18258 | 913 | N913W | C | 1.1e-05 | 8.3e-06 | 1.4e-05 | 1.110E-05 | 2.851E-06 | 8.300E-06 | 1.400E-05 |
| 18259 | 913 | N913Y | C | 8.1e-06 | 9.7e-06 | 1.6e-05 | 1.127E-05 | 4.177E-06 | 8.100E-06 | 1.600E-05 |
| 18260 | 913 | N913V | C | 1.1e-05 | 1.7e-05 | 1.6e-05 | 1.467E-05 | 3.215E-06 | 1.100E-05 | 1.700E-05 |
| 18261 | 914 | T914A | C | 1.7e-05 | 1.7e-05 | 8.7e-06 | 1.423E-05 | 4.792E-06 | 8.700E-06 | 1.700E-05 |
| 18262 | 914 | T914R | C | 1.6e-05 | 1.2e-05 | 1.6e-05 | 1.467E-05 | 2.309E-06 | 1.200E-05 | 1.600E-05 |
| 18263 | 914 | T914N | C | 8.7e-06 | 1.1e-05 | 9.4e-06 | 9.700E-06 | 1.179E-06 | 8.700E-06 | 1.100E-05 |
| 18264 | 914 | T914D | C | 1.1e-05 | 6.6e-06 | 1.1e-05 | 9.533E-06 | 2.540E-06 | 6.600E-06 | 1.100E-05 |
| 18265 | 914 | T914C | C | 1e-05 | 1.1e-05 | 1.5e-05 | 1.200E-05 | 2.646E-06 | 1.000E-05 | 1.500E-05 |
| 18266 | 914 | T914E | C | 1.1e-05 | 1e-05 | 1.2e-05 | 1.100E-05 | 1.000E-06 | 1.000E-05 | 1.200E-05 |
| 18267 | 914 | T914Q | C | 9.1e-06 | 1.7e-05 | 1.2e-05 | 1.270E-05 | 3.996E-06 | 9.100E-06 | 1.700E-05 |
| 18268 | 914 | T914G | C | 1e-05 | 1.3e-05 | 1e-05 | 1.100E-05 | 1.732E-06 | 1.000E-05 | 1.300E-05 |
| 18269 | 914 | T914H | C | 1.6e-05 | 1.1e-05 | 1.4e-05 | 1.367E-05 | 2.517E-06 | 1.100E-05 | 1.600E-05 |
| 18270 | 914 | T914I | C | 9.4e-06 | 1.5e-05 | 1.1e-05 | 1.180E-05 | 2.884E-06 | 9.400E-06 | 1.500E-05 |
| 18271 | 914 | T914L | C | 1.3e-05 | 1.8e-05 | 1.1e-05 | 1.400E-05 | 3.606E-06 | 1.100E-05 | 1.800E-05 |
| 18272 | 914 | T914K | C | 1.1e-05 | 1.3e-05 | 1.4e-05 | 1.267E-05 | 1.528E-06 | 1.100E-05 | 1.400E-05 |
| 18273 | 914 | T914M | C | 1.7e-05 | 1e-05 | 1.3e-05 | 1.333E-05 | 3.512E-06 | 1.000E-05 | 1.700E-05 |
| 18274 | 914 | T914F | C | 1.4e-05 | 1.3e-05 | 1.2e-05 | 1.300E-05 | 1.000E-06 | 1.200E-05 | 1.400E-05 |

| | | | | | | | | | | |
|---|---|---|---|---|---|---|---|---|---|---|
| 18275 | 914 | T914P | C | 1.1e-05 | 1.3e-05 | 1.5e-05 | 1.300E-05 | 2.000E-06 | 1.100E-05 | 1.500E-05 |
| 18276 | 914 | T914S | C | 1.3e-05 | 1.3e-05 | 2.3e-05 | 1.633E-05 | 5.774E-06 | 1.300E-05 | 2.300E-05 |
| 18277 | 914 | T914T | C | 1.2e-05 | 1.1e-05 | 1.4e-05 | 1.233E-05 | 1.528E-06 | 1.100E-05 | 1.400E-05 |
| 18278 | 914 | T914W | C | 6e-06 | 1e-05 | 9.6e-06 | 8.533E-06 | 2.203E-06 | 6.000E-06 | 1.000E-05 |
| 18279 | 914 | T914Y | C | 1.5e-05 | 1.5e-05 | 9.5e-06 | 1.317E-05 | 3.175E-06 | 9.500E-06 | 1.500E-05 |
| 18280 | 914 | T914V | C | 8.7e-06 | 9.3e-06 | 1.3e-05 | 1.033E-05 | 2.329E-06 | 8.700E-06 | 1.300E-05 |
| 18281 | 915 | L915A | C | 8.7e-06 | 1.7e-05 | 1.1e-05 | 1.223E-05 | 4.285E-06 | 8.700E-06 | 1.700E-05 |
| 18282 | 915 | L915R | C | 1.5e-05 | 1.4e-05 | 9.5e-06 | 1.283E-05 | 2.930E-06 | 9.500E-06 | 1.500E-05 |
| 18283 | 915 | L915N | C | 1.2e-05 | 1.2e-05 | 8.8e-06 | 1.093E-05 | 1.848E-06 | 8.800E-06 | 1.200E-05 |
| 18284 | 915 | L915D | C | 8.6e-06 | 1.7e-05 | 5.4e-06 | 1.033E-05 | 5.991E-06 | 5.400E-06 | 1.700E-05 |
| 18285 | 915 | L915C | C | 1.1e-05 | 1.4e-05 | 1.1e-05 | 1.200E-05 | 1.732E-06 | 1.100E-05 | 1.400E-05 |
| 18286 | 915 | L915E | C | 1.5e-05 | 8.3e-06 | 1e-05 | 1.110E-05 | 3.483E-06 | 8.300E-06 | 1.500E-05 |
| 18287 | 915 | L915Q | C | 1.6e-05 | 1.4e-05 | 1.4e-05 | 1.467E-05 | 1.155E-06 | 1.400E-05 | 1.600E-05 |
| 18288 | 915 | L915G | C | 1.6e-05 | 1.1e-05 | 1.4e-05 | 1.367E-05 | 2.517E-06 | 1.100E-05 | 1.600E-05 |
| 18289 | 915 | L915H | C | 1.2e-05 | 1.4e-05 | 1.1e-05 | 1.233E-05 | 1.528E-06 | 1.100E-05 | 1.400E-05 |
| 18290 | 915 | L915I | C | 9.2e-06 | 1.7e-05 | 1.6e-05 | 1.407E-05 | 4.244E-06 | 9.200E-06 | 1.700E-05 |
| 18291 | 915 | L915L | C | 1.4e-05 | 1.5e-05 | 1.5e-05 | 1.467E-05 | 5.774E-07 | 1.400E-05 | 1.500E-05 |
| 18292 | 915 | L915K | C | 1.3e-05 | 1.5e-05 | 1.1e-05 | 1.300E-05 | 2.000E-06 | 1.100E-05 | 1.500E-05 |
| 18293 | 915 | L915M | C | 1.1e-05 | 1e-05 | 1.1e-05 | 1.067E-05 | 5.774E-07 | 1.000E-05 | 1.100E-05 |
| 18294 | 915 | L915F | C | 1.7e-05 | 1.1e-05 | 1.7e-05 | 1.500E-05 | 3.464E-06 | 1.100E-05 | 1.700E-05 |
| 18295 | 915 | L915P | C | 1.1e-05 | 1.1e-05 | 1.5e-05 | 1.233E-05 | 2.309E-06 | 1.100E-05 | 1.500E-05 |
| 18296 | 915 | L915S | C | 1e-05 | 9.4e-06 | 1.2e-05 | 1.047E-05 | 1.361E-06 | 9.400E-06 | 1.200E-05 |
| 18297 | 915 | L915T | C | 1.3e-05 | 1.5e-05 | 1.4e-05 | 1.400E-05 | 1.000E-06 | 1.300E-05 | 1.500E-05 |
| 18298 | 915 | L915W | C | 1.3e-05 | 1.6e-05 | 9.3e-06 | 1.277E-05 | 3.356E-06 | 9.300E-06 | 1.600E-05 |
| 18299 | 915 | L915Y | C | 1.4e-05 | 1.2e-05 | 1.6e-05 | 1.400E-05 | 2.000E-06 | 1.200E-05 | 1.600E-05 |
| 18300 | 915 | L915V | C | 1.5e-05 | 1e-05 | 9.5e-06 | 1.150E-05 | 3.041E-06 | 9.500E-06 | 1.500E-05 |
| 18301 | 916 | Y916A | C | 1.4e-05 | 1.9e-05 | 1.7e-05 | 1.667E-05 | 2.517E-06 | 1.400E-05 | 1.900E-05 |
| 18302 | 916 | Y916R | C | 1.4e-05 | 1.6e-05 | 1.6e-05 | 1.533E-05 | 1.155E-06 | 1.400E-05 | 1.600E-05 |
| 18303 | 916 | Y916N | C | 1.4e-05 | 1e-05 | 1.4e-05 | 1.267E-05 | 2.309E-06 | 1.000E-05 | 1.400E-05 |
| 18304 | 916 | Y916D | C | 9.2e-06 | 1.5e-05 | 1e-05 | 1.140E-05 | 3.143E-06 | 9.200E-06 | 1.500E-05 |
| 18305 | 916 | Y916C | C | 8.5e-06 | 1.4e-05 | 9.4e-06 | 1.063E-05 | 2.950E-06 | 8.500E-06 | 1.400E-05 |
| 18306 | 916 | Y916E | C | 9.9e-06 | 1.5e-05 | 1.7e-05 | 1.397E-05 | 3.661E-06 | 9.900E-06 | 1.700E-05 |
| 18307 | 916 | Y916Q | C | 1.6e-05 | 1.1e-05 | 1.2e-05 | 1.300E-05 | 2.646E-06 | 1.100E-05 | 1.600E-05 |
| 18308 | 916 | Y916G | C | 1.6e-05 | 1.3e-05 | 1.5e-05 | 1.467E-05 | 1.528E-06 | 1.300E-05 | 1.600E-05 |
| 18309 | 916 | Y916H | C | 1e-05 | 1.2e-05 | 1.5e-05 | 1.233E-05 | 2.517E-06 | 1.000E-05 | 1.500E-05 |
| 18310 | 916 | Y916I | C | 1.5e-05 | 1.6e-05 | 1.1e-05 | 1.400E-05 | 2.646E-06 | 1.100E-05 | 1.600E-05 |
| 18311 | 916 | Y916L | C | 1.5e-05 | 1.4e-05 | 1.6e-05 | 1.500E-05 | 1.000E-06 | 1.400E-05 | 1.600E-05 |
| 18312 | 916 | Y916K | C | 1.6e-05 | 1.7e-05 | 1.3e-05 | 1.533E-05 | 2.082E-06 | 1.300E-05 | 1.700E-05 |
| 18313 | 916 | Y916M | C | 1.5e-05 | 9.6e-06 | 1.7e-05 | 1.387E-05 | 3.828E-06 | 9.600E-06 | 1.700E-05 |
| 18314 | 916 | Y916F | C | 1.4e-05 | 1.5e-05 | 1.7e-05 | 1.533E-05 | 1.528E-06 | 1.400E-05 | 1.700E-05 |
| 18315 | 916 | Y916P | C | 1e-05 | 1.2e-05 | 1.6e-05 | 1.267E-05 | 3.055E-06 | 1.000E-05 | 1.600E-05 |
| 18316 | 916 | Y916S | C | 9.5e-06 | 1.1e-05 | 1.3e-05 | 1.117E-05 | 1.756E-06 | 9.500E-06 | 1.300E-05 |
| 18317 | 916 | Y916T | C | 1.5e-05 | 8.6e-06 | 1.3e-05 | 1.220E-05 | 3.274E-06 | 8.600E-06 | 1.500E-05 |
| 18318 | 916 | Y916W | C | 1.5e-05 | 1.1e-05 | 1.6e-05 | 1.400E-05 | 2.646E-06 | 1.100E-05 | 1.600E-05 |
| 18319 | 916 | Y916Y | C | 1.2e-05 | 1e-05 | 1.5e-05 | 1.233E-05 | 2.517E-06 | 1.000E-05 | 1.500E-05 |
| 18320 | 916 | Y916V | C | 1.1e-05 | 1.5e-05 | 1e-05 | 1.200E-05 | 2.646E-06 | 1.000E-05 | 1.500E-05 |
| 18321 | 917 | L917A | C | 1.6e-05 | 1.5e-05 | 1.1e-05 | 1.400E-05 | 2.646E-06 | 1.100E-05 | 1.600E-05 |
| 18322 | 917 | L917R | C | 1.6e-05 | 1.3e-05 | 1.4e-05 | 1.433E-05 | 1.528E-06 | 1.300E-05 | 1.600E-05 |
| 18323 | 917 | L917N | C | 1.1e-05 | 1.7e-05 | 1.1e-05 | 1.300E-05 | 3.464E-06 | 1.100E-05 | 1.700E-05 |
| 18324 | 917 | L917D | C | 1e-05 | 1e-05 | 1.5e-05 | 1.167E-05 | 2.887E-06 | 1.000E-05 | 1.500E-05 |
| 18325 | 917 | L917C | C | 1.6e-05 | 1e-05 | 1.5e-05 | 1.367E-05 | 3.215E-06 | 1.000E-05 | 1.600E-05 |
| 18326 | 917 | L917E | C | 1e-05 | 1e-05 | 1.1e-05 | 1.033E-05 | 5.774E-07 | 1.000E-05 | 1.100E-05 |
| 18327 | 917 | L917Q | C | 1.2e-05 | 8e-06 | 8.5e-06 | 9.500E-06 | 2.179E-06 | 8.000E-06 | 1.200E-05 |
| 18328 | 917 | L917G | C | 1.5e-05 | 1.5e-05 | 1.4e-05 | 1.467E-05 | 5.774E-07 | 1.400E-05 | 1.500E-05 |
| 18329 | 917 | L917H | C | 1.3e-05 | 1.4e-05 | 1.1e-05 | 1.267E-05 | 1.528E-06 | 1.100E-05 | 1.400E-05 |
| 18330 | 917 | L917I | C | 1e-05 | 1.5e-05 | 1.5e-05 | 1.333E-05 | 2.887E-06 | 1.000E-05 | 1.500E-05 |
| 18331 | 917 | L917L | C | 1e-05 | 1.2e-05 | 9.6e-06 | 1.053E-05 | 1.286E-06 | 9.600E-06 | 1.200E-05 |
| 18332 | 917 | L917K | C | 1.2e-05 | 1.1e-05 | 8.1e-06 | 1.037E-05 | 2.026E-06 | 8.100E-06 | 1.200E-05 |
| 18333 | 917 | L917M | C | 8.7e-06 | 8.7e-06 | 1.5e-05 | 1.080E-05 | 3.637E-06 | 8.700E-06 | 1.500E-05 |
| 18334 | 917 | L917F | C | 1e-05 | 1.2e-05 | 1.1e-05 | 1.100E-05 | 1.000E-06 | 1.000E-05 | 1.200E-05 |
| 18335 | 917 | L917P | C | 1.7e-05 | 1e-05 | 1.5e-05 | 1.400E-05 | 3.606E-06 | 1.000E-05 | 1.700E-05 |
| 18336 | 917 | L917S | C | 1.1e-05 | 1.4e-05 | 1e-05 | 1.167E-05 | 2.082E-06 | 1.000E-05 | 1.400E-05 |
| 18337 | 917 | L917T | C | 1.5e-05 | 1.6e-05 | 1.1e-05 | 1.400E-05 | 2.646E-06 | 1.100E-05 | 1.600E-05 |
| 18338 | 917 | L917W | C | 1.1e-05 | 1.2e-05 | 1.4e-05 | 1.233E-05 | 1.528E-06 | 1.100E-05 | 1.400E-05 |
| 18339 | 917 | L917Y | C | 1.5e-05 | 9.6e-06 | 8.7e-06 | 1.110E-05 | 3.407E-06 | 8.700E-06 | 1.500E-05 |
| 18340 | 917 | L917V | C | 1.1e-05 | 9.8e-06 | 1.4e-05 | 1.160E-05 | 2.163E-06 | 9.800E-06 | 1.400E-05 |
| 18341 | 918 | Q918A | C | 1.2e-05 | 7.6e-06 | 1.5e-05 | 1.153E-05 | 3.722E-06 | 7.600E-06 | 1.500E-05 |
| 18342 | 918 | Q918R | C | 1.2e-05 | 8.5e-06 | 1.6e-05 | 1.217E-05 | 3.753E-06 | 8.500E-06 | 1.600E-05 |
| 18343 | 918 | Q918N | C | 9.1e-06 | 9.7e-06 | 1.2e-05 | 1.027E-05 | 1.531E-06 | 9.100E-06 | 1.200E-05 |
| 18344 | 918 | Q918D | C | 9.4e-06 | 1.5e-05 | 9.5e-06 | 1.130E-05 | 3.205E-06 | 9.400E-06 | 1.500E-05 |
| 18345 | 918 | Q918C | C | 1.5e-05 | 1.2e-05 | 1.2e-05 | 1.300E-05 | 1.732E-06 | 1.200E-05 | 1.500E-05 |
| 18346 | 918 | Q918E | C | 1.4e-05 | 1.3e-05 | 1.1e-05 | 1.267E-05 | 1.528E-06 | 1.100E-05 | 1.400E-05 |
| 18347 | 918 | Q918Q | C | 2e-05 | 1.3e-05 | 1e-05 | 1.433E-05 | 5.132E-06 | 1.000E-05 | 2.000E-05 |
| 18348 | 918 | Q918G | C | 8e-06 | 1.5e-05 | 1.4e-05 | 1.233E-05 | 3.786E-06 | 8.000E-06 | 1.500E-05 |
| 18349 | 918 | Q918H | C | 1e-05 | 1.7e-05 | 9.7e-06 | 1.223E-05 | 4.131E-06 | 9.700E-06 | 1.700E-05 |
| 18350 | 918 | Q918I | C | 1.2e-05 | 1.5e-05 | 1.5e-05 | 1.400E-05 | 1.732E-06 | 1.200E-05 | 1.500E-05 |
| 18351 | 918 | Q918L | C | 1.1e-05 | 1e-05 | 1.4e-05 | 1.167E-05 | 2.082E-06 | 1.000E-05 | 1.400E-05 |
| 18352 | 918 | Q918K | C | 1e-05 | 1e-05 | 1.3e-05 | 1.100E-05 | 1.732E-06 | 1.000E-05 | 1.300E-05 |
| 18353 | 918 | Q918M | C | 1.7e-05 | 1.4e-05 | 1.8e-05 | 1.633E-05 | 2.082E-06 | 1.400E-05 | 1.800E-05 |
| 18354 | 918 | Q918F | C | 8.8e-06 | 1.5e-05 | 1.5e-05 | 1.293E-05 | 3.580E-06 | 8.800E-06 | 1.500E-05 |
| 18355 | 918 | Q918P | C | 1.5e-05 | 1.3e-05 | 1.1e-05 | 1.300E-05 | 2.000E-06 | 1.100E-05 | 1.500E-05 |
| 18356 | 918 | Q918S | C | 9.6e-06 | 1.2e-05 | 1.4e-05 | 1.187E-05 | 2.203E-06 | 9.600E-06 | 1.400E-05 |
| 18357 | 918 | Q918T | C | 1.4e-05 | 1.3e-05 | 1.5e-05 | 1.400E-05 | 1.000E-06 | 1.300E-05 | 1.500E-05 |
| 18358 | 918 | Q918W | C | 1.5e-05 | 1.4e-05 | 1.2e-05 | 1.367E-05 | 1.528E-06 | 1.200E-05 | 1.500E-05 |
| 18359 | 918 | Q918Y | C | 1.4e-05 | 8.2e-06 | 1.5e-05 | 1.240E-05 | 3.672E-06 | 8.200E-06 | 1.500E-05 |

| | | | | | | | | | | |
|---|---|---|---|---|---|---|---|---|---|---|
| 18360 | 918 | Q918V | C | 1.5e-05 | 1.5e-05 | 9.8e-06 | 1.327E-05 | 3.002E-06 | 9.800E-06 | 1.500E-05 |
| 18361 | 919 | M919A | C | 1.5e-05 | 1.6e-05 | 1.4e-05 | 1.500E-05 | 1.000E-06 | 1.400E-05 | 1.600E-05 |
| 18362 | 919 | M919R | C | 9.2e-06 | 1.1e-05 | 8.5e-06 | 9.567E-06 | 1.290E-06 | 8.500E-06 | 1.100E-05 |
| 18363 | 919 | M919N | C | 1.2e-05 | 8.2e-06 | 6.5e-06 | 8.900E-06 | 2.816E-06 | 6.500E-06 | 1.200E-05 |
| 18364 | 919 | M919D | C | 1.2e-05 | 1.1e-05 | 1.8e-05 | 1.367E-05 | 3.786E-06 | 1.100E-05 | 1.800E-05 |
| 18365 | 919 | M919C | C | 1.6e-05 | 1.5e-05 | 1.3e-05 | 1.467E-05 | 1.528E-06 | 1.300E-05 | 1.600E-05 |
| 18366 | 919 | M919E | C | 1.1e-05 | 1.7e-05 | 8.9e-06 | 1.230E-05 | 4.204E-06 | 8.900E-06 | 1.700E-05 |
| 18367 | 919 | M919Q | C | 7.9e-06 | 1.3e-05 | 1.4e-05 | 1.163E-05 | 3.272E-06 | 7.900E-06 | 1.400E-05 |
| 18368 | 919 | M919G | C | 1e-05 | 1.6e-05 | 1.7e-05 | 1.433E-05 | 3.786E-06 | 1.000E-05 | 1.700E-05 |
| 18369 | 919 | M919H | C | 1.5e-05 | 1e-05 | 9.5e-06 | 1.150E-05 | 3.041E-06 | 9.500E-06 | 1.500E-05 |
| 18370 | 919 | M919I | C | 1.5e-05 | 1.2e-05 | 1.4e-05 | 1.367E-05 | 1.528E-06 | 1.200E-05 | 1.500E-05 |
| 18371 | 919 | M919L | C | 9e-06 | 7.8e-06 | 1.6e-05 | 1.093E-05 | 4.429E-06 | 7.800E-06 | 1.600E-05 |
| 18372 | 919 | M919K | C | 1.2e-05 | 1.6e-05 | 1.5e-05 | 1.433E-05 | 2.082E-06 | 1.200E-05 | 1.600E-05 |
| 18373 | 919 | M919M | C | 1.8e-05 | 1.1e-05 | 1.1e-05 | 1.333E-05 | 4.041E-06 | 1.100E-05 | 1.800E-05 |
| 18374 | 919 | M919F | C | 1e-05 | 1.6e-05 | 1.5e-05 | 1.367E-05 | 3.215E-06 | 1.000E-05 | 1.600E-05 |
| 18375 | 919 | M919P | C | 1.5e-05 | 1.7e-05 | 1.2e-05 | 1.467E-05 | 2.517E-06 | 1.200E-05 | 1.700E-05 |
| 18376 | 919 | M919S | C | 1.4e-05 | 1.7e-05 | 1e-05 | 1.367E-05 | 3.512E-06 | 1.000E-05 | 1.700E-05 |
| 18377 | 919 | M919T | C | 1.3e-05 | 1.5e-05 | 9.5e-06 | 1.250E-05 | 2.784E-06 | 9.500E-06 | 1.500E-05 |
| 18378 | 919 | M919W | C | 9.6e-06 | 1e-05 | 1.3e-05 | 1.087E-05 | 1.858E-06 | 9.600E-06 | 1.300E-05 |
| 18379 | 919 | M919Y | C | 1.4e-05 | 8.3e-06 | 9.3e-06 | 1.053E-05 | 3.044E-06 | 8.300E-06 | 1.400E-05 |
| 18380 | 919 | M919V | C | 8.8e-06 | 1.2e-05 | 1.4e-05 | 1.160E-05 | 2.623E-06 | 8.800E-06 | 1.400E-05 |
| 18381 | 920 | N920A | C | 1.4e-05 | 9.5e-06 | 1.3e-05 | 1.217E-05 | 2.363E-06 | 9.500E-06 | 1.400E-05 |
| 18382 | 920 | N920R | C | 1.1e-05 | 1.2e-05 | 8.9e-06 | 1.063E-05 | 1.582E-06 | 8.900E-06 | 1.200E-05 |
| 18383 | 920 | N920N | C | 9.6e-06 | 1.6e-05 | 8.7e-06 | 1.143E-05 | 3.980E-06 | 8.700E-06 | 1.600E-05 |
| 18384 | 920 | N920D | C | 1.6e-05 | 1.4e-05 | 1.1e-05 | 1.367E-05 | 2.517E-06 | 1.100E-05 | 1.600E-05 |
| 18385 | 920 | N920C | C | 1.6e-05 | 1.3e-05 | 8.9e-06 | 1.263E-05 | 3.564E-06 | 8.900E-06 | 1.600E-05 |
| 18386 | 920 | N920E | C | 1.2e-05 | 1.6e-05 | 9.6e-06 | 1.253E-05 | 3.233E-06 | 9.600E-06 | 1.600E-05 |
| 18387 | 920 | N920Q | C | 1.1e-05 | 1.5e-05 | 1.5e-05 | 1.367E-05 | 2.309E-06 | 1.100E-05 | 1.500E-05 |
| 18388 | 920 | N920G | C | 1.6e-05 | 1.6e-05 | 1.1e-05 | 1.433E-05 | 2.887E-06 | 1.100E-05 | 1.600E-05 |
| 18389 | 920 | N920H | C | 1.1e-05 | 1e-05 | 1.5e-05 | 1.200E-05 | 2.646E-06 | 1.000E-05 | 1.500E-05 |
| 18390 | 920 | N920I | C | 8e-06 | 1.1e-05 | 1.3e-05 | 1.067E-05 | 2.517E-06 | 8.000E-06 | 1.300E-05 |
| 18391 | 920 | N920L | C | 1.1e-05 | 1.1e-05 | 1.2e-05 | 1.133E-05 | 5.774E-07 | 1.100E-05 | 1.200E-05 |
| 18392 | 920 | N920K | C | 1.3e-05 | 1.2e-05 | 1.3e-05 | 1.267E-05 | 5.774E-07 | 1.200E-05 | 1.300E-05 |
| 18393 | 920 | N920M | C | 1.7e-05 | 1.4e-05 | 8.1e-06 | 1.303E-05 | 4.528E-06 | 8.100E-06 | 1.700E-05 |
| 18394 | 920 | N920F | C | 1.5e-05 | 1.2e-05 | 1.2e-05 | 1.300E-05 | 1.732E-06 | 1.200E-05 | 1.500E-05 |
| 18395 | 920 | N920P | C | 1.4e-05 | 1e-05 | 1.7e-05 | 1.367E-05 | 3.512E-06 | 1.000E-05 | 1.700E-05 |
| 18396 | 920 | N920S | C | 1.6e-05 | 1.5e-05 | 1.4e-05 | 1.500E-05 | 1.000E-06 | 1.400E-05 | 1.600E-05 |
| 18397 | 920 | N920T | C | 8.9e-06 | 1.2e-05 | 1.1e-05 | 1.063E-05 | 1.582E-06 | 8.900E-06 | 1.200E-05 |
| 18398 | 920 | N920W | C | 8.7e-06 | 1.1e-05 | 1.5e-05 | 1.157E-05 | 3.188E-06 | 8.700E-06 | 1.500E-05 |
| 18399 | 920 | N920Y | C | 9.7e-06 | 1.4e-05 | 1.6e-05 | 1.323E-05 | 3.219E-06 | 9.700E-06 | 1.600E-05 |
| 18400 | 920 | N920V | C | 9.1e-06 | 1.4e-05 | 1.5e-05 | 1.270E-05 | 3.158E-06 | 9.100E-06 | 1.500E-05 |
| 18401 | 921 | S921A | C | 9.6e-06 | 1.5e-05 | 1.7e-05 | 1.387E-05 | 3.828E-06 | 9.600E-06 | 1.700E-05 |
| 18402 | 921 | S921R | C | 1.8e-05 | 1.7e-05 | 1.1e-05 | 1.533E-05 | 3.786E-06 | 1.100E-05 | 1.800E-05 |
| 18403 | 921 | S921N | C | 8.8e-06 | 1.1e-05 | 1.7e-05 | 1.227E-05 | 4.244E-06 | 8.800E-06 | 1.700E-05 |
| 18404 | 921 | S921D | C | 1e-05 | 1.6e-05 | 1.7e-05 | 1.433E-05 | 3.786E-06 | 1.000E-05 | 1.700E-05 |
| 18405 | 921 | S921C | C | 1.5e-05 | 1.2e-05 | 1.4e-05 | 1.367E-05 | 1.528E-06 | 1.200E-05 | 1.500E-05 |
| 18406 | 921 | S921E | C | 1e-05 | 2.3e-05 | 1.4e-05 | 1.567E-05 | 6.658E-06 | 1.000E-05 | 2.300E-05 |
| 18407 | 921 | S921Q | C | 1.8e-05 | 1.5e-05 | 1.7e-05 | 1.667E-05 | 1.528E-06 | 1.500E-05 | 1.800E-05 |
| 18408 | 921 | S921G | C | 8.6e-06 | 1.3e-05 | 1.5e-05 | 1.220E-05 | 3.274E-06 | 8.600E-06 | 1.500E-05 |
| 18409 | 921 | S921H | C | 1.6e-05 | 1.6e-05 | 1.6e-05 | 1.600E-05 | 0.000E+00 | 1.600E-05 | 1.600E-05 |
| 18410 | 921 | S921I | C | 1.4e-05 | 1.4e-05 | 1.6e-05 | 1.467E-05 | 1.155E-06 | 1.400E-05 | 1.600E-05 |
| 18411 | 921 | S921L | C | 1.4e-05 | 1.4e-05 | 1.4e-05 | 1.400E-05 | 0.000E+00 | 1.400E-05 | 1.400E-05 |
| 18412 | 921 | S921K | C | 1.5e-05 | 9.2e-06 | 1.5e-05 | 1.307E-05 | 3.349E-06 | 9.200E-06 | 1.500E-05 |
| 18413 | 921 | S921M | C | 9.9e-06 | 1.6e-05 | 8.9e-06 | 1.160E-05 | 3.843E-06 | 8.900E-06 | 1.600E-05 |
| 18414 | 921 | S921F | C | 1.3e-05 | 1.1e-05 | 1.1e-05 | 1.167E-05 | 1.155E-06 | 1.100E-05 | 1.300E-05 |
| 18415 | 921 | S921P | C | 1e-05 | 1.6e-05 | 9.2e-06 | 1.173E-05 | 3.717E-06 | 9.200E-06 | 1.600E-05 |
| 18416 | 921 | S921S | C | 7.5e-06 | 1.1e-05 | 1.7e-05 | 1.183E-05 | 4.805E-06 | 7.500E-06 | 1.700E-05 |
| 18417 | 921 | S921T | C | 1.1e-05 | 1.5e-05 | 1.1e-05 | 1.233E-05 | 2.309E-06 | 1.100E-05 | 1.500E-05 |
| 18418 | 921 | S921W | C | 1.6e-05 | 9.5e-06 | 9.2e-06 | 1.157E-05 | 3.842E-06 | 9.200E-06 | 1.600E-05 |
| 18419 | 921 | S921Y | C | 1.5e-05 | 9.8e-06 | 1.1e-05 | 1.193E-05 | 2.723E-06 | 9.800E-06 | 1.500E-05 |
| 18420 | 921 | S921V | C | 9.2e-06 | 1.7e-05 | 1.2e-05 | 1.273E-05 | 3.951E-06 | 9.200E-06 | 1.700E-05 |
| 18421 | 922 | L922A | C | 9.5e-06 | 1.5e-05 | 8e-06 | 1.083E-05 | 3.686E-06 | 8.000E-06 | 1.500E-05 |
| 18422 | 922 | L922R | C | 1.4e-05 | 1.6e-05 | 1.2e-05 | 1.400E-05 | 2.000E-06 | 1.200E-05 | 1.600E-05 |
| 18423 | 922 | L922N | C | 1.4e-05 | 1.7e-05 | 1.5e-05 | 1.533E-05 | 1.528E-06 | 1.400E-05 | 1.700E-05 |
| 18424 | 922 | L922D | C | 1e-05 | 1.6e-05 | 9.9e-06 | 1.197E-05 | 3.493E-06 | 9.900E-06 | 1.600E-05 |
| 18425 | 922 | L922C | C | 9.3e-06 | 1.4e-05 | 1.5e-05 | 1.277E-05 | 3.044E-06 | 9.300E-06 | 1.500E-05 |
| 18426 | 922 | L922E | C | 9.9e-06 | 1.6e-05 | 1.2e-05 | 1.263E-05 | 3.099E-06 | 9.900E-06 | 1.600E-05 |
| 18427 | 922 | L922Q | C | 1.6e-05 | 1.2e-05 | 1.6e-05 | 1.467E-05 | 2.309E-06 | 1.200E-05 | 1.600E-05 |
| 18428 | 922 | L922G | C | 1.5e-05 | 1.5e-05 | 1.6e-05 | 1.533E-05 | 5.774E-07 | 1.500E-05 | 1.600E-05 |
| 18429 | 922 | L922H | C | 1.4e-05 | 1e-05 | 1.4e-05 | 1.267E-05 | 2.309E-06 | 1.000E-05 | 1.400E-05 |
| 18430 | 922 | L922I | C | 1e-05 | 1.2e-05 | 1.3e-05 | 1.167E-05 | 1.528E-06 | 1.000E-05 | 1.300E-05 |
| 18431 | 922 | L922L | C | 8.1e-06 | 7.9e-06 | 1.5e-05 | 1.033E-05 | 4.043E-06 | 7.900E-06 | 1.500E-05 |
| 18432 | 922 | L922K | C | 1e-05 | 1e-05 | 1.4e-05 | 1.133E-05 | 2.309E-06 | 1.000E-05 | 1.400E-05 |
| 18433 | 922 | L922M | C | 8.7e-06 | 8.5e-06 | 1.1e-05 | 9.400E-06 | 1.389E-06 | 8.500E-06 | 1.100E-05 |
| 18434 | 922 | L922F | C | 1.2e-05 | 9.8e-06 | 9.9e-06 | 1.057E-05 | 1.242E-06 | 9.800E-06 | 1.200E-05 |
| 18435 | 922 | L922P | C | 1e-05 | 1.5e-05 | 8.9e-06 | 1.130E-05 | 3.251E-06 | 8.900E-06 | 1.500E-05 |
| 18436 | 922 | L922S | C | 1.3e-05 | 1.1e-05 | 1.5e-05 | 1.300E-05 | 2.000E-06 | 1.100E-05 | 1.500E-05 |
| 18437 | 922 | L922T | C | 1.4e-05 | 1e-05 | 1.1e-05 | 1.167E-05 | 2.082E-06 | 1.000E-05 | 1.400E-05 |
| 18438 | 922 | L922W | C | 1.4e-05 | 1.4e-05 | 1.2e-05 | 1.333E-05 | 1.155E-06 | 1.200E-05 | 1.400E-05 |
| 18439 | 922 | L922Y | C | 1.4e-05 | 1.4e-05 | 1e-05 | 1.267E-05 | 2.309E-06 | 1.000E-05 | 1.400E-05 |
| 18440 | 922 | L922V | C | 1.4e-05 | 1.5e-05 | 8.1e-06 | 1.237E-05 | 3.729E-06 | 8.100E-06 | 1.500E-05 |
| 18441 | 923 | R923A | C | 1.2e-05 | 1.1e-05 | 1.5e-05 | 1.267E-05 | 2.082E-06 | 1.100E-05 | 1.500E-05 |
| 18442 | 923 | R923R | C | 1.6e-05 | 9.1e-06 | 1e-05 | 1.170E-05 | 3.751E-06 | 9.100E-06 | 1.600E-05 |
| 18443 | 923 | R923N | C | 1.1e-05 | 9.7e-06 | 1.4e-05 | 1.157E-05 | 2.205E-06 | 9.700E-06 | 1.400E-05 |
| 18444 | 923 | R923D | C | 1.2e-05 | 6e-06 | 1.6e-05 | 1.133E-05 | 5.033E-06 | 6.000E-06 | 1.600E-05 |

| | | | | | | | | | | |
|---|---|---|---|---|---|---|---|---|---|---|
| 18445 | 923 | R923C | C | 1.8e-05 | 9.1e-06 | 1e-05 | 1.237E-05 | 4.899E-06 | 9.100E-06 | 1.800E-05 |
| 18446 | 923 | R923E | C | 1.6e-05 | 1.6e-05 | 1.1e-05 | 1.433E-05 | 2.887E-06 | 1.100E-05 | 1.600E-05 |
| 18447 | 923 | R923Q | C | 1.4e-05 | 1.4e-05 | 9.4e-06 | 1.247E-05 | 2.656E-06 | 9.400E-06 | 1.400E-05 |
| 18448 | 923 | R923G | C | 9e-06 | 1.5e-05 | 1.3e-05 | 1.233E-05 | 3.055E-06 | 9.000E-06 | 1.500E-05 |
| 18449 | 923 | R923H | C | 1.5e-05 | 1.4e-05 | 1.5e-05 | 1.467E-05 | 5.774E-07 | 1.400E-05 | 1.500E-05 |
| 18450 | 923 | R923I | C | 1.5e-05 | 1.4e-05 | 1e-05 | 1.300E-05 | 2.646E-06 | 1.000E-05 | 1.500E-05 |
| 18451 | 923 | R923L | C | 1.2e-05 | 1.3e-05 | 1.5e-05 | 1.333E-05 | 1.528E-06 | 1.200E-05 | 1.500E-05 |
| 18452 | 923 | R923K | C | 1.5e-05 | 1.5e-05 | 1.5e-05 | 1.500E-05 | 0.000E+00 | 1.500E-05 | 1.500E-05 |
| 18453 | 923 | R923M | C | 1.2e-05 | 1.5e-05 | 1e-05 | 1.233E-05 | 2.517E-06 | 1.000E-05 | 1.500E-05 |
| 18454 | 923 | R923F | C | 1.4e-05 | 9.7e-06 | 1e-05 | 1.123E-05 | 2.401E-06 | 9.700E-06 | 1.400E-05 |
| 18455 | 923 | R923P | C | 1.1e-05 | 1.1e-05 | 9.9e-06 | 1.063E-05 | 6.351E-07 | 9.900E-06 | 1.100E-05 |
| 18456 | 923 | R923S | C | 8.6e-06 | 1.2e-05 | 1e-05 | 1.020E-05 | 1.709E-06 | 8.600E-06 | 1.200E-05 |
| 18457 | 923 | R923T | C | 1.6e-05 | 1.6e-05 | 9.5e-06 | 1.383E-05 | 3.753E-06 | 9.500E-06 | 1.600E-05 |
| 18458 | 923 | R923W | C | 8.8e-06 | 9.6e-06 | 9.1e-06 | 9.167E-06 | 4.041E-07 | 8.800E-06 | 9.600E-06 |
| 18459 | 923 | R923Y | C | 7.4e-06 | 9.9e-06 | 1.5e-05 | 1.077E-05 | 3.873E-06 | 7.400E-06 | 1.500E-05 |
| 18460 | 923 | R923V | C | 8.3e-06 | 9.9e-06 | 1.6e-05 | 1.140E-05 | 4.063E-06 | 8.300E-06 | 1.600E-05 |
| 18461 | 924 | A924A | C | 1.3e-05 | 1.5e-05 | 1.4e-05 | 1.400E-05 | 1.000E-06 | 1.300E-05 | 1.500E-05 |
| 18462 | 924 | A924R | C | 1.6e-05 | 1.2e-05 | 1.1e-05 | 1.300E-05 | 2.646E-06 | 1.100E-05 | 1.600E-05 |
| 18463 | 924 | A924N | C | 1.2e-05 | 1.4e-05 | 1.4e-05 | 1.333E-05 | 1.155E-06 | 1.200E-05 | 1.400E-05 |
| 18464 | 924 | A924D | C | 1.4e-05 | 8.2e-06 | 1.5e-05 | 1.240E-05 | 3.672E-06 | 8.200E-06 | 1.500E-05 |
| 18465 | 924 | A924C | C | 1.5e-05 | 6.9e-06 | 9.2e-06 | 1.037E-05 | 4.174E-06 | 6.900E-06 | 1.500E-05 |
| 18466 | 924 | A924E | C | 1.3e-05 | 1.4e-05 | 1.2e-05 | 1.300E-05 | 1.000E-06 | 1.200E-05 | 1.400E-05 |
| 18467 | 924 | A924Q | C | 1.2e-05 | 1.4e-05 | 7.5e-06 | 1.117E-05 | 3.329E-06 | 7.500E-06 | 1.400E-05 |
| 18468 | 924 | A924G | C | 1.5e-05 | 1.6e-05 | 1.1e-05 | 1.400E-05 | 2.646E-06 | 1.100E-05 | 1.600E-05 |
| 18469 | 924 | A924H | C | 1.4e-05 | 9.5e-06 | 7.5e-06 | 1.033E-05 | 3.329E-06 | 7.500E-06 | 1.400E-05 |
| 18470 | 924 | A924I | C | 1.6e-05 | 1.5e-05 | 1.4e-05 | 1.500E-05 | 1.000E-06 | 1.400E-05 | 1.600E-05 |
| 18471 | 924 | A924L | C | 1.3e-05 | 1.6e-05 | 1e-05 | 1.300E-05 | 3.000E-06 | 1.000E-05 | 1.600E-05 |
| 18472 | 924 | A924K | C | 1.1e-05 | 1.1e-05 | 7.6e-06 | 9.867E-06 | 1.963E-06 | 7.600E-06 | 1.100E-05 |
| 18473 | 924 | A924M | C | 9.5e-06 | 1.2e-05 | 1.1e-05 | 1.083E-05 | 1.258E-06 | 9.500E-06 | 1.200E-05 |
| 18474 | 924 | A924F | C | 1.4e-05 | 9.4e-06 | 1.1e-05 | 1.147E-05 | 2.335E-06 | 9.400E-06 | 1.400E-05 |
| 18475 | 924 | A924P | C | 1.4e-05 | 1.1e-05 | 1.2e-05 | 1.233E-05 | 1.528E-06 | 1.100E-05 | 1.400E-05 |
| 18476 | 924 | A924S | C | 1.5e-05 | 1.1e-05 | 1.4e-05 | 1.333E-05 | 2.082E-06 | 1.100E-05 | 1.500E-05 |
| 18477 | 924 | A924T | C | 1.6e-05 | 1.5e-05 | 6.6e-06 | 1.253E-05 | 5.163E-06 | 6.600E-06 | 1.600E-05 |
| 18478 | 924 | A924W | C | 1.1e-05 | 1.6e-05 | 1.5e-05 | 1.400E-05 | 2.646E-06 | 1.100E-05 | 1.600E-05 |
| 18479 | 924 | A924Y | C | 1.3e-05 | 8.6e-06 | 8.4e-06 | 1.000E-05 | 2.600E-06 | 8.400E-06 | 1.300E-05 |
| 18480 | 924 | A924V | C | 1.4e-05 | 9.6e-06 | 1.1e-05 | 1.153E-05 | 2.248E-06 | 9.600E-06 | 1.400E-05 |
| 18481 | 925 | E925A | C | 9e-06 | 1.5e-05 | 1.4e-05 | 1.267E-05 | 3.215E-06 | 9.000E-06 | 1.500E-05 |
| 18482 | 925 | E925R | C | 8e-06 | 1.4e-05 | 1.5e-05 | 1.233E-05 | 3.786E-06 | 8.000E-06 | 1.500E-05 |
| 18483 | 925 | E925N | C | 1.1e-05 | 9.8e-06 | 1.7e-05 | 1.260E-05 | 3.857E-06 | 9.800E-06 | 1.700E-05 |
| 18484 | 925 | E925D | C | 1.3e-05 | 1.2e-05 | 1.5e-05 | 1.333E-05 | 1.528E-06 | 1.200E-05 | 1.500E-05 |
| 18485 | 925 | E925C | C | 1.6e-05 | 1.5e-05 | 1.3e-05 | 1.467E-05 | 1.528E-06 | 1.300E-05 | 1.600E-05 |
| 18486 | 925 | E925E | C | 1.4e-05 | 1.1e-05 | 1.1e-05 | 1.200E-05 | 1.732E-06 | 1.100E-05 | 1.400E-05 |
| 18487 | 925 | E925Q | C | 8.8e-06 | 1.3e-05 | 9e-06 | 1.027E-05 | 2.369E-06 | 8.800E-06 | 1.300E-05 |
| 18488 | 925 | E925G | C | 8.9e-06 | 1.5e-05 | 1.4e-05 | 1.263E-05 | 3.272E-06 | 8.900E-06 | 1.500E-05 |
| 18489 | 925 | E925H | C | 1.3e-05 | 1e-05 | 1.6e-05 | 1.300E-05 | 3.000E-06 | 1.000E-05 | 1.600E-05 |
| 18490 | 925 | E925I | C | 2.5e-05 | 1.3e-05 | 1e-05 | 1.600E-05 | 7.937E-06 | 1.000E-05 | 2.500E-05 |
| 18491 | 925 | E925L | C | 1.1e-05 | 1.1e-05 | 1.1e-05 | 1.100E-05 | 2.075E-21 | 1.100E-05 | 1.100E-05 |
| 18492 | 925 | E925K | C | 1.6e-05 | 1.4e-05 | 9.9e-06 | 1.330E-05 | 3.110E-06 | 9.900E-06 | 1.600E-05 |
| 18493 | 925 | E925M | C | 9.8e-06 | 1.5e-05 | 1.5e-05 | 1.327E-05 | 3.002E-06 | 9.800E-06 | 1.500E-05 |
| 18494 | 925 | E925F | C | 1.1e-05 | 1.5e-05 | 1.3e-05 | 1.300E-05 | 2.000E-06 | 1.100E-05 | 1.500E-05 |
| 18495 | 925 | E925P | C | 1.5e-05 | 1.6e-05 | 1.6e-05 | 1.567E-05 | 5.774E-07 | 1.500E-05 | 1.600E-05 |
| 18496 | 925 | E925S | C | 8.7e-06 | 1.6e-05 | 1.4e-05 | 1.290E-05 | 3.772E-06 | 8.700E-06 | 1.600E-05 |
| 18497 | 925 | E925T | C | 1.8e-05 | 1.4e-05 | 1.6e-05 | 1.600E-05 | 2.000E-06 | 1.400E-05 | 1.800E-05 |
| 18498 | 925 | E925W | C | 9.7e-06 | 1.7e-05 | 1.4e-05 | 1.357E-05 | 3.669E-06 | 9.700E-06 | 1.700E-05 |
| 18499 | 925 | E925Y | C | 1.3e-05 | 1e-05 | 1.5e-05 | 1.267E-05 | 2.517E-06 | 1.000E-05 | 1.500E-05 |
| 18500 | 925 | E925V | C | 1.1e-05 | 1.1e-05 | 9.8e-06 | 1.060E-05 | 6.928E-07 | 9.800E-06 | 1.100E-05 |
| 18501 | 926 | D926A | C | 1.1e-05 | 1.1e-05 | 1.6e-05 | 1.267E-05 | 2.887E-06 | 1.100E-05 | 1.600E-05 |
| 18502 | 926 | D926R | C | 1.5e-05 | 1e-05 | 1.8e-05 | 1.433E-05 | 4.041E-06 | 1.000E-05 | 1.800E-05 |
| 18503 | 926 | D926N | C | 9.6e-06 | 1.1e-05 | 1.4e-05 | 1.153E-05 | 2.248E-06 | 9.600E-06 | 1.400E-05 |
| 18504 | 926 | D926D | C | 1.1e-05 | 1e-05 | 9.1e-06 | 1.003E-05 | 9.504E-07 | 9.100E-06 | 1.100E-05 |
| 18505 | 926 | D926C | C | 1.4e-05 | 1e-05 | 9.9e-06 | 1.130E-05 | 2.339E-06 | 9.900E-06 | 1.400E-05 |
| 18506 | 926 | D926E | C | 1.4e-05 | 1e-05 | 1.4e-05 | 1.267E-05 | 2.309E-06 | 1.000E-05 | 1.400E-05 |
| 18507 | 926 | D926Q | C | 1.5e-05 | 6.4e-06 | 1.1e-05 | 1.080E-05 | 4.303E-06 | 6.400E-06 | 1.500E-05 |
| 18508 | 926 | D926G | C | 1.6e-05 | 1.2e-05 | 8.5e-06 | 1.217E-05 | 3.753E-06 | 8.500E-06 | 1.600E-05 |
| 18509 | 926 | D926H | C | 1.5e-05 | 6.8e-06 | 1.1e-05 | 1.093E-05 | 4.100E-06 | 6.800E-06 | 1.500E-05 |
| 18510 | 926 | D926I | C | 1e-05 | 1.1e-05 | 1.3e-05 | 1.133E-05 | 1.528E-06 | 1.000E-05 | 1.300E-05 |
| 18511 | 926 | D926L | C | 9.8e-06 | 1e-05 | 1e-05 | 9.933E-06 | 1.155E-07 | 9.800E-06 | 1.000E-05 |
| 18512 | 926 | D926K | C | 1.5e-05 | 1.3e-05 | 1.2e-05 | 1.333E-05 | 1.528E-06 | 1.200E-05 | 1.500E-05 |
| 18513 | 926 | D926M | C | 1e-05 | 6.8e-06 | 1e-05 | 8.933E-06 | 1.848E-06 | 6.800E-06 | 1.000E-05 |
| 18514 | 926 | D926F | C | 1.2e-05 | 8.7e-06 | 1.2e-05 | 1.090E-05 | 1.905E-06 | 8.700E-06 | 1.200E-05 |
| 18515 | 926 | D926P | C | 1.6e-05 | 1e-05 | 1e-05 | 1.200E-05 | 3.464E-06 | 1.000E-05 | 1.600E-05 |
| 18516 | 926 | D926S | C | 1.1e-05 | 1.3e-05 | 1.4e-05 | 1.267E-05 | 1.528E-06 | 1.100E-05 | 1.400E-05 |
| 18517 | 926 | D926T | C | 1.1e-05 | 1.5e-05 | 1.3e-05 | 1.300E-05 | 2.000E-06 | 1.100E-05 | 1.500E-05 |
| 18518 | 926 | D926W | C | 2.2e-05 | 1.1e-05 | 1.3e-05 | 1.533E-05 | 5.859E-06 | 1.100E-05 | 2.200E-05 |
| 18519 | 926 | D926Y | C | 1.4e-05 | 9.9e-06 | 1.2e-05 | 1.197E-05 | 2.050E-06 | 9.900E-06 | 1.400E-05 |
| 18520 | 926 | D926V | C | 1.1e-05 | 1.4e-05 | 1.1e-05 | 1.200E-05 | 1.732E-06 | 1.100E-05 | 1.400E-05 |
| 18521 | 927 | T927A | C | 1.3e-05 | 1.1e-05 | 1.5e-05 | 1.300E-05 | 2.000E-06 | 1.100E-05 | 1.500E-05 |
| 18522 | 927 | T927R | C | 8.8e-06 | 9.7e-06 | 9.7e-06 | 9.400E-06 | 5.196E-07 | 8.800E-06 | 9.700E-06 |
| 18523 | 927 | T927N | C | 8.6e-06 | 1.6e-05 | 1.4e-05 | 1.287E-05 | 3.828E-06 | 8.600E-06 | 1.600E-05 |
| 18524 | 927 | T927D | C | 1.5e-05 | 1.2e-05 | 1.8e-05 | 1.500E-05 | 3.000E-06 | 1.200E-05 | 1.800E-05 |
| 18525 | 927 | T927C | C | 1.5e-05 | 1.5e-05 | 8.4e-06 | 1.280E-05 | 3.811E-06 | 8.400E-06 | 1.500E-05 |
| 18526 | 927 | T927E | C | 1e-05 | 1.4e-05 | 1.1e-05 | 1.167E-05 | 2.082E-06 | 1.000E-05 | 1.400E-05 |
| 18527 | 927 | T927Q | C | 9.6e-06 | 1.2e-05 | 1.4e-05 | 1.187E-05 | 2.203E-06 | 9.600E-06 | 1.400E-05 |
| 18528 | 927 | T927G | C | 1.1e-05 | 9.4e-06 | 1.1e-05 | 1.047E-05 | 9.238E-07 | 9.400E-06 | 1.100E-05 |
| 18529 | 927 | T927H | C | 1.5e-05 | 1.2e-05 | 1.4e-05 | 1.367E-05 | 1.528E-06 | 1.200E-05 | 1.500E-05 |

| | | | | | | | | | | |
|---|---|---|---|---|---|---|---|---|---|---|
| 18530 | 927 | T927I | C | 1.6e-05 | 1.6e-05 | 1.6e-05 | 1.600E-05 | 0.000E+00 | 1.600E-05 | 1.600E-05 |
| 18531 | 927 | T927L | C | 8.9e-06 | 1e-05 | 1.1e-05 | 9.967E-06 | 1.050E-06 | 8.900E-06 | 1.100E-05 |
| 18532 | 927 | T927K | C | 1.5e-05 | 7.3e-06 | 1.5e-05 | 1.243E-05 | 4.446E-06 | 7.300E-06 | 1.500E-05 |
| 18533 | 927 | T927M | C | 9.9e-06 | 1.7e-05 | 1.1e-05 | 1.263E-05 | 3.821E-06 | 9.900E-06 | 1.700E-05 |
| 18534 | 927 | T927F | C | 1.1e-05 | 1e-05 | 1.5e-05 | 1.200E-05 | 2.646E-06 | 1.000E-05 | 1.500E-05 |
| 18535 | 927 | T927P | C | 1.4e-05 | 1.6e-05 | 7.2e-06 | 1.240E-05 | 4.613E-06 | 7.200E-06 | 1.600E-05 |
| 18536 | 927 | T927S | C | 1e-05 | 1.3e-05 | 9.9e-06 | 1.097E-05 | 1.762E-06 | 9.900E-06 | 1.300E-05 |
| 18537 | 927 | T927T | C | 1.4e-05 | 1.5e-05 | 9.8e-06 | 1.293E-05 | 2.759E-06 | 9.800E-06 | 1.500E-05 |
| 18538 | 927 | T927W | C | 1.1e-05 | 1.4e-05 | 1e-05 | 1.167E-05 | 2.082E-06 | 1.000E-05 | 1.400E-05 |
| 18539 | 927 | T927Y | C | 1.6e-05 | 1.2e-05 | 1.1e-05 | 1.300E-05 | 2.646E-06 | 1.100E-05 | 1.600E-05 |
| 18540 | 927 | T927V | C | 9.6e-06 | 1.5e-05 | 1.6e-05 | 1.353E-05 | 3.443E-06 | 9.600E-06 | 1.600E-05 |
| 18541 | 928 | A928A | C | 9.9e-06 | 1.6e-05 | 1.3e-05 | 1.297E-05 | 3.050E-06 | 9.900E-06 | 1.600E-05 |
| 18542 | 928 | A928R | C | 8.6e-06 | 1.3e-05 | 1.4e-05 | 1.187E-05 | 2.873E-06 | 8.600E-06 | 1.400E-05 |
| 18543 | 928 | A928N | C | 9.4e-06 | 9.3e-06 | 1.3e-05 | 1.057E-05 | 2.108E-06 | 9.300E-06 | 1.300E-05 |
| 18544 | 928 | A928D | C | 1.4e-05 | 1.1e-05 | 1.5e-05 | 1.333E-05 | 2.082E-06 | 1.100E-05 | 1.500E-05 |
| 18545 | 928 | A928C | C | 9.7e-06 | 1.7e-05 | 1.7e-05 | 1.457E-05 | 4.215E-06 | 9.700E-06 | 1.700E-05 |
| 18546 | 928 | A928E | C | 1.4e-05 | 1.7e-05 | 1e-05 | 1.367E-05 | 3.512E-06 | 1.000E-05 | 1.700E-05 |
| 18547 | 928 | A928Q | C | 9.8e-06 | 2.5e-05 | 1.6e-05 | 1.693E-05 | 7.643E-06 | 9.800E-06 | 2.500E-05 |
| 18548 | 928 | A928G | C | 1.6e-05 | 1.2e-05 | 1.4e-05 | 1.400E-05 | 2.000E-06 | 1.200E-05 | 1.600E-05 |
| 18549 | 928 | A928H | C | 1.2e-05 | 1.1e-05 | 9.9e-06 | 1.097E-05 | 1.050E-06 | 9.900E-06 | 1.200E-05 |
| 18550 | 928 | A928I | C | 1.6e-05 | 8.7e-06 | 1.3e-05 | 1.257E-05 | 3.669E-06 | 8.700E-06 | 1.600E-05 |
| 18551 | 928 | A928L | C | 1e-05 | 1.2e-05 | 9.9e-06 | 1.063E-05 | 1.185E-06 | 9.900E-06 | 1.200E-05 |
| 18552 | 928 | A928K | C | 1.3e-05 | 8.8e-06 | 1.4e-05 | 1.193E-05 | 2.759E-06 | 8.800E-06 | 1.400E-05 |
| 18553 | 928 | A928M | C | 1.1e-05 | 1.3e-05 | 1.5e-05 | 1.300E-05 | 2.000E-06 | 1.100E-05 | 1.500E-05 |
| 18554 | 928 | A928F | C | 1.5e-05 | 1.5e-05 | 1.1e-05 | 1.367E-05 | 2.309E-06 | 1.100E-05 | 1.500E-05 |
| 18555 | 928 | A928P | C | 1.6e-05 | 9.5e-06 | 1e-05 | 1.183E-05 | 3.617E-06 | 9.500E-06 | 1.600E-05 |
| 18556 | 928 | A928S | C | 9.8e-06 | 1.6e-05 | 1.3e-05 | 1.293E-05 | 3.101E-06 | 9.800E-06 | 1.600E-05 |
| 18557 | 928 | A928T | C | 1.4e-05 | 1.1e-05 | 1.5e-05 | 1.333E-05 | 2.082E-06 | 1.100E-05 | 1.500E-05 |
| 18558 | 928 | A928W | C | 1.2e-05 | 1.6e-05 | 1.4e-05 | 1.400E-05 | 2.000E-06 | 1.200E-05 | 1.600E-05 |
| 18559 | 928 | A928Y | C | 1.5e-05 | 1.4e-05 | 1.6e-05 | 1.500E-05 | 1.000E-06 | 1.400E-05 | 1.600E-05 |
| 18560 | 928 | A928V | C | 1.4e-05 | 1.1e-05 | 1.5e-05 | 1.333E-05 | 2.082E-06 | 1.100E-05 | 1.500E-05 |
| 18561 | 929 | V929A | C | 1.5e-05 | 1.6e-05 | 1e-05 | 1.367E-05 | 3.215E-06 | 1.000E-05 | 1.600E-05 |
| 18562 | 929 | V929R | C | 1.1e-05 | 1.3e-05 | 8.3e-06 | 1.077E-05 | 2.359E-06 | 8.300E-06 | 1.300E-05 |
| 18563 | 929 | V929N | C | 1.1e-05 | 1.6e-05 | 1e-05 | 1.233E-05 | 3.215E-06 | 1.000E-05 | 1.600E-05 |
| 18564 | 929 | V929D | C | 1e-05 | 1.5e-05 | 8.2e-06 | 1.107E-05 | 3.523E-06 | 8.200E-06 | 1.500E-05 |
| 18565 | 929 | V929C | C | 1.2e-05 | 1.1e-05 | 9.2e-06 | 1.073E-05 | 1.419E-06 | 9.200E-06 | 1.200E-05 |
| 18566 | 929 | V929E | C | 1.5e-05 | 1.5e-05 | 1.4e-05 | 1.467E-05 | 5.774E-07 | 1.400E-05 | 1.500E-05 |
| 18567 | 929 | V929Q | C | 1.6e-05 | 8e-06 | 1.5e-05 | 1.300E-05 | 4.359E-06 | 8.000E-06 | 1.600E-05 |
| 18568 | 929 | V929G | C | 9.8e-06 | 1.3e-05 | 9.4e-06 | 1.073E-05 | 1.973E-06 | 9.400E-06 | 1.300E-05 |
| 18569 | 929 | V929H | C | 1.1e-05 | 1.1e-05 | 1e-05 | 1.067E-05 | 5.774E-07 | 1.000E-05 | 1.100E-05 |
| 18570 | 929 | V929I | C | 1.4e-05 | 1.6e-05 | 1e-05 | 1.333E-05 | 3.055E-06 | 1.000E-05 | 1.600E-05 |
| 18571 | 929 | V929L | C | 1.8e-05 | 1.3e-05 | 1.3e-05 | 1.467E-05 | 2.887E-06 | 1.300E-05 | 1.800E-05 |
| 18572 | 929 | V929K | C | 1.4e-05 | 1.6e-05 | 8.9e-06 | 1.297E-05 | 3.661E-06 | 8.900E-06 | 1.600E-05 |
| 18573 | 929 | V929M | C | 1.5e-05 | 1.2e-05 | 1.2e-05 | 1.300E-05 | 1.732E-06 | 1.200E-05 | 1.500E-05 |
| 18574 | 929 | V929F | C | 8.7e-06 | 1.5e-05 | 1.5e-05 | 1.290E-05 | 3.637E-06 | 8.700E-06 | 1.500E-05 |
| 18575 | 929 | V929P | C | 9.3e-06 | 8.9e-06 | 1.5e-05 | 1.107E-05 | 3.412E-06 | 8.900E-06 | 1.500E-05 |
| 18576 | 929 | V929S | C | 1.1e-05 | 9.8e-06 | 9.8e-06 | 1.020E-05 | 6.928E-07 | 9.800E-06 | 1.100E-05 |
| 18577 | 929 | V929T | C | 8.9e-06 | 1.6e-05 | 1.3e-05 | 1.263E-05 | 3.564E-06 | 8.900E-06 | 1.600E-05 |
| 18578 | 929 | V929W | C | 1.4e-05 | 7.4e-06 | 1.5e-05 | 1.213E-05 | 4.130E-06 | 7.400E-06 | 1.500E-05 |
| 18579 | 929 | V929Y | C | 1.1e-05 | 1.6e-05 | 9.6e-06 | 1.220E-05 | 3.365E-06 | 9.600E-06 | 1.600E-05 |
| 18580 | 929 | V929V | C | 1e-05 | 1.2e-05 | 1.5e-05 | 1.233E-05 | 2.517E-06 | 1.000E-05 | 1.500E-05 |
| 18581 | 930 | Y930A | C | 9.9e-06 | 1.2e-05 | 9.7e-06 | 1.053E-05 | 1.274E-06 | 9.700E-06 | 1.200E-05 |
| 18582 | 930 | Y930R | C | 8.5e-06 | 1.5e-05 | 1.4e-05 | 1.250E-05 | 3.500E-06 | 8.500E-06 | 1.500E-05 |
| 18583 | 930 | Y930N | C | 9.9e-06 | 1.1e-05 | 9.1e-06 | 1.000E-05 | 9.539E-07 | 9.100E-06 | 1.100E-05 |
| 18584 | 930 | Y930D | C | 1.1e-05 | 1.7e-05 | 1.2e-05 | 1.333E-05 | 3.215E-06 | 1.100E-05 | 1.700E-05 |
| 18585 | 930 | Y930C | C | 1.7e-05 | 1.5e-05 | 1.2e-05 | 1.467E-05 | 2.517E-06 | 1.200E-05 | 1.700E-05 |
| 18586 | 930 | Y930E | C | 1.8e-05 | 1e-05 | 9.6e-06 | 1.253E-05 | 4.738E-06 | 9.600E-06 | 1.800E-05 |
| 18587 | 930 | Y930Q | C | 1.5e-05 | 9.5e-06 | 5.8e-06 | 1.010E-05 | 4.629E-06 | 5.800E-06 | 1.500E-05 |
| 18588 | 930 | Y930G | C | 1.7e-05 | 7.7e-06 | 1.4e-05 | 1.290E-05 | 4.747E-06 | 7.700E-06 | 1.700E-05 |
| 18589 | 930 | Y930H | C | 1.5e-05 | 1.5e-05 | 8.6e-06 | 1.287E-05 | 3.695E-06 | 8.600E-06 | 1.500E-05 |
| 18590 | 930 | Y930I | C | 1.4e-05 | 9.2e-06 | 1.7e-05 | 1.340E-05 | 3.934E-06 | 9.200E-06 | 1.700E-05 |
| 18591 | 930 | Y930L | C | 1e-05 | 1.3e-05 | 1.8e-05 | 1.367E-05 | 4.041E-06 | 1.000E-05 | 1.800E-05 |
| 18592 | 930 | Y930K | C | 1.6e-05 | 9.7e-06 | 1e-05 | 1.190E-05 | 3.554E-06 | 9.700E-06 | 1.600E-05 |
| 18593 | 930 | Y930M | C | 1.8e-05 | 1e-05 | 1.1e-05 | 1.300E-05 | 4.359E-06 | 1.000E-05 | 1.800E-05 |
| 18594 | 930 | Y930F | C | 1.5e-05 | 1.5e-05 | 1.1e-05 | 1.367E-05 | 2.309E-06 | 1.100E-05 | 1.500E-05 |
| 18595 | 930 | Y930P | C | 1.6e-05 | 1e-05 | 1.7e-05 | 1.433E-05 | 3.786E-06 | 1.000E-05 | 1.700E-05 |
| 18596 | 930 | Y930S | C | 1.1e-05 | 1.3e-05 | 1.6e-05 | 1.333E-05 | 2.517E-06 | 1.100E-05 | 1.600E-05 |
| 18597 | 930 | Y930T | C | 1.5e-05 | 1.1e-05 | 1.5e-05 | 1.367E-05 | 2.309E-06 | 1.100E-05 | 1.500E-05 |
| 18598 | 930 | Y930W | C | 1.7e-05 | 1.3e-05 | 9.8e-06 | 1.327E-05 | 3.607E-06 | 9.800E-06 | 1.700E-05 |
| 18599 | 930 | Y930Y | C | 1.6e-05 | 1.4e-05 | 1.5e-05 | 1.500E-05 | 1.000E-06 | 1.400E-05 | 1.600E-05 |
| 18600 | 930 | Y930V | C | 1.3e-05 | 1.6e-05 | 7.6e-06 | 1.220E-05 | 4.257E-06 | 7.600E-06 | 1.600E-05 |
| 18601 | 931 | Y931A | C | 1e-05 | 1.2e-05 | 1.6e-05 | 1.267E-05 | 3.055E-06 | 1.000E-05 | 1.600E-05 |
| 18602 | 931 | Y931R | C | 1.5e-05 | 1.5e-05 | 1.6e-05 | 1.533E-05 | 5.774E-07 | 1.500E-05 | 1.600E-05 |
| 18603 | 931 | Y931N | C | 1.1e-05 | 1.6e-05 | 1.6e-05 | 1.433E-05 | 2.887E-06 | 1.100E-05 | 1.600E-05 |
| 18604 | 931 | Y931D | C | 9.2e-06 | 1.3e-05 | 1.6e-05 | 1.273E-05 | 3.408E-06 | 9.200E-06 | 1.600E-05 |
| 18605 | 931 | Y931C | C | 1.2e-05 | 1.1e-05 | 1.7e-05 | 1.333E-05 | 3.215E-06 | 1.100E-05 | 1.700E-05 |
| 18606 | 931 | Y931E | C | 1.1e-05 | 1.5e-05 | 1.7e-05 | 1.433E-05 | 3.055E-06 | 1.100E-05 | 1.700E-05 |
| 18607 | 931 | Y931Q | C | 1.6e-05 | 1e-05 | 1.1e-05 | 1.233E-05 | 3.215E-06 | 1.000E-05 | 1.600E-05 |
| 18608 | 931 | Y931G | C | 1.2e-05 | 8.5e-06 | 1.5e-05 | 1.183E-05 | 3.253E-06 | 8.500E-06 | 1.500E-05 |
| 18609 | 931 | Y931H | C | 1.6e-05 | 1.6e-05 | 1.7e-05 | 1.633E-05 | 5.774E-07 | 1.600E-05 | 1.700E-05 |
| 18610 | 931 | Y931I | C | 1.6e-05 | 1.7e-05 | 1e-05 | 1.433E-05 | 3.786E-06 | 1.000E-05 | 1.700E-05 |
| 18611 | 931 | Y931L | C | 8e-06 | 1.1e-05 | 1e-05 | 9.667E-06 | 1.528E-06 | 8.000E-06 | 1.100E-05 |
| 18612 | 931 | Y931K | C | 1.1e-05 | 9e-06 | 1e-05 | 1.000E-05 | 1.000E-06 | 9.000E-06 | 1.100E-05 |
| 18613 | 931 | Y931M | C | 9.9e-06 | 1.2e-05 | 1.3e-05 | 1.163E-05 | 1.582E-06 | 9.900E-06 | 1.300E-05 |
| 18614 | 931 | Y931F | C | 1.8e-05 | 2.4e-05 | 1.4e-05 | 1.867E-05 | 5.033E-06 | 1.400E-05 | 2.400E-05 |

| | | | | | | | | | | |
|---|---|---|---|---|---|---|---|---|---|---|
| 18615 | 931 | Y931P | C | 1.5e-05 | 1.6e-05 | 1.2e-05 | 1.433E-05 | 2.082E-06 | 1.200E-05 | 1.600E-05 |
| 18616 | 931 | Y931S | C | 1.6e-05 | 1.6e-05 | 1e-05 | 1.400E-05 | 3.464E-06 | 1.000E-05 | 1.600E-05 |
| 18617 | 931 | Y931T | C | 1.6e-05 | 9.2e-06 | 1.3e-05 | 1.273E-05 | 3.408E-06 | 9.200E-06 | 1.600E-05 |
| 18618 | 931 | Y931W | C | 1e-05 | 1.1e-05 | 1.6e-05 | 1.233E-05 | 3.215E-06 | 1.000E-05 | 1.600E-05 |
| 18619 | 931 | Y931Y | C | 1.7e-05 | 9.5e-06 | 1.5e-05 | 1.383E-05 | 3.884E-06 | 9.500E-06 | 1.700E-05 |
| 18620 | 931 | Y931V | C | 1.6e-05 | 1.5e-05 | 1.2e-05 | 1.433E-05 | 2.082E-06 | 1.200E-05 | 1.600E-05 |
| 18621 | 932 | C932A | C | 1.3e-05 | 1.4e-05 | 1.3e-05 | 1.333E-05 | 5.774E-07 | 1.300E-05 | 1.400E-05 |
| 18622 | 932 | C932R | C | 1.1e-05 | 1.4e-05 | 1.6e-05 | 1.367E-05 | 2.517E-06 | 1.100E-05 | 1.600E-05 |
| 18623 | 932 | C932N | C | 1.2e-05 | 1e-05 | 1.3e-05 | 1.167E-05 | 1.528E-06 | 1.000E-05 | 1.300E-05 |
| 18624 | 932 | C932D | C | 1.5e-05 | 1.4e-05 | 1.6e-05 | 1.500E-05 | 1.000E-06 | 1.400E-05 | 1.600E-05 |
| 18625 | 932 | C932C | C | 1.4e-05 | 8.1e-06 | 8.8e-06 | 1.030E-05 | 3.223E-06 | 8.100E-06 | 1.400E-05 |
| 18626 | 932 | C932E | C | 1.1e-05 | 1e-05 | 1.4e-05 | 1.167E-05 | 2.082E-06 | 1.000E-05 | 1.400E-05 |
| 18627 | 932 | C932Q | C | 1.4e-05 | 2e-05 | 1.4e-05 | 1.600E-05 | 3.464E-06 | 1.400E-05 | 2.000E-05 |
| 18628 | 932 | C932G | C | 1.4e-05 | 8.2e-06 | 1.5e-05 | 1.240E-05 | 3.672E-06 | 8.200E-06 | 1.500E-05 |
| 18629 | 932 | C932H | C | 1.4e-05 | 8.8e-06 | 1.6e-05 | 1.293E-05 | 3.717E-06 | 8.800E-06 | 1.600E-05 |
| 18630 | 932 | C932I | C | 1.6e-05 | 1.2e-05 | 8e-06 | 1.200E-05 | 4.000E-06 | 8.000E-06 | 1.600E-05 |
| 18631 | 932 | C932L | C | 1.6e-05 | 9e-06 | 1.1e-05 | 1.200E-05 | 3.606E-06 | 9.000E-06 | 1.600E-05 |
| 18632 | 932 | C932K | C | 1.4e-05 | 1e-05 | 1.4e-05 | 1.267E-05 | 2.309E-06 | 1.000E-05 | 1.400E-05 |
| 18633 | 932 | C932M | C | 1.5e-05 | 1.7e-05 | 1.5e-05 | 1.567E-05 | 1.155E-06 | 1.500E-05 | 1.700E-05 |
| 18634 | 932 | C932F | C | 6.8e-06 | 8.8e-06 | 1.5e-05 | 1.020E-05 | 4.276E-06 | 6.800E-06 | 1.500E-05 |
| 18635 | 932 | C932P | C | 1.5e-05 | 1.6e-05 | 1e-05 | 1.367E-05 | 3.215E-06 | 1.000E-05 | 1.600E-05 |
| 18636 | 932 | C932S | C | 7.4e-06 | 1.5e-05 | 1.5e-05 | 1.247E-05 | 4.388E-06 | 7.400E-06 | 1.500E-05 |
| 18637 | 932 | C932T | C | 1.4e-05 | 9.7e-06 | 1e-05 | 1.123E-05 | 2.401E-06 | 9.700E-06 | 1.400E-05 |
| 18638 | 932 | C932W | C | 1.1e-05 | 1.6e-05 | 8.8e-06 | 1.193E-05 | 3.690E-06 | 8.800E-06 | 1.600E-05 |
| 18639 | 932 | C932Y | C | 8.8e-06 | 1e-05 | 3.8e-06 | 7.533E-06 | 3.288E-06 | 3.800E-06 | 1.000E-05 |
| 18640 | 932 | C932V | C | 1.2e-05 | 1.1e-05 | 1.5e-05 | 1.267E-05 | 2.082E-06 | 1.100E-05 | 1.500E-05 |
| 18641 | 933 | A933A | C | 1e-05 | 1.5e-05 | 1.1e-05 | 1.200E-05 | 2.646E-06 | 1.000E-05 | 1.500E-05 |
| 18642 | 933 | A933R | C | 1.1e-05 | 1e-05 | 1.2e-05 | 1.100E-05 | 1.000E-06 | 1.000E-05 | 1.200E-05 |
| 18643 | 933 | A933N | C | 1.7e-05 | 1.6e-05 | 1.3e-05 | 1.533E-05 | 2.082E-06 | 1.300E-05 | 1.700E-05 |
| 18644 | 933 | A933D | C | 1.5e-05 | 1.5e-05 | 1e-05 | 1.333E-05 | 2.887E-06 | 1.000E-05 | 1.500E-05 |
| 18645 | 933 | A933C | C | 1.1e-05 | 1.2e-05 | 1.5e-05 | 1.267E-05 | 2.082E-06 | 1.100E-05 | 1.500E-05 |
| 18646 | 933 | A933E | C | 1.5e-05 | 1.5e-05 | 1.1e-05 | 1.367E-05 | 2.309E-06 | 1.100E-05 | 1.500E-05 |
| 18647 | 933 | A933Q | C | 1e-05 | 1.3e-05 | 1.5e-05 | 1.267E-05 | 2.517E-06 | 1.000E-05 | 1.500E-05 |
| 18648 | 933 | A933G | C | 9.4e-06 | 1.2e-05 | 1.1e-05 | 1.080E-05 | 1.311E-06 | 9.400E-06 | 1.200E-05 |
| 18649 | 933 | A933H | C | 1e-05 | 1.4e-05 | 1.1e-05 | 1.167E-05 | 2.082E-06 | 1.000E-05 | 1.400E-05 |
| 18650 | 933 | A933I | C | 1e-05 | 1.5e-05 | 1.8e-05 | 1.433E-05 | 4.041E-06 | 1.000E-05 | 1.800E-05 |
| 18651 | 933 | A933L | C | 1.5e-05 | 1.3e-05 | 1.4e-05 | 1.400E-05 | 1.000E-06 | 1.300E-05 | 1.500E-05 |
| 18652 | 933 | A933K | C | 1.4e-05 | 1.1e-05 | 1e-05 | 1.167E-05 | 2.082E-06 | 1.000E-05 | 1.400E-05 |
| 18653 | 933 | A933M | C | 1.1e-05 | 1e-05 | 1.7e-05 | 1.267E-05 | 3.786E-06 | 1.000E-05 | 1.700E-05 |
| 18654 | 933 | A933F | C | 1.4e-05 | 1.5e-05 | 1.5e-05 | 1.467E-05 | 5.774E-07 | 1.400E-05 | 1.500E-05 |
| 18655 | 933 | A933P | C | 1.6e-05 | 1.1e-05 | 9.5e-06 | 1.217E-05 | 3.403E-06 | 9.500E-06 | 1.600E-05 |
| 18656 | 933 | A933S | C | 1.1e-05 | 1.5e-05 | 1.3e-05 | 1.300E-05 | 2.000E-06 | 1.100E-05 | 1.500E-05 |
| 18657 | 933 | A933T | C | 2e-05 | 1.4e-05 | 8.8e-06 | 1.427E-05 | 5.605E-06 | 8.800E-06 | 2.000E-05 |
| 18658 | 933 | A933W | C | 9.8e-06 | 1.6e-05 | 1.1e-05 | 1.227E-05 | 3.288E-06 | 9.800E-06 | 1.600E-05 |
| 18659 | 933 | A933Y | C | 2.5e-05 | 1e-05 | 1.3e-05 | 1.600E-05 | 7.937E-06 | 1.000E-05 | 2.500E-05 |
| 18660 | 933 | A933V | C | 1.4e-05 | 1.1e-05 | 9.6e-06 | 1.153E-05 | 2.248E-06 | 9.600E-06 | 1.400E-05 |
| 18661 | 934 | R934A | C | 1.2e-05 | 1.5e-05 | 1.5e-05 | 1.400E-05 | 1.732E-06 | 1.200E-05 | 1.500E-05 |
| 18662 | 934 | R934R | C | 1.4e-05 | 1.8e-05 | 1.7e-05 | 1.633E-05 | 2.082E-06 | 1.400E-05 | 1.800E-05 |
| 18663 | 934 | R934N | C | 9.3e-06 | 1.4e-05 | 1.5e-05 | 1.277E-05 | 3.044E-06 | 9.300E-06 | 1.500E-05 |
| 18664 | 934 | R934D | C | 9.3e-06 | 1.3e-05 | 1.6e-05 | 1.277E-05 | 3.356E-06 | 9.300E-06 | 1.600E-05 |
| 18665 | 934 | R934C | C | 1.6e-05 | 1.2e-05 | 1.1e-05 | 1.300E-05 | 2.646E-06 | 1.100E-05 | 1.600E-05 |
| 18666 | 934 | R934E | C | 6.7e-06 | 1e-05 | 1.1e-05 | 9.233E-06 | 2.250E-06 | 6.700E-06 | 1.100E-05 |
| 18667 | 934 | R934Q | C | 1.5e-05 | 1.1e-05 | 1.4e-05 | 1.333E-05 | 2.082E-06 | 1.100E-05 | 1.500E-05 |
| 18668 | 934 | R934G | C | 1.5e-05 | 1.3e-05 | 1e-05 | 1.267E-05 | 2.517E-06 | 1.000E-05 | 1.500E-05 |
| 18669 | 934 | R934H | C | 1.3e-05 | 1.1e-05 | 1.5e-05 | 1.300E-05 | 2.000E-06 | 1.100E-05 | 1.500E-05 |
| 18670 | 934 | R934I | C | 1.5e-05 | 1.2e-05 | 1.7e-05 | 1.467E-05 | 2.517E-06 | 1.200E-05 | 1.700E-05 |
| 18671 | 934 | R934L | C | 1.3e-05 | 1.1e-05 | 1e-05 | 1.133E-05 | 1.528E-06 | 1.000E-05 | 1.300E-05 |
| 18672 | 934 | R934K | C | 9.3e-06 | 1.1e-05 | 7.4e-06 | 9.233E-06 | 1.801E-06 | 7.400E-06 | 1.100E-05 |
| 18673 | 934 | R934M | C | 1.4e-05 | 1.1e-05 | 1.6e-05 | 1.367E-05 | 2.517E-06 | 1.100E-05 | 1.600E-05 |
| 18674 | 934 | R934F | C | 1.3e-05 | 1.1e-05 | 1e-05 | 1.133E-05 | 1.528E-06 | 1.000E-05 | 1.300E-05 |
| 18675 | 934 | R934P | C | 1.2e-05 | 1.3e-05 | 1.2e-05 | 1.233E-05 | 5.774E-07 | 1.200E-05 | 1.300E-05 |
| 18676 | 934 | R934S | C | 1.1e-05 | 9.6e-06 | 1.5e-05 | 1.187E-05 | 2.802E-06 | 9.600E-06 | 1.500E-05 |
| 18677 | 934 | R934T | C | 8.2e-06 | 1.5e-05 | 7.8e-06 | 1.033E-05 | 4.046E-06 | 7.800E-06 | 1.500E-05 |
| 18678 | 934 | R934W | C | 1.3e-05 | 1e-05 | 1.3e-05 | 1.200E-05 | 1.732E-06 | 1.000E-05 | 1.300E-05 |
| 18679 | 934 | R934Y | C | 1.2e-05 | 9.5e-06 | 7e-06 | 9.500E-06 | 2.500E-06 | 7.000E-06 | 1.200E-05 |
| 18680 | 934 | R934V | C | 1e-05 | 1.3e-05 | 1.1e-05 | 1.133E-05 | 1.528E-06 | 1.000E-05 | 1.300E-05 |
| 18681 | 935 | N935A | C | 9.9e-06 | 1.6e-05 | 1.6e-05 | 1.397E-05 | 3.522E-06 | 9.900E-06 | 1.600E-05 |
| 18682 | 935 | N935R | C | 1.5e-05 | 1.2e-05 | 1.4e-05 | 1.367E-05 | 1.528E-06 | 1.200E-05 | 1.500E-05 |
| 18683 | 935 | N935N | C | 1.4e-05 | 1e-05 | 1.5e-05 | 1.300E-05 | 2.646E-06 | 1.000E-05 | 1.500E-05 |
| 18684 | 935 | N935D | C | 1.5e-05 | 1.6e-05 | 1e-05 | 1.367E-05 | 3.215E-06 | 1.000E-05 | 1.600E-05 |
| 18685 | 935 | N935C | C | 1.5e-05 | 1.6e-05 | 1.7e-05 | 1.600E-05 | 1.000E-06 | 1.500E-05 | 1.700E-05 |
| 18686 | 935 | N935E | C | 1.4e-05 | 1.1e-05 | 1.1e-05 | 1.200E-05 | 1.732E-06 | 1.100E-05 | 1.400E-05 |
| 18687 | 935 | N935Q | C | 1.4e-05 | 1.1e-05 | 1.1e-05 | 1.200E-05 | 1.732E-06 | 1.100E-05 | 1.400E-05 |
| 18688 | 935 | N935G | C | 1.5e-05 | 2.2e-05 | 1.7e-05 | 1.800E-05 | 3.606E-06 | 1.500E-05 | 2.200E-05 |
| 18689 | 935 | N935H | C | 1.5e-05 | 1.6e-05 | 9.9e-06 | 1.363E-05 | 3.272E-06 | 9.900E-06 | 1.600E-05 |
| 18690 | 935 | N935I | C | 1.7e-05 | 6.7e-06 | 1e-05 | 1.123E-05 | 5.260E-06 | 6.700E-06 | 1.700E-05 |
| 18691 | 935 | N935L | C | 1.6e-05 | 1.5e-05 | 1.1e-05 | 1.400E-05 | 2.646E-06 | 1.100E-05 | 1.600E-05 |
| 18692 | 935 | N935K | C | 1.4e-05 | 1.5e-05 | 1.3e-05 | 1.400E-05 | 1.000E-06 | 1.300E-05 | 1.500E-05 |
| 18693 | 935 | N935M | C | 1.1e-05 | 1.6e-05 | 9.6e-06 | 1.220E-05 | 3.365E-06 | 9.600E-06 | 1.600E-05 |
| 18694 | 935 | N935F | C | 1.2e-05 | 1.1e-05 | 1.3e-05 | 1.200E-05 | 1.000E-06 | 1.100E-05 | 1.300E-05 |
| 18695 | 935 | N935P | C | 1.5e-05 | 1.5e-05 | 1.6e-05 | 1.533E-05 | 5.774E-07 | 1.500E-05 | 1.600E-05 |
| 18696 | 935 | N935S | C | 9.9e-06 | 8.5e-06 | 1.6e-05 | 1.147E-05 | 3.988E-06 | 8.500E-06 | 1.600E-05 |
| 18697 | 935 | N935T | C | 1.3e-05 | 1.1e-05 | 1.5e-05 | 1.300E-05 | 2.000E-06 | 1.100E-05 | 1.500E-05 |
| 18698 | 935 | N935W | C | 1.5e-05 | 1.5e-05 | 9.2e-06 | 1.307E-05 | 3.349E-06 | 9.200E-06 | 1.500E-05 |
| 18699 | 935 | N935Y | C | 9.9e-06 | 1.3e-05 | 1.1e-05 | 1.130E-05 | 1.572E-06 | 9.900E-06 | 1.300E-05 |

| | | | | | | | | | | |
|---|---|---|---|---|---|---|---|---|---|---|
| 18700 | 935 | N935V | C | 1.1e-05 | 1.3e-05 | 1.3e-05 | 1.233E-05 | 1.155E-06 | 1.100E-05 | 1.300E-05 |
| 18701 | 936 | L936A | C | 1.2e-05 | 1.1e-05 | 1.5e-05 | 1.267E-05 | 2.082E-06 | 1.100E-05 | 1.500E-05 |
| 18702 | 936 | L936R | C | 1.4e-05 | 1.1e-05 | 1.5e-05 | 1.333E-05 | 2.082E-06 | 1.100E-05 | 1.500E-05 |
| 18703 | 936 | L936N | C | 1e-05 | 1.4e-05 | 1.5e-05 | 1.300E-05 | 2.646E-06 | 1.000E-05 | 1.500E-05 |
| 18704 | 936 | L936D | C | 1.3e-05 | 1.4e-05 | 1e-05 | 1.233E-05 | 2.082E-06 | 1.000E-05 | 1.400E-05 |
| 18705 | 936 | L936C | C | 1e-05 | 1e-05 | 1.4e-05 | 1.133E-05 | 2.309E-06 | 1.000E-05 | 1.400E-05 |
| 18706 | 936 | L936E | C | 1.1e-05 | 1.1e-05 | 1.1e-05 | 1.100E-05 | 2.075E-21 | 1.100E-05 | 1.100E-05 |
| 18707 | 936 | L936Q | C | 1.3e-05 | 1e-05 | 1e-05 | 1.100E-05 | 1.732E-06 | 1.000E-05 | 1.300E-05 |
| 18708 | 936 | L936G | C | 9.4e-06 | 1.6e-05 | 1.5e-05 | 1.347E-05 | 3.557E-06 | 9.400E-06 | 1.600E-05 |
| 18709 | 936 | L936H | C | 9.7e-06 | 1.5e-05 | 1.5e-05 | 1.323E-05 | 3.060E-06 | 9.700E-06 | 1.500E-05 |
| 18710 | 936 | L936I | C | 1.4e-05 | 1e-05 | 1.6e-05 | 1.333E-05 | 3.055E-06 | 1.000E-05 | 1.600E-05 |
| 18711 | 936 | L936L | C | 1.4e-05 | 1.2e-05 | 9.5e-06 | 1.183E-05 | 2.255E-06 | 9.500E-06 | 1.400E-05 |
| 18712 | 936 | L936K | C | 1.6e-05 | 1.3e-05 | 1.1e-05 | 1.333E-05 | 2.517E-06 | 1.100E-05 | 1.600E-05 |
| 18713 | 936 | L936M | C | 1.5e-05 | 1e-05 | 1.6e-05 | 1.367E-05 | 3.215E-06 | 1.000E-05 | 1.600E-05 |
| 18714 | 936 | L936F | C | 1.4e-05 | 1.4e-05 | 1e-05 | 1.267E-05 | 2.309E-06 | 1.000E-05 | 1.400E-05 |
| 18715 | 936 | L936P | C | 1.6e-05 | 1.5e-05 | 1.1e-05 | 1.400E-05 | 2.646E-06 | 1.100E-05 | 1.600E-05 |
| 18716 | 936 | L936S | C | 9.6e-06 | 9.7e-06 | 1.6e-05 | 1.177E-05 | 3.667E-06 | 9.600E-06 | 1.600E-05 |
| 18717 | 936 | L936T | C | 1.5e-05 | 1.1e-05 | 1.6e-05 | 1.400E-05 | 2.646E-06 | 1.100E-05 | 1.600E-05 |
| 18718 | 936 | L936W | C | 1.9e-05 | 1.7e-05 | 1.3e-05 | 1.633E-05 | 3.055E-06 | 1.300E-05 | 1.900E-05 |
| 18719 | 936 | L936Y | C | 9.5e-06 | 1.6e-05 | 1.6e-05 | 1.383E-05 | 3.753E-06 | 9.500E-06 | 1.600E-05 |
| 18720 | 936 | L936V | C | 1.1e-05 | 1.7e-05 | 1.6e-05 | 1.467E-05 | 3.215E-06 | 1.100E-05 | 1.700E-05 |
| 18721 | 937 | G937A | C | 1.4e-05 | 8.8e-06 | 1.6e-05 | 1.293E-05 | 3.717E-06 | 8.800E-06 | 1.600E-05 |
| 18722 | 937 | G937R | C | 1.6e-05 | 1.6e-05 | 9.2e-06 | 1.373E-05 | 3.926E-06 | 9.200E-06 | 1.600E-05 |
| 18723 | 937 | G937N | C | 1.5e-05 | 1.5e-05 | 1e-05 | 1.333E-05 | 2.887E-06 | 1.000E-05 | 1.500E-05 |
| 18724 | 937 | G937D | C | 1e-05 | 9.6e-06 | 1e-05 | 9.867E-06 | 2.309E-07 | 9.600E-06 | 1.000E-05 |
| 18725 | 937 | G937C | C | 1.3e-05 | 1.4e-05 | 1.4e-05 | 1.367E-05 | 5.774E-07 | 1.300E-05 | 1.400E-05 |
| 18726 | 937 | G937E | C | 1.6e-05 | 1.5e-05 | 6.5e-06 | 1.250E-05 | 5.220E-06 | 6.500E-06 | 1.600E-05 |
| 18727 | 937 | G937Q | C | 1.6e-05 | 1.5e-05 | 1.6e-05 | 1.567E-05 | 5.774E-07 | 1.500E-05 | 1.600E-05 |
| 18728 | 937 | G937G | C | 9.1e-06 | 9.4e-06 | 9.7e-06 | 9.400E-06 | 3.000E-07 | 9.100E-06 | 9.700E-06 |
| 18729 | 937 | G937H | C | 1.4e-05 | 1.4e-05 | 8.9e-06 | 1.230E-05 | 2.944E-06 | 8.900E-06 | 1.400E-05 |
| 18730 | 937 | G937I | C | 1.3e-05 | 1.1e-05 | 1.5e-05 | 1.300E-05 | 2.000E-06 | 1.100E-05 | 1.500E-05 |
| 18731 | 937 | G937L | C | 1.6e-05 | 1.6e-05 | 1e-05 | 1.400E-05 | 3.464E-06 | 1.000E-05 | 1.600E-05 |
| 18732 | 937 | G937K | C | 1.6e-05 | 1.1e-05 | 1.1e-05 | 1.267E-05 | 2.887E-06 | 1.100E-05 | 1.600E-05 |
| 18733 | 937 | G937M | C | 1.1e-05 | 1.6e-05 | 1.1e-05 | 1.267E-05 | 2.887E-06 | 1.100E-05 | 1.600E-05 |
| 18734 | 937 | G937F | C | 1.5e-05 | 1.4e-05 | 1.2e-05 | 1.367E-05 | 1.528E-06 | 1.200E-05 | 1.500E-05 |
| 18735 | 937 | G937P | C | 9.6e-06 | 1.7e-05 | 1.5e-05 | 1.387E-05 | 3.828E-06 | 9.600E-06 | 1.700E-05 |
| 18736 | 937 | G937S | C | 1.1e-05 | 1.4e-05 | 1.1e-05 | 1.200E-05 | 1.732E-06 | 1.100E-05 | 1.400E-05 |
| 18737 | 937 | G937T | C | 6.9e-06 | 1e-05 | 1.1e-05 | 9.300E-06 | 2.138E-06 | 6.900E-06 | 1.100E-05 |
| 18738 | 937 | G937W | C | 1.1e-05 | 9.1e-06 | 1.6e-05 | 1.203E-05 | 3.564E-06 | 9.100E-06 | 1.600E-05 |
| 18739 | 937 | G937Y | C | 7.7e-06 | 1.4e-05 | 7.6e-06 | 9.767E-06 | 3.667E-06 | 7.600E-06 | 1.400E-05 |
| 18740 | 937 | G937V | C | 1.6e-05 | 1.7e-05 | 1.4e-05 | 1.567E-05 | 1.528E-06 | 1.400E-05 | 1.700E-05 |
| 18741 | 938 | P938A | C | 1.5e-05 | 1.5e-05 | 1.6e-05 | 1.533E-05 | 5.774E-07 | 1.500E-05 | 1.600E-05 |
| 18742 | 938 | P938R | C | 1.6e-05 | 1.5e-05 | 2.1e-05 | 1.733E-05 | 3.215E-06 | 1.500E-05 | 2.100E-05 |
| 18743 | 938 | P938N | C | 1.2e-05 | 9.1e-06 | 9.9e-06 | 1.033E-05 | 1.498E-06 | 9.100E-06 | 1.200E-05 |
| 18744 | 938 | P938D | C | 1.4e-05 | 1.5e-05 | 1.2e-05 | 1.367E-05 | 1.528E-06 | 1.200E-05 | 1.500E-05 |
| 18745 | 938 | P938C | C | 1.1e-05 | 9.6e-06 | 1.2e-05 | 1.087E-05 | 1.206E-06 | 9.600E-06 | 1.200E-05 |
| 18746 | 938 | P938E | C | 1.1e-05 | 9e-06 | 9.7e-06 | 9.900E-06 | 1.015E-06 | 9.000E-06 | 1.100E-05 |
| 18747 | 938 | P938Q | C | 9.3e-06 | 1.4e-05 | 1.6e-05 | 1.310E-05 | 3.439E-06 | 9.300E-06 | 1.600E-05 |
| 18748 | 938 | P938G | C | 1.7e-05 | 1.5e-05 | 1.4e-05 | 1.533E-05 | 1.528E-06 | 1.400E-05 | 1.700E-05 |
| 18749 | 938 | P938H | C | 8.4e-06 | 1.6e-05 | 7.7e-06 | 1.070E-05 | 4.603E-06 | 7.700E-06 | 1.600E-05 |
| 18750 | 938 | P938I | C | 1.3e-05 | 1.6e-05 | 1.1e-05 | 1.333E-05 | 2.517E-06 | 1.100E-05 | 1.600E-05 |
| 18751 | 938 | P938L | C | 9.4e-06 | 1.1e-05 | 1e-05 | 1.013E-05 | 8.083E-07 | 9.400E-06 | 1.100E-05 |
| 18752 | 938 | P938K | C | 1.2e-05 | 1.3e-05 | 9.2e-06 | 1.140E-05 | 1.970E-06 | 9.200E-06 | 1.300E-05 |
| 18753 | 938 | P938M | C | 1e-05 | 1.4e-05 | 1.8e-05 | 1.400E-05 | 4.000E-06 | 1.000E-05 | 1.800E-05 |
| 18754 | 938 | P938F | C | 8e-06 | 1.4e-05 | 1.3e-05 | 1.167E-05 | 3.215E-06 | 8.000E-06 | 1.400E-05 |
| 18755 | 938 | P938P | C | 1.5e-05 | 1.1e-05 | 9.2e-06 | 1.173E-05 | 2.969E-06 | 9.200E-06 | 1.500E-05 |
| 18756 | 938 | P938S | C | 1.2e-05 | 1e-05 | 9.7e-06 | 1.057E-05 | 1.250E-06 | 9.700E-06 | 1.200E-05 |
| 18757 | 938 | P938T | C | 1.1e-05 | 1.4e-05 | 1.2e-05 | 1.233E-05 | 1.528E-06 | 1.100E-05 | 1.400E-05 |
| 18758 | 938 | P938W | C | 1.5e-05 | 1.4e-05 | 1.5e-05 | 1.467E-05 | 5.774E-07 | 1.400E-05 | 1.500E-05 |
| 18759 | 938 | P938Y | C | 1.3e-05 | 1.5e-05 | 9.7e-06 | 1.257E-05 | 2.676E-06 | 9.700E-06 | 1.500E-05 |
| 18760 | 938 | P938V | C | 1.6e-05 | 9.1e-06 | 8.9e-06 | 1.133E-05 | 4.043E-06 | 8.900E-06 | 1.600E-05 |
| 18761 | 939 | S939A | C | 1.5e-05 | 1e-05 | 1.6e-05 | 1.367E-05 | 3.215E-06 | 1.000E-05 | 1.600E-05 |
| 18762 | 939 | S939R | C | 1.6e-05 | 1.5e-05 | 9.9e-06 | 1.363E-05 | 3.272E-06 | 9.900E-06 | 1.600E-05 |
| 18763 | 939 | S939N | C | 1e-05 | 1.1e-05 | 9.4e-06 | 1.013E-05 | 8.083E-07 | 9.400E-06 | 1.100E-05 |
| 18764 | 939 | S939D | C | 1.1e-05 | 1e-05 | 1.1e-05 | 1.067E-05 | 5.774E-07 | 1.000E-05 | 1.100E-05 |
| 18765 | 939 | S939C | C | 1.7e-05 | 1.7e-05 | 9.5e-06 | 1.450E-05 | 4.330E-06 | 9.500E-06 | 1.700E-05 |
| 18766 | 939 | S939E | C | 1.1e-05 | 1.2e-05 | 8.6e-06 | 1.053E-05 | 1.747E-06 | 8.600E-06 | 1.200E-05 |
| 18767 | 939 | S939Q | C | 1.1e-05 | 1.6e-05 | 8.5e-06 | 1.183E-05 | 3.819E-06 | 8.500E-06 | 1.600E-05 |
| 18768 | 939 | S939G | C | 1.1e-05 | 1.5e-05 | 1.5e-05 | 1.367E-05 | 2.309E-06 | 1.100E-05 | 1.500E-05 |
| 18769 | 939 | S939H | C | 9.5e-06 | 9.7e-06 | 1e-05 | 9.733E-06 | 2.517E-07 | 9.500E-06 | 1.000E-05 |
| 18770 | 939 | S939I | C | 1.1e-05 | 1.5e-05 | 1.5e-05 | 1.367E-05 | 2.309E-06 | 1.100E-05 | 1.500E-05 |
| 18771 | 939 | S939L | C | 1.6e-05 | 1.8e-05 | 1.5e-05 | 1.633E-05 | 1.528E-06 | 1.500E-05 | 1.800E-05 |
| 18772 | 939 | S939K | C | 1.4e-05 | 1e-05 | 1.2e-05 | 1.200E-05 | 2.000E-06 | 1.000E-05 | 1.400E-05 |
| 18773 | 939 | S939M | C | 1.8e-05 | 1.7e-05 | 1.6e-05 | 1.700E-05 | 1.000E-06 | 1.600E-05 | 1.800E-05 |
| 18774 | 939 | S939F | C | 1.2e-05 | 8.6e-06 | 1.6e-05 | 1.220E-05 | 3.704E-06 | 8.600E-06 | 1.600E-05 |
| 18775 | 939 | S939P | C | 9.9e-06 | 1.5e-05 | 1.5e-05 | 1.330E-05 | 2.944E-06 | 9.900E-06 | 1.500E-05 |
| 18776 | 939 | S939S | C | 1.6e-05 | 1.2e-05 | 1.1e-05 | 1.300E-05 | 2.646E-06 | 1.100E-05 | 1.600E-05 |
| 18777 | 939 | S939T | C | 1.5e-05 | 1.6e-05 | 1.5e-05 | 1.533E-05 | 5.774E-07 | 1.500E-05 | 1.600E-05 |
| 18778 | 939 | S939W | C | 1.4e-05 | 1.4e-05 | 1.2e-05 | 1.333E-05 | 1.155E-06 | 1.200E-05 | 1.400E-05 |
| 18779 | 939 | S939Y | C | 1.1e-05 | 1.5e-05 | 1.5e-05 | 1.367E-05 | 2.309E-06 | 1.100E-05 | 1.500E-05 |
| 18780 | 939 | S939V | C | 8.6e-06 | 1.6e-05 | 1.5e-05 | 1.320E-05 | 4.015E-06 | 8.600E-06 | 1.600E-05 |
| 18781 | 940 | F940A | C | 8e-06 | 1.2e-05 | 1.2e-05 | 1.067E-05 | 2.309E-06 | 8.000E-06 | 1.200E-05 |
| 18782 | 940 | F940R | C | 1.2e-05 | 1.4e-05 | 7.4e-06 | 1.113E-05 | 3.384E-06 | 7.400E-06 | 1.400E-05 |
| 18783 | 940 | F940N | C | 1.5e-05 | 1.2e-05 | 1.4e-05 | 1.367E-05 | 1.528E-06 | 1.200E-05 | 1.500E-05 |
| 18784 | 940 | F940D | C | 1.6e-05 | 1.1e-05 | 1.1e-05 | 1.267E-05 | 2.887E-06 | 1.100E-05 | 1.600E-05 |

| | | | | | | | | | | |
|---|---|---|---|---|---|---|---|---|---|---|
| 18785 | 940 | F940C | C | 1.4e-05 | 1.6e-05 | 1.1e-05 | 1.367E-05 | 2.517E-06 | 1.100E-05 | 1.600E-05 |
| 18786 | 940 | F940E | C | 9.8e-06 | 1e-05 | 1.5e-05 | 1.160E-05 | 2.946E-06 | 9.800E-06 | 1.500E-05 |
| 18787 | 940 | F940Q | C | 9.6e-06 | 1e-05 | 7.6e-06 | 9.067E-06 | 1.286E-06 | 7.600E-06 | 1.000E-05 |
| 18788 | 940 | F940G | C | 1.4e-05 | 1.7e-05 | 1.5e-05 | 1.533E-05 | 1.528E-06 | 1.400E-05 | 1.700E-05 |
| 18789 | 940 | F940H | C | 1.5e-05 | 1.3e-05 | 1.5e-05 | 1.433E-05 | 1.155E-06 | 1.300E-05 | 1.500E-05 |
| 18790 | 940 | F940I | C | 1.6e-05 | 9.9e-06 | 1.2e-05 | 1.263E-05 | 3.099E-06 | 9.900E-06 | 1.600E-05 |
| 18791 | 940 | F940L | C | 8.5e-06 | 9.7e-06 | 1.6e-05 | 1.140E-05 | 4.029E-06 | 8.500E-06 | 1.600E-05 |
| 18792 | 940 | F940K | C | 2.2e-05 | 1e-05 | 1.5e-05 | 1.567E-05 | 6.028E-06 | 1.000E-05 | 2.200E-05 |
| 18793 | 940 | F940M | C | 1.5e-05 | 1e-05 | 8e-06 | 1.100E-05 | 3.606E-06 | 8.000E-06 | 1.500E-05 |
| 18794 | 940 | F940F | C | 1.3e-05 | 8e-06 | 1.5e-05 | 1.200E-05 | 3.606E-06 | 8.000E-06 | 1.500E-05 |
| 18795 | 940 | F940P | C | 9.7e-06 | 1.1e-05 | 1.6e-05 | 1.223E-05 | 3.326E-06 | 9.700E-06 | 1.600E-05 |
| 18796 | 940 | F940S | C | 1.3e-05 | 1e-05 | 1.1e-05 | 1.133E-05 | 1.528E-06 | 1.000E-05 | 1.300E-05 |
| 18797 | 940 | F940T | C | 1.1e-05 | 6.7e-06 | 1.5e-05 | 1.090E-05 | 4.151E-06 | 6.700E-06 | 1.500E-05 |
| 18798 | 940 | F940W | C | 1.4e-05 | 1.4e-05 | 1.4e-05 | 1.400E-05 | 0.000E+00 | 1.400E-05 | 1.400E-05 |
| 18799 | 940 | F940Y | C | 9.5e-06 | 1.1e-05 | 6.1e-06 | 8.867E-06 | 2.511E-06 | 6.100E-06 | 1.100E-05 |
| 18800 | 940 | F940V | C | 1.3e-05 | 1.6e-05 | 1.5e-05 | 1.467E-05 | 1.528E-06 | 1.300E-05 | 1.600E-05 |
| 18801 | 941 | Y941A | C | 1.7e-05 | 1.3e-05 | 1.6e-05 | 1.533E-05 | 2.082E-06 | 1.300E-05 | 1.700E-05 |
| 18802 | 941 | Y941R | C | 1.4e-05 | 1.2e-05 | 1.4e-05 | 1.333E-05 | 1.155E-06 | 1.200E-05 | 1.400E-05 |
| 18803 | 941 | Y941N | C | 1.1e-05 | 1.5e-05 | 1.6e-05 | 1.400E-05 | 2.646E-06 | 1.100E-05 | 1.600E-05 |
| 18804 | 941 | Y941D | C | 1.7e-05 | 1.5e-05 | 7.9e-06 | 1.330E-05 | 4.782E-06 | 7.900E-06 | 1.700E-05 |
| 18805 | 941 | Y941C | C | 1.6e-05 | 1.5e-05 | 1.1e-05 | 1.400E-05 | 2.646E-06 | 1.100E-05 | 1.600E-05 |
| 18806 | 941 | Y941E | C | 1.6e-05 | 1.6e-05 | 9.3e-06 | 1.377E-05 | 3.868E-06 | 9.300E-06 | 1.600E-05 |
| 18807 | 941 | Y941Q | C | 1.5e-05 | 1.7e-05 | 1.4e-05 | 1.533E-05 | 1.528E-06 | 1.400E-05 | 1.700E-05 |
| 18808 | 941 | Y941G | C | 1.6e-05 | 1.2e-05 | 1.5e-05 | 1.433E-05 | 2.082E-06 | 1.200E-05 | 1.600E-05 |
| 18809 | 941 | Y941H | C | 9.9e-06 | 1.2e-05 | 1.5e-05 | 1.230E-05 | 2.563E-06 | 9.900E-06 | 1.500E-05 |
| 18810 | 941 | Y941I | C | 1.7e-05 | 1e-05 | 1.1e-05 | 1.267E-05 | 3.786E-06 | 1.000E-05 | 1.700E-05 |
| 18811 | 941 | Y941L | C | 1.5e-05 | 1.5e-05 | 1.5e-05 | 1.500E-05 | 0.000E+00 | 1.500E-05 | 1.500E-05 |
| 18812 | 941 | Y941K | C | 1.6e-05 | 1.6e-05 | 1.6e-05 | 1.600E-05 | 0.000E+00 | 1.600E-05 | 1.600E-05 |
| 18813 | 941 | Y941M | C | 1.4e-05 | 1e-05 | 9.6e-06 | 1.120E-05 | 2.433E-06 | 9.600E-06 | 1.400E-05 |
| 18814 | 941 | Y941F | C | 1.9e-05 | 1.5e-05 | 1e-05 | 1.467E-05 | 4.509E-06 | 1.000E-05 | 1.900E-05 |
| 18815 | 941 | Y941P | C | 1.6e-05 | 1.4e-05 | 1.2e-05 | 1.400E-05 | 2.000E-06 | 1.200E-05 | 1.600E-05 |
| 18816 | 941 | Y941S | C | 1.4e-05 | 1.1e-05 | 1.2e-05 | 1.233E-05 | 1.528E-06 | 1.100E-05 | 1.400E-05 |
| 18817 | 941 | Y941T | C | 2.2e-05 | 1.7e-05 | 1.4e-05 | 1.767E-05 | 4.041E-06 | 1.400E-05 | 2.200E-05 |
| 18818 | 941 | Y941W | C | 1.3e-05 | 1.6e-05 | 1.4e-05 | 1.433E-05 | 1.528E-06 | 1.300E-05 | 1.600E-05 |
| 18819 | 941 | Y941Y | C | 1.2e-05 | 1.5e-05 | 8.5e-06 | 1.183E-05 | 3.253E-06 | 8.500E-06 | 1.500E-05 |
| 18820 | 941 | Y941V | C | 1.1e-05 | 1.1e-05 | 1.6e-05 | 1.267E-05 | 2.887E-06 | 1.100E-05 | 1.600E-05 |
| 18821 | 942 | F942A | C | 1.5e-05 | 1e-05 | 9.5e-06 | 1.150E-05 | 3.041E-06 | 9.500E-06 | 1.500E-05 |
| 18822 | 942 | F942R | C | 1.4e-05 | 1e-05 | 1.2e-05 | 1.200E-05 | 2.000E-06 | 1.000E-05 | 1.400E-05 |
| 18823 | 942 | F942N | C | 1.5e-05 | 1.6e-05 | 9.5e-06 | 1.350E-05 | 3.500E-06 | 9.500E-06 | 1.600E-05 |
| 18824 | 942 | F942D | C | 9.2e-06 | 1.1e-05 | 1.6e-05 | 1.207E-05 | 3.523E-06 | 9.200E-06 | 1.600E-05 |
| 18825 | 942 | F942C | C | 1.5e-05 | 9.8e-06 | 9.5e-06 | 1.143E-05 | 3.092E-06 | 9.500E-06 | 1.500E-05 |
| 18826 | 942 | F942E | C | 1.2e-05 | 1.5e-05 | 2e-05 | 1.567E-05 | 4.041E-06 | 1.200E-05 | 2.000E-05 |
| 18827 | 942 | F942Q | C | 1.3e-05 | 1.4e-05 | 9.9e-06 | 1.230E-05 | 2.138E-06 | 9.900E-06 | 1.400E-05 |
| 18828 | 942 | F942G | C | 1.5e-05 | 1.4e-05 | 2.2e-05 | 1.700E-05 | 4.359E-06 | 1.400E-05 | 2.200E-05 |
| 18829 | 942 | F942H | C | 9.6e-06 | 1.4e-05 | 7.8e-06 | 1.047E-05 | 3.190E-06 | 7.800E-06 | 1.400E-05 |
| 18830 | 942 | F942I | C | 1.3e-05 | 1.5e-05 | 1.4e-05 | 1.400E-05 | 1.000E-06 | 1.300E-05 | 1.500E-05 |
| 18831 | 942 | F942L | C | 1.3e-05 | 1e-05 | 1.6e-05 | 1.300E-05 | 3.000E-06 | 1.000E-05 | 1.600E-05 |
| 18832 | 942 | F942K | C | 9.9e-06 | 1.1e-05 | 1.5e-05 | 1.197E-05 | 2.684E-06 | 9.900E-06 | 1.500E-05 |
| 18833 | 942 | F942M | C | 1.3e-05 | 1.5e-05 | 1.5e-05 | 1.433E-05 | 1.155E-06 | 1.300E-05 | 1.500E-05 |
| 18834 | 942 | F942F | C | 1.5e-05 | 1e-05 | 1e-05 | 1.167E-05 | 2.887E-06 | 1.000E-05 | 1.500E-05 |
| 18835 | 942 | F942P | C | 8.7e-06 | 1.7e-05 | 1.5e-05 | 1.357E-05 | 4.332E-06 | 8.700E-06 | 1.700E-05 |
| 18836 | 942 | F942S | C | 1e-05 | 1.5e-05 | 1.3e-05 | 1.267E-05 | 2.517E-06 | 1.000E-05 | 1.500E-05 |
| 18837 | 942 | F942T | C | 9.1e-06 | 1.1e-05 | 7.9e-06 | 9.333E-06 | 1.563E-06 | 7.900E-06 | 1.100E-05 |
| 18838 | 942 | F942W | C | 8.6e-06 | 1.5e-05 | 1.3e-05 | 1.220E-05 | 3.274E-06 | 8.600E-06 | 1.500E-05 |
| 18839 | 942 | F942Y | C | 1.3e-05 | 1.2e-05 | 1e-05 | 1.167E-05 | 1.528E-06 | 1.000E-05 | 1.300E-05 |
| 18840 | 942 | F942V | C | 8.9e-06 | 1.1e-05 | 1.5e-05 | 1.163E-05 | 3.099E-06 | 8.900E-06 | 1.500E-05 |
| 18841 | 943 | D943A | C | 1.4e-05 | 1.5e-05 | 1.5e-05 | 1.467E-05 | 5.774E-07 | 1.400E-05 | 1.500E-05 |
| 18842 | 943 | D943R | C | 1e-05 | 1.6e-05 | 1.5e-05 | 1.367E-05 | 3.215E-06 | 1.000E-05 | 1.600E-05 |
| 18843 | 943 | D943N | C | 1.5e-05 | 1.7e-05 | 9.5e-06 | 1.383E-05 | 3.884E-06 | 9.500E-06 | 1.700E-05 |
| 18844 | 943 | D943D | C | 1.5e-05 | 1e-05 | 5.6e-06 | 1.020E-05 | 4.703E-06 | 5.600E-06 | 1.500E-05 |
| 18845 | 943 | D943C | C | 1e-05 | 1.2e-05 | 1.3e-05 | 1.167E-05 | 1.528E-06 | 1.000E-05 | 1.300E-05 |
| 18846 | 943 | D943E | C | 1.4e-05 | 1.1e-05 | 1.6e-05 | 1.367E-05 | 2.517E-06 | 1.100E-05 | 1.600E-05 |
| 18847 | 943 | D943Q | C | 9.5e-06 | 1.4e-05 | 1e-05 | 1.117E-05 | 2.466E-06 | 9.500E-06 | 1.400E-05 |
| 18848 | 943 | D943G | C | 1.1e-05 | 1.2e-05 | 1.5e-05 | 1.267E-05 | 2.082E-06 | 1.100E-05 | 1.500E-05 |
| 18849 | 943 | D943H | C | 1e-05 | 1e-05 | 1.4e-05 | 1.133E-05 | 2.309E-06 | 1.000E-05 | 1.400E-05 |
| 18850 | 943 | D943I | C | 1.1e-05 | 1.5e-05 | 1.3e-05 | 1.300E-05 | 2.000E-06 | 1.100E-05 | 1.500E-05 |
| 18851 | 943 | D943L | C | 1.8e-05 | 9.6e-06 | 1.1e-05 | 1.287E-05 | 4.500E-06 | 9.600E-06 | 1.800E-05 |
| 18852 | 943 | D943K | C | 1.5e-05 | 1.1e-05 | 1.2e-05 | 1.267E-05 | 2.082E-06 | 1.100E-05 | 1.500E-05 |
| 18853 | 943 | D943M | C | 1.5e-05 | 1.4e-05 | 1.5e-05 | 1.467E-05 | 5.774E-07 | 1.400E-05 | 1.500E-05 |
| 18854 | 943 | D943F | C | 1.3e-05 | 9.8e-06 | 1e-05 | 1.093E-05 | 1.793E-06 | 9.800E-06 | 1.300E-05 |
| 18855 | 943 | D943P | C | 1.1e-05 | 1.1e-05 | 9.8e-06 | 1.060E-05 | 6.928E-07 | 9.800E-06 | 1.100E-05 |
| 18856 | 943 | D943S | C | 1e-05 | 9.8e-06 | 9.7e-06 | 9.833E-06 | 1.528E-07 | 9.700E-06 | 1.000E-05 |
| 18857 | 943 | D943T | C | 7.2e-06 | 7.9e-06 | 9.6e-06 | 8.233E-06 | 1.234E-06 | 7.200E-06 | 9.600E-06 |
| 18858 | 943 | D943W | C | 1.4e-05 | 1.5e-05 | 1e-05 | 1.300E-05 | 2.646E-06 | 1.000E-05 | 1.500E-05 |
| 18859 | 943 | D943Y | C | 1.5e-05 | 9.7e-06 | 1.7e-05 | 1.390E-05 | 3.772E-06 | 9.700E-06 | 1.700E-05 |
| 18860 | 943 | D943V | C | 9.5e-06 | 1.1e-05 | 1.4e-05 | 1.150E-05 | 2.291E-06 | 9.500E-06 | 1.400E-05 |
| 18861 | 944 | Y944A | C | 1.6e-05 | 1.6e-05 | 1e-05 | 1.400E-05 | 3.464E-06 | 1.000E-05 | 1.600E-05 |
| 18862 | 944 | Y944R | C | 1.4e-05 | 1.2e-05 | 1.5e-05 | 1.367E-05 | 1.528E-06 | 1.200E-05 | 1.500E-05 |
| 18863 | 944 | Y944N | C | 1.4e-05 | 1.6e-05 | 1.2e-05 | 1.400E-05 | 2.000E-06 | 1.200E-05 | 1.600E-05 |
| 18864 | 944 | Y944D | C | 1.4e-05 | 1.6e-05 | 1.2e-05 | 1.400E-05 | 2.000E-06 | 1.200E-05 | 1.600E-05 |
| 18865 | 944 | Y944C | C | 1.1e-05 | 9.2e-06 | 1e-05 | 1.007E-05 | 9.018E-07 | 9.200E-06 | 1.100E-05 |
| 18866 | 944 | Y944E | C | 8.7e-06 | 1e-05 | 1e-05 | 9.567E-06 | 7.506E-07 | 8.700E-06 | 1.000E-05 |
| 18867 | 944 | Y944Q | C | 1.7e-05 | 1.5e-05 | 1.3e-05 | 1.500E-05 | 2.000E-06 | 1.300E-05 | 1.700E-05 |
| 18868 | 944 | Y944G | C | 8.8e-06 | 9.3e-06 | 1.2e-05 | 1.003E-05 | 1.721E-06 | 8.800E-06 | 1.200E-05 |
| 18869 | 944 | Y944H | C | 1e-05 | 1.2e-05 | 1e-05 | 1.067E-05 | 1.155E-06 | 1.000E-05 | 1.200E-05 |

| | | | | | | | | | | |
|---|---|---|---|---|---|---|---|---|---|---|
| 18870 | 944 | Y944I | C | 1.1e-05 | 1.5e-05 | 1.6e-05 | 1.400E-05 | 2.646E-06 | 1.100E-05 | 1.600E-05 |
| 18871 | 944 | Y944L | C | 1.6e-05 | 1.2e-05 | 1.6e-05 | 1.467E-05 | 2.309E-06 | 1.200E-05 | 1.600E-05 |
| 18872 | 944 | Y944K | C | 1e-05 | 1.7e-05 | 1.6e-05 | 1.433E-05 | 3.786E-06 | 1.000E-05 | 1.700E-05 |
| 18873 | 944 | Y944M | C | 8.4e-06 | 9.9e-06 | 1.6e-05 | 1.143E-05 | 4.025E-06 | 8.400E-06 | 1.600E-05 |
| 18874 | 944 | Y944F | C | 1e-05 | 1.1e-05 | 1.3e-05 | 1.133E-05 | 1.528E-06 | 1.000E-05 | 1.300E-05 |
| 18875 | 944 | Y944P | C | 1.5e-05 | 1.2e-05 | 1.5e-05 | 1.400E-05 | 1.732E-06 | 1.200E-05 | 1.500E-05 |
| 18876 | 944 | Y944S | C | 1.1e-05 | 1.5e-05 | 1.2e-05 | 1.267E-05 | 2.082E-06 | 1.100E-05 | 1.500E-05 |
| 18877 | 944 | Y944T | C | 1.5e-05 | 1.5e-05 | 1.4e-05 | 1.467E-05 | 5.774E-07 | 1.400E-05 | 1.500E-05 |
| 18878 | 944 | Y944W | C | 1.5e-05 | 1.6e-05 | 1.5e-05 | 1.533E-05 | 5.774E-07 | 1.500E-05 | 1.600E-05 |
| 18879 | 944 | Y944Y | C | 1.1e-05 | 1.2e-05 | 1.7e-05 | 1.333E-05 | 3.215E-06 | 1.100E-05 | 1.700E-05 |
| 18880 | 944 | Y944V | C | 1.6e-05 | 1.7e-05 | 1.5e-05 | 1.600E-05 | 1.000E-06 | 1.500E-05 | 1.700E-05 |
| 18881 | 945 | W945A | C | 1.7e-05 | 6.6e-06 | 8.5e-06 | 1.070E-05 | 5.538E-06 | 6.600E-06 | 1.700E-05 |
| 18882 | 945 | W945R | C | 9.2e-06 | 1.4e-05 | 1.6e-05 | 1.307E-05 | 3.495E-06 | 9.200E-06 | 1.600E-05 |
| 18883 | 945 | W945N | C | 1e-05 | 1.4e-05 | 1.5e-05 | 1.300E-05 | 2.646E-06 | 1.000E-05 | 1.500E-05 |
| 18884 | 945 | W945D | C | 1.1e-05 | 1.6e-05 | 1.2e-05 | 1.300E-05 | 2.646E-06 | 1.100E-05 | 1.600E-05 |
| 18885 | 945 | W945C | C | 1e-05 | 8.1e-06 | 7.2e-06 | 8.433E-06 | 1.429E-06 | 7.200E-06 | 1.000E-05 |
| 18886 | 945 | W945E | C | 1.7e-05 | 9.5e-06 | 1.1e-05 | 1.250E-05 | 3.969E-06 | 9.500E-06 | 1.700E-05 |
| 18887 | 945 | W945Q | C | 1.2e-05 | 1.1e-05 | 9.8e-06 | 1.093E-05 | 1.102E-06 | 9.800E-06 | 1.200E-05 |
| 18888 | 945 | W945G | C | 1e-05 | 1e-05 | 1.4e-05 | 1.133E-05 | 2.309E-06 | 1.000E-05 | 1.400E-05 |
| 18889 | 945 | W945H | C | 1e-05 | 2.1e-05 | 1.5e-05 | 1.533E-05 | 5.508E-06 | 1.000E-05 | 2.100E-05 |
| 18890 | 945 | W945I | C | 1.7e-05 | 1.1e-05 | 1.5e-05 | 1.433E-05 | 3.055E-06 | 1.100E-05 | 1.700E-05 |
| 18891 | 945 | W945L | C | 1.2e-05 | 1.6e-05 | 1.1e-05 | 1.300E-05 | 2.646E-06 | 1.100E-05 | 1.600E-05 |
| 18892 | 945 | W945K | C | 1.6e-05 | 1e-05 | 1.5e-05 | 1.367E-05 | 3.215E-06 | 1.000E-05 | 1.600E-05 |
| 18893 | 945 | W945M | C | 1.1e-05 | 1e-05 | 1.5e-05 | 1.200E-05 | 2.646E-06 | 1.000E-05 | 1.500E-05 |
| 18894 | 945 | W945F | C | 1.1e-05 | 1.1e-05 | 9.8e-06 | 1.060E-05 | 6.928E-07 | 9.800E-06 | 1.100E-05 |
| 18895 | 945 | W945P | C | 1.1e-05 | 1.1e-05 | 1.8e-05 | 1.333E-05 | 4.041E-06 | 1.100E-05 | 1.800E-05 |
| 18896 | 945 | W945S | C | 1.5e-05 | 1.6e-05 | 6.3e-06 | 1.243E-05 | 5.335E-06 | 6.300E-06 | 1.600E-05 |
| 18897 | 945 | W945T | C | 1.1e-05 | 1.4e-05 | 1.4e-05 | 1.300E-05 | 1.732E-06 | 1.100E-05 | 1.400E-05 |
| 18898 | 945 | W945W | C | 9.3e-06 | 9.4e-06 | 1.4e-05 | 1.090E-05 | 2.685E-06 | 9.300E-06 | 1.400E-05 |
| 18899 | 945 | W945Y | C | 9.9e-06 | 1.5e-05 | 1.2e-05 | 1.230E-05 | 2.563E-06 | 9.900E-06 | 1.500E-05 |
| 18900 | 945 | W945V | C | 1.5e-05 | 1.6e-05 | 1.3e-05 | 1.467E-05 | 1.528E-06 | 1.300E-05 | 1.600E-05 |
| 18901 | 946 | G946A | C | 1.5e-05 | 9.7e-06 | 1.1e-05 | 1.190E-05 | 2.762E-06 | 9.700E-06 | 1.500E-05 |
| 18902 | 946 | G946R | C | 1.5e-05 | 1.1e-05 | 1.6e-05 | 1.400E-05 | 2.646E-06 | 1.100E-05 | 1.600E-05 |
| 18903 | 946 | G946N | C | 1.4e-05 | 1.1e-05 | 1.1e-05 | 1.200E-05 | 1.732E-06 | 1.100E-05 | 1.400E-05 |
| 18904 | 946 | G946D | C | 1e-05 | 1.6e-05 | 1.4e-05 | 1.333E-05 | 3.055E-06 | 1.000E-05 | 1.600E-05 |
| 18905 | 946 | G946C | C | 1.4e-05 | 1e-05 | 1.4e-05 | 1.267E-05 | 2.309E-06 | 1.000E-05 | 1.400E-05 |
| 18906 | 946 | G946E | C | 1.5e-05 | 1.5e-05 | 1e-05 | 1.333E-05 | 2.887E-06 | 1.000E-05 | 1.500E-05 |
| 18907 | 946 | G946Q | C | 1.3e-05 | 1.4e-05 | 1.6e-05 | 1.433E-05 | 1.528E-06 | 1.300E-05 | 1.600E-05 |
| 18908 | 946 | G946G | C | 1.6e-05 | 8.7e-06 | 1.5e-05 | 1.323E-05 | 3.958E-06 | 8.700E-06 | 1.600E-05 |
| 18909 | 946 | G946H | C | 1.3e-05 | 1.4e-05 | 9.8e-06 | 1.227E-05 | 2.194E-06 | 9.800E-06 | 1.400E-05 |
| 18910 | 946 | G946I | C | 1.5e-05 | 1.4e-05 | 1.1e-05 | 1.333E-05 | 2.082E-06 | 1.100E-05 | 1.500E-05 |
| 18911 | 946 | G946L | C | 1.5e-05 | 1.4e-05 | 1.6e-05 | 1.500E-05 | 1.000E-06 | 1.400E-05 | 1.600E-05 |
| 18912 | 946 | G946K | C | 9.3e-06 | 9.8e-06 | 9.9e-06 | 9.667E-06 | 3.215E-07 | 9.300E-06 | 9.900E-06 |
| 18913 | 946 | G946M | C | 1.4e-05 | 1.1e-05 | 1.4e-05 | 1.300E-05 | 1.732E-06 | 1.100E-05 | 1.400E-05 |
| 18914 | 946 | G946F | C | 1.5e-05 | 1.5e-05 | 8.3e-06 | 1.277E-05 | 3.868E-06 | 8.300E-06 | 1.500E-05 |
| 18915 | 946 | G946P | C | 1.5e-05 | 8.2e-06 | 1.5e-05 | 1.273E-05 | 3.926E-06 | 8.200E-06 | 1.500E-05 |
| 18916 | 946 | G946S | C | 9.8e-06 | 1e-05 | 1.2e-05 | 1.060E-05 | 1.217E-06 | 9.800E-06 | 1.200E-05 |
| 18917 | 946 | G946T | C | 1.1e-05 | 6.7e-06 | 1e-05 | 9.233E-06 | 2.250E-06 | 6.700E-06 | 1.100E-05 |
| 18918 | 946 | G946W | C | 1e-05 | 1.1e-05 | 1.3e-05 | 1.133E-05 | 1.528E-06 | 1.000E-05 | 1.300E-05 |
| 18919 | 946 | G946Y | C | 1.4e-05 | 9.6e-06 | 1.1e-05 | 1.153E-05 | 2.248E-06 | 9.600E-06 | 1.400E-05 |
| 18920 | 946 | G946V | C | 1.6e-05 | 1.1e-05 | 1.7e-05 | 1.467E-05 | 3.215E-06 | 1.100E-05 | 1.700E-05 |
| 18921 | 947 | Q947A | C | 1.1e-05 | 1.2e-05 | 8.4e-06 | 1.047E-05 | 1.858E-06 | 8.400E-06 | 1.200E-05 |
| 18922 | 947 | Q947R | C | 1.1e-05 | 1.7e-05 | 1.4e-05 | 1.400E-05 | 3.000E-06 | 1.100E-05 | 1.700E-05 |
| 18923 | 947 | Q947N | C | 7.2e-06 | 1.1e-05 | 9.3e-06 | 9.167E-06 | 1.904E-06 | 7.200E-06 | 1.100E-05 |
| 18924 | 947 | Q947D | C | 1.2e-05 | 1.6e-05 | 9.2e-06 | 1.240E-05 | 3.418E-06 | 9.200E-06 | 1.600E-05 |
| 18925 | 947 | Q947C | C | 1.8e-05 | 1.5e-05 | 9.4e-06 | 1.413E-05 | 4.365E-06 | 9.400E-06 | 1.800E-05 |
| 18926 | 947 | Q947E | C | 1.1e-05 | 1.1e-05 | 1.6e-05 | 1.267E-05 | 2.887E-06 | 1.100E-05 | 1.600E-05 |
| 18927 | 947 | Q947Q | C | 1e-05 | 7.7e-06 | 1.1e-05 | 9.567E-06 | 1.692E-06 | 7.700E-06 | 1.100E-05 |
| 18928 | 947 | Q947G | C | 1.8e-05 | 1.1e-05 | 1e-05 | 1.300E-05 | 4.359E-06 | 1.000E-05 | 1.800E-05 |
| 18929 | 947 | Q947H | C | 1.7e-05 | 2.3e-05 | 1.7e-05 | 1.900E-05 | 3.464E-06 | 1.700E-05 | 2.300E-05 |
| 18930 | 947 | Q947I | C | 1.1e-05 | 1.3e-05 | 1.2e-05 | 1.200E-05 | 1.000E-06 | 1.100E-05 | 1.300E-05 |
| 18931 | 947 | Q947L | C | 1.6e-05 | 1.7e-05 | 1.1e-05 | 1.467E-05 | 3.215E-06 | 1.100E-05 | 1.700E-05 |
| 18932 | 947 | Q947K | C | 1e-05 | 9.3e-06 | 8.8e-06 | 9.367E-06 | 6.028E-07 | 8.800E-06 | 1.000E-05 |
| 18933 | 947 | Q947M | C | 1.2e-05 | 1.2e-05 | 1.3e-05 | 1.233E-05 | 5.774E-07 | 1.200E-05 | 1.300E-05 |
| 18934 | 947 | Q947F | C | 1.2e-05 | 6.6e-06 | 8.8e-06 | 9.133E-06 | 2.715E-06 | 6.600E-06 | 1.200E-05 |
| 18935 | 947 | Q947P | C | 1.1e-05 | 1.1e-05 | 1.2e-05 | 1.133E-05 | 5.774E-07 | 1.100E-05 | 1.200E-05 |
| 18936 | 947 | Q947S | C | 9.2e-06 | 1e-05 | 1.5e-05 | 1.140E-05 | 3.143E-06 | 9.200E-06 | 1.500E-05 |
| 18937 | 947 | Q947T | C | 1e-05 | 1e-05 | 1.7e-05 | 1.233E-05 | 4.041E-06 | 1.000E-05 | 1.700E-05 |
| 18938 | 947 | Q947W | C | 9.1e-06 | 1e-05 | 1.1e-05 | 1.003E-05 | 9.504E-07 | 9.100E-06 | 1.100E-05 |
| 18939 | 947 | Q947Y | C | 1.1e-05 | 1.1e-05 | 1.5e-05 | 1.233E-05 | 2.309E-06 | 1.100E-05 | 1.500E-05 |
| 18940 | 947 | Q947V | C | 1.4e-05 | 1e-05 | 1.1e-05 | 1.167E-05 | 2.082E-06 | 1.000E-05 | 1.400E-05 |
| 18941 | 948 | G948A | C | 1.3e-05 | 1.3e-05 | 1.4e-05 | 1.333E-05 | 5.774E-07 | 1.300E-05 | 1.400E-05 |
| 18942 | 948 | G948R | C | 1.1e-05 | 9.7e-06 | 1.4e-05 | 1.157E-05 | 2.205E-06 | 9.700E-06 | 1.400E-05 |
| 18943 | 948 | G948N | C | 1.5e-05 | 9.1e-06 | 1.4e-05 | 1.270E-05 | 3.158E-06 | 9.100E-06 | 1.500E-05 |
| 18944 | 948 | G948D | C | 1e-05 | 1.3e-05 | 1.5e-05 | 1.267E-05 | 2.517E-06 | 1.000E-05 | 1.500E-05 |
| 18945 | 948 | G948C | C | 1.4e-05 | 1.1e-05 | 9.7e-06 | 1.157E-05 | 2.205E-06 | 9.700E-06 | 1.400E-05 |
| 18946 | 948 | G948E | C | 1.5e-05 | 9.8e-06 | 1.6e-05 | 1.360E-05 | 3.329E-06 | 9.800E-06 | 1.600E-05 |
| 18947 | 948 | G948Q | C | 1e-05 | 8.5e-06 | 1e-05 | 9.500E-06 | 8.660E-07 | 8.500E-06 | 1.000E-05 |
| 18948 | 948 | G948G | C | 1.5e-05 | 1.4e-05 | 9e-06 | 1.267E-05 | 3.215E-06 | 9.000E-06 | 1.500E-05 |
| 18949 | 948 | G948H | C | 1.3e-05 | 1.3e-05 | 9.9e-06 | 1.197E-05 | 1.790E-06 | 9.900E-06 | 1.300E-05 |
| 18950 | 948 | G948I | C | 1.4e-05 | 1.6e-05 | 9.6e-06 | 1.320E-05 | 3.274E-06 | 9.600E-06 | 1.600E-05 |
| 18951 | 948 | G948L | C | 1.1e-05 | 1.4e-05 | 1.1e-05 | 1.200E-05 | 1.732E-06 | 1.100E-05 | 1.400E-05 |
| 18952 | 948 | G948K | C | 1.1e-05 | 1.6e-05 | 9.3e-06 | 1.210E-05 | 3.483E-06 | 9.300E-06 | 1.600E-05 |
| 18953 | 948 | G948M | C | 1.4e-05 | 1e-05 | 1.6e-05 | 1.333E-05 | 3.055E-06 | 1.000E-05 | 1.600E-05 |
| 18954 | 948 | G948F | C | 1.4e-05 | 1.6e-05 | 1.5e-05 | 1.500E-05 | 1.000E-06 | 1.400E-05 | 1.600E-05 |

| | | | | | | | | | | |
|---|---|---|---|---|---|---|---|---|---|---|
| 18955 | 948 | G948P | C | 1.5e-05 | 1.5e-05 | 1.6e-05 | 1.533E-05 | 5.774E-07 | 1.500E-05 | 1.600E-05 |
| 18956 | 948 | G948S | C | 2.1e-05 | 1.1e-05 | 1.2e-05 | 1.467E-05 | 5.508E-06 | 1.100E-05 | 2.100E-05 |
| 18957 | 948 | G948T | C | 1.4e-05 | 8.3e-06 | 7.7e-06 | 1.000E-05 | 3.477E-06 | 7.700E-06 | 1.400E-05 |
| 18958 | 948 | G948W | C | 1.1e-05 | 1.5e-05 | 2.9e-05 | 1.833E-05 | 9.452E-06 | 1.100E-05 | 2.900E-05 |
| 18959 | 948 | G948Y | C | 1.1e-05 | 1.5e-05 | 1.5e-05 | 1.367E-05 | 2.309E-06 | 1.100E-05 | 1.500E-05 |
| 18960 | 948 | G948V | C | 9.7e-06 | 1.5e-05 | 1.8e-05 | 1.423E-05 | 4.203E-06 | 9.700E-06 | 1.800E-05 |
| 18961 | 949 | T949A | C | 1.6e-05 | 1.4e-05 | 1.2e-05 | 1.400E-05 | 2.000E-06 | 1.200E-05 | 1.600E-05 |
| 18962 | 949 | T949R | C | 1.6e-05 | 1.6e-05 | 9.9e-06 | 1.397E-05 | 3.522E-06 | 9.900E-06 | 1.600E-05 |
| 18963 | 949 | T949N | C | 1.4e-05 | 1.3e-05 | 1.2e-05 | 1.300E-05 | 1.000E-06 | 1.200E-05 | 1.400E-05 |
| 18964 | 949 | T949D | C | 1.5e-05 | 1.1e-05 | 1.3e-05 | 1.300E-05 | 2.000E-06 | 1.100E-05 | 1.500E-05 |
| 18965 | 949 | T949C | C | 1.6e-05 | 9.7e-06 | 7.4e-06 | 1.103E-05 | 4.452E-06 | 7.400E-06 | 1.600E-05 |
| 18966 | 949 | T949E | C | 1e-05 | 1.2e-05 | 1.3e-05 | 1.167E-05 | 1.528E-06 | 1.000E-05 | 1.300E-05 |
| 18967 | 949 | T949Q | C | 1e-05 | 1e-05 | 1.5e-05 | 1.167E-05 | 2.887E-06 | 1.000E-05 | 1.500E-05 |
| 18968 | 949 | T949G | C | 1.5e-05 | 1.4e-05 | 1.6e-05 | 1.500E-05 | 1.000E-06 | 1.400E-05 | 1.600E-05 |
| 18969 | 949 | T949H | C | 1e-05 | 1.4e-05 | 1.5e-05 | 1.300E-05 | 2.646E-06 | 1.000E-05 | 1.500E-05 |
| 18970 | 949 | T949I | C | 9.6e-06 | 1.1e-05 | 1.6e-05 | 1.220E-05 | 3.365E-06 | 9.600E-06 | 1.600E-05 |
| 18971 | 949 | T949L | C | 1.1e-05 | 1.6e-05 | 1.4e-05 | 1.367E-05 | 2.517E-06 | 1.100E-05 | 1.600E-05 |
| 18972 | 949 | T949K | C | 1.2e-05 | 1.5e-05 | 1e-05 | 1.233E-05 | 2.517E-06 | 1.000E-05 | 1.500E-05 |
| 18973 | 949 | T949M | C | 1.6e-05 | 1.1e-05 | 1.6e-05 | 1.433E-05 | 2.887E-06 | 1.100E-05 | 1.600E-05 |
| 18974 | 949 | T949F | C | 1.8e-05 | 1.5e-05 | 1e-05 | 1.433E-05 | 4.041E-06 | 1.000E-05 | 1.800E-05 |
| 18975 | 949 | T949P | C | 9.4e-06 | 1.3e-05 | 1.6e-05 | 1.280E-05 | 3.305E-06 | 9.400E-06 | 1.600E-05 |
| 18976 | 949 | T949S | C | 1.6e-05 | 1.7e-05 | 1.2e-05 | 1.500E-05 | 2.646E-06 | 1.200E-05 | 1.700E-05 |
| 18977 | 949 | T949T | C | 1.6e-05 | 1.2e-05 | 1.6e-05 | 1.467E-05 | 2.309E-06 | 1.200E-05 | 1.600E-05 |
| 18978 | 949 | T949W | C | 8.7e-06 | 1.2e-05 | 7.5e-06 | 9.400E-06 | 2.330E-06 | 7.500E-06 | 1.200E-05 |
| 18979 | 949 | T949Y | C | 1.1e-05 | 1.1e-05 | 1.1e-05 | 1.100E-05 | 2.075E-21 | 1.100E-05 | 1.100E-05 |
| 18980 | 949 | T949V | C | 1.7e-05 | 1.1e-05 | 1.5e-05 | 1.433E-05 | 3.055E-06 | 1.100E-05 | 1.700E-05 |
| 18981 | 950 | L950A | C | 1.6e-05 | 1.4e-05 | 1.1e-05 | 1.367E-05 | 2.517E-06 | 1.100E-05 | 1.600E-05 |
| 18982 | 950 | L950R | C | 1.5e-05 | 1.4e-05 | 1.4e-05 | 1.433E-05 | 5.774E-07 | 1.400E-05 | 1.500E-05 |
| 18983 | 950 | L950N | C | 1.4e-05 | 1e-05 | 8.9e-06 | 1.097E-05 | 2.684E-06 | 8.900E-06 | 1.400E-05 |
| 18984 | 950 | L950D | C | 8.2e-06 | 1.1e-05 | 1.1e-05 | 1.007E-05 | 1.617E-06 | 8.200E-06 | 1.100E-05 |
| 18985 | 950 | L950C | C | 8.4e-06 | 7.3e-06 | 1.2e-05 | 9.233E-06 | 2.458E-06 | 7.300E-06 | 1.200E-05 |
| 18986 | 950 | L950E | C | 1.5e-05 | 1.9e-05 | 1.1e-05 | 1.500E-05 | 4.000E-06 | 1.100E-05 | 1.900E-05 |
| 18987 | 950 | L950Q | C | 9.5e-06 | 1.6e-05 | 1.1e-05 | 1.217E-05 | 3.403E-06 | 9.500E-06 | 1.600E-05 |
| 18988 | 950 | L950G | C | 1.6e-05 | 1.3e-05 | 1.5e-05 | 1.467E-05 | 1.528E-06 | 1.300E-05 | 1.600E-05 |
| 18989 | 950 | L950H | C | 1.3e-05 | 1.4e-05 | 6.4e-06 | 1.113E-05 | 4.130E-06 | 6.400E-06 | 1.400E-05 |
| 18990 | 950 | L950I | C | 1.4e-05 | 1.2e-05 | 1.4e-05 | 1.333E-05 | 1.155E-06 | 1.200E-05 | 1.400E-05 |
| 18991 | 950 | L950L | C | 1.2e-05 | 1.5e-05 | 1.1e-05 | 1.267E-05 | 2.082E-06 | 1.100E-05 | 1.500E-05 |
| 18992 | 950 | L950K | C | 1.6e-05 | 1.5e-05 | 1.8e-05 | 1.633E-05 | 1.528E-06 | 1.500E-05 | 1.800E-05 |
| 18993 | 950 | L950M | C | 1.5e-05 | 1.3e-05 | 1.2e-05 | 1.333E-05 | 1.528E-06 | 1.200E-05 | 1.500E-05 |
| 18994 | 950 | L950F | C | 1.7e-05 | 1.4e-05 | 1.1e-05 | 1.400E-05 | 3.000E-06 | 1.100E-05 | 1.700E-05 |
| 18995 | 950 | L950P | C | 1.8e-05 | 1e-05 | 1.7e-05 | 1.500E-05 | 4.359E-06 | 1.000E-05 | 1.800E-05 |
| 18996 | 950 | L950S | C | 1.4e-05 | 1.5e-05 | 1.2e-05 | 1.367E-05 | 1.528E-06 | 1.200E-05 | 1.500E-05 |
| 18997 | 950 | L950T | C | 1.4e-05 | 8.7e-06 | 1.3e-05 | 1.190E-05 | 2.816E-06 | 8.700E-06 | 1.400E-05 |
| 18998 | 950 | L950W | C | 1.2e-05 | 1.6e-05 | 1.6e-05 | 1.467E-05 | 2.309E-06 | 1.200E-05 | 1.600E-05 |
| 18999 | 950 | L950Y | C | 1e-05 | 1.3e-05 | 1.1e-05 | 1.133E-05 | 1.528E-06 | 1.000E-05 | 1.300E-05 |
| 19000 | 950 | L950V | C | 1.3e-05 | 1.6e-05 | 2.5e-05 | 1.800E-05 | 6.245E-06 | 1.300E-05 | 2.500E-05 |
| 19001 | 951 | V951A | C | 1.4e-05 | 1.5e-05 | 1.8e-05 | 1.567E-05 | 2.082E-06 | 1.400E-05 | 1.800E-05 |
| 19002 | 951 | V951R | C | 1e-05 | 1.1e-05 | 9.6e-06 | 1.020E-05 | 7.211E-07 | 9.600E-06 | 1.100E-05 |
| 19003 | 951 | V951N | C | 1.1e-05 | 1.3e-05 | 1.4e-05 | 1.267E-05 | 1.528E-06 | 1.100E-05 | 1.400E-05 |
| 19004 | 951 | V951D | C | 1.3e-05 | 1.2e-05 | 7.4e-06 | 1.080E-05 | 2.987E-06 | 7.400E-06 | 1.300E-05 |
| 19005 | 951 | V951C | C | 1.5e-05 | 1.5e-05 | 9.2e-06 | 1.307E-05 | 3.349E-06 | 9.200E-06 | 1.500E-05 |
| 19006 | 951 | V951E | C | 9.8e-06 | 9.7e-06 | 1.5e-05 | 1.150E-05 | 3.032E-06 | 9.700E-06 | 1.500E-05 |
| 19007 | 951 | V951Q | C | 1.4e-05 | 1.5e-05 | 7.5e-06 | 1.217E-05 | 4.072E-06 | 7.500E-06 | 1.500E-05 |
| 19008 | 951 | V951G | C | 1.5e-05 | 8.3e-06 | 1.7e-05 | 1.343E-05 | 4.557E-06 | 8.300E-06 | 1.700E-05 |
| 19009 | 951 | V951H | C | 1.1e-05 | 1.8e-05 | 1.4e-05 | 1.433E-05 | 3.512E-06 | 1.100E-05 | 1.800E-05 |
| 19010 | 951 | V951I | C | 9.5e-06 | 1.5e-05 | 1.3e-05 | 1.250E-05 | 2.784E-06 | 9.500E-06 | 1.500E-05 |
| 19011 | 951 | V951L | C | 1.5e-05 | 1.1e-05 | 1.1e-05 | 1.233E-05 | 2.309E-06 | 1.100E-05 | 1.500E-05 |
| 19012 | 951 | V951K | C | 1e-05 | 1.5e-05 | 1.4e-05 | 1.300E-05 | 2.646E-06 | 1.000E-05 | 1.500E-05 |
| 19013 | 951 | V951M | C | 1e-05 | 1.3e-05 | 1.1e-05 | 1.133E-05 | 1.528E-06 | 1.000E-05 | 1.300E-05 |
| 19014 | 951 | V951F | C | 1.1e-05 | 8.9e-06 | 1.1e-05 | 1.030E-05 | 1.212E-06 | 8.900E-06 | 1.100E-05 |
| 19015 | 951 | V951P | C | 6.2e-06 | 9.8e-06 | 1.6e-05 | 1.067E-05 | 4.957E-06 | 6.200E-06 | 1.600E-05 |
| 19016 | 951 | V951S | C | 1.4e-05 | 1.6e-05 | 1.4e-05 | 1.467E-05 | 1.155E-06 | 1.400E-05 | 1.600E-05 |
| 19017 | 951 | V951T | C | 1e-05 | 1.6e-05 | 1.1e-05 | 1.233E-05 | 3.215E-06 | 1.000E-05 | 1.600E-05 |
| 19018 | 951 | V951W | C | 1.4e-05 | 1.2e-05 | 1.1e-05 | 1.233E-05 | 1.528E-06 | 1.100E-05 | 1.400E-05 |
| 19019 | 951 | V951Y | C | 1.2e-05 | 1.4e-05 | 1.7e-05 | 1.433E-05 | 2.517E-06 | 1.200E-05 | 1.700E-05 |
| 19020 | 951 | V951V | C | 1.6e-05 | 1.6e-05 | 1.2e-05 | 1.467E-05 | 2.309E-06 | 1.200E-05 | 1.600E-05 |
| 19021 | 952 | T952A | C | 9.6e-06 | 1.1e-05 | 2.2e-05 | 1.420E-05 | 6.791E-06 | 9.600E-06 | 2.200E-05 |
| 19022 | 952 | T952R | C | 7.6e-06 | 1e-05 | 1.6e-05 | 1.120E-05 | 4.327E-06 | 7.600E-06 | 1.600E-05 |
| 19023 | 952 | T952N | C | 9.4e-06 | 1.6e-05 | 1.5e-05 | 1.347E-05 | 3.557E-06 | 9.400E-06 | 1.600E-05 |
| 19024 | 952 | T952D | C | 1.1e-05 | 1.8e-05 | 1.6e-05 | 1.500E-05 | 3.606E-06 | 1.100E-05 | 1.800E-05 |
| 19025 | 952 | T952C | C | 9.4e-06 | 1.1e-05 | 1.5e-05 | 1.180E-05 | 2.884E-06 | 9.400E-06 | 1.500E-05 |
| 19026 | 952 | T952E | C | 1e-05 | 1.6e-05 | 1e-05 | 1.200E-05 | 3.464E-06 | 1.000E-05 | 1.600E-05 |
| 19027 | 952 | T952Q | C | 1.3e-05 | 1.3e-05 | 1.5e-05 | 1.367E-05 | 1.155E-06 | 1.300E-05 | 1.500E-05 |
| 19028 | 952 | T952G | C | 1.1e-05 | 9.5e-06 | 1.1e-05 | 1.050E-05 | 8.660E-07 | 9.500E-06 | 1.100E-05 |
| 19029 | 952 | T952H | C | 9.3e-06 | 1e-05 | 1.5e-05 | 1.143E-05 | 3.109E-06 | 9.300E-06 | 1.500E-05 |
| 19030 | 952 | T952I | C | 1.7e-05 | 1.1e-05 | 9.4e-06 | 1.247E-05 | 4.007E-06 | 9.400E-06 | 1.700E-05 |
| 19031 | 952 | T952L | C | 1.1e-05 | 1.4e-05 | 1.5e-05 | 1.333E-05 | 2.082E-06 | 1.100E-05 | 1.500E-05 |
| 19032 | 952 | T952K | C | 1.5e-05 | 9.6e-06 | 1.1e-05 | 1.187E-05 | 2.802E-06 | 9.600E-06 | 1.500E-05 |
| 19033 | 952 | T952M | C | 1.7e-05 | 1.4e-05 | 1.6e-05 | 1.567E-05 | 1.528E-06 | 1.400E-05 | 1.700E-05 |
| 19034 | 952 | T952F | C | 1.4e-05 | 1.1e-05 | 1.5e-05 | 1.333E-05 | 2.082E-06 | 1.100E-05 | 1.500E-05 |
| 19035 | 952 | T952P | C | 1e-05 | 1.5e-05 | 1.1e-05 | 1.200E-05 | 2.646E-06 | 1.000E-05 | 1.500E-05 |
| 19036 | 952 | T952S | C | 1.1e-05 | 1.2e-05 | 1e-05 | 1.100E-05 | 1.000E-06 | 1.000E-05 | 1.200E-05 |
| 19037 | 952 | T952T | C | 1e-05 | 1.9e-05 | 1.1e-05 | 1.333E-05 | 4.933E-06 | 1.000E-05 | 1.900E-05 |
| 19038 | 952 | T952W | C | 1.6e-05 | 1.5e-05 | 1.2e-05 | 1.433E-05 | 2.082E-06 | 1.200E-05 | 1.600E-05 |
| 19039 | 952 | T952Y | C | 1.4e-05 | 1.6e-05 | 1.3e-05 | 1.433E-05 | 1.528E-06 | 1.300E-05 | 1.600E-05 |

| | | | | | | | | | | |
|---|---|---|---|---|---|---|---|---|---|---|
| 19040 | 952 | T952V | C | 1.5e-05 | 1.1e-05 | 1.4e-05 | 1.333E-05 | 2.082E-06 | 1.100E-05 | 1.500E-05 |
| 19041 | 953 | V953A | C | 1.5e-05 | 1.7e-05 | 1.5e-05 | 1.567E-05 | 1.155E-06 | 1.500E-05 | 1.700E-05 |
| 19042 | 953 | V953R | C | 8.5e-06 | 1.5e-05 | 1.5e-05 | 1.283E-05 | 3.753E-06 | 8.500E-06 | 1.500E-05 |
| 19043 | 953 | V953N | C | 1.5e-05 | 1e-05 | 8.9e-06 | 1.130E-05 | 3.251E-06 | 8.900E-06 | 1.500E-05 |
| 19044 | 953 | V953D | C | 1.5e-05 | 1e-05 | 1.6e-05 | 1.367E-05 | 3.215E-06 | 1.000E-05 | 1.600E-05 |
| 19045 | 953 | V953C | C | 1.3e-05 | 8.2e-06 | 1.1e-05 | 1.073E-05 | 2.411E-06 | 8.200E-06 | 1.300E-05 |
| 19046 | 953 | V953E | C | 1.1e-05 | 1.1e-05 | 1.2e-05 | 1.133E-05 | 5.774E-07 | 1.100E-05 | 1.200E-05 |
| 19047 | 953 | V953Q | C | 1.4e-05 | 1.1e-05 | 1.4e-05 | 1.300E-05 | 1.732E-06 | 1.100E-05 | 1.400E-05 |
| 19048 | 953 | V953G | C | 1.6e-05 | 1.2e-05 | 1.4e-05 | 1.400E-05 | 2.000E-06 | 1.200E-05 | 1.600E-05 |
| 19049 | 953 | V953H | C | 1.5e-05 | 1.4e-05 | 1.4e-05 | 1.433E-05 | 5.774E-07 | 1.400E-05 | 1.500E-05 |
| 19050 | 953 | V953I | C | 1.1e-05 | 1.2e-05 | 1.3e-05 | 1.200E-05 | 1.000E-06 | 1.100E-05 | 1.300E-05 |
| 19051 | 953 | V953L | C | 1.1e-05 | 1.3e-05 | 1.1e-05 | 1.167E-05 | 1.155E-06 | 1.100E-05 | 1.300E-05 |
| 19052 | 953 | V953K | C | 8.6e-06 | 1e-05 | 1.4e-05 | 1.087E-05 | 2.802E-06 | 8.600E-06 | 1.400E-05 |
| 19053 | 953 | V953M | C | 8.9e-06 | 1e-05 | 1.4e-05 | 1.097E-05 | 2.684E-06 | 8.900E-06 | 1.400E-05 |
| 19054 | 953 | V953F | C | 1.5e-05 | 9.9e-06 | 1.1e-05 | 1.197E-05 | 2.684E-06 | 9.900E-06 | 1.500E-05 |
| 19055 | 953 | V953P | C | 1.8e-05 | 9.4e-06 | 7.5e-06 | 1.163E-05 | 5.595E-06 | 7.500E-06 | 1.800E-05 |
| 19056 | 953 | V953S | C | 7.5e-06 | 1.1e-05 | 1e-05 | 9.500E-06 | 1.803E-06 | 7.500E-06 | 1.100E-05 |
| 19057 | 953 | V953T | C | 9.4e-06 | 1.3e-05 | 1.6e-05 | 1.280E-05 | 3.305E-06 | 9.400E-06 | 1.600E-05 |
| 19058 | 953 | V953W | C | 1.2e-05 | 2.3e-05 | 1.5e-05 | 1.667E-05 | 5.686E-06 | 1.200E-05 | 2.300E-05 |
| 19059 | 953 | V953Y | C | 1.4e-05 | 1.2e-05 | 9.8e-06 | 1.193E-05 | 2.101E-06 | 9.800E-06 | 1.400E-05 |
| 19060 | 953 | V953V | C | 1e-05 | 9.9e-06 | 1.6e-05 | 1.197E-05 | 3.493E-06 | 9.900E-06 | 1.600E-05 |
| 19061 | 954 | S954A | C | 1.4e-05 | 1.2e-05 | 1.6e-05 | 1.400E-05 | 2.000E-06 | 1.200E-05 | 1.600E-05 |
| 19062 | 954 | S954R | C | 1.1e-05 | 1.1e-05 | 1e-05 | 1.067E-05 | 5.774E-07 | 1.000E-05 | 1.100E-05 |
| 19063 | 954 | S954N | C | 1.6e-05 | 1.6e-05 | 1.1e-05 | 1.433E-05 | 2.887E-06 | 1.100E-05 | 1.600E-05 |
| 19064 | 954 | S954D | C | 9.2e-06 | 1.6e-05 | 1e-05 | 1.173E-05 | 3.717E-06 | 9.200E-06 | 1.600E-05 |
| 19065 | 954 | S954C | C | 8.9e-06 | 1.2e-05 | 1e-05 | 1.030E-05 | 1.572E-06 | 8.900E-06 | 1.200E-05 |
| 19066 | 954 | S954E | C | 1.7e-05 | 2.3e-05 | 1.6e-05 | 1.867E-05 | 3.786E-06 | 1.600E-05 | 2.300E-05 |
| 19067 | 954 | S954Q | C | 1.6e-05 | 1.5e-05 | 8.3e-06 | 1.310E-05 | 4.187E-06 | 8.300E-06 | 1.600E-05 |
| 19068 | 954 | S954G | C | 8.4e-06 | 1.2e-05 | 1.1e-05 | 1.047E-05 | 1.858E-06 | 8.400E-06 | 1.200E-05 |
| 19069 | 954 | S954H | C | 1.3e-05 | 1.2e-05 | 1.5e-05 | 1.333E-05 | 1.528E-06 | 1.200E-05 | 1.500E-05 |
| 19070 | 954 | S954I | C | 1.6e-05 | 8.7e-06 | 1.6e-05 | 1.357E-05 | 4.215E-06 | 8.700E-06 | 1.600E-05 |
| 19071 | 954 | S954L | C | 1.6e-05 | 1.4e-05 | 1.1e-05 | 1.367E-05 | 2.517E-06 | 1.100E-05 | 1.600E-05 |
| 19072 | 954 | S954K | C | 1.4e-05 | 1.4e-05 | 9.4e-06 | 1.247E-05 | 2.656E-06 | 9.400E-06 | 1.400E-05 |
| 19073 | 954 | S954M | C | 1.5e-05 | 1.5e-05 | 1.5e-05 | 1.500E-05 | 0.000E+00 | 1.500E-05 | 1.500E-05 |
| 19074 | 954 | S954F | C | 1.5e-05 | 8.7e-06 | 7.8e-06 | 1.050E-05 | 3.923E-06 | 7.800E-06 | 1.500E-05 |
| 19075 | 954 | S954P | C | 1.6e-05 | 1.8e-05 | 1e-05 | 1.467E-05 | 4.163E-06 | 1.000E-05 | 1.800E-05 |
| 19076 | 954 | S954S | C | 1.1e-05 | 9.2e-06 | 1.5e-05 | 1.173E-05 | 2.969E-06 | 9.200E-06 | 1.500E-05 |
| 19077 | 954 | S954T | C | 1.4e-05 | 1e-05 | 1.5e-05 | 1.300E-05 | 2.646E-06 | 1.000E-05 | 1.500E-05 |
| 19078 | 954 | S954W | C | 1.3e-05 | 1.5e-05 | 1.5e-05 | 1.433E-05 | 1.155E-06 | 1.300E-05 | 1.500E-05 |
| 19079 | 954 | S954Y | C | 9.9e-06 | 1.1e-05 | 1.6e-05 | 1.230E-05 | 3.251E-06 | 9.900E-06 | 1.600E-05 |
| 19080 | 954 | S954V | C | 1.6e-05 | 1.1e-05 | 1.7e-05 | 1.467E-05 | 3.215E-06 | 1.100E-05 | 1.700E-05 |
| 19081 | 955 | S955A | C | 1.5e-05 | 1.6e-05 | 1.5e-05 | 1.533E-05 | 5.774E-07 | 1.500E-05 | 1.600E-05 |
| 19082 | 955 | S955R | C | 1.5e-05 | 1.1e-05 | 1.2e-05 | 1.267E-05 | 2.082E-06 | 1.100E-05 | 1.500E-05 |
| 19083 | 955 | S955N | C | 1.5e-05 | 1.5e-05 | 1.1e-05 | 1.367E-05 | 2.309E-06 | 1.100E-05 | 1.500E-05 |
| 19084 | 955 | S955D | C | 1.5e-05 | 1.6e-05 | 6.7e-06 | 1.257E-05 | 5.105E-06 | 6.700E-06 | 1.600E-05 |
| 19085 | 955 | S955C | C | 1.2e-05 | 1.4e-05 | 1e-05 | 1.200E-05 | 2.000E-06 | 1.000E-05 | 1.400E-05 |
| 19086 | 955 | S955E | C | 1.5e-05 | 1.1e-05 | 1.4e-05 | 1.333E-05 | 2.082E-06 | 1.100E-05 | 1.500E-05 |
| 19087 | 955 | S955Q | C | 9.9e-06 | 1.6e-05 | 6.6e-06 | 1.083E-05 | 4.769E-06 | 6.600E-06 | 1.600E-05 |
| 19088 | 955 | S955G | C | 1.6e-05 | 1.6e-05 | 9.1e-06 | 1.370E-05 | 3.984E-06 | 9.100E-06 | 1.600E-05 |
| 19089 | 955 | S955H | C | 1.3e-05 | 1.2e-05 | 1.5e-05 | 1.333E-05 | 1.528E-06 | 1.200E-05 | 1.500E-05 |
| 19090 | 955 | S955I | C | 1.5e-05 | 1.1e-05 | 1.7e-05 | 1.433E-05 | 3.055E-06 | 1.100E-05 | 1.700E-05 |
| 19091 | 955 | S955L | C | 1.5e-05 | 8.2e-06 | 2e-05 | 1.440E-05 | 5.923E-06 | 8.200E-06 | 2.000E-05 |
| 19092 | 955 | S955K | C | 1.2e-05 | 1.4e-05 | 8.6e-06 | 1.153E-05 | 2.730E-06 | 8.600E-06 | 1.400E-05 |
| 19093 | 955 | S955M | C | 1e-05 | 1.4e-05 | 1.1e-05 | 1.167E-05 | 2.082E-06 | 1.000E-05 | 1.400E-05 |
| 19094 | 955 | S955F | C | 7.4e-06 | 1.3e-05 | 1.2e-05 | 1.080E-05 | 2.987E-06 | 7.400E-06 | 1.300E-05 |
| 19095 | 955 | S955P | C | 1.6e-05 | 1.1e-05 | 1.1e-05 | 1.267E-05 | 2.887E-06 | 1.100E-05 | 1.600E-05 |
| 19096 | 955 | S955S | C | 7.4e-06 | 9.4e-06 | 1.4e-05 | 1.027E-05 | 3.384E-06 | 7.400E-06 | 1.400E-05 |
| 19097 | 955 | S955T | C | 1.2e-05 | 1.5e-05 | 1.5e-05 | 1.400E-05 | 1.732E-06 | 1.200E-05 | 1.500E-05 |
| 19098 | 955 | S955W | C | 1.6e-05 | 1.5e-05 | 1.1e-05 | 1.400E-05 | 2.646E-06 | 1.100E-05 | 1.600E-05 |
| 19099 | 955 | S955Y | C | 1.2e-05 | 1e-05 | 1.4e-05 | 1.200E-05 | 2.000E-06 | 1.000E-05 | 1.400E-05 |
| 19100 | 955 | S955V | C | 1.2e-05 | 1.1e-05 | 1.6e-05 | 1.300E-05 | 2.646E-06 | 1.100E-05 | 1.600E-05 |
| 19101 | 956 | A956A | C | 1.4e-05 | 8.9e-06 | 1e-05 | 1.097E-05 | 2.684E-06 | 8.900E-06 | 1.400E-05 |
| 19102 | 956 | A956R | C | 1.3e-05 | 1.1e-05 | 1.1e-05 | 1.167E-05 | 1.155E-06 | 1.100E-05 | 1.300E-05 |
| 19103 | 956 | A956N | C | 1.3e-05 | 8.9e-06 | 9.2e-06 | 1.037E-05 | 2.285E-06 | 8.900E-06 | 1.300E-05 |
| 19104 | 956 | A956D | C | 1.5e-05 | 8.4e-06 | 1.5e-05 | 1.280E-05 | 3.811E-06 | 8.400E-06 | 1.500E-05 |
| 19105 | 956 | A956C | C | 1.1e-05 | 9.1e-06 | 1.6e-05 | 1.203E-05 | 3.564E-06 | 9.100E-06 | 1.600E-05 |
| 19106 | 956 | A956E | C | 1.2e-05 | 8.8e-06 | 7.8e-06 | 9.533E-06 | 2.194E-06 | 7.800E-06 | 1.200E-05 |
| 19107 | 956 | A956Q | C | 1.7e-05 | 1.1e-05 | 1.4e-05 | 1.400E-05 | 3.000E-06 | 1.100E-05 | 1.700E-05 |
| 19108 | 956 | A956G | C | 1.3e-05 | 1.4e-05 | 1.1e-05 | 1.267E-05 | 1.528E-06 | 1.100E-05 | 1.400E-05 |
| 19109 | 956 | A956H | C | 1.5e-05 | 1e-05 | 1.5e-05 | 1.333E-05 | 2.887E-06 | 1.000E-05 | 1.500E-05 |
| 19110 | 956 | A956I | C | 1.2e-05 | 1.2e-05 | 1.1e-05 | 1.167E-05 | 5.774E-07 | 1.100E-05 | 1.200E-05 |
| 19111 | 956 | A956L | C | 1.2e-05 | 8.7e-06 | 1.1e-05 | 1.057E-05 | 1.692E-06 | 8.700E-06 | 1.200E-05 |
| 19112 | 956 | A956K | C | 1.5e-05 | 9e-06 | 2.2e-05 | 1.533E-05 | 6.506E-06 | 9.000E-06 | 2.200E-05 |
| 19113 | 956 | A956M | C | 1.4e-05 | 1.5e-05 | 1.5e-05 | 1.467E-05 | 5.774E-07 | 1.400E-05 | 1.500E-05 |
| 19114 | 956 | A956F | C | 1.1e-05 | 1.5e-05 | 1.1e-05 | 1.233E-05 | 2.309E-06 | 1.100E-05 | 1.500E-05 |
| 19115 | 956 | A956P | C | 2.7e-05 | 1.4e-05 | 1e-05 | 1.700E-05 | 8.888E-06 | 1.000E-05 | 2.700E-05 |
| 19116 | 956 | A956S | C | 1.4e-05 | 1.1e-05 | 9.3e-06 | 1.143E-05 | 2.380E-06 | 9.300E-06 | 1.400E-05 |
| 19117 | 956 | A956T | C | 1.6e-05 | 1.5e-05 | 1.4e-05 | 1.500E-05 | 1.000E-06 | 1.400E-05 | 1.600E-05 |
| 19118 | 956 | A956W | C | 1.4e-05 | 9.7e-06 | 1.2e-05 | 1.190E-05 | 2.152E-06 | 9.700E-06 | 1.400E-05 |
| 19119 | 956 | A956Y | C | 9.4e-06 | 1.5e-05 | 1.1e-05 | 1.180E-05 | 2.884E-06 | 9.400E-06 | 1.500E-05 |
| 19120 | 956 | A956V | C | 1e-05 | 1.1e-05 | 1.3e-05 | 1.133E-05 | 1.528E-06 | 1.000E-05 | 1.300E-05 |
| 19121 | 957 | S957A | C | 1e-05 | 1.7e-05 | 1.2e-05 | 1.300E-05 | 3.606E-06 | 1.000E-05 | 1.700E-05 |
| 19122 | 957 | S957R | C | 9.7e-06 | 1.5e-05 | 1.6e-05 | 1.357E-05 | 3.386E-06 | 9.700E-06 | 1.600E-05 |
| 19123 | 957 | S957N | C | 1.5e-05 | 1.1e-05 | 1e-05 | 1.200E-05 | 2.646E-06 | 1.000E-05 | 1.500E-05 |
| 19124 | 957 | S957D | C | 1.1e-05 | 1.5e-05 | 8.9e-06 | 1.163E-05 | 3.099E-06 | 8.900E-06 | 1.500E-05 |

| | | | | | | | | | | |
|---|---|---|---|---|---|---|---|---|---|---|
| 19125 | 957 | S957C | C | 1.5e-05 | 1.4e-05 | 9.4e-06 | 1.280E-05 | 2.987E-06 | 9.400E-06 | 1.500E-05 |
| 19126 | 957 | S957E | C | 1e-05 | 1.3e-05 | 1.5e-05 | 1.267E-05 | 2.517E-06 | 1.000E-05 | 1.500E-05 |
| 19127 | 957 | S957Q | C | 1e-05 | 1.1e-05 | 1.7e-05 | 1.267E-05 | 3.786E-06 | 1.000E-05 | 1.700E-05 |
| 19128 | 957 | S957G | C | 1.5e-05 | 1e-05 | 1.7e-05 | 1.400E-05 | 3.606E-06 | 1.000E-05 | 1.700E-05 |
| 19129 | 957 | S957H | C | 1.1e-05 | 1.7e-05 | 1.4e-05 | 1.400E-05 | 3.000E-06 | 1.100E-05 | 1.700E-05 |
| 19130 | 957 | S957I | C | 1.8e-05 | 1.6e-05 | 1.1e-05 | 1.500E-05 | 3.606E-06 | 1.100E-05 | 1.800E-05 |
| 19131 | 957 | S957L | C | 1.7e-05 | 1.6e-05 | 1e-05 | 1.433E-05 | 3.786E-06 | 1.000E-05 | 1.700E-05 |
| 19132 | 957 | S957K | C | 1.6e-05 | 8.6e-06 | 9.5e-06 | 1.137E-05 | 4.038E-06 | 8.600E-06 | 1.600E-05 |
| 19133 | 957 | S957M | C | 9.7e-06 | 1.3e-05 | 1.6e-05 | 1.290E-05 | 3.151E-06 | 9.700E-06 | 1.600E-05 |
| 19134 | 957 | S957F | C | 1.5e-05 | 1e-05 | 8.7e-06 | 1.123E-05 | 3.326E-06 | 8.700E-06 | 1.500E-05 |
| 19135 | 957 | S957P | C | 1e-05 | 1.6e-05 | 1.6e-05 | 1.400E-05 | 3.464E-06 | 1.000E-05 | 1.600E-05 |
| 19136 | 957 | S957S | C | 1.6e-05 | 1.4e-05 | 1.4e-05 | 1.467E-05 | 1.155E-06 | 1.400E-05 | 1.600E-05 |
| 19137 | 957 | S957T | C | 1e-05 | 1.5e-05 | 8.5e-06 | 1.117E-05 | 3.403E-06 | 8.500E-06 | 1.500E-05 |
| 19138 | 957 | S957W | C | 1.4e-05 | 1.5e-05 | 1.5e-05 | 1.467E-05 | 5.774E-07 | 1.400E-05 | 1.500E-05 |
| 19139 | 957 | S957Y | C | 9.9e-06 | 8.9e-06 | 1.2e-05 | 1.027E-05 | 1.582E-06 | 8.900E-06 | 1.200E-05 |
| 19140 | 957 | S957V | C | 1.5e-05 | 1.6e-05 | 1.1e-05 | 1.400E-05 | 2.646E-06 | 1.100E-05 | 1.600E-05 |
| 19141 | 958 | T958A | C | 9.3e-06 | 1e-05 | 1e-05 | 9.767E-06 | 4.041E-07 | 9.300E-06 | 1.000E-05 |
| 19142 | 958 | T958R | C | 1.5e-05 | 1.1e-05 | 1.5e-05 | 1.367E-05 | 2.309E-06 | 1.100E-05 | 1.500E-05 |
| 19143 | 958 | T958N | C | 1.6e-05 | 8.9e-06 | 1.7e-05 | 1.397E-05 | 4.416E-06 | 8.900E-06 | 1.700E-05 |
| 19144 | 958 | T958D | C | 1.5e-05 | 1.4e-05 | 1e-05 | 1.300E-05 | 2.646E-06 | 1.000E-05 | 1.500E-05 |
| 19145 | 958 | T958C | C | 1.5e-05 | 8.7e-06 | 1.1e-05 | 1.157E-05 | 3.188E-06 | 8.700E-06 | 1.500E-05 |
| 19146 | 958 | T958E | C | 1e-05 | 1.1e-05 | 1.5e-05 | 1.200E-05 | 2.646E-06 | 1.000E-05 | 1.500E-05 |
| 19147 | 958 | T958Q | C | 9.9e-06 | 1.5e-05 | 9.3e-06 | 1.140E-05 | 3.132E-06 | 9.300E-06 | 1.500E-05 |
| 19148 | 958 | T958G | C | 1.2e-05 | 1.5e-05 | 1.6e-05 | 1.433E-05 | 2.082E-06 | 1.200E-05 | 1.600E-05 |
| 19149 | 958 | T958H | C | 1.1e-05 | 1.6e-05 | 9.9e-06 | 1.230E-05 | 3.251E-06 | 9.900E-06 | 1.600E-05 |
| 19150 | 958 | T958I | C | 1.4e-05 | 1.1e-05 | 1.1e-05 | 1.200E-05 | 1.732E-06 | 1.100E-05 | 1.400E-05 |
| 19151 | 958 | T958L | C | 1.6e-05 | 1.7e-05 | 1.5e-05 | 1.600E-05 | 1.000E-06 | 1.500E-05 | 1.700E-05 |
| 19152 | 958 | T958K | C | 7.8e-06 | 1.3e-05 | 1.6e-05 | 1.227E-05 | 4.149E-06 | 7.800E-06 | 1.600E-05 |
| 19153 | 958 | T958M | C | 1.6e-05 | 1e-05 | 1.5e-05 | 1.367E-05 | 3.215E-06 | 1.000E-05 | 1.600E-05 |
| 19154 | 958 | T958F | C | 1.3e-05 | 1.6e-05 | 1.3e-05 | 1.400E-05 | 1.732E-06 | 1.300E-05 | 1.600E-05 |
| 19155 | 958 | T958P | C | 1.1e-05 | 8.6e-06 | 1.5e-05 | 1.153E-05 | 3.233E-06 | 8.600E-06 | 1.500E-05 |
| 19156 | 958 | T958S | C | 1e-05 | 1.4e-05 | 1e-05 | 1.133E-05 | 2.309E-06 | 1.000E-05 | 1.400E-05 |
| 19157 | 958 | T958T | C | 1.1e-05 | 1.5e-05 | 9.1e-06 | 1.170E-05 | 3.012E-06 | 9.100E-06 | 1.500E-05 |
| 19158 | 958 | T958W | C | 1.4e-05 | 1.5e-05 | 1.1e-05 | 1.333E-05 | 2.082E-06 | 1.100E-05 | 1.500E-05 |
| 19159 | 958 | T958Y | C | 1.1e-05 | 1.4e-05 | 1.1e-05 | 1.200E-05 | 1.732E-06 | 1.100E-05 | 1.400E-05 |
| 19160 | 958 | T958V | C | 1.7e-05 | 1.6e-05 | 1.1e-05 | 1.467E-05 | 3.215E-06 | 1.100E-05 | 1.700E-05 |
| 19161 | 959 | K959A | C | 1.4e-05 | 1.4e-05 | 1.1e-05 | 1.300E-05 | 1.732E-06 | 1.100E-05 | 1.400E-05 |
| 19162 | 959 | K959R | C | 1e-05 | 1.1e-05 | 9.8e-06 | 1.027E-05 | 6.429E-07 | 9.800E-06 | 1.100E-05 |
| 19163 | 959 | K959N | C | 1.5e-05 | 1.5e-05 | 1.4e-05 | 1.467E-05 | 5.774E-07 | 1.400E-05 | 1.500E-05 |
| 19164 | 959 | K959D | C | 1.3e-05 | 1.5e-05 | 1.6e-05 | 1.467E-05 | 1.528E-06 | 1.300E-05 | 1.600E-05 |
| 19165 | 959 | K959C | C | 1.3e-05 | 1.3e-05 | 1.1e-05 | 1.233E-05 | 1.155E-06 | 1.100E-05 | 1.300E-05 |
| 19166 | 959 | K959E | C | 1.1e-05 | 1.3e-05 | 7e-06 | 1.033E-05 | 3.055E-06 | 7.000E-06 | 1.300E-05 |
| 19167 | 959 | K959Q | C | 9.9e-06 | 9.6e-06 | 1e-05 | 9.833E-06 | 2.082E-07 | 9.600E-06 | 1.000E-05 |
| 19168 | 959 | K959G | C | 1.4e-05 | 1.2e-05 | 8.9e-06 | 1.163E-05 | 2.570E-06 | 8.900E-06 | 1.400E-05 |
| 19169 | 959 | K959H | C | 9.1e-06 | 1.4e-05 | 9.8e-06 | 1.097E-05 | 2.650E-06 | 9.100E-06 | 1.400E-05 |
| 19170 | 959 | K959I | C | 1.3e-05 | 1.2e-05 | 1.2e-05 | 1.233E-05 | 5.774E-07 | 1.200E-05 | 1.300E-05 |
| 19171 | 959 | K959L | C | 1.1e-05 | 1.1e-05 | 9.9e-06 | 1.063E-05 | 6.351E-07 | 9.900E-06 | 1.100E-05 |
| 19172 | 959 | K959K | C | 1.2e-05 | 1.6e-05 | 1.2e-05 | 1.333E-05 | 2.309E-06 | 1.200E-05 | 1.600E-05 |
| 19173 | 959 | K959M | C | 8.4e-06 | 1.2e-05 | 2.8e-05 | 1.613E-05 | 1.043E-05 | 8.400E-06 | 2.800E-05 |
| 19174 | 959 | K959F | C | 1.5e-05 | 1.7e-05 | 2.6e-05 | 1.933E-05 | 5.859E-06 | 1.500E-05 | 2.600E-05 |
| 19175 | 959 | K959P | C | 1.2e-05 | 1.1e-05 | 1.5e-05 | 1.267E-05 | 2.082E-06 | 1.100E-05 | 1.500E-05 |
| 19176 | 959 | K959S | C | 1.4e-05 | 1e-05 | 1.5e-05 | 1.300E-05 | 2.646E-06 | 1.000E-05 | 1.500E-05 |
| 19177 | 959 | K959T | C | 1e-05 | 1e-05 | 9.8e-06 | 9.933E-06 | 1.155E-07 | 9.800E-06 | 1.000E-05 |
| 19178 | 959 | K959W | C | 1.6e-05 | 1.1e-05 | 1.5e-05 | 1.400E-05 | 2.646E-06 | 1.100E-05 | 1.600E-05 |
| 19179 | 959 | K959Y | C | 1.5e-05 | 1.3e-05 | 1e-05 | 1.267E-05 | 2.517E-06 | 1.000E-05 | 1.500E-05 |
| 19180 | 959 | K959V | C | 1.4e-05 | 1.2e-05 | 1.6e-05 | 1.400E-05 | 2.000E-06 | 1.200E-05 | 1.600E-05 |
| 19181 | 960 | G960A | C | 8.6e-06 | 8.9e-06 | 1.7e-05 | 1.150E-05 | 4.766E-06 | 8.600E-06 | 1.700E-05 |
| 19182 | 960 | G960R | C | 1.2e-05 | 1.3e-05 | 9.9e-06 | 1.163E-05 | 1.582E-06 | 9.900E-06 | 1.300E-05 |
| 19183 | 960 | G960N | C | 8.1e-06 | 1.4e-05 | 6.5e-06 | 9.533E-06 | 3.950E-06 | 6.500E-06 | 1.400E-05 |
| 19184 | 960 | G960D | C | 1e-05 | 1.5e-05 | 1.5e-05 | 1.333E-05 | 2.887E-06 | 1.000E-05 | 1.500E-05 |
| 19185 | 960 | G960C | C | 1.4e-05 | 1.2e-05 | 1.6e-05 | 1.400E-05 | 2.000E-06 | 1.200E-05 | 1.600E-05 |
| 19186 | 960 | G960E | C | 1.1e-05 | 1.4e-05 | 1.3e-05 | 1.267E-05 | 1.528E-06 | 1.100E-05 | 1.400E-05 |
| 19187 | 960 | G960Q | C | 1.3e-05 | 1.4e-05 | 1.5e-05 | 1.400E-05 | 1.000E-06 | 1.300E-05 | 1.500E-05 |
| 19188 | 960 | G960G | C | 1.3e-05 | 1.3e-05 | 1.4e-05 | 1.333E-05 | 5.774E-07 | 1.300E-05 | 1.400E-05 |
| 19189 | 960 | G960H | C | 9.1e-06 | 1.1e-05 | 1.5e-05 | 1.170E-05 | 3.012E-06 | 9.100E-06 | 1.500E-05 |
| 19190 | 960 | G960I | C | 1.4e-05 | 1.4e-05 | 1.1e-05 | 1.300E-05 | 1.732E-06 | 1.100E-05 | 1.400E-05 |
| 19191 | 960 | G960L | C | 1.9e-05 | 1.7e-05 | 1.1e-05 | 1.567E-05 | 4.163E-06 | 1.100E-05 | 1.900E-05 |
| 19192 | 960 | G960K | C | 9.2e-06 | 1.1e-05 | 1.4e-05 | 1.140E-05 | 2.425E-06 | 9.200E-06 | 1.400E-05 |
| 19193 | 960 | G960M | C | 1.5e-05 | 1.6e-05 | 9.2e-06 | 1.340E-05 | 3.672E-06 | 9.200E-06 | 1.600E-05 |
| 19194 | 960 | G960F | C | 1.1e-05 | 9.9e-06 | 9e-06 | 9.967E-06 | 1.002E-06 | 9.000E-06 | 1.100E-05 |
| 19195 | 960 | G960P | C | 9.7e-06 | 9.7e-06 | 1.3e-05 | 1.080E-05 | 1.905E-06 | 9.700E-06 | 1.300E-05 |
| 19196 | 960 | G960S | C | 9.1e-06 | 2.2e-05 | 1.4e-05 | 1.503E-05 | 6.512E-06 | 9.100E-06 | 2.200E-05 |
| 19197 | 960 | G960T | C | 1e-05 | 1.4e-05 | 9.9e-06 | 1.130E-05 | 2.339E-06 | 9.900E-06 | 1.400E-05 |
| 19198 | 960 | G960W | C | 1.5e-05 | 9.5e-06 | 1e-05 | 1.150E-05 | 3.041E-06 | 9.500E-06 | 1.500E-05 |
| 19199 | 960 | G960Y | C | 1.3e-05 | 1.5e-05 | 1.4e-05 | 1.400E-05 | 1.000E-06 | 1.300E-05 | 1.500E-05 |
| 19200 | 960 | G960V | C | 1e-05 | 1e-05 | 1.4e-05 | 1.133E-05 | 2.309E-06 | 1.000E-05 | 1.400E-05 |
| 19201 | 961 | P961A | C | 1.5e-05 | 1.1e-05 | 1.7e-05 | 1.433E-05 | 3.055E-06 | 1.100E-05 | 1.700E-05 |
| 19202 | 961 | P961R | C | 8.8e-06 | 1e-05 | 1e-05 | 9.600E-06 | 6.928E-07 | 8.800E-06 | 1.000E-05 |
| 19203 | 961 | P961N | C | 1.5e-05 | 1.4e-05 | 1e-05 | 1.300E-05 | 2.646E-06 | 1.000E-05 | 1.500E-05 |
| 19204 | 961 | P961D | C | 1.5e-05 | 1.1e-05 | 7.4e-06 | 1.113E-05 | 3.802E-06 | 7.400E-06 | 1.500E-05 |
| 19205 | 961 | P961C | C | 1.1e-05 | 1.4e-05 | 1.5e-05 | 1.333E-05 | 2.082E-06 | 1.100E-05 | 1.500E-05 |
| 19206 | 961 | P961E | C | 1.6e-05 | 1.7e-05 | 1.6e-05 | 1.633E-05 | 5.774E-07 | 1.600E-05 | 1.700E-05 |
| 19207 | 961 | P961Q | C | 1.4e-05 | 1.2e-05 | 1.5e-05 | 1.367E-05 | 1.528E-06 | 1.200E-05 | 1.500E-05 |
| 19208 | 961 | P961G | C | 1e-05 | 1.1e-05 | 9.8e-06 | 1.027E-05 | 6.429E-07 | 9.800E-06 | 1.100E-05 |
| 19209 | 961 | P961H | C | 1.4e-05 | 8.6e-06 | 1.1e-05 | 1.120E-05 | 2.706E-06 | 8.600E-06 | 1.400E-05 |

| | | | | | | | | | | |
|---|---|---|---|---|---|---|---|---|---|---|
| 19210 | 961 | P961I | C | 9.8e-06 | 1.1e-05 | 7.8e-06 | 9.533E-06 | 1.617E-06 | 7.800E-06 | 1.100E-05 |
| 19211 | 961 | P961L | C | 1.5e-05 | 9.6e-06 | 1.4e-05 | 1.287E-05 | 2.873E-06 | 9.600E-06 | 1.500E-05 |
| 19212 | 961 | P961K | C | 1e-05 | 8.1e-06 | 1.3e-05 | 1.037E-05 | 2.470E-06 | 8.100E-06 | 1.300E-05 |
| 19213 | 961 | P961M | C | 1.6e-05 | 1.7e-05 | 1.5e-05 | 1.600E-05 | 1.000E-06 | 1.500E-05 | 1.700E-05 |
| 19214 | 961 | P961F | C | 1.4e-05 | 9.9e-06 | 1.6e-05 | 1.330E-05 | 3.110E-06 | 9.900E-06 | 1.600E-05 |
| 19215 | 961 | P961P | C | 1.4e-05 | 9.2e-06 | 1.1e-05 | 1.140E-05 | 2.425E-06 | 9.200E-06 | 1.400E-05 |
| 19216 | 961 | P961S | C | 1.4e-05 | 1.5e-05 | 1.5e-05 | 1.467E-05 | 5.774E-07 | 1.400E-05 | 1.500E-05 |
| 19217 | 961 | P961T | C | 1e-05 | 1.1e-05 | 9.1e-06 | 1.003E-05 | 9.504E-07 | 9.100E-06 | 1.100E-05 |
| 19218 | 961 | P961W | C | 1.5e-05 | 1.5e-05 | 1.5e-05 | 1.500E-05 | 0.000E+00 | 1.500E-05 | 1.500E-05 |
| 19219 | 961 | P961Y | C | 1.5e-05 | 1.5e-05 | 1.5e-05 | 1.500E-05 | 0.000E+00 | 1.500E-05 | 1.500E-05 |
| 19220 | 961 | P961V | C | 9.1e-06 | 1.7e-05 | 9.8e-06 | 1.197E-05 | 4.373E-06 | 9.100E-06 | 1.700E-05 |
| 19221 | 962 | S962A | C | 1.1e-05 | 1.6e-05 | 1.6e-05 | 1.433E-05 | 2.887E-06 | 1.100E-05 | 1.600E-05 |
| 19222 | 962 | S962R | C | 1.3e-05 | 1.1e-05 | 1.3e-05 | 1.233E-05 | 1.155E-06 | 1.100E-05 | 1.300E-05 |
| 19223 | 962 | S962N | C | 1.3e-05 | 1e-05 | 1.1e-05 | 1.133E-05 | 1.528E-06 | 1.000E-05 | 1.300E-05 |
| 19224 | 962 | S962D | C | 1.6e-05 | 1.5e-05 | 1.3e-05 | 1.467E-05 | 1.528E-06 | 1.300E-05 | 1.600E-05 |
| 19225 | 962 | S962C | C | 1.2e-05 | 1.5e-05 | 1e-05 | 1.233E-05 | 2.517E-06 | 1.000E-05 | 1.500E-05 |
| 19226 | 962 | S962E | C | 1.4e-05 | 1.1e-05 | 9.9e-06 | 1.163E-05 | 2.122E-06 | 9.900E-06 | 1.400E-05 |
| 19227 | 962 | S962Q | C | 9.7e-06 | 1.5e-05 | 9e-06 | 1.123E-05 | 3.281E-06 | 9.000E-06 | 1.500E-05 |
| 19228 | 962 | S962G | C | 7e-06 | 1.4e-05 | 1.1e-05 | 1.067E-05 | 3.512E-06 | 7.000E-06 | 1.400E-05 |
| 19229 | 962 | S962H | C | 1.7e-05 | 1.5e-05 | 1e-05 | 1.400E-05 | 3.606E-06 | 1.000E-05 | 1.700E-05 |
| 19230 | 962 | S962I | C | 1.2e-05 | 1e-05 | 1.1e-05 | 1.100E-05 | 1.000E-06 | 1.000E-05 | 1.200E-05 |
| 19231 | 962 | S962L | C | 1.2e-05 | 1.5e-05 | 9.4e-06 | 1.213E-05 | 2.802E-06 | 9.400E-06 | 1.500E-05 |
| 19232 | 962 | S962K | C | 1.1e-05 | 1.7e-05 | 1.7e-05 | 1.500E-05 | 3.464E-06 | 1.100E-05 | 1.700E-05 |
| 19233 | 962 | S962M | C | 1.7e-05 | 1.3e-05 | 1.1e-05 | 1.367E-05 | 3.055E-06 | 1.100E-05 | 1.700E-05 |
| 19234 | 962 | S962F | C | 1.5e-05 | 1.2e-05 | 1.1e-05 | 1.267E-05 | 2.082E-06 | 1.100E-05 | 1.500E-05 |
| 19235 | 962 | S962P | C | 8.8e-06 | 1e-05 | 1e-05 | 9.600E-06 | 6.928E-07 | 8.800E-06 | 1.000E-05 |
| 19236 | 962 | S962S | C | 1.4e-05 | 1.2e-05 | 1.6e-05 | 1.400E-05 | 2.000E-06 | 1.200E-05 | 1.600E-05 |
| 19237 | 962 | S962T | C | 1e-05 | 1.1e-05 | 1.5e-05 | 1.200E-05 | 2.646E-06 | 1.000E-05 | 1.500E-05 |
| 19238 | 962 | S962W | C | 1.5e-05 | 1.3e-05 | 1.7e-05 | 1.500E-05 | 2.000E-06 | 1.300E-05 | 1.700E-05 |
| 19239 | 962 | S962Y | C | 8.4e-06 | 8.5e-06 | 1e-05 | 8.967E-06 | 8.963E-07 | 8.400E-06 | 1.000E-05 |
| 19240 | 962 | S962V | C | 1.1e-05 | 1.7e-05 | 1.5e-05 | 1.433E-05 | 3.055E-06 | 1.100E-05 | 1.700E-05 |
| 19241 | 963 | V963A | C | 1.5e-05 | 1.4e-05 | 1.4e-05 | 1.433E-05 | 5.774E-07 | 1.400E-05 | 1.500E-05 |
| 19242 | 963 | V963R | C | 1.1e-05 | 1.6e-05 | 1.3e-05 | 1.333E-05 | 2.517E-06 | 1.100E-05 | 1.600E-05 |
| 19243 | 963 | V963N | C | 8.4e-06 | 1.5e-05 | 1.5e-05 | 1.280E-05 | 3.811E-06 | 8.400E-06 | 1.500E-05 |
| 19244 | 963 | V963D | C | 1.1e-05 | 9.5e-06 | 1.4e-05 | 1.150E-05 | 2.291E-06 | 9.500E-06 | 1.400E-05 |
| 19245 | 963 | V963C | C | 1.4e-05 | 1.6e-05 | 8.3e-06 | 1.277E-05 | 3.995E-06 | 8.300E-06 | 1.600E-05 |
| 19246 | 963 | V963E | C | 9.3e-06 | 1.5e-05 | 1.6e-05 | 1.343E-05 | 3.614E-06 | 9.300E-06 | 1.600E-05 |
| 19247 | 963 | V963Q | C | 1e-05 | 9.4e-06 | 8.1e-06 | 9.167E-06 | 9.713E-07 | 8.100E-06 | 1.000E-05 |
| 19248 | 963 | V963G | C | 1.6e-05 | 1.5e-05 | 1.6e-05 | 1.567E-05 | 5.774E-07 | 1.500E-05 | 1.600E-05 |
| 19249 | 963 | V963H | C | 1.1e-05 | 1.1e-05 | 6.8e-06 | 9.600E-06 | 2.425E-06 | 6.800E-06 | 1.100E-05 |
| 19250 | 963 | V963I | C | 1.4e-05 | 1.3e-05 | 8.9e-06 | 1.197E-05 | 2.702E-06 | 8.900E-06 | 1.400E-05 |
| 19251 | 963 | V963L | C | 1.6e-05 | 9.3e-06 | 1.3e-05 | 1.277E-05 | 3.356E-06 | 9.300E-06 | 1.600E-05 |
| 19252 | 963 | V963K | C | 1.3e-05 | 1e-05 | 1.6e-05 | 1.300E-05 | 3.000E-06 | 1.000E-05 | 1.600E-05 |
| 19253 | 963 | V963M | C | 1.2e-05 | 1e-05 | 1e-05 | 1.067E-05 | 1.155E-06 | 1.000E-05 | 1.200E-05 |
| 19254 | 963 | V963F | C | 1.5e-05 | 1.3e-05 | 1e-05 | 1.267E-05 | 2.517E-06 | 1.000E-05 | 1.500E-05 |
| 19255 | 963 | V963P | C | 1.5e-05 | 1.2e-05 | 1.1e-05 | 1.267E-05 | 2.082E-06 | 1.100E-05 | 1.500E-05 |
| 19256 | 963 | V963S | C | 1.4e-05 | 1.7e-05 | 1.6e-05 | 1.567E-05 | 1.528E-06 | 1.400E-05 | 1.700E-05 |
| 19257 | 963 | V963T | C | 1.5e-05 | 1.1e-05 | 1.3e-05 | 1.300E-05 | 2.000E-06 | 1.100E-05 | 1.500E-05 |
| 19258 | 963 | V963W | C | 9.8e-06 | 1.3e-05 | 1.3e-05 | 1.193E-05 | 1.848E-06 | 9.800E-06 | 1.300E-05 |
| 19259 | 963 | V963Y | C | 9.3e-06 | 1.1e-05 | 8.2e-06 | 9.500E-06 | 1.411E-06 | 8.200E-06 | 1.100E-05 |
| 19260 | 963 | V963V | C | 1.5e-05 | 9.9e-06 | 1.6e-05 | 1.363E-05 | 3.272E-06 | 9.900E-06 | 1.600E-05 |
| 19261 | 964 | F964A | C | 1.1e-05 | 1.1e-05 | 1.1e-05 | 1.100E-05 | 2.075E-21 | 1.100E-05 | 1.100E-05 |
| 19262 | 964 | F964R | C | 1.7e-05 | 6.2e-06 | 1.4e-05 | 1.240E-05 | 5.575E-06 | 6.200E-06 | 1.700E-05 |
| 19263 | 964 | F964N | C | 1.1e-05 | 1.5e-05 | 1e-05 | 1.200E-05 | 2.646E-06 | 1.000E-05 | 1.500E-05 |
| 19264 | 964 | F964D | C | 1.5e-05 | 1.4e-05 | 1e-05 | 1.300E-05 | 2.646E-06 | 1.000E-05 | 1.500E-05 |
| 19265 | 964 | F964C | C | 1.4e-05 | 1.4e-05 | 9.1e-06 | 1.237E-05 | 2.829E-06 | 9.100E-06 | 1.400E-05 |
| 19266 | 964 | F964E | C | 1.5e-05 | 1.4e-05 | 1.7e-05 | 1.533E-05 | 1.528E-06 | 1.400E-05 | 1.700E-05 |
| 19267 | 964 | F964Q | C | 1.4e-05 | 1.3e-05 | 1.4e-05 | 1.367E-05 | 5.774E-07 | 1.300E-05 | 1.400E-05 |
| 19268 | 964 | F964G | C | 1.5e-05 | 1.5e-05 | 1.5e-05 | 1.500E-05 | 0.000E+00 | 1.500E-05 | 1.500E-05 |
| 19269 | 964 | F964H | C | 1.2e-05 | 7.9e-06 | 1.5e-05 | 1.163E-05 | 3.564E-06 | 7.900E-06 | 1.500E-05 |
| 19270 | 964 | F964I | C | 1.4e-05 | 1.6e-05 | 8.1e-06 | 1.270E-05 | 4.107E-06 | 8.100E-06 | 1.600E-05 |
| 19271 | 964 | F964L | C | 8.4e-06 | 1.1e-05 | 1.4e-05 | 1.113E-05 | 2.802E-06 | 8.400E-06 | 1.400E-05 |
| 19272 | 964 | F964K | C | 1.4e-05 | 1.1e-05 | 1.5e-05 | 1.333E-05 | 2.082E-06 | 1.100E-05 | 1.500E-05 |
| 19273 | 964 | F964M | C | 1.1e-05 | 1.3e-05 | 1.7e-05 | 1.367E-05 | 3.055E-06 | 1.100E-05 | 1.700E-05 |
| 19274 | 964 | F964F | C | 1.6e-05 | 1.1e-05 | 1.5e-05 | 1.400E-05 | 2.646E-06 | 1.100E-05 | 1.600E-05 |
| 19275 | 964 | F964P | C | 8.5e-06 | 1e-05 | 9.5e-06 | 9.333E-06 | 7.638E-07 | 8.500E-06 | 1.000E-05 |
| 19276 | 964 | F964S | C | 1.2e-05 | 1.1e-05 | 1.5e-05 | 1.267E-05 | 2.082E-06 | 1.100E-05 | 1.500E-05 |
| 19277 | 964 | F964T | C | 1.4e-05 | 1.3e-05 | 1e-05 | 1.233E-05 | 2.082E-06 | 1.000E-05 | 1.400E-05 |
| 19278 | 964 | F964W | C | 1.3e-05 | 1.3e-05 | 1.4e-05 | 1.333E-05 | 5.774E-07 | 1.300E-05 | 1.400E-05 |
| 19279 | 964 | F964Y | C | 1.5e-05 | 9.7e-06 | 1.4e-05 | 1.290E-05 | 2.816E-06 | 9.700E-06 | 1.500E-05 |
| 19280 | 964 | F964V | C | 1e-05 | 1.5e-05 | 1.5e-05 | 1.333E-05 | 2.887E-06 | 1.000E-05 | 1.500E-05 |
| 19281 | 965 | P965A | C | 1.6e-05 | 1.1e-05 | 1.2e-05 | 1.300E-05 | 2.646E-06 | 1.100E-05 | 1.600E-05 |
| 19282 | 965 | P965R | C | 1.3e-05 | 9.5e-06 | 1.1e-05 | 1.117E-05 | 1.756E-06 | 9.500E-06 | 1.300E-05 |
| 19283 | 965 | P965N | C | 1.5e-05 | 1.4e-05 | 1.5e-05 | 1.467E-05 | 5.774E-07 | 1.400E-05 | 1.500E-05 |
| 19284 | 965 | P965D | C | 1.5e-05 | 1.3e-05 | 1.5e-05 | 1.433E-05 | 1.155E-06 | 1.300E-05 | 1.500E-05 |
| 19285 | 965 | P965C | C | 1.6e-05 | 7.4e-06 | 1.5e-05 | 1.280E-05 | 4.703E-06 | 7.400E-06 | 1.600E-05 |
| 19286 | 965 | P965E | C | 1.2e-05 | 1.8e-05 | 1.3e-05 | 1.433E-05 | 3.215E-06 | 1.200E-05 | 1.800E-05 |
| 19287 | 965 | P965Q | C | 9.9e-06 | 1e-05 | 1.5e-05 | 1.163E-05 | 2.916E-06 | 9.900E-06 | 1.500E-05 |
| 19288 | 965 | P965G | C | 1.2e-05 | 9.3e-06 | 9.6e-06 | 1.030E-05 | 1.480E-06 | 9.300E-06 | 1.200E-05 |
| 19289 | 965 | P965H | C | 1.4e-05 | 1.4e-05 | 1.4e-05 | 1.400E-05 | 0.000E+00 | 1.400E-05 | 1.400E-05 |
| 19290 | 965 | P965I | C | 1.1e-05 | 1.4e-05 | 1.3e-05 | 1.267E-05 | 1.528E-06 | 1.100E-05 | 1.400E-05 |
| 19291 | 965 | P965L | C | 1.2e-05 | 1.8e-05 | 1.6e-05 | 1.533E-05 | 3.055E-06 | 1.200E-05 | 1.800E-05 |
| 19292 | 965 | P965K | C | 1.1e-05 | 1.4e-05 | 1.7e-05 | 1.400E-05 | 3.000E-06 | 1.100E-05 | 1.700E-05 |
| 19293 | 965 | P965M | C | 1.1e-05 | 1.4e-05 | 1.4e-05 | 1.300E-05 | 1.732E-06 | 1.100E-05 | 1.400E-05 |
| 19294 | 965 | P965F | C | 1.6e-05 | 1.5e-05 | 8.8e-06 | 1.327E-05 | 3.900E-06 | 8.800E-06 | 1.600E-05 |

| | | | | | | | | | | |
|---|---|---|---|---|---|---|---|---|---|---|
| 19295 | 965 | P965P | C | 1e-05 | 1.7e-05 | 1.2e-05 | 1.300E-05 | 3.606E-06 | 1.000E-05 | 1.700E-05 |
| 19296 | 965 | P965S | C | 1.3e-05 | 1.5e-05 | 1.5e-05 | 1.433E-05 | 1.155E-06 | 1.300E-05 | 1.500E-05 |
| 19297 | 965 | P965T | C | 1.4e-05 | 1.6e-05 | 1.5e-05 | 1.500E-05 | 1.000E-06 | 1.400E-05 | 1.600E-05 |
| 19298 | 965 | P965W | C | 1.1e-05 | 1.4e-05 | 1.5e-05 | 1.333E-05 | 2.082E-06 | 1.100E-05 | 1.500E-05 |
| 19299 | 965 | P965Y | C | 1e-05 | 9.8e-06 | 9e-06 | 9.600E-06 | 5.292E-07 | 9.000E-06 | 1.000E-05 |
| 19300 | 965 | P965V | C | 1.5e-05 | 1e-05 | 1.6e-05 | 1.367E-05 | 3.215E-06 | 1.000E-05 | 1.600E-05 |
| 19301 | 966 | L966A | C | 1.6e-05 | 1.5e-05 | 1.6e-05 | 1.567E-05 | 5.774E-07 | 1.500E-05 | 1.600E-05 |
| 19302 | 966 | L966R | C | 1.3e-05 | 1.4e-05 | 1.3e-05 | 1.333E-05 | 5.774E-07 | 1.300E-05 | 1.400E-05 |
| 19303 | 966 | L966N | C | 1.2e-05 | 1.5e-05 | 9.3e-06 | 1.210E-05 | 2.851E-06 | 9.300E-06 | 1.500E-05 |
| 19304 | 966 | L966D | C | 1.4e-05 | 9.9e-06 | 1.5e-05 | 1.297E-05 | 2.702E-06 | 9.900E-06 | 1.500E-05 |
| 19305 | 966 | L966C | C | 1.4e-05 | 9.9e-06 | 1e-05 | 1.130E-05 | 2.339E-06 | 9.900E-06 | 1.400E-05 |
| 19306 | 966 | L966E | C | 1.3e-05 | 1.1e-05 | 1.3e-05 | 1.233E-05 | 1.155E-06 | 1.100E-05 | 1.300E-05 |
| 19307 | 966 | L966Q | C | 8.7e-06 | 1.5e-05 | 1.5e-05 | 1.290E-05 | 3.637E-06 | 8.700E-06 | 1.500E-05 |
| 19308 | 966 | L966G | C | 1.5e-05 | 1.5e-05 | 1.5e-05 | 1.500E-05 | 0.000E+00 | 1.500E-05 | 1.500E-05 |
| 19309 | 966 | L966H | C | 1.4e-05 | 9.7e-06 | 1.4e-05 | 1.257E-05 | 2.483E-06 | 9.700E-06 | 1.400E-05 |
| 19310 | 966 | L966I | C | 1.6e-05 | 1.6e-05 | 8.7e-06 | 1.357E-05 | 4.215E-06 | 8.700E-06 | 1.600E-05 |
| 19311 | 966 | L966L | C | 9.3e-06 | 1.6e-05 | 1.6e-05 | 1.377E-05 | 3.868E-06 | 9.300E-06 | 1.600E-05 |
| 19312 | 966 | L966K | C | 1.5e-05 | 9.9e-06 | 1.5e-05 | 1.330E-05 | 2.944E-06 | 9.900E-06 | 1.500E-05 |
| 19313 | 966 | L966M | C | 1.1e-05 | 1.7e-05 | 1e-05 | 1.267E-05 | 3.786E-06 | 1.000E-05 | 1.700E-05 |
| 19314 | 966 | L966F | C | 1e-05 | 1.4e-05 | 1.7e-05 | 1.367E-05 | 3.512E-06 | 1.000E-05 | 1.700E-05 |
| 19315 | 966 | L966P | C | 1.3e-05 | 1.8e-05 | 1.2e-05 | 1.433E-05 | 3.215E-06 | 1.200E-05 | 1.800E-05 |
| 19316 | 966 | L966S | C | 1.2e-05 | 8.8e-06 | 1.6e-05 | 1.227E-05 | 3.607E-06 | 8.800E-06 | 1.600E-05 |
| 19317 | 966 | L966T | C | 1.1e-05 | 1.6e-05 | 1.5e-05 | 1.400E-05 | 2.646E-06 | 1.100E-05 | 1.600E-05 |
| 19318 | 966 | L966W | C | 1.5e-05 | 1.4e-05 | 1.6e-05 | 1.500E-05 | 1.000E-06 | 1.400E-05 | 1.600E-05 |
| 19319 | 966 | L966Y | C | 1.3e-05 | 1.1e-05 | 8.6e-06 | 1.087E-05 | 2.203E-06 | 8.600E-06 | 1.300E-05 |
| 19320 | 966 | L966V | C | 1.3e-05 | 9.5e-06 | 1.6e-05 | 1.283E-05 | 3.253E-06 | 9.500E-06 | 1.600E-05 |
| 19321 | 967 | A967A | C | 1.2e-05 | 1.1e-05 | 1.5e-05 | 1.267E-05 | 2.082E-06 | 1.100E-05 | 1.500E-05 |
| 19322 | 967 | A967R | C | 1e-05 | 1.1e-05 | 8.1e-06 | 9.700E-06 | 1.473E-06 | 8.100E-06 | 1.100E-05 |
| 19323 | 967 | A967N | C | 1e-05 | 8.1e-06 | 1.5e-05 | 1.103E-05 | 3.564E-06 | 8.100E-06 | 1.500E-05 |
| 19324 | 967 | A967D | C | 1.4e-05 | 1.5e-05 | 8.5e-06 | 1.250E-05 | 3.500E-06 | 8.500E-06 | 1.500E-05 |
| 19325 | 967 | A967C | C | 1.4e-05 | 1.2e-05 | 1.6e-05 | 1.400E-05 | 2.000E-06 | 1.200E-05 | 1.600E-05 |
| 19326 | 967 | A967E | C | 1.1e-05 | 1.5e-05 | 1.1e-05 | 1.233E-05 | 2.309E-06 | 1.100E-05 | 1.500E-05 |
| 19327 | 967 | A967Q | C | 6.8e-06 | 6.5e-06 | 1e-05 | 7.767E-06 | 1.940E-06 | 6.500E-06 | 1.000E-05 |
| 19328 | 967 | A967G | C | 1.9e-05 | 8.2e-06 | 1e-05 | 1.240E-05 | 5.786E-06 | 8.200E-06 | 1.900E-05 |
| 19329 | 967 | A967H | C | 1.3e-05 | 1.5e-05 | 1.4e-05 | 1.400E-05 | 1.000E-06 | 1.300E-05 | 1.500E-05 |
| 19330 | 967 | A967I | C | 1.7e-05 | 1.5e-05 | 1e-05 | 1.400E-05 | 3.606E-06 | 1.000E-05 | 1.700E-05 |
| 19331 | 967 | A967L | C | 1.6e-05 | 1.6e-05 | 1.5e-05 | 1.567E-05 | 5.774E-07 | 1.500E-05 | 1.600E-05 |
| 19332 | 967 | A967K | C | 1.4e-05 | 1.5e-05 | 8.4e-06 | 1.247E-05 | 3.557E-06 | 8.400E-06 | 1.500E-05 |
| 19333 | 967 | A967M | C | 1.2e-05 | 1.7e-05 | 1.1e-05 | 1.333E-05 | 3.215E-06 | 1.100E-05 | 1.700E-05 |
| 19334 | 967 | A967F | C | 1e-05 | 1.2e-05 | 9.3e-06 | 1.043E-05 | 1.401E-06 | 9.300E-06 | 1.200E-05 |
| 19335 | 967 | A967P | C | 1.5e-05 | 1.1e-05 | 1.4e-05 | 1.333E-05 | 2.082E-06 | 1.100E-05 | 1.500E-05 |
| 19336 | 967 | A967S | C | 1.5e-05 | 1.3e-05 | 1.1e-05 | 1.300E-05 | 2.000E-06 | 1.100E-05 | 1.500E-05 |
| 19337 | 967 | A967T | C | 1.5e-05 | 1.5e-05 | 1.1e-05 | 1.367E-05 | 2.309E-06 | 1.100E-05 | 1.500E-05 |
| 19338 | 967 | A967W | C | 1.2e-05 | 9.6e-06 | 9.6e-06 | 1.040E-05 | 1.386E-06 | 9.600E-06 | 1.200E-05 |
| 19339 | 967 | A967Y | C | 1e-05 | 1.4e-05 | 1e-05 | 1.133E-05 | 2.309E-06 | 1.000E-05 | 1.400E-05 |
| 19340 | 967 | A967V | C | 1e-05 | 1.4e-05 | 1.4e-05 | 1.267E-05 | 2.309E-06 | 1.000E-05 | 1.400E-05 |
| 19341 | 968 | P968A | C | 1.8e-05 | 1.1e-05 | 1.5e-05 | 1.467E-05 | 3.512E-06 | 1.100E-05 | 1.800E-05 |
| 19342 | 968 | P968R | C | 1e-05 | 1e-05 | 1.6e-05 | 1.200E-05 | 3.464E-06 | 1.000E-05 | 1.600E-05 |
| 19343 | 968 | P968N | C | 8.8e-06 | 9.8e-06 | 1.2e-05 | 1.020E-05 | 1.637E-06 | 8.800E-06 | 1.200E-05 |
| 19344 | 968 | P968D | C | 1.1e-05 | 1.2e-05 | 1.5e-05 | 1.267E-05 | 2.082E-06 | 1.100E-05 | 1.500E-05 |
| 19345 | 968 | P968C | C | 1.4e-05 | 1.6e-05 | 8.9e-06 | 1.297E-05 | 3.661E-06 | 8.900E-06 | 1.600E-05 |
| 19346 | 968 | P968E | C | 1.1e-05 | 1.4e-05 | 1.3e-05 | 1.267E-05 | 1.528E-06 | 1.100E-05 | 1.400E-05 |
| 19347 | 968 | P968Q | C | 1.7e-05 | 1.5e-05 | 1.5e-05 | 1.567E-05 | 1.155E-06 | 1.500E-05 | 1.700E-05 |
| 19348 | 968 | P968G | C | 1.5e-05 | 1e-05 | 1e-05 | 1.167E-05 | 2.887E-06 | 1.000E-05 | 1.500E-05 |
| 19349 | 968 | P968H | C | 1.4e-05 | 1.5e-05 | 1.4e-05 | 1.433E-05 | 5.774E-07 | 1.400E-05 | 1.500E-05 |
| 19350 | 968 | P968I | C | 1.6e-05 | 1.1e-05 | 1.5e-05 | 1.400E-05 | 2.646E-06 | 1.100E-05 | 1.600E-05 |
| 19351 | 968 | P968L | C | 9.7e-06 | 1e-05 | 1.4e-05 | 1.123E-05 | 2.401E-06 | 9.700E-06 | 1.400E-05 |
| 19352 | 968 | P968K | C | 1.1e-05 | 1.4e-05 | 1.1e-05 | 1.200E-05 | 1.732E-06 | 1.100E-05 | 1.400E-05 |
| 19353 | 968 | P968M | C | 1.4e-05 | 1e-05 | 1.7e-05 | 1.367E-05 | 3.512E-06 | 1.000E-05 | 1.700E-05 |
| 19354 | 968 | P968F | C | 1.5e-05 | 1.6e-05 | 2.9e-05 | 2.000E-05 | 7.810E-06 | 1.500E-05 | 2.900E-05 |
| 19355 | 968 | P968P | C | 1.5e-05 | 1.1e-05 | 9.9e-06 | 1.197E-05 | 2.684E-06 | 9.900E-06 | 1.500E-05 |
| 19356 | 968 | P968S | C | 1e-05 | 1.4e-05 | 1.6e-05 | 1.333E-05 | 3.055E-06 | 1.000E-05 | 1.600E-05 |
| 19357 | 968 | P968T | C | 1.2e-05 | 1.4e-05 | 1.4e-05 | 1.333E-05 | 1.155E-06 | 1.200E-05 | 1.400E-05 |
| 19358 | 968 | P968W | C | 1.1e-05 | 9.9e-06 | 1.5e-05 | 1.197E-05 | 2.684E-06 | 9.900E-06 | 1.500E-05 |
| 19359 | 968 | P968Y | C | 1e-05 | 9e-06 | 1.3e-05 | 1.067E-05 | 2.082E-06 | 9.000E-06 | 1.300E-05 |
| 19360 | 968 | P968V | C | 9.6e-06 | 1.6e-05 | 1.4e-05 | 1.320E-05 | 3.274E-06 | 9.600E-06 | 1.600E-05 |
| 19361 | 969 | S969A | C | 9.8e-06 | 1.1e-05 | 1.1e-05 | 1.060E-05 | 6.928E-07 | 9.800E-06 | 1.100E-05 |
| 19362 | 969 | S969R | C | 1.6e-05 | 6.8e-06 | 1.6e-05 | 1.293E-05 | 5.312E-06 | 6.800E-06 | 1.600E-05 |
| 19363 | 969 | S969N | C | 1.1e-05 | 9.1e-06 | 1.5e-05 | 1.170E-05 | 3.012E-06 | 9.100E-06 | 1.500E-05 |
| 19364 | 969 | S969D | C | 1.5e-05 | 1.1e-05 | 1.3e-05 | 1.300E-05 | 2.000E-06 | 1.100E-05 | 1.500E-05 |
| 19365 | 969 | S969C | C | 1.5e-05 | 1.6e-05 | 1.5e-05 | 1.533E-05 | 5.774E-07 | 1.500E-05 | 1.600E-05 |
| 19366 | 969 | S969E | C | 1.6e-05 | 1.5e-05 | 7.8e-06 | 1.293E-05 | 4.474E-06 | 7.800E-06 | 1.600E-05 |
| 19367 | 969 | S969Q | C | 1.6e-05 | 1.4e-05 | 1.7e-05 | 1.567E-05 | 1.528E-06 | 1.400E-05 | 1.700E-05 |
| 19368 | 969 | S969G | C | 1.1e-05 | 1.8e-05 | 8.4e-06 | 1.247E-05 | 4.965E-06 | 8.400E-06 | 1.800E-05 |
| 19369 | 969 | S969H | C | 1.2e-05 | 1.6e-05 | 1.6e-05 | 1.467E-05 | 2.309E-06 | 1.200E-05 | 1.600E-05 |
| 19370 | 969 | S969I | C | 1.1e-05 | 9.8e-06 | 1.1e-05 | 1.060E-05 | 6.928E-07 | 9.800E-06 | 1.100E-05 |
| 19371 | 969 | S969L | C | 1e-05 | 1.9e-05 | 1.1e-05 | 1.333E-05 | 4.933E-06 | 1.000E-05 | 1.900E-05 |
| 19372 | 969 | S969K | C | 1.7e-05 | 1.6e-05 | 1.5e-05 | 1.600E-05 | 1.000E-06 | 1.500E-05 | 1.700E-05 |
| 19373 | 969 | S969M | C | 1.4e-05 | 1.6e-05 | 1.2e-05 | 1.400E-05 | 2.000E-06 | 1.200E-05 | 1.600E-05 |
| 19374 | 969 | S969F | C | 1.6e-05 | 1.6e-05 | 1.4e-05 | 1.533E-05 | 1.155E-06 | 1.400E-05 | 1.600E-05 |
| 19375 | 969 | S969P | C | 8.9e-06 | 1.4e-05 | 1e-05 | 1.097E-05 | 2.684E-06 | 8.900E-06 | 1.400E-05 |
| 19376 | 969 | S969S | C | 1.1e-05 | 1.1e-05 | 1.1e-05 | 1.100E-05 | 2.075E-21 | 1.100E-05 | 1.100E-05 |
| 19377 | 969 | S969T | C | 1.4e-05 | 1.3e-05 | 1.1e-05 | 1.267E-05 | 1.528E-06 | 1.100E-05 | 1.400E-05 |
| 19378 | 969 | S969W | C | 1.5e-05 | 1.2e-05 | 1.4e-05 | 1.367E-05 | 1.528E-06 | 1.200E-05 | 1.500E-05 |
| 19379 | 969 | S969Y | C | 1.6e-05 | 1.2e-05 | 1.6e-05 | 1.467E-05 | 2.309E-06 | 1.200E-05 | 1.600E-05 |

| | | | | | | | | | | |
|---|---|---|---|---|---|---|---|---|---|---|
| 19380 | 969 | S969V | C | 1.1e-05 | 1e-05 | 1.4e-05 | 1.167E-05 | 2.082E-06 | 1.000E-05 | 1.400E-05 |
| 19381 | 970 | S970A | C | 1.5e-05 | 7.5e-06 | 2.2e-05 | 1.483E-05 | 7.251E-06 | 7.500E-06 | 2.200E-05 |
| 19382 | 970 | S970R | C | 1.3e-05 | 1.5e-05 | 1.3e-05 | 1.367E-05 | 1.155E-06 | 1.300E-05 | 1.500E-05 |
| 19383 | 970 | S970N | C | 1.5e-05 | 1.5e-05 | 1.6e-05 | 1.533E-05 | 5.774E-07 | 1.500E-05 | 1.600E-05 |
| 19384 | 970 | S970D | C | 9.9e-06 | 7.2e-06 | 9.7e-06 | 8.933E-06 | 1.504E-06 | 7.200E-06 | 9.900E-06 |
| 19385 | 970 | S970C | C | 1.1e-05 | 1.4e-05 | 1.5e-05 | 1.333E-05 | 2.082E-06 | 1.100E-05 | 1.500E-05 |
| 19386 | 970 | S970E | C | 1.5e-05 | 1.7e-05 | 1.5e-05 | 1.567E-05 | 1.155E-06 | 1.500E-05 | 1.700E-05 |
| 19387 | 970 | S970Q | C | 1.7e-05 | 9.5e-06 | 1.6e-05 | 1.417E-05 | 4.072E-06 | 9.500E-06 | 1.700E-05 |
| 19388 | 970 | S970G | C | 1e-05 | 1e-05 | 1.1e-05 | 1.033E-05 | 5.774E-07 | 1.000E-05 | 1.100E-05 |
| 19389 | 970 | S970H | C | 9.7e-06 | 1.4e-05 | 1e-05 | 1.123E-05 | 2.401E-06 | 9.700E-06 | 1.400E-05 |
| 19390 | 970 | S970I | C | 8e-06 | 1.4e-05 | 1.1e-05 | 1.100E-05 | 3.000E-06 | 8.000E-06 | 1.400E-05 |
| 19391 | 970 | S970L | C | 1.1e-05 | 1.6e-05 | 1.1e-05 | 1.267E-05 | 2.887E-06 | 1.100E-05 | 1.600E-05 |
| 19392 | 970 | S970K | C | 1.1e-05 | 1.6e-05 | 1.6e-05 | 1.433E-05 | 2.887E-06 | 1.100E-05 | 1.600E-05 |
| 19393 | 970 | S970M | C | 1.6e-05 | 9.6e-06 | 1.2e-05 | 1.253E-05 | 3.233E-06 | 9.600E-06 | 1.600E-05 |
| 19394 | 970 | S970F | C | 1.6e-05 | 1.5e-05 | 1.7e-05 | 1.600E-05 | 1.000E-06 | 1.500E-05 | 1.700E-05 |
| 19395 | 970 | S970P | C | 1.4e-05 | 7.8e-06 | 2.3e-05 | 1.493E-05 | 7.643E-06 | 7.800E-06 | 2.300E-05 |
| 19396 | 970 | S970S | C | 1.1e-05 | 1.2e-05 | 9.7e-06 | 1.090E-05 | 1.153E-06 | 9.700E-06 | 1.200E-05 |
| 19397 | 970 | S970T | C | 1.2e-05 | 1.6e-05 | 1.2e-05 | 1.333E-05 | 2.309E-06 | 1.200E-05 | 1.600E-05 |
| 19398 | 970 | S970W | C | 1.1e-05 | 1.2e-05 | 1.6e-05 | 1.300E-05 | 2.646E-06 | 1.100E-05 | 1.600E-05 |
| 19399 | 970 | S970Y | C | 1.6e-05 | 1.3e-05 | 1.4e-05 | 1.433E-05 | 1.528E-06 | 1.300E-05 | 1.600E-05 |
| 19400 | 970 | S970V | C | 1.3e-05 | 9.8e-06 | 1.6e-05 | 1.293E-05 | 3.101E-06 | 9.800E-06 | 1.600E-05 |
| 19401 | 971 | K971A | C | 1.7e-05 | 1.8e-05 | 1.7e-05 | 1.733E-05 | 5.774E-07 | 1.700E-05 | 1.800E-05 |
| 19402 | 971 | K971R | C | 1.1e-05 | 1.6e-05 | 1e-05 | 1.233E-05 | 3.215E-06 | 1.000E-05 | 1.600E-05 |
| 19403 | 971 | K971N | C | 1.4e-05 | 9.6e-06 | 1.5e-05 | 1.287E-05 | 2.873E-06 | 9.600E-06 | 1.500E-05 |
| 19404 | 971 | K971D | C | 1.4e-05 | 8.5e-06 | 1.5e-05 | 1.250E-05 | 3.500E-06 | 8.500E-06 | 1.500E-05 |
| 19405 | 971 | K971C | C | 1.7e-05 | 8.6e-06 | 1.4e-05 | 1.320E-05 | 4.257E-06 | 8.600E-06 | 1.700E-05 |
| 19406 | 971 | K971E | C | 1.4e-05 | 1.5e-05 | 1.6e-05 | 1.500E-05 | 1.000E-06 | 1.400E-05 | 1.600E-05 |
| 19407 | 971 | K971Q | C | 1e-05 | 9.6e-06 | 1.5e-05 | 1.153E-05 | 3.009E-06 | 9.600E-06 | 1.500E-05 |
| 19408 | 971 | K971G | C | 1.2e-05 | 1.5e-05 | 1.7e-05 | 1.467E-05 | 2.517E-06 | 1.200E-05 | 1.700E-05 |
| 19409 | 971 | K971H | C | 1.2e-05 | 8.4e-06 | 1.3e-05 | 1.113E-05 | 2.419E-06 | 8.400E-06 | 1.300E-05 |
| 19410 | 971 | K971I | C | 1.2e-05 | 1.2e-05 | 1.3e-05 | 1.233E-05 | 5.774E-07 | 1.200E-05 | 1.300E-05 |
| 19411 | 971 | K971L | C | 1.5e-05 | 1.6e-05 | 1.4e-05 | 1.500E-05 | 1.000E-06 | 1.400E-05 | 1.600E-05 |
| 19412 | 971 | K971K | C | 9.1e-06 | 1.3e-05 | 1.4e-05 | 1.203E-05 | 2.589E-06 | 9.100E-06 | 1.400E-05 |
| 19413 | 971 | K971M | C | 1.5e-05 | 1.1e-05 | 1.6e-05 | 1.400E-05 | 2.646E-06 | 1.100E-05 | 1.600E-05 |
| 19414 | 971 | K971F | C | 1.5e-05 | 1.6e-05 | 1.4e-05 | 1.500E-05 | 1.000E-06 | 1.400E-05 | 1.600E-05 |
| 19415 | 971 | K971P | C | 1.4e-05 | 1.7e-05 | 1.5e-05 | 1.533E-05 | 1.528E-06 | 1.400E-05 | 1.700E-05 |
| 19416 | 971 | K971S | C | 1e-05 | 1.1e-05 | 1.1e-05 | 1.067E-05 | 5.774E-07 | 1.000E-05 | 1.100E-05 |
| 19417 | 971 | K971T | C | 8.4e-06 | 1.3e-05 | 1.1e-05 | 1.080E-05 | 2.307E-06 | 8.400E-06 | 1.300E-05 |
| 19418 | 971 | K971W | C | 9.9e-06 | 1.3e-05 | 1.1e-05 | 1.130E-05 | 1.572E-06 | 9.900E-06 | 1.300E-05 |
| 19419 | 971 | K971Y | C | 1.5e-05 | 1.4e-05 | 1.5e-05 | 1.467E-05 | 5.774E-07 | 1.400E-05 | 1.500E-05 |
| 19420 | 971 | K971V | C | 1.5e-05 | 9.4e-06 | 1.5e-05 | 1.313E-05 | 3.233E-06 | 9.400E-06 | 1.500E-05 |
| 19421 | 972 | S972A | C | 1.1e-05 | 1.4e-05 | 1.5e-05 | 1.333E-05 | 2.082E-06 | 1.100E-05 | 1.500E-05 |
| 19422 | 972 | S972R | C | 1.6e-05 | 1.4e-05 | 1.5e-05 | 1.500E-05 | 1.000E-06 | 1.400E-05 | 1.600E-05 |
| 19423 | 972 | S972N | C | 9.7e-06 | 1.5e-05 | 1.4e-05 | 1.290E-05 | 2.816E-06 | 9.700E-06 | 1.500E-05 |
| 19424 | 972 | S972D | C | 1.5e-05 | 1.7e-05 | 1.4e-05 | 1.533E-05 | 1.528E-06 | 1.400E-05 | 1.700E-05 |
| 19425 | 972 | S972C | C | 1.5e-05 | 1e-05 | 1.6e-05 | 1.367E-05 | 3.215E-06 | 1.000E-05 | 1.600E-05 |
| 19426 | 972 | S972E | C | 1.5e-05 | 1.4e-05 | 1.6e-05 | 1.500E-05 | 1.000E-06 | 1.400E-05 | 1.600E-05 |
| 19427 | 972 | S972Q | C | 1.2e-05 | 1.4e-05 | 8.8e-06 | 1.160E-05 | 2.623E-06 | 8.800E-06 | 1.400E-05 |
| 19428 | 972 | S972G | C | 1.5e-05 | 1.3e-05 | 1.7e-05 | 1.500E-05 | 2.000E-06 | 1.300E-05 | 1.700E-05 |
| 19429 | 972 | S972H | C | 1.3e-05 | 5.6e-06 | 1.4e-05 | 1.087E-05 | 4.588E-06 | 5.600E-06 | 1.400E-05 |
| 19430 | 972 | S972I | C | 1.4e-05 | 1.4e-05 | 1.6e-05 | 1.467E-05 | 1.155E-06 | 1.400E-05 | 1.600E-05 |
| 19431 | 972 | S972L | C | 1.3e-05 | 1.1e-05 | 1.1e-05 | 1.167E-05 | 1.155E-06 | 1.100E-05 | 1.300E-05 |
| 19432 | 972 | S972K | C | 1.2e-05 | 1.6e-05 | 1.2e-05 | 1.333E-05 | 2.309E-06 | 1.200E-05 | 1.600E-05 |
| 19433 | 972 | S972M | C | 9.3e-06 | 1.1e-05 | 1.1e-05 | 1.043E-05 | 9.815E-07 | 9.300E-06 | 1.100E-05 |
| 19434 | 972 | S972F | C | 1e-05 | 1.6e-05 | 8.5e-06 | 1.150E-05 | 3.969E-06 | 8.500E-06 | 1.600E-05 |
| 19435 | 972 | S972P | C | 1e-05 | 1.5e-05 | 1e-05 | 1.167E-05 | 2.887E-06 | 1.000E-05 | 1.500E-05 |
| 19436 | 972 | S972S | C | 1.3e-05 | 1e-05 | 1.6e-05 | 1.300E-05 | 3.000E-06 | 1.000E-05 | 1.600E-05 |
| 19437 | 972 | S972T | C | 1.5e-05 | 1.2e-05 | 1.3e-05 | 1.333E-05 | 1.528E-06 | 1.200E-05 | 1.500E-05 |
| 19438 | 972 | S972W | C | 9.9e-06 | 1.1e-05 | 1.1e-05 | 1.063E-05 | 6.351E-07 | 9.900E-06 | 1.100E-05 |
| 19439 | 972 | S972Y | C | 1.1e-05 | 1.3e-05 | 1e-05 | 1.133E-05 | 1.528E-06 | 1.000E-05 | 1.300E-05 |
| 19440 | 972 | S972V | C | 1.5e-05 | 1.1e-05 | 1.4e-05 | 1.333E-05 | 2.082E-06 | 1.100E-05 | 1.500E-05 |
| 19441 | 973 | T973A | C | 1.6e-05 | 1.5e-05 | 1.1e-05 | 1.400E-05 | 2.646E-06 | 1.100E-05 | 1.600E-05 |
| 19442 | 973 | T973R | C | 9.2e-06 | 9.4e-06 | 1.3e-05 | 1.053E-05 | 2.139E-06 | 9.200E-06 | 1.300E-05 |
| 19443 | 973 | T973N | C | 1.5e-05 | 1.5e-05 | 1.1e-05 | 1.367E-05 | 2.309E-06 | 1.100E-05 | 1.500E-05 |
| 19444 | 973 | T973D | C | 1.2e-05 | 9.7e-06 | 1.7e-05 | 1.290E-05 | 3.732E-06 | 9.700E-06 | 1.700E-05 |
| 19445 | 973 | T973C | C | 1.1e-05 | 1.7e-05 | 1.5e-05 | 1.433E-05 | 3.055E-06 | 1.100E-05 | 1.700E-05 |
| 19446 | 973 | T973E | C | 1.5e-05 | 1.1e-05 | 1.4e-05 | 1.333E-05 | 2.082E-06 | 1.100E-05 | 1.500E-05 |
| 19447 | 973 | T973Q | C | 8.9e-06 | 1.5e-05 | 9.7e-06 | 1.120E-05 | 3.315E-06 | 8.900E-06 | 1.500E-05 |
| 19448 | 973 | T973G | C | 8.6e-06 | 1.7e-05 | 8.8e-06 | 1.147E-05 | 4.793E-06 | 8.600E-06 | 1.700E-05 |
| 19449 | 973 | T973H | C | 8.5e-06 | 1.5e-05 | 1.3e-05 | 1.217E-05 | 3.329E-06 | 8.500E-06 | 1.500E-05 |
| 19450 | 973 | T973I | C | 1.6e-05 | 8.5e-06 | 1.3e-05 | 1.250E-05 | 3.775E-06 | 8.500E-06 | 1.600E-05 |
| 19451 | 973 | T973L | C | 1.5e-05 | 1.1e-05 | 1.6e-05 | 1.400E-05 | 2.646E-06 | 1.100E-05 | 1.600E-05 |
| 19452 | 973 | T973K | C | 1.4e-05 | 1.7e-05 | 1.2e-05 | 1.433E-05 | 2.517E-06 | 1.200E-05 | 1.700E-05 |
| 19453 | 973 | T973M | C | 1.6e-05 | 1e-05 | 9.5e-06 | 1.183E-05 | 3.617E-06 | 9.500E-06 | 1.600E-05 |
| 19454 | 973 | T973F | C | 9.6e-06 | 1.1e-05 | 1.4e-05 | 1.153E-05 | 2.248E-06 | 9.600E-06 | 1.400E-05 |
| 19455 | 973 | T973P | C | 6.1e-06 | 8.9e-06 | 1.2e-05 | 9.000E-06 | 2.951E-06 | 6.100E-06 | 1.200E-05 |
| 19456 | 973 | T973S | C | 1.5e-05 | 1.3e-05 | 1.3e-05 | 1.367E-05 | 1.155E-06 | 1.300E-05 | 1.500E-05 |
| 19457 | 973 | T973T | C | 1.4e-05 | 1.6e-05 | 9.5e-06 | 1.317E-05 | 3.329E-06 | 9.500E-06 | 1.600E-05 |
| 19458 | 973 | T973W | C | 1.5e-05 | 1.5e-05 | 1.1e-05 | 1.367E-05 | 2.309E-06 | 1.100E-05 | 1.500E-05 |
| 19459 | 973 | T973Y | C | 1e-05 | 1.5e-05 | 9.4e-06 | 1.147E-05 | 3.075E-06 | 9.400E-06 | 1.500E-05 |
| 19460 | 973 | T973V | C | 1.8e-05 | 1.1e-05 | 1.1e-05 | 1.333E-05 | 4.041E-06 | 1.100E-05 | 1.800E-05 |
| 19461 | 974 | S974A | C | 9.7e-06 | 1.7e-05 | 1.7e-05 | 1.457E-05 | 4.215E-06 | 9.700E-06 | 1.700E-05 |
| 19462 | 974 | S974R | C | 1.4e-05 | 1.6e-05 | 7.8e-06 | 1.260E-05 | 4.276E-06 | 7.800E-06 | 1.600E-05 |
| 19463 | 974 | S974N | C | 1.6e-05 | 1.6e-05 | 2e-05 | 1.733E-05 | 2.309E-06 | 1.600E-05 | 2.000E-05 |
| 19464 | 974 | S974D | C | 1.2e-05 | 1.5e-05 | 1.4e-05 | 1.367E-05 | 1.528E-06 | 1.200E-05 | 1.500E-05 |

| | | | | | | | | | | |
|---|---|---|---|---|---|---|---|---|---|---|
| 19465 | 974 | S974C | C | 1e-05 | 1.5e-05 | 1.1e-05 | 1.200E-05 | 2.646E-06 | 1.000E-05 | 1.500E-05 |
| 19466 | 974 | S974E | C | 1e-05 | 1.6e-05 | 1.5e-05 | 1.367E-05 | 3.215E-06 | 1.000E-05 | 1.600E-05 |
| 19467 | 974 | S974Q | C | 1.5e-05 | 1.5e-05 | 1.7e-05 | 1.567E-05 | 1.155E-06 | 1.500E-05 | 1.700E-05 |
| 19468 | 974 | S974G | C | 1.6e-05 | 1.6e-05 | 1.5e-05 | 1.567E-05 | 5.774E-07 | 1.500E-05 | 1.600E-05 |
| 19469 | 974 | S974H | C | 1.1e-05 | 1.3e-05 | 1.5e-05 | 1.300E-05 | 2.000E-06 | 1.100E-05 | 1.500E-05 |
| 19470 | 974 | S974I | C | 9.2e-06 | 1.5e-05 | 1.1e-05 | 1.173E-05 | 2.969E-06 | 9.200E-06 | 1.500E-05 |
| 19471 | 974 | S974L | C | 1.1e-05 | 1.4e-05 | 1.1e-05 | 1.200E-05 | 1.732E-06 | 1.100E-05 | 1.400E-05 |
| 19472 | 974 | S974K | C | 1.1e-05 | 1.5e-05 | 1e-05 | 1.200E-05 | 2.646E-06 | 1.000E-05 | 1.500E-05 |
| 19473 | 974 | S974M | C | 1.6e-05 | 9.6e-06 | 1.5e-05 | 1.353E-05 | 3.443E-06 | 9.600E-06 | 1.600E-05 |
| 19474 | 974 | S974F | C | 8.9e-06 | 1.2e-05 | 9.7e-06 | 1.020E-05 | 1.609E-06 | 8.900E-06 | 1.200E-05 |
| 19475 | 974 | S974P | C | 1.6e-05 | 1.5e-05 | 1.5e-05 | 1.533E-05 | 5.774E-07 | 1.500E-05 | 1.600E-05 |
| 19476 | 974 | S974S | C | 1.1e-05 | 1.6e-05 | 1.4e-05 | 1.367E-05 | 2.517E-06 | 1.100E-05 | 1.600E-05 |
| 19477 | 974 | S974T | C | 9.5e-06 | 1.6e-05 | 1e-05 | 1.183E-05 | 3.617E-06 | 9.500E-06 | 1.600E-05 |
| 19478 | 974 | S974W | C | 1.5e-05 | 9.7e-06 | 1.5e-05 | 1.323E-05 | 3.060E-06 | 9.700E-06 | 1.500E-05 |
| 19479 | 974 | S974Y | C | 1.7e-05 | 1.5e-05 | 1.4e-05 | 1.533E-05 | 1.528E-06 | 1.400E-05 | 1.700E-05 |
| 19480 | 974 | S974V | C | 1.5e-05 | 1.7e-05 | 1.2e-05 | 1.467E-05 | 2.517E-06 | 1.200E-05 | 1.700E-05 |
| 19481 | 975 | G975A | C | 1.5e-05 | 1.5e-05 | 8.2e-06 | 1.273E-05 | 3.926E-06 | 8.200E-06 | 1.500E-05 |
| 19482 | 975 | G975R | C | 1.1e-05 | 1.1e-05 | 1.1e-05 | 1.100E-05 | 2.075E-21 | 1.100E-05 | 1.100E-05 |
| 19483 | 975 | G975N | C | 1.5e-05 | 1.5e-05 | 1.1e-05 | 1.367E-05 | 2.309E-06 | 1.100E-05 | 1.500E-05 |
| 19484 | 975 | G975D | C | 1.5e-05 | 1.3e-05 | 1.5e-05 | 1.433E-05 | 1.155E-06 | 1.300E-05 | 1.500E-05 |
| 19485 | 975 | G975C | C | 1e-05 | 6.7e-06 | 1.7e-05 | 1.123E-05 | 5.260E-06 | 6.700E-06 | 1.700E-05 |
| 19486 | 975 | G975E | C | 1.3e-05 | 8.9e-06 | 1.3e-05 | 1.163E-05 | 2.367E-06 | 8.900E-06 | 1.300E-05 |
| 19487 | 975 | G975Q | C | 1.1e-05 | 1.3e-05 | 1e-05 | 1.133E-05 | 1.528E-06 | 1.000E-05 | 1.300E-05 |
| 19488 | 975 | G975G | C | 1.4e-05 | 1e-05 | 1.1e-05 | 1.167E-05 | 2.082E-06 | 1.000E-05 | 1.400E-05 |
| 19489 | 975 | G975H | C | 1.5e-05 | 9.7e-06 | 9.3e-06 | 1.133E-05 | 3.182E-06 | 9.300E-06 | 1.500E-05 |
| 19490 | 975 | G975I | C | 1.5e-05 | 1.3e-05 | 1.5e-05 | 1.433E-05 | 1.155E-06 | 1.300E-05 | 1.500E-05 |
| 19491 | 975 | G975L | C | 1.6e-05 | 1.4e-05 | 1.4e-05 | 1.467E-05 | 1.155E-06 | 1.400E-05 | 1.600E-05 |
| 19492 | 975 | G975K | C | 1.4e-05 | 1.2e-05 | 1.3e-05 | 1.300E-05 | 1.000E-06 | 1.200E-05 | 1.400E-05 |
| 19493 | 975 | G975M | C | 1.4e-05 | 1.7e-05 | 1.5e-05 | 1.533E-05 | 1.528E-06 | 1.400E-05 | 1.700E-05 |
| 19494 | 975 | G975F | C | 1.5e-05 | 1e-05 | 9.4e-06 | 1.147E-05 | 3.075E-06 | 9.400E-06 | 1.500E-05 |
| 19495 | 975 | G975P | C | 1.6e-05 | 1.1e-05 | 1e-05 | 1.233E-05 | 3.215E-06 | 1.000E-05 | 1.600E-05 |
| 19496 | 975 | G975S | C | 1.1e-05 | 1.4e-05 | 1.6e-05 | 1.367E-05 | 2.517E-06 | 1.100E-05 | 1.600E-05 |
| 19497 | 975 | G975T | C | 1.5e-05 | 1.3e-05 | 1e-05 | 1.267E-05 | 2.517E-06 | 1.000E-05 | 1.500E-05 |
| 19498 | 975 | G975W | C | 1.5e-05 | 9.9e-06 | 1.7e-05 | 1.397E-05 | 3.661E-06 | 9.900E-06 | 1.700E-05 |
| 19499 | 975 | G975Y | C | 1.1e-05 | 1e-05 | 1.6e-05 | 1.233E-05 | 3.215E-06 | 1.000E-05 | 1.600E-05 |
| 19500 | 975 | G975V | C | 1.5e-05 | 1.1e-05 | 1.6e-05 | 1.400E-05 | 2.646E-06 | 1.100E-05 | 1.600E-05 |
| 19501 | 976 | G976A | C | 1e-05 | 1.2e-05 | 1.5e-05 | 1.233E-05 | 2.517E-06 | 1.000E-05 | 1.500E-05 |
| 19502 | 976 | G976R | C | 1.7e-05 | 1.3e-05 | 1.6e-05 | 1.533E-05 | 2.082E-06 | 1.300E-05 | 1.700E-05 |
| 19503 | 976 | G976N | C | 9.4e-06 | 1.3e-05 | 1.2e-05 | 1.147E-05 | 1.858E-06 | 9.400E-06 | 1.300E-05 |
| 19504 | 976 | G976D | C | 1.6e-05 | 1.1e-05 | 1.1e-05 | 1.267E-05 | 2.887E-06 | 1.100E-05 | 1.600E-05 |
| 19505 | 976 | G976C | C | 1.3e-05 | 1.1e-05 | 1.6e-05 | 1.333E-05 | 2.517E-06 | 1.100E-05 | 1.600E-05 |
| 19506 | 976 | G976E | C | 1.3e-05 | 1.3e-05 | 1e-05 | 1.200E-05 | 1.732E-06 | 1.000E-05 | 1.300E-05 |
| 19507 | 976 | G976Q | C | 1.6e-05 | 9.6e-06 | 1.6e-05 | 1.387E-05 | 3.695E-06 | 9.600E-06 | 1.600E-05 |
| 19508 | 976 | G976G | C | 1.6e-05 | 1.3e-05 | 1.3e-05 | 1.400E-05 | 1.732E-06 | 1.300E-05 | 1.600E-05 |
| 19509 | 976 | G976H | C | 1.4e-05 | 1.6e-05 | 8e-06 | 1.267E-05 | 4.163E-06 | 8.000E-06 | 1.600E-05 |
| 19510 | 976 | G976I | C | 1.1e-05 | 9.6e-06 | 1e-05 | 1.020E-05 | 7.211E-07 | 9.600E-06 | 1.100E-05 |
| 19511 | 976 | G976L | C | 1.5e-05 | 8.5e-06 | 1.2e-05 | 1.183E-05 | 3.253E-06 | 8.500E-06 | 1.500E-05 |
| 19512 | 976 | G976K | C | 1.4e-05 | 1.5e-05 | 1.1e-05 | 1.333E-05 | 2.082E-06 | 1.100E-05 | 1.500E-05 |
| 19513 | 976 | G976M | C | 1.4e-05 | 1.2e-05 | 9.2e-06 | 1.173E-05 | 2.411E-06 | 9.200E-06 | 1.400E-05 |
| 19514 | 976 | G976F | C | 9.9e-06 | 1.6e-05 | 9.5e-06 | 1.180E-05 | 3.643E-06 | 9.500E-06 | 1.600E-05 |
| 19515 | 976 | G976P | C | 1e-05 | 2.6e-05 | 1.5e-05 | 1.700E-05 | 8.185E-06 | 1.000E-05 | 2.600E-05 |
| 19516 | 976 | G976S | C | 1.6e-05 | 1.2e-05 | 1.5e-05 | 1.433E-05 | 2.082E-06 | 1.200E-05 | 1.600E-05 |
| 19517 | 976 | G976T | C | 1.1e-05 | 8.3e-06 | 1.5e-05 | 1.143E-05 | 3.371E-06 | 8.300E-06 | 1.500E-05 |
| 19518 | 976 | G976W | C | 1.4e-05 | 1.3e-05 | 9.3e-06 | 1.210E-05 | 2.476E-06 | 9.300E-06 | 1.400E-05 |
| 19519 | 976 | G976Y | C | 1.1e-05 | 1.6e-05 | 1.4e-05 | 1.367E-05 | 2.517E-06 | 1.100E-05 | 1.600E-05 |
| 19520 | 976 | G976V | C | 9.9e-06 | 9.4e-06 | 1.5e-05 | 1.143E-05 | 3.099E-06 | 9.400E-06 | 1.500E-05 |
| 19521 | 977 | T977A | C | 1.5e-05 | 1.1e-05 | 9.8e-06 | 1.193E-05 | 2.723E-06 | 9.800E-06 | 1.500E-05 |
| 19522 | 977 | T977R | C | 1.6e-05 | 1.6e-05 | 1.1e-05 | 1.433E-05 | 2.887E-06 | 1.100E-05 | 1.600E-05 |
| 19523 | 977 | T977N | C | 1.1e-05 | 1.6e-05 | 1.5e-05 | 1.400E-05 | 2.646E-06 | 1.100E-05 | 1.600E-05 |
| 19524 | 977 | T977D | C | 1.1e-05 | 7.9e-06 | 1e-05 | 9.633E-06 | 1.582E-06 | 7.900E-06 | 1.100E-05 |
| 19525 | 977 | T977C | C | 1.5e-05 | 9.9e-06 | 8.4e-06 | 1.110E-05 | 3.460E-06 | 8.400E-06 | 1.500E-05 |
| 19526 | 977 | T977E | C | 9.6e-06 | 9e-06 | 1.5e-05 | 1.120E-05 | 3.305E-06 | 9.000E-06 | 1.500E-05 |
| 19527 | 977 | T977Q | C | 1.4e-05 | 1.3e-05 | 1.4e-05 | 1.367E-05 | 5.774E-07 | 1.300E-05 | 1.400E-05 |
| 19528 | 977 | T977G | C | 9e-06 | 1.5e-05 | 1.5e-05 | 1.300E-05 | 3.464E-06 | 9.000E-06 | 1.500E-05 |
| 19529 | 977 | T977H | C | 1.4e-05 | 1.4e-05 | 1.1e-05 | 1.300E-05 | 1.732E-06 | 1.100E-05 | 1.400E-05 |
| 19530 | 977 | T977I | C | 1.6e-05 | 1.6e-05 | 1.8e-05 | 1.667E-05 | 1.155E-06 | 1.600E-05 | 1.800E-05 |
| 19531 | 977 | T977L | C | 1.1e-05 | 1.6e-05 | 1.6e-05 | 1.433E-05 | 2.887E-06 | 1.100E-05 | 1.600E-05 |
| 19532 | 977 | T977K | C | 1.3e-05 | 8.7e-06 | 1.2e-05 | 1.123E-05 | 2.250E-06 | 8.700E-06 | 1.300E-05 |
| 19533 | 977 | T977M | C | 1.5e-05 | 1.3e-05 | 1e-05 | 1.267E-05 | 2.517E-06 | 1.000E-05 | 1.500E-05 |
| 19534 | 977 | T977F | C | 9.4e-06 | 1.5e-05 | 1.6e-05 | 1.347E-05 | 3.557E-06 | 9.400E-06 | 1.600E-05 |
| 19535 | 977 | T977P | C | 9.6e-06 | 1.1e-05 | 1.1e-05 | 1.053E-05 | 8.083E-07 | 9.600E-06 | 1.100E-05 |
| 19536 | 977 | T977S | C | 1.4e-05 | 1.5e-05 | 1.6e-05 | 1.500E-05 | 1.000E-06 | 1.400E-05 | 1.600E-05 |
| 19537 | 977 | T977T | C | 1.7e-05 | 1.4e-05 | 1.5e-05 | 1.533E-05 | 1.528E-06 | 1.400E-05 | 1.700E-05 |
| 19538 | 977 | T977W | C | 1.1e-05 | 7.8e-06 | 1.3e-05 | 1.060E-05 | 2.623E-06 | 7.800E-06 | 1.300E-05 |
| 19539 | 977 | T977Y | C | 9.8e-06 | 1.6e-05 | 9e-06 | 1.160E-05 | 3.831E-06 | 9.000E-06 | 1.600E-05 |
| 19540 | 977 | T977V | C | 1.5e-05 | 1.7e-05 | 1.3e-05 | 1.500E-05 | 2.000E-06 | 1.300E-05 | 1.700E-05 |
| 19541 | 978 | A978A | C | 8.4e-06 | 1.1e-05 | 1.6e-05 | 1.180E-05 | 3.863E-06 | 8.400E-06 | 1.600E-05 |
| 19542 | 978 | A978R | C | 9.6e-06 | 1.5e-05 | 1.3e-05 | 1.253E-05 | 2.730E-06 | 9.600E-06 | 1.500E-05 |
| 19543 | 978 | A978N | C | 9.3e-06 | 1.5e-05 | 1.6e-05 | 1.343E-05 | 3.614E-06 | 9.300E-06 | 1.600E-05 |
| 19544 | 978 | A978D | C | 1.6e-05 | 1.1e-05 | 9.6e-06 | 1.220E-05 | 3.365E-06 | 9.600E-06 | 1.600E-05 |
| 19545 | 978 | A978C | C | 5.8e-06 | 1.1e-05 | 1.7e-05 | 1.127E-05 | 5.605E-06 | 5.800E-06 | 1.700E-05 |
| 19546 | 978 | A978E | C | 1.1e-05 | 8.2e-06 | 1.4e-05 | 1.107E-05 | 2.901E-06 | 8.200E-06 | 1.400E-05 |
| 19547 | 978 | A978Q | C | 1.3e-05 | 1.1e-05 | 1.6e-05 | 1.333E-05 | 2.517E-06 | 1.100E-05 | 1.600E-05 |
| 19548 | 978 | A978G | C | 1.7e-05 | 9.5e-06 | 1.6e-05 | 1.417E-05 | 4.072E-06 | 9.500E-06 | 1.700E-05 |
| 19549 | 978 | A978H | C | 1.6e-05 | 1.6e-05 | 1.1e-05 | 1.433E-05 | 2.887E-06 | 1.100E-05 | 1.600E-05 |

| | | | | | | | | | | |
|---|---|---|---|---|---|---|---|---|---|---|
| 19550 | 978 | A978I | C | 1.2e-05 | 8.9e-06 | 1.4e-05 | 1.163E-05 | 2.570E-06 | 8.900E-06 | 1.400E-05 |
| 19551 | 978 | A978L | C | 1e-05 | 1.6e-05 | 1.5e-05 | 1.367E-05 | 3.215E-06 | 1.000E-05 | 1.600E-05 |
| 19552 | 978 | A978K | C | 1.5e-05 | 9.1e-06 | 1.5e-05 | 1.303E-05 | 3.406E-06 | 9.100E-06 | 1.500E-05 |
| 19553 | 978 | A978M | C | 1.2e-05 | 1.5e-05 | 1.3e-05 | 1.333E-05 | 1.528E-06 | 1.200E-05 | 1.500E-05 |
| 19554 | 978 | A978F | C | 1.1e-05 | 9.5e-06 | 9.2e-06 | 9.900E-06 | 9.644E-07 | 9.200E-06 | 1.100E-05 |
| 19555 | 978 | A978P | C | 1.1e-05 | 8.3e-06 | 1e-05 | 9.767E-06 | 1.365E-06 | 8.300E-06 | 1.100E-05 |
| 19556 | 978 | A978S | C | 1.7e-05 | 1.1e-05 | 9.6e-06 | 1.253E-05 | 3.931E-06 | 9.600E-06 | 1.700E-05 |
| 19557 | 978 | A978T | C | 1.1e-05 | 8.3e-06 | 1.3e-05 | 1.077E-05 | 2.359E-06 | 8.300E-06 | 1.300E-05 |
| 19558 | 978 | A978W | C | 1e-05 | 1.4e-05 | 1.5e-05 | 1.300E-05 | 2.646E-06 | 1.000E-05 | 1.500E-05 |
| 19559 | 978 | A978Y | C | 1.2e-05 | 1e-05 | 1.6e-05 | 1.267E-05 | 3.055E-06 | 1.000E-05 | 1.600E-05 |
| 19560 | 978 | A978V | C | 8.8e-06 | 1.1e-05 | 9.9e-06 | 9.900E-06 | 1.100E-06 | 8.800E-06 | 1.100E-05 |
| 19561 | 979 | A979A | C | 1.6e-05 | 1.1e-05 | 1.2e-05 | 1.300E-05 | 2.646E-06 | 1.100E-05 | 1.600E-05 |
| 19562 | 979 | A979R | C | 1.5e-05 | 1.4e-05 | 1.6e-05 | 1.500E-05 | 1.000E-06 | 1.400E-05 | 1.600E-05 |
| 19563 | 979 | A979N | C | 8.9e-06 | 1e-05 | 1.4e-05 | 1.097E-05 | 2.684E-06 | 8.900E-06 | 1.400E-05 |
| 19564 | 979 | A979D | C | 1.5e-05 | 1e-05 | 1e-05 | 1.167E-05 | 2.887E-06 | 1.000E-05 | 1.500E-05 |
| 19565 | 979 | A979C | C | 9.8e-06 | 1.4e-05 | 1.5e-05 | 1.293E-05 | 2.759E-06 | 9.800E-06 | 1.500E-05 |
| 19566 | 979 | A979E | C | 1.4e-05 | 1e-05 | 1.2e-05 | 1.200E-05 | 2.000E-06 | 1.000E-05 | 1.400E-05 |
| 19567 | 979 | A979Q | C | 1.1e-05 | 1.5e-05 | 9.5e-06 | 1.183E-05 | 2.843E-06 | 9.500E-06 | 1.500E-05 |
| 19568 | 979 | A979G | C | 9.7e-06 | 1.1e-05 | 1.1e-05 | 1.057E-05 | 7.506E-07 | 9.700E-06 | 1.100E-05 |
| 19569 | 979 | A979H | C | 1.3e-05 | 1.9e-05 | 1.5e-05 | 1.567E-05 | 3.055E-06 | 1.300E-05 | 1.900E-05 |
| 19570 | 979 | A979I | C | 1.6e-05 | 1.2e-05 | 1.6e-05 | 1.467E-05 | 2.309E-06 | 1.200E-05 | 1.600E-05 |
| 19571 | 979 | A979L | C | 1.6e-05 | 1.4e-05 | 2e-05 | 1.667E-05 | 3.055E-06 | 1.400E-05 | 2.000E-05 |
| 19572 | 979 | A979K | C | 1.4e-05 | 1.4e-05 | 8.5e-06 | 1.217E-05 | 3.175E-06 | 8.500E-06 | 1.400E-05 |
| 19573 | 979 | A979M | C | 1.5e-05 | 1.7e-05 | 1.5e-05 | 1.567E-05 | 1.155E-06 | 1.500E-05 | 1.700E-05 |
| 19574 | 979 | A979F | C | 7.9e-06 | 1.3e-05 | 1.5e-05 | 1.197E-05 | 3.661E-06 | 7.900E-06 | 1.500E-05 |
| 19575 | 979 | A979P | C | 1.1e-05 | 6.8e-06 | 9.1e-06 | 8.967E-06 | 2.103E-06 | 6.800E-06 | 1.100E-05 |
| 19576 | 979 | A979S | C | 7.7e-06 | 1.3e-05 | 1.2e-05 | 1.090E-05 | 2.816E-06 | 7.700E-06 | 1.300E-05 |
| 19577 | 979 | A979T | C | 1.4e-05 | 9.3e-06 | 8.3e-06 | 1.053E-05 | 3.044E-06 | 8.300E-06 | 1.400E-05 |
| 19578 | 979 | A979W | C | 9.6e-06 | 1.2e-05 | 1.5e-05 | 1.220E-05 | 2.706E-06 | 9.600E-06 | 1.500E-05 |
| 19579 | 979 | A979Y | C | 9.2e-06 | 1.5e-05 | 6.5e-06 | 1.023E-05 | 4.343E-06 | 6.500E-06 | 1.500E-05 |
| 19580 | 979 | A979V | C | 1e-05 | 7.3e-06 | 1.7e-05 | 1.143E-05 | 5.006E-06 | 7.300E-06 | 1.700E-05 |
| 19581 | 980 | L980A | C | 8.4e-06 | 1.1e-05 | 1.5e-05 | 1.147E-05 | 3.325E-06 | 8.400E-06 | 1.500E-05 |
| 19582 | 980 | L980R | C | 1.2e-05 | 9.4e-06 | 2e-05 | 1.380E-05 | 5.524E-06 | 9.400E-06 | 2.000E-05 |
| 19583 | 980 | L980N | C | 1.3e-05 | 1.7e-05 | 1.2e-05 | 1.400E-05 | 2.646E-06 | 1.200E-05 | 1.700E-05 |
| 19584 | 980 | L980D | C | 1.6e-05 | 1.5e-05 | 1.5e-05 | 1.533E-05 | 5.774E-07 | 1.500E-05 | 1.600E-05 |
| 19585 | 980 | L980C | C | 1.1e-05 | 9.4e-06 | 1.4e-05 | 1.147E-05 | 2.335E-06 | 9.400E-06 | 1.400E-05 |
| 19586 | 980 | L980E | C | 1.1e-05 | 1.6e-05 | 1.5e-05 | 1.400E-05 | 2.646E-06 | 1.100E-05 | 1.600E-05 |
| 19587 | 980 | L980Q | C | 1.5e-05 | 1.4e-05 | 9.6e-06 | 1.287E-05 | 2.873E-06 | 9.600E-06 | 1.500E-05 |
| 19588 | 980 | L980G | C | 1.6e-05 | 1.4e-05 | 1.5e-05 | 1.500E-05 | 1.000E-06 | 1.400E-05 | 1.600E-05 |
| 19589 | 980 | L980H | C | 1.5e-05 | 1.1e-05 | 1.5e-05 | 1.367E-05 | 2.309E-06 | 1.100E-05 | 1.500E-05 |
| 19590 | 980 | L980I | C | 1.6e-05 | 1.4e-05 | 1.1e-05 | 1.367E-05 | 2.517E-06 | 1.100E-05 | 1.600E-05 |
| 19591 | 980 | L980L | C | 1.4e-05 | 8.9e-06 | 1.6e-05 | 1.297E-05 | 3.661E-06 | 8.900E-06 | 1.600E-05 |
| 19592 | 980 | L980K | C | 1e-05 | 1.6e-05 | 1.8e-05 | 1.467E-05 | 4.163E-06 | 1.000E-05 | 1.800E-05 |
| 19593 | 980 | L980M | C | 1.4e-05 | 1.4e-05 | 1e-05 | 1.267E-05 | 2.309E-06 | 1.000E-05 | 1.400E-05 |
| 19594 | 980 | L980F | C | 1.4e-05 | 1.5e-05 | 1.4e-05 | 1.433E-05 | 5.774E-07 | 1.400E-05 | 1.500E-05 |
| 19595 | 980 | L980P | C | 1.1e-05 | 1.6e-05 | 7.7e-06 | 1.157E-05 | 4.179E-06 | 7.700E-06 | 1.600E-05 |
| 19596 | 980 | L980S | C | 5.9e-06 | 1.7e-05 | 1.3e-05 | 1.197E-05 | 5.622E-06 | 5.900E-06 | 1.700E-05 |
| 19597 | 980 | L980T | C | 1.6e-05 | 1.4e-05 | 1.5e-05 | 1.500E-05 | 1.000E-06 | 1.400E-05 | 1.600E-05 |
| 19598 | 980 | L980W | C | 1.6e-05 | 1e-05 | 6.6e-06 | 1.087E-05 | 4.760E-06 | 6.600E-06 | 1.600E-05 |
| 19599 | 980 | L980Y | C | 9.3e-06 | 1.3e-05 | 1.5e-05 | 1.243E-05 | 2.892E-06 | 9.300E-06 | 1.500E-05 |
| 19600 | 980 | L980V | C | 1.6e-05 | 1.6e-05 | 1.7e-05 | 1.633E-05 | 5.774E-07 | 1.600E-05 | 1.700E-05 |
| 19601 | 981 | G981A | C | 1.4e-05 | 1e-05 | 1.1e-05 | 1.167E-05 | 2.082E-06 | 1.000E-05 | 1.400E-05 |
| 19602 | 981 | G981R | C | 1.4e-05 | 1.1e-05 | 9.7e-06 | 1.157E-05 | 2.205E-06 | 9.700E-06 | 1.400E-05 |
| 19603 | 981 | G981N | C | 1.5e-05 | 1.2e-05 | 1.4e-05 | 1.367E-05 | 1.528E-06 | 1.200E-05 | 1.500E-05 |
| 19604 | 981 | G981D | C | 1.4e-05 | 1.6e-05 | 1.5e-05 | 1.500E-05 | 1.000E-06 | 1.400E-05 | 1.600E-05 |
| 19605 | 981 | G981C | C | 1.6e-05 | 1.5e-05 | 8.6e-06 | 1.320E-05 | 4.015E-06 | 8.600E-06 | 1.600E-05 |
| 19606 | 981 | G981E | C | 1.4e-05 | 9.5e-06 | 1.5e-05 | 1.283E-05 | 2.930E-06 | 9.500E-06 | 1.500E-05 |
| 19607 | 981 | G981Q | C | 1.1e-05 | 1.4e-05 | 1.1e-05 | 1.200E-05 | 1.732E-06 | 1.100E-05 | 1.400E-05 |
| 19608 | 981 | G981G | C | 1.1e-05 | 1.5e-05 | 1.2e-05 | 1.267E-05 | 2.082E-06 | 1.100E-05 | 1.500E-05 |
| 19609 | 981 | G981H | C | 1.6e-05 | 1e-05 | 1.7e-05 | 1.433E-05 | 3.786E-06 | 1.000E-05 | 1.700E-05 |
| 19610 | 981 | G981I | C | 1.5e-05 | 1.6e-05 | 1.3e-05 | 1.467E-05 | 1.528E-06 | 1.300E-05 | 1.600E-05 |
| 19611 | 981 | G981L | C | 9.7e-06 | 1.2e-05 | 1.5e-05 | 1.223E-05 | 2.658E-06 | 9.700E-06 | 1.500E-05 |
| 19612 | 981 | G981K | C | 1.4e-05 | 8.3e-06 | 1.9e-05 | 1.377E-05 | 5.354E-06 | 8.300E-06 | 1.900E-05 |
| 19613 | 981 | G981M | C | 1.6e-05 | 1.1e-05 | 1.5e-05 | 1.400E-05 | 2.646E-06 | 1.100E-05 | 1.600E-05 |
| 19614 | 981 | G981F | C | 1.6e-05 | 1.3e-05 | 1.4e-05 | 1.433E-05 | 1.528E-06 | 1.300E-05 | 1.600E-05 |
| 19615 | 981 | G981P | C | 8.8e-06 | 9.5e-06 | 1.3e-05 | 1.043E-05 | 2.250E-06 | 8.800E-06 | 1.300E-05 |
| 19616 | 981 | G981S | C | 1.5e-05 | 1e-05 | 1.5e-05 | 1.333E-05 | 2.887E-06 | 1.000E-05 | 1.500E-05 |
| 19617 | 981 | G981T | C | 1.3e-05 | 1.1e-05 | 9.2e-06 | 1.107E-05 | 1.901E-06 | 9.200E-06 | 1.300E-05 |
| 19618 | 981 | G981W | C | 1.2e-05 | 1.5e-05 | 1.4e-05 | 1.367E-05 | 1.528E-06 | 1.200E-05 | 1.500E-05 |
| 19619 | 981 | G981Y | C | 1.7e-05 | 1.6e-05 | 9.1e-06 | 1.403E-05 | 4.302E-06 | 9.100E-06 | 1.700E-05 |
| 19620 | 981 | G981V | C | 1.1e-05 | 1.6e-05 | 1.5e-05 | 1.400E-05 | 2.646E-06 | 1.100E-05 | 1.600E-05 |
| 19621 | 982 | C982A | C | 1e-05 | 1.5e-05 | 1.6e-05 | 1.367E-05 | 3.215E-06 | 1.000E-05 | 1.600E-05 |
| 19622 | 982 | C982R | C | 1.5e-05 | 9.5e-06 | 1.4e-05 | 1.283E-05 | 2.930E-06 | 9.500E-06 | 1.500E-05 |
| 19623 | 982 | C982N | C | 1.5e-05 | 1.4e-05 | 9.5e-06 | 1.283E-05 | 2.930E-06 | 9.500E-06 | 1.500E-05 |
| 19624 | 982 | C982D | C | 1.1e-05 | 1.2e-05 | 1.5e-05 | 1.267E-05 | 2.082E-06 | 1.100E-05 | 1.500E-05 |
| 19625 | 982 | C982C | C | 1.1e-05 | 1.6e-05 | 1.4e-05 | 1.367E-05 | 2.517E-06 | 1.100E-05 | 1.600E-05 |
| 19626 | 982 | C982E | C | 1.5e-05 | 1.7e-05 | 1.6e-05 | 1.600E-05 | 1.000E-06 | 1.500E-05 | 1.700E-05 |
| 19627 | 982 | C982Q | C | 1.6e-05 | 6.1e-06 | 9.9e-06 | 1.067E-05 | 4.994E-06 | 6.100E-06 | 1.600E-05 |
| 19628 | 982 | C982G | C | 1.6e-05 | 9e-06 | 1.3e-05 | 1.267E-05 | 3.512E-06 | 9.000E-06 | 1.600E-05 |
| 19629 | 982 | C982H | C | 1.1e-05 | 1e-05 | 1.2e-05 | 1.100E-05 | 1.000E-06 | 1.000E-05 | 1.200E-05 |
| 19630 | 982 | C982I | C | 9.8e-06 | 1.3e-05 | 9.7e-06 | 1.083E-05 | 1.877E-06 | 9.700E-06 | 1.300E-05 |
| 19631 | 982 | C982L | C | 1.5e-05 | 1e-05 | 1.6e-05 | 1.367E-05 | 3.215E-06 | 1.000E-05 | 1.600E-05 |
| 19632 | 982 | C982K | C | 1.1e-05 | 1.5e-05 | 1.1e-05 | 1.233E-05 | 2.309E-06 | 1.100E-05 | 1.500E-05 |
| 19633 | 982 | C982M | C | 1.7e-05 | 1e-05 | 1.1e-05 | 1.267E-05 | 3.786E-06 | 1.000E-05 | 1.700E-05 |
| 19634 | 982 | C982F | C | 1.4e-05 | 1e-05 | 1e-05 | 1.133E-05 | 2.309E-06 | 1.000E-05 | 1.400E-05 |

| | | | | | | | | | | |
|---|---|---|---|---|---|---|---|---|---|---|
| 19635 | 982 | C982P | C | 1.5e-05 | 9.3e-06 | 1.5e-05 | 1.310E-05 | 3.291E-06 | 9.300E-06 | 1.500E-05 |
| 19636 | 982 | C982S | C | 1.6e-05 | 1.2e-05 | 1.3e-05 | 1.367E-05 | 2.082E-06 | 1.200E-05 | 1.600E-05 |
| 19637 | 982 | C982T | C | 8.4e-06 | 1.1e-05 | 7.1e-06 | 8.833E-06 | 1.986E-06 | 7.100E-06 | 1.100E-05 |
| 19638 | 982 | C982W | C | 8.8e-06 | 1.4e-05 | 1.1e-05 | 1.127E-05 | 2.610E-06 | 8.800E-06 | 1.400E-05 |
| 19639 | 982 | C982Y | C | 1e-05 | 1e-05 | 1.6e-05 | 1.200E-05 | 3.464E-06 | 1.000E-05 | 1.600E-05 |
| 19640 | 982 | C982V | C | 1.5e-05 | 8.3e-06 | 1.4e-05 | 1.243E-05 | 3.614E-06 | 8.300E-06 | 1.500E-05 |
| 19641 | 983 | L983A | C | 1e-05 | 1.6e-05 | 1.5e-05 | 1.367E-05 | 3.215E-06 | 1.000E-05 | 1.600E-05 |
| 19642 | 983 | L983R | C | 1.1e-05 | 1.4e-05 | 1e-05 | 1.167E-05 | 2.082E-06 | 1.000E-05 | 1.400E-05 |
| 19643 | 983 | L983N | C | 1.5e-05 | 1e-05 | 8.8e-06 | 1.127E-05 | 3.288E-06 | 8.800E-06 | 1.500E-05 |
| 19644 | 983 | L983D | C | 1.5e-05 | 1.3e-05 | 5.7e-06 | 1.123E-05 | 4.895E-06 | 5.700E-06 | 1.500E-05 |
| 19645 | 983 | L983C | C | 1.7e-05 | 1.5e-05 | 1.1e-05 | 1.433E-05 | 3.055E-06 | 1.100E-05 | 1.700E-05 |
| 19646 | 983 | L983E | C | 1e-05 | 8.3e-06 | 1.4e-05 | 1.077E-05 | 2.926E-06 | 8.300E-06 | 1.400E-05 |
| 19647 | 983 | L983Q | C | 1.5e-05 | 1.3e-05 | 1.2e-05 | 1.333E-05 | 1.528E-06 | 1.200E-05 | 1.500E-05 |
| 19648 | 983 | L983G | C | 1e-05 | 1.5e-05 | 1.6e-05 | 1.367E-05 | 3.215E-06 | 1.000E-05 | 1.600E-05 |
| 19649 | 983 | L983H | C | 1.5e-05 | 1.1e-05 | 1.5e-05 | 1.367E-05 | 2.309E-06 | 1.100E-05 | 1.500E-05 |
| 19650 | 983 | L983I | C | 1.1e-05 | 1.5e-05 | 1.6e-05 | 1.400E-05 | 2.646E-06 | 1.100E-05 | 1.600E-05 |
| 19651 | 983 | L983L | C | 1.5e-05 | 1.6e-05 | 1.5e-05 | 1.533E-05 | 5.774E-07 | 1.500E-05 | 1.600E-05 |
| 19652 | 983 | L983K | C | 1e-05 | 1.4e-05 | 1.5e-05 | 1.300E-05 | 2.646E-06 | 1.000E-05 | 1.500E-05 |
| 19653 | 983 | L983M | C | 1.4e-05 | 1.4e-05 | 1.4e-05 | 1.400E-05 | 0.000E+00 | 1.400E-05 | 1.400E-05 |
| 19654 | 983 | L983F | C | 1.1e-05 | 1.2e-05 | 1.3e-05 | 1.200E-05 | 1.000E-06 | 1.100E-05 | 1.300E-05 |
| 19655 | 983 | L983P | C | 9.4e-06 | 1.4e-05 | 1.2e-05 | 1.180E-05 | 2.307E-06 | 9.400E-06 | 1.400E-05 |
| 19656 | 983 | L983S | C | 1.6e-05 | 1.4e-05 | 9.9e-06 | 1.330E-05 | 3.110E-06 | 9.900E-06 | 1.600E-05 |
| 19657 | 983 | L983T | C | 1.5e-05 | 1.5e-05 | 9.7e-06 | 1.323E-05 | 3.060E-06 | 9.700E-06 | 1.500E-05 |
| 19658 | 983 | L983W | C | 9.8e-06 | 9.2e-06 | 9.6e-06 | 9.533E-06 | 3.055E-07 | 9.200E-06 | 9.800E-06 |
| 19659 | 983 | L983Y | C | 1.4e-05 | 8.4e-06 | 1.1e-05 | 1.113E-05 | 2.802E-06 | 8.400E-06 | 1.400E-05 |
| 19660 | 983 | L983V | C | 1.6e-05 | 1.2e-05 | 1.6e-05 | 1.467E-05 | 2.309E-06 | 1.200E-05 | 1.600E-05 |
| 19661 | 984 | V984A | C | 1.3e-05 | 1.2e-05 | 1e-05 | 1.167E-05 | 1.528E-06 | 1.000E-05 | 1.300E-05 |
| 19662 | 984 | V984R | C | 9.5e-06 | 1.4e-05 | 1.1e-05 | 1.150E-05 | 2.291E-06 | 9.500E-06 | 1.400E-05 |
| 19663 | 984 | V984N | C | 8.6e-06 | 1.4e-05 | 1.5e-05 | 1.253E-05 | 3.443E-06 | 8.600E-06 | 1.500E-05 |
| 19664 | 984 | V984D | C | 1.4e-05 | 1.1e-05 | 1.7e-05 | 1.400E-05 | 3.000E-06 | 1.100E-05 | 1.700E-05 |
| 19665 | 984 | V984C | C | 8.4e-06 | 1e-05 | 1.6e-05 | 1.147E-05 | 4.007E-06 | 8.400E-06 | 1.600E-05 |
| 19666 | 984 | V984E | C | 9.5e-06 | 1.1e-05 | 1.4e-05 | 1.150E-05 | 2.291E-06 | 9.500E-06 | 1.400E-05 |
| 19667 | 984 | V984Q | C | 1.1e-05 | 1.1e-05 | 1.4e-05 | 1.200E-05 | 1.732E-06 | 1.100E-05 | 1.400E-05 |
| 19668 | 984 | V984G | C | 1.2e-05 | 1.6e-05 | 1.1e-05 | 1.300E-05 | 2.646E-06 | 1.100E-05 | 1.600E-05 |
| 19669 | 984 | V984H | C | 1.5e-05 | 8.6e-06 | 8.2e-06 | 1.060E-05 | 3.816E-06 | 8.200E-06 | 1.500E-05 |
| 19670 | 984 | V984I | C | 9.6e-06 | 1.4e-05 | 5.7e-06 | 9.767E-06 | 4.153E-06 | 5.700E-06 | 1.400E-05 |
| 19671 | 984 | V984L | C | 9.6e-06 | 9.7e-06 | 1.5e-05 | 1.143E-05 | 3.089E-06 | 9.600E-06 | 1.500E-05 |
| 19672 | 984 | V984K | C | 1.5e-05 | 1.7e-05 | 1.5e-05 | 1.567E-05 | 1.155E-06 | 1.500E-05 | 1.700E-05 |
| 19673 | 984 | V984M | C | 1.5e-05 | 1.1e-05 | 1.3e-05 | 1.300E-05 | 2.000E-06 | 1.100E-05 | 1.500E-05 |
| 19674 | 984 | V984F | C | 1.6e-05 | 1.1e-05 | 1.3e-05 | 1.333E-05 | 2.517E-06 | 1.100E-05 | 1.600E-05 |
| 19675 | 984 | V984P | C | 1.7e-05 | 9.4e-06 | 8.4e-06 | 1.160E-05 | 4.703E-06 | 8.400E-06 | 1.700E-05 |
| 19676 | 984 | V984S | C | 9.3e-06 | 1.3e-05 | 1.3e-05 | 1.177E-05 | 2.136E-06 | 9.300E-06 | 1.300E-05 |
| 19677 | 984 | V984T | C | 1.2e-05 | 1.3e-05 | 1.2e-05 | 1.233E-05 | 5.774E-07 | 1.200E-05 | 1.300E-05 |
| 19678 | 984 | V984W | C | 1.7e-05 | 1.5e-05 | 1.7e-05 | 1.633E-05 | 1.155E-06 | 1.500E-05 | 1.700E-05 |
| 19679 | 984 | V984Y | C | 1.8e-05 | 8.7e-06 | 1e-05 | 1.223E-05 | 5.036E-06 | 8.700E-06 | 1.800E-05 |
| 19680 | 984 | V984V | C | 6.6e-06 | 1.5e-05 | 1.5e-05 | 1.220E-05 | 4.850E-06 | 6.600E-06 | 1.500E-05 |
| 19681 | 985 | K985A | C | 1.1e-05 | 1.5e-05 | 1.3e-05 | 1.300E-05 | 2.000E-06 | 1.100E-05 | 1.500E-05 |
| 19682 | 985 | K985R | C | 9.8e-06 | 1.4e-05 | 1.1e-05 | 1.160E-05 | 2.163E-06 | 9.800E-06 | 1.400E-05 |
| 19683 | 985 | K985N | C | 1.7e-05 | 1.5e-05 | 8.8e-06 | 1.360E-05 | 4.276E-06 | 8.800E-06 | 1.700E-05 |
| 19684 | 985 | K985D | C | 1.3e-05 | 1.1e-05 | 1.5e-05 | 1.300E-05 | 2.000E-06 | 1.100E-05 | 1.500E-05 |
| 19685 | 985 | K985C | C | 5.7e-06 | 1.1e-05 | 1.6e-05 | 1.090E-05 | 5.151E-06 | 5.700E-06 | 1.600E-05 |
| 19686 | 985 | K985E | C | 1e-05 | 1.4e-05 | 1.1e-05 | 1.167E-05 | 2.082E-06 | 1.000E-05 | 1.400E-05 |
| 19687 | 985 | K985Q | C | 1.4e-05 | 1.5e-05 | 8.9e-06 | 1.263E-05 | 3.272E-06 | 8.900E-06 | 1.500E-05 |
| 19688 | 985 | K985G | C | 1.6e-05 | 1.4e-05 | 7.6e-06 | 1.253E-05 | 4.388E-06 | 7.600E-06 | 1.600E-05 |
| 19689 | 985 | K985H | C | 1e-05 | 1.4e-05 | 6.4e-06 | 1.013E-05 | 3.802E-06 | 6.400E-06 | 1.400E-05 |
| 19690 | 985 | K985I | C | 1.2e-05 | 9.8e-06 | 1.3e-05 | 1.160E-05 | 1.637E-06 | 9.800E-06 | 1.300E-05 |
| 19691 | 985 | K985L | C | 1.7e-05 | 7.8e-06 | 1.6e-05 | 1.360E-05 | 5.048E-06 | 7.800E-06 | 1.700E-05 |
| 19692 | 985 | K985K | C | 1.6e-05 | 1.4e-05 | 1.3e-05 | 1.433E-05 | 1.528E-06 | 1.300E-05 | 1.600E-05 |
| 19693 | 985 | K985M | C | 1.4e-05 | 1.6e-05 | 1.1e-05 | 1.367E-05 | 2.517E-06 | 1.100E-05 | 1.600E-05 |
| 19694 | 985 | K985F | C | 1.5e-05 | 1.6e-05 | 1.7e-05 | 1.600E-05 | 1.000E-06 | 1.500E-05 | 1.700E-05 |
| 19695 | 985 | K985P | C | 9.1e-06 | 7.3e-06 | 1.6e-05 | 1.080E-05 | 4.592E-06 | 7.300E-06 | 1.600E-05 |
| 19696 | 985 | K985S | C | 9.2e-06 | 1.6e-05 | 9e-06 | 1.140E-05 | 3.985E-06 | 9.000E-06 | 1.600E-05 |
| 19697 | 985 | K985T | C | 1e-05 | 9.1e-06 | 1e-05 | 9.700E-06 | 5.196E-07 | 9.100E-06 | 1.000E-05 |
| 19698 | 985 | K985W | C | 1e-05 | 1.5e-05 | 1.3e-05 | 1.267E-05 | 2.517E-06 | 1.000E-05 | 1.500E-05 |
| 19699 | 985 | K985Y | C | 1.4e-05 | 8.5e-06 | 1.5e-05 | 1.250E-05 | 3.500E-06 | 8.500E-06 | 1.500E-05 |
| 19700 | 985 | K985V | C | 1e-05 | 1e-05 | 1.7e-05 | 1.233E-05 | 4.041E-06 | 1.000E-05 | 1.700E-05 |
| 19701 | 986 | D986A | C | 1.1e-05 | 1.1e-05 | 1.1e-05 | 1.100E-05 | 2.075E-21 | 1.100E-05 | 1.100E-05 |
| 19702 | 986 | D986R | C | 1.2e-05 | 1.2e-05 | 8.8e-06 | 1.093E-05 | 1.848E-06 | 8.800E-06 | 1.200E-05 |
| 19703 | 986 | D986N | C | 1.4e-05 | 1.5e-05 | 9.1e-06 | 1.270E-05 | 3.158E-06 | 9.100E-06 | 1.500E-05 |
| 19704 | 986 | D986D | C | 1e-05 | 1.5e-05 | 2.3e-05 | 1.600E-05 | 6.557E-06 | 1.000E-05 | 2.300E-05 |
| 19705 | 986 | D986C | C | 1.5e-05 | 8.8e-06 | 1.2e-05 | 1.193E-05 | 3.101E-06 | 8.800E-06 | 1.500E-05 |
| 19706 | 986 | D986E | C | 1e-05 | 1.6e-05 | 1.9e-05 | 1.500E-05 | 4.583E-06 | 1.000E-05 | 1.900E-05 |
| 19707 | 986 | D986Q | C | 1.3e-05 | 1.2e-05 | 8.2e-06 | 1.107E-05 | 2.532E-06 | 8.200E-06 | 1.300E-05 |
| 19708 | 986 | D986G | C | 1.4e-05 | 1.2e-05 | 1.4e-05 | 1.333E-05 | 1.155E-06 | 1.200E-05 | 1.400E-05 |
| 19709 | 986 | D986H | C | 1.5e-05 | 8.8e-06 | 1.5e-05 | 1.293E-05 | 3.580E-06 | 8.800E-06 | 1.500E-05 |
| 19710 | 986 | D986I | C | 1.2e-05 | 1.2e-05 | 1.3e-05 | 1.233E-05 | 5.774E-07 | 1.200E-05 | 1.300E-05 |
| 19711 | 986 | D986L | C | 1.3e-05 | 1.5e-05 | 1e-05 | 1.267E-05 | 2.517E-06 | 1.000E-05 | 1.500E-05 |
| 19712 | 986 | D986K | C | 1.5e-05 | 1.5e-05 | 1.6e-05 | 1.533E-05 | 5.774E-07 | 1.500E-05 | 1.600E-05 |
| 19713 | 986 | D986M | C | 1e-05 | 1.4e-05 | 9.6e-06 | 1.120E-05 | 2.433E-06 | 9.600E-06 | 1.400E-05 |
| 19714 | 986 | D986F | C | 1.1e-05 | 1e-05 | 1.3e-05 | 1.133E-05 | 1.528E-06 | 1.000E-05 | 1.300E-05 |
| 19715 | 986 | D986P | C | 1.5e-05 | 1.4e-05 | 9.9e-06 | 1.297E-05 | 2.702E-06 | 9.900E-06 | 1.500E-05 |
| 19716 | 986 | D986S | C | 1.1e-05 | 1.5e-05 | 9.2e-06 | 1.173E-05 | 2.969E-06 | 9.200E-06 | 1.500E-05 |
| 19717 | 986 | D986T | C | 1.5e-05 | 1.4e-05 | 1.1e-05 | 1.333E-05 | 2.082E-06 | 1.100E-05 | 1.500E-05 |
| 19718 | 986 | D986W | C | 1.4e-05 | 1.4e-05 | 1.2e-05 | 1.333E-05 | 1.155E-06 | 1.200E-05 | 1.400E-05 |
| 19719 | 986 | D986Y | C | 1.1e-05 | 1.4e-05 | 1.4e-05 | 1.300E-05 | 1.732E-06 | 1.100E-05 | 1.400E-05 |

| | | | | | | | | | | |
|---|---|---|---|---|---|---|---|---|---|---|
| 19720 | 986 | D986V | C | 1.5e-05 | 9.1e-06 | 9.4e-06 | 1.117E-05 | 3.323E-06 | 9.100E-06 | 1.500E-05 |
| 19721 | 987 | Y987A | C | 8.5e-06 | 1.6e-05 | 8e-06 | 1.083E-05 | 4.481E-06 | 8.000E-06 | 1.600E-05 |
| 19722 | 987 | Y987R | C | 1.5e-05 | 1.6e-05 | 1.5e-05 | 1.533E-05 | 5.774E-07 | 1.500E-05 | 1.600E-05 |
| 19723 | 987 | Y987N | C | 1.5e-05 | 1.2e-05 | 1e-05 | 1.233E-05 | 2.517E-06 | 1.000E-05 | 1.500E-05 |
| 19724 | 987 | Y987D | C | 9e-06 | 1.2e-05 | 1.3e-05 | 1.133E-05 | 2.082E-06 | 9.000E-06 | 1.300E-05 |
| 19725 | 987 | Y987C | C | 1.5e-05 | 1.3e-05 | 1.1e-05 | 1.300E-05 | 2.000E-06 | 1.100E-05 | 1.500E-05 |
| 19726 | 987 | Y987E | C | 9.1e-06 | 1.5e-05 | 1.8e-05 | 1.403E-05 | 4.528E-06 | 9.100E-06 | 1.800E-05 |
| 19727 | 987 | Y987Q | C | 1.6e-05 | 1.6e-05 | 9.8e-06 | 1.393E-05 | 3.580E-06 | 9.800E-06 | 1.600E-05 |
| 19728 | 987 | Y987G | C | 1.6e-05 | 1.4e-05 | 1.5e-05 | 1.500E-05 | 1.000E-06 | 1.400E-05 | 1.600E-05 |
| 19729 | 987 | Y987H | C | 1.4e-05 | 1.5e-05 | 9.2e-06 | 1.273E-05 | 3.101E-06 | 9.200E-06 | 1.500E-05 |
| 19730 | 987 | Y987I | C | 1.4e-05 | 1.1e-05 | 1.4e-05 | 1.300E-05 | 1.732E-06 | 1.100E-05 | 1.400E-05 |
| 19731 | 987 | Y987L | C | 1.4e-05 | 1.5e-05 | 1.5e-05 | 1.467E-05 | 5.774E-07 | 1.400E-05 | 1.500E-05 |
| 19732 | 987 | Y987K | C | 9.3e-06 | 1.5e-05 | 1.7e-05 | 1.377E-05 | 3.995E-06 | 9.300E-06 | 1.700E-05 |
| 19733 | 987 | Y987M | C | 1.8e-05 | 1.1e-05 | 1.6e-05 | 1.500E-05 | 3.606E-06 | 1.100E-05 | 1.800E-05 |
| 19734 | 987 | Y987F | C | 1.6e-05 | 1.2e-05 | 8.9e-06 | 1.230E-05 | 3.559E-06 | 8.900E-06 | 1.600E-05 |
| 19735 | 987 | Y987P | C | 1.5e-05 | 1.1e-05 | 1.2e-05 | 1.267E-05 | 2.082E-06 | 1.100E-05 | 1.500E-05 |
| 19736 | 987 | Y987S | C | 1.5e-05 | 1.4e-05 | 8.3e-06 | 1.243E-05 | 3.614E-06 | 8.300E-06 | 1.500E-05 |
| 19737 | 987 | Y987T | C | 1.6e-05 | 1.1e-05 | 1e-05 | 1.233E-05 | 3.215E-06 | 1.000E-05 | 1.600E-05 |
| 19738 | 987 | Y987W | C | 1.4e-05 | 1.3e-05 | 1.1e-05 | 1.267E-05 | 1.528E-06 | 1.100E-05 | 1.400E-05 |
| 19739 | 987 | Y987Y | C | 9.3e-06 | 1.5e-05 | 1.6e-05 | 1.343E-05 | 3.614E-06 | 9.300E-06 | 1.600E-05 |
| 19740 | 987 | Y987V | C | 1e-05 | 1e-05 | 1.5e-05 | 1.167E-05 | 2.887E-06 | 1.000E-05 | 1.500E-05 |
| 19741 | 988 | F988A | C | 1.1e-05 | 1.7e-05 | 1e-05 | 1.267E-05 | 3.786E-06 | 1.000E-05 | 1.700E-05 |
| 19742 | 988 | F988R | C | 1e-05 | 9.3e-06 | 1.4e-05 | 1.110E-05 | 2.536E-06 | 9.300E-06 | 1.400E-05 |
| 19743 | 988 | F988N | C | 1.2e-05 | 9.7e-06 | 1.2e-05 | 1.123E-05 | 1.328E-06 | 9.700E-06 | 1.200E-05 |
| 19744 | 988 | F988D | C | 1e-05 | 6.9e-06 | 9.2e-06 | 8.700E-06 | 1.609E-06 | 6.900E-06 | 1.000E-05 |
| 19745 | 988 | F988C | C | 9.1e-06 | 1.4e-05 | 7.7e-06 | 1.027E-05 | 3.308E-06 | 7.700E-06 | 1.400E-05 |
| 19746 | 988 | F988E | C | 1.2e-05 | 1.4e-05 | 8.6e-06 | 1.153E-05 | 2.730E-06 | 8.600E-06 | 1.400E-05 |
| 19747 | 988 | F988Q | C | 1e-05 | 8.3e-06 | 1.5e-05 | 1.110E-05 | 3.483E-06 | 8.300E-06 | 1.500E-05 |
| 19748 | 988 | F988G | C | 1.6e-05 | 1e-05 | 1e-05 | 1.200E-05 | 3.464E-06 | 1.000E-05 | 1.600E-05 |
| 19749 | 988 | F988H | C | 1.1e-05 | 1.5e-05 | 9.3e-06 | 1.177E-05 | 2.926E-06 | 9.300E-06 | 1.500E-05 |
| 19750 | 988 | F988I | C | 1.5e-05 | 9.9e-06 | 1.3e-05 | 1.263E-05 | 2.570E-06 | 9.900E-06 | 1.500E-05 |
| 19751 | 988 | F988L | C | 1.1e-05 | 1.6e-05 | 1e-05 | 1.233E-05 | 3.215E-06 | 1.000E-05 | 1.600E-05 |
| 19752 | 988 | F988K | C | 9.5e-06 | 1.1e-05 | 1.5e-05 | 1.183E-05 | 2.843E-06 | 9.500E-06 | 1.500E-05 |
| 19753 | 988 | F988M | C | 1.1e-05 | 1e-05 | 1.5e-05 | 1.200E-05 | 2.646E-06 | 1.000E-05 | 1.500E-05 |
| 19754 | 988 | F988F | C | 1.5e-05 | 6.8e-06 | 1.2e-05 | 1.127E-05 | 4.149E-06 | 6.800E-06 | 1.500E-05 |
| 19755 | 988 | F988P | C | 9.7e-06 | 1.4e-05 | 1.5e-05 | 1.290E-05 | 2.816E-06 | 9.700E-06 | 1.500E-05 |
| 19756 | 988 | F988S | C | 1.6e-05 | 1.4e-05 | 1.1e-05 | 1.367E-05 | 2.517E-06 | 1.100E-05 | 1.600E-05 |
| 19757 | 988 | F988T | C | 1.4e-05 | 9e-06 | 1.4e-05 | 1.233E-05 | 2.887E-06 | 9.000E-06 | 1.400E-05 |
| 19758 | 988 | F988W | C | 9.8e-06 | 2.2e-05 | 1.3e-05 | 1.493E-05 | 6.326E-06 | 9.800E-06 | 2.200E-05 |
| 19759 | 988 | F988Y | C | 1.5e-05 | 1.4e-05 | 1.4e-05 | 1.433E-05 | 5.774E-07 | 1.400E-05 | 1.500E-05 |
| 19760 | 988 | F988V | C | 9.4e-06 | 1.1e-05 | 1.7e-05 | 1.247E-05 | 4.007E-06 | 9.400E-06 | 1.700E-05 |
| 19761 | 989 | P989A | C | 1.5e-05 | 1.6e-05 | 9.6e-06 | 1.353E-05 | 3.443E-06 | 9.600E-06 | 1.600E-05 |
| 19762 | 989 | P989R | C | 1.7e-05 | 1.7e-05 | 1.3e-05 | 1.567E-05 | 2.309E-06 | 1.300E-05 | 1.700E-05 |
| 19763 | 989 | P989N | C | 9.8e-06 | 1.1e-05 | 1e-05 | 1.027E-05 | 6.429E-07 | 9.800E-06 | 1.100E-05 |
| 19764 | 989 | P989D | C | 1.1e-05 | 1.5e-05 | 1e-05 | 1.200E-05 | 2.646E-06 | 1.000E-05 | 1.500E-05 |
| 19765 | 989 | P989C | C | 1.2e-05 | 1.6e-05 | 1e-05 | 1.267E-05 | 3.055E-06 | 1.000E-05 | 1.600E-05 |
| 19766 | 989 | P989E | C | 1.9e-05 | 1.2e-05 | 1.5e-05 | 1.533E-05 | 3.512E-06 | 1.200E-05 | 1.900E-05 |
| 19767 | 989 | P989Q | C | 1.6e-05 | 1.4e-05 | 1.5e-05 | 1.500E-05 | 1.000E-06 | 1.400E-05 | 1.600E-05 |
| 19768 | 989 | P989G | C | 1.3e-05 | 1.1e-05 | 8.7e-06 | 1.090E-05 | 2.152E-06 | 8.700E-06 | 1.300E-05 |
| 19769 | 989 | P989H | C | 1.5e-05 | 8.2e-06 | 1.4e-05 | 1.240E-05 | 3.672E-06 | 8.200E-06 | 1.500E-05 |
| 19770 | 989 | P989I | C | 1.4e-05 | 1.6e-05 | 9.5e-06 | 1.317E-05 | 3.329E-06 | 9.500E-06 | 1.600E-05 |
| 19771 | 989 | P989L | C | 1.6e-05 | 1.6e-05 | 1e-05 | 1.400E-05 | 3.464E-06 | 1.000E-05 | 1.600E-05 |
| 19772 | 989 | P989K | C | 1.5e-05 | 1.1e-05 | 1.1e-05 | 1.233E-05 | 2.309E-06 | 1.100E-05 | 1.500E-05 |
| 19773 | 989 | P989M | C | 8.6e-06 | 9.8e-06 | 1.3e-05 | 1.047E-05 | 2.274E-06 | 8.600E-06 | 1.300E-05 |
| 19774 | 989 | P989F | C | 9.1e-06 | 9.9e-06 | 7.1e-06 | 8.700E-06 | 1.442E-06 | 7.100E-06 | 9.900E-06 |
| 19775 | 989 | P989P | C | 1.6e-05 | 8.7e-06 | 1.1e-05 | 1.190E-05 | 3.732E-06 | 8.700E-06 | 1.600E-05 |
| 19776 | 989 | P989S | C | 1.3e-05 | 1e-05 | 9.2e-06 | 1.073E-05 | 2.003E-06 | 9.200E-06 | 1.300E-05 |
| 19777 | 989 | P989T | C | 1.1e-05 | 8.5e-06 | 1.5e-05 | 1.150E-05 | 3.279E-06 | 8.500E-06 | 1.500E-05 |
| 19778 | 989 | P989W | C | 1.6e-05 | 9.7e-06 | 9.9e-06 | 1.187E-05 | 3.581E-06 | 9.700E-06 | 1.600E-05 |
| 19779 | 989 | P989Y | C | 1.5e-05 | 9.2e-06 | 7.9e-06 | 1.070E-05 | 3.780E-06 | 7.900E-06 | 1.500E-05 |
| 19780 | 989 | P989V | C | 1.6e-05 | 1.6e-05 | 1.5e-05 | 1.567E-05 | 5.774E-07 | 1.500E-05 | 1.600E-05 |
| 19781 | 990 | E990A | C | 1.5e-05 | 1.1e-05 | 9.7e-06 | 1.190E-05 | 2.762E-06 | 9.700E-06 | 1.500E-05 |
| 19782 | 990 | E990R | C | 1e-05 | 1.7e-05 | 7.6e-06 | 1.153E-05 | 4.884E-06 | 7.600E-06 | 1.700E-05 |
| 19783 | 990 | E990N | C | 1.3e-05 | 1.3e-05 | 1.4e-05 | 1.333E-05 | 5.774E-07 | 1.300E-05 | 1.400E-05 |
| 19784 | 990 | E990D | C | 1.4e-05 | 1.3e-05 | 1.5e-05 | 1.400E-05 | 1.000E-06 | 1.300E-05 | 1.500E-05 |
| 19785 | 990 | E990C | C | 1.1e-05 | 1.5e-05 | 1.4e-05 | 1.333E-05 | 2.082E-06 | 1.100E-05 | 1.500E-05 |
| 19786 | 990 | E990E | C | 1.1e-05 | 1.4e-05 | 9.6e-06 | 1.153E-05 | 2.248E-06 | 9.600E-06 | 1.400E-05 |
| 19787 | 990 | E990Q | C | 9.6e-06 | 1.4e-05 | 1.6e-05 | 1.320E-05 | 3.274E-06 | 9.600E-06 | 1.600E-05 |
| 19788 | 990 | E990G | C | 9.6e-06 | 1.1e-05 | 8.9e-06 | 9.833E-06 | 1.069E-06 | 8.900E-06 | 1.100E-05 |
| 19789 | 990 | E990H | C | 9.5e-06 | 8.7e-06 | 1.5e-05 | 1.107E-05 | 3.430E-06 | 8.700E-06 | 1.500E-05 |
| 19790 | 990 | E990I | C | 1e-05 | 8e-06 | 1.1e-05 | 9.667E-06 | 1.528E-06 | 8.000E-06 | 1.100E-05 |
| 19791 | 990 | E990L | C | 1e-05 | 1.8e-05 | 1.5e-05 | 1.433E-05 | 4.041E-06 | 1.000E-05 | 1.800E-05 |
| 19792 | 990 | E990K | C | 1.5e-05 | 1.1e-05 | 1.1e-05 | 1.233E-05 | 2.309E-06 | 1.100E-05 | 1.500E-05 |
| 19793 | 990 | E990M | C | 1.5e-05 | 1.5e-05 | 1.1e-05 | 1.367E-05 | 2.309E-06 | 1.100E-05 | 1.500E-05 |
| 19794 | 990 | E990F | C | 6.9e-06 | 1.6e-05 | 7e-06 | 9.967E-06 | 5.225E-06 | 6.900E-06 | 1.600E-05 |
| 19795 | 990 | E990P | C | 5.1e-06 | 1.1e-05 | 1.2e-05 | 9.367E-06 | 3.729E-06 | 5.100E-06 | 1.200E-05 |
| 19796 | 990 | E990S | C | 1.4e-05 | 1.1e-05 | 1.4e-05 | 1.300E-05 | 1.732E-06 | 1.100E-05 | 1.400E-05 |
| 19797 | 990 | E990T | C | 9.9e-06 | 9.6e-06 | 9.5e-06 | 9.667E-06 | 2.082E-07 | 9.500E-06 | 9.900E-06 |
| 19798 | 990 | E990W | C | 8.7e-06 | 9.9e-06 | 1.2e-05 | 1.020E-05 | 1.670E-06 | 8.700E-06 | 1.200E-05 |
| 19799 | 990 | E990Y | C | 1.5e-05 | 7.5e-06 | 1.6e-05 | 1.283E-05 | 4.646E-06 | 7.500E-06 | 1.600E-05 |
| 19800 | 990 | E990V | C | 7.1e-06 | 9.8e-06 | 1.7e-05 | 1.130E-05 | 5.118E-06 | 7.100E-06 | 1.700E-05 |
| 19801 | 991 | P991A | C | 1.2e-05 | 1.3e-05 | 1.4e-05 | 1.300E-05 | 1.000E-06 | 1.200E-05 | 1.400E-05 |
| 19802 | 991 | P991R | C | 1.6e-05 | 1.1e-05 | 9.3e-06 | 1.210E-05 | 3.483E-06 | 9.300E-06 | 1.600E-05 |
| 19803 | 991 | P991N | C | 9.5e-06 | 1.6e-05 | 1.1e-05 | 1.217E-05 | 3.403E-06 | 9.500E-06 | 1.600E-05 |
| 19804 | 991 | P991D | C | 1.2e-05 | 1e-05 | 1.3e-05 | 1.167E-05 | 1.528E-06 | 1.000E-05 | 1.300E-05 |

| | | | | | | | | | | |
|---|---|---|---|---|---|---|---|---|---|---|
| 19805 | 991 | P991C | C | 1.2e-05 | 1e-05 | 1e-05 | 1.067E-05 | 1.155E-06 | 1.000E-05 | 1.200E-05 |
| 19806 | 991 | P991E | C | 1.3e-05 | 9.9e-06 | 9.5e-06 | 1.080E-05 | 1.916E-06 | 9.500E-06 | 1.300E-05 |
| 19807 | 991 | P991Q | C | 8.1e-06 | 1.1e-05 | 1.9e-05 | 1.270E-05 | 5.645E-06 | 8.100E-06 | 1.900E-05 |
| 19808 | 991 | P991G | C | 1.3e-05 | 9.2e-06 | 8.6e-06 | 1.027E-05 | 2.386E-06 | 8.600E-06 | 1.300E-05 |
| 19809 | 991 | P991H | C | 1e-05 | 9.6e-06 | 9.7e-06 | 9.767E-06 | 2.082E-07 | 9.600E-06 | 1.000E-05 |
| 19810 | 991 | P991I | C | 1.7e-05 | 1.5e-05 | 1.3e-05 | 1.500E-05 | 2.000E-06 | 1.300E-05 | 1.700E-05 |
| 19811 | 991 | P991L | C | 1.3e-05 | 1.6e-05 | 1.2e-05 | 1.367E-05 | 2.082E-06 | 1.200E-05 | 1.600E-05 |
| 19812 | 991 | P991K | C | 1.5e-05 | 1e-05 | 1.4e-05 | 1.300E-05 | 2.646E-06 | 1.000E-05 | 1.500E-05 |
| 19813 | 991 | P991M | C | 9.6e-06 | 1.4e-05 | 1.6e-05 | 1.320E-05 | 3.274E-06 | 9.600E-06 | 1.600E-05 |
| 19814 | 991 | P991F | C | 1.4e-05 | 1e-05 | 1.5e-05 | 1.300E-05 | 2.646E-06 | 1.000E-05 | 1.500E-05 |
| 19815 | 991 | P991P | C | 8.2e-06 | 1e-05 | 1.4e-05 | 1.073E-05 | 2.969E-06 | 8.200E-06 | 1.400E-05 |
| 19816 | 991 | P991S | C | 9.7e-06 | 9.1e-06 | 1.6e-05 | 1.160E-05 | 3.822E-06 | 9.100E-06 | 1.600E-05 |
| 19817 | 991 | P991T | C | 1.5e-05 | 1.5e-05 | 1.6e-05 | 1.533E-05 | 5.774E-07 | 1.500E-05 | 1.600E-05 |
| 19818 | 991 | P991W | C | 1.4e-05 | 1.4e-05 | 1e-05 | 1.267E-05 | 2.309E-06 | 1.000E-05 | 1.400E-05 |
| 19819 | 991 | P991Y | C | 1.1e-05 | 1.4e-05 | 1.5e-05 | 1.333E-05 | 2.082E-06 | 1.100E-05 | 1.500E-05 |
| 19820 | 991 | P991V | C | 1.1e-05 | 1.6e-05 | 8.9e-06 | 1.197E-05 | 3.647E-06 | 8.900E-06 | 1.600E-05 |
| 19821 | 992 | V992A | C | 9.6e-06 | 1.5e-05 | 1.8e-05 | 1.420E-05 | 4.257E-06 | 9.600E-06 | 1.800E-05 |
| 19822 | 992 | V992R | C | 1e-05 | 1.5e-05 | 1e-05 | 1.167E-05 | 2.887E-06 | 1.000E-05 | 1.500E-05 |
| 19823 | 992 | V992N | C | 1.3e-05 | 8.8e-06 | 1.4e-05 | 1.193E-05 | 2.759E-06 | 8.800E-06 | 1.400E-05 |
| 19824 | 992 | V992D | C | 1.5e-05 | 1e-05 | 1.2e-05 | 1.233E-05 | 2.517E-06 | 1.000E-05 | 1.500E-05 |
| 19825 | 992 | V992C | C | 1.1e-05 | 1.3e-05 | 1.3e-05 | 1.233E-05 | 1.155E-06 | 1.100E-05 | 1.300E-05 |
| 19826 | 992 | V992E | C | 1.3e-05 | 1.1e-05 | 1.4e-05 | 1.267E-05 | 1.528E-06 | 1.100E-05 | 1.400E-05 |
| 19827 | 992 | V992Q | C | 1.3e-05 | 1.3e-05 | 1.4e-05 | 1.333E-05 | 5.774E-07 | 1.300E-05 | 1.400E-05 |
| 19828 | 992 | V992G | C | 1e-05 | 1.5e-05 | 1.2e-05 | 1.233E-05 | 2.517E-06 | 1.000E-05 | 1.500E-05 |
| 19829 | 992 | V992H | C | 1e-05 | 1.5e-05 | 1.6e-05 | 1.367E-05 | 3.215E-06 | 1.000E-05 | 1.600E-05 |
| 19830 | 992 | V992I | C | 8.1e-06 | 1.5e-05 | 8.8e-06 | 1.063E-05 | 3.798E-06 | 8.100E-06 | 1.500E-05 |
| 19831 | 992 | V992L | C | 1.4e-05 | 1.4e-05 | 6.6e-06 | 1.153E-05 | 4.272E-06 | 6.600E-06 | 1.400E-05 |
| 19832 | 992 | V992K | C | 1.6e-05 | 1.3e-05 | 1.2e-05 | 1.367E-05 | 2.082E-06 | 1.200E-05 | 1.600E-05 |
| 19833 | 992 | V992M | C | 1.3e-05 | 1.5e-05 | 1.7e-05 | 1.500E-05 | 2.000E-06 | 1.300E-05 | 1.700E-05 |
| 19834 | 992 | V992F | C | 1.1e-05 | 1.2e-05 | 1.7e-05 | 1.333E-05 | 3.215E-06 | 1.100E-05 | 1.700E-05 |
| 19835 | 992 | V992P | C | 9.9e-06 | 1.6e-05 | 9.3e-06 | 1.173E-05 | 3.707E-06 | 9.300E-06 | 1.600E-05 |
| 19836 | 992 | V992S | C | 1.6e-05 | 1.6e-05 | 1.5e-05 | 1.567E-05 | 5.774E-07 | 1.500E-05 | 1.600E-05 |
| 19837 | 992 | V992T | C | 9e-06 | 1.4e-05 | 7.9e-06 | 1.030E-05 | 3.251E-06 | 7.900E-06 | 1.400E-05 |
| 19838 | 992 | V992W | C | 9.9e-06 | 1.4e-05 | 8.9e-06 | 1.093E-05 | 2.702E-06 | 8.900E-06 | 1.400E-05 |
| 19839 | 992 | V992Y | C | 9.3e-06 | 1e-05 | 1.5e-05 | 1.143E-05 | 3.109E-06 | 9.300E-06 | 1.500E-05 |
| 19840 | 992 | V992V | C | 1.2e-05 | 1e-05 | 1.5e-05 | 1.233E-05 | 2.517E-06 | 1.000E-05 | 1.500E-05 |
| 19841 | 993 | T993A | C | 1e-05 | 1.9e-05 | 9.5e-06 | 1.283E-05 | 5.346E-06 | 9.500E-06 | 1.900E-05 |
| 19842 | 993 | T993R | C | 1.1e-05 | 1.6e-05 | 1.1e-05 | 1.267E-05 | 2.887E-06 | 1.100E-05 | 1.600E-05 |
| 19843 | 993 | T993N | C | 1.5e-05 | 1.3e-05 | 9.6e-06 | 1.253E-05 | 2.730E-06 | 9.600E-06 | 1.500E-05 |
| 19844 | 993 | T993D | C | 1.6e-05 | 1e-05 | 1.4e-05 | 1.333E-05 | 3.055E-06 | 1.000E-05 | 1.600E-05 |
| 19845 | 993 | T993C | C | 9.6e-06 | 1.4e-05 | 1.5e-05 | 1.287E-05 | 2.873E-06 | 9.600E-06 | 1.500E-05 |
| 19846 | 993 | T993E | C | 1.5e-05 | 2.2e-05 | 1.2e-05 | 1.633E-05 | 5.132E-06 | 1.200E-05 | 2.200E-05 |
| 19847 | 993 | T993Q | C | 1.4e-05 | 1.2e-05 | 1.3e-05 | 1.300E-05 | 1.000E-06 | 1.200E-05 | 1.400E-05 |
| 19848 | 993 | T993G | C | 9.8e-06 | 1.1e-05 | 1e-05 | 1.027E-05 | 6.429E-07 | 9.800E-06 | 1.100E-05 |
| 19849 | 993 | T993H | C | 1e-05 | 1.5e-05 | 1.3e-05 | 1.267E-05 | 2.517E-06 | 1.000E-05 | 1.500E-05 |
| 19850 | 993 | T993I | C | 1.5e-05 | 1.5e-05 | 1.1e-05 | 1.367E-05 | 2.309E-06 | 1.100E-05 | 1.500E-05 |
| 19851 | 993 | T993L | C | 1.1e-05 | 1.4e-05 | 1.4e-05 | 1.300E-05 | 1.732E-06 | 1.100E-05 | 1.400E-05 |
| 19852 | 993 | T993K | C | 1.4e-05 | 1.5e-05 | 1.4e-05 | 1.433E-05 | 5.774E-07 | 1.400E-05 | 1.500E-05 |
| 19853 | 993 | T993M | C | 1.8e-05 | 1.6e-05 | 1.2e-05 | 1.533E-05 | 3.055E-06 | 1.200E-05 | 1.800E-05 |
| 19854 | 993 | T993F | C | 9.5e-06 | 1.1e-05 | 1.4e-05 | 1.150E-05 | 2.291E-06 | 9.500E-06 | 1.400E-05 |
| 19855 | 993 | T993P | C | 1.2e-05 | 1.4e-05 | 9.8e-06 | 1.193E-05 | 2.101E-06 | 9.800E-06 | 1.400E-05 |
| 19856 | 993 | T993S | C | 1e-05 | 9.5e-06 | 1.4e-05 | 1.117E-05 | 2.466E-06 | 9.500E-06 | 1.400E-05 |
| 19857 | 993 | T993T | C | 1.3e-05 | 1.5e-05 | 1.4e-05 | 1.400E-05 | 1.000E-06 | 1.300E-05 | 1.500E-05 |
| 19858 | 993 | T993W | C | 9.8e-06 | 1e-05 | 1e-05 | 9.933E-06 | 1.155E-07 | 9.800E-06 | 1.000E-05 |
| 19859 | 993 | T993Y | C | 1.1e-05 | 1.6e-05 | 8.9e-06 | 1.197E-05 | 3.647E-06 | 8.900E-06 | 1.600E-05 |
| 19860 | 993 | T993V | C | 1.4e-05 | 1.5e-05 | 7.8e-06 | 1.227E-05 | 3.900E-06 | 7.800E-06 | 1.500E-05 |
| 19861 | 994 | V994A | C | 1.5e-05 | 1.6e-05 | 1e-05 | 1.367E-05 | 3.215E-06 | 1.000E-05 | 1.600E-05 |
| 19862 | 994 | V994R | C | 1.4e-05 | 9e-06 | 1.5e-05 | 1.267E-05 | 3.215E-06 | 9.000E-06 | 1.500E-05 |
| 19863 | 994 | V994N | C | 1.1e-05 | 1.5e-05 | 1.3e-05 | 1.300E-05 | 2.000E-06 | 1.100E-05 | 1.500E-05 |
| 19864 | 994 | V994D | C | 1.5e-05 | 8.4e-06 | 1.4e-05 | 1.247E-05 | 3.557E-06 | 8.400E-06 | 1.500E-05 |
| 19865 | 994 | V994C | C | 9.7e-06 | 1.4e-05 | 1e-05 | 1.123E-05 | 2.401E-06 | 9.700E-06 | 1.400E-05 |
| 19866 | 994 | V994E | C | 1.5e-05 | 1.5e-05 | 1.1e-05 | 1.367E-05 | 2.309E-06 | 1.100E-05 | 1.500E-05 |
| 19867 | 994 | V994Q | C | 1.1e-05 | 1.4e-05 | 9.4e-06 | 1.147E-05 | 2.335E-06 | 9.400E-06 | 1.400E-05 |
| 19868 | 994 | V994G | C | 9.5e-06 | 1.1e-05 | 1.1e-05 | 1.050E-05 | 8.660E-07 | 9.500E-06 | 1.100E-05 |
| 19869 | 994 | V994H | C | 9.1e-06 | 1.3e-05 | 9.2e-06 | 1.043E-05 | 2.223E-06 | 9.100E-06 | 1.300E-05 |
| 19870 | 994 | V994I | C | 1.4e-05 | 1.4e-05 | 1.1e-05 | 1.300E-05 | 1.732E-06 | 1.100E-05 | 1.400E-05 |
| 19871 | 994 | V994L | C | 1.7e-05 | 1.5e-05 | 1.5e-05 | 1.567E-05 | 1.155E-06 | 1.500E-05 | 1.700E-05 |
| 19872 | 994 | V994K | C | 1.1e-05 | 9e-06 | 1.2e-05 | 1.067E-05 | 1.528E-06 | 9.000E-06 | 1.200E-05 |
| 19873 | 994 | V994M | C | 1.6e-05 | 1.5e-05 | 1.5e-05 | 1.533E-05 | 5.774E-07 | 1.500E-05 | 1.600E-05 |
| 19874 | 994 | V994F | C | 9.5e-06 | 1e-05 | 1e-05 | 9.833E-06 | 2.887E-07 | 9.500E-06 | 1.000E-05 |
| 19875 | 994 | V994P | C | 9.6e-06 | 1e-05 | 1.4e-05 | 1.120E-05 | 2.433E-06 | 9.600E-06 | 1.400E-05 |
| 19876 | 994 | V994S | C | 5.8e-06 | 1.3e-05 | 1.3e-05 | 1.060E-05 | 4.157E-06 | 5.800E-06 | 1.300E-05 |
| 19877 | 994 | V994T | C | 1e-05 | 1.1e-05 | 1.3e-05 | 1.133E-05 | 1.528E-06 | 1.000E-05 | 1.300E-05 |
| 19878 | 994 | V994W | C | 1.3e-05 | 1.6e-05 | 1.5e-05 | 1.467E-05 | 1.528E-06 | 1.300E-05 | 1.600E-05 |
| 19879 | 994 | V994Y | C | 1.1e-05 | 1.4e-05 | 1.4e-05 | 1.300E-05 | 1.732E-06 | 1.100E-05 | 1.400E-05 |
| 19880 | 994 | V994V | C | 1.4e-05 | 1.6e-05 | 1.6e-05 | 1.533E-05 | 1.155E-06 | 1.400E-05 | 1.600E-05 |
| 19881 | 995 | S995A | C | 1.1e-05 | 1.7e-05 | 1.5e-05 | 1.433E-05 | 3.055E-06 | 1.100E-05 | 1.700E-05 |
| 19882 | 995 | S995R | C | 1.5e-05 | 8.5e-06 | 1e-05 | 1.117E-05 | 3.403E-06 | 8.500E-06 | 1.500E-05 |
| 19883 | 995 | S995N | C | 8.1e-06 | 1.2e-05 | 1.4e-05 | 1.137E-05 | 3.001E-06 | 8.100E-06 | 1.400E-05 |
| 19884 | 995 | S995D | C | 1e-05 | 1e-05 | 1.2e-05 | 1.067E-05 | 1.155E-06 | 1.000E-05 | 1.200E-05 |
| 19885 | 995 | S995C | C | 1.6e-05 | 1.5e-05 | 1.1e-05 | 1.400E-05 | 2.646E-06 | 1.100E-05 | 1.600E-05 |
| 19886 | 995 | S995E | C | 1e-05 | 1.4e-05 | 1.3e-05 | 1.233E-05 | 2.082E-06 | 1.000E-05 | 1.400E-05 |
| 19887 | 995 | S995Q | C | 1.7e-05 | 1.3e-05 | 1e-05 | 1.333E-05 | 3.512E-06 | 1.000E-05 | 1.700E-05 |
| 19888 | 995 | S995G | C | 1.1e-05 | 8.4e-06 | 1.3e-05 | 1.080E-05 | 2.307E-06 | 8.400E-06 | 1.300E-05 |
| 19889 | 995 | S995H | C | 1.4e-05 | 1.2e-05 | 1e-05 | 1.200E-05 | 2.000E-06 | 1.000E-05 | 1.400E-05 |

| | | | | | | | | | | |
|---|---|---|---|---|---|---|---|---|---|---|
| 19890 | 995 | S995I | C | 9.2e-06 | 1e-05 | 1.4e-05 | 1.107E-05 | 2.572E-06 | 9.200E-06 | 1.400E-05 |
| 19891 | 995 | S995L | C | 1e-05 | 1.3e-05 | 1.5e-05 | 1.267E-05 | 2.517E-06 | 1.000E-05 | 1.500E-05 |
| 19892 | 995 | S995K | C | 1e-05 | 1.1e-05 | 1.3e-05 | 1.133E-05 | 1.528E-06 | 1.000E-05 | 1.300E-05 |
| 19893 | 995 | S995M | C | 1.5e-05 | 1.5e-05 | 1.1e-05 | 1.367E-05 | 2.309E-06 | 1.100E-05 | 1.500E-05 |
| 19894 | 995 | S995F | C | 1.6e-05 | 1e-05 | 1.5e-05 | 1.367E-05 | 3.215E-06 | 1.000E-05 | 1.600E-05 |
| 19895 | 995 | S995P | C | 1.5e-05 | 1.9e-05 | 1.4e-05 | 1.600E-05 | 2.646E-06 | 1.400E-05 | 1.900E-05 |
| 19896 | 995 | S995S | C | 1.6e-05 | 1.4e-05 | 1.9e-05 | 1.633E-05 | 2.517E-06 | 1.400E-05 | 1.900E-05 |
| 19897 | 995 | S995T | C | 1.6e-05 | 1.5e-05 | 1.6e-05 | 1.567E-05 | 5.774E-07 | 1.500E-05 | 1.600E-05 |
| 19898 | 995 | S995W | C | 9.1e-06 | 9.8e-06 | 1e-05 | 9.633E-06 | 4.726E-07 | 9.100E-06 | 1.000E-05 |
| 19899 | 995 | S995Y | C | 1.7e-05 | 1.2e-05 | 1.1e-05 | 1.333E-05 | 3.215E-06 | 1.100E-05 | 1.700E-05 |
| 19900 | 995 | S995V | C | 8.5e-06 | 1.4e-05 | 1.1e-05 | 1.117E-05 | 2.754E-06 | 8.500E-06 | 1.400E-05 |
| 19901 | 996 | W996A | C | 1.1e-05 | 9e-06 | 1.7e-05 | 1.233E-05 | 4.163E-06 | 9.000E-06 | 1.700E-05 |
| 19902 | 996 | W996R | C | 1.1e-05 | 1.2e-05 | 9.5e-06 | 1.083E-05 | 1.258E-06 | 9.500E-06 | 1.200E-05 |
| 19903 | 996 | W996N | C | 1.6e-05 | 1.4e-05 | 1.1e-05 | 1.367E-05 | 2.517E-06 | 1.100E-05 | 1.600E-05 |
| 19904 | 996 | W996D | C | 9.9e-06 | 1.5e-05 | 9.9e-06 | 1.160E-05 | 2.944E-06 | 9.900E-06 | 1.500E-05 |
| 19905 | 996 | W996C | C | 1.6e-05 | 1e-05 | 1.5e-05 | 1.367E-05 | 3.215E-06 | 1.000E-05 | 1.600E-05 |
| 19906 | 996 | W996E | C | 1e-05 | 1.1e-05 | 1.4e-05 | 1.167E-05 | 2.082E-06 | 1.000E-05 | 1.400E-05 |
| 19907 | 996 | W996Q | C | 1.3e-05 | 1.5e-05 | 1.4e-05 | 1.400E-05 | 1.000E-06 | 1.300E-05 | 1.500E-05 |
| 19908 | 996 | W996G | C | 1.2e-05 | 8.1e-06 | 9.7e-06 | 9.933E-06 | 1.960E-06 | 8.100E-06 | 1.200E-05 |
| 19909 | 996 | W996H | C | 1e-05 | 1.5e-05 | 1e-05 | 1.167E-05 | 2.887E-06 | 1.000E-05 | 1.500E-05 |
| 19910 | 996 | W996I | C | 1.4e-05 | 1.5e-05 | 1.1e-05 | 1.333E-05 | 2.082E-06 | 1.100E-05 | 1.500E-05 |
| 19911 | 996 | W996L | C | 7.6e-06 | 9.4e-06 | 1.6e-05 | 1.100E-05 | 4.423E-06 | 7.600E-06 | 1.600E-05 |
| 19912 | 996 | W996K | C | 1.3e-05 | 1.1e-05 | 9.8e-06 | 1.127E-05 | 1.617E-06 | 9.800E-06 | 1.300E-05 |
| 19913 | 996 | W996M | C | 1.6e-05 | 1.1e-05 | 8.9e-06 | 1.197E-05 | 3.647E-06 | 8.900E-06 | 1.600E-05 |
| 19914 | 996 | W996F | C | 1.1e-05 | 1.2e-05 | 1.6e-05 | 1.300E-05 | 2.646E-06 | 1.100E-05 | 1.600E-05 |
| 19915 | 996 | W996P | C | 1.1e-05 | 1.5e-05 | 8.4e-06 | 1.147E-05 | 3.325E-06 | 8.400E-06 | 1.500E-05 |
| 19916 | 996 | W996S | C | 1.3e-05 | 1.4e-05 | 9.4e-06 | 1.213E-05 | 2.419E-06 | 9.400E-06 | 1.400E-05 |
| 19917 | 996 | W996T | C | 1.5e-05 | 9.7e-06 | 1.4e-05 | 1.290E-05 | 2.816E-06 | 9.700E-06 | 1.500E-05 |
| 19918 | 996 | W996W | C | 1.6e-05 | 1.1e-05 | 1.1e-05 | 1.267E-05 | 2.887E-06 | 1.100E-05 | 1.600E-05 |
| 19919 | 996 | W996Y | C | 1.5e-05 | 1.2e-05 | 1.4e-05 | 1.367E-05 | 1.528E-06 | 1.200E-05 | 1.500E-05 |
| 19920 | 996 | W996V | C | 8e-06 | 8.9e-06 | 1.3e-05 | 9.967E-06 | 2.665E-06 | 8.000E-06 | 1.300E-05 |
| 19921 | 997 | N997A | C | 1.6e-05 | 1.4e-05 | 1.7e-05 | 1.567E-05 | 1.528E-06 | 1.400E-05 | 1.700E-05 |
| 19922 | 997 | N997R | C | 1.6e-05 | 1.3e-05 | 6.5e-06 | 1.183E-05 | 4.856E-06 | 6.500E-06 | 1.600E-05 |
| 19923 | 997 | N997N | C | 1.6e-05 | 9e-06 | 1e-05 | 1.167E-05 | 3.786E-06 | 9.000E-06 | 1.600E-05 |
| 19924 | 997 | N997D | C | 1.1e-05 | 1.4e-05 | 1.4e-05 | 1.300E-05 | 1.732E-06 | 1.100E-05 | 1.400E-05 |
| 19925 | 997 | N997C | C | 1.1e-05 | 1.2e-05 | 1.1e-05 | 1.133E-05 | 5.774E-07 | 1.100E-05 | 1.200E-05 |
| 19926 | 997 | N997E | C | 1.4e-05 | 1.6e-05 | 1.2e-05 | 1.400E-05 | 2.000E-06 | 1.200E-05 | 1.600E-05 |
| 19927 | 997 | N997Q | C | 8.9e-06 | 9.6e-06 | 1.1e-05 | 9.833E-06 | 1.069E-06 | 8.900E-06 | 1.100E-05 |
| 19928 | 997 | N997G | C | 1.5e-05 | 1.1e-05 | 1.5e-05 | 1.367E-05 | 2.309E-06 | 1.100E-05 | 1.500E-05 |
| 19929 | 997 | N997H | C | 1.2e-05 | 1.6e-05 | 1.6e-05 | 1.467E-05 | 2.309E-06 | 1.200E-05 | 1.600E-05 |
| 19930 | 997 | N997I | C | 1e-05 | 7.5e-06 | 1.3e-05 | 1.017E-05 | 2.754E-06 | 7.500E-06 | 1.300E-05 |
| 19931 | 997 | N997L | C | 1.2e-05 | 1.2e-05 | 1.1e-05 | 1.167E-05 | 5.774E-07 | 1.100E-05 | 1.200E-05 |
| 19932 | 997 | N997K | C | 1.6e-05 | 1.2e-05 | 1.9e-05 | 1.567E-05 | 3.512E-06 | 1.200E-05 | 1.900E-05 |
| 19933 | 997 | N997M | C | 1.1e-05 | 9.7e-06 | 8.5e-06 | 9.733E-06 | 1.250E-06 | 8.500E-06 | 1.100E-05 |
| 19934 | 997 | N997F | C | 9.3e-06 | 1.7e-05 | 1.4e-05 | 1.343E-05 | 3.881E-06 | 9.300E-06 | 1.700E-05 |
| 19935 | 997 | N997P | C | 9.6e-06 | 1e-05 | 1.2e-05 | 1.053E-05 | 1.286E-06 | 9.600E-06 | 1.200E-05 |
| 19936 | 997 | N997S | C | 1.1e-05 | 1.6e-05 | 1.6e-05 | 1.433E-05 | 2.887E-06 | 1.100E-05 | 1.600E-05 |
| 19937 | 997 | N997T | C | 1e-05 | 1.4e-05 | 1.5e-05 | 1.300E-05 | 2.646E-06 | 1.000E-05 | 1.500E-05 |
| 19938 | 997 | N997W | C | 1.6e-05 | 1.6e-05 | 1e-05 | 1.400E-05 | 3.464E-06 | 1.000E-05 | 1.600E-05 |
| 19939 | 997 | N997Y | C | 1.2e-05 | 9.5e-06 | 9.8e-06 | 1.043E-05 | 1.365E-06 | 9.500E-06 | 1.200E-05 |
| 19940 | 997 | N997V | C | 1.1e-05 | 1.1e-05 | 1.6e-05 | 1.267E-05 | 2.887E-06 | 1.100E-05 | 1.600E-05 |
| 19941 | 998 | S998A | C | 1.3e-05 | 1e-05 | 1e-05 | 1.100E-05 | 1.732E-06 | 1.000E-05 | 1.300E-05 |
| 19942 | 998 | S998R | C | 1.3e-05 | 1.2e-05 | 1.1e-05 | 1.200E-05 | 1.000E-06 | 1.100E-05 | 1.300E-05 |
| 19943 | 998 | S998N | C | 9.8e-06 | 1.6e-05 | 1.4e-05 | 1.327E-05 | 3.164E-06 | 9.800E-06 | 1.600E-05 |
| 19944 | 998 | S998D | C | 1.5e-05 | 1e-05 | 1.5e-05 | 1.333E-05 | 2.887E-06 | 1.000E-05 | 1.500E-05 |
| 19945 | 998 | S998C | C | 1.1e-05 | 9.8e-06 | 1.5e-05 | 1.193E-05 | 2.723E-06 | 9.800E-06 | 1.500E-05 |
| 19946 | 998 | S998E | C | 1e-05 | 7.1e-06 | 1.7e-05 | 1.137E-05 | 5.090E-06 | 7.100E-06 | 1.700E-05 |
| 19947 | 998 | S998Q | C | 1.4e-05 | 1.3e-05 | 1.3e-05 | 1.333E-05 | 5.774E-07 | 1.300E-05 | 1.400E-05 |
| 19948 | 998 | S998G | C | 1.8e-05 | 9.7e-06 | 1e-05 | 1.257E-05 | 4.708E-06 | 9.700E-06 | 1.800E-05 |
| 19949 | 998 | S998H | C | 1.6e-05 | 8.4e-06 | 1.2e-05 | 1.213E-05 | 3.802E-06 | 8.400E-06 | 1.600E-05 |
| 19950 | 998 | S998I | C | 1.6e-05 | 1.4e-05 | 9.4e-06 | 1.313E-05 | 3.384E-06 | 9.400E-06 | 1.600E-05 |
| 19951 | 998 | S998L | C | 1.2e-05 | 1.7e-05 | 1.5e-05 | 1.467E-05 | 2.517E-06 | 1.200E-05 | 1.700E-05 |
| 19952 | 998 | S998K | C | 1.5e-05 | 8.7e-06 | 1e-05 | 1.123E-05 | 3.326E-06 | 8.700E-06 | 1.500E-05 |
| 19953 | 998 | S998M | C | 1.6e-05 | 1.1e-05 | 1.7e-05 | 1.467E-05 | 3.215E-06 | 1.100E-05 | 1.700E-05 |
| 19954 | 998 | S998F | C | 1.4e-05 | 1.2e-05 | 1.4e-05 | 1.333E-05 | 1.155E-06 | 1.200E-05 | 1.400E-05 |
| 19955 | 998 | S998P | C | 1.4e-05 | 1.2e-05 | 1.5e-05 | 1.367E-05 | 1.528E-06 | 1.200E-05 | 1.500E-05 |
| 19956 | 998 | S998S | C | 9.9e-06 | 1.6e-05 | 1.4e-05 | 1.330E-05 | 3.110E-06 | 9.900E-06 | 1.600E-05 |
| 19957 | 998 | S998T | C | 1.1e-05 | 1.4e-05 | 1.4e-05 | 1.300E-05 | 1.732E-06 | 1.100E-05 | 1.400E-05 |
| 19958 | 998 | S998W | C | 1.1e-05 | 8.4e-06 | 1e-05 | 9.800E-06 | 1.311E-06 | 8.400E-06 | 1.100E-05 |
| 19959 | 998 | S998Y | C | 1.1e-05 | 1e-05 | 1.1e-05 | 1.067E-05 | 5.774E-07 | 1.000E-05 | 1.100E-05 |
| 19960 | 998 | S998V | C | 1.8e-05 | 1.5e-05 | 1.4e-05 | 1.567E-05 | 2.082E-06 | 1.400E-05 | 1.800E-05 |
| 19961 | 999 | G999A | C | 1.7e-05 | 1.3e-05 | 9.3e-06 | 1.310E-05 | 3.851E-06 | 9.300E-06 | 1.700E-05 |
| 19962 | 999 | G999R | C | 1.6e-05 | 9.3e-06 | 9.2e-06 | 1.150E-05 | 3.897E-06 | 9.200E-06 | 1.600E-05 |
| 19963 | 999 | G999N | C | 1.4e-05 | 9.4e-06 | 1.3e-05 | 1.213E-05 | 2.419E-06 | 9.400E-06 | 1.400E-05 |
| 19964 | 999 | G999D | C | 1.1e-05 | 1.5e-05 | 8.5e-06 | 1.150E-05 | 3.279E-06 | 8.500E-06 | 1.500E-05 |
| 19965 | 999 | G999C | C | 1.4e-05 | 1.4e-05 | 1.6e-05 | 1.467E-05 | 1.155E-06 | 1.400E-05 | 1.600E-05 |
| 19966 | 999 | G999E | C | 1.1e-05 | 1e-05 | 1.4e-05 | 1.167E-05 | 2.082E-06 | 1.000E-05 | 1.400E-05 |
| 19967 | 999 | G999Q | C | 1.3e-05 | 1.4e-05 | 1.4e-05 | 1.367E-05 | 5.774E-07 | 1.300E-05 | 1.400E-05 |
| 19968 | 999 | G999G | C | 8.9e-06 | 1.1e-05 | 1.8e-05 | 1.263E-05 | 4.765E-06 | 8.900E-06 | 1.800E-05 |
| 19969 | 999 | G999H | C | 1.4e-05 | 1.3e-05 | 1.3e-05 | 1.333E-05 | 5.774E-07 | 1.300E-05 | 1.400E-05 |
| 19970 | 999 | G999I | C | 8.5e-06 | 1e-05 | 1.5e-05 | 1.117E-05 | 3.403E-06 | 8.500E-06 | 1.500E-05 |
| 19971 | 999 | G999L | C | 1.4e-05 | 1e-05 | 1.6e-05 | 1.333E-05 | 3.055E-06 | 1.000E-05 | 1.600E-05 |
| 19972 | 999 | G999K | C | 1.7e-05 | 1e-05 | 1.1e-05 | 1.267E-05 | 3.786E-06 | 1.000E-05 | 1.700E-05 |
| 19973 | 999 | G999M | C | 1.5e-05 | 9.6e-06 | 1.5e-05 | 1.320E-05 | 3.118E-06 | 9.600E-06 | 1.500E-05 |
| 19974 | 999 | G999F | C | 1.4e-05 | 1.6e-05 | 1.4e-05 | 1.467E-05 | 1.155E-06 | 1.400E-05 | 1.600E-05 |

| | | | | | | | | | | |
|---|---|---|---|---|---|---|---|---|---|---|
| 19975 | 999 | G999P | C | 1e-05 | 1.5e-05 | 9.3e-06 | 1.143E-05 | 3.109E-06 | 9.300E-06 | 1.500E-05 |
| 19976 | 999 | G999S | C | 1.5e-05 | 1.2e-05 | 9.2e-06 | 1.207E-05 | 2.901E-06 | 9.200E-06 | 1.500E-05 |
| 19977 | 999 | G999T | C | 1.6e-05 | 9.6e-06 | 1.5e-05 | 1.353E-05 | 3.443E-06 | 9.600E-06 | 1.600E-05 |
| 19978 | 999 | G999W | C | 1.6e-05 | 1.6e-05 | 1.5e-05 | 1.567E-05 | 5.774E-07 | 1.500E-05 | 1.600E-05 |
| 19979 | 999 | G999Y | C | 1.2e-05 | 1.7e-05 | 1.4e-05 | 1.433E-05 | 2.517E-06 | 1.200E-05 | 1.700E-05 |
| 19980 | 999 | G999V | C | 8.5e-06 | 1.4e-05 | 1.4e-05 | 1.217E-05 | 3.175E-06 | 8.500E-06 | 1.400E-05 |
| 19981 | 1000 | A1000A | C | 1.5e-05 | 1.4e-05 | 1.4e-05 | 1.433E-05 | 5.774E-07 | 1.400E-05 | 1.500E-05 |
| 19982 | 1000 | A1000R | C | 1.6e-05 | 1.3e-05 | 9.8e-06 | 1.293E-05 | 3.101E-06 | 9.800E-06 | 1.600E-05 |
| 19983 | 1000 | A1000N | C | 1.5e-05 | 8.1e-06 | 1.5e-05 | 1.270E-05 | 3.984E-06 | 8.100E-06 | 1.500E-05 |
| 19984 | 1000 | A1000D | C | 1.1e-05 | 1.5e-05 | 8.3e-06 | 1.143E-05 | 3.371E-06 | 8.300E-06 | 1.500E-05 |
| 19985 | 1000 | A1000C | C | 9.7e-06 | 8.3e-06 | 1.4e-05 | 1.067E-05 | 2.970E-06 | 8.300E-06 | 1.400E-05 |
| 19986 | 1000 | A1000E | C | 1.6e-05 | 1e-05 | 1.3e-05 | 1.300E-05 | 3.000E-06 | 1.000E-05 | 1.600E-05 |
| 19987 | 1000 | A1000Q | C | 9.8e-06 | 1.5e-05 | 1.1e-05 | 1.193E-05 | 2.723E-06 | 9.800E-06 | 1.500E-05 |
| 19988 | 1000 | A1000G | C | 1.7e-05 | 1.5e-05 | 1.5e-05 | 1.567E-05 | 1.155E-06 | 1.500E-05 | 1.700E-05 |
| 19989 | 1000 | A1000H | C | 1.7e-05 | 8.6e-06 | 1.6e-05 | 1.387E-05 | 4.588E-06 | 8.600E-06 | 1.700E-05 |
| 19990 | 1000 | A1000I | C | 9.4e-06 | 1.6e-05 | 1.5e-05 | 1.347E-05 | 3.557E-06 | 9.400E-06 | 1.600E-05 |
| 19991 | 1000 | A1000L | C | 1.8e-05 | 1.2e-05 | 1.5e-05 | 1.500E-05 | 3.000E-06 | 1.200E-05 | 1.800E-05 |
| 19992 | 1000 | A1000K | C | 1.2e-05 | 1.1e-05 | 1e-05 | 1.100E-05 | 1.000E-06 | 1.000E-05 | 1.200E-05 |
| 19993 | 1000 | A1000M | C | 1e-05 | 7.2e-06 | 1.6e-05 | 1.107E-05 | 4.496E-06 | 7.200E-06 | 1.600E-05 |
| 19994 | 1000 | A1000F | C | 1.1e-05 | 1.1e-05 | 1e-05 | 1.067E-05 | 5.774E-07 | 1.000E-05 | 1.100E-05 |
| 19995 | 1000 | A1000P | C | 1.2e-05 | 1.3e-05 | 1.5e-05 | 1.333E-05 | 1.528E-06 | 1.200E-05 | 1.500E-05 |
| 19996 | 1000 | A1000S | C | 1.6e-05 | 1.5e-05 | 1.3e-05 | 1.467E-05 | 1.528E-06 | 1.300E-05 | 1.600E-05 |
| 19997 | 1000 | A1000T | C | 1.8e-05 | 1.5e-05 | 1.3e-05 | 1.533E-05 | 2.517E-06 | 1.300E-05 | 1.800E-05 |
| 19998 | 1000 | A1000W | C | 9.7e-06 | 1.4e-05 | 1.1e-05 | 1.157E-05 | 2.205E-06 | 9.700E-06 | 1.400E-05 |
| 19999 | 1000 | A1000Y | C | 9.2e-06 | 1.3e-05 | 1.5e-05 | 1.240E-05 | 2.946E-06 | 9.200E-06 | 1.500E-05 |
| 20000 | 1000 | A1000V | C | 1.3e-05 | 1.2e-05 | 1.1e-05 | 1.200E-05 | 1.000E-06 | 1.100E-05 | 1.300E-05 |
| 20001 | 1001 | L1001A | C | 1.4e-05 | 1.4e-05 | 1.3e-05 | 1.367E-05 | 5.774E-07 | 1.300E-05 | 1.400E-05 |
| 20002 | 1001 | L1001R | C | 1.1e-05 | 1.6e-05 | 9.9e-06 | 1.230E-05 | 3.251E-06 | 9.900E-06 | 1.600E-05 |
| 20003 | 1001 | L1001N | C | 1e-05 | 1.1e-05 | 1.2e-05 | 1.100E-05 | 1.000E-06 | 1.000E-05 | 1.200E-05 |
| 20004 | 1001 | L1001D | C | 1.6e-05 | 1e-05 | 1.3e-05 | 1.300E-05 | 3.000E-06 | 1.000E-05 | 1.600E-05 |
| 20005 | 1001 | L1001C | C | 1.5e-05 | 1.2e-05 | 1.5e-05 | 1.400E-05 | 1.732E-06 | 1.200E-05 | 1.500E-05 |
| 20006 | 1001 | L1001E | C | 1.2e-05 | 1.4e-05 | 1.6e-05 | 1.400E-05 | 2.000E-06 | 1.200E-05 | 1.600E-05 |
| 20007 | 1001 | L1001Q | C | 1.1e-05 | 1.6e-05 | 1.2e-05 | 1.300E-05 | 2.646E-06 | 1.100E-05 | 1.600E-05 |
| 20008 | 1001 | L1001G | C | 1.4e-05 | 1e-05 | 1.1e-05 | 1.167E-05 | 2.082E-06 | 1.000E-05 | 1.400E-05 |
| 20009 | 1001 | L1001H | C | 7.9e-06 | 9.6e-06 | 9.8e-06 | 9.100E-06 | 1.044E-06 | 7.900E-06 | 9.800E-06 |
| 20010 | 1001 | L1001I | C | 1.4e-05 | 1.4e-05 | 1.2e-05 | 1.333E-05 | 1.155E-06 | 1.200E-05 | 1.400E-05 |
| 20011 | 1001 | L1001L | C | 1.1e-05 | 9e-06 | 1.4e-05 | 1.133E-05 | 2.517E-06 | 9.000E-06 | 1.400E-05 |
| 20012 | 1001 | L1001K | C | 1.5e-05 | 9.8e-06 | 9.4e-06 | 1.140E-05 | 3.124E-06 | 9.400E-06 | 1.500E-05 |
| 20013 | 1001 | L1001M | C | 1.1e-05 | 1.4e-05 | 1e-05 | 1.167E-05 | 2.082E-06 | 1.000E-05 | 1.400E-05 |
| 20014 | 1001 | L1001F | C | 1e-05 | 1.1e-05 | 9.9e-06 | 1.030E-05 | 6.083E-07 | 9.900E-06 | 1.100E-05 |
| 20015 | 1001 | L1001P | C | 9.5e-06 | 9.4e-06 | 1.3e-05 | 1.063E-05 | 2.050E-06 | 9.400E-06 | 1.300E-05 |
| 20016 | 1001 | L1001S | C | 1.7e-05 | 1.1e-05 | 1.1e-05 | 1.300E-05 | 3.464E-06 | 1.100E-05 | 1.700E-05 |
| 20017 | 1001 | L1001T | C | 1.3e-05 | 1.4e-05 | 1.2e-05 | 1.300E-05 | 1.000E-06 | 1.200E-05 | 1.400E-05 |
| 20018 | 1001 | L1001W | C | 1e-05 | 1.6e-05 | 1.1e-05 | 1.233E-05 | 3.215E-06 | 1.000E-05 | 1.600E-05 |
| 20019 | 1001 | L1001Y | C | 1.1e-05 | 1.4e-05 | 1e-05 | 1.167E-05 | 2.082E-06 | 1.000E-05 | 1.400E-05 |
| 20020 | 1001 | L1001V | C | 1.6e-05 | 1.1e-05 | 1.5e-05 | 1.400E-05 | 2.646E-06 | 1.100E-05 | 1.600E-05 |
| 20021 | 1002 | T1002A | C | 1.5e-05 | 1.6e-05 | 1e-05 | 1.367E-05 | 3.215E-06 | 1.000E-05 | 1.600E-05 |
| 20022 | 1002 | T1002R | C | 1.5e-05 | 1.1e-05 | 1.3e-05 | 1.300E-05 | 2.000E-06 | 1.100E-05 | 1.500E-05 |
| 20023 | 1002 | T1002N | C | 9.2e-06 | 1.6e-05 | 1.4e-05 | 1.307E-05 | 3.495E-06 | 9.200E-06 | 1.600E-05 |
| 20024 | 1002 | T1002D | C | 1.1e-05 | 1.1e-05 | 1.6e-05 | 1.267E-05 | 2.887E-06 | 1.100E-05 | 1.600E-05 |
| 20025 | 1002 | T1002C | C | 1.1e-05 | 1e-05 | 1.1e-05 | 1.067E-05 | 5.774E-07 | 1.000E-05 | 1.100E-05 |
| 20026 | 1002 | T1002E | C | 1.6e-05 | 1.4e-05 | 1.4e-05 | 1.467E-05 | 1.155E-06 | 1.400E-05 | 1.600E-05 |
| 20027 | 1002 | T1002Q | C | 1.6e-05 | 1.2e-05 | 1e-05 | 1.267E-05 | 3.055E-06 | 1.000E-05 | 1.600E-05 |
| 20028 | 1002 | T1002G | C | 9e-06 | 8.7e-06 | 1.6e-05 | 1.123E-05 | 4.131E-06 | 8.700E-06 | 1.600E-05 |
| 20029 | 1002 | T1002H | C | 1e-05 | 1.3e-05 | 1.5e-05 | 1.267E-05 | 2.517E-06 | 1.000E-05 | 1.500E-05 |
| 20030 | 1002 | T1002I | C | 1.1e-05 | 1.5e-05 | 1.1e-05 | 1.233E-05 | 2.309E-06 | 1.100E-05 | 1.500E-05 |
| 20031 | 1002 | T1002L | C | 1.5e-05 | 1.7e-05 | 1.2e-05 | 1.467E-05 | 2.517E-06 | 1.200E-05 | 1.700E-05 |
| 20032 | 1002 | T1002K | C | 1.6e-05 | 1.1e-05 | 1.5e-05 | 1.400E-05 | 2.646E-06 | 1.100E-05 | 1.600E-05 |
| 20033 | 1002 | T1002M | C | 9e-06 | 1.5e-05 | 1.7e-05 | 1.367E-05 | 4.163E-06 | 9.000E-06 | 1.700E-05 |
| 20034 | 1002 | T1002F | C | 9.8e-06 | 1.6e-05 | 9.7e-06 | 1.183E-05 | 3.609E-06 | 9.700E-06 | 1.600E-05 |
| 20035 | 1002 | T1002P | C | 1.8e-05 | 1.4e-05 | 1.6e-05 | 1.600E-05 | 2.000E-06 | 1.400E-05 | 1.800E-05 |
| 20036 | 1002 | T1002S | C | 1.5e-05 | 1.6e-05 | 1.2e-05 | 1.433E-05 | 2.082E-06 | 1.200E-05 | 1.600E-05 |
| 20037 | 1002 | T1002T | C | 1.5e-05 | 1.4e-05 | 1.4e-05 | 1.433E-05 | 5.774E-07 | 1.400E-05 | 1.500E-05 |
| 20038 | 1002 | T1002W | C | 1.1e-05 | 7.5e-06 | 8.7e-06 | 9.067E-06 | 1.779E-06 | 7.500E-06 | 1.100E-05 |
| 20039 | 1002 | T1002Y | C | 9.5e-06 | 1.6e-05 | 9.3e-06 | 1.160E-05 | 3.812E-06 | 9.300E-06 | 1.600E-05 |
| 20040 | 1002 | T1002V | C | 7.5e-06 | 1.1e-05 | 1e-05 | 9.500E-06 | 1.803E-06 | 7.500E-06 | 1.100E-05 |
| 20041 | 1003 | S1003A | C | 1.1e-05 | 1e-05 | 1.6e-05 | 1.233E-05 | 3.215E-06 | 1.000E-05 | 1.600E-05 |
| 20042 | 1003 | S1003R | C | 1.5e-05 | 1.3e-05 | 1.2e-05 | 1.333E-05 | 1.528E-06 | 1.200E-05 | 1.500E-05 |
| 20043 | 1003 | S1003N | C | 1.6e-05 | 1e-05 | 1.1e-05 | 1.233E-05 | 3.215E-06 | 1.000E-05 | 1.600E-05 |
| 20044 | 1003 | S1003D | C | 1.7e-05 | 1.1e-05 | 9.5e-06 | 1.250E-05 | 3.969E-06 | 9.500E-06 | 1.700E-05 |
| 20045 | 1003 | S1003C | C | 1.6e-05 | 1.1e-05 | 8.7e-06 | 1.190E-05 | 3.732E-06 | 8.700E-06 | 1.600E-05 |
| 20046 | 1003 | S1003E | C | 7.8e-06 | 1e-05 | 1.4e-05 | 1.060E-05 | 3.143E-06 | 7.800E-06 | 1.400E-05 |
| 20047 | 1003 | S1003Q | C | 9e-06 | 1.8e-05 | 1.2e-05 | 1.300E-05 | 4.583E-06 | 9.000E-06 | 1.800E-05 |
| 20048 | 1003 | S1003G | C | 9.1e-06 | 1.2e-05 | 6.6e-06 | 9.233E-06 | 2.702E-06 | 6.600E-06 | 1.200E-05 |
| 20049 | 1003 | S1003H | C | 7.3e-06 | 1.7e-05 | 9.7e-06 | 1.133E-05 | 5.052E-06 | 7.300E-06 | 1.700E-05 |
| 20050 | 1003 | S1003I | C | 1.6e-05 | 1.3e-05 | 1.7e-05 | 1.533E-05 | 2.082E-06 | 1.300E-05 | 1.700E-05 |
| 20051 | 1003 | S1003L | C | 1.1e-05 | 1.6e-05 | 1.3e-05 | 1.333E-05 | 2.517E-06 | 1.100E-05 | 1.600E-05 |
| 20052 | 1003 | S1003K | C | 1.1e-05 | 1.6e-05 | 1.6e-05 | 1.433E-05 | 2.887E-06 | 1.100E-05 | 1.600E-05 |
| 20053 | 1003 | S1003M | C | 9.4e-06 | 1.5e-05 | 1.5e-05 | 1.313E-05 | 3.233E-06 | 9.400E-06 | 1.500E-05 |
| 20054 | 1003 | S1003F | C | 1.6e-05 | 1.6e-05 | 9e-06 | 1.367E-05 | 4.041E-06 | 9.000E-06 | 1.600E-05 |
| 20055 | 1003 | S1003P | C | 1e-05 | 1.1e-05 | 9e-06 | 1.000E-05 | 1.000E-06 | 9.000E-06 | 1.100E-05 |
| 20056 | 1003 | S1003S | C | 1.4e-05 | 9.5e-06 | 1.1e-05 | 1.150E-05 | 2.291E-06 | 9.500E-06 | 1.400E-05 |
| 20057 | 1003 | S1003T | C | 7.1e-06 | 1.6e-05 | 1.1e-05 | 1.137E-05 | 4.461E-06 | 7.100E-06 | 1.600E-05 |
| 20058 | 1003 | S1003W | C | 1e-05 | 9.2e-06 | 1.3e-05 | 1.073E-05 | 2.003E-06 | 9.200E-06 | 1.300E-05 |
| 20059 | 1003 | S1003Y | C | 1.6e-05 | 1.4e-05 | 1.1e-05 | 1.367E-05 | 2.517E-06 | 1.100E-05 | 1.600E-05 |

| | | | | | | | | | | |
|---|---|---|---|---|---|---|---|---|---|---|
| 20060 | 1003 | S1003V | C | 8.7e-06 | 1.6e-05 | 9.3e-06 | 1.133E-05 | 4.053E-06 | 8.700E-06 | 1.600E-05 |
| 20061 | 1004 | G1004A | C | 1e-05 | 1.5e-05 | 1.1e-05 | 1.200E-05 | 2.646E-06 | 1.000E-05 | 1.500E-05 |
| 20062 | 1004 | G1004R | C | 8.9e-06 | 1.5e-05 | 1.1e-05 | 1.163E-05 | 3.099E-06 | 8.900E-06 | 1.500E-05 |
| 20063 | 1004 | G1004N | C | 1.1e-05 | 1.4e-05 | 1.2e-05 | 1.233E-05 | 1.528E-06 | 1.100E-05 | 1.400E-05 |
| 20064 | 1004 | G1004D | C | 1.5e-05 | 1e-05 | 9.7e-06 | 1.157E-05 | 2.977E-06 | 9.700E-06 | 1.500E-05 |
| 20065 | 1004 | G1004C | C | 8.7e-06 | 1.4e-05 | 1e-05 | 1.090E-05 | 2.762E-06 | 8.700E-06 | 1.400E-05 |
| 20066 | 1004 | G1004E | C | 1e-05 | 9.5e-06 | 1e-05 | 9.833E-06 | 2.887E-07 | 9.500E-06 | 1.000E-05 |
| 20067 | 1004 | G1004Q | C | 1e-05 | 1.4e-05 | 1.5e-05 | 1.300E-05 | 2.646E-06 | 1.000E-05 | 1.500E-05 |
| 20068 | 1004 | G1004G | C | 1.2e-05 | 1e-05 | 1.4e-05 | 1.200E-05 | 2.000E-06 | 1.000E-05 | 1.400E-05 |
| 20069 | 1004 | G1004H | C | 1.7e-05 | 8.2e-06 | 8.5e-06 | 1.123E-05 | 4.996E-06 | 8.200E-06 | 1.700E-05 |
| 20070 | 1004 | G1004I | C | 9.9e-06 | 1.1e-05 | 9.2e-06 | 1.003E-05 | 9.074E-07 | 9.200E-06 | 1.100E-05 |
| 20071 | 1004 | G1004L | C | 1.3e-05 | 1e-05 | 1.1e-05 | 1.133E-05 | 1.528E-06 | 1.000E-05 | 1.300E-05 |
| 20072 | 1004 | G1004K | C | 1.3e-05 | 8.9e-06 | 1.6e-05 | 1.263E-05 | 3.564E-06 | 8.900E-06 | 1.600E-05 |
| 20073 | 1004 | G1004M | C | 1.5e-05 | 1.7e-05 | 9e-06 | 1.367E-05 | 4.163E-06 | 9.000E-06 | 1.700E-05 |
| 20074 | 1004 | G1004F | C | 1.3e-05 | 1.1e-05 | 1.5e-05 | 1.300E-05 | 2.000E-06 | 1.100E-05 | 1.500E-05 |
| 20075 | 1004 | G1004P | C | 1.6e-05 | 1.1e-05 | 1.1e-05 | 1.267E-05 | 2.887E-06 | 1.100E-05 | 1.600E-05 |
| 20076 | 1004 | G1004S | C | 8.6e-06 | 1.6e-05 | 8.8e-06 | 1.113E-05 | 4.216E-06 | 8.600E-06 | 1.600E-05 |
| 20077 | 1004 | G1004T | C | 8.8e-06 | 1.5e-05 | 1.5e-05 | 1.293E-05 | 3.580E-06 | 8.800E-06 | 1.500E-05 |
| 20078 | 1004 | G1004W | C | 1.1e-05 | 1.3e-05 | 1.5e-05 | 1.300E-05 | 2.000E-06 | 1.100E-05 | 1.500E-05 |
| 20079 | 1004 | G1004Y | C | 1.5e-05 | 1.5e-05 | 9.1e-06 | 1.303E-05 | 3.406E-06 | 9.100E-06 | 1.500E-05 |
| 20080 | 1004 | G1004V | C | 9.7e-06 | 9.5e-06 | 8.2e-06 | 9.133E-06 | 8.145E-07 | 8.200E-06 | 9.700E-06 |
| 20081 | 1005 | V1005A | C | 1.2e-05 | 1.5e-05 | 1.6e-05 | 1.433E-05 | 2.082E-06 | 1.200E-05 | 1.600E-05 |
| 20082 | 1005 | V1005R | C | 1.5e-05 | 1.4e-05 | 1.5e-05 | 1.467E-05 | 5.774E-07 | 1.400E-05 | 1.500E-05 |
| 20083 | 1005 | V1005N | C | 1.5e-05 | 1.6e-05 | 1.6e-05 | 1.567E-05 | 5.774E-07 | 1.500E-05 | 1.600E-05 |
| 20084 | 1005 | V1005D | C | 1.4e-05 | 1.6e-05 | 1.3e-05 | 1.433E-05 | 1.528E-06 | 1.300E-05 | 1.600E-05 |
| 20085 | 1005 | V1005C | C | 1.4e-05 | 8.9e-06 | 1.3e-05 | 1.197E-05 | 2.702E-06 | 8.900E-06 | 1.400E-05 |
| 20086 | 1005 | V1005E | C | 1.4e-05 | 1.4e-05 | 1e-05 | 1.267E-05 | 2.309E-06 | 1.000E-05 | 1.400E-05 |
| 20087 | 1005 | V1005Q | C | 1.5e-05 | 1.3e-05 | 9.8e-06 | 1.260E-05 | 2.623E-06 | 9.800E-06 | 1.500E-05 |
| 20088 | 1005 | V1005G | C | 1.5e-05 | 1.6e-05 | 1e-05 | 1.367E-05 | 3.215E-06 | 1.000E-05 | 1.600E-05 |
| 20089 | 1005 | V1005H | C | 9.6e-06 | 1.6e-05 | 7.8e-06 | 1.113E-05 | 4.310E-06 | 7.800E-06 | 1.600E-05 |
| 20090 | 1005 | V1005I | C | 1.1e-05 | 9.6e-06 | 1.8e-05 | 1.287E-05 | 4.500E-06 | 9.600E-06 | 1.800E-05 |
| 20091 | 1005 | V1005L | C | 1.5e-05 | 1.2e-05 | 1e-05 | 1.233E-05 | 2.517E-06 | 1.000E-05 | 1.500E-05 |
| 20092 | 1005 | V1005K | C | 8.1e-06 | 1.2e-05 | 1e-05 | 1.003E-05 | 1.950E-06 | 8.100E-06 | 1.200E-05 |
| 20093 | 1005 | V1005M | C | 1.1e-05 | 1.4e-05 | 1.1e-05 | 1.200E-05 | 1.732E-06 | 1.100E-05 | 1.400E-05 |
| 20094 | 1005 | V1005F | C | 1.1e-05 | 1.1e-05 | 1.2e-05 | 1.133E-05 | 5.774E-07 | 1.100E-05 | 1.200E-05 |
| 20095 | 1005 | V1005P | C | 1.6e-05 | 1.6e-05 | 1.6e-05 | 1.600E-05 | 0.000E+00 | 1.600E-05 | 1.600E-05 |
| 20096 | 1005 | V1005S | C | 1e-05 | 1e-05 | 9.6e-06 | 9.867E-06 | 2.309E-07 | 9.600E-06 | 1.000E-05 |
| 20097 | 1005 | V1005T | C | 1.2e-05 | 1.2e-05 | 1.2e-05 | 1.200E-05 | 0.000E+00 | 1.200E-05 | 1.200E-05 |
| 20098 | 1005 | V1005W | C | 1.6e-05 | 9.3e-06 | 9.9e-06 | 1.173E-05 | 3.707E-06 | 9.300E-06 | 1.600E-05 |
| 20099 | 1005 | V1005Y | C | 9e-06 | 7e-06 | 1e-05 | 8.667E-06 | 1.528E-06 | 7.000E-06 | 1.000E-05 |
| 20100 | 1005 | V1005V | C | 1.7e-05 | 1.5e-05 | 1.6e-05 | 1.600E-05 | 1.000E-06 | 1.500E-05 | 1.700E-05 |
| 20101 | 1006 | H1006A | C | 1.6e-05 | 1.7e-05 | 1.1e-05 | 1.467E-05 | 3.215E-06 | 1.100E-05 | 1.700E-05 |
| 20102 | 1006 | H1006R | C | 1.7e-05 | 1.3e-05 | 1.4e-05 | 1.467E-05 | 2.082E-06 | 1.300E-05 | 1.700E-05 |
| 20103 | 1006 | H1006N | C | 9.7e-06 | 1.5e-05 | 8.9e-06 | 1.120E-05 | 3.315E-06 | 8.900E-06 | 1.500E-05 |
| 20104 | 1006 | H1006D | C | 1.5e-05 | 1.6e-05 | 1.5e-05 | 1.533E-05 | 5.774E-07 | 1.500E-05 | 1.600E-05 |
| 20105 | 1006 | H1006C | C | 1e-05 | 9.9e-06 | 1.3e-05 | 1.097E-05 | 1.762E-06 | 9.900E-06 | 1.300E-05 |
| 20106 | 1006 | H1006E | C | 8.5e-06 | 1.6e-05 | 1e-05 | 1.150E-05 | 3.969E-06 | 8.500E-06 | 1.600E-05 |
| 20107 | 1006 | H1006Q | C | 1.5e-05 | 1e-05 | 1.5e-05 | 1.333E-05 | 2.887E-06 | 1.000E-05 | 1.500E-05 |
| 20108 | 1006 | H1006G | C | 1.3e-05 | 1.8e-05 | 1e-05 | 1.367E-05 | 4.041E-06 | 1.000E-05 | 1.800E-05 |
| 20109 | 1006 | H1006H | C | 1.6e-05 | 1.4e-05 | 1.3e-05 | 1.433E-05 | 1.528E-06 | 1.300E-05 | 1.600E-05 |
| 20110 | 1006 | H1006I | C | 1.6e-05 | 1.2e-05 | 9.3e-06 | 1.243E-05 | 3.371E-06 | 9.300E-06 | 1.600E-05 |
| 20111 | 1006 | H1006L | C | 1.6e-05 | 9.6e-06 | 1.1e-05 | 1.220E-05 | 3.365E-06 | 9.600E-06 | 1.600E-05 |
| 20112 | 1006 | H1006K | C | 7.7e-06 | 1.5e-05 | 1.3e-05 | 1.190E-05 | 3.772E-06 | 7.700E-06 | 1.500E-05 |
| 20113 | 1006 | H1006M | C | 1.1e-05 | 1.1e-05 | 1.5e-05 | 1.233E-05 | 2.309E-06 | 1.100E-05 | 1.500E-05 |
| 20114 | 1006 | H1006F | C | 1.5e-05 | 1.5e-05 | 1.5e-05 | 1.500E-05 | 0.000E+00 | 1.500E-05 | 1.500E-05 |
| 20115 | 1006 | H1006P | C | 1.6e-05 | 8.9e-06 | 1.4e-05 | 1.297E-05 | 3.661E-06 | 8.900E-06 | 1.600E-05 |
| 20116 | 1006 | H1006S | C | 1.4e-05 | 1e-05 | 9.6e-06 | 1.120E-05 | 2.433E-06 | 9.600E-06 | 1.400E-05 |
| 20117 | 1006 | H1006T | C | 1.4e-05 | 9.9e-06 | 8.5e-06 | 1.080E-05 | 2.858E-06 | 8.500E-06 | 1.400E-05 |
| 20118 | 1006 | H1006W | C | 1.6e-05 | 1.5e-05 | 1.3e-05 | 1.467E-05 | 1.528E-06 | 1.300E-05 | 1.600E-05 |
| 20119 | 1006 | H1006Y | C | 1.5e-05 | 1.3e-05 | 1.5e-05 | 1.433E-05 | 1.155E-06 | 1.300E-05 | 1.500E-05 |
| 20120 | 1006 | H1006V | C | 1.4e-05 | 1.1e-05 | 1.5e-05 | 1.333E-05 | 2.082E-06 | 1.100E-05 | 1.500E-05 |
| 20121 | 1007 | T1007A | C | 1.5e-05 | 7.5e-06 | 1.4e-05 | 1.217E-05 | 4.072E-06 | 7.500E-06 | 1.500E-05 |
| 20122 | 1007 | T1007R | C | 1.1e-05 | 1e-05 | 1.3e-05 | 1.133E-05 | 1.528E-06 | 1.000E-05 | 1.300E-05 |
| 20123 | 1007 | T1007N | C | 1.1e-05 | 1.4e-05 | 1.7e-05 | 1.400E-05 | 3.000E-06 | 1.100E-05 | 1.700E-05 |
| 20124 | 1007 | T1007D | C | 1.1e-05 | 1.4e-05 | 1.1e-05 | 1.200E-05 | 1.732E-06 | 1.100E-05 | 1.400E-05 |
| 20125 | 1007 | T1007C | C | 8.9e-06 | 1.5e-05 | 1e-05 | 1.130E-05 | 3.251E-06 | 8.900E-06 | 1.500E-05 |
| 20126 | 1007 | T1007E | C | 1.1e-05 | 1.7e-05 | 9.8e-06 | 1.260E-05 | 3.857E-06 | 9.800E-06 | 1.700E-05 |
| 20127 | 1007 | T1007Q | C | 1.4e-05 | 1.1e-05 | 1e-05 | 1.167E-05 | 2.082E-06 | 1.000E-05 | 1.400E-05 |
| 20128 | 1007 | T1007G | C | 1.3e-05 | 1.5e-05 | 1.7e-05 | 1.500E-05 | 2.000E-06 | 1.300E-05 | 1.700E-05 |
| 20129 | 1007 | T1007H | C | 1.6e-05 | 9.9e-06 | 6.6e-06 | 1.083E-05 | 4.769E-06 | 6.600E-06 | 1.600E-05 |
| 20130 | 1007 | T1007I | C | 1.4e-05 | 8.9e-06 | 2e-05 | 1.430E-05 | 5.556E-06 | 8.900E-06 | 2.000E-05 |
| 20131 | 1007 | T1007L | C | 1.1e-05 | 1.1e-05 | 1e-05 | 1.067E-05 | 5.774E-07 | 1.000E-05 | 1.100E-05 |
| 20132 | 1007 | T1007K | C | 1.2e-05 | 1.5e-05 | 1.1e-05 | 1.267E-05 | 2.082E-06 | 1.100E-05 | 1.500E-05 |
| 20133 | 1007 | T1007M | C | 7.4e-06 | 9.8e-06 | 9.4e-06 | 8.867E-06 | 1.286E-06 | 7.400E-06 | 9.800E-06 |
| 20134 | 1007 | T1007F | C | 1.3e-05 | 1.6e-05 | 1.5e-05 | 1.467E-05 | 1.528E-06 | 1.300E-05 | 1.600E-05 |
| 20135 | 1007 | T1007P | C | 1.1e-05 | 1.7e-05 | 1.1e-05 | 1.300E-05 | 3.464E-06 | 1.100E-05 | 1.700E-05 |
| 20136 | 1007 | T1007S | C | 1.3e-05 | 9.1e-06 | 1e-05 | 1.070E-05 | 2.042E-06 | 9.100E-06 | 1.300E-05 |
| 20137 | 1007 | T1007T | C | 1.6e-05 | 1.5e-05 | 1.4e-05 | 1.500E-05 | 1.000E-06 | 1.400E-05 | 1.600E-05 |
| 20138 | 1007 | T1007W | C | 1.7e-05 | 1.1e-05 | 1e-05 | 1.267E-05 | 3.786E-06 | 1.000E-05 | 1.700E-05 |
| 20139 | 1007 | T1007Y | C | 1.5e-05 | 1.5e-05 | 1.1e-05 | 1.367E-05 | 2.309E-06 | 1.100E-05 | 1.500E-05 |
| 20140 | 1007 | T1007V | C | 1.1e-05 | 1.6e-05 | 1.1e-05 | 1.267E-05 | 2.887E-06 | 1.100E-05 | 1.600E-05 |
| 20141 | 1008 | F1008A | C | 1e-05 | 9.5e-06 | 1.7e-05 | 1.217E-05 | 4.193E-06 | 9.500E-06 | 1.700E-05 |
| 20142 | 1008 | F1008R | C | 1.5e-05 | 1.6e-05 | 1.5e-05 | 1.533E-05 | 5.774E-07 | 1.500E-05 | 1.600E-05 |
| 20143 | 1008 | F1008N | C | 1.3e-05 | 1.1e-05 | 1.3e-05 | 1.233E-05 | 1.155E-06 | 1.100E-05 | 1.300E-05 |
| 20144 | 1008 | F1008D | C | 1.8e-05 | 1.5e-05 | 1e-05 | 1.433E-05 | 4.041E-06 | 1.000E-05 | 1.800E-05 |

| | | | | | | | | | | |
|---|---|---|---|---|---|---|---|---|---|---|
| 20145 | 1008 | F1008C | C | 1.3e-05 | 1.6e-05 | 9.9e-06 | 1.297E-05 | 3.050E-06 | 9.900E-06 | 1.600E-05 |
| 20146 | 1008 | F1008E | C | 1.1e-05 | 9e-06 | 1.1e-05 | 1.033E-05 | 1.155E-06 | 9.000E-06 | 1.100E-05 |
| 20147 | 1008 | F1008Q | C | 1e-05 | 1.1e-05 | 1.1e-05 | 1.067E-05 | 5.774E-07 | 1.000E-05 | 1.100E-05 |
| 20148 | 1008 | F1008G | C | 8.8e-06 | 2.7e-05 | 1.5e-05 | 1.693E-05 | 9.253E-06 | 8.800E-06 | 2.700E-05 |
| 20149 | 1008 | F1008H | C | 1.5e-05 | 1.5e-05 | 9.2e-06 | 1.307E-05 | 3.349E-06 | 9.200E-06 | 1.500E-05 |
| 20150 | 1008 | F1008I | C | 1.6e-05 | 1e-05 | 1.5e-05 | 1.367E-05 | 3.215E-06 | 1.000E-05 | 1.600E-05 |
| 20151 | 1008 | F1008L | C | 8.6e-06 | 1e-05 | 9.8e-06 | 9.467E-06 | 7.572E-07 | 8.600E-06 | 1.000E-05 |
| 20152 | 1008 | F1008K | C | 1.8e-05 | 8.7e-06 | 1.5e-05 | 1.390E-05 | 4.747E-06 | 8.700E-06 | 1.800E-05 |
| 20153 | 1008 | F1008M | C | 1.1e-05 | 1.6e-05 | 1.1e-05 | 1.267E-05 | 2.887E-06 | 1.100E-05 | 1.600E-05 |
| 20154 | 1008 | F1008F | C | 1e-05 | 6e-06 | 1.6e-05 | 1.067E-05 | 5.033E-06 | 6.000E-06 | 1.600E-05 |
| 20155 | 1008 | F1008P | C | 1.3e-05 | 1e-05 | 1.6e-05 | 1.300E-05 | 3.000E-06 | 1.000E-05 | 1.600E-05 |
| 20156 | 1008 | F1008S | C | 1.5e-05 | 1.3e-05 | 1.4e-05 | 1.400E-05 | 1.000E-06 | 1.300E-05 | 1.500E-05 |
| 20157 | 1008 | F1008T | C | 1.7e-05 | 1.6e-05 | 9.4e-06 | 1.413E-05 | 4.130E-06 | 9.400E-06 | 1.700E-05 |
| 20158 | 1008 | F1008W | C | 1.3e-05 | 1.4e-05 | 9.7e-06 | 1.223E-05 | 2.250E-06 | 9.700E-06 | 1.400E-05 |
| 20159 | 1008 | F1008Y | C | 9.4e-06 | 1.5e-05 | 1.3e-05 | 1.247E-05 | 2.838E-06 | 9.400E-06 | 1.500E-05 |
| 20160 | 1008 | F1008V | C | 1.8e-05 | 1.5e-05 | 8.6e-06 | 1.387E-05 | 4.801E-06 | 8.600E-06 | 1.800E-05 |
| 20161 | 1009 | P1009A | C | 1e-05 | 1.4e-05 | 1.1e-05 | 1.167E-05 | 2.082E-06 | 1.000E-05 | 1.400E-05 |
| 20162 | 1009 | P1009R | C | 1.5e-05 | 1.5e-05 | 1.4e-05 | 1.467E-05 | 5.774E-07 | 1.400E-05 | 1.500E-05 |
| 20163 | 1009 | P1009N | C | 1.3e-05 | 9.3e-06 | 1.6e-05 | 1.277E-05 | 3.356E-06 | 9.300E-06 | 1.600E-05 |
| 20164 | 1009 | P1009D | C | 1e-05 | 1e-05 | 1.4e-05 | 1.133E-05 | 2.309E-06 | 1.000E-05 | 1.400E-05 |
| 20165 | 1009 | P1009C | C | 1.6e-05 | 1e-05 | 9.3e-06 | 1.177E-05 | 3.683E-06 | 9.300E-06 | 1.600E-05 |
| 20166 | 1009 | P1009E | C | 1.1e-05 | 1.5e-05 | 1.4e-05 | 1.333E-05 | 2.082E-06 | 1.100E-05 | 1.500E-05 |
| 20167 | 1009 | P1009Q | C | 1.5e-05 | 1.4e-05 | 1.1e-05 | 1.333E-05 | 2.082E-06 | 1.100E-05 | 1.500E-05 |
| 20168 | 1009 | P1009G | C | 1.5e-05 | 9.3e-06 | 9.3e-06 | 1.120E-05 | 3.291E-06 | 9.300E-06 | 1.500E-05 |
| 20169 | 1009 | P1009H | C | 1e-05 | 1.4e-05 | 9.3e-06 | 1.110E-05 | 2.536E-06 | 9.300E-06 | 1.400E-05 |
| 20170 | 1009 | P1009I | C | 1.4e-05 | 1.5e-05 | 1.1e-05 | 1.333E-05 | 2.082E-06 | 1.100E-05 | 1.500E-05 |
| 20171 | 1009 | P1009L | C | 1e-05 | 1.5e-05 | 1.1e-05 | 1.200E-05 | 2.646E-06 | 1.000E-05 | 1.500E-05 |
| 20172 | 1009 | P1009K | C | 1.5e-05 | 1.1e-05 | 1.4e-05 | 1.333E-05 | 2.082E-06 | 1.100E-05 | 1.500E-05 |
| 20173 | 1009 | P1009M | C | 9.2e-06 | 9.5e-06 | 1.6e-05 | 1.157E-05 | 3.842E-06 | 9.200E-06 | 1.600E-05 |
| 20174 | 1009 | P1009F | C | 1.5e-05 | 1.6e-05 | 1.3e-05 | 1.467E-05 | 1.528E-06 | 1.300E-05 | 1.600E-05 |
| 20175 | 1009 | P1009P | C | 1.5e-05 | 9.8e-06 | 1.3e-05 | 1.260E-05 | 2.623E-06 | 9.800E-06 | 1.500E-05 |
| 20176 | 1009 | P1009S | C | 1.6e-05 | 1.5e-05 | 1.1e-05 | 1.400E-05 | 2.646E-06 | 1.100E-05 | 1.600E-05 |
| 20177 | 1009 | P1009T | C | 1.7e-05 | 1.1e-05 | 1.2e-05 | 1.333E-05 | 3.215E-06 | 1.100E-05 | 1.700E-05 |
| 20178 | 1009 | P1009W | C | 1.4e-05 | 1.6e-05 | 1.4e-05 | 1.467E-05 | 1.155E-06 | 1.400E-05 | 1.600E-05 |
| 20179 | 1009 | P1009Y | C | 1e-05 | 1.6e-05 | 1.3e-05 | 1.300E-05 | 3.000E-06 | 1.000E-05 | 1.600E-05 |
| 20180 | 1009 | P1009V | C | 1e-05 | 1.7e-05 | 9e-06 | 1.200E-05 | 4.359E-06 | 9.000E-06 | 1.700E-05 |
| 20181 | 1010 | A1010A | C | 1.2e-05 | 6.2e-06 | 1.7e-05 | 1.173E-05 | 5.405E-06 | 6.200E-06 | 1.700E-05 |
| 20182 | 1010 | A1010R | C | 1.5e-05 | 1.2e-05 | 1.2e-05 | 1.300E-05 | 1.732E-06 | 1.200E-05 | 1.500E-05 |
| 20183 | 1010 | A1010N | C | 1.4e-05 | 9.6e-06 | 1.3e-05 | 1.220E-05 | 2.307E-06 | 9.600E-06 | 1.400E-05 |
| 20184 | 1010 | A1010D | C | 1.1e-05 | 1.7e-05 | 9.9e-06 | 1.263E-05 | 3.821E-06 | 9.900E-06 | 1.700E-05 |
| 20185 | 1010 | A1010C | C | 1.6e-05 | 1.4e-05 | 1.5e-05 | 1.500E-05 | 1.000E-06 | 1.400E-05 | 1.600E-05 |
| 20186 | 1010 | A1010E | C | 1.6e-05 | 1.2e-05 | 1.1e-05 | 1.300E-05 | 2.646E-06 | 1.100E-05 | 1.600E-05 |
| 20187 | 1010 | A1010Q | C | 1.5e-05 | 1.2e-05 | 9.9e-06 | 1.230E-05 | 2.563E-06 | 9.900E-06 | 1.500E-05 |
| 20188 | 1010 | A1010G | C | 1e-05 | 1.6e-05 | 1.5e-05 | 1.367E-05 | 3.215E-06 | 1.000E-05 | 1.600E-05 |
| 20189 | 1010 | A1010H | C | 1.3e-05 | 1.3e-05 | 1.6e-05 | 1.400E-05 | 1.732E-06 | 1.300E-05 | 1.600E-05 |
| 20190 | 1010 | A1010I | C | 1.5e-05 | 1.3e-05 | 1.3e-05 | 1.367E-05 | 1.155E-06 | 1.300E-05 | 1.500E-05 |
| 20191 | 1010 | A1010L | C | 1.1e-05 | 8.8e-06 | 1.3e-05 | 1.093E-05 | 2.101E-06 | 8.800E-06 | 1.300E-05 |
| 20192 | 1010 | A1010K | C | 2.5e-05 | 1.7e-05 | 1.4e-05 | 1.867E-05 | 5.686E-06 | 1.400E-05 | 2.500E-05 |
| 20193 | 1010 | A1010M | C | 1.3e-05 | 1.3e-05 | 1.4e-05 | 1.333E-05 | 5.774E-07 | 1.300E-05 | 1.400E-05 |
| 20194 | 1010 | A1010F | C | 1.1e-05 | 1.4e-05 | 1.2e-05 | 1.233E-05 | 1.528E-06 | 1.100E-05 | 1.400E-05 |
| 20195 | 1010 | A1010P | C | 1.4e-05 | 1.1e-05 | 1e-05 | 1.167E-05 | 2.082E-06 | 1.000E-05 | 1.400E-05 |
| 20196 | 1010 | A1010S | C | 9.6e-06 | 1.5e-05 | 1.5e-05 | 1.320E-05 | 3.118E-06 | 9.600E-06 | 1.500E-05 |
| 20197 | 1010 | A1010T | C | 1.6e-05 | 1.1e-05 | 1.1e-05 | 1.267E-05 | 2.887E-06 | 1.100E-05 | 1.600E-05 |
| 20198 | 1010 | A1010W | C | 1.3e-05 | 1.5e-05 | 8.8e-06 | 1.227E-05 | 3.164E-06 | 8.800E-06 | 1.500E-05 |
| 20199 | 1010 | A1010Y | C | 1.5e-05 | 7.9e-06 | 1.5e-05 | 1.263E-05 | 4.099E-06 | 7.900E-06 | 1.500E-05 |
| 20200 | 1010 | A1010V | C | 1.4e-05 | 1.4e-05 | 1.5e-05 | 1.433E-05 | 5.774E-07 | 1.400E-05 | 1.500E-05 |
| 20201 | 1011 | V1011A | C | 9.9e-06 | 1.5e-05 | 1.4e-05 | 1.297E-05 | 2.702E-06 | 9.900E-06 | 1.500E-05 |
| 20202 | 1011 | V1011R | C | 1.4e-05 | 1.6e-05 | 1.3e-05 | 1.433E-05 | 1.528E-06 | 1.300E-05 | 1.600E-05 |
| 20203 | 1011 | V1011N | C | 1.7e-05 | 8.6e-06 | 1.9e-05 | 1.487E-05 | 5.518E-06 | 8.600E-06 | 1.900E-05 |
| 20204 | 1011 | V1011D | C | 1.3e-05 | 1.4e-05 | 1.5e-05 | 1.400E-05 | 1.000E-06 | 1.300E-05 | 1.500E-05 |
| 20205 | 1011 | V1011C | C | 1.4e-05 | 1.2e-05 | 7e-06 | 1.100E-05 | 3.606E-06 | 7.000E-06 | 1.400E-05 |
| 20206 | 1011 | V1011E | C | 1.5e-05 | 1.3e-05 | 1.5e-05 | 1.433E-05 | 1.155E-06 | 1.300E-05 | 1.500E-05 |
| 20207 | 1011 | V1011Q | C | 1.2e-05 | 1.1e-05 | 1.5e-05 | 1.267E-05 | 2.082E-06 | 1.100E-05 | 1.500E-05 |
| 20208 | 1011 | V1011G | C | 1.6e-05 | 9.7e-06 | 1.1e-05 | 1.223E-05 | 3.326E-06 | 9.700E-06 | 1.600E-05 |
| 20209 | 1011 | V1011H | C | 1.3e-05 | 1e-05 | 9.5e-06 | 1.083E-05 | 1.893E-06 | 9.500E-06 | 1.300E-05 |
| 20210 | 1011 | V1011I | C | 1.4e-05 | 1.5e-05 | 1e-05 | 1.300E-05 | 2.646E-06 | 1.000E-05 | 1.500E-05 |
| 20211 | 1011 | V1011L | C | 7.6e-06 | 1.5e-05 | 9.2e-06 | 1.060E-05 | 3.894E-06 | 7.600E-06 | 1.500E-05 |
| 20212 | 1011 | V1011K | C | 1.3e-05 | 9.7e-06 | 1.4e-05 | 1.223E-05 | 2.250E-06 | 9.700E-06 | 1.400E-05 |
| 20213 | 1011 | V1011M | C | 1e-05 | 9.4e-06 | 1e-05 | 9.800E-06 | 3.464E-07 | 9.400E-06 | 1.000E-05 |
| 20214 | 1011 | V1011F | C | 1.6e-05 | 1.1e-05 | 1.6e-05 | 1.433E-05 | 2.887E-06 | 1.100E-05 | 1.600E-05 |
| 20215 | 1011 | V1011P | C | 1.1e-05 | 1.5e-05 | 1.4e-05 | 1.333E-05 | 2.082E-06 | 1.100E-05 | 1.500E-05 |
| 20216 | 1011 | V1011S | C | 9.5e-06 | 1.1e-05 | 1e-05 | 1.017E-05 | 7.638E-07 | 9.500E-06 | 1.100E-05 |
| 20217 | 1011 | V1011T | C | 9.3e-06 | 1.4e-05 | 1.4e-05 | 1.243E-05 | 2.714E-06 | 9.300E-06 | 1.400E-05 |
| 20218 | 1011 | V1011W | C | 1.1e-05 | 1.7e-05 | 1.3e-05 | 1.367E-05 | 3.055E-06 | 1.100E-05 | 1.700E-05 |
| 20219 | 1011 | V1011Y | C | 1.6e-05 | 1.1e-05 | 1.1e-05 | 1.267E-05 | 2.887E-06 | 1.100E-05 | 1.600E-05 |
| 20220 | 1011 | V1011V | C | 1.1e-05 | 1.5e-05 | 1.3e-05 | 1.300E-05 | 2.000E-06 | 1.100E-05 | 1.500E-05 |
| 20221 | 1012 | L1012A | C | 1e-05 | 1.4e-05 | 1.5e-05 | 1.300E-05 | 2.646E-06 | 1.000E-05 | 1.500E-05 |
| 20222 | 1012 | L1012R | C | 1.1e-05 | 1.5e-05 | 1.1e-05 | 1.233E-05 | 2.309E-06 | 1.100E-05 | 1.500E-05 |
| 20223 | 1012 | L1012N | C | 1.1e-05 | 8.6e-06 | 1.6e-05 | 1.187E-05 | 3.775E-06 | 8.600E-06 | 1.600E-05 |
| 20224 | 1012 | L1012D | C | 1.6e-05 | 1e-05 | 1e-05 | 1.200E-05 | 3.464E-06 | 1.000E-05 | 1.600E-05 |
| 20225 | 1012 | L1012C | C | 1.5e-05 | 1.5e-05 | 8.8e-06 | 1.293E-05 | 3.580E-06 | 8.800E-06 | 1.500E-05 |
| 20226 | 1012 | L1012E | C | 1.6e-05 | 1.5e-05 | 1.9e-05 | 1.667E-05 | 2.082E-06 | 1.500E-05 | 1.900E-05 |
| 20227 | 1012 | L1012Q | C | 1e-05 | 1.5e-05 | 1e-05 | 1.167E-05 | 2.887E-06 | 1.000E-05 | 1.500E-05 |
| 20228 | 1012 | L1012G | C | 1.4e-05 | 1.4e-05 | 1e-05 | 1.267E-05 | 2.309E-06 | 1.000E-05 | 1.400E-05 |
| 20229 | 1012 | L1012H | C | 1.6e-05 | 1.1e-05 | 1.4e-05 | 1.367E-05 | 2.517E-06 | 1.100E-05 | 1.600E-05 |

| | | | | | | | | | | |
|---|---|---|---|---|---|---|---|---|---|---|
| 20230 | 1012 | L1012I | C | 1.5e-05 | 1.3e-05 | 1e-05 | 1.267E-05 | 2.517E-06 | 1.000E-05 | 1.500E-05 |
| 20231 | 1012 | L1012L | C | 1.5e-05 | 1.5e-05 | 1.1e-05 | 1.367E-05 | 2.309E-06 | 1.100E-05 | 1.500E-05 |
| 20232 | 1012 | L1012K | C | 1.1e-05 | 1.7e-05 | 1.2e-05 | 1.333E-05 | 3.215E-06 | 1.100E-05 | 1.700E-05 |
| 20233 | 1012 | L1012M | C | 1e-05 | 1.5e-05 | 1.5e-05 | 1.333E-05 | 2.887E-06 | 1.000E-05 | 1.500E-05 |
| 20234 | 1012 | L1012F | C | 1.1e-05 | 5.2e-06 | 1.3e-05 | 9.733E-06 | 4.051E-06 | 5.200E-06 | 1.300E-05 |
| 20235 | 1012 | L1012P | C | 1e-05 | 1.4e-05 | 1.4e-05 | 1.267E-05 | 2.309E-06 | 1.000E-05 | 1.400E-05 |
| 20236 | 1012 | L1012S | C | 9.4e-06 | 1e-05 | 1.3e-05 | 1.080E-05 | 1.929E-06 | 9.400E-06 | 1.300E-05 |
| 20237 | 1012 | L1012T | C | 1.2e-05 | 9.6e-06 | 1.5e-05 | 1.220E-05 | 2.706E-06 | 9.600E-06 | 1.500E-05 |
| 20238 | 1012 | L1012W | C | 8.1e-06 | 1.3e-05 | 8.4e-06 | 9.833E-06 | 2.747E-06 | 8.100E-06 | 1.300E-05 |
| 20239 | 1012 | L1012Y | C | 8.9e-06 | 1.6e-05 | 1.5e-05 | 1.330E-05 | 3.843E-06 | 8.900E-06 | 1.600E-05 |
| 20240 | 1012 | L1012V | C | 9.1e-06 | 1e-05 | 1e-05 | 9.700E-06 | 5.196E-07 | 9.100E-06 | 1.000E-05 |
| 20241 | 1013 | Q1013A | C | 8.7e-06 | 7.2e-06 | 1.1e-05 | 8.967E-06 | 1.914E-06 | 7.200E-06 | 1.100E-05 |
| 20242 | 1013 | Q1013R | C | 1.4e-05 | 1.4e-05 | 1.3e-05 | 1.367E-05 | 5.774E-07 | 1.300E-05 | 1.400E-05 |
| 20243 | 1013 | Q1013N | C | 1.6e-05 | 1.4e-05 | 1.1e-05 | 1.367E-05 | 2.517E-06 | 1.100E-05 | 1.600E-05 |
| 20244 | 1013 | Q1013D | C | 1.4e-05 | 1e-05 | 1.1e-05 | 1.167E-05 | 2.082E-06 | 1.000E-05 | 1.400E-05 |
| 20245 | 1013 | Q1013C | C | 9e-06 | 1.4e-05 | 1.5e-05 | 1.267E-05 | 3.215E-06 | 9.000E-06 | 1.500E-05 |
| 20246 | 1013 | Q1013E | C | 1.4e-05 | 1.5e-05 | 1.5e-05 | 1.467E-05 | 5.774E-07 | 1.400E-05 | 1.500E-05 |
| 20247 | 1013 | Q1013Q | C | 1.3e-05 | 1.1e-05 | 1.4e-05 | 1.267E-05 | 1.528E-06 | 1.100E-05 | 1.400E-05 |
| 20248 | 1013 | Q1013G | C | 1.5e-05 | 1.4e-05 | 1e-05 | 1.300E-05 | 2.646E-06 | 1.000E-05 | 1.500E-05 |
| 20249 | 1013 | Q1013H | C | 9.6e-06 | 1.3e-05 | 8.4e-06 | 1.033E-05 | 2.386E-06 | 8.400E-06 | 1.300E-05 |
| 20250 | 1013 | Q1013I | C | 1.3e-05 | 1.5e-05 | 1.2e-05 | 1.333E-05 | 1.528E-06 | 1.200E-05 | 1.500E-05 |
| 20251 | 1013 | Q1013L | C | 1e-05 | 1.7e-05 | 1.4e-05 | 1.367E-05 | 3.512E-06 | 1.000E-05 | 1.700E-05 |
| 20252 | 1013 | Q1013K | C | 1.3e-05 | 1.6e-05 | 1e-05 | 1.300E-05 | 3.000E-06 | 1.000E-05 | 1.600E-05 |
| 20253 | 1013 | Q1013M | C | 9.7e-06 | 1.5e-05 | 8.3e-06 | 1.100E-05 | 3.534E-06 | 8.300E-06 | 1.500E-05 |
| 20254 | 1013 | Q1013F | C | 1.3e-05 | 8.6e-06 | 9.4e-06 | 1.033E-05 | 2.344E-06 | 8.600E-06 | 1.300E-05 |
| 20255 | 1013 | Q1013P | C | 8.2e-06 | 1e-05 | 9.4e-06 | 9.200E-06 | 9.165E-07 | 8.200E-06 | 1.000E-05 |
| 20256 | 1013 | Q1013S | C | 1.4e-05 | 1.5e-05 | 9.1e-06 | 1.270E-05 | 3.158E-06 | 9.100E-06 | 1.500E-05 |
| 20257 | 1013 | Q1013T | C | 9.7e-06 | 1.1e-05 | 1.5e-05 | 1.190E-05 | 2.762E-06 | 9.700E-06 | 1.500E-05 |
| 20258 | 1013 | Q1013W | C | 1.4e-05 | 1.1e-05 | 1.2e-05 | 1.233E-05 | 1.528E-06 | 1.100E-05 | 1.400E-05 |
| 20259 | 1013 | Q1013Y | C | 1.1e-05 | 1.5e-05 | 9.5e-06 | 1.183E-05 | 2.843E-06 | 9.500E-06 | 1.500E-05 |
| 20260 | 1013 | Q1013V | C | 9.2e-06 | 1.5e-05 | 9.1e-06 | 1.110E-05 | 3.378E-06 | 9.100E-06 | 1.500E-05 |
| 20261 | 1014 | S1014A | C | 1.4e-05 | 1.1e-05 | 1.5e-05 | 1.333E-05 | 2.082E-06 | 1.100E-05 | 1.500E-05 |
| 20262 | 1014 | S1014R | C | 1.5e-05 | 1.1e-05 | 1.1e-05 | 1.233E-05 | 2.309E-06 | 1.100E-05 | 1.500E-05 |
| 20263 | 1014 | S1014N | C | 9.1e-06 | 1.3e-05 | 1.1e-05 | 1.103E-05 | 1.950E-06 | 9.100E-06 | 1.300E-05 |
| 20264 | 1014 | S1014D | C | 1e-05 | 1.6e-05 | 1.6e-05 | 1.400E-05 | 3.464E-06 | 1.000E-05 | 1.600E-05 |
| 20265 | 1014 | S1014C | C | 1.7e-05 | 1.5e-05 | 9.4e-06 | 1.380E-05 | 3.940E-06 | 9.400E-06 | 1.700E-05 |
| 20266 | 1014 | S1014E | C | 1.6e-05 | 1.1e-05 | 1.2e-05 | 1.300E-05 | 2.646E-06 | 1.100E-05 | 1.600E-05 |
| 20267 | 1014 | S1014Q | C | 1.3e-05 | 1.6e-05 | 1.5e-05 | 1.467E-05 | 1.528E-06 | 1.300E-05 | 1.600E-05 |
| 20268 | 1014 | S1014G | C | 1.1e-05 | 1.3e-05 | 1.3e-05 | 1.233E-05 | 1.155E-06 | 1.100E-05 | 1.300E-05 |
| 20269 | 1014 | S1014H | C | 1.5e-05 | 1e-05 | 1.6e-05 | 1.367E-05 | 3.215E-06 | 1.000E-05 | 1.600E-05 |
| 20270 | 1014 | S1014I | C | 1.2e-05 | 1.1e-05 | 1.2e-05 | 1.167E-05 | 5.774E-07 | 1.100E-05 | 1.200E-05 |
| 20271 | 1014 | S1014L | C | 1.4e-05 | 8.4e-06 | 1.2e-05 | 1.147E-05 | 2.838E-06 | 8.400E-06 | 1.400E-05 |
| 20272 | 1014 | S1014K | C | 1.5e-05 | 1.2e-05 | 1.5e-05 | 1.400E-05 | 1.732E-06 | 1.200E-05 | 1.500E-05 |
| 20273 | 1014 | S1014M | C | 8.9e-06 | 1.5e-05 | 1.7e-05 | 1.363E-05 | 4.219E-06 | 8.900E-06 | 1.700E-05 |
| 20274 | 1014 | S1014F | C | 1.7e-05 | 1.9e-05 | 1e-05 | 1.533E-05 | 4.726E-06 | 1.000E-05 | 1.900E-05 |
| 20275 | 1014 | S1014P | C | 1.3e-05 | 1.1e-05 | 7.4e-06 | 1.047E-05 | 2.838E-06 | 7.400E-06 | 1.300E-05 |
| 20276 | 1014 | S1014S | C | 1.5e-05 | 1.4e-05 | 1.2e-05 | 1.367E-05 | 1.528E-06 | 1.200E-05 | 1.500E-05 |
| 20277 | 1014 | S1014T | C | 1e-05 | 1.6e-05 | 1.6e-05 | 1.400E-05 | 3.464E-06 | 1.000E-05 | 1.600E-05 |
| 20278 | 1014 | S1014W | C | 9.1e-06 | 1.2e-05 | 9.6e-06 | 1.023E-05 | 1.550E-06 | 9.100E-06 | 1.200E-05 |
| 20279 | 1014 | S1014Y | C | 1.2e-05 | 1.4e-05 | 1.5e-05 | 1.367E-05 | 1.528E-06 | 1.200E-05 | 1.500E-05 |
| 20280 | 1014 | S1014V | C | 1.1e-05 | 1.5e-05 | 1.4e-05 | 1.333E-05 | 2.082E-06 | 1.100E-05 | 1.500E-05 |
| 20281 | 1015 | S1015A | C | 1.4e-05 | 1.2e-05 | 1.5e-05 | 1.367E-05 | 1.528E-06 | 1.200E-05 | 1.500E-05 |
| 20282 | 1015 | S1015R | C | 9.2e-06 | 1.1e-05 | 1.2e-05 | 1.073E-05 | 1.419E-06 | 9.200E-06 | 1.200E-05 |
| 20283 | 1015 | S1015N | C | 1.5e-05 | 1.1e-05 | 1.4e-05 | 1.333E-05 | 2.082E-06 | 1.100E-05 | 1.500E-05 |
| 20284 | 1015 | S1015D | C | 8.7e-06 | 1e-05 | 9.6e-06 | 9.433E-06 | 6.658E-07 | 8.700E-06 | 1.000E-05 |
| 20285 | 1015 | S1015C | C | 1.6e-05 | 1.4e-05 | 1.6e-05 | 1.533E-05 | 1.155E-06 | 1.400E-05 | 1.600E-05 |
| 20286 | 1015 | S1015E | C | 1.3e-05 | 1.4e-05 | 1.4e-05 | 1.367E-05 | 5.774E-07 | 1.300E-05 | 1.400E-05 |
| 20287 | 1015 | S1015Q | C | 1e-05 | 8.9e-06 | 1.6e-05 | 1.163E-05 | 3.821E-06 | 8.900E-06 | 1.600E-05 |
| 20288 | 1015 | S1015G | C | 1.5e-05 | 1.6e-05 | 1.5e-05 | 1.533E-05 | 5.774E-07 | 1.500E-05 | 1.600E-05 |
| 20289 | 1015 | S1015H | C | 1.3e-05 | 1.6e-05 | 1.8e-05 | 1.567E-05 | 2.517E-06 | 1.300E-05 | 1.800E-05 |
| 20290 | 1015 | S1015I | C | 1.4e-05 | 1e-05 | 1.8e-05 | 1.400E-05 | 4.000E-06 | 1.000E-05 | 1.800E-05 |
| 20291 | 1015 | S1015L | C | 1.4e-05 | 1e-05 | 1.2e-05 | 1.200E-05 | 2.000E-06 | 1.000E-05 | 1.400E-05 |
| 20292 | 1015 | S1015K | C | 1.4e-05 | 1.1e-05 | 1.6e-05 | 1.367E-05 | 2.517E-06 | 1.100E-05 | 1.600E-05 |
| 20293 | 1015 | S1015M | C | 1.5e-05 | 1.5e-05 | 1e-05 | 1.333E-05 | 2.887E-06 | 1.000E-05 | 1.500E-05 |
| 20294 | 1015 | S1015F | C | 1.5e-05 | 1.6e-05 | 1.5e-05 | 1.533E-05 | 5.774E-07 | 1.500E-05 | 1.600E-05 |
| 20295 | 1015 | S1015P | C | 1.4e-05 | 1.5e-05 | 1.3e-05 | 1.400E-05 | 1.000E-06 | 1.300E-05 | 1.500E-05 |
| 20296 | 1015 | S1015S | C | 1e-05 | 1.5e-05 | 1.5e-05 | 1.333E-05 | 2.887E-06 | 1.000E-05 | 1.500E-05 |
| 20297 | 1015 | S1015T | C | 1e-05 | 1e-05 | 9.6e-06 | 9.867E-06 | 2.309E-07 | 9.600E-06 | 1.000E-05 |
| 20298 | 1015 | S1015W | C | 1.6e-05 | 1.4e-05 | 2.2e-05 | 1.733E-05 | 4.163E-06 | 1.400E-05 | 2.200E-05 |
| 20299 | 1015 | S1015Y | C | 1.1e-05 | 1.5e-05 | 9.6e-06 | 1.187E-05 | 2.802E-06 | 9.600E-06 | 1.500E-05 |
| 20300 | 1015 | S1015V | C | 1.1e-05 | 1.7e-05 | 1.1e-05 | 1.300E-05 | 3.464E-06 | 1.100E-05 | 1.700E-05 |
| 20301 | 1016 | G1016A | C | 1.5e-05 | 1.5e-05 | 2.5e-05 | 1.833E-05 | 5.774E-06 | 1.500E-05 | 2.500E-05 |
| 20302 | 1016 | G1016R | C | 1.5e-05 | 1.5e-05 | 1.5e-05 | 1.500E-05 | 0.000E+00 | 1.500E-05 | 1.500E-05 |
| 20303 | 1016 | G1016N | C | 1.1e-05 | 1e-05 | 1.1e-05 | 1.067E-05 | 5.774E-07 | 1.000E-05 | 1.100E-05 |
| 20304 | 1016 | G1016D | C | 1.1e-05 | 1e-05 | 1.7e-05 | 1.267E-05 | 3.786E-06 | 1.000E-05 | 1.700E-05 |
| 20305 | 1016 | G1016C | C | 1.4e-05 | 1.1e-05 | 1e-05 | 1.167E-05 | 2.082E-06 | 1.000E-05 | 1.400E-05 |
| 20306 | 1016 | G1016E | C | 9.9e-06 | 1.5e-05 | 1.7e-05 | 1.397E-05 | 3.661E-06 | 9.900E-06 | 1.700E-05 |
| 20307 | 1016 | G1016Q | C | 1.5e-05 | 9.8e-06 | 1e-05 | 1.160E-05 | 2.946E-06 | 9.800E-06 | 1.500E-05 |
| 20308 | 1016 | G1016G | C | 1.2e-05 | 1e-05 | 1.5e-05 | 1.233E-05 | 2.517E-06 | 1.000E-05 | 1.500E-05 |
| 20309 | 1016 | G1016H | C | 1e-05 | 1e-05 | 1.4e-05 | 1.133E-05 | 2.309E-06 | 1.000E-05 | 1.400E-05 |
| 20310 | 1016 | G1016I | C | 1.5e-05 | 9.6e-06 | 1.3e-05 | 1.253E-05 | 2.730E-06 | 9.600E-06 | 1.500E-05 |
| 20311 | 1016 | G1016L | C | 1.6e-05 | 1.2e-05 | 1.5e-05 | 1.433E-05 | 2.082E-06 | 1.200E-05 | 1.600E-05 |
| 20312 | 1016 | G1016K | C | 9.1e-06 | 1.3e-05 | 1.5e-05 | 1.237E-05 | 3.001E-06 | 9.100E-06 | 1.500E-05 |
| 20313 | 1016 | G1016M | C | 1.4e-05 | 1.2e-05 | 9.8e-06 | 1.193E-05 | 2.101E-06 | 9.800E-06 | 1.400E-05 |
| 20314 | 1016 | G1016F | C | 1.4e-05 | 1.1e-05 | 9.3e-06 | 1.143E-05 | 2.380E-06 | 9.300E-06 | 1.400E-05 |

| | | | | | | | | | | |
|---|---|---|---|---|---|---|---|---|---|---|
| 20315 | 1016 | G1016P | C | 1.2e-05 | 1.6e-05 | 1.1e-05 | 1.300E-05 | 2.646E-06 | 1.100E-05 | 1.600E-05 |
| 20316 | 1016 | G1016S | C | 1e-05 | 9.7e-06 | 1.4e-05 | 1.123E-05 | 2.401E-06 | 9.700E-06 | 1.400E-05 |
| 20317 | 1016 | G1016T | C | 1.7e-05 | 9.8e-06 | 1.1e-05 | 1.260E-05 | 3.857E-06 | 9.800E-06 | 1.700E-05 |
| 20318 | 1016 | G1016W | C | 1.5e-05 | 1e-05 | 9.4e-06 | 1.147E-05 | 3.075E-06 | 9.400E-06 | 1.500E-05 |
| 20319 | 1016 | G1016Y | C | 1.4e-05 | 1.3e-05 | 1.1e-05 | 1.267E-05 | 1.528E-06 | 1.100E-05 | 1.400E-05 |
| 20320 | 1016 | G1016V | C | 1.5e-05 | 1.1e-05 | 9.8e-06 | 1.193E-05 | 2.723E-06 | 9.800E-06 | 1.500E-05 |
| 20321 | 1017 | L1017A | C | 1e-05 | 9.6e-06 | 1.6e-05 | 1.187E-05 | 3.585E-06 | 9.600E-06 | 1.600E-05 |
| 20322 | 1017 | L1017R | C | 1e-05 | 1.8e-05 | 1.1e-05 | 1.300E-05 | 4.359E-06 | 1.000E-05 | 1.800E-05 |
| 20323 | 1017 | L1017N | C | 9.9e-06 | 1.2e-05 | 8.2e-06 | 1.003E-05 | 1.904E-06 | 8.200E-06 | 1.200E-05 |
| 20324 | 1017 | L1017D | C | 1e-05 | 1.1e-05 | 1.5e-05 | 1.200E-05 | 2.646E-06 | 1.000E-05 | 1.500E-05 |
| 20325 | 1017 | L1017C | C | 8.9e-06 | 9e-06 | 1.1e-05 | 9.633E-06 | 1.185E-06 | 8.900E-06 | 1.100E-05 |
| 20326 | 1017 | L1017E | C | 1.3e-05 | 1e-05 | 1.6e-05 | 1.300E-05 | 3.000E-06 | 1.000E-05 | 1.600E-05 |
| 20327 | 1017 | L1017Q | C | 9.6e-06 | 1e-05 | 1.6e-05 | 1.187E-05 | 3.585E-06 | 9.600E-06 | 1.600E-05 |
| 20328 | 1017 | L1017G | C | 1.3e-05 | 1e-05 | 1.5e-05 | 1.267E-05 | 2.517E-06 | 1.000E-05 | 1.500E-05 |
| 20329 | 1017 | L1017H | C | 1e-05 | 9.9e-06 | 1.5e-05 | 1.163E-05 | 2.916E-06 | 9.900E-06 | 1.500E-05 |
| 20330 | 1017 | L1017I | C | 1.5e-05 | 1e-05 | 1.1e-05 | 1.200E-05 | 2.646E-06 | 1.000E-05 | 1.500E-05 |
| 20331 | 1017 | L1017L | C | 1.3e-05 | 1.6e-05 | 1.6e-05 | 1.500E-05 | 1.732E-06 | 1.300E-05 | 1.600E-05 |
| 20332 | 1017 | L1017K | C | 1e-05 | 1.5e-05 | 9.6e-06 | 1.153E-05 | 3.009E-06 | 9.600E-06 | 1.500E-05 |
| 20333 | 1017 | L1017M | C | 1.2e-05 | 8.6e-06 | 1.2e-05 | 1.087E-05 | 1.963E-06 | 8.600E-06 | 1.200E-05 |
| 20334 | 1017 | L1017F | C | 1.5e-05 | 1.2e-05 | 9.4e-06 | 1.213E-05 | 2.802E-06 | 9.400E-06 | 1.500E-05 |
| 20335 | 1017 | L1017P | C | 1.1e-05 | 1.6e-05 | 8.1e-06 | 1.170E-05 | 3.996E-06 | 8.100E-06 | 1.600E-05 |
| 20336 | 1017 | L1017S | C | 1.1e-05 | 9.1e-06 | 1.4e-05 | 1.137E-05 | 2.470E-06 | 9.100E-06 | 1.400E-05 |
| 20337 | 1017 | L1017T | C | 1e-05 | 1.4e-05 | 8.8e-06 | 1.093E-05 | 2.723E-06 | 8.800E-06 | 1.400E-05 |
| 20338 | 1017 | L1017W | C | 1.6e-05 | 1.6e-05 | 1.4e-05 | 1.533E-05 | 1.155E-06 | 1.400E-05 | 1.600E-05 |
| 20339 | 1017 | L1017Y | C | 1.1e-05 | 1.5e-05 | 1.5e-05 | 1.367E-05 | 2.309E-06 | 1.100E-05 | 1.500E-05 |
| 20340 | 1017 | L1017V | C | 8.7e-06 | 9.5e-06 | 9.1e-06 | 9.100E-06 | 4.000E-07 | 8.700E-06 | 9.500E-06 |
| 20341 | 1018 | Y1018A | C | 9.1e-06 | 1.7e-05 | 1.1e-05 | 1.237E-05 | 4.124E-06 | 9.100E-06 | 1.700E-05 |
| 20342 | 1018 | Y1018R | C | 1.6e-05 | 1.1e-05 | 1e-05 | 1.233E-05 | 3.215E-06 | 1.000E-05 | 1.600E-05 |
| 20343 | 1018 | Y1018N | C | 1.6e-05 | 9.8e-06 | 1.2e-05 | 1.260E-05 | 3.143E-06 | 9.800E-06 | 1.600E-05 |
| 20344 | 1018 | Y1018D | C | 1.6e-05 | 1.8e-05 | 8e-06 | 1.400E-05 | 5.292E-06 | 8.000E-06 | 1.800E-05 |
| 20345 | 1018 | Y1018C | C | 1.5e-05 | 1.1e-05 | 9.2e-06 | 1.173E-05 | 2.969E-06 | 9.200E-06 | 1.500E-05 |
| 20346 | 1018 | Y1018E | C | 1.1e-05 | 1.1e-05 | 1.1e-05 | 1.100E-05 | 2.075E-21 | 1.100E-05 | 1.100E-05 |
| 20347 | 1018 | Y1018Q | C | 9.7e-06 | 1e-05 | 8.7e-06 | 9.467E-06 | 6.807E-07 | 8.700E-06 | 1.000E-05 |
| 20348 | 1018 | Y1018G | C | 1.3e-05 | 1.5e-05 | 1.6e-05 | 1.467E-05 | 1.528E-06 | 1.300E-05 | 1.600E-05 |
| 20349 | 1018 | Y1018H | C | 1.1e-05 | 9.8e-06 | 1.5e-05 | 1.193E-05 | 2.723E-06 | 9.800E-06 | 1.500E-05 |
| 20350 | 1018 | Y1018I | C | 1.2e-05 | 1.5e-05 | 1.1e-05 | 1.267E-05 | 2.082E-06 | 1.100E-05 | 1.500E-05 |
| 20351 | 1018 | Y1018L | C | 8.5e-06 | 1.1e-05 | 1.4e-05 | 1.117E-05 | 2.754E-06 | 8.500E-06 | 1.400E-05 |
| 20352 | 1018 | Y1018K | C | 1.1e-05 | 1.5e-05 | 1.1e-05 | 1.233E-05 | 2.309E-06 | 1.100E-05 | 1.500E-05 |
| 20353 | 1018 | Y1018M | C | 9.7e-06 | 1.1e-05 | 1.4e-05 | 1.157E-05 | 2.205E-06 | 9.700E-06 | 1.400E-05 |
| 20354 | 1018 | Y1018F | C | 1.2e-05 | 1.6e-05 | 8.5e-06 | 1.217E-05 | 3.753E-06 | 8.500E-06 | 1.600E-05 |
| 20355 | 1018 | Y1018P | C | 1e-05 | 1.1e-05 | 1.3e-05 | 1.133E-05 | 1.528E-06 | 1.000E-05 | 1.300E-05 |
| 20356 | 1018 | Y1018S | C | 9.3e-06 | 9.4e-06 | 1.5e-05 | 1.123E-05 | 3.262E-06 | 9.300E-06 | 1.500E-05 |
| 20357 | 1018 | Y1018T | C | 1e-05 | 1.5e-05 | 1.5e-05 | 1.333E-05 | 2.887E-06 | 1.000E-05 | 1.500E-05 |
| 20358 | 1018 | Y1018W | C | 9.7e-06 | 1e-05 | 9.3e-06 | 9.667E-06 | 3.512E-07 | 9.300E-06 | 1.000E-05 |
| 20359 | 1018 | Y1018Y | C | 1.4e-05 | 1e-05 | 9.8e-06 | 1.127E-05 | 2.369E-06 | 9.800E-06 | 1.400E-05 |
| 20360 | 1018 | Y1018V | C | 1.5e-05 | 1.5e-05 | 8.8e-06 | 1.293E-05 | 3.580E-06 | 8.800E-06 | 1.500E-05 |
| 20361 | 1019 | S1019A | C | 1.5e-05 | 1.7e-05 | 1.3e-05 | 1.500E-05 | 2.000E-06 | 1.300E-05 | 1.700E-05 |
| 20362 | 1019 | S1019R | C | 9.9e-06 | 8.3e-06 | 1.1e-05 | 9.733E-06 | 1.358E-06 | 8.300E-06 | 1.100E-05 |
| 20363 | 1019 | S1019N | C | 1.4e-05 | 1.1e-05 | 1e-05 | 1.167E-05 | 2.082E-06 | 1.000E-05 | 1.400E-05 |
| 20364 | 1019 | S1019D | C | 8.7e-06 | 8.8e-06 | 1.9e-05 | 1.217E-05 | 5.918E-06 | 8.700E-06 | 1.900E-05 |
| 20365 | 1019 | S1019C | C | 1.1e-05 | 1.5e-05 | 1.7e-05 | 1.433E-05 | 3.055E-06 | 1.100E-05 | 1.700E-05 |
| 20366 | 1019 | S1019E | C | 1.1e-05 | 9.6e-06 | 1.2e-05 | 1.087E-05 | 1.206E-06 | 9.600E-06 | 1.200E-05 |
| 20367 | 1019 | S1019Q | C | 1.5e-05 | 8.7e-06 | 1.4e-05 | 1.257E-05 | 3.386E-06 | 8.700E-06 | 1.500E-05 |
| 20368 | 1019 | S1019G | C | 1.5e-05 | 1.2e-05 | 1.6e-05 | 1.433E-05 | 2.082E-06 | 1.200E-05 | 1.600E-05 |
| 20369 | 1019 | S1019H | C | 9.3e-06 | 1.4e-05 | 1.4e-05 | 1.243E-05 | 2.714E-06 | 9.300E-06 | 1.400E-05 |
| 20370 | 1019 | S1019I | C | 8.7e-06 | 1.2e-05 | 9.4e-06 | 1.003E-05 | 1.739E-06 | 8.700E-06 | 1.200E-05 |
| 20371 | 1019 | S1019L | C | 9.2e-06 | 9.2e-06 | 1.5e-05 | 1.113E-05 | 3.349E-06 | 9.200E-06 | 1.500E-05 |
| 20372 | 1019 | S1019K | C | 1.1e-05 | 1e-05 | 1.6e-05 | 1.233E-05 | 3.215E-06 | 1.000E-05 | 1.600E-05 |
| 20373 | 1019 | S1019M | C | 1.4e-05 | 1.1e-05 | 1.1e-05 | 1.200E-05 | 1.732E-06 | 1.100E-05 | 1.400E-05 |
| 20374 | 1019 | S1019F | C | 1.5e-05 | 1.3e-05 | 1.1e-05 | 1.300E-05 | 2.000E-06 | 1.100E-05 | 1.500E-05 |
| 20375 | 1019 | S1019P | C | 1.4e-05 | 1.5e-05 | 1.1e-05 | 1.333E-05 | 2.082E-06 | 1.100E-05 | 1.500E-05 |
| 20376 | 1019 | S1019S | C | 1.6e-05 | 1.3e-05 | 1.2e-05 | 1.367E-05 | 2.082E-06 | 1.200E-05 | 1.600E-05 |
| 20377 | 1019 | S1019T | C | 1e-05 | 9.6e-06 | 8.4e-06 | 9.333E-06 | 8.327E-07 | 8.400E-06 | 1.000E-05 |
| 20378 | 1019 | S1019W | C | 1.2e-05 | 1.1e-05 | 1.5e-05 | 1.267E-05 | 2.082E-06 | 1.100E-05 | 1.500E-05 |
| 20379 | 1019 | S1019Y | C | 1.5e-05 | 1.5e-05 | 1.1e-05 | 1.367E-05 | 2.309E-06 | 1.100E-05 | 1.500E-05 |
| 20380 | 1019 | S1019V | C | 1e-05 | 1.1e-05 | 1.2e-05 | 1.100E-05 | 1.000E-06 | 1.000E-05 | 1.200E-05 |
| 20381 | 1020 | L1020A | C | 1.4e-05 | 1e-05 | 1e-05 | 1.133E-05 | 2.309E-06 | 1.000E-05 | 1.400E-05 |
| 20382 | 1020 | L1020R | C | 1.6e-05 | 1.7e-05 | 9e-06 | 1.400E-05 | 4.359E-06 | 9.000E-06 | 1.700E-05 |
| 20383 | 1020 | L1020N | C | 1.2e-05 | 1.1e-05 | 7.7e-06 | 1.023E-05 | 2.250E-06 | 7.700E-06 | 1.200E-05 |
| 20384 | 1020 | L1020D | C | 1.7e-05 | 1e-05 | 1.2e-05 | 1.300E-05 | 3.606E-06 | 1.000E-05 | 1.700E-05 |
| 20385 | 1020 | L1020C | C | 8.5e-06 | 1.4e-05 | 1.2e-05 | 1.150E-05 | 2.784E-06 | 8.500E-06 | 1.400E-05 |
| 20386 | 1020 | L1020E | C | 1.1e-05 | 1.6e-05 | 7.2e-06 | 1.140E-05 | 4.414E-06 | 7.200E-06 | 1.600E-05 |
| 20387 | 1020 | L1020Q | C | 1e-05 | 1.3e-05 | 1.2e-05 | 1.167E-05 | 1.528E-06 | 1.000E-05 | 1.300E-05 |
| 20388 | 1020 | L1020G | C | 9.9e-06 | 1.6e-05 | 1.1e-05 | 1.230E-05 | 3.251E-06 | 9.900E-06 | 1.600E-05 |
| 20389 | 1020 | L1020H | C | 1.3e-05 | 1.3e-05 | 1.5e-05 | 1.367E-05 | 1.155E-06 | 1.300E-05 | 1.500E-05 |
| 20390 | 1020 | L1020I | C | 1.6e-05 | 1.3e-05 | 9.6e-06 | 1.287E-05 | 3.202E-06 | 9.600E-06 | 1.600E-05 |
| 20391 | 1020 | L1020L | C | 8.6e-06 | 1e-05 | 9.6e-06 | 9.400E-06 | 7.211E-07 | 8.600E-06 | 1.000E-05 |
| 20392 | 1020 | L1020K | C | 1.7e-05 | 1.4e-05 | 1.1e-05 | 1.400E-05 | 3.000E-06 | 1.100E-05 | 1.700E-05 |
| 20393 | 1020 | L1020M | C | 1e-05 | 1.5e-05 | 1.6e-05 | 1.367E-05 | 3.215E-06 | 1.000E-05 | 1.600E-05 |
| 20394 | 1020 | L1020F | C | 1.4e-05 | 1.6e-05 | 1.1e-05 | 1.367E-05 | 2.517E-06 | 1.100E-05 | 1.600E-05 |
| 20395 | 1020 | L1020P | C | 1.3e-05 | 1.1e-05 | 1.2e-05 | 1.200E-05 | 1.000E-06 | 1.100E-05 | 1.300E-05 |
| 20396 | 1020 | L1020S | C | 9.6e-06 | 1.5e-05 | 1e-05 | 1.153E-05 | 3.009E-06 | 9.600E-06 | 1.500E-05 |
| 20397 | 1020 | L1020T | C | 1.3e-05 | 1.4e-05 | 1.7e-05 | 1.467E-05 | 2.082E-06 | 1.300E-05 | 1.700E-05 |
| 20398 | 1020 | L1020W | C | 1e-05 | 1.3e-05 | 1.5e-05 | 1.267E-05 | 2.517E-06 | 1.000E-05 | 1.500E-05 |
| 20399 | 1020 | L1020Y | C | 1.1e-05 | 1.1e-05 | 1e-05 | 1.067E-05 | 5.774E-07 | 1.000E-05 | 1.100E-05 |

| | | | | | | | | | | |
|---|---|---|---|---|---|---|---|---|---|---|
| 20400 | 1020 | L1020V | C | 1.4e-05 | 1.1e-05 | 1.4e-05 | 1.300E-05 | 1.732E-06 | 1.100E-05 | 1.400E-05 |
| 20401 | 1021 | S1021A | C | 1.8e-05 | 1.5e-05 | 1.5e-05 | 1.600E-05 | 1.732E-06 | 1.500E-05 | 1.800E-05 |
| 20402 | 1021 | S1021R | C | 1.1e-05 | 1.6e-05 | 1.5e-05 | 1.400E-05 | 2.646E-06 | 1.100E-05 | 1.600E-05 |
| 20403 | 1021 | S1021N | C | 1.2e-05 | 1.5e-05 | 1.5e-05 | 1.400E-05 | 1.732E-06 | 1.200E-05 | 1.500E-05 |
| 20404 | 1021 | S1021D | C | 1e-05 | 1.5e-05 | 1.1e-05 | 1.200E-05 | 2.646E-06 | 1.000E-05 | 1.500E-05 |
| 20405 | 1021 | S1021C | C | 6.7e-06 | 1.2e-05 | 1.7e-05 | 1.190E-05 | 5.151E-06 | 6.700E-06 | 1.700E-05 |
| 20406 | 1021 | S1021E | C | 1.4e-05 | 1.1e-05 | 1.4e-05 | 1.300E-05 | 1.732E-06 | 1.100E-05 | 1.400E-05 |
| 20407 | 1021 | S1021Q | C | 9.9e-06 | 9.8e-06 | 1.5e-05 | 1.157E-05 | 2.974E-06 | 9.800E-06 | 1.500E-05 |
| 20408 | 1021 | S1021G | C | 1.3e-05 | 1.4e-05 | 1.1e-05 | 1.267E-05 | 1.528E-06 | 1.100E-05 | 1.400E-05 |
| 20409 | 1021 | S1021H | C | 1e-05 | 1.1e-05 | 1.6e-05 | 1.233E-05 | 3.215E-06 | 1.000E-05 | 1.600E-05 |
| 20410 | 1021 | S1021I | C | 1.1e-05 | 8e-06 | 1.4e-05 | 1.100E-05 | 3.000E-06 | 8.000E-06 | 1.400E-05 |
| 20411 | 1021 | S1021L | C | 1.1e-05 | 1.6e-05 | 1.3e-05 | 1.333E-05 | 2.517E-06 | 1.100E-05 | 1.600E-05 |
| 20412 | 1021 | S1021K | C | 1.5e-05 | 9.9e-06 | 1.6e-05 | 1.363E-05 | 3.272E-06 | 9.900E-06 | 1.600E-05 |
| 20413 | 1021 | S1021M | C | 9.4e-06 | 1.2e-05 | 1.2e-05 | 1.113E-05 | 1.501E-06 | 9.400E-06 | 1.200E-05 |
| 20414 | 1021 | S1021F | C | 8.8e-06 | 1.5e-05 | 9.2e-06 | 1.100E-05 | 3.470E-06 | 8.800E-06 | 1.500E-05 |
| 20415 | 1021 | S1021P | C | 6.6e-06 | 1e-05 | 1e-05 | 8.867E-06 | 1.963E-06 | 6.600E-06 | 1.000E-05 |
| 20416 | 1021 | S1021S | C | 1.1e-05 | 1.6e-05 | 1.1e-05 | 1.267E-05 | 2.887E-06 | 1.100E-05 | 1.600E-05 |
| 20417 | 1021 | S1021T | C | 1.3e-05 | 1.4e-05 | 7.8e-06 | 1.160E-05 | 3.329E-06 | 7.800E-06 | 1.400E-05 |
| 20418 | 1021 | S1021W | C | 1.5e-05 | 1.4e-05 | 1.1e-05 | 1.333E-05 | 2.082E-06 | 1.100E-05 | 1.500E-05 |
| 20419 | 1021 | S1021Y | C | 1.5e-05 | 1.7e-05 | 1.1e-05 | 1.433E-05 | 3.055E-06 | 1.100E-05 | 1.700E-05 |
| 20420 | 1021 | S1021V | C | 1e-05 | 1.6e-05 | 1.6e-05 | 1.400E-05 | 3.464E-06 | 1.000E-05 | 1.600E-05 |
| 20421 | 1022 | S1022A | C | 9.9e-06 | 8.9e-06 | 1.6e-05 | 1.160E-05 | 3.843E-06 | 8.900E-06 | 1.600E-05 |
| 20422 | 1022 | S1022R | C | 1.5e-05 | 1.1e-05 | 1.4e-05 | 1.333E-05 | 2.082E-06 | 1.100E-05 | 1.500E-05 |
| 20423 | 1022 | S1022N | C | 1.2e-05 | 1.5e-05 | 8.1e-06 | 1.170E-05 | 3.460E-06 | 8.100E-06 | 1.500E-05 |
| 20424 | 1022 | S1022D | C | 1.4e-05 | 1.5e-05 | 1.3e-05 | 1.400E-05 | 1.000E-06 | 1.300E-05 | 1.500E-05 |
| 20425 | 1022 | S1022C | C | 1.6e-05 | 1.6e-05 | 1.6e-05 | 1.600E-05 | 0.000E+00 | 1.600E-05 | 1.600E-05 |
| 20426 | 1022 | S1022E | C | 1e-05 | 9.9e-06 | 6.8e-06 | 8.900E-06 | 1.819E-06 | 6.800E-06 | 1.000E-05 |
| 20427 | 1022 | S1022Q | C | 1e-05 | 1.1e-05 | 8.7e-06 | 9.900E-06 | 1.153E-06 | 8.700E-06 | 1.100E-05 |
| 20428 | 1022 | S1022G | C | 1.1e-05 | 1.6e-05 | 1e-05 | 1.233E-05 | 3.215E-06 | 1.000E-05 | 1.600E-05 |
| 20429 | 1022 | S1022H | C | 1.6e-05 | 1.5e-05 | 1.3e-05 | 1.467E-05 | 1.528E-06 | 1.300E-05 | 1.600E-05 |
| 20430 | 1022 | S1022I | C | 1.1e-05 | 1.5e-05 | 9.3e-06 | 1.177E-05 | 2.926E-06 | 9.300E-06 | 1.500E-05 |
| 20431 | 1022 | S1022L | C | 1.5e-05 | 1.5e-05 | 1.5e-05 | 1.500E-05 | 0.000E+00 | 1.500E-05 | 1.500E-05 |
| 20432 | 1022 | S1022K | C | 1.1e-05 | 1.3e-05 | 1.5e-05 | 1.300E-05 | 2.000E-06 | 1.100E-05 | 1.500E-05 |
| 20433 | 1022 | S1022M | C | 1.2e-05 | 1.4e-05 | 1.2e-05 | 1.267E-05 | 1.155E-06 | 1.200E-05 | 1.400E-05 |
| 20434 | 1022 | S1022F | C | 1.4e-05 | 1.6e-05 | 1.5e-05 | 1.500E-05 | 1.000E-06 | 1.400E-05 | 1.600E-05 |
| 20435 | 1022 | S1022P | C | 1.3e-05 | 1.5e-05 | 1e-05 | 1.267E-05 | 2.517E-06 | 1.000E-05 | 1.500E-05 |
| 20436 | 1022 | S1022S | C | 1.6e-05 | 1.5e-05 | 1.1e-05 | 1.400E-05 | 2.646E-06 | 1.100E-05 | 1.600E-05 |
| 20437 | 1022 | S1022T | C | 1.7e-05 | 1.3e-05 | 1.1e-05 | 1.367E-05 | 3.055E-06 | 1.100E-05 | 1.700E-05 |
| 20438 | 1022 | S1022W | C | 1.2e-05 | 1.1e-05 | 1.2e-05 | 1.167E-05 | 5.774E-07 | 1.100E-05 | 1.200E-05 |
| 20439 | 1022 | S1022Y | C | 1.1e-05 | 9.3e-06 | 1.3e-05 | 1.110E-05 | 1.852E-06 | 9.300E-06 | 1.300E-05 |
| 20440 | 1022 | S1022V | C | 1.5e-05 | 1.6e-05 | 1.6e-05 | 1.567E-05 | 5.774E-07 | 1.500E-05 | 1.600E-05 |
| 20441 | 1023 | V1023A | C | 1.1e-05 | 1.1e-05 | 1.1e-05 | 1.100E-05 | 2.075E-21 | 1.100E-05 | 1.100E-05 |
| 20442 | 1023 | V1023R | C | 1.4e-05 | 1.5e-05 | 1.1e-05 | 1.333E-05 | 2.082E-06 | 1.100E-05 | 1.500E-05 |
| 20443 | 1023 | V1023N | C | 9.6e-06 | 1.1e-05 | 1.6e-05 | 1.220E-05 | 3.365E-06 | 9.600E-06 | 1.600E-05 |
| 20444 | 1023 | V1023D | C | 1.5e-05 | 8.3e-06 | 1.1e-05 | 1.143E-05 | 3.371E-06 | 8.300E-06 | 1.500E-05 |
| 20445 | 1023 | V1023C | C | 1.6e-05 | 1.5e-05 | 1.4e-05 | 1.500E-05 | 1.000E-06 | 1.400E-05 | 1.600E-05 |
| 20446 | 1023 | V1023E | C | 1.5e-05 | 8.4e-06 | 2.7e-05 | 1.680E-05 | 9.430E-06 | 8.400E-06 | 2.700E-05 |
| 20447 | 1023 | V1023Q | C | 1.1e-05 | 1.4e-05 | 1.6e-05 | 1.367E-05 | 2.517E-06 | 1.100E-05 | 1.600E-05 |
| 20448 | 1023 | V1023G | C | 1.1e-05 | 9.6e-06 | 1.6e-05 | 1.220E-05 | 3.365E-06 | 9.600E-06 | 1.600E-05 |
| 20449 | 1023 | V1023H | C | 7.4e-06 | 1.3e-05 | 1.1e-05 | 1.047E-05 | 2.838E-06 | 7.400E-06 | 1.300E-05 |
| 20450 | 1023 | V1023I | C | 1.5e-05 | 1.7e-05 | 1.5e-05 | 1.567E-05 | 1.155E-06 | 1.500E-05 | 1.700E-05 |
| 20451 | 1023 | V1023L | C | 1.2e-05 | 1.2e-05 | 1.4e-05 | 1.267E-05 | 1.155E-06 | 1.200E-05 | 1.400E-05 |
| 20452 | 1023 | V1023K | C | 1.4e-05 | 1.5e-05 | 8.9e-06 | 1.263E-05 | 3.272E-06 | 8.900E-06 | 1.500E-05 |
| 20453 | 1023 | V1023M | C | 1.7e-05 | 1.4e-05 | 9.3e-06 | 1.343E-05 | 3.881E-06 | 9.300E-06 | 1.700E-05 |
| 20454 | 1023 | V1023F | C | 1e-05 | 9.7e-06 | 8.8e-06 | 9.500E-06 | 6.245E-07 | 8.800E-06 | 1.000E-05 |
| 20455 | 1023 | V1023P | C | 1e-05 | 1.4e-05 | 1.5e-05 | 1.300E-05 | 2.646E-06 | 1.000E-05 | 1.500E-05 |
| 20456 | 1023 | V1023S | C | 1.4e-05 | 1.1e-05 | 1.3e-05 | 1.267E-05 | 1.528E-06 | 1.100E-05 | 1.400E-05 |
| 20457 | 1023 | V1023T | C | 1.2e-05 | 1.6e-05 | 8e-06 | 1.200E-05 | 4.000E-06 | 8.000E-06 | 1.600E-05 |
| 20458 | 1023 | V1023W | C | 1.2e-05 | 1.5e-05 | 1.4e-05 | 1.367E-05 | 1.528E-06 | 1.200E-05 | 1.500E-05 |
| 20459 | 1023 | V1023Y | C | 1.5e-05 | 1e-05 | 1.2e-05 | 1.233E-05 | 2.517E-06 | 1.000E-05 | 1.500E-05 |
| 20460 | 1023 | V1023V | C | 9.7e-06 | 1.1e-05 | 1.4e-05 | 1.157E-05 | 2.205E-06 | 9.700E-06 | 1.400E-05 |
| 20461 | 1024 | V1024A | C | 1.4e-05 | 1.7e-05 | 9.9e-06 | 1.363E-05 | 3.564E-06 | 9.900E-06 | 1.700E-05 |
| 20462 | 1024 | V1024R | C | 1e-05 | 1.1e-05 | 1.4e-05 | 1.167E-05 | 2.082E-06 | 1.000E-05 | 1.400E-05 |
| 20463 | 1024 | V1024N | C | 1.3e-05 | 1.5e-05 | 1.1e-05 | 1.300E-05 | 2.000E-06 | 1.100E-05 | 1.500E-05 |
| 20464 | 1024 | V1024D | C | 1.4e-05 | 9e-06 | 1.1e-05 | 1.133E-05 | 2.517E-06 | 9.000E-06 | 1.400E-05 |
| 20465 | 1024 | V1024C | C | 1e-05 | 1e-05 | 1.5e-05 | 1.167E-05 | 2.887E-06 | 1.000E-05 | 1.500E-05 |
| 20466 | 1024 | V1024E | C | 1e-05 | 9.6e-06 | 1.1e-05 | 1.020E-05 | 7.211E-07 | 9.600E-06 | 1.100E-05 |
| 20467 | 1024 | V1024Q | C | 1.6e-05 | 1.5e-05 | 1.5e-05 | 1.533E-05 | 5.774E-07 | 1.500E-05 | 1.600E-05 |
| 20468 | 1024 | V1024G | C | 1.6e-05 | 1.6e-05 | 1.1e-05 | 1.433E-05 | 2.887E-06 | 1.100E-05 | 1.600E-05 |
| 20469 | 1024 | V1024H | C | 1.3e-05 | 9.6e-06 | 1.6e-05 | 1.287E-05 | 3.202E-06 | 9.600E-06 | 1.600E-05 |
| 20470 | 1024 | V1024I | C | 1.4e-05 | 1e-05 | 1.5e-05 | 1.300E-05 | 2.646E-06 | 1.000E-05 | 1.500E-05 |
| 20471 | 1024 | V1024L | C | 1.5e-05 | 3.4e-05 | 1e-05 | 1.967E-05 | 1.266E-05 | 1.000E-05 | 3.400E-05 |
| 20472 | 1024 | V1024K | C | 1e-05 | 1.4e-05 | 1.1e-05 | 1.167E-05 | 2.082E-06 | 1.000E-05 | 1.400E-05 |
| 20473 | 1024 | V1024M | C | 1.6e-05 | 6.1e-06 | 1.4e-05 | 1.203E-05 | 5.235E-06 | 6.100E-06 | 1.600E-05 |
| 20474 | 1024 | V1024F | C | 1e-05 | 1.4e-05 | 8.7e-06 | 1.090E-05 | 2.762E-06 | 8.700E-06 | 1.400E-05 |
| 20475 | 1024 | V1024P | C | 9.1e-06 | 1.1e-05 | 1.4e-05 | 1.137E-05 | 2.470E-06 | 9.100E-06 | 1.400E-05 |
| 20476 | 1024 | V1024S | C | 9.2e-06 | 1.2e-05 | 8.1e-06 | 9.767E-06 | 2.011E-06 | 8.100E-06 | 1.200E-05 |
| 20477 | 1024 | V1024T | C | 1.1e-05 | 1.3e-05 | 1.6e-05 | 1.333E-05 | 2.517E-06 | 1.100E-05 | 1.600E-05 |
| 20478 | 1024 | V1024W | C | 1.3e-05 | 1.2e-05 | 1.6e-05 | 1.367E-05 | 2.082E-06 | 1.200E-05 | 1.600E-05 |
| 20479 | 1024 | V1024Y | C | 1.5e-05 | 1.6e-05 | 1.3e-05 | 1.467E-05 | 1.528E-06 | 1.300E-05 | 1.600E-05 |
| 20480 | 1024 | V1024V | C | 9.8e-06 | 1.4e-05 | 1e-05 | 1.127E-05 | 2.369E-06 | 9.800E-06 | 1.400E-05 |
| 20481 | 1025 | T1025A | C | 1.2e-05 | 1.1e-05 | 1.3e-05 | 1.200E-05 | 1.000E-06 | 1.100E-05 | 1.300E-05 |
| 20482 | 1025 | T1025R | C | 1.5e-05 | 1.6e-05 | 9.9e-06 | 1.363E-05 | 3.272E-06 | 9.900E-06 | 1.600E-05 |
| 20483 | 1025 | T1025N | C | 1.4e-05 | 1.9e-05 | 1.4e-05 | 1.567E-05 | 2.887E-06 | 1.400E-05 | 1.900E-05 |
| 20484 | 1025 | T1025D | C | 1.4e-05 | 4.4e-05 | 1.6e-05 | 2.467E-05 | 1.677E-05 | 1.400E-05 | 4.400E-05 |

| | | | | | | | | | | |
|---|---|---|---|---|---|---|---|---|---|---|
| 20485 | 1025 | T1025C | C | 8.9e-06 | 7.6e-06 | 1.1e-05 | 9.167E-06 | 1.716E-06 | 7.600E-06 | 1.100E-05 |
| 20486 | 1025 | T1025E | C | 1.6e-05 | 1.6e-05 | 1.1e-05 | 1.433E-05 | 2.887E-06 | 1.100E-05 | 1.600E-05 |
| 20487 | 1025 | T1025Q | C | 1.4e-05 | 1.3e-05 | 1.1e-05 | 1.267E-05 | 1.528E-06 | 1.100E-05 | 1.400E-05 |
| 20488 | 1025 | T1025G | C | 1.1e-05 | 1.5e-05 | 1.2e-05 | 1.267E-05 | 2.082E-06 | 1.100E-05 | 1.500E-05 |
| 20489 | 1025 | T1025H | C | 9.5e-06 | 1.6e-05 | 1.1e-05 | 1.217E-05 | 3.403E-06 | 9.500E-06 | 1.600E-05 |
| 20490 | 1025 | T1025I | C | 1e-05 | 1.1e-05 | 1.6e-05 | 1.233E-05 | 3.215E-06 | 1.000E-05 | 1.600E-05 |
| 20491 | 1025 | T1025L | C | 1.4e-05 | 1.5e-05 | 1.2e-05 | 1.367E-05 | 1.528E-06 | 1.200E-05 | 1.500E-05 |
| 20492 | 1025 | T1025K | C | 1.4e-05 | 1.7e-05 | 1.3e-05 | 1.467E-05 | 2.082E-06 | 1.300E-05 | 1.700E-05 |
| 20493 | 1025 | T1025M | C | 1.6e-05 | 9.9e-06 | 1.5e-05 | 1.363E-05 | 3.272E-06 | 9.900E-06 | 1.600E-05 |
| 20494 | 1025 | T1025F | C | 1.5e-05 | 1.1e-05 | 1.6e-05 | 1.400E-05 | 2.646E-06 | 1.100E-05 | 1.600E-05 |
| 20495 | 1025 | T1025P | C | 1.3e-05 | 1.7e-05 | 8.9e-06 | 1.297E-05 | 4.050E-06 | 8.900E-06 | 1.700E-05 |
| 20496 | 1025 | T1025S | C | 1.4e-05 | 1.1e-05 | 1e-05 | 1.167E-05 | 2.082E-06 | 1.000E-05 | 1.400E-05 |
| 20497 | 1025 | T1025T | C | 1.5e-05 | 1e-05 | 1.2e-05 | 1.233E-05 | 2.517E-06 | 1.000E-05 | 1.500E-05 |
| 20498 | 1025 | T1025W | C | 9.4e-06 | 1.2e-05 | 1e-05 | 1.047E-05 | 1.361E-06 | 9.400E-06 | 1.200E-05 |
| 20499 | 1025 | T1025Y | C | 1.5e-05 | 1.4e-05 | 1.3e-05 | 1.400E-05 | 1.000E-06 | 1.300E-05 | 1.500E-05 |
| 20500 | 1025 | T1025V | C | 9.5e-06 | 9.9e-06 | 1.7e-05 | 1.213E-05 | 4.219E-06 | 9.500E-06 | 1.700E-05 |
| 20501 | 1026 | V1026A | C | 2e-05 | 1.4e-05 | 9.5e-06 | 1.450E-05 | 5.268E-06 | 9.500E-06 | 2.000E-05 |
| 20502 | 1026 | V1026R | C | 1.5e-05 | 1.1e-05 | 1.5e-05 | 1.367E-05 | 2.309E-06 | 1.100E-05 | 1.500E-05 |
| 20503 | 1026 | V1026N | C | 7.4e-06 | 1.4e-05 | 9.8e-06 | 1.040E-05 | 3.341E-06 | 7.400E-06 | 1.400E-05 |
| 20504 | 1026 | V1026D | C | 1.1e-05 | 1.1e-05 | 9.3e-06 | 1.043E-05 | 9.815E-07 | 9.300E-06 | 1.100E-05 |
| 20505 | 1026 | V1026C | C | 1.5e-05 | 9.3e-06 | 1.4e-05 | 1.277E-05 | 3.044E-06 | 9.300E-06 | 1.500E-05 |
| 20506 | 1026 | V1026E | C | 1.5e-05 | 8.8e-06 | 1.6e-05 | 1.327E-05 | 3.900E-06 | 8.800E-06 | 1.600E-05 |
| 20507 | 1026 | V1026Q | C | 1.3e-05 | 1.1e-05 | 1.2e-05 | 1.200E-05 | 1.000E-06 | 1.100E-05 | 1.300E-05 |
| 20508 | 1026 | V1026G | C | 1.5e-05 | 1.6e-05 | 1.3e-05 | 1.467E-05 | 1.528E-06 | 1.300E-05 | 1.600E-05 |
| 20509 | 1026 | V1026H | C | 9.9e-06 | 1.6e-05 | 7.9e-06 | 1.127E-05 | 4.219E-06 | 7.900E-06 | 1.600E-05 |
| 20510 | 1026 | V1026I | C | 9.6e-06 | 9.2e-06 | 1.3e-05 | 1.060E-05 | 2.088E-06 | 9.200E-06 | 1.300E-05 |
| 20511 | 1026 | V1026L | C | 1.7e-05 | 1.3e-05 | 7.6e-06 | 1.253E-05 | 4.717E-06 | 7.600E-06 | 1.700E-05 |
| 20512 | 1026 | V1026K | C | 9.4e-06 | 1.1e-05 | 1.1e-05 | 1.047E-05 | 9.238E-07 | 9.400E-06 | 1.100E-05 |
| 20513 | 1026 | V1026M | C | 1.5e-05 | 1e-05 | 1.8e-05 | 1.433E-05 | 4.041E-06 | 1.000E-05 | 1.800E-05 |
| 20514 | 1026 | V1026F | C | 1.6e-05 | 1.5e-05 | 1.4e-05 | 1.500E-05 | 1.000E-06 | 1.400E-05 | 1.600E-05 |
| 20515 | 1026 | V1026P | C | 1.7e-05 | 1.1e-05 | 9.6e-06 | 1.253E-05 | 3.931E-06 | 9.600E-06 | 1.700E-05 |
| 20516 | 1026 | V1026S | C | 1.6e-05 | 1.1e-05 | 9e-06 | 1.200E-05 | 3.606E-06 | 9.000E-06 | 1.600E-05 |
| 20517 | 1026 | V1026T | C | 1.5e-05 | 1.1e-05 | 1.5e-05 | 1.367E-05 | 2.309E-06 | 1.100E-05 | 1.500E-05 |
| 20518 | 1026 | V1026W | C | 1.1e-05 | 1.3e-05 | 7.6e-06 | 1.053E-05 | 2.730E-06 | 7.600E-06 | 1.300E-05 |
| 20519 | 1026 | V1026Y | C | 1.7e-05 | 1.7e-05 | 1.3e-05 | 1.567E-05 | 2.309E-06 | 1.300E-05 | 1.700E-05 |
| 20520 | 1026 | V1026V | C | 8e-06 | 1.2e-05 | 1.6e-05 | 1.200E-05 | 4.000E-06 | 8.000E-06 | 1.600E-05 |
| 20521 | 1027 | P1027A | C | 1.7e-05 | 1.1e-05 | 1.6e-05 | 1.467E-05 | 3.215E-06 | 1.100E-05 | 1.700E-05 |
| 20522 | 1027 | P1027R | C | 1e-05 | 1.2e-05 | 5.8e-06 | 9.267E-06 | 3.164E-06 | 5.800E-06 | 1.200E-05 |
| 20523 | 1027 | P1027N | C | 1.6e-05 | 1.5e-05 | 1.5e-05 | 1.533E-05 | 5.774E-07 | 1.500E-05 | 1.600E-05 |
| 20524 | 1027 | P1027D | C | 1.4e-05 | 1.1e-05 | 1.5e-05 | 1.333E-05 | 2.082E-06 | 1.100E-05 | 1.500E-05 |
| 20525 | 1027 | P1027C | C | 1.2e-05 | 1.3e-05 | 1.2e-05 | 1.233E-05 | 5.774E-07 | 1.200E-05 | 1.300E-05 |
| 20526 | 1027 | P1027E | C | 1.5e-05 | 1.5e-05 | 1.3e-05 | 1.433E-05 | 1.155E-06 | 1.300E-05 | 1.500E-05 |
| 20527 | 1027 | P1027Q | C | 1.6e-05 | 1.5e-05 | 1.1e-05 | 1.400E-05 | 2.646E-06 | 1.100E-05 | 1.600E-05 |
| 20528 | 1027 | P1027G | C | 1.6e-05 | 1e-05 | 9.3e-06 | 1.177E-05 | 3.683E-06 | 9.300E-06 | 1.600E-05 |
| 20529 | 1027 | P1027H | C | 7.3e-06 | 1.1e-05 | 1.3e-05 | 1.043E-05 | 2.892E-06 | 7.300E-06 | 1.300E-05 |
| 20530 | 1027 | P1027I | C | 1.5e-05 | 9.4e-06 | 1.4e-05 | 1.280E-05 | 2.987E-06 | 9.400E-06 | 1.500E-05 |
| 20531 | 1027 | P1027L | C | 1.6e-05 | 1.4e-05 | 1.3e-05 | 1.433E-05 | 1.528E-06 | 1.300E-05 | 1.600E-05 |
| 20532 | 1027 | P1027K | C | 1e-05 | 1.1e-05 | 1.4e-05 | 1.167E-05 | 2.082E-06 | 1.000E-05 | 1.400E-05 |
| 20533 | 1027 | P1027M | C | 1e-05 | 1.5e-05 | 1.6e-05 | 1.367E-05 | 3.215E-06 | 1.000E-05 | 1.600E-05 |
| 20534 | 1027 | P1027F | C | 1.5e-05 | 1.5e-05 | 1.5e-05 | 1.500E-05 | 0.000E+00 | 1.500E-05 | 1.500E-05 |
| 20535 | 1027 | P1027P | C | 1e-05 | 1e-05 | 1.1e-05 | 1.033E-05 | 5.774E-07 | 1.000E-05 | 1.100E-05 |
| 20536 | 1027 | P1027S | C | 1.6e-05 | 1.3e-05 | 1.7e-05 | 1.533E-05 | 2.082E-06 | 1.300E-05 | 1.700E-05 |
| 20537 | 1027 | P1027T | C | 1e-05 | 9.6e-06 | 1.1e-05 | 1.020E-05 | 7.211E-07 | 9.600E-06 | 1.100E-05 |
| 20538 | 1027 | P1027W | C | 1.1e-05 | 9.8e-06 | 1e-05 | 1.027E-05 | 6.429E-07 | 9.800E-06 | 1.100E-05 |
| 20539 | 1027 | P1027Y | C | 9.5e-06 | 1.3e-05 | 1.6e-05 | 1.283E-05 | 3.253E-06 | 9.500E-06 | 1.600E-05 |
| 20540 | 1027 | P1027V | C | 1.4e-05 | 1.5e-05 | 9.6e-06 | 1.287E-05 | 2.873E-06 | 9.600E-06 | 1.500E-05 |
| 20541 | 1028 | S1028A | C | 9.5e-06 | 1.5e-05 | 1.5e-05 | 1.317E-05 | 3.175E-06 | 9.500E-06 | 1.500E-05 |
| 20542 | 1028 | S1028R | C | 1.1e-05 | 1.4e-05 | 1.1e-05 | 1.200E-05 | 1.732E-06 | 1.100E-05 | 1.400E-05 |
| 20543 | 1028 | S1028N | C | 1.6e-05 | 8.4e-06 | 1.5e-05 | 1.313E-05 | 4.130E-06 | 8.400E-06 | 1.600E-05 |
| 20544 | 1028 | S1028D | C | 1.6e-05 | 7.6e-06 | 1.5e-05 | 1.287E-05 | 4.588E-06 | 7.600E-06 | 1.600E-05 |
| 20545 | 1028 | S1028C | C | 1.4e-05 | 8.2e-06 | 1.2e-05 | 1.140E-05 | 2.946E-06 | 8.200E-06 | 1.400E-05 |
| 20546 | 1028 | S1028E | C | 1.5e-05 | 1.6e-05 | 1e-05 | 1.367E-05 | 3.215E-06 | 1.000E-05 | 1.600E-05 |
| 20547 | 1028 | S1028Q | C | 1.4e-05 | 8.9e-06 | 1.1e-05 | 1.130E-05 | 2.563E-06 | 8.900E-06 | 1.400E-05 |
| 20548 | 1028 | S1028G | C | 9.6e-06 | 7.9e-06 | 1.2e-05 | 9.833E-06 | 2.060E-06 | 7.900E-06 | 1.200E-05 |
| 20549 | 1028 | S1028H | C | 1.6e-05 | 1.6e-05 | 1e-05 | 1.400E-05 | 3.464E-06 | 1.000E-05 | 1.600E-05 |
| 20550 | 1028 | S1028I | C | 1.5e-05 | 1.5e-05 | 1.6e-05 | 1.533E-05 | 5.774E-07 | 1.500E-05 | 1.600E-05 |
| 20551 | 1028 | S1028L | C | 1.4e-05 | 1.8e-05 | 1.1e-05 | 1.433E-05 | 3.512E-06 | 1.100E-05 | 1.800E-05 |
| 20552 | 1028 | S1028K | C | 1.1e-05 | 1.6e-05 | 1.2e-05 | 1.300E-05 | 2.646E-06 | 1.100E-05 | 1.600E-05 |
| 20553 | 1028 | S1028M | C | 1e-05 | 1.6e-05 | 1.6e-05 | 1.400E-05 | 3.464E-06 | 1.000E-05 | 1.600E-05 |
| 20554 | 1028 | S1028F | C | 1.4e-05 | 1.5e-05 | 1.6e-05 | 1.500E-05 | 1.000E-06 | 1.400E-05 | 1.600E-05 |
| 20555 | 1028 | S1028P | C | 1.6e-05 | 1.1e-05 | 1.1e-05 | 1.267E-05 | 2.887E-06 | 1.100E-05 | 1.600E-05 |
| 20556 | 1028 | S1028S | C | 1.6e-05 | 9.8e-06 | 1.5e-05 | 1.360E-05 | 3.329E-06 | 9.800E-06 | 1.600E-05 |
| 20557 | 1028 | S1028T | C | 8.8e-06 | 1.4e-05 | 1.5e-05 | 1.260E-05 | 3.329E-06 | 8.800E-06 | 1.500E-05 |
| 20558 | 1028 | S1028W | C | 1.6e-05 | 1.8e-05 | 1.1e-05 | 1.500E-05 | 3.606E-06 | 1.100E-05 | 1.800E-05 |
| 20559 | 1028 | S1028Y | C | 1.2e-05 | 1.6e-05 | 1.4e-05 | 1.400E-05 | 2.000E-06 | 1.200E-05 | 1.600E-05 |
| 20560 | 1028 | S1028V | C | 1e-05 | 7.6e-06 | 1.2e-05 | 9.867E-06 | 2.203E-06 | 7.600E-06 | 1.200E-05 |
| 20561 | 1029 | S1029A | C | 1.1e-05 | 9e-06 | 8.7e-06 | 9.567E-06 | 1.250E-06 | 8.700E-06 | 1.100E-05 |
| 20562 | 1029 | S1029R | C | 1.5e-05 | 1.4e-05 | 1.4e-05 | 1.433E-05 | 5.774E-07 | 1.400E-05 | 1.500E-05 |
| 20563 | 1029 | S1029N | C | 8.6e-06 | 1.2e-05 | 1.5e-05 | 1.187E-05 | 3.202E-06 | 8.600E-06 | 1.500E-05 |
| 20564 | 1029 | S1029D | C | 1.5e-05 | 1.7e-05 | 1.1e-05 | 1.433E-05 | 3.055E-06 | 1.100E-05 | 1.700E-05 |
| 20565 | 1029 | S1029C | C | 1.5e-05 | 1.6e-05 | 1.4e-05 | 1.500E-05 | 1.000E-06 | 1.400E-05 | 1.600E-05 |
| 20566 | 1029 | S1029E | C | 8.3e-06 | 1.5e-05 | 1e-05 | 1.110E-05 | 3.483E-06 | 8.300E-06 | 1.500E-05 |
| 20567 | 1029 | S1029Q | C | 1.5e-05 | 1.2e-05 | 1.5e-05 | 1.400E-05 | 1.732E-06 | 1.200E-05 | 1.500E-05 |
| 20568 | 1029 | S1029G | C | 1.1e-05 | 1.7e-05 | 1.6e-05 | 1.467E-05 | 3.215E-06 | 1.100E-05 | 1.700E-05 |
| 20569 | 1029 | S1029H | C | 1.5e-05 | 1.5e-05 | 9.6e-06 | 1.320E-05 | 3.118E-06 | 9.600E-06 | 1.500E-05 |

| | | | | | | | | | | |
|---|---|---|---|---|---|---|---|---|---|---|
| 20570 | 1029 | S1029I | C | 1.3e-05 | 1e-05 | 1.6e-05 | 1.300E-05 | 3.000E-06 | 1.000E-05 | 1.600E-05 |
| 20571 | 1029 | S1029L | C | 1.5e-05 | 1.1e-05 | 1.4e-05 | 1.333E-05 | 2.082E-06 | 1.100E-05 | 1.500E-05 |
| 20572 | 1029 | S1029K | C | 1.5e-05 | 1.5e-05 | 1.3e-05 | 1.433E-05 | 1.155E-06 | 1.300E-05 | 1.500E-05 |
| 20573 | 1029 | S1029M | C | 1.1e-05 | 1.2e-05 | 1.6e-05 | 1.300E-05 | 2.646E-06 | 1.100E-05 | 1.600E-05 |
| 20574 | 1029 | S1029F | C | 1.1e-05 | 1.5e-05 | 7.9e-06 | 1.130E-05 | 3.559E-06 | 7.900E-06 | 1.500E-05 |
| 20575 | 1029 | S1029P | C | 9.6e-06 | 1.6e-05 | 9.3e-06 | 1.163E-05 | 3.785E-06 | 9.300E-06 | 1.600E-05 |
| 20576 | 1029 | S1029S | C | 1.6e-05 | 1.5e-05 | 1.5e-05 | 1.533E-05 | 5.774E-07 | 1.500E-05 | 1.600E-05 |
| 20577 | 1029 | S1029T | C | 1.5e-05 | 1.1e-05 | 1e-05 | 1.200E-05 | 2.646E-06 | 1.000E-05 | 1.500E-05 |
| 20578 | 1029 | S1029W | C | 9.1e-06 | 1.5e-05 | 1.2e-05 | 1.203E-05 | 2.950E-06 | 9.100E-06 | 1.500E-05 |
| 20579 | 1029 | S1029Y | C | 9.3e-06 | 1e-05 | 9.3e-06 | 9.533E-06 | 4.041E-07 | 9.300E-06 | 1.000E-05 |
| 20580 | 1029 | S1029V | C | 1.7e-05 | 1.4e-05 | 9.7e-06 | 1.357E-05 | 3.669E-06 | 9.700E-06 | 1.700E-05 |
| 20581 | 1030 | S1030A | C | 1.6e-05 | 1.7e-05 | 1e-05 | 1.433E-05 | 3.786E-06 | 1.000E-05 | 1.700E-05 |
| 20582 | 1030 | S1030R | C | 1.6e-05 | 1.5e-05 | 1.5e-05 | 1.533E-05 | 5.774E-07 | 1.500E-05 | 1.600E-05 |
| 20583 | 1030 | S1030N | C | 1.2e-05 | 1e-05 | 1.6e-05 | 1.267E-05 | 3.055E-06 | 1.000E-05 | 1.600E-05 |
| 20584 | 1030 | S1030D | C | 8.8e-06 | 1.5e-05 | 1.5e-05 | 1.293E-05 | 3.580E-06 | 8.800E-06 | 1.500E-05 |
| 20585 | 1030 | S1030C | C | 1.7e-05 | 1.4e-05 | 9.4e-06 | 1.347E-05 | 3.828E-06 | 9.400E-06 | 1.700E-05 |
| 20586 | 1030 | S1030E | C | 1.8e-05 | 1.1e-05 | 1.4e-05 | 1.433E-05 | 3.512E-06 | 1.100E-05 | 1.800E-05 |
| 20587 | 1030 | S1030Q | C | 9e-06 | 1.5e-05 | 1.6e-05 | 1.333E-05 | 3.786E-06 | 9.000E-06 | 1.600E-05 |
| 20588 | 1030 | S1030G | C | 1.4e-05 | 1.6e-05 | 1.6e-05 | 1.533E-05 | 1.155E-06 | 1.400E-05 | 1.600E-05 |
| 20589 | 1030 | S1030H | C | 1.5e-05 | 1.7e-05 | 1e-05 | 1.400E-05 | 3.606E-06 | 1.000E-05 | 1.700E-05 |
| 20590 | 1030 | S1030I | C | 1.4e-05 | 1.1e-05 | 1.5e-05 | 1.333E-05 | 2.082E-06 | 1.100E-05 | 1.500E-05 |
| 20591 | 1030 | S1030L | C | 1.2e-05 | 1.3e-05 | 1e-05 | 1.167E-05 | 1.528E-06 | 1.000E-05 | 1.300E-05 |
| 20592 | 1030 | S1030K | C | 1.1e-05 | 7.2e-06 | 1.4e-05 | 1.073E-05 | 3.408E-06 | 7.200E-06 | 1.400E-05 |
| 20593 | 1030 | S1030M | C | 1.1e-05 | 1.5e-05 | 1.1e-05 | 1.233E-05 | 2.309E-06 | 1.100E-05 | 1.500E-05 |
| 20594 | 1030 | S1030F | C | 1.2e-05 | 1.4e-05 | 1.5e-05 | 1.367E-05 | 1.528E-06 | 1.200E-05 | 1.500E-05 |
| 20595 | 1030 | S1030P | C | 1.3e-05 | 1.8e-05 | 1.2e-05 | 1.433E-05 | 3.215E-06 | 1.200E-05 | 1.800E-05 |
| 20596 | 1030 | S1030S | C | 1e-05 | 1.2e-05 | 8.8e-06 | 1.027E-05 | 1.617E-06 | 8.800E-06 | 1.200E-05 |
| 20597 | 1030 | S1030T | C | 8.6e-06 | 1.6e-05 | 1e-05 | 1.153E-05 | 3.931E-06 | 8.600E-06 | 1.600E-05 |
| 20598 | 1030 | S1030W | C | 1.2e-05 | 1.1e-05 | 1.4e-05 | 1.233E-05 | 1.528E-06 | 1.100E-05 | 1.400E-05 |
| 20599 | 1030 | S1030Y | C | 1.3e-05 | 1.8e-05 | 1.6e-05 | 1.567E-05 | 2.517E-06 | 1.300E-05 | 1.800E-05 |
| 20600 | 1030 | S1030V | C | 1e-05 | 1.5e-05 | 1.3e-05 | 1.267E-05 | 2.517E-06 | 1.000E-05 | 1.500E-05 |
| 20601 | 1031 | L1031A | C | 1e-05 | 1.3e-05 | 1.5e-05 | 1.267E-05 | 2.517E-06 | 1.000E-05 | 1.500E-05 |
| 20602 | 1031 | L1031R | C | 9.2e-06 | 1.6e-05 | 9.2e-06 | 1.147E-05 | 3.926E-06 | 9.200E-06 | 1.600E-05 |
| 20603 | 1031 | L1031N | C | 1.4e-05 | 1.7e-05 | 1.5e-05 | 1.533E-05 | 1.528E-06 | 1.400E-05 | 1.700E-05 |
| 20604 | 1031 | L1031D | C | 7.9e-06 | 1e-05 | 1.6e-05 | 1.130E-05 | 4.204E-06 | 7.900E-06 | 1.600E-05 |
| 20605 | 1031 | L1031C | C | 1.2e-05 | 1.6e-05 | 9.7e-06 | 1.257E-05 | 3.188E-06 | 9.700E-06 | 1.600E-05 |
| 20606 | 1031 | L1031E | C | 1.1e-05 | 8.5e-06 | 1.7e-05 | 1.217E-05 | 4.368E-06 | 8.500E-06 | 1.700E-05 |
| 20607 | 1031 | L1031Q | C | 9.7e-06 | 9.9e-06 | 1e-05 | 9.867E-06 | 1.528E-07 | 9.700E-06 | 1.000E-05 |
| 20608 | 1031 | L1031G | C | 1.5e-05 | 1.5e-05 | 1.4e-05 | 1.467E-05 | 5.774E-07 | 1.400E-05 | 1.500E-05 |
| 20609 | 1031 | L1031H | C | 9.9e-06 | 9.2e-06 | 1.5e-05 | 1.137E-05 | 3.166E-06 | 9.200E-06 | 1.500E-05 |
| 20610 | 1031 | L1031I | C | 1e-05 | 1.5e-05 | 1.1e-05 | 1.200E-05 | 2.646E-06 | 1.000E-05 | 1.500E-05 |
| 20611 | 1031 | L1031L | C | 1.2e-05 | 9.2e-06 | 9.9e-06 | 1.037E-05 | 1.457E-06 | 9.200E-06 | 1.200E-05 |
| 20612 | 1031 | L1031K | C | 1.5e-05 | 2.2e-05 | 1.6e-05 | 1.767E-05 | 3.786E-06 | 1.500E-05 | 2.200E-05 |
| 20613 | 1031 | L1031M | C | 1.6e-05 | 1.6e-05 | 1.5e-05 | 1.567E-05 | 5.774E-07 | 1.500E-05 | 1.600E-05 |
| 20614 | 1031 | L1031F | C | 1.4e-05 | 1.5e-05 | 1.4e-05 | 1.433E-05 | 5.774E-07 | 1.400E-05 | 1.500E-05 |
| 20615 | 1031 | L1031P | C | 1.2e-05 | 7.5e-06 | 1.2e-05 | 1.050E-05 | 2.598E-06 | 7.500E-06 | 1.200E-05 |
| 20616 | 1031 | L1031S | C | 1.3e-05 | 1.5e-05 | 7.7e-06 | 1.190E-05 | 3.772E-06 | 7.700E-06 | 1.500E-05 |
| 20617 | 1031 | L1031T | C | 1.1e-05 | 9.9e-06 | 1e-05 | 1.030E-05 | 6.083E-07 | 9.900E-06 | 1.100E-05 |
| 20618 | 1031 | L1031W | C | 1.5e-05 | 9.8e-06 | 1.3e-05 | 1.260E-05 | 2.623E-06 | 9.800E-06 | 1.500E-05 |
| 20619 | 1031 | L1031Y | C | 1e-05 | 1.6e-05 | 1e-05 | 1.200E-05 | 3.464E-06 | 1.000E-05 | 1.600E-05 |
| 20620 | 1031 | L1031V | C | 1.5e-05 | 8.8e-06 | 9.5e-06 | 1.110E-05 | 3.396E-06 | 8.800E-06 | 1.500E-05 |
| 20621 | 1032 | G1032A | C | 1.1e-05 | 1e-05 | 1.1e-05 | 1.067E-05 | 5.774E-07 | 1.000E-05 | 1.100E-05 |
| 20622 | 1032 | G1032R | C | 1e-05 | 1.6e-05 | 1e-05 | 1.200E-05 | 3.464E-06 | 1.000E-05 | 1.600E-05 |
| 20623 | 1032 | G1032N | C | 1.3e-05 | 1.4e-05 | 1.4e-05 | 1.367E-05 | 5.774E-07 | 1.300E-05 | 1.400E-05 |
| 20624 | 1032 | G1032D | C | 1.1e-05 | 1.4e-05 | 1.5e-05 | 1.333E-05 | 2.082E-06 | 1.100E-05 | 1.500E-05 |
| 20625 | 1032 | G1032C | C | 1.2e-05 | 8.7e-06 | 1.2e-05 | 1.090E-05 | 1.905E-06 | 8.700E-06 | 1.200E-05 |
| 20626 | 1032 | G1032E | C | 1.4e-05 | 1.3e-05 | 1e-05 | 1.233E-05 | 2.082E-06 | 1.000E-05 | 1.400E-05 |
| 20627 | 1032 | G1032Q | C | 1.3e-05 | 1.5e-05 | 1.1e-05 | 1.300E-05 | 2.000E-06 | 1.100E-05 | 1.500E-05 |
| 20628 | 1032 | G1032G | C | 1.3e-05 | 1.5e-05 | 1.1e-05 | 1.300E-05 | 2.000E-06 | 1.100E-05 | 1.500E-05 |
| 20629 | 1032 | G1032H | C | 1.5e-05 | 1.5e-05 | 1.4e-05 | 1.467E-05 | 5.774E-07 | 1.400E-05 | 1.500E-05 |
| 20630 | 1032 | G1032I | C | 1e-05 | 1.1e-05 | 1.3e-05 | 1.133E-05 | 1.528E-06 | 1.000E-05 | 1.300E-05 |
| 20631 | 1032 | G1032L | C | 1.5e-05 | 1.7e-05 | 8.3e-06 | 1.343E-05 | 4.557E-06 | 8.300E-06 | 1.700E-05 |
| 20632 | 1032 | G1032K | C | 1.1e-05 | 9.7e-06 | 1.4e-05 | 1.157E-05 | 2.205E-06 | 9.700E-06 | 1.400E-05 |
| 20633 | 1032 | G1032M | C | 1.4e-05 | 7.4e-06 | 1.1e-05 | 1.080E-05 | 3.305E-06 | 7.400E-06 | 1.400E-05 |
| 20634 | 1032 | G1032F | C | 1.5e-05 | 1.1e-05 | 1.5e-05 | 1.367E-05 | 2.309E-06 | 1.100E-05 | 1.500E-05 |
| 20635 | 1032 | G1032P | C | 1.5e-05 | 1.1e-05 | 1.5e-05 | 1.367E-05 | 2.309E-06 | 1.100E-05 | 1.500E-05 |
| 20636 | 1032 | G1032S | C | 1.2e-05 | 1.1e-05 | 1.2e-05 | 1.167E-05 | 5.774E-07 | 1.100E-05 | 1.200E-05 |
| 20637 | 1032 | G1032T | C | 1.3e-05 | 9.9e-06 | 1.8e-05 | 1.363E-05 | 4.087E-06 | 9.900E-06 | 1.800E-05 |
| 20638 | 1032 | G1032W | C | 9.8e-06 | 1.7e-05 | 1.5e-05 | 1.393E-05 | 3.717E-06 | 9.800E-06 | 1.700E-05 |
| 20639 | 1032 | G1032Y | C | 1.2e-05 | 1.5e-05 | 9.4e-06 | 1.213E-05 | 2.802E-06 | 9.400E-06 | 1.500E-05 |
| 20640 | 1032 | G1032V | C | 1.2e-05 | 1.5e-05 | 1.3e-05 | 1.333E-05 | 1.528E-06 | 1.200E-05 | 1.500E-05 |
| 20641 | 1033 | T1033A | C | 1e-05 | 1.4e-05 | 1.7e-05 | 1.367E-05 | 3.512E-06 | 1.000E-05 | 1.700E-05 |
| 20642 | 1033 | T1033R | C | 9e-06 | 1.5e-05 | 1.6e-05 | 1.333E-05 | 3.786E-06 | 9.000E-06 | 1.600E-05 |
| 20643 | 1033 | T1033N | C | 1.5e-05 | 1.3e-05 | 1.6e-05 | 1.467E-05 | 1.528E-06 | 1.300E-05 | 1.600E-05 |
| 20644 | 1033 | T1033D | C | 1.2e-05 | 1.5e-05 | 1.4e-05 | 1.367E-05 | 1.528E-06 | 1.200E-05 | 1.500E-05 |
| 20645 | 1033 | T1033C | C | 1.3e-05 | 1.1e-05 | 1.5e-05 | 1.300E-05 | 2.000E-06 | 1.100E-05 | 1.500E-05 |
| 20646 | 1033 | T1033E | C | 7.8e-06 | 1.6e-05 | 1.2e-05 | 1.193E-05 | 4.100E-06 | 7.800E-06 | 1.600E-05 |
| 20647 | 1033 | T1033Q | C | 1.3e-05 | 1.3e-05 | 1.3e-05 | 1.300E-05 | 0.000E+00 | 1.300E-05 | 1.300E-05 |
| 20648 | 1033 | T1033G | C | 1.5e-05 | 1.2e-05 | 1.8e-05 | 1.500E-05 | 3.000E-06 | 1.200E-05 | 1.800E-05 |
| 20649 | 1033 | T1033H | C | 1.5e-05 | 8.2e-06 | 1.5e-05 | 1.273E-05 | 3.926E-06 | 8.200E-06 | 1.500E-05 |
| 20650 | 1033 | T1033I | C | 1.6e-05 | 7.1e-06 | 1.3e-05 | 1.203E-05 | 4.528E-06 | 7.100E-06 | 1.600E-05 |
| 20651 | 1033 | T1033L | C | 1.1e-05 | 1e-05 | 1.4e-05 | 1.167E-05 | 2.082E-06 | 1.000E-05 | 1.400E-05 |
| 20652 | 1033 | T1033K | C | 1.5e-05 | 1.1e-05 | 1.3e-05 | 1.300E-05 | 2.000E-06 | 1.100E-05 | 1.500E-05 |
| 20653 | 1033 | T1033M | C | 1.6e-05 | 1.2e-05 | 1.6e-05 | 1.467E-05 | 2.309E-06 | 1.200E-05 | 1.600E-05 |
| 20654 | 1033 | T1033F | C | 1.5e-05 | 1.1e-05 | 9.8e-06 | 1.193E-05 | 2.723E-06 | 9.800E-06 | 1.500E-05 |

| | | | | | | | | | | |
|---|---|---|---|---|---|---|---|---|---|---|
| 20655 | 1033 | T1033P | C | 8.9e-06 | 1.2e-05 | 1.1e-05 | 1.063E-05 | 1.582E-06 | 8.900E-06 | 1.200E-05 |
| 20656 | 1033 | T1033S | C | 1.5e-05 | 1.6e-05 | 1.2e-05 | 1.433E-05 | 2.082E-06 | 1.200E-05 | 1.600E-05 |
| 20657 | 1033 | T1033T | C | 1.3e-05 | 1e-05 | 1.5e-05 | 1.267E-05 | 2.517E-06 | 1.000E-05 | 1.500E-05 |
| 20658 | 1033 | T1033W | C | 1.2e-05 | 1.6e-05 | 1.4e-05 | 1.400E-05 | 2.000E-06 | 1.200E-05 | 1.600E-05 |
| 20659 | 1033 | T1033Y | C | 1.5e-05 | 1.5e-05 | 1.5e-05 | 1.500E-05 | 0.000E+00 | 1.500E-05 | 1.500E-05 |
| 20660 | 1033 | T1033V | C | 1.2e-05 | 1e-05 | 1.6e-05 | 1.267E-05 | 3.055E-06 | 1.000E-05 | 1.600E-05 |
| 20661 | 1034 | Q1034A | C | 1.2e-05 | 1.2e-05 | 9.6e-06 | 1.120E-05 | 1.386E-06 | 9.600E-06 | 1.200E-05 |
| 20662 | 1034 | Q1034R | C | 9.9e-06 | 1.5e-05 | 1.5e-05 | 1.330E-05 | 2.944E-06 | 9.900E-06 | 1.500E-05 |
| 20663 | 1034 | Q1034N | C | 1.2e-05 | 8.2e-06 | 1.6e-05 | 1.207E-05 | 3.900E-06 | 8.200E-06 | 1.600E-05 |
| 20664 | 1034 | Q1034D | C | 1.1e-05 | 1.2e-05 | 2.4e-05 | 1.567E-05 | 7.234E-06 | 1.100E-05 | 2.400E-05 |
| 20665 | 1034 | Q1034C | C | 1e-05 | 1.1e-05 | 8.9e-06 | 9.967E-06 | 1.050E-06 | 8.900E-06 | 1.100E-05 |
| 20666 | 1034 | Q1034E | C | 8.9e-06 | 1.6e-05 | 1.2e-05 | 1.230E-05 | 3.559E-06 | 8.900E-06 | 1.600E-05 |
| 20667 | 1034 | Q1034Q | C | 1e-05 | 9.7e-06 | 1.5e-05 | 1.157E-05 | 2.977E-06 | 9.700E-06 | 1.500E-05 |
| 20668 | 1034 | Q1034G | C | 1.4e-05 | 1.8e-05 | 1.2e-05 | 1.467E-05 | 3.055E-06 | 1.200E-05 | 1.800E-05 |
| 20669 | 1034 | Q1034H | C | 1e-05 | 1.5e-05 | 1.4e-05 | 1.300E-05 | 2.646E-06 | 1.000E-05 | 1.500E-05 |
| 20670 | 1034 | Q1034I | C | 1.2e-05 | 1.8e-05 | 1.5e-05 | 1.500E-05 | 3.000E-06 | 1.200E-05 | 1.800E-05 |
| 20671 | 1034 | Q1034L | C | 1.7e-05 | 1.3e-05 | 8.7e-06 | 1.290E-05 | 4.151E-06 | 8.700E-06 | 1.700E-05 |
| 20672 | 1034 | Q1034K | C | 1.5e-05 | 1.1e-05 | 1.5e-05 | 1.367E-05 | 2.309E-06 | 1.100E-05 | 1.500E-05 |
| 20673 | 1034 | Q1034M | C | 9.6e-06 | 1.2e-05 | 1.6e-05 | 1.253E-05 | 3.233E-06 | 9.600E-06 | 1.600E-05 |
| 20674 | 1034 | Q1034F | C | 1.3e-05 | 1.3e-05 | 1.2e-05 | 1.267E-05 | 5.774E-07 | 1.200E-05 | 1.300E-05 |
| 20675 | 1034 | Q1034P | C | 1.1e-05 | 1.6e-05 | 1.8e-05 | 1.500E-05 | 3.606E-06 | 1.100E-05 | 1.800E-05 |
| 20676 | 1034 | Q1034S | C | 1.6e-05 | 1.4e-05 | 1.1e-05 | 1.367E-05 | 2.517E-06 | 1.100E-05 | 1.600E-05 |
| 20677 | 1034 | Q1034T | C | 1.3e-05 | 1.5e-05 | 1.1e-05 | 1.300E-05 | 2.000E-06 | 1.100E-05 | 1.500E-05 |
| 20678 | 1034 | Q1034W | C | 1.4e-05 | 1e-05 | 1.3e-05 | 1.233E-05 | 2.082E-06 | 1.000E-05 | 1.400E-05 |
| 20679 | 1034 | Q1034Y | C | 1.1e-05 | 6.4e-06 | 1.4e-05 | 1.047E-05 | 3.828E-06 | 6.400E-06 | 1.400E-05 |
| 20680 | 1034 | Q1034V | C | 1.5e-05 | 1.1e-05 | 1.5e-05 | 1.367E-05 | 2.309E-06 | 1.100E-05 | 1.500E-05 |
| 20681 | 1035 | T1035A | C | 1.5e-05 | 1.5e-05 | 1.5e-05 | 1.500E-05 | 0.000E+00 | 1.500E-05 | 1.500E-05 |
| 20682 | 1035 | T1035R | C | 1.8e-05 | 9.5e-06 | 1.5e-05 | 1.417E-05 | 4.311E-06 | 9.500E-06 | 1.800E-05 |
| 20683 | 1035 | T1035N | C | 1.6e-05 | 1.2e-05 | 1.1e-05 | 1.300E-05 | 2.646E-06 | 1.100E-05 | 1.600E-05 |
| 20684 | 1035 | T1035D | C | 8.6e-06 | 1.5e-05 | 1e-05 | 1.120E-05 | 3.365E-06 | 8.600E-06 | 1.500E-05 |
| 20685 | 1035 | T1035C | C | 1.4e-05 | 1.3e-05 | 9.5e-06 | 1.217E-05 | 2.363E-06 | 9.500E-06 | 1.400E-05 |
| 20686 | 1035 | T1035E | C | 8.2e-06 | 1.1e-05 | 7e-06 | 8.733E-06 | 2.053E-06 | 7.000E-06 | 1.100E-05 |
| 20687 | 1035 | T1035Q | C | 1.3e-05 | 1.1e-05 | 1.6e-05 | 1.333E-05 | 2.517E-06 | 1.100E-05 | 1.600E-05 |
| 20688 | 1035 | T1035G | C | 1.5e-05 | 1.6e-05 | 1.9e-05 | 1.667E-05 | 2.082E-06 | 1.500E-05 | 1.900E-05 |
| 20689 | 1035 | T1035H | C | 1.1e-05 | 1.3e-05 | 1.7e-05 | 1.367E-05 | 3.055E-06 | 1.100E-05 | 1.700E-05 |
| 20690 | 1035 | T1035I | C | 1.5e-05 | 1.3e-05 | 1.7e-05 | 1.500E-05 | 2.000E-06 | 1.300E-05 | 1.700E-05 |
| 20691 | 1035 | T1035L | C | 6.1e-06 | 1.6e-05 | 9.9e-06 | 1.067E-05 | 4.994E-06 | 6.100E-06 | 1.600E-05 |
| 20692 | 1035 | T1035K | C | 1.5e-05 | 1.7e-05 | 1.3e-05 | 1.500E-05 | 2.000E-06 | 1.300E-05 | 1.700E-05 |
| 20693 | 1035 | T1035M | C | 1.7e-05 | 1e-05 | 1.2e-05 | 1.300E-05 | 3.606E-06 | 1.000E-05 | 1.700E-05 |
| 20694 | 1035 | T1035F | C | 1.3e-05 | 8.6e-06 | 1.4e-05 | 1.187E-05 | 2.873E-06 | 8.600E-06 | 1.400E-05 |
| 20695 | 1035 | T1035P | C | 1.6e-05 | 1.5e-05 | 1.6e-05 | 1.567E-05 | 5.774E-07 | 1.500E-05 | 1.600E-05 |
| 20696 | 1035 | T1035S | C | 1e-05 | 1.5e-05 | 1.5e-05 | 1.333E-05 | 2.887E-06 | 1.000E-05 | 1.500E-05 |
| 20697 | 1035 | T1035T | C | 1.6e-05 | 1.5e-05 | 1.4e-05 | 1.500E-05 | 1.000E-06 | 1.400E-05 | 1.600E-05 |
| 20698 | 1035 | T1035W | C | 1.6e-05 | 1e-05 | 1.4e-05 | 1.333E-05 | 3.055E-06 | 1.000E-05 | 1.600E-05 |
| 20699 | 1035 | T1035Y | C | 9.3e-06 | 8.6e-06 | 1.1e-05 | 9.633E-06 | 1.234E-06 | 8.600E-06 | 1.100E-05 |
| 20700 | 1035 | T1035V | C | 1.2e-05 | 1.7e-05 | 9.8e-06 | 1.293E-05 | 3.690E-06 | 9.800E-06 | 1.700E-05 |
| 20701 | 1036 | Y1036A | C | 1e-05 | 9.7e-06 | 1.5e-05 | 1.157E-05 | 2.977E-06 | 9.700E-06 | 1.500E-05 |
| 20702 | 1036 | Y1036R | C | 8.2e-06 | 1.5e-05 | 1.2e-05 | 1.173E-05 | 3.408E-06 | 8.200E-06 | 1.500E-05 |
| 20703 | 1036 | Y1036N | C | 1.7e-05 | 7.2e-06 | 1.1e-05 | 1.173E-05 | 4.941E-06 | 7.200E-06 | 1.700E-05 |
| 20704 | 1036 | Y1036D | C | 1.2e-05 | 1e-05 | 1.3e-05 | 1.167E-05 | 1.528E-06 | 1.000E-05 | 1.300E-05 |
| 20705 | 1036 | Y1036C | C | 8.9e-06 | 1.6e-05 | 1.6e-05 | 1.363E-05 | 4.099E-06 | 8.900E-06 | 1.600E-05 |
| 20706 | 1036 | Y1036E | C | 1e-05 | 1.3e-05 | 1.6e-05 | 1.300E-05 | 3.000E-06 | 1.000E-05 | 1.600E-05 |
| 20707 | 1036 | Y1036Q | C | 1.7e-05 | 9.1e-06 | 1.6e-05 | 1.403E-05 | 4.302E-06 | 9.100E-06 | 1.700E-05 |
| 20708 | 1036 | Y1036G | C | 9e-06 | 1e-05 | 1.4e-05 | 1.100E-05 | 2.646E-06 | 9.000E-06 | 1.400E-05 |
| 20709 | 1036 | Y1036H | C | 1.2e-05 | 1.1e-05 | 9.3e-06 | 1.077E-05 | 1.365E-06 | 9.300E-06 | 1.200E-05 |
| 20710 | 1036 | Y1036I | C | 1.4e-05 | 1.6e-05 | 1e-05 | 1.333E-05 | 3.055E-06 | 1.000E-05 | 1.600E-05 |
| 20711 | 1036 | Y1036L | C | 9.2e-06 | 1.7e-05 | 1.6e-05 | 1.407E-05 | 4.244E-06 | 9.200E-06 | 1.700E-05 |
| 20712 | 1036 | Y1036K | C | 1.1e-05 | 1.3e-05 | 9.7e-06 | 1.123E-05 | 1.662E-06 | 9.700E-06 | 1.300E-05 |
| 20713 | 1036 | Y1036M | C | 1.1e-05 | 1.3e-05 | 1.2e-05 | 1.200E-05 | 1.000E-06 | 1.100E-05 | 1.300E-05 |
| 20714 | 1036 | Y1036F | C | 1.5e-05 | 1e-05 | 1.3e-05 | 1.267E-05 | 2.517E-06 | 1.000E-05 | 1.500E-05 |
| 20715 | 1036 | Y1036P | C | 9.2e-06 | 1.5e-05 | 1.6e-05 | 1.340E-05 | 3.672E-06 | 9.200E-06 | 1.600E-05 |
| 20716 | 1036 | Y1036S | C | 1.6e-05 | 1.6e-05 | 1.4e-05 | 1.533E-05 | 1.155E-06 | 1.400E-05 | 1.600E-05 |
| 20717 | 1036 | Y1036T | C | 1.5e-05 | 1.3e-05 | 1.7e-05 | 1.500E-05 | 2.000E-06 | 1.300E-05 | 1.700E-05 |
| 20718 | 1036 | Y1036W | C | 1.5e-05 | 1.5e-05 | 1.5e-05 | 1.500E-05 | 0.000E+00 | 1.500E-05 | 1.500E-05 |
| 20719 | 1036 | Y1036Y | C | 1.7e-05 | 1.4e-05 | 1e-05 | 1.367E-05 | 3.512E-06 | 1.000E-05 | 1.700E-05 |
| 20720 | 1036 | Y1036V | C | 1.4e-05 | 1.3e-05 | 1.4e-05 | 1.367E-05 | 5.774E-07 | 1.300E-05 | 1.400E-05 |
| 20721 | 1037 | I1037A | C | 1.2e-05 | 1e-05 | 9.9e-06 | 1.063E-05 | 1.185E-06 | 9.900E-06 | 1.200E-05 |
| 20722 | 1037 | I1037R | C | 1.1e-05 | 1.2e-05 | 1.1e-05 | 1.133E-05 | 5.774E-07 | 1.100E-05 | 1.200E-05 |
| 20723 | 1037 | I1037N | C | 8.8e-06 | 1.8e-05 | 1.4e-05 | 1.360E-05 | 4.613E-06 | 8.800E-06 | 1.800E-05 |
| 20724 | 1037 | I1037D | C | 1.4e-05 | 1.6e-05 | 1.4e-05 | 1.467E-05 | 1.155E-06 | 1.400E-05 | 1.600E-05 |
| 20725 | 1037 | I1037C | C | 1.1e-05 | 1.5e-05 | 1.2e-05 | 1.267E-05 | 2.082E-06 | 1.100E-05 | 1.500E-05 |
| 20726 | 1037 | I1037E | C | 1.4e-05 | 1.5e-05 | 9.5e-06 | 1.283E-05 | 2.930E-06 | 9.500E-06 | 1.500E-05 |
| 20727 | 1037 | I1037Q | C | 2.3e-05 | 1.5e-05 | 1.4e-05 | 1.733E-05 | 4.933E-06 | 1.400E-05 | 2.300E-05 |
| 20728 | 1037 | I1037G | C | 1e-05 | 1.1e-05 | 9.6e-06 | 1.020E-05 | 7.211E-07 | 9.600E-06 | 1.100E-05 |
| 20729 | 1037 | I1037H | C | 9.3e-06 | 1.6e-05 | 1e-05 | 1.177E-05 | 3.683E-06 | 9.300E-06 | 1.600E-05 |
| 20730 | 1037 | I1037I | C | 1.1e-05 | 1.3e-05 | 1.5e-05 | 1.300E-05 | 2.000E-06 | 1.100E-05 | 1.500E-05 |
| 20731 | 1037 | I1037L | C | 1.5e-05 | 1e-05 | 1.5e-05 | 1.333E-05 | 2.887E-06 | 1.000E-05 | 1.500E-05 |
| 20732 | 1037 | I1037K | C | 9.9e-06 | 8.6e-06 | 1.6e-05 | 1.150E-05 | 3.951E-06 | 8.600E-06 | 1.600E-05 |
| 20733 | 1037 | I1037M | C | 8.3e-06 | 1.5e-05 | 1.5e-05 | 1.277E-05 | 3.868E-06 | 8.300E-06 | 1.500E-05 |
| 20734 | 1037 | I1037F | C | 9.1e-06 | 1.7e-05 | 1.5e-05 | 1.370E-05 | 4.107E-06 | 9.100E-06 | 1.700E-05 |
| 20735 | 1037 | I1037P | C | 1.1e-05 | 1.6e-05 | 1.2e-05 | 1.300E-05 | 2.646E-06 | 1.100E-05 | 1.600E-05 |
| 20736 | 1037 | I1037S | C | 1.3e-05 | 1.1e-05 | 1.1e-05 | 1.167E-05 | 1.155E-06 | 1.100E-05 | 1.300E-05 |
| 20737 | 1037 | I1037T | C | 1e-05 | 1.6e-05 | 9.1e-06 | 1.170E-05 | 3.751E-06 | 9.100E-06 | 1.600E-05 |
| 20738 | 1037 | I1037W | C | 1e-05 | 9.9e-06 | 1.4e-05 | 1.130E-05 | 2.339E-06 | 9.900E-06 | 1.400E-05 |
| 20739 | 1037 | I1037Y | C | 1.4e-05 | 1.3e-05 | 1.5e-05 | 1.400E-05 | 1.000E-06 | 1.300E-05 | 1.500E-05 |

| | | | | | | | | | | |
|---|---|---|---|---|---|---|---|---|---|---|
| 20740 | 1037 | I1037V | C | 1.6e-05 | 1.2e-05 | 1e-05 | 1.267E-05 | 3.055E-06 | 1.000E-05 | 1.600E-05 |
| 20741 | 1038 | C1038A | C | 1.4e-05 | 1.3e-05 | 1.2e-05 | 1.300E-05 | 1.000E-06 | 1.200E-05 | 1.400E-05 |
| 20742 | 1038 | C1038R | C | 1.5e-05 | 1.4e-05 | 1.4e-05 | 1.433E-05 | 5.774E-07 | 1.400E-05 | 1.500E-05 |
| 20743 | 1038 | C1038N | C | 1.3e-05 | 1.6e-05 | 1.1e-05 | 1.333E-05 | 2.517E-06 | 1.100E-05 | 1.600E-05 |
| 20744 | 1038 | C1038D | C | 1e-05 | 1.3e-05 | 8.7e-06 | 1.057E-05 | 2.205E-06 | 8.700E-06 | 1.300E-05 |
| 20745 | 1038 | C1038C | C | 1.7e-05 | 1e-05 | 1.5e-05 | 1.400E-05 | 3.606E-06 | 1.000E-05 | 1.700E-05 |
| 20746 | 1038 | C1038E | C | 1.6e-05 | 1.1e-05 | 8.6e-06 | 1.187E-05 | 3.775E-06 | 8.600E-06 | 1.600E-05 |
| 20747 | 1038 | C1038Q | C | 9.2e-06 | 1e-05 | 1.1e-05 | 1.007E-05 | 9.018E-07 | 9.200E-06 | 1.100E-05 |
| 20748 | 1038 | C1038G | C | 1.1e-05 | 1.1e-05 | 1.7e-05 | 1.300E-05 | 3.464E-06 | 1.100E-05 | 1.700E-05 |
| 20749 | 1038 | C1038H | C | 1.6e-05 | 1.1e-05 | 9.8e-06 | 1.227E-05 | 3.288E-06 | 9.800E-06 | 1.600E-05 |
| 20750 | 1038 | C1038I | C | 8.7e-06 | 9.5e-06 | 1.1e-05 | 9.733E-06 | 1.168E-06 | 8.700E-06 | 1.100E-05 |
| 20751 | 1038 | C1038L | C | 1.4e-05 | 1.1e-05 | 9.7e-06 | 1.157E-05 | 2.205E-06 | 9.700E-06 | 1.400E-05 |
| 20752 | 1038 | C1038K | C | 1.3e-05 | 1.1e-05 | 9.3e-06 | 1.110E-05 | 1.852E-06 | 9.300E-06 | 1.300E-05 |
| 20753 | 1038 | C1038M | C | 1.6e-05 | 1.3e-05 | 9.9e-06 | 1.297E-05 | 3.050E-06 | 9.900E-06 | 1.600E-05 |
| 20754 | 1038 | C1038F | C | 1.5e-05 | 8e-06 | 8.3e-06 | 1.043E-05 | 3.958E-06 | 8.000E-06 | 1.500E-05 |
| 20755 | 1038 | C1038P | C | 9.6e-06 | 1.4e-05 | 1.5e-05 | 1.287E-05 | 2.873E-06 | 9.600E-06 | 1.500E-05 |
| 20756 | 1038 | C1038S | C | 1.6e-05 | 1e-05 | 1.1e-05 | 1.233E-05 | 3.215E-06 | 1.000E-05 | 1.600E-05 |
| 20757 | 1038 | C1038T | C | 1.6e-05 | 1.5e-05 | 1.6e-05 | 1.567E-05 | 5.774E-07 | 1.500E-05 | 1.600E-05 |
| 20758 | 1038 | C1038W | C | 1.7e-05 | 1.3e-05 | 9.9e-06 | 1.330E-05 | 3.559E-06 | 9.900E-06 | 1.700E-05 |
| 20759 | 1038 | C1038Y | C | 9.2e-06 | 1.5e-05 | 1e-05 | 1.140E-05 | 3.143E-06 | 9.200E-06 | 1.500E-05 |
| 20760 | 1038 | C1038V | C | 1.5e-05 | 1.4e-05 | 1.5e-05 | 1.467E-05 | 5.774E-07 | 1.400E-05 | 1.500E-05 |
| 20761 | 1039 | N1039A | C | 1.1e-05 | 1.5e-05 | 1.7e-05 | 1.433E-05 | 3.055E-06 | 1.100E-05 | 1.700E-05 |
| 20762 | 1039 | N1039R | C | 7.8e-06 | 1.7e-05 | 1.6e-05 | 1.360E-05 | 5.048E-06 | 7.800E-06 | 1.700E-05 |
| 20763 | 1039 | N1039N | C | 2.3e-05 | 1e-05 | 1.1e-05 | 1.467E-05 | 7.234E-06 | 1.000E-05 | 2.300E-05 |
| 20764 | 1039 | N1039D | C | 1.3e-05 | 1.6e-05 | 1.2e-05 | 1.367E-05 | 2.082E-06 | 1.200E-05 | 1.600E-05 |
| 20765 | 1039 | N1039C | C | 1e-05 | 9.8e-06 | 1.5e-05 | 1.160E-05 | 2.946E-06 | 9.800E-06 | 1.500E-05 |
| 20766 | 1039 | N1039E | C | 1.5e-05 | 1.2e-05 | 9e-06 | 1.200E-05 | 3.000E-06 | 9.000E-06 | 1.500E-05 |
| 20767 | 1039 | N1039Q | C | 1.1e-05 | 1.4e-05 | 1.5e-05 | 1.333E-05 | 2.082E-06 | 1.100E-05 | 1.500E-05 |
| 20768 | 1039 | N1039G | C | 9.3e-06 | 1.7e-05 | 1.1e-05 | 1.243E-05 | 4.045E-06 | 9.300E-06 | 1.700E-05 |
| 20769 | 1039 | N1039H | C | 9.3e-06 | 1.5e-05 | 1.1e-05 | 1.177E-05 | 2.926E-06 | 9.300E-06 | 1.500E-05 |
| 20770 | 1039 | N1039I | C | 1.5e-05 | 1.4e-05 | 9e-06 | 1.267E-05 | 3.215E-06 | 9.000E-06 | 1.500E-05 |
| 20771 | 1039 | N1039L | C | 1.4e-05 | 1.4e-05 | 1.8e-05 | 1.533E-05 | 2.309E-06 | 1.400E-05 | 1.800E-05 |
| 20772 | 1039 | N1039K | C | 1.2e-05 | 1.2e-05 | 1.6e-05 | 1.333E-05 | 2.309E-06 | 1.200E-05 | 1.600E-05 |
| 20773 | 1039 | N1039M | C | 1.5e-05 | 1.4e-05 | 1.6e-05 | 1.500E-05 | 1.000E-06 | 1.400E-05 | 1.600E-05 |
| 20774 | 1039 | N1039F | C | 1.3e-05 | 9.7e-06 | 8.2e-06 | 1.030E-05 | 2.456E-06 | 8.200E-06 | 1.300E-05 |
| 20775 | 1039 | N1039P | C | 1.5e-05 | 1.1e-05 | 8.2e-06 | 1.140E-05 | 3.418E-06 | 8.200E-06 | 1.500E-05 |
| 20776 | 1039 | N1039S | C | 1e-05 | 1.7e-05 | 1.1e-05 | 1.267E-05 | 3.786E-06 | 1.000E-05 | 1.700E-05 |
| 20777 | 1039 | N1039T | C | 1.6e-05 | 9.8e-06 | 1.5e-05 | 1.360E-05 | 3.329E-06 | 9.800E-06 | 1.600E-05 |
| 20778 | 1039 | N1039W | C | 9.3e-06 | 1.2e-05 | 1.6e-05 | 1.243E-05 | 3.371E-06 | 9.300E-06 | 1.600E-05 |
| 20779 | 1039 | N1039Y | C | 1.6e-05 | 1.5e-05 | 1.6e-05 | 1.567E-05 | 5.774E-07 | 1.500E-05 | 1.600E-05 |
| 20780 | 1039 | N1039V | C | 1.1e-05 | 1.4e-05 | 1.5e-05 | 1.333E-05 | 2.082E-06 | 1.100E-05 | 1.500E-05 |
| 20781 | 1040 | V1040A | C | 1.6e-05 | 1.1e-05 | 1.4e-05 | 1.367E-05 | 2.517E-06 | 1.100E-05 | 1.600E-05 |
| 20782 | 1040 | V1040R | C | 1.5e-05 | 1.1e-05 | 9e-06 | 1.167E-05 | 3.055E-06 | 9.000E-06 | 1.500E-05 |
| 20783 | 1040 | V1040N | C | 1.1e-05 | 9.6e-06 | 1.6e-05 | 1.220E-05 | 3.365E-06 | 9.600E-06 | 1.600E-05 |
| 20784 | 1040 | V1040D | C | 8.8e-06 | 1.5e-05 | 1.4e-05 | 1.260E-05 | 3.329E-06 | 8.800E-06 | 1.500E-05 |
| 20785 | 1040 | V1040C | C | 1.5e-05 | 1e-05 | 1e-05 | 1.167E-05 | 2.887E-06 | 1.000E-05 | 1.500E-05 |
| 20786 | 1040 | V1040E | C | 1.3e-05 | 1e-05 | 1.4e-05 | 1.233E-05 | 2.082E-06 | 1.000E-05 | 1.400E-05 |
| 20787 | 1040 | V1040Q | C | 1.4e-05 | 1.1e-05 | 1.3e-05 | 1.267E-05 | 1.528E-06 | 1.100E-05 | 1.400E-05 |
| 20788 | 1040 | V1040G | C | 8.8e-06 | 1.6e-05 | 1.4e-05 | 1.293E-05 | 3.717E-06 | 8.800E-06 | 1.600E-05 |
| 20789 | 1040 | V1040H | C | 1.5e-05 | 1.5e-05 | 8.8e-06 | 1.293E-05 | 3.580E-06 | 8.800E-06 | 1.500E-05 |
| 20790 | 1040 | V1040I | C | 1.7e-05 | 1.5e-05 | 1.6e-05 | 1.600E-05 | 1.000E-06 | 1.500E-05 | 1.700E-05 |
| 20791 | 1040 | V1040L | C | 1.3e-05 | 1.2e-05 | 6.9e-06 | 1.063E-05 | 3.272E-06 | 6.900E-06 | 1.300E-05 |
| 20792 | 1040 | V1040K | C | 1.5e-05 | 1.7e-05 | 1.5e-05 | 1.567E-05 | 1.155E-06 | 1.500E-05 | 1.700E-05 |
| 20793 | 1040 | V1040M | C | 1.5e-05 | 1.4e-05 | 1.1e-05 | 1.333E-05 | 2.082E-06 | 1.100E-05 | 1.500E-05 |
| 20794 | 1040 | V1040F | C | 9.6e-06 | 9.6e-06 | 1e-05 | 9.733E-06 | 2.309E-07 | 9.600E-06 | 1.000E-05 |
| 20795 | 1040 | V1040P | C | 1.2e-05 | 1.4e-05 | 1.4e-05 | 1.333E-05 | 1.155E-06 | 1.200E-05 | 1.400E-05 |
| 20796 | 1040 | V1040S | C | 8.1e-06 | 9.1e-06 | 1.1e-05 | 9.400E-06 | 1.473E-06 | 8.100E-06 | 1.100E-05 |
| 20797 | 1040 | V1040T | C | 1.6e-05 | 1.2e-05 | 1.5e-05 | 1.433E-05 | 2.082E-06 | 1.200E-05 | 1.600E-05 |
| 20798 | 1040 | V1040W | C | 9e-06 | 8.9e-06 | 1.4e-05 | 1.063E-05 | 2.916E-06 | 8.900E-06 | 1.400E-05 |
| 20799 | 1040 | V1040Y | C | 1.6e-05 | 1.2e-05 | 1.1e-05 | 1.300E-05 | 2.646E-06 | 1.100E-05 | 1.600E-05 |
| 20800 | 1040 | V1040V | C | 7.7e-06 | 1.7e-05 | 1.3e-05 | 1.257E-05 | 4.665E-06 | 7.700E-06 | 1.700E-05 |
| 20801 | 1041 | N1041A | C | 8.2e-06 | 1.1e-05 | 1.6e-05 | 1.173E-05 | 3.951E-06 | 8.200E-06 | 1.600E-05 |
| 20802 | 1041 | N1041R | C | 9.9e-06 | 1.6e-05 | 1.6e-05 | 1.397E-05 | 3.522E-06 | 9.900E-06 | 1.600E-05 |
| 20803 | 1041 | N1041N | C | 1.6e-05 | 8.5e-06 | 1.6e-05 | 1.350E-05 | 4.330E-06 | 8.500E-06 | 1.600E-05 |
| 20804 | 1041 | N1041D | C | 1.5e-05 | 1e-05 | 1.5e-05 | 1.333E-05 | 2.887E-06 | 1.000E-05 | 1.500E-05 |
| 20805 | 1041 | N1041C | C | 9.3e-06 | 1.5e-05 | 1.2e-05 | 1.210E-05 | 2.851E-06 | 9.300E-06 | 1.500E-05 |
| 20806 | 1041 | N1041E | C | 1.1e-05 | 7.5e-06 | 1.2e-05 | 1.017E-05 | 2.363E-06 | 7.500E-06 | 1.200E-05 |
| 20807 | 1041 | N1041Q | C | 1.5e-05 | 1.1e-05 | 1e-05 | 1.200E-05 | 2.646E-06 | 1.000E-05 | 1.500E-05 |
| 20808 | 1041 | N1041G | C | 1.1e-05 | 1.3e-05 | 1.5e-05 | 1.300E-05 | 2.000E-06 | 1.100E-05 | 1.500E-05 |
| 20809 | 1041 | N1041H | C | 1.5e-05 | 1.1e-05 | 9.1e-06 | 1.170E-05 | 3.012E-06 | 9.100E-06 | 1.500E-05 |
| 20810 | 1041 | N1041I | C | 1.3e-05 | 1.6e-05 | 1.4e-05 | 1.433E-05 | 1.528E-06 | 1.300E-05 | 1.600E-05 |
| 20811 | 1041 | N1041L | C | 1.6e-05 | 1.1e-05 | 1.4e-05 | 1.367E-05 | 2.517E-06 | 1.100E-05 | 1.600E-05 |
| 20812 | 1041 | N1041K | C | 1e-05 | 1.1e-05 | 1.4e-05 | 1.167E-05 | 2.082E-06 | 1.000E-05 | 1.400E-05 |
| 20813 | 1041 | N1041M | C | 1.5e-05 | 1.5e-05 | 1.3e-05 | 1.433E-05 | 1.155E-06 | 1.300E-05 | 1.500E-05 |
| 20814 | 1041 | N1041F | C | 1e-05 | 1.5e-05 | 1.5e-05 | 1.333E-05 | 2.887E-06 | 1.000E-05 | 1.500E-05 |
| 20815 | 1041 | N1041P | C | 8.3e-06 | 1.1e-05 | 1.6e-05 | 1.177E-05 | 3.907E-06 | 8.300E-06 | 1.600E-05 |
| 20816 | 1041 | N1041S | C | 1e-05 | 1.7e-05 | 1.4e-05 | 1.367E-05 | 3.512E-06 | 1.000E-05 | 1.700E-05 |
| 20817 | 1041 | N1041T | C | 1.4e-05 | 1.1e-05 | 1.4e-05 | 1.300E-05 | 1.732E-06 | 1.100E-05 | 1.400E-05 |
| 20818 | 1041 | N1041W | C | 1.5e-05 | 1.1e-05 | 1.4e-05 | 1.333E-05 | 2.082E-06 | 1.100E-05 | 1.500E-05 |
| 20819 | 1041 | N1041Y | C | 1.1e-05 | 1.6e-05 | 1.2e-05 | 1.300E-05 | 2.646E-06 | 1.100E-05 | 1.600E-05 |
| 20820 | 1041 | N1041V | C | 1.1e-05 | 1.6e-05 | 1.6e-05 | 1.433E-05 | 2.887E-06 | 1.100E-05 | 1.600E-05 |
| 20821 | 1042 | H1042A | C | 1.6e-05 | 1.4e-05 | 1.3e-05 | 1.433E-05 | 1.528E-06 | 1.300E-05 | 1.600E-05 |
| 20822 | 1042 | H1042R | C | 1e-05 | 1.4e-05 | 7.2e-06 | 1.040E-05 | 3.418E-06 | 7.200E-06 | 1.400E-05 |
| 20823 | 1042 | H1042N | C | 1e-05 | 8.2e-06 | 1.6e-05 | 1.140E-05 | 4.084E-06 | 8.200E-06 | 1.600E-05 |
| 20824 | 1042 | H1042D | C | 9e-06 | 1.4e-05 | 1.1e-05 | 1.133E-05 | 2.517E-06 | 9.000E-06 | 1.400E-05 |

| | | | | | | | | | | |
|---|---|---|---|---|---|---|---|---|---|---|
| 20825 | 1042 | H1042C | C | 5.9e-06 | 1.4e-05 | 8.2e-06 | 9.367E-06 | 4.174E-06 | 5.900E-06 | 1.400E-05 |
| 20826 | 1042 | H1042E | C | 1.6e-05 | 1e-05 | 1e-05 | 1.200E-05 | 3.464E-06 | 1.000E-05 | 1.600E-05 |
| 20827 | 1042 | H1042Q | C | 1.5e-05 | 1.6e-05 | 1e-05 | 1.367E-05 | 3.215E-06 | 1.000E-05 | 1.600E-05 |
| 20828 | 1042 | H1042G | C | 1.3e-05 | 1.1e-05 | 1.1e-05 | 1.167E-05 | 1.155E-06 | 1.100E-05 | 1.300E-05 |
| 20829 | 1042 | H1042H | C | 1.7e-05 | 1.6e-05 | 1.5e-05 | 1.600E-05 | 1.000E-06 | 1.500E-05 | 1.700E-05 |
| 20830 | 1042 | H1042I | C | 1.2e-05 | 9e-06 | 1.6e-05 | 1.233E-05 | 3.512E-06 | 9.000E-06 | 1.600E-05 |
| 20831 | 1042 | H1042L | C | 1.3e-05 | 1.4e-05 | 1.2e-05 | 1.300E-05 | 1.000E-06 | 1.200E-05 | 1.400E-05 |
| 20832 | 1042 | H1042K | C | 1.2e-05 | 1.4e-05 | 1.5e-05 | 1.367E-05 | 1.528E-06 | 1.200E-05 | 1.500E-05 |
| 20833 | 1042 | H1042M | C | 8.1e-06 | 1.5e-05 | 1.6e-05 | 1.303E-05 | 4.302E-06 | 8.100E-06 | 1.600E-05 |
| 20834 | 1042 | H1042F | C | 1.5e-05 | 1.5e-05 | 1.2e-05 | 1.400E-05 | 1.732E-06 | 1.200E-05 | 1.500E-05 |
| 20835 | 1042 | H1042P | C | 1.5e-05 | 1.2e-05 | 1.4e-05 | 1.367E-05 | 1.528E-06 | 1.200E-05 | 1.500E-05 |
| 20836 | 1042 | H1042S | C | 5.7e-06 | 1.6e-05 | 1e-05 | 1.057E-05 | 5.173E-06 | 5.700E-06 | 1.600E-05 |
| 20837 | 1042 | H1042T | C | 1.7e-05 | 9.6e-06 | 1.2e-05 | 1.287E-05 | 3.775E-06 | 9.600E-06 | 1.700E-05 |
| 20838 | 1042 | H1042W | C | 1.6e-05 | 9.7e-06 | 9.8e-06 | 1.183E-05 | 3.609E-06 | 9.700E-06 | 1.600E-05 |
| 20839 | 1042 | H1042Y | C | 1.5e-05 | 9.3e-06 | 1.6e-05 | 1.343E-05 | 3.614E-06 | 9.300E-06 | 1.600E-05 |
| 20840 | 1042 | H1042V | C | 1e-05 | 1.1e-05 | 1e-05 | 1.033E-05 | 5.774E-07 | 1.000E-05 | 1.100E-05 |
| 20841 | 1043 | K1043A | C | 1.1e-05 | 1.1e-05 | 1.2e-05 | 1.133E-05 | 5.774E-07 | 1.100E-05 | 1.200E-05 |
| 20842 | 1043 | K1043R | C | 1.6e-05 | 1.4e-05 | 1.3e-05 | 1.433E-05 | 1.528E-06 | 1.300E-05 | 1.600E-05 |
| 20843 | 1043 | K1043N | C | 9.7e-06 | 9.9e-06 | 3.1e-05 | 1.687E-05 | 1.224E-05 | 9.700E-06 | 3.100E-05 |
| 20844 | 1043 | K1043D | C | 8.1e-06 | 1.4e-05 | 1.7e-05 | 1.303E-05 | 4.528E-06 | 8.100E-06 | 1.700E-05 |
| 20845 | 1043 | K1043C | C | 1.1e-05 | 1e-05 | 1.5e-05 | 1.200E-05 | 2.646E-06 | 1.000E-05 | 1.500E-05 |
| 20846 | 1043 | K1043E | C | 1.4e-05 | 9.9e-06 | 1.5e-05 | 1.297E-05 | 2.702E-06 | 9.900E-06 | 1.500E-05 |
| 20847 | 1043 | K1043Q | C | 1.1e-05 | 1.5e-05 | 1.4e-05 | 1.333E-05 | 2.082E-06 | 1.100E-05 | 1.500E-05 |
| 20848 | 1043 | K1043G | C | 1.3e-05 | 8.1e-06 | 1.5e-05 | 1.203E-05 | 3.550E-06 | 8.100E-06 | 1.500E-05 |
| 20849 | 1043 | K1043H | C | 1e-05 | 1.4e-05 | 1.5e-05 | 1.300E-05 | 2.646E-06 | 1.000E-05 | 1.500E-05 |
| 20850 | 1043 | K1043I | C | 1.5e-05 | 1.1e-05 | 1.3e-05 | 1.300E-05 | 2.000E-06 | 1.100E-05 | 1.500E-05 |
| 20851 | 1043 | K1043L | C | 1.1e-05 | 1.5e-05 | 1e-05 | 1.200E-05 | 2.646E-06 | 1.000E-05 | 1.500E-05 |
| 20852 | 1043 | K1043K | C | 1.4e-05 | 9.9e-06 | 1e-05 | 1.130E-05 | 2.339E-06 | 9.900E-06 | 1.400E-05 |
| 20853 | 1043 | K1043M | C | 1.1e-05 | 1.3e-05 | 9.8e-06 | 1.127E-05 | 1.617E-06 | 9.800E-06 | 1.300E-05 |
| 20854 | 1043 | K1043F | C | 9.8e-06 | 1.5e-05 | 1e-05 | 1.160E-05 | 2.946E-06 | 9.800E-06 | 1.500E-05 |
| 20855 | 1043 | K1043P | C | 1.3e-05 | 1.6e-05 | 1.2e-05 | 1.367E-05 | 2.082E-06 | 1.200E-05 | 1.600E-05 |
| 20856 | 1043 | K1043S | C | 1.6e-05 | 1.5e-05 | 1.2e-05 | 1.433E-05 | 2.082E-06 | 1.200E-05 | 1.600E-05 |
| 20857 | 1043 | K1043T | C | 1.1e-05 | 1.1e-05 | 1.3e-05 | 1.167E-05 | 1.155E-06 | 1.100E-05 | 1.300E-05 |
| 20858 | 1043 | K1043W | C | 1.2e-05 | 1.2e-05 | 1.5e-05 | 1.300E-05 | 1.732E-06 | 1.200E-05 | 1.500E-05 |
| 20859 | 1043 | K1043Y | C | 1.1e-05 | 1.5e-05 | 1.5e-05 | 1.367E-05 | 2.309E-06 | 1.100E-05 | 1.500E-05 |
| 20860 | 1043 | K1043V | C | 1e-05 | 1.4e-05 | 1.6e-05 | 1.333E-05 | 3.055E-06 | 1.000E-05 | 1.600E-05 |
| 20861 | 1044 | P1044A | C | 1.5e-05 | 1.5e-05 | 1.1e-05 | 1.367E-05 | 2.309E-06 | 1.100E-05 | 1.500E-05 |
| 20862 | 1044 | P1044R | C | 9.5e-06 | 1e-05 | 9.2e-06 | 9.567E-06 | 4.041E-07 | 9.200E-06 | 1.000E-05 |
| 20863 | 1044 | P1044N | C | 1.1e-05 | 1e-05 | 9.5e-06 | 1.017E-05 | 7.638E-07 | 9.500E-06 | 1.100E-05 |
| 20864 | 1044 | P1044D | C | 1.2e-05 | 1.5e-05 | 1.6e-05 | 1.433E-05 | 2.082E-06 | 1.200E-05 | 1.600E-05 |
| 20865 | 1044 | P1044C | C | 1.5e-05 | 1e-05 | 1.4e-05 | 1.300E-05 | 2.646E-06 | 1.000E-05 | 1.500E-05 |
| 20866 | 1044 | P1044E | C | 1.1e-05 | 1.5e-05 | 1.7e-05 | 1.433E-05 | 3.055E-06 | 1.100E-05 | 1.700E-05 |
| 20867 | 1044 | P1044Q | C | 9.9e-06 | 1.2e-05 | 1.3e-05 | 1.163E-05 | 1.582E-06 | 9.900E-06 | 1.300E-05 |
| 20868 | 1044 | P1044G | C | 1.3e-05 | 1.5e-05 | 1.2e-05 | 1.333E-05 | 1.528E-06 | 1.200E-05 | 1.500E-05 |
| 20869 | 1044 | P1044H | C | 1.5e-05 | 1e-05 | 1e-05 | 1.167E-05 | 2.887E-06 | 1.000E-05 | 1.500E-05 |
| 20870 | 1044 | P1044I | C | 1e-05 | 1.5e-05 | 1.5e-05 | 1.333E-05 | 2.887E-06 | 1.000E-05 | 1.500E-05 |
| 20871 | 1044 | P1044L | C | 9.8e-06 | 1.3e-05 | 1.4e-05 | 1.227E-05 | 2.194E-06 | 9.800E-06 | 1.400E-05 |
| 20872 | 1044 | P1044K | C | 1.5e-05 | 9.7e-06 | 8.8e-06 | 1.117E-05 | 3.350E-06 | 8.800E-06 | 1.500E-05 |
| 20873 | 1044 | P1044M | C | 6.4e-06 | 8.9e-06 | 1.5e-05 | 1.010E-05 | 4.424E-06 | 6.400E-06 | 1.500E-05 |
| 20874 | 1044 | P1044F | C | 1e-05 | 1.5e-05 | 1.9e-05 | 1.467E-05 | 4.509E-06 | 1.000E-05 | 1.900E-05 |
| 20875 | 1044 | P1044P | C | 8.9e-06 | 1.2e-05 | 1.4e-05 | 1.163E-05 | 2.570E-06 | 8.900E-06 | 1.400E-05 |
| 20876 | 1044 | P1044S | C | 1.6e-05 | 1.2e-05 | 1.2e-05 | 1.333E-05 | 2.309E-06 | 1.200E-05 | 1.600E-05 |
| 20877 | 1044 | P1044T | C | 7e-06 | 1.6e-05 | 8.6e-06 | 1.053E-05 | 4.801E-06 | 7.000E-06 | 1.600E-05 |
| 20878 | 1044 | P1044W | C | 1.6e-05 | 8.6e-06 | 1e-05 | 1.153E-05 | 3.931E-06 | 8.600E-06 | 1.600E-05 |
| 20879 | 1044 | P1044Y | C | 1e-05 | 1.4e-05 | 7.5e-06 | 1.050E-05 | 3.279E-06 | 7.500E-06 | 1.400E-05 |
| 20880 | 1044 | P1044V | C | 8.2e-06 | 1.2e-05 | 1.6e-05 | 1.207E-05 | 3.900E-06 | 8.200E-06 | 1.600E-05 |
| 20881 | 1045 | S1045A | C | 1.5e-05 | 1.1e-05 | 1.1e-05 | 1.233E-05 | 2.309E-06 | 1.100E-05 | 1.500E-05 |
| 20882 | 1045 | S1045R | C | 1.1e-05 | 1.4e-05 | 1.5e-05 | 1.333E-05 | 2.082E-06 | 1.100E-05 | 1.500E-05 |
| 20883 | 1045 | S1045N | C | 1.2e-05 | 1.6e-05 | 1.1e-05 | 1.300E-05 | 2.646E-06 | 1.100E-05 | 1.600E-05 |
| 20884 | 1045 | S1045D | C | 1.5e-05 | 1.1e-05 | 1.5e-05 | 1.367E-05 | 2.309E-06 | 1.100E-05 | 1.500E-05 |
| 20885 | 1045 | S1045C | C | 1.1e-05 | 1.3e-05 | 1e-05 | 1.133E-05 | 1.528E-06 | 1.000E-05 | 1.300E-05 |
| 20886 | 1045 | S1045E | C | 1e-05 | 1.4e-05 | 1.3e-05 | 1.233E-05 | 2.082E-06 | 1.000E-05 | 1.400E-05 |
| 20887 | 1045 | S1045Q | C | 1.7e-05 | 1.3e-05 | 1.1e-05 | 1.367E-05 | 3.055E-06 | 1.100E-05 | 1.700E-05 |
| 20888 | 1045 | S1045G | C | 9.7e-06 | 1.1e-05 | 1.5e-05 | 1.190E-05 | 2.762E-06 | 9.700E-06 | 1.500E-05 |
| 20889 | 1045 | S1045H | C | 1.6e-05 | 1.1e-05 | 9.1e-06 | 1.203E-05 | 3.564E-06 | 9.100E-06 | 1.600E-05 |
| 20890 | 1045 | S1045I | C | 1.3e-05 | 1.6e-05 | 1.3e-05 | 1.400E-05 | 1.732E-06 | 1.300E-05 | 1.600E-05 |
| 20891 | 1045 | S1045L | C | 1.1e-05 | 1.5e-05 | 1.7e-05 | 1.433E-05 | 3.055E-06 | 1.100E-05 | 1.700E-05 |
| 20892 | 1045 | S1045K | C | 1.1e-05 | 9.7e-06 | 1.5e-05 | 1.190E-05 | 2.762E-06 | 9.700E-06 | 1.500E-05 |
| 20893 | 1045 | S1045M | C | 9.7e-06 | 1.5e-05 | 9.9e-06 | 1.153E-05 | 3.004E-06 | 9.700E-06 | 1.500E-05 |
| 20894 | 1045 | S1045F | C | 9.2e-06 | 9.7e-06 | 1.1e-05 | 9.967E-06 | 9.292E-07 | 9.200E-06 | 1.100E-05 |
| 20895 | 1045 | S1045P | C | 1.5e-05 | 1.5e-05 | 1.2e-05 | 1.400E-05 | 1.732E-06 | 1.200E-05 | 1.500E-05 |
| 20896 | 1045 | S1045S | C | 1.6e-05 | 1.1e-05 | 1.5e-05 | 1.400E-05 | 2.646E-06 | 1.100E-05 | 1.600E-05 |
| 20897 | 1045 | S1045T | C | 1.6e-05 | 7.3e-06 | 1.1e-05 | 1.143E-05 | 4.366E-06 | 7.300E-06 | 1.600E-05 |
| 20898 | 1045 | S1045W | C | 9.8e-06 | 9.4e-06 | 1e-05 | 9.733E-06 | 3.055E-07 | 9.400E-06 | 1.000E-05 |
| 20899 | 1045 | S1045Y | C | 1.1e-05 | 1.2e-05 | 1.6e-05 | 1.300E-05 | 2.646E-06 | 1.100E-05 | 1.600E-05 |
| 20900 | 1045 | S1045V | C | 1.5e-05 | 1e-05 | 1.3e-05 | 1.267E-05 | 2.517E-06 | 1.000E-05 | 1.500E-05 |
| 20901 | 1046 | N1046A | C | 1.5e-05 | 1.5e-05 | 1.1e-05 | 1.367E-05 | 2.309E-06 | 1.100E-05 | 1.500E-05 |
| 20902 | 1046 | N1046R | C | 1e-05 | 1.4e-05 | 1.5e-05 | 1.300E-05 | 2.646E-06 | 1.000E-05 | 1.500E-05 |
| 20903 | 1046 | N1046N | C | 7.2e-06 | 1.7e-05 | 1.5e-05 | 1.307E-05 | 5.178E-06 | 7.200E-06 | 1.700E-05 |
| 20904 | 1046 | N1046D | C | 1.1e-05 | 1.1e-05 | 1.5e-05 | 1.233E-05 | 2.309E-06 | 1.100E-05 | 1.500E-05 |
| 20905 | 1046 | N1046C | C | 2.7e-05 | 1.1e-05 | 1.6e-05 | 1.800E-05 | 8.185E-06 | 1.100E-05 | 2.700E-05 |
| 20906 | 1046 | N1046E | C | 1.5e-05 | 1.1e-05 | 9.8e-06 | 1.193E-05 | 2.723E-06 | 9.800E-06 | 1.500E-05 |
| 20907 | 1046 | N1046Q | C | 1.5e-05 | 8e-06 | 1.2e-05 | 1.167E-05 | 3.512E-06 | 8.000E-06 | 1.500E-05 |
| 20908 | 1046 | N1046G | C | 1.3e-05 | 1.6e-05 | 1.1e-05 | 1.333E-05 | 2.517E-06 | 1.100E-05 | 1.600E-05 |
| 20909 | 1046 | N1046H | C | 2.2e-05 | 1.1e-05 | 1.6e-05 | 1.633E-05 | 5.508E-06 | 1.100E-05 | 2.200E-05 |

| | | | | | | | | | | |
|---|---|---|---|---|---|---|---|---|---|---|
| 20910 | 1046 | N1046I | C | 1.7e-05 | 1.5e-05 | 1.1e-05 | 1.433E-05 | 3.055E-06 | 1.100E-05 | 1.700E-05 |
| 20911 | 1046 | N1046L | C | 1.5e-05 | 1.6e-05 | 1.6e-05 | 1.567E-05 | 5.774E-07 | 1.500E-05 | 1.600E-05 |
| 20912 | 1046 | N1046K | C | 1.4e-05 | 1.7e-05 | 8.4e-06 | 1.313E-05 | 4.365E-06 | 8.400E-06 | 1.700E-05 |
| 20913 | 1046 | N1046M | C | 1.5e-05 | 9.6e-06 | 1.1e-05 | 1.187E-05 | 2.802E-06 | 9.600E-06 | 1.500E-05 |
| 20914 | 1046 | N1046F | C | 1.8e-05 | 1.4e-05 | 8e-06 | 1.333E-05 | 5.033E-06 | 8.000E-06 | 1.800E-05 |
| 20915 | 1046 | N1046P | C | 9.5e-06 | 9.6e-06 | 1.6e-05 | 1.170E-05 | 3.724E-06 | 9.500E-06 | 1.600E-05 |
| 20916 | 1046 | N1046S | C | 1.1e-05 | 1.4e-05 | 9.5e-06 | 1.150E-05 | 2.291E-06 | 9.500E-06 | 1.400E-05 |
| 20917 | 1046 | N1046T | C | 1.4e-05 | 1.1e-05 | 1.4e-05 | 1.300E-05 | 1.732E-06 | 1.100E-05 | 1.400E-05 |
| 20918 | 1046 | N1046W | C | 1.7e-05 | 9.8e-06 | 9.3e-06 | 1.203E-05 | 4.309E-06 | 9.300E-06 | 1.700E-05 |
| 20919 | 1046 | N1046Y | C | 1.5e-05 | 8.1e-06 | 1.4e-05 | 1.237E-05 | 3.729E-06 | 8.100E-06 | 1.500E-05 |
| 20920 | 1046 | N1046V | C | 1.4e-05 | 1.1e-05 | 1.3e-05 | 1.267E-05 | 1.528E-06 | 1.100E-05 | 1.400E-05 |
| 20921 | 1047 | T1047A | C | 1.6e-05 | 9.6e-06 | 1.6e-05 | 1.387E-05 | 3.695E-06 | 9.600E-06 | 1.600E-05 |
| 20922 | 1047 | T1047R | C | 7.9e-06 | 1.2e-05 | 1.6e-05 | 1.197E-05 | 4.050E-06 | 7.900E-06 | 1.600E-05 |
| 20923 | 1047 | T1047N | C | 1.7e-05 | 9.5e-06 | 1.3e-05 | 1.317E-05 | 3.753E-06 | 9.500E-06 | 1.700E-05 |
| 20924 | 1047 | T1047D | C | 1.1e-05 | 1.1e-05 | 1.5e-05 | 1.233E-05 | 2.309E-06 | 1.100E-05 | 1.500E-05 |
| 20925 | 1047 | T1047C | C | 8.6e-06 | 1e-05 | 1.1e-05 | 9.867E-06 | 1.206E-06 | 8.600E-06 | 1.100E-05 |
| 20926 | 1047 | T1047E | C | 1e-05 | 1.4e-05 | 8.2e-06 | 1.073E-05 | 2.969E-06 | 8.200E-06 | 1.400E-05 |
| 20927 | 1047 | T1047Q | C | 1.1e-05 | 4.6e-06 | 1.3e-05 | 9.533E-06 | 4.388E-06 | 4.600E-06 | 1.300E-05 |
| 20928 | 1047 | T1047G | C | 9.6e-06 | 1.4e-05 | 1.4e-05 | 1.253E-05 | 2.540E-06 | 9.600E-06 | 1.400E-05 |
| 20929 | 1047 | T1047H | C | 6.4e-06 | 9.2e-06 | 1.1e-05 | 8.867E-06 | 2.318E-06 | 6.400E-06 | 1.100E-05 |
| 20930 | 1047 | T1047I | C | 1.1e-05 | 1.7e-05 | 9.3e-06 | 1.243E-05 | 4.045E-06 | 9.300E-06 | 1.700E-05 |
| 20931 | 1047 | T1047L | C | 1.2e-05 | 1.4e-05 | 1.4e-05 | 1.333E-05 | 1.155E-06 | 1.200E-05 | 1.400E-05 |
| 20932 | 1047 | T1047K | C | 9.6e-06 | 1.5e-05 | 9.3e-06 | 1.130E-05 | 3.208E-06 | 9.300E-06 | 1.500E-05 |
| 20933 | 1047 | T1047M | C | 9.3e-06 | 9.6e-06 | 1.6e-05 | 1.163E-05 | 3.785E-06 | 9.300E-06 | 1.600E-05 |
| 20934 | 1047 | T1047F | C | 9.6e-06 | 9.5e-06 | 1.1e-05 | 1.003E-05 | 8.386E-07 | 9.500E-06 | 1.100E-05 |
| 20935 | 1047 | T1047P | C | 1.6e-05 | 9.7e-06 | 1.1e-05 | 1.223E-05 | 3.326E-06 | 9.700E-06 | 1.600E-05 |
| 20936 | 1047 | T1047S | C | 1.1e-05 | 1.2e-05 | 6.6e-06 | 9.867E-06 | 2.873E-06 | 6.600E-06 | 1.200E-05 |
| 20937 | 1047 | T1047T | C | 8.9e-06 | 1.5e-05 | 1.2e-05 | 1.197E-05 | 3.050E-06 | 8.900E-06 | 1.500E-05 |
| 20938 | 1047 | T1047W | C | 1.5e-05 | 1.1e-05 | 1.4e-05 | 1.333E-05 | 2.082E-06 | 1.100E-05 | 1.500E-05 |
| 20939 | 1047 | T1047Y | C | 1.1e-05 | 1.1e-05 | 1.1e-05 | 1.100E-05 | 2.075E-21 | 1.100E-05 | 1.100E-05 |
| 20940 | 1047 | T1047V | C | 1e-05 | 1.5e-05 | 8.9e-06 | 1.130E-05 | 3.251E-06 | 8.900E-06 | 1.500E-05 |
| 20941 | 1048 | K1048A | C | 1.1e-05 | 8.1e-06 | 8.7e-06 | 9.267E-06 | 1.531E-06 | 8.100E-06 | 1.100E-05 |
| 20942 | 1048 | K1048R | C | 1e-05 | 1.1e-05 | 1.7e-05 | 1.267E-05 | 3.786E-06 | 1.000E-05 | 1.700E-05 |
| 20943 | 1048 | K1048N | C | 1.4e-05 | 1.2e-05 | 1.7e-05 | 1.433E-05 | 2.517E-06 | 1.200E-05 | 1.700E-05 |
| 20944 | 1048 | K1048D | C | 1.3e-05 | 1.3e-05 | 1.1e-05 | 1.233E-05 | 1.155E-06 | 1.100E-05 | 1.300E-05 |
| 20945 | 1048 | K1048C | C | 2.7e-05 | 1.6e-05 | 1.1e-05 | 1.800E-05 | 8.185E-06 | 1.100E-05 | 2.700E-05 |
| 20946 | 1048 | K1048E | C | 1.1e-05 | 1.6e-05 | 1.5e-05 | 1.400E-05 | 2.646E-06 | 1.100E-05 | 1.600E-05 |
| 20947 | 1048 | K1048Q | C | 1.3e-05 | 1.5e-05 | 1.4e-05 | 1.400E-05 | 1.000E-06 | 1.300E-05 | 1.500E-05 |
| 20948 | 1048 | K1048G | C | 1.1e-05 | 1.5e-05 | 1.5e-05 | 1.367E-05 | 2.309E-06 | 1.100E-05 | 1.500E-05 |
| 20949 | 1048 | K1048H | C | 1.4e-05 | 1.7e-05 | 6.8e-06 | 1.260E-05 | 5.242E-06 | 6.800E-06 | 1.700E-05 |
| 20950 | 1048 | K1048I | C | 1.1e-05 | 1.7e-05 | 1.6e-05 | 1.467E-05 | 3.215E-06 | 1.100E-05 | 1.700E-05 |
| 20951 | 1048 | K1048L | C | 1e-05 | 1.5e-05 | 1.6e-05 | 1.367E-05 | 3.215E-06 | 1.000E-05 | 1.600E-05 |
| 20952 | 1048 | K1048K | C | 9.8e-06 | 1e-05 | 1.2e-05 | 1.060E-05 | 1.217E-06 | 9.800E-06 | 1.200E-05 |
| 20953 | 1048 | K1048M | C | 1.6e-05 | 1.6e-05 | 1.7e-05 | 1.633E-05 | 5.774E-07 | 1.600E-05 | 1.700E-05 |
| 20954 | 1048 | K1048F | C | 9.5e-06 | 1.4e-05 | 1.1e-05 | 1.150E-05 | 2.291E-06 | 9.500E-06 | 1.400E-05 |
| 20955 | 1048 | K1048P | C | 1.1e-05 | 1.2e-05 | 1e-05 | 1.100E-05 | 1.000E-06 | 1.000E-05 | 1.200E-05 |
| 20956 | 1048 | K1048S | C | 1.5e-05 | 1.7e-05 | 1.6e-05 | 1.600E-05 | 1.000E-06 | 1.500E-05 | 1.700E-05 |
| 20957 | 1048 | K1048T | C | 9.8e-06 | 1.5e-05 | 1.1e-05 | 1.193E-05 | 2.723E-06 | 9.800E-06 | 1.500E-05 |
| 20958 | 1048 | K1048W | C | 7.8e-06 | 1.5e-05 | 1.5e-05 | 1.260E-05 | 4.157E-06 | 7.800E-06 | 1.500E-05 |
| 20959 | 1048 | K1048Y | C | 1.4e-05 | 1e-05 | 1.2e-05 | 1.200E-05 | 2.000E-06 | 1.000E-05 | 1.400E-05 |
| 20960 | 1048 | K1048V | C | 1.4e-05 | 1.1e-05 | 1.5e-05 | 1.333E-05 | 2.082E-06 | 1.100E-05 | 1.500E-05 |
| 20961 | 1049 | V1049A | C | 1e-05 | 1.7e-05 | 1.6e-05 | 1.433E-05 | 3.786E-06 | 1.000E-05 | 1.700E-05 |
| 20962 | 1049 | V1049R | C | 1.3e-05 | 1.4e-05 | 1.2e-05 | 1.300E-05 | 1.000E-06 | 1.200E-05 | 1.400E-05 |
| 20963 | 1049 | V1049N | C | 8.6e-06 | 1.4e-05 | 1e-05 | 1.087E-05 | 2.802E-06 | 8.600E-06 | 1.400E-05 |
| 20964 | 1049 | V1049D | C | 1.1e-05 | 9.6e-06 | 1.6e-05 | 1.220E-05 | 3.365E-06 | 9.600E-06 | 1.600E-05 |
| 20965 | 1049 | V1049C | C | 1.2e-05 | 9.6e-06 | 1.4e-05 | 1.187E-05 | 2.203E-06 | 9.600E-06 | 1.400E-05 |
| 20966 | 1049 | V1049E | C | 9.9e-06 | 1.4e-05 | 1.3e-05 | 1.230E-05 | 2.138E-06 | 9.900E-06 | 1.400E-05 |
| 20967 | 1049 | V1049Q | C | 1.5e-05 | 1.4e-05 | 1e-05 | 1.300E-05 | 2.646E-06 | 1.000E-05 | 1.500E-05 |
| 20968 | 1049 | V1049G | C | 1.7e-05 | 1.4e-05 | 1.3e-05 | 1.467E-05 | 2.082E-06 | 1.300E-05 | 1.700E-05 |
| 20969 | 1049 | V1049H | C | 1e-05 | 1.1e-05 | 1.5e-05 | 1.200E-05 | 2.646E-06 | 1.000E-05 | 1.500E-05 |
| 20970 | 1049 | V1049I | C | 1e-05 | 1e-05 | 1e-05 | 1.000E-05 | 0.000E+00 | 1.000E-05 | 1.000E-05 |
| 20971 | 1049 | V1049L | C | 1e-05 | 9.6e-06 | 1e-05 | 9.867E-06 | 2.309E-07 | 9.600E-06 | 1.000E-05 |
| 20972 | 1049 | V1049K | C | 1.5e-05 | 1.3e-05 | 1.4e-05 | 1.400E-05 | 1.000E-06 | 1.300E-05 | 1.500E-05 |
| 20973 | 1049 | V1049M | C | 1.4e-05 | 1.4e-05 | 1.1e-05 | 1.300E-05 | 1.732E-06 | 1.100E-05 | 1.400E-05 |
| 20974 | 1049 | V1049F | C | 1e-05 | 7.5e-06 | 1.3e-05 | 1.017E-05 | 2.754E-06 | 7.500E-06 | 1.300E-05 |
| 20975 | 1049 | V1049P | C | 1.5e-05 | 1.5e-05 | 1e-05 | 1.333E-05 | 2.887E-06 | 1.000E-05 | 1.500E-05 |
| 20976 | 1049 | V1049S | C | 9.8e-06 | 9.1e-06 | 1.3e-05 | 1.063E-05 | 2.079E-06 | 9.100E-06 | 1.300E-05 |
| 20977 | 1049 | V1049T | C | 1.5e-05 | 1.1e-05 | 1e-05 | 1.200E-05 | 2.646E-06 | 1.000E-05 | 1.500E-05 |
| 20978 | 1049 | V1049W | C | 1.6e-05 | 1.2e-05 | 1.3e-05 | 1.367E-05 | 2.082E-06 | 1.200E-05 | 1.600E-05 |
| 20979 | 1049 | V1049Y | C | 1.4e-05 | 1e-05 | 1.6e-05 | 1.333E-05 | 3.055E-06 | 1.000E-05 | 1.600E-05 |
| 20980 | 1049 | V1049V | C | 1.2e-05 | 1.5e-05 | 1.5e-05 | 1.400E-05 | 1.732E-06 | 1.200E-05 | 1.500E-05 |
| 20981 | 1050 | D1050A | C | 1.1e-05 | 1.1e-05 | 1.2e-05 | 1.133E-05 | 5.774E-07 | 1.100E-05 | 1.200E-05 |
| 20982 | 1050 | D1050R | C | 1.6e-05 | 1.6e-05 | 1.4e-05 | 1.533E-05 | 1.155E-06 | 1.400E-05 | 1.600E-05 |
| 20983 | 1050 | D1050N | C | 1.6e-05 | 1.4e-05 | 7.2e-06 | 1.240E-05 | 4.613E-06 | 7.200E-06 | 1.600E-05 |
| 20984 | 1050 | D1050D | C | 1.5e-05 | 1.6e-05 | 1.5e-05 | 1.533E-05 | 5.774E-07 | 1.500E-05 | 1.600E-05 |
| 20985 | 1050 | D1050C | C | 9.3e-06 | 1.6e-05 | 1.4e-05 | 1.310E-05 | 3.439E-06 | 9.300E-06 | 1.600E-05 |
| 20986 | 1050 | D1050E | C | 1.4e-05 | 1.7e-05 | 1.1e-05 | 1.400E-05 | 3.000E-06 | 1.100E-05 | 1.700E-05 |
| 20987 | 1050 | D1050Q | C | 1.1e-05 | 8e-06 | 9.9e-06 | 9.633E-06 | 1.518E-06 | 8.000E-06 | 1.100E-05 |
| 20988 | 1050 | D1050G | C | 1.3e-05 | 1.5e-05 | 9.5e-06 | 1.250E-05 | 2.784E-06 | 9.500E-06 | 1.500E-05 |
| 20989 | 1050 | D1050H | C | 1.6e-05 | 1e-05 | 9.8e-06 | 1.193E-05 | 3.523E-06 | 9.800E-06 | 1.600E-05 |
| 20990 | 1050 | D1050I | C | 1.7e-05 | 8.8e-06 | 1.1e-05 | 1.227E-05 | 4.244E-06 | 8.800E-06 | 1.700E-05 |
| 20991 | 1050 | D1050L | C | 1.2e-05 | 1.5e-05 | 1.3e-05 | 1.333E-05 | 1.528E-06 | 1.200E-05 | 1.500E-05 |
| 20992 | 1050 | D1050K | C | 1.2e-05 | 1.5e-05 | 8e-06 | 1.167E-05 | 3.512E-06 | 8.000E-06 | 1.500E-05 |
| 20993 | 1050 | D1050M | C | 1.5e-05 | 1e-05 | 1.5e-05 | 1.333E-05 | 2.887E-06 | 1.000E-05 | 1.500E-05 |
| 20994 | 1050 | D1050F | C | 8.9e-06 | 1.5e-05 | 1.3e-05 | 1.230E-05 | 3.110E-06 | 8.900E-06 | 1.500E-05 |

| | | | | | | | | | | |
|---|---|---|---|---|---|---|---|---|---|---|
| 20995 | 1050 | D1050P | C | 1.6e-05 | 1.5e-05 | 1.5e-05 | 1.533E-05 | 5.774E-07 | 1.500E-05 | 1.600E-05 |
| 20996 | 1050 | D1050S | C | 1.7e-05 | 1.6e-05 | 8.9e-06 | 1.397E-05 | 4.416E-06 | 8.900E-06 | 1.700E-05 |
| 20997 | 1050 | D1050T | C | 9.7e-06 | 1e-05 | 1.5e-05 | 1.157E-05 | 2.977E-06 | 9.700E-06 | 1.500E-05 |
| 20998 | 1050 | D1050W | C | 9.9e-06 | 1e-05 | 1.4e-05 | 1.130E-05 | 2.339E-06 | 9.900E-06 | 1.400E-05 |
| 20999 | 1050 | D1050Y | C | 1.2e-05 | 1.5e-05 | 1.1e-05 | 1.267E-05 | 2.082E-06 | 1.100E-05 | 1.500E-05 |
| 21000 | 1050 | D1050V | C | 1.7e-05 | 1e-05 | 1.1e-05 | 1.267E-05 | 3.786E-06 | 1.000E-05 | 1.700E-05 |
| 21001 | 1051 | K1051A | C | 1.6e-05 | 1.3e-05 | 1.5e-05 | 1.467E-05 | 1.528E-06 | 1.300E-05 | 1.600E-05 |
| 21002 | 1051 | K1051R | C | 1.6e-05 | 1.2e-05 | 1.2e-05 | 1.333E-05 | 2.309E-06 | 1.200E-05 | 1.600E-05 |
| 21003 | 1051 | K1051N | C | 1e-05 | 1.4e-05 | 1.1e-05 | 1.167E-05 | 2.082E-06 | 1.000E-05 | 1.400E-05 |
| 21004 | 1051 | K1051D | C | 1.5e-05 | 1.3e-05 | 1.1e-05 | 1.300E-05 | 2.000E-06 | 1.100E-05 | 1.500E-05 |
| 21005 | 1051 | K1051C | C | 1.2e-05 | 1.4e-05 | 1.3e-05 | 1.300E-05 | 1.000E-06 | 1.200E-05 | 1.400E-05 |
| 21006 | 1051 | K1051E | C | 8.6e-06 | 1.5e-05 | 1.3e-05 | 1.220E-05 | 3.274E-06 | 8.600E-06 | 1.500E-05 |
| 21007 | 1051 | K1051Q | C | 9.9e-06 | 1.4e-05 | 9.6e-06 | 1.117E-05 | 2.458E-06 | 9.600E-06 | 1.400E-05 |
| 21008 | 1051 | K1051G | C | 7.1e-06 | 1.1e-05 | 8e-06 | 8.700E-06 | 2.042E-06 | 7.100E-06 | 1.100E-05 |
| 21009 | 1051 | K1051H | C | 1.5e-05 | 1e-05 | 1.4e-05 | 1.300E-05 | 2.646E-06 | 1.000E-05 | 1.500E-05 |
| 21010 | 1051 | K1051I | C | 1.5e-05 | 1.4e-05 | 1.6e-05 | 1.500E-05 | 1.000E-06 | 1.400E-05 | 1.600E-05 |
| 21011 | 1051 | K1051L | C | 9.9e-06 | 1.5e-05 | 1.3e-05 | 1.263E-05 | 2.570E-06 | 9.900E-06 | 1.500E-05 |
| 21012 | 1051 | K1051K | C | 1.1e-05 | 1e-05 | 1e-05 | 1.033E-05 | 5.774E-07 | 1.000E-05 | 1.100E-05 |
| 21013 | 1051 | K1051M | C | 1.5e-05 | 1.4e-05 | 8.7e-06 | 1.257E-05 | 3.386E-06 | 8.700E-06 | 1.500E-05 |
| 21014 | 1051 | K1051F | C | 1.1e-05 | 1.5e-05 | 1.4e-05 | 1.333E-05 | 2.082E-06 | 1.100E-05 | 1.500E-05 |
| 21015 | 1051 | K1051P | C | 1.2e-05 | 1.1e-05 | 1e-05 | 1.100E-05 | 1.000E-06 | 1.000E-05 | 1.200E-05 |
| 21016 | 1051 | K1051S | C | 1.4e-05 | 1.6e-05 | 1.3e-05 | 1.433E-05 | 1.528E-06 | 1.300E-05 | 1.600E-05 |
| 21017 | 1051 | K1051T | C | 1.1e-05 | 1.5e-05 | 1.4e-05 | 1.333E-05 | 2.082E-06 | 1.100E-05 | 1.500E-05 |
| 21018 | 1051 | K1051W | C | 9.1e-06 | 1.4e-05 | 1.2e-05 | 1.170E-05 | 2.464E-06 | 9.100E-06 | 1.400E-05 |
| 21019 | 1051 | K1051Y | C | 1.4e-05 | 9.4e-06 | 1.6e-05 | 1.313E-05 | 3.384E-06 | 9.400E-06 | 1.600E-05 |
| 21020 | 1051 | K1051V | C | 1.6e-05 | 1.7e-05 | 1.6e-05 | 1.633E-05 | 5.774E-07 | 1.600E-05 | 1.700E-05 |
| 21021 | 1052 | K1052A | C | 1.2e-05 | 8.6e-06 | 1.5e-05 | 1.187E-05 | 3.202E-06 | 8.600E-06 | 1.500E-05 |
| 21022 | 1052 | K1052R | C | 1.4e-05 | 1.2e-05 | 1.5e-05 | 1.367E-05 | 1.528E-06 | 1.200E-05 | 1.500E-05 |
| 21023 | 1052 | K1052N | C | 1.5e-05 | 9.8e-06 | 1.4e-05 | 1.293E-05 | 2.759E-06 | 9.800E-06 | 1.500E-05 |
| 21024 | 1052 | K1052D | C | 1e-05 | 1.1e-05 | 9.7e-06 | 1.023E-05 | 6.807E-07 | 9.700E-06 | 1.100E-05 |
| 21025 | 1052 | K1052C | C | 9.9e-06 | 1.5e-05 | 1.5e-05 | 1.330E-05 | 2.944E-06 | 9.900E-06 | 1.500E-05 |
| 21026 | 1052 | K1052E | C | 1e-05 | 9.1e-06 | 1.5e-05 | 1.137E-05 | 3.179E-06 | 9.100E-06 | 1.500E-05 |
| 21027 | 1052 | K1052Q | C | 1e-05 | 1.4e-05 | 1.4e-05 | 1.267E-05 | 2.309E-06 | 1.000E-05 | 1.400E-05 |
| 21028 | 1052 | K1052G | C | 1e-05 | 1.3e-05 | 1.2e-05 | 1.167E-05 | 1.528E-06 | 1.000E-05 | 1.300E-05 |
| 21029 | 1052 | K1052H | C | 7.5e-06 | 1.5e-05 | 1.5e-05 | 1.250E-05 | 4.330E-06 | 7.500E-06 | 1.500E-05 |
| 21030 | 1052 | K1052I | C | 1.5e-05 | 1.1e-05 | 1e-05 | 1.200E-05 | 2.646E-06 | 1.000E-05 | 1.500E-05 |
| 21031 | 1052 | K1052L | C | 1.6e-05 | 1.3e-05 | 1.3e-05 | 1.400E-05 | 1.732E-06 | 1.300E-05 | 1.600E-05 |
| 21032 | 1052 | K1052K | C | 9.2e-06 | 1.4e-05 | 1.6e-05 | 1.307E-05 | 3.495E-06 | 9.200E-06 | 1.600E-05 |
| 21033 | 1052 | K1052M | C | 1.4e-05 | 1.1e-05 | 1.8e-05 | 1.433E-05 | 3.512E-06 | 1.100E-05 | 1.800E-05 |
| 21034 | 1052 | K1052F | C | 1.1e-05 | 1.6e-05 | 1e-05 | 1.233E-05 | 3.215E-06 | 1.000E-05 | 1.600E-05 |
| 21035 | 1052 | K1052P | C | 9.3e-06 | 9.2e-06 | 1.5e-05 | 1.117E-05 | 3.320E-06 | 9.200E-06 | 1.500E-05 |
| 21036 | 1052 | K1052S | C | 1.5e-05 | 1.8e-05 | 1.1e-05 | 1.467E-05 | 3.512E-06 | 1.100E-05 | 1.800E-05 |
| 21037 | 1052 | K1052T | C | 1.3e-05 | 9.6e-06 | 1.1e-05 | 1.120E-05 | 1.709E-06 | 9.600E-06 | 1.300E-05 |
| 21038 | 1052 | K1052W | C | 1.1e-05 | 9.6e-06 | 1.3e-05 | 1.120E-05 | 1.709E-06 | 9.600E-06 | 1.300E-05 |
| 21039 | 1052 | K1052Y | C | 1.7e-05 | 1.1e-05 | 1.4e-05 | 1.400E-05 | 3.000E-06 | 1.100E-05 | 1.700E-05 |
| 21040 | 1052 | K1052V | C | 1.6e-05 | 1e-05 | 1.6e-05 | 1.400E-05 | 3.464E-06 | 1.000E-05 | 1.600E-05 |
| 21041 | 1053 | V1053A | C | 1.1e-05 | 1.4e-05 | 1.2e-05 | 1.233E-05 | 1.528E-06 | 1.100E-05 | 1.400E-05 |
| 21042 | 1053 | V1053R | C | 1.4e-05 | 1.5e-05 | 1.5e-05 | 1.467E-05 | 5.774E-07 | 1.400E-05 | 1.500E-05 |
| 21043 | 1053 | V1053N | C | 1.6e-05 | 1e-05 | 1e-05 | 1.200E-05 | 3.464E-06 | 1.000E-05 | 1.600E-05 |
| 21044 | 1053 | V1053D | C | 6.9e-06 | 1.4e-05 | 1.4e-05 | 1.163E-05 | 4.099E-06 | 6.900E-06 | 1.400E-05 |
| 21045 | 1053 | V1053C | C | 1.6e-05 | 8.9e-06 | 1e-05 | 1.163E-05 | 3.821E-06 | 8.900E-06 | 1.600E-05 |
| 21046 | 1053 | V1053E | C | 1.4e-05 | 9.4e-06 | 1.4e-05 | 1.247E-05 | 2.656E-06 | 9.400E-06 | 1.400E-05 |
| 21047 | 1053 | V1053Q | C | 9.4e-06 | 1.4e-05 | 1.5e-05 | 1.280E-05 | 2.987E-06 | 9.400E-06 | 1.500E-05 |
| 21048 | 1053 | V1053G | C | 9.7e-06 | 9.4e-06 | 1.2e-05 | 1.037E-05 | 1.422E-06 | 9.400E-06 | 1.200E-05 |
| 21049 | 1053 | V1053H | C | 8.4e-06 | 1.9e-05 | 1.4e-05 | 1.380E-05 | 5.303E-06 | 8.400E-06 | 1.900E-05 |
| 21050 | 1053 | V1053I | C | 1.3e-05 | 1.1e-05 | 1.7e-05 | 1.367E-05 | 3.055E-06 | 1.100E-05 | 1.700E-05 |
| 21051 | 1053 | V1053L | C | 1.5e-05 | 1.5e-05 | 1.1e-05 | 1.367E-05 | 2.309E-06 | 1.100E-05 | 1.500E-05 |
| 21052 | 1053 | V1053K | C | 1.6e-05 | 6.8e-06 | 1.1e-05 | 1.127E-05 | 4.606E-06 | 6.800E-06 | 1.600E-05 |
| 21053 | 1053 | V1053M | C | 1.1e-05 | 1e-05 | 9.3e-06 | 1.010E-05 | 8.544E-07 | 9.300E-06 | 1.100E-05 |
| 21054 | 1053 | V1053F | C | 1.3e-05 | 1.5e-05 | 7.9e-06 | 1.197E-05 | 3.661E-06 | 7.900E-06 | 1.500E-05 |
| 21055 | 1053 | V1053P | C | 1.5e-05 | 1.5e-05 | 1.4e-05 | 1.467E-05 | 5.774E-07 | 1.400E-05 | 1.500E-05 |
| 21056 | 1053 | V1053S | C | 1.3e-05 | 9.8e-06 | 1.1e-05 | 1.127E-05 | 1.617E-06 | 9.800E-06 | 1.300E-05 |
| 21057 | 1053 | V1053T | C | 1.4e-05 | 1.6e-05 | 1.5e-05 | 1.500E-05 | 1.000E-06 | 1.400E-05 | 1.600E-05 |
| 21058 | 1053 | V1053W | C | 1.3e-05 | 1e-05 | 1.5e-05 | 1.267E-05 | 2.517E-06 | 1.000E-05 | 1.500E-05 |
| 21059 | 1053 | V1053Y | C | 1.1e-05 | 1.4e-05 | 9.4e-06 | 1.147E-05 | 2.335E-06 | 9.400E-06 | 1.400E-05 |
| 21060 | 1053 | V1053V | C | 1e-05 | 1.1e-05 | 1.2e-05 | 1.100E-05 | 1.000E-06 | 1.000E-05 | 1.200E-05 |
| 21061 | 1054 | E1054A | C | 1.5e-05 | 1.7e-05 | 1.6e-05 | 1.600E-05 | 1.000E-06 | 1.500E-05 | 1.700E-05 |
| 21062 | 1054 | E1054R | C | 1.5e-05 | 1.5e-05 | 1.6e-05 | 1.533E-05 | 5.774E-07 | 1.500E-05 | 1.600E-05 |
| 21063 | 1054 | E1054N | C | 1.1e-05 | 1.4e-05 | 1.1e-05 | 1.200E-05 | 1.732E-06 | 1.100E-05 | 1.400E-05 |
| 21064 | 1054 | E1054D | C | 1.7e-05 | 9.8e-06 | 9.8e-06 | 1.220E-05 | 4.157E-06 | 9.800E-06 | 1.700E-05 |
| 21065 | 1054 | E1054C | C | 1.2e-05 | 9e-06 | 1.3e-05 | 1.133E-05 | 2.082E-06 | 9.000E-06 | 1.300E-05 |
| 21066 | 1054 | E1054E | C | 9.4e-06 | 9.3e-06 | 1.5e-05 | 1.123E-05 | 3.262E-06 | 9.300E-06 | 1.500E-05 |
| 21067 | 1054 | E1054Q | C | 9.9e-06 | 1.7e-05 | 1.5e-05 | 1.397E-05 | 3.661E-06 | 9.900E-06 | 1.700E-05 |
| 21068 | 1054 | E1054G | C | 1.1e-05 | 1e-05 | 1.8e-05 | 1.300E-05 | 4.359E-06 | 1.000E-05 | 1.800E-05 |
| 21069 | 1054 | E1054H | C | 1.4e-05 | 1.5e-05 | 1.1e-05 | 1.333E-05 | 2.082E-06 | 1.100E-05 | 1.500E-05 |
| 21070 | 1054 | E1054I | C | 1.1e-05 | 9.9e-06 | 1.6e-05 | 1.230E-05 | 3.251E-06 | 9.900E-06 | 1.600E-05 |
| 21071 | 1054 | E1054L | C | 1.1e-05 | 1.1e-05 | 1.3e-05 | 1.167E-05 | 1.155E-06 | 1.100E-05 | 1.300E-05 |
| 21072 | 1054 | E1054K | C | 1.5e-05 | 1.6e-05 | 1.4e-05 | 1.500E-05 | 1.000E-06 | 1.400E-05 | 1.600E-05 |
| 21073 | 1054 | E1054M | C | 1.5e-05 | 9.7e-06 | 1.5e-05 | 1.323E-05 | 3.060E-06 | 9.700E-06 | 1.500E-05 |
| 21074 | 1054 | E1054F | C | 1.6e-05 | 1.6e-05 | 1e-05 | 1.400E-05 | 3.464E-06 | 1.000E-05 | 1.600E-05 |
| 21075 | 1054 | E1054P | C | 1.2e-05 | 8.6e-06 | 1.5e-05 | 1.187E-05 | 3.202E-06 | 8.600E-06 | 1.500E-05 |
| 21076 | 1054 | E1054S | C | 1.2e-05 | 1.1e-05 | 1.2e-05 | 1.167E-05 | 5.774E-07 | 1.100E-05 | 1.200E-05 |
| 21077 | 1054 | E1054T | C | 9.8e-06 | 9.1e-06 | 1.6e-05 | 1.163E-05 | 3.798E-06 | 9.100E-06 | 1.600E-05 |
| 21078 | 1054 | E1054W | C | 9.5e-06 | 1.4e-05 | 1.4e-05 | 1.250E-05 | 2.598E-06 | 9.500E-06 | 1.400E-05 |
| 21079 | 1054 | E1054Y | C | 1.6e-05 | 8.2e-06 | 8.9e-06 | 1.103E-05 | 4.315E-06 | 8.200E-06 | 1.600E-05 |

| | | | | | | | | | | |
|---|---|---|---|---|---|---|---|---|---|---|
| 21080 | 1054 | E1054V | C | 1.5e-05 | 1.6e-05 | 1.5e-05 | 1.533E-05 | 5.774E-07 | 1.500E-05 | 1.600E-05 |
| 21081 | 1055 | P1055A | C | 1.5e-05 | 1.6e-05 | 1.1e-05 | 1.400E-05 | 2.646E-06 | 1.100E-05 | 1.600E-05 |
| 21082 | 1055 | P1055R | C | 1.2e-05 | 1.6e-05 | 1.5e-05 | 1.433E-05 | 2.082E-06 | 1.200E-05 | 1.600E-05 |
| 21083 | 1055 | P1055N | C | 1.5e-05 | 1e-05 | 8.7e-06 | 1.123E-05 | 3.326E-06 | 8.700E-06 | 1.500E-05 |
| 21084 | 1055 | P1055D | C | 1.7e-05 | 1.2e-05 | 9.7e-06 | 1.290E-05 | 3.732E-06 | 9.700E-06 | 1.700E-05 |
| 21085 | 1055 | P1055C | C | 1.4e-05 | 8.8e-06 | 1.3e-05 | 1.193E-05 | 2.759E-06 | 8.800E-06 | 1.400E-05 |
| 21086 | 1055 | P1055E | C | 1.5e-05 | 7.9e-06 | 1.1e-05 | 1.130E-05 | 3.559E-06 | 7.900E-06 | 1.500E-05 |
| 21087 | 1055 | P1055Q | C | 1.7e-05 | 1.1e-05 | 1.4e-05 | 1.400E-05 | 3.000E-06 | 1.100E-05 | 1.700E-05 |
| 21088 | 1055 | P1055G | C | 1.1e-05 | 1.3e-05 | 9.2e-06 | 1.107E-05 | 1.901E-06 | 9.200E-06 | 1.300E-05 |
| 21089 | 1055 | P1055H | C | 8.2e-06 | 1.4e-05 | 1e-05 | 1.073E-05 | 2.969E-06 | 8.200E-06 | 1.400E-05 |
| 21090 | 1055 | P1055I | C | 1.3e-05 | 1.6e-05 | 1e-05 | 1.300E-05 | 3.000E-06 | 1.000E-05 | 1.600E-05 |
| 21091 | 1055 | P1055L | C | 1.4e-05 | 1e-05 | 1.1e-05 | 1.167E-05 | 2.082E-06 | 1.000E-05 | 1.400E-05 |
| 21092 | 1055 | P1055K | C | 1e-05 | 9.4e-06 | 1.5e-05 | 1.147E-05 | 3.075E-06 | 9.400E-06 | 1.500E-05 |
| 21093 | 1055 | P1055M | C | 9.7e-06 | 1e-05 | 8.9e-06 | 9.533E-06 | 5.686E-07 | 8.900E-06 | 1.000E-05 |
| 21094 | 1055 | P1055F | C | 1.5e-05 | 1.5e-05 | 1.2e-05 | 1.400E-05 | 1.732E-06 | 1.200E-05 | 1.500E-05 |
| 21095 | 1055 | P1055P | C | 1.5e-05 | 8.5e-06 | 1.7e-05 | 1.350E-05 | 4.444E-06 | 8.500E-06 | 1.700E-05 |
| 21096 | 1055 | P1055S | C | 1.4e-05 | 1.2e-05 | 8.4e-06 | 1.147E-05 | 2.838E-06 | 8.400E-06 | 1.400E-05 |
| 21097 | 1055 | P1055T | C | 1.5e-05 | 1e-05 | 1.3e-05 | 1.267E-05 | 2.517E-06 | 1.000E-05 | 1.500E-05 |
| 21098 | 1055 | P1055W | C | 1.3e-05 | 1e-05 | 1e-05 | 1.100E-05 | 1.732E-06 | 1.000E-05 | 1.300E-05 |
| 21099 | 1055 | P1055Y | C | 8.9e-06 | 1e-05 | 1.1e-05 | 9.967E-06 | 1.050E-06 | 8.900E-06 | 1.100E-05 |
| 21100 | 1055 | P1055V | C | 1.1e-05 | 1.1e-05 | 1e-05 | 1.067E-05 | 5.774E-07 | 1.000E-05 | 1.100E-05 |
| 21101 | 1056 | K1056A | C | 1.1e-05 | 1.6e-05 | 1.4e-05 | 1.367E-05 | 2.517E-06 | 1.100E-05 | 1.600E-05 |
| 21102 | 1056 | K1056R | C | 1.5e-05 | 9.5e-06 | 1.6e-05 | 1.350E-05 | 3.500E-06 | 9.500E-06 | 1.600E-05 |
| 21103 | 1056 | K1056N | C | 1.6e-05 | 1.3e-05 | 1.1e-05 | 1.333E-05 | 2.517E-06 | 1.100E-05 | 1.600E-05 |
| 21104 | 1056 | K1056D | C | 1.1e-05 | 9.4e-06 | 9.1e-06 | 9.833E-06 | 1.021E-06 | 9.100E-06 | 1.100E-05 |
| 21105 | 1056 | K1056C | C | 1.5e-05 | 9.9e-06 | 1.7e-05 | 1.397E-05 | 3.661E-06 | 9.900E-06 | 1.700E-05 |
| 21106 | 1056 | K1056E | C | 1.5e-05 | 1.4e-05 | 1e-05 | 1.300E-05 | 2.646E-06 | 1.000E-05 | 1.500E-05 |
| 21107 | 1056 | K1056Q | C | 8.2e-06 | 1.4e-05 | 1.1e-05 | 1.107E-05 | 2.901E-06 | 8.200E-06 | 1.400E-05 |
| 21108 | 1056 | K1056G | C | 1.5e-05 | 9.5e-06 | 1.5e-05 | 1.317E-05 | 3.175E-06 | 9.500E-06 | 1.500E-05 |
| 21109 | 1056 | K1056H | C | 1.5e-05 | 1.1e-05 | 1.5e-05 | 1.367E-05 | 2.309E-06 | 1.100E-05 | 1.500E-05 |
| 21110 | 1056 | K1056I | C | 1.5e-05 | 1.5e-05 | 1e-05 | 1.333E-05 | 2.887E-06 | 1.000E-05 | 1.500E-05 |
| 21111 | 1056 | K1056L | C | 8.8e-06 | 1.5e-05 | 8e-06 | 1.060E-05 | 3.831E-06 | 8.000E-06 | 1.500E-05 |
| 21112 | 1056 | K1056K | C | 7.9e-06 | 1.2e-05 | 9.3e-06 | 9.733E-06 | 2.084E-06 | 7.900E-06 | 1.200E-05 |
| 21113 | 1056 | K1056M | C | 1.6e-05 | 1.1e-05 | 9.3e-06 | 1.210E-05 | 3.483E-06 | 9.300E-06 | 1.600E-05 |
| 21114 | 1056 | K1056F | C | 1.5e-05 | 1.5e-05 | 1.4e-05 | 1.467E-05 | 5.774E-07 | 1.400E-05 | 1.500E-05 |
| 21115 | 1056 | K1056P | C | 1.1e-05 | 1e-05 | 7.4e-06 | 9.467E-06 | 1.858E-06 | 7.400E-06 | 1.100E-05 |
| 21116 | 1056 | K1056S | C | 1.7e-05 | 1.5e-05 | 1.5e-05 | 1.567E-05 | 1.155E-06 | 1.500E-05 | 1.700E-05 |
| 21117 | 1056 | K1056T | C | 1.1e-05 | 1.5e-05 | 1.5e-05 | 1.367E-05 | 2.309E-06 | 1.100E-05 | 1.500E-05 |
| 21118 | 1056 | K1056W | C | 1.1e-05 | 1.1e-05 | 1.2e-05 | 1.133E-05 | 5.774E-07 | 1.100E-05 | 1.200E-05 |
| 21119 | 1056 | K1056Y | C | 1e-05 | 1.3e-05 | 9.8e-06 | 1.093E-05 | 1.793E-06 | 9.800E-06 | 1.300E-05 |
| 21120 | 1056 | K1056V | C | 1.6e-05 | 9.5e-06 | 8.9e-06 | 1.147E-05 | 3.937E-06 | 8.900E-06 | 1.600E-05 |
| 21121 | 1057 | S1057A | C | 8.9e-06 | 1.6e-05 | 1.3e-05 | 1.263E-05 | 3.564E-06 | 8.900E-06 | 1.600E-05 |
| 21122 | 1057 | S1057R | C | 1.2e-05 | 8.4e-06 | 1.2e-05 | 1.080E-05 | 2.078E-06 | 8.400E-06 | 1.200E-05 |
| 21123 | 1057 | S1057N | C | 1.1e-05 | 1.4e-05 | 1.1e-05 | 1.200E-05 | 1.732E-06 | 1.100E-05 | 1.400E-05 |
| 21124 | 1057 | S1057D | C | 1.2e-05 | 1.2e-05 | 1.6e-05 | 1.333E-05 | 2.309E-06 | 1.200E-05 | 1.600E-05 |
| 21125 | 1057 | S1057C | C | 8.9e-06 | 1.5e-05 | 1e-05 | 1.130E-05 | 3.251E-06 | 8.900E-06 | 1.500E-05 |
| 21126 | 1057 | S1057E | C | 9.6e-06 | 1.3e-05 | 1.3e-05 | 1.187E-05 | 1.963E-06 | 9.600E-06 | 1.300E-05 |
| 21127 | 1057 | S1057Q | C | 8e-06 | 9.6e-06 | 1e-05 | 9.200E-06 | 1.058E-06 | 8.000E-06 | 1.000E-05 |
| 21128 | 1057 | S1057G | C | 1e-05 | 1.5e-05 | 1.1e-05 | 1.200E-05 | 2.646E-06 | 1.000E-05 | 1.500E-05 |
| 21129 | 1057 | S1057H | C | 1.6e-05 | 1.2e-05 | 1.5e-05 | 1.433E-05 | 2.082E-06 | 1.200E-05 | 1.600E-05 |
| 21130 | 1057 | S1057I | C | 1e-05 | 1.4e-05 | 1.4e-05 | 1.267E-05 | 2.309E-06 | 1.000E-05 | 1.400E-05 |
| 21131 | 1057 | S1057L | C | 1.1e-05 | 1.7e-05 | 8.7e-06 | 1.223E-05 | 4.285E-06 | 8.700E-06 | 1.700E-05 |
| 21132 | 1057 | S1057K | C | 1.6e-05 | 1.6e-05 | 1.1e-05 | 1.433E-05 | 2.887E-06 | 1.100E-05 | 1.600E-05 |
| 21133 | 1057 | S1057M | C | 8.8e-06 | 1.1e-05 | 1e-05 | 9.933E-06 | 1.102E-06 | 8.800E-06 | 1.100E-05 |
| 21134 | 1057 | S1057F | C | 1.5e-05 | 1.5e-05 | 1.6e-05 | 1.533E-05 | 5.774E-07 | 1.500E-05 | 1.600E-05 |
| 21135 | 1057 | S1057P | C | 1.1e-05 | 1.4e-05 | 1.2e-05 | 1.233E-05 | 1.528E-06 | 1.100E-05 | 1.400E-05 |
| 21136 | 1057 | S1057S | C | 8.8e-06 | 9.5e-06 | 1.7e-05 | 1.177E-05 | 4.546E-06 | 8.800E-06 | 1.700E-05 |
| 21137 | 1057 | S1057T | C | 1.4e-05 | 1.4e-05 | 1.3e-05 | 1.367E-05 | 5.774E-07 | 1.300E-05 | 1.400E-05 |
| 21138 | 1057 | S1057W | C | 1.5e-05 | 1.1e-05 | 1.4e-05 | 1.333E-05 | 2.082E-06 | 1.100E-05 | 1.500E-05 |
| 21139 | 1057 | S1057Y | C | 1e-05 | 1.4e-05 | 1.6e-05 | 1.333E-05 | 3.055E-06 | 1.000E-05 | 1.600E-05 |
| 21140 | 1057 | S1057V | C | 1.6e-05 | 1.5e-05 | 8.5e-06 | 1.317E-05 | 4.072E-06 | 8.500E-06 | 1.600E-05 |
| 21141 | 1058 | C1058A | C | 1.2e-05 | 1.1e-05 | 1.4e-05 | 1.233E-05 | 1.528E-06 | 1.100E-05 | 1.400E-05 |
| 21142 | 1058 | C1058R | C | 1.1e-05 | 1e-05 | 1.6e-05 | 1.233E-05 | 3.215E-06 | 1.000E-05 | 1.600E-05 |
| 21143 | 1058 | C1058N | C | 1.5e-05 | 1.5e-05 | 9.5e-06 | 1.317E-05 | 3.175E-06 | 9.500E-06 | 1.500E-05 |
| 21144 | 1058 | C1058D | C | 1.1e-05 | 1.6e-05 | 1.4e-05 | 1.367E-05 | 2.517E-06 | 1.100E-05 | 1.600E-05 |
| 21145 | 1058 | C1058C | C | 1.3e-05 | 1.1e-05 | 9.9e-06 | 1.130E-05 | 1.572E-06 | 9.900E-06 | 1.300E-05 |
| 21146 | 1058 | C1058E | C | 1.1e-05 | 9e-06 | 1.1e-05 | 1.033E-05 | 1.155E-06 | 9.000E-06 | 1.100E-05 |
| 21147 | 1058 | C1058Q | C | 1.3e-05 | 1.8e-05 | 1.5e-05 | 1.533E-05 | 2.517E-06 | 1.300E-05 | 1.800E-05 |
| 21148 | 1058 | C1058G | C | 1.6e-05 | 1.3e-05 | 1.5e-05 | 1.467E-05 | 1.528E-06 | 1.300E-05 | 1.600E-05 |
| 21149 | 1058 | C1058H | C | 1.2e-05 | 9.8e-06 | 1.3e-05 | 1.160E-05 | 1.637E-06 | 9.800E-06 | 1.300E-05 |
| 21150 | 1058 | C1058I | C | 1.6e-05 | 1e-05 | 1.3e-05 | 1.300E-05 | 3.000E-06 | 1.000E-05 | 1.600E-05 |
| 21151 | 1058 | C1058L | C | 9.1e-06 | 1.5e-05 | 1.2e-05 | 1.203E-05 | 2.950E-06 | 9.100E-06 | 1.500E-05 |
| 21152 | 1058 | C1058K | C | 1.4e-05 | 9.9e-06 | 1.3e-05 | 1.230E-05 | 2.138E-06 | 9.900E-06 | 1.400E-05 |
| 21153 | 1058 | C1058M | C | 1.1e-05 | 1.8e-05 | 1.7e-05 | 1.533E-05 | 3.786E-06 | 1.100E-05 | 1.800E-05 |
| 21154 | 1058 | C1058F | C | 1.7e-05 | 1.1e-05 | 1.4e-05 | 1.400E-05 | 3.000E-06 | 1.100E-05 | 1.700E-05 |
| 21155 | 1058 | C1058P | C | 1.5e-05 | 1.4e-05 | 8.6e-06 | 1.253E-05 | 3.443E-06 | 8.600E-06 | 1.500E-05 |
| 21156 | 1058 | C1058S | C | 1.7e-05 | 1.5e-05 | 1.5e-05 | 1.567E-05 | 1.155E-06 | 1.500E-05 | 1.700E-05 |
| 21157 | 1058 | C1058T | C | 1.5e-05 | 1.5e-05 | 9.6e-06 | 1.320E-05 | 3.118E-06 | 9.600E-06 | 1.500E-05 |
| 21158 | 1058 | C1058W | C | 1.4e-05 | 1.4e-05 | 1.5e-05 | 1.433E-05 | 5.774E-07 | 1.400E-05 | 1.500E-05 |
| 21159 | 1058 | C1058Y | C | 1.1e-05 | 9.8e-06 | 1.2e-05 | 1.093E-05 | 1.102E-06 | 9.800E-06 | 1.200E-05 |
| 21160 | 1058 | C1058V | C | 1.5e-05 | 1.5e-05 | 1.2e-05 | 1.400E-05 | 1.732E-06 | 1.200E-05 | 1.500E-05 |
| 21161 | 1059 | D1059A | D | 1.4e-05 | 1e-05 | 1.5e-05 | 1.300E-05 | 2.646E-06 | 1.000E-05 | 1.500E-05 |
| 21162 | 1059 | D1059R | D | 1.3e-05 | 1.1e-05 | 9.9e-06 | 1.130E-05 | 1.572E-06 | 9.900E-06 | 1.300E-05 |
| 21163 | 1059 | D1059N | D | 1.4e-05 | 1.3e-05 | 1.5e-05 | 1.400E-05 | 1.000E-06 | 1.300E-05 | 1.500E-05 |
| 21164 | 1059 | D1059D | D | 9.6e-06 | 1.6e-05 | 1.6e-05 | 1.387E-05 | 3.695E-06 | 9.600E-06 | 1.600E-05 |

| | | | | | | | | | | |
|---|---|---|---|---|---|---|---|---|---|---|
| 21165 | 1059 | D1059C | D | 1.2e-05 | 1.4e-05 | 1.1e-05 | 1.233E-05 | 1.528E-06 | 1.100E-05 | 1.400E-05 |
| 21166 | 1059 | D1059E | D | 9.6e-06 | 1.4e-05 | 1e-05 | 1.120E-05 | 2.433E-06 | 9.600E-06 | 1.400E-05 |
| 21167 | 1059 | D1059Q | D | 1.5e-05 | 1.4e-05 | 1.4e-05 | 1.433E-05 | 5.774E-07 | 1.400E-05 | 1.500E-05 |
| 21168 | 1059 | D1059G | D | 1.3e-05 | 1.6e-05 | 1.5e-05 | 1.467E-05 | 1.528E-06 | 1.300E-05 | 1.600E-05 |
| 21169 | 1059 | D1059H | D | 8.7e-06 | 9.2e-06 | 1.1e-05 | 9.633E-06 | 1.210E-06 | 8.700E-06 | 1.100E-05 |
| 21170 | 1059 | D1059I | D | 1.5e-05 | 1e-05 | 1.4e-05 | 1.300E-05 | 2.646E-06 | 1.000E-05 | 1.500E-05 |
| 21171 | 1059 | D1059L | D | 1e-05 | 1.7e-05 | 7.8e-06 | 1.160E-05 | 4.804E-06 | 7.800E-06 | 1.700E-05 |
| 21172 | 1059 | D1059K | D | 6.6e-06 | 1.1e-05 | 1.5e-05 | 1.087E-05 | 4.202E-06 | 6.600E-06 | 1.500E-05 |
| 21173 | 1059 | D1059M | D | 1.5e-05 | 1.7e-05 | 1.2e-05 | 1.467E-05 | 2.517E-06 | 1.200E-05 | 1.700E-05 |
| 21174 | 1059 | D1059F | D | 1.3e-05 | 8.2e-06 | 1.8e-05 | 1.307E-05 | 4.900E-06 | 8.200E-06 | 1.800E-05 |
| 21175 | 1059 | D1059P | D | 9.4e-06 | 1.7e-05 | 1.2e-05 | 1.280E-05 | 3.863E-06 | 9.400E-06 | 1.700E-05 |
| 21176 | 1059 | D1059S | D | 1.7e-05 | 1.2e-05 | 1.3e-05 | 1.400E-05 | 2.646E-06 | 1.200E-05 | 1.700E-05 |
| 21177 | 1059 | D1059T | D | 9.3e-06 | 8.4e-06 | 1.5e-05 | 1.090E-05 | 3.579E-06 | 8.400E-06 | 1.500E-05 |
| 21178 | 1059 | D1059W | D | 1.6e-05 | 1.2e-05 | 1.3e-05 | 1.367E-05 | 2.082E-06 | 1.200E-05 | 1.600E-05 |
| 21179 | 1059 | D1059Y | D | 1.3e-05 | 9.3e-06 | 1.4e-05 | 1.210E-05 | 2.476E-06 | 9.300E-06 | 1.400E-05 |
| 21180 | 1059 | D1059V | D | 9.6e-06 | 1.7e-05 | 1.3e-05 | 1.320E-05 | 3.704E-06 | 9.600E-06 | 1.700E-05 |
| 21181 | 1060 | I1060A | D | 1e-05 | 1.1e-05 | 1.5e-05 | 1.200E-05 | 2.646E-06 | 1.000E-05 | 1.500E-05 |
| 21182 | 1060 | I1060R | D | 1.5e-05 | 1.5e-05 | 1.3e-05 | 1.433E-05 | 1.155E-06 | 1.300E-05 | 1.500E-05 |
| 21183 | 1060 | I1060N | D | 1.6e-05 | 9.5e-06 | 1.1e-05 | 1.217E-05 | 3.403E-06 | 9.500E-06 | 1.600E-05 |
| 21184 | 1060 | I1060D | D | 8.5e-06 | 9.9e-06 | 1.6e-05 | 1.147E-05 | 3.988E-06 | 8.500E-06 | 1.600E-05 |
| 21185 | 1060 | I1060C | D | 9.3e-06 | 1.7e-05 | 1.3e-05 | 1.310E-05 | 3.851E-06 | 9.300E-06 | 1.700E-05 |
| 21186 | 1060 | I1060E | D | 1.5e-05 | 1.1e-05 | 1.5e-05 | 1.367E-05 | 2.309E-06 | 1.100E-05 | 1.500E-05 |
| 21187 | 1060 | I1060Q | D | 1.5e-05 | 1.5e-05 | 1.1e-05 | 1.367E-05 | 2.309E-06 | 1.100E-05 | 1.500E-05 |
| 21188 | 1060 | I1060G | D | 9.8e-06 | 1.2e-05 | 1.5e-05 | 1.227E-05 | 2.610E-06 | 9.800E-06 | 1.500E-05 |
| 21189 | 1060 | I1060H | D | 1.6e-05 | 1.3e-05 | 1e-05 | 1.300E-05 | 3.000E-06 | 1.000E-05 | 1.600E-05 |
| 21190 | 1060 | I1060I | D | 1.2e-05 | 1.5e-05 | 1.5e-05 | 1.400E-05 | 1.732E-06 | 1.200E-05 | 1.500E-05 |
| 21191 | 1060 | I1060L | D | 1.4e-05 | 1.8e-05 | 1.5e-05 | 1.567E-05 | 2.082E-06 | 1.400E-05 | 1.800E-05 |
| 21192 | 1060 | I1060K | D | 9.2e-06 | 1.1e-05 | 1.1e-05 | 1.040E-05 | 1.039E-06 | 9.200E-06 | 1.100E-05 |
| 21193 | 1060 | I1060M | D | 1e-05 | 1.4e-05 | 1.1e-05 | 1.167E-05 | 2.082E-06 | 1.000E-05 | 1.400E-05 |
| 21194 | 1060 | I1060F | D | 1e-05 | 8.3e-06 | 9.4e-06 | 9.233E-06 | 8.622E-07 | 8.300E-06 | 1.000E-05 |
| 21195 | 1060 | I1060P | D | 9.6e-06 | 1.6e-05 | 8.1e-06 | 1.123E-05 | 4.196E-06 | 8.100E-06 | 1.600E-05 |
| 21196 | 1060 | I1060S | D | 1.5e-05 | 1.6e-05 | 8.1e-06 | 1.303E-05 | 4.302E-06 | 8.100E-06 | 1.600E-05 |
| 21197 | 1060 | I1060T | D | 1.5e-05 | 1.6e-05 | 1.7e-05 | 1.600E-05 | 1.000E-06 | 1.500E-05 | 1.700E-05 |
| 21198 | 1060 | I1060W | D | 1.1e-05 | 9.7e-06 | 1.3e-05 | 1.123E-05 | 1.662E-06 | 9.700E-06 | 1.300E-05 |
| 21199 | 1060 | I1060Y | D | 1.6e-05 | 1.4e-05 | 1.5e-05 | 1.500E-05 | 1.000E-06 | 1.400E-05 | 1.600E-05 |
| 21200 | 1060 | I1060V | D | 1e-05 | 1.4e-05 | 1.4e-05 | 1.267E-05 | 2.309E-06 | 1.000E-05 | 1.400E-05 |
| 21201 | 1061 | Q1061A | D | 1.4e-05 | 1.5e-05 | 8.8e-06 | 1.260E-05 | 3.329E-06 | 8.800E-06 | 1.500E-05 |
| 21202 | 1061 | Q1061R | D | 1.5e-05 | 1.6e-05 | 1.1e-05 | 1.400E-05 | 2.646E-06 | 1.100E-05 | 1.600E-05 |
| 21203 | 1061 | Q1061N | D | 7.4e-06 | 1.5e-05 | 1.5e-05 | 1.247E-05 | 4.388E-06 | 7.400E-06 | 1.500E-05 |
| 21204 | 1061 | Q1061D | D | 1.3e-05 | 1.1e-05 | 1.7e-05 | 1.367E-05 | 3.055E-06 | 1.100E-05 | 1.700E-05 |
| 21205 | 1061 | Q1061C | D | 1.5e-05 | 9.8e-06 | 1.5e-05 | 1.327E-05 | 3.002E-06 | 9.800E-06 | 1.500E-05 |
| 21206 | 1061 | Q1061E | D | 1.4e-05 | 1.4e-05 | 1.6e-05 | 1.467E-05 | 1.155E-06 | 1.400E-05 | 1.600E-05 |
| 21207 | 1061 | Q1061Q | D | 1.1e-05 | 1e-05 | 8.7e-06 | 9.900E-06 | 1.153E-06 | 8.700E-06 | 1.100E-05 |
| 21208 | 1061 | Q1061G | D | 1.8e-05 | 1.2e-05 | 1.1e-05 | 1.367E-05 | 3.786E-06 | 1.100E-05 | 1.800E-05 |
| 21209 | 1061 | Q1061H | D | 1.4e-05 | 9.7e-06 | 1.5e-05 | 1.290E-05 | 2.816E-06 | 9.700E-06 | 1.500E-05 |
| 21210 | 1061 | Q1061I | D | 1.4e-05 | 1.6e-05 | 9.7e-06 | 1.323E-05 | 3.219E-06 | 9.700E-06 | 1.600E-05 |
| 21211 | 1061 | Q1061L | D | 1e-05 | 1.2e-05 | 1.5e-05 | 1.233E-05 | 2.517E-06 | 1.000E-05 | 1.500E-05 |
| 21212 | 1061 | Q1061K | D | 1e-05 | 1.4e-05 | 1.4e-05 | 1.267E-05 | 2.309E-06 | 1.000E-05 | 1.400E-05 |
| 21213 | 1061 | Q1061M | D | 1e-05 | 1.6e-05 | 1.2e-05 | 1.267E-05 | 3.055E-06 | 1.000E-05 | 1.600E-05 |
| 21214 | 1061 | Q1061F | D | 1.3e-05 | 1.7e-05 | 1.6e-05 | 1.533E-05 | 2.082E-06 | 1.300E-05 | 1.700E-05 |
| 21215 | 1061 | Q1061P | D | 1.5e-05 | 9.9e-06 | 8.6e-06 | 1.117E-05 | 3.383E-06 | 8.600E-06 | 1.500E-05 |
| 21216 | 1061 | Q1061S | D | 1.2e-05 | 1.3e-05 | 1.4e-05 | 1.300E-05 | 1.000E-06 | 1.200E-05 | 1.400E-05 |
| 21217 | 1061 | Q1061T | D | 1.1e-05 | 9.3e-06 | 1.6e-05 | 1.210E-05 | 3.483E-06 | 9.300E-06 | 1.600E-05 |
| 21218 | 1061 | Q1061W | D | 1.5e-05 | 1.1e-05 | 1.1e-05 | 1.233E-05 | 2.309E-06 | 1.100E-05 | 1.500E-05 |
| 21219 | 1061 | Q1061Y | D | 1.2e-05 | 1e-05 | 1.5e-05 | 1.233E-05 | 2.517E-06 | 1.000E-05 | 1.500E-05 |
| 21220 | 1061 | Q1061V | D | 1.7e-05 | 1.3e-05 | 1.4e-05 | 1.467E-05 | 2.082E-06 | 1.300E-05 | 1.700E-05 |
| 21221 | 1062 | M1062A | D | 1.6e-05 | 1e-05 | 9.9e-06 | 1.197E-05 | 3.493E-06 | 9.900E-06 | 1.600E-05 |
| 21222 | 1062 | M1062R | D | 1.3e-05 | 9.4e-06 | 9.3e-06 | 1.057E-05 | 2.108E-06 | 9.300E-06 | 1.300E-05 |
| 21223 | 1062 | M1062N | D | 1.2e-05 | 1.6e-05 | 8.8e-06 | 1.227E-05 | 3.607E-06 | 8.800E-06 | 1.600E-05 |
| 21224 | 1062 | M1062D | D | 1.5e-05 | 1.6e-05 | 1.1e-05 | 1.400E-05 | 2.646E-06 | 1.100E-05 | 1.600E-05 |
| 21225 | 1062 | M1062C | D | 9.3e-06 | 7.4e-06 | 7.7e-06 | 8.133E-06 | 1.021E-06 | 7.400E-06 | 9.300E-06 |
| 21226 | 1062 | M1062E | D | 1.1e-05 | 1e-05 | 1.1e-05 | 1.067E-05 | 5.774E-07 | 1.000E-05 | 1.100E-05 |
| 21227 | 1062 | M1062Q | D | 9.7e-06 | 1.3e-05 | 1.4e-05 | 1.223E-05 | 2.250E-06 | 9.700E-06 | 1.400E-05 |
| 21228 | 1062 | M1062G | D | 1.5e-05 | 1.3e-05 | 1e-05 | 1.267E-05 | 2.517E-06 | 1.000E-05 | 1.500E-05 |
| 21229 | 1062 | M1062H | D | 1.4e-05 | 1e-05 | 8.9e-06 | 1.097E-05 | 2.684E-06 | 8.900E-06 | 1.400E-05 |
| 21230 | 1062 | M1062I | D | 1.1e-05 | 1.5e-05 | 9.7e-06 | 1.190E-05 | 2.762E-06 | 9.700E-06 | 1.500E-05 |
| 21231 | 1062 | M1062L | D | 1.4e-05 | 1.3e-05 | 1.3e-05 | 1.333E-05 | 5.774E-07 | 1.300E-05 | 1.400E-05 |
| 21232 | 1062 | M1062K | D | 8.9e-06 | 1.7e-05 | 1e-05 | 1.197E-05 | 4.394E-06 | 8.900E-06 | 1.700E-05 |
| 21233 | 1062 | M1062M | D | 1.4e-05 | 1.4e-05 | 1.1e-05 | 1.300E-05 | 1.732E-06 | 1.100E-05 | 1.400E-05 |
| 21234 | 1062 | M1062F | D | 1.2e-05 | 1.6e-05 | 7.6e-06 | 1.187E-05 | 4.202E-06 | 7.600E-06 | 1.600E-05 |
| 21235 | 1062 | M1062P | D | 1.6e-05 | 9.6e-06 | 1.6e-05 | 1.387E-05 | 3.695E-06 | 9.600E-06 | 1.600E-05 |
| 21236 | 1062 | M1062S | D | 1e-05 | 9.2e-06 | 1.1e-05 | 1.007E-05 | 9.018E-07 | 9.200E-06 | 1.100E-05 |
| 21237 | 1062 | M1062T | D | 1.3e-05 | 1.3e-05 | 1.3e-05 | 1.300E-05 | 0.000E+00 | 1.300E-05 | 1.300E-05 |
| 21238 | 1062 | M1062W | D | 8.9e-06 | 1e-05 | 9.3e-06 | 9.400E-06 | 5.568E-07 | 8.900E-06 | 1.000E-05 |
| 21239 | 1062 | M1062Y | D | 1.6e-05 | 1e-05 | 9.7e-06 | 1.190E-05 | 3.554E-06 | 9.700E-06 | 1.600E-05 |
| 21240 | 1062 | M1062V | D | 8.8e-06 | 1.5e-05 | 1.4e-05 | 1.260E-05 | 3.329E-06 | 8.800E-06 | 1.500E-05 |
| 21241 | 1063 | T1063A | D | 1.7e-05 | 1.6e-05 | 1.2e-05 | 1.500E-05 | 2.646E-06 | 1.200E-05 | 1.700E-05 |
| 21242 | 1063 | T1063R | D | 1.6e-05 | 1.1e-05 | 1.4e-05 | 1.367E-05 | 2.517E-06 | 1.100E-05 | 1.600E-05 |
| 21243 | 1063 | T1063N | D | 9.9e-06 | 7.2e-06 | 1.4e-05 | 1.037E-05 | 3.424E-06 | 7.200E-06 | 1.400E-05 |
| 21244 | 1063 | T1063D | D | 1.4e-05 | 1.7e-05 | 1.6e-05 | 1.567E-05 | 1.528E-06 | 1.400E-05 | 1.700E-05 |
| 21245 | 1063 | T1063C | D | 9.6e-06 | 1.5e-05 | 1.3e-05 | 1.253E-05 | 2.730E-06 | 9.600E-06 | 1.500E-05 |
| 21246 | 1063 | T1063E | D | 9.8e-06 | 1.4e-05 | 1.6e-05 | 1.327E-05 | 3.164E-06 | 9.800E-06 | 1.600E-05 |
| 21247 | 1063 | T1063Q | D | 9.6e-06 | 1.5e-05 | 1.6e-05 | 1.353E-05 | 3.443E-06 | 9.600E-06 | 1.600E-05 |
| 21248 | 1063 | T1063G | D | 1.2e-05 | 1e-05 | 1.6e-05 | 1.267E-05 | 3.055E-06 | 1.000E-05 | 1.600E-05 |
| 21249 | 1063 | T1063H | D | 2.5e-05 | 1.1e-05 | 1e-05 | 1.533E-05 | 8.386E-06 | 1.000E-05 | 2.500E-05 |

| | | | | | | | | | | |
|---|---|---|---|---|---|---|---|---|---|---|
| 21250 | 1063 | T1063I | D | 1.6e-05 | 8.9e-06 | 1.5e-05 | 1.330E-05 | 3.843E-06 | 8.900E-06 | 1.600E-05 |
| 21251 | 1063 | T1063L | D | 7.2e-06 | 1.5e-05 | 1.6e-05 | 1.273E-05 | 4.818E-06 | 7.200E-06 | 1.600E-05 |
| 21252 | 1063 | T1063K | D | 1.7e-05 | 1.4e-05 | 1.1e-05 | 1.400E-05 | 3.000E-06 | 1.100E-05 | 1.700E-05 |
| 21253 | 1063 | T1063M | D | 1.5e-05 | 1.6e-05 | 1.5e-05 | 1.533E-05 | 5.774E-07 | 1.500E-05 | 1.600E-05 |
| 21254 | 1063 | T1063F | D | 1.6e-05 | 1.6e-05 | 1.1e-05 | 1.433E-05 | 2.887E-06 | 1.100E-05 | 1.600E-05 |
| 21255 | 1063 | T1063P | D | 7.7e-06 | 1.4e-05 | 1.5e-05 | 1.223E-05 | 3.958E-06 | 7.700E-06 | 1.500E-05 |
| 21256 | 1063 | T1063S | D | 1.2e-05 | 1.3e-05 | 1.4e-05 | 1.300E-05 | 1.000E-06 | 1.200E-05 | 1.400E-05 |
| 21257 | 1063 | T1063T | D | 1.6e-05 | 1.1e-05 | 1.1e-05 | 1.267E-05 | 2.887E-06 | 1.100E-05 | 1.600E-05 |
| 21258 | 1063 | T1063W | D | 8.4e-06 | 1e-05 | 1e-05 | 9.467E-06 | 9.238E-07 | 8.400E-06 | 1.000E-05 |
| 21259 | 1063 | T1063Y | D | 1.5e-05 | 1.5e-05 | 9.1e-06 | 1.303E-05 | 3.406E-06 | 9.100E-06 | 1.500E-05 |
| 21260 | 1063 | T1063V | D | 1.7e-05 | 2.3e-05 | 1e-05 | 1.667E-05 | 6.506E-06 | 1.000E-05 | 2.300E-05 |
| 21261 | 1064 | Q1064A | D | 1.3e-05 | 7.3e-06 | 1.3e-05 | 1.110E-05 | 3.291E-06 | 7.300E-06 | 1.300E-05 |
| 21262 | 1064 | Q1064R | D | 1e-05 | 1.3e-05 | 1.6e-05 | 1.300E-05 | 3.000E-06 | 1.000E-05 | 1.600E-05 |
| 21263 | 1064 | Q1064N | D | 1.5e-05 | 1.7e-05 | 1.4e-05 | 1.533E-05 | 1.528E-06 | 1.400E-05 | 1.700E-05 |
| 21264 | 1064 | Q1064D | D | 1.5e-05 | 1.6e-05 | 1.1e-05 | 1.400E-05 | 2.646E-06 | 1.100E-05 | 1.600E-05 |
| 21265 | 1064 | Q1064C | D | 1e-05 | 1.5e-05 | 1.1e-05 | 1.200E-05 | 2.646E-06 | 1.000E-05 | 1.500E-05 |
| 21266 | 1064 | Q1064E | D | 9.8e-06 | 6.9e-06 | 9.9e-06 | 8.867E-06 | 1.704E-06 | 6.900E-06 | 9.900E-06 |
| 21267 | 1064 | Q1064Q | D | 1.6e-05 | 1.6e-05 | 1e-05 | 1.400E-05 | 3.464E-06 | 1.000E-05 | 1.600E-05 |
| 21268 | 1064 | Q1064G | D | 1.1e-05 | 1.6e-05 | 1.4e-05 | 1.367E-05 | 2.517E-06 | 1.100E-05 | 1.600E-05 |
| 21269 | 1064 | Q1064H | D | 1.4e-05 | 1.2e-05 | 1.5e-05 | 1.367E-05 | 1.528E-06 | 1.200E-05 | 1.500E-05 |
| 21270 | 1064 | Q1064I | D | 1.4e-05 | 1.5e-05 | 1.3e-05 | 1.400E-05 | 1.000E-06 | 1.300E-05 | 1.500E-05 |
| 21271 | 1064 | Q1064L | D | 1.2e-05 | 1.4e-05 | 1.6e-05 | 1.400E-05 | 2.000E-06 | 1.200E-05 | 1.600E-05 |
| 21272 | 1064 | Q1064K | D | 8.7e-06 | 1.1e-05 | 1.6e-05 | 1.190E-05 | 3.732E-06 | 8.700E-06 | 1.600E-05 |
| 21273 | 1064 | Q1064M | D | 9.4e-06 | 1.7e-05 | 1.3e-05 | 1.313E-05 | 3.802E-06 | 9.400E-06 | 1.700E-05 |
| 21274 | 1064 | Q1064F | D | 1.7e-05 | 1.6e-05 | 1.5e-05 | 1.600E-05 | 1.000E-06 | 1.500E-05 | 1.700E-05 |
| 21275 | 1064 | Q1064P | D | 1.3e-05 | 9.9e-06 | 1.4e-05 | 1.230E-05 | 2.138E-06 | 9.900E-06 | 1.400E-05 |
| 21276 | 1064 | Q1064S | D | 1.5e-05 | 1.2e-05 | 1.3e-05 | 1.333E-05 | 1.528E-06 | 1.200E-05 | 1.500E-05 |
| 21277 | 1064 | Q1064T | D | 1e-05 | 1e-05 | 6.4e-06 | 8.800E-06 | 2.078E-06 | 6.400E-06 | 1.000E-05 |
| 21278 | 1064 | Q1064W | D | 1.4e-05 | 1.4e-05 | 1.5e-05 | 1.433E-05 | 5.774E-07 | 1.400E-05 | 1.500E-05 |
| 21279 | 1064 | Q1064Y | D | 1.6e-05 | 1.3e-05 | 1.5e-05 | 1.467E-05 | 1.528E-06 | 1.300E-05 | 1.600E-05 |
| 21280 | 1064 | Q1064V | D | 1.2e-05 | 1.1e-05 | 1.4e-05 | 1.233E-05 | 1.528E-06 | 1.100E-05 | 1.400E-05 |
| 21281 | 1065 | S1065A | D | 1.6e-05 | 1.7e-05 | 1.4e-05 | 1.567E-05 | 1.528E-06 | 1.400E-05 | 1.700E-05 |
| 21282 | 1065 | S1065R | D | 9.7e-06 | 1.5e-05 | 1.2e-05 | 1.223E-05 | 2.658E-06 | 9.700E-06 | 1.500E-05 |
| 21283 | 1065 | S1065N | D | 1.8e-05 | 1.4e-05 | 1.4e-05 | 1.533E-05 | 2.309E-06 | 1.400E-05 | 1.800E-05 |
| 21284 | 1065 | S1065D | D | 8.9e-06 | 1.5e-05 | 1.6e-05 | 1.330E-05 | 3.843E-06 | 8.900E-06 | 1.600E-05 |
| 21285 | 1065 | S1065C | D | 1.2e-05 | 2.2e-05 | 1.1e-05 | 1.500E-05 | 6.083E-06 | 1.100E-05 | 2.200E-05 |
| 21286 | 1065 | S1065E | D | 9.7e-06 | 1.5e-05 | 8.9e-06 | 1.120E-05 | 3.315E-06 | 8.900E-06 | 1.500E-05 |
| 21287 | 1065 | S1065Q | D | 1.6e-05 | 8.1e-06 | 9.5e-06 | 1.120E-05 | 4.215E-06 | 8.100E-06 | 1.600E-05 |
| 21288 | 1065 | S1065G | D | 9.6e-06 | 1.2e-05 | 1.5e-05 | 1.220E-05 | 2.706E-06 | 9.600E-06 | 1.500E-05 |
| 21289 | 1065 | S1065H | D | 1.4e-05 | 1.1e-05 | 1.1e-05 | 1.200E-05 | 1.732E-06 | 1.100E-05 | 1.400E-05 |
| 21290 | 1065 | S1065I | D | 1.2e-05 | 1.6e-05 | 1.5e-05 | 1.433E-05 | 2.082E-06 | 1.200E-05 | 1.600E-05 |
| 21291 | 1065 | S1065L | D | 1.8e-05 | 1.6e-05 | 9.8e-06 | 1.460E-05 | 4.276E-06 | 9.800E-06 | 1.800E-05 |
| 21292 | 1065 | S1065K | D | 1.6e-05 | 1.6e-05 | 1.7e-05 | 1.633E-05 | 5.774E-07 | 1.600E-05 | 1.700E-05 |
| 21293 | 1065 | S1065M | D | 1e-05 | 1.1e-05 | 1.4e-05 | 1.167E-05 | 2.082E-06 | 1.000E-05 | 1.400E-05 |
| 21294 | 1065 | S1065F | D | 9e-06 | 1.1e-05 | 1.5e-05 | 1.167E-05 | 3.055E-06 | 9.000E-06 | 1.500E-05 |
| 21295 | 1065 | S1065P | D | 9.3e-06 | 8.1e-06 | 1.6e-05 | 1.113E-05 | 4.257E-06 | 8.100E-06 | 1.600E-05 |
| 21296 | 1065 | S1065S | D | 7.9e-06 | 9.9e-06 | 1e-05 | 9.267E-06 | 1.185E-06 | 7.900E-06 | 1.000E-05 |
| 21297 | 1065 | S1065T | D | 1e-05 | 1.3e-05 | 8.8e-06 | 1.060E-05 | 2.163E-06 | 8.800E-06 | 1.300E-05 |
| 21298 | 1065 | S1065W | D | 1.6e-05 | 1.8e-05 | 1.6e-05 | 1.667E-05 | 1.155E-06 | 1.600E-05 | 1.800E-05 |
| 21299 | 1065 | S1065Y | D | 9.4e-06 | 1.2e-05 | 1e-05 | 1.047E-05 | 1.361E-06 | 9.400E-06 | 1.200E-05 |
| 21300 | 1065 | S1065V | D | 1.2e-05 | 2e-05 | 1.1e-05 | 1.433E-05 | 4.933E-06 | 1.100E-05 | 2.000E-05 |
| 21301 | 1066 | P1066A | D | 1.4e-05 | 1.6e-05 | 1.6e-05 | 1.533E-05 | 1.155E-06 | 1.400E-05 | 1.600E-05 |
| 21302 | 1066 | P1066R | D | 1.6e-05 | 1.6e-05 | 1.1e-05 | 1.433E-05 | 2.887E-06 | 1.100E-05 | 1.600E-05 |
| 21303 | 1066 | P1066N | D | 1.4e-05 | 1.1e-05 | 1.4e-05 | 1.300E-05 | 1.732E-06 | 1.100E-05 | 1.400E-05 |
| 21304 | 1066 | P1066D | D | 1.6e-05 | 1e-05 | 1.5e-05 | 1.367E-05 | 3.215E-06 | 1.000E-05 | 1.600E-05 |
| 21305 | 1066 | P1066C | D | 1.4e-05 | 1.5e-05 | 9.8e-06 | 1.293E-05 | 2.759E-06 | 9.800E-06 | 1.500E-05 |
| 21306 | 1066 | P1066E | D | 1.5e-05 | 1.5e-05 | 1.2e-05 | 1.400E-05 | 1.732E-06 | 1.200E-05 | 1.500E-05 |
| 21307 | 1066 | P1066Q | D | 7.5e-06 | 1.5e-05 | 1.5e-05 | 1.250E-05 | 4.330E-06 | 7.500E-06 | 1.500E-05 |
| 21308 | 1066 | P1066G | D | 1.1e-05 | 9.4e-06 | 8.3e-06 | 9.567E-06 | 1.358E-06 | 8.300E-06 | 1.100E-05 |
| 21309 | 1066 | P1066H | D | 1.4e-05 | 1.5e-05 | 1.4e-05 | 1.433E-05 | 5.774E-07 | 1.400E-05 | 1.500E-05 |
| 21310 | 1066 | P1066I | D | 1.4e-05 | 1.2e-05 | 1.5e-05 | 1.367E-05 | 1.528E-06 | 1.200E-05 | 1.500E-05 |
| 21311 | 1066 | P1066L | D | 1e-05 | 1.6e-05 | 1.5e-05 | 1.367E-05 | 3.215E-06 | 1.000E-05 | 1.600E-05 |
| 21312 | 1066 | P1066K | D | 1.6e-05 | 1.5e-05 | 7.7e-06 | 1.290E-05 | 4.531E-06 | 7.700E-06 | 1.600E-05 |
| 21313 | 1066 | P1066M | D | 1.5e-05 | 1.1e-05 | 1.1e-05 | 1.233E-05 | 2.309E-06 | 1.100E-05 | 1.500E-05 |
| 21314 | 1066 | P1066F | D | 9.3e-06 | 9.6e-06 | 9.5e-06 | 9.467E-06 | 1.528E-07 | 9.300E-06 | 9.600E-06 |
| 21315 | 1066 | P1066P | D | 1.3e-05 | 2.7e-05 | 9.6e-06 | 1.653E-05 | 9.222E-06 | 9.600E-06 | 2.700E-05 |
| 21316 | 1066 | P1066S | D | 1.1e-05 | 1.5e-05 | 1.1e-05 | 1.233E-05 | 2.309E-06 | 1.100E-05 | 1.500E-05 |
| 21317 | 1066 | P1066T | D | 1.5e-05 | 9.7e-06 | 1.5e-05 | 1.323E-05 | 3.060E-06 | 9.700E-06 | 1.500E-05 |
| 21318 | 1066 | P1066W | D | 1.3e-05 | 1.5e-05 | 8.3e-06 | 1.210E-05 | 3.439E-06 | 8.300E-06 | 1.500E-05 |
| 21319 | 1066 | P1066Y | D | 1.1e-05 | 1.1e-05 | 1.1e-05 | 1.100E-05 | 2.075E-21 | 1.100E-05 | 1.100E-05 |
| 21320 | 1066 | P1066V | D | 1.2e-05 | 1.7e-05 | 1.5e-05 | 1.467E-05 | 2.517E-06 | 1.200E-05 | 1.700E-05 |
| 21321 | 1067 | S1067A | D | 1.7e-05 | 1.9e-05 | 1.2e-05 | 1.600E-05 | 3.606E-06 | 1.200E-05 | 1.900E-05 |
| 21322 | 1067 | S1067R | D | 6.7e-06 | 1.5e-05 | 1.1e-05 | 1.090E-05 | 4.151E-06 | 6.700E-06 | 1.500E-05 |
| 21323 | 1067 | S1067N | D | 1.4e-05 | 1.4e-05 | 9.7e-06 | 1.257E-05 | 2.483E-06 | 9.700E-06 | 1.400E-05 |
| 21324 | 1067 | S1067D | D | 1.2e-05 | 1.2e-05 | 2.6e-05 | 1.667E-05 | 8.083E-06 | 1.200E-05 | 2.600E-05 |
| 21325 | 1067 | S1067C | D | 1.2e-05 | 9.6e-06 | 1.6e-05 | 1.253E-05 | 3.233E-06 | 9.600E-06 | 1.600E-05 |
| 21326 | 1067 | S1067E | D | 1.6e-05 | 1e-05 | 1.1e-05 | 1.233E-05 | 3.215E-06 | 1.000E-05 | 1.600E-05 |
| 21327 | 1067 | S1067Q | D | 1.7e-05 | 1.5e-05 | 1.2e-05 | 1.467E-05 | 2.517E-06 | 1.200E-05 | 1.700E-05 |
| 21328 | 1067 | S1067G | D | 9.2e-06 | 1.1e-05 | 9.7e-06 | 9.967E-06 | 9.292E-07 | 9.200E-06 | 1.100E-05 |
| 21329 | 1067 | S1067H | D | 1.3e-05 | 1.5e-05 | 7.9e-06 | 1.197E-05 | 3.661E-06 | 7.900E-06 | 1.500E-05 |
| 21330 | 1067 | S1067I | D | 7.6e-06 | 1.3e-05 | 1.3e-05 | 1.120E-05 | 3.118E-06 | 7.600E-06 | 1.300E-05 |
| 21331 | 1067 | S1067L | D | 8e-06 | 1.6e-05 | 1.5e-05 | 1.300E-05 | 4.359E-06 | 8.000E-06 | 1.600E-05 |
| 21332 | 1067 | S1067K | D | 1.5e-05 | 1.7e-05 | 1.5e-05 | 1.567E-05 | 1.155E-06 | 1.500E-05 | 1.700E-05 |
| 21333 | 1067 | S1067M | D | 9.9e-06 | 1.4e-05 | 1.4e-05 | 1.263E-05 | 2.367E-06 | 9.900E-06 | 1.400E-05 |
| 21334 | 1067 | S1067F | D | 1.6e-05 | 1.2e-05 | 1.5e-05 | 1.433E-05 | 2.082E-06 | 1.200E-05 | 1.600E-05 |

| | | | | | | | | | | |
|---|---|---|---|---|---|---|---|---|---|---|
| 21335 | 1067 | S1067P | D | 1.4e-05 | 1.2e-05 | 1.6e-05 | 1.400E-05 | 2.000E-06 | 1.200E-05 | 1.600E-05 |
| 21336 | 1067 | S1067S | D | 1.3e-05 | 1.5e-05 | 1.7e-05 | 1.500E-05 | 2.000E-06 | 1.300E-05 | 1.700E-05 |
| 21337 | 1067 | S1067T | D | 1.4e-05 | 1.5e-05 | 1.6e-05 | 1.500E-05 | 1.000E-06 | 1.400E-05 | 1.600E-05 |
| 21338 | 1067 | S1067W | D | 1.1e-05 | 1.6e-05 | 1.1e-05 | 1.267E-05 | 2.887E-06 | 1.100E-05 | 1.600E-05 |
| 21339 | 1067 | S1067Y | D | 9.7e-06 | 1.6e-05 | 9.2e-06 | 1.163E-05 | 3.790E-06 | 9.200E-06 | 1.600E-05 |
| 21340 | 1067 | S1067V | D | 1.7e-05 | 1.1e-05 | 1.1e-05 | 1.300E-05 | 3.464E-06 | 1.100E-05 | 1.700E-05 |
| 21341 | 1068 | S1068A | D | 1.5e-05 | 1.4e-05 | 1.6e-05 | 1.500E-05 | 1.000E-06 | 1.400E-05 | 1.600E-05 |
| 21342 | 1068 | S1068R | D | 1.6e-05 | 1.3e-05 | 1.4e-05 | 1.433E-05 | 1.528E-06 | 1.300E-05 | 1.600E-05 |
| 21343 | 1068 | S1068N | D | 1.5e-05 | 1.5e-05 | 1.1e-05 | 1.367E-05 | 2.309E-06 | 1.100E-05 | 1.500E-05 |
| 21344 | 1068 | S1068D | D | 1.7e-05 | 1.1e-05 | 1.1e-05 | 1.300E-05 | 3.464E-06 | 1.100E-05 | 1.700E-05 |
| 21345 | 1068 | S1068C | D | 1e-05 | 1.5e-05 | 1.4e-05 | 1.300E-05 | 2.646E-06 | 1.000E-05 | 1.500E-05 |
| 21346 | 1068 | S1068E | D | 1.3e-05 | 1.6e-05 | 1.1e-05 | 1.333E-05 | 2.517E-06 | 1.100E-05 | 1.600E-05 |
| 21347 | 1068 | S1068Q | D | 9.1e-06 | 1.2e-05 | 1.5e-05 | 1.203E-05 | 2.950E-06 | 9.100E-06 | 1.500E-05 |
| 21348 | 1068 | S1068G | D | 1.7e-05 | 1.3e-05 | 1.4e-05 | 1.467E-05 | 2.082E-06 | 1.300E-05 | 1.700E-05 |
| 21349 | 1068 | S1068H | D | 1.4e-05 | 1.1e-05 | 2.7e-05 | 1.733E-05 | 8.505E-06 | 1.100E-05 | 2.700E-05 |
| 21350 | 1068 | S1068I | D | 1.6e-05 | 1.1e-05 | 1.6e-05 | 1.433E-05 | 2.887E-06 | 1.100E-05 | 1.600E-05 |
| 21351 | 1068 | S1068L | D | 1e-05 | 1.1e-05 | 1.1e-05 | 1.067E-05 | 5.774E-07 | 1.000E-05 | 1.100E-05 |
| 21352 | 1068 | S1068K | D | 1.5e-05 | 8.2e-06 | 1.7e-05 | 1.340E-05 | 4.613E-06 | 8.200E-06 | 1.700E-05 |
| 21353 | 1068 | S1068M | D | 1.6e-05 | 8.3e-06 | 1.5e-05 | 1.310E-05 | 4.187E-06 | 8.300E-06 | 1.600E-05 |
| 21354 | 1068 | S1068F | D | 1.5e-05 | 1e-05 | 1.8e-05 | 1.433E-05 | 4.041E-06 | 1.000E-05 | 1.800E-05 |
| 21355 | 1068 | S1068P | D | 1e-05 | 7.2e-06 | 1.5e-05 | 1.073E-05 | 3.951E-06 | 7.200E-06 | 1.500E-05 |
| 21356 | 1068 | S1068S | D | 1.5e-05 | 1.5e-05 | 1.6e-05 | 1.533E-05 | 5.774E-07 | 1.500E-05 | 1.600E-05 |
| 21357 | 1068 | S1068T | D | 1.1e-05 | 1.5e-05 | 1.1e-05 | 1.233E-05 | 2.309E-06 | 1.100E-05 | 1.500E-05 |
| 21358 | 1068 | S1068W | D | 1.5e-05 | 1.4e-05 | 1.4e-05 | 1.433E-05 | 5.774E-07 | 1.400E-05 | 1.500E-05 |
| 21359 | 1068 | S1068Y | D | 1e-05 | 9.8e-06 | 1.5e-05 | 1.160E-05 | 2.946E-06 | 9.800E-06 | 1.500E-05 |
| 21360 | 1068 | S1068V | D | 8.3e-06 | 1.4e-05 | 1.1e-05 | 1.110E-05 | 2.851E-06 | 8.300E-06 | 1.400E-05 |
| 21361 | 1069 | L1069A | D | 8.4e-06 | 1.6e-05 | 1.4e-05 | 1.280E-05 | 3.940E-06 | 8.400E-06 | 1.600E-05 |
| 21362 | 1069 | L1069R | D | 8.6e-06 | 9e-06 | 1.5e-05 | 1.087E-05 | 3.585E-06 | 8.600E-06 | 1.500E-05 |
| 21363 | 1069 | L1069N | D | 1.2e-05 | 1e-05 | 1.6e-05 | 1.267E-05 | 3.055E-06 | 1.000E-05 | 1.600E-05 |
| 21364 | 1069 | L1069D | D | 1.6e-05 | 1.5e-05 | 1.6e-05 | 1.567E-05 | 5.774E-07 | 1.500E-05 | 1.600E-05 |
| 21365 | 1069 | L1069C | D | 1.6e-05 | 9.4e-06 | 1.4e-05 | 1.313E-05 | 3.384E-06 | 9.400E-06 | 1.600E-05 |
| 21366 | 1069 | L1069E | D | 1.4e-05 | 1e-05 | 8.6e-06 | 1.087E-05 | 2.802E-06 | 8.600E-06 | 1.400E-05 |
| 21367 | 1069 | L1069Q | D | 1e-05 | 9.5e-06 | 8.7e-06 | 9.400E-06 | 6.557E-07 | 8.700E-06 | 1.000E-05 |
| 21368 | 1069 | L1069G | D | 1.5e-05 | 1.6e-05 | 1e-05 | 1.367E-05 | 3.215E-06 | 1.000E-05 | 1.600E-05 |
| 21369 | 1069 | L1069H | D | 1.4e-05 | 1.5e-05 | 1.3e-05 | 1.400E-05 | 1.000E-06 | 1.300E-05 | 1.500E-05 |
| 21370 | 1069 | L1069I | D | 1.4e-05 | 9.8e-06 | 1.6e-05 | 1.327E-05 | 3.164E-06 | 9.800E-06 | 1.600E-05 |
| 21371 | 1069 | L1069L | D | 1e-05 | 7.2e-06 | 1.1e-05 | 9.400E-06 | 1.970E-06 | 7.200E-06 | 1.100E-05 |
| 21372 | 1069 | L1069K | D | 2.5e-05 | 1.5e-05 | 1.4e-05 | 1.800E-05 | 6.083E-06 | 1.400E-05 | 2.500E-05 |
| 21373 | 1069 | L1069M | D | 1.1e-05 | 1e-05 | 1.4e-05 | 1.167E-05 | 2.082E-06 | 1.000E-05 | 1.400E-05 |
| 21374 | 1069 | L1069F | D | 1.1e-05 | 1.4e-05 | 1.1e-05 | 1.200E-05 | 1.732E-06 | 1.100E-05 | 1.400E-05 |
| 21375 | 1069 | L1069P | D | 1.5e-05 | 1e-05 | 1.4e-05 | 1.300E-05 | 2.646E-06 | 1.000E-05 | 1.500E-05 |
| 21376 | 1069 | L1069S | D | 1.3e-05 | 1.3e-05 | 1.1e-05 | 1.233E-05 | 1.155E-06 | 1.100E-05 | 1.300E-05 |
| 21377 | 1069 | L1069T | D | 1.4e-05 | 1.3e-05 | 9.2e-06 | 1.207E-05 | 2.532E-06 | 9.200E-06 | 1.400E-05 |
| 21378 | 1069 | L1069W | D | 1.4e-05 | 1.7e-05 | 1e-05 | 1.367E-05 | 3.512E-06 | 1.000E-05 | 1.700E-05 |
| 21379 | 1069 | L1069Y | D | 1.2e-05 | 1.1e-05 | 1.5e-05 | 1.267E-05 | 2.082E-06 | 1.100E-05 | 1.500E-05 |
| 21380 | 1069 | L1069V | D | 1.3e-05 | 1.1e-05 | 8.9e-06 | 1.097E-05 | 2.050E-06 | 8.900E-06 | 1.300E-05 |
| 21381 | 1070 | S1070A | D | 9.8e-06 | 9.1e-06 | 1.3e-05 | 1.063E-05 | 2.079E-06 | 9.100E-06 | 1.300E-05 |
| 21382 | 1070 | S1070R | D | 1e-05 | 1e-05 | 1.5e-05 | 1.167E-05 | 2.887E-06 | 1.000E-05 | 1.500E-05 |
| 21383 | 1070 | S1070N | D | 1.6e-05 | 1.4e-05 | 1.6e-05 | 1.533E-05 | 1.155E-06 | 1.400E-05 | 1.600E-05 |
| 21384 | 1070 | S1070D | D | 9.1e-06 | 1e-05 | 1.6e-05 | 1.170E-05 | 3.751E-06 | 9.100E-06 | 1.600E-05 |
| 21385 | 1070 | S1070C | D | 8.9e-06 | 1.3e-05 | 1e-05 | 1.063E-05 | 2.122E-06 | 8.900E-06 | 1.300E-05 |
| 21386 | 1070 | S1070E | D | 1.4e-05 | 1.6e-05 | 1.1e-05 | 1.367E-05 | 2.517E-06 | 1.100E-05 | 1.600E-05 |
| 21387 | 1070 | S1070Q | D | 1.6e-05 | 1.7e-05 | 1.5e-05 | 1.600E-05 | 1.000E-06 | 1.500E-05 | 1.700E-05 |
| 21388 | 1070 | S1070G | D | 1.3e-05 | 1.5e-05 | 1.3e-05 | 1.367E-05 | 1.155E-06 | 1.300E-05 | 1.500E-05 |
| 21389 | 1070 | S1070H | D | 8.6e-06 | 1e-05 | 1e-05 | 9.533E-06 | 8.083E-07 | 8.600E-06 | 1.000E-05 |
| 21390 | 1070 | S1070I | D | 8.9e-06 | 1e-05 | 1.3e-05 | 1.063E-05 | 2.122E-06 | 8.900E-06 | 1.300E-05 |
| 21391 | 1070 | S1070L | D | 1.7e-05 | 1.2e-05 | 7.6e-06 | 1.220E-05 | 4.703E-06 | 7.600E-06 | 1.700E-05 |
| 21392 | 1070 | S1070K | D | 1.7e-05 | 1.4e-05 | 1.6e-05 | 1.567E-05 | 1.528E-06 | 1.400E-05 | 1.700E-05 |
| 21393 | 1070 | S1070M | D | 6.8e-06 | 1.2e-05 | 1.6e-05 | 1.160E-05 | 4.613E-06 | 6.800E-06 | 1.600E-05 |
| 21394 | 1070 | S1070F | D | 1.5e-05 | 9.8e-06 | 1.6e-05 | 1.360E-05 | 3.329E-06 | 9.800E-06 | 1.600E-05 |
| 21395 | 1070 | S1070P | D | 1.6e-05 | 1.5e-05 | 1.3e-05 | 1.467E-05 | 1.528E-06 | 1.300E-05 | 1.600E-05 |
| 21396 | 1070 | S1070S | D | 1.4e-05 | 1.1e-05 | 1.1e-05 | 1.200E-05 | 1.732E-06 | 1.100E-05 | 1.400E-05 |
| 21397 | 1070 | S1070T | D | 1e-05 | 1.1e-05 | 1.4e-05 | 1.167E-05 | 2.082E-06 | 1.000E-05 | 1.400E-05 |
| 21398 | 1070 | S1070W | D | 7.5e-06 | 1.1e-05 | 1.1e-05 | 9.833E-06 | 2.021E-06 | 7.500E-06 | 1.100E-05 |
| 21399 | 1070 | S1070Y | D | 1.4e-05 | 1.5e-05 | 1.1e-05 | 1.333E-05 | 2.082E-06 | 1.100E-05 | 1.500E-05 |
| 21400 | 1070 | S1070V | D | 9.4e-06 | 1.4e-05 | 1.3e-05 | 1.213E-05 | 2.419E-06 | 9.400E-06 | 1.400E-05 |
| 21401 | 1071 | A1071A | D | 1e-05 | 1.5e-05 | 9.9e-06 | 1.163E-05 | 2.916E-06 | 9.900E-06 | 1.500E-05 |
| 21402 | 1071 | A1071R | D | 1.6e-05 | 8.3e-06 | 1.6e-05 | 1.343E-05 | 4.446E-06 | 8.300E-06 | 1.600E-05 |
| 21403 | 1071 | A1071N | D | 1.4e-05 | 1.5e-05 | 1.4e-05 | 1.433E-05 | 5.774E-07 | 1.400E-05 | 1.500E-05 |
| 21404 | 1071 | A1071D | D | 1.4e-05 | 1.6e-05 | 1e-05 | 1.333E-05 | 3.055E-06 | 1.000E-05 | 1.600E-05 |
| 21405 | 1071 | A1071C | D | 1.4e-05 | 9.7e-06 | 9.7e-06 | 1.113E-05 | 2.483E-06 | 9.700E-06 | 1.400E-05 |
| 21406 | 1071 | A1071E | D | 1.5e-05 | 1.4e-05 | 1.3e-05 | 1.400E-05 | 1.000E-06 | 1.300E-05 | 1.500E-05 |
| 21407 | 1071 | A1071Q | D | 1.2e-05 | 1.5e-05 | 1.4e-05 | 1.367E-05 | 1.528E-06 | 1.200E-05 | 1.500E-05 |
| 21408 | 1071 | A1071G | D | 1.7e-05 | 1e-05 | 9.3e-06 | 1.210E-05 | 4.258E-06 | 9.300E-06 | 1.700E-05 |
| 21409 | 1071 | A1071H | D | 8.2e-06 | 1.5e-05 | 9.5e-06 | 1.090E-05 | 3.610E-06 | 8.200E-06 | 1.500E-05 |
| 21410 | 1071 | A1071I | D | 1.5e-05 | 1.1e-05 | 9.8e-06 | 1.193E-05 | 2.723E-06 | 9.800E-06 | 1.500E-05 |
| 21411 | 1071 | A1071L | D | 1.5e-05 | 1.6e-05 | 1.6e-05 | 1.567E-05 | 5.774E-07 | 1.500E-05 | 1.600E-05 |
| 21412 | 1071 | A1071K | D | 1.8e-05 | 1.6e-05 | 1e-05 | 1.467E-05 | 4.163E-06 | 1.000E-05 | 1.800E-05 |
| 21413 | 1071 | A1071M | D | 1.6e-05 | 1.1e-05 | 1.2e-05 | 1.300E-05 | 2.646E-06 | 1.100E-05 | 1.600E-05 |
| 21414 | 1071 | A1071F | D | 1.3e-05 | 1.7e-05 | 9.8e-06 | 1.327E-05 | 3.607E-06 | 9.800E-06 | 1.700E-05 |
| 21415 | 1071 | A1071P | D | 1.4e-05 | 9.4e-06 | 1e-05 | 1.113E-05 | 2.501E-06 | 9.400E-06 | 1.400E-05 |
| 21416 | 1071 | A1071S | D | 1.1e-05 | 1.4e-05 | 1.1e-05 | 1.200E-05 | 1.732E-06 | 1.100E-05 | 1.400E-05 |
| 21417 | 1071 | A1071T | D | 1.1e-05 | 1.5e-05 | 1.5e-05 | 1.367E-05 | 2.309E-06 | 1.100E-05 | 1.500E-05 |
| 21418 | 1071 | A1071W | D | 1.3e-05 | 9.6e-06 | 1.4e-05 | 1.220E-05 | 2.307E-06 | 9.600E-06 | 1.400E-05 |
| 21419 | 1071 | A1071Y | D | 1.4e-05 | 1e-05 | 1.1e-05 | 1.167E-05 | 2.082E-06 | 1.000E-05 | 1.400E-05 |

| | | | | | | | | | | |
|---|---|---|---|---|---|---|---|---|---|---|
| 21420 | 1071 | A1071V | D | 9.4e-06 | 9.6e-06 | 1.1e-05 | 1.000E-05 | 8.718E-07 | 9.400E-06 | 1.100E-05 |
| 21421 | 1072 | S1072A | D | 1.6e-05 | 1.5e-05 | 1.5e-05 | 1.533E-05 | 5.774E-07 | 1.500E-05 | 1.600E-05 |
| 21422 | 1072 | S1072R | D | 8.6e-06 | 1.4e-05 | 1.1e-05 | 1.120E-05 | 2.706E-06 | 8.600E-06 | 1.400E-05 |
| 21423 | 1072 | S1072N | D | 9.9e-06 | 1.1e-05 | 1.3e-05 | 1.130E-05 | 1.572E-06 | 9.900E-06 | 1.300E-05 |
| 21424 | 1072 | S1072D | D | 1.1e-05 | 9.9e-06 | 1e-05 | 1.030E-05 | 6.083E-07 | 9.900E-06 | 1.100E-05 |
| 21425 | 1072 | S1072C | D | 1.5e-05 | 1.5e-05 | 1.5e-05 | 1.500E-05 | 0.000E+00 | 1.500E-05 | 1.500E-05 |
| 21426 | 1072 | S1072E | D | 1.6e-05 | 1.6e-05 | 1.1e-05 | 1.433E-05 | 2.887E-06 | 1.100E-05 | 1.600E-05 |
| 21427 | 1072 | S1072Q | D | 9.9e-06 | 9.7e-06 | 1.4e-05 | 1.120E-05 | 2.427E-06 | 9.700E-06 | 1.400E-05 |
| 21428 | 1072 | S1072G | D | 1.9e-05 | 1.4e-05 | 1.2e-05 | 1.500E-05 | 3.606E-06 | 1.200E-05 | 1.900E-05 |
| 21429 | 1072 | S1072H | D | 1.5e-05 | 1.5e-05 | 1.2e-05 | 1.400E-05 | 1.732E-06 | 1.200E-05 | 1.500E-05 |
| 21430 | 1072 | S1072I | D | 1e-05 | 1e-05 | 1.4e-05 | 1.133E-05 | 2.309E-06 | 1.000E-05 | 1.400E-05 |
| 21431 | 1072 | S1072L | D | 8.2e-06 | 1.3e-05 | 1.6e-05 | 1.240E-05 | 3.934E-06 | 8.200E-06 | 1.600E-05 |
| 21432 | 1072 | S1072K | D | 9e-06 | 9.8e-06 | 1.1e-05 | 9.933E-06 | 1.007E-06 | 9.000E-06 | 1.100E-05 |
| 21433 | 1072 | S1072M | D | 9.5e-06 | 1.8e-05 | 1.4e-05 | 1.383E-05 | 4.252E-06 | 9.500E-06 | 1.800E-05 |
| 21434 | 1072 | S1072F | D | 1.5e-05 | 1.1e-05 | 1e-05 | 1.200E-05 | 2.646E-06 | 1.000E-05 | 1.500E-05 |
| 21435 | 1072 | S1072P | D | 1e-05 | 1.1e-05 | 9.9e-06 | 1.030E-05 | 6.083E-07 | 9.900E-06 | 1.100E-05 |
| 21436 | 1072 | S1072S | D | 1.5e-05 | 1.5e-05 | 1.6e-05 | 1.533E-05 | 5.774E-07 | 1.500E-05 | 1.600E-05 |
| 21437 | 1072 | S1072T | D | 1.4e-05 | 8.5e-06 | 1.6e-05 | 1.283E-05 | 3.884E-06 | 8.500E-06 | 1.600E-05 |
| 21438 | 1072 | S1072W | D | 1.1e-05 | 1.6e-05 | 1.5e-05 | 1.400E-05 | 2.646E-06 | 1.100E-05 | 1.600E-05 |
| 21439 | 1072 | S1072Y | D | 1.6e-05 | 1.3e-05 | 1.5e-05 | 1.467E-05 | 1.528E-06 | 1.300E-05 | 1.600E-05 |
| 21440 | 1072 | S1072V | D | 1.5e-05 | 1.1e-05 | 1.5e-05 | 1.367E-05 | 2.309E-06 | 1.100E-05 | 1.500E-05 |
| 21441 | 1073 | V1073A | D | 1.5e-05 | 2.1e-05 | 1e-05 | 1.533E-05 | 5.508E-06 | 1.000E-05 | 2.100E-05 |
| 21442 | 1073 | V1073R | D | 1.1e-05 | 1.1e-05 | 1.4e-05 | 1.200E-05 | 1.732E-06 | 1.100E-05 | 1.400E-05 |
| 21443 | 1073 | V1073N | D | 9.8e-06 | 9.3e-06 | 1.2e-05 | 1.037E-05 | 1.436E-06 | 9.300E-06 | 1.200E-05 |
| 21444 | 1073 | V1073D | D | 1.5e-05 | 1.2e-05 | 1.3e-05 | 1.333E-05 | 1.528E-06 | 1.200E-05 | 1.500E-05 |
| 21445 | 1073 | V1073C | D | 1.3e-05 | 1.4e-05 | 1.4e-05 | 1.367E-05 | 5.774E-07 | 1.300E-05 | 1.400E-05 |
| 21446 | 1073 | V1073E | D | 1.3e-05 | 1.5e-05 | 9.4e-06 | 1.247E-05 | 2.838E-06 | 9.400E-06 | 1.500E-05 |
| 21447 | 1073 | V1073Q | D | 1.4e-05 | 1.3e-05 | 9.3e-06 | 1.210E-05 | 2.476E-06 | 9.300E-06 | 1.400E-05 |
| 21448 | 1073 | V1073G | D | 1.3e-05 | 7.2e-06 | 8.6e-06 | 9.600E-06 | 3.027E-06 | 7.200E-06 | 1.300E-05 |
| 21449 | 1073 | V1073H | D | 1.1e-05 | 1.6e-05 | 1.4e-05 | 1.367E-05 | 2.517E-06 | 1.100E-05 | 1.600E-05 |
| 21450 | 1073 | V1073I | D | 1.5e-05 | 8.7e-06 | 1.1e-05 | 1.157E-05 | 3.188E-06 | 8.700E-06 | 1.500E-05 |
| 21451 | 1073 | V1073L | D | 1.1e-05 | 1.6e-05 | 1.5e-05 | 1.400E-05 | 2.646E-06 | 1.100E-05 | 1.600E-05 |
| 21452 | 1073 | V1073K | D | 8.6e-06 | 1.4e-05 | 1.1e-05 | 1.120E-05 | 2.706E-06 | 8.600E-06 | 1.400E-05 |
| 21453 | 1073 | V1073M | D | 1.3e-05 | 1.4e-05 | 6.6e-06 | 1.120E-05 | 4.015E-06 | 6.600E-06 | 1.400E-05 |
| 21454 | 1073 | V1073F | D | 1.2e-05 | 8.9e-06 | 1.5e-05 | 1.197E-05 | 3.050E-06 | 8.900E-06 | 1.500E-05 |
| 21455 | 1073 | V1073P | D | 1.5e-05 | 1.5e-05 | 1.7e-05 | 1.567E-05 | 1.155E-06 | 1.500E-05 | 1.700E-05 |
| 21456 | 1073 | V1073S | D | 1.1e-05 | 1.7e-05 | 8.3e-06 | 1.210E-05 | 4.453E-06 | 8.300E-06 | 1.700E-05 |
| 21457 | 1073 | V1073T | D | 1.5e-05 | 1.3e-05 | 1.4e-05 | 1.400E-05 | 1.000E-06 | 1.300E-05 | 1.500E-05 |
| 21458 | 1073 | V1073W | D | 1.4e-05 | 1.6e-05 | 6.5e-06 | 1.217E-05 | 5.008E-06 | 6.500E-06 | 1.600E-05 |
| 21459 | 1073 | V1073Y | D | 9.5e-06 | 1.4e-05 | 1e-05 | 1.117E-05 | 2.466E-06 | 9.500E-06 | 1.400E-05 |
| 21460 | 1073 | V1073V | D | 1e-05 | 1.5e-05 | 1.1e-05 | 1.200E-05 | 2.646E-06 | 1.000E-05 | 1.500E-05 |
| 21461 | 1074 | G1074A | D | 1.2e-05 | 1.2e-05 | 1.5e-05 | 1.300E-05 | 1.732E-06 | 1.200E-05 | 1.500E-05 |
| 21462 | 1074 | G1074R | D | 9.8e-06 | 1.1e-05 | 1.3e-05 | 1.127E-05 | 1.617E-06 | 9.800E-06 | 1.300E-05 |
| 21463 | 1074 | G1074N | D | 1.6e-05 | 1.2e-05 | 1.6e-05 | 1.467E-05 | 2.309E-06 | 1.200E-05 | 1.600E-05 |
| 21464 | 1074 | G1074D | D | 1.4e-05 | 1.6e-05 | 1.5e-05 | 1.500E-05 | 1.000E-06 | 1.400E-05 | 1.600E-05 |
| 21465 | 1074 | G1074C | D | 8.2e-06 | 1e-05 | 8.1e-06 | 8.767E-06 | 1.069E-06 | 8.100E-06 | 1.000E-05 |
| 21466 | 1074 | G1074E | D | 1e-05 | 1.3e-05 | 1.6e-05 | 1.300E-05 | 3.000E-06 | 1.000E-05 | 1.600E-05 |
| 21467 | 1074 | G1074Q | D | 9.3e-06 | 1.5e-05 | 1.6e-05 | 1.343E-05 | 3.614E-06 | 9.300E-06 | 1.600E-05 |
| 21468 | 1074 | G1074G | D | 1.5e-05 | 1.5e-05 | 1.1e-05 | 1.367E-05 | 2.309E-06 | 1.100E-05 | 1.500E-05 |
| 21469 | 1074 | G1074H | D | 7.9e-06 | 1.2e-05 | 1.4e-05 | 1.130E-05 | 3.110E-06 | 7.900E-06 | 1.400E-05 |
| 21470 | 1074 | G1074I | D | 1.5e-05 | 9.5e-06 | 1.8e-05 | 1.417E-05 | 4.311E-06 | 9.500E-06 | 1.800E-05 |
| 21471 | 1074 | G1074L | D | 9.2e-06 | 1.3e-05 | 1.5e-05 | 1.240E-05 | 2.946E-06 | 9.200E-06 | 1.500E-05 |
| 21472 | 1074 | G1074K | D | 7.8e-06 | 7.7e-06 | 1.1e-05 | 8.833E-06 | 1.877E-06 | 7.700E-06 | 1.100E-05 |
| 21473 | 1074 | G1074M | D | 1.1e-05 | 1.6e-05 | 9.9e-06 | 1.230E-05 | 3.251E-06 | 9.900E-06 | 1.600E-05 |
| 21474 | 1074 | G1074F | D | 7.6e-06 | 1.4e-05 | 9.9e-06 | 1.050E-05 | 3.242E-06 | 7.600E-06 | 1.400E-05 |
| 21475 | 1074 | G1074P | D | 8.5e-06 | 1.2e-05 | 1.2e-05 | 1.083E-05 | 2.021E-06 | 8.500E-06 | 1.200E-05 |
| 21476 | 1074 | G1074S | D | 1.1e-05 | 1.3e-05 | 1e-05 | 1.133E-05 | 1.528E-06 | 1.000E-05 | 1.300E-05 |
| 21477 | 1074 | G1074T | D | 1.6e-05 | 1.4e-05 | 1.5e-05 | 1.500E-05 | 1.000E-06 | 1.400E-05 | 1.600E-05 |
| 21478 | 1074 | G1074W | D | 9.8e-06 | 9.4e-06 | 1.3e-05 | 1.073E-05 | 1.973E-06 | 9.400E-06 | 1.300E-05 |
| 21479 | 1074 | G1074Y | D | 1e-05 | 1.3e-05 | 1.3e-05 | 1.200E-05 | 1.732E-06 | 1.000E-05 | 1.300E-05 |
| 21480 | 1074 | G1074V | D | 1.5e-05 | 1e-05 | 1.1e-05 | 1.200E-05 | 2.646E-06 | 1.000E-05 | 1.500E-05 |
| 21481 | 1075 | D1075A | D | 1.6e-05 | 1.4e-05 | 8.8e-06 | 1.293E-05 | 3.717E-06 | 8.800E-06 | 1.600E-05 |
| 21482 | 1075 | D1075R | D | 1.5e-05 | 1.2e-05 | 9.7e-06 | 1.223E-05 | 2.658E-06 | 9.700E-06 | 1.500E-05 |
| 21483 | 1075 | D1075N | D | 1.3e-05 | 1.4e-05 | 1.5e-05 | 1.400E-05 | 1.000E-06 | 1.300E-05 | 1.500E-05 |
| 21484 | 1075 | D1075D | D | 1.7e-05 | 1.7e-05 | 1.1e-05 | 1.500E-05 | 3.464E-06 | 1.100E-05 | 1.700E-05 |
| 21485 | 1075 | D1075C | D | 1.4e-05 | 9e-06 | 8.9e-06 | 1.063E-05 | 2.916E-06 | 8.900E-06 | 1.400E-05 |
| 21486 | 1075 | D1075E | D | 1e-05 | 1.3e-05 | 1.5e-05 | 1.267E-05 | 2.517E-06 | 1.000E-05 | 1.500E-05 |
| 21487 | 1075 | D1075Q | D | 1.1e-05 | 1e-05 | 1.5e-05 | 1.200E-05 | 2.646E-06 | 1.000E-05 | 1.500E-05 |
| 21488 | 1075 | D1075G | D | 1e-05 | 8.8e-06 | 1.4e-05 | 1.093E-05 | 2.723E-06 | 8.800E-06 | 1.400E-05 |
| 21489 | 1075 | D1075H | D | 9.1e-06 | 1.3e-05 | 1.4e-05 | 1.203E-05 | 2.589E-06 | 9.100E-06 | 1.400E-05 |
| 21490 | 1075 | D1075I | D | 1.5e-05 | 1.1e-05 | 1.3e-05 | 1.300E-05 | 2.000E-06 | 1.100E-05 | 1.500E-05 |
| 21491 | 1075 | D1075L | D | 1.4e-05 | 1.6e-05 | 1.1e-05 | 1.367E-05 | 2.517E-06 | 1.100E-05 | 1.600E-05 |
| 21492 | 1075 | D1075K | D | 1.5e-05 | 1.4e-05 | 9.8e-06 | 1.293E-05 | 2.759E-06 | 9.800E-06 | 1.500E-05 |
| 21493 | 1075 | D1075M | D | 1.5e-05 | 1.4e-05 | 1.5e-05 | 1.467E-05 | 5.774E-07 | 1.400E-05 | 1.500E-05 |
| 21494 | 1075 | D1075F | D | 1e-05 | 9.6e-06 | 1.5e-05 | 1.153E-05 | 3.009E-06 | 9.600E-06 | 1.500E-05 |
| 21495 | 1075 | D1075P | D | 1.1e-05 | 1.5e-05 | 1.3e-05 | 1.300E-05 | 2.000E-06 | 1.100E-05 | 1.500E-05 |
| 21496 | 1075 | D1075S | D | 1.2e-05 | 1.4e-05 | 1.3e-05 | 1.300E-05 | 1.000E-06 | 1.200E-05 | 1.400E-05 |
| 21497 | 1075 | D1075T | D | 1e-05 | 1.5e-05 | 9.3e-06 | 1.143E-05 | 3.109E-06 | 9.300E-06 | 1.500E-05 |
| 21498 | 1075 | D1075W | D | 8.6e-06 | 9.9e-06 | 9.2e-06 | 9.233E-06 | 6.506E-07 | 8.600E-06 | 9.900E-06 |
| 21499 | 1075 | D1075Y | D | 1.5e-05 | 1.4e-05 | 1.4e-05 | 1.433E-05 | 5.774E-07 | 1.400E-05 | 1.500E-05 |
| 21500 | 1075 | D1075V | D | 9.9e-06 | 1.1e-05 | 1e-05 | 1.030E-05 | 6.083E-07 | 9.900E-06 | 1.100E-05 |
| 21501 | 1076 | R1076A | D | 1.1e-05 | 1.6e-05 | 9.3e-06 | 1.210E-05 | 3.483E-06 | 9.300E-06 | 1.600E-05 |
| 21502 | 1076 | R1076R | D | 9.3e-06 | 1e-05 | 9.1e-06 | 9.467E-06 | 4.726E-07 | 9.100E-06 | 1.000E-05 |
| 21503 | 1076 | R1076N | D | 1.5e-05 | 1.1e-05 | 1.6e-05 | 1.400E-05 | 2.646E-06 | 1.100E-05 | 1.600E-05 |
| 21504 | 1076 | R1076D | D | 1.6e-05 | 1.3e-05 | 1.1e-05 | 1.333E-05 | 2.517E-06 | 1.100E-05 | 1.600E-05 |

| | | | | | | | | | | |
|---|---|---|---|---|---|---|---|---|---|---|
| 21505 | 1076 | R1076C | D | 1.4e-05 | 9.4e-06 | 9.3e-06 | 1.090E-05 | 2.685E-06 | 9.300E-06 | 1.400E-05 |
| 21506 | 1076 | R1076E | D | 1.3e-05 | 1.4e-05 | 1.1e-05 | 1.267E-05 | 1.528E-06 | 1.100E-05 | 1.400E-05 |
| 21507 | 1076 | R1076Q | D | 1.2e-05 | 1.4e-05 | 1.4e-05 | 1.333E-05 | 1.155E-06 | 1.200E-05 | 1.400E-05 |
| 21508 | 1076 | R1076G | D | 1.5e-05 | 1.5e-05 | 1e-05 | 1.333E-05 | 2.887E-06 | 1.000E-05 | 1.500E-05 |
| 21509 | 1076 | R1076H | D | 8.5e-06 | 1.4e-05 | 1.6e-05 | 1.283E-05 | 3.884E-06 | 8.500E-06 | 1.600E-05 |
| 21510 | 1076 | R1076I | D | 9.6e-06 | 1.4e-05 | 1.4e-05 | 1.253E-05 | 2.540E-06 | 9.600E-06 | 1.400E-05 |
| 21511 | 1076 | R1076L | D | 9.1e-06 | 8.8e-06 | 1.4e-05 | 1.063E-05 | 2.919E-06 | 8.800E-06 | 1.400E-05 |
| 21512 | 1076 | R1076K | D | 1.2e-05 | 1.7e-05 | 1.4e-05 | 1.433E-05 | 2.517E-06 | 1.200E-05 | 1.700E-05 |
| 21513 | 1076 | R1076M | D | 1.5e-05 | 1.4e-05 | 1.3e-05 | 1.400E-05 | 1.000E-06 | 1.300E-05 | 1.500E-05 |
| 21514 | 1076 | R1076F | D | 1.6e-05 | 1.4e-05 | 1.1e-05 | 1.367E-05 | 2.517E-06 | 1.100E-05 | 1.600E-05 |
| 21515 | 1076 | R1076P | D | 8.3e-06 | 1.3e-05 | 1.8e-05 | 1.310E-05 | 4.851E-06 | 8.300E-06 | 1.800E-05 |
| 21516 | 1076 | R1076S | D | 1.5e-05 | 1.1e-05 | 1.7e-05 | 1.433E-05 | 3.055E-06 | 1.100E-05 | 1.700E-05 |
| 21517 | 1076 | R1076T | D | 1.4e-05 | 1.5e-05 | 1.5e-05 | 1.467E-05 | 5.774E-07 | 1.400E-05 | 1.500E-05 |
| 21518 | 1076 | R1076W | D | 1.1e-05 | 1.1e-05 | 9.9e-06 | 1.063E-05 | 6.351E-07 | 9.900E-06 | 1.100E-05 |
| 21519 | 1076 | R1076Y | D | 1.1e-05 | 1.3e-05 | 1.2e-05 | 1.200E-05 | 1.000E-06 | 1.100E-05 | 1.300E-05 |
| 21520 | 1076 | R1076V | D | 1.6e-05 | 1.2e-05 | 1.1e-05 | 1.300E-05 | 2.646E-06 | 1.100E-05 | 1.600E-05 |
| 21521 | 1077 | V1077A | D | 9.7e-06 | 2.1e-05 | 1.5e-05 | 1.523E-05 | 5.654E-06 | 9.700E-06 | 2.100E-05 |
| 21522 | 1077 | V1077R | D | 1.4e-05 | 1e-05 | 1.7e-05 | 1.367E-05 | 3.512E-06 | 1.000E-05 | 1.700E-05 |
| 21523 | 1077 | V1077N | D | 1.6e-05 | 1.6e-05 | 1.1e-05 | 1.433E-05 | 2.887E-06 | 1.100E-05 | 1.600E-05 |
| 21524 | 1077 | V1077D | D | 9.5e-06 | 1.1e-05 | 1.1e-05 | 1.050E-05 | 8.660E-07 | 9.500E-06 | 1.100E-05 |
| 21525 | 1077 | V1077C | D | 1.4e-05 | 1.5e-05 | 8.7e-06 | 1.257E-05 | 3.386E-06 | 8.700E-06 | 1.500E-05 |
| 21526 | 1077 | V1077E | D | 9.8e-06 | 1.4e-05 | 1.6e-05 | 1.327E-05 | 3.164E-06 | 9.800E-06 | 1.600E-05 |
| 21527 | 1077 | V1077Q | D | 1.3e-05 | 1.3e-05 | 1.3e-05 | 1.300E-05 | 0.000E+00 | 1.300E-05 | 1.300E-05 |
| 21528 | 1077 | V1077G | D | 9.6e-06 | 1.2e-05 | 1e-05 | 1.053E-05 | 1.286E-06 | 9.600E-06 | 1.200E-05 |
| 21529 | 1077 | V1077H | D | 9e-06 | 1.6e-05 | 1e-05 | 1.167E-05 | 3.786E-06 | 9.000E-06 | 1.600E-05 |
| 21530 | 1077 | V1077I | D | 9.5e-06 | 1.5e-05 | 1.5e-05 | 1.317E-05 | 3.175E-06 | 9.500E-06 | 1.500E-05 |
| 21531 | 1077 | V1077L | D | 1.1e-05 | 1.2e-05 | 1e-05 | 1.100E-05 | 1.000E-06 | 1.000E-05 | 1.200E-05 |
| 21532 | 1077 | V1077K | D | 8e-06 | 9.5e-06 | 1.3e-05 | 1.017E-05 | 2.566E-06 | 8.000E-06 | 1.300E-05 |
| 21533 | 1077 | V1077M | D | 1.6e-05 | 1.6e-05 | 1.5e-05 | 1.567E-05 | 5.774E-07 | 1.500E-05 | 1.600E-05 |
| 21534 | 1077 | V1077F | D | 9.8e-06 | 1e-05 | 7.2e-06 | 9.000E-06 | 1.562E-06 | 7.200E-06 | 1.000E-05 |
| 21535 | 1077 | V1077P | D | 1.6e-05 | 1.6e-05 | 1e-05 | 1.400E-05 | 3.464E-06 | 1.000E-05 | 1.600E-05 |
| 21536 | 1077 | V1077S | D | 1.1e-05 | 1.5e-05 | 1.5e-05 | 1.367E-05 | 2.309E-06 | 1.100E-05 | 1.500E-05 |
| 21537 | 1077 | V1077T | D | 1.6e-05 | 1.5e-05 | 1.4e-05 | 1.500E-05 | 1.000E-06 | 1.400E-05 | 1.600E-05 |
| 21538 | 1077 | V1077W | D | 1.4e-05 | 1e-05 | 1.1e-05 | 1.167E-05 | 2.082E-06 | 1.000E-05 | 1.400E-05 |
| 21539 | 1077 | V1077Y | D | 1.4e-05 | 9.5e-06 | 1.5e-05 | 1.283E-05 | 2.930E-06 | 9.500E-06 | 1.500E-05 |
| 21540 | 1077 | V1077V | D | 1.1e-05 | 1.7e-05 | 8.8e-06 | 1.227E-05 | 4.244E-06 | 8.800E-06 | 1.700E-05 |
| 21541 | 1078 | T1078A | D | 1.4e-05 | 1.1e-05 | 1.7e-05 | 1.400E-05 | 3.000E-06 | 1.100E-05 | 1.700E-05 |
| 21542 | 1078 | T1078R | D | 1.2e-05 | 1.1e-05 | 1.6e-05 | 1.300E-05 | 2.646E-06 | 1.100E-05 | 1.600E-05 |
| 21543 | 1078 | T1078N | D | 9e-06 | 1e-05 | 1.6e-05 | 1.167E-05 | 3.786E-06 | 9.000E-06 | 1.600E-05 |
| 21544 | 1078 | T1078D | D | 1.8e-05 | 1.4e-05 | 1.4e-05 | 1.533E-05 | 2.309E-06 | 1.400E-05 | 1.800E-05 |
| 21545 | 1078 | T1078C | D | 9.2e-06 | 1.3e-05 | 1.3e-05 | 1.173E-05 | 2.194E-06 | 9.200E-06 | 1.300E-05 |
| 21546 | 1078 | T1078E | D | 1.4e-05 | 1.1e-05 | 1e-05 | 1.167E-05 | 2.082E-06 | 1.000E-05 | 1.400E-05 |
| 21547 | 1078 | T1078Q | D | 1.6e-05 | 9.6e-06 | 1.1e-05 | 1.220E-05 | 3.365E-06 | 9.600E-06 | 1.600E-05 |
| 21548 | 1078 | T1078G | D | 1.2e-05 | 1.5e-05 | 1.3e-05 | 1.333E-05 | 1.528E-06 | 1.200E-05 | 1.500E-05 |
| 21549 | 1078 | T1078H | D | 1.5e-05 | 1.4e-05 | 8.7e-06 | 1.257E-05 | 3.386E-06 | 8.700E-06 | 1.500E-05 |
| 21550 | 1078 | T1078I | D | 1.5e-05 | 1.5e-05 | 1.5e-05 | 1.500E-05 | 0.000E+00 | 1.500E-05 | 1.500E-05 |
| 21551 | 1078 | T1078L | D | 1.5e-05 | 1.7e-05 | 1.8e-05 | 1.667E-05 | 1.528E-06 | 1.500E-05 | 1.800E-05 |
| 21552 | 1078 | T1078K | D | 1.2e-05 | 1.3e-05 | 1.3e-05 | 1.267E-05 | 5.774E-07 | 1.200E-05 | 1.300E-05 |
| 21553 | 1078 | T1078M | D | 1.3e-05 | 1.4e-05 | 1.7e-05 | 1.467E-05 | 2.082E-06 | 1.300E-05 | 1.700E-05 |
| 21554 | 1078 | T1078F | D | 1.4e-05 | 9.1e-06 | 9e-06 | 1.070E-05 | 2.858E-06 | 9.000E-06 | 1.400E-05 |
| 21555 | 1078 | T1078P | D | 1.6e-05 | 1.1e-05 | 1.2e-05 | 1.300E-05 | 2.646E-06 | 1.100E-05 | 1.600E-05 |
| 21556 | 1078 | T1078S | D | 1.1e-05 | 1.5e-05 | 1.2e-05 | 1.267E-05 | 2.082E-06 | 1.100E-05 | 1.500E-05 |
| 21557 | 1078 | T1078T | D | 1.6e-05 | 1e-05 | 1.8e-05 | 1.467E-05 | 4.163E-06 | 1.000E-05 | 1.800E-05 |
| 21558 | 1078 | T1078W | D | 1.5e-05 | 1.2e-05 | 1.3e-05 | 1.333E-05 | 1.528E-06 | 1.200E-05 | 1.500E-05 |
| 21559 | 1078 | T1078Y | D | 1.6e-05 | 1e-05 | 1.6e-05 | 1.400E-05 | 3.464E-06 | 1.000E-05 | 1.600E-05 |
| 21560 | 1078 | T1078V | D | 9.4e-06 | 1e-05 | 1.2e-05 | 1.047E-05 | 1.361E-06 | 9.400E-06 | 1.200E-05 |
| 21561 | 1079 | I1079A | D | 1.1e-05 | 1.4e-05 | 9e-06 | 1.133E-05 | 2.517E-06 | 9.000E-06 | 1.400E-05 |
| 21562 | 1079 | I1079R | D | 1.7e-05 | 1e-05 | 1.1e-05 | 1.267E-05 | 3.786E-06 | 1.000E-05 | 1.700E-05 |
| 21563 | 1079 | I1079N | D | 1.6e-05 | 1.1e-05 | 1.1e-05 | 1.267E-05 | 2.887E-06 | 1.100E-05 | 1.600E-05 |
| 21564 | 1079 | I1079D | D | 1.4e-05 | 7.5e-06 | 1e-05 | 1.050E-05 | 3.279E-06 | 7.500E-06 | 1.400E-05 |
| 21565 | 1079 | I1079C | D | 1.5e-05 | 1.1e-05 | 1.5e-05 | 1.367E-05 | 2.309E-06 | 1.100E-05 | 1.500E-05 |
| 21566 | 1079 | I1079E | D | 8.7e-06 | 9.4e-06 | 1.1e-05 | 9.700E-06 | 1.179E-06 | 8.700E-06 | 1.100E-05 |
| 21567 | 1079 | I1079Q | D | 1.3e-05 | 1.1e-05 | 1.6e-05 | 1.333E-05 | 2.517E-06 | 1.100E-05 | 1.600E-05 |
| 21568 | 1079 | I1079G | D | 1.4e-05 | 1.5e-05 | 1.1e-05 | 1.333E-05 | 2.082E-06 | 1.100E-05 | 1.500E-05 |
| 21569 | 1079 | I1079H | D | 1.3e-05 | 1.7e-05 | 1.3e-05 | 1.433E-05 | 2.309E-06 | 1.300E-05 | 1.700E-05 |
| 21570 | 1079 | I1079I | D | 1e-05 | 1.6e-05 | 1.5e-05 | 1.367E-05 | 3.215E-06 | 1.000E-05 | 1.600E-05 |
| 21571 | 1079 | I1079L | D | 1.2e-05 | 9.6e-06 | 1e-05 | 1.053E-05 | 1.286E-06 | 9.600E-06 | 1.200E-05 |
| 21572 | 1079 | I1079K | D | 1.6e-05 | 1.4e-05 | 1.6e-05 | 1.533E-05 | 1.155E-06 | 1.400E-05 | 1.600E-05 |
| 21573 | 1079 | I1079M | D | 8.8e-06 | 1.4e-05 | 9.5e-06 | 1.077E-05 | 2.822E-06 | 8.800E-06 | 1.400E-05 |
| 21574 | 1079 | I1079F | D | 1.6e-05 | 9.7e-06 | 1.4e-05 | 1.323E-05 | 3.219E-06 | 9.700E-06 | 1.600E-05 |
| 21575 | 1079 | I1079P | D | 1.7e-05 | 1.5e-05 | 9.1e-06 | 1.370E-05 | 4.107E-06 | 9.100E-06 | 1.700E-05 |
| 21576 | 1079 | I1079S | D | 1.4e-05 | 1.4e-05 | 1.4e-05 | 1.400E-05 | 0.000E+00 | 1.400E-05 | 1.400E-05 |
| 21577 | 1079 | I1079T | D | 1.2e-05 | 1.5e-05 | 1.3e-05 | 1.333E-05 | 1.528E-06 | 1.200E-05 | 1.500E-05 |
| 21578 | 1079 | I1079W | D | 1.3e-05 | 1.2e-05 | 1.3e-05 | 1.267E-05 | 5.774E-07 | 1.200E-05 | 1.300E-05 |
| 21579 | 1079 | I1079Y | D | 9.2e-06 | 1.4e-05 | 1.6e-05 | 1.307E-05 | 3.495E-06 | 9.200E-06 | 1.600E-05 |
| 21580 | 1079 | I1079V | D | 1.5e-05 | 1.4e-05 | 9.4e-06 | 1.280E-05 | 2.987E-06 | 9.400E-06 | 1.500E-05 |
| 21581 | 1080 | T1080A | D | 1.1e-05 | 1.3e-05 | 1.6e-05 | 1.333E-05 | 2.517E-06 | 1.100E-05 | 1.600E-05 |
| 21582 | 1080 | T1080R | D | 1.2e-05 | 1.2e-05 | 1.2e-05 | 1.200E-05 | 0.000E+00 | 1.200E-05 | 1.200E-05 |
| 21583 | 1080 | T1080N | D | 1.1e-05 | 9.4e-06 | 1.3e-05 | 1.113E-05 | 1.804E-06 | 9.400E-06 | 1.300E-05 |
| 21584 | 1080 | T1080D | D | 1.6e-05 | 8.7e-06 | 2e-05 | 1.490E-05 | 5.730E-06 | 8.700E-06 | 2.000E-05 |
| 21585 | 1080 | T1080C | D | 1.5e-05 | 1.6e-05 | 1.5e-05 | 1.533E-05 | 5.774E-07 | 1.500E-05 | 1.600E-05 |
| 21586 | 1080 | T1080E | D | 8.9e-06 | 1.3e-05 | 1.5e-05 | 1.230E-05 | 3.110E-06 | 8.900E-06 | 1.500E-05 |
| 21587 | 1080 | T1080Q | D | 1.5e-05 | 1.4e-05 | 1.1e-05 | 1.333E-05 | 2.082E-06 | 1.100E-05 | 1.500E-05 |
| 21588 | 1080 | T1080G | D | 9.5e-06 | 1.3e-05 | 8.4e-06 | 1.030E-05 | 2.402E-06 | 8.400E-06 | 1.300E-05 |
| 21589 | 1080 | T1080H | D | 1.8e-05 | 1.2e-05 | 1.5e-05 | 1.500E-05 | 3.000E-06 | 1.200E-05 | 1.800E-05 |

| | | | | | | | | | | |
|---|---|---|---|---|---|---|---|---|---|---|
| 21590 | 1080 | T1080I | D | 1.5e-05 | 1.6e-05 | 1.6e-05 | 1.567E-05 | 5.774E-07 | 1.500E-05 | 1.600E-05 |
| 21591 | 1080 | T1080L | D | 1.7e-05 | 1.1e-05 | 1.1e-05 | 1.300E-05 | 3.464E-06 | 1.100E-05 | 1.700E-05 |
| 21592 | 1080 | T1080K | D | 9e-06 | 1.5e-05 | 1.1e-05 | 1.167E-05 | 3.055E-06 | 9.000E-06 | 1.500E-05 |
| 21593 | 1080 | T1080M | D | 9.8e-06 | 9.9e-06 | 1.1e-05 | 1.023E-05 | 6.658E-07 | 9.800E-06 | 1.100E-05 |
| 21594 | 1080 | T1080F | D | 1.6e-05 | 1.3e-05 | 1.6e-05 | 1.500E-05 | 1.732E-06 | 1.300E-05 | 1.600E-05 |
| 21595 | 1080 | T1080P | D | 1e-05 | 8e-06 | 1e-05 | 9.333E-06 | 1.155E-06 | 8.000E-06 | 1.000E-05 |
| 21596 | 1080 | T1080S | D | 1.7e-05 | 1.2e-05 | 1.5e-05 | 1.467E-05 | 2.517E-06 | 1.200E-05 | 1.700E-05 |
| 21597 | 1080 | T1080T | D | 1.6e-05 | 9.7e-06 | 1.6e-05 | 1.390E-05 | 3.637E-06 | 9.700E-06 | 1.600E-05 |
| 21598 | 1080 | T1080W | D | 1.6e-05 | 1.5e-05 | 1.6e-05 | 1.567E-05 | 5.774E-07 | 1.500E-05 | 1.600E-05 |
| 21599 | 1080 | T1080Y | D | 9.4e-06 | 1.5e-05 | 1.1e-05 | 1.180E-05 | 2.884E-06 | 9.400E-06 | 1.500E-05 |
| 21600 | 1080 | T1080V | D | 1.1e-05 | 1.1e-05 | 1.5e-05 | 1.233E-05 | 2.309E-06 | 1.100E-05 | 1.500E-05 |
| 21601 | 1081 | C1081A | D | 1.2e-05 | 1.1e-05 | 1.5e-05 | 1.267E-05 | 2.082E-06 | 1.100E-05 | 1.500E-05 |
| 21602 | 1081 | C1081R | D | 1.5e-05 | 1.3e-05 | 1e-05 | 1.267E-05 | 2.517E-06 | 1.000E-05 | 1.500E-05 |
| 21603 | 1081 | C1081N | D | 8e-06 | 1.5e-05 | 1.4e-05 | 1.233E-05 | 3.786E-06 | 8.000E-06 | 1.500E-05 |
| 21604 | 1081 | C1081D | D | 1.6e-05 | 1.6e-05 | 8e-06 | 1.333E-05 | 4.619E-06 | 8.000E-06 | 1.600E-05 |
| 21605 | 1081 | C1081C | D | 9.6e-06 | 9.9e-06 | 1.6e-05 | 1.183E-05 | 3.612E-06 | 9.600E-06 | 1.600E-05 |
| 21606 | 1081 | C1081E | D | 1.4e-05 | 1.6e-05 | 1.1e-05 | 1.367E-05 | 2.517E-06 | 1.100E-05 | 1.600E-05 |
| 21607 | 1081 | C1081Q | D | 1.2e-05 | 1.8e-05 | 1.6e-05 | 1.533E-05 | 3.055E-06 | 1.200E-05 | 1.800E-05 |
| 21608 | 1081 | C1081G | D | 1.6e-05 | 1.5e-05 | 1.5e-05 | 1.533E-05 | 5.774E-07 | 1.500E-05 | 1.600E-05 |
| 21609 | 1081 | C1081H | D | 1.6e-05 | 1e-05 | 1e-05 | 1.200E-05 | 3.464E-06 | 1.000E-05 | 1.600E-05 |
| 21610 | 1081 | C1081I | D | 1.4e-05 | 1e-05 | 9.7e-06 | 1.123E-05 | 2.401E-06 | 9.700E-06 | 1.400E-05 |
| 21611 | 1081 | C1081L | D | 1.2e-05 | 9.4e-06 | 1.6e-05 | 1.247E-05 | 3.325E-06 | 9.400E-06 | 1.600E-05 |
| 21612 | 1081 | C1081K | D | 1.3e-05 | 1.1e-05 | 1.3e-05 | 1.233E-05 | 1.155E-06 | 1.100E-05 | 1.300E-05 |
| 21613 | 1081 | C1081M | D | 1.5e-05 | 1.1e-05 | 9.2e-06 | 1.173E-05 | 2.969E-06 | 9.200E-06 | 1.500E-05 |
| 21614 | 1081 | C1081F | D | 9.4e-06 | 1.3e-05 | 1.4e-05 | 1.213E-05 | 2.419E-06 | 9.400E-06 | 1.400E-05 |
| 21615 | 1081 | C1081P | D | 1.6e-05 | 1e-05 | 9.6e-06 | 1.187E-05 | 3.585E-06 | 9.600E-06 | 1.600E-05 |
| 21616 | 1081 | C1081S | D | 1.5e-05 | 1.5e-05 | 1.5e-05 | 1.500E-05 | 0.000E+00 | 1.500E-05 | 1.500E-05 |
| 21617 | 1081 | C1081T | D | 1.1e-05 | 7.1e-06 | 1e-05 | 9.367E-06 | 2.026E-06 | 7.100E-06 | 1.100E-05 |
| 21618 | 1081 | C1081W | D | 8.7e-06 | 9.4e-06 | 1.1e-05 | 9.700E-06 | 1.179E-06 | 8.700E-06 | 1.100E-05 |
| 21619 | 1081 | C1081Y | D | 1.3e-05 | 9.7e-06 | 1.4e-05 | 1.223E-05 | 2.250E-06 | 9.700E-06 | 1.400E-05 |
| 21620 | 1081 | C1081V | D | 1.5e-05 | 1e-05 | 1.3e-05 | 1.267E-05 | 2.517E-06 | 1.000E-05 | 1.500E-05 |
| 21621 | 1082 | R1082A | D | 1.4e-05 | 1.2e-05 | 1e-05 | 1.200E-05 | 2.000E-06 | 1.000E-05 | 1.400E-05 |
| 21622 | 1082 | R1082R | D | 1.7e-05 | 8.7e-06 | 1.5e-05 | 1.357E-05 | 4.332E-06 | 8.700E-06 | 1.700E-05 |
| 21623 | 1082 | R1082N | D | 1e-05 | 1.6e-05 | 1e-05 | 1.200E-05 | 3.464E-06 | 1.000E-05 | 1.600E-05 |
| 21624 | 1082 | R1082D | D | 1.1e-05 | 8.9e-06 | 1.5e-05 | 1.163E-05 | 3.099E-06 | 8.900E-06 | 1.500E-05 |
| 21625 | 1082 | R1082C | D | 1.5e-05 | 1.4e-05 | 1.4e-05 | 1.433E-05 | 5.774E-07 | 1.400E-05 | 1.500E-05 |
| 21626 | 1082 | R1082E | D | 9.1e-06 | 1.2e-05 | 1.4e-05 | 1.170E-05 | 2.464E-06 | 9.100E-06 | 1.400E-05 |
| 21627 | 1082 | R1082Q | D | 1.5e-05 | 1.6e-05 | 9.7e-06 | 1.357E-05 | 3.386E-06 | 9.700E-06 | 1.600E-05 |
| 21628 | 1082 | R1082G | D | 1e-05 | 1.5e-05 | 1.5e-05 | 1.333E-05 | 2.887E-06 | 1.000E-05 | 1.500E-05 |
| 21629 | 1082 | R1082H | D | 1.4e-05 | 1.5e-05 | 1.3e-05 | 1.400E-05 | 1.000E-06 | 1.300E-05 | 1.500E-05 |
| 21630 | 1082 | R1082I | D | 1.4e-05 | 1.6e-05 | 1.3e-05 | 1.433E-05 | 1.528E-06 | 1.300E-05 | 1.600E-05 |
| 21631 | 1082 | R1082L | D | 1.5e-05 | 9.6e-06 | 1.1e-05 | 1.187E-05 | 2.802E-06 | 9.600E-06 | 1.500E-05 |
| 21632 | 1082 | R1082K | D | 1.5e-05 | 1.1e-05 | 1.2e-05 | 1.267E-05 | 2.082E-06 | 1.100E-05 | 1.500E-05 |
| 21633 | 1082 | R1082M | D | 1.4e-05 | 1.5e-05 | 1.1e-05 | 1.333E-05 | 2.082E-06 | 1.100E-05 | 1.500E-05 |
| 21634 | 1082 | R1082F | D | 1.5e-05 | 8.4e-06 | 1.1e-05 | 1.147E-05 | 3.325E-06 | 8.400E-06 | 1.500E-05 |
| 21635 | 1082 | R1082P | D | 1.2e-05 | 7.8e-06 | 1.2e-05 | 1.060E-05 | 2.425E-06 | 7.800E-06 | 1.200E-05 |
| 21636 | 1082 | R1082S | D | 1.7e-05 | 1.4e-05 | 1.3e-05 | 1.467E-05 | 2.082E-06 | 1.300E-05 | 1.700E-05 |
| 21637 | 1082 | R1082T | D | 1e-05 | 1.3e-05 | 1.3e-05 | 1.200E-05 | 1.732E-06 | 1.000E-05 | 1.300E-05 |
| 21638 | 1082 | R1082W | D | 1.4e-05 | 1.1e-05 | 1.5e-05 | 1.333E-05 | 2.082E-06 | 1.100E-05 | 1.500E-05 |
| 21639 | 1082 | R1082Y | D | 1.6e-05 | 1.5e-05 | 9e-06 | 1.333E-05 | 3.786E-06 | 9.000E-06 | 1.600E-05 |
| 21640 | 1082 | R1082V | D | 1.4e-05 | 1.4e-05 | 1.1e-05 | 1.300E-05 | 1.732E-06 | 1.100E-05 | 1.400E-05 |
| 21641 | 1083 | A1083A | D | 1.1e-05 | 1e-05 | 1e-05 | 1.033E-05 | 5.774E-07 | 1.000E-05 | 1.100E-05 |
| 21642 | 1083 | A1083R | D | 1.4e-05 | 1.6e-05 | 1.4e-05 | 1.467E-05 | 1.155E-06 | 1.400E-05 | 1.600E-05 |
| 21643 | 1083 | A1083N | D | 8.9e-06 | 1e-05 | 9.9e-06 | 9.600E-06 | 6.083E-07 | 8.900E-06 | 1.000E-05 |
| 21644 | 1083 | A1083D | D | 1.1e-05 | 1.5e-05 | 1.5e-05 | 1.367E-05 | 2.309E-06 | 1.100E-05 | 1.500E-05 |
| 21645 | 1083 | A1083C | D | 8.9e-06 | 1.4e-05 | 1.5e-05 | 1.263E-05 | 3.272E-06 | 8.900E-06 | 1.500E-05 |
| 21646 | 1083 | A1083E | D | 1.5e-05 | 1.2e-05 | 1.6e-05 | 1.433E-05 | 2.082E-06 | 1.200E-05 | 1.600E-05 |
| 21647 | 1083 | A1083Q | D | 9.7e-06 | 1.3e-05 | 1.4e-05 | 1.223E-05 | 2.250E-06 | 9.700E-06 | 1.400E-05 |
| 21648 | 1083 | A1083G | D | 1.1e-05 | 1.4e-05 | 1.3e-05 | 1.267E-05 | 1.528E-06 | 1.100E-05 | 1.400E-05 |
| 21649 | 1083 | A1083H | D | 1.1e-05 | 6.9e-06 | 1.4e-05 | 1.063E-05 | 3.564E-06 | 6.900E-06 | 1.400E-05 |
| 21650 | 1083 | A1083I | D | 9.8e-06 | 1.5e-05 | 1.1e-05 | 1.193E-05 | 2.723E-06 | 9.800E-06 | 1.500E-05 |
| 21651 | 1083 | A1083L | D | 1.7e-05 | 1.5e-05 | 1.3e-05 | 1.500E-05 | 2.000E-06 | 1.300E-05 | 1.700E-05 |
| 21652 | 1083 | A1083K | D | 1.4e-05 | 1.6e-05 | 1.3e-05 | 1.433E-05 | 1.528E-06 | 1.300E-05 | 1.600E-05 |
| 21653 | 1083 | A1083M | D | 1e-05 | 1.6e-05 | 7.9e-06 | 1.130E-05 | 4.204E-06 | 7.900E-06 | 1.600E-05 |
| 21654 | 1083 | A1083F | D | 1.5e-05 | 1.5e-05 | 1.3e-05 | 1.433E-05 | 1.155E-06 | 1.300E-05 | 1.500E-05 |
| 21655 | 1083 | A1083P | D | 9.9e-06 | 1e-05 | 1.6e-05 | 1.197E-05 | 3.493E-06 | 9.900E-06 | 1.600E-05 |
| 21656 | 1083 | A1083S | D | 1.5e-05 | 1.4e-05 | 1e-05 | 1.300E-05 | 2.646E-06 | 1.000E-05 | 1.500E-05 |
| 21657 | 1083 | A1083T | D | 1e-05 | 1.1e-05 | 1.4e-05 | 1.167E-05 | 2.082E-06 | 1.000E-05 | 1.400E-05 |
| 21658 | 1083 | A1083W | D | 8e-06 | 9.6e-06 | 1.3e-05 | 1.020E-05 | 2.553E-06 | 8.000E-06 | 1.300E-05 |
| 21659 | 1083 | A1083Y | D | 1.2e-05 | 1.1e-05 | 1e-05 | 1.100E-05 | 1.000E-06 | 1.000E-05 | 1.200E-05 |
| 21660 | 1083 | A1083V | D | 1.3e-05 | 1.4e-05 | 1.6e-05 | 1.433E-05 | 1.528E-06 | 1.300E-05 | 1.600E-05 |
| 21661 | 1084 | S1084A | D | 1.6e-05 | 1e-05 | 1.9e-05 | 1.500E-05 | 4.583E-06 | 1.000E-05 | 1.900E-05 |
| 21662 | 1084 | S1084R | D | 1.2e-05 | 7.2e-06 | 1.1e-05 | 1.007E-05 | 2.532E-06 | 7.200E-06 | 1.200E-05 |
| 21663 | 1084 | S1084N | D | 1.5e-05 | 1.2e-05 | 1.5e-05 | 1.400E-05 | 1.732E-06 | 1.200E-05 | 1.500E-05 |
| 21664 | 1084 | S1084D | D | 8.4e-06 | 8.4e-06 | 1.6e-05 | 1.093E-05 | 4.388E-06 | 8.400E-06 | 1.600E-05 |
| 21665 | 1084 | S1084C | D | 1e-05 | 1.1e-05 | 1e-05 | 1.033E-05 | 5.774E-07 | 1.000E-05 | 1.100E-05 |
| 21666 | 1084 | S1084E | D | 1.1e-05 | 1.7e-05 | 1.1e-05 | 1.300E-05 | 3.464E-06 | 1.100E-05 | 1.700E-05 |
| 21667 | 1084 | S1084Q | D | 1.5e-05 | 1e-05 | 1.1e-05 | 1.200E-05 | 2.646E-06 | 1.000E-05 | 1.500E-05 |
| 21668 | 1084 | S1084G | D | 1.1e-05 | 1.1e-05 | 9.1e-06 | 1.037E-05 | 1.097E-06 | 9.100E-06 | 1.100E-05 |
| 21669 | 1084 | S1084H | D | 1.7e-05 | 1.6e-05 | 1.1e-05 | 1.467E-05 | 3.215E-06 | 1.100E-05 | 1.700E-05 |
| 21670 | 1084 | S1084I | D | 1.7e-05 | 1.7e-05 | 1.3e-05 | 1.567E-05 | 2.309E-06 | 1.300E-05 | 1.700E-05 |
| 21671 | 1084 | S1084L | D | 1.2e-05 | 1.3e-05 | 1.4e-05 | 1.300E-05 | 1.000E-06 | 1.200E-05 | 1.400E-05 |
| 21672 | 1084 | S1084K | D | 1.6e-05 | 1.5e-05 | 1.5e-05 | 1.533E-05 | 5.774E-07 | 1.500E-05 | 1.600E-05 |
| 21673 | 1084 | S1084M | D | 2.2e-05 | 1.5e-05 | 1.6e-05 | 1.767E-05 | 3.786E-06 | 1.500E-05 | 2.200E-05 |
| 21674 | 1084 | S1084F | D | 1.5e-05 | 1.4e-05 | 1.5e-05 | 1.467E-05 | 5.774E-07 | 1.400E-05 | 1.500E-05 |

| | | | | | | | | | | |
|---|---|---|---|---|---|---|---|---|---|---|
| 21675 | 1084 | S1084P | D | 1.1e-05 | 1.3e-05 | 1.6e-05 | 1.333E-05 | 2.517E-06 | 1.100E-05 | 1.600E-05 |
| 21676 | 1084 | S1084S | D | 1.4e-05 | 1.7e-05 | 1.5e-05 | 1.533E-05 | 1.528E-06 | 1.400E-05 | 1.700E-05 |
| 21677 | 1084 | S1084T | D | 2.4e-05 | 1.3e-05 | 1.4e-05 | 1.700E-05 | 6.083E-06 | 1.300E-05 | 2.400E-05 |
| 21678 | 1084 | S1084W | D | 1.6e-05 | 1.4e-05 | 1e-05 | 1.333E-05 | 3.055E-06 | 1.000E-05 | 1.600E-05 |
| 21679 | 1084 | S1084Y | D | 1.5e-05 | 1.5e-05 | 1.8e-05 | 1.600E-05 | 1.732E-06 | 1.500E-05 | 1.800E-05 |
| 21680 | 1084 | S1084V | D | 1.5e-05 | 9.4e-06 | 1.6e-05 | 1.347E-05 | 3.557E-06 | 9.400E-06 | 1.600E-05 |
| 21681 | 1085 | Q1085A | D | 8.1e-06 | 1.1e-05 | 7.5e-06 | 8.867E-06 | 1.872E-06 | 7.500E-06 | 1.100E-05 |
| 21682 | 1085 | Q1085R | D | 1.3e-05 | 1.3e-05 | 1.4e-05 | 1.333E-05 | 5.774E-07 | 1.300E-05 | 1.400E-05 |
| 21683 | 1085 | Q1085N | D | 9.7e-06 | 1.4e-05 | 1.3e-05 | 1.223E-05 | 2.250E-06 | 9.700E-06 | 1.400E-05 |
| 21684 | 1085 | Q1085D | D | 1.8e-05 | 1.1e-05 | 1.7e-05 | 1.533E-05 | 3.786E-06 | 1.100E-05 | 1.800E-05 |
| 21685 | 1085 | Q1085C | D | 1.5e-05 | 1.1e-05 | 1.2e-05 | 1.267E-05 | 2.082E-06 | 1.100E-05 | 1.500E-05 |
| 21686 | 1085 | Q1085E | D | 9.3e-06 | 1.4e-05 | 1.8e-05 | 1.377E-05 | 4.355E-06 | 9.300E-06 | 1.800E-05 |
| 21687 | 1085 | Q1085Q | D | 1.1e-05 | 1.5e-05 | 1.1e-05 | 1.233E-05 | 2.309E-06 | 1.100E-05 | 1.500E-05 |
| 21688 | 1085 | Q1085G | D | 1.1e-05 | 1.4e-05 | 1.2e-05 | 1.233E-05 | 1.528E-06 | 1.100E-05 | 1.400E-05 |
| 21689 | 1085 | Q1085H | D | 8.9e-06 | 1.5e-05 | 1.4e-05 | 1.263E-05 | 3.272E-06 | 8.900E-06 | 1.500E-05 |
| 21690 | 1085 | Q1085I | D | 1.6e-05 | 9.9e-06 | 1e-05 | 1.197E-05 | 3.493E-06 | 9.900E-06 | 1.600E-05 |
| 21691 | 1085 | Q1085L | D | 1.4e-05 | 1.6e-05 | 1.4e-05 | 1.467E-05 | 1.155E-06 | 1.400E-05 | 1.600E-05 |
| 21692 | 1085 | Q1085K | D | 1.5e-05 | 1e-05 | 1.1e-05 | 1.200E-05 | 2.646E-06 | 1.000E-05 | 1.500E-05 |
| 21693 | 1085 | Q1085M | D | 1.6e-05 | 1.3e-05 | 1.5e-05 | 1.467E-05 | 1.528E-06 | 1.300E-05 | 1.600E-05 |
| 21694 | 1085 | Q1085F | D | 1.5e-05 | 1.1e-05 | 9.1e-06 | 1.170E-05 | 3.012E-06 | 9.100E-06 | 1.500E-05 |
| 21695 | 1085 | Q1085P | D | 1.5e-05 | 1.2e-05 | 9.8e-06 | 1.227E-05 | 2.610E-06 | 9.800E-06 | 1.500E-05 |
| 21696 | 1085 | Q1085S | D | 9.7e-06 | 1.3e-05 | 1.2e-05 | 1.157E-05 | 1.692E-06 | 9.700E-06 | 1.300E-05 |
| 21697 | 1085 | Q1085T | D | 1.1e-05 | 1e-05 | 1.5e-05 | 1.200E-05 | 2.646E-06 | 1.000E-05 | 1.500E-05 |
| 21698 | 1085 | Q1085W | D | 1.5e-05 | 1.6e-05 | 1.1e-05 | 1.400E-05 | 2.646E-06 | 1.100E-05 | 1.600E-05 |
| 21699 | 1085 | Q1085Y | D | 9.5e-06 | 9.2e-06 | 1.1e-05 | 9.900E-06 | 9.644E-07 | 9.200E-06 | 1.100E-05 |
| 21700 | 1085 | Q1085V | D | 1.7e-05 | 1.7e-05 | 1.6e-05 | 1.667E-05 | 5.774E-07 | 1.600E-05 | 1.700E-05 |
| 21701 | 1086 | D1086A | D | 1.4e-05 | 1.4e-05 | 1.5e-05 | 1.433E-05 | 5.774E-07 | 1.400E-05 | 1.500E-05 |
| 21702 | 1086 | D1086R | D | 1.6e-05 | 1.6e-05 | 9.6e-06 | 1.387E-05 | 3.695E-06 | 9.600E-06 | 1.600E-05 |
| 21703 | 1086 | D1086N | D | 1.1e-05 | 1.5e-05 | 1.6e-05 | 1.400E-05 | 2.646E-06 | 1.100E-05 | 1.600E-05 |
| 21704 | 1086 | D1086D | D | 1.1e-05 | 9.8e-06 | 1.6e-05 | 1.227E-05 | 3.288E-06 | 9.800E-06 | 1.600E-05 |
| 21705 | 1086 | D1086C | D | 1.5e-05 | 1e-05 | 1.1e-05 | 1.200E-05 | 2.646E-06 | 1.000E-05 | 1.500E-05 |
| 21706 | 1086 | D1086E | D | 9.2e-06 | 9.4e-06 | 1.5e-05 | 1.120E-05 | 3.292E-06 | 9.200E-06 | 1.500E-05 |
| 21707 | 1086 | D1086Q | D | 1.1e-05 | 1.5e-05 | 1e-05 | 1.200E-05 | 2.646E-06 | 1.000E-05 | 1.500E-05 |
| 21708 | 1086 | D1086G | D | 1.2e-05 | 9.2e-06 | 1e-05 | 1.040E-05 | 1.442E-06 | 9.200E-06 | 1.200E-05 |
| 21709 | 1086 | D1086H | D | 1.4e-05 | 1.6e-05 | 1.1e-05 | 1.367E-05 | 2.517E-06 | 1.100E-05 | 1.600E-05 |
| 21710 | 1086 | D1086I | D | 1.1e-05 | 9.6e-06 | 1.5e-05 | 1.187E-05 | 2.802E-06 | 9.600E-06 | 1.500E-05 |
| 21711 | 1086 | D1086L | D | 1.5e-05 | 1.7e-05 | 1.5e-05 | 1.567E-05 | 1.155E-06 | 1.500E-05 | 1.700E-05 |
| 21712 | 1086 | D1086K | D | 1e-05 | 1.3e-05 | 1.5e-05 | 1.267E-05 | 2.517E-06 | 1.000E-05 | 1.500E-05 |
| 21713 | 1086 | D1086M | D | 1.2e-05 | 1.5e-05 | 1.4e-05 | 1.367E-05 | 1.528E-06 | 1.200E-05 | 1.500E-05 |
| 21714 | 1086 | D1086F | D | 1.5e-05 | 1.4e-05 | 1.1e-05 | 1.333E-05 | 2.082E-06 | 1.100E-05 | 1.500E-05 |
| 21715 | 1086 | D1086P | D | 1.6e-05 | 1.1e-05 | 1.3e-05 | 1.333E-05 | 2.517E-06 | 1.100E-05 | 1.600E-05 |
| 21716 | 1086 | D1086S | D | 1.1e-05 | 1.2e-05 | 9.6e-06 | 1.087E-05 | 1.206E-06 | 9.600E-06 | 1.200E-05 |
| 21717 | 1086 | D1086T | D | 7.2e-06 | 1.5e-05 | 1.2e-05 | 1.140E-05 | 3.934E-06 | 7.200E-06 | 1.500E-05 |
| 21718 | 1086 | D1086W | D | 1e-05 | 1.5e-05 | 1.4e-05 | 1.300E-05 | 2.646E-06 | 1.000E-05 | 1.500E-05 |
| 21719 | 1086 | D1086Y | D | 1.4e-05 | 1.3e-05 | 1.1e-05 | 1.267E-05 | 1.528E-06 | 1.100E-05 | 1.400E-05 |
| 21720 | 1086 | D1086V | D | 1.4e-05 | 1.2e-05 | 1.4e-05 | 1.333E-05 | 1.155E-06 | 1.200E-05 | 1.400E-05 |
| 21721 | 1087 | V1087A | D | 1.1e-05 | 1.7e-05 | 1.5e-05 | 1.433E-05 | 3.055E-06 | 1.100E-05 | 1.700E-05 |
| 21722 | 1087 | V1087R | D | 1.7e-05 | 1e-05 | 1e-05 | 1.233E-05 | 4.041E-06 | 1.000E-05 | 1.700E-05 |
| 21723 | 1087 | V1087N | D | 1.5e-05 | 1e-05 | 1.5e-05 | 1.333E-05 | 2.887E-06 | 1.000E-05 | 1.500E-05 |
| 21724 | 1087 | V1087D | D | 1.5e-05 | 1e-05 | 1.1e-05 | 1.200E-05 | 2.646E-06 | 1.000E-05 | 1.500E-05 |
| 21725 | 1087 | V1087C | D | 1.4e-05 | 1.4e-05 | 1e-05 | 1.267E-05 | 2.309E-06 | 1.000E-05 | 1.400E-05 |
| 21726 | 1087 | V1087E | D | 1.4e-05 | 1.5e-05 | 1.3e-05 | 1.400E-05 | 1.000E-06 | 1.300E-05 | 1.500E-05 |
| 21727 | 1087 | V1087Q | D | 1.2e-05 | 1.3e-05 | 1.2e-05 | 1.233E-05 | 5.774E-07 | 1.200E-05 | 1.300E-05 |
| 21728 | 1087 | V1087G | D | 1.7e-05 | 1.1e-05 | 2.6e-05 | 1.800E-05 | 7.550E-06 | 1.100E-05 | 2.600E-05 |
| 21729 | 1087 | V1087H | D | 1.4e-05 | 8.3e-06 | 6.9e-06 | 9.733E-06 | 3.761E-06 | 6.900E-06 | 1.400E-05 |
| 21730 | 1087 | V1087I | D | 1.4e-05 | 1.1e-05 | 1.6e-05 | 1.367E-05 | 2.517E-06 | 1.100E-05 | 1.600E-05 |
| 21731 | 1087 | V1087L | D | 1e-05 | 9.2e-06 | 9.7e-06 | 9.633E-06 | 4.041E-07 | 9.200E-06 | 1.000E-05 |
| 21732 | 1087 | V1087K | D | 1.4e-05 | 9.4e-06 | 8.5e-06 | 1.063E-05 | 2.950E-06 | 8.500E-06 | 1.400E-05 |
| 21733 | 1087 | V1087M | D | 1e-05 | 1.5e-05 | 1.5e-05 | 1.333E-05 | 2.887E-06 | 1.000E-05 | 1.500E-05 |
| 21734 | 1087 | V1087F | D | 1.4e-05 | 1e-05 | 8.8e-06 | 1.093E-05 | 2.723E-06 | 8.800E-06 | 1.400E-05 |
| 21735 | 1087 | V1087P | D | 1e-05 | 1.5e-05 | 1.2e-05 | 1.233E-05 | 2.517E-06 | 1.000E-05 | 1.500E-05 |
| 21736 | 1087 | V1087S | D | 9.9e-06 | 9.3e-06 | 9e-06 | 9.400E-06 | 4.583E-07 | 9.000E-06 | 9.900E-06 |
| 21737 | 1087 | V1087T | D | 8.5e-06 | 1.6e-05 | 8.8e-06 | 1.110E-05 | 4.246E-06 | 8.500E-06 | 1.600E-05 |
| 21738 | 1087 | V1087W | D | 1.4e-05 | 1.5e-05 | 1.6e-05 | 1.500E-05 | 1.000E-06 | 1.400E-05 | 1.600E-05 |
| 21739 | 1087 | V1087Y | D | 1.5e-05 | 1.6e-05 | 1.4e-05 | 1.500E-05 | 1.000E-06 | 1.400E-05 | 1.600E-05 |
| 21740 | 1087 | V1087V | D | 1.4e-05 | 1.6e-05 | 9.3e-06 | 1.310E-05 | 3.439E-06 | 9.300E-06 | 1.600E-05 |
| 21741 | 1088 | N1088A | D | 1.5e-05 | 1.6e-05 | 1.3e-05 | 1.467E-05 | 1.528E-06 | 1.300E-05 | 1.600E-05 |
| 21742 | 1088 | N1088R | D | 1e-05 | 1e-05 | 1.1e-05 | 1.033E-05 | 5.774E-07 | 1.000E-05 | 1.100E-05 |
| 21743 | 1088 | N1088N | D | 1e-05 | 1.6e-05 | 1.4e-05 | 1.333E-05 | 3.055E-06 | 1.000E-05 | 1.600E-05 |
| 21744 | 1088 | N1088D | D | 1.7e-05 | 1.5e-05 | 1.1e-05 | 1.433E-05 | 3.055E-06 | 1.100E-05 | 1.700E-05 |
| 21745 | 1088 | N1088C | D | 1.5e-05 | 1.1e-05 | 1.4e-05 | 1.333E-05 | 2.082E-06 | 1.100E-05 | 1.500E-05 |
| 21746 | 1088 | N1088E | D | 1e-05 | 1.3e-05 | 1.5e-05 | 1.267E-05 | 2.517E-06 | 1.000E-05 | 1.500E-05 |
| 21747 | 1088 | N1088Q | D | 9.4e-06 | 1.4e-05 | 1.6e-05 | 1.313E-05 | 3.384E-06 | 9.400E-06 | 1.600E-05 |
| 21748 | 1088 | N1088G | D | 9.2e-06 | 1.7e-05 | 1.7e-05 | 1.440E-05 | 4.503E-06 | 9.200E-06 | 1.700E-05 |
| 21749 | 1088 | N1088H | D | 1.5e-05 | 1e-05 | 1.6e-05 | 1.367E-05 | 3.215E-06 | 1.000E-05 | 1.600E-05 |
| 21750 | 1088 | N1088I | D | 1.2e-05 | 1.2e-05 | 1.6e-05 | 1.333E-05 | 2.309E-06 | 1.200E-05 | 1.600E-05 |
| 21751 | 1088 | N1088L | D | 1.3e-05 | 1.5e-05 | 8.7e-06 | 1.223E-05 | 3.219E-06 | 8.700E-06 | 1.500E-05 |
| 21752 | 1088 | N1088K | D | 1.3e-05 | 1.3e-05 | 1.4e-05 | 1.333E-05 | 5.774E-07 | 1.300E-05 | 1.400E-05 |
| 21753 | 1088 | N1088M | D | 1.5e-05 | 1.6e-05 | 1.1e-05 | 1.400E-05 | 2.646E-06 | 1.100E-05 | 1.600E-05 |
| 21754 | 1088 | N1088F | D | 1.5e-05 | 1.3e-05 | 1.5e-05 | 1.433E-05 | 1.155E-06 | 1.300E-05 | 1.500E-05 |
| 21755 | 1088 | N1088P | D | 1.1e-05 | 1.1e-05 | 1.1e-05 | 1.100E-05 | 2.075E-21 | 1.100E-05 | 1.100E-05 |
| 21756 | 1088 | N1088S | D | 1.1e-05 | 1.5e-05 | 1.5e-05 | 1.367E-05 | 2.309E-06 | 1.100E-05 | 1.500E-05 |
| 21757 | 1088 | N1088T | D | 1.1e-05 | 1.3e-05 | 1.5e-05 | 1.300E-05 | 2.000E-06 | 1.100E-05 | 1.500E-05 |
| 21758 | 1088 | N1088W | D | 7.2e-06 | 1e-05 | 1.1e-05 | 9.400E-06 | 1.970E-06 | 7.200E-06 | 1.100E-05 |
| 21759 | 1088 | N1088Y | D | 1.2e-05 | 1.4e-05 | 1.5e-05 | 1.367E-05 | 1.528E-06 | 1.200E-05 | 1.500E-05 |

| | | | | | | | | | | |
|---|---|---|---|---|---|---|---|---|---|---|
| 21760 | 1088 | N1088V | D | 1.6e-05 | 1.6e-05 | 1.7e-05 | 1.633E-05 | 5.774E-07 | 1.600E-05 | 1.700E-05 |
| 21761 | 1089 | T1089A | D | 1.6e-05 | 1.5e-05 | 1.5e-05 | 1.533E-05 | 5.774E-07 | 1.500E-05 | 1.600E-05 |
| 21762 | 1089 | T1089R | D | 1e-05 | 4.5e-06 | 1.4e-05 | 9.500E-06 | 4.770E-06 | 4.500E-06 | 1.400E-05 |
| 21763 | 1089 | T1089N | D | 1.7e-05 | 1.5e-05 | 1.4e-05 | 1.533E-05 | 1.528E-06 | 1.400E-05 | 1.700E-05 |
| 21764 | 1089 | T1089D | D | 1.4e-05 | 1.1e-05 | 6.5e-06 | 1.050E-05 | 3.775E-06 | 6.500E-06 | 1.400E-05 |
| 21765 | 1089 | T1089C | D | 1.1e-05 | 1.1e-05 | 1.1e-05 | 1.100E-05 | 2.075E-21 | 1.100E-05 | 1.100E-05 |
| 21766 | 1089 | T1089E | D | 1.6e-05 | 1e-05 | 8.7e-06 | 1.157E-05 | 3.894E-06 | 8.700E-06 | 1.600E-05 |
| 21767 | 1089 | T1089Q | D | 1.1e-05 | 1.3e-05 | 1.4e-05 | 1.267E-05 | 1.528E-06 | 1.100E-05 | 1.400E-05 |
| 21768 | 1089 | T1089G | D | 1.1e-05 | 1.1e-05 | 1.7e-05 | 1.300E-05 | 3.464E-06 | 1.100E-05 | 1.700E-05 |
| 21769 | 1089 | T1089H | D | 1e-05 | 1.1e-05 | 9e-06 | 1.000E-05 | 1.000E-06 | 9.000E-06 | 1.100E-05 |
| 21770 | 1089 | T1089I | D | 1.1e-05 | 1.1e-05 | 2.2e-05 | 1.467E-05 | 6.351E-06 | 1.100E-05 | 2.200E-05 |
| 21771 | 1089 | T1089L | D | 1.1e-05 | 1.1e-05 | 1.5e-05 | 1.233E-05 | 2.309E-06 | 1.100E-05 | 1.500E-05 |
| 21772 | 1089 | T1089K | D | 1e-05 | 1.5e-05 | 1.5e-05 | 1.333E-05 | 2.887E-06 | 1.000E-05 | 1.500E-05 |
| 21773 | 1089 | T1089M | D | 9.1e-06 | 1.6e-05 | 1.3e-05 | 1.270E-05 | 3.460E-06 | 9.100E-06 | 1.600E-05 |
| 21774 | 1089 | T1089F | D | 9.8e-06 | 9.8e-06 | 9.9e-06 | 9.833E-06 | 5.774E-08 | 9.800E-06 | 9.900E-06 |
| 21775 | 1089 | T1089P | D | 1.6e-05 | 1.3e-05 | 1.5e-05 | 1.467E-05 | 1.528E-06 | 1.300E-05 | 1.600E-05 |
| 21776 | 1089 | T1089S | D | 1.3e-05 | 9.6e-06 | 1.5e-05 | 1.253E-05 | 2.730E-06 | 9.600E-06 | 1.500E-05 |
| 21777 | 1089 | T1089T | D | 1.6e-05 | 9.5e-06 | 1.3e-05 | 1.283E-05 | 3.253E-06 | 9.500E-06 | 1.600E-05 |
| 21778 | 1089 | T1089W | D | 1.1e-05 | 1.2e-05 | 8.7e-06 | 1.057E-05 | 1.692E-06 | 8.700E-06 | 1.200E-05 |
| 21779 | 1089 | T1089Y | D | 1.3e-05 | 1.1e-05 | 9.7e-06 | 1.123E-05 | 1.662E-06 | 9.700E-06 | 1.300E-05 |
| 21780 | 1089 | T1089V | D | 1.4e-05 | 1.3e-05 | 1.4e-05 | 1.367E-05 | 5.774E-07 | 1.300E-05 | 1.400E-05 |
| 21781 | 1090 | A1090A | D | 9.8e-06 | 1.2e-05 | 9e-06 | 1.027E-05 | 1.553E-06 | 9.000E-06 | 1.200E-05 |
| 21782 | 1090 | A1090R | D | 1.4e-05 | 1.5e-05 | 1.1e-05 | 1.333E-05 | 2.082E-06 | 1.100E-05 | 1.500E-05 |
| 21783 | 1090 | A1090N | D | 1.4e-05 | 1.4e-05 | 9.3e-06 | 1.243E-05 | 2.714E-06 | 9.300E-06 | 1.400E-05 |
| 21784 | 1090 | A1090D | D | 1.1e-05 | 9.7e-06 | 1.2e-05 | 1.090E-05 | 1.153E-06 | 9.700E-06 | 1.200E-05 |
| 21785 | 1090 | A1090C | D | 1.5e-05 | 1.6e-05 | 1.6e-05 | 1.567E-05 | 5.774E-07 | 1.500E-05 | 1.600E-05 |
| 21786 | 1090 | A1090E | D | 1.1e-05 | 8.2e-06 | 1.3e-05 | 1.073E-05 | 2.411E-06 | 8.200E-06 | 1.300E-05 |
| 21787 | 1090 | A1090Q | D | 1.6e-05 | 1.6e-05 | 1.1e-05 | 1.433E-05 | 2.887E-06 | 1.100E-05 | 1.600E-05 |
| 21788 | 1090 | A1090G | D | 1.6e-05 | 1.3e-05 | 1.5e-05 | 1.467E-05 | 1.528E-06 | 1.300E-05 | 1.600E-05 |
| 21789 | 1090 | A1090H | D | 1.6e-05 | 1.4e-05 | 8.9e-06 | 1.297E-05 | 3.661E-06 | 8.900E-06 | 1.600E-05 |
| 21790 | 1090 | A1090I | D | 9.9e-06 | 1.5e-05 | 1.5e-05 | 1.330E-05 | 2.944E-06 | 9.900E-06 | 1.500E-05 |
| 21791 | 1090 | A1090L | D | 1.4e-05 | 1.5e-05 | 1.5e-05 | 1.467E-05 | 5.774E-07 | 1.400E-05 | 1.500E-05 |
| 21792 | 1090 | A1090K | D | 1.6e-05 | 1.4e-05 | 1.3e-05 | 1.433E-05 | 1.528E-06 | 1.300E-05 | 1.600E-05 |
| 21793 | 1090 | A1090M | D | 1.4e-05 | 1.1e-05 | 1.6e-05 | 1.367E-05 | 2.517E-06 | 1.100E-05 | 1.600E-05 |
| 21794 | 1090 | A1090F | D | 1.6e-05 | 1.5e-05 | 8.4e-06 | 1.313E-05 | 4.130E-06 | 8.400E-06 | 1.600E-05 |
| 21795 | 1090 | A1090P | D | 1e-05 | 1.5e-05 | 1.6e-05 | 1.367E-05 | 3.215E-06 | 1.000E-05 | 1.600E-05 |
| 21796 | 1090 | A1090S | D | 1.1e-05 | 1e-05 | 1.7e-05 | 1.267E-05 | 3.786E-06 | 1.000E-05 | 1.700E-05 |
| 21797 | 1090 | A1090T | D | 1.5e-05 | 1.5e-05 | 1.6e-05 | 1.533E-05 | 5.774E-07 | 1.500E-05 | 1.600E-05 |
| 21798 | 1090 | A1090W | D | 1.4e-05 | 1.3e-05 | 1.1e-05 | 1.267E-05 | 1.528E-06 | 1.100E-05 | 1.400E-05 |
| 21799 | 1090 | A1090Y | D | 1.3e-05 | 1.6e-05 | 1.6e-05 | 1.500E-05 | 1.732E-06 | 1.300E-05 | 1.600E-05 |
| 21800 | 1090 | A1090V | D | 1.4e-05 | 1.4e-05 | 1.7e-05 | 1.500E-05 | 1.732E-06 | 1.400E-05 | 1.700E-05 |
| 21801 | 1091 | V1091A | D | 7.3e-06 | 1.5e-05 | 1.5e-05 | 1.243E-05 | 4.446E-06 | 7.300E-06 | 1.500E-05 |
| 21802 | 1091 | V1091R | D | 1e-05 | 1.6e-05 | 1.6e-05 | 1.400E-05 | 3.464E-06 | 1.000E-05 | 1.600E-05 |
| 21803 | 1091 | V1091N | D | 1.3e-05 | 1.4e-05 | 1e-05 | 1.233E-05 | 2.082E-06 | 1.000E-05 | 1.400E-05 |
| 21804 | 1091 | V1091D | D | 1.6e-05 | 1.3e-05 | 1.5e-05 | 1.467E-05 | 1.528E-06 | 1.300E-05 | 1.600E-05 |
| 21805 | 1091 | V1091C | D | 9e-06 | 1.4e-05 | 1.2e-05 | 1.167E-05 | 2.517E-06 | 9.000E-06 | 1.400E-05 |
| 21806 | 1091 | V1091E | D | 9.6e-06 | 8.2e-06 | 1.5e-05 | 1.093E-05 | 3.591E-06 | 8.200E-06 | 1.500E-05 |
| 21807 | 1091 | V1091Q | D | 1.2e-05 | 8.1e-06 | 1.6e-05 | 1.203E-05 | 3.950E-06 | 8.100E-06 | 1.600E-05 |
| 21808 | 1091 | V1091G | D | 9e-06 | 9.9e-06 | 1.1e-05 | 9.967E-06 | 1.002E-06 | 9.000E-06 | 1.100E-05 |
| 21809 | 1091 | V1091H | D | 1.4e-05 | 1.5e-05 | 8.4e-06 | 1.247E-05 | 3.557E-06 | 8.400E-06 | 1.500E-05 |
| 21810 | 1091 | V1091I | D | 1.7e-05 | 1.1e-05 | 8.3e-06 | 1.210E-05 | 4.453E-06 | 8.300E-06 | 1.700E-05 |
| 21811 | 1091 | V1091L | D | 1e-05 | 1.5e-05 | 1.4e-05 | 1.300E-05 | 2.646E-06 | 1.000E-05 | 1.500E-05 |
| 21812 | 1091 | V1091K | D | 1e-05 | 1.5e-05 | 9.8e-06 | 1.160E-05 | 2.946E-06 | 9.800E-06 | 1.500E-05 |
| 21813 | 1091 | V1091M | D | 1.5e-05 | 1.6e-05 | 1.3e-05 | 1.467E-05 | 1.528E-06 | 1.300E-05 | 1.600E-05 |
| 21814 | 1091 | V1091F | D | 1.6e-05 | 1.5e-05 | 1.5e-05 | 1.533E-05 | 5.774E-07 | 1.500E-05 | 1.600E-05 |
| 21815 | 1091 | V1091P | D | 1.4e-05 | 1.4e-05 | 1.7e-05 | 1.500E-05 | 1.732E-06 | 1.400E-05 | 1.700E-05 |
| 21816 | 1091 | V1091S | D | 1.1e-05 | 9.3e-06 | 1.3e-05 | 1.110E-05 | 1.852E-06 | 9.300E-06 | 1.300E-05 |
| 21817 | 1091 | V1091T | D | 1e-05 | 1.3e-05 | 1.5e-05 | 1.267E-05 | 2.517E-06 | 1.000E-05 | 1.500E-05 |
| 21818 | 1091 | V1091W | D | 1.1e-05 | 1e-05 | 9.4e-06 | 1.013E-05 | 8.083E-07 | 9.400E-06 | 1.100E-05 |
| 21819 | 1091 | V1091Y | D | 1.5e-05 | 1.6e-05 | 9.1e-06 | 1.337E-05 | 3.729E-06 | 9.100E-06 | 1.600E-05 |
| 21820 | 1091 | V1091V | D | 1.2e-05 | 1e-05 | 1.1e-05 | 1.100E-05 | 1.000E-06 | 1.000E-05 | 1.200E-05 |
| 21821 | 1092 | A1092A | D | 1.4e-05 | 1.2e-05 | 1e-05 | 1.200E-05 | 2.000E-06 | 1.000E-05 | 1.400E-05 |
| 21822 | 1092 | A1092R | D | 8.6e-06 | 1e-05 | 1.6e-05 | 1.153E-05 | 3.931E-06 | 8.600E-06 | 1.600E-05 |
| 21823 | 1092 | A1092N | D | 8.7e-06 | 2.6e-05 | 1.1e-05 | 1.523E-05 | 9.395E-06 | 8.700E-06 | 2.600E-05 |
| 21824 | 1092 | A1092D | D | 1.4e-05 | 1.1e-05 | 1.3e-05 | 1.267E-05 | 1.528E-06 | 1.100E-05 | 1.400E-05 |
| 21825 | 1092 | A1092C | D | 1.4e-05 | 1e-05 | 9.1e-06 | 1.103E-05 | 2.608E-06 | 9.100E-06 | 1.400E-05 |
| 21826 | 1092 | A1092E | D | 1.7e-05 | 1.1e-05 | 1.6e-05 | 1.467E-05 | 3.215E-06 | 1.100E-05 | 1.700E-05 |
| 21827 | 1092 | A1092Q | D | 1.5e-05 | 1.1e-05 | 1.6e-05 | 1.400E-05 | 2.646E-06 | 1.100E-05 | 1.600E-05 |
| 21828 | 1092 | A1092G | D | 1.1e-05 | 1.5e-05 | 1.2e-05 | 1.267E-05 | 2.082E-06 | 1.100E-05 | 1.500E-05 |
| 21829 | 1092 | A1092H | D | 1.5e-05 | 1.5e-05 | 1.6e-05 | 1.533E-05 | 5.774E-07 | 1.500E-05 | 1.600E-05 |
| 21830 | 1092 | A1092I | D | 1.5e-05 | 1.6e-05 | 9.6e-06 | 1.353E-05 | 3.443E-06 | 9.600E-06 | 1.600E-05 |
| 21831 | 1092 | A1092L | D | 1.1e-05 | 1.7e-05 | 1.4e-05 | 1.400E-05 | 3.000E-06 | 1.100E-05 | 1.700E-05 |
| 21832 | 1092 | A1092K | D | 9.3e-06 | 1.5e-05 | 1.4e-05 | 1.277E-05 | 3.044E-06 | 9.300E-06 | 1.500E-05 |
| 21833 | 1092 | A1092M | D | 1.3e-05 | 9.8e-06 | 1.1e-05 | 1.127E-05 | 1.617E-06 | 9.800E-06 | 1.300E-05 |
| 21834 | 1092 | A1092F | D | 1.4e-05 | 1.6e-05 | 1.9e-05 | 1.633E-05 | 2.517E-06 | 1.400E-05 | 1.900E-05 |
| 21835 | 1092 | A1092P | D | 1.4e-05 | 1.6e-05 | 1.3e-05 | 1.433E-05 | 1.528E-06 | 1.300E-05 | 1.600E-05 |
| 21836 | 1092 | A1092S | D | 1.4e-05 | 9.8e-06 | 1.1e-05 | 1.160E-05 | 2.163E-06 | 9.800E-06 | 1.400E-05 |
| 21837 | 1092 | A1092T | D | 1.4e-05 | 1.5e-05 | 1.5e-05 | 1.467E-05 | 5.774E-07 | 1.400E-05 | 1.500E-05 |
| 21838 | 1092 | A1092W | D | 1.7e-05 | 1e-05 | 1.1e-05 | 1.267E-05 | 3.786E-06 | 1.000E-05 | 1.700E-05 |
| 21839 | 1092 | A1092Y | D | 1.4e-05 | 5.7e-06 | 1.1e-05 | 1.023E-05 | 4.203E-06 | 5.700E-06 | 1.400E-05 |
| 21840 | 1092 | A1092V | D | 1.3e-05 | 1.5e-05 | 1.2e-05 | 1.333E-05 | 1.528E-06 | 1.200E-05 | 1.500E-05 |
| 21841 | 1093 | W1093A | D | 1.6e-05 | 3.4e-05 | 1.1e-05 | 2.033E-05 | 1.210E-05 | 1.100E-05 | 3.400E-05 |
| 21842 | 1093 | W1093R | D | 9.1e-06 | 1.3e-05 | 1.3e-05 | 1.170E-05 | 2.252E-06 | 9.100E-06 | 1.300E-05 |
| 21843 | 1093 | W1093N | D | 6e-06 | 1.6e-05 | 1e-05 | 1.067E-05 | 5.033E-06 | 6.000E-06 | 1.600E-05 |
| 21844 | 1093 | W1093D | D | 1.1e-05 | 1.5e-05 | 1.4e-05 | 1.333E-05 | 2.082E-06 | 1.100E-05 | 1.500E-05 |

| | | | | | | | | | | |
|---|---|---|---|---|---|---|---|---|---|---|
| 21845 | 1093 | W1093C | D | 1.4e-05 | 1.5e-05 | 1.5e-05 | 1.467E-05 | 5.774E-07 | 1.400E-05 | 1.500E-05 |
| 21846 | 1093 | W1093E | D | 1.1e-05 | 1.6e-05 | 1.7e-05 | 1.467E-05 | 3.215E-06 | 1.100E-05 | 1.700E-05 |
| 21847 | 1093 | W1093Q | D | 1.1e-05 | 1.1e-05 | 1.2e-05 | 1.133E-05 | 5.774E-07 | 1.100E-05 | 1.200E-05 |
| 21848 | 1093 | W1093G | D | 1.6e-05 | 1.4e-05 | 1.3e-05 | 1.433E-05 | 1.528E-06 | 1.300E-05 | 1.600E-05 |
| 21849 | 1093 | W1093H | D | 1.1e-05 | 1.1e-05 | 1.5e-05 | 1.233E-05 | 2.309E-06 | 1.100E-05 | 1.500E-05 |
| 21850 | 1093 | W1093I | D | 7.6e-06 | 1.3e-05 | 1.6e-05 | 1.220E-05 | 4.257E-06 | 7.600E-06 | 1.600E-05 |
| 21851 | 1093 | W1093L | D | 1.5e-05 | 9.9e-06 | 1.4e-05 | 1.297E-05 | 2.702E-06 | 9.900E-06 | 1.500E-05 |
| 21852 | 1093 | W1093K | D | 1.9e-05 | 9.1e-06 | 1.1e-05 | 1.303E-05 | 5.254E-06 | 9.100E-06 | 1.900E-05 |
| 21853 | 1093 | W1093M | D | 1.6e-05 | 1.2e-05 | 1.7e-05 | 1.500E-05 | 2.646E-06 | 1.200E-05 | 1.700E-05 |
| 21854 | 1093 | W1093F | D | 1.5e-05 | 1.5e-05 | 1.5e-05 | 1.500E-05 | 0.000E+00 | 1.500E-05 | 1.500E-05 |
| 21855 | 1093 | W1093P | D | 1.5e-05 | 1.1e-05 | 1.5e-05 | 1.367E-05 | 2.309E-06 | 1.100E-05 | 1.500E-05 |
| 21856 | 1093 | W1093S | D | 1e-05 | 1.1e-05 | 1.7e-05 | 1.267E-05 | 3.786E-06 | 1.000E-05 | 1.700E-05 |
| 21857 | 1093 | W1093T | D | 1.5e-05 | 1.5e-05 | 1.2e-05 | 1.400E-05 | 1.732E-06 | 1.200E-05 | 1.500E-05 |
| 21858 | 1093 | W1093W | D | 1e-05 | 1.5e-05 | 1e-05 | 1.167E-05 | 2.887E-06 | 1.000E-05 | 1.500E-05 |
| 21859 | 1093 | W1093Y | D | 1.6e-05 | 1.2e-05 | 1.1e-05 | 1.300E-05 | 2.646E-06 | 1.100E-05 | 1.600E-05 |
| 21860 | 1093 | W1093V | D | 1.3e-05 | 1.6e-05 | 1.6e-05 | 1.500E-05 | 1.732E-06 | 1.300E-05 | 1.600E-05 |
| 21861 | 1094 | Y1094A | D | 1.5e-05 | 1.6e-05 | 1.4e-05 | 1.500E-05 | 1.000E-06 | 1.400E-05 | 1.600E-05 |
| 21862 | 1094 | Y1094R | D | 9.9e-06 | 1.2e-05 | 1.2e-05 | 1.130E-05 | 1.212E-06 | 9.900E-06 | 1.200E-05 |
| 21863 | 1094 | Y1094N | D | 1.6e-05 | 3.6e-05 | 1.3e-05 | 2.167E-05 | 1.250E-05 | 1.300E-05 | 3.600E-05 |
| 21864 | 1094 | Y1094D | D | 1e-05 | 1.1e-05 | 1.2e-05 | 1.100E-05 | 1.000E-06 | 1.000E-05 | 1.200E-05 |
| 21865 | 1094 | Y1094C | D | 1.5e-05 | 1.8e-05 | 9.1e-06 | 1.403E-05 | 4.528E-06 | 9.100E-06 | 1.800E-05 |
| 21866 | 1094 | Y1094E | D | 1.4e-05 | 1.1e-05 | 1.6e-05 | 1.367E-05 | 2.517E-06 | 1.100E-05 | 1.600E-05 |
| 21867 | 1094 | Y1094Q | D | 1.1e-05 | 1.1e-05 | 8.7e-06 | 1.023E-05 | 1.328E-06 | 8.700E-06 | 1.100E-05 |
| 21868 | 1094 | Y1094G | D | 1.4e-05 | 1.6e-05 | 1.5e-05 | 1.500E-05 | 1.000E-06 | 1.400E-05 | 1.600E-05 |
| 21869 | 1094 | Y1094H | D | 1.5e-05 | 1.1e-05 | 1.5e-05 | 1.367E-05 | 2.309E-06 | 1.100E-05 | 1.500E-05 |
| 21870 | 1094 | Y1094I | D | 9.1e-06 | 1.7e-05 | 9.2e-06 | 1.177E-05 | 4.532E-06 | 9.100E-06 | 1.700E-05 |
| 21871 | 1094 | Y1094L | D | 1.1e-05 | 1.1e-05 | 1.8e-05 | 1.333E-05 | 4.041E-06 | 1.100E-05 | 1.800E-05 |
| 21872 | 1094 | Y1094K | D | 1.6e-05 | 9.3e-06 | 1.4e-05 | 1.310E-05 | 3.439E-06 | 9.300E-06 | 1.600E-05 |
| 21873 | 1094 | Y1094M | D | 9.7e-06 | 1.1e-05 | 1e-05 | 1.023E-05 | 6.807E-07 | 9.700E-06 | 1.100E-05 |
| 21874 | 1094 | Y1094F | D | 1.4e-05 | 8.7e-06 | 9.7e-06 | 1.080E-05 | 2.816E-06 | 8.700E-06 | 1.400E-05 |
| 21875 | 1094 | Y1094P | D | 1.2e-05 | 1.1e-05 | 1.5e-05 | 1.267E-05 | 2.082E-06 | 1.100E-05 | 1.500E-05 |
| 21876 | 1094 | Y1094S | D | 1.4e-05 | 9.1e-06 | 1.6e-05 | 1.303E-05 | 3.550E-06 | 9.100E-06 | 1.600E-05 |
| 21877 | 1094 | Y1094T | D | 9.5e-06 | 1.5e-05 | 8.1e-06 | 1.087E-05 | 3.647E-06 | 8.100E-06 | 1.500E-05 |
| 21878 | 1094 | Y1094W | D | 1.1e-05 | 8.6e-06 | 1.6e-05 | 1.187E-05 | 3.775E-06 | 8.600E-06 | 1.600E-05 |
| 21879 | 1094 | Y1094Y | D | 1.3e-05 | 9.6e-06 | 1.3e-05 | 1.187E-05 | 1.963E-06 | 9.600E-06 | 1.300E-05 |
| 21880 | 1094 | Y1094V | D | 1.2e-05 | 1.6e-05 | 1.6e-05 | 1.467E-05 | 2.309E-06 | 1.200E-05 | 1.600E-05 |
| 21881 | 1095 | Q1095A | D | 1.4e-05 | 7.4e-06 | 1e-05 | 1.047E-05 | 3.325E-06 | 7.400E-06 | 1.400E-05 |
| 21882 | 1095 | Q1095R | D | 9.3e-06 | 1.4e-05 | 1.6e-05 | 1.310E-05 | 3.439E-06 | 9.300E-06 | 1.600E-05 |
| 21883 | 1095 | Q1095N | D | 1.7e-05 | 1.6e-05 | 1.1e-05 | 1.467E-05 | 3.215E-06 | 1.100E-05 | 1.700E-05 |
| 21884 | 1095 | Q1095D | D | 1.1e-05 | 1.6e-05 | 9.6e-06 | 1.220E-05 | 3.365E-06 | 9.600E-06 | 1.600E-05 |
| 21885 | 1095 | Q1095C | D | 1e-05 | 1.6e-05 | 1.1e-05 | 1.233E-05 | 3.215E-06 | 1.000E-05 | 1.600E-05 |
| 21886 | 1095 | Q1095E | D | 1e-05 | 1.7e-05 | 1.5e-05 | 1.400E-05 | 3.606E-06 | 1.000E-05 | 1.700E-05 |
| 21887 | 1095 | Q1095Q | D | 8e-06 | 1.6e-05 | 1.3e-05 | 1.233E-05 | 4.041E-06 | 8.000E-06 | 1.600E-05 |
| 21888 | 1095 | Q1095G | D | 8.6e-06 | 1e-05 | 8.3e-06 | 8.967E-06 | 9.074E-07 | 8.300E-06 | 1.000E-05 |
| 21889 | 1095 | Q1095H | D | 1.8e-05 | 1.2e-05 | 1.1e-05 | 1.367E-05 | 3.786E-06 | 1.100E-05 | 1.800E-05 |
| 21890 | 1095 | Q1095I | D | 1.5e-05 | 1.1e-05 | 1.5e-05 | 1.367E-05 | 2.309E-06 | 1.100E-05 | 1.500E-05 |
| 21891 | 1095 | Q1095L | D | 1.1e-05 | 8.4e-06 | 7.2e-06 | 8.867E-06 | 1.943E-06 | 7.200E-06 | 1.100E-05 |
| 21892 | 1095 | Q1095K | D | 9.8e-06 | 1.6e-05 | 1.4e-05 | 1.327E-05 | 3.164E-06 | 9.800E-06 | 1.600E-05 |
| 21893 | 1095 | Q1095M | D | 1.4e-05 | 1.6e-05 | 9e-06 | 1.300E-05 | 3.606E-06 | 9.000E-06 | 1.600E-05 |
| 21894 | 1095 | Q1095F | D | 1.4e-05 | 1.1e-05 | 1e-05 | 1.167E-05 | 2.082E-06 | 1.000E-05 | 1.400E-05 |
| 21895 | 1095 | Q1095P | D | 1.6e-05 | 1.2e-05 | 1.8e-05 | 1.533E-05 | 3.055E-06 | 1.200E-05 | 1.800E-05 |
| 21896 | 1095 | Q1095S | D | 1.1e-05 | 1.1e-05 | 1.8e-05 | 1.333E-05 | 4.041E-06 | 1.100E-05 | 1.800E-05 |
| 21897 | 1095 | Q1095T | D | 1.3e-05 | 1.2e-05 | 1.5e-05 | 1.333E-05 | 1.528E-06 | 1.200E-05 | 1.500E-05 |
| 21898 | 1095 | Q1095W | D | 1.1e-05 | 1.3e-05 | 1.3e-05 | 1.233E-05 | 1.155E-06 | 1.100E-05 | 1.300E-05 |
| 21899 | 1095 | Q1095Y | D | 1.6e-05 | 1.2e-05 | 1.6e-05 | 1.467E-05 | 2.309E-06 | 1.200E-05 | 1.600E-05 |
| 21900 | 1095 | Q1095V | D | 1.6e-05 | 1.7e-05 | 1.2e-05 | 1.500E-05 | 2.646E-06 | 1.200E-05 | 1.700E-05 |
| 21901 | 1096 | Q1096A | D | 1.1e-05 | 1.5e-05 | 9.8e-06 | 1.193E-05 | 2.723E-06 | 9.800E-06 | 1.500E-05 |
| 21902 | 1096 | Q1096R | D | 1.1e-05 | 1.1e-05 | 1.6e-05 | 1.267E-05 | 2.887E-06 | 1.100E-05 | 1.600E-05 |
| 21903 | 1096 | Q1096N | D | 7.5e-06 | 1.3e-05 | 1.4e-05 | 1.150E-05 | 3.500E-06 | 7.500E-06 | 1.400E-05 |
| 21904 | 1096 | Q1096D | D | 1.5e-05 | 1.1e-05 | 1.5e-05 | 1.367E-05 | 2.309E-06 | 1.100E-05 | 1.500E-05 |
| 21905 | 1096 | Q1096C | D | 1.4e-05 | 1.4e-05 | 1.6e-05 | 1.467E-05 | 1.155E-06 | 1.400E-05 | 1.600E-05 |
| 21906 | 1096 | Q1096E | D | 1.3e-05 | 1.6e-05 | 1.3e-05 | 1.400E-05 | 1.732E-06 | 1.300E-05 | 1.600E-05 |
| 21907 | 1096 | Q1096Q | D | 1.4e-05 | 9.8e-06 | 1.6e-05 | 1.327E-05 | 3.164E-06 | 9.800E-06 | 1.600E-05 |
| 21908 | 1096 | Q1096G | D | 1.4e-05 | 9.7e-06 | 8.3e-06 | 1.067E-05 | 2.970E-06 | 8.300E-06 | 1.400E-05 |
| 21909 | 1096 | Q1096H | D | 9.6e-06 | 8.8e-06 | 1e-05 | 9.467E-06 | 6.110E-07 | 8.800E-06 | 1.000E-05 |
| 21910 | 1096 | Q1096I | D | 1.6e-05 | 7.6e-06 | 1.6e-05 | 1.320E-05 | 4.850E-06 | 7.600E-06 | 1.600E-05 |
| 21911 | 1096 | Q1096L | D | 1.7e-05 | 1.5e-05 | 1.7e-05 | 1.633E-05 | 1.155E-06 | 1.500E-05 | 1.700E-05 |
| 21912 | 1096 | Q1096K | D | 1.1e-05 | 1.4e-05 | 1e-05 | 1.167E-05 | 2.082E-06 | 1.000E-05 | 1.400E-05 |
| 21913 | 1096 | Q1096M | D | 1.4e-05 | 1.7e-05 | 1.7e-05 | 1.600E-05 | 1.732E-06 | 1.400E-05 | 1.700E-05 |
| 21914 | 1096 | Q1096F | D | 1.1e-05 | 1.1e-05 | 1.7e-05 | 1.300E-05 | 3.464E-06 | 1.100E-05 | 1.700E-05 |
| 21915 | 1096 | Q1096P | D | 1.5e-05 | 1.6e-05 | 1.2e-05 | 1.433E-05 | 2.082E-06 | 1.200E-05 | 1.600E-05 |
| 21916 | 1096 | Q1096S | D | 1.2e-05 | 1.1e-05 | 1.5e-05 | 1.267E-05 | 2.082E-06 | 1.100E-05 | 1.500E-05 |
| 21917 | 1096 | Q1096T | D | 1.1e-05 | 1e-05 | 9.4e-06 | 1.013E-05 | 8.083E-07 | 9.400E-06 | 1.100E-05 |
| 21918 | 1096 | Q1096W | D | 1.2e-05 | 1.5e-05 | 8.2e-06 | 1.173E-05 | 3.408E-06 | 8.200E-06 | 1.500E-05 |
| 21919 | 1096 | Q1096Y | D | 1e-05 | 1.6e-05 | 1.7e-05 | 1.433E-05 | 3.786E-06 | 1.000E-05 | 1.700E-05 |
| 21920 | 1096 | Q1096V | D | 1.2e-05 | 1e-05 | 1.5e-05 | 1.233E-05 | 2.517E-06 | 1.000E-05 | 1.500E-05 |
| 21921 | 1097 | K1097A | D | 1.5e-05 | 1.2e-05 | 1.4e-05 | 1.367E-05 | 1.528E-06 | 1.200E-05 | 1.500E-05 |
| 21922 | 1097 | K1097R | D | 1.4e-05 | 1.4e-05 | 1.6e-05 | 1.467E-05 | 1.155E-06 | 1.400E-05 | 1.600E-05 |
| 21923 | 1097 | K1097N | D | 1.2e-05 | 1.4e-05 | 1.4e-05 | 1.333E-05 | 1.155E-06 | 1.200E-05 | 1.400E-05 |
| 21924 | 1097 | K1097D | D | 1.2e-05 | 3e-05 | 6.6e-06 | 1.620E-05 | 1.225E-05 | 6.600E-06 | 3.000E-05 |
| 21925 | 1097 | K1097C | D | 8.6e-06 | 1.5e-05 | 1.5e-05 | 1.287E-05 | 3.695E-06 | 8.600E-06 | 1.500E-05 |
| 21926 | 1097 | K1097E | D | 1e-05 | 1.5e-05 | 1.3e-05 | 1.267E-05 | 2.517E-06 | 1.000E-05 | 1.500E-05 |
| 21927 | 1097 | K1097Q | D | 9e-06 | 1.5e-05 | 9.7e-06 | 1.123E-05 | 3.281E-06 | 9.000E-06 | 1.500E-05 |
| 21928 | 1097 | K1097G | D | 1e-05 | 1.6e-05 | 9.2e-06 | 1.173E-05 | 3.717E-06 | 9.200E-06 | 1.600E-05 |
| 21929 | 1097 | K1097H | D | 8.9e-06 | 1.4e-05 | 1.4e-05 | 1.230E-05 | 2.944E-06 | 8.900E-06 | 1.400E-05 |

| | | | | | | | | | | |
|---|---|---|---|---|---|---|---|---|---|---|
| 21930 | 1097 | K1097I | D | 1.5e-05 | 1.4e-05 | 1.4e-05 | 1.433E-05 | 5.774E-07 | 1.400E-05 | 1.500E-05 |
| 21931 | 1097 | K1097L | D | 1.1e-05 | 1.5e-05 | 1.1e-05 | 1.233E-05 | 2.309E-06 | 1.100E-05 | 1.500E-05 |
| 21932 | 1097 | K1097K | D | 1e-05 | 1.5e-05 | 1.5e-05 | 1.333E-05 | 2.887E-06 | 1.000E-05 | 1.500E-05 |
| 21933 | 1097 | K1097M | D | 8.7e-06 | 1.4e-05 | 1.4e-05 | 1.223E-05 | 3.060E-06 | 8.700E-06 | 1.400E-05 |
| 21934 | 1097 | K1097F | D | 1.5e-05 | 1e-05 | 1.5e-05 | 1.333E-05 | 2.887E-06 | 1.000E-05 | 1.500E-05 |
| 21935 | 1097 | K1097P | D | 1.1e-05 | 1.1e-05 | 1.1e-05 | 1.100E-05 | 2.075E-21 | 1.100E-05 | 1.100E-05 |
| 21936 | 1097 | K1097S | D | 1.3e-05 | 1.7e-05 | 1.1e-05 | 1.367E-05 | 3.055E-06 | 1.100E-05 | 1.700E-05 |
| 21937 | 1097 | K1097T | D | 1.5e-05 | 1.1e-05 | 1.6e-05 | 1.400E-05 | 2.646E-06 | 1.100E-05 | 1.600E-05 |
| 21938 | 1097 | K1097W | D | 9.7e-06 | 1.6e-05 | 1.7e-05 | 1.423E-05 | 3.958E-06 | 9.700E-06 | 1.700E-05 |
| 21939 | 1097 | K1097Y | D | 1.1e-05 | 1.2e-05 | 1.2e-05 | 1.167E-05 | 5.774E-07 | 1.100E-05 | 1.200E-05 |
| 21940 | 1097 | K1097V | D | 9.8e-06 | 1.6e-05 | 1.5e-05 | 1.360E-05 | 3.329E-06 | 9.800E-06 | 1.600E-05 |
| 21941 | 1098 | P1098A | D | 1.1e-05 | 9.5e-06 | 1.4e-05 | 1.150E-05 | 2.291E-06 | 9.500E-06 | 1.400E-05 |
| 21942 | 1098 | P1098R | D | 1.1e-05 | 1.5e-05 | 1.3e-05 | 1.300E-05 | 2.000E-06 | 1.100E-05 | 1.500E-05 |
| 21943 | 1098 | P1098N | D | 1.2e-05 | 1.4e-05 | 1.4e-05 | 1.333E-05 | 1.155E-06 | 1.200E-05 | 1.400E-05 |
| 21944 | 1098 | P1098D | D | 1.7e-05 | 1.5e-05 | 9.1e-06 | 1.370E-05 | 4.107E-06 | 9.100E-06 | 1.700E-05 |
| 21945 | 1098 | P1098C | D | 1.5e-05 | 9.1e-06 | 8.7e-06 | 1.093E-05 | 3.528E-06 | 8.700E-06 | 1.500E-05 |
| 21946 | 1098 | P1098E | D | 1.5e-05 | 8.4e-06 | 1.6e-05 | 1.313E-05 | 4.130E-06 | 8.400E-06 | 1.600E-05 |
| 21947 | 1098 | P1098Q | D | 1e-05 | 1.5e-05 | 9.3e-06 | 1.143E-05 | 3.109E-06 | 9.300E-06 | 1.500E-05 |
| 21948 | 1098 | P1098G | D | 1.5e-05 | 1.7e-05 | 1.1e-05 | 1.433E-05 | 3.055E-06 | 1.100E-05 | 1.700E-05 |
| 21949 | 1098 | P1098H | D | 9.9e-06 | 1.3e-05 | 1.1e-05 | 1.130E-05 | 1.572E-06 | 9.900E-06 | 1.300E-05 |
| 21950 | 1098 | P1098I | D | 9.6e-06 | 1.4e-05 | 9.1e-06 | 1.090E-05 | 2.696E-06 | 9.100E-06 | 1.400E-05 |
| 21951 | 1098 | P1098L | D | 1e-05 | 1.6e-05 | 1e-05 | 1.200E-05 | 3.464E-06 | 1.000E-05 | 1.600E-05 |
| 21952 | 1098 | P1098K | D | 1.3e-05 | 9.2e-06 | 1.5e-05 | 1.240E-05 | 2.946E-06 | 9.200E-06 | 1.500E-05 |
| 21953 | 1098 | P1098M | D | 1.4e-05 | 1.7e-05 | 1.3e-05 | 1.467E-05 | 2.082E-06 | 1.300E-05 | 1.700E-05 |
| 21954 | 1098 | P1098F | D | 1.2e-05 | 1.3e-05 | 1.7e-05 | 1.400E-05 | 2.646E-06 | 1.200E-05 | 1.700E-05 |
| 21955 | 1098 | P1098P | D | 1.1e-05 | 1.5e-05 | 1.7e-05 | 1.433E-05 | 3.055E-06 | 1.100E-05 | 1.700E-05 |
| 21956 | 1098 | P1098S | D | 1e-05 | 1.3e-05 | 1.1e-05 | 1.133E-05 | 1.528E-06 | 1.000E-05 | 1.300E-05 |
| 21957 | 1098 | P1098T | D | 9e-06 | 9.8e-06 | 1e-05 | 9.600E-06 | 5.292E-07 | 9.000E-06 | 1.000E-05 |
| 21958 | 1098 | P1098W | D | 1.5e-05 | 1.4e-05 | 9.8e-06 | 1.293E-05 | 2.759E-06 | 9.800E-06 | 1.500E-05 |
| 21959 | 1098 | P1098Y | D | 1.5e-05 | 1.1e-05 | 1.4e-05 | 1.333E-05 | 2.082E-06 | 1.100E-05 | 1.500E-05 |
| 21960 | 1098 | P1098V | D | 9.2e-06 | 1e-05 | 1.8e-05 | 1.240E-05 | 4.866E-06 | 9.200E-06 | 1.800E-05 |
| 21961 | 1099 | G1099A | D | 1.1e-05 | 1.5e-05 | 1.4e-05 | 1.333E-05 | 2.082E-06 | 1.100E-05 | 1.500E-05 |
| 21962 | 1099 | G1099R | D | 1.1e-05 | 7.8e-06 | 1.6e-05 | 1.160E-05 | 4.133E-06 | 7.800E-06 | 1.600E-05 |
| 21963 | 1099 | G1099N | D | 9.6e-06 | 1.2e-05 | 1.5e-05 | 1.220E-05 | 2.706E-06 | 9.600E-06 | 1.500E-05 |
| 21964 | 1099 | G1099D | D | 1.5e-05 | 1.4e-05 | 1.5e-05 | 1.467E-05 | 5.774E-07 | 1.400E-05 | 1.500E-05 |
| 21965 | 1099 | G1099C | D | 1.6e-05 | 1.6e-05 | 1.1e-05 | 1.433E-05 | 2.887E-06 | 1.100E-05 | 1.600E-05 |
| 21966 | 1099 | G1099E | D | 1.6e-05 | 9.8e-06 | 1.2e-05 | 1.260E-05 | 3.143E-06 | 9.800E-06 | 1.600E-05 |
| 21967 | 1099 | G1099Q | D | 1.6e-05 | 1e-05 | 1e-05 | 1.200E-05 | 3.464E-06 | 1.000E-05 | 1.600E-05 |
| 21968 | 1099 | G1099G | D | 1.3e-05 | 1.6e-05 | 1.7e-05 | 1.533E-05 | 2.082E-06 | 1.300E-05 | 1.700E-05 |
| 21969 | 1099 | G1099H | D | 9.4e-06 | 1.6e-05 | 1.1e-05 | 1.213E-05 | 3.443E-06 | 9.400E-06 | 1.600E-05 |
| 21970 | 1099 | G1099I | D | 1.1e-05 | 9.5e-06 | 1e-05 | 1.017E-05 | 7.638E-07 | 9.500E-06 | 1.100E-05 |
| 21971 | 1099 | G1099L | D | 1.2e-05 | 1e-05 | 1.1e-05 | 1.100E-05 | 1.000E-06 | 1.000E-05 | 1.200E-05 |
| 21972 | 1099 | G1099K | D | 6e-06 | 1.6e-05 | 1.1e-05 | 1.100E-05 | 5.000E-06 | 6.000E-06 | 1.600E-05 |
| 21973 | 1099 | G1099M | D | 1.1e-05 | 6.2e-06 | 9.2e-06 | 8.800E-06 | 2.425E-06 | 6.200E-06 | 1.100E-05 |
| 21974 | 1099 | G1099F | D | 8.4e-06 | 1.5e-05 | 8.9e-06 | 1.077E-05 | 3.675E-06 | 8.400E-06 | 1.500E-05 |
| 21975 | 1099 | G1099P | D | 9.7e-06 | 1.5e-05 | 1.5e-05 | 1.323E-05 | 3.060E-06 | 9.700E-06 | 1.500E-05 |
| 21976 | 1099 | G1099S | D | 8.4e-06 | 1.4e-05 | 1.3e-05 | 1.180E-05 | 2.987E-06 | 8.400E-06 | 1.400E-05 |
| 21977 | 1099 | G1099T | D | 1.2e-05 | 8e-06 | 1.6e-05 | 1.200E-05 | 4.000E-06 | 8.000E-06 | 1.600E-05 |
| 21978 | 1099 | G1099W | D | 1.3e-05 | 1.5e-05 | 1.4e-05 | 1.400E-05 | 1.000E-06 | 1.300E-05 | 1.500E-05 |
| 21979 | 1099 | G1099Y | D | 1.4e-05 | 1.1e-05 | 1e-05 | 1.167E-05 | 2.082E-06 | 1.000E-05 | 1.400E-05 |
| 21980 | 1099 | G1099V | D | 1.5e-05 | 9.6e-06 | 1e-05 | 1.153E-05 | 3.009E-06 | 9.600E-06 | 1.500E-05 |
| 21981 | 1100 | K1100A | D | 1.7e-05 | 1e-05 | 1.3e-05 | 1.333E-05 | 3.512E-06 | 1.000E-05 | 1.700E-05 |
| 21982 | 1100 | K1100R | D | 1.4e-05 | 1e-05 | 1.5e-05 | 1.300E-05 | 2.646E-06 | 1.000E-05 | 1.500E-05 |
| 21983 | 1100 | K1100N | D | 1.1e-05 | 1.4e-05 | 1.1e-05 | 1.200E-05 | 1.732E-06 | 1.100E-05 | 1.400E-05 |
| 21984 | 1100 | K1100D | D | 9.9e-06 | 1.7e-05 | 1.4e-05 | 1.363E-05 | 3.564E-06 | 9.900E-06 | 1.700E-05 |
| 21985 | 1100 | K1100C | D | 8.2e-06 | 1.5e-05 | 1.1e-05 | 1.140E-05 | 3.418E-06 | 8.200E-06 | 1.500E-05 |
| 21986 | 1100 | K1100E | D | 1.3e-05 | 9.9e-06 | 9e-06 | 1.063E-05 | 2.098E-06 | 9.000E-06 | 1.300E-05 |
| 21987 | 1100 | K1100Q | D | 1.3e-05 | 1.5e-05 | 1.4e-05 | 1.400E-05 | 1.000E-06 | 1.300E-05 | 1.500E-05 |
| 21988 | 1100 | K1100G | D | 9.6e-06 | 7.5e-06 | 9.4e-06 | 8.833E-06 | 1.159E-06 | 7.500E-06 | 9.600E-06 |
| 21989 | 1100 | K1100H | D | 1.4e-05 | 1.2e-05 | 9.9e-06 | 1.197E-05 | 2.050E-06 | 9.900E-06 | 1.400E-05 |
| 21990 | 1100 | K1100I | D | 7.3e-06 | 8.7e-06 | 1.5e-05 | 1.033E-05 | 4.102E-06 | 7.300E-06 | 1.500E-05 |
| 21991 | 1100 | K1100L | D | 7e-06 | 1.6e-05 | 1.5e-05 | 1.267E-05 | 4.933E-06 | 7.000E-06 | 1.600E-05 |
| 21992 | 1100 | K1100K | D | 1.4e-05 | 1e-05 | 1.6e-05 | 1.333E-05 | 3.055E-06 | 1.000E-05 | 1.600E-05 |
| 21993 | 1100 | K1100M | D | 1.6e-05 | 9.7e-06 | 1e-05 | 1.190E-05 | 3.554E-06 | 9.700E-06 | 1.600E-05 |
| 21994 | 1100 | K1100F | D | 1.5e-05 | 1.5e-05 | 1.6e-05 | 1.533E-05 | 5.774E-07 | 1.500E-05 | 1.600E-05 |
| 21995 | 1100 | K1100P | D | 1.5e-05 | 1.7e-05 | 1e-05 | 1.400E-05 | 3.606E-06 | 1.000E-05 | 1.700E-05 |
| 21996 | 1100 | K1100S | D | 1.1e-05 | 8.9e-06 | 9.2e-06 | 9.700E-06 | 1.136E-06 | 8.900E-06 | 1.100E-05 |
| 21997 | 1100 | K1100T | D | 1.4e-05 | 1.8e-05 | 1.3e-05 | 1.500E-05 | 2.646E-06 | 1.300E-05 | 1.800E-05 |
| 21998 | 1100 | K1100W | D | 1.5e-05 | 1.2e-05 | 7e-06 | 1.133E-05 | 4.041E-06 | 7.000E-06 | 1.500E-05 |
| 21999 | 1100 | K1100Y | D | 1.5e-05 | 1.3e-05 | 1.3e-05 | 1.367E-05 | 1.155E-06 | 1.300E-05 | 1.500E-05 |
| 22000 | 1100 | K1100V | D | 1.6e-05 | 1.6e-05 | 1.5e-05 | 1.567E-05 | 5.774E-07 | 1.500E-05 | 1.600E-05 |
| 22001 | 1101 | A1101A | D | 9.8e-06 | 1.1e-05 | 1.5e-05 | 1.193E-05 | 2.723E-06 | 9.800E-06 | 1.500E-05 |
| 22002 | 1101 | A1101R | D | 1.6e-05 | 1e-05 | 1.7e-05 | 1.433E-05 | 3.786E-06 | 1.000E-05 | 1.700E-05 |
| 22003 | 1101 | A1101N | D | 1.4e-05 | 1.1e-05 | 1.5e-05 | 1.333E-05 | 2.082E-06 | 1.100E-05 | 1.500E-05 |
| 22004 | 1101 | A1101D | D | 1.5e-05 | 1.6e-05 | 1.5e-05 | 1.533E-05 | 5.774E-07 | 1.500E-05 | 1.600E-05 |
| 22005 | 1101 | A1101C | D | 1.2e-05 | 9.9e-06 | 1.5e-05 | 1.230E-05 | 2.563E-06 | 9.900E-06 | 1.500E-05 |
| 22006 | 1101 | A1101E | D | 1.1e-05 | 1.5e-05 | 1e-05 | 1.200E-05 | 2.646E-06 | 1.000E-05 | 1.500E-05 |
| 22007 | 1101 | A1101Q | D | 1.5e-05 | 1.7e-05 | 9.7e-06 | 1.390E-05 | 3.772E-06 | 9.700E-06 | 1.700E-05 |
| 22008 | 1101 | A1101G | D | 2.2e-05 | 1.1e-05 | 1.5e-05 | 1.600E-05 | 5.568E-06 | 1.100E-05 | 2.200E-05 |
| 22009 | 1101 | A1101H | D | 7.8e-06 | 9.3e-06 | 1e-05 | 9.033E-06 | 1.124E-06 | 7.800E-06 | 1.000E-05 |
| 22010 | 1101 | A1101I | D | 1.6e-05 | 9.6e-06 | 9.1e-06 | 1.157E-05 | 3.848E-06 | 9.100E-06 | 1.600E-05 |
| 22011 | 1101 | A1101L | D | 1.4e-05 | 1.3e-05 | 1.6e-05 | 1.433E-05 | 1.528E-06 | 1.300E-05 | 1.600E-05 |
| 22012 | 1101 | A1101K | D | 1.2e-05 | 1.7e-05 | 1.5e-05 | 1.467E-05 | 2.517E-06 | 1.200E-05 | 1.700E-05 |
| 22013 | 1101 | A1101M | D | 9.3e-06 | 1.2e-05 | 1.4e-05 | 1.177E-05 | 2.359E-06 | 9.300E-06 | 1.400E-05 |
| 22014 | 1101 | A1101F | D | 1.5e-05 | 1.7e-05 | 1.5e-05 | 1.567E-05 | 1.155E-06 | 1.500E-05 | 1.700E-05 |

| | | | | | | | | | | |
|---|---|---|---|---|---|---|---|---|---|---|
| 22015 | 1101 | A1101P | D | 1.1e-05 | 1.4e-05 | 9.4e-06 | 1.147E-05 | 2.335E-06 | 9.400E-06 | 1.400E-05 |
| 22016 | 1101 | A1101S | D | 1.6e-05 | 9.4e-06 | 1.1e-05 | 1.213E-05 | 3.443E-06 | 9.400E-06 | 1.600E-05 |
| 22017 | 1101 | A1101T | D | 1.5e-05 | 1.4e-05 | 1.5e-05 | 1.467E-05 | 5.774E-07 | 1.400E-05 | 1.500E-05 |
| 22018 | 1101 | A1101W | D | 1.5e-05 | 1.4e-05 | 1.4e-05 | 1.433E-05 | 5.774E-07 | 1.400E-05 | 1.500E-05 |
| 22019 | 1101 | A1101Y | D | 1.5e-05 | 1.4e-05 | 1.4e-05 | 1.433E-05 | 5.774E-07 | 1.400E-05 | 1.500E-05 |
| 22020 | 1101 | A1101V | D | 1.3e-05 | 1.6e-05 | 1e-05 | 1.300E-05 | 3.000E-06 | 1.000E-05 | 1.600E-05 |
| 22021 | 1102 | P1102A | D | 1.6e-05 | 1.6e-05 | 1.5e-05 | 1.567E-05 | 5.774E-07 | 1.500E-05 | 1.600E-05 |
| 22022 | 1102 | P1102R | D | 1e-05 | 1e-05 | 1.4e-05 | 1.133E-05 | 2.309E-06 | 1.000E-05 | 1.400E-05 |
| 22023 | 1102 | P1102N | D | 1.3e-05 | 1.6e-05 | 1.1e-05 | 1.333E-05 | 2.517E-06 | 1.100E-05 | 1.600E-05 |
| 22024 | 1102 | P1102D | D | 9.8e-06 | 9.2e-06 | 1.6e-05 | 1.167E-05 | 3.765E-06 | 9.200E-06 | 1.600E-05 |
| 22025 | 1102 | P1102C | D | 1.1e-05 | 1.7e-05 | 1e-05 | 1.267E-05 | 3.786E-06 | 1.000E-05 | 1.700E-05 |
| 22026 | 1102 | P1102E | D | 1.1e-05 | 1.5e-05 | 1.3e-05 | 1.300E-05 | 2.000E-06 | 1.100E-05 | 1.500E-05 |
| 22027 | 1102 | P1102Q | D | 9.4e-06 | 1.2e-05 | 8.4e-06 | 9.933E-06 | 1.858E-06 | 8.400E-06 | 1.200E-05 |
| 22028 | 1102 | P1102G | D | 1.1e-05 | 1.3e-05 | 1.8e-05 | 1.400E-05 | 3.606E-06 | 1.100E-05 | 1.800E-05 |
| 22029 | 1102 | P1102H | D | 1.4e-05 | 9.8e-06 | 1.5e-05 | 1.293E-05 | 2.759E-06 | 9.800E-06 | 1.500E-05 |
| 22030 | 1102 | P1102I | D | 1.5e-05 | 1.2e-05 | 1.4e-05 | 1.367E-05 | 1.528E-06 | 1.200E-05 | 1.500E-05 |
| 22031 | 1102 | P1102L | D | 1.5e-05 | 1e-05 | 1.4e-05 | 1.300E-05 | 2.646E-06 | 1.000E-05 | 1.500E-05 |
| 22032 | 1102 | P1102K | D | 1e-05 | 1.5e-05 | 1.1e-05 | 1.200E-05 | 2.646E-06 | 1.000E-05 | 1.500E-05 |
| 22033 | 1102 | P1102M | D | 1.2e-05 | 1.4e-05 | 1e-05 | 1.200E-05 | 2.000E-06 | 1.000E-05 | 1.400E-05 |
| 22034 | 1102 | P1102F | D | 8.2e-06 | 1.3e-05 | 1.6e-05 | 1.240E-05 | 3.934E-06 | 8.200E-06 | 1.600E-05 |
| 22035 | 1102 | P1102P | D | 1e-05 | 1.4e-05 | 9.3e-06 | 1.110E-05 | 2.536E-06 | 9.300E-06 | 1.400E-05 |
| 22036 | 1102 | P1102S | D | 1.4e-05 | 1.3e-05 | 1.3e-05 | 1.333E-05 | 5.774E-07 | 1.300E-05 | 1.400E-05 |
| 22037 | 1102 | P1102T | D | 1.1e-05 | 1e-05 | 1.4e-05 | 1.167E-05 | 2.082E-06 | 1.000E-05 | 1.400E-05 |
| 22038 | 1102 | P1102W | D | 1.6e-05 | 1.4e-05 | 1.1e-05 | 1.367E-05 | 2.517E-06 | 1.100E-05 | 1.600E-05 |
| 22039 | 1102 | P1102Y | D | 1.1e-05 | 9.7e-06 | 1.3e-05 | 1.123E-05 | 1.662E-06 | 9.700E-06 | 1.300E-05 |
| 22040 | 1102 | P1102V | D | 1.2e-05 | 1.2e-05 | 8.9e-06 | 1.097E-05 | 1.790E-06 | 8.900E-06 | 1.200E-05 |
| 22041 | 1103 | K1103A | D | 1.5e-05 | 1.6e-05 | 1.5e-05 | 1.533E-05 | 5.774E-07 | 1.500E-05 | 1.600E-05 |
| 22042 | 1103 | K1103R | D | 1.3e-05 | 1.6e-05 | 1.5e-05 | 1.467E-05 | 1.528E-06 | 1.300E-05 | 1.600E-05 |
| 22043 | 1103 | K1103N | D | 1.4e-05 | 1e-05 | 9.3e-06 | 1.110E-05 | 2.536E-06 | 9.300E-06 | 1.400E-05 |
| 22044 | 1103 | K1103D | D | 1.4e-05 | 1.4e-05 | 1.3e-05 | 1.367E-05 | 5.774E-07 | 1.300E-05 | 1.400E-05 |
| 22045 | 1103 | K1103C | D | 8.4e-06 | 9e-06 | 6.8e-06 | 8.067E-06 | 1.137E-06 | 6.800E-06 | 9.000E-06 |
| 22046 | 1103 | K1103E | D | 1.1e-05 | 1.1e-05 | 1.1e-05 | 1.100E-05 | 2.075E-21 | 1.100E-05 | 1.100E-05 |
| 22047 | 1103 | K1103Q | D | 1.1e-05 | 1e-05 | 1.2e-05 | 1.100E-05 | 1.000E-06 | 1.000E-05 | 1.200E-05 |
| 22048 | 1103 | K1103G | D | 1.6e-05 | 1.6e-05 | 1.6e-05 | 1.600E-05 | 0.000E+00 | 1.600E-05 | 1.600E-05 |
| 22049 | 1103 | K1103H | D | 9.9e-06 | 1.5e-05 | 1.6e-05 | 1.363E-05 | 3.272E-06 | 9.900E-06 | 1.600E-05 |
| 22050 | 1103 | K1103I | D | 1.4e-05 | 1.5e-05 | 1e-05 | 1.300E-05 | 2.646E-06 | 1.000E-05 | 1.500E-05 |
| 22051 | 1103 | K1103L | D | 1.6e-05 | 1.1e-05 | 1.7e-05 | 1.467E-05 | 3.215E-06 | 1.100E-05 | 1.700E-05 |
| 22052 | 1103 | K1103K | D | 2e-05 | 1.4e-05 | 1e-05 | 1.467E-05 | 5.033E-06 | 1.000E-05 | 2.000E-05 |
| 22053 | 1103 | K1103M | D | 1.5e-05 | 1e-05 | 1.3e-05 | 1.267E-05 | 2.517E-06 | 1.000E-05 | 1.500E-05 |
| 22054 | 1103 | K1103F | D | 2e-05 | 1.1e-05 | 1.6e-05 | 1.567E-05 | 4.509E-06 | 1.100E-05 | 2.000E-05 |
| 22055 | 1103 | K1103P | D | 1.5e-05 | 1.3e-05 | 1.7e-05 | 1.500E-05 | 2.000E-06 | 1.300E-05 | 1.700E-05 |
| 22056 | 1103 | K1103S | D | 1.6e-05 | 1.2e-05 | 1.7e-05 | 1.500E-05 | 2.646E-06 | 1.200E-05 | 1.700E-05 |
| 22057 | 1103 | K1103T | D | 1.4e-05 | 1e-05 | 1.8e-05 | 1.400E-05 | 4.000E-06 | 1.000E-05 | 1.800E-05 |
| 22058 | 1103 | K1103W | D | 1.7e-05 | 1.5e-05 | 1.1e-05 | 1.433E-05 | 3.055E-06 | 1.100E-05 | 1.700E-05 |
| 22059 | 1103 | K1103Y | D | 1e-05 | 1.3e-05 | 1.3e-05 | 1.200E-05 | 1.732E-06 | 1.000E-05 | 1.300E-05 |
| 22060 | 1103 | K1103V | D | 1.5e-05 | 1.5e-05 | 1.5e-05 | 1.500E-05 | 0.000E+00 | 1.500E-05 | 1.500E-05 |
| 22061 | 1104 | L1104A | D | 8.6e-06 | 5.6e-06 | 1.6e-05 | 1.007E-05 | 5.353E-06 | 5.600E-06 | 1.600E-05 |
| 22062 | 1104 | L1104R | D | 1.5e-05 | 8.2e-06 | 1.5e-05 | 1.273E-05 | 3.926E-06 | 8.200E-06 | 1.500E-05 |
| 22063 | 1104 | L1104N | D | 1.2e-05 | 1.4e-05 | 1e-05 | 1.200E-05 | 2.000E-06 | 1.000E-05 | 1.400E-05 |
| 22064 | 1104 | L1104D | D | 1e-05 | 1.3e-05 | 8.4e-06 | 1.047E-05 | 2.335E-06 | 8.400E-06 | 1.300E-05 |
| 22065 | 1104 | L1104C | D | 8.7e-06 | 1.1e-05 | 1.6e-05 | 1.190E-05 | 3.732E-06 | 8.700E-06 | 1.600E-05 |
| 22066 | 1104 | L1104E | D | 1.1e-05 | 1.4e-05 | 1.1e-05 | 1.200E-05 | 1.732E-06 | 1.100E-05 | 1.400E-05 |
| 22067 | 1104 | L1104Q | D | 1.2e-05 | 9.1e-06 | 1.7e-05 | 1.270E-05 | 3.996E-06 | 9.100E-06 | 1.700E-05 |
| 22068 | 1104 | L1104G | D | 1.2e-05 | 9.7e-06 | 1.1e-05 | 1.090E-05 | 1.153E-06 | 9.700E-06 | 1.200E-05 |
| 22069 | 1104 | L1104H | D | 9.5e-06 | 1.3e-05 | 1.4e-05 | 1.217E-05 | 2.363E-06 | 9.500E-06 | 1.400E-05 |
| 22070 | 1104 | L1104I | D | 1.1e-05 | 1.1e-05 | 1e-05 | 1.067E-05 | 5.774E-07 | 1.000E-05 | 1.100E-05 |
| 22071 | 1104 | L1104L | D | 1.4e-05 | 1.6e-05 | 1.1e-05 | 1.367E-05 | 2.517E-06 | 1.100E-05 | 1.600E-05 |
| 22072 | 1104 | L1104K | D | 9.6e-06 | 1.5e-05 | 1.1e-05 | 1.187E-05 | 2.802E-06 | 9.600E-06 | 1.500E-05 |
| 22073 | 1104 | L1104M | D | 2e-05 | 1.5e-05 | 1.1e-05 | 1.533E-05 | 4.509E-06 | 1.100E-05 | 2.000E-05 |
| 22074 | 1104 | L1104F | D | 1.4e-05 | 1.3e-05 | 1e-05 | 1.233E-05 | 2.082E-06 | 1.000E-05 | 1.400E-05 |
| 22075 | 1104 | L1104P | D | 1.3e-05 | 9.1e-06 | 9e-06 | 1.037E-05 | 2.281E-06 | 9.000E-06 | 1.300E-05 |
| 22076 | 1104 | L1104S | D | 6.4e-06 | 1.4e-05 | 1.4e-05 | 1.147E-05 | 4.388E-06 | 6.400E-06 | 1.400E-05 |
| 22077 | 1104 | L1104T | D | 1.5e-05 | 1e-05 | 8.7e-06 | 1.123E-05 | 3.326E-06 | 8.700E-06 | 1.500E-05 |
| 22078 | 1104 | L1104W | D | 1e-05 | 1.5e-05 | 1.1e-05 | 1.200E-05 | 2.646E-06 | 1.000E-05 | 1.500E-05 |
| 22079 | 1104 | L1104Y | D | 1.3e-05 | 1.4e-05 | 9e-06 | 1.200E-05 | 2.646E-06 | 9.000E-06 | 1.400E-05 |
| 22080 | 1104 | L1104V | D | 1.4e-05 | 9.4e-06 | 1.4e-05 | 1.247E-05 | 2.656E-06 | 9.400E-06 | 1.400E-05 |
| 22081 | 1105 | L1105A | D | 8.7e-06 | 1.6e-05 | 1.3e-05 | 1.257E-05 | 3.669E-06 | 8.700E-06 | 1.600E-05 |
| 22082 | 1105 | L1105R | D | 1.5e-05 | 9.9e-06 | 1.1e-05 | 1.197E-05 | 2.684E-06 | 9.900E-06 | 1.500E-05 |
| 22083 | 1105 | L1105N | D | 8.5e-06 | 1.5e-05 | 1.1e-05 | 1.150E-05 | 3.279E-06 | 8.500E-06 | 1.500E-05 |
| 22084 | 1105 | L1105D | D | 1e-05 | 1.1e-05 | 1.2e-05 | 1.100E-05 | 1.000E-06 | 1.000E-05 | 1.200E-05 |
| 22085 | 1105 | L1105C | D | 1.7e-05 | 8.8e-06 | 8.5e-06 | 1.143E-05 | 4.823E-06 | 8.500E-06 | 1.700E-05 |
| 22086 | 1105 | L1105E | D | 8.1e-06 | 8.9e-06 | 9.9e-06 | 8.967E-06 | 9.018E-07 | 8.100E-06 | 9.900E-06 |
| 22087 | 1105 | L1105Q | D | 1.3e-05 | 1e-05 | 1.4e-05 | 1.233E-05 | 2.082E-06 | 1.000E-05 | 1.400E-05 |
| 22088 | 1105 | L1105G | D | 9.7e-06 | 1.1e-05 | 1.4e-05 | 1.157E-05 | 2.205E-06 | 9.700E-06 | 1.400E-05 |
| 22089 | 1105 | L1105H | D | 9.8e-06 | 1.6e-05 | 1.6e-05 | 1.393E-05 | 3.580E-06 | 9.800E-06 | 1.600E-05 |
| 22090 | 1105 | L1105I | D | 1.1e-05 | 9.1e-06 | 1.1e-05 | 1.037E-05 | 1.097E-06 | 9.100E-06 | 1.100E-05 |
| 22091 | 1105 | L1105L | D | 9.5e-06 | 1.6e-05 | 1.3e-05 | 1.283E-05 | 3.253E-06 | 9.500E-06 | 1.600E-05 |
| 22092 | 1105 | L1105K | D | 1.3e-05 | 1.6e-05 | 1.4e-05 | 1.433E-05 | 1.528E-06 | 1.300E-05 | 1.600E-05 |
| 22093 | 1105 | L1105M | D | 1.6e-05 | 1.4e-05 | 9.6e-06 | 1.320E-05 | 3.274E-06 | 9.600E-06 | 1.600E-05 |
| 22094 | 1105 | L1105F | D | 1.1e-05 | 1.4e-05 | 1.6e-05 | 1.367E-05 | 2.517E-06 | 1.100E-05 | 1.600E-05 |
| 22095 | 1105 | L1105P | D | 1e-05 | 1.4e-05 | 1.1e-05 | 1.167E-05 | 2.082E-06 | 1.000E-05 | 1.400E-05 |
| 22096 | 1105 | L1105S | D | 1.4e-05 | 1.3e-05 | 9.9e-06 | 1.230E-05 | 2.138E-06 | 9.900E-06 | 1.400E-05 |
| 22097 | 1105 | L1105T | D | 1.5e-05 | 1.6e-05 | 1e-05 | 1.367E-05 | 3.215E-06 | 1.000E-05 | 1.600E-05 |
| 22098 | 1105 | L1105W | D | 1.4e-05 | 9.7e-06 | 1.6e-05 | 1.323E-05 | 3.219E-06 | 9.700E-06 | 1.600E-05 |
| 22099 | 1105 | L1105Y | D | 1.6e-05 | 8.7e-06 | 8.4e-06 | 1.103E-05 | 4.304E-06 | 8.400E-06 | 1.600E-05 |

| | | | | | | | | | | |
|---|---|---|---|---|---|---|---|---|---|---|
| 22100 | 1105 | L1105V | D | 1.5e-05 | 1.5e-05 | 1.3e-05 | 1.433E-05 | 1.155E-06 | 1.300E-05 | 1.500E-05 |
| 22101 | 1106 | I1106A | D | 1.5e-05 | 1e-05 | 1e-05 | 1.167E-05 | 2.887E-06 | 1.000E-05 | 1.500E-05 |
| 22102 | 1106 | I1106R | D | 9e-06 | 1.6e-05 | 1.6e-05 | 1.367E-05 | 4.041E-06 | 9.000E-06 | 1.600E-05 |
| 22103 | 1106 | I1106N | D | 1.1e-05 | 1.6e-05 | 1.2e-05 | 1.300E-05 | 2.646E-06 | 1.100E-05 | 1.600E-05 |
| 22104 | 1106 | I1106D | D | 1.1e-05 | 9.6e-06 | 1e-05 | 1.020E-05 | 7.211E-07 | 9.600E-06 | 1.100E-05 |
| 22105 | 1106 | I1106C | D | 9.3e-06 | 1.3e-05 | 1.5e-05 | 1.243E-05 | 2.892E-06 | 9.300E-06 | 1.500E-05 |
| 22106 | 1106 | I1106E | D | 9.9e-06 | 1.4e-05 | 1.5e-05 | 1.297E-05 | 2.702E-06 | 9.900E-06 | 1.500E-05 |
| 22107 | 1106 | I1106Q | D | 1.2e-05 | 1.2e-05 | 1.4e-05 | 1.267E-05 | 1.155E-06 | 1.200E-05 | 1.400E-05 |
| 22108 | 1106 | I1106G | D | 1.4e-05 | 1.4e-05 | 1.5e-05 | 1.433E-05 | 5.774E-07 | 1.400E-05 | 1.500E-05 |
| 22109 | 1106 | I1106H | D | 1.8e-05 | 1.7e-05 | 1.5e-05 | 1.667E-05 | 1.528E-06 | 1.500E-05 | 1.800E-05 |
| 22110 | 1106 | I1106I | D | 1.5e-05 | 1.4e-05 | 1.4e-05 | 1.433E-05 | 5.774E-07 | 1.400E-05 | 1.500E-05 |
| 22111 | 1106 | I1106L | D | 9.6e-06 | 1.6e-05 | 1.2e-05 | 1.253E-05 | 3.233E-06 | 9.600E-06 | 1.600E-05 |
| 22112 | 1106 | I1106K | D | 1.5e-05 | 1.6e-05 | 1.2e-05 | 1.433E-05 | 2.082E-06 | 1.200E-05 | 1.600E-05 |
| 22113 | 1106 | I1106M | D | 1.3e-05 | 1e-05 | 1.5e-05 | 1.267E-05 | 2.517E-06 | 1.000E-05 | 1.500E-05 |
| 22114 | 1106 | I1106F | D | 1.8e-05 | 1.2e-05 | 9.9e-06 | 1.330E-05 | 4.204E-06 | 9.900E-06 | 1.800E-05 |
| 22115 | 1106 | I1106P | D | 9e-06 | 1.1e-05 | 9.8e-06 | 9.933E-06 | 1.007E-06 | 9.000E-06 | 1.100E-05 |
| 22116 | 1106 | I1106S | D | 1.6e-05 | 1.7e-05 | 1.5e-05 | 1.600E-05 | 1.000E-06 | 1.500E-05 | 1.700E-05 |
| 22117 | 1106 | I1106T | D | 8.8e-06 | 9.5e-06 | 1.3e-05 | 1.043E-05 | 2.250E-06 | 8.800E-06 | 1.300E-05 |
| 22118 | 1106 | I1106W | D | 1.4e-05 | 1.5e-05 | 1.1e-05 | 1.333E-05 | 2.082E-06 | 1.100E-05 | 1.500E-05 |
| 22119 | 1106 | I1106Y | D | 1.6e-05 | 8.3e-06 | 1.5e-05 | 1.310E-05 | 4.187E-06 | 8.300E-06 | 1.600E-05 |
| 22120 | 1106 | I1106V | D | 1.6e-05 | 9.6e-06 | 1.5e-05 | 1.353E-05 | 3.443E-06 | 9.600E-06 | 1.600E-05 |
| 22121 | 1107 | Y1107A | D | 1.6e-05 | 1.5e-05 | 1e-05 | 1.367E-05 | 3.215E-06 | 1.000E-05 | 1.600E-05 |
| 22122 | 1107 | Y1107R | D | 1.5e-05 | 1.1e-05 | 1.4e-05 | 1.333E-05 | 2.082E-06 | 1.100E-05 | 1.500E-05 |
| 22123 | 1107 | Y1107N | D | 1.5e-05 | 1.1e-05 | 1.5e-05 | 1.367E-05 | 2.309E-06 | 1.100E-05 | 1.500E-05 |
| 22124 | 1107 | Y1107D | D | 1.1e-05 | 1.4e-05 | 1.1e-05 | 1.200E-05 | 1.732E-06 | 1.100E-05 | 1.400E-05 |
| 22125 | 1107 | Y1107C | D | 8.6e-06 | 1.5e-05 | 1.1e-05 | 1.153E-05 | 3.233E-06 | 8.600E-06 | 1.500E-05 |
| 22126 | 1107 | Y1107E | D | 1e-05 | 1.6e-05 | 1.5e-05 | 1.367E-05 | 3.215E-06 | 1.000E-05 | 1.600E-05 |
| 22127 | 1107 | Y1107Q | D | 9.7e-06 | 1.3e-05 | 1.5e-05 | 1.257E-05 | 2.676E-06 | 9.700E-06 | 1.500E-05 |
| 22128 | 1107 | Y1107G | D | 1.1e-05 | 1.6e-05 | 1.6e-05 | 1.433E-05 | 2.887E-06 | 1.100E-05 | 1.600E-05 |
| 22129 | 1107 | Y1107H | D | 1.1e-05 | 1.2e-05 | 1.2e-05 | 1.167E-05 | 5.774E-07 | 1.100E-05 | 1.200E-05 |
| 22130 | 1107 | Y1107I | D | 1.6e-05 | 1.5e-05 | 1.6e-05 | 1.567E-05 | 5.774E-07 | 1.500E-05 | 1.600E-05 |
| 22131 | 1107 | Y1107L | D | 1.4e-05 | 8.6e-06 | 1.3e-05 | 1.187E-05 | 2.873E-06 | 8.600E-06 | 1.400E-05 |
| 22132 | 1107 | Y1107K | D | 1.6e-05 | 1.5e-05 | 1e-05 | 1.367E-05 | 3.215E-06 | 1.000E-05 | 1.600E-05 |
| 22133 | 1107 | Y1107M | D | 1.1e-05 | 1.3e-05 | 9.3e-06 | 1.110E-05 | 1.852E-06 | 9.300E-06 | 1.300E-05 |
| 22134 | 1107 | Y1107F | D | 1.6e-05 | 1.4e-05 | 1.2e-05 | 1.400E-05 | 2.000E-06 | 1.200E-05 | 1.600E-05 |
| 22135 | 1107 | Y1107P | D | 1.6e-05 | 1.1e-05 | 1.6e-05 | 1.433E-05 | 2.887E-06 | 1.100E-05 | 1.600E-05 |
| 22136 | 1107 | Y1107S | D | 1e-05 | 1.4e-05 | 1.1e-05 | 1.167E-05 | 2.082E-06 | 1.000E-05 | 1.400E-05 |
| 22137 | 1107 | Y1107T | D | 1.4e-05 | 1.6e-05 | 1.1e-05 | 1.367E-05 | 2.517E-06 | 1.100E-05 | 1.600E-05 |
| 22138 | 1107 | Y1107W | D | 9.2e-06 | 1.4e-05 | 1.4e-05 | 1.240E-05 | 2.771E-06 | 9.200E-06 | 1.400E-05 |
| 22139 | 1107 | Y1107Y | D | 1.4e-05 | 1.3e-05 | 1.5e-05 | 1.400E-05 | 1.000E-06 | 1.300E-05 | 1.500E-05 |
| 22140 | 1107 | Y1107V | D | 1.1e-05 | 8.8e-06 | 1.5e-05 | 1.160E-05 | 3.143E-06 | 8.800E-06 | 1.500E-05 |
| 22141 | 1108 | S1108A | D | 1.7e-05 | 1.6e-05 | 1.4e-05 | 1.567E-05 | 1.528E-06 | 1.400E-05 | 1.700E-05 |
| 22142 | 1108 | S1108R | D | 1.4e-05 | 1e-05 | 1.4e-05 | 1.267E-05 | 2.309E-06 | 1.000E-05 | 1.400E-05 |
| 22143 | 1108 | S1108N | D | 1.5e-05 | 8.3e-06 | 9.6e-06 | 1.097E-05 | 3.553E-06 | 8.300E-06 | 1.500E-05 |
| 22144 | 1108 | S1108D | D | 1.1e-05 | 1.1e-05 | 1.1e-05 | 1.100E-05 | 2.075E-21 | 1.100E-05 | 1.100E-05 |
| 22145 | 1108 | S1108C | D | 9.7e-06 | 8.3e-06 | 1.5e-05 | 1.100E-05 | 3.534E-06 | 8.300E-06 | 1.500E-05 |
| 22146 | 1108 | S1108E | D | 1.1e-05 | 1.6e-05 | 9.9e-06 | 1.230E-05 | 3.251E-06 | 9.900E-06 | 1.600E-05 |
| 22147 | 1108 | S1108Q | D | 1.4e-05 | 1.2e-05 | 9.5e-06 | 1.183E-05 | 2.255E-06 | 9.500E-06 | 1.400E-05 |
| 22148 | 1108 | S1108G | D | 1.7e-05 | 1.3e-05 | 1.2e-05 | 1.400E-05 | 2.646E-06 | 1.200E-05 | 1.700E-05 |
| 22149 | 1108 | S1108H | D | 1.5e-05 | 8.3e-06 | 1.5e-05 | 1.277E-05 | 3.868E-06 | 8.300E-06 | 1.500E-05 |
| 22150 | 1108 | S1108I | D | 1e-05 | 1.2e-05 | 1.3e-05 | 1.167E-05 | 1.528E-06 | 1.000E-05 | 1.300E-05 |
| 22151 | 1108 | S1108L | D | 1.2e-05 | 1.7e-05 | 1.3e-05 | 1.400E-05 | 2.646E-06 | 1.200E-05 | 1.700E-05 |
| 22152 | 1108 | S1108K | D | 9.9e-06 | 1.5e-05 | 1.4e-05 | 1.297E-05 | 2.702E-06 | 9.900E-06 | 1.500E-05 |
| 22153 | 1108 | S1108M | D | 1e-05 | 1.2e-05 | 2.2e-05 | 1.467E-05 | 6.429E-06 | 1.000E-05 | 2.200E-05 |
| 22154 | 1108 | S1108F | D | 1.2e-05 | 1.6e-05 | 1.3e-05 | 1.367E-05 | 2.082E-06 | 1.200E-05 | 1.600E-05 |
| 22155 | 1108 | S1108P | D | 1e-05 | 1.6e-05 | 1.5e-05 | 1.367E-05 | 3.215E-06 | 1.000E-05 | 1.600E-05 |
| 22156 | 1108 | S1108S | D | 1e-05 | 1.6e-05 | 1.1e-05 | 1.233E-05 | 3.215E-06 | 1.000E-05 | 1.600E-05 |
| 22157 | 1108 | S1108T | D | 1.6e-05 | 1.1e-05 | 1.1e-05 | 1.267E-05 | 2.887E-06 | 1.100E-05 | 1.600E-05 |
| 22158 | 1108 | S1108W | D | 1.5e-05 | 8.1e-06 | 8.7e-06 | 1.060E-05 | 3.822E-06 | 8.100E-06 | 1.500E-05 |
| 22159 | 1108 | S1108Y | D | 1.5e-05 | 1.3e-05 | 1.5e-05 | 1.433E-05 | 1.155E-06 | 1.300E-05 | 1.500E-05 |
| 22160 | 1108 | S1108V | D | 1.6e-05 | 1.6e-05 | 1.6e-05 | 1.600E-05 | 0.000E+00 | 1.600E-05 | 1.600E-05 |
| 22161 | 1109 | A1109A | D | 1.1e-05 | 1.4e-05 | 1.6e-05 | 1.367E-05 | 2.517E-06 | 1.100E-05 | 1.600E-05 |
| 22162 | 1109 | A1109R | D | 1e-05 | 1.1e-05 | 1.3e-05 | 1.133E-05 | 1.528E-06 | 1.000E-05 | 1.300E-05 |
| 22163 | 1109 | A1109N | D | 1.5e-05 | 9.4e-06 | 1.7e-05 | 1.380E-05 | 3.940E-06 | 9.400E-06 | 1.700E-05 |
| 22164 | 1109 | A1109D | D | 1e-05 | 8.4e-06 | 1.5e-05 | 1.113E-05 | 3.443E-06 | 8.400E-06 | 1.500E-05 |
| 22165 | 1109 | A1109C | D | 8.9e-06 | 1.1e-05 | 9.7e-06 | 9.867E-06 | 1.060E-06 | 8.900E-06 | 1.100E-05 |
| 22166 | 1109 | A1109E | D | 1.3e-05 | 1e-05 | 1.2e-05 | 1.167E-05 | 1.528E-06 | 1.000E-05 | 1.300E-05 |
| 22167 | 1109 | A1109Q | D | 1.5e-05 | 1.4e-05 | 1.1e-05 | 1.333E-05 | 2.082E-06 | 1.100E-05 | 1.500E-05 |
| 22168 | 1109 | A1109G | D | 9.1e-06 | 9.3e-06 | 1.5e-05 | 1.113E-05 | 3.350E-06 | 9.100E-06 | 1.500E-05 |
| 22169 | 1109 | A1109H | D | 1e-05 | 8.6e-06 | 1e-05 | 9.533E-06 | 8.083E-07 | 8.600E-06 | 1.000E-05 |
| 22170 | 1109 | A1109I | D | 8.5e-06 | 1.6e-05 | 1.4e-05 | 1.283E-05 | 3.884E-06 | 8.500E-06 | 1.600E-05 |
| 22171 | 1109 | A1109L | D | 9.8e-06 | 1.6e-05 | 1.5e-05 | 1.360E-05 | 3.329E-06 | 9.800E-06 | 1.600E-05 |
| 22172 | 1109 | A1109K | D | 1.1e-05 | 8.4e-06 | 1.1e-05 | 1.013E-05 | 1.501E-06 | 8.400E-06 | 1.100E-05 |
| 22173 | 1109 | A1109M | D | 1.1e-05 | 1.6e-05 | 1.4e-05 | 1.367E-05 | 2.517E-06 | 1.100E-05 | 1.600E-05 |
| 22174 | 1109 | A1109F | D | 1.4e-05 | 1.1e-05 | 9.5e-06 | 1.150E-05 | 2.291E-06 | 9.500E-06 | 1.400E-05 |
| 22175 | 1109 | A1109P | D | 1.4e-05 | 1e-05 | 1.1e-05 | 1.167E-05 | 2.082E-06 | 1.000E-05 | 1.400E-05 |
| 22176 | 1109 | A1109S | D | 1.8e-05 | 1.5e-05 | 1.1e-05 | 1.467E-05 | 3.512E-06 | 1.100E-05 | 1.800E-05 |
| 22177 | 1109 | A1109T | D | 1.1e-05 | 1e-05 | 1.5e-05 | 1.200E-05 | 2.646E-06 | 1.000E-05 | 1.500E-05 |
| 22178 | 1109 | A1109W | D | 1e-05 | 1.4e-05 | 1.1e-05 | 1.167E-05 | 2.082E-06 | 1.000E-05 | 1.400E-05 |
| 22179 | 1109 | A1109Y | D | 1.4e-05 | 1.5e-05 | 1.4e-05 | 1.433E-05 | 5.774E-07 | 1.400E-05 | 1.500E-05 |
| 22180 | 1109 | A1109V | D | 1.2e-05 | 1.5e-05 | 1.5e-05 | 1.400E-05 | 1.732E-06 | 1.200E-05 | 1.500E-05 |
| 22181 | 1110 | S1110A | D | 2e-05 | 1.5e-05 | 1.1e-05 | 1.533E-05 | 4.509E-06 | 1.100E-05 | 2.000E-05 |
| 22182 | 1110 | S1110R | D | 8.8e-06 | 1.3e-05 | 1.3e-05 | 1.160E-05 | 2.425E-06 | 8.800E-06 | 1.300E-05 |
| 22183 | 1110 | S1110N | D | 1.4e-05 | 1.4e-05 | 1.2e-05 | 1.333E-05 | 1.155E-06 | 1.200E-05 | 1.400E-05 |
| 22184 | 1110 | S1110D | D | 2.6e-05 | 1.1e-05 | 9.8e-06 | 1.560E-05 | 9.027E-06 | 9.800E-06 | 2.600E-05 |

| | | | | | | | | | | |
|---|---|---|---|---|---|---|---|---|---|---|
| 22185 | 1110 | S1110C | D | 1.4e-05 | 1e-05 | 6.9e-06 | 1.030E-05 | 3.559E-06 | 6.900E-06 | 1.400E-05 |
| 22186 | 1110 | S1110E | D | 1.6e-05 | 1.1e-05 | 2.2e-05 | 1.633E-05 | 5.508E-06 | 1.100E-05 | 2.200E-05 |
| 22187 | 1110 | S1110Q | D | 1.3e-05 | 1.5e-05 | 1.1e-05 | 1.300E-05 | 2.000E-06 | 1.100E-05 | 1.500E-05 |
| 22188 | 1110 | S1110G | D | 1.5e-05 | 1e-05 | 9.9e-06 | 1.163E-05 | 2.916E-06 | 9.900E-06 | 1.500E-05 |
| 22189 | 1110 | S1110H | D | 1.2e-05 | 1.6e-05 | 1.3e-05 | 1.367E-05 | 2.082E-06 | 1.200E-05 | 1.600E-05 |
| 22190 | 1110 | S1110I | D | 1.4e-05 | 1e-05 | 6.8e-06 | 1.027E-05 | 3.607E-06 | 6.800E-06 | 1.400E-05 |
| 22191 | 1110 | S1110L | D | 1.6e-05 | 1.1e-05 | 1.5e-05 | 1.400E-05 | 2.646E-06 | 1.100E-05 | 1.600E-05 |
| 22192 | 1110 | S1110K | D | 1e-05 | 1.4e-05 | 1.4e-05 | 1.267E-05 | 2.309E-06 | 1.000E-05 | 1.400E-05 |
| 22193 | 1110 | S1110M | D | 1.4e-05 | 1e-05 | 1.1e-05 | 1.167E-05 | 2.082E-06 | 1.000E-05 | 1.400E-05 |
| 22194 | 1110 | S1110F | D | 9.5e-06 | 1.3e-05 | 1.6e-05 | 1.283E-05 | 3.253E-06 | 9.500E-06 | 1.600E-05 |
| 22195 | 1110 | S1110P | D | 1.2e-05 | 1.5e-05 | 1.4e-05 | 1.367E-05 | 1.528E-06 | 1.200E-05 | 1.500E-05 |
| 22196 | 1110 | S1110S | D | 1.7e-05 | 1.1e-05 | 1.1e-05 | 1.300E-05 | 3.464E-06 | 1.100E-05 | 1.700E-05 |
| 22197 | 1110 | S1110T | D | 1.6e-05 | 1.4e-05 | 9.7e-06 | 1.323E-05 | 3.219E-06 | 9.700E-06 | 1.600E-05 |
| 22198 | 1110 | S1110W | D | 1.6e-05 | 1e-05 | 1e-05 | 1.200E-05 | 3.464E-06 | 1.000E-05 | 1.600E-05 |
| 22199 | 1110 | S1110Y | D | 1e-05 | 8.2e-06 | 1.6e-05 | 1.140E-05 | 4.084E-06 | 8.200E-06 | 1.600E-05 |
| 22200 | 1110 | S1110V | D | 1.2e-05 | 1e-05 | 1.1e-05 | 1.100E-05 | 1.000E-06 | 1.000E-05 | 1.200E-05 |
| 22201 | 1111 | F1111A | D | 1.2e-05 | 1.5e-05 | 1.4e-05 | 1.367E-05 | 1.528E-06 | 1.200E-05 | 1.500E-05 |
| 22202 | 1111 | F1111R | D | 1.5e-05 | 9.6e-06 | 1.3e-05 | 1.253E-05 | 2.730E-06 | 9.600E-06 | 1.500E-05 |
| 22203 | 1111 | F1111N | D | 1.6e-05 | 1.4e-05 | 8.8e-06 | 1.293E-05 | 3.717E-06 | 8.800E-06 | 1.600E-05 |
| 22204 | 1111 | F1111D | D | 9.9e-06 | 1.6e-05 | 1.5e-05 | 1.363E-05 | 3.272E-06 | 9.900E-06 | 1.600E-05 |
| 22205 | 1111 | F1111C | D | 8.9e-06 | 1.7e-05 | 1.5e-05 | 1.363E-05 | 4.219E-06 | 8.900E-06 | 1.700E-05 |
| 22206 | 1111 | F1111E | D | 8.1e-06 | 1.3e-05 | 2e-05 | 1.370E-05 | 5.981E-06 | 8.100E-06 | 2.000E-05 |
| 22207 | 1111 | F1111Q | D | 1e-05 | 1.4e-05 | 1.1e-05 | 1.167E-05 | 2.082E-06 | 1.000E-05 | 1.400E-05 |
| 22208 | 1111 | F1111G | D | 1.5e-05 | 7.9e-06 | 1.7e-05 | 1.330E-05 | 4.782E-06 | 7.900E-06 | 1.700E-05 |
| 22209 | 1111 | F1111H | D | 1.2e-05 | 1.6e-05 | 1.5e-05 | 1.433E-05 | 2.082E-06 | 1.200E-05 | 1.600E-05 |
| 22210 | 1111 | F1111I | D | 1.5e-05 | 1.2e-05 | 9.5e-06 | 1.217E-05 | 2.754E-06 | 9.500E-06 | 1.500E-05 |
| 22211 | 1111 | F1111L | D | 1.4e-05 | 1e-05 | 1.4e-05 | 1.267E-05 | 2.309E-06 | 1.000E-05 | 1.400E-05 |
| 22212 | 1111 | F1111K | D | 1.1e-05 | 1.6e-05 | 1.1e-05 | 1.267E-05 | 2.887E-06 | 1.100E-05 | 1.600E-05 |
| 22213 | 1111 | F1111M | D | 1.2e-05 | 1.3e-05 | 9.7e-06 | 1.157E-05 | 1.692E-06 | 9.700E-06 | 1.300E-05 |
| 22214 | 1111 | F1111F | D | 1.6e-05 | 1.5e-05 | 8.6e-06 | 1.320E-05 | 4.015E-06 | 8.600E-06 | 1.600E-05 |
| 22215 | 1111 | F1111P | D | 1.5e-05 | 1.7e-05 | 1.1e-05 | 1.433E-05 | 3.055E-06 | 1.100E-05 | 1.700E-05 |
| 22216 | 1111 | F1111S | D | 1.6e-05 | 1.2e-05 | 8.7e-06 | 1.223E-05 | 3.656E-06 | 8.700E-06 | 1.600E-05 |
| 22217 | 1111 | F1111T | D | 9.7e-06 | 8.9e-06 | 1.4e-05 | 1.087E-05 | 2.743E-06 | 8.900E-06 | 1.400E-05 |
| 22218 | 1111 | F1111W | D | 1.4e-05 | 1.4e-05 | 1.1e-05 | 1.300E-05 | 1.732E-06 | 1.100E-05 | 1.400E-05 |
| 22219 | 1111 | F1111Y | D | 1.6e-05 | 1.3e-05 | 9.3e-06 | 1.277E-05 | 3.356E-06 | 9.300E-06 | 1.600E-05 |
| 22220 | 1111 | F1111V | D | 9.6e-06 | 1.1e-05 | 1.3e-05 | 1.120E-05 | 1.709E-06 | 9.600E-06 | 1.300E-05 |
| 22221 | 1112 | L1112A | D | 1.5e-05 | 1.1e-05 | 1.5e-05 | 1.367E-05 | 2.309E-06 | 1.100E-05 | 1.500E-05 |
| 22222 | 1112 | L1112R | D | 1.3e-05 | 1.7e-05 | 1.7e-05 | 1.567E-05 | 2.309E-06 | 1.300E-05 | 1.700E-05 |
| 22223 | 1112 | L1112N | D | 1.2e-05 | 1.3e-05 | 1.4e-05 | 1.300E-05 | 1.000E-06 | 1.200E-05 | 1.400E-05 |
| 22224 | 1112 | L1112D | D | 9.3e-06 | 9.6e-06 | 1.1e-05 | 9.967E-06 | 9.074E-07 | 9.300E-06 | 1.100E-05 |
| 22225 | 1112 | L1112C | D | 1.5e-05 | 1.1e-05 | 1.7e-05 | 1.433E-05 | 3.055E-06 | 1.100E-05 | 1.700E-05 |
| 22226 | 1112 | L1112E | D | 1.6e-05 | 9.1e-06 | 1e-05 | 1.170E-05 | 3.751E-06 | 9.100E-06 | 1.600E-05 |
| 22227 | 1112 | L1112Q | D | 1.5e-05 | 8.3e-06 | 1.3e-05 | 1.210E-05 | 3.439E-06 | 8.300E-06 | 1.500E-05 |
| 22228 | 1112 | L1112G | D | 1.5e-05 | 1.3e-05 | 1.6e-05 | 1.467E-05 | 1.528E-06 | 1.300E-05 | 1.600E-05 |
| 22229 | 1112 | L1112H | D | 1.4e-05 | 1.2e-05 | 1.3e-05 | 1.300E-05 | 1.000E-06 | 1.200E-05 | 1.400E-05 |
| 22230 | 1112 | L1112I | D | 1.4e-05 | 1.5e-05 | 1.4e-05 | 1.433E-05 | 5.774E-07 | 1.400E-05 | 1.500E-05 |
| 22231 | 1112 | L1112L | D | 1.2e-05 | 1.2e-05 | 1.5e-05 | 1.300E-05 | 1.732E-06 | 1.200E-05 | 1.500E-05 |
| 22232 | 1112 | L1112K | D | 1.5e-05 | 1.4e-05 | 1e-05 | 1.300E-05 | 2.646E-06 | 1.000E-05 | 1.500E-05 |
| 22233 | 1112 | L1112M | D | 1.5e-05 | 1.5e-05 | 1.3e-05 | 1.433E-05 | 1.155E-06 | 1.300E-05 | 1.500E-05 |
| 22234 | 1112 | L1112F | D | 9.8e-06 | 1.6e-05 | 1.7e-05 | 1.427E-05 | 3.900E-06 | 9.800E-06 | 1.700E-05 |
| 22235 | 1112 | L1112P | D | 9.8e-06 | 1.5e-05 | 1.2e-05 | 1.227E-05 | 2.610E-06 | 9.800E-06 | 1.500E-05 |
| 22236 | 1112 | L1112S | D | 1.1e-05 | 1.6e-05 | 1e-05 | 1.233E-05 | 3.215E-06 | 1.000E-05 | 1.600E-05 |
| 22237 | 1112 | L1112T | D | 1.6e-05 | 1.4e-05 | 1.6e-05 | 1.533E-05 | 1.155E-06 | 1.400E-05 | 1.600E-05 |
| 22238 | 1112 | L1112W | D | 1.1e-05 | 1e-05 | 1.4e-05 | 1.167E-05 | 2.082E-06 | 1.000E-05 | 1.400E-05 |
| 22239 | 1112 | L1112Y | D | 1.2e-05 | 1.1e-05 | 1.3e-05 | 1.200E-05 | 1.000E-06 | 1.100E-05 | 1.300E-05 |
| 22240 | 1112 | L1112V | D | 1.4e-05 | 1.4e-05 | 1.4e-05 | 1.400E-05 | 0.000E+00 | 1.400E-05 | 1.400E-05 |
| 22241 | 1113 | Y1113A | D | 1.5e-05 | 1.5e-05 | 1.4e-05 | 1.467E-05 | 5.774E-07 | 1.400E-05 | 1.500E-05 |
| 22242 | 1113 | Y1113R | D | 1.3e-05 | 1.3e-05 | 1.2e-05 | 1.267E-05 | 5.774E-07 | 1.200E-05 | 1.300E-05 |
| 22243 | 1113 | Y1113N | D | 6.6e-06 | 1.7e-05 | 1.6e-05 | 1.320E-05 | 5.738E-06 | 6.600E-06 | 1.700E-05 |
| 22244 | 1113 | Y1113D | D | 1.1e-05 | 1.1e-05 | 1e-05 | 1.067E-05 | 5.774E-07 | 1.000E-05 | 1.100E-05 |
| 22245 | 1113 | Y1113C | D | 1.5e-05 | 9.8e-06 | 1.1e-05 | 1.193E-05 | 2.723E-06 | 9.800E-06 | 1.500E-05 |
| 22246 | 1113 | Y1113E | D | 1.4e-05 | 1e-05 | 1.2e-05 | 1.200E-05 | 2.000E-06 | 1.000E-05 | 1.400E-05 |
| 22247 | 1113 | Y1113Q | D | 1.4e-05 | 1e-05 | 1.5e-05 | 1.300E-05 | 2.646E-06 | 1.000E-05 | 1.500E-05 |
| 22248 | 1113 | Y1113G | D | 1.8e-05 | 1.2e-05 | 9.1e-06 | 1.303E-05 | 4.539E-06 | 9.100E-06 | 1.800E-05 |
| 22249 | 1113 | Y1113H | D | 1.7e-05 | 1e-05 | 1.4e-05 | 1.367E-05 | 3.512E-06 | 1.000E-05 | 1.700E-05 |
| 22250 | 1113 | Y1113I | D | 1.5e-05 | 1.6e-05 | 1.5e-05 | 1.533E-05 | 5.774E-07 | 1.500E-05 | 1.600E-05 |
| 22251 | 1113 | Y1113L | D | 1.5e-05 | 1.4e-05 | 1.2e-05 | 1.367E-05 | 1.528E-06 | 1.200E-05 | 1.500E-05 |
| 22252 | 1113 | Y1113K | D | 1.4e-05 | 9.1e-06 | 1.3e-05 | 1.203E-05 | 2.589E-06 | 9.100E-06 | 1.400E-05 |
| 22253 | 1113 | Y1113M | D | 1.1e-05 | 1.6e-05 | 9.7e-06 | 1.223E-05 | 3.326E-06 | 9.700E-06 | 1.600E-05 |
| 22254 | 1113 | Y1113F | D | 1.3e-05 | 1.4e-05 | 1.6e-05 | 1.433E-05 | 1.528E-06 | 1.300E-05 | 1.600E-05 |
| 22255 | 1113 | Y1113P | D | 1.7e-05 | 1.4e-05 | 9.7e-06 | 1.357E-05 | 3.669E-06 | 9.700E-06 | 1.700E-05 |
| 22256 | 1113 | Y1113S | D | 1.4e-05 | 1.1e-05 | 1.3e-05 | 1.267E-05 | 1.528E-06 | 1.100E-05 | 1.400E-05 |
| 22257 | 1113 | Y1113T | D | 1.5e-05 | 1e-05 | 1.5e-05 | 1.333E-05 | 2.887E-06 | 1.000E-05 | 1.500E-05 |
| 22258 | 1113 | Y1113W | D | 1.5e-05 | 1e-05 | 5.9e-06 | 1.030E-05 | 4.557E-06 | 5.900E-06 | 1.500E-05 |
| 22259 | 1113 | Y1113Y | D | 1.2e-05 | 1.2e-05 | 1.1e-05 | 1.167E-05 | 5.774E-07 | 1.100E-05 | 1.200E-05 |
| 22260 | 1113 | Y1113V | D | 1.8e-05 | 1.5e-05 | 1.5e-05 | 1.600E-05 | 1.732E-06 | 1.500E-05 | 1.800E-05 |
| 22261 | 1114 | S1114A | D | 1.6e-05 | 9.1e-06 | 1.3e-05 | 1.270E-05 | 3.460E-06 | 9.100E-06 | 1.600E-05 |
| 22262 | 1114 | S1114R | D | 1.2e-05 | 1.4e-05 | 1.1e-05 | 1.233E-05 | 1.528E-06 | 1.100E-05 | 1.400E-05 |
| 22263 | 1114 | S1114N | D | 1.1e-05 | 1.4e-05 | 1.5e-05 | 1.333E-05 | 2.082E-06 | 1.100E-05 | 1.500E-05 |
| 22264 | 1114 | S1114D | D | 1.7e-05 | 1.4e-05 | 1.4e-05 | 1.500E-05 | 1.732E-06 | 1.400E-05 | 1.700E-05 |
| 22265 | 1114 | S1114C | D | 1.4e-05 | 1.4e-05 | 1.4e-05 | 1.400E-05 | 0.000E+00 | 1.400E-05 | 1.400E-05 |
| 22266 | 1114 | S1114E | D | 1.5e-05 | 1.5e-05 | 1.5e-05 | 1.500E-05 | 0.000E+00 | 1.500E-05 | 1.500E-05 |
| 22267 | 1114 | S1114Q | D | 1.7e-05 | 1.5e-05 | 1.7e-05 | 1.633E-05 | 1.155E-06 | 1.500E-05 | 1.700E-05 |
| 22268 | 1114 | S1114G | D | 9.5e-06 | 1.4e-05 | 1.7e-05 | 1.350E-05 | 3.775E-06 | 9.500E-06 | 1.700E-05 |
| 22269 | 1114 | S1114H | D | 9.5e-06 | 1.1e-05 | 1.2e-05 | 1.083E-05 | 1.258E-06 | 9.500E-06 | 1.200E-05 |

| | | | | | | | | | | |
|---|---|---|---|---|---|---|---|---|---|---|
| 22270 | 1114 | S1114I | D | 8.5e-06 | 1.1e-05 | 1.1e-05 | 1.017E-05 | 1.443E-06 | 8.500E-06 | 1.100E-05 |
| 22271 | 1114 | S1114L | D | 1.1e-05 | 9.1e-06 | 1.4e-05 | 1.137E-05 | 2.470E-06 | 9.100E-06 | 1.400E-05 |
| 22272 | 1114 | S1114K | D | 1.6e-05 | 1.7e-05 | 1e-05 | 1.433E-05 | 3.786E-06 | 1.000E-05 | 1.700E-05 |
| 22273 | 1114 | S1114M | D | 8.8e-06 | 1.7e-05 | 1.1e-05 | 1.227E-05 | 4.244E-06 | 8.800E-06 | 1.700E-05 |
| 22274 | 1114 | S1114F | D | 1e-05 | 1.5e-05 | 6.4e-06 | 1.047E-05 | 4.319E-06 | 6.400E-06 | 1.500E-05 |
| 22275 | 1114 | S1114P | D | 1.2e-05 | 1.1e-05 | 1.5e-05 | 1.267E-05 | 2.082E-06 | 1.100E-05 | 1.500E-05 |
| 22276 | 1114 | S1114S | D | 1e-05 | 8.7e-06 | 1.6e-05 | 1.157E-05 | 3.894E-06 | 8.700E-06 | 1.600E-05 |
| 22277 | 1114 | S1114T | D | 1.1e-05 | 1.4e-05 | 1.3e-05 | 1.267E-05 | 1.528E-06 | 1.100E-05 | 1.400E-05 |
| 22278 | 1114 | S1114W | D | 9.1e-06 | 1.4e-05 | 1.7e-05 | 1.337E-05 | 3.988E-06 | 9.100E-06 | 1.700E-05 |
| 22279 | 1114 | S1114Y | D | 1.1e-05 | 1.5e-05 | 1.4e-05 | 1.333E-05 | 2.082E-06 | 1.100E-05 | 1.500E-05 |
| 22280 | 1114 | S1114V | D | 1.1e-05 | 1.6e-05 | 1.5e-05 | 1.400E-05 | 2.646E-06 | 1.100E-05 | 1.600E-05 |
| 22281 | 1115 | G1115A | D | 1.7e-05 | 1e-05 | 1.1e-05 | 1.267E-05 | 3.786E-06 | 1.000E-05 | 1.700E-05 |
| 22282 | 1115 | G1115R | D | 1.4e-05 | 1.5e-05 | 1.1e-05 | 1.333E-05 | 2.082E-06 | 1.100E-05 | 1.500E-05 |
| 22283 | 1115 | G1115N | D | 1.5e-05 | 7.6e-06 | 1.2e-05 | 1.153E-05 | 3.722E-06 | 7.600E-06 | 1.500E-05 |
| 22284 | 1115 | G1115D | D | 1.1e-05 | 1.2e-05 | 1.1e-05 | 1.133E-05 | 5.774E-07 | 1.100E-05 | 1.200E-05 |
| 22285 | 1115 | G1115C | D | 1.4e-05 | 1.4e-05 | 1.3e-05 | 1.367E-05 | 5.774E-07 | 1.300E-05 | 1.400E-05 |
| 22286 | 1115 | G1115E | D | 1.6e-05 | 1.6e-05 | 1.7e-05 | 1.633E-05 | 5.774E-07 | 1.600E-05 | 1.700E-05 |
| 22287 | 1115 | G1115Q | D | 1.2e-05 | 9e-06 | 8.8e-06 | 9.933E-06 | 1.793E-06 | 8.800E-06 | 1.200E-05 |
| 22288 | 1115 | G1115G | D | 1e-05 | 1.1e-05 | 1.7e-05 | 1.267E-05 | 3.786E-06 | 1.000E-05 | 1.700E-05 |
| 22289 | 1115 | G1115H | D | 1e-05 | 1e-05 | 1.6e-05 | 1.200E-05 | 3.464E-06 | 1.000E-05 | 1.600E-05 |
| 22290 | 1115 | G1115I | D | 1.5e-05 | 1.5e-05 | 1e-05 | 1.333E-05 | 2.887E-06 | 1.000E-05 | 1.500E-05 |
| 22291 | 1115 | G1115L | D | 1.2e-05 | 1.6e-05 | 1e-05 | 1.267E-05 | 3.055E-06 | 1.000E-05 | 1.600E-05 |
| 22292 | 1115 | G1115K | D | 1.9e-05 | 1.1e-05 | 1.6e-05 | 1.533E-05 | 4.041E-06 | 1.100E-05 | 1.900E-05 |
| 22293 | 1115 | G1115M | D | 1.5e-05 | 1.4e-05 | 1.6e-05 | 1.500E-05 | 1.000E-06 | 1.400E-05 | 1.600E-05 |
| 22294 | 1115 | G1115F | D | 1.6e-05 | 9.9e-06 | 1.1e-05 | 1.230E-05 | 3.251E-06 | 9.900E-06 | 1.600E-05 |
| 22295 | 1115 | G1115P | D | 1.6e-05 | 1.1e-05 | 1.6e-05 | 1.433E-05 | 2.887E-06 | 1.100E-05 | 1.600E-05 |
| 22296 | 1115 | G1115S | D | 1.6e-05 | 1.5e-05 | 1.6e-05 | 1.567E-05 | 5.774E-07 | 1.500E-05 | 1.600E-05 |
| 22297 | 1115 | G1115T | D | 9.8e-06 | 1.5e-05 | 9.1e-06 | 1.130E-05 | 3.223E-06 | 9.100E-06 | 1.500E-05 |
| 22298 | 1115 | G1115W | D | 1.6e-05 | 1.1e-05 | 1e-05 | 1.233E-05 | 3.215E-06 | 1.000E-05 | 1.600E-05 |
| 22299 | 1115 | G1115Y | D | 1.4e-05 | 1e-05 | 1.4e-05 | 1.267E-05 | 2.309E-06 | 1.000E-05 | 1.400E-05 |
| 22300 | 1115 | G1115V | D | 1.1e-05 | 1.3e-05 | 1.6e-05 | 1.333E-05 | 2.517E-06 | 1.100E-05 | 1.600E-05 |
| 22301 | 1116 | V1116A | D | 1.1e-05 | 9.8e-06 | 1.1e-05 | 1.060E-05 | 6.928E-07 | 9.800E-06 | 1.100E-05 |
| 22302 | 1116 | V1116R | D | 1.1e-05 | 8.8e-06 | 1.1e-05 | 1.027E-05 | 1.270E-06 | 8.800E-06 | 1.100E-05 |
| 22303 | 1116 | V1116N | D | 1.5e-05 | 1.5e-05 | 1.4e-05 | 1.467E-05 | 5.774E-07 | 1.400E-05 | 1.500E-05 |
| 22304 | 1116 | V1116D | D | 1.1e-05 | 9.5e-06 | 1.5e-05 | 1.183E-05 | 2.843E-06 | 9.500E-06 | 1.500E-05 |
| 22305 | 1116 | V1116C | D | 1.4e-05 | 1.5e-05 | 1.5e-05 | 1.467E-05 | 5.774E-07 | 1.400E-05 | 1.500E-05 |
| 22306 | 1116 | V1116E | D | 1e-05 | 1.6e-05 | 1.6e-05 | 1.400E-05 | 3.464E-06 | 1.000E-05 | 1.600E-05 |
| 22307 | 1116 | V1116Q | D | 1.1e-05 | 1.1e-05 | 1.2e-05 | 1.133E-05 | 5.774E-07 | 1.100E-05 | 1.200E-05 |
| 22308 | 1116 | V1116G | D | 1.2e-05 | 1.1e-05 | 1.7e-05 | 1.333E-05 | 3.215E-06 | 1.100E-05 | 1.700E-05 |
| 22309 | 1116 | V1116H | D | 1.6e-05 | 1.2e-05 | 1.9e-05 | 1.567E-05 | 3.512E-06 | 1.200E-05 | 1.900E-05 |
| 22310 | 1116 | V1116I | D | 1.4e-05 | 1.4e-05 | 9.8e-06 | 1.260E-05 | 2.425E-06 | 9.800E-06 | 1.400E-05 |
| 22311 | 1116 | V1116L | D | 1e-05 | 1e-05 | 1.5e-05 | 1.167E-05 | 2.887E-06 | 1.000E-05 | 1.500E-05 |
| 22312 | 1116 | V1116K | D | 1.4e-05 | 1.6e-05 | 1.5e-05 | 1.500E-05 | 1.000E-06 | 1.400E-05 | 1.600E-05 |
| 22313 | 1116 | V1116M | D | 1.1e-05 | 1.5e-05 | 1.4e-05 | 1.333E-05 | 2.082E-06 | 1.100E-05 | 1.500E-05 |
| 22314 | 1116 | V1116F | D | 1e-05 | 1.1e-05 | 9.2e-06 | 1.007E-05 | 9.018E-07 | 9.200E-06 | 1.100E-05 |
| 22315 | 1116 | V1116P | D | 1.6e-05 | 1.4e-05 | 1.2e-05 | 1.400E-05 | 2.000E-06 | 1.200E-05 | 1.600E-05 |
| 22316 | 1116 | V1116S | D | 1.4e-05 | 1.3e-05 | 1.5e-05 | 1.400E-05 | 1.000E-06 | 1.300E-05 | 1.500E-05 |
| 22317 | 1116 | V1116T | D | 1e-05 | 1.4e-05 | 1.4e-05 | 1.267E-05 | 2.309E-06 | 1.000E-05 | 1.400E-05 |
| 22318 | 1116 | V1116W | D | 8.9e-06 | 1.2e-05 | 1.5e-05 | 1.197E-05 | 3.050E-06 | 8.900E-06 | 1.500E-05 |
| 22319 | 1116 | V1116Y | D | 1.3e-05 | 1.2e-05 | 7.8e-06 | 1.093E-05 | 2.759E-06 | 7.800E-06 | 1.300E-05 |
| 22320 | 1116 | V1116V | D | 1.7e-05 | 1.6e-05 | 9.3e-06 | 1.410E-05 | 4.187E-06 | 9.300E-06 | 1.700E-05 |
| 22321 | 1117 | P1117A | D | 1.5e-05 | 1.1e-05 | 1.1e-05 | 1.233E-05 | 2.309E-06 | 1.100E-05 | 1.500E-05 |
| 22322 | 1117 | P1117R | D | 1.5e-05 | 1.2e-05 | 1.6e-05 | 1.433E-05 | 2.082E-06 | 1.200E-05 | 1.600E-05 |
| 22323 | 1117 | P1117N | D | 1.5e-05 | 1e-05 | 9.7e-06 | 1.157E-05 | 2.977E-06 | 9.700E-06 | 1.500E-05 |
| 22324 | 1117 | P1117D | D | 1.1e-05 | 1.4e-05 | 9.8e-06 | 1.160E-05 | 2.163E-06 | 9.800E-06 | 1.400E-05 |
| 22325 | 1117 | P1117C | D | 1.5e-05 | 1.6e-05 | 1e-05 | 1.367E-05 | 3.215E-06 | 1.000E-05 | 1.600E-05 |
| 22326 | 1117 | P1117E | D | 1.1e-05 | 1e-05 | 1.4e-05 | 1.167E-05 | 2.082E-06 | 1.000E-05 | 1.400E-05 |
| 22327 | 1117 | P1117Q | D | 1.6e-05 | 1.6e-05 | 1e-05 | 1.400E-05 | 3.464E-06 | 1.000E-05 | 1.600E-05 |
| 22328 | 1117 | P1117G | D | 1.5e-05 | 9e-06 | 1.3e-05 | 1.233E-05 | 3.055E-06 | 9.000E-06 | 1.500E-05 |
| 22329 | 1117 | P1117H | D | 1.1e-05 | 1.4e-05 | 1.5e-05 | 1.333E-05 | 2.082E-06 | 1.100E-05 | 1.500E-05 |
| 22330 | 1117 | P1117I | D | 1.5e-05 | 1.5e-05 | 1.3e-05 | 1.433E-05 | 1.155E-06 | 1.300E-05 | 1.500E-05 |
| 22331 | 1117 | P1117L | D | 1.8e-05 | 1.5e-05 | 1e-05 | 1.433E-05 | 4.041E-06 | 1.000E-05 | 1.800E-05 |
| 22332 | 1117 | P1117K | D | 1.4e-05 | 1.4e-05 | 8.8e-06 | 1.227E-05 | 3.002E-06 | 8.800E-06 | 1.400E-05 |
| 22333 | 1117 | P1117M | D | 9.4e-06 | 1.4e-05 | 1.5e-05 | 1.280E-05 | 2.987E-06 | 9.400E-06 | 1.500E-05 |
| 22334 | 1117 | P1117F | D | 1.6e-05 | 1.5e-05 | 1.6e-05 | 1.567E-05 | 5.774E-07 | 1.500E-05 | 1.600E-05 |
| 22335 | 1117 | P1117P | D | 1.4e-05 | 1.2e-05 | 9.7e-06 | 1.190E-05 | 2.152E-06 | 9.700E-06 | 1.400E-05 |
| 22336 | 1117 | P1117S | D | 9.9e-06 | 1.1e-05 | 1.5e-05 | 1.197E-05 | 2.684E-06 | 9.900E-06 | 1.500E-05 |
| 22337 | 1117 | P1117T | D | 1.5e-05 | 1.1e-05 | 1.5e-05 | 1.367E-05 | 2.309E-06 | 1.100E-05 | 1.500E-05 |
| 22338 | 1117 | P1117W | D | 6.8e-06 | 1.1e-05 | 1.6e-05 | 1.127E-05 | 4.606E-06 | 6.800E-06 | 1.600E-05 |
| 22339 | 1117 | P1117Y | D | 1.2e-05 | 1e-05 | 1e-05 | 1.067E-05 | 1.155E-06 | 1.000E-05 | 1.200E-05 |
| 22340 | 1117 | P1117V | D | 1.1e-05 | 1.5e-05 | 1.6e-05 | 1.400E-05 | 2.646E-06 | 1.100E-05 | 1.600E-05 |
| 22341 | 1118 | S1118A | D | 1e-05 | 1.4e-05 | 1.4e-05 | 1.267E-05 | 2.309E-06 | 1.000E-05 | 1.400E-05 |
| 22342 | 1118 | S1118R | D | 1.1e-05 | 8.7e-06 | 9e-06 | 9.567E-06 | 1.250E-06 | 8.700E-06 | 1.100E-05 |
| 22343 | 1118 | S1118N | D | 1.1e-05 | 9.1e-06 | 1.4e-05 | 1.137E-05 | 2.470E-06 | 9.100E-06 | 1.400E-05 |
| 22344 | 1118 | S1118D | D | 8.2e-06 | 1.1e-05 | 1.1e-05 | 1.007E-05 | 1.617E-06 | 8.200E-06 | 1.100E-05 |
| 22345 | 1118 | S1118C | D | 1.5e-05 | 1e-05 | 9.6e-06 | 1.153E-05 | 3.009E-06 | 9.600E-06 | 1.500E-05 |
| 22346 | 1118 | S1118E | D | 1e-05 | 7.8e-06 | 1.5e-05 | 1.093E-05 | 3.690E-06 | 7.800E-06 | 1.500E-05 |
| 22347 | 1118 | S1118Q | D | 1.4e-05 | 9.2e-06 | 1.6e-05 | 1.307E-05 | 3.495E-06 | 9.200E-06 | 1.600E-05 |
| 22348 | 1118 | S1118G | D | 1.6e-05 | 1.1e-05 | 1.1e-05 | 1.267E-05 | 2.887E-06 | 1.100E-05 | 1.600E-05 |
| 22349 | 1118 | S1118H | D | 1.5e-05 | 9.2e-06 | 9.9e-06 | 1.137E-05 | 3.166E-06 | 9.200E-06 | 1.500E-05 |
| 22350 | 1118 | S1118I | D | 1.2e-05 | 1e-05 | 2.2e-05 | 1.467E-05 | 6.429E-06 | 1.000E-05 | 2.200E-05 |
| 22351 | 1118 | S1118L | D | 1.3e-05 | 1.2e-05 | 1.3e-05 | 1.267E-05 | 5.774E-07 | 1.200E-05 | 1.300E-05 |
| 22352 | 1118 | S1118K | D | 8.8e-06 | 1e-05 | 1.4e-05 | 1.093E-05 | 2.723E-06 | 8.800E-06 | 1.400E-05 |
| 22353 | 1118 | S1118M | D | 1.6e-05 | 1.5e-05 | 1.6e-05 | 1.567E-05 | 5.774E-07 | 1.500E-05 | 1.600E-05 |
| 22354 | 1118 | S1118F | D | 1.2e-05 | 1.3e-05 | 9.3e-06 | 1.143E-05 | 1.914E-06 | 9.300E-06 | 1.300E-05 |

| | | | | | | | | | | |
|---|---|---|---|---|---|---|---|---|---|---|
| 22355 | 1118 | S1118P | D | 1.3e-05 | 1.2e-05 | 1.2e-05 | 1.233E-05 | 5.774E-07 | 1.200E-05 | 1.300E-05 |
| 22356 | 1118 | S1118S | D | 1.5e-05 | 1.1e-05 | 1e-05 | 1.200E-05 | 2.646E-06 | 1.000E-05 | 1.500E-05 |
| 22357 | 1118 | S1118T | D | 1.5e-05 | 1.4e-05 | 1.1e-05 | 1.333E-05 | 2.082E-06 | 1.100E-05 | 1.500E-05 |
| 22358 | 1118 | S1118W | D | 1.1e-05 | 1.2e-05 | 1.3e-05 | 1.200E-05 | 1.000E-06 | 1.100E-05 | 1.300E-05 |
| 22359 | 1118 | S1118Y | D | 1e-05 | 1.4e-05 | 9e-06 | 1.100E-05 | 2.646E-06 | 9.000E-06 | 1.400E-05 |
| 22360 | 1118 | S1118V | D | 1.5e-05 | 1.6e-05 | 1.7e-05 | 1.600E-05 | 1.000E-06 | 1.500E-05 | 1.700E-05 |
| 22361 | 1119 | R1119A | D | 1.1e-05 | 7e-06 | 1.6e-05 | 1.133E-05 | 4.509E-06 | 7.000E-06 | 1.600E-05 |
| 22362 | 1119 | R1119R | D | 1.1e-05 | 1.6e-05 | 1.6e-05 | 1.433E-05 | 2.887E-06 | 1.100E-05 | 1.600E-05 |
| 22363 | 1119 | R1119N | D | 1.5e-05 | 1.5e-05 | 9.2e-06 | 1.307E-05 | 3.349E-06 | 9.200E-06 | 1.500E-05 |
| 22364 | 1119 | R1119D | D | 1.6e-05 | 1.6e-05 | 9.1e-06 | 1.370E-05 | 3.984E-06 | 9.100E-06 | 1.600E-05 |
| 22365 | 1119 | R1119C | D | 1.5e-05 | 1.5e-05 | 1.1e-05 | 1.367E-05 | 2.309E-06 | 1.100E-05 | 1.500E-05 |
| 22366 | 1119 | R1119E | D | 7.4e-06 | 1e-05 | 1.5e-05 | 1.080E-05 | 3.863E-06 | 7.400E-06 | 1.500E-05 |
| 22367 | 1119 | R1119Q | D | 1.4e-05 | 1.5e-05 | 1e-05 | 1.300E-05 | 2.646E-06 | 1.000E-05 | 1.500E-05 |
| 22368 | 1119 | R1119G | D | 1e-05 | 1.6e-05 | 1.1e-05 | 1.233E-05 | 3.215E-06 | 1.000E-05 | 1.600E-05 |
| 22369 | 1119 | R1119H | D | 8.6e-06 | 8.8e-06 | 1.4e-05 | 1.047E-05 | 3.062E-06 | 8.600E-06 | 1.400E-05 |
| 22370 | 1119 | R1119I | D | 6.3e-06 | 1.2e-05 | 2.2e-05 | 1.343E-05 | 7.948E-06 | 6.300E-06 | 2.200E-05 |
| 22371 | 1119 | R1119L | D | 1.4e-05 | 1.5e-05 | 1.1e-05 | 1.333E-05 | 2.082E-06 | 1.100E-05 | 1.500E-05 |
| 22372 | 1119 | R1119K | D | 1.5e-05 | 1.4e-05 | 1e-05 | 1.300E-05 | 2.646E-06 | 1.000E-05 | 1.500E-05 |
| 22373 | 1119 | R1119M | D | 1.3e-05 | 1.1e-05 | 1.4e-05 | 1.267E-05 | 1.528E-06 | 1.100E-05 | 1.400E-05 |
| 22374 | 1119 | R1119F | D | 1.6e-05 | 1.5e-05 | 1.1e-05 | 1.400E-05 | 2.646E-06 | 1.100E-05 | 1.600E-05 |
| 22375 | 1119 | R1119P | D | 1.1e-05 | 1.5e-05 | 1.7e-05 | 1.433E-05 | 3.055E-06 | 1.100E-05 | 1.700E-05 |
| 22376 | 1119 | R1119S | D | 1.1e-05 | 1e-05 | 1.1e-05 | 1.067E-05 | 5.774E-07 | 1.000E-05 | 1.100E-05 |
| 22377 | 1119 | R1119T | D | 1e-05 | 1.3e-05 | 1.5e-05 | 1.267E-05 | 2.517E-06 | 1.000E-05 | 1.500E-05 |
| 22378 | 1119 | R1119W | D | 9.6e-06 | 1.6e-05 | 1.5e-05 | 1.353E-05 | 3.443E-06 | 9.600E-06 | 1.600E-05 |
| 22379 | 1119 | R1119Y | D | 1.7e-05 | 7.1e-06 | 1.3e-05 | 1.237E-05 | 4.980E-06 | 7.100E-06 | 1.700E-05 |
| 22380 | 1119 | R1119V | D | 1.3e-05 | 9.5e-06 | 1e-05 | 1.083E-05 | 1.893E-06 | 9.500E-06 | 1.300E-05 |
| 22381 | 1120 | F1120A | D | 1.5e-05 | 1e-05 | 1.5e-05 | 1.333E-05 | 2.887E-06 | 1.000E-05 | 1.500E-05 |
| 22382 | 1120 | F1120R | D | 9.9e-06 | 1.5e-05 | 1.6e-05 | 1.363E-05 | 3.272E-06 | 9.900E-06 | 1.600E-05 |
| 22383 | 1120 | F1120N | D | 1.1e-05 | 1e-05 | 1.5e-05 | 1.200E-05 | 2.646E-06 | 1.000E-05 | 1.500E-05 |
| 22384 | 1120 | F1120D | D | 1.1e-05 | 1e-05 | 1.3e-05 | 1.133E-05 | 1.528E-06 | 1.000E-05 | 1.300E-05 |
| 22385 | 1120 | F1120C | D | 1.1e-05 | 1.5e-05 | 1.7e-05 | 1.433E-05 | 3.055E-06 | 1.100E-05 | 1.700E-05 |
| 22386 | 1120 | F1120E | D | 1.5e-05 | 1.6e-05 | 1.5e-05 | 1.533E-05 | 5.774E-07 | 1.500E-05 | 1.600E-05 |
| 22387 | 1120 | F1120Q | D | 1.7e-05 | 1.4e-05 | 1.6e-05 | 1.567E-05 | 1.528E-06 | 1.400E-05 | 1.700E-05 |
| 22388 | 1120 | F1120G | D | 1.5e-05 | 3.9e-05 | 1.5e-05 | 2.300E-05 | 1.386E-05 | 1.500E-05 | 3.900E-05 |
| 22389 | 1120 | F1120H | D | 2.1e-05 | 1.3e-05 | 1.3e-05 | 1.567E-05 | 4.619E-06 | 1.300E-05 | 2.100E-05 |
| 22390 | 1120 | F1120I | D | 1.1e-05 | 1.5e-05 | 1.4e-05 | 1.333E-05 | 2.082E-06 | 1.100E-05 | 1.500E-05 |
| 22391 | 1120 | F1120L | D | 1.5e-05 | 1e-05 | 1.5e-05 | 1.333E-05 | 2.887E-06 | 1.000E-05 | 1.500E-05 |
| 22392 | 1120 | F1120K | D | 1e-05 | 7.5e-06 | 9.3e-06 | 8.933E-06 | 1.290E-06 | 7.500E-06 | 1.000E-05 |
| 22393 | 1120 | F1120M | D | 1.7e-05 | 9.9e-06 | 1.5e-05 | 1.397E-05 | 3.661E-06 | 9.900E-06 | 1.700E-05 |
| 22394 | 1120 | F1120F | D | 1.5e-05 | 1e-05 | 1.4e-05 | 1.300E-05 | 2.646E-06 | 1.000E-05 | 1.500E-05 |
| 22395 | 1120 | F1120P | D | 1.3e-05 | 1.2e-05 | 1.6e-05 | 1.367E-05 | 2.082E-06 | 1.200E-05 | 1.600E-05 |
| 22396 | 1120 | F1120S | D | 8e-06 | 9.8e-06 | 6.2e-06 | 8.000E-06 | 1.800E-06 | 6.200E-06 | 9.800E-06 |
| 22397 | 1120 | F1120T | D | 8.5e-06 | 1.6e-05 | 1.1e-05 | 1.183E-05 | 3.819E-06 | 8.500E-06 | 1.600E-05 |
| 22398 | 1120 | F1120W | D | 9.1e-06 | 1.3e-05 | 1e-05 | 1.070E-05 | 2.042E-06 | 9.100E-06 | 1.300E-05 |
| 22399 | 1120 | F1120Y | D | 1.1e-05 | 1.5e-05 | 9.1e-06 | 1.170E-05 | 3.012E-06 | 9.100E-06 | 1.500E-05 |
| 22400 | 1120 | F1120V | D | 1.2e-05 | 1.4e-05 | 1.3e-05 | 1.300E-05 | 1.000E-06 | 1.200E-05 | 1.400E-05 |
| 22401 | 1121 | S1121A | D | 9e-06 | 1.7e-05 | 1.8e-05 | 1.467E-05 | 4.933E-06 | 9.000E-06 | 1.800E-05 |
| 22402 | 1121 | S1121R | D | 1.5e-05 | 1.5e-05 | 1.7e-05 | 1.567E-05 | 1.155E-06 | 1.500E-05 | 1.700E-05 |
| 22403 | 1121 | S1121N | D | 1.1e-05 | 9.7e-06 | 1.3e-05 | 1.123E-05 | 1.662E-06 | 9.700E-06 | 1.300E-05 |
| 22404 | 1121 | S1121D | D | 9.3e-06 | 1.3e-05 | 1.2e-05 | 1.143E-05 | 1.914E-06 | 9.300E-06 | 1.300E-05 |
| 22405 | 1121 | S1121C | D | 1.5e-05 | 1.3e-05 | 1e-05 | 1.267E-05 | 2.517E-06 | 1.000E-05 | 1.500E-05 |
| 22406 | 1121 | S1121E | D | 1.2e-05 | 1.9e-05 | 1.1e-05 | 1.400E-05 | 4.359E-06 | 1.100E-05 | 1.900E-05 |
| 22407 | 1121 | S1121Q | D | 1e-05 | 1.3e-05 | 1e-05 | 1.100E-05 | 1.732E-06 | 1.000E-05 | 1.300E-05 |
| 22408 | 1121 | S1121G | D | 9.9e-06 | 1.7e-05 | 8.8e-06 | 1.190E-05 | 4.451E-06 | 8.800E-06 | 1.700E-05 |
| 22409 | 1121 | S1121H | D | 1.1e-05 | 1.6e-05 | 9.4e-06 | 1.213E-05 | 3.443E-06 | 9.400E-06 | 1.600E-05 |
| 22410 | 1121 | S1121I | D | 1.1e-05 | 1.6e-05 | 1.7e-05 | 1.467E-05 | 3.215E-06 | 1.100E-05 | 1.700E-05 |
| 22411 | 1121 | S1121L | D | 1.2e-05 | 1.5e-05 | 1.1e-05 | 1.267E-05 | 2.082E-06 | 1.100E-05 | 1.500E-05 |
| 22412 | 1121 | S1121K | D | 1.3e-05 | 1.4e-05 | 1.5e-05 | 1.400E-05 | 1.000E-06 | 1.300E-05 | 1.500E-05 |
| 22413 | 1121 | S1121M | D | 1.7e-05 | 1.3e-05 | 1.5e-05 | 1.500E-05 | 2.000E-06 | 1.300E-05 | 1.700E-05 |
| 22414 | 1121 | S1121F | D | 1.6e-05 | 1.2e-05 | 1.8e-05 | 1.533E-05 | 3.055E-06 | 1.200E-05 | 1.800E-05 |
| 22415 | 1121 | S1121P | D | 1.5e-05 | 1.1e-05 | 1.1e-05 | 1.233E-05 | 2.309E-06 | 1.100E-05 | 1.500E-05 |
| 22416 | 1121 | S1121S | D | 1.8e-05 | 1.1e-05 | 1.4e-05 | 1.433E-05 | 3.512E-06 | 1.100E-05 | 1.800E-05 |
| 22417 | 1121 | S1121T | D | 1.3e-05 | 9.8e-06 | 1.7e-05 | 1.327E-05 | 3.607E-06 | 9.800E-06 | 1.700E-05 |
| 22418 | 1121 | S1121W | D | 1.1e-05 | 9.1e-06 | 1.5e-05 | 1.170E-05 | 3.012E-06 | 9.100E-06 | 1.500E-05 |
| 22419 | 1121 | S1121Y | D | 1.5e-05 | 1.4e-05 | 1.1e-05 | 1.333E-05 | 2.082E-06 | 1.100E-05 | 1.500E-05 |
| 22420 | 1121 | S1121V | D | 1e-05 | 1.1e-05 | 9.1e-06 | 1.003E-05 | 9.504E-07 | 9.100E-06 | 1.100E-05 |
| 22421 | 1122 | G1122A | D | 1.7e-05 | 6.9e-06 | 1.1e-05 | 1.163E-05 | 5.080E-06 | 6.900E-06 | 1.700E-05 |
| 22422 | 1122 | G1122R | D | 1.3e-05 | 1.1e-05 | 1.5e-05 | 1.300E-05 | 2.000E-06 | 1.100E-05 | 1.500E-05 |
| 22423 | 1122 | G1122N | D | 1.3e-05 | 9.4e-06 | 1.1e-05 | 1.113E-05 | 1.804E-06 | 9.400E-06 | 1.300E-05 |
| 22424 | 1122 | G1122D | D | 1.3e-05 | 1.1e-05 | 1.5e-05 | 1.300E-05 | 2.000E-06 | 1.100E-05 | 1.500E-05 |
| 22425 | 1122 | G1122C | D | 1.4e-05 | 1.5e-05 | 1.4e-05 | 1.433E-05 | 5.774E-07 | 1.400E-05 | 1.500E-05 |
| 22426 | 1122 | G1122E | D | 1.5e-05 | 1.5e-05 | 1.4e-05 | 1.467E-05 | 5.774E-07 | 1.400E-05 | 1.500E-05 |
| 22427 | 1122 | G1122Q | D | 9.6e-06 | 9.7e-06 | 1.5e-05 | 1.143E-05 | 3.089E-06 | 9.600E-06 | 1.500E-05 |
| 22428 | 1122 | G1122G | D | 7.3e-06 | 9.9e-06 | 1.1e-05 | 9.400E-06 | 1.900E-06 | 7.300E-06 | 1.100E-05 |
| 22429 | 1122 | G1122H | D | 8.6e-06 | 1.6e-05 | 1e-05 | 1.153E-05 | 3.931E-06 | 8.600E-06 | 1.600E-05 |
| 22430 | 1122 | G1122I | D | 9.5e-06 | 1e-05 | 1.1e-05 | 1.017E-05 | 7.638E-07 | 9.500E-06 | 1.100E-05 |
| 22431 | 1122 | G1122L | D | 1.6e-05 | 1.5e-05 | 1e-05 | 1.367E-05 | 3.215E-06 | 1.000E-05 | 1.600E-05 |
| 22432 | 1122 | G1122K | D | 1.4e-05 | 1.3e-05 | 1.7e-05 | 1.467E-05 | 2.082E-06 | 1.300E-05 | 1.700E-05 |
| 22433 | 1122 | G1122M | D | 1.5e-05 | 1.7e-05 | 1.1e-05 | 1.433E-05 | 3.055E-06 | 1.100E-05 | 1.700E-05 |
| 22434 | 1122 | G1122F | D | 1.1e-05 | 9e-06 | 1.1e-05 | 1.033E-05 | 1.155E-06 | 9.000E-06 | 1.100E-05 |
| 22435 | 1122 | G1122P | D | 1.1e-05 | 1.5e-05 | 1.6e-05 | 1.400E-05 | 2.646E-06 | 1.100E-05 | 1.600E-05 |
| 22436 | 1122 | G1122S | D | 9e-06 | 1.4e-05 | 1.5e-05 | 1.267E-05 | 3.215E-06 | 9.000E-06 | 1.500E-05 |
| 22437 | 1122 | G1122T | D | 1.4e-05 | 1e-05 | 1.2e-05 | 1.200E-05 | 2.000E-06 | 1.000E-05 | 1.400E-05 |
| 22438 | 1122 | G1122W | D | 1.4e-05 | 1.7e-05 | 1.2e-05 | 1.433E-05 | 2.517E-06 | 1.200E-05 | 1.700E-05 |
| 22439 | 1122 | G1122Y | D | 1.5e-05 | 1.9e-05 | 9.3e-06 | 1.443E-05 | 4.875E-06 | 9.300E-06 | 1.900E-05 |

| | | | | | | | | | | |
|---|---|---|---|---|---|---|---|---|---|---|
| 22440 | 1122 | G1122V | D | 8.7e-06 | 1.8e-05 | 1.7e-05 | 1.457E-05 | 5.105E-06 | 8.700E-06 | 1.800E-05 |
| 22441 | 1123 | S1123A | D | 1.4e-05 | 9.7e-06 | 1.5e-05 | 1.290E-05 | 2.816E-06 | 9.700E-06 | 1.500E-05 |
| 22442 | 1123 | S1123R | D | 6.9e-06 | 1e-05 | 1.1e-05 | 9.300E-06 | 2.138E-06 | 6.900E-06 | 1.100E-05 |
| 22443 | 1123 | S1123N | D | 1e-05 | 8.1e-06 | 1.4e-05 | 1.070E-05 | 3.012E-06 | 8.100E-06 | 1.400E-05 |
| 22444 | 1123 | S1123D | D | 1.2e-05 | 1.2e-05 | 9.5e-06 | 1.117E-05 | 1.443E-06 | 9.500E-06 | 1.200E-05 |
| 22445 | 1123 | S1123C | D | 1e-05 | 1e-05 | 8.3e-06 | 9.433E-06 | 9.815E-07 | 8.300E-06 | 1.000E-05 |
| 22446 | 1123 | S1123E | D | 1.2e-05 | 1.1e-05 | 2.7e-05 | 1.667E-05 | 8.963E-06 | 1.100E-05 | 2.700E-05 |
| 22447 | 1123 | S1123Q | D | 8.2e-06 | 1.2e-05 | 9.1e-06 | 9.767E-06 | 1.986E-06 | 8.200E-06 | 1.200E-05 |
| 22448 | 1123 | S1123G | D | 1.4e-05 | 1.6e-05 | 6.6e-06 | 1.220E-05 | 4.952E-06 | 6.600E-06 | 1.600E-05 |
| 22449 | 1123 | S1123H | D | 1.5e-05 | 1.4e-05 | 1.4e-05 | 1.433E-05 | 5.774E-07 | 1.400E-05 | 1.500E-05 |
| 22450 | 1123 | S1123I | D | 1.5e-05 | 9.7e-06 | 1.2e-05 | 1.223E-05 | 2.658E-06 | 9.700E-06 | 1.500E-05 |
| 22451 | 1123 | S1123L | D | 1.1e-05 | 1.5e-05 | 1.1e-05 | 1.233E-05 | 2.309E-06 | 1.100E-05 | 1.500E-05 |
| 22452 | 1123 | S1123K | D | 1.1e-05 | 1e-05 | 1.7e-05 | 1.267E-05 | 3.786E-06 | 1.000E-05 | 1.700E-05 |
| 22453 | 1123 | S1123M | D | 1.4e-05 | 1.4e-05 | 1.7e-05 | 1.500E-05 | 1.732E-06 | 1.400E-05 | 1.700E-05 |
| 22454 | 1123 | S1123F | D | 1.6e-05 | 9.2e-06 | 1.5e-05 | 1.340E-05 | 3.672E-06 | 9.200E-06 | 1.600E-05 |
| 22455 | 1123 | S1123P | D | 1.2e-05 | 9.7e-06 | 1.5e-05 | 1.223E-05 | 2.658E-06 | 9.700E-06 | 1.500E-05 |
| 22456 | 1123 | S1123S | D | 1.7e-05 | 1.1e-05 | 1.5e-05 | 1.433E-05 | 3.055E-06 | 1.100E-05 | 1.700E-05 |
| 22457 | 1123 | S1123T | D | 8.3e-06 | 1.5e-05 | 7.8e-06 | 1.037E-05 | 4.020E-06 | 7.800E-06 | 1.500E-05 |
| 22458 | 1123 | S1123W | D | 8.5e-06 | 1.3e-05 | 9.6e-06 | 1.037E-05 | 2.346E-06 | 8.500E-06 | 1.300E-05 |
| 22459 | 1123 | S1123Y | D | 9.1e-06 | 1.5e-05 | 1.6e-05 | 1.337E-05 | 3.729E-06 | 9.100E-06 | 1.600E-05 |
| 22460 | 1123 | S1123V | D | 1.4e-05 | 1.4e-05 | 1.3e-05 | 1.367E-05 | 5.774E-07 | 1.300E-05 | 1.400E-05 |
| 22461 | 1124 | R1124A | D | 1.8e-05 | 1.3e-05 | 1.4e-05 | 1.500E-05 | 2.646E-06 | 1.300E-05 | 1.800E-05 |
| 22462 | 1124 | R1124R | D | 1.7e-05 | 1.1e-05 | 9.2e-06 | 1.240E-05 | 4.084E-06 | 9.200E-06 | 1.700E-05 |
| 22463 | 1124 | R1124N | D | 1.6e-05 | 1.5e-05 | 1.2e-05 | 1.433E-05 | 2.082E-06 | 1.200E-05 | 1.600E-05 |
| 22464 | 1124 | R1124D | D | 1e-05 | 1.6e-05 | 1.4e-05 | 1.333E-05 | 3.055E-06 | 1.000E-05 | 1.600E-05 |
| 22465 | 1124 | R1124C | D | 1.1e-05 | 1.4e-05 | 1.2e-05 | 1.233E-05 | 1.528E-06 | 1.100E-05 | 1.400E-05 |
| 22466 | 1124 | R1124E | D | 1.6e-05 | 1.1e-05 | 9.9e-06 | 1.230E-05 | 3.251E-06 | 9.900E-06 | 1.600E-05 |
| 22467 | 1124 | R1124Q | D | 1.5e-05 | 1.5e-05 | 1.1e-05 | 1.367E-05 | 2.309E-06 | 1.100E-05 | 1.500E-05 |
| 22468 | 1124 | R1124G | D | 1.5e-05 | 1.5e-05 | 1.5e-05 | 1.500E-05 | 0.000E+00 | 1.500E-05 | 1.500E-05 |
| 22469 | 1124 | R1124H | D | 1.2e-05 | 1.1e-05 | 8.7e-06 | 1.057E-05 | 1.692E-06 | 8.700E-06 | 1.200E-05 |
| 22470 | 1124 | R1124I | D | 1.1e-05 | 1.6e-05 | 1.4e-05 | 1.367E-05 | 2.517E-06 | 1.100E-05 | 1.600E-05 |
| 22471 | 1124 | R1124L | D | 9.9e-06 | 1.7e-05 | 1.6e-05 | 1.430E-05 | 3.843E-06 | 9.900E-06 | 1.700E-05 |
| 22472 | 1124 | R1124K | D | 1.6e-05 | 1.1e-05 | 1.6e-05 | 1.433E-05 | 2.887E-06 | 1.100E-05 | 1.600E-05 |
| 22473 | 1124 | R1124M | D | 9.3e-06 | 9.8e-06 | 1.4e-05 | 1.103E-05 | 2.581E-06 | 9.300E-06 | 1.400E-05 |
| 22474 | 1124 | R1124F | D | 1e-05 | 1.6e-05 | 1.4e-05 | 1.333E-05 | 3.055E-06 | 1.000E-05 | 1.600E-05 |
| 22475 | 1124 | R1124P | D | 1.6e-05 | 1e-05 | 1.1e-05 | 1.233E-05 | 3.215E-06 | 1.000E-05 | 1.600E-05 |
| 22476 | 1124 | R1124S | D | 1.5e-05 | 1.4e-05 | 9.6e-06 | 1.287E-05 | 2.873E-06 | 9.600E-06 | 1.500E-05 |
| 22477 | 1124 | R1124T | D | 1.1e-05 | 1e-05 | 1.6e-05 | 1.233E-05 | 3.215E-06 | 1.000E-05 | 1.600E-05 |
| 22478 | 1124 | R1124W | D | 1.6e-05 | 1.1e-05 | 1.4e-05 | 1.367E-05 | 2.517E-06 | 1.100E-05 | 1.600E-05 |
| 22479 | 1124 | R1124Y | D | 1.1e-05 | 1.5e-05 | 9.5e-06 | 1.183E-05 | 2.843E-06 | 9.500E-06 | 1.500E-05 |
| 22480 | 1124 | R1124V | D | 1.1e-05 | 1.1e-05 | 1.1e-05 | 1.100E-05 | 2.075E-21 | 1.100E-05 | 1.100E-05 |
| 22481 | 1125 | S1125A | D | 9.5e-06 | 1.5e-05 | 1.5e-05 | 1.317E-05 | 3.175E-06 | 9.500E-06 | 1.500E-05 |
| 22482 | 1125 | S1125R | D | 1.6e-05 | 2.4e-05 | 1e-05 | 1.667E-05 | 7.024E-06 | 1.000E-05 | 2.400E-05 |
| 22483 | 1125 | S1125N | D | 1.5e-05 | 1.4e-05 | 1.4e-05 | 1.433E-05 | 5.774E-07 | 1.400E-05 | 1.500E-05 |
| 22484 | 1125 | S1125D | D | 1.1e-05 | 1.4e-05 | 9e-06 | 1.133E-05 | 2.517E-06 | 9.000E-06 | 1.400E-05 |
| 22485 | 1125 | S1125C | D | 1.6e-05 | 1e-05 | 1.5e-05 | 1.367E-05 | 3.215E-06 | 1.000E-05 | 1.600E-05 |
| 22486 | 1125 | S1125E | D | 1.1e-05 | 1.5e-05 | 1e-05 | 1.200E-05 | 2.646E-06 | 1.000E-05 | 1.500E-05 |
| 22487 | 1125 | S1125Q | D | 1.5e-05 | 5.9e-06 | 1.1e-05 | 1.063E-05 | 4.561E-06 | 5.900E-06 | 1.500E-05 |
| 22488 | 1125 | S1125G | D | 1.2e-05 | 1.1e-05 | 1.7e-05 | 1.333E-05 | 3.215E-06 | 1.100E-05 | 1.700E-05 |
| 22489 | 1125 | S1125H | D | 9.8e-06 | 9.6e-06 | 9e-06 | 9.467E-06 | 4.163E-07 | 9.000E-06 | 9.800E-06 |
| 22490 | 1125 | S1125I | D | 1.3e-05 | 1e-05 | 9.5e-06 | 1.083E-05 | 1.893E-06 | 9.500E-06 | 1.300E-05 |
| 22491 | 1125 | S1125L | D | 1.6e-05 | 1.4e-05 | 1.2e-05 | 1.400E-05 | 2.000E-06 | 1.200E-05 | 1.600E-05 |
| 22492 | 1125 | S1125K | D | 1.4e-05 | 7.1e-06 | 1.1e-05 | 1.070E-05 | 3.460E-06 | 7.100E-06 | 1.400E-05 |
| 22493 | 1125 | S1125M | D | 1.4e-05 | 1.1e-05 | 1.1e-05 | 1.200E-05 | 1.732E-06 | 1.100E-05 | 1.400E-05 |
| 22494 | 1125 | S1125F | D | 1e-05 | 1.1e-05 | 1.1e-05 | 1.067E-05 | 5.774E-07 | 1.000E-05 | 1.100E-05 |
| 22495 | 1125 | S1125P | D | 7.9e-06 | 1.5e-05 | 1.5e-05 | 1.263E-05 | 4.099E-06 | 7.900E-06 | 1.500E-05 |
| 22496 | 1125 | S1125S | D | 1.5e-05 | 1.6e-05 | 1.3e-05 | 1.467E-05 | 1.528E-06 | 1.300E-05 | 1.600E-05 |
| 22497 | 1125 | S1125T | D | 1e-05 | 1.5e-05 | 1.1e-05 | 1.200E-05 | 2.646E-06 | 1.000E-05 | 1.500E-05 |
| 22498 | 1125 | S1125W | D | 8.3e-06 | 1.5e-05 | 1.5e-05 | 1.277E-05 | 3.868E-06 | 8.300E-06 | 1.500E-05 |
| 22499 | 1125 | S1125Y | D | 1.2e-05 | 1.4e-05 | 1.1e-05 | 1.233E-05 | 1.528E-06 | 1.100E-05 | 1.400E-05 |
| 22500 | 1125 | S1125V | D | 1.3e-05 | 9.7e-06 | 1.6e-05 | 1.290E-05 | 3.151E-06 | 9.700E-06 | 1.600E-05 |
| 22501 | 1126 | G1126A | D | 9.8e-06 | 1.1e-05 | 1.5e-05 | 1.193E-05 | 2.723E-06 | 9.800E-06 | 1.500E-05 |
| 22502 | 1126 | G1126R | D | 9.2e-06 | 1.3e-05 | 8.3e-06 | 1.017E-05 | 2.495E-06 | 8.300E-06 | 1.300E-05 |
| 22503 | 1126 | G1126N | D | 1.6e-05 | 9.3e-06 | 1.6e-05 | 1.377E-05 | 3.868E-06 | 9.300E-06 | 1.600E-05 |
| 22504 | 1126 | G1126D | D | 1.4e-05 | 1.5e-05 | 1.6e-05 | 1.500E-05 | 1.000E-06 | 1.400E-05 | 1.600E-05 |
| 22505 | 1126 | G1126C | D | 1.6e-05 | 1.5e-05 | 1e-05 | 1.367E-05 | 3.215E-06 | 1.000E-05 | 1.600E-05 |
| 22506 | 1126 | G1126E | D | 1e-05 | 9e-06 | 1.3e-05 | 1.067E-05 | 2.082E-06 | 9.000E-06 | 1.300E-05 |
| 22507 | 1126 | G1126Q | D | 1.4e-05 | 1e-05 | 1.5e-05 | 1.300E-05 | 2.646E-06 | 1.000E-05 | 1.500E-05 |
| 22508 | 1126 | G1126G | D | 1.3e-05 | 1.6e-05 | 1.5e-05 | 1.467E-05 | 1.528E-06 | 1.300E-05 | 1.600E-05 |
| 22509 | 1126 | G1126H | D | 8.6e-06 | 1.6e-05 | 1e-05 | 1.153E-05 | 3.931E-06 | 8.600E-06 | 1.600E-05 |
| 22510 | 1126 | G1126I | D | 1.5e-05 | 1.5e-05 | 1.2e-05 | 1.400E-05 | 1.732E-06 | 1.200E-05 | 1.500E-05 |
| 22511 | 1126 | G1126L | D | 1.4e-05 | 1.6e-05 | 1.7e-05 | 1.567E-05 | 1.528E-06 | 1.400E-05 | 1.700E-05 |
| 22512 | 1126 | G1126K | D | 1e-05 | 1.1e-05 | 1e-05 | 1.033E-05 | 5.774E-07 | 1.000E-05 | 1.100E-05 |
| 22513 | 1126 | G1126M | D | 1.6e-05 | 1.4e-05 | 1.6e-05 | 1.533E-05 | 1.155E-06 | 1.400E-05 | 1.600E-05 |
| 22514 | 1126 | G1126F | D | 1.4e-05 | 1.4e-05 | 9.7e-06 | 1.257E-05 | 2.483E-06 | 9.700E-06 | 1.400E-05 |
| 22515 | 1126 | G1126P | D | 9.9e-06 | 1.6e-05 | 8.8e-06 | 1.157E-05 | 3.879E-06 | 8.800E-06 | 1.600E-05 |
| 22516 | 1126 | G1126S | D | 1.4e-05 | 1e-05 | 1.5e-05 | 1.300E-05 | 2.646E-06 | 1.000E-05 | 1.500E-05 |
| 22517 | 1126 | G1126T | D | 1.5e-05 | 1.2e-05 | 1.5e-05 | 1.400E-05 | 1.732E-06 | 1.200E-05 | 1.500E-05 |
| 22518 | 1126 | G1126W | D | 1e-05 | 9.5e-06 | 1.4e-05 | 1.117E-05 | 2.466E-06 | 9.500E-06 | 1.400E-05 |
| 22519 | 1126 | G1126Y | D | 1.6e-05 | 1.5e-05 | 1.5e-05 | 1.533E-05 | 5.774E-07 | 1.500E-05 | 1.600E-05 |
| 22520 | 1126 | G1126V | D | 1.6e-05 | 1.2e-05 | 1.4e-05 | 1.400E-05 | 2.000E-06 | 1.200E-05 | 1.600E-05 |
| 22521 | 1127 | T1127A | D | 8.7e-06 | 1.4e-05 | 1.8e-05 | 1.357E-05 | 4.665E-06 | 8.700E-06 | 1.800E-05 |
| 22522 | 1127 | T1127R | D | 1e-05 | 1.1e-05 | 1.3e-05 | 1.133E-05 | 1.528E-06 | 1.000E-05 | 1.300E-05 |
| 22523 | 1127 | T1127N | D | 1.1e-05 | 1.1e-05 | 1.5e-05 | 1.233E-05 | 2.309E-06 | 1.100E-05 | 1.500E-05 |
| 22524 | 1127 | T1127D | D | 9.9e-06 | 1.7e-05 | 8.5e-06 | 1.180E-05 | 4.557E-06 | 8.500E-06 | 1.700E-05 |

| | | | | | | | | | | |
|---|---|---|---|---|---|---|---|---|---|---|
| 22525 | 1127 | T1127C | D | 1.6e-05 | 1.6e-05 | 1e-05 | 1.400E-05 | 3.464E-06 | 1.000E-05 | 1.600E-05 |
| 22526 | 1127 | T1127E | D | 1e-05 | 1.7e-05 | 1.6e-05 | 1.433E-05 | 3.786E-06 | 1.000E-05 | 1.700E-05 |
| 22527 | 1127 | T1127Q | D | 1.4e-05 | 1e-05 | 9.8e-06 | 1.127E-05 | 2.369E-06 | 9.800E-06 | 1.400E-05 |
| 22528 | 1127 | T1127G | D | 1.5e-05 | 1.7e-05 | 1.6e-05 | 1.600E-05 | 1.000E-06 | 1.500E-05 | 1.700E-05 |
| 22529 | 1127 | T1127H | D | 9.1e-06 | 1e-05 | 1e-05 | 9.700E-06 | 5.196E-07 | 9.100E-06 | 1.000E-05 |
| 22530 | 1127 | T1127I | D | 1.4e-05 | 1.1e-05 | 1.1e-05 | 1.200E-05 | 1.732E-06 | 1.100E-05 | 1.400E-05 |
| 22531 | 1127 | T1127L | D | 9.3e-06 | 1.4e-05 | 1e-05 | 1.110E-05 | 2.536E-06 | 9.300E-06 | 1.400E-05 |
| 22532 | 1127 | T1127K | D | 1e-05 | 1.6e-05 | 1.3e-05 | 1.300E-05 | 3.000E-06 | 1.000E-05 | 1.600E-05 |
| 22533 | 1127 | T1127M | D | 1.1e-05 | 1.7e-05 | 1.1e-05 | 1.300E-05 | 3.464E-06 | 1.100E-05 | 1.700E-05 |
| 22534 | 1127 | T1127F | D | 1.1e-05 | 1.5e-05 | 1.7e-05 | 1.433E-05 | 3.055E-06 | 1.100E-05 | 1.700E-05 |
| 22535 | 1127 | T1127P | D | 1.2e-05 | 1.8e-05 | 1.4e-05 | 1.467E-05 | 3.055E-06 | 1.200E-05 | 1.800E-05 |
| 22536 | 1127 | T1127S | D | 1.1e-05 | 1.5e-05 | 1.6e-05 | 1.400E-05 | 2.646E-06 | 1.100E-05 | 1.600E-05 |
| 22537 | 1127 | T1127T | D | 8.3e-06 | 1e-05 | 1.4e-05 | 1.077E-05 | 2.926E-06 | 8.300E-06 | 1.400E-05 |
| 22538 | 1127 | T1127W | D | 1.2e-05 | 1.3e-05 | 1.5e-05 | 1.333E-05 | 1.528E-06 | 1.200E-05 | 1.500E-05 |
| 22539 | 1127 | T1127Y | D | 1.1e-05 | 1.2e-05 | 1.1e-05 | 1.133E-05 | 5.774E-07 | 1.100E-05 | 1.200E-05 |
| 22540 | 1127 | T1127V | D | 1e-05 | 1.6e-05 | 1.6e-05 | 1.400E-05 | 3.464E-06 | 1.000E-05 | 1.600E-05 |
| 22541 | 1128 | D1128A | D | 1.5e-05 | 8.9e-06 | 1.4e-05 | 1.263E-05 | 3.272E-06 | 8.900E-06 | 1.500E-05 |
| 22542 | 1128 | D1128R | D | 1.2e-05 | 9.5e-06 | 1.4e-05 | 1.183E-05 | 2.255E-06 | 9.500E-06 | 1.400E-05 |
| 22543 | 1128 | D1128N | D | 1.1e-05 | 1.3e-05 | 1.2e-05 | 1.200E-05 | 1.000E-06 | 1.100E-05 | 1.300E-05 |
| 22544 | 1128 | D1128D | D | 1e-05 | 1.3e-05 | 1.4e-05 | 1.233E-05 | 2.082E-06 | 1.000E-05 | 1.400E-05 |
| 22545 | 1128 | D1128C | D | 1.1e-05 | 1.4e-05 | 1.6e-05 | 1.367E-05 | 2.517E-06 | 1.100E-05 | 1.600E-05 |
| 22546 | 1128 | D1128E | D | 1.4e-05 | 1.5e-05 | 1.4e-05 | 1.433E-05 | 5.774E-07 | 1.400E-05 | 1.500E-05 |
| 22547 | 1128 | D1128Q | D | 1e-05 | 1.7e-05 | 8.3e-06 | 1.177E-05 | 4.611E-06 | 8.300E-06 | 1.700E-05 |
| 22548 | 1128 | D1128G | D | 9.1e-06 | 1.2e-05 | 1.5e-05 | 1.203E-05 | 2.950E-06 | 9.100E-06 | 1.500E-05 |
| 22549 | 1128 | D1128H | D | 1.7e-05 | 1.4e-05 | 9.7e-06 | 1.357E-05 | 3.669E-06 | 9.700E-06 | 1.700E-05 |
| 22550 | 1128 | D1128I | D | 1.4e-05 | 1e-05 | 1.7e-05 | 1.367E-05 | 3.512E-06 | 1.000E-05 | 1.700E-05 |
| 22551 | 1128 | D1128L | D | 1.6e-05 | 1.6e-05 | 9.3e-06 | 1.377E-05 | 3.868E-06 | 9.300E-06 | 1.600E-05 |
| 22552 | 1128 | D1128K | D | 1.8e-05 | 1.6e-05 | 1.1e-05 | 1.500E-05 | 3.606E-06 | 1.100E-05 | 1.800E-05 |
| 22553 | 1128 | D1128M | D | 1.8e-05 | 1.7e-05 | 1.4e-05 | 1.633E-05 | 2.082E-06 | 1.400E-05 | 1.800E-05 |
| 22554 | 1128 | D1128F | D | 9.9e-06 | 1.1e-05 | 1e-05 | 1.030E-05 | 6.083E-07 | 9.900E-06 | 1.100E-05 |
| 22555 | 1128 | D1128P | D | 1.4e-05 | 1e-05 | 1.1e-05 | 1.167E-05 | 2.082E-06 | 1.000E-05 | 1.400E-05 |
| 22556 | 1128 | D1128S | D | 1.6e-05 | 1.3e-05 | 1.5e-05 | 1.467E-05 | 1.528E-06 | 1.300E-05 | 1.600E-05 |
| 22557 | 1128 | D1128T | D | 1.1e-05 | 1.5e-05 | 1.6e-05 | 1.400E-05 | 2.646E-06 | 1.100E-05 | 1.600E-05 |
| 22558 | 1128 | D1128W | D | 1.1e-05 | 1.3e-05 | 1.5e-05 | 1.300E-05 | 2.000E-06 | 1.100E-05 | 1.500E-05 |
| 22559 | 1128 | D1128Y | D | 1.7e-05 | 1.4e-05 | 1.4e-05 | 1.500E-05 | 1.732E-06 | 1.400E-05 | 1.700E-05 |
| 22560 | 1128 | D1128V | D | 1.4e-05 | 1.2e-05 | 9.7e-06 | 1.190E-05 | 2.152E-06 | 9.700E-06 | 1.400E-05 |
| 22561 | 1129 | F1129A | D | 1.2e-05 | 8.3e-06 | 1.1e-05 | 1.043E-05 | 1.914E-06 | 8.300E-06 | 1.200E-05 |
| 22562 | 1129 | F1129R | D | 1.3e-05 | 1.4e-05 | 1.5e-05 | 1.400E-05 | 1.000E-06 | 1.300E-05 | 1.500E-05 |
| 22563 | 1129 | F1129N | D | 1.5e-05 | 1.1e-05 | 9.5e-06 | 1.183E-05 | 2.843E-06 | 9.500E-06 | 1.500E-05 |
| 22564 | 1129 | F1129D | D | 1.5e-05 | 2.6e-05 | 1.7e-05 | 1.933E-05 | 5.859E-06 | 1.500E-05 | 2.600E-05 |
| 22565 | 1129 | F1129C | D | 1.4e-05 | 1.4e-05 | 1.5e-05 | 1.433E-05 | 5.774E-07 | 1.400E-05 | 1.500E-05 |
| 22566 | 1129 | F1129E | D | 1.5e-05 | 1.5e-05 | 8.1e-06 | 1.270E-05 | 3.984E-06 | 8.100E-06 | 1.500E-05 |
| 22567 | 1129 | F1129Q | D | 8.2e-06 | 9.5e-06 | 1e-05 | 9.233E-06 | 9.292E-07 | 8.200E-06 | 1.000E-05 |
| 22568 | 1129 | F1129G | D | 1.7e-05 | 1.1e-05 | 1.1e-05 | 1.300E-05 | 3.464E-06 | 1.100E-05 | 1.700E-05 |
| 22569 | 1129 | F1129H | D | 8.6e-06 | 1.5e-05 | 9.3e-06 | 1.097E-05 | 3.510E-06 | 8.600E-06 | 1.500E-05 |
| 22570 | 1129 | F1129I | D | 1.6e-05 | 1.3e-05 | 1.1e-05 | 1.333E-05 | 2.517E-06 | 1.100E-05 | 1.600E-05 |
| 22571 | 1129 | F1129L | D | 5.8e-06 | 1.5e-05 | 1.1e-05 | 1.060E-05 | 4.613E-06 | 5.800E-06 | 1.500E-05 |
| 22572 | 1129 | F1129K | D | 1.3e-05 | 1.2e-05 | 1.4e-05 | 1.300E-05 | 1.000E-06 | 1.200E-05 | 1.400E-05 |
| 22573 | 1129 | F1129M | D | 1.4e-05 | 1.5e-05 | 8.6e-06 | 1.253E-05 | 3.443E-06 | 8.600E-06 | 1.500E-05 |
| 22574 | 1129 | F1129F | D | 1.6e-05 | 8.4e-06 | 1.4e-05 | 1.280E-05 | 3.940E-06 | 8.400E-06 | 1.600E-05 |
| 22575 | 1129 | F1129P | D | 1.1e-05 | 9.9e-06 | 1.1e-05 | 1.063E-05 | 6.351E-07 | 9.900E-06 | 1.100E-05 |
| 22576 | 1129 | F1129S | D | 1.5e-05 | 1e-05 | 1.5e-05 | 1.333E-05 | 2.887E-06 | 1.000E-05 | 1.500E-05 |
| 22577 | 1129 | F1129T | D | 1.4e-05 | 1.7e-05 | 1.5e-05 | 1.533E-05 | 1.528E-06 | 1.400E-05 | 1.700E-05 |
| 22578 | 1129 | F1129W | D | 1.5e-05 | 1.5e-05 | 1.5e-05 | 1.500E-05 | 0.000E+00 | 1.500E-05 | 1.500E-05 |
| 22579 | 1129 | F1129Y | D | 1.4e-05 | 9.1e-06 | 2.3e-05 | 1.537E-05 | 7.050E-06 | 9.100E-06 | 2.300E-05 |
| 22580 | 1129 | F1129V | D | 1.5e-05 | 1.3e-05 | 1.7e-05 | 1.500E-05 | 2.000E-06 | 1.300E-05 | 1.700E-05 |
| 22581 | 1130 | T1130A | D | 1.1e-05 | 1e-05 | 1.3e-05 | 1.133E-05 | 1.528E-06 | 1.000E-05 | 1.300E-05 |
| 22582 | 1130 | T1130R | D | 1.6e-05 | 1.1e-05 | 8.9e-06 | 1.197E-05 | 3.647E-06 | 8.900E-06 | 1.600E-05 |
| 22583 | 1130 | T1130N | D | 1.4e-05 | 1.2e-05 | 1.5e-05 | 1.367E-05 | 1.528E-06 | 1.200E-05 | 1.500E-05 |
| 22584 | 1130 | T1130D | D | 1.6e-05 | 1.1e-05 | 1e-05 | 1.233E-05 | 3.215E-06 | 1.000E-05 | 1.600E-05 |
| 22585 | 1130 | T1130C | D | 1e-05 | 1.2e-05 | 8.9e-06 | 1.030E-05 | 1.572E-06 | 8.900E-06 | 1.200E-05 |
| 22586 | 1130 | T1130E | D | 1.6e-05 | 1.3e-05 | 1.4e-05 | 1.433E-05 | 1.528E-06 | 1.300E-05 | 1.600E-05 |
| 22587 | 1130 | T1130Q | D | 1.1e-05 | 1.4e-05 | 2.3e-05 | 1.600E-05 | 6.245E-06 | 1.100E-05 | 2.300E-05 |
| 22588 | 1130 | T1130G | D | 1.4e-05 | 1.1e-05 | 1.4e-05 | 1.300E-05 | 1.732E-06 | 1.100E-05 | 1.400E-05 |
| 22589 | 1130 | T1130H | D | 1.5e-05 | 9.5e-06 | 1.1e-05 | 1.183E-05 | 2.843E-06 | 9.500E-06 | 1.500E-05 |
| 22590 | 1130 | T1130I | D | 1.7e-05 | 1.5e-05 | 1.1e-05 | 1.433E-05 | 3.055E-06 | 1.100E-05 | 1.700E-05 |
| 22591 | 1130 | T1130L | D | 8.7e-06 | 1.5e-05 | 1.5e-05 | 1.290E-05 | 3.637E-06 | 8.700E-06 | 1.500E-05 |
| 22592 | 1130 | T1130K | D | 1.8e-05 | 1.1e-05 | 1.5e-05 | 1.467E-05 | 3.512E-06 | 1.100E-05 | 1.800E-05 |
| 22593 | 1130 | T1130M | D | 1e-05 | 1.6e-05 | 1.3e-05 | 1.300E-05 | 3.000E-06 | 1.000E-05 | 1.600E-05 |
| 22594 | 1130 | T1130F | D | 1.6e-05 | 1.5e-05 | 1.6e-05 | 1.567E-05 | 5.774E-07 | 1.500E-05 | 1.600E-05 |
| 22595 | 1130 | T1130P | D | 1.4e-05 | 1.1e-05 | 9.9e-06 | 1.163E-05 | 2.122E-06 | 9.900E-06 | 1.400E-05 |
| 22596 | 1130 | T1130S | D | 1.5e-05 | 1.2e-05 | 1.5e-05 | 1.400E-05 | 1.732E-06 | 1.200E-05 | 1.500E-05 |
| 22597 | 1130 | T1130T | D | 1.5e-05 | 1e-05 | 1e-05 | 1.167E-05 | 2.887E-06 | 1.000E-05 | 1.500E-05 |
| 22598 | 1130 | T1130W | D | 1.1e-05 | 9.6e-06 | 1.3e-05 | 1.120E-05 | 1.709E-06 | 9.600E-06 | 1.300E-05 |
| 22599 | 1130 | T1130Y | D | 1.5e-05 | 9.5e-06 | 1.4e-05 | 1.283E-05 | 2.930E-06 | 9.500E-06 | 1.500E-05 |
| 22600 | 1130 | T1130V | D | 1.2e-05 | 1.7e-05 | 1.4e-05 | 1.433E-05 | 2.517E-06 | 1.200E-05 | 1.700E-05 |
| 22601 | 1131 | L1131A | D | 1.1e-05 | 1.5e-05 | 1e-05 | 1.200E-05 | 2.646E-06 | 1.000E-05 | 1.500E-05 |
| 22602 | 1131 | L1131R | D | 1.1e-05 | 1.7e-05 | 1.4e-05 | 1.400E-05 | 3.000E-06 | 1.100E-05 | 1.700E-05 |
| 22603 | 1131 | L1131N | D | 1.3e-05 | 1.6e-05 | 1.3e-05 | 1.400E-05 | 1.732E-06 | 1.300E-05 | 1.600E-05 |
| 22604 | 1131 | L1131D | D | 6.5e-06 | 1.5e-05 | 1.5e-05 | 1.217E-05 | 4.907E-06 | 6.500E-06 | 1.500E-05 |
| 22605 | 1131 | L1131C | D | 1.7e-05 | 1.5e-05 | 1.1e-05 | 1.433E-05 | 3.055E-06 | 1.100E-05 | 1.700E-05 |
| 22606 | 1131 | L1131E | D | 1.5e-05 | 1.2e-05 | 9e-06 | 1.200E-05 | 3.000E-06 | 9.000E-06 | 1.500E-05 |
| 22607 | 1131 | L1131Q | D | 1.4e-05 | 1.4e-05 | 1.4e-05 | 1.400E-05 | 0.000E+00 | 1.400E-05 | 1.400E-05 |
| 22608 | 1131 | L1131G | D | 1.5e-05 | 1.6e-05 | 1.1e-05 | 1.400E-05 | 2.646E-06 | 1.100E-05 | 1.600E-05 |
| 22609 | 1131 | L1131H | D | 1.7e-05 | 1e-05 | 9.9e-06 | 1.230E-05 | 4.071E-06 | 9.900E-06 | 1.700E-05 |

| | | | | | | | | | | |
|---|---|---|---|---|---|---|---|---|---|---|
| 22610 | 1131 | L1131I | D | 1.4e-05 | 9.7e-06 | 7.7e-06 | 1.047E-05 | 3.219E-06 | 7.700E-06 | 1.400E-05 |
| 22611 | 1131 | L1131L | D | 1e-05 | 1.3e-05 | 1e-05 | 1.100E-05 | 1.732E-06 | 1.000E-05 | 1.300E-05 |
| 22612 | 1131 | L1131K | D | 1.5e-05 | 1.5e-05 | 1.5e-05 | 1.500E-05 | 0.000E+00 | 1.500E-05 | 1.500E-05 |
| 22613 | 1131 | L1131M | D | 1.3e-05 | 1.6e-05 | 9.8e-06 | 1.293E-05 | 3.101E-06 | 9.800E-06 | 1.600E-05 |
| 22614 | 1131 | L1131F | D | 1.5e-05 | 1.6e-05 | 1.6e-05 | 1.567E-05 | 5.774E-07 | 1.500E-05 | 1.600E-05 |
| 22615 | 1131 | L1131P | D | 1.2e-05 | 1.3e-05 | 1.3e-05 | 1.267E-05 | 5.774E-07 | 1.200E-05 | 1.300E-05 |
| 22616 | 1131 | L1131S | D | 1.1e-05 | 9.6e-06 | 1.5e-05 | 1.187E-05 | 2.802E-06 | 9.600E-06 | 1.500E-05 |
| 22617 | 1131 | L1131T | D | 8.7e-06 | 9.9e-06 | 1.5e-05 | 1.120E-05 | 3.345E-06 | 8.700E-06 | 1.500E-05 |
| 22618 | 1131 | L1131W | D | 1.6e-05 | 1.6e-05 | 1.6e-05 | 1.600E-05 | 0.000E+00 | 1.600E-05 | 1.600E-05 |
| 22619 | 1131 | L1131Y | D | 1.2e-05 | 1.5e-05 | 1.1e-05 | 1.267E-05 | 2.082E-06 | 1.100E-05 | 1.500E-05 |
| 22620 | 1131 | L1131V | D | 1.4e-05 | 1.4e-05 | 9.5e-06 | 1.250E-05 | 2.598E-06 | 9.500E-06 | 1.400E-05 |
| 22621 | 1132 | T1132A | D | 1.2e-05 | 1.1e-05 | 1.1e-05 | 1.133E-05 | 5.774E-07 | 1.100E-05 | 1.200E-05 |
| 22622 | 1132 | T1132R | D | 1.6e-05 | 1.1e-05 | 1.4e-05 | 1.367E-05 | 2.517E-06 | 1.100E-05 | 1.600E-05 |
| 22623 | 1132 | T1132N | D | 1.1e-05 | 1.6e-05 | 2.1e-05 | 1.600E-05 | 5.000E-06 | 1.100E-05 | 2.100E-05 |
| 22624 | 1132 | T1132D | D | 9.7e-06 | 1.1e-05 | 1.4e-05 | 1.157E-05 | 2.205E-06 | 9.700E-06 | 1.400E-05 |
| 22625 | 1132 | T1132C | D | 1.5e-05 | 1.6e-05 | 1.5e-05 | 1.533E-05 | 5.774E-07 | 1.500E-05 | 1.600E-05 |
| 22626 | 1132 | T1132E | D | 9.3e-06 | 1.3e-05 | 1.6e-05 | 1.277E-05 | 3.356E-06 | 9.300E-06 | 1.600E-05 |
| 22627 | 1132 | T1132Q | D | 1.5e-05 | 1.3e-05 | 1.5e-05 | 1.433E-05 | 1.155E-06 | 1.300E-05 | 1.500E-05 |
| 22628 | 1132 | T1132G | D | 1.1e-05 | 1.7e-05 | 1e-05 | 1.267E-05 | 3.786E-06 | 1.000E-05 | 1.700E-05 |
| 22629 | 1132 | T1132H | D | 9.6e-06 | 9.4e-06 | 2.7e-05 | 1.533E-05 | 1.010E-05 | 9.400E-06 | 2.700E-05 |
| 22630 | 1132 | T1132I | D | 1.2e-05 | 6.2e-06 | 8.5e-06 | 8.900E-06 | 2.921E-06 | 6.200E-06 | 1.200E-05 |
| 22631 | 1132 | T1132L | D | 1.2e-05 | 1.1e-05 | 1.5e-05 | 1.267E-05 | 2.082E-06 | 1.100E-05 | 1.500E-05 |
| 22632 | 1132 | T1132K | D | 1.5e-05 | 9.8e-06 | 9.7e-06 | 1.150E-05 | 3.032E-06 | 9.700E-06 | 1.500E-05 |
| 22633 | 1132 | T1132M | D | 1.6e-05 | 1.4e-05 | 1.5e-05 | 1.500E-05 | 1.000E-06 | 1.400E-05 | 1.600E-05 |
| 22634 | 1132 | T1132F | D | 1.2e-05 | 1.5e-05 | 9.6e-06 | 1.220E-05 | 2.706E-06 | 9.600E-06 | 1.500E-05 |
| 22635 | 1132 | T1132P | D | 9.4e-06 | 1.4e-05 | 1.5e-05 | 1.280E-05 | 2.987E-06 | 9.400E-06 | 1.500E-05 |
| 22636 | 1132 | T1132S | D | 8.5e-06 | 1e-05 | 1.7e-05 | 1.183E-05 | 4.537E-06 | 8.500E-06 | 1.700E-05 |
| 22637 | 1132 | T1132T | D | 1.1e-05 | 1.5e-05 | 1.3e-05 | 1.300E-05 | 2.000E-06 | 1.100E-05 | 1.500E-05 |
| 22638 | 1132 | T1132W | D | 1.2e-05 | 1.3e-05 | 1.3e-05 | 1.267E-05 | 5.774E-07 | 1.200E-05 | 1.300E-05 |
| 22639 | 1132 | T1132Y | D | 1.1e-05 | 8.5e-06 | 1.2e-05 | 1.050E-05 | 1.803E-06 | 8.500E-06 | 1.200E-05 |
| 22640 | 1132 | T1132V | D | 1.5e-05 | 1.1e-05 | 1.5e-05 | 1.367E-05 | 2.309E-06 | 1.100E-05 | 1.500E-05 |
| 22641 | 1133 | I1133A | D | 9.2e-06 | 2.6e-05 | 9.8e-06 | 1.500E-05 | 9.531E-06 | 9.200E-06 | 2.600E-05 |
| 22642 | 1133 | I1133R | D | 1e-05 | 1.2e-05 | 1.3e-05 | 1.167E-05 | 1.528E-06 | 1.000E-05 | 1.300E-05 |
| 22643 | 1133 | I1133N | D | 1.3e-05 | 7.8e-06 | 1.5e-05 | 1.193E-05 | 3.717E-06 | 7.800E-06 | 1.500E-05 |
| 22644 | 1133 | I1133D | D | 1e-05 | 1.6e-05 | 1.4e-05 | 1.333E-05 | 3.055E-06 | 1.000E-05 | 1.600E-05 |
| 22645 | 1133 | I1133C | D | 1.2e-05 | 1.3e-05 | 2.2e-05 | 1.567E-05 | 5.508E-06 | 1.200E-05 | 2.200E-05 |
| 22646 | 1133 | I1133E | D | 9.8e-06 | 1.6e-05 | 1e-05 | 1.193E-05 | 3.523E-06 | 9.800E-06 | 1.600E-05 |
| 22647 | 1133 | I1133Q | D | 9.3e-06 | 1.1e-05 | 1.4e-05 | 1.143E-05 | 2.380E-06 | 9.300E-06 | 1.400E-05 |
| 22648 | 1133 | I1133G | D | 8.3e-06 | 9.8e-06 | 1.5e-05 | 1.103E-05 | 3.516E-06 | 8.300E-06 | 1.500E-05 |
| 22649 | 1133 | I1133H | D | 1.4e-05 | 1e-05 | 1.1e-05 | 1.167E-05 | 2.082E-06 | 1.000E-05 | 1.400E-05 |
| 22650 | 1133 | I1133I | D | 8.7e-06 | 1.5e-05 | 1.7e-05 | 1.357E-05 | 4.332E-06 | 8.700E-06 | 1.700E-05 |
| 22651 | 1133 | I1133L | D | 1.6e-05 | 1.6e-05 | 1.2e-05 | 1.467E-05 | 2.309E-06 | 1.200E-05 | 1.600E-05 |
| 22652 | 1133 | I1133K | D | 1.5e-05 | 9.2e-06 | 1.5e-05 | 1.307E-05 | 3.349E-06 | 9.200E-06 | 1.500E-05 |
| 22653 | 1133 | I1133M | D | 2.5e-05 | 9.9e-06 | 1e-05 | 1.497E-05 | 8.689E-06 | 9.900E-06 | 2.500E-05 |
| 22654 | 1133 | I1133F | D | 2.1e-05 | 1.4e-05 | 1.6e-05 | 1.700E-05 | 3.606E-06 | 1.400E-05 | 2.100E-05 |
| 22655 | 1133 | I1133P | D | 1.1e-05 | 9.7e-06 | 1.1e-05 | 1.057E-05 | 7.506E-07 | 9.700E-06 | 1.100E-05 |
| 22656 | 1133 | I1133S | D | 8.9e-06 | 1.5e-05 | 1.5e-05 | 1.297E-05 | 3.522E-06 | 8.900E-06 | 1.500E-05 |
| 22657 | 1133 | I1133T | D | 1e-05 | 1e-05 | 1.4e-05 | 1.133E-05 | 2.309E-06 | 1.000E-05 | 1.400E-05 |
| 22658 | 1133 | I1133W | D | 1.1e-05 | 9.8e-06 | 1.2e-05 | 1.093E-05 | 1.102E-06 | 9.800E-06 | 1.200E-05 |
| 22659 | 1133 | I1133Y | D | 1.2e-05 | 8.2e-06 | 1.5e-05 | 1.173E-05 | 3.408E-06 | 8.200E-06 | 1.500E-05 |
| 22660 | 1133 | I1133V | D | 1.6e-05 | 1.1e-05 | 1.3e-05 | 1.333E-05 | 2.517E-06 | 1.100E-05 | 1.600E-05 |
| 22661 | 1134 | S1134A | D | 1.6e-05 | 1.1e-05 | 1.4e-05 | 1.367E-05 | 2.517E-06 | 1.100E-05 | 1.600E-05 |
| 22662 | 1134 | S1134R | D | 1.5e-05 | 9.9e-06 | 9.7e-06 | 1.153E-05 | 3.004E-06 | 9.700E-06 | 1.500E-05 |
| 22663 | 1134 | S1134N | D | 1.6e-05 | 1.3e-05 | 1.4e-05 | 1.433E-05 | 1.528E-06 | 1.300E-05 | 1.600E-05 |
| 22664 | 1134 | S1134D | D | 9.6e-06 | 1.6e-05 | 1.5e-05 | 1.353E-05 | 3.443E-06 | 9.600E-06 | 1.600E-05 |
| 22665 | 1134 | S1134C | D | 1.7e-05 | 1.1e-05 | 1.2e-05 | 1.333E-05 | 3.215E-06 | 1.100E-05 | 1.700E-05 |
| 22666 | 1134 | S1134E | D | 1e-05 | 1.4e-05 | 1.5e-05 | 1.300E-05 | 2.646E-06 | 1.000E-05 | 1.500E-05 |
| 22667 | 1134 | S1134Q | D | 1.5e-05 | 1.6e-05 | 1.6e-05 | 1.567E-05 | 5.774E-07 | 1.500E-05 | 1.600E-05 |
| 22668 | 1134 | S1134G | D | 1.6e-05 | 1.5e-05 | 1.5e-05 | 1.533E-05 | 5.774E-07 | 1.500E-05 | 1.600E-05 |
| 22669 | 1134 | S1134H | D | 9.4e-06 | 1.6e-05 | 1.1e-05 | 1.213E-05 | 3.443E-06 | 9.400E-06 | 1.600E-05 |
| 22670 | 1134 | S1134I | D | 1.4e-05 | 1.4e-05 | 1.6e-05 | 1.467E-05 | 1.155E-06 | 1.400E-05 | 1.600E-05 |
| 22671 | 1134 | S1134L | D | 1.5e-05 | 8.8e-06 | 9.4e-06 | 1.107E-05 | 3.420E-06 | 8.800E-06 | 1.500E-05 |
| 22672 | 1134 | S1134K | D | 8.3e-06 | 1.3e-05 | 8.7e-06 | 1.000E-05 | 2.606E-06 | 8.300E-06 | 1.300E-05 |
| 22673 | 1134 | S1134M | D | 8.5e-06 | 1.4e-05 | 1.4e-05 | 1.217E-05 | 3.175E-06 | 8.500E-06 | 1.400E-05 |
| 22674 | 1134 | S1134F | D | 7.8e-06 | 1.6e-05 | 1.5e-05 | 1.293E-05 | 4.474E-06 | 7.800E-06 | 1.600E-05 |
| 22675 | 1134 | S1134P | D | 9.3e-06 | 8.4e-06 | 1.7e-05 | 1.157E-05 | 4.727E-06 | 8.400E-06 | 1.700E-05 |
| 22676 | 1134 | S1134S | D | 1e-05 | 1.6e-05 | 1.4e-05 | 1.333E-05 | 3.055E-06 | 1.000E-05 | 1.600E-05 |
| 22677 | 1134 | S1134T | D | 1.1e-05 | 1.3e-05 | 9.7e-06 | 1.123E-05 | 1.662E-06 | 9.700E-06 | 1.300E-05 |
| 22678 | 1134 | S1134W | D | 1.5e-05 | 1.7e-05 | 1.3e-05 | 1.500E-05 | 2.000E-06 | 1.300E-05 | 1.700E-05 |
| 22679 | 1134 | S1134Y | D | 1.2e-05 | 1.4e-05 | 9.4e-06 | 1.180E-05 | 2.307E-06 | 9.400E-06 | 1.400E-05 |
| 22680 | 1134 | S1134V | D | 1.1e-05 | 9.1e-06 | 1.6e-05 | 1.203E-05 | 3.564E-06 | 9.100E-06 | 1.600E-05 |
| 22681 | 1135 | S1135A | D | 1.5e-05 | 1.4e-05 | 1.1e-05 | 1.333E-05 | 2.082E-06 | 1.100E-05 | 1.500E-05 |
| 22682 | 1135 | S1135R | D | 1.1e-05 | 1.4e-05 | 1.1e-05 | 1.200E-05 | 1.732E-06 | 1.100E-05 | 1.400E-05 |
| 22683 | 1135 | S1135N | D | 1.4e-05 | 1e-05 | 1.5e-05 | 1.300E-05 | 2.646E-06 | 1.000E-05 | 1.500E-05 |
| 22684 | 1135 | S1135D | D | 1.1e-05 | 1.6e-05 | 1.2e-05 | 1.300E-05 | 2.646E-06 | 1.100E-05 | 1.600E-05 |
| 22685 | 1135 | S1135C | D | 1.5e-05 | 1.6e-05 | 9.3e-06 | 1.343E-05 | 3.614E-06 | 9.300E-06 | 1.600E-05 |
| 22686 | 1135 | S1135E | D | 1e-05 | 1.5e-05 | 1.6e-05 | 1.367E-05 | 3.215E-06 | 1.000E-05 | 1.600E-05 |
| 22687 | 1135 | S1135Q | D | 1.2e-05 | 1.6e-05 | 1.1e-05 | 1.300E-05 | 2.646E-06 | 1.100E-05 | 1.600E-05 |
| 22688 | 1135 | S1135G | D | 1.4e-05 | 1.1e-05 | 1e-05 | 1.167E-05 | 2.082E-06 | 1.000E-05 | 1.400E-05 |
| 22689 | 1135 | S1135H | D | 1.4e-05 | 1.1e-05 | 1.4e-05 | 1.300E-05 | 1.732E-06 | 1.100E-05 | 1.400E-05 |
| 22690 | 1135 | S1135I | D | 1e-05 | 1.4e-05 | 9.4e-06 | 1.113E-05 | 2.501E-06 | 9.400E-06 | 1.400E-05 |
| 22691 | 1135 | S1135L | D | 8e-06 | 1.1e-05 | 1.7e-05 | 1.200E-05 | 4.583E-06 | 8.000E-06 | 1.700E-05 |
| 22692 | 1135 | S1135K | D | 1.1e-05 | 1.8e-05 | 1.1e-05 | 1.333E-05 | 4.041E-06 | 1.100E-05 | 1.800E-05 |
| 22693 | 1135 | S1135M | D | 1.3e-05 | 1e-05 | 1.5e-05 | 1.267E-05 | 2.517E-06 | 1.000E-05 | 1.500E-05 |
| 22694 | 1135 | S1135F | D | 1.5e-05 | 1.5e-05 | 1.3e-05 | 1.433E-05 | 1.155E-06 | 1.300E-05 | 1.500E-05 |

| | | | | | | | | | | |
|---|---|---|---|---|---|---|---|---|---|---|
| 22695 | 1135 | S1135P | D | 1.6e-05 | 1.1e-05 | 1.5e-05 | 1.400E-05 | 2.646E-06 | 1.100E-05 | 1.600E-05 |
| 22696 | 1135 | S1135S | D | 9.9e-06 | 1.5e-05 | 1.5e-05 | 1.330E-05 | 2.944E-06 | 9.900E-06 | 1.500E-05 |
| 22697 | 1135 | S1135T | D | 1.1e-05 | 1.4e-05 | 1.1e-05 | 1.200E-05 | 1.732E-06 | 1.100E-05 | 1.400E-05 |
| 22698 | 1135 | S1135W | D | 1.7e-05 | 8.9e-06 | 1.2e-05 | 1.263E-05 | 4.087E-06 | 8.900E-06 | 1.700E-05 |
| 22699 | 1135 | S1135Y | D | 1.1e-05 | 1.4e-05 | 1.2e-05 | 1.233E-05 | 1.528E-06 | 1.100E-05 | 1.400E-05 |
| 22700 | 1135 | S1135V | D | 9.6e-06 | 1.3e-05 | 1.4e-05 | 1.220E-05 | 2.307E-06 | 9.600E-06 | 1.400E-05 |
| 22701 | 1136 | L1136A | D | 1.2e-05 | 3e-05 | 1.4e-05 | 1.867E-05 | 9.866E-06 | 1.200E-05 | 3.000E-05 |
| 22702 | 1136 | L1136R | D | 1.2e-05 | 8.2e-06 | 1.1e-05 | 1.040E-05 | 1.970E-06 | 8.200E-06 | 1.200E-05 |
| 22703 | 1136 | L1136N | D | 1.5e-05 | 1.3e-05 | 9.4e-06 | 1.247E-05 | 2.838E-06 | 9.400E-06 | 1.500E-05 |
| 22704 | 1136 | L1136D | D | 1.3e-05 | 1.4e-05 | 1.2e-05 | 1.300E-05 | 1.000E-06 | 1.200E-05 | 1.400E-05 |
| 22705 | 1136 | L1136C | D | 1.5e-05 | 1.3e-05 | 1.3e-05 | 1.367E-05 | 1.155E-06 | 1.300E-05 | 1.500E-05 |
| 22706 | 1136 | L1136E | D | 1.4e-05 | 9.6e-06 | 1.7e-05 | 1.353E-05 | 3.722E-06 | 9.600E-06 | 1.700E-05 |
| 22707 | 1136 | L1136Q | D | 9.9e-06 | 1.6e-05 | 1.2e-05 | 1.263E-05 | 3.099E-06 | 9.900E-06 | 1.600E-05 |
| 22708 | 1136 | L1136G | D | 1.4e-05 | 9.9e-06 | 1.4e-05 | 1.263E-05 | 2.367E-06 | 9.900E-06 | 1.400E-05 |
| 22709 | 1136 | L1136H | D | 1.5e-05 | 1.5e-05 | 1e-05 | 1.333E-05 | 2.887E-06 | 1.000E-05 | 1.500E-05 |
| 22710 | 1136 | L1136I | D | 1.5e-05 | 8.6e-06 | 1.5e-05 | 1.287E-05 | 3.695E-06 | 8.600E-06 | 1.500E-05 |
| 22711 | 1136 | L1136L | D | 1.5e-05 | 9.1e-06 | 1.1e-05 | 1.170E-05 | 3.012E-06 | 9.100E-06 | 1.500E-05 |
| 22712 | 1136 | L1136K | D | 1.7e-05 | 1.1e-05 | 1.2e-05 | 1.333E-05 | 3.215E-06 | 1.100E-05 | 1.700E-05 |
| 22713 | 1136 | L1136M | D | 1.9e-05 | 9.2e-06 | 1.2e-05 | 1.340E-05 | 5.048E-06 | 9.200E-06 | 1.900E-05 |
| 22714 | 1136 | L1136F | D | 1.7e-05 | 9.8e-06 | 1.5e-05 | 1.393E-05 | 3.717E-06 | 9.800E-06 | 1.700E-05 |
| 22715 | 1136 | L1136P | D | 9.7e-06 | 1.3e-05 | 1.4e-05 | 1.223E-05 | 2.250E-06 | 9.700E-06 | 1.400E-05 |
| 22716 | 1136 | L1136S | D | 1.2e-05 | 1.6e-05 | 1.3e-05 | 1.367E-05 | 2.082E-06 | 1.200E-05 | 1.600E-05 |
| 22717 | 1136 | L1136T | D | 1.5e-05 | 1.3e-05 | 1.5e-05 | 1.433E-05 | 1.155E-06 | 1.300E-05 | 1.500E-05 |
| 22718 | 1136 | L1136W | D | 1.6e-05 | 1.3e-05 | 1.5e-05 | 1.467E-05 | 1.528E-06 | 1.300E-05 | 1.600E-05 |
| 22719 | 1136 | L1136Y | D | 1.1e-05 | 1.1e-05 | 1.4e-05 | 1.200E-05 | 1.732E-06 | 1.100E-05 | 1.400E-05 |
| 22720 | 1136 | L1136V | D | 1.5e-05 | 1.7e-05 | 1.6e-05 | 1.600E-05 | 1.000E-06 | 1.500E-05 | 1.700E-05 |
| 22721 | 1137 | Q1137A | D | 1.6e-05 | 1.4e-05 | 1.5e-05 | 1.500E-05 | 1.000E-06 | 1.400E-05 | 1.600E-05 |
| 22722 | 1137 | Q1137R | D | 1.1e-05 | 1.5e-05 | 1.1e-05 | 1.233E-05 | 2.309E-06 | 1.100E-05 | 1.500E-05 |
| 22723 | 1137 | Q1137N | D | 1.1e-05 | 1.7e-05 | 1.1e-05 | 1.300E-05 | 3.464E-06 | 1.100E-05 | 1.700E-05 |
| 22724 | 1137 | Q1137D | D | 1.6e-05 | 1.4e-05 | 1.1e-05 | 1.367E-05 | 2.517E-06 | 1.100E-05 | 1.600E-05 |
| 22725 | 1137 | Q1137C | D | 1.1e-05 | 8.5e-06 | 1.2e-05 | 1.050E-05 | 1.803E-06 | 8.500E-06 | 1.200E-05 |
| 22726 | 1137 | Q1137E | D | 1.8e-05 | 1.6e-05 | 1.1e-05 | 1.500E-05 | 3.606E-06 | 1.100E-05 | 1.800E-05 |
| 22727 | 1137 | Q1137Q | D | 1.7e-05 | 1.1e-05 | 1e-05 | 1.267E-05 | 3.786E-06 | 1.000E-05 | 1.700E-05 |
| 22728 | 1137 | Q1137G | D | 1.3e-05 | 1.4e-05 | 1.7e-05 | 1.467E-05 | 2.082E-06 | 1.300E-05 | 1.700E-05 |
| 22729 | 1137 | Q1137H | D | 1.5e-05 | 1e-05 | 1.1e-05 | 1.200E-05 | 2.646E-06 | 1.000E-05 | 1.500E-05 |
| 22730 | 1137 | Q1137I | D | 1.1e-05 | 1e-05 | 9.5e-06 | 1.017E-05 | 7.638E-07 | 9.500E-06 | 1.100E-05 |
| 22731 | 1137 | Q1137L | D | 1.2e-05 | 2.9e-05 | 1.6e-05 | 1.900E-05 | 8.888E-06 | 1.200E-05 | 2.900E-05 |
| 22732 | 1137 | Q1137K | D | 1.4e-05 | 1.4e-05 | 8.6e-06 | 1.220E-05 | 3.118E-06 | 8.600E-06 | 1.400E-05 |
| 22733 | 1137 | Q1137M | D | 1e-05 | 9.5e-06 | 1.1e-05 | 1.017E-05 | 7.638E-07 | 9.500E-06 | 1.100E-05 |
| 22734 | 1137 | Q1137F | D | 1e-05 | 1.3e-05 | 1.5e-05 | 1.267E-05 | 2.517E-06 | 1.000E-05 | 1.500E-05 |
| 22735 | 1137 | Q1137P | D | 1.2e-05 | 9.6e-06 | 1.6e-05 | 1.253E-05 | 3.233E-06 | 9.600E-06 | 1.600E-05 |
| 22736 | 1137 | Q1137S | D | 1.4e-05 | 1.2e-05 | 8.6e-06 | 1.153E-05 | 2.730E-06 | 8.600E-06 | 1.400E-05 |
| 22737 | 1137 | Q1137T | D | 1.4e-05 | 1.2e-05 | 1.1e-05 | 1.233E-05 | 1.528E-06 | 1.100E-05 | 1.400E-05 |
| 22738 | 1137 | Q1137W | D | 1.5e-05 | 1e-05 | 1.1e-05 | 1.200E-05 | 2.646E-06 | 1.000E-05 | 1.500E-05 |
| 22739 | 1137 | Q1137Y | D | 1.5e-05 | 1.1e-05 | 1.1e-05 | 1.233E-05 | 2.309E-06 | 1.100E-05 | 1.500E-05 |
| 22740 | 1137 | Q1137V | D | 1.1e-05 | 1.4e-05 | 1.4e-05 | 1.300E-05 | 1.732E-06 | 1.100E-05 | 1.400E-05 |
| 22741 | 1138 | P1138A | D | 1.1e-05 | 1e-05 | 1.5e-05 | 1.200E-05 | 2.646E-06 | 1.000E-05 | 1.500E-05 |
| 22742 | 1138 | P1138R | D | 1.5e-05 | 1.3e-05 | 8.9e-06 | 1.230E-05 | 3.110E-06 | 8.900E-06 | 1.500E-05 |
| 22743 | 1138 | P1138N | D | 1.3e-05 | 1.4e-05 | 1.2e-05 | 1.300E-05 | 1.000E-06 | 1.200E-05 | 1.400E-05 |
| 22744 | 1138 | P1138D | D | 1.5e-05 | 1.5e-05 | 1.5e-05 | 1.500E-05 | 0.000E+00 | 1.500E-05 | 1.500E-05 |
| 22745 | 1138 | P1138C | D | 1.4e-05 | 1.3e-05 | 1.2e-05 | 1.300E-05 | 1.000E-06 | 1.200E-05 | 1.400E-05 |
| 22746 | 1138 | P1138E | D | 1e-05 | 1.2e-05 | 1.4e-05 | 1.200E-05 | 2.000E-06 | 1.000E-05 | 1.400E-05 |
| 22747 | 1138 | P1138Q | D | 1.1e-05 | 9.5e-06 | 9.8e-06 | 1.010E-05 | 7.937E-07 | 9.500E-06 | 1.100E-05 |
| 22748 | 1138 | P1138G | D | 1.5e-05 | 1.5e-05 | 1.4e-05 | 1.467E-05 | 5.774E-07 | 1.400E-05 | 1.500E-05 |
| 22749 | 1138 | P1138H | D | 1e-05 | 1.6e-05 | 1.4e-05 | 1.333E-05 | 3.055E-06 | 1.000E-05 | 1.600E-05 |
| 22750 | 1138 | P1138I | D | 1.2e-05 | 1.5e-05 | 1.5e-05 | 1.400E-05 | 1.732E-06 | 1.200E-05 | 1.500E-05 |
| 22751 | 1138 | P1138L | D | 1.4e-05 | 1.6e-05 | 1.5e-05 | 1.500E-05 | 1.000E-06 | 1.400E-05 | 1.600E-05 |
| 22752 | 1138 | P1138K | D | 1.1e-05 | 1.1e-05 | 1e-05 | 1.067E-05 | 5.774E-07 | 1.000E-05 | 1.100E-05 |
| 22753 | 1138 | P1138M | D | 1e-05 | 9.1e-06 | 9.2e-06 | 9.433E-06 | 4.933E-07 | 9.100E-06 | 1.000E-05 |
| 22754 | 1138 | P1138F | D | 1.1e-05 | 1e-05 | 1e-05 | 1.033E-05 | 5.774E-07 | 1.000E-05 | 1.100E-05 |
| 22755 | 1138 | P1138P | D | 1.3e-05 | 1.1e-05 | 1.6e-05 | 1.333E-05 | 2.517E-06 | 1.100E-05 | 1.600E-05 |
| 22756 | 1138 | P1138S | D | 1.2e-05 | 9.5e-06 | 8.8e-06 | 1.010E-05 | 1.682E-06 | 8.800E-06 | 1.200E-05 |
| 22757 | 1138 | P1138T | D | 1.4e-05 | 1e-05 | 1.1e-05 | 1.167E-05 | 2.082E-06 | 1.000E-05 | 1.400E-05 |
| 22758 | 1138 | P1138W | D | 9.9e-06 | 1.6e-05 | 1.6e-05 | 1.397E-05 | 3.522E-06 | 9.900E-06 | 1.600E-05 |
| 22759 | 1138 | P1138Y | D | 1.7e-05 | 8.2e-06 | 1.7e-05 | 1.407E-05 | 5.081E-06 | 8.200E-06 | 1.700E-05 |
| 22760 | 1138 | P1138V | D | 1.5e-05 | 1.1e-05 | 1.4e-05 | 1.333E-05 | 2.082E-06 | 1.100E-05 | 1.500E-05 |
| 22761 | 1139 | E1139A | D | 2.2e-05 | 1.1e-05 | 1.6e-05 | 1.633E-05 | 5.508E-06 | 1.100E-05 | 2.200E-05 |
| 22762 | 1139 | E1139R | D | 7.8e-06 | 1.7e-05 | 9e-06 | 1.127E-05 | 5.001E-06 | 7.800E-06 | 1.700E-05 |
| 22763 | 1139 | E1139N | D | 8.7e-06 | 1e-05 | 1.6e-05 | 1.157E-05 | 3.894E-06 | 8.700E-06 | 1.600E-05 |
| 22764 | 1139 | E1139D | D | 9e-06 | 1.5e-05 | 8.6e-06 | 1.087E-05 | 3.585E-06 | 8.600E-06 | 1.500E-05 |
| 22765 | 1139 | E1139C | D | 1.4e-05 | 1.6e-05 | 1.4e-05 | 1.467E-05 | 1.155E-06 | 1.400E-05 | 1.600E-05 |
| 22766 | 1139 | E1139E | D | 1.5e-05 | 8.9e-06 | 1.2e-05 | 1.197E-05 | 3.050E-06 | 8.900E-06 | 1.500E-05 |
| 22767 | 1139 | E1139Q | D | 9.5e-06 | 1e-05 | 1.1e-05 | 1.017E-05 | 7.638E-07 | 9.500E-06 | 1.100E-05 |
| 22768 | 1139 | E1139G | D | 1.5e-05 | 1.2e-05 | 8.9e-06 | 1.197E-05 | 3.050E-06 | 8.900E-06 | 1.500E-05 |
| 22769 | 1139 | E1139H | D | 1.5e-05 | 5.6e-06 | 1.5e-05 | 1.187E-05 | 5.427E-06 | 5.600E-06 | 1.500E-05 |
| 22770 | 1139 | E1139I | D | 1.1e-05 | 1.1e-05 | 1.6e-05 | 1.267E-05 | 2.887E-06 | 1.100E-05 | 1.600E-05 |
| 22771 | 1139 | E1139L | D | 1.8e-05 | 1.5e-05 | 1.5e-05 | 1.600E-05 | 1.732E-06 | 1.500E-05 | 1.800E-05 |
| 22772 | 1139 | E1139K | D | 1.2e-05 | 9.7e-06 | 1.1e-05 | 1.090E-05 | 1.153E-06 | 9.700E-06 | 1.200E-05 |
| 22773 | 1139 | E1139M | D | 9.4e-06 | 7.9e-06 | 1.3e-05 | 1.010E-05 | 2.621E-06 | 7.900E-06 | 1.300E-05 |
| 22774 | 1139 | E1139F | D | 1.7e-05 | 1.7e-05 | 1.1e-05 | 1.500E-05 | 3.464E-06 | 1.100E-05 | 1.700E-05 |
| 22775 | 1139 | E1139P | D | 1.3e-05 | 1.6e-05 | 1.2e-05 | 1.367E-05 | 2.082E-06 | 1.200E-05 | 1.600E-05 |
| 22776 | 1139 | E1139S | D | 1.3e-05 | 1.1e-05 | 9.1e-06 | 1.103E-05 | 1.950E-06 | 9.100E-06 | 1.300E-05 |
| 22777 | 1139 | E1139T | D | 1.4e-05 | 1.3e-05 | 1.2e-05 | 1.300E-05 | 1.000E-06 | 1.200E-05 | 1.400E-05 |
| 22778 | 1139 | E1139W | D | 1.2e-05 | 1.2e-05 | 1.1e-05 | 1.167E-05 | 5.774E-07 | 1.100E-05 | 1.200E-05 |
| 22779 | 1139 | E1139Y | D | 1.3e-05 | 1.5e-05 | 1.4e-05 | 1.400E-05 | 1.000E-06 | 1.300E-05 | 1.500E-05 |

| | | | | | | | | | | |
|---|---|---|---|---|---|---|---|---|---|---|
| 22780 | 1139 | E1139V | D | 1e-05 | 1.4e-05 | 7.3e-06 | 1.043E-05 | 3.371E-06 | 7.300E-06 | 1.400E-05 |
| 22781 | 1140 | D1140A | D | 1.1e-05 | 1.4e-05 | 1.1e-05 | 1.200E-05 | 1.732E-06 | 1.100E-05 | 1.400E-05 |
| 22782 | 1140 | D1140R | D | 1.3e-05 | 1e-05 | 1.5e-05 | 1.267E-05 | 2.517E-06 | 1.000E-05 | 1.500E-05 |
| 22783 | 1140 | D1140N | D | 1.4e-05 | 1.1e-05 | 1.5e-05 | 1.333E-05 | 2.082E-06 | 1.100E-05 | 1.500E-05 |
| 22784 | 1140 | D1140D | D | 1.4e-05 | 1e-05 | 1.4e-05 | 1.267E-05 | 2.309E-06 | 1.000E-05 | 1.400E-05 |
| 22785 | 1140 | D1140C | D | 9.8e-06 | 1.6e-05 | 1.1e-05 | 1.227E-05 | 3.288E-06 | 9.800E-06 | 1.600E-05 |
| 22786 | 1140 | D1140E | D | 1.4e-05 | 1.4e-05 | 1.7e-05 | 1.500E-05 | 1.732E-06 | 1.400E-05 | 1.700E-05 |
| 22787 | 1140 | D1140Q | D | 1.1e-05 | 1.5e-05 | 1e-05 | 1.200E-05 | 2.646E-06 | 1.000E-05 | 1.500E-05 |
| 22788 | 1140 | D1140G | D | 9.8e-06 | 1.1e-05 | 9.3e-06 | 1.003E-05 | 8.737E-07 | 9.300E-06 | 1.100E-05 |
| 22789 | 1140 | D1140H | D | 1.1e-05 | 1e-05 | 1.1e-05 | 1.067E-05 | 5.774E-07 | 1.000E-05 | 1.100E-05 |
| 22790 | 1140 | D1140I | D | 1.4e-05 | 1.1e-05 | 1.3e-05 | 1.267E-05 | 1.528E-06 | 1.100E-05 | 1.400E-05 |
| 22791 | 1140 | D1140L | D | 1.6e-05 | 1.2e-05 | 1.4e-05 | 1.400E-05 | 2.000E-06 | 1.200E-05 | 1.600E-05 |
| 22792 | 1140 | D1140K | D | 8.8e-06 | 1.5e-05 | 1.1e-05 | 1.160E-05 | 3.143E-06 | 8.800E-06 | 1.500E-05 |
| 22793 | 1140 | D1140M | D | 1e-05 | 9.9e-06 | 1.2e-05 | 1.063E-05 | 1.185E-06 | 9.900E-06 | 1.200E-05 |
| 22794 | 1140 | D1140F | D | 1.6e-05 | 1.6e-05 | 1.2e-05 | 1.467E-05 | 2.309E-06 | 1.200E-05 | 1.600E-05 |
| 22795 | 1140 | D1140P | D | 1.5e-05 | 1.4e-05 | 1.2e-05 | 1.367E-05 | 1.528E-06 | 1.200E-05 | 1.500E-05 |
| 22796 | 1140 | D1140S | D | 1.1e-05 | 1.1e-05 | 1.1e-05 | 1.100E-05 | 2.075E-21 | 1.100E-05 | 1.100E-05 |
| 22797 | 1140 | D1140T | D | 1.3e-05 | 1.6e-05 | 1.3e-05 | 1.400E-05 | 1.732E-06 | 1.300E-05 | 1.600E-05 |
| 22798 | 1140 | D1140W | D | 9.4e-06 | 9.4e-06 | 1.5e-05 | 1.127E-05 | 3.233E-06 | 9.400E-06 | 1.500E-05 |
| 22799 | 1140 | D1140Y | D | 1.6e-05 | 9.8e-06 | 1.4e-05 | 1.327E-05 | 3.164E-06 | 9.800E-06 | 1.600E-05 |
| 22800 | 1140 | D1140V | D | 1.5e-05 | 1.1e-05 | 1.5e-05 | 1.367E-05 | 2.309E-06 | 1.100E-05 | 1.500E-05 |
| 22801 | 1141 | F1141A | D | 1e-05 | 1.1e-05 | 1.4e-05 | 1.167E-05 | 2.082E-06 | 1.000E-05 | 1.400E-05 |
| 22802 | 1141 | F1141R | D | 1.4e-05 | 8.7e-06 | 1.6e-05 | 1.290E-05 | 3.772E-06 | 8.700E-06 | 1.600E-05 |
| 22803 | 1141 | F1141N | D | 9.5e-06 | 9.3e-06 | 8.8e-06 | 9.200E-06 | 3.606E-07 | 8.800E-06 | 9.500E-06 |
| 22804 | 1141 | F1141D | D | 1.2e-05 | 1.7e-05 | 1.6e-05 | 1.500E-05 | 2.646E-06 | 1.200E-05 | 1.700E-05 |
| 22805 | 1141 | F1141C | D | 1.4e-05 | 1.1e-05 | 1.4e-05 | 1.300E-05 | 1.732E-06 | 1.100E-05 | 1.400E-05 |
| 22806 | 1141 | F1141E | D | 1.3e-05 | 1.8e-05 | 1.5e-05 | 1.533E-05 | 2.517E-06 | 1.300E-05 | 1.800E-05 |
| 22807 | 1141 | F1141Q | D | 1e-05 | 1e-05 | 1.3e-05 | 1.100E-05 | 1.732E-06 | 1.000E-05 | 1.300E-05 |
| 22808 | 1141 | F1141G | D | 1.4e-05 | 8.9e-06 | 1.3e-05 | 1.197E-05 | 2.702E-06 | 8.900E-06 | 1.400E-05 |
| 22809 | 1141 | F1141H | D | 1e-05 | 1.1e-05 | 1.6e-05 | 1.233E-05 | 3.215E-06 | 1.000E-05 | 1.600E-05 |
| 22810 | 1141 | F1141I | D | 1.5e-05 | 1.3e-05 | 1.2e-05 | 1.333E-05 | 1.528E-06 | 1.200E-05 | 1.500E-05 |
| 22811 | 1141 | F1141L | D | 8.5e-06 | 1.1e-05 | 1.3e-05 | 1.083E-05 | 2.255E-06 | 8.500E-06 | 1.300E-05 |
| 22812 | 1141 | F1141K | D | 8.8e-06 | 1.6e-05 | 1.1e-05 | 1.193E-05 | 3.690E-06 | 8.800E-06 | 1.600E-05 |
| 22813 | 1141 | F1141M | D | 1e-05 | 1.4e-05 | 1.6e-05 | 1.333E-05 | 3.055E-06 | 1.000E-05 | 1.600E-05 |
| 22814 | 1141 | F1141F | D | 1.7e-05 | 1.7e-05 | 9.5e-06 | 1.450E-05 | 4.330E-06 | 9.500E-06 | 1.700E-05 |
| 22815 | 1141 | F1141P | D | 1e-05 | 1.1e-05 | 1.7e-05 | 1.267E-05 | 3.786E-06 | 1.000E-05 | 1.700E-05 |
| 22816 | 1141 | F1141S | D | 1.4e-05 | 1.4e-05 | 1.5e-05 | 1.433E-05 | 5.774E-07 | 1.400E-05 | 1.500E-05 |
| 22817 | 1141 | F1141T | D | 1.5e-05 | 1.4e-05 | 1.6e-05 | 1.500E-05 | 1.000E-06 | 1.400E-05 | 1.600E-05 |
| 22818 | 1141 | F1141W | D | 9.2e-06 | 1.5e-05 | 1.1e-05 | 1.173E-05 | 2.969E-06 | 9.200E-06 | 1.500E-05 |
| 22819 | 1141 | F1141Y | D | 1.5e-05 | 1.7e-05 | 1.3e-05 | 1.500E-05 | 2.000E-06 | 1.300E-05 | 1.700E-05 |
| 22820 | 1141 | F1141V | D | 1.4e-05 | 1.6e-05 | 1.7e-05 | 1.567E-05 | 1.528E-06 | 1.400E-05 | 1.700E-05 |
| 22821 | 1142 | A1142A | D | 1.4e-05 | 1.4e-05 | 1e-05 | 1.267E-05 | 2.309E-06 | 1.000E-05 | 1.400E-05 |
| 22822 | 1142 | A1142R | D | 1.4e-05 | 1.6e-05 | 1.5e-05 | 1.500E-05 | 1.000E-06 | 1.400E-05 | 1.600E-05 |
| 22823 | 1142 | A1142N | D | 1.4e-05 | 1.1e-05 | 1.4e-05 | 1.300E-05 | 1.732E-06 | 1.100E-05 | 1.400E-05 |
| 22824 | 1142 | A1142D | D | 1.2e-05 | 1.6e-05 | 1e-05 | 1.267E-05 | 3.055E-06 | 1.000E-05 | 1.600E-05 |
| 22825 | 1142 | A1142C | D | 1.5e-05 | 1.5e-05 | 1e-05 | 1.333E-05 | 2.887E-06 | 1.000E-05 | 1.500E-05 |
| 22826 | 1142 | A1142E | D | 9.2e-06 | 1e-05 | 1.7e-05 | 1.207E-05 | 4.291E-06 | 9.200E-06 | 1.700E-05 |
| 22827 | 1142 | A1142Q | D | 1.3e-05 | 1e-05 | 1.5e-05 | 1.267E-05 | 2.517E-06 | 1.000E-05 | 1.500E-05 |
| 22828 | 1142 | A1142G | D | 9.8e-06 | 1e-05 | 1.5e-05 | 1.160E-05 | 2.946E-06 | 9.800E-06 | 1.500E-05 |
| 22829 | 1142 | A1142H | D | 1.4e-05 | 1e-05 | 1.6e-05 | 1.333E-05 | 3.055E-06 | 1.000E-05 | 1.600E-05 |
| 22830 | 1142 | A1142I | D | 1.6e-05 | 1.8e-05 | 1e-05 | 1.467E-05 | 4.163E-06 | 1.000E-05 | 1.800E-05 |
| 22831 | 1142 | A1142L | D | 1.1e-05 | 1.6e-05 | 1.4e-05 | 1.367E-05 | 2.517E-06 | 1.100E-05 | 1.600E-05 |
| 22832 | 1142 | A1142K | D | 1.5e-05 | 1.4e-05 | 1.1e-05 | 1.333E-05 | 2.082E-06 | 1.100E-05 | 1.500E-05 |
| 22833 | 1142 | A1142M | D | 1.9e-05 | 1.7e-05 | 1.6e-05 | 1.733E-05 | 1.528E-06 | 1.600E-05 | 1.900E-05 |
| 22834 | 1142 | A1142F | D | 1.6e-05 | 1.6e-05 | 9.8e-06 | 1.393E-05 | 3.580E-06 | 9.800E-06 | 1.600E-05 |
| 22835 | 1142 | A1142P | D | 8.2e-06 | 9.9e-06 | 1.5e-05 | 1.103E-05 | 3.539E-06 | 8.200E-06 | 1.500E-05 |
| 22836 | 1142 | A1142S | D | 9.8e-06 | 1.3e-05 | 1e-05 | 1.093E-05 | 1.793E-06 | 9.800E-06 | 1.300E-05 |
| 22837 | 1142 | A1142T | D | 1.7e-05 | 1.6e-05 | 1.7e-05 | 1.667E-05 | 5.774E-07 | 1.600E-05 | 1.700E-05 |
| 22838 | 1142 | A1142W | D | 1.4e-05 | 1.1e-05 | 1.5e-05 | 1.333E-05 | 2.082E-06 | 1.100E-05 | 1.500E-05 |
| 22839 | 1142 | A1142Y | D | 9.8e-06 | 1.2e-05 | 1.3e-05 | 1.160E-05 | 1.637E-06 | 9.800E-06 | 1.300E-05 |
| 22840 | 1142 | A1142V | D | 1.1e-05 | 1.5e-05 | 1.4e-05 | 1.333E-05 | 2.082E-06 | 1.100E-05 | 1.500E-05 |
| 22841 | 1143 | T1143A | D | 1.1e-05 | 1.2e-05 | 1.5e-05 | 1.267E-05 | 2.082E-06 | 1.100E-05 | 1.500E-05 |
| 22842 | 1143 | T1143R | D | 9.1e-06 | 1.5e-05 | 1.1e-05 | 1.170E-05 | 3.012E-06 | 9.100E-06 | 1.500E-05 |
| 22843 | 1143 | T1143N | D | 7.3e-06 | 1.6e-05 | 1.7e-05 | 1.343E-05 | 5.335E-06 | 7.300E-06 | 1.700E-05 |
| 22844 | 1143 | T1143D | D | 1.6e-05 | 1.6e-05 | 1e-05 | 1.400E-05 | 3.464E-06 | 1.000E-05 | 1.600E-05 |
| 22845 | 1143 | T1143C | D | 1.5e-05 | 1.3e-05 | 1.5e-05 | 1.433E-05 | 1.155E-06 | 1.300E-05 | 1.500E-05 |
| 22846 | 1143 | T1143E | D | 1.5e-05 | 1.2e-05 | 1.4e-05 | 1.367E-05 | 1.528E-06 | 1.200E-05 | 1.500E-05 |
| 22847 | 1143 | T1143Q | D | 1.2e-05 | 1.3e-05 | 1.6e-05 | 1.367E-05 | 2.082E-06 | 1.200E-05 | 1.600E-05 |
| 22848 | 1143 | T1143G | D | 1.5e-05 | 1.4e-05 | 1.5e-05 | 1.467E-05 | 5.774E-07 | 1.400E-05 | 1.500E-05 |
| 22849 | 1143 | T1143H | D | 7.4e-06 | 1.6e-05 | 1e-05 | 1.113E-05 | 4.411E-06 | 7.400E-06 | 1.600E-05 |
| 22850 | 1143 | T1143I | D | 1.6e-05 | 1e-05 | 1e-05 | 1.200E-05 | 3.464E-06 | 1.000E-05 | 1.600E-05 |
| 22851 | 1143 | T1143L | D | 1.4e-05 | 5.2e-06 | 1.6e-05 | 1.173E-05 | 5.746E-06 | 5.200E-06 | 1.600E-05 |
| 22852 | 1143 | T1143K | D | 1e-05 | 1.2e-05 | 1.5e-05 | 1.233E-05 | 2.517E-06 | 1.000E-05 | 1.500E-05 |
| 22853 | 1143 | T1143M | D | 1.9e-05 | 1.6e-05 | 1.5e-05 | 1.667E-05 | 2.082E-06 | 1.500E-05 | 1.900E-05 |
| 22854 | 1143 | T1143F | D | 1.5e-05 | 1.7e-05 | 1.3e-05 | 1.500E-05 | 2.000E-06 | 1.300E-05 | 1.700E-05 |
| 22855 | 1143 | T1143P | D | 1.6e-05 | 8.6e-06 | 1e-05 | 1.153E-05 | 3.931E-06 | 8.600E-06 | 1.600E-05 |
| 22856 | 1143 | T1143S | D | 1.5e-05 | 8.8e-06 | 1.6e-05 | 1.327E-05 | 3.900E-06 | 8.800E-06 | 1.600E-05 |
| 22857 | 1143 | T1143T | D | 1.5e-05 | 1.5e-05 | 1.1e-05 | 1.367E-05 | 2.309E-06 | 1.100E-05 | 1.500E-05 |
| 22858 | 1143 | T1143W | D | 1.7e-05 | 9.9e-06 | 1.4e-05 | 1.363E-05 | 3.564E-06 | 9.900E-06 | 1.700E-05 |
| 22859 | 1143 | T1143Y | D | 1.4e-05 | 1.3e-05 | 1.1e-05 | 1.267E-05 | 1.528E-06 | 1.100E-05 | 1.400E-05 |
| 22860 | 1143 | T1143V | D | 1e-05 | 1.3e-05 | 9.9e-06 | 1.097E-05 | 1.762E-06 | 9.900E-06 | 1.300E-05 |
| 22861 | 1144 | Y1144A | D | 1.2e-05 | 1.6e-05 | 1.5e-05 | 1.433E-05 | 2.082E-06 | 1.200E-05 | 1.600E-05 |
| 22862 | 1144 | Y1144R | D | 1.2e-05 | 1.5e-05 | 1.7e-05 | 1.467E-05 | 2.517E-06 | 1.200E-05 | 1.700E-05 |
| 22863 | 1144 | Y1144N | D | 1.2e-05 | 1.3e-05 | 1.1e-05 | 1.200E-05 | 1.000E-06 | 1.100E-05 | 1.300E-05 |
| 22864 | 1144 | Y1144D | D | 1e-05 | 9.7e-06 | 1.2e-05 | 1.057E-05 | 1.250E-06 | 9.700E-06 | 1.200E-05 |

| | | | | | | | | | | |
|---|---|---|---|---|---|---|---|---|---|---|
| 22865 | 1144 | Y1144C | D | 1.5e-05 | 1e-05 | 2.3e-05 | 1.600E-05 | 6.557E-06 | 1.000E-05 | 2.300E-05 |
| 22866 | 1144 | Y1144E | D | 9.4e-06 | 1.5e-05 | 1.6e-05 | 1.347E-05 | 3.557E-06 | 9.400E-06 | 1.600E-05 |
| 22867 | 1144 | Y1144Q | D | 1.5e-05 | 1.1e-05 | 1.2e-05 | 1.267E-05 | 2.082E-06 | 1.100E-05 | 1.500E-05 |
| 22868 | 1144 | Y1144G | D | 1e-05 | 1.7e-05 | 1.1e-05 | 1.267E-05 | 3.786E-06 | 1.000E-05 | 1.700E-05 |
| 22869 | 1144 | Y1144H | D | 1.1e-05 | 1.5e-05 | 1.1e-05 | 1.233E-05 | 2.309E-06 | 1.100E-05 | 1.500E-05 |
| 22870 | 1144 | Y1144I | D | 1.3e-05 | 9.3e-06 | 1.6e-05 | 1.277E-05 | 3.356E-06 | 9.300E-06 | 1.600E-05 |
| 22871 | 1144 | Y1144L | D | 1.6e-05 | 1.1e-05 | 7.5e-06 | 1.150E-05 | 4.272E-06 | 7.500E-06 | 1.600E-05 |
| 22872 | 1144 | Y1144K | D | 8.3e-06 | 1e-05 | 1.5e-05 | 1.110E-05 | 3.483E-06 | 8.300E-06 | 1.500E-05 |
| 22873 | 1144 | Y1144M | D | 1.5e-05 | 1.6e-05 | 1.5e-05 | 1.533E-05 | 5.774E-07 | 1.500E-05 | 1.600E-05 |
| 22874 | 1144 | Y1144F | D | 1.4e-05 | 8.8e-06 | 1.6e-05 | 1.293E-05 | 3.717E-06 | 8.800E-06 | 1.600E-05 |
| 22875 | 1144 | Y1144P | D | 1.5e-05 | 1.1e-05 | 1.4e-05 | 1.333E-05 | 2.082E-06 | 1.100E-05 | 1.500E-05 |
| 22876 | 1144 | Y1144S | D | 1.7e-05 | 1.7e-05 | 1.1e-05 | 1.500E-05 | 3.464E-06 | 1.100E-05 | 1.700E-05 |
| 22877 | 1144 | Y1144T | D | 1.6e-05 | 9e-06 | 1.6e-05 | 1.367E-05 | 4.041E-06 | 9.000E-06 | 1.600E-05 |
| 22878 | 1144 | Y1144W | D | 1.4e-05 | 1.5e-05 | 1.3e-05 | 1.400E-05 | 1.000E-06 | 1.300E-05 | 1.500E-05 |
| 22879 | 1144 | Y1144Y | D | 1e-05 | 1.5e-05 | 1.5e-05 | 1.333E-05 | 2.887E-06 | 1.000E-05 | 1.500E-05 |
| 22880 | 1144 | Y1144V | D | 1.4e-05 | 1.2e-05 | 1.5e-05 | 1.367E-05 | 1.528E-06 | 1.200E-05 | 1.500E-05 |
| 22881 | 1145 | Y1145A | D | 1.7e-05 | 1.1e-05 | 1.5e-05 | 1.433E-05 | 3.055E-06 | 1.100E-05 | 1.700E-05 |
| 22882 | 1145 | Y1145R | D | 1.7e-05 | 9e-06 | 1.5e-05 | 1.367E-05 | 4.163E-06 | 9.000E-06 | 1.700E-05 |
| 22883 | 1145 | Y1145N | D | 1.5e-05 | 1.5e-05 | 1.4e-05 | 1.467E-05 | 5.774E-07 | 1.400E-05 | 1.500E-05 |
| 22884 | 1145 | Y1145D | D | 1.4e-05 | 1.5e-05 | 1e-05 | 1.300E-05 | 2.646E-06 | 1.000E-05 | 1.500E-05 |
| 22885 | 1145 | Y1145C | D | 9.3e-06 | 9.7e-06 | 1.4e-05 | 1.100E-05 | 2.606E-06 | 9.300E-06 | 1.400E-05 |
| 22886 | 1145 | Y1145E | D | 9.8e-06 | 9.1e-06 | 9.6e-06 | 9.500E-06 | 3.606E-07 | 9.100E-06 | 9.800E-06 |
| 22887 | 1145 | Y1145Q | D | 9.4e-06 | 1.5e-05 | 1.6e-05 | 1.347E-05 | 3.557E-06 | 9.400E-06 | 1.600E-05 |
| 22888 | 1145 | Y1145G | D | 1.2e-05 | 1.4e-05 | 1.1e-05 | 1.233E-05 | 1.528E-06 | 1.100E-05 | 1.400E-05 |
| 22889 | 1145 | Y1145H | D | 1.3e-05 | 9.2e-06 | 1.5e-05 | 1.240E-05 | 2.946E-06 | 9.200E-06 | 1.500E-05 |
| 22890 | 1145 | Y1145I | D | 1.4e-05 | 1.6e-05 | 1e-05 | 1.333E-05 | 3.055E-06 | 1.000E-05 | 1.600E-05 |
| 22891 | 1145 | Y1145L | D | 1e-05 | 9.3e-06 | 1.5e-05 | 1.143E-05 | 3.109E-06 | 9.300E-06 | 1.500E-05 |
| 22892 | 1145 | Y1145K | D | 7.3e-06 | 1.5e-05 | 1.2e-05 | 1.143E-05 | 3.881E-06 | 7.300E-06 | 1.500E-05 |
| 22893 | 1145 | Y1145M | D | 9e-06 | 1.4e-05 | 1.7e-05 | 1.333E-05 | 4.041E-06 | 9.000E-06 | 1.700E-05 |
| 22894 | 1145 | Y1145F | D | 1.1e-05 | 1.3e-05 | 8e-06 | 1.067E-05 | 2.517E-06 | 8.000E-06 | 1.300E-05 |
| 22895 | 1145 | Y1145P | D | 9.8e-06 | 1.1e-05 | 1.3e-05 | 1.127E-05 | 1.617E-06 | 9.800E-06 | 1.300E-05 |
| 22896 | 1145 | Y1145S | D | 1.1e-05 | 8.1e-06 | 1.1e-05 | 1.003E-05 | 1.674E-06 | 8.100E-06 | 1.100E-05 |
| 22897 | 1145 | Y1145T | D | 1e-05 | 8.9e-06 | 9.6e-06 | 9.500E-06 | 5.568E-07 | 8.900E-06 | 1.000E-05 |
| 22898 | 1145 | Y1145W | D | 1.2e-05 | 1.1e-05 | 1.5e-05 | 1.267E-05 | 2.082E-06 | 1.100E-05 | 1.500E-05 |
| 22899 | 1145 | Y1145Y | D | 8.7e-06 | 1.7e-05 | 1.4e-05 | 1.323E-05 | 4.203E-06 | 8.700E-06 | 1.700E-05 |
| 22900 | 1145 | Y1145V | D | 1.1e-05 | 1e-05 | 1.7e-05 | 1.267E-05 | 3.786E-06 | 1.000E-05 | 1.700E-05 |
| 22901 | 1146 | C1146A | D | 1.1e-05 | 1.2e-05 | 1.6e-05 | 1.300E-05 | 2.646E-06 | 1.100E-05 | 1.600E-05 |
| 22902 | 1146 | C1146R | D | 1.2e-05 | 1e-05 | 1.6e-05 | 1.267E-05 | 3.055E-06 | 1.000E-05 | 1.600E-05 |
| 22903 | 1146 | C1146N | D | 1.4e-05 | 9e-06 | 1.4e-05 | 1.233E-05 | 2.887E-06 | 9.000E-06 | 1.400E-05 |
| 22904 | 1146 | C1146D | D | 8.4e-06 | 1.1e-05 | 1.5e-05 | 1.147E-05 | 3.325E-06 | 8.400E-06 | 1.500E-05 |
| 22905 | 1146 | C1146C | D | 9.3e-06 | 1.2e-05 | 1.6e-05 | 1.243E-05 | 3.371E-06 | 9.300E-06 | 1.600E-05 |
| 22906 | 1146 | C1146E | D | 1.6e-05 | 1.1e-05 | 1.7e-05 | 1.467E-05 | 3.215E-06 | 1.100E-05 | 1.700E-05 |
| 22907 | 1146 | C1146Q | D | 1.1e-05 | 1.6e-05 | 9.1e-06 | 1.203E-05 | 3.564E-06 | 9.100E-06 | 1.600E-05 |
| 22908 | 1146 | C1146G | D | 9.7e-06 | 1e-05 | 1e-05 | 9.900E-06 | 1.732E-07 | 9.700E-06 | 1.000E-05 |
| 22909 | 1146 | C1146H | D | 7.2e-06 | 1.3e-05 | 1e-05 | 1.007E-05 | 2.901E-06 | 7.200E-06 | 1.300E-05 |
| 22910 | 1146 | C1146I | D | 1.2e-05 | 1.4e-05 | 1e-05 | 1.200E-05 | 2.000E-06 | 1.000E-05 | 1.400E-05 |
| 22911 | 1146 | C1146L | D | 1.1e-05 | 1.4e-05 | 9.9e-06 | 1.163E-05 | 2.122E-06 | 9.900E-06 | 1.400E-05 |
| 22912 | 1146 | C1146K | D | 1.1e-05 | 9.6e-06 | 1e-05 | 1.020E-05 | 7.211E-07 | 9.600E-06 | 1.100E-05 |
| 22913 | 1146 | C1146M | D | 1.3e-05 | 1.2e-05 | 9.3e-06 | 1.143E-05 | 1.914E-06 | 9.300E-06 | 1.300E-05 |
| 22914 | 1146 | C1146F | D | 1.5e-05 | 1.6e-05 | 1.6e-05 | 1.567E-05 | 5.774E-07 | 1.500E-05 | 1.600E-05 |
| 22915 | 1146 | C1146P | D | 1.4e-05 | 1.4e-05 | 1e-05 | 1.267E-05 | 2.309E-06 | 1.000E-05 | 1.400E-05 |
| 22916 | 1146 | C1146S | D | 1.2e-05 | 8.8e-06 | 1.4e-05 | 1.160E-05 | 2.623E-06 | 8.800E-06 | 1.400E-05 |
| 22917 | 1146 | C1146T | D | 1.3e-05 | 1.6e-05 | 1.2e-05 | 1.367E-05 | 2.082E-06 | 1.200E-05 | 1.600E-05 |
| 22918 | 1146 | C1146W | D | 8.3e-06 | 1.5e-05 | 1.5e-05 | 1.277E-05 | 3.868E-06 | 8.300E-06 | 1.500E-05 |
| 22919 | 1146 | C1146Y | D | 1.1e-05 | 1.2e-05 | 1.2e-05 | 1.167E-05 | 5.774E-07 | 1.100E-05 | 1.200E-05 |
| 22920 | 1146 | C1146V | D | 1.5e-05 | 1.2e-05 | 1e-05 | 1.233E-05 | 2.517E-06 | 1.000E-05 | 1.500E-05 |
| 22921 | 1147 | Q1147A | D | 1.4e-05 | 9e-06 | 1.7e-05 | 1.333E-05 | 4.041E-06 | 9.000E-06 | 1.700E-05 |
| 22922 | 1147 | Q1147R | D | 1.5e-05 | 1.7e-05 | 1.1e-05 | 1.433E-05 | 3.055E-06 | 1.100E-05 | 1.700E-05 |
| 22923 | 1147 | Q1147N | D | 1.3e-05 | 9.2e-06 | 1.7e-05 | 1.307E-05 | 3.900E-06 | 9.200E-06 | 1.700E-05 |
| 22924 | 1147 | Q1147D | D | 1.3e-05 | 1.1e-05 | 1.5e-05 | 1.300E-05 | 2.000E-06 | 1.100E-05 | 1.500E-05 |
| 22925 | 1147 | Q1147C | D | 1.5e-05 | 1.4e-05 | 1e-05 | 1.300E-05 | 2.646E-06 | 1.000E-05 | 1.500E-05 |
| 22926 | 1147 | Q1147E | D | 1.2e-05 | 9.8e-06 | 1.3e-05 | 1.160E-05 | 1.637E-06 | 9.800E-06 | 1.300E-05 |
| 22927 | 1147 | Q1147Q | D | 1.6e-05 | 8.5e-06 | 1.1e-05 | 1.183E-05 | 3.819E-06 | 8.500E-06 | 1.600E-05 |
| 22928 | 1147 | Q1147G | D | 1.7e-05 | 1.1e-05 | 1.2e-05 | 1.333E-05 | 3.215E-06 | 1.100E-05 | 1.700E-05 |
| 22929 | 1147 | Q1147H | D | 8.8e-06 | 1.3e-05 | 1.2e-05 | 1.127E-05 | 2.194E-06 | 8.800E-06 | 1.300E-05 |
| 22930 | 1147 | Q1147I | D | 1.5e-05 | 1.1e-05 | 1.4e-05 | 1.333E-05 | 2.082E-06 | 1.100E-05 | 1.500E-05 |
| 22931 | 1147 | Q1147L | D | 1.5e-05 | 1.2e-05 | 1.6e-05 | 1.433E-05 | 2.082E-06 | 1.200E-05 | 1.600E-05 |
| 22932 | 1147 | Q1147K | D | 8.9e-06 | 1.5e-05 | 1.5e-05 | 1.297E-05 | 3.522E-06 | 8.900E-06 | 1.500E-05 |
| 22933 | 1147 | Q1147M | D | 1.1e-05 | 1.5e-05 | 1.6e-05 | 1.400E-05 | 2.646E-06 | 1.100E-05 | 1.600E-05 |
| 22934 | 1147 | Q1147F | D | 1e-05 | 1e-05 | 1.5e-05 | 1.167E-05 | 2.887E-06 | 1.000E-05 | 1.500E-05 |
| 22935 | 1147 | Q1147P | D | 9.3e-06 | 1.3e-05 | 1.5e-05 | 1.243E-05 | 2.892E-06 | 9.300E-06 | 1.500E-05 |
| 22936 | 1147 | Q1147S | D | 1.5e-05 | 1.4e-05 | 1.5e-05 | 1.467E-05 | 5.774E-07 | 1.400E-05 | 1.500E-05 |
| 22937 | 1147 | Q1147T | D | 2.6e-05 | 1.2e-05 | 9.6e-06 | 1.587E-05 | 8.857E-06 | 9.600E-06 | 2.600E-05 |
| 22938 | 1147 | Q1147W | D | 1.1e-05 | 8.9e-06 | 1.6e-05 | 1.197E-05 | 3.647E-06 | 8.900E-06 | 1.600E-05 |
| 22939 | 1147 | Q1147Y | D | 8.8e-06 | 1e-05 | 8.1e-06 | 8.967E-06 | 9.609E-07 | 8.100E-06 | 1.000E-05 |
| 22940 | 1147 | Q1147V | D | 1.7e-05 | 1.3e-05 | 1e-05 | 1.333E-05 | 3.512E-06 | 1.000E-05 | 1.700E-05 |
| 22941 | 1148 | Q1148A | D | 1e-05 | 7.7e-06 | 2.8e-05 | 1.523E-05 | 1.112E-05 | 7.700E-06 | 2.800E-05 |
| 22942 | 1148 | Q1148R | D | 1.3e-05 | 1.2e-05 | 1.1e-05 | 1.200E-05 | 1.000E-06 | 1.100E-05 | 1.300E-05 |
| 22943 | 1148 | Q1148N | D | 1.5e-05 | 1.9e-05 | 1.5e-05 | 1.633E-05 | 2.309E-06 | 1.500E-05 | 1.900E-05 |
| 22944 | 1148 | Q1148D | D | 8.8e-06 | 1.6e-05 | 1.6e-05 | 1.360E-05 | 4.157E-06 | 8.800E-06 | 1.600E-05 |
| 22945 | 1148 | Q1148C | D | 1.4e-05 | 1.4e-05 | 1.1e-05 | 1.300E-05 | 1.732E-06 | 1.100E-05 | 1.400E-05 |
| 22946 | 1148 | Q1148E | D | 1.1e-05 | 1.1e-05 | 9.8e-06 | 1.060E-05 | 6.928E-07 | 9.800E-06 | 1.100E-05 |
| 22947 | 1148 | Q1148Q | D | 8.7e-06 | 8.7e-06 | 1.5e-05 | 1.080E-05 | 3.637E-06 | 8.700E-06 | 1.500E-05 |
| 22948 | 1148 | Q1148G | D | 1.5e-05 | 1.5e-05 | 1.1e-05 | 1.367E-05 | 2.309E-06 | 1.100E-05 | 1.500E-05 |
| 22949 | 1148 | Q1148H | D | 9.7e-06 | 1.2e-05 | 1.3e-05 | 1.157E-05 | 1.692E-06 | 9.700E-06 | 1.300E-05 |

| | | | | | | | | | | |
|---|---|---|---|---|---|---|---|---|---|---|
| 22950 | 1148 | Q1148I | D | 1.5e-05 | 9.2e-06 | 1.5e-05 | 1.307E-05 | 3.349E-06 | 9.200E-06 | 1.500E-05 |
| 22951 | 1148 | Q1148L | D | 1.6e-05 | 9.9e-06 | 1.4e-05 | 1.330E-05 | 3.110E-06 | 9.900E-06 | 1.600E-05 |
| 22952 | 1148 | Q1148K | D | 9.3e-06 | 8.8e-06 | 9.3e-06 | 9.133E-06 | 2.887E-07 | 8.800E-06 | 9.300E-06 |
| 22953 | 1148 | Q1148M | D | 1e-05 | 9.4e-06 | 1.6e-05 | 1.180E-05 | 3.650E-06 | 9.400E-06 | 1.600E-05 |
| 22954 | 1148 | Q1148F | D | 1.5e-05 | 1.3e-05 | 1e-05 | 1.267E-05 | 2.517E-06 | 1.000E-05 | 1.500E-05 |
| 22955 | 1148 | Q1148P | D | 9.8e-06 | 1.6e-05 | 9.7e-06 | 1.183E-05 | 3.609E-06 | 9.700E-06 | 1.600E-05 |
| 22956 | 1148 | Q1148S | D | 1.2e-05 | 1.5e-05 | 1.2e-05 | 1.300E-05 | 1.732E-06 | 1.200E-05 | 1.500E-05 |
| 22957 | 1148 | Q1148T | D | 1.4e-05 | 1.1e-05 | 9.7e-06 | 1.157E-05 | 2.205E-06 | 9.700E-06 | 1.400E-05 |
| 22958 | 1148 | Q1148W | D | 1e-05 | 1.1e-05 | 1.4e-05 | 1.167E-05 | 2.082E-06 | 1.000E-05 | 1.400E-05 |
| 22959 | 1148 | Q1148Y | D | 8.5e-06 | 1.4e-05 | 1.2e-05 | 1.150E-05 | 2.784E-06 | 8.500E-06 | 1.400E-05 |
| 22960 | 1148 | Q1148V | D | 1e-05 | 1.5e-05 | 1.1e-05 | 1.200E-05 | 2.646E-06 | 1.000E-05 | 1.500E-05 |
| 22961 | 1149 | H1149A | D | 1.6e-05 | 1.1e-05 | 1.4e-05 | 1.367E-05 | 2.517E-06 | 1.100E-05 | 1.600E-05 |
| 22962 | 1149 | H1149R | D | 1.5e-05 | 1.6e-05 | 6.5e-06 | 1.250E-05 | 5.220E-06 | 6.500E-06 | 1.600E-05 |
| 22963 | 1149 | H1149N | D | 1.7e-05 | 1.5e-05 | 1.1e-05 | 1.433E-05 | 3.055E-06 | 1.100E-05 | 1.700E-05 |
| 22964 | 1149 | H1149D | D | 1.1e-05 | 9.8e-06 | 1.6e-05 | 1.227E-05 | 3.288E-06 | 9.800E-06 | 1.600E-05 |
| 22965 | 1149 | H1149C | D | 1e-05 | 1.5e-05 | 1.3e-05 | 1.267E-05 | 2.517E-06 | 1.000E-05 | 1.500E-05 |
| 22966 | 1149 | H1149E | D | 1.6e-05 | 1.7e-05 | 1.6e-05 | 1.633E-05 | 5.774E-07 | 1.600E-05 | 1.700E-05 |
| 22967 | 1149 | H1149Q | D | 1.4e-05 | 1.3e-05 | 1.5e-05 | 1.400E-05 | 1.000E-06 | 1.300E-05 | 1.500E-05 |
| 22968 | 1149 | H1149G | D | 9.7e-06 | 1.5e-05 | 1e-05 | 1.157E-05 | 2.977E-06 | 9.700E-06 | 1.500E-05 |
| 22969 | 1149 | H1149H | D | 2.3e-05 | 2.4e-05 | 1.6e-05 | 2.100E-05 | 4.359E-06 | 1.600E-05 | 2.400E-05 |
| 22970 | 1149 | H1149I | D | 1.3e-05 | 1.1e-05 | 1.7e-05 | 1.367E-05 | 3.055E-06 | 1.100E-05 | 1.700E-05 |
| 22971 | 1149 | H1149L | D | 1.3e-05 | 1.5e-05 | 7.7e-06 | 1.190E-05 | 3.772E-06 | 7.700E-06 | 1.500E-05 |
| 22972 | 1149 | H1149K | D | 5.4e-06 | 1e-05 | 9.9e-06 | 8.433E-06 | 2.627E-06 | 5.400E-06 | 1.000E-05 |
| 22973 | 1149 | H1149M | D | 1.4e-05 | 1.1e-05 | 1.6e-05 | 1.367E-05 | 2.517E-06 | 1.100E-05 | 1.600E-05 |
| 22974 | 1149 | H1149F | D | 1.6e-05 | 1.3e-05 | 1.2e-05 | 1.367E-05 | 2.082E-06 | 1.200E-05 | 1.600E-05 |
| 22975 | 1149 | H1149P | D | 1.7e-05 | 8.5e-06 | 1.6e-05 | 1.383E-05 | 4.646E-06 | 8.500E-06 | 1.700E-05 |
| 22976 | 1149 | H1149S | D | 1.5e-05 | 1.6e-05 | 1.5e-05 | 1.533E-05 | 5.774E-07 | 1.500E-05 | 1.600E-05 |
| 22977 | 1149 | H1149T | D | 1e-05 | 1.5e-05 | 1.3e-05 | 1.267E-05 | 2.517E-06 | 1.000E-05 | 1.500E-05 |
| 22978 | 1149 | H1149W | D | 1.4e-05 | 7.1e-06 | 1.2e-05 | 1.103E-05 | 3.550E-06 | 7.100E-06 | 1.400E-05 |
| 22979 | 1149 | H1149Y | D | 1.5e-05 | 9.5e-06 | 1e-05 | 1.150E-05 | 3.041E-06 | 9.500E-06 | 1.500E-05 |
| 22980 | 1149 | H1149V | D | 1.1e-05 | 9.6e-06 | 1.7e-05 | 1.253E-05 | 3.931E-06 | 9.600E-06 | 1.700E-05 |
| 22981 | 1150 | Y1150A | D | 9.2e-06 | 1.3e-05 | 1.4e-05 | 1.207E-05 | 2.532E-06 | 9.200E-06 | 1.400E-05 |
| 22982 | 1150 | Y1150R | D | 1.4e-05 | 9.3e-06 | 1.1e-05 | 1.143E-05 | 2.380E-06 | 9.300E-06 | 1.400E-05 |
| 22983 | 1150 | Y1150N | D | 1.2e-05 | 1.6e-05 | 1.1e-05 | 1.300E-05 | 2.646E-06 | 1.100E-05 | 1.600E-05 |
| 22984 | 1150 | Y1150D | D | 1.5e-05 | 1.2e-05 | 1.5e-05 | 1.400E-05 | 1.732E-06 | 1.200E-05 | 1.500E-05 |
| 22985 | 1150 | Y1150C | D | 1.2e-05 | 1.6e-05 | 8.9e-06 | 1.230E-05 | 3.559E-06 | 8.900E-06 | 1.600E-05 |
| 22986 | 1150 | Y1150E | D | 2.4e-05 | 1.6e-05 | 1.5e-05 | 1.833E-05 | 4.933E-06 | 1.500E-05 | 2.400E-05 |
| 22987 | 1150 | Y1150Q | D | 9.2e-06 | 1.6e-05 | 1e-05 | 1.173E-05 | 3.717E-06 | 9.200E-06 | 1.600E-05 |
| 22988 | 1150 | Y1150G | D | 1.7e-05 | 1.6e-05 | 1.5e-05 | 1.600E-05 | 1.000E-06 | 1.500E-05 | 1.700E-05 |
| 22989 | 1150 | Y1150H | D | 9.4e-06 | 1.7e-05 | 1.5e-05 | 1.380E-05 | 3.940E-06 | 9.400E-06 | 1.700E-05 |
| 22990 | 1150 | Y1150I | D | 1.6e-05 | 1.4e-05 | 1e-05 | 1.333E-05 | 3.055E-06 | 1.000E-05 | 1.600E-05 |
| 22991 | 1150 | Y1150L | D | 1e-05 | 1.3e-05 | 1.3e-05 | 1.200E-05 | 1.732E-06 | 1.000E-05 | 1.300E-05 |
| 22992 | 1150 | Y1150K | D | 1.2e-05 | 1.6e-05 | 1.2e-05 | 1.333E-05 | 2.309E-06 | 1.200E-05 | 1.600E-05 |
| 22993 | 1150 | Y1150M | D | 1e-05 | 1.6e-05 | 1.6e-05 | 1.400E-05 | 3.464E-06 | 1.000E-05 | 1.600E-05 |
| 22994 | 1150 | Y1150F | D | 1.3e-05 | 1.6e-05 | 8.1e-06 | 1.237E-05 | 3.988E-06 | 8.100E-06 | 1.600E-05 |
| 22995 | 1150 | Y1150P | D | 1.3e-05 | 1.5e-05 | 9.9e-06 | 1.263E-05 | 2.570E-06 | 9.900E-06 | 1.500E-05 |
| 22996 | 1150 | Y1150S | D | 1.6e-05 | 1e-05 | 1.1e-05 | 1.233E-05 | 3.215E-06 | 1.000E-05 | 1.600E-05 |
| 22997 | 1150 | Y1150T | D | 1.6e-05 | 8.4e-06 | 1e-05 | 1.147E-05 | 4.007E-06 | 8.400E-06 | 1.600E-05 |
| 22998 | 1150 | Y1150W | D | 1.1e-05 | 1.6e-05 | 1.6e-05 | 1.433E-05 | 2.887E-06 | 1.100E-05 | 1.600E-05 |
| 22999 | 1150 | Y1150Y | D | 9e-06 | 1.5e-05 | 1.4e-05 | 1.267E-05 | 3.215E-06 | 9.000E-06 | 1.500E-05 |
| 23000 | 1150 | Y1150V | D | 1.4e-05 | 1.3e-05 | 6.3e-06 | 1.110E-05 | 4.187E-06 | 6.300E-06 | 1.400E-05 |
| 23001 | 1151 | T1151A | D | 1.5e-05 | 1.5e-05 | 1.1e-05 | 1.367E-05 | 2.309E-06 | 1.100E-05 | 1.500E-05 |
| 23002 | 1151 | T1151R | D | 1.4e-05 | 1.4e-05 | 1e-05 | 1.267E-05 | 2.309E-06 | 1.000E-05 | 1.400E-05 |
| 23003 | 1151 | T1151N | D | 1.4e-05 | 1e-05 | 1.2e-05 | 1.200E-05 | 2.000E-06 | 1.000E-05 | 1.400E-05 |
| 23004 | 1151 | T1151D | D | 1.5e-05 | 1.5e-05 | 1.1e-05 | 1.367E-05 | 2.309E-06 | 1.100E-05 | 1.500E-05 |
| 23005 | 1151 | T1151C | D | 1.4e-05 | 1.6e-05 | 9.5e-06 | 1.317E-05 | 3.329E-06 | 9.500E-06 | 1.600E-05 |
| 23006 | 1151 | T1151E | D | 1e-05 | 1.5e-05 | 8e-06 | 1.100E-05 | 3.606E-06 | 8.000E-06 | 1.500E-05 |
| 23007 | 1151 | T1151Q | D | 1.4e-05 | 1.3e-05 | 1.3e-05 | 1.333E-05 | 5.774E-07 | 1.300E-05 | 1.400E-05 |
| 23008 | 1151 | T1151G | D | 1.6e-05 | 1.5e-05 | 1.2e-05 | 1.433E-05 | 2.082E-06 | 1.200E-05 | 1.600E-05 |
| 23009 | 1151 | T1151H | D | 1.4e-05 | 1.4e-05 | 1e-05 | 1.267E-05 | 2.309E-06 | 1.000E-05 | 1.400E-05 |
| 23010 | 1151 | T1151I | D | 1.3e-05 | 1.6e-05 | 1.6e-05 | 1.500E-05 | 1.732E-06 | 1.300E-05 | 1.600E-05 |
| 23011 | 1151 | T1151L | D | 1.1e-05 | 1.1e-05 | 1.3e-05 | 1.167E-05 | 1.155E-06 | 1.100E-05 | 1.300E-05 |
| 23012 | 1151 | T1151K | D | 1.6e-05 | 1.1e-05 | 1.1e-05 | 1.267E-05 | 2.887E-06 | 1.100E-05 | 1.600E-05 |
| 23013 | 1151 | T1151M | D | 6e-06 | 1.2e-05 | 1.1e-05 | 9.667E-06 | 3.215E-06 | 6.000E-06 | 1.200E-05 |
| 23014 | 1151 | T1151F | D | 1.2e-05 | 1.5e-05 | 1.7e-05 | 1.467E-05 | 2.517E-06 | 1.200E-05 | 1.700E-05 |
| 23015 | 1151 | T1151P | D | 1.3e-05 | 1.3e-05 | 1.1e-05 | 1.233E-05 | 1.155E-06 | 1.100E-05 | 1.300E-05 |
| 23016 | 1151 | T1151S | D | 1.4e-05 | 1.7e-05 | 1.6e-05 | 1.567E-05 | 1.528E-06 | 1.400E-05 | 1.700E-05 |
| 23017 | 1151 | T1151T | D | 1.3e-05 | 1.5e-05 | 1.5e-05 | 1.433E-05 | 1.155E-06 | 1.300E-05 | 1.500E-05 |
| 23018 | 1151 | T1151W | D | 1.5e-05 | 1.7e-05 | 1.5e-05 | 1.567E-05 | 1.155E-06 | 1.500E-05 | 1.700E-05 |
| 23019 | 1151 | T1151Y | D | 9.8e-06 | 9.4e-06 | 1.3e-05 | 1.073E-05 | 1.973E-06 | 9.400E-06 | 1.300E-05 |
| 23020 | 1151 | T1151V | D | 1.1e-05 | 8.2e-06 | 1.4e-05 | 1.107E-05 | 2.901E-06 | 8.200E-06 | 1.400E-05 |
| 23021 | 1152 | T1152A | D | 1.7e-05 | 9.8e-06 | 1.4e-05 | 1.360E-05 | 3.617E-06 | 9.800E-06 | 1.700E-05 |
| 23022 | 1152 | T1152R | D | 1.4e-05 | 1.5e-05 | 1.4e-05 | 1.433E-05 | 5.774E-07 | 1.400E-05 | 1.500E-05 |
| 23023 | 1152 | T1152N | D | 1.2e-05 | 1.1e-05 | 9.5e-06 | 1.083E-05 | 1.258E-06 | 9.500E-06 | 1.200E-05 |
| 23024 | 1152 | T1152D | D | 1.5e-05 | 1.8e-05 | 1.1e-05 | 1.467E-05 | 3.512E-06 | 1.100E-05 | 1.800E-05 |
| 23025 | 1152 | T1152C | D | 1.2e-05 | 9.5e-06 | 1.3e-05 | 1.150E-05 | 1.803E-06 | 9.500E-06 | 1.300E-05 |
| 23026 | 1152 | T1152E | D | 1.1e-05 | 1.1e-05 | 1.6e-05 | 1.267E-05 | 2.887E-06 | 1.100E-05 | 1.600E-05 |
| 23027 | 1152 | T1152Q | D | 1.3e-05 | 1.4e-05 | 1.1e-05 | 1.267E-05 | 1.528E-06 | 1.100E-05 | 1.400E-05 |
| 23028 | 1152 | T1152G | D | 1.7e-05 | 1.1e-05 | 1.1e-05 | 1.300E-05 | 3.464E-06 | 1.100E-05 | 1.700E-05 |
| 23029 | 1152 | T1152H | D | 1.1e-05 | 1.5e-05 | 1.6e-05 | 1.400E-05 | 2.646E-06 | 1.100E-05 | 1.600E-05 |
| 23030 | 1152 | T1152I | D | 1.1e-05 | 9e-06 | 1.3e-05 | 1.100E-05 | 2.000E-06 | 9.000E-06 | 1.300E-05 |
| 23031 | 1152 | T1152L | D | 7.7e-06 | 1.1e-05 | 9.5e-06 | 9.400E-06 | 1.652E-06 | 7.700E-06 | 1.100E-05 |
| 23032 | 1152 | T1152K | D | 1.6e-05 | 1.4e-05 | 1.4e-05 | 1.467E-05 | 1.155E-06 | 1.400E-05 | 1.600E-05 |
| 23033 | 1152 | T1152M | D | 1.5e-05 | 1.5e-05 | 7.6e-06 | 1.253E-05 | 4.272E-06 | 7.600E-06 | 1.500E-05 |
| 23034 | 1152 | T1152F | D | 1.1e-05 | 1.4e-05 | 9.5e-06 | 1.150E-05 | 2.291E-06 | 9.500E-06 | 1.400E-05 |

| | | | | | | | | | | |
|---|---|---|---|---|---|---|---|---|---|---|
| 23035 | 1152 | T1152P | D | 1.3e-05 | 9.2e-06 | 1.1e-05 | 1.107E-05 | 1.901E-06 | 9.200E-06 | 1.300E-05 |
| 23036 | 1152 | T1152S | D | 9.8e-06 | 1.1e-05 | 1.1e-05 | 1.060E-05 | 6.928E-07 | 9.800E-06 | 1.100E-05 |
| 23037 | 1152 | T1152T | D | 1.5e-05 | 1.1e-05 | 1.4e-05 | 1.333E-05 | 2.082E-06 | 1.100E-05 | 1.500E-05 |
| 23038 | 1152 | T1152W | D | 1.3e-05 | 1.1e-05 | 1.4e-05 | 1.267E-05 | 1.528E-06 | 1.100E-05 | 1.400E-05 |
| 23039 | 1152 | T1152Y | D | 1.5e-05 | 1e-05 | 1.2e-05 | 1.233E-05 | 2.517E-06 | 1.000E-05 | 1.500E-05 |
| 23040 | 1152 | T1152V | D | 1.1e-05 | 1.5e-05 | 1.2e-05 | 1.267E-05 | 2.082E-06 | 1.100E-05 | 1.500E-05 |
| 23041 | 1153 | P1153A | D | 1.3e-05 | 1.2e-05 | 1.1e-05 | 1.200E-05 | 1.000E-06 | 1.100E-05 | 1.300E-05 |
| 23042 | 1153 | P1153R | D | 1.6e-05 | 1.4e-05 | 1.4e-05 | 1.467E-05 | 1.155E-06 | 1.400E-05 | 1.600E-05 |
| 23043 | 1153 | P1153N | D | 1.5e-05 | 1e-05 | 1.6e-05 | 1.367E-05 | 3.215E-06 | 1.000E-05 | 1.600E-05 |
| 23044 | 1153 | P1153D | D | 1.5e-05 | 1.1e-05 | 9.9e-06 | 1.197E-05 | 2.684E-06 | 9.900E-06 | 1.500E-05 |
| 23045 | 1153 | P1153C | D | 1.3e-05 | 1.3e-05 | 1.1e-05 | 1.233E-05 | 1.155E-06 | 1.100E-05 | 1.300E-05 |
| 23046 | 1153 | P1153E | D | 1.1e-05 | 1.6e-05 | 1.5e-05 | 1.400E-05 | 2.646E-06 | 1.100E-05 | 1.600E-05 |
| 23047 | 1153 | P1153Q | D | 1.3e-05 | 1.5e-05 | 1.1e-05 | 1.300E-05 | 2.000E-06 | 1.100E-05 | 1.500E-05 |
| 23048 | 1153 | P1153G | D | 1.4e-05 | 9.8e-06 | 9.3e-06 | 1.103E-05 | 2.581E-06 | 9.300E-06 | 1.400E-05 |
| 23049 | 1153 | P1153H | D | 9.8e-06 | 1.1e-05 | 1.2e-05 | 1.093E-05 | 1.102E-06 | 9.800E-06 | 1.200E-05 |
| 23050 | 1153 | P1153I | D | 1.3e-05 | 1.7e-05 | 1e-05 | 1.333E-05 | 3.512E-06 | 1.000E-05 | 1.700E-05 |
| 23051 | 1153 | P1153L | D | 1.5e-05 | 1.6e-05 | 1.3e-05 | 1.467E-05 | 1.528E-06 | 1.300E-05 | 1.600E-05 |
| 23052 | 1153 | P1153K | D | 1.4e-05 | 1.5e-05 | 1.6e-05 | 1.500E-05 | 1.000E-06 | 1.400E-05 | 1.600E-05 |
| 23053 | 1153 | P1153M | D | 1.7e-05 | 1.4e-05 | 1.4e-05 | 1.500E-05 | 1.732E-06 | 1.400E-05 | 1.700E-05 |
| 23054 | 1153 | P1153F | D | 1.5e-05 | 1.5e-05 | 1.5e-05 | 1.500E-05 | 0.000E+00 | 1.500E-05 | 1.500E-05 |
| 23055 | 1153 | P1153P | D | 1.6e-05 | 1.4e-05 | 1.5e-05 | 1.500E-05 | 1.000E-06 | 1.400E-05 | 1.600E-05 |
| 23056 | 1153 | P1153S | D | 9.2e-06 | 1.4e-05 | 1.3e-05 | 1.207E-05 | 2.532E-06 | 9.200E-06 | 1.400E-05 |
| 23057 | 1153 | P1153T | D | 1.2e-05 | 8.4e-06 | 1.6e-05 | 1.213E-05 | 3.802E-06 | 8.400E-06 | 1.600E-05 |
| 23058 | 1153 | P1153W | D | 1.4e-05 | 1e-05 | 9.3e-06 | 1.110E-05 | 2.536E-06 | 9.300E-06 | 1.400E-05 |
| 23059 | 1153 | P1153Y | D | 1e-05 | 1.4e-05 | 9e-06 | 1.100E-05 | 2.646E-06 | 9.000E-06 | 1.400E-05 |
| 23060 | 1153 | P1153V | D | 1.3e-05 | 1.1e-05 | 1.5e-05 | 1.300E-05 | 2.000E-06 | 1.100E-05 | 1.500E-05 |
| 23061 | 1154 | P1154A | D | 1.3e-05 | 1e-05 | 1.2e-05 | 1.167E-05 | 1.528E-06 | 1.000E-05 | 1.300E-05 |
| 23062 | 1154 | P1154R | D | 1e-05 | 1.5e-05 | 1e-05 | 1.167E-05 | 2.887E-06 | 1.000E-05 | 1.500E-05 |
| 23063 | 1154 | P1154N | D | 1.1e-05 | 1.1e-05 | 8.2e-06 | 1.007E-05 | 1.617E-06 | 8.200E-06 | 1.100E-05 |
| 23064 | 1154 | P1154D | D | 9.5e-06 | 1.5e-05 | 1e-05 | 1.150E-05 | 3.041E-06 | 9.500E-06 | 1.500E-05 |
| 23065 | 1154 | P1154C | D | 1.7e-05 | 9.3e-06 | 1.5e-05 | 1.377E-05 | 3.995E-06 | 9.300E-06 | 1.700E-05 |
| 23066 | 1154 | P1154E | D | 8.9e-06 | 1.5e-05 | 1.6e-05 | 1.330E-05 | 3.843E-06 | 8.900E-06 | 1.600E-05 |
| 23067 | 1154 | P1154Q | D | 1.2e-05 | 1.3e-05 | 1.1e-05 | 1.200E-05 | 1.000E-06 | 1.100E-05 | 1.300E-05 |
| 23068 | 1154 | P1154G | D | 1.2e-05 | 1.5e-05 | 8.4e-06 | 1.180E-05 | 3.305E-06 | 8.400E-06 | 1.500E-05 |
| 23069 | 1154 | P1154H | D | 1.4e-05 | 3.4e-05 | 1.4e-05 | 2.067E-05 | 1.155E-05 | 1.400E-05 | 3.400E-05 |
| 23070 | 1154 | P1154I | D | 1.4e-05 | 1.1e-05 | 1.3e-05 | 1.267E-05 | 1.528E-06 | 1.100E-05 | 1.400E-05 |
| 23071 | 1154 | P1154L | D | 8.7e-06 | 9.6e-06 | 1.7e-05 | 1.177E-05 | 4.554E-06 | 8.700E-06 | 1.700E-05 |
| 23072 | 1154 | P1154K | D | 1.1e-05 | 9.2e-06 | 1.3e-05 | 1.107E-05 | 1.901E-06 | 9.200E-06 | 1.300E-05 |
| 23073 | 1154 | P1154M | D | 1.6e-05 | 1.4e-05 | 9.5e-06 | 1.317E-05 | 3.329E-06 | 9.500E-06 | 1.600E-05 |
| 23074 | 1154 | P1154F | D | 1.1e-05 | 1.1e-05 | 1.1e-05 | 1.100E-05 | 2.075E-21 | 1.100E-05 | 1.100E-05 |
| 23075 | 1154 | P1154P | D | 1e-05 | 7.5e-06 | 8.5e-06 | 8.667E-06 | 1.258E-06 | 7.500E-06 | 1.000E-05 |
| 23076 | 1154 | P1154S | D | 7.8e-06 | 1.4e-05 | 9.9e-06 | 1.057E-05 | 3.153E-06 | 7.800E-06 | 1.400E-05 |
| 23077 | 1154 | P1154T | D | 1.1e-05 | 1.1e-05 | 1.5e-05 | 1.233E-05 | 2.309E-06 | 1.100E-05 | 1.500E-05 |
| 23078 | 1154 | P1154W | D | 9.4e-06 | 1.6e-05 | 1.5e-05 | 1.347E-05 | 3.557E-06 | 9.400E-06 | 1.600E-05 |
| 23079 | 1154 | P1154Y | D | 1e-05 | 2.5e-05 | 1e-05 | 1.500E-05 | 8.660E-06 | 1.000E-05 | 2.500E-05 |
| 23080 | 1154 | P1154V | D | 1e-05 | 9.5e-06 | 1.2e-05 | 1.050E-05 | 1.323E-06 | 9.500E-06 | 1.200E-05 |
| 23081 | 1155 | T1155A | D | 8.3e-06 | 1.7e-05 | 1e-05 | 1.177E-05 | 4.611E-06 | 8.300E-06 | 1.700E-05 |
| 23082 | 1155 | T1155R | D | 1.5e-05 | 1.6e-05 | 1.7e-05 | 1.600E-05 | 1.000E-06 | 1.500E-05 | 1.700E-05 |
| 23083 | 1155 | T1155N | D | 1.1e-05 | 1.4e-05 | 9.9e-06 | 1.163E-05 | 2.122E-06 | 9.900E-06 | 1.400E-05 |
| 23084 | 1155 | T1155D | D | 9e-06 | 1.5e-05 | 1.2e-05 | 1.200E-05 | 3.000E-06 | 9.000E-06 | 1.500E-05 |
| 23085 | 1155 | T1155C | D | 1.4e-05 | 1.1e-05 | 1.3e-05 | 1.267E-05 | 1.528E-06 | 1.100E-05 | 1.400E-05 |
| 23086 | 1155 | T1155E | D | 1e-05 | 1.4e-05 | 9.6e-06 | 1.120E-05 | 2.433E-06 | 9.600E-06 | 1.400E-05 |
| 23087 | 1155 | T1155Q | D | 1.3e-05 | 1.3e-05 | 1.2e-05 | 1.267E-05 | 5.774E-07 | 1.200E-05 | 1.300E-05 |
| 23088 | 1155 | T1155G | D | 1.7e-05 | 1.7e-05 | 1.2e-05 | 1.533E-05 | 2.887E-06 | 1.200E-05 | 1.700E-05 |
| 23089 | 1155 | T1155H | D | 3e-05 | 1.1e-05 | 1.6e-05 | 1.900E-05 | 9.849E-06 | 1.100E-05 | 3.000E-05 |
| 23090 | 1155 | T1155I | D | 1e-05 | 1.1e-05 | 5.5e-06 | 8.833E-06 | 2.930E-06 | 5.500E-06 | 1.100E-05 |
| 23091 | 1155 | T1155L | D | 1.3e-05 | 1.3e-05 | 1.6e-05 | 1.400E-05 | 1.732E-06 | 1.300E-05 | 1.600E-05 |
| 23092 | 1155 | T1155K | D | 8.6e-06 | 1.5e-05 | 1.5e-05 | 1.287E-05 | 3.695E-06 | 8.600E-06 | 1.500E-05 |
| 23093 | 1155 | T1155M | D | 1.6e-05 | 1.4e-05 | 1.6e-05 | 1.533E-05 | 1.155E-06 | 1.400E-05 | 1.600E-05 |
| 23094 | 1155 | T1155F | D | 1.1e-05 | 1.5e-05 | 9e-06 | 1.167E-05 | 3.055E-06 | 9.000E-06 | 1.500E-05 |
| 23095 | 1155 | T1155P | D | 1.1e-05 | 1.4e-05 | 1e-05 | 1.167E-05 | 2.082E-06 | 1.000E-05 | 1.400E-05 |
| 23096 | 1155 | T1155S | D | 1e-05 | 1.6e-05 | 8.6e-06 | 1.153E-05 | 3.931E-06 | 8.600E-06 | 1.600E-05 |
| 23097 | 1155 | T1155T | D | 1.5e-05 | 7.4e-06 | 1.3e-05 | 1.180E-05 | 3.940E-06 | 7.400E-06 | 1.500E-05 |
| 23098 | 1155 | T1155W | D | 1.2e-05 | 1.1e-05 | 1.5e-05 | 1.267E-05 | 2.082E-06 | 1.100E-05 | 1.500E-05 |
| 23099 | 1155 | T1155Y | D | 1.2e-05 | 1.2e-05 | 1e-05 | 1.133E-05 | 1.155E-06 | 1.000E-05 | 1.200E-05 |
| 23100 | 1155 | T1155V | D | 1.5e-05 | 1.2e-05 | 1.7e-05 | 1.467E-05 | 2.517E-06 | 1.200E-05 | 1.700E-05 |
| 23101 | 1156 | F1156A | D | 1.2e-05 | 1.1e-05 | 7.1e-06 | 1.003E-05 | 2.589E-06 | 7.100E-06 | 1.200E-05 |
| 23102 | 1156 | F1156R | D | 1.1e-05 | 1.6e-05 | 1.4e-05 | 1.367E-05 | 2.517E-06 | 1.100E-05 | 1.600E-05 |
| 23103 | 1156 | F1156N | D | 1.3e-05 | 1.3e-05 | 1e-05 | 1.200E-05 | 1.732E-06 | 1.000E-05 | 1.300E-05 |
| 23104 | 1156 | F1156D | D | 1e-05 | 1.6e-05 | 8.5e-06 | 1.150E-05 | 3.969E-06 | 8.500E-06 | 1.600E-05 |
| 23105 | 1156 | F1156C | D | 1.1e-05 | 1.4e-05 | 1.2e-05 | 1.233E-05 | 1.528E-06 | 1.100E-05 | 1.400E-05 |
| 23106 | 1156 | F1156E | D | 1.2e-05 | 1.6e-05 | 8.4e-06 | 1.213E-05 | 3.802E-06 | 8.400E-06 | 1.600E-05 |
| 23107 | 1156 | F1156Q | D | 1e-05 | 9e-06 | 1.3e-05 | 1.067E-05 | 2.082E-06 | 9.000E-06 | 1.300E-05 |
| 23108 | 1156 | F1156G | D | 1.2e-05 | 1.5e-05 | 1.4e-05 | 1.367E-05 | 1.528E-06 | 1.200E-05 | 1.500E-05 |
| 23109 | 1156 | F1156H | D | 1.7e-05 | 1.1e-05 | 1.5e-05 | 1.433E-05 | 3.055E-06 | 1.100E-05 | 1.700E-05 |
| 23110 | 1156 | F1156I | D | 1.6e-05 | 1.4e-05 | 1.5e-05 | 1.500E-05 | 1.000E-06 | 1.400E-05 | 1.600E-05 |
| 23111 | 1156 | F1156L | D | 7.5e-06 | 1.3e-05 | 1e-05 | 1.017E-05 | 2.754E-06 | 7.500E-06 | 1.300E-05 |
| 23112 | 1156 | F1156K | D | 1.4e-05 | 1e-05 | 9.7e-06 | 1.123E-05 | 2.401E-06 | 9.700E-06 | 1.400E-05 |
| 23113 | 1156 | F1156M | D | 1.6e-05 | 1.2e-05 | 7.9e-06 | 1.197E-05 | 4.050E-06 | 7.900E-06 | 1.600E-05 |
| 23114 | 1156 | F1156F | D | 1.4e-05 | 1.5e-05 | 1.4e-05 | 1.433E-05 | 5.774E-07 | 1.400E-05 | 1.500E-05 |
| 23115 | 1156 | F1156P | D | 7.1e-06 | 1.5e-05 | 1.7e-05 | 1.303E-05 | 5.235E-06 | 7.100E-06 | 1.700E-05 |
| 23116 | 1156 | F1156S | D | 8.8e-06 | 1.5e-05 | 1.2e-05 | 1.193E-05 | 3.101E-06 | 8.800E-06 | 1.500E-05 |
| 23117 | 1156 | F1156T | D | 1.4e-05 | 1.3e-05 | 8.5e-06 | 1.183E-05 | 2.930E-06 | 8.500E-06 | 1.400E-05 |
| 23118 | 1156 | F1156W | D | 9.1e-06 | 1.7e-05 | 9.6e-06 | 1.190E-05 | 4.424E-06 | 9.100E-06 | 1.700E-05 |
| 23119 | 1156 | F1156Y | D | 1.5e-05 | 1.5e-05 | 1.1e-05 | 1.367E-05 | 2.309E-06 | 1.100E-05 | 1.500E-05 |

| | | | | | | | | | | |
|---|---|---|---|---|---|---|---|---|---|---|
| 23120 | 1156 | F1156V | D | 1.5e-05 | 1.5e-05 | 1.5e-05 | 1.500E-05 | 0.000E+00 | 1.500E-05 | 1.500E-05 |
| 23121 | 1157 | G1157A | D | 1.7e-05 | 1.3e-05 | 1.4e-05 | 1.467E-05 | 2.082E-06 | 1.300E-05 | 1.700E-05 |
| 23122 | 1157 | G1157R | D | 9.9e-06 | 9.5e-06 | 9.6e-06 | 9.667E-06 | 2.082E-07 | 9.500E-06 | 9.900E-06 |
| 23123 | 1157 | G1157N | D | 1.2e-05 | 1.6e-05 | 1.2e-05 | 1.333E-05 | 2.309E-06 | 1.200E-05 | 1.600E-05 |
| 23124 | 1157 | G1157D | D | 1.6e-05 | 9.9e-06 | 1.2e-05 | 1.263E-05 | 3.099E-06 | 9.900E-06 | 1.600E-05 |
| 23125 | 1157 | G1157C | D | 1.4e-05 | 1.5e-05 | 1.7e-05 | 1.533E-05 | 1.528E-06 | 1.400E-05 | 1.700E-05 |
| 23126 | 1157 | G1157E | D | 9.8e-06 | 7.6e-06 | 1.6e-05 | 1.113E-05 | 4.356E-06 | 7.600E-06 | 1.600E-05 |
| 23127 | 1157 | G1157Q | D | 1.1e-05 | 1.3e-05 | 1e-05 | 1.133E-05 | 1.528E-06 | 1.000E-05 | 1.300E-05 |
| 23128 | 1157 | G1157G | D | 1.1e-05 | 1.3e-05 | 1e-05 | 1.133E-05 | 1.528E-06 | 1.000E-05 | 1.300E-05 |
| 23129 | 1157 | G1157H | D | 1.4e-05 | 1.5e-05 | 1.1e-05 | 1.333E-05 | 2.082E-06 | 1.100E-05 | 1.500E-05 |
| 23130 | 1157 | G1157I | D | 1e-05 | 1.1e-05 | 1.4e-05 | 1.167E-05 | 2.082E-06 | 1.000E-05 | 1.400E-05 |
| 23131 | 1157 | G1157L | D | 1.1e-05 | 1.6e-05 | 1.4e-05 | 1.367E-05 | 2.517E-06 | 1.100E-05 | 1.600E-05 |
| 23132 | 1157 | G1157K | D | 1.4e-05 | 8.3e-06 | 1.4e-05 | 1.210E-05 | 3.291E-06 | 8.300E-06 | 1.400E-05 |
| 23133 | 1157 | G1157M | D | 1.3e-05 | 8.3e-06 | 1.5e-05 | 1.210E-05 | 3.439E-06 | 8.300E-06 | 1.500E-05 |
| 23134 | 1157 | G1157F | D | 1.5e-05 | 1e-05 | 9.7e-06 | 1.157E-05 | 2.977E-06 | 9.700E-06 | 1.500E-05 |
| 23135 | 1157 | G1157P | D | 1.5e-05 | 1.5e-05 | 1.4e-05 | 1.467E-05 | 5.774E-07 | 1.400E-05 | 1.500E-05 |
| 23136 | 1157 | G1157S | D | 2.5e-05 | 9.4e-06 | 1.6e-05 | 1.680E-05 | 7.831E-06 | 9.400E-06 | 2.500E-05 |
| 23137 | 1157 | G1157T | D | 1e-05 | 1e-05 | 8.5e-06 | 9.500E-06 | 8.660E-07 | 8.500E-06 | 1.000E-05 |
| 23138 | 1157 | G1157W | D | 1.5e-05 | 1e-05 | 1.7e-05 | 1.400E-05 | 3.606E-06 | 1.000E-05 | 1.700E-05 |
| 23139 | 1157 | G1157Y | D | 1e-05 | 1.1e-05 | 1.1e-05 | 1.067E-05 | 5.774E-07 | 1.000E-05 | 1.100E-05 |
| 23140 | 1157 | G1157V | D | 7.4e-06 | 1.3e-05 | 1.4e-05 | 1.147E-05 | 3.557E-06 | 7.400E-06 | 1.400E-05 |
| 23141 | 1158 | Q1158A | D | 8.7e-06 | 1.1e-05 | 1.9e-05 | 1.290E-05 | 5.406E-06 | 8.700E-06 | 1.900E-05 |
| 23142 | 1158 | Q1158R | D | 1.1e-05 | 1.2e-05 | 1.1e-05 | 1.133E-05 | 5.774E-07 | 1.100E-05 | 1.200E-05 |
| 23143 | 1158 | Q1158N | D | 1.5e-05 | 1.4e-05 | 8.7e-06 | 1.257E-05 | 3.386E-06 | 8.700E-06 | 1.500E-05 |
| 23144 | 1158 | Q1158D | D | 1.5e-05 | 9.6e-06 | 1.4e-05 | 1.287E-05 | 2.873E-06 | 9.600E-06 | 1.500E-05 |
| 23145 | 1158 | Q1158C | D | 1.5e-05 | 1.5e-05 | 1.3e-05 | 1.433E-05 | 1.155E-06 | 1.300E-05 | 1.500E-05 |
| 23146 | 1158 | Q1158E | D | 1.3e-05 | 1.2e-05 | 1.1e-05 | 1.200E-05 | 1.000E-06 | 1.100E-05 | 1.300E-05 |
| 23147 | 1158 | Q1158Q | D | 1.6e-05 | 1.5e-05 | 1e-05 | 1.367E-05 | 3.215E-06 | 1.000E-05 | 1.600E-05 |
| 23148 | 1158 | Q1158G | D | 1.6e-05 | 1.7e-05 | 1.5e-05 | 1.600E-05 | 1.000E-06 | 1.500E-05 | 1.700E-05 |
| 23149 | 1158 | Q1158H | D | 1.1e-05 | 9.7e-06 | 1.4e-05 | 1.157E-05 | 2.205E-06 | 9.700E-06 | 1.400E-05 |
| 23150 | 1158 | Q1158I | D | 1.6e-05 | 1.4e-05 | 1.6e-05 | 1.533E-05 | 1.155E-06 | 1.400E-05 | 1.600E-05 |
| 23151 | 1158 | Q1158L | D | 1e-05 | 1.5e-05 | 1.6e-05 | 1.367E-05 | 3.215E-06 | 1.000E-05 | 1.600E-05 |
| 23152 | 1158 | Q1158K | D | 1.3e-05 | 1.1e-05 | 1.6e-05 | 1.333E-05 | 2.517E-06 | 1.100E-05 | 1.600E-05 |
| 23153 | 1158 | Q1158M | D | 1.5e-05 | 2.4e-05 | 1.1e-05 | 1.667E-05 | 6.658E-06 | 1.100E-05 | 2.400E-05 |
| 23154 | 1158 | Q1158F | D | 1.5e-05 | 1.2e-05 | 1.4e-05 | 1.367E-05 | 1.528E-06 | 1.200E-05 | 1.500E-05 |
| 23155 | 1158 | Q1158P | D | 1.7e-05 | 1.2e-05 | 1.5e-05 | 1.467E-05 | 2.517E-06 | 1.200E-05 | 1.700E-05 |
| 23156 | 1158 | Q1158S | D | 1.2e-05 | 1.5e-05 | 1.6e-05 | 1.433E-05 | 2.082E-06 | 1.200E-05 | 1.600E-05 |
| 23157 | 1158 | Q1158T | D | 1.5e-05 | 1e-05 | 1.5e-05 | 1.333E-05 | 2.887E-06 | 1.000E-05 | 1.500E-05 |
| 23158 | 1158 | Q1158W | D | 1.3e-05 | 1.4e-05 | 1.5e-05 | 1.400E-05 | 1.000E-06 | 1.300E-05 | 1.500E-05 |
| 23159 | 1158 | Q1158Y | D | 1.4e-05 | 1.4e-05 | 9.4e-06 | 1.247E-05 | 2.656E-06 | 9.400E-06 | 1.400E-05 |
| 23160 | 1158 | Q1158V | D | 1.7e-05 | 1.1e-05 | 1e-05 | 1.267E-05 | 3.786E-06 | 1.000E-05 | 1.700E-05 |
| 23161 | 1159 | G1159A | D | 1.4e-05 | 1.4e-05 | 1.7e-05 | 1.500E-05 | 1.732E-06 | 1.400E-05 | 1.700E-05 |
| 23162 | 1159 | G1159R | D | 1.5e-05 | 1.2e-05 | 1.5e-05 | 1.400E-05 | 1.732E-06 | 1.200E-05 | 1.500E-05 |
| 23163 | 1159 | G1159N | D | 1.2e-05 | 9.7e-06 | 1.7e-05 | 1.290E-05 | 3.732E-06 | 9.700E-06 | 1.700E-05 |
| 23164 | 1159 | G1159D | D | 1.4e-05 | 1.7e-05 | 1e-05 | 1.367E-05 | 3.512E-06 | 1.000E-05 | 1.700E-05 |
| 23165 | 1159 | G1159C | D | 2.1e-05 | 1.1e-05 | 1e-05 | 1.400E-05 | 6.083E-06 | 1.000E-05 | 2.100E-05 |
| 23166 | 1159 | G1159E | D | 1e-05 | 1.2e-05 | 1.3e-05 | 1.167E-05 | 1.528E-06 | 1.000E-05 | 1.300E-05 |
| 23167 | 1159 | G1159Q | D | 1.4e-05 | 1.4e-05 | 2e-05 | 1.600E-05 | 3.464E-06 | 1.400E-05 | 2.000E-05 |
| 23168 | 1159 | G1159G | D | 1.1e-05 | 1.6e-05 | 1.6e-05 | 1.433E-05 | 2.887E-06 | 1.100E-05 | 1.600E-05 |
| 23169 | 1159 | G1159H | D | 1.5e-05 | 1e-05 | 6.2e-06 | 1.040E-05 | 4.414E-06 | 6.200E-06 | 1.500E-05 |
| 23170 | 1159 | G1159I | D | 1.4e-05 | 1.4e-05 | 1.6e-05 | 1.467E-05 | 1.155E-06 | 1.400E-05 | 1.600E-05 |
| 23171 | 1159 | G1159L | D | 1.7e-05 | 1e-05 | 1.5e-05 | 1.400E-05 | 3.606E-06 | 1.000E-05 | 1.700E-05 |
| 23172 | 1159 | G1159K | D | 1.6e-05 | 1.2e-05 | 1.1e-05 | 1.300E-05 | 2.646E-06 | 1.100E-05 | 1.600E-05 |
| 23173 | 1159 | G1159M | D | 8.5e-06 | 1.6e-05 | 1.1e-05 | 1.183E-05 | 3.819E-06 | 8.500E-06 | 1.600E-05 |
| 23174 | 1159 | G1159F | D | 1.5e-05 | 1.3e-05 | 1.2e-05 | 1.333E-05 | 1.528E-06 | 1.200E-05 | 1.500E-05 |
| 23175 | 1159 | G1159P | D | 1.5e-05 | 1.6e-05 | 1e-05 | 1.367E-05 | 3.215E-06 | 1.000E-05 | 1.600E-05 |
| 23176 | 1159 | G1159S | D | 2.4e-05 | 1.3e-05 | 7.8e-06 | 1.493E-05 | 8.271E-06 | 7.800E-06 | 2.400E-05 |
| 23177 | 1159 | G1159T | D | 1.6e-05 | 8.8e-06 | 9.2e-06 | 1.133E-05 | 4.046E-06 | 8.800E-06 | 1.600E-05 |
| 23178 | 1159 | G1159W | D | 9.4e-06 | 1.4e-05 | 1.5e-05 | 1.280E-05 | 2.987E-06 | 9.400E-06 | 1.500E-05 |
| 23179 | 1159 | G1159Y | D | 7.5e-06 | 1.5e-05 | 1.5e-05 | 1.250E-05 | 4.330E-06 | 7.500E-06 | 1.500E-05 |
| 23180 | 1159 | G1159V | D | 1.1e-05 | 8.9e-06 | 8.7e-06 | 9.533E-06 | 1.274E-06 | 8.700E-06 | 1.100E-05 |
| 23181 | 1160 | T1160A | D | 6.6e-06 | 1.6e-05 | 1.5e-05 | 1.253E-05 | 5.163E-06 | 6.600E-06 | 1.600E-05 |
| 23182 | 1160 | T1160R | D | 1.4e-05 | 1.2e-05 | 9.7e-06 | 1.190E-05 | 2.152E-06 | 9.700E-06 | 1.400E-05 |
| 23183 | 1160 | T1160N | D | 1.6e-05 | 1.6e-05 | 1.5e-05 | 1.567E-05 | 5.774E-07 | 1.500E-05 | 1.600E-05 |
| 23184 | 1160 | T1160D | D | 9.6e-06 | 1.1e-05 | 1e-05 | 1.020E-05 | 7.211E-07 | 9.600E-06 | 1.100E-05 |
| 23185 | 1160 | T1160C | D | 9.9e-06 | 1e-05 | 6.9e-06 | 8.933E-06 | 1.762E-06 | 6.900E-06 | 1.000E-05 |
| 23186 | 1160 | T1160E | D | 1.5e-05 | 1.5e-05 | 9.8e-06 | 1.327E-05 | 3.002E-06 | 9.800E-06 | 1.500E-05 |
| 23187 | 1160 | T1160Q | D | 1.3e-05 | 1.4e-05 | 1.5e-05 | 1.400E-05 | 1.000E-06 | 1.300E-05 | 1.500E-05 |
| 23188 | 1160 | T1160G | D | 1.2e-05 | 9.5e-06 | 1.3e-05 | 1.150E-05 | 1.803E-06 | 9.500E-06 | 1.300E-05 |
| 23189 | 1160 | T1160H | D | 1.6e-05 | 8.7e-06 | 1.5e-05 | 1.323E-05 | 3.958E-06 | 8.700E-06 | 1.600E-05 |
| 23190 | 1160 | T1160I | D | 1.8e-05 | 1.1e-05 | 1.5e-05 | 1.467E-05 | 3.512E-06 | 1.100E-05 | 1.800E-05 |
| 23191 | 1160 | T1160L | D | 1e-05 | 1.2e-05 | 1.6e-05 | 1.267E-05 | 3.055E-06 | 1.000E-05 | 1.600E-05 |
| 23192 | 1160 | T1160K | D | 1.6e-05 | 1e-05 | 1.1e-05 | 1.233E-05 | 3.215E-06 | 1.000E-05 | 1.600E-05 |
| 23193 | 1160 | T1160M | D | 1e-05 | 1.5e-05 | 1.6e-05 | 1.367E-05 | 3.215E-06 | 1.000E-05 | 1.600E-05 |
| 23194 | 1160 | T1160F | D | 1.1e-05 | 1.5e-05 | 1.4e-05 | 1.333E-05 | 2.082E-06 | 1.100E-05 | 1.500E-05 |
| 23195 | 1160 | T1160P | D | 1.5e-05 | 1.5e-05 | 1.5e-05 | 1.500E-05 | 0.000E+00 | 1.500E-05 | 1.500E-05 |
| 23196 | 1160 | T1160S | D | 8.7e-06 | 1.1e-05 | 7.3e-06 | 9.000E-06 | 1.868E-06 | 7.300E-06 | 1.100E-05 |
| 23197 | 1160 | T1160T | D | 1.2e-05 | 1.3e-05 | 1.1e-05 | 1.200E-05 | 1.000E-06 | 1.100E-05 | 1.300E-05 |
| 23198 | 1160 | T1160W | D | 1.8e-05 | 1.3e-05 | 1.3e-05 | 1.467E-05 | 2.887E-06 | 1.300E-05 | 1.800E-05 |
| 23199 | 1160 | T1160Y | D | 1e-05 | 9.9e-06 | 9.3e-06 | 9.733E-06 | 3.786E-07 | 9.300E-06 | 1.000E-05 |
| 23200 | 1160 | T1160V | D | 1.2e-05 | 1.8e-05 | 8.5e-06 | 1.283E-05 | 4.805E-06 | 8.500E-06 | 1.800E-05 |
| 23201 | 1161 | K1161A | D | 9.2e-06 | 1.5e-05 | 1.4e-05 | 1.273E-05 | 3.101E-06 | 9.200E-06 | 1.500E-05 |
| 23202 | 1161 | K1161R | D | 1.4e-05 | 1.3e-05 | 1.5e-05 | 1.400E-05 | 1.000E-06 | 1.300E-05 | 1.500E-05 |
| 23203 | 1161 | K1161N | D | 1e-05 | 1.4e-05 | 1.3e-05 | 1.233E-05 | 2.082E-06 | 1.000E-05 | 1.400E-05 |
| 23204 | 1161 | K1161D | D | 1.3e-05 | 1.1e-05 | 9.6e-06 | 1.120E-05 | 1.709E-06 | 9.600E-06 | 1.300E-05 |

| | | | | | | | | | | |
|---|---|---|---|---|---|---|---|---|---|---|
| 23205 | 1161 | K1161C | D | 8.7e-06 | 1.1e-05 | 1.5e-05 | 1.157E-05 | 3.188E-06 | 8.700E-06 | 1.500E-05 |
| 23206 | 1161 | K1161E | D | 1.5e-05 | 1.5e-05 | 8e-06 | 1.267E-05 | 4.041E-06 | 8.000E-06 | 1.500E-05 |
| 23207 | 1161 | K1161Q | D | 9.7e-06 | 1.4e-05 | 1.4e-05 | 1.257E-05 | 2.483E-06 | 9.700E-06 | 1.400E-05 |
| 23208 | 1161 | K1161G | D | 1.6e-05 | 1e-05 | 8.1e-06 | 1.137E-05 | 4.124E-06 | 8.100E-06 | 1.600E-05 |
| 23209 | 1161 | K1161H | D | 8.7e-06 | 1.5e-05 | 1.7e-05 | 1.357E-05 | 4.332E-06 | 8.700E-06 | 1.700E-05 |
| 23210 | 1161 | K1161I | D | 1.5e-05 | 1.3e-05 | 1.3e-05 | 1.367E-05 | 1.155E-06 | 1.300E-05 | 1.500E-05 |
| 23211 | 1161 | K1161L | D | 1.8e-05 | 1.7e-05 | 9e-06 | 1.467E-05 | 4.933E-06 | 9.000E-06 | 1.800E-05 |
| 23212 | 1161 | K1161K | D | 9.4e-06 | 1.4e-05 | 1.1e-05 | 1.147E-05 | 2.335E-06 | 9.400E-06 | 1.400E-05 |
| 23213 | 1161 | K1161M | D | 1.5e-05 | 1.6e-05 | 1.4e-05 | 1.500E-05 | 1.000E-06 | 1.400E-05 | 1.600E-05 |
| 23214 | 1161 | K1161F | D | 1.2e-05 | 9.3e-06 | 1.1e-05 | 1.077E-05 | 1.365E-06 | 9.300E-06 | 1.200E-05 |
| 23215 | 1161 | K1161P | D | 1.1e-05 | 8.8e-06 | 1.4e-05 | 1.127E-05 | 2.610E-06 | 8.800E-06 | 1.400E-05 |
| 23216 | 1161 | K1161S | D | 1.4e-05 | 9.8e-06 | 1.6e-05 | 1.327E-05 | 3.164E-06 | 9.800E-06 | 1.600E-05 |
| 23217 | 1161 | K1161T | D | 8.4e-06 | 9.8e-06 | 1.1e-05 | 9.733E-06 | 1.301E-06 | 8.400E-06 | 1.100E-05 |
| 23218 | 1161 | K1161W | D | 1.6e-05 | 1.4e-05 | 1.4e-05 | 1.467E-05 | 1.155E-06 | 1.400E-05 | 1.600E-05 |
| 23219 | 1161 | K1161Y | D | 1.2e-05 | 9.8e-06 | 1.2e-05 | 1.127E-05 | 1.270E-06 | 9.800E-06 | 1.200E-05 |
| 23220 | 1161 | K1161V | D | 1.5e-05 | 9.5e-06 | 7.5e-06 | 1.067E-05 | 3.884E-06 | 7.500E-06 | 1.500E-05 |
| 23221 | 1162 | V1162A | D | 1.3e-05 | 1.4e-05 | 1.6e-05 | 1.433E-05 | 1.528E-06 | 1.300E-05 | 1.600E-05 |
| 23222 | 1162 | V1162R | D | 1.6e-05 | 9.7e-06 | 1.4e-05 | 1.323E-05 | 3.219E-06 | 9.700E-06 | 1.600E-05 |
| 23223 | 1162 | V1162N | D | 1.2e-05 | 1.5e-05 | 1.3e-05 | 1.333E-05 | 1.528E-06 | 1.200E-05 | 1.500E-05 |
| 23224 | 1162 | V1162D | D | 9.8e-06 | 1.5e-05 | 1.5e-05 | 1.327E-05 | 3.002E-06 | 9.800E-06 | 1.500E-05 |
| 23225 | 1162 | V1162C | D | 9.5e-06 | 9.8e-06 | 1.2e-05 | 1.043E-05 | 1.365E-06 | 9.500E-06 | 1.200E-05 |
| 23226 | 1162 | V1162E | D | 1.4e-05 | 1.5e-05 | 9.4e-06 | 1.280E-05 | 2.987E-06 | 9.400E-06 | 1.500E-05 |
| 23227 | 1162 | V1162Q | D | 9.2e-06 | 1.3e-05 | 1.4e-05 | 1.207E-05 | 2.532E-06 | 9.200E-06 | 1.400E-05 |
| 23228 | 1162 | V1162G | D | 1.3e-05 | 1.1e-05 | 1.5e-05 | 1.300E-05 | 2.000E-06 | 1.100E-05 | 1.500E-05 |
| 23229 | 1162 | V1162H | D | 8.9e-06 | 1.5e-05 | 9.9e-06 | 1.127E-05 | 3.272E-06 | 8.900E-06 | 1.500E-05 |
| 23230 | 1162 | V1162I | D | 1.5e-05 | 9.3e-06 | 1.5e-05 | 1.310E-05 | 3.291E-06 | 9.300E-06 | 1.500E-05 |
| 23231 | 1162 | V1162L | D | 1e-05 | 1.5e-05 | 1.2e-05 | 1.233E-05 | 2.517E-06 | 1.000E-05 | 1.500E-05 |
| 23232 | 1162 | V1162K | D | 1.2e-05 | 8.5e-06 | 1.6e-05 | 1.217E-05 | 3.753E-06 | 8.500E-06 | 1.600E-05 |
| 23233 | 1162 | V1162M | D | 1.1e-05 | 9.6e-06 | 1.2e-05 | 1.087E-05 | 1.206E-06 | 9.600E-06 | 1.200E-05 |
| 23234 | 1162 | V1162F | D | 1.6e-05 | 1.6e-05 | 9.6e-06 | 1.387E-05 | 3.695E-06 | 9.600E-06 | 1.600E-05 |
| 23235 | 1162 | V1162P | D | 1e-05 | 1.6e-05 | 1.7e-05 | 1.433E-05 | 3.786E-06 | 1.000E-05 | 1.700E-05 |
| 23236 | 1162 | V1162S | D | 1.3e-05 | 1.1e-05 | 8.4e-06 | 1.080E-05 | 2.307E-06 | 8.400E-06 | 1.300E-05 |
| 23237 | 1162 | V1162T | D | 9.3e-06 | 1.4e-05 | 1.4e-05 | 1.243E-05 | 2.714E-06 | 9.300E-06 | 1.400E-05 |
| 23238 | 1162 | V1162W | D | 1.5e-05 | 9.3e-06 | 1e-05 | 1.143E-05 | 3.109E-06 | 9.300E-06 | 1.500E-05 |
| 23239 | 1162 | V1162Y | D | 8e-06 | 1.5e-05 | 1e-05 | 1.100E-05 | 3.606E-06 | 8.000E-06 | 1.500E-05 |
| 23240 | 1162 | V1162V | D | 1.2e-05 | 9.9e-06 | 1.5e-05 | 1.230E-05 | 2.563E-06 | 9.900E-06 | 1.500E-05 |
| 23241 | 1163 | E1163A | D | 1.4e-05 | 1.3e-05 | 1.3e-05 | 1.333E-05 | 5.774E-07 | 1.300E-05 | 1.400E-05 |
| 23242 | 1163 | E1163R | D | 1.4e-05 | 1.2e-05 | 1.4e-05 | 1.333E-05 | 1.155E-06 | 1.200E-05 | 1.400E-05 |
| 23243 | 1163 | E1163N | D | 1.1e-05 | 1.6e-05 | 1.4e-05 | 1.367E-05 | 2.517E-06 | 1.100E-05 | 1.600E-05 |
| 23244 | 1163 | E1163D | D | 9.8e-06 | 1.5e-05 | 8.6e-06 | 1.113E-05 | 3.402E-06 | 8.600E-06 | 1.500E-05 |
| 23245 | 1163 | E1163C | D | 9.3e-06 | 1.3e-05 | 9e-06 | 1.043E-05 | 2.228E-06 | 9.000E-06 | 1.300E-05 |
| 23246 | 1163 | E1163E | D | 8.5e-06 | 1.5e-05 | 1.5e-05 | 1.283E-05 | 3.753E-06 | 8.500E-06 | 1.500E-05 |
| 23247 | 1163 | E1163Q | D | 1.6e-05 | 9.1e-06 | 1e-05 | 1.170E-05 | 3.751E-06 | 9.100E-06 | 1.600E-05 |
| 23248 | 1163 | E1163G | D | 1.5e-05 | 2.1e-05 | 1.2e-05 | 1.600E-05 | 4.583E-06 | 1.200E-05 | 2.100E-05 |
| 23249 | 1163 | E1163H | D | 1.5e-05 | 1.5e-05 | 1.1e-05 | 1.367E-05 | 2.309E-06 | 1.100E-05 | 1.500E-05 |
| 23250 | 1163 | E1163I | D | 1.6e-05 | 1.4e-05 | 1.3e-05 | 1.433E-05 | 1.528E-06 | 1.300E-05 | 1.600E-05 |
| 23251 | 1163 | E1163L | D | 1.5e-05 | 1.5e-05 | 1.2e-05 | 1.400E-05 | 1.732E-06 | 1.200E-05 | 1.500E-05 |
| 23252 | 1163 | E1163K | D | 1.6e-05 | 8.7e-06 | 1.2e-05 | 1.223E-05 | 3.656E-06 | 8.700E-06 | 1.600E-05 |
| 23253 | 1163 | E1163M | D | 1.1e-05 | 1.3e-05 | 1.6e-05 | 1.333E-05 | 2.517E-06 | 1.100E-05 | 1.600E-05 |
| 23254 | 1163 | E1163F | D | 1.5e-05 | 1.4e-05 | 9.1e-06 | 1.270E-05 | 3.158E-06 | 9.100E-06 | 1.500E-05 |
| 23255 | 1163 | E1163P | D | 1e-05 | 1.2e-05 | 1.6e-05 | 1.267E-05 | 3.055E-06 | 1.000E-05 | 1.600E-05 |
| 23256 | 1163 | E1163S | D | 1.2e-05 | 6.2e-06 | 1.3e-05 | 1.040E-05 | 3.672E-06 | 6.200E-06 | 1.300E-05 |
| 23257 | 1163 | E1163T | D | 1.3e-05 | 1.4e-05 | 1.3e-05 | 1.333E-05 | 5.774E-07 | 1.300E-05 | 1.400E-05 |
| 23258 | 1163 | E1163W | D | 1.5e-05 | 5.9e-06 | 8.8e-06 | 9.900E-06 | 4.649E-06 | 5.900E-06 | 1.500E-05 |
| 23259 | 1163 | E1163Y | D | 9.4e-06 | 1.5e-05 | 9.4e-06 | 1.127E-05 | 3.233E-06 | 9.400E-06 | 1.500E-05 |
| 23260 | 1163 | E1163V | D | 1.1e-05 | 1.6e-05 | 1e-05 | 1.233E-05 | 3.215E-06 | 1.000E-05 | 1.600E-05 |
| 23261 | 1164 | I1164A | D | 1.4e-05 | 1.3e-05 | 1.3e-05 | 1.333E-05 | 5.774E-07 | 1.300E-05 | 1.400E-05 |
| 23262 | 1164 | I1164R | D | 1.5e-05 | 1.1e-05 | 1e-05 | 1.200E-05 | 2.646E-06 | 1.000E-05 | 1.500E-05 |
| 23263 | 1164 | I1164N | D | 1.4e-05 | 2.5e-05 | 1.6e-05 | 1.833E-05 | 5.859E-06 | 1.400E-05 | 2.500E-05 |
| 23264 | 1164 | I1164D | D | 1e-05 | 1.1e-05 | 1e-05 | 1.033E-05 | 5.774E-07 | 1.000E-05 | 1.100E-05 |
| 23265 | 1164 | I1164C | D | 8.5e-06 | 9.1e-06 | 1.5e-05 | 1.087E-05 | 3.592E-06 | 8.500E-06 | 1.500E-05 |
| 23266 | 1164 | I1164E | D | 8.2e-06 | 1.4e-05 | 1.1e-05 | 1.107E-05 | 2.901E-06 | 8.200E-06 | 1.400E-05 |
| 23267 | 1164 | I1164Q | D | 1e-05 | 1.2e-05 | 8.8e-06 | 1.027E-05 | 1.617E-06 | 8.800E-06 | 1.200E-05 |
| 23268 | 1164 | I1164G | D | 1.1e-05 | 9.8e-06 | 1.4e-05 | 1.160E-05 | 2.163E-06 | 9.800E-06 | 1.400E-05 |
| 23269 | 1164 | I1164H | D | 1e-05 | 9e-06 | 1.6e-05 | 1.167E-05 | 3.786E-06 | 9.000E-06 | 1.600E-05 |
| 23270 | 1164 | I1164I | D | 1.5e-05 | 9.3e-06 | 1e-05 | 1.143E-05 | 3.109E-06 | 9.300E-06 | 1.500E-05 |
| 23271 | 1164 | I1164L | D | 1e-05 | 1e-05 | 1.5e-05 | 1.167E-05 | 2.887E-06 | 1.000E-05 | 1.500E-05 |
| 23272 | 1164 | I1164K | D | 9.8e-06 | 8.3e-06 | 1.4e-05 | 1.070E-05 | 2.955E-06 | 8.300E-06 | 1.400E-05 |
| 23273 | 1164 | I1164M | D | 1.3e-05 | 1.6e-05 | 1.5e-05 | 1.467E-05 | 1.528E-06 | 1.300E-05 | 1.600E-05 |
| 23274 | 1164 | I1164F | D | 1.7e-05 | 1.1e-05 | 1.6e-05 | 1.467E-05 | 3.215E-06 | 1.100E-05 | 1.700E-05 |
| 23275 | 1164 | I1164P | D | 1.7e-05 | 1.2e-05 | 1.5e-05 | 1.467E-05 | 2.517E-06 | 1.200E-05 | 1.700E-05 |
| 23276 | 1164 | I1164S | D | 3.3e-05 | 8.5e-06 | 1.6e-05 | 1.917E-05 | 1.255E-05 | 8.500E-06 | 3.300E-05 |
| 23277 | 1164 | I1164T | D | 1.1e-05 | 1.6e-05 | 1.4e-05 | 1.367E-05 | 2.517E-06 | 1.100E-05 | 1.600E-05 |
| 23278 | 1164 | I1164W | D | 1e-05 | 1.4e-05 | 1e-05 | 1.133E-05 | 2.309E-06 | 1.000E-05 | 1.400E-05 |
| 23279 | 1164 | I1164Y | D | 1.3e-05 | 8.2e-06 | 1.1e-05 | 1.073E-05 | 2.411E-06 | 8.200E-06 | 1.300E-05 |
| 23280 | 1164 | I1164V | D | 1.3e-05 | 1.6e-05 | 9.3e-06 | 1.277E-05 | 3.356E-06 | 9.300E-06 | 1.600E-05 |
| 23281 | 1165 | K1165A | D | 1.3e-05 | 1.5e-05 | 1.7e-05 | 1.500E-05 | 2.000E-06 | 1.300E-05 | 1.700E-05 |
| 23282 | 1165 | K1165R | D | 1.4e-05 | 1.5e-05 | 1.1e-05 | 1.333E-05 | 2.082E-06 | 1.100E-05 | 1.500E-05 |
| 23283 | 1165 | K1165N | D | 1.4e-05 | 1.6e-05 | 9.6e-06 | 1.320E-05 | 3.274E-06 | 9.600E-06 | 1.600E-05 |
| 23284 | 1165 | K1165D | D | 1.5e-05 | 8.5e-06 | 9.4e-06 | 1.097E-05 | 3.522E-06 | 8.500E-06 | 1.500E-05 |
| 23285 | 1165 | K1165C | D | 8.7e-06 | 1.4e-05 | 1e-05 | 1.090E-05 | 2.762E-06 | 8.700E-06 | 1.400E-05 |
| 23286 | 1165 | K1165E | D | 1.5e-05 | 1.1e-05 | 1.1e-05 | 1.233E-05 | 2.309E-06 | 1.100E-05 | 1.500E-05 |
| 23287 | 1165 | K1165Q | D | 1.2e-05 | 1.5e-05 | 1.5e-05 | 1.400E-05 | 1.732E-06 | 1.200E-05 | 1.500E-05 |
| 23288 | 1165 | K1165G | D | 1.5e-05 | 1.6e-05 | 9e-06 | 1.333E-05 | 3.786E-06 | 9.000E-06 | 1.600E-05 |
| 23289 | 1165 | K1165H | D | 1.7e-05 | 1.4e-05 | 1.5e-05 | 1.533E-05 | 1.528E-06 | 1.400E-05 | 1.700E-05 |

| | | | | | | | | | | |
|---|---|---|---|---|---|---|---|---|---|---|
| 23290 | 1165 | K1165I | D | 1.5e-05 | 1.9e-05 | 9.3e-06 | 1.443E-05 | 4.875E-06 | 9.300E-06 | 1.900E-05 |
| 23291 | 1165 | K1165L | D | 1.1e-05 | 1.7e-05 | 1.3e-05 | 1.367E-05 | 3.055E-06 | 1.100E-05 | 1.700E-05 |
| 23292 | 1165 | K1165K | D | 1.1e-05 | 1.6e-05 | 1.2e-05 | 1.300E-05 | 2.646E-06 | 1.100E-05 | 1.600E-05 |
| 23293 | 1165 | K1165M | D | 1.6e-05 | 1.8e-05 | 9.6e-06 | 1.453E-05 | 4.388E-06 | 9.600E-06 | 1.800E-05 |
| 23294 | 1165 | K1165F | D | 2.2e-05 | 1.2e-05 | 1.4e-05 | 1.600E-05 | 5.292E-06 | 1.200E-05 | 2.200E-05 |
| 23295 | 1165 | K1165P | D | 1e-05 | 1.8e-05 | 1.6e-05 | 1.467E-05 | 4.163E-06 | 1.000E-05 | 1.800E-05 |
| 23296 | 1165 | K1165S | D | 1.5e-05 | 1e-05 | 1.7e-05 | 1.400E-05 | 3.606E-06 | 1.000E-05 | 1.700E-05 |
| 23297 | 1165 | K1165T | D | 1e-05 | 1.1e-05 | 1.4e-05 | 1.167E-05 | 2.082E-06 | 1.000E-05 | 1.400E-05 |
| 23298 | 1165 | K1165W | D | 1e-05 | 1.5e-05 | 1.3e-05 | 1.267E-05 | 2.517E-06 | 1.000E-05 | 1.500E-05 |
| 23299 | 1165 | K1165Y | D | 8.5e-06 | 1.5e-05 | 1.3e-05 | 1.217E-05 | 3.329E-06 | 8.500E-06 | 1.500E-05 |
| 23300 | 1165 | K1165V | D | 1.5e-05 | 1.3e-05 | 1.2e-05 | 1.333E-05 | 1.528E-06 | 1.200E-05 | 1.500E-05 |
| 23301 | 1166 | R1166A | D | 1.5e-05 | 1.1e-05 | 1.1e-05 | 1.233E-05 | 2.309E-06 | 1.100E-05 | 1.500E-05 |
| 23302 | 1166 | R1166R | D | 1e-05 | 1.5e-05 | 8.8e-06 | 1.127E-05 | 3.288E-06 | 8.800E-06 | 1.500E-05 |
| 23303 | 1166 | R1166N | D | 8.7e-06 | 1e-05 | 8.9e-06 | 9.200E-06 | 7.000E-07 | 8.700E-06 | 1.000E-05 |
| 23304 | 1166 | R1166D | D | 9.8e-06 | 9.7e-06 | 1.5e-05 | 1.150E-05 | 3.032E-06 | 9.700E-06 | 1.500E-05 |
| 23305 | 1166 | R1166C | D | 1.1e-05 | 9.1e-06 | 1e-05 | 1.003E-05 | 9.504E-07 | 9.100E-06 | 1.100E-05 |
| 23306 | 1166 | R1166E | D | 1e-05 | 1.2e-05 | 1.5e-05 | 1.233E-05 | 2.517E-06 | 1.000E-05 | 1.500E-05 |
| 23307 | 1166 | R1166Q | D | 1.1e-05 | 8.8e-06 | 8.6e-06 | 9.467E-06 | 1.332E-06 | 8.600E-06 | 1.100E-05 |
| 23308 | 1166 | R1166G | D | 1.1e-05 | 1.2e-05 | 1.4e-05 | 1.233E-05 | 1.528E-06 | 1.100E-05 | 1.400E-05 |
| 23309 | 1166 | R1166H | D | 8.4e-06 | 1.2e-05 | 1e-05 | 1.013E-05 | 1.804E-06 | 8.400E-06 | 1.200E-05 |
| 23310 | 1166 | R1166I | D | 1.5e-05 | 1.6e-05 | 1.4e-05 | 1.500E-05 | 1.000E-06 | 1.400E-05 | 1.600E-05 |
| 23311 | 1166 | R1166L | D | 9.8e-06 | 1e-05 | 1.1e-05 | 1.027E-05 | 6.429E-07 | 9.800E-06 | 1.100E-05 |
| 23312 | 1166 | R1166K | D | 1e-05 | 1.3e-05 | 1.6e-05 | 1.300E-05 | 3.000E-06 | 1.000E-05 | 1.600E-05 |
| 23313 | 1166 | R1166M | D | 8.9e-06 | 1.2e-05 | 1.6e-05 | 1.230E-05 | 3.559E-06 | 8.900E-06 | 1.600E-05 |
| 23314 | 1166 | R1166F | D | 1.1e-05 | 1e-05 | 8.4e-06 | 9.800E-06 | 1.311E-06 | 8.400E-06 | 1.100E-05 |
| 23315 | 1166 | R1166P | D | 1.4e-05 | 1.5e-05 | 1.5e-05 | 1.467E-05 | 5.774E-07 | 1.400E-05 | 1.500E-05 |
| 23316 | 1166 | R1166S | D | 1.4e-05 | 1e-05 | 1.5e-05 | 1.300E-05 | 2.646E-06 | 1.000E-05 | 1.500E-05 |
| 23317 | 1166 | R1166T | D | 1.5e-05 | 1e-05 | 1.6e-05 | 1.367E-05 | 3.215E-06 | 1.000E-05 | 1.600E-05 |
| 23318 | 1166 | R1166W | D | 1.4e-05 | 9e-06 | 8.4e-06 | 1.047E-05 | 3.075E-06 | 8.400E-06 | 1.400E-05 |
| 23319 | 1166 | R1166Y | D | 1.3e-05 | 1.1e-05 | 2.1e-05 | 1.500E-05 | 5.292E-06 | 1.100E-05 | 2.100E-05 |
| 23320 | 1166 | R1166V | D | 1.6e-05 | 1.6e-05 | 1.1e-05 | 1.433E-05 | 2.887E-06 | 1.100E-05 | 1.600E-05 |
| 23321 | 1167 | T1167A | D | 1.4e-05 | 1.3e-05 | 1.1e-05 | 1.267E-05 | 1.528E-06 | 1.100E-05 | 1.400E-05 |
| 23322 | 1167 | T1167R | D | 1.1e-05 | 1.3e-05 | 1e-05 | 1.133E-05 | 1.528E-06 | 1.000E-05 | 1.300E-05 |
| 23323 | 1167 | T1167N | D | 1.4e-05 | 9e-06 | 1.6e-05 | 1.300E-05 | 3.606E-06 | 9.000E-06 | 1.600E-05 |
| 23324 | 1167 | T1167D | D | 8.1e-06 | 1.5e-05 | 1.5e-05 | 1.270E-05 | 3.984E-06 | 8.100E-06 | 1.500E-05 |
| 23325 | 1167 | T1167C | D | 1.4e-05 | 1.5e-05 | 1.1e-05 | 1.333E-05 | 2.082E-06 | 1.100E-05 | 1.500E-05 |
| 23326 | 1167 | T1167E | D | 1.6e-05 | 1.3e-05 | 1.1e-05 | 1.333E-05 | 2.517E-06 | 1.100E-05 | 1.600E-05 |
| 23327 | 1167 | T1167Q | D | 1e-05 | 8.7e-06 | 8e-06 | 8.900E-06 | 1.015E-06 | 8.000E-06 | 1.000E-05 |
| 23328 | 1167 | T1167G | D | 1.7e-05 | 1.2e-05 | 1.4e-05 | 1.433E-05 | 2.517E-06 | 1.200E-05 | 1.700E-05 |
| 23329 | 1167 | T1167H | D | 9.8e-06 | 1e-05 | 1.1e-05 | 1.027E-05 | 6.429E-07 | 9.800E-06 | 1.100E-05 |
| 23330 | 1167 | T1167I | D | 1.6e-05 | 1.5e-05 | 1.5e-05 | 1.533E-05 | 5.774E-07 | 1.500E-05 | 1.600E-05 |
| 23331 | 1167 | T1167L | D | 1.5e-05 | 9.9e-06 | 1.5e-05 | 1.330E-05 | 2.944E-06 | 9.900E-06 | 1.500E-05 |
| 23332 | 1167 | T1167K | D | 1.2e-05 | 8.6e-06 | 1.1e-05 | 1.053E-05 | 1.747E-06 | 8.600E-06 | 1.200E-05 |
| 23333 | 1167 | T1167M | D | 1.6e-05 | 1.5e-05 | 1.7e-05 | 1.600E-05 | 1.000E-06 | 1.500E-05 | 1.700E-05 |
| 23334 | 1167 | T1167F | D | 1.5e-05 | 1.1e-05 | 1.7e-05 | 1.433E-05 | 3.055E-06 | 1.100E-05 | 1.700E-05 |
| 23335 | 1167 | T1167P | D | 1e-05 | 1.4e-05 | 8.8e-06 | 1.093E-05 | 2.723E-06 | 8.800E-06 | 1.400E-05 |
| 23336 | 1167 | T1167S | D | 1.3e-05 | 1.1e-05 | 1.7e-05 | 1.367E-05 | 3.055E-06 | 1.100E-05 | 1.700E-05 |
| 23337 | 1167 | T1167T | D | 1.5e-05 | 1.1e-05 | 9.5e-06 | 1.183E-05 | 2.843E-06 | 9.500E-06 | 1.500E-05 |
| 23338 | 1167 | T1167W | D | 1.1e-05 | 1.2e-05 | 1.5e-05 | 1.267E-05 | 2.082E-06 | 1.100E-05 | 1.500E-05 |
| 23339 | 1167 | T1167Y | D | 1e-05 | 1.4e-05 | 1.1e-05 | 1.167E-05 | 2.082E-06 | 1.000E-05 | 1.400E-05 |
| 23340 | 1167 | T1167V | D | 1.4e-05 | 1.5e-05 | 1.1e-05 | 1.333E-05 | 2.082E-06 | 1.100E-05 | 1.500E-05 |
| 23341 | 1168 | V1168A | D | 9.1e-06 | 1.8e-05 | 1.7e-05 | 1.470E-05 | 4.875E-06 | 9.100E-06 | 1.800E-05 |
| 23342 | 1168 | V1168R | D | 1.1e-05 | 1.1e-05 | 1.6e-05 | 1.267E-05 | 2.887E-06 | 1.100E-05 | 1.600E-05 |
| 23343 | 1168 | V1168N | D | 1.1e-05 | 1.2e-05 | 9.8e-06 | 1.093E-05 | 1.102E-06 | 9.800E-06 | 1.200E-05 |
| 23344 | 1168 | V1168D | D | 1.7e-05 | 9.6e-06 | 1e-05 | 1.220E-05 | 4.162E-06 | 9.600E-06 | 1.700E-05 |
| 23345 | 1168 | V1168C | D | 1.3e-05 | 1.5e-05 | 1.1e-05 | 1.300E-05 | 2.000E-06 | 1.100E-05 | 1.500E-05 |
| 23346 | 1168 | V1168E | D | 9.5e-06 | 1.6e-05 | 1.4e-05 | 1.317E-05 | 3.329E-06 | 9.500E-06 | 1.600E-05 |
| 23347 | 1168 | V1168Q | D | 1.2e-05 | 1.2e-05 | 1.1e-05 | 1.167E-05 | 5.774E-07 | 1.100E-05 | 1.200E-05 |
| 23348 | 1168 | V1168G | D | 9.5e-06 | 1.1e-05 | 1.1e-05 | 1.050E-05 | 8.660E-07 | 9.500E-06 | 1.100E-05 |
| 23349 | 1168 | V1168H | D | 1e-05 | 9.7e-06 | 1.5e-05 | 1.157E-05 | 2.977E-06 | 9.700E-06 | 1.500E-05 |
| 23350 | 1168 | V1168I | D | 1.5e-05 | 8.1e-06 | 1.1e-05 | 1.137E-05 | 3.465E-06 | 8.100E-06 | 1.500E-05 |
| 23351 | 1168 | V1168L | D | 8.1e-06 | 1.5e-05 | 1e-05 | 1.103E-05 | 3.564E-06 | 8.100E-06 | 1.500E-05 |
| 23352 | 1168 | V1168K | D | 1.3e-05 | 1.4e-05 | 9.6e-06 | 1.220E-05 | 2.307E-06 | 9.600E-06 | 1.400E-05 |
| 23353 | 1168 | V1168M | D | 1.1e-05 | 1.5e-05 | 1e-05 | 1.200E-05 | 2.646E-06 | 1.000E-05 | 1.500E-05 |
| 23354 | 1168 | V1168F | D | 1.1e-05 | 1.5e-05 | 1.3e-05 | 1.300E-05 | 2.000E-06 | 1.100E-05 | 1.500E-05 |
| 23355 | 1168 | V1168P | D | 1.5e-05 | 1.2e-05 | 6.9e-06 | 1.130E-05 | 4.095E-06 | 6.900E-06 | 1.500E-05 |
| 23356 | 1168 | V1168S | D | 1e-05 | 1.5e-05 | 9.5e-06 | 1.150E-05 | 3.041E-06 | 9.500E-06 | 1.500E-05 |
| 23357 | 1168 | V1168T | D | 6.5e-06 | 9.5e-06 | 1.2e-05 | 9.333E-06 | 2.754E-06 | 6.500E-06 | 1.200E-05 |
| 23358 | 1168 | V1168W | D | 1.6e-05 | 1.8e-05 | 7.9e-06 | 1.397E-05 | 5.348E-06 | 7.900E-06 | 1.800E-05 |
| 23359 | 1168 | V1168Y | D | 2.2e-05 | 1.4e-05 | 8.9e-06 | 1.497E-05 | 6.603E-06 | 8.900E-06 | 2.200E-05 |
| 23360 | 1168 | V1168V | D | 6.7e-06 | 1.4e-05 | 1.4e-05 | 1.157E-05 | 4.215E-06 | 6.700E-06 | 1.400E-05 |
| 23361 | 1169 | A1169A | D | 1.3e-05 | 1.5e-05 | 1.5e-05 | 1.433E-05 | 1.155E-06 | 1.300E-05 | 1.500E-05 |
| 23362 | 1169 | A1169R | D | 1.2e-05 | 1.6e-05 | 1.7e-05 | 1.500E-05 | 2.646E-06 | 1.200E-05 | 1.700E-05 |
| 23363 | 1169 | A1169N | D | 1.4e-05 | 1.2e-05 | 1.4e-05 | 1.333E-05 | 1.155E-06 | 1.200E-05 | 1.400E-05 |
| 23364 | 1169 | A1169D | D | 1.1e-05 | 1.2e-05 | 1.4e-05 | 1.233E-05 | 1.528E-06 | 1.100E-05 | 1.400E-05 |
| 23365 | 1169 | A1169C | D | 1.7e-05 | 1.4e-05 | 1.6e-05 | 1.567E-05 | 1.528E-06 | 1.400E-05 | 1.700E-05 |
| 23366 | 1169 | A1169E | D | 1.5e-05 | 9.1e-06 | 1.6e-05 | 1.337E-05 | 3.729E-06 | 9.100E-06 | 1.600E-05 |
| 23367 | 1169 | A1169Q | D | 1.2e-05 | 9.6e-06 | 1.6e-05 | 1.253E-05 | 3.233E-06 | 9.600E-06 | 1.600E-05 |
| 23368 | 1169 | A1169G | D | 1.3e-05 | 1.4e-05 | 1.2e-05 | 1.300E-05 | 1.000E-06 | 1.200E-05 | 1.400E-05 |
| 23369 | 1169 | A1169H | D | 9.5e-06 | 1.5e-05 | 1.5e-05 | 1.317E-05 | 3.175E-06 | 9.500E-06 | 1.500E-05 |
| 23370 | 1169 | A1169I | D | 1.3e-05 | 1.6e-05 | 9.7e-06 | 1.290E-05 | 3.151E-06 | 9.700E-06 | 1.600E-05 |
| 23371 | 1169 | A1169L | D | 1.1e-05 | 9.9e-06 | 1e-05 | 1.030E-05 | 6.083E-07 | 9.900E-06 | 1.100E-05 |
| 23372 | 1169 | A1169K | D | 1.5e-05 | 1.3e-05 | 1.1e-05 | 1.300E-05 | 2.000E-06 | 1.100E-05 | 1.500E-05 |
| 23373 | 1169 | A1169M | D | 1.2e-05 | 1.3e-05 | 1e-05 | 1.167E-05 | 1.528E-06 | 1.000E-05 | 1.300E-05 |
| 23374 | 1169 | A1169F | D | 1.6e-05 | 1.6e-05 | 9.4e-06 | 1.380E-05 | 3.811E-06 | 9.400E-06 | 1.600E-05 |

| | | | | | | | | | | |
|---|---|---|---|---|---|---|---|---|---|---|
| 23375 | 1169 | A1169P | D | 1e-05 | 1.6e-05 | 1.5e-05 | 1.367E-05 | 3.215E-06 | 1.000E-05 | 1.600E-05 |
| 23376 | 1169 | A1169S | D | 7.9e-06 | 8.9e-06 | 8.7e-06 | 8.500E-06 | 5.292E-07 | 7.900E-06 | 8.900E-06 |
| 23377 | 1169 | A1169T | D | 1.4e-05 | 1.3e-05 | 1.1e-05 | 1.267E-05 | 1.528E-06 | 1.100E-05 | 1.400E-05 |
| 23378 | 1169 | A1169W | D | 1.5e-05 | 9.6e-06 | 1.6e-05 | 1.353E-05 | 3.443E-06 | 9.600E-06 | 1.600E-05 |
| 23379 | 1169 | A1169Y | D | 1.5e-05 | 1.3e-05 | 1.2e-05 | 1.333E-05 | 1.528E-06 | 1.200E-05 | 1.500E-05 |
| 23380 | 1169 | A1169V | D | 1.1e-05 | 1.6e-05 | 2e-05 | 1.567E-05 | 4.509E-06 | 1.100E-05 | 2.000E-05 |
| 23381 | 1170 | A1170A | D | 9.6e-06 | 1e-05 | 1.2e-05 | 1.053E-05 | 1.286E-06 | 9.600E-06 | 1.200E-05 |
| 23382 | 1170 | A1170R | D | 1.7e-05 | 1.1e-05 | 1.3e-05 | 1.367E-05 | 3.055E-06 | 1.100E-05 | 1.700E-05 |
| 23383 | 1170 | A1170N | D | 9e-06 | 1.1e-05 | 8.2e-06 | 9.400E-06 | 1.442E-06 | 8.200E-06 | 1.100E-05 |
| 23384 | 1170 | A1170D | D | 1.1e-05 | 1.2e-05 | 1.1e-05 | 1.133E-05 | 5.774E-07 | 1.100E-05 | 1.200E-05 |
| 23385 | 1170 | A1170C | D | 8.7e-06 | 1.5e-05 | 1.4e-05 | 1.257E-05 | 3.386E-06 | 8.700E-06 | 1.500E-05 |
| 23386 | 1170 | A1170E | D | 9.2e-06 | 1.6e-05 | 7.1e-06 | 1.077E-05 | 4.652E-06 | 7.100E-06 | 1.600E-05 |
| 23387 | 1170 | A1170Q | D | 1.5e-05 | 1.2e-05 | 1.5e-05 | 1.400E-05 | 1.732E-06 | 1.200E-05 | 1.500E-05 |
| 23388 | 1170 | A1170G | D | 1.6e-05 | 9.9e-06 | 1e-05 | 1.197E-05 | 3.493E-06 | 9.900E-06 | 1.600E-05 |
| 23389 | 1170 | A1170H | D | 1.5e-05 | 1.5e-05 | 1.1e-05 | 1.367E-05 | 2.309E-06 | 1.100E-05 | 1.500E-05 |
| 23390 | 1170 | A1170I | D | 9.7e-06 | 1.2e-05 | 1.2e-05 | 1.123E-05 | 1.328E-06 | 9.700E-06 | 1.200E-05 |
| 23391 | 1170 | A1170L | D | 8.2e-06 | 1.1e-05 | 1.5e-05 | 1.140E-05 | 3.418E-06 | 8.200E-06 | 1.500E-05 |
| 23392 | 1170 | A1170K | D | 1.5e-05 | 1.6e-05 | 1.1e-05 | 1.400E-05 | 2.646E-06 | 1.100E-05 | 1.600E-05 |
| 23393 | 1170 | A1170M | D | 9.8e-06 | 1.5e-05 | 1.2e-05 | 1.227E-05 | 2.610E-06 | 9.800E-06 | 1.500E-05 |
| 23394 | 1170 | A1170F | D | 1.6e-05 | 1e-05 | 1.2e-05 | 1.267E-05 | 3.055E-06 | 1.000E-05 | 1.600E-05 |
| 23395 | 1170 | A1170P | D | 1.6e-05 | 9.1e-06 | 1.5e-05 | 1.337E-05 | 3.729E-06 | 9.100E-06 | 1.600E-05 |
| 23396 | 1170 | A1170S | D | 8.1e-06 | 1.1e-05 | 1.1e-05 | 1.003E-05 | 1.674E-06 | 8.100E-06 | 1.100E-05 |
| 23397 | 1170 | A1170T | D | 1.3e-05 | 9.5e-06 | 8.7e-06 | 1.040E-05 | 2.287E-06 | 8.700E-06 | 1.300E-05 |
| 23398 | 1170 | A1170W | D | 1.4e-05 | 1.1e-05 | 9.6e-06 | 1.153E-05 | 2.248E-06 | 9.600E-06 | 1.400E-05 |
| 23399 | 1170 | A1170Y | D | 9.7e-06 | 1.4e-05 | 9.6e-06 | 1.110E-05 | 2.512E-06 | 9.600E-06 | 1.400E-05 |
| 23400 | 1170 | A1170V | D | 9.6e-06 | 1.3e-05 | 1.2e-05 | 1.153E-05 | 1.747E-06 | 9.600E-06 | 1.300E-05 |
| 23401 | 1171 | P1171A | D | 1.5e-05 | 9.3e-06 | 1.3e-05 | 1.243E-05 | 2.892E-06 | 9.300E-06 | 1.500E-05 |
| 23402 | 1171 | P1171R | D | 8.5e-06 | 1.4e-05 | 8.6e-06 | 1.037E-05 | 3.147E-06 | 8.500E-06 | 1.400E-05 |
| 23403 | 1171 | P1171N | D | 9.7e-06 | 9.8e-06 | 1.3e-05 | 1.083E-05 | 1.877E-06 | 9.700E-06 | 1.300E-05 |
| 23404 | 1171 | P1171D | D | 1.7e-05 | 1.1e-05 | 1.6e-05 | 1.467E-05 | 3.215E-06 | 1.100E-05 | 1.700E-05 |
| 23405 | 1171 | P1171C | D | 1.3e-05 | 1.7e-05 | 9.8e-06 | 1.327E-05 | 3.607E-06 | 9.800E-06 | 1.700E-05 |
| 23406 | 1171 | P1171E | D | 1.1e-05 | 1.5e-05 | 9.6e-06 | 1.187E-05 | 2.802E-06 | 9.600E-06 | 1.500E-05 |
| 23407 | 1171 | P1171Q | D | 1.5e-05 | 1.6e-05 | 8.6e-06 | 1.320E-05 | 4.015E-06 | 8.600E-06 | 1.600E-05 |
| 23408 | 1171 | P1171G | D | 1.5e-05 | 1e-05 | 1.1e-05 | 1.200E-05 | 2.646E-06 | 1.000E-05 | 1.500E-05 |
| 23409 | 1171 | P1171H | D | 1.7e-05 | 1.5e-05 | 1.4e-05 | 1.533E-05 | 1.528E-06 | 1.400E-05 | 1.700E-05 |
| 23410 | 1171 | P1171I | D | 1.6e-05 | 1.6e-05 | 1.4e-05 | 1.533E-05 | 1.155E-06 | 1.400E-05 | 1.600E-05 |
| 23411 | 1171 | P1171L | D | 1.1e-05 | 1.4e-05 | 1.1e-05 | 1.200E-05 | 1.732E-06 | 1.100E-05 | 1.400E-05 |
| 23412 | 1171 | P1171K | D | 1.8e-05 | 1.4e-05 | 9.9e-06 | 1.397E-05 | 4.050E-06 | 9.900E-06 | 1.800E-05 |
| 23413 | 1171 | P1171M | D | 1.5e-05 | 1.5e-05 | 8.6e-06 | 1.287E-05 | 3.695E-06 | 8.600E-06 | 1.500E-05 |
| 23414 | 1171 | P1171F | D | 1.5e-05 | 9e-06 | 1.1e-05 | 1.167E-05 | 3.055E-06 | 9.000E-06 | 1.500E-05 |
| 23415 | 1171 | P1171P | D | 9.9e-06 | 1.5e-05 | 9.4e-06 | 1.143E-05 | 3.099E-06 | 9.400E-06 | 1.500E-05 |
| 23416 | 1171 | P1171S | D | 9.8e-06 | 1.4e-05 | 1.6e-05 | 1.327E-05 | 3.164E-06 | 9.800E-06 | 1.600E-05 |
| 23417 | 1171 | P1171T | D | 1.6e-05 | 1.2e-05 | 8e-06 | 1.200E-05 | 4.000E-06 | 8.000E-06 | 1.600E-05 |
| 23418 | 1171 | P1171W | D | 1.2e-05 | 1.3e-05 | 1.9e-05 | 1.467E-05 | 3.786E-06 | 1.200E-05 | 1.900E-05 |
| 23419 | 1171 | P1171Y | D | 9e-06 | 1.3e-05 | 1.4e-05 | 1.200E-05 | 2.646E-06 | 9.000E-06 | 1.400E-05 |
| 23420 | 1171 | P1171V | D | 1.5e-05 | 1.5e-05 | 1.5e-05 | 1.500E-05 | 0.000E+00 | 1.500E-05 | 1.500E-05 |
| 23421 | 1172 | S1172A | D | 1.1e-05 | 9.2e-06 | 1.6e-05 | 1.207E-05 | 3.523E-06 | 9.200E-06 | 1.600E-05 |
| 23422 | 1172 | S1172R | D | 1e-05 | 1.6e-05 | 1e-05 | 1.200E-05 | 3.464E-06 | 1.000E-05 | 1.600E-05 |
| 23423 | 1172 | S1172N | D | 1.2e-05 | 2.5e-05 | 1.3e-05 | 1.667E-05 | 7.234E-06 | 1.200E-05 | 2.500E-05 |
| 23424 | 1172 | S1172D | D | 2.4e-05 | 1e-05 | 1.1e-05 | 1.500E-05 | 7.810E-06 | 1.000E-05 | 2.400E-05 |
| 23425 | 1172 | S1172C | D | 1.4e-05 | 1.5e-05 | 8.4e-06 | 1.247E-05 | 3.557E-06 | 8.400E-06 | 1.500E-05 |
| 23426 | 1172 | S1172E | D | 1.2e-05 | 1.6e-05 | 9.9e-06 | 1.263E-05 | 3.099E-06 | 9.900E-06 | 1.600E-05 |
| 23427 | 1172 | S1172Q | D | 9.3e-06 | 1.5e-05 | 1.1e-05 | 1.177E-05 | 2.926E-06 | 9.300E-06 | 1.500E-05 |
| 23428 | 1172 | S1172G | D | 1.3e-05 | 1.5e-05 | 1.2e-05 | 1.333E-05 | 1.528E-06 | 1.200E-05 | 1.500E-05 |
| 23429 | 1172 | S1172H | D | 9.2e-06 | 1.6e-05 | 1.6e-05 | 1.373E-05 | 3.926E-06 | 9.200E-06 | 1.600E-05 |
| 23430 | 1172 | S1172I | D | 1.1e-05 | 1.1e-05 | 1.8e-05 | 1.333E-05 | 4.041E-06 | 1.100E-05 | 1.800E-05 |
| 23431 | 1172 | S1172L | D | 1e-05 | 1e-05 | 1.4e-05 | 1.133E-05 | 2.309E-06 | 1.000E-05 | 1.400E-05 |
| 23432 | 1172 | S1172K | D | 1.2e-05 | 1e-05 | 1.1e-05 | 1.100E-05 | 1.000E-06 | 1.000E-05 | 1.200E-05 |
| 23433 | 1172 | S1172M | D | 1e-05 | 1.6e-05 | 1.5e-05 | 1.367E-05 | 3.215E-06 | 1.000E-05 | 1.600E-05 |
| 23434 | 1172 | S1172F | D | 8.1e-06 | 1.6e-05 | 1.1e-05 | 1.170E-05 | 3.996E-06 | 8.100E-06 | 1.600E-05 |
| 23435 | 1172 | S1172P | D | 1.4e-05 | 1.7e-05 | 1.6e-05 | 1.567E-05 | 1.528E-06 | 1.400E-05 | 1.700E-05 |
| 23436 | 1172 | S1172S | D | 1.6e-05 | 1.4e-05 | 1.5e-05 | 1.500E-05 | 1.000E-06 | 1.400E-05 | 1.600E-05 |
| 23437 | 1172 | S1172T | D | 1.4e-05 | 9.2e-06 | 1.2e-05 | 1.173E-05 | 2.411E-06 | 9.200E-06 | 1.400E-05 |
| 23438 | 1172 | S1172W | D | 1e-05 | 1.2e-05 | 1e-05 | 1.067E-05 | 1.155E-06 | 1.000E-05 | 1.200E-05 |
| 23439 | 1172 | S1172Y | D | 1.4e-05 | 1.1e-05 | 1.1e-05 | 1.200E-05 | 1.732E-06 | 1.100E-05 | 1.400E-05 |
| 23440 | 1172 | S1172V | D | 9.8e-06 | 1e-05 | 1.1e-05 | 1.027E-05 | 6.429E-07 | 9.800E-06 | 1.100E-05 |
| 23441 | 1173 | V1173A | D | 9.5e-06 | 1e-05 | 7.1e-06 | 8.867E-06 | 1.550E-06 | 7.100E-06 | 1.000E-05 |
| 23442 | 1173 | V1173R | D | 1.1e-05 | 9.7e-06 | 1.5e-05 | 1.190E-05 | 2.762E-06 | 9.700E-06 | 1.500E-05 |
| 23443 | 1173 | V1173N | D | 1e-05 | 1.1e-05 | 1.1e-05 | 1.067E-05 | 5.774E-07 | 1.000E-05 | 1.100E-05 |
| 23444 | 1173 | V1173D | D | 1.4e-05 | 1.2e-05 | 6.9e-06 | 1.097E-05 | 3.661E-06 | 6.900E-06 | 1.400E-05 |
| 23445 | 1173 | V1173C | D | 8.5e-06 | 1.4e-05 | 1.5e-05 | 1.250E-05 | 3.500E-06 | 8.500E-06 | 1.500E-05 |
| 23446 | 1173 | V1173E | D | 1.6e-05 | 1.5e-05 | 1.1e-05 | 1.400E-05 | 2.646E-06 | 1.100E-05 | 1.600E-05 |
| 23447 | 1173 | V1173Q | D | 1e-05 | 1.6e-05 | 8.8e-06 | 1.160E-05 | 3.857E-06 | 8.800E-06 | 1.600E-05 |
| 23448 | 1173 | V1173G | D | 1.2e-05 | 1.3e-05 | 1.4e-05 | 1.300E-05 | 1.000E-06 | 1.200E-05 | 1.400E-05 |
| 23449 | 1173 | V1173H | D | 8.1e-06 | 1.5e-05 | 1.5e-05 | 1.270E-05 | 3.984E-06 | 8.100E-06 | 1.500E-05 |
| 23450 | 1173 | V1173I | D | 1.2e-05 | 1.4e-05 | 1.2e-05 | 1.267E-05 | 1.155E-06 | 1.200E-05 | 1.400E-05 |
| 23451 | 1173 | V1173L | D | 1.6e-05 | 1.1e-05 | 2.2e-05 | 1.633E-05 | 5.508E-06 | 1.100E-05 | 2.200E-05 |
| 23452 | 1173 | V1173K | D | 1.5e-05 | 1.7e-05 | 8e-06 | 1.333E-05 | 4.726E-06 | 8.000E-06 | 1.700E-05 |
| 23453 | 1173 | V1173M | D | 1.3e-05 | 5.6e-06 | 8.5e-06 | 9.033E-06 | 3.729E-06 | 5.600E-06 | 1.300E-05 |
| 23454 | 1173 | V1173F | D | 1.1e-05 | 1.6e-05 | 1.7e-05 | 1.467E-05 | 3.215E-06 | 1.100E-05 | 1.700E-05 |
| 23455 | 1173 | V1173P | D | 1.2e-05 | 1e-05 | 1.1e-05 | 1.100E-05 | 1.000E-06 | 1.000E-05 | 1.200E-05 |
| 23456 | 1173 | V1173S | D | 1.4e-05 | 1.6e-05 | 1.1e-05 | 1.367E-05 | 2.517E-06 | 1.100E-05 | 1.600E-05 |
| 23457 | 1173 | V1173T | D | 1e-05 | 1.4e-05 | 1.4e-05 | 1.267E-05 | 2.309E-06 | 1.000E-05 | 1.400E-05 |
| 23458 | 1173 | V1173W | D | 1.5e-05 | 9.4e-06 | 1.1e-05 | 1.180E-05 | 2.884E-06 | 9.400E-06 | 1.500E-05 |
| 23459 | 1173 | V1173Y | D | 8.1e-06 | 1.4e-05 | 1.4e-05 | 1.203E-05 | 3.406E-06 | 8.100E-06 | 1.400E-05 |

| | | | | | | | | | | |
|---|---|---|---|---|---|---|---|---|---|---|
| 23460 | 1173 | V1173V | D | 1.1e-05 | 1.4e-05 | 1e-05 | 1.167E-05 | 2.082E-06 | 1.000E-05 | 1.400E-05 |
| 23461 | 1174 | F1174A | D | 1.9e-05 | 1.1e-05 | 1.1e-05 | 1.367E-05 | 4.619E-06 | 1.100E-05 | 1.900E-05 |
| 23462 | 1174 | F1174R | D | 9.1e-06 | 1.9e-05 | 1e-05 | 1.270E-05 | 5.474E-06 | 9.100E-06 | 1.900E-05 |
| 23463 | 1174 | F1174N | D | 7.8e-06 | 1.5e-05 | 1.6e-05 | 1.293E-05 | 4.474E-06 | 7.800E-06 | 1.600E-05 |
| 23464 | 1174 | F1174D | D | 1.6e-05 | 1.3e-05 | 1.6e-05 | 1.500E-05 | 1.732E-06 | 1.300E-05 | 1.600E-05 |
| 23465 | 1174 | F1174C | D | 8.7e-06 | 1e-05 | 1e-05 | 9.567E-06 | 7.506E-07 | 8.700E-06 | 1.000E-05 |
| 23466 | 1174 | F1174E | D | 1e-05 | 1.4e-05 | 1.5e-05 | 1.300E-05 | 2.646E-06 | 1.000E-05 | 1.500E-05 |
| 23467 | 1174 | F1174Q | D | 6.5e-06 | 1.7e-05 | 1.2e-05 | 1.183E-05 | 5.252E-06 | 6.500E-06 | 1.700E-05 |
| 23468 | 1174 | F1174G | D | 9.9e-06 | 1.1e-05 | 1.1e-05 | 1.063E-05 | 6.351E-07 | 9.900E-06 | 1.100E-05 |
| 23469 | 1174 | F1174H | D | 1.6e-05 | 1e-05 | 1.5e-05 | 1.367E-05 | 3.215E-06 | 1.000E-05 | 1.600E-05 |
| 23470 | 1174 | F1174I | D | 1.2e-05 | 1e-05 | 1.7e-05 | 1.300E-05 | 3.606E-06 | 1.000E-05 | 1.700E-05 |
| 23471 | 1174 | F1174L | D | 1.6e-05 | 9.4e-06 | 1.4e-05 | 1.313E-05 | 3.384E-06 | 9.400E-06 | 1.600E-05 |
| 23472 | 1174 | F1174K | D | 1e-05 | 1.8e-05 | 1.5e-05 | 1.433E-05 | 4.041E-06 | 1.000E-05 | 1.800E-05 |
| 23473 | 1174 | F1174M | D | 1e-05 | 8.6e-06 | 9.5e-06 | 9.367E-06 | 7.095E-07 | 8.600E-06 | 1.000E-05 |
| 23474 | 1174 | F1174F | D | 1.7e-05 | 1.6e-05 | 1.1e-05 | 1.467E-05 | 3.215E-06 | 1.100E-05 | 1.700E-05 |
| 23475 | 1174 | F1174P | D | 1.4e-05 | 1.1e-05 | 1e-05 | 1.167E-05 | 2.082E-06 | 1.000E-05 | 1.400E-05 |
| 23476 | 1174 | F1174S | D | 1.3e-05 | 1.6e-05 | 1.1e-05 | 1.333E-05 | 2.517E-06 | 1.100E-05 | 1.600E-05 |
| 23477 | 1174 | F1174T | D | 1.2e-05 | 1.4e-05 | 1.4e-05 | 1.333E-05 | 1.155E-06 | 1.200E-05 | 1.400E-05 |
| 23478 | 1174 | F1174W | D | 1.4e-05 | 1.1e-05 | 1.1e-05 | 1.200E-05 | 1.732E-06 | 1.100E-05 | 1.400E-05 |
| 23479 | 1174 | F1174Y | D | 1.6e-05 | 1.4e-05 | 1.4e-05 | 1.467E-05 | 1.155E-06 | 1.400E-05 | 1.600E-05 |
| 23480 | 1174 | F1174V | D | 1.8e-05 | 1.4e-05 | 1e-05 | 1.400E-05 | 4.000E-06 | 1.000E-05 | 1.800E-05 |
| 23481 | 1175 | I1175A | D | 1e-05 | 1.5e-05 | 1.7e-05 | 1.400E-05 | 3.606E-06 | 1.000E-05 | 1.700E-05 |
| 23482 | 1175 | I1175R | D | 1.4e-05 | 1.3e-05 | 9.7e-06 | 1.223E-05 | 2.250E-06 | 9.700E-06 | 1.400E-05 |
| 23483 | 1175 | I1175N | D | 1.8e-05 | 1.3e-05 | 1e-05 | 1.367E-05 | 4.041E-06 | 1.000E-05 | 1.800E-05 |
| 23484 | 1175 | I1175D | D | 1e-05 | 1.1e-05 | 1.6e-05 | 1.233E-05 | 3.215E-06 | 1.000E-05 | 1.600E-05 |
| 23485 | 1175 | I1175C | D | 1.1e-05 | 1.7e-05 | 1e-05 | 1.267E-05 | 3.786E-06 | 1.000E-05 | 1.700E-05 |
| 23486 | 1175 | I1175E | D | 1.1e-05 | 1.1e-05 | 1.5e-05 | 1.233E-05 | 2.309E-06 | 1.100E-05 | 1.500E-05 |
| 23487 | 1175 | I1175Q | D | 1.4e-05 | 9.6e-06 | 1.4e-05 | 1.253E-05 | 2.540E-06 | 9.600E-06 | 1.400E-05 |
| 23488 | 1175 | I1175G | D | 1.1e-05 | 1e-05 | 1.3e-05 | 1.133E-05 | 1.528E-06 | 1.000E-05 | 1.300E-05 |
| 23489 | 1175 | I1175H | D | 1.5e-05 | 9.2e-06 | 1.5e-05 | 1.307E-05 | 3.349E-06 | 9.200E-06 | 1.500E-05 |
| 23490 | 1175 | I1175I | D | 1.1e-05 | 1.5e-05 | 1.1e-05 | 1.233E-05 | 2.309E-06 | 1.100E-05 | 1.500E-05 |
| 23491 | 1175 | I1175L | D | 9.4e-06 | 1.4e-05 | 1.2e-05 | 1.180E-05 | 2.307E-06 | 9.400E-06 | 1.400E-05 |
| 23492 | 1175 | I1175K | D | 1.3e-05 | 9.9e-06 | 1.3e-05 | 1.197E-05 | 1.790E-06 | 9.900E-06 | 1.300E-05 |
| 23493 | 1175 | I1175M | D | 1.2e-05 | 1.4e-05 | 1e-05 | 1.200E-05 | 2.000E-06 | 1.000E-05 | 1.400E-05 |
| 23494 | 1175 | I1175F | D | 9.5e-06 | 1.4e-05 | 9.4e-06 | 1.097E-05 | 2.627E-06 | 9.400E-06 | 1.400E-05 |
| 23495 | 1175 | I1175P | D | 1.1e-05 | 1.6e-05 | 1.1e-05 | 1.267E-05 | 2.887E-06 | 1.100E-05 | 1.600E-05 |
| 23496 | 1175 | I1175S | D | 7.1e-06 | 1.4e-05 | 1e-05 | 1.037E-05 | 3.465E-06 | 7.100E-06 | 1.400E-05 |
| 23497 | 1175 | I1175T | D | 1.6e-05 | 1.6e-05 | 1.4e-05 | 1.533E-05 | 1.155E-06 | 1.400E-05 | 1.600E-05 |
| 23498 | 1175 | I1175W | D | 9.4e-06 | 1.1e-05 | 1e-05 | 1.013E-05 | 8.083E-07 | 9.400E-06 | 1.100E-05 |
| 23499 | 1175 | I1175Y | D | 1.5e-05 | 1.6e-05 | 1.4e-05 | 1.500E-05 | 1.000E-06 | 1.400E-05 | 1.600E-05 |
| 23500 | 1175 | I1175V | D | 9.1e-06 | 1e-05 | 1.3e-05 | 1.070E-05 | 2.042E-06 | 9.100E-06 | 1.300E-05 |
| 23501 | 1176 | F1176A | D | 1.1e-05 | 1.1e-05 | 1.5e-05 | 1.233E-05 | 2.309E-06 | 1.100E-05 | 1.500E-05 |
| 23502 | 1176 | F1176R | D | 9.4e-06 | 9.7e-06 | 1.6e-05 | 1.170E-05 | 3.727E-06 | 9.400E-06 | 1.600E-05 |
| 23503 | 1176 | F1176N | D | 7.2e-06 | 1.5e-05 | 1.5e-05 | 1.240E-05 | 4.503E-06 | 7.200E-06 | 1.500E-05 |
| 23504 | 1176 | F1176D | D | 2e-05 | 1.8e-05 | 1.6e-05 | 1.800E-05 | 2.000E-06 | 1.600E-05 | 2.000E-05 |
| 23505 | 1176 | F1176C | D | 1.5e-05 | 1.3e-05 | 1.5e-05 | 1.433E-05 | 1.155E-06 | 1.300E-05 | 1.500E-05 |
| 23506 | 1176 | F1176E | D | 1e-05 | 1.5e-05 | 8.9e-06 | 1.130E-05 | 3.251E-06 | 8.900E-06 | 1.500E-05 |
| 23507 | 1176 | F1176Q | D | 1.5e-05 | 9.1e-06 | 1.4e-05 | 1.270E-05 | 3.158E-06 | 9.100E-06 | 1.500E-05 |
| 23508 | 1176 | F1176G | D | 1.5e-05 | 1.6e-05 | 1.5e-05 | 1.533E-05 | 5.774E-07 | 1.500E-05 | 1.600E-05 |
| 23509 | 1176 | F1176H | D | 9.8e-06 | 1.1e-05 | 1.5e-05 | 1.193E-05 | 2.723E-06 | 9.800E-06 | 1.500E-05 |
| 23510 | 1176 | F1176I | D | 1.2e-05 | 1.3e-05 | 1.5e-05 | 1.333E-05 | 1.528E-06 | 1.200E-05 | 1.500E-05 |
| 23511 | 1176 | F1176L | D | 8.8e-06 | 9.6e-06 | 1.2e-05 | 1.013E-05 | 1.665E-06 | 8.800E-06 | 1.200E-05 |
| 23512 | 1176 | F1176K | D | 9.8e-06 | 1.4e-05 | 1.4e-05 | 1.260E-05 | 2.425E-06 | 9.800E-06 | 1.400E-05 |
| 23513 | 1176 | F1176M | D | 1e-05 | 9.6e-06 | 1.4e-05 | 1.120E-05 | 2.433E-06 | 9.600E-06 | 1.400E-05 |
| 23514 | 1176 | F1176F | D | 1.4e-05 | 1.5e-05 | 1.6e-05 | 1.500E-05 | 1.000E-06 | 1.400E-05 | 1.600E-05 |
| 23515 | 1176 | F1176P | D | 1e-05 | 2.9e-05 | 1.4e-05 | 1.767E-05 | 1.002E-05 | 1.000E-05 | 2.900E-05 |
| 23516 | 1176 | F1176S | D | 1.8e-05 | 9.7e-06 | 1.6e-05 | 1.457E-05 | 4.332E-06 | 9.700E-06 | 1.800E-05 |
| 23517 | 1176 | F1176T | D | 1.5e-05 | 1.5e-05 | 9.4e-06 | 1.313E-05 | 3.233E-06 | 9.400E-06 | 1.500E-05 |
| 23518 | 1176 | F1176W | D | 9.7e-06 | 1.7e-05 | 1.4e-05 | 1.357E-05 | 3.669E-06 | 9.700E-06 | 1.700E-05 |
| 23519 | 1176 | F1176Y | D | 1.3e-05 | 8.2e-06 | 1.5e-05 | 1.207E-05 | 3.495E-06 | 8.200E-06 | 1.500E-05 |
| 23520 | 1176 | F1176V | D | 1.1e-05 | 1.6e-05 | 1.4e-05 | 1.367E-05 | 2.517E-06 | 1.100E-05 | 1.600E-05 |
| 23521 | 1177 | P1177A | D | 1.4e-05 | 1.6e-05 | 1.1e-05 | 1.367E-05 | 2.517E-06 | 1.100E-05 | 1.600E-05 |
| 23522 | 1177 | P1177R | D | 8.8e-06 | 8.5e-06 | 8.5e-06 | 8.600E-06 | 1.732E-07 | 8.500E-06 | 8.800E-06 |
| 23523 | 1177 | P1177N | D | 1.4e-05 | 1.1e-05 | 9.5e-06 | 1.150E-05 | 2.291E-06 | 9.500E-06 | 1.400E-05 |
| 23524 | 1177 | P1177D | D | 1.5e-05 | 9.9e-06 | 1.5e-05 | 1.330E-05 | 2.944E-06 | 9.900E-06 | 1.500E-05 |
| 23525 | 1177 | P1177C | D | 9.6e-06 | 1.5e-05 | 1.3e-05 | 1.253E-05 | 2.730E-06 | 9.600E-06 | 1.500E-05 |
| 23526 | 1177 | P1177E | D | 1.3e-05 | 6.8e-06 | 1.1e-05 | 1.027E-05 | 3.164E-06 | 6.800E-06 | 1.300E-05 |
| 23527 | 1177 | P1177Q | D | 9.7e-06 | 1.4e-05 | 1.7e-05 | 1.357E-05 | 3.669E-06 | 9.700E-06 | 1.700E-05 |
| 23528 | 1177 | P1177G | D | 1.4e-05 | 1.3e-05 | 1.2e-05 | 1.300E-05 | 1.000E-06 | 1.200E-05 | 1.400E-05 |
| 23529 | 1177 | P1177H | D | 1.2e-05 | 1.4e-05 | 1.1e-05 | 1.233E-05 | 1.528E-06 | 1.100E-05 | 1.400E-05 |
| 23530 | 1177 | P1177I | D | 1.4e-05 | 1.1e-05 | 1.3e-05 | 1.267E-05 | 1.528E-06 | 1.100E-05 | 1.400E-05 |
| 23531 | 1177 | P1177L | D | 1e-05 | 9.6e-06 | 1.6e-05 | 1.187E-05 | 3.585E-06 | 9.600E-06 | 1.600E-05 |
| 23532 | 1177 | P1177K | D | 1.2e-05 | 1.1e-05 | 1.2e-05 | 1.167E-05 | 5.774E-07 | 1.100E-05 | 1.200E-05 |
| 23533 | 1177 | P1177M | D | 1.5e-05 | 1.1e-05 | 1.6e-05 | 1.400E-05 | 2.646E-06 | 1.100E-05 | 1.600E-05 |
| 23534 | 1177 | P1177F | D | 8.9e-06 | 1.1e-05 | 9.6e-06 | 9.833E-06 | 1.069E-06 | 8.900E-06 | 1.100E-05 |
| 23535 | 1177 | P1177P | D | 1e-05 | 1.1e-05 | 1.1e-05 | 1.067E-05 | 5.774E-07 | 1.000E-05 | 1.100E-05 |
| 23536 | 1177 | P1177S | D | 9.2e-06 | 1.5e-05 | 9.5e-06 | 1.123E-05 | 3.265E-06 | 9.200E-06 | 1.500E-05 |
| 23537 | 1177 | P1177T | D | 1.1e-05 | 1.6e-05 | 1e-05 | 1.233E-05 | 3.215E-06 | 1.000E-05 | 1.600E-05 |
| 23538 | 1177 | P1177W | D | 1e-05 | 8.2e-06 | 1.3e-05 | 1.040E-05 | 2.425E-06 | 8.200E-06 | 1.300E-05 |
| 23539 | 1177 | P1177Y | D | 1.4e-05 | 1.4e-05 | 1.4e-05 | 1.400E-05 | 0.000E+00 | 1.400E-05 | 1.400E-05 |
| 23540 | 1177 | P1177V | D | 1.5e-05 | 1.5e-05 | 1.6e-05 | 1.533E-05 | 5.774E-07 | 1.500E-05 | 1.600E-05 |
| 23541 | 1178 | P1178A | D | 1.2e-05 | 1.1e-05 | 1.3e-05 | 1.200E-05 | 1.000E-06 | 1.100E-05 | 1.300E-05 |
| 23542 | 1178 | P1178R | D | 1.7e-05 | 1.6e-05 | 9.2e-06 | 1.407E-05 | 4.244E-06 | 9.200E-06 | 1.700E-05 |
| 23543 | 1178 | P1178N | D | 1.3e-05 | 1.5e-05 | 9.8e-06 | 1.260E-05 | 2.623E-06 | 9.800E-06 | 1.500E-05 |
| 23544 | 1178 | P1178D | D | 1.3e-05 | 1.6e-05 | 8.8e-06 | 1.260E-05 | 3.617E-06 | 8.800E-06 | 1.600E-05 |

| | | | | | | | | | | |
|---|---|---|---|---|---|---|---|---|---|---|
| 23545 | 1178 | P1178C | D | 9.4e-06 | 1.4e-05 | 1.6e-05 | 1.313E-05 | 3.384E-06 | 9.400E-06 | 1.600E-05 |
| 23546 | 1178 | P1178E | D | 1.3e-05 | 1e-05 | 7.3e-06 | 1.010E-05 | 2.851E-06 | 7.300E-06 | 1.300E-05 |
| 23547 | 1178 | P1178Q | D | 1.2e-05 | 1.2e-05 | 1e-05 | 1.133E-05 | 1.155E-06 | 1.000E-05 | 1.200E-05 |
| 23548 | 1178 | P1178G | D | 1.7e-05 | 1.5e-05 | 1.4e-05 | 1.533E-05 | 1.528E-06 | 1.400E-05 | 1.700E-05 |
| 23549 | 1178 | P1178H | D | 1e-05 | 8.7e-06 | 8.8e-06 | 9.167E-06 | 7.234E-07 | 8.700E-06 | 1.000E-05 |
| 23550 | 1178 | P1178I | D | 1.7e-05 | 1.5e-05 | 1e-05 | 1.400E-05 | 3.606E-06 | 1.000E-05 | 1.700E-05 |
| 23551 | 1178 | P1178L | D | 1.4e-05 | 1.1e-05 | 7.7e-06 | 1.090E-05 | 3.151E-06 | 7.700E-06 | 1.400E-05 |
| 23552 | 1178 | P1178K | D | 1.5e-05 | 9.9e-06 | 1.6e-05 | 1.363E-05 | 3.272E-06 | 9.900E-06 | 1.600E-05 |
| 23553 | 1178 | P1178M | D | 1.5e-05 | 9.8e-06 | 1.5e-05 | 1.327E-05 | 3.002E-06 | 9.800E-06 | 1.500E-05 |
| 23554 | 1178 | P1178F | D | 1.5e-05 | 1.5e-05 | 1.5e-05 | 1.500E-05 | 0.000E+00 | 1.500E-05 | 1.500E-05 |
| 23555 | 1178 | P1178P | D | 1.1e-05 | 1e-05 | 1.6e-05 | 1.233E-05 | 3.215E-06 | 1.000E-05 | 1.600E-05 |
| 23556 | 1178 | P1178S | D | 1.4e-05 | 7.8e-06 | 2.4e-05 | 1.527E-05 | 8.174E-06 | 7.800E-06 | 2.400E-05 |
| 23557 | 1178 | P1178T | D | 1.6e-05 | 1.4e-05 | 1e-05 | 1.333E-05 | 3.055E-06 | 1.000E-05 | 1.600E-05 |
| 23558 | 1178 | P1178W | D | 1.6e-05 | 9.9e-06 | 9.8e-06 | 1.190E-05 | 3.551E-06 | 9.800E-06 | 1.600E-05 |
| 23559 | 1178 | P1178Y | D | 1.1e-05 | 7.1e-06 | 1e-05 | 9.367E-06 | 2.026E-06 | 7.100E-06 | 1.100E-05 |
| 23560 | 1178 | P1178V | D | 9.8e-06 | 1.6e-05 | 9.9e-06 | 1.190E-05 | 3.551E-06 | 9.800E-06 | 1.600E-05 |
| 23561 | 1179 | S1179A | D | 1.6e-05 | 1.4e-05 | 1.3e-05 | 1.433E-05 | 1.528E-06 | 1.300E-05 | 1.600E-05 |
| 23562 | 1179 | S1179R | D | 1.1e-05 | 1.6e-05 | 1.6e-05 | 1.433E-05 | 2.887E-06 | 1.100E-05 | 1.600E-05 |
| 23563 | 1179 | S1179N | D | 1.1e-05 | 1.1e-05 | 1.2e-05 | 1.133E-05 | 5.774E-07 | 1.100E-05 | 1.200E-05 |
| 23564 | 1179 | S1179D | D | 1.5e-05 | 1.2e-05 | 1e-05 | 1.233E-05 | 2.517E-06 | 1.000E-05 | 1.500E-05 |
| 23565 | 1179 | S1179C | D | 1.4e-05 | 1.6e-05 | 8.5e-06 | 1.283E-05 | 3.884E-06 | 8.500E-06 | 1.600E-05 |
| 23566 | 1179 | S1179E | D | 1.1e-05 | 8.5e-06 | 9e-06 | 9.500E-06 | 1.323E-06 | 8.500E-06 | 1.100E-05 |
| 23567 | 1179 | S1179Q | D | 1e-05 | 1e-05 | 9.6e-06 | 9.867E-06 | 2.309E-07 | 9.600E-06 | 1.000E-05 |
| 23568 | 1179 | S1179G | D | 1.3e-05 | 1.5e-05 | 1.5e-05 | 1.433E-05 | 1.155E-06 | 1.300E-05 | 1.500E-05 |
| 23569 | 1179 | S1179H | D | 1.6e-05 | 1.6e-05 | 1.1e-05 | 1.433E-05 | 2.887E-06 | 1.100E-05 | 1.600E-05 |
| 23570 | 1179 | S1179I | D | 1e-05 | 1.5e-05 | 1.4e-05 | 1.300E-05 | 2.646E-06 | 1.000E-05 | 1.500E-05 |
| 23571 | 1179 | S1179L | D | 1e-05 | 1.7e-05 | 1.9e-05 | 1.533E-05 | 4.726E-06 | 1.000E-05 | 1.900E-05 |
| 23572 | 1179 | S1179K | D | 1.5e-05 | 1.1e-05 | 1.4e-05 | 1.333E-05 | 2.082E-06 | 1.100E-05 | 1.500E-05 |
| 23573 | 1179 | S1179M | D | 1e-05 | 1.1e-05 | 8.7e-06 | 9.900E-06 | 1.153E-06 | 8.700E-06 | 1.100E-05 |
| 23574 | 1179 | S1179F | D | 1.6e-05 | 1e-05 | 1.1e-05 | 1.233E-05 | 3.215E-06 | 1.000E-05 | 1.600E-05 |
| 23575 | 1179 | S1179P | D | 9.7e-06 | 1.5e-05 | 9e-06 | 1.123E-05 | 3.281E-06 | 9.000E-06 | 1.500E-05 |
| 23576 | 1179 | S1179S | D | 9.7e-06 | 1.5e-05 | 1.7e-05 | 1.390E-05 | 3.772E-06 | 9.700E-06 | 1.700E-05 |
| 23577 | 1179 | S1179T | D | 1.4e-05 | 1.3e-05 | 1.3e-05 | 1.333E-05 | 5.774E-07 | 1.300E-05 | 1.400E-05 |
| 23578 | 1179 | S1179W | D | 1.5e-05 | 1e-05 | 9.6e-06 | 1.153E-05 | 3.009E-06 | 9.600E-06 | 1.500E-05 |
| 23579 | 1179 | S1179Y | D | 9.1e-06 | 1.6e-05 | 1.4e-05 | 1.303E-05 | 3.550E-06 | 9.100E-06 | 1.600E-05 |
| 23580 | 1179 | S1179V | D | 1.5e-05 | 1.6e-05 | 1.1e-05 | 1.400E-05 | 2.646E-06 | 1.100E-05 | 1.600E-05 |
| 23581 | 1180 | D1180A | D | 9.4e-06 | 1.6e-05 | 1e-05 | 1.180E-05 | 3.650E-06 | 9.400E-06 | 1.600E-05 |
| 23582 | 1180 | D1180R | D | 1e-05 | 1.2e-05 | 1.4e-05 | 1.200E-05 | 2.000E-06 | 1.000E-05 | 1.400E-05 |
| 23583 | 1180 | D1180N | D | 1.6e-05 | 9.9e-06 | 1e-05 | 1.197E-05 | 3.493E-06 | 9.900E-06 | 1.600E-05 |
| 23584 | 1180 | D1180D | D | 1.9e-05 | 8.6e-06 | 1.5e-05 | 1.420E-05 | 5.246E-06 | 8.600E-06 | 1.900E-05 |
| 23585 | 1180 | D1180C | D | 1.3e-05 | 2.6e-05 | 1.5e-05 | 1.800E-05 | 7.000E-06 | 1.300E-05 | 2.600E-05 |
| 23586 | 1180 | D1180E | D | 1.5e-05 | 1.5e-05 | 8.4e-06 | 1.280E-05 | 3.811E-06 | 8.400E-06 | 1.500E-05 |
| 23587 | 1180 | D1180Q | D | 8.9e-06 | 1e-05 | 2.3e-05 | 1.397E-05 | 7.842E-06 | 8.900E-06 | 2.300E-05 |
| 23588 | 1180 | D1180G | D | 1.4e-05 | 9.6e-06 | 1.8e-05 | 1.387E-05 | 4.202E-06 | 9.600E-06 | 1.800E-05 |
| 23589 | 1180 | D1180H | D | 1.3e-05 | 8.3e-06 | 9.8e-06 | 1.037E-05 | 2.401E-06 | 8.300E-06 | 1.300E-05 |
| 23590 | 1180 | D1180I | D | 1.6e-05 | 1.1e-05 | 1.4e-05 | 1.367E-05 | 2.517E-06 | 1.100E-05 | 1.600E-05 |
| 23591 | 1180 | D1180L | D | 1.6e-05 | 1.6e-05 | 1.6e-05 | 1.600E-05 | 0.000E+00 | 1.600E-05 | 1.600E-05 |
| 23592 | 1180 | D1180K | D | 1.6e-05 | 1.3e-05 | 1.7e-05 | 1.533E-05 | 2.082E-06 | 1.300E-05 | 1.700E-05 |
| 23593 | 1180 | D1180M | D | 1.3e-05 | 1e-05 | 1.2e-05 | 1.167E-05 | 1.528E-06 | 1.000E-05 | 1.300E-05 |
| 23594 | 1180 | D1180F | D | 1.4e-05 | 1.5e-05 | 1.3e-05 | 1.400E-05 | 1.000E-06 | 1.300E-05 | 1.500E-05 |
| 23595 | 1180 | D1180P | D | 9.5e-06 | 1.5e-05 | 1.1e-05 | 1.183E-05 | 2.843E-06 | 9.500E-06 | 1.500E-05 |
| 23596 | 1180 | D1180S | D | 1.6e-05 | 1e-05 | 1.4e-05 | 1.333E-05 | 3.055E-06 | 1.000E-05 | 1.600E-05 |
| 23597 | 1180 | D1180T | D | 1.3e-05 | 9.8e-06 | 1.4e-05 | 1.227E-05 | 2.194E-06 | 9.800E-06 | 1.400E-05 |
| 23598 | 1180 | D1180W | D | 9e-06 | 1.5e-05 | 1.5e-05 | 1.300E-05 | 3.464E-06 | 9.000E-06 | 1.500E-05 |
| 23599 | 1180 | D1180Y | D | 9.8e-06 | 1.5e-05 | 1.6e-05 | 1.360E-05 | 3.329E-06 | 9.800E-06 | 1.600E-05 |
| 23600 | 1180 | D1180V | D | 1e-05 | 1.8e-05 | 1.6e-05 | 1.467E-05 | 4.163E-06 | 1.000E-05 | 1.800E-05 |
| 23601 | 1181 | E1181A | D | 1.7e-05 | 8.9e-06 | 8.8e-06 | 1.157E-05 | 4.706E-06 | 8.800E-06 | 1.700E-05 |
| 23602 | 1181 | E1181R | D | 9.5e-06 | 7.6e-06 | 1.5e-05 | 1.070E-05 | 3.843E-06 | 7.600E-06 | 1.500E-05 |
| 23603 | 1181 | E1181N | D | 9.9e-06 | 1.6e-05 | 1.5e-05 | 1.363E-05 | 3.272E-06 | 9.900E-06 | 1.600E-05 |
| 23604 | 1181 | E1181D | D | 1e-05 | 1.5e-05 | 9.3e-06 | 1.143E-05 | 3.109E-06 | 9.300E-06 | 1.500E-05 |
| 23605 | 1181 | E1181C | D | 9.9e-06 | 1.5e-05 | 9.8e-06 | 1.157E-05 | 2.974E-06 | 9.800E-06 | 1.500E-05 |
| 23606 | 1181 | E1181E | D | 1.7e-05 | 1.2e-05 | 1.2e-05 | 1.367E-05 | 2.887E-06 | 1.200E-05 | 1.700E-05 |
| 23607 | 1181 | E1181Q | D | 1.5e-05 | 1.1e-05 | 1e-05 | 1.200E-05 | 2.646E-06 | 1.000E-05 | 1.500E-05 |
| 23608 | 1181 | E1181G | D | 1.5e-05 | 1.4e-05 | 1.5e-05 | 1.467E-05 | 5.774E-07 | 1.400E-05 | 1.500E-05 |
| 23609 | 1181 | E1181H | D | 1.5e-05 | 1.1e-05 | 1.2e-05 | 1.267E-05 | 2.082E-06 | 1.100E-05 | 1.500E-05 |
| 23610 | 1181 | E1181I | D | 1.1e-05 | 1.3e-05 | 1.6e-05 | 1.333E-05 | 2.517E-06 | 1.100E-05 | 1.600E-05 |
| 23611 | 1181 | E1181L | D | 9.6e-06 | 8.2e-06 | 1.3e-05 | 1.027E-05 | 2.468E-06 | 8.200E-06 | 1.300E-05 |
| 23612 | 1181 | E1181K | D | 1.1e-05 | 1.5e-05 | 1.1e-05 | 1.233E-05 | 2.309E-06 | 1.100E-05 | 1.500E-05 |
| 23613 | 1181 | E1181M | D | 1.1e-05 | 1.5e-05 | 1e-05 | 1.200E-05 | 2.646E-06 | 1.000E-05 | 1.500E-05 |
| 23614 | 1181 | E1181F | D | 1.8e-05 | 1.1e-05 | 1.1e-05 | 1.333E-05 | 4.041E-06 | 1.100E-05 | 1.800E-05 |
| 23615 | 1181 | E1181P | D | 1e-05 | 1.5e-05 | 1.1e-05 | 1.200E-05 | 2.646E-06 | 1.000E-05 | 1.500E-05 |
| 23616 | 1181 | E1181S | D | 1.1e-05 | 9.8e-06 | 1.5e-05 | 1.193E-05 | 2.723E-06 | 9.800E-06 | 1.500E-05 |
| 23617 | 1181 | E1181T | D | 2.2e-05 | 9.3e-06 | 1.3e-05 | 1.477E-05 | 6.532E-06 | 9.300E-06 | 2.200E-05 |
| 23618 | 1181 | E1181W | D | 1.4e-05 | 1.3e-05 | 1.1e-05 | 1.267E-05 | 1.528E-06 | 1.100E-05 | 1.400E-05 |
| 23619 | 1181 | E1181Y | D | 1.6e-05 | 1.4e-05 | 1.4e-05 | 1.467E-05 | 1.155E-06 | 1.400E-05 | 1.600E-05 |
| 23620 | 1181 | E1181V | D | 1.4e-05 | 1.3e-05 | 1.4e-05 | 1.367E-05 | 5.774E-07 | 1.300E-05 | 1.400E-05 |
| 23621 | 1182 | Q1182A | D | 1.5e-05 | 1.5e-05 | 1.7e-05 | 1.567E-05 | 1.155E-06 | 1.500E-05 | 1.700E-05 |
| 23622 | 1182 | Q1182R | D | 1.4e-05 | 1e-05 | 9.3e-06 | 1.110E-05 | 2.536E-06 | 9.300E-06 | 1.400E-05 |
| 23623 | 1182 | Q1182N | D | 1e-05 | 1e-05 | 9.5e-06 | 9.833E-06 | 2.887E-07 | 9.500E-06 | 1.000E-05 |
| 23624 | 1182 | Q1182D | D | 1.3e-05 | 1.2e-05 | 1.3e-05 | 1.267E-05 | 5.774E-07 | 1.200E-05 | 1.300E-05 |
| 23625 | 1182 | Q1182C | D | 1e-05 | 2.1e-05 | 1.4e-05 | 1.500E-05 | 5.568E-06 | 1.000E-05 | 2.100E-05 |
| 23626 | 1182 | Q1182E | D | 2.1e-05 | 1.5e-05 | 1.9e-05 | 1.833E-05 | 3.055E-06 | 1.500E-05 | 2.100E-05 |
| 23627 | 1182 | Q1182Q | D | 1e-05 | 1.1e-05 | 1e-05 | 1.033E-05 | 5.774E-07 | 1.000E-05 | 1.100E-05 |
| 23628 | 1182 | Q1182G | D | 1.2e-05 | 9.9e-06 | 1e-05 | 1.063E-05 | 1.185E-06 | 9.900E-06 | 1.200E-05 |
| 23629 | 1182 | Q1182H | D | 1.5e-05 | 9.5e-06 | 7.3e-06 | 1.060E-05 | 3.966E-06 | 7.300E-06 | 1.500E-05 |

| | | | | | | | | | | |
|---|---|---|---|---|---|---|---|---|---|---|
| 23630 | 1182 | Q1182I | D | 1e-05 | 1e-05 | 1.4e-05 | 1.133E-05 | 2.309E-06 | 1.000E-05 | 1.400E-05 |
| 23631 | 1182 | Q1182L | D | 1.7e-05 | 1.1e-05 | 1.2e-05 | 1.333E-05 | 3.215E-06 | 1.100E-05 | 1.700E-05 |
| 23632 | 1182 | Q1182K | D | 1.4e-05 | 1.5e-05 | 9.8e-06 | 1.293E-05 | 2.759E-06 | 9.800E-06 | 1.500E-05 |
| 23633 | 1182 | Q1182M | D | 1.5e-05 | 1.8e-05 | 1.5e-05 | 1.600E-05 | 1.732E-06 | 1.500E-05 | 1.800E-05 |
| 23634 | 1182 | Q1182F | D | 1.4e-05 | 1.3e-05 | 1.6e-05 | 1.433E-05 | 1.528E-06 | 1.300E-05 | 1.600E-05 |
| 23635 | 1182 | Q1182P | D | 1.3e-05 | 1.4e-05 | 9.3e-06 | 1.210E-05 | 2.476E-06 | 9.300E-06 | 1.400E-05 |
| 23636 | 1182 | Q1182S | D | 1.6e-05 | 1.6e-05 | 1.3e-05 | 1.500E-05 | 1.732E-06 | 1.300E-05 | 1.600E-05 |
| 23637 | 1182 | Q1182T | D | 1.5e-05 | 1.4e-05 | 1.1e-05 | 1.333E-05 | 2.082E-06 | 1.100E-05 | 1.500E-05 |
| 23638 | 1182 | Q1182W | D | 1.3e-05 | 1.7e-05 | 1e-05 | 1.333E-05 | 3.512E-06 | 1.000E-05 | 1.700E-05 |
| 23639 | 1182 | Q1182Y | D | 1.5e-05 | 1.2e-05 | 8.8e-06 | 1.193E-05 | 3.101E-06 | 8.800E-06 | 1.500E-05 |
| 23640 | 1182 | Q1182V | D | 1.1e-05 | 1.3e-05 | 1.3e-05 | 1.233E-05 | 1.155E-06 | 1.100E-05 | 1.300E-05 |
| 23641 | 1183 | L1183A | D | 1.2e-05 | 9.9e-06 | 1.4e-05 | 1.197E-05 | 2.050E-06 | 9.900E-06 | 1.400E-05 |
| 23642 | 1183 | L1183R | D | 1.4e-05 | 1.5e-05 | 9e-06 | 1.267E-05 | 3.215E-06 | 9.000E-06 | 1.500E-05 |
| 23643 | 1183 | L1183N | D | 1.3e-05 | 1.1e-05 | 1.4e-05 | 1.267E-05 | 1.528E-06 | 1.100E-05 | 1.400E-05 |
| 23644 | 1183 | L1183D | D | 1.6e-05 | 7.9e-06 | 1.4e-05 | 1.263E-05 | 4.219E-06 | 7.900E-06 | 1.600E-05 |
| 23645 | 1183 | L1183C | D | 1.4e-05 | 1.4e-05 | 1.7e-05 | 1.500E-05 | 1.732E-06 | 1.400E-05 | 1.700E-05 |
| 23646 | 1183 | L1183E | D | 1.4e-05 | 9.8e-06 | 9.1e-06 | 1.097E-05 | 2.650E-06 | 9.100E-06 | 1.400E-05 |
| 23647 | 1183 | L1183Q | D | 1.3e-05 | 1.6e-05 | 1.1e-05 | 1.333E-05 | 2.517E-06 | 1.100E-05 | 1.600E-05 |
| 23648 | 1183 | L1183G | D | 9.7e-06 | 1e-05 | 1e-05 | 9.900E-06 | 1.732E-07 | 9.700E-06 | 1.000E-05 |
| 23649 | 1183 | L1183H | D | 1.1e-05 | 1e-05 | 1.7e-05 | 1.267E-05 | 3.786E-06 | 1.000E-05 | 1.700E-05 |
| 23650 | 1183 | L1183I | D | 1.4e-05 | 1.5e-05 | 1.6e-05 | 1.500E-05 | 1.000E-06 | 1.400E-05 | 1.600E-05 |
| 23651 | 1183 | L1183L | D | 1.3e-05 | 1.1e-05 | 1.2e-05 | 1.200E-05 | 1.000E-06 | 1.100E-05 | 1.300E-05 |
| 23652 | 1183 | L1183K | D | 1.4e-05 | 1.4e-05 | 1.4e-05 | 1.400E-05 | 0.000E+00 | 1.400E-05 | 1.400E-05 |
| 23653 | 1183 | L1183M | D | 1.4e-05 | 1.4e-05 | 1.1e-05 | 1.300E-05 | 1.732E-06 | 1.100E-05 | 1.400E-05 |
| 23654 | 1183 | L1183F | D | 1.2e-05 | 1.1e-05 | 1.1e-05 | 1.133E-05 | 5.774E-07 | 1.100E-05 | 1.200E-05 |
| 23655 | 1183 | L1183P | D | 1.2e-05 | 9.6e-06 | 1.5e-05 | 1.220E-05 | 2.706E-06 | 9.600E-06 | 1.500E-05 |
| 23656 | 1183 | L1183S | D | 1.5e-05 | 1e-05 | 1.4e-05 | 1.300E-05 | 2.646E-06 | 1.000E-05 | 1.500E-05 |
| 23657 | 1183 | L1183T | D | 1.6e-05 | 1.6e-05 | 1.2e-05 | 1.467E-05 | 2.309E-06 | 1.200E-05 | 1.600E-05 |
| 23658 | 1183 | L1183W | D | 9.5e-06 | 1e-05 | 8.6e-06 | 9.367E-06 | 7.095E-07 | 8.600E-06 | 1.000E-05 |
| 23659 | 1183 | L1183Y | D | 1.6e-05 | 1.3e-05 | 1.5e-05 | 1.467E-05 | 1.528E-06 | 1.300E-05 | 1.600E-05 |
| 23660 | 1183 | L1183V | D | 1.3e-05 | 1.6e-05 | 1.6e-05 | 1.500E-05 | 1.732E-06 | 1.300E-05 | 1.600E-05 |
| 23661 | 1184 | K1184A | D | 1.6e-05 | 1.6e-05 | 1.3e-05 | 1.500E-05 | 1.732E-06 | 1.300E-05 | 1.600E-05 |
| 23662 | 1184 | K1184R | D | 1e-05 | 1.6e-05 | 1.3e-05 | 1.300E-05 | 3.000E-06 | 1.000E-05 | 1.600E-05 |
| 23663 | 1184 | K1184N | D | 1.4e-05 | 9.9e-06 | 1.4e-05 | 1.263E-05 | 2.367E-06 | 9.900E-06 | 1.400E-05 |
| 23664 | 1184 | K1184D | D | 7.3e-06 | 1.1e-05 | 1e-05 | 9.433E-06 | 1.914E-06 | 7.300E-06 | 1.100E-05 |
| 23665 | 1184 | K1184C | D | 6.8e-06 | 8.2e-06 | 1.5e-05 | 1.000E-05 | 4.386E-06 | 6.800E-06 | 1.500E-05 |
| 23666 | 1184 | K1184E | D | 1.2e-05 | 1.6e-05 | 1.6e-05 | 1.467E-05 | 2.309E-06 | 1.200E-05 | 1.600E-05 |
| 23667 | 1184 | K1184Q | D | 1.7e-05 | 1.4e-05 | 1.1e-05 | 1.400E-05 | 3.000E-06 | 1.100E-05 | 1.700E-05 |
| 23668 | 1184 | K1184G | D | 1.5e-05 | 1.3e-05 | 1.1e-05 | 1.300E-05 | 2.000E-06 | 1.100E-05 | 1.500E-05 |
| 23669 | 1184 | K1184H | D | 1.4e-05 | 8.8e-06 | 1.6e-05 | 1.293E-05 | 3.717E-06 | 8.800E-06 | 1.600E-05 |
| 23670 | 1184 | K1184I | D | 1.2e-05 | 1.1e-05 | 1.2e-05 | 1.167E-05 | 5.774E-07 | 1.100E-05 | 1.200E-05 |
| 23671 | 1184 | K1184L | D | 1.1e-05 | 8.6e-06 | 1.5e-05 | 1.153E-05 | 3.233E-06 | 8.600E-06 | 1.500E-05 |
| 23672 | 1184 | K1184K | D | 1e-05 | 1.1e-05 | 1.2e-05 | 1.100E-05 | 1.000E-06 | 1.000E-05 | 1.200E-05 |
| 23673 | 1184 | K1184M | D | 1e-05 | 8.1e-06 | 1.6e-05 | 1.137E-05 | 4.124E-06 | 8.100E-06 | 1.600E-05 |
| 23674 | 1184 | K1184F | D | 1.6e-05 | 1.7e-05 | 1.3e-05 | 1.533E-05 | 2.082E-06 | 1.300E-05 | 1.700E-05 |
| 23675 | 1184 | K1184P | D | 1.7e-05 | 1.6e-05 | 1.5e-05 | 1.600E-05 | 1.000E-06 | 1.500E-05 | 1.700E-05 |
| 23676 | 1184 | K1184S | D | 1.6e-05 | 1e-05 | 1.5e-05 | 1.367E-05 | 3.215E-06 | 1.000E-05 | 1.600E-05 |
| 23677 | 1184 | K1184T | D | 4.4e-05 | 1.5e-05 | 6e-06 | 2.167E-05 | 1.986E-05 | 6.000E-06 | 4.400E-05 |
| 23678 | 1184 | K1184W | D | 1.6e-05 | 1.6e-05 | 9.9e-06 | 1.397E-05 | 3.522E-06 | 9.900E-06 | 1.600E-05 |
| 23679 | 1184 | K1184Y | D | 1e-05 | 1.4e-05 | 1.5e-05 | 1.300E-05 | 2.646E-06 | 1.000E-05 | 1.500E-05 |
| 23680 | 1184 | K1184V | D | 1.5e-05 | 1.1e-05 | 9.9e-06 | 1.197E-05 | 2.684E-06 | 9.900E-06 | 1.500E-05 |
| 23681 | 1185 | S1185A | D | 1.4e-05 | 1.5e-05 | 1.3e-05 | 1.400E-05 | 1.000E-06 | 1.300E-05 | 1.500E-05 |
| 23682 | 1185 | S1185R | D | 9e-06 | 1.3e-05 | 1.1e-05 | 1.100E-05 | 2.000E-06 | 9.000E-06 | 1.300E-05 |
| 23683 | 1185 | S1185N | D | 1.5e-05 | 1.4e-05 | 1.1e-05 | 1.333E-05 | 2.082E-06 | 1.100E-05 | 1.500E-05 |
| 23684 | 1185 | S1185D | D | 1.5e-05 | 1.7e-05 | 1.3e-05 | 1.500E-05 | 2.000E-06 | 1.300E-05 | 1.700E-05 |
| 23685 | 1185 | S1185C | D | 1.5e-05 | 1.2e-05 | 1.4e-05 | 1.367E-05 | 1.528E-06 | 1.200E-05 | 1.500E-05 |
| 23686 | 1185 | S1185E | D | 1.5e-05 | 1.5e-05 | 1.5e-05 | 1.500E-05 | 0.000E+00 | 1.500E-05 | 1.500E-05 |
| 23687 | 1185 | S1185Q | D | 1e-05 | 9.3e-06 | 1.3e-05 | 1.077E-05 | 1.966E-06 | 9.300E-06 | 1.300E-05 |
| 23688 | 1185 | S1185G | D | 1.5e-05 | 1.5e-05 | 1.1e-05 | 1.367E-05 | 2.309E-06 | 1.100E-05 | 1.500E-05 |
| 23689 | 1185 | S1185H | D | 1.3e-05 | 1.4e-05 | 1.1e-05 | 1.267E-05 | 1.528E-06 | 1.100E-05 | 1.400E-05 |
| 23690 | 1185 | S1185I | D | 1.6e-05 | 1.2e-05 | 1.6e-05 | 1.467E-05 | 2.309E-06 | 1.200E-05 | 1.600E-05 |
| 23691 | 1185 | S1185L | D | 9.7e-06 | 1.2e-05 | 8.4e-06 | 1.003E-05 | 1.823E-06 | 8.400E-06 | 1.200E-05 |
| 23692 | 1185 | S1185K | D | 1e-05 | 1.5e-05 | 1.7e-05 | 1.400E-05 | 3.606E-06 | 1.000E-05 | 1.700E-05 |
| 23693 | 1185 | S1185M | D | 1.5e-05 | 1.1e-05 | 1.1e-05 | 1.233E-05 | 2.309E-06 | 1.100E-05 | 1.500E-05 |
| 23694 | 1185 | S1185F | D | 1.5e-05 | 1.5e-05 | 1.3e-05 | 1.433E-05 | 1.155E-06 | 1.300E-05 | 1.500E-05 |
| 23695 | 1185 | S1185P | D | 1.5e-05 | 9.7e-06 | 9.8e-06 | 1.150E-05 | 3.032E-06 | 9.700E-06 | 1.500E-05 |
| 23696 | 1185 | S1185S | D | 1.4e-05 | 1.1e-05 | 9.9e-06 | 1.163E-05 | 2.122E-06 | 9.900E-06 | 1.400E-05 |
| 23697 | 1185 | S1185T | D | 8.5e-06 | 9.8e-06 | 1.6e-05 | 1.143E-05 | 4.008E-06 | 8.500E-06 | 1.600E-05 |
| 23698 | 1185 | S1185W | D | 1.2e-05 | 9.8e-06 | 1.5e-05 | 1.227E-05 | 2.610E-06 | 9.800E-06 | 1.500E-05 |
| 23699 | 1185 | S1185Y | D | 6.9e-06 | 1.2e-05 | 9.5e-06 | 9.467E-06 | 2.550E-06 | 6.900E-06 | 1.200E-05 |
| 23700 | 1185 | S1185V | D | 1.4e-05 | 1.2e-05 | 1.6e-05 | 1.400E-05 | 2.000E-06 | 1.200E-05 | 1.600E-05 |
| 23701 | 1186 | G1186A | D | 1.2e-05 | 9.7e-06 | 9.7e-06 | 1.047E-05 | 1.328E-06 | 9.700E-06 | 1.200E-05 |
| 23702 | 1186 | G1186R | D | 1.4e-05 | 1.5e-05 | 1e-05 | 1.300E-05 | 2.646E-06 | 1.000E-05 | 1.500E-05 |
| 23703 | 1186 | G1186N | D | 1.5e-05 | 1.5e-05 | 1.4e-05 | 1.467E-05 | 5.774E-07 | 1.400E-05 | 1.500E-05 |
| 23704 | 1186 | G1186D | D | 9.2e-06 | 8.5e-06 | 1.5e-05 | 1.090E-05 | 3.568E-06 | 8.500E-06 | 1.500E-05 |
| 23705 | 1186 | G1186C | D | 1.1e-05 | 1e-05 | 1.1e-05 | 1.067E-05 | 5.774E-07 | 1.000E-05 | 1.100E-05 |
| 23706 | 1186 | G1186E | D | 1.1e-05 | 8.7e-06 | 2.3e-05 | 1.423E-05 | 7.679E-06 | 8.700E-06 | 2.300E-05 |
| 23707 | 1186 | G1186Q | D | 8.4e-06 | 8.3e-06 | 1.4e-05 | 1.023E-05 | 3.262E-06 | 8.300E-06 | 1.400E-05 |
| 23708 | 1186 | G1186G | D | 1.5e-05 | 1.2e-05 | 1.7e-05 | 1.467E-05 | 2.517E-06 | 1.200E-05 | 1.700E-05 |
| 23709 | 1186 | G1186H | D | 1.3e-05 | 1.1e-05 | 1.4e-05 | 1.267E-05 | 1.528E-06 | 1.100E-05 | 1.400E-05 |
| 23710 | 1186 | G1186I | D | 1.5e-05 | 1.5e-05 | 1e-05 | 1.333E-05 | 2.887E-06 | 1.000E-05 | 1.500E-05 |
| 23711 | 1186 | G1186L | D | 1.2e-05 | 1.1e-05 | 7.1e-06 | 1.003E-05 | 2.589E-06 | 7.100E-06 | 1.200E-05 |
| 23712 | 1186 | G1186K | D | 1.5e-05 | 9.5e-06 | 9e-06 | 1.117E-05 | 3.329E-06 | 9.000E-06 | 1.500E-05 |
| 23713 | 1186 | G1186M | D | 1.1e-05 | 1.3e-05 | 1.5e-05 | 1.300E-05 | 2.000E-06 | 1.100E-05 | 1.500E-05 |
| 23714 | 1186 | G1186F | D | 1.1e-05 | 1.2e-05 | 1.6e-05 | 1.300E-05 | 2.646E-06 | 1.100E-05 | 1.600E-05 |

| | | | | | | | | | | |
|---|---|---|---|---|---|---|---|---|---|---|
| 23715 | 1186 | G1186P | D | 1e-05 | 1.7e-05 | 1.6e-05 | 1.433E-05 | 3.786E-06 | 1.000E-05 | 1.700E-05 |
| 23716 | 1186 | G1186S | D | 1e-05 | 1.5e-05 | 1.2e-05 | 1.233E-05 | 2.517E-06 | 1.000E-05 | 1.500E-05 |
| 23717 | 1186 | G1186T | D | 1.2e-05 | 8.2e-06 | 1.4e-05 | 1.140E-05 | 2.946E-06 | 8.200E-06 | 1.400E-05 |
| 23718 | 1186 | G1186W | D | 1.5e-05 | 1.4e-05 | 1.4e-05 | 1.433E-05 | 5.774E-07 | 1.400E-05 | 1.500E-05 |
| 23719 | 1186 | G1186Y | D | 1.5e-05 | 1.7e-05 | 9.1e-06 | 1.370E-05 | 4.107E-06 | 9.100E-06 | 1.700E-05 |
| 23720 | 1186 | G1186V | D | 1.4e-05 | 1.6e-05 | 1.5e-05 | 1.500E-05 | 1.000E-06 | 1.400E-05 | 1.600E-05 |
| 23721 | 1187 | T1187A | D | 1.1e-05 | 1.1e-05 | 1e-05 | 1.067E-05 | 5.774E-07 | 1.000E-05 | 1.100E-05 |
| 23722 | 1187 | T1187R | D | 1.6e-05 | 1.5e-05 | 1e-05 | 1.367E-05 | 3.215E-06 | 1.000E-05 | 1.600E-05 |
| 23723 | 1187 | T1187N | D | 1.1e-05 | 1.5e-05 | 1e-05 | 1.200E-05 | 2.646E-06 | 1.000E-05 | 1.500E-05 |
| 23724 | 1187 | T1187D | D | 1.9e-05 | 1e-05 | 1.5e-05 | 1.467E-05 | 4.509E-06 | 1.000E-05 | 1.900E-05 |
| 23725 | 1187 | T1187C | D | 1.5e-05 | 1.7e-05 | 1.6e-05 | 1.600E-05 | 1.000E-06 | 1.500E-05 | 1.700E-05 |
| 23726 | 1187 | T1187E | D | 1.5e-05 | 1.5e-05 | 1.5e-05 | 1.500E-05 | 0.000E+00 | 1.500E-05 | 1.500E-05 |
| 23727 | 1187 | T1187Q | D | 1e-05 | 1.1e-05 | 1.2e-05 | 1.100E-05 | 1.000E-06 | 1.000E-05 | 1.200E-05 |
| 23728 | 1187 | T1187G | D | 1.5e-05 | 1.5e-05 | 1.4e-05 | 1.467E-05 | 5.774E-07 | 1.400E-05 | 1.500E-05 |
| 23729 | 1187 | T1187H | D | 1.6e-05 | 1.4e-05 | 1.1e-05 | 1.367E-05 | 2.517E-06 | 1.100E-05 | 1.600E-05 |
| 23730 | 1187 | T1187I | D | 1.4e-05 | 8.8e-06 | 1.6e-05 | 1.293E-05 | 3.717E-06 | 8.800E-06 | 1.600E-05 |
| 23731 | 1187 | T1187L | D | 1.7e-05 | 1.7e-05 | 1.3e-05 | 1.567E-05 | 2.309E-06 | 1.300E-05 | 1.700E-05 |
| 23732 | 1187 | T1187K | D | 1.2e-05 | 1.1e-05 | 9.6e-06 | 1.087E-05 | 1.206E-06 | 9.600E-06 | 1.200E-05 |
| 23733 | 1187 | T1187M | D | 1.5e-05 | 1.7e-05 | 1.3e-05 | 1.500E-05 | 2.000E-06 | 1.300E-05 | 1.700E-05 |
| 23734 | 1187 | T1187F | D | 6.5e-06 | 1.5e-05 | 1.6e-05 | 1.250E-05 | 5.220E-06 | 6.500E-06 | 1.600E-05 |
| 23735 | 1187 | T1187P | D | 1.5e-05 | 1.2e-05 | 1.5e-05 | 1.400E-05 | 1.732E-06 | 1.200E-05 | 1.500E-05 |
| 23736 | 1187 | T1187S | D | 1.7e-05 | 1.5e-05 | 1.5e-05 | 1.567E-05 | 1.155E-06 | 1.500E-05 | 1.700E-05 |
| 23737 | 1187 | T1187T | D | 1.3e-05 | 1.3e-05 | 1.6e-05 | 1.400E-05 | 1.732E-06 | 1.300E-05 | 1.600E-05 |
| 23738 | 1187 | T1187W | D | 9.8e-06 | 9.5e-06 | 1e-05 | 9.767E-06 | 2.517E-07 | 9.500E-06 | 1.000E-05 |
| 23739 | 1187 | T1187Y | D | 1.5e-05 | 1.3e-05 | 1.4e-05 | 1.400E-05 | 1.000E-06 | 1.300E-05 | 1.500E-05 |
| 23740 | 1187 | T1187V | D | 9.7e-06 | 1.5e-05 | 1.1e-05 | 1.190E-05 | 2.762E-06 | 9.700E-06 | 1.500E-05 |
| 23741 | 1188 | A1188A | D | 1.3e-05 | 1.3e-05 | 1.1e-05 | 1.233E-05 | 1.155E-06 | 1.100E-05 | 1.300E-05 |
| 23742 | 1188 | A1188R | D | 9.4e-06 | 1.7e-05 | 1e-05 | 1.213E-05 | 4.225E-06 | 9.400E-06 | 1.700E-05 |
| 23743 | 1188 | A1188N | D | 8.7e-06 | 1.4e-05 | 1.1e-05 | 1.123E-05 | 2.658E-06 | 8.700E-06 | 1.400E-05 |
| 23744 | 1188 | A1188D | D | 6.9e-06 | 8.5e-06 | 1e-05 | 8.467E-06 | 1.550E-06 | 6.900E-06 | 1.000E-05 |
| 23745 | 1188 | A1188C | D | 1.5e-05 | 1.1e-05 | 9.9e-06 | 1.197E-05 | 2.684E-06 | 9.900E-06 | 1.500E-05 |
| 23746 | 1188 | A1188E | D | 9.2e-06 | 9.9e-06 | 9.7e-06 | 9.600E-06 | 3.606E-07 | 9.200E-06 | 9.900E-06 |
| 23747 | 1188 | A1188Q | D | 9.2e-06 | 1.5e-05 | 1.5e-05 | 1.307E-05 | 3.349E-06 | 9.200E-06 | 1.500E-05 |
| 23748 | 1188 | A1188G | D | 1.4e-05 | 1.1e-05 | 1.6e-05 | 1.367E-05 | 2.517E-06 | 1.100E-05 | 1.600E-05 |
| 23749 | 1188 | A1188H | D | 1.5e-05 | 1.5e-05 | 1.2e-05 | 1.400E-05 | 1.732E-06 | 1.200E-05 | 1.500E-05 |
| 23750 | 1188 | A1188I | D | 1.5e-05 | 1.2e-05 | 1.3e-05 | 1.333E-05 | 1.528E-06 | 1.200E-05 | 1.500E-05 |
| 23751 | 1188 | A1188L | D | 9.1e-06 | 1.5e-05 | 9.9e-06 | 1.133E-05 | 3.201E-06 | 9.100E-06 | 1.500E-05 |
| 23752 | 1188 | A1188K | D | 1.4e-05 | 1.3e-05 | 1.1e-05 | 1.267E-05 | 1.528E-06 | 1.100E-05 | 1.400E-05 |
| 23753 | 1188 | A1188M | D | 1e-05 | 1.1e-05 | 7.3e-06 | 9.433E-06 | 1.914E-06 | 7.300E-06 | 1.100E-05 |
| 23754 | 1188 | A1188F | D | 1.1e-05 | 9.3e-06 | 1.5e-05 | 1.177E-05 | 2.926E-06 | 9.300E-06 | 1.500E-05 |
| 23755 | 1188 | A1188P | D | 1.5e-05 | 1e-05 | 1.5e-05 | 1.333E-05 | 2.887E-06 | 1.000E-05 | 1.500E-05 |
| 23756 | 1188 | A1188S | D | 9.3e-06 | 1.3e-05 | 1.4e-05 | 1.210E-05 | 2.476E-06 | 9.300E-06 | 1.400E-05 |
| 23757 | 1188 | A1188T | D | 1.3e-05 | 1.6e-05 | 1.1e-05 | 1.333E-05 | 2.517E-06 | 1.100E-05 | 1.600E-05 |
| 23758 | 1188 | A1188W | D | 1.3e-05 | 1.5e-05 | 9.8e-06 | 1.260E-05 | 2.623E-06 | 9.800E-06 | 1.500E-05 |
| 23759 | 1188 | A1188Y | D | 1.6e-05 | 1.2e-05 | 1.2e-05 | 1.333E-05 | 2.309E-06 | 1.200E-05 | 1.600E-05 |
| 23760 | 1188 | A1188V | D | 1.3e-05 | 1.6e-05 | 1.5e-05 | 1.467E-05 | 1.528E-06 | 1.300E-05 | 1.600E-05 |
| 23761 | 1189 | S1189A | D | 1.1e-05 | 1.2e-05 | 1.2e-05 | 1.167E-05 | 5.774E-07 | 1.100E-05 | 1.200E-05 |
| 23762 | 1189 | S1189R | D | 1.1e-05 | 1.6e-05 | 1e-05 | 1.233E-05 | 3.215E-06 | 1.000E-05 | 1.600E-05 |
| 23763 | 1189 | S1189N | D | 8.8e-06 | 6.5e-06 | 1.5e-05 | 1.010E-05 | 4.397E-06 | 6.500E-06 | 1.500E-05 |
| 23764 | 1189 | S1189D | D | 1.1e-05 | 1.5e-05 | 1.4e-05 | 1.333E-05 | 2.082E-06 | 1.100E-05 | 1.500E-05 |
| 23765 | 1189 | S1189C | D | 1.1e-05 | 1.2e-05 | 1.3e-05 | 1.200E-05 | 1.000E-06 | 1.100E-05 | 1.300E-05 |
| 23766 | 1189 | S1189E | D | 1.5e-05 | 1.5e-05 | 1.1e-05 | 1.367E-05 | 2.309E-06 | 1.100E-05 | 1.500E-05 |
| 23767 | 1189 | S1189Q | D | 1.6e-05 | 1.2e-05 | 9.5e-06 | 1.250E-05 | 3.279E-06 | 9.500E-06 | 1.600E-05 |
| 23768 | 1189 | S1189G | D | 1.4e-05 | 1.3e-05 | 1.7e-05 | 1.467E-05 | 2.082E-06 | 1.300E-05 | 1.700E-05 |
| 23769 | 1189 | S1189H | D | 1.4e-05 | 9e-06 | 8.7e-06 | 1.057E-05 | 2.977E-06 | 8.700E-06 | 1.400E-05 |
| 23770 | 1189 | S1189I | D | 1e-05 | 1.3e-05 | 9.8e-06 | 1.093E-05 | 1.793E-06 | 9.800E-06 | 1.300E-05 |
| 23771 | 1189 | S1189L | D | 1e-05 | 1.1e-05 | 1.5e-05 | 1.200E-05 | 2.646E-06 | 1.000E-05 | 1.500E-05 |
| 23772 | 1189 | S1189K | D | 1e-05 | 1.7e-05 | 1.2e-05 | 1.300E-05 | 3.606E-06 | 1.000E-05 | 1.700E-05 |
| 23773 | 1189 | S1189M | D | 1.4e-05 | 1.5e-05 | 1.6e-05 | 1.500E-05 | 1.000E-06 | 1.400E-05 | 1.600E-05 |
| 23774 | 1189 | S1189F | D | 1.3e-05 | 5.7e-06 | 1.5e-05 | 1.123E-05 | 4.895E-06 | 5.700E-06 | 1.500E-05 |
| 23775 | 1189 | S1189P | D | 8.9e-06 | 1.6e-05 | 3.5e-05 | 1.997E-05 | 1.349E-05 | 8.900E-06 | 3.500E-05 |
| 23776 | 1189 | S1189S | D | 1.4e-05 | 1.5e-05 | 1e-05 | 1.300E-05 | 2.646E-06 | 1.000E-05 | 1.500E-05 |
| 23777 | 1189 | S1189T | D | 2.1e-05 | 1.6e-05 | 1.5e-05 | 1.733E-05 | 3.215E-06 | 1.500E-05 | 2.100E-05 |
| 23778 | 1189 | S1189W | D | 1.1e-05 | 1.1e-05 | 1.1e-05 | 1.100E-05 | 2.075E-21 | 1.100E-05 | 1.100E-05 |
| 23779 | 1189 | S1189Y | D | 1.3e-05 | 1.3e-05 | 9.4e-06 | 1.180E-05 | 2.078E-06 | 9.400E-06 | 1.300E-05 |
| 23780 | 1189 | S1189V | D | 1.5e-05 | 1.5e-05 | 1e-05 | 1.333E-05 | 2.887E-06 | 1.000E-05 | 1.500E-05 |
| 23781 | 1190 | V1190A | D | 1.6e-05 | 1.4e-05 | 1.1e-05 | 1.367E-05 | 2.517E-06 | 1.100E-05 | 1.600E-05 |
| 23782 | 1190 | V1190R | D | 1.5e-05 | 1.7e-05 | 1.1e-05 | 1.433E-05 | 3.055E-06 | 1.100E-05 | 1.700E-05 |
| 23783 | 1190 | V1190N | D | 1e-05 | 1.4e-05 | 1.1e-05 | 1.167E-05 | 2.082E-06 | 1.000E-05 | 1.400E-05 |
| 23784 | 1190 | V1190D | D | 1.4e-05 | 1e-05 | 9.8e-06 | 1.127E-05 | 2.369E-06 | 9.800E-06 | 1.400E-05 |
| 23785 | 1190 | V1190C | D | 1.1e-05 | 7.8e-06 | 1.4e-05 | 1.093E-05 | 3.101E-06 | 7.800E-06 | 1.400E-05 |
| 23786 | 1190 | V1190E | D | 9.3e-06 | 1.4e-05 | 1e-05 | 1.110E-05 | 2.536E-06 | 9.300E-06 | 1.400E-05 |
| 23787 | 1190 | V1190Q | D | 1.4e-05 | 1.5e-05 | 1.6e-05 | 1.500E-05 | 1.000E-06 | 1.400E-05 | 1.600E-05 |
| 23788 | 1190 | V1190G | D | 1.5e-05 | 1.1e-05 | 1.1e-05 | 1.233E-05 | 2.309E-06 | 1.100E-05 | 1.500E-05 |
| 23789 | 1190 | V1190H | D | 1.1e-05 | 1.4e-05 | 1.6e-05 | 1.367E-05 | 2.517E-06 | 1.100E-05 | 1.600E-05 |
| 23790 | 1190 | V1190I | D | 1.4e-05 | 1e-05 | 1.7e-05 | 1.367E-05 | 3.512E-06 | 1.000E-05 | 1.700E-05 |
| 23791 | 1190 | V1190L | D | 1.6e-05 | 1.6e-05 | 8.6e-06 | 1.353E-05 | 4.272E-06 | 8.600E-06 | 1.600E-05 |
| 23792 | 1190 | V1190K | D | 1.1e-05 | 1e-05 | 1.5e-05 | 1.200E-05 | 2.646E-06 | 1.000E-05 | 1.500E-05 |
| 23793 | 1190 | V1190M | D | 8.3e-06 | 8.8e-06 | 1.5e-05 | 1.070E-05 | 3.732E-06 | 8.300E-06 | 1.500E-05 |
| 23794 | 1190 | V1190F | D | 1.5e-05 | 1.4e-05 | 1.1e-05 | 1.333E-05 | 2.082E-06 | 1.100E-05 | 1.500E-05 |
| 23795 | 1190 | V1190P | D | 1e-05 | 1.5e-05 | 1.6e-05 | 1.367E-05 | 3.215E-06 | 1.000E-05 | 1.600E-05 |
| 23796 | 1190 | V1190S | D | 9.9e-06 | 1.1e-05 | 1.1e-05 | 1.063E-05 | 6.351E-07 | 9.900E-06 | 1.100E-05 |
| 23797 | 1190 | V1190T | D | 1.4e-05 | 1.3e-05 | 1.1e-05 | 1.267E-05 | 1.528E-06 | 1.100E-05 | 1.400E-05 |
| 23798 | 1190 | V1190W | D | 1.4e-05 | 1e-05 | 1.3e-05 | 1.233E-05 | 2.082E-06 | 1.000E-05 | 1.400E-05 |
| 23799 | 1190 | V1190Y | D | 1.1e-05 | 1.5e-05 | 1.1e-05 | 1.233E-05 | 2.309E-06 | 1.100E-05 | 1.500E-05 |

| | | | | | | | | | | |
|---|---|---|---|---|---|---|---|---|---|---|
| 23800 | 1190 | V1190V | D | 9.4e-06 | 1e-05 | 1.7e-05 | 1.213E-05 | 4.225E-06 | 9.400E-06 | 1.700E-05 |
| 23801 | 1191 | V1191A | D | 1.6e-05 | 1.4e-05 | 1.4e-05 | 1.467E-05 | 1.155E-06 | 1.400E-05 | 1.600E-05 |
| 23802 | 1191 | V1191R | D | 1.6e-05 | 1.5e-05 | 9.7e-06 | 1.357E-05 | 3.386E-06 | 9.700E-06 | 1.600E-05 |
| 23803 | 1191 | V1191N | D | 8.7e-06 | 1.1e-05 | 1.3e-05 | 1.090E-05 | 2.152E-06 | 8.700E-06 | 1.300E-05 |
| 23804 | 1191 | V1191D | D | 1.5e-05 | 1.1e-05 | 1.7e-05 | 1.433E-05 | 3.055E-06 | 1.100E-05 | 1.700E-05 |
| 23805 | 1191 | V1191C | D | 1.5e-05 | 9.6e-06 | 1.5e-05 | 1.320E-05 | 3.118E-06 | 9.600E-06 | 1.500E-05 |
| 23806 | 1191 | V1191E | D | 1.2e-05 | 1.4e-05 | 9.6e-06 | 1.187E-05 | 2.203E-06 | 9.600E-06 | 1.400E-05 |
| 23807 | 1191 | V1191Q | D | 1.5e-05 | 8.3e-06 | 1.4e-05 | 1.243E-05 | 3.614E-06 | 8.300E-06 | 1.500E-05 |
| 23808 | 1191 | V1191G | D | 1.1e-05 | 9.7e-06 | 1.5e-05 | 1.190E-05 | 2.762E-06 | 9.700E-06 | 1.500E-05 |
| 23809 | 1191 | V1191H | D | 1.1e-05 | 1.5e-05 | 1.5e-05 | 1.367E-05 | 2.309E-06 | 1.100E-05 | 1.500E-05 |
| 23810 | 1191 | V1191I | D | 1.4e-05 | 1.5e-05 | 9.4e-06 | 1.280E-05 | 2.987E-06 | 9.400E-06 | 1.500E-05 |
| 23811 | 1191 | V1191L | D | 1.1e-05 | 1.3e-05 | 1.2e-05 | 1.200E-05 | 1.000E-06 | 1.100E-05 | 1.300E-05 |
| 23812 | 1191 | V1191K | D | 1.4e-05 | 1.5e-05 | 1.3e-05 | 1.400E-05 | 1.000E-06 | 1.300E-05 | 1.500E-05 |
| 23813 | 1191 | V1191M | D | 1.7e-05 | 1e-05 | 1e-05 | 1.233E-05 | 4.041E-06 | 1.000E-05 | 1.700E-05 |
| 23814 | 1191 | V1191F | D | 1.7e-05 | 1e-05 | 1.5e-05 | 1.400E-05 | 3.606E-06 | 1.000E-05 | 1.700E-05 |
| 23815 | 1191 | V1191P | D | 1.4e-05 | 1.5e-05 | 1.4e-05 | 1.433E-05 | 5.774E-07 | 1.400E-05 | 1.500E-05 |
| 23816 | 1191 | V1191S | D | 1.6e-05 | 1.1e-05 | 9.8e-06 | 1.227E-05 | 3.288E-06 | 9.800E-06 | 1.600E-05 |
| 23817 | 1191 | V1191T | D | 9.5e-06 | 8.8e-06 | 1.1e-05 | 9.767E-06 | 1.124E-06 | 8.800E-06 | 1.100E-05 |
| 23818 | 1191 | V1191W | D | 8.9e-06 | 1.5e-05 | 9.7e-06 | 1.120E-05 | 3.315E-06 | 8.900E-06 | 1.500E-05 |
| 23819 | 1191 | V1191Y | D | 9.6e-06 | 1e-05 | 1.6e-05 | 1.187E-05 | 3.585E-06 | 9.600E-06 | 1.600E-05 |
| 23820 | 1191 | V1191V | D | 1.5e-05 | 9.9e-06 | 1.1e-05 | 1.197E-05 | 2.684E-06 | 9.900E-06 | 1.500E-05 |
| 23821 | 1192 | C1192A | D | 1.7e-05 | 9.4e-06 | 1.5e-05 | 1.380E-05 | 3.940E-06 | 9.400E-06 | 1.700E-05 |
| 23822 | 1192 | C1192R | D | 1.3e-05 | 1.4e-05 | 1.1e-05 | 1.267E-05 | 1.528E-06 | 1.100E-05 | 1.400E-05 |
| 23823 | 1192 | C1192N | D | 1.2e-05 | 1.1e-05 | 1e-05 | 1.100E-05 | 1.000E-06 | 1.000E-05 | 1.200E-05 |
| 23824 | 1192 | C1192D | D | 9.9e-06 | 1.5e-05 | 1.6e-05 | 1.363E-05 | 3.272E-06 | 9.900E-06 | 1.600E-05 |
| 23825 | 1192 | C1192C | D | 1.7e-05 | 1.6e-05 | 1.5e-05 | 1.600E-05 | 1.000E-06 | 1.500E-05 | 1.700E-05 |
| 23826 | 1192 | C1192E | D | 7.8e-06 | 9.1e-06 | 1.4e-05 | 1.030E-05 | 3.270E-06 | 7.800E-06 | 1.400E-05 |
| 23827 | 1192 | C1192Q | D | 1.2e-05 | 1e-05 | 1.2e-05 | 1.133E-05 | 1.155E-06 | 1.000E-05 | 1.200E-05 |
| 23828 | 1192 | C1192G | D | 1e-05 | 1.7e-05 | 1.1e-05 | 1.267E-05 | 3.786E-06 | 1.000E-05 | 1.700E-05 |
| 23829 | 1192 | C1192H | D | 1.6e-05 | 1.5e-05 | 1.7e-05 | 1.600E-05 | 1.000E-06 | 1.500E-05 | 1.700E-05 |
| 23830 | 1192 | C1192I | D | 1.7e-05 | 1.4e-05 | 1.2e-05 | 1.433E-05 | 2.517E-06 | 1.200E-05 | 1.700E-05 |
| 23831 | 1192 | C1192L | D | 1.2e-05 | 1.4e-05 | 9.6e-06 | 1.187E-05 | 2.203E-06 | 9.600E-06 | 1.400E-05 |
| 23832 | 1192 | C1192K | D | 1.1e-05 | 1e-05 | 8.4e-06 | 9.800E-06 | 1.311E-06 | 8.400E-06 | 1.100E-05 |
| 23833 | 1192 | C1192M | D | 1.1e-05 | 1.7e-05 | 1.5e-05 | 1.433E-05 | 3.055E-06 | 1.100E-05 | 1.700E-05 |
| 23834 | 1192 | C1192F | D | 1.6e-05 | 1.3e-05 | 1e-05 | 1.300E-05 | 3.000E-06 | 1.000E-05 | 1.600E-05 |
| 23835 | 1192 | C1192P | D | 1.5e-05 | 9.8e-06 | 1.1e-05 | 1.193E-05 | 2.723E-06 | 9.800E-06 | 1.500E-05 |
| 23836 | 1192 | C1192S | D | 9.8e-06 | 1.5e-05 | 1.3e-05 | 1.260E-05 | 2.623E-06 | 9.800E-06 | 1.500E-05 |
| 23837 | 1192 | C1192T | D | 1.4e-05 | 1.6e-05 | 9.6e-06 | 1.320E-05 | 3.274E-06 | 9.600E-06 | 1.600E-05 |
| 23838 | 1192 | C1192W | D | 1.2e-05 | 1.6e-05 | 9.6e-06 | 1.253E-05 | 3.233E-06 | 9.600E-06 | 1.600E-05 |
| 23839 | 1192 | C1192Y | D | 1.2e-05 | 1.4e-05 | 1.1e-05 | 1.233E-05 | 1.528E-06 | 1.100E-05 | 1.400E-05 |
| 23840 | 1192 | C1192V | D | 1.4e-05 | 1.8e-05 | 1.1e-05 | 1.433E-05 | 3.512E-06 | 1.100E-05 | 1.800E-05 |
| 23841 | 1193 | L1193A | D | 1.5e-05 | 1.5e-05 | 1.4e-05 | 1.467E-05 | 5.774E-07 | 1.400E-05 | 1.500E-05 |
| 23842 | 1193 | L1193R | D | 9.9e-06 | 9.7e-06 | 1.7e-05 | 1.220E-05 | 4.158E-06 | 9.700E-06 | 1.700E-05 |
| 23843 | 1193 | L1193N | D | 1.5e-05 | 1.4e-05 | 1.4e-05 | 1.433E-05 | 5.774E-07 | 1.400E-05 | 1.500E-05 |
| 23844 | 1193 | L1193D | D | 8.3e-06 | 1.5e-05 | 1.1e-05 | 1.143E-05 | 3.371E-06 | 8.300E-06 | 1.500E-05 |
| 23845 | 1193 | L1193C | D | 1.5e-05 | 9.6e-06 | 1.4e-05 | 1.287E-05 | 2.873E-06 | 9.600E-06 | 1.500E-05 |
| 23846 | 1193 | L1193E | D | 1.1e-05 | 1.1e-05 | 1.5e-05 | 1.233E-05 | 2.309E-06 | 1.100E-05 | 1.500E-05 |
| 23847 | 1193 | L1193Q | D | 1e-05 | 1.5e-05 | 1.4e-05 | 1.300E-05 | 2.646E-06 | 1.000E-05 | 1.500E-05 |
| 23848 | 1193 | L1193G | D | 9.7e-06 | 1.4e-05 | 1.4e-05 | 1.257E-05 | 2.483E-06 | 9.700E-06 | 1.400E-05 |
| 23849 | 1193 | L1193H | D | 1.4e-05 | 9e-06 | 8.3e-06 | 1.043E-05 | 3.109E-06 | 8.300E-06 | 1.400E-05 |
| 23850 | 1193 | L1193I | D | 1.7e-05 | 1.1e-05 | 1.4e-05 | 1.400E-05 | 3.000E-06 | 1.100E-05 | 1.700E-05 |
| 23851 | 1193 | L1193L | D | 1.5e-05 | 1.2e-05 | 1.4e-05 | 1.367E-05 | 1.528E-06 | 1.200E-05 | 1.500E-05 |
| 23852 | 1193 | L1193K | D | 1.4e-05 | 1.5e-05 | 1.1e-05 | 1.333E-05 | 2.082E-06 | 1.100E-05 | 1.500E-05 |
| 23853 | 1193 | L1193M | D | 1.6e-05 | 1.6e-05 | 8e-06 | 1.333E-05 | 4.619E-06 | 8.000E-06 | 1.600E-05 |
| 23854 | 1193 | L1193F | D | 9.6e-06 | 1.1e-05 | 1.5e-05 | 1.187E-05 | 2.802E-06 | 9.600E-06 | 1.500E-05 |
| 23855 | 1193 | L1193P | D | 1.5e-05 | 1e-05 | 1.5e-05 | 1.333E-05 | 2.887E-06 | 1.000E-05 | 1.500E-05 |
| 23856 | 1193 | L1193S | D | 1.7e-05 | 1.8e-05 | 1.1e-05 | 1.533E-05 | 3.786E-06 | 1.100E-05 | 1.800E-05 |
| 23857 | 1193 | L1193T | D | 9.5e-06 | 1.1e-05 | 9.8e-06 | 1.010E-05 | 7.937E-07 | 9.500E-06 | 1.100E-05 |
| 23858 | 1193 | L1193W | D | 1.6e-05 | 1.1e-05 | 1.5e-05 | 1.400E-05 | 2.646E-06 | 1.100E-05 | 1.600E-05 |
| 23859 | 1193 | L1193Y | D | 1.5e-05 | 1.7e-05 | 1.6e-05 | 1.600E-05 | 1.000E-06 | 1.500E-05 | 1.700E-05 |
| 23860 | 1193 | L1193V | D | 7.9e-06 | 1.4e-05 | 1.6e-05 | 1.263E-05 | 4.219E-06 | 7.900E-06 | 1.600E-05 |
| 23861 | 1194 | L1194A | D | 1.7e-05 | 8.8e-06 | 1.5e-05 | 1.360E-05 | 4.276E-06 | 8.800E-06 | 1.700E-05 |
| 23862 | 1194 | L1194R | D | 1.7e-05 | 1.2e-05 | 1.5e-05 | 1.467E-05 | 2.517E-06 | 1.200E-05 | 1.700E-05 |
| 23863 | 1194 | L1194N | D | 1.6e-05 | 8.9e-06 | 8.3e-06 | 1.107E-05 | 4.283E-06 | 8.300E-06 | 1.600E-05 |
| 23864 | 1194 | L1194D | D | 1.1e-05 | 1.4e-05 | 1.4e-05 | 1.300E-05 | 1.732E-06 | 1.100E-05 | 1.400E-05 |
| 23865 | 1194 | L1194C | D | 1e-05 | 1.5e-05 | 1.4e-05 | 1.300E-05 | 2.646E-06 | 1.000E-05 | 1.500E-05 |
| 23866 | 1194 | L1194E | D | 1.1e-05 | 1.1e-05 | 1.4e-05 | 1.200E-05 | 1.732E-06 | 1.100E-05 | 1.400E-05 |
| 23867 | 1194 | L1194Q | D | 1.4e-05 | 1.2e-05 | 1.1e-05 | 1.233E-05 | 1.528E-06 | 1.100E-05 | 1.400E-05 |
| 23868 | 1194 | L1194G | D | 1.2e-05 | 1.5e-05 | 1.5e-05 | 1.400E-05 | 1.732E-06 | 1.200E-05 | 1.500E-05 |
| 23869 | 1194 | L1194H | D | 1.1e-05 | 1.5e-05 | 1.6e-05 | 1.400E-05 | 2.646E-06 | 1.100E-05 | 1.600E-05 |
| 23870 | 1194 | L1194I | D | 9.6e-06 | 1.6e-05 | 1.5e-05 | 1.353E-05 | 3.443E-06 | 9.600E-06 | 1.600E-05 |
| 23871 | 1194 | L1194L | D | 1.4e-05 | 1.7e-05 | 1.4e-05 | 1.500E-05 | 1.732E-06 | 1.400E-05 | 1.700E-05 |
| 23872 | 1194 | L1194K | D | 1.6e-05 | 1.6e-05 | 1.6e-05 | 1.600E-05 | 0.000E+00 | 1.600E-05 | 1.600E-05 |
| 23873 | 1194 | L1194M | D | 1.5e-05 | 1.7e-05 | 1.2e-05 | 1.467E-05 | 2.517E-06 | 1.200E-05 | 1.700E-05 |
| 23874 | 1194 | L1194F | D | 1.6e-05 | 1.3e-05 | 1.5e-05 | 1.467E-05 | 1.528E-06 | 1.300E-05 | 1.600E-05 |
| 23875 | 1194 | L1194P | D | 1.1e-05 | 1e-05 | 1.8e-05 | 1.300E-05 | 4.359E-06 | 1.000E-05 | 1.800E-05 |
| 23876 | 1194 | L1194S | D | 1.5e-05 | 1.5e-05 | 1.6e-05 | 1.533E-05 | 5.774E-07 | 1.500E-05 | 1.600E-05 |
| 23877 | 1194 | L1194T | D | 1.4e-05 | 1e-05 | 9.9e-06 | 1.130E-05 | 2.339E-06 | 9.900E-06 | 1.400E-05 |
| 23878 | 1194 | L1194W | D | 1.6e-05 | 1.5e-05 | 1.2e-05 | 1.433E-05 | 2.082E-06 | 1.200E-05 | 1.600E-05 |
| 23879 | 1194 | L1194Y | D | 1.5e-05 | 1.5e-05 | 1.4e-05 | 1.467E-05 | 5.774E-07 | 1.400E-05 | 1.500E-05 |
| 23880 | 1194 | L1194V | D | 1.7e-05 | 1.1e-05 | 1.7e-05 | 1.500E-05 | 3.464E-06 | 1.100E-05 | 1.700E-05 |
| 23881 | 1195 | N1195A | D | 1.6e-05 | 1.2e-05 | 1.6e-05 | 1.467E-05 | 2.309E-06 | 1.200E-05 | 1.600E-05 |
| 23882 | 1195 | N1195R | D | 1.6e-05 | 1.7e-05 | 1.6e-05 | 1.633E-05 | 5.774E-07 | 1.600E-05 | 1.700E-05 |
| 23883 | 1195 | N1195N | D | 1.7e-05 | 1.1e-05 | 1e-05 | 1.267E-05 | 3.786E-06 | 1.000E-05 | 1.700E-05 |
| 23884 | 1195 | N1195D | D | 1.7e-05 | 1.5e-05 | 1.7e-05 | 1.633E-05 | 1.155E-06 | 1.500E-05 | 1.700E-05 |

| | | | | | | | | | | |
|---|---|---|---|---|---|---|---|---|---|---|
| 23885 | 1195 | N1195C | D | 1.5e-05 | 1.6e-05 | 1.4e-05 | 1.500E-05 | 1.000E-06 | 1.400E-05 | 1.600E-05 |
| 23886 | 1195 | N1195E | D | 1e-05 | 1e-05 | 1.4e-05 | 1.133E-05 | 2.309E-06 | 1.000E-05 | 1.400E-05 |
| 23887 | 1195 | N1195Q | D | 1.4e-05 | 1.2e-05 | 1.6e-05 | 1.400E-05 | 2.000E-06 | 1.200E-05 | 1.600E-05 |
| 23888 | 1195 | N1195G | D | 1.1e-05 | 1.4e-05 | 1.1e-05 | 1.200E-05 | 1.732E-06 | 1.100E-05 | 1.400E-05 |
| 23889 | 1195 | N1195H | D | 9.7e-06 | 7.8e-06 | 1.1e-05 | 9.500E-06 | 1.609E-06 | 7.800E-06 | 1.100E-05 |
| 23890 | 1195 | N1195I | D | 1.6e-05 | 8.5e-06 | 1.4e-05 | 1.283E-05 | 3.884E-06 | 8.500E-06 | 1.600E-05 |
| 23891 | 1195 | N1195L | D | 1e-05 | 1.8e-05 | 1.1e-05 | 1.300E-05 | 4.359E-06 | 1.000E-05 | 1.800E-05 |
| 23892 | 1195 | N1195K | D | 1.1e-05 | 1e-05 | 1.4e-05 | 1.167E-05 | 2.082E-06 | 1.000E-05 | 1.400E-05 |
| 23893 | 1195 | N1195M | D | 9e-06 | 1.5e-05 | 1.5e-05 | 1.300E-05 | 3.464E-06 | 9.000E-06 | 1.500E-05 |
| 23894 | 1195 | N1195F | D | 1.1e-05 | 1.6e-05 | 1.7e-05 | 1.467E-05 | 3.215E-06 | 1.100E-05 | 1.700E-05 |
| 23895 | 1195 | N1195P | D | 9e-06 | 1.5e-05 | 9e-06 | 1.100E-05 | 3.464E-06 | 9.000E-06 | 1.500E-05 |
| 23896 | 1195 | N1195S | D | 8.7e-06 | 1.9e-05 | 9.9e-06 | 1.253E-05 | 5.632E-06 | 8.700E-06 | 1.900E-05 |
| 23897 | 1195 | N1195T | D | 1.1e-05 | 1e-05 | 1.6e-05 | 1.233E-05 | 3.215E-06 | 1.000E-05 | 1.600E-05 |
| 23898 | 1195 | N1195W | D | 1.3e-05 | 8.5e-06 | 1.1e-05 | 1.083E-05 | 2.255E-06 | 8.500E-06 | 1.300E-05 |
| 23899 | 1195 | N1195Y | D | 1.2e-05 | 1.5e-05 | 1.8e-05 | 1.500E-05 | 3.000E-06 | 1.200E-05 | 1.800E-05 |
| 23900 | 1195 | N1195V | D | 1e-05 | 1.3e-05 | 1.5e-05 | 1.267E-05 | 2.517E-06 | 1.000E-05 | 1.500E-05 |
| 23901 | 1196 | N1196A | D | 1.4e-05 | 1e-05 | 1.6e-05 | 1.333E-05 | 3.055E-06 | 1.000E-05 | 1.600E-05 |
| 23902 | 1196 | N1196R | D | 1.3e-05 | 7.1e-06 | 1.7e-05 | 1.237E-05 | 4.980E-06 | 7.100E-06 | 1.700E-05 |
| 23903 | 1196 | N1196N | D | 1e-05 | 1.7e-05 | 1.4e-05 | 1.367E-05 | 3.512E-06 | 1.000E-05 | 1.700E-05 |
| 23904 | 1196 | N1196D | D | 1.1e-05 | 1.5e-05 | 1.5e-05 | 1.367E-05 | 2.309E-06 | 1.100E-05 | 1.500E-05 |
| 23905 | 1196 | N1196C | D | 1.5e-05 | 9.4e-06 | 1.4e-05 | 1.280E-05 | 2.987E-06 | 9.400E-06 | 1.500E-05 |
| 23906 | 1196 | N1196E | D | 1.4e-05 | 8.2e-06 | 1.6e-05 | 1.273E-05 | 4.051E-06 | 8.200E-06 | 1.600E-05 |
| 23907 | 1196 | N1196Q | D | 1.5e-05 | 1e-05 | 1.1e-05 | 1.200E-05 | 2.646E-06 | 1.000E-05 | 1.500E-05 |
| 23908 | 1196 | N1196G | D | 1e-05 | 1.4e-05 | 1.1e-05 | 1.167E-05 | 2.082E-06 | 1.000E-05 | 1.400E-05 |
| 23909 | 1196 | N1196H | D | 1.6e-05 | 1.2e-05 | 1.6e-05 | 1.467E-05 | 2.309E-06 | 1.200E-05 | 1.600E-05 |
| 23910 | 1196 | N1196I | D | 1.6e-05 | 1.1e-05 | 1.1e-05 | 1.267E-05 | 2.887E-06 | 1.100E-05 | 1.600E-05 |
| 23911 | 1196 | N1196L | D | 7.1e-06 | 1.3e-05 | 1.4e-05 | 1.137E-05 | 3.729E-06 | 7.100E-06 | 1.400E-05 |
| 23912 | 1196 | N1196K | D | 1.4e-05 | 1.6e-05 | 1.1e-05 | 1.367E-05 | 2.517E-06 | 1.100E-05 | 1.600E-05 |
| 23913 | 1196 | N1196M | D | 1.4e-05 | 1e-05 | 1.6e-05 | 1.333E-05 | 3.055E-06 | 1.000E-05 | 1.600E-05 |
| 23914 | 1196 | N1196F | D | 1.6e-05 | 9.2e-06 | 9.8e-06 | 1.167E-05 | 3.765E-06 | 9.200E-06 | 1.600E-05 |
| 23915 | 1196 | N1196P | D | 1.1e-05 | 1.1e-05 | 1e-05 | 1.067E-05 | 5.774E-07 | 1.000E-05 | 1.100E-05 |
| 23916 | 1196 | N1196S | D | 9.3e-06 | 1.3e-05 | 1e-05 | 1.077E-05 | 1.966E-06 | 9.300E-06 | 1.300E-05 |
| 23917 | 1196 | N1196T | D | 8.9e-06 | 1.7e-05 | 1.6e-05 | 1.397E-05 | 4.416E-06 | 8.900E-06 | 1.700E-05 |
| 23918 | 1196 | N1196W | D | 1.8e-05 | 1.5e-05 | 8.7e-06 | 1.390E-05 | 4.747E-06 | 8.700E-06 | 1.800E-05 |
| 23919 | 1196 | N1196Y | D | 1.5e-05 | 1e-05 | 1.4e-05 | 1.300E-05 | 2.646E-06 | 1.000E-05 | 1.500E-05 |
| 23920 | 1196 | N1196V | D | 9.9e-06 | 1.1e-05 | 1.4e-05 | 1.163E-05 | 2.122E-06 | 9.900E-06 | 1.400E-05 |
| 23921 | 1197 | F1197A | D | 1.6e-05 | 1.1e-05 | 1.1e-05 | 1.267E-05 | 2.887E-06 | 1.100E-05 | 1.600E-05 |
| 23922 | 1197 | F1197R | D | 9.5e-06 | 1.4e-05 | 1.4e-05 | 1.250E-05 | 2.598E-06 | 9.500E-06 | 1.400E-05 |
| 23923 | 1197 | F1197N | D | 1.2e-05 | 1.4e-05 | 1.6e-05 | 1.400E-05 | 2.000E-06 | 1.200E-05 | 1.600E-05 |
| 23924 | 1197 | F1197D | D | 1.6e-05 | 1e-05 | 1.1e-05 | 1.233E-05 | 3.215E-06 | 1.000E-05 | 1.600E-05 |
| 23925 | 1197 | F1197C | D | 1.4e-05 | 1.1e-05 | 1.2e-05 | 1.233E-05 | 1.528E-06 | 1.100E-05 | 1.400E-05 |
| 23926 | 1197 | F1197E | D | 1.4e-05 | 1e-05 | 1.3e-05 | 1.233E-05 | 2.082E-06 | 1.000E-05 | 1.400E-05 |
| 23927 | 1197 | F1197Q | D | 8e-06 | 1.6e-05 | 1.4e-05 | 1.267E-05 | 4.163E-06 | 8.000E-06 | 1.600E-05 |
| 23928 | 1197 | F1197G | D | 1.6e-05 | 8.4e-06 | 1.1e-05 | 1.180E-05 | 3.863E-06 | 8.400E-06 | 1.600E-05 |
| 23929 | 1197 | F1197H | D | 9.9e-06 | 1.1e-05 | 1.4e-05 | 1.163E-05 | 2.122E-06 | 9.900E-06 | 1.400E-05 |
| 23930 | 1197 | F1197I | D | 1.4e-05 | 1.5e-05 | 1.1e-05 | 1.333E-05 | 2.082E-06 | 1.100E-05 | 1.500E-05 |
| 23931 | 1197 | F1197L | D | 1.2e-05 | 8.9e-06 | 1.5e-05 | 1.197E-05 | 3.050E-06 | 8.900E-06 | 1.500E-05 |
| 23932 | 1197 | F1197K | D | 9.9e-06 | 1.4e-05 | 1.4e-05 | 1.263E-05 | 2.367E-06 | 9.900E-06 | 1.400E-05 |
| 23933 | 1197 | F1197M | D | 1.4e-05 | 1.7e-05 | 1.4e-05 | 1.500E-05 | 1.732E-06 | 1.400E-05 | 1.700E-05 |
| 23934 | 1197 | F1197F | D | 1.2e-05 | 1.3e-05 | 1.4e-05 | 1.300E-05 | 1.000E-06 | 1.200E-05 | 1.400E-05 |
| 23935 | 1197 | F1197P | D | 1.2e-05 | 9e-06 | 8.6e-06 | 9.867E-06 | 1.858E-06 | 8.600E-06 | 1.200E-05 |
| 23936 | 1197 | F1197S | D | 1.5e-05 | 1.3e-05 | 1.3e-05 | 1.367E-05 | 1.155E-06 | 1.300E-05 | 1.500E-05 |
| 23937 | 1197 | F1197T | D | 1e-05 | 1.4e-05 | 1.5e-05 | 1.300E-05 | 2.646E-06 | 1.000E-05 | 1.500E-05 |
| 23938 | 1197 | F1197W | D | 1e-05 | 1.7e-05 | 1.6e-05 | 1.433E-05 | 3.786E-06 | 1.000E-05 | 1.700E-05 |
| 23939 | 1197 | F1197Y | D | 1.2e-05 | 1.2e-05 | 7.1e-06 | 1.037E-05 | 2.829E-06 | 7.100E-06 | 1.200E-05 |
| 23940 | 1197 | F1197V | D | 9.7e-06 | 1.3e-05 | 1.5e-05 | 1.257E-05 | 2.676E-06 | 9.700E-06 | 1.500E-05 |
| 23941 | 1198 | Y1198A | D | 1e-05 | 1.6e-05 | 1.3e-05 | 1.300E-05 | 3.000E-06 | 1.000E-05 | 1.600E-05 |
| 23942 | 1198 | Y1198R | D | 1.4e-05 | 9.4e-06 | 1.7e-05 | 1.347E-05 | 3.828E-06 | 9.400E-06 | 1.700E-05 |
| 23943 | 1198 | Y1198N | D | 1e-05 | 1.6e-05 | 8.8e-06 | 1.160E-05 | 3.857E-06 | 8.800E-06 | 1.600E-05 |
| 23944 | 1198 | Y1198D | D | 1.5e-05 | 1.2e-05 | 1e-05 | 1.233E-05 | 2.517E-06 | 1.000E-05 | 1.500E-05 |
| 23945 | 1198 | Y1198C | D | 1.1e-05 | 1e-05 | 1.5e-05 | 1.200E-05 | 2.646E-06 | 1.000E-05 | 1.500E-05 |
| 23946 | 1198 | Y1198E | D | 1.4e-05 | 9.2e-06 | 1.4e-05 | 1.240E-05 | 2.771E-06 | 9.200E-06 | 1.400E-05 |
| 23947 | 1198 | Y1198Q | D | 1.4e-05 | 1.5e-05 | 1e-05 | 1.300E-05 | 2.646E-06 | 1.000E-05 | 1.500E-05 |
| 23948 | 1198 | Y1198G | D | 1e-05 | 1.4e-05 | 1.5e-05 | 1.300E-05 | 2.646E-06 | 1.000E-05 | 1.500E-05 |
| 23949 | 1198 | Y1198H | D | 9.1e-06 | 1.5e-05 | 1.4e-05 | 1.270E-05 | 3.158E-06 | 9.100E-06 | 1.500E-05 |
| 23950 | 1198 | Y1198I | D | 1.3e-05 | 1.5e-05 | 8.2e-06 | 1.207E-05 | 3.495E-06 | 8.200E-06 | 1.500E-05 |
| 23951 | 1198 | Y1198L | D | 1.5e-05 | 1.2e-05 | 1.5e-05 | 1.400E-05 | 1.732E-06 | 1.200E-05 | 1.500E-05 |
| 23952 | 1198 | Y1198K | D | 1.4e-05 | 1.5e-05 | 9.7e-06 | 1.290E-05 | 2.816E-06 | 9.700E-06 | 1.500E-05 |
| 23953 | 1198 | Y1198M | D | 9.6e-06 | 1.8e-05 | 1.7e-05 | 1.487E-05 | 4.588E-06 | 9.600E-06 | 1.800E-05 |
| 23954 | 1198 | Y1198F | D | 1.5e-05 | 1.4e-05 | 7.8e-06 | 1.227E-05 | 3.900E-06 | 7.800E-06 | 1.500E-05 |
| 23955 | 1198 | Y1198P | D | 9.7e-06 | 1.4e-05 | 1.3e-05 | 1.223E-05 | 2.250E-06 | 9.700E-06 | 1.400E-05 |
| 23956 | 1198 | Y1198S | D | 1.1e-05 | 1.4e-05 | 1e-05 | 1.167E-05 | 2.082E-06 | 1.000E-05 | 1.400E-05 |
| 23957 | 1198 | Y1198T | D | 1.5e-05 | 1.6e-05 | 1e-05 | 1.367E-05 | 3.215E-06 | 1.000E-05 | 1.600E-05 |
| 23958 | 1198 | Y1198W | D | 1.3e-05 | 1.1e-05 | 1e-05 | 1.133E-05 | 1.528E-06 | 1.000E-05 | 1.300E-05 |
| 23959 | 1198 | Y1198Y | D | 1.8e-05 | 1.4e-05 | 1.1e-05 | 1.433E-05 | 3.512E-06 | 1.100E-05 | 1.800E-05 |
| 23960 | 1198 | Y1198V | D | 1.4e-05 | 1.1e-05 | 1.3e-05 | 1.267E-05 | 1.528E-06 | 1.100E-05 | 1.400E-05 |
| 23961 | 1199 | P1199A | D | 1.4e-05 | 1.5e-05 | 2.6e-05 | 1.833E-05 | 6.658E-06 | 1.400E-05 | 2.600E-05 |
| 23962 | 1199 | P1199R | D | 1.5e-05 | 1.7e-05 | 1.2e-05 | 1.467E-05 | 2.517E-06 | 1.200E-05 | 1.700E-05 |
| 23963 | 1199 | P1199N | D | 7.6e-06 | 1.2e-05 | 1.5e-05 | 1.153E-05 | 3.722E-06 | 7.600E-06 | 1.500E-05 |
| 23964 | 1199 | P1199D | D | 1.1e-05 | 1.5e-05 | 1.2e-05 | 1.267E-05 | 2.082E-06 | 1.100E-05 | 1.500E-05 |
| 23965 | 1199 | P1199C | D | 9.6e-06 | 1.1e-05 | 1.3e-05 | 1.120E-05 | 1.709E-06 | 9.600E-06 | 1.300E-05 |
| 23966 | 1199 | P1199E | D | 1e-05 | 1.1e-05 | 1.5e-05 | 1.200E-05 | 2.646E-06 | 1.000E-05 | 1.500E-05 |
| 23967 | 1199 | P1199Q | D | 1e-05 | 1.6e-05 | 1.5e-05 | 1.367E-05 | 3.215E-06 | 1.000E-05 | 1.600E-05 |
| 23968 | 1199 | P1199G | D | 1.5e-05 | 1.5e-05 | 1.4e-05 | 1.467E-05 | 5.774E-07 | 1.400E-05 | 1.500E-05 |
| 23969 | 1199 | P1199H | D | 1.4e-05 | 1.4e-05 | 1.3e-05 | 1.367E-05 | 5.774E-07 | 1.300E-05 | 1.400E-05 |

| | | | | | | | | | | |
|---|---|---|---|---|---|---|---|---|---|---|
| 23970 | 1199 | P1199I | D | 8.3e-06 | 1.1e-05 | 1.5e-05 | 1.143E-05 | 3.371E-06 | 8.300E-06 | 1.500E-05 |
| 23971 | 1199 | P1199L | D | 1.2e-05 | 1.6e-05 | 1.4e-05 | 1.400E-05 | 2.000E-06 | 1.200E-05 | 1.600E-05 |
| 23972 | 1199 | P1199K | D | 1.5e-05 | 1.6e-05 | 1.6e-05 | 1.567E-05 | 5.774E-07 | 1.500E-05 | 1.600E-05 |
| 23973 | 1199 | P1199M | D | 1.1e-05 | 2.2e-05 | 1.5e-05 | 1.600E-05 | 5.568E-06 | 1.100E-05 | 2.200E-05 |
| 23974 | 1199 | P1199F | D | 1.1e-05 | 8.8e-06 | 1.6e-05 | 1.193E-05 | 3.690E-06 | 8.800E-06 | 1.600E-05 |
| 23975 | 1199 | P1199P | D | 7.9e-06 | 1.4e-05 | 1.2e-05 | 1.130E-05 | 3.110E-06 | 7.900E-06 | 1.400E-05 |
| 23976 | 1199 | P1199S | D | 8.3e-06 | 1.6e-05 | 9.3e-06 | 1.120E-05 | 4.187E-06 | 8.300E-06 | 1.600E-05 |
| 23977 | 1199 | P1199T | D | 8.2e-06 | 1.3e-05 | 1.2e-05 | 1.107E-05 | 2.532E-06 | 8.200E-06 | 1.300E-05 |
| 23978 | 1199 | P1199W | D | 1.4e-05 | 1.1e-05 | 1.4e-05 | 1.300E-05 | 1.732E-06 | 1.100E-05 | 1.400E-05 |
| 23979 | 1199 | P1199Y | D | 7.4e-06 | 1.1e-05 | 1.1e-05 | 9.800E-06 | 2.078E-06 | 7.400E-06 | 1.100E-05 |
| 23980 | 1199 | P1199V | D | 1.1e-05 | 1.1e-05 | 1.8e-05 | 1.333E-05 | 4.041E-06 | 1.100E-05 | 1.800E-05 |
| 23981 | 1200 | R1200A | D | 1.2e-05 | 1.5e-05 | 1.2e-05 | 1.300E-05 | 1.732E-06 | 1.200E-05 | 1.500E-05 |
| 23982 | 1200 | R1200R | D | 9.1e-06 | 1.7e-05 | 1.5e-05 | 1.370E-05 | 4.107E-06 | 9.100E-06 | 1.700E-05 |
| 23983 | 1200 | R1200N | D | 1.6e-05 | 1.5e-05 | 1.4e-05 | 1.500E-05 | 1.000E-06 | 1.400E-05 | 1.600E-05 |
| 23984 | 1200 | R1200D | D | 1.3e-05 | 1.1e-05 | 1.6e-05 | 1.333E-05 | 2.517E-06 | 1.100E-05 | 1.600E-05 |
| 23985 | 1200 | R1200C | D | 1e-05 | 1e-05 | 2.3e-05 | 1.433E-05 | 7.506E-06 | 1.000E-05 | 2.300E-05 |
| 23986 | 1200 | R1200E | D | 1.5e-05 | 9.1e-06 | 9.4e-06 | 1.117E-05 | 3.323E-06 | 9.100E-06 | 1.500E-05 |
| 23987 | 1200 | R1200Q | D | 1e-05 | 9.7e-06 | 1.1e-05 | 1.023E-05 | 6.807E-07 | 9.700E-06 | 1.100E-05 |
| 23988 | 1200 | R1200G | D | 1.5e-05 | 1.5e-05 | 1.3e-05 | 1.433E-05 | 1.155E-06 | 1.300E-05 | 1.500E-05 |
| 23989 | 1200 | R1200H | D | 8.5e-06 | 1.2e-05 | 1.5e-05 | 1.183E-05 | 3.253E-06 | 8.500E-06 | 1.500E-05 |
| 23990 | 1200 | R1200I | D | 1.6e-05 | 1.6e-05 | 1.1e-05 | 1.433E-05 | 2.887E-06 | 1.100E-05 | 1.600E-05 |
| 23991 | 1200 | R1200L | D | 1.3e-05 | 1.6e-05 | 1.5e-05 | 1.467E-05 | 1.528E-06 | 1.300E-05 | 1.600E-05 |
| 23992 | 1200 | R1200K | D | 1.2e-05 | 1.3e-05 | 9e-06 | 1.133E-05 | 2.082E-06 | 9.000E-06 | 1.300E-05 |
| 23993 | 1200 | R1200M | D | 1.5e-05 | 1e-05 | 1.3e-05 | 1.267E-05 | 2.517E-06 | 1.000E-05 | 1.500E-05 |
| 23994 | 1200 | R1200F | D | 1.5e-05 | 1.6e-05 | 1.5e-05 | 1.533E-05 | 5.774E-07 | 1.500E-05 | 1.600E-05 |
| 23995 | 1200 | R1200P | D | 1.1e-05 | 8.6e-06 | 8.3e-06 | 9.300E-06 | 1.480E-06 | 8.300E-06 | 1.100E-05 |
| 23996 | 1200 | R1200S | D | 1.1e-05 | 9.3e-06 | 1.1e-05 | 1.043E-05 | 9.815E-07 | 9.300E-06 | 1.100E-05 |
| 23997 | 1200 | R1200T | D | 1.2e-05 | 1e-05 | 1.4e-05 | 1.200E-05 | 2.000E-06 | 1.000E-05 | 1.400E-05 |
| 23998 | 1200 | R1200W | D | 1.6e-05 | 1.2e-05 | 1.5e-05 | 1.433E-05 | 2.082E-06 | 1.200E-05 | 1.600E-05 |
| 23999 | 1200 | R1200Y | D | 1.8e-05 | 1.3e-05 | 1.6e-05 | 1.567E-05 | 2.517E-06 | 1.300E-05 | 1.800E-05 |
| 24000 | 1200 | R1200V | D | 1e-05 | 1.1e-05 | 1.5e-05 | 1.200E-05 | 2.646E-06 | 1.000E-05 | 1.500E-05 |
| 24001 | 1201 | E1201A | D | 1.1e-05 | 1.6e-05 | 1.6e-05 | 1.433E-05 | 2.887E-06 | 1.100E-05 | 1.600E-05 |
| 24002 | 1201 | E1201R | D | 1.6e-05 | 1.5e-05 | 1e-05 | 1.367E-05 | 3.215E-06 | 1.000E-05 | 1.600E-05 |
| 24003 | 1201 | E1201N | D | 1.1e-05 | 1.1e-05 | 9.6e-06 | 1.053E-05 | 8.083E-07 | 9.600E-06 | 1.100E-05 |
| 24004 | 1201 | E1201D | D | 9e-06 | 1.4e-05 | 1.3e-05 | 1.200E-05 | 2.646E-06 | 9.000E-06 | 1.400E-05 |
| 24005 | 1201 | E1201C | D | 1.5e-05 | 1e-05 | 1.6e-05 | 1.367E-05 | 3.215E-06 | 1.000E-05 | 1.600E-05 |
| 24006 | 1201 | E1201E | D | 1.1e-05 | 9.8e-06 | 1.7e-05 | 1.260E-05 | 3.857E-06 | 9.800E-06 | 1.700E-05 |
| 24007 | 1201 | E1201Q | D | 1.4e-05 | 2.3e-05 | 1.1e-05 | 1.600E-05 | 6.245E-06 | 1.100E-05 | 2.300E-05 |
| 24008 | 1201 | E1201G | D | 1.3e-05 | 1.3e-05 | 9.6e-06 | 1.187E-05 | 1.963E-06 | 9.600E-06 | 1.300E-05 |
| 24009 | 1201 | E1201H | D | 1.4e-05 | 1e-05 | 1.6e-05 | 1.333E-05 | 3.055E-06 | 1.000E-05 | 1.600E-05 |
| 24010 | 1201 | E1201I | D | 1e-05 | 1.5e-05 | 1.1e-05 | 1.200E-05 | 2.646E-06 | 1.000E-05 | 1.500E-05 |
| 24011 | 1201 | E1201L | D | 1.9e-05 | 1e-05 | 1.1e-05 | 1.333E-05 | 4.933E-06 | 1.000E-05 | 1.900E-05 |
| 24012 | 1201 | E1201K | D | 1.6e-05 | 1.2e-05 | 1.3e-05 | 1.367E-05 | 2.082E-06 | 1.200E-05 | 1.600E-05 |
| 24013 | 1201 | E1201M | D | 1.4e-05 | 9.6e-06 | 1.5e-05 | 1.287E-05 | 2.873E-06 | 9.600E-06 | 1.500E-05 |
| 24014 | 1201 | E1201F | D | 1.5e-05 | 1.3e-05 | 1.3e-05 | 1.367E-05 | 1.155E-06 | 1.300E-05 | 1.500E-05 |
| 24015 | 1201 | E1201P | D | 1.4e-05 | 1.6e-05 | 1.1e-05 | 1.367E-05 | 2.517E-06 | 1.100E-05 | 1.600E-05 |
| 24016 | 1201 | E1201S | D | 8.8e-06 | 1.4e-05 | 1.6e-05 | 1.293E-05 | 3.717E-06 | 8.800E-06 | 1.600E-05 |
| 24017 | 1201 | E1201T | D | 1.5e-05 | 1.5e-05 | 9.6e-06 | 1.320E-05 | 3.118E-06 | 9.600E-06 | 1.500E-05 |
| 24018 | 1201 | E1201W | D | 1e-05 | 1.2e-05 | 1e-05 | 1.067E-05 | 1.155E-06 | 1.000E-05 | 1.200E-05 |
| 24019 | 1201 | E1201Y | D | 1.1e-05 | 1.5e-05 | 1.5e-05 | 1.367E-05 | 2.309E-06 | 1.100E-05 | 1.500E-05 |
| 24020 | 1201 | E1201V | D | 9.8e-06 | 1.2e-05 | 1.5e-05 | 1.227E-05 | 2.610E-06 | 9.800E-06 | 1.500E-05 |
| 24021 | 1202 | A1202A | D | 1.5e-05 | 1.1e-05 | 1.1e-05 | 1.233E-05 | 2.309E-06 | 1.100E-05 | 1.500E-05 |
| 24022 | 1202 | A1202R | D | 1e-05 | 1.6e-05 | 1.4e-05 | 1.333E-05 | 3.055E-06 | 1.000E-05 | 1.600E-05 |
| 24023 | 1202 | A1202N | D | 1.2e-05 | 1.1e-05 | 1.5e-05 | 1.267E-05 | 2.082E-06 | 1.100E-05 | 1.500E-05 |
| 24024 | 1202 | A1202D | D | 1e-05 | 1.3e-05 | 1.1e-05 | 1.133E-05 | 1.528E-06 | 1.000E-05 | 1.300E-05 |
| 24025 | 1202 | A1202C | D | 1.4e-05 | 1.7e-05 | 1.4e-05 | 1.500E-05 | 1.732E-06 | 1.400E-05 | 1.700E-05 |
| 24026 | 1202 | A1202E | D | 1e-05 | 7.4e-06 | 1.1e-05 | 9.467E-06 | 1.858E-06 | 7.400E-06 | 1.100E-05 |
| 24027 | 1202 | A1202Q | D | 1.4e-05 | 1.5e-05 | 1.1e-05 | 1.333E-05 | 2.082E-06 | 1.100E-05 | 1.500E-05 |
| 24028 | 1202 | A1202G | D | 1.4e-05 | 1.6e-05 | 1.4e-05 | 1.467E-05 | 1.155E-06 | 1.400E-05 | 1.600E-05 |
| 24029 | 1202 | A1202H | D | 1.7e-05 | 8e-06 | 1e-05 | 1.167E-05 | 4.726E-06 | 8.000E-06 | 1.700E-05 |
| 24030 | 1202 | A1202I | D | 9.3e-06 | 1e-05 | 1.7e-05 | 1.210E-05 | 4.258E-06 | 9.300E-06 | 1.700E-05 |
| 24031 | 1202 | A1202L | D | 1.4e-05 | 1.2e-05 | 1.1e-05 | 1.233E-05 | 1.528E-06 | 1.100E-05 | 1.400E-05 |
| 24032 | 1202 | A1202K | D | 1.5e-05 | 1e-05 | 1.5e-05 | 1.333E-05 | 2.887E-06 | 1.000E-05 | 1.500E-05 |
| 24033 | 1202 | A1202M | D | 1.3e-05 | 7.4e-06 | 9.2e-06 | 9.867E-06 | 2.859E-06 | 7.400E-06 | 1.300E-05 |
| 24034 | 1202 | A1202F | D | 1.5e-05 | 9.8e-06 | 9.3e-06 | 1.137E-05 | 3.156E-06 | 9.300E-06 | 1.500E-05 |
| 24035 | 1202 | A1202P | D | 1.1e-05 | 1.6e-05 | 1.1e-05 | 1.267E-05 | 2.887E-06 | 1.100E-05 | 1.600E-05 |
| 24036 | 1202 | A1202S | D | 1.5e-05 | 1.3e-05 | 1.6e-05 | 1.467E-05 | 1.528E-06 | 1.300E-05 | 1.600E-05 |
| 24037 | 1202 | A1202T | D | 1e-05 | 9.7e-06 | 5.9e-06 | 8.533E-06 | 2.285E-06 | 5.900E-06 | 1.000E-05 |
| 24038 | 1202 | A1202W | D | 1.2e-05 | 9.9e-06 | 1.6e-05 | 1.263E-05 | 3.099E-06 | 9.900E-06 | 1.600E-05 |
| 24039 | 1202 | A1202Y | D | 1.4e-05 | 9.4e-06 | 1.2e-05 | 1.180E-05 | 2.307E-06 | 9.400E-06 | 1.400E-05 |
| 24040 | 1202 | A1202V | D | 1.5e-05 | 1.5e-05 | 8.5e-06 | 1.283E-05 | 3.753E-06 | 8.500E-06 | 1.500E-05 |
| 24041 | 1203 | K1203A | D | 2e-05 | 9.8e-06 | 1.1e-05 | 1.360E-05 | 5.575E-06 | 9.800E-06 | 2.000E-05 |
| 24042 | 1203 | K1203R | D | 1.1e-05 | 1.5e-05 | 1.5e-05 | 1.367E-05 | 2.309E-06 | 1.100E-05 | 1.500E-05 |
| 24043 | 1203 | K1203N | D | 1.3e-05 | 9e-06 | 7.8e-06 | 9.933E-06 | 2.723E-06 | 7.800E-06 | 1.300E-05 |
| 24044 | 1203 | K1203D | D | 1.4e-05 | 1.6e-05 | 6.7e-06 | 1.223E-05 | 4.895E-06 | 6.700E-06 | 1.600E-05 |
| 24045 | 1203 | K1203C | D | 1.5e-05 | 1.2e-05 | 1.1e-05 | 1.267E-05 | 2.082E-06 | 1.100E-05 | 1.500E-05 |
| 24046 | 1203 | K1203E | D | 1.8e-05 | 1.1e-05 | 1.7e-05 | 1.533E-05 | 3.786E-06 | 1.100E-05 | 1.800E-05 |
| 24047 | 1203 | K1203Q | D | 1.1e-05 | 1.2e-05 | 9.4e-06 | 1.080E-05 | 1.311E-06 | 9.400E-06 | 1.200E-05 |
| 24048 | 1203 | K1203G | D | 1.4e-05 | 1e-05 | 1.4e-05 | 1.267E-05 | 2.309E-06 | 1.000E-05 | 1.400E-05 |
| 24049 | 1203 | K1203H | D | 1e-05 | 1.4e-05 | 1.4e-05 | 1.267E-05 | 2.309E-06 | 1.000E-05 | 1.400E-05 |
| 24050 | 1203 | K1203I | D | 1e-05 | 1e-05 | 1.2e-05 | 1.067E-05 | 1.155E-06 | 1.000E-05 | 1.200E-05 |
| 24051 | 1203 | K1203L | D | 1.1e-05 | 1.6e-05 | 1.5e-05 | 1.400E-05 | 2.646E-06 | 1.100E-05 | 1.600E-05 |
| 24052 | 1203 | K1203K | D | 9.9e-06 | 1.3e-05 | 8.3e-06 | 1.040E-05 | 2.390E-06 | 8.300E-06 | 1.300E-05 |
| 24053 | 1203 | K1203M | D | 9.3e-06 | 1.4e-05 | 1.3e-05 | 1.210E-05 | 2.476E-06 | 9.300E-06 | 1.400E-05 |
| 24054 | 1203 | K1203F | D | 1.1e-05 | 1.4e-05 | 9.8e-06 | 1.160E-05 | 2.163E-06 | 9.800E-06 | 1.400E-05 |

| | | | | | | | | | | |
|---|---|---|---|---|---|---|---|---|---|---|
| 24055 | 1203 | K1203P | D | 1e-05 | 1.6e-05 | 1.1e-05 | 1.233E-05 | 3.215E-06 | 1.000E-05 | 1.600E-05 |
| 24056 | 1203 | K1203S | D | 1e-05 | 1.3e-05 | 9.2e-06 | 1.073E-05 | 2.003E-06 | 9.200E-06 | 1.300E-05 |
| 24057 | 1203 | K1203T | D | 1.1e-05 | 9.2e-06 | 9.5e-06 | 9.900E-06 | 9.644E-07 | 9.200E-06 | 1.100E-05 |
| 24058 | 1203 | K1203W | D | 1.5e-05 | 8.9e-06 | 1.5e-05 | 1.297E-05 | 3.522E-06 | 8.900E-06 | 1.500E-05 |
| 24059 | 1203 | K1203Y | D | 1.4e-05 | 1.3e-05 | 1.1e-05 | 1.267E-05 | 1.528E-06 | 1.100E-05 | 1.400E-05 |
| 24060 | 1203 | K1203V | D | 1e-05 | 1.6e-05 | 1.4e-05 | 1.333E-05 | 3.055E-06 | 1.000E-05 | 1.600E-05 |
| 24061 | 1204 | V1204A | D | 1.6e-05 | 1.4e-05 | 1.7e-05 | 1.567E-05 | 1.528E-06 | 1.400E-05 | 1.700E-05 |
| 24062 | 1204 | V1204R | D | 1.3e-05 | 1.6e-05 | 1.5e-05 | 1.467E-05 | 1.528E-06 | 1.300E-05 | 1.600E-05 |
| 24063 | 1204 | V1204N | D | 1.4e-05 | 1.4e-05 | 1.6e-05 | 1.467E-05 | 1.155E-06 | 1.400E-05 | 1.600E-05 |
| 24064 | 1204 | V1204D | D | 1.2e-05 | 1.3e-05 | 1.7e-05 | 1.400E-05 | 2.646E-06 | 1.200E-05 | 1.700E-05 |
| 24065 | 1204 | V1204C | D | 1.4e-05 | 1.4e-05 | 1.1e-05 | 1.300E-05 | 1.732E-06 | 1.100E-05 | 1.400E-05 |
| 24066 | 1204 | V1204E | D | 8.6e-06 | 1.4e-05 | 1.2e-05 | 1.153E-05 | 2.730E-06 | 8.600E-06 | 1.400E-05 |
| 24067 | 1204 | V1204Q | D | 1.3e-05 | 1.5e-05 | 9e-06 | 1.233E-05 | 3.055E-06 | 9.000E-06 | 1.500E-05 |
| 24068 | 1204 | V1204G | D | 1.4e-05 | 1.5e-05 | 1.5e-05 | 1.467E-05 | 5.774E-07 | 1.400E-05 | 1.500E-05 |
| 24069 | 1204 | V1204H | D | 1e-05 | 1.6e-05 | 6.6e-06 | 1.087E-05 | 4.760E-06 | 6.600E-06 | 1.600E-05 |
| 24070 | 1204 | V1204I | D | 1.2e-05 | 7.9e-06 | 1.2e-05 | 1.063E-05 | 2.367E-06 | 7.900E-06 | 1.200E-05 |
| 24071 | 1204 | V1204L | D | 1.5e-05 | 1.4e-05 | 1e-05 | 1.300E-05 | 2.646E-06 | 1.000E-05 | 1.500E-05 |
| 24072 | 1204 | V1204K | D | 1.1e-05 | 1.2e-05 | 1.1e-05 | 1.133E-05 | 5.774E-07 | 1.100E-05 | 1.200E-05 |
| 24073 | 1204 | V1204M | D | 9.7e-06 | 1.3e-05 | 1.6e-05 | 1.290E-05 | 3.151E-06 | 9.700E-06 | 1.600E-05 |
| 24074 | 1204 | V1204F | D | 1e-05 | 9.8e-06 | 8.2e-06 | 9.333E-06 | 9.866E-07 | 8.200E-06 | 1.000E-05 |
| 24075 | 1204 | V1204P | D | 1.1e-05 | 1.4e-05 | 1.1e-05 | 1.200E-05 | 1.732E-06 | 1.100E-05 | 1.400E-05 |
| 24076 | 1204 | V1204S | D | 9.5e-06 | 1e-05 | 1.4e-05 | 1.117E-05 | 2.466E-06 | 9.500E-06 | 1.400E-05 |
| 24077 | 1204 | V1204T | D | 1.4e-05 | 1.5e-05 | 9.5e-06 | 1.283E-05 | 2.930E-06 | 9.500E-06 | 1.500E-05 |
| 24078 | 1204 | V1204W | D | 1.4e-05 | 1e-05 | 9.6e-06 | 1.120E-05 | 2.433E-06 | 9.600E-06 | 1.400E-05 |
| 24079 | 1204 | V1204Y | D | 5.8e-06 | 1e-05 | 1.3e-05 | 9.600E-06 | 3.617E-06 | 5.800E-06 | 1.300E-05 |
| 24080 | 1204 | V1204V | D | 1.1e-05 | 1.6e-05 | 1.6e-05 | 1.433E-05 | 2.887E-06 | 1.100E-05 | 1.600E-05 |
| 24081 | 1205 | Q1205A | D | 1.7e-05 | 1.7e-05 | 9.4e-06 | 1.447E-05 | 4.388E-06 | 9.400E-06 | 1.700E-05 |
| 24082 | 1205 | Q1205R | D | 1.6e-05 | 1.3e-05 | 1.6e-05 | 1.500E-05 | 1.732E-06 | 1.300E-05 | 1.600E-05 |
| 24083 | 1205 | Q1205N | D | 1.3e-05 | 1.5e-05 | 9.2e-06 | 1.240E-05 | 2.946E-06 | 9.200E-06 | 1.500E-05 |
| 24084 | 1205 | Q1205D | D | 1.5e-05 | 1.4e-05 | 1.6e-05 | 1.500E-05 | 1.000E-06 | 1.400E-05 | 1.600E-05 |
| 24085 | 1205 | Q1205C | D | 9.1e-06 | 1.4e-05 | 1.2e-05 | 1.170E-05 | 2.464E-06 | 9.100E-06 | 1.400E-05 |
| 24086 | 1205 | Q1205E | D | 1.1e-05 | 1.6e-05 | 1.1e-05 | 1.267E-05 | 2.887E-06 | 1.100E-05 | 1.600E-05 |
| 24087 | 1205 | Q1205Q | D | 9.6e-06 | 1.5e-05 | 1.5e-05 | 1.320E-05 | 3.118E-06 | 9.600E-06 | 1.500E-05 |
| 24088 | 1205 | Q1205G | D | 1.3e-05 | 1.5e-05 | 1.6e-05 | 1.467E-05 | 1.528E-06 | 1.300E-05 | 1.600E-05 |
| 24089 | 1205 | Q1205H | D | 1.1e-05 | 1.2e-05 | 1.2e-05 | 1.167E-05 | 5.774E-07 | 1.100E-05 | 1.200E-05 |
| 24090 | 1205 | Q1205I | D | 1.7e-05 | 1.4e-05 | 1.1e-05 | 1.400E-05 | 3.000E-06 | 1.100E-05 | 1.700E-05 |
| 24091 | 1205 | Q1205L | D | 1.4e-05 | 1.5e-05 | 1.7e-05 | 1.533E-05 | 1.528E-06 | 1.400E-05 | 1.700E-05 |
| 24092 | 1205 | Q1205K | D | 1.6e-05 | 1.4e-05 | 1.7e-05 | 1.567E-05 | 1.528E-06 | 1.400E-05 | 1.700E-05 |
| 24093 | 1205 | Q1205M | D | 1.1e-05 | 1.7e-05 | 1.8e-05 | 1.533E-05 | 3.786E-06 | 1.100E-05 | 1.800E-05 |
| 24094 | 1205 | Q1205F | D | 9.3e-06 | 1e-05 | 1.4e-05 | 1.110E-05 | 2.536E-06 | 9.300E-06 | 1.400E-05 |
| 24095 | 1205 | Q1205P | D | 1.5e-05 | 1.5e-05 | 1.5e-05 | 1.500E-05 | 0.000E+00 | 1.500E-05 | 1.500E-05 |
| 24096 | 1205 | Q1205S | D | 1.3e-05 | 1.3e-05 | 1.2e-05 | 1.267E-05 | 5.774E-07 | 1.200E-05 | 1.300E-05 |
| 24097 | 1205 | Q1205T | D | 1.4e-05 | 1.1e-05 | 1.4e-05 | 1.300E-05 | 1.732E-06 | 1.100E-05 | 1.400E-05 |
| 24098 | 1205 | Q1205W | D | 1.1e-05 | 1e-05 | 1e-05 | 1.033E-05 | 5.774E-07 | 1.000E-05 | 1.100E-05 |
| 24099 | 1205 | Q1205Y | D | 1.5e-05 | 1.6e-05 | 1.6e-05 | 1.567E-05 | 5.774E-07 | 1.500E-05 | 1.600E-05 |
| 24100 | 1205 | Q1205V | D | 1.3e-05 | 1.6e-05 | 1.6e-05 | 1.500E-05 | 1.732E-06 | 1.300E-05 | 1.600E-05 |
| 24101 | 1206 | W1206A | D | 1.3e-05 | 8.6e-06 | 1.3e-05 | 1.153E-05 | 2.540E-06 | 8.600E-06 | 1.300E-05 |
| 24102 | 1206 | W1206R | D | 1e-05 | 6.8e-06 | 1.6e-05 | 1.093E-05 | 4.670E-06 | 6.800E-06 | 1.600E-05 |
| 24103 | 1206 | W1206N | D | 1.3e-05 | 1.4e-05 | 1.1e-05 | 1.267E-05 | 1.528E-06 | 1.100E-05 | 1.400E-05 |
| 24104 | 1206 | W1206D | D | 1.5e-05 | 1.5e-05 | 1.1e-05 | 1.367E-05 | 2.309E-06 | 1.100E-05 | 1.500E-05 |
| 24105 | 1206 | W1206C | D | 1.5e-05 | 1.1e-05 | 1.2e-05 | 1.267E-05 | 2.082E-06 | 1.100E-05 | 1.500E-05 |
| 24106 | 1206 | W1206E | D | 1.6e-05 | 9.1e-06 | 9.6e-06 | 1.157E-05 | 3.848E-06 | 9.100E-06 | 1.600E-05 |
| 24107 | 1206 | W1206Q | D | 1.2e-05 | 9.3e-06 | 1.5e-05 | 1.210E-05 | 2.851E-06 | 9.300E-06 | 1.500E-05 |
| 24108 | 1206 | W1206G | D | 1.6e-05 | 1.4e-05 | 1.4e-05 | 1.467E-05 | 1.155E-06 | 1.400E-05 | 1.600E-05 |
| 24109 | 1206 | W1206H | D | 1.2e-05 | 1.5e-05 | 1.4e-05 | 1.367E-05 | 1.528E-06 | 1.200E-05 | 1.500E-05 |
| 24110 | 1206 | W1206I | D | 1.3e-05 | 1.6e-05 | 1.1e-05 | 1.333E-05 | 2.517E-06 | 1.100E-05 | 1.600E-05 |
| 24111 | 1206 | W1206L | D | 1.5e-05 | 9.9e-06 | 1.4e-05 | 1.297E-05 | 2.702E-06 | 9.900E-06 | 1.500E-05 |
| 24112 | 1206 | W1206K | D | 1.1e-05 | 1.7e-05 | 1.4e-05 | 1.400E-05 | 3.000E-06 | 1.100E-05 | 1.700E-05 |
| 24113 | 1206 | W1206M | D | 1.5e-05 | 1.6e-05 | 9.4e-06 | 1.347E-05 | 3.557E-06 | 9.400E-06 | 1.600E-05 |
| 24114 | 1206 | W1206F | D | 7e-06 | 5.8e-06 | 1.5e-05 | 9.267E-06 | 5.001E-06 | 5.800E-06 | 1.500E-05 |
| 24115 | 1206 | W1206P | D | 8.6e-06 | 1.4e-05 | 1.6e-05 | 1.287E-05 | 3.828E-06 | 8.600E-06 | 1.600E-05 |
| 24116 | 1206 | W1206S | D | 1.4e-05 | 8.2e-06 | 1e-05 | 1.073E-05 | 2.969E-06 | 8.200E-06 | 1.400E-05 |
| 24117 | 1206 | W1206T | D | 1.2e-05 | 1.6e-05 | 1.2e-05 | 1.333E-05 | 2.309E-06 | 1.200E-05 | 1.600E-05 |
| 24118 | 1206 | W1206W | D | 8.9e-06 | 1e-05 | 1e-05 | 9.633E-06 | 6.351E-07 | 8.900E-06 | 1.000E-05 |
| 24119 | 1206 | W1206Y | D | 1.5e-05 | 8.7e-06 | 6.5e-06 | 1.007E-05 | 4.412E-06 | 6.500E-06 | 1.500E-05 |
| 24120 | 1206 | W1206V | D | 1.1e-05 | 1.6e-05 | 1.4e-05 | 1.367E-05 | 2.517E-06 | 1.100E-05 | 1.600E-05 |
| 24121 | 1207 | K1207A | D | 1.2e-05 | 1.4e-05 | 1.4e-05 | 1.333E-05 | 1.155E-06 | 1.200E-05 | 1.400E-05 |
| 24122 | 1207 | K1207R | D | 1.4e-05 | 8.1e-06 | 1.3e-05 | 1.170E-05 | 3.158E-06 | 8.100E-06 | 1.400E-05 |
| 24123 | 1207 | K1207N | D | 1.1e-05 | 1.6e-05 | 1.1e-05 | 1.267E-05 | 2.887E-06 | 1.100E-05 | 1.600E-05 |
| 24124 | 1207 | K1207D | D | 2.1e-05 | 1.6e-05 | 1.6e-05 | 1.767E-05 | 2.887E-06 | 1.600E-05 | 2.100E-05 |
| 24125 | 1207 | K1207C | D | 1.5e-05 | 1.1e-05 | 1.5e-05 | 1.367E-05 | 2.309E-06 | 1.100E-05 | 1.500E-05 |
| 24126 | 1207 | K1207E | D | 8.9e-06 | 1e-05 | 1.5e-05 | 1.130E-05 | 3.251E-06 | 8.900E-06 | 1.500E-05 |
| 24127 | 1207 | K1207Q | D | 1e-05 | 1.6e-05 | 8.9e-06 | 1.163E-05 | 3.821E-06 | 8.900E-06 | 1.600E-05 |
| 24128 | 1207 | K1207G | D | 1.6e-05 | 1e-05 | 9.8e-06 | 1.193E-05 | 3.523E-06 | 9.800E-06 | 1.600E-05 |
| 24129 | 1207 | K1207H | D | 1.6e-05 | 1.1e-05 | 1.3e-05 | 1.333E-05 | 2.517E-06 | 1.100E-05 | 1.600E-05 |
| 24130 | 1207 | K1207I | D | 1.1e-05 | 1.3e-05 | 9.8e-06 | 1.127E-05 | 1.617E-06 | 9.800E-06 | 1.300E-05 |
| 24131 | 1207 | K1207L | D | 1.7e-05 | 1.1e-05 | 9.7e-06 | 1.257E-05 | 3.894E-06 | 9.700E-06 | 1.700E-05 |
| 24132 | 1207 | K1207K | D | 1.4e-05 | 1.6e-05 | 9.9e-06 | 1.330E-05 | 3.110E-06 | 9.900E-06 | 1.600E-05 |
| 24133 | 1207 | K1207M | D | 1.2e-05 | 1.7e-05 | 8.8e-06 | 1.260E-05 | 4.133E-06 | 8.800E-06 | 1.700E-05 |
| 24134 | 1207 | K1207F | D | 1.3e-05 | 1.2e-05 | 9e-06 | 1.133E-05 | 2.082E-06 | 9.000E-06 | 1.300E-05 |
| 24135 | 1207 | K1207P | D | 1e-05 | 1.1e-05 | 1.1e-05 | 1.067E-05 | 5.774E-07 | 1.000E-05 | 1.100E-05 |
| 24136 | 1207 | K1207S | D | 1.3e-05 | 1.5e-05 | 1.2e-05 | 1.333E-05 | 1.528E-06 | 1.200E-05 | 1.500E-05 |
| 24137 | 1207 | K1207T | D | 1.2e-05 | 1e-05 | 1e-05 | 1.067E-05 | 1.155E-06 | 1.000E-05 | 1.200E-05 |
| 24138 | 1207 | K1207W | D | 1.6e-05 | 9.3e-06 | 1.9e-05 | 1.477E-05 | 4.966E-06 | 9.300E-06 | 1.900E-05 |
| 24139 | 1207 | K1207Y | D | 1.4e-05 | 9.9e-06 | 9.9e-06 | 1.127E-05 | 2.367E-06 | 9.900E-06 | 1.400E-05 |

| | | | | | | | | | | |
|---|---|---|---|---|---|---|---|---|---|---|
| 24140 | 1207 | K1207V | D | 1.6e-05 | 9.5e-06 | 1e-05 | 1.183E-05 | 3.617E-06 | 9.500E-06 | 1.600E-05 |
| 24141 | 1208 | V1208A | D | 1.3e-05 | 1.2e-05 | 1.6e-05 | 1.367E-05 | 2.082E-06 | 1.200E-05 | 1.600E-05 |
| 24142 | 1208 | V1208R | D | 1.1e-05 | 1e-05 | 1.1e-05 | 1.067E-05 | 5.774E-07 | 1.000E-05 | 1.100E-05 |
| 24143 | 1208 | V1208N | D | 1.2e-05 | 1.7e-05 | 1e-05 | 1.300E-05 | 3.606E-06 | 1.000E-05 | 1.700E-05 |
| 24144 | 1208 | V1208D | D | 1.9e-05 | 9.4e-06 | 1.7e-05 | 1.513E-05 | 5.065E-06 | 9.400E-06 | 1.900E-05 |
| 24145 | 1208 | V1208C | D | 1e-05 | 1.4e-05 | 1.2e-05 | 1.200E-05 | 2.000E-06 | 1.000E-05 | 1.400E-05 |
| 24146 | 1208 | V1208E | D | 1.5e-05 | 1.5e-05 | 1.3e-05 | 1.433E-05 | 1.155E-06 | 1.300E-05 | 1.500E-05 |
| 24147 | 1208 | V1208Q | D | 1.4e-05 | 1.1e-05 | 1.7e-05 | 1.400E-05 | 3.000E-06 | 1.100E-05 | 1.700E-05 |
| 24148 | 1208 | V1208G | D | 1.1e-05 | 1e-05 | 1.8e-05 | 1.300E-05 | 4.359E-06 | 1.000E-05 | 1.800E-05 |
| 24149 | 1208 | V1208H | D | 1.4e-05 | 1.6e-05 | 1e-05 | 1.333E-05 | 3.055E-06 | 1.000E-05 | 1.600E-05 |
| 24150 | 1208 | V1208I | D | 1.4e-05 | 1e-05 | 1.5e-05 | 1.300E-05 | 2.646E-06 | 1.000E-05 | 1.500E-05 |
| 24151 | 1208 | V1208L | D | 1.1e-05 | 1.5e-05 | 1.5e-05 | 1.367E-05 | 2.309E-06 | 1.100E-05 | 1.500E-05 |
| 24152 | 1208 | V1208K | D | 1.5e-05 | 1.6e-05 | 9.3e-06 | 1.343E-05 | 3.614E-06 | 9.300E-06 | 1.600E-05 |
| 24153 | 1208 | V1208M | D | 1.6e-05 | 1.3e-05 | 1.5e-05 | 1.467E-05 | 1.528E-06 | 1.300E-05 | 1.600E-05 |
| 24154 | 1208 | V1208F | D | 1.1e-05 | 1.3e-05 | 1.5e-05 | 1.300E-05 | 2.000E-06 | 1.100E-05 | 1.500E-05 |
| 24155 | 1208 | V1208P | D | 1.2e-05 | 1.6e-05 | 1e-05 | 1.267E-05 | 3.055E-06 | 1.000E-05 | 1.600E-05 |
| 24156 | 1208 | V1208S | D | 1.5e-05 | 1e-05 | 1.1e-05 | 1.200E-05 | 2.646E-06 | 1.000E-05 | 1.500E-05 |
| 24157 | 1208 | V1208T | D | 9.8e-06 | 1.7e-05 | 9.9e-06 | 1.223E-05 | 4.128E-06 | 9.800E-06 | 1.700E-05 |
| 24158 | 1208 | V1208W | D | 1.5e-05 | 1.6e-05 | 1.4e-05 | 1.500E-05 | 1.000E-06 | 1.400E-05 | 1.600E-05 |
| 24159 | 1208 | V1208Y | D | 1.1e-05 | 1.3e-05 | 1.6e-05 | 1.333E-05 | 2.517E-06 | 1.100E-05 | 1.600E-05 |
| 24160 | 1208 | V1208V | D | 1e-05 | 1.6e-05 | 1.5e-05 | 1.367E-05 | 3.215E-06 | 1.000E-05 | 1.600E-05 |
| 24161 | 1209 | D1209A | D | 1.4e-05 | 1.2e-05 | 1.3e-05 | 1.300E-05 | 1.000E-06 | 1.200E-05 | 1.400E-05 |
| 24162 | 1209 | D1209R | D | 1e-05 | 9e-06 | 1.1e-05 | 1.000E-05 | 1.000E-06 | 9.000E-06 | 1.100E-05 |
| 24163 | 1209 | D1209N | D | 7.9e-06 | 9.2e-06 | 1.6e-05 | 1.103E-05 | 4.350E-06 | 7.900E-06 | 1.600E-05 |
| 24164 | 1209 | D1209D | D | 2.5e-05 | 1.5e-05 | 1.5e-05 | 1.833E-05 | 5.774E-06 | 1.500E-05 | 2.500E-05 |
| 24165 | 1209 | D1209C | D | 1.4e-05 | 9.8e-06 | 1.4e-05 | 1.260E-05 | 2.425E-06 | 9.800E-06 | 1.400E-05 |
| 24166 | 1209 | D1209E | D | 1.5e-05 | 1.5e-05 | 1e-05 | 1.333E-05 | 2.887E-06 | 1.000E-05 | 1.500E-05 |
| 24167 | 1209 | D1209Q | D | 1.3e-05 | 1e-05 | 1.1e-05 | 1.133E-05 | 1.528E-06 | 1.000E-05 | 1.300E-05 |
| 24168 | 1209 | D1209G | D | 1.3e-05 | 1.3e-05 | 8.3e-06 | 1.143E-05 | 2.714E-06 | 8.300E-06 | 1.300E-05 |
| 24169 | 1209 | D1209H | D | 7.6e-06 | 1.1e-05 | 1.4e-05 | 1.087E-05 | 3.202E-06 | 7.600E-06 | 1.400E-05 |
| 24170 | 1209 | D1209I | D | 1.4e-05 | 1e-05 | 1.1e-05 | 1.167E-05 | 2.082E-06 | 1.000E-05 | 1.400E-05 |
| 24171 | 1209 | D1209L | D | 1.5e-05 | 1e-05 | 1.4e-05 | 1.300E-05 | 2.646E-06 | 1.000E-05 | 1.500E-05 |
| 24172 | 1209 | D1209K | D | 1e-05 | 1.1e-05 | 1.6e-05 | 1.233E-05 | 3.215E-06 | 1.000E-05 | 1.600E-05 |
| 24173 | 1209 | D1209M | D | 1.1e-05 | 1.3e-05 | 1.1e-05 | 1.167E-05 | 1.155E-06 | 1.100E-05 | 1.300E-05 |
| 24174 | 1209 | D1209F | D | 1.4e-05 | 1.1e-05 | 1.2e-05 | 1.233E-05 | 1.528E-06 | 1.100E-05 | 1.400E-05 |
| 24175 | 1209 | D1209P | D | 1.4e-05 | 1.2e-05 | 1.1e-05 | 1.233E-05 | 1.528E-06 | 1.100E-05 | 1.400E-05 |
| 24176 | 1209 | D1209S | D | 9.9e-06 | 1.2e-05 | 9.9e-06 | 1.060E-05 | 1.212E-06 | 9.900E-06 | 1.200E-05 |
| 24177 | 1209 | D1209T | D | 1.1e-05 | 9.7e-06 | 1.5e-05 | 1.190E-05 | 2.762E-06 | 9.700E-06 | 1.500E-05 |
| 24178 | 1209 | D1209W | D | 1.1e-05 | 9.8e-06 | 1.4e-05 | 1.160E-05 | 2.163E-06 | 9.800E-06 | 1.400E-05 |
| 24179 | 1209 | D1209Y | D | 1e-05 | 1.2e-05 | 1e-05 | 1.067E-05 | 1.155E-06 | 1.000E-05 | 1.200E-05 |
| 24180 | 1209 | D1209V | D | 1.3e-05 | 1e-05 | 1.4e-05 | 1.233E-05 | 2.082E-06 | 1.000E-05 | 1.400E-05 |
| 24181 | 1210 | N1210A | D | 1.1e-05 | 9.6e-06 | 1.1e-05 | 1.053E-05 | 8.083E-07 | 9.600E-06 | 1.100E-05 |
| 24182 | 1210 | N1210R | D | 1.4e-05 | 1.1e-05 | 1.6e-05 | 1.367E-05 | 2.517E-06 | 1.100E-05 | 1.600E-05 |
| 24183 | 1210 | N1210N | D | 1.5e-05 | 1.6e-05 | 1.7e-05 | 1.600E-05 | 1.000E-06 | 1.500E-05 | 1.700E-05 |
| 24184 | 1210 | N1210D | D | 1.2e-05 | 1.6e-05 | 1.5e-05 | 1.433E-05 | 2.082E-06 | 1.200E-05 | 1.600E-05 |
| 24185 | 1210 | N1210C | D | 1.5e-05 | 1e-05 | 1.1e-05 | 1.200E-05 | 2.646E-06 | 1.000E-05 | 1.500E-05 |
| 24186 | 1210 | N1210E | D | 1.1e-05 | 9.9e-06 | 1.1e-05 | 1.063E-05 | 6.351E-07 | 9.900E-06 | 1.100E-05 |
| 24187 | 1210 | N1210Q | D | 1.1e-05 | 1.6e-05 | 1.1e-05 | 1.267E-05 | 2.887E-06 | 1.100E-05 | 1.600E-05 |
| 24188 | 1210 | N1210G | D | 1.8e-05 | 1.6e-05 | 1e-05 | 1.467E-05 | 4.163E-06 | 1.000E-05 | 1.800E-05 |
| 24189 | 1210 | N1210H | D | 1.5e-05 | 1.7e-05 | 1.6e-05 | 1.600E-05 | 1.000E-06 | 1.500E-05 | 1.700E-05 |
| 24190 | 1210 | N1210I | D | 1.6e-05 | 1.5e-05 | 1.5e-05 | 1.533E-05 | 5.774E-07 | 1.500E-05 | 1.600E-05 |
| 24191 | 1210 | N1210L | D | 1.1e-05 | 1.4e-05 | 1.5e-05 | 1.333E-05 | 2.082E-06 | 1.100E-05 | 1.500E-05 |
| 24192 | 1210 | N1210K | D | 1.3e-05 | 1.1e-05 | 1.5e-05 | 1.300E-05 | 2.000E-06 | 1.100E-05 | 1.500E-05 |
| 24193 | 1210 | N1210M | D | 1.5e-05 | 8.9e-06 | 1.5e-05 | 1.297E-05 | 3.522E-06 | 8.900E-06 | 1.500E-05 |
| 24194 | 1210 | N1210F | D | 7.4e-06 | 1.5e-05 | 1.7e-05 | 1.313E-05 | 5.065E-06 | 7.400E-06 | 1.700E-05 |
| 24195 | 1210 | N1210P | D | 1.2e-05 | 1.5e-05 | 1.6e-05 | 1.433E-05 | 2.082E-06 | 1.200E-05 | 1.600E-05 |
| 24196 | 1210 | N1210S | D | 9.4e-06 | 1.5e-05 | 1.4e-05 | 1.280E-05 | 2.987E-06 | 9.400E-06 | 1.500E-05 |
| 24197 | 1210 | N1210T | D | 1.5e-05 | 1.6e-05 | 1.4e-05 | 1.500E-05 | 1.000E-06 | 1.400E-05 | 1.600E-05 |
| 24198 | 1210 | N1210W | D | 1.6e-05 | 1.1e-05 | 1.5e-05 | 1.400E-05 | 2.646E-06 | 1.100E-05 | 1.600E-05 |
| 24199 | 1210 | N1210Y | D | 1.6e-05 | 2.2e-05 | 9.7e-06 | 1.590E-05 | 6.151E-06 | 9.700E-06 | 2.200E-05 |
| 24200 | 1210 | N1210V | D | 1.6e-05 | 1.6e-05 | 1.5e-05 | 1.567E-05 | 5.774E-07 | 1.500E-05 | 1.600E-05 |
| 24201 | 1211 | A1211A | D | 1.4e-05 | 1.5e-05 | 1.7e-05 | 1.533E-05 | 1.528E-06 | 1.400E-05 | 1.700E-05 |
| 24202 | 1211 | A1211R | D | 1.5e-05 | 1e-05 | 1.7e-05 | 1.400E-05 | 3.606E-06 | 1.000E-05 | 1.700E-05 |
| 24203 | 1211 | A1211N | D | 1.4e-05 | 1.1e-05 | 9.2e-06 | 1.140E-05 | 2.425E-06 | 9.200E-06 | 1.400E-05 |
| 24204 | 1211 | A1211D | D | 1.6e-05 | 1.4e-05 | 1.1e-05 | 1.367E-05 | 2.517E-06 | 1.100E-05 | 1.600E-05 |
| 24205 | 1211 | A1211C | D | 1.1e-05 | 1.3e-05 | 1.4e-05 | 1.267E-05 | 1.528E-06 | 1.100E-05 | 1.400E-05 |
| 24206 | 1211 | A1211E | D | 1.1e-05 | 9.2e-06 | 1e-05 | 1.007E-05 | 9.018E-07 | 9.200E-06 | 1.100E-05 |
| 24207 | 1211 | A1211Q | D | 1.5e-05 | 1e-05 | 1.4e-05 | 1.300E-05 | 2.646E-06 | 1.000E-05 | 1.500E-05 |
| 24208 | 1211 | A1211G | D | 9.7e-06 | 9.6e-06 | 1.6e-05 | 1.177E-05 | 3.667E-06 | 9.600E-06 | 1.600E-05 |
| 24209 | 1211 | A1211H | D | 1.6e-05 | 1.7e-05 | 1.4e-05 | 1.567E-05 | 1.528E-06 | 1.400E-05 | 1.700E-05 |
| 24210 | 1211 | A1211I | D | 6.2e-06 | 1e-05 | 1e-05 | 8.733E-06 | 2.194E-06 | 6.200E-06 | 1.000E-05 |
| 24211 | 1211 | A1211L | D | 1.5e-05 | 1.5e-05 | 1.4e-05 | 1.467E-05 | 5.774E-07 | 1.400E-05 | 1.500E-05 |
| 24212 | 1211 | A1211K | D | 1.2e-05 | 1.4e-05 | 1.1e-05 | 1.233E-05 | 1.528E-06 | 1.100E-05 | 1.400E-05 |
| 24213 | 1211 | A1211M | D | 9.1e-06 | 9.2e-06 | 9.7e-06 | 9.333E-06 | 3.215E-07 | 9.100E-06 | 9.700E-06 |
| 24214 | 1211 | A1211F | D | 9e-06 | 1.3e-05 | 1.4e-05 | 1.200E-05 | 2.646E-06 | 9.000E-06 | 1.400E-05 |
| 24215 | 1211 | A1211P | D | 1.6e-05 | 1e-05 | 1.6e-05 | 1.400E-05 | 3.464E-06 | 1.000E-05 | 1.600E-05 |
| 24216 | 1211 | A1211S | D | 1.1e-05 | 1.4e-05 | 1.6e-05 | 1.367E-05 | 2.517E-06 | 1.100E-05 | 1.600E-05 |
| 24217 | 1211 | A1211T | D | 9.9e-06 | 1.2e-05 | 1.5e-05 | 1.230E-05 | 2.563E-06 | 9.900E-06 | 1.500E-05 |
| 24218 | 1211 | A1211W | D | 1.5e-05 | 1.5e-05 | 1.4e-05 | 1.467E-05 | 5.774E-07 | 1.400E-05 | 1.500E-05 |
| 24219 | 1211 | A1211Y | D | 8.3e-06 | 1.5e-05 | 1.4e-05 | 1.243E-05 | 3.614E-06 | 8.300E-06 | 1.500E-05 |
| 24220 | 1211 | A1211V | D | 1.4e-05 | 1.6e-05 | 5.8e-06 | 1.193E-05 | 5.405E-06 | 5.800E-06 | 1.600E-05 |
| 24221 | 1212 | L1212A | D | 1.2e-05 | 1.2e-05 | 1e-05 | 1.133E-05 | 1.155E-06 | 1.000E-05 | 1.200E-05 |
| 24222 | 1212 | L1212R | D | 9.3e-06 | 1e-05 | 1.2e-05 | 1.043E-05 | 1.401E-06 | 9.300E-06 | 1.200E-05 |
| 24223 | 1212 | L1212N | D | 1.4e-05 | 1.4e-05 | 1.5e-05 | 1.433E-05 | 5.774E-07 | 1.400E-05 | 1.500E-05 |
| 24224 | 1212 | L1212D | D | 9.1e-06 | 1.3e-05 | 1.3e-05 | 1.170E-05 | 2.252E-06 | 9.100E-06 | 1.300E-05 |

| | | | | | | | | | | |
|---|---|---|---|---|---|---|---|---|---|---|
| 24225 | 1212 | L1212C | D | 1.5e-05 | 9.4e-06 | 1e-05 | 1.147E-05 | 3.075E-06 | 9.400E-06 | 1.500E-05 |
| 24226 | 1212 | L1212E | D | 1.3e-05 | 1.1e-05 | 8.9e-06 | 1.097E-05 | 2.050E-06 | 8.900E-06 | 1.300E-05 |
| 24227 | 1212 | L1212Q | D | 1e-05 | 1.5e-05 | 1.5e-05 | 1.333E-05 | 2.887E-06 | 1.000E-05 | 1.500E-05 |
| 24228 | 1212 | L1212G | D | 1.2e-05 | 1.1e-05 | 9.9e-06 | 1.097E-05 | 1.050E-06 | 9.900E-06 | 1.200E-05 |
| 24229 | 1212 | L1212H | D | 1.4e-05 | 1.5e-05 | 1.4e-05 | 1.433E-05 | 5.774E-07 | 1.400E-05 | 1.500E-05 |
| 24230 | 1212 | L1212I | D | 9.7e-06 | 1.4e-05 | 1e-05 | 1.123E-05 | 2.401E-06 | 9.700E-06 | 1.400E-05 |
| 24231 | 1212 | L1212L | D | 1.5e-05 | 1.7e-05 | 9.7e-06 | 1.390E-05 | 3.772E-06 | 9.700E-06 | 1.700E-05 |
| 24232 | 1212 | L1212K | D | 1.1e-05 | 1.8e-05 | 9.4e-06 | 1.280E-05 | 4.574E-06 | 9.400E-06 | 1.800E-05 |
| 24233 | 1212 | L1212M | D | 8.8e-06 | 1.6e-05 | 1e-05 | 1.160E-05 | 3.857E-06 | 8.800E-06 | 1.600E-05 |
| 24234 | 1212 | L1212F | D | 1.6e-05 | 1.1e-05 | 1.2e-05 | 1.300E-05 | 2.646E-06 | 1.100E-05 | 1.600E-05 |
| 24235 | 1212 | L1212P | D | 1.1e-05 | 1.4e-05 | 1e-05 | 1.167E-05 | 2.082E-06 | 1.000E-05 | 1.400E-05 |
| 24236 | 1212 | L1212S | D | 9.2e-06 | 1e-05 | 1e-05 | 9.733E-06 | 4.619E-07 | 9.200E-06 | 1.000E-05 |
| 24237 | 1212 | L1212T | D | 1.5e-05 | 9.8e-06 | 1.7e-05 | 1.393E-05 | 3.717E-06 | 9.800E-06 | 1.700E-05 |
| 24238 | 1212 | L1212W | D | 1.5e-05 | 1.3e-05 | 1.1e-05 | 1.300E-05 | 2.000E-06 | 1.100E-05 | 1.500E-05 |
| 24239 | 1212 | L1212Y | D | 1.5e-05 | 1.4e-05 | 1e-05 | 1.300E-05 | 2.646E-06 | 1.000E-05 | 1.500E-05 |
| 24240 | 1212 | L1212V | D | 1.7e-05 | 1.5e-05 | 1.1e-05 | 1.433E-05 | 3.055E-06 | 1.100E-05 | 1.700E-05 |
| 24241 | 1213 | Q1213A | D | 1.1e-05 | 9.9e-06 | 1.1e-05 | 1.063E-05 | 6.351E-07 | 9.900E-06 | 1.100E-05 |
| 24242 | 1213 | Q1213R | D | 8.7e-06 | 1.1e-05 | 1.4e-05 | 1.123E-05 | 2.658E-06 | 8.700E-06 | 1.400E-05 |
| 24243 | 1213 | Q1213N | D | 1.3e-05 | 1.1e-05 | 1.2e-05 | 1.200E-05 | 1.000E-06 | 1.100E-05 | 1.300E-05 |
| 24244 | 1213 | Q1213D | D | 1.1e-05 | 1e-05 | 1e-05 | 1.033E-05 | 5.774E-07 | 1.000E-05 | 1.100E-05 |
| 24245 | 1213 | Q1213C | D | 1.5e-05 | 9.7e-06 | 1.4e-05 | 1.290E-05 | 2.816E-06 | 9.700E-06 | 1.500E-05 |
| 24246 | 1213 | Q1213E | D | 1.7e-05 | 1.5e-05 | 1.4e-05 | 1.533E-05 | 1.528E-06 | 1.400E-05 | 1.700E-05 |
| 24247 | 1213 | Q1213Q | D | 1.1e-05 | 6.4e-06 | 1.6e-05 | 1.113E-05 | 4.801E-06 | 6.400E-06 | 1.600E-05 |
| 24248 | 1213 | Q1213G | D | 9.8e-06 | 1.5e-05 | 1.5e-05 | 1.327E-05 | 3.002E-06 | 9.800E-06 | 1.500E-05 |
| 24249 | 1213 | Q1213H | D | 1.7e-05 | 1.5e-05 | 1.1e-05 | 1.433E-05 | 3.055E-06 | 1.100E-05 | 1.700E-05 |
| 24250 | 1213 | Q1213I | D | 1.2e-05 | 1.3e-05 | 1.4e-05 | 1.300E-05 | 1.000E-06 | 1.200E-05 | 1.400E-05 |
| 24251 | 1213 | Q1213L | D | 1.4e-05 | 9.4e-06 | 9.8e-06 | 1.107E-05 | 2.548E-06 | 9.400E-06 | 1.400E-05 |
| 24252 | 1213 | Q1213K | D | 9.9e-06 | 7.4e-06 | 1.6e-05 | 1.110E-05 | 4.424E-06 | 7.400E-06 | 1.600E-05 |
| 24253 | 1213 | Q1213M | D | 1.5e-05 | 1.6e-05 | 1.1e-05 | 1.400E-05 | 2.646E-06 | 1.100E-05 | 1.600E-05 |
| 24254 | 1213 | Q1213F | D | 1.2e-05 | 8.3e-06 | 9.8e-06 | 1.003E-05 | 1.861E-06 | 8.300E-06 | 1.200E-05 |
| 24255 | 1213 | Q1213P | D | 1.3e-05 | 7.9e-06 | 7.2e-06 | 9.367E-06 | 3.166E-06 | 7.200E-06 | 1.300E-05 |
| 24256 | 1213 | Q1213S | D | 9.8e-06 | 1.6e-05 | 9.7e-06 | 1.183E-05 | 3.609E-06 | 9.700E-06 | 1.600E-05 |
| 24257 | 1213 | Q1213T | D | 1.1e-05 | 1.5e-05 | 1.5e-05 | 1.367E-05 | 2.309E-06 | 1.100E-05 | 1.500E-05 |
| 24258 | 1213 | Q1213W | D | 1.4e-05 | 1.4e-05 | 1.3e-05 | 1.367E-05 | 5.774E-07 | 1.300E-05 | 1.400E-05 |
| 24259 | 1213 | Q1213Y | D | 1e-05 | 1.7e-05 | 1.6e-05 | 1.433E-05 | 3.786E-06 | 1.000E-05 | 1.700E-05 |
| 24260 | 1213 | Q1213V | D | 1.6e-05 | 1.3e-05 | 1.3e-05 | 1.400E-05 | 1.732E-06 | 1.300E-05 | 1.600E-05 |
| 24261 | 1214 | S1214A | D | 1.8e-05 | 1.7e-05 | 1.1e-05 | 1.533E-05 | 3.786E-06 | 1.100E-05 | 1.800E-05 |
| 24262 | 1214 | S1214R | D | 1e-05 | 1.6e-05 | 1.7e-05 | 1.433E-05 | 3.786E-06 | 1.000E-05 | 1.700E-05 |
| 24263 | 1214 | S1214N | D | 1.5e-05 | 1.2e-05 | 1.5e-05 | 1.400E-05 | 1.732E-06 | 1.200E-05 | 1.500E-05 |
| 24264 | 1214 | S1214D | D | 1.5e-05 | 1e-05 | 1.6e-05 | 1.367E-05 | 3.215E-06 | 1.000E-05 | 1.600E-05 |
| 24265 | 1214 | S1214C | D | 1e-05 | 1.5e-05 | 1.3e-05 | 1.267E-05 | 2.517E-06 | 1.000E-05 | 1.500E-05 |
| 24266 | 1214 | S1214E | D | 1.8e-05 | 1.7e-05 | 1.6e-05 | 1.700E-05 | 1.000E-06 | 1.600E-05 | 1.800E-05 |
| 24267 | 1214 | S1214Q | D | 1.1e-05 | 1.1e-05 | 1.4e-05 | 1.200E-05 | 1.732E-06 | 1.100E-05 | 1.400E-05 |
| 24268 | 1214 | S1214G | D | 1e-05 | 1.6e-05 | 1.2e-05 | 1.267E-05 | 3.055E-06 | 1.000E-05 | 1.600E-05 |
| 24269 | 1214 | S1214H | D | 1.6e-05 | 1.4e-05 | 1.5e-05 | 1.500E-05 | 1.000E-06 | 1.400E-05 | 1.600E-05 |
| 24270 | 1214 | S1214I | D | 1.5e-05 | 1.7e-05 | 1.4e-05 | 1.533E-05 | 1.528E-06 | 1.400E-05 | 1.700E-05 |
| 24271 | 1214 | S1214L | D | 1.8e-05 | 2.8e-05 | 1.6e-05 | 2.067E-05 | 6.429E-06 | 1.600E-05 | 2.800E-05 |
| 24272 | 1214 | S1214K | D | 1.1e-05 | 1.6e-05 | 1.6e-05 | 1.433E-05 | 2.887E-06 | 1.100E-05 | 1.600E-05 |
| 24273 | 1214 | S1214M | D | 1.7e-05 | 1.3e-05 | 1.5e-05 | 1.500E-05 | 2.000E-06 | 1.300E-05 | 1.700E-05 |
| 24274 | 1214 | S1214F | D | 1.5e-05 | 1e-05 | 1.2e-05 | 1.233E-05 | 2.517E-06 | 1.000E-05 | 1.500E-05 |
| 24275 | 1214 | S1214P | D | 1.1e-05 | 1.5e-05 | 1.5e-05 | 1.367E-05 | 2.309E-06 | 1.100E-05 | 1.500E-05 |
| 24276 | 1214 | S1214S | D | 1.2e-05 | 1.7e-05 | 8.5e-06 | 1.250E-05 | 4.272E-06 | 8.500E-06 | 1.700E-05 |
| 24277 | 1214 | S1214T | D | 9e-06 | 1e-05 | 1.1e-05 | 1.000E-05 | 1.000E-06 | 9.000E-06 | 1.100E-05 |
| 24278 | 1214 | S1214W | D | 1.5e-05 | 1.3e-05 | 8.3e-06 | 1.210E-05 | 3.439E-06 | 8.300E-06 | 1.500E-05 |
| 24279 | 1214 | S1214Y | D | 1.1e-05 | 1.4e-05 | 1.1e-05 | 1.200E-05 | 1.732E-06 | 1.100E-05 | 1.400E-05 |
| 24280 | 1214 | S1214V | D | 1.1e-05 | 1.7e-05 | 1.5e-05 | 1.433E-05 | 3.055E-06 | 1.100E-05 | 1.700E-05 |
| 24281 | 1215 | G1215A | D | 1.5e-05 | 8.9e-06 | 1.7e-05 | 1.363E-05 | 4.219E-06 | 8.900E-06 | 1.700E-05 |
| 24282 | 1215 | G1215R | D | 1.5e-05 | 1.5e-05 | 1.4e-05 | 1.467E-05 | 5.774E-07 | 1.400E-05 | 1.500E-05 |
| 24283 | 1215 | G1215N | D | 1.4e-05 | 1.4e-05 | 8.9e-06 | 1.230E-05 | 2.944E-06 | 8.900E-06 | 1.400E-05 |
| 24284 | 1215 | G1215D | D | 1.2e-05 | 1.5e-05 | 1.5e-05 | 1.400E-05 | 1.732E-06 | 1.200E-05 | 1.500E-05 |
| 24285 | 1215 | G1215C | D | 1.6e-05 | 1.3e-05 | 1.6e-05 | 1.500E-05 | 1.732E-06 | 1.300E-05 | 1.600E-05 |
| 24286 | 1215 | G1215E | D | 1.6e-05 | 1.8e-05 | 1.2e-05 | 1.533E-05 | 3.055E-06 | 1.200E-05 | 1.800E-05 |
| 24287 | 1215 | G1215Q | D | 8.4e-06 | 1.4e-05 | 1.6e-05 | 1.280E-05 | 3.940E-06 | 8.400E-06 | 1.600E-05 |
| 24288 | 1215 | G1215G | D | 1.1e-05 | 1.1e-05 | 1.5e-05 | 1.233E-05 | 2.309E-06 | 1.100E-05 | 1.500E-05 |
| 24289 | 1215 | G1215H | D | 7e-06 | 9.7e-06 | 1.1e-05 | 9.233E-06 | 2.040E-06 | 7.000E-06 | 1.100E-05 |
| 24290 | 1215 | G1215I | D | 1.1e-05 | 1.5e-05 | 6.2e-06 | 1.073E-05 | 4.406E-06 | 6.200E-06 | 1.500E-05 |
| 24291 | 1215 | G1215L | D | 1.1e-05 | 9e-06 | 7.4e-06 | 9.133E-06 | 1.804E-06 | 7.400E-06 | 1.100E-05 |
| 24292 | 1215 | G1215K | D | 9.7e-06 | 1.5e-05 | 1.6e-05 | 1.357E-05 | 3.386E-06 | 9.700E-06 | 1.600E-05 |
| 24293 | 1215 | G1215M | D | 9.9e-06 | 1.3e-05 | 9.9e-06 | 1.093E-05 | 1.790E-06 | 9.900E-06 | 1.300E-05 |
| 24294 | 1215 | G1215F | D | 1.3e-05 | 1.1e-05 | 1.1e-05 | 1.167E-05 | 1.155E-06 | 1.100E-05 | 1.300E-05 |
| 24295 | 1215 | G1215P | D | 9e-06 | 1.1e-05 | 1.6e-05 | 1.200E-05 | 3.606E-06 | 9.000E-06 | 1.600E-05 |
| 24296 | 1215 | G1215S | D | 1.6e-05 | 1.4e-05 | 1.4e-05 | 1.467E-05 | 1.155E-06 | 1.400E-05 | 1.600E-05 |
| 24297 | 1215 | G1215T | D | 1.1e-05 | 9.5e-06 | 1.3e-05 | 1.117E-05 | 1.756E-06 | 9.500E-06 | 1.300E-05 |
| 24298 | 1215 | G1215W | D | 1.4e-05 | 1.4e-05 | 8.5e-06 | 1.217E-05 | 3.175E-06 | 8.500E-06 | 1.400E-05 |
| 24299 | 1215 | G1215Y | D | 1.2e-05 | 1.4e-05 | 1e-05 | 1.200E-05 | 2.000E-06 | 1.000E-05 | 1.400E-05 |
| 24300 | 1215 | G1215V | D | 1.2e-05 | 1.4e-05 | 1.5e-05 | 1.367E-05 | 1.528E-06 | 1.200E-05 | 1.500E-05 |
| 24301 | 1216 | N1216A | D | 1.2e-05 | 1.5e-05 | 8.9e-06 | 1.197E-05 | 3.050E-06 | 8.900E-06 | 1.500E-05 |
| 24302 | 1216 | N1216R | D | 8.3e-06 | 2.2e-05 | 1.7e-05 | 1.577E-05 | 6.933E-06 | 8.300E-06 | 2.200E-05 |
| 24303 | 1216 | N1216N | D | 1.3e-05 | 1.5e-05 | 9.1e-06 | 1.237E-05 | 3.001E-06 | 9.100E-06 | 1.500E-05 |
| 24304 | 1216 | N1216D | D | 1.7e-05 | 1.7e-05 | 1.4e-05 | 1.600E-05 | 1.732E-06 | 1.400E-05 | 1.700E-05 |
| 24305 | 1216 | N1216C | D | 1.6e-05 | 1e-05 | 1.5e-05 | 1.367E-05 | 3.215E-06 | 1.000E-05 | 1.600E-05 |
| 24306 | 1216 | N1216E | D | 1.7e-05 | 1.5e-05 | 1.6e-05 | 1.600E-05 | 1.000E-06 | 1.500E-05 | 1.700E-05 |
| 24307 | 1216 | N1216Q | D | 1.1e-05 | 1.4e-05 | 1.5e-05 | 1.333E-05 | 2.082E-06 | 1.100E-05 | 1.500E-05 |
| 24308 | 1216 | N1216G | D | 1.4e-05 | 1.1e-05 | 1.5e-05 | 1.333E-05 | 2.082E-06 | 1.100E-05 | 1.500E-05 |
| 24309 | 1216 | N1216H | D | 1.6e-05 | 1.3e-05 | 1.6e-05 | 1.500E-05 | 1.732E-06 | 1.300E-05 | 1.600E-05 |

| | | | | | | | | | | |
|---|---|---|---|---|---|---|---|---|---|---|
| 24310 | 1216 | N1216I | D | 1.5e-05 | 9.9e-06 | 6.4e-06 | 1.043E-05 | 4.325E-06 | 6.400E-06 | 1.500E-05 |
| 24311 | 1216 | N1216L | D | 9.1e-06 | 1.7e-05 | 1.1e-05 | 1.237E-05 | 4.124E-06 | 9.100E-06 | 1.700E-05 |
| 24312 | 1216 | N1216K | D | 1e-05 | 1.1e-05 | 1.5e-05 | 1.200E-05 | 2.646E-06 | 1.000E-05 | 1.500E-05 |
| 24313 | 1216 | N1216M | D | 1.6e-05 | 1.1e-05 | 1.4e-05 | 1.367E-05 | 2.517E-06 | 1.100E-05 | 1.600E-05 |
| 24314 | 1216 | N1216F | D | 1.4e-05 | 1.8e-05 | 1.6e-05 | 1.600E-05 | 2.000E-06 | 1.400E-05 | 1.800E-05 |
| 24315 | 1216 | N1216P | D | 1.4e-05 | 1e-05 | 1e-05 | 1.133E-05 | 2.309E-06 | 1.000E-05 | 1.400E-05 |
| 24316 | 1216 | N1216S | D | 8.7e-06 | 9e-06 | 1.7e-05 | 1.157E-05 | 4.708E-06 | 8.700E-06 | 1.700E-05 |
| 24317 | 1216 | N1216T | D | 1.1e-05 | 1.5e-05 | 9e-06 | 1.167E-05 | 3.055E-06 | 9.000E-06 | 1.500E-05 |
| 24318 | 1216 | N1216W | D | 1.4e-05 | 1.1e-05 | 1e-05 | 1.167E-05 | 2.082E-06 | 1.000E-05 | 1.400E-05 |
| 24319 | 1216 | N1216Y | D | 1.2e-05 | 9.7e-06 | 1.4e-05 | 1.190E-05 | 2.152E-06 | 9.700E-06 | 1.400E-05 |
| 24320 | 1216 | N1216V | D | 1.5e-05 | 1.5e-05 | 1.4e-05 | 1.467E-05 | 5.774E-07 | 1.400E-05 | 1.500E-05 |
| 24321 | 1217 | S1217A | D | 1.6e-05 | 1.7e-05 | 1.4e-05 | 1.567E-05 | 1.528E-06 | 1.400E-05 | 1.700E-05 |
| 24322 | 1217 | S1217R | D | 1.1e-05 | 1.5e-05 | 1.6e-05 | 1.400E-05 | 2.646E-06 | 1.100E-05 | 1.600E-05 |
| 24323 | 1217 | S1217N | D | 1.5e-05 | 1.4e-05 | 1.4e-05 | 1.433E-05 | 5.774E-07 | 1.400E-05 | 1.500E-05 |
| 24324 | 1217 | S1217D | D | 6.7e-06 | 1.2e-05 | 1.4e-05 | 1.090E-05 | 3.772E-06 | 6.700E-06 | 1.400E-05 |
| 24325 | 1217 | S1217C | D | 1.7e-05 | 1.5e-05 | 1.5e-05 | 1.567E-05 | 1.155E-06 | 1.500E-05 | 1.700E-05 |
| 24326 | 1217 | S1217E | D | 1.6e-05 | 1.1e-05 | 9.8e-06 | 1.227E-05 | 3.288E-06 | 9.800E-06 | 1.600E-05 |
| 24327 | 1217 | S1217Q | D | 9.6e-06 | 1e-05 | 1.6e-05 | 1.187E-05 | 3.585E-06 | 9.600E-06 | 1.600E-05 |
| 24328 | 1217 | S1217G | D | 1.5e-05 | 1.1e-05 | 9.5e-06 | 1.183E-05 | 2.843E-06 | 9.500E-06 | 1.500E-05 |
| 24329 | 1217 | S1217H | D | 1.6e-05 | 1.2e-05 | 1.3e-05 | 1.367E-05 | 2.082E-06 | 1.200E-05 | 1.600E-05 |
| 24330 | 1217 | S1217I | D | 1.7e-05 | 1.5e-05 | 1.5e-05 | 1.567E-05 | 1.155E-06 | 1.500E-05 | 1.700E-05 |
| 24331 | 1217 | S1217L | D | 1.6e-05 | 9.9e-06 | 1e-05 | 1.197E-05 | 3.493E-06 | 9.900E-06 | 1.600E-05 |
| 24332 | 1217 | S1217K | D | 9.6e-06 | 1.4e-05 | 1e-05 | 1.120E-05 | 2.433E-06 | 9.600E-06 | 1.400E-05 |
| 24333 | 1217 | S1217M | D | 9.5e-06 | 1.6e-05 | 9.2e-06 | 1.157E-05 | 3.842E-06 | 9.200E-06 | 1.600E-05 |
| 24334 | 1217 | S1217F | D | 1.6e-05 | 9.1e-06 | 1.6e-05 | 1.370E-05 | 3.984E-06 | 9.100E-06 | 1.600E-05 |
| 24335 | 1217 | S1217P | D | 1e-05 | 1.3e-05 | 1.3e-05 | 1.200E-05 | 1.732E-06 | 1.000E-05 | 1.300E-05 |
| 24336 | 1217 | S1217S | D | 1e-05 | 9.9e-06 | 1.1e-05 | 1.030E-05 | 6.083E-07 | 9.900E-06 | 1.100E-05 |
| 24337 | 1217 | S1217T | D | 1.5e-05 | 1.3e-05 | 1.7e-05 | 1.500E-05 | 2.000E-06 | 1.300E-05 | 1.700E-05 |
| 24338 | 1217 | S1217W | D | 1.6e-05 | 1.2e-05 | 1e-05 | 1.267E-05 | 3.055E-06 | 1.000E-05 | 1.600E-05 |
| 24339 | 1217 | S1217Y | D | 1.1e-05 | 1e-05 | 1.1e-05 | 1.067E-05 | 5.774E-07 | 1.000E-05 | 1.100E-05 |
| 24340 | 1217 | S1217V | D | 1.1e-05 | 1.4e-05 | 1.7e-05 | 1.400E-05 | 3.000E-06 | 1.100E-05 | 1.700E-05 |
| 24341 | 1218 | Q1218A | D | 1.4e-05 | 1.5e-05 | 9.5e-06 | 1.283E-05 | 2.930E-06 | 9.500E-06 | 1.500E-05 |
| 24342 | 1218 | Q1218R | D | 1.5e-05 | 1.4e-05 | 1.1e-05 | 1.333E-05 | 2.082E-06 | 1.100E-05 | 1.500E-05 |
| 24343 | 1218 | Q1218N | D | 1.6e-05 | 1.4e-05 | 1.7e-05 | 1.567E-05 | 1.528E-06 | 1.400E-05 | 1.700E-05 |
| 24344 | 1218 | Q1218D | D | 1.6e-05 | 1.3e-05 | 1.1e-05 | 1.333E-05 | 2.517E-06 | 1.100E-05 | 1.600E-05 |
| 24345 | 1218 | Q1218C | D | 1.3e-05 | 1.7e-05 | 9.1e-06 | 1.303E-05 | 3.950E-06 | 9.100E-06 | 1.700E-05 |
| 24346 | 1218 | Q1218E | D | 1.4e-05 | 1.1e-05 | 1.5e-05 | 1.333E-05 | 2.082E-06 | 1.100E-05 | 1.500E-05 |
| 24347 | 1218 | Q1218Q | D | 1.5e-05 | 1e-05 | 1.4e-05 | 1.300E-05 | 2.646E-06 | 1.000E-05 | 1.500E-05 |
| 24348 | 1218 | Q1218G | D | 1e-05 | 1.6e-05 | 1.6e-05 | 1.400E-05 | 3.464E-06 | 1.000E-05 | 1.600E-05 |
| 24349 | 1218 | Q1218H | D | 1.6e-05 | 1.1e-05 | 1.1e-05 | 1.267E-05 | 2.887E-06 | 1.100E-05 | 1.600E-05 |
| 24350 | 1218 | Q1218I | D | 9.8e-06 | 1.3e-05 | 1.6e-05 | 1.293E-05 | 3.101E-06 | 9.800E-06 | 1.600E-05 |
| 24351 | 1218 | Q1218L | D | 7.7e-06 | 1.5e-05 | 1.6e-05 | 1.290E-05 | 4.531E-06 | 7.700E-06 | 1.600E-05 |
| 24352 | 1218 | Q1218K | D | 1e-05 | 1.1e-05 | 1.6e-05 | 1.233E-05 | 3.215E-06 | 1.000E-05 | 1.600E-05 |
| 24353 | 1218 | Q1218M | D | 1.7e-05 | 1.5e-05 | 1.6e-05 | 1.600E-05 | 1.000E-06 | 1.500E-05 | 1.700E-05 |
| 24354 | 1218 | Q1218F | D | 1.5e-05 | 1.1e-05 | 8.7e-06 | 1.157E-05 | 3.188E-06 | 8.700E-06 | 1.500E-05 |
| 24355 | 1218 | Q1218P | D | 9.9e-06 | 1e-05 | 1e-05 | 9.967E-06 | 5.774E-08 | 9.900E-06 | 1.000E-05 |
| 24356 | 1218 | Q1218S | D | 1.6e-05 | 1.3e-05 | 1.5e-05 | 1.467E-05 | 1.528E-06 | 1.300E-05 | 1.600E-05 |
| 24357 | 1218 | Q1218T | D | 1.5e-05 | 1.2e-05 | 1.1e-05 | 1.267E-05 | 2.082E-06 | 1.100E-05 | 1.500E-05 |
| 24358 | 1218 | Q1218W | D | 9.7e-06 | 1.5e-05 | 1.3e-05 | 1.257E-05 | 2.676E-06 | 9.700E-06 | 1.500E-05 |
| 24359 | 1218 | Q1218Y | D | 9.5e-06 | 1.4e-05 | 7.1e-06 | 1.020E-05 | 3.503E-06 | 7.100E-06 | 1.400E-05 |
| 24360 | 1218 | Q1218V | D | 2.4e-05 | 1.6e-05 | 1.3e-05 | 1.767E-05 | 5.686E-06 | 1.300E-05 | 2.400E-05 |
| 24361 | 1219 | E1219A | D | 1.8e-05 | 1.1e-05 | 1.4e-05 | 1.433E-05 | 3.512E-06 | 1.100E-05 | 1.800E-05 |
| 24362 | 1219 | E1219R | D | 1e-05 | 1e-05 | 1e-05 | 1.000E-05 | 0.000E+00 | 1.000E-05 | 1.000E-05 |
| 24363 | 1219 | E1219N | D | 1.2e-05 | 1.1e-05 | 1.7e-05 | 1.333E-05 | 3.215E-06 | 1.100E-05 | 1.700E-05 |
| 24364 | 1219 | E1219D | D | 1.4e-05 | 1.5e-05 | 1e-05 | 1.300E-05 | 2.646E-06 | 1.000E-05 | 1.500E-05 |
| 24365 | 1219 | E1219C | D | 9.6e-06 | 1.6e-05 | 1.3e-05 | 1.287E-05 | 3.202E-06 | 9.600E-06 | 1.600E-05 |
| 24366 | 1219 | E1219E | D | 8.1e-06 | 1e-05 | 1.3e-05 | 1.037E-05 | 2.470E-06 | 8.100E-06 | 1.300E-05 |
| 24367 | 1219 | E1219Q | D | 1.2e-05 | 1.6e-05 | 1.3e-05 | 1.367E-05 | 2.082E-06 | 1.200E-05 | 1.600E-05 |
| 24368 | 1219 | E1219G | D | 1.1e-05 | 6.7e-06 | 9.4e-06 | 9.033E-06 | 2.173E-06 | 6.700E-06 | 1.100E-05 |
| 24369 | 1219 | E1219H | D | 8.8e-06 | 9.6e-06 | 1.5e-05 | 1.113E-05 | 3.372E-06 | 8.800E-06 | 1.500E-05 |
| 24370 | 1219 | E1219I | D | 1.1e-05 | 1.1e-05 | 1.3e-05 | 1.167E-05 | 1.155E-06 | 1.100E-05 | 1.300E-05 |
| 24371 | 1219 | E1219L | D | 1.1e-05 | 1.5e-05 | 1.1e-05 | 1.233E-05 | 2.309E-06 | 1.100E-05 | 1.500E-05 |
| 24372 | 1219 | E1219K | D | 1.7e-05 | 1.3e-05 | 1.5e-05 | 1.500E-05 | 2.000E-06 | 1.300E-05 | 1.700E-05 |
| 24373 | 1219 | E1219M | D | 9.9e-06 | 1.5e-05 | 1.5e-05 | 1.330E-05 | 2.944E-06 | 9.900E-06 | 1.500E-05 |
| 24374 | 1219 | E1219F | D | 1.1e-05 | 1.5e-05 | 1.4e-05 | 1.333E-05 | 2.082E-06 | 1.100E-05 | 1.500E-05 |
| 24375 | 1219 | E1219P | D | 1.3e-05 | 1.2e-05 | 1.2e-05 | 1.233E-05 | 5.774E-07 | 1.200E-05 | 1.300E-05 |
| 24376 | 1219 | E1219S | D | 1e-05 | 1e-05 | 1.5e-05 | 1.167E-05 | 2.887E-06 | 1.000E-05 | 1.500E-05 |
| 24377 | 1219 | E1219T | D | 1.3e-05 | 1.4e-05 | 1.4e-05 | 1.367E-05 | 5.774E-07 | 1.300E-05 | 1.400E-05 |
| 24378 | 1219 | E1219W | D | 1.5e-05 | 1.1e-05 | 1.3e-05 | 1.300E-05 | 2.000E-06 | 1.100E-05 | 1.500E-05 |
| 24379 | 1219 | E1219Y | D | 1.4e-05 | 1.5e-05 | 1.5e-05 | 1.467E-05 | 5.774E-07 | 1.400E-05 | 1.500E-05 |
| 24380 | 1219 | E1219V | D | 1.4e-05 | 1.1e-05 | 1.5e-05 | 1.333E-05 | 2.082E-06 | 1.100E-05 | 1.500E-05 |
| 24381 | 1220 | S1220A | D | 1.1e-05 | 1e-05 | 1e-05 | 1.033E-05 | 5.774E-07 | 1.000E-05 | 1.100E-05 |
| 24382 | 1220 | S1220R | D | 1.8e-05 | 1.1e-05 | 1.7e-05 | 1.533E-05 | 3.786E-06 | 1.100E-05 | 1.800E-05 |
| 24383 | 1220 | S1220N | D | 1.6e-05 | 1.3e-05 | 9.9e-06 | 1.297E-05 | 3.050E-06 | 9.900E-06 | 1.600E-05 |
| 24384 | 1220 | S1220D | D | 8.9e-06 | 1.2e-05 | 1.1e-05 | 1.063E-05 | 1.582E-06 | 8.900E-06 | 1.200E-05 |
| 24385 | 1220 | S1220C | D | 1.4e-05 | 9.9e-06 | 7.6e-06 | 1.050E-05 | 3.242E-06 | 7.600E-06 | 1.400E-05 |
| 24386 | 1220 | S1220E | D | 1.6e-05 | 1.4e-05 | 1.1e-05 | 1.367E-05 | 2.517E-06 | 1.100E-05 | 1.600E-05 |
| 24387 | 1220 | S1220Q | D | 1.5e-05 | 1.1e-05 | 1.2e-05 | 1.267E-05 | 2.082E-06 | 1.100E-05 | 1.500E-05 |
| 24388 | 1220 | S1220G | D | 1.7e-05 | 1e-05 | 1.1e-05 | 1.267E-05 | 3.786E-06 | 1.000E-05 | 1.700E-05 |
| 24389 | 1220 | S1220H | D | 1.3e-05 | 1.2e-05 | 1.1e-05 | 1.200E-05 | 1.000E-06 | 1.100E-05 | 1.300E-05 |
| 24390 | 1220 | S1220I | D | 1.1e-05 | 1.3e-05 | 1.7e-05 | 1.367E-05 | 3.055E-06 | 1.100E-05 | 1.700E-05 |
| 24391 | 1220 | S1220L | D | 1.5e-05 | 1e-05 | 1.2e-05 | 1.233E-05 | 2.517E-06 | 1.000E-05 | 1.500E-05 |
| 24392 | 1220 | S1220K | D | 1.1e-05 | 1.4e-05 | 1.7e-05 | 1.400E-05 | 3.000E-06 | 1.100E-05 | 1.700E-05 |
| 24393 | 1220 | S1220M | D | 1.1e-05 | 1.6e-05 | 1.6e-05 | 1.433E-05 | 2.887E-06 | 1.100E-05 | 1.600E-05 |
| 24394 | 1220 | S1220F | D | 1.4e-05 | 1.1e-05 | 1.1e-05 | 1.200E-05 | 1.732E-06 | 1.100E-05 | 1.400E-05 |

| | | | | | | | | | | |
|---|---|---|---|---|---|---|---|---|---|---|
| 24395 | 1220 | S1220P | D | 1.7e-05 | 1.4e-05 | 1.7e-05 | 1.600E-05 | 1.732E-06 | 1.400E-05 | 1.700E-05 |
| 24396 | 1220 | S1220S | D | 9.3e-06 | 9.6e-06 | 1.4e-05 | 1.097E-05 | 2.631E-06 | 9.300E-06 | 1.400E-05 |
| 24397 | 1220 | S1220T | D | 9.6e-06 | 1.4e-05 | 1.4e-05 | 1.253E-05 | 2.540E-06 | 9.600E-06 | 1.400E-05 |
| 24398 | 1220 | S1220W | D | 1.5e-05 | 1.1e-05 | 1.5e-05 | 1.367E-05 | 2.309E-06 | 1.100E-05 | 1.500E-05 |
| 24399 | 1220 | S1220Y | D | 1.1e-05 | 1.4e-05 | 9.2e-06 | 1.140E-05 | 2.425E-06 | 9.200E-06 | 1.400E-05 |
| 24400 | 1220 | S1220V | D | 1.1e-05 | 1.6e-05 | 1.5e-05 | 1.400E-05 | 2.646E-06 | 1.100E-05 | 1.600E-05 |
| 24401 | 1221 | V1221A | D | 9.6e-06 | 1.2e-05 | 1e-05 | 1.053E-05 | 1.286E-06 | 9.600E-06 | 1.200E-05 |
| 24402 | 1221 | V1221R | D | 1.6e-05 | 8.1e-06 | 1.7e-05 | 1.370E-05 | 4.875E-06 | 8.100E-06 | 1.700E-05 |
| 24403 | 1221 | V1221N | D | 1.6e-05 | 1.3e-05 | 1.6e-05 | 1.500E-05 | 1.732E-06 | 1.300E-05 | 1.600E-05 |
| 24404 | 1221 | V1221D | D | 1.1e-05 | 1.1e-05 | 1.5e-05 | 1.233E-05 | 2.309E-06 | 1.100E-05 | 1.500E-05 |
| 24405 | 1221 | V1221C | D | 1.4e-05 | 1.4e-05 | 1e-05 | 1.267E-05 | 2.309E-06 | 1.000E-05 | 1.400E-05 |
| 24406 | 1221 | V1221E | D | 7.9e-06 | 1.3e-05 | 1.4e-05 | 1.163E-05 | 3.272E-06 | 7.900E-06 | 1.400E-05 |
| 24407 | 1221 | V1221Q | D | 1.4e-05 | 1e-05 | 1.2e-05 | 1.200E-05 | 2.000E-06 | 1.000E-05 | 1.400E-05 |
| 24408 | 1221 | V1221G | D | 9.6e-06 | 1.6e-05 | 1e-05 | 1.187E-05 | 3.585E-06 | 9.600E-06 | 1.600E-05 |
| 24409 | 1221 | V1221H | D | 8e-06 | 1.1e-05 | 9.6e-06 | 9.533E-06 | 1.501E-06 | 8.000E-06 | 1.100E-05 |
| 24410 | 1221 | V1221I | D | 1.1e-05 | 1.2e-05 | 1.4e-05 | 1.233E-05 | 1.528E-06 | 1.100E-05 | 1.400E-05 |
| 24411 | 1221 | V1221L | D | 1.6e-05 | 1.6e-05 | 9.8e-06 | 1.393E-05 | 3.580E-06 | 9.800E-06 | 1.600E-05 |
| 24412 | 1221 | V1221K | D | 1e-05 | 1.5e-05 | 1.2e-05 | 1.233E-05 | 2.517E-06 | 1.000E-05 | 1.500E-05 |
| 24413 | 1221 | V1221M | D | 1.5e-05 | 1.5e-05 | 1.4e-05 | 1.467E-05 | 5.774E-07 | 1.400E-05 | 1.500E-05 |
| 24414 | 1221 | V1221F | D | 1.3e-05 | 1e-05 | 1.7e-05 | 1.333E-05 | 3.512E-06 | 1.000E-05 | 1.700E-05 |
| 24415 | 1221 | V1221P | D | 1.9e-05 | 1.5e-05 | 1.4e-05 | 1.600E-05 | 2.646E-06 | 1.400E-05 | 1.900E-05 |
| 24416 | 1221 | V1221S | D | 1.1e-05 | 1.5e-05 | 1e-05 | 1.200E-05 | 2.646E-06 | 1.000E-05 | 1.500E-05 |
| 24417 | 1221 | V1221T | D | 1.1e-05 | 8.5e-06 | 1.4e-05 | 1.117E-05 | 2.754E-06 | 8.500E-06 | 1.400E-05 |
| 24418 | 1221 | V1221W | D | 1.4e-05 | 1.4e-05 | 1.5e-05 | 1.433E-05 | 5.774E-07 | 1.400E-05 | 1.500E-05 |
| 24419 | 1221 | V1221Y | D | 8.8e-06 | 1.5e-05 | 1.5e-05 | 1.293E-05 | 3.580E-06 | 8.800E-06 | 1.500E-05 |
| 24420 | 1221 | V1221V | D | 1.5e-05 | 1.4e-05 | 1.6e-05 | 1.500E-05 | 1.000E-06 | 1.400E-05 | 1.600E-05 |
| 24421 | 1222 | T1222A | D | 1.4e-05 | 1.2e-05 | 1.1e-05 | 1.233E-05 | 1.528E-06 | 1.100E-05 | 1.400E-05 |
| 24422 | 1222 | T1222R | D | 1.3e-05 | 1.1e-05 | 1.5e-05 | 1.300E-05 | 2.000E-06 | 1.100E-05 | 1.500E-05 |
| 24423 | 1222 | T1222N | D | 1.6e-05 | 1e-05 | 1e-05 | 1.200E-05 | 3.464E-06 | 1.000E-05 | 1.600E-05 |
| 24424 | 1222 | T1222D | D | 1.1e-05 | 1.8e-05 | 9.7e-06 | 1.290E-05 | 4.464E-06 | 9.700E-06 | 1.800E-05 |
| 24425 | 1222 | T1222C | D | 1.1e-05 | 1.4e-05 | 1e-05 | 1.167E-05 | 2.082E-06 | 1.000E-05 | 1.400E-05 |
| 24426 | 1222 | T1222E | D | 1.7e-05 | 1.1e-05 | 1.2e-05 | 1.333E-05 | 3.215E-06 | 1.100E-05 | 1.700E-05 |
| 24427 | 1222 | T1222Q | D | 8.2e-06 | 9.9e-06 | 1.2e-05 | 1.003E-05 | 1.904E-06 | 8.200E-06 | 1.200E-05 |
| 24428 | 1222 | T1222G | D | 1.5e-05 | 1.3e-05 | 1.7e-05 | 1.500E-05 | 2.000E-06 | 1.300E-05 | 1.700E-05 |
| 24429 | 1222 | T1222H | D | 1.5e-05 | 1e-05 | 1.2e-05 | 1.233E-05 | 2.517E-06 | 1.000E-05 | 1.500E-05 |
| 24430 | 1222 | T1222I | D | 1.1e-05 | 1.6e-05 | 1.5e-05 | 1.400E-05 | 2.646E-06 | 1.100E-05 | 1.600E-05 |
| 24431 | 1222 | T1222L | D | 1.8e-05 | 1.7e-05 | 1.5e-05 | 1.667E-05 | 1.528E-06 | 1.500E-05 | 1.800E-05 |
| 24432 | 1222 | T1222K | D | 1.7e-05 | 1.5e-05 | 1.3e-05 | 1.500E-05 | 2.000E-06 | 1.300E-05 | 1.700E-05 |
| 24433 | 1222 | T1222M | D | 1.5e-05 | 1e-05 | 1.1e-05 | 1.200E-05 | 2.646E-06 | 1.000E-05 | 1.500E-05 |
| 24434 | 1222 | T1222F | D | 1.6e-05 | 1.5e-05 | 1.4e-05 | 1.500E-05 | 1.000E-06 | 1.400E-05 | 1.600E-05 |
| 24435 | 1222 | T1222P | D | 1.2e-05 | 9.6e-06 | 1.7e-05 | 1.287E-05 | 3.775E-06 | 9.600E-06 | 1.700E-05 |
| 24436 | 1222 | T1222S | D | 1e-05 | 1.1e-05 | 1.3e-05 | 1.133E-05 | 1.528E-06 | 1.000E-05 | 1.300E-05 |
| 24437 | 1222 | T1222T | D | 9.5e-06 | 8.3e-06 | 1.7e-05 | 1.160E-05 | 4.715E-06 | 8.300E-06 | 1.700E-05 |
| 24438 | 1222 | T1222W | D | 1.6e-05 | 1.6e-05 | 1.1e-05 | 1.433E-05 | 2.887E-06 | 1.100E-05 | 1.600E-05 |
| 24439 | 1222 | T1222Y | D | 1e-05 | 1.3e-05 | 8.7e-06 | 1.057E-05 | 2.205E-06 | 8.700E-06 | 1.300E-05 |
| 24440 | 1222 | T1222V | D | 1.1e-05 | 1.2e-05 | 1.5e-05 | 1.267E-05 | 2.082E-06 | 1.100E-05 | 1.500E-05 |
| 24441 | 1223 | E1223A | D | 1e-05 | 9.1e-06 | 1.4e-05 | 1.103E-05 | 2.608E-06 | 9.100E-06 | 1.400E-05 |
| 24442 | 1223 | E1223R | D | 1e-05 | 1.1e-05 | 1.5e-05 | 1.200E-05 | 2.646E-06 | 1.000E-05 | 1.500E-05 |
| 24443 | 1223 | E1223N | D | 1.7e-05 | 1.6e-05 | 1.4e-05 | 1.567E-05 | 1.528E-06 | 1.400E-05 | 1.700E-05 |
| 24444 | 1223 | E1223D | D | 1e-05 | 8.6e-06 | 9.9e-06 | 9.500E-06 | 7.810E-07 | 8.600E-06 | 1.000E-05 |
| 24445 | 1223 | E1223C | D | 1.4e-05 | 1.2e-05 | 1.4e-05 | 1.333E-05 | 1.155E-06 | 1.200E-05 | 1.400E-05 |
| 24446 | 1223 | E1223E | D | 1.2e-05 | 1.5e-05 | 1.2e-05 | 1.300E-05 | 1.732E-06 | 1.200E-05 | 1.500E-05 |
| 24447 | 1223 | E1223Q | D | 1.6e-05 | 1.3e-05 | 1.1e-05 | 1.333E-05 | 2.517E-06 | 1.100E-05 | 1.600E-05 |
| 24448 | 1223 | E1223G | D | 1e-05 | 1e-05 | 1.3e-05 | 1.100E-05 | 1.732E-06 | 1.000E-05 | 1.300E-05 |
| 24449 | 1223 | E1223H | D | 1.2e-05 | 1.5e-05 | 1.3e-05 | 1.333E-05 | 1.528E-06 | 1.200E-05 | 1.500E-05 |
| 24450 | 1223 | E1223I | D | 1.2e-05 | 1.2e-05 | 1.3e-05 | 1.233E-05 | 5.774E-07 | 1.200E-05 | 1.300E-05 |
| 24451 | 1223 | E1223L | D | 1.2e-05 | 1e-05 | 1.1e-05 | 1.100E-05 | 1.000E-06 | 1.000E-05 | 1.200E-05 |
| 24452 | 1223 | E1223K | D | 1e-05 | 1.2e-05 | 8e-06 | 1.000E-05 | 2.000E-06 | 8.000E-06 | 1.200E-05 |
| 24453 | 1223 | E1223M | D | 1.6e-05 | 1.3e-05 | 1.6e-05 | 1.500E-05 | 1.732E-06 | 1.300E-05 | 1.600E-05 |
| 24454 | 1223 | E1223F | D | 1.3e-05 | 1.6e-05 | 1.5e-05 | 1.467E-05 | 1.528E-06 | 1.300E-05 | 1.600E-05 |
| 24455 | 1223 | E1223P | D | 6.1e-06 | 9.6e-06 | 1.6e-05 | 1.057E-05 | 5.020E-06 | 6.100E-06 | 1.600E-05 |
| 24456 | 1223 | E1223S | D | 9.2e-06 | 1.1e-05 | 8.3e-06 | 9.500E-06 | 1.375E-06 | 8.300E-06 | 1.100E-05 |
| 24457 | 1223 | E1223T | D | 1.3e-05 | 1.1e-05 | 1.6e-05 | 1.333E-05 | 2.517E-06 | 1.100E-05 | 1.600E-05 |
| 24458 | 1223 | E1223W | D | 1.1e-05 | 9.6e-06 | 1.5e-05 | 1.187E-05 | 2.802E-06 | 9.600E-06 | 1.500E-05 |
| 24459 | 1223 | E1223Y | D | 1.5e-05 | 1.7e-05 | 1.4e-05 | 1.533E-05 | 1.528E-06 | 1.400E-05 | 1.700E-05 |
| 24460 | 1223 | E1223V | D | 1.1e-05 | 1.4e-05 | 1.5e-05 | 1.333E-05 | 2.082E-06 | 1.100E-05 | 1.500E-05 |
| 24461 | 1224 | Q1224A | D | 9.8e-06 | 1.6e-05 | 1.6e-05 | 1.393E-05 | 3.580E-06 | 9.800E-06 | 1.600E-05 |
| 24462 | 1224 | Q1224R | D | 1.1e-05 | 1e-05 | 1.3e-05 | 1.133E-05 | 1.528E-06 | 1.000E-05 | 1.300E-05 |
| 24463 | 1224 | Q1224N | D | 1.2e-05 | 1.6e-05 | 1.6e-05 | 1.467E-05 | 2.309E-06 | 1.200E-05 | 1.600E-05 |
| 24464 | 1224 | Q1224D | D | 1.5e-05 | 8.7e-06 | 1e-05 | 1.123E-05 | 3.326E-06 | 8.700E-06 | 1.500E-05 |
| 24465 | 1224 | Q1224C | D | 9.7e-06 | 1.4e-05 | 8.8e-06 | 1.083E-05 | 2.779E-06 | 8.800E-06 | 1.400E-05 |
| 24466 | 1224 | Q1224E | D | 1.5e-05 | 9.5e-06 | 1.7e-05 | 1.383E-05 | 3.884E-06 | 9.500E-06 | 1.700E-05 |
| 24467 | 1224 | Q1224Q | D | 1.3e-05 | 1.5e-05 | 1e-05 | 1.267E-05 | 2.517E-06 | 1.000E-05 | 1.500E-05 |
| 24468 | 1224 | Q1224G | D | 1.4e-05 | 8e-06 | 1.6e-05 | 1.267E-05 | 4.163E-06 | 8.000E-06 | 1.600E-05 |
| 24469 | 1224 | Q1224H | D | 1.5e-05 | 1.2e-05 | 1.4e-05 | 1.367E-05 | 1.528E-06 | 1.200E-05 | 1.500E-05 |
| 24470 | 1224 | Q1224I | D | 1.7e-05 | 1.6e-05 | 1.6e-05 | 1.633E-05 | 5.774E-07 | 1.600E-05 | 1.700E-05 |
| 24471 | 1224 | Q1224L | D | 1.3e-05 | 1.5e-05 | 9.4e-06 | 1.247E-05 | 2.838E-06 | 9.400E-06 | 1.500E-05 |
| 24472 | 1224 | Q1224K | D | 1.7e-05 | 1.3e-05 | 9.3e-06 | 1.310E-05 | 3.851E-06 | 9.300E-06 | 1.700E-05 |
| 24473 | 1224 | Q1224M | D | 8.9e-06 | 1.6e-05 | 1.1e-05 | 1.197E-05 | 3.647E-06 | 8.900E-06 | 1.600E-05 |
| 24474 | 1224 | Q1224F | D | 1.7e-05 | 1.2e-05 | 1.6e-05 | 1.500E-05 | 2.646E-06 | 1.200E-05 | 1.700E-05 |
| 24475 | 1224 | Q1224P | D | 1.6e-05 | 1.6e-05 | 7.6e-06 | 1.320E-05 | 4.850E-06 | 7.600E-06 | 1.600E-05 |
| 24476 | 1224 | Q1224S | D | 9.7e-06 | 9.9e-06 | 9.9e-06 | 9.833E-06 | 1.155E-07 | 9.700E-06 | 9.900E-06 |
| 24477 | 1224 | Q1224T | D | 1.5e-05 | 1.1e-05 | 9.8e-06 | 1.193E-05 | 2.723E-06 | 9.800E-06 | 1.500E-05 |
| 24478 | 1224 | Q1224W | D | 1.5e-05 | 1.6e-05 | 1.5e-05 | 1.533E-05 | 5.774E-07 | 1.500E-05 | 1.600E-05 |
| 24479 | 1224 | Q1224Y | D | 1.5e-05 | 1.4e-05 | 1.2e-05 | 1.367E-05 | 1.528E-06 | 1.200E-05 | 1.500E-05 |

| | | | | | | | | | | |
|---|---|---|---|---|---|---|---|---|---|---|
| 24480 | 1224 | Q1224V | D | 1.1e-05 | 1.6e-05 | 1.1e-05 | 1.267E-05 | 2.887E-06 | 1.100E-05 | 1.600E-05 |
| 24481 | 1225 | D1225A | D | 1.5e-05 | 1.6e-05 | 9.1e-06 | 1.337E-05 | 3.729E-06 | 9.100E-06 | 1.600E-05 |
| 24482 | 1225 | D1225R | D | 1.5e-05 | 1e-05 | 1.6e-05 | 1.367E-05 | 3.215E-06 | 1.000E-05 | 1.600E-05 |
| 24483 | 1225 | D1225N | D | 1.5e-05 | 9.9e-06 | 1.4e-05 | 1.297E-05 | 2.702E-06 | 9.900E-06 | 1.500E-05 |
| 24484 | 1225 | D1225D | D | 1e-05 | 1.5e-05 | 1.1e-05 | 1.200E-05 | 2.646E-06 | 1.000E-05 | 1.500E-05 |
| 24485 | 1225 | D1225C | D | 1.6e-05 | 1.7e-05 | 9.5e-06 | 1.417E-05 | 4.072E-06 | 9.500E-06 | 1.700E-05 |
| 24486 | 1225 | D1225E | D | 1.5e-05 | 9.2e-06 | 1.1e-05 | 1.173E-05 | 2.969E-06 | 9.200E-06 | 1.500E-05 |
| 24487 | 1225 | D1225Q | D | 1.6e-05 | 1.2e-05 | 9.9e-06 | 1.263E-05 | 3.099E-06 | 9.900E-06 | 1.600E-05 |
| 24488 | 1225 | D1225G | D | 9e-06 | 1.1e-05 | 8.6e-06 | 9.533E-06 | 1.286E-06 | 8.600E-06 | 1.100E-05 |
| 24489 | 1225 | D1225H | D | 1e-05 | 1.2e-05 | 1.1e-05 | 1.100E-05 | 1.000E-06 | 1.000E-05 | 1.200E-05 |
| 24490 | 1225 | D1225I | D | 1.7e-05 | 1.4e-05 | 1.6e-05 | 1.567E-05 | 1.528E-06 | 1.400E-05 | 1.700E-05 |
| 24491 | 1225 | D1225L | D | 1.4e-05 | 1.3e-05 | 1.3e-05 | 1.333E-05 | 5.774E-07 | 1.300E-05 | 1.400E-05 |
| 24492 | 1225 | D1225K | D | 1.5e-05 | 1.6e-05 | 9.3e-06 | 1.343E-05 | 3.614E-06 | 9.300E-06 | 1.600E-05 |
| 24493 | 1225 | D1225M | D | 1.3e-05 | 1.1e-05 | 1.5e-05 | 1.300E-05 | 2.000E-06 | 1.100E-05 | 1.500E-05 |
| 24494 | 1225 | D1225F | D | 1.1e-05 | 1.4e-05 | 1.5e-05 | 1.333E-05 | 2.082E-06 | 1.100E-05 | 1.500E-05 |
| 24495 | 1225 | D1225P | D | 9.2e-06 | 1.5e-05 | 1.7e-05 | 1.373E-05 | 4.051E-06 | 9.200E-06 | 1.700E-05 |
| 24496 | 1225 | D1225S | D | 1.5e-05 | 9.7e-06 | 1.1e-05 | 1.190E-05 | 2.762E-06 | 9.700E-06 | 1.500E-05 |
| 24497 | 1225 | D1225T | D | 1.1e-05 | 1.6e-05 | 1.5e-05 | 1.400E-05 | 2.646E-06 | 1.100E-05 | 1.600E-05 |
| 24498 | 1225 | D1225W | D | 1.6e-05 | 1.4e-05 | 1.6e-05 | 1.533E-05 | 1.155E-06 | 1.400E-05 | 1.600E-05 |
| 24499 | 1225 | D1225Y | D | 9.8e-06 | 9.8e-06 | 8.9e-06 | 9.500E-06 | 5.196E-07 | 8.900E-06 | 9.800E-06 |
| 24500 | 1225 | D1225V | D | 1.4e-05 | 8.4e-06 | 1.1e-05 | 1.113E-05 | 2.802E-06 | 8.400E-06 | 1.400E-05 |
| 24501 | 1226 | S1226A | D | 1.3e-05 | 1.7e-05 | 1.8e-05 | 1.600E-05 | 2.646E-06 | 1.300E-05 | 1.800E-05 |
| 24502 | 1226 | S1226R | D | 1.5e-05 | 1.5e-05 | 1.4e-05 | 1.467E-05 | 5.774E-07 | 1.400E-05 | 1.500E-05 |
| 24503 | 1226 | S1226N | D | 9.1e-06 | 1.5e-05 | 1.4e-05 | 1.270E-05 | 3.158E-06 | 9.100E-06 | 1.500E-05 |
| 24504 | 1226 | S1226D | D | 1.7e-05 | 1e-05 | 2.4e-05 | 1.700E-05 | 7.000E-06 | 1.000E-05 | 2.400E-05 |
| 24505 | 1226 | S1226C | D | 1e-05 | 1.5e-05 | 1.4e-05 | 1.300E-05 | 2.646E-06 | 1.000E-05 | 1.500E-05 |
| 24506 | 1226 | S1226E | D | 1.1e-05 | 1.2e-05 | 1.4e-05 | 1.233E-05 | 1.528E-06 | 1.100E-05 | 1.400E-05 |
| 24507 | 1226 | S1226Q | D | 1.7e-05 | 1.3e-05 | 1.2e-05 | 1.400E-05 | 2.646E-06 | 1.200E-05 | 1.700E-05 |
| 24508 | 1226 | S1226G | D | 9.6e-06 | 8.5e-06 | 1e-05 | 9.367E-06 | 7.767E-07 | 8.500E-06 | 1.000E-05 |
| 24509 | 1226 | S1226H | D | 1.4e-05 | 1.4e-05 | 1.7e-05 | 1.500E-05 | 1.732E-06 | 1.400E-05 | 1.700E-05 |
| 24510 | 1226 | S1226I | D | 1.6e-05 | 1.5e-05 | 9.7e-06 | 1.357E-05 | 3.386E-06 | 9.700E-06 | 1.600E-05 |
| 24511 | 1226 | S1226L | D | 1.2e-05 | 1.6e-05 | 1.1e-05 | 1.300E-05 | 2.646E-06 | 1.100E-05 | 1.600E-05 |
| 24512 | 1226 | S1226K | D | 8.3e-06 | 1.1e-05 | 9.8e-06 | 9.700E-06 | 1.353E-06 | 8.300E-06 | 1.100E-05 |
| 24513 | 1226 | S1226M | D | 1.6e-05 | 9.7e-06 | 1.6e-05 | 1.390E-05 | 3.637E-06 | 9.700E-06 | 1.600E-05 |
| 24514 | 1226 | S1226F | D | 1.5e-05 | 1.4e-05 | 1.5e-05 | 1.467E-05 | 5.774E-07 | 1.400E-05 | 1.500E-05 |
| 24515 | 1226 | S1226P | D | 1.1e-05 | 1.6e-05 | 1.6e-05 | 1.433E-05 | 2.887E-06 | 1.100E-05 | 1.600E-05 |
| 24516 | 1226 | S1226S | D | 1.4e-05 | 1.3e-05 | 9.8e-06 | 1.227E-05 | 2.194E-06 | 9.800E-06 | 1.400E-05 |
| 24517 | 1226 | S1226T | D | 1.6e-05 | 1.4e-05 | 2.6e-05 | 1.867E-05 | 6.429E-06 | 1.400E-05 | 2.600E-05 |
| 24518 | 1226 | S1226W | D | 1.6e-05 | 1e-05 | 1.5e-05 | 1.367E-05 | 3.215E-06 | 1.000E-05 | 1.600E-05 |
| 24519 | 1226 | S1226Y | D | 1.6e-05 | 1.4e-05 | 1.7e-05 | 1.567E-05 | 1.528E-06 | 1.400E-05 | 1.700E-05 |
| 24520 | 1226 | S1226V | D | 1.6e-05 | 1.7e-05 | 1.2e-05 | 1.500E-05 | 2.646E-06 | 1.200E-05 | 1.700E-05 |
| 24521 | 1227 | K1227A | D | 1e-05 | 1.4e-05 | 1.4e-05 | 1.267E-05 | 2.309E-06 | 1.000E-05 | 1.400E-05 |
| 24522 | 1227 | K1227R | D | 1.3e-05 | 1.5e-05 | 1.1e-05 | 1.300E-05 | 2.000E-06 | 1.100E-05 | 1.500E-05 |
| 24523 | 1227 | K1227N | D | 9.5e-06 | 1.5e-05 | 9.7e-06 | 1.140E-05 | 3.119E-06 | 9.500E-06 | 1.500E-05 |
| 24524 | 1227 | K1227D | D | 9.1e-06 | 1.3e-05 | 1.2e-05 | 1.137E-05 | 2.026E-06 | 9.100E-06 | 1.300E-05 |
| 24525 | 1227 | K1227C | D | 1.5e-05 | 1.4e-05 | 1.1e-05 | 1.333E-05 | 2.082E-06 | 1.100E-05 | 1.500E-05 |
| 24526 | 1227 | K1227E | D | 1.1e-05 | 1.4e-05 | 1.6e-05 | 1.367E-05 | 2.517E-06 | 1.100E-05 | 1.600E-05 |
| 24527 | 1227 | K1227Q | D | 1.7e-05 | 1.1e-05 | 1.5e-05 | 1.433E-05 | 3.055E-06 | 1.100E-05 | 1.700E-05 |
| 24528 | 1227 | K1227G | D | 1.5e-05 | 1.4e-05 | 1.3e-05 | 1.400E-05 | 1.000E-06 | 1.300E-05 | 1.500E-05 |
| 24529 | 1227 | K1227H | D | 8.6e-06 | 1.5e-05 | 1.7e-05 | 1.353E-05 | 4.388E-06 | 8.600E-06 | 1.700E-05 |
| 24530 | 1227 | K1227I | D | 1.4e-05 | 1e-05 | 1.2e-05 | 1.200E-05 | 2.000E-06 | 1.000E-05 | 1.400E-05 |
| 24531 | 1227 | K1227L | D | 1.5e-05 | 1.6e-05 | 1.1e-05 | 1.400E-05 | 2.646E-06 | 1.100E-05 | 1.600E-05 |
| 24532 | 1227 | K1227K | D | 1.1e-05 | 9.4e-06 | 8.7e-06 | 9.700E-06 | 1.179E-06 | 8.700E-06 | 1.100E-05 |
| 24533 | 1227 | K1227M | D | 1.5e-05 | 1.7e-05 | 9.6e-06 | 1.387E-05 | 3.828E-06 | 9.600E-06 | 1.700E-05 |
| 24534 | 1227 | K1227F | D | 1.2e-05 | 8e-06 | 9.3e-06 | 9.767E-06 | 2.040E-06 | 8.000E-06 | 1.200E-05 |
| 24535 | 1227 | K1227P | D | 1.4e-05 | 1e-05 | 1.3e-05 | 1.233E-05 | 2.082E-06 | 1.000E-05 | 1.400E-05 |
| 24536 | 1227 | K1227S | D | 1.2e-05 | 9.4e-06 | 1e-05 | 1.047E-05 | 1.361E-06 | 9.400E-06 | 1.200E-05 |
| 24537 | 1227 | K1227T | D | 1.5e-05 | 1.7e-05 | 9.5e-06 | 1.383E-05 | 3.884E-06 | 9.500E-06 | 1.700E-05 |
| 24538 | 1227 | K1227W | D | 9.8e-06 | 1.1e-05 | 1.2e-05 | 1.093E-05 | 1.102E-06 | 9.800E-06 | 1.200E-05 |
| 24539 | 1227 | K1227Y | D | 9.1e-06 | 1e-05 | 1e-05 | 9.700E-06 | 5.196E-07 | 9.100E-06 | 1.000E-05 |
| 24540 | 1227 | K1227V | D | 1.7e-05 | 1.5e-05 | 7.3e-06 | 1.310E-05 | 5.122E-06 | 7.300E-06 | 1.700E-05 |
| 24541 | 1228 | D1228A | D | 1.4e-05 | 1.5e-05 | 1.8e-05 | 1.567E-05 | 2.082E-06 | 1.400E-05 | 1.800E-05 |
| 24542 | 1228 | D1228R | D | 1.5e-05 | 1.6e-05 | 2.9e-05 | 2.000E-05 | 7.810E-06 | 1.500E-05 | 2.900E-05 |
| 24543 | 1228 | D1228N | D | 1.5e-05 | 9.7e-06 | 1.1e-05 | 1.190E-05 | 2.762E-06 | 9.700E-06 | 1.500E-05 |
| 24544 | 1228 | D1228D | D | 1.2e-05 | 9.9e-06 | 1.6e-05 | 1.263E-05 | 3.099E-06 | 9.900E-06 | 1.600E-05 |
| 24545 | 1228 | D1228C | D | 1.1e-05 | 1.5e-05 | 1e-05 | 1.200E-05 | 2.646E-06 | 1.000E-05 | 1.500E-05 |
| 24546 | 1228 | D1228E | D | 1.6e-05 | 9.8e-06 | 1.4e-05 | 1.327E-05 | 3.164E-06 | 9.800E-06 | 1.600E-05 |
| 24547 | 1228 | D1228Q | D | 9.2e-06 | 1.2e-05 | 1e-05 | 1.040E-05 | 1.442E-06 | 9.200E-06 | 1.200E-05 |
| 24548 | 1228 | D1228G | D | 1.5e-05 | 1.1e-05 | 1.1e-05 | 1.233E-05 | 2.309E-06 | 1.100E-05 | 1.500E-05 |
| 24549 | 1228 | D1228H | D | 1.5e-05 | 1.5e-05 | 1.3e-05 | 1.433E-05 | 1.155E-06 | 1.300E-05 | 1.500E-05 |
| 24550 | 1228 | D1228I | D | 6.8e-06 | 1e-05 | 1.4e-05 | 1.027E-05 | 3.607E-06 | 6.800E-06 | 1.400E-05 |
| 24551 | 1228 | D1228L | D | 1.5e-05 | 1.4e-05 | 1.3e-05 | 1.400E-05 | 1.000E-06 | 1.300E-05 | 1.500E-05 |
| 24552 | 1228 | D1228K | D | 1.5e-05 | 1.6e-05 | 1.6e-05 | 1.567E-05 | 5.774E-07 | 1.500E-05 | 1.600E-05 |
| 24553 | 1228 | D1228M | D | 1.6e-05 | 1.6e-05 | 1e-05 | 1.400E-05 | 3.464E-06 | 1.000E-05 | 1.600E-05 |
| 24554 | 1228 | D1228F | D | 1e-05 | 9.9e-06 | 6.3e-06 | 8.733E-06 | 2.108E-06 | 6.300E-06 | 1.000E-05 |
| 24555 | 1228 | D1228P | D | 1.1e-05 | 1e-05 | 1.1e-05 | 1.067E-05 | 5.774E-07 | 1.000E-05 | 1.100E-05 |
| 24556 | 1228 | D1228S | D | 1.5e-05 | 1.5e-05 | 1e-05 | 1.333E-05 | 2.887E-06 | 1.000E-05 | 1.500E-05 |
| 24557 | 1228 | D1228T | D | 1e-05 | 1.6e-05 | 1e-05 | 1.200E-05 | 3.464E-06 | 1.000E-05 | 1.600E-05 |
| 24558 | 1228 | D1228W | D | 1.4e-05 | 1.6e-05 | 1.4e-05 | 1.467E-05 | 1.155E-06 | 1.400E-05 | 1.600E-05 |
| 24559 | 1228 | D1228Y | D | 1.1e-05 | 1e-05 | 1.5e-05 | 1.200E-05 | 2.646E-06 | 1.000E-05 | 1.500E-05 |
| 24560 | 1228 | D1228V | D | 1.1e-05 | 1.4e-05 | 1.1e-05 | 1.200E-05 | 1.732E-06 | 1.100E-05 | 1.400E-05 |
| 24561 | 1229 | S1229A | D | 1.2e-05 | 1.2e-05 | 1.1e-05 | 1.167E-05 | 5.774E-07 | 1.100E-05 | 1.200E-05 |
| 24562 | 1229 | S1229R | D | 1.1e-05 | 1.2e-05 | 1.4e-05 | 1.233E-05 | 1.528E-06 | 1.100E-05 | 1.400E-05 |
| 24563 | 1229 | S1229N | D | 1.5e-05 | 8e-06 | 1.1e-05 | 1.133E-05 | 3.512E-06 | 8.000E-06 | 1.500E-05 |
| 24564 | 1229 | S1229D | D | 1.3e-05 | 1.5e-05 | 8.4e-06 | 1.213E-05 | 3.384E-06 | 8.400E-06 | 1.500E-05 |

| | | | | | | | | | | |
|---|---|---|---|---|---|---|---|---|---|---|
| 24565 | 1229 | S1229C | D | 1.4e-05 | 1.3e-05 | 9.8e-06 | 1.227E-05 | 2.194E-06 | 9.800E-06 | 1.400E-05 |
| 24566 | 1229 | S1229E | D | 1.6e-05 | 9.3e-06 | 1.1e-05 | 1.210E-05 | 3.483E-06 | 9.300E-06 | 1.600E-05 |
| 24567 | 1229 | S1229Q | D | 1.4e-05 | 1.6e-05 | 1.5e-05 | 1.500E-05 | 1.000E-06 | 1.400E-05 | 1.600E-05 |
| 24568 | 1229 | S1229G | D | 1.5e-05 | 1.1e-05 | 1.4e-05 | 1.333E-05 | 2.082E-06 | 1.100E-05 | 1.500E-05 |
| 24569 | 1229 | S1229H | D | 1.2e-05 | 1.1e-05 | 1.4e-05 | 1.233E-05 | 1.528E-06 | 1.100E-05 | 1.400E-05 |
| 24570 | 1229 | S1229I | D | 1.4e-05 | 1.1e-05 | 1.5e-05 | 1.333E-05 | 2.082E-06 | 1.100E-05 | 1.500E-05 |
| 24571 | 1229 | S1229L | D | 1e-05 | 1e-05 | 1.5e-05 | 1.167E-05 | 2.887E-06 | 1.000E-05 | 1.500E-05 |
| 24572 | 1229 | S1229K | D | 1.4e-05 | 1.6e-05 | 1.1e-05 | 1.367E-05 | 2.517E-06 | 1.100E-05 | 1.600E-05 |
| 24573 | 1229 | S1229M | D | 1.1e-05 | 1.1e-05 | 1.2e-05 | 1.133E-05 | 5.774E-07 | 1.100E-05 | 1.200E-05 |
| 24574 | 1229 | S1229F | D | 1.5e-05 | 1.1e-05 | 1.1e-05 | 1.233E-05 | 2.309E-06 | 1.100E-05 | 1.500E-05 |
| 24575 | 1229 | S1229P | D | 1.6e-05 | 1.3e-05 | 1.5e-05 | 1.467E-05 | 1.528E-06 | 1.300E-05 | 1.600E-05 |
| 24576 | 1229 | S1229S | D | 1.7e-05 | 1.6e-05 | 1.4e-05 | 1.567E-05 | 1.528E-06 | 1.400E-05 | 1.700E-05 |
| 24577 | 1229 | S1229T | D | 1.7e-05 | 1.5e-05 | 1.1e-05 | 1.433E-05 | 3.055E-06 | 1.100E-05 | 1.700E-05 |
| 24578 | 1229 | S1229W | D | 1.4e-05 | 1.6e-05 | 1.5e-05 | 1.500E-05 | 1.000E-06 | 1.400E-05 | 1.600E-05 |
| 24579 | 1229 | S1229Y | D | 1.7e-05 | 1.4e-05 | 9.3e-06 | 1.343E-05 | 3.881E-06 | 9.300E-06 | 1.700E-05 |
| 24580 | 1229 | S1229V | D | 1.1e-05 | 1.3e-05 | 1.5e-05 | 1.300E-05 | 2.000E-06 | 1.100E-05 | 1.500E-05 |
| 24581 | 1230 | T1230A | D | 1.5e-05 | 1.2e-05 | 1.4e-05 | 1.367E-05 | 1.528E-06 | 1.200E-05 | 1.500E-05 |
| 24582 | 1230 | T1230R | D | 9.5e-06 | 1.6e-05 | 1.5e-05 | 1.350E-05 | 3.500E-06 | 9.500E-06 | 1.600E-05 |
| 24583 | 1230 | T1230N | D | 1.1e-05 | 1.5e-05 | 1.6e-05 | 1.400E-05 | 2.646E-06 | 1.100E-05 | 1.600E-05 |
| 24584 | 1230 | T1230D | D | 1e-05 | 1.4e-05 | 1.1e-05 | 1.167E-05 | 2.082E-06 | 1.000E-05 | 1.400E-05 |
| 24585 | 1230 | T1230C | D | 1.1e-05 | 1.2e-05 | 1.6e-05 | 1.300E-05 | 2.646E-06 | 1.100E-05 | 1.600E-05 |
| 24586 | 1230 | T1230E | D | 1.5e-05 | 1.1e-05 | 1.5e-05 | 1.367E-05 | 2.309E-06 | 1.100E-05 | 1.500E-05 |
| 24587 | 1230 | T1230Q | D | 1.6e-05 | 1.1e-05 | 9.9e-06 | 1.230E-05 | 3.251E-06 | 9.900E-06 | 1.600E-05 |
| 24588 | 1230 | T1230G | D | 1.1e-05 | 1.4e-05 | 1.2e-05 | 1.233E-05 | 1.528E-06 | 1.100E-05 | 1.400E-05 |
| 24589 | 1230 | T1230H | D | 1.4e-05 | 1.5e-05 | 1.5e-05 | 1.467E-05 | 5.774E-07 | 1.400E-05 | 1.500E-05 |
| 24590 | 1230 | T1230I | D | 1.6e-05 | 1.1e-05 | 1.7e-05 | 1.467E-05 | 3.215E-06 | 1.100E-05 | 1.700E-05 |
| 24591 | 1230 | T1230L | D | 1.2e-05 | 1.6e-05 | 1.6e-05 | 1.467E-05 | 2.309E-06 | 1.200E-05 | 1.600E-05 |
| 24592 | 1230 | T1230K | D | 1e-05 | 1.5e-05 | 1.1e-05 | 1.200E-05 | 2.646E-06 | 1.000E-05 | 1.500E-05 |
| 24593 | 1230 | T1230M | D | 1.5e-05 | 1.5e-05 | 8.6e-06 | 1.287E-05 | 3.695E-06 | 8.600E-06 | 1.500E-05 |
| 24594 | 1230 | T1230F | D | 2.3e-05 | 1.5e-05 | 1.2e-05 | 1.667E-05 | 5.686E-06 | 1.200E-05 | 2.300E-05 |
| 24595 | 1230 | T1230P | D | 1.1e-05 | 8.6e-06 | 1.1e-05 | 1.020E-05 | 1.386E-06 | 8.600E-06 | 1.100E-05 |
| 24596 | 1230 | T1230S | D | 1e-05 | 1.1e-05 | 1.8e-05 | 1.300E-05 | 4.359E-06 | 1.000E-05 | 1.800E-05 |
| 24597 | 1230 | T1230T | D | 1.4e-05 | 1.1e-05 | 1.1e-05 | 1.200E-05 | 1.732E-06 | 1.100E-05 | 1.400E-05 |
| 24598 | 1230 | T1230W | D | 9.7e-06 | 8.8e-06 | 7.7e-06 | 8.733E-06 | 1.002E-06 | 7.700E-06 | 9.700E-06 |
| 24599 | 1230 | T1230Y | D | 8.9e-06 | 9.8e-06 | 1.6e-05 | 1.157E-05 | 3.866E-06 | 8.900E-06 | 1.600E-05 |
| 24600 | 1230 | T1230V | D | 1.5e-05 | 9.2e-06 | 1.1e-05 | 1.173E-05 | 2.969E-06 | 9.200E-06 | 1.500E-05 |
| 24601 | 1231 | Y1231A | D | 1.1e-05 | 1.5e-05 | 9.8e-06 | 1.193E-05 | 2.723E-06 | 9.800E-06 | 1.500E-05 |
| 24602 | 1231 | Y1231R | D | 1.6e-05 | 1.5e-05 | 1.5e-05 | 1.533E-05 | 5.774E-07 | 1.500E-05 | 1.600E-05 |
| 24603 | 1231 | Y1231N | D | 1e-05 | 1.3e-05 | 7.5e-06 | 1.017E-05 | 2.754E-06 | 7.500E-06 | 1.300E-05 |
| 24604 | 1231 | Y1231D | D | 9.2e-06 | 1.5e-05 | 1.1e-05 | 1.173E-05 | 2.969E-06 | 9.200E-06 | 1.500E-05 |
| 24605 | 1231 | Y1231C | D | 9.8e-06 | 1.4e-05 | 1.1e-05 | 1.160E-05 | 2.163E-06 | 9.800E-06 | 1.400E-05 |
| 24606 | 1231 | Y1231E | D | 1.6e-05 | 1.4e-05 | 1.4e-05 | 1.467E-05 | 1.155E-06 | 1.400E-05 | 1.600E-05 |
| 24607 | 1231 | Y1231Q | D | 1.5e-05 | 1.3e-05 | 9.1e-06 | 1.237E-05 | 3.001E-06 | 9.100E-06 | 1.500E-05 |
| 24608 | 1231 | Y1231G | D | 1.1e-05 | 1.2e-05 | 1.5e-05 | 1.267E-05 | 2.082E-06 | 1.100E-05 | 1.500E-05 |
| 24609 | 1231 | Y1231H | D | 8.9e-06 | 8.9e-06 | 7.9e-06 | 8.567E-06 | 5.774E-07 | 7.900E-06 | 8.900E-06 |
| 24610 | 1231 | Y1231I | D | 1.5e-05 | 1.4e-05 | 1.2e-05 | 1.367E-05 | 1.528E-06 | 1.200E-05 | 1.500E-05 |
| 24611 | 1231 | Y1231L | D | 1.1e-05 | 7e-06 | 9.6e-06 | 9.200E-06 | 2.030E-06 | 7.000E-06 | 1.100E-05 |
| 24612 | 1231 | Y1231K | D | 1e-05 | 2.8e-05 | 1e-05 | 1.600E-05 | 1.039E-05 | 1.000E-05 | 2.800E-05 |
| 24613 | 1231 | Y1231M | D | 8e-06 | 1e-05 | 1.3e-05 | 1.033E-05 | 2.517E-06 | 8.000E-06 | 1.300E-05 |
| 24614 | 1231 | Y1231F | D | 1.5e-05 | 1e-05 | 1.6e-05 | 1.367E-05 | 3.215E-06 | 1.000E-05 | 1.600E-05 |
| 24615 | 1231 | Y1231P | D | 1.3e-05 | 1.6e-05 | 1.6e-05 | 1.500E-05 | 1.732E-06 | 1.300E-05 | 1.600E-05 |
| 24616 | 1231 | Y1231S | D | 1.5e-05 | 1.4e-05 | 1e-05 | 1.300E-05 | 2.646E-06 | 1.000E-05 | 1.500E-05 |
| 24617 | 1231 | Y1231T | D | 1.3e-05 | 1.5e-05 | 1e-05 | 1.267E-05 | 2.517E-06 | 1.000E-05 | 1.500E-05 |
| 24618 | 1231 | Y1231W | D | 1.1e-05 | 1.2e-05 | 1.6e-05 | 1.300E-05 | 2.646E-06 | 1.100E-05 | 1.600E-05 |
| 24619 | 1231 | Y1231Y | D | 1.4e-05 | 1.5e-05 | 1.7e-05 | 1.533E-05 | 1.528E-06 | 1.400E-05 | 1.700E-05 |
| 24620 | 1231 | Y1231V | D | 1.7e-05 | 1.3e-05 | 9.6e-06 | 1.320E-05 | 3.704E-06 | 9.600E-06 | 1.700E-05 |
| 24621 | 1232 | S1232A | D | 1e-05 | 1.5e-05 | 1.6e-05 | 1.367E-05 | 3.215E-06 | 1.000E-05 | 1.600E-05 |
| 24622 | 1232 | S1232R | D | 1.1e-05 | 1.6e-05 | 1.1e-05 | 1.267E-05 | 2.887E-06 | 1.100E-05 | 1.600E-05 |
| 24623 | 1232 | S1232N | D | 1.1e-05 | 1.2e-05 | 1.5e-05 | 1.267E-05 | 2.082E-06 | 1.100E-05 | 1.500E-05 |
| 24624 | 1232 | S1232D | D | 1.1e-05 | 1.1e-05 | 9.6e-06 | 1.053E-05 | 8.083E-07 | 9.600E-06 | 1.100E-05 |
| 24625 | 1232 | S1232C | D | 1.4e-05 | 1.6e-05 | 1.1e-05 | 1.367E-05 | 2.517E-06 | 1.100E-05 | 1.600E-05 |
| 24626 | 1232 | S1232E | D | 8.8e-06 | 1.5e-05 | 1.1e-05 | 1.160E-05 | 3.143E-06 | 8.800E-06 | 1.500E-05 |
| 24627 | 1232 | S1232Q | D | 1.4e-05 | 9.2e-06 | 1.5e-05 | 1.273E-05 | 3.101E-06 | 9.200E-06 | 1.500E-05 |
| 24628 | 1232 | S1232G | D | 1e-05 | 1.5e-05 | 1.6e-05 | 1.367E-05 | 3.215E-06 | 1.000E-05 | 1.600E-05 |
| 24629 | 1232 | S1232H | D | 1.5e-05 | 1.3e-05 | 9.9e-06 | 1.263E-05 | 2.570E-06 | 9.900E-06 | 1.500E-05 |
| 24630 | 1232 | S1232I | D | 1.5e-05 | 1.4e-05 | 1.2e-05 | 1.367E-05 | 1.528E-06 | 1.200E-05 | 1.500E-05 |
| 24631 | 1232 | S1232L | D | 1.1e-05 | 1.7e-05 | 1.6e-05 | 1.467E-05 | 3.215E-06 | 1.100E-05 | 1.700E-05 |
| 24632 | 1232 | S1232K | D | 1.3e-05 | 1.6e-05 | 9.6e-06 | 1.287E-05 | 3.202E-06 | 9.600E-06 | 1.600E-05 |
| 24633 | 1232 | S1232M | D | 1.4e-05 | 9.9e-06 | 1.4e-05 | 1.263E-05 | 2.367E-06 | 9.900E-06 | 1.400E-05 |
| 24634 | 1232 | S1232F | D | 1.5e-05 | 1.5e-05 | 1.5e-05 | 1.500E-05 | 0.000E+00 | 1.500E-05 | 1.500E-05 |
| 24635 | 1232 | S1232P | D | 1.1e-05 | 1.3e-05 | 9.9e-06 | 1.130E-05 | 1.572E-06 | 9.900E-06 | 1.300E-05 |
| 24636 | 1232 | S1232S | D | 1.3e-05 | 1.1e-05 | 9.8e-06 | 1.127E-05 | 1.617E-06 | 9.800E-06 | 1.300E-05 |
| 24637 | 1232 | S1232T | D | 1.1e-05 | 1e-05 | 1.4e-05 | 1.167E-05 | 2.082E-06 | 1.000E-05 | 1.400E-05 |
| 24638 | 1232 | S1232W | D | 9.8e-06 | 9.8e-06 | 1.3e-05 | 1.087E-05 | 1.848E-06 | 9.800E-06 | 1.300E-05 |
| 24639 | 1232 | S1232Y | D | 1e-05 | 1e-05 | 9e-06 | 9.667E-06 | 5.774E-07 | 9.000E-06 | 1.000E-05 |
| 24640 | 1232 | S1232V | D | 1.6e-05 | 8.5e-06 | 1.4e-05 | 1.283E-05 | 3.884E-06 | 8.500E-06 | 1.600E-05 |
| 24641 | 1233 | L1233A | D | 1.4e-05 | 1.5e-05 | 1.3e-05 | 1.400E-05 | 1.000E-06 | 1.300E-05 | 1.500E-05 |
| 24642 | 1233 | L1233R | D | 9.3e-06 | 1.1e-05 | 1.1e-05 | 1.043E-05 | 9.815E-07 | 9.300E-06 | 1.100E-05 |
| 24643 | 1233 | L1233N | D | 1.4e-05 | 1.4e-05 | 9.6e-06 | 1.253E-05 | 2.540E-06 | 9.600E-06 | 1.400E-05 |
| 24644 | 1233 | L1233D | D | 1.3e-05 | 1.4e-05 | 8.8e-06 | 1.193E-05 | 2.759E-06 | 8.800E-06 | 1.400E-05 |
| 24645 | 1233 | L1233C | D | 8.2e-06 | 8.5e-06 | 8.9e-06 | 8.533E-06 | 3.512E-07 | 8.200E-06 | 8.900E-06 |
| 24646 | 1233 | L1233E | D | 1.2e-05 | 1.2e-05 | 1.1e-05 | 1.167E-05 | 5.774E-07 | 1.100E-05 | 1.200E-05 |
| 24647 | 1233 | L1233Q | D | 1.4e-05 | 1.6e-05 | 1.6e-05 | 1.533E-05 | 1.155E-06 | 1.400E-05 | 1.600E-05 |
| 24648 | 1233 | L1233G | D | 1.3e-05 | 1.6e-05 | 1.5e-05 | 1.467E-05 | 1.528E-06 | 1.300E-05 | 1.600E-05 |
| 24649 | 1233 | L1233H | D | 1.6e-05 | 1.5e-05 | 8.3e-06 | 1.310E-05 | 4.187E-06 | 8.300E-06 | 1.600E-05 |

| | | | | | | | | | | |
|---|---|---|---|---|---|---|---|---|---|---|
| 24650 | 1233 | L1233I | D | 1.5e-05 | 9.6e-06 | 1.1e-05 | 1.187E-05 | 2.802E-06 | 9.600E-06 | 1.500E-05 |
| 24651 | 1233 | L1233L | D | 1e-05 | 1.3e-05 | 1.3e-05 | 1.200E-05 | 1.732E-06 | 1.000E-05 | 1.300E-05 |
| 24652 | 1233 | L1233K | D | 1.1e-05 | 1.1e-05 | 1.4e-05 | 1.200E-05 | 1.732E-06 | 1.100E-05 | 1.400E-05 |
| 24653 | 1233 | L1233M | D | 1.1e-05 | 1.6e-05 | 9.3e-06 | 1.210E-05 | 3.483E-06 | 9.300E-06 | 1.600E-05 |
| 24654 | 1233 | L1233F | D | 1.4e-05 | 1.1e-05 | 1.4e-05 | 1.300E-05 | 1.732E-06 | 1.100E-05 | 1.400E-05 |
| 24655 | 1233 | L1233P | D | 1.3e-05 | 1.4e-05 | 9.5e-06 | 1.217E-05 | 2.363E-06 | 9.500E-06 | 1.400E-05 |
| 24656 | 1233 | L1233S | D | 1.6e-05 | 1.4e-05 | 1.5e-05 | 1.500E-05 | 1.000E-06 | 1.400E-05 | 1.600E-05 |
| 24657 | 1233 | L1233T | D | 1.3e-05 | 1.6e-05 | 1.4e-05 | 1.433E-05 | 1.528E-06 | 1.300E-05 | 1.600E-05 |
| 24658 | 1233 | L1233W | D | 1.3e-05 | 1.4e-05 | 1.4e-05 | 1.367E-05 | 5.774E-07 | 1.300E-05 | 1.400E-05 |
| 24659 | 1233 | L1233Y | D | 9.1e-06 | 9.4e-06 | 9.7e-06 | 9.400E-06 | 3.000E-07 | 9.100E-06 | 9.700E-06 |
| 24660 | 1233 | L1233V | D | 1.2e-05 | 1.1e-05 | 1.1e-05 | 1.133E-05 | 5.774E-07 | 1.100E-05 | 1.200E-05 |
| 24661 | 1234 | S1234A | D | 1.1e-05 | 1.4e-05 | 2.4e-05 | 1.633E-05 | 6.807E-06 | 1.100E-05 | 2.400E-05 |
| 24662 | 1234 | S1234R | D | 1.1e-05 | 1.6e-05 | 1e-05 | 1.233E-05 | 3.215E-06 | 1.000E-05 | 1.600E-05 |
| 24663 | 1234 | S1234N | D | 1.6e-05 | 1.2e-05 | 1.5e-05 | 1.433E-05 | 2.082E-06 | 1.200E-05 | 1.600E-05 |
| 24664 | 1234 | S1234D | D | 1e-05 | 1.1e-05 | 9.7e-06 | 1.023E-05 | 6.807E-07 | 9.700E-06 | 1.100E-05 |
| 24665 | 1234 | S1234C | D | 1.5e-05 | 1.4e-05 | 1.5e-05 | 1.467E-05 | 5.774E-07 | 1.400E-05 | 1.500E-05 |
| 24666 | 1234 | S1234E | D | 1e-05 | 6.2e-06 | 1e-05 | 8.733E-06 | 2.194E-06 | 6.200E-06 | 1.000E-05 |
| 24667 | 1234 | S1234Q | D | 1.3e-05 | 1.6e-05 | 1.6e-05 | 1.500E-05 | 1.732E-06 | 1.300E-05 | 1.600E-05 |
| 24668 | 1234 | S1234G | D | 1.6e-05 | 1.1e-05 | 1.3e-05 | 1.333E-05 | 2.517E-06 | 1.100E-05 | 1.600E-05 |
| 24669 | 1234 | S1234H | D | 8.6e-06 | 1.2e-05 | 1.2e-05 | 1.087E-05 | 1.963E-06 | 8.600E-06 | 1.200E-05 |
| 24670 | 1234 | S1234I | D | 1.1e-05 | 8.3e-06 | 1.5e-05 | 1.143E-05 | 3.371E-06 | 8.300E-06 | 1.500E-05 |
| 24671 | 1234 | S1234L | D | 1.3e-05 | 1.5e-05 | 1.7e-05 | 1.500E-05 | 2.000E-06 | 1.300E-05 | 1.700E-05 |
| 24672 | 1234 | S1234K | D | 9.9e-06 | 1.4e-05 | 1.5e-05 | 1.297E-05 | 2.702E-06 | 9.900E-06 | 1.500E-05 |
| 24673 | 1234 | S1234M | D | 1.4e-05 | 1.5e-05 | 1.4e-05 | 1.433E-05 | 5.774E-07 | 1.400E-05 | 1.500E-05 |
| 24674 | 1234 | S1234F | D | 1.2e-05 | 1.5e-05 | 6.2e-06 | 1.107E-05 | 4.474E-06 | 6.200E-06 | 1.500E-05 |
| 24675 | 1234 | S1234P | D | 1.7e-05 | 1.5e-05 | 9e-06 | 1.367E-05 | 4.163E-06 | 9.000E-06 | 1.700E-05 |
| 24676 | 1234 | S1234S | D | 1.5e-05 | 1.4e-05 | 1.4e-05 | 1.433E-05 | 5.774E-07 | 1.400E-05 | 1.500E-05 |
| 24677 | 1234 | S1234T | D | 1.5e-05 | 2.5e-05 | 1.5e-05 | 1.833E-05 | 5.774E-06 | 1.500E-05 | 2.500E-05 |
| 24678 | 1234 | S1234W | D | 1.6e-05 | 1e-05 | 1.4e-05 | 1.333E-05 | 3.055E-06 | 1.000E-05 | 1.600E-05 |
| 24679 | 1234 | S1234Y | D | 1.5e-05 | 1.5e-05 | 1.2e-05 | 1.400E-05 | 1.732E-06 | 1.200E-05 | 1.500E-05 |
| 24680 | 1234 | S1234V | D | 6.7e-06 | 1.5e-05 | 1.6e-05 | 1.257E-05 | 5.105E-06 | 6.700E-06 | 1.600E-05 |
| 24681 | 1235 | S1235A | D | 1.1e-05 | 1.7e-05 | 1.4e-05 | 1.400E-05 | 3.000E-06 | 1.100E-05 | 1.700E-05 |
| 24682 | 1235 | S1235R | D | 7e-06 | 1.1e-05 | 1.5e-05 | 1.100E-05 | 4.000E-06 | 7.000E-06 | 1.500E-05 |
| 24683 | 1235 | S1235N | D | 9.7e-06 | 9.1e-06 | 1.1e-05 | 9.933E-06 | 9.713E-07 | 9.100E-06 | 1.100E-05 |
| 24684 | 1235 | S1235D | D | 9.4e-06 | 1.5e-05 | 1.4e-05 | 1.280E-05 | 2.987E-06 | 9.400E-06 | 1.500E-05 |
| 24685 | 1235 | S1235C | D | 1.3e-05 | 1.5e-05 | 1.6e-05 | 1.467E-05 | 1.528E-06 | 1.300E-05 | 1.600E-05 |
| 24686 | 1235 | S1235E | D | 1.2e-05 | 1.3e-05 | 1.4e-05 | 1.300E-05 | 1.000E-06 | 1.200E-05 | 1.400E-05 |
| 24687 | 1235 | S1235Q | D | 8.2e-06 | 1.5e-05 | 1.1e-05 | 1.140E-05 | 3.418E-06 | 8.200E-06 | 1.500E-05 |
| 24688 | 1235 | S1235G | D | 1.5e-05 | 1.1e-05 | 1.1e-05 | 1.233E-05 | 2.309E-06 | 1.100E-05 | 1.500E-05 |
| 24689 | 1235 | S1235H | D | 9e-06 | 1e-05 | 1.5e-05 | 1.133E-05 | 3.215E-06 | 9.000E-06 | 1.500E-05 |
| 24690 | 1235 | S1235I | D | 1.2e-05 | 8.8e-06 | 1.7e-05 | 1.260E-05 | 4.133E-06 | 8.800E-06 | 1.700E-05 |
| 24691 | 1235 | S1235L | D | 1.5e-05 | 1.5e-05 | 1.3e-05 | 1.433E-05 | 1.155E-06 | 1.300E-05 | 1.500E-05 |
| 24692 | 1235 | S1235K | D | 1e-05 | 1.6e-05 | 1.2e-05 | 1.267E-05 | 3.055E-06 | 1.000E-05 | 1.600E-05 |
| 24693 | 1235 | S1235M | D | 1.4e-05 | 1.4e-05 | 1.1e-05 | 1.300E-05 | 1.732E-06 | 1.100E-05 | 1.400E-05 |
| 24694 | 1235 | S1235F | D | 1.4e-05 | 1.4e-05 | 7.7e-06 | 1.190E-05 | 3.637E-06 | 7.700E-06 | 1.400E-05 |
| 24695 | 1235 | S1235P | D | 1.6e-05 | 1.6e-05 | 1e-05 | 1.400E-05 | 3.464E-06 | 1.000E-05 | 1.600E-05 |
| 24696 | 1235 | S1235S | D | 1.4e-05 | 9.8e-06 | 9.4e-06 | 1.107E-05 | 2.548E-06 | 9.400E-06 | 1.400E-05 |
| 24697 | 1235 | S1235T | D | 1.7e-05 | 1.3e-05 | 1.6e-05 | 1.533E-05 | 2.082E-06 | 1.300E-05 | 1.700E-05 |
| 24698 | 1235 | S1235W | D | 1.5e-05 | 1.6e-05 | 1.4e-05 | 1.500E-05 | 1.000E-06 | 1.400E-05 | 1.600E-05 |
| 24699 | 1235 | S1235Y | D | 1.1e-05 | 1.5e-05 | 7.2e-06 | 1.107E-05 | 3.900E-06 | 7.200E-06 | 1.500E-05 |
| 24700 | 1235 | S1235V | D | 1e-05 | 1.2e-05 | 1e-05 | 1.067E-05 | 1.155E-06 | 1.000E-05 | 1.200E-05 |
| 24701 | 1236 | T1236A | D | 1.5e-05 | 9.2e-06 | 1e-05 | 1.140E-05 | 3.143E-06 | 9.200E-06 | 1.500E-05 |
| 24702 | 1236 | T1236R | D | 1.1e-05 | 1.2e-05 | 1.4e-05 | 1.233E-05 | 1.528E-06 | 1.100E-05 | 1.400E-05 |
| 24703 | 1236 | T1236N | D | 9.8e-06 | 1.6e-05 | 8.6e-06 | 1.147E-05 | 3.972E-06 | 8.600E-06 | 1.600E-05 |
| 24704 | 1236 | T1236D | D | 1.1e-05 | 1.5e-05 | 1.6e-05 | 1.400E-05 | 2.646E-06 | 1.100E-05 | 1.600E-05 |
| 24705 | 1236 | T1236C | D | 9.1e-06 | 1.5e-05 | 1.5e-05 | 1.303E-05 | 3.406E-06 | 9.100E-06 | 1.500E-05 |
| 24706 | 1236 | T1236E | D | 1.7e-05 | 7.4e-06 | 8.1e-06 | 1.083E-05 | 5.352E-06 | 7.400E-06 | 1.700E-05 |
| 24707 | 1236 | T1236Q | D | 1.1e-05 | 1.6e-05 | 1.4e-05 | 1.367E-05 | 2.517E-06 | 1.100E-05 | 1.600E-05 |
| 24708 | 1236 | T1236G | D | 9.1e-06 | 1.4e-05 | 1.6e-05 | 1.303E-05 | 3.550E-06 | 9.100E-06 | 1.600E-05 |
| 24709 | 1236 | T1236H | D | 1e-05 | 1.1e-05 | 1.4e-05 | 1.167E-05 | 2.082E-06 | 1.000E-05 | 1.400E-05 |
| 24710 | 1236 | T1236I | D | 1.6e-05 | 1.1e-05 | 1.4e-05 | 1.367E-05 | 2.517E-06 | 1.100E-05 | 1.600E-05 |
| 24711 | 1236 | T1236L | D | 9.7e-06 | 1.2e-05 | 9.2e-06 | 1.030E-05 | 1.493E-06 | 9.200E-06 | 1.200E-05 |
| 24712 | 1236 | T1236K | D | 1.2e-05 | 1.7e-05 | 1.1e-05 | 1.333E-05 | 3.215E-06 | 1.100E-05 | 1.700E-05 |
| 24713 | 1236 | T1236M | D | 9.8e-06 | 1.5e-05 | 1.5e-05 | 1.327E-05 | 3.002E-06 | 9.800E-06 | 1.500E-05 |
| 24714 | 1236 | T1236F | D | 1e-05 | 9.5e-06 | 1.6e-05 | 1.183E-05 | 3.617E-06 | 9.500E-06 | 1.600E-05 |
| 24715 | 1236 | T1236P | D | 8.8e-06 | 8.9e-06 | 1.1e-05 | 9.567E-06 | 1.242E-06 | 8.800E-06 | 1.100E-05 |
| 24716 | 1236 | T1236S | D | 1e-05 | 1.1e-05 | 1.7e-05 | 1.267E-05 | 3.786E-06 | 1.000E-05 | 1.700E-05 |
| 24717 | 1236 | T1236T | D | 1.6e-05 | 1e-05 | 1.7e-05 | 1.433E-05 | 3.786E-06 | 1.000E-05 | 1.700E-05 |
| 24718 | 1236 | T1236W | D | 1.1e-05 | 1.5e-05 | 1.5e-05 | 1.367E-05 | 2.309E-06 | 1.100E-05 | 1.500E-05 |
| 24719 | 1236 | T1236Y | D | 1.5e-05 | 9.6e-06 | 1.4e-05 | 1.287E-05 | 2.873E-06 | 9.600E-06 | 1.500E-05 |
| 24720 | 1236 | T1236V | D | 1.6e-05 | 1.6e-05 | 1.3e-05 | 1.500E-05 | 1.732E-06 | 1.300E-05 | 1.600E-05 |
| 24721 | 1237 | L1237A | D | 1.7e-05 | 9.4e-06 | 1.2e-05 | 1.280E-05 | 3.863E-06 | 9.400E-06 | 1.700E-05 |
| 24722 | 1237 | L1237R | D | 1.7e-05 | 1.2e-05 | 9.1e-06 | 1.270E-05 | 3.996E-06 | 9.100E-06 | 1.700E-05 |
| 24723 | 1237 | L1237N | D | 9.5e-06 | 1.1e-05 | 1e-05 | 1.017E-05 | 7.638E-07 | 9.500E-06 | 1.100E-05 |
| 24724 | 1237 | L1237D | D | 1.2e-05 | 1.4e-05 | 1e-05 | 1.200E-05 | 2.000E-06 | 1.000E-05 | 1.400E-05 |
| 24725 | 1237 | L1237C | D | 1.1e-05 | 6.7e-06 | 8.5e-06 | 8.733E-06 | 2.159E-06 | 6.700E-06 | 1.100E-05 |
| 24726 | 1237 | L1237E | D | 1.2e-05 | 9.3e-06 | 8.2e-06 | 9.833E-06 | 1.955E-06 | 8.200E-06 | 1.200E-05 |
| 24727 | 1237 | L1237Q | D | 1.7e-05 | 1.6e-05 | 1.4e-05 | 1.567E-05 | 1.528E-06 | 1.400E-05 | 1.700E-05 |
| 24728 | 1237 | L1237G | D | 1.3e-05 | 1.5e-05 | 1.6e-05 | 1.467E-05 | 1.528E-06 | 1.300E-05 | 1.600E-05 |
| 24729 | 1237 | L1237H | D | 1.2e-05 | 9.9e-06 | 1.4e-05 | 1.197E-05 | 2.050E-06 | 9.900E-06 | 1.400E-05 |
| 24730 | 1237 | L1237I | D | 1e-05 | 1e-05 | 1.1e-05 | 1.033E-05 | 5.774E-07 | 1.000E-05 | 1.100E-05 |
| 24731 | 1237 | L1237L | D | 8.2e-06 | 1.3e-05 | 1.6e-05 | 1.240E-05 | 3.934E-06 | 8.200E-06 | 1.600E-05 |
| 24732 | 1237 | L1237K | D | 7.9e-06 | 1.4e-05 | 1.7e-05 | 1.297E-05 | 4.637E-06 | 7.900E-06 | 1.700E-05 |
| 24733 | 1237 | L1237M | D | 8.5e-06 | 1.2e-05 | 1.5e-05 | 1.183E-05 | 3.253E-06 | 8.500E-06 | 1.500E-05 |
| 24734 | 1237 | L1237F | D | 1.4e-05 | 1.9e-05 | 9e-06 | 1.400E-05 | 5.000E-06 | 9.000E-06 | 1.900E-05 |

| | | | | | | | | | | |
|---|---|---|---|---|---|---|---|---|---|---|
| 24735 | 1237 | L1237P | D | 1e-05 | 9.8e-06 | 1.6e-05 | 1.193E-05 | 3.523E-06 | 9.800E-06 | 1.600E-05 |
| 24736 | 1237 | L1237S | D | 1.3e-05 | 1e-05 | 1.9e-05 | 1.400E-05 | 4.583E-06 | 1.000E-05 | 1.900E-05 |
| 24737 | 1237 | L1237T | D | 1.6e-05 | 1.1e-05 | 1.3e-05 | 1.333E-05 | 2.517E-06 | 1.100E-05 | 1.600E-05 |
| 24738 | 1237 | L1237W | D | 1.2e-05 | 1.4e-05 | 8.8e-06 | 1.160E-05 | 2.623E-06 | 8.800E-06 | 1.400E-05 |
| 24739 | 1237 | L1237Y | D | 9.6e-06 | 9.8e-06 | 9.9e-06 | 9.767E-06 | 1.528E-07 | 9.600E-06 | 9.900E-06 |
| 24740 | 1237 | L1237V | D | 1.3e-05 | 1.5e-05 | 1.1e-05 | 1.300E-05 | 2.000E-06 | 1.100E-05 | 1.500E-05 |
| 24741 | 1238 | T1238A | D | 8.5e-06 | 1.9e-05 | 1.6e-05 | 1.450E-05 | 5.408E-06 | 8.500E-06 | 1.900E-05 |
| 24742 | 1238 | T1238R | D | 9.4e-06 | 1.3e-05 | 8.9e-06 | 1.043E-05 | 2.237E-06 | 8.900E-06 | 1.300E-05 |
| 24743 | 1238 | T1238N | D | 1.4e-05 | 1.1e-05 | 1.4e-05 | 1.300E-05 | 1.732E-06 | 1.100E-05 | 1.400E-05 |
| 24744 | 1238 | T1238D | D | 9.3e-06 | 9.8e-06 | 1.4e-05 | 1.103E-05 | 2.581E-06 | 9.300E-06 | 1.400E-05 |
| 24745 | 1238 | T1238C | D | 1.7e-05 | 8.8e-06 | 1.4e-05 | 1.327E-05 | 4.149E-06 | 8.800E-06 | 1.700E-05 |
| 24746 | 1238 | T1238E | D | 1.6e-05 | 1.5e-05 | 1.3e-05 | 1.467E-05 | 1.528E-06 | 1.300E-05 | 1.600E-05 |
| 24747 | 1238 | T1238Q | D | 1.6e-05 | 1.6e-05 | 1.7e-05 | 1.633E-05 | 5.774E-07 | 1.600E-05 | 1.700E-05 |
| 24748 | 1238 | T1238G | D | 1.7e-05 | 1.3e-05 | 1.7e-05 | 1.567E-05 | 2.309E-06 | 1.300E-05 | 1.700E-05 |
| 24749 | 1238 | T1238H | D | 9.5e-06 | 1.4e-05 | 1e-05 | 1.117E-05 | 2.466E-06 | 9.500E-06 | 1.400E-05 |
| 24750 | 1238 | T1238I | D | 8.2e-06 | 1.1e-05 | 1.6e-05 | 1.173E-05 | 3.951E-06 | 8.200E-06 | 1.600E-05 |
| 24751 | 1238 | T1238L | D | 1e-05 | 1.4e-05 | 1.3e-05 | 1.233E-05 | 2.082E-06 | 1.000E-05 | 1.400E-05 |
| 24752 | 1238 | T1238K | D | 8.2e-06 | 1.5e-05 | 9e-06 | 1.073E-05 | 3.717E-06 | 8.200E-06 | 1.500E-05 |
| 24753 | 1238 | T1238M | D | 1.3e-05 | 1.4e-05 | 1.8e-05 | 1.500E-05 | 2.646E-06 | 1.300E-05 | 1.800E-05 |
| 24754 | 1238 | T1238F | D | 1.1e-05 | 9e-06 | 1e-05 | 1.000E-05 | 1.000E-06 | 9.000E-06 | 1.100E-05 |
| 24755 | 1238 | T1238P | D | 1.4e-05 | 1.4e-05 | 1.7e-05 | 1.500E-05 | 1.732E-06 | 1.400E-05 | 1.700E-05 |
| 24756 | 1238 | T1238S | D | 9.6e-06 | 9.8e-06 | 1.5e-05 | 1.147E-05 | 3.062E-06 | 9.600E-06 | 1.500E-05 |
| 24757 | 1238 | T1238T | D | 1.2e-05 | 1.3e-05 | 1.5e-05 | 1.333E-05 | 1.528E-06 | 1.200E-05 | 1.500E-05 |
| 24758 | 1238 | T1238W | D | 1.1e-05 | 7.6e-06 | 1.5e-05 | 1.120E-05 | 3.704E-06 | 7.600E-06 | 1.500E-05 |
| 24759 | 1238 | T1238Y | D | 1.6e-05 | 1.5e-05 | 1.1e-05 | 1.400E-05 | 2.646E-06 | 1.100E-05 | 1.600E-05 |
| 24760 | 1238 | T1238V | D | 1.7e-05 | 1.6e-05 | 9.5e-06 | 1.417E-05 | 4.072E-06 | 9.500E-06 | 1.700E-05 |
| 24761 | 1239 | L1239A | D | 1.5e-05 | 1.4e-05 | 9.3e-06 | 1.277E-05 | 3.044E-06 | 9.300E-06 | 1.500E-05 |
| 24762 | 1239 | L1239R | D | 8.6e-06 | 1.6e-05 | 1.2e-05 | 1.220E-05 | 3.704E-06 | 8.600E-06 | 1.600E-05 |
| 24763 | 1239 | L1239N | D | 1.3e-05 | 1e-05 | 1e-05 | 1.100E-05 | 1.732E-06 | 1.000E-05 | 1.300E-05 |
| 24764 | 1239 | L1239D | D | 1.5e-05 | 9.8e-06 | 1.6e-05 | 1.360E-05 | 3.329E-06 | 9.800E-06 | 1.600E-05 |
| 24765 | 1239 | L1239C | D | 1.2e-05 | 1e-05 | 1e-05 | 1.067E-05 | 1.155E-06 | 1.000E-05 | 1.200E-05 |
| 24766 | 1239 | L1239E | D | 1.1e-05 | 1e-05 | 1.4e-05 | 1.167E-05 | 2.082E-06 | 1.000E-05 | 1.400E-05 |
| 24767 | 1239 | L1239Q | D | 1.4e-05 | 1.1e-05 | 1.4e-05 | 1.300E-05 | 1.732E-06 | 1.100E-05 | 1.400E-05 |
| 24768 | 1239 | L1239G | D | 1.7e-05 | 1.5e-05 | 1.6e-05 | 1.600E-05 | 1.000E-06 | 1.500E-05 | 1.700E-05 |
| 24769 | 1239 | L1239H | D | 1.3e-05 | 1.4e-05 | 1e-05 | 1.233E-05 | 2.082E-06 | 1.000E-05 | 1.400E-05 |
| 24770 | 1239 | L1239I | D | 1.4e-05 | 8.4e-06 | 8.9e-06 | 1.043E-05 | 3.099E-06 | 8.400E-06 | 1.400E-05 |
| 24771 | 1239 | L1239L | D | 1.4e-05 | 9.4e-06 | 1.2e-05 | 1.180E-05 | 2.307E-06 | 9.400E-06 | 1.400E-05 |
| 24772 | 1239 | L1239K | D | 1.5e-05 | 9.2e-06 | 5.8e-06 | 1.000E-05 | 4.652E-06 | 5.800E-06 | 1.500E-05 |
| 24773 | 1239 | L1239M | D | 1e-05 | 1.5e-05 | 7.5e-06 | 1.083E-05 | 3.819E-06 | 7.500E-06 | 1.500E-05 |
| 24774 | 1239 | L1239F | D | 1.4e-05 | 1.7e-05 | 1e-05 | 1.367E-05 | 3.512E-06 | 1.000E-05 | 1.700E-05 |
| 24775 | 1239 | L1239P | D | 1.1e-05 | 1.6e-05 | 1.6e-05 | 1.433E-05 | 2.887E-06 | 1.100E-05 | 1.600E-05 |
| 24776 | 1239 | L1239S | D | 1.1e-05 | 9.5e-06 | 1.4e-05 | 1.150E-05 | 2.291E-06 | 9.500E-06 | 1.400E-05 |
| 24777 | 1239 | L1239T | D | 1e-05 | 1.1e-05 | 1.3e-05 | 1.133E-05 | 1.528E-06 | 1.000E-05 | 1.300E-05 |
| 24778 | 1239 | L1239W | D | 1.5e-05 | 1.6e-05 | 1.1e-05 | 1.400E-05 | 2.646E-06 | 1.100E-05 | 1.600E-05 |
| 24779 | 1239 | L1239Y | D | 1.3e-05 | 1e-05 | 9.5e-06 | 1.083E-05 | 1.893E-06 | 9.500E-06 | 1.300E-05 |
| 24780 | 1239 | L1239V | D | 1.3e-05 | 1.6e-05 | 1.6e-05 | 1.500E-05 | 1.732E-06 | 1.300E-05 | 1.600E-05 |
| 24781 | 1240 | S1240A | D | 1.1e-05 | 1.1e-05 | 1.2e-05 | 1.133E-05 | 5.774E-07 | 1.100E-05 | 1.200E-05 |
| 24782 | 1240 | S1240R | D | 9.2e-06 | 1.1e-05 | 9.1e-06 | 9.767E-06 | 1.069E-06 | 9.100E-06 | 1.100E-05 |
| 24783 | 1240 | S1240N | D | 8.3e-06 | 1.5e-05 | 7.9e-06 | 1.040E-05 | 3.989E-06 | 7.900E-06 | 1.500E-05 |
| 24784 | 1240 | S1240D | D | 1.4e-05 | 1.4e-05 | 1.4e-05 | 1.400E-05 | 0.000E+00 | 1.400E-05 | 1.400E-05 |
| 24785 | 1240 | S1240C | D | 1.4e-05 | 1.1e-05 | 1.1e-05 | 1.200E-05 | 1.732E-06 | 1.100E-05 | 1.400E-05 |
| 24786 | 1240 | S1240E | D | 1.1e-05 | 1.3e-05 | 1.6e-05 | 1.333E-05 | 2.517E-06 | 1.100E-05 | 1.600E-05 |
| 24787 | 1240 | S1240Q | D | 1.1e-05 | 1.1e-05 | 1.6e-05 | 1.267E-05 | 2.887E-06 | 1.100E-05 | 1.600E-05 |
| 24788 | 1240 | S1240G | D | 1e-05 | 8.2e-06 | 1.4e-05 | 1.073E-05 | 2.969E-06 | 8.200E-06 | 1.400E-05 |
| 24789 | 1240 | S1240H | D | 9.2e-06 | 9.5e-06 | 1.4e-05 | 1.090E-05 | 2.689E-06 | 9.200E-06 | 1.400E-05 |
| 24790 | 1240 | S1240I | D | 1.6e-05 | 1.5e-05 | 1.2e-05 | 1.433E-05 | 2.082E-06 | 1.200E-05 | 1.600E-05 |
| 24791 | 1240 | S1240L | D | 1.4e-05 | 1.7e-05 | 9.7e-06 | 1.357E-05 | 3.669E-06 | 9.700E-06 | 1.700E-05 |
| 24792 | 1240 | S1240K | D | 1.3e-05 | 1.8e-05 | 9.6e-06 | 1.353E-05 | 4.225E-06 | 9.600E-06 | 1.800E-05 |
| 24793 | 1240 | S1240M | D | 1.5e-05 | 1.6e-05 | 1.1e-05 | 1.400E-05 | 2.646E-06 | 1.100E-05 | 1.600E-05 |
| 24794 | 1240 | S1240F | D | 1.1e-05 | 1.2e-05 | 9.4e-06 | 1.080E-05 | 1.311E-06 | 9.400E-06 | 1.200E-05 |
| 24795 | 1240 | S1240P | D | 9.9e-06 | 1e-05 | 1.2e-05 | 1.063E-05 | 1.185E-06 | 9.900E-06 | 1.200E-05 |
| 24796 | 1240 | S1240S | D | 9.7e-06 | 1.7e-05 | 8.9e-06 | 1.187E-05 | 4.464E-06 | 8.900E-06 | 1.700E-05 |
| 24797 | 1240 | S1240T | D | 1e-05 | 9.5e-06 | 1.1e-05 | 1.017E-05 | 7.638E-07 | 9.500E-06 | 1.100E-05 |
| 24798 | 1240 | S1240W | D | 6.5e-06 | 9.8e-06 | 1.2e-05 | 9.433E-06 | 2.768E-06 | 6.500E-06 | 1.200E-05 |
| 24799 | 1240 | S1240Y | D | 1.1e-05 | 1.3e-05 | 1.2e-05 | 1.200E-05 | 1.000E-06 | 1.100E-05 | 1.300E-05 |
| 24800 | 1240 | S1240V | D | 1.1e-05 | 1.7e-05 | 1.3e-05 | 1.367E-05 | 3.055E-06 | 1.100E-05 | 1.700E-05 |
| 24801 | 1241 | K1241A | D | 1.2e-05 | 1.4e-05 | 9.8e-06 | 1.193E-05 | 2.101E-06 | 9.800E-06 | 1.400E-05 |
| 24802 | 1241 | K1241R | D | 9.6e-06 | 1.3e-05 | 1.3e-05 | 1.187E-05 | 1.963E-06 | 9.600E-06 | 1.300E-05 |
| 24803 | 1241 | K1241N | D | 1.5e-05 | 1.4e-05 | 1.5e-05 | 1.467E-05 | 5.774E-07 | 1.400E-05 | 1.500E-05 |
| 24804 | 1241 | K1241D | D | 9.4e-06 | 1e-05 | 1.5e-05 | 1.147E-05 | 3.075E-06 | 9.400E-06 | 1.500E-05 |
| 24805 | 1241 | K1241C | D | 1.5e-05 | 1.5e-05 | 1.5e-05 | 1.500E-05 | 0.000E+00 | 1.500E-05 | 1.500E-05 |
| 24806 | 1241 | K1241E | D | 1e-05 | 9.8e-06 | 1.4e-05 | 1.127E-05 | 2.369E-06 | 9.800E-06 | 1.400E-05 |
| 24807 | 1241 | K1241Q | D | 1.5e-05 | 9.9e-06 | 1.5e-05 | 1.330E-05 | 2.944E-06 | 9.900E-06 | 1.500E-05 |
| 24808 | 1241 | K1241G | D | 1.6e-05 | 1.7e-05 | 1.4e-05 | 1.567E-05 | 1.528E-06 | 1.400E-05 | 1.700E-05 |
| 24809 | 1241 | K1241H | D | 9.8e-06 | 1.5e-05 | 1.1e-05 | 1.193E-05 | 2.723E-06 | 9.800E-06 | 1.500E-05 |
| 24810 | 1241 | K1241I | D | 1.4e-05 | 1e-05 | 1e-05 | 1.133E-05 | 2.309E-06 | 1.000E-05 | 1.400E-05 |
| 24811 | 1241 | K1241L | D | 1.1e-05 | 1.6e-05 | 1.2e-05 | 1.300E-05 | 2.646E-06 | 1.100E-05 | 1.600E-05 |
| 24812 | 1241 | K1241K | D | 8.4e-06 | 1.5e-05 | 8.7e-06 | 1.070E-05 | 3.727E-06 | 8.400E-06 | 1.500E-05 |
| 24813 | 1241 | K1241M | D | 1.7e-05 | 1.5e-05 | 1.3e-05 | 1.500E-05 | 2.000E-06 | 1.300E-05 | 1.700E-05 |
| 24814 | 1241 | K1241F | D | 1.4e-05 | 1.6e-05 | 9.9e-06 | 1.330E-05 | 3.110E-06 | 9.900E-06 | 1.600E-05 |
| 24815 | 1241 | K1241P | D | 9.4e-06 | 1.2e-05 | 9.4e-06 | 1.027E-05 | 1.501E-06 | 9.400E-06 | 1.200E-05 |
| 24816 | 1241 | K1241S | D | 1.5e-05 | 1.4e-05 | 1.3e-05 | 1.400E-05 | 1.000E-06 | 1.300E-05 | 1.500E-05 |
| 24817 | 1241 | K1241T | D | 9.5e-06 | 1e-05 | 1.5e-05 | 1.150E-05 | 3.041E-06 | 9.500E-06 | 1.500E-05 |
| 24818 | 1241 | K1241W | D | 1.2e-05 | 9.1e-06 | 1.5e-05 | 1.203E-05 | 2.950E-06 | 9.100E-06 | 1.500E-05 |
| 24819 | 1241 | K1241Y | D | 1.4e-05 | 1.3e-05 | 8.9e-06 | 1.197E-05 | 2.702E-06 | 8.900E-06 | 1.400E-05 |

| | | | | | | | | | | |
|---|---|---|---|---|---|---|---|---|---|---|
| 24820 | 1241 | K1241V | D | 1.2e-05 | 1.4e-05 | 9.7e-06 | 1.190E-05 | 2.152E-06 | 9.700E-06 | 1.400E-05 |
| 24821 | 1242 | A1242A | D | 1.4e-05 | 1.6e-05 | 1.5e-05 | 1.500E-05 | 1.000E-06 | 1.400E-05 | 1.600E-05 |
| 24822 | 1242 | A1242R | D | 8.5e-06 | 1.1e-05 | 1.1e-05 | 1.017E-05 | 1.443E-06 | 8.500E-06 | 1.100E-05 |
| 24823 | 1242 | A1242N | D | 1.5e-05 | 8.1e-06 | 9.4e-06 | 1.083E-05 | 3.667E-06 | 8.100E-06 | 1.500E-05 |
| 24824 | 1242 | A1242D | D | 1.6e-05 | 1.1e-05 | 1.5e-05 | 1.400E-05 | 2.646E-06 | 1.100E-05 | 1.600E-05 |
| 24825 | 1242 | A1242C | D | 1.3e-05 | 1.1e-05 | 1e-05 | 1.133E-05 | 1.528E-06 | 1.000E-05 | 1.300E-05 |
| 24826 | 1242 | A1242E | D | 9.7e-06 | 1.2e-05 | 1.4e-05 | 1.190E-05 | 2.152E-06 | 9.700E-06 | 1.400E-05 |
| 24827 | 1242 | A1242Q | D | 1.4e-05 | 9.3e-06 | 1.7e-05 | 1.343E-05 | 3.881E-06 | 9.300E-06 | 1.700E-05 |
| 24828 | 1242 | A1242G | D | 1.4e-05 | 1.7e-05 | 1e-05 | 1.367E-05 | 3.512E-06 | 1.000E-05 | 1.700E-05 |
| 24829 | 1242 | A1242H | D | 1.6e-05 | 1.3e-05 | 1.5e-05 | 1.467E-05 | 1.528E-06 | 1.300E-05 | 1.600E-05 |
| 24830 | 1242 | A1242I | D | 8.8e-06 | 6.9e-06 | 3.8e-05 | 1.790E-05 | 1.743E-05 | 6.900E-06 | 3.800E-05 |
| 24831 | 1242 | A1242L | D | 8.6e-06 | 1.4e-05 | 1.5e-05 | 1.253E-05 | 3.443E-06 | 8.600E-06 | 1.500E-05 |
| 24832 | 1242 | A1242K | D | 1.7e-05 | 9.3e-06 | 1.5e-05 | 1.377E-05 | 3.995E-06 | 9.300E-06 | 1.700E-05 |
| 24833 | 1242 | A1242M | D | 1.5e-05 | 1.6e-05 | 1e-05 | 1.367E-05 | 3.215E-06 | 1.000E-05 | 1.600E-05 |
| 24834 | 1242 | A1242F | D | 1.2e-05 | 1.2e-05 | 1.6e-05 | 1.333E-05 | 2.309E-06 | 1.200E-05 | 1.600E-05 |
| 24835 | 1242 | A1242P | D | 1.6e-05 | 1.4e-05 | 1.6e-05 | 1.533E-05 | 1.155E-06 | 1.400E-05 | 1.600E-05 |
| 24836 | 1242 | A1242S | D | 1.5e-05 | 1.5e-05 | 1.1e-05 | 1.367E-05 | 2.309E-06 | 1.100E-05 | 1.500E-05 |
| 24837 | 1242 | A1242T | D | 1.4e-05 | 1.6e-05 | 1.4e-05 | 1.467E-05 | 1.155E-06 | 1.400E-05 | 1.600E-05 |
| 24838 | 1242 | A1242W | D | 1.6e-05 | 1.5e-05 | 1e-05 | 1.367E-05 | 3.215E-06 | 1.000E-05 | 1.600E-05 |
| 24839 | 1242 | A1242Y | D | 1.5e-05 | 1.3e-05 | 1e-05 | 1.267E-05 | 2.517E-06 | 1.000E-05 | 1.500E-05 |
| 24840 | 1242 | A1242V | D | 1.6e-05 | 1.7e-05 | 1.1e-05 | 1.467E-05 | 3.215E-06 | 1.100E-05 | 1.700E-05 |
| 24841 | 1243 | D1243A | D | 1.4e-05 | 1.5e-05 | 1.3e-05 | 1.400E-05 | 1.000E-06 | 1.300E-05 | 1.500E-05 |
| 24842 | 1243 | D1243R | D | 1.1e-05 | 1.6e-05 | 1.1e-05 | 1.267E-05 | 2.887E-06 | 1.100E-05 | 1.600E-05 |
| 24843 | 1243 | D1243N | D | 8.1e-06 | 1.2e-05 | 1.4e-05 | 1.137E-05 | 3.001E-06 | 8.100E-06 | 1.400E-05 |
| 24844 | 1243 | D1243D | D | 1.3e-05 | 1.6e-05 | 1.3e-05 | 1.400E-05 | 1.732E-06 | 1.300E-05 | 1.600E-05 |
| 24845 | 1243 | D1243C | D | 1.1e-05 | 1.5e-05 | 9.6e-06 | 1.187E-05 | 2.802E-06 | 9.600E-06 | 1.500E-05 |
| 24846 | 1243 | D1243E | D | 1.1e-05 | 1.7e-05 | 1.4e-05 | 1.400E-05 | 3.000E-06 | 1.100E-05 | 1.700E-05 |
| 24847 | 1243 | D1243Q | D | 1.1e-05 | 1.3e-05 | 1.4e-05 | 1.267E-05 | 1.528E-06 | 1.100E-05 | 1.400E-05 |
| 24848 | 1243 | D1243G | D | 1e-05 | 2.2e-05 | 1.5e-05 | 1.567E-05 | 6.028E-06 | 1.000E-05 | 2.200E-05 |
| 24849 | 1243 | D1243H | D | 1.7e-05 | 8.6e-06 | 1.6e-05 | 1.387E-05 | 4.588E-06 | 8.600E-06 | 1.700E-05 |
| 24850 | 1243 | D1243I | D | 1.1e-05 | 1.4e-05 | 1.5e-05 | 1.333E-05 | 2.082E-06 | 1.100E-05 | 1.500E-05 |
| 24851 | 1243 | D1243L | D | 1.5e-05 | 1.1e-05 | 1.2e-05 | 1.267E-05 | 2.082E-06 | 1.100E-05 | 1.500E-05 |
| 24852 | 1243 | D1243K | D | 1e-05 | 1.5e-05 | 1e-05 | 1.167E-05 | 2.887E-06 | 1.000E-05 | 1.500E-05 |
| 24853 | 1243 | D1243M | D | 1.8e-05 | 1.4e-05 | 1.2e-05 | 1.467E-05 | 3.055E-06 | 1.200E-05 | 1.800E-05 |
| 24854 | 1243 | D1243F | D | 9.4e-06 | 1.1e-05 | 1e-05 | 1.013E-05 | 8.083E-07 | 9.400E-06 | 1.100E-05 |
| 24855 | 1243 | D1243P | D | 1.6e-05 | 8.7e-06 | 1.5e-05 | 1.323E-05 | 3.958E-06 | 8.700E-06 | 1.600E-05 |
| 24856 | 1243 | D1243S | D | 8.7e-06 | 1.5e-05 | 1.4e-05 | 1.257E-05 | 3.386E-06 | 8.700E-06 | 1.500E-05 |
| 24857 | 1243 | D1243T | D | 1.1e-05 | 1.3e-05 | 1.5e-05 | 1.300E-05 | 2.000E-06 | 1.100E-05 | 1.500E-05 |
| 24858 | 1243 | D1243W | D | 9.9e-06 | 1.3e-05 | 1e-05 | 1.097E-05 | 1.762E-06 | 9.900E-06 | 1.300E-05 |
| 24859 | 1243 | D1243Y | D | 1.4e-05 | 1.3e-05 | 1.3e-05 | 1.333E-05 | 5.774E-07 | 1.300E-05 | 1.400E-05 |
| 24860 | 1243 | D1243V | D | 1.4e-05 | 1.1e-05 | 8.3e-06 | 1.110E-05 | 2.851E-06 | 8.300E-06 | 1.400E-05 |
| 24861 | 1244 | Y1244A | D | 1.5e-05 | 1.1e-05 | 1.1e-05 | 1.233E-05 | 2.309E-06 | 1.100E-05 | 1.500E-05 |
| 24862 | 1244 | Y1244R | D | 1.5e-05 | 1.3e-05 | 1e-05 | 1.267E-05 | 2.517E-06 | 1.000E-05 | 1.500E-05 |
| 24863 | 1244 | Y1244N | D | 1.3e-05 | 1.1e-05 | 1.5e-05 | 1.300E-05 | 2.000E-06 | 1.100E-05 | 1.500E-05 |
| 24864 | 1244 | Y1244D | D | 1.6e-05 | 8.9e-06 | 9.3e-06 | 1.140E-05 | 3.989E-06 | 8.900E-06 | 1.600E-05 |
| 24865 | 1244 | Y1244C | D | 1.5e-05 | 8.9e-06 | 1.6e-05 | 1.330E-05 | 3.843E-06 | 8.900E-06 | 1.600E-05 |
| 24866 | 1244 | Y1244E | D | 1.1e-05 | 1.3e-05 | 1.4e-05 | 1.267E-05 | 1.528E-06 | 1.100E-05 | 1.400E-05 |
| 24867 | 1244 | Y1244Q | D | 1.6e-05 | 1.1e-05 | 1.4e-05 | 1.367E-05 | 2.517E-06 | 1.100E-05 | 1.600E-05 |
| 24868 | 1244 | Y1244G | D | 1.3e-05 | 1.4e-05 | 9.9e-06 | 1.230E-05 | 2.138E-06 | 9.900E-06 | 1.400E-05 |
| 24869 | 1244 | Y1244H | D | 9.8e-06 | 9e-06 | 1.6e-05 | 1.160E-05 | 3.831E-06 | 9.000E-06 | 1.600E-05 |
| 24870 | 1244 | Y1244I | D | 1.5e-05 | 1.4e-05 | 9.6e-06 | 1.287E-05 | 2.873E-06 | 9.600E-06 | 1.500E-05 |
| 24871 | 1244 | Y1244L | D | 1.6e-05 | 1e-05 | 1.6e-05 | 1.400E-05 | 3.464E-06 | 1.000E-05 | 1.600E-05 |
| 24872 | 1244 | Y1244K | D | 1.7e-05 | 1e-05 | 1e-05 | 1.233E-05 | 4.041E-06 | 1.000E-05 | 1.700E-05 |
| 24873 | 1244 | Y1244M | D | 1.5e-05 | 9.7e-06 | 1.1e-05 | 1.190E-05 | 2.762E-06 | 9.700E-06 | 1.500E-05 |
| 24874 | 1244 | Y1244F | D | 1.1e-05 | 1.3e-05 | 1.3e-05 | 1.233E-05 | 1.155E-06 | 1.100E-05 | 1.300E-05 |
| 24875 | 1244 | Y1244P | D | 1.4e-05 | 1.7e-05 | 1.4e-05 | 1.500E-05 | 1.732E-06 | 1.400E-05 | 1.700E-05 |
| 24876 | 1244 | Y1244S | D | 1.8e-05 | 1e-05 | 1.4e-05 | 1.400E-05 | 4.000E-06 | 1.000E-05 | 1.800E-05 |
| 24877 | 1244 | Y1244T | D | 1.4e-05 | 9.3e-06 | 1e-05 | 1.110E-05 | 2.536E-06 | 9.300E-06 | 1.400E-05 |
| 24878 | 1244 | Y1244W | D | 1.1e-05 | 1.5e-05 | 1.1e-05 | 1.233E-05 | 2.309E-06 | 1.100E-05 | 1.500E-05 |
| 24879 | 1244 | Y1244Y | D | 1.5e-05 | 1.2e-05 | 9.8e-06 | 1.227E-05 | 2.610E-06 | 9.800E-06 | 1.500E-05 |
| 24880 | 1244 | Y1244V | D | 1.3e-05 | 1.5e-05 | 1.1e-05 | 1.300E-05 | 2.000E-06 | 1.100E-05 | 1.500E-05 |
| 24881 | 1245 | E1245A | D | 1e-05 | 9.3e-06 | 1.5e-05 | 1.143E-05 | 3.109E-06 | 9.300E-06 | 1.500E-05 |
| 24882 | 1245 | E1245R | D | 1.3e-05 | 1e-05 | 1.7e-05 | 1.333E-05 | 3.512E-06 | 1.000E-05 | 1.700E-05 |
| 24883 | 1245 | E1245N | D | 1.4e-05 | 9e-06 | 1.3e-05 | 1.200E-05 | 2.646E-06 | 9.000E-06 | 1.400E-05 |
| 24884 | 1245 | E1245D | D | 1.9e-05 | 1.4e-05 | 1.6e-05 | 1.633E-05 | 2.517E-06 | 1.400E-05 | 1.900E-05 |
| 24885 | 1245 | E1245C | D | 1e-05 | 1.2e-05 | 1.6e-05 | 1.267E-05 | 3.055E-06 | 1.000E-05 | 1.600E-05 |
| 24886 | 1245 | E1245E | D | 1.4e-05 | 1.1e-05 | 1.5e-05 | 1.333E-05 | 2.082E-06 | 1.100E-05 | 1.500E-05 |
| 24887 | 1245 | E1245Q | D | 9.3e-06 | 1e-05 | 1e-05 | 9.767E-06 | 4.041E-07 | 9.300E-06 | 1.000E-05 |
| 24888 | 1245 | E1245G | D | 1.4e-05 | 1e-05 | 1.6e-05 | 1.333E-05 | 3.055E-06 | 1.000E-05 | 1.600E-05 |
| 24889 | 1245 | E1245H | D | 1.6e-05 | 1.7e-05 | 1.3e-05 | 1.533E-05 | 2.082E-06 | 1.300E-05 | 1.700E-05 |
| 24890 | 1245 | E1245I | D | 1.4e-05 | 1.1e-05 | 1e-05 | 1.167E-05 | 2.082E-06 | 1.000E-05 | 1.400E-05 |
| 24891 | 1245 | E1245L | D | 1.1e-05 | 1.4e-05 | 9.1e-06 | 1.137E-05 | 2.470E-06 | 9.100E-06 | 1.400E-05 |
| 24892 | 1245 | E1245K | D | 1.5e-05 | 1.2e-05 | 1.6e-05 | 1.433E-05 | 2.082E-06 | 1.200E-05 | 1.600E-05 |
| 24893 | 1245 | E1245M | D | 1.6e-05 | 1.6e-05 | 1.1e-05 | 1.433E-05 | 2.887E-06 | 1.100E-05 | 1.600E-05 |
| 24894 | 1245 | E1245F | D | 1.1e-05 | 1.4e-05 | 1e-05 | 1.167E-05 | 2.082E-06 | 1.000E-05 | 1.400E-05 |
| 24895 | 1245 | E1245P | D | 1.5e-05 | 1.6e-05 | 1.1e-05 | 1.400E-05 | 2.646E-06 | 1.100E-05 | 1.600E-05 |
| 24896 | 1245 | E1245S | D | 1.5e-05 | 1.3e-05 | 1.4e-05 | 1.400E-05 | 1.000E-06 | 1.300E-05 | 1.500E-05 |
| 24897 | 1245 | E1245T | D | 1.1e-05 | 1.5e-05 | 1.2e-05 | 1.267E-05 | 2.082E-06 | 1.100E-05 | 1.500E-05 |
| 24898 | 1245 | E1245W | D | 1.1e-05 | 9.5e-06 | 1.4e-05 | 1.150E-05 | 2.291E-06 | 9.500E-06 | 1.400E-05 |
| 24899 | 1245 | E1245Y | D | 1.6e-05 | 7e-06 | 1.4e-05 | 1.233E-05 | 4.726E-06 | 7.000E-06 | 1.600E-05 |
| 24900 | 1245 | E1245V | D | 1.4e-05 | 1.4e-05 | 1.1e-05 | 1.300E-05 | 1.732E-06 | 1.100E-05 | 1.400E-05 |
| 24901 | 1246 | K1246A | D | 8.6e-06 | 1.5e-05 | 1.5e-05 | 1.287E-05 | 3.695E-06 | 8.600E-06 | 1.500E-05 |
| 24902 | 1246 | K1246R | D | 1.5e-05 | 9.7e-06 | 1.1e-05 | 1.190E-05 | 2.762E-06 | 9.700E-06 | 1.500E-05 |
| 24903 | 1246 | K1246N | D | 9.5e-06 | 1.3e-05 | 1.5e-05 | 1.250E-05 | 2.784E-06 | 9.500E-06 | 1.500E-05 |
| 24904 | 1246 | K1246D | D | 1.5e-05 | 1e-05 | 1.7e-05 | 1.400E-05 | 3.606E-06 | 1.000E-05 | 1.700E-05 |

| | | | | | | | | | | |
|---|---|---|---|---|---|---|---|---|---|---|
| 24905 | 1246 | K1246C | D | 9.9e-06 | 1.3e-05 | 1.3e-05 | 1.197E-05 | 1.790E-06 | 9.900E-06 | 1.300E-05 |
| 24906 | 1246 | K1246E | D | 1.1e-05 | 1.6e-05 | 1.4e-05 | 1.367E-05 | 2.517E-06 | 1.100E-05 | 1.600E-05 |
| 24907 | 1246 | K1246Q | D | 1.5e-05 | 1.3e-05 | 1.1e-05 | 1.300E-05 | 2.000E-06 | 1.100E-05 | 1.500E-05 |
| 24908 | 1246 | K1246G | D | 1.1e-05 | 1.5e-05 | 1.6e-05 | 1.400E-05 | 2.646E-06 | 1.100E-05 | 1.600E-05 |
| 24909 | 1246 | K1246H | D | 1.5e-05 | 1.4e-05 | 9e-06 | 1.267E-05 | 3.215E-06 | 9.000E-06 | 1.500E-05 |
| 24910 | 1246 | K1246I | D | 1.5e-05 | 1.3e-05 | 1.5e-05 | 1.433E-05 | 1.155E-06 | 1.300E-05 | 1.500E-05 |
| 24911 | 1246 | K1246L | D | 1e-05 | 8.4e-06 | 1.7e-05 | 1.180E-05 | 4.574E-06 | 8.400E-06 | 1.700E-05 |
| 24912 | 1246 | K1246K | D | 1.4e-05 | 1.6e-05 | 1.1e-05 | 1.367E-05 | 2.517E-06 | 1.100E-05 | 1.600E-05 |
| 24913 | 1246 | K1246M | D | 8.3e-06 | 9.7e-06 | 1.5e-05 | 1.100E-05 | 3.534E-06 | 8.300E-06 | 1.500E-05 |
| 24914 | 1246 | K1246F | D | 9.6e-06 | 1.4e-05 | 8.8e-06 | 1.080E-05 | 2.800E-06 | 8.800E-06 | 1.400E-05 |
| 24915 | 1246 | K1246P | D | 6.2e-06 | 1.3e-05 | 1.5e-05 | 1.140E-05 | 4.613E-06 | 6.200E-06 | 1.500E-05 |
| 24916 | 1246 | K1246S | D | 1.4e-05 | 1e-05 | 1.1e-05 | 1.167E-05 | 2.082E-06 | 1.000E-05 | 1.400E-05 |
| 24917 | 1246 | K1246T | D | 1.5e-05 | 1e-05 | 1e-05 | 1.167E-05 | 2.887E-06 | 1.000E-05 | 1.500E-05 |
| 24918 | 1246 | K1246W | D | 1.5e-05 | 1.5e-05 | 1e-05 | 1.333E-05 | 2.887E-06 | 1.000E-05 | 1.500E-05 |
| 24919 | 1246 | K1246Y | D | 1.1e-05 | 1.5e-05 | 1.6e-05 | 1.400E-05 | 2.646E-06 | 1.100E-05 | 1.600E-05 |
| 24920 | 1246 | K1246V | D | 1.5e-05 | 1.1e-05 | 1.5e-05 | 1.367E-05 | 2.309E-06 | 1.100E-05 | 1.500E-05 |
| 24921 | 1247 | H1247A | D | 1e-05 | 1.1e-05 | 1.2e-05 | 1.100E-05 | 1.000E-06 | 1.000E-05 | 1.200E-05 |
| 24922 | 1247 | H1247R | D | 1.3e-05 | 1.7e-05 | 1.4e-05 | 1.467E-05 | 2.082E-06 | 1.300E-05 | 1.700E-05 |
| 24923 | 1247 | H1247N | D | 1.5e-05 | 1.1e-05 | 7.8e-06 | 1.127E-05 | 3.607E-06 | 7.800E-06 | 1.500E-05 |
| 24924 | 1247 | H1247D | D | 1.5e-05 | 1.6e-05 | 1.4e-05 | 1.500E-05 | 1.000E-06 | 1.400E-05 | 1.600E-05 |
| 24925 | 1247 | H1247C | D | 1e-05 | 8.8e-06 | 1.4e-05 | 1.093E-05 | 2.723E-06 | 8.800E-06 | 1.400E-05 |
| 24926 | 1247 | H1247E | D | 1.6e-05 | 1.4e-05 | 8.8e-06 | 1.293E-05 | 3.717E-06 | 8.800E-06 | 1.600E-05 |
| 24927 | 1247 | H1247Q | D | 1.3e-05 | 1.6e-05 | 1.5e-05 | 1.467E-05 | 1.528E-06 | 1.300E-05 | 1.600E-05 |
| 24928 | 1247 | H1247G | D | 1.5e-05 | 1.1e-05 | 1.1e-05 | 1.233E-05 | 2.309E-06 | 1.100E-05 | 1.500E-05 |
| 24929 | 1247 | H1247H | D | 1.4e-05 | 8.8e-06 | 1.5e-05 | 1.260E-05 | 3.329E-06 | 8.800E-06 | 1.500E-05 |
| 24930 | 1247 | H1247I | D | 9.3e-06 | 1.1e-05 | 1.4e-05 | 1.143E-05 | 2.380E-06 | 9.300E-06 | 1.400E-05 |
| 24931 | 1247 | H1247L | D | 1.2e-05 | 1.1e-05 | 1.1e-05 | 1.133E-05 | 5.774E-07 | 1.100E-05 | 1.200E-05 |
| 24932 | 1247 | H1247K | D | 9.7e-06 | 1.4e-05 | 1.3e-05 | 1.223E-05 | 2.250E-06 | 9.700E-06 | 1.400E-05 |
| 24933 | 1247 | H1247M | D | 1.3e-05 | 1.1e-05 | 1.2e-05 | 1.200E-05 | 1.000E-06 | 1.100E-05 | 1.300E-05 |
| 24934 | 1247 | H1247F | D | 1e-05 | 9.2e-06 | 1.7e-05 | 1.207E-05 | 4.291E-06 | 9.200E-06 | 1.700E-05 |
| 24935 | 1247 | H1247P | D | 1.5e-05 | 1.3e-05 | 8.2e-06 | 1.207E-05 | 3.495E-06 | 8.200E-06 | 1.500E-05 |
| 24936 | 1247 | H1247S | D | 1.1e-05 | 1.8e-05 | 1.5e-05 | 1.467E-05 | 3.512E-06 | 1.100E-05 | 1.800E-05 |
| 24937 | 1247 | H1247T | D | 1.5e-05 | 1.1e-05 | 1.5e-05 | 1.367E-05 | 2.309E-06 | 1.100E-05 | 1.500E-05 |
| 24938 | 1247 | H1247W | D | 1.4e-05 | 1.6e-05 | 1.1e-05 | 1.367E-05 | 2.517E-06 | 1.100E-05 | 1.600E-05 |
| 24939 | 1247 | H1247Y | D | 1.5e-05 | 1.4e-05 | 1.4e-05 | 1.433E-05 | 5.774E-07 | 1.400E-05 | 1.500E-05 |
| 24940 | 1247 | H1247V | D | 8.7e-06 | 8.7e-06 | 9.2e-06 | 8.867E-06 | 2.887E-07 | 8.700E-06 | 9.200E-06 |
| 24941 | 1248 | K1248A | D | 1.1e-05 | 1.5e-05 | 1e-05 | 1.200E-05 | 2.646E-06 | 1.000E-05 | 1.500E-05 |
| 24942 | 1248 | K1248R | D | 1.5e-05 | 8.8e-06 | 9.8e-06 | 1.120E-05 | 3.329E-06 | 8.800E-06 | 1.500E-05 |
| 24943 | 1248 | K1248N | D | 1.1e-05 | 1.4e-05 | 1.4e-05 | 1.300E-05 | 1.732E-06 | 1.100E-05 | 1.400E-05 |
| 24944 | 1248 | K1248D | D | 1.4e-05 | 1.4e-05 | 1.2e-05 | 1.333E-05 | 1.155E-06 | 1.200E-05 | 1.400E-05 |
| 24945 | 1248 | K1248C | D | 1.6e-05 | 1.1e-05 | 1.4e-05 | 1.367E-05 | 2.517E-06 | 1.100E-05 | 1.600E-05 |
| 24946 | 1248 | K1248E | D | 1.4e-05 | 1.6e-05 | 1e-05 | 1.333E-05 | 3.055E-06 | 1.000E-05 | 1.600E-05 |
| 24947 | 1248 | K1248Q | D | 1.3e-05 | 1.3e-05 | 1.6e-05 | 1.400E-05 | 1.732E-06 | 1.300E-05 | 1.600E-05 |
| 24948 | 1248 | K1248G | D | 1.2e-05 | 1.6e-05 | 1e-05 | 1.267E-05 | 3.055E-06 | 1.000E-05 | 1.600E-05 |
| 24949 | 1248 | K1248H | D | 1e-05 | 8.8e-06 | 1.2e-05 | 1.027E-05 | 1.617E-06 | 8.800E-06 | 1.200E-05 |
| 24950 | 1248 | K1248I | D | 1.7e-05 | 9.1e-06 | 1.1e-05 | 1.237E-05 | 4.124E-06 | 9.100E-06 | 1.700E-05 |
| 24951 | 1248 | K1248L | D | 1.6e-05 | 1e-05 | 1.2e-05 | 1.267E-05 | 3.055E-06 | 1.000E-05 | 1.600E-05 |
| 24952 | 1248 | K1248K | D | 1e-05 | 1.4e-05 | 1e-05 | 1.133E-05 | 2.309E-06 | 1.000E-05 | 1.400E-05 |
| 24953 | 1248 | K1248M | D | 1.1e-05 | 1.5e-05 | 1e-05 | 1.200E-05 | 2.646E-06 | 1.000E-05 | 1.500E-05 |
| 24954 | 1248 | K1248F | D | 1.6e-05 | 1.3e-05 | 1e-05 | 1.300E-05 | 3.000E-06 | 1.000E-05 | 1.600E-05 |
| 24955 | 1248 | K1248P | D | 8.7e-06 | 1e-05 | 1.6e-05 | 1.157E-05 | 3.894E-06 | 8.700E-06 | 1.600E-05 |
| 24956 | 1248 | K1248S | D | 9.1e-06 | 9.7e-06 | 1.4e-05 | 1.093E-05 | 2.673E-06 | 9.100E-06 | 1.400E-05 |
| 24957 | 1248 | K1248T | D | 1.3e-05 | 1.4e-05 | 9.6e-06 | 1.220E-05 | 2.307E-06 | 9.600E-06 | 1.400E-05 |
| 24958 | 1248 | K1248W | D | 1.2e-05 | 1.7e-05 | 1.5e-05 | 1.467E-05 | 2.517E-06 | 1.200E-05 | 1.700E-05 |
| 24959 | 1248 | K1248Y | D | 9.9e-06 | 9.9e-06 | 1.4e-05 | 1.127E-05 | 2.367E-06 | 9.900E-06 | 1.400E-05 |
| 24960 | 1248 | K1248V | D | 1.6e-05 | 1.8e-05 | 3.1e-05 | 2.167E-05 | 8.145E-06 | 1.600E-05 | 3.100E-05 |
| 24961 | 1249 | V1249A | D | 9.8e-06 | 1.2e-05 | 1.2e-05 | 1.127E-05 | 1.270E-06 | 9.800E-06 | 1.200E-05 |
| 24962 | 1249 | V1249R | D | 9.7e-06 | 1.5e-05 | 1.4e-05 | 1.290E-05 | 2.816E-06 | 9.700E-06 | 1.500E-05 |
| 24963 | 1249 | V1249N | D | 1e-05 | 1.5e-05 | 1e-05 | 1.167E-05 | 2.887E-06 | 1.000E-05 | 1.500E-05 |
| 24964 | 1249 | V1249D | D | 1.1e-05 | 1.5e-05 | 1.4e-05 | 1.333E-05 | 2.082E-06 | 1.100E-05 | 1.500E-05 |
| 24965 | 1249 | V1249C | D | 1.4e-05 | 9.9e-06 | 9.5e-06 | 1.113E-05 | 2.491E-06 | 9.500E-06 | 1.400E-05 |
| 24966 | 1249 | V1249E | D | 9.5e-06 | 1.1e-05 | 1e-05 | 1.017E-05 | 7.638E-07 | 9.500E-06 | 1.100E-05 |
| 24967 | 1249 | V1249Q | D | 1e-05 | 1.1e-05 | 1.5e-05 | 1.200E-05 | 2.646E-06 | 1.000E-05 | 1.500E-05 |
| 24968 | 1249 | V1249G | D | 1.4e-05 | 1.5e-05 | 1.5e-05 | 1.467E-05 | 5.774E-07 | 1.400E-05 | 1.500E-05 |
| 24969 | 1249 | V1249H | D | 1.6e-05 | 1.3e-05 | 1.5e-05 | 1.467E-05 | 1.528E-06 | 1.300E-05 | 1.600E-05 |
| 24970 | 1249 | V1249I | D | 1.7e-05 | 7.7e-06 | 1.2e-05 | 1.223E-05 | 4.654E-06 | 7.700E-06 | 1.700E-05 |
| 24971 | 1249 | V1249L | D | 1.1e-05 | 9.7e-06 | 5.7e-06 | 8.800E-06 | 2.762E-06 | 5.700E-06 | 1.100E-05 |
| 24972 | 1249 | V1249K | D | 1.6e-05 | 7.4e-06 | 1.3e-05 | 1.213E-05 | 4.365E-06 | 7.400E-06 | 1.600E-05 |
| 24973 | 1249 | V1249M | D | 1.9e-05 | 1.3e-05 | 1.6e-05 | 1.600E-05 | 3.000E-06 | 1.300E-05 | 1.900E-05 |
| 24974 | 1249 | V1249F | D | 1.1e-05 | 1.2e-05 | 6.7e-06 | 9.900E-06 | 2.816E-06 | 6.700E-06 | 1.200E-05 |
| 24975 | 1249 | V1249P | D | 9.4e-06 | 1.6e-05 | 1.1e-05 | 1.213E-05 | 3.443E-06 | 9.400E-06 | 1.600E-05 |
| 24976 | 1249 | V1249S | D | 1.4e-05 | 1.3e-05 | 1.5e-05 | 1.400E-05 | 1.000E-06 | 1.300E-05 | 1.500E-05 |
| 24977 | 1249 | V1249T | D | 1.7e-05 | 1e-05 | 1.2e-05 | 1.300E-05 | 3.606E-06 | 1.000E-05 | 1.700E-05 |
| 24978 | 1249 | V1249W | D | 1.4e-05 | 1.1e-05 | 1.4e-05 | 1.300E-05 | 1.732E-06 | 1.100E-05 | 1.400E-05 |
| 24979 | 1249 | V1249Y | D | 1.4e-05 | 1.6e-05 | 1.3e-05 | 1.433E-05 | 1.528E-06 | 1.300E-05 | 1.600E-05 |
| 24980 | 1249 | V1249V | D | 1.4e-05 | 1.6e-05 | 1.3e-05 | 1.433E-05 | 1.528E-06 | 1.300E-05 | 1.600E-05 |
| 24981 | 1250 | Y1250A | D | 1.3e-05 | 1e-05 | 1.1e-05 | 1.133E-05 | 1.528E-06 | 1.000E-05 | 1.300E-05 |
| 24982 | 1250 | Y1250R | D | 1.6e-05 | 1.4e-05 | 1.3e-05 | 1.433E-05 | 1.528E-06 | 1.300E-05 | 1.600E-05 |
| 24983 | 1250 | Y1250N | D | 1.5e-05 | 1.5e-05 | 1.1e-05 | 1.367E-05 | 2.309E-06 | 1.100E-05 | 1.500E-05 |
| 24984 | 1250 | Y1250D | D | 9.7e-06 | 8.9e-06 | 1.1e-05 | 9.867E-06 | 1.060E-06 | 8.900E-06 | 1.100E-05 |
| 24985 | 1250 | Y1250C | D | 1.4e-05 | 1.8e-05 | 8.7e-06 | 1.357E-05 | 4.665E-06 | 8.700E-06 | 1.800E-05 |
| 24986 | 1250 | Y1250E | D | 9.5e-06 | 1.2e-05 | 1.1e-05 | 1.083E-05 | 1.258E-06 | 9.500E-06 | 1.200E-05 |
| 24987 | 1250 | Y1250Q | D | 1.3e-05 | 1.1e-05 | 1.1e-05 | 1.167E-05 | 1.155E-06 | 1.100E-05 | 1.300E-05 |
| 24988 | 1250 | Y1250G | D | 1.2e-05 | 8.1e-06 | 1.1e-05 | 1.037E-05 | 2.026E-06 | 8.100E-06 | 1.200E-05 |
| 24989 | 1250 | Y1250H | D | 1.1e-05 | 1.5e-05 | 8.8e-06 | 1.160E-05 | 3.143E-06 | 8.800E-06 | 1.500E-05 |

| | | | | | | | | | | |
|---|---|---|---|---|---|---|---|---|---|---|
| 24990 | 1250 | Y1250I | D | 1.5e-05 | 1.4e-05 | 1.2e-05 | 1.367E-05 | 1.528E-06 | 1.200E-05 | 1.500E-05 |
| 24991 | 1250 | Y1250L | D | 1.6e-05 | 1.4e-05 | 9.9e-06 | 1.330E-05 | 3.110E-06 | 9.900E-06 | 1.600E-05 |
| 24992 | 1250 | Y1250K | D | 1.2e-05 | 1.6e-05 | 1.4e-05 | 1.400E-05 | 2.000E-06 | 1.200E-05 | 1.600E-05 |
| 24993 | 1250 | Y1250M | D | 1.1e-05 | 1.2e-05 | 1e-05 | 1.100E-05 | 1.000E-06 | 1.000E-05 | 1.200E-05 |
| 24994 | 1250 | Y1250F | D | 1.1e-05 | 1.1e-05 | 1.7e-05 | 1.300E-05 | 3.464E-06 | 1.100E-05 | 1.700E-05 |
| 24995 | 1250 | Y1250P | D | 9.9e-06 | 1.5e-05 | 1e-05 | 1.163E-05 | 2.916E-06 | 9.900E-06 | 1.500E-05 |
| 24996 | 1250 | Y1250S | D | 6.7e-06 | 1.5e-05 | 1.6e-05 | 1.257E-05 | 5.105E-06 | 6.700E-06 | 1.600E-05 |
| 24997 | 1250 | Y1250T | D | 1.5e-05 | 1.7e-05 | 1.2e-05 | 1.467E-05 | 2.517E-06 | 1.200E-05 | 1.700E-05 |
| 24998 | 1250 | Y1250W | D | 1.5e-05 | 9.9e-06 | 1.1e-05 | 1.197E-05 | 2.684E-06 | 9.900E-06 | 1.500E-05 |
| 24999 | 1250 | Y1250Y | D | 1.3e-05 | 1e-05 | 1.5e-05 | 1.267E-05 | 2.517E-06 | 1.000E-05 | 1.500E-05 |
| 25000 | 1250 | Y1250V | D | 1.5e-05 | 1.5e-05 | 1.5e-05 | 1.500E-05 | 0.000E+00 | 1.500E-05 | 1.500E-05 |
| 25001 | 1251 | A1251A | D | 1.1e-05 | 1.4e-05 | 1.6e-05 | 1.367E-05 | 2.517E-06 | 1.100E-05 | 1.600E-05 |
| 25002 | 1251 | A1251R | D | 1.1e-05 | 1e-05 | 1.2e-05 | 1.100E-05 | 1.000E-06 | 1.000E-05 | 1.200E-05 |
| 25003 | 1251 | A1251N | D | 1.5e-05 | 1.6e-05 | 1.5e-05 | 1.533E-05 | 5.774E-07 | 1.500E-05 | 1.600E-05 |
| 25004 | 1251 | A1251D | D | 1.6e-05 | 1.6e-05 | 1.2e-05 | 1.467E-05 | 2.309E-06 | 1.200E-05 | 1.600E-05 |
| 25005 | 1251 | A1251C | D | 2e-05 | 1.2e-05 | 8.1e-06 | 1.337E-05 | 6.067E-06 | 8.100E-06 | 2.000E-05 |
| 25006 | 1251 | A1251E | D | 1.6e-05 | 9.4e-06 | 1.5e-05 | 1.347E-05 | 3.557E-06 | 9.400E-06 | 1.600E-05 |
| 25007 | 1251 | A1251Q | D | 1.6e-05 | 1.3e-05 | 1.5e-05 | 1.467E-05 | 1.528E-06 | 1.300E-05 | 1.600E-05 |
| 25008 | 1251 | A1251G | D | 9.9e-06 | 1.1e-05 | 1.1e-05 | 1.063E-05 | 6.351E-07 | 9.900E-06 | 1.100E-05 |
| 25009 | 1251 | A1251H | D | 1.5e-05 | 1.7e-05 | 1.6e-05 | 1.600E-05 | 1.000E-06 | 1.500E-05 | 1.700E-05 |
| 25010 | 1251 | A1251I | D | 8.6e-06 | 1.4e-05 | 1.6e-05 | 1.287E-05 | 3.828E-06 | 8.600E-06 | 1.600E-05 |
| 25011 | 1251 | A1251L | D | 1.7e-05 | 1.4e-05 | 1.6e-05 | 1.567E-05 | 1.528E-06 | 1.400E-05 | 1.700E-05 |
| 25012 | 1251 | A1251K | D | 1.2e-05 | 1.4e-05 | 1e-05 | 1.200E-05 | 2.000E-06 | 1.000E-05 | 1.400E-05 |
| 25013 | 1251 | A1251M | D | 1.3e-05 | 1.6e-05 | 1.5e-05 | 1.467E-05 | 1.528E-06 | 1.300E-05 | 1.600E-05 |
| 25014 | 1251 | A1251F | D | 1.1e-05 | 1.4e-05 | 9.5e-06 | 1.150E-05 | 2.291E-06 | 9.500E-06 | 1.400E-05 |
| 25015 | 1251 | A1251P | D | 1.5e-05 | 1.1e-05 | 1.4e-05 | 1.333E-05 | 2.082E-06 | 1.100E-05 | 1.500E-05 |
| 25016 | 1251 | A1251S | D | 1.3e-05 | 1.2e-05 | 1.6e-05 | 1.367E-05 | 2.082E-06 | 1.200E-05 | 1.600E-05 |
| 25017 | 1251 | A1251T | D | 1.3e-05 | 1.1e-05 | 1.6e-05 | 1.333E-05 | 2.517E-06 | 1.100E-05 | 1.600E-05 |
| 25018 | 1251 | A1251W | D | 1.5e-05 | 1.1e-05 | 1.5e-05 | 1.367E-05 | 2.309E-06 | 1.100E-05 | 1.500E-05 |
| 25019 | 1251 | A1251Y | D | 1.2e-05 | 9.9e-06 | 9.5e-06 | 1.047E-05 | 1.343E-06 | 9.500E-06 | 1.200E-05 |
| 25020 | 1251 | A1251V | D | 9.9e-06 | 1.5e-05 | 1.7e-05 | 1.397E-05 | 3.661E-06 | 9.900E-06 | 1.700E-05 |
| 25021 | 1252 | C1252A | D | 1.5e-05 | 1.1e-05 | 1.1e-05 | 1.233E-05 | 2.309E-06 | 1.100E-05 | 1.500E-05 |
| 25022 | 1252 | C1252R | D | 9.1e-06 | 1.5e-05 | 1.1e-05 | 1.170E-05 | 3.012E-06 | 9.100E-06 | 1.500E-05 |
| 25023 | 1252 | C1252N | D | 1.4e-05 | 1e-05 | 9.8e-06 | 1.127E-05 | 2.369E-06 | 9.800E-06 | 1.400E-05 |
| 25024 | 1252 | C1252D | D | 1.4e-05 | 1.2e-05 | 1.6e-05 | 1.400E-05 | 2.000E-06 | 1.200E-05 | 1.600E-05 |
| 25025 | 1252 | C1252C | D | 1.6e-05 | 1.6e-05 | 1.3e-05 | 1.500E-05 | 1.732E-06 | 1.300E-05 | 1.600E-05 |
| 25026 | 1252 | C1252E | D | 1.1e-05 | 1e-05 | 1.1e-05 | 1.067E-05 | 5.774E-07 | 1.000E-05 | 1.100E-05 |
| 25027 | 1252 | C1252Q | D | 1.4e-05 | 1.6e-05 | 1.6e-05 | 1.533E-05 | 1.155E-06 | 1.400E-05 | 1.600E-05 |
| 25028 | 1252 | C1252G | D | 1.5e-05 | 1.6e-05 | 1.1e-05 | 1.400E-05 | 2.646E-06 | 1.100E-05 | 1.600E-05 |
| 25029 | 1252 | C1252H | D | 1.3e-05 | 1.2e-05 | 1.8e-05 | 1.433E-05 | 3.215E-06 | 1.200E-05 | 1.800E-05 |
| 25030 | 1252 | C1252I | D | 1.7e-05 | 1.6e-05 | 1e-05 | 1.433E-05 | 3.786E-06 | 1.000E-05 | 1.700E-05 |
| 25031 | 1252 | C1252L | D | 1.1e-05 | 1.6e-05 | 9.8e-06 | 1.227E-05 | 3.288E-06 | 9.800E-06 | 1.600E-05 |
| 25032 | 1252 | C1252K | D | 1.1e-05 | 1.6e-05 | 1.7e-05 | 1.467E-05 | 3.215E-06 | 1.100E-05 | 1.700E-05 |
| 25033 | 1252 | C1252M | D | 1.5e-05 | 9.6e-06 | 1.5e-05 | 1.320E-05 | 3.118E-06 | 9.600E-06 | 1.500E-05 |
| 25034 | 1252 | C1252F | D | 9.2e-06 | 1.5e-05 | 1.7e-05 | 1.373E-05 | 4.051E-06 | 9.200E-06 | 1.700E-05 |
| 25035 | 1252 | C1252P | D | 1.5e-05 | 1.2e-05 | 1.5e-05 | 1.400E-05 | 1.732E-06 | 1.200E-05 | 1.500E-05 |
| 25036 | 1252 | C1252S | D | 1.6e-05 | 8.5e-06 | 1.3e-05 | 1.250E-05 | 3.775E-06 | 8.500E-06 | 1.600E-05 |
| 25037 | 1252 | C1252T | D | 1.1e-05 | 1.6e-05 | 1.5e-05 | 1.400E-05 | 2.646E-06 | 1.100E-05 | 1.600E-05 |
| 25038 | 1252 | C1252W | D | 1.3e-05 | 1.5e-05 | 1.1e-05 | 1.300E-05 | 2.000E-06 | 1.100E-05 | 1.500E-05 |
| 25039 | 1252 | C1252Y | D | 7.8e-06 | 1.2e-05 | 9.3e-06 | 9.700E-06 | 2.128E-06 | 7.800E-06 | 1.200E-05 |
| 25040 | 1252 | C1252V | D | 1.5e-05 | 1.5e-05 | 1.6e-05 | 1.533E-05 | 5.774E-07 | 1.500E-05 | 1.600E-05 |
| 25041 | 1253 | E1253A | D | 1.5e-05 | 1.5e-05 | 1e-05 | 1.333E-05 | 2.887E-06 | 1.000E-05 | 1.500E-05 |
| 25042 | 1253 | E1253R | D | 1.6e-05 | 1.6e-05 | 1.5e-05 | 1.567E-05 | 5.774E-07 | 1.500E-05 | 1.600E-05 |
| 25043 | 1253 | E1253N | D | 1.6e-05 | 1e-05 | 1.5e-05 | 1.367E-05 | 3.215E-06 | 1.000E-05 | 1.600E-05 |
| 25044 | 1253 | E1253D | D | 1.1e-05 | 1.5e-05 | 1.6e-05 | 1.400E-05 | 2.646E-06 | 1.100E-05 | 1.600E-05 |
| 25045 | 1253 | E1253C | D | 9e-06 | 1e-05 | 1.3e-05 | 1.067E-05 | 2.082E-06 | 9.000E-06 | 1.300E-05 |
| 25046 | 1253 | E1253E | D | 1.6e-05 | 1.3e-05 | 8.1e-06 | 1.237E-05 | 3.988E-06 | 8.100E-06 | 1.600E-05 |
| 25047 | 1253 | E1253Q | D | 1.4e-05 | 1.1e-05 | 1.5e-05 | 1.333E-05 | 2.082E-06 | 1.100E-05 | 1.500E-05 |
| 25048 | 1253 | E1253G | D | 1.3e-05 | 1.3e-05 | 1.2e-05 | 1.267E-05 | 5.774E-07 | 1.200E-05 | 1.300E-05 |
| 25049 | 1253 | E1253H | D | 8.7e-06 | 8.1e-06 | 9.7e-06 | 8.833E-06 | 8.083E-07 | 8.100E-06 | 9.700E-06 |
| 25050 | 1253 | E1253I | D | 1.6e-05 | 9.8e-06 | 1.1e-05 | 1.227E-05 | 3.288E-06 | 9.800E-06 | 1.600E-05 |
| 25051 | 1253 | E1253L | D | 1e-05 | 8.5e-06 | 9.9e-06 | 9.467E-06 | 8.386E-07 | 8.500E-06 | 1.000E-05 |
| 25052 | 1253 | E1253K | D | 1.2e-05 | 9.1e-06 | 1.5e-05 | 1.203E-05 | 2.950E-06 | 9.100E-06 | 1.500E-05 |
| 25053 | 1253 | E1253M | D | 1e-05 | 1.4e-05 | 1.4e-05 | 1.267E-05 | 2.309E-06 | 1.000E-05 | 1.400E-05 |
| 25054 | 1253 | E1253F | D | 1.5e-05 | 1.5e-05 | 1.1e-05 | 1.367E-05 | 2.309E-06 | 1.100E-05 | 1.500E-05 |
| 25055 | 1253 | E1253P | D | 1.1e-05 | 1e-05 | 1.5e-05 | 1.200E-05 | 2.646E-06 | 1.000E-05 | 1.500E-05 |
| 25056 | 1253 | E1253S | D | 1e-05 | 1.5e-05 | 1.5e-05 | 1.333E-05 | 2.887E-06 | 1.000E-05 | 1.500E-05 |
| 25057 | 1253 | E1253T | D | 9.3e-06 | 1.4e-05 | 1.4e-05 | 1.243E-05 | 2.714E-06 | 9.300E-06 | 1.400E-05 |
| 25058 | 1253 | E1253W | D | 1.5e-05 | 1.1e-05 | 9.2e-06 | 1.173E-05 | 2.969E-06 | 9.200E-06 | 1.500E-05 |
| 25059 | 1253 | E1253Y | D | 9.7e-06 | 9.6e-06 | 1e-05 | 9.767E-06 | 2.082E-07 | 9.600E-06 | 1.000E-05 |
| 25060 | 1253 | E1253V | D | 1.5e-05 | 1.2e-05 | 1.3e-05 | 1.333E-05 | 1.528E-06 | 1.200E-05 | 1.500E-05 |
| 25061 | 1254 | V1254A | D | 1e-05 | 1.6e-05 | 9.4e-06 | 1.180E-05 | 3.650E-06 | 9.400E-06 | 1.600E-05 |
| 25062 | 1254 | V1254R | D | 7.4e-06 | 1.5e-05 | 1.5e-05 | 1.247E-05 | 4.388E-06 | 7.400E-06 | 1.500E-05 |
| 25063 | 1254 | V1254N | D | 1.2e-05 | 1.4e-05 | 2e-05 | 1.533E-05 | 4.163E-06 | 1.200E-05 | 2.000E-05 |
| 25064 | 1254 | V1254D | D | 1.4e-05 | 1.6e-05 | 1.7e-05 | 1.567E-05 | 1.528E-06 | 1.400E-05 | 1.700E-05 |
| 25065 | 1254 | V1254C | D | 1.1e-05 | 7.9e-06 | 1.5e-05 | 1.130E-05 | 3.559E-06 | 7.900E-06 | 1.500E-05 |
| 25066 | 1254 | V1254E | D | 9.1e-06 | 1.2e-05 | 1.2e-05 | 1.103E-05 | 1.674E-06 | 9.100E-06 | 1.200E-05 |
| 25067 | 1254 | V1254Q | D | 1.5e-05 | 8.1e-06 | 1.4e-05 | 1.237E-05 | 3.729E-06 | 8.100E-06 | 1.500E-05 |
| 25068 | 1254 | V1254G | D | 1.5e-05 | 1.8e-05 | 1.6e-05 | 1.633E-05 | 1.528E-06 | 1.500E-05 | 1.800E-05 |
| 25069 | 1254 | V1254H | D | 9.3e-06 | 9.5e-06 | 1.4e-05 | 1.093E-05 | 2.658E-06 | 9.300E-06 | 1.400E-05 |
| 25070 | 1254 | V1254I | D | 1.3e-05 | 1.4e-05 | 1e-05 | 1.233E-05 | 2.082E-06 | 1.000E-05 | 1.400E-05 |
| 25071 | 1254 | V1254L | D | 9.9e-06 | 1.1e-05 | 1.6e-05 | 1.230E-05 | 3.251E-06 | 9.900E-06 | 1.600E-05 |
| 25072 | 1254 | V1254K | D | 9.4e-06 | 1.3e-05 | 1e-05 | 1.080E-05 | 1.929E-06 | 9.400E-06 | 1.300E-05 |
| 25073 | 1254 | V1254M | D | 1.3e-05 | 1.5e-05 | 1.3e-05 | 1.367E-05 | 1.155E-06 | 1.300E-05 | 1.500E-05 |
| 25074 | 1254 | V1254F | D | 1e-05 | 1.4e-05 | 1.6e-05 | 1.333E-05 | 3.055E-06 | 1.000E-05 | 1.600E-05 |

| | | | | | | | | | | |
|---|---|---|---|---|---|---|---|---|---|---|
| 25075 | 1254 | V1254P | D | 1.1e-05 | 1.1e-05 | 1.4e-05 | 1.200E-05 | 1.732E-06 | 1.100E-05 | 1.400E-05 |
| 25076 | 1254 | V1254S | D | 1.1e-05 | 1.5e-05 | 1.6e-05 | 1.400E-05 | 2.646E-06 | 1.100E-05 | 1.600E-05 |
| 25077 | 1254 | V1254T | D | 9.4e-06 | 1.5e-05 | 1.4e-05 | 1.280E-05 | 2.987E-06 | 9.400E-06 | 1.500E-05 |
| 25078 | 1254 | V1254W | D | 1.2e-05 | 1.2e-05 | 1.6e-05 | 1.333E-05 | 2.309E-06 | 1.200E-05 | 1.600E-05 |
| 25079 | 1254 | V1254Y | D | 8.4e-06 | 9.2e-06 | 1.1e-05 | 9.533E-06 | 1.332E-06 | 8.400E-06 | 1.100E-05 |
| 25080 | 1254 | V1254V | D | 1.3e-05 | 1.3e-05 | 1.4e-05 | 1.333E-05 | 5.774E-07 | 1.300E-05 | 1.400E-05 |
| 25081 | 1255 | T1255A | D | 1.6e-05 | 1.6e-05 | 1e-05 | 1.400E-05 | 3.464E-06 | 1.000E-05 | 1.600E-05 |
| 25082 | 1255 | T1255R | D | 9.6e-06 | 1.4e-05 | 1.4e-05 | 1.253E-05 | 2.540E-06 | 9.600E-06 | 1.400E-05 |
| 25083 | 1255 | T1255N | D | 1.7e-05 | 1.3e-05 | 1.5e-05 | 1.500E-05 | 2.000E-06 | 1.300E-05 | 1.700E-05 |
| 25084 | 1255 | T1255D | D | 1.3e-05 | 1.5e-05 | 1.5e-05 | 1.433E-05 | 1.155E-06 | 1.300E-05 | 1.500E-05 |
| 25085 | 1255 | T1255C | D | 1.3e-05 | 1e-05 | 1.1e-05 | 1.133E-05 | 1.528E-06 | 1.000E-05 | 1.300E-05 |
| 25086 | 1255 | T1255E | D | 1.4e-05 | 1.2e-05 | 1.8e-05 | 1.467E-05 | 3.055E-06 | 1.200E-05 | 1.800E-05 |
| 25087 | 1255 | T1255Q | D | 1.3e-05 | 1.3e-05 | 9.7e-06 | 1.190E-05 | 1.905E-06 | 9.700E-06 | 1.300E-05 |
| 25088 | 1255 | T1255G | D | 1.1e-05 | 8.5e-06 | 1.5e-05 | 1.150E-05 | 3.279E-06 | 8.500E-06 | 1.500E-05 |
| 25089 | 1255 | T1255H | D | 1.8e-05 | 9.6e-06 | 1.4e-05 | 1.387E-05 | 4.202E-06 | 9.600E-06 | 1.800E-05 |
| 25090 | 1255 | T1255I | D | 1.7e-05 | 1.6e-05 | 1.1e-05 | 1.467E-05 | 3.215E-06 | 1.100E-05 | 1.700E-05 |
| 25091 | 1255 | T1255L | D | 9.3e-06 | 9.5e-06 | 1.3e-05 | 1.060E-05 | 2.081E-06 | 9.300E-06 | 1.300E-05 |
| 25092 | 1255 | T1255K | D | 1.3e-05 | 7.8e-06 | 1.2e-05 | 1.093E-05 | 2.759E-06 | 7.800E-06 | 1.300E-05 |
| 25093 | 1255 | T1255M | D | 9.6e-06 | 1.6e-05 | 1.2e-05 | 1.253E-05 | 3.233E-06 | 9.600E-06 | 1.600E-05 |
| 25094 | 1255 | T1255F | D | 9.1e-06 | 1.4e-05 | 1.1e-05 | 1.137E-05 | 2.470E-06 | 9.100E-06 | 1.400E-05 |
| 25095 | 1255 | T1255P | D | 1.2e-05 | 1.5e-05 | 1e-05 | 1.233E-05 | 2.517E-06 | 1.000E-05 | 1.500E-05 |
| 25096 | 1255 | T1255S | D | 1.2e-05 | 1.5e-05 | 9.9e-06 | 1.230E-05 | 2.563E-06 | 9.900E-06 | 1.500E-05 |
| 25097 | 1255 | T1255T | D | 1.4e-05 | 1.5e-05 | 1e-05 | 1.300E-05 | 2.646E-06 | 1.000E-05 | 1.500E-05 |
| 25098 | 1255 | T1255W | D | 1.5e-05 | 1.1e-05 | 8.2e-06 | 1.140E-05 | 3.418E-06 | 8.200E-06 | 1.500E-05 |
| 25099 | 1255 | T1255Y | D | 9.2e-06 | 1.3e-05 | 9.6e-06 | 1.060E-05 | 2.088E-06 | 9.200E-06 | 1.300E-05 |
| 25100 | 1255 | T1255V | D | 1.2e-05 | 1.6e-05 | 1.4e-05 | 1.400E-05 | 2.000E-06 | 1.200E-05 | 1.600E-05 |
| 25101 | 1256 | H1256A | D | 8.8e-06 | 1.6e-05 | 1.1e-05 | 1.193E-05 | 3.690E-06 | 8.800E-06 | 1.600E-05 |
| 25102 | 1256 | H1256R | D | 9e-06 | 1e-05 | 9.5e-06 | 9.500E-06 | 5.000E-07 | 9.000E-06 | 1.000E-05 |
| 25103 | 1256 | H1256N | D | 1.5e-05 | 1.4e-05 | 1.5e-05 | 1.467E-05 | 5.774E-07 | 1.400E-05 | 1.500E-05 |
| 25104 | 1256 | H1256D | D | 1.5e-05 | 1.1e-05 | 8.6e-06 | 1.153E-05 | 3.233E-06 | 8.600E-06 | 1.500E-05 |
| 25105 | 1256 | H1256C | D | 1.4e-05 | 1.4e-05 | 1.6e-05 | 1.467E-05 | 1.155E-06 | 1.400E-05 | 1.600E-05 |
| 25106 | 1256 | H1256E | D | 1.6e-05 | 1.1e-05 | 1.5e-05 | 1.400E-05 | 2.646E-06 | 1.100E-05 | 1.600E-05 |
| 25107 | 1256 | H1256Q | D | 1.4e-05 | 1.1e-05 | 1e-05 | 1.167E-05 | 2.082E-06 | 1.000E-05 | 1.400E-05 |
| 25108 | 1256 | H1256G | D | 1.7e-05 | 9.7e-06 | 9.4e-06 | 1.203E-05 | 4.304E-06 | 9.400E-06 | 1.700E-05 |
| 25109 | 1256 | H1256H | D | 1.5e-05 | 8.9e-06 | 1.3e-05 | 1.230E-05 | 3.110E-06 | 8.900E-06 | 1.500E-05 |
| 25110 | 1256 | H1256I | D | 1.4e-05 | 1.4e-05 | 1.1e-05 | 1.300E-05 | 1.732E-06 | 1.100E-05 | 1.400E-05 |
| 25111 | 1256 | H1256L | D | 1.5e-05 | 1.2e-05 | 1.6e-05 | 1.433E-05 | 2.082E-06 | 1.200E-05 | 1.600E-05 |
| 25112 | 1256 | H1256K | D | 1.4e-05 | 1.3e-05 | 1.1e-05 | 1.267E-05 | 1.528E-06 | 1.100E-05 | 1.400E-05 |
| 25113 | 1256 | H1256M | D | 8e-06 | 1.7e-05 | 1e-05 | 1.167E-05 | 4.726E-06 | 8.000E-06 | 1.700E-05 |
| 25114 | 1256 | H1256F | D | 1.3e-05 | 1.5e-05 | 1.7e-05 | 1.500E-05 | 2.000E-06 | 1.300E-05 | 1.700E-05 |
| 25115 | 1256 | H1256P | D | 1.6e-05 | 1.5e-05 | 1.8e-05 | 1.633E-05 | 1.528E-06 | 1.500E-05 | 1.800E-05 |
| 25116 | 1256 | H1256S | D | 1.5e-05 | 1.4e-05 | 1.1e-05 | 1.333E-05 | 2.082E-06 | 1.100E-05 | 1.500E-05 |
| 25117 | 1256 | H1256T | D | 1e-05 | 1.1e-05 | 1.4e-05 | 1.167E-05 | 2.082E-06 | 1.000E-05 | 1.400E-05 |
| 25118 | 1256 | H1256W | D | 1.3e-05 | 1.4e-05 | 1.6e-05 | 1.433E-05 | 1.528E-06 | 1.300E-05 | 1.600E-05 |
| 25119 | 1256 | H1256Y | D | 7.8e-06 | 1.1e-05 | 9.5e-06 | 9.433E-06 | 1.601E-06 | 7.800E-06 | 1.100E-05 |
| 25120 | 1256 | H1256V | D | 1.5e-05 | 1.2e-05 | 1.5e-05 | 1.400E-05 | 1.732E-06 | 1.200E-05 | 1.500E-05 |
| 25121 | 1257 | Q1257A | D | 1.7e-05 | 1.5e-05 | 1.2e-05 | 1.467E-05 | 2.517E-06 | 1.200E-05 | 1.700E-05 |
| 25122 | 1257 | Q1257R | D | 1e-05 | 1.3e-05 | 9.1e-06 | 1.070E-05 | 2.042E-06 | 9.100E-06 | 1.300E-05 |
| 25123 | 1257 | Q1257N | D | 1.1e-05 | 1.6e-05 | 1.6e-05 | 1.433E-05 | 2.887E-06 | 1.100E-05 | 1.600E-05 |
| 25124 | 1257 | Q1257D | D | 1.3e-05 | 1e-05 | 1e-05 | 1.100E-05 | 1.732E-06 | 1.000E-05 | 1.300E-05 |
| 25125 | 1257 | Q1257C | D | 1.2e-05 | 7.5e-06 | 1.1e-05 | 1.017E-05 | 2.363E-06 | 7.500E-06 | 1.200E-05 |
| 25126 | 1257 | Q1257E | D | 1.5e-05 | 1.5e-05 | 8.4e-06 | 1.280E-05 | 3.811E-06 | 8.400E-06 | 1.500E-05 |
| 25127 | 1257 | Q1257Q | D | 1.6e-05 | 1.3e-05 | 1.5e-05 | 1.467E-05 | 1.528E-06 | 1.300E-05 | 1.600E-05 |
| 25128 | 1257 | Q1257G | D | 1.5e-05 | 2.2e-05 | 1.5e-05 | 1.733E-05 | 4.041E-06 | 1.500E-05 | 2.200E-05 |
| 25129 | 1257 | Q1257H | D | 1.5e-05 | 1.6e-05 | 1.3e-05 | 1.467E-05 | 1.528E-06 | 1.300E-05 | 1.600E-05 |
| 25130 | 1257 | Q1257I | D | 1.6e-05 | 1e-05 | 1e-05 | 1.200E-05 | 3.464E-06 | 1.000E-05 | 1.600E-05 |
| 25131 | 1257 | Q1257L | D | 8.4e-06 | 1e-05 | 1.6e-05 | 1.147E-05 | 4.007E-06 | 8.400E-06 | 1.600E-05 |
| 25132 | 1257 | Q1257K | D | 1.4e-05 | 1.1e-05 | 1.1e-05 | 1.200E-05 | 1.732E-06 | 1.100E-05 | 1.400E-05 |
| 25133 | 1257 | Q1257M | D | 1.4e-05 | 1.1e-05 | 1.1e-05 | 1.200E-05 | 1.732E-06 | 1.100E-05 | 1.400E-05 |
| 25134 | 1257 | Q1257F | D | 1.2e-05 | 1.2e-05 | 1.2e-05 | 1.200E-05 | 0.000E+00 | 1.200E-05 | 1.200E-05 |
| 25135 | 1257 | Q1257P | D | 1.7e-05 | 1.6e-05 | 1.4e-05 | 1.567E-05 | 1.528E-06 | 1.400E-05 | 1.700E-05 |
| 25136 | 1257 | Q1257S | D | 1.4e-05 | 1.4e-05 | 1.1e-05 | 1.300E-05 | 1.732E-06 | 1.100E-05 | 1.400E-05 |
| 25137 | 1257 | Q1257T | D | 1.1e-05 | 1.5e-05 | 1.5e-05 | 1.367E-05 | 2.309E-06 | 1.100E-05 | 1.500E-05 |
| 25138 | 1257 | Q1257W | D | 1.5e-05 | 1.5e-05 | 1.6e-05 | 1.533E-05 | 5.774E-07 | 1.500E-05 | 1.600E-05 |
| 25139 | 1257 | Q1257Y | D | 8.7e-06 | 8.9e-06 | 1e-05 | 9.200E-06 | 7.000E-07 | 8.700E-06 | 1.000E-05 |
| 25140 | 1257 | Q1257V | D | 1.6e-05 | 1.5e-05 | 1.2e-05 | 1.433E-05 | 2.082E-06 | 1.200E-05 | 1.600E-05 |
| 25141 | 1258 | G1258A | D | 1.6e-05 | 1.3e-05 | 1.2e-05 | 1.367E-05 | 2.082E-06 | 1.200E-05 | 1.600E-05 |
| 25142 | 1258 | G1258R | D | 1.4e-05 | 1.3e-05 | 1e-05 | 1.233E-05 | 2.082E-06 | 1.000E-05 | 1.400E-05 |
| 25143 | 1258 | G1258N | D | 8.5e-06 | 1.4e-05 | 1e-05 | 1.083E-05 | 2.843E-06 | 8.500E-06 | 1.400E-05 |
| 25144 | 1258 | G1258D | D | 8.5e-06 | 1.1e-05 | 1e-05 | 9.833E-06 | 1.258E-06 | 8.500E-06 | 1.100E-05 |
| 25145 | 1258 | G1258C | D | 1.4e-05 | 9.6e-06 | 1.2e-05 | 1.187E-05 | 2.203E-06 | 9.600E-06 | 1.400E-05 |
| 25146 | 1258 | G1258E | D | 1.1e-05 | 1.3e-05 | 2.5e-05 | 1.633E-05 | 7.572E-06 | 1.100E-05 | 2.500E-05 |
| 25147 | 1258 | G1258Q | D | 1.5e-05 | 1.4e-05 | 1.3e-05 | 1.400E-05 | 1.000E-06 | 1.300E-05 | 1.500E-05 |
| 25148 | 1258 | G1258G | D | 1.5e-05 | 1.2e-05 | 1.5e-05 | 1.400E-05 | 1.732E-06 | 1.200E-05 | 1.500E-05 |
| 25149 | 1258 | G1258H | D | 9.7e-06 | 1.2e-05 | 8.8e-06 | 1.017E-05 | 1.650E-06 | 8.800E-06 | 1.200E-05 |
| 25150 | 1258 | G1258I | D | 1.4e-05 | 1.3e-05 | 1.5e-05 | 1.400E-05 | 1.000E-06 | 1.300E-05 | 1.500E-05 |
| 25151 | 1258 | G1258L | D | 1.1e-05 | 1.5e-05 | 9.6e-06 | 1.187E-05 | 2.802E-06 | 9.600E-06 | 1.500E-05 |
| 25152 | 1258 | G1258K | D | 1.5e-05 | 1.2e-05 | 1.5e-05 | 1.400E-05 | 1.732E-06 | 1.200E-05 | 1.500E-05 |
| 25153 | 1258 | G1258M | D | 8.1e-06 | 1.1e-05 | 1.1e-05 | 1.003E-05 | 1.674E-06 | 8.100E-06 | 1.100E-05 |
| 25154 | 1258 | G1258F | D | 6.8e-06 | 1.4e-05 | 1.5e-05 | 1.193E-05 | 4.474E-06 | 6.800E-06 | 1.500E-05 |
| 25155 | 1258 | G1258P | D | 1e-05 | 1.5e-05 | 1.4e-05 | 1.300E-05 | 2.646E-06 | 1.000E-05 | 1.500E-05 |
| 25156 | 1258 | G1258S | D | 1e-05 | 1.4e-05 | 1.5e-05 | 1.300E-05 | 2.646E-06 | 1.000E-05 | 1.500E-05 |
| 25157 | 1258 | G1258T | D | 1.4e-05 | 1.3e-05 | 1.5e-05 | 1.400E-05 | 1.000E-06 | 1.300E-05 | 1.500E-05 |
| 25158 | 1258 | G1258W | D | 1.3e-05 | 1.2e-05 | 1.1e-05 | 1.200E-05 | 1.000E-06 | 1.100E-05 | 1.300E-05 |
| 25159 | 1258 | G1258Y | D | 9.7e-06 | 1.5e-05 | 8.8e-06 | 1.117E-05 | 3.350E-06 | 8.800E-06 | 1.500E-05 |

| | | | | | | | | | | |
|---|---|---|---|---|---|---|---|---|---|---|
| 25160 | 1258 | G1258V | D | 9.4e-06 | 1.4e-05 | 1.1e-05 | 1.147E-05 | 2.335E-06 | 9.400E-06 | 1.400E-05 |
| 25161 | 1259 | L1259A | D | 1.4e-05 | 1.1e-05 | 1.1e-05 | 1.200E-05 | 1.732E-06 | 1.100E-05 | 1.400E-05 |
| 25162 | 1259 | L1259R | D | 9.4e-06 | 1.5e-05 | 9.1e-06 | 1.117E-05 | 3.323E-06 | 9.100E-06 | 1.500E-05 |
| 25163 | 1259 | L1259N | D | 1.5e-05 | 1.5e-05 | 1.6e-05 | 1.533E-05 | 5.774E-07 | 1.500E-05 | 1.600E-05 |
| 25164 | 1259 | L1259D | D | 1.6e-05 | 7.2e-06 | 1.2e-05 | 1.173E-05 | 4.406E-06 | 7.200E-06 | 1.600E-05 |
| 25165 | 1259 | L1259C | D | 1.5e-05 | 1.1e-05 | 1.5e-05 | 1.367E-05 | 2.309E-06 | 1.100E-05 | 1.500E-05 |
| 25166 | 1259 | L1259E | D | 9.2e-06 | 1.1e-05 | 1.3e-05 | 1.107E-05 | 1.901E-06 | 9.200E-06 | 1.300E-05 |
| 25167 | 1259 | L1259Q | D | 8.1e-06 | 9.9e-06 | 1.1e-05 | 9.667E-06 | 1.464E-06 | 8.100E-06 | 1.100E-05 |
| 25168 | 1259 | L1259G | D | 1e-05 | 1.5e-05 | 1.2e-05 | 1.233E-05 | 2.517E-06 | 1.000E-05 | 1.500E-05 |
| 25169 | 1259 | L1259H | D | 1.4e-05 | 1.6e-05 | 1.5e-05 | 1.500E-05 | 1.000E-06 | 1.400E-05 | 1.600E-05 |
| 25170 | 1259 | L1259I | D | 1.3e-05 | 1e-05 | 1.6e-05 | 1.300E-05 | 3.000E-06 | 1.000E-05 | 1.600E-05 |
| 25171 | 1259 | L1259L | D | 1e-05 | 1.6e-05 | 1.1e-05 | 1.233E-05 | 3.215E-06 | 1.000E-05 | 1.600E-05 |
| 25172 | 1259 | L1259K | D | 9.9e-06 | 1.1e-05 | 1.5e-05 | 1.197E-05 | 2.684E-06 | 9.900E-06 | 1.500E-05 |
| 25173 | 1259 | L1259M | D | 1.3e-05 | 1.7e-05 | 1.5e-05 | 1.500E-05 | 2.000E-06 | 1.300E-05 | 1.700E-05 |
| 25174 | 1259 | L1259F | D | 1.1e-05 | 1.1e-05 | 1.6e-05 | 1.267E-05 | 2.887E-06 | 1.100E-05 | 1.600E-05 |
| 25175 | 1259 | L1259P | D | 1.5e-05 | 9e-06 | 1.5e-05 | 1.300E-05 | 3.464E-06 | 9.000E-06 | 1.500E-05 |
| 25176 | 1259 | L1259S | D | 1.8e-05 | 1.5e-05 | 1.3e-05 | 1.533E-05 | 2.517E-06 | 1.300E-05 | 1.800E-05 |
| 25177 | 1259 | L1259T | D | 1.4e-05 | 1.6e-05 | 1.6e-05 | 1.533E-05 | 1.155E-06 | 1.400E-05 | 1.600E-05 |
| 25178 | 1259 | L1259W | D | 7.9e-06 | 1.5e-05 | 1.6e-05 | 1.297E-05 | 4.416E-06 | 7.900E-06 | 1.600E-05 |
| 25179 | 1259 | L1259Y | D | 1.3e-05 | 1.1e-05 | 1.2e-05 | 1.200E-05 | 1.000E-06 | 1.100E-05 | 1.300E-05 |
| 25180 | 1259 | L1259V | D | 1.6e-05 | 1.2e-05 | 8.9e-06 | 1.230E-05 | 3.559E-06 | 8.900E-06 | 1.600E-05 |
| 25181 | 1260 | S1260A | D | 1.8e-05 | 1.3e-05 | 1.4e-05 | 1.500E-05 | 2.646E-06 | 1.300E-05 | 1.800E-05 |
| 25182 | 1260 | S1260R | D | 1.6e-05 | 9.8e-06 | 1.1e-05 | 1.227E-05 | 3.288E-06 | 9.800E-06 | 1.600E-05 |
| 25183 | 1260 | S1260N | D | 1.4e-05 | 1.4e-05 | 1e-05 | 1.267E-05 | 2.309E-06 | 1.000E-05 | 1.400E-05 |
| 25184 | 1260 | S1260D | D | 1.4e-05 | 1e-05 | 1.3e-05 | 1.233E-05 | 2.082E-06 | 1.000E-05 | 1.400E-05 |
| 25185 | 1260 | S1260C | D | 1.5e-05 | 8.9e-06 | 7.6e-06 | 1.050E-05 | 3.951E-06 | 7.600E-06 | 1.500E-05 |
| 25186 | 1260 | S1260E | D | 1.6e-05 | 1.3e-05 | 1.6e-05 | 1.500E-05 | 1.732E-06 | 1.300E-05 | 1.600E-05 |
| 25187 | 1260 | S1260Q | D | 1.3e-05 | 1.4e-05 | 1.5e-05 | 1.400E-05 | 1.000E-06 | 1.300E-05 | 1.500E-05 |
| 25188 | 1260 | S1260G | D | 1.3e-05 | 1.5e-05 | 6.4e-06 | 1.147E-05 | 4.500E-06 | 6.400E-06 | 1.500E-05 |
| 25189 | 1260 | S1260H | D | 1.3e-05 | 9.8e-06 | 1.1e-05 | 1.127E-05 | 1.617E-06 | 9.800E-06 | 1.300E-05 |
| 25190 | 1260 | S1260I | D | 1.1e-05 | 1.8e-05 | 1.3e-05 | 1.400E-05 | 3.606E-06 | 1.100E-05 | 1.800E-05 |
| 25191 | 1260 | S1260L | D | 1e-05 | 1.1e-05 | 1.1e-05 | 1.067E-05 | 5.774E-07 | 1.000E-05 | 1.100E-05 |
| 25192 | 1260 | S1260K | D | 1.6e-05 | 1.5e-05 | 1.6e-05 | 1.567E-05 | 5.774E-07 | 1.500E-05 | 1.600E-05 |
| 25193 | 1260 | S1260M | D | 1.2e-05 | 1.5e-05 | 1.4e-05 | 1.367E-05 | 1.528E-06 | 1.200E-05 | 1.500E-05 |
| 25194 | 1260 | S1260F | D | 1e-05 | 1.1e-05 | 1.1e-05 | 1.067E-05 | 5.774E-07 | 1.000E-05 | 1.100E-05 |
| 25195 | 1260 | S1260P | D | 1.5e-05 | 1.3e-05 | 1.1e-05 | 1.300E-05 | 2.000E-06 | 1.100E-05 | 1.500E-05 |
| 25196 | 1260 | S1260S | D | 1.2e-05 | 1.5e-05 | 1.4e-05 | 1.367E-05 | 1.528E-06 | 1.200E-05 | 1.500E-05 |
| 25197 | 1260 | S1260T | D | 1.1e-05 | 1e-05 | 2.3e-05 | 1.467E-05 | 7.234E-06 | 1.000E-05 | 2.300E-05 |
| 25198 | 1260 | S1260W | D | 1.6e-05 | 1.8e-05 | 1.5e-05 | 1.633E-05 | 1.528E-06 | 1.500E-05 | 1.800E-05 |
| 25199 | 1260 | S1260Y | D | 8.7e-06 | 8.9e-06 | 8.6e-06 | 8.733E-06 | 1.528E-07 | 8.600E-06 | 8.900E-06 |
| 25200 | 1260 | S1260V | D | 1.4e-05 | 1.1e-05 | 9.9e-06 | 1.163E-05 | 2.122E-06 | 9.900E-06 | 1.400E-05 |
| 25201 | 1261 | S1261A | D | 1.5e-05 | 1e-05 | 1.6e-05 | 1.367E-05 | 3.215E-06 | 1.000E-05 | 1.600E-05 |
| 25202 | 1261 | S1261R | D | 8.9e-06 | 9.3e-06 | 1.6e-05 | 1.140E-05 | 3.989E-06 | 8.900E-06 | 1.600E-05 |
| 25203 | 1261 | S1261N | D | 1e-05 | 1.4e-05 | 1.1e-05 | 1.167E-05 | 2.082E-06 | 1.000E-05 | 1.400E-05 |
| 25204 | 1261 | S1261D | D | 1.5e-05 | 1.4e-05 | 1.7e-05 | 1.533E-05 | 1.528E-06 | 1.400E-05 | 1.700E-05 |
| 25205 | 1261 | S1261C | D | 1.4e-05 | 1.3e-05 | 1e-05 | 1.233E-05 | 2.082E-06 | 1.000E-05 | 1.400E-05 |
| 25206 | 1261 | S1261E | D | 1.1e-05 | 1.4e-05 | 1.7e-05 | 1.400E-05 | 3.000E-06 | 1.100E-05 | 1.700E-05 |
| 25207 | 1261 | S1261Q | D | 9.9e-06 | 1.4e-05 | 1.2e-05 | 1.197E-05 | 2.050E-06 | 9.900E-06 | 1.400E-05 |
| 25208 | 1261 | S1261G | D | 9e-06 | 1.6e-05 | 1.1e-05 | 1.200E-05 | 3.606E-06 | 9.000E-06 | 1.600E-05 |
| 25209 | 1261 | S1261H | D | 1.3e-05 | 1.6e-05 | 1e-05 | 1.300E-05 | 3.000E-06 | 1.000E-05 | 1.600E-05 |
| 25210 | 1261 | S1261I | D | 1.1e-05 | 8.8e-06 | 1.4e-05 | 1.127E-05 | 2.610E-06 | 8.800E-06 | 1.400E-05 |
| 25211 | 1261 | S1261L | D | 1e-05 | 1.7e-05 | 1.2e-05 | 1.300E-05 | 3.606E-06 | 1.000E-05 | 1.700E-05 |
| 25212 | 1261 | S1261K | D | 1.6e-05 | 7.8e-06 | 1.6e-05 | 1.327E-05 | 4.734E-06 | 7.800E-06 | 1.600E-05 |
| 25213 | 1261 | S1261M | D | 8.8e-06 | 1.1e-05 | 1.3e-05 | 1.093E-05 | 2.101E-06 | 8.800E-06 | 1.300E-05 |
| 25214 | 1261 | S1261F | D | 1.1e-05 | 1.2e-05 | 1.1e-05 | 1.133E-05 | 5.774E-07 | 1.100E-05 | 1.200E-05 |
| 25215 | 1261 | S1261P | D | 1.7e-05 | 1.8e-05 | 1.6e-05 | 1.700E-05 | 1.000E-06 | 1.600E-05 | 1.800E-05 |
| 25216 | 1261 | S1261S | D | 1.1e-05 | 1.5e-05 | 1.1e-05 | 1.233E-05 | 2.309E-06 | 1.100E-05 | 1.500E-05 |
| 25217 | 1261 | S1261T | D | 1.4e-05 | 1.4e-05 | 1.2e-05 | 1.333E-05 | 1.155E-06 | 1.200E-05 | 1.400E-05 |
| 25218 | 1261 | S1261W | D | 1.6e-05 | 1.5e-05 | 1.5e-05 | 1.533E-05 | 5.774E-07 | 1.500E-05 | 1.600E-05 |
| 25219 | 1261 | S1261Y | D | 1.3e-05 | 1.5e-05 | 1.2e-05 | 1.333E-05 | 1.528E-06 | 1.200E-05 | 1.500E-05 |
| 25220 | 1261 | S1261V | D | 1.4e-05 | 1.2e-05 | 1.3e-05 | 1.300E-05 | 1.000E-06 | 1.200E-05 | 1.400E-05 |
| 25221 | 1262 | P1262A | D | 1.3e-05 | 1e-05 | 9.1e-06 | 1.070E-05 | 2.042E-06 | 9.100E-06 | 1.300E-05 |
| 25222 | 1262 | P1262R | D | 1e-05 | 1.4e-05 | 1.1e-05 | 1.167E-05 | 2.082E-06 | 1.000E-05 | 1.400E-05 |
| 25223 | 1262 | P1262N | D | 1.4e-05 | 9.7e-06 | 1.5e-05 | 1.290E-05 | 2.816E-06 | 9.700E-06 | 1.500E-05 |
| 25224 | 1262 | P1262D | D | 1.6e-05 | 1.3e-05 | 1.5e-05 | 1.467E-05 | 1.528E-06 | 1.300E-05 | 1.600E-05 |
| 25225 | 1262 | P1262C | D | 1e-05 | 1.2e-05 | 8.3e-06 | 1.010E-05 | 1.852E-06 | 8.300E-06 | 1.200E-05 |
| 25226 | 1262 | P1262E | D | 1.3e-05 | 2e-05 | 1.5e-05 | 1.600E-05 | 3.606E-06 | 1.300E-05 | 2.000E-05 |
| 25227 | 1262 | P1262Q | D | 9.5e-06 | 1e-05 | 1.4e-05 | 1.117E-05 | 2.466E-06 | 9.500E-06 | 1.400E-05 |
| 25228 | 1262 | P1262G | D | 9.7e-06 | 1.2e-05 | 1.1e-05 | 1.090E-05 | 1.153E-06 | 9.700E-06 | 1.200E-05 |
| 25229 | 1262 | P1262H | D | 1.1e-05 | 9.5e-06 | 1.1e-05 | 1.050E-05 | 8.660E-07 | 9.500E-06 | 1.100E-05 |
| 25230 | 1262 | P1262I | D | 1.1e-05 | 1.2e-05 | 1.6e-05 | 1.300E-05 | 2.646E-06 | 1.100E-05 | 1.600E-05 |
| 25231 | 1262 | P1262L | D | 1e-05 | 1.5e-05 | 1.6e-05 | 1.367E-05 | 3.215E-06 | 1.000E-05 | 1.600E-05 |
| 25232 | 1262 | P1262K | D | 1.6e-05 | 1.1e-05 | 1.4e-05 | 1.367E-05 | 2.517E-06 | 1.100E-05 | 1.600E-05 |
| 25233 | 1262 | P1262M | D | 1.4e-05 | 1.4e-05 | 1e-05 | 1.267E-05 | 2.309E-06 | 1.000E-05 | 1.400E-05 |
| 25234 | 1262 | P1262F | D | 1.5e-05 | 1.2e-05 | 9.9e-06 | 1.230E-05 | 2.563E-06 | 9.900E-06 | 1.500E-05 |
| 25235 | 1262 | P1262P | D | 1.4e-05 | 1.3e-05 | 1.1e-05 | 1.267E-05 | 1.528E-06 | 1.100E-05 | 1.400E-05 |
| 25236 | 1262 | P1262S | D | 1.9e-05 | 8.6e-06 | 1.2e-05 | 1.320E-05 | 5.303E-06 | 8.600E-06 | 1.900E-05 |
| 25237 | 1262 | P1262T | D | 1.6e-05 | 1.3e-05 | 1.5e-05 | 1.467E-05 | 1.528E-06 | 1.300E-05 | 1.600E-05 |
| 25238 | 1262 | P1262W | D | 1e-05 | 9.7e-06 | 1.5e-05 | 1.157E-05 | 2.977E-06 | 9.700E-06 | 1.500E-05 |
| 25239 | 1262 | P1262Y | D | 1.4e-05 | 1.6e-05 | 1.4e-05 | 1.467E-05 | 1.155E-06 | 1.400E-05 | 1.600E-05 |
| 25240 | 1262 | P1262V | D | 1.7e-05 | 1.5e-05 | 1.2e-05 | 1.467E-05 | 2.517E-06 | 1.200E-05 | 1.700E-05 |
| 25241 | 1263 | V1263A | D | 1.5e-05 | 1.3e-05 | 9.5e-06 | 1.250E-05 | 2.784E-06 | 9.500E-06 | 1.500E-05 |
| 25242 | 1263 | V1263R | D | 1.6e-05 | 1.2e-05 | 1.2e-05 | 1.333E-05 | 2.309E-06 | 1.200E-05 | 1.600E-05 |
| 25243 | 1263 | V1263N | D | 1.5e-05 | 1.2e-05 | 1.6e-05 | 1.433E-05 | 2.082E-06 | 1.200E-05 | 1.600E-05 |
| 25244 | 1263 | V1263D | D | 1.5e-05 | 1.1e-05 | 1.4e-05 | 1.333E-05 | 2.082E-06 | 1.100E-05 | 1.500E-05 |

| | | | | | | | | | | |
|---|---|---|---|---|---|---|---|---|---|---|
| 25245 | 1263 | V1263C | D | 8.9e-06 | 8.7e-06 | 1.5e-05 | 1.087E-05 | 3.581E-06 | 8.700E-06 | 1.500E-05 |
| 25246 | 1263 | V1263E | D | 1.5e-05 | 1.1e-05 | 1.5e-05 | 1.367E-05 | 2.309E-06 | 1.100E-05 | 1.500E-05 |
| 25247 | 1263 | V1263Q | D | 1e-05 | 1.5e-05 | 1.5e-05 | 1.333E-05 | 2.887E-06 | 1.000E-05 | 1.500E-05 |
| 25248 | 1263 | V1263G | D | 1.5e-05 | 1e-05 | 1.6e-05 | 1.367E-05 | 3.215E-06 | 1.000E-05 | 1.600E-05 |
| 25249 | 1263 | V1263H | D | 1.3e-05 | 1.5e-05 | 1.6e-05 | 1.467E-05 | 1.528E-06 | 1.300E-05 | 1.600E-05 |
| 25250 | 1263 | V1263I | D | 1.4e-05 | 1.5e-05 | 1.6e-05 | 1.500E-05 | 1.000E-06 | 1.400E-05 | 1.600E-05 |
| 25251 | 1263 | V1263L | D | 8.5e-06 | 1.6e-05 | 8.8e-06 | 1.110E-05 | 4.246E-06 | 8.500E-06 | 1.600E-05 |
| 25252 | 1263 | V1263K | D | 9.1e-06 | 1.5e-05 | 1.6e-05 | 1.337E-05 | 3.729E-06 | 9.100E-06 | 1.600E-05 |
| 25253 | 1263 | V1263M | D | 1.4e-05 | 1.4e-05 | 1.5e-05 | 1.433E-05 | 5.774E-07 | 1.400E-05 | 1.500E-05 |
| 25254 | 1263 | V1263F | D | 1e-05 | 1.4e-05 | 1e-05 | 1.133E-05 | 2.309E-06 | 1.000E-05 | 1.400E-05 |
| 25255 | 1263 | V1263P | D | 9.9e-06 | 1.7e-05 | 8.9e-06 | 1.193E-05 | 4.416E-06 | 8.900E-06 | 1.700E-05 |
| 25256 | 1263 | V1263S | D | 1.5e-05 | 1e-05 | 1.5e-05 | 1.333E-05 | 2.887E-06 | 1.000E-05 | 1.500E-05 |
| 25257 | 1263 | V1263T | D | 1.1e-05 | 9.3e-06 | 1.5e-05 | 1.177E-05 | 2.926E-06 | 9.300E-06 | 1.500E-05 |
| 25258 | 1263 | V1263W | D | 1.4e-05 | 1.1e-05 | 1.1e-05 | 1.200E-05 | 1.732E-06 | 1.100E-05 | 1.400E-05 |
| 25259 | 1263 | V1263Y | D | 1.2e-05 | 9.7e-06 | 8.7e-06 | 1.013E-05 | 1.692E-06 | 8.700E-06 | 1.200E-05 |
| 25260 | 1263 | V1263V | D | 1.1e-05 | 1.5e-05 | 1.5e-05 | 1.367E-05 | 2.309E-06 | 1.100E-05 | 1.500E-05 |
| 25261 | 1264 | T1264A | D | 1.5e-05 | 1.7e-05 | 1.6e-05 | 1.600E-05 | 1.000E-06 | 1.500E-05 | 1.700E-05 |
| 25262 | 1264 | T1264R | D | 8.1e-06 | 8.8e-06 | 8.4e-06 | 8.433E-06 | 3.512E-07 | 8.100E-06 | 8.800E-06 |
| 25263 | 1264 | T1264N | D | 9.8e-06 | 1.1e-05 | 9.3e-06 | 1.003E-05 | 8.737E-07 | 9.300E-06 | 1.100E-05 |
| 25264 | 1264 | T1264D | D | 1.4e-05 | 1.7e-05 | 1.8e-05 | 1.633E-05 | 2.082E-06 | 1.400E-05 | 1.800E-05 |
| 25265 | 1264 | T1264C | D | 9.6e-06 | 1.5e-05 | 1.1e-05 | 1.187E-05 | 2.802E-06 | 9.600E-06 | 1.500E-05 |
| 25266 | 1264 | T1264E | D | 1.4e-05 | 9.3e-06 | 9.7e-06 | 1.100E-05 | 2.606E-06 | 9.300E-06 | 1.400E-05 |
| 25267 | 1264 | T1264Q | D | 8.2e-06 | 1.5e-05 | 1.1e-05 | 1.140E-05 | 3.418E-06 | 8.200E-06 | 1.500E-05 |
| 25268 | 1264 | T1264G | D | 1.1e-05 | 1.7e-05 | 1.5e-05 | 1.433E-05 | 3.055E-06 | 1.100E-05 | 1.700E-05 |
| 25269 | 1264 | T1264H | D | 1.5e-05 | 1.5e-05 | 7.3e-06 | 1.243E-05 | 4.446E-06 | 7.300E-06 | 1.500E-05 |
| 25270 | 1264 | T1264I | D | 1e-05 | 1.7e-05 | 1.6e-05 | 1.433E-05 | 3.786E-06 | 1.000E-05 | 1.700E-05 |
| 25271 | 1264 | T1264L | D | 1.6e-05 | 1.2e-05 | 9.1e-06 | 1.237E-05 | 3.465E-06 | 9.100E-06 | 1.600E-05 |
| 25272 | 1264 | T1264K | D | 1.3e-05 | 1.1e-05 | 1.5e-05 | 1.300E-05 | 2.000E-06 | 1.100E-05 | 1.500E-05 |
| 25273 | 1264 | T1264M | D | 1.8e-05 | 1.6e-05 | 1.6e-05 | 1.667E-05 | 1.155E-06 | 1.600E-05 | 1.800E-05 |
| 25274 | 1264 | T1264F | D | 1e-05 | 1.1e-05 | 1.3e-05 | 1.133E-05 | 1.528E-06 | 1.000E-05 | 1.300E-05 |
| 25275 | 1264 | T1264P | D | 1.4e-05 | 1.4e-05 | 1.6e-05 | 1.467E-05 | 1.155E-06 | 1.400E-05 | 1.600E-05 |
| 25276 | 1264 | T1264S | D | 1.1e-05 | 1.4e-05 | 1.2e-05 | 1.233E-05 | 1.528E-06 | 1.100E-05 | 1.400E-05 |
| 25277 | 1264 | T1264T | D | 1.5e-05 | 7.3e-06 | 1.1e-05 | 1.110E-05 | 3.851E-06 | 7.300E-06 | 1.500E-05 |
| 25278 | 1264 | T1264W | D | 1.3e-05 | 1.1e-05 | 1.6e-05 | 1.333E-05 | 2.517E-06 | 1.100E-05 | 1.600E-05 |
| 25279 | 1264 | T1264Y | D | 1.1e-05 | 1.6e-05 | 1.8e-05 | 1.500E-05 | 3.606E-06 | 1.100E-05 | 1.800E-05 |
| 25280 | 1264 | T1264V | D | 1.5e-05 | 1.6e-05 | 1e-05 | 1.367E-05 | 3.215E-06 | 1.000E-05 | 1.600E-05 |
| 25281 | 1265 | K1265A | D | 1.2e-05 | 9.7e-06 | 1.1e-05 | 1.090E-05 | 1.153E-06 | 9.700E-06 | 1.200E-05 |
| 25282 | 1265 | K1265R | D | 1.1e-05 | 1e-05 | 1.6e-05 | 1.233E-05 | 3.215E-06 | 1.000E-05 | 1.600E-05 |
| 25283 | 1265 | K1265N | D | 1.4e-05 | 1.5e-05 | 1.1e-05 | 1.333E-05 | 2.082E-06 | 1.100E-05 | 1.500E-05 |
| 25284 | 1265 | K1265D | D | 1e-05 | 9.3e-06 | 1.1e-05 | 1.010E-05 | 8.544E-07 | 9.300E-06 | 1.100E-05 |
| 25285 | 1265 | K1265C | D | 1.3e-05 | 1.5e-05 | 1.4e-05 | 1.400E-05 | 1.000E-06 | 1.300E-05 | 1.500E-05 |
| 25286 | 1265 | K1265E | D | 1e-05 | 1.6e-05 | 9.6e-06 | 1.187E-05 | 3.585E-06 | 9.600E-06 | 1.600E-05 |
| 25287 | 1265 | K1265Q | D | 1e-05 | 1e-05 | 1.6e-05 | 1.200E-05 | 3.464E-06 | 1.000E-05 | 1.600E-05 |
| 25288 | 1265 | K1265G | D | 1.3e-05 | 8.2e-06 | 1.4e-05 | 1.173E-05 | 3.101E-06 | 8.200E-06 | 1.400E-05 |
| 25289 | 1265 | K1265H | D | 1.6e-05 | 1.2e-05 | 2.5e-05 | 1.767E-05 | 6.658E-06 | 1.200E-05 | 2.500E-05 |
| 25290 | 1265 | K1265I | D | 1.2e-05 | 1e-05 | 1.4e-05 | 1.200E-05 | 2.000E-06 | 1.000E-05 | 1.400E-05 |
| 25291 | 1265 | K1265L | D | 1.1e-05 | 1.5e-05 | 1.5e-05 | 1.367E-05 | 2.309E-06 | 1.100E-05 | 1.500E-05 |
| 25292 | 1265 | K1265K | D | 1.1e-05 | 1e-05 | 1.1e-05 | 1.067E-05 | 5.774E-07 | 1.000E-05 | 1.100E-05 |
| 25293 | 1265 | K1265M | D | 1.1e-05 | 1.6e-05 | 9.3e-06 | 1.210E-05 | 3.483E-06 | 9.300E-06 | 1.600E-05 |
| 25294 | 1265 | K1265F | D | 1.4e-05 | 1.6e-05 | 1.1e-05 | 1.367E-05 | 2.517E-06 | 1.100E-05 | 1.600E-05 |
| 25295 | 1265 | K1265P | D | 1.6e-05 | 1.6e-05 | 1.2e-05 | 1.467E-05 | 2.309E-06 | 1.200E-05 | 1.600E-05 |
| 25296 | 1265 | K1265S | D | 1.3e-05 | 1.2e-05 | 1.6e-05 | 1.367E-05 | 2.082E-06 | 1.200E-05 | 1.600E-05 |
| 25297 | 1265 | K1265T | D | 1.3e-05 | 1.7e-05 | 1.1e-05 | 1.367E-05 | 3.055E-06 | 1.100E-05 | 1.700E-05 |
| 25298 | 1265 | K1265W | D | 1.5e-05 | 1.5e-05 | 1.3e-05 | 1.433E-05 | 1.155E-06 | 1.300E-05 | 1.500E-05 |
| 25299 | 1265 | K1265Y | D | 1.1e-05 | 1.5e-05 | 1.6e-05 | 1.400E-05 | 2.646E-06 | 1.100E-05 | 1.600E-05 |
| 25300 | 1265 | K1265V | D | 1.6e-05 | 2.1e-05 | 9.8e-06 | 1.560E-05 | 5.611E-06 | 9.800E-06 | 2.100E-05 |
| 25301 | 1266 | S1266A | D | 1.5e-05 | 1e-05 | 1.6e-05 | 1.367E-05 | 3.215E-06 | 1.000E-05 | 1.600E-05 |
| 25302 | 1266 | S1266R | D | 7.4e-06 | 1e-05 | 1.2e-05 | 9.800E-06 | 2.307E-06 | 7.400E-06 | 1.200E-05 |
| 25303 | 1266 | S1266N | D | 1e-05 | 1.5e-05 | 1.4e-05 | 1.300E-05 | 2.646E-06 | 1.000E-05 | 1.500E-05 |
| 25304 | 1266 | S1266D | D | 1.1e-05 | 1.5e-05 | 1.6e-05 | 1.400E-05 | 2.646E-06 | 1.100E-05 | 1.600E-05 |
| 25305 | 1266 | S1266C | D | 1.5e-05 | 1.5e-05 | 1.2e-05 | 1.400E-05 | 1.732E-06 | 1.200E-05 | 1.500E-05 |
| 25306 | 1266 | S1266E | D | 1e-05 | 1.4e-05 | 1.6e-05 | 1.333E-05 | 3.055E-06 | 1.000E-05 | 1.600E-05 |
| 25307 | 1266 | S1266Q | D | 1.2e-05 | 1e-05 | 1.6e-05 | 1.267E-05 | 3.055E-06 | 1.000E-05 | 1.600E-05 |
| 25308 | 1266 | S1266G | D | 1.1e-05 | 1.7e-05 | 1e-05 | 1.267E-05 | 3.786E-06 | 1.000E-05 | 1.700E-05 |
| 25309 | 1266 | S1266H | D | 1e-05 | 1.4e-05 | 1e-05 | 1.133E-05 | 2.309E-06 | 1.000E-05 | 1.400E-05 |
| 25310 | 1266 | S1266I | D | 1.2e-05 | 9.7e-06 | 1e-05 | 1.057E-05 | 1.250E-06 | 9.700E-06 | 1.200E-05 |
| 25311 | 1266 | S1266L | D | 1.1e-05 | 1.2e-05 | 9e-06 | 1.067E-05 | 1.528E-06 | 9.000E-06 | 1.200E-05 |
| 25312 | 1266 | S1266K | D | 1.6e-05 | 1.4e-05 | 1.5e-05 | 1.500E-05 | 1.000E-06 | 1.400E-05 | 1.600E-05 |
| 25313 | 1266 | S1266M | D | 1.6e-05 | 1.4e-05 | 1.4e-05 | 1.467E-05 | 1.155E-06 | 1.400E-05 | 1.600E-05 |
| 25314 | 1266 | S1266F | D | 1.1e-05 | 1.6e-05 | 1.6e-05 | 1.433E-05 | 2.887E-06 | 1.100E-05 | 1.600E-05 |
| 25315 | 1266 | S1266P | D | 1.1e-05 | 1.6e-05 | 1.4e-05 | 1.367E-05 | 2.517E-06 | 1.100E-05 | 1.600E-05 |
| 25316 | 1266 | S1266S | D | 1.4e-05 | 9.3e-06 | 1.4e-05 | 1.243E-05 | 2.714E-06 | 9.300E-06 | 1.400E-05 |
| 25317 | 1266 | S1266T | D | 1.5e-05 | 1.1e-05 | 1e-05 | 1.200E-05 | 2.646E-06 | 1.000E-05 | 1.500E-05 |
| 25318 | 1266 | S1266W | D | 1.3e-05 | 1.6e-05 | 1.1e-05 | 1.333E-05 | 2.517E-06 | 1.100E-05 | 1.600E-05 |
| 25319 | 1266 | S1266Y | D | 6.9e-06 | 1.3e-05 | 9.4e-06 | 9.767E-06 | 3.066E-06 | 6.900E-06 | 1.300E-05 |
| 25320 | 1266 | S1266V | D | 8.6e-06 | 9.5e-06 | 1.5e-05 | 1.103E-05 | 3.465E-06 | 8.600E-06 | 1.500E-05 |
| 25321 | 1267 | F1267A | D | 9.4e-06 | 1.3e-05 | 1.3e-05 | 1.180E-05 | 2.078E-06 | 9.400E-06 | 1.300E-05 |
| 25322 | 1267 | F1267R | D | 1e-05 | 1.5e-05 | 1.3e-05 | 1.267E-05 | 2.517E-06 | 1.000E-05 | 1.500E-05 |
| 25323 | 1267 | F1267N | D | 1.1e-05 | 1.3e-05 | 1.2e-05 | 1.200E-05 | 1.000E-06 | 1.100E-05 | 1.300E-05 |
| 25324 | 1267 | F1267D | D | 9.8e-06 | 1.5e-05 | 1e-05 | 1.160E-05 | 2.946E-06 | 9.800E-06 | 1.500E-05 |
| 25325 | 1267 | F1267C | D | 1.4e-05 | 1.2e-05 | 1.3e-05 | 1.300E-05 | 1.000E-06 | 1.200E-05 | 1.400E-05 |
| 25326 | 1267 | F1267E | D | 1.1e-05 | 1.6e-05 | 1.5e-05 | 1.400E-05 | 2.646E-06 | 1.100E-05 | 1.600E-05 |
| 25327 | 1267 | F1267Q | D | 8.5e-06 | 1.9e-05 | 1.3e-05 | 1.350E-05 | 5.268E-06 | 8.500E-06 | 1.900E-05 |
| 25328 | 1267 | F1267G | D | 1e-05 | 8.8e-06 | 1.5e-05 | 1.127E-05 | 3.288E-06 | 8.800E-06 | 1.500E-05 |
| 25329 | 1267 | F1267H | D | 1.4e-05 | 1.5e-05 | 1.5e-05 | 1.467E-05 | 5.774E-07 | 1.400E-05 | 1.500E-05 |

| | | | | | | | | | | |
|---|---|---|---|---|---|---|---|---|---|---|
| 25330 | 1267 | F1267I | D | 1.5e-05 | 1.4e-05 | 1.1e-05 | 1.333E-05 | 2.082E-06 | 1.100E-05 | 1.500E-05 |
| 25331 | 1267 | F1267L | D | 1.1e-05 | 1e-05 | 1.5e-05 | 1.200E-05 | 2.646E-06 | 1.000E-05 | 1.500E-05 |
| 25332 | 1267 | F1267K | D | 1.1e-05 | 1.4e-05 | 1.1e-05 | 1.200E-05 | 1.732E-06 | 1.100E-05 | 1.400E-05 |
| 25333 | 1267 | F1267M | D | 1.4e-05 | 1.1e-05 | 1.3e-05 | 1.267E-05 | 1.528E-06 | 1.100E-05 | 1.400E-05 |
| 25334 | 1267 | F1267F | D | 1.5e-05 | 1.6e-05 | 1.1e-05 | 1.400E-05 | 2.646E-06 | 1.100E-05 | 1.600E-05 |
| 25335 | 1267 | F1267P | D | 8.2e-06 | 1e-05 | 1.3e-05 | 1.040E-05 | 2.425E-06 | 8.200E-06 | 1.300E-05 |
| 25336 | 1267 | F1267S | D | 9.6e-06 | 9.8e-06 | 1.2e-05 | 1.047E-05 | 1.332E-06 | 9.600E-06 | 1.200E-05 |
| 25337 | 1267 | F1267T | D | 1.4e-05 | 1.2e-05 | 1.1e-05 | 1.233E-05 | 1.528E-06 | 1.100E-05 | 1.400E-05 |
| 25338 | 1267 | F1267W | D | 8.4e-06 | 1.5e-05 | 1.6e-05 | 1.313E-05 | 4.130E-06 | 8.400E-06 | 1.600E-05 |
| 25339 | 1267 | F1267Y | D | 7.5e-06 | 7e-06 | 1.1e-05 | 8.500E-06 | 2.179E-06 | 7.000E-06 | 1.100E-05 |
| 25340 | 1267 | F1267V | D | 1.4e-05 | 9.9e-06 | 1.5e-05 | 1.297E-05 | 2.702E-06 | 9.900E-06 | 1.500E-05 |
| 25341 | 1268 | N1268A | D | 1.6e-05 | 9.7e-06 | 6.9e-06 | 1.087E-05 | 4.661E-06 | 6.900E-06 | 1.600E-05 |
| 25342 | 1268 | N1268R | D | 1.6e-05 | 1e-05 | 1.7e-05 | 1.433E-05 | 3.786E-06 | 1.000E-05 | 1.700E-05 |
| 25343 | 1268 | N1268N | D | 1.3e-05 | 9.1e-06 | 1e-05 | 1.070E-05 | 2.042E-06 | 9.100E-06 | 1.300E-05 |
| 25344 | 1268 | N1268D | D | 1e-05 | 1.5e-05 | 1.6e-05 | 1.367E-05 | 3.215E-06 | 1.000E-05 | 1.600E-05 |
| 25345 | 1268 | N1268C | D | 8.3e-06 | 1.5e-05 | 9.4e-06 | 1.090E-05 | 3.593E-06 | 8.300E-06 | 1.500E-05 |
| 25346 | 1268 | N1268E | D | 1.5e-05 | 1.1e-05 | 1.4e-05 | 1.333E-05 | 2.082E-06 | 1.100E-05 | 1.500E-05 |
| 25347 | 1268 | N1268Q | D | 1.4e-05 | 1.6e-05 | 1.4e-05 | 1.467E-05 | 1.155E-06 | 1.400E-05 | 1.600E-05 |
| 25348 | 1268 | N1268G | D | 1e-05 | 1.5e-05 | 1.5e-05 | 1.333E-05 | 2.887E-06 | 1.000E-05 | 1.500E-05 |
| 25349 | 1268 | N1268H | D | 1.1e-05 | 1.2e-05 | 1.5e-05 | 1.267E-05 | 2.082E-06 | 1.100E-05 | 1.500E-05 |
| 25350 | 1268 | N1268I | D | 1.3e-05 | 1e-05 | 1.4e-05 | 1.233E-05 | 2.082E-06 | 1.000E-05 | 1.400E-05 |
| 25351 | 1268 | N1268L | D | 1.1e-05 | 1.6e-05 | 1.1e-05 | 1.267E-05 | 2.887E-06 | 1.100E-05 | 1.600E-05 |
| 25352 | 1268 | N1268K | D | 1.3e-05 | 1.1e-05 | 1.3e-05 | 1.233E-05 | 1.155E-06 | 1.100E-05 | 1.300E-05 |
| 25353 | 1268 | N1268M | D | 1.1e-05 | 1.2e-05 | 1.4e-05 | 1.233E-05 | 1.528E-06 | 1.100E-05 | 1.400E-05 |
| 25354 | 1268 | N1268F | D | 1.1e-05 | 1.2e-05 | 9.6e-06 | 1.087E-05 | 1.206E-06 | 9.600E-06 | 1.200E-05 |
| 25355 | 1268 | N1268P | D | 1.1e-05 | 9.1e-06 | 1.3e-05 | 1.103E-05 | 1.950E-06 | 9.100E-06 | 1.300E-05 |
| 25356 | 1268 | N1268S | D | 1.3e-05 | 1.5e-05 | 1.6e-05 | 1.467E-05 | 1.528E-06 | 1.300E-05 | 1.600E-05 |
| 25357 | 1268 | N1268T | D | 1.6e-05 | 8.4e-06 | 9.6e-06 | 1.133E-05 | 4.086E-06 | 8.400E-06 | 1.600E-05 |
| 25358 | 1268 | N1268W | D | 1.1e-05 | 9.8e-06 | 1.6e-05 | 1.227E-05 | 3.288E-06 | 9.800E-06 | 1.600E-05 |
| 25359 | 1268 | N1268Y | D | 1e-05 | 8.5e-06 | 1.3e-05 | 1.050E-05 | 2.291E-06 | 8.500E-06 | 1.300E-05 |
| 25360 | 1268 | N1268V | D | 1e-05 | 1.7e-05 | 1.2e-05 | 1.300E-05 | 3.606E-06 | 1.000E-05 | 1.700E-05 |
| 25361 | 1269 | R1269A | D | 8.6e-06 | 1.4e-05 | 1.1e-05 | 1.120E-05 | 2.706E-06 | 8.600E-06 | 1.400E-05 |
| 25362 | 1269 | R1269R | D | 1.1e-05 | 1.5e-05 | 1.5e-05 | 1.367E-05 | 2.309E-06 | 1.100E-05 | 1.500E-05 |
| 25363 | 1269 | R1269N | D | 1e-05 | 1.5e-05 | 1.2e-05 | 1.233E-05 | 2.517E-06 | 1.000E-05 | 1.500E-05 |
| 25364 | 1269 | R1269D | D | 1.2e-05 | 1.1e-05 | 1.4e-05 | 1.233E-05 | 1.528E-06 | 1.100E-05 | 1.400E-05 |
| 25365 | 1269 | R1269C | D | 1.1e-05 | 1.5e-05 | 1.4e-05 | 1.333E-05 | 2.082E-06 | 1.100E-05 | 1.500E-05 |
| 25366 | 1269 | R1269E | D | 9.2e-06 | 9.8e-06 | 1.7e-05 | 1.200E-05 | 4.341E-06 | 9.200E-06 | 1.700E-05 |
| 25367 | 1269 | R1269Q | D | 9.9e-06 | 1.3e-05 | 7.8e-06 | 1.023E-05 | 2.616E-06 | 7.800E-06 | 1.300E-05 |
| 25368 | 1269 | R1269G | D | 1.1e-05 | 9.3e-06 | 1.3e-05 | 1.110E-05 | 1.852E-06 | 9.300E-06 | 1.300E-05 |
| 25369 | 1269 | R1269H | D | 1.4e-05 | 8e-06 | 7.6e-06 | 9.867E-06 | 3.585E-06 | 7.600E-06 | 1.400E-05 |
| 25370 | 1269 | R1269I | D | 1.7e-05 | 1.4e-05 | 1.6e-05 | 1.567E-05 | 1.528E-06 | 1.400E-05 | 1.700E-05 |
| 25371 | 1269 | R1269L | D | 1.5e-05 | 1.4e-05 | 8.8e-06 | 1.260E-05 | 3.329E-06 | 8.800E-06 | 1.500E-05 |
| 25372 | 1269 | R1269K | D | 1.5e-05 | 1.4e-05 | 9.8e-06 | 1.293E-05 | 2.759E-06 | 9.800E-06 | 1.500E-05 |
| 25373 | 1269 | R1269M | D | 1.4e-05 | 8.4e-06 | 1.1e-05 | 1.113E-05 | 2.802E-06 | 8.400E-06 | 1.400E-05 |
| 25374 | 1269 | R1269F | D | 1.5e-05 | 7.7e-06 | 1.2e-05 | 1.157E-05 | 3.669E-06 | 7.700E-06 | 1.500E-05 |
| 25375 | 1269 | R1269P | D | 1.1e-05 | 1.3e-05 | 1.7e-05 | 1.367E-05 | 3.055E-06 | 1.100E-05 | 1.700E-05 |
| 25376 | 1269 | R1269S | D | 1.5e-05 | 1e-05 | 1.5e-05 | 1.333E-05 | 2.887E-06 | 1.000E-05 | 1.500E-05 |
| 25377 | 1269 | R1269T | D | 1.5e-05 | 1.5e-05 | 1.6e-05 | 1.533E-05 | 5.774E-07 | 1.500E-05 | 1.600E-05 |
| 25378 | 1269 | R1269W | D | 1.1e-05 | 1.3e-05 | 1.2e-05 | 1.200E-05 | 1.000E-06 | 1.100E-05 | 1.300E-05 |
| 25379 | 1269 | R1269Y | D | 1.4e-05 | 1.5e-05 | 1.6e-05 | 1.500E-05 | 1.000E-06 | 1.400E-05 | 1.600E-05 |
| 25380 | 1269 | R1269V | D | 1.2e-05 | 5.3e-06 | 1.3e-05 | 1.010E-05 | 4.187E-06 | 5.300E-06 | 1.300E-05 |
| 25381 | 1270 | G1270A | D | 1.3e-05 | 1.2e-05 | 1.3e-05 | 1.267E-05 | 5.774E-07 | 1.200E-05 | 1.300E-05 |
| 25382 | 1270 | G1270R | D | 1.3e-05 | 9.3e-06 | 1.5e-05 | 1.243E-05 | 2.892E-06 | 9.300E-06 | 1.500E-05 |
| 25383 | 1270 | G1270N | D | 8.7e-06 | 1.5e-05 | 1.5e-05 | 1.290E-05 | 3.637E-06 | 8.700E-06 | 1.500E-05 |
| 25384 | 1270 | G1270D | D | 1.7e-05 | 1.6e-05 | 6.6e-06 | 1.320E-05 | 5.738E-06 | 6.600E-06 | 1.700E-05 |
| 25385 | 1270 | G1270C | D | 1e-05 | 1.5e-05 | 1.1e-05 | 1.200E-05 | 2.646E-06 | 1.000E-05 | 1.500E-05 |
| 25386 | 1270 | G1270E | D | 8.7e-06 | 9.8e-06 | 8.4e-06 | 8.967E-06 | 7.371E-07 | 8.400E-06 | 9.800E-06 |
| 25387 | 1270 | G1270Q | D | 1.1e-05 | 1.5e-05 | 1.7e-05 | 1.433E-05 | 3.055E-06 | 1.100E-05 | 1.700E-05 |
| 25388 | 1270 | G1270G | D | 1.7e-05 | 1.5e-05 | 1e-05 | 1.400E-05 | 3.606E-06 | 1.000E-05 | 1.700E-05 |
| 25389 | 1270 | G1270H | D | 8.8e-06 | 1.4e-05 | 1.4e-05 | 1.227E-05 | 3.002E-06 | 8.800E-06 | 1.400E-05 |
| 25390 | 1270 | G1270I | D | 1.4e-05 | 1.6e-05 | 1e-05 | 1.333E-05 | 3.055E-06 | 1.000E-05 | 1.600E-05 |
| 25391 | 1270 | G1270L | D | 1.5e-05 | 1.4e-05 | 1.1e-05 | 1.333E-05 | 2.082E-06 | 1.100E-05 | 1.500E-05 |
| 25392 | 1270 | G1270K | D | 1.3e-05 | 1.3e-05 | 1.7e-05 | 1.433E-05 | 2.309E-06 | 1.300E-05 | 1.700E-05 |
| 25393 | 1270 | G1270M | D | 7.7e-06 | 1.6e-05 | 1.5e-05 | 1.290E-05 | 4.531E-06 | 7.700E-06 | 1.600E-05 |
| 25394 | 1270 | G1270F | D | 1.6e-05 | 1.2e-05 | 1.6e-05 | 1.467E-05 | 2.309E-06 | 1.200E-05 | 1.600E-05 |
| 25395 | 1270 | G1270P | D | 1e-05 | 9.8e-06 | 8e-06 | 9.267E-06 | 1.102E-06 | 8.000E-06 | 1.000E-05 |
| 25396 | 1270 | G1270S | D | 1.3e-05 | 6.3e-06 | 8.3e-06 | 9.200E-06 | 3.439E-06 | 6.300E-06 | 1.300E-05 |
| 25397 | 1270 | G1270T | D | 1.3e-05 | 1.5e-05 | 1.5e-05 | 1.433E-05 | 1.155E-06 | 1.300E-05 | 1.500E-05 |
| 25398 | 1270 | G1270W | D | 9.5e-06 | 1.4e-05 | 1e-05 | 1.117E-05 | 2.466E-06 | 9.500E-06 | 1.400E-05 |
| 25399 | 1270 | G1270Y | D | 1.5e-05 | 1.2e-05 | 1.2e-05 | 1.300E-05 | 1.732E-06 | 1.200E-05 | 1.500E-05 |
| 25400 | 1270 | G1270V | D | 1.7e-05 | 1.5e-05 | 1.1e-05 | 1.433E-05 | 3.055E-06 | 1.100E-05 | 1.700E-05 |
| 25401 | 1271 | E1271A | D | 1.1e-05 | 1.4e-05 | 1.6e-05 | 1.367E-05 | 2.517E-06 | 1.100E-05 | 1.600E-05 |
| 25402 | 1271 | E1271R | D | 1.3e-05 | 1.3e-05 | 1.6e-05 | 1.400E-05 | 1.732E-06 | 1.300E-05 | 1.600E-05 |
| 25403 | 1271 | E1271N | D | 9.1e-06 | 8.4e-06 | 1e-05 | 9.167E-06 | 8.021E-07 | 8.400E-06 | 1.000E-05 |
| 25404 | 1271 | E1271D | D | 1.5e-05 | 1.5e-05 | 9.3e-06 | 1.310E-05 | 3.291E-06 | 9.300E-06 | 1.500E-05 |
| 25405 | 1271 | E1271C | D | 1.4e-05 | 1.5e-05 | 1.4e-05 | 1.433E-05 | 5.774E-07 | 1.400E-05 | 1.500E-05 |
| 25406 | 1271 | E1271E | D | 1.4e-05 | 1e-05 | 1.2e-05 | 1.200E-05 | 2.000E-06 | 1.000E-05 | 1.400E-05 |
| 25407 | 1271 | E1271Q | D | 1.4e-05 | 1.3e-05 | 1.6e-05 | 1.433E-05 | 1.528E-06 | 1.300E-05 | 1.600E-05 |
| 25408 | 1271 | E1271G | D | 1.6e-05 | 1.5e-05 | 1.4e-05 | 1.500E-05 | 1.000E-06 | 1.400E-05 | 1.600E-05 |
| 25409 | 1271 | E1271H | D | 1.4e-05 | 1.5e-05 | 1.6e-05 | 1.500E-05 | 1.000E-06 | 1.400E-05 | 1.600E-05 |
| 25410 | 1271 | E1271I | D | 8e-06 | 1.4e-05 | 1.1e-05 | 1.100E-05 | 3.000E-06 | 8.000E-06 | 1.400E-05 |
| 25411 | 1271 | E1271L | D | 8.9e-06 | 1.6e-05 | 1.6e-05 | 1.363E-05 | 4.099E-06 | 8.900E-06 | 1.600E-05 |
| 25412 | 1271 | E1271K | D | 1.2e-05 | 1e-05 | 1.5e-05 | 1.233E-05 | 2.517E-06 | 1.000E-05 | 1.500E-05 |
| 25413 | 1271 | E1271M | D | 1.5e-05 | 1.1e-05 | 1.4e-05 | 1.333E-05 | 2.082E-06 | 1.100E-05 | 1.500E-05 |
| 25414 | 1271 | E1271F | D | 8.7e-06 | 1.5e-05 | 8.9e-06 | 1.087E-05 | 3.581E-06 | 8.700E-06 | 1.500E-05 |

| | | | | | | | | | | |
|---|---|---|---|---|---|---|---|---|---|---|
| 25415 | 1271 | E1271P | D | 1.6e-05 | 1.1e-05 | 1.5e-05 | 1.400E-05 | 2.646E-06 | 1.100E-05 | 1.600E-05 |
| 25416 | 1271 | E1271S | D | 1.4e-05 | 1e-05 | 8.6e-06 | 1.087E-05 | 2.802E-06 | 8.600E-06 | 1.400E-05 |
| 25417 | 1271 | E1271T | D | 1e-05 | 9.9e-06 | 1.2e-05 | 1.063E-05 | 1.185E-06 | 9.900E-06 | 1.200E-05 |
| 25418 | 1271 | E1271W | D | 1.2e-05 | 1.3e-05 | 1e-05 | 1.167E-05 | 1.528E-06 | 1.000E-05 | 1.300E-05 |
| 25419 | 1271 | E1271Y | D | 1.1e-05 | 1e-05 | 1.4e-05 | 1.167E-05 | 2.082E-06 | 1.000E-05 | 1.400E-05 |
| 25420 | 1271 | E1271V | D | 1e-05 | 9.2e-06 | 1.5e-05 | 1.140E-05 | 3.143E-06 | 9.200E-06 | 1.500E-05 |
| 25421 | 1272 | C1272A | D | 1e-05 | 1.4e-05 | 1.7e-05 | 1.367E-05 | 3.512E-06 | 1.000E-05 | 1.700E-05 |
| 25422 | 1272 | C1272R | D | 1.2e-05 | 1.5e-05 | 1.1e-05 | 1.267E-05 | 2.082E-06 | 1.100E-05 | 1.500E-05 |
| 25423 | 1272 | C1272N | D | 1.3e-05 | 9.5e-06 | 2.1e-05 | 1.450E-05 | 5.895E-06 | 9.500E-06 | 2.100E-05 |
| 25424 | 1272 | C1272D | D | 1.7e-05 | 1.7e-05 | 1.6e-05 | 1.667E-05 | 5.774E-07 | 1.600E-05 | 1.700E-05 |
| 25425 | 1272 | C1272C | D | 1e-05 | 9.6e-06 | 1.7e-05 | 1.220E-05 | 4.162E-06 | 9.600E-06 | 1.700E-05 |
| 25426 | 1272 | C1272E | D | 1.5e-05 | 8.8e-06 | 1.1e-05 | 1.160E-05 | 3.143E-06 | 8.800E-06 | 1.500E-05 |
| 25427 | 1272 | C1272Q | D | 1.3e-05 | 1.5e-05 | 9.7e-06 | 1.257E-05 | 2.676E-06 | 9.700E-06 | 1.500E-05 |
| 25428 | 1272 | C1272G | D | 1.3e-05 | 9e-06 | 1e-05 | 1.067E-05 | 2.082E-06 | 9.000E-06 | 1.300E-05 |
| 25429 | 1272 | C1272H | D | 1e-05 | 1.6e-05 | 9.1e-06 | 1.170E-05 | 3.751E-06 | 9.100E-06 | 1.600E-05 |
| 25430 | 1272 | C1272I | D | 9.6e-06 | 1.2e-05 | 1e-05 | 1.053E-05 | 1.286E-06 | 9.600E-06 | 1.200E-05 |
| 25431 | 1272 | C1272L | D | 9.5e-06 | 1.8e-05 | 9.7e-06 | 1.240E-05 | 4.851E-06 | 9.500E-06 | 1.800E-05 |
| 25432 | 1272 | C1272K | D | 8.4e-06 | 1.5e-05 | 1e-05 | 1.113E-05 | 3.443E-06 | 8.400E-06 | 1.500E-05 |
| 25433 | 1272 | C1272M | D | 1.7e-05 | 1.4e-05 | 9.9e-06 | 1.363E-05 | 3.564E-06 | 9.900E-06 | 1.700E-05 |
| 25434 | 1272 | C1272F | D | 1.5e-05 | 1.2e-05 | 1.6e-05 | 1.433E-05 | 2.082E-06 | 1.200E-05 | 1.600E-05 |
| 25435 | 1272 | C1272P | D | 9.2e-06 | 1.6e-05 | 1.5e-05 | 1.340E-05 | 3.672E-06 | 9.200E-06 | 1.600E-05 |
| 25436 | 1272 | C1272S | D | 1.6e-05 | 9.8e-06 | 1.5e-05 | 1.360E-05 | 3.329E-06 | 9.800E-06 | 1.600E-05 |
| 25437 | 1272 | C1272T | D | 9.2e-06 | 1.5e-05 | 9.2e-06 | 1.113E-05 | 3.349E-06 | 9.200E-06 | 1.500E-05 |
| 25438 | 1272 | C1272W | D | 1e-05 | 1.4e-05 | 1e-05 | 1.133E-05 | 2.309E-06 | 1.000E-05 | 1.400E-05 |
| 25439 | 1272 | C1272Y | D | 1.6e-05 | 1.7e-05 | 1.8e-05 | 1.700E-05 | 1.000E-06 | 1.600E-05 | 1.800E-05 |
| 25440 | 1272 | C1272V | D | 1.1e-05 | 1e-05 | 1.3e-05 | 1.133E-05 | 1.528E-06 | 1.000E-05 | 1.300E-05 |
| 25441 | 1273 | E1273A | E | 1.4e-05 | 1e-05 | 8.7e-06 | 1.090E-05 | 2.762E-06 | 8.700E-06 | 1.400E-05 |
| 25442 | 1273 | E1273R | E | 9.9e-06 | 1.5e-05 | 1.4e-05 | 1.297E-05 | 2.702E-06 | 9.900E-06 | 1.500E-05 |
| 25443 | 1273 | E1273N | E | 1.6e-05 | 1.4e-05 | 1.2e-05 | 1.400E-05 | 2.000E-06 | 1.200E-05 | 1.600E-05 |
| 25444 | 1273 | E1273D | E | 1.5e-05 | 1e-05 | 1e-05 | 1.167E-05 | 2.887E-06 | 1.000E-05 | 1.500E-05 |
| 25445 | 1273 | E1273C | E | 1.6e-05 | 1.4e-05 | 1.4e-05 | 1.467E-05 | 1.155E-06 | 1.400E-05 | 1.600E-05 |
| 25446 | 1273 | E1273E | E | 1.5e-05 | 1.5e-05 | 1.6e-05 | 1.533E-05 | 5.774E-07 | 1.500E-05 | 1.600E-05 |
| 25447 | 1273 | E1273Q | E | 1.2e-05 | 2.1e-05 | 1.7e-05 | 1.667E-05 | 4.509E-06 | 1.200E-05 | 2.100E-05 |
| 25448 | 1273 | E1273G | E | 1.3e-05 | 1.6e-05 | 1.5e-05 | 1.467E-05 | 1.528E-06 | 1.300E-05 | 1.600E-05 |
| 25449 | 1273 | E1273H | E | 8.1e-06 | 1.5e-05 | 1.5e-05 | 1.270E-05 | 3.984E-06 | 8.100E-06 | 1.500E-05 |
| 25450 | 1273 | E1273I | E | 1.4e-05 | 1.1e-05 | 9.9e-06 | 1.163E-05 | 2.122E-06 | 9.900E-06 | 1.400E-05 |
| 25451 | 1273 | E1273L | E | 9.3e-06 | 1.7e-05 | 1.3e-05 | 1.310E-05 | 3.851E-06 | 9.300E-06 | 1.700E-05 |
| 25452 | 1273 | E1273K | E | 8.9e-06 | 9.7e-06 | 1.4e-05 | 1.087E-05 | 2.743E-06 | 8.900E-06 | 1.400E-05 |
| 25453 | 1273 | E1273M | E | 1.1e-05 | 1.7e-05 | 1.1e-05 | 1.300E-05 | 3.464E-06 | 1.100E-05 | 1.700E-05 |
| 25454 | 1273 | E1273F | E | 1.6e-05 | 1e-05 | 1.5e-05 | 1.367E-05 | 3.215E-06 | 1.000E-05 | 1.600E-05 |
| 25455 | 1273 | E1273P | E | 1.5e-05 | 9.6e-06 | 1.1e-05 | 1.187E-05 | 2.802E-06 | 9.600E-06 | 1.500E-05 |
| 25456 | 1273 | E1273S | E | 1.6e-05 | 1.4e-05 | 7.1e-06 | 1.237E-05 | 4.669E-06 | 7.100E-06 | 1.600E-05 |
| 25457 | 1273 | E1273T | E | 1.1e-05 | 1.3e-05 | 1e-05 | 1.133E-05 | 1.528E-06 | 1.000E-05 | 1.300E-05 |
| 25458 | 1273 | E1273W | E | 1.2e-05 | 2.3e-05 | 1.7e-05 | 1.733E-05 | 5.508E-06 | 1.200E-05 | 2.300E-05 |
| 25459 | 1273 | E1273Y | E | 1.5e-05 | 1.5e-05 | 1.2e-05 | 1.400E-05 | 1.732E-06 | 1.200E-05 | 1.500E-05 |
| 25460 | 1273 | E1273V | E | 9.5e-06 | 1.6e-05 | 1.2e-05 | 1.250E-05 | 3.279E-06 | 9.500E-06 | 1.600E-05 |
| 25461 | 1274 | V1274A | E | 9.2e-06 | 1e-05 | 1.5e-05 | 1.140E-05 | 3.143E-06 | 9.200E-06 | 1.500E-05 |
| 25462 | 1274 | V1274R | E | 1.5e-05 | 1.2e-05 | 1.1e-05 | 1.267E-05 | 2.082E-06 | 1.100E-05 | 1.500E-05 |
| 25463 | 1274 | V1274N | E | 1.5e-05 | 1.5e-05 | 1.5e-05 | 1.500E-05 | 0.000E+00 | 1.500E-05 | 1.500E-05 |
| 25464 | 1274 | V1274D | E | 1.4e-05 | 1.4e-05 | 1.1e-05 | 1.300E-05 | 1.732E-06 | 1.100E-05 | 1.400E-05 |
| 25465 | 1274 | V1274C | E | 1.1e-05 | 1e-05 | 1.4e-05 | 1.167E-05 | 2.082E-06 | 1.000E-05 | 1.400E-05 |
| 25466 | 1274 | V1274E | E | 8.4e-06 | 1.1e-05 | 1.1e-05 | 1.013E-05 | 1.501E-06 | 8.400E-06 | 1.100E-05 |
| 25467 | 1274 | V1274Q | E | 1.4e-05 | 1.4e-05 | 9.1e-06 | 1.237E-05 | 2.829E-06 | 9.100E-06 | 1.400E-05 |
| 25468 | 1274 | V1274G | E | 1.6e-05 | 1.1e-05 | 1.6e-05 | 1.433E-05 | 2.887E-06 | 1.100E-05 | 1.600E-05 |
| 25469 | 1274 | V1274H | E | 8.8e-06 | 1.4e-05 | 1.6e-05 | 1.293E-05 | 3.717E-06 | 8.800E-06 | 1.600E-05 |
| 25470 | 1274 | V1274I | E | 1.1e-05 | 1e-05 | 1.4e-05 | 1.167E-05 | 2.082E-06 | 1.000E-05 | 1.400E-05 |
| 25471 | 1274 | V1274L | E | 9.2e-06 | 9.7e-06 | 1e-05 | 9.633E-06 | 4.041E-07 | 9.200E-06 | 1.000E-05 |
| 25472 | 1274 | V1274K | E | 7.3e-06 | 1.1e-05 | 1.1e-05 | 9.767E-06 | 2.136E-06 | 7.300E-06 | 1.100E-05 |
| 25473 | 1274 | V1274M | E | 1.3e-05 | 1.5e-05 | 1.7e-05 | 1.500E-05 | 2.000E-06 | 1.300E-05 | 1.700E-05 |
| 25474 | 1274 | V1274F | E | 7.8e-06 | 1.5e-05 | 1.5e-05 | 1.260E-05 | 4.157E-06 | 7.800E-06 | 1.500E-05 |
| 25475 | 1274 | V1274P | E | 1.3e-05 | 1.6e-05 | 1.6e-05 | 1.500E-05 | 1.732E-06 | 1.300E-05 | 1.600E-05 |
| 25476 | 1274 | V1274S | E | 1.6e-05 | 1.5e-05 | 1.2e-05 | 1.433E-05 | 2.082E-06 | 1.200E-05 | 1.600E-05 |
| 25477 | 1274 | V1274T | E | 1.4e-05 | 1.5e-05 | 8.7e-06 | 1.257E-05 | 3.386E-06 | 8.700E-06 | 1.500E-05 |
| 25478 | 1274 | V1274W | E | 1.2e-05 | 1.6e-05 | 1.4e-05 | 1.400E-05 | 2.000E-06 | 1.200E-05 | 1.600E-05 |
| 25479 | 1274 | V1274Y | E | 1.5e-05 | 1.2e-05 | 1e-05 | 1.233E-05 | 2.517E-06 | 1.000E-05 | 1.500E-05 |
| 25480 | 1274 | V1274V | E | 1.1e-05 | 1.5e-05 | 1.5e-05 | 1.367E-05 | 2.309E-06 | 1.100E-05 | 1.500E-05 |
| 25481 | 1275 | Q1275A | E | 1.1e-05 | 1.5e-05 | 8e-06 | 1.133E-05 | 3.512E-06 | 8.000E-06 | 1.500E-05 |
| 25482 | 1275 | Q1275R | E | 1.5e-05 | 1.1e-05 | 1.6e-05 | 1.400E-05 | 2.646E-06 | 1.100E-05 | 1.600E-05 |
| 25483 | 1275 | Q1275N | E | 7.8e-06 | 1.6e-05 | 1e-05 | 1.127E-05 | 4.244E-06 | 7.800E-06 | 1.600E-05 |
| 25484 | 1275 | Q1275D | E | 1e-05 | 1.5e-05 | 1.5e-05 | 1.333E-05 | 2.887E-06 | 1.000E-05 | 1.500E-05 |
| 25485 | 1275 | Q1275C | E | 1.4e-05 | 1.4e-05 | 1.4e-05 | 1.400E-05 | 0.000E+00 | 1.400E-05 | 1.400E-05 |
| 25486 | 1275 | Q1275E | E | 9.4e-06 | 1.9e-05 | 1.3e-05 | 1.380E-05 | 4.850E-06 | 9.400E-06 | 1.900E-05 |
| 25487 | 1275 | Q1275Q | E | 9.9e-06 | 1.4e-05 | 1.1e-05 | 1.163E-05 | 2.122E-06 | 9.900E-06 | 1.400E-05 |
| 25488 | 1275 | Q1275G | E | 1.7e-05 | 1.6e-05 | 6.3e-06 | 1.310E-05 | 5.910E-06 | 6.300E-06 | 1.700E-05 |
| 25489 | 1275 | Q1275H | E | 1.1e-05 | 1.1e-05 | 1.5e-05 | 1.233E-05 | 2.309E-06 | 1.100E-05 | 1.500E-05 |
| 25490 | 1275 | Q1275I | E | 1.1e-05 | 1.1e-05 | 1.5e-05 | 1.233E-05 | 2.309E-06 | 1.100E-05 | 1.500E-05 |
| 25491 | 1275 | Q1275L | E | 1.3e-05 | 1.1e-05 | 8.7e-06 | 1.090E-05 | 2.152E-06 | 8.700E-06 | 1.300E-05 |
| 25492 | 1275 | Q1275K | E | 1.1e-05 | 1.5e-05 | 1.2e-05 | 1.267E-05 | 2.082E-06 | 1.100E-05 | 1.500E-05 |
| 25493 | 1275 | Q1275M | E | 1.6e-05 | 8.3e-06 | 1.6e-05 | 1.343E-05 | 4.446E-06 | 8.300E-06 | 1.600E-05 |
| 25494 | 1275 | Q1275F | E | 1.1e-05 | 1.7e-05 | 9.7e-06 | 1.257E-05 | 3.894E-06 | 9.700E-06 | 1.700E-05 |
| 25495 | 1275 | Q1275P | E | 1e-05 | 1e-05 | 1.7e-05 | 1.233E-05 | 4.041E-06 | 1.000E-05 | 1.700E-05 |
| 25496 | 1275 | Q1275S | E | 1.5e-05 | 1.5e-05 | 1.3e-05 | 1.433E-05 | 1.155E-06 | 1.300E-05 | 1.500E-05 |
| 25497 | 1275 | Q1275T | E | 1.6e-05 | 1.6e-05 | 9.9e-06 | 1.397E-05 | 3.522E-06 | 9.900E-06 | 1.600E-05 |
| 25498 | 1275 | Q1275W | E | 1.5e-05 | 1.5e-05 | 1.4e-05 | 1.467E-05 | 5.774E-07 | 1.400E-05 | 1.500E-05 |
| 25499 | 1275 | Q1275Y | E | 1e-05 | 1.1e-05 | 1.6e-05 | 1.233E-05 | 3.215E-06 | 1.000E-05 | 1.600E-05 |

| | | | | | | | | | | |
|---|---|---|---|---|---|---|---|---|---|---|
| 25500 | 1275 | Q1275V | E | 1.5e-05 | 1.4e-05 | 1.8e-05 | 1.567E-05 | 2.082E-06 | 1.400E-05 | 1.800E-05 |
| 25501 | 1276 | L1276A | E | 1.4e-05 | 1.2e-05 | 1.1e-05 | 1.233E-05 | 1.528E-06 | 1.100E-05 | 1.400E-05 |
| 25502 | 1276 | L1276R | E | 1.2e-05 | 1.4e-05 | 1.6e-05 | 1.400E-05 | 2.000E-06 | 1.200E-05 | 1.600E-05 |
| 25503 | 1276 | L1276N | E | 8.4e-06 | 1.3e-05 | 9.8e-06 | 1.040E-05 | 2.358E-06 | 8.400E-06 | 1.300E-05 |
| 25504 | 1276 | L1276D | E | 1.6e-05 | 9.4e-06 | 1.6e-05 | 1.380E-05 | 3.811E-06 | 9.400E-06 | 1.600E-05 |
| 25505 | 1276 | L1276C | E | 1.2e-05 | 1.6e-05 | 1e-05 | 1.267E-05 | 3.055E-06 | 1.000E-05 | 1.600E-05 |
| 25506 | 1276 | L1276E | E | 9.8e-06 | 1.7e-05 | 1.6e-05 | 1.427E-05 | 3.900E-06 | 9.800E-06 | 1.700E-05 |
| 25507 | 1276 | L1276Q | E | 8.6e-06 | 1.6e-05 | 1.7e-05 | 1.387E-05 | 4.588E-06 | 8.600E-06 | 1.700E-05 |
| 25508 | 1276 | L1276G | E | 1.1e-05 | 1.3e-05 | 1.2e-05 | 1.200E-05 | 1.000E-06 | 1.100E-05 | 1.300E-05 |
| 25509 | 1276 | L1276H | E | 1.1e-05 | 1.3e-05 | 8.9e-06 | 1.097E-05 | 2.050E-06 | 8.900E-06 | 1.300E-05 |
| 25510 | 1276 | L1276I | E | 1.7e-05 | 1.6e-05 | 1.1e-05 | 1.467E-05 | 3.215E-06 | 1.100E-05 | 1.700E-05 |
| 25511 | 1276 | L1276L | E | 1.3e-05 | 1.1e-05 | 1.4e-05 | 1.267E-05 | 1.528E-06 | 1.100E-05 | 1.400E-05 |
| 25512 | 1276 | L1276K | E | 1.3e-05 | 1.6e-05 | 1.3e-05 | 1.400E-05 | 1.732E-06 | 1.300E-05 | 1.600E-05 |
| 25513 | 1276 | L1276M | E | 9.9e-06 | 1.5e-05 | 1.3e-05 | 1.263E-05 | 2.570E-06 | 9.900E-06 | 1.500E-05 |
| 25514 | 1276 | L1276F | E | 8.9e-06 | 1.5e-05 | 9.1e-06 | 1.100E-05 | 3.466E-06 | 8.900E-06 | 1.500E-05 |
| 25515 | 1276 | L1276P | E | 1.3e-05 | 1.4e-05 | 1.5e-05 | 1.400E-05 | 1.000E-06 | 1.300E-05 | 1.500E-05 |
| 25516 | 1276 | L1276S | E | 9e-06 | 1e-05 | 9.1e-06 | 9.367E-06 | 5.508E-07 | 9.000E-06 | 1.000E-05 |
| 25517 | 1276 | L1276T | E | 1.2e-05 | 1.4e-05 | 1.3e-05 | 1.300E-05 | 1.000E-06 | 1.200E-05 | 1.400E-05 |
| 25518 | 1276 | L1276W | E | 8.1e-06 | 1.7e-05 | 9.5e-06 | 1.153E-05 | 4.786E-06 | 8.100E-06 | 1.700E-05 |
| 25519 | 1276 | L1276Y | E | 1.5e-05 | 9.2e-06 | 1.4e-05 | 1.273E-05 | 3.101E-06 | 9.200E-06 | 1.500E-05 |
| 25520 | 1276 | L1276V | E | 1e-05 | 1e-05 | 1e-05 | 1.000E-05 | 0.000E+00 | 1.000E-05 | 1.000E-05 |
| 25521 | 1277 | V1277A | E | 1.5e-05 | 1.1e-05 | 1e-05 | 1.200E-05 | 2.646E-06 | 1.000E-05 | 1.500E-05 |
| 25522 | 1277 | V1277R | E | 1.6e-05 | 1e-05 | 1.5e-05 | 1.367E-05 | 3.215E-06 | 1.000E-05 | 1.600E-05 |
| 25523 | 1277 | V1277N | E | 1.2e-05 | 1e-05 | 1.1e-05 | 1.100E-05 | 1.000E-06 | 1.000E-05 | 1.200E-05 |
| 25524 | 1277 | V1277D | E | 1.5e-05 | 1.4e-05 | 9.7e-06 | 1.290E-05 | 2.816E-06 | 9.700E-06 | 1.500E-05 |
| 25525 | 1277 | V1277C | E | 1.4e-05 | 1.2e-05 | 1.4e-05 | 1.333E-05 | 1.155E-06 | 1.200E-05 | 1.400E-05 |
| 25526 | 1277 | V1277E | E | 1.6e-05 | 1.3e-05 | 1.1e-05 | 1.333E-05 | 2.517E-06 | 1.100E-05 | 1.600E-05 |
| 25527 | 1277 | V1277Q | E | 1e-05 | 1.5e-05 | 1.3e-05 | 1.267E-05 | 2.517E-06 | 1.000E-05 | 1.500E-05 |
| 25528 | 1277 | V1277G | E | 8.7e-06 | 9.4e-06 | 1.5e-05 | 1.103E-05 | 3.453E-06 | 8.700E-06 | 1.500E-05 |
| 25529 | 1277 | V1277H | E | 1.2e-05 | 9.5e-06 | 8.3e-06 | 9.933E-06 | 1.888E-06 | 8.300E-06 | 1.200E-05 |
| 25530 | 1277 | V1277I | E | 8.4e-06 | 5.6e-06 | 1.2e-05 | 8.667E-06 | 3.208E-06 | 5.600E-06 | 1.200E-05 |
| 25531 | 1277 | V1277L | E | 7.9e-06 | 1.5e-05 | 1e-05 | 1.097E-05 | 3.647E-06 | 7.900E-06 | 1.500E-05 |
| 25532 | 1277 | V1277K | E | 1.1e-05 | 1.1e-05 | 9e-06 | 1.033E-05 | 1.155E-06 | 9.000E-06 | 1.100E-05 |
| 25533 | 1277 | V1277M | E | 1.6e-05 | 1e-05 | 1.4e-05 | 1.333E-05 | 3.055E-06 | 1.000E-05 | 1.600E-05 |
| 25534 | 1277 | V1277F | E | 1.5e-05 | 1.4e-05 | 1.1e-05 | 1.333E-05 | 2.082E-06 | 1.100E-05 | 1.500E-05 |
| 25535 | 1277 | V1277P | E | 1.4e-05 | 1e-05 | 9.7e-06 | 1.123E-05 | 2.401E-06 | 9.700E-06 | 1.400E-05 |
| 25536 | 1277 | V1277S | E | 9.3e-06 | 1.5e-05 | 1.2e-05 | 1.210E-05 | 2.851E-06 | 9.300E-06 | 1.500E-05 |
| 25537 | 1277 | V1277T | E | 1.5e-05 | 1.4e-05 | 1.3e-05 | 1.400E-05 | 1.000E-06 | 1.300E-05 | 1.500E-05 |
| 25538 | 1277 | V1277W | E | 1.6e-05 | 3.4e-05 | 1.4e-05 | 2.133E-05 | 1.102E-05 | 1.400E-05 | 3.400E-05 |
| 25539 | 1277 | V1277Y | E | 9.5e-06 | 1.5e-05 | 1e-05 | 1.150E-05 | 3.041E-06 | 9.500E-06 | 1.500E-05 |
| 25540 | 1277 | V1277V | E | 1.4e-05 | 8.4e-06 | 1.2e-05 | 1.147E-05 | 2.838E-06 | 8.400E-06 | 1.400E-05 |
| 25541 | 1278 | E1278A | E | 1.1e-05 | 1.8e-05 | 1.4e-05 | 1.433E-05 | 3.512E-06 | 1.100E-05 | 1.800E-05 |
| 25542 | 1278 | E1278R | E | 1.6e-05 | 1e-05 | 9.3e-06 | 1.177E-05 | 3.683E-06 | 9.300E-06 | 1.600E-05 |
| 25543 | 1278 | E1278N | E | 1.1e-05 | 1.4e-05 | 1.6e-05 | 1.367E-05 | 2.517E-06 | 1.100E-05 | 1.600E-05 |
| 25544 | 1278 | E1278D | E | 1.7e-05 | 9.4e-06 | 9.8e-06 | 1.207E-05 | 4.277E-06 | 9.400E-06 | 1.700E-05 |
| 25545 | 1278 | E1278C | E | 1.2e-05 | 1.3e-05 | 1.5e-05 | 1.333E-05 | 1.528E-06 | 1.200E-05 | 1.500E-05 |
| 25546 | 1278 | E1278E | E | 1.5e-05 | 1.6e-05 | 1.5e-05 | 1.533E-05 | 5.774E-07 | 1.500E-05 | 1.600E-05 |
| 25547 | 1278 | E1278Q | E | 7.7e-06 | 9.8e-06 | 1e-05 | 9.167E-06 | 1.274E-06 | 7.700E-06 | 1.000E-05 |
| 25548 | 1278 | E1278G | E | 1.4e-05 | 1.7e-05 | 1.5e-05 | 1.533E-05 | 1.528E-06 | 1.400E-05 | 1.700E-05 |
| 25549 | 1278 | E1278H | E | 1.6e-05 | 1.4e-05 | 1.6e-05 | 1.533E-05 | 1.155E-06 | 1.400E-05 | 1.600E-05 |
| 25550 | 1278 | E1278I | E | 8.8e-06 | 1.7e-05 | 1.1e-05 | 1.227E-05 | 4.244E-06 | 8.800E-06 | 1.700E-05 |
| 25551 | 1278 | E1278L | E | 1.6e-05 | 1.1e-05 | 1.5e-05 | 1.400E-05 | 2.646E-06 | 1.100E-05 | 1.600E-05 |
| 25552 | 1278 | E1278K | E | 8e-06 | 8.4e-06 | 1.3e-05 | 9.800E-06 | 2.778E-06 | 8.000E-06 | 1.300E-05 |
| 25553 | 1278 | E1278M | E | 8.3e-06 | 8.9e-06 | 1.5e-05 | 1.073E-05 | 3.707E-06 | 8.300E-06 | 1.500E-05 |
| 25554 | 1278 | E1278F | E | 1.1e-05 | 1e-05 | 1.5e-05 | 1.200E-05 | 2.646E-06 | 1.000E-05 | 1.500E-05 |
| 25555 | 1278 | E1278P | E | 9.6e-06 | 1.4e-05 | 1.4e-05 | 1.253E-05 | 2.540E-06 | 9.600E-06 | 1.400E-05 |
| 25556 | 1278 | E1278S | E | 1e-05 | 1.6e-05 | 1e-05 | 1.200E-05 | 3.464E-06 | 1.000E-05 | 1.600E-05 |
| 25557 | 1278 | E1278T | E | 1.1e-05 | 1e-05 | 1.3e-05 | 1.133E-05 | 1.528E-06 | 1.000E-05 | 1.300E-05 |
| 25558 | 1278 | E1278W | E | 1.3e-05 | 1.6e-05 | 1.7e-05 | 1.533E-05 | 2.082E-06 | 1.300E-05 | 1.700E-05 |
| 25559 | 1278 | E1278Y | E | 1.7e-05 | 1.7e-05 | 1.6e-05 | 1.667E-05 | 5.774E-07 | 1.600E-05 | 1.700E-05 |
| 25560 | 1278 | E1278V | E | 1.6e-05 | 9.1e-06 | 1e-05 | 1.170E-05 | 3.751E-06 | 9.100E-06 | 1.600E-05 |
| 25561 | 1279 | S1279A | E | 9.9e-06 | 1.6e-05 | 1.3e-05 | 1.297E-05 | 3.050E-06 | 9.900E-06 | 1.600E-05 |
| 25562 | 1279 | S1279R | E | 1.5e-05 | 1.6e-05 | 1.1e-05 | 1.400E-05 | 2.646E-06 | 1.100E-05 | 1.600E-05 |
| 25563 | 1279 | S1279N | E | 1.9e-05 | 7.5e-06 | 1.5e-05 | 1.383E-05 | 5.838E-06 | 7.500E-06 | 1.900E-05 |
| 25564 | 1279 | S1279D | E | 1.4e-05 | 1.1e-05 | 9.4e-06 | 1.147E-05 | 2.335E-06 | 9.400E-06 | 1.400E-05 |
| 25565 | 1279 | S1279C | E | 1.4e-05 | 1.4e-05 | 1.5e-05 | 1.433E-05 | 5.774E-07 | 1.400E-05 | 1.500E-05 |
| 25566 | 1279 | S1279E | E | 1.4e-05 | 1.6e-05 | 1.1e-05 | 1.367E-05 | 2.517E-06 | 1.100E-05 | 1.600E-05 |
| 25567 | 1279 | S1279Q | E | 1.1e-05 | 9.4e-06 | 1.4e-05 | 1.147E-05 | 2.335E-06 | 9.400E-06 | 1.400E-05 |
| 25568 | 1279 | S1279G | E | 1.2e-05 | 1.5e-05 | 9.5e-06 | 1.217E-05 | 2.754E-06 | 9.500E-06 | 1.500E-05 |
| 25569 | 1279 | S1279H | E | 1.1e-05 | 9.6e-06 | 1.4e-05 | 1.153E-05 | 2.248E-06 | 9.600E-06 | 1.400E-05 |
| 25570 | 1279 | S1279I | E | 1.6e-05 | 7.1e-06 | 9e-06 | 1.070E-05 | 4.687E-06 | 7.100E-06 | 1.600E-05 |
| 25571 | 1279 | S1279L | E | 1.1e-05 | 9.2e-06 | 1.2e-05 | 1.073E-05 | 1.419E-06 | 9.200E-06 | 1.200E-05 |
| 25572 | 1279 | S1279K | E | 1.3e-05 | 1.2e-05 | 1.1e-05 | 1.200E-05 | 1.000E-06 | 1.100E-05 | 1.300E-05 |
| 25573 | 1279 | S1279M | E | 9.9e-06 | 1.4e-05 | 1.2e-05 | 1.197E-05 | 2.050E-06 | 9.900E-06 | 1.400E-05 |
| 25574 | 1279 | S1279F | E | 1.4e-05 | 1.6e-05 | 1.7e-05 | 1.567E-05 | 1.528E-06 | 1.400E-05 | 1.700E-05 |
| 25575 | 1279 | S1279P | E | 1.5e-05 | 2.1e-05 | 1.5e-05 | 1.700E-05 | 3.464E-06 | 1.500E-05 | 2.100E-05 |
| 25576 | 1279 | S1279S | E | 1e-05 | 1.1e-05 | 1.1e-05 | 1.067E-05 | 5.774E-07 | 1.000E-05 | 1.100E-05 |
| 25577 | 1279 | S1279T | E | 1e-05 | 9.9e-06 | 1.1e-05 | 1.030E-05 | 6.083E-07 | 9.900E-06 | 1.100E-05 |
| 25578 | 1279 | S1279W | E | 9.5e-06 | 1.7e-05 | 1.5e-05 | 1.383E-05 | 3.884E-06 | 9.500E-06 | 1.700E-05 |
| 25579 | 1279 | S1279Y | E | 9.7e-06 | 1.6e-05 | 1.4e-05 | 1.323E-05 | 3.219E-06 | 9.700E-06 | 1.600E-05 |
| 25580 | 1279 | S1279V | E | 1.7e-05 | 1.6e-05 | 1.1e-05 | 1.467E-05 | 3.215E-06 | 1.100E-05 | 1.700E-05 |
| 25581 | 1280 | G1280A | E | 1.2e-05 | 1.4e-05 | 1.7e-05 | 1.433E-05 | 2.517E-06 | 1.200E-05 | 1.700E-05 |
| 25582 | 1280 | G1280R | E | 1.4e-05 | 1.1e-05 | 1.1e-05 | 1.200E-05 | 1.732E-06 | 1.100E-05 | 1.400E-05 |
| 25583 | 1280 | G1280N | E | 1e-05 | 1.4e-05 | 1.6e-05 | 1.333E-05 | 3.055E-06 | 1.000E-05 | 1.600E-05 |
| 25584 | 1280 | G1280D | E | 8.8e-06 | 1.4e-05 | 9.1e-06 | 1.063E-05 | 2.919E-06 | 8.800E-06 | 1.400E-05 |

| | | | | | | | | | | |
|---|---|---|---|---|---|---|---|---|---|---|
| 25585 | 1280 | G1280C | E | 1.5e-05 | 1.5e-05 | 9.5e-06 | 1.317E-05 | 3.175E-06 | 9.500E-06 | 1.500E-05 |
| 25586 | 1280 | G1280E | E | 8.4e-06 | 8.8e-06 | 7.4e-06 | 8.200E-06 | 7.211E-07 | 7.400E-06 | 8.800E-06 |
| 25587 | 1280 | G1280Q | E | 1.4e-05 | 1.1e-05 | 1.5e-05 | 1.333E-05 | 2.082E-06 | 1.100E-05 | 1.500E-05 |
| 25588 | 1280 | G1280G | E | 1.1e-05 | 1.1e-05 | 9.5e-06 | 1.050E-05 | 8.660E-07 | 9.500E-06 | 1.100E-05 |
| 25589 | 1280 | G1280H | E | 1.6e-05 | 1.4e-05 | 1.5e-05 | 1.500E-05 | 1.000E-06 | 1.400E-05 | 1.600E-05 |
| 25590 | 1280 | G1280I | E | 8.4e-06 | 1.4e-05 | 1.4e-05 | 1.213E-05 | 3.233E-06 | 8.400E-06 | 1.400E-05 |
| 25591 | 1280 | G1280L | E | 1.5e-05 | 9.3e-06 | 1.5e-05 | 1.310E-05 | 3.291E-06 | 9.300E-06 | 1.500E-05 |
| 25592 | 1280 | G1280K | E | 1.2e-05 | 1.4e-05 | 9.3e-06 | 1.177E-05 | 2.359E-06 | 9.300E-06 | 1.400E-05 |
| 25593 | 1280 | G1280M | E | 1.7e-05 | 1.1e-05 | 1.6e-05 | 1.467E-05 | 3.215E-06 | 1.100E-05 | 1.700E-05 |
| 25594 | 1280 | G1280F | E | 9e-06 | 9.6e-06 | 1.6e-05 | 1.153E-05 | 3.880E-06 | 9.000E-06 | 1.600E-05 |
| 25595 | 1280 | G1280P | E | 1.3e-05 | 1.3e-05 | 8.3e-06 | 1.143E-05 | 2.714E-06 | 8.300E-06 | 1.300E-05 |
| 25596 | 1280 | G1280S | E | 1.4e-05 | 1.4e-05 | 1.3e-05 | 1.367E-05 | 5.774E-07 | 1.300E-05 | 1.400E-05 |
| 25597 | 1280 | G1280T | E | 1.5e-05 | 1.6e-05 | 1.4e-05 | 1.500E-05 | 1.000E-06 | 1.400E-05 | 1.600E-05 |
| 25598 | 1280 | G1280W | E | 1.5e-05 | 1.6e-05 | 9e-06 | 1.333E-05 | 3.786E-06 | 9.000E-06 | 1.600E-05 |
| 25599 | 1280 | G1280Y | E | 1.1e-05 | 9.4e-06 | 9.3e-06 | 9.900E-06 | 9.539E-07 | 9.300E-06 | 1.100E-05 |
| 25600 | 1280 | G1280V | E | 1.6e-05 | 1e-05 | 1.3e-05 | 1.300E-05 | 3.000E-06 | 1.000E-05 | 1.600E-05 |
| 25601 | 1281 | G1281A | E | 1.4e-05 | 1.1e-05 | 1.6e-05 | 1.367E-05 | 2.517E-06 | 1.100E-05 | 1.600E-05 |
| 25602 | 1281 | G1281R | E | 1.5e-05 | 1.4e-05 | 8.3e-06 | 1.243E-05 | 3.614E-06 | 8.300E-06 | 1.500E-05 |
| 25603 | 1281 | G1281N | E | 1.6e-05 | 1e-05 | 8.5e-06 | 1.150E-05 | 3.969E-06 | 8.500E-06 | 1.600E-05 |
| 25604 | 1281 | G1281D | E | 1.4e-05 | 1.1e-05 | 1.5e-05 | 1.333E-05 | 2.082E-06 | 1.100E-05 | 1.500E-05 |
| 25605 | 1281 | G1281C | E | 9.9e-06 | 9.6e-06 | 1.2e-05 | 1.050E-05 | 1.308E-06 | 9.600E-06 | 1.200E-05 |
| 25606 | 1281 | G1281E | E | 9.9e-06 | 1.4e-05 | 1e-05 | 1.130E-05 | 2.339E-06 | 9.900E-06 | 1.400E-05 |
| 25607 | 1281 | G1281Q | E | 8.7e-06 | 8.4e-06 | 9.3e-06 | 8.800E-06 | 4.583E-07 | 8.400E-06 | 9.300E-06 |
| 25608 | 1281 | G1281G | E | 8.2e-06 | 1e-05 | 9.9e-06 | 9.367E-06 | 1.012E-06 | 8.200E-06 | 1.000E-05 |
| 25609 | 1281 | G1281H | E | 1.4e-05 | 1.3e-05 | 1.4e-05 | 1.367E-05 | 5.774E-07 | 1.300E-05 | 1.400E-05 |
| 25610 | 1281 | G1281I | E | 1e-05 | 1e-05 | 1.5e-05 | 1.167E-05 | 2.887E-06 | 1.000E-05 | 1.500E-05 |
| 25611 | 1281 | G1281L | E | 9.7e-06 | 1.1e-05 | 1e-05 | 1.023E-05 | 6.807E-07 | 9.700E-06 | 1.100E-05 |
| 25612 | 1281 | G1281K | E | 1.2e-05 | 1.1e-05 | 9.5e-06 | 1.083E-05 | 1.258E-06 | 9.500E-06 | 1.200E-05 |
| 25613 | 1281 | G1281M | E | 1.3e-05 | 7.2e-06 | 1.2e-05 | 1.073E-05 | 3.101E-06 | 7.200E-06 | 1.300E-05 |
| 25614 | 1281 | G1281F | E | 1.1e-05 | 1.6e-05 | 1.2e-05 | 1.300E-05 | 2.646E-06 | 1.100E-05 | 1.600E-05 |
| 25615 | 1281 | G1281P | E | 1.7e-05 | 1e-05 | 7.9e-06 | 1.163E-05 | 4.765E-06 | 7.900E-06 | 1.700E-05 |
| 25616 | 1281 | G1281S | E | 1.4e-05 | 1.4e-05 | 9.1e-06 | 1.237E-05 | 2.829E-06 | 9.100E-06 | 1.400E-05 |
| 25617 | 1281 | G1281T | E | 1.4e-05 | 8.7e-06 | 1.5e-05 | 1.257E-05 | 3.386E-06 | 8.700E-06 | 1.500E-05 |
| 25618 | 1281 | G1281W | E | 8e-06 | 1.6e-05 | 1e-05 | 1.133E-05 | 4.163E-06 | 8.000E-06 | 1.600E-05 |
| 25619 | 1281 | G1281Y | E | 1.1e-05 | 1.3e-05 | 1.5e-05 | 1.300E-05 | 2.000E-06 | 1.100E-05 | 1.500E-05 |
| 25620 | 1281 | G1281V | E | 1.3e-05 | 1.1e-05 | 1.1e-05 | 1.167E-05 | 1.155E-06 | 1.100E-05 | 1.300E-05 |
| 25621 | 1282 | G1282A | E | 1.2e-05 | 1.5e-05 | 1.7e-05 | 1.467E-05 | 2.517E-06 | 1.200E-05 | 1.700E-05 |
| 25622 | 1282 | G1282R | E | 1.4e-05 | 1.6e-05 | 6e-06 | 1.200E-05 | 5.292E-06 | 6.000E-06 | 1.600E-05 |
| 25623 | 1282 | G1282N | E | 1.2e-05 | 1.4e-05 | 1.5e-05 | 1.367E-05 | 1.528E-06 | 1.200E-05 | 1.500E-05 |
| 25624 | 1282 | G1282D | E | 1.6e-05 | 8.6e-06 | 1.4e-05 | 1.287E-05 | 3.828E-06 | 8.600E-06 | 1.600E-05 |
| 25625 | 1282 | G1282C | E | 9.5e-06 | 1.5e-05 | 1.5e-05 | 1.317E-05 | 3.175E-06 | 9.500E-06 | 1.500E-05 |
| 25626 | 1282 | G1282E | E | 1.5e-05 | 1.1e-05 | 1.4e-05 | 1.333E-05 | 2.082E-06 | 1.100E-05 | 1.500E-05 |
| 25627 | 1282 | G1282Q | E | 1e-05 | 1.2e-05 | 1.4e-05 | 1.200E-05 | 2.000E-06 | 1.000E-05 | 1.400E-05 |
| 25628 | 1282 | G1282G | E | 1.4e-05 | 1.1e-05 | 1.2e-05 | 1.233E-05 | 1.528E-06 | 1.100E-05 | 1.400E-05 |
| 25629 | 1282 | G1282H | E | 7.9e-06 | 1.5e-05 | 1.1e-05 | 1.130E-05 | 3.559E-06 | 7.900E-06 | 1.500E-05 |
| 25630 | 1282 | G1282I | E | 1e-05 | 1.6e-05 | 1.6e-05 | 1.400E-05 | 3.464E-06 | 1.000E-05 | 1.600E-05 |
| 25631 | 1282 | G1282L | E | 1.5e-05 | 1.2e-05 | 1.5e-05 | 1.400E-05 | 1.732E-06 | 1.200E-05 | 1.500E-05 |
| 25632 | 1282 | G1282K | E | 1.4e-05 | 1.1e-05 | 1.1e-05 | 1.200E-05 | 1.732E-06 | 1.100E-05 | 1.400E-05 |
| 25633 | 1282 | G1282M | E | 1e-05 | 1.7e-05 | 8.5e-06 | 1.183E-05 | 4.537E-06 | 8.500E-06 | 1.700E-05 |
| 25634 | 1282 | G1282F | E | 9.8e-06 | 1.5e-05 | 1.7e-05 | 1.393E-05 | 3.717E-06 | 9.800E-06 | 1.700E-05 |
| 25635 | 1282 | G1282P | E | 1.5e-05 | 1.4e-05 | 1.5e-05 | 1.467E-05 | 5.774E-07 | 1.400E-05 | 1.500E-05 |
| 25636 | 1282 | G1282S | E | 1.5e-05 | 1.2e-05 | 1.4e-05 | 1.367E-05 | 1.528E-06 | 1.200E-05 | 1.500E-05 |
| 25637 | 1282 | G1282T | E | 1.5e-05 | 1e-05 | 1.5e-05 | 1.333E-05 | 2.887E-06 | 1.000E-05 | 1.500E-05 |
| 25638 | 1282 | G1282W | E | 9.9e-06 | 1.5e-05 | 1.5e-05 | 1.330E-05 | 2.944E-06 | 9.900E-06 | 1.500E-05 |
| 25639 | 1282 | G1282Y | E | 9.7e-06 | 1.3e-05 | 1.1e-05 | 1.123E-05 | 1.662E-06 | 9.700E-06 | 1.300E-05 |
| 25640 | 1282 | G1282V | E | 6.4e-06 | 1.2e-05 | 1.1e-05 | 9.800E-06 | 2.987E-06 | 6.400E-06 | 1.200E-05 |
| 25641 | 1283 | L1283A | E | 1.6e-05 | 1.1e-05 | 1.4e-05 | 1.367E-05 | 2.517E-06 | 1.100E-05 | 1.600E-05 |
| 25642 | 1283 | L1283R | E | 1.6e-05 | 8.6e-06 | 1.8e-05 | 1.420E-05 | 4.952E-06 | 8.600E-06 | 1.800E-05 |
| 25643 | 1283 | L1283N | E | 1.1e-05 | 1.3e-05 | 1e-05 | 1.133E-05 | 1.528E-06 | 1.000E-05 | 1.300E-05 |
| 25644 | 1283 | L1283D | E | 1.4e-05 | 9.7e-06 | 8.2e-06 | 1.063E-05 | 3.011E-06 | 8.200E-06 | 1.400E-05 |
| 25645 | 1283 | L1283C | E | 9.2e-06 | 1.6e-05 | 1.4e-05 | 1.307E-05 | 3.495E-06 | 9.200E-06 | 1.600E-05 |
| 25646 | 1283 | L1283E | E | 1.4e-05 | 8.7e-06 | 1.6e-05 | 1.290E-05 | 3.772E-06 | 8.700E-06 | 1.600E-05 |
| 25647 | 1283 | L1283Q | E | 1.5e-05 | 1.5e-05 | 9.5e-06 | 1.317E-05 | 3.175E-06 | 9.500E-06 | 1.500E-05 |
| 25648 | 1283 | L1283G | E | 1.6e-05 | 8.8e-06 | 1.5e-05 | 1.327E-05 | 3.900E-06 | 8.800E-06 | 1.600E-05 |
| 25649 | 1283 | L1283H | E | 7.6e-06 | 1.7e-05 | 1.5e-05 | 1.320E-05 | 4.952E-06 | 7.600E-06 | 1.700E-05 |
| 25650 | 1283 | L1283I | E | 8e-06 | 1.5e-05 | 1.3e-05 | 1.200E-05 | 3.606E-06 | 8.000E-06 | 1.500E-05 |
| 25651 | 1283 | L1283L | E | 1.4e-05 | 5.9e-06 | 1.4e-05 | 1.130E-05 | 4.677E-06 | 5.900E-06 | 1.400E-05 |
| 25652 | 1283 | L1283K | E | 1.4e-05 | 1.1e-05 | 1e-05 | 1.167E-05 | 2.082E-06 | 1.000E-05 | 1.400E-05 |
| 25653 | 1283 | L1283M | E | 1.9e-05 | 1.4e-05 | 1e-05 | 1.433E-05 | 4.509E-06 | 1.000E-05 | 1.900E-05 |
| 25654 | 1283 | L1283F | E | 1.5e-05 | 1.5e-05 | 1.4e-05 | 1.467E-05 | 5.774E-07 | 1.400E-05 | 1.500E-05 |
| 25655 | 1283 | L1283P | E | 1.1e-05 | 1.5e-05 | 9.6e-06 | 1.187E-05 | 2.802E-06 | 9.600E-06 | 1.500E-05 |
| 25656 | 1283 | L1283S | E | 1.4e-05 | 1.1e-05 | 1e-05 | 1.167E-05 | 2.082E-06 | 1.000E-05 | 1.400E-05 |
| 25657 | 1283 | L1283T | E | 1e-05 | 1.5e-05 | 9.2e-06 | 1.140E-05 | 3.143E-06 | 9.200E-06 | 1.500E-05 |
| 25658 | 1283 | L1283W | E | 1e-05 | 1.1e-05 | 1.1e-05 | 1.067E-05 | 5.774E-07 | 1.000E-05 | 1.100E-05 |
| 25659 | 1283 | L1283Y | E | 1.4e-05 | 1.4e-05 | 1.3e-05 | 1.367E-05 | 5.774E-07 | 1.300E-05 | 1.400E-05 |
| 25660 | 1283 | L1283V | E | 1e-05 | 1.4e-05 | 1.1e-05 | 1.167E-05 | 2.082E-06 | 1.000E-05 | 1.400E-05 |
| 25661 | 1284 | V1284A | E | 1.5e-05 | 1.7e-05 | 1.1e-05 | 1.433E-05 | 3.055E-06 | 1.100E-05 | 1.700E-05 |
| 25662 | 1284 | V1284R | E | 9.6e-06 | 1.2e-05 | 9.4e-06 | 1.033E-05 | 1.447E-06 | 9.400E-06 | 1.200E-05 |
| 25663 | 1284 | V1284N | E | 1.7e-05 | 9.7e-06 | 9.6e-06 | 1.210E-05 | 4.244E-06 | 9.600E-06 | 1.700E-05 |
| 25664 | 1284 | V1284D | E | 1.4e-05 | 1.1e-05 | 9.9e-06 | 1.163E-05 | 2.122E-06 | 9.900E-06 | 1.400E-05 |
| 25665 | 1284 | V1284C | E | 1.1e-05 | 1.1e-05 | 1e-05 | 1.067E-05 | 5.774E-07 | 1.000E-05 | 1.100E-05 |
| 25666 | 1284 | V1284E | E | 1e-05 | 8.2e-06 | 1.3e-05 | 1.040E-05 | 2.425E-06 | 8.200E-06 | 1.300E-05 |
| 25667 | 1284 | V1284Q | E | 1.4e-05 | 1.4e-05 | 1.4e-05 | 1.400E-05 | 0.000E+00 | 1.400E-05 | 1.400E-05 |
| 25668 | 1284 | V1284G | E | 1.3e-05 | 1.5e-05 | 9e-06 | 1.233E-05 | 3.055E-06 | 9.000E-06 | 1.500E-05 |
| 25669 | 1284 | V1284H | E | 1e-05 | 1e-05 | 1.5e-05 | 1.167E-05 | 2.887E-06 | 1.000E-05 | 1.500E-05 |

| | | | | | | | | | | |
|---|---|---|---|---|---|---|---|---|---|---|
| 25670 | 1284 | V1284I | E | 1.1e-05 | 1e-05 | 1.5e-05 | 1.200E-05 | 2.646E-06 | 1.000E-05 | 1.500E-05 |
| 25671 | 1284 | V1284L | E | 1.3e-05 | 1.4e-05 | 9.8e-06 | 1.227E-05 | 2.194E-06 | 9.800E-06 | 1.400E-05 |
| 25672 | 1284 | V1284K | E | 9.7e-06 | 1.6e-05 | 9e-06 | 1.157E-05 | 3.855E-06 | 9.000E-06 | 1.600E-05 |
| 25673 | 1284 | V1284M | E | 1.3e-05 | 1.7e-05 | 1.4e-05 | 1.467E-05 | 2.082E-06 | 1.300E-05 | 1.700E-05 |
| 25674 | 1284 | V1284F | E | 8.1e-06 | 1e-05 | 1.2e-05 | 1.003E-05 | 1.950E-06 | 8.100E-06 | 1.200E-05 |
| 25675 | 1284 | V1284P | E | 1.7e-05 | 1.2e-05 | 9.8e-06 | 1.293E-05 | 3.690E-06 | 9.800E-06 | 1.700E-05 |
| 25676 | 1284 | V1284S | E | 1.5e-05 | 1.5e-05 | 1e-05 | 1.333E-05 | 2.887E-06 | 1.000E-05 | 1.500E-05 |
| 25677 | 1284 | V1284T | E | 1e-05 | 1.5e-05 | 1.6e-05 | 1.367E-05 | 3.215E-06 | 1.000E-05 | 1.600E-05 |
| 25678 | 1284 | V1284W | E | 1.5e-05 | 1.6e-05 | 1.1e-05 | 1.400E-05 | 2.646E-06 | 1.100E-05 | 1.600E-05 |
| 25679 | 1284 | V1284Y | E | 1.6e-05 | 1.3e-05 | 7.5e-06 | 1.217E-05 | 4.311E-06 | 7.500E-06 | 1.600E-05 |
| 25680 | 1284 | V1284V | E | 9.2e-06 | 1.3e-05 | 9.6e-06 | 1.060E-05 | 2.088E-06 | 9.200E-06 | 1.300E-05 |
| 25681 | 1285 | Q1285A | E | 1e-05 | 1.5e-05 | 1.2e-05 | 1.233E-05 | 2.517E-06 | 1.000E-05 | 1.500E-05 |
| 25682 | 1285 | Q1285R | E | 1.6e-05 | 1.6e-05 | 1.3e-05 | 1.500E-05 | 1.732E-06 | 1.300E-05 | 1.600E-05 |
| 25683 | 1285 | Q1285N | E | 9.1e-06 | 9.9e-06 | 1.7e-05 | 1.200E-05 | 4.349E-06 | 9.100E-06 | 1.700E-05 |
| 25684 | 1285 | Q1285D | E | 1.5e-05 | 7.9e-06 | 1.2e-05 | 1.163E-05 | 3.564E-06 | 7.900E-06 | 1.500E-05 |
| 25685 | 1285 | Q1285C | E | 1.4e-05 | 1.6e-05 | 9.2e-06 | 1.307E-05 | 3.495E-06 | 9.200E-06 | 1.600E-05 |
| 25686 | 1285 | Q1285E | E | 1.9e-05 | 8.2e-06 | 1.6e-05 | 1.440E-05 | 5.575E-06 | 8.200E-06 | 1.900E-05 |
| 25687 | 1285 | Q1285Q | E | 1.4e-05 | 1.3e-05 | 9.3e-06 | 1.210E-05 | 2.476E-06 | 9.300E-06 | 1.400E-05 |
| 25688 | 1285 | Q1285G | E | 1e-05 | 1e-05 | 1e-05 | 1.000E-05 | 0.000E+00 | 1.000E-05 | 1.000E-05 |
| 25689 | 1285 | Q1285H | E | 1e-05 | 1.3e-05 | 1.5e-05 | 1.267E-05 | 2.517E-06 | 1.000E-05 | 1.500E-05 |
| 25690 | 1285 | Q1285I | E | 1.5e-05 | 1.7e-05 | 1.7e-05 | 1.633E-05 | 1.155E-06 | 1.500E-05 | 1.700E-05 |
| 25691 | 1285 | Q1285L | E | 1.4e-05 | 1.1e-05 | 1.3e-05 | 1.267E-05 | 1.528E-06 | 1.100E-05 | 1.400E-05 |
| 25692 | 1285 | Q1285K | E | 1.1e-05 | 1e-05 | 1.4e-05 | 1.167E-05 | 2.082E-06 | 1.000E-05 | 1.400E-05 |
| 25693 | 1285 | Q1285M | E | 1.2e-05 | 1.6e-05 | 1.4e-05 | 1.400E-05 | 2.000E-06 | 1.200E-05 | 1.600E-05 |
| 25694 | 1285 | Q1285F | E | 1.3e-05 | 1.4e-05 | 1.6e-05 | 1.433E-05 | 1.528E-06 | 1.300E-05 | 1.600E-05 |
| 25695 | 1285 | Q1285P | E | 1.2e-05 | 1.5e-05 | 1.7e-05 | 1.467E-05 | 2.517E-06 | 1.200E-05 | 1.700E-05 |
| 25696 | 1285 | Q1285S | E | 1.6e-05 | 1e-05 | 1.7e-05 | 1.433E-05 | 3.786E-06 | 1.000E-05 | 1.700E-05 |
| 25697 | 1285 | Q1285T | E | 9.2e-06 | 1.5e-05 | 1.2e-05 | 1.207E-05 | 2.901E-06 | 9.200E-06 | 1.500E-05 |
| 25698 | 1285 | Q1285W | E | 1.3e-05 | 9.6e-06 | 1e-05 | 1.087E-05 | 1.858E-06 | 9.600E-06 | 1.300E-05 |
| 25699 | 1285 | Q1285Y | E | 9.8e-06 | 1.3e-05 | 1e-05 | 1.093E-05 | 1.793E-06 | 9.800E-06 | 1.300E-05 |
| 25700 | 1285 | Q1285V | E | 1.5e-05 | 1.1e-05 | 1.6e-05 | 1.400E-05 | 2.646E-06 | 1.100E-05 | 1.600E-05 |
| 25701 | 1286 | P1286A | E | 1.2e-05 | 1e-05 | 1.4e-05 | 1.200E-05 | 2.000E-06 | 1.000E-05 | 1.400E-05 |
| 25702 | 1286 | P1286R | E | 9.8e-06 | 1.5e-05 | 1.3e-05 | 1.260E-05 | 2.623E-06 | 9.800E-06 | 1.500E-05 |
| 25703 | 1286 | P1286N | E | 8.5e-06 | 1.2e-05 | 1.3e-05 | 1.117E-05 | 2.363E-06 | 8.500E-06 | 1.300E-05 |
| 25704 | 1286 | P1286D | E | 1.4e-05 | 1.3e-05 | 8.7e-06 | 1.190E-05 | 2.816E-06 | 8.700E-06 | 1.400E-05 |
| 25705 | 1286 | P1286C | E | 1.2e-05 | 1.3e-05 | 1.4e-05 | 1.300E-05 | 1.000E-06 | 1.200E-05 | 1.400E-05 |
| 25706 | 1286 | P1286E | E | 1.2e-05 | 1.6e-05 | 9.7e-06 | 1.257E-05 | 3.188E-06 | 9.700E-06 | 1.600E-05 |
| 25707 | 1286 | P1286Q | E | 1.4e-05 | 1.3e-05 | 1.3e-05 | 1.333E-05 | 5.774E-07 | 1.300E-05 | 1.400E-05 |
| 25708 | 1286 | P1286G | E | 1.2e-05 | 1.6e-05 | 1.5e-05 | 1.433E-05 | 2.082E-06 | 1.200E-05 | 1.600E-05 |
| 25709 | 1286 | P1286H | E | 1e-05 | 8.7e-06 | 1.5e-05 | 1.123E-05 | 3.326E-06 | 8.700E-06 | 1.500E-05 |
| 25710 | 1286 | P1286I | E | 8.6e-06 | 2e-05 | 1.8e-05 | 1.553E-05 | 6.087E-06 | 8.600E-06 | 2.000E-05 |
| 25711 | 1286 | P1286L | E | 1.5e-05 | 1.2e-05 | 1.1e-05 | 1.267E-05 | 2.082E-06 | 1.100E-05 | 1.500E-05 |
| 25712 | 1286 | P1286K | E | 1.1e-05 | 1e-05 | 9.5e-06 | 1.017E-05 | 7.638E-07 | 9.500E-06 | 1.100E-05 |
| 25713 | 1286 | P1286M | E | 9.5e-06 | 1e-05 | 1e-05 | 9.833E-06 | 2.887E-07 | 9.500E-06 | 1.000E-05 |
| 25714 | 1286 | P1286F | E | 1.6e-05 | 1e-05 | 9.4e-06 | 1.180E-05 | 3.650E-06 | 9.400E-06 | 1.600E-05 |
| 25715 | 1286 | P1286P | E | 1e-05 | 9.9e-06 | 9.2e-06 | 9.700E-06 | 4.359E-07 | 9.200E-06 | 1.000E-05 |
| 25716 | 1286 | P1286S | E | 1.6e-05 | 1.3e-05 | 1.5e-05 | 1.467E-05 | 1.528E-06 | 1.300E-05 | 1.600E-05 |
| 25717 | 1286 | P1286T | E | 1.6e-05 | 8.2e-06 | 9.1e-06 | 1.110E-05 | 4.267E-06 | 8.200E-06 | 1.600E-05 |
| 25718 | 1286 | P1286W | E | 9.9e-06 | 1.5e-05 | 1.5e-05 | 1.330E-05 | 2.944E-06 | 9.900E-06 | 1.500E-05 |
| 25719 | 1286 | P1286Y | E | 9.9e-06 | 1.1e-05 | 1.5e-05 | 1.197E-05 | 2.684E-06 | 9.900E-06 | 1.500E-05 |
| 25720 | 1286 | P1286V | E | 1.5e-05 | 1.5e-05 | 9.9e-06 | 1.330E-05 | 2.944E-06 | 9.900E-06 | 1.500E-05 |
| 25721 | 1287 | G1287A | E | 1.4e-05 | 1.1e-05 | 1.1e-05 | 1.200E-05 | 1.732E-06 | 1.100E-05 | 1.400E-05 |
| 25722 | 1287 | G1287R | E | 1.3e-05 | 1.6e-05 | 1.4e-05 | 1.433E-05 | 1.528E-06 | 1.300E-05 | 1.600E-05 |
| 25723 | 1287 | G1287N | E | 1.1e-05 | 1.2e-05 | 8.9e-06 | 1.063E-05 | 1.582E-06 | 8.900E-06 | 1.200E-05 |
| 25724 | 1287 | G1287D | E | 9.9e-06 | 1.6e-05 | 1.4e-05 | 1.330E-05 | 3.110E-06 | 9.900E-06 | 1.600E-05 |
| 25725 | 1287 | G1287C | E | 1e-05 | 1.1e-05 | 1.3e-05 | 1.133E-05 | 1.528E-06 | 1.000E-05 | 1.300E-05 |
| 25726 | 1287 | G1287E | E | 1.7e-05 | 1.5e-05 | 1.5e-05 | 1.567E-05 | 1.155E-06 | 1.500E-05 | 1.700E-05 |
| 25727 | 1287 | G1287Q | E | 9.4e-06 | 5.9e-06 | 9.5e-06 | 8.267E-06 | 2.050E-06 | 5.900E-06 | 9.500E-06 |
| 25728 | 1287 | G1287G | E | 1.5e-05 | 1.6e-05 | 1.3e-05 | 1.467E-05 | 1.528E-06 | 1.300E-05 | 1.600E-05 |
| 25729 | 1287 | G1287H | E | 1.4e-05 | 1.2e-05 | 9e-06 | 1.167E-05 | 2.517E-06 | 9.000E-06 | 1.400E-05 |
| 25730 | 1287 | G1287I | E | 9.1e-06 | 1.3e-05 | 1.4e-05 | 1.203E-05 | 2.589E-06 | 9.100E-06 | 1.400E-05 |
| 25731 | 1287 | G1287L | E | 1.1e-05 | 1.4e-05 | 1e-05 | 1.167E-05 | 2.082E-06 | 1.000E-05 | 1.400E-05 |
| 25732 | 1287 | G1287K | E | 8.8e-06 | 1.4e-05 | 1.6e-05 | 1.293E-05 | 3.717E-06 | 8.800E-06 | 1.600E-05 |
| 25733 | 1287 | G1287M | E | 1e-05 | 9.7e-06 | 1.4e-05 | 1.123E-05 | 2.401E-06 | 9.700E-06 | 1.400E-05 |
| 25734 | 1287 | G1287F | E | 1.4e-05 | 1.1e-05 | 1e-05 | 1.167E-05 | 2.082E-06 | 1.000E-05 | 1.400E-05 |
| 25735 | 1287 | G1287P | E | 1.3e-05 | 9.8e-06 | 1.6e-05 | 1.293E-05 | 3.101E-06 | 9.800E-06 | 1.600E-05 |
| 25736 | 1287 | G1287S | E | 1.1e-05 | 9.2e-06 | 1.5e-05 | 1.173E-05 | 2.969E-06 | 9.200E-06 | 1.500E-05 |
| 25737 | 1287 | G1287T | E | 1.3e-05 | 1.6e-05 | 1.1e-05 | 1.333E-05 | 2.517E-06 | 1.100E-05 | 1.600E-05 |
| 25738 | 1287 | G1287W | E | 1.6e-05 | 1.5e-05 | 1.1e-05 | 1.400E-05 | 2.646E-06 | 1.100E-05 | 1.600E-05 |
| 25739 | 1287 | G1287Y | E | 9.9e-06 | 1.1e-05 | 1e-05 | 1.030E-05 | 6.083E-07 | 9.900E-06 | 1.100E-05 |
| 25740 | 1287 | G1287V | E | 1.5e-05 | 7.5e-06 | 8e-06 | 1.017E-05 | 4.193E-06 | 7.500E-06 | 1.500E-05 |
| 25741 | 1288 | G1288A | E | 1.5e-05 | 1.7e-05 | 9.3e-06 | 1.377E-05 | 3.995E-06 | 9.300E-06 | 1.700E-05 |
| 25742 | 1288 | G1288R | E | 1.6e-05 | 1e-05 | 1.5e-05 | 1.367E-05 | 3.215E-06 | 1.000E-05 | 1.600E-05 |
| 25743 | 1288 | G1288N | E | 8.3e-06 | 9.1e-06 | 1.7e-05 | 1.147E-05 | 4.809E-06 | 8.300E-06 | 1.700E-05 |
| 25744 | 1288 | G1288D | E | 1.6e-05 | 1.2e-05 | 1.1e-05 | 1.300E-05 | 2.646E-06 | 1.100E-05 | 1.600E-05 |
| 25745 | 1288 | G1288C | E | 1.5e-05 | 1e-05 | 1.5e-05 | 1.333E-05 | 2.887E-06 | 1.000E-05 | 1.500E-05 |
| 25746 | 1288 | G1288E | E | 1.4e-05 | 1.3e-05 | 9.8e-06 | 1.227E-05 | 2.194E-06 | 9.800E-06 | 1.400E-05 |
| 25747 | 1288 | G1288Q | E | 1.4e-05 | 8.5e-06 | 1.6e-05 | 1.283E-05 | 3.884E-06 | 8.500E-06 | 1.600E-05 |
| 25748 | 1288 | G1288G | E | 9.6e-06 | 1.3e-05 | 1.4e-05 | 1.220E-05 | 2.307E-06 | 9.600E-06 | 1.400E-05 |
| 25749 | 1288 | G1288H | E | 9.7e-06 | 1.3e-05 | 8e-06 | 1.023E-05 | 2.542E-06 | 8.000E-06 | 1.300E-05 |
| 25750 | 1288 | G1288I | E | 1.5e-05 | 1.1e-05 | 1.6e-05 | 1.400E-05 | 2.646E-06 | 1.100E-05 | 1.600E-05 |
| 25751 | 1288 | G1288L | E | 1.7e-05 | 9.4e-06 | 9.5e-06 | 1.197E-05 | 4.359E-06 | 9.400E-06 | 1.700E-05 |
| 25752 | 1288 | G1288K | E | 9.8e-06 | 1.3e-05 | 1.6e-05 | 1.293E-05 | 3.101E-06 | 9.800E-06 | 1.600E-05 |
| 25753 | 1288 | G1288M | E | 1.3e-05 | 1.7e-05 | 1.7e-05 | 1.567E-05 | 2.309E-06 | 1.300E-05 | 1.700E-05 |
| 25754 | 1288 | G1288F | E | 8.4e-06 | 1.3e-05 | 9.1e-06 | 1.017E-05 | 2.479E-06 | 8.400E-06 | 1.300E-05 |

| | | | | | | | | | | |
|---|---|---|---|---|---|---|---|---|---|---|
| 25755 | 1288 | G1288P | E | 1.5e-05 | 1.1e-05 | 1.6e-05 | 1.400E-05 | 2.646E-06 | 1.100E-05 | 1.600E-05 |
| 25756 | 1288 | G1288S | E | 1.5e-05 | 1.1e-05 | 1.4e-05 | 1.333E-05 | 2.082E-06 | 1.100E-05 | 1.500E-05 |
| 25757 | 1288 | G1288T | E | 1.5e-05 | 1.6e-05 | 1e-05 | 1.367E-05 | 3.215E-06 | 1.000E-05 | 1.600E-05 |
| 25758 | 1288 | G1288W | E | 1.5e-05 | 1.2e-05 | 1.1e-05 | 1.267E-05 | 2.082E-06 | 1.100E-05 | 1.500E-05 |
| 25759 | 1288 | G1288Y | E | 1.2e-05 | 9.9e-06 | 1.7e-05 | 1.297E-05 | 3.647E-06 | 9.900E-06 | 1.700E-05 |
| 25760 | 1288 | G1288V | E | 1e-05 | 1e-05 | 1.3e-05 | 1.100E-05 | 1.732E-06 | 1.000E-05 | 1.300E-05 |
| 25761 | 1289 | S1289A | E | 1.5e-05 | 1e-05 | 1.4e-05 | 1.300E-05 | 2.646E-06 | 1.000E-05 | 1.500E-05 |
| 25762 | 1289 | S1289R | E | 1.7e-05 | 1.8e-05 | 1.6e-05 | 1.700E-05 | 1.000E-06 | 1.600E-05 | 1.800E-05 |
| 25763 | 1289 | S1289N | E | 1e-05 | 1.6e-05 | 1.2e-05 | 1.267E-05 | 3.055E-06 | 1.000E-05 | 1.600E-05 |
| 25764 | 1289 | S1289D | E | 1.6e-05 | 1.1e-05 | 1.5e-05 | 1.400E-05 | 2.646E-06 | 1.100E-05 | 1.600E-05 |
| 25765 | 1289 | S1289C | E | 1.4e-05 | 1.6e-05 | 1.1e-05 | 1.367E-05 | 2.517E-06 | 1.100E-05 | 1.600E-05 |
| 25766 | 1289 | S1289E | E | 1.1e-05 | 1.5e-05 | 1e-05 | 1.200E-05 | 2.646E-06 | 1.000E-05 | 1.500E-05 |
| 25767 | 1289 | S1289Q | E | 8.7e-06 | 1.6e-05 | 1.5e-05 | 1.323E-05 | 3.958E-06 | 8.700E-06 | 1.600E-05 |
| 25768 | 1289 | S1289G | E | 9.6e-06 | 1e-05 | 1.5e-05 | 1.153E-05 | 3.009E-06 | 9.600E-06 | 1.500E-05 |
| 25769 | 1289 | S1289H | E | 1.4e-05 | 9.9e-06 | 1.1e-05 | 1.163E-05 | 2.122E-06 | 9.900E-06 | 1.400E-05 |
| 25770 | 1289 | S1289I | E | 1.3e-05 | 1.5e-05 | 1.2e-05 | 1.333E-05 | 1.528E-06 | 1.200E-05 | 1.500E-05 |
| 25771 | 1289 | S1289L | E | 1.7e-05 | 1.7e-05 | 1.6e-05 | 1.667E-05 | 5.774E-07 | 1.600E-05 | 1.700E-05 |
| 25772 | 1289 | S1289K | E | 1.6e-05 | 1.6e-05 | 1.8e-05 | 1.667E-05 | 1.155E-06 | 1.600E-05 | 1.800E-05 |
| 25773 | 1289 | S1289M | E | 1.1e-05 | 1.1e-05 | 1.6e-05 | 1.267E-05 | 2.887E-06 | 1.100E-05 | 1.600E-05 |
| 25774 | 1289 | S1289F | E | 1.6e-05 | 1.1e-05 | 7.6e-06 | 1.153E-05 | 4.225E-06 | 7.600E-06 | 1.600E-05 |
| 25775 | 1289 | S1289P | E | 9.5e-06 | 1.5e-05 | 1.1e-05 | 1.183E-05 | 2.843E-06 | 9.500E-06 | 1.500E-05 |
| 25776 | 1289 | S1289S | E | 8.4e-06 | 1e-05 | 1.2e-05 | 1.013E-05 | 1.804E-06 | 8.400E-06 | 1.200E-05 |
| 25777 | 1289 | S1289T | E | 1.2e-05 | 9.4e-06 | 9.9e-06 | 1.043E-05 | 1.380E-06 | 9.400E-06 | 1.200E-05 |
| 25778 | 1289 | S1289W | E | 1.3e-05 | 1.6e-05 | 1.6e-05 | 1.500E-05 | 1.732E-06 | 1.300E-05 | 1.600E-05 |
| 25779 | 1289 | S1289Y | E | 9.1e-06 | 1.4e-05 | 1.3e-05 | 1.203E-05 | 2.589E-06 | 9.100E-06 | 1.400E-05 |
| 25780 | 1289 | S1289V | E | 1.5e-05 | 1.1e-05 | 1.1e-05 | 1.233E-05 | 2.309E-06 | 1.100E-05 | 1.500E-05 |
| 25781 | 1290 | L1290A | E | 1.5e-05 | 1.6e-05 | 1.2e-05 | 1.433E-05 | 2.082E-06 | 1.200E-05 | 1.600E-05 |
| 25782 | 1290 | L1290R | E | 1e-05 | 8.2e-06 | 1.2e-05 | 1.007E-05 | 1.901E-06 | 8.200E-06 | 1.200E-05 |
| 25783 | 1290 | L1290N | E | 1.1e-05 | 1.5e-05 | 1.3e-05 | 1.300E-05 | 2.000E-06 | 1.100E-05 | 1.500E-05 |
| 25784 | 1290 | L1290D | E | 8.2e-06 | 1.5e-05 | 9.4e-06 | 1.087E-05 | 3.630E-06 | 8.200E-06 | 1.500E-05 |
| 25785 | 1290 | L1290C | E | 1.2e-05 | 1.7e-05 | 1.3e-05 | 1.400E-05 | 2.646E-06 | 1.200E-05 | 1.700E-05 |
| 25786 | 1290 | L1290E | E | 1.6e-05 | 1.1e-05 | 7.7e-06 | 1.157E-05 | 4.179E-06 | 7.700E-06 | 1.600E-05 |
| 25787 | 1290 | L1290Q | E | 1.5e-05 | 9.5e-06 | 1.1e-05 | 1.183E-05 | 2.843E-06 | 9.500E-06 | 1.500E-05 |
| 25788 | 1290 | L1290G | E | 7.8e-06 | 1.1e-05 | 1.5e-05 | 1.127E-05 | 3.607E-06 | 7.800E-06 | 1.500E-05 |
| 25789 | 1290 | L1290H | E | 1.4e-05 | 8.5e-06 | 9e-06 | 1.050E-05 | 3.041E-06 | 8.500E-06 | 1.400E-05 |
| 25790 | 1290 | L1290I | E | 1.5e-05 | 9.7e-06 | 9.5e-06 | 1.140E-05 | 3.119E-06 | 9.500E-06 | 1.500E-05 |
| 25791 | 1290 | L1290L | E | 8.5e-06 | 9.7e-06 | 1.5e-05 | 1.107E-05 | 3.459E-06 | 8.500E-06 | 1.500E-05 |
| 25792 | 1290 | L1290K | E | 1.5e-05 | 1.4e-05 | 1e-05 | 1.300E-05 | 2.646E-06 | 1.000E-05 | 1.500E-05 |
| 25793 | 1290 | L1290M | E | 1.6e-05 | 1.6e-05 | 1.1e-05 | 1.433E-05 | 2.887E-06 | 1.100E-05 | 1.600E-05 |
| 25794 | 1290 | L1290F | E | 1.5e-05 | 1.2e-05 | 1.5e-05 | 1.400E-05 | 1.732E-06 | 1.200E-05 | 1.500E-05 |
| 25795 | 1290 | L1290P | E | 1e-05 | 1.4e-05 | 1.2e-05 | 1.200E-05 | 2.000E-06 | 1.000E-05 | 1.400E-05 |
| 25796 | 1290 | L1290S | E | 1.4e-05 | 1.5e-05 | 1.5e-05 | 1.467E-05 | 5.774E-07 | 1.400E-05 | 1.500E-05 |
| 25797 | 1290 | L1290T | E | 1.5e-05 | 1.3e-05 | 1.3e-05 | 1.367E-05 | 1.155E-06 | 1.300E-05 | 1.500E-05 |
| 25798 | 1290 | L1290W | E | 1.6e-05 | 1.6e-05 | 9.8e-06 | 1.393E-05 | 3.580E-06 | 9.800E-06 | 1.600E-05 |
| 25799 | 1290 | L1290Y | E | 1.5e-05 | 8.4e-06 | 1.7e-05 | 1.347E-05 | 4.500E-06 | 8.400E-06 | 1.700E-05 |
| 25800 | 1290 | L1290V | E | 9.6e-06 | 1.4e-05 | 1.6e-05 | 1.320E-05 | 3.274E-06 | 9.600E-06 | 1.600E-05 |
| 25801 | 1291 | R1291A | E | 1.6e-05 | 1.6e-05 | 9.4e-06 | 1.380E-05 | 3.811E-06 | 9.400E-06 | 1.600E-05 |
| 25802 | 1291 | R1291R | E | 1.1e-05 | 1.5e-05 | 1.5e-05 | 1.367E-05 | 2.309E-06 | 1.100E-05 | 1.500E-05 |
| 25803 | 1291 | R1291N | E | 1.6e-05 | 8.5e-06 | 1.4e-05 | 1.283E-05 | 3.884E-06 | 8.500E-06 | 1.600E-05 |
| 25804 | 1291 | R1291D | E | 9.6e-06 | 1.5e-05 | 1.5e-05 | 1.320E-05 | 3.118E-06 | 9.600E-06 | 1.500E-05 |
| 25805 | 1291 | R1291C | E | 1.3e-05 | 1.5e-05 | 1.5e-05 | 1.433E-05 | 1.155E-06 | 1.300E-05 | 1.500E-05 |
| 25806 | 1291 | R1291E | E | 8.5e-06 | 8.7e-06 | 9.8e-06 | 9.000E-06 | 7.000E-07 | 8.500E-06 | 9.800E-06 |
| 25807 | 1291 | R1291Q | E | 1.4e-05 | 1.2e-05 | 7.9e-06 | 1.130E-05 | 3.110E-06 | 7.900E-06 | 1.400E-05 |
| 25808 | 1291 | R1291G | E | 1.4e-05 | 1.4e-05 | 1.1e-05 | 1.300E-05 | 1.732E-06 | 1.100E-05 | 1.400E-05 |
| 25809 | 1291 | R1291H | E | 1.2e-05 | 1.3e-05 | 1.4e-05 | 1.300E-05 | 1.000E-06 | 1.200E-05 | 1.400E-05 |
| 25810 | 1291 | R1291I | E | 1.6e-05 | 1.5e-05 | 1e-05 | 1.367E-05 | 3.215E-06 | 1.000E-05 | 1.600E-05 |
| 25811 | 1291 | R1291L | E | 1.4e-05 | 1.7e-05 | 1.1e-05 | 1.400E-05 | 3.000E-06 | 1.100E-05 | 1.700E-05 |
| 25812 | 1291 | R1291K | E | 9.9e-06 | 1.4e-05 | 1.6e-05 | 1.330E-05 | 3.110E-06 | 9.900E-06 | 1.600E-05 |
| 25813 | 1291 | R1291M | E | 1.1e-05 | 1.7e-05 | 1e-05 | 1.267E-05 | 3.786E-06 | 1.000E-05 | 1.700E-05 |
| 25814 | 1291 | R1291F | E | 1e-05 | 1.6e-05 | 1.3e-05 | 1.300E-05 | 3.000E-06 | 1.000E-05 | 1.600E-05 |
| 25815 | 1291 | R1291P | E | 1.1e-05 | 1e-05 | 1.4e-05 | 1.167E-05 | 2.082E-06 | 1.000E-05 | 1.400E-05 |
| 25816 | 1291 | R1291S | E | 1e-05 | 1.6e-05 | 1.5e-05 | 1.367E-05 | 3.215E-06 | 1.000E-05 | 1.600E-05 |
| 25817 | 1291 | R1291T | E | 7.2e-06 | 1.1e-05 | 1.4e-05 | 1.073E-05 | 3.408E-06 | 7.200E-06 | 1.400E-05 |
| 25818 | 1291 | R1291W | E | 1.5e-05 | 1.4e-05 | 1.1e-05 | 1.333E-05 | 2.082E-06 | 1.100E-05 | 1.500E-05 |
| 25819 | 1291 | R1291Y | E | 1e-05 | 7.9e-06 | 1.1e-05 | 9.633E-06 | 1.582E-06 | 7.900E-06 | 1.100E-05 |
| 25820 | 1291 | R1291V | E | 1.5e-05 | 1.1e-05 | 1.2e-05 | 1.267E-05 | 2.082E-06 | 1.100E-05 | 1.500E-05 |
| 25821 | 1292 | L1292A | E | 1.5e-05 | 1e-05 | 1e-05 | 1.167E-05 | 2.887E-06 | 1.000E-05 | 1.500E-05 |
| 25822 | 1292 | L1292R | E | 1.4e-05 | 1e-05 | 1.1e-05 | 1.167E-05 | 2.082E-06 | 1.000E-05 | 1.400E-05 |
| 25823 | 1292 | L1292N | E | 9.7e-06 | 1.3e-05 | 1.4e-05 | 1.223E-05 | 2.250E-06 | 9.700E-06 | 1.400E-05 |
| 25824 | 1292 | L1292D | E | 1.5e-05 | 1.6e-05 | 1.2e-05 | 1.433E-05 | 2.082E-06 | 1.200E-05 | 1.600E-05 |
| 25825 | 1292 | L1292C | E | 1.1e-05 | 1.3e-05 | 1.3e-05 | 1.233E-05 | 1.155E-06 | 1.100E-05 | 1.300E-05 |
| 25826 | 1292 | L1292E | E | 1.5e-05 | 9.8e-06 | 1.4e-05 | 1.293E-05 | 2.759E-06 | 9.800E-06 | 1.500E-05 |
| 25827 | 1292 | L1292Q | E | 7.9e-06 | 1.6e-05 | 1e-05 | 1.130E-05 | 4.204E-06 | 7.900E-06 | 1.600E-05 |
| 25828 | 1292 | L1292G | E | 1.6e-05 | 1.6e-05 | 1.5e-05 | 1.567E-05 | 5.774E-07 | 1.500E-05 | 1.600E-05 |
| 25829 | 1292 | L1292H | E | 7.8e-06 | 1.1e-05 | 1.6e-05 | 1.160E-05 | 4.133E-06 | 7.800E-06 | 1.600E-05 |
| 25830 | 1292 | L1292I | E | 1.7e-05 | 1e-05 | 1e-05 | 1.233E-05 | 4.041E-06 | 1.000E-05 | 1.700E-05 |
| 25831 | 1292 | L1292L | E | 9.5e-06 | 1.7e-05 | 1.3e-05 | 1.317E-05 | 3.753E-06 | 9.500E-06 | 1.700E-05 |
| 25832 | 1292 | L1292K | E | 1.5e-05 | 1.4e-05 | 1e-05 | 1.300E-05 | 2.646E-06 | 1.000E-05 | 1.500E-05 |
| 25833 | 1292 | L1292M | E | 1.4e-05 | 1.4e-05 | 8.2e-06 | 1.207E-05 | 3.349E-06 | 8.200E-06 | 1.400E-05 |
| 25834 | 1292 | L1292F | E | 1.5e-05 | 9.4e-06 | 1.5e-05 | 1.313E-05 | 3.233E-06 | 9.400E-06 | 1.500E-05 |
| 25835 | 1292 | L1292P | E | 1.5e-05 | 1.1e-05 | 1.5e-05 | 1.367E-05 | 2.309E-06 | 1.100E-05 | 1.500E-05 |
| 25836 | 1292 | L1292S | E | 1.4e-05 | 1.4e-05 | 1.3e-05 | 1.367E-05 | 5.774E-07 | 1.300E-05 | 1.400E-05 |
| 25837 | 1292 | L1292T | E | 1.6e-05 | 1.5e-05 | 8.5e-06 | 1.317E-05 | 4.072E-06 | 8.500E-06 | 1.600E-05 |
| 25838 | 1292 | L1292W | E | 1.1e-05 | 1.6e-05 | 1.4e-05 | 1.367E-05 | 2.517E-06 | 1.100E-05 | 1.600E-05 |
| 25839 | 1292 | L1292Y | E | 1.5e-05 | 9.7e-06 | 1.4e-05 | 1.290E-05 | 2.816E-06 | 9.700E-06 | 1.500E-05 |

| | | | | | | | | | | |
|---|---|---|---|---|---|---|---|---|---|---|
| 25840 | 1292 | L1292V | E | 1e-05 | 1.3e-05 | 1.2e-05 | 1.167E-05 | 1.528E-06 | 1.000E-05 | 1.300E-05 |
| 25841 | 1293 | S1293A | E | 1.3e-05 | 8e-06 | 1.2e-05 | 1.100E-05 | 2.646E-06 | 8.000E-06 | 1.300E-05 |
| 25842 | 1293 | S1293R | E | 8.3e-06 | 1.3e-05 | 1.6e-05 | 1.243E-05 | 3.881E-06 | 8.300E-06 | 1.600E-05 |
| 25843 | 1293 | S1293N | E | 1.2e-05 | 1e-05 | 6.8e-06 | 9.600E-06 | 2.623E-06 | 6.800E-06 | 1.200E-05 |
| 25844 | 1293 | S1293D | E | 9.7e-06 | 1.7e-05 | 1e-05 | 1.223E-05 | 4.131E-06 | 9.700E-06 | 1.700E-05 |
| 25845 | 1293 | S1293C | E | 1.5e-05 | 8.4e-06 | 1.1e-05 | 1.147E-05 | 3.325E-06 | 8.400E-06 | 1.500E-05 |
| 25846 | 1293 | S1293E | E | 1.8e-05 | 1.1e-05 | 1e-05 | 1.300E-05 | 4.359E-06 | 1.000E-05 | 1.800E-05 |
| 25847 | 1293 | S1293Q | E | 1.3e-05 | 1.4e-05 | 9.5e-06 | 1.217E-05 | 2.363E-06 | 9.500E-06 | 1.400E-05 |
| 25848 | 1293 | S1293G | E | 1.7e-05 | 1.1e-05 | 1.3e-05 | 1.367E-05 | 3.055E-06 | 1.100E-05 | 1.700E-05 |
| 25849 | 1293 | S1293H | E | 1.4e-05 | 1.6e-05 | 1.6e-05 | 1.533E-05 | 1.155E-06 | 1.400E-05 | 1.600E-05 |
| 25850 | 1293 | S1293I | E | 1.4e-05 | 1.6e-05 | 1.1e-05 | 1.367E-05 | 2.517E-06 | 1.100E-05 | 1.600E-05 |
| 25851 | 1293 | S1293L | E | 1.5e-05 | 1.3e-05 | 1.1e-05 | 1.300E-05 | 2.000E-06 | 1.100E-05 | 1.500E-05 |
| 25852 | 1293 | S1293K | E | 1.3e-05 | 1.4e-05 | 1.1e-05 | 1.267E-05 | 1.528E-06 | 1.100E-05 | 1.400E-05 |
| 25853 | 1293 | S1293M | E | 1.7e-05 | 1.2e-05 | 1.7e-05 | 1.533E-05 | 2.887E-06 | 1.200E-05 | 1.700E-05 |
| 25854 | 1293 | S1293F | E | 1.9e-05 | 1.4e-05 | 1.5e-05 | 1.600E-05 | 2.646E-06 | 1.400E-05 | 1.900E-05 |
| 25855 | 1293 | S1293P | E | 1.2e-05 | 9.1e-06 | 1.6e-05 | 1.237E-05 | 3.465E-06 | 9.100E-06 | 1.600E-05 |
| 25856 | 1293 | S1293S | E | 9.6e-06 | 1.5e-05 | 1.4e-05 | 1.287E-05 | 2.873E-06 | 9.600E-06 | 1.500E-05 |
| 25857 | 1293 | S1293T | E | 1e-05 | 1.4e-05 | 1.6e-05 | 1.333E-05 | 3.055E-06 | 1.000E-05 | 1.600E-05 |
| 25858 | 1293 | S1293W | E | 1.3e-05 | 1.4e-05 | 1e-05 | 1.233E-05 | 2.082E-06 | 1.000E-05 | 1.400E-05 |
| 25859 | 1293 | S1293Y | E | 1.6e-05 | 1e-05 | 1.5e-05 | 1.367E-05 | 3.215E-06 | 1.000E-05 | 1.600E-05 |
| 25860 | 1293 | S1293V | E | 1.1e-05 | 1e-05 | 9.8e-06 | 1.027E-05 | 6.429E-07 | 9.800E-06 | 1.100E-05 |
| 25861 | 1294 | C1294A | E | 1.6e-05 | 9.4e-06 | 1.5e-05 | 1.347E-05 | 3.557E-06 | 9.400E-06 | 1.600E-05 |
| 25862 | 1294 | C1294R | E | 1.3e-05 | 9.3e-06 | 1.4e-05 | 1.210E-05 | 2.476E-06 | 9.300E-06 | 1.400E-05 |
| 25863 | 1294 | C1294N | E | 7.2e-06 | 1.6e-05 | 1.1e-05 | 1.140E-05 | 4.414E-06 | 7.200E-06 | 1.600E-05 |
| 25864 | 1294 | C1294D | E | 1.6e-05 | 1.1e-05 | 1.5e-05 | 1.400E-05 | 2.646E-06 | 1.100E-05 | 1.600E-05 |
| 25865 | 1294 | C1294C | E | 1.6e-05 | 1.6e-05 | 1.1e-05 | 1.433E-05 | 2.887E-06 | 1.100E-05 | 1.600E-05 |
| 25866 | 1294 | C1294E | E | 1.4e-05 | 1.4e-05 | 1.5e-05 | 1.433E-05 | 5.774E-07 | 1.400E-05 | 1.500E-05 |
| 25867 | 1294 | C1294Q | E | 6.6e-06 | 1.2e-05 | 1.6e-05 | 1.153E-05 | 4.717E-06 | 6.600E-06 | 1.600E-05 |
| 25868 | 1294 | C1294G | E | 2.7e-05 | 1.4e-05 | 1.5e-05 | 1.867E-05 | 7.234E-06 | 1.400E-05 | 2.700E-05 |
| 25869 | 1294 | C1294H | E | 1.4e-05 | 1.6e-05 | 1.4e-05 | 1.467E-05 | 1.155E-06 | 1.400E-05 | 1.600E-05 |
| 25870 | 1294 | C1294I | E | 1.4e-05 | 1.2e-05 | 1.1e-05 | 1.233E-05 | 1.528E-06 | 1.100E-05 | 1.400E-05 |
| 25871 | 1294 | C1294L | E | 1.3e-05 | 1.2e-05 | 9.7e-06 | 1.157E-05 | 1.692E-06 | 9.700E-06 | 1.300E-05 |
| 25872 | 1294 | C1294K | E | 9.3e-06 | 1.2e-05 | 1.4e-05 | 1.177E-05 | 2.359E-06 | 9.300E-06 | 1.400E-05 |
| 25873 | 1294 | C1294M | E | 1e-05 | 9.3e-06 | 1.5e-05 | 1.143E-05 | 3.109E-06 | 9.300E-06 | 1.500E-05 |
| 25874 | 1294 | C1294F | E | 1.4e-05 | 1.6e-05 | 1.2e-05 | 1.400E-05 | 2.000E-06 | 1.200E-05 | 1.600E-05 |
| 25875 | 1294 | C1294P | E | 1e-05 | 7.6e-06 | 1.5e-05 | 1.087E-05 | 3.775E-06 | 7.600E-06 | 1.500E-05 |
| 25876 | 1294 | C1294S | E | 9.3e-06 | 1.4e-05 | 9.2e-06 | 1.083E-05 | 2.743E-06 | 9.200E-06 | 1.400E-05 |
| 25877 | 1294 | C1294T | E | 1.4e-05 | 1.5e-05 | 1.2e-05 | 1.367E-05 | 1.528E-06 | 1.200E-05 | 1.500E-05 |
| 25878 | 1294 | C1294W | E | 1.7e-05 | 1.3e-05 | 9.1e-06 | 1.303E-05 | 3.950E-06 | 9.100E-06 | 1.700E-05 |
| 25879 | 1294 | C1294Y | E | 9.9e-06 | 8.2e-06 | 1.2e-05 | 1.003E-05 | 1.904E-06 | 8.200E-06 | 1.200E-05 |
| 25880 | 1294 | C1294V | E | 1.6e-05 | 1.5e-05 | 1.5e-05 | 1.533E-05 | 5.774E-07 | 1.500E-05 | 1.600E-05 |
| 25881 | 1295 | A1295A | E | 1.5e-05 | 1.6e-05 | 1.5e-05 | 1.533E-05 | 5.774E-07 | 1.500E-05 | 1.600E-05 |
| 25882 | 1295 | A1295R | E | 8.6e-06 | 8.6e-06 | 1.1e-05 | 9.400E-06 | 1.386E-06 | 8.600E-06 | 1.100E-05 |
| 25883 | 1295 | A1295N | E | 1.4e-05 | 1.6e-05 | 1.1e-05 | 1.367E-05 | 2.517E-06 | 1.100E-05 | 1.600E-05 |
| 25884 | 1295 | A1295D | E | 1.1e-05 | 1.4e-05 | 7.2e-06 | 1.073E-05 | 3.408E-06 | 7.200E-06 | 1.400E-05 |
| 25885 | 1295 | A1295C | E | 1e-05 | 1.4e-05 | 1e-05 | 1.133E-05 | 2.309E-06 | 1.000E-05 | 1.400E-05 |
| 25886 | 1295 | A1295E | E | 6.7e-06 | 1.5e-05 | 1.2e-05 | 1.123E-05 | 4.203E-06 | 6.700E-06 | 1.500E-05 |
| 25887 | 1295 | A1295Q | E | 9.9e-06 | 1.3e-05 | 1.3e-05 | 1.197E-05 | 1.790E-06 | 9.900E-06 | 1.300E-05 |
| 25888 | 1295 | A1295G | E | 1.7e-05 | 1.4e-05 | 9.2e-06 | 1.340E-05 | 3.934E-06 | 9.200E-06 | 1.700E-05 |
| 25889 | 1295 | A1295H | E | 1.1e-05 | 1.2e-05 | 1.6e-05 | 1.300E-05 | 2.646E-06 | 1.100E-05 | 1.600E-05 |
| 25890 | 1295 | A1295I | E | 1e-05 | 1.1e-05 | 9.5e-06 | 1.017E-05 | 7.638E-07 | 9.500E-06 | 1.100E-05 |
| 25891 | 1295 | A1295L | E | 1.5e-05 | 1.3e-05 | 1.4e-05 | 1.400E-05 | 1.000E-06 | 1.300E-05 | 1.500E-05 |
| 25892 | 1295 | A1295K | E | 1.2e-05 | 1.4e-05 | 9.8e-06 | 1.193E-05 | 2.101E-06 | 9.800E-06 | 1.400E-05 |
| 25893 | 1295 | A1295M | E | 1.3e-05 | 2e-05 | 1.3e-05 | 1.533E-05 | 4.041E-06 | 1.300E-05 | 2.000E-05 |
| 25894 | 1295 | A1295F | E | 1.6e-05 | 9.4e-06 | 1.5e-05 | 1.347E-05 | 3.557E-06 | 9.400E-06 | 1.600E-05 |
| 25895 | 1295 | A1295P | E | 1e-05 | 1.1e-05 | 9.7e-06 | 1.023E-05 | 6.807E-07 | 9.700E-06 | 1.100E-05 |
| 25896 | 1295 | A1295S | E | 1.3e-05 | 1.3e-05 | 9.3e-06 | 1.177E-05 | 2.136E-06 | 9.300E-06 | 1.300E-05 |
| 25897 | 1295 | A1295T | E | 7.4e-06 | 1.5e-05 | 1.5e-05 | 1.247E-05 | 4.388E-06 | 7.400E-06 | 1.500E-05 |
| 25898 | 1295 | A1295W | E | 6.1e-06 | 1.2e-05 | 1.5e-05 | 1.103E-05 | 4.528E-06 | 6.100E-06 | 1.500E-05 |
| 25899 | 1295 | A1295Y | E | 1.4e-05 | 1.2e-05 | 9e-06 | 1.167E-05 | 2.517E-06 | 9.000E-06 | 1.400E-05 |
| 25900 | 1295 | A1295V | E | 9.7e-06 | 9.5e-06 | 1.3e-05 | 1.073E-05 | 1.966E-06 | 9.500E-06 | 1.300E-05 |
| 25901 | 1296 | A1296A | E | 9.4e-06 | 1.5e-05 | 1.3e-05 | 1.247E-05 | 2.838E-06 | 9.400E-06 | 1.500E-05 |
| 25902 | 1296 | A1296R | E | 8.6e-06 | 1e-05 | 1.1e-05 | 9.867E-06 | 1.206E-06 | 8.600E-06 | 1.100E-05 |
| 25903 | 1296 | A1296N | E | 1.5e-05 | 1e-05 | 1.1e-05 | 1.200E-05 | 2.646E-06 | 1.000E-05 | 1.500E-05 |
| 25904 | 1296 | A1296D | E | 9.8e-06 | 1.4e-05 | 1.5e-05 | 1.293E-05 | 2.759E-06 | 9.800E-06 | 1.500E-05 |
| 25905 | 1296 | A1296C | E | 1.4e-05 | 1.5e-05 | 1.4e-05 | 1.433E-05 | 5.774E-07 | 1.400E-05 | 1.500E-05 |
| 25906 | 1296 | A1296E | E | 9.2e-06 | 1.6e-05 | 1.1e-05 | 1.207E-05 | 3.523E-06 | 9.200E-06 | 1.600E-05 |
| 25907 | 1296 | A1296Q | E | 1.4e-05 | 1.1e-05 | 1.1e-05 | 1.200E-05 | 1.732E-06 | 1.100E-05 | 1.400E-05 |
| 25908 | 1296 | A1296G | E | 1.6e-05 | 1.2e-05 | 9.9e-06 | 1.263E-05 | 3.099E-06 | 9.900E-06 | 1.600E-05 |
| 25909 | 1296 | A1296H | E | 1.2e-05 | 8.5e-06 | 1.5e-05 | 1.183E-05 | 3.253E-06 | 8.500E-06 | 1.500E-05 |
| 25910 | 1296 | A1296I | E | 1.2e-05 | 1.7e-05 | 1.6e-05 | 1.500E-05 | 2.646E-06 | 1.200E-05 | 1.700E-05 |
| 25911 | 1296 | A1296L | E | 9.1e-06 | 1.5e-05 | 8.8e-06 | 1.097E-05 | 3.496E-06 | 8.800E-06 | 1.500E-05 |
| 25912 | 1296 | A1296K | E | 1.4e-05 | 1.5e-05 | 1.7e-05 | 1.533E-05 | 1.528E-06 | 1.400E-05 | 1.700E-05 |
| 25913 | 1296 | A1296M | E | 8.7e-06 | 1.5e-05 | 1.2e-05 | 1.190E-05 | 3.151E-06 | 8.700E-06 | 1.500E-05 |
| 25914 | 1296 | A1296F | E | 1.1e-05 | 1.6e-05 | 9.8e-06 | 1.227E-05 | 3.288E-06 | 9.800E-06 | 1.600E-05 |
| 25915 | 1296 | A1296P | E | 9e-06 | 9.7e-06 | 1.6e-05 | 1.157E-05 | 3.855E-06 | 9.000E-06 | 1.600E-05 |
| 25916 | 1296 | A1296S | E | 1.5e-05 | 1.5e-05 | 1.3e-05 | 1.433E-05 | 1.155E-06 | 1.300E-05 | 1.500E-05 |
| 25917 | 1296 | A1296T | E | 1.6e-05 | 7.9e-06 | 1.7e-05 | 1.363E-05 | 4.990E-06 | 7.900E-06 | 1.700E-05 |
| 25918 | 1296 | A1296W | E | 1.1e-05 | 9.5e-06 | 1.6e-05 | 1.217E-05 | 3.403E-06 | 9.500E-06 | 1.600E-05 |
| 25919 | 1296 | A1296Y | E | 1.3e-05 | 1.3e-05 | 1.1e-05 | 1.233E-05 | 1.155E-06 | 1.100E-05 | 1.300E-05 |
| 25920 | 1296 | A1296V | E | 8.7e-06 | 1.5e-05 | 1.6e-05 | 1.323E-05 | 3.958E-06 | 8.700E-06 | 1.600E-05 |
| 25921 | 1297 | S1297A | E | 1.6e-05 | 1.2e-05 | 1.3e-05 | 1.367E-05 | 2.082E-06 | 1.200E-05 | 1.600E-05 |
| 25922 | 1297 | S1297R | E | 9.9e-06 | 1.6e-05 | 1e-05 | 1.197E-05 | 3.493E-06 | 9.900E-06 | 1.600E-05 |
| 25923 | 1297 | S1297N | E | 1.5e-05 | 1.4e-05 | 9.9e-06 | 1.297E-05 | 2.702E-06 | 9.900E-06 | 1.500E-05 |
| 25924 | 1297 | S1297D | E | 1.5e-05 | 1.5e-05 | 1.7e-05 | 1.567E-05 | 1.155E-06 | 1.500E-05 | 1.700E-05 |

| | | | | | | | | | | |
|---|---|---|---|---|---|---|---|---|---|---|
| 25925 | 1297 | S1297C | E | 1.6e-05 | 1.7e-05 | 1.4e-05 | 1.567E-05 | 1.528E-06 | 1.400E-05 | 1.700E-05 |
| 25926 | 1297 | S1297E | E | 1.2e-05 | 1.1e-05 | 8.3e-06 | 1.043E-05 | 1.914E-06 | 8.300E-06 | 1.200E-05 |
| 25927 | 1297 | S1297Q | E | 1.3e-05 | 1.1e-05 | 1.2e-05 | 1.200E-05 | 1.000E-06 | 1.100E-05 | 1.300E-05 |
| 25928 | 1297 | S1297G | E | 6.3e-06 | 1.6e-05 | 1.2e-05 | 1.143E-05 | 4.875E-06 | 6.300E-06 | 1.600E-05 |
| 25929 | 1297 | S1297H | E | 1e-05 | 1.1e-05 | 1.1e-05 | 1.067E-05 | 5.774E-07 | 1.000E-05 | 1.100E-05 |
| 25930 | 1297 | S1297I | E | 1.1e-05 | 1.4e-05 | 1.7e-05 | 1.400E-05 | 3.000E-06 | 1.100E-05 | 1.700E-05 |
| 25931 | 1297 | S1297L | E | 8.9e-06 | 9.9e-06 | 1.6e-05 | 1.160E-05 | 3.843E-06 | 8.900E-06 | 1.600E-05 |
| 25932 | 1297 | S1297K | E | 1.2e-05 | 1.4e-05 | 5.5e-06 | 1.050E-05 | 4.444E-06 | 5.500E-06 | 1.400E-05 |
| 25933 | 1297 | S1297M | E | 1.5e-05 | 1.1e-05 | 1.5e-05 | 1.367E-05 | 2.309E-06 | 1.100E-05 | 1.500E-05 |
| 25934 | 1297 | S1297F | E | 9.5e-06 | 1.6e-05 | 7.9e-06 | 1.113E-05 | 4.290E-06 | 7.900E-06 | 1.600E-05 |
| 25935 | 1297 | S1297P | E | 9.6e-06 | 1e-05 | 1.2e-05 | 1.053E-05 | 1.286E-06 | 9.600E-06 | 1.200E-05 |
| 25936 | 1297 | S1297S | E | 9.8e-06 | 1.5e-05 | 1.1e-05 | 1.193E-05 | 2.723E-06 | 9.800E-06 | 1.500E-05 |
| 25937 | 1297 | S1297T | E | 7.3e-06 | 1e-05 | 1e-05 | 9.100E-06 | 1.559E-06 | 7.300E-06 | 1.000E-05 |
| 25938 | 1297 | S1297W | E | 1.1e-05 | 1.4e-05 | 1.4e-05 | 1.300E-05 | 1.732E-06 | 1.100E-05 | 1.400E-05 |
| 25939 | 1297 | S1297Y | E | 1.1e-05 | 1.2e-05 | 1.4e-05 | 1.233E-05 | 1.528E-06 | 1.100E-05 | 1.400E-05 |
| 25940 | 1297 | S1297V | E | 1e-05 | 1.1e-05 | 1.4e-05 | 1.167E-05 | 2.082E-06 | 1.000E-05 | 1.400E-05 |
| 25941 | 1298 | G1298A | E | 1.5e-05 | 9.3e-06 | 1.5e-05 | 1.310E-05 | 3.291E-06 | 9.300E-06 | 1.500E-05 |
| 25942 | 1298 | G1298R | E | 1.7e-05 | 9.4e-06 | 1.1e-05 | 1.247E-05 | 4.007E-06 | 9.400E-06 | 1.700E-05 |
| 25943 | 1298 | G1298N | E | 1.6e-05 | 1.5e-05 | 9.9e-06 | 1.363E-05 | 3.272E-06 | 9.900E-06 | 1.600E-05 |
| 25944 | 1298 | G1298D | E | 1.6e-05 | 9.8e-06 | 1.5e-05 | 1.360E-05 | 3.329E-06 | 9.800E-06 | 1.600E-05 |
| 25945 | 1298 | G1298C | E | 1.5e-05 | 1.4e-05 | 1.3e-05 | 1.400E-05 | 1.000E-06 | 1.300E-05 | 1.500E-05 |
| 25946 | 1298 | G1298E | E | 1.6e-05 | 1e-05 | 1.5e-05 | 1.367E-05 | 3.215E-06 | 1.000E-05 | 1.600E-05 |
| 25947 | 1298 | G1298Q | E | 1.5e-05 | 1.1e-05 | 1.4e-05 | 1.333E-05 | 2.082E-06 | 1.100E-05 | 1.500E-05 |
| 25948 | 1298 | G1298G | E | 1.1e-05 | 1.1e-05 | 1e-05 | 1.067E-05 | 5.774E-07 | 1.000E-05 | 1.100E-05 |
| 25949 | 1298 | G1298H | E | 1.4e-05 | 1e-05 | 6.9e-06 | 1.030E-05 | 3.559E-06 | 6.900E-06 | 1.400E-05 |
| 25950 | 1298 | G1298I | E | 1.5e-05 | 1.7e-05 | 1.3e-05 | 1.500E-05 | 2.000E-06 | 1.300E-05 | 1.700E-05 |
| 25951 | 1298 | G1298L | E | 1.6e-05 | 1e-05 | 1.2e-05 | 1.267E-05 | 3.055E-06 | 1.000E-05 | 1.600E-05 |
| 25952 | 1298 | G1298K | E | 9.7e-06 | 9.2e-06 | 1.5e-05 | 1.130E-05 | 3.214E-06 | 9.200E-06 | 1.500E-05 |
| 25953 | 1298 | G1298M | E | 1.4e-05 | 1.4e-05 | 1.5e-05 | 1.433E-05 | 5.774E-07 | 1.400E-05 | 1.500E-05 |
| 25954 | 1298 | G1298F | E | 9.5e-06 | 2.1e-05 | 1e-05 | 1.350E-05 | 6.500E-06 | 9.500E-06 | 2.100E-05 |
| 25955 | 1298 | G1298P | E | 1.4e-05 | 1.4e-05 | 1.1e-05 | 1.300E-05 | 1.732E-06 | 1.100E-05 | 1.400E-05 |
| 25956 | 1298 | G1298S | E | 1.1e-05 | 1.4e-05 | 1.2e-05 | 1.233E-05 | 1.528E-06 | 1.100E-05 | 1.400E-05 |
| 25957 | 1298 | G1298T | E | 1.2e-05 | 1.1e-05 | 1.5e-05 | 1.267E-05 | 2.082E-06 | 1.100E-05 | 1.500E-05 |
| 25958 | 1298 | G1298W | E | 1.4e-05 | 7.1e-06 | 1.3e-05 | 1.137E-05 | 3.729E-06 | 7.100E-06 | 1.400E-05 |
| 25959 | 1298 | G1298Y | E | 9.7e-06 | 9.8e-06 | 1.4e-05 | 1.117E-05 | 2.454E-06 | 9.700E-06 | 1.400E-05 |
| 25960 | 1298 | G1298V | E | 8.3e-06 | 1.5e-05 | 1.2e-05 | 1.177E-05 | 3.356E-06 | 8.300E-06 | 1.500E-05 |
| 25961 | 1299 | F1299A | E | 1.1e-05 | 1.5e-05 | 2e-05 | 1.533E-05 | 4.509E-06 | 1.100E-05 | 2.000E-05 |
| 25962 | 1299 | F1299R | E | 1.3e-05 | 1.4e-05 | 1.2e-05 | 1.300E-05 | 1.000E-06 | 1.200E-05 | 1.400E-05 |
| 25963 | 1299 | F1299N | E | 8.4e-06 | 8.5e-06 | 1.7e-05 | 1.130E-05 | 4.937E-06 | 8.400E-06 | 1.700E-05 |
| 25964 | 1299 | F1299D | E | 1.1e-05 | 1.1e-05 | 1.5e-05 | 1.233E-05 | 2.309E-06 | 1.100E-05 | 1.500E-05 |
| 25965 | 1299 | F1299C | E | 8.8e-06 | 1.1e-05 | 1.4e-05 | 1.127E-05 | 2.610E-06 | 8.800E-06 | 1.400E-05 |
| 25966 | 1299 | F1299E | E | 1.1e-05 | 1.3e-05 | 1.5e-05 | 1.300E-05 | 2.000E-06 | 1.100E-05 | 1.500E-05 |
| 25967 | 1299 | F1299Q | E | 1.4e-05 | 1e-05 | 7.5e-06 | 1.050E-05 | 3.279E-06 | 7.500E-06 | 1.400E-05 |
| 25968 | 1299 | F1299G | E | 1.3e-05 | 9.4e-06 | 1.5e-05 | 1.247E-05 | 2.838E-06 | 9.400E-06 | 1.500E-05 |
| 25969 | 1299 | F1299H | E | 1.6e-05 | 1e-05 | 1e-05 | 1.200E-05 | 3.464E-06 | 1.000E-05 | 1.600E-05 |
| 25970 | 1299 | F1299I | E | 1.2e-05 | 1.1e-05 | 9e-06 | 1.067E-05 | 1.528E-06 | 9.000E-06 | 1.200E-05 |
| 25971 | 1299 | F1299L | E | 1.1e-05 | 7.1e-06 | 1.5e-05 | 1.103E-05 | 3.950E-06 | 7.100E-06 | 1.500E-05 |
| 25972 | 1299 | F1299K | E | 9.6e-06 | 1.5e-05 | 1.5e-05 | 1.320E-05 | 3.118E-06 | 9.600E-06 | 1.500E-05 |
| 25973 | 1299 | F1299M | E | 1.5e-05 | 9.2e-06 | 1.1e-05 | 1.173E-05 | 2.969E-06 | 9.200E-06 | 1.500E-05 |
| 25974 | 1299 | F1299F | E | 1.6e-05 | 1.1e-05 | 9.7e-06 | 1.223E-05 | 3.326E-06 | 9.700E-06 | 1.600E-05 |
| 25975 | 1299 | F1299P | E | 1.1e-05 | 1.1e-05 | 1.2e-05 | 1.133E-05 | 5.774E-07 | 1.100E-05 | 1.200E-05 |
| 25976 | 1299 | F1299S | E | 1.5e-05 | 1.1e-05 | 1e-05 | 1.200E-05 | 2.646E-06 | 1.000E-05 | 1.500E-05 |
| 25977 | 1299 | F1299T | E | 1.5e-05 | 8.7e-06 | 1.3e-05 | 1.223E-05 | 3.219E-06 | 8.700E-06 | 1.500E-05 |
| 25978 | 1299 | F1299W | E | 1.1e-05 | 1.3e-05 | 1.5e-05 | 1.300E-05 | 2.000E-06 | 1.100E-05 | 1.500E-05 |
| 25979 | 1299 | F1299Y | E | 1e-05 | 9.9e-06 | 1.6e-05 | 1.197E-05 | 3.493E-06 | 9.900E-06 | 1.600E-05 |
| 25980 | 1299 | F1299V | E | 1.5e-05 | 1.4e-05 | 1.4e-05 | 1.433E-05 | 5.774E-07 | 1.400E-05 | 1.500E-05 |
| 25981 | 1300 | N1300A | E | 8.3e-06 | 1.6e-05 | 1.4e-05 | 1.277E-05 | 3.995E-06 | 8.300E-06 | 1.600E-05 |
| 25982 | 1300 | N1300R | E | 1.5e-05 | 1.1e-05 | 1.6e-05 | 1.400E-05 | 2.646E-06 | 1.100E-05 | 1.600E-05 |
| 25983 | 1300 | N1300N | E | 1.3e-05 | 1.1e-05 | 1.4e-05 | 1.267E-05 | 1.528E-06 | 1.100E-05 | 1.400E-05 |
| 25984 | 1300 | N1300D | E | 1e-05 | 1.7e-05 | 1.3e-05 | 1.333E-05 | 3.512E-06 | 1.000E-05 | 1.700E-05 |
| 25985 | 1300 | N1300C | E | 1.5e-05 | 1.1e-05 | 1e-05 | 1.200E-05 | 2.646E-06 | 1.000E-05 | 1.500E-05 |
| 25986 | 1300 | N1300E | E | 1.6e-05 | 1e-05 | 1.4e-05 | 1.333E-05 | 3.055E-06 | 1.000E-05 | 1.600E-05 |
| 25987 | 1300 | N1300Q | E | 1.5e-05 | 1.1e-05 | 1.4e-05 | 1.333E-05 | 2.082E-06 | 1.100E-05 | 1.500E-05 |
| 25988 | 1300 | N1300G | E | 1.4e-05 | 1.1e-05 | 1.5e-05 | 1.333E-05 | 2.082E-06 | 1.100E-05 | 1.500E-05 |
| 25989 | 1300 | N1300H | E | 1.5e-05 | 8.5e-06 | 1.4e-05 | 1.250E-05 | 3.500E-06 | 8.500E-06 | 1.500E-05 |
| 25990 | 1300 | N1300I | E | 1.4e-05 | 1e-05 | 8.7e-06 | 1.090E-05 | 2.762E-06 | 8.700E-06 | 1.400E-05 |
| 25991 | 1300 | N1300L | E | 1.1e-05 | 1.7e-05 | 1.3e-05 | 1.367E-05 | 3.055E-06 | 1.100E-05 | 1.700E-05 |
| 25992 | 1300 | N1300K | E | 9.2e-06 | 9.4e-06 | 1.4e-05 | 1.087E-05 | 2.715E-06 | 9.200E-06 | 1.400E-05 |
| 25993 | 1300 | N1300M | E | 1.5e-05 | 1.2e-05 | 1.5e-05 | 1.400E-05 | 1.732E-06 | 1.200E-05 | 1.500E-05 |
| 25994 | 1300 | N1300F | E | 1.1e-05 | 1.3e-05 | 1.1e-05 | 1.167E-05 | 1.155E-06 | 1.100E-05 | 1.300E-05 |
| 25995 | 1300 | N1300P | E | 1.4e-05 | 1.3e-05 | 1.6e-05 | 1.433E-05 | 1.528E-06 | 1.300E-05 | 1.600E-05 |
| 25996 | 1300 | N1300S | E | 1.6e-05 | 1.5e-05 | 9.1e-06 | 1.337E-05 | 3.729E-06 | 9.100E-06 | 1.600E-05 |
| 25997 | 1300 | N1300T | E | 9.3e-06 | 1.7e-05 | 1.5e-05 | 1.377E-05 | 3.995E-06 | 9.300E-06 | 1.700E-05 |
| 25998 | 1300 | N1300W | E | 1.1e-05 | 1.5e-05 | 1.2e-05 | 1.267E-05 | 2.082E-06 | 1.100E-05 | 1.500E-05 |
| 25999 | 1300 | N1300Y | E | 9.6e-06 | 9e-06 | 1.3e-05 | 1.053E-05 | 2.157E-06 | 9.000E-06 | 1.300E-05 |
| 26000 | 1300 | N1300V | E | 1e-05 | 1.7e-05 | 1.5e-05 | 1.400E-05 | 3.606E-06 | 1.000E-05 | 1.700E-05 |
| 26001 | 1301 | I1301A | E | 1.5e-05 | 1.6e-05 | 1.6e-05 | 1.567E-05 | 5.774E-07 | 1.500E-05 | 1.600E-05 |
| 26002 | 1301 | I1301R | E | 1.1e-05 | 1.4e-05 | 1.4e-05 | 1.300E-05 | 1.732E-06 | 1.100E-05 | 1.400E-05 |
| 26003 | 1301 | I1301N | E | 9.9e-06 | 1.4e-05 | 9.9e-06 | 1.127E-05 | 2.367E-06 | 9.900E-06 | 1.400E-05 |
| 26004 | 1301 | I1301D | E | 1.4e-05 | 1.5e-05 | 1e-05 | 1.300E-05 | 2.646E-06 | 1.000E-05 | 1.500E-05 |
| 26005 | 1301 | I1301C | E | 1.1e-05 | 1.6e-05 | 1.2e-05 | 1.300E-05 | 2.646E-06 | 1.100E-05 | 1.600E-05 |
| 26006 | 1301 | I1301E | E | 7.7e-06 | 1e-05 | 1.3e-05 | 1.023E-05 | 2.658E-06 | 7.700E-06 | 1.300E-05 |
| 26007 | 1301 | I1301Q | E | 9.5e-06 | 1.5e-05 | 1.6e-05 | 1.350E-05 | 3.500E-06 | 9.500E-06 | 1.600E-05 |
| 26008 | 1301 | I1301G | E | 1.1e-05 | 1.4e-05 | 1.6e-05 | 1.367E-05 | 2.517E-06 | 1.100E-05 | 1.600E-05 |
| 26009 | 1301 | I1301H | E | 1.1e-05 | 1e-05 | 9.8e-06 | 1.027E-05 | 6.429E-07 | 9.800E-06 | 1.100E-05 |

| | | | | | | | | | | |
|---|---|---|---|---|---|---|---|---|---|---|
| 26010 | 1301 | I1301I | E | 1.3e-05 | 1.4e-05 | 6.2e-06 | 1.107E-05 | 4.244E-06 | 6.200E-06 | 1.400E-05 |
| 26011 | 1301 | I1301L | E | 1.4e-05 | 1e-05 | 1.6e-05 | 1.333E-05 | 3.055E-06 | 1.000E-05 | 1.600E-05 |
| 26012 | 1301 | I1301K | E | 9.5e-06 | 1.7e-05 | 8.7e-06 | 1.173E-05 | 4.579E-06 | 8.700E-06 | 1.700E-05 |
| 26013 | 1301 | I1301M | E | 1.2e-05 | 9.9e-06 | 8.5e-06 | 1.013E-05 | 1.762E-06 | 8.500E-06 | 1.200E-05 |
| 26014 | 1301 | I1301F | E | 9.4e-06 | 1.1e-05 | 1.3e-05 | 1.113E-05 | 1.804E-06 | 9.400E-06 | 1.300E-05 |
| 26015 | 1301 | I1301P | E | 1.6e-05 | 1.6e-05 | 1.5e-05 | 1.567E-05 | 5.774E-07 | 1.500E-05 | 1.600E-05 |
| 26016 | 1301 | I1301S | E | 7.3e-06 | 9.6e-06 | 9.1e-06 | 8.667E-06 | 1.210E-06 | 7.300E-06 | 9.600E-06 |
| 26017 | 1301 | I1301T | E | 1.6e-05 | 1.6e-05 | 1.6e-05 | 1.600E-05 | 0.000E+00 | 1.600E-05 | 1.600E-05 |
| 26018 | 1301 | I1301W | E | 8.9e-06 | 1.5e-05 | 1.1e-05 | 1.163E-05 | 3.099E-06 | 8.900E-06 | 1.500E-05 |
| 26019 | 1301 | I1301Y | E | 1.6e-05 | 1.4e-05 | 1.6e-05 | 1.533E-05 | 1.155E-06 | 1.400E-05 | 1.600E-05 |
| 26020 | 1301 | I1301V | E | 1.5e-05 | 1.1e-05 | 1.5e-05 | 1.367E-05 | 2.309E-06 | 1.100E-05 | 1.500E-05 |
| 26021 | 1302 | K1302A | E | 1.5e-05 | 9.9e-06 | 1e-05 | 1.163E-05 | 2.916E-06 | 9.900E-06 | 1.500E-05 |
| 26022 | 1302 | K1302R | E | 1.1e-05 | 9e-06 | 9e-06 | 9.667E-06 | 1.155E-06 | 9.000E-06 | 1.100E-05 |
| 26023 | 1302 | K1302N | E | 9.7e-06 | 1.4e-05 | 1.1e-05 | 1.157E-05 | 2.205E-06 | 9.700E-06 | 1.400E-05 |
| 26024 | 1302 | K1302D | E | 1.1e-05 | 1.7e-05 | 1.6e-05 | 1.467E-05 | 3.215E-06 | 1.100E-05 | 1.700E-05 |
| 26025 | 1302 | K1302C | E | 1.5e-05 | 9.2e-06 | 1.2e-05 | 1.207E-05 | 2.901E-06 | 9.200E-06 | 1.500E-05 |
| 26026 | 1302 | K1302E | E | 1.6e-05 | 1.4e-05 | 1e-05 | 1.333E-05 | 3.055E-06 | 1.000E-05 | 1.600E-05 |
| 26027 | 1302 | K1302Q | E | 1.6e-05 | 1e-05 | 1.5e-05 | 1.367E-05 | 3.215E-06 | 1.000E-05 | 1.600E-05 |
| 26028 | 1302 | K1302G | E | 1.6e-05 | 1.5e-05 | 1.6e-05 | 1.567E-05 | 5.774E-07 | 1.500E-05 | 1.600E-05 |
| 26029 | 1302 | K1302H | E | 9.1e-06 | 3.1e-05 | 1.6e-05 | 1.870E-05 | 1.120E-05 | 9.100E-06 | 3.100E-05 |
| 26030 | 1302 | K1302I | E | 9.5e-06 | 1.5e-05 | 9e-06 | 1.117E-05 | 3.329E-06 | 9.000E-06 | 1.500E-05 |
| 26031 | 1302 | K1302L | E | 9.8e-06 | 1.2e-05 | 8.9e-06 | 1.023E-05 | 1.595E-06 | 8.900E-06 | 1.200E-05 |
| 26032 | 1302 | K1302K | E | 1.5e-05 | 9.5e-06 | 1e-05 | 1.150E-05 | 3.041E-06 | 9.500E-06 | 1.500E-05 |
| 26033 | 1302 | K1302M | E | 1.5e-05 | 1.5e-05 | 9.8e-06 | 1.327E-05 | 3.002E-06 | 9.800E-06 | 1.500E-05 |
| 26034 | 1302 | K1302F | E | 1.7e-05 | 9.3e-06 | 1.1e-05 | 1.243E-05 | 4.045E-06 | 9.300E-06 | 1.700E-05 |
| 26035 | 1302 | K1302P | E | 6.6e-06 | 1.4e-05 | 1.6e-05 | 1.220E-05 | 4.952E-06 | 6.600E-06 | 1.600E-05 |
| 26036 | 1302 | K1302S | E | 1.5e-05 | 1.5e-05 | 1.4e-05 | 1.467E-05 | 5.774E-07 | 1.400E-05 | 1.500E-05 |
| 26037 | 1302 | K1302T | E | 1.1e-05 | 1.6e-05 | 1.1e-05 | 1.267E-05 | 2.887E-06 | 1.100E-05 | 1.600E-05 |
| 26038 | 1302 | K1302W | E | 1.4e-05 | 1e-05 | 1.5e-05 | 1.300E-05 | 2.646E-06 | 1.000E-05 | 1.500E-05 |
| 26039 | 1302 | K1302Y | E | 1.1e-05 | 1.1e-05 | 1.6e-05 | 1.267E-05 | 2.887E-06 | 1.100E-05 | 1.600E-05 |
| 26040 | 1302 | K1302V | E | 1.4e-05 | 9.8e-06 | 1.6e-05 | 1.327E-05 | 3.164E-06 | 9.800E-06 | 1.600E-05 |
| 26041 | 1303 | D1303A | E | 1.3e-05 | 1.1e-05 | 9.8e-06 | 1.127E-05 | 1.617E-06 | 9.800E-06 | 1.300E-05 |
| 26042 | 1303 | D1303R | E | 9.4e-06 | 9e-06 | 2e-05 | 1.280E-05 | 6.239E-06 | 9.000E-06 | 2.000E-05 |
| 26043 | 1303 | D1303N | E | 1.5e-05 | 9e-06 | 8.7e-06 | 1.090E-05 | 3.554E-06 | 8.700E-06 | 1.500E-05 |
| 26044 | 1303 | D1303D | E | 9.6e-06 | 1.1e-05 | 1.1e-05 | 1.053E-05 | 8.083E-07 | 9.600E-06 | 1.100E-05 |
| 26045 | 1303 | D1303C | E | 9.5e-06 | 7.9e-06 | 1.2e-05 | 9.800E-06 | 2.066E-06 | 7.900E-06 | 1.200E-05 |
| 26046 | 1303 | D1303E | E | 1.4e-05 | 9.1e-06 | 1e-05 | 1.103E-05 | 2.608E-06 | 9.100E-06 | 1.400E-05 |
| 26047 | 1303 | D1303Q | E | 1.4e-05 | 1.4e-05 | 1.1e-05 | 1.300E-05 | 1.732E-06 | 1.100E-05 | 1.400E-05 |
| 26048 | 1303 | D1303G | E | 1.2e-05 | 1.4e-05 | 1.2e-05 | 1.267E-05 | 1.155E-06 | 1.200E-05 | 1.400E-05 |
| 26049 | 1303 | D1303H | E | 8.5e-06 | 1.5e-05 | 1.2e-05 | 1.183E-05 | 3.253E-06 | 8.500E-06 | 1.500E-05 |
| 26050 | 1303 | D1303I | E | 1.4e-05 | 1.4e-05 | 1.1e-05 | 1.300E-05 | 1.732E-06 | 1.100E-05 | 1.400E-05 |
| 26051 | 1303 | D1303L | E | 1.2e-05 | 1.6e-05 | 1e-05 | 1.267E-05 | 3.055E-06 | 1.000E-05 | 1.600E-05 |
| 26052 | 1303 | D1303K | E | 9.1e-06 | 1.1e-05 | 9.7e-06 | 9.933E-06 | 9.713E-07 | 9.100E-06 | 1.100E-05 |
| 26053 | 1303 | D1303M | E | 1.6e-05 | 1.2e-05 | 9e-06 | 1.233E-05 | 3.512E-06 | 9.000E-06 | 1.600E-05 |
| 26054 | 1303 | D1303F | E | 1.4e-05 | 1.1e-05 | 1.2e-05 | 1.233E-05 | 1.528E-06 | 1.100E-05 | 1.400E-05 |
| 26055 | 1303 | D1303P | E | 7.5e-06 | 1.6e-05 | 1.6e-05 | 1.317E-05 | 4.907E-06 | 7.500E-06 | 1.600E-05 |
| 26056 | 1303 | D1303S | E | 1.4e-05 | 1e-05 | 8.6e-06 | 1.087E-05 | 2.802E-06 | 8.600E-06 | 1.400E-05 |
| 26057 | 1303 | D1303T | E | 1.4e-05 | 1.4e-05 | 1.6e-05 | 1.467E-05 | 1.155E-06 | 1.400E-05 | 1.600E-05 |
| 26058 | 1303 | D1303W | E | 7.4e-06 | 1e-05 | 1.4e-05 | 1.047E-05 | 3.325E-06 | 7.400E-06 | 1.400E-05 |
| 26059 | 1303 | D1303Y | E | 1.4e-05 | 1.7e-05 | 1.6e-05 | 1.567E-05 | 1.528E-06 | 1.400E-05 | 1.700E-05 |
| 26060 | 1303 | D1303V | E | 1.4e-05 | 1.2e-05 | 1.3e-05 | 1.300E-05 | 1.000E-06 | 1.200E-05 | 1.400E-05 |
| 26061 | 1304 | T1304A | E | 1.5e-05 | 1.5e-05 | 9.4e-06 | 1.313E-05 | 3.233E-06 | 9.400E-06 | 1.500E-05 |
| 26062 | 1304 | T1304R | E | 1.1e-05 | 1.5e-05 | 1.2e-05 | 1.267E-05 | 2.082E-06 | 1.100E-05 | 1.500E-05 |
| 26063 | 1304 | T1304N | E | 1.5e-05 | 1.6e-05 | 9.4e-06 | 1.347E-05 | 3.557E-06 | 9.400E-06 | 1.600E-05 |
| 26064 | 1304 | T1304D | E | 1.5e-05 | 1.2e-05 | 1.6e-05 | 1.433E-05 | 2.082E-06 | 1.200E-05 | 1.600E-05 |
| 26065 | 1304 | T1304C | E | 1.7e-05 | 1.4e-05 | 2e-05 | 1.700E-05 | 3.000E-06 | 1.400E-05 | 2.000E-05 |
| 26066 | 1304 | T1304E | E | 9.7e-06 | 1e-05 | 1e-05 | 9.900E-06 | 1.732E-07 | 9.700E-06 | 1.000E-05 |
| 26067 | 1304 | T1304Q | E | 9e-06 | 1e-05 | 1.6e-05 | 1.167E-05 | 3.786E-06 | 9.000E-06 | 1.600E-05 |
| 26068 | 1304 | T1304G | E | 1.6e-05 | 1.6e-05 | 1.6e-05 | 1.600E-05 | 0.000E+00 | 1.600E-05 | 1.600E-05 |
| 26069 | 1304 | T1304H | E | 1.1e-05 | 8.9e-06 | 9.4e-06 | 9.767E-06 | 1.097E-06 | 8.900E-06 | 1.100E-05 |
| 26070 | 1304 | T1304I | E | 1.6e-05 | 9.1e-06 | 1.5e-05 | 1.337E-05 | 3.729E-06 | 9.100E-06 | 1.600E-05 |
| 26071 | 1304 | T1304L | E | 1e-05 | 1.4e-05 | 1.1e-05 | 1.167E-05 | 2.082E-06 | 1.000E-05 | 1.400E-05 |
| 26072 | 1304 | T1304K | E | 1.6e-05 | 1e-05 | 1.3e-05 | 1.300E-05 | 3.000E-06 | 1.000E-05 | 1.600E-05 |
| 26073 | 1304 | T1304M | E | 1.5e-05 | 1e-05 | 1.1e-05 | 1.200E-05 | 2.646E-06 | 1.000E-05 | 1.500E-05 |
| 26074 | 1304 | T1304F | E | 1.5e-05 | 1.2e-05 | 1.6e-05 | 1.433E-05 | 2.082E-06 | 1.200E-05 | 1.600E-05 |
| 26075 | 1304 | T1304P | E | 1.3e-05 | 8.7e-06 | 1.3e-05 | 1.157E-05 | 2.483E-06 | 8.700E-06 | 1.300E-05 |
| 26076 | 1304 | T1304S | E | 1.1e-05 | 1.5e-05 | 1.2e-05 | 1.267E-05 | 2.082E-06 | 1.100E-05 | 1.500E-05 |
| 26077 | 1304 | T1304T | E | 1.3e-05 | 1.5e-05 | 8.7e-06 | 1.223E-05 | 3.219E-06 | 8.700E-06 | 1.500E-05 |
| 26078 | 1304 | T1304W | E | 8.6e-06 | 1.1e-05 | 1.3e-05 | 1.087E-05 | 2.203E-06 | 8.600E-06 | 1.300E-05 |
| 26079 | 1304 | T1304Y | E | 1.1e-05 | 1.5e-05 | 1.2e-05 | 1.267E-05 | 2.082E-06 | 1.100E-05 | 1.500E-05 |
| 26080 | 1304 | T1304V | E | 1.6e-05 | 1.4e-05 | 1.5e-05 | 1.500E-05 | 1.000E-06 | 1.400E-05 | 1.600E-05 |
| 26081 | 1305 | Y1305A | E | 1e-05 | 1.8e-05 | 1e-05 | 1.267E-05 | 4.619E-06 | 1.000E-05 | 1.800E-05 |
| 26082 | 1305 | Y1305R | E | 1.6e-05 | 9.4e-06 | 1e-05 | 1.180E-05 | 3.650E-06 | 9.400E-06 | 1.600E-05 |
| 26083 | 1305 | Y1305N | E | 8.7e-06 | 1.5e-05 | 1.4e-05 | 1.257E-05 | 3.386E-06 | 8.700E-06 | 1.500E-05 |
| 26084 | 1305 | Y1305D | E | 8.8e-06 | 1.3e-05 | 1e-05 | 1.060E-05 | 2.163E-06 | 8.800E-06 | 1.300E-05 |
| 26085 | 1305 | Y1305C | E | 1.6e-05 | 8.4e-06 | 9.6e-06 | 1.133E-05 | 4.086E-06 | 8.400E-06 | 1.600E-05 |
| 26086 | 1305 | Y1305E | E | 1.1e-05 | 1.6e-05 | 1.5e-05 | 1.400E-05 | 2.646E-06 | 1.100E-05 | 1.600E-05 |
| 26087 | 1305 | Y1305Q | E | 1.5e-05 | 1.1e-05 | 1.5e-05 | 1.367E-05 | 2.309E-06 | 1.100E-05 | 1.500E-05 |
| 26088 | 1305 | Y1305G | E | 1e-05 | 1.7e-05 | 1.5e-05 | 1.400E-05 | 3.606E-06 | 1.000E-05 | 1.700E-05 |
| 26089 | 1305 | Y1305H | E | 1.5e-05 | 1.5e-05 | 1.1e-05 | 1.367E-05 | 2.309E-06 | 1.100E-05 | 1.500E-05 |
| 26090 | 1305 | Y1305I | E | 1.5e-05 | 7.4e-06 | 1e-05 | 1.080E-05 | 3.863E-06 | 7.400E-06 | 1.500E-05 |
| 26091 | 1305 | Y1305L | E | 1e-05 | 1.6e-05 | 1.1e-05 | 1.233E-05 | 3.215E-06 | 1.000E-05 | 1.600E-05 |
| 26092 | 1305 | Y1305K | E | 1.5e-05 | 1.5e-05 | 1.5e-05 | 1.500E-05 | 0.000E+00 | 1.500E-05 | 1.500E-05 |
| 26093 | 1305 | Y1305M | E | 1.1e-05 | 1.6e-05 | 1.3e-05 | 1.333E-05 | 2.517E-06 | 1.100E-05 | 1.600E-05 |
| 26094 | 1305 | Y1305F | E | 1.4e-05 | 9.2e-06 | 1.5e-05 | 1.273E-05 | 3.101E-06 | 9.200E-06 | 1.500E-05 |

| | | | | | | | | | | |
|---|---|---|---|---|---|---|---|---|---|---|
| 26095 | 1305 | Y1305P | E | 1.3e-05 | 1.1e-05 | 1.5e-05 | 1.300E-05 | 2.000E-06 | 1.100E-05 | 1.500E-05 |
| 26096 | 1305 | Y1305S | E | 1.5e-05 | 1e-05 | 1.2e-05 | 1.233E-05 | 2.517E-06 | 1.000E-05 | 1.500E-05 |
| 26097 | 1305 | Y1305T | E | 1.5e-05 | 1.6e-05 | 1.4e-05 | 1.500E-05 | 1.000E-06 | 1.400E-05 | 1.600E-05 |
| 26098 | 1305 | Y1305W | E | 1.1e-05 | 9.3e-06 | 1.5e-05 | 1.177E-05 | 2.926E-06 | 9.300E-06 | 1.500E-05 |
| 26099 | 1305 | Y1305Y | E | 1.4e-05 | 1.6e-05 | 1.7e-05 | 1.567E-05 | 1.528E-06 | 1.400E-05 | 1.700E-05 |
| 26100 | 1305 | Y1305V | E | 1.8e-05 | 1.4e-05 | 1.7e-05 | 1.633E-05 | 2.082E-06 | 1.400E-05 | 1.800E-05 |
| 26101 | 1306 | I1306A | E | 1.6e-05 | 7.7e-06 | 7.4e-06 | 1.037E-05 | 4.881E-06 | 7.400E-06 | 1.600E-05 |
| 26102 | 1306 | I1306R | E | 8e-06 | 1.3e-05 | 1.6e-05 | 1.233E-05 | 4.041E-06 | 8.000E-06 | 1.600E-05 |
| 26103 | 1306 | I1306N | E | 1.6e-05 | 8.5e-06 | 1e-05 | 1.150E-05 | 3.969E-06 | 8.500E-06 | 1.600E-05 |
| 26104 | 1306 | I1306D | E | 1e-05 | 1.3e-05 | 1.6e-05 | 1.300E-05 | 3.000E-06 | 1.000E-05 | 1.600E-05 |
| 26105 | 1306 | I1306C | E | 8.5e-06 | 1e-05 | 1.1e-05 | 9.833E-06 | 1.258E-06 | 8.500E-06 | 1.100E-05 |
| 26106 | 1306 | I1306E | E | 1.5e-05 | 1.1e-05 | 1.4e-05 | 1.333E-05 | 2.082E-06 | 1.100E-05 | 1.500E-05 |
| 26107 | 1306 | I1306Q | E | 8.1e-06 | 9e-06 | 8.9e-06 | 8.667E-06 | 4.933E-07 | 8.100E-06 | 9.000E-06 |
| 26108 | 1306 | I1306G | E | 1.5e-05 | 1.2e-05 | 1.6e-05 | 1.433E-05 | 2.082E-06 | 1.200E-05 | 1.600E-05 |
| 26109 | 1306 | I1306H | E | 1.1e-05 | 1.5e-05 | 7.9e-06 | 1.130E-05 | 3.559E-06 | 7.900E-06 | 1.500E-05 |
| 26110 | 1306 | I1306I | E | 1e-05 | 1.5e-05 | 1.2e-05 | 1.233E-05 | 2.517E-06 | 1.000E-05 | 1.500E-05 |
| 26111 | 1306 | I1306L | E | 1.3e-05 | 1.1e-05 | 1.5e-05 | 1.300E-05 | 2.000E-06 | 1.100E-05 | 1.500E-05 |
| 26112 | 1306 | I1306K | E | 1.1e-05 | 1e-05 | 1.6e-05 | 1.233E-05 | 3.215E-06 | 1.000E-05 | 1.600E-05 |
| 26113 | 1306 | I1306M | E | 1.1e-05 | 1.1e-05 | 1.1e-05 | 1.100E-05 | 2.075E-21 | 1.100E-05 | 1.100E-05 |
| 26114 | 1306 | I1306F | E | 1.3e-05 | 9.9e-06 | 1.7e-05 | 1.330E-05 | 3.559E-06 | 9.900E-06 | 1.700E-05 |
| 26115 | 1306 | I1306P | E | 1.5e-05 | 1.2e-05 | 9.6e-06 | 1.220E-05 | 2.706E-06 | 9.600E-06 | 1.500E-05 |
| 26116 | 1306 | I1306S | E | 1.4e-05 | 9.3e-06 | 1.1e-05 | 1.143E-05 | 2.380E-06 | 9.300E-06 | 1.400E-05 |
| 26117 | 1306 | I1306T | E | 1e-05 | 9.8e-06 | 1.5e-05 | 1.160E-05 | 2.946E-06 | 9.800E-06 | 1.500E-05 |
| 26118 | 1306 | I1306W | E | 1e-05 | 1.5e-05 | 1.1e-05 | 1.200E-05 | 2.646E-06 | 1.000E-05 | 1.500E-05 |
| 26119 | 1306 | I1306Y | E | 1.3e-05 | 1.6e-05 | 1e-05 | 1.300E-05 | 3.000E-06 | 1.000E-05 | 1.600E-05 |
| 26120 | 1306 | I1306V | E | 1.5e-05 | 1.1e-05 | 9.6e-06 | 1.187E-05 | 2.802E-06 | 9.600E-06 | 1.500E-05 |
| 26121 | 1307 | H1307A | E | 8.9e-06 | 1.1e-05 | 1.2e-05 | 1.063E-05 | 1.582E-06 | 8.900E-06 | 1.200E-05 |
| 26122 | 1307 | H1307R | E | 1.3e-05 | 1e-05 | 1.3e-05 | 1.200E-05 | 1.732E-06 | 1.000E-05 | 1.300E-05 |
| 26123 | 1307 | H1307N | E | 1.5e-05 | 1.6e-05 | 1.1e-05 | 1.400E-05 | 2.646E-06 | 1.100E-05 | 1.600E-05 |
| 26124 | 1307 | H1307D | E | 1.1e-05 | 1.4e-05 | 1.2e-05 | 1.233E-05 | 1.528E-06 | 1.100E-05 | 1.400E-05 |
| 26125 | 1307 | H1307C | E | 1.2e-05 | 1.5e-05 | 9.7e-06 | 1.223E-05 | 2.658E-06 | 9.700E-06 | 1.500E-05 |
| 26126 | 1307 | H1307E | E | 1.1e-05 | 1.6e-05 | 1.1e-05 | 1.267E-05 | 2.887E-06 | 1.100E-05 | 1.600E-05 |
| 26127 | 1307 | H1307Q | E | 9.7e-06 | 1e-05 | 1.2e-05 | 1.057E-05 | 1.250E-06 | 9.700E-06 | 1.200E-05 |
| 26128 | 1307 | H1307G | E | 1.7e-05 | 1.5e-05 | 1.7e-05 | 1.633E-05 | 1.155E-06 | 1.500E-05 | 1.700E-05 |
| 26129 | 1307 | H1307H | E | 1.1e-05 | 1.5e-05 | 1e-05 | 1.200E-05 | 2.646E-06 | 1.000E-05 | 1.500E-05 |
| 26130 | 1307 | H1307I | E | 1.6e-05 | 1.6e-05 | 1.4e-05 | 1.533E-05 | 1.155E-06 | 1.400E-05 | 1.600E-05 |
| 26131 | 1307 | H1307L | E | 1.5e-05 | 1.5e-05 | 1e-05 | 1.333E-05 | 2.887E-06 | 1.000E-05 | 1.500E-05 |
| 26132 | 1307 | H1307K | E | 1.5e-05 | 1e-05 | 1.5e-05 | 1.333E-05 | 2.887E-06 | 1.000E-05 | 1.500E-05 |
| 26133 | 1307 | H1307M | E | 1.6e-05 | 1.2e-05 | 1.8e-05 | 1.533E-05 | 3.055E-06 | 1.200E-05 | 1.800E-05 |
| 26134 | 1307 | H1307F | E | 1.4e-05 | 1.1e-05 | 1.6e-05 | 1.367E-05 | 2.517E-06 | 1.100E-05 | 1.600E-05 |
| 26135 | 1307 | H1307P | E | 9.1e-06 | 7.9e-06 | 1.5e-05 | 1.067E-05 | 3.800E-06 | 7.900E-06 | 1.500E-05 |
| 26136 | 1307 | H1307S | E | 1.3e-05 | 1.2e-05 | 1.5e-05 | 1.333E-05 | 1.528E-06 | 1.200E-05 | 1.500E-05 |
| 26137 | 1307 | H1307T | E | 9.1e-06 | 9.9e-06 | 9e-06 | 9.333E-06 | 4.933E-07 | 9.000E-06 | 9.900E-06 |
| 26138 | 1307 | H1307W | E | 1.3e-05 | 9.6e-06 | 1.1e-05 | 1.120E-05 | 1.709E-06 | 9.600E-06 | 1.300E-05 |
| 26139 | 1307 | H1307Y | E | 1.1e-05 | 1.1e-05 | 1.5e-05 | 1.233E-05 | 2.309E-06 | 1.100E-05 | 1.500E-05 |
| 26140 | 1307 | H1307V | E | 1.1e-05 | 1.5e-05 | 1.3e-05 | 1.300E-05 | 2.000E-06 | 1.100E-05 | 1.500E-05 |
| 26141 | 1308 | W1308A | E | 1.4e-05 | 1.3e-05 | 1.7e-05 | 1.467E-05 | 2.082E-06 | 1.300E-05 | 1.700E-05 |
| 26142 | 1308 | W1308R | E | 1.4e-05 | 9.8e-06 | 1.1e-05 | 1.160E-05 | 2.163E-06 | 9.800E-06 | 1.400E-05 |
| 26143 | 1308 | W1308N | E | 9.5e-06 | 1e-05 | 1.6e-05 | 1.183E-05 | 3.617E-06 | 9.500E-06 | 1.600E-05 |
| 26144 | 1308 | W1308D | E | 1.4e-05 | 1.6e-05 | 1.6e-05 | 1.533E-05 | 1.155E-06 | 1.400E-05 | 1.600E-05 |
| 26145 | 1308 | W1308C | E | 1e-05 | 1.5e-05 | 1.1e-05 | 1.200E-05 | 2.646E-06 | 1.000E-05 | 1.500E-05 |
| 26146 | 1308 | W1308E | E | 1.2e-05 | 1.1e-05 | 1.3e-05 | 1.200E-05 | 1.000E-06 | 1.100E-05 | 1.300E-05 |
| 26147 | 1308 | W1308Q | E | 8.5e-06 | 1.1e-05 | 1.5e-05 | 1.150E-05 | 3.279E-06 | 8.500E-06 | 1.500E-05 |
| 26148 | 1308 | W1308G | E | 1.5e-05 | 8.9e-06 | 1.4e-05 | 1.263E-05 | 3.272E-06 | 8.900E-06 | 1.500E-05 |
| 26149 | 1308 | W1308H | E | 1.1e-05 | 1e-05 | 9.6e-06 | 1.020E-05 | 7.211E-07 | 9.600E-06 | 1.100E-05 |
| 26150 | 1308 | W1308I | E | 1.5e-05 | 1.3e-05 | 9.7e-06 | 1.257E-05 | 2.676E-06 | 9.700E-06 | 1.500E-05 |
| 26151 | 1308 | W1308L | E | 1.2e-05 | 1.5e-05 | 1.4e-05 | 1.367E-05 | 1.528E-06 | 1.200E-05 | 1.500E-05 |
| 26152 | 1308 | W1308K | E | 1.4e-05 | 1e-05 | 1e-05 | 1.133E-05 | 2.309E-06 | 1.000E-05 | 1.400E-05 |
| 26153 | 1308 | W1308M | E | 1.4e-05 | 9.6e-06 | 1.5e-05 | 1.287E-05 | 2.873E-06 | 9.600E-06 | 1.500E-05 |
| 26154 | 1308 | W1308F | E | 1.1e-05 | 8.3e-06 | 1.2e-05 | 1.043E-05 | 1.914E-06 | 8.300E-06 | 1.200E-05 |
| 26155 | 1308 | W1308P | E | 1.3e-05 | 1.5e-05 | 1.3e-05 | 1.367E-05 | 1.155E-06 | 1.300E-05 | 1.500E-05 |
| 26156 | 1308 | W1308S | E | 1e-05 | 8.4e-06 | 1.6e-05 | 1.147E-05 | 4.007E-06 | 8.400E-06 | 1.600E-05 |
| 26157 | 1308 | W1308T | E | 8.6e-06 | 1.4e-05 | 1.1e-05 | 1.120E-05 | 2.706E-06 | 8.600E-06 | 1.400E-05 |
| 26158 | 1308 | W1308W | E | 1.5e-05 | 9.9e-06 | 1e-05 | 1.163E-05 | 2.916E-06 | 9.900E-06 | 1.500E-05 |
| 26159 | 1308 | W1308Y | E | 1.3e-05 | 1.5e-05 | 1.6e-05 | 1.467E-05 | 1.528E-06 | 1.300E-05 | 1.600E-05 |
| 26160 | 1308 | W1308V | E | 1.1e-05 | 1.4e-05 | 1.6e-05 | 1.367E-05 | 2.517E-06 | 1.100E-05 | 1.600E-05 |
| 26161 | 1309 | V1309A | E | 1.9e-05 | 1.6e-05 | 1.4e-05 | 1.633E-05 | 2.517E-06 | 1.400E-05 | 1.900E-05 |
| 26162 | 1309 | V1309R | E | 1.7e-05 | 1e-05 | 1.4e-05 | 1.367E-05 | 3.512E-06 | 1.000E-05 | 1.700E-05 |
| 26163 | 1309 | V1309N | E | 1.3e-05 | 1.2e-05 | 1.7e-05 | 1.400E-05 | 2.646E-06 | 1.200E-05 | 1.700E-05 |
| 26164 | 1309 | V1309D | E | 1.4e-05 | 1.4e-05 | 1.4e-05 | 1.400E-05 | 0.000E+00 | 1.400E-05 | 1.400E-05 |
| 26165 | 1309 | V1309C | E | 1.2e-05 | 1.4e-05 | 1e-05 | 1.200E-05 | 2.000E-06 | 1.000E-05 | 1.400E-05 |
| 26166 | 1309 | V1309E | E | 1.6e-05 | 1.4e-05 | 9.7e-06 | 1.323E-05 | 3.219E-06 | 9.700E-06 | 1.600E-05 |
| 26167 | 1309 | V1309Q | E | 9.7e-06 | 1.1e-05 | 9.2e-06 | 9.967E-06 | 9.292E-07 | 9.200E-06 | 1.100E-05 |
| 26168 | 1309 | V1309G | E | 1.4e-05 | 7.8e-06 | 1.6e-05 | 1.260E-05 | 4.276E-06 | 7.800E-06 | 1.600E-05 |
| 26169 | 1309 | V1309H | E | 1.4e-05 | 1.5e-05 | 1.1e-05 | 1.333E-05 | 2.082E-06 | 1.100E-05 | 1.500E-05 |
| 26170 | 1309 | V1309I | E | 8.1e-06 | 1.4e-05 | 1.5e-05 | 1.237E-05 | 3.729E-06 | 8.100E-06 | 1.500E-05 |
| 26171 | 1309 | V1309L | E | 1.7e-05 | 1e-05 | 1.4e-05 | 1.367E-05 | 3.512E-06 | 1.000E-05 | 1.700E-05 |
| 26172 | 1309 | V1309K | E | 1.2e-05 | 1e-05 | 8.5e-06 | 1.017E-05 | 1.756E-06 | 8.500E-06 | 1.200E-05 |
| 26173 | 1309 | V1309M | E | 1.3e-05 | 9.6e-06 | 1.1e-05 | 1.120E-05 | 1.709E-06 | 9.600E-06 | 1.300E-05 |
| 26174 | 1309 | V1309F | E | 1.4e-05 | 1.2e-05 | 9.5e-06 | 1.183E-05 | 2.255E-06 | 9.500E-06 | 1.400E-05 |
| 26175 | 1309 | V1309P | E | 4.4e-06 | 1.6e-05 | 1.1e-05 | 1.047E-05 | 5.818E-06 | 4.400E-06 | 1.600E-05 |
| 26176 | 1309 | V1309S | E | 1.6e-05 | 1.5e-05 | 1.5e-05 | 1.533E-05 | 5.774E-07 | 1.500E-05 | 1.600E-05 |
| 26177 | 1309 | V1309T | E | 9.2e-06 | 1.5e-05 | 1.6e-05 | 1.340E-05 | 3.672E-06 | 9.200E-06 | 1.600E-05 |
| 26178 | 1309 | V1309W | E | 1.3e-05 | 1.3e-05 | 1e-05 | 1.200E-05 | 1.732E-06 | 1.000E-05 | 1.300E-05 |
| 26179 | 1309 | V1309Y | E | 1.4e-05 | 1e-05 | 1.3e-05 | 1.233E-05 | 2.082E-06 | 1.000E-05 | 1.400E-05 |

| | | | | | | | | | | |
|---|---|---|---|---|---|---|---|---|---|---|
| 26180 | 1309 | V1309V | E | 9.1e-06 | 1e-05 | 1.1e-05 | 1.003E-05 | 9.504E-07 | 9.100E-06 | 1.100E-05 |
| 26181 | 1310 | R1310A | E | 8.6e-06 | 1.2e-05 | 9.2e-06 | 9.933E-06 | 1.815E-06 | 8.600E-06 | 1.200E-05 |
| 26182 | 1310 | R1310R | E | 9.5e-06 | 1.1e-05 | 1.5e-05 | 1.183E-05 | 2.843E-06 | 9.500E-06 | 1.500E-05 |
| 26183 | 1310 | R1310N | E | 8.2e-06 | 1.4e-05 | 1e-05 | 1.073E-05 | 2.969E-06 | 8.200E-06 | 1.400E-05 |
| 26184 | 1310 | R1310D | E | 9.8e-06 | 1.7e-05 | 8.9e-06 | 1.190E-05 | 4.440E-06 | 8.900E-06 | 1.700E-05 |
| 26185 | 1310 | R1310C | E | 1.6e-05 | 1.1e-05 | 1.5e-05 | 1.400E-05 | 2.646E-06 | 1.100E-05 | 1.600E-05 |
| 26186 | 1310 | R1310E | E | 1.5e-05 | 1.6e-05 | 1e-05 | 1.367E-05 | 3.215E-06 | 1.000E-05 | 1.600E-05 |
| 26187 | 1310 | R1310Q | E | 1.1e-05 | 1.2e-05 | 9e-06 | 1.067E-05 | 1.528E-06 | 9.000E-06 | 1.200E-05 |
| 26188 | 1310 | R1310G | E | 1.4e-05 | 8.4e-06 | 1.6e-05 | 1.280E-05 | 3.940E-06 | 8.400E-06 | 1.600E-05 |
| 26189 | 1310 | R1310H | E | 1.5e-05 | 1.1e-05 | 1.3e-05 | 1.300E-05 | 2.000E-06 | 1.100E-05 | 1.500E-05 |
| 26190 | 1310 | R1310I | E | 1.3e-05 | 1.7e-05 | 1.6e-05 | 1.533E-05 | 2.082E-06 | 1.300E-05 | 1.700E-05 |
| 26191 | 1310 | R1310L | E | 1.3e-05 | 1.3e-05 | 1e-05 | 1.200E-05 | 1.732E-06 | 1.000E-05 | 1.300E-05 |
| 26192 | 1310 | R1310K | E | 1.1e-05 | 1.5e-05 | 1.1e-05 | 1.233E-05 | 2.309E-06 | 1.100E-05 | 1.500E-05 |
| 26193 | 1310 | R1310M | E | 1.7e-05 | 1.1e-05 | 1.5e-05 | 1.433E-05 | 3.055E-06 | 1.100E-05 | 1.700E-05 |
| 26194 | 1310 | R1310F | E | 9.6e-06 | 1e-05 | 1.6e-05 | 1.187E-05 | 3.585E-06 | 9.600E-06 | 1.600E-05 |
| 26195 | 1310 | R1310P | E | 1.5e-05 | 1.6e-05 | 1.4e-05 | 1.500E-05 | 1.000E-06 | 1.400E-05 | 1.600E-05 |
| 26196 | 1310 | R1310S | E | 1.5e-05 | 1e-05 | 1.4e-05 | 1.300E-05 | 2.646E-06 | 1.000E-05 | 1.500E-05 |
| 26197 | 1310 | R1310T | E | 1.5e-05 | 1e-05 | 1.2e-05 | 1.233E-05 | 2.517E-06 | 1.000E-05 | 1.500E-05 |
| 26198 | 1310 | R1310W | E | 1e-05 | 9.1e-06 | 1.8e-05 | 1.237E-05 | 4.899E-06 | 9.100E-06 | 1.800E-05 |
| 26199 | 1310 | R1310Y | E | 1.1e-05 | 1.2e-05 | 1.1e-05 | 1.133E-05 | 5.774E-07 | 1.100E-05 | 1.200E-05 |
| 26200 | 1310 | R1310V | E | 1.5e-05 | 1.5e-05 | 1.4e-05 | 1.467E-05 | 5.774E-07 | 1.400E-05 | 1.500E-05 |
| 26201 | 1311 | Q1311A | E | 1.2e-05 | 1.4e-05 | 1.1e-05 | 1.233E-05 | 1.528E-06 | 1.100E-05 | 1.400E-05 |
| 26202 | 1311 | Q1311R | E | 1.7e-05 | 1.4e-05 | 8.6e-06 | 1.320E-05 | 4.257E-06 | 8.600E-06 | 1.700E-05 |
| 26203 | 1311 | Q1311N | E | 1.7e-05 | 9.8e-06 | 1.4e-05 | 1.360E-05 | 3.617E-06 | 9.800E-06 | 1.700E-05 |
| 26204 | 1311 | Q1311D | E | 1.3e-05 | 1.5e-05 | 1.1e-05 | 1.300E-05 | 2.000E-06 | 1.100E-05 | 1.500E-05 |
| 26205 | 1311 | Q1311C | E | 1.1e-05 | 8e-06 | 1.1e-05 | 1.000E-05 | 1.732E-06 | 8.000E-06 | 1.100E-05 |
| 26206 | 1311 | Q1311E | E | 1.6e-05 | 1.4e-05 | 1.3e-05 | 1.433E-05 | 1.528E-06 | 1.300E-05 | 1.600E-05 |
| 26207 | 1311 | Q1311Q | E | 1.7e-05 | 9.1e-06 | 8.2e-06 | 1.143E-05 | 4.842E-06 | 8.200E-06 | 1.700E-05 |
| 26208 | 1311 | Q1311G | E | 1.1e-05 | 1.4e-05 | 1.4e-05 | 1.300E-05 | 1.732E-06 | 1.100E-05 | 1.400E-05 |
| 26209 | 1311 | Q1311H | E | 1.4e-05 | 1.4e-05 | 9.9e-06 | 1.263E-05 | 2.367E-06 | 9.900E-06 | 1.400E-05 |
| 26210 | 1311 | Q1311I | E | 1.5e-05 | 1.5e-05 | 1.4e-05 | 1.467E-05 | 5.774E-07 | 1.400E-05 | 1.500E-05 |
| 26211 | 1311 | Q1311L | E | 1.9e-05 | 1.8e-05 | 1.4e-05 | 1.700E-05 | 2.646E-06 | 1.400E-05 | 1.900E-05 |
| 26212 | 1311 | Q1311K | E | 8.9e-06 | 1.5e-05 | 1.6e-05 | 1.330E-05 | 3.843E-06 | 8.900E-06 | 1.600E-05 |
| 26213 | 1311 | Q1311M | E | 1.1e-05 | 1.3e-05 | 1.3e-05 | 1.233E-05 | 1.155E-06 | 1.100E-05 | 1.300E-05 |
| 26214 | 1311 | Q1311F | E | 1.6e-05 | 1.5e-05 | 1.7e-05 | 1.600E-05 | 1.000E-06 | 1.500E-05 | 1.700E-05 |
| 26215 | 1311 | Q1311P | E | 1e-05 | 9.3e-06 | 1.5e-05 | 1.143E-05 | 3.109E-06 | 9.300E-06 | 1.500E-05 |
| 26216 | 1311 | Q1311S | E | 2.3e-05 | 1.1e-05 | 1.8e-05 | 1.733E-05 | 6.028E-06 | 1.100E-05 | 2.300E-05 |
| 26217 | 1311 | Q1311T | E | 1e-05 | 1.5e-05 | 1.7e-05 | 1.400E-05 | 3.606E-06 | 1.000E-05 | 1.700E-05 |
| 26218 | 1311 | Q1311W | E | 1.5e-05 | 1.1e-05 | 1.5e-05 | 1.367E-05 | 2.309E-06 | 1.100E-05 | 1.500E-05 |
| 26219 | 1311 | Q1311Y | E | 1.6e-05 | 1e-05 | 1.7e-05 | 1.433E-05 | 3.786E-06 | 1.000E-05 | 1.700E-05 |
| 26220 | 1311 | Q1311V | E | 1.5e-05 | 8.3e-06 | 1e-05 | 1.110E-05 | 3.483E-06 | 8.300E-06 | 1.500E-05 |
| 26221 | 1312 | A1312A | E | 9.8e-06 | 1.2e-05 | 1e-05 | 1.060E-05 | 1.217E-06 | 9.800E-06 | 1.200E-05 |
| 26222 | 1312 | A1312R | E | 1.5e-05 | 1.5e-05 | 1.6e-05 | 1.533E-05 | 5.774E-07 | 1.500E-05 | 1.600E-05 |
| 26223 | 1312 | A1312N | E | 9.6e-06 | 1.2e-05 | 9.3e-06 | 1.030E-05 | 1.480E-06 | 9.300E-06 | 1.200E-05 |
| 26224 | 1312 | A1312D | E | 1.3e-05 | 1.5e-05 | 1e-05 | 1.267E-05 | 2.517E-06 | 1.000E-05 | 1.500E-05 |
| 26225 | 1312 | A1312C | E | 1.5e-05 | 1.1e-05 | 1.2e-05 | 1.267E-05 | 2.082E-06 | 1.100E-05 | 1.500E-05 |
| 26226 | 1312 | A1312E | E | 1.6e-05 | 1.5e-05 | 2.3e-05 | 1.800E-05 | 4.359E-06 | 1.500E-05 | 2.300E-05 |
| 26227 | 1312 | A1312Q | E | 1.5e-05 | 1.3e-05 | 1.3e-05 | 1.367E-05 | 1.155E-06 | 1.300E-05 | 1.500E-05 |
| 26228 | 1312 | A1312G | E | 9.5e-06 | 8.7e-06 | 1.3e-05 | 1.040E-05 | 2.287E-06 | 8.700E-06 | 1.300E-05 |
| 26229 | 1312 | A1312H | E | 9.9e-06 | 1.4e-05 | 1.1e-05 | 1.163E-05 | 2.122E-06 | 9.900E-06 | 1.400E-05 |
| 26230 | 1312 | A1312I | E | 1.1e-05 | 1.6e-05 | 1.5e-05 | 1.400E-05 | 2.646E-06 | 1.100E-05 | 1.600E-05 |
| 26231 | 1312 | A1312L | E | 1.1e-05 | 1e-05 | 1.5e-05 | 1.200E-05 | 2.646E-06 | 1.000E-05 | 1.500E-05 |
| 26232 | 1312 | A1312K | E | 1.6e-05 | 1e-05 | 1e-05 | 1.200E-05 | 3.464E-06 | 1.000E-05 | 1.600E-05 |
| 26233 | 1312 | A1312M | E | 1e-05 | 1.5e-05 | 1.5e-05 | 1.333E-05 | 2.887E-06 | 1.000E-05 | 1.500E-05 |
| 26234 | 1312 | A1312F | E | 1.6e-05 | 8.9e-06 | 1.6e-05 | 1.363E-05 | 4.099E-06 | 8.900E-06 | 1.600E-05 |
| 26235 | 1312 | A1312P | E | 1.4e-05 | 9.4e-06 | 1.5e-05 | 1.280E-05 | 2.987E-06 | 9.400E-06 | 1.500E-05 |
| 26236 | 1312 | A1312S | E | 8.8e-06 | 1.5e-05 | 1.7e-05 | 1.360E-05 | 4.276E-06 | 8.800E-06 | 1.700E-05 |
| 26237 | 1312 | A1312T | E | 1.5e-05 | 1.4e-05 | 8.7e-06 | 1.257E-05 | 3.386E-06 | 8.700E-06 | 1.500E-05 |
| 26238 | 1312 | A1312W | E | 8.6e-06 | 1.4e-05 | 1.5e-05 | 1.253E-05 | 3.443E-06 | 8.600E-06 | 1.500E-05 |
| 26239 | 1312 | A1312Y | E | 1.5e-05 | 7.8e-06 | 9.7e-06 | 1.083E-05 | 3.731E-06 | 7.800E-06 | 1.500E-05 |
| 26240 | 1312 | A1312V | E | 1.5e-05 | 1e-05 | 1.2e-05 | 1.233E-05 | 2.517E-06 | 1.000E-05 | 1.500E-05 |
| 26241 | 1313 | P1313A | E | 1.4e-05 | 8.9e-06 | 1e-05 | 1.097E-05 | 2.684E-06 | 8.900E-06 | 1.400E-05 |
| 26242 | 1313 | P1313R | E | 1.3e-05 | 1.4e-05 | 1.2e-05 | 1.300E-05 | 1.000E-06 | 1.200E-05 | 1.400E-05 |
| 26243 | 1313 | P1313N | E | 8.8e-06 | 1.4e-05 | 1.7e-05 | 1.327E-05 | 4.149E-06 | 8.800E-06 | 1.700E-05 |
| 26244 | 1313 | P1313D | E | 1.6e-05 | 1.6e-05 | 9.6e-06 | 1.387E-05 | 3.695E-06 | 9.600E-06 | 1.600E-05 |
| 26245 | 1313 | P1313C | E | 1.7e-05 | 8.6e-06 | 1e-05 | 1.187E-05 | 4.500E-06 | 8.600E-06 | 1.700E-05 |
| 26246 | 1313 | P1313E | E | 1.4e-05 | 1.4e-05 | 1.3e-05 | 1.367E-05 | 5.774E-07 | 1.300E-05 | 1.400E-05 |
| 26247 | 1313 | P1313Q | E | 1.1e-05 | 1e-05 | 1.4e-05 | 1.167E-05 | 2.082E-06 | 1.000E-05 | 1.400E-05 |
| 26248 | 1313 | P1313G | E | 1.6e-05 | 1e-05 | 1.1e-05 | 1.233E-05 | 3.215E-06 | 1.000E-05 | 1.600E-05 |
| 26249 | 1313 | P1313H | E | 1.6e-05 | 1.2e-05 | 1.6e-05 | 1.467E-05 | 2.309E-06 | 1.200E-05 | 1.600E-05 |
| 26250 | 1313 | P1313I | E | 1.2e-05 | 1e-05 | 1.6e-05 | 1.267E-05 | 3.055E-06 | 1.000E-05 | 1.600E-05 |
| 26251 | 1313 | P1313L | E | 1.6e-05 | 1.7e-05 | 1.1e-05 | 1.467E-05 | 3.215E-06 | 1.100E-05 | 1.700E-05 |
| 26252 | 1313 | P1313K | E | 1.3e-05 | 1.6e-05 | 1.3e-05 | 1.400E-05 | 1.732E-06 | 1.300E-05 | 1.600E-05 |
| 26253 | 1313 | P1313M | E | 1.5e-05 | 1.1e-05 | 1.4e-05 | 1.333E-05 | 2.082E-06 | 1.100E-05 | 1.500E-05 |
| 26254 | 1313 | P1313F | E | 9.5e-06 | 1.5e-05 | 1.1e-05 | 1.183E-05 | 2.843E-06 | 9.500E-06 | 1.500E-05 |
| 26255 | 1313 | P1313P | E | 1.2e-05 | 1.5e-05 | 1.5e-05 | 1.400E-05 | 1.732E-06 | 1.200E-05 | 1.500E-05 |
| 26256 | 1313 | P1313S | E | 1.1e-05 | 1.6e-05 | 6.5e-06 | 1.117E-05 | 4.752E-06 | 6.500E-06 | 1.600E-05 |
| 26257 | 1313 | P1313T | E | 8.4e-06 | 1.5e-05 | 1.4e-05 | 1.247E-05 | 3.557E-06 | 8.400E-06 | 1.500E-05 |
| 26258 | 1313 | P1313W | E | 2.2e-05 | 1.3e-05 | 1.6e-05 | 1.700E-05 | 4.583E-06 | 1.300E-05 | 2.200E-05 |
| 26259 | 1313 | P1313Y | E | 1.6e-05 | 9.5e-06 | 1.4e-05 | 1.317E-05 | 3.329E-06 | 9.500E-06 | 1.600E-05 |
| 26260 | 1313 | P1313V | E | 1e-05 | 1.7e-05 | 1.6e-05 | 1.433E-05 | 3.786E-06 | 1.000E-05 | 1.700E-05 |
| 26261 | 1314 | G1314A | E | 1.6e-05 | 9.9e-06 | 1.7e-05 | 1.430E-05 | 3.843E-06 | 9.900E-06 | 1.700E-05 |
| 26262 | 1314 | G1314R | E | 1.3e-05 | 1.5e-05 | 9.3e-06 | 1.243E-05 | 2.892E-06 | 9.300E-06 | 1.500E-05 |
| 26263 | 1314 | G1314N | E | 1.2e-05 | 1.5e-05 | 1.4e-05 | 1.367E-05 | 1.528E-06 | 1.200E-05 | 1.500E-05 |
| 26264 | 1314 | G1314D | E | 1.1e-05 | 1.5e-05 | 1.2e-05 | 1.267E-05 | 2.082E-06 | 1.100E-05 | 1.500E-05 |

| | | | | | | | | | | |
|---|---|---|---|---|---|---|---|---|---|---|
| 26265 | 1314 | G1314C | E | 1.5e-05 | 1.3e-05 | 8.2e-06 | 1.207E-05 | 3.495E-06 | 8.200E-06 | 1.500E-05 |
| 26266 | 1314 | G1314E | E | 1e-05 | 1.2e-05 | 1.6e-05 | 1.267E-05 | 3.055E-06 | 1.000E-05 | 1.600E-05 |
| 26267 | 1314 | G1314Q | E | 1.5e-05 | 1.4e-05 | 9.9e-06 | 1.297E-05 | 2.702E-06 | 9.900E-06 | 1.500E-05 |
| 26268 | 1314 | G1314G | E | 1e-05 | 1.5e-05 | 1.7e-05 | 1.400E-05 | 3.606E-06 | 1.000E-05 | 1.700E-05 |
| 26269 | 1314 | G1314H | E | 1.4e-05 | 9.6e-06 | 1.4e-05 | 1.253E-05 | 2.540E-06 | 9.600E-06 | 1.400E-05 |
| 26270 | 1314 | G1314I | E | 1.5e-05 | 1e-05 | 1.2e-05 | 1.233E-05 | 2.517E-06 | 1.000E-05 | 1.500E-05 |
| 26271 | 1314 | G1314L | E | 1.6e-05 | 1.4e-05 | 1e-05 | 1.333E-05 | 3.055E-06 | 1.000E-05 | 1.600E-05 |
| 26272 | 1314 | G1314K | E | 1.3e-05 | 9e-06 | 1.5e-05 | 1.233E-05 | 3.055E-06 | 9.000E-06 | 1.500E-05 |
| 26273 | 1314 | G1314M | E | 1.5e-05 | 1.4e-05 | 1.5e-05 | 1.467E-05 | 5.774E-07 | 1.400E-05 | 1.500E-05 |
| 26274 | 1314 | G1314F | E | 1.6e-05 | 1.4e-05 | 1.2e-05 | 1.400E-05 | 2.000E-06 | 1.200E-05 | 1.600E-05 |
| 26275 | 1314 | G1314P | E | 1.5e-05 | 1e-05 | 1.7e-05 | 1.400E-05 | 3.606E-06 | 1.000E-05 | 1.700E-05 |
| 26276 | 1314 | G1314S | E | 1.4e-05 | 1e-05 | 1.3e-05 | 1.233E-05 | 2.082E-06 | 1.000E-05 | 1.400E-05 |
| 26277 | 1314 | G1314T | E | 1.4e-05 | 1e-05 | 1.5e-05 | 1.300E-05 | 2.646E-06 | 1.000E-05 | 1.500E-05 |
| 26278 | 1314 | G1314W | E | 1.1e-05 | 8.8e-06 | 1.1e-05 | 1.027E-05 | 1.270E-06 | 8.800E-06 | 1.100E-05 |
| 26279 | 1314 | G1314Y | E | 1.3e-05 | 1.4e-05 | 1.4e-05 | 1.367E-05 | 5.774E-07 | 1.300E-05 | 1.400E-05 |
| 26280 | 1314 | G1314V | E | 1.5e-05 | 1.2e-05 | 1.3e-05 | 1.333E-05 | 1.528E-06 | 1.200E-05 | 1.500E-05 |
| 26281 | 1315 | K1315A | E | 1.5e-05 | 1e-05 | 1.1e-05 | 1.200E-05 | 2.646E-06 | 1.000E-05 | 1.500E-05 |
| 26282 | 1315 | K1315R | E | 1.6e-05 | 9.9e-06 | 9.9e-06 | 1.193E-05 | 3.522E-06 | 9.900E-06 | 1.600E-05 |
| 26283 | 1315 | K1315N | E | 1.1e-05 | 1.7e-05 | 1.6e-05 | 1.467E-05 | 3.215E-06 | 1.100E-05 | 1.700E-05 |
| 26284 | 1315 | K1315D | E | 1.6e-05 | 1.1e-05 | 6.9e-06 | 1.130E-05 | 4.557E-06 | 6.900E-06 | 1.600E-05 |
| 26285 | 1315 | K1315C | E | 1.5e-05 | 1.4e-05 | 1.4e-05 | 1.433E-05 | 5.774E-07 | 1.400E-05 | 1.500E-05 |
| 26286 | 1315 | K1315E | E | 1.5e-05 | 1.6e-05 | 1.1e-05 | 1.400E-05 | 2.646E-06 | 1.100E-05 | 1.600E-05 |
| 26287 | 1315 | K1315Q | E | 1.1e-05 | 1e-05 | 1e-05 | 1.033E-05 | 5.774E-07 | 1.000E-05 | 1.100E-05 |
| 26288 | 1315 | K1315G | E | 1.5e-05 | 9.6e-06 | 1.2e-05 | 1.220E-05 | 2.706E-06 | 9.600E-06 | 1.500E-05 |
| 26289 | 1315 | K1315H | E | 1.4e-05 | 9.6e-06 | 1.5e-05 | 1.287E-05 | 2.873E-06 | 9.600E-06 | 1.500E-05 |
| 26290 | 1315 | K1315I | E | 1.7e-05 | 1.7e-05 | 1.4e-05 | 1.600E-05 | 1.732E-06 | 1.400E-05 | 1.700E-05 |
| 26291 | 1315 | K1315L | E | 1.6e-05 | 1.1e-05 | 1.1e-05 | 1.267E-05 | 2.887E-06 | 1.100E-05 | 1.600E-05 |
| 26292 | 1315 | K1315K | E | 1.1e-05 | 1.5e-05 | 1.5e-05 | 1.367E-05 | 2.309E-06 | 1.100E-05 | 1.500E-05 |
| 26293 | 1315 | K1315M | E | 1.6e-05 | 1.4e-05 | 7.5e-06 | 1.250E-05 | 4.444E-06 | 7.500E-06 | 1.600E-05 |
| 26294 | 1315 | K1315F | E | 1.4e-05 | 1.6e-05 | 1.1e-05 | 1.367E-05 | 2.517E-06 | 1.100E-05 | 1.600E-05 |
| 26295 | 1315 | K1315P | E | 1.6e-05 | 1.5e-05 | 1.1e-05 | 1.400E-05 | 2.646E-06 | 1.100E-05 | 1.600E-05 |
| 26296 | 1315 | K1315S | E | 1.2e-05 | 1.4e-05 | 1.5e-05 | 1.367E-05 | 1.528E-06 | 1.200E-05 | 1.500E-05 |
| 26297 | 1315 | K1315T | E | 8.4e-06 | 1.8e-05 | 1e-05 | 1.213E-05 | 5.143E-06 | 8.400E-06 | 1.800E-05 |
| 26298 | 1315 | K1315W | E | 8.3e-06 | 6.3e-06 | 1.5e-05 | 9.867E-06 | 4.557E-06 | 6.300E-06 | 1.500E-05 |
| 26299 | 1315 | K1315Y | E | 1e-05 | 9.8e-06 | 1e-05 | 9.933E-06 | 1.155E-07 | 9.800E-06 | 1.000E-05 |
| 26300 | 1315 | K1315V | E | 1.2e-05 | 1.2e-05 | 1e-05 | 1.133E-05 | 1.155E-06 | 1.000E-05 | 1.200E-05 |
| 26301 | 1316 | G1316A | E | 1.4e-05 | 9.3e-06 | 1.6e-05 | 1.310E-05 | 3.439E-06 | 9.300E-06 | 1.600E-05 |
| 26302 | 1316 | G1316R | E | 1.3e-05 | 1.5e-05 | 1.5e-05 | 1.433E-05 | 1.155E-06 | 1.300E-05 | 1.500E-05 |
| 26303 | 1316 | G1316N | E | 1.4e-05 | 1.1e-05 | 1.4e-05 | 1.300E-05 | 1.732E-06 | 1.100E-05 | 1.400E-05 |
| 26304 | 1316 | G1316D | E | 1.5e-05 | 1.1e-05 | 1.4e-05 | 1.333E-05 | 2.082E-06 | 1.100E-05 | 1.500E-05 |
| 26305 | 1316 | G1316C | E | 1.5e-05 | 1e-05 | 1.5e-05 | 1.333E-05 | 2.887E-06 | 1.000E-05 | 1.500E-05 |
| 26306 | 1316 | G1316E | E | 2.4e-05 | 1.6e-05 | 1.4e-05 | 1.800E-05 | 5.292E-06 | 1.400E-05 | 2.400E-05 |
| 26307 | 1316 | G1316Q | E | 1.6e-05 | 1.3e-05 | 9.8e-06 | 1.293E-05 | 3.101E-06 | 9.800E-06 | 1.600E-05 |
| 26308 | 1316 | G1316G | E | 1.3e-05 | 1.2e-05 | 8.5e-06 | 1.117E-05 | 2.363E-06 | 8.500E-06 | 1.300E-05 |
| 26309 | 1316 | G1316H | E | 1e-05 | 1.4e-05 | 1e-05 | 1.133E-05 | 2.309E-06 | 1.000E-05 | 1.400E-05 |
| 26310 | 1316 | G1316I | E | 1e-05 | 9.8e-06 | 1.7e-05 | 1.227E-05 | 4.100E-06 | 9.800E-06 | 1.700E-05 |
| 26311 | 1316 | G1316L | E | 1.5e-05 | 8.5e-06 | 9.7e-06 | 1.107E-05 | 3.459E-06 | 8.500E-06 | 1.500E-05 |
| 26312 | 1316 | G1316K | E | 9.7e-06 | 1.3e-05 | 9.8e-06 | 1.083E-05 | 1.877E-06 | 9.700E-06 | 1.300E-05 |
| 26313 | 1316 | G1316M | E | 1.3e-05 | 1.4e-05 | 1.6e-05 | 1.433E-05 | 1.528E-06 | 1.300E-05 | 1.600E-05 |
| 26314 | 1316 | G1316F | E | 1.1e-05 | 1e-05 | 1.3e-05 | 1.133E-05 | 1.528E-06 | 1.000E-05 | 1.300E-05 |
| 26315 | 1316 | G1316P | E | 1.4e-05 | 2.3e-05 | 1.6e-05 | 1.767E-05 | 4.726E-06 | 1.400E-05 | 2.300E-05 |
| 26316 | 1316 | G1316S | E | 1.5e-05 | 8.6e-06 | 1.5e-05 | 1.287E-05 | 3.695E-06 | 8.600E-06 | 1.500E-05 |
| 26317 | 1316 | G1316T | E | 1.3e-05 | 8.2e-06 | 1.1e-05 | 1.073E-05 | 2.411E-06 | 8.200E-06 | 1.300E-05 |
| 26318 | 1316 | G1316W | E | 1.3e-05 | 8.6e-06 | 9.9e-06 | 1.050E-05 | 2.261E-06 | 8.600E-06 | 1.300E-05 |
| 26319 | 1316 | G1316Y | E | 1.3e-05 | 1.3e-05 | 1.4e-05 | 1.333E-05 | 5.774E-07 | 1.300E-05 | 1.400E-05 |
| 26320 | 1316 | G1316V | E | 6.9e-06 | 1e-05 | 9.9e-06 | 8.933E-06 | 1.762E-06 | 6.900E-06 | 1.000E-05 |
| 26321 | 1317 | L1317A | E | 1.1e-05 | 1.6e-05 | 1.5e-05 | 1.400E-05 | 2.646E-06 | 1.100E-05 | 1.600E-05 |
| 26322 | 1317 | L1317R | E | 1.8e-05 | 1.2e-05 | 1.4e-05 | 1.467E-05 | 3.055E-06 | 1.200E-05 | 1.800E-05 |
| 26323 | 1317 | L1317N | E | 1.1e-05 | 1.4e-05 | 9.1e-06 | 1.137E-05 | 2.470E-06 | 9.100E-06 | 1.400E-05 |
| 26324 | 1317 | L1317D | E | 1.3e-05 | 1.5e-05 | 9.4e-06 | 1.247E-05 | 2.838E-06 | 9.400E-06 | 1.500E-05 |
| 26325 | 1317 | L1317C | E | 1.4e-05 | 7.8e-06 | 1.6e-05 | 1.260E-05 | 4.276E-06 | 7.800E-06 | 1.600E-05 |
| 26326 | 1317 | L1317E | E | 1.3e-05 | 1e-05 | 1.2e-05 | 1.167E-05 | 1.528E-06 | 1.000E-05 | 1.300E-05 |
| 26327 | 1317 | L1317Q | E | 1.4e-05 | 1.4e-05 | 8.1e-06 | 1.203E-05 | 3.406E-06 | 8.100E-06 | 1.400E-05 |
| 26328 | 1317 | L1317G | E | 1.6e-05 | 1.4e-05 | 1.5e-05 | 1.500E-05 | 1.000E-06 | 1.400E-05 | 1.600E-05 |
| 26329 | 1317 | L1317H | E | 7.9e-06 | 1.5e-05 | 1.1e-05 | 1.130E-05 | 3.559E-06 | 7.900E-06 | 1.500E-05 |
| 26330 | 1317 | L1317I | E | 8.8e-06 | 1.5e-05 | 1.4e-05 | 1.260E-05 | 3.329E-06 | 8.800E-06 | 1.500E-05 |
| 26331 | 1317 | L1317L | E | 1.6e-05 | 1.1e-05 | 1.5e-05 | 1.400E-05 | 2.646E-06 | 1.100E-05 | 1.600E-05 |
| 26332 | 1317 | L1317K | E | 1e-05 | 8.8e-06 | 1e-05 | 9.600E-06 | 6.928E-07 | 8.800E-06 | 1.000E-05 |
| 26333 | 1317 | L1317M | E | 1.8e-05 | 1.2e-05 | 1.4e-05 | 1.467E-05 | 3.055E-06 | 1.200E-05 | 1.800E-05 |
| 26334 | 1317 | L1317F | E | 1.5e-05 | 1.7e-05 | 1.5e-05 | 1.567E-05 | 1.155E-06 | 1.500E-05 | 1.700E-05 |
| 26335 | 1317 | L1317P | E | 7.8e-06 | 1.5e-05 | 1.4e-05 | 1.227E-05 | 3.900E-06 | 7.800E-06 | 1.500E-05 |
| 26336 | 1317 | L1317S | E | 8.8e-06 | 1.3e-05 | 1.4e-05 | 1.193E-05 | 2.759E-06 | 8.800E-06 | 1.400E-05 |
| 26337 | 1317 | L1317T | E | 1e-05 | 1.1e-05 | 1.5e-05 | 1.200E-05 | 2.646E-06 | 1.000E-05 | 1.500E-05 |
| 26338 | 1317 | L1317W | E | 1.5e-05 | 1.5e-05 | 1.5e-05 | 1.500E-05 | 0.000E+00 | 1.500E-05 | 1.500E-05 |
| 26339 | 1317 | L1317Y | E | 1.2e-05 | 1.2e-05 | 1.5e-05 | 1.300E-05 | 1.732E-06 | 1.200E-05 | 1.500E-05 |
| 26340 | 1317 | L1317V | E | 1.7e-05 | 8.4e-06 | 9.7e-06 | 1.170E-05 | 4.636E-06 | 8.400E-06 | 1.700E-05 |
| 26341 | 1318 | E1318A | E | 1.6e-05 | 1.4e-05 | 7.9e-06 | 1.263E-05 | 4.219E-06 | 7.900E-06 | 1.600E-05 |
| 26342 | 1318 | E1318R | E | 9e-06 | 1.4e-05 | 1.5e-05 | 1.267E-05 | 3.215E-06 | 9.000E-06 | 1.500E-05 |
| 26343 | 1318 | E1318N | E | 9.4e-06 | 1.3e-05 | 1.5e-05 | 1.247E-05 | 2.838E-06 | 9.400E-06 | 1.500E-05 |
| 26344 | 1318 | E1318D | E | 8.3e-06 | 1.6e-05 | 1.5e-05 | 1.310E-05 | 4.187E-06 | 8.300E-06 | 1.600E-05 |
| 26345 | 1318 | E1318C | E | 1.4e-05 | 1.3e-05 | 1e-05 | 1.233E-05 | 2.082E-06 | 1.000E-05 | 1.400E-05 |
| 26346 | 1318 | E1318E | E | 1.4e-05 | 1.5e-05 | 1.6e-05 | 1.500E-05 | 1.000E-06 | 1.400E-05 | 1.600E-05 |
| 26347 | 1318 | E1318Q | E | 1.5e-05 | 1.4e-05 | 1e-05 | 1.300E-05 | 2.646E-06 | 1.000E-05 | 1.500E-05 |
| 26348 | 1318 | E1318G | E | 1.3e-05 | 1.7e-05 | 1.1e-05 | 1.367E-05 | 3.055E-06 | 1.100E-05 | 1.700E-05 |
| 26349 | 1318 | E1318H | E | 1.7e-05 | 9.7e-06 | 1.3e-05 | 1.323E-05 | 3.656E-06 | 9.700E-06 | 1.700E-05 |

| | | | | | | | | | | |
|---|---|---|---|---|---|---|---|---|---|---|
| 26350 | 1318 | E1318I | E | 1.4e-05 | 9.4e-06 | 1.1e-05 | 1.147E-05 | 2.335E-06 | 9.400E-06 | 1.400E-05 |
| 26351 | 1318 | E1318L | E | 1.1e-05 | 1.6e-05 | 1.3e-05 | 1.333E-05 | 2.517E-06 | 1.100E-05 | 1.600E-05 |
| 26352 | 1318 | E1318K | E | 1.7e-05 | 1.4e-05 | 1.4e-05 | 1.500E-05 | 1.732E-06 | 1.400E-05 | 1.700E-05 |
| 26353 | 1318 | E1318M | E | 1.6e-05 | 1.6e-05 | 1e-05 | 1.400E-05 | 3.464E-06 | 1.000E-05 | 1.600E-05 |
| 26354 | 1318 | E1318F | E | 1.5e-05 | 1.6e-05 | 1.5e-05 | 1.533E-05 | 5.774E-07 | 1.500E-05 | 1.600E-05 |
| 26355 | 1318 | E1318P | E | 7e-06 | 1.1e-05 | 8.3e-06 | 8.767E-06 | 2.040E-06 | 7.000E-06 | 1.100E-05 |
| 26356 | 1318 | E1318S | E | 1.4e-05 | 1.4e-05 | 1.6e-05 | 1.467E-05 | 1.155E-06 | 1.400E-05 | 1.600E-05 |
| 26357 | 1318 | E1318T | E | 1.6e-05 | 9.6e-06 | 1e-05 | 1.187E-05 | 3.585E-06 | 9.600E-06 | 1.600E-05 |
| 26358 | 1318 | E1318W | E | 1.1e-05 | 1.4e-05 | 1.4e-05 | 1.300E-05 | 1.732E-06 | 1.100E-05 | 1.400E-05 |
| 26359 | 1318 | E1318Y | E | 1.5e-05 | 1.5e-05 | 9.1e-06 | 1.303E-05 | 3.406E-06 | 9.100E-06 | 1.500E-05 |
| 26360 | 1318 | E1318V | E | 1.1e-05 | 1e-05 | 1.5e-05 | 1.200E-05 | 2.646E-06 | 1.000E-05 | 1.500E-05 |
| 26361 | 1319 | W1319A | E | 1.4e-05 | 1.4e-05 | 1e-05 | 1.267E-05 | 2.309E-06 | 1.000E-05 | 1.400E-05 |
| 26362 | 1319 | W1319R | E | 1.5e-05 | 1.6e-05 | 1.2e-05 | 1.433E-05 | 2.082E-06 | 1.200E-05 | 1.600E-05 |
| 26363 | 1319 | W1319N | E | 1.5e-05 | 1.6e-05 | 1.4e-05 | 1.500E-05 | 1.000E-06 | 1.400E-05 | 1.600E-05 |
| 26364 | 1319 | W1319D | E | 1.4e-05 | 9.2e-06 | 1.5e-05 | 1.273E-05 | 3.101E-06 | 9.200E-06 | 1.500E-05 |
| 26365 | 1319 | W1319C | E | 1e-05 | 1.4e-05 | 1.5e-05 | 1.300E-05 | 2.646E-06 | 1.000E-05 | 1.500E-05 |
| 26366 | 1319 | W1319E | E | 1.1e-05 | 1.6e-05 | 1.1e-05 | 1.267E-05 | 2.887E-06 | 1.100E-05 | 1.600E-05 |
| 26367 | 1319 | W1319Q | E | 1.2e-05 | 1.5e-05 | 8.5e-06 | 1.183E-05 | 3.253E-06 | 8.500E-06 | 1.500E-05 |
| 26368 | 1319 | W1319G | E | 1.3e-05 | 1.5e-05 | 1e-05 | 1.267E-05 | 2.517E-06 | 1.000E-05 | 1.500E-05 |
| 26369 | 1319 | W1319H | E | 9.7e-06 | 1.4e-05 | 1.5e-05 | 1.290E-05 | 2.816E-06 | 9.700E-06 | 1.500E-05 |
| 26370 | 1319 | W1319I | E | 1.4e-05 | 1.5e-05 | 1.3e-05 | 1.400E-05 | 1.000E-06 | 1.300E-05 | 1.500E-05 |
| 26371 | 1319 | W1319L | E | 1.5e-05 | 1e-05 | 1.1e-05 | 1.200E-05 | 2.646E-06 | 1.000E-05 | 1.500E-05 |
| 26372 | 1319 | W1319K | E | 1.5e-05 | 6.7e-06 | 1.1e-05 | 1.090E-05 | 4.151E-06 | 6.700E-06 | 1.500E-05 |
| 26373 | 1319 | W1319M | E | 1.4e-05 | 1.2e-05 | 1.1e-05 | 1.233E-05 | 1.528E-06 | 1.100E-05 | 1.400E-05 |
| 26374 | 1319 | W1319F | E | 9.9e-06 | 1.4e-05 | 1.3e-05 | 1.230E-05 | 2.138E-06 | 9.900E-06 | 1.400E-05 |
| 26375 | 1319 | W1319P | E | 1.5e-05 | 1.6e-05 | 1.7e-05 | 1.600E-05 | 1.000E-06 | 1.500E-05 | 1.700E-05 |
| 26376 | 1319 | W1319S | E | 9.7e-06 | 1.6e-05 | 1.5e-05 | 1.357E-05 | 3.386E-06 | 9.700E-06 | 1.600E-05 |
| 26377 | 1319 | W1319T | E | 1.3e-05 | 1.5e-05 | 1.3e-05 | 1.367E-05 | 1.155E-06 | 1.300E-05 | 1.500E-05 |
| 26378 | 1319 | W1319W | E | 1.1e-05 | 1e-05 | 1.4e-05 | 1.167E-05 | 2.082E-06 | 1.000E-05 | 1.400E-05 |
| 26379 | 1319 | W1319Y | E | 1.1e-05 | 1.4e-05 | 1.4e-05 | 1.300E-05 | 1.732E-06 | 1.100E-05 | 1.400E-05 |
| 26380 | 1319 | W1319V | E | 1.4e-05 | 1.7e-05 | 1.6e-05 | 1.567E-05 | 1.528E-06 | 1.400E-05 | 1.700E-05 |
| 26381 | 1320 | V1320A | E | 1.4e-05 | 1.6e-05 | 1e-05 | 1.333E-05 | 3.055E-06 | 1.000E-05 | 1.600E-05 |
| 26382 | 1320 | V1320R | E | 1.1e-05 | 1.3e-05 | 1.5e-05 | 1.300E-05 | 2.000E-06 | 1.100E-05 | 1.500E-05 |
| 26383 | 1320 | V1320N | E | 1.2e-05 | 1.4e-05 | 1.7e-05 | 1.433E-05 | 2.517E-06 | 1.200E-05 | 1.700E-05 |
| 26384 | 1320 | V1320D | E | 1.1e-05 | 1.1e-05 | 1.6e-05 | 1.267E-05 | 2.887E-06 | 1.100E-05 | 1.600E-05 |
| 26385 | 1320 | V1320C | E | 9.7e-06 | 1.2e-05 | 1.3e-05 | 1.157E-05 | 1.692E-06 | 9.700E-06 | 1.300E-05 |
| 26386 | 1320 | V1320E | E | 1.5e-05 | 1.4e-05 | 1.1e-05 | 1.333E-05 | 2.082E-06 | 1.100E-05 | 1.500E-05 |
| 26387 | 1320 | V1320Q | E | 1e-05 | 1.1e-05 | 1.3e-05 | 1.133E-05 | 1.528E-06 | 1.000E-05 | 1.300E-05 |
| 26388 | 1320 | V1320G | E | 1.6e-05 | 1.2e-05 | 1.6e-05 | 1.467E-05 | 2.309E-06 | 1.200E-05 | 1.600E-05 |
| 26389 | 1320 | V1320H | E | 6e-06 | 9.1e-06 | 1.4e-05 | 9.700E-06 | 4.034E-06 | 6.000E-06 | 1.400E-05 |
| 26390 | 1320 | V1320I | E | 1e-05 | 1.7e-05 | 1.7e-05 | 1.467E-05 | 4.041E-06 | 1.000E-05 | 1.700E-05 |
| 26391 | 1320 | V1320L | E | 1.5e-05 | 9.5e-06 | 1.2e-05 | 1.217E-05 | 2.754E-06 | 9.500E-06 | 1.500E-05 |
| 26392 | 1320 | V1320K | E | 9.4e-06 | 1.1e-05 | 9.9e-06 | 1.010E-05 | 8.185E-07 | 9.400E-06 | 1.100E-05 |
| 26393 | 1320 | V1320M | E | 1.5e-05 | 9.9e-06 | 1.4e-05 | 1.297E-05 | 2.702E-06 | 9.900E-06 | 1.500E-05 |
| 26394 | 1320 | V1320F | E | 1.4e-05 | 1.6e-05 | 8.9e-06 | 1.297E-05 | 3.661E-06 | 8.900E-06 | 1.600E-05 |
| 26395 | 1320 | V1320P | E | 1.3e-05 | 1.1e-05 | 1.5e-05 | 1.300E-05 | 2.000E-06 | 1.100E-05 | 1.500E-05 |
| 26396 | 1320 | V1320S | E | 1e-05 | 1.5e-05 | 1.5e-05 | 1.333E-05 | 2.887E-06 | 1.000E-05 | 1.500E-05 |
| 26397 | 1320 | V1320T | E | 1.3e-05 | 1.1e-05 | 9.3e-06 | 1.110E-05 | 1.852E-06 | 9.300E-06 | 1.300E-05 |
| 26398 | 1320 | V1320W | E | 1.3e-05 | 6.6e-06 | 1.6e-05 | 1.187E-05 | 4.801E-06 | 6.600E-06 | 1.600E-05 |
| 26399 | 1320 | V1320Y | E | 1.1e-05 | 1.4e-05 | 9.1e-06 | 1.137E-05 | 2.470E-06 | 9.100E-06 | 1.400E-05 |
| 26400 | 1320 | V1320V | E | 1.4e-05 | 9.7e-06 | 1.5e-05 | 1.290E-05 | 2.816E-06 | 9.700E-06 | 1.500E-05 |
| 26401 | 1321 | A1321A | E | 1.4e-05 | 9.8e-06 | 9e-06 | 1.093E-05 | 2.686E-06 | 9.000E-06 | 1.400E-05 |
| 26402 | 1321 | A1321R | E | 9.7e-06 | 1.5e-05 | 1.1e-05 | 1.190E-05 | 2.762E-06 | 9.700E-06 | 1.500E-05 |
| 26403 | 1321 | A1321N | E | 1.2e-05 | 1.6e-05 | 1.7e-05 | 1.500E-05 | 2.646E-06 | 1.200E-05 | 1.700E-05 |
| 26404 | 1321 | A1321D | E | 1.4e-05 | 1.6e-05 | 1.5e-05 | 1.500E-05 | 1.000E-06 | 1.400E-05 | 1.600E-05 |
| 26405 | 1321 | A1321C | E | 1.5e-05 | 8e-06 | 1.6e-05 | 1.300E-05 | 4.359E-06 | 8.000E-06 | 1.600E-05 |
| 26406 | 1321 | A1321E | E | 1.1e-05 | 1.1e-05 | 7.9e-06 | 9.967E-06 | 1.790E-06 | 7.900E-06 | 1.100E-05 |
| 26407 | 1321 | A1321Q | E | 1e-05 | 1.1e-05 | 1.1e-05 | 1.067E-05 | 5.774E-07 | 1.000E-05 | 1.100E-05 |
| 26408 | 1321 | A1321G | E | 1.2e-05 | 8.6e-06 | 1.1e-05 | 1.053E-05 | 1.747E-06 | 8.600E-06 | 1.200E-05 |
| 26409 | 1321 | A1321H | E | 1.1e-05 | 1.4e-05 | 1.5e-05 | 1.333E-05 | 2.082E-06 | 1.100E-05 | 1.500E-05 |
| 26410 | 1321 | A1321I | E | 1.2e-05 | 1e-05 | 1.5e-05 | 1.233E-05 | 2.517E-06 | 1.000E-05 | 1.500E-05 |
| 26411 | 1321 | A1321L | E | 7.7e-06 | 1.4e-05 | 1.1e-05 | 1.090E-05 | 3.151E-06 | 7.700E-06 | 1.400E-05 |
| 26412 | 1321 | A1321K | E | 1e-05 | 1e-05 | 9.6e-06 | 9.867E-06 | 2.309E-07 | 9.600E-06 | 1.000E-05 |
| 26413 | 1321 | A1321M | E | 1.5e-05 | 1.5e-05 | 1.5e-05 | 1.500E-05 | 0.000E+00 | 1.500E-05 | 1.500E-05 |
| 26414 | 1321 | A1321F | E | 1.8e-05 | 1.4e-05 | 8.6e-06 | 1.353E-05 | 4.717E-06 | 8.600E-06 | 1.800E-05 |
| 26415 | 1321 | A1321P | E | 9.9e-06 | 9.6e-06 | 1.4e-05 | 1.117E-05 | 2.458E-06 | 9.600E-06 | 1.400E-05 |
| 26416 | 1321 | A1321S | E | 1e-05 | 1.4e-05 | 1.2e-05 | 1.200E-05 | 2.000E-06 | 1.000E-05 | 1.400E-05 |
| 26417 | 1321 | A1321T | E | 1.3e-05 | 8e-06 | 1.4e-05 | 1.167E-05 | 3.215E-06 | 8.000E-06 | 1.400E-05 |
| 26418 | 1321 | A1321W | E | 9.4e-06 | 1.1e-05 | 1.6e-05 | 1.213E-05 | 3.443E-06 | 9.400E-06 | 1.600E-05 |
| 26419 | 1321 | A1321Y | E | 1e-05 | 1.2e-05 | 9.6e-06 | 1.053E-05 | 1.286E-06 | 9.600E-06 | 1.200E-05 |
| 26420 | 1321 | A1321V | E | 1.6e-05 | 1.5e-05 | 1.5e-05 | 1.533E-05 | 5.774E-07 | 1.500E-05 | 1.600E-05 |
| 26421 | 1322 | R1322A | E | 1.7e-05 | 1e-05 | 1.6e-05 | 1.433E-05 | 3.786E-06 | 1.000E-05 | 1.700E-05 |
| 26422 | 1322 | R1322R | E | 2.2e-05 | 1.2e-05 | 1.2e-05 | 1.533E-05 | 5.774E-06 | 1.200E-05 | 2.200E-05 |
| 26423 | 1322 | R1322N | E | 1.3e-05 | 1.1e-05 | 1.5e-05 | 1.300E-05 | 2.000E-06 | 1.100E-05 | 1.500E-05 |
| 26424 | 1322 | R1322D | E | 1.1e-05 | 1.1e-05 | 1.6e-05 | 1.267E-05 | 2.887E-06 | 1.100E-05 | 1.600E-05 |
| 26425 | 1322 | R1322C | E | 1.2e-05 | 1.4e-05 | 9.7e-06 | 1.190E-05 | 2.152E-06 | 9.700E-06 | 1.400E-05 |
| 26426 | 1322 | R1322E | E | 1.2e-05 | 8.2e-06 | 1.1e-05 | 1.040E-05 | 1.970E-06 | 8.200E-06 | 1.200E-05 |
| 26427 | 1322 | R1322Q | E | 9.2e-06 | 1.7e-05 | 1.4e-05 | 1.340E-05 | 3.934E-06 | 9.200E-06 | 1.700E-05 |
| 26428 | 1322 | R1322G | E | 1.5e-05 | 9.5e-06 | 1.5e-05 | 1.317E-05 | 3.175E-06 | 9.500E-06 | 1.500E-05 |
| 26429 | 1322 | R1322H | E | 1.5e-05 | 1.4e-05 | 1.3e-05 | 1.400E-05 | 1.000E-06 | 1.300E-05 | 1.500E-05 |
| 26430 | 1322 | R1322I | E | 1.4e-05 | 1.4e-05 | 9.3e-06 | 1.243E-05 | 2.714E-06 | 9.300E-06 | 1.400E-05 |
| 26431 | 1322 | R1322L | E | 1.3e-05 | 1e-05 | 1e-05 | 1.100E-05 | 1.732E-06 | 1.000E-05 | 1.300E-05 |
| 26432 | 1322 | R1322K | E | 1.8e-05 | 1.5e-05 | 8.5e-06 | 1.383E-05 | 4.856E-06 | 8.500E-06 | 1.800E-05 |
| 26433 | 1322 | R1322M | E | 1.8e-05 | 1.7e-05 | 1.1e-05 | 1.533E-05 | 3.786E-06 | 1.100E-05 | 1.800E-05 |
| 26434 | 1322 | R1322F | E | 1e-05 | 1.3e-05 | 1.1e-05 | 1.133E-05 | 1.528E-06 | 1.000E-05 | 1.300E-05 |

| | | | | | | | | | | |
|---|---|---|---|---|---|---|---|---|---|---|
| 26435 | 1322 | R1322P | E | 9.9e-06 | 1.1e-05 | 1.4e-05 | 1.163E-05 | 2.122E-06 | 9.900E-06 | 1.400E-05 |
| 26436 | 1322 | R1322S | E | 1.5e-05 | 9.3e-06 | 1.3e-05 | 1.243E-05 | 2.892E-06 | 9.300E-06 | 1.500E-05 |
| 26437 | 1322 | R1322T | E | 1.6e-05 | 9.5e-06 | 1e-05 | 1.183E-05 | 3.617E-06 | 9.500E-06 | 1.600E-05 |
| 26438 | 1322 | R1322W | E | 1e-05 | 1.5e-05 | 1.6e-05 | 1.367E-05 | 3.215E-06 | 1.000E-05 | 1.600E-05 |
| 26439 | 1322 | R1322Y | E | 1.1e-05 | 1.4e-05 | 1.5e-05 | 1.333E-05 | 2.082E-06 | 1.100E-05 | 1.500E-05 |
| 26440 | 1322 | R1322V | E | 9.9e-06 | 1.5e-05 | 1.3e-05 | 1.263E-05 | 2.570E-06 | 9.900E-06 | 1.500E-05 |
| 26441 | 1323 | I1323A | E | 1.7e-05 | 1.8e-05 | 1.6e-05 | 1.700E-05 | 1.000E-06 | 1.600E-05 | 1.800E-05 |
| 26442 | 1323 | I1323R | E | 1.1e-05 | 1.1e-05 | 1.6e-05 | 1.267E-05 | 2.887E-06 | 1.100E-05 | 1.600E-05 |
| 26443 | 1323 | I1323N | E | 1.5e-05 | 1.4e-05 | 1e-05 | 1.300E-05 | 2.646E-06 | 1.000E-05 | 1.500E-05 |
| 26444 | 1323 | I1323D | E | 1.4e-05 | 2.5e-05 | 1.5e-05 | 1.800E-05 | 6.083E-06 | 1.400E-05 | 2.500E-05 |
| 26445 | 1323 | I1323C | E | 1.5e-05 | 1.1e-05 | 1.4e-05 | 1.333E-05 | 2.082E-06 | 1.100E-05 | 1.500E-05 |
| 26446 | 1323 | I1323E | E | 1e-05 | 1.6e-05 | 1.1e-05 | 1.233E-05 | 3.215E-06 | 1.000E-05 | 1.600E-05 |
| 26447 | 1323 | I1323Q | E | 1e-05 | 1.2e-05 | 1.6e-05 | 1.267E-05 | 3.055E-06 | 1.000E-05 | 1.600E-05 |
| 26448 | 1323 | I1323G | E | 1.6e-05 | 1.4e-05 | 1.6e-05 | 1.533E-05 | 1.155E-06 | 1.400E-05 | 1.600E-05 |
| 26449 | 1323 | I1323H | E | 1.6e-05 | 1.4e-05 | 1e-05 | 1.333E-05 | 3.055E-06 | 1.000E-05 | 1.600E-05 |
| 26450 | 1323 | I1323I | E | 1.5e-05 | 1.5e-05 | 1.1e-05 | 1.367E-05 | 2.309E-06 | 1.100E-05 | 1.500E-05 |
| 26451 | 1323 | I1323L | E | 1e-05 | 1.6e-05 | 1.4e-05 | 1.333E-05 | 3.055E-06 | 1.000E-05 | 1.600E-05 |
| 26452 | 1323 | I1323K | E | 1.1e-05 | 1.5e-05 | 1.4e-05 | 1.333E-05 | 2.082E-06 | 1.100E-05 | 1.500E-05 |
| 26453 | 1323 | I1323M | E | 1e-05 | 9.9e-06 | 1.6e-05 | 1.197E-05 | 3.493E-06 | 9.900E-06 | 1.600E-05 |
| 26454 | 1323 | I1323F | E | 1.6e-05 | 1.5e-05 | 1e-05 | 1.367E-05 | 3.215E-06 | 1.000E-05 | 1.600E-05 |
| 26455 | 1323 | I1323P | E | 1.1e-05 | 1.1e-05 | 1.2e-05 | 1.133E-05 | 5.774E-07 | 1.100E-05 | 1.200E-05 |
| 26456 | 1323 | I1323S | E | 9.4e-06 | 1.5e-05 | 1.4e-05 | 1.280E-05 | 2.987E-06 | 9.400E-06 | 1.500E-05 |
| 26457 | 1323 | I1323T | E | 9.6e-06 | 1.7e-05 | 1.7e-05 | 1.453E-05 | 4.272E-06 | 9.600E-06 | 1.700E-05 |
| 26458 | 1323 | I1323W | E | 1.4e-05 | 1.4e-05 | 1.6e-05 | 1.467E-05 | 1.155E-06 | 1.400E-05 | 1.600E-05 |
| 26459 | 1323 | I1323Y | E | 1.3e-05 | 1.4e-05 | 1.2e-05 | 1.300E-05 | 1.000E-06 | 1.200E-05 | 1.400E-05 |
| 26460 | 1323 | I1323V | E | 1.1e-05 | 1e-05 | 1.6e-05 | 1.233E-05 | 3.215E-06 | 1.000E-05 | 1.600E-05 |
| 26461 | 1324 | Y1324A | E | 1.1e-05 | 1.5e-05 | 1.3e-05 | 1.300E-05 | 2.000E-06 | 1.100E-05 | 1.500E-05 |
| 26462 | 1324 | Y1324R | E | 1.5e-05 | 1.4e-05 | 1.5e-05 | 1.467E-05 | 5.774E-07 | 1.400E-05 | 1.500E-05 |
| 26463 | 1324 | Y1324N | E | 9.7e-06 | 1.5e-05 | 1.7e-05 | 1.390E-05 | 3.772E-06 | 9.700E-06 | 1.700E-05 |
| 26464 | 1324 | Y1324D | E | 1.5e-05 | 1.2e-05 | 1.7e-05 | 1.467E-05 | 2.517E-06 | 1.200E-05 | 1.700E-05 |
| 26465 | 1324 | Y1324C | E | 1.8e-05 | 1.6e-05 | 1.6e-05 | 1.667E-05 | 1.155E-06 | 1.600E-05 | 1.800E-05 |
| 26466 | 1324 | Y1324E | E | 1.4e-05 | 1.3e-05 | 1.5e-05 | 1.400E-05 | 1.000E-06 | 1.300E-05 | 1.500E-05 |
| 26467 | 1324 | Y1324Q | E | 1.2e-05 | 1.6e-05 | 1.6e-05 | 1.467E-05 | 2.309E-06 | 1.200E-05 | 1.600E-05 |
| 26468 | 1324 | Y1324G | E | 1.4e-05 | 1.2e-05 | 1e-05 | 1.200E-05 | 2.000E-06 | 1.000E-05 | 1.400E-05 |
| 26469 | 1324 | Y1324H | E | 1.5e-05 | 9.7e-06 | 1.8e-05 | 1.423E-05 | 4.203E-06 | 9.700E-06 | 1.800E-05 |
| 26470 | 1324 | Y1324I | E | 1.7e-05 | 1.4e-05 | 1.8e-05 | 1.633E-05 | 2.082E-06 | 1.400E-05 | 1.800E-05 |
| 26471 | 1324 | Y1324L | E | 9.6e-06 | 1.5e-05 | 1.1e-05 | 1.187E-05 | 2.802E-06 | 9.600E-06 | 1.500E-05 |
| 26472 | 1324 | Y1324K | E | 1.4e-05 | 1.3e-05 | 8.3e-06 | 1.177E-05 | 3.044E-06 | 8.300E-06 | 1.400E-05 |
| 26473 | 1324 | Y1324M | E | 1.2e-05 | 1.2e-05 | 1.4e-05 | 1.267E-05 | 1.155E-06 | 1.200E-05 | 1.400E-05 |
| 26474 | 1324 | Y1324F | E | 1.2e-05 | 1e-05 | 1.5e-05 | 1.233E-05 | 2.517E-06 | 1.000E-05 | 1.500E-05 |
| 26475 | 1324 | Y1324P | E | 1.4e-05 | 1.2e-05 | 1.1e-05 | 1.233E-05 | 1.528E-06 | 1.100E-05 | 1.400E-05 |
| 26476 | 1324 | Y1324S | E | 1.1e-05 | 1.3e-05 | 1e-05 | 1.133E-05 | 1.528E-06 | 1.000E-05 | 1.300E-05 |
| 26477 | 1324 | Y1324T | E | 1.4e-05 | 1.3e-05 | 1e-05 | 1.233E-05 | 2.082E-06 | 1.000E-05 | 1.400E-05 |
| 26478 | 1324 | Y1324W | E | 1.5e-05 | 1.3e-05 | 1e-05 | 1.267E-05 | 2.517E-06 | 1.000E-05 | 1.500E-05 |
| 26479 | 1324 | Y1324Y | E | 1.4e-05 | 1.5e-05 | 1.4e-05 | 1.433E-05 | 5.774E-07 | 1.400E-05 | 1.500E-05 |
| 26480 | 1324 | Y1324V | E | 1.1e-05 | 1.2e-05 | 1.5e-05 | 1.267E-05 | 2.082E-06 | 1.100E-05 | 1.500E-05 |
| 26481 | 1325 | P1325A | E | 1e-05 | 9.3e-06 | 1.6e-05 | 1.177E-05 | 3.683E-06 | 9.300E-06 | 1.600E-05 |
| 26482 | 1325 | P1325R | E | 1.6e-05 | 2.5e-05 | 1.6e-05 | 1.900E-05 | 5.196E-06 | 1.600E-05 | 2.500E-05 |
| 26483 | 1325 | P1325N | E | 1.1e-05 | 1.6e-05 | 1.6e-05 | 1.433E-05 | 2.887E-06 | 1.100E-05 | 1.600E-05 |
| 26484 | 1325 | P1325D | E | 1.4e-05 | 1.4e-05 | 1.5e-05 | 1.433E-05 | 5.774E-07 | 1.400E-05 | 1.500E-05 |
| 26485 | 1325 | P1325C | E | 1.4e-05 | 1e-05 | 1e-05 | 1.133E-05 | 2.309E-06 | 1.000E-05 | 1.400E-05 |
| 26486 | 1325 | P1325E | E | 1.7e-05 | 1.1e-05 | 1e-05 | 1.267E-05 | 3.786E-06 | 1.000E-05 | 1.700E-05 |
| 26487 | 1325 | P1325Q | E | 1.4e-05 | 1.5e-05 | 1.3e-05 | 1.400E-05 | 1.000E-06 | 1.300E-05 | 1.500E-05 |
| 26488 | 1325 | P1325G | E | 1.4e-05 | 1e-05 | 1.7e-05 | 1.367E-05 | 3.512E-06 | 1.000E-05 | 1.700E-05 |
| 26489 | 1325 | P1325H | E | 9.3e-06 | 1.4e-05 | 1.6e-05 | 1.310E-05 | 3.439E-06 | 9.300E-06 | 1.600E-05 |
| 26490 | 1325 | P1325I | E | 1.4e-05 | 1.1e-05 | 1.3e-05 | 1.267E-05 | 1.528E-06 | 1.100E-05 | 1.400E-05 |
| 26491 | 1325 | P1325L | E | 1.4e-05 | 1.6e-05 | 1.7e-05 | 1.567E-05 | 1.528E-06 | 1.400E-05 | 1.700E-05 |
| 26492 | 1325 | P1325K | E | 7.4e-06 | 1.3e-05 | 1.6e-05 | 1.213E-05 | 4.365E-06 | 7.400E-06 | 1.600E-05 |
| 26493 | 1325 | P1325M | E | 1.1e-05 | 1.2e-05 | 1.5e-05 | 1.267E-05 | 2.082E-06 | 1.100E-05 | 1.500E-05 |
| 26494 | 1325 | P1325F | E | 9.1e-06 | 1.1e-05 | 1.4e-05 | 1.137E-05 | 2.470E-06 | 9.100E-06 | 1.400E-05 |
| 26495 | 1325 | P1325P | E | 1.4e-05 | 8e-06 | 1e-05 | 1.067E-05 | 3.055E-06 | 8.000E-06 | 1.400E-05 |
| 26496 | 1325 | P1325S | E | 7.5e-06 | 1.1e-05 | 1.5e-05 | 1.117E-05 | 3.753E-06 | 7.500E-06 | 1.500E-05 |
| 26497 | 1325 | P1325T | E | 1e-05 | 8.3e-06 | 1.6e-05 | 1.143E-05 | 4.045E-06 | 8.300E-06 | 1.600E-05 |
| 26498 | 1325 | P1325W | E | 1.4e-05 | 1e-05 | 1.4e-05 | 1.267E-05 | 2.309E-06 | 1.000E-05 | 1.400E-05 |
| 26499 | 1325 | P1325Y | E | 1.4e-05 | 1.6e-05 | 1.5e-05 | 1.500E-05 | 1.000E-06 | 1.400E-05 | 1.600E-05 |
| 26500 | 1325 | P1325V | E | 1.5e-05 | 1.5e-05 | 1.3e-05 | 1.433E-05 | 1.155E-06 | 1.300E-05 | 1.500E-05 |
| 26501 | 1326 | T1326A | E | 1.3e-05 | 1e-05 | 1.5e-05 | 1.267E-05 | 2.517E-06 | 1.000E-05 | 1.500E-05 |
| 26502 | 1326 | T1326R | E | 1.5e-05 | 1.6e-05 | 9e-06 | 1.333E-05 | 3.786E-06 | 9.000E-06 | 1.600E-05 |
| 26503 | 1326 | T1326N | E | 1e-05 | 9.7e-06 | 1.1e-05 | 1.023E-05 | 6.807E-07 | 9.700E-06 | 1.100E-05 |
| 26504 | 1326 | T1326D | E | 1.3e-05 | 1.4e-05 | 1.6e-05 | 1.433E-05 | 1.528E-06 | 1.300E-05 | 1.600E-05 |
| 26505 | 1326 | T1326C | E | 1e-05 | 1.6e-05 | 9.3e-06 | 1.177E-05 | 3.683E-06 | 9.300E-06 | 1.600E-05 |
| 26506 | 1326 | T1326E | E | 1.4e-05 | 1.3e-05 | 1.7e-05 | 1.467E-05 | 2.082E-06 | 1.300E-05 | 1.700E-05 |
| 26507 | 1326 | T1326Q | E | 1.1e-05 | 1.5e-05 | 1.6e-05 | 1.400E-05 | 2.646E-06 | 1.100E-05 | 1.600E-05 |
| 26508 | 1326 | T1326G | E | 1.6e-05 | 9.6e-06 | 1e-05 | 1.187E-05 | 3.585E-06 | 9.600E-06 | 1.600E-05 |
| 26509 | 1326 | T1326H | E | 1.1e-05 | 9.1e-06 | 9.2e-06 | 9.767E-06 | 1.069E-06 | 9.100E-06 | 1.100E-05 |
| 26510 | 1326 | T1326I | E | 1.6e-05 | 1.2e-05 | 1.6e-05 | 1.467E-05 | 2.309E-06 | 1.200E-05 | 1.600E-05 |
| 26511 | 1326 | T1326L | E | 8.9e-06 | 1.5e-05 | 1.1e-05 | 1.163E-05 | 3.099E-06 | 8.900E-06 | 1.500E-05 |
| 26512 | 1326 | T1326K | E | 1.3e-05 | 1.5e-05 | 1e-05 | 1.267E-05 | 2.517E-06 | 1.000E-05 | 1.500E-05 |
| 26513 | 1326 | T1326M | E | 1.5e-05 | 9.9e-06 | 9.8e-06 | 1.157E-05 | 2.974E-06 | 9.800E-06 | 1.500E-05 |
| 26514 | 1326 | T1326F | E | 9.7e-06 | 1.8e-05 | 1.3e-05 | 1.357E-05 | 4.179E-06 | 9.700E-06 | 1.800E-05 |
| 26515 | 1326 | T1326P | E | 2.3e-05 | 1e-05 | 1.1e-05 | 1.467E-05 | 7.234E-06 | 1.000E-05 | 2.300E-05 |
| 26516 | 1326 | T1326S | E | 1.4e-05 | 1.1e-05 | 1.4e-05 | 1.300E-05 | 1.732E-06 | 1.100E-05 | 1.400E-05 |
| 26517 | 1326 | T1326T | E | 1e-05 | 1.4e-05 | 1.1e-05 | 1.167E-05 | 2.082E-06 | 1.000E-05 | 1.400E-05 |
| 26518 | 1326 | T1326W | E | 1.3e-05 | 1.5e-05 | 1.2e-05 | 1.333E-05 | 1.528E-06 | 1.200E-05 | 1.500E-05 |
| 26519 | 1326 | T1326Y | E | 9.4e-06 | 9.9e-06 | 1.5e-05 | 1.143E-05 | 3.099E-06 | 9.400E-06 | 1.500E-05 |

| | | | | | | | | | | |
|---|---|---|---|---|---|---|---|---|---|---|
| 26520 | 1326 | T1326V | E | 1.4e-05 | 1.5e-05 | 1.2e-05 | 1.367E-05 | 1.528E-06 | 1.200E-05 | 1.500E-05 |
| 26521 | 1327 | N1327A | E | 8.5e-06 | 1.5e-05 | 1.5e-05 | 1.283E-05 | 3.753E-06 | 8.500E-06 | 1.500E-05 |
| 26522 | 1327 | N1327R | E | 1.6e-05 | 1.6e-05 | 1.1e-05 | 1.433E-05 | 2.887E-06 | 1.100E-05 | 1.600E-05 |
| 26523 | 1327 | N1327N | E | 1.4e-05 | 1.6e-05 | 1.5e-05 | 1.500E-05 | 1.000E-06 | 1.400E-05 | 1.600E-05 |
| 26524 | 1327 | N1327D | E | 1.7e-05 | 1.7e-05 | 1.1e-05 | 1.500E-05 | 3.464E-06 | 1.100E-05 | 1.700E-05 |
| 26525 | 1327 | N1327C | E | 1.4e-05 | 1.5e-05 | 1e-05 | 1.300E-05 | 2.646E-06 | 1.000E-05 | 1.500E-05 |
| 26526 | 1327 | N1327E | E | 1.1e-05 | 1.5e-05 | 1.6e-05 | 1.400E-05 | 2.646E-06 | 1.100E-05 | 1.600E-05 |
| 26527 | 1327 | N1327Q | E | 8.7e-06 | 8.4e-06 | 1.1e-05 | 9.367E-06 | 1.422E-06 | 8.400E-06 | 1.100E-05 |
| 26528 | 1327 | N1327G | E | 1.1e-05 | 1.3e-05 | 1.4e-05 | 1.267E-05 | 1.528E-06 | 1.100E-05 | 1.400E-05 |
| 26529 | 1327 | N1327H | E | 1.5e-05 | 1e-05 | 1.4e-05 | 1.300E-05 | 2.646E-06 | 1.000E-05 | 1.500E-05 |
| 26530 | 1327 | N1327I | E | 1.3e-05 | 1.3e-05 | 1.5e-05 | 1.367E-05 | 1.155E-06 | 1.300E-05 | 1.500E-05 |
| 26531 | 1327 | N1327L | E | 1.4e-05 | 9.7e-06 | 1.5e-05 | 1.290E-05 | 2.816E-06 | 9.700E-06 | 1.500E-05 |
| 26532 | 1327 | N1327K | E | 9.9e-06 | 1.1e-05 | 1.7e-05 | 1.263E-05 | 3.821E-06 | 9.900E-06 | 1.700E-05 |
| 26533 | 1327 | N1327M | E | 9.2e-06 | 8.4e-06 | 1.5e-05 | 1.087E-05 | 3.602E-06 | 8.400E-06 | 1.500E-05 |
| 26534 | 1327 | N1327F | E | 1.4e-05 | 1e-05 | 1.1e-05 | 1.167E-05 | 2.082E-06 | 1.000E-05 | 1.400E-05 |
| 26535 | 1327 | N1327P | E | 1.5e-05 | 1.3e-05 | 2.6e-05 | 1.800E-05 | 7.000E-06 | 1.300E-05 | 2.600E-05 |
| 26536 | 1327 | N1327S | E | 1e-05 | 1.6e-05 | 1.5e-05 | 1.367E-05 | 3.215E-06 | 1.000E-05 | 1.600E-05 |
| 26537 | 1327 | N1327T | E | 6.4e-06 | 1e-05 | 1.5e-05 | 1.047E-05 | 4.319E-06 | 6.400E-06 | 1.500E-05 |
| 26538 | 1327 | N1327W | E | 1.5e-05 | 1.5e-05 | 1e-05 | 1.333E-05 | 2.887E-06 | 1.000E-05 | 1.500E-05 |
| 26539 | 1327 | N1327Y | E | 1.5e-05 | 1.1e-05 | 1.4e-05 | 1.333E-05 | 2.082E-06 | 1.100E-05 | 1.500E-05 |
| 26540 | 1327 | N1327V | E | 1.4e-05 | 1.3e-05 | 8.9e-06 | 1.197E-05 | 2.702E-06 | 8.900E-06 | 1.400E-05 |
| 26541 | 1328 | G1328A | E | 1.1e-05 | 8.3e-06 | 2.7e-05 | 1.543E-05 | 1.011E-05 | 8.300E-06 | 2.700E-05 |
| 26542 | 1328 | G1328R | E | 1.3e-05 | 1.6e-05 | 1.7e-05 | 1.533E-05 | 2.082E-06 | 1.300E-05 | 1.700E-05 |
| 26543 | 1328 | G1328N | E | 1.4e-05 | 1.6e-05 | 1.5e-05 | 1.500E-05 | 1.000E-06 | 1.400E-05 | 1.600E-05 |
| 26544 | 1328 | G1328D | E | 1.5e-05 | 1.2e-05 | 1.5e-05 | 1.400E-05 | 1.732E-06 | 1.200E-05 | 1.500E-05 |
| 26545 | 1328 | G1328C | E | 1.5e-05 | 1.5e-05 | 9.2e-06 | 1.307E-05 | 3.349E-06 | 9.200E-06 | 1.500E-05 |
| 26546 | 1328 | G1328E | E | 1.1e-05 | 9.5e-06 | 1e-05 | 1.017E-05 | 7.638E-07 | 9.500E-06 | 1.100E-05 |
| 26547 | 1328 | G1328Q | E | 9.3e-06 | 1.5e-05 | 1.5e-05 | 1.310E-05 | 3.291E-06 | 9.300E-06 | 1.500E-05 |
| 26548 | 1328 | G1328G | E | 8.4e-06 | 1.5e-05 | 9.9e-06 | 1.110E-05 | 3.460E-06 | 8.400E-06 | 1.500E-05 |
| 26549 | 1328 | G1328H | E | 1.6e-05 | 1.6e-05 | 1.3e-05 | 1.500E-05 | 1.732E-06 | 1.300E-05 | 1.600E-05 |
| 26550 | 1328 | G1328I | E | 8.4e-06 | 1.4e-05 | 1.6e-05 | 1.280E-05 | 3.940E-06 | 8.400E-06 | 1.600E-05 |
| 26551 | 1328 | G1328L | E | 1.2e-05 | 1.5e-05 | 1.6e-05 | 1.433E-05 | 2.082E-06 | 1.200E-05 | 1.600E-05 |
| 26552 | 1328 | G1328K | E | 1.5e-05 | 1.3e-05 | 9.5e-06 | 1.250E-05 | 2.784E-06 | 9.500E-06 | 1.500E-05 |
| 26553 | 1328 | G1328M | E | 1.5e-05 | 1.2e-05 | 9.7e-06 | 1.223E-05 | 2.658E-06 | 9.700E-06 | 1.500E-05 |
| 26554 | 1328 | G1328F | E | 1.2e-05 | 1.4e-05 | 6.9e-06 | 1.097E-05 | 3.661E-06 | 6.900E-06 | 1.400E-05 |
| 26555 | 1328 | G1328P | E | 1e-05 | 1.7e-05 | 1.6e-05 | 1.433E-05 | 3.786E-06 | 1.000E-05 | 1.700E-05 |
| 26556 | 1328 | G1328S | E | 1.4e-05 | 1.2e-05 | 1e-05 | 1.200E-05 | 2.000E-06 | 1.000E-05 | 1.400E-05 |
| 26557 | 1328 | G1328T | E | 1.6e-05 | 1.7e-05 | 9.4e-06 | 1.413E-05 | 4.130E-06 | 9.400E-06 | 1.700E-05 |
| 26558 | 1328 | G1328W | E | 1.3e-05 | 1.2e-05 | 1.4e-05 | 1.300E-05 | 1.000E-06 | 1.200E-05 | 1.400E-05 |
| 26559 | 1328 | G1328Y | E | 8.8e-06 | 1.3e-05 | 1.5e-05 | 1.227E-05 | 3.164E-06 | 8.800E-06 | 1.500E-05 |
| 26560 | 1328 | G1328V | E | 1e-05 | 1.4e-05 | 2.3e-05 | 1.567E-05 | 6.658E-06 | 1.000E-05 | 2.300E-05 |
| 26561 | 1329 | Y1329A | E | 1.6e-05 | 1e-05 | 1.2e-05 | 1.267E-05 | 3.055E-06 | 1.000E-05 | 1.600E-05 |
| 26562 | 1329 | Y1329R | E | 1.5e-05 | 1.2e-05 | 1.2e-05 | 1.300E-05 | 1.732E-06 | 1.200E-05 | 1.500E-05 |
| 26563 | 1329 | Y1329N | E | 1.3e-05 | 1.7e-05 | 1.2e-05 | 1.400E-05 | 2.646E-06 | 1.200E-05 | 1.700E-05 |
| 26564 | 1329 | Y1329D | E | 9.7e-06 | 1.5e-05 | 7.2e-06 | 1.063E-05 | 3.983E-06 | 7.200E-06 | 1.500E-05 |
| 26565 | 1329 | Y1329C | E | 1.3e-05 | 1.4e-05 | 1.5e-05 | 1.400E-05 | 1.000E-06 | 1.300E-05 | 1.500E-05 |
| 26566 | 1329 | Y1329E | E | 1.7e-05 | 1e-05 | 9.1e-06 | 1.203E-05 | 4.325E-06 | 9.100E-06 | 1.700E-05 |
| 26567 | 1329 | Y1329Q | E | 1.4e-05 | 1.5e-05 | 1.4e-05 | 1.433E-05 | 5.774E-07 | 1.400E-05 | 1.500E-05 |
| 26568 | 1329 | Y1329G | E | 1.1e-05 | 1.1e-05 | 1.5e-05 | 1.233E-05 | 2.309E-06 | 1.100E-05 | 1.500E-05 |
| 26569 | 1329 | Y1329H | E | 1.3e-05 | 8.6e-06 | 1.6e-05 | 1.253E-05 | 3.722E-06 | 8.600E-06 | 1.600E-05 |
| 26570 | 1329 | Y1329I | E | 1.2e-05 | 1e-05 | 1.2e-05 | 1.133E-05 | 1.155E-06 | 1.000E-05 | 1.200E-05 |
| 26571 | 1329 | Y1329L | E | 1.2e-05 | 1.6e-05 | 1.6e-05 | 1.467E-05 | 2.309E-06 | 1.200E-05 | 1.600E-05 |
| 26572 | 1329 | Y1329K | E | 1.1e-05 | 8.6e-06 | 1.2e-05 | 1.053E-05 | 1.747E-06 | 8.600E-06 | 1.200E-05 |
| 26573 | 1329 | Y1329M | E | 1.4e-05 | 1e-05 | 9.5e-06 | 1.117E-05 | 2.466E-06 | 9.500E-06 | 1.400E-05 |
| 26574 | 1329 | Y1329F | E | 7.8e-06 | 1.6e-05 | 1.5e-05 | 1.293E-05 | 4.474E-06 | 7.800E-06 | 1.600E-05 |
| 26575 | 1329 | Y1329P | E | 1e-05 | 7e-06 | 1.5e-05 | 1.067E-05 | 4.041E-06 | 7.000E-06 | 1.500E-05 |
| 26576 | 1329 | Y1329S | E | 1.1e-05 | 9.9e-06 | 9.7e-06 | 1.020E-05 | 7.000E-07 | 9.700E-06 | 1.100E-05 |
| 26577 | 1329 | Y1329T | E | 1.6e-05 | 1.3e-05 | 1.6e-05 | 1.500E-05 | 1.732E-06 | 1.300E-05 | 1.600E-05 |
| 26578 | 1329 | Y1329W | E | 1.1e-05 | 1.5e-05 | 1e-05 | 1.200E-05 | 2.646E-06 | 1.000E-05 | 1.500E-05 |
| 26579 | 1329 | Y1329Y | E | 1.4e-05 | 1.5e-05 | 1.4e-05 | 1.433E-05 | 5.774E-07 | 1.400E-05 | 1.500E-05 |
| 26580 | 1329 | Y1329V | E | 1.3e-05 | 1e-05 | 9.8e-06 | 1.093E-05 | 1.793E-06 | 9.800E-06 | 1.300E-05 |
| 26581 | 1330 | T1330A | E | 1e-05 | 1.3e-05 | 1.4e-05 | 1.233E-05 | 2.082E-06 | 1.000E-05 | 1.400E-05 |
| 26582 | 1330 | T1330R | E | 1.4e-05 | 9.9e-06 | 1.3e-05 | 1.230E-05 | 2.138E-06 | 9.900E-06 | 1.400E-05 |
| 26583 | 1330 | T1330N | E | 1.5e-05 | 1.1e-05 | 1.5e-05 | 1.367E-05 | 2.309E-06 | 1.100E-05 | 1.500E-05 |
| 26584 | 1330 | T1330D | E | 1.3e-05 | 1.3e-05 | 9.4e-06 | 1.180E-05 | 2.078E-06 | 9.400E-06 | 1.300E-05 |
| 26585 | 1330 | T1330C | E | 1.3e-05 | 1.3e-05 | 1.7e-05 | 1.433E-05 | 2.309E-06 | 1.300E-05 | 1.700E-05 |
| 26586 | 1330 | T1330E | E | 1.2e-05 | 1e-05 | 1.5e-05 | 1.233E-05 | 2.517E-06 | 1.000E-05 | 1.500E-05 |
| 26587 | 1330 | T1330Q | E | 1e-05 | 1.4e-05 | 1.2e-05 | 1.200E-05 | 2.000E-06 | 1.000E-05 | 1.400E-05 |
| 26588 | 1330 | T1330G | E | 7.6e-06 | 1.6e-05 | 1.1e-05 | 1.153E-05 | 4.225E-06 | 7.600E-06 | 1.600E-05 |
| 26589 | 1330 | T1330H | E | 1.5e-05 | 8.7e-06 | 9.8e-06 | 1.117E-05 | 3.365E-06 | 8.700E-06 | 1.500E-05 |
| 26590 | 1330 | T1330I | E | 1.5e-05 | 9.1e-06 | 1.1e-05 | 1.170E-05 | 3.012E-06 | 9.100E-06 | 1.500E-05 |
| 26591 | 1330 | T1330L | E | 1.2e-05 | 1.5e-05 | 1.1e-05 | 1.267E-05 | 2.082E-06 | 1.100E-05 | 1.500E-05 |
| 26592 | 1330 | T1330K | E | 1.1e-05 | 1e-05 | 2.2e-05 | 1.433E-05 | 6.658E-06 | 1.000E-05 | 2.200E-05 |
| 26593 | 1330 | T1330M | E | 1.1e-05 | 1.5e-05 | 1e-05 | 1.200E-05 | 2.646E-06 | 1.000E-05 | 1.500E-05 |
| 26594 | 1330 | T1330F | E | 9.3e-06 | 1.2e-05 | 1.9e-05 | 1.343E-05 | 5.006E-06 | 9.300E-06 | 1.900E-05 |
| 26595 | 1330 | T1330P | E | 1e-05 | 1.6e-05 | 1.5e-05 | 1.367E-05 | 3.215E-06 | 1.000E-05 | 1.600E-05 |
| 26596 | 1330 | T1330S | E | 1.1e-05 | 1.5e-05 | 1.4e-05 | 1.333E-05 | 2.082E-06 | 1.100E-05 | 1.500E-05 |
| 26597 | 1330 | T1330T | E | 9.1e-06 | 1.4e-05 | 9.6e-06 | 1.090E-05 | 2.696E-06 | 9.100E-06 | 1.400E-05 |
| 26598 | 1330 | T1330W | E | 1e-05 | 1.1e-05 | 8.5e-06 | 9.833E-06 | 1.258E-06 | 8.500E-06 | 1.100E-05 |
| 26599 | 1330 | T1330Y | E | 1.7e-05 | 1.6e-05 | 1.6e-05 | 1.633E-05 | 5.774E-07 | 1.600E-05 | 1.700E-05 |
| 26600 | 1330 | T1330V | E | 1.4e-05 | 1.1e-05 | 1.3e-05 | 1.267E-05 | 1.528E-06 | 1.100E-05 | 1.400E-05 |
| 26601 | 1331 | R1331A | E | 1.2e-05 | 1.5e-05 | 1.6e-05 | 1.433E-05 | 2.082E-06 | 1.200E-05 | 1.600E-05 |
| 26602 | 1331 | R1331R | E | 1.1e-05 | 9.8e-06 | 1.5e-05 | 1.193E-05 | 2.723E-06 | 9.800E-06 | 1.500E-05 |
| 26603 | 1331 | R1331N | E | 1.6e-05 | 1e-05 | 1.4e-05 | 1.333E-05 | 3.055E-06 | 1.000E-05 | 1.600E-05 |
| 26604 | 1331 | R1331D | E | 1e-05 | 1.6e-05 | 1.5e-05 | 1.367E-05 | 3.215E-06 | 1.000E-05 | 1.600E-05 |

| | | | | | | | | | | |
|---|---|---|---|---|---|---|---|---|---|---|
| 26605 | 1331 | R1331C | E | 9e-06 | 1e-05 | 1.5e-05 | 1.133E-05 | 3.215E-06 | 9.000E-06 | 1.500E-05 |
| 26606 | 1331 | R1331E | E | 8.4e-06 | 9.5e-06 | 1e-05 | 9.300E-06 | 8.185E-07 | 8.400E-06 | 1.000E-05 |
| 26607 | 1331 | R1331Q | E | 1.5e-05 | 1.1e-05 | 9.5e-06 | 1.183E-05 | 2.843E-06 | 9.500E-06 | 1.500E-05 |
| 26608 | 1331 | R1331G | E | 1.5e-05 | 1.5e-05 | 7.7e-06 | 1.257E-05 | 4.215E-06 | 7.700E-06 | 1.500E-05 |
| 26609 | 1331 | R1331H | E | 1.2e-05 | 1e-05 | 1.7e-05 | 1.300E-05 | 3.606E-06 | 1.000E-05 | 1.700E-05 |
| 26610 | 1331 | R1331I | E | 1.3e-05 | 1.2e-05 | 1.6e-05 | 1.367E-05 | 2.082E-06 | 1.200E-05 | 1.600E-05 |
| 26611 | 1331 | R1331L | E | 1.8e-05 | 1.6e-05 | 1.2e-05 | 1.533E-05 | 3.055E-06 | 1.200E-05 | 1.800E-05 |
| 26612 | 1331 | R1331K | E | 8.5e-06 | 8.9e-06 | 1.5e-05 | 1.080E-05 | 3.643E-06 | 8.500E-06 | 1.500E-05 |
| 26613 | 1331 | R1331M | E | 1.6e-05 | 1.6e-05 | 2.2e-05 | 1.800E-05 | 3.464E-06 | 1.600E-05 | 2.200E-05 |
| 26614 | 1331 | R1331F | E | 1e-05 | 9.8e-06 | 1e-05 | 9.933E-06 | 1.155E-07 | 9.800E-06 | 1.000E-05 |
| 26615 | 1331 | R1331P | E | 1.2e-05 | 1.6e-05 | 1.8e-05 | 1.533E-05 | 3.055E-06 | 1.200E-05 | 1.800E-05 |
| 26616 | 1331 | R1331S | E | 1.3e-05 | 9.8e-06 | 9.8e-06 | 1.087E-05 | 1.848E-06 | 9.800E-06 | 1.300E-05 |
| 26617 | 1331 | R1331T | E | 1.2e-05 | 1.1e-05 | 1.3e-05 | 1.200E-05 | 1.000E-06 | 1.100E-05 | 1.300E-05 |
| 26618 | 1331 | R1331W | E | 1.5e-05 | 1.4e-05 | 1.1e-05 | 1.333E-05 | 2.082E-06 | 1.100E-05 | 1.500E-05 |
| 26619 | 1331 | R1331Y | E | 1.1e-05 | 1.5e-05 | 1.5e-05 | 1.367E-05 | 2.309E-06 | 1.100E-05 | 1.500E-05 |
| 26620 | 1331 | R1331V | E | 1.2e-05 | 1.5e-05 | 1.2e-05 | 1.300E-05 | 1.732E-06 | 1.200E-05 | 1.500E-05 |
| 26621 | 1332 | Y1332A | E | 1.2e-05 | 9.7e-06 | 1.6e-05 | 1.257E-05 | 3.188E-06 | 9.700E-06 | 1.600E-05 |
| 26622 | 1332 | Y1332R | E | 1.3e-05 | 1.5e-05 | 1.7e-05 | 1.500E-05 | 2.000E-06 | 1.300E-05 | 1.700E-05 |
| 26623 | 1332 | Y1332N | E | 1.7e-05 | 1.8e-05 | 1.3e-05 | 1.600E-05 | 2.646E-06 | 1.300E-05 | 1.800E-05 |
| 26624 | 1332 | Y1332D | E | 1.6e-05 | 1.5e-05 | 1.5e-05 | 1.533E-05 | 5.774E-07 | 1.500E-05 | 1.600E-05 |
| 26625 | 1332 | Y1332C | E | 8.8e-06 | 1.1e-05 | 1.3e-05 | 1.093E-05 | 2.101E-06 | 8.800E-06 | 1.300E-05 |
| 26626 | 1332 | Y1332E | E | 1.3e-05 | 1.6e-05 | 9.7e-06 | 1.290E-05 | 3.151E-06 | 9.700E-06 | 1.600E-05 |
| 26627 | 1332 | Y1332Q | E | 1.4e-05 | 1.1e-05 | 1e-05 | 1.167E-05 | 2.082E-06 | 1.000E-05 | 1.400E-05 |
| 26628 | 1332 | Y1332G | E | 1.2e-05 | 6.5e-06 | 1.3e-05 | 1.050E-05 | 3.500E-06 | 6.500E-06 | 1.300E-05 |
| 26629 | 1332 | Y1332H | E | 9.1e-06 | 1.5e-05 | 1.2e-05 | 1.203E-05 | 2.950E-06 | 9.100E-06 | 1.500E-05 |
| 26630 | 1332 | Y1332I | E | 1.5e-05 | 1.5e-05 | 1.1e-05 | 1.367E-05 | 2.309E-06 | 1.100E-05 | 1.500E-05 |
| 26631 | 1332 | Y1332L | E | 1.8e-05 | 9.4e-06 | 7.5e-06 | 1.163E-05 | 5.595E-06 | 7.500E-06 | 1.800E-05 |
| 26632 | 1332 | Y1332K | E | 1.5e-05 | 1.1e-05 | 9.1e-06 | 1.170E-05 | 3.012E-06 | 9.100E-06 | 1.500E-05 |
| 26633 | 1332 | Y1332M | E | 1.6e-05 | 1.6e-05 | 1.3e-05 | 1.500E-05 | 1.732E-06 | 1.300E-05 | 1.600E-05 |
| 26634 | 1332 | Y1332F | E | 1.2e-05 | 8.8e-06 | 1.1e-05 | 1.060E-05 | 1.637E-06 | 8.800E-06 | 1.200E-05 |
| 26635 | 1332 | Y1332P | E | 1.9e-05 | 1.1e-05 | 1.5e-05 | 1.500E-05 | 4.000E-06 | 1.100E-05 | 1.900E-05 |
| 26636 | 1332 | Y1332S | E | 1.5e-05 | 1.1e-05 | 1e-05 | 1.200E-05 | 2.646E-06 | 1.000E-05 | 1.500E-05 |
| 26637 | 1332 | Y1332T | E | 1.7e-05 | 8.5e-06 | 8.8e-06 | 1.143E-05 | 4.823E-06 | 8.500E-06 | 1.700E-05 |
| 26638 | 1332 | Y1332W | E | 1.6e-05 | 1e-05 | 1.5e-05 | 1.367E-05 | 3.215E-06 | 1.000E-05 | 1.600E-05 |
| 26639 | 1332 | Y1332Y | E | 1.6e-05 | 9.8e-06 | 1e-05 | 1.193E-05 | 3.523E-06 | 9.800E-06 | 1.600E-05 |
| 26640 | 1332 | Y1332V | E | 1.1e-05 | 1.4e-05 | 1.1e-05 | 1.200E-05 | 1.732E-06 | 1.100E-05 | 1.400E-05 |
| 26641 | 1333 | A1333A | E | 1.5e-05 | 1e-05 | 1e-05 | 1.167E-05 | 2.887E-06 | 1.000E-05 | 1.500E-05 |
| 26642 | 1333 | A1333R | E | 9.3e-06 | 1.3e-05 | 1.1e-05 | 1.110E-05 | 1.852E-06 | 9.300E-06 | 1.300E-05 |
| 26643 | 1333 | A1333N | E | 1.2e-05 | 1.6e-05 | 9.1e-06 | 1.237E-05 | 3.465E-06 | 9.100E-06 | 1.600E-05 |
| 26644 | 1333 | A1333D | E | 1.4e-05 | 1.4e-05 | 9.4e-06 | 1.247E-05 | 2.656E-06 | 9.400E-06 | 1.400E-05 |
| 26645 | 1333 | A1333C | E | 1.3e-05 | 1.2e-05 | 1e-05 | 1.167E-05 | 1.528E-06 | 1.000E-05 | 1.300E-05 |
| 26646 | 1333 | A1333E | E | 1.1e-05 | 1.5e-05 | 1.5e-05 | 1.367E-05 | 2.309E-06 | 1.100E-05 | 1.500E-05 |
| 26647 | 1333 | A1333Q | E | 9.4e-06 | 1.7e-05 | 1.1e-05 | 1.247E-05 | 4.007E-06 | 9.400E-06 | 1.700E-05 |
| 26648 | 1333 | A1333G | E | 1.4e-05 | 1.1e-05 | 1.7e-05 | 1.400E-05 | 3.000E-06 | 1.100E-05 | 1.700E-05 |
| 26649 | 1333 | A1333H | E | 1e-05 | 1.7e-05 | 1e-05 | 1.233E-05 | 4.041E-06 | 1.000E-05 | 1.700E-05 |
| 26650 | 1333 | A1333I | E | 1.1e-05 | 1.1e-05 | 1.8e-05 | 1.333E-05 | 4.041E-06 | 1.100E-05 | 1.800E-05 |
| 26651 | 1333 | A1333L | E | 1.4e-05 | 1.5e-05 | 1.1e-05 | 1.333E-05 | 2.082E-06 | 1.100E-05 | 1.500E-05 |
| 26652 | 1333 | A1333K | E | 1e-05 | 1.4e-05 | 1.5e-05 | 1.300E-05 | 2.646E-06 | 1.000E-05 | 1.500E-05 |
| 26653 | 1333 | A1333M | E | 1.4e-05 | 8.3e-06 | 1.1e-05 | 1.110E-05 | 2.851E-06 | 8.300E-06 | 1.400E-05 |
| 26654 | 1333 | A1333F | E | 1.1e-05 | 1.4e-05 | 1.4e-05 | 1.300E-05 | 1.732E-06 | 1.100E-05 | 1.400E-05 |
| 26655 | 1333 | A1333P | E | 1.4e-05 | 1.5e-05 | 1.5e-05 | 1.467E-05 | 5.774E-07 | 1.400E-05 | 1.500E-05 |
| 26656 | 1333 | A1333S | E | 1.5e-05 | 1.4e-05 | 1.6e-05 | 1.500E-05 | 1.000E-06 | 1.400E-05 | 1.600E-05 |
| 26657 | 1333 | A1333T | E | 1.2e-05 | 1e-05 | 1.4e-05 | 1.200E-05 | 2.000E-06 | 1.000E-05 | 1.400E-05 |
| 26658 | 1333 | A1333W | E | 1.5e-05 | 1.1e-05 | 1.1e-05 | 1.233E-05 | 2.309E-06 | 1.100E-05 | 1.500E-05 |
| 26659 | 1333 | A1333Y | E | 1.2e-05 | 9.2e-06 | 1.5e-05 | 1.207E-05 | 2.901E-06 | 9.200E-06 | 1.500E-05 |
| 26660 | 1333 | A1333V | E | 1.1e-05 | 1e-05 | 1.7e-05 | 1.267E-05 | 3.786E-06 | 1.000E-05 | 1.700E-05 |
| 26661 | 1334 | D1334A | E | 9.9e-06 | 9.7e-06 | 9.9e-06 | 9.833E-06 | 1.155E-07 | 9.700E-06 | 9.900E-06 |
| 26662 | 1334 | D1334R | E | 1e-05 | 1.1e-05 | 1.3e-05 | 1.133E-05 | 1.528E-06 | 1.000E-05 | 1.300E-05 |
| 26663 | 1334 | D1334N | E | 1.3e-05 | 2.9e-05 | 1.6e-05 | 1.933E-05 | 8.505E-06 | 1.300E-05 | 2.900E-05 |
| 26664 | 1334 | D1334D | E | 1e-05 | 1.5e-05 | 1.6e-05 | 1.367E-05 | 3.215E-06 | 1.000E-05 | 1.600E-05 |
| 26665 | 1334 | D1334C | E | 1e-05 | 9.6e-06 | 1.6e-05 | 1.187E-05 | 3.585E-06 | 9.600E-06 | 1.600E-05 |
| 26666 | 1334 | D1334E | E | 1e-05 | 8.1e-06 | 1.1e-05 | 9.700E-06 | 1.473E-06 | 8.100E-06 | 1.100E-05 |
| 26667 | 1334 | D1334Q | E | 1.4e-05 | 1.7e-05 | 9.7e-06 | 1.357E-05 | 3.669E-06 | 9.700E-06 | 1.700E-05 |
| 26668 | 1334 | D1334G | E | 8.4e-06 | 1.1e-05 | 1.3e-05 | 1.080E-05 | 2.307E-06 | 8.400E-06 | 1.300E-05 |
| 26669 | 1334 | D1334H | E | 9.3e-06 | 1.5e-05 | 1.5e-05 | 1.310E-05 | 3.291E-06 | 9.300E-06 | 1.500E-05 |
| 26670 | 1334 | D1334I | E | 1.7e-05 | 9.2e-06 | 1.1e-05 | 1.240E-05 | 4.084E-06 | 9.200E-06 | 1.700E-05 |
| 26671 | 1334 | D1334L | E | 1e-05 | 1.7e-05 | 1e-05 | 1.233E-05 | 4.041E-06 | 1.000E-05 | 1.700E-05 |
| 26672 | 1334 | D1334K | E | 1.6e-05 | 1.6e-05 | 1.3e-05 | 1.500E-05 | 1.732E-06 | 1.300E-05 | 1.600E-05 |
| 26673 | 1334 | D1334M | E | 1e-05 | 1.2e-05 | 1.5e-05 | 1.233E-05 | 2.517E-06 | 1.000E-05 | 1.500E-05 |
| 26674 | 1334 | D1334F | E | 1.5e-05 | 1.6e-05 | 1.5e-05 | 1.533E-05 | 5.774E-07 | 1.500E-05 | 1.600E-05 |
| 26675 | 1334 | D1334P | E | 1e-05 | 1.8e-05 | 1.5e-05 | 1.433E-05 | 4.041E-06 | 1.000E-05 | 1.800E-05 |
| 26676 | 1334 | D1334S | E | 1.4e-05 | 6.9e-06 | 8.5e-06 | 9.800E-06 | 3.724E-06 | 6.900E-06 | 1.400E-05 |
| 26677 | 1334 | D1334T | E | 1e-05 | 1.4e-05 | 1.2e-05 | 1.200E-05 | 2.000E-06 | 1.000E-05 | 1.400E-05 |
| 26678 | 1334 | D1334W | E | 1e-05 | 1.3e-05 | 1.6e-05 | 1.300E-05 | 3.000E-06 | 1.000E-05 | 1.600E-05 |
| 26679 | 1334 | D1334Y | E | 1.5e-05 | 1.2e-05 | 1e-05 | 1.233E-05 | 2.517E-06 | 1.000E-05 | 1.500E-05 |
| 26680 | 1334 | D1334V | E | 1.6e-05 | 1.1e-05 | 1.2e-05 | 1.300E-05 | 2.646E-06 | 1.100E-05 | 1.600E-05 |
| 26681 | 1335 | S1335A | E | 1.5e-05 | 1.5e-05 | 1e-05 | 1.333E-05 | 2.887E-06 | 1.000E-05 | 1.500E-05 |
| 26682 | 1335 | S1335R | E | 8.8e-06 | 1.5e-05 | 1.6e-05 | 1.327E-05 | 3.900E-06 | 8.800E-06 | 1.600E-05 |
| 26683 | 1335 | S1335N | E | 1.6e-05 | 8.6e-06 | 1.2e-05 | 1.220E-05 | 3.704E-06 | 8.600E-06 | 1.600E-05 |
| 26684 | 1335 | S1335D | E | 9.1e-06 | 1.1e-05 | 1.1e-05 | 1.037E-05 | 1.097E-06 | 9.100E-06 | 1.100E-05 |
| 26685 | 1335 | S1335C | E | 1.1e-05 | 1.6e-05 | 1.6e-05 | 1.433E-05 | 2.887E-06 | 1.100E-05 | 1.600E-05 |
| 26686 | 1335 | S1335E | E | 1.4e-05 | 1.5e-05 | 1.1e-05 | 1.333E-05 | 2.082E-06 | 1.100E-05 | 1.500E-05 |
| 26687 | 1335 | S1335Q | E | 1e-05 | 1.1e-05 | 1.5e-05 | 1.200E-05 | 2.646E-06 | 1.000E-05 | 1.500E-05 |
| 26688 | 1335 | S1335G | E | 1.1e-05 | 1.2e-05 | 1.6e-05 | 1.300E-05 | 2.646E-06 | 1.100E-05 | 1.600E-05 |
| 26689 | 1335 | S1335H | E | 1.3e-05 | 1.4e-05 | 1.3e-05 | 1.333E-05 | 5.774E-07 | 1.300E-05 | 1.400E-05 |

| | | | | | | | | | | |
|---|---|---|---|---|---|---|---|---|---|---|
| 26690 | 1335 | S1335I | E | 1.5e-05 | 9.9e-06 | 1.7e-05 | 1.397E-05 | 3.661E-06 | 9.900E-06 | 1.700E-05 |
| 26691 | 1335 | S1335L | E | 1.8e-05 | 6.5e-06 | 1.1e-05 | 1.183E-05 | 5.795E-06 | 6.500E-06 | 1.800E-05 |
| 26692 | 1335 | S1335K | E | 1.1e-05 | 1.6e-05 | 1.5e-05 | 1.400E-05 | 2.646E-06 | 1.100E-05 | 1.600E-05 |
| 26693 | 1335 | S1335M | E | 1.3e-05 | 1e-05 | 1.5e-05 | 1.267E-05 | 2.517E-06 | 1.000E-05 | 1.500E-05 |
| 26694 | 1335 | S1335F | E | 1.8e-05 | 1.1e-05 | 1.5e-05 | 1.467E-05 | 3.512E-06 | 1.100E-05 | 1.800E-05 |
| 26695 | 1335 | S1335P | E | 1.5e-05 | 1.4e-05 | 9.6e-06 | 1.287E-05 | 2.873E-06 | 9.600E-06 | 1.500E-05 |
| 26696 | 1335 | S1335S | E | 1.5e-05 | 1.2e-05 | 1.6e-05 | 1.433E-05 | 2.082E-06 | 1.200E-05 | 1.600E-05 |
| 26697 | 1335 | S1335T | E | 1.4e-05 | 1.1e-05 | 9.7e-06 | 1.157E-05 | 2.205E-06 | 9.700E-06 | 1.400E-05 |
| 26698 | 1335 | S1335W | E | 9.1e-06 | 1.4e-05 | 9.6e-06 | 1.090E-05 | 2.696E-06 | 9.100E-06 | 1.400E-05 |
| 26699 | 1335 | S1335Y | E | 1.5e-05 | 1.6e-05 | 1.2e-05 | 1.433E-05 | 2.082E-06 | 1.200E-05 | 1.600E-05 |
| 26700 | 1335 | S1335V | E | 1.4e-05 | 8.9e-06 | 1.3e-05 | 1.197E-05 | 2.702E-06 | 8.900E-06 | 1.400E-05 |
| 26701 | 1336 | V1336A | E | 8.6e-06 | 9.7e-06 | 1.5e-05 | 1.110E-05 | 3.422E-06 | 8.600E-06 | 1.500E-05 |
| 26702 | 1336 | V1336R | E | 1.2e-05 | 1.5e-05 | 1.6e-05 | 1.433E-05 | 2.082E-06 | 1.200E-05 | 1.600E-05 |
| 26703 | 1336 | V1336N | E | 1.5e-05 | 1.4e-05 | 1.2e-05 | 1.367E-05 | 1.528E-06 | 1.200E-05 | 1.500E-05 |
| 26704 | 1336 | V1336D | E | 1.4e-05 | 1.3e-05 | 8.7e-06 | 1.190E-05 | 2.816E-06 | 8.700E-06 | 1.400E-05 |
| 26705 | 1336 | V1336C | E | 9.3e-06 | 1.2e-05 | 1.5e-05 | 1.210E-05 | 2.851E-06 | 9.300E-06 | 1.500E-05 |
| 26706 | 1336 | V1336E | E | 1.4e-05 | 1.5e-05 | 1.3e-05 | 1.400E-05 | 1.000E-06 | 1.300E-05 | 1.500E-05 |
| 26707 | 1336 | V1336Q | E | 1.5e-05 | 1.2e-05 | 1.4e-05 | 1.367E-05 | 1.528E-06 | 1.200E-05 | 1.500E-05 |
| 26708 | 1336 | V1336G | E | 1e-05 | 1.1e-05 | 3.6e-05 | 1.900E-05 | 1.473E-05 | 1.000E-05 | 3.600E-05 |
| 26709 | 1336 | V1336H | E | 1.2e-05 | 1.5e-05 | 1.3e-05 | 1.333E-05 | 1.528E-06 | 1.200E-05 | 1.500E-05 |
| 26710 | 1336 | V1336I | E | 8.3e-06 | 1.7e-05 | 1.1e-05 | 1.210E-05 | 4.453E-06 | 8.300E-06 | 1.700E-05 |
| 26711 | 1336 | V1336L | E | 8.6e-06 | 9.6e-06 | 1.8e-05 | 1.207E-05 | 5.163E-06 | 8.600E-06 | 1.800E-05 |
| 26712 | 1336 | V1336K | E | 9.9e-06 | 1.5e-05 | 1.2e-05 | 1.230E-05 | 2.563E-06 | 9.900E-06 | 1.500E-05 |
| 26713 | 1336 | V1336M | E | 1.5e-05 | 1.1e-05 | 9.6e-06 | 1.187E-05 | 2.802E-06 | 9.600E-06 | 1.500E-05 |
| 26714 | 1336 | V1336F | E | 1.6e-05 | 1e-05 | 1.5e-05 | 1.367E-05 | 3.215E-06 | 1.000E-05 | 1.600E-05 |
| 26715 | 1336 | V1336P | E | 1.4e-05 | 1.5e-05 | 1.1e-05 | 1.333E-05 | 2.082E-06 | 1.100E-05 | 1.500E-05 |
| 26716 | 1336 | V1336S | E | 1.2e-05 | 1.1e-05 | 9.5e-06 | 1.083E-05 | 1.258E-06 | 9.500E-06 | 1.200E-05 |
| 26717 | 1336 | V1336T | E | 1.1e-05 | 1.4e-05 | 8.5e-06 | 1.117E-05 | 2.754E-06 | 8.500E-06 | 1.400E-05 |
| 26718 | 1336 | V1336W | E | 1.5e-05 | 1e-05 | 1.5e-05 | 1.333E-05 | 2.887E-06 | 1.000E-05 | 1.500E-05 |
| 26719 | 1336 | V1336Y | E | 1e-05 | 8.4e-06 | 9.7e-06 | 9.367E-06 | 8.505E-07 | 8.400E-06 | 1.000E-05 |
| 26720 | 1336 | V1336V | E | 1.5e-05 | 1.6e-05 | 1.4e-05 | 1.500E-05 | 1.000E-06 | 1.400E-05 | 1.600E-05 |
| 26721 | 1337 | K1337A | E | 1.1e-05 | 1.4e-05 | 1.1e-05 | 1.200E-05 | 1.732E-06 | 1.100E-05 | 1.400E-05 |
| 26722 | 1337 | K1337R | E | 1.2e-05 | 1e-05 | 1.9e-05 | 1.367E-05 | 4.726E-06 | 1.000E-05 | 1.900E-05 |
| 26723 | 1337 | K1337N | E | 1.6e-05 | 1e-05 | 1.6e-05 | 1.400E-05 | 3.464E-06 | 1.000E-05 | 1.600E-05 |
| 26724 | 1337 | K1337D | E | 1e-05 | 9.7e-06 | 1.1e-05 | 1.023E-05 | 6.807E-07 | 9.700E-06 | 1.100E-05 |
| 26725 | 1337 | K1337C | E | 1.4e-05 | 1.1e-05 | 1.4e-05 | 1.300E-05 | 1.732E-06 | 1.100E-05 | 1.400E-05 |
| 26726 | 1337 | K1337E | E | 1.4e-05 | 1e-05 | 1.1e-05 | 1.167E-05 | 2.082E-06 | 1.000E-05 | 1.400E-05 |
| 26727 | 1337 | K1337Q | E | 9.2e-06 | 1.2e-05 | 1.5e-05 | 1.207E-05 | 2.901E-06 | 9.200E-06 | 1.500E-05 |
| 26728 | 1337 | K1337G | E | 1.4e-05 | 1.7e-05 | 1.7e-05 | 1.600E-05 | 1.732E-06 | 1.400E-05 | 1.700E-05 |
| 26729 | 1337 | K1337H | E | 1.3e-05 | 9.6e-06 | 1e-05 | 1.087E-05 | 1.858E-06 | 9.600E-06 | 1.300E-05 |
| 26730 | 1337 | K1337I | E | 1.5e-05 | 1.2e-05 | 1.9e-05 | 1.533E-05 | 3.512E-06 | 1.200E-05 | 1.900E-05 |
| 26731 | 1337 | K1337L | E | 1.1e-05 | 1.5e-05 | 1.1e-05 | 1.233E-05 | 2.309E-06 | 1.100E-05 | 1.500E-05 |
| 26732 | 1337 | K1337K | E | 9.6e-06 | 1.1e-05 | 1.6e-05 | 1.220E-05 | 3.365E-06 | 9.600E-06 | 1.600E-05 |
| 26733 | 1337 | K1337M | E | 1.4e-05 | 1.5e-05 | 1.7e-05 | 1.533E-05 | 1.528E-06 | 1.400E-05 | 1.700E-05 |
| 26734 | 1337 | K1337F | E | 9.7e-06 | 1e-05 | 1.1e-05 | 1.023E-05 | 6.807E-07 | 9.700E-06 | 1.100E-05 |
| 26735 | 1337 | K1337P | E | 1e-05 | 1.7e-05 | 1.6e-05 | 1.433E-05 | 3.786E-06 | 1.000E-05 | 1.700E-05 |
| 26736 | 1337 | K1337S | E | 9.9e-06 | 1.5e-05 | 1.4e-05 | 1.297E-05 | 2.702E-06 | 9.900E-06 | 1.500E-05 |
| 26737 | 1337 | K1337T | E | 9e-06 | 1.1e-05 | 1.7e-05 | 1.233E-05 | 4.163E-06 | 9.000E-06 | 1.700E-05 |
| 26738 | 1337 | K1337W | E | 1.6e-05 | 1.5e-05 | 1.6e-05 | 1.567E-05 | 5.774E-07 | 1.500E-05 | 1.600E-05 |
| 26739 | 1337 | K1337Y | E | 9.8e-06 | 9.1e-06 | 1e-05 | 9.633E-06 | 4.726E-07 | 9.100E-06 | 1.000E-05 |
| 26740 | 1337 | K1337V | E | 9.9e-06 | 8.7e-06 | 8.4e-06 | 9.000E-06 | 7.937E-07 | 8.400E-06 | 9.900E-06 |
| 26741 | 1338 | G1338A | E | 1.7e-05 | 1.1e-05 | 1.7e-05 | 1.500E-05 | 3.464E-06 | 1.100E-05 | 1.700E-05 |
| 26742 | 1338 | G1338R | E | 1.5e-05 | 1.5e-05 | 1.6e-05 | 1.533E-05 | 5.774E-07 | 1.500E-05 | 1.600E-05 |
| 26743 | 1338 | G1338N | E | 1.4e-05 | 1.2e-05 | 1.4e-05 | 1.333E-05 | 1.155E-06 | 1.200E-05 | 1.400E-05 |
| 26744 | 1338 | G1338D | E | 1.1e-05 | 8.9e-06 | 1.2e-05 | 1.063E-05 | 1.582E-06 | 8.900E-06 | 1.200E-05 |
| 26745 | 1338 | G1338C | E | 9.2e-06 | 1.1e-05 | 1.6e-05 | 1.207E-05 | 3.523E-06 | 9.200E-06 | 1.600E-05 |
| 26746 | 1338 | G1338E | E | 1.1e-05 | 9.9e-06 | 1.6e-05 | 1.230E-05 | 3.251E-06 | 9.900E-06 | 1.600E-05 |
| 26747 | 1338 | G1338Q | E | 1.5e-05 | 1.2e-05 | 1.5e-05 | 1.400E-05 | 1.732E-06 | 1.200E-05 | 1.500E-05 |
| 26748 | 1338 | G1338G | E | 1.1e-05 | 1.1e-05 | 9.5e-06 | 1.050E-05 | 8.660E-07 | 9.500E-06 | 1.100E-05 |
| 26749 | 1338 | G1338H | E | 1.3e-05 | 1.3e-05 | 1.4e-05 | 1.333E-05 | 5.774E-07 | 1.300E-05 | 1.400E-05 |
| 26750 | 1338 | G1338I | E | 1.4e-05 | 1.1e-05 | 1.1e-05 | 1.200E-05 | 1.732E-06 | 1.100E-05 | 1.400E-05 |
| 26751 | 1338 | G1338L | E | 1.1e-05 | 1.5e-05 | 1.6e-05 | 1.400E-05 | 2.646E-06 | 1.100E-05 | 1.600E-05 |
| 26752 | 1338 | G1338K | E | 1e-05 | 1.2e-05 | 9.5e-06 | 1.050E-05 | 1.323E-06 | 9.500E-06 | 1.200E-05 |
| 26753 | 1338 | G1338M | E | 1.1e-05 | 1.7e-05 | 1.7e-05 | 1.500E-05 | 3.464E-06 | 1.100E-05 | 1.700E-05 |
| 26754 | 1338 | G1338F | E | 1.1e-05 | 1.5e-05 | 1.1e-05 | 1.233E-05 | 2.309E-06 | 1.100E-05 | 1.500E-05 |
| 26755 | 1338 | G1338P | E | 1e-05 | 1.5e-05 | 1.4e-05 | 1.300E-05 | 2.646E-06 | 1.000E-05 | 1.500E-05 |
| 26756 | 1338 | G1338S | E | 1.5e-05 | 1.5e-05 | 1.5e-05 | 1.500E-05 | 0.000E+00 | 1.500E-05 | 1.500E-05 |
| 26757 | 1338 | G1338T | E | 9.7e-06 | 1e-05 | 1.5e-05 | 1.157E-05 | 2.977E-06 | 9.700E-06 | 1.500E-05 |
| 26758 | 1338 | G1338W | E | 1.4e-05 | 1.5e-05 | 1.5e-05 | 1.467E-05 | 5.774E-07 | 1.400E-05 | 1.500E-05 |
| 26759 | 1338 | G1338Y | E | 1.7e-05 | 1.4e-05 | 1.3e-05 | 1.467E-05 | 2.082E-06 | 1.300E-05 | 1.700E-05 |
| 26760 | 1338 | G1338V | E | 1.1e-05 | 1.6e-05 | 1.5e-05 | 1.400E-05 | 2.646E-06 | 1.100E-05 | 1.600E-05 |
| 26761 | 1339 | R1339A | E | 1.5e-05 | 1.1e-05 | 1.5e-05 | 1.367E-05 | 2.309E-06 | 1.100E-05 | 1.500E-05 |
| 26762 | 1339 | R1339R | E | 5.6e-06 | 9.8e-06 | 1.6e-05 | 1.047E-05 | 5.232E-06 | 5.600E-06 | 1.600E-05 |
| 26763 | 1339 | R1339N | E | 1.4e-05 | 9.6e-06 | 9.7e-06 | 1.110E-05 | 2.512E-06 | 9.600E-06 | 1.400E-05 |
| 26764 | 1339 | R1339D | E | 1.4e-05 | 1.7e-05 | 1.6e-05 | 1.567E-05 | 1.528E-06 | 1.400E-05 | 1.700E-05 |
| 26765 | 1339 | R1339C | E | 1.1e-05 | 1.1e-05 | 8.9e-06 | 1.030E-05 | 1.212E-06 | 8.900E-06 | 1.100E-05 |
| 26766 | 1339 | R1339E | E | 1.6e-05 | 1.5e-05 | 1e-05 | 1.367E-05 | 3.215E-06 | 1.000E-05 | 1.600E-05 |
| 26767 | 1339 | R1339Q | E | 1.2e-05 | 1.6e-05 | 1.4e-05 | 1.400E-05 | 2.000E-06 | 1.200E-05 | 1.600E-05 |
| 26768 | 1339 | R1339G | E | 1.5e-05 | 1.5e-05 | 1.5e-05 | 1.500E-05 | 0.000E+00 | 1.500E-05 | 1.500E-05 |
| 26769 | 1339 | R1339H | E | 7.4e-06 | 1e-05 | 1.5e-05 | 1.080E-05 | 3.863E-06 | 7.400E-06 | 1.500E-05 |
| 26770 | 1339 | R1339I | E | 1e-05 | 1.8e-05 | 1.5e-05 | 1.433E-05 | 4.041E-06 | 1.000E-05 | 1.800E-05 |
| 26771 | 1339 | R1339L | E | 9.7e-06 | 1.1e-05 | 1.5e-05 | 1.190E-05 | 2.762E-06 | 9.700E-06 | 1.500E-05 |
| 26772 | 1339 | R1339K | E | 1e-05 | 1e-05 | 1.5e-05 | 1.167E-05 | 2.887E-06 | 1.000E-05 | 1.500E-05 |
| 26773 | 1339 | R1339M | E | 8.1e-06 | 1.4e-05 | 9.4e-06 | 1.050E-05 | 3.100E-06 | 8.100E-06 | 1.400E-05 |
| 26774 | 1339 | R1339F | E | 1.5e-05 | 9.5e-06 | 1e-05 | 1.150E-05 | 3.041E-06 | 9.500E-06 | 1.500E-05 |

| | | | | | | | | | | |
|---|---|---|---|---|---|---|---|---|---|---|
| 26775 | 1339 | R1339P | E | 1.6e-05 | 9.9e-06 | 1.1e-05 | 1.230E-05 | 3.251E-06 | 9.900E-06 | 1.600E-05 |
| 26776 | 1339 | R1339S | E | 1.6e-05 | 1e-05 | 1e-05 | 1.200E-05 | 3.464E-06 | 1.000E-05 | 1.600E-05 |
| 26777 | 1339 | R1339T | E | 1.4e-05 | 1.4e-05 | 1.1e-05 | 1.300E-05 | 1.732E-06 | 1.100E-05 | 1.400E-05 |
| 26778 | 1339 | R1339W | E | 1.5e-05 | 1.4e-05 | 1.3e-05 | 1.400E-05 | 1.000E-06 | 1.300E-05 | 1.500E-05 |
| 26779 | 1339 | R1339Y | E | 1.5e-05 | 1e-05 | 1e-05 | 1.167E-05 | 2.887E-06 | 1.000E-05 | 1.500E-05 |
| 26780 | 1339 | R1339V | E | 1.1e-05 | 1.5e-05 | 1.3e-05 | 1.300E-05 | 2.000E-06 | 1.100E-05 | 1.500E-05 |
| 26781 | 1340 | F1340A | E | 1.4e-05 | 8.7e-06 | 1.1e-05 | 1.123E-05 | 2.658E-06 | 8.700E-06 | 1.400E-05 |
| 26782 | 1340 | F1340R | E | 9.6e-06 | 1.1e-05 | 9.7e-06 | 1.010E-05 | 7.810E-07 | 9.600E-06 | 1.100E-05 |
| 26783 | 1340 | F1340N | E | 9.3e-06 | 1.4e-05 | 1e-05 | 1.110E-05 | 2.536E-06 | 9.300E-06 | 1.400E-05 |
| 26784 | 1340 | F1340D | E | 1.5e-05 | 1.6e-05 | 1.5e-05 | 1.533E-05 | 5.774E-07 | 1.500E-05 | 1.600E-05 |
| 26785 | 1340 | F1340C | E | 1.4e-05 | 1.4e-05 | 9.7e-06 | 1.257E-05 | 2.483E-06 | 9.700E-06 | 1.400E-05 |
| 26786 | 1340 | F1340E | E | 1e-05 | 1.6e-05 | 1.7e-05 | 1.433E-05 | 3.786E-06 | 1.000E-05 | 1.700E-05 |
| 26787 | 1340 | F1340Q | E | 9.4e-06 | 9.1e-06 | 8.9e-06 | 9.133E-06 | 2.517E-07 | 8.900E-06 | 9.400E-06 |
| 26788 | 1340 | F1340G | E | 1.3e-05 | 1.3e-05 | 1e-05 | 1.200E-05 | 1.732E-06 | 1.000E-05 | 1.300E-05 |
| 26789 | 1340 | F1340H | E | 1.5e-05 | 1.1e-05 | 1.5e-05 | 1.367E-05 | 2.309E-06 | 1.100E-05 | 1.500E-05 |
| 26790 | 1340 | F1340I | E | 7.8e-06 | 1.4e-05 | 1.1e-05 | 1.093E-05 | 3.101E-06 | 7.800E-06 | 1.400E-05 |
| 26791 | 1340 | F1340L | E | 9.4e-06 | 1.4e-05 | 1.6e-05 | 1.313E-05 | 3.384E-06 | 9.400E-06 | 1.600E-05 |
| 26792 | 1340 | F1340K | E | 1.3e-05 | 1.5e-05 | 1.5e-05 | 1.433E-05 | 1.155E-06 | 1.300E-05 | 1.500E-05 |
| 26793 | 1340 | F1340M | E | 1.2e-05 | 1.5e-05 | 1.6e-05 | 1.433E-05 | 2.082E-06 | 1.200E-05 | 1.600E-05 |
| 26794 | 1340 | F1340F | E | 1e-05 | 1.3e-05 | 1e-05 | 1.100E-05 | 1.732E-06 | 1.000E-05 | 1.300E-05 |
| 26795 | 1340 | F1340P | E | 1.6e-05 | 1.4e-05 | 1.3e-05 | 1.433E-05 | 1.528E-06 | 1.300E-05 | 1.600E-05 |
| 26796 | 1340 | F1340S | E | 1.7e-05 | 1.6e-05 | 1.4e-05 | 1.567E-05 | 1.528E-06 | 1.400E-05 | 1.700E-05 |
| 26797 | 1340 | F1340T | E | 1.3e-05 | 1.1e-05 | 9.9e-06 | 1.130E-05 | 1.572E-06 | 9.900E-06 | 1.300E-05 |
| 26798 | 1340 | F1340W | E | 1.6e-05 | 1.6e-05 | 1.5e-05 | 1.567E-05 | 5.774E-07 | 1.500E-05 | 1.600E-05 |
| 26799 | 1340 | F1340Y | E | 7.9e-06 | 1.6e-05 | 1.2e-05 | 1.197E-05 | 4.050E-06 | 7.900E-06 | 1.600E-05 |
| 26800 | 1340 | F1340V | E | 1.1e-05 | 1.2e-05 | 9.1e-06 | 1.070E-05 | 1.473E-06 | 9.100E-06 | 1.200E-05 |
| 26801 | 1341 | T1341A | E | 1.5e-05 | 1e-05 | 1.7e-05 | 1.400E-05 | 3.606E-06 | 1.000E-05 | 1.700E-05 |
| 26802 | 1341 | T1341R | E | 1.5e-05 | 1.4e-05 | 7.3e-06 | 1.210E-05 | 4.187E-06 | 7.300E-06 | 1.500E-05 |
| 26803 | 1341 | T1341N | E | 1.5e-05 | 1.4e-05 | 1.1e-05 | 1.333E-05 | 2.082E-06 | 1.100E-05 | 1.500E-05 |
| 26804 | 1341 | T1341D | E | 1e-05 | 1.6e-05 | 1.2e-05 | 1.267E-05 | 3.055E-06 | 1.000E-05 | 1.600E-05 |
| 26805 | 1341 | T1341C | E | 1.1e-05 | 1.7e-05 | 9.7e-06 | 1.257E-05 | 3.894E-06 | 9.700E-06 | 1.700E-05 |
| 26806 | 1341 | T1341E | E | 1.5e-05 | 1.4e-05 | 1.5e-05 | 1.467E-05 | 5.774E-07 | 1.400E-05 | 1.500E-05 |
| 26807 | 1341 | T1341Q | E | 1.1e-05 | 1.3e-05 | 1e-05 | 1.133E-05 | 1.528E-06 | 1.000E-05 | 1.300E-05 |
| 26808 | 1341 | T1341G | E | 1.6e-05 | 1.4e-05 | 1.4e-05 | 1.467E-05 | 1.155E-06 | 1.400E-05 | 1.600E-05 |
| 26809 | 1341 | T1341H | E | 1.2e-05 | 8e-06 | 1.3e-05 | 1.100E-05 | 2.646E-06 | 8.000E-06 | 1.300E-05 |
| 26810 | 1341 | T1341I | E | 9.4e-06 | 1.1e-05 | 1.2e-05 | 1.080E-05 | 1.311E-06 | 9.400E-06 | 1.200E-05 |
| 26811 | 1341 | T1341L | E | 1.1e-05 | 1.1e-05 | 1.6e-05 | 1.267E-05 | 2.887E-06 | 1.100E-05 | 1.600E-05 |
| 26812 | 1341 | T1341K | E | 9.2e-06 | 1.1e-05 | 9.8e-06 | 1.000E-05 | 9.165E-07 | 9.200E-06 | 1.100E-05 |
| 26813 | 1341 | T1341M | E | 1.1e-05 | 1.1e-05 | 1.5e-05 | 1.233E-05 | 2.309E-06 | 1.100E-05 | 1.500E-05 |
| 26814 | 1341 | T1341F | E | 1.7e-05 | 1.3e-05 | 1.8e-05 | 1.600E-05 | 2.646E-06 | 1.300E-05 | 1.800E-05 |
| 26815 | 1341 | T1341P | E | 1.8e-05 | 1.5e-05 | 1e-05 | 1.433E-05 | 4.041E-06 | 1.000E-05 | 1.800E-05 |
| 26816 | 1341 | T1341S | E | 1.6e-05 | 1.5e-05 | 9e-06 | 1.333E-05 | 3.786E-06 | 9.000E-06 | 1.600E-05 |
| 26817 | 1341 | T1341T | E | 1.7e-05 | 1e-05 | 1.1e-05 | 1.267E-05 | 3.786E-06 | 1.000E-05 | 1.700E-05 |
| 26818 | 1341 | T1341W | E | 1.4e-05 | 1.5e-05 | 1.2e-05 | 1.367E-05 | 1.528E-06 | 1.200E-05 | 1.500E-05 |
| 26819 | 1341 | T1341Y | E | 1.5e-05 | 1.4e-05 | 1.2e-05 | 1.367E-05 | 1.528E-06 | 1.200E-05 | 1.500E-05 |
| 26820 | 1341 | T1341V | E | 1.6e-05 | 1.6e-05 | 8.2e-06 | 1.340E-05 | 4.503E-06 | 8.200E-06 | 1.600E-05 |
| 26821 | 1342 | I1342A | E | 1e-05 | 8.7e-06 | 1.8e-05 | 1.223E-05 | 5.036E-06 | 8.700E-06 | 1.800E-05 |
| 26822 | 1342 | I1342R | E | 1.2e-05 | 1.3e-05 | 1.6e-05 | 1.367E-05 | 2.082E-06 | 1.200E-05 | 1.600E-05 |
| 26823 | 1342 | I1342N | E | 1.5e-05 | 1e-05 | 1.6e-05 | 1.367E-05 | 3.215E-06 | 1.000E-05 | 1.600E-05 |
| 26824 | 1342 | I1342D | E | 1.4e-05 | 1.5e-05 | 1.3e-05 | 1.400E-05 | 1.000E-06 | 1.300E-05 | 1.500E-05 |
| 26825 | 1342 | I1342C | E | 9.5e-06 | 1.4e-05 | 8.6e-06 | 1.070E-05 | 2.893E-06 | 8.600E-06 | 1.400E-05 |
| 26826 | 1342 | I1342E | E | 1.1e-05 | 1.3e-05 | 1.6e-05 | 1.333E-05 | 2.517E-06 | 1.100E-05 | 1.600E-05 |
| 26827 | 1342 | I1342Q | E | 1.4e-05 | 8.9e-06 | 1.1e-05 | 1.130E-05 | 2.563E-06 | 8.900E-06 | 1.400E-05 |
| 26828 | 1342 | I1342G | E | 1.6e-05 | 1.5e-05 | 1.3e-05 | 1.467E-05 | 1.528E-06 | 1.300E-05 | 1.600E-05 |
| 26829 | 1342 | I1342H | E | 1.5e-05 | 1.4e-05 | 1.4e-05 | 1.433E-05 | 5.774E-07 | 1.400E-05 | 1.500E-05 |
| 26830 | 1342 | I1342I | E | 1.7e-05 | 1.5e-05 | 1.5e-05 | 1.567E-05 | 1.155E-06 | 1.500E-05 | 1.700E-05 |
| 26831 | 1342 | I1342L | E | 1.1e-05 | 1e-05 | 1.1e-05 | 1.067E-05 | 5.774E-07 | 1.000E-05 | 1.100E-05 |
| 26832 | 1342 | I1342K | E | 1.6e-05 | 1.2e-05 | 1e-05 | 1.267E-05 | 3.055E-06 | 1.000E-05 | 1.600E-05 |
| 26833 | 1342 | I1342M | E | 1.6e-05 | 1.2e-05 | 8.5e-06 | 1.217E-05 | 3.753E-06 | 8.500E-06 | 1.600E-05 |
| 26834 | 1342 | I1342F | E | 1.7e-05 | 1.3e-05 | 1.5e-05 | 1.500E-05 | 2.000E-06 | 1.300E-05 | 1.700E-05 |
| 26835 | 1342 | I1342P | E | 1.2e-05 | 1.4e-05 | 1.5e-05 | 1.367E-05 | 1.528E-06 | 1.200E-05 | 1.500E-05 |
| 26836 | 1342 | I1342S | E | 9.3e-06 | 1.4e-05 | 1.2e-05 | 1.177E-05 | 2.359E-06 | 9.300E-06 | 1.400E-05 |
| 26837 | 1342 | I1342T | E | 1e-05 | 1.6e-05 | 1.1e-05 | 1.233E-05 | 3.215E-06 | 1.000E-05 | 1.600E-05 |
| 26838 | 1342 | I1342W | E | 8.6e-06 | 1.2e-05 | 1.6e-05 | 1.220E-05 | 3.704E-06 | 8.600E-06 | 1.600E-05 |
| 26839 | 1342 | I1342Y | E | 1.4e-05 | 9.3e-06 | 1.7e-05 | 1.343E-05 | 3.881E-06 | 9.300E-06 | 1.700E-05 |
| 26840 | 1342 | I1342V | E | 1.1e-05 | 1.3e-05 | 1.3e-05 | 1.233E-05 | 1.155E-06 | 1.100E-05 | 1.300E-05 |
| 26841 | 1343 | S1343A | E | 6.5e-06 | 9.3e-06 | 8.6e-06 | 8.133E-06 | 1.457E-06 | 6.500E-06 | 9.300E-06 |
| 26842 | 1343 | S1343R | E | 1.3e-05 | 1.5e-05 | 1.5e-05 | 1.433E-05 | 1.155E-06 | 1.300E-05 | 1.500E-05 |
| 26843 | 1343 | S1343N | E | 1.1e-05 | 1.2e-05 | 1.1e-05 | 1.133E-05 | 5.774E-07 | 1.100E-05 | 1.200E-05 |
| 26844 | 1343 | S1343D | E | 1.4e-05 | 1.6e-05 | 1e-05 | 1.333E-05 | 3.055E-06 | 1.000E-05 | 1.600E-05 |
| 26845 | 1343 | S1343C | E | 9.4e-06 | 1.3e-05 | 1e-05 | 1.080E-05 | 1.929E-06 | 9.400E-06 | 1.300E-05 |
| 26846 | 1343 | S1343E | E | 1.3e-05 | 1.4e-05 | 1e-05 | 1.233E-05 | 2.082E-06 | 1.000E-05 | 1.400E-05 |
| 26847 | 1343 | S1343Q | E | 9.8e-06 | 7.5e-06 | 1.5e-05 | 1.077E-05 | 3.842E-06 | 7.500E-06 | 1.500E-05 |
| 26848 | 1343 | S1343G | E | 1.5e-05 | 1.6e-05 | 9.1e-06 | 1.337E-05 | 3.729E-06 | 9.100E-06 | 1.600E-05 |
| 26849 | 1343 | S1343H | E | 1.1e-05 | 1.4e-05 | 1.4e-05 | 1.300E-05 | 1.732E-06 | 1.100E-05 | 1.400E-05 |
| 26850 | 1343 | S1343I | E | 1.2e-05 | 1.1e-05 | 1.2e-05 | 1.167E-05 | 5.774E-07 | 1.100E-05 | 1.200E-05 |
| 26851 | 1343 | S1343L | E | 1.8e-05 | 1.6e-05 | 1.7e-05 | 1.700E-05 | 1.000E-06 | 1.600E-05 | 1.800E-05 |
| 26852 | 1343 | S1343K | E | 1.6e-05 | 1.1e-05 | 1.1e-05 | 1.267E-05 | 2.887E-06 | 1.100E-05 | 1.600E-05 |
| 26853 | 1343 | S1343M | E | 1.4e-05 | 1.1e-05 | 1.4e-05 | 1.300E-05 | 1.732E-06 | 1.100E-05 | 1.400E-05 |
| 26854 | 1343 | S1343F | E | 1.5e-05 | 1.3e-05 | 8.3e-06 | 1.210E-05 | 3.439E-06 | 8.300E-06 | 1.500E-05 |
| 26855 | 1343 | S1343P | E | 1e-05 | 1.7e-05 | 1.1e-05 | 1.267E-05 | 3.786E-06 | 1.000E-05 | 1.700E-05 |
| 26856 | 1343 | S1343S | E | 1.4e-05 | 1.5e-05 | 1.4e-05 | 1.433E-05 | 5.774E-07 | 1.400E-05 | 1.500E-05 |
| 26857 | 1343 | S1343T | E | 1.4e-05 | 1e-05 | 9.6e-06 | 1.120E-05 | 2.433E-06 | 9.600E-06 | 1.400E-05 |
| 26858 | 1343 | S1343W | E | 9e-06 | 1e-05 | 1.6e-05 | 1.167E-05 | 3.786E-06 | 9.000E-06 | 1.600E-05 |
| 26859 | 1343 | S1343Y | E | 1.4e-05 | 1.5e-05 | 9e-06 | 1.267E-05 | 3.215E-06 | 9.000E-06 | 1.500E-05 |

| | | | | | | | | | | |
|---|---|---|---|---|---|---|---|---|---|---|
| 26860 | 1343 | S1343V | E | 1.6e-05 | 1.3e-05 | 1.6e-05 | 1.500E-05 | 1.732E-06 | 1.300E-05 | 1.600E-05 |
| 26861 | 1344 | A1344A | E | 8.7e-06 | 1.6e-05 | 1.3e-05 | 1.257E-05 | 3.669E-06 | 8.700E-06 | 1.600E-05 |
| 26862 | 1344 | A1344R | E | 1e-05 | 1e-05 | 1.5e-05 | 1.167E-05 | 2.887E-06 | 1.000E-05 | 1.500E-05 |
| 26863 | 1344 | A1344N | E | 1.3e-05 | 1.4e-05 | 1.1e-05 | 1.267E-05 | 1.528E-06 | 1.100E-05 | 1.400E-05 |
| 26864 | 1344 | A1344D | E | 1.3e-05 | 9.2e-06 | 1.5e-05 | 1.240E-05 | 2.946E-06 | 9.200E-06 | 1.500E-05 |
| 26865 | 1344 | A1344C | E | 1.2e-05 | 1.8e-05 | 1.5e-05 | 1.500E-05 | 3.000E-06 | 1.200E-05 | 1.800E-05 |
| 26866 | 1344 | A1344E | E | 1e-05 | 1.1e-05 | 1e-05 | 1.033E-05 | 5.774E-07 | 1.000E-05 | 1.100E-05 |
| 26867 | 1344 | A1344Q | E | 9.5e-06 | 8.7e-06 | 1.4e-05 | 1.073E-05 | 2.857E-06 | 8.700E-06 | 1.400E-05 |
| 26868 | 1344 | A1344G | E | 1.1e-05 | 1.8e-05 | 1.1e-05 | 1.333E-05 | 4.041E-06 | 1.100E-05 | 1.800E-05 |
| 26869 | 1344 | A1344H | E | 1.6e-05 | 1.4e-05 | 1.4e-05 | 1.467E-05 | 1.155E-06 | 1.400E-05 | 1.600E-05 |
| 26870 | 1344 | A1344I | E | 1.6e-05 | 1.4e-05 | 1.4e-05 | 1.467E-05 | 1.155E-06 | 1.400E-05 | 1.600E-05 |
| 26871 | 1344 | A1344L | E | 9.5e-06 | 1.6e-05 | 1.5e-05 | 1.350E-05 | 3.500E-06 | 9.500E-06 | 1.600E-05 |
| 26872 | 1344 | A1344K | E | 1e-05 | 1.4e-05 | 1.3e-05 | 1.233E-05 | 2.082E-06 | 1.000E-05 | 1.400E-05 |
| 26873 | 1344 | A1344M | E | 1.4e-05 | 1.5e-05 | 1.6e-05 | 1.500E-05 | 1.000E-06 | 1.400E-05 | 1.600E-05 |
| 26874 | 1344 | A1344F | E | 1e-05 | 1.5e-05 | 1.5e-05 | 1.333E-05 | 2.887E-06 | 1.000E-05 | 1.500E-05 |
| 26875 | 1344 | A1344P | E | 1.3e-05 | 1.1e-05 | 1.5e-05 | 1.300E-05 | 2.000E-06 | 1.100E-05 | 1.500E-05 |
| 26876 | 1344 | A1344S | E | 1.5e-05 | 9.3e-06 | 1.6e-05 | 1.343E-05 | 3.614E-06 | 9.300E-06 | 1.600E-05 |
| 26877 | 1344 | A1344T | E | 5e-06 | 9.6e-06 | 1.3e-05 | 9.200E-06 | 4.015E-06 | 5.000E-06 | 1.300E-05 |
| 26878 | 1344 | A1344W | E | 1.4e-05 | 8.9e-06 | 1.2e-05 | 1.163E-05 | 2.570E-06 | 8.900E-06 | 1.400E-05 |
| 26879 | 1344 | A1344Y | E | 8.9e-06 | 1.4e-05 | 1e-05 | 1.097E-05 | 2.684E-06 | 8.900E-06 | 1.400E-05 |
| 26880 | 1344 | A1344V | E | 1.5e-05 | 1.1e-05 | 1.4e-05 | 1.333E-05 | 2.082E-06 | 1.100E-05 | 1.500E-05 |
| 26881 | 1345 | D1345A | E | 1.3e-05 | 1.1e-05 | 1.6e-05 | 1.333E-05 | 2.517E-06 | 1.100E-05 | 1.600E-05 |
| 26882 | 1345 | D1345R | E | 7.9e-06 | 1.4e-05 | 1.1e-05 | 1.097E-05 | 3.050E-06 | 7.900E-06 | 1.400E-05 |
| 26883 | 1345 | D1345N | E | 1.5e-05 | 1.4e-05 | 1.5e-05 | 1.467E-05 | 5.774E-07 | 1.400E-05 | 1.500E-05 |
| 26884 | 1345 | D1345D | E | 1.5e-05 | 1.7e-05 | 1.1e-05 | 1.433E-05 | 3.055E-06 | 1.100E-05 | 1.700E-05 |
| 26885 | 1345 | D1345C | E | 1e-05 | 8.5e-06 | 1e-05 | 9.500E-06 | 8.660E-07 | 8.500E-06 | 1.000E-05 |
| 26886 | 1345 | D1345E | E | 1.1e-05 | 1.1e-05 | 1.6e-05 | 1.267E-05 | 2.887E-06 | 1.100E-05 | 1.600E-05 |
| 26887 | 1345 | D1345Q | E | 1.6e-05 | 1.2e-05 | 1e-05 | 1.267E-05 | 3.055E-06 | 1.000E-05 | 1.600E-05 |
| 26888 | 1345 | D1345G | E | 1.5e-05 | 1.1e-05 | 1e-05 | 1.200E-05 | 2.646E-06 | 1.000E-05 | 1.500E-05 |
| 26889 | 1345 | D1345H | E | 7e-06 | 1.5e-05 | 1.6e-05 | 1.267E-05 | 4.933E-06 | 7.000E-06 | 1.600E-05 |
| 26890 | 1345 | D1345I | E | 1.6e-05 | 1e-05 | 9e-06 | 1.167E-05 | 3.786E-06 | 9.000E-06 | 1.600E-05 |
| 26891 | 1345 | D1345L | E | 9.1e-06 | 1.1e-05 | 1.1e-05 | 1.037E-05 | 1.097E-06 | 9.100E-06 | 1.100E-05 |
| 26892 | 1345 | D1345K | E | 1.6e-05 | 1.7e-05 | 1.7e-05 | 1.667E-05 | 5.774E-07 | 1.600E-05 | 1.700E-05 |
| 26893 | 1345 | D1345M | E | 1.3e-05 | 8.3e-06 | 1.5e-05 | 1.210E-05 | 3.439E-06 | 8.300E-06 | 1.500E-05 |
| 26894 | 1345 | D1345F | E | 1.6e-05 | 9e-06 | 9.3e-06 | 1.143E-05 | 3.958E-06 | 9.000E-06 | 1.600E-05 |
| 26895 | 1345 | D1345P | E | 1.5e-05 | 1.5e-05 | 1.5e-05 | 1.500E-05 | 0.000E+00 | 1.500E-05 | 1.500E-05 |
| 26896 | 1345 | D1345S | E | 9.9e-06 | 1.4e-05 | 1e-05 | 1.130E-05 | 2.339E-06 | 9.900E-06 | 1.400E-05 |
| 26897 | 1345 | D1345T | E | 1.4e-05 | 8.4e-06 | 2.5e-05 | 1.580E-05 | 8.445E-06 | 8.400E-06 | 2.500E-05 |
| 26898 | 1345 | D1345W | E | 1.1e-05 | 1.4e-05 | 1.5e-05 | 1.333E-05 | 2.082E-06 | 1.100E-05 | 1.500E-05 |
| 26899 | 1345 | D1345Y | E | 1.3e-05 | 1e-05 | 1e-05 | 1.100E-05 | 1.732E-06 | 1.000E-05 | 1.300E-05 |
| 26900 | 1345 | D1345V | E | 6.4e-06 | 1e-05 | 1.4e-05 | 1.013E-05 | 3.802E-06 | 6.400E-06 | 1.400E-05 |
| 26901 | 1346 | T1346A | E | 1.5e-05 | 1.1e-05 | 8.7e-06 | 1.157E-05 | 3.188E-06 | 8.700E-06 | 1.500E-05 |
| 26902 | 1346 | T1346R | E | 1.2e-05 | 1.3e-05 | 9.4e-06 | 1.147E-05 | 1.858E-06 | 9.400E-06 | 1.300E-05 |
| 26903 | 1346 | T1346N | E | 9.9e-06 | 1.5e-05 | 1e-05 | 1.163E-05 | 2.916E-06 | 9.900E-06 | 1.500E-05 |
| 26904 | 1346 | T1346D | E | 9.9e-06 | 1.7e-05 | 1e-05 | 1.230E-05 | 4.071E-06 | 9.900E-06 | 1.700E-05 |
| 26905 | 1346 | T1346C | E | 9.2e-06 | 1.6e-05 | 1e-05 | 1.173E-05 | 3.717E-06 | 9.200E-06 | 1.600E-05 |
| 26906 | 1346 | T1346E | E | 1.6e-05 | 1.4e-05 | 1.4e-05 | 1.467E-05 | 1.155E-06 | 1.400E-05 | 1.600E-05 |
| 26907 | 1346 | T1346Q | E | 7.2e-06 | 9.6e-06 | 1.6e-05 | 1.093E-05 | 4.549E-06 | 7.200E-06 | 1.600E-05 |
| 26908 | 1346 | T1346G | E | 1.6e-05 | 1.7e-05 | 1.6e-05 | 1.633E-05 | 5.774E-07 | 1.600E-05 | 1.700E-05 |
| 26909 | 1346 | T1346H | E | 1.9e-05 | 1e-05 | 1.6e-05 | 1.500E-05 | 4.583E-06 | 1.000E-05 | 1.900E-05 |
| 26910 | 1346 | T1346I | E | 1e-05 | 1.1e-05 | 1.2e-05 | 1.100E-05 | 1.000E-06 | 1.000E-05 | 1.200E-05 |
| 26911 | 1346 | T1346L | E | 1.1e-05 | 1.1e-05 | 1.7e-05 | 1.300E-05 | 3.464E-06 | 1.100E-05 | 1.700E-05 |
| 26912 | 1346 | T1346K | E | 1.5e-05 | 1.5e-05 | 1.5e-05 | 1.500E-05 | 0.000E+00 | 1.500E-05 | 1.500E-05 |
| 26913 | 1346 | T1346M | E | 1.2e-05 | 1.1e-05 | 1.6e-05 | 1.300E-05 | 2.646E-06 | 1.100E-05 | 1.600E-05 |
| 26914 | 1346 | T1346F | E | 1.4e-05 | 1.5e-05 | 1.4e-05 | 1.433E-05 | 5.774E-07 | 1.400E-05 | 1.500E-05 |
| 26915 | 1346 | T1346P | E | 1.1e-05 | 8.7e-06 | 7.4e-06 | 9.033E-06 | 1.823E-06 | 7.400E-06 | 1.100E-05 |
| 26916 | 1346 | T1346S | E | 8.4e-06 | 1.5e-05 | 1.2e-05 | 1.180E-05 | 3.305E-06 | 8.400E-06 | 1.500E-05 |
| 26917 | 1346 | T1346T | E | 9.2e-06 | 1.5e-05 | 1.5e-05 | 1.307E-05 | 3.349E-06 | 9.200E-06 | 1.500E-05 |
| 26918 | 1346 | T1346W | E | 9.2e-06 | 1.2e-05 | 1.3e-05 | 1.140E-05 | 1.970E-06 | 9.200E-06 | 1.300E-05 |
| 26919 | 1346 | T1346Y | E | 8.3e-06 | 1e-05 | 1.1e-05 | 9.767E-06 | 1.365E-06 | 8.300E-06 | 1.100E-05 |
| 26920 | 1346 | T1346V | E | 1.4e-05 | 1.4e-05 | 1.5e-05 | 1.433E-05 | 5.774E-07 | 1.400E-05 | 1.500E-05 |
| 26921 | 1347 | S1347A | E | 1.1e-05 | 1.1e-05 | 1.8e-05 | 1.333E-05 | 4.041E-06 | 1.100E-05 | 1.800E-05 |
| 26922 | 1347 | S1347R | E | 1.5e-05 | 1.5e-05 | 1.1e-05 | 1.367E-05 | 2.309E-06 | 1.100E-05 | 1.500E-05 |
| 26923 | 1347 | S1347N | E | 8.2e-06 | 9.9e-06 | 1.2e-05 | 1.003E-05 | 1.904E-06 | 8.200E-06 | 1.200E-05 |
| 26924 | 1347 | S1347D | E | 1.1e-05 | 1.1e-05 | 1.5e-05 | 1.233E-05 | 2.309E-06 | 1.100E-05 | 1.500E-05 |
| 26925 | 1347 | S1347C | E | 9.5e-06 | 1.5e-05 | 1.1e-05 | 1.183E-05 | 2.843E-06 | 9.500E-06 | 1.500E-05 |
| 26926 | 1347 | S1347E | E | 1.4e-05 | 9.1e-06 | 1.5e-05 | 1.270E-05 | 3.158E-06 | 9.100E-06 | 1.500E-05 |
| 26927 | 1347 | S1347Q | E | 1.6e-05 | 9.9e-06 | 1.7e-05 | 1.430E-05 | 3.843E-06 | 9.900E-06 | 1.700E-05 |
| 26928 | 1347 | S1347G | E | 1.5e-05 | 1.6e-05 | 1.5e-05 | 1.533E-05 | 5.774E-07 | 1.500E-05 | 1.600E-05 |
| 26929 | 1347 | S1347H | E | 1.1e-05 | 1e-05 | 9.5e-06 | 1.017E-05 | 7.638E-07 | 9.500E-06 | 1.100E-05 |
| 26930 | 1347 | S1347I | E | 1.6e-05 | 7.7e-06 | 1.5e-05 | 1.290E-05 | 4.531E-06 | 7.700E-06 | 1.600E-05 |
| 26931 | 1347 | S1347L | E | 1.2e-05 | 9.9e-06 | 8.3e-06 | 1.007E-05 | 1.856E-06 | 8.300E-06 | 1.200E-05 |
| 26932 | 1347 | S1347K | E | 1.6e-05 | 1.1e-05 | 1.7e-05 | 1.467E-05 | 3.215E-06 | 1.100E-05 | 1.700E-05 |
| 26933 | 1347 | S1347M | E | 1.5e-05 | 1.6e-05 | 1.1e-05 | 1.400E-05 | 2.646E-06 | 1.100E-05 | 1.600E-05 |
| 26934 | 1347 | S1347F | E | 1.5e-05 | 1.7e-05 | 1.1e-05 | 1.433E-05 | 3.055E-06 | 1.100E-05 | 1.700E-05 |
| 26935 | 1347 | S1347P | E | 1.1e-05 | 1.2e-05 | 1.1e-05 | 1.133E-05 | 5.774E-07 | 1.100E-05 | 1.200E-05 |
| 26936 | 1347 | S1347S | E | 1.1e-05 | 1.4e-05 | 9.9e-06 | 1.163E-05 | 2.122E-06 | 9.900E-06 | 1.400E-05 |
| 26937 | 1347 | S1347T | E | 1.4e-05 | 1.5e-05 | 9.1e-06 | 1.270E-05 | 3.158E-06 | 9.100E-06 | 1.500E-05 |
| 26938 | 1347 | S1347W | E | 1.6e-05 | 1.4e-05 | 9.2e-06 | 1.307E-05 | 3.495E-06 | 9.200E-06 | 1.600E-05 |
| 26939 | 1347 | S1347Y | E | 9.2e-06 | 9.4e-06 | 1.5e-05 | 1.120E-05 | 3.292E-06 | 9.200E-06 | 1.500E-05 |
| 26940 | 1347 | S1347V | E | 1e-05 | 9.3e-06 | 1e-05 | 9.767E-06 | 4.041E-07 | 9.300E-06 | 1.000E-05 |
| 26941 | 1348 | K1348A | E | 1.6e-05 | 1.7e-05 | 1.1e-05 | 1.467E-05 | 3.215E-06 | 1.100E-05 | 1.700E-05 |
| 26942 | 1348 | K1348R | E | 1.1e-05 | 1e-05 | 1.5e-05 | 1.200E-05 | 2.646E-06 | 1.000E-05 | 1.500E-05 |
| 26943 | 1348 | K1348N | E | 1.6e-05 | 9.8e-06 | 1.4e-05 | 1.327E-05 | 3.164E-06 | 9.800E-06 | 1.600E-05 |
| 26944 | 1348 | K1348D | E | 1.1e-05 | 1.4e-05 | 8.1e-06 | 1.103E-05 | 2.950E-06 | 8.100E-06 | 1.400E-05 |

| | | | | | | | | | | |
|---|---|---|---|---|---|---|---|---|---|---|
| 26945 | 1348 | K1348C | E | 1.3e-05 | 1.4e-05 | 1e-05 | 1.233E-05 | 2.082E-06 | 1.000E-05 | 1.400E-05 |
| 26946 | 1348 | K1348E | E | 1.1e-05 | 1.3e-05 | 1.5e-05 | 1.300E-05 | 2.000E-06 | 1.100E-05 | 1.500E-05 |
| 26947 | 1348 | K1348Q | E | 1.6e-05 | 1.4e-05 | 9.7e-06 | 1.323E-05 | 3.219E-06 | 9.700E-06 | 1.600E-05 |
| 26948 | 1348 | K1348G | E | 1.3e-05 | 8.7e-06 | 9.5e-06 | 1.040E-05 | 2.287E-06 | 8.700E-06 | 1.300E-05 |
| 26949 | 1348 | K1348H | E | 1.4e-05 | 1.5e-05 | 1.1e-05 | 1.333E-05 | 2.082E-06 | 1.100E-05 | 1.500E-05 |
| 26950 | 1348 | K1348I | E | 1.1e-05 | 1.5e-05 | 1.5e-05 | 1.367E-05 | 2.309E-06 | 1.100E-05 | 1.500E-05 |
| 26951 | 1348 | K1348L | E | 1e-05 | 1.5e-05 | 1e-05 | 1.167E-05 | 2.887E-06 | 1.000E-05 | 1.500E-05 |
| 26952 | 1348 | K1348K | E | 1.5e-05 | 1e-05 | 8.7e-06 | 1.123E-05 | 3.326E-06 | 8.700E-06 | 1.500E-05 |
| 26953 | 1348 | K1348M | E | 1.2e-05 | 9.4e-06 | 1.7e-05 | 1.280E-05 | 3.863E-06 | 9.400E-06 | 1.700E-05 |
| 26954 | 1348 | K1348F | E | 1.6e-05 | 1.4e-05 | 1.6e-05 | 1.533E-05 | 1.155E-06 | 1.400E-05 | 1.600E-05 |
| 26955 | 1348 | K1348P | E | 1e-05 | 9.4e-06 | 1.5e-05 | 1.147E-05 | 3.075E-06 | 9.400E-06 | 1.500E-05 |
| 26956 | 1348 | K1348S | E | 1.6e-05 | 1.1e-05 | 9.9e-06 | 1.230E-05 | 3.251E-06 | 9.900E-06 | 1.600E-05 |
| 26957 | 1348 | K1348T | E | 1.6e-05 | 1.5e-05 | 1.6e-05 | 1.567E-05 | 5.774E-07 | 1.500E-05 | 1.600E-05 |
| 26958 | 1348 | K1348W | E | 8.5e-06 | 1.5e-05 | 1.1e-05 | 1.150E-05 | 3.279E-06 | 8.500E-06 | 1.500E-05 |
| 26959 | 1348 | K1348Y | E | 1e-05 | 1.5e-05 | 1.6e-05 | 1.367E-05 | 3.215E-06 | 1.000E-05 | 1.600E-05 |
| 26960 | 1348 | K1348V | E | 1.1e-05 | 1.6e-05 | 1.1e-05 | 1.267E-05 | 2.887E-06 | 1.100E-05 | 1.600E-05 |
| 26961 | 1349 | N1349A | E | 1.1e-05 | 1.5e-05 | 1.7e-05 | 1.433E-05 | 3.055E-06 | 1.100E-05 | 1.700E-05 |
| 26962 | 1349 | N1349R | E | 1.7e-05 | 1.4e-05 | 1.2e-05 | 1.433E-05 | 2.517E-06 | 1.200E-05 | 1.700E-05 |
| 26963 | 1349 | N1349N | E | 1.2e-05 | 1.7e-05 | 9.9e-06 | 1.297E-05 | 3.647E-06 | 9.900E-06 | 1.700E-05 |
| 26964 | 1349 | N1349D | E | 8e-06 | 1.1e-05 | 1.1e-05 | 1.000E-05 | 1.732E-06 | 8.000E-06 | 1.100E-05 |
| 26965 | 1349 | N1349C | E | 1.5e-05 | 1.1e-05 | 1.5e-05 | 1.367E-05 | 2.309E-06 | 1.100E-05 | 1.500E-05 |
| 26966 | 1349 | N1349E | E | 1e-05 | 1.1e-05 | 1.6e-05 | 1.233E-05 | 3.215E-06 | 1.000E-05 | 1.600E-05 |
| 26967 | 1349 | N1349Q | E | 1.4e-05 | 1e-05 | 2.3e-05 | 1.567E-05 | 6.658E-06 | 1.000E-05 | 2.300E-05 |
| 26968 | 1349 | N1349G | E | 1.5e-05 | 1.5e-05 | 1.5e-05 | 1.500E-05 | 0.000E+00 | 1.500E-05 | 1.500E-05 |
| 26969 | 1349 | N1349H | E | 1.2e-05 | 1.7e-05 | 9.5e-06 | 1.283E-05 | 3.819E-06 | 9.500E-06 | 1.700E-05 |
| 26970 | 1349 | N1349I | E | 8.8e-06 | 1.6e-05 | 1.7e-05 | 1.393E-05 | 4.474E-06 | 8.800E-06 | 1.700E-05 |
| 26971 | 1349 | N1349L | E | 1.3e-05 | 1.4e-05 | 1.1e-05 | 1.267E-05 | 1.528E-06 | 1.100E-05 | 1.400E-05 |
| 26972 | 1349 | N1349K | E | 1.6e-05 | 1.3e-05 | 1.1e-05 | 1.333E-05 | 2.517E-06 | 1.100E-05 | 1.600E-05 |
| 26973 | 1349 | N1349M | E | 1.1e-05 | 8.8e-06 | 9.8e-06 | 9.867E-06 | 1.102E-06 | 8.800E-06 | 1.100E-05 |
| 26974 | 1349 | N1349F | E | 1.7e-05 | 9.8e-06 | 7.4e-06 | 1.140E-05 | 4.996E-06 | 7.400E-06 | 1.700E-05 |
| 26975 | 1349 | N1349P | E | 1.6e-05 | 9.4e-06 | 1.1e-05 | 1.213E-05 | 3.443E-06 | 9.400E-06 | 1.600E-05 |
| 26976 | 1349 | N1349S | E | 1.4e-05 | 9.9e-06 | 1.6e-05 | 1.330E-05 | 3.110E-06 | 9.900E-06 | 1.600E-05 |
| 26977 | 1349 | N1349T | E | 1.4e-05 | 1.7e-05 | 1.6e-05 | 1.567E-05 | 1.528E-06 | 1.400E-05 | 1.700E-05 |
| 26978 | 1349 | N1349W | E | 1.4e-05 | 1.4e-05 | 1.1e-05 | 1.300E-05 | 1.732E-06 | 1.100E-05 | 1.400E-05 |
| 26979 | 1349 | N1349Y | E | 1.6e-05 | 1.7e-05 | 1.4e-05 | 1.567E-05 | 1.528E-06 | 1.400E-05 | 1.700E-05 |
| 26980 | 1349 | N1349V | E | 1.6e-05 | 1.3e-05 | 1.4e-05 | 1.433E-05 | 1.528E-06 | 1.300E-05 | 1.600E-05 |
| 26981 | 1350 | T1350A | E | 1.7e-05 | 1.2e-05 | 1.1e-05 | 1.333E-05 | 3.215E-06 | 1.100E-05 | 1.700E-05 |
| 26982 | 1350 | T1350R | E | 7.5e-06 | 1.3e-05 | 9.6e-06 | 1.003E-05 | 2.775E-06 | 7.500E-06 | 1.300E-05 |
| 26983 | 1350 | T1350N | E | 8.8e-06 | 1.9e-05 | 5.7e-06 | 1.117E-05 | 6.959E-06 | 5.700E-06 | 1.900E-05 |
| 26984 | 1350 | T1350D | E | 8.7e-06 | 1.1e-05 | 8.9e-06 | 9.533E-06 | 1.274E-06 | 8.700E-06 | 1.100E-05 |
| 26985 | 1350 | T1350C | E | 1.4e-05 | 1.6e-05 | 1.5e-05 | 1.500E-05 | 1.000E-06 | 1.400E-05 | 1.600E-05 |
| 26986 | 1350 | T1350E | E | 1.6e-05 | 1e-05 | 7.4e-06 | 1.113E-05 | 4.411E-06 | 7.400E-06 | 1.600E-05 |
| 26987 | 1350 | T1350Q | E | 1.7e-05 | 1.1e-05 | 1.6e-05 | 1.467E-05 | 3.215E-06 | 1.100E-05 | 1.700E-05 |
| 26988 | 1350 | T1350G | E | 8.8e-06 | 1.7e-05 | 8.7e-06 | 1.150E-05 | 4.763E-06 | 8.700E-06 | 1.700E-05 |
| 26989 | 1350 | T1350H | E | 1.1e-05 | 1.5e-05 | 1.3e-05 | 1.300E-05 | 2.000E-06 | 1.100E-05 | 1.500E-05 |
| 26990 | 1350 | T1350I | E | 8.6e-06 | 9.1e-06 | 1.5e-05 | 1.090E-05 | 3.559E-06 | 8.600E-06 | 1.500E-05 |
| 26991 | 1350 | T1350L | E | 1.5e-05 | 1e-05 | 1.2e-05 | 1.233E-05 | 2.517E-06 | 1.000E-05 | 1.500E-05 |
| 26992 | 1350 | T1350K | E | 1e-05 | 1.2e-05 | 1e-05 | 1.067E-05 | 1.155E-06 | 1.000E-05 | 1.200E-05 |
| 26993 | 1350 | T1350M | E | 1.4e-05 | 1.1e-05 | 1.7e-05 | 1.400E-05 | 3.000E-06 | 1.100E-05 | 1.700E-05 |
| 26994 | 1350 | T1350F | E | 1.2e-05 | 1.1e-05 | 1.1e-05 | 1.133E-05 | 5.774E-07 | 1.100E-05 | 1.200E-05 |
| 26995 | 1350 | T1350P | E | 1.5e-05 | 1.3e-05 | 1.3e-05 | 1.367E-05 | 1.155E-06 | 1.300E-05 | 1.500E-05 |
| 26996 | 1350 | T1350S | E | 1e-05 | 1.2e-05 | 9.4e-06 | 1.047E-05 | 1.361E-06 | 9.400E-06 | 1.200E-05 |
| 26997 | 1350 | T1350T | E | 1.6e-05 | 7.2e-06 | 1.6e-05 | 1.307E-05 | 5.081E-06 | 7.200E-06 | 1.600E-05 |
| 26998 | 1350 | T1350W | E | 1.5e-05 | 1.7e-05 | 9.5e-06 | 1.383E-05 | 3.884E-06 | 9.500E-06 | 1.700E-05 |
| 26999 | 1350 | T1350Y | E | 1e-05 | 1.7e-05 | 1.5e-05 | 1.400E-05 | 3.606E-06 | 1.000E-05 | 1.700E-05 |
| 27000 | 1350 | T1350V | E | 1.6e-05 | 1.6e-05 | 4.2e-05 | 2.467E-05 | 1.501E-05 | 1.600E-05 | 4.200E-05 |
| 27001 | 1351 | A1351A | E | 1e-05 | 1.1e-05 | 9.1e-06 | 1.003E-05 | 9.504E-07 | 9.100E-06 | 1.100E-05 |
| 27002 | 1351 | A1351R | E | 9.5e-06 | 1.1e-05 | 1.1e-05 | 1.050E-05 | 8.660E-07 | 9.500E-06 | 1.100E-05 |
| 27003 | 1351 | A1351N | E | 1.1e-05 | 1e-05 | 1.6e-05 | 1.233E-05 | 3.215E-06 | 1.000E-05 | 1.600E-05 |
| 27004 | 1351 | A1351D | E | 1.6e-05 | 1.1e-05 | 1.5e-05 | 1.400E-05 | 2.646E-06 | 1.100E-05 | 1.600E-05 |
| 27005 | 1351 | A1351C | E | 1.4e-05 | 1.4e-05 | 1.4e-05 | 1.400E-05 | 0.000E+00 | 1.400E-05 | 1.400E-05 |
| 27006 | 1351 | A1351E | E | 1.5e-05 | 1.4e-05 | 1.6e-05 | 1.500E-05 | 1.000E-06 | 1.400E-05 | 1.600E-05 |
| 27007 | 1351 | A1351Q | E | 1.6e-05 | 1.2e-05 | 1.4e-05 | 1.400E-05 | 2.000E-06 | 1.200E-05 | 1.600E-05 |
| 27008 | 1351 | A1351G | E | 9.8e-06 | 1.5e-05 | 1.7e-05 | 1.393E-05 | 3.717E-06 | 9.800E-06 | 1.700E-05 |
| 27009 | 1351 | A1351H | E | 9.7e-06 | 1.1e-05 | 1.3e-05 | 1.123E-05 | 1.662E-06 | 9.700E-06 | 1.300E-05 |
| 27010 | 1351 | A1351I | E | 9.7e-06 | 1.5e-05 | 1.5e-05 | 1.323E-05 | 3.060E-06 | 9.700E-06 | 1.500E-05 |
| 27011 | 1351 | A1351L | E | 9.3e-06 | 1e-05 | 1.3e-05 | 1.077E-05 | 1.966E-06 | 9.300E-06 | 1.300E-05 |
| 27012 | 1351 | A1351K | E | 1.4e-05 | 1e-05 | 1.1e-05 | 1.167E-05 | 2.082E-06 | 1.000E-05 | 1.400E-05 |
| 27013 | 1351 | A1351M | E | 1.7e-05 | 1.6e-05 | 1e-05 | 1.433E-05 | 3.786E-06 | 1.000E-05 | 1.700E-05 |
| 27014 | 1351 | A1351F | E | 1.7e-05 | 1.6e-05 | 1.2e-05 | 1.500E-05 | 2.646E-06 | 1.200E-05 | 1.700E-05 |
| 27015 | 1351 | A1351P | E | 1.5e-05 | 1.1e-05 | 1.1e-05 | 1.233E-05 | 2.309E-06 | 1.100E-05 | 1.500E-05 |
| 27016 | 1351 | A1351S | E | 8.8e-06 | 1.2e-05 | 1.1e-05 | 1.060E-05 | 1.637E-06 | 8.800E-06 | 1.200E-05 |
| 27017 | 1351 | A1351T | E | 1.1e-05 | 9.3e-06 | 1.5e-05 | 1.177E-05 | 2.926E-06 | 9.300E-06 | 1.500E-05 |
| 27018 | 1351 | A1351W | E | 9e-06 | 1e-05 | 8.5e-06 | 9.167E-06 | 7.638E-07 | 8.500E-06 | 1.000E-05 |
| 27019 | 1351 | A1351Y | E | 1.5e-05 | 1.2e-05 | 1.1e-05 | 1.267E-05 | 2.082E-06 | 1.100E-05 | 1.500E-05 |
| 27020 | 1351 | A1351V | E | 1.4e-05 | 1.5e-05 | 1.2e-05 | 1.367E-05 | 1.528E-06 | 1.200E-05 | 1.500E-05 |
| 27021 | 1352 | Y1352A | E | 1.1e-05 | 1.7e-05 | 9.9e-06 | 1.263E-05 | 3.821E-06 | 9.900E-06 | 1.700E-05 |
| 27022 | 1352 | Y1352R | E | 1.7e-05 | 1.7e-05 | 1e-05 | 1.467E-05 | 4.041E-06 | 1.000E-05 | 1.700E-05 |
| 27023 | 1352 | Y1352N | E | 1.5e-05 | 1.5e-05 | 1.3e-05 | 1.433E-05 | 1.155E-06 | 1.300E-05 | 1.500E-05 |
| 27024 | 1352 | Y1352D | E | 1.6e-05 | 1.4e-05 | 1.1e-05 | 1.367E-05 | 2.517E-06 | 1.100E-05 | 1.600E-05 |
| 27025 | 1352 | Y1352C | E | 1.6e-05 | 1.4e-05 | 1.5e-05 | 1.500E-05 | 1.000E-06 | 1.400E-05 | 1.600E-05 |
| 27026 | 1352 | Y1352E | E | 1e-05 | 1.6e-05 | 9e-06 | 1.167E-05 | 3.786E-06 | 9.000E-06 | 1.600E-05 |
| 27027 | 1352 | Y1352Q | E | 1.5e-05 | 1.2e-05 | 1.1e-05 | 1.267E-05 | 2.082E-06 | 1.100E-05 | 1.500E-05 |
| 27028 | 1352 | Y1352G | E | 1.1e-05 | 2.7e-05 | 1.3e-05 | 1.700E-05 | 8.718E-06 | 1.100E-05 | 2.700E-05 |
| 27029 | 1352 | Y1352H | E | 1.2e-05 | 1.3e-05 | 1.5e-05 | 1.333E-05 | 1.528E-06 | 1.200E-05 | 1.500E-05 |

| | | | | | | | | | | |
|---|---|---|---|---|---|---|---|---|---|---|
| 27030 | 1352 | Y1352I | E | 1.4e-05 | 1.6e-05 | 1.6e-05 | 1.533E-05 | 1.155E-06 | 1.400E-05 | 1.600E-05 |
| 27031 | 1352 | Y1352L | E | 1.6e-05 | 1.1e-05 | 9.7e-06 | 1.223E-05 | 3.326E-06 | 9.700E-06 | 1.600E-05 |
| 27032 | 1352 | Y1352K | E | 1.6e-05 | 1.3e-05 | 1.6e-05 | 1.500E-05 | 1.732E-06 | 1.300E-05 | 1.600E-05 |
| 27033 | 1352 | Y1352M | E | 9.2e-06 | 1e-05 | 9.6e-06 | 9.600E-06 | 4.000E-07 | 9.200E-06 | 1.000E-05 |
| 27034 | 1352 | Y1352F | E | 1.4e-05 | 1.1e-05 | 1.3e-05 | 1.267E-05 | 1.528E-06 | 1.100E-05 | 1.400E-05 |
| 27035 | 1352 | Y1352P | E | 1.2e-05 | 1e-05 | 1.5e-05 | 1.233E-05 | 2.517E-06 | 1.000E-05 | 1.500E-05 |
| 27036 | 1352 | Y1352S | E | 1.1e-05 | 1.5e-05 | 1.6e-05 | 1.400E-05 | 2.646E-06 | 1.100E-05 | 1.600E-05 |
| 27037 | 1352 | Y1352T | E | 1e-05 | 1.6e-05 | 1e-05 | 1.200E-05 | 3.464E-06 | 1.000E-05 | 1.600E-05 |
| 27038 | 1352 | Y1352W | E | 1.4e-05 | 1.2e-05 | 1.7e-05 | 1.433E-05 | 2.517E-06 | 1.200E-05 | 1.700E-05 |
| 27039 | 1352 | Y1352Y | E | 1.6e-05 | 1.4e-05 | 1.5e-05 | 1.500E-05 | 1.000E-06 | 1.400E-05 | 1.600E-05 |
| 27040 | 1352 | Y1352V | E | 1.6e-05 | 1.6e-05 | 1.5e-05 | 1.567E-05 | 5.774E-07 | 1.500E-05 | 1.600E-05 |
| 27041 | 1353 | L1353A | E | 9.1e-06 | 1e-05 | 1e-05 | 9.700E-06 | 5.196E-07 | 9.100E-06 | 1.000E-05 |
| 27042 | 1353 | L1353R | E | 1.4e-05 | 1.4e-05 | 1.4e-05 | 1.400E-05 | 0.000E+00 | 1.400E-05 | 1.400E-05 |
| 27043 | 1353 | L1353N | E | 1.1e-05 | 1.7e-05 | 1.5e-05 | 1.433E-05 | 3.055E-06 | 1.100E-05 | 1.700E-05 |
| 27044 | 1353 | L1353D | E | 1.6e-05 | 8.8e-06 | 1.5e-05 | 1.327E-05 | 3.900E-06 | 8.800E-06 | 1.600E-05 |
| 27045 | 1353 | L1353C | E | 1e-05 | 1.7e-05 | 9.6e-06 | 1.220E-05 | 4.162E-06 | 9.600E-06 | 1.700E-05 |
| 27046 | 1353 | L1353E | E | 1.1e-05 | 1.1e-05 | 1.4e-05 | 1.200E-05 | 1.732E-06 | 1.100E-05 | 1.400E-05 |
| 27047 | 1353 | L1353Q | E | 1.3e-05 | 9.3e-06 | 1.3e-05 | 1.177E-05 | 2.136E-06 | 9.300E-06 | 1.300E-05 |
| 27048 | 1353 | L1353G | E | 1.4e-05 | 1.5e-05 | 1.6e-05 | 1.500E-05 | 1.000E-06 | 1.400E-05 | 1.600E-05 |
| 27049 | 1353 | L1353H | E | 1.2e-05 | 1.1e-05 | 1.6e-05 | 1.300E-05 | 2.646E-06 | 1.100E-05 | 1.600E-05 |
| 27050 | 1353 | L1353I | E | 9.6e-06 | 8.9e-06 | 1.8e-05 | 1.217E-05 | 5.064E-06 | 8.900E-06 | 1.800E-05 |
| 27051 | 1353 | L1353L | E | 1.2e-05 | 1.5e-05 | 1.1e-05 | 1.267E-05 | 2.082E-06 | 1.100E-05 | 1.500E-05 |
| 27052 | 1353 | L1353K | E | 5.9e-06 | 1.2e-05 | 9.7e-06 | 9.200E-06 | 3.081E-06 | 5.900E-06 | 1.200E-05 |
| 27053 | 1353 | L1353M | E | 1.7e-05 | 1.4e-05 | 1.3e-05 | 1.467E-05 | 2.082E-06 | 1.300E-05 | 1.700E-05 |
| 27054 | 1353 | L1353F | E | 1.2e-05 | 1.5e-05 | 1.2e-05 | 1.300E-05 | 1.732E-06 | 1.200E-05 | 1.500E-05 |
| 27055 | 1353 | L1353P | E | 9e-06 | 1.1e-05 | 8e-06 | 9.333E-06 | 1.528E-06 | 8.000E-06 | 1.100E-05 |
| 27056 | 1353 | L1353S | E | 1.5e-05 | 8.8e-06 | 8.8e-06 | 1.087E-05 | 3.580E-06 | 8.800E-06 | 1.500E-05 |
| 27057 | 1353 | L1353T | E | 1.6e-05 | 1.5e-05 | 2.1e-05 | 1.733E-05 | 3.215E-06 | 1.500E-05 | 2.100E-05 |
| 27058 | 1353 | L1353W | E | 1.1e-05 | 1.3e-05 | 9e-06 | 1.100E-05 | 2.000E-06 | 9.000E-06 | 1.300E-05 |
| 27059 | 1353 | L1353Y | E | 1.4e-05 | 1.4e-05 | 9.6e-06 | 1.253E-05 | 2.540E-06 | 9.600E-06 | 1.400E-05 |
| 27060 | 1353 | L1353V | E | 1.6e-05 | 1.5e-05 | 1.5e-05 | 1.533E-05 | 5.774E-07 | 1.500E-05 | 1.600E-05 |
| 27061 | 1354 | Q1354A | E | 1.7e-05 | 1.7e-05 | 1.3e-05 | 1.567E-05 | 2.309E-06 | 1.300E-05 | 1.700E-05 |
| 27062 | 1354 | Q1354R | E | 9.9e-06 | 9.3e-06 | 1.5e-05 | 1.140E-05 | 3.132E-06 | 9.300E-06 | 1.500E-05 |
| 27063 | 1354 | Q1354N | E | 1.5e-05 | 1e-05 | 1.1e-05 | 1.200E-05 | 2.646E-06 | 1.000E-05 | 1.500E-05 |
| 27064 | 1354 | Q1354D | E | 1.5e-05 | 1.5e-05 | 1e-05 | 1.333E-05 | 2.887E-06 | 1.000E-05 | 1.500E-05 |
| 27065 | 1354 | Q1354C | E | 2.2e-05 | 1.4e-05 | 9.6e-06 | 1.520E-05 | 6.286E-06 | 9.600E-06 | 2.200E-05 |
| 27066 | 1354 | Q1354E | E | 1.7e-05 | 1e-05 | 1.4e-05 | 1.367E-05 | 3.512E-06 | 1.000E-05 | 1.700E-05 |
| 27067 | 1354 | Q1354Q | E | 8.8e-06 | 1.3e-05 | 9.8e-06 | 1.053E-05 | 2.194E-06 | 8.800E-06 | 1.300E-05 |
| 27068 | 1354 | Q1354G | E | 1.7e-05 | 1.5e-05 | 1.4e-05 | 1.533E-05 | 1.528E-06 | 1.400E-05 | 1.700E-05 |
| 27069 | 1354 | Q1354H | E | 1.1e-05 | 1.1e-05 | 1.1e-05 | 1.100E-05 | 2.075E-21 | 1.100E-05 | 1.100E-05 |
| 27070 | 1354 | Q1354I | E | 1.7e-05 | 9.8e-06 | 1.1e-05 | 1.260E-05 | 3.857E-06 | 9.800E-06 | 1.700E-05 |
| 27071 | 1354 | Q1354L | E | 1e-05 | 1e-05 | 1.5e-05 | 1.167E-05 | 2.887E-06 | 1.000E-05 | 1.500E-05 |
| 27072 | 1354 | Q1354K | E | 1.2e-05 | 1.5e-05 | 1e-05 | 1.233E-05 | 2.517E-06 | 1.000E-05 | 1.500E-05 |
| 27073 | 1354 | Q1354M | E | 1.6e-05 | 1.5e-05 | 1.6e-05 | 1.567E-05 | 5.774E-07 | 1.500E-05 | 1.600E-05 |
| 27074 | 1354 | Q1354F | E | 1.2e-05 | 1.2e-05 | 9.3e-06 | 1.110E-05 | 1.559E-06 | 9.300E-06 | 1.200E-05 |
| 27075 | 1354 | Q1354P | E | 1.1e-05 | 1.2e-05 | 1.1e-05 | 1.133E-05 | 5.774E-07 | 1.100E-05 | 1.200E-05 |
| 27076 | 1354 | Q1354S | E | 1.1e-05 | 1e-05 | 1.5e-05 | 1.200E-05 | 2.646E-06 | 1.000E-05 | 1.500E-05 |
| 27077 | 1354 | Q1354T | E | 1.1e-05 | 1.4e-05 | 1.4e-05 | 1.300E-05 | 1.732E-06 | 1.100E-05 | 1.400E-05 |
| 27078 | 1354 | Q1354W | E | 1.5e-05 | 1.4e-05 | 1e-05 | 1.300E-05 | 2.646E-06 | 1.000E-05 | 1.500E-05 |
| 27079 | 1354 | Q1354Y | E | 1.3e-05 | 8.7e-06 | 1.4e-05 | 1.190E-05 | 2.816E-06 | 8.700E-06 | 1.400E-05 |
| 27080 | 1354 | Q1354V | E | 1.2e-05 | 1.5e-05 | 1.4e-05 | 1.367E-05 | 1.528E-06 | 1.200E-05 | 1.500E-05 |
| 27081 | 1355 | M1355A | E | 1.5e-05 | 1.1e-05 | 1.3e-05 | 1.300E-05 | 2.000E-06 | 1.100E-05 | 1.500E-05 |
| 27082 | 1355 | M1355R | E | 1.3e-05 | 1.2e-05 | 1.6e-05 | 1.367E-05 | 2.082E-06 | 1.200E-05 | 1.600E-05 |
| 27083 | 1355 | M1355N | E | 1.2e-05 | 1.1e-05 | 1.4e-05 | 1.233E-05 | 1.528E-06 | 1.100E-05 | 1.400E-05 |
| 27084 | 1355 | M1355D | E | 1.4e-05 | 1.2e-05 | 1.5e-05 | 1.367E-05 | 1.528E-06 | 1.200E-05 | 1.500E-05 |
| 27085 | 1355 | M1355C | E | 1.3e-05 | 9.3e-06 | 1.5e-05 | 1.243E-05 | 2.892E-06 | 9.300E-06 | 1.500E-05 |
| 27086 | 1355 | M1355E | E | 1.8e-05 | 1.7e-05 | 9.8e-06 | 1.493E-05 | 4.474E-06 | 9.800E-06 | 1.800E-05 |
| 27087 | 1355 | M1355Q | E | 1.4e-05 | 1.4e-05 | 9.5e-06 | 1.250E-05 | 2.598E-06 | 9.500E-06 | 1.400E-05 |
| 27088 | 1355 | M1355G | E | 1.4e-05 | 1.2e-05 | 1.6e-05 | 1.400E-05 | 2.000E-06 | 1.200E-05 | 1.600E-05 |
| 27089 | 1355 | M1355H | E | 1.1e-05 | 1.6e-05 | 1.6e-05 | 1.433E-05 | 2.887E-06 | 1.100E-05 | 1.600E-05 |
| 27090 | 1355 | M1355I | E | 1.4e-05 | 8.4e-06 | 1.5e-05 | 1.247E-05 | 3.557E-06 | 8.400E-06 | 1.500E-05 |
| 27091 | 1355 | M1355L | E | 1.7e-05 | 1.6e-05 | 1.6e-05 | 1.633E-05 | 5.774E-07 | 1.600E-05 | 1.700E-05 |
| 27092 | 1355 | M1355K | E | 1.2e-05 | 1.6e-05 | 1.4e-05 | 1.400E-05 | 2.000E-06 | 1.200E-05 | 1.600E-05 |
| 27093 | 1355 | M1355M | E | 9.2e-06 | 8.4e-06 | 1e-05 | 9.200E-06 | 8.000E-07 | 8.400E-06 | 1.000E-05 |
| 27094 | 1355 | M1355F | E | 1.6e-05 | 1.5e-05 | 8.3e-06 | 1.310E-05 | 4.187E-06 | 8.300E-06 | 1.600E-05 |
| 27095 | 1355 | M1355P | E | 1.5e-05 | 1e-05 | 1.3e-05 | 1.267E-05 | 2.517E-06 | 1.000E-05 | 1.500E-05 |
| 27096 | 1355 | M1355S | E | 1.5e-05 | 1.5e-05 | 8.8e-06 | 1.293E-05 | 3.580E-06 | 8.800E-06 | 1.500E-05 |
| 27097 | 1355 | M1355T | E | 1.1e-05 | 1.6e-05 | 1.4e-05 | 1.367E-05 | 2.517E-06 | 1.100E-05 | 1.600E-05 |
| 27098 | 1355 | M1355W | E | 1.7e-05 | 1.6e-05 | 8.2e-06 | 1.373E-05 | 4.818E-06 | 8.200E-06 | 1.700E-05 |
| 27099 | 1355 | M1355Y | E | 1.5e-05 | 1.3e-05 | 1.1e-05 | 1.300E-05 | 2.000E-06 | 1.100E-05 | 1.500E-05 |
| 27100 | 1355 | M1355V | E | 1e-05 | 1e-05 | 1.6e-05 | 1.200E-05 | 3.464E-06 | 1.000E-05 | 1.600E-05 |
| 27101 | 1356 | N1356A | E | 1.3e-05 | 1.1e-05 | 1.6e-05 | 1.333E-05 | 2.517E-06 | 1.100E-05 | 1.600E-05 |
| 27102 | 1356 | N1356R | E | 1.3e-05 | 1.5e-05 | 1.6e-05 | 1.467E-05 | 1.528E-06 | 1.300E-05 | 1.600E-05 |
| 27103 | 1356 | N1356N | E | 1.1e-05 | 1.1e-05 | 7.7e-06 | 9.900E-06 | 1.905E-06 | 7.700E-06 | 1.100E-05 |
| 27104 | 1356 | N1356D | E | 2.7e-05 | 1.2e-05 | 1.4e-05 | 1.767E-05 | 8.145E-06 | 1.200E-05 | 2.700E-05 |
| 27105 | 1356 | N1356C | E | 1.6e-05 | 1.1e-05 | 1.6e-05 | 1.433E-05 | 2.887E-06 | 1.100E-05 | 1.600E-05 |
| 27106 | 1356 | N1356E | E | 1.2e-05 | 1.6e-05 | 1.1e-05 | 1.300E-05 | 2.646E-06 | 1.100E-05 | 1.600E-05 |
| 27107 | 1356 | N1356Q | E | 8.5e-06 | 1.2e-05 | 6.6e-06 | 9.033E-06 | 2.739E-06 | 6.600E-06 | 1.200E-05 |
| 27108 | 1356 | N1356G | E | 1.6e-05 | 7.7e-06 | 1.1e-05 | 1.157E-05 | 4.179E-06 | 7.700E-06 | 1.600E-05 |
| 27109 | 1356 | N1356H | E | 1.3e-05 | 1.6e-05 | 1e-05 | 1.300E-05 | 3.000E-06 | 1.000E-05 | 1.600E-05 |
| 27110 | 1356 | N1356I | E | 1.3e-05 | 1.3e-05 | 1.2e-05 | 1.267E-05 | 5.774E-07 | 1.200E-05 | 1.300E-05 |
| 27111 | 1356 | N1356L | E | 1.1e-05 | 1.6e-05 | 1.2e-05 | 1.300E-05 | 2.646E-06 | 1.100E-05 | 1.600E-05 |
| 27112 | 1356 | N1356K | E | 1.6e-05 | 1.1e-05 | 9.7e-06 | 1.223E-05 | 3.326E-06 | 9.700E-06 | 1.600E-05 |
| 27113 | 1356 | N1356M | E | 1.4e-05 | 1.1e-05 | 9.7e-06 | 1.157E-05 | 2.205E-06 | 9.700E-06 | 1.400E-05 |
| 27114 | 1356 | N1356F | E | 1.5e-05 | 9.7e-06 | 1.5e-05 | 1.323E-05 | 3.060E-06 | 9.700E-06 | 1.500E-05 |

| | | | | | | | | | | |
|---|---|---|---|---|---|---|---|---|---|---|
| 27115 | 1356 | N1356P | E | 8.2e-06 | 8.2e-06 | 1.1e-05 | 9.133E-06 | 1.617E-06 | 8.200E-06 | 1.100E-05 |
| 27116 | 1356 | N1356S | E | 6.8e-06 | 1.6e-05 | 9.5e-06 | 1.077E-05 | 4.729E-06 | 6.800E-06 | 1.600E-05 |
| 27117 | 1356 | N1356T | E | 1.6e-05 | 7.9e-06 | 1.6e-05 | 1.330E-05 | 4.677E-06 | 7.900E-06 | 1.600E-05 |
| 27118 | 1356 | N1356W | E | 9.8e-06 | 1.4e-05 | 1.5e-05 | 1.293E-05 | 2.759E-06 | 9.800E-06 | 1.500E-05 |
| 27119 | 1356 | N1356Y | E | 9.5e-06 | 9.8e-06 | 1.7e-05 | 1.210E-05 | 4.246E-06 | 9.500E-06 | 1.700E-05 |
| 27120 | 1356 | N1356V | E | 9.5e-06 | 1.7e-05 | 1.1e-05 | 1.250E-05 | 3.969E-06 | 9.500E-06 | 1.700E-05 |
| 27121 | 1357 | S1357A | E | 1.1e-05 | 8.4e-06 | 8.6e-06 | 9.333E-06 | 1.447E-06 | 8.400E-06 | 1.100E-05 |
| 27122 | 1357 | S1357R | E | 1.5e-05 | 9.7e-06 | 1.6e-05 | 1.357E-05 | 3.386E-06 | 9.700E-06 | 1.600E-05 |
| 27123 | 1357 | S1357N | E | 9.5e-06 | 1.4e-05 | 1.1e-05 | 1.150E-05 | 2.291E-06 | 9.500E-06 | 1.400E-05 |
| 27124 | 1357 | S1357D | E | 1e-05 | 1.5e-05 | 1.7e-05 | 1.400E-05 | 3.606E-06 | 1.000E-05 | 1.700E-05 |
| 27125 | 1357 | S1357C | E | 1e-05 | 1.1e-05 | 8.1e-06 | 9.700E-06 | 1.473E-06 | 8.100E-06 | 1.100E-05 |
| 27126 | 1357 | S1357E | E | 1.4e-05 | 1.6e-05 | 1.4e-05 | 1.467E-05 | 1.155E-06 | 1.400E-05 | 1.600E-05 |
| 27127 | 1357 | S1357Q | E | 1.5e-05 | 8.8e-06 | 1.5e-05 | 1.293E-05 | 3.580E-06 | 8.800E-06 | 1.500E-05 |
| 27128 | 1357 | S1357G | E | 1.1e-05 | 3e-05 | 1.7e-05 | 1.933E-05 | 9.713E-06 | 1.100E-05 | 3.000E-05 |
| 27129 | 1357 | S1357H | E | 1.2e-05 | 1.6e-05 | 1.6e-05 | 1.467E-05 | 2.309E-06 | 1.200E-05 | 1.600E-05 |
| 27130 | 1357 | S1357I | E | 1.5e-05 | 9.7e-06 | 1.7e-05 | 1.390E-05 | 3.772E-06 | 9.700E-06 | 1.700E-05 |
| 27131 | 1357 | S1357L | E | 1.6e-05 | 1.6e-05 | 1.4e-05 | 1.533E-05 | 1.155E-06 | 1.400E-05 | 1.600E-05 |
| 27132 | 1357 | S1357K | E | 1.3e-05 | 1.5e-05 | 1.8e-05 | 1.533E-05 | 2.517E-06 | 1.300E-05 | 1.800E-05 |
| 27133 | 1357 | S1357M | E | 8.8e-06 | 1.6e-05 | 1.6e-05 | 1.360E-05 | 4.157E-06 | 8.800E-06 | 1.600E-05 |
| 27134 | 1357 | S1357F | E | 1.2e-05 | 9.9e-06 | 1.1e-05 | 1.097E-05 | 1.050E-06 | 9.900E-06 | 1.200E-05 |
| 27135 | 1357 | S1357P | E | 1.1e-05 | 1.5e-05 | 1.6e-05 | 1.400E-05 | 2.646E-06 | 1.100E-05 | 1.600E-05 |
| 27136 | 1357 | S1357S | E | 1.5e-05 | 1.6e-05 | 9.1e-06 | 1.337E-05 | 3.729E-06 | 9.100E-06 | 1.600E-05 |
| 27137 | 1357 | S1357T | E | 1.5e-05 | 1.5e-05 | 1.1e-05 | 1.367E-05 | 2.309E-06 | 1.100E-05 | 1.500E-05 |
| 27138 | 1357 | S1357W | E | 9.3e-06 | 1.3e-05 | 1.7e-05 | 1.310E-05 | 3.851E-06 | 9.300E-06 | 1.700E-05 |
| 27139 | 1357 | S1357Y | E | 1.2e-05 | 1.1e-05 | 1.1e-05 | 1.133E-05 | 5.774E-07 | 1.100E-05 | 1.200E-05 |
| 27140 | 1357 | S1357V | E | 1.5e-05 | 1.4e-05 | 1.1e-05 | 1.333E-05 | 2.082E-06 | 1.100E-05 | 1.500E-05 |
| 27141 | 1358 | L1358A | E | 1.3e-05 | 1.1e-05 | 1.4e-05 | 1.267E-05 | 1.528E-06 | 1.100E-05 | 1.400E-05 |
| 27142 | 1358 | L1358R | E | 1.6e-05 | 2e-05 | 1.2e-05 | 1.600E-05 | 4.000E-06 | 1.200E-05 | 2.000E-05 |
| 27143 | 1358 | L1358N | E | 1.5e-05 | 1.3e-05 | 1.3e-05 | 1.367E-05 | 1.155E-06 | 1.300E-05 | 1.500E-05 |
| 27144 | 1358 | L1358D | E | 1.1e-05 | 1.3e-05 | 1.1e-05 | 1.167E-05 | 1.155E-06 | 1.100E-05 | 1.300E-05 |
| 27145 | 1358 | L1358C | E | 9.7e-06 | 1.6e-05 | 7.7e-06 | 1.113E-05 | 4.332E-06 | 7.700E-06 | 1.600E-05 |
| 27146 | 1358 | L1358E | E | 9e-06 | 1.5e-05 | 1.3e-05 | 1.233E-05 | 3.055E-06 | 9.000E-06 | 1.500E-05 |
| 27147 | 1358 | L1358Q | E | 1.4e-05 | 1.1e-05 | 8.6e-06 | 1.120E-05 | 2.706E-06 | 8.600E-06 | 1.400E-05 |
| 27148 | 1358 | L1358G | E | 1.5e-05 | 1.5e-05 | 8.8e-06 | 1.293E-05 | 3.580E-06 | 8.800E-06 | 1.500E-05 |
| 27149 | 1358 | L1358H | E | 1.4e-05 | 1.4e-05 | 1.2e-05 | 1.333E-05 | 1.155E-06 | 1.200E-05 | 1.400E-05 |
| 27150 | 1358 | L1358I | E | 1.6e-05 | 1.6e-05 | 1.6e-05 | 1.600E-05 | 0.000E+00 | 1.600E-05 | 1.600E-05 |
| 27151 | 1358 | L1358L | E | 1.1e-05 | 1.4e-05 | 1.3e-05 | 1.267E-05 | 1.528E-06 | 1.100E-05 | 1.400E-05 |
| 27152 | 1358 | L1358K | E | 9.8e-06 | 1.5e-05 | 8.8e-06 | 1.120E-05 | 3.329E-06 | 8.800E-06 | 1.500E-05 |
| 27153 | 1358 | L1358M | E | 1.5e-05 | 9.6e-06 | 1.3e-05 | 1.253E-05 | 2.730E-06 | 9.600E-06 | 1.500E-05 |
| 27154 | 1358 | L1358F | E | 1.1e-05 | 1.3e-05 | 1.4e-05 | 1.267E-05 | 1.528E-06 | 1.100E-05 | 1.400E-05 |
| 27155 | 1358 | L1358P | E | 1e-05 | 8.9e-06 | 1.2e-05 | 1.030E-05 | 1.572E-06 | 8.900E-06 | 1.200E-05 |
| 27156 | 1358 | L1358S | E | 1.1e-05 | 1.4e-05 | 8.4e-06 | 1.113E-05 | 2.802E-06 | 8.400E-06 | 1.400E-05 |
| 27157 | 1358 | L1358T | E | 1.5e-05 | 1.6e-05 | 1.4e-05 | 1.500E-05 | 1.000E-06 | 1.400E-05 | 1.600E-05 |
| 27158 | 1358 | L1358W | E | 1.6e-05 | 9.1e-06 | 1e-05 | 1.170E-05 | 3.751E-06 | 9.100E-06 | 1.600E-05 |
| 27159 | 1358 | L1358Y | E | 9.4e-06 | 1.7e-05 | 1.6e-05 | 1.413E-05 | 4.130E-06 | 9.400E-06 | 1.700E-05 |
| 27160 | 1358 | L1358V | E | 1e-05 | 1.4e-05 | 1.1e-05 | 1.167E-05 | 2.082E-06 | 1.000E-05 | 1.400E-05 |
| 27161 | 1359 | R1359A | E | 1e-05 | 9.8e-06 | 1.7e-05 | 1.227E-05 | 4.100E-06 | 9.800E-06 | 1.700E-05 |
| 27162 | 1359 | R1359R | E | 1.1e-05 | 9.6e-06 | 1.4e-05 | 1.153E-05 | 2.248E-06 | 9.600E-06 | 1.400E-05 |
| 27163 | 1359 | R1359N | E | 9.3e-06 | 1.6e-05 | 1.4e-05 | 1.310E-05 | 3.439E-06 | 9.300E-06 | 1.600E-05 |
| 27164 | 1359 | R1359D | E | 1.1e-05 | 9.6e-06 | 1.5e-05 | 1.187E-05 | 2.802E-06 | 9.600E-06 | 1.500E-05 |
| 27165 | 1359 | R1359C | E | 1e-05 | 1.4e-05 | 8.8e-06 | 1.093E-05 | 2.723E-06 | 8.800E-06 | 1.400E-05 |
| 27166 | 1359 | R1359E | E | 1.2e-05 | 1.3e-05 | 1.7e-05 | 1.400E-05 | 2.646E-06 | 1.200E-05 | 1.700E-05 |
| 27167 | 1359 | R1359Q | E | 1.1e-05 | 1.1e-05 | 8.5e-06 | 1.017E-05 | 1.443E-06 | 8.500E-06 | 1.100E-05 |
| 27168 | 1359 | R1359G | E | 6.8e-06 | 1.4e-05 | 1.5e-05 | 1.193E-05 | 4.474E-06 | 6.800E-06 | 1.500E-05 |
| 27169 | 1359 | R1359H | E | 9.3e-06 | 1e-05 | 8.9e-06 | 9.400E-06 | 5.568E-07 | 8.900E-06 | 1.000E-05 |
| 27170 | 1359 | R1359I | E | 1.5e-05 | 1.5e-05 | 1.5e-05 | 1.500E-05 | 0.000E+00 | 1.500E-05 | 1.500E-05 |
| 27171 | 1359 | R1359L | E | 3e-05 | 1e-05 | 1.6e-05 | 1.867E-05 | 1.026E-05 | 1.000E-05 | 3.000E-05 |
| 27172 | 1359 | R1359K | E | 1.1e-05 | 1.5e-05 | 1e-05 | 1.200E-05 | 2.646E-06 | 1.000E-05 | 1.500E-05 |
| 27173 | 1359 | R1359M | E | 1.4e-05 | 9e-06 | 1e-05 | 1.100E-05 | 2.646E-06 | 9.000E-06 | 1.400E-05 |
| 27174 | 1359 | R1359F | E | 1.5e-05 | 1.5e-05 | 1.1e-05 | 1.367E-05 | 2.309E-06 | 1.100E-05 | 1.500E-05 |
| 27175 | 1359 | R1359P | E | 9.3e-06 | 1.6e-05 | 1.5e-05 | 1.343E-05 | 3.614E-06 | 9.300E-06 | 1.600E-05 |
| 27176 | 1359 | R1359S | E | 1.4e-05 | 1.2e-05 | 1.5e-05 | 1.367E-05 | 1.528E-06 | 1.200E-05 | 1.500E-05 |
| 27177 | 1359 | R1359T | E | 1e-05 | 1.4e-05 | 9.6e-06 | 1.120E-05 | 2.433E-06 | 9.600E-06 | 1.400E-05 |
| 27178 | 1359 | R1359W | E | 1e-05 | 1.3e-05 | 1.5e-05 | 1.267E-05 | 2.517E-06 | 1.000E-05 | 1.500E-05 |
| 27179 | 1359 | R1359Y | E | 1.5e-05 | 1.4e-05 | 1e-05 | 1.300E-05 | 2.646E-06 | 1.000E-05 | 1.500E-05 |
| 27180 | 1359 | R1359V | E | 7.6e-06 | 1.1e-05 | 1.6e-05 | 1.153E-05 | 4.225E-06 | 7.600E-06 | 1.600E-05 |
| 27181 | 1360 | A1360A | E | 1.4e-05 | 1.2e-05 | 1.2e-05 | 1.267E-05 | 1.155E-06 | 1.200E-05 | 1.400E-05 |
| 27182 | 1360 | A1360R | E | 1.2e-05 | 1.3e-05 | 1.6e-05 | 1.367E-05 | 2.082E-06 | 1.200E-05 | 1.600E-05 |
| 27183 | 1360 | A1360N | E | 1.5e-05 | 1.1e-05 | 1.6e-05 | 1.400E-05 | 2.646E-06 | 1.100E-05 | 1.600E-05 |
| 27184 | 1360 | A1360D | E | 1.4e-05 | 1.5e-05 | 1.6e-05 | 1.500E-05 | 1.000E-06 | 1.400E-05 | 1.600E-05 |
| 27185 | 1360 | A1360C | E | 1.6e-05 | 1.5e-05 | 1e-05 | 1.367E-05 | 3.215E-06 | 1.000E-05 | 1.600E-05 |
| 27186 | 1360 | A1360E | E | 1.2e-05 | 1.5e-05 | 1e-05 | 1.233E-05 | 2.517E-06 | 1.000E-05 | 1.500E-05 |
| 27187 | 1360 | A1360Q | E | 1.1e-05 | 1.5e-05 | 7.8e-06 | 1.127E-05 | 3.607E-06 | 7.800E-06 | 1.500E-05 |
| 27188 | 1360 | A1360G | E | 1.2e-05 | 1.1e-05 | 1.7e-05 | 1.333E-05 | 3.215E-06 | 1.100E-05 | 1.700E-05 |
| 27189 | 1360 | A1360H | E | 1.6e-05 | 1.5e-05 | 1.5e-05 | 1.533E-05 | 5.774E-07 | 1.500E-05 | 1.600E-05 |
| 27190 | 1360 | A1360I | E | 1.7e-05 | 1.1e-05 | 1.6e-05 | 1.467E-05 | 3.215E-06 | 1.100E-05 | 1.700E-05 |
| 27191 | 1360 | A1360L | E | 1.5e-05 | 1.5e-05 | 1.5e-05 | 1.500E-05 | 0.000E+00 | 1.500E-05 | 1.500E-05 |
| 27192 | 1360 | A1360K | E | 1e-05 | 1.3e-05 | 1.5e-05 | 1.267E-05 | 2.517E-06 | 1.000E-05 | 1.500E-05 |
| 27193 | 1360 | A1360M | E | 9.2e-06 | 1.2e-05 | 1.5e-05 | 1.207E-05 | 2.901E-06 | 9.200E-06 | 1.500E-05 |
| 27194 | 1360 | A1360F | E | 1.3e-05 | 1.4e-05 | 1.1e-05 | 1.267E-05 | 1.528E-06 | 1.100E-05 | 1.400E-05 |
| 27195 | 1360 | A1360P | E | 6.9e-06 | 1.5e-05 | 1.5e-05 | 1.230E-05 | 4.677E-06 | 6.900E-06 | 1.500E-05 |
| 27196 | 1360 | A1360S | E | 8.2e-06 | 9.6e-06 | 1.4e-05 | 1.060E-05 | 3.027E-06 | 8.200E-06 | 1.400E-05 |
| 27197 | 1360 | A1360T | E | 9.9e-06 | 1e-05 | 1.8e-05 | 1.263E-05 | 4.648E-06 | 9.900E-06 | 1.800E-05 |
| 27198 | 1360 | A1360W | E | 1.4e-05 | 1.1e-05 | 8.3e-06 | 1.110E-05 | 2.851E-06 | 8.300E-06 | 1.400E-05 |
| 27199 | 1360 | A1360Y | E | 9.3e-06 | 1.4e-05 | 1e-05 | 1.110E-05 | 2.536E-06 | 9.300E-06 | 1.400E-05 |

| | | | | | | | | | | |
|---|---|---|---|---|---|---|---|---|---|---|
| 27200 | 1360 | A1360V | E | 9.7e-06 | 1.7e-05 | 1.6e-05 | 1.423E-05 | 3.958E-06 | 9.700E-06 | 1.700E-05 |
| 27201 | 1361 | E1361A | E | 8.7e-06 | 1.5e-05 | 1.2e-05 | 1.190E-05 | 3.151E-06 | 8.700E-06 | 1.500E-05 |
| 27202 | 1361 | E1361R | E | 8.1e-06 | 8.4e-06 | 1.6e-05 | 1.083E-05 | 4.477E-06 | 8.100E-06 | 1.600E-05 |
| 27203 | 1361 | E1361N | E | 1.2e-05 | 9.9e-06 | 1.3e-05 | 1.163E-05 | 1.582E-06 | 9.900E-06 | 1.300E-05 |
| 27204 | 1361 | E1361D | E | 8.8e-06 | 1e-05 | 9.9e-06 | 9.567E-06 | 6.658E-07 | 8.800E-06 | 1.000E-05 |
| 27205 | 1361 | E1361C | E | 1.1e-05 | 9e-06 | 9.3e-06 | 9.767E-06 | 1.079E-06 | 9.000E-06 | 1.100E-05 |
| 27206 | 1361 | E1361E | E | 1.7e-05 | 9.7e-06 | 9.9e-06 | 1.220E-05 | 4.158E-06 | 9.700E-06 | 1.700E-05 |
| 27207 | 1361 | E1361Q | E | 1.6e-05 | 1.4e-05 | 1.5e-05 | 1.500E-05 | 1.000E-06 | 1.400E-05 | 1.600E-05 |
| 27208 | 1361 | E1361G | E | 1.1e-05 | 1e-05 | 1.7e-05 | 1.267E-05 | 3.786E-06 | 1.000E-05 | 1.700E-05 |
| 27209 | 1361 | E1361H | E | 1.1e-05 | 1.3e-05 | 1.6e-05 | 1.333E-05 | 2.517E-06 | 1.100E-05 | 1.600E-05 |
| 27210 | 1361 | E1361I | E | 1.7e-05 | 1.6e-05 | 1e-05 | 1.433E-05 | 3.786E-06 | 1.000E-05 | 1.700E-05 |
| 27211 | 1361 | E1361L | E | 1.1e-05 | 8.7e-06 | 1.6e-05 | 1.190E-05 | 3.732E-06 | 8.700E-06 | 1.600E-05 |
| 27212 | 1361 | E1361K | E | 1.6e-05 | 1.5e-05 | 2.2e-05 | 1.767E-05 | 3.786E-06 | 1.500E-05 | 2.200E-05 |
| 27213 | 1361 | E1361M | E | 1.1e-05 | 1.1e-05 | 1.4e-05 | 1.200E-05 | 1.732E-06 | 1.100E-05 | 1.400E-05 |
| 27214 | 1361 | E1361F | E | 1.1e-05 | 1.6e-05 | 1.2e-05 | 1.300E-05 | 2.646E-06 | 1.100E-05 | 1.600E-05 |
| 27215 | 1361 | E1361P | E | 1e-05 | 9.4e-06 | 8.9e-06 | 9.433E-06 | 5.508E-07 | 8.900E-06 | 1.000E-05 |
| 27216 | 1361 | E1361S | E | 9.8e-06 | 1.5e-05 | 1e-05 | 1.160E-05 | 2.946E-06 | 9.800E-06 | 1.500E-05 |
| 27217 | 1361 | E1361T | E | 1.2e-05 | 9.9e-06 | 9.2e-06 | 1.037E-05 | 1.457E-06 | 9.200E-06 | 1.200E-05 |
| 27218 | 1361 | E1361W | E | 1.1e-05 | 1.3e-05 | 1.5e-05 | 1.300E-05 | 2.000E-06 | 1.100E-05 | 1.500E-05 |
| 27219 | 1361 | E1361Y | E | 1.3e-05 | 1.1e-05 | 9.1e-06 | 1.103E-05 | 1.950E-06 | 9.100E-06 | 1.300E-05 |
| 27220 | 1361 | E1361V | E | 1.6e-05 | 1.5e-05 | 1.5e-05 | 1.533E-05 | 5.774E-07 | 1.500E-05 | 1.600E-05 |
| 27221 | 1362 | D1362A | E | 1e-05 | 1e-05 | 8.4e-06 | 9.467E-06 | 9.238E-07 | 8.400E-06 | 1.000E-05 |
| 27222 | 1362 | D1362R | E | 1.1e-05 | 1.5e-05 | 1.1e-05 | 1.233E-05 | 2.309E-06 | 1.100E-05 | 1.500E-05 |
| 27223 | 1362 | D1362N | E | 8.5e-06 | 1.3e-05 | 1.2e-05 | 1.117E-05 | 2.363E-06 | 8.500E-06 | 1.300E-05 |
| 27224 | 1362 | D1362D | E | 1.5e-05 | 1.3e-05 | 9.8e-06 | 1.260E-05 | 2.623E-06 | 9.800E-06 | 1.500E-05 |
| 27225 | 1362 | D1362C | E | 1.6e-05 | 1.7e-05 | 1.5e-05 | 1.600E-05 | 1.000E-06 | 1.500E-05 | 1.700E-05 |
| 27226 | 1362 | D1362E | E | 1.3e-05 | 1.6e-05 | 1.6e-05 | 1.500E-05 | 1.732E-06 | 1.300E-05 | 1.600E-05 |
| 27227 | 1362 | D1362Q | E | 9.4e-06 | 1.3e-05 | 1.3e-05 | 1.180E-05 | 2.078E-06 | 9.400E-06 | 1.300E-05 |
| 27228 | 1362 | D1362G | E | 9.6e-06 | 1.3e-05 | 1.5e-05 | 1.253E-05 | 2.730E-06 | 9.600E-06 | 1.500E-05 |
| 27229 | 1362 | D1362H | E | 1.6e-05 | 1.2e-05 | 1.4e-05 | 1.400E-05 | 2.000E-06 | 1.200E-05 | 1.600E-05 |
| 27230 | 1362 | D1362I | E | 1.5e-05 | 1.4e-05 | 9.3e-06 | 1.277E-05 | 3.044E-06 | 9.300E-06 | 1.500E-05 |
| 27231 | 1362 | D1362L | E | 1.8e-05 | 1.4e-05 | 1.4e-05 | 1.533E-05 | 2.309E-06 | 1.400E-05 | 1.800E-05 |
| 27232 | 1362 | D1362K | E | 1.1e-05 | 7.4e-06 | 1.4e-05 | 1.080E-05 | 3.305E-06 | 7.400E-06 | 1.400E-05 |
| 27233 | 1362 | D1362M | E | 1.1e-05 | 1.6e-05 | 1.3e-05 | 1.333E-05 | 2.517E-06 | 1.100E-05 | 1.600E-05 |
| 27234 | 1362 | D1362F | E | 1.3e-05 | 1.6e-05 | 1.6e-05 | 1.500E-05 | 1.732E-06 | 1.300E-05 | 1.600E-05 |
| 27235 | 1362 | D1362P | E | 1.5e-05 | 8.2e-06 | 1.4e-05 | 1.240E-05 | 3.672E-06 | 8.200E-06 | 1.500E-05 |
| 27236 | 1362 | D1362S | E | 1.5e-05 | 1.6e-05 | 1.5e-05 | 1.533E-05 | 5.774E-07 | 1.500E-05 | 1.600E-05 |
| 27237 | 1362 | D1362T | E | 2.9e-05 | 1.3e-05 | 8.6e-06 | 1.687E-05 | 1.074E-05 | 8.600E-06 | 2.900E-05 |
| 27238 | 1362 | D1362W | E | 1.5e-05 | 9.1e-06 | 1.2e-05 | 1.203E-05 | 2.950E-06 | 9.100E-06 | 1.500E-05 |
| 27239 | 1362 | D1362Y | E | 1e-05 | 1.1e-05 | 9.9e-06 | 1.030E-05 | 6.083E-07 | 9.900E-06 | 1.100E-05 |
| 27240 | 1362 | D1362V | E | 1.4e-05 | 1.6e-05 | 1.4e-05 | 1.467E-05 | 1.155E-06 | 1.400E-05 | 1.600E-05 |
| 27241 | 1363 | T1363A | E | 1.5e-05 | 1.7e-05 | 1.5e-05 | 1.567E-05 | 1.155E-06 | 1.500E-05 | 1.700E-05 |
| 27242 | 1363 | T1363R | E | 1.2e-05 | 1.6e-05 | 1.4e-05 | 1.400E-05 | 2.000E-06 | 1.200E-05 | 1.600E-05 |
| 27243 | 1363 | T1363N | E | 1.7e-05 | 9.6e-06 | 1.6e-05 | 1.420E-05 | 4.015E-06 | 9.600E-06 | 1.700E-05 |
| 27244 | 1363 | T1363D | E | 9.6e-06 | 1.5e-05 | 1.8e-05 | 1.420E-05 | 4.257E-06 | 9.600E-06 | 1.800E-05 |
| 27245 | 1363 | T1363C | E | 1e-05 | 9.7e-06 | 1.2e-05 | 1.057E-05 | 1.250E-06 | 9.700E-06 | 1.200E-05 |
| 27246 | 1363 | T1363E | E | 1.6e-05 | 1.1e-05 | 1.7e-05 | 1.467E-05 | 3.215E-06 | 1.100E-05 | 1.700E-05 |
| 27247 | 1363 | T1363Q | E | 1.5e-05 | 1.5e-05 | 1.5e-05 | 1.500E-05 | 0.000E+00 | 1.500E-05 | 1.500E-05 |
| 27248 | 1363 | T1363G | E | 1.6e-05 | 1.5e-05 | 1.4e-05 | 1.500E-05 | 1.000E-06 | 1.400E-05 | 1.600E-05 |
| 27249 | 1363 | T1363H | E | 1.1e-05 | 1.5e-05 | 1e-05 | 1.200E-05 | 2.646E-06 | 1.000E-05 | 1.500E-05 |
| 27250 | 1363 | T1363I | E | 1.2e-05 | 1.3e-05 | 1.1e-05 | 1.200E-05 | 1.000E-06 | 1.100E-05 | 1.300E-05 |
| 27251 | 1363 | T1363L | E | 9.9e-06 | 9.2e-06 | 1.5e-05 | 1.137E-05 | 3.166E-06 | 9.200E-06 | 1.500E-05 |
| 27252 | 1363 | T1363K | E | 1.1e-05 | 1.3e-05 | 1.6e-05 | 1.333E-05 | 2.517E-06 | 1.100E-05 | 1.600E-05 |
| 27253 | 1363 | T1363M | E | 1.5e-05 | 1.7e-05 | 1.3e-05 | 1.500E-05 | 2.000E-06 | 1.300E-05 | 1.700E-05 |
| 27254 | 1363 | T1363F | E | 1.5e-05 | 1.5e-05 | 1.5e-05 | 1.500E-05 | 0.000E+00 | 1.500E-05 | 1.500E-05 |
| 27255 | 1363 | T1363P | E | 1.1e-05 | 9.8e-06 | 1e-05 | 1.027E-05 | 6.429E-07 | 9.800E-06 | 1.100E-05 |
| 27256 | 1363 | T1363S | E | 1e-05 | 1.3e-05 | 1.6e-05 | 1.300E-05 | 3.000E-06 | 1.000E-05 | 1.600E-05 |
| 27257 | 1363 | T1363T | E | 8.4e-06 | 1.5e-05 | 1.6e-05 | 1.313E-05 | 4.130E-06 | 8.400E-06 | 1.600E-05 |
| 27258 | 1363 | T1363W | E | 9.3e-06 | 1.1e-05 | 1.3e-05 | 1.110E-05 | 1.852E-06 | 9.300E-06 | 1.300E-05 |
| 27259 | 1363 | T1363Y | E | 1.1e-05 | 1.3e-05 | 1.3e-05 | 1.233E-05 | 1.155E-06 | 1.100E-05 | 1.300E-05 |
| 27260 | 1363 | T1363V | E | 1.1e-05 | 1.7e-05 | 8.2e-06 | 1.207E-05 | 4.496E-06 | 8.200E-06 | 1.700E-05 |
| 27261 | 1364 | A1364A | E | 1.5e-05 | 1e-05 | 1e-05 | 1.167E-05 | 2.887E-06 | 1.000E-05 | 1.500E-05 |
| 27262 | 1364 | A1364R | E | 1e-05 | 1.6e-05 | 1.6e-05 | 1.400E-05 | 3.464E-06 | 1.000E-05 | 1.600E-05 |
| 27263 | 1364 | A1364N | E | 1.1e-05 | 1.6e-05 | 1.3e-05 | 1.333E-05 | 2.517E-06 | 1.100E-05 | 1.600E-05 |
| 27264 | 1364 | A1364D | E | 1.6e-05 | 1e-05 | 1.2e-05 | 1.267E-05 | 3.055E-06 | 1.000E-05 | 1.600E-05 |
| 27265 | 1364 | A1364C | E | 1.4e-05 | 9.1e-06 | 1e-05 | 1.103E-05 | 2.608E-06 | 9.100E-06 | 1.400E-05 |
| 27266 | 1364 | A1364E | E | 1.3e-05 | 1e-05 | 1.8e-05 | 1.367E-05 | 4.041E-06 | 1.000E-05 | 1.800E-05 |
| 27267 | 1364 | A1364Q | E | 9.2e-06 | 1.5e-05 | 1.4e-05 | 1.273E-05 | 3.101E-06 | 9.200E-06 | 1.500E-05 |
| 27268 | 1364 | A1364G | E | 1.4e-05 | 9.7e-06 | 7.4e-06 | 1.037E-05 | 3.350E-06 | 7.400E-06 | 1.400E-05 |
| 27269 | 1364 | A1364H | E | 1.4e-05 | 1e-05 | 1.5e-05 | 1.300E-05 | 2.646E-06 | 1.000E-05 | 1.500E-05 |
| 27270 | 1364 | A1364I | E | 1.3e-05 | 1.6e-05 | 1.1e-05 | 1.333E-05 | 2.517E-06 | 1.100E-05 | 1.600E-05 |
| 27271 | 1364 | A1364L | E | 1.4e-05 | 1.5e-05 | 1.6e-05 | 1.500E-05 | 1.000E-06 | 1.400E-05 | 1.600E-05 |
| 27272 | 1364 | A1364K | E | 1.5e-05 | 1.5e-05 | 1.4e-05 | 1.467E-05 | 5.774E-07 | 1.400E-05 | 1.500E-05 |
| 27273 | 1364 | A1364M | E | 1.6e-05 | 9.3e-06 | 1e-05 | 1.177E-05 | 3.683E-06 | 9.300E-06 | 1.600E-05 |
| 27274 | 1364 | A1364F | E | 1.2e-05 | 6.5e-06 | 8.7e-06 | 9.067E-06 | 2.768E-06 | 6.500E-06 | 1.200E-05 |
| 27275 | 1364 | A1364P | E | 1.6e-05 | 1.4e-05 | 9.7e-06 | 1.323E-05 | 3.219E-06 | 9.700E-06 | 1.600E-05 |
| 27276 | 1364 | A1364S | E | 1.6e-05 | 1.1e-05 | 1.1e-05 | 1.267E-05 | 2.887E-06 | 1.100E-05 | 1.600E-05 |
| 27277 | 1364 | A1364T | E | 1.6e-05 | 1.7e-05 | 1.2e-05 | 1.500E-05 | 2.646E-06 | 1.200E-05 | 1.700E-05 |
| 27278 | 1364 | A1364W | E | 1e-05 | 2e-05 | 1.5e-05 | 1.500E-05 | 5.000E-06 | 1.000E-05 | 2.000E-05 |
| 27279 | 1364 | A1364Y | E | 1.6e-05 | 1.4e-05 | 1e-05 | 1.333E-05 | 3.055E-06 | 1.000E-05 | 1.600E-05 |
| 27280 | 1364 | A1364V | E | 1e-05 | 7.7e-06 | 1.1e-05 | 9.567E-06 | 1.692E-06 | 7.700E-06 | 1.100E-05 |
| 27281 | 1365 | V1365A | E | 1.4e-05 | 1.1e-05 | 1.3e-05 | 1.267E-05 | 1.528E-06 | 1.100E-05 | 1.400E-05 |
| 27282 | 1365 | V1365R | E | 1.4e-05 | 1.4e-05 | 1.3e-05 | 1.367E-05 | 5.774E-07 | 1.300E-05 | 1.400E-05 |
| 27283 | 1365 | V1365N | E | 9.7e-06 | 9.7e-06 | 1.1e-05 | 1.013E-05 | 7.506E-07 | 9.700E-06 | 1.100E-05 |
| 27284 | 1365 | V1365D | E | 1.1e-05 | 6.9e-06 | 1.4e-05 | 1.063E-05 | 3.564E-06 | 6.900E-06 | 1.400E-05 |

| | | | | | | | | | | |
|---|---|---|---|---|---|---|---|---|---|---|
| 27285 | 1365 | V1365C | E | 1.6e-05 | 1.1e-05 | 1.4e-05 | 1.367E-05 | 2.517E-06 | 1.100E-05 | 1.600E-05 |
| 27286 | 1365 | V1365E | E | 1.5e-05 | 8.5e-06 | 8.1e-06 | 1.053E-05 | 3.873E-06 | 8.100E-06 | 1.500E-05 |
| 27287 | 1365 | V1365Q | E | 9.8e-06 | 9.6e-06 | 9.9e-06 | 9.767E-06 | 1.528E-07 | 9.600E-06 | 9.900E-06 |
| 27288 | 1365 | V1365G | E | 9.7e-06 | 1.1e-05 | 1.2e-05 | 1.090E-05 | 1.153E-06 | 9.700E-06 | 1.200E-05 |
| 27289 | 1365 | V1365H | E | 1.5e-05 | 1e-05 | 1.5e-05 | 1.333E-05 | 2.887E-06 | 1.000E-05 | 1.500E-05 |
| 27290 | 1365 | V1365I | E | 1.1e-05 | 1.2e-05 | 1.3e-05 | 1.200E-05 | 1.000E-06 | 1.100E-05 | 1.300E-05 |
| 27291 | 1365 | V1365L | E | 1.1e-05 | 1.3e-05 | 1.1e-05 | 1.167E-05 | 1.155E-06 | 1.100E-05 | 1.300E-05 |
| 27292 | 1365 | V1365K | E | 1.5e-05 | 1.6e-05 | 1.5e-05 | 1.533E-05 | 5.774E-07 | 1.500E-05 | 1.600E-05 |
| 27293 | 1365 | V1365M | E | 1.5e-05 | 1.4e-05 | 1.3e-05 | 1.400E-05 | 1.000E-06 | 1.300E-05 | 1.500E-05 |
| 27294 | 1365 | V1365F | E | 1e-05 | 1.8e-05 | 1.6e-05 | 1.467E-05 | 4.163E-06 | 1.000E-05 | 1.800E-05 |
| 27295 | 1365 | V1365P | E | 1.7e-05 | 1.2e-05 | 9.3e-06 | 1.277E-05 | 3.907E-06 | 9.300E-06 | 1.700E-05 |
| 27296 | 1365 | V1365S | E | 8.9e-06 | 1.4e-05 | 1.4e-05 | 1.230E-05 | 2.944E-06 | 8.900E-06 | 1.400E-05 |
| 27297 | 1365 | V1365T | E | 1.2e-05 | 1.4e-05 | 1.4e-05 | 1.333E-05 | 1.155E-06 | 1.200E-05 | 1.400E-05 |
| 27298 | 1365 | V1365W | E | 1.5e-05 | 9.8e-06 | 9.2e-06 | 1.133E-05 | 3.190E-06 | 9.200E-06 | 1.500E-05 |
| 27299 | 1365 | V1365Y | E | 1.4e-05 | 1.4e-05 | 1.1e-05 | 1.300E-05 | 1.732E-06 | 1.100E-05 | 1.400E-05 |
| 27300 | 1365 | V1365V | E | 1.5e-05 | 1.5e-05 | 1.5e-05 | 1.500E-05 | 0.000E+00 | 1.500E-05 | 1.500E-05 |
| 27301 | 1366 | Y1366A | E | 1.7e-05 | 1.8e-05 | 1.7e-05 | 1.733E-05 | 5.774E-07 | 1.700E-05 | 1.800E-05 |
| 27302 | 1366 | Y1366R | E | 1.2e-05 | 1.4e-05 | 1.5e-05 | 1.367E-05 | 1.528E-06 | 1.200E-05 | 1.500E-05 |
| 27303 | 1366 | Y1366N | E | 1.1e-05 | 1.6e-05 | 1.3e-05 | 1.333E-05 | 2.517E-06 | 1.100E-05 | 1.600E-05 |
| 27304 | 1366 | Y1366D | E | 1.1e-05 | 9.9e-06 | 1.2e-05 | 1.097E-05 | 1.050E-06 | 9.900E-06 | 1.200E-05 |
| 27305 | 1366 | Y1366C | E | 1.6e-05 | 1.6e-05 | 1.6e-05 | 1.600E-05 | 0.000E+00 | 1.600E-05 | 1.600E-05 |
| 27306 | 1366 | Y1366E | E | 1.2e-05 | 1.5e-05 | 1.5e-05 | 1.400E-05 | 1.732E-06 | 1.200E-05 | 1.500E-05 |
| 27307 | 1366 | Y1366Q | E | 1.5e-05 | 1.6e-05 | 1.1e-05 | 1.400E-05 | 2.646E-06 | 1.100E-05 | 1.600E-05 |
| 27308 | 1366 | Y1366G | E | 1.9e-05 | 1.9e-05 | 1.7e-05 | 1.833E-05 | 1.155E-06 | 1.700E-05 | 1.900E-05 |
| 27309 | 1366 | Y1366H | E | 9.5e-06 | 1.6e-05 | 1e-05 | 1.183E-05 | 3.617E-06 | 9.500E-06 | 1.600E-05 |
| 27310 | 1366 | Y1366I | E | 1.4e-05 | 1.6e-05 | 1e-05 | 1.333E-05 | 3.055E-06 | 1.000E-05 | 1.600E-05 |
| 27311 | 1366 | Y1366L | E | 1.5e-05 | 1.8e-05 | 9.3e-06 | 1.410E-05 | 4.419E-06 | 9.300E-06 | 1.800E-05 |
| 27312 | 1366 | Y1366K | E | 1.4e-05 | 1.3e-05 | 1.1e-05 | 1.267E-05 | 1.528E-06 | 1.100E-05 | 1.400E-05 |
| 27313 | 1366 | Y1366M | E | 1.2e-05 | 1.7e-05 | 1.4e-05 | 1.433E-05 | 2.517E-06 | 1.200E-05 | 1.700E-05 |
| 27314 | 1366 | Y1366F | E | 1.2e-05 | 1.1e-05 | 1.4e-05 | 1.233E-05 | 1.528E-06 | 1.100E-05 | 1.400E-05 |
| 27315 | 1366 | Y1366P | E | 1.6e-05 | 9e-06 | 1.7e-05 | 1.400E-05 | 4.359E-06 | 9.000E-06 | 1.700E-05 |
| 27316 | 1366 | Y1366S | E | 1.6e-05 | 8e-06 | 1e-05 | 1.133E-05 | 4.163E-06 | 8.000E-06 | 1.600E-05 |
| 27317 | 1366 | Y1366T | E | 1.1e-05 | 1.5e-05 | 1.1e-05 | 1.233E-05 | 2.309E-06 | 1.100E-05 | 1.500E-05 |
| 27318 | 1366 | Y1366W | E | 9e-06 | 1e-05 | 1.3e-05 | 1.067E-05 | 2.082E-06 | 9.000E-06 | 1.300E-05 |
| 27319 | 1366 | Y1366Y | E | 1.5e-05 | 1.3e-05 | 1.6e-05 | 1.467E-05 | 1.528E-06 | 1.300E-05 | 1.600E-05 |
| 27320 | 1366 | Y1366V | E | 9.7e-06 | 1.9e-05 | 1.1e-05 | 1.323E-05 | 5.036E-06 | 9.700E-06 | 1.900E-05 |
| 27321 | 1367 | Y1367A | E | 1.5e-05 | 1.1e-05 | 8.4e-06 | 1.147E-05 | 3.325E-06 | 8.400E-06 | 1.500E-05 |
| 27322 | 1367 | Y1367R | E | 1e-05 | 1.3e-05 | 9.6e-06 | 1.087E-05 | 1.858E-06 | 9.600E-06 | 1.300E-05 |
| 27323 | 1367 | Y1367N | E | 1.5e-05 | 1.1e-05 | 1.2e-05 | 1.267E-05 | 2.082E-06 | 1.100E-05 | 1.500E-05 |
| 27324 | 1367 | Y1367D | E | 1.1e-05 | 1.6e-05 | 9.8e-06 | 1.227E-05 | 3.288E-06 | 9.800E-06 | 1.600E-05 |
| 27325 | 1367 | Y1367C | E | 1.1e-05 | 9.8e-06 | 1.5e-05 | 1.193E-05 | 2.723E-06 | 9.800E-06 | 1.500E-05 |
| 27326 | 1367 | Y1367E | E | 1.8e-05 | 1.3e-05 | 1.2e-05 | 1.433E-05 | 3.215E-06 | 1.200E-05 | 1.800E-05 |
| 27327 | 1367 | Y1367Q | E | 1.5e-05 | 1.4e-05 | 1.1e-05 | 1.333E-05 | 2.082E-06 | 1.100E-05 | 1.500E-05 |
| 27328 | 1367 | Y1367G | E | 1.5e-05 | 9.1e-06 | 1.1e-05 | 1.170E-05 | 3.012E-06 | 9.100E-06 | 1.500E-05 |
| 27329 | 1367 | Y1367H | E | 1.7e-05 | 1.4e-05 | 1.3e-05 | 1.467E-05 | 2.082E-06 | 1.300E-05 | 1.700E-05 |
| 27330 | 1367 | Y1367I | E | 1.1e-05 | 1.2e-05 | 1.7e-05 | 1.333E-05 | 3.215E-06 | 1.100E-05 | 1.700E-05 |
| 27331 | 1367 | Y1367L | E | 8.1e-06 | 1.6e-05 | 7.1e-06 | 1.040E-05 | 4.875E-06 | 7.100E-06 | 1.600E-05 |
| 27332 | 1367 | Y1367K | E | 1e-05 | 1.2e-05 | 1.8e-05 | 1.333E-05 | 4.163E-06 | 1.000E-05 | 1.800E-05 |
| 27333 | 1367 | Y1367M | E | 1.5e-05 | 1.4e-05 | 1.4e-05 | 1.433E-05 | 5.774E-07 | 1.400E-05 | 1.500E-05 |
| 27334 | 1367 | Y1367F | E | 1e-05 | 1.6e-05 | 1.1e-05 | 1.233E-05 | 3.215E-06 | 1.000E-05 | 1.600E-05 |
| 27335 | 1367 | Y1367P | E | 1.6e-05 | 1.5e-05 | 1.3e-05 | 1.467E-05 | 1.528E-06 | 1.300E-05 | 1.600E-05 |
| 27336 | 1367 | Y1367S | E | 1.4e-05 | 9.9e-06 | 1.6e-05 | 1.330E-05 | 3.110E-06 | 9.900E-06 | 1.600E-05 |
| 27337 | 1367 | Y1367T | E | 1.2e-05 | 1.6e-05 | 8.9e-06 | 1.230E-05 | 3.559E-06 | 8.900E-06 | 1.600E-05 |
| 27338 | 1367 | Y1367W | E | 1.6e-05 | 1.4e-05 | 1.1e-05 | 1.367E-05 | 2.517E-06 | 1.100E-05 | 1.600E-05 |
| 27339 | 1367 | Y1367Y | E | 1.1e-05 | 1.3e-05 | 1.5e-05 | 1.300E-05 | 2.000E-06 | 1.100E-05 | 1.500E-05 |
| 27340 | 1367 | Y1367V | E | 1.4e-05 | 1.2e-05 | 1.4e-05 | 1.333E-05 | 1.155E-06 | 1.200E-05 | 1.400E-05 |
| 27341 | 1368 | C1368A | E | 1e-05 | 1e-05 | 1.5e-05 | 1.167E-05 | 2.887E-06 | 1.000E-05 | 1.500E-05 |
| 27342 | 1368 | C1368R | E | 1.2e-05 | 9.7e-06 | 1.5e-05 | 1.223E-05 | 2.658E-06 | 9.700E-06 | 1.500E-05 |
| 27343 | 1368 | C1368N | E | 1.5e-05 | 1e-05 | 9.8e-06 | 1.160E-05 | 2.946E-06 | 9.800E-06 | 1.500E-05 |
| 27344 | 1368 | C1368D | E | 1.4e-05 | 1.5e-05 | 1.7e-05 | 1.533E-05 | 1.528E-06 | 1.400E-05 | 1.700E-05 |
| 27345 | 1368 | C1368C | E | 9.5e-06 | 3.4e-05 | 1.5e-05 | 1.950E-05 | 1.285E-05 | 9.500E-06 | 3.400E-05 |
| 27346 | 1368 | C1368E | E | 1e-05 | 6.6e-06 | 1.6e-05 | 1.087E-05 | 4.760E-06 | 6.600E-06 | 1.600E-05 |
| 27347 | 1368 | C1368Q | E | 9.6e-06 | 1.5e-05 | 8.3e-06 | 1.097E-05 | 3.553E-06 | 8.300E-06 | 1.500E-05 |
| 27348 | 1368 | C1368G | E | 9.6e-06 | 1.5e-05 | 1.4e-05 | 1.287E-05 | 2.873E-06 | 9.600E-06 | 1.500E-05 |
| 27349 | 1368 | C1368H | E | 1.4e-05 | 1.1e-05 | 1e-05 | 1.167E-05 | 2.082E-06 | 1.000E-05 | 1.400E-05 |
| 27350 | 1368 | C1368I | E | 1e-05 | 1.4e-05 | 1e-05 | 1.133E-05 | 2.309E-06 | 1.000E-05 | 1.400E-05 |
| 27351 | 1368 | C1368L | E | 1.1e-05 | 1.5e-05 | 9e-06 | 1.167E-05 | 3.055E-06 | 9.000E-06 | 1.500E-05 |
| 27352 | 1368 | C1368K | E | 1e-05 | 1.4e-05 | 1.6e-05 | 1.333E-05 | 3.055E-06 | 1.000E-05 | 1.600E-05 |
| 27353 | 1368 | C1368M | E | 1e-05 | 1.5e-05 | 1e-05 | 1.167E-05 | 2.887E-06 | 1.000E-05 | 1.500E-05 |
| 27354 | 1368 | C1368F | E | 1.5e-05 | 1.1e-05 | 8.9e-06 | 1.163E-05 | 3.099E-06 | 8.900E-06 | 1.500E-05 |
| 27355 | 1368 | C1368P | E | 1.1e-05 | 9.6e-06 | 1e-05 | 1.020E-05 | 7.211E-07 | 9.600E-06 | 1.100E-05 |
| 27356 | 1368 | C1368S | E | 1e-05 | 1.3e-05 | 1.6e-05 | 1.300E-05 | 3.000E-06 | 1.000E-05 | 1.600E-05 |
| 27357 | 1368 | C1368T | E | 9.2e-06 | 1e-05 | 1.3e-05 | 1.073E-05 | 2.003E-06 | 9.200E-06 | 1.300E-05 |
| 27358 | 1368 | C1368W | E | 1.6e-05 | 1.6e-05 | 1e-05 | 1.400E-05 | 3.464E-06 | 1.000E-05 | 1.600E-05 |
| 27359 | 1368 | C1368Y | E | 1e-05 | 1.5e-05 | 8.6e-06 | 1.120E-05 | 3.365E-06 | 8.600E-06 | 1.500E-05 |
| 27360 | 1368 | C1368V | E | 1.2e-05 | 8.5e-06 | 1.5e-05 | 1.183E-05 | 3.253E-06 | 8.500E-06 | 1.500E-05 |
| 27361 | 1369 | S1369A | E | 1.6e-05 | 1.4e-05 | 9.1e-06 | 1.303E-05 | 3.550E-06 | 9.100E-06 | 1.600E-05 |
| 27362 | 1369 | S1369R | E | 1.1e-05 | 7.8e-06 | 1e-05 | 9.600E-06 | 1.637E-06 | 7.800E-06 | 1.100E-05 |
| 27363 | 1369 | S1369N | E | 9.8e-06 | 8.8e-06 | 1.5e-05 | 1.120E-05 | 3.329E-06 | 8.800E-06 | 1.500E-05 |
| 27364 | 1369 | S1369D | E | 1.2e-05 | 6.7e-06 | 1.1e-05 | 9.900E-06 | 2.816E-06 | 6.700E-06 | 1.200E-05 |
| 27365 | 1369 | S1369C | E | 1e-05 | 1.5e-05 | 1.1e-05 | 1.200E-05 | 2.646E-06 | 1.000E-05 | 1.500E-05 |
| 27366 | 1369 | S1369E | E | 1.1e-05 | 6.9e-06 | 9.9e-06 | 9.267E-06 | 2.122E-06 | 6.900E-06 | 1.100E-05 |
| 27367 | 1369 | S1369Q | E | 9.5e-06 | 1.4e-05 | 1.2e-05 | 1.183E-05 | 2.255E-06 | 9.500E-06 | 1.400E-05 |
| 27368 | 1369 | S1369G | E | 9.2e-06 | 1.4e-05 | 1.2e-05 | 1.173E-05 | 2.411E-06 | 9.200E-06 | 1.400E-05 |
| 27369 | 1369 | S1369H | E | 8.1e-06 | 1e-05 | 1e-05 | 9.367E-06 | 1.097E-06 | 8.100E-06 | 1.000E-05 |

| | | | | | | | | | | |
|---|---|---|---|---|---|---|---|---|---|---|
| 27370 | 1369 | S1369I | E | 1.5e-05 | 1.1e-05 | 8.8e-06 | 1.160E-05 | 3.143E-06 | 8.800E-06 | 1.500E-05 |
| 27371 | 1369 | S1369L | E | 1.2e-05 | 1.5e-05 | 1.2e-05 | 1.300E-05 | 1.732E-06 | 1.200E-05 | 1.500E-05 |
| 27372 | 1369 | S1369K | E | 1.1e-05 | 1.1e-05 | 1.1e-05 | 1.100E-05 | 2.075E-21 | 1.100E-05 | 1.100E-05 |
| 27373 | 1369 | S1369M | E | 1.1e-05 | 1.6e-05 | 1.4e-05 | 1.367E-05 | 2.517E-06 | 1.100E-05 | 1.600E-05 |
| 27374 | 1369 | S1369F | E | 1.1e-05 | 1e-05 | 1.4e-05 | 1.167E-05 | 2.082E-06 | 1.000E-05 | 1.400E-05 |
| 27375 | 1369 | S1369P | E | 1.3e-05 | 1e-05 | 1.5e-05 | 1.267E-05 | 2.517E-06 | 1.000E-05 | 1.500E-05 |
| 27376 | 1369 | S1369S | E | 1.2e-05 | 1e-05 | 1.3e-05 | 1.167E-05 | 1.528E-06 | 1.000E-05 | 1.300E-05 |
| 27377 | 1369 | S1369T | E | 1.6e-05 | 1e-05 | 2.1e-05 | 1.567E-05 | 5.508E-06 | 1.000E-05 | 2.100E-05 |
| 27378 | 1369 | S1369W | E | 1.6e-05 | 1.5e-05 | 1.6e-05 | 1.567E-05 | 5.774E-07 | 1.500E-05 | 1.600E-05 |
| 27379 | 1369 | S1369Y | E | 1.6e-05 | 1.3e-05 | 1.1e-05 | 1.333E-05 | 2.517E-06 | 1.100E-05 | 1.600E-05 |
| 27380 | 1369 | S1369V | E | 1e-05 | 1.6e-05 | 1.4e-05 | 1.333E-05 | 3.055E-06 | 1.000E-05 | 1.600E-05 |
| 27381 | 1370 | R1370A | E | 1.6e-05 | 9.4e-06 | 1.5e-05 | 1.347E-05 | 3.557E-06 | 9.400E-06 | 1.600E-05 |
| 27382 | 1370 | R1370R | E | 1e-05 | 2e-05 | 6.7e-06 | 1.223E-05 | 6.926E-06 | 6.700E-06 | 2.000E-05 |
| 27383 | 1370 | R1370N | E | 1.7e-05 | 1.7e-05 | 1.1e-05 | 1.500E-05 | 3.464E-06 | 1.100E-05 | 1.700E-05 |
| 27384 | 1370 | R1370D | E | 1e-05 | 9.9e-06 | 1.2e-05 | 1.063E-05 | 1.185E-06 | 9.900E-06 | 1.200E-05 |
| 27385 | 1370 | R1370C | E | 8.7e-06 | 1.3e-05 | 1.1e-05 | 1.090E-05 | 2.152E-06 | 8.700E-06 | 1.300E-05 |
| 27386 | 1370 | R1370E | E | 1.5e-05 | 1.6e-05 | 1.2e-05 | 1.433E-05 | 2.082E-06 | 1.200E-05 | 1.600E-05 |
| 27387 | 1370 | R1370Q | E | 1e-05 | 1.3e-05 | 8.4e-06 | 1.047E-05 | 2.335E-06 | 8.400E-06 | 1.300E-05 |
| 27388 | 1370 | R1370G | E | 1.3e-05 | 1e-05 | 1.6e-05 | 1.300E-05 | 3.000E-06 | 1.000E-05 | 1.600E-05 |
| 27389 | 1370 | R1370H | E | 1.1e-05 | 8.7e-06 | 8.4e-06 | 9.367E-06 | 1.422E-06 | 8.400E-06 | 1.100E-05 |
| 27390 | 1370 | R1370I | E | 1.2e-05 | 1.7e-05 | 1.4e-05 | 1.433E-05 | 2.517E-06 | 1.200E-05 | 1.700E-05 |
| 27391 | 1370 | R1370L | E | 1.3e-05 | 1.3e-05 | 1.6e-05 | 1.400E-05 | 1.732E-06 | 1.300E-05 | 1.600E-05 |
| 27392 | 1370 | R1370K | E | 1.1e-05 | 1.1e-05 | 2e-05 | 1.400E-05 | 5.196E-06 | 1.100E-05 | 2.000E-05 |
| 27393 | 1370 | R1370M | E | 1.4e-05 | 1.4e-05 | 1.4e-05 | 1.400E-05 | 0.000E+00 | 1.400E-05 | 1.400E-05 |
| 27394 | 1370 | R1370F | E | 1.2e-05 | 1e-05 | 1.6e-05 | 1.267E-05 | 3.055E-06 | 1.000E-05 | 1.600E-05 |
| 27395 | 1370 | R1370P | E | 2e-05 | 9.6e-06 | 1.3e-05 | 1.420E-05 | 5.303E-06 | 9.600E-06 | 2.000E-05 |
| 27396 | 1370 | R1370S | E | 1.2e-05 | 1.5e-05 | 1.4e-05 | 1.367E-05 | 1.528E-06 | 1.200E-05 | 1.500E-05 |
| 27397 | 1370 | R1370T | E | 1.6e-05 | 1e-05 | 1.1e-05 | 1.233E-05 | 3.215E-06 | 1.000E-05 | 1.600E-05 |
| 27398 | 1370 | R1370W | E | 7.7e-06 | 9.9e-06 | 1.2e-05 | 9.867E-06 | 2.150E-06 | 7.700E-06 | 1.200E-05 |
| 27399 | 1370 | R1370Y | E | 1.5e-05 | 1.6e-05 | 1e-05 | 1.367E-05 | 3.215E-06 | 1.000E-05 | 1.600E-05 |
| 27400 | 1370 | R1370V | E | 1.1e-05 | 9.6e-06 | 1.4e-05 | 1.153E-05 | 2.248E-06 | 9.600E-06 | 1.400E-05 |
| 27401 | 1371 | W1371A | E | 9.7e-06 | 1.7e-05 | 8e-06 | 1.157E-05 | 4.782E-06 | 8.000E-06 | 1.700E-05 |
| 27402 | 1371 | W1371R | E | 1.5e-05 | 8.9e-06 | 1.6e-05 | 1.330E-05 | 3.843E-06 | 8.900E-06 | 1.600E-05 |
| 27403 | 1371 | W1371N | E | 1.3e-05 | 1.6e-05 | 9.2e-06 | 1.273E-05 | 3.408E-06 | 9.200E-06 | 1.600E-05 |
| 27404 | 1371 | W1371D | E | 2.1e-05 | 1.5e-05 | 1.6e-05 | 1.733E-05 | 3.215E-06 | 1.500E-05 | 2.100E-05 |
| 27405 | 1371 | W1371C | E | 1.7e-05 | 8.7e-06 | 8.7e-06 | 1.147E-05 | 4.792E-06 | 8.700E-06 | 1.700E-05 |
| 27406 | 1371 | W1371E | E | 1.6e-05 | 1.5e-05 | 1.7e-05 | 1.600E-05 | 1.000E-06 | 1.500E-05 | 1.700E-05 |
| 27407 | 1371 | W1371Q | E | 1.5e-05 | 1.3e-05 | 1.5e-05 | 1.433E-05 | 1.155E-06 | 1.300E-05 | 1.500E-05 |
| 27408 | 1371 | W1371G | E | 1e-05 | 9.4e-06 | 1.7e-05 | 1.213E-05 | 4.225E-06 | 9.400E-06 | 1.700E-05 |
| 27409 | 1371 | W1371H | E | 1.2e-05 | 9.5e-06 | 1e-05 | 1.050E-05 | 1.323E-06 | 9.500E-06 | 1.200E-05 |
| 27410 | 1371 | W1371I | E | 1.6e-05 | 1.1e-05 | 1.6e-05 | 1.433E-05 | 2.887E-06 | 1.100E-05 | 1.600E-05 |
| 27411 | 1371 | W1371L | E | 1.2e-05 | 1.6e-05 | 7.6e-06 | 1.187E-05 | 4.202E-06 | 7.600E-06 | 1.600E-05 |
| 27412 | 1371 | W1371K | E | 1.5e-05 | 9.5e-06 | 1.1e-05 | 1.183E-05 | 2.843E-06 | 9.500E-06 | 1.500E-05 |
| 27413 | 1371 | W1371M | E | 1.6e-05 | 1.6e-05 | 1.1e-05 | 1.433E-05 | 2.887E-06 | 1.100E-05 | 1.600E-05 |
| 27414 | 1371 | W1371F | E | 8.7e-06 | 1.5e-05 | 1.5e-05 | 1.290E-05 | 3.637E-06 | 8.700E-06 | 1.500E-05 |
| 27415 | 1371 | W1371P | E | 1.1e-05 | 1.1e-05 | 8.6e-06 | 1.020E-05 | 1.386E-06 | 8.600E-06 | 1.100E-05 |
| 27416 | 1371 | W1371S | E | 9e-06 | 1.7e-05 | 9.1e-06 | 1.170E-05 | 4.590E-06 | 9.000E-06 | 1.700E-05 |
| 27417 | 1371 | W1371T | E | 1.7e-05 | 1.4e-05 | 1.5e-05 | 1.533E-05 | 1.528E-06 | 1.400E-05 | 1.700E-05 |
| 27418 | 1371 | W1371W | E | 1.4e-05 | 1.1e-05 | 1.5e-05 | 1.333E-05 | 2.082E-06 | 1.100E-05 | 1.500E-05 |
| 27419 | 1371 | W1371Y | E | 1e-05 | 1.1e-05 | 1.3e-05 | 1.133E-05 | 1.528E-06 | 1.000E-05 | 1.300E-05 |
| 27420 | 1371 | W1371V | E | 1.3e-05 | 1.7e-05 | 1.5e-05 | 1.500E-05 | 2.000E-06 | 1.300E-05 | 1.700E-05 |
| 27421 | 1372 | G1372A | E | 6.3e-06 | 1e-05 | 1.1e-05 | 9.100E-06 | 2.476E-06 | 6.300E-06 | 1.100E-05 |
| 27422 | 1372 | G1372R | E | 1.1e-05 | 1.4e-05 | 1.6e-05 | 1.367E-05 | 2.517E-06 | 1.100E-05 | 1.600E-05 |
| 27423 | 1372 | G1372N | E | 1.7e-05 | 1.5e-05 | 8.7e-06 | 1.357E-05 | 4.332E-06 | 8.700E-06 | 1.700E-05 |
| 27424 | 1372 | G1372D | E | 1.4e-05 | 8.4e-06 | 9.7e-06 | 1.070E-05 | 2.931E-06 | 8.400E-06 | 1.400E-05 |
| 27425 | 1372 | G1372C | E | 8.4e-06 | 1.2e-05 | 1.6e-05 | 1.213E-05 | 3.802E-06 | 8.400E-06 | 1.600E-05 |
| 27426 | 1372 | G1372E | E | 1e-05 | 9.7e-06 | 9.6e-06 | 9.767E-06 | 2.082E-07 | 9.600E-06 | 1.000E-05 |
| 27427 | 1372 | G1372Q | E | 1.4e-05 | 1e-05 | 9.9e-06 | 1.130E-05 | 2.339E-06 | 9.900E-06 | 1.400E-05 |
| 27428 | 1372 | G1372G | E | 1.3e-05 | 1e-05 | 1.2e-05 | 1.167E-05 | 1.528E-06 | 1.000E-05 | 1.300E-05 |
| 27429 | 1372 | G1372H | E | 1.4e-05 | 1.6e-05 | 9.2e-06 | 1.307E-05 | 3.495E-06 | 9.200E-06 | 1.600E-05 |
| 27430 | 1372 | G1372I | E | 1e-05 | 1e-05 | 1.4e-05 | 1.133E-05 | 2.309E-06 | 1.000E-05 | 1.400E-05 |
| 27431 | 1372 | G1372L | E | 1.1e-05 | 6.7e-06 | 1e-05 | 9.233E-06 | 2.250E-06 | 6.700E-06 | 1.100E-05 |
| 27432 | 1372 | G1372K | E | 1.3e-05 | 1.5e-05 | 1e-05 | 1.267E-05 | 2.517E-06 | 1.000E-05 | 1.500E-05 |
| 27433 | 1372 | G1372M | E | 1.5e-05 | 1.6e-05 | 1.1e-05 | 1.400E-05 | 2.646E-06 | 1.100E-05 | 1.600E-05 |
| 27434 | 1372 | G1372F | E | 1.1e-05 | 1.1e-05 | 1.1e-05 | 1.100E-05 | 2.075E-21 | 1.100E-05 | 1.100E-05 |
| 27435 | 1372 | G1372P | E | 9.8e-06 | 1.4e-05 | 1.3e-05 | 1.227E-05 | 2.194E-06 | 9.800E-06 | 1.400E-05 |
| 27436 | 1372 | G1372S | E | 1.3e-05 | 1.6e-05 | 1e-05 | 1.300E-05 | 3.000E-06 | 1.000E-05 | 1.600E-05 |
| 27437 | 1372 | G1372T | E | 1.4e-05 | 1.3e-05 | 9e-06 | 1.200E-05 | 2.646E-06 | 9.000E-06 | 1.400E-05 |
| 27438 | 1372 | G1372W | E | 1.4e-05 | 1e-05 | 1.2e-05 | 1.200E-05 | 2.000E-06 | 1.000E-05 | 1.400E-05 |
| 27439 | 1372 | G1372Y | E | 9.8e-06 | 1.5e-05 | 1.6e-05 | 1.360E-05 | 3.329E-06 | 9.800E-06 | 1.600E-05 |
| 27440 | 1372 | G1372V | E | 1e-05 | 8.4e-06 | 1.5e-05 | 1.113E-05 | 3.443E-06 | 8.400E-06 | 1.500E-05 |
| 27441 | 1373 | G1373A | E | 1.5e-05 | 1.6e-05 | 9.7e-06 | 1.357E-05 | 3.386E-06 | 9.700E-06 | 1.600E-05 |
| 27442 | 1373 | G1373R | E | 1.3e-05 | 1.6e-05 | 1.3e-05 | 1.400E-05 | 1.732E-06 | 1.300E-05 | 1.600E-05 |
| 27443 | 1373 | G1373N | E | 1.5e-05 | 8.2e-06 | 7.3e-06 | 1.017E-05 | 4.210E-06 | 7.300E-06 | 1.500E-05 |
| 27444 | 1373 | G1373D | E | 1.6e-05 | 1.2e-05 | 9.8e-06 | 1.260E-05 | 3.143E-06 | 9.800E-06 | 1.600E-05 |
| 27445 | 1373 | G1373C | E | 9e-06 | 1.6e-05 | 1.7e-05 | 1.400E-05 | 4.359E-06 | 9.000E-06 | 1.700E-05 |
| 27446 | 1373 | G1373E | E | 8.8e-06 | 9.3e-06 | 1.7e-05 | 1.170E-05 | 4.597E-06 | 8.800E-06 | 1.700E-05 |
| 27447 | 1373 | G1373Q | E | 1.1e-05 | 1.3e-05 | 1.3e-05 | 1.233E-05 | 1.155E-06 | 1.100E-05 | 1.300E-05 |
| 27448 | 1373 | G1373G | E | 1.1e-05 | 1e-05 | 1.6e-05 | 1.233E-05 | 3.215E-06 | 1.000E-05 | 1.600E-05 |
| 27449 | 1373 | G1373H | E | 1.4e-05 | 1.5e-05 | 1e-05 | 1.300E-05 | 2.646E-06 | 1.000E-05 | 1.500E-05 |
| 27450 | 1373 | G1373I | E | 1e-05 | 1.1e-05 | 1.3e-05 | 1.133E-05 | 1.528E-06 | 1.000E-05 | 1.300E-05 |
| 27451 | 1373 | G1373L | E | 9.8e-06 | 1.2e-05 | 2.7e-05 | 1.627E-05 | 9.360E-06 | 9.800E-06 | 2.700E-05 |
| 27452 | 1373 | G1373K | E | 1.1e-05 | 8.7e-06 | 1.4e-05 | 1.123E-05 | 2.658E-06 | 8.700E-06 | 1.400E-05 |
| 27453 | 1373 | G1373M | E | 1.5e-05 | 1.5e-05 | 1.4e-05 | 1.467E-05 | 5.774E-07 | 1.400E-05 | 1.500E-05 |
| 27454 | 1373 | G1373F | E | 1.5e-05 | 1.4e-05 | 1.4e-05 | 1.433E-05 | 5.774E-07 | 1.400E-05 | 1.500E-05 |

| | | | | | | | | | | |
|---|---|---|---|---|---|---|---|---|---|---|
| 27455 | 1373 | G1373P | E | 1.2e-05 | 1.5e-05 | 8.8e-06 | 1.193E-05 | 3.101E-06 | 8.800E-06 | 1.500E-05 |
| 27456 | 1373 | G1373S | E | 1.4e-05 | 1.1e-05 | 1.4e-05 | 1.300E-05 | 1.732E-06 | 1.100E-05 | 1.400E-05 |
| 27457 | 1373 | G1373T | E | 1.6e-05 | 1.1e-05 | 8.8e-06 | 1.193E-05 | 3.690E-06 | 8.800E-06 | 1.600E-05 |
| 27458 | 1373 | G1373W | E | 1.3e-05 | 1.6e-05 | 1.1e-05 | 1.333E-05 | 2.517E-06 | 1.100E-05 | 1.600E-05 |
| 27459 | 1373 | G1373Y | E | 1.5e-05 | 1.3e-05 | 1e-05 | 1.267E-05 | 2.517E-06 | 1.000E-05 | 1.500E-05 |
| 27460 | 1373 | G1373V | E | 1.3e-05 | 1.7e-05 | 8e-06 | 1.267E-05 | 4.509E-06 | 8.000E-06 | 1.700E-05 |
| 27461 | 1374 | D1374A | E | 9.8e-06 | 1e-05 | 1.7e-05 | 1.227E-05 | 4.100E-06 | 9.800E-06 | 1.700E-05 |
| 27462 | 1374 | D1374R | E | 1e-05 | 2e-05 | 1.7e-05 | 1.567E-05 | 5.132E-06 | 1.000E-05 | 2.000E-05 |
| 27463 | 1374 | D1374N | E | 1.5e-05 | 1.2e-05 | 1.2e-05 | 1.300E-05 | 1.732E-06 | 1.200E-05 | 1.500E-05 |
| 27464 | 1374 | D1374D | E | 1e-05 | 1.4e-05 | 1.5e-05 | 1.300E-05 | 2.646E-06 | 1.000E-05 | 1.500E-05 |
| 27465 | 1374 | D1374C | E | 8.7e-06 | 1.2e-05 | 9.8e-06 | 1.017E-05 | 1.680E-06 | 8.700E-06 | 1.200E-05 |
| 27466 | 1374 | D1374E | E | 1.5e-05 | 1.8e-05 | 9.7e-06 | 1.423E-05 | 4.203E-06 | 9.700E-06 | 1.800E-05 |
| 27467 | 1374 | D1374Q | E | 9.7e-06 | 1.4e-05 | 1.1e-05 | 1.157E-05 | 2.205E-06 | 9.700E-06 | 1.400E-05 |
| 27468 | 1374 | D1374G | E | 9.1e-06 | 9.9e-06 | 1.6e-05 | 1.167E-05 | 3.774E-06 | 9.100E-06 | 1.600E-05 |
| 27469 | 1374 | D1374H | E | 8.7e-06 | 1e-05 | 1.1e-05 | 9.900E-06 | 1.153E-06 | 8.700E-06 | 1.100E-05 |
| 27470 | 1374 | D1374I | E | 1.4e-05 | 1.8e-05 | 1.4e-05 | 1.533E-05 | 2.309E-06 | 1.400E-05 | 1.800E-05 |
| 27471 | 1374 | D1374L | E | 7e-06 | 1.3e-05 | 9.6e-06 | 9.867E-06 | 3.009E-06 | 7.000E-06 | 1.300E-05 |
| 27472 | 1374 | D1374K | E | 1.4e-05 | 1.3e-05 | 1.5e-05 | 1.400E-05 | 1.000E-06 | 1.300E-05 | 1.500E-05 |
| 27473 | 1374 | D1374M | E | 1.6e-05 | 1e-05 | 1.6e-05 | 1.400E-05 | 3.464E-06 | 1.000E-05 | 1.600E-05 |
| 27474 | 1374 | D1374F | E | 1.4e-05 | 1.6e-05 | 1.1e-05 | 1.367E-05 | 2.517E-06 | 1.100E-05 | 1.600E-05 |
| 27475 | 1374 | D1374P | E | 1.1e-05 | 1.4e-05 | 1.5e-05 | 1.333E-05 | 2.082E-06 | 1.100E-05 | 1.500E-05 |
| 27476 | 1374 | D1374S | E | 1.1e-05 | 1.6e-05 | 1.1e-05 | 1.267E-05 | 2.887E-06 | 1.100E-05 | 1.600E-05 |
| 27477 | 1374 | D1374T | E | 1.7e-05 | 1.6e-05 | 1.6e-05 | 1.633E-05 | 5.774E-07 | 1.600E-05 | 1.700E-05 |
| 27478 | 1374 | D1374W | E | 1.5e-05 | 1.2e-05 | 1.6e-05 | 1.433E-05 | 2.082E-06 | 1.200E-05 | 1.600E-05 |
| 27479 | 1374 | D1374Y | E | 1.1e-05 | 9.6e-06 | 1.6e-05 | 1.220E-05 | 3.365E-06 | 9.600E-06 | 1.600E-05 |
| 27480 | 1374 | D1374V | E | 1.1e-05 | 1.2e-05 | 1.1e-05 | 1.133E-05 | 5.774E-07 | 1.100E-05 | 1.200E-05 |
| 27481 | 1375 | G1375A | E | 1.1e-05 | 1.1e-05 | 1e-05 | 1.067E-05 | 5.774E-07 | 1.000E-05 | 1.100E-05 |
| 27482 | 1375 | G1375R | E | 1.1e-05 | 1.3e-05 | 1.5e-05 | 1.300E-05 | 2.000E-06 | 1.100E-05 | 1.500E-05 |
| 27483 | 1375 | G1375N | E | 1.4e-05 | 1.7e-05 | 1e-05 | 1.367E-05 | 3.512E-06 | 1.000E-05 | 1.700E-05 |
| 27484 | 1375 | G1375D | E | 1.3e-05 | 9e-06 | 1.7e-05 | 1.300E-05 | 4.000E-06 | 9.000E-06 | 1.700E-05 |
| 27485 | 1375 | G1375C | E | 1e-05 | 8.4e-06 | 1e-05 | 9.467E-06 | 9.238E-07 | 8.400E-06 | 1.000E-05 |
| 27486 | 1375 | G1375E | E | 1.3e-05 | 9.7e-06 | 9.4e-06 | 1.070E-05 | 1.997E-06 | 9.400E-06 | 1.300E-05 |
| 27487 | 1375 | G1375Q | E | 1e-05 | 9.2e-06 | 1.2e-05 | 1.040E-05 | 1.442E-06 | 9.200E-06 | 1.200E-05 |
| 27488 | 1375 | G1375G | E | 9.9e-06 | 1.5e-05 | 1.2e-05 | 1.230E-05 | 2.563E-06 | 9.900E-06 | 1.500E-05 |
| 27489 | 1375 | G1375H | E | 9.9e-06 | 1.4e-05 | 1e-05 | 1.130E-05 | 2.339E-06 | 9.900E-06 | 1.400E-05 |
| 27490 | 1375 | G1375I | E | 1e-05 | 1.5e-05 | 1.5e-05 | 1.333E-05 | 2.887E-06 | 1.000E-05 | 1.500E-05 |
| 27491 | 1375 | G1375L | E | 1.5e-05 | 1.8e-05 | 6.3e-06 | 1.310E-05 | 6.077E-06 | 6.300E-06 | 1.800E-05 |
| 27492 | 1375 | G1375K | E | 1.6e-05 | 1.5e-05 | 1.5e-05 | 1.533E-05 | 5.774E-07 | 1.500E-05 | 1.600E-05 |
| 27493 | 1375 | G1375M | E | 1.6e-05 | 1e-05 | 9.6e-06 | 1.187E-05 | 3.585E-06 | 9.600E-06 | 1.600E-05 |
| 27494 | 1375 | G1375F | E | 1.4e-05 | 1.4e-05 | 1.5e-05 | 1.433E-05 | 5.774E-07 | 1.400E-05 | 1.500E-05 |
| 27495 | 1375 | G1375P | E | 1.3e-05 | 1e-05 | 8.9e-06 | 1.063E-05 | 2.122E-06 | 8.900E-06 | 1.300E-05 |
| 27496 | 1375 | G1375S | E | 1.4e-05 | 9.5e-06 | 1.2e-05 | 1.183E-05 | 2.255E-06 | 9.500E-06 | 1.400E-05 |
| 27497 | 1375 | G1375T | E | 1.3e-05 | 1.3e-05 | 1.5e-05 | 1.367E-05 | 1.155E-06 | 1.300E-05 | 1.500E-05 |
| 27498 | 1375 | G1375W | E | 8.3e-06 | 1.5e-05 | 9.8e-06 | 1.103E-05 | 3.516E-06 | 8.300E-06 | 1.500E-05 |
| 27499 | 1375 | G1375Y | E | 1.5e-05 | 8.9e-06 | 1.2e-05 | 1.197E-05 | 3.050E-06 | 8.900E-06 | 1.500E-05 |
| 27500 | 1375 | G1375V | E | 1.3e-05 | 1.5e-05 | 8.7e-06 | 1.223E-05 | 3.219E-06 | 8.700E-06 | 1.500E-05 |
| 27501 | 1376 | F1376A | E | 8.9e-06 | 1.2e-05 | 1.6e-05 | 1.230E-05 | 3.559E-06 | 8.900E-06 | 1.600E-05 |
| 27502 | 1376 | F1376R | E | 1.4e-05 | 1.5e-05 | 1.3e-05 | 1.400E-05 | 1.000E-06 | 1.300E-05 | 1.500E-05 |
| 27503 | 1376 | F1376N | E | 1.2e-05 | 1.5e-05 | 1.2e-05 | 1.300E-05 | 1.732E-06 | 1.200E-05 | 1.500E-05 |
| 27504 | 1376 | F1376D | E | 9.5e-06 | 1.4e-05 | 1.2e-05 | 1.183E-05 | 2.255E-06 | 9.500E-06 | 1.400E-05 |
| 27505 | 1376 | F1376C | E | 7.4e-06 | 1e-05 | 1.1e-05 | 9.467E-06 | 1.858E-06 | 7.400E-06 | 1.100E-05 |
| 27506 | 1376 | F1376E | E | 1.1e-05 | 1.7e-05 | 9.9e-06 | 1.263E-05 | 3.821E-06 | 9.900E-06 | 1.700E-05 |
| 27507 | 1376 | F1376Q | E | 1e-05 | 1.3e-05 | 1.1e-05 | 1.133E-05 | 1.528E-06 | 1.000E-05 | 1.300E-05 |
| 27508 | 1376 | F1376G | E | 2.2e-05 | 1.1e-05 | 1.4e-05 | 1.567E-05 | 5.686E-06 | 1.100E-05 | 2.200E-05 |
| 27509 | 1376 | F1376H | E | 1.5e-05 | 1.1e-05 | 1.3e-05 | 1.300E-05 | 2.000E-06 | 1.100E-05 | 1.500E-05 |
| 27510 | 1376 | F1376I | E | 1.4e-05 | 1.5e-05 | 1.4e-05 | 1.433E-05 | 5.774E-07 | 1.400E-05 | 1.500E-05 |
| 27511 | 1376 | F1376L | E | 1e-05 | 1.3e-05 | 1.1e-05 | 1.133E-05 | 1.528E-06 | 1.000E-05 | 1.300E-05 |
| 27512 | 1376 | F1376K | E | 1.5e-05 | 1.5e-05 | 1.5e-05 | 1.500E-05 | 0.000E+00 | 1.500E-05 | 1.500E-05 |
| 27513 | 1376 | F1376M | E | 9.4e-06 | 9.9e-06 | 1e-05 | 9.767E-06 | 3.215E-07 | 9.400E-06 | 1.000E-05 |
| 27514 | 1376 | F1376F | E | 1.4e-05 | 1.5e-05 | 1.1e-05 | 1.333E-05 | 2.082E-06 | 1.100E-05 | 1.500E-05 |
| 27515 | 1376 | F1376P | E | 1.2e-05 | 1.3e-05 | 1.5e-05 | 1.333E-05 | 1.528E-06 | 1.200E-05 | 1.500E-05 |
| 27516 | 1376 | F1376S | E | 1.6e-05 | 9.8e-06 | 1.1e-05 | 1.227E-05 | 3.288E-06 | 9.800E-06 | 1.600E-05 |
| 27517 | 1376 | F1376T | E | 1.4e-05 | 1e-05 | 1.2e-05 | 1.200E-05 | 2.000E-06 | 1.000E-05 | 1.400E-05 |
| 27518 | 1376 | F1376W | E | 1.4e-05 | 1.2e-05 | 1.1e-05 | 1.233E-05 | 1.528E-06 | 1.100E-05 | 1.400E-05 |
| 27519 | 1376 | F1376Y | E | 8.8e-06 | 9.7e-06 | 1.4e-05 | 1.083E-05 | 2.779E-06 | 8.800E-06 | 1.400E-05 |
| 27520 | 1376 | F1376V | E | 1.1e-05 | 2.5e-05 | 1e-05 | 1.533E-05 | 8.386E-06 | 1.000E-05 | 2.500E-05 |
| 27521 | 1377 | Y1377A | E | 7.1e-06 | 1.6e-05 | 9.3e-06 | 1.080E-05 | 4.636E-06 | 7.100E-06 | 1.600E-05 |
| 27522 | 1377 | Y1377R | E | 1.6e-05 | 1.2e-05 | 2.2e-05 | 1.667E-05 | 5.033E-06 | 1.200E-05 | 2.200E-05 |
| 27523 | 1377 | Y1377N | E | 9.6e-06 | 1e-05 | 1.3e-05 | 1.087E-05 | 1.858E-06 | 9.600E-06 | 1.300E-05 |
| 27524 | 1377 | Y1377D | E | 9e-06 | 8.8e-06 | 1.1e-05 | 9.600E-06 | 1.217E-06 | 8.800E-06 | 1.100E-05 |
| 27525 | 1377 | Y1377C | E | 1.5e-05 | 1.4e-05 | 1.2e-05 | 1.367E-05 | 1.528E-06 | 1.200E-05 | 1.500E-05 |
| 27526 | 1377 | Y1377E | E | 1.6e-05 | 1.5e-05 | 1.1e-05 | 1.400E-05 | 2.646E-06 | 1.100E-05 | 1.600E-05 |
| 27527 | 1377 | Y1377Q | E | 1.5e-05 | 9.5e-06 | 9.9e-06 | 1.147E-05 | 3.066E-06 | 9.500E-06 | 1.500E-05 |
| 27528 | 1377 | Y1377G | E | 1.5e-05 | 9.9e-06 | 1e-05 | 1.163E-05 | 2.916E-06 | 9.900E-06 | 1.500E-05 |
| 27529 | 1377 | Y1377H | E | 1.5e-05 | 1.2e-05 | 1.7e-05 | 1.467E-05 | 2.517E-06 | 1.200E-05 | 1.700E-05 |
| 27530 | 1377 | Y1377I | E | 8.9e-06 | 1e-05 | 9.6e-06 | 9.500E-06 | 5.568E-07 | 8.900E-06 | 1.000E-05 |
| 27531 | 1377 | Y1377L | E | 1.8e-05 | 9.4e-06 | 1.4e-05 | 1.380E-05 | 4.303E-06 | 9.400E-06 | 1.800E-05 |
| 27532 | 1377 | Y1377K | E | 8.5e-06 | 1.6e-05 | 1.5e-05 | 1.317E-05 | 4.072E-06 | 8.500E-06 | 1.600E-05 |
| 27533 | 1377 | Y1377M | E | 1.1e-05 | 1.7e-05 | 9e-06 | 1.233E-05 | 4.163E-06 | 9.000E-06 | 1.700E-05 |
| 27534 | 1377 | Y1377F | E | 8.6e-06 | 1.4e-05 | 1.1e-05 | 1.120E-05 | 2.706E-06 | 8.600E-06 | 1.400E-05 |
| 27535 | 1377 | Y1377P | E | 1.1e-05 | 1.2e-05 | 1.5e-05 | 1.267E-05 | 2.082E-06 | 1.100E-05 | 1.500E-05 |
| 27536 | 1377 | Y1377S | E | 1e-05 | 1.1e-05 | 1e-05 | 1.033E-05 | 5.774E-07 | 1.000E-05 | 1.100E-05 |
| 27537 | 1377 | Y1377T | E | 1.4e-05 | 9.7e-06 | 1.5e-05 | 1.290E-05 | 2.816E-06 | 9.700E-06 | 1.500E-05 |
| 27538 | 1377 | Y1377W | E | 1.3e-05 | 1.6e-05 | 1.3e-05 | 1.400E-05 | 1.732E-06 | 1.300E-05 | 1.600E-05 |
| 27539 | 1377 | Y1377Y | E | 1.4e-05 | 1.6e-05 | 1e-05 | 1.333E-05 | 3.055E-06 | 1.000E-05 | 1.600E-05 |

| | | | | | | | | | | |
|---|---|---|---|---|---|---|---|---|---|---|
| 27540 | 1377 | Y1377V | E | 1.8e-05 | 9.5e-06 | 9.1e-06 | 1.220E-05 | 5.027E-06 | 9.100E-06 | 1.800E-05 |
| 27541 | 1378 | A1378A | E | 1.5e-05 | 1.5e-05 | 9.2e-06 | 1.307E-05 | 3.349E-06 | 9.200E-06 | 1.500E-05 |
| 27542 | 1378 | A1378R | E | 1.1e-05 | 1.3e-05 | 1.1e-05 | 1.167E-05 | 1.155E-06 | 1.100E-05 | 1.300E-05 |
| 27543 | 1378 | A1378N | E | 1.4e-05 | 9.7e-06 | 1.6e-05 | 1.323E-05 | 3.219E-06 | 9.700E-06 | 1.600E-05 |
| 27544 | 1378 | A1378D | E | 1.2e-05 | 7.2e-06 | 9.5e-06 | 9.567E-06 | 2.401E-06 | 7.200E-06 | 1.200E-05 |
| 27545 | 1378 | A1378C | E | 1.5e-05 | 7.6e-06 | 8.1e-06 | 1.023E-05 | 4.136E-06 | 7.600E-06 | 1.500E-05 |
| 27546 | 1378 | A1378E | E | 1.5e-05 | 1.4e-05 | 1.4e-05 | 1.433E-05 | 5.774E-07 | 1.400E-05 | 1.500E-05 |
| 27547 | 1378 | A1378Q | E | 1.6e-05 | 8.9e-06 | 1.6e-05 | 1.363E-05 | 4.099E-06 | 8.900E-06 | 1.600E-05 |
| 27548 | 1378 | A1378G | E | 1.1e-05 | 9.8e-06 | 1.1e-05 | 1.060E-05 | 6.928E-07 | 9.800E-06 | 1.100E-05 |
| 27549 | 1378 | A1378H | E | 1.7e-05 | 8.9e-06 | 8.6e-06 | 1.150E-05 | 4.766E-06 | 8.600E-06 | 1.700E-05 |
| 27550 | 1378 | A1378I | E | 1e-05 | 2.6e-05 | 1.5e-05 | 1.700E-05 | 8.185E-06 | 1.000E-05 | 2.600E-05 |
| 27551 | 1378 | A1378L | E | 1.4e-05 | 1e-05 | 1.1e-05 | 1.167E-05 | 2.082E-06 | 1.000E-05 | 1.400E-05 |
| 27552 | 1378 | A1378K | E | 1.5e-05 | 8.8e-06 | 1.4e-05 | 1.260E-05 | 3.329E-06 | 8.800E-06 | 1.500E-05 |
| 27553 | 1378 | A1378M | E | 1.7e-05 | 1.5e-05 | 1.5e-05 | 1.567E-05 | 1.155E-06 | 1.500E-05 | 1.700E-05 |
| 27554 | 1378 | A1378F | E | 1.1e-05 | 9.5e-06 | 1.1e-05 | 1.050E-05 | 8.660E-07 | 9.500E-06 | 1.100E-05 |
| 27555 | 1378 | A1378P | E | 1.3e-05 | 1.5e-05 | 1.2e-05 | 1.333E-05 | 1.528E-06 | 1.200E-05 | 1.500E-05 |
| 27556 | 1378 | A1378S | E | 1.5e-05 | 1e-05 | 1.5e-05 | 1.333E-05 | 2.887E-06 | 1.000E-05 | 1.500E-05 |
| 27557 | 1378 | A1378T | E | 1.6e-05 | 1.4e-05 | 1.4e-05 | 1.467E-05 | 1.155E-06 | 1.400E-05 | 1.600E-05 |
| 27558 | 1378 | A1378W | E | 1.4e-05 | 1.6e-05 | 1.5e-05 | 1.500E-05 | 1.000E-06 | 1.400E-05 | 1.600E-05 |
| 27559 | 1378 | A1378Y | E | 1.6e-05 | 8.2e-06 | 1.4e-05 | 1.273E-05 | 4.051E-06 | 8.200E-06 | 1.600E-05 |
| 27560 | 1378 | A1378V | E | 1.3e-05 | 1.5e-05 | 1e-05 | 1.267E-05 | 2.517E-06 | 1.000E-05 | 1.500E-05 |
| 27561 | 1379 | M1379A | E | 1.1e-05 | 1.5e-05 | 8.6e-06 | 1.153E-05 | 3.233E-06 | 8.600E-06 | 1.500E-05 |
| 27562 | 1379 | M1379R | E | 1.5e-05 | 1.1e-05 | 1.7e-05 | 1.433E-05 | 3.055E-06 | 1.100E-05 | 1.700E-05 |
| 27563 | 1379 | M1379N | E | 1.6e-05 | 7.8e-06 | 1.1e-05 | 1.160E-05 | 4.133E-06 | 7.800E-06 | 1.600E-05 |
| 27564 | 1379 | M1379D | E | 9.6e-06 | 8.7e-06 | 8.7e-06 | 9.000E-06 | 5.196E-07 | 8.700E-06 | 9.600E-06 |
| 27565 | 1379 | M1379C | E | 1.1e-05 | 1.4e-05 | 1.5e-05 | 1.333E-05 | 2.082E-06 | 1.100E-05 | 1.500E-05 |
| 27566 | 1379 | M1379E | E | 1.4e-05 | 1.6e-05 | 1.3e-05 | 1.433E-05 | 1.528E-06 | 1.300E-05 | 1.600E-05 |
| 27567 | 1379 | M1379Q | E | 1.2e-05 | 1.8e-05 | 8e-06 | 1.267E-05 | 5.033E-06 | 8.000E-06 | 1.800E-05 |
| 27568 | 1379 | M1379G | E | 8.3e-06 | 1.1e-05 | 1.4e-05 | 1.110E-05 | 2.851E-06 | 8.300E-06 | 1.400E-05 |
| 27569 | 1379 | M1379H | E | 9.4e-06 | 1e-05 | 1.2e-05 | 1.047E-05 | 1.361E-06 | 9.400E-06 | 1.200E-05 |
| 27570 | 1379 | M1379I | E | 1.5e-05 | 1.6e-05 | 1.1e-05 | 1.400E-05 | 2.646E-06 | 1.100E-05 | 1.600E-05 |
| 27571 | 1379 | M1379L | E | 1e-05 | 8.5e-06 | 1e-05 | 9.500E-06 | 8.660E-07 | 8.500E-06 | 1.000E-05 |
| 27572 | 1379 | M1379K | E | 1.4e-05 | 1.1e-05 | 1e-05 | 1.167E-05 | 2.082E-06 | 1.000E-05 | 1.400E-05 |
| 27573 | 1379 | M1379M | E | 1.5e-05 | 1.3e-05 | 1.3e-05 | 1.367E-05 | 1.155E-06 | 1.300E-05 | 1.500E-05 |
| 27574 | 1379 | M1379F | E | 1.1e-05 | 1e-05 | 1.5e-05 | 1.200E-05 | 2.646E-06 | 1.000E-05 | 1.500E-05 |
| 27575 | 1379 | M1379P | E | 1.6e-05 | 1.3e-05 | 8.9e-06 | 1.263E-05 | 3.564E-06 | 8.900E-06 | 1.600E-05 |
| 27576 | 1379 | M1379S | E | 1.1e-05 | 1.5e-05 | 1.6e-05 | 1.400E-05 | 2.646E-06 | 1.100E-05 | 1.600E-05 |
| 27577 | 1379 | M1379T | E | 1.5e-05 | 1.3e-05 | 8.6e-06 | 1.220E-05 | 3.274E-06 | 8.600E-06 | 1.500E-05 |
| 27578 | 1379 | M1379W | E | 1.2e-05 | 1.6e-05 | 1.6e-05 | 1.467E-05 | 2.309E-06 | 1.200E-05 | 1.600E-05 |
| 27579 | 1379 | M1379Y | E | 9e-06 | 1.7e-05 | 1.5e-05 | 1.367E-05 | 4.163E-06 | 9.000E-06 | 1.700E-05 |
| 27580 | 1379 | M1379V | E | 1.4e-05 | 1.4e-05 | 1.4e-05 | 1.400E-05 | 0.000E+00 | 1.400E-05 | 1.400E-05 |
| 27581 | 1380 | D1380A | E | 1.5e-05 | 1.5e-05 | 8.7e-06 | 1.290E-05 | 3.637E-06 | 8.700E-06 | 1.500E-05 |
| 27582 | 1380 | D1380R | E | 1.5e-05 | 1.7e-05 | 1e-05 | 1.400E-05 | 3.606E-06 | 1.000E-05 | 1.700E-05 |
| 27583 | 1380 | D1380N | E | 1.7e-05 | 1.3e-05 | 9.8e-06 | 1.327E-05 | 3.607E-06 | 9.800E-06 | 1.700E-05 |
| 27584 | 1380 | D1380D | E | 1.1e-05 | 1.5e-05 | 1.6e-05 | 1.400E-05 | 2.646E-06 | 1.100E-05 | 1.600E-05 |
| 27585 | 1380 | D1380C | E | 1.6e-05 | 1.1e-05 | 1.3e-05 | 1.333E-05 | 2.517E-06 | 1.100E-05 | 1.600E-05 |
| 27586 | 1380 | D1380E | E | 1.6e-05 | 1.5e-05 | 9.8e-06 | 1.360E-05 | 3.329E-06 | 9.800E-06 | 1.600E-05 |
| 27587 | 1380 | D1380Q | E | 1.1e-05 | 8.3e-06 | 1.4e-05 | 1.110E-05 | 2.851E-06 | 8.300E-06 | 1.400E-05 |
| 27588 | 1380 | D1380G | E | 1.1e-05 | 1.6e-05 | 1.5e-05 | 1.400E-05 | 2.646E-06 | 1.100E-05 | 1.600E-05 |
| 27589 | 1380 | D1380H | E | 1.5e-05 | 1.4e-05 | 1.5e-05 | 1.467E-05 | 5.774E-07 | 1.400E-05 | 1.500E-05 |
| 27590 | 1380 | D1380I | E | 1e-05 | 1.5e-05 | 1.6e-05 | 1.367E-05 | 3.215E-06 | 1.000E-05 | 1.600E-05 |
| 27591 | 1380 | D1380L | E | 1.4e-05 | 9.4e-06 | 1.5e-05 | 1.280E-05 | 2.987E-06 | 9.400E-06 | 1.500E-05 |
| 27592 | 1380 | D1380K | E | 1.1e-05 | 1.5e-05 | 1e-05 | 1.200E-05 | 2.646E-06 | 1.000E-05 | 1.500E-05 |
| 27593 | 1380 | D1380M | E | 8.9e-06 | 1.1e-05 | 1.1e-05 | 1.030E-05 | 1.212E-06 | 8.900E-06 | 1.100E-05 |
| 27594 | 1380 | D1380F | E | 9.7e-06 | 1.5e-05 | 1.3e-05 | 1.257E-05 | 2.676E-06 | 9.700E-06 | 1.500E-05 |
| 27595 | 1380 | D1380P | E | 1e-05 | 1.5e-05 | 1.6e-05 | 1.367E-05 | 3.215E-06 | 1.000E-05 | 1.600E-05 |
| 27596 | 1380 | D1380S | E | 1.3e-05 | 1.2e-05 | 8.8e-06 | 1.127E-05 | 2.194E-06 | 8.800E-06 | 1.300E-05 |
| 27597 | 1380 | D1380T | E | 1.5e-05 | 1.3e-05 | 1e-05 | 1.267E-05 | 2.517E-06 | 1.000E-05 | 1.500E-05 |
| 27598 | 1380 | D1380W | E | 1.5e-05 | 1.1e-05 | 1.4e-05 | 1.333E-05 | 2.082E-06 | 1.100E-05 | 1.500E-05 |
| 27599 | 1380 | D1380Y | E | 9.3e-06 | 1e-05 | 1.6e-05 | 1.177E-05 | 3.683E-06 | 9.300E-06 | 1.600E-05 |
| 27600 | 1380 | D1380V | E | 1.4e-05 | 9.8e-06 | 1.1e-05 | 1.160E-05 | 2.163E-06 | 9.800E-06 | 1.400E-05 |
| 27601 | 1381 | Y1381A | E | 1.1e-05 | 2e-05 | 1.4e-05 | 1.500E-05 | 4.583E-06 | 1.100E-05 | 2.000E-05 |
| 27602 | 1381 | Y1381R | E | 1.5e-05 | 1.6e-05 | 1.1e-05 | 1.400E-05 | 2.646E-06 | 1.100E-05 | 1.600E-05 |
| 27603 | 1381 | Y1381N | E | 1.1e-05 | 1.1e-05 | 1e-05 | 1.067E-05 | 5.774E-07 | 1.000E-05 | 1.100E-05 |
| 27604 | 1381 | Y1381D | E | 9.5e-06 | 1.7e-05 | 1.5e-05 | 1.383E-05 | 3.884E-06 | 9.500E-06 | 1.700E-05 |
| 27605 | 1381 | Y1381C | E | 7.8e-06 | 1.6e-05 | 1e-05 | 1.127E-05 | 4.244E-06 | 7.800E-06 | 1.600E-05 |
| 27606 | 1381 | Y1381E | E | 2e-05 | 1.6e-05 | 1.2e-05 | 1.600E-05 | 4.000E-06 | 1.200E-05 | 2.000E-05 |
| 27607 | 1381 | Y1381Q | E | 1.1e-05 | 1e-05 | 1.5e-05 | 1.200E-05 | 2.646E-06 | 1.000E-05 | 1.500E-05 |
| 27608 | 1381 | Y1381G | E | 1.6e-05 | 1.1e-05 | 1.6e-05 | 1.433E-05 | 2.887E-06 | 1.100E-05 | 1.600E-05 |
| 27609 | 1381 | Y1381H | E | 1.3e-05 | 1.2e-05 | 1.3e-05 | 1.267E-05 | 5.774E-07 | 1.200E-05 | 1.300E-05 |
| 27610 | 1381 | Y1381I | E | 1.7e-05 | 1.4e-05 | 1.1e-05 | 1.400E-05 | 3.000E-06 | 1.100E-05 | 1.700E-05 |
| 27611 | 1381 | Y1381L | E | 1.7e-05 | 1e-05 | 1.3e-05 | 1.333E-05 | 3.512E-06 | 1.000E-05 | 1.700E-05 |
| 27612 | 1381 | Y1381K | E | 1.1e-05 | 1.7e-05 | 1.9e-05 | 1.567E-05 | 4.163E-06 | 1.100E-05 | 1.900E-05 |
| 27613 | 1381 | Y1381M | E | 9.7e-06 | 1.1e-05 | 7.3e-06 | 9.333E-06 | 1.877E-06 | 7.300E-06 | 1.100E-05 |
| 27614 | 1381 | Y1381F | E | 1e-05 | 1.3e-05 | 1.4e-05 | 1.233E-05 | 2.082E-06 | 1.000E-05 | 1.400E-05 |
| 27615 | 1381 | Y1381P | E | 1e-05 | 9.9e-06 | 1.4e-05 | 1.130E-05 | 2.339E-06 | 9.900E-06 | 1.400E-05 |
| 27616 | 1381 | Y1381S | E | 1.3e-05 | 7.2e-06 | 1.3e-05 | 1.107E-05 | 3.349E-06 | 7.200E-06 | 1.300E-05 |
| 27617 | 1381 | Y1381T | E | 1.4e-05 | 1e-05 | 1.6e-05 | 1.333E-05 | 3.055E-06 | 1.000E-05 | 1.600E-05 |
| 27618 | 1381 | Y1381W | E | 1.3e-05 | 1.4e-05 | 1.3e-05 | 1.333E-05 | 5.774E-07 | 1.300E-05 | 1.400E-05 |
| 27619 | 1381 | Y1381Y | E | 1.4e-05 | 1e-05 | 1.4e-05 | 1.267E-05 | 2.309E-06 | 1.000E-05 | 1.400E-05 |
| 27620 | 1381 | Y1381V | E | 1.2e-05 | 1.4e-05 | 1.1e-05 | 1.233E-05 | 1.528E-06 | 1.100E-05 | 1.400E-05 |
| 27621 | 1382 | W1382A | E | 1e-05 | 7.5e-06 | 9.9e-06 | 9.133E-06 | 1.415E-06 | 7.500E-06 | 1.000E-05 |
| 27622 | 1382 | W1382R | E | 9.6e-06 | 9.6e-06 | 1.6e-05 | 1.173E-05 | 3.695E-06 | 9.600E-06 | 1.600E-05 |
| 27623 | 1382 | W1382N | E | 1.2e-05 | 9.1e-06 | 1.4e-05 | 1.170E-05 | 2.464E-06 | 9.100E-06 | 1.400E-05 |
| 27624 | 1382 | W1382D | E | 1.4e-05 | 1.8e-05 | 1.5e-05 | 1.567E-05 | 2.082E-06 | 1.400E-05 | 1.800E-05 |

| | | | | | | | | | | |
|---|---|---|---|---|---|---|---|---|---|---|
| 27625 | 1382 | W1382C | E | 1.4e-05 | 1.2e-05 | 1.3e-05 | 1.300E-05 | 1.000E-06 | 1.200E-05 | 1.400E-05 |
| 27626 | 1382 | W1382E | E | 1.3e-05 | 1.4e-05 | 1.1e-05 | 1.267E-05 | 1.528E-06 | 1.100E-05 | 1.400E-05 |
| 27627 | 1382 | W1382Q | E | 1.6e-05 | 1.6e-05 | 1.1e-05 | 1.433E-05 | 2.887E-06 | 1.100E-05 | 1.600E-05 |
| 27628 | 1382 | W1382G | E | 1.5e-05 | 1.6e-05 | 1e-05 | 1.367E-05 | 3.215E-06 | 1.000E-05 | 1.600E-05 |
| 27629 | 1382 | W1382H | E | 1.5e-05 | 1.1e-05 | 1.3e-05 | 1.300E-05 | 2.000E-06 | 1.100E-05 | 1.500E-05 |
| 27630 | 1382 | W1382I | E | 1.1e-05 | 1.6e-05 | 1.1e-05 | 1.267E-05 | 2.887E-06 | 1.100E-05 | 1.600E-05 |
| 27631 | 1382 | W1382L | E | 1.7e-05 | 9e-06 | 1.5e-05 | 1.367E-05 | 4.163E-06 | 9.000E-06 | 1.700E-05 |
| 27632 | 1382 | W1382K | E | 1.4e-05 | 1.5e-05 | 1.6e-05 | 1.500E-05 | 1.000E-06 | 1.400E-05 | 1.600E-05 |
| 27633 | 1382 | W1382M | E | 9.8e-06 | 1.8e-05 | 1.5e-05 | 1.427E-05 | 4.149E-06 | 9.800E-06 | 1.800E-05 |
| 27634 | 1382 | W1382F | E | 1.6e-05 | 1.9e-05 | 1.1e-05 | 1.533E-05 | 4.041E-06 | 1.100E-05 | 1.900E-05 |
| 27635 | 1382 | W1382P | E | 1.4e-05 | 1.2e-05 | 1.2e-05 | 1.267E-05 | 1.155E-06 | 1.200E-05 | 1.400E-05 |
| 27636 | 1382 | W1382S | E | 8.6e-06 | 1.5e-05 | 1.4e-05 | 1.253E-05 | 3.443E-06 | 8.600E-06 | 1.500E-05 |
| 27637 | 1382 | W1382T | E | 1.3e-05 | 1.8e-05 | 1.6e-05 | 1.567E-05 | 2.517E-06 | 1.300E-05 | 1.800E-05 |
| 27638 | 1382 | W1382W | E | 1.4e-05 | 7.5e-06 | 1.2e-05 | 1.117E-05 | 3.329E-06 | 7.500E-06 | 1.400E-05 |
| 27639 | 1382 | W1382Y | E | 8.2e-06 | 1.1e-05 | 1.5e-05 | 1.140E-05 | 3.418E-06 | 8.200E-06 | 1.500E-05 |
| 27640 | 1382 | W1382V | E | 1.5e-05 | 1.5e-05 | 7.9e-06 | 1.263E-05 | 4.099E-06 | 7.900E-06 | 1.500E-05 |
| 27641 | 1383 | G1383A | E | 8.8e-06 | 1e-05 | 8.7e-06 | 9.167E-06 | 7.234E-07 | 8.700E-06 | 1.000E-05 |
| 27642 | 1383 | G1383R | E | 1.5e-05 | 1.5e-05 | 1.4e-05 | 1.467E-05 | 5.774E-07 | 1.400E-05 | 1.500E-05 |
| 27643 | 1383 | G1383N | E | 7.6e-06 | 1.5e-05 | 1.5e-05 | 1.253E-05 | 4.272E-06 | 7.600E-06 | 1.500E-05 |
| 27644 | 1383 | G1383D | E | 1.7e-05 | 1.1e-05 | 1.1e-05 | 1.300E-05 | 3.464E-06 | 1.100E-05 | 1.700E-05 |
| 27645 | 1383 | G1383C | E | 1.3e-05 | 1.2e-05 | 1.1e-05 | 1.200E-05 | 1.000E-06 | 1.100E-05 | 1.300E-05 |
| 27646 | 1383 | G1383E | E | 1.5e-05 | 1.1e-05 | 1.4e-05 | 1.333E-05 | 2.082E-06 | 1.100E-05 | 1.500E-05 |
| 27647 | 1383 | G1383Q | E | 1.1e-05 | 1.2e-05 | 1.4e-05 | 1.233E-05 | 1.528E-06 | 1.100E-05 | 1.400E-05 |
| 27648 | 1383 | G1383G | E | 9e-06 | 1.1e-05 | 1e-05 | 1.000E-05 | 1.000E-06 | 9.000E-06 | 1.100E-05 |
| 27649 | 1383 | G1383H | E | 9.3e-06 | 1.1e-05 | 1.6e-05 | 1.210E-05 | 3.483E-06 | 9.300E-06 | 1.600E-05 |
| 27650 | 1383 | G1383I | E | 6.5e-06 | 1.5e-05 | 1.4e-05 | 1.183E-05 | 4.646E-06 | 6.500E-06 | 1.500E-05 |
| 27651 | 1383 | G1383L | E | 9.2e-06 | 9.6e-06 | 1.5e-05 | 1.127E-05 | 3.239E-06 | 9.200E-06 | 1.500E-05 |
| 27652 | 1383 | G1383K | E | 1.5e-05 | 1.4e-05 | 1.4e-05 | 1.433E-05 | 5.774E-07 | 1.400E-05 | 1.500E-05 |
| 27653 | 1383 | G1383M | E | 1.2e-05 | 1.1e-05 | 1.1e-05 | 1.133E-05 | 5.774E-07 | 1.100E-05 | 1.200E-05 |
| 27654 | 1383 | G1383F | E | 1.5e-05 | 1.1e-05 | 1.4e-05 | 1.333E-05 | 2.082E-06 | 1.100E-05 | 1.500E-05 |
| 27655 | 1383 | G1383P | E | 1e-05 | 1.5e-05 | 1e-05 | 1.167E-05 | 2.887E-06 | 1.000E-05 | 1.500E-05 |
| 27656 | 1383 | G1383S | E | 1.6e-05 | 1.6e-05 | 1.6e-05 | 1.600E-05 | 0.000E+00 | 1.600E-05 | 1.600E-05 |
| 27657 | 1383 | G1383T | E | 1.5e-05 | 1.4e-05 | 9.7e-06 | 1.290E-05 | 2.816E-06 | 9.700E-06 | 1.500E-05 |
| 27658 | 1383 | G1383W | E | 1.5e-05 | 1.5e-05 | 8.5e-06 | 1.283E-05 | 3.753E-06 | 8.500E-06 | 1.500E-05 |
| 27659 | 1383 | G1383Y | E | 1.4e-05 | 1.1e-05 | 1.6e-05 | 1.367E-05 | 2.517E-06 | 1.100E-05 | 1.600E-05 |
| 27660 | 1383 | G1383V | E | 1.6e-05 | 1.5e-05 | 1.9e-05 | 1.667E-05 | 2.082E-06 | 1.500E-05 | 1.900E-05 |
| 27661 | 1384 | Q1384A | E | 1.2e-05 | 1.5e-05 | 9.9e-06 | 1.230E-05 | 2.563E-06 | 9.900E-06 | 1.500E-05 |
| 27662 | 1384 | Q1384R | E | 1.5e-05 | 1.2e-05 | 1.6e-05 | 1.433E-05 | 2.082E-06 | 1.200E-05 | 1.600E-05 |
| 27663 | 1384 | Q1384N | E | 1.2e-05 | 1.6e-05 | 1.4e-05 | 1.400E-05 | 2.000E-06 | 1.200E-05 | 1.600E-05 |
| 27664 | 1384 | Q1384D | E | 1.1e-05 | 1.1e-05 | 1.2e-05 | 1.133E-05 | 5.774E-07 | 1.100E-05 | 1.200E-05 |
| 27665 | 1384 | Q1384C | E | 1.3e-05 | 1.5e-05 | 1.3e-05 | 1.367E-05 | 1.155E-06 | 1.300E-05 | 1.500E-05 |
| 27666 | 1384 | Q1384E | E | 8.6e-06 | 1e-05 | 1.1e-05 | 9.867E-06 | 1.206E-06 | 8.600E-06 | 1.100E-05 |
| 27667 | 1384 | Q1384Q | E | 8.8e-06 | 1.1e-05 | 1.8e-05 | 1.260E-05 | 4.804E-06 | 8.800E-06 | 1.800E-05 |
| 27668 | 1384 | Q1384G | E | 1.2e-05 | 1.2e-05 | 1.3e-05 | 1.233E-05 | 5.774E-07 | 1.200E-05 | 1.300E-05 |
| 27669 | 1384 | Q1384H | E | 1.5e-05 | 1.6e-05 | 8.9e-06 | 1.330E-05 | 3.843E-06 | 8.900E-06 | 1.600E-05 |
| 27670 | 1384 | Q1384I | E | 1.1e-05 | 1.2e-05 | 1.6e-05 | 1.300E-05 | 2.646E-06 | 1.100E-05 | 1.600E-05 |
| 27671 | 1384 | Q1384L | E | 1.6e-05 | 1.4e-05 | 1e-05 | 1.333E-05 | 3.055E-06 | 1.000E-05 | 1.600E-05 |
| 27672 | 1384 | Q1384K | E | 1.8e-05 | 9.4e-06 | 1.6e-05 | 1.447E-05 | 4.500E-06 | 9.400E-06 | 1.800E-05 |
| 27673 | 1384 | Q1384M | E | 1.6e-05 | 1.5e-05 | 1.4e-05 | 1.500E-05 | 1.000E-06 | 1.400E-05 | 1.600E-05 |
| 27674 | 1384 | Q1384F | E | 1.9e-05 | 1.4e-05 | 9.4e-06 | 1.413E-05 | 4.801E-06 | 9.400E-06 | 1.900E-05 |
| 27675 | 1384 | Q1384P | E | 8.5e-06 | 1e-05 | 1.7e-05 | 1.183E-05 | 4.537E-06 | 8.500E-06 | 1.700E-05 |
| 27676 | 1384 | Q1384S | E | 1.6e-05 | 1.4e-05 | 1.6e-05 | 1.533E-05 | 1.155E-06 | 1.400E-05 | 1.600E-05 |
| 27677 | 1384 | Q1384T | E | 1.3e-05 | 1e-05 | 1.6e-05 | 1.300E-05 | 3.000E-06 | 1.000E-05 | 1.600E-05 |
| 27678 | 1384 | Q1384W | E | 1.3e-05 | 1.2e-05 | 1.1e-05 | 1.200E-05 | 1.000E-06 | 1.100E-05 | 1.300E-05 |
| 27679 | 1384 | Q1384Y | E | 1.3e-05 | 1.5e-05 | 9.8e-06 | 1.260E-05 | 2.623E-06 | 9.800E-06 | 1.500E-05 |
| 27680 | 1384 | Q1384V | E | 1.5e-05 | 1.1e-05 | 1.6e-05 | 1.400E-05 | 2.646E-06 | 1.100E-05 | 1.600E-05 |
| 27681 | 1385 | G1385A | E | 9.4e-06 | 7.3e-06 | 1.5e-05 | 1.057E-05 | 3.980E-06 | 7.300E-06 | 1.500E-05 |
| 27682 | 1385 | G1385R | E | 6.5e-06 | 1.1e-05 | 1.6e-05 | 1.117E-05 | 4.752E-06 | 6.500E-06 | 1.600E-05 |
| 27683 | 1385 | G1385N | E | 1.3e-05 | 1.4e-05 | 1e-05 | 1.233E-05 | 2.082E-06 | 1.000E-05 | 1.400E-05 |
| 27684 | 1385 | G1385D | E | 1e-05 | 1.6e-05 | 1.1e-05 | 1.233E-05 | 3.215E-06 | 1.000E-05 | 1.600E-05 |
| 27685 | 1385 | G1385C | E | 9.2e-06 | 8.6e-06 | 1.6e-05 | 1.127E-05 | 4.110E-06 | 8.600E-06 | 1.600E-05 |
| 27686 | 1385 | G1385E | E | 9.4e-06 | 1.5e-05 | 9.9e-06 | 1.143E-05 | 3.099E-06 | 9.400E-06 | 1.500E-05 |
| 27687 | 1385 | G1385Q | E | 1.1e-05 | 1.3e-05 | 1.1e-05 | 1.167E-05 | 1.155E-06 | 1.100E-05 | 1.300E-05 |
| 27688 | 1385 | G1385G | E | 1e-05 | 1.1e-05 | 1.2e-05 | 1.100E-05 | 1.000E-06 | 1.000E-05 | 1.200E-05 |
| 27689 | 1385 | G1385H | E | 1.5e-05 | 1.5e-05 | 1.5e-05 | 1.500E-05 | 0.000E+00 | 1.500E-05 | 1.500E-05 |
| 27690 | 1385 | G1385I | E | 1.2e-05 | 1.4e-05 | 1e-05 | 1.200E-05 | 2.000E-06 | 1.000E-05 | 1.400E-05 |
| 27691 | 1385 | G1385L | E | 1e-05 | 1.6e-05 | 1.5e-05 | 1.367E-05 | 3.215E-06 | 1.000E-05 | 1.600E-05 |
| 27692 | 1385 | G1385K | E | 8.5e-06 | 1.5e-05 | 8.9e-06 | 1.080E-05 | 3.643E-06 | 8.500E-06 | 1.500E-05 |
| 27693 | 1385 | G1385M | E | 1.4e-05 | 1.2e-05 | 1.5e-05 | 1.367E-05 | 1.528E-06 | 1.200E-05 | 1.500E-05 |
| 27694 | 1385 | G1385F | E | 1e-05 | 1.5e-05 | 1.3e-05 | 1.267E-05 | 2.517E-06 | 1.000E-05 | 1.500E-05 |
| 27695 | 1385 | G1385P | E | 1.4e-05 | 1.7e-05 | 9.6e-06 | 1.353E-05 | 3.722E-06 | 9.600E-06 | 1.700E-05 |
| 27696 | 1385 | G1385S | E | 1.4e-05 | 1.4e-05 | 1.3e-05 | 1.367E-05 | 5.774E-07 | 1.300E-05 | 1.400E-05 |
| 27697 | 1385 | G1385T | E | 1.1e-05 | 9.7e-06 | 1e-05 | 1.023E-05 | 6.807E-07 | 9.700E-06 | 1.100E-05 |
| 27698 | 1385 | G1385W | E | 1.2e-05 | 1.5e-05 | 1e-05 | 1.233E-05 | 2.517E-06 | 1.000E-05 | 1.500E-05 |
| 27699 | 1385 | G1385Y | E | 1.4e-05 | 1.4e-05 | 1.6e-05 | 1.467E-05 | 1.155E-06 | 1.400E-05 | 1.600E-05 |
| 27700 | 1385 | G1385V | E | 1e-05 | 1.2e-05 | 1.4e-05 | 1.200E-05 | 2.000E-06 | 1.000E-05 | 1.400E-05 |
| 27701 | 1386 | T1386A | E | 1.5e-05 | 1.6e-05 | 6.1e-06 | 1.237E-05 | 5.450E-06 | 6.100E-06 | 1.600E-05 |
| 27702 | 1386 | T1386R | E | 1.4e-05 | 1e-05 | 1.2e-05 | 1.200E-05 | 2.000E-06 | 1.000E-05 | 1.400E-05 |
| 27703 | 1386 | T1386N | E | 1e-05 | 1.8e-05 | 9.3e-06 | 1.243E-05 | 4.834E-06 | 9.300E-06 | 1.800E-05 |
| 27704 | 1386 | T1386D | E | 1.6e-05 | 9.2e-06 | 1.2e-05 | 1.240E-05 | 3.418E-06 | 9.200E-06 | 1.600E-05 |
| 27705 | 1386 | T1386C | E | 1.2e-05 | 6.5e-06 | 1.8e-05 | 1.217E-05 | 5.752E-06 | 6.500E-06 | 1.800E-05 |
| 27706 | 1386 | T1386E | E | 1.5e-05 | 1e-05 | 1.7e-05 | 1.400E-05 | 3.606E-06 | 1.000E-05 | 1.700E-05 |
| 27707 | 1386 | T1386Q | E | 1.2e-05 | 1.1e-05 | 1.7e-05 | 1.333E-05 | 3.215E-06 | 1.100E-05 | 1.700E-05 |
| 27708 | 1386 | T1386G | E | 1.1e-05 | 1.1e-05 | 1.4e-05 | 1.200E-05 | 1.732E-06 | 1.100E-05 | 1.400E-05 |
| 27709 | 1386 | T1386H | E | 1.5e-05 | 8.3e-06 | 9.6e-06 | 1.097E-05 | 3.553E-06 | 8.300E-06 | 1.500E-05 |

| | | | | | | | | | | |
|---|---|---|---|---|---|---|---|---|---|---|
| 27710 | 1386 | T1386I | E | 1.7e-05 | 1.7e-05 | 1.6e-05 | 1.667E-05 | 5.774E-07 | 1.600E-05 | 1.700E-05 |
| 27711 | 1386 | T1386L | E | 1.5e-05 | 1.1e-05 | 9.6e-06 | 1.187E-05 | 2.802E-06 | 9.600E-06 | 1.500E-05 |
| 27712 | 1386 | T1386K | E | 1.5e-05 | 9.5e-06 | 1.5e-05 | 1.317E-05 | 3.175E-06 | 9.500E-06 | 1.500E-05 |
| 27713 | 1386 | T1386M | E | 7.4e-06 | 9.9e-06 | 1.9e-05 | 1.210E-05 | 6.105E-06 | 7.400E-06 | 1.900E-05 |
| 27714 | 1386 | T1386F | E | 1.6e-05 | 1.1e-05 | 1.3e-05 | 1.333E-05 | 2.517E-06 | 1.100E-05 | 1.600E-05 |
| 27715 | 1386 | T1386P | E | 1.5e-05 | 1.5e-05 | 1.3e-05 | 1.433E-05 | 1.155E-06 | 1.300E-05 | 1.500E-05 |
| 27716 | 1386 | T1386S | E | 9.4e-06 | 1.1e-05 | 1e-05 | 1.013E-05 | 8.083E-07 | 9.400E-06 | 1.100E-05 |
| 27717 | 1386 | T1386T | E | 8.4e-06 | 1.1e-05 | 9.1e-06 | 9.500E-06 | 1.345E-06 | 8.400E-06 | 1.100E-05 |
| 27718 | 1386 | T1386W | E | 1.1e-05 | 1.4e-05 | 1.3e-05 | 1.267E-05 | 1.528E-06 | 1.100E-05 | 1.400E-05 |
| 27719 | 1386 | T1386Y | E | 1e-05 | 1.1e-05 | 9.4e-06 | 1.013E-05 | 8.083E-07 | 9.400E-06 | 1.100E-05 |
| 27720 | 1386 | T1386V | E | 1.6e-05 | 1e-05 | 9.6e-06 | 1.187E-05 | 3.585E-06 | 9.600E-06 | 1.600E-05 |
| 27721 | 1387 | L1387A | E | 1.7e-05 | 1e-05 | 1.2e-05 | 1.300E-05 | 3.606E-06 | 1.000E-05 | 1.700E-05 |
| 27722 | 1387 | L1387R | E | 1.7e-05 | 1e-05 | 9.4e-06 | 1.213E-05 | 4.225E-06 | 9.400E-06 | 1.700E-05 |
| 27723 | 1387 | L1387N | E | 9.3e-06 | 8.4e-06 | 1.5e-05 | 1.090E-05 | 3.579E-06 | 8.400E-06 | 1.500E-05 |
| 27724 | 1387 | L1387D | E | 9.9e-06 | 9.7e-06 | 1e-05 | 9.867E-06 | 1.528E-07 | 9.700E-06 | 1.000E-05 |
| 27725 | 1387 | L1387C | E | 9.6e-06 | 1.4e-05 | 1.2e-05 | 1.187E-05 | 2.203E-06 | 9.600E-06 | 1.400E-05 |
| 27726 | 1387 | L1387E | E | 1.5e-05 | 9.7e-06 | 1.5e-05 | 1.323E-05 | 3.060E-06 | 9.700E-06 | 1.500E-05 |
| 27727 | 1387 | L1387Q | E | 1.7e-05 | 1.4e-05 | 1.3e-05 | 1.467E-05 | 2.082E-06 | 1.300E-05 | 1.700E-05 |
| 27728 | 1387 | L1387G | E | 9.4e-06 | 8.5e-06 | 1.5e-05 | 1.097E-05 | 3.522E-06 | 8.500E-06 | 1.500E-05 |
| 27729 | 1387 | L1387H | E | 9.8e-06 | 9e-06 | 1.1e-05 | 9.933E-06 | 1.007E-06 | 9.000E-06 | 1.100E-05 |
| 27730 | 1387 | L1387I | E | 1.1e-05 | 1.6e-05 | 8e-06 | 1.167E-05 | 4.041E-06 | 8.000E-06 | 1.600E-05 |
| 27731 | 1387 | L1387L | E | 1.1e-05 | 1.2e-05 | 8.2e-06 | 1.040E-05 | 1.970E-06 | 8.200E-06 | 1.200E-05 |
| 27732 | 1387 | L1387K | E | 1.5e-05 | 9.1e-06 | 1.6e-05 | 1.337E-05 | 3.729E-06 | 9.100E-06 | 1.600E-05 |
| 27733 | 1387 | L1387M | E | 9.3e-06 | 1.5e-05 | 9.4e-06 | 1.123E-05 | 3.262E-06 | 9.300E-06 | 1.500E-05 |
| 27734 | 1387 | L1387F | E | 1e-05 | 1.4e-05 | 1.6e-05 | 1.333E-05 | 3.055E-06 | 1.000E-05 | 1.600E-05 |
| 27735 | 1387 | L1387P | E | 1.5e-05 | 1e-05 | 1.5e-05 | 1.333E-05 | 2.887E-06 | 1.000E-05 | 1.500E-05 |
| 27736 | 1387 | L1387S | E | 8.7e-06 | 1.1e-05 | 1.4e-05 | 1.123E-05 | 2.658E-06 | 8.700E-06 | 1.400E-05 |
| 27737 | 1387 | L1387T | E | 1.5e-05 | 1.6e-05 | 1.7e-05 | 1.600E-05 | 1.000E-06 | 1.500E-05 | 1.700E-05 |
| 27738 | 1387 | L1387W | E | 1.5e-05 | 1.5e-05 | 8.7e-06 | 1.290E-05 | 3.637E-06 | 8.700E-06 | 1.500E-05 |
| 27739 | 1387 | L1387Y | E | 1.5e-05 | 1.5e-05 | 1.2e-05 | 1.400E-05 | 1.732E-06 | 1.200E-05 | 1.500E-05 |
| 27740 | 1387 | L1387V | E | 1e-05 | 1.3e-05 | 9.3e-06 | 1.077E-05 | 1.966E-06 | 9.300E-06 | 1.300E-05 |
| 27741 | 1388 | V1388A | E | 1e-05 | 1.5e-05 | 1.3e-05 | 1.267E-05 | 2.517E-06 | 1.000E-05 | 1.500E-05 |
| 27742 | 1388 | V1388R | E | 1.6e-05 | 1.2e-05 | 1e-05 | 1.267E-05 | 3.055E-06 | 1.000E-05 | 1.600E-05 |
| 27743 | 1388 | V1388N | E | 1.6e-05 | 1.5e-05 | 1.4e-05 | 1.500E-05 | 1.000E-06 | 1.400E-05 | 1.600E-05 |
| 27744 | 1388 | V1388D | E | 1.3e-05 | 1.4e-05 | 8.9e-06 | 1.197E-05 | 2.702E-06 | 8.900E-06 | 1.400E-05 |
| 27745 | 1388 | V1388C | E | 9.9e-06 | 1e-05 | 1.4e-05 | 1.130E-05 | 2.339E-06 | 9.900E-06 | 1.400E-05 |
| 27746 | 1388 | V1388E | E | 1.2e-05 | 1.4e-05 | 1.3e-05 | 1.300E-05 | 1.000E-06 | 1.200E-05 | 1.400E-05 |
| 27747 | 1388 | V1388Q | E | 1.4e-05 | 1.1e-05 | 1.1e-05 | 1.200E-05 | 1.732E-06 | 1.100E-05 | 1.400E-05 |
| 27748 | 1388 | V1388G | E | 9e-06 | 1.7e-05 | 1.1e-05 | 1.233E-05 | 4.163E-06 | 9.000E-06 | 1.700E-05 |
| 27749 | 1388 | V1388H | E | 1.4e-05 | 1.5e-05 | 1.1e-05 | 1.333E-05 | 2.082E-06 | 1.100E-05 | 1.500E-05 |
| 27750 | 1388 | V1388I | E | 1.6e-05 | 1.6e-05 | 1.6e-05 | 1.600E-05 | 0.000E+00 | 1.600E-05 | 1.600E-05 |
| 27751 | 1388 | V1388L | E | 9.9e-06 | 1.4e-05 | 1e-05 | 1.130E-05 | 2.339E-06 | 9.900E-06 | 1.400E-05 |
| 27752 | 1388 | V1388K | E | 1.7e-05 | 1e-05 | 1.5e-05 | 1.400E-05 | 3.606E-06 | 1.000E-05 | 1.700E-05 |
| 27753 | 1388 | V1388M | E | 1.5e-05 | 1.4e-05 | 1.1e-05 | 1.333E-05 | 2.082E-06 | 1.100E-05 | 1.500E-05 |
| 27754 | 1388 | V1388F | E | 5.8e-06 | 1e-05 | 1.3e-05 | 9.600E-06 | 3.617E-06 | 5.800E-06 | 1.300E-05 |
| 27755 | 1388 | V1388P | E | 1.5e-05 | 9.7e-06 | 1.3e-05 | 1.257E-05 | 2.676E-06 | 9.700E-06 | 1.500E-05 |
| 27756 | 1388 | V1388S | E | 1.4e-05 | 1.5e-05 | 1.5e-05 | 1.467E-05 | 5.774E-07 | 1.400E-05 | 1.500E-05 |
| 27757 | 1388 | V1388T | E | 9.2e-06 | 1.6e-05 | 8.9e-06 | 1.137E-05 | 4.015E-06 | 8.900E-06 | 1.600E-05 |
| 27758 | 1388 | V1388W | E | 1.2e-05 | 1.2e-05 | 1.5e-05 | 1.300E-05 | 1.732E-06 | 1.200E-05 | 1.500E-05 |
| 27759 | 1388 | V1388Y | E | 1.1e-05 | 1.1e-05 | 1.1e-05 | 1.100E-05 | 2.075E-21 | 1.100E-05 | 1.100E-05 |
| 27760 | 1388 | V1388V | E | 1.3e-05 | 1e-05 | 7.5e-06 | 1.017E-05 | 2.754E-06 | 7.500E-06 | 1.300E-05 |
| 27761 | 1389 | T1389A | E | 1.4e-05 | 1.3e-05 | 1e-05 | 1.233E-05 | 2.082E-06 | 1.000E-05 | 1.400E-05 |
| 27762 | 1389 | T1389R | E | 1.6e-05 | 1.6e-05 | 8.2e-06 | 1.340E-05 | 4.503E-06 | 8.200E-06 | 1.600E-05 |
| 27763 | 1389 | T1389N | E | 9.1e-06 | 1.7e-05 | 1.1e-05 | 1.237E-05 | 4.124E-06 | 9.100E-06 | 1.700E-05 |
| 27764 | 1389 | T1389D | E | 1.6e-05 | 1.5e-05 | 1.4e-05 | 1.500E-05 | 1.000E-06 | 1.400E-05 | 1.600E-05 |
| 27765 | 1389 | T1389C | E | 1.3e-05 | 1e-05 | 1.4e-05 | 1.233E-05 | 2.082E-06 | 1.000E-05 | 1.400E-05 |
| 27766 | 1389 | T1389E | E | 1.5e-05 | 1.1e-05 | 1.4e-05 | 1.333E-05 | 2.082E-06 | 1.100E-05 | 1.500E-05 |
| 27767 | 1389 | T1389Q | E | 1.7e-05 | 1.6e-05 | 9.7e-06 | 1.423E-05 | 3.958E-06 | 9.700E-06 | 1.700E-05 |
| 27768 | 1389 | T1389G | E | 1.2e-05 | 1.1e-05 | 1.5e-05 | 1.267E-05 | 2.082E-06 | 1.100E-05 | 1.500E-05 |
| 27769 | 1389 | T1389H | E | 1.1e-05 | 8.9e-06 | 1.4e-05 | 1.130E-05 | 2.563E-06 | 8.900E-06 | 1.400E-05 |
| 27770 | 1389 | T1389I | E | 1.4e-05 | 1.4e-05 | 1.5e-05 | 1.433E-05 | 5.774E-07 | 1.400E-05 | 1.500E-05 |
| 27771 | 1389 | T1389L | E | 1.6e-05 | 1.7e-05 | 1.5e-05 | 1.600E-05 | 1.000E-06 | 1.500E-05 | 1.700E-05 |
| 27772 | 1389 | T1389K | E | 1.4e-05 | 1.6e-05 | 1.4e-05 | 1.467E-05 | 1.155E-06 | 1.400E-05 | 1.600E-05 |
| 27773 | 1389 | T1389M | E | 1.5e-05 | 1e-05 | 1.1e-05 | 1.200E-05 | 2.646E-06 | 1.000E-05 | 1.500E-05 |
| 27774 | 1389 | T1389F | E | 8.4e-06 | 1.1e-05 | 1.5e-05 | 1.147E-05 | 3.325E-06 | 8.400E-06 | 1.500E-05 |
| 27775 | 1389 | T1389P | E | 1.6e-05 | 1.4e-05 | 1.4e-05 | 1.467E-05 | 1.155E-06 | 1.400E-05 | 1.600E-05 |
| 27776 | 1389 | T1389S | E | 1e-05 | 1.1e-05 | 1.5e-05 | 1.200E-05 | 2.646E-06 | 1.000E-05 | 1.500E-05 |
| 27777 | 1389 | T1389T | E | 1.6e-05 | 1.3e-05 | 1.3e-05 | 1.400E-05 | 1.732E-06 | 1.300E-05 | 1.600E-05 |
| 27778 | 1389 | T1389W | E | 1e-05 | 1e-05 | 1.5e-05 | 1.167E-05 | 2.887E-06 | 1.000E-05 | 1.500E-05 |
| 27779 | 1389 | T1389Y | E | 9.2e-06 | 1.2e-05 | 1e-05 | 1.040E-05 | 1.442E-06 | 9.200E-06 | 1.200E-05 |
| 27780 | 1389 | T1389V | E | 1.1e-05 | 1.5e-05 | 1e-05 | 1.200E-05 | 2.646E-06 | 1.000E-05 | 1.500E-05 |
| 27781 | 1390 | V1390A | E | 9.4e-06 | 1.2e-05 | 1.7e-05 | 1.280E-05 | 3.863E-06 | 9.400E-06 | 1.700E-05 |
| 27782 | 1390 | V1390R | E | 9.6e-06 | 1.5e-05 | 1.4e-05 | 1.287E-05 | 2.873E-06 | 9.600E-06 | 1.500E-05 |
| 27783 | 1390 | V1390N | E | 1.6e-05 | 1.5e-05 | 1.6e-05 | 1.567E-05 | 5.774E-07 | 1.500E-05 | 1.600E-05 |
| 27784 | 1390 | V1390D | E | 1e-05 | 9.1e-06 | 1.5e-05 | 1.137E-05 | 3.179E-06 | 9.100E-06 | 1.500E-05 |
| 27785 | 1390 | V1390C | E | 1.5e-05 | 1e-05 | 1.1e-05 | 1.200E-05 | 2.646E-06 | 1.000E-05 | 1.500E-05 |
| 27786 | 1390 | V1390E | E | 1.2e-05 | 1.5e-05 | 8.8e-06 | 1.193E-05 | 3.101E-06 | 8.800E-06 | 1.500E-05 |
| 27787 | 1390 | V1390Q | E | 1.3e-05 | 1.1e-05 | 1.6e-05 | 1.333E-05 | 2.517E-06 | 1.100E-05 | 1.600E-05 |
| 27788 | 1390 | V1390G | E | 1.1e-05 | 1.5e-05 | 8.4e-06 | 1.147E-05 | 3.325E-06 | 8.400E-06 | 1.500E-05 |
| 27789 | 1390 | V1390H | E | 1.5e-05 | 1.1e-05 | 1.1e-05 | 1.233E-05 | 2.309E-06 | 1.100E-05 | 1.500E-05 |
| 27790 | 1390 | V1390I | E | 9.9e-06 | 1.4e-05 | 1.7e-05 | 1.363E-05 | 3.564E-06 | 9.900E-06 | 1.700E-05 |
| 27791 | 1390 | V1390L | E | 1.1e-05 | 8.2e-06 | 1.6e-05 | 1.173E-05 | 3.951E-06 | 8.200E-06 | 1.600E-05 |
| 27792 | 1390 | V1390K | E | 1.1e-05 | 9.3e-06 | 1.5e-05 | 1.177E-05 | 2.926E-06 | 9.300E-06 | 1.500E-05 |
| 27793 | 1390 | V1390M | E | 1.5e-05 | 1.7e-05 | 1.2e-05 | 1.467E-05 | 2.517E-06 | 1.200E-05 | 1.700E-05 |
| 27794 | 1390 | V1390F | E | 8.3e-06 | 1.5e-05 | 8.7e-06 | 1.067E-05 | 3.758E-06 | 8.300E-06 | 1.500E-05 |

| | | | | | | | | | | |
|---|---|---|---|---|---|---|---|---|---|---|
| 27795 | 1390 | V1390P | E | 5.8e-06 | 1.4e-05 | 1.4e-05 | 1.127E-05 | 4.734E-06 | 5.800E-06 | 1.400E-05 |
| 27796 | 1390 | V1390S | E | 1.4e-05 | 1.6e-05 | 1.1e-05 | 1.367E-05 | 2.517E-06 | 1.100E-05 | 1.600E-05 |
| 27797 | 1390 | V1390T | E | 9.5e-06 | 1e-05 | 1.4e-05 | 1.117E-05 | 2.466E-06 | 9.500E-06 | 1.400E-05 |
| 27798 | 1390 | V1390W | E | 1.5e-05 | 9.7e-06 | 1.3e-05 | 1.257E-05 | 2.676E-06 | 9.700E-06 | 1.500E-05 |
| 27799 | 1390 | V1390Y | E | 1.5e-05 | 1.1e-05 | 9.7e-06 | 1.190E-05 | 2.762E-06 | 9.700E-06 | 1.500E-05 |
| 27800 | 1390 | V1390V | E | 1e-05 | 1.6e-05 | 1.6e-05 | 1.400E-05 | 3.464E-06 | 1.000E-05 | 1.600E-05 |
| 27801 | 1391 | S1391A | E | 1.2e-05 | 7.6e-06 | 1.5e-05 | 1.153E-05 | 3.722E-06 | 7.600E-06 | 1.500E-05 |
| 27802 | 1391 | S1391R | E | 1.5e-05 | 8e-06 | 1.3e-05 | 1.200E-05 | 3.606E-06 | 8.000E-06 | 1.500E-05 |
| 27803 | 1391 | S1391N | E | 1.5e-05 | 1.6e-05 | 8.1e-06 | 1.303E-05 | 4.302E-06 | 8.100E-06 | 1.600E-05 |
| 27804 | 1391 | S1391D | E | 1.1e-05 | 1.3e-05 | 1.6e-05 | 1.333E-05 | 2.517E-06 | 1.100E-05 | 1.600E-05 |
| 27805 | 1391 | S1391C | E | 1.2e-05 | 1.5e-05 | 1.4e-05 | 1.367E-05 | 1.528E-06 | 1.200E-05 | 1.500E-05 |
| 27806 | 1391 | S1391E | E | 1e-05 | 1.1e-05 | 1.4e-05 | 1.167E-05 | 2.082E-06 | 1.000E-05 | 1.400E-05 |
| 27807 | 1391 | S1391Q | E | 1.1e-05 | 1.5e-05 | 1.5e-05 | 1.367E-05 | 2.309E-06 | 1.100E-05 | 1.500E-05 |
| 27808 | 1391 | S1391G | E | 1.7e-05 | 1.1e-05 | 1.1e-05 | 1.300E-05 | 3.464E-06 | 1.100E-05 | 1.700E-05 |
| 27809 | 1391 | S1391H | E | 1.1e-05 | 1.3e-05 | 1.2e-05 | 1.200E-05 | 1.000E-06 | 1.100E-05 | 1.300E-05 |
| 27810 | 1391 | S1391I | E | 1e-05 | 1.1e-05 | 1.5e-05 | 1.200E-05 | 2.646E-06 | 1.000E-05 | 1.500E-05 |
| 27811 | 1391 | S1391L | E | 1.4e-05 | 1.1e-05 | 9.9e-06 | 1.163E-05 | 2.122E-06 | 9.900E-06 | 1.400E-05 |
| 27812 | 1391 | S1391K | E | 9.8e-06 | 1.6e-05 | 1e-05 | 1.193E-05 | 3.523E-06 | 9.800E-06 | 1.600E-05 |
| 27813 | 1391 | S1391M | E | 1e-05 | 1.2e-05 | 9.7e-06 | 1.057E-05 | 1.250E-06 | 9.700E-06 | 1.200E-05 |
| 27814 | 1391 | S1391F | E | 1.6e-05 | 1.3e-05 | 1.1e-05 | 1.333E-05 | 2.517E-06 | 1.100E-05 | 1.600E-05 |
| 27815 | 1391 | S1391P | E | 1.4e-05 | 8.2e-06 | 1.6e-05 | 1.273E-05 | 4.051E-06 | 8.200E-06 | 1.600E-05 |
| 27816 | 1391 | S1391S | E | 1.2e-05 | 1.2e-05 | 9.1e-06 | 1.103E-05 | 1.674E-06 | 9.100E-06 | 1.200E-05 |
| 27817 | 1391 | S1391T | E | 1.4e-05 | 1.1e-05 | 1.6e-05 | 1.367E-05 | 2.517E-06 | 1.100E-05 | 1.600E-05 |
| 27818 | 1391 | S1391W | E | 1.5e-05 | 1.1e-05 | 9.4e-06 | 1.180E-05 | 2.884E-06 | 9.400E-06 | 1.500E-05 |
| 27819 | 1391 | S1391Y | E | 1.5e-05 | 1.2e-05 | 1.7e-05 | 1.467E-05 | 2.517E-06 | 1.200E-05 | 1.700E-05 |
| 27820 | 1391 | S1391V | E | 1e-05 | 1.1e-05 | 1.1e-05 | 1.067E-05 | 5.774E-07 | 1.000E-05 | 1.100E-05 |
| 27821 | 1392 | S1392A | E | 1.6e-05 | 1.3e-05 | 1.6e-05 | 1.500E-05 | 1.732E-06 | 1.300E-05 | 1.600E-05 |
| 27822 | 1392 | S1392R | E | 1.4e-05 | 1.4e-05 | 1.1e-05 | 1.300E-05 | 1.732E-06 | 1.100E-05 | 1.400E-05 |
| 27823 | 1392 | S1392N | E | 1.2e-05 | 1.4e-05 | 1.6e-05 | 1.400E-05 | 2.000E-06 | 1.200E-05 | 1.600E-05 |
| 27824 | 1392 | S1392D | E | 1.4e-05 | 9.2e-06 | 1.5e-05 | 1.273E-05 | 3.101E-06 | 9.200E-06 | 1.500E-05 |
| 27825 | 1392 | S1392C | E | 8.7e-06 | 1e-05 | 8.4e-06 | 9.033E-06 | 8.505E-07 | 8.400E-06 | 1.000E-05 |
| 27826 | 1392 | S1392E | E | 9e-06 | 1.6e-05 | 9e-06 | 1.133E-05 | 4.041E-06 | 9.000E-06 | 1.600E-05 |
| 27827 | 1392 | S1392Q | E | 9.5e-06 | 1e-05 | 1e-05 | 9.833E-06 | 2.887E-07 | 9.500E-06 | 1.000E-05 |
| 27828 | 1392 | S1392G | E | 1.6e-05 | 1.6e-05 | 1.4e-05 | 1.533E-05 | 1.155E-06 | 1.400E-05 | 1.600E-05 |
| 27829 | 1392 | S1392H | E | 9.8e-06 | 1.1e-05 | 1.1e-05 | 1.060E-05 | 6.928E-07 | 9.800E-06 | 1.100E-05 |
| 27830 | 1392 | S1392I | E | 1.7e-05 | 1.5e-05 | 1.4e-05 | 1.533E-05 | 1.528E-06 | 1.400E-05 | 1.700E-05 |
| 27831 | 1392 | S1392L | E | 1.3e-05 | 1.5e-05 | 1e-05 | 1.267E-05 | 2.517E-06 | 1.000E-05 | 1.500E-05 |
| 27832 | 1392 | S1392K | E | 9.6e-06 | 1.5e-05 | 9.2e-06 | 1.127E-05 | 3.239E-06 | 9.200E-06 | 1.500E-05 |
| 27833 | 1392 | S1392M | E | 1e-05 | 1.3e-05 | 1.5e-05 | 1.267E-05 | 2.517E-06 | 1.000E-05 | 1.500E-05 |
| 27834 | 1392 | S1392F | E | 1.3e-05 | 1.1e-05 | 1.2e-05 | 1.200E-05 | 1.000E-06 | 1.100E-05 | 1.300E-05 |
| 27835 | 1392 | S1392P | E | 1.4e-05 | 1.6e-05 | 1.6e-05 | 1.533E-05 | 1.155E-06 | 1.400E-05 | 1.600E-05 |
| 27836 | 1392 | S1392S | E | 1.2e-05 | 1.3e-05 | 1.3e-05 | 1.267E-05 | 5.774E-07 | 1.200E-05 | 1.300E-05 |
| 27837 | 1392 | S1392T | E | 1.6e-05 | 5.9e-06 | 1e-05 | 1.063E-05 | 5.080E-06 | 5.900E-06 | 1.600E-05 |
| 27838 | 1392 | S1392W | E | 1.7e-05 | 1.5e-05 | 1.4e-05 | 1.533E-05 | 1.528E-06 | 1.400E-05 | 1.700E-05 |
| 27839 | 1392 | S1392Y | E | 1.4e-05 | 1.5e-05 | 9.6e-06 | 1.287E-05 | 2.873E-06 | 9.600E-06 | 1.500E-05 |
| 27840 | 1392 | S1392V | E | 8.4e-06 | 1.5e-05 | 1.5e-05 | 1.280E-05 | 3.811E-06 | 8.400E-06 | 1.500E-05 |
| 27841 | 1393 | A1393A | E | 1e-05 | 7e-06 | 1.2e-05 | 9.667E-06 | 2.517E-06 | 7.000E-06 | 1.200E-05 |
| 27842 | 1393 | A1393R | E | 1.3e-05 | 1.5e-05 | 1e-05 | 1.267E-05 | 2.517E-06 | 1.000E-05 | 1.500E-05 |
| 27843 | 1393 | A1393N | E | 1.4e-05 | 1.6e-05 | 9e-06 | 1.300E-05 | 3.606E-06 | 9.000E-06 | 1.600E-05 |
| 27844 | 1393 | A1393D | E | 8.2e-06 | 1e-05 | 1.6e-05 | 1.140E-05 | 4.084E-06 | 8.200E-06 | 1.600E-05 |
| 27845 | 1393 | A1393C | E | 1e-05 | 1.1e-05 | 1e-05 | 1.033E-05 | 5.774E-07 | 1.000E-05 | 1.100E-05 |
| 27846 | 1393 | A1393E | E | 9.5e-06 | 1.5e-05 | 1.4e-05 | 1.283E-05 | 2.930E-06 | 9.500E-06 | 1.500E-05 |
| 27847 | 1393 | A1393Q | E | 6.3e-06 | 1.1e-05 | 1.6e-05 | 1.110E-05 | 4.851E-06 | 6.300E-06 | 1.600E-05 |
| 27848 | 1393 | A1393G | E | 1.6e-05 | 1.1e-05 | 1.1e-05 | 1.267E-05 | 2.887E-06 | 1.100E-05 | 1.600E-05 |
| 27849 | 1393 | A1393H | E | 1.8e-05 | 1.1e-05 | 1e-05 | 1.300E-05 | 4.359E-06 | 1.000E-05 | 1.800E-05 |
| 27850 | 1393 | A1393I | E | 1e-05 | 1.3e-05 | 1e-05 | 1.100E-05 | 1.732E-06 | 1.000E-05 | 1.300E-05 |
| 27851 | 1393 | A1393L | E | 1.3e-05 | 1.2e-05 | 1.1e-05 | 1.200E-05 | 1.000E-06 | 1.100E-05 | 1.300E-05 |
| 27852 | 1393 | A1393K | E | 1.1e-05 | 1.1e-05 | 1.6e-05 | 1.267E-05 | 2.887E-06 | 1.100E-05 | 1.600E-05 |
| 27853 | 1393 | A1393M | E | 1.5e-05 | 8.4e-06 | 1.6e-05 | 1.313E-05 | 4.130E-06 | 8.400E-06 | 1.600E-05 |
| 27854 | 1393 | A1393F | E | 1.3e-05 | 1.6e-05 | 1.3e-05 | 1.400E-05 | 1.732E-06 | 1.300E-05 | 1.600E-05 |
| 27855 | 1393 | A1393P | E | 1.2e-05 | 1e-05 | 1.4e-05 | 1.200E-05 | 2.000E-06 | 1.000E-05 | 1.400E-05 |
| 27856 | 1393 | A1393S | E | 1.1e-05 | 1.3e-05 | 2.2e-05 | 1.533E-05 | 5.859E-06 | 1.100E-05 | 2.200E-05 |
| 27857 | 1393 | A1393T | E | 1e-05 | 1.4e-05 | 1.5e-05 | 1.300E-05 | 2.646E-06 | 1.000E-05 | 1.500E-05 |
| 27858 | 1393 | A1393W | E | 1.6e-05 | 1.1e-05 | 1.4e-05 | 1.367E-05 | 2.517E-06 | 1.100E-05 | 1.600E-05 |
| 27859 | 1393 | A1393Y | E | 1.2e-05 | 1.3e-05 | 1.3e-05 | 1.267E-05 | 5.774E-07 | 1.200E-05 | 1.300E-05 |
| 27860 | 1393 | A1393V | E | 9.4e-06 | 1.6e-05 | 1e-05 | 1.180E-05 | 3.650E-06 | 9.400E-06 | 1.600E-05 |
| 27861 | 1394 | S1394A | E | 1.4e-05 | 9.1e-06 | 1.1e-05 | 1.137E-05 | 2.470E-06 | 9.100E-06 | 1.400E-05 |
| 27862 | 1394 | S1394R | E | 1.6e-05 | 9.7e-06 | 1.5e-05 | 1.357E-05 | 3.386E-06 | 9.700E-06 | 1.600E-05 |
| 27863 | 1394 | S1394N | E | 1.6e-05 | 9e-06 | 8.9e-06 | 1.130E-05 | 4.071E-06 | 8.900E-06 | 1.600E-05 |
| 27864 | 1394 | S1394D | E | 1.5e-05 | 1.4e-05 | 1.1e-05 | 1.333E-05 | 2.082E-06 | 1.100E-05 | 1.500E-05 |
| 27865 | 1394 | S1394C | E | 1.2e-05 | 1.3e-05 | 1.5e-05 | 1.333E-05 | 1.528E-06 | 1.200E-05 | 1.500E-05 |
| 27866 | 1394 | S1394E | E | 1.5e-05 | 1.5e-05 | 1.7e-05 | 1.567E-05 | 1.155E-06 | 1.500E-05 | 1.700E-05 |
| 27867 | 1394 | S1394Q | E | 1e-05 | 1.1e-05 | 1e-05 | 1.033E-05 | 5.774E-07 | 1.000E-05 | 1.100E-05 |
| 27868 | 1394 | S1394G | E | 1.1e-05 | 1.1e-05 | 1e-05 | 1.067E-05 | 5.774E-07 | 1.000E-05 | 1.100E-05 |
| 27869 | 1394 | S1394H | E | 1e-05 | 1.6e-05 | 1.5e-05 | 1.367E-05 | 3.215E-06 | 1.000E-05 | 1.600E-05 |
| 27870 | 1394 | S1394I | E | 1.7e-05 | 1.1e-05 | 1.2e-05 | 1.333E-05 | 3.215E-06 | 1.100E-05 | 1.700E-05 |
| 27871 | 1394 | S1394L | E | 1.6e-05 | 1.2e-05 | 1.5e-05 | 1.433E-05 | 2.082E-06 | 1.200E-05 | 1.600E-05 |
| 27872 | 1394 | S1394K | E | 1.5e-05 | 1.2e-05 | 1.1e-05 | 1.267E-05 | 2.082E-06 | 1.100E-05 | 1.500E-05 |
| 27873 | 1394 | S1394M | E | 9.6e-06 | 1.1e-05 | 1.4e-05 | 1.153E-05 | 2.248E-06 | 9.600E-06 | 1.400E-05 |
| 27874 | 1394 | S1394F | E | 1.7e-05 | 7.7e-06 | 1.4e-05 | 1.290E-05 | 4.747E-06 | 7.700E-06 | 1.700E-05 |
| 27875 | 1394 | S1394P | E | 1.6e-05 | 1.1e-05 | 9.7e-06 | 1.223E-05 | 3.326E-06 | 9.700E-06 | 1.600E-05 |
| 27876 | 1394 | S1394S | E | 9.7e-06 | 1.1e-05 | 1.6e-05 | 1.223E-05 | 3.326E-06 | 9.700E-06 | 1.600E-05 |
| 27877 | 1394 | S1394T | E | 1.1e-05 | 1.4e-05 | 9.7e-06 | 1.157E-05 | 2.205E-06 | 9.700E-06 | 1.400E-05 |
| 27878 | 1394 | S1394W | E | 1.1e-05 | 9.7e-06 | 1.1e-05 | 1.057E-05 | 7.506E-07 | 9.700E-06 | 1.100E-05 |
| 27879 | 1394 | S1394Y | E | 1e-05 | 1.1e-05 | 1.5e-05 | 1.200E-05 | 2.646E-06 | 1.000E-05 | 1.500E-05 |

| | | | | | | | | | | |
|---|---|---|---|---|---|---|---|---|---|---|
| 27880 | 1394 | S1394V | E | 1.1e-05 | 9.5e-06 | 9.7e-06 | 1.007E-05 | 8.145E-07 | 9.500E-06 | 1.100E-05 |
| 27881 | 1395 | T1395A | E | 1.1e-05 | 1.3e-05 | 9.8e-06 | 1.127E-05 | 1.617E-06 | 9.800E-06 | 1.300E-05 |
| 27882 | 1395 | T1395R | E | 1.6e-05 | 1.1e-05 | 1.6e-05 | 1.433E-05 | 2.887E-06 | 1.100E-05 | 1.600E-05 |
| 27883 | 1395 | T1395N | E | 1.5e-05 | 1.5e-05 | 1.1e-05 | 1.367E-05 | 2.309E-06 | 1.100E-05 | 1.500E-05 |
| 27884 | 1395 | T1395D | E | 1.1e-05 | 1.4e-05 | 1.4e-05 | 1.300E-05 | 1.732E-06 | 1.100E-05 | 1.400E-05 |
| 27885 | 1395 | T1395C | E | 1.5e-05 | 1e-05 | 1.4e-05 | 1.300E-05 | 2.646E-06 | 1.000E-05 | 1.500E-05 |
| 27886 | 1395 | T1395E | E | 1.7e-05 | 1.7e-05 | 1.5e-05 | 1.633E-05 | 1.155E-06 | 1.500E-05 | 1.700E-05 |
| 27887 | 1395 | T1395Q | E | 1e-05 | 1.4e-05 | 1e-05 | 1.133E-05 | 2.309E-06 | 1.000E-05 | 1.400E-05 |
| 27888 | 1395 | T1395G | E | 1.1e-05 | 2.4e-05 | 1.3e-05 | 1.600E-05 | 7.000E-06 | 1.100E-05 | 2.400E-05 |
| 27889 | 1395 | T1395H | E | 1.6e-05 | 1.8e-05 | 1.5e-05 | 1.633E-05 | 1.528E-06 | 1.500E-05 | 1.800E-05 |
| 27890 | 1395 | T1395I | E | 1.5e-05 | 1.4e-05 | 1e-05 | 1.300E-05 | 2.646E-06 | 1.000E-05 | 1.500E-05 |
| 27891 | 1395 | T1395L | E | 2.1e-05 | 1.6e-05 | 1.5e-05 | 1.733E-05 | 3.215E-06 | 1.500E-05 | 2.100E-05 |
| 27892 | 1395 | T1395K | E | 9.8e-06 | 1.2e-05 | 1.5e-05 | 1.227E-05 | 2.610E-06 | 9.800E-06 | 1.500E-05 |
| 27893 | 1395 | T1395M | E | 1.2e-05 | 1.2e-05 | 8.7e-06 | 1.090E-05 | 1.905E-06 | 8.700E-06 | 1.200E-05 |
| 27894 | 1395 | T1395F | E | 1.3e-05 | 1.1e-05 | 1.5e-05 | 1.300E-05 | 2.000E-06 | 1.100E-05 | 1.500E-05 |
| 27895 | 1395 | T1395P | E | 1.2e-05 | 1.6e-05 | 1.2e-05 | 1.333E-05 | 2.309E-06 | 1.200E-05 | 1.600E-05 |
| 27896 | 1395 | T1395S | E | 1.4e-05 | 9.3e-06 | 1e-05 | 1.110E-05 | 2.536E-06 | 9.300E-06 | 1.400E-05 |
| 27897 | 1395 | T1395T | E | 1e-05 | 1.5e-05 | 1.1e-05 | 1.200E-05 | 2.646E-06 | 1.000E-05 | 1.500E-05 |
| 27898 | 1395 | T1395W | E | 9.6e-06 | 8.5e-06 | 1.1e-05 | 9.700E-06 | 1.253E-06 | 8.500E-06 | 1.100E-05 |
| 27899 | 1395 | T1395Y | E | 1.4e-05 | 1.4e-05 | 1.5e-05 | 1.433E-05 | 5.774E-07 | 1.400E-05 | 1.500E-05 |
| 27900 | 1395 | T1395V | E | 1e-05 | 1.5e-05 | 1.4e-05 | 1.300E-05 | 2.646E-06 | 1.000E-05 | 1.500E-05 |
| 27901 | 1396 | K1396A | E | 1.5e-05 | 1.5e-05 | 1.6e-05 | 1.533E-05 | 5.774E-07 | 1.500E-05 | 1.600E-05 |
| 27902 | 1396 | K1396R | E | 1.1e-05 | 1.5e-05 | 1.4e-05 | 1.333E-05 | 2.082E-06 | 1.100E-05 | 1.500E-05 |
| 27903 | 1396 | K1396N | E | 9.2e-06 | 1e-05 | 1.3e-05 | 1.073E-05 | 2.003E-06 | 9.200E-06 | 1.300E-05 |
| 27904 | 1396 | K1396D | E | 7.1e-06 | 1.1e-05 | 1.7e-05 | 1.170E-05 | 4.987E-06 | 7.100E-06 | 1.700E-05 |
| 27905 | 1396 | K1396C | E | 1.5e-05 | 1.1e-05 | 1.6e-05 | 1.400E-05 | 2.646E-06 | 1.100E-05 | 1.600E-05 |
| 27906 | 1396 | K1396E | E | 1.5e-05 | 1.6e-05 | 1e-05 | 1.367E-05 | 3.215E-06 | 1.000E-05 | 1.600E-05 |
| 27907 | 1396 | K1396Q | E | 1.7e-05 | 8.8e-06 | 1.6e-05 | 1.393E-05 | 4.474E-06 | 8.800E-06 | 1.700E-05 |
| 27908 | 1396 | K1396G | E | 9.3e-06 | 1.3e-05 | 1.2e-05 | 1.143E-05 | 1.914E-06 | 9.300E-06 | 1.300E-05 |
| 27909 | 1396 | K1396H | E | 1.4e-05 | 1.5e-05 | 7.9e-06 | 1.230E-05 | 3.843E-06 | 7.900E-06 | 1.500E-05 |
| 27910 | 1396 | K1396I | E | 1.3e-05 | 1.5e-05 | 1e-05 | 1.267E-05 | 2.517E-06 | 1.000E-05 | 1.500E-05 |
| 27911 | 1396 | K1396L | E | 1.4e-05 | 1.1e-05 | 1.7e-05 | 1.400E-05 | 3.000E-06 | 1.100E-05 | 1.700E-05 |
| 27912 | 1396 | K1396K | E | 1.1e-05 | 1.1e-05 | 1.1e-05 | 1.100E-05 | 2.075E-21 | 1.100E-05 | 1.100E-05 |
| 27913 | 1396 | K1396M | E | 1e-05 | 1.5e-05 | 1.4e-05 | 1.300E-05 | 2.646E-06 | 1.000E-05 | 1.500E-05 |
| 27914 | 1396 | K1396F | E | 9.3e-06 | 1.5e-05 | 1.4e-05 | 1.277E-05 | 3.044E-06 | 9.300E-06 | 1.500E-05 |
| 27915 | 1396 | K1396P | E | 9.4e-06 | 1.1e-05 | 1.1e-05 | 1.047E-05 | 9.238E-07 | 9.400E-06 | 1.100E-05 |
| 27916 | 1396 | K1396S | E | 1e-05 | 1.5e-05 | 1.7e-05 | 1.400E-05 | 3.606E-06 | 1.000E-05 | 1.700E-05 |
| 27917 | 1396 | K1396T | E | 1.5e-05 | 1.1e-05 | 1e-05 | 1.200E-05 | 2.646E-06 | 1.000E-05 | 1.500E-05 |
| 27918 | 1396 | K1396W | E | 1e-05 | 8.3e-06 | 1.1e-05 | 9.767E-06 | 1.365E-06 | 8.300E-06 | 1.100E-05 |
| 27919 | 1396 | K1396Y | E | 1.3e-05 | 1.5e-05 | 1.5e-05 | 1.433E-05 | 1.155E-06 | 1.300E-05 | 1.500E-05 |
| 27920 | 1396 | K1396V | E | 1e-05 | 1.3e-05 | 1.6e-05 | 1.300E-05 | 3.000E-06 | 1.000E-05 | 1.600E-05 |
| 27921 | 1397 | G1397A | E | 1e-05 | 1.3e-05 | 1.4e-05 | 1.233E-05 | 2.082E-06 | 1.000E-05 | 1.400E-05 |
| 27922 | 1397 | G1397R | E | 1.6e-05 | 9.7e-06 | 1e-05 | 1.190E-05 | 3.554E-06 | 9.700E-06 | 1.600E-05 |
| 27923 | 1397 | G1397N | E | 1.6e-05 | 1e-05 | 1e-05 | 1.200E-05 | 3.464E-06 | 1.000E-05 | 1.600E-05 |
| 27924 | 1397 | G1397D | E | 1.4e-05 | 1.5e-05 | 1.1e-05 | 1.333E-05 | 2.082E-06 | 1.100E-05 | 1.500E-05 |
| 27925 | 1397 | G1397C | E | 1.5e-05 | 1.5e-05 | 1.4e-05 | 1.467E-05 | 5.774E-07 | 1.400E-05 | 1.500E-05 |
| 27926 | 1397 | G1397E | E | 9e-06 | 8.7e-06 | 9.4e-06 | 9.033E-06 | 3.512E-07 | 8.700E-06 | 9.400E-06 |
| 27927 | 1397 | G1397Q | E | 1e-05 | 1.4e-05 | 1e-05 | 1.133E-05 | 2.309E-06 | 1.000E-05 | 1.400E-05 |
| 27928 | 1397 | G1397G | E | 1.6e-05 | 1.3e-05 | 1.5e-05 | 1.467E-05 | 1.528E-06 | 1.300E-05 | 1.600E-05 |
| 27929 | 1397 | G1397H | E | 9.6e-06 | 9.7e-06 | 1.1e-05 | 1.010E-05 | 7.810E-07 | 9.600E-06 | 1.100E-05 |
| 27930 | 1397 | G1397I | E | 1.4e-05 | 1.1e-05 | 1.1e-05 | 1.200E-05 | 1.732E-06 | 1.100E-05 | 1.400E-05 |
| 27931 | 1397 | G1397L | E | 1.2e-05 | 1.7e-05 | 1.3e-05 | 1.400E-05 | 2.646E-06 | 1.200E-05 | 1.700E-05 |
| 27932 | 1397 | G1397K | E | 1.5e-05 | 1.4e-05 | 1.5e-05 | 1.467E-05 | 5.774E-07 | 1.400E-05 | 1.500E-05 |
| 27933 | 1397 | G1397M | E | 1.2e-05 | 1.1e-05 | 9.8e-06 | 1.093E-05 | 1.102E-06 | 9.800E-06 | 1.200E-05 |
| 27934 | 1397 | G1397F | E | 1.1e-05 | 9.6e-06 | 1.2e-05 | 1.087E-05 | 1.206E-06 | 9.600E-06 | 1.200E-05 |
| 27935 | 1397 | G1397P | E | 1.6e-05 | 1.6e-05 | 9.9e-06 | 1.397E-05 | 3.522E-06 | 9.900E-06 | 1.600E-05 |
| 27936 | 1397 | G1397S | E | 8.6e-06 | 1.5e-05 | 1.4e-05 | 1.253E-05 | 3.443E-06 | 8.600E-06 | 1.500E-05 |
| 27937 | 1397 | G1397T | E | 1.5e-05 | 8.4e-06 | 1.5e-05 | 1.280E-05 | 3.811E-06 | 8.400E-06 | 1.500E-05 |
| 27938 | 1397 | G1397W | E | 1.6e-05 | 1.7e-05 | 1.1e-05 | 1.467E-05 | 3.215E-06 | 1.100E-05 | 1.700E-05 |
| 27939 | 1397 | G1397Y | E | 1.1e-05 | 1.5e-05 | 1.5e-05 | 1.367E-05 | 2.309E-06 | 1.100E-05 | 1.500E-05 |
| 27940 | 1397 | G1397V | E | 1.6e-05 | 9.8e-06 | 1.2e-05 | 1.260E-05 | 3.143E-06 | 9.800E-06 | 1.600E-05 |
| 27941 | 1398 | P1398A | E | 1e-05 | 1.2e-05 | 1.1e-05 | 1.100E-05 | 1.000E-06 | 1.000E-05 | 1.200E-05 |
| 27942 | 1398 | P1398R | E | 2.3e-05 | 1.5e-05 | 8.5e-06 | 1.550E-05 | 7.263E-06 | 8.500E-06 | 2.300E-05 |
| 27943 | 1398 | P1398N | E | 1e-05 | 1.6e-05 | 1.1e-05 | 1.233E-05 | 3.215E-06 | 1.000E-05 | 1.600E-05 |
| 27944 | 1398 | P1398D | E | 1.2e-05 | 1.4e-05 | 1.3e-05 | 1.300E-05 | 1.000E-06 | 1.200E-05 | 1.400E-05 |
| 27945 | 1398 | P1398C | E | 1.5e-05 | 5.9e-06 | 7.1e-06 | 9.333E-06 | 4.944E-06 | 5.900E-06 | 1.500E-05 |
| 27946 | 1398 | P1398E | E | 1.6e-05 | 1.4e-05 | 1.6e-05 | 1.533E-05 | 1.155E-06 | 1.400E-05 | 1.600E-05 |
| 27947 | 1398 | P1398Q | E | 7.7e-06 | 1.6e-05 | 1e-05 | 1.123E-05 | 4.285E-06 | 7.700E-06 | 1.600E-05 |
| 27948 | 1398 | P1398G | E | 1.3e-05 | 1.1e-05 | 1.6e-05 | 1.333E-05 | 2.517E-06 | 1.100E-05 | 1.600E-05 |
| 27949 | 1398 | P1398H | E | 1.4e-05 | 1.5e-05 | 1.3e-05 | 1.400E-05 | 1.000E-06 | 1.300E-05 | 1.500E-05 |
| 27950 | 1398 | P1398I | E | 1.5e-05 | 1.4e-05 | 1.4e-05 | 1.433E-05 | 5.774E-07 | 1.400E-05 | 1.500E-05 |
| 27951 | 1398 | P1398L | E | 1.6e-05 | 1.5e-05 | 1.4e-05 | 1.500E-05 | 1.000E-06 | 1.400E-05 | 1.600E-05 |
| 27952 | 1398 | P1398K | E | 1.5e-05 | 1e-05 | 1.4e-05 | 1.300E-05 | 2.646E-06 | 1.000E-05 | 1.500E-05 |
| 27953 | 1398 | P1398M | E | 9.1e-06 | 1.6e-05 | 1.4e-05 | 1.303E-05 | 3.550E-06 | 9.100E-06 | 1.600E-05 |
| 27954 | 1398 | P1398F | E | 9.8e-06 | 1.6e-05 | 1.2e-05 | 1.260E-05 | 3.143E-06 | 9.800E-06 | 1.600E-05 |
| 27955 | 1398 | P1398P | E | 1e-05 | 1e-05 | 1.6e-05 | 1.200E-05 | 3.464E-06 | 1.000E-05 | 1.600E-05 |
| 27956 | 1398 | P1398S | E | 1.6e-05 | 9.6e-06 | 1.3e-05 | 1.287E-05 | 3.202E-06 | 9.600E-06 | 1.600E-05 |
| 27957 | 1398 | P1398T | E | 9e-06 | 1.5e-05 | 1.3e-05 | 1.233E-05 | 3.055E-06 | 9.000E-06 | 1.500E-05 |
| 27958 | 1398 | P1398W | E | 8.8e-06 | 1.4e-05 | 1.6e-05 | 1.293E-05 | 3.717E-06 | 8.800E-06 | 1.600E-05 |
| 27959 | 1398 | P1398Y | E | 1.3e-05 | 1.5e-05 | 1.4e-05 | 1.400E-05 | 1.000E-06 | 1.300E-05 | 1.500E-05 |
| 27960 | 1398 | P1398V | E | 9.7e-06 | 1.4e-05 | 1.4e-05 | 1.257E-05 | 2.483E-06 | 9.700E-06 | 1.400E-05 |
| 27961 | 1399 | S1399A | E | 1.4e-05 | 1.7e-05 | 1.6e-05 | 1.567E-05 | 1.528E-06 | 1.400E-05 | 1.700E-05 |
| 27962 | 1399 | S1399R | E | 1.5e-05 | 1.8e-05 | 1.5e-05 | 1.600E-05 | 1.732E-06 | 1.500E-05 | 1.800E-05 |
| 27963 | 1399 | S1399N | E | 1.6e-05 | 1.5e-05 | 1.5e-05 | 1.533E-05 | 5.774E-07 | 1.500E-05 | 1.600E-05 |
| 27964 | 1399 | S1399D | E | 1.6e-05 | 1.5e-05 | 1.2e-05 | 1.433E-05 | 2.082E-06 | 1.200E-05 | 1.600E-05 |

| | | | | | | | | | | |
|---|---|---|---|---|---|---|---|---|---|---|
| 27965 | 1399 | S1399C | E | 1.1e-05 | 9.6e-06 | 1.4e-05 | 1.153E-05 | 2.248E-06 | 9.600E-06 | 1.400E-05 |
| 27966 | 1399 | S1399E | E | 1.5e-05 | 1.1e-05 | 1.3e-05 | 1.300E-05 | 2.000E-06 | 1.100E-05 | 1.500E-05 |
| 27967 | 1399 | S1399Q | E | 1.2e-05 | 1.5e-05 | 1.7e-05 | 1.467E-05 | 2.517E-06 | 1.200E-05 | 1.700E-05 |
| 27968 | 1399 | S1399G | E | 1.7e-05 | 1.4e-05 | 1.4e-05 | 1.500E-05 | 1.732E-06 | 1.400E-05 | 1.700E-05 |
| 27969 | 1399 | S1399H | E | 1.5e-05 | 1.1e-05 | 1.3e-05 | 1.300E-05 | 2.000E-06 | 1.100E-05 | 1.500E-05 |
| 27970 | 1399 | S1399I | E | 1.2e-05 | 1.6e-05 | 1e-05 | 1.267E-05 | 3.055E-06 | 1.000E-05 | 1.600E-05 |
| 27971 | 1399 | S1399L | E | 1.1e-05 | 1.1e-05 | 1.1e-05 | 1.100E-05 | 2.075E-21 | 1.100E-05 | 1.100E-05 |
| 27972 | 1399 | S1399K | E | 1.1e-05 | 1.4e-05 | 1.6e-05 | 1.367E-05 | 2.517E-06 | 1.100E-05 | 1.600E-05 |
| 27973 | 1399 | S1399M | E | 1.1e-05 | 8.9e-06 | 1.1e-05 | 1.030E-05 | 1.212E-06 | 8.900E-06 | 1.100E-05 |
| 27974 | 1399 | S1399F | E | 1.1e-05 | 1.6e-05 | 1.6e-05 | 1.433E-05 | 2.887E-06 | 1.100E-05 | 1.600E-05 |
| 27975 | 1399 | S1399P | E | 1e-05 | 1.3e-05 | 1.5e-05 | 1.267E-05 | 2.517E-06 | 1.000E-05 | 1.500E-05 |
| 27976 | 1399 | S1399S | E | 1.6e-05 | 9.1e-06 | 8.4e-06 | 1.117E-05 | 4.200E-06 | 8.400E-06 | 1.600E-05 |
| 27977 | 1399 | S1399T | E | 1.3e-05 | 1e-05 | 1.7e-05 | 1.333E-05 | 3.512E-06 | 1.000E-05 | 1.700E-05 |
| 27978 | 1399 | S1399W | E | 9.3e-06 | 9.7e-06 | 1.3e-05 | 1.067E-05 | 2.031E-06 | 9.300E-06 | 1.300E-05 |
| 27979 | 1399 | S1399Y | E | 1.3e-05 | 1.6e-05 | 1.5e-05 | 1.467E-05 | 1.528E-06 | 1.300E-05 | 1.600E-05 |
| 27980 | 1399 | S1399V | E | 1.6e-05 | 1.6e-05 | 8.5e-06 | 1.350E-05 | 4.330E-06 | 8.500E-06 | 1.600E-05 |
| 27981 | 1400 | V1400A | E | 1.7e-05 | 1.5e-05 | 1.5e-05 | 1.567E-05 | 1.155E-06 | 1.500E-05 | 1.700E-05 |
| 27982 | 1400 | V1400R | E | 1.1e-05 | 1.5e-05 | 1.7e-05 | 1.433E-05 | 3.055E-06 | 1.100E-05 | 1.700E-05 |
| 27983 | 1400 | V1400N | E | 1.4e-05 | 1.6e-05 | 1.5e-05 | 1.500E-05 | 1.000E-06 | 1.400E-05 | 1.600E-05 |
| 27984 | 1400 | V1400D | E | 9.4e-06 | 9.6e-06 | 1.5e-05 | 1.133E-05 | 3.177E-06 | 9.400E-06 | 1.500E-05 |
| 27985 | 1400 | V1400C | E | 1.5e-05 | 1e-05 | 1.5e-05 | 1.333E-05 | 2.887E-06 | 1.000E-05 | 1.500E-05 |
| 27986 | 1400 | V1400E | E | 1.6e-05 | 1e-05 | 1.7e-05 | 1.433E-05 | 3.786E-06 | 1.000E-05 | 1.700E-05 |
| 27987 | 1400 | V1400Q | E | 1.2e-05 | 1.3e-05 | 1.5e-05 | 1.333E-05 | 1.528E-06 | 1.200E-05 | 1.500E-05 |
| 27988 | 1400 | V1400G | E | 1.6e-05 | 1.6e-05 | 9.5e-06 | 1.383E-05 | 3.753E-06 | 9.500E-06 | 1.600E-05 |
| 27989 | 1400 | V1400H | E | 1.6e-05 | 1.4e-05 | 1.1e-05 | 1.367E-05 | 2.517E-06 | 1.100E-05 | 1.600E-05 |
| 27990 | 1400 | V1400I | E | 1.6e-05 | 1.6e-05 | 1.5e-05 | 1.567E-05 | 5.774E-07 | 1.500E-05 | 1.600E-05 |
| 27991 | 1400 | V1400L | E | 1.5e-05 | 1.6e-05 | 9.4e-06 | 1.347E-05 | 3.557E-06 | 9.400E-06 | 1.600E-05 |
| 27992 | 1400 | V1400K | E | 5.5e-06 | 1.1e-05 | 1.3e-05 | 9.833E-06 | 3.884E-06 | 5.500E-06 | 1.300E-05 |
| 27993 | 1400 | V1400M | E | 9.9e-06 | 1e-05 | 1e-05 | 9.967E-06 | 5.774E-08 | 9.900E-06 | 1.000E-05 |
| 27994 | 1400 | V1400F | E | 1.3e-05 | 1.5e-05 | 1.1e-05 | 1.300E-05 | 2.000E-06 | 1.100E-05 | 1.500E-05 |
| 27995 | 1400 | V1400P | E | 1.1e-05 | 1.2e-05 | 1.6e-05 | 1.300E-05 | 2.646E-06 | 1.100E-05 | 1.600E-05 |
| 27996 | 1400 | V1400S | E | 8.8e-06 | 1.4e-05 | 1.4e-05 | 1.227E-05 | 3.002E-06 | 8.800E-06 | 1.400E-05 |
| 27997 | 1400 | V1400T | E | 1.5e-05 | 1.5e-05 | 1.6e-05 | 1.533E-05 | 5.774E-07 | 1.500E-05 | 1.600E-05 |
| 27998 | 1400 | V1400W | E | 2.6e-05 | 1.3e-05 | 1e-05 | 1.633E-05 | 8.505E-06 | 1.000E-05 | 2.600E-05 |
| 27999 | 1400 | V1400Y | E | 1.5e-05 | 1e-05 | 1.1e-05 | 1.200E-05 | 2.646E-06 | 1.000E-05 | 1.500E-05 |
| 28000 | 1400 | V1400V | E | 1.6e-05 | 1.2e-05 | 1.4e-05 | 1.400E-05 | 2.000E-06 | 1.200E-05 | 1.600E-05 |
| 28001 | 1401 | F1401A | E | 1.1e-05 | 1.3e-05 | 1.5e-05 | 1.300E-05 | 2.000E-06 | 1.100E-05 | 1.500E-05 |
| 28002 | 1401 | F1401R | E | 1.5e-05 | 1.7e-05 | 1.5e-05 | 1.567E-05 | 1.155E-06 | 1.500E-05 | 1.700E-05 |
| 28003 | 1401 | F1401N | E | 1.6e-05 | 9.2e-06 | 1.1e-05 | 1.207E-05 | 3.523E-06 | 9.200E-06 | 1.600E-05 |
| 28004 | 1401 | F1401D | E | 1.5e-05 | 1.1e-05 | 9.8e-06 | 1.193E-05 | 2.723E-06 | 9.800E-06 | 1.500E-05 |
| 28005 | 1401 | F1401C | E | 1.5e-05 | 1.5e-05 | 1.4e-05 | 1.467E-05 | 5.774E-07 | 1.400E-05 | 1.500E-05 |
| 28006 | 1401 | F1401E | E | 1.1e-05 | 1.1e-05 | 7.3e-06 | 9.767E-06 | 2.136E-06 | 7.300E-06 | 1.100E-05 |
| 28007 | 1401 | F1401Q | E | 1.1e-05 | 1e-05 | 9.9e-06 | 1.030E-05 | 6.083E-07 | 9.900E-06 | 1.100E-05 |
| 28008 | 1401 | F1401G | E | 8e-06 | 1.2e-05 | 1.6e-05 | 1.200E-05 | 4.000E-06 | 8.000E-06 | 1.600E-05 |
| 28009 | 1401 | F1401H | E | 1.5e-05 | 1.5e-05 | 1.5e-05 | 1.500E-05 | 0.000E+00 | 1.500E-05 | 1.500E-05 |
| 28010 | 1401 | F1401I | E | 1.3e-05 | 1.1e-05 | 9.7e-06 | 1.123E-05 | 1.662E-06 | 9.700E-06 | 1.300E-05 |
| 28011 | 1401 | F1401L | E | 1e-05 | 1.1e-05 | 1.4e-05 | 1.167E-05 | 2.082E-06 | 1.000E-05 | 1.400E-05 |
| 28012 | 1401 | F1401K | E | 1.3e-05 | 8.3e-06 | 1.6e-05 | 1.243E-05 | 3.881E-06 | 8.300E-06 | 1.600E-05 |
| 28013 | 1401 | F1401M | E | 1.3e-05 | 1.4e-05 | 1.2e-05 | 1.300E-05 | 1.000E-06 | 1.200E-05 | 1.400E-05 |
| 28014 | 1401 | F1401F | E | 1.6e-05 | 1.3e-05 | 1.6e-05 | 1.500E-05 | 1.732E-06 | 1.300E-05 | 1.600E-05 |
| 28015 | 1401 | F1401P | E | 1.4e-05 | 1.1e-05 | 1.1e-05 | 1.200E-05 | 1.732E-06 | 1.100E-05 | 1.400E-05 |
| 28016 | 1401 | F1401S | E | 1.1e-05 | 8.6e-06 | 1.1e-05 | 1.020E-05 | 1.386E-06 | 8.600E-06 | 1.100E-05 |
| 28017 | 1401 | F1401T | E | 1.5e-05 | 8.6e-06 | 9.5e-06 | 1.103E-05 | 3.465E-06 | 8.600E-06 | 1.500E-05 |
| 28018 | 1401 | F1401W | E | 1.4e-05 | 8.9e-06 | 1.3e-05 | 1.197E-05 | 2.702E-06 | 8.900E-06 | 1.400E-05 |
| 28019 | 1401 | F1401Y | E | 1.1e-05 | 8.9e-06 | 1e-05 | 9.967E-06 | 1.050E-06 | 8.900E-06 | 1.100E-05 |
| 28020 | 1401 | F1401V | E | 1.6e-05 | 1.5e-05 | 1.3e-05 | 1.467E-05 | 1.528E-06 | 1.300E-05 | 1.600E-05 |
| 28021 | 1402 | P1402A | E | 1.5e-05 | 1.1e-05 | 8.1e-06 | 1.137E-05 | 3.465E-06 | 8.100E-06 | 1.500E-05 |
| 28022 | 1402 | P1402R | E | 1.4e-05 | 1.2e-05 | 1e-05 | 1.200E-05 | 2.000E-06 | 1.000E-05 | 1.400E-05 |
| 28023 | 1402 | P1402N | E | 1.1e-05 | 1e-05 | 1e-05 | 1.033E-05 | 5.774E-07 | 1.000E-05 | 1.100E-05 |
| 28024 | 1402 | P1402D | E | 1.4e-05 | 9.2e-06 | 1.4e-05 | 1.240E-05 | 2.771E-06 | 9.200E-06 | 1.400E-05 |
| 28025 | 1402 | P1402C | E | 1.3e-05 | 1.2e-05 | 8.7e-06 | 1.123E-05 | 2.250E-06 | 8.700E-06 | 1.300E-05 |
| 28026 | 1402 | P1402E | E | 1.4e-05 | 1.3e-05 | 1.5e-05 | 1.400E-05 | 1.000E-06 | 1.300E-05 | 1.500E-05 |
| 28027 | 1402 | P1402Q | E | 8.7e-06 | 1.4e-05 | 1.1e-05 | 1.123E-05 | 2.658E-06 | 8.700E-06 | 1.400E-05 |
| 28028 | 1402 | P1402G | E | 1.5e-05 | 1.4e-05 | 1.1e-05 | 1.333E-05 | 2.082E-06 | 1.100E-05 | 1.500E-05 |
| 28029 | 1402 | P1402H | E | 1e-05 | 1.5e-05 | 1.1e-05 | 1.200E-05 | 2.646E-06 | 1.000E-05 | 1.500E-05 |
| 28030 | 1402 | P1402I | E | 1.4e-05 | 1.5e-05 | 1.5e-05 | 1.467E-05 | 5.774E-07 | 1.400E-05 | 1.500E-05 |
| 28031 | 1402 | P1402L | E | 1.5e-05 | 1.5e-05 | 1.5e-05 | 1.500E-05 | 0.000E+00 | 1.500E-05 | 1.500E-05 |
| 28032 | 1402 | P1402K | E | 9.6e-06 | 8.3e-06 | 1.5e-05 | 1.097E-05 | 3.553E-06 | 8.300E-06 | 1.500E-05 |
| 28033 | 1402 | P1402M | E | 9.7e-06 | 1.6e-05 | 1e-05 | 1.190E-05 | 3.554E-06 | 9.700E-06 | 1.600E-05 |
| 28034 | 1402 | P1402F | E | 9.3e-06 | 1.6e-05 | 1.5e-05 | 1.343E-05 | 3.614E-06 | 9.300E-06 | 1.600E-05 |
| 28035 | 1402 | P1402P | E | 1e-05 | 1.4e-05 | 9.4e-06 | 1.113E-05 | 2.501E-06 | 9.400E-06 | 1.400E-05 |
| 28036 | 1402 | P1402S | E | 1.5e-05 | 1.4e-05 | 6e-06 | 1.167E-05 | 4.933E-06 | 6.000E-06 | 1.500E-05 |
| 28037 | 1402 | P1402T | E | 1.4e-05 | 1.1e-05 | 1.4e-05 | 1.300E-05 | 1.732E-06 | 1.100E-05 | 1.400E-05 |
| 28038 | 1402 | P1402W | E | 1.2e-05 | 1.1e-05 | 9e-06 | 1.067E-05 | 1.528E-06 | 9.000E-06 | 1.200E-05 |
| 28039 | 1402 | P1402Y | E | 1.2e-05 | 1.3e-05 | 1.4e-05 | 1.300E-05 | 1.000E-06 | 1.200E-05 | 1.400E-05 |
| 28040 | 1402 | P1402V | E | 1.6e-05 | 9.7e-06 | 1.5e-05 | 1.357E-05 | 3.386E-06 | 9.700E-06 | 1.600E-05 |
| 28041 | 1403 | L1403A | E | 9e-06 | 1.3e-05 | 1.3e-05 | 1.167E-05 | 2.309E-06 | 9.000E-06 | 1.300E-05 |
| 28042 | 1403 | L1403R | E | 8.8e-06 | 1.4e-05 | 1.1e-05 | 1.127E-05 | 2.610E-06 | 8.800E-06 | 1.400E-05 |
| 28043 | 1403 | L1403N | E | 1.6e-05 | 1.1e-05 | 1.5e-05 | 1.400E-05 | 2.646E-06 | 1.100E-05 | 1.600E-05 |
| 28044 | 1403 | L1403D | E | 1.3e-05 | 1.6e-05 | 1.1e-05 | 1.333E-05 | 2.517E-06 | 1.100E-05 | 1.600E-05 |
| 28045 | 1403 | L1403C | E | 1.4e-05 | 1.5e-05 | 1.1e-05 | 1.333E-05 | 2.082E-06 | 1.100E-05 | 1.500E-05 |
| 28046 | 1403 | L1403E | E | 1.1e-05 | 1.5e-05 | 1.2e-05 | 1.267E-05 | 2.082E-06 | 1.100E-05 | 1.500E-05 |
| 28047 | 1403 | L1403Q | E | 1e-05 | 8.6e-06 | 1.5e-05 | 1.120E-05 | 3.365E-06 | 8.600E-06 | 1.500E-05 |
| 28048 | 1403 | L1403G | E | 1.6e-05 | 1e-05 | 9.1e-06 | 1.170E-05 | 3.751E-06 | 9.100E-06 | 1.600E-05 |
| 28049 | 1403 | L1403H | E | 1.4e-05 | 1.6e-05 | 9.1e-06 | 1.303E-05 | 3.550E-06 | 9.100E-06 | 1.600E-05 |

| | | | | | | | | | | |
|---|---|---|---|---|---|---|---|---|---|---|
| 28050 | 1403 | L1403I | E | 8.4e-06 | 1.6e-05 | 1.5e-05 | 1.313E-05 | 4.130E-06 | 8.400E-06 | 1.600E-05 |
| 28051 | 1403 | L1403L | E | 1.3e-05 | 1.5e-05 | 1.5e-05 | 1.433E-05 | 1.155E-06 | 1.300E-05 | 1.500E-05 |
| 28052 | 1403 | L1403K | E | 1.1e-05 | 1.6e-05 | 1.6e-05 | 1.433E-05 | 2.887E-06 | 1.100E-05 | 1.600E-05 |
| 28053 | 1403 | L1403M | E | 1.5e-05 | 1.4e-05 | 1.5e-05 | 1.467E-05 | 5.774E-07 | 1.400E-05 | 1.500E-05 |
| 28054 | 1403 | L1403F | E | 1.5e-05 | 1.6e-05 | 1.4e-05 | 1.500E-05 | 1.000E-06 | 1.400E-05 | 1.600E-05 |
| 28055 | 1403 | L1403P | E | 1.4e-05 | 1.5e-05 | 1.5e-05 | 1.467E-05 | 5.774E-07 | 1.400E-05 | 1.500E-05 |
| 28056 | 1403 | L1403S | E | 1e-05 | 1.1e-05 | 8.3e-06 | 9.767E-06 | 1.365E-06 | 8.300E-06 | 1.100E-05 |
| 28057 | 1403 | L1403T | E | 1e-05 | 1.4e-05 | 8.7e-06 | 1.090E-05 | 2.762E-06 | 8.700E-06 | 1.400E-05 |
| 28058 | 1403 | L1403W | E | 1.1e-05 | 1.1e-05 | 1.7e-05 | 1.300E-05 | 3.464E-06 | 1.100E-05 | 1.700E-05 |
| 28059 | 1403 | L1403Y | E | 1e-05 | 1.5e-05 | 1.6e-05 | 1.367E-05 | 3.215E-06 | 1.000E-05 | 1.600E-05 |
| 28060 | 1403 | L1403V | E | 1.5e-05 | 1.7e-05 | 1e-05 | 1.400E-05 | 3.606E-06 | 1.000E-05 | 1.700E-05 |
| 28061 | 1404 | A1404A | E | 1.2e-05 | 1.2e-05 | 1.5e-05 | 1.300E-05 | 1.732E-06 | 1.200E-05 | 1.500E-05 |
| 28062 | 1404 | A1404R | E | 1.5e-05 | 1e-05 | 1.5e-05 | 1.333E-05 | 2.887E-06 | 1.000E-05 | 1.500E-05 |
| 28063 | 1404 | A1404N | E | 1e-05 | 1e-05 | 1.5e-05 | 1.167E-05 | 2.887E-06 | 1.000E-05 | 1.500E-05 |
| 28064 | 1404 | A1404D | E | 1.7e-05 | 1.1e-05 | 1.6e-05 | 1.467E-05 | 3.215E-06 | 1.100E-05 | 1.700E-05 |
| 28065 | 1404 | A1404C | E | 1.4e-05 | 1.2e-05 | 1e-05 | 1.200E-05 | 2.000E-06 | 1.000E-05 | 1.400E-05 |
| 28066 | 1404 | A1404E | E | 1.4e-05 | 1.5e-05 | 1.7e-05 | 1.533E-05 | 1.528E-06 | 1.400E-05 | 1.700E-05 |
| 28067 | 1404 | A1404Q | E | 1.4e-05 | 1.7e-05 | 1e-05 | 1.367E-05 | 3.512E-06 | 1.000E-05 | 1.700E-05 |
| 28068 | 1404 | A1404G | E | 1e-05 | 8.9e-06 | 1.4e-05 | 1.097E-05 | 2.684E-06 | 8.900E-06 | 1.400E-05 |
| 28069 | 1404 | A1404H | E | 1.5e-05 | 1.4e-05 | 1.3e-05 | 1.400E-05 | 1.000E-06 | 1.300E-05 | 1.500E-05 |
| 28070 | 1404 | A1404I | E | 1.4e-05 | 8.8e-06 | 8.6e-06 | 1.047E-05 | 3.062E-06 | 8.600E-06 | 1.400E-05 |
| 28071 | 1404 | A1404L | E | 1.2e-05 | 9e-06 | 1.1e-05 | 1.067E-05 | 1.528E-06 | 9.000E-06 | 1.200E-05 |
| 28072 | 1404 | A1404K | E | 1.3e-05 | 9.5e-06 | 1.5e-05 | 1.250E-05 | 2.784E-06 | 9.500E-06 | 1.500E-05 |
| 28073 | 1404 | A1404M | E | 1.6e-05 | 1.4e-05 | 9e-06 | 1.300E-05 | 3.606E-06 | 9.000E-06 | 1.600E-05 |
| 28074 | 1404 | A1404F | E | 1.5e-05 | 9e-06 | 1.1e-05 | 1.167E-05 | 3.055E-06 | 9.000E-06 | 1.500E-05 |
| 28075 | 1404 | A1404P | E | 1.5e-05 | 1.9e-05 | 1.2e-05 | 1.533E-05 | 3.512E-06 | 1.200E-05 | 1.900E-05 |
| 28076 | 1404 | A1404S | E | 1.5e-05 | 1.4e-05 | 1e-05 | 1.300E-05 | 2.646E-06 | 1.000E-05 | 1.500E-05 |
| 28077 | 1404 | A1404T | E | 1.4e-05 | 1e-05 | 1.2e-05 | 1.200E-05 | 2.000E-06 | 1.000E-05 | 1.400E-05 |
| 28078 | 1404 | A1404W | E | 1.3e-05 | 1e-05 | 1.4e-05 | 1.233E-05 | 2.082E-06 | 1.000E-05 | 1.400E-05 |
| 28079 | 1404 | A1404Y | E | 1.3e-05 | 1.1e-05 | 1.5e-05 | 1.300E-05 | 2.000E-06 | 1.100E-05 | 1.500E-05 |
| 28080 | 1404 | A1404V | E | 1.1e-05 | 9.8e-06 | 1.1e-05 | 1.060E-05 | 6.928E-07 | 9.800E-06 | 1.100E-05 |
| 28081 | 1405 | P1405A | E | 7.8e-06 | 8.7e-06 | 7.9e-06 | 8.133E-06 | 4.933E-07 | 7.800E-06 | 8.700E-06 |
| 28082 | 1405 | P1405R | E | 9.8e-06 | 1.6e-05 | 8.9e-06 | 1.157E-05 | 3.866E-06 | 8.900E-06 | 1.600E-05 |
| 28083 | 1405 | P1405N | E | 9.5e-06 | 9.4e-06 | 1.6e-05 | 1.163E-05 | 3.782E-06 | 9.400E-06 | 1.600E-05 |
| 28084 | 1405 | P1405D | E | 1.2e-05 | 1.5e-05 | 1.5e-05 | 1.400E-05 | 1.732E-06 | 1.200E-05 | 1.500E-05 |
| 28085 | 1405 | P1405C | E | 1.4e-05 | 1.5e-05 | 9.2e-06 | 1.273E-05 | 3.101E-06 | 9.200E-06 | 1.500E-05 |
| 28086 | 1405 | P1405E | E | 1.6e-05 | 1.3e-05 | 1.5e-05 | 1.467E-05 | 1.528E-06 | 1.300E-05 | 1.600E-05 |
| 28087 | 1405 | P1405Q | E | 1e-05 | 1.4e-05 | 8.8e-06 | 1.093E-05 | 2.723E-06 | 8.800E-06 | 1.400E-05 |
| 28088 | 1405 | P1405G | E | 1.4e-05 | 1e-05 | 1.6e-05 | 1.333E-05 | 3.055E-06 | 1.000E-05 | 1.600E-05 |
| 28089 | 1405 | P1405H | E | 7.4e-06 | 8.4e-06 | 1.6e-05 | 1.060E-05 | 4.703E-06 | 7.400E-06 | 1.600E-05 |
| 28090 | 1405 | P1405I | E | 9.3e-06 | 1.1e-05 | 1.6e-05 | 1.210E-05 | 3.483E-06 | 9.300E-06 | 1.600E-05 |
| 28091 | 1405 | P1405L | E | 1.5e-05 | 1.5e-05 | 1.6e-05 | 1.533E-05 | 5.774E-07 | 1.500E-05 | 1.600E-05 |
| 28092 | 1405 | P1405K | E | 1.5e-05 | 9.3e-06 | 9.5e-06 | 1.127E-05 | 3.235E-06 | 9.300E-06 | 1.500E-05 |
| 28093 | 1405 | P1405M | E | 1.6e-05 | 8.1e-06 | 1.6e-05 | 1.337E-05 | 4.561E-06 | 8.100E-06 | 1.600E-05 |
| 28094 | 1405 | P1405F | E | 1.6e-05 | 1.4e-05 | 9.2e-06 | 1.307E-05 | 3.495E-06 | 9.200E-06 | 1.600E-05 |
| 28095 | 1405 | P1405P | E | 1.5e-05 | 1.5e-05 | 1e-05 | 1.333E-05 | 2.887E-06 | 1.000E-05 | 1.500E-05 |
| 28096 | 1405 | P1405S | E | 1.5e-05 | 9.7e-06 | 1.3e-05 | 1.257E-05 | 2.676E-06 | 9.700E-06 | 1.500E-05 |
| 28097 | 1405 | P1405T | E | 1.3e-05 | 1.5e-05 | 1e-05 | 1.267E-05 | 2.517E-06 | 1.000E-05 | 1.500E-05 |
| 28098 | 1405 | P1405W | E | 9.1e-06 | 1.5e-05 | 9.3e-06 | 1.113E-05 | 3.350E-06 | 9.100E-06 | 1.500E-05 |
| 28099 | 1405 | P1405Y | E | 1.3e-05 | 1.5e-05 | 7.7e-06 | 1.190E-05 | 3.772E-06 | 7.700E-06 | 1.500E-05 |
| 28100 | 1405 | P1405V | E | 1.2e-05 | 1.7e-05 | 1.5e-05 | 1.467E-05 | 2.517E-06 | 1.200E-05 | 1.700E-05 |
| 28101 | 1406 | S1406A | E | 9.3e-06 | 1.7e-05 | 8.3e-06 | 1.153E-05 | 4.761E-06 | 8.300E-06 | 1.700E-05 |
| 28102 | 1406 | S1406R | E | 1.3e-05 | 1.5e-05 | 1.3e-05 | 1.367E-05 | 1.155E-06 | 1.300E-05 | 1.500E-05 |
| 28103 | 1406 | S1406N | E | 1.3e-05 | 1.6e-05 | 1.5e-05 | 1.467E-05 | 1.528E-06 | 1.300E-05 | 1.600E-05 |
| 28104 | 1406 | S1406D | E | 1.4e-05 | 1.6e-05 | 9.1e-06 | 1.303E-05 | 3.550E-06 | 9.100E-06 | 1.600E-05 |
| 28105 | 1406 | S1406C | E | 1.6e-05 | 1.5e-05 | 1.4e-05 | 1.500E-05 | 1.000E-06 | 1.400E-05 | 1.600E-05 |
| 28106 | 1406 | S1406E | E | 1.1e-05 | 9.6e-06 | 1.5e-05 | 1.187E-05 | 2.802E-06 | 9.600E-06 | 1.500E-05 |
| 28107 | 1406 | S1406Q | E | 1.5e-05 | 1.5e-05 | 1.5e-05 | 1.500E-05 | 0.000E+00 | 1.500E-05 | 1.500E-05 |
| 28108 | 1406 | S1406G | E | 1.7e-05 | 1.3e-05 | 1.5e-05 | 1.500E-05 | 2.000E-06 | 1.300E-05 | 1.700E-05 |
| 28109 | 1406 | S1406H | E | 1.2e-05 | 1.3e-05 | 1.5e-05 | 1.333E-05 | 1.528E-06 | 1.200E-05 | 1.500E-05 |
| 28110 | 1406 | S1406I | E | 1e-05 | 1.2e-05 | 1e-05 | 1.067E-05 | 1.155E-06 | 1.000E-05 | 1.200E-05 |
| 28111 | 1406 | S1406L | E | 9.3e-06 | 1.4e-05 | 1.5e-05 | 1.277E-05 | 3.044E-06 | 9.300E-06 | 1.500E-05 |
| 28112 | 1406 | S1406K | E | 1.6e-05 | 1.6e-05 | 1e-05 | 1.400E-05 | 3.464E-06 | 1.000E-05 | 1.600E-05 |
| 28113 | 1406 | S1406M | E | 1.6e-05 | 1.6e-05 | 1.6e-05 | 1.600E-05 | 0.000E+00 | 1.600E-05 | 1.600E-05 |
| 28114 | 1406 | S1406F | E | 8.8e-06 | 1.6e-05 | 2.3e-05 | 1.593E-05 | 7.100E-06 | 8.800E-06 | 2.300E-05 |
| 28115 | 1406 | S1406P | E | 9.8e-06 | 7.5e-06 | 1.5e-05 | 1.077E-05 | 3.842E-06 | 7.500E-06 | 1.500E-05 |
| 28116 | 1406 | S1406S | E | 8.9e-06 | 1.1e-05 | 9e-06 | 9.633E-06 | 1.185E-06 | 8.900E-06 | 1.100E-05 |
| 28117 | 1406 | S1406T | E | 1.3e-05 | 1.2e-05 | 1.6e-05 | 1.367E-05 | 2.082E-06 | 1.200E-05 | 1.600E-05 |
| 28118 | 1406 | S1406W | E | 1.3e-05 | 1.3e-05 | 1.6e-05 | 1.400E-05 | 1.732E-06 | 1.300E-05 | 1.600E-05 |
| 28119 | 1406 | S1406Y | E | 1.6e-05 | 1.5e-05 | 1.1e-05 | 1.400E-05 | 2.646E-06 | 1.100E-05 | 1.600E-05 |
| 28120 | 1406 | S1406V | E | 1.3e-05 | 1.7e-05 | 6.2e-06 | 1.207E-05 | 5.460E-06 | 6.200E-06 | 1.700E-05 |
| 28121 | 1407 | S1407A | E | 1.6e-05 | 9.3e-06 | 7.9e-06 | 1.107E-05 | 4.329E-06 | 7.900E-06 | 1.600E-05 |
| 28122 | 1407 | S1407R | E | 1.2e-05 | 1.3e-05 | 1.6e-05 | 1.367E-05 | 2.082E-06 | 1.200E-05 | 1.600E-05 |
| 28123 | 1407 | S1407N | E | 1.5e-05 | 2.4e-05 | 1.6e-05 | 1.833E-05 | 4.933E-06 | 1.500E-05 | 2.400E-05 |
| 28124 | 1407 | S1407D | E | 1e-05 | 1.5e-05 | 1.6e-05 | 1.367E-05 | 3.215E-06 | 1.000E-05 | 1.600E-05 |
| 28125 | 1407 | S1407C | E | 1.6e-05 | 1.1e-05 | 1.1e-05 | 1.267E-05 | 2.887E-06 | 1.100E-05 | 1.600E-05 |
| 28126 | 1407 | S1407E | E | 1.8e-05 | 1.3e-05 | 1.5e-05 | 1.533E-05 | 2.517E-06 | 1.300E-05 | 1.800E-05 |
| 28127 | 1407 | S1407Q | E | 1.5e-05 | 1.2e-05 | 1.4e-05 | 1.367E-05 | 1.528E-06 | 1.200E-05 | 1.500E-05 |
| 28128 | 1407 | S1407G | E | 1.1e-05 | 1.6e-05 | 1.5e-05 | 1.400E-05 | 2.646E-06 | 1.100E-05 | 1.600E-05 |
| 28129 | 1407 | S1407H | E | 9.4e-06 | 1.4e-05 | 1.4e-05 | 1.247E-05 | 2.656E-06 | 9.400E-06 | 1.400E-05 |
| 28130 | 1407 | S1407I | E | 1.5e-05 | 1.6e-05 | 8.6e-06 | 1.320E-05 | 4.015E-06 | 8.600E-06 | 1.600E-05 |
| 28131 | 1407 | S1407L | E | 1.7e-05 | 9.8e-06 | 9.9e-06 | 1.223E-05 | 4.128E-06 | 9.800E-06 | 1.700E-05 |
| 28132 | 1407 | S1407K | E | 1.7e-05 | 1.6e-05 | 1.5e-05 | 1.600E-05 | 1.000E-06 | 1.500E-05 | 1.700E-05 |
| 28133 | 1407 | S1407M | E | 1.4e-05 | 9.8e-06 | 1.5e-05 | 1.293E-05 | 2.759E-06 | 9.800E-06 | 1.500E-05 |
| 28134 | 1407 | S1407F | E | 1.7e-05 | 9.6e-06 | 1.5e-05 | 1.387E-05 | 3.828E-06 | 9.600E-06 | 1.700E-05 |

| | | | | | | | | | | |
|---|---|---|---|---|---|---|---|---|---|---|
| 28135 | 1407 | S1407P | E | 1.1e-05 | 1.6e-05 | 1.5e-05 | 1.400E-05 | 2.646E-06 | 1.100E-05 | 1.600E-05 |
| 28136 | 1407 | S1407S | E | 1.1e-05 | 1.5e-05 | 1.1e-05 | 1.233E-05 | 2.309E-06 | 1.100E-05 | 1.500E-05 |
| 28137 | 1407 | S1407T | E | 1.6e-05 | 1e-05 | 1.1e-05 | 1.233E-05 | 3.215E-06 | 1.000E-05 | 1.600E-05 |
| 28138 | 1407 | S1407W | E | 9e-06 | 1.1e-05 | 1.6e-05 | 1.200E-05 | 3.606E-06 | 9.000E-06 | 1.600E-05 |
| 28139 | 1407 | S1407Y | E | 1.2e-05 | 1.7e-05 | 1e-05 | 1.300E-05 | 3.606E-06 | 1.000E-05 | 1.700E-05 |
| 28140 | 1407 | S1407V | E | 9.5e-06 | 1.5e-05 | 1e-05 | 1.150E-05 | 3.041E-06 | 9.500E-06 | 1.500E-05 |
| 28141 | 1408 | K1408A | E | 1e-05 | 1e-05 | 6.2e-06 | 8.733E-06 | 2.194E-06 | 6.200E-06 | 1.000E-05 |
| 28142 | 1408 | K1408R | E | 1.2e-05 | 1.4e-05 | 9.8e-06 | 1.193E-05 | 2.101E-06 | 9.800E-06 | 1.400E-05 |
| 28143 | 1408 | K1408N | E | 9.4e-06 | 1.5e-05 | 9.3e-06 | 1.123E-05 | 3.262E-06 | 9.300E-06 | 1.500E-05 |
| 28144 | 1408 | K1408D | E | 1.1e-05 | 7.6e-06 | 1e-05 | 9.533E-06 | 1.747E-06 | 7.600E-06 | 1.100E-05 |
| 28145 | 1408 | K1408C | E | 1.1e-05 | 1.2e-05 | 1e-05 | 1.100E-05 | 1.000E-06 | 1.000E-05 | 1.200E-05 |
| 28146 | 1408 | K1408E | E | 1.4e-05 | 1.4e-05 | 1.3e-05 | 1.367E-05 | 5.774E-07 | 1.300E-05 | 1.400E-05 |
| 28147 | 1408 | K1408Q | E | 1.2e-05 | 8.9e-06 | 1.2e-05 | 1.097E-05 | 1.790E-06 | 8.900E-06 | 1.200E-05 |
| 28148 | 1408 | K1408G | E | 9.3e-06 | 1.1e-05 | 9.5e-06 | 9.933E-06 | 9.292E-07 | 9.300E-06 | 1.100E-05 |
| 28149 | 1408 | K1408H | E | 1.6e-05 | 1.5e-05 | 1.6e-05 | 1.567E-05 | 5.774E-07 | 1.500E-05 | 1.600E-05 |
| 28150 | 1408 | K1408I | E | 9.4e-06 | 1.5e-05 | 1.1e-05 | 1.180E-05 | 2.884E-06 | 9.400E-06 | 1.500E-05 |
| 28151 | 1408 | K1408L | E | 3.3e-05 | 1.3e-05 | 1.8e-05 | 2.133E-05 | 1.041E-05 | 1.300E-05 | 3.300E-05 |
| 28152 | 1408 | K1408K | E | 1e-05 | 9.7e-06 | 1.5e-05 | 1.157E-05 | 2.977E-06 | 9.700E-06 | 1.500E-05 |
| 28153 | 1408 | K1408M | E | 6.7e-06 | 1.6e-05 | 1.5e-05 | 1.257E-05 | 5.105E-06 | 6.700E-06 | 1.600E-05 |
| 28154 | 1408 | K1408F | E | 1.7e-05 | 1.6e-05 | 6.1e-06 | 1.303E-05 | 6.025E-06 | 6.100E-06 | 1.700E-05 |
| 28155 | 1408 | K1408P | E | 1.2e-05 | 1.5e-05 | 1.1e-05 | 1.267E-05 | 2.082E-06 | 1.100E-05 | 1.500E-05 |
| 28156 | 1408 | K1408S | E | 1.1e-05 | 9.9e-06 | 1.4e-05 | 1.163E-05 | 2.122E-06 | 9.900E-06 | 1.400E-05 |
| 28157 | 1408 | K1408T | E | 1.5e-05 | 1.1e-05 | 1.7e-05 | 1.433E-05 | 3.055E-06 | 1.100E-05 | 1.700E-05 |
| 28158 | 1408 | K1408W | E | 1.6e-05 | 1.4e-05 | 1.1e-05 | 1.367E-05 | 2.517E-06 | 1.100E-05 | 1.600E-05 |
| 28159 | 1408 | K1408Y | E | 8.3e-06 | 9.3e-06 | 1e-05 | 9.200E-06 | 8.544E-07 | 8.300E-06 | 1.000E-05 |
| 28160 | 1408 | K1408V | E | 1.5e-05 | 9.7e-06 | 1.3e-05 | 1.257E-05 | 2.676E-06 | 9.700E-06 | 1.500E-05 |
| 28161 | 1409 | S1409A | E | 1e-05 | 1e-05 | 1.6e-05 | 1.200E-05 | 3.464E-06 | 1.000E-05 | 1.600E-05 |
| 28162 | 1409 | S1409R | E | 1.7e-05 | 1.6e-05 | 1.4e-05 | 1.567E-05 | 1.528E-06 | 1.400E-05 | 1.700E-05 |
| 28163 | 1409 | S1409N | E | 9.1e-06 | 1.1e-05 | 1.1e-05 | 1.037E-05 | 1.097E-06 | 9.100E-06 | 1.100E-05 |
| 28164 | 1409 | S1409D | E | 8.1e-06 | 1.4e-05 | 8.6e-06 | 1.023E-05 | 3.272E-06 | 8.100E-06 | 1.400E-05 |
| 28165 | 1409 | S1409C | E | 1.6e-05 | 1.5e-05 | 1.7e-05 | 1.600E-05 | 1.000E-06 | 1.500E-05 | 1.700E-05 |
| 28166 | 1409 | S1409E | E | 1.1e-05 | 1.7e-05 | 8.9e-06 | 1.230E-05 | 4.204E-06 | 8.900E-06 | 1.700E-05 |
| 28167 | 1409 | S1409Q | E | 1e-05 | 1e-05 | 1.1e-05 | 1.033E-05 | 5.774E-07 | 1.000E-05 | 1.100E-05 |
| 28168 | 1409 | S1409G | E | 1.5e-05 | 1.5e-05 | 1.7e-05 | 1.567E-05 | 1.155E-06 | 1.500E-05 | 1.700E-05 |
| 28169 | 1409 | S1409H | E | 1.4e-05 | 1.8e-05 | 1.6e-05 | 1.600E-05 | 2.000E-06 | 1.400E-05 | 1.800E-05 |
| 28170 | 1409 | S1409I | E | 1.1e-05 | 1.1e-05 | 1.5e-05 | 1.233E-05 | 2.309E-06 | 1.100E-05 | 1.500E-05 |
| 28171 | 1409 | S1409L | E | 1.3e-05 | 1.2e-05 | 1e-05 | 1.167E-05 | 1.528E-06 | 1.000E-05 | 1.300E-05 |
| 28172 | 1409 | S1409K | E | 9.4e-06 | 1e-05 | 1.9e-05 | 1.280E-05 | 5.378E-06 | 9.400E-06 | 1.900E-05 |
| 28173 | 1409 | S1409M | E | 1.5e-05 | 1.5e-05 | 1.3e-05 | 1.433E-05 | 1.155E-06 | 1.300E-05 | 1.500E-05 |
| 28174 | 1409 | S1409F | E | 1.5e-05 | 1e-05 | 1.7e-05 | 1.400E-05 | 3.606E-06 | 1.000E-05 | 1.700E-05 |
| 28175 | 1409 | S1409P | E | 9.2e-06 | 1.6e-05 | 1.1e-05 | 1.207E-05 | 3.523E-06 | 9.200E-06 | 1.600E-05 |
| 28176 | 1409 | S1409S | E | 1.5e-05 | 1e-05 | 1.1e-05 | 1.200E-05 | 2.646E-06 | 1.000E-05 | 1.500E-05 |
| 28177 | 1409 | S1409T | E | 1.3e-05 | 1.1e-05 | 1e-05 | 1.133E-05 | 1.528E-06 | 1.000E-05 | 1.300E-05 |
| 28178 | 1409 | S1409W | E | 1.2e-05 | 9.8e-06 | 1.5e-05 | 1.227E-05 | 2.610E-06 | 9.800E-06 | 1.500E-05 |
| 28179 | 1409 | S1409Y | E | 1.4e-05 | 7.4e-06 | 1.5e-05 | 1.213E-05 | 4.130E-06 | 7.400E-06 | 1.500E-05 |
| 28180 | 1409 | S1409V | E | 6.8e-06 | 1.7e-05 | 1.6e-05 | 1.327E-05 | 5.623E-06 | 6.800E-06 | 1.700E-05 |
| 28181 | 1410 | T1410A | E | 1.2e-05 | 1.4e-05 | 1e-05 | 1.200E-05 | 2.000E-06 | 1.000E-05 | 1.400E-05 |
| 28182 | 1410 | T1410R | E | 1.5e-05 | 1e-05 | 1e-05 | 1.167E-05 | 2.887E-06 | 1.000E-05 | 1.500E-05 |
| 28183 | 1410 | T1410N | E | 1.5e-05 | 1.6e-05 | 9.5e-06 | 1.350E-05 | 3.500E-06 | 9.500E-06 | 1.600E-05 |
| 28184 | 1410 | T1410D | E | 1.4e-05 | 8.7e-06 | 1.1e-05 | 1.123E-05 | 2.658E-06 | 8.700E-06 | 1.400E-05 |
| 28185 | 1410 | T1410C | E | 9.9e-06 | 1.1e-05 | 1.1e-05 | 1.063E-05 | 6.351E-07 | 9.900E-06 | 1.100E-05 |
| 28186 | 1410 | T1410E | E | 1e-05 | 9.9e-06 | 9.6e-06 | 9.833E-06 | 2.082E-07 | 9.600E-06 | 1.000E-05 |
| 28187 | 1410 | T1410Q | E | 1.5e-05 | 1.4e-05 | 1.4e-05 | 1.433E-05 | 5.774E-07 | 1.400E-05 | 1.500E-05 |
| 28188 | 1410 | T1410G | E | 9.1e-06 | 8.4e-06 | 6.9e-06 | 8.133E-06 | 1.124E-06 | 6.900E-06 | 9.100E-06 |
| 28189 | 1410 | T1410H | E | 1.2e-05 | 1.4e-05 | 9.2e-06 | 1.173E-05 | 2.411E-06 | 9.200E-06 | 1.400E-05 |
| 28190 | 1410 | T1410I | E | 1e-05 | 1.6e-05 | 1.6e-05 | 1.400E-05 | 3.464E-06 | 1.000E-05 | 1.600E-05 |
| 28191 | 1410 | T1410L | E | 1.5e-05 | 1.3e-05 | 8.8e-06 | 1.227E-05 | 3.164E-06 | 8.800E-06 | 1.500E-05 |
| 28192 | 1410 | T1410K | E | 1.3e-05 | 1.1e-05 | 1.4e-05 | 1.267E-05 | 1.528E-06 | 1.100E-05 | 1.400E-05 |
| 28193 | 1410 | T1410M | E | 1.6e-05 | 1.1e-05 | 1.5e-05 | 1.400E-05 | 2.646E-06 | 1.100E-05 | 1.600E-05 |
| 28194 | 1410 | T1410F | E | 1e-05 | 1.1e-05 | 1.5e-05 | 1.200E-05 | 2.646E-06 | 1.000E-05 | 1.500E-05 |
| 28195 | 1410 | T1410P | E | 1.1e-05 | 1.1e-05 | 1.5e-05 | 1.233E-05 | 2.309E-06 | 1.100E-05 | 1.500E-05 |
| 28196 | 1410 | T1410S | E | 1.5e-05 | 1.4e-05 | 1.6e-05 | 1.500E-05 | 1.000E-06 | 1.400E-05 | 1.600E-05 |
| 28197 | 1410 | T1410T | E | 1.3e-05 | 1.5e-05 | 8.1e-06 | 1.203E-05 | 3.550E-06 | 8.100E-06 | 1.500E-05 |
| 28198 | 1410 | T1410W | E | 1.2e-05 | 1.3e-05 | 1.5e-05 | 1.333E-05 | 1.528E-06 | 1.200E-05 | 1.500E-05 |
| 28199 | 1410 | T1410Y | E | 9.5e-06 | 1.5e-05 | 1.7e-05 | 1.383E-05 | 3.884E-06 | 9.500E-06 | 1.700E-05 |
| 28200 | 1410 | T1410V | E | 1.5e-05 | 1.8e-05 | 1.2e-05 | 1.500E-05 | 3.000E-06 | 1.200E-05 | 1.800E-05 |
| 28201 | 1411 | S1411A | E | 1.4e-05 | 1.1e-05 | 1.6e-05 | 1.367E-05 | 2.517E-06 | 1.100E-05 | 1.600E-05 |
| 28202 | 1411 | S1411R | E | 1.6e-05 | 9.1e-06 | 1.7e-05 | 1.403E-05 | 4.302E-06 | 9.100E-06 | 1.700E-05 |
| 28203 | 1411 | S1411N | E | 1.2e-05 | 1.6e-05 | 1.7e-05 | 1.500E-05 | 2.646E-06 | 1.200E-05 | 1.700E-05 |
| 28204 | 1411 | S1411D | E | 1.4e-05 | 1.4e-05 | 1.1e-05 | 1.300E-05 | 1.732E-06 | 1.100E-05 | 1.400E-05 |
| 28205 | 1411 | S1411C | E | 9.8e-06 | 9.3e-06 | 1.1e-05 | 1.003E-05 | 8.737E-07 | 9.300E-06 | 1.100E-05 |
| 28206 | 1411 | S1411E | E | 1.2e-05 | 1.3e-05 | 1.2e-05 | 1.233E-05 | 5.774E-07 | 1.200E-05 | 1.300E-05 |
| 28207 | 1411 | S1411Q | E | 1.2e-05 | 1.4e-05 | 1e-05 | 1.200E-05 | 2.000E-06 | 1.000E-05 | 1.400E-05 |
| 28208 | 1411 | S1411G | E | 1e-05 | 1.4e-05 | 1.6e-05 | 1.333E-05 | 3.055E-06 | 1.000E-05 | 1.600E-05 |
| 28209 | 1411 | S1411H | E | 1.1e-05 | 1.4e-05 | 1.1e-05 | 1.200E-05 | 1.732E-06 | 1.100E-05 | 1.400E-05 |
| 28210 | 1411 | S1411I | E | 1.1e-05 | 1.5e-05 | 8.9e-06 | 1.163E-05 | 3.099E-06 | 8.900E-06 | 1.500E-05 |
| 28211 | 1411 | S1411L | E | 1.1e-05 | 1.4e-05 | 1.1e-05 | 1.200E-05 | 1.732E-06 | 1.100E-05 | 1.400E-05 |
| 28212 | 1411 | S1411K | E | 9.7e-06 | 1.7e-05 | 3.1e-05 | 1.923E-05 | 1.082E-05 | 9.700E-06 | 3.100E-05 |
| 28213 | 1411 | S1411M | E | 9.2e-06 | 8.6e-06 | 1.4e-05 | 1.060E-05 | 2.960E-06 | 8.600E-06 | 1.400E-05 |
| 28214 | 1411 | S1411F | E | 1.5e-05 | 9e-06 | 1.4e-05 | 1.267E-05 | 3.215E-06 | 9.000E-06 | 1.500E-05 |
| 28215 | 1411 | S1411P | E | 9.5e-06 | 1e-05 | 1.5e-05 | 1.150E-05 | 3.041E-06 | 9.500E-06 | 1.500E-05 |
| 28216 | 1411 | S1411S | E | 1.6e-05 | 1.7e-05 | 1.6e-05 | 1.633E-05 | 5.774E-07 | 1.600E-05 | 1.700E-05 |
| 28217 | 1411 | S1411T | E | 1.3e-05 | 1e-05 | 1.4e-05 | 1.233E-05 | 2.082E-06 | 1.000E-05 | 1.400E-05 |
| 28218 | 1411 | S1411W | E | 1e-05 | 1.3e-05 | 1.7e-05 | 1.333E-05 | 3.512E-06 | 1.000E-05 | 1.700E-05 |
| 28219 | 1411 | S1411Y | E | 1.1e-05 | 1.1e-05 | 9.8e-06 | 1.060E-05 | 6.928E-07 | 9.800E-06 | 1.100E-05 |

| | | | | | | | | | | |
|---|---|---|---|---|---|---|---|---|---|---|
| 28220 | 1411 | S1411V | E | 1.3e-05 | 1.2e-05 | 8.4e-06 | 1.113E-05 | 2.419E-06 | 8.400E-06 | 1.300E-05 |
| 28221 | 1412 | G1412A | E | 1.5e-05 | 8.9e-06 | 1.6e-05 | 1.330E-05 | 3.843E-06 | 8.900E-06 | 1.600E-05 |
| 28222 | 1412 | G1412R | E | 1.5e-05 | 1.5e-05 | 1.1e-05 | 1.367E-05 | 2.309E-06 | 1.100E-05 | 1.500E-05 |
| 28223 | 1412 | G1412N | E | 9.4e-06 | 1.4e-05 | 9.4e-06 | 1.093E-05 | 2.656E-06 | 9.400E-06 | 1.400E-05 |
| 28224 | 1412 | G1412D | E | 1.5e-05 | 9.3e-06 | 8.9e-06 | 1.107E-05 | 3.412E-06 | 8.900E-06 | 1.500E-05 |
| 28225 | 1412 | G1412C | E | 1.4e-05 | 1.1e-05 | 1.1e-05 | 1.200E-05 | 1.732E-06 | 1.100E-05 | 1.400E-05 |
| 28226 | 1412 | G1412E | E | 1.6e-05 | 1.7e-05 | 1.3e-05 | 1.533E-05 | 2.082E-06 | 1.300E-05 | 1.700E-05 |
| 28227 | 1412 | G1412Q | E | 1.8e-05 | 1e-05 | 9.3e-06 | 1.243E-05 | 4.834E-06 | 9.300E-06 | 1.800E-05 |
| 28228 | 1412 | G1412G | E | 1.5e-05 | 1.1e-05 | 1.6e-05 | 1.400E-05 | 2.646E-06 | 1.100E-05 | 1.600E-05 |
| 28229 | 1412 | G1412H | E | 1.5e-05 | 9.8e-06 | 9.6e-06 | 1.147E-05 | 3.062E-06 | 9.600E-06 | 1.500E-05 |
| 28230 | 1412 | G1412I | E | 1e-05 | 9.7e-06 | 1.2e-05 | 1.057E-05 | 1.250E-06 | 9.700E-06 | 1.200E-05 |
| 28231 | 1412 | G1412L | E | 1.5e-05 | 1.5e-05 | 1.2e-05 | 1.400E-05 | 1.732E-06 | 1.200E-05 | 1.500E-05 |
| 28232 | 1412 | G1412K | E | 1.3e-05 | 1e-05 | 1.4e-05 | 1.233E-05 | 2.082E-06 | 1.000E-05 | 1.400E-05 |
| 28233 | 1412 | G1412M | E | 1.1e-05 | 1.3e-05 | 1.4e-05 | 1.267E-05 | 1.528E-06 | 1.100E-05 | 1.400E-05 |
| 28234 | 1412 | G1412F | E | 1e-05 | 1.5e-05 | 7.6e-06 | 1.087E-05 | 3.775E-06 | 7.600E-06 | 1.500E-05 |
| 28235 | 1412 | G1412P | E | 1.5e-05 | 1.4e-05 | 1.5e-05 | 1.467E-05 | 5.774E-07 | 1.400E-05 | 1.500E-05 |
| 28236 | 1412 | G1412S | E | 1.3e-05 | 1.1e-05 | 8.3e-06 | 1.077E-05 | 2.359E-06 | 8.300E-06 | 1.300E-05 |
| 28237 | 1412 | G1412T | E | 1e-05 | 1.1e-05 | 1.5e-05 | 1.200E-05 | 2.646E-06 | 1.000E-05 | 1.500E-05 |
| 28238 | 1412 | G1412W | E | 1e-05 | 9.1e-06 | 1.5e-05 | 1.137E-05 | 3.179E-06 | 9.100E-06 | 1.500E-05 |
| 28239 | 1412 | G1412Y | E | 1.3e-05 | 1.4e-05 | 1.2e-05 | 1.300E-05 | 1.000E-06 | 1.200E-05 | 1.400E-05 |
| 28240 | 1412 | G1412V | E | 8.2e-06 | 1.1e-05 | 1.4e-05 | 1.107E-05 | 2.901E-06 | 8.200E-06 | 1.400E-05 |
| 28241 | 1413 | G1413A | E | 9.9e-06 | 1.6e-05 | 1.3e-05 | 1.297E-05 | 3.050E-06 | 9.900E-06 | 1.600E-05 |
| 28242 | 1413 | G1413R | E | 1.5e-05 | 1.6e-05 | 1.5e-05 | 1.533E-05 | 5.774E-07 | 1.500E-05 | 1.600E-05 |
| 28243 | 1413 | G1413N | E | 1.4e-05 | 1.5e-05 | 1.6e-05 | 1.500E-05 | 1.000E-06 | 1.400E-05 | 1.600E-05 |
| 28244 | 1413 | G1413D | E | 1.8e-05 | 1.6e-05 | 1.5e-05 | 1.633E-05 | 1.528E-06 | 1.500E-05 | 1.800E-05 |
| 28245 | 1413 | G1413C | E | 1e-05 | 1.4e-05 | 1.4e-05 | 1.267E-05 | 2.309E-06 | 1.000E-05 | 1.400E-05 |
| 28246 | 1413 | G1413E | E | 1.5e-05 | 1.3e-05 | 9.5e-06 | 1.250E-05 | 2.784E-06 | 9.500E-06 | 1.500E-05 |
| 28247 | 1413 | G1413Q | E | 1.5e-05 | 8.1e-06 | 8.7e-06 | 1.060E-05 | 3.822E-06 | 8.100E-06 | 1.500E-05 |
| 28248 | 1413 | G1413G | E | 1.2e-05 | 1e-05 | 1.5e-05 | 1.233E-05 | 2.517E-06 | 1.000E-05 | 1.500E-05 |
| 28249 | 1413 | G1413H | E | 1.4e-05 | 9.3e-06 | 1.5e-05 | 1.277E-05 | 3.044E-06 | 9.300E-06 | 1.500E-05 |
| 28250 | 1413 | G1413I | E | 9.9e-06 | 1e-05 | 1.6e-05 | 1.197E-05 | 3.493E-06 | 9.900E-06 | 1.600E-05 |
| 28251 | 1413 | G1413L | E | 1.4e-05 | 1.5e-05 | 1e-05 | 1.300E-05 | 2.646E-06 | 1.000E-05 | 1.500E-05 |
| 28252 | 1413 | G1413K | E | 1.1e-05 | 1.7e-05 | 9.9e-06 | 1.263E-05 | 3.821E-06 | 9.900E-06 | 1.700E-05 |
| 28253 | 1413 | G1413M | E | 9.2e-06 | 1.5e-05 | 1.4e-05 | 1.273E-05 | 3.101E-06 | 9.200E-06 | 1.500E-05 |
| 28254 | 1413 | G1413F | E | 8.9e-06 | 1.4e-05 | 1.5e-05 | 1.263E-05 | 3.272E-06 | 8.900E-06 | 1.500E-05 |
| 28255 | 1413 | G1413P | E | 1.1e-05 | 1.4e-05 | 1.2e-05 | 1.233E-05 | 1.528E-06 | 1.100E-05 | 1.400E-05 |
| 28256 | 1413 | G1413S | E | 1.6e-05 | 1.5e-05 | 8.4e-06 | 1.313E-05 | 4.130E-06 | 8.400E-06 | 1.600E-05 |
| 28257 | 1413 | G1413T | E | 1e-05 | 1.4e-05 | 1.4e-05 | 1.267E-05 | 2.309E-06 | 1.000E-05 | 1.400E-05 |
| 28258 | 1413 | G1413W | E | 1.3e-05 | 1.1e-05 | 1.2e-05 | 1.200E-05 | 1.000E-06 | 1.100E-05 | 1.300E-05 |
| 28259 | 1413 | G1413Y | E | 1.3e-05 | 1.2e-05 | 1e-05 | 1.167E-05 | 1.528E-06 | 1.000E-05 | 1.300E-05 |
| 28260 | 1413 | G1413V | E | 1.5e-05 | 1.5e-05 | 1.5e-05 | 1.500E-05 | 0.000E+00 | 1.500E-05 | 1.500E-05 |
| 28261 | 1414 | T1414A | E | 1.5e-05 | 8.5e-06 | 1.4e-05 | 1.250E-05 | 3.500E-06 | 8.500E-06 | 1.500E-05 |
| 28262 | 1414 | T1414R | E | 1.3e-05 | 1.5e-05 | 1.4e-05 | 1.400E-05 | 1.000E-06 | 1.300E-05 | 1.500E-05 |
| 28263 | 1414 | T1414N | E | 9.6e-06 | 1.5e-05 | 1.1e-05 | 1.187E-05 | 2.802E-06 | 9.600E-06 | 1.500E-05 |
| 28264 | 1414 | T1414D | E | 1e-05 | 1.6e-05 | 9.3e-06 | 1.177E-05 | 3.683E-06 | 9.300E-06 | 1.600E-05 |
| 28265 | 1414 | T1414C | E | 1.1e-05 | 6.4e-06 | 1.2e-05 | 9.800E-06 | 2.987E-06 | 6.400E-06 | 1.200E-05 |
| 28266 | 1414 | T1414E | E | 1e-05 | 1.7e-05 | 1e-05 | 1.233E-05 | 4.041E-06 | 1.000E-05 | 1.700E-05 |
| 28267 | 1414 | T1414Q | E | 7.4e-06 | 9.8e-06 | 1.1e-05 | 9.400E-06 | 1.833E-06 | 7.400E-06 | 1.100E-05 |
| 28268 | 1414 | T1414G | E | 1.8e-05 | 1e-05 | 1.6e-05 | 1.467E-05 | 4.163E-06 | 1.000E-05 | 1.800E-05 |
| 28269 | 1414 | T1414H | E | 2.2e-05 | 1e-05 | 1.6e-05 | 1.600E-05 | 6.000E-06 | 1.000E-05 | 2.200E-05 |
| 28270 | 1414 | T1414I | E | 1.1e-05 | 1.3e-05 | 1.4e-05 | 1.267E-05 | 1.528E-06 | 1.100E-05 | 1.400E-05 |
| 28271 | 1414 | T1414L | E | 6.1e-06 | 1.2e-05 | 1.5e-05 | 1.103E-05 | 4.528E-06 | 6.100E-06 | 1.500E-05 |
| 28272 | 1414 | T1414K | E | 1.2e-05 | 8.7e-06 | 9.1e-06 | 9.933E-06 | 1.801E-06 | 8.700E-06 | 1.200E-05 |
| 28273 | 1414 | T1414M | E | 1.1e-05 | 1e-05 | 1.4e-05 | 1.167E-05 | 2.082E-06 | 1.000E-05 | 1.400E-05 |
| 28274 | 1414 | T1414F | E | 1.3e-05 | 1.1e-05 | 1e-05 | 1.133E-05 | 1.528E-06 | 1.000E-05 | 1.300E-05 |
| 28275 | 1414 | T1414P | E | 9.8e-06 | 1.8e-05 | 1.1e-05 | 1.293E-05 | 4.429E-06 | 9.800E-06 | 1.800E-05 |
| 28276 | 1414 | T1414S | E | 1.2e-05 | 9e-06 | 1.3e-05 | 1.133E-05 | 2.082E-06 | 9.000E-06 | 1.300E-05 |
| 28277 | 1414 | T1414T | E | 1e-05 | 1.3e-05 | 1.5e-05 | 1.267E-05 | 2.517E-06 | 1.000E-05 | 1.500E-05 |
| 28278 | 1414 | T1414W | E | 1.3e-05 | 8.6e-06 | 1.4e-05 | 1.187E-05 | 2.873E-06 | 8.600E-06 | 1.400E-05 |
| 28279 | 1414 | T1414Y | E | 1.1e-05 | 1.4e-05 | 1.6e-05 | 1.367E-05 | 2.517E-06 | 1.100E-05 | 1.600E-05 |
| 28280 | 1414 | T1414V | E | 9.8e-06 | 1.1e-05 | 1.6e-05 | 1.227E-05 | 3.288E-06 | 9.800E-06 | 1.600E-05 |
| 28281 | 1415 | A1415A | E | 1.4e-05 | 1e-05 | 1e-05 | 1.133E-05 | 2.309E-06 | 1.000E-05 | 1.400E-05 |
| 28282 | 1415 | A1415R | E | 9.8e-06 | 1.1e-05 | 1.4e-05 | 1.160E-05 | 2.163E-06 | 9.800E-06 | 1.400E-05 |
| 28283 | 1415 | A1415N | E | 1.1e-05 | 1.5e-05 | 1.6e-05 | 1.400E-05 | 2.646E-06 | 1.100E-05 | 1.600E-05 |
| 28284 | 1415 | A1415D | E | 7.4e-06 | 3.6e-05 | 1e-05 | 1.780E-05 | 1.582E-05 | 7.400E-06 | 3.600E-05 |
| 28285 | 1415 | A1415C | E | 1.4e-05 | 8e-06 | 1.6e-05 | 1.267E-05 | 4.163E-06 | 8.000E-06 | 1.600E-05 |
| 28286 | 1415 | A1415E | E | 1e-05 | 1e-05 | 1.3e-05 | 1.100E-05 | 1.732E-06 | 1.000E-05 | 1.300E-05 |
| 28287 | 1415 | A1415Q | E | 8.4e-06 | 9.7e-06 | 9.4e-06 | 9.167E-06 | 6.807E-07 | 8.400E-06 | 9.700E-06 |
| 28288 | 1415 | A1415G | E | 1.5e-05 | 1.1e-05 | 1.1e-05 | 1.233E-05 | 2.309E-06 | 1.100E-05 | 1.500E-05 |
| 28289 | 1415 | A1415H | E | 1.4e-05 | 1.4e-05 | 1.6e-05 | 1.467E-05 | 1.155E-06 | 1.400E-05 | 1.600E-05 |
| 28290 | 1415 | A1415I | E | 1.6e-05 | 6.6e-06 | 8.9e-06 | 1.050E-05 | 4.900E-06 | 6.600E-06 | 1.600E-05 |
| 28291 | 1415 | A1415L | E | 1.6e-05 | 8.8e-06 | 1.5e-05 | 1.327E-05 | 3.900E-06 | 8.800E-06 | 1.600E-05 |
| 28292 | 1415 | A1415K | E | 1.3e-05 | 1.1e-05 | 1e-05 | 1.133E-05 | 1.528E-06 | 1.000E-05 | 1.300E-05 |
| 28293 | 1415 | A1415M | E | 1.2e-05 | 1e-05 | 1.1e-05 | 1.100E-05 | 1.000E-06 | 1.000E-05 | 1.200E-05 |
| 28294 | 1415 | A1415F | E | 1.3e-05 | 8.8e-06 | 1.5e-05 | 1.227E-05 | 3.164E-06 | 8.800E-06 | 1.500E-05 |
| 28295 | 1415 | A1415P | E | 6.4e-06 | 1.5e-05 | 1e-05 | 1.047E-05 | 4.319E-06 | 6.400E-06 | 1.500E-05 |
| 28296 | 1415 | A1415S | E | 1.4e-05 | 1.3e-05 | 1e-05 | 1.233E-05 | 2.082E-06 | 1.000E-05 | 1.400E-05 |
| 28297 | 1415 | A1415T | E | 1.2e-05 | 1.6e-05 | 1.5e-05 | 1.433E-05 | 2.082E-06 | 1.200E-05 | 1.600E-05 |
| 28298 | 1415 | A1415W | E | 1.5e-05 | 1.4e-05 | 1.5e-05 | 1.467E-05 | 5.774E-07 | 1.400E-05 | 1.500E-05 |
| 28299 | 1415 | A1415Y | E | 1.1e-05 | 1.5e-05 | 1.4e-05 | 1.333E-05 | 2.082E-06 | 1.100E-05 | 1.500E-05 |
| 28300 | 1415 | A1415V | E | 1.4e-05 | 1.5e-05 | 1.3e-05 | 1.400E-05 | 1.000E-06 | 1.300E-05 | 1.500E-05 |
| 28301 | 1416 | A1416A | E | 1.3e-05 | 1.5e-05 | 9.1e-06 | 1.237E-05 | 3.001E-06 | 9.100E-06 | 1.500E-05 |
| 28302 | 1416 | A1416R | E | 9.2e-06 | 1.5e-05 | 1.2e-05 | 1.207E-05 | 2.901E-06 | 9.200E-06 | 1.500E-05 |
| 28303 | 1416 | A1416N | E | 1.4e-05 | 7.8e-06 | 1.4e-05 | 1.193E-05 | 3.580E-06 | 7.800E-06 | 1.400E-05 |
| 28304 | 1416 | A1416D | E | 1.1e-05 | 1.3e-05 | 1.4e-05 | 1.267E-05 | 1.528E-06 | 1.100E-05 | 1.400E-05 |

| | | | | | | | | | | |
|---|---|---|---|---|---|---|---|---|---|---|
| 28305 | 1416 | A1416C | E | 1.3e-05 | 1.5e-05 | 1.8e-05 | 1.533E-05 | 2.517E-06 | 1.300E-05 | 1.800E-05 |
| 28306 | 1416 | A1416E | E | 1.2e-05 | 1.6e-05 | 1.5e-05 | 1.433E-05 | 2.082E-06 | 1.200E-05 | 1.600E-05 |
| 28307 | 1416 | A1416Q | E | 1.1e-05 | 1e-05 | 1.4e-05 | 1.167E-05 | 2.082E-06 | 1.000E-05 | 1.400E-05 |
| 28308 | 1416 | A1416G | E | 1.4e-05 | 1e-05 | 1.1e-05 | 1.167E-05 | 2.082E-06 | 1.000E-05 | 1.400E-05 |
| 28309 | 1416 | A1416H | E | 1.2e-05 | 9.5e-06 | 1.1e-05 | 1.083E-05 | 1.258E-06 | 9.500E-06 | 1.200E-05 |
| 28310 | 1416 | A1416I | E | 1.3e-05 | 1.5e-05 | 1.1e-05 | 1.300E-05 | 2.000E-06 | 1.100E-05 | 1.500E-05 |
| 28311 | 1416 | A1416L | E | 1.3e-05 | 1.6e-05 | 8.7e-06 | 1.257E-05 | 3.669E-06 | 8.700E-06 | 1.600E-05 |
| 28312 | 1416 | A1416K | E | 1e-05 | 1.5e-05 | 1.1e-05 | 1.200E-05 | 2.646E-06 | 1.000E-05 | 1.500E-05 |
| 28313 | 1416 | A1416M | E | 1.4e-05 | 1.6e-05 | 1.2e-05 | 1.400E-05 | 2.000E-06 | 1.200E-05 | 1.600E-05 |
| 28314 | 1416 | A1416F | E | 1.2e-05 | 8.2e-06 | 1.1e-05 | 1.040E-05 | 1.970E-06 | 8.200E-06 | 1.200E-05 |
| 28315 | 1416 | A1416P | E | 1.5e-05 | 9.8e-06 | 1.4e-05 | 1.293E-05 | 2.759E-06 | 9.800E-06 | 1.500E-05 |
| 28316 | 1416 | A1416S | E | 1.6e-05 | 1e-05 | 1.6e-05 | 1.400E-05 | 3.464E-06 | 1.000E-05 | 1.600E-05 |
| 28317 | 1416 | A1416T | E | 1.4e-05 | 9.6e-06 | 9.6e-06 | 1.107E-05 | 2.540E-06 | 9.600E-06 | 1.400E-05 |
| 28318 | 1416 | A1416W | E | 1.7e-05 | 9.1e-06 | 8.6e-06 | 1.157E-05 | 4.712E-06 | 8.600E-06 | 1.700E-05 |
| 28319 | 1416 | A1416Y | E | 1.1e-05 | 1.5e-05 | 1e-05 | 1.200E-05 | 2.646E-06 | 1.000E-05 | 1.500E-05 |
| 28320 | 1416 | A1416V | E | 8.7e-06 | 1.6e-05 | 1.1e-05 | 1.190E-05 | 3.732E-06 | 8.700E-06 | 1.600E-05 |
| 28321 | 1417 | L1417A | E | 1.6e-05 | 1.5e-05 | 1.6e-05 | 1.567E-05 | 5.774E-07 | 1.500E-05 | 1.600E-05 |
| 28322 | 1417 | L1417R | E | 7.9e-06 | 1.3e-05 | 1.5e-05 | 1.197E-05 | 3.661E-06 | 7.900E-06 | 1.500E-05 |
| 28323 | 1417 | L1417N | E | 1.5e-05 | 1e-05 | 1.4e-05 | 1.300E-05 | 2.646E-06 | 1.000E-05 | 1.500E-05 |
| 28324 | 1417 | L1417D | E | 1.7e-05 | 9.5e-06 | 1.1e-05 | 1.250E-05 | 3.969E-06 | 9.500E-06 | 1.700E-05 |
| 28325 | 1417 | L1417C | E | 1.7e-05 | 1.6e-05 | 1.7e-05 | 1.667E-05 | 5.774E-07 | 1.600E-05 | 1.700E-05 |
| 28326 | 1417 | L1417E | E | 1.1e-05 | 1.3e-05 | 1.4e-05 | 1.267E-05 | 1.528E-06 | 1.100E-05 | 1.400E-05 |
| 28327 | 1417 | L1417Q | E | 1e-05 | 1.1e-05 | 1.3e-05 | 1.133E-05 | 1.528E-06 | 1.000E-05 | 1.300E-05 |
| 28328 | 1417 | L1417G | E | 1.6e-05 | 1.6e-05 | 9.7e-06 | 1.390E-05 | 3.637E-06 | 9.700E-06 | 1.600E-05 |
| 28329 | 1417 | L1417H | E | 7.7e-06 | 1.5e-05 | 1.4e-05 | 1.223E-05 | 3.958E-06 | 7.700E-06 | 1.500E-05 |
| 28330 | 1417 | L1417I | E | 1.1e-05 | 9.8e-06 | 1e-05 | 1.027E-05 | 6.429E-07 | 9.800E-06 | 1.100E-05 |
| 28331 | 1417 | L1417L | E | 9.1e-06 | 1.5e-05 | 1.3e-05 | 1.237E-05 | 3.001E-06 | 9.100E-06 | 1.500E-05 |
| 28332 | 1417 | L1417K | E | 1.9e-05 | 9.7e-06 | 1.3e-05 | 1.390E-05 | 4.715E-06 | 9.700E-06 | 1.900E-05 |
| 28333 | 1417 | L1417M | E | 9.7e-06 | 1.1e-05 | 1.4e-05 | 1.157E-05 | 2.205E-06 | 9.700E-06 | 1.400E-05 |
| 28334 | 1417 | L1417F | E | 1.5e-05 | 8.8e-06 | 1e-05 | 1.127E-05 | 3.288E-06 | 8.800E-06 | 1.500E-05 |
| 28335 | 1417 | L1417P | E | 1.5e-05 | 1.3e-05 | 1.4e-05 | 1.400E-05 | 1.000E-06 | 1.300E-05 | 1.500E-05 |
| 28336 | 1417 | L1417S | E | 1.5e-05 | 1e-05 | 1.1e-05 | 1.200E-05 | 2.646E-06 | 1.000E-05 | 1.500E-05 |
| 28337 | 1417 | L1417T | E | 1.5e-05 | 1.2e-05 | 9.1e-06 | 1.203E-05 | 2.950E-06 | 9.100E-06 | 1.500E-05 |
| 28338 | 1417 | L1417W | E | 1.4e-05 | 1.6e-05 | 1e-05 | 1.333E-05 | 3.055E-06 | 1.000E-05 | 1.600E-05 |
| 28339 | 1417 | L1417Y | E | 1.4e-05 | 1.2e-05 | 1.4e-05 | 1.333E-05 | 1.155E-06 | 1.200E-05 | 1.400E-05 |
| 28340 | 1417 | L1417V | E | 1.2e-05 | 1.5e-05 | 1.5e-05 | 1.400E-05 | 1.732E-06 | 1.200E-05 | 1.500E-05 |
| 28341 | 1418 | G1418A | E | 1.4e-05 | 1e-05 | 7.6e-06 | 1.053E-05 | 3.233E-06 | 7.600E-06 | 1.400E-05 |
| 28342 | 1418 | G1418R | E | 9.8e-06 | 9.4e-06 | 1e-05 | 9.733E-06 | 3.055E-07 | 9.400E-06 | 1.000E-05 |
| 28343 | 1418 | G1418N | E | 1e-05 | 1.7e-05 | 1.5e-05 | 1.400E-05 | 3.606E-06 | 1.000E-05 | 1.700E-05 |
| 28344 | 1418 | G1418D | E | 1.5e-05 | 1.4e-05 | 1.5e-05 | 1.467E-05 | 5.774E-07 | 1.400E-05 | 1.500E-05 |
| 28345 | 1418 | G1418C | E | 9e-06 | 8.8e-06 | 9.8e-06 | 9.200E-06 | 5.292E-07 | 8.800E-06 | 9.800E-06 |
| 28346 | 1418 | G1418E | E | 8.9e-06 | 1.2e-05 | 1e-05 | 1.030E-05 | 1.572E-06 | 8.900E-06 | 1.200E-05 |
| 28347 | 1418 | G1418Q | E | 1.5e-05 | 1.5e-05 | 1.4e-05 | 1.467E-05 | 5.774E-07 | 1.400E-05 | 1.500E-05 |
| 28348 | 1418 | G1418G | E | 1.6e-05 | 1.4e-05 | 1e-05 | 1.333E-05 | 3.055E-06 | 1.000E-05 | 1.600E-05 |
| 28349 | 1418 | G1418H | E | 1.3e-05 | 1.6e-05 | 1.1e-05 | 1.333E-05 | 2.517E-06 | 1.100E-05 | 1.600E-05 |
| 28350 | 1418 | G1418I | E | 1.6e-05 | 9.6e-06 | 1.1e-05 | 1.220E-05 | 3.365E-06 | 9.600E-06 | 1.600E-05 |
| 28351 | 1418 | G1418L | E | 1.4e-05 | 1.4e-05 | 1e-05 | 1.267E-05 | 2.309E-06 | 1.000E-05 | 1.400E-05 |
| 28352 | 1418 | G1418K | E | 1.5e-05 | 1.1e-05 | 8.1e-06 | 1.137E-05 | 3.465E-06 | 8.100E-06 | 1.500E-05 |
| 28353 | 1418 | G1418M | E | 1e-05 | 1.4e-05 | 9.5e-06 | 1.117E-05 | 2.466E-06 | 9.500E-06 | 1.400E-05 |
| 28354 | 1418 | G1418F | E | 1.6e-05 | 1.6e-05 | 1.7e-05 | 1.633E-05 | 5.774E-07 | 1.600E-05 | 1.700E-05 |
| 28355 | 1418 | G1418P | E | 8.5e-06 | 1.1e-05 | 1.3e-05 | 1.083E-05 | 2.255E-06 | 8.500E-06 | 1.300E-05 |
| 28356 | 1418 | G1418S | E | 1.6e-05 | 1.1e-05 | 1.5e-05 | 1.400E-05 | 2.646E-06 | 1.100E-05 | 1.600E-05 |
| 28357 | 1418 | G1418T | E | 1.5e-05 | 1.1e-05 | 1.6e-05 | 1.400E-05 | 2.646E-06 | 1.100E-05 | 1.600E-05 |
| 28358 | 1418 | G1418W | E | 1.1e-05 | 1.3e-05 | 1.2e-05 | 1.200E-05 | 1.000E-06 | 1.100E-05 | 1.300E-05 |
| 28359 | 1418 | G1418Y | E | 1.1e-05 | 1.6e-05 | 1.1e-05 | 1.267E-05 | 2.887E-06 | 1.100E-05 | 1.600E-05 |
| 28360 | 1418 | G1418V | E | 1.1e-05 | 1.3e-05 | 1.8e-05 | 1.400E-05 | 3.606E-06 | 1.100E-05 | 1.800E-05 |
| 28361 | 1419 | C1419A | E | 1.7e-05 | 9.2e-06 | 1.5e-05 | 1.373E-05 | 4.051E-06 | 9.200E-06 | 1.700E-05 |
| 28362 | 1419 | C1419R | E | 1.2e-05 | 2.2e-05 | 1.6e-05 | 1.667E-05 | 5.033E-06 | 1.200E-05 | 2.200E-05 |
| 28363 | 1419 | C1419N | E | 1.5e-05 | 1.4e-05 | 1.7e-05 | 1.533E-05 | 1.528E-06 | 1.400E-05 | 1.700E-05 |
| 28364 | 1419 | C1419D | E | 9.8e-06 | 1.5e-05 | 1.5e-05 | 1.327E-05 | 3.002E-06 | 9.800E-06 | 1.500E-05 |
| 28365 | 1419 | C1419C | E | 1.6e-05 | 1.6e-05 | 1.6e-05 | 1.600E-05 | 0.000E+00 | 1.600E-05 | 1.600E-05 |
| 28366 | 1419 | C1419E | E | 1.1e-05 | 1.6e-05 | 1.1e-05 | 1.267E-05 | 2.887E-06 | 1.100E-05 | 1.600E-05 |
| 28367 | 1419 | C1419Q | E | 1.3e-05 | 1.1e-05 | 1e-05 | 1.133E-05 | 1.528E-06 | 1.000E-05 | 1.300E-05 |
| 28368 | 1419 | C1419G | E | 1.1e-05 | 1.4e-05 | 1.6e-05 | 1.367E-05 | 2.517E-06 | 1.100E-05 | 1.600E-05 |
| 28369 | 1419 | C1419H | E | 1.4e-05 | 1.5e-05 | 1.4e-05 | 1.433E-05 | 5.774E-07 | 1.400E-05 | 1.500E-05 |
| 28370 | 1419 | C1419I | E | 1.4e-05 | 1.4e-05 | 1.6e-05 | 1.467E-05 | 1.155E-06 | 1.400E-05 | 1.600E-05 |
| 28371 | 1419 | C1419L | E | 1.1e-05 | 9.7e-06 | 1.2e-05 | 1.090E-05 | 1.153E-06 | 9.700E-06 | 1.200E-05 |
| 28372 | 1419 | C1419K | E | 1.1e-05 | 1.4e-05 | 1.2e-05 | 1.233E-05 | 1.528E-06 | 1.100E-05 | 1.400E-05 |
| 28373 | 1419 | C1419M | E | 1.5e-05 | 9.4e-06 | 1.4e-05 | 1.280E-05 | 2.987E-06 | 9.400E-06 | 1.500E-05 |
| 28374 | 1419 | C1419F | E | 4.8e-05 | 1.6e-05 | 8.8e-06 | 2.427E-05 | 2.087E-05 | 8.800E-06 | 4.800E-05 |
| 28375 | 1419 | C1419P | E | 1.6e-05 | 1.1e-05 | 1.6e-05 | 1.433E-05 | 2.887E-06 | 1.100E-05 | 1.600E-05 |
| 28376 | 1419 | C1419S | E | 1.6e-05 | 1.7e-05 | 7.9e-06 | 1.363E-05 | 4.990E-06 | 7.900E-06 | 1.700E-05 |
| 28377 | 1419 | C1419T | E | 1.3e-05 | 1.4e-05 | 1.5e-05 | 1.400E-05 | 1.000E-06 | 1.300E-05 | 1.500E-05 |
| 28378 | 1419 | C1419W | E | 1.7e-05 | 1.6e-05 | 1.3e-05 | 1.533E-05 | 2.082E-06 | 1.300E-05 | 1.700E-05 |
| 28379 | 1419 | C1419Y | E | 1.3e-05 | 1.6e-05 | 9.9e-06 | 1.297E-05 | 3.050E-06 | 9.900E-06 | 1.600E-05 |
| 28380 | 1419 | C1419V | E | 1.5e-05 | 1.4e-05 | 9.2e-06 | 1.273E-05 | 3.101E-06 | 9.200E-06 | 1.500E-05 |
| 28381 | 1420 | L1420A | E | 1.6e-05 | 1.5e-05 | 1.1e-05 | 1.400E-05 | 2.646E-06 | 1.100E-05 | 1.600E-05 |
| 28382 | 1420 | L1420R | E | 9.5e-06 | 1.4e-05 | 1.6e-05 | 1.317E-05 | 3.329E-06 | 9.500E-06 | 1.600E-05 |
| 28383 | 1420 | L1420N | E | 1.4e-05 | 1.5e-05 | 1.1e-05 | 1.333E-05 | 2.082E-06 | 1.100E-05 | 1.500E-05 |
| 28384 | 1420 | L1420D | E | 1e-05 | 1.6e-05 | 1e-05 | 1.200E-05 | 3.464E-06 | 1.000E-05 | 1.600E-05 |
| 28385 | 1420 | L1420C | E | 1.5e-05 | 1.1e-05 | 7.7e-06 | 1.123E-05 | 3.656E-06 | 7.700E-06 | 1.500E-05 |
| 28386 | 1420 | L1420E | E | 1.5e-05 | 1.4e-05 | 7.2e-06 | 1.207E-05 | 4.244E-06 | 7.200E-06 | 1.500E-05 |
| 28387 | 1420 | L1420Q | E | 1.1e-05 | 1.3e-05 | 1.6e-05 | 1.333E-05 | 2.517E-06 | 1.100E-05 | 1.600E-05 |
| 28388 | 1420 | L1420G | E | 1.2e-05 | 1.4e-05 | 1.4e-05 | 1.333E-05 | 1.155E-06 | 1.200E-05 | 1.400E-05 |
| 28389 | 1420 | L1420H | E | 1.2e-05 | 1.4e-05 | 1.6e-05 | 1.400E-05 | 2.000E-06 | 1.200E-05 | 1.600E-05 |

| | | | | | | | | | | |
|---|---|---|---|---|---|---|---|---|---|---|
| 28390 | 1420 | L1420I | E | 2.1e-05 | 1.5e-05 | 1.4e-05 | 1.667E-05 | 3.786E-06 | 1.400E-05 | 2.100E-05 |
| 28391 | 1420 | L1420L | E | 1.7e-05 | 1.6e-05 | 1.7e-05 | 1.667E-05 | 5.774E-07 | 1.600E-05 | 1.700E-05 |
| 28392 | 1420 | L1420K | E | 1.3e-05 | 1.6e-05 | 1.7e-05 | 1.533E-05 | 2.082E-06 | 1.300E-05 | 1.700E-05 |
| 28393 | 1420 | L1420M | E | 1.6e-05 | 8.8e-06 | 1.6e-05 | 1.360E-05 | 4.157E-06 | 8.800E-06 | 1.600E-05 |
| 28394 | 1420 | L1420F | E | 1.1e-05 | 1.3e-05 | 1.1e-05 | 1.167E-05 | 1.155E-06 | 1.100E-05 | 1.300E-05 |
| 28395 | 1420 | L1420P | E | 9.6e-06 | 1.5e-05 | 1.1e-05 | 1.187E-05 | 2.802E-06 | 9.600E-06 | 1.500E-05 |
| 28396 | 1420 | L1420S | E | 1.4e-05 | 8.8e-06 | 8.7e-06 | 1.050E-05 | 3.032E-06 | 8.700E-06 | 1.400E-05 |
| 28397 | 1420 | L1420T | E | 1.3e-05 | 1.5e-05 | 1.5e-05 | 1.433E-05 | 1.155E-06 | 1.300E-05 | 1.500E-05 |
| 28398 | 1420 | L1420W | E | 1e-05 | 9e-06 | 9.9e-06 | 9.633E-06 | 5.508E-07 | 9.000E-06 | 1.000E-05 |
| 28399 | 1420 | L1420Y | E | 1.5e-05 | 1.2e-05 | 1.6e-05 | 1.433E-05 | 2.082E-06 | 1.200E-05 | 1.600E-05 |
| 28400 | 1420 | L1420V | E | 1.1e-05 | 1.2e-05 | 8.8e-06 | 1.060E-05 | 1.637E-06 | 8.800E-06 | 1.200E-05 |
| 28401 | 1421 | V1421A | E | 9.9e-06 | 1.1e-05 | 1e-05 | 1.030E-05 | 6.083E-07 | 9.900E-06 | 1.100E-05 |
| 28402 | 1421 | V1421R | E | 1.7e-05 | 1.3e-05 | 1.4e-05 | 1.467E-05 | 2.082E-06 | 1.300E-05 | 1.700E-05 |
| 28403 | 1421 | V1421N | E | 1.6e-05 | 2.2e-05 | 1.6e-05 | 1.800E-05 | 3.464E-06 | 1.600E-05 | 2.200E-05 |
| 28404 | 1421 | V1421D | E | 9.8e-06 | 1.5e-05 | 1.2e-05 | 1.227E-05 | 2.610E-06 | 9.800E-06 | 1.500E-05 |
| 28405 | 1421 | V1421C | E | 1e-05 | 1.4e-05 | 1.1e-05 | 1.167E-05 | 2.082E-06 | 1.000E-05 | 1.400E-05 |
| 28406 | 1421 | V1421E | E | 1.2e-05 | 1.1e-05 | 1.6e-05 | 1.300E-05 | 2.646E-06 | 1.100E-05 | 1.600E-05 |
| 28407 | 1421 | V1421Q | E | 1.1e-05 | 9.5e-06 | 9.9e-06 | 1.013E-05 | 7.767E-07 | 9.500E-06 | 1.100E-05 |
| 28408 | 1421 | V1421G | E | 7.4e-06 | 1.5e-05 | 1.7e-05 | 1.313E-05 | 5.065E-06 | 7.400E-06 | 1.700E-05 |
| 28409 | 1421 | V1421H | E | 9.8e-06 | 1.4e-05 | 8.4e-06 | 1.073E-05 | 2.914E-06 | 8.400E-06 | 1.400E-05 |
| 28410 | 1421 | V1421I | E | 1.2e-05 | 1.6e-05 | 9e-06 | 1.233E-05 | 3.512E-06 | 9.000E-06 | 1.600E-05 |
| 28411 | 1421 | V1421L | E | 1.1e-05 | 1.5e-05 | 1.6e-05 | 1.400E-05 | 2.646E-06 | 1.100E-05 | 1.600E-05 |
| 28412 | 1421 | V1421K | E | 1e-05 | 1.5e-05 | 1.3e-05 | 1.267E-05 | 2.517E-06 | 1.000E-05 | 1.500E-05 |
| 28413 | 1421 | V1421M | E | 9.9e-06 | 1.1e-05 | 1.7e-05 | 1.263E-05 | 3.821E-06 | 9.900E-06 | 1.700E-05 |
| 28414 | 1421 | V1421F | E | 1.1e-05 | 1.4e-05 | 1.2e-05 | 1.233E-05 | 1.528E-06 | 1.100E-05 | 1.400E-05 |
| 28415 | 1421 | V1421P | E | 1.1e-05 | 9.2e-06 | 1.4e-05 | 1.140E-05 | 2.425E-06 | 9.200E-06 | 1.400E-05 |
| 28416 | 1421 | V1421S | E | 1.6e-05 | 1.5e-05 | 1.6e-05 | 1.567E-05 | 5.774E-07 | 1.500E-05 | 1.600E-05 |
| 28417 | 1421 | V1421T | E | 1.4e-05 | 1.4e-05 | 1.5e-05 | 1.433E-05 | 5.774E-07 | 1.400E-05 | 1.500E-05 |
| 28418 | 1421 | V1421W | E | 7.3e-06 | 1.1e-05 | 9.9e-06 | 9.400E-06 | 1.900E-06 | 7.300E-06 | 1.100E-05 |
| 28419 | 1421 | V1421Y | E | 1.4e-05 | 1.7e-05 | 1.5e-05 | 1.533E-05 | 1.528E-06 | 1.400E-05 | 1.700E-05 |
| 28420 | 1421 | V1421V | E | 8.9e-06 | 1.4e-05 | 1.7e-05 | 1.330E-05 | 4.095E-06 | 8.900E-06 | 1.700E-05 |
| 28421 | 1422 | K1422A | E | 9.5e-06 | 1e-05 | 1.5e-05 | 1.150E-05 | 3.041E-06 | 9.500E-06 | 1.500E-05 |
| 28422 | 1422 | K1422R | E | 1.5e-05 | 1.8e-05 | 9.7e-06 | 1.423E-05 | 4.203E-06 | 9.700E-06 | 1.800E-05 |
| 28423 | 1422 | K1422N | E | 9.5e-06 | 1.4e-05 | 1.6e-05 | 1.317E-05 | 3.329E-06 | 9.500E-06 | 1.600E-05 |
| 28424 | 1422 | K1422D | E | 1.3e-05 | 1.4e-05 | 1.4e-05 | 1.367E-05 | 5.774E-07 | 1.300E-05 | 1.400E-05 |
| 28425 | 1422 | K1422C | E | 1.1e-05 | 1.5e-05 | 1.1e-05 | 1.233E-05 | 2.309E-06 | 1.100E-05 | 1.500E-05 |
| 28426 | 1422 | K1422E | E | 1.4e-05 | 1e-05 | 1.3e-05 | 1.233E-05 | 2.082E-06 | 1.000E-05 | 1.400E-05 |
| 28427 | 1422 | K1422Q | E | 1.6e-05 | 1.4e-05 | 8.8e-06 | 1.293E-05 | 3.717E-06 | 8.800E-06 | 1.600E-05 |
| 28428 | 1422 | K1422G | E | 1.3e-05 | 1.5e-05 | 1.1e-05 | 1.300E-05 | 2.000E-06 | 1.100E-05 | 1.500E-05 |
| 28429 | 1422 | K1422H | E | 3.1e-05 | 1.4e-05 | 1.1e-05 | 1.867E-05 | 1.079E-05 | 1.100E-05 | 3.100E-05 |
| 28430 | 1422 | K1422I | E | 1.3e-05 | 1.3e-05 | 1.1e-05 | 1.233E-05 | 1.155E-06 | 1.100E-05 | 1.300E-05 |
| 28431 | 1422 | K1422L | E | 1.7e-05 | 1.4e-05 | 1.7e-05 | 1.600E-05 | 1.732E-06 | 1.400E-05 | 1.700E-05 |
| 28432 | 1422 | K1422K | E | 1.4e-05 | 9.3e-06 | 1.3e-05 | 1.210E-05 | 2.476E-06 | 9.300E-06 | 1.400E-05 |
| 28433 | 1422 | K1422M | E | 1.2e-05 | 9.6e-06 | 9.7e-06 | 1.043E-05 | 1.358E-06 | 9.600E-06 | 1.200E-05 |
| 28434 | 1422 | K1422F | E | 1.4e-05 | 9.8e-06 | 1.4e-05 | 1.260E-05 | 2.425E-06 | 9.800E-06 | 1.400E-05 |
| 28435 | 1422 | K1422P | E | 8.3e-06 | 1.5e-05 | 1.1e-05 | 1.143E-05 | 3.371E-06 | 8.300E-06 | 1.500E-05 |
| 28436 | 1422 | K1422S | E | 1.5e-05 | 1e-05 | 1.4e-05 | 1.300E-05 | 2.646E-06 | 1.000E-05 | 1.500E-05 |
| 28437 | 1422 | K1422T | E | 9.1e-06 | 8.1e-06 | 8.8e-06 | 8.667E-06 | 5.132E-07 | 8.100E-06 | 9.100E-06 |
| 28438 | 1422 | K1422W | E | 1e-05 | 9.6e-06 | 9.8e-06 | 9.800E-06 | 2.000E-07 | 9.600E-06 | 1.000E-05 |
| 28439 | 1422 | K1422Y | E | 1.6e-05 | 1.3e-05 | 1.5e-05 | 1.467E-05 | 1.528E-06 | 1.300E-05 | 1.600E-05 |
| 28440 | 1422 | K1422V | E | 1.5e-05 | 1.4e-05 | 1.1e-05 | 1.333E-05 | 2.082E-06 | 1.100E-05 | 1.500E-05 |
| 28441 | 1423 | D1423A | E | 1.4e-05 | 1.1e-05 | 1.5e-05 | 1.333E-05 | 2.082E-06 | 1.100E-05 | 1.500E-05 |
| 28442 | 1423 | D1423R | E | 1.3e-05 | 1.5e-05 | 1.2e-05 | 1.333E-05 | 1.528E-06 | 1.200E-05 | 1.500E-05 |
| 28443 | 1423 | D1423N | E | 9.4e-06 | 1.5e-05 | 9.6e-06 | 1.133E-05 | 3.177E-06 | 9.400E-06 | 1.500E-05 |
| 28444 | 1423 | D1423D | E | 1.2e-05 | 1.5e-05 | 1.5e-05 | 1.400E-05 | 1.732E-06 | 1.200E-05 | 1.500E-05 |
| 28445 | 1423 | D1423C | E | 1.1e-05 | 1.5e-05 | 1.1e-05 | 1.233E-05 | 2.309E-06 | 1.100E-05 | 1.500E-05 |
| 28446 | 1423 | D1423E | E | 1.3e-05 | 8.4e-06 | 1.6e-05 | 1.247E-05 | 3.828E-06 | 8.400E-06 | 1.600E-05 |
| 28447 | 1423 | D1423Q | E | 1.4e-05 | 2.8e-05 | 1.2e-05 | 1.800E-05 | 8.718E-06 | 1.200E-05 | 2.800E-05 |
| 28448 | 1423 | D1423G | E | 8.5e-06 | 1.4e-05 | 1.1e-05 | 1.117E-05 | 2.754E-06 | 8.500E-06 | 1.400E-05 |
| 28449 | 1423 | D1423H | E | 1.1e-05 | 1.7e-05 | 1.2e-05 | 1.333E-05 | 3.215E-06 | 1.100E-05 | 1.700E-05 |
| 28450 | 1423 | D1423I | E | 1.5e-05 | 1.2e-05 | 1e-05 | 1.233E-05 | 2.517E-06 | 1.000E-05 | 1.500E-05 |
| 28451 | 1423 | D1423L | E | 1.1e-05 | 1.7e-05 | 1.6e-05 | 1.467E-05 | 3.215E-06 | 1.100E-05 | 1.700E-05 |
| 28452 | 1423 | D1423K | E | 9.6e-06 | 1.4e-05 | 9.9e-06 | 1.117E-05 | 2.458E-06 | 9.600E-06 | 1.400E-05 |
| 28453 | 1423 | D1423M | E | 1.9e-05 | 1.3e-05 | 1.5e-05 | 1.567E-05 | 3.055E-06 | 1.300E-05 | 1.900E-05 |
| 28454 | 1423 | D1423F | E | 1.4e-05 | 1.3e-05 | 1.1e-05 | 1.267E-05 | 1.528E-06 | 1.100E-05 | 1.400E-05 |
| 28455 | 1423 | D1423P | E | 1.4e-05 | 1.8e-05 | 1.5e-05 | 1.567E-05 | 2.082E-06 | 1.400E-05 | 1.800E-05 |
| 28456 | 1423 | D1423S | E | 7.4e-06 | 1.1e-05 | 1.4e-05 | 1.080E-05 | 3.305E-06 | 7.400E-06 | 1.400E-05 |
| 28457 | 1423 | D1423T | E | 1.5e-05 | 1.6e-05 | 1.4e-05 | 1.500E-05 | 1.000E-06 | 1.400E-05 | 1.600E-05 |
| 28458 | 1423 | D1423W | E | 1.4e-05 | 1.5e-05 | 1.2e-05 | 1.367E-05 | 1.528E-06 | 1.200E-05 | 1.500E-05 |
| 28459 | 1423 | D1423Y | E | 1.5e-05 | 9.4e-06 | 1.6e-05 | 1.347E-05 | 3.557E-06 | 9.400E-06 | 1.600E-05 |
| 28460 | 1423 | D1423V | E | 1.4e-05 | 1.4e-05 | 1.6e-05 | 1.467E-05 | 1.155E-06 | 1.400E-05 | 1.600E-05 |
| 28461 | 1424 | Y1424A | E | 1.5e-05 | 1.6e-05 | 1.6e-05 | 1.567E-05 | 5.774E-07 | 1.500E-05 | 1.600E-05 |
| 28462 | 1424 | Y1424R | E | 1.4e-05 | 1.1e-05 | 1.4e-05 | 1.300E-05 | 1.732E-06 | 1.100E-05 | 1.400E-05 |
| 28463 | 1424 | Y1424N | E | 7e-06 | 1.4e-05 | 1.6e-05 | 1.233E-05 | 4.726E-06 | 7.000E-06 | 1.600E-05 |
| 28464 | 1424 | Y1424D | E | 1.4e-05 | 9.9e-06 | 1.5e-05 | 1.297E-05 | 2.702E-06 | 9.900E-06 | 1.500E-05 |
| 28465 | 1424 | Y1424C | E | 1.4e-05 | 9.7e-06 | 1.5e-05 | 1.290E-05 | 2.816E-06 | 9.700E-06 | 1.500E-05 |
| 28466 | 1424 | Y1424E | E | 1e-05 | 1.2e-05 | 1.4e-05 | 1.200E-05 | 2.000E-06 | 1.000E-05 | 1.400E-05 |
| 28467 | 1424 | Y1424Q | E | 9.8e-06 | 1.7e-05 | 1.5e-05 | 1.393E-05 | 3.717E-06 | 9.800E-06 | 1.700E-05 |
| 28468 | 1424 | Y1424G | E | 1.5e-05 | 9.6e-06 | 1.2e-05 | 1.220E-05 | 2.706E-06 | 9.600E-06 | 1.500E-05 |
| 28469 | 1424 | Y1424H | E | 1.4e-05 | 1.6e-05 | 1.5e-05 | 1.500E-05 | 1.000E-06 | 1.400E-05 | 1.600E-05 |
| 28470 | 1424 | Y1424I | E | 1.1e-05 | 1.5e-05 | 1.5e-05 | 1.367E-05 | 2.309E-06 | 1.100E-05 | 1.500E-05 |
| 28471 | 1424 | Y1424L | E | 1.5e-05 | 1.1e-05 | 1.4e-05 | 1.333E-05 | 2.082E-06 | 1.100E-05 | 1.500E-05 |
| 28472 | 1424 | Y1424K | E | 1.2e-05 | 1.4e-05 | 1e-05 | 1.200E-05 | 2.000E-06 | 1.000E-05 | 1.400E-05 |
| 28473 | 1424 | Y1424M | E | 1.4e-05 | 1.1e-05 | 1.4e-05 | 1.300E-05 | 1.732E-06 | 1.100E-05 | 1.400E-05 |
| 28474 | 1424 | Y1424F | E | 1.7e-05 | 1.6e-05 | 1e-05 | 1.433E-05 | 3.786E-06 | 1.000E-05 | 1.700E-05 |

| | | | | | | | | | | |
|---|---|---|---|---|---|---|---|---|---|---|
| 28475 | 1424 | Y1424P | E | 1.5e-05 | 1.2e-05 | 2.1e-05 | 1.600E-05 | 4.583E-06 | 1.200E-05 | 2.100E-05 |
| 28476 | 1424 | Y1424S | E | 1.1e-05 | 1.4e-05 | 1.3e-05 | 1.267E-05 | 1.528E-06 | 1.100E-05 | 1.400E-05 |
| 28477 | 1424 | Y1424T | E | 1e-05 | 1.3e-05 | 1.8e-05 | 1.367E-05 | 4.041E-06 | 1.000E-05 | 1.800E-05 |
| 28478 | 1424 | Y1424W | E | 1.5e-05 | 8.6e-06 | 1.4e-05 | 1.253E-05 | 3.443E-06 | 8.600E-06 | 1.500E-05 |
| 28479 | 1424 | Y1424Y | E | 1.4e-05 | 1.4e-05 | 1e-05 | 1.267E-05 | 2.309E-06 | 1.000E-05 | 1.400E-05 |
| 28480 | 1424 | Y1424V | E | 1.4e-05 | 1.3e-05 | 1.5e-05 | 1.400E-05 | 1.000E-06 | 1.300E-05 | 1.500E-05 |
| 28481 | 1425 | F1425A | E | 9.9e-06 | 1.4e-05 | 1.4e-05 | 1.263E-05 | 2.367E-06 | 9.900E-06 | 1.400E-05 |
| 28482 | 1425 | F1425R | E | 8.5e-06 | 8.3e-06 | 1.5e-05 | 1.060E-05 | 3.812E-06 | 8.300E-06 | 1.500E-05 |
| 28483 | 1425 | F1425N | E | 1.1e-05 | 8.8e-06 | 1.6e-05 | 1.193E-05 | 3.690E-06 | 8.800E-06 | 1.600E-05 |
| 28484 | 1425 | F1425D | E | 1.5e-05 | 1.4e-05 | 1.5e-05 | 1.467E-05 | 5.774E-07 | 1.400E-05 | 1.500E-05 |
| 28485 | 1425 | F1425C | E | 8.3e-06 | 1.1e-05 | 1.6e-05 | 1.177E-05 | 3.907E-06 | 8.300E-06 | 1.600E-05 |
| 28486 | 1425 | F1425E | E | 1.4e-05 | 1e-05 | 1.6e-05 | 1.333E-05 | 3.055E-06 | 1.000E-05 | 1.600E-05 |
| 28487 | 1425 | F1425Q | E | 1.3e-05 | 1.5e-05 | 7.8e-06 | 1.193E-05 | 3.717E-06 | 7.800E-06 | 1.500E-05 |
| 28488 | 1425 | F1425G | E | 1.7e-05 | 1.5e-05 | 1e-05 | 1.400E-05 | 3.606E-06 | 1.000E-05 | 1.700E-05 |
| 28489 | 1425 | F1425H | E | 1e-05 | 1.5e-05 | 1.1e-05 | 1.200E-05 | 2.646E-06 | 1.000E-05 | 1.500E-05 |
| 28490 | 1425 | F1425I | E | 1.6e-05 | 1.5e-05 | 1.5e-05 | 1.533E-05 | 5.774E-07 | 1.500E-05 | 1.600E-05 |
| 28491 | 1425 | F1425L | E | 1.3e-05 | 1.4e-05 | 1.1e-05 | 1.267E-05 | 1.528E-06 | 1.100E-05 | 1.400E-05 |
| 28492 | 1425 | F1425K | E | 1.2e-05 | 1e-05 | 8.6e-06 | 1.020E-05 | 1.709E-06 | 8.600E-06 | 1.200E-05 |
| 28493 | 1425 | F1425M | E | 8.3e-06 | 1.6e-05 | 1e-05 | 1.143E-05 | 4.045E-06 | 8.300E-06 | 1.600E-05 |
| 28494 | 1425 | F1425F | E | 1.2e-05 | 1.6e-05 | 1e-05 | 1.267E-05 | 3.055E-06 | 1.000E-05 | 1.600E-05 |
| 28495 | 1425 | F1425P | E | 1.6e-05 | 1.6e-05 | 1.1e-05 | 1.433E-05 | 2.887E-06 | 1.100E-05 | 1.600E-05 |
| 28496 | 1425 | F1425S | E | 7.8e-06 | 1.3e-05 | 1.1e-05 | 1.060E-05 | 2.623E-06 | 7.800E-06 | 1.300E-05 |
| 28497 | 1425 | F1425T | E | 1.5e-05 | 1.5e-05 | 1e-05 | 1.333E-05 | 2.887E-06 | 1.000E-05 | 1.500E-05 |
| 28498 | 1425 | F1425W | E | 1.3e-05 | 1.1e-05 | 1.4e-05 | 1.267E-05 | 1.528E-06 | 1.100E-05 | 1.400E-05 |
| 28499 | 1425 | F1425Y | E | 7.6e-06 | 1.1e-05 | 1.7e-05 | 1.187E-05 | 4.760E-06 | 7.600E-06 | 1.700E-05 |
| 28500 | 1425 | F1425V | E | 1.4e-05 | 1.1e-05 | 1.3e-05 | 1.267E-05 | 1.528E-06 | 1.100E-05 | 1.400E-05 |
| 28501 | 1426 | P1426A | E | 1.6e-05 | 1.1e-05 | 1.6e-05 | 1.433E-05 | 2.887E-06 | 1.100E-05 | 1.600E-05 |
| 28502 | 1426 | P1426R | E | 1.2e-05 | 1.1e-05 | 1.1e-05 | 1.133E-05 | 5.774E-07 | 1.100E-05 | 1.200E-05 |
| 28503 | 1426 | P1426N | E | 9.6e-06 | 1.6e-05 | 1.5e-05 | 1.353E-05 | 3.443E-06 | 9.600E-06 | 1.600E-05 |
| 28504 | 1426 | P1426D | E | 1.5e-05 | 1.5e-05 | 1.2e-05 | 1.400E-05 | 1.732E-06 | 1.200E-05 | 1.500E-05 |
| 28505 | 1426 | P1426C | E | 1.5e-05 | 9.7e-06 | 1.4e-05 | 1.290E-05 | 2.816E-06 | 9.700E-06 | 1.500E-05 |
| 28506 | 1426 | P1426E | E | 1.6e-05 | 9.5e-06 | 1.2e-05 | 1.250E-05 | 3.279E-06 | 9.500E-06 | 1.600E-05 |
| 28507 | 1426 | P1426Q | E | 1.3e-05 | 9.5e-06 | 1.5e-05 | 1.250E-05 | 2.784E-06 | 9.500E-06 | 1.500E-05 |
| 28508 | 1426 | P1426G | E | 1.3e-05 | 1.1e-05 | 1e-05 | 1.133E-05 | 1.528E-06 | 1.000E-05 | 1.300E-05 |
| 28509 | 1426 | P1426H | E | 1.7e-05 | 8.4e-06 | 9e-06 | 1.147E-05 | 4.801E-06 | 8.400E-06 | 1.700E-05 |
| 28510 | 1426 | P1426I | E | 1.5e-05 | 1.5e-05 | 1.8e-05 | 1.600E-05 | 1.732E-06 | 1.500E-05 | 1.800E-05 |
| 28511 | 1426 | P1426L | E | 1.1e-05 | 1.5e-05 | 1e-05 | 1.200E-05 | 2.646E-06 | 1.000E-05 | 1.500E-05 |
| 28512 | 1426 | P1426K | E | 1.4e-05 | 1.4e-05 | 1.4e-05 | 1.400E-05 | 0.000E+00 | 1.400E-05 | 1.400E-05 |
| 28513 | 1426 | P1426M | E | 1e-05 | 1.5e-05 | 1.3e-05 | 1.267E-05 | 2.517E-06 | 1.000E-05 | 1.500E-05 |
| 28514 | 1426 | P1426F | E | 1.2e-05 | 1.5e-05 | 1.3e-05 | 1.333E-05 | 1.528E-06 | 1.200E-05 | 1.500E-05 |
| 28515 | 1426 | P1426P | E | 1.1e-05 | 1.4e-05 | 1.5e-05 | 1.333E-05 | 2.082E-06 | 1.100E-05 | 1.500E-05 |
| 28516 | 1426 | P1426S | E | 1.5e-05 | 1.4e-05 | 1.1e-05 | 1.333E-05 | 2.082E-06 | 1.100E-05 | 1.500E-05 |
| 28517 | 1426 | P1426T | E | 1.5e-05 | 9.7e-06 | 1.6e-05 | 1.357E-05 | 3.386E-06 | 9.700E-06 | 1.600E-05 |
| 28518 | 1426 | P1426W | E | 1.1e-05 | 1.4e-05 | 9.3e-06 | 1.143E-05 | 2.380E-06 | 9.300E-06 | 1.400E-05 |
| 28519 | 1426 | P1426Y | E | 1e-05 | 1.5e-05 | 1.5e-05 | 1.333E-05 | 2.887E-06 | 1.000E-05 | 1.500E-05 |
| 28520 | 1426 | P1426V | E | 1.1e-05 | 1.2e-05 | 1e-05 | 1.100E-05 | 1.000E-06 | 1.000E-05 | 1.200E-05 |
| 28521 | 1427 | E1427A | E | 1e-05 | 1.3e-05 | 1.9e-05 | 1.400E-05 | 4.583E-06 | 1.000E-05 | 1.900E-05 |
| 28522 | 1427 | E1427R | E | 9.8e-06 | 2.5e-05 | 1.5e-05 | 1.660E-05 | 7.725E-06 | 9.800E-06 | 2.500E-05 |
| 28523 | 1427 | E1427N | E | 9e-06 | 1.5e-05 | 8.8e-06 | 1.093E-05 | 3.523E-06 | 8.800E-06 | 1.500E-05 |
| 28524 | 1427 | E1427D | E | 9.6e-06 | 9e-06 | 7.7e-06 | 8.767E-06 | 9.713E-07 | 7.700E-06 | 9.600E-06 |
| 28525 | 1427 | E1427C | E | 1.5e-05 | 8.3e-06 | 1.2e-05 | 1.177E-05 | 3.356E-06 | 8.300E-06 | 1.500E-05 |
| 28526 | 1427 | E1427E | E | 1.1e-05 | 1.5e-05 | 1.5e-05 | 1.367E-05 | 2.309E-06 | 1.100E-05 | 1.500E-05 |
| 28527 | 1427 | E1427Q | E | 1.6e-05 | 1.2e-05 | 9.4e-06 | 1.247E-05 | 3.325E-06 | 9.400E-06 | 1.600E-05 |
| 28528 | 1427 | E1427G | E | 8.8e-06 | 1.7e-05 | 1.5e-05 | 1.360E-05 | 4.276E-06 | 8.800E-06 | 1.700E-05 |
| 28529 | 1427 | E1427H | E | 8.7e-06 | 1.2e-05 | 7.9e-06 | 9.533E-06 | 2.173E-06 | 7.900E-06 | 1.200E-05 |
| 28530 | 1427 | E1427I | E | 1.5e-05 | 5.5e-06 | 1.5e-05 | 1.183E-05 | 5.485E-06 | 5.500E-06 | 1.500E-05 |
| 28531 | 1427 | E1427L | E | 9.5e-06 | 1.1e-05 | 1.9e-05 | 1.317E-05 | 5.107E-06 | 9.500E-06 | 1.900E-05 |
| 28532 | 1427 | E1427K | E | 1.5e-05 | 8.8e-06 | 9.1e-06 | 1.097E-05 | 3.496E-06 | 8.800E-06 | 1.500E-05 |
| 28533 | 1427 | E1427M | E | 1.4e-05 | 1.8e-05 | 1.5e-05 | 1.567E-05 | 2.082E-06 | 1.400E-05 | 1.800E-05 |
| 28534 | 1427 | E1427F | E | 1.4e-05 | 1.7e-05 | 1.2e-05 | 1.433E-05 | 2.517E-06 | 1.200E-05 | 1.700E-05 |
| 28535 | 1427 | E1427P | E | 1.7e-05 | 1.5e-05 | 1.5e-05 | 1.567E-05 | 1.155E-06 | 1.500E-05 | 1.700E-05 |
| 28536 | 1427 | E1427S | E | 1.3e-05 | 1.6e-05 | 1e-05 | 1.300E-05 | 3.000E-06 | 1.000E-05 | 1.600E-05 |
| 28537 | 1427 | E1427T | E | 1.3e-05 | 9.5e-06 | 1e-05 | 1.083E-05 | 1.893E-06 | 9.500E-06 | 1.300E-05 |
| 28538 | 1427 | E1427W | E | 1.4e-05 | 1.4e-05 | 1e-05 | 1.267E-05 | 2.309E-06 | 1.000E-05 | 1.400E-05 |
| 28539 | 1427 | E1427Y | E | 1.5e-05 | 9.3e-06 | 8.5e-06 | 1.093E-05 | 3.544E-06 | 8.500E-06 | 1.500E-05 |
| 28540 | 1427 | E1427V | E | 1.3e-05 | 1.1e-05 | 1.4e-05 | 1.267E-05 | 1.528E-06 | 1.100E-05 | 1.400E-05 |
| 28541 | 1428 | P1428A | E | 1e-05 | 1.5e-05 | 1.6e-05 | 1.367E-05 | 3.215E-06 | 1.000E-05 | 1.600E-05 |
| 28542 | 1428 | P1428R | E | 1.4e-05 | 8.2e-06 | 1.5e-05 | 1.240E-05 | 3.672E-06 | 8.200E-06 | 1.500E-05 |
| 28543 | 1428 | P1428N | E | 1.5e-05 | 1.4e-05 | 1.5e-05 | 1.467E-05 | 5.774E-07 | 1.400E-05 | 1.500E-05 |
| 28544 | 1428 | P1428D | E | 1.5e-05 | 9.4e-06 | 1.4e-05 | 1.280E-05 | 2.987E-06 | 9.400E-06 | 1.500E-05 |
| 28545 | 1428 | P1428C | E | 1.5e-05 | 1.4e-05 | 1.2e-05 | 1.367E-05 | 1.528E-06 | 1.200E-05 | 1.500E-05 |
| 28546 | 1428 | P1428E | E | 8.3e-06 | 9.2e-06 | 1.5e-05 | 1.083E-05 | 3.636E-06 | 8.300E-06 | 1.500E-05 |
| 28547 | 1428 | P1428Q | E | 8.9e-06 | 9.4e-06 | 1e-05 | 9.433E-06 | 5.508E-07 | 8.900E-06 | 1.000E-05 |
| 28548 | 1428 | P1428G | E | 1e-05 | 1e-05 | 1.5e-05 | 1.167E-05 | 2.887E-06 | 1.000E-05 | 1.500E-05 |
| 28549 | 1428 | P1428H | E | 1.1e-05 | 1.5e-05 | 1.4e-05 | 1.333E-05 | 2.082E-06 | 1.100E-05 | 1.500E-05 |
| 28550 | 1428 | P1428I | E | 8.4e-06 | 1.2e-05 | 1.1e-05 | 1.047E-05 | 1.858E-06 | 8.400E-06 | 1.200E-05 |
| 28551 | 1428 | P1428L | E | 9e-06 | 9.9e-06 | 1e-05 | 9.633E-06 | 5.508E-07 | 9.000E-06 | 1.000E-05 |
| 28552 | 1428 | P1428K | E | 1.7e-05 | 8.9e-06 | 1.1e-05 | 1.230E-05 | 4.204E-06 | 8.900E-06 | 1.700E-05 |
| 28553 | 1428 | P1428M | E | 1.6e-05 | 9.6e-06 | 1.5e-05 | 1.353E-05 | 3.443E-06 | 9.600E-06 | 1.600E-05 |
| 28554 | 1428 | P1428F | E | 1.1e-05 | 1.5e-05 | 8.5e-06 | 1.150E-05 | 3.279E-06 | 8.500E-06 | 1.500E-05 |
| 28555 | 1428 | P1428P | E | 1.1e-05 | 1.4e-05 | 1.2e-05 | 1.233E-05 | 1.528E-06 | 1.100E-05 | 1.400E-05 |
| 28556 | 1428 | P1428S | E | 8.6e-06 | 6.8e-06 | 1.5e-05 | 1.013E-05 | 4.310E-06 | 6.800E-06 | 1.500E-05 |
| 28557 | 1428 | P1428T | E | 5.7e-06 | 1.7e-05 | 8.9e-06 | 1.053E-05 | 5.824E-06 | 5.700E-06 | 1.700E-05 |
| 28558 | 1428 | P1428W | E | 1.5e-05 | 1.7e-05 | 1.3e-05 | 1.500E-05 | 2.000E-06 | 1.300E-05 | 1.700E-05 |
| 28559 | 1428 | P1428Y | E | 9.9e-06 | 1.4e-05 | 1.3e-05 | 1.230E-05 | 2.138E-06 | 9.900E-06 | 1.400E-05 |

| | | | | | | | | | | |
|---|---|---|---|---|---|---|---|---|---|---|
| 28560 | 1428 | P1428V | E | 7.8e-06 | 1.5e-05 | 1.3e-05 | 1.193E-05 | 3.717E-06 | 7.800E-06 | 1.500E-05 |
| 28561 | 1429 | V1429A | E | 9.1e-06 | 1.1e-05 | 1.5e-05 | 1.170E-05 | 3.012E-06 | 9.100E-06 | 1.500E-05 |
| 28562 | 1429 | V1429R | E | 1.5e-05 | 1.2e-05 | 1.5e-05 | 1.400E-05 | 1.732E-06 | 1.200E-05 | 1.500E-05 |
| 28563 | 1429 | V1429N | E | 1.1e-05 | 1.5e-05 | 1e-05 | 1.200E-05 | 2.646E-06 | 1.000E-05 | 1.500E-05 |
| 28564 | 1429 | V1429D | E | 1.5e-05 | 1.3e-05 | 1.4e-05 | 1.400E-05 | 1.000E-06 | 1.300E-05 | 1.500E-05 |
| 28565 | 1429 | V1429C | E | 1.4e-05 | 1.4e-05 | 1.3e-05 | 1.367E-05 | 5.774E-07 | 1.300E-05 | 1.400E-05 |
| 28566 | 1429 | V1429E | E | 1.5e-05 | 1.5e-05 | 1.7e-05 | 1.567E-05 | 1.155E-06 | 1.500E-05 | 1.700E-05 |
| 28567 | 1429 | V1429Q | E | 1.4e-05 | 1.1e-05 | 1.2e-05 | 1.233E-05 | 1.528E-06 | 1.100E-05 | 1.400E-05 |
| 28568 | 1429 | V1429G | E | 1.1e-05 | 1.1e-05 | 1.3e-05 | 1.167E-05 | 1.155E-06 | 1.100E-05 | 1.300E-05 |
| 28569 | 1429 | V1429H | E | 1.4e-05 | 1.3e-05 | 1.2e-05 | 1.300E-05 | 1.000E-06 | 1.200E-05 | 1.400E-05 |
| 28570 | 1429 | V1429I | E | 3.2e-05 | 1.4e-05 | 1.6e-05 | 2.067E-05 | 9.866E-06 | 1.400E-05 | 3.200E-05 |
| 28571 | 1429 | V1429L | E | 1.1e-05 | 1e-05 | 1.5e-05 | 1.200E-05 | 2.646E-06 | 1.000E-05 | 1.500E-05 |
| 28572 | 1429 | V1429K | E | 1.5e-05 | 1.6e-05 | 1.5e-05 | 1.533E-05 | 5.774E-07 | 1.500E-05 | 1.600E-05 |
| 28573 | 1429 | V1429M | E | 1.6e-05 | 1e-05 | 1.2e-05 | 1.267E-05 | 3.055E-06 | 1.000E-05 | 1.600E-05 |
| 28574 | 1429 | V1429F | E | 9.2e-06 | 8.7e-06 | 1.4e-05 | 1.063E-05 | 2.926E-06 | 8.700E-06 | 1.400E-05 |
| 28575 | 1429 | V1429P | E | 9.3e-06 | 1.6e-05 | 8.1e-06 | 1.113E-05 | 4.257E-06 | 8.100E-06 | 1.600E-05 |
| 28576 | 1429 | V1429S | E | 1.4e-05 | 1.5e-05 | 1.1e-05 | 1.333E-05 | 2.082E-06 | 1.100E-05 | 1.500E-05 |
| 28577 | 1429 | V1429T | E | 1e-05 | 1.4e-05 | 1.4e-05 | 1.267E-05 | 2.309E-06 | 1.000E-05 | 1.400E-05 |
| 28578 | 1429 | V1429W | E | 1.6e-05 | 1.4e-05 | 1e-05 | 1.333E-05 | 3.055E-06 | 1.000E-05 | 1.600E-05 |
| 28579 | 1429 | V1429Y | E | 1.5e-05 | 1.6e-05 | 1.6e-05 | 1.567E-05 | 5.774E-07 | 1.500E-05 | 1.600E-05 |
| 28580 | 1429 | V1429V | E | 1.4e-05 | 1.9e-05 | 1e-05 | 1.433E-05 | 4.509E-06 | 1.000E-05 | 1.900E-05 |
| 28581 | 1430 | T1430A | E | 1.6e-05 | 1.6e-05 | 1.4e-05 | 1.533E-05 | 1.155E-06 | 1.400E-05 | 1.600E-05 |
| 28582 | 1430 | T1430R | E | 1.4e-05 | 1.6e-05 | 1.1e-05 | 1.367E-05 | 2.517E-06 | 1.100E-05 | 1.600E-05 |
| 28583 | 1430 | T1430N | E | 1e-05 | 1.3e-05 | 9.8e-06 | 1.093E-05 | 1.793E-06 | 9.800E-06 | 1.300E-05 |
| 28584 | 1430 | T1430D | E | 1e-05 | 1.1e-05 | 1.7e-05 | 1.267E-05 | 3.786E-06 | 1.000E-05 | 1.700E-05 |
| 28585 | 1430 | T1430C | E | 1.7e-05 | 1.1e-05 | 9.3e-06 | 1.243E-05 | 4.045E-06 | 9.300E-06 | 1.700E-05 |
| 28586 | 1430 | T1430E | E | 1.7e-05 | 1.5e-05 | 1.1e-05 | 1.433E-05 | 3.055E-06 | 1.100E-05 | 1.700E-05 |
| 28587 | 1430 | T1430Q | E | 9.5e-06 | 1.3e-05 | 8.5e-06 | 1.033E-05 | 2.363E-06 | 8.500E-06 | 1.300E-05 |
| 28588 | 1430 | T1430G | E | 1.7e-05 | 1.3e-05 | 1.2e-05 | 1.400E-05 | 2.646E-06 | 1.200E-05 | 1.700E-05 |
| 28589 | 1430 | T1430H | E | 7.9e-06 | 1.1e-05 | 1.3e-05 | 1.063E-05 | 2.570E-06 | 7.900E-06 | 1.300E-05 |
| 28590 | 1430 | T1430I | E | 1.6e-05 | 1.6e-05 | 1.4e-05 | 1.533E-05 | 1.155E-06 | 1.400E-05 | 1.600E-05 |
| 28591 | 1430 | T1430L | E | 1.6e-05 | 1.5e-05 | 1.6e-05 | 1.567E-05 | 5.774E-07 | 1.500E-05 | 1.600E-05 |
| 28592 | 1430 | T1430K | E | 1.2e-05 | 1.1e-05 | 1.2e-05 | 1.167E-05 | 5.774E-07 | 1.100E-05 | 1.200E-05 |
| 28593 | 1430 | T1430M | E | 1e-05 | 7.9e-06 | 1.6e-05 | 1.130E-05 | 4.204E-06 | 7.900E-06 | 1.600E-05 |
| 28594 | 1430 | T1430F | E | 1.5e-05 | 9.5e-06 | 1.1e-05 | 1.183E-05 | 2.843E-06 | 9.500E-06 | 1.500E-05 |
| 28595 | 1430 | T1430P | E | 9.2e-06 | 1.6e-05 | 1.6e-05 | 1.373E-05 | 3.926E-06 | 9.200E-06 | 1.600E-05 |
| 28596 | 1430 | T1430S | E | 1e-05 | 1.6e-05 | 1.2e-05 | 1.267E-05 | 3.055E-06 | 1.000E-05 | 1.600E-05 |
| 28597 | 1430 | T1430T | E | 1.5e-05 | 1.2e-05 | 9.4e-06 | 1.213E-05 | 2.802E-06 | 9.400E-06 | 1.500E-05 |
| 28598 | 1430 | T1430W | E | 1e-05 | 1.3e-05 | 1.5e-05 | 1.267E-05 | 2.517E-06 | 1.000E-05 | 1.500E-05 |
| 28599 | 1430 | T1430Y | E | 1.4e-05 | 1.4e-05 | 8.4e-06 | 1.213E-05 | 3.233E-06 | 8.400E-06 | 1.400E-05 |
| 28600 | 1430 | T1430V | E | 9.3e-06 | 1.5e-05 | 1.6e-05 | 1.343E-05 | 3.614E-06 | 9.300E-06 | 1.600E-05 |
| 28601 | 1431 | V1431A | E | 1.3e-05 | 1.6e-05 | 1e-05 | 1.300E-05 | 3.000E-06 | 1.000E-05 | 1.600E-05 |
| 28602 | 1431 | V1431R | E | 1.6e-05 | 1.4e-05 | 1.7e-05 | 1.567E-05 | 1.528E-06 | 1.400E-05 | 1.700E-05 |
| 28603 | 1431 | V1431N | E | 1.6e-05 | 1.4e-05 | 1.1e-05 | 1.367E-05 | 2.517E-06 | 1.100E-05 | 1.600E-05 |
| 28604 | 1431 | V1431D | E | 1.4e-05 | 1.7e-05 | 1.5e-05 | 1.533E-05 | 1.528E-06 | 1.400E-05 | 1.700E-05 |
| 28605 | 1431 | V1431C | E | 1.4e-05 | 1e-05 | 1.5e-05 | 1.300E-05 | 2.646E-06 | 1.000E-05 | 1.500E-05 |
| 28606 | 1431 | V1431E | E | 1.3e-05 | 1e-05 | 1.3e-05 | 1.200E-05 | 1.732E-06 | 1.000E-05 | 1.300E-05 |
| 28607 | 1431 | V1431Q | E | 1.6e-05 | 1.1e-05 | 1.5e-05 | 1.400E-05 | 2.646E-06 | 1.100E-05 | 1.600E-05 |
| 28608 | 1431 | V1431G | E | 1.5e-05 | 1.6e-05 | 1.4e-05 | 1.500E-05 | 1.000E-06 | 1.400E-05 | 1.600E-05 |
| 28609 | 1431 | V1431H | E | 1e-05 | 1e-05 | 1.3e-05 | 1.100E-05 | 1.732E-06 | 1.000E-05 | 1.300E-05 |
| 28610 | 1431 | V1431I | E | 1.1e-05 | 1.6e-05 | 9.8e-06 | 1.227E-05 | 3.288E-06 | 9.800E-06 | 1.600E-05 |
| 28611 | 1431 | V1431L | E | 9.4e-06 | 1.4e-05 | 8.7e-06 | 1.070E-05 | 2.879E-06 | 8.700E-06 | 1.400E-05 |
| 28612 | 1431 | V1431K | E | 1.5e-05 | 1.6e-05 | 1.1e-05 | 1.400E-05 | 2.646E-06 | 1.100E-05 | 1.600E-05 |
| 28613 | 1431 | V1431M | E | 1e-05 | 1.4e-05 | 8.4e-06 | 1.080E-05 | 2.884E-06 | 8.400E-06 | 1.400E-05 |
| 28614 | 1431 | V1431F | E | 9.9e-06 | 1.5e-05 | 1.3e-05 | 1.263E-05 | 2.570E-06 | 9.900E-06 | 1.500E-05 |
| 28615 | 1431 | V1431P | E | 8.4e-06 | 1.5e-05 | 1.5e-05 | 1.280E-05 | 3.811E-06 | 8.400E-06 | 1.500E-05 |
| 28616 | 1431 | V1431S | E | 1.6e-05 | 9.5e-06 | 1.5e-05 | 1.350E-05 | 3.500E-06 | 9.500E-06 | 1.600E-05 |
| 28617 | 1431 | V1431T | E | 9.2e-06 | 1.2e-05 | 9.9e-06 | 1.037E-05 | 1.457E-06 | 9.200E-06 | 1.200E-05 |
| 28618 | 1431 | V1431W | E | 1.3e-05 | 7.5e-06 | 1e-05 | 1.017E-05 | 2.754E-06 | 7.500E-06 | 1.300E-05 |
| 28619 | 1431 | V1431Y | E | 1.5e-05 | 1.1e-05 | 1.1e-05 | 1.233E-05 | 2.309E-06 | 1.100E-05 | 1.500E-05 |
| 28620 | 1431 | V1431V | E | 2.2e-05 | 1.3e-05 | 1.2e-05 | 1.567E-05 | 5.508E-06 | 1.200E-05 | 2.200E-05 |
| 28621 | 1432 | S1432A | E | 1.5e-05 | 1.1e-05 | 1.4e-05 | 1.333E-05 | 2.082E-06 | 1.100E-05 | 1.500E-05 |
| 28622 | 1432 | S1432R | E | 1.6e-05 | 1e-05 | 1.7e-05 | 1.433E-05 | 3.786E-06 | 1.000E-05 | 1.700E-05 |
| 28623 | 1432 | S1432N | E | 1.4e-05 | 1.5e-05 | 1.2e-05 | 1.367E-05 | 1.528E-06 | 1.200E-05 | 1.500E-05 |
| 28624 | 1432 | S1432D | E | 1.3e-05 | 1.6e-05 | 1.2e-05 | 1.367E-05 | 2.082E-06 | 1.200E-05 | 1.600E-05 |
| 28625 | 1432 | S1432C | E | 1.2e-05 | 1.6e-05 | 1e-05 | 1.267E-05 | 3.055E-06 | 1.000E-05 | 1.600E-05 |
| 28626 | 1432 | S1432E | E | 9.9e-06 | 8.6e-06 | 1e-05 | 9.500E-06 | 7.810E-07 | 8.600E-06 | 1.000E-05 |
| 28627 | 1432 | S1432Q | E | 1.3e-05 | 9.6e-06 | 1.5e-05 | 1.253E-05 | 2.730E-06 | 9.600E-06 | 1.500E-05 |
| 28628 | 1432 | S1432G | E | 1.1e-05 | 1.3e-05 | 1.1e-05 | 1.167E-05 | 1.155E-06 | 1.100E-05 | 1.300E-05 |
| 28629 | 1432 | S1432H | E | 1.6e-05 | 1.7e-05 | 1.5e-05 | 1.600E-05 | 1.000E-06 | 1.500E-05 | 1.700E-05 |
| 28630 | 1432 | S1432I | E | 1.2e-05 | 1.1e-05 | 1.2e-05 | 1.167E-05 | 5.774E-07 | 1.100E-05 | 1.200E-05 |
| 28631 | 1432 | S1432L | E | 1.3e-05 | 1.3e-05 | 1.1e-05 | 1.233E-05 | 1.155E-06 | 1.100E-05 | 1.300E-05 |
| 28632 | 1432 | S1432K | E | 9.2e-06 | 9.8e-06 | 1.3e-05 | 1.067E-05 | 2.043E-06 | 9.200E-06 | 1.300E-05 |
| 28633 | 1432 | S1432M | E | 1.4e-05 | 1.5e-05 | 1.1e-05 | 1.333E-05 | 2.082E-06 | 1.100E-05 | 1.500E-05 |
| 28634 | 1432 | S1432F | E | 1.5e-05 | 1.5e-05 | 1.1e-05 | 1.367E-05 | 2.309E-06 | 1.100E-05 | 1.500E-05 |
| 28635 | 1432 | S1432P | E | 1.1e-05 | 1.1e-05 | 1.6e-05 | 1.267E-05 | 2.887E-06 | 1.100E-05 | 1.600E-05 |
| 28636 | 1432 | S1432S | E | 1.4e-05 | 8.2e-06 | 1e-05 | 1.073E-05 | 2.969E-06 | 8.200E-06 | 1.400E-05 |
| 28637 | 1432 | S1432T | E | 1.6e-05 | 1.6e-05 | 1.1e-05 | 1.433E-05 | 2.887E-06 | 1.100E-05 | 1.600E-05 |
| 28638 | 1432 | S1432W | E | 1.3e-05 | 1.4e-05 | 1.3e-05 | 1.333E-05 | 5.774E-07 | 1.300E-05 | 1.400E-05 |
| 28639 | 1432 | S1432Y | E | 1.3e-05 | 9e-06 | 9.3e-06 | 1.043E-05 | 2.228E-06 | 9.000E-06 | 1.300E-05 |
| 28640 | 1432 | S1432V | E | 1.3e-05 | 1.6e-05 | 1e-05 | 1.300E-05 | 3.000E-06 | 1.000E-05 | 1.600E-05 |
| 28641 | 1433 | W1433A | E | 1.2e-05 | 1.2e-05 | 1.5e-05 | 1.300E-05 | 1.732E-06 | 1.200E-05 | 1.500E-05 |
| 28642 | 1433 | W1433R | E | 1.4e-05 | 1e-05 | 1.3e-05 | 1.233E-05 | 2.082E-06 | 1.000E-05 | 1.400E-05 |
| 28643 | 1433 | W1433N | E | 1.1e-05 | 9e-06 | 9.5e-06 | 9.833E-06 | 1.041E-06 | 9.000E-06 | 1.100E-05 |
| 28644 | 1433 | W1433D | E | 1.6e-05 | 1.5e-05 | 1.6e-05 | 1.567E-05 | 5.774E-07 | 1.500E-05 | 1.600E-05 |

| | | | | | | | | | | |
|---|---|---|---|---|---|---|---|---|---|---|
| 28645 | 1433 | W1433C | E | 1.3e-05 | 1e-05 | 1.2e-05 | 1.167E-05 | 1.528E-06 | 1.000E-05 | 1.300E-05 |
| 28646 | 1433 | W1433E | E | 1.1e-05 | 9.4e-06 | 1.6e-05 | 1.213E-05 | 3.443E-06 | 9.400E-06 | 1.600E-05 |
| 28647 | 1433 | W1433Q | E | 5.9e-06 | 1.8e-05 | 7.9e-06 | 1.060E-05 | 6.486E-06 | 5.900E-06 | 1.800E-05 |
| 28648 | 1433 | W1433G | E | 8.8e-06 | 1.6e-05 | 9.4e-06 | 1.140E-05 | 3.995E-06 | 8.800E-06 | 1.600E-05 |
| 28649 | 1433 | W1433H | E | 1e-05 | 8.7e-06 | 1.1e-05 | 9.900E-06 | 1.153E-06 | 8.700E-06 | 1.100E-05 |
| 28650 | 1433 | W1433I | E | 1.6e-05 | 1.6e-05 | 1.6e-05 | 1.600E-05 | 0.000E+00 | 1.600E-05 | 1.600E-05 |
| 28651 | 1433 | W1433L | E | 1.5e-05 | 1.5e-05 | 1.6e-05 | 1.533E-05 | 5.774E-07 | 1.500E-05 | 1.600E-05 |
| 28652 | 1433 | W1433K | E | 1.1e-05 | 1.1e-05 | 9.8e-06 | 1.060E-05 | 6.928E-07 | 9.800E-06 | 1.100E-05 |
| 28653 | 1433 | W1433M | E | 1e-05 | 1.1e-05 | 1.6e-05 | 1.233E-05 | 3.215E-06 | 1.000E-05 | 1.600E-05 |
| 28654 | 1433 | W1433F | E | 1.1e-05 | 1.4e-05 | 1.5e-05 | 1.333E-05 | 2.082E-06 | 1.100E-05 | 1.500E-05 |
| 28655 | 1433 | W1433P | E | 1.2e-05 | 1.6e-05 | 8.6e-06 | 1.220E-05 | 3.704E-06 | 8.600E-06 | 1.600E-05 |
| 28656 | 1433 | W1433S | E | 1.3e-05 | 1.6e-05 | 1.6e-05 | 1.500E-05 | 1.732E-06 | 1.300E-05 | 1.600E-05 |
| 28657 | 1433 | W1433T | E | 1.1e-05 | 1.6e-05 | 1.1e-05 | 1.267E-05 | 2.887E-06 | 1.100E-05 | 1.600E-05 |
| 28658 | 1433 | W1433W | E | 1.7e-05 | 9.7e-06 | 1.8e-05 | 1.490E-05 | 4.531E-06 | 9.700E-06 | 1.800E-05 |
| 28659 | 1433 | W1433Y | E | 1.6e-05 | 1.1e-05 | 1.3e-05 | 1.333E-05 | 2.517E-06 | 1.100E-05 | 1.600E-05 |
| 28660 | 1433 | W1433V | E | 1.1e-05 | 1.5e-05 | 1.5e-05 | 1.367E-05 | 2.309E-06 | 1.100E-05 | 1.500E-05 |
| 28661 | 1434 | N1434A | E | 1.7e-05 | 1.1e-05 | 1.1e-05 | 1.300E-05 | 3.464E-06 | 1.100E-05 | 1.700E-05 |
| 28662 | 1434 | N1434R | E | 1.2e-05 | 1.2e-05 | 1.3e-05 | 1.233E-05 | 5.774E-07 | 1.200E-05 | 1.300E-05 |
| 28663 | 1434 | N1434N | E | 1.5e-05 | 1.5e-05 | 9.1e-06 | 1.303E-05 | 3.406E-06 | 9.100E-06 | 1.500E-05 |
| 28664 | 1434 | N1434D | E | 1.4e-05 | 1.4e-05 | 1.2e-05 | 1.333E-05 | 1.155E-06 | 1.200E-05 | 1.400E-05 |
| 28665 | 1434 | N1434C | E | 9.3e-06 | 1.6e-05 | 1.2e-05 | 1.243E-05 | 3.371E-06 | 9.300E-06 | 1.600E-05 |
| 28666 | 1434 | N1434E | E | 1.1e-05 | 1.8e-05 | 1.8e-05 | 1.567E-05 | 4.041E-06 | 1.100E-05 | 1.800E-05 |
| 28667 | 1434 | N1434Q | E | 1.2e-05 | 1.4e-05 | 1.5e-05 | 1.367E-05 | 1.528E-06 | 1.200E-05 | 1.500E-05 |
| 28668 | 1434 | N1434G | E | 1.6e-05 | 1.3e-05 | 1.4e-05 | 1.433E-05 | 1.528E-06 | 1.300E-05 | 1.600E-05 |
| 28669 | 1434 | N1434H | E | 1.2e-05 | 1.3e-05 | 1.1e-05 | 1.200E-05 | 1.000E-06 | 1.100E-05 | 1.300E-05 |
| 28670 | 1434 | N1434I | E | 1.7e-05 | 9.8e-06 | 1.5e-05 | 1.393E-05 | 3.717E-06 | 9.800E-06 | 1.700E-05 |
| 28671 | 1434 | N1434L | E | 1.6e-05 | 1.6e-05 | 1.4e-05 | 1.533E-05 | 1.155E-06 | 1.400E-05 | 1.600E-05 |
| 28672 | 1434 | N1434K | E | 1.1e-05 | 1.1e-05 | 1.4e-05 | 1.200E-05 | 1.732E-06 | 1.100E-05 | 1.400E-05 |
| 28673 | 1434 | N1434M | E | 1.5e-05 | 1.3e-05 | 1.1e-05 | 1.300E-05 | 2.000E-06 | 1.100E-05 | 1.500E-05 |
| 28674 | 1434 | N1434F | E | 1.3e-05 | 9.3e-06 | 8e-06 | 1.010E-05 | 2.594E-06 | 8.000E-06 | 1.300E-05 |
| 28675 | 1434 | N1434P | E | 1.6e-05 | 8.5e-06 | 1.2e-05 | 1.217E-05 | 3.753E-06 | 8.500E-06 | 1.600E-05 |
| 28676 | 1434 | N1434S | E | 1.4e-05 | 1.6e-05 | 1.1e-05 | 1.367E-05 | 2.517E-06 | 1.100E-05 | 1.600E-05 |
| 28677 | 1434 | N1434T | E | 1.1e-05 | 9.9e-06 | 1.4e-05 | 1.163E-05 | 2.122E-06 | 9.900E-06 | 1.400E-05 |
| 28678 | 1434 | N1434W | E | 1.4e-05 | 1.6e-05 | 1.2e-05 | 1.400E-05 | 2.000E-06 | 1.200E-05 | 1.600E-05 |
| 28679 | 1434 | N1434Y | E | 1.5e-05 | 1.4e-05 | 1.1e-05 | 1.333E-05 | 2.082E-06 | 1.100E-05 | 1.500E-05 |
| 28680 | 1434 | N1434V | E | 1.1e-05 | 1e-05 | 1.2e-05 | 1.100E-05 | 1.000E-06 | 1.000E-05 | 1.200E-05 |
| 28681 | 1435 | S1435A | E | 1.6e-05 | 1e-05 | 1.1e-05 | 1.233E-05 | 3.215E-06 | 1.000E-05 | 1.600E-05 |
| 28682 | 1435 | S1435R | E | 1.7e-05 | 1.1e-05 | 1.5e-05 | 1.433E-05 | 3.055E-06 | 1.100E-05 | 1.700E-05 |
| 28683 | 1435 | S1435N | E | 1.2e-05 | 1.1e-05 | 1.2e-05 | 1.167E-05 | 5.774E-07 | 1.100E-05 | 1.200E-05 |
| 28684 | 1435 | S1435D | E | 1.5e-05 | 1.5e-05 | 1.3e-05 | 1.433E-05 | 1.155E-06 | 1.300E-05 | 1.500E-05 |
| 28685 | 1435 | S1435C | E | 1.2e-05 | 9.3e-06 | 9.3e-06 | 1.020E-05 | 1.559E-06 | 9.300E-06 | 1.200E-05 |
| 28686 | 1435 | S1435E | E | 1e-05 | 9.6e-06 | 9.8e-06 | 9.800E-06 | 2.000E-07 | 9.600E-06 | 1.000E-05 |
| 28687 | 1435 | S1435Q | E | 9.8e-06 | 1.5e-05 | 1.4e-05 | 1.293E-05 | 2.759E-06 | 9.800E-06 | 1.500E-05 |
| 28688 | 1435 | S1435G | E | 1e-05 | 1.1e-05 | 1.5e-05 | 1.200E-05 | 2.646E-06 | 1.000E-05 | 1.500E-05 |
| 28689 | 1435 | S1435H | E | 3.3e-05 | 1.5e-05 | 1e-05 | 1.933E-05 | 1.210E-05 | 1.000E-05 | 3.300E-05 |
| 28690 | 1435 | S1435I | E | 1.7e-05 | 1.5e-05 | 1.1e-05 | 1.433E-05 | 3.055E-06 | 1.100E-05 | 1.700E-05 |
| 28691 | 1435 | S1435L | E | 9.5e-06 | 1.6e-05 | 1.1e-05 | 1.217E-05 | 3.403E-06 | 9.500E-06 | 1.600E-05 |
| 28692 | 1435 | S1435K | E | 1.1e-05 | 1.5e-05 | 1.5e-05 | 1.367E-05 | 2.309E-06 | 1.100E-05 | 1.500E-05 |
| 28693 | 1435 | S1435M | E | 1.1e-05 | 1e-05 | 1.7e-05 | 1.267E-05 | 3.786E-06 | 1.000E-05 | 1.700E-05 |
| 28694 | 1435 | S1435F | E | 1.3e-05 | 1e-05 | 7.6e-06 | 1.020E-05 | 2.706E-06 | 7.600E-06 | 1.300E-05 |
| 28695 | 1435 | S1435P | E | 1.5e-05 | 1.7e-05 | 1.8e-05 | 1.667E-05 | 1.528E-06 | 1.500E-05 | 1.800E-05 |
| 28696 | 1435 | S1435S | E | 1e-05 | 1.3e-05 | 1.5e-05 | 1.267E-05 | 2.517E-06 | 1.000E-05 | 1.500E-05 |
| 28697 | 1435 | S1435T | E | 1.2e-05 | 1.4e-05 | 1e-05 | 1.200E-05 | 2.000E-06 | 1.000E-05 | 1.400E-05 |
| 28698 | 1435 | S1435W | E | 1.2e-05 | 1.2e-05 | 1e-05 | 1.133E-05 | 1.155E-06 | 1.000E-05 | 1.200E-05 |
| 28699 | 1435 | S1435Y | E | 1.1e-05 | 9.9e-06 | 9.6e-06 | 1.017E-05 | 7.371E-07 | 9.600E-06 | 1.100E-05 |
| 28700 | 1435 | S1435V | E | 1.4e-05 | 1.6e-05 | 1.6e-05 | 1.533E-05 | 1.155E-06 | 1.400E-05 | 1.600E-05 |
| 28701 | 1436 | G1436A | E | 9.7e-06 | 1.6e-05 | 1e-05 | 1.190E-05 | 3.554E-06 | 9.700E-06 | 1.600E-05 |
| 28702 | 1436 | G1436R | E | 1.4e-05 | 1e-05 | 1.3e-05 | 1.233E-05 | 2.082E-06 | 1.000E-05 | 1.400E-05 |
| 28703 | 1436 | G1436N | E | 9.7e-06 | 1.6e-05 | 1.1e-05 | 1.223E-05 | 3.326E-06 | 9.700E-06 | 1.600E-05 |
| 28704 | 1436 | G1436D | E | 1.5e-05 | 1.1e-05 | 9.6e-06 | 1.187E-05 | 2.802E-06 | 9.600E-06 | 1.500E-05 |
| 28705 | 1436 | G1436C | E | 1.5e-05 | 1.5e-05 | 1.5e-05 | 1.500E-05 | 0.000E+00 | 1.500E-05 | 1.500E-05 |
| 28706 | 1436 | G1436E | E | 1e-05 | 1.5e-05 | 9.6e-06 | 1.153E-05 | 3.009E-06 | 9.600E-06 | 1.500E-05 |
| 28707 | 1436 | G1436Q | E | 1.5e-05 | 8.4e-06 | 1.5e-05 | 1.280E-05 | 3.811E-06 | 8.400E-06 | 1.500E-05 |
| 28708 | 1436 | G1436G | E | 1.7e-05 | 1.4e-05 | 1.4e-05 | 1.500E-05 | 1.732E-06 | 1.400E-05 | 1.700E-05 |
| 28709 | 1436 | G1436H | E | 1.4e-05 | 1e-05 | 1e-05 | 1.133E-05 | 2.309E-06 | 1.000E-05 | 1.400E-05 |
| 28710 | 1436 | G1436I | E | 1.3e-05 | 1e-05 | 1e-05 | 1.100E-05 | 1.732E-06 | 1.000E-05 | 1.300E-05 |
| 28711 | 1436 | G1436L | E | 9e-06 | 1.5e-05 | 1.2e-05 | 1.200E-05 | 3.000E-06 | 9.000E-06 | 1.500E-05 |
| 28712 | 1436 | G1436K | E | 1.2e-05 | 8.1e-06 | 1.2e-05 | 1.070E-05 | 2.252E-06 | 8.100E-06 | 1.200E-05 |
| 28713 | 1436 | G1436M | E | 1.4e-05 | 1.3e-05 | 1.4e-05 | 1.367E-05 | 5.774E-07 | 1.300E-05 | 1.400E-05 |
| 28714 | 1436 | G1436F | E | 1.4e-05 | 1.7e-05 | 1.2e-05 | 1.433E-05 | 2.517E-06 | 1.200E-05 | 1.700E-05 |
| 28715 | 1436 | G1436P | E | 1.6e-05 | 1.1e-05 | 1.5e-05 | 1.400E-05 | 2.646E-06 | 1.100E-05 | 1.600E-05 |
| 28716 | 1436 | G1436S | E | 1e-05 | 1.1e-05 | 1e-05 | 1.033E-05 | 5.774E-07 | 1.000E-05 | 1.100E-05 |
| 28717 | 1436 | G1436T | E | 1.5e-05 | 1.3e-05 | 1.1e-05 | 1.300E-05 | 2.000E-06 | 1.100E-05 | 1.500E-05 |
| 28718 | 1436 | G1436W | E | 1.6e-05 | 1.2e-05 | 1.2e-05 | 1.333E-05 | 2.309E-06 | 1.200E-05 | 1.600E-05 |
| 28719 | 1436 | G1436Y | E | 1e-05 | 1.4e-05 | 8.5e-06 | 1.083E-05 | 2.843E-06 | 8.500E-06 | 1.400E-05 |
| 28720 | 1436 | G1436V | E | 1.6e-05 | 1.1e-05 | 1e-05 | 1.233E-05 | 3.215E-06 | 1.000E-05 | 1.600E-05 |
| 28721 | 1437 | A1437A | E | 1.6e-05 | 1.6e-05 | 1.4e-05 | 1.533E-05 | 1.155E-06 | 1.400E-05 | 1.600E-05 |
| 28722 | 1437 | A1437R | E | 1e-05 | 1.5e-05 | 1.1e-05 | 1.200E-05 | 2.646E-06 | 1.000E-05 | 1.500E-05 |
| 28723 | 1437 | A1437N | E | 1.2e-05 | 1.5e-05 | 1.3e-05 | 1.333E-05 | 1.528E-06 | 1.200E-05 | 1.500E-05 |
| 28724 | 1437 | A1437D | E | 1.4e-05 | 1.4e-05 | 6.5e-06 | 1.150E-05 | 4.330E-06 | 6.500E-06 | 1.400E-05 |
| 28725 | 1437 | A1437C | E | 1.6e-05 | 1.7e-05 | 1.5e-05 | 1.600E-05 | 1.000E-06 | 1.500E-05 | 1.700E-05 |
| 28726 | 1437 | A1437E | E | 1.1e-05 | 1.5e-05 | 8.1e-06 | 1.137E-05 | 3.465E-06 | 8.100E-06 | 1.500E-05 |
| 28727 | 1437 | A1437Q | E | 1.4e-05 | 1.5e-05 | 9.8e-06 | 1.293E-05 | 2.759E-06 | 9.800E-06 | 1.500E-05 |
| 28728 | 1437 | A1437G | E | 1.4e-05 | 1.4e-05 | 1.1e-05 | 1.300E-05 | 1.732E-06 | 1.100E-05 | 1.400E-05 |
| 28729 | 1437 | A1437H | E | 1.5e-05 | 1.7e-05 | 1.5e-05 | 1.567E-05 | 1.155E-06 | 1.500E-05 | 1.700E-05 |

| | | | | | | | | | | |
|---|---|---|---|---|---|---|---|---|---|---|
| 28730 | 1437 | A1437I | E | 1e-05 | 9.1e-06 | 1.6e-05 | 1.170E-05 | 3.751E-06 | 9.100E-06 | 1.600E-05 |
| 28731 | 1437 | A1437L | E | 1.5e-05 | 1.6e-05 | 1e-05 | 1.367E-05 | 3.215E-06 | 1.000E-05 | 1.600E-05 |
| 28732 | 1437 | A1437K | E | 1.5e-05 | 1e-05 | 1.4e-05 | 1.300E-05 | 2.646E-06 | 1.000E-05 | 1.500E-05 |
| 28733 | 1437 | A1437M | E | 8.4e-06 | 1.5e-05 | 9.2e-06 | 1.087E-05 | 3.602E-06 | 8.400E-06 | 1.500E-05 |
| 28734 | 1437 | A1437F | E | 1.4e-05 | 9.6e-06 | 1e-05 | 1.120E-05 | 2.433E-06 | 9.600E-06 | 1.400E-05 |
| 28735 | 1437 | A1437P | E | 1e-05 | 1.7e-05 | 1.5e-05 | 1.400E-05 | 3.606E-06 | 1.000E-05 | 1.700E-05 |
| 28736 | 1437 | A1437S | E | 1.4e-05 | 1.6e-05 | 8.8e-06 | 1.293E-05 | 3.717E-06 | 8.800E-06 | 1.600E-05 |
| 28737 | 1437 | A1437T | E | 1.3e-05 | 8.4e-06 | 1.5e-05 | 1.213E-05 | 3.384E-06 | 8.400E-06 | 1.500E-05 |
| 28738 | 1437 | A1437W | E | 1.3e-05 | 9.3e-06 | 1.6e-05 | 1.277E-05 | 3.356E-06 | 9.300E-06 | 1.600E-05 |
| 28739 | 1437 | A1437Y | E | 9.4e-06 | 1e-05 | 1.3e-05 | 1.080E-05 | 1.929E-06 | 9.400E-06 | 1.300E-05 |
| 28740 | 1437 | A1437V | E | 1.3e-05 | 9.1e-06 | 9.6e-06 | 1.057E-05 | 2.122E-06 | 9.100E-06 | 1.300E-05 |
| 28741 | 1438 | L1438A | E | 1.5e-05 | 9.5e-06 | 1.1e-05 | 1.183E-05 | 2.843E-06 | 9.500E-06 | 1.500E-05 |
| 28742 | 1438 | L1438R | E | 1.6e-05 | 1.3e-05 | 1.5e-05 | 1.467E-05 | 1.528E-06 | 1.300E-05 | 1.600E-05 |
| 28743 | 1438 | L1438N | E | 1.2e-05 | 1.5e-05 | 1.2e-05 | 1.300E-05 | 1.732E-06 | 1.200E-05 | 1.500E-05 |
| 28744 | 1438 | L1438D | E | 8.9e-06 | 1.3e-05 | 1.4e-05 | 1.197E-05 | 2.702E-06 | 8.900E-06 | 1.400E-05 |
| 28745 | 1438 | L1438C | E | 1.6e-05 | 1.5e-05 | 1.5e-05 | 1.533E-05 | 5.774E-07 | 1.500E-05 | 1.600E-05 |
| 28746 | 1438 | L1438E | E | 8.1e-06 | 1.3e-05 | 1.4e-05 | 1.170E-05 | 3.158E-06 | 8.100E-06 | 1.400E-05 |
| 28747 | 1438 | L1438Q | E | 1.1e-05 | 1.5e-05 | 9.2e-06 | 1.173E-05 | 2.969E-06 | 9.200E-06 | 1.500E-05 |
| 28748 | 1438 | L1438G | E | 1.3e-05 | 1e-05 | 1.5e-05 | 1.267E-05 | 2.517E-06 | 1.000E-05 | 1.500E-05 |
| 28749 | 1438 | L1438H | E | 2.1e-05 | 1.5e-05 | 1.2e-05 | 1.600E-05 | 4.583E-06 | 1.200E-05 | 2.100E-05 |
| 28750 | 1438 | L1438I | E | 1.7e-05 | 1.5e-05 | 1.7e-05 | 1.633E-05 | 1.155E-06 | 1.500E-05 | 1.700E-05 |
| 28751 | 1438 | L1438L | E | 1.7e-05 | 1e-05 | 9.9e-06 | 1.230E-05 | 4.071E-06 | 9.900E-06 | 1.700E-05 |
| 28752 | 1438 | L1438K | E | 1.4e-05 | 1.7e-05 | 1.5e-05 | 1.533E-05 | 1.528E-06 | 1.400E-05 | 1.700E-05 |
| 28753 | 1438 | L1438M | E | 1.5e-05 | 1.3e-05 | 1.4e-05 | 1.400E-05 | 1.000E-06 | 1.300E-05 | 1.500E-05 |
| 28754 | 1438 | L1438F | E | 1.5e-05 | 1e-05 | 1.5e-05 | 1.333E-05 | 2.887E-06 | 1.000E-05 | 1.500E-05 |
| 28755 | 1438 | L1438P | E | 1.5e-05 | 1.5e-05 | 1e-05 | 1.333E-05 | 2.887E-06 | 1.000E-05 | 1.500E-05 |
| 28756 | 1438 | L1438S | E | 1.5e-05 | 1.7e-05 | 1.5e-05 | 1.567E-05 | 1.155E-06 | 1.500E-05 | 1.700E-05 |
| 28757 | 1438 | L1438T | E | 1.3e-05 | 1e-05 | 1.6e-05 | 1.300E-05 | 3.000E-06 | 1.000E-05 | 1.600E-05 |
| 28758 | 1438 | L1438W | E | 1.3e-05 | 9.9e-06 | 1.2e-05 | 1.163E-05 | 1.582E-06 | 9.900E-06 | 1.300E-05 |
| 28759 | 1438 | L1438Y | E | 1.4e-05 | 1.5e-05 | 1.1e-05 | 1.333E-05 | 2.082E-06 | 1.100E-05 | 1.500E-05 |
| 28760 | 1438 | L1438V | E | 7.5e-06 | 1.5e-05 | 8.9e-06 | 1.047E-05 | 3.988E-06 | 7.500E-06 | 1.500E-05 |
| 28761 | 1439 | T1439A | E | 1e-05 | 1.4e-05 | 1.7e-05 | 1.367E-05 | 3.512E-06 | 1.000E-05 | 1.700E-05 |
| 28762 | 1439 | T1439R | E | 1.2e-05 | 7.6e-06 | 1.4e-05 | 1.120E-05 | 3.274E-06 | 7.600E-06 | 1.400E-05 |
| 28763 | 1439 | T1439N | E | 9.5e-06 | 1.5e-05 | 1.5e-05 | 1.317E-05 | 3.175E-06 | 9.500E-06 | 1.500E-05 |
| 28764 | 1439 | T1439D | E | 1.2e-05 | 1.7e-05 | 1.6e-05 | 1.500E-05 | 2.646E-06 | 1.200E-05 | 1.700E-05 |
| 28765 | 1439 | T1439C | E | 1e-05 | 1.1e-05 | 9e-06 | 1.000E-05 | 1.000E-06 | 9.000E-06 | 1.100E-05 |
| 28766 | 1439 | T1439E | E | 1.5e-05 | 1.6e-05 | 1.2e-05 | 1.433E-05 | 2.082E-06 | 1.200E-05 | 1.600E-05 |
| 28767 | 1439 | T1439Q | E | 1.4e-05 | 1.5e-05 | 1.1e-05 | 1.333E-05 | 2.082E-06 | 1.100E-05 | 1.500E-05 |
| 28768 | 1439 | T1439G | E | 1.1e-05 | 1.5e-05 | 1.1e-05 | 1.233E-05 | 2.309E-06 | 1.100E-05 | 1.500E-05 |
| 28769 | 1439 | T1439H | E | 1.1e-05 | 1e-05 | 9.9e-06 | 1.030E-05 | 6.083E-07 | 9.900E-06 | 1.100E-05 |
| 28770 | 1439 | T1439I | E | 1.7e-05 | 1e-05 | 1.2e-05 | 1.300E-05 | 3.606E-06 | 1.000E-05 | 1.700E-05 |
| 28771 | 1439 | T1439L | E | 1.6e-05 | 8.4e-06 | 1.6e-05 | 1.347E-05 | 4.388E-06 | 8.400E-06 | 1.600E-05 |
| 28772 | 1439 | T1439K | E | 1.3e-05 | 1.5e-05 | 1.3e-05 | 1.367E-05 | 1.155E-06 | 1.300E-05 | 1.500E-05 |
| 28773 | 1439 | T1439M | E | 7.8e-06 | 1.7e-05 | 9.8e-06 | 1.153E-05 | 4.839E-06 | 7.800E-06 | 1.700E-05 |
| 28774 | 1439 | T1439F | E | 1.3e-05 | 1e-05 | 1.3e-05 | 1.200E-05 | 1.732E-06 | 1.000E-05 | 1.300E-05 |
| 28775 | 1439 | T1439P | E | 1.6e-05 | 8.9e-06 | 1.6e-05 | 1.363E-05 | 4.099E-06 | 8.900E-06 | 1.600E-05 |
| 28776 | 1439 | T1439S | E | 1.1e-05 | 1.2e-05 | 1.3e-05 | 1.200E-05 | 1.000E-06 | 1.100E-05 | 1.300E-05 |
| 28777 | 1439 | T1439T | E | 1.5e-05 | 1.1e-05 | 1.7e-05 | 1.433E-05 | 3.055E-06 | 1.100E-05 | 1.700E-05 |
| 28778 | 1439 | T1439W | E | 1.1e-05 | 1.5e-05 | 1e-05 | 1.200E-05 | 2.646E-06 | 1.000E-05 | 1.500E-05 |
| 28779 | 1439 | T1439Y | E | 8.2e-06 | 1.5e-05 | 1e-05 | 1.107E-05 | 3.523E-06 | 8.200E-06 | 1.500E-05 |
| 28780 | 1439 | T1439V | E | 1.2e-05 | 9.9e-06 | 1.7e-05 | 1.297E-05 | 3.647E-06 | 9.900E-06 | 1.700E-05 |
| 28781 | 1440 | S1440A | E | 1.1e-05 | 1.6e-05 | 1.6e-05 | 1.433E-05 | 2.887E-06 | 1.100E-05 | 1.600E-05 |
| 28782 | 1440 | S1440R | E | 1.7e-05 | 7.7e-06 | 1.2e-05 | 1.223E-05 | 4.654E-06 | 7.700E-06 | 1.700E-05 |
| 28783 | 1440 | S1440N | E | 1.5e-05 | 1.6e-05 | 1.1e-05 | 1.400E-05 | 2.646E-06 | 1.100E-05 | 1.600E-05 |
| 28784 | 1440 | S1440D | E | 1e-05 | 1.7e-05 | 1.4e-05 | 1.367E-05 | 3.512E-06 | 1.000E-05 | 1.700E-05 |
| 28785 | 1440 | S1440C | E | 7.7e-06 | 1.5e-05 | 1e-05 | 1.090E-05 | 3.732E-06 | 7.700E-06 | 1.500E-05 |
| 28786 | 1440 | S1440E | E | 1.6e-05 | 1.6e-05 | 1.1e-05 | 1.433E-05 | 2.887E-06 | 1.100E-05 | 1.600E-05 |
| 28787 | 1440 | S1440Q | E | 1.3e-05 | 1.5e-05 | 1.4e-05 | 1.400E-05 | 1.000E-06 | 1.300E-05 | 1.500E-05 |
| 28788 | 1440 | S1440G | E | 1.6e-05 | 1.2e-05 | 9.9e-06 | 1.263E-05 | 3.099E-06 | 9.900E-06 | 1.600E-05 |
| 28789 | 1440 | S1440H | E | 1.1e-05 | 7.9e-06 | 1.6e-05 | 1.163E-05 | 4.087E-06 | 7.900E-06 | 1.600E-05 |
| 28790 | 1440 | S1440I | E | 1.3e-05 | 1.7e-05 | 1.6e-05 | 1.533E-05 | 2.082E-06 | 1.300E-05 | 1.700E-05 |
| 28791 | 1440 | S1440L | E | 1.5e-05 | 1.5e-05 | 1.8e-05 | 1.600E-05 | 1.732E-06 | 1.500E-05 | 1.800E-05 |
| 28792 | 1440 | S1440K | E | 1.5e-05 | 1.1e-05 | 1.6e-05 | 1.400E-05 | 2.646E-06 | 1.100E-05 | 1.600E-05 |
| 28793 | 1440 | S1440M | E | 1e-05 | 1.6e-05 | 1.5e-05 | 1.367E-05 | 3.215E-06 | 1.000E-05 | 1.600E-05 |
| 28794 | 1440 | S1440F | E | 1.6e-05 | 9.4e-06 | 1.6e-05 | 1.380E-05 | 3.811E-06 | 9.400E-06 | 1.600E-05 |
| 28795 | 1440 | S1440P | E | 1e-05 | 1.3e-05 | 1.5e-05 | 1.267E-05 | 2.517E-06 | 1.000E-05 | 1.500E-05 |
| 28796 | 1440 | S1440S | E | 1e-05 | 1.5e-05 | 1.1e-05 | 1.200E-05 | 2.646E-06 | 1.000E-05 | 1.500E-05 |
| 28797 | 1440 | S1440T | E | 9.5e-06 | 1.2e-05 | 1.2e-05 | 1.117E-05 | 1.443E-06 | 9.500E-06 | 1.200E-05 |
| 28798 | 1440 | S1440W | E | 1.3e-05 | 1.7e-05 | 1.5e-05 | 1.500E-05 | 2.000E-06 | 1.300E-05 | 1.700E-05 |
| 28799 | 1440 | S1440Y | E | 8.2e-06 | 1.5e-05 | 1.3e-05 | 1.207E-05 | 3.495E-06 | 8.200E-06 | 1.500E-05 |
| 28800 | 1440 | S1440V | E | 1.1e-05 | 1.1e-05 | 1.6e-05 | 1.267E-05 | 2.887E-06 | 1.100E-05 | 1.600E-05 |
| 28801 | 1441 | G1441A | E | 1.1e-05 | 1.5e-05 | 1.4e-05 | 1.333E-05 | 2.082E-06 | 1.100E-05 | 1.500E-05 |
| 28802 | 1441 | G1441R | E | 1.6e-05 | 2e-05 | 1.1e-05 | 1.567E-05 | 4.509E-06 | 1.100E-05 | 2.000E-05 |
| 28803 | 1441 | G1441N | E | 1.5e-05 | 9.8e-06 | 1.6e-05 | 1.360E-05 | 3.329E-06 | 9.800E-06 | 1.600E-05 |
| 28804 | 1441 | G1441D | E | 9.7e-06 | 1.4e-05 | 1e-05 | 1.123E-05 | 2.401E-06 | 9.700E-06 | 1.400E-05 |
| 28805 | 1441 | G1441C | E | 1.4e-05 | 1.5e-05 | 7.7e-06 | 1.223E-05 | 3.958E-06 | 7.700E-06 | 1.500E-05 |
| 28806 | 1441 | G1441E | E | 8.2e-06 | 1.6e-05 | 1.5e-05 | 1.307E-05 | 4.244E-06 | 8.200E-06 | 1.600E-05 |
| 28807 | 1441 | G1441Q | E | 1.5e-05 | 1.7e-05 | 6.5e-06 | 1.283E-05 | 5.575E-06 | 6.500E-06 | 1.700E-05 |
| 28808 | 1441 | G1441G | E | 1.3e-05 | 7.7e-06 | 1.3e-05 | 1.123E-05 | 3.060E-06 | 7.700E-06 | 1.300E-05 |
| 28809 | 1441 | G1441H | E | 1e-05 | 1.1e-05 | 1.1e-05 | 1.067E-05 | 5.774E-07 | 1.000E-05 | 1.100E-05 |
| 28810 | 1441 | G1441I | E | 1.6e-05 | 1.5e-05 | 1.7e-05 | 1.600E-05 | 1.000E-06 | 1.500E-05 | 1.700E-05 |
| 28811 | 1441 | G1441L | E | 1.1e-05 | 1.5e-05 | 1.5e-05 | 1.367E-05 | 2.309E-06 | 1.100E-05 | 1.500E-05 |
| 28812 | 1441 | G1441K | E | 1.1e-05 | 6.7e-06 | 8.6e-06 | 8.767E-06 | 2.155E-06 | 6.700E-06 | 1.100E-05 |
| 28813 | 1441 | G1441M | E | 8.4e-06 | 1.6e-05 | 1.7e-05 | 1.380E-05 | 4.703E-06 | 8.400E-06 | 1.700E-05 |
| 28814 | 1441 | G1441F | E | 1.5e-05 | 1.1e-05 | 1.6e-05 | 1.400E-05 | 2.646E-06 | 1.100E-05 | 1.600E-05 |

| | | | | | | | | | | |
|---|---|---|---|---|---|---|---|---|---|---|
| 28815 | 1441 | G1441P | E | 1.3e-05 | 1.5e-05 | 1.5e-05 | 1.433E-05 | 1.155E-06 | 1.300E-05 | 1.500E-05 |
| 28816 | 1441 | G1441S | E | 1.5e-05 | 1.3e-05 | 1.1e-05 | 1.300E-05 | 2.000E-06 | 1.100E-05 | 1.500E-05 |
| 28817 | 1441 | G1441T | E | 1.6e-05 | 1.3e-05 | 9.9e-06 | 1.297E-05 | 3.050E-06 | 9.900E-06 | 1.600E-05 |
| 28818 | 1441 | G1441W | E | 9.9e-06 | 1.4e-05 | 1.1e-05 | 1.163E-05 | 2.122E-06 | 9.900E-06 | 1.400E-05 |
| 28819 | 1441 | G1441Y | E | 9.5e-06 | 9.3e-06 | 1.6e-05 | 1.160E-05 | 3.812E-06 | 9.300E-06 | 1.600E-05 |
| 28820 | 1441 | G1441V | E | 1.4e-05 | 1.5e-05 | 1.4e-05 | 1.433E-05 | 5.774E-07 | 1.400E-05 | 1.500E-05 |
| 28821 | 1442 | V1442A | E | 1.5e-05 | 1.6e-05 | 1.1e-05 | 1.400E-05 | 2.646E-06 | 1.100E-05 | 1.600E-05 |
| 28822 | 1442 | V1442R | E | 1.4e-05 | 1.3e-05 | 1.5e-05 | 1.400E-05 | 1.000E-06 | 1.300E-05 | 1.500E-05 |
| 28823 | 1442 | V1442N | E | 1e-05 | 1.5e-05 | 9.5e-06 | 1.150E-05 | 3.041E-06 | 9.500E-06 | 1.500E-05 |
| 28824 | 1442 | V1442D | E | 1.4e-05 | 1.2e-05 | 8.9e-06 | 1.163E-05 | 2.570E-06 | 8.900E-06 | 1.400E-05 |
| 28825 | 1442 | V1442C | E | 6.9e-06 | 8.4e-06 | 1.4e-05 | 9.767E-06 | 3.742E-06 | 6.900E-06 | 1.400E-05 |
| 28826 | 1442 | V1442E | E | 1.1e-05 | 1.5e-05 | 1.4e-05 | 1.333E-05 | 2.082E-06 | 1.100E-05 | 1.500E-05 |
| 28827 | 1442 | V1442Q | E | 1.6e-05 | 1.5e-05 | 1.1e-05 | 1.400E-05 | 2.646E-06 | 1.100E-05 | 1.600E-05 |
| 28828 | 1442 | V1442G | E | 1.6e-05 | 1.5e-05 | 1.4e-05 | 1.500E-05 | 1.000E-06 | 1.400E-05 | 1.600E-05 |
| 28829 | 1442 | V1442H | E | 1e-05 | 9.4e-06 | 1.3e-05 | 1.080E-05 | 1.929E-06 | 9.400E-06 | 1.300E-05 |
| 28830 | 1442 | V1442I | E | 1.5e-05 | 1.4e-05 | 1.6e-05 | 1.500E-05 | 1.000E-06 | 1.400E-05 | 1.600E-05 |
| 28831 | 1442 | V1442L | E | 1.2e-05 | 1.6e-05 | 1.5e-05 | 1.433E-05 | 2.082E-06 | 1.200E-05 | 1.600E-05 |
| 28832 | 1442 | V1442K | E | 1.5e-05 | 1.6e-05 | 9.2e-06 | 1.340E-05 | 3.672E-06 | 9.200E-06 | 1.600E-05 |
| 28833 | 1442 | V1442M | E | 1.3e-05 | 1.1e-05 | 1.5e-05 | 1.300E-05 | 2.000E-06 | 1.100E-05 | 1.500E-05 |
| 28834 | 1442 | V1442F | E | 1.4e-05 | 9.4e-06 | 1.7e-05 | 1.347E-05 | 3.828E-06 | 9.400E-06 | 1.700E-05 |
| 28835 | 1442 | V1442P | E | 1.6e-05 | 1.1e-05 | 1.4e-05 | 1.367E-05 | 2.517E-06 | 1.100E-05 | 1.600E-05 |
| 28836 | 1442 | V1442S | E | 1.4e-05 | 1.5e-05 | 1.2e-05 | 1.367E-05 | 1.528E-06 | 1.200E-05 | 1.500E-05 |
| 28837 | 1442 | V1442T | E | 1.4e-05 | 1.5e-05 | 1.7e-05 | 1.533E-05 | 1.528E-06 | 1.400E-05 | 1.700E-05 |
| 28838 | 1442 | V1442W | E | 1.5e-05 | 1.1e-05 | 1.5e-05 | 1.367E-05 | 2.309E-06 | 1.100E-05 | 1.500E-05 |
| 28839 | 1442 | V1442Y | E | 1.6e-05 | 7.8e-06 | 1.6e-05 | 1.327E-05 | 4.734E-06 | 7.800E-06 | 1.600E-05 |
| 28840 | 1442 | V1442V | E | 9.7e-06 | 1.5e-05 | 7.2e-06 | 1.063E-05 | 3.983E-06 | 7.200E-06 | 1.500E-05 |
| 28841 | 1443 | H1443A | E | 1.4e-05 | 9.4e-06 | 1.2e-05 | 1.180E-05 | 2.307E-06 | 9.400E-06 | 1.400E-05 |
| 28842 | 1443 | H1443R | E | 1e-05 | 1.5e-05 | 1e-05 | 1.167E-05 | 2.887E-06 | 1.000E-05 | 1.500E-05 |
| 28843 | 1443 | H1443N | E | 1.5e-05 | 1.5e-05 | 8.8e-06 | 1.293E-05 | 3.580E-06 | 8.800E-06 | 1.500E-05 |
| 28844 | 1443 | H1443D | E | 1e-05 | 9.7e-06 | 1.5e-05 | 1.157E-05 | 2.977E-06 | 9.700E-06 | 1.500E-05 |
| 28845 | 1443 | H1443C | E | 1.5e-05 | 8.7e-06 | 6.5e-06 | 1.007E-05 | 4.412E-06 | 6.500E-06 | 1.500E-05 |
| 28846 | 1443 | H1443E | E | 1.1e-05 | 1.7e-05 | 1.5e-05 | 1.433E-05 | 3.055E-06 | 1.100E-05 | 1.700E-05 |
| 28847 | 1443 | H1443Q | E | 1.5e-05 | 1.2e-05 | 1.1e-05 | 1.267E-05 | 2.082E-06 | 1.100E-05 | 1.500E-05 |
| 28848 | 1443 | H1443G | E | 1.4e-05 | 1.6e-05 | 1.6e-05 | 1.533E-05 | 1.155E-06 | 1.400E-05 | 1.600E-05 |
| 28849 | 1443 | H1443H | E | 1.5e-05 | 1.7e-05 | 1.3e-05 | 1.500E-05 | 2.000E-06 | 1.300E-05 | 1.700E-05 |
| 28850 | 1443 | H1443I | E | 1.6e-05 | 6.6e-06 | 1.8e-05 | 1.353E-05 | 6.087E-06 | 6.600E-06 | 1.800E-05 |
| 28851 | 1443 | H1443L | E | 1.7e-05 | 1.6e-05 | 8.1e-06 | 1.370E-05 | 4.875E-06 | 8.100E-06 | 1.700E-05 |
| 28852 | 1443 | H1443K | E | 1.6e-05 | 1.1e-05 | 1e-05 | 1.233E-05 | 3.215E-06 | 1.000E-05 | 1.600E-05 |
| 28853 | 1443 | H1443M | E | 1.4e-05 | 1.4e-05 | 1.1e-05 | 1.300E-05 | 1.732E-06 | 1.100E-05 | 1.400E-05 |
| 28854 | 1443 | H1443F | E | 1.1e-05 | 8.4e-06 | 1.6e-05 | 1.180E-05 | 3.863E-06 | 8.400E-06 | 1.600E-05 |
| 28855 | 1443 | H1443P | E | 1.6e-05 | 1.4e-05 | 1.7e-05 | 1.567E-05 | 1.528E-06 | 1.400E-05 | 1.700E-05 |
| 28856 | 1443 | H1443S | E | 8.9e-06 | 1e-05 | 1.4e-05 | 1.097E-05 | 2.684E-06 | 8.900E-06 | 1.400E-05 |
| 28857 | 1443 | H1443T | E | 9.8e-06 | 1e-05 | 1e-05 | 9.933E-06 | 1.155E-07 | 9.800E-06 | 1.000E-05 |
| 28858 | 1443 | H1443W | E | 1.1e-05 | 1.4e-05 | 8.2e-06 | 1.107E-05 | 2.901E-06 | 8.200E-06 | 1.400E-05 |
| 28859 | 1443 | H1443Y | E | 1e-05 | 1.3e-05 | 1.6e-05 | 1.300E-05 | 3.000E-06 | 1.000E-05 | 1.600E-05 |
| 28860 | 1443 | H1443V | E | 7.2e-06 | 1.1e-05 | 1.1e-05 | 9.733E-06 | 2.194E-06 | 7.200E-06 | 1.100E-05 |
| 28861 | 1444 | T1444A | E | 1.1e-05 | 1.4e-05 | 9.8e-06 | 1.160E-05 | 2.163E-06 | 9.800E-06 | 1.400E-05 |
| 28862 | 1444 | T1444R | E | 1.4e-05 | 1.1e-05 | 1.7e-05 | 1.400E-05 | 3.000E-06 | 1.100E-05 | 1.700E-05 |
| 28863 | 1444 | T1444N | E | 1e-05 | 8.6e-06 | 1e-05 | 9.533E-06 | 8.083E-07 | 8.600E-06 | 1.000E-05 |
| 28864 | 1444 | T1444D | E | 7e-06 | 9.6e-06 | 1.2e-05 | 9.533E-06 | 2.501E-06 | 7.000E-06 | 1.200E-05 |
| 28865 | 1444 | T1444C | E | 1.4e-05 | 8.5e-06 | 9.9e-06 | 1.080E-05 | 2.858E-06 | 8.500E-06 | 1.400E-05 |
| 28866 | 1444 | T1444E | E | 1e-05 | 1.7e-05 | 1.4e-05 | 1.367E-05 | 3.512E-06 | 1.000E-05 | 1.700E-05 |
| 28867 | 1444 | T1444Q | E | 7.3e-06 | 1e-05 | 1.3e-05 | 1.010E-05 | 2.851E-06 | 7.300E-06 | 1.300E-05 |
| 28868 | 1444 | T1444G | E | 1.7e-05 | 1.3e-05 | 1.1e-05 | 1.367E-05 | 3.055E-06 | 1.100E-05 | 1.700E-05 |
| 28869 | 1444 | T1444H | E | 1.4e-05 | 1.5e-05 | 1e-05 | 1.300E-05 | 2.646E-06 | 1.000E-05 | 1.500E-05 |
| 28870 | 1444 | T1444I | E | 1.1e-05 | 1.6e-05 | 1.6e-05 | 1.433E-05 | 2.887E-06 | 1.100E-05 | 1.600E-05 |
| 28871 | 1444 | T1444L | E | 1.3e-05 | 1.6e-05 | 1.5e-05 | 1.467E-05 | 1.528E-06 | 1.300E-05 | 1.600E-05 |
| 28872 | 1444 | T1444K | E | 1.5e-05 | 1e-05 | 9.5e-06 | 1.150E-05 | 3.041E-06 | 9.500E-06 | 1.500E-05 |
| 28873 | 1444 | T1444M | E | 1.6e-05 | 9.2e-06 | 1.5e-05 | 1.340E-05 | 3.672E-06 | 9.200E-06 | 1.600E-05 |
| 28874 | 1444 | T1444F | E | 1.1e-05 | 1e-05 | 1.5e-05 | 1.200E-05 | 2.646E-06 | 1.000E-05 | 1.500E-05 |
| 28875 | 1444 | T1444P | E | 1.5e-05 | 1.6e-05 | 8.9e-06 | 1.330E-05 | 3.843E-06 | 8.900E-06 | 1.600E-05 |
| 28876 | 1444 | T1444S | E | 1.6e-05 | 1.7e-05 | 1.1e-05 | 1.467E-05 | 3.215E-06 | 1.100E-05 | 1.700E-05 |
| 28877 | 1444 | T1444T | E | 1.5e-05 | 9.6e-06 | 1.5e-05 | 1.320E-05 | 3.118E-06 | 9.600E-06 | 1.500E-05 |
| 28878 | 1444 | T1444W | E | 1.8e-05 | 8.1e-06 | 6.5e-06 | 1.087E-05 | 6.229E-06 | 6.500E-06 | 1.800E-05 |
| 28879 | 1444 | T1444Y | E | 1.1e-05 | 1.4e-05 | 1.7e-05 | 1.400E-05 | 3.000E-06 | 1.100E-05 | 1.700E-05 |
| 28880 | 1444 | T1444V | E | 1.5e-05 | 9.1e-06 | 1.7e-05 | 1.370E-05 | 4.107E-06 | 9.100E-06 | 1.700E-05 |
| 28881 | 1445 | F1445A | E | 1.6e-05 | 1.1e-05 | 9.9e-06 | 1.230E-05 | 3.251E-06 | 9.900E-06 | 1.600E-05 |
| 28882 | 1445 | F1445R | E | 1.1e-05 | 1.4e-05 | 1.5e-05 | 1.333E-05 | 2.082E-06 | 1.100E-05 | 1.500E-05 |
| 28883 | 1445 | F1445N | E | 9.5e-06 | 1.5e-05 | 1.1e-05 | 1.183E-05 | 2.843E-06 | 9.500E-06 | 1.500E-05 |
| 28884 | 1445 | F1445D | E | 1.4e-05 | 1.5e-05 | 9.5e-06 | 1.283E-05 | 2.930E-06 | 9.500E-06 | 1.500E-05 |
| 28885 | 1445 | F1445C | E | 8.3e-06 | 9.5e-06 | 1.4e-05 | 1.060E-05 | 3.005E-06 | 8.300E-06 | 1.400E-05 |
| 28886 | 1445 | F1445E | E | 9.6e-06 | 1.5e-05 | 1e-05 | 1.153E-05 | 3.009E-06 | 9.600E-06 | 1.500E-05 |
| 28887 | 1445 | F1445Q | E | 1e-05 | 1.5e-05 | 1e-05 | 1.167E-05 | 2.887E-06 | 1.000E-05 | 1.500E-05 |
| 28888 | 1445 | F1445G | E | 1.1e-05 | 1.7e-05 | 1.3e-05 | 1.367E-05 | 3.055E-06 | 1.100E-05 | 1.700E-05 |
| 28889 | 1445 | F1445H | E | 1.5e-05 | 1.1e-05 | 1.4e-05 | 1.333E-05 | 2.082E-06 | 1.100E-05 | 1.500E-05 |
| 28890 | 1445 | F1445I | E | 1e-05 | 9.9e-06 | 1.2e-05 | 1.063E-05 | 1.185E-06 | 9.900E-06 | 1.200E-05 |
| 28891 | 1445 | F1445L | E | 1.6e-05 | 1.7e-05 | 1.4e-05 | 1.567E-05 | 1.528E-06 | 1.400E-05 | 1.700E-05 |
| 28892 | 1445 | F1445K | E | 1.3e-05 | 1.2e-05 | 1.4e-05 | 1.300E-05 | 1.000E-06 | 1.200E-05 | 1.400E-05 |
| 28893 | 1445 | F1445M | E | 1.3e-05 | 1.5e-05 | 8.5e-06 | 1.217E-05 | 3.329E-06 | 8.500E-06 | 1.500E-05 |
| 28894 | 1445 | F1445F | E | 1.5e-05 | 1.2e-05 | 1.5e-05 | 1.400E-05 | 1.732E-06 | 1.200E-05 | 1.500E-05 |
| 28895 | 1445 | F1445P | E | 1e-05 | 1e-05 | 1.1e-05 | 1.033E-05 | 5.774E-07 | 1.000E-05 | 1.100E-05 |
| 28896 | 1445 | F1445S | E | 1.4e-05 | 8.9e-06 | 1.5e-05 | 1.263E-05 | 3.272E-06 | 8.900E-06 | 1.500E-05 |
| 28897 | 1445 | F1445T | E | 1.6e-05 | 1.6e-05 | 1.4e-05 | 1.533E-05 | 1.155E-06 | 1.400E-05 | 1.600E-05 |
| 28898 | 1445 | F1445W | E | 1e-05 | 1.1e-05 | 1e-05 | 1.033E-05 | 5.774E-07 | 1.000E-05 | 1.100E-05 |
| 28899 | 1445 | F1445Y | E | 1.5e-05 | 1.5e-05 | 1.3e-05 | 1.433E-05 | 1.155E-06 | 1.300E-05 | 1.500E-05 |

| | | | | | | | | | | |
|---|---|---|---|---|---|---|---|---|---|---|
| 28900 | 1445 | F1445V | E | 1.5e-05 | 1.4e-05 | 1e-05 | 1.300E-05 | 2.646E-06 | 1.000E-05 | 1.500E-05 |
| 28901 | 1446 | P1446A | E | 1.6e-05 | 1.3e-05 | 1.6e-05 | 1.500E-05 | 1.732E-06 | 1.300E-05 | 1.600E-05 |
| 28902 | 1446 | P1446R | E | 9e-06 | 1e-05 | 2.4e-05 | 1.433E-05 | 8.386E-06 | 9.000E-06 | 2.400E-05 |
| 28903 | 1446 | P1446N | E | 1.4e-05 | 1e-05 | 9.8e-06 | 1.127E-05 | 2.369E-06 | 9.800E-06 | 1.400E-05 |
| 28904 | 1446 | P1446D | E | 1.1e-05 | 1.5e-05 | 1.5e-05 | 1.367E-05 | 2.309E-06 | 1.100E-05 | 1.500E-05 |
| 28905 | 1446 | P1446C | E | 1e-05 | 8.1e-06 | 9.9e-06 | 9.333E-06 | 1.069E-06 | 8.100E-06 | 1.000E-05 |
| 28906 | 1446 | P1446E | E | 1.4e-05 | 1.6e-05 | 1.2e-05 | 1.400E-05 | 2.000E-06 | 1.200E-05 | 1.600E-05 |
| 28907 | 1446 | P1446Q | E | 1.5e-05 | 6.4e-06 | 1e-05 | 1.047E-05 | 4.319E-06 | 6.400E-06 | 1.500E-05 |
| 28908 | 1446 | P1446G | E | 8.4e-06 | 1e-05 | 1.1e-05 | 9.800E-06 | 1.311E-06 | 8.400E-06 | 1.100E-05 |
| 28909 | 1446 | P1446H | E | 1.4e-05 | 9.9e-06 | 9e-06 | 1.097E-05 | 2.665E-06 | 9.000E-06 | 1.400E-05 |
| 28910 | 1446 | P1446I | E | 1.4e-05 | 9.9e-06 | 1.6e-05 | 1.330E-05 | 3.110E-06 | 9.900E-06 | 1.600E-05 |
| 28911 | 1446 | P1446L | E | 9.1e-06 | 8.6e-06 | 1.3e-05 | 1.023E-05 | 2.409E-06 | 8.600E-06 | 1.300E-05 |
| 28912 | 1446 | P1446K | E | 1.7e-05 | 1.4e-05 | 1.1e-05 | 1.400E-05 | 3.000E-06 | 1.100E-05 | 1.700E-05 |
| 28913 | 1446 | P1446M | E | 1.4e-05 | 1.5e-05 | 1e-05 | 1.300E-05 | 2.646E-06 | 1.000E-05 | 1.500E-05 |
| 28914 | 1446 | P1446F | E | 1.6e-05 | 1.5e-05 | 1.5e-05 | 1.533E-05 | 5.774E-07 | 1.500E-05 | 1.600E-05 |
| 28915 | 1446 | P1446P | E | 1.8e-05 | 1e-05 | 1.8e-05 | 1.533E-05 | 4.619E-06 | 1.000E-05 | 1.800E-05 |
| 28916 | 1446 | P1446S | E | 1.5e-05 | 1.1e-05 | 1.3e-05 | 1.300E-05 | 2.000E-06 | 1.100E-05 | 1.500E-05 |
| 28917 | 1446 | P1446T | E | 8e-06 | 8.4e-06 | 1.5e-05 | 1.047E-05 | 3.931E-06 | 8.000E-06 | 1.500E-05 |
| 28918 | 1446 | P1446W | E | 9.7e-06 | 1e-05 | 1.5e-05 | 1.157E-05 | 2.977E-06 | 9.700E-06 | 1.500E-05 |
| 28919 | 1446 | P1446Y | E | 1.1e-05 | 1.4e-05 | 1.5e-05 | 1.333E-05 | 2.082E-06 | 1.100E-05 | 1.500E-05 |
| 28920 | 1446 | P1446V | E | 1.5e-05 | 1.1e-05 | 1.4e-05 | 1.333E-05 | 2.082E-06 | 1.100E-05 | 1.500E-05 |
| 28921 | 1447 | A1447A | E | 1.6e-05 | 1.6e-05 | 1e-05 | 1.400E-05 | 3.464E-06 | 1.000E-05 | 1.600E-05 |
| 28922 | 1447 | A1447R | E | 8.7e-06 | 1.5e-05 | 9e-06 | 1.090E-05 | 3.554E-06 | 8.700E-06 | 1.500E-05 |
| 28923 | 1447 | A1447N | E | 8.7e-06 | 1.4e-05 | 1.1e-05 | 1.123E-05 | 2.658E-06 | 8.700E-06 | 1.400E-05 |
| 28924 | 1447 | A1447D | E | 9.1e-06 | 1.5e-05 | 1.6e-05 | 1.337E-05 | 3.729E-06 | 9.100E-06 | 1.600E-05 |
| 28925 | 1447 | A1447C | E | 1.6e-05 | 1.4e-05 | 1.1e-05 | 1.367E-05 | 2.517E-06 | 1.100E-05 | 1.600E-05 |
| 28926 | 1447 | A1447E | E | 1.4e-05 | 1.1e-05 | 1.5e-05 | 1.333E-05 | 2.082E-06 | 1.100E-05 | 1.500E-05 |
| 28927 | 1447 | A1447Q | E | 7.6e-06 | 1.4e-05 | 1.6e-05 | 1.253E-05 | 4.388E-06 | 7.600E-06 | 1.600E-05 |
| 28928 | 1447 | A1447G | E | 1.1e-05 | 1.6e-05 | 1.7e-05 | 1.467E-05 | 3.215E-06 | 1.100E-05 | 1.700E-05 |
| 28929 | 1447 | A1447H | E | 1.1e-05 | 1.7e-05 | 1.5e-05 | 1.433E-05 | 3.055E-06 | 1.100E-05 | 1.700E-05 |
| 28930 | 1447 | A1447I | E | 1.3e-05 | 1e-05 | 1.1e-05 | 1.133E-05 | 1.528E-06 | 1.000E-05 | 1.300E-05 |
| 28931 | 1447 | A1447L | E | 1.6e-05 | 1.5e-05 | 1.6e-05 | 1.567E-05 | 5.774E-07 | 1.500E-05 | 1.600E-05 |
| 28932 | 1447 | A1447K | E | 1.5e-05 | 9.7e-06 | 9.8e-06 | 1.150E-05 | 3.032E-06 | 9.700E-06 | 1.500E-05 |
| 28933 | 1447 | A1447M | E | 1.3e-05 | 1.1e-05 | 1.5e-05 | 1.300E-05 | 2.000E-06 | 1.100E-05 | 1.500E-05 |
| 28934 | 1447 | A1447F | E | 1.5e-05 | 1.5e-05 | 1.1e-05 | 1.367E-05 | 2.309E-06 | 1.100E-05 | 1.500E-05 |
| 28935 | 1447 | A1447P | E | 1.6e-05 | 1.1e-05 | 1.2e-05 | 1.300E-05 | 2.646E-06 | 1.100E-05 | 1.600E-05 |
| 28936 | 1447 | A1447S | E | 1.4e-05 | 1.3e-05 | 1.1e-05 | 1.267E-05 | 1.528E-06 | 1.100E-05 | 1.400E-05 |
| 28937 | 1447 | A1447T | E | 1.1e-05 | 9e-06 | 1.2e-05 | 1.067E-05 | 1.528E-06 | 9.000E-06 | 1.200E-05 |
| 28938 | 1447 | A1447W | E | 1.6e-05 | 9.3e-06 | 1.4e-05 | 1.310E-05 | 3.439E-06 | 9.300E-06 | 1.600E-05 |
| 28939 | 1447 | A1447Y | E | 1.4e-05 | 1.4e-05 | 2.4e-05 | 1.733E-05 | 5.774E-06 | 1.400E-05 | 2.400E-05 |
| 28940 | 1447 | A1447V | E | 1.3e-05 | 1.5e-05 | 1.5e-05 | 1.433E-05 | 1.155E-06 | 1.300E-05 | 1.500E-05 |
| 28941 | 1448 | V1448A | E | 1.4e-05 | 1e-05 | 1.3e-05 | 1.233E-05 | 2.082E-06 | 1.000E-05 | 1.400E-05 |
| 28942 | 1448 | V1448R | E | 9.6e-06 | 1.9e-05 | 1.1e-05 | 1.320E-05 | 5.071E-06 | 9.600E-06 | 1.900E-05 |
| 28943 | 1448 | V1448N | E | 1.4e-05 | 9.1e-06 | 1.5e-05 | 1.270E-05 | 3.158E-06 | 9.100E-06 | 1.500E-05 |
| 28944 | 1448 | V1448D | E | 1.5e-05 | 1.1e-05 | 1.5e-05 | 1.367E-05 | 2.309E-06 | 1.100E-05 | 1.500E-05 |
| 28945 | 1448 | V1448C | E | 2.1e-05 | 1.4e-05 | 1.3e-05 | 1.600E-05 | 4.359E-06 | 1.300E-05 | 2.100E-05 |
| 28946 | 1448 | V1448E | E | 8.9e-06 | 1.5e-05 | 1.4e-05 | 1.263E-05 | 3.272E-06 | 8.900E-06 | 1.500E-05 |
| 28947 | 1448 | V1448Q | E | 1.3e-05 | 1.2e-05 | 1.3e-05 | 1.267E-05 | 5.774E-07 | 1.200E-05 | 1.300E-05 |
| 28948 | 1448 | V1448G | E | 1.1e-05 | 1.1e-05 | 1.6e-05 | 1.267E-05 | 2.887E-06 | 1.100E-05 | 1.600E-05 |
| 28949 | 1448 | V1448H | E | 1e-05 | 1.5e-05 | 9.4e-06 | 1.147E-05 | 3.075E-06 | 9.400E-06 | 1.500E-05 |
| 28950 | 1448 | V1448I | E | 1.3e-05 | 1.1e-05 | 1.5e-05 | 1.300E-05 | 2.000E-06 | 1.100E-05 | 1.500E-05 |
| 28951 | 1448 | V1448L | E | 1e-05 | 1.5e-05 | 1.5e-05 | 1.333E-05 | 2.887E-06 | 1.000E-05 | 1.500E-05 |
| 28952 | 1448 | V1448K | E | 9.7e-06 | 1.5e-05 | 1.3e-05 | 1.257E-05 | 2.676E-06 | 9.700E-06 | 1.500E-05 |
| 28953 | 1448 | V1448M | E | 1.1e-05 | 9.9e-06 | 1.5e-05 | 1.197E-05 | 2.684E-06 | 9.900E-06 | 1.500E-05 |
| 28954 | 1448 | V1448F | E | 1.5e-05 | 1.4e-05 | 1.2e-05 | 1.367E-05 | 1.528E-06 | 1.200E-05 | 1.500E-05 |
| 28955 | 1448 | V1448P | E | 1e-05 | 1.6e-05 | 1.3e-05 | 1.300E-05 | 3.000E-06 | 1.000E-05 | 1.600E-05 |
| 28956 | 1448 | V1448S | E | 1.1e-05 | 1.5e-05 | 1e-05 | 1.200E-05 | 2.646E-06 | 1.000E-05 | 1.500E-05 |
| 28957 | 1448 | V1448T | E | 1.4e-05 | 1.7e-05 | 1e-05 | 1.367E-05 | 3.512E-06 | 1.000E-05 | 1.700E-05 |
| 28958 | 1448 | V1448W | E | 9.9e-06 | 1.1e-05 | 1.5e-05 | 1.197E-05 | 2.684E-06 | 9.900E-06 | 1.500E-05 |
| 28959 | 1448 | V1448Y | E | 9.9e-06 | 8.3e-06 | 8e-06 | 8.733E-06 | 1.021E-06 | 8.000E-06 | 9.900E-06 |
| 28960 | 1448 | V1448V | E | 1.5e-05 | 1.5e-05 | 1.6e-05 | 1.533E-05 | 5.774E-07 | 1.500E-05 | 1.600E-05 |
| 28961 | 1449 | L1449A | E | 1.7e-05 | 1.4e-05 | 1.6e-05 | 1.567E-05 | 1.528E-06 | 1.400E-05 | 1.700E-05 |
| 28962 | 1449 | L1449R | E | 1.1e-05 | 1.5e-05 | 1.1e-05 | 1.233E-05 | 2.309E-06 | 1.100E-05 | 1.500E-05 |
| 28963 | 1449 | L1449N | E | 1.2e-05 | 1.6e-05 | 1.1e-05 | 1.300E-05 | 2.646E-06 | 1.100E-05 | 1.600E-05 |
| 28964 | 1449 | L1449D | E | 9.7e-06 | 9.8e-06 | 1.4e-05 | 1.117E-05 | 2.454E-06 | 9.700E-06 | 1.400E-05 |
| 28965 | 1449 | L1449C | E | 8.5e-06 | 1.4e-05 | 8.6e-06 | 1.037E-05 | 3.147E-06 | 8.500E-06 | 1.400E-05 |
| 28966 | 1449 | L1449E | E | 1e-05 | 1.5e-05 | 1.1e-05 | 1.200E-05 | 2.646E-06 | 1.000E-05 | 1.500E-05 |
| 28967 | 1449 | L1449Q | E | 1e-05 | 9.7e-06 | 9.4e-06 | 9.700E-06 | 3.000E-07 | 9.400E-06 | 1.000E-05 |
| 28968 | 1449 | L1449G | E | 1.6e-05 | 1.5e-05 | 1e-05 | 1.367E-05 | 3.215E-06 | 1.000E-05 | 1.600E-05 |
| 28969 | 1449 | L1449H | E | 1.7e-05 | 9.7e-06 | 1.5e-05 | 1.390E-05 | 3.772E-06 | 9.700E-06 | 1.700E-05 |
| 28970 | 1449 | L1449I | E | 1.4e-05 | 1e-05 | 1e-05 | 1.133E-05 | 2.309E-06 | 1.000E-05 | 1.400E-05 |
| 28971 | 1449 | L1449L | E | 1.6e-05 | 1.5e-05 | 8.9e-06 | 1.330E-05 | 3.843E-06 | 8.900E-06 | 1.600E-05 |
| 28972 | 1449 | L1449K | E | 1.7e-05 | 1.7e-05 | 1.6e-05 | 1.667E-05 | 5.774E-07 | 1.600E-05 | 1.700E-05 |
| 28973 | 1449 | L1449M | E | 1.3e-05 | 1.5e-05 | 1.5e-05 | 1.433E-05 | 1.155E-06 | 1.300E-05 | 1.500E-05 |
| 28974 | 1449 | L1449F | E | 1.4e-05 | 8.4e-06 | 1.3e-05 | 1.180E-05 | 2.987E-06 | 8.400E-06 | 1.400E-05 |
| 28975 | 1449 | L1449P | E | 1.1e-05 | 1e-05 | 1.4e-05 | 1.167E-05 | 2.082E-06 | 1.000E-05 | 1.400E-05 |
| 28976 | 1449 | L1449S | E | 1.5e-05 | 1.5e-05 | 8.3e-06 | 1.277E-05 | 3.868E-06 | 8.300E-06 | 1.500E-05 |
| 28977 | 1449 | L1449T | E | 9e-06 | 1.4e-05 | 1e-05 | 1.100E-05 | 2.646E-06 | 9.000E-06 | 1.400E-05 |
| 28978 | 1449 | L1449W | E | 1.3e-05 | 8.8e-06 | 1.4e-05 | 1.193E-05 | 2.759E-06 | 8.800E-06 | 1.400E-05 |
| 28979 | 1449 | L1449Y | E | 1e-05 | 1e-05 | 1.5e-05 | 1.167E-05 | 2.887E-06 | 1.000E-05 | 1.500E-05 |
| 28980 | 1449 | L1449V | E | 1.7e-05 | 1.6e-05 | 1.3e-05 | 1.533E-05 | 2.082E-06 | 1.300E-05 | 1.700E-05 |
| 28981 | 1450 | Q1450A | E | 1.5e-05 | 1.2e-05 | 8.9e-06 | 1.197E-05 | 3.050E-06 | 8.900E-06 | 1.500E-05 |
| 28982 | 1450 | Q1450R | E | 1.7e-05 | 1.8e-05 | 9.9e-06 | 1.497E-05 | 4.416E-06 | 9.900E-06 | 1.800E-05 |
| 28983 | 1450 | Q1450N | E | 9e-06 | 1.6e-05 | 1.1e-05 | 1.200E-05 | 3.606E-06 | 9.000E-06 | 1.600E-05 |
| 28984 | 1450 | Q1450D | E | 9.6e-06 | 1.8e-05 | 1.6e-05 | 1.453E-05 | 4.388E-06 | 9.600E-06 | 1.800E-05 |

| | | | | | | | | | | |
|---|---|---|---|---|---|---|---|---|---|---|
| 28985 | 1450 | Q1450C | E | 1.2e-05 | 1.1e-05 | 1.5e-05 | 1.267E-05 | 2.082E-06 | 1.100E-05 | 1.500E-05 |
| 28986 | 1450 | Q1450E | E | 1.4e-05 | 1.5e-05 | 1.1e-05 | 1.333E-05 | 2.082E-06 | 1.100E-05 | 1.500E-05 |
| 28987 | 1450 | Q1450Q | E | 8.7e-06 | 1.1e-05 | 1.2e-05 | 1.057E-05 | 1.692E-06 | 8.700E-06 | 1.200E-05 |
| 28988 | 1450 | Q1450G | E | 1.1e-05 | 1.6e-05 | 1.1e-05 | 1.267E-05 | 2.887E-06 | 1.100E-05 | 1.600E-05 |
| 28989 | 1450 | Q1450H | E | 1.4e-05 | 1.5e-05 | 1.6e-05 | 1.500E-05 | 1.000E-06 | 1.400E-05 | 1.600E-05 |
| 28990 | 1450 | Q1450I | E | 1.6e-05 | 1.6e-05 | 1.5e-05 | 1.567E-05 | 5.774E-07 | 1.500E-05 | 1.600E-05 |
| 28991 | 1450 | Q1450L | E | 1.6e-05 | 1.3e-05 | 1.6e-05 | 1.500E-05 | 1.732E-06 | 1.300E-05 | 1.600E-05 |
| 28992 | 1450 | Q1450K | E | 1.5e-05 | 1e-05 | 1.5e-05 | 1.333E-05 | 2.887E-06 | 1.000E-05 | 1.500E-05 |
| 28993 | 1450 | Q1450M | E | 1.1e-05 | 1.3e-05 | 1.5e-05 | 1.300E-05 | 2.000E-06 | 1.100E-05 | 1.500E-05 |
| 28994 | 1450 | Q1450F | E | 1.2e-05 | 1.2e-05 | 1.5e-05 | 1.300E-05 | 1.732E-06 | 1.200E-05 | 1.500E-05 |
| 28995 | 1450 | Q1450P | E | 1.5e-05 | 1.5e-05 | 8.8e-06 | 1.293E-05 | 3.580E-06 | 8.800E-06 | 1.500E-05 |
| 28996 | 1450 | Q1450S | E | 1.4e-05 | 1.4e-05 | 1.1e-05 | 1.300E-05 | 1.732E-06 | 1.100E-05 | 1.400E-05 |
| 28997 | 1450 | Q1450T | E | 1.6e-05 | 1.5e-05 | 1.6e-05 | 1.567E-05 | 5.774E-07 | 1.500E-05 | 1.600E-05 |
| 28998 | 1450 | Q1450W | E | 8.5e-06 | 1.3e-05 | 1.4e-05 | 1.183E-05 | 2.930E-06 | 8.500E-06 | 1.400E-05 |
| 28999 | 1450 | Q1450Y | E | 1.5e-05 | 1.5e-05 | 1.1e-05 | 1.367E-05 | 2.309E-06 | 1.100E-05 | 1.500E-05 |
| 29000 | 1450 | Q1450V | E | 1.1e-05 | 1.5e-05 | 1.1e-05 | 1.233E-05 | 2.309E-06 | 1.100E-05 | 1.500E-05 |
| 29001 | 1451 | S1451A | E | 1.1e-05 | 1.2e-05 | 9.8e-06 | 1.093E-05 | 1.102E-06 | 9.800E-06 | 1.200E-05 |
| 29002 | 1451 | S1451R | E | 1.2e-05 | 1.1e-05 | 1.1e-05 | 1.133E-05 | 5.774E-07 | 1.100E-05 | 1.200E-05 |
| 29003 | 1451 | S1451N | E | 1.1e-05 | 1.1e-05 | 1.4e-05 | 1.200E-05 | 1.732E-06 | 1.100E-05 | 1.400E-05 |
| 29004 | 1451 | S1451D | E | 1.3e-05 | 1.5e-05 | 9.8e-06 | 1.260E-05 | 2.623E-06 | 9.800E-06 | 1.500E-05 |
| 29005 | 1451 | S1451C | E | 1.3e-05 | 1.1e-05 | 1e-05 | 1.133E-05 | 1.528E-06 | 1.000E-05 | 1.300E-05 |
| 29006 | 1451 | S1451E | E | 1.1e-05 | 1.5e-05 | 1.7e-05 | 1.433E-05 | 3.055E-06 | 1.100E-05 | 1.700E-05 |
| 29007 | 1451 | S1451Q | E | 1.2e-05 | 8.3e-06 | 1.6e-05 | 1.210E-05 | 3.851E-06 | 8.300E-06 | 1.600E-05 |
| 29008 | 1451 | S1451G | E | 1.1e-05 | 1.1e-05 | 1.8e-05 | 1.333E-05 | 4.041E-06 | 1.100E-05 | 1.800E-05 |
| 29009 | 1451 | S1451H | E | 1.4e-05 | 8.8e-06 | 1.5e-05 | 1.260E-05 | 3.329E-06 | 8.800E-06 | 1.500E-05 |
| 29010 | 1451 | S1451I | E | 1.1e-05 | 1.3e-05 | 1.1e-05 | 1.167E-05 | 1.155E-06 | 1.100E-05 | 1.300E-05 |
| 29011 | 1451 | S1451L | E | 9.8e-06 | 1.5e-05 | 1.1e-05 | 1.193E-05 | 2.723E-06 | 9.800E-06 | 1.500E-05 |
| 29012 | 1451 | S1451K | E | 9e-06 | 1.7e-05 | 1.5e-05 | 1.367E-05 | 4.163E-06 | 9.000E-06 | 1.700E-05 |
| 29013 | 1451 | S1451M | E | 1.5e-05 | 1.6e-05 | 9.8e-06 | 1.360E-05 | 3.329E-06 | 9.800E-06 | 1.600E-05 |
| 29014 | 1451 | S1451F | E | 1e-05 | 9.6e-06 | 1e-05 | 9.867E-06 | 2.309E-07 | 9.600E-06 | 1.000E-05 |
| 29015 | 1451 | S1451P | E | 9.3e-06 | 1.6e-05 | 1.5e-05 | 1.343E-05 | 3.614E-06 | 9.300E-06 | 1.600E-05 |
| 29016 | 1451 | S1451S | E | 1.7e-05 | 1.1e-05 | 1.2e-05 | 1.333E-05 | 3.215E-06 | 1.100E-05 | 1.700E-05 |
| 29017 | 1451 | S1451T | E | 1e-05 | 1.5e-05 | 1.1e-05 | 1.200E-05 | 2.646E-06 | 1.000E-05 | 1.500E-05 |
| 29018 | 1451 | S1451W | E | 1.7e-05 | 1.6e-05 | 6.7e-06 | 1.323E-05 | 5.680E-06 | 6.700E-06 | 1.700E-05 |
| 29019 | 1451 | S1451Y | E | 1.4e-05 | 1.7e-05 | 1.1e-05 | 1.400E-05 | 3.000E-06 | 1.100E-05 | 1.700E-05 |
| 29020 | 1451 | S1451V | E | 1.7e-05 | 1e-05 | 1.4e-05 | 1.367E-05 | 3.512E-06 | 1.000E-05 | 1.700E-05 |
| 29021 | 1452 | S1452A | E | 1.4e-05 | 1.2e-05 | 1.4e-05 | 1.333E-05 | 1.155E-06 | 1.200E-05 | 1.400E-05 |
| 29022 | 1452 | S1452R | E | 1.6e-05 | 1.7e-05 | 1e-05 | 1.433E-05 | 3.786E-06 | 1.000E-05 | 1.700E-05 |
| 29023 | 1452 | S1452N | E | 1.3e-05 | 1.3e-05 | 1.6e-05 | 1.400E-05 | 1.732E-06 | 1.300E-05 | 1.600E-05 |
| 29024 | 1452 | S1452D | E | 1.6e-05 | 1.6e-05 | 1.2e-05 | 1.467E-05 | 2.309E-06 | 1.200E-05 | 1.600E-05 |
| 29025 | 1452 | S1452C | E | 1.2e-05 | 1.1e-05 | 1e-05 | 1.100E-05 | 1.000E-06 | 1.000E-05 | 1.200E-05 |
| 29026 | 1452 | S1452E | E | 8.6e-06 | 1.4e-05 | 1.4e-05 | 1.220E-05 | 3.118E-06 | 8.600E-06 | 1.400E-05 |
| 29027 | 1452 | S1452Q | E | 1.5e-05 | 9.9e-06 | 9.9e-06 | 1.160E-05 | 2.944E-06 | 9.900E-06 | 1.500E-05 |
| 29028 | 1452 | S1452G | E | 1.1e-05 | 1.8e-05 | 1e-05 | 1.300E-05 | 4.359E-06 | 1.000E-05 | 1.800E-05 |
| 29029 | 1452 | S1452H | E | 1.1e-05 | 1.6e-05 | 1.6e-05 | 1.433E-05 | 2.887E-06 | 1.100E-05 | 1.600E-05 |
| 29030 | 1452 | S1452I | E | 1.4e-05 | 1.7e-05 | 1.6e-05 | 1.567E-05 | 1.528E-06 | 1.400E-05 | 1.700E-05 |
| 29031 | 1452 | S1452L | E | 1.4e-05 | 1.4e-05 | 1e-05 | 1.267E-05 | 2.309E-06 | 1.000E-05 | 1.400E-05 |
| 29032 | 1452 | S1452K | E | 1.3e-05 | 1e-05 | 1.1e-05 | 1.133E-05 | 1.528E-06 | 1.000E-05 | 1.300E-05 |
| 29033 | 1452 | S1452M | E | 1.4e-05 | 1.2e-05 | 1.4e-05 | 1.333E-05 | 1.155E-06 | 1.200E-05 | 1.400E-05 |
| 29034 | 1452 | S1452F | E | 1.4e-05 | 1.2e-05 | 1.5e-05 | 1.367E-05 | 1.528E-06 | 1.200E-05 | 1.500E-05 |
| 29035 | 1452 | S1452P | E | 1.1e-05 | 1.4e-05 | 1.6e-05 | 1.367E-05 | 2.517E-06 | 1.100E-05 | 1.600E-05 |
| 29036 | 1452 | S1452S | E | 1e-05 | 1.7e-05 | 1.3e-05 | 1.333E-05 | 3.512E-06 | 1.000E-05 | 1.700E-05 |
| 29037 | 1452 | S1452T | E | 9.8e-06 | 1.7e-05 | 1.3e-05 | 1.327E-05 | 3.607E-06 | 9.800E-06 | 1.700E-05 |
| 29038 | 1452 | S1452W | E | 1.6e-05 | 1.1e-05 | 8.6e-06 | 1.187E-05 | 3.775E-06 | 8.600E-06 | 1.600E-05 |
| 29039 | 1452 | S1452Y | E | 9.8e-06 | 1.4e-05 | 1.3e-05 | 1.227E-05 | 2.194E-06 | 9.800E-06 | 1.400E-05 |
| 29040 | 1452 | S1452V | E | 1.5e-05 | 1.2e-05 | 8.2e-06 | 1.173E-05 | 3.408E-06 | 8.200E-06 | 1.500E-05 |
| 29041 | 1453 | G1453A | E | 9.9e-06 | 1.5e-05 | 1.4e-05 | 1.297E-05 | 2.702E-06 | 9.900E-06 | 1.500E-05 |
| 29042 | 1453 | G1453R | E | 1.5e-05 | 1.5e-05 | 1.6e-05 | 1.533E-05 | 5.774E-07 | 1.500E-05 | 1.600E-05 |
| 29043 | 1453 | G1453N | E | 1e-05 | 8.6e-06 | 1.5e-05 | 1.120E-05 | 3.365E-06 | 8.600E-06 | 1.500E-05 |
| 29044 | 1453 | G1453D | E | 1.5e-05 | 1.2e-05 | 8e-06 | 1.167E-05 | 3.512E-06 | 8.000E-06 | 1.500E-05 |
| 29045 | 1453 | G1453C | E | 1.1e-05 | 9.4e-06 | 1.5e-05 | 1.180E-05 | 2.884E-06 | 9.400E-06 | 1.500E-05 |
| 29046 | 1453 | G1453E | E | 1.3e-05 | 9.6e-06 | 1.5e-05 | 1.253E-05 | 2.730E-06 | 9.600E-06 | 1.500E-05 |
| 29047 | 1453 | G1453Q | E | 1.3e-05 | 1.8e-05 | 1.6e-05 | 1.567E-05 | 2.517E-06 | 1.300E-05 | 1.800E-05 |
| 29048 | 1453 | G1453G | E | 1.2e-05 | 9.9e-06 | 1.7e-05 | 1.297E-05 | 3.647E-06 | 9.900E-06 | 1.700E-05 |
| 29049 | 1453 | G1453H | E | 1.5e-05 | 1.2e-05 | 1.2e-05 | 1.300E-05 | 1.732E-06 | 1.200E-05 | 1.500E-05 |
| 29050 | 1453 | G1453I | E | 1.1e-05 | 1e-05 | 1.5e-05 | 1.200E-05 | 2.646E-06 | 1.000E-05 | 1.500E-05 |
| 29051 | 1453 | G1453L | E | 1.5e-05 | 1.4e-05 | 8.4e-06 | 1.247E-05 | 3.557E-06 | 8.400E-06 | 1.500E-05 |
| 29052 | 1453 | G1453K | E | 1.4e-05 | 1.5e-05 | 1.3e-05 | 1.400E-05 | 1.000E-06 | 1.300E-05 | 1.500E-05 |
| 29053 | 1453 | G1453M | E | 8.6e-06 | 1.4e-05 | 1.5e-05 | 1.253E-05 | 3.443E-06 | 8.600E-06 | 1.500E-05 |
| 29054 | 1453 | G1453F | E | 9.2e-06 | 1.2e-05 | 1.5e-05 | 1.207E-05 | 2.901E-06 | 9.200E-06 | 1.500E-05 |
| 29055 | 1453 | G1453P | E | 9.8e-06 | 1.4e-05 | 1.6e-05 | 1.327E-05 | 3.164E-06 | 9.800E-06 | 1.600E-05 |
| 29056 | 1453 | G1453S | E | 6.5e-06 | 1.3e-05 | 1.2e-05 | 1.050E-05 | 3.500E-06 | 6.500E-06 | 1.300E-05 |
| 29057 | 1453 | G1453T | E | 1.2e-05 | 5.7e-06 | 1.5e-05 | 1.090E-05 | 4.747E-06 | 5.700E-06 | 1.500E-05 |
| 29058 | 1453 | G1453W | E | 1.1e-05 | 1.6e-05 | 9.4e-06 | 1.213E-05 | 3.443E-06 | 9.400E-06 | 1.600E-05 |
| 29059 | 1453 | G1453Y | E | 1.4e-05 | 1.5e-05 | 1.6e-05 | 1.500E-05 | 1.000E-06 | 1.400E-05 | 1.600E-05 |
| 29060 | 1453 | G1453V | E | 1.2e-05 | 1.3e-05 | 1.2e-05 | 1.233E-05 | 5.774E-07 | 1.200E-05 | 1.300E-05 |
| 29061 | 1454 | L1454A | E | 1e-05 | 1.4e-05 | 9.6e-06 | 1.120E-05 | 2.433E-06 | 9.600E-06 | 1.400E-05 |
| 29062 | 1454 | L1454R | E | 1.9e-05 | 1.4e-05 | 1.6e-05 | 1.633E-05 | 2.517E-06 | 1.400E-05 | 1.900E-05 |
| 29063 | 1454 | L1454N | E | 1.5e-05 | 1.4e-05 | 1.2e-05 | 1.367E-05 | 1.528E-06 | 1.200E-05 | 1.500E-05 |
| 29064 | 1454 | L1454D | E | 1.2e-05 | 1e-05 | 1e-05 | 1.067E-05 | 1.155E-06 | 1.000E-05 | 1.200E-05 |
| 29065 | 1454 | L1454C | E | 1.6e-05 | 1.6e-05 | 1.3e-05 | 1.500E-05 | 1.732E-06 | 1.300E-05 | 1.600E-05 |
| 29066 | 1454 | L1454E | E | 9.8e-06 | 8.3e-06 | 1.3e-05 | 1.037E-05 | 2.401E-06 | 8.300E-06 | 1.300E-05 |
| 29067 | 1454 | L1454Q | E | 8.6e-06 | 1.5e-05 | 9.3e-06 | 1.097E-05 | 3.510E-06 | 8.600E-06 | 1.500E-05 |
| 29068 | 1454 | L1454G | E | 3.9e-05 | 1.2e-05 | 1.3e-05 | 2.133E-05 | 1.531E-05 | 1.200E-05 | 3.900E-05 |
| 29069 | 1454 | L1454H | E | 1.5e-05 | 1.2e-05 | 1.2e-05 | 1.300E-05 | 1.732E-06 | 1.200E-05 | 1.500E-05 |

| | | | | | | | | | | |
|---|---|---|---|---|---|---|---|---|---|---|
| 29070 | 1454 | L1454I | E | 1.1e-05 | 9.7e-06 | 1.5e-05 | 1.190E-05 | 2.762E-06 | 9.700E-06 | 1.500E-05 |
| 29071 | 1454 | L1454L | E | 1e-05 | 8.6e-06 | 1.7e-05 | 1.187E-05 | 4.500E-06 | 8.600E-06 | 1.700E-05 |
| 29072 | 1454 | L1454K | E | 1.4e-05 | 1.3e-05 | 1.4e-05 | 1.367E-05 | 5.774E-07 | 1.300E-05 | 1.400E-05 |
| 29073 | 1454 | L1454M | E | 1e-05 | 1.6e-05 | 8.7e-06 | 1.157E-05 | 3.894E-06 | 8.700E-06 | 1.600E-05 |
| 29074 | 1454 | L1454F | E | 1.4e-05 | 1.6e-05 | 8.4e-06 | 1.280E-05 | 3.940E-06 | 8.400E-06 | 1.600E-05 |
| 29075 | 1454 | L1454P | E | 1e-05 | 1.6e-05 | 1.2e-05 | 1.267E-05 | 3.055E-06 | 1.000E-05 | 1.600E-05 |
| 29076 | 1454 | L1454S | E | 1.4e-05 | 1.4e-05 | 1.1e-05 | 1.300E-05 | 1.732E-06 | 1.100E-05 | 1.400E-05 |
| 29077 | 1454 | L1454T | E | 1.1e-05 | 9.6e-06 | 1.5e-05 | 1.187E-05 | 2.802E-06 | 9.600E-06 | 1.500E-05 |
| 29078 | 1454 | L1454W | E | 1.6e-05 | 1e-05 | 1.3e-05 | 1.300E-05 | 3.000E-06 | 1.000E-05 | 1.600E-05 |
| 29079 | 1454 | L1454Y | E | 1.5e-05 | 1.3e-05 | 1.5e-05 | 1.433E-05 | 1.155E-06 | 1.300E-05 | 1.500E-05 |
| 29080 | 1454 | L1454V | E | 8.2e-06 | 1e-05 | 1.6e-05 | 1.140E-05 | 4.084E-06 | 8.200E-06 | 1.600E-05 |
| 29081 | 1455 | Y1455A | E | 9.8e-06 | 1.5e-05 | 1.5e-05 | 1.327E-05 | 3.002E-06 | 9.800E-06 | 1.500E-05 |
| 29082 | 1455 | Y1455R | E | 1.1e-05 | 8.8e-06 | 1.8e-05 | 1.260E-05 | 4.804E-06 | 8.800E-06 | 1.800E-05 |
| 29083 | 1455 | Y1455N | E | 9.3e-06 | 1.1e-05 | 1.5e-05 | 1.177E-05 | 2.926E-06 | 9.300E-06 | 1.500E-05 |
| 29084 | 1455 | Y1455D | E | 9.2e-06 | 1.5e-05 | 1.4e-05 | 1.273E-05 | 3.101E-06 | 9.200E-06 | 1.500E-05 |
| 29085 | 1455 | Y1455C | E | 8.7e-06 | 1.4e-05 | 1.4e-05 | 1.223E-05 | 3.060E-06 | 8.700E-06 | 1.400E-05 |
| 29086 | 1455 | Y1455E | E | 1.6e-05 | 1.3e-05 | 6.9e-06 | 1.197E-05 | 4.637E-06 | 6.900E-06 | 1.600E-05 |
| 29087 | 1455 | Y1455Q | E | 1.1e-05 | 1.5e-05 | 9e-06 | 1.167E-05 | 3.055E-06 | 9.000E-06 | 1.500E-05 |
| 29088 | 1455 | Y1455G | E | 1.5e-05 | 1.5e-05 | 1.1e-05 | 1.367E-05 | 2.309E-06 | 1.100E-05 | 1.500E-05 |
| 29089 | 1455 | Y1455H | E | 9.7e-06 | 1.4e-05 | 1.4e-05 | 1.257E-05 | 2.483E-06 | 9.700E-06 | 1.400E-05 |
| 29090 | 1455 | Y1455I | E | 7.4e-06 | 1.7e-05 | 1.8e-05 | 1.413E-05 | 5.853E-06 | 7.400E-06 | 1.800E-05 |
| 29091 | 1455 | Y1455L | E | 1.4e-05 | 1.3e-05 | 1.5e-05 | 1.400E-05 | 1.000E-06 | 1.300E-05 | 1.500E-05 |
| 29092 | 1455 | Y1455K | E | 1.2e-05 | 1.7e-05 | 1.1e-05 | 1.333E-05 | 3.215E-06 | 1.100E-05 | 1.700E-05 |
| 29093 | 1455 | Y1455M | E | 1.1e-05 | 9.9e-06 | 1.5e-05 | 1.197E-05 | 2.684E-06 | 9.900E-06 | 1.500E-05 |
| 29094 | 1455 | Y1455F | E | 1.5e-05 | 1.6e-05 | 1.5e-05 | 1.533E-05 | 5.774E-07 | 1.500E-05 | 1.600E-05 |
| 29095 | 1455 | Y1455P | E | 1.5e-05 | 1.5e-05 | 1e-05 | 1.333E-05 | 2.887E-06 | 1.000E-05 | 1.500E-05 |
| 29096 | 1455 | Y1455S | E | 1.3e-05 | 7.5e-06 | 1.1e-05 | 1.050E-05 | 2.784E-06 | 7.500E-06 | 1.300E-05 |
| 29097 | 1455 | Y1455T | E | 9.9e-06 | 1e-05 | 1.4e-05 | 1.130E-05 | 2.339E-06 | 9.900E-06 | 1.400E-05 |
| 29098 | 1455 | Y1455W | E | 1.7e-05 | 9.9e-06 | 8.6e-06 | 1.183E-05 | 4.521E-06 | 8.600E-06 | 1.700E-05 |
| 29099 | 1455 | Y1455Y | E | 8.2e-06 | 1e-05 | 1.1e-05 | 9.733E-06 | 1.419E-06 | 8.200E-06 | 1.100E-05 |
| 29100 | 1455 | Y1455V | E | 1.7e-05 | 1.6e-05 | 1e-05 | 1.433E-05 | 3.786E-06 | 1.000E-05 | 1.700E-05 |
| 29101 | 1456 | S1456A | E | 1.4e-05 | 1.1e-05 | 1.5e-05 | 1.333E-05 | 2.082E-06 | 1.100E-05 | 1.500E-05 |
| 29102 | 1456 | S1456R | E | 1.1e-05 | 1.6e-05 | 1.4e-05 | 1.367E-05 | 2.517E-06 | 1.100E-05 | 1.600E-05 |
| 29103 | 1456 | S1456N | E | 1.5e-05 | 1e-05 | 1.5e-05 | 1.333E-05 | 2.887E-06 | 1.000E-05 | 1.500E-05 |
| 29104 | 1456 | S1456D | E | 1.6e-05 | 1.5e-05 | 1.1e-05 | 1.400E-05 | 2.646E-06 | 1.100E-05 | 1.600E-05 |
| 29105 | 1456 | S1456C | E | 1.5e-05 | 1.1e-05 | 1.3e-05 | 1.300E-05 | 2.000E-06 | 1.100E-05 | 1.500E-05 |
| 29106 | 1456 | S1456E | E | 1.6e-05 | 7.2e-06 | 1.6e-05 | 1.307E-05 | 5.081E-06 | 7.200E-06 | 1.600E-05 |
| 29107 | 1456 | S1456Q | E | 9e-06 | 1.5e-05 | 1.6e-05 | 1.333E-05 | 3.786E-06 | 9.000E-06 | 1.600E-05 |
| 29108 | 1456 | S1456G | E | 8.7e-06 | 1.7e-05 | 1.8e-05 | 1.457E-05 | 5.105E-06 | 8.700E-06 | 1.800E-05 |
| 29109 | 1456 | S1456H | E | 1.6e-05 | 1.4e-05 | 1.2e-05 | 1.400E-05 | 2.000E-06 | 1.200E-05 | 1.600E-05 |
| 29110 | 1456 | S1456I | E | 1.1e-05 | 1.6e-05 | 1e-05 | 1.233E-05 | 3.215E-06 | 1.000E-05 | 1.600E-05 |
| 29111 | 1456 | S1456L | E | 1.1e-05 | 9.6e-06 | 1.4e-05 | 1.153E-05 | 2.248E-06 | 9.600E-06 | 1.400E-05 |
| 29112 | 1456 | S1456K | E | 1.6e-05 | 1.4e-05 | 1e-05 | 1.333E-05 | 3.055E-06 | 1.000E-05 | 1.600E-05 |
| 29113 | 1456 | S1456M | E | 1.6e-05 | 9.8e-06 | 1.1e-05 | 1.227E-05 | 3.288E-06 | 9.800E-06 | 1.600E-05 |
| 29114 | 1456 | S1456F | E | 8.3e-06 | 1.3e-05 | 1.6e-05 | 1.243E-05 | 3.881E-06 | 8.300E-06 | 1.600E-05 |
| 29115 | 1456 | S1456P | E | 9.8e-06 | 1.4e-05 | 1.1e-05 | 1.160E-05 | 2.163E-06 | 9.800E-06 | 1.400E-05 |
| 29116 | 1456 | S1456S | E | 1.1e-05 | 1.4e-05 | 1.5e-05 | 1.333E-05 | 2.082E-06 | 1.100E-05 | 1.500E-05 |
| 29117 | 1456 | S1456T | E | 1.4e-05 | 1.3e-05 | 1.1e-05 | 1.267E-05 | 1.528E-06 | 1.100E-05 | 1.400E-05 |
| 29118 | 1456 | S1456W | E | 9.7e-06 | 1.2e-05 | 1.3e-05 | 1.157E-05 | 1.692E-06 | 9.700E-06 | 1.300E-05 |
| 29119 | 1456 | S1456Y | E | 1.1e-05 | 1e-05 | 9.9e-06 | 1.030E-05 | 6.083E-07 | 9.900E-06 | 1.100E-05 |
| 29120 | 1456 | S1456V | E | 1.4e-05 | 1.8e-05 | 1.1e-05 | 1.433E-05 | 3.512E-06 | 1.100E-05 | 1.800E-05 |
| 29121 | 1457 | L1457A | E | 1.1e-05 | 1.5e-05 | 1.5e-05 | 1.367E-05 | 2.309E-06 | 1.100E-05 | 1.500E-05 |
| 29122 | 1457 | L1457R | E | 1.5e-05 | 1.2e-05 | 1.6e-05 | 1.433E-05 | 2.082E-06 | 1.200E-05 | 1.600E-05 |
| 29123 | 1457 | L1457N | E | 1.1e-05 | 9.4e-06 | 1.4e-05 | 1.147E-05 | 2.335E-06 | 9.400E-06 | 1.400E-05 |
| 29124 | 1457 | L1457D | E | 9.6e-06 | 1.2e-05 | 1.1e-05 | 1.087E-05 | 1.206E-06 | 9.600E-06 | 1.200E-05 |
| 29125 | 1457 | L1457C | E | 1e-05 | 1e-05 | 1.4e-05 | 1.133E-05 | 2.309E-06 | 1.000E-05 | 1.400E-05 |
| 29126 | 1457 | L1457E | E | 1.4e-05 | 1e-05 | 1.5e-05 | 1.300E-05 | 2.646E-06 | 1.000E-05 | 1.500E-05 |
| 29127 | 1457 | L1457Q | E | 9.8e-06 | 1e-05 | 1.4e-05 | 1.127E-05 | 2.369E-06 | 9.800E-06 | 1.400E-05 |
| 29128 | 1457 | L1457G | E | 1.5e-05 | 8.3e-06 | 1.5e-05 | 1.277E-05 | 3.868E-06 | 8.300E-06 | 1.500E-05 |
| 29129 | 1457 | L1457H | E | 1.6e-05 | 9.3e-06 | 9.8e-06 | 1.170E-05 | 3.732E-06 | 9.300E-06 | 1.600E-05 |
| 29130 | 1457 | L1457I | E | 1.6e-05 | 1.6e-05 | 1.2e-05 | 1.467E-05 | 2.309E-06 | 1.200E-05 | 1.600E-05 |
| 29131 | 1457 | L1457L | E | 1.5e-05 | 9.1e-06 | 4.9e-06 | 9.667E-06 | 5.074E-06 | 4.900E-06 | 1.500E-05 |
| 29132 | 1457 | L1457K | E | 1.1e-05 | 9.5e-06 | 1.3e-05 | 1.117E-05 | 1.756E-06 | 9.500E-06 | 1.300E-05 |
| 29133 | 1457 | L1457M | E | 1.6e-05 | 1.1e-05 | 1e-05 | 1.233E-05 | 3.215E-06 | 1.000E-05 | 1.600E-05 |
| 29134 | 1457 | L1457F | E | 1.1e-05 | 1e-05 | 1.7e-05 | 1.267E-05 | 3.786E-06 | 1.000E-05 | 1.700E-05 |
| 29135 | 1457 | L1457P | E | 1.5e-05 | 1.4e-05 | 8.5e-06 | 1.250E-05 | 3.500E-06 | 8.500E-06 | 1.500E-05 |
| 29136 | 1457 | L1457S | E | 1.5e-05 | 1.6e-05 | 1.5e-05 | 1.533E-05 | 5.774E-07 | 1.500E-05 | 1.600E-05 |
| 29137 | 1457 | L1457T | E | 1.4e-05 | 1.5e-05 | 1.5e-05 | 1.467E-05 | 5.774E-07 | 1.400E-05 | 1.500E-05 |
| 29138 | 1457 | L1457W | E | 1.4e-05 | 1.4e-05 | 1.1e-05 | 1.300E-05 | 1.732E-06 | 1.100E-05 | 1.400E-05 |
| 29139 | 1457 | L1457Y | E | 1.6e-05 | 1.6e-05 | 1.1e-05 | 1.433E-05 | 2.887E-06 | 1.100E-05 | 1.600E-05 |
| 29140 | 1457 | L1457V | E | 1.3e-05 | 2.1e-05 | 1.8e-05 | 1.733E-05 | 4.041E-06 | 1.300E-05 | 2.100E-05 |
| 29141 | 1458 | S1458A | E | 1.1e-05 | 1.3e-05 | 1.7e-05 | 1.367E-05 | 3.055E-06 | 1.100E-05 | 1.700E-05 |
| 29142 | 1458 | S1458R | E | 1.5e-05 | 1.6e-05 | 1.5e-05 | 1.533E-05 | 5.774E-07 | 1.500E-05 | 1.600E-05 |
| 29143 | 1458 | S1458N | E | 1.1e-05 | 1.4e-05 | 6.5e-06 | 1.050E-05 | 3.775E-06 | 6.500E-06 | 1.400E-05 |
| 29144 | 1458 | S1458D | E | 1.9e-05 | 1.1e-05 | 1.1e-05 | 1.367E-05 | 4.619E-06 | 1.100E-05 | 1.900E-05 |
| 29145 | 1458 | S1458C | E | 7.5e-06 | 1.3e-05 | 1.1e-05 | 1.050E-05 | 2.784E-06 | 7.500E-06 | 1.300E-05 |
| 29146 | 1458 | S1458E | E | 1.2e-05 | 1.4e-05 | 1e-05 | 1.200E-05 | 2.000E-06 | 1.000E-05 | 1.400E-05 |
| 29147 | 1458 | S1458Q | E | 1.3e-05 | 1.1e-05 | 1e-05 | 1.133E-05 | 1.528E-06 | 1.000E-05 | 1.300E-05 |
| 29148 | 1458 | S1458G | E | 1e-05 | 1.5e-05 | 9.8e-06 | 1.160E-05 | 2.946E-06 | 9.800E-06 | 1.500E-05 |
| 29149 | 1458 | S1458H | E | 9.3e-06 | 1.5e-05 | 1.2e-05 | 1.210E-05 | 2.851E-06 | 9.300E-06 | 1.500E-05 |
| 29150 | 1458 | S1458I | E | 1.5e-05 | 1.4e-05 | 1.5e-05 | 1.467E-05 | 5.774E-07 | 1.400E-05 | 1.500E-05 |
| 29151 | 1458 | S1458L | E | 1.4e-05 | 1.1e-05 | 1.6e-05 | 1.367E-05 | 2.517E-06 | 1.100E-05 | 1.600E-05 |
| 29152 | 1458 | S1458K | E | 1.9e-05 | 1.3e-05 | 1e-05 | 1.400E-05 | 4.583E-06 | 1.000E-05 | 1.900E-05 |
| 29153 | 1458 | S1458M | E | 1e-05 | 1.3e-05 | 1.2e-05 | 1.167E-05 | 1.528E-06 | 1.000E-05 | 1.300E-05 |
| 29154 | 1458 | S1458F | E | 1.9e-05 | 1e-05 | 1.3e-05 | 1.400E-05 | 4.583E-06 | 1.000E-05 | 1.900E-05 |

| | | | | | | | | | | |
|---|---|---|---|---|---|---|---|---|---|---|
| 29155 | 1458 | S1458P | E | 1.6e-05 | 9.5e-06 | 1.6e-05 | 1.383E-05 | 3.753E-06 | 9.500E-06 | 1.600E-05 |
| 29156 | 1458 | S1458S | E | 1.7e-05 | 1.4e-05 | 1.7e-05 | 1.600E-05 | 1.732E-06 | 1.400E-05 | 1.700E-05 |
| 29157 | 1458 | S1458T | E | 7.6e-06 | 1e-05 | 1e-05 | 9.200E-06 | 1.386E-06 | 7.600E-06 | 1.000E-05 |
| 29158 | 1458 | S1458W | E | 9.4e-06 | 1.4e-05 | 6.9e-06 | 1.010E-05 | 3.601E-06 | 6.900E-06 | 1.400E-05 |
| 29159 | 1458 | S1458Y | E | 1.5e-05 | 1.2e-05 | 1e-05 | 1.233E-05 | 2.517E-06 | 1.000E-05 | 1.500E-05 |
| 29160 | 1458 | S1458V | E | 9.9e-06 | 9.9e-06 | 1.5e-05 | 1.160E-05 | 2.944E-06 | 9.900E-06 | 1.500E-05 |
| 29161 | 1459 | S1459A | E | 6.9e-06 | 1.1e-05 | 1.3e-05 | 1.030E-05 | 3.110E-06 | 6.900E-06 | 1.300E-05 |
| 29162 | 1459 | S1459R | E | 1.1e-05 | 9.8e-06 | 1e-05 | 1.027E-05 | 6.429E-07 | 9.800E-06 | 1.100E-05 |
| 29163 | 1459 | S1459N | E | 1.1e-05 | 1.1e-05 | 1.5e-05 | 1.233E-05 | 2.309E-06 | 1.100E-05 | 1.500E-05 |
| 29164 | 1459 | S1459D | E | 1.6e-05 | 1.5e-05 | 6.7e-06 | 1.257E-05 | 5.105E-06 | 6.700E-06 | 1.600E-05 |
| 29165 | 1459 | S1459C | E | 1.7e-05 | 1.6e-05 | 1.1e-05 | 1.467E-05 | 3.215E-06 | 1.100E-05 | 1.700E-05 |
| 29166 | 1459 | S1459E | E | 1.4e-05 | 8.9e-06 | 9.2e-06 | 1.070E-05 | 2.862E-06 | 8.900E-06 | 1.400E-05 |
| 29167 | 1459 | S1459Q | E | 1e-05 | 1.4e-05 | 1.6e-05 | 1.333E-05 | 3.055E-06 | 1.000E-05 | 1.600E-05 |
| 29168 | 1459 | S1459G | E | 7.9e-06 | 9.8e-06 | 1.2e-05 | 9.900E-06 | 2.052E-06 | 7.900E-06 | 1.200E-05 |
| 29169 | 1459 | S1459H | E | 1.5e-05 | 1.3e-05 | 1.1e-05 | 1.300E-05 | 2.000E-06 | 1.100E-05 | 1.500E-05 |
| 29170 | 1459 | S1459I | E | 1e-05 | 1.4e-05 | 1.2e-05 | 1.200E-05 | 2.000E-06 | 1.000E-05 | 1.400E-05 |
| 29171 | 1459 | S1459L | E | 1e-05 | 1.5e-05 | 1.4e-05 | 1.300E-05 | 2.646E-06 | 1.000E-05 | 1.500E-05 |
| 29172 | 1459 | S1459K | E | 9.8e-06 | 1.6e-05 | 1.5e-05 | 1.360E-05 | 3.329E-06 | 9.800E-06 | 1.600E-05 |
| 29173 | 1459 | S1459M | E | 1.8e-05 | 1.1e-05 | 1.5e-05 | 1.467E-05 | 3.512E-06 | 1.100E-05 | 1.800E-05 |
| 29174 | 1459 | S1459F | E | 1.4e-05 | 1.4e-05 | 9.5e-06 | 1.250E-05 | 2.598E-06 | 9.500E-06 | 1.400E-05 |
| 29175 | 1459 | S1459P | E | 1.2e-05 | 1e-05 | 1.4e-05 | 1.200E-05 | 2.000E-06 | 1.000E-05 | 1.400E-05 |
| 29176 | 1459 | S1459S | E | 1.6e-05 | 1.1e-05 | 1.1e-05 | 1.267E-05 | 2.887E-06 | 1.100E-05 | 1.600E-05 |
| 29177 | 1459 | S1459T | E | 9.3e-06 | 1.3e-05 | 1.5e-05 | 1.243E-05 | 2.892E-06 | 9.300E-06 | 1.500E-05 |
| 29178 | 1459 | S1459W | E | 8.9e-06 | 7.1e-06 | 1.1e-05 | 9.000E-06 | 1.952E-06 | 7.100E-06 | 1.100E-05 |
| 29179 | 1459 | S1459Y | E | 1e-05 | 1.8e-05 | 9e-06 | 1.233E-05 | 4.933E-06 | 9.000E-06 | 1.800E-05 |
| 29180 | 1459 | S1459V | E | 2.2e-05 | 1.4e-05 | 9.2e-06 | 1.507E-05 | 6.466E-06 | 9.200E-06 | 2.200E-05 |
| 29181 | 1460 | V1460A | E | 1.1e-05 | 1.2e-05 | 1.6e-05 | 1.300E-05 | 2.646E-06 | 1.100E-05 | 1.600E-05 |
| 29182 | 1460 | V1460R | E | 1.1e-05 | 1.5e-05 | 9.4e-06 | 1.180E-05 | 2.884E-06 | 9.400E-06 | 1.500E-05 |
| 29183 | 1460 | V1460N | E | 8.5e-06 | 1.4e-05 | 1.1e-05 | 1.117E-05 | 2.754E-06 | 8.500E-06 | 1.400E-05 |
| 29184 | 1460 | V1460D | E | 1.1e-05 | 1.5e-05 | 1.3e-05 | 1.300E-05 | 2.000E-06 | 1.100E-05 | 1.500E-05 |
| 29185 | 1460 | V1460C | E | 1.1e-05 | 1.5e-05 | 1.6e-05 | 1.400E-05 | 2.646E-06 | 1.100E-05 | 1.600E-05 |
| 29186 | 1460 | V1460E | E | 1.7e-05 | 1.5e-05 | 8.6e-06 | 1.353E-05 | 4.388E-06 | 8.600E-06 | 1.700E-05 |
| 29187 | 1460 | V1460Q | E | 1.5e-05 | 9.7e-06 | 1.5e-05 | 1.323E-05 | 3.060E-06 | 9.700E-06 | 1.500E-05 |
| 29188 | 1460 | V1460G | E | 1.6e-05 | 1e-05 | 7.7e-06 | 1.123E-05 | 4.285E-06 | 7.700E-06 | 1.600E-05 |
| 29189 | 1460 | V1460H | E | 1.3e-05 | 1.2e-05 | 1e-05 | 1.167E-05 | 1.528E-06 | 1.000E-05 | 1.300E-05 |
| 29190 | 1460 | V1460I | E | 1e-05 | 1.4e-05 | 1.3e-05 | 1.233E-05 | 2.082E-06 | 1.000E-05 | 1.400E-05 |
| 29191 | 1460 | V1460L | E | 1.5e-05 | 1.5e-05 | 9.6e-06 | 1.320E-05 | 3.118E-06 | 9.600E-06 | 1.500E-05 |
| 29192 | 1460 | V1460K | E | 1.1e-05 | 1.1e-05 | 1.5e-05 | 1.233E-05 | 2.309E-06 | 1.100E-05 | 1.500E-05 |
| 29193 | 1460 | V1460M | E | 1.5e-05 | 9.9e-06 | 1.5e-05 | 1.330E-05 | 2.944E-06 | 9.900E-06 | 1.500E-05 |
| 29194 | 1460 | V1460F | E | 1.4e-05 | 9.7e-06 | 1e-05 | 1.123E-05 | 2.401E-06 | 9.700E-06 | 1.400E-05 |
| 29195 | 1460 | V1460P | E | 1.1e-05 | 1.4e-05 | 1.6e-05 | 1.367E-05 | 2.517E-06 | 1.100E-05 | 1.600E-05 |
| 29196 | 1460 | V1460S | E | 1.4e-05 | 1.3e-05 | 1.3e-05 | 1.333E-05 | 5.774E-07 | 1.300E-05 | 1.400E-05 |
| 29197 | 1460 | V1460T | E | 1.2e-05 | 1e-05 | 1.3e-05 | 1.167E-05 | 1.528E-06 | 1.000E-05 | 1.300E-05 |
| 29198 | 1460 | V1460W | E | 1.4e-05 | 1.1e-05 | 1.7e-05 | 1.400E-05 | 3.000E-06 | 1.100E-05 | 1.700E-05 |
| 29199 | 1460 | V1460Y | E | 1.4e-05 | 9.5e-06 | 1.7e-05 | 1.350E-05 | 3.775E-06 | 9.500E-06 | 1.700E-05 |
| 29200 | 1460 | V1460V | E | 1.1e-05 | 9.4e-06 | 1e-05 | 1.013E-05 | 8.083E-07 | 9.400E-06 | 1.100E-05 |
| 29201 | 1461 | V1461A | E | 1.6e-05 | 1e-05 | 1.6e-05 | 1.400E-05 | 3.464E-06 | 1.000E-05 | 1.600E-05 |
| 29202 | 1461 | V1461R | E | 8.7e-06 | 1.2e-05 | 1.2e-05 | 1.090E-05 | 1.905E-06 | 8.700E-06 | 1.200E-05 |
| 29203 | 1461 | V1461N | E | 1.2e-05 | 1.1e-05 | 1.5e-05 | 1.267E-05 | 2.082E-06 | 1.100E-05 | 1.500E-05 |
| 29204 | 1461 | V1461D | E | 1e-05 | 1.6e-05 | 9e-06 | 1.167E-05 | 3.786E-06 | 9.000E-06 | 1.600E-05 |
| 29205 | 1461 | V1461C | E | 9.7e-06 | 1.1e-05 | 1.7e-05 | 1.257E-05 | 3.894E-06 | 9.700E-06 | 1.700E-05 |
| 29206 | 1461 | V1461E | E | 1.4e-05 | 9.7e-06 | 9.7e-06 | 1.113E-05 | 2.483E-06 | 9.700E-06 | 1.400E-05 |
| 29207 | 1461 | V1461Q | E | 9.2e-06 | 1.5e-05 | 9.8e-06 | 1.133E-05 | 3.190E-06 | 9.200E-06 | 1.500E-05 |
| 29208 | 1461 | V1461G | E | 1e-05 | 1.4e-05 | 1.1e-05 | 1.167E-05 | 2.082E-06 | 1.000E-05 | 1.400E-05 |
| 29209 | 1461 | V1461H | E | 1.4e-05 | 2.3e-05 | 1e-05 | 1.567E-05 | 6.658E-06 | 1.000E-05 | 2.300E-05 |
| 29210 | 1461 | V1461I | E | 1.6e-05 | 1.4e-05 | 1e-05 | 1.333E-05 | 3.055E-06 | 1.000E-05 | 1.600E-05 |
| 29211 | 1461 | V1461L | E | 1.4e-05 | 1e-05 | 1.1e-05 | 1.167E-05 | 2.082E-06 | 1.000E-05 | 1.400E-05 |
| 29212 | 1461 | V1461K | E | 1.5e-05 | 1.1e-05 | 1.5e-05 | 1.367E-05 | 2.309E-06 | 1.100E-05 | 1.500E-05 |
| 29213 | 1461 | V1461M | E | 1.3e-05 | 1.2e-05 | 9.3e-06 | 1.143E-05 | 1.914E-06 | 9.300E-06 | 1.300E-05 |
| 29214 | 1461 | V1461F | E | 9.7e-06 | 1.6e-05 | 1.1e-05 | 1.223E-05 | 3.326E-06 | 9.700E-06 | 1.600E-05 |
| 29215 | 1461 | V1461P | E | 1.3e-05 | 1.4e-05 | 7.9e-06 | 1.163E-05 | 3.272E-06 | 7.900E-06 | 1.400E-05 |
| 29216 | 1461 | V1461S | E | 1.4e-05 | 2.7e-05 | 1.3e-05 | 1.800E-05 | 7.810E-06 | 1.300E-05 | 2.700E-05 |
| 29217 | 1461 | V1461T | E | 1.2e-05 | 1.1e-05 | 1.4e-05 | 1.233E-05 | 1.528E-06 | 1.100E-05 | 1.400E-05 |
| 29218 | 1461 | V1461W | E | 1.4e-05 | 9.2e-06 | 1.3e-05 | 1.207E-05 | 2.532E-06 | 9.200E-06 | 1.400E-05 |
| 29219 | 1461 | V1461Y | E | 1.1e-05 | 1.3e-05 | 1.1e-05 | 1.167E-05 | 1.155E-06 | 1.100E-05 | 1.300E-05 |
| 29220 | 1461 | V1461V | E | 1e-05 | 1.5e-05 | 1.1e-05 | 1.200E-05 | 2.646E-06 | 1.000E-05 | 1.500E-05 |
| 29221 | 1462 | T1462A | E | 1.6e-05 | 1.2e-05 | 1.3e-05 | 1.367E-05 | 2.082E-06 | 1.200E-05 | 1.600E-05 |
| 29222 | 1462 | T1462R | E | 1.2e-05 | 1e-05 | 1.3e-05 | 1.167E-05 | 1.528E-06 | 1.000E-05 | 1.300E-05 |
| 29223 | 1462 | T1462N | E | 2.4e-05 | 1.5e-05 | 9.3e-06 | 1.610E-05 | 7.411E-06 | 9.300E-06 | 2.400E-05 |
| 29224 | 1462 | T1462D | E | 1.5e-05 | 1.3e-05 | 1.7e-05 | 1.500E-05 | 2.000E-06 | 1.300E-05 | 1.700E-05 |
| 29225 | 1462 | T1462C | E | 1.5e-05 | 8.7e-06 | 1.5e-05 | 1.290E-05 | 3.637E-06 | 8.700E-06 | 1.500E-05 |
| 29226 | 1462 | T1462E | E | 1.7e-05 | 1.6e-05 | 1.7e-05 | 1.667E-05 | 5.774E-07 | 1.600E-05 | 1.700E-05 |
| 29227 | 1462 | T1462Q | E | 1.5e-05 | 1.5e-05 | 1.6e-05 | 1.533E-05 | 5.774E-07 | 1.500E-05 | 1.600E-05 |
| 29228 | 1462 | T1462G | E | 1.1e-05 | 1.7e-05 | 1.2e-05 | 1.333E-05 | 3.215E-06 | 1.100E-05 | 1.700E-05 |
| 29229 | 1462 | T1462H | E | 1.6e-05 | 1.4e-05 | 1.6e-05 | 1.533E-05 | 1.155E-06 | 1.400E-05 | 1.600E-05 |
| 29230 | 1462 | T1462I | E | 1.5e-05 | 1.2e-05 | 1.1e-05 | 1.267E-05 | 2.082E-06 | 1.100E-05 | 1.500E-05 |
| 29231 | 1462 | T1462L | E | 1.7e-05 | 1.2e-05 | 1.1e-05 | 1.333E-05 | 3.215E-06 | 1.100E-05 | 1.700E-05 |
| 29232 | 1462 | T1462K | E | 1.7e-05 | 1.6e-05 | 9.3e-06 | 1.410E-05 | 4.187E-06 | 9.300E-06 | 1.700E-05 |
| 29233 | 1462 | T1462M | E | 1.1e-05 | 1.2e-05 | 1.4e-05 | 1.233E-05 | 1.528E-06 | 1.100E-05 | 1.400E-05 |
| 29234 | 1462 | T1462F | E | 1.4e-05 | 7.8e-06 | 1.7e-05 | 1.293E-05 | 4.692E-06 | 7.800E-06 | 1.700E-05 |
| 29235 | 1462 | T1462P | E | 1.6e-05 | 1.7e-05 | 1.4e-05 | 1.567E-05 | 1.528E-06 | 1.400E-05 | 1.700E-05 |
| 29236 | 1462 | T1462S | E | 1.7e-05 | 1.5e-05 | 1.3e-05 | 1.500E-05 | 2.000E-06 | 1.300E-05 | 1.700E-05 |
| 29237 | 1462 | T1462T | E | 1.3e-05 | 2e-05 | 1e-05 | 1.433E-05 | 5.132E-06 | 1.000E-05 | 2.000E-05 |
| 29238 | 1462 | T1462W | E | 1.4e-05 | 1.5e-05 | 1.5e-05 | 1.467E-05 | 5.774E-07 | 1.400E-05 | 1.500E-05 |
| 29239 | 1462 | T1462Y | E | 8.6e-06 | 1.1e-05 | 1.3e-05 | 1.087E-05 | 2.203E-06 | 8.600E-06 | 1.300E-05 |

| | | | | | | | | | | |
|---|---|---|---|---|---|---|---|---|---|---|
| 29240 | 1462 | T1462V | E | 9.7e-06 | 1.3e-05 | 1.5e-05 | 1.257E-05 | 2.676E-06 | 9.700E-06 | 1.500E-05 |
| 29241 | 1463 | V1463A | E | 1.2e-05 | 1.1e-05 | 1.4e-05 | 1.233E-05 | 1.528E-06 | 1.100E-05 | 1.400E-05 |
| 29242 | 1463 | V1463R | E | 9.9e-06 | 1.1e-05 | 1.4e-05 | 1.163E-05 | 2.122E-06 | 9.900E-06 | 1.400E-05 |
| 29243 | 1463 | V1463N | E | 1.5e-05 | 1.6e-05 | 1.4e-05 | 1.500E-05 | 1.000E-06 | 1.400E-05 | 1.600E-05 |
| 29244 | 1463 | V1463D | E | 1.2e-05 | 1.3e-05 | 1.6e-05 | 1.367E-05 | 2.082E-06 | 1.200E-05 | 1.600E-05 |
| 29245 | 1463 | V1463C | E | 9.7e-06 | 9.7e-06 | 1e-05 | 9.800E-06 | 1.732E-07 | 9.700E-06 | 1.000E-05 |
| 29246 | 1463 | V1463E | E | 8.3e-06 | 1.5e-05 | 9.4e-06 | 1.090E-05 | 3.593E-06 | 8.300E-06 | 1.500E-05 |
| 29247 | 1463 | V1463Q | E | 8.9e-06 | 1.3e-05 | 1.6e-05 | 1.263E-05 | 3.564E-06 | 8.900E-06 | 1.600E-05 |
| 29248 | 1463 | V1463G | E | 9.5e-06 | 1.1e-05 | 1e-05 | 1.017E-05 | 7.638E-07 | 9.500E-06 | 1.100E-05 |
| 29249 | 1463 | V1463H | E | 1.2e-05 | 1.5e-05 | 8.5e-06 | 1.183E-05 | 3.253E-06 | 8.500E-06 | 1.500E-05 |
| 29250 | 1463 | V1463I | E | 1.6e-05 | 1.3e-05 | 9.4e-06 | 1.280E-05 | 3.305E-06 | 9.400E-06 | 1.600E-05 |
| 29251 | 1463 | V1463L | E | 1.4e-05 | 1.3e-05 | 1.1e-05 | 1.267E-05 | 1.528E-06 | 1.100E-05 | 1.400E-05 |
| 29252 | 1463 | V1463K | E | 1.4e-05 | 1.6e-05 | 1.3e-05 | 1.433E-05 | 1.528E-06 | 1.300E-05 | 1.600E-05 |
| 29253 | 1463 | V1463M | E | 1.2e-05 | 1.1e-05 | 1.5e-05 | 1.267E-05 | 2.082E-06 | 1.100E-05 | 1.500E-05 |
| 29254 | 1463 | V1463F | E | 1.6e-05 | 1.7e-05 | 1.4e-05 | 1.567E-05 | 1.528E-06 | 1.400E-05 | 1.700E-05 |
| 29255 | 1463 | V1463P | E | 7.2e-06 | 1.6e-05 | 9.8e-06 | 1.100E-05 | 4.521E-06 | 7.200E-06 | 1.600E-05 |
| 29256 | 1463 | V1463S | E | 1.4e-05 | 1.5e-05 | 1.4e-05 | 1.433E-05 | 5.774E-07 | 1.400E-05 | 1.500E-05 |
| 29257 | 1463 | V1463T | E | 1.1e-05 | 8.2e-06 | 1.5e-05 | 1.140E-05 | 3.418E-06 | 8.200E-06 | 1.500E-05 |
| 29258 | 1463 | V1463W | E | 1.5e-05 | 9.9e-06 | 1e-05 | 1.163E-05 | 2.916E-06 | 9.900E-06 | 1.500E-05 |
| 29259 | 1463 | V1463Y | E | 1.3e-05 | 1.5e-05 | 9.8e-06 | 1.260E-05 | 2.623E-06 | 9.800E-06 | 1.500E-05 |
| 29260 | 1463 | V1463V | E | 1.6e-05 | 1.1e-05 | 1e-05 | 1.233E-05 | 3.215E-06 | 1.000E-05 | 1.600E-05 |
| 29261 | 1464 | P1464A | E | 1.1e-05 | 1.5e-05 | 9.8e-06 | 1.193E-05 | 2.723E-06 | 9.800E-06 | 1.500E-05 |
| 29262 | 1464 | P1464R | E | 8.2e-06 | 1.1e-05 | 1.1e-05 | 1.007E-05 | 1.617E-06 | 8.200E-06 | 1.100E-05 |
| 29263 | 1464 | P1464N | E | 1.4e-05 | 1.4e-05 | 1.6e-05 | 1.467E-05 | 1.155E-06 | 1.400E-05 | 1.600E-05 |
| 29264 | 1464 | P1464D | E | 1e-05 | 1.6e-05 | 1.6e-05 | 1.400E-05 | 3.464E-06 | 1.000E-05 | 1.600E-05 |
| 29265 | 1464 | P1464C | E | 1e-05 | 1.3e-05 | 9.8e-06 | 1.093E-05 | 1.793E-06 | 9.800E-06 | 1.300E-05 |
| 29266 | 1464 | P1464E | E | 8.3e-06 | 1.1e-05 | 8.2e-06 | 9.167E-06 | 1.589E-06 | 8.200E-06 | 1.100E-05 |
| 29267 | 1464 | P1464Q | E | 8.3e-06 | 1.2e-05 | 1.3e-05 | 1.110E-05 | 2.476E-06 | 8.300E-06 | 1.300E-05 |
| 29268 | 1464 | P1464G | E | 1.6e-05 | 1e-05 | 1.1e-05 | 1.233E-05 | 3.215E-06 | 1.000E-05 | 1.600E-05 |
| 29269 | 1464 | P1464H | E | 8.5e-06 | 1.4e-05 | 1.6e-05 | 1.283E-05 | 3.884E-06 | 8.500E-06 | 1.600E-05 |
| 29270 | 1464 | P1464I | E | 1e-05 | 1.7e-05 | 1.5e-05 | 1.400E-05 | 3.606E-06 | 1.000E-05 | 1.700E-05 |
| 29271 | 1464 | P1464L | E | 1.1e-05 | 1.4e-05 | 9.9e-06 | 1.163E-05 | 2.122E-06 | 9.900E-06 | 1.400E-05 |
| 29272 | 1464 | P1464K | E | 1.7e-05 | 1.5e-05 | 1.2e-05 | 1.467E-05 | 2.517E-06 | 1.200E-05 | 1.700E-05 |
| 29273 | 1464 | P1464M | E | 1.4e-05 | 1.2e-05 | 1.3e-05 | 1.300E-05 | 1.000E-06 | 1.200E-05 | 1.400E-05 |
| 29274 | 1464 | P1464F | E | 1.4e-05 | 1.6e-05 | 1.6e-05 | 1.533E-05 | 1.155E-06 | 1.400E-05 | 1.600E-05 |
| 29275 | 1464 | P1464P | E | 1.1e-05 | 1.5e-05 | 1.5e-05 | 1.367E-05 | 2.309E-06 | 1.100E-05 | 1.500E-05 |
| 29276 | 1464 | P1464S | E | 1e-05 | 1.5e-05 | 9.5e-06 | 1.150E-05 | 3.041E-06 | 9.500E-06 | 1.500E-05 |
| 29277 | 1464 | P1464T | E | 1.6e-05 | 1.4e-05 | 1.5e-05 | 1.500E-05 | 1.000E-06 | 1.400E-05 | 1.600E-05 |
| 29278 | 1464 | P1464W | E | 1.2e-05 | 1e-05 | 1.1e-05 | 1.100E-05 | 1.000E-06 | 1.000E-05 | 1.200E-05 |
| 29279 | 1464 | P1464Y | E | 9.7e-06 | 1e-05 | 1.4e-05 | 1.123E-05 | 2.401E-06 | 9.700E-06 | 1.400E-05 |
| 29280 | 1464 | P1464V | E | 1.7e-05 | 1.3e-05 | 9e-06 | 1.300E-05 | 4.000E-06 | 9.000E-06 | 1.700E-05 |
| 29281 | 1465 | S1465A | E | 1.6e-05 | 1.2e-05 | 1.4e-05 | 1.400E-05 | 2.000E-06 | 1.200E-05 | 1.600E-05 |
| 29282 | 1465 | S1465R | E | 1.5e-05 | 9.4e-06 | 1.9e-05 | 1.447E-05 | 4.822E-06 | 9.400E-06 | 1.900E-05 |
| 29283 | 1465 | S1465N | E | 9.6e-06 | 8e-06 | 1e-05 | 9.200E-06 | 1.058E-06 | 8.000E-06 | 1.000E-05 |
| 29284 | 1465 | S1465D | E | 1.1e-05 | 1.6e-05 | 1.5e-05 | 1.400E-05 | 2.646E-06 | 1.100E-05 | 1.600E-05 |
| 29285 | 1465 | S1465C | E | 1.5e-05 | 1.5e-05 | 1e-05 | 1.333E-05 | 2.887E-06 | 1.000E-05 | 1.500E-05 |
| 29286 | 1465 | S1465E | E | 1.5e-05 | 1.2e-05 | 1.1e-05 | 1.267E-05 | 2.082E-06 | 1.100E-05 | 1.500E-05 |
| 29287 | 1465 | S1465Q | E | 1.6e-05 | 1e-05 | 1.1e-05 | 1.233E-05 | 3.215E-06 | 1.000E-05 | 1.600E-05 |
| 29288 | 1465 | S1465G | E | 1.5e-05 | 1.1e-05 | 1.5e-05 | 1.367E-05 | 2.309E-06 | 1.100E-05 | 1.500E-05 |
| 29289 | 1465 | S1465H | E | 1e-05 | 1e-05 | 9.1e-06 | 9.700E-06 | 5.196E-07 | 9.100E-06 | 1.000E-05 |
| 29290 | 1465 | S1465I | E | 1.4e-05 | 8.5e-06 | 1.6e-05 | 1.283E-05 | 3.884E-06 | 8.500E-06 | 1.600E-05 |
| 29291 | 1465 | S1465L | E | 1.5e-05 | 1.2e-05 | 1.5e-05 | 1.400E-05 | 1.732E-06 | 1.200E-05 | 1.500E-05 |
| 29292 | 1465 | S1465K | E | 7.5e-06 | 1.4e-05 | 1.1e-05 | 1.083E-05 | 3.253E-06 | 7.500E-06 | 1.400E-05 |
| 29293 | 1465 | S1465M | E | 8.8e-06 | 1.1e-05 | 9e-06 | 9.600E-06 | 1.217E-06 | 8.800E-06 | 1.100E-05 |
| 29294 | 1465 | S1465F | E | 9e-06 | 1.3e-05 | 1.2e-05 | 1.133E-05 | 2.082E-06 | 9.000E-06 | 1.300E-05 |
| 29295 | 1465 | S1465P | E | 7.2e-06 | 1.1e-05 | 1e-05 | 9.400E-06 | 1.970E-06 | 7.200E-06 | 1.100E-05 |
| 29296 | 1465 | S1465S | E | 1.5e-05 | 8.9e-06 | 1e-05 | 1.130E-05 | 3.251E-06 | 8.900E-06 | 1.500E-05 |
| 29297 | 1465 | S1465T | E | 1e-05 | 1.3e-05 | 8.2e-06 | 1.040E-05 | 2.425E-06 | 8.200E-06 | 1.300E-05 |
| 29298 | 1465 | S1465W | E | 1.4e-05 | 1.3e-05 | 1.5e-05 | 1.400E-05 | 1.000E-06 | 1.300E-05 | 1.500E-05 |
| 29299 | 1465 | S1465Y | E | 1.4e-05 | 1e-05 | 1.5e-05 | 1.300E-05 | 2.646E-06 | 1.000E-05 | 1.500E-05 |
| 29300 | 1465 | S1465V | E | 1.2e-05 | 1.6e-05 | 1.6e-05 | 1.467E-05 | 2.309E-06 | 1.200E-05 | 1.600E-05 |
| 29301 | 1466 | S1466A | E | 1.5e-05 | 1.7e-05 | 1.3e-05 | 1.500E-05 | 2.000E-06 | 1.300E-05 | 1.700E-05 |
| 29302 | 1466 | S1466R | E | 1.5e-05 | 1e-05 | 1.6e-05 | 1.367E-05 | 3.215E-06 | 1.000E-05 | 1.600E-05 |
| 29303 | 1466 | S1466N | E | 1.5e-05 | 1.4e-05 | 1.4e-05 | 1.433E-05 | 5.774E-07 | 1.400E-05 | 1.500E-05 |
| 29304 | 1466 | S1466D | E | 9.4e-06 | 1.6e-05 | 1.4e-05 | 1.313E-05 | 3.384E-06 | 9.400E-06 | 1.600E-05 |
| 29305 | 1466 | S1466C | E | 6.9e-06 | 1.5e-05 | 1.1e-05 | 1.097E-05 | 4.050E-06 | 6.900E-06 | 1.500E-05 |
| 29306 | 1466 | S1466E | E | 1.6e-05 | 8.2e-06 | 1.5e-05 | 1.307E-05 | 4.244E-06 | 8.200E-06 | 1.600E-05 |
| 29307 | 1466 | S1466Q | E | 1e-05 | 1.4e-05 | 1.4e-05 | 1.267E-05 | 2.309E-06 | 1.000E-05 | 1.400E-05 |
| 29308 | 1466 | S1466G | E | 1.4e-05 | 1.7e-05 | 1.1e-05 | 1.400E-05 | 3.000E-06 | 1.100E-05 | 1.700E-05 |
| 29309 | 1466 | S1466H | E | 9.4e-06 | 9.4e-06 | 9.8e-06 | 9.533E-06 | 2.309E-07 | 9.400E-06 | 9.800E-06 |
| 29310 | 1466 | S1466I | E | 1.6e-05 | 2e-05 | 9.4e-06 | 1.513E-05 | 5.353E-06 | 9.400E-06 | 2.000E-05 |
| 29311 | 1466 | S1466L | E | 1.6e-05 | 9.4e-06 | 1.6e-05 | 1.380E-05 | 3.811E-06 | 9.400E-06 | 1.600E-05 |
| 29312 | 1466 | S1466K | E | 1.1e-05 | 1.6e-05 | 1.3e-05 | 1.333E-05 | 2.517E-06 | 1.100E-05 | 1.600E-05 |
| 29313 | 1466 | S1466M | E | 1.6e-05 | 1.2e-05 | 1.1e-05 | 1.300E-05 | 2.646E-06 | 1.100E-05 | 1.600E-05 |
| 29314 | 1466 | S1466F | E | 1e-05 | 1.1e-05 | 1.4e-05 | 1.167E-05 | 2.082E-06 | 1.000E-05 | 1.400E-05 |
| 29315 | 1466 | S1466P | E | 1.7e-05 | 1.6e-05 | 1.7e-05 | 1.667E-05 | 5.774E-07 | 1.600E-05 | 1.700E-05 |
| 29316 | 1466 | S1466S | E | 1.4e-05 | 1.1e-05 | 1.1e-05 | 1.200E-05 | 1.732E-06 | 1.100E-05 | 1.400E-05 |
| 29317 | 1466 | S1466T | E | 1.5e-05 | 9.3e-06 | 1.5e-05 | 1.310E-05 | 3.291E-06 | 9.300E-06 | 1.500E-05 |
| 29318 | 1466 | S1466W | E | 1e-05 | 2e-05 | 1.6e-05 | 1.533E-05 | 5.033E-06 | 1.000E-05 | 2.000E-05 |
| 29319 | 1466 | S1466Y | E | 1.6e-05 | 1.6e-05 | 8.2e-06 | 1.340E-05 | 4.503E-06 | 8.200E-06 | 1.600E-05 |
| 29320 | 1466 | S1466V | E | 8e-06 | 1.1e-05 | 1.6e-05 | 1.167E-05 | 4.041E-06 | 8.000E-06 | 1.600E-05 |
| 29321 | 1467 | S1467A | E | 1.3e-05 | 7.3e-06 | 1.3e-05 | 1.110E-05 | 3.291E-06 | 7.300E-06 | 1.300E-05 |
| 29322 | 1467 | S1467R | E | 9.6e-06 | 8.9e-06 | 1.6e-05 | 1.150E-05 | 3.913E-06 | 8.900E-06 | 1.600E-05 |
| 29323 | 1467 | S1467N | E | 1.3e-05 | 1.4e-05 | 1.5e-05 | 1.400E-05 | 1.000E-06 | 1.300E-05 | 1.500E-05 |
| 29324 | 1467 | S1467D | E | 9.8e-06 | 1e-05 | 1.4e-05 | 1.127E-05 | 2.369E-06 | 9.800E-06 | 1.400E-05 |

| | | | | | | | | | | |
|---|---|---|---|---|---|---|---|---|---|---|
| 29325 | 1467 | S1467C | E | 1.1e-05 | 1.6e-05 | 1.2e-05 | 1.300E-05 | 2.646E-06 | 1.100E-05 | 1.600E-05 |
| 29326 | 1467 | S1467E | E | 1.6e-05 | 1e-05 | 1.1e-05 | 1.233E-05 | 3.215E-06 | 1.000E-05 | 1.600E-05 |
| 29327 | 1467 | S1467Q | E | 1.4e-05 | 6.3e-06 | 1.5e-05 | 1.177E-05 | 4.761E-06 | 6.300E-06 | 1.500E-05 |
| 29328 | 1467 | S1467G | E | 8.9e-06 | 1.5e-05 | 1.6e-05 | 1.330E-05 | 3.843E-06 | 8.900E-06 | 1.600E-05 |
| 29329 | 1467 | S1467H | E | 2.2e-05 | 1.5e-05 | 1.3e-05 | 1.667E-05 | 4.726E-06 | 1.300E-05 | 2.200E-05 |
| 29330 | 1467 | S1467I | E | 1.6e-05 | 1.5e-05 | 1.4e-05 | 1.500E-05 | 1.000E-06 | 1.400E-05 | 1.600E-05 |
| 29331 | 1467 | S1467L | E | 1.5e-05 | 1.4e-05 | 1.3e-05 | 1.400E-05 | 1.000E-06 | 1.300E-05 | 1.500E-05 |
| 29332 | 1467 | S1467K | E | 9.3e-06 | 1.5e-05 | 1.2e-05 | 1.210E-05 | 2.851E-06 | 9.300E-06 | 1.500E-05 |
| 29333 | 1467 | S1467M | E | 1e-05 | 6.5e-06 | 1.4e-05 | 1.017E-05 | 3.753E-06 | 6.500E-06 | 1.400E-05 |
| 29334 | 1467 | S1467F | E | 1.5e-05 | 1e-05 | 1.1e-05 | 1.200E-05 | 2.646E-06 | 1.000E-05 | 1.500E-05 |
| 29335 | 1467 | S1467P | E | 2.2e-05 | 1.1e-05 | 1.3e-05 | 1.533E-05 | 5.859E-06 | 1.100E-05 | 2.200E-05 |
| 29336 | 1467 | S1467S | E | 1.6e-05 | 9.2e-06 | 8.6e-06 | 1.127E-05 | 4.110E-06 | 8.600E-06 | 1.600E-05 |
| 29337 | 1467 | S1467T | E | 1.4e-05 | 1.3e-05 | 1.5e-05 | 1.400E-05 | 1.000E-06 | 1.300E-05 | 1.500E-05 |
| 29338 | 1467 | S1467W | E | 1.5e-05 | 1.4e-05 | 1.1e-05 | 1.333E-05 | 2.082E-06 | 1.100E-05 | 1.500E-05 |
| 29339 | 1467 | S1467Y | E | 1.1e-05 | 9.6e-06 | 1.4e-05 | 1.153E-05 | 2.248E-06 | 9.600E-06 | 1.400E-05 |
| 29340 | 1467 | S1467V | E | 1.4e-05 | 9.3e-06 | 1.6e-05 | 1.310E-05 | 3.439E-06 | 9.300E-06 | 1.600E-05 |
| 29341 | 1468 | L1468A | E | 1.7e-05 | 1.2e-05 | 1.5e-05 | 1.467E-05 | 2.517E-06 | 1.200E-05 | 1.700E-05 |
| 29342 | 1468 | L1468R | E | 1.1e-05 | 1.1e-05 | 1.4e-05 | 1.200E-05 | 1.732E-06 | 1.100E-05 | 1.400E-05 |
| 29343 | 1468 | L1468N | E | 1e-05 | 1.1e-05 | 1e-05 | 1.033E-05 | 5.774E-07 | 1.000E-05 | 1.100E-05 |
| 29344 | 1468 | L1468D | E | 1.1e-05 | 1.1e-05 | 1e-05 | 1.067E-05 | 5.774E-07 | 1.000E-05 | 1.100E-05 |
| 29345 | 1468 | L1468C | E | 1.4e-05 | 1.8e-05 | 1.6e-05 | 1.600E-05 | 2.000E-06 | 1.400E-05 | 1.800E-05 |
| 29346 | 1468 | L1468E | E | 1.6e-05 | 1.1e-05 | 1.4e-05 | 1.367E-05 | 2.517E-06 | 1.100E-05 | 1.600E-05 |
| 29347 | 1468 | L1468Q | E | 1.5e-05 | 1e-05 | 1.5e-05 | 1.333E-05 | 2.887E-06 | 1.000E-05 | 1.500E-05 |
| 29348 | 1468 | L1468G | E | 9.4e-06 | 1.1e-05 | 1.1e-05 | 1.047E-05 | 9.238E-07 | 9.400E-06 | 1.100E-05 |
| 29349 | 1468 | L1468H | E | 1.6e-05 | 8.4e-06 | 1.4e-05 | 1.280E-05 | 3.940E-06 | 8.400E-06 | 1.600E-05 |
| 29350 | 1468 | L1468I | E | 1.6e-05 | 1.1e-05 | 1.2e-05 | 1.300E-05 | 2.646E-06 | 1.100E-05 | 1.600E-05 |
| 29351 | 1468 | L1468L | E | 1.6e-05 | 1.3e-05 | 1.3e-05 | 1.400E-05 | 1.732E-06 | 1.300E-05 | 1.600E-05 |
| 29352 | 1468 | L1468K | E | 1.1e-05 | 9.9e-06 | 1.5e-05 | 1.197E-05 | 2.684E-06 | 9.900E-06 | 1.500E-05 |
| 29353 | 1468 | L1468M | E | 1.5e-05 | 1.1e-05 | 1e-05 | 1.200E-05 | 2.646E-06 | 1.000E-05 | 1.500E-05 |
| 29354 | 1468 | L1468F | E | 1.5e-05 | 9.9e-06 | 1.4e-05 | 1.297E-05 | 2.702E-06 | 9.900E-06 | 1.500E-05 |
| 29355 | 1468 | L1468P | E | 1.1e-05 | 1.1e-05 | 7.5e-06 | 9.833E-06 | 2.021E-06 | 7.500E-06 | 1.100E-05 |
| 29356 | 1468 | L1468S | E | 9.6e-06 | 1.6e-05 | 1.3e-05 | 1.287E-05 | 3.202E-06 | 9.600E-06 | 1.600E-05 |
| 29357 | 1468 | L1468T | E | 1.5e-05 | 1.6e-05 | 8.5e-06 | 1.317E-05 | 4.072E-06 | 8.500E-06 | 1.600E-05 |
| 29358 | 1468 | L1468W | E | 1.4e-05 | 1.3e-05 | 1.4e-05 | 1.367E-05 | 5.774E-07 | 1.300E-05 | 1.400E-05 |
| 29359 | 1468 | L1468Y | E | 1.1e-05 | 1e-05 | 9.1e-06 | 1.003E-05 | 9.504E-07 | 9.100E-06 | 1.100E-05 |
| 29360 | 1468 | L1468V | E | 1.4e-05 | 1.5e-05 | 1.6e-05 | 1.500E-05 | 1.000E-06 | 1.400E-05 | 1.600E-05 |
| 29361 | 1469 | G1469A | E | 1.5e-05 | 1.5e-05 | 1e-05 | 1.333E-05 | 2.887E-06 | 1.000E-05 | 1.500E-05 |
| 29362 | 1469 | G1469R | E | 1.5e-05 | 9.6e-06 | 1e-05 | 1.153E-05 | 3.009E-06 | 9.600E-06 | 1.500E-05 |
| 29363 | 1469 | G1469N | E | 9.3e-06 | 1.1e-05 | 1.3e-05 | 1.110E-05 | 1.852E-06 | 9.300E-06 | 1.300E-05 |
| 29364 | 1469 | G1469D | E | 1e-05 | 1.5e-05 | 1e-05 | 1.167E-05 | 2.887E-06 | 1.000E-05 | 1.500E-05 |
| 29365 | 1469 | G1469C | E | 1.4e-05 | 1.4e-05 | 2.3e-05 | 1.700E-05 | 5.196E-06 | 1.400E-05 | 2.300E-05 |
| 29366 | 1469 | G1469E | E | 1.5e-05 | 1.7e-05 | 1.5e-05 | 1.567E-05 | 1.155E-06 | 1.500E-05 | 1.700E-05 |
| 29367 | 1469 | G1469Q | E | 2.3e-05 | 8.8e-06 | 1e-05 | 1.393E-05 | 7.875E-06 | 8.800E-06 | 2.300E-05 |
| 29368 | 1469 | G1469G | E | 1.3e-05 | 9.9e-06 | 7.6e-06 | 1.017E-05 | 2.710E-06 | 7.600E-06 | 1.300E-05 |
| 29369 | 1469 | G1469H | E | 1.5e-05 | 1.6e-05 | 1.4e-05 | 1.500E-05 | 1.000E-06 | 1.400E-05 | 1.600E-05 |
| 29370 | 1469 | G1469I | E | 9.8e-06 | 1e-05 | 1e-05 | 9.933E-06 | 1.155E-07 | 9.800E-06 | 1.000E-05 |
| 29371 | 1469 | G1469L | E | 9.9e-06 | 1.3e-05 | 1e-05 | 1.097E-05 | 1.762E-06 | 9.900E-06 | 1.300E-05 |
| 29372 | 1469 | G1469K | E | 1.4e-05 | 1.2e-05 | 9.3e-06 | 1.177E-05 | 2.359E-06 | 9.300E-06 | 1.400E-05 |
| 29373 | 1469 | G1469M | E | 1e-05 | 1.4e-05 | 1.4e-05 | 1.267E-05 | 2.309E-06 | 1.000E-05 | 1.400E-05 |
| 29374 | 1469 | G1469F | E | 1.5e-05 | 1.5e-05 | 1.4e-05 | 1.467E-05 | 5.774E-07 | 1.400E-05 | 1.500E-05 |
| 29375 | 1469 | G1469P | E | 1.7e-05 | 1.1e-05 | 1.1e-05 | 1.300E-05 | 3.464E-06 | 1.100E-05 | 1.700E-05 |
| 29376 | 1469 | G1469S | E | 8.5e-06 | 1.7e-05 | 1.1e-05 | 1.217E-05 | 4.368E-06 | 8.500E-06 | 1.700E-05 |
| 29377 | 1469 | G1469T | E | 9.6e-06 | 1.3e-05 | 1.1e-05 | 1.120E-05 | 1.709E-06 | 9.600E-06 | 1.300E-05 |
| 29378 | 1469 | G1469W | E | 1.1e-05 | 1.8e-05 | 9.7e-06 | 1.290E-05 | 4.464E-06 | 9.700E-06 | 1.800E-05 |
| 29379 | 1469 | G1469Y | E | 1.4e-05 | 1.5e-05 | 1.6e-05 | 1.500E-05 | 1.000E-06 | 1.400E-05 | 1.600E-05 |
| 29380 | 1469 | G1469V | E | 1.7e-05 | 9.3e-06 | 9.9e-06 | 1.207E-05 | 4.283E-06 | 9.300E-06 | 1.700E-05 |
| 29381 | 1470 | T1470A | E | 1.6e-05 | 1.1e-05 | 1.3e-05 | 1.333E-05 | 2.517E-06 | 1.100E-05 | 1.600E-05 |
| 29382 | 1470 | T1470R | E | 1.5e-05 | 1.4e-05 | 1e-05 | 1.300E-05 | 2.646E-06 | 1.000E-05 | 1.500E-05 |
| 29383 | 1470 | T1470N | E | 1.4e-05 | 9.2e-06 | 1e-05 | 1.107E-05 | 2.572E-06 | 9.200E-06 | 1.400E-05 |
| 29384 | 1470 | T1470D | E | 1.5e-05 | 8.7e-06 | 1e-05 | 1.123E-05 | 3.326E-06 | 8.700E-06 | 1.500E-05 |
| 29385 | 1470 | T1470C | E | 1e-05 | 1.5e-05 | 1.6e-05 | 1.367E-05 | 3.215E-06 | 1.000E-05 | 1.600E-05 |
| 29386 | 1470 | T1470E | E | 1.2e-05 | 1e-05 | 1.5e-05 | 1.233E-05 | 2.517E-06 | 1.000E-05 | 1.500E-05 |
| 29387 | 1470 | T1470Q | E | 1.7e-05 | 1.7e-05 | 9.4e-06 | 1.447E-05 | 4.388E-06 | 9.400E-06 | 1.700E-05 |
| 29388 | 1470 | T1470G | E | 1e-05 | 1.3e-05 | 1.1e-05 | 1.133E-05 | 1.528E-06 | 1.000E-05 | 1.300E-05 |
| 29389 | 1470 | T1470H | E | 1.6e-05 | 1.5e-05 | 1e-05 | 1.367E-05 | 3.215E-06 | 1.000E-05 | 1.600E-05 |
| 29390 | 1470 | T1470I | E | 9.6e-06 | 1.4e-05 | 1.4e-05 | 1.253E-05 | 2.540E-06 | 9.600E-06 | 1.400E-05 |
| 29391 | 1470 | T1470L | E | 1.7e-05 | 1.6e-05 | 1.2e-05 | 1.500E-05 | 2.646E-06 | 1.200E-05 | 1.700E-05 |
| 29392 | 1470 | T1470K | E | 1.5e-05 | 1.3e-05 | 1.2e-05 | 1.333E-05 | 1.528E-06 | 1.200E-05 | 1.500E-05 |
| 29393 | 1470 | T1470M | E | 1.6e-05 | 1.5e-05 | 1.3e-05 | 1.467E-05 | 1.528E-06 | 1.300E-05 | 1.600E-05 |
| 29394 | 1470 | T1470F | E | 1.7e-05 | 1.4e-05 | 1.5e-05 | 1.533E-05 | 1.528E-06 | 1.400E-05 | 1.700E-05 |
| 29395 | 1470 | T1470P | E | 1.1e-05 | 1.3e-05 | 1.2e-05 | 1.200E-05 | 1.000E-06 | 1.100E-05 | 1.300E-05 |
| 29396 | 1470 | T1470S | E | 1.1e-05 | 1e-05 | 1.6e-05 | 1.233E-05 | 3.215E-06 | 1.000E-05 | 1.600E-05 |
| 29397 | 1470 | T1470T | E | 1.5e-05 | 1e-05 | 1e-05 | 1.167E-05 | 2.887E-06 | 1.000E-05 | 1.500E-05 |
| 29398 | 1470 | T1470W | E | 1.1e-05 | 1.2e-05 | 7.2e-06 | 1.007E-05 | 2.532E-06 | 7.200E-06 | 1.200E-05 |
| 29399 | 1470 | T1470Y | E | 1.5e-05 | 1.6e-05 | 1.2e-05 | 1.433E-05 | 2.082E-06 | 1.200E-05 | 1.600E-05 |
| 29400 | 1470 | T1470V | E | 1.6e-05 | 1.5e-05 | 1.3e-05 | 1.467E-05 | 1.528E-06 | 1.300E-05 | 1.600E-05 |
| 29401 | 1471 | Q1471A | E | 1.6e-05 | 1.5e-05 | 1.6e-05 | 1.567E-05 | 5.774E-07 | 1.500E-05 | 1.600E-05 |
| 29402 | 1471 | Q1471R | E | 1.4e-05 | 1.2e-05 | 1e-05 | 1.200E-05 | 2.000E-06 | 1.000E-05 | 1.400E-05 |
| 29403 | 1471 | Q1471N | E | 1e-05 | 9.3e-06 | 1.1e-05 | 1.010E-05 | 8.544E-07 | 9.300E-06 | 1.100E-05 |
| 29404 | 1471 | Q1471D | E | 9.7e-06 | 1.4e-05 | 2.3e-05 | 1.557E-05 | 6.787E-06 | 9.700E-06 | 2.300E-05 |
| 29405 | 1471 | Q1471C | E | 1.2e-05 | 1e-05 | 1.3e-05 | 1.167E-05 | 1.528E-06 | 1.000E-05 | 1.300E-05 |
| 29406 | 1471 | Q1471E | E | 1.1e-05 | 9e-06 | 1.1e-05 | 1.033E-05 | 1.155E-06 | 9.000E-06 | 1.100E-05 |
| 29407 | 1471 | Q1471Q | E | 1.7e-05 | 9.2e-06 | 1.5e-05 | 1.373E-05 | 4.051E-06 | 9.200E-06 | 1.700E-05 |
| 29408 | 1471 | Q1471G | E | 1.5e-05 | 1.5e-05 | 1.5e-05 | 1.500E-05 | 0.000E+00 | 1.500E-05 | 1.500E-05 |
| 29409 | 1471 | Q1471H | E | 1.5e-05 | 1.7e-05 | 1.5e-05 | 1.567E-05 | 1.155E-06 | 1.500E-05 | 1.700E-05 |

| | | | | | | | | | | |
|---|---|---|---|---|---|---|---|---|---|---|
| 29410 | 1471 | Q1471I | E | 7.8e-06 | 1.5e-05 | 1.3e-05 | 1.193E-05 | 3.717E-06 | 7.800E-06 | 1.500E-05 |
| 29411 | 1471 | Q1471L | E | 1.1e-05 | 1.5e-05 | 1.2e-05 | 1.267E-05 | 2.082E-06 | 1.100E-05 | 1.500E-05 |
| 29412 | 1471 | Q1471K | E | 1.1e-05 | 1.5e-05 | 1.5e-05 | 1.367E-05 | 2.309E-06 | 1.100E-05 | 1.500E-05 |
| 29413 | 1471 | Q1471M | E | 1.7e-05 | 1.1e-05 | 6.4e-06 | 1.147E-05 | 5.315E-06 | 6.400E-06 | 1.700E-05 |
| 29414 | 1471 | Q1471F | E | 1.7e-05 | 1.2e-05 | 1.6e-05 | 1.500E-05 | 2.646E-06 | 1.200E-05 | 1.700E-05 |
| 29415 | 1471 | Q1471P | E | 1.4e-05 | 1.6e-05 | 9.1e-06 | 1.303E-05 | 3.550E-06 | 9.100E-06 | 1.600E-05 |
| 29416 | 1471 | Q1471S | E | 9.3e-06 | 1.5e-05 | 1.5e-05 | 1.310E-05 | 3.291E-06 | 9.300E-06 | 1.500E-05 |
| 29417 | 1471 | Q1471T | E | 1.4e-05 | 1.7e-05 | 1.1e-05 | 1.400E-05 | 3.000E-06 | 1.100E-05 | 1.700E-05 |
| 29418 | 1471 | Q1471W | E | 1.4e-05 | 1.5e-05 | 1.3e-05 | 1.400E-05 | 1.000E-06 | 1.300E-05 | 1.500E-05 |
| 29419 | 1471 | Q1471Y | E | 1.6e-05 | 8.4e-06 | 1.5e-05 | 1.313E-05 | 4.130E-06 | 8.400E-06 | 1.600E-05 |
| 29420 | 1471 | Q1471V | E | 1.3e-05 | 1.1e-05 | 1.6e-05 | 1.333E-05 | 2.517E-06 | 1.100E-05 | 1.600E-05 |
| 29421 | 1472 | T1472A | E | 1.6e-05 | 2.1e-05 | 9.2e-06 | 1.540E-05 | 5.923E-06 | 9.200E-06 | 2.100E-05 |
| 29422 | 1472 | T1472R | E | 1.1e-05 | 1.2e-05 | 1.5e-05 | 1.267E-05 | 2.082E-06 | 1.100E-05 | 1.500E-05 |
| 29423 | 1472 | T1472N | E | 1e-05 | 7.2e-06 | 8.2e-06 | 8.467E-06 | 1.419E-06 | 7.200E-06 | 1.000E-05 |
| 29424 | 1472 | T1472D | E | 1.5e-05 | 1e-05 | 9.6e-06 | 1.153E-05 | 3.009E-06 | 9.600E-06 | 1.500E-05 |
| 29425 | 1472 | T1472C | E | 6.8e-06 | 1.4e-05 | 1.5e-05 | 1.193E-05 | 4.474E-06 | 6.800E-06 | 1.500E-05 |
| 29426 | 1472 | T1472E | E | 1.6e-05 | 9.8e-06 | 1.6e-05 | 1.393E-05 | 3.580E-06 | 9.800E-06 | 1.600E-05 |
| 29427 | 1472 | T1472Q | E | 1.1e-05 | 1.2e-05 | 1e-05 | 1.100E-05 | 1.000E-06 | 1.000E-05 | 1.200E-05 |
| 29428 | 1472 | T1472G | E | 1.5e-05 | 1.4e-05 | 1.5e-05 | 1.467E-05 | 5.774E-07 | 1.400E-05 | 1.500E-05 |
| 29429 | 1472 | T1472H | E | 9.5e-06 | 1e-05 | 1.1e-05 | 1.017E-05 | 7.638E-07 | 9.500E-06 | 1.100E-05 |
| 29430 | 1472 | T1472I | E | 1.1e-05 | 1.5e-05 | 8.7e-06 | 1.157E-05 | 3.188E-06 | 8.700E-06 | 1.500E-05 |
| 29431 | 1472 | T1472L | E | 1.6e-05 | 1.3e-05 | 1.3e-05 | 1.400E-05 | 1.732E-06 | 1.300E-05 | 1.600E-05 |
| 29432 | 1472 | T1472K | E | 1.5e-05 | 1.6e-05 | 1.6e-05 | 1.567E-05 | 5.774E-07 | 1.500E-05 | 1.600E-05 |
| 29433 | 1472 | T1472M | E | 1.2e-05 | 9.5e-06 | 9.8e-06 | 1.043E-05 | 1.365E-06 | 9.500E-06 | 1.200E-05 |
| 29434 | 1472 | T1472F | E | 1.5e-05 | 8.5e-06 | 9.7e-06 | 1.107E-05 | 3.459E-06 | 8.500E-06 | 1.500E-05 |
| 29435 | 1472 | T1472P | E | 1.5e-05 | 1.1e-05 | 1e-05 | 1.200E-05 | 2.646E-06 | 1.000E-05 | 1.500E-05 |
| 29436 | 1472 | T1472S | E | 1.1e-05 | 1.6e-05 | 1.1e-05 | 1.267E-05 | 2.887E-06 | 1.100E-05 | 1.600E-05 |
| 29437 | 1472 | T1472T | E | 1.6e-05 | 1.3e-05 | 1.1e-05 | 1.333E-05 | 2.517E-06 | 1.100E-05 | 1.600E-05 |
| 29438 | 1472 | T1472W | E | 1.7e-05 | 1.1e-05 | 1.4e-05 | 1.400E-05 | 3.000E-06 | 1.100E-05 | 1.700E-05 |
| 29439 | 1472 | T1472Y | E | 1.1e-05 | 1.6e-05 | 1.2e-05 | 1.300E-05 | 2.646E-06 | 1.100E-05 | 1.600E-05 |
| 29440 | 1472 | T1472V | E | 1.5e-05 | 1.3e-05 | 1.5e-05 | 1.433E-05 | 1.155E-06 | 1.300E-05 | 1.500E-05 |
| 29441 | 1473 | Y1473A | E | 1.7e-05 | 1e-05 | 1.4e-05 | 1.367E-05 | 3.512E-06 | 1.000E-05 | 1.700E-05 |
| 29442 | 1473 | Y1473R | E | 1e-05 | 1.2e-05 | 1.2e-05 | 1.133E-05 | 1.155E-06 | 1.000E-05 | 1.200E-05 |
| 29443 | 1473 | Y1473N | E | 8.6e-06 | 9.2e-06 | 1.3e-05 | 1.027E-05 | 2.386E-06 | 8.600E-06 | 1.300E-05 |
| 29444 | 1473 | Y1473D | E | 1.4e-05 | 1.5e-05 | 1.5e-05 | 1.467E-05 | 5.774E-07 | 1.400E-05 | 1.500E-05 |
| 29445 | 1473 | Y1473C | E | 1.5e-05 | 8.8e-06 | 1.4e-05 | 1.260E-05 | 3.329E-06 | 8.800E-06 | 1.500E-05 |
| 29446 | 1473 | Y1473E | E | 1.5e-05 | 1.7e-05 | 1.1e-05 | 1.433E-05 | 3.055E-06 | 1.100E-05 | 1.700E-05 |
| 29447 | 1473 | Y1473Q | E | 1.8e-05 | 8.8e-06 | 1.6e-05 | 1.427E-05 | 4.839E-06 | 8.800E-06 | 1.800E-05 |
| 29448 | 1473 | Y1473G | E | 1.6e-05 | 1.6e-05 | 9.2e-06 | 1.373E-05 | 3.926E-06 | 9.200E-06 | 1.600E-05 |
| 29449 | 1473 | Y1473H | E | 1.6e-05 | 1.5e-05 | 1.1e-05 | 1.400E-05 | 2.646E-06 | 1.100E-05 | 1.600E-05 |
| 29450 | 1473 | Y1473I | E | 1.2e-05 | 8.3e-06 | 1.7e-05 | 1.243E-05 | 4.366E-06 | 8.300E-06 | 1.700E-05 |
| 29451 | 1473 | Y1473L | E | 7.9e-06 | 8.7e-06 | 1.6e-05 | 1.087E-05 | 4.464E-06 | 7.900E-06 | 1.600E-05 |
| 29452 | 1473 | Y1473K | E | 1.6e-05 | 1.6e-05 | 1.6e-05 | 1.600E-05 | 0.000E+00 | 1.600E-05 | 1.600E-05 |
| 29453 | 1473 | Y1473M | E | 1.1e-05 | 1.1e-05 | 1.7e-05 | 1.300E-05 | 3.464E-06 | 1.100E-05 | 1.700E-05 |
| 29454 | 1473 | Y1473F | E | 1.1e-05 | 1.1e-05 | 9.5e-06 | 1.050E-05 | 8.660E-07 | 9.500E-06 | 1.100E-05 |
| 29455 | 1473 | Y1473P | E | 8.3e-06 | 1.1e-05 | 1.4e-05 | 1.110E-05 | 2.851E-06 | 8.300E-06 | 1.400E-05 |
| 29456 | 1473 | Y1473S | E | 1.5e-05 | 1e-05 | 9.2e-06 | 1.140E-05 | 3.143E-06 | 9.200E-06 | 1.500E-05 |
| 29457 | 1473 | Y1473T | E | 9.2e-06 | 1.1e-05 | 9.8e-06 | 1.000E-05 | 9.165E-07 | 9.200E-06 | 1.100E-05 |
| 29458 | 1473 | Y1473W | E | 1.5e-05 | 9.1e-06 | 8.3e-06 | 1.080E-05 | 3.659E-06 | 8.300E-06 | 1.500E-05 |
| 29459 | 1473 | Y1473Y | E | 1.1e-05 | 1.5e-05 | 9.2e-06 | 1.173E-05 | 2.969E-06 | 9.200E-06 | 1.500E-05 |
| 29460 | 1473 | Y1473V | E | 1.4e-05 | 1.3e-05 | 1.5e-05 | 1.400E-05 | 1.000E-06 | 1.300E-05 | 1.500E-05 |
| 29461 | 1474 | I1474A | E | 1.3e-05 | 1e-05 | 1.2e-05 | 1.167E-05 | 1.528E-06 | 1.000E-05 | 1.300E-05 |
| 29462 | 1474 | I1474R | E | 9.8e-06 | 8.8e-06 | 1.3e-05 | 1.053E-05 | 2.194E-06 | 8.800E-06 | 1.300E-05 |
| 29463 | 1474 | I1474N | E | 1.1e-05 | 1.8e-05 | 9.2e-06 | 1.273E-05 | 4.649E-06 | 9.200E-06 | 1.800E-05 |
| 29464 | 1474 | I1474D | E | 5.9e-06 | 1e-05 | 1.6e-05 | 1.063E-05 | 5.080E-06 | 5.900E-06 | 1.600E-05 |
| 29465 | 1474 | I1474C | E | 1.4e-05 | 1.4e-05 | 1e-05 | 1.267E-05 | 2.309E-06 | 1.000E-05 | 1.400E-05 |
| 29466 | 1474 | I1474E | E | 1.4e-05 | 1.1e-05 | 9.4e-06 | 1.147E-05 | 2.335E-06 | 9.400E-06 | 1.400E-05 |
| 29467 | 1474 | I1474Q | E | 1.5e-05 | 1.6e-05 | 1.4e-05 | 1.500E-05 | 1.000E-06 | 1.400E-05 | 1.600E-05 |
| 29468 | 1474 | I1474G | E | 1.5e-05 | 1.4e-05 | 1e-05 | 1.300E-05 | 2.646E-06 | 1.000E-05 | 1.500E-05 |
| 29469 | 1474 | I1474H | E | 1.3e-05 | 1.4e-05 | 1.6e-05 | 1.433E-05 | 1.528E-06 | 1.300E-05 | 1.600E-05 |
| 29470 | 1474 | I1474I | E | 7.1e-06 | 1.5e-05 | 1.5e-05 | 1.237E-05 | 4.561E-06 | 7.100E-06 | 1.500E-05 |
| 29471 | 1474 | I1474L | E | 1.5e-05 | 1.6e-05 | 1.3e-05 | 1.467E-05 | 1.528E-06 | 1.300E-05 | 1.600E-05 |
| 29472 | 1474 | I1474K | E | 1.9e-05 | 9.6e-06 | 9e-06 | 1.253E-05 | 5.608E-06 | 9.000E-06 | 1.900E-05 |
| 29473 | 1474 | I1474M | E | 1.5e-05 | 8.6e-06 | 1.6e-05 | 1.320E-05 | 4.015E-06 | 8.600E-06 | 1.600E-05 |
| 29474 | 1474 | I1474F | E | 1.2e-05 | 9.6e-06 | 1.1e-05 | 1.087E-05 | 1.206E-06 | 9.600E-06 | 1.200E-05 |
| 29475 | 1474 | I1474P | E | 1.7e-05 | 1.5e-05 | 9.9e-06 | 1.397E-05 | 3.661E-06 | 9.900E-06 | 1.700E-05 |
| 29476 | 1474 | I1474S | E | 1.1e-05 | 1.1e-05 | 1e-05 | 1.067E-05 | 5.774E-07 | 1.000E-05 | 1.100E-05 |
| 29477 | 1474 | I1474T | E | 9.6e-06 | 1.2e-05 | 8.9e-06 | 1.017E-05 | 1.626E-06 | 8.900E-06 | 1.200E-05 |
| 29478 | 1474 | I1474W | E | 1.1e-05 | 1.5e-05 | 9.3e-06 | 1.177E-05 | 2.926E-06 | 9.300E-06 | 1.500E-05 |
| 29479 | 1474 | I1474Y | E | 8.7e-06 | 1.5e-05 | 1.2e-05 | 1.190E-05 | 3.151E-06 | 8.700E-06 | 1.500E-05 |
| 29480 | 1474 | I1474V | E | 1.4e-05 | 1.6e-05 | 1.1e-05 | 1.367E-05 | 2.517E-06 | 1.100E-05 | 1.600E-05 |
| 29481 | 1475 | C1475A | E | 1.5e-05 | 1e-05 | 1.1e-05 | 1.200E-05 | 2.646E-06 | 1.000E-05 | 1.500E-05 |
| 29482 | 1475 | C1475R | E | 9.7e-06 | 1e-05 | 1.1e-05 | 1.023E-05 | 6.807E-07 | 9.700E-06 | 1.100E-05 |
| 29483 | 1475 | C1475N | E | 1.3e-05 | 1.4e-05 | 1.4e-05 | 1.367E-05 | 5.774E-07 | 1.300E-05 | 1.400E-05 |
| 29484 | 1475 | C1475D | E | 1.1e-05 | 1.4e-05 | 1.5e-05 | 1.333E-05 | 2.082E-06 | 1.100E-05 | 1.500E-05 |
| 29485 | 1475 | C1475C | E | 1.5e-05 | 1.5e-05 | 1.1e-05 | 1.367E-05 | 2.309E-06 | 1.100E-05 | 1.500E-05 |
| 29486 | 1475 | C1475E | E | 1.4e-05 | 9.8e-06 | 1.2e-05 | 1.193E-05 | 2.101E-06 | 9.800E-06 | 1.400E-05 |
| 29487 | 1475 | C1475Q | E | 1e-05 | 1e-05 | 1.5e-05 | 1.167E-05 | 2.887E-06 | 1.000E-05 | 1.500E-05 |
| 29488 | 1475 | C1475G | E | 1.5e-05 | 1.5e-05 | 1e-05 | 1.333E-05 | 2.887E-06 | 1.000E-05 | 1.500E-05 |
| 29489 | 1475 | C1475H | E | 9.9e-06 | 1.5e-05 | 1.6e-05 | 1.363E-05 | 3.272E-06 | 9.900E-06 | 1.600E-05 |
| 29490 | 1475 | C1475I | E | 1e-05 | 1.2e-05 | 9.1e-06 | 1.037E-05 | 1.484E-06 | 9.100E-06 | 1.200E-05 |
| 29491 | 1475 | C1475L | E | 1.2e-05 | 1.1e-05 | 1.5e-05 | 1.267E-05 | 2.082E-06 | 1.100E-05 | 1.500E-05 |
| 29492 | 1475 | C1475K | E | 1.2e-05 | 1.4e-05 | 1.6e-05 | 1.400E-05 | 2.000E-06 | 1.200E-05 | 1.600E-05 |
| 29493 | 1475 | C1475M | E | 1.6e-05 | 1.5e-05 | 1.1e-05 | 1.400E-05 | 2.646E-06 | 1.100E-05 | 1.600E-05 |
| 29494 | 1475 | C1475F | E | 1.2e-05 | 1.3e-05 | 1.6e-05 | 1.367E-05 | 2.082E-06 | 1.200E-05 | 1.600E-05 |

| | | | | | | | | | | |
|---|---|---|---|---|---|---|---|---|---|---|
| 29495 | 1475 | C1475P | E | 1e-05 | 1.5e-05 | 1.3e-05 | 1.267E-05 | 2.517E-06 | 1.000E-05 | 1.500E-05 |
| 29496 | 1475 | C1475S | E | 1.5e-05 | 1e-05 | 7.8e-06 | 1.093E-05 | 3.690E-06 | 7.800E-06 | 1.500E-05 |
| 29497 | 1475 | C1475T | E | 1.6e-05 | 1.5e-05 | 1.1e-05 | 1.400E-05 | 2.646E-06 | 1.100E-05 | 1.600E-05 |
| 29498 | 1475 | C1475W | E | 1e-05 | 1.1e-05 | 2.2e-05 | 1.433E-05 | 6.658E-06 | 1.000E-05 | 2.200E-05 |
| 29499 | 1475 | C1475Y | E | 1.5e-05 | 1.5e-05 | 1.5e-05 | 1.500E-05 | 0.000E+00 | 1.500E-05 | 1.500E-05 |
| 29500 | 1475 | C1475V | E | 1.5e-05 | 9e-06 | 1.6e-05 | 1.333E-05 | 3.786E-06 | 9.000E-06 | 1.600E-05 |
| 29501 | 1476 | N1476A | E | 1.4e-05 | 1.2e-05 | 1.1e-05 | 1.233E-05 | 1.528E-06 | 1.100E-05 | 1.400E-05 |
| 29502 | 1476 | N1476R | E | 1e-05 | 1.8e-05 | 1.6e-05 | 1.467E-05 | 4.163E-06 | 1.000E-05 | 1.800E-05 |
| 29503 | 1476 | N1476N | E | 1.1e-05 | 1.4e-05 | 1.6e-05 | 1.367E-05 | 2.517E-06 | 1.100E-05 | 1.600E-05 |
| 29504 | 1476 | N1476D | E | 1.5e-05 | 9.1e-06 | 1.4e-05 | 1.270E-05 | 3.158E-06 | 9.100E-06 | 1.500E-05 |
| 29505 | 1476 | N1476C | E | 1.7e-05 | 1.1e-05 | 1.4e-05 | 1.400E-05 | 3.000E-06 | 1.100E-05 | 1.700E-05 |
| 29506 | 1476 | N1476E | E | 1.4e-05 | 1e-05 | 1.5e-05 | 1.300E-05 | 2.646E-06 | 1.000E-05 | 1.500E-05 |
| 29507 | 1476 | N1476Q | E | 1e-05 | 9.2e-06 | 1.1e-05 | 1.007E-05 | 9.018E-07 | 9.200E-06 | 1.100E-05 |
| 29508 | 1476 | N1476G | E | 1.3e-05 | 1.5e-05 | 1.1e-05 | 1.300E-05 | 2.000E-06 | 1.100E-05 | 1.500E-05 |
| 29509 | 1476 | N1476H | E | 8.6e-06 | 1.1e-05 | 9.2e-06 | 9.600E-06 | 1.249E-06 | 8.600E-06 | 1.100E-05 |
| 29510 | 1476 | N1476I | E | 1.5e-05 | 1.1e-05 | 1.1e-05 | 1.233E-05 | 2.309E-06 | 1.100E-05 | 1.500E-05 |
| 29511 | 1476 | N1476L | E | 1.1e-05 | 1.6e-05 | 1.1e-05 | 1.267E-05 | 2.887E-06 | 1.100E-05 | 1.600E-05 |
| 29512 | 1476 | N1476K | E | 1.6e-05 | 1.3e-05 | 1.7e-05 | 1.533E-05 | 2.082E-06 | 1.300E-05 | 1.700E-05 |
| 29513 | 1476 | N1476M | E | 1.2e-05 | 1.4e-05 | 1.2e-05 | 1.267E-05 | 1.155E-06 | 1.200E-05 | 1.400E-05 |
| 29514 | 1476 | N1476F | E | 1.6e-05 | 1.6e-05 | 1.7e-05 | 1.633E-05 | 5.774E-07 | 1.600E-05 | 1.700E-05 |
| 29515 | 1476 | N1476P | E | 1.6e-05 | 1.4e-05 | 1.2e-05 | 1.400E-05 | 2.000E-06 | 1.200E-05 | 1.600E-05 |
| 29516 | 1476 | N1476S | E | 1.5e-05 | 1.5e-05 | 1.6e-05 | 1.533E-05 | 5.774E-07 | 1.500E-05 | 1.600E-05 |
| 29517 | 1476 | N1476T | E | 1.8e-05 | 1.1e-05 | 1.5e-05 | 1.467E-05 | 3.512E-06 | 1.100E-05 | 1.800E-05 |
| 29518 | 1476 | N1476W | E | 1.3e-05 | 1.6e-05 | 1e-05 | 1.300E-05 | 3.000E-06 | 1.000E-05 | 1.600E-05 |
| 29519 | 1476 | N1476Y | E | 1.4e-05 | 1e-05 | 1.1e-05 | 1.167E-05 | 2.082E-06 | 1.000E-05 | 1.400E-05 |
| 29520 | 1476 | N1476V | E | 1.7e-05 | 1.6e-05 | 1.2e-05 | 1.500E-05 | 2.646E-06 | 1.200E-05 | 1.700E-05 |
| 29521 | 1477 | V1477A | E | 1.6e-05 | 1.1e-05 | 1.3e-05 | 1.333E-05 | 2.517E-06 | 1.100E-05 | 1.600E-05 |
| 29522 | 1477 | V1477R | E | 1.4e-05 | 1e-05 | 7.8e-06 | 1.060E-05 | 3.143E-06 | 7.800E-06 | 1.400E-05 |
| 29523 | 1477 | V1477N | E | 1.6e-05 | 1.5e-05 | 1.5e-05 | 1.533E-05 | 5.774E-07 | 1.500E-05 | 1.600E-05 |
| 29524 | 1477 | V1477D | E | 1.6e-05 | 1.5e-05 | 1.5e-05 | 1.533E-05 | 5.774E-07 | 1.500E-05 | 1.600E-05 |
| 29525 | 1477 | V1477C | E | 1.4e-05 | 1.6e-05 | 1.5e-05 | 1.500E-05 | 1.000E-06 | 1.400E-05 | 1.600E-05 |
| 29526 | 1477 | V1477E | E | 1.3e-05 | 1.1e-05 | 1.6e-05 | 1.333E-05 | 2.517E-06 | 1.100E-05 | 1.600E-05 |
| 29527 | 1477 | V1477Q | E | 1.2e-05 | 8.6e-06 | 1.2e-05 | 1.087E-05 | 1.963E-06 | 8.600E-06 | 1.200E-05 |
| 29528 | 1477 | V1477G | E | 1.5e-05 | 1.4e-05 | 1.4e-05 | 1.433E-05 | 5.774E-07 | 1.400E-05 | 1.500E-05 |
| 29529 | 1477 | V1477H | E | 1.3e-05 | 8.3e-06 | 1.3e-05 | 1.143E-05 | 2.714E-06 | 8.300E-06 | 1.300E-05 |
| 29530 | 1477 | V1477I | E | 1.2e-05 | 1.7e-05 | 1.8e-05 | 1.567E-05 | 3.215E-06 | 1.200E-05 | 1.800E-05 |
| 29531 | 1477 | V1477L | E | 1.2e-05 | 1.1e-05 | 9.1e-06 | 1.070E-05 | 1.473E-06 | 9.100E-06 | 1.200E-05 |
| 29532 | 1477 | V1477K | E | 1e-05 | 1.4e-05 | 1.1e-05 | 1.167E-05 | 2.082E-06 | 1.000E-05 | 1.400E-05 |
| 29533 | 1477 | V1477M | E | 1.4e-05 | 1.1e-05 | 1.7e-05 | 1.400E-05 | 3.000E-06 | 1.100E-05 | 1.700E-05 |
| 29534 | 1477 | V1477F | E | 1.5e-05 | 2.9e-05 | 1.5e-05 | 1.967E-05 | 8.083E-06 | 1.500E-05 | 2.900E-05 |
| 29535 | 1477 | V1477P | E | 1.2e-05 | 1e-05 | 1.6e-05 | 1.267E-05 | 3.055E-06 | 1.000E-05 | 1.600E-05 |
| 29536 | 1477 | V1477S | E | 1.4e-05 | 1.7e-05 | 1.4e-05 | 1.500E-05 | 1.732E-06 | 1.400E-05 | 1.700E-05 |
| 29537 | 1477 | V1477T | E | 9.4e-06 | 1.6e-05 | 1.6e-05 | 1.380E-05 | 3.811E-06 | 9.400E-06 | 1.600E-05 |
| 29538 | 1477 | V1477W | E | 1.1e-05 | 1.2e-05 | 1e-05 | 1.100E-05 | 1.000E-06 | 1.000E-05 | 1.200E-05 |
| 29539 | 1477 | V1477Y | E | 1.4e-05 | 1.3e-05 | 1.1e-05 | 1.267E-05 | 1.528E-06 | 1.100E-05 | 1.400E-05 |
| 29540 | 1477 | V1477V | E | 1.6e-05 | 1.6e-05 | 1.1e-05 | 1.433E-05 | 2.887E-06 | 1.100E-05 | 1.600E-05 |
| 29541 | 1478 | N1478A | E | 1.7e-05 | 1.2e-05 | 1.7e-05 | 1.533E-05 | 2.887E-06 | 1.200E-05 | 1.700E-05 |
| 29542 | 1478 | N1478R | E | 6.9e-06 | 7.4e-06 | 1.5e-05 | 9.767E-06 | 4.539E-06 | 6.900E-06 | 1.500E-05 |
| 29543 | 1478 | N1478N | E | 8e-06 | 9e-06 | 1.6e-05 | 1.100E-05 | 4.359E-06 | 8.000E-06 | 1.600E-05 |
| 29544 | 1478 | N1478D | E | 1.3e-05 | 8.3e-06 | 1.4e-05 | 1.177E-05 | 3.044E-06 | 8.300E-06 | 1.400E-05 |
| 29545 | 1478 | N1478C | E | 1.4e-05 | 1.3e-05 | 1.1e-05 | 1.267E-05 | 1.528E-06 | 1.100E-05 | 1.400E-05 |
| 29546 | 1478 | N1478E | E | 1.1e-05 | 1.1e-05 | 1.2e-05 | 1.133E-05 | 5.774E-07 | 1.100E-05 | 1.200E-05 |
| 29547 | 1478 | N1478Q | E | 1.5e-05 | 1.5e-05 | 1e-05 | 1.333E-05 | 2.887E-06 | 1.000E-05 | 1.500E-05 |
| 29548 | 1478 | N1478G | E | 1.4e-05 | 1.3e-05 | 1e-05 | 1.233E-05 | 2.082E-06 | 1.000E-05 | 1.400E-05 |
| 29549 | 1478 | N1478H | E | 1e-05 | 8.8e-06 | 9.7e-06 | 9.500E-06 | 6.245E-07 | 8.800E-06 | 1.000E-05 |
| 29550 | 1478 | N1478I | E | 1e-05 | 1.5e-05 | 1.2e-05 | 1.233E-05 | 2.517E-06 | 1.000E-05 | 1.500E-05 |
| 29551 | 1478 | N1478L | E | 9.5e-06 | 1.1e-05 | 1.2e-05 | 1.083E-05 | 1.258E-06 | 9.500E-06 | 1.200E-05 |
| 29552 | 1478 | N1478K | E | 1.6e-05 | 1.1e-05 | 1.2e-05 | 1.300E-05 | 2.646E-06 | 1.100E-05 | 1.600E-05 |
| 29553 | 1478 | N1478M | E | 1e-05 | 1.3e-05 | 1.6e-05 | 1.300E-05 | 3.000E-06 | 1.000E-05 | 1.600E-05 |
| 29554 | 1478 | N1478F | E | 1.6e-05 | 1.3e-05 | 8.9e-06 | 1.263E-05 | 3.564E-06 | 8.900E-06 | 1.600E-05 |
| 29555 | 1478 | N1478P | E | 1e-05 | 1.3e-05 | 1.3e-05 | 1.200E-05 | 1.732E-06 | 1.000E-05 | 1.300E-05 |
| 29556 | 1478 | N1478S | E | 1.1e-05 | 1.6e-05 | 1.1e-05 | 1.267E-05 | 2.887E-06 | 1.100E-05 | 1.600E-05 |
| 29557 | 1478 | N1478T | E | 1e-05 | 2.5e-05 | 9.7e-06 | 1.490E-05 | 8.748E-06 | 9.700E-06 | 2.500E-05 |
| 29558 | 1478 | N1478W | E | 1.7e-05 | 1.6e-05 | 9.2e-06 | 1.407E-05 | 4.244E-06 | 9.200E-06 | 1.700E-05 |
| 29559 | 1478 | N1478Y | E | 1.5e-05 | 1.2e-05 | 1e-05 | 1.233E-05 | 2.517E-06 | 1.000E-05 | 1.500E-05 |
| 29560 | 1478 | N1478V | E | 1.1e-05 | 1.2e-05 | 1.6e-05 | 1.300E-05 | 2.646E-06 | 1.100E-05 | 1.600E-05 |
| 29561 | 1479 | H1479A | E | 9.9e-06 | 1e-05 | 1.4e-05 | 1.130E-05 | 2.339E-06 | 9.900E-06 | 1.400E-05 |
| 29562 | 1479 | H1479R | E | 8.7e-06 | 1.4e-05 | 1.1e-05 | 1.123E-05 | 2.658E-06 | 8.700E-06 | 1.400E-05 |
| 29563 | 1479 | H1479N | E | 1.3e-05 | 1.3e-05 | 1.2e-05 | 1.267E-05 | 5.774E-07 | 1.200E-05 | 1.300E-05 |
| 29564 | 1479 | H1479D | E | 1.1e-05 | 1.6e-05 | 1e-05 | 1.233E-05 | 3.215E-06 | 1.000E-05 | 1.600E-05 |
| 29565 | 1479 | H1479C | E | 1.5e-05 | 1.5e-05 | 1.5e-05 | 1.500E-05 | 0.000E+00 | 1.500E-05 | 1.500E-05 |
| 29566 | 1479 | H1479E | E | 1.6e-05 | 1.1e-05 | 1.6e-05 | 1.433E-05 | 2.887E-06 | 1.100E-05 | 1.600E-05 |
| 29567 | 1479 | H1479Q | E | 1.4e-05 | 1.2e-05 | 1e-05 | 1.200E-05 | 2.000E-06 | 1.000E-05 | 1.400E-05 |
| 29568 | 1479 | H1479G | E | 9.5e-06 | 1.4e-05 | 9.4e-06 | 1.097E-05 | 2.627E-06 | 9.400E-06 | 1.400E-05 |
| 29569 | 1479 | H1479H | E | 9.4e-06 | 9.5e-06 | 1.3e-05 | 1.063E-05 | 2.050E-06 | 9.400E-06 | 1.300E-05 |
| 29570 | 1479 | H1479I | E | 1.5e-05 | 1.1e-05 | 1.4e-05 | 1.333E-05 | 2.082E-06 | 1.100E-05 | 1.500E-05 |
| 29571 | 1479 | H1479L | E | 1.4e-05 | 8.5e-06 | 1.7e-05 | 1.317E-05 | 4.311E-06 | 8.500E-06 | 1.700E-05 |
| 29572 | 1479 | H1479K | E | 1.4e-05 | 1.4e-05 | 9.9e-06 | 1.263E-05 | 2.367E-06 | 9.900E-06 | 1.400E-05 |
| 29573 | 1479 | H1479M | E | 1.6e-05 | 1.5e-05 | 1.5e-05 | 1.533E-05 | 5.774E-07 | 1.500E-05 | 1.600E-05 |
| 29574 | 1479 | H1479F | E | 1.6e-05 | 1e-05 | 1.3e-05 | 1.300E-05 | 3.000E-06 | 1.000E-05 | 1.600E-05 |
| 29575 | 1479 | H1479P | E | 1.4e-05 | 1.2e-05 | 1.6e-05 | 1.400E-05 | 2.000E-06 | 1.200E-05 | 1.600E-05 |
| 29576 | 1479 | H1479S | E | 1.4e-05 | 9.6e-06 | 9e-06 | 1.087E-05 | 2.730E-06 | 9.000E-06 | 1.400E-05 |
| 29577 | 1479 | H1479T | E | 9.9e-06 | 1.5e-05 | 1.1e-05 | 1.197E-05 | 2.684E-06 | 9.900E-06 | 1.500E-05 |
| 29578 | 1479 | H1479W | E | 1.6e-05 | 1.5e-05 | 1.6e-05 | 1.567E-05 | 5.774E-07 | 1.500E-05 | 1.600E-05 |
| 29579 | 1479 | H1479Y | E | 1.1e-05 | 1.5e-05 | 1.4e-05 | 1.333E-05 | 2.082E-06 | 1.100E-05 | 1.500E-05 |

| | | | | | | | | | | |
|---|---|---|---|---|---|---|---|---|---|---|
| 29580 | 1479 | H1479V | E | 1.5e-05 | 8.7e-06 | 1.1e-05 | 1.157E-05 | 3.188E-06 | 8.700E-06 | 1.500E-05 |
| 29581 | 1480 | K1480A | E | 1.5e-05 | 1.5e-05 | 1.2e-05 | 1.400E-05 | 1.732E-06 | 1.200E-05 | 1.500E-05 |
| 29582 | 1480 | K1480R | E | 1.7e-05 | 1.5e-05 | 1.1e-05 | 1.433E-05 | 3.055E-06 | 1.100E-05 | 1.700E-05 |
| 29583 | 1480 | K1480N | E | 9.4e-06 | 1e-05 | 1.5e-05 | 1.147E-05 | 3.075E-06 | 9.400E-06 | 1.500E-05 |
| 29584 | 1480 | K1480D | E | 8.3e-06 | 1e-05 | 1.6e-05 | 1.143E-05 | 4.045E-06 | 8.300E-06 | 1.600E-05 |
| 29585 | 1480 | K1480C | E | 1.7e-05 | 9.8e-06 | 1.4e-05 | 1.360E-05 | 3.617E-06 | 9.800E-06 | 1.700E-05 |
| 29586 | 1480 | K1480E | E | 9.8e-06 | 1.5e-05 | 9.6e-06 | 1.147E-05 | 3.062E-06 | 9.600E-06 | 1.500E-05 |
| 29587 | 1480 | K1480Q | E | 1e-05 | 9.3e-06 | 1.4e-05 | 1.110E-05 | 2.536E-06 | 9.300E-06 | 1.400E-05 |
| 29588 | 1480 | K1480G | E | 1.7e-05 | 1.6e-05 | 1.1e-05 | 1.467E-05 | 3.215E-06 | 1.100E-05 | 1.700E-05 |
| 29589 | 1480 | K1480H | E | 1.1e-05 | 1.1e-05 | 1.2e-05 | 1.133E-05 | 5.774E-07 | 1.100E-05 | 1.200E-05 |
| 29590 | 1480 | K1480I | E | 9.2e-06 | 1.5e-05 | 1.1e-05 | 1.173E-05 | 2.969E-06 | 9.200E-06 | 1.500E-05 |
| 29591 | 1480 | K1480L | E | 1.2e-05 | 1e-05 | 1.2e-05 | 1.133E-05 | 1.155E-06 | 1.000E-05 | 1.200E-05 |
| 29592 | 1480 | K1480K | E | 1.1e-05 | 1.6e-05 | 1.5e-05 | 1.400E-05 | 2.646E-06 | 1.100E-05 | 1.600E-05 |
| 29593 | 1480 | K1480M | E | 1e-05 | 1.6e-05 | 1.1e-05 | 1.233E-05 | 3.215E-06 | 1.000E-05 | 1.600E-05 |
| 29594 | 1480 | K1480F | E | 1.5e-05 | 1.1e-05 | 1.6e-05 | 1.400E-05 | 2.646E-06 | 1.100E-05 | 1.600E-05 |
| 29595 | 1480 | K1480P | E | 1.2e-05 | 9.4e-06 | 1.1e-05 | 1.080E-05 | 1.311E-06 | 9.400E-06 | 1.200E-05 |
| 29596 | 1480 | K1480S | E | 9.5e-06 | 1.4e-05 | 1e-05 | 1.117E-05 | 2.466E-06 | 9.500E-06 | 1.400E-05 |
| 29597 | 1480 | K1480T | E | 8.8e-06 | 8.3e-06 | 1.5e-05 | 1.070E-05 | 3.732E-06 | 8.300E-06 | 1.500E-05 |
| 29598 | 1480 | K1480W | E | 1.6e-05 | 1.5e-05 | 1.4e-05 | 1.500E-05 | 1.000E-06 | 1.400E-05 | 1.600E-05 |
| 29599 | 1480 | K1480Y | E | 1.5e-05 | 1.3e-05 | 1.4e-05 | 1.400E-05 | 1.000E-06 | 1.300E-05 | 1.500E-05 |
| 29600 | 1480 | K1480V | E | 1.4e-05 | 1.5e-05 | 9.4e-06 | 1.280E-05 | 2.987E-06 | 9.400E-06 | 1.500E-05 |
| 29601 | 1481 | P1481A | E | 9.8e-06 | 1.6e-05 | 1.4e-05 | 1.327E-05 | 3.164E-06 | 9.800E-06 | 1.600E-05 |
| 29602 | 1481 | P1481R | E | 1.6e-05 | 8.4e-06 | 1.6e-05 | 1.347E-05 | 4.388E-06 | 8.400E-06 | 1.600E-05 |
| 29603 | 1481 | P1481N | E | 2.1e-05 | 1.5e-05 | 1e-05 | 1.533E-05 | 5.508E-06 | 1.000E-05 | 2.100E-05 |
| 29604 | 1481 | P1481D | E | 1.3e-05 | 1.3e-05 | 9.1e-06 | 1.170E-05 | 2.252E-06 | 9.100E-06 | 1.300E-05 |
| 29605 | 1481 | P1481C | E | 1e-05 | 1.5e-05 | 9e-06 | 1.133E-05 | 3.215E-06 | 9.000E-06 | 1.500E-05 |
| 29606 | 1481 | P1481E | E | 1.2e-05 | 1.1e-05 | 6e-06 | 9.667E-06 | 3.215E-06 | 6.000E-06 | 1.200E-05 |
| 29607 | 1481 | P1481Q | E | 1.8e-05 | 1e-05 | 1.5e-05 | 1.433E-05 | 4.041E-06 | 1.000E-05 | 1.800E-05 |
| 29608 | 1481 | P1481G | E | 9.8e-06 | 9.7e-06 | 1.3e-05 | 1.083E-05 | 1.877E-06 | 9.700E-06 | 1.300E-05 |
| 29609 | 1481 | P1481H | E | 1.5e-05 | 1.6e-05 | 1.6e-05 | 1.567E-05 | 5.774E-07 | 1.500E-05 | 1.600E-05 |
| 29610 | 1481 | P1481I | E | 1e-05 | 1.5e-05 | 1.5e-05 | 1.333E-05 | 2.887E-06 | 1.000E-05 | 1.500E-05 |
| 29611 | 1481 | P1481L | E | 1.5e-05 | 1.2e-05 | 1.4e-05 | 1.367E-05 | 1.528E-06 | 1.200E-05 | 1.500E-05 |
| 29612 | 1481 | P1481K | E | 1e-05 | 1.6e-05 | 1.1e-05 | 1.233E-05 | 3.215E-06 | 1.000E-05 | 1.600E-05 |
| 29613 | 1481 | P1481M | E | 1.5e-05 | 1.6e-05 | 1e-05 | 1.367E-05 | 3.215E-06 | 1.000E-05 | 1.600E-05 |
| 29614 | 1481 | P1481F | E | 1.3e-05 | 1.4e-05 | 1.2e-05 | 1.300E-05 | 1.000E-06 | 1.200E-05 | 1.400E-05 |
| 29615 | 1481 | P1481P | E | 9.8e-06 | 1.2e-05 | 1.1e-05 | 1.093E-05 | 1.102E-06 | 9.800E-06 | 1.200E-05 |
| 29616 | 1481 | P1481S | E | 1.4e-05 | 1.1e-05 | 1.1e-05 | 1.200E-05 | 1.732E-06 | 1.100E-05 | 1.400E-05 |
| 29617 | 1481 | P1481T | E | 7.7e-06 | 1.2e-05 | 1.1e-05 | 1.023E-05 | 2.250E-06 | 7.700E-06 | 1.200E-05 |
| 29618 | 1481 | P1481W | E | 1e-05 | 1e-05 | 1.6e-05 | 1.200E-05 | 3.464E-06 | 1.000E-05 | 1.600E-05 |
| 29619 | 1481 | P1481Y | E | 1.4e-05 | 1.4e-05 | 1.6e-05 | 1.467E-05 | 1.155E-06 | 1.400E-05 | 1.600E-05 |
| 29620 | 1481 | P1481V | E | 9e-06 | 8.4e-06 | 1.8e-05 | 1.180E-05 | 5.378E-06 | 8.400E-06 | 1.800E-05 |
| 29621 | 1482 | S1482A | E | 1.4e-05 | 1e-05 | 1.1e-05 | 1.167E-05 | 2.082E-06 | 1.000E-05 | 1.400E-05 |
| 29622 | 1482 | S1482R | E | 8.4e-06 | 1.3e-05 | 1e-05 | 1.047E-05 | 2.335E-06 | 8.400E-06 | 1.300E-05 |
| 29623 | 1482 | S1482N | E | 1e-05 | 1.6e-05 | 1.1e-05 | 1.233E-05 | 3.215E-06 | 1.000E-05 | 1.600E-05 |
| 29624 | 1482 | S1482D | E | 1.5e-05 | 1e-05 | 1.1e-05 | 1.200E-05 | 2.646E-06 | 1.000E-05 | 1.500E-05 |
| 29625 | 1482 | S1482C | E | 1.4e-05 | 1.1e-05 | 1.3e-05 | 1.267E-05 | 1.528E-06 | 1.100E-05 | 1.400E-05 |
| 29626 | 1482 | S1482E | E | 1.6e-05 | 8.9e-06 | 1.6e-05 | 1.363E-05 | 4.099E-06 | 8.900E-06 | 1.600E-05 |
| 29627 | 1482 | S1482Q | E | 1e-05 | 1e-05 | 1.5e-05 | 1.167E-05 | 2.887E-06 | 1.000E-05 | 1.500E-05 |
| 29628 | 1482 | S1482G | E | 1.5e-05 | 1.2e-05 | 1.2e-05 | 1.300E-05 | 1.732E-06 | 1.200E-05 | 1.500E-05 |
| 29629 | 1482 | S1482H | E | 9e-06 | 1.3e-05 | 1.4e-05 | 1.200E-05 | 2.646E-06 | 9.000E-06 | 1.400E-05 |
| 29630 | 1482 | S1482I | E | 1.5e-05 | 1e-05 | 1.4e-05 | 1.300E-05 | 2.646E-06 | 1.000E-05 | 1.500E-05 |
| 29631 | 1482 | S1482L | E | 1e-05 | 1.5e-05 | 9.8e-06 | 1.160E-05 | 2.946E-06 | 9.800E-06 | 1.500E-05 |
| 29632 | 1482 | S1482K | E | 1.4e-05 | 1.1e-05 | 1.1e-05 | 1.200E-05 | 1.732E-06 | 1.100E-05 | 1.400E-05 |
| 29633 | 1482 | S1482M | E | 1.7e-05 | 1.1e-05 | 1.1e-05 | 1.300E-05 | 3.464E-06 | 1.100E-05 | 1.700E-05 |
| 29634 | 1482 | S1482F | E | 1.3e-05 | 1.5e-05 | 9.7e-06 | 1.257E-05 | 2.676E-06 | 9.700E-06 | 1.500E-05 |
| 29635 | 1482 | S1482P | E | 1.2e-05 | 9.3e-06 | 1.8e-05 | 1.310E-05 | 4.453E-06 | 9.300E-06 | 1.800E-05 |
| 29636 | 1482 | S1482S | E | 6.4e-06 | 2.1e-05 | 1e-05 | 1.247E-05 | 7.606E-06 | 6.400E-06 | 2.100E-05 |
| 29637 | 1482 | S1482T | E | 1.1e-05 | 1.5e-05 | 1.5e-05 | 1.367E-05 | 2.309E-06 | 1.100E-05 | 1.500E-05 |
| 29638 | 1482 | S1482W | E | 9.4e-06 | 1.6e-05 | 1.2e-05 | 1.247E-05 | 3.325E-06 | 9.400E-06 | 1.600E-05 |
| 29639 | 1482 | S1482Y | E | 1.4e-05 | 1.1e-05 | 1.5e-05 | 1.333E-05 | 2.082E-06 | 1.100E-05 | 1.500E-05 |
| 29640 | 1482 | S1482V | E | 1e-05 | 1e-05 | 1.6e-05 | 1.200E-05 | 3.464E-06 | 1.000E-05 | 1.600E-05 |
| 29641 | 1483 | N1483A | E | 1e-05 | 1.3e-05 | 1.5e-05 | 1.267E-05 | 2.517E-06 | 1.000E-05 | 1.500E-05 |
| 29642 | 1483 | N1483R | E | 1.5e-05 | 9.4e-06 | 9.9e-06 | 1.143E-05 | 3.099E-06 | 9.400E-06 | 1.500E-05 |
| 29643 | 1483 | N1483N | E | 1.1e-05 | 1.2e-05 | 1.5e-05 | 1.267E-05 | 2.082E-06 | 1.100E-05 | 1.500E-05 |
| 29644 | 1483 | N1483D | E | 1.7e-05 | 1.2e-05 | 1.5e-05 | 1.467E-05 | 2.517E-06 | 1.200E-05 | 1.700E-05 |
| 29645 | 1483 | N1483C | E | 9e-06 | 1.5e-05 | 1.2e-05 | 1.200E-05 | 3.000E-06 | 9.000E-06 | 1.500E-05 |
| 29646 | 1483 | N1483E | E | 1.1e-05 | 1.4e-05 | 1.1e-05 | 1.200E-05 | 1.732E-06 | 1.100E-05 | 1.400E-05 |
| 29647 | 1483 | N1483Q | E | 1.5e-05 | 1.6e-05 | 1.5e-05 | 1.533E-05 | 5.774E-07 | 1.500E-05 | 1.600E-05 |
| 29648 | 1483 | N1483G | E | 1.1e-05 | 1.6e-05 | 9.6e-06 | 1.220E-05 | 3.365E-06 | 9.600E-06 | 1.600E-05 |
| 29649 | 1483 | N1483H | E | 1.5e-05 | 8.8e-06 | 1.6e-05 | 1.327E-05 | 3.900E-06 | 8.800E-06 | 1.600E-05 |
| 29650 | 1483 | N1483I | E | 1.2e-05 | 1.4e-05 | 1.2e-05 | 1.267E-05 | 1.155E-06 | 1.200E-05 | 1.400E-05 |
| 29651 | 1483 | N1483L | E | 1.5e-05 | 1.3e-05 | 1.5e-05 | 1.433E-05 | 1.155E-06 | 1.300E-05 | 1.500E-05 |
| 29652 | 1483 | N1483K | E | 1.5e-05 | 1.6e-05 | 9.7e-06 | 1.357E-05 | 3.386E-06 | 9.700E-06 | 1.600E-05 |
| 29653 | 1483 | N1483M | E | 1.5e-05 | 1.5e-05 | 1.6e-05 | 1.533E-05 | 5.774E-07 | 1.500E-05 | 1.600E-05 |
| 29654 | 1483 | N1483F | E | 1.5e-05 | 1.6e-05 | 1.5e-05 | 1.533E-05 | 5.774E-07 | 1.500E-05 | 1.600E-05 |
| 29655 | 1483 | N1483P | E | 1e-05 | 1e-05 | 1.2e-05 | 1.067E-05 | 1.155E-06 | 1.000E-05 | 1.200E-05 |
| 29656 | 1483 | N1483S | E | 1.5e-05 | 8.8e-06 | 1.1e-05 | 1.160E-05 | 3.143E-06 | 8.800E-06 | 1.500E-05 |
| 29657 | 1483 | N1483T | E | 1.1e-05 | 1.6e-05 | 1.4e-05 | 1.367E-05 | 2.517E-06 | 1.100E-05 | 1.600E-05 |
| 29658 | 1483 | N1483W | E | 1.7e-05 | 1.6e-05 | 1.2e-05 | 1.500E-05 | 2.646E-06 | 1.200E-05 | 1.700E-05 |
| 29659 | 1483 | N1483Y | E | 1.5e-05 | 1e-05 | 1.4e-05 | 1.300E-05 | 2.646E-06 | 1.000E-05 | 1.500E-05 |
| 29660 | 1483 | N1483V | E | 1e-05 | 1.1e-05 | 1.6e-05 | 1.233E-05 | 3.215E-06 | 1.000E-05 | 1.600E-05 |
| 29661 | 1484 | T1484A | E | 1.7e-05 | 1.6e-05 | 1.5e-05 | 1.600E-05 | 1.000E-06 | 1.500E-05 | 1.700E-05 |
| 29662 | 1484 | T1484R | E | 1.6e-05 | 8.3e-06 | 1.4e-05 | 1.277E-05 | 3.995E-06 | 8.300E-06 | 1.600E-05 |
| 29663 | 1484 | T1484N | E | 1.4e-05 | 1.4e-05 | 1e-05 | 1.267E-05 | 2.309E-06 | 1.000E-05 | 1.400E-05 |
| 29664 | 1484 | T1484D | E | 9.7e-06 | 1.1e-05 | 1.7e-05 | 1.257E-05 | 3.894E-06 | 9.700E-06 | 1.700E-05 |

| | | | | | | | | | | |
|---|---|---|---|---|---|---|---|---|---|---|
| 29665 | 1484 | T1484C | E | 1.6e-05 | 1.1e-05 | 1e-05 | 1.233E-05 | 3.215E-06 | 1.000E-05 | 1.600E-05 |
| 29666 | 1484 | T1484E | E | 1.6e-05 | 1.4e-05 | 1e-05 | 1.333E-05 | 3.055E-06 | 1.000E-05 | 1.600E-05 |
| 29667 | 1484 | T1484Q | E | 1.3e-05 | 1.5e-05 | 1.6e-05 | 1.467E-05 | 1.528E-06 | 1.300E-05 | 1.600E-05 |
| 29668 | 1484 | T1484G | E | 1.2e-05 | 1.8e-05 | 1.6e-05 | 1.533E-05 | 3.055E-06 | 1.200E-05 | 1.800E-05 |
| 29669 | 1484 | T1484H | E | 1.6e-05 | 9.9e-06 | 1.4e-05 | 1.330E-05 | 3.110E-06 | 9.900E-06 | 1.600E-05 |
| 29670 | 1484 | T1484I | E | 1e-05 | 1.1e-05 | 1e-05 | 1.033E-05 | 5.774E-07 | 1.000E-05 | 1.100E-05 |
| 29671 | 1484 | T1484L | E | 7.2e-06 | 1.5e-05 | 1.3e-05 | 1.173E-05 | 4.051E-06 | 7.200E-06 | 1.500E-05 |
| 29672 | 1484 | T1484K | E | 1.8e-05 | 1.5e-05 | 1.1e-05 | 1.467E-05 | 3.512E-06 | 1.100E-05 | 1.800E-05 |
| 29673 | 1484 | T1484M | E | 1.7e-05 | 1.1e-05 | 1.5e-05 | 1.433E-05 | 3.055E-06 | 1.100E-05 | 1.700E-05 |
| 29674 | 1484 | T1484F | E | 1.6e-05 | 1.6e-05 | 9.1e-06 | 1.370E-05 | 3.984E-06 | 9.100E-06 | 1.600E-05 |
| 29675 | 1484 | T1484P | E | 1.6e-05 | 1.6e-05 | 1e-05 | 1.400E-05 | 3.464E-06 | 1.000E-05 | 1.600E-05 |
| 29676 | 1484 | T1484S | E | 1.7e-05 | 1.1e-05 | 1.8e-05 | 1.533E-05 | 3.786E-06 | 1.100E-05 | 1.800E-05 |
| 29677 | 1484 | T1484T | E | 1.2e-05 | 1.8e-05 | 1.5e-05 | 1.500E-05 | 3.000E-06 | 1.200E-05 | 1.800E-05 |
| 29678 | 1484 | T1484W | E | 1.5e-05 | 1.5e-05 | 1.3e-05 | 1.433E-05 | 1.155E-06 | 1.300E-05 | 1.500E-05 |
| 29679 | 1484 | T1484Y | E | 1.5e-05 | 8.3e-06 | 1.6e-05 | 1.310E-05 | 4.187E-06 | 8.300E-06 | 1.600E-05 |
| 29680 | 1484 | T1484V | E | 9.9e-06 | 1.4e-05 | 7e-06 | 1.030E-05 | 3.517E-06 | 7.000E-06 | 1.400E-05 |
| 29681 | 1485 | K1485A | E | 1.5e-05 | 9.3e-06 | 1e-05 | 1.143E-05 | 3.109E-06 | 9.300E-06 | 1.500E-05 |
| 29682 | 1485 | K1485R | E | 1.1e-05 | 1.6e-05 | 1.2e-05 | 1.300E-05 | 2.646E-06 | 1.100E-05 | 1.600E-05 |
| 29683 | 1485 | K1485N | E | 1e-05 | 8.6e-06 | 9.8e-06 | 9.467E-06 | 7.572E-07 | 8.600E-06 | 1.000E-05 |
| 29684 | 1485 | K1485D | E | 2.1e-05 | 8.7e-06 | 1.5e-05 | 1.490E-05 | 6.151E-06 | 8.700E-06 | 2.100E-05 |
| 29685 | 1485 | K1485C | E | 1.9e-05 | 1.1e-05 | 1.4e-05 | 1.467E-05 | 4.041E-06 | 1.100E-05 | 1.900E-05 |
| 29686 | 1485 | K1485E | E | 9.3e-06 | 1.1e-05 | 8.6e-06 | 9.633E-06 | 1.234E-06 | 8.600E-06 | 1.100E-05 |
| 29687 | 1485 | K1485Q | E | 1.5e-05 | 9.3e-06 | 9.7e-06 | 1.133E-05 | 3.182E-06 | 9.300E-06 | 1.500E-05 |
| 29688 | 1485 | K1485G | E | 1.2e-05 | 9.4e-06 | 8.4e-06 | 9.933E-06 | 1.858E-06 | 8.400E-06 | 1.200E-05 |
| 29689 | 1485 | K1485H | E | 9.9e-06 | 1.5e-05 | 1e-05 | 1.163E-05 | 2.916E-06 | 9.900E-06 | 1.500E-05 |
| 29690 | 1485 | K1485I | E | 1.6e-05 | 1.1e-05 | 9.7e-06 | 1.223E-05 | 3.326E-06 | 9.700E-06 | 1.600E-05 |
| 29691 | 1485 | K1485L | E | 1.1e-05 | 1e-05 | 1.9e-05 | 1.333E-05 | 4.933E-06 | 1.000E-05 | 1.900E-05 |
| 29692 | 1485 | K1485K | E | 1.1e-05 | 1.5e-05 | 1.5e-05 | 1.367E-05 | 2.309E-06 | 1.100E-05 | 1.500E-05 |
| 29693 | 1485 | K1485M | E | 1.6e-05 | 1.5e-05 | 1e-05 | 1.367E-05 | 3.215E-06 | 1.000E-05 | 1.600E-05 |
| 29694 | 1485 | K1485F | E | 1.5e-05 | 1.9e-05 | 8.3e-06 | 1.410E-05 | 5.406E-06 | 8.300E-06 | 1.900E-05 |
| 29695 | 1485 | K1485P | E | 1.1e-05 | 1.5e-05 | 1.3e-05 | 1.300E-05 | 2.000E-06 | 1.100E-05 | 1.500E-05 |
| 29696 | 1485 | K1485S | E | 1e-05 | 9.7e-06 | 1.1e-05 | 1.023E-05 | 6.807E-07 | 9.700E-06 | 1.100E-05 |
| 29697 | 1485 | K1485T | E | 1.9e-05 | 8.5e-06 | 1.4e-05 | 1.383E-05 | 5.252E-06 | 8.500E-06 | 1.900E-05 |
| 29698 | 1485 | K1485W | E | 1.5e-05 | 1.1e-05 | 8.8e-06 | 1.160E-05 | 3.143E-06 | 8.800E-06 | 1.500E-05 |
| 29699 | 1485 | K1485Y | E | 9.7e-06 | 1.3e-05 | 1.6e-05 | 1.290E-05 | 3.151E-06 | 9.700E-06 | 1.600E-05 |
| 29700 | 1485 | K1485V | E | 1.2e-05 | 1.3e-05 | 1.6e-05 | 1.367E-05 | 2.082E-06 | 1.200E-05 | 1.600E-05 |
| 29701 | 1486 | V1486A | E | 9.9e-06 | 1.4e-05 | 9.7e-06 | 1.120E-05 | 2.427E-06 | 9.700E-06 | 1.400E-05 |
| 29702 | 1486 | V1486R | E | 1e-05 | 1.3e-05 | 1.7e-05 | 1.333E-05 | 3.512E-06 | 1.000E-05 | 1.700E-05 |
| 29703 | 1486 | V1486N | E | 1e-05 | 1.5e-05 | 8.4e-06 | 1.113E-05 | 3.443E-06 | 8.400E-06 | 1.500E-05 |
| 29704 | 1486 | V1486D | E | 1.5e-05 | 1.6e-05 | 1.1e-05 | 1.400E-05 | 2.646E-06 | 1.100E-05 | 1.600E-05 |
| 29705 | 1486 | V1486C | E | 9.8e-06 | 1.6e-05 | 1.3e-05 | 1.293E-05 | 3.101E-06 | 9.800E-06 | 1.600E-05 |
| 29706 | 1486 | V1486E | E | 1.3e-05 | 1.1e-05 | 1.1e-05 | 1.167E-05 | 1.155E-06 | 1.100E-05 | 1.300E-05 |
| 29707 | 1486 | V1486Q | E | 1e-05 | 1.6e-05 | 1.5e-05 | 1.367E-05 | 3.215E-06 | 1.000E-05 | 1.600E-05 |
| 29708 | 1486 | V1486G | E | 1e-05 | 1.2e-05 | 1.5e-05 | 1.233E-05 | 2.517E-06 | 1.000E-05 | 1.500E-05 |
| 29709 | 1486 | V1486H | E | 1.4e-05 | 1.5e-05 | 9.8e-06 | 1.293E-05 | 2.759E-06 | 9.800E-06 | 1.500E-05 |
| 29710 | 1486 | V1486I | E | 1.4e-05 | 8e-06 | 1.2e-05 | 1.133E-05 | 3.055E-06 | 8.000E-06 | 1.400E-05 |
| 29711 | 1486 | V1486L | E | 9.8e-06 | 1.5e-05 | 1.4e-05 | 1.293E-05 | 2.759E-06 | 9.800E-06 | 1.500E-05 |
| 29712 | 1486 | V1486K | E | 1.5e-05 | 1.2e-05 | 1.1e-05 | 1.267E-05 | 2.082E-06 | 1.100E-05 | 1.500E-05 |
| 29713 | 1486 | V1486M | E | 1e-05 | 1.2e-05 | 1.4e-05 | 1.200E-05 | 2.000E-06 | 1.000E-05 | 1.400E-05 |
| 29714 | 1486 | V1486F | E | 1.2e-05 | 1e-05 | 1.1e-05 | 1.100E-05 | 1.000E-06 | 1.000E-05 | 1.200E-05 |
| 29715 | 1486 | V1486P | E | 1e-05 | 8.7e-06 | 1.5e-05 | 1.123E-05 | 3.326E-06 | 8.700E-06 | 1.500E-05 |
| 29716 | 1486 | V1486S | E | 1.4e-05 | 1.3e-05 | 1.6e-05 | 1.433E-05 | 1.528E-06 | 1.300E-05 | 1.600E-05 |
| 29717 | 1486 | V1486T | E | 2.2e-05 | 1.5e-05 | 1.5e-05 | 1.733E-05 | 4.041E-06 | 1.500E-05 | 2.200E-05 |
| 29718 | 1486 | V1486W | E | 9.8e-06 | 1.4e-05 | 2.3e-05 | 1.560E-05 | 6.744E-06 | 9.800E-06 | 2.300E-05 |
| 29719 | 1486 | V1486Y | E | 1e-05 | 9e-06 | 1.3e-05 | 1.067E-05 | 2.082E-06 | 9.000E-06 | 1.300E-05 |
| 29720 | 1486 | V1486V | E | 1.4e-05 | 1.6e-05 | 1.1e-05 | 1.367E-05 | 2.517E-06 | 1.100E-05 | 1.600E-05 |
| 29721 | 1487 | D1487A | E | 9.9e-06 | 1.3e-05 | 1.4e-05 | 1.230E-05 | 2.138E-06 | 9.900E-06 | 1.400E-05 |
| 29722 | 1487 | D1487R | E | 1.5e-05 | 1.2e-05 | 1.1e-05 | 1.267E-05 | 2.082E-06 | 1.100E-05 | 1.500E-05 |
| 29723 | 1487 | D1487N | E | 1e-05 | 1e-05 | 1.6e-05 | 1.200E-05 | 3.464E-06 | 1.000E-05 | 1.600E-05 |
| 29724 | 1487 | D1487D | E | 9.4e-06 | 1.2e-05 | 1.1e-05 | 1.080E-05 | 1.311E-06 | 9.400E-06 | 1.200E-05 |
| 29725 | 1487 | D1487C | E | 1.5e-05 | 8.8e-06 | 8.6e-06 | 1.080E-05 | 3.639E-06 | 8.600E-06 | 1.500E-05 |
| 29726 | 1487 | D1487E | E | 8e-06 | 1.6e-05 | 1.6e-05 | 1.333E-05 | 4.619E-06 | 8.000E-06 | 1.600E-05 |
| 29727 | 1487 | D1487Q | E | 9.1e-06 | 1.4e-05 | 8.1e-06 | 1.040E-05 | 3.158E-06 | 8.100E-06 | 1.400E-05 |
| 29728 | 1487 | D1487G | E | 1.1e-05 | 1.3e-05 | 8e-06 | 1.067E-05 | 2.517E-06 | 8.000E-06 | 1.300E-05 |
| 29729 | 1487 | D1487H | E | 1.4e-05 | 1.5e-05 | 1.4e-05 | 1.433E-05 | 5.774E-07 | 1.400E-05 | 1.500E-05 |
| 29730 | 1487 | D1487I | E | 1.1e-05 | 1.2e-05 | 9.7e-06 | 1.090E-05 | 1.153E-06 | 9.700E-06 | 1.200E-05 |
| 29731 | 1487 | D1487L | E | 1.4e-05 | 1.3e-05 | 1.5e-05 | 1.400E-05 | 1.000E-06 | 1.300E-05 | 1.500E-05 |
| 29732 | 1487 | D1487K | E | 1.5e-05 | 1.2e-05 | 1e-05 | 1.233E-05 | 2.517E-06 | 1.000E-05 | 1.500E-05 |
| 29733 | 1487 | D1487M | E | 1.1e-05 | 1.6e-05 | 7e-06 | 1.133E-05 | 4.509E-06 | 7.000E-06 | 1.600E-05 |
| 29734 | 1487 | D1487F | E | 9.8e-06 | 1.7e-05 | 1.4e-05 | 1.360E-05 | 3.617E-06 | 9.800E-06 | 1.700E-05 |
| 29735 | 1487 | D1487P | E | 1.5e-05 | 1.5e-05 | 2.3e-05 | 1.767E-05 | 4.619E-06 | 1.500E-05 | 2.300E-05 |
| 29736 | 1487 | D1487S | E | 9.9e-06 | 1.2e-05 | 1.1e-05 | 1.097E-05 | 1.050E-06 | 9.900E-06 | 1.200E-05 |
| 29737 | 1487 | D1487T | E | 1.1e-05 | 1e-05 | 1.3e-05 | 1.133E-05 | 1.528E-06 | 1.000E-05 | 1.300E-05 |
| 29738 | 1487 | D1487W | E | 1e-05 | 1.6e-05 | 1.5e-05 | 1.367E-05 | 3.215E-06 | 1.000E-05 | 1.600E-05 |
| 29739 | 1487 | D1487Y | E | 1.5e-05 | 1.1e-05 | 6.5e-06 | 1.083E-05 | 4.252E-06 | 6.500E-06 | 1.500E-05 |
| 29740 | 1487 | D1487V | E | 1.7e-05 | 1.4e-05 | 1.4e-05 | 1.500E-05 | 1.732E-06 | 1.400E-05 | 1.700E-05 |
| 29741 | 1488 | K1488A | E | 9.1e-06 | 1e-05 | 1.4e-05 | 1.103E-05 | 2.608E-06 | 9.100E-06 | 1.400E-05 |
| 29742 | 1488 | K1488R | E | 1.6e-05 | 1.4e-05 | 1.3e-05 | 1.433E-05 | 1.528E-06 | 1.300E-05 | 1.600E-05 |
| 29743 | 1488 | K1488N | E | 9.9e-06 | 1.1e-05 | 1.4e-05 | 1.163E-05 | 2.122E-06 | 9.900E-06 | 1.400E-05 |
| 29744 | 1488 | K1488D | E | 6.7e-06 | 1.2e-05 | 1.1e-05 | 9.900E-06 | 2.816E-06 | 6.700E-06 | 1.200E-05 |
| 29745 | 1488 | K1488C | E | 1.4e-05 | 1.5e-05 | 1.4e-05 | 1.433E-05 | 5.774E-07 | 1.400E-05 | 1.500E-05 |
| 29746 | 1488 | K1488E | E | 1.2e-05 | 1.2e-05 | 1.3e-05 | 1.233E-05 | 5.774E-07 | 1.200E-05 | 1.300E-05 |
| 29747 | 1488 | K1488Q | E | 1.5e-05 | 1.5e-05 | 1.1e-05 | 1.367E-05 | 2.309E-06 | 1.100E-05 | 1.500E-05 |
| 29748 | 1488 | K1488G | E | 9.9e-06 | 1.3e-05 | 1.3e-05 | 1.197E-05 | 1.790E-06 | 9.900E-06 | 1.300E-05 |
| 29749 | 1488 | K1488H | E | 1.3e-05 | 1.4e-05 | 1.2e-05 | 1.300E-05 | 1.000E-06 | 1.200E-05 | 1.400E-05 |

| | | | | | | | | | | |
|---|---|---|---|---|---|---|---|---|---|---|
| 29750 | 1488 | K1488I | E | 9.1e-06 | 1.2e-05 | 1.6e-05 | 1.237E-05 | 3.465E-06 | 9.100E-06 | 1.600E-05 |
| 29751 | 1488 | K1488L | E | 1.6e-05 | 1.7e-05 | 1.1e-05 | 1.467E-05 | 3.215E-06 | 1.100E-05 | 1.700E-05 |
| 29752 | 1488 | K1488K | E | 1.3e-05 | 1.3e-05 | 1.6e-05 | 1.400E-05 | 1.732E-06 | 1.300E-05 | 1.600E-05 |
| 29753 | 1488 | K1488M | E | 1e-05 | 1.4e-05 | 9.6e-06 | 1.120E-05 | 2.433E-06 | 9.600E-06 | 1.400E-05 |
| 29754 | 1488 | K1488F | E | 1.5e-05 | 1.5e-05 | 1.4e-05 | 1.467E-05 | 5.774E-07 | 1.400E-05 | 1.500E-05 |
| 29755 | 1488 | K1488P | E | 1e-05 | 1.2e-05 | 7.1e-06 | 9.700E-06 | 2.464E-06 | 7.100E-06 | 1.200E-05 |
| 29756 | 1488 | K1488S | E | 1.5e-05 | 1e-05 | 1.5e-05 | 1.333E-05 | 2.887E-06 | 1.000E-05 | 1.500E-05 |
| 29757 | 1488 | K1488T | E | 1.4e-05 | 1.4e-05 | 1.4e-05 | 1.400E-05 | 0.000E+00 | 1.400E-05 | 1.400E-05 |
| 29758 | 1488 | K1488W | E | 9.9e-06 | 1e-05 | 9e-06 | 9.633E-06 | 5.508E-07 | 9.000E-06 | 1.000E-05 |
| 29759 | 1488 | K1488Y | E | 1.4e-05 | 1.6e-05 | 9e-06 | 1.300E-05 | 3.606E-06 | 9.000E-06 | 1.600E-05 |
| 29760 | 1488 | K1488V | E | 1.5e-05 | 1.5e-05 | 1.3e-05 | 1.433E-05 | 1.155E-06 | 1.300E-05 | 1.500E-05 |
| 29761 | 1489 | K1489A | E | 9.5e-06 | 1.5e-05 | 1.6e-05 | 1.350E-05 | 3.500E-06 | 9.500E-06 | 1.600E-05 |
| 29762 | 1489 | K1489R | E | 9.5e-06 | 1.1e-05 | 1e-05 | 1.017E-05 | 7.638E-07 | 9.500E-06 | 1.100E-05 |
| 29763 | 1489 | K1489N | E | 1.3e-05 | 1.3e-05 | 1.3e-05 | 1.300E-05 | 0.000E+00 | 1.300E-05 | 1.300E-05 |
| 29764 | 1489 | K1489D | E | 8.6e-06 | 1.7e-05 | 1.1e-05 | 1.220E-05 | 4.327E-06 | 8.600E-06 | 1.700E-05 |
| 29765 | 1489 | K1489C | E | 1.3e-05 | 1.1e-05 | 1.1e-05 | 1.167E-05 | 1.155E-06 | 1.100E-05 | 1.300E-05 |
| 29766 | 1489 | K1489E | E | 1e-05 | 9.8e-06 | 9.9e-06 | 9.900E-06 | 1.000E-07 | 9.800E-06 | 1.000E-05 |
| 29767 | 1489 | K1489Q | E | 1.6e-05 | 9.4e-06 | 1.4e-05 | 1.313E-05 | 3.384E-06 | 9.400E-06 | 1.600E-05 |
| 29768 | 1489 | K1489G | E | 1.3e-05 | 1.6e-05 | 9.1e-06 | 1.270E-05 | 3.460E-06 | 9.100E-06 | 1.600E-05 |
| 29769 | 1489 | K1489H | E | 1e-05 | 1.2e-05 | 1.3e-05 | 1.167E-05 | 1.528E-06 | 1.000E-05 | 1.300E-05 |
| 29770 | 1489 | K1489I | E | 1e-05 | 1.1e-05 | 1.7e-05 | 1.267E-05 | 3.786E-06 | 1.000E-05 | 1.700E-05 |
| 29771 | 1489 | K1489L | E | 1.5e-05 | 9.3e-06 | 1.2e-05 | 1.210E-05 | 2.851E-06 | 9.300E-06 | 1.500E-05 |
| 29772 | 1489 | K1489K | E | 9.9e-06 | 1.1e-05 | 1.4e-05 | 1.163E-05 | 2.122E-06 | 9.900E-06 | 1.400E-05 |
| 29773 | 1489 | K1489M | E | 1.7e-05 | 1.5e-05 | 1.1e-05 | 1.433E-05 | 3.055E-06 | 1.100E-05 | 1.700E-05 |
| 29774 | 1489 | K1489F | E | 1.5e-05 | 8.8e-06 | 1.6e-05 | 1.327E-05 | 3.900E-06 | 8.800E-06 | 1.600E-05 |
| 29775 | 1489 | K1489P | E | 1.6e-05 | 1.1e-05 | 9.6e-06 | 1.220E-05 | 3.365E-06 | 9.600E-06 | 1.600E-05 |
| 29776 | 1489 | K1489S | E | 9.6e-06 | 9.7e-06 | 1.4e-05 | 1.110E-05 | 2.512E-06 | 9.600E-06 | 1.400E-05 |
| 29777 | 1489 | K1489T | E | 1.6e-05 | 1.7e-05 | 1e-05 | 1.433E-05 | 3.786E-06 | 1.000E-05 | 1.700E-05 |
| 29778 | 1489 | K1489W | E | 8.2e-06 | 1.1e-05 | 1.5e-05 | 1.140E-05 | 3.418E-06 | 8.200E-06 | 1.500E-05 |
| 29779 | 1489 | K1489Y | E | 1.1e-05 | 8.2e-06 | 1.6e-05 | 1.173E-05 | 3.951E-06 | 8.200E-06 | 1.600E-05 |
| 29780 | 1489 | K1489V | E | 9.8e-06 | 1.5e-05 | 1.1e-05 | 1.193E-05 | 2.723E-06 | 9.800E-06 | 1.500E-05 |
| 29781 | 1490 | V1490A | E | 1.1e-05 | 1.1e-05 | 9.4e-06 | 1.047E-05 | 9.238E-07 | 9.400E-06 | 1.100E-05 |
| 29782 | 1490 | V1490R | E | 1.1e-05 | 1.4e-05 | 1e-05 | 1.167E-05 | 2.082E-06 | 1.000E-05 | 1.400E-05 |
| 29783 | 1490 | V1490N | E | 1.4e-05 | 1.2e-05 | 1.6e-05 | 1.400E-05 | 2.000E-06 | 1.200E-05 | 1.600E-05 |
| 29784 | 1490 | V1490D | E | 1.3e-05 | 1.5e-05 | 1.5e-05 | 1.433E-05 | 1.155E-06 | 1.300E-05 | 1.500E-05 |
| 29785 | 1490 | V1490C | E | 9.8e-06 | 1.5e-05 | 1.4e-05 | 1.293E-05 | 2.759E-06 | 9.800E-06 | 1.500E-05 |
| 29786 | 1490 | V1490E | E | 1.6e-05 | 1.4e-05 | 1.5e-05 | 1.500E-05 | 1.000E-06 | 1.400E-05 | 1.600E-05 |
| 29787 | 1490 | V1490Q | E | 1e-05 | 1.5e-05 | 1.4e-05 | 1.300E-05 | 2.646E-06 | 1.000E-05 | 1.500E-05 |
| 29788 | 1490 | V1490G | E | 1.5e-05 | 1.4e-05 | 1.6e-05 | 1.500E-05 | 1.000E-06 | 1.400E-05 | 1.600E-05 |
| 29789 | 1490 | V1490H | E | 1.1e-05 | 1.2e-05 | 1.6e-05 | 1.300E-05 | 2.646E-06 | 1.100E-05 | 1.600E-05 |
| 29790 | 1490 | V1490I | E | 1.5e-05 | 7.2e-06 | 1.2e-05 | 1.140E-05 | 3.934E-06 | 7.200E-06 | 1.500E-05 |
| 29791 | 1490 | V1490L | E | 1.4e-05 | 1.4e-05 | 1.1e-05 | 1.300E-05 | 1.732E-06 | 1.100E-05 | 1.400E-05 |
| 29792 | 1490 | V1490K | E | 1.2e-05 | 1e-05 | 9e-06 | 1.033E-05 | 1.528E-06 | 9.000E-06 | 1.200E-05 |
| 29793 | 1490 | V1490M | E | 1.4e-05 | 8.3e-06 | 1.5e-05 | 1.243E-05 | 3.614E-06 | 8.300E-06 | 1.500E-05 |
| 29794 | 1490 | V1490F | E | 1.5e-05 | 1.1e-05 | 1.5e-05 | 1.367E-05 | 2.309E-06 | 1.100E-05 | 1.500E-05 |
| 29795 | 1490 | V1490P | E | 1.3e-05 | 1.1e-05 | 1.1e-05 | 1.167E-05 | 1.155E-06 | 1.100E-05 | 1.300E-05 |
| 29796 | 1490 | V1490S | E | 1.5e-05 | 1.1e-05 | 1.4e-05 | 1.333E-05 | 2.082E-06 | 1.100E-05 | 1.500E-05 |
| 29797 | 1490 | V1490T | E | 1.4e-05 | 1.1e-05 | 1.3e-05 | 1.267E-05 | 1.528E-06 | 1.100E-05 | 1.400E-05 |
| 29798 | 1490 | V1490W | E | 1.5e-05 | 1e-05 | 1.5e-05 | 1.333E-05 | 2.887E-06 | 1.000E-05 | 1.500E-05 |
| 29799 | 1490 | V1490Y | E | 9.8e-06 | 1.3e-05 | 1e-05 | 1.093E-05 | 1.793E-06 | 9.800E-06 | 1.300E-05 |
| 29800 | 1490 | V1490V | E | 1.5e-05 | 1.1e-05 | 1.1e-05 | 1.233E-05 | 2.309E-06 | 1.100E-05 | 1.500E-05 |
| 29801 | 1491 | E1491A | E | 1.1e-05 | 1.6e-05 | 1.1e-05 | 1.267E-05 | 2.887E-06 | 1.100E-05 | 1.600E-05 |
| 29802 | 1491 | E1491R | E | 1.5e-05 | 1.5e-05 | 1.4e-05 | 1.467E-05 | 5.774E-07 | 1.400E-05 | 1.500E-05 |
| 29803 | 1491 | E1491N | E | 1.6e-05 | 7.9e-06 | 1.3e-05 | 1.230E-05 | 4.095E-06 | 7.900E-06 | 1.600E-05 |
| 29804 | 1491 | E1491D | E | 1.1e-05 | 1.3e-05 | 6.6e-06 | 1.020E-05 | 3.274E-06 | 6.600E-06 | 1.300E-05 |
| 29805 | 1491 | E1491C | E | 1.5e-05 | 1.5e-05 | 8.8e-06 | 1.293E-05 | 3.580E-06 | 8.800E-06 | 1.500E-05 |
| 29806 | 1491 | E1491E | E | 9.2e-06 | 1.7e-05 | 1.4e-05 | 1.340E-05 | 3.934E-06 | 9.200E-06 | 1.700E-05 |
| 29807 | 1491 | E1491Q | E | 1.5e-05 | 1.7e-05 | 1.4e-05 | 1.533E-05 | 1.528E-06 | 1.400E-05 | 1.700E-05 |
| 29808 | 1491 | E1491G | E | 1.5e-05 | 1.6e-05 | 1.1e-05 | 1.400E-05 | 2.646E-06 | 1.100E-05 | 1.600E-05 |
| 29809 | 1491 | E1491H | E | 1.4e-05 | 1.4e-05 | 1e-05 | 1.267E-05 | 2.309E-06 | 1.000E-05 | 1.400E-05 |
| 29810 | 1491 | E1491I | E | 1.4e-05 | 1.1e-05 | 1.7e-05 | 1.400E-05 | 3.000E-06 | 1.100E-05 | 1.700E-05 |
| 29811 | 1491 | E1491L | E | 1.4e-05 | 1e-05 | 1.4e-05 | 1.267E-05 | 2.309E-06 | 1.000E-05 | 1.400E-05 |
| 29812 | 1491 | E1491K | E | 1.6e-05 | 1.5e-05 | 1.3e-05 | 1.467E-05 | 1.528E-06 | 1.300E-05 | 1.600E-05 |
| 29813 | 1491 | E1491M | E | 1.5e-05 | 1e-05 | 9.9e-06 | 1.163E-05 | 2.916E-06 | 9.900E-06 | 1.500E-05 |
| 29814 | 1491 | E1491F | E | 1e-05 | 1.1e-05 | 1e-05 | 1.033E-05 | 5.774E-07 | 1.000E-05 | 1.100E-05 |
| 29815 | 1491 | E1491P | E | 1.2e-05 | 6.8e-06 | 1.2e-05 | 1.027E-05 | 3.002E-06 | 6.800E-06 | 1.200E-05 |
| 29816 | 1491 | E1491S | E | 1.6e-05 | 1.3e-05 | 1.3e-05 | 1.400E-05 | 1.732E-06 | 1.300E-05 | 1.600E-05 |
| 29817 | 1491 | E1491T | E | 8.4e-06 | 1.1e-05 | 1.4e-05 | 1.113E-05 | 2.802E-06 | 8.400E-06 | 1.400E-05 |
| 29818 | 1491 | E1491W | E | 1.5e-05 | 1e-05 | 1.5e-05 | 1.333E-05 | 2.887E-06 | 1.000E-05 | 1.500E-05 |
| 29819 | 1491 | E1491Y | E | 1.1e-05 | 1.1e-05 | 1e-05 | 1.067E-05 | 5.774E-07 | 1.000E-05 | 1.100E-05 |
| 29820 | 1491 | E1491V | E | 1.5e-05 | 8.8e-06 | 1.4e-05 | 1.260E-05 | 3.329E-06 | 8.800E-06 | 1.500E-05 |
| 29821 | 1492 | P1492A | E | 1.3e-05 | 1.2e-05 | 1.5e-05 | 1.333E-05 | 1.528E-06 | 1.200E-05 | 1.500E-05 |
| 29822 | 1492 | P1492R | E | 1.2e-05 | 1.6e-05 | 1.6e-05 | 1.467E-05 | 2.309E-06 | 1.200E-05 | 1.600E-05 |
| 29823 | 1492 | P1492N | E | 1.7e-05 | 1.1e-05 | 1.1e-05 | 1.300E-05 | 3.464E-06 | 1.100E-05 | 1.700E-05 |
| 29824 | 1492 | P1492D | E | 1.1e-05 | 1.1e-05 | 1.2e-05 | 1.133E-05 | 5.774E-07 | 1.100E-05 | 1.200E-05 |
| 29825 | 1492 | P1492C | E | 1.4e-05 | 1.4e-05 | 1e-05 | 1.267E-05 | 2.309E-06 | 1.000E-05 | 1.400E-05 |
| 29826 | 1492 | P1492E | E | 1.5e-05 | 1.3e-05 | 9.9e-06 | 1.263E-05 | 2.570E-06 | 9.900E-06 | 1.500E-05 |
| 29827 | 1492 | P1492Q | E | 9e-06 | 1.1e-05 | 1.5e-05 | 1.167E-05 | 3.055E-06 | 9.000E-06 | 1.500E-05 |
| 29828 | 1492 | P1492G | E | 1.4e-05 | 1.7e-05 | 1.3e-05 | 1.467E-05 | 2.082E-06 | 1.300E-05 | 1.700E-05 |
| 29829 | 1492 | P1492H | E | 1.1e-05 | 1.1e-05 | 1.6e-05 | 1.267E-05 | 2.887E-06 | 1.100E-05 | 1.600E-05 |
| 29830 | 1492 | P1492I | E | 1.4e-05 | 1.4e-05 | 9.9e-06 | 1.263E-05 | 2.367E-06 | 9.900E-06 | 1.400E-05 |
| 29831 | 1492 | P1492L | E | 8.8e-06 | 1.3e-05 | 8.8e-06 | 1.020E-05 | 2.425E-06 | 8.800E-06 | 1.300E-05 |
| 29832 | 1492 | P1492K | E | 1.4e-05 | 1e-05 | 1.5e-05 | 1.300E-05 | 2.646E-06 | 1.000E-05 | 1.500E-05 |
| 29833 | 1492 | P1492M | E | 1.4e-05 | 9.9e-06 | 1.2e-05 | 1.197E-05 | 2.050E-06 | 9.900E-06 | 1.400E-05 |
| 29834 | 1492 | P1492F | E | 1.1e-05 | 9.6e-06 | 1.6e-05 | 1.220E-05 | 3.365E-06 | 9.600E-06 | 1.600E-05 |

| | | | | | | | | | | |
|---|---|---|---|---|---|---|---|---|---|---|
| 29835 | 1492 | P1492P | E | 1.5e-05 | 1e-05 | 1e-05 | 1.167E-05 | 2.887E-06 | 1.000E-05 | 1.500E-05 |
| 29836 | 1492 | P1492S | E | 9.9e-06 | 2e-05 | 1.4e-05 | 1.463E-05 | 5.080E-06 | 9.900E-06 | 2.000E-05 |
| 29837 | 1492 | P1492T | E | 1.3e-05 | 1e-05 | 1.2e-05 | 1.167E-05 | 1.528E-06 | 1.000E-05 | 1.300E-05 |
| 29838 | 1492 | P1492W | E | 1.5e-05 | 1.6e-05 | 1.6e-05 | 1.567E-05 | 5.774E-07 | 1.500E-05 | 1.600E-05 |
| 29839 | 1492 | P1492Y | E | 1.1e-05 | 1.1e-05 | 9.2e-06 | 1.040E-05 | 1.039E-06 | 9.200E-06 | 1.100E-05 |
| 29840 | 1492 | P1492V | E | 1.2e-05 | 1.3e-05 | 1.2e-05 | 1.233E-05 | 5.774E-07 | 1.200E-05 | 1.300E-05 |

Table 3: A scalable Her2-Trastuzumab-Pertuzumab binding affinity landscape. In this table, **ResID** represents the amino acid residue number of PDB entry 6OGE, while **Muta** represents the site-specific missense mutation introduced into the 1492 amino acid residues in PDB entry 6OGE, **Min**, **Max**, **Mean** and **Std** represent the minimum, the maximum, the average and the standard deviation of the $K_d$ values calculated using Prodigy, respectively.

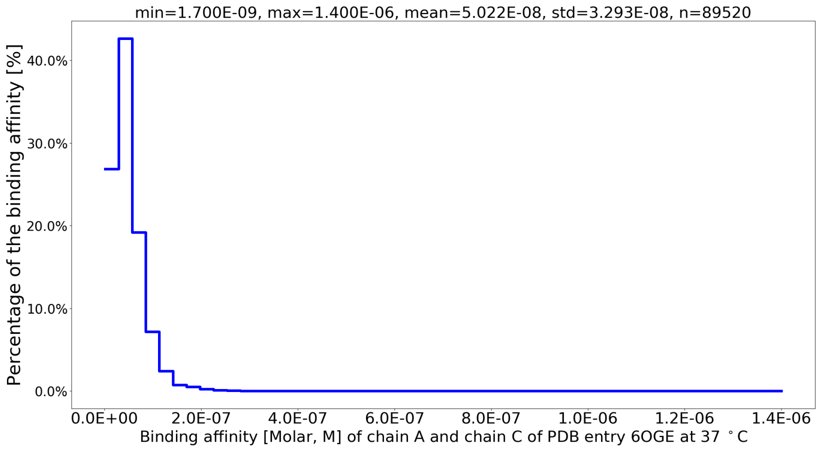


Figure 2: A histogram of the distribution pattern of the Her2-Trastuzumab-Pertuzumab binding affinities between chain A and chain C (listed in Table 1) of PDB entry 6OGE with one site-specific missense mutation.

| No. | ResID | Muta | ChainID | $K_{d1}$ (M) | $K_{d2}$ (M) | $K_{d3}$ (M) | Mean (M) | Std (M) | Min (M) | Max (M) |
|---|---|---|---|---|---|---|---|---|---|---|
| 1 | 1 | T1A | A | 8.2e-08 | 8.2e-08 | 8.2e-08 | 8.200E-08 | 0.000E+00 | 8.200E-08 | 8.200E-08 |
| 2 | 1 | T1R | A | 5.8e-08 | 5.8e-08 | 5.8e-08 | 5.800E-08 | 8.105E-24 | 5.800E-08 | 5.800E-08 |
| 3 | 1 | T1N | A | 6.5e-08 | 6.5e-08 | 6.5e-08 | 6.500E-08 | 0.000E+00 | 6.500E-08 | 6.500E-08 |
| 4 | 1 | T1D | A | 2e-08 | 2e-08 | 2e-08 | 2.000E-08 | 4.052E-24 | 2.000E-08 | 2.000E-08 |
| 5 | 1 | T1C | A | 2.2e-08 | 2.2e-08 | 2.2e-08 | 2.200E-08 | 0.000E+00 | 2.200E-08 | 2.200E-08 |
| 6 | 1 | T1E | A | 8.6e-08 | 8.6e-08 | 8.6e-08 | 8.600E-08 | 0.000E+00 | 8.600E-08 | 8.600E-08 |
| 7 | 1 | T1Q | A | 5.4e-08 | 5.4e-08 | 5.4e-08 | 5.400E-08 | 0.000E+00 | 5.400E-08 | 5.400E-08 |
| 8 | 1 | T1G | A | 7.3e-08 | 7.3e-08 | 7.3e-08 | 7.300E-08 | 0.000E+00 | 7.300E-08 | 7.300E-08 |
| 9 | 1 | T1H | A | 7.9e-08 | 7.9e-08 | 7.9e-08 | 7.900E-08 | 0.000E+00 | 7.900E-08 | 7.900E-08 |
| 10 | 1 | T1I | A | 8e-08 | 8e-08 | 8e-08 | 8.000E-08 | 1.621E-23 | 8.000E-08 | 8.000E-08 |
| 11 | 1 | T1L | A | 4.2e-08 | 4.2e-08 | 4.2e-08 | 4.200E-08 | 0.000E+00 | 4.200E-08 | 4.200E-08 |
| 12 | 1 | T1K | A | 9.4e-08 | 9.4e-08 | 9.4e-08 | 9.400E-08 | 1.621E-23 | 9.400E-08 | 9.400E-08 |
| 13 | 1 | T1M | A | 2.6e-08 | 2.6e-08 | 2.6e-08 | 2.600E-08 | 4.052E-24 | 2.600E-08 | 2.600E-08 |
| 14 | 1 | T1F | A | 2.4e-08 | 2.4e-08 | 2.4e-08 | 2.400E-08 | 0.000E+00 | 2.400E-08 | 2.400E-08 |
| 15 | 1 | T1P | A | 1.7e-07 | 1.7e-07 | 1.7e-07 | 1.700E-07 | 0.000E+00 | 1.700E-07 | 1.700E-07 |
| 16 | 1 | T1S | A | 2.2e-08 | 2.2e-08 | 2.2e-08 | 2.200E-08 | 0.000E+00 | 2.200E-08 | 2.200E-08 |
| 17 | 1 | T1T | A | 5.3e-08 | 5.3e-08 | 5.3e-08 | 5.300E-08 | 8.105E-24 | 5.300E-08 | 5.300E-08 |
| 18 | 1 | T1W | A | 6.6e-08 | 6.6e-08 | 6.6e-08 | 6.600E-08 | 0.000E+00 | 6.600E-08 | 6.600E-08 |
| 19 | 1 | T1Y | A | 1.6e-07 | 1.6e-07 | 1.6e-07 | 1.600E-07 | 3.242E-23 | 1.600E-07 | 1.600E-07 |
| 20 | 1 | T1V | A | 9e-08 | 9e-08 | 9e-08 | 9.000E-08 | 0.000E+00 | 9.000E-08 | 9.000E-08 |
| 21 | 2 | Q2A | A | 4.1e-08 | 4.1e-08 | 4.1e-08 | 4.100E-08 | 0.000E+00 | 4.100E-08 | 4.100E-08 |
| 22 | 2 | Q2R | A | 8.9e-08 | 8.9e-08 | 8.9e-08 | 8.900E-08 | 0.000E+00 | 8.900E-08 | 8.900E-08 |
| 23 | 2 | Q2N | A | 8.7e-09 | 8.7e-09 | 8.7e-09 | 8.700E-09 | 0.000E+00 | 8.700E-09 | 8.700E-09 |
| 24 | 2 | Q2D | A | 2.3e-08 | 2.3e-08 | 2.3e-08 | 2.300E-08 | 4.052E-24 | 2.300E-08 | 2.300E-08 |
| 25 | 2 | Q2C | A | 7.7e-08 | 2.2e-08 | 2.2e-08 | 4.033E-08 | 3.175E-08 | 2.200E-08 | 7.700E-08 |
| 26 | 2 | Q2E | A | 4.1e-08 | 3.2e-08 | 4.4e-08 | 3.900E-08 | 6.245E-09 | 3.200E-08 | 4.400E-08 |
| 27 | 2 | Q2Q | A | 1.7e-08 | 7.1e-08 | 8.1e-09 | 3.203E-08 | 3.404E-08 | 8.100E-09 | 7.100E-08 |
| 28 | 2 | Q2G | A | 1.1e-07 | 3.5e-08 | 1.1e-07 | 8.500E-08 | 4.330E-08 | 3.500E-08 | 1.100E-07 |
| 29 | 2 | Q2H | A | 1.1e-07 | 6.6e-08 | 6.6e-08 | 8.067E-08 | 2.540E-08 | 6.600E-08 | 1.100E-07 |
| 30 | 2 | Q2I | A | 3.7e-08 | 1e-08 | 3.7e-08 | 2.800E-08 | 1.559E-08 | 1.000E-08 | 3.700E-08 |
| 31 | 2 | Q2L | A | 1.7e-08 | 4.9e-08 | 1.7e-08 | 2.767E-08 | 1.848E-08 | 1.700E-08 | 4.900E-08 |
| 32 | 2 | Q2K | A | 3.3e-08 | 3.3e-08 | 8.3e-08 | 4.967E-08 | 2.887E-08 | 3.300E-08 | 8.300E-08 |
| 33 | 2 | Q2M | A | 1.3e-07 | 1.3e-07 | 5e-08 | 1.033E-07 | 4.619E-08 | 5.000E-08 | 1.300E-07 |
| 34 | 2 | Q2F | A | 1.9e-08 | 3.4e-08 | 1.9e-08 | 2.400E-08 | 8.660E-09 | 1.900E-08 | 3.400E-08 |
| 35 | 2 | Q2P | A | 6.6e-08 | 8.3e-08 | 6.6e-08 | 7.167E-08 | 9.815E-09 | 6.600E-08 | 8.300E-08 |
| 36 | 2 | Q2S | A | 9e-08 | 9e-08 | 9e-08 | 9.000E-08 | 0.000E+00 | 9.000E-08 | 9.000E-08 |
| 37 | 2 | Q2T | A | 7.5e-08 | 4.7e-08 | 4.3e-08 | 5.500E-08 | 1.744E-08 | 4.300E-08 | 7.500E-08 |
| 38 | 2 | Q2W | A | 5e-08 | 1.8e-08 | 6.1e-08 | 4.300E-08 | 2.234E-08 | 1.800E-08 | 6.100E-08 |
| 39 | 2 | Q2Y | A | 3.2e-08 | 2.1e-08 | 2.8e-08 | 2.700E-08 | 5.568E-09 | 2.100E-08 | 3.200E-08 |
| 40 | 2 | Q2V | A | 3.1e-08 | 1.6e-07 | 1.6e-07 | 1.170E-07 | 7.448E-08 | 3.100E-08 | 1.600E-07 |
| 41 | 3 | V3A | A | 1.4e-07 | 1.4e-07 | 3.1e-08 | 1.037E-07 | 6.293E-08 | 3.100E-08 | 1.400E-07 |
| 42 | 3 | V3R | A | 1.8e-08 | 3.9e-08 | 3.9e-08 | 3.200E-08 | 1.212E-08 | 1.800E-08 | 3.900E-08 |
| 43 | 3 | V3N | A | 5.5e-08 | 5.5e-08 | 5.5e-08 | 5.500E-08 | 0.000E+00 | 5.500E-08 | 5.500E-08 |
| 44 | 3 | V3D | A | 4e-08 | 4e-08 | 4e-08 | 4.000E-08 | 8.105E-24 | 4.000E-08 | 4.000E-08 |
| 45 | 3 | V3C | A | 8.1e-08 | 8.1e-08 | 3.4e-08 | 6.533E-08 | 2.714E-08 | 3.400E-08 | 8.100E-08 |
| 46 | 3 | V3E | A | 7.9e-08 | 7.9e-08 | 2.6e-08 | 6.133E-08 | 3.060E-08 | 2.600E-08 | 7.900E-08 |
| 47 | 3 | V3Q | A | 4e-08 | 4e-08 | 7.6e-08 | 5.200E-08 | 2.078E-08 | 4.000E-08 | 7.600E-08 |
| 48 | 3 | V3G | A | 2.7e-08 | 2.7e-08 | 2.7e-08 | 2.700E-08 | 0.000E+00 | 2.700E-08 | 2.700E-08 |
| 49 | 3 | V3H | A | 5e-08 | 7.3e-08 | 6.9e-08 | 6.400E-08 | 1.229E-08 | 5.000E-08 | 7.300E-08 |
| 50 | 3 | V3I | A | 9.5e-08 | 2.9e-08 | 9.5e-08 | 7.300E-08 | 3.811E-08 | 2.900E-08 | 9.500E-08 |
| 51 | 3 | V3L | A | 4.9e-08 | 4.9e-08 | 4e-08 | 4.600E-08 | 5.196E-09 | 4.000E-08 | 4.900E-08 |
| 52 | 3 | V3K | A | 3.6e-08 | 2.9e-08 | 2.8e-08 | 3.100E-08 | 4.359E-09 | 2.800E-08 | 3.600E-08 |
| 53 | 3 | V3M | A | 1.6e-07 | 3e-08 | 3e-08 | 7.333E-08 | 7.506E-08 | 3.000E-08 | 1.600E-07 |
| 54 | 3 | V3F | A | 2.5e-08 | 4.6e-08 | 5.6e-08 | 4.233E-08 | 1.582E-08 | 2.500E-08 | 5.600E-08 |
| 55 | 3 | V3P | A | 4.6e-08 | 1.9e-08 | 1.9e-08 | 2.800E-08 | 1.559E-08 | 1.900E-08 | 4.600E-08 |
| 56 | 3 | V3S | A | 9.2e-09 | 9.2e-09 | 7.9e-08 | 3.247E-08 | 4.030E-08 | 9.200E-09 | 7.900E-08 |
| 57 | 3 | V3T | A | 8.4e-08 | 6.7e-08 | 3.6e-08 | 6.233E-08 | 2.434E-08 | 3.600E-08 | 8.400E-08 |
| 58 | 3 | V3W | A | 5.4e-08 | 5.4e-08 | 3.4e-08 | 4.733E-08 | 1.155E-08 | 3.400E-08 | 5.400E-08 |
| 59 | 3 | V3Y | A | 2.9e-08 | 7.1e-08 | 1.8e-08 | 3.933E-08 | 2.797E-08 | 1.800E-08 | 7.100E-08 |
| 60 | 3 | V3V | A | 1.1e-08 | 1.4e-08 | 5.9e-08 | 2.800E-08 | 2.689E-08 | 1.100E-08 | 5.900E-08 |
| 61 | 4 | C4A | A | 2.7e-08 | 1.1e-07 | 1.4e-08 | 5.033E-08 | 5.208E-08 | 1.400E-08 | 1.100E-07 |
| 62 | 4 | C4R | A | 1.1e-08 | 6.6e-08 | 3.2e-08 | 3.633E-08 | 2.775E-08 | 1.100E-08 | 6.600E-08 |
| 63 | 4 | C4N | A | 4.4e-08 | 2.7e-08 | 2e-08 | 3.033E-08 | 1.234E-08 | 2.000E-08 | 4.400E-08 |
| 64 | 4 | C4D | A | 9.5e-09 | 1.1e-07 | 4.9e-08 | 5.617E-08 | 5.063E-08 | 9.500E-09 | 1.100E-07 |
| 65 | 4 | C4C | A | 1.2e-07 | 1.2e-07 | 1.2e-07 | 1.200E-07 | 0.000E+00 | 1.200E-07 | 1.200E-07 |
| 66 | 4 | C4E | A | 4.8e-08 | 4.5e-08 | 4.3e-08 | 4.533E-08 | 2.517E-09 | 4.300E-08 | 4.800E-08 |
| 67 | 4 | C4Q | A | 5.5e-08 | 8.4e-08 | 1.8e-07 | 1.063E-07 | 6.542E-08 | 5.500E-08 | 1.800E-07 |
| 68 | 4 | C4G | A | 7e-08 | 6.9e-08 | 4.5e-08 | 6.133E-08 | 1.415E-08 | 4.500E-08 | 7.000E-08 |
| 69 | 4 | C4H | A | 5.9e-08 | 1.6e-08 | 4.5e-08 | 4.000E-08 | 2.193E-08 | 1.600E-08 | 5.900E-08 |
| 70 | 4 | C4I | A | 3.1e-08 | 3.1e-08 | 2.3e-08 | 2.833E-08 | 4.619E-09 | 2.300E-08 | 3.100E-08 |
| 71 | 4 | C4L | A | 4.3e-08 | 3.4e-08 | 1.6e-08 | 3.100E-08 | 1.375E-08 | 1.600E-08 | 4.300E-08 |
| 72 | 4 | C4K | A | 9.4e-08 | 9.4e-08 | 4.6e-08 | 7.800E-08 | 2.771E-08 | 4.600E-08 | 9.400E-08 |
| 73 | 4 | C4M | A | 1.2e-07 | 2.1e-08 | 5.2e-08 | 6.433E-08 | 5.064E-08 | 2.100E-08 | 1.200E-07 |
| 74 | 4 | C4F | A | 4.8e-08 | 2.8e-08 | 1.7e-08 | 3.100E-08 | 1.572E-08 | 1.700E-08 | 4.800E-08 |
| 75 | 4 | C4P | A | 5.3e-08 | 5.5e-08 | 1.2e-07 | 7.600E-08 | 3.812E-08 | 5.300E-08 | 1.200E-07 |
| 76 | 4 | C4S | A | 5.2e-08 | 3.2e-08 | 7.1e-08 | 5.167E-08 | 1.950E-08 | 3.200E-08 | 7.100E-08 |
| 77 | 4 | C4T | A | 7.1e-08 | 4.6e-08 | 6.6e-08 | 6.100E-08 | 1.323E-08 | 4.600E-08 | 7.100E-08 |
| 78 | 4 | C4W | A | 6.4e-08 | 1.1e-07 | 3e-08 | 6.800E-08 | 4.015E-08 | 3.000E-08 | 1.100E-07 |
| 79 | 4 | C4Y | A | 5.8e-08 | 3.2e-08 | 4.3e-08 | 4.433E-08 | 1.305E-08 | 3.200E-08 | 5.800E-08 |
| 80 | 4 | C4V | A | 2.4e-08 | 5.8e-08 | 3.1e-08 | 3.767E-08 | 1.795E-08 | 2.400E-08 | 5.800E-08 |
| 81 | 5 | T5A | A | 5e-08 | 7.5e-08 | 4e-08 | 5.500E-08 | 1.803E-08 | 4.000E-08 | 7.500E-08 |
| 82 | 5 | T5R | A | 4.8e-08 | 1.9e-08 | 3.6e-08 | 3.433E-08 | 1.457E-08 | 1.900E-08 | 4.800E-08 |
| 83 | 5 | T5N | A | 3.6e-08 | 1.9e-08 | 6.3e-08 | 3.933E-08 | 2.219E-08 | 1.900E-08 | 6.300E-08 |
| 84 | 5 | T5D | A | 3.1e-08 | 1.6e-08 | 1.6e-08 | 2.100E-08 | 8.660E-09 | 1.600E-08 | 3.100E-08 |

| | | | | | | | | | | |
|---|---|---|---|---|---|---|---|---|---|---|
| 85 | 5 | T5C | A | 3e-08 | 1.7e-08 | 1.3e-08 | 2.000E-08 | 8.888E-09 | 1.300E-08 | 3.000E-08 |
| 86 | 5 | T5E | A | 3e-08 | 7.5e-08 | 1.8e-08 | 4.100E-08 | 3.005E-08 | 1.800E-08 | 7.500E-08 |
| 87 | 5 | T5Q | A | 4.3e-08 | 2.2e-08 | 4.8e-08 | 3.767E-08 | 1.380E-08 | 2.200E-08 | 4.800E-08 |
| 88 | 5 | T5G | A | 3.6e-08 | 3.5e-08 | 7e-08 | 4.700E-08 | 1.992E-08 | 3.500E-08 | 7.000E-08 |
| 89 | 5 | T5H | A | 1.4e-08 | 5.4e-08 | 4.2e-08 | 3.667E-08 | 2.053E-08 | 1.400E-08 | 5.400E-08 |
| 90 | 5 | T5I | A | 9.1e-08 | 2.2e-08 | 1.5e-07 | 8.767E-08 | 6.407E-08 | 2.200E-08 | 1.500E-07 |
| 91 | 5 | T5L | A | 3.2e-08 | 3.2e-08 | 6.5e-08 | 4.300E-08 | 1.905E-08 | 3.200E-08 | 6.500E-08 |
| 92 | 5 | T5K | A | 5.9e-08 | 3e-08 | 3.2e-08 | 4.033E-08 | 1.620E-08 | 3.000E-08 | 5.900E-08 |
| 93 | 5 | T5M | A | 1.7e-08 | 2.2e-08 | 4.7e-08 | 2.867E-08 | 1.607E-08 | 1.700E-08 | 4.700E-08 |
| 94 | 5 | T5F | A | 7.7e-08 | 1.8e-08 | 1.8e-08 | 3.767E-08 | 3.406E-08 | 1.800E-08 | 7.700E-08 |
| 95 | 5 | T5P | A | 8.3e-08 | 3.3e-08 | 3.5e-08 | 5.033E-08 | 2.831E-08 | 3.300E-08 | 8.300E-08 |
| 96 | 5 | T5S | A | 8.4e-08 | 2.5e-08 | 3.8e-08 | 4.900E-08 | 3.100E-08 | 2.500E-08 | 8.400E-08 |
| 97 | 5 | T5T | A | 7.3e-08 | 5.5e-08 | 6.6e-08 | 6.467E-08 | 9.074E-09 | 5.500E-08 | 7.300E-08 |
| 98 | 5 | T5W | A | 2.3e-08 | 6.7e-08 | 1.7e-07 | 8.667E-08 | 7.545E-08 | 2.300E-08 | 1.700E-07 |
| 99 | 5 | T5Y | A | 3.3e-08 | 4.5e-08 | 3.6e-08 | 3.800E-08 | 6.245E-09 | 3.300E-08 | 4.500E-08 |
| 100 | 5 | T5V | A | 5.8e-08 | 3.7e-08 | 1.8e-08 | 3.767E-08 | 2.001E-08 | 1.800E-08 | 5.800E-08 |
| 101 | 6 | G6A | A | 8.8e-08 | 2.2e-08 | 2.9e-08 | 4.633E-08 | 3.625E-08 | 2.200E-08 | 8.800E-08 |
| 102 | 6 | G6R | A | 1.1e-07 | 4.9e-08 | 2.5e-08 | 6.133E-08 | 4.382E-08 | 2.500E-08 | 1.100E-07 |
| 103 | 6 | G6N | A | 4.2e-08 | 5.9e-08 | 1.5e-08 | 3.867E-08 | 2.219E-08 | 1.500E-08 | 5.900E-08 |
| 104 | 6 | G6D | A | 1.1e-08 | 6.1e-08 | 2.1e-08 | 3.100E-08 | 2.646E-08 | 1.100E-08 | 6.100E-08 |
| 105 | 6 | G6C | A | 1.8e-08 | 4e-08 | 4.4e-08 | 3.400E-08 | 1.400E-08 | 1.800E-08 | 4.400E-08 |
| 106 | 6 | G6E | A | 2.3e-08 | 3.7e-08 | 5.2e-08 | 3.733E-08 | 1.450E-08 | 2.300E-08 | 5.200E-08 |
| 107 | 6 | G6Q | A | 3.6e-08 | 6.8e-08 | 3.9e-08 | 4.767E-08 | 1.767E-08 | 3.600E-08 | 6.800E-08 |
| 108 | 6 | G6G | A | 2.6e-08 | 7.1e-08 | 3.7e-08 | 4.467E-08 | 2.346E-08 | 2.600E-08 | 7.100E-08 |
| 109 | 6 | G6H | A | 4.1e-08 | 2.4e-08 | 3.5e-08 | 3.333E-08 | 8.622E-09 | 2.400E-08 | 4.100E-08 |
| 110 | 6 | G6I | A | 7.1e-08 | 5.5e-08 | 6.1e-08 | 6.233E-08 | 8.083E-09 | 5.500E-08 | 7.100E-08 |
| 111 | 6 | G6L | A | 5.1e-08 | 3.4e-08 | 9.5e-08 | 6.000E-08 | 3.148E-08 | 3.400E-08 | 9.500E-08 |
| 112 | 6 | G6K | A | 2.4e-08 | 7e-08 | 1.4e-07 | 7.800E-08 | 5.841E-08 | 2.400E-08 | 1.400E-07 |
| 113 | 6 | G6M | A | 6.2e-08 | 9.1e-08 | 7.6e-08 | 7.633E-08 | 1.450E-08 | 6.200E-08 | 9.100E-08 |
| 114 | 6 | G6F | A | 3.8e-08 | 1.4e-08 | 3e-08 | 2.733E-08 | 1.222E-08 | 1.400E-08 | 3.800E-08 |
| 115 | 6 | G6P | A | 7.4e-08 | 3.9e-08 | 2.3e-08 | 4.533E-08 | 2.608E-08 | 2.300E-08 | 7.400E-08 |
| 116 | 6 | G6S | A | 4.1e-08 | 8.1e-08 | 5.6e-08 | 5.933E-08 | 2.021E-08 | 4.100E-08 | 8.100E-08 |
| 117 | 6 | G6T | A | 6e-08 | 3e-08 | 8e-08 | 5.667E-08 | 2.517E-08 | 3.000E-08 | 8.000E-08 |
| 118 | 6 | G6W | A | 4.2e-08 | 5e-08 | 4e-08 | 4.400E-08 | 5.292E-09 | 4.000E-08 | 5.000E-08 |
| 119 | 6 | G6Y | A | 4.5e-08 | 8.2e-08 | 4.4e-08 | 5.700E-08 | 2.166E-08 | 4.400E-08 | 8.200E-08 |
| 120 | 6 | G6V | A | 5.4e-08 | 8.2e-08 | 2.6e-08 | 5.400E-08 | 2.800E-08 | 2.600E-08 | 8.200E-08 |
| 121 | 7 | T7A | A | 4.4e-08 | 1.9e-08 | 2.2e-08 | 2.833E-08 | 1.365E-08 | 1.900E-08 | 4.400E-08 |
| 122 | 7 | T7R | A | 2.7e-08 | 6.3e-08 | 7.1e-08 | 5.367E-08 | 2.344E-08 | 2.700E-08 | 7.100E-08 |
| 123 | 7 | T7N | A | 3.1e-08 | 9.9e-08 | 5.9e-08 | 6.300E-08 | 3.418E-08 | 3.100E-08 | 9.900E-08 |
| 124 | 7 | T7D | A | 1.2e-07 | 6.9e-08 | 3.6e-08 | 7.500E-08 | 4.232E-08 | 3.600E-08 | 1.200E-07 |
| 125 | 7 | T7C | A | 5.3e-08 | 9e-08 | 8.4e-08 | 7.567E-08 | 1.986E-08 | 5.300E-08 | 9.000E-08 |
| 126 | 7 | T7E | A | 4.1e-08 | 2.7e-08 | 4.1e-08 | 3.633E-08 | 8.083E-09 | 2.700E-08 | 4.100E-08 |
| 127 | 7 | T7Q | A | 3e-08 | 5.8e-08 | 6.3e-08 | 5.033E-08 | 1.779E-08 | 3.000E-08 | 6.300E-08 |
| 128 | 7 | T7G | A | 1.8e-08 | 1.2e-07 | 1e-07 | 7.933E-08 | 5.405E-08 | 1.800E-08 | 1.200E-07 |
| 129 | 7 | T7H | A | 3.4e-08 | 3.9e-08 | 3.9e-08 | 3.733E-08 | 2.887E-09 | 3.400E-08 | 3.900E-08 |
| 130 | 7 | T7I | A | 6.1e-08 | 5e-08 | 7.6e-08 | 6.233E-08 | 1.305E-08 | 5.000E-08 | 7.600E-08 |
| 131 | 7 | T7L | A | 4e-08 | 2.2e-08 | 1.7e-08 | 2.633E-08 | 1.210E-08 | 1.700E-08 | 4.000E-08 |
| 132 | 7 | T7K | A | 3.1e-08 | 4.4e-08 | 3.9e-08 | 3.800E-08 | 6.557E-09 | 3.100E-08 | 4.400E-08 |
| 133 | 7 | T7M | A | 5.5e-08 | 3.1e-08 | 4.4e-08 | 4.333E-08 | 1.201E-08 | 3.100E-08 | 5.500E-08 |
| 134 | 7 | T7F | A | 1.5e-08 | 1.6e-08 | 7.7e-08 | 3.600E-08 | 3.551E-08 | 1.500E-08 | 7.700E-08 |
| 135 | 7 | T7P | A | 5.2e-08 | 4.4e-08 | 4.3e-08 | 4.633E-08 | 4.933E-09 | 4.300E-08 | 5.200E-08 |
| 136 | 7 | T7S | A | 3.6e-08 | 1.1e-08 | 8.9e-08 | 4.533E-08 | 3.983E-08 | 1.100E-08 | 8.900E-08 |
| 137 | 7 | T7T | A | 4.2e-08 | 3e-07 | 5e-08 | 1.307E-07 | 1.467E-07 | 4.200E-08 | 3.000E-07 |
| 138 | 7 | T7W | A | 2.5e-08 | 7.6e-08 | 3.7e-08 | 4.600E-08 | 2.666E-08 | 2.500E-08 | 7.600E-08 |
| 139 | 7 | T7Y | A | 4.7e-08 | 9.2e-08 | 3.6e-08 | 5.833E-08 | 2.967E-08 | 3.600E-08 | 9.200E-08 |
| 140 | 7 | T7V | A | 4.4e-08 | 1.1e-07 | 2.9e-08 | 6.100E-08 | 4.309E-08 | 2.900E-08 | 1.100E-07 |
| 141 | 8 | D8A | A | 2.8e-08 | 6e-08 | 3.3e-08 | 4.033E-08 | 1.721E-08 | 2.800E-08 | 6.000E-08 |
| 142 | 8 | D8R | A | 6.9e-08 | 4.9e-08 | 4.9e-08 | 5.567E-08 | 1.155E-08 | 4.900E-08 | 6.900E-08 |
| 143 | 8 | D8N | A | 4.5e-08 | 9.5e-09 | 2.9e-08 | 2.783E-08 | 1.778E-08 | 9.500E-09 | 4.500E-08 |
| 144 | 8 | D8D | A | 7.2e-08 | 4.9e-08 | 7.2e-08 | 6.433E-08 | 1.328E-08 | 4.900E-08 | 7.200E-08 |
| 145 | 8 | D8C | A | 1.6e-08 | 4.5e-08 | 2.3e-08 | 2.800E-08 | 1.513E-08 | 1.600E-08 | 4.500E-08 |
| 146 | 8 | D8E | A | 1e-07 | 3.2e-08 | 1.4e-07 | 9.067E-08 | 5.460E-08 | 3.200E-08 | 1.400E-07 |
| 147 | 8 | D8Q | A | 4.8e-08 | 9.5e-08 | 2.2e-08 | 5.500E-08 | 3.700E-08 | 2.200E-08 | 9.500E-08 |
| 148 | 8 | D8G | A | 2.7e-08 | 4.5e-08 | 3.7e-08 | 3.633E-08 | 9.018E-09 | 2.700E-08 | 4.500E-08 |
| 149 | 8 | D8H | A | 3.9e-08 | 3.9e-08 | 3.1e-08 | 3.633E-08 | 4.619E-09 | 3.100E-08 | 3.900E-08 |
| 150 | 8 | D8I | A | 4.6e-08 | 5.5e-08 | 6.5e-08 | 5.533E-08 | 9.504E-09 | 4.600E-08 | 6.500E-08 |
| 151 | 8 | D8L | A | 2e-08 | 3.5e-08 | 7.5e-09 | 2.083E-08 | 1.377E-08 | 7.500E-09 | 3.500E-08 |
| 152 | 8 | D8K | A | 3.8e-08 | 1.8e-08 | 2e-08 | 2.533E-08 | 1.102E-08 | 1.800E-08 | 3.800E-08 |
| 153 | 8 | D8M | A | 2e-08 | 3.5e-08 | 4.1e-08 | 3.200E-08 | 1.082E-08 | 2.000E-08 | 4.100E-08 |
| 154 | 8 | D8F | A | 5.9e-08 | 4.7e-08 | 4.9e-08 | 5.167E-08 | 6.429E-09 | 4.700E-08 | 5.900E-08 |
| 155 | 8 | D8P | A | 5.5e-08 | 1.1e-07 | 3.2e-08 | 6.567E-08 | 4.008E-08 | 3.200E-08 | 1.100E-07 |
| 156 | 8 | D8S | A | 4.5e-08 | 2.3e-08 | 5.2e-08 | 4.000E-08 | 1.513E-08 | 2.300E-08 | 5.200E-08 |
| 157 | 8 | D8T | A | 2.5e-08 | 4.6e-08 | 8.7e-08 | 5.267E-08 | 3.153E-08 | 2.500E-08 | 8.700E-08 |
| 158 | 8 | D8W | A | 1e-07 | 3.7e-08 | 2.5e-08 | 5.400E-08 | 4.029E-08 | 2.500E-08 | 1.000E-07 |
| 159 | 8 | D8Y | A | 1.2e-07 | 1.4e-08 | 6.9e-08 | 6.767E-08 | 5.301E-08 | 1.400E-08 | 1.200E-07 |
| 160 | 8 | D8V | A | 2.3e-08 | 1.4e-08 | 4.4e-08 | 2.700E-08 | 1.539E-08 | 1.400E-08 | 4.400E-08 |
| 161 | 9 | M9A | A | 4.5e-08 | 5.1e-08 | 4e-08 | 4.533E-08 | 5.508E-09 | 4.000E-08 | 5.100E-08 |
| 162 | 9 | M9R | A | 4.7e-08 | 6.8e-08 | 8.5e-09 | 4.117E-08 | 3.018E-08 | 8.500E-09 | 6.800E-08 |
| 163 | 9 | M9N | A | 3.2e-08 | 4.5e-08 | 4.6e-08 | 4.100E-08 | 7.810E-09 | 3.200E-08 | 4.600E-08 |
| 164 | 9 | M9D | A | 8.4e-08 | 1.7e-08 | 5.4e-08 | 5.167E-08 | 3.356E-08 | 1.700E-08 | 8.400E-08 |
| 165 | 9 | M9C | A | 4.8e-08 | 2.6e-08 | 2.8e-08 | 3.400E-08 | 1.217E-08 | 2.600E-08 | 4.800E-08 |
| 166 | 9 | M9E | A | 5.8e-08 | 5e-08 | 5.3e-08 | 5.367E-08 | 4.041E-09 | 5.000E-08 | 5.800E-08 |
| 167 | 9 | M9Q | A | 2.3e-08 | 1.2e-08 | 1.1e-07 | 4.833E-08 | 5.369E-08 | 1.200E-08 | 1.100E-07 |
| 168 | 9 | M9G | A | 4.4e-08 | 4.5e-08 | 1.7e-08 | 3.533E-08 | 1.589E-08 | 1.700E-08 | 4.500E-08 |
| 169 | 9 | M9H | A | 5.9e-08 | 5.2e-08 | 6.1e-08 | 5.733E-08 | 4.726E-09 | 5.200E-08 | 6.100E-08 |

| | | | | | | | | | | |
|---|---|---|---|---|---|---|---|---|---|---|
| 170 | 9 | M9I | A | 6.1e-08 | 3e-08 | 3.1e-08 | 4.067E-08 | 1.762E-08 | 3.000E-08 | 6.100E-08 |
| 171 | 9 | M9L | A | 2.9e-08 | 2e-08 | 6.2e-08 | 3.700E-08 | 2.211E-08 | 2.000E-08 | 6.200E-08 |
| 172 | 9 | M9K | A | 6.7e-08 | 6.6e-08 | 2.1e-08 | 5.133E-08 | 2.627E-08 | 2.100E-08 | 6.700E-08 |
| 173 | 9 | M9M | A | 9.5e-08 | 5.6e-08 | 7.2e-08 | 7.433E-08 | 1.960E-08 | 5.600E-08 | 9.500E-08 |
| 174 | 9 | M9F | A | 5.5e-08 | 7.4e-08 | 4.9e-08 | 5.933E-08 | 1.305E-08 | 4.900E-08 | 7.400E-08 |
| 175 | 9 | M9P | A | 2e-08 | 4.4e-08 | 4.9e-08 | 3.767E-08 | 1.550E-08 | 2.000E-08 | 4.900E-08 |
| 176 | 9 | M9S | A | 2.9e-08 | 2.6e-08 | 3.1e-08 | 2.867E-08 | 2.517E-09 | 2.600E-08 | 3.100E-08 |
| 177 | 9 | M9T | A | 9.4e-08 | 7.6e-09 | 4.4e-08 | 4.853E-08 | 4.338E-08 | 7.600E-09 | 9.400E-08 |
| 178 | 9 | M9W | A | 5e-08 | 8.7e-08 | 4.9e-08 | 6.200E-08 | 2.166E-08 | 4.900E-08 | 8.700E-08 |
| 179 | 9 | M9Y | A | 5e-08 | 4.9e-08 | 1e-07 | 6.633E-08 | 2.916E-08 | 4.900E-08 | 1.000E-07 |
| 180 | 9 | M9V | A | 2e-08 | 7.9e-08 | 3.1e-08 | 4.333E-08 | 3.137E-08 | 2.000E-08 | 7.900E-08 |
| 181 | 10 | K10A | A | 1.6e-08 | 1e-07 | 7.9e-08 | 6.500E-08 | 4.371E-08 | 1.600E-08 | 1.000E-07 |
| 182 | 10 | K10R | A | 1.7e-08 | 9.7e-08 | 5.1e-08 | 5.500E-08 | 4.015E-08 | 1.700E-08 | 9.700E-08 |
| 183 | 10 | K10N | A | 1.7e-08 | 6.3e-08 | 5.2e-08 | 4.400E-08 | 2.402E-08 | 1.700E-08 | 6.300E-08 |
| 184 | 10 | K10D | A | 1.4e-08 | 5.3e-08 | 2.2e-08 | 2.967E-08 | 2.060E-08 | 1.400E-08 | 5.300E-08 |
| 185 | 10 | K10C | A | 3.9e-08 | 2.2e-08 | 2.4e-08 | 2.833E-08 | 9.292E-09 | 2.200E-08 | 3.900E-08 |
| 186 | 10 | K10E | A | 2.4e-08 | 2.9e-08 | 1.7e-08 | 2.333E-08 | 6.028E-09 | 1.700E-08 | 2.900E-08 |
| 187 | 10 | K10Q | A | 7.6e-08 | 8.9e-08 | 4.6e-08 | 7.033E-08 | 2.205E-08 | 4.600E-08 | 8.900E-08 |
| 188 | 10 | K10G | A | 3.4e-08 | 2e-08 | 4.8e-08 | 3.400E-08 | 1.400E-08 | 2.000E-08 | 4.800E-08 |
| 189 | 10 | K10H | A | 3e-08 | 3.4e-08 | 9.7e-08 | 5.367E-08 | 3.758E-08 | 3.000E-08 | 9.700E-08 |
| 190 | 10 | K10I | A | 1.3e-08 | 1.7e-08 | 3.8e-08 | 2.267E-08 | 1.343E-08 | 1.300E-08 | 3.800E-08 |
| 191 | 10 | K10L | A | 1e-08 | 4.1e-08 | 3.1e-08 | 2.733E-08 | 1.582E-08 | 1.000E-08 | 4.100E-08 |
| 192 | 10 | K10K | A | 6.6e-08 | 5.6e-08 | 2.5e-08 | 4.900E-08 | 2.138E-08 | 2.500E-08 | 6.600E-08 |
| 193 | 10 | K10M | A | 3e-08 | 6.3e-08 | 4.1e-08 | 4.467E-08 | 1.680E-08 | 3.000E-08 | 6.300E-08 |
| 194 | 10 | K10F | A | 1.9e-08 | 1.5e-08 | 4.2e-08 | 2.533E-08 | 1.457E-08 | 1.500E-08 | 4.200E-08 |
| 195 | 10 | K10P | A | 9.6e-08 | 2.9e-08 | 3.2e-08 | 5.233E-08 | 3.785E-08 | 2.900E-08 | 9.600E-08 |
| 196 | 10 | K10S | A | 5.8e-08 | 1e-07 | 2.8e-08 | 6.200E-08 | 3.617E-08 | 2.800E-08 | 1.000E-07 |
| 197 | 10 | K10T | A | 6.4e-08 | 4.8e-08 | 3.9e-08 | 5.033E-08 | 1.266E-08 | 3.900E-08 | 6.400E-08 |
| 198 | 10 | K10W | A | 6.5e-08 | 2.1e-08 | 4.1e-08 | 4.233E-08 | 2.203E-08 | 2.100E-08 | 6.500E-08 |
| 199 | 10 | K10Y | A | 2.5e-08 | 1.8e-08 | 3.8e-08 | 2.700E-08 | 1.015E-08 | 1.800E-08 | 3.800E-08 |
| 200 | 10 | K10V | A | 5.3e-08 | 2.1e-08 | 3.5e-08 | 3.633E-08 | 1.604E-08 | 2.100E-08 | 5.300E-08 |
| 201 | 11 | L11A | A | 7.4e-08 | 1.6e-08 | 3.5e-08 | 4.167E-08 | 2.957E-08 | 1.600E-08 | 7.400E-08 |
| 202 | 11 | L11R | A | 1.1e-08 | 1.1e-07 | 1.1e-07 | 7.700E-08 | 5.716E-08 | 1.100E-08 | 1.100E-07 |
| 203 | 11 | L11N | A | 5.9e-08 | 4.3e-08 | 3.5e-08 | 4.567E-08 | 1.222E-08 | 3.500E-08 | 5.900E-08 |
| 204 | 11 | L11D | A | 1.5e-08 | 2.4e-08 | 1.6e-08 | 1.833E-08 | 4.933E-09 | 1.500E-08 | 2.400E-08 |
| 205 | 11 | L11C | A | 1.8e-08 | 1.9e-08 | 6.5e-08 | 3.400E-08 | 2.685E-08 | 1.800E-08 | 6.500E-08 |
| 206 | 11 | L11E | A | 6.1e-08 | 8.3e-08 | 6.7e-08 | 7.033E-08 | 1.137E-08 | 6.100E-08 | 8.300E-08 |
| 207 | 11 | L11Q | A | 3.2e-08 | 5.6e-08 | 5.2e-08 | 4.667E-08 | 1.286E-08 | 3.200E-08 | 5.600E-08 |
| 208 | 11 | L11G | A | 5.9e-08 | 5.9e-08 | 5.7e-08 | 5.833E-08 | 1.155E-09 | 5.700E-08 | 5.900E-08 |
| 209 | 11 | L11H | A | 2.5e-08 | 1.8e-08 | 2.5e-08 | 2.267E-08 | 4.041E-09 | 1.800E-08 | 2.500E-08 |
| 210 | 11 | L11I | A | 3.4e-08 | 3.9e-08 | 2.1e-08 | 3.133E-08 | 9.292E-09 | 2.100E-08 | 3.900E-08 |
| 211 | 11 | L11L | A | 2.3e-08 | 8.6e-08 | 2.6e-08 | 4.500E-08 | 3.554E-08 | 2.300E-08 | 8.600E-08 |
| 212 | 11 | L11K | A | 2.4e-08 | 4.8e-08 | 9.4e-08 | 5.533E-08 | 3.557E-08 | 2.400E-08 | 9.400E-08 |
| 213 | 11 | L11M | A | 8.5e-08 | 3.5e-08 | 2.2e-08 | 4.733E-08 | 3.326E-08 | 2.200E-08 | 8.500E-08 |
| 214 | 11 | L11F | A | 4.5e-08 | 1.3e-08 | 3.3e-08 | 3.033E-08 | 1.617E-08 | 1.300E-08 | 4.500E-08 |
| 215 | 11 | L11P | A | 2.6e-08 | 5.7e-08 | 4.9e-08 | 4.400E-08 | 1.609E-08 | 2.600E-08 | 5.700E-08 |
| 216 | 11 | L11S | A | 2.3e-08 | 8e-08 | 6.2e-08 | 5.500E-08 | 2.914E-08 | 2.300E-08 | 8.000E-08 |
| 217 | 11 | L11T | A | 6.4e-08 | 2e-08 | 6.3e-08 | 4.900E-08 | 2.512E-08 | 2.000E-08 | 6.400E-08 |
| 218 | 11 | L11W | A | 3.1e-08 | 1.2e-07 | 3.6e-08 | 6.233E-08 | 5.000E-08 | 3.100E-08 | 1.200E-07 |
| 219 | 11 | L11Y | A | 2.4e-08 | 9.7e-08 | 6.1e-08 | 6.067E-08 | 3.650E-08 | 2.400E-08 | 9.700E-08 |
| 220 | 11 | L11V | A | 1.5e-08 | 3.2e-08 | 3.3e-08 | 2.667E-08 | 1.012E-08 | 1.500E-08 | 3.300E-08 |
| 221 | 12 | R12A | A | 2.4e-08 | 1.5e-08 | 5.5e-08 | 3.133E-08 | 2.098E-08 | 1.500E-08 | 5.500E-08 |
| 222 | 12 | R12R | A | 2.5e-08 | 4.5e-08 | 1.7e-08 | 2.900E-08 | 1.442E-08 | 1.700E-08 | 4.500E-08 |
| 223 | 12 | R12N | A | 8.6e-08 | 2e-08 | 1e-07 | 6.867E-08 | 4.272E-08 | 2.000E-08 | 1.000E-07 |
| 224 | 12 | R12D | A | 4.1e-08 | 2.4e-08 | 4.4e-08 | 3.633E-08 | 1.079E-08 | 2.400E-08 | 4.400E-08 |
| 225 | 12 | R12C | A | 3.6e-08 | 7.2e-08 | 2.5e-08 | 4.433E-08 | 2.458E-08 | 2.500E-08 | 7.200E-08 |
| 226 | 12 | R12E | A | 2.4e-08 | 5.6e-08 | 2.8e-08 | 3.600E-08 | 1.744E-08 | 2.400E-08 | 5.600E-08 |
| 227 | 12 | R12Q | A | 6.4e-08 | 8.1e-08 | 7.2e-08 | 7.233E-08 | 8.505E-09 | 6.400E-08 | 8.100E-08 |
| 228 | 12 | R12G | A | 4e-08 | 4.8e-08 | 1.6e-08 | 3.467E-08 | 1.665E-08 | 1.600E-08 | 4.800E-08 |
| 229 | 12 | R12H | A | 2.5e-08 | 7.5e-08 | 3.9e-08 | 4.633E-08 | 2.579E-08 | 2.500E-08 | 7.500E-08 |
| 230 | 12 | R12I | A | 2.7e-08 | 2.6e-08 | 4.1e-08 | 3.133E-08 | 8.386E-09 | 2.600E-08 | 4.100E-08 |
| 231 | 12 | R12L | A | 1.4e-08 | 1e-07 | 6e-08 | 5.800E-08 | 4.303E-08 | 1.400E-08 | 1.000E-07 |
| 232 | 12 | R12K | A | 3.9e-08 | 1.7e-07 | 7.8e-08 | 9.567E-08 | 6.726E-08 | 3.900E-08 | 1.700E-07 |
| 233 | 12 | R12M | A | 2.4e-08 | 1.1e-08 | 3.3e-08 | 2.267E-08 | 1.106E-08 | 1.100E-08 | 3.300E-08 |
| 234 | 12 | R12F | A | 2.7e-08 | 4.5e-08 | 1.9e-08 | 3.033E-08 | 1.332E-08 | 1.900E-08 | 4.500E-08 |
| 235 | 12 | R12P | A | 6.9e-08 | 2.5e-08 | 7.6e-08 | 5.667E-08 | 2.765E-08 | 2.500E-08 | 7.600E-08 |
| 236 | 12 | R12S | A | 5.1e-08 | 2.5e-08 | 4.9e-08 | 4.167E-08 | 1.447E-08 | 2.500E-08 | 5.100E-08 |
| 237 | 12 | R12T | A | 6.7e-08 | 5.3e-08 | 1.1e-07 | 7.667E-08 | 2.970E-08 | 5.300E-08 | 1.100E-07 |
| 238 | 12 | R12W | A | 6.8e-08 | 4.3e-08 | 4.7e-08 | 5.267E-08 | 1.343E-08 | 4.300E-08 | 6.800E-08 |
| 239 | 12 | R12Y | A | 4.9e-08 | 2.5e-08 | 8.7e-08 | 5.367E-08 | 3.126E-08 | 2.500E-08 | 8.700E-08 |
| 240 | 12 | R12V | A | 1.4e-08 | 6.7e-08 | 3.6e-08 | 3.900E-08 | 2.663E-08 | 1.400E-08 | 6.700E-08 |
| 241 | 13 | L13A | A | 3e-08 | 2.6e-08 | 6.2e-08 | 3.933E-08 | 1.973E-08 | 2.600E-08 | 6.200E-08 |
| 242 | 13 | L13R | A | 1.1e-07 | 1.9e-08 | 3.1e-08 | 5.333E-08 | 4.944E-08 | 1.900E-08 | 1.100E-07 |
| 243 | 13 | L13N | A | 3.1e-08 | 5.1e-08 | 2e-08 | 3.400E-08 | 1.572E-08 | 2.000E-08 | 5.100E-08 |
| 244 | 13 | L13D | A | 3.7e-08 | 9.1e-08 | 6.8e-08 | 6.533E-08 | 2.710E-08 | 3.700E-08 | 9.100E-08 |
| 245 | 13 | L13C | A | 5.9e-08 | 3.1e-08 | 2.5e-08 | 3.833E-08 | 1.815E-08 | 2.500E-08 | 5.900E-08 |
| 246 | 13 | L13E | A | 2.7e-08 | 3.5e-08 | 3.4e-08 | 3.200E-08 | 4.359E-09 | 2.700E-08 | 3.500E-08 |
| 247 | 13 | L13Q | A | 5.5e-08 | 4.6e-08 | 1.3e-07 | 7.700E-08 | 4.612E-08 | 4.600E-08 | 1.300E-07 |
| 248 | 13 | L13G | A | 2.7e-08 | 2.1e-08 | 2.5e-08 | 2.433E-08 | 3.055E-09 | 2.100E-08 | 2.700E-08 |
| 249 | 13 | L13H | A | 3.3e-08 | 4.8e-08 | 9.4e-08 | 5.833E-08 | 3.179E-08 | 3.300E-08 | 9.400E-08 |
| 250 | 13 | L13I | A | 5.8e-08 | 4.7e-08 | 5.3e-08 | 5.267E-08 | 5.508E-09 | 4.700E-08 | 5.800E-08 |
| 251 | 13 | L13L | A | 6.4e-08 | 4.5e-08 | 6.2e-08 | 5.700E-08 | 1.044E-08 | 4.500E-08 | 6.400E-08 |
| 252 | 13 | L13K | A | 8.6e-08 | 3.7e-08 | 4.4e-08 | 5.567E-08 | 2.650E-08 | 3.700E-08 | 8.600E-08 |
| 253 | 13 | L13M | A | 3.2e-08 | 1.8e-08 | 7.7e-08 | 4.233E-08 | 3.083E-08 | 1.800E-08 | 7.700E-08 |
| 254 | 13 | L13F | A | 2.6e-08 | 7.7e-08 | 7.4e-08 | 5.900E-08 | 2.862E-08 | 2.600E-08 | 7.700E-08 |

| | | | | | | | | | | |
|---|---|---|---|---|---|---|---|---|---|---|
| 255 | 13 | L13P | A | 2.8e-08 | 4.3e-08 | 5.7e-08 | 4.267E-08 | 1.450E-08 | 2.800E-08 | 5.700E-08 |
| 256 | 13 | L13S | A | 4.6e-08 | 4.9e-08 | 3.7e-08 | 4.400E-08 | 6.245E-09 | 3.700E-08 | 4.900E-08 |
| 257 | 13 | L13T | A | 4.1e-08 | 1.3e-07 | 2e-08 | 6.367E-08 | 5.840E-08 | 2.000E-08 | 1.300E-07 |
| 258 | 13 | L13W | A | 6e-08 | 2.5e-08 | 3.4e-08 | 3.967E-08 | 1.818E-08 | 2.500E-08 | 6.000E-08 |
| 259 | 13 | L13Y | A | 5.1e-08 | 3.7e-08 | 7e-08 | 5.267E-08 | 1.656E-08 | 3.700E-08 | 7.000E-08 |
| 260 | 13 | L13V | A | 2.3e-08 | 7.9e-08 | 1.2e-08 | 3.800E-08 | 3.593E-08 | 1.200E-08 | 7.900E-08 |
| 261 | 14 | P14A | A | 9.9e-08 | 2.3e-08 | 4.6e-08 | 5.600E-08 | 3.897E-08 | 2.300E-08 | 9.900E-08 |
| 262 | 14 | P14R | A | 3.5e-08 | 6.4e-08 | 3e-08 | 4.300E-08 | 1.836E-08 | 3.000E-08 | 6.400E-08 |
| 263 | 14 | P14N | A | 6.8e-08 | 9.6e-08 | 1e-07 | 8.800E-08 | 1.744E-08 | 6.800E-08 | 1.000E-07 |
| 264 | 14 | P14D | A | 2.2e-08 | 2e-08 | 1e-07 | 4.733E-08 | 4.562E-08 | 2.000E-08 | 1.000E-07 |
| 265 | 14 | P14C | A | 4e-08 | 4.7e-08 | 1.7e-08 | 3.467E-08 | 1.570E-08 | 1.700E-08 | 4.700E-08 |
| 266 | 14 | P14E | A | 5.3e-08 | 5.2e-08 | 3.3e-08 | 4.600E-08 | 1.127E-08 | 3.300E-08 | 5.300E-08 |
| 267 | 14 | P14Q | A | 1.4e-08 | 3.5e-08 | 2.5e-08 | 2.467E-08 | 1.050E-08 | 1.400E-08 | 3.500E-08 |
| 268 | 14 | P14G | A | 3.2e-08 | 2.5e-08 | 1.1e-07 | 5.567E-08 | 4.718E-08 | 2.500E-08 | 1.100E-07 |
| 269 | 14 | P14H | A | 3.8e-08 | 1.4e-07 | 5.3e-08 | 7.700E-08 | 5.507E-08 | 3.800E-08 | 1.400E-07 |
| 270 | 14 | P14I | A | 8.9e-08 | 6.2e-08 | 4.4e-08 | 6.500E-08 | 2.265E-08 | 4.400E-08 | 8.900E-08 |
| 271 | 14 | P14L | A | 2.4e-08 | 5.8e-08 | 9.6e-08 | 5.933E-08 | 3.602E-08 | 2.400E-08 | 9.600E-08 |
| 272 | 14 | P14K | A | 8.3e-09 | 7.4e-08 | 3.3e-08 | 3.843E-08 | 3.319E-08 | 8.300E-09 | 7.400E-08 |
| 273 | 14 | P14M | A | 1.2e-07 | 6.6e-08 | 2.7e-08 | 7.100E-08 | 4.670E-08 | 2.700E-08 | 1.200E-07 |
| 274 | 14 | P14F | A | 3.4e-08 | 2.9e-08 | 5.2e-08 | 3.833E-08 | 1.210E-08 | 2.900E-08 | 5.200E-08 |
| 275 | 14 | P14P | A | 3.6e-08 | 5.1e-08 | 5e-08 | 4.567E-08 | 8.386E-09 | 3.600E-08 | 5.100E-08 |
| 276 | 14 | P14S | A | 1.7e-08 | 2.8e-08 | 1.7e-08 | 2.067E-08 | 6.351E-09 | 1.700E-08 | 2.800E-08 |
| 277 | 14 | P14T | A | 3.8e-09 | 3e-08 | 4e-08 | 2.460E-08 | 1.869E-08 | 3.800E-09 | 4.000E-08 |
| 278 | 14 | P14W | A | 5.1e-08 | 1.9e-08 | 4.9e-08 | 3.967E-08 | 1.793E-08 | 1.900E-08 | 5.100E-08 |
| 279 | 14 | P14Y | A | 3e-08 | 2.7e-08 | 6e-08 | 3.900E-08 | 1.825E-08 | 2.700E-08 | 6.000E-08 |
| 280 | 14 | P14V | A | 2.9e-08 | 6.6e-08 | 6.6e-08 | 5.367E-08 | 2.136E-08 | 2.900E-08 | 6.600E-08 |
| 281 | 15 | A15A | A | 2.6e-08 | 8.6e-08 | 6.7e-08 | 5.967E-08 | 3.066E-08 | 2.600E-08 | 8.600E-08 |
| 282 | 15 | A15R | A | 2.3e-08 | 5.7e-08 | 7.1e-08 | 5.033E-08 | 2.468E-08 | 2.300E-08 | 7.100E-08 |
| 283 | 15 | A15N | A | 1.1e-07 | 4.9e-08 | 2.7e-08 | 6.200E-08 | 4.300E-08 | 2.700E-08 | 1.100E-07 |
| 284 | 15 | A15D | A | 2.4e-08 | 1.8e-08 | 4.7e-08 | 2.967E-08 | 1.531E-08 | 1.800E-08 | 4.700E-08 |
| 285 | 15 | A15C | A | 2.9e-08 | 9.3e-08 | 6e-08 | 6.067E-08 | 3.201E-08 | 2.900E-08 | 9.300E-08 |
| 286 | 15 | A15E | A | 3.8e-08 | 3.1e-08 | 1.4e-07 | 6.967E-08 | 6.101E-08 | 3.100E-08 | 1.400E-07 |
| 287 | 15 | A15Q | A | 4.5e-07 | 3.7e-08 | 8.2e-08 | 1.897E-07 | 2.266E-07 | 3.700E-08 | 4.500E-07 |
| 288 | 15 | A15G | A | 6.6e-08 | 2.2e-08 | 5.6e-08 | 4.800E-08 | 2.307E-08 | 2.200E-08 | 6.600E-08 |
| 289 | 15 | A15H | A | 6.3e-08 | 4.1e-08 | 3.2e-08 | 4.533E-08 | 1.595E-08 | 3.200E-08 | 6.300E-08 |
| 290 | 15 | A15I | A | 2.3e-08 | 2.6e-08 | 5.5e-08 | 3.467E-08 | 1.767E-08 | 2.300E-08 | 5.500E-08 |
| 291 | 15 | A15L | A | 7.6e-08 | 9.7e-09 | 1.3e-08 | 3.290E-08 | 3.736E-08 | 9.700E-09 | 7.600E-08 |
| 292 | 15 | A15K | A | 2.7e-08 | 3.5e-08 | 1.7e-07 | 7.733E-08 | 8.035E-08 | 2.700E-08 | 1.700E-07 |
| 293 | 15 | A15M | A | 4e-08 | 3.2e-08 | 8.2e-08 | 5.133E-08 | 2.686E-08 | 3.200E-08 | 8.200E-08 |
| 294 | 15 | A15F | A | 2.1e-08 | 1.5e-08 | 5.8e-08 | 3.133E-08 | 2.329E-08 | 1.500E-08 | 5.800E-08 |
| 295 | 15 | A15P | A | 1.4e-07 | 3e-08 | 9.5e-08 | 8.833E-08 | 5.530E-08 | 3.000E-08 | 1.400E-07 |
| 296 | 15 | A15S | A | 4.9e-08 | 3.4e-08 | 3.4e-08 | 3.900E-08 | 8.660E-09 | 3.400E-08 | 4.900E-08 |
| 297 | 15 | A15T | A | 1e-07 | 3.7e-08 | 3.5e-08 | 5.733E-08 | 3.696E-08 | 3.500E-08 | 1.000E-07 |
| 298 | 15 | A15W | A | 5.7e-08 | 2.1e-08 | 2.7e-08 | 3.500E-08 | 1.929E-08 | 2.100E-08 | 5.700E-08 |
| 299 | 15 | A15Y | A | 1e-07 | 1.3e-08 | 4.2e-08 | 5.167E-08 | 4.430E-08 | 1.300E-08 | 1.000E-07 |
| 300 | 15 | A15V | A | 5.6e-08 | 9.3e-08 | 6.6e-08 | 7.167E-08 | 1.914E-08 | 5.600E-08 | 9.300E-08 |
| 301 | 16 | S16A | A | 2e-08 | 1.2e-07 | 1.2e-07 | 8.667E-08 | 5.774E-08 | 2.000E-08 | 1.200E-07 |
| 302 | 16 | S16R | A | 4.7e-08 | 4.1e-08 | 5.6e-08 | 4.800E-08 | 7.550E-09 | 4.100E-08 | 5.600E-08 |
| 303 | 16 | S16N | A | 6e-08 | 1.3e-07 | 9e-08 | 9.333E-08 | 3.512E-08 | 6.000E-08 | 1.300E-07 |
| 304 | 16 | S16D | A | 5.6e-08 | 1.1e-08 | 5.2e-08 | 3.967E-08 | 2.491E-08 | 1.100E-08 | 5.600E-08 |
| 305 | 16 | S16C | A | 4.3e-08 | 7.4e-08 | 4.2e-08 | 5.300E-08 | 1.819E-08 | 4.200E-08 | 7.400E-08 |
| 306 | 16 | S16E | A | 2.9e-08 | 2.2e-08 | 4e-08 | 3.033E-08 | 9.074E-09 | 2.200E-08 | 4.000E-08 |
| 307 | 16 | S16Q | A | 7.4e-08 | 1e-07 | 9.5e-08 | 8.967E-08 | 1.380E-08 | 7.400E-08 | 1.000E-07 |
| 308 | 16 | S16G | A | 5.9e-08 | 7.4e-08 | 4.9e-08 | 6.067E-08 | 1.258E-08 | 4.900E-08 | 7.400E-08 |
| 309 | 16 | S16H | A | 2.4e-08 | 1.4e-08 | 3.6e-08 | 2.467E-08 | 1.102E-08 | 1.400E-08 | 3.600E-08 |
| 310 | 16 | S16I | A | 3.8e-08 | 6.1e-08 | 3.4e-08 | 4.433E-08 | 1.457E-08 | 3.400E-08 | 6.100E-08 |
| 311 | 16 | S16L | A | 4.4e-08 | 2.6e-08 | 2.6e-08 | 3.200E-08 | 1.039E-08 | 2.600E-08 | 4.400E-08 |
| 312 | 16 | S16K | A | 2.7e-08 | 5.9e-08 | 5.7e-08 | 4.767E-08 | 1.793E-08 | 2.700E-08 | 5.900E-08 |
| 313 | 16 | S16M | A | 4e-07 | 2e-08 | 8.8e-08 | 1.693E-07 | 2.026E-07 | 2.000E-08 | 4.000E-07 |
| 314 | 16 | S16F | A | 9.4e-08 | 8.1e-08 | 3.5e-08 | 7.000E-08 | 3.100E-08 | 3.500E-08 | 9.400E-08 |
| 315 | 16 | S16P | A | 4.2e-08 | 8.8e-08 | 3.4e-08 | 5.467E-08 | 2.914E-08 | 3.400E-08 | 8.800E-08 |
| 316 | 16 | S16S | A | 1.4e-07 | 3.2e-08 | 5.6e-08 | 7.600E-08 | 5.671E-08 | 3.200E-08 | 1.400E-07 |
| 317 | 16 | S16T | A | 2.8e-08 | 6.2e-08 | 5e-08 | 4.667E-08 | 1.724E-08 | 2.800E-08 | 6.200E-08 |
| 318 | 16 | S16W | A | 6.5e-08 | 3.6e-08 | 3.4e-08 | 4.500E-08 | 1.735E-08 | 3.400E-08 | 6.500E-08 |
| 319 | 16 | S16Y | A | 7.4e-08 | 4.6e-08 | 2.9e-08 | 4.967E-08 | 2.272E-08 | 2.900E-08 | 7.400E-08 |
| 320 | 16 | S16V | A | 4e-08 | 1.2e-07 | 1.2e-08 | 5.733E-08 | 5.605E-08 | 1.200E-08 | 1.200E-07 |
| 321 | 17 | P17A | A | 1.3e-08 | 4.4e-08 | 9.8e-08 | 5.167E-08 | 4.302E-08 | 1.300E-08 | 9.800E-08 |
| 322 | 17 | P17R | A | 4.6e-08 | 5.3e-08 | 7.7e-08 | 5.867E-08 | 1.626E-08 | 4.600E-08 | 7.700E-08 |
| 323 | 17 | P17N | A | 2e-08 | 5.7e-08 | 2.1e-08 | 3.267E-08 | 2.108E-08 | 2.000E-08 | 5.700E-08 |
| 324 | 17 | P17D | A | 3.1e-08 | 4.9e-08 | 2.1e-08 | 3.367E-08 | 1.419E-08 | 2.100E-08 | 4.900E-08 |
| 325 | 17 | P17C | A | 7e-08 | 2.9e-08 | 2.3e-08 | 4.067E-08 | 2.558E-08 | 2.300E-08 | 7.000E-08 |
| 326 | 17 | P17E | A | 8.1e-08 | 5.8e-08 | 3e-08 | 5.633E-08 | 2.554E-08 | 3.000E-08 | 8.100E-08 |
| 327 | 17 | P17Q | A | 2.9e-08 | 4.4e-08 | 1.6e-07 | 7.767E-08 | 7.170E-08 | 2.900E-08 | 1.600E-07 |
| 328 | 17 | P17G | A | 5e-08 | 2.8e-08 | 3.6e-08 | 3.800E-08 | 1.114E-08 | 2.800E-08 | 5.000E-08 |
| 329 | 17 | P17H | A | 4.9e-08 | 1.5e-07 | 2.7e-08 | 7.533E-08 | 6.559E-08 | 2.700E-08 | 1.500E-07 |
| 330 | 17 | P17I | A | 5.1e-08 | 2.6e-08 | 1.4e-08 | 3.033E-08 | 1.888E-08 | 1.400E-08 | 5.100E-08 |
| 331 | 17 | P17L | A | 1.1e-08 | 1.3e-08 | 3.2e-08 | 1.867E-08 | 1.159E-08 | 1.100E-08 | 3.200E-08 |
| 332 | 17 | P17K | A | 1.3e-07 | 5.6e-08 | 2.8e-08 | 7.133E-08 | 5.270E-08 | 2.800E-08 | 1.300E-07 |
| 333 | 17 | P17M | A | 1.1e-08 | 3.9e-08 | 3.7e-08 | 2.900E-08 | 1.562E-08 | 1.100E-08 | 3.900E-08 |
| 334 | 17 | P17F | A | 6.4e-08 | 2.4e-08 | 1.2e-07 | 6.933E-08 | 4.822E-08 | 2.400E-08 | 1.200E-07 |
| 335 | 17 | P17P | A | 2e-07 | 2.7e-08 | 3.9e-08 | 8.867E-08 | 9.660E-08 | 2.700E-08 | 2.000E-07 |
| 336 | 17 | P17S | A | 1.5e-08 | 2.9e-08 | 3e-08 | 2.467E-08 | 8.386E-09 | 1.500E-08 | 3.000E-08 |
| 337 | 17 | P17T | A | 1.6e-08 | 8.7e-08 | 2.2e-08 | 4.167E-08 | 3.937E-08 | 1.600E-08 | 8.700E-08 |
| 338 | 17 | P17W | A | 2.5e-08 | 2.4e-08 | 6.2e-09 | 1.840E-08 | 1.058E-08 | 6.200E-09 | 2.500E-08 |
| 339 | 17 | P17Y | A | 9.3e-08 | 4.8e-08 | 1.3e-07 | 9.033E-08 | 4.106E-08 | 4.800E-08 | 1.300E-07 |

| | | | | | | | | | | |
|---|---|---|---|---|---|---|---|---|---|---|
| 340 | 17 | P17V | A | 5.2e-08 | 5.6e-08 | 1.6e-08 | 4.133E-08 | 2.203E-08 | 1.600E-08 | 5.600E-08 |
| 341 | 18 | E18A | A | 3.8e-08 | 8.7e-08 | 4.7e-08 | 5.733E-08 | 2.608E-08 | 3.800E-08 | 8.700E-08 |
| 342 | 18 | E18R | A | 3e-08 | 6.3e-08 | 4.4e-08 | 4.567E-08 | 1.656E-08 | 3.000E-08 | 6.300E-08 |
| 343 | 18 | E18N | A | 4.4e-08 | 4.3e-08 | 2.3e-08 | 3.667E-08 | 1.185E-08 | 2.300E-08 | 4.400E-08 |
| 344 | 18 | E18D | A | 2.1e-08 | 6.3e-08 | 4.9e-08 | 4.433E-08 | 2.139E-08 | 2.100E-08 | 6.300E-08 |
| 345 | 18 | E18C | A | 3.4e-08 | 5.6e-08 | 7.6e-08 | 5.533E-08 | 2.101E-08 | 3.400E-08 | 7.600E-08 |
| 346 | 18 | E18E | A | 4e-08 | 2.5e-08 | 2.4e-08 | 2.967E-08 | 8.963E-09 | 2.400E-08 | 4.000E-08 |
| 347 | 18 | E18Q | A | 1.6e-08 | 3.8e-08 | 3.6e-08 | 3.000E-08 | 1.217E-08 | 1.600E-08 | 3.800E-08 |
| 348 | 18 | E18G | A | 1e-07 | 7.8e-08 | 3e-08 | 6.933E-08 | 3.580E-08 | 3.000E-08 | 1.000E-07 |
| 349 | 18 | E18H | A | 6.8e-08 | 3e-08 | 8.2e-08 | 6.000E-08 | 2.691E-08 | 3.000E-08 | 8.200E-08 |
| 350 | 18 | E18I | A | 2.6e-08 | 4.6e-08 | 8.6e-08 | 5.267E-08 | 3.055E-08 | 2.600E-08 | 8.600E-08 |
| 351 | 18 | E18L | A | 7.8e-08 | 1.6e-08 | 3.6e-08 | 4.333E-08 | 3.164E-08 | 1.600E-08 | 7.800E-08 |
| 352 | 18 | E18K | A | 2e-08 | 9e-09 | 6.1e-08 | 3.000E-08 | 2.740E-08 | 9.000E-09 | 6.100E-08 |
| 353 | 18 | E18M | A | 1.1e-07 | 3.1e-08 | 9.2e-08 | 7.767E-08 | 4.140E-08 | 3.100E-08 | 1.100E-07 |
| 354 | 18 | E18F | A | 3.8e-08 | 4.8e-08 | 2.9e-08 | 3.833E-08 | 9.504E-09 | 2.900E-08 | 4.800E-08 |
| 355 | 18 | E18P | A | 5.5e-08 | 4.1e-08 | 6.5e-08 | 5.367E-08 | 1.206E-08 | 4.100E-08 | 6.500E-08 |
| 356 | 18 | E18S | A | 3.1e-08 | 2.9e-08 | 4.1e-08 | 3.367E-08 | 6.429E-09 | 2.900E-08 | 4.100E-08 |
| 357 | 18 | E18T | A | 2e-08 | 4.6e-08 | 3.5e-08 | 3.367E-08 | 1.305E-08 | 2.000E-08 | 4.600E-08 |
| 358 | 18 | E18W | A | 1.2e-07 | 7.7e-08 | 6.8e-08 | 8.833E-08 | 2.779E-08 | 6.800E-08 | 1.200E-07 |
| 359 | 18 | E18Y | A | 1.6e-08 | 2.5e-08 | 6.9e-08 | 3.667E-08 | 2.836E-08 | 1.600E-08 | 6.900E-08 |
| 360 | 18 | E18V | A | 1.9e-08 | 3e-08 | 4.1e-08 | 3.000E-08 | 1.100E-08 | 1.900E-08 | 4.100E-08 |
| 361 | 19 | T19A | A | 3.2e-08 | 4.9e-08 | 1.3e-07 | 7.033E-08 | 5.237E-08 | 3.200E-08 | 1.300E-07 |
| 362 | 19 | T19R | A | 8.4e-08 | 2.6e-08 | 3.1e-08 | 4.700E-08 | 3.214E-08 | 2.600E-08 | 8.400E-08 |
| 363 | 19 | T19N | A | 5.9e-08 | 4e-08 | 2.9e-08 | 4.267E-08 | 1.518E-08 | 2.900E-08 | 5.900E-08 |
| 364 | 19 | T19D | A | 2.5e-08 | 6e-08 | 3.2e-08 | 3.900E-08 | 1.852E-08 | 2.500E-08 | 6.000E-08 |
| 365 | 19 | T19C | A | 1.2e-07 | 7.9e-08 | 7e-08 | 8.967E-08 | 2.665E-08 | 7.000E-08 | 1.200E-07 |
| 366 | 19 | T19E | A | 4.3e-08 | 6e-08 | 5.5e-08 | 5.267E-08 | 8.737E-09 | 4.300E-08 | 6.000E-08 |
| 367 | 19 | T19Q | A | 6e-08 | 6.2e-08 | 7.5e-08 | 6.567E-08 | 8.145E-09 | 6.000E-08 | 7.500E-08 |
| 368 | 19 | T19G | A | 2.4e-08 | 5e-08 | 4.4e-08 | 3.933E-08 | 1.361E-08 | 2.400E-08 | 5.000E-08 |
| 369 | 19 | T19H | A | 4.4e-08 | 4e-08 | 3.4e-08 | 3.933E-08 | 5.033E-09 | 3.400E-08 | 4.400E-08 |
| 370 | 19 | T19I | A | 2.1e-08 | 6.7e-08 | 5.2e-08 | 4.667E-08 | 2.346E-08 | 2.100E-08 | 6.700E-08 |
| 371 | 19 | T19L | A | 8.2e-08 | 2.6e-07 | 4e-08 | 1.273E-07 | 1.168E-07 | 4.000E-08 | 2.600E-07 |
| 372 | 19 | T19K | A | 3.6e-08 | 1.4e-07 | 4.7e-08 | 7.433E-08 | 5.713E-08 | 3.600E-08 | 1.400E-07 |
| 373 | 19 | T19M | A | 4.8e-08 | 2.5e-08 | 2e-08 | 3.100E-08 | 1.493E-08 | 2.000E-08 | 4.800E-08 |
| 374 | 19 | T19F | A | 1.3e-08 | 5e-08 | 5e-08 | 3.767E-08 | 2.136E-08 | 1.300E-08 | 5.000E-08 |
| 375 | 19 | T19P | A | 6.8e-08 | 2e-08 | 5.6e-08 | 4.800E-08 | 2.498E-08 | 2.000E-08 | 6.800E-08 |
| 376 | 19 | T19S | A | 1.2e-08 | 1.5e-08 | 4.1e-08 | 2.267E-08 | 1.595E-08 | 1.200E-08 | 4.100E-08 |
| 377 | 19 | T19T | A | 7e-08 | 4.7e-08 | 3.4e-08 | 5.033E-08 | 1.823E-08 | 3.400E-08 | 7.000E-08 |
| 378 | 19 | T19W | A | 4.1e-08 | 1.1e-07 | 4.4e-08 | 6.500E-08 | 3.900E-08 | 4.100E-08 | 1.100E-07 |
| 379 | 19 | T19Y | A | 3.1e-08 | 2.8e-08 | 3.3e-08 | 3.067E-08 | 2.517E-09 | 2.800E-08 | 3.300E-08 |
| 380 | 19 | T19V | A | 4.4e-08 | 2.6e-08 | 1.6e-08 | 2.867E-08 | 1.419E-08 | 1.600E-08 | 4.400E-08 |
| 381 | 20 | H20A | A | 6.9e-08 | 5.3e-08 | 1.1e-07 | 7.733E-08 | 2.940E-08 | 5.300E-08 | 1.100E-07 |
| 382 | 20 | H20R | A | 4.4e-08 | 6.3e-08 | 8.2e-08 | 6.300E-08 | 1.900E-08 | 4.400E-08 | 8.200E-08 |
| 383 | 20 | H20N | A | 5.3e-08 | 2.3e-08 | 4.1e-08 | 3.900E-08 | 1.510E-08 | 2.300E-08 | 5.300E-08 |
| 384 | 20 | H20D | A | 9.6e-08 | 3.3e-08 | 4.5e-08 | 5.800E-08 | 3.345E-08 | 3.300E-08 | 9.600E-08 |
| 385 | 20 | H20C | A | 2.9e-08 | 4.6e-08 | 3.3e-08 | 3.600E-08 | 8.888E-09 | 2.900E-08 | 4.600E-08 |
| 386 | 20 | H20E | A | 3.9e-08 | 5.9e-08 | 9.4e-09 | 3.580E-08 | 2.495E-08 | 9.400E-09 | 5.900E-08 |
| 387 | 20 | H20Q | A | 3.5e-08 | 4.9e-08 | 5.8e-08 | 4.733E-08 | 1.159E-08 | 3.500E-08 | 5.800E-08 |
| 388 | 20 | H20G | A | 8.6e-08 | 3.2e-08 | 2.4e-08 | 4.733E-08 | 3.372E-08 | 2.400E-08 | 8.600E-08 |
| 389 | 20 | H20H | A | 3.2e-08 | 6.1e-08 | 2.9e-08 | 4.067E-08 | 1.767E-08 | 2.900E-08 | 6.100E-08 |
| 390 | 20 | H20I | A | 8.2e-08 | 3.8e-08 | 7.5e-08 | 6.500E-08 | 2.364E-08 | 3.800E-08 | 8.200E-08 |
| 391 | 20 | H20L | A | 3.7e-08 | 1.1e-07 | 3.8e-08 | 6.167E-08 | 4.186E-08 | 3.700E-08 | 1.100E-07 |
| 392 | 20 | H20K | A | 2.8e-08 | 1.1e-08 | 2.8e-08 | 2.233E-08 | 9.815E-09 | 1.100E-08 | 2.800E-08 |
| 393 | 20 | H20M | A | 5.5e-08 | 4e-08 | 4e-08 | 4.500E-08 | 8.660E-09 | 4.000E-08 | 5.500E-08 |
| 394 | 20 | H20F | A | 2.9e-08 | 3.8e-08 | 4.7e-08 | 3.800E-08 | 9.000E-09 | 2.900E-08 | 4.700E-08 |
| 395 | 20 | H20P | A | 4.6e-08 | 2.8e-08 | 4.9e-08 | 4.100E-08 | 1.136E-08 | 2.800E-08 | 4.900E-08 |
| 396 | 20 | H20S | A | 3e-08 | 4.8e-08 | 4.5e-08 | 4.100E-08 | 9.644E-09 | 3.000E-08 | 4.800E-08 |
| 397 | 20 | H20T | A | 7.4e-08 | 4.9e-08 | 2e-08 | 4.767E-08 | 2.702E-08 | 2.000E-08 | 7.400E-08 |
| 398 | 20 | H20W | A | 2.5e-08 | 5.7e-08 | 1.1e-07 | 6.400E-08 | 4.293E-08 | 2.500E-08 | 1.100E-07 |
| 399 | 20 | H20Y | A | 4.2e-08 | 3.7e-08 | 1.2e-07 | 6.633E-08 | 4.654E-08 | 3.700E-08 | 1.200E-07 |
| 400 | 20 | H20V | A | 4.7e-08 | 4.4e-08 | 5.5e-08 | 4.867E-08 | 5.686E-09 | 4.400E-08 | 5.500E-08 |
| 401 | 21 | L21A | A | 4.9e-08 | 1.1e-07 | 4.6e-08 | 6.833E-08 | 3.612E-08 | 4.600E-08 | 1.100E-07 |
| 402 | 21 | L21R | A | 3.1e-08 | 6.1e-08 | 3.7e-08 | 4.300E-08 | 1.587E-08 | 3.100E-08 | 6.100E-08 |
| 403 | 21 | L21N | A | 2e-08 | 2.5e-08 | 5.4e-08 | 3.300E-08 | 1.836E-08 | 2.000E-08 | 5.400E-08 |
| 404 | 21 | L21D | A | 5.1e-08 | 5.4e-08 | 2.4e-08 | 4.300E-08 | 1.652E-08 | 2.400E-08 | 5.400E-08 |
| 405 | 21 | L21C | A | 4.9e-08 | 1.9e-08 | 9e-08 | 5.267E-08 | 3.564E-08 | 1.900E-08 | 9.000E-08 |
| 406 | 21 | L21E | A | 5e-08 | 2.1e-08 | 1e-08 | 2.700E-08 | 2.066E-08 | 1.000E-08 | 5.000E-08 |
| 407 | 21 | L21Q | A | 3.7e-08 | 2e-08 | 5.9e-08 | 3.867E-08 | 1.955E-08 | 2.000E-08 | 5.900E-08 |
| 408 | 21 | L21G | A | 5.9e-08 | 2.4e-08 | 5.2e-08 | 4.500E-08 | 1.852E-08 | 2.400E-08 | 5.900E-08 |
| 409 | 21 | L21H | A | 6.9e-08 | 3e-08 | 8.8e-08 | 6.233E-08 | 2.957E-08 | 3.000E-08 | 8.800E-08 |
| 410 | 21 | L21I | A | 1.8e-08 | 2.4e-08 | 1.1e-07 | 5.067E-08 | 5.147E-08 | 1.800E-08 | 1.100E-07 |
| 411 | 21 | L21L | A | 3.3e-08 | 3.2e-08 | 2.5e-08 | 3.000E-08 | 4.359E-09 | 2.500E-08 | 3.300E-08 |
| 412 | 21 | L21K | A | 7e-08 | 3.9e-08 | 2.6e-08 | 4.500E-08 | 2.261E-08 | 2.600E-08 | 7.000E-08 |
| 413 | 21 | L21M | A | 3.7e-08 | 6e-08 | 4.8e-08 | 4.833E-08 | 1.150E-08 | 3.700E-08 | 6.000E-08 |
| 414 | 21 | L21F | A | 7.7e-08 | 9.9e-08 | 3.8e-08 | 7.133E-08 | 3.089E-08 | 3.800E-08 | 9.900E-08 |
| 415 | 21 | L21P | A | 7.2e-08 | 3.3e-08 | 3.7e-08 | 4.733E-08 | 2.146E-08 | 3.300E-08 | 7.200E-08 |
| 416 | 21 | L21S | A | 2.7e-08 | 5.3e-08 | 3.5e-08 | 3.833E-08 | 1.332E-08 | 2.700E-08 | 5.300E-08 |
| 417 | 21 | L21T | A | 1.6e-08 | 7.2e-08 | 5.8e-08 | 4.867E-08 | 2.914E-08 | 1.600E-08 | 7.200E-08 |
| 418 | 21 | L21W | A | 3.3e-08 | 6.8e-08 | 7.7e-08 | 5.933E-08 | 2.325E-08 | 3.300E-08 | 7.700E-08 |
| 419 | 21 | L21Y | A | 4.8e-08 | 1e-07 | 2.6e-08 | 5.800E-08 | 3.800E-08 | 2.600E-08 | 1.000E-07 |
| 420 | 21 | L21V | A | 2.1e-08 | 3e-08 | 3.2e-08 | 2.767E-08 | 5.859E-09 | 2.100E-08 | 3.200E-08 |
| 421 | 22 | D22A | A | 4.8e-08 | 3.1e-08 | 1e-07 | 5.967E-08 | 3.595E-08 | 3.100E-08 | 1.000E-07 |
| 422 | 22 | D22R | A | 5.4e-08 | 9.8e-08 | 2.4e-08 | 5.867E-08 | 3.722E-08 | 2.400E-08 | 9.800E-08 |
| 423 | 22 | D22N | A | 3.5e-08 | 7.9e-08 | 9.5e-08 | 6.967E-08 | 3.107E-08 | 3.500E-08 | 9.500E-08 |
| 424 | 22 | D22D | A | 8.6e-08 | 1.2e-07 | 1.8e-08 | 7.467E-08 | 5.194E-08 | 1.800E-08 | 1.200E-07 |

| | | | | | | | | | | |
|---|---|---|---|---|---|---|---|---|---|---|
| 425 | 22 | D22C | A | 5.2e-08 | 3.4e-08 | 4.8e-08 | 4.467E-08 | 9.452E-09 | 3.400E-08 | 5.200E-08 |
| 426 | 22 | D22E | A | 9.4e-08 | 3.4e-08 | 2.1e-08 | 4.967E-08 | 3.894E-08 | 2.100E-08 | 9.400E-08 |
| 427 | 22 | D22Q | A | 6.9e-08 | 5.1e-08 | 1.3e-07 | 8.333E-08 | 4.140E-08 | 5.100E-08 | 1.300E-07 |
| 428 | 22 | D22G | A | 1.1e-08 | 4.5e-08 | 2.8e-08 | 2.800E-08 | 1.700E-08 | 1.100E-08 | 4.500E-08 |
| 429 | 22 | D22H | A | 3.4e-08 | 3.5e-08 | 3e-08 | 3.300E-08 | 2.646E-09 | 3.000E-08 | 3.500E-08 |
| 430 | 22 | D22I | A | 2.6e-08 | 6.1e-08 | 1e-07 | 6.233E-08 | 3.702E-08 | 2.600E-08 | 1.000E-07 |
| 431 | 22 | D22L | A | 3.5e-08 | 8.3e-08 | 4e-08 | 5.267E-08 | 2.639E-08 | 3.500E-08 | 8.300E-08 |
| 432 | 22 | D22K | A | 6.5e-08 | 2.3e-08 | 4.7e-08 | 4.500E-08 | 2.107E-08 | 2.300E-08 | 6.500E-08 |
| 433 | 22 | D22M | A | 1.6e-08 | 8e-08 | 4e-08 | 4.533E-08 | 3.233E-08 | 1.600E-08 | 8.000E-08 |
| 434 | 22 | D22F | A | 3.4e-08 | 2.3e-08 | 2.4e-08 | 2.700E-08 | 6.083E-09 | 2.300E-08 | 3.400E-08 |
| 435 | 22 | D22P | A | 2.8e-08 | 4.4e-08 | 5.1e-08 | 4.100E-08 | 1.179E-08 | 2.800E-08 | 5.100E-08 |
| 436 | 22 | D22S | A | 3.9e-08 | 3.7e-08 | 3.3e-08 | 3.633E-08 | 3.055E-09 | 3.300E-08 | 3.900E-08 |
| 437 | 22 | D22T | A | 3.1e-08 | 4.9e-08 | 1.8e-08 | 3.267E-08 | 1.557E-08 | 1.800E-08 | 4.900E-08 |
| 438 | 22 | D22W | A | 1.4e-07 | 5.1e-08 | 4e-08 | 7.700E-08 | 5.484E-08 | 4.000E-08 | 1.400E-07 |
| 439 | 22 | D22Y | A | 1e-07 | 8.2e-08 | 8.5e-08 | 8.900E-08 | 9.644E-09 | 8.200E-08 | 1.000E-07 |
| 440 | 22 | D22V | A | 7.6e-08 | 4e-08 | 2e-08 | 4.533E-08 | 2.838E-08 | 2.000E-08 | 7.600E-08 |
| 441 | 23 | M23A | A | 5.4e-08 | 4.3e-08 | 5.8e-08 | 5.167E-08 | 7.767E-09 | 4.300E-08 | 5.800E-08 |
| 442 | 23 | M23R | A | 7.6e-08 | 6e-08 | 8.8e-08 | 7.467E-08 | 1.405E-08 | 6.000E-08 | 8.800E-08 |
| 443 | 23 | M23N | A | 8.9e-08 | 9.4e-08 | 2.9e-08 | 7.067E-08 | 3.617E-08 | 2.900E-08 | 9.400E-08 |
| 444 | 23 | M23D | A | 2.2e-08 | 1.6e-08 | 4.5e-08 | 2.767E-08 | 1.531E-08 | 1.600E-08 | 4.500E-08 |
| 445 | 23 | M23C | A | 4.2e-08 | 4e-08 | 5.9e-08 | 4.700E-08 | 1.044E-08 | 4.000E-08 | 5.900E-08 |
| 446 | 23 | M23E | A | 2e-08 | 5.8e-08 | 8.2e-08 | 5.333E-08 | 3.126E-08 | 2.000E-08 | 8.200E-08 |
| 447 | 23 | M23Q | A | 7.1e-08 | 7e-08 | 7.9e-08 | 7.333E-08 | 4.933E-09 | 7.000E-08 | 7.900E-08 |
| 448 | 23 | M23G | A | 6.1e-08 | 5.3e-08 | 3.2e-08 | 4.867E-08 | 1.498E-08 | 3.200E-08 | 6.100E-08 |
| 449 | 23 | M23H | A | 8e-08 | 5.7e-08 | 1e-07 | 7.900E-08 | 2.152E-08 | 5.700E-08 | 1.000E-07 |
| 450 | 23 | M23I | A | 2.9e-08 | 1.9e-07 | 2.5e-08 | 8.133E-08 | 9.413E-08 | 2.500E-08 | 1.900E-07 |
| 451 | 23 | M23L | A | 1.7e-07 | 6.1e-08 | 1.9e-08 | 8.333E-08 | 7.794E-08 | 1.900E-08 | 1.700E-07 |
| 452 | 23 | M23K | A | 4.9e-08 | 2.6e-08 | 3.7e-08 | 3.733E-08 | 1.150E-08 | 2.600E-08 | 4.900E-08 |
| 453 | 23 | M23M | A | 9.6e-08 | 5.4e-08 | 1.1e-07 | 8.667E-08 | 2.914E-08 | 5.400E-08 | 1.100E-07 |
| 454 | 23 | M23F | A | 3.8e-08 | 2.6e-08 | 2.8e-08 | 3.067E-08 | 6.429E-09 | 2.600E-08 | 3.800E-08 |
| 455 | 23 | M23P | A | 3.2e-08 | 1.2e-07 | 3.4e-08 | 6.200E-08 | 5.024E-08 | 3.200E-08 | 1.200E-07 |
| 456 | 23 | M23S | A | 6.5e-08 | 3.2e-08 | 7.5e-08 | 5.733E-08 | 2.250E-08 | 3.200E-08 | 7.500E-08 |
| 457 | 23 | M23T | A | 3.6e-08 | 8.5e-08 | 6.1e-08 | 6.067E-08 | 2.450E-08 | 3.600E-08 | 8.500E-08 |
| 458 | 23 | M23W | A | 4.7e-08 | 3.1e-08 | 7e-08 | 4.933E-08 | 1.960E-08 | 3.100E-08 | 7.000E-08 |
| 459 | 23 | M23Y | A | 4.2e-08 | 2.1e-08 | 9.7e-08 | 5.333E-08 | 3.925E-08 | 2.100E-08 | 9.700E-08 |
| 460 | 23 | M23V | A | 2.1e-08 | 1.2e-07 | 7.6e-08 | 7.233E-08 | 4.960E-08 | 2.100E-08 | 1.200E-07 |
| 461 | 24 | L24A | A | 1e-07 | 1.1e-08 | 1.5e-07 | 8.700E-08 | 7.041E-08 | 1.100E-08 | 1.500E-07 |
| 462 | 24 | L24R | A | 7.1e-08 | 2.2e-08 | 5.9e-08 | 5.067E-08 | 2.554E-08 | 2.200E-08 | 7.100E-08 |
| 463 | 24 | L24N | A | 7.3e-08 | 5.5e-08 | 1.1e-07 | 7.933E-08 | 2.804E-08 | 5.500E-08 | 1.100E-07 |
| 464 | 24 | L24D | A | 5.8e-08 | 6.3e-08 | 6.5e-08 | 6.200E-08 | 3.606E-09 | 5.800E-08 | 6.500E-08 |
| 465 | 24 | L24C | A | 6.4e-08 | 5.3e-08 | 1.5e-07 | 8.900E-08 | 5.311E-08 | 5.300E-08 | 1.500E-07 |
| 466 | 24 | L24E | A | 2.9e-08 | 4.8e-08 | 6.6e-08 | 4.767E-08 | 1.850E-08 | 2.900E-08 | 6.600E-08 |
| 467 | 24 | L24Q | A | 4.4e-08 | 5.5e-08 | 2.9e-08 | 4.267E-08 | 1.305E-08 | 2.900E-08 | 5.500E-08 |
| 468 | 24 | L24G | A | 4.9e-08 | 1.9e-08 | 2.5e-08 | 3.100E-08 | 1.587E-08 | 1.900E-08 | 4.900E-08 |
| 469 | 24 | L24H | A | 1.1e-08 | 4.7e-08 | 6.2e-08 | 4.000E-08 | 2.621E-08 | 1.100E-08 | 6.200E-08 |
| 470 | 24 | L24I | A | 3.5e-08 | 5.1e-08 | 3.7e-08 | 4.100E-08 | 8.718E-09 | 3.500E-08 | 5.100E-08 |
| 471 | 24 | L24L | A | 5.9e-08 | 6.5e-08 | 5.2e-08 | 5.867E-08 | 6.506E-09 | 5.200E-08 | 6.500E-08 |
| 472 | 24 | L24K | A | 2.6e-08 | 4.1e-08 | 3.6e-08 | 3.433E-08 | 7.638E-09 | 2.600E-08 | 4.100E-08 |
| 473 | 24 | L24M | A | 5e-08 | 3.2e-08 | 3.1e-08 | 3.767E-08 | 1.069E-08 | 3.100E-08 | 5.000E-08 |
| 474 | 24 | L24F | A | 2.8e-08 | 6.4e-08 | 4.6e-08 | 4.600E-08 | 1.800E-08 | 2.800E-08 | 6.400E-08 |
| 475 | 24 | L24P | A | 2.9e-08 | 6.8e-08 | 1.1e-07 | 6.900E-08 | 4.051E-08 | 2.900E-08 | 1.100E-07 |
| 476 | 24 | L24S | A | 3.7e-08 | 1.4e-08 | 2.4e-08 | 2.500E-08 | 1.153E-08 | 1.400E-08 | 3.700E-08 |
| 477 | 24 | L24T | A | 1.2e-07 | 6.9e-08 | 1.9e-08 | 6.933E-08 | 5.050E-08 | 1.900E-08 | 1.200E-07 |
| 478 | 24 | L24W | A | 5.3e-08 | 3.6e-08 | 3.5e-08 | 4.133E-08 | 1.012E-08 | 3.500E-08 | 5.300E-08 |
| 479 | 24 | L24Y | A | 5.3e-08 | 1.6e-08 | 3.8e-08 | 3.567E-08 | 1.861E-08 | 1.600E-08 | 5.300E-08 |
| 480 | 24 | L24V | A | 6.2e-08 | 4.1e-08 | 3.2e-08 | 4.500E-08 | 1.539E-08 | 3.200E-08 | 6.200E-08 |
| 481 | 25 | R25A | A | 3.3e-08 | 7.1e-08 | 4.4e-08 | 4.933E-08 | 1.955E-08 | 3.300E-08 | 7.100E-08 |
| 482 | 25 | R25R | A | 1.1e-08 | 5.6e-08 | 3.5e-08 | 3.400E-08 | 2.252E-08 | 1.100E-08 | 5.600E-08 |
| 483 | 25 | R25N | A | 7.9e-08 | 1.6e-08 | 3.8e-08 | 4.433E-08 | 3.197E-08 | 1.600E-08 | 7.900E-08 |
| 484 | 25 | R25D | A | 5.6e-08 | 2.1e-07 | 2.9e-08 | 9.833E-08 | 9.764E-08 | 2.900E-08 | 2.100E-07 |
| 485 | 25 | R25C | A | 8.4e-08 | 1.1e-07 | 5.2e-08 | 8.200E-08 | 2.905E-08 | 5.200E-08 | 1.100E-07 |
| 486 | 25 | R25E | A | 5.4e-08 | 6.8e-08 | 3.3e-08 | 5.167E-08 | 1.762E-08 | 3.300E-08 | 6.800E-08 |
| 487 | 25 | R25Q | A | 2e-08 | 7.1e-08 | 2.7e-08 | 3.933E-08 | 2.765E-08 | 2.000E-08 | 7.100E-08 |
| 488 | 25 | R25G | A | 2.3e-08 | 9.1e-08 | 8.7e-08 | 6.700E-08 | 3.816E-08 | 2.300E-08 | 9.100E-08 |
| 489 | 25 | R25H | A | 3.5e-08 | 4e-08 | 3.4e-08 | 3.633E-08 | 3.215E-09 | 3.400E-08 | 4.000E-08 |
| 490 | 25 | R25I | A | 7.1e-08 | 1.9e-08 | 5.5e-08 | 4.833E-08 | 2.663E-08 | 1.900E-08 | 7.100E-08 |
| 491 | 25 | R25L | A | 3.5e-08 | 4.7e-08 | 7.7e-08 | 5.300E-08 | 2.163E-08 | 3.500E-08 | 7.700E-08 |
| 492 | 25 | R25K | A | 6.7e-08 | 9.1e-08 | 4e-08 | 6.600E-08 | 2.551E-08 | 4.000E-08 | 9.100E-08 |
| 493 | 25 | R25M | A | 3.3e-08 | 2.7e-08 | 4.3e-08 | 3.433E-08 | 8.083E-09 | 2.700E-08 | 4.300E-08 |
| 494 | 25 | R25F | A | 1.1e-08 | 5e-08 | 6.6e-09 | 2.253E-08 | 2.389E-08 | 6.600E-09 | 5.000E-08 |
| 495 | 25 | R25P | A | 4e-08 | 4.3e-08 | 1.4e-08 | 3.233E-08 | 1.595E-08 | 1.400E-08 | 4.300E-08 |
| 496 | 25 | R25S | A | 8.5e-08 | 6.6e-08 | 3.9e-08 | 6.333E-08 | 2.312E-08 | 3.900E-08 | 8.500E-08 |
| 497 | 25 | R25T | A | 7.6e-08 | 4.3e-08 | 6.2e-08 | 6.033E-08 | 1.656E-08 | 4.300E-08 | 7.600E-08 |
| 498 | 25 | R25W | A | 9.4e-08 | 1e-08 | 2.2e-08 | 4.200E-08 | 4.543E-08 | 1.000E-08 | 9.400E-08 |
| 499 | 25 | R25Y | A | 1.4e-08 | 6e-08 | 5.9e-08 | 4.433E-08 | 2.627E-08 | 1.400E-08 | 6.000E-08 |
| 500 | 25 | R25V | A | 2e-08 | 1.4e-08 | 4.7e-08 | 2.700E-08 | 1.758E-08 | 1.400E-08 | 4.700E-08 |
| 501 | 26 | H26A | A | 6.3e-08 | 3.7e-08 | 3.9e-08 | 4.633E-08 | 1.447E-08 | 3.700E-08 | 6.300E-08 |
| 502 | 26 | H26R | A | 6.1e-08 | 2.3e-08 | 5.3e-08 | 4.567E-08 | 2.003E-08 | 2.300E-08 | 6.100E-08 |
| 503 | 26 | H26N | A | 8e-08 | 2.4e-08 | 4.6e-08 | 5.000E-08 | 2.821E-08 | 2.400E-08 | 8.000E-08 |
| 504 | 26 | H26D | A | 1.6e-08 | 5.3e-08 | 5.8e-08 | 4.233E-08 | 2.294E-08 | 1.600E-08 | 5.800E-08 |
| 505 | 26 | H26C | A | 1.1e-07 | 2.1e-08 | 6.8e-08 | 6.633E-08 | 4.452E-08 | 2.100E-08 | 1.100E-07 |
| 506 | 26 | H26E | A | 3.8e-08 | 3.8e-08 | 3.4e-08 | 3.667E-08 | 2.309E-09 | 3.400E-08 | 3.800E-08 |
| 507 | 26 | H26Q | A | 5e-08 | 6.5e-08 | 8.8e-08 | 6.767E-08 | 1.914E-08 | 5.000E-08 | 8.800E-08 |
| 508 | 26 | H26G | A | 2.3e-08 | 6.1e-08 | 4.5e-08 | 4.300E-08 | 1.908E-08 | 2.300E-08 | 6.100E-08 |
| 509 | 26 | H26H | A | 7.8e-08 | 6.6e-08 | 1.4e-07 | 9.467E-08 | 3.972E-08 | 6.600E-08 | 1.400E-07 |

| | | | | | | | | | | |
|---|---|---|---|---|---|---|---|---|---|---|
| 510 | 26 | H26I | A | 4.7e-08 | 8e-08 | 4.1e-08 | 5.600E-08 | 2.100E-08 | 4.100E-08 | 8.000E-08 |
| 511 | 26 | H26L | A | 6.8e-08 | 6e-08 | 3.2e-08 | 5.333E-08 | 1.890E-08 | 3.200E-08 | 6.800E-08 |
| 512 | 26 | H26K | A | 1.8e-07 | 4.9e-08 | 2.1e-08 | 8.333E-08 | 8.488E-08 | 2.100E-08 | 1.800E-07 |
| 513 | 26 | H26M | A | 3.9e-08 | 5.1e-08 | 4.6e-08 | 4.533E-08 | 6.028E-09 | 3.900E-08 | 5.100E-08 |
| 514 | 26 | H26F | A | 3.7e-08 | 1.5e-08 | 3.5e-08 | 2.900E-08 | 1.217E-08 | 1.500E-08 | 3.700E-08 |
| 515 | 26 | H26P | A | 6.1e-08 | 8.7e-08 | 5.8e-08 | 6.867E-08 | 1.595E-08 | 5.800E-08 | 8.700E-08 |
| 516 | 26 | H26S | A | 5.2e-08 | 8e-08 | 1.4e-08 | 4.867E-08 | 3.313E-08 | 1.400E-08 | 8.000E-08 |
| 517 | 26 | H26T | A | 4.7e-08 | 1.8e-08 | 4.4e-08 | 3.633E-08 | 1.595E-08 | 1.800E-08 | 4.700E-08 |
| 518 | 26 | H26W | A | 3.4e-08 | 7.9e-08 | 5e-08 | 5.433E-08 | 2.281E-08 | 3.400E-08 | 7.900E-08 |
| 519 | 26 | H26Y | A | 3.3e-08 | 1.1e-07 | 1.7e-08 | 5.333E-08 | 4.972E-08 | 1.700E-08 | 1.100E-07 |
| 520 | 26 | H26V | A | 6.3e-08 | 3.6e-08 | 1.4e-08 | 3.767E-08 | 2.454E-08 | 1.400E-08 | 6.300E-08 |
| 521 | 27 | L27A | A | 3.4e-08 | 1e-07 | 3.8e-08 | 5.733E-08 | 3.700E-08 | 3.400E-08 | 1.000E-07 |
| 522 | 27 | L27R | A | 3.1e-08 | 2.3e-08 | 3.8e-08 | 3.067E-08 | 7.506E-09 | 2.300E-08 | 3.800E-08 |
| 523 | 27 | L27N | A | 1e-07 | 2.4e-08 | 2.1e-08 | 4.833E-08 | 4.477E-08 | 2.100E-08 | 1.000E-07 |
| 524 | 27 | L27D | A | 2.6e-08 | 3.8e-08 | 5.3e-08 | 3.900E-08 | 1.353E-08 | 2.600E-08 | 5.300E-08 |
| 525 | 27 | L27C | A | 3e-08 | 5.2e-08 | 4.1e-08 | 4.100E-08 | 1.100E-08 | 3.000E-08 | 5.200E-08 |
| 526 | 27 | L27E | A | 4.1e-08 | 6.3e-08 | 9.2e-08 | 6.533E-08 | 2.558E-08 | 4.100E-08 | 9.200E-08 |
| 527 | 27 | L27Q | A | 1.7e-07 | 3e-08 | 2.5e-08 | 7.500E-08 | 8.231E-08 | 2.500E-08 | 1.700E-07 |
| 528 | 27 | L27G | A | 5e-08 | 2.6e-08 | 7.5e-08 | 5.033E-08 | 2.450E-08 | 2.600E-08 | 7.500E-08 |
| 529 | 27 | L27H | A | 5.4e-08 | 1.5e-07 | 8.4e-08 | 9.600E-08 | 4.911E-08 | 5.400E-08 | 1.500E-07 |
| 530 | 27 | L27I | A | 4e-08 | 5.5e-08 | 1.7e-08 | 3.733E-08 | 1.914E-08 | 1.700E-08 | 5.500E-08 |
| 531 | 27 | L27L | A | 1.2e-07 | 4.8e-08 | 2.6e-08 | 6.467E-08 | 4.917E-08 | 2.600E-08 | 1.200E-07 |
| 532 | 27 | L27K | A | 1.6e-08 | 5.6e-08 | 2.1e-08 | 3.100E-08 | 2.179E-08 | 1.600E-08 | 5.600E-08 |
| 533 | 27 | L27M | A | 2.3e-08 | 1.2e-08 | 3.1e-08 | 2.200E-08 | 9.539E-09 | 1.200E-08 | 3.100E-08 |
| 534 | 27 | L27F | A | 2.7e-08 | 3e-08 | 1.8e-08 | 2.500E-08 | 6.245E-09 | 1.800E-08 | 3.000E-08 |
| 535 | 27 | L27P | A | 3.6e-08 | 2.4e-08 | 5.5e-08 | 3.833E-08 | 1.563E-08 | 2.400E-08 | 5.500E-08 |
| 536 | 27 | L27S | A | 1e-07 | 4.2e-08 | 4.4e-08 | 6.200E-08 | 3.292E-08 | 4.200E-08 | 1.000E-07 |
| 537 | 27 | L27T | A | 7e-08 | 9.8e-09 | 1.1e-07 | 6.327E-08 | 5.044E-08 | 9.800E-09 | 1.100E-07 |
| 538 | 27 | L27W | A | 3e-08 | 3.8e-08 | 1.4e-07 | 6.933E-08 | 6.133E-08 | 3.000E-08 | 1.400E-07 |
| 539 | 27 | L27Y | A | 2.2e-08 | 4.3e-08 | 7.3e-08 | 4.600E-08 | 2.563E-08 | 2.200E-08 | 7.300E-08 |
| 540 | 27 | L27V | A | 2.8e-08 | 3e-08 | 3e-08 | 2.933E-08 | 1.155E-09 | 2.800E-08 | 3.000E-08 |
| 541 | 28 | Y28A | A | 1.1e-07 | 1.1e-07 | 2.9e-08 | 8.300E-08 | 4.677E-08 | 2.900E-08 | 1.100E-07 |
| 542 | 28 | Y28R | A | 1.8e-08 | 1.2e-08 | 2.9e-08 | 1.967E-08 | 8.622E-09 | 1.200E-08 | 2.900E-08 |
| 543 | 28 | Y28N | A | 9.7e-09 | 3.8e-08 | 4.6e-08 | 3.123E-08 | 1.907E-08 | 9.700E-09 | 4.600E-08 |
| 544 | 28 | Y28D | A | 1.3e-07 | 1.2e-08 | 3.5e-08 | 5.900E-08 | 6.255E-08 | 1.200E-08 | 1.300E-07 |
| 545 | 28 | Y28C | A | 1.7e-08 | 3.8e-08 | 2.7e-08 | 2.733E-08 | 1.050E-08 | 1.700E-08 | 3.800E-08 |
| 546 | 28 | Y28E | A | 9.3e-08 | 7.3e-08 | 6.6e-08 | 7.733E-08 | 1.401E-08 | 6.600E-08 | 9.300E-08 |
| 547 | 28 | Y28Q | A | 2.3e-08 | 5.5e-08 | 5.6e-08 | 4.467E-08 | 1.877E-08 | 2.300E-08 | 5.600E-08 |
| 548 | 28 | Y28G | A | 3.9e-08 | 5.8e-08 | 1.2e-07 | 7.233E-08 | 4.236E-08 | 3.900E-08 | 1.200E-07 |
| 549 | 28 | Y28H | A | 5.1e-08 | 2.8e-08 | 7.6e-08 | 5.167E-08 | 2.401E-08 | 2.800E-08 | 7.600E-08 |
| 550 | 28 | Y28I | A | 1.3e-07 | 1.1e-07 | 4.2e-08 | 9.400E-08 | 4.613E-08 | 4.200E-08 | 1.300E-07 |
| 551 | 28 | Y28L | A | 2e-08 | 3.3e-08 | 1.5e-08 | 2.267E-08 | 9.292E-09 | 1.500E-08 | 3.300E-08 |
| 552 | 28 | Y28K | A | 7e-08 | 1.9e-08 | 4.2e-08 | 4.367E-08 | 2.554E-08 | 1.900E-08 | 7.000E-08 |
| 553 | 28 | Y28M | A | 1.6e-08 | 3.9e-08 | 2e-08 | 2.500E-08 | 1.229E-08 | 1.600E-08 | 3.900E-08 |
| 554 | 28 | Y28F | A | 2.5e-08 | 3.7e-08 | 1.3e-08 | 2.500E-08 | 1.200E-08 | 1.300E-08 | 3.700E-08 |
| 555 | 28 | Y28P | A | 3.4e-08 | 7e-08 | 7.6e-08 | 6.000E-08 | 2.272E-08 | 3.400E-08 | 7.600E-08 |
| 556 | 28 | Y28S | A | 2.2e-08 | 5.2e-08 | 1.7e-08 | 3.033E-08 | 1.893E-08 | 1.700E-08 | 5.200E-08 |
| 557 | 28 | Y28T | A | 4.9e-08 | 8.1e-08 | 3.9e-08 | 5.633E-08 | 2.194E-08 | 3.900E-08 | 8.100E-08 |
| 558 | 28 | Y28W | A | 4e-08 | 4.7e-08 | 6.6e-08 | 5.100E-08 | 1.345E-08 | 4.000E-08 | 6.600E-08 |
| 559 | 28 | Y28Y | A | 6e-08 | 3.4e-08 | 6.1e-08 | 5.167E-08 | 1.531E-08 | 3.400E-08 | 6.100E-08 |
| 560 | 28 | Y28V | A | 2.7e-08 | 2.8e-08 | 2.6e-08 | 2.700E-08 | 1.000E-09 | 2.600E-08 | 2.800E-08 |
| 561 | 29 | Q29A | A | 8.9e-08 | 2.5e-08 | 3.9e-08 | 5.100E-08 | 3.365E-08 | 2.500E-08 | 8.900E-08 |
| 562 | 29 | Q29R | A | 1.4e-08 | 2.8e-08 | 5.4e-08 | 3.200E-08 | 2.030E-08 | 1.400E-08 | 5.400E-08 |
| 563 | 29 | Q29N | A | 6.7e-08 | 4.3e-08 | 2.6e-08 | 4.533E-08 | 2.060E-08 | 2.600E-08 | 6.700E-08 |
| 564 | 29 | Q29D | A | 5.2e-08 | 5.8e-08 | 9.3e-08 | 6.767E-08 | 2.214E-08 | 5.200E-08 | 9.300E-08 |
| 565 | 29 | Q29C | A | 1.5e-08 | 4.1e-08 | 2.3e-08 | 2.633E-08 | 1.332E-08 | 1.500E-08 | 4.100E-08 |
| 566 | 29 | Q29E | A | 7.4e-08 | 6.3e-08 | 1.2e-08 | 4.967E-08 | 3.308E-08 | 1.200E-08 | 7.400E-08 |
| 567 | 29 | Q29Q | A | 4.9e-08 | 3e-08 | 5.4e-08 | 4.433E-08 | 1.266E-08 | 3.000E-08 | 5.400E-08 |
| 568 | 29 | Q29G | A | 7.9e-08 | 2.9e-08 | 5.3e-08 | 5.367E-08 | 2.501E-08 | 2.900E-08 | 7.900E-08 |
| 569 | 29 | Q29H | A | 4.2e-08 | 6e-08 | 4.7e-08 | 4.967E-08 | 9.292E-09 | 4.200E-08 | 6.000E-08 |
| 570 | 29 | Q29I | A | 1.8e-08 | 6.3e-08 | 3e-08 | 3.700E-08 | 2.330E-08 | 1.800E-08 | 6.300E-08 |
| 571 | 29 | Q29L | A | 3.1e-08 | 3.7e-08 | 3.7e-08 | 3.500E-08 | 3.464E-09 | 3.100E-08 | 3.700E-08 |
| 572 | 29 | Q29K | A | 2e-08 | 3.4e-08 | 2e-08 | 2.467E-08 | 8.083E-09 | 2.000E-08 | 3.400E-08 |
| 573 | 29 | Q29M | A | 5.8e-08 | 9.6e-08 | 7.5e-08 | 7.633E-08 | 1.904E-08 | 5.800E-08 | 9.600E-08 |
| 574 | 29 | Q29F | A | 5.7e-08 | 1.7e-08 | 5.6e-08 | 4.333E-08 | 2.281E-08 | 1.700E-08 | 5.700E-08 |
| 575 | 29 | Q29P | A | 1.5e-08 | 4.5e-08 | 1.2e-07 | 6.000E-08 | 5.408E-08 | 1.500E-08 | 1.200E-07 |
| 576 | 29 | Q29S | A | 8.4e-08 | 4.6e-08 | 2.6e-08 | 5.200E-08 | 2.946E-08 | 2.600E-08 | 8.400E-08 |
| 577 | 29 | Q29T | A | 4.7e-08 | 4.7e-08 | 2.5e-08 | 3.967E-08 | 1.270E-08 | 2.500E-08 | 4.700E-08 |
| 578 | 29 | Q29W | A | 3.3e-08 | 1.2e-08 | 1.8e-08 | 2.100E-08 | 1.082E-08 | 1.200E-08 | 3.300E-08 |
| 579 | 29 | Q29Y | A | 8.4e-08 | 2.4e-08 | 5.5e-08 | 5.433E-08 | 3.001E-08 | 2.400E-08 | 8.400E-08 |
| 580 | 29 | Q29V | A | 3.7e-08 | 5.3e-08 | 3.6e-08 | 4.200E-08 | 9.539E-09 | 3.600E-08 | 5.300E-08 |
| 581 | 30 | G30A | A | 1.7e-08 | 4.2e-08 | 4.4e-08 | 3.433E-08 | 1.504E-08 | 1.700E-08 | 4.400E-08 |
| 582 | 30 | G30R | A | 5.3e-08 | 3e-08 | 3.1e-08 | 3.800E-08 | 1.300E-08 | 3.000E-08 | 5.300E-08 |
| 583 | 30 | G30N | A | 5.8e-08 | 2.2e-08 | 5.4e-08 | 4.467E-08 | 1.973E-08 | 2.200E-08 | 5.800E-08 |
| 584 | 30 | G30D | A | 9.4e-08 | 1.2e-07 | 2.7e-08 | 8.033E-08 | 4.798E-08 | 2.700E-08 | 1.200E-07 |
| 585 | 30 | G30C | A | 2e-07 | 4.9e-08 | 2.3e-08 | 9.067E-08 | 9.557E-08 | 2.300E-08 | 2.000E-07 |
| 586 | 30 | G30E | A | 2.8e-08 | 5.5e-08 | 2.3e-08 | 3.533E-08 | 1.721E-08 | 2.300E-08 | 5.500E-08 |
| 587 | 30 | G30Q | A | 4.3e-08 | 8.4e-09 | 1.8e-08 | 2.313E-08 | 1.786E-08 | 8.400E-09 | 4.300E-08 |
| 588 | 30 | G30G | A | 9.1e-08 | 4e-08 | 7.3e-08 | 6.800E-08 | 2.587E-08 | 4.000E-08 | 9.100E-08 |
| 589 | 30 | G30H | A | 5.5e-08 | 4.9e-09 | 2.1e-08 | 2.697E-08 | 2.558E-08 | 4.900E-09 | 5.500E-08 |
| 590 | 30 | G30I | A | 1.8e-08 | 5.8e-08 | 8.8e-08 | 5.467E-08 | 3.512E-08 | 1.800E-08 | 8.800E-08 |
| 591 | 30 | G30L | A | 2.3e-08 | 4.3e-08 | 2.6e-08 | 3.067E-08 | 1.079E-08 | 2.300E-08 | 4.300E-08 |
| 592 | 30 | G30K | A | 6e-08 | 6.8e-08 | 2e-08 | 4.933E-08 | 2.572E-08 | 2.000E-08 | 6.800E-08 |
| 593 | 30 | G30M | A | 3.4e-08 | 5.6e-08 | 7.2e-08 | 5.400E-08 | 1.908E-08 | 3.400E-08 | 7.200E-08 |
| 594 | 30 | G30F | A | 4.5e-08 | 4.6e-08 | 8.8e-08 | 5.967E-08 | 2.454E-08 | 4.500E-08 | 8.800E-08 |

| | | | | | | | | | | |
|---|---|---|---|---|---|---|---|---|---|---|
| 595 | 30 | G30P | A | 5.6e-08 | 4.9e-08 | 3.2e-08 | 4.567E-08 | 1.234E-08 | 3.200E-08 | 5.600E-08 |
| 596 | 30 | G30S | A | 1.5e-07 | 5e-08 | 3.4e-08 | 7.800E-08 | 6.286E-08 | 3.400E-08 | 1.500E-07 |
| 597 | 30 | G30T | A | 6.1e-08 | 3.4e-08 | 3.3e-08 | 4.267E-08 | 1.589E-08 | 3.300E-08 | 6.100E-08 |
| 598 | 30 | G30W | A | 2.3e-08 | 5.5e-08 | 3.1e-08 | 3.633E-08 | 1.665E-08 | 2.300E-08 | 5.500E-08 |
| 599 | 30 | G30Y | A | 3.7e-08 | 6.2e-08 | 1.3e-08 | 3.733E-08 | 2.450E-08 | 1.300E-08 | 6.200E-08 |
| 600 | 30 | G30V | A | 4.1e-08 | 1.5e-08 | 5.6e-08 | 3.733E-08 | 2.074E-08 | 1.500E-08 | 5.600E-08 |
| 601 | 31 | C31A | A | 5e-08 | 4.8e-08 | 5.2e-08 | 5.000E-08 | 2.000E-09 | 4.800E-08 | 5.200E-08 |
| 602 | 31 | C31R | A | 3e-07 | 5.1e-08 | 1.5e-08 | 1.220E-07 | 1.552E-07 | 1.500E-08 | 3.000E-07 |
| 603 | 31 | C31N | A | 8.1e-08 | 8.8e-08 | 8.3e-08 | 8.400E-08 | 3.606E-09 | 8.100E-08 | 8.800E-08 |
| 604 | 31 | C31D | A | 1.1e-07 | 7.7e-08 | 8.6e-08 | 9.100E-08 | 1.706E-08 | 7.700E-08 | 1.100E-07 |
| 605 | 31 | C31C | A | 5.6e-08 | 7.2e-08 | 8.3e-08 | 7.033E-08 | 1.358E-08 | 5.600E-08 | 8.300E-08 |
| 606 | 31 | C31E | A | 6.1e-08 | 5.2e-08 | 2.5e-08 | 4.600E-08 | 1.873E-08 | 2.500E-08 | 6.100E-08 |
| 607 | 31 | C31Q | A | 2.3e-08 | 7.5e-08 | 4.3e-08 | 4.700E-08 | 2.623E-08 | 2.300E-08 | 7.500E-08 |
| 608 | 31 | C31G | A | 1.1e-07 | 1.1e-07 | 1.6e-08 | 7.867E-08 | 5.427E-08 | 1.600E-08 | 1.100E-07 |
| 609 | 31 | C31H | A | 1.4e-08 | 4.1e-08 | 7.7e-08 | 4.400E-08 | 3.161E-08 | 1.400E-08 | 7.700E-08 |
| 610 | 31 | C31I | A | 7.5e-08 | 9.9e-08 | 2.1e-08 | 6.500E-08 | 3.995E-08 | 2.100E-08 | 9.900E-08 |
| 611 | 31 | C31L | A | 5.9e-08 | 9.1e-08 | 2.5e-08 | 5.833E-08 | 3.301E-08 | 2.500E-08 | 9.100E-08 |
| 612 | 31 | C31K | A | 4.1e-08 | 1e-07 | 6.5e-08 | 6.867E-08 | 2.967E-08 | 4.100E-08 | 1.000E-07 |
| 613 | 31 | C31M | A | 2.9e-08 | 4.3e-08 | 6.9e-08 | 4.700E-08 | 2.030E-08 | 2.900E-08 | 6.900E-08 |
| 614 | 31 | C31F | A | 5.6e-08 | 1e-07 | 5.3e-08 | 6.967E-08 | 2.631E-08 | 5.300E-08 | 1.000E-07 |
| 615 | 31 | C31P | A | 4e-08 | 3.3e-08 | 4.8e-08 | 4.033E-08 | 7.506E-09 | 3.300E-08 | 4.800E-08 |
| 616 | 31 | C31S | A | 1.2e-07 | 3.9e-08 | 2.6e-08 | 6.167E-08 | 5.093E-08 | 2.600E-08 | 1.200E-07 |
| 617 | 31 | C31T | A | 2.7e-08 | 1.9e-08 | 6.8e-08 | 3.800E-08 | 2.629E-08 | 1.900E-08 | 6.800E-08 |
| 618 | 31 | C31W | A | 9.5e-08 | 6.3e-08 | 3.3e-08 | 6.367E-08 | 3.101E-08 | 3.300E-08 | 9.500E-08 |
| 619 | 31 | C31Y | A | 6.8e-08 | 5.2e-08 | 5.9e-08 | 5.967E-08 | 8.021E-09 | 5.200E-08 | 6.800E-08 |
| 620 | 31 | C31V | A | 3.2e-08 | 3.4e-08 | 2.3e-08 | 2.967E-08 | 5.859E-09 | 2.300E-08 | 3.400E-08 |
| 621 | 32 | Q32A | A | 1.1e-07 | 3.1e-08 | 1.3e-07 | 9.033E-08 | 5.235E-08 | 3.100E-08 | 1.300E-07 |
| 622 | 32 | Q32R | A | 1.2e-07 | 7.6e-08 | 3.4e-08 | 7.667E-08 | 4.300E-08 | 3.400E-08 | 1.200E-07 |
| 623 | 32 | Q32N | A | 4.1e-08 | 7.1e-08 | 2.3e-08 | 4.500E-08 | 2.425E-08 | 2.300E-08 | 7.100E-08 |
| 624 | 32 | Q32D | A | 1.8e-07 | 3e-08 | 1.2e-07 | 1.100E-07 | 7.550E-08 | 3.000E-08 | 1.800E-07 |
| 625 | 32 | Q32C | A | 9.5e-08 | 6.3e-08 | 3.1e-08 | 6.300E-08 | 3.200E-08 | 3.100E-08 | 9.500E-08 |
| 626 | 32 | Q32E | A | 2.6e-08 | 5e-08 | 4.2e-08 | 3.933E-08 | 1.222E-08 | 2.600E-08 | 5.000E-08 |
| 627 | 32 | Q32Q | A | 3.6e-08 | 4.6e-08 | 8.1e-08 | 5.433E-08 | 2.363E-08 | 3.600E-08 | 8.100E-08 |
| 628 | 32 | Q32G | A | 9.5e-08 | 3.5e-08 | 5.6e-08 | 6.200E-08 | 3.045E-08 | 3.500E-08 | 9.500E-08 |
| 629 | 32 | Q32H | A | 5.4e-08 | 6.7e-08 | 3.4e-08 | 5.167E-08 | 1.662E-08 | 3.400E-08 | 6.700E-08 |
| 630 | 32 | Q32I | A | 3.4e-08 | 2.9e-08 | 5.3e-08 | 3.867E-08 | 1.266E-08 | 2.900E-08 | 5.300E-08 |
| 631 | 32 | Q32L | A | 5.4e-08 | 2.7e-08 | 5.4e-08 | 4.500E-08 | 1.559E-08 | 2.700E-08 | 5.400E-08 |
| 632 | 32 | Q32K | A | 1e-07 | 1.4e-07 | 3.1e-08 | 9.033E-08 | 5.514E-08 | 3.100E-08 | 1.400E-07 |
| 633 | 32 | Q32M | A | 6.8e-08 | 1.3e-07 | 5.4e-08 | 8.400E-08 | 4.045E-08 | 5.400E-08 | 1.300E-07 |
| 634 | 32 | Q32F | A | 1.8e-08 | 2.1e-08 | 1.8e-08 | 1.900E-08 | 1.732E-09 | 1.800E-08 | 2.100E-08 |
| 635 | 32 | Q32P | A | 4.3e-08 | 3.4e-08 | 4.8e-08 | 4.167E-08 | 7.095E-09 | 3.400E-08 | 4.800E-08 |
| 636 | 32 | Q32S | A | 2.2e-08 | 2.4e-08 | 5.1e-08 | 3.233E-08 | 1.620E-08 | 2.200E-08 | 5.100E-08 |
| 637 | 32 | Q32T | A | 2e-08 | 6e-08 | 7.1e-09 | 2.903E-08 | 2.758E-08 | 7.100E-09 | 6.000E-08 |
| 638 | 32 | Q32W | A | 5.7e-08 | 2.6e-08 | 3.9e-08 | 4.067E-08 | 1.557E-08 | 2.600E-08 | 5.700E-08 |
| 639 | 32 | Q32Y | A | 2.6e-08 | 4.1e-08 | 4.2e-08 | 3.633E-08 | 8.963E-09 | 2.600E-08 | 4.200E-08 |
| 640 | 32 | Q32V | A | 4.6e-08 | 3.3e-08 | 4.7e-08 | 4.200E-08 | 7.810E-09 | 3.300E-08 | 4.700E-08 |
| 641 | 33 | V33A | A | 1.9e-08 | 2.2e-08 | 2.9e-08 | 2.333E-08 | 5.132E-09 | 1.900E-08 | 2.900E-08 |
| 642 | 33 | V33R | A | 2.8e-08 | 9.1e-08 | 4.5e-08 | 5.467E-08 | 3.259E-08 | 2.800E-08 | 9.100E-08 |
| 643 | 33 | V33N | A | 2.4e-08 | 8.7e-08 | 5.9e-08 | 5.667E-08 | 3.156E-08 | 2.400E-08 | 8.700E-08 |
| 644 | 33 | V33D | A | 5.9e-08 | 5.5e-08 | 9.2e-08 | 6.867E-08 | 2.031E-08 | 5.500E-08 | 9.200E-08 |
| 645 | 33 | V33C | A | 1.4e-08 | 5.7e-08 | 3e-08 | 3.367E-08 | 2.173E-08 | 1.400E-08 | 5.700E-08 |
| 646 | 33 | V33E | A | 5.4e-08 | 1.7e-08 | 3e-08 | 3.367E-08 | 1.877E-08 | 1.700E-08 | 5.400E-08 |
| 647 | 33 | V33Q | A | 7.3e-08 | 1.8e-08 | 6e-08 | 5.033E-08 | 2.875E-08 | 1.800E-08 | 7.300E-08 |
| 648 | 33 | V33G | A | 7.3e-08 | 2.2e-08 | 8.6e-08 | 6.033E-08 | 3.383E-08 | 2.200E-08 | 8.600E-08 |
| 649 | 33 | V33H | A | 4.8e-08 | 2.4e-08 | 3.5e-08 | 3.567E-08 | 1.201E-08 | 2.400E-08 | 4.800E-08 |
| 650 | 33 | V33I | A | 2.3e-08 | 2.5e-08 | 8.6e-08 | 4.467E-08 | 3.581E-08 | 2.300E-08 | 8.600E-08 |
| 651 | 33 | V33L | A | 3.4e-08 | 5.1e-08 | 6.9e-08 | 5.133E-08 | 1.750E-08 | 3.400E-08 | 6.900E-08 |
| 652 | 33 | V33K | A | 9.7e-08 | 2.3e-08 | 3.2e-08 | 5.067E-08 | 4.038E-08 | 2.300E-08 | 9.700E-08 |
| 653 | 33 | V33M | A | 6e-08 | 3.8e-08 | 2.5e-08 | 4.100E-08 | 1.769E-08 | 2.500E-08 | 6.000E-08 |
| 654 | 33 | V33F | A | 2e-08 | 2.5e-08 | 7.3e-08 | 3.933E-08 | 2.926E-08 | 2.000E-08 | 7.300E-08 |
| 655 | 33 | V33P | A | 1.5e-07 | 2.6e-08 | 2.9e-08 | 6.833E-08 | 7.074E-08 | 2.600E-08 | 1.500E-07 |
| 656 | 33 | V33S | A | 6.1e-08 | 1.5e-08 | 1.9e-08 | 3.167E-08 | 2.548E-08 | 1.500E-08 | 6.100E-08 |
| 657 | 33 | V33T | A | 2.4e-08 | 3.6e-08 | 7.9e-08 | 4.633E-08 | 2.892E-08 | 2.400E-08 | 7.900E-08 |
| 658 | 33 | V33W | A | 4.7e-08 | 2.6e-08 | 3.5e-08 | 3.600E-08 | 1.054E-08 | 2.600E-08 | 4.700E-08 |
| 659 | 33 | V33Y | A | 2.8e-08 | 5.7e-08 | 2.4e-08 | 3.633E-08 | 1.801E-08 | 2.400E-08 | 5.700E-08 |
| 660 | 33 | V33V | A | 8.8e-08 | 5.9e-08 | 2.4e-08 | 5.700E-08 | 3.205E-08 | 2.400E-08 | 8.800E-08 |
| 661 | 34 | V34A | A | 6.9e-08 | 1.8e-08 | 3.3e-08 | 4.000E-08 | 2.621E-08 | 1.800E-08 | 6.900E-08 |
| 662 | 34 | V34R | A | 3.4e-08 | 2.7e-08 | 1.7e-07 | 7.700E-08 | 8.062E-08 | 2.700E-08 | 1.700E-07 |
| 663 | 34 | V34N | A | 2.6e-08 | 7.7e-08 | 2.8e-08 | 4.367E-08 | 2.888E-08 | 2.600E-08 | 7.700E-08 |
| 664 | 34 | V34D | A | 5.4e-08 | 6.8e-08 | 4.6e-08 | 5.600E-08 | 1.114E-08 | 4.600E-08 | 6.800E-08 |
| 665 | 34 | V34C | A | 2.9e-08 | 3.8e-08 | 4.3e-08 | 3.667E-08 | 7.095E-09 | 2.900E-08 | 4.300E-08 |
| 666 | 34 | V34E | A | 3.4e-08 | 6.4e-08 | 3.8e-08 | 4.533E-08 | 1.629E-08 | 3.400E-08 | 6.400E-08 |
| 667 | 34 | V34Q | A | 4.5e-08 | 2.1e-08 | 3.9e-08 | 3.500E-08 | 1.249E-08 | 2.100E-08 | 4.500E-08 |
| 668 | 34 | V34G | A | 3.8e-08 | 3.6e-08 | 4.3e-08 | 3.900E-08 | 3.606E-09 | 3.600E-08 | 4.300E-08 |
| 669 | 34 | V34H | A | 5.6e-08 | 3.5e-08 | 1.8e-08 | 3.633E-08 | 1.904E-08 | 1.800E-08 | 5.600E-08 |
| 670 | 34 | V34I | A | 4.1e-08 | 5.9e-08 | 2.7e-08 | 4.233E-08 | 1.604E-08 | 2.700E-08 | 5.900E-08 |
| 671 | 34 | V34L | A | 5.2e-08 | 3.3e-08 | 2.9e-08 | 3.800E-08 | 1.229E-08 | 2.900E-08 | 5.200E-08 |
| 672 | 34 | V34K | A | 1.7e-08 | 6.5e-08 | 5e-08 | 4.400E-08 | 2.456E-08 | 1.700E-08 | 6.500E-08 |
| 673 | 34 | V34M | A | 7.5e-08 | 8.4e-08 | 5.2e-08 | 7.033E-08 | 1.650E-08 | 5.200E-08 | 8.400E-08 |
| 674 | 34 | V34F | A | 2.5e-08 | 2e-08 | 5.3e-08 | 3.267E-08 | 1.779E-08 | 2.000E-08 | 5.300E-08 |
| 675 | 34 | V34P | A | 1.3e-07 | 5.4e-08 | 3e-08 | 7.133E-08 | 5.220E-08 | 3.000E-08 | 1.300E-07 |
| 676 | 34 | V34S | A | 1.8e-08 | 5.1e-08 | 1.6e-08 | 2.833E-08 | 1.966E-08 | 1.600E-08 | 5.100E-08 |
| 677 | 34 | V34T | A | 3e-08 | 8e-08 | 1.3e-07 | 8.000E-08 | 5.000E-08 | 3.000E-08 | 1.300E-07 |
| 678 | 34 | V34W | A | 4e-08 | 5e-08 | 7.5e-08 | 5.500E-08 | 1.803E-08 | 4.000E-08 | 7.500E-08 |
| 679 | 34 | V34Y | A | 4.4e-08 | 1.4e-07 | 6.2e-08 | 8.200E-08 | 5.103E-08 | 4.400E-08 | 1.400E-07 |

| | | | | | | | | | | |
|---|---|---|---|---|---|---|---|---|---|---|
| 680 | 34 | V34V | A | 6.8e-08 | 2.9e-08 | 2.5e-08 | 4.067E-08 | 2.376E-08 | 2.500E-08 | 6.800E-08 |
| 681 | 35 | Q35A | A | 4.7e-08 | 9.2e-08 | 6.4e-08 | 6.767E-08 | 2.272E-08 | 4.700E-08 | 9.200E-08 |
| 682 | 35 | Q35R | A | 3.6e-08 | 4e-08 | 7.4e-08 | 5.000E-08 | 2.088E-08 | 3.600E-08 | 7.400E-08 |
| 683 | 35 | Q35N | A | 2.4e-08 | 5.2e-08 | 3.2e-08 | 3.600E-08 | 1.442E-08 | 2.400E-08 | 5.200E-08 |
| 684 | 35 | Q35D | A | 4.4e-08 | 2.9e-08 | 2.1e-08 | 3.133E-08 | 1.168E-08 | 2.100E-08 | 4.400E-08 |
| 685 | 35 | Q35C | A | 4.5e-08 | 6.8e-08 | 4.1e-08 | 5.133E-08 | 1.457E-08 | 4.100E-08 | 6.800E-08 |
| 686 | 35 | Q35E | A | 7e-08 | 9.6e-08 | 5.7e-08 | 7.433E-08 | 1.986E-08 | 5.700E-08 | 9.600E-08 |
| 687 | 35 | Q35Q | A | 6e-08 | 3.2e-08 | 1.4e-08 | 3.533E-08 | 2.318E-08 | 1.400E-08 | 6.000E-08 |
| 688 | 35 | Q35G | A | 4.3e-08 | 2.5e-08 | 5.4e-08 | 4.067E-08 | 1.464E-08 | 2.500E-08 | 5.400E-08 |
| 689 | 35 | Q35H | A | 3.5e-08 | 5.5e-08 | 5.7e-08 | 4.900E-08 | 1.217E-08 | 3.500E-08 | 5.700E-08 |
| 690 | 35 | Q35I | A | 1.1e-07 | 5.4e-08 | 6e-08 | 7.467E-08 | 3.075E-08 | 5.400E-08 | 1.100E-07 |
| 691 | 35 | Q35L | A | 5.6e-08 | 6.2e-08 | 4e-08 | 5.267E-08 | 1.137E-08 | 4.000E-08 | 6.200E-08 |
| 692 | 35 | Q35K | A | 2.7e-08 | 2.1e-08 | 6.2e-08 | 3.667E-08 | 2.214E-08 | 2.100E-08 | 6.200E-08 |
| 693 | 35 | Q35M | A | 8e-08 | 5e-08 | 4.9e-08 | 5.967E-08 | 1.762E-08 | 4.900E-08 | 8.000E-08 |
| 694 | 35 | Q35F | A | 1.3e-08 | 2.7e-08 | 8.3e-08 | 4.100E-08 | 3.704E-08 | 1.300E-08 | 8.300E-08 |
| 695 | 35 | Q35P | A | 2.6e-08 | 3.4e-08 | 2.2e-08 | 2.733E-08 | 6.110E-09 | 2.200E-08 | 3.400E-08 |
| 696 | 35 | Q35S | A | 9.6e-08 | 2.6e-08 | 2e-08 | 4.733E-08 | 4.225E-08 | 2.000E-08 | 9.600E-08 |
| 697 | 35 | Q35T | A | 7.4e-08 | 2.5e-08 | 1e-07 | 6.633E-08 | 3.808E-08 | 2.500E-08 | 1.000E-07 |
| 698 | 35 | Q35W | A | 3.2e-08 | 7.6e-08 | 4.2e-08 | 5.000E-08 | 2.307E-08 | 3.200E-08 | 7.600E-08 |
| 699 | 35 | Q35Y | A | 5.8e-08 | 5.8e-08 | 3.6e-08 | 5.067E-08 | 1.270E-08 | 3.600E-08 | 5.800E-08 |
| 700 | 35 | Q35V | A | 5.7e-08 | 7.6e-08 | 4e-08 | 5.767E-08 | 1.801E-08 | 4.000E-08 | 7.600E-08 |
| 701 | 36 | G36A | A | 3.9e-08 | 2.6e-08 | 2e-08 | 2.833E-08 | 9.713E-09 | 2.000E-08 | 3.900E-08 |
| 702 | 36 | G36R | A | 3.9e-08 | 1.3e-08 | 4.7e-08 | 3.300E-08 | 1.778E-08 | 1.300E-08 | 4.700E-08 |
| 703 | 36 | G36N | A | 3.1e-08 | 7.5e-08 | 1.5e-08 | 4.033E-08 | 3.107E-08 | 1.500E-08 | 7.500E-08 |
| 704 | 36 | G36D | A | 4.4e-08 | 3.7e-08 | 5.7e-08 | 4.600E-08 | 1.015E-08 | 3.700E-08 | 5.700E-08 |
| 705 | 36 | G36C | A | 2.9e-08 | 7.6e-08 | 3.4e-08 | 4.633E-08 | 2.581E-08 | 2.900E-08 | 7.600E-08 |
| 706 | 36 | G36E | A | 5.1e-08 | 4e-08 | 3.4e-08 | 4.167E-08 | 8.622E-09 | 3.400E-08 | 5.100E-08 |
| 707 | 36 | G36Q | A | 2.3e-08 | 1.5e-08 | 2.2e-08 | 2.000E-08 | 4.359E-09 | 1.500E-08 | 2.300E-08 |
| 708 | 36 | G36G | A | 5.4e-08 | 5.5e-08 | 1.5e-08 | 4.133E-08 | 2.281E-08 | 1.500E-08 | 5.500E-08 |
| 709 | 36 | G36H | A | 4.4e-08 | 2.8e-08 | 5.6e-08 | 4.267E-08 | 1.405E-08 | 2.800E-08 | 5.600E-08 |
| 710 | 36 | G36I | A | 1.8e-07 | 6.1e-08 | 2.1e-08 | 8.733E-08 | 8.271E-08 | 2.100E-08 | 1.800E-07 |
| 711 | 36 | G36L | A | 1e-07 | 6.8e-08 | 4.6e-08 | 7.133E-08 | 2.715E-08 | 4.600E-08 | 1.000E-07 |
| 712 | 36 | G36K | A | 1.3e-07 | 3.7e-08 | 6e-08 | 7.567E-08 | 4.844E-08 | 3.700E-08 | 1.300E-07 |
| 713 | 36 | G36M | A | 1.5e-08 | 5.5e-08 | 3.2e-08 | 3.400E-08 | 2.007E-08 | 1.500E-08 | 5.500E-08 |
| 714 | 36 | G36F | A | 5.9e-08 | 3e-08 | 3.7e-08 | 4.200E-08 | 1.513E-08 | 3.000E-08 | 5.900E-08 |
| 715 | 36 | G36P | A | 1.8e-07 | 4.3e-08 | 2.6e-08 | 8.300E-08 | 8.443E-08 | 2.600E-08 | 1.800E-07 |
| 716 | 36 | G36S | A | 4e-08 | 4.6e-08 | 3.5e-08 | 4.033E-08 | 5.508E-09 | 3.500E-08 | 4.600E-08 |
| 717 | 36 | G36T | A | 4e-08 | 1.5e-07 | 4.5e-08 | 7.833E-08 | 6.212E-08 | 4.000E-08 | 1.500E-07 |
| 718 | 36 | G36W | A | 3.2e-08 | 2.4e-08 | 5e-08 | 3.533E-08 | 1.332E-08 | 2.400E-08 | 5.000E-08 |
| 719 | 36 | G36Y | A | 4e-08 | 2.1e-08 | 8.1e-08 | 4.733E-08 | 3.066E-08 | 2.100E-08 | 8.100E-08 |
| 720 | 36 | G36V | A | 3.5e-08 | 3.3e-08 | 8.5e-08 | 5.100E-08 | 2.946E-08 | 3.300E-08 | 8.500E-08 |
| 721 | 37 | N37A | A | 7.3e-08 | 5.1e-08 | 1.7e-08 | 4.700E-08 | 2.821E-08 | 1.700E-08 | 7.300E-08 |
| 722 | 37 | N37R | A | 5.9e-08 | 4.7e-08 | 5.1e-08 | 5.233E-08 | 6.110E-09 | 4.700E-08 | 5.900E-08 |
| 723 | 37 | N37N | A | 4.2e-08 | 1.7e-08 | 8.2e-08 | 4.700E-08 | 3.279E-08 | 1.700E-08 | 8.200E-08 |
| 724 | 37 | N37D | A | 4.9e-08 | 6.9e-08 | 2e-08 | 4.600E-08 | 2.464E-08 | 2.000E-08 | 6.900E-08 |
| 725 | 37 | N37C | A | 5.2e-08 | 1.1e-07 | 3.2e-08 | 6.467E-08 | 4.051E-08 | 3.200E-08 | 1.100E-07 |
| 726 | 37 | N37E | A | 3e-08 | 5.1e-08 | 5.1e-08 | 4.400E-08 | 1.212E-08 | 3.000E-08 | 5.100E-08 |
| 727 | 37 | N37Q | A | 7.7e-08 | 4.2e-08 | 1.2e-07 | 7.967E-08 | 3.907E-08 | 4.200E-08 | 1.200E-07 |
| 728 | 37 | N37G | A | 1.6e-08 | 3.9e-08 | 3.6e-08 | 3.033E-08 | 1.250E-08 | 1.600E-08 | 3.900E-08 |
| 729 | 37 | N37H | A | 2.4e-08 | 8e-08 | 1.4e-07 | 8.133E-08 | 5.801E-08 | 2.400E-08 | 1.400E-07 |
| 730 | 37 | N37I | A | 4.8e-08 | 1.7e-08 | 3.8e-08 | 3.433E-08 | 1.582E-08 | 1.700E-08 | 4.800E-08 |
| 731 | 37 | N37L | A | 3.7e-08 | 2.4e-08 | 3.6e-08 | 3.233E-08 | 7.234E-09 | 2.400E-08 | 3.700E-08 |
| 732 | 37 | N37K | A | 4.2e-08 | 5.7e-08 | 2e-08 | 3.967E-08 | 1.861E-08 | 2.000E-08 | 5.700E-08 |
| 733 | 37 | N37M | A | 3.5e-08 | 1.8e-08 | 5.2e-08 | 3.500E-08 | 1.700E-08 | 1.800E-08 | 5.200E-08 |
| 734 | 37 | N37F | A | 4.1e-08 | 1.9e-08 | 8.9e-08 | 4.967E-08 | 3.580E-08 | 1.900E-08 | 8.900E-08 |
| 735 | 37 | N37P | A | 2.2e-08 | 1.1e-07 | 4.5e-08 | 5.900E-08 | 4.564E-08 | 2.200E-08 | 1.100E-07 |
| 736 | 37 | N37S | A | 5e-08 | 3.4e-08 | 1.3e-08 | 3.233E-08 | 1.856E-08 | 1.300E-08 | 5.000E-08 |
| 737 | 37 | N37T | A | 4.9e-08 | 5.9e-08 | 3e-08 | 4.600E-08 | 1.473E-08 | 3.000E-08 | 5.900E-08 |
| 738 | 37 | N37W | A | 5.3e-08 | 1.1e-08 | 2e-08 | 2.800E-08 | 2.211E-08 | 1.100E-08 | 5.300E-08 |
| 739 | 37 | N37Y | A | 1.8e-08 | 3.3e-08 | 3.9e-08 | 3.000E-08 | 1.082E-08 | 1.800E-08 | 3.900E-08 |
| 740 | 37 | N37V | A | 5.5e-08 | 4.4e-08 | 4.2e-08 | 4.700E-08 | 7.000E-09 | 4.200E-08 | 5.500E-08 |
| 741 | 38 | L38A | A | 3.9e-08 | 2.3e-08 | 5.4e-08 | 3.867E-08 | 1.550E-08 | 2.300E-08 | 5.400E-08 |
| 742 | 38 | L38R | A | 6.9e-08 | 3.3e-08 | 2.2e-08 | 4.133E-08 | 2.458E-08 | 2.200E-08 | 6.900E-08 |
| 743 | 38 | L38N | A | 3.5e-08 | 6.5e-08 | 3.9e-08 | 4.633E-08 | 1.629E-08 | 3.500E-08 | 6.500E-08 |
| 744 | 38 | L38D | A | 3.6e-08 | 8.2e-08 | 4.3e-08 | 5.367E-08 | 2.479E-08 | 3.600E-08 | 8.200E-08 |
| 745 | 38 | L38C | A | 5.6e-08 | 6.3e-08 | 3.1e-08 | 5.000E-08 | 1.682E-08 | 3.100E-08 | 6.300E-08 |
| 746 | 38 | L38E | A | 4.4e-08 | 1.3e-08 | 6.7e-08 | 4.133E-08 | 2.710E-08 | 1.300E-08 | 6.700E-08 |
| 747 | 38 | L38Q | A | 9.1e-08 | 5.4e-08 | 8.5e-08 | 7.667E-08 | 1.986E-08 | 5.400E-08 | 9.100E-08 |
| 748 | 38 | L38G | A | 4.5e-08 | 2.4e-08 | 9.9e-09 | 2.630E-08 | 1.766E-08 | 9.900E-09 | 4.500E-08 |
| 749 | 38 | L38H | A | 1.1e-07 | 4.6e-08 | 2.3e-08 | 5.967E-08 | 4.508E-08 | 2.300E-08 | 1.100E-07 |
| 750 | 38 | L38I | A | 4e-08 | 4e-08 | 2.9e-08 | 3.633E-08 | 6.351E-09 | 2.900E-08 | 4.000E-08 |
| 751 | 38 | L38L | A | 3.1e-08 | 8.8e-08 | 1.1e-07 | 7.633E-08 | 4.077E-08 | 3.100E-08 | 1.100E-07 |
| 752 | 38 | L38K | A | 1.3e-08 | 2.1e-07 | 3.5e-08 | 8.600E-08 | 1.079E-07 | 1.300E-08 | 2.100E-07 |
| 753 | 38 | L38M | A | 4.8e-08 | 5.6e-08 | 4.8e-08 | 5.067E-08 | 4.619E-09 | 4.800E-08 | 5.600E-08 |
| 754 | 38 | L38F | A | 2.2e-08 | 2.1e-08 | 2.2e-08 | 2.167E-08 | 5.774E-10 | 2.100E-08 | 2.200E-08 |
| 755 | 38 | L38P | A | 1.1e-07 | 4.1e-08 | 1.7e-08 | 5.600E-08 | 4.828E-08 | 1.700E-08 | 1.100E-07 |
| 756 | 38 | L38S | A | 7.1e-08 | 1.5e-08 | 9.7e-08 | 6.100E-08 | 4.190E-08 | 1.500E-08 | 9.700E-08 |
| 757 | 38 | L38T | A | 5.7e-08 | 4.6e-08 | 3e-08 | 4.433E-08 | 1.358E-08 | 3.000E-08 | 5.700E-08 |
| 758 | 38 | L38W | A | 9.1e-08 | 5.3e-08 | 3.2e-08 | 5.867E-08 | 2.991E-08 | 3.200E-08 | 9.100E-08 |
| 759 | 38 | L38Y | A | 3.7e-08 | 6.9e-08 | 9.1e-08 | 6.567E-08 | 2.715E-08 | 3.700E-08 | 9.100E-08 |
| 760 | 38 | L38V | A | 5.7e-08 | 5.7e-08 | 4.9e-08 | 5.433E-08 | 4.619E-09 | 4.900E-08 | 5.700E-08 |
| 761 | 39 | E39A | A | 4.1e-08 | 3.4e-08 | 2.6e-08 | 3.367E-08 | 7.506E-09 | 2.600E-08 | 4.100E-08 |
| 762 | 39 | E39R | A | 2e-08 | 7.8e-08 | 6.9e-08 | 5.567E-08 | 3.121E-08 | 2.000E-08 | 7.800E-08 |
| 763 | 39 | E39N | A | 2.5e-08 | 5e-08 | 6.9e-08 | 4.800E-08 | 2.207E-08 | 2.500E-08 | 6.900E-08 |
| 764 | 39 | E39D | A | 8.3e-08 | 2.7e-08 | 4.7e-08 | 5.233E-08 | 2.838E-08 | 2.700E-08 | 8.300E-08 |

| | | | | | | | | | | |
|---|---|---|---|---|---|---|---|---|---|---|
| 765 | 39 | E39C | A | 1.8e-08 | 4.8e-08 | 3.1e-08 | 3.233E-08 | 1.504E-08 | 1.800E-08 | 4.800E-08 |
| 766 | 39 | E39E | A | 4e-08 | 1.4e-08 | 4.1e-08 | 3.167E-08 | 1.531E-08 | 1.400E-08 | 4.100E-08 |
| 767 | 39 | E39Q | A | 1.8e-08 | 3.3e-08 | 5.1e-08 | 3.400E-08 | 1.652E-08 | 1.800E-08 | 5.100E-08 |
| 768 | 39 | E39G | A | 6.7e-08 | 3.8e-08 | 4.8e-08 | 5.100E-08 | 1.473E-08 | 3.800E-08 | 6.700E-08 |
| 769 | 39 | E39H | A | 9.2e-08 | 6.8e-08 | 6.5e-08 | 7.500E-08 | 1.480E-08 | 6.500E-08 | 9.200E-08 |
| 770 | 39 | E39I | A | 4.6e-08 | 2e-08 | 2.8e-08 | 3.133E-08 | 1.332E-08 | 2.000E-08 | 4.600E-08 |
| 771 | 39 | E39L | A | 4.1e-08 | 2.5e-08 | 7.9e-08 | 4.833E-08 | 2.774E-08 | 2.500E-08 | 7.900E-08 |
| 772 | 39 | E39K | A | 8.7e-08 | 2.6e-08 | 2.2e-08 | 4.500E-08 | 3.643E-08 | 2.200E-08 | 8.700E-08 |
| 773 | 39 | E39M | A | 2.6e-08 | 2.7e-08 | 7e-08 | 4.100E-08 | 2.512E-08 | 2.600E-08 | 7.000E-08 |
| 774 | 39 | E39F | A | 6.9e-08 | 9.7e-08 | 3.1e-08 | 6.567E-08 | 3.313E-08 | 3.100E-08 | 9.700E-08 |
| 775 | 39 | E39P | A | 3.6e-08 | 3.3e-08 | 1.5e-08 | 2.800E-08 | 1.136E-08 | 1.500E-08 | 3.600E-08 |
| 776 | 39 | E39S | A | 4.1e-08 | 5.6e-08 | 7.1e-08 | 5.600E-08 | 1.500E-08 | 4.100E-08 | 7.100E-08 |
| 777 | 39 | E39T | A | 2.5e-08 | 3.9e-08 | 2.4e-08 | 2.933E-08 | 8.386E-09 | 2.400E-08 | 3.900E-08 |
| 778 | 39 | E39W | A | 7.1e-08 | 2.4e-08 | 3.4e-08 | 4.300E-08 | 2.476E-08 | 2.400E-08 | 7.100E-08 |
| 779 | 39 | E39Y | A | 7.3e-08 | 1.5e-08 | 6.8e-08 | 5.200E-08 | 3.214E-08 | 1.500E-08 | 7.300E-08 |
| 780 | 39 | E39V | A | 4e-08 | 2.5e-08 | 1.7e-08 | 2.733E-08 | 1.168E-08 | 1.700E-08 | 4.000E-08 |
| 781 | 40 | L40A | A | 1.6e-08 | 2.6e-08 | 4.3e-08 | 2.833E-08 | 1.365E-08 | 1.600E-08 | 4.300E-08 |
| 782 | 40 | L40R | A | 6.5e-08 | 9.4e-09 | 2.5e-08 | 3.313E-08 | 2.868E-08 | 9.400E-09 | 6.500E-08 |
| 783 | 40 | L40N | A | 8.7e-09 | 2.6e-08 | 5.7e-08 | 3.057E-08 | 2.447E-08 | 8.700E-09 | 5.700E-08 |
| 784 | 40 | L40D | A | 5.9e-08 | 5.8e-08 | 5.3e-08 | 5.667E-08 | 3.215E-09 | 5.300E-08 | 5.900E-08 |
| 785 | 40 | L40C | A | 3.1e-08 | 5.2e-08 | 4.6e-08 | 4.300E-08 | 1.082E-08 | 3.100E-08 | 5.200E-08 |
| 786 | 40 | L40E | A | 2.9e-08 | 3.1e-08 | 2.8e-08 | 2.933E-08 | 1.528E-09 | 2.800E-08 | 3.100E-08 |
| 787 | 40 | L40Q | A | 3.4e-08 | 3.5e-08 | 4.5e-08 | 3.800E-08 | 6.083E-09 | 3.400E-08 | 4.500E-08 |
| 788 | 40 | L40G | A | 3.2e-08 | 4.6e-08 | 2.6e-08 | 3.467E-08 | 1.026E-08 | 2.600E-08 | 4.600E-08 |
| 789 | 40 | L40H | A | 2e-08 | 4.1e-08 | 6.9e-08 | 4.333E-08 | 2.458E-08 | 2.000E-08 | 6.900E-08 |
| 790 | 40 | L40I | A | 1.4e-08 | 6.4e-08 | 6e-08 | 4.600E-08 | 2.778E-08 | 1.400E-08 | 6.400E-08 |
| 791 | 40 | L40L | A | 3.3e-08 | 5.7e-08 | 7.1e-08 | 5.367E-08 | 1.922E-08 | 3.300E-08 | 7.100E-08 |
| 792 | 40 | L40K | A | 1.6e-08 | 8.5e-08 | 3.9e-08 | 4.667E-08 | 3.513E-08 | 1.600E-08 | 8.500E-08 |
| 793 | 40 | L40M | A | 1.6e-08 | 1.4e-08 | 2e-08 | 1.667E-08 | 3.055E-09 | 1.400E-08 | 2.000E-08 |
| 794 | 40 | L40F | A | 3.9e-08 | 3.5e-08 | 2.5e-08 | 3.300E-08 | 7.211E-09 | 2.500E-08 | 3.900E-08 |
| 795 | 40 | L40P | A | 3.3e-08 | 3.7e-08 | 1.2e-08 | 2.733E-08 | 1.343E-08 | 1.200E-08 | 3.700E-08 |
| 796 | 40 | L40S | A | 4.4e-08 | 5.1e-08 | 2.2e-08 | 3.900E-08 | 1.513E-08 | 2.200E-08 | 5.100E-08 |
| 797 | 40 | L40T | A | 8.3e-08 | 8.4e-08 | 2.7e-08 | 6.467E-08 | 3.262E-08 | 2.700E-08 | 8.400E-08 |
| 798 | 40 | L40W | A | 1.3e-07 | 5.1e-08 | 1e-07 | 9.367E-08 | 3.988E-08 | 5.100E-08 | 1.300E-07 |
| 799 | 40 | L40Y | A | 5.7e-08 | 5.4e-08 | 4.2e-08 | 5.100E-08 | 7.937E-09 | 4.200E-08 | 5.700E-08 |
| 800 | 40 | L40V | A | 4.4e-08 | 5e-08 | 5.4e-08 | 4.933E-08 | 5.033E-09 | 4.400E-08 | 5.400E-08 |
| 801 | 41 | T41A | A | 4.5e-08 | 2.9e-08 | 6.4e-08 | 4.600E-08 | 1.752E-08 | 2.900E-08 | 6.400E-08 |
| 802 | 41 | T41R | A | 4.3e-08 | 5.4e-08 | 2e-07 | 9.900E-08 | 8.764E-08 | 4.300E-08 | 2.000E-07 |
| 803 | 41 | T41N | A | 2.8e-08 | 9.2e-09 | 3e-08 | 2.240E-08 | 1.148E-08 | 9.200E-09 | 3.000E-08 |
| 804 | 41 | T41D | A | 4.8e-08 | 5.1e-08 | 3.7e-08 | 4.533E-08 | 7.371E-09 | 3.700E-08 | 5.100E-08 |
| 805 | 41 | T41C | A | 2.6e-08 | 2.9e-08 | 6.3e-08 | 3.933E-08 | 2.055E-08 | 2.600E-08 | 6.300E-08 |
| 806 | 41 | T41E | A | 5e-08 | 2.9e-08 | 5.8e-08 | 4.567E-08 | 1.498E-08 | 2.900E-08 | 5.800E-08 |
| 807 | 41 | T41Q | A | 9.7e-08 | 5.4e-08 | 7.3e-08 | 7.467E-08 | 2.155E-08 | 5.400E-08 | 9.700E-08 |
| 808 | 41 | T41G | A | 7.7e-08 | 7.6e-08 | 1e-07 | 8.433E-08 | 1.358E-08 | 7.600E-08 | 1.000E-07 |
| 809 | 41 | T41H | A | 4.9e-08 | 6.7e-08 | 4.7e-08 | 5.433E-08 | 1.102E-08 | 4.700E-08 | 6.700E-08 |
| 810 | 41 | T41I | A | 3e-08 | 4.2e-08 | 3.8e-08 | 3.667E-08 | 6.110E-09 | 3.000E-08 | 4.200E-08 |
| 811 | 41 | T41L | A | 4.2e-08 | 8.5e-08 | 4.9e-08 | 5.867E-08 | 2.307E-08 | 4.200E-08 | 8.500E-08 |
| 812 | 41 | T41K | A | 4.6e-08 | 3.1e-08 | 8.4e-08 | 5.367E-08 | 2.732E-08 | 3.100E-08 | 8.400E-08 |
| 813 | 41 | T41M | A | 5.3e-08 | 8.2e-08 | 2.4e-08 | 5.300E-08 | 2.900E-08 | 2.400E-08 | 8.200E-08 |
| 814 | 41 | T41F | A | 8.3e-08 | 1.3e-07 | 2.3e-08 | 7.867E-08 | 5.363E-08 | 2.300E-08 | 1.300E-07 |
| 815 | 41 | T41P | A | 8.7e-08 | 5.8e-08 | 4.4e-08 | 6.300E-08 | 2.193E-08 | 4.400E-08 | 8.700E-08 |
| 816 | 41 | T41S | A | 3.4e-08 | 2.8e-08 | 2.5e-08 | 2.900E-08 | 4.583E-09 | 2.500E-08 | 3.400E-08 |
| 817 | 41 | T41T | A | 3.6e-08 | 8.1e-08 | 4.1e-08 | 5.267E-08 | 2.466E-08 | 3.600E-08 | 8.100E-08 |
| 818 | 41 | T41W | A | 6.9e-08 | 5.2e-08 | 4.1e-08 | 5.400E-08 | 1.411E-08 | 4.100E-08 | 6.900E-08 |
| 819 | 41 | T41Y | A | 4.6e-08 | 2.4e-08 | 6.8e-08 | 4.600E-08 | 2.200E-08 | 2.400E-08 | 6.800E-08 |
| 820 | 41 | T41V | A | 3.7e-08 | 5e-08 | 1.4e-08 | 3.367E-08 | 1.823E-08 | 1.400E-08 | 5.000E-08 |
| 821 | 42 | Y42A | A | 2.3e-08 | 4e-08 | 6.9e-08 | 4.400E-08 | 2.326E-08 | 2.300E-08 | 6.900E-08 |
| 822 | 42 | Y42R | A | 8.7e-08 | 1.6e-07 | 5.3e-08 | 1.000E-07 | 5.467E-08 | 5.300E-08 | 1.600E-07 |
| 823 | 42 | Y42N | A | 1.6e-08 | 9.7e-08 | 4.4e-08 | 5.233E-08 | 4.114E-08 | 1.600E-08 | 9.700E-08 |
| 824 | 42 | Y42D | A | 6.4e-08 | 2.6e-08 | 4.4e-08 | 4.467E-08 | 1.901E-08 | 2.600E-08 | 6.400E-08 |
| 825 | 42 | Y42C | A | 4e-08 | 1.4e-08 | 6e-08 | 3.800E-08 | 2.307E-08 | 1.400E-08 | 6.000E-08 |
| 826 | 42 | Y42E | A | 4.9e-08 | 9.2e-08 | 5.5e-08 | 6.533E-08 | 2.329E-08 | 4.900E-08 | 9.200E-08 |
| 827 | 42 | Y42Q | A | 4.4e-08 | 1.6e-08 | 3e-08 | 3.000E-08 | 1.400E-08 | 1.600E-08 | 4.400E-08 |
| 828 | 42 | Y42G | A | 7.8e-08 | 1.3e-07 | 6e-08 | 8.933E-08 | 3.635E-08 | 6.000E-08 | 1.300E-07 |
| 829 | 42 | Y42H | A | 3.5e-08 | 4.3e-08 | 5.9e-08 | 4.567E-08 | 1.222E-08 | 3.500E-08 | 5.900E-08 |
| 830 | 42 | Y42I | A | 7.1e-08 | 5.6e-08 | 2.1e-08 | 4.933E-08 | 2.566E-08 | 2.100E-08 | 7.100E-08 |
| 831 | 42 | Y42L | A | 9.5e-08 | 5.2e-08 | 7.4e-08 | 7.367E-08 | 2.150E-08 | 5.200E-08 | 9.500E-08 |
| 832 | 42 | Y42K | A | 1e-07 | 1.4e-08 | 3.7e-08 | 5.033E-08 | 4.452E-08 | 1.400E-08 | 1.000E-07 |
| 833 | 42 | Y42M | A | 1.4e-07 | 8.6e-08 | 7.8e-08 | 1.013E-07 | 3.372E-08 | 7.800E-08 | 1.400E-07 |
| 834 | 42 | Y42F | A | 1.4e-08 | 2e-08 | 4.4e-08 | 2.600E-08 | 1.587E-08 | 1.400E-08 | 4.400E-08 |
| 835 | 42 | Y42P | A | 6.2e-08 | 8.2e-08 | 2.6e-08 | 5.667E-08 | 2.838E-08 | 2.600E-08 | 8.200E-08 |
| 836 | 42 | Y42S | A | 5.4e-08 | 3.6e-08 | 9.8e-08 | 6.267E-08 | 3.190E-08 | 3.600E-08 | 9.800E-08 |
| 837 | 42 | Y42T | A | 4e-08 | 5.3e-08 | 4.7e-08 | 4.667E-08 | 6.506E-09 | 4.000E-08 | 5.300E-08 |
| 838 | 42 | Y42W | A | 6.5e-08 | 1.7e-08 | 9.9e-08 | 6.033E-08 | 4.120E-08 | 1.700E-08 | 9.900E-08 |
| 839 | 42 | Y42Y | A | 3.2e-08 | 1.2e-07 | 8.9e-08 | 8.033E-08 | 4.464E-08 | 3.200E-08 | 1.200E-07 |
| 840 | 42 | Y42V | A | 5.9e-08 | 1.3e-07 | 5.2e-08 | 8.033E-08 | 4.315E-08 | 5.200E-08 | 1.300E-07 |
| 841 | 43 | L43A | A | 4.7e-08 | 7.4e-08 | 1e-07 | 7.367E-08 | 2.650E-08 | 4.700E-08 | 1.000E-07 |
| 842 | 43 | L43R | A | 2.2e-08 | 2.2e-08 | 8.5e-08 | 4.300E-08 | 3.637E-08 | 2.200E-08 | 8.500E-08 |
| 843 | 43 | L43N | A | 2.8e-08 | 8.4e-08 | 3.3e-08 | 4.833E-08 | 3.099E-08 | 2.800E-08 | 8.400E-08 |
| 844 | 43 | L43D | A | 1e-07 | 3.9e-08 | 6.3e-08 | 6.733E-08 | 3.073E-08 | 3.900E-08 | 1.000E-07 |
| 845 | 43 | L43C | A | 1.4e-08 | 1.9e-08 | 3.2e-08 | 2.167E-08 | 9.292E-09 | 1.400E-08 | 3.200E-08 |
| 846 | 43 | L43E | A | 3.5e-08 | 7e-08 | 2.5e-08 | 4.333E-08 | 2.363E-08 | 2.500E-08 | 7.000E-08 |
| 847 | 43 | L43Q | A | 1.1e-07 | 2.8e-08 | 2.5e-08 | 5.433E-08 | 4.823E-08 | 2.500E-08 | 1.100E-07 |
| 848 | 43 | L43G | A | 3.1e-08 | 3.6e-08 | 2.7e-08 | 3.133E-08 | 4.509E-09 | 2.700E-08 | 3.600E-08 |
| 849 | 43 | L43H | A | 5.3e-08 | 9.2e-08 | 5.4e-08 | 6.633E-08 | 2.223E-08 | 5.300E-08 | 9.200E-08 |

| | | | | | | | | | | |
|---|---|---|---|---|---|---|---|---|---|---|
| 850 | 43 | L43I | A | 5.2e-08 | 3.3e-08 | 5.9e-08 | 4.800E-08 | 1.345E-08 | 3.300E-08 | 5.900E-08 |
| 851 | 43 | L43L | A | 3.9e-08 | 2e-08 | 3.4e-08 | 3.100E-08 | 9.849E-09 | 2.000E-08 | 3.900E-08 |
| 852 | 43 | L43K | A | 8.6e-08 | 2.1e-08 | 4.5e-08 | 5.067E-08 | 3.287E-08 | 2.100E-08 | 8.600E-08 |
| 853 | 43 | L43M | A | 6.9e-08 | 7.7e-08 | 1.2e-08 | 5.267E-08 | 3.544E-08 | 1.200E-08 | 7.700E-08 |
| 854 | 43 | L43F | A | 4.1e-08 | 5.1e-08 | 4.1e-08 | 4.433E-08 | 5.774E-09 | 4.100E-08 | 5.100E-08 |
| 855 | 43 | L43P | A | 4.8e-08 | 3.4e-08 | 7.2e-08 | 5.133E-08 | 1.922E-08 | 3.400E-08 | 7.200E-08 |
| 856 | 43 | L43S | A | 8e-08 | 6.1e-08 | 3.4e-08 | 5.833E-08 | 2.312E-08 | 3.400E-08 | 8.000E-08 |
| 857 | 43 | L43T | A | 5.1e-08 | 4.5e-08 | 2.7e-08 | 4.100E-08 | 1.249E-08 | 2.700E-08 | 5.100E-08 |
| 858 | 43 | L43W | A | 1e-07 | 1.1e-08 | 3.8e-08 | 4.967E-08 | 4.563E-08 | 1.100E-08 | 1.000E-07 |
| 859 | 43 | L43Y | A | 5.4e-08 | 1.7e-08 | 5.8e-08 | 4.300E-08 | 2.261E-08 | 1.700E-08 | 5.800E-08 |
| 860 | 43 | L43V | A | 4.8e-08 | 4.6e-08 | 1.7e-07 | 8.800E-08 | 7.102E-08 | 4.600E-08 | 1.700E-07 |
| 861 | 44 | P44A | A | 4e-08 | 2.1e-08 | 3.9e-08 | 3.333E-08 | 1.069E-08 | 2.100E-08 | 4.000E-08 |
| 862 | 44 | P44R | A | 5e-08 | 5.6e-08 | 4.5e-08 | 5.033E-08 | 5.508E-09 | 4.500E-08 | 5.600E-08 |
| 863 | 44 | P44N | A | 5.2e-08 | 3.6e-08 | 3.5e-08 | 4.100E-08 | 9.539E-09 | 3.500E-08 | 5.200E-08 |
| 864 | 44 | P44D | A | 2.6e-08 | 6.6e-08 | 3.3e-08 | 4.167E-08 | 2.136E-08 | 2.600E-08 | 6.600E-08 |
| 865 | 44 | P44C | A | 5.8e-08 | 5e-08 | 3.6e-08 | 4.800E-08 | 1.114E-08 | 3.600E-08 | 5.800E-08 |
| 866 | 44 | P44E | A | 5.7e-08 | 6.1e-08 | 3.3e-08 | 5.033E-08 | 1.514E-08 | 3.300E-08 | 6.100E-08 |
| 867 | 44 | P44Q | A | 6.1e-08 | 2e-08 | 6.3e-08 | 4.800E-08 | 2.427E-08 | 2.000E-08 | 6.300E-08 |
| 868 | 44 | P44G | A | 6.2e-08 | 8.2e-08 | 2.2e-08 | 5.533E-08 | 3.055E-08 | 2.200E-08 | 8.200E-08 |
| 869 | 44 | P44H | A | 5e-08 | 3.4e-08 | 3.5e-08 | 3.967E-08 | 8.963E-09 | 3.400E-08 | 5.000E-08 |
| 870 | 44 | P44I | A | 4.7e-08 | 4.3e-08 | 7.3e-08 | 5.433E-08 | 1.629E-08 | 4.300E-08 | 7.300E-08 |
| 871 | 44 | P44L | A | 1.7e-08 | 4.5e-08 | 4.6e-08 | 3.600E-08 | 1.646E-08 | 1.700E-08 | 4.600E-08 |
| 872 | 44 | P44K | A | 4.6e-08 | 4e-08 | 1.9e-08 | 3.500E-08 | 1.418E-08 | 1.900E-08 | 4.600E-08 |
| 873 | 44 | P44M | A | 2.4e-08 | 1.1e-07 | 4.8e-08 | 6.067E-08 | 4.438E-08 | 2.400E-08 | 1.100E-07 |
| 874 | 44 | P44F | A | 4.7e-08 | 4.6e-08 | 5.6e-08 | 4.967E-08 | 5.508E-09 | 4.600E-08 | 5.600E-08 |
| 875 | 44 | P44P | A | 5.5e-08 | 3.4e-08 | 3.5e-08 | 4.133E-08 | 1.185E-08 | 3.400E-08 | 5.500E-08 |
| 876 | 44 | P44S | A | 3.5e-08 | 5.8e-08 | 6.6e-08 | 5.300E-08 | 1.609E-08 | 3.500E-08 | 6.600E-08 |
| 877 | 44 | P44T | A | 6.4e-08 | 5.2e-08 | 3.5e-08 | 5.033E-08 | 1.457E-08 | 3.500E-08 | 6.400E-08 |
| 878 | 44 | P44W | A | 2.8e-08 | 2.9e-08 | 4.2e-08 | 3.300E-08 | 7.810E-09 | 2.800E-08 | 4.200E-08 |
| 879 | 44 | P44Y | A | 1.6e-08 | 1.7e-08 | 8.2e-08 | 3.833E-08 | 3.782E-08 | 1.600E-08 | 8.200E-08 |
| 880 | 44 | P44V | A | 4.8e-08 | 3.7e-08 | 4e-08 | 4.167E-08 | 5.686E-09 | 3.700E-08 | 4.800E-08 |
| 881 | 45 | T45A | A | 2.6e-08 | 6.9e-08 | 1.3e-07 | 7.500E-08 | 5.226E-08 | 2.600E-08 | 1.300E-07 |
| 882 | 45 | T45R | A | 6.3e-08 | 9.3e-08 | 8.4e-08 | 8.000E-08 | 1.539E-08 | 6.300E-08 | 9.300E-08 |
| 883 | 45 | T45N | A | 4.6e-08 | 3.1e-08 | 3.8e-08 | 3.833E-08 | 7.506E-09 | 3.100E-08 | 4.600E-08 |
| 884 | 45 | T45D | A | 3.7e-08 | 5.7e-08 | 6.2e-08 | 5.200E-08 | 1.323E-08 | 3.700E-08 | 6.200E-08 |
| 885 | 45 | T45C | A | 6.9e-09 | 5.6e-08 | 5.9e-08 | 4.063E-08 | 2.925E-08 | 6.900E-09 | 5.900E-08 |
| 886 | 45 | T45E | A | 3.2e-08 | 8.7e-08 | 3.5e-08 | 5.133E-08 | 3.092E-08 | 3.200E-08 | 8.700E-08 |
| 887 | 45 | T45Q | A | 4.8e-08 | 5.1e-08 | 3.5e-08 | 4.467E-08 | 8.505E-09 | 3.500E-08 | 5.100E-08 |
| 888 | 45 | T45G | A | 4.4e-08 | 4.8e-08 | 4.6e-08 | 4.600E-08 | 2.000E-09 | 4.400E-08 | 4.800E-08 |
| 889 | 45 | T45H | A | 6.5e-08 | 1.1e-07 | 3.1e-08 | 6.867E-08 | 3.963E-08 | 3.100E-08 | 1.100E-07 |
| 890 | 45 | T45I | A | 5.7e-08 | 5.1e-08 | 1.3e-08 | 4.033E-08 | 2.386E-08 | 1.300E-08 | 5.700E-08 |
| 891 | 45 | T45L | A | 3.4e-08 | 4.9e-08 | 1.2e-07 | 6.767E-08 | 4.594E-08 | 3.400E-08 | 1.200E-07 |
| 892 | 45 | T45K | A | 3.9e-08 | 6.2e-08 | 3.7e-08 | 4.600E-08 | 1.389E-08 | 3.700E-08 | 6.200E-08 |
| 893 | 45 | T45M | A | 6.3e-08 | 5.5e-08 | 2.5e-08 | 4.767E-08 | 2.003E-08 | 2.500E-08 | 6.300E-08 |
| 894 | 45 | T45F | A | 2.8e-08 | 2.2e-08 | 3.5e-08 | 2.833E-08 | 6.506E-09 | 2.200E-08 | 3.500E-08 |
| 895 | 45 | T45P | A | 4.3e-08 | 6.1e-08 | 4.1e-08 | 4.833E-08 | 1.102E-08 | 4.100E-08 | 6.100E-08 |
| 896 | 45 | T45S | A | 5.1e-08 | 6.5e-08 | 1.9e-08 | 4.500E-08 | 2.358E-08 | 1.900E-08 | 6.500E-08 |
| 897 | 45 | T45T | A | 5e-08 | 5.2e-08 | 1.5e-08 | 3.900E-08 | 2.081E-08 | 1.500E-08 | 5.200E-08 |
| 898 | 45 | T45W | A | 8.5e-08 | 1.7e-08 | 4.7e-08 | 4.967E-08 | 3.408E-08 | 1.700E-08 | 8.500E-08 |
| 899 | 45 | T45Y | A | 4.8e-08 | 6.9e-08 | 2.9e-08 | 4.867E-08 | 2.001E-08 | 2.900E-08 | 6.900E-08 |
| 900 | 45 | T45V | A | 4.6e-08 | 2.6e-07 | 2.8e-08 | 1.113E-07 | 1.291E-07 | 2.800E-08 | 2.600E-07 |
| 901 | 46 | N46A | A | 6e-08 | 5.7e-08 | 8.3e-08 | 6.667E-08 | 1.422E-08 | 5.700E-08 | 8.300E-08 |
| 902 | 46 | N46R | A | 2.7e-08 | 9.7e-08 | 1.9e-08 | 4.767E-08 | 4.291E-08 | 1.900E-08 | 9.700E-08 |
| 903 | 46 | N46N | A | 2.1e-08 | 3.3e-08 | 5.7e-08 | 3.700E-08 | 1.833E-08 | 2.100E-08 | 5.700E-08 |
| 904 | 46 | N46D | A | 1.7e-08 | 2.6e-08 | 7.1e-08 | 3.800E-08 | 2.893E-08 | 1.700E-08 | 7.100E-08 |
| 905 | 46 | N46C | A | 5.9e-08 | 4.6e-08 | 8.6e-08 | 6.367E-08 | 2.040E-08 | 4.600E-08 | 8.600E-08 |
| 906 | 46 | N46E | A | 2.6e-08 | 5.2e-08 | 4.2e-08 | 4.000E-08 | 1.311E-08 | 2.600E-08 | 5.200E-08 |
| 907 | 46 | N46Q | A | 3.6e-08 | 6.1e-08 | 2.8e-08 | 4.167E-08 | 1.721E-08 | 2.800E-08 | 6.100E-08 |
| 908 | 46 | N46G | A | 4.7e-08 | 3.6e-08 | 1.1e-08 | 3.133E-08 | 1.845E-08 | 1.100E-08 | 4.700E-08 |
| 909 | 46 | N46H | A | 3.7e-08 | 1.2e-08 | 9e-08 | 4.633E-08 | 3.983E-08 | 1.200E-08 | 9.000E-08 |
| 910 | 46 | N46I | A | 3.8e-08 | 4.4e-08 | 3.2e-08 | 3.800E-08 | 6.000E-09 | 3.200E-08 | 4.400E-08 |
| 911 | 46 | N46L | A | 1.4e-08 | 4.3e-08 | 3e-08 | 2.900E-08 | 1.453E-08 | 1.400E-08 | 4.300E-08 |
| 912 | 46 | N46K | A | 2.9e-08 | 4.8e-08 | 1.9e-08 | 3.200E-08 | 1.473E-08 | 1.900E-08 | 4.800E-08 |
| 913 | 46 | N46M | A | 5.9e-08 | 4.1e-08 | 5.7e-08 | 5.233E-08 | 9.866E-09 | 4.100E-08 | 5.900E-08 |
| 914 | 46 | N46F | A | 6.8e-08 | 3.8e-08 | 2.5e-08 | 4.367E-08 | 2.205E-08 | 2.500E-08 | 6.800E-08 |
| 915 | 46 | N46P | A | 4.9e-08 | 3.2e-08 | 3.2e-08 | 3.767E-08 | 9.815E-09 | 3.200E-08 | 4.900E-08 |
| 916 | 46 | N46S | A | 5.5e-08 | 6.9e-08 | 2.8e-08 | 5.067E-08 | 2.084E-08 | 2.800E-08 | 6.900E-08 |
| 917 | 46 | N46T | A | 5.2e-08 | 3.1e-08 | 4.6e-08 | 4.300E-08 | 1.082E-08 | 3.100E-08 | 5.200E-08 |
| 918 | 46 | N46W | A | 4.8e-08 | 2.4e-08 | 2.3e-08 | 3.167E-08 | 1.415E-08 | 2.300E-08 | 4.800E-08 |
| 919 | 46 | N46Y | A | 6.1e-08 | 7.3e-08 | 7.6e-08 | 7.000E-08 | 7.937E-09 | 6.100E-08 | 7.600E-08 |
| 920 | 46 | N46V | A | 7.3e-08 | 1.2e-07 | 2.9e-08 | 7.400E-08 | 4.551E-08 | 2.900E-08 | 1.200E-07 |
| 921 | 47 | A47A | A | 1.2e-08 | 5.1e-08 | 1.2e-07 | 6.100E-08 | 5.469E-08 | 1.200E-08 | 1.200E-07 |
| 922 | 47 | A47R | A | 9e-08 | 4e-08 | 2.9e-08 | 5.300E-08 | 3.251E-08 | 2.900E-08 | 9.000E-08 |
| 923 | 47 | A47N | A | 4.3e-08 | 3.8e-08 | 2.8e-08 | 3.633E-08 | 7.638E-09 | 2.800E-08 | 4.300E-08 |
| 924 | 47 | A47D | A | 2e-08 | 4.6e-08 | 4.9e-08 | 3.833E-08 | 1.595E-08 | 2.000E-08 | 4.900E-08 |
| 925 | 47 | A47C | A | 4.7e-08 | 5.1e-08 | 7.8e-08 | 5.867E-08 | 1.686E-08 | 4.700E-08 | 7.800E-08 |
| 926 | 47 | A47E | A | 1.2e-07 | 8.5e-08 | 2.9e-08 | 7.800E-08 | 4.590E-08 | 2.900E-08 | 1.200E-07 |
| 927 | 47 | A47Q | A | 6.2e-08 | 2.3e-08 | 3.8e-08 | 4.100E-08 | 1.967E-08 | 2.300E-08 | 6.200E-08 |
| 928 | 47 | A47G | A | 3.7e-08 | 8.8e-08 | 2.5e-08 | 5.000E-08 | 3.345E-08 | 2.500E-08 | 8.800E-08 |
| 929 | 47 | A47H | A | 5.4e-08 | 4.9e-08 | 4e-08 | 4.767E-08 | 7.095E-09 | 4.000E-08 | 5.400E-08 |
| 930 | 47 | A47I | A | 1e-07 | 2.5e-08 | 7.3e-08 | 6.600E-08 | 3.799E-08 | 2.500E-08 | 1.000E-07 |
| 931 | 47 | A47L | A | 4e-08 | 3.1e-08 | 3.2e-08 | 3.433E-08 | 4.933E-09 | 3.100E-08 | 4.000E-08 |
| 932 | 47 | A47K | A | 1.5e-08 | 2.1e-08 | 4.5e-08 | 2.700E-08 | 1.587E-08 | 1.500E-08 | 4.500E-08 |
| 933 | 47 | A47M | A | 7.9e-08 | 5.1e-08 | 9.2e-08 | 7.400E-08 | 2.095E-08 | 5.100E-08 | 9.200E-08 |
| 934 | 47 | A47F | A | 1.7e-08 | 3.6e-08 | 4.7e-08 | 3.333E-08 | 1.518E-08 | 1.700E-08 | 4.700E-08 |

| | | | | | | | | | | |
|---|---|---|---|---|---|---|---|---|---|---|
| 935 | 47 | A47P | A | 8.6e-08 | 6.2e-08 | 8.1e-08 | 7.633E-08 | 1.266E-08 | 6.200E-08 | 8.600E-08 |
| 936 | 47 | A47S | A | 2.4e-08 | 9.8e-08 | 2.6e-08 | 4.933E-08 | 4.216E-08 | 2.400E-08 | 9.800E-08 |
| 937 | 47 | A47T | A | 1.2e-08 | 4e-08 | 1.2e-07 | 5.733E-08 | 5.605E-08 | 1.200E-08 | 1.200E-07 |
| 938 | 47 | A47W | A | 4.2e-08 | 6.4e-08 | 3.5e-08 | 4.700E-08 | 1.513E-08 | 3.500E-08 | 6.400E-08 |
| 939 | 47 | A47Y | A | 2.4e-08 | 8.1e-08 | 9.4e-08 | 6.633E-08 | 3.723E-08 | 2.400E-08 | 9.400E-08 |
| 940 | 47 | A47V | A | 3.7e-08 | 2e-08 | 7e-08 | 4.233E-08 | 2.542E-08 | 2.000E-08 | 7.000E-08 |
| 941 | 48 | S48A | A | 4.3e-08 | 7.1e-08 | 5.2e-08 | 5.533E-08 | 1.429E-08 | 4.300E-08 | 7.100E-08 |
| 942 | 48 | S48R | A | 2.5e-08 | 6e-08 | 2e-08 | 3.500E-08 | 2.179E-08 | 2.000E-08 | 6.000E-08 |
| 943 | 48 | S48N | A | 3.5e-08 | 7.1e-08 | 7.5e-08 | 6.033E-08 | 2.203E-08 | 3.500E-08 | 7.500E-08 |
| 944 | 48 | S48D | A | 3.2e-08 | 4.9e-08 | 5e-08 | 4.367E-08 | 1.012E-08 | 3.200E-08 | 5.000E-08 |
| 945 | 48 | S48C | A | 1e-07 | 5.1e-08 | 6.6e-08 | 7.233E-08 | 2.511E-08 | 5.100E-08 | 1.000E-07 |
| 946 | 48 | S48E | A | 7.5e-08 | 6.8e-08 | 7.1e-08 | 7.133E-08 | 3.512E-09 | 6.800E-08 | 7.500E-08 |
| 947 | 48 | S48Q | A | 1.7e-08 | 7.4e-08 | 5.5e-08 | 4.867E-08 | 2.902E-08 | 1.700E-08 | 7.400E-08 |
| 948 | 48 | S48G | A | 1.9e-07 | 5e-08 | 5.3e-08 | 9.767E-08 | 7.998E-08 | 5.000E-08 | 1.900E-07 |
| 949 | 48 | S48H | A | 3.6e-08 | 1.3e-07 | 2.1e-08 | 6.233E-08 | 5.908E-08 | 2.100E-08 | 1.300E-07 |
| 950 | 48 | S48I | A | 4e-08 | 3.8e-08 | 7.3e-08 | 5.033E-08 | 1.966E-08 | 3.800E-08 | 7.300E-08 |
| 951 | 48 | S48L | A | 3.4e-08 | 4.4e-08 | 1.1e-07 | 6.267E-08 | 4.130E-08 | 3.400E-08 | 1.100E-07 |
| 952 | 48 | S48K | A | 5.6e-08 | 2e-08 | 5.2e-08 | 4.267E-08 | 1.973E-08 | 2.000E-08 | 5.600E-08 |
| 953 | 48 | S48M | A | 1.3e-07 | 2.9e-08 | 4.4e-08 | 6.767E-08 | 5.450E-08 | 2.900E-08 | 1.300E-07 |
| 954 | 48 | S48F | A | 1.5e-08 | 5e-08 | 7e-08 | 4.500E-08 | 2.784E-08 | 1.500E-08 | 7.000E-08 |
| 955 | 48 | S48P | A | 1.2e-08 | 2e-08 | 2.2e-08 | 1.800E-08 | 5.292E-09 | 1.200E-08 | 2.200E-08 |
| 956 | 48 | S48S | A | 1.4e-08 | 5.6e-08 | 3.5e-08 | 3.500E-08 | 2.100E-08 | 1.400E-08 | 5.600E-08 |
| 957 | 48 | S48T | A | 2.3e-08 | 5.6e-08 | 3.7e-08 | 3.867E-08 | 1.656E-08 | 2.300E-08 | 5.600E-08 |
| 958 | 48 | S48W | A | 7.5e-08 | 1.9e-08 | 6.7e-08 | 5.367E-08 | 3.029E-08 | 1.900E-08 | 7.500E-08 |
| 959 | 48 | S48Y | A | 6.3e-08 | 4.1e-08 | 1.5e-08 | 3.967E-08 | 2.403E-08 | 1.500E-08 | 6.300E-08 |
| 960 | 48 | S48V | A | 1.2e-07 | 4.1e-08 | 4.8e-08 | 6.967E-08 | 4.373E-08 | 4.100E-08 | 1.200E-07 |
| 961 | 49 | L49A | A | 4.8e-08 | 1.1e-08 | 2.1e-08 | 2.667E-08 | 1.914E-08 | 1.100E-08 | 4.800E-08 |
| 962 | 49 | L49R | A | 4.6e-08 | 4.2e-08 | 6e-08 | 4.933E-08 | 9.452E-09 | 4.200E-08 | 6.000E-08 |
| 963 | 49 | L49N | A | 8.2e-08 | 5.3e-08 | 5.9e-08 | 6.467E-08 | 1.531E-08 | 5.300E-08 | 8.200E-08 |
| 964 | 49 | L49D | A | 1.5e-08 | 3.9e-08 | 5.8e-08 | 3.733E-08 | 2.155E-08 | 1.500E-08 | 5.800E-08 |
| 965 | 49 | L49C | A | 1.5e-08 | 8e-08 | 2e-08 | 3.833E-08 | 3.617E-08 | 1.500E-08 | 8.000E-08 |
| 966 | 49 | L49E | A | 2.6e-08 | 6.2e-08 | 7.5e-08 | 5.433E-08 | 2.538E-08 | 2.600E-08 | 7.500E-08 |
| 967 | 49 | L49Q | A | 7e-09 | 4.1e-08 | 2.6e-08 | 2.467E-08 | 1.704E-08 | 7.000E-09 | 4.100E-08 |
| 968 | 49 | L49G | A | 3e-08 | 2.9e-08 | 1.1e-07 | 5.633E-08 | 4.648E-08 | 2.900E-08 | 1.100E-07 |
| 969 | 49 | L49H | A | 2.8e-08 | 2.9e-08 | 3.6e-08 | 3.100E-08 | 4.359E-09 | 2.800E-08 | 3.600E-08 |
| 970 | 49 | L49I | A | 5.9e-08 | 6.4e-08 | 3e-08 | 5.100E-08 | 1.836E-08 | 3.000E-08 | 6.400E-08 |
| 971 | 49 | L49L | A | 1.5e-07 | 2.7e-08 | 2.8e-08 | 6.833E-08 | 7.073E-08 | 2.700E-08 | 1.500E-07 |
| 972 | 49 | L49K | A | 8.9e-08 | 3.3e-08 | 4.1e-08 | 5.433E-08 | 3.029E-08 | 3.300E-08 | 8.900E-08 |
| 973 | 49 | L49M | A | 2.5e-08 | 6.3e-08 | 2.4e-08 | 3.733E-08 | 2.223E-08 | 2.400E-08 | 6.300E-08 |
| 974 | 49 | L49F | A | 4.7e-08 | 2.5e-08 | 3e-08 | 3.400E-08 | 1.153E-08 | 2.500E-08 | 4.700E-08 |
| 975 | 49 | L49P | A | 1.8e-08 | 4.6e-08 | 4.2e-08 | 3.533E-08 | 1.514E-08 | 1.800E-08 | 4.600E-08 |
| 976 | 49 | L49S | A | 5.2e-08 | 8.5e-08 | 8.2e-08 | 7.300E-08 | 1.825E-08 | 5.200E-08 | 8.500E-08 |
| 977 | 49 | L49T | A | 5.8e-08 | 4.6e-08 | 5.3e-08 | 5.233E-08 | 6.028E-09 | 4.600E-08 | 5.800E-08 |
| 978 | 49 | L49W | A | 4.9e-08 | 3.6e-08 | 4.3e-08 | 4.267E-08 | 6.506E-09 | 3.600E-08 | 4.900E-08 |
| 979 | 49 | L49Y | A | 5.1e-08 | 8.6e-08 | 1.5e-07 | 9.567E-08 | 5.020E-08 | 5.100E-08 | 1.500E-07 |
| 980 | 49 | L49V | A | 3.2e-08 | 5.2e-08 | 6.1e-08 | 4.833E-08 | 1.484E-08 | 3.200E-08 | 6.100E-08 |
| 981 | 50 | S50A | A | 5.1e-08 | 3.4e-08 | 2.2e-08 | 3.567E-08 | 1.457E-08 | 2.200E-08 | 5.100E-08 |
| 982 | 50 | S50R | A | 2.5e-08 | 2.5e-08 | 8.9e-08 | 4.633E-08 | 3.695E-08 | 2.500E-08 | 8.900E-08 |
| 983 | 50 | S50N | A | 1.8e-08 | 2.6e-08 | 1.1e-07 | 5.133E-08 | 5.096E-08 | 1.800E-08 | 1.100E-07 |
| 984 | 50 | S50D | A | 2.5e-08 | 4.7e-08 | 2.4e-08 | 3.200E-08 | 1.300E-08 | 2.400E-08 | 4.700E-08 |
| 985 | 50 | S50C | A | 2.2e-08 | 2.3e-08 | 1.5e-08 | 2.000E-08 | 4.359E-09 | 1.500E-08 | 2.300E-08 |
| 986 | 50 | S50E | A | 2.5e-08 | 2.1e-08 | 1.2e-07 | 5.533E-08 | 5.604E-08 | 2.100E-08 | 1.200E-07 |
| 987 | 50 | S50Q | A | 1.6e-08 | 2.3e-08 | 1.5e-08 | 1.800E-08 | 4.359E-09 | 1.500E-08 | 2.300E-08 |
| 988 | 50 | S50G | A | 9.7e-09 | 1.4e-07 | 5.6e-08 | 6.857E-08 | 6.605E-08 | 9.700E-09 | 1.400E-07 |
| 989 | 50 | S50H | A | 2.1e-08 | 1.3e-08 | 4.4e-08 | 2.600E-08 | 1.609E-08 | 1.300E-08 | 4.400E-08 |
| 990 | 50 | S50I | A | 3.4e-08 | 6.7e-08 | 2.7e-07 | 1.237E-07 | 1.278E-07 | 3.400E-08 | 2.700E-07 |
| 991 | 50 | S50L | A | 2.6e-08 | 3.6e-08 | 4.2e-08 | 3.467E-08 | 8.083E-09 | 2.600E-08 | 4.200E-08 |
| 992 | 50 | S50K | A | 8.1e-08 | 4.3e-08 | 2.2e-08 | 4.867E-08 | 2.991E-08 | 2.200E-08 | 8.100E-08 |
| 993 | 50 | S50M | A | 2.3e-08 | 2.9e-08 | 1.2e-07 | 5.733E-08 | 5.435E-08 | 2.300E-08 | 1.200E-07 |
| 994 | 50 | S50F | A | 2.7e-08 | 2.1e-08 | 4.2e-08 | 3.000E-08 | 1.082E-08 | 2.100E-08 | 4.200E-08 |
| 995 | 50 | S50P | A | 6.2e-08 | 9.5e-08 | 7.7e-08 | 7.800E-08 | 1.652E-08 | 6.200E-08 | 9.500E-08 |
| 996 | 50 | S50S | A | 2.3e-08 | 1.3e-08 | 1.4e-07 | 5.867E-08 | 7.061E-08 | 1.300E-08 | 1.400E-07 |
| 997 | 50 | S50T | A | 1.6e-08 | 7.3e-08 | 4.8e-08 | 4.567E-08 | 2.857E-08 | 1.600E-08 | 7.300E-08 |
| 998 | 50 | S50W | A | 5.9e-08 | 4.2e-08 | 2.4e-08 | 4.167E-08 | 1.750E-08 | 2.400E-08 | 5.900E-08 |
| 999 | 50 | S50Y | A | 1.3e-07 | 1.3e-08 | 5.5e-08 | 6.600E-08 | 5.927E-08 | 1.300E-08 | 1.300E-07 |
| 1000 | 50 | S50V | A | 7.4e-08 | 6.1e-08 | 2.4e-08 | 5.300E-08 | 2.594E-08 | 2.400E-08 | 7.400E-08 |
| 1001 | 51 | F51A | A | 4.9e-08 | 2.4e-08 | 2.5e-08 | 3.267E-08 | 1.415E-08 | 2.400E-08 | 4.900E-08 |
| 1002 | 51 | F51R | A | 6.8e-08 | 9.4e-08 | 4.2e-08 | 6.800E-08 | 2.600E-08 | 4.200E-08 | 9.400E-08 |
| 1003 | 51 | F51N | A | 3.1e-08 | 2.6e-08 | 3.7e-08 | 3.133E-08 | 5.508E-09 | 2.600E-08 | 3.700E-08 |
| 1004 | 51 | F51D | A | 6e-08 | 3.3e-08 | 1.4e-07 | 7.767E-08 | 5.564E-08 | 3.300E-08 | 1.400E-07 |
| 1005 | 51 | F51C | A | 4.8e-08 | 1e-07 | 2.9e-08 | 5.900E-08 | 3.676E-08 | 2.900E-08 | 1.000E-07 |
| 1006 | 51 | F51E | A | 4.2e-08 | 2.4e-08 | 2.6e-08 | 3.067E-08 | 9.866E-09 | 2.400E-08 | 4.200E-08 |
| 1007 | 51 | F51Q | A | 3.4e-08 | 3.1e-08 | 1.6e-07 | 7.500E-08 | 7.363E-08 | 3.100E-08 | 1.600E-07 |
| 1008 | 51 | F51G | A | 5.2e-08 | 5.3e-08 | 2.3e-08 | 4.267E-08 | 1.704E-08 | 2.300E-08 | 5.300E-08 |
| 1009 | 51 | F51H | A | 1.1e-08 | 1.8e-08 | 6e-08 | 2.967E-08 | 2.650E-08 | 1.100E-08 | 6.000E-08 |
| 1010 | 51 | F51I | A | 2.7e-08 | 7.5e-08 | 4.2e-08 | 4.800E-08 | 2.456E-08 | 2.700E-08 | 7.500E-08 |
| 1011 | 51 | F51L | A | 3.9e-08 | 1.7e-08 | 1.1e-07 | 5.533E-08 | 4.860E-08 | 1.700E-08 | 1.100E-07 |
| 1012 | 51 | F51K | A | 4.2e-08 | 9.9e-08 | 3.2e-08 | 5.767E-08 | 3.614E-08 | 3.200E-08 | 9.900E-08 |
| 1013 | 51 | F51M | A | 4.8e-08 | 4.5e-08 | 2.3e-07 | 1.077E-07 | 1.060E-07 | 4.500E-08 | 2.300E-07 |
| 1014 | 51 | F51F | A | 1e-07 | 2.2e-08 | 1.8e-08 | 4.667E-08 | 4.623E-08 | 1.800E-08 | 1.000E-07 |
| 1015 | 51 | F51P | A | 3.8e-08 | 7.2e-08 | 4.7e-08 | 5.233E-08 | 1.762E-08 | 3.800E-08 | 7.200E-08 |
| 1016 | 51 | F51S | A | 1.7e-08 | 3.2e-08 | 6.8e-08 | 3.900E-08 | 2.621E-08 | 1.700E-08 | 6.800E-08 |
| 1017 | 51 | F51T | A | 7.3e-08 | 4.3e-08 | 7.6e-09 | 4.120E-08 | 3.274E-08 | 7.600E-09 | 7.300E-08 |
| 1018 | 51 | F51W | A | 4.7e-08 | 5.8e-08 | 1.1e-07 | 7.167E-08 | 3.365E-08 | 4.700E-08 | 1.100E-07 |
| 1019 | 51 | F51Y | A | 4.3e-08 | 7.1e-08 | 2.7e-08 | 4.700E-08 | 2.227E-08 | 2.700E-08 | 7.100E-08 |

| | | | | | | | | | | |
|---|---|---|---|---|---|---|---|---|---|---|
| 1020 | 51 | F51V | A | 5.1e-08 | 4.4e-08 | 1.9e-08 | 3.800E-08 | 1.682E-08 | 1.900E-08 | 5.100E-08 |
| 1021 | 52 | L52A | A | 4.1e-08 | 6.1e-08 | 5.7e-08 | 5.300E-08 | 1.058E-08 | 4.100E-08 | 6.100E-08 |
| 1022 | 52 | L52R | A | 5.1e-08 | 1.6e-07 | 2.6e-08 | 7.900E-08 | 7.125E-08 | 2.600E-08 | 1.600E-07 |
| 1023 | 52 | L52N | A | 2.2e-08 | 5.3e-08 | 1.8e-08 | 3.100E-08 | 1.916E-08 | 1.800E-08 | 5.300E-08 |
| 1024 | 52 | L52D | A | 1.1e-07 | 5e-08 | 6.3e-08 | 7.433E-08 | 3.156E-08 | 5.000E-08 | 1.100E-07 |
| 1025 | 52 | L52C | A | 4.9e-08 | 4.3e-08 | 1e-07 | 6.400E-08 | 3.132E-08 | 4.300E-08 | 1.000E-07 |
| 1026 | 52 | L52E | A | 1.2e-08 | 2.4e-08 | 1.9e-08 | 1.833E-08 | 6.028E-09 | 1.200E-08 | 2.400E-08 |
| 1027 | 52 | L52Q | A | 2.1e-08 | 7.3e-08 | 4e-08 | 4.467E-08 | 2.631E-08 | 2.100E-08 | 7.300E-08 |
| 1028 | 52 | L52G | A | 4.3e-08 | 6.9e-08 | 2.8e-08 | 4.667E-08 | 2.074E-08 | 2.800E-08 | 6.900E-08 |
| 1029 | 52 | L52H | A | 2e-07 | 6.1e-08 | 5.1e-08 | 1.040E-07 | 8.329E-08 | 5.100E-08 | 2.000E-07 |
| 1030 | 52 | L52I | A | 3.4e-08 | 1e-07 | 4.2e-08 | 5.867E-08 | 3.602E-08 | 3.400E-08 | 1.000E-07 |
| 1031 | 52 | L52L | A | 3.4e-08 | 4.9e-08 | 2.6e-08 | 3.633E-08 | 1.168E-08 | 2.600E-08 | 4.900E-08 |
| 1032 | 52 | L52K | A | 2.7e-08 | 3.9e-08 | 2.5e-08 | 3.033E-08 | 7.572E-09 | 2.500E-08 | 3.900E-08 |
| 1033 | 52 | L52M | A | 2e-08 | 1.4e-07 | 5.5e-08 | 7.167E-08 | 6.171E-08 | 2.000E-08 | 1.400E-07 |
| 1034 | 52 | L52F | A | 3.7e-08 | 4.6e-08 | 2.5e-08 | 3.600E-08 | 1.054E-08 | 2.500E-08 | 4.600E-08 |
| 1035 | 52 | L52P | A | 3.5e-08 | 3.5e-08 | 2.8e-08 | 3.267E-08 | 4.041E-09 | 2.800E-08 | 3.500E-08 |
| 1036 | 52 | L52S | A | 8.4e-09 | 4e-08 | 3.9e-08 | 2.913E-08 | 1.796E-08 | 8.400E-09 | 4.000E-08 |
| 1037 | 52 | L52T | A | 2.6e-08 | 6.5e-08 | 1.9e-08 | 3.667E-08 | 2.479E-08 | 1.900E-08 | 6.500E-08 |
| 1038 | 52 | L52W | A | 6.3e-08 | 6.6e-08 | 9.3e-08 | 7.400E-08 | 1.652E-08 | 6.300E-08 | 9.300E-08 |
| 1039 | 52 | L52Y | A | 3.6e-08 | 5.3e-08 | 2.3e-08 | 3.733E-08 | 1.504E-08 | 2.300E-08 | 5.300E-08 |
| 1040 | 52 | L52V | A | 1.4e-08 | 2e-08 | 4.5e-08 | 2.633E-08 | 1.644E-08 | 1.400E-08 | 4.500E-08 |
| 1041 | 53 | Q53A | A | 3.3e-08 | 5.4e-08 | 3e-08 | 3.900E-08 | 1.308E-08 | 3.000E-08 | 5.400E-08 |
| 1042 | 53 | Q53R | A | 7e-08 | 7.8e-08 | 1.7e-08 | 5.500E-08 | 3.315E-08 | 1.700E-08 | 7.800E-08 |
| 1043 | 53 | Q53N | A | 3.7e-08 | 6.1e-08 | 3.3e-08 | 4.367E-08 | 1.514E-08 | 3.300E-08 | 6.100E-08 |
| 1044 | 53 | Q53D | A | 1.9e-08 | 7.3e-08 | 3.3e-08 | 4.167E-08 | 2.802E-08 | 1.900E-08 | 7.300E-08 |
| 1045 | 53 | Q53C | A | 2.1e-08 | 4.7e-08 | 7.9e-08 | 4.900E-08 | 2.905E-08 | 2.100E-08 | 7.900E-08 |
| 1046 | 53 | Q53E | A | 1.4e-07 | 3e-08 | 1.1e-08 | 6.033E-08 | 6.964E-08 | 1.100E-08 | 1.400E-07 |
| 1047 | 53 | Q53Q | A | 1.1e-08 | 2.9e-08 | 7.6e-08 | 3.867E-08 | 3.356E-08 | 1.100E-08 | 7.600E-08 |
| 1048 | 53 | Q53G | A | 8.9e-08 | 6.3e-08 | 3.7e-08 | 6.300E-08 | 2.600E-08 | 3.700E-08 | 8.900E-08 |
| 1049 | 53 | Q53H | A | 3.7e-08 | 3.9e-08 | 7.7e-08 | 5.100E-08 | 2.254E-08 | 3.700E-08 | 7.700E-08 |
| 1050 | 53 | Q53I | A | 3.2e-08 | 7.7e-09 | 8.9e-08 | 4.290E-08 | 4.173E-08 | 7.700E-09 | 8.900E-08 |
| 1051 | 53 | Q53L | A | 7.4e-08 | 4e-08 | 6.6e-08 | 6.000E-08 | 1.778E-08 | 4.000E-08 | 7.400E-08 |
| 1052 | 53 | Q53K | A | 1.3e-08 | 6.6e-08 | 2.2e-08 | 3.367E-08 | 2.836E-08 | 1.300E-08 | 6.600E-08 |
| 1053 | 53 | Q53M | A | 6.2e-08 | 1.6e-08 | 5.8e-08 | 4.533E-08 | 2.548E-08 | 1.600E-08 | 6.200E-08 |
| 1054 | 53 | Q53F | A | 3.7e-08 | 4.6e-08 | 5.4e-08 | 4.567E-08 | 8.505E-09 | 3.700E-08 | 5.400E-08 |
| 1055 | 53 | Q53P | A | 2.5e-08 | 7.5e-08 | 3.2e-08 | 4.400E-08 | 2.707E-08 | 2.500E-08 | 7.500E-08 |
| 1056 | 53 | Q53S | A | 5.6e-08 | 7.8e-09 | 4.4e-08 | 3.593E-08 | 2.509E-08 | 7.800E-09 | 5.600E-08 |
| 1057 | 53 | Q53T | A | 2.2e-07 | 4.2e-08 | 2.3e-08 | 9.500E-08 | 1.087E-07 | 2.300E-08 | 2.200E-07 |
| 1058 | 53 | Q53W | A | 5.8e-08 | 2.5e-08 | 7.7e-08 | 5.333E-08 | 2.631E-08 | 2.500E-08 | 7.700E-08 |
| 1059 | 53 | Q53Y | A | 9.2e-08 | 1.1e-07 | 3.3e-08 | 7.833E-08 | 4.028E-08 | 3.300E-08 | 1.100E-07 |
| 1060 | 53 | Q53V | A | 3.1e-08 | 3.5e-08 | 3.8e-08 | 3.467E-08 | 3.512E-09 | 3.100E-08 | 3.800E-08 |
| 1061 | 54 | D54A | A | 2.1e-08 | 2.1e-08 | 1.2e-07 | 5.400E-08 | 5.716E-08 | 2.100E-08 | 1.200E-07 |
| 1062 | 54 | D54R | A | 7e-08 | 9.4e-08 | 5.2e-08 | 7.200E-08 | 2.107E-08 | 5.200E-08 | 9.400E-08 |
| 1063 | 54 | D54N | A | 1.3e-07 | 2.1e-07 | 4.3e-08 | 1.277E-07 | 8.352E-08 | 4.300E-08 | 2.100E-07 |
| 1064 | 54 | D54D | A | 2.8e-08 | 3.1e-08 | 9.9e-08 | 5.267E-08 | 4.015E-08 | 2.800E-08 | 9.900E-08 |
| 1065 | 54 | D54C | A | 8.5e-08 | 5.7e-08 | 2.3e-08 | 5.500E-08 | 3.105E-08 | 2.300E-08 | 8.500E-08 |
| 1066 | 54 | D54E | A | 4.2e-08 | 7.9e-08 | 8e-08 | 6.700E-08 | 2.166E-08 | 4.200E-08 | 8.000E-08 |
| 1067 | 54 | D54Q | A | 7.4e-08 | 2.5e-08 | 7.5e-09 | 3.550E-08 | 3.447E-08 | 7.500E-09 | 7.400E-08 |
| 1068 | 54 | D54G | A | 7.6e-08 | 2.7e-08 | 6.2e-08 | 5.500E-08 | 2.524E-08 | 2.700E-08 | 7.600E-08 |
| 1069 | 54 | D54H | A | 3.2e-08 | 1.3e-08 | 4.4e-08 | 2.967E-08 | 1.563E-08 | 1.300E-08 | 4.400E-08 |
| 1070 | 54 | D54I | A | 1.9e-08 | 6.2e-08 | 1.1e-07 | 6.367E-08 | 4.552E-08 | 1.900E-08 | 1.100E-07 |
| 1071 | 54 | D54L | A | 5.3e-08 | 3.3e-08 | 1.6e-08 | 3.400E-08 | 1.852E-08 | 1.600E-08 | 5.300E-08 |
| 1072 | 54 | D54K | A | 1.4e-08 | 3.3e-08 | 1.8e-08 | 2.167E-08 | 1.002E-08 | 1.400E-08 | 3.300E-08 |
| 1073 | 54 | D54M | A | 4.8e-08 | 2e-08 | 4.1e-08 | 3.633E-08 | 1.457E-08 | 2.000E-08 | 4.800E-08 |
| 1074 | 54 | D54F | A | 2.2e-08 | 1.2e-07 | 9e-08 | 7.733E-08 | 5.021E-08 | 2.200E-08 | 1.200E-07 |
| 1075 | 54 | D54P | A | 1.5e-08 | 5.4e-08 | 7.9e-08 | 4.933E-08 | 3.225E-08 | 1.500E-08 | 7.900E-08 |
| 1076 | 54 | D54S | A | 8.6e-08 | 1.9e-08 | 8.9e-08 | 6.467E-08 | 3.958E-08 | 1.900E-08 | 8.900E-08 |
| 1077 | 54 | D54T | A | 2.6e-08 | 6.6e-08 | 6e-08 | 5.067E-08 | 2.157E-08 | 2.600E-08 | 6.600E-08 |
| 1078 | 54 | D54W | A | 4.4e-08 | 5.6e-08 | 3.6e-08 | 4.533E-08 | 1.007E-08 | 3.600E-08 | 5.600E-08 |
| 1079 | 54 | D54Y | A | 5.5e-08 | 4e-08 | 3.9e-08 | 4.467E-08 | 8.963E-09 | 3.900E-08 | 5.500E-08 |
| 1080 | 54 | D54V | A | 5.2e-08 | 6.1e-08 | 9.6e-08 | 6.967E-08 | 2.325E-08 | 5.200E-08 | 9.600E-08 |
| 1081 | 55 | I55A | A | 3.1e-08 | 6.6e-08 | 6.4e-08 | 5.367E-08 | 1.966E-08 | 3.100E-08 | 6.600E-08 |
| 1082 | 55 | I55R | A | 6.2e-08 | 3.3e-08 | 2.1e-08 | 3.867E-08 | 2.108E-08 | 2.100E-08 | 6.200E-08 |
| 1083 | 55 | I55N | A | 6.2e-08 | 5.3e-08 | 8.5e-08 | 6.667E-08 | 1.650E-08 | 5.300E-08 | 8.500E-08 |
| 1084 | 55 | I55D | A | 4.1e-08 | 3.7e-08 | 3.8e-08 | 3.867E-08 | 2.082E-09 | 3.700E-08 | 4.100E-08 |
| 1085 | 55 | I55C | A | 4.4e-08 | 3.9e-08 | 4.5e-08 | 4.267E-08 | 3.215E-09 | 3.900E-08 | 4.500E-08 |
| 1086 | 55 | I55E | A | 3.4e-08 | 5.8e-08 | 1.4e-07 | 7.733E-08 | 5.558E-08 | 3.400E-08 | 1.400E-07 |
| 1087 | 55 | I55Q | A | 4.7e-08 | 2.9e-08 | 8.5e-08 | 5.367E-08 | 2.859E-08 | 2.900E-08 | 8.500E-08 |
| 1088 | 55 | I55G | A | 8.3e-08 | 1.2e-08 | 5.8e-08 | 5.100E-08 | 3.601E-08 | 1.200E-08 | 8.300E-08 |
| 1089 | 55 | I55H | A | 3.3e-08 | 3.2e-08 | 8.5e-08 | 5.000E-08 | 3.032E-08 | 3.200E-08 | 8.500E-08 |
| 1090 | 55 | I55I | A | 4.5e-08 | 1.1e-07 | 5.4e-08 | 6.967E-08 | 3.522E-08 | 4.500E-08 | 1.100E-07 |
| 1091 | 55 | I55L | A | 3.1e-08 | 1.3e-08 | 3e-08 | 2.467E-08 | 1.012E-08 | 1.300E-08 | 3.100E-08 |
| 1092 | 55 | I55K | A | 5.2e-08 | 3.9e-08 | 8.7e-08 | 5.933E-08 | 2.483E-08 | 3.900E-08 | 8.700E-08 |
| 1093 | 55 | I55M | A | 4.4e-08 | 6e-08 | 3.2e-08 | 4.533E-08 | 1.405E-08 | 3.200E-08 | 6.000E-08 |
| 1094 | 55 | I55F | A | 4e-08 | 5e-08 | 2.4e-08 | 3.800E-08 | 1.311E-08 | 2.400E-08 | 5.000E-08 |
| 1095 | 55 | I55P | A | 6.8e-08 | 6.4e-08 | 7e-08 | 6.733E-08 | 3.055E-09 | 6.400E-08 | 7.000E-08 |
| 1096 | 55 | I55S | A | 1.1e-07 | 7.2e-08 | 4.5e-08 | 7.567E-08 | 3.265E-08 | 4.500E-08 | 1.100E-07 |
| 1097 | 55 | I55T | A | 3.3e-08 | 1.9e-08 | 1.2e-08 | 2.133E-08 | 1.069E-08 | 1.200E-08 | 3.300E-08 |
| 1098 | 55 | I55W | A | 4.4e-08 | 2.7e-08 | 3e-08 | 3.367E-08 | 9.074E-09 | 2.700E-08 | 4.400E-08 |
| 1099 | 55 | I55Y | A | 2.4e-08 | 2.7e-08 | 5.3e-08 | 3.467E-08 | 1.595E-08 | 2.400E-08 | 5.300E-08 |
| 1100 | 55 | I55V | A | 2.2e-08 | 3.9e-08 | 2.2e-08 | 2.767E-08 | 9.815E-09 | 2.200E-08 | 3.900E-08 |
| 1101 | 56 | Q56A | A | 5.8e-08 | 2e-08 | 4.4e-08 | 4.067E-08 | 1.922E-08 | 2.000E-08 | 5.800E-08 |
| 1102 | 56 | Q56R | A | 1.2e-07 | 8.8e-08 | 3.1e-08 | 7.967E-08 | 4.508E-08 | 3.100E-08 | 1.200E-07 |
| 1103 | 56 | Q56N | A | 6.7e-08 | 3.8e-08 | 3.2e-08 | 4.567E-08 | 1.872E-08 | 3.200E-08 | 6.700E-08 |
| 1104 | 56 | Q56D | A | 7.7e-08 | 3.8e-08 | 1.3e-07 | 8.167E-08 | 4.618E-08 | 3.800E-08 | 1.300E-07 |

| | | | | | | | | | | |
|---|---|---|---|---|---|---|---|---|---|---|
| 1105 | 56 | Q56C | A | 3.7e-08 | 3.9e-08 | 3.4e-08 | 3.667E-08 | 2.517E-09 | 3.400E-08 | 3.900E-08 |
| 1106 | 56 | Q56E | A | 1.8e-08 | 3.7e-08 | 4.8e-08 | 3.433E-08 | 1.518E-08 | 1.800E-08 | 4.800E-08 |
| 1107 | 56 | Q56Q | A | 3.2e-08 | 5.8e-08 | 2.6e-08 | 3.867E-08 | 1.701E-08 | 2.600E-08 | 5.800E-08 |
| 1108 | 56 | Q56G | A | 3.5e-08 | 5.4e-08 | 7.8e-08 | 5.567E-08 | 2.155E-08 | 3.500E-08 | 7.800E-08 |
| 1109 | 56 | Q56H | A | 4.8e-08 | 3.8e-08 | 8.1e-08 | 5.567E-08 | 2.250E-08 | 3.800E-08 | 8.100E-08 |
| 1110 | 56 | Q56I | A | 6.6e-08 | 4.9e-08 | 2.7e-08 | 4.733E-08 | 1.955E-08 | 2.700E-08 | 6.600E-08 |
| 1111 | 56 | Q56L | A | 8.5e-09 | 4.2e-08 | 1.2e-07 | 5.683E-08 | 5.721E-08 | 8.500E-09 | 1.200E-07 |
| 1112 | 56 | Q56K | A | 4.5e-08 | 7e-08 | 6e-08 | 5.833E-08 | 1.258E-08 | 4.500E-08 | 7.000E-08 |
| 1113 | 56 | Q56M | A | 4.8e-08 | 9e-08 | 3.2e-08 | 5.667E-08 | 2.996E-08 | 3.200E-08 | 9.000E-08 |
| 1114 | 56 | Q56F | A | 3.3e-08 | 2.2e-08 | 4.4e-08 | 3.300E-08 | 1.100E-08 | 2.200E-08 | 4.400E-08 |
| 1115 | 56 | Q56P | A | 1.7e-08 | 2.2e-08 | 4.7e-08 | 2.867E-08 | 1.607E-08 | 1.700E-08 | 4.700E-08 |
| 1116 | 56 | Q56S | A | 3.9e-08 | 2.1e-08 | 3.2e-08 | 3.067E-08 | 9.074E-09 | 2.100E-08 | 3.900E-08 |
| 1117 | 56 | Q56T | A | 4.8e-08 | 1.1e-07 | 1.3e-07 | 9.600E-08 | 4.276E-08 | 4.800E-08 | 1.300E-07 |
| 1118 | 56 | Q56W | A | 2.9e-08 | 7.8e-08 | 6e-08 | 5.567E-08 | 2.479E-08 | 2.900E-08 | 7.800E-08 |
| 1119 | 56 | Q56Y | A | 3.1e-08 | 8.7e-08 | 3.7e-08 | 5.167E-08 | 3.075E-08 | 3.100E-08 | 8.700E-08 |
| 1120 | 56 | Q56V | A | 4.1e-08 | 2.5e-08 | 3.4e-08 | 3.333E-08 | 8.021E-09 | 2.500E-08 | 4.100E-08 |
| 1121 | 57 | E57A | A | 9.4e-08 | 2.7e-08 | 3.5e-08 | 5.200E-08 | 3.659E-08 | 2.700E-08 | 9.400E-08 |
| 1122 | 57 | E57R | A | 7.5e-08 | 5.4e-08 | 1.3e-07 | 8.633E-08 | 3.925E-08 | 5.400E-08 | 1.300E-07 |
| 1123 | 57 | E57N | A | 3.1e-08 | 3.6e-08 | 3.6e-08 | 3.433E-08 | 2.887E-09 | 3.100E-08 | 3.600E-08 |
| 1124 | 57 | E57D | A | 3.1e-08 | 2.2e-08 | 5.8e-08 | 3.700E-08 | 1.873E-08 | 2.200E-08 | 5.800E-08 |
| 1125 | 57 | E57C | A | 6.4e-08 | 4.3e-08 | 1.5e-07 | 8.567E-08 | 5.670E-08 | 4.300E-08 | 1.500E-07 |
| 1126 | 57 | E57E | A | 3.5e-08 | 5.4e-08 | 7.1e-08 | 5.333E-08 | 1.801E-08 | 3.500E-08 | 7.100E-08 |
| 1127 | 57 | E57Q | A | 6.3e-08 | 7.7e-08 | 4.3e-08 | 6.100E-08 | 1.709E-08 | 4.300E-08 | 7.700E-08 |
| 1128 | 57 | E57G | A | 1.4e-08 | 2.8e-08 | 1.4e-07 | 6.067E-08 | 6.906E-08 | 1.400E-08 | 1.400E-07 |
| 1129 | 57 | E57H | A | 3.1e-08 | 8.5e-08 | 4.1e-08 | 5.233E-08 | 2.873E-08 | 3.100E-08 | 8.500E-08 |
| 1130 | 57 | E57I | A | 2.5e-08 | 5.1e-08 | 3.2e-08 | 3.600E-08 | 1.345E-08 | 2.500E-08 | 5.100E-08 |
| 1131 | 57 | E57L | A | 3.4e-08 | 5.9e-08 | 2e-08 | 3.767E-08 | 1.976E-08 | 2.000E-08 | 5.900E-08 |
| 1132 | 57 | E57K | A | 1.2e-07 | 2.5e-08 | 6.8e-08 | 7.100E-08 | 4.757E-08 | 2.500E-08 | 1.200E-07 |
| 1133 | 57 | E57M | A | 6.9e-08 | 3.3e-08 | 8.2e-08 | 6.133E-08 | 2.538E-08 | 3.300E-08 | 8.200E-08 |
| 1134 | 57 | E57F | A | 2.5e-08 | 3.2e-08 | 8.7e-08 | 4.800E-08 | 3.396E-08 | 2.500E-08 | 8.700E-08 |
| 1135 | 57 | E57P | A | 3.4e-08 | 4.4e-08 | 4.9e-08 | 4.233E-08 | 7.638E-09 | 3.400E-08 | 4.900E-08 |
| 1136 | 57 | E57S | A | 3e-08 | 7.1e-08 | 3.7e-08 | 4.600E-08 | 2.193E-08 | 3.000E-08 | 7.100E-08 |
| 1137 | 57 | E57T | A | 5.1e-08 | 1.3e-08 | 7.4e-08 | 4.600E-08 | 3.081E-08 | 1.300E-08 | 7.400E-08 |
| 1138 | 57 | E57W | A | 1.4e-08 | 3.4e-08 | 2.8e-08 | 2.533E-08 | 1.026E-08 | 1.400E-08 | 3.400E-08 |
| 1139 | 57 | E57Y | A | 1.9e-08 | 4.6e-08 | 3.8e-08 | 3.433E-08 | 1.387E-08 | 1.900E-08 | 4.600E-08 |
| 1140 | 57 | E57V | A | 4.9e-08 | 4.1e-08 | 4.3e-08 | 4.433E-08 | 4.163E-09 | 4.100E-08 | 4.900E-08 |
| 1141 | 58 | V58A | A | 5.1e-08 | 2.5e-08 | 1.2e-07 | 6.533E-08 | 4.910E-08 | 2.500E-08 | 1.200E-07 |
| 1142 | 58 | V58R | A | 4.9e-08 | 3.9e-08 | 3e-08 | 3.933E-08 | 9.504E-09 | 3.000E-08 | 4.900E-08 |
| 1143 | 58 | V58N | A | 4.7e-08 | 4.6e-08 | 4.4e-08 | 4.567E-08 | 1.528E-09 | 4.400E-08 | 4.700E-08 |
| 1144 | 58 | V58D | A | 2.2e-08 | 9.9e-09 | 4.9e-08 | 2.697E-08 | 2.002E-08 | 9.900E-09 | 4.900E-08 |
| 1145 | 58 | V58C | A | 8.3e-08 | 3.9e-08 | 4.8e-08 | 5.667E-08 | 2.325E-08 | 3.900E-08 | 8.300E-08 |
| 1146 | 58 | V58E | A | 8.6e-08 | 2.6e-08 | 3.6e-08 | 4.933E-08 | 3.215E-08 | 2.600E-08 | 8.600E-08 |
| 1147 | 58 | V58Q | A | 4.4e-08 | 5.4e-08 | 7.1e-08 | 5.633E-08 | 1.365E-08 | 4.400E-08 | 7.100E-08 |
| 1148 | 58 | V58G | A | 9.4e-08 | 1.3e-07 | 2.2e-08 | 8.200E-08 | 5.499E-08 | 2.200E-08 | 1.300E-07 |
| 1149 | 58 | V58H | A | 3.1e-08 | 4.1e-08 | 3.9e-08 | 3.700E-08 | 5.292E-09 | 3.100E-08 | 4.100E-08 |
| 1150 | 58 | V58I | A | 8.7e-08 | 5.9e-08 | 3.3e-08 | 5.967E-08 | 2.701E-08 | 3.300E-08 | 8.700E-08 |
| 1151 | 58 | V58L | A | 6.2e-08 | 4.3e-08 | 7.3e-08 | 5.933E-08 | 1.518E-08 | 4.300E-08 | 7.300E-08 |
| 1152 | 58 | V58K | A | 2.9e-08 | 5.9e-08 | 4e-08 | 4.267E-08 | 1.518E-08 | 2.900E-08 | 5.900E-08 |
| 1153 | 58 | V58M | A | 2.2e-08 | 6.8e-08 | 6.8e-09 | 3.227E-08 | 3.187E-08 | 6.800E-09 | 6.800E-08 |
| 1154 | 58 | V58F | A | 3.9e-08 | 3e-08 | 1.2e-08 | 2.700E-08 | 1.375E-08 | 1.200E-08 | 3.900E-08 |
| 1155 | 58 | V58P | A | 7.6e-08 | 1.5e-08 | 2.6e-08 | 3.900E-08 | 3.251E-08 | 1.500E-08 | 7.600E-08 |
| 1156 | 58 | V58S | A | 4.6e-08 | 8e-08 | 6.4e-08 | 6.333E-08 | 1.701E-08 | 4.600E-08 | 8.000E-08 |
| 1157 | 58 | V58T | A | 4.3e-08 | 4.1e-08 | 2e-08 | 3.467E-08 | 1.274E-08 | 2.000E-08 | 4.300E-08 |
| 1158 | 58 | V58W | A | 3.7e-08 | 7.1e-08 | 6.8e-08 | 5.867E-08 | 1.882E-08 | 3.700E-08 | 7.100E-08 |
| 1159 | 58 | V58Y | A | 4.9e-08 | 2.9e-08 | 2.9e-08 | 3.567E-08 | 1.155E-08 | 2.900E-08 | 4.900E-08 |
| 1160 | 58 | V58V | A | 8.3e-08 | 5.2e-08 | 6.2e-08 | 6.567E-08 | 1.582E-08 | 5.200E-08 | 8.300E-08 |
| 1161 | 59 | Q59A | A | 5.6e-08 | 1.9e-08 | 3.1e-08 | 3.533E-08 | 1.888E-08 | 1.900E-08 | 5.600E-08 |
| 1162 | 59 | Q59R | A | 2.9e-08 | 4.6e-08 | 4.1e-08 | 3.867E-08 | 8.737E-09 | 2.900E-08 | 4.600E-08 |
| 1163 | 59 | Q59N | A | 6.7e-08 | 2.2e-08 | 5.5e-08 | 4.800E-08 | 2.330E-08 | 2.200E-08 | 6.700E-08 |
| 1164 | 59 | Q59D | A | 1.9e-08 | 1.7e-08 | 4.8e-08 | 2.800E-08 | 1.735E-08 | 1.700E-08 | 4.800E-08 |
| 1165 | 59 | Q59C | A | 2.5e-08 | 7.3e-08 | 3.1e-08 | 4.300E-08 | 2.615E-08 | 2.500E-08 | 7.300E-08 |
| 1166 | 59 | Q59E | A | 4.6e-08 | 1.8e-08 | 4e-08 | 3.467E-08 | 1.474E-08 | 1.800E-08 | 4.600E-08 |
| 1167 | 59 | Q59Q | A | 5.6e-08 | 2.2e-08 | 3.3e-08 | 3.700E-08 | 1.735E-08 | 2.200E-08 | 5.600E-08 |
| 1168 | 59 | Q59G | A | 3e-08 | 8.1e-08 | 5.7e-08 | 5.600E-08 | 2.551E-08 | 3.000E-08 | 8.100E-08 |
| 1169 | 59 | Q59H | A | 8.9e-08 | 9.1e-08 | 5.5e-08 | 7.833E-08 | 2.023E-08 | 5.500E-08 | 9.100E-08 |
| 1170 | 59 | Q59I | A | 1.8e-08 | 1.1e-07 | 5.3e-08 | 6.033E-08 | 4.644E-08 | 1.800E-08 | 1.100E-07 |
| 1171 | 59 | Q59L | A | 3.8e-08 | 3e-08 | 3.7e-08 | 3.500E-08 | 4.359E-09 | 3.000E-08 | 3.800E-08 |
| 1172 | 59 | Q59K | A | 3.6e-08 | 5.4e-08 | 1.9e-08 | 3.633E-08 | 1.750E-08 | 1.900E-08 | 5.400E-08 |
| 1173 | 59 | Q59M | A | 5.1e-08 | 4.7e-08 | 4.9e-08 | 4.900E-08 | 2.000E-09 | 4.700E-08 | 5.100E-08 |
| 1174 | 59 | Q59F | A | 4.9e-08 | 3.9e-08 | 3.1e-08 | 3.967E-08 | 9.018E-09 | 3.100E-08 | 4.900E-08 |
| 1175 | 59 | Q59P | A | 5.1e-08 | 5.7e-08 | 3.5e-08 | 4.767E-08 | 1.137E-08 | 3.500E-08 | 5.700E-08 |
| 1176 | 59 | Q59S | A | 1.2e-08 | 7.1e-08 | 2.5e-08 | 3.600E-08 | 3.100E-08 | 1.200E-08 | 7.100E-08 |
| 1177 | 59 | Q59T | A | 5.7e-08 | 5.2e-08 | 4.6e-08 | 5.167E-08 | 5.508E-09 | 4.600E-08 | 5.700E-08 |
| 1178 | 59 | Q59W | A | 3.8e-08 | 8.1e-09 | 3.3e-08 | 2.637E-08 | 1.602E-08 | 8.100E-09 | 3.800E-08 |
| 1179 | 59 | Q59Y | A | 5.9e-08 | 1.7e-08 | 4e-08 | 3.867E-08 | 2.103E-08 | 1.700E-08 | 5.900E-08 |
| 1180 | 59 | Q59V | A | 4.2e-08 | 3.1e-08 | 1.6e-07 | 7.767E-08 | 7.151E-08 | 3.100E-08 | 1.600E-07 |
| 1181 | 60 | G60A | A | 7.2e-09 | 4.8e-08 | 2.7e-08 | 2.740E-08 | 2.040E-08 | 7.200E-09 | 4.800E-08 |
| 1182 | 60 | G60R | A | 9.7e-08 | 3.8e-08 | 1.1e-07 | 8.167E-08 | 3.837E-08 | 3.800E-08 | 1.100E-07 |
| 1183 | 60 | G60N | A | 5.2e-08 | 4.7e-08 | 2.2e-08 | 4.033E-08 | 1.607E-08 | 2.200E-08 | 5.200E-08 |
| 1184 | 60 | G60D | A | 2.5e-08 | 2.3e-08 | 1.5e-07 | 6.600E-08 | 7.275E-08 | 2.300E-08 | 1.500E-07 |
| 1185 | 60 | G60C | A | 1.5e-08 | 3.7e-08 | 5.1e-08 | 3.433E-08 | 1.815E-08 | 1.500E-08 | 5.100E-08 |
| 1186 | 60 | G60E | A | 2.4e-08 | 9.3e-08 | 2.3e-08 | 4.667E-08 | 4.013E-08 | 2.300E-08 | 9.300E-08 |
| 1187 | 60 | G60Q | A | 9.3e-09 | 3.6e-08 | 5.8e-08 | 3.443E-08 | 2.439E-08 | 9.300E-09 | 5.800E-08 |
| 1188 | 60 | G60G | A | 6.4e-08 | 7e-08 | 3.9e-08 | 5.767E-08 | 1.644E-08 | 3.900E-08 | 7.000E-08 |
| 1189 | 60 | G60H | A | 3.6e-08 | 1.1e-07 | 3.3e-08 | 5.967E-08 | 4.362E-08 | 3.300E-08 | 1.100E-07 |

| | | | | | | | | | | |
|---|---|---|---|---|---|---|---|---|---|---|
| 1190 | 60 | G60I | A | 9.2e-08 | 7.2e-08 | 5.6e-08 | 7.333E-08 | 1.804E-08 | 5.600E-08 | 9.200E-08 |
| 1191 | 60 | G60L | A | 3.8e-08 | 2e-07 | 1.5e-08 | 8.433E-08 | 1.008E-07 | 1.500E-08 | 2.000E-07 |
| 1192 | 60 | G60K | A | 2.8e-08 | 1.3e-07 | 1.9e-08 | 5.900E-08 | 6.165E-08 | 1.900E-08 | 1.300E-07 |
| 1193 | 60 | G60M | A | 2.6e-08 | 2.3e-08 | 4.4e-08 | 3.100E-08 | 1.136E-08 | 2.300E-08 | 4.400E-08 |
| 1194 | 60 | G60F | A | 7.8e-09 | 3.6e-08 | 1.9e-08 | 2.093E-08 | 1.420E-08 | 7.800E-09 | 3.600E-08 |
| 1195 | 60 | G60P | A | 6.1e-08 | 1.3e-07 | 2.7e-08 | 7.267E-08 | 5.248E-08 | 2.700E-08 | 1.300E-07 |
| 1196 | 60 | G60S | A | 2.7e-08 | 4.3e-08 | 7.4e-08 | 4.800E-08 | 2.390E-08 | 2.700E-08 | 7.400E-08 |
| 1197 | 60 | G60T | A | 3.7e-08 | 6.7e-08 | 2.5e-08 | 4.300E-08 | 2.163E-08 | 2.500E-08 | 6.700E-08 |
| 1198 | 60 | G60W | A | 4.1e-08 | 1.4e-07 | 5.8e-08 | 7.967E-08 | 5.294E-08 | 4.100E-08 | 1.400E-07 |
| 1199 | 60 | G60Y | A | 1.5e-08 | 1.6e-08 | 4.3e-08 | 2.467E-08 | 1.589E-08 | 1.500E-08 | 4.300E-08 |
| 1200 | 60 | G60V | A | 3.4e-08 | 6.2e-08 | 5.3e-08 | 4.967E-08 | 1.429E-08 | 3.400E-08 | 6.200E-08 |
| 1201 | 61 | Y61A | A | 4.5e-08 | 4.4e-08 | 1.9e-08 | 3.600E-08 | 1.473E-08 | 1.900E-08 | 4.500E-08 |
| 1202 | 61 | Y61R | A | 3.3e-08 | 9.1e-08 | 3.2e-08 | 5.200E-08 | 3.378E-08 | 3.200E-08 | 9.100E-08 |
| 1203 | 61 | Y61N | A | 2.2e-08 | 4.6e-08 | 8.5e-08 | 5.100E-08 | 3.180E-08 | 2.200E-08 | 8.500E-08 |
| 1204 | 61 | Y61D | A | 1.4e-07 | 2.6e-08 | 7e-08 | 7.867E-08 | 5.749E-08 | 2.600E-08 | 1.400E-07 |
| 1205 | 61 | Y61C | A | 3e-08 | 1.1e-07 | 3.3e-08 | 5.767E-08 | 4.535E-08 | 3.000E-08 | 1.100E-07 |
| 1206 | 61 | Y61E | A | 5.5e-08 | 3.7e-08 | 1.9e-08 | 3.700E-08 | 1.800E-08 | 1.900E-08 | 5.500E-08 |
| 1207 | 61 | Y61Q | A | 4.4e-08 | 6.6e-08 | 5.1e-08 | 5.367E-08 | 1.124E-08 | 4.400E-08 | 6.600E-08 |
| 1208 | 61 | Y61G | A | 3.3e-08 | 2.9e-08 | 5.2e-08 | 3.800E-08 | 1.229E-08 | 2.900E-08 | 5.200E-08 |
| 1209 | 61 | Y61H | A | 1.1e-07 | 7.5e-08 | 7.5e-08 | 8.667E-08 | 2.021E-08 | 7.500E-08 | 1.100E-07 |
| 1210 | 61 | Y61I | A | 4.7e-08 | 5.2e-08 | 5.6e-08 | 5.167E-08 | 4.509E-09 | 4.700E-08 | 5.600E-08 |
| 1211 | 61 | Y61L | A | 6.6e-08 | 5.5e-08 | 2.4e-08 | 4.833E-08 | 2.178E-08 | 2.400E-08 | 6.600E-08 |
| 1212 | 61 | Y61K | A | 4.3e-08 | 5.4e-08 | 5.3e-08 | 5.000E-08 | 6.083E-09 | 4.300E-08 | 5.400E-08 |
| 1213 | 61 | Y61M | A | 3.4e-08 | 5.7e-08 | 2.5e-08 | 3.867E-08 | 1.650E-08 | 2.500E-08 | 5.700E-08 |
| 1214 | 61 | Y61F | A | 5.6e-08 | 1.2e-07 | 2.4e-08 | 6.667E-08 | 4.888E-08 | 2.400E-08 | 1.200E-07 |
| 1215 | 61 | Y61P | A | 7.5e-08 | 5e-08 | 3.3e-08 | 5.267E-08 | 2.113E-08 | 3.300E-08 | 7.500E-08 |
| 1216 | 61 | Y61S | A | 3.7e-08 | 3.3e-08 | 9e-08 | 5.333E-08 | 3.182E-08 | 3.300E-08 | 9.000E-08 |
| 1217 | 61 | Y61T | A | 6.7e-08 | 2e-08 | 5.5e-08 | 4.733E-08 | 2.442E-08 | 2.000E-08 | 6.700E-08 |
| 1218 | 61 | Y61W | A | 2.2e-08 | 2e-08 | 4.9e-08 | 3.033E-08 | 1.620E-08 | 2.000E-08 | 4.900E-08 |
| 1219 | 61 | Y61Y | A | 6.2e-08 | 7.2e-08 | 3.3e-08 | 5.567E-08 | 2.026E-08 | 3.300E-08 | 7.200E-08 |
| 1220 | 61 | Y61V | A | 7.4e-08 | 1.7e-08 | 1.1e-07 | 6.700E-08 | 4.689E-08 | 1.700E-08 | 1.100E-07 |
| 1221 | 62 | V62A | A | 5.1e-08 | 2.4e-08 | 5.2e-08 | 4.233E-08 | 1.589E-08 | 2.400E-08 | 5.200E-08 |
| 1222 | 62 | V62R | A | 1.9e-08 | 1.3e-07 | 6.3e-08 | 7.067E-08 | 5.590E-08 | 1.900E-08 | 1.300E-07 |
| 1223 | 62 | V62N | A | 4.4e-08 | 2.1e-07 | 4.4e-08 | 9.933E-08 | 9.584E-08 | 4.400E-08 | 2.100E-07 |
| 1224 | 62 | V62D | A | 2e-08 | 9.2e-08 | 4.8e-08 | 5.333E-08 | 3.630E-08 | 2.000E-08 | 9.200E-08 |
| 1225 | 62 | V62C | A | 4.5e-08 | 3.4e-08 | 3.8e-08 | 3.900E-08 | 5.568E-09 | 3.400E-08 | 4.500E-08 |
| 1226 | 62 | V62E | A | 2.3e-08 | 2.3e-08 | 2.5e-08 | 2.367E-08 | 1.155E-09 | 2.300E-08 | 2.500E-08 |
| 1227 | 62 | V62Q | A | 5e-08 | 3.1e-08 | 5.2e-08 | 4.433E-08 | 1.159E-08 | 3.100E-08 | 5.200E-08 |
| 1228 | 62 | V62G | A | 4.5e-08 | 6.6e-08 | 2.7e-08 | 4.600E-08 | 1.952E-08 | 2.700E-08 | 6.600E-08 |
| 1229 | 62 | V62H | A | 3.4e-08 | 2.1e-08 | 2.2e-08 | 2.567E-08 | 7.234E-09 | 2.100E-08 | 3.400E-08 |
| 1230 | 62 | V62I | A | 2.5e-08 | 2e-08 | 4.7e-08 | 3.067E-08 | 1.436E-08 | 2.000E-08 | 4.700E-08 |
| 1231 | 62 | V62L | A | 4.9e-08 | 2.9e-08 | 4.9e-08 | 4.233E-08 | 1.155E-08 | 2.900E-08 | 4.900E-08 |
| 1232 | 62 | V62K | A | 3.2e-08 | 9.4e-08 | 1.6e-08 | 4.733E-08 | 4.120E-08 | 1.600E-08 | 9.400E-08 |
| 1233 | 62 | V62M | A | 3.3e-08 | 8.8e-08 | 2.3e-08 | 4.800E-08 | 3.500E-08 | 2.300E-08 | 8.800E-08 |
| 1234 | 62 | V62F | A | 3e-08 | 1.2e-07 | 3.3e-08 | 6.100E-08 | 5.112E-08 | 3.000E-08 | 1.200E-07 |
| 1235 | 62 | V62P | A | 6.3e-08 | 6e-08 | 6.3e-08 | 6.200E-08 | 1.732E-09 | 6.000E-08 | 6.300E-08 |
| 1236 | 62 | V62S | A | 4e-08 | 3e-08 | 9.5e-08 | 5.500E-08 | 3.500E-08 | 3.000E-08 | 9.500E-08 |
| 1237 | 62 | V62T | A | 2.8e-08 | 6.2e-08 | 5.4e-08 | 4.800E-08 | 1.778E-08 | 2.800E-08 | 6.200E-08 |
| 1238 | 62 | V62W | A | 9.6e-09 | 2.5e-08 | 3.2e-08 | 2.220E-08 | 1.146E-08 | 9.600E-09 | 3.200E-08 |
| 1239 | 62 | V62Y | A | 3.3e-08 | 2.5e-08 | 5.7e-08 | 3.833E-08 | 1.665E-08 | 2.500E-08 | 5.700E-08 |
| 1240 | 62 | V62V | A | 5.8e-08 | 2.6e-08 | 1.2e-07 | 6.800E-08 | 4.779E-08 | 2.600E-08 | 1.200E-07 |
| 1241 | 63 | L63A | A | 2.8e-08 | 1.6e-08 | 7.2e-08 | 3.867E-08 | 2.948E-08 | 1.600E-08 | 7.200E-08 |
| 1242 | 63 | L63R | A | 1.7e-07 | 6.5e-08 | 7e-08 | 1.017E-07 | 5.923E-08 | 6.500E-08 | 1.700E-07 |
| 1243 | 63 | L63N | A | 6e-08 | 4.3e-08 | 3.8e-08 | 4.700E-08 | 1.153E-08 | 3.800E-08 | 6.000E-08 |
| 1244 | 63 | L63D | A | 5.6e-08 | 5.6e-08 | 9e-08 | 6.733E-08 | 1.963E-08 | 5.600E-08 | 9.000E-08 |
| 1245 | 63 | L63C | A | 3e-08 | 7.8e-08 | 1.9e-08 | 4.233E-08 | 3.137E-08 | 1.900E-08 | 7.800E-08 |
| 1246 | 63 | L63E | A | 1e-07 | 4.1e-08 | 2.8e-08 | 5.633E-08 | 3.837E-08 | 2.800E-08 | 1.000E-07 |
| 1247 | 63 | L63Q | A | 8.8e-08 | 4.6e-08 | 3e-08 | 5.467E-08 | 2.996E-08 | 3.000E-08 | 8.800E-08 |
| 1248 | 63 | L63G | A | 9.4e-09 | 2e-08 | 4.5e-08 | 2.480E-08 | 1.828E-08 | 9.400E-09 | 4.500E-08 |
| 1249 | 63 | L63H | A | 8.6e-08 | 8.7e-08 | 3.4e-08 | 6.900E-08 | 3.032E-08 | 3.400E-08 | 8.700E-08 |
| 1250 | 63 | L63I | A | 2.3e-08 | 5.7e-08 | 5.8e-08 | 4.600E-08 | 1.992E-08 | 2.300E-08 | 5.800E-08 |
| 1251 | 63 | L63L | A | 9.5e-08 | 5.1e-08 | 8.3e-08 | 7.633E-08 | 2.274E-08 | 5.100E-08 | 9.500E-08 |
| 1252 | 63 | L63K | A | 2.9e-08 | 4.2e-08 | 5.2e-08 | 4.100E-08 | 1.153E-08 | 2.900E-08 | 5.200E-08 |
| 1253 | 63 | L63M | A | 4.6e-08 | 1.1e-07 | 4.7e-08 | 6.767E-08 | 3.667E-08 | 4.600E-08 | 1.100E-07 |
| 1254 | 63 | L63F | A | 9.4e-08 | 6.3e-08 | 3.4e-08 | 6.367E-08 | 3.001E-08 | 3.400E-08 | 9.400E-08 |
| 1255 | 63 | L63P | A | 5.1e-08 | 6.8e-08 | 4.6e-08 | 5.500E-08 | 1.153E-08 | 4.600E-08 | 6.800E-08 |
| 1256 | 63 | L63S | A | 4.3e-08 | 9.6e-08 | 2.6e-08 | 5.500E-08 | 3.651E-08 | 2.600E-08 | 9.600E-08 |
| 1257 | 63 | L63T | A | 5.6e-08 | 2.6e-08 | 5.9e-08 | 4.700E-08 | 1.825E-08 | 2.600E-08 | 5.900E-08 |
| 1258 | 63 | L63W | A | 6.6e-08 | 3.2e-08 | 6.2e-08 | 5.333E-08 | 1.858E-08 | 3.200E-08 | 6.600E-08 |
| 1259 | 63 | L63Y | A | 4.6e-08 | 5.7e-08 | 1.8e-08 | 4.033E-08 | 2.011E-08 | 1.800E-08 | 5.700E-08 |
| 1260 | 63 | L63V | A | 6e-08 | 2.5e-08 | 2.8e-08 | 3.767E-08 | 1.940E-08 | 2.500E-08 | 6.000E-08 |
| 1261 | 64 | I64A | A | 6.9e-08 | 3.2e-08 | 6.7e-08 | 5.600E-08 | 2.081E-08 | 3.200E-08 | 6.900E-08 |
| 1262 | 64 | I64R | A | 2.9e-08 | 7.3e-08 | 5.8e-08 | 5.333E-08 | 2.237E-08 | 2.900E-08 | 7.300E-08 |
| 1263 | 64 | I64N | A | 3.7e-08 | 7.4e-08 | 3.7e-08 | 4.933E-08 | 2.136E-08 | 3.700E-08 | 7.400E-08 |
| 1264 | 64 | I64D | A | 6.5e-08 | 4e-08 | 8.4e-08 | 6.300E-08 | 2.207E-08 | 4.000E-08 | 8.400E-08 |
| 1265 | 64 | I64C | A | 5.2e-08 | 1.1e-07 | 5.8e-08 | 7.333E-08 | 3.190E-08 | 5.200E-08 | 1.100E-07 |
| 1266 | 64 | I64E | A | 4.9e-08 | 7.3e-08 | 3.7e-08 | 5.300E-08 | 1.833E-08 | 3.700E-08 | 7.300E-08 |
| 1267 | 64 | I64Q | A | 4.7e-08 | 5.2e-08 | 9.1e-08 | 6.333E-08 | 2.409E-08 | 4.700E-08 | 9.100E-08 |
| 1268 | 64 | I64G | A | 2.8e-08 | 5.3e-08 | 3.5e-08 | 3.867E-08 | 1.290E-08 | 2.800E-08 | 5.300E-08 |
| 1269 | 64 | I64H | A | 4.4e-08 | 9.2e-08 | 8.2e-08 | 7.267E-08 | 2.532E-08 | 4.400E-08 | 9.200E-08 |
| 1270 | 64 | I64I | A | 1.1e-07 | 4.9e-08 | 5.7e-08 | 7.200E-08 | 3.315E-08 | 4.900E-08 | 1.100E-07 |
| 1271 | 64 | I64L | A | 3.5e-08 | 1.1e-08 | 5.5e-08 | 3.367E-08 | 2.203E-08 | 1.100E-08 | 5.500E-08 |
| 1272 | 64 | I64K | A | 5.5e-08 | 8.4e-08 | 2.7e-08 | 5.533E-08 | 2.850E-08 | 2.700E-08 | 8.400E-08 |
| 1273 | 64 | I64M | A | 4.7e-08 | 2.7e-08 | 6.5e-08 | 4.633E-08 | 1.901E-08 | 2.700E-08 | 6.500E-08 |
| 1274 | 64 | I64F | A | 3.9e-08 | 4.9e-08 | 2.4e-08 | 3.733E-08 | 1.258E-08 | 2.400E-08 | 4.900E-08 |

| | | | | | | | | | | |
|---|---|---|---|---|---|---|---|---|---|---|
| 1275 | 64 | I64P | A | 2.1e-08 | 3.4e-08 | 9.3e-08 | 4.933E-08 | 3.837E-08 | 2.100E-08 | 9.300E-08 |
| 1276 | 64 | I64S | A | 1.4e-08 | 7.6e-08 | 6.5e-08 | 5.167E-08 | 3.308E-08 | 1.400E-08 | 7.600E-08 |
| 1277 | 64 | I64T | A | 3.3e-08 | 6.1e-08 | 3.6e-08 | 4.333E-08 | 1.537E-08 | 3.300E-08 | 6.100E-08 |
| 1278 | 64 | I64W | A | 3.1e-08 | 6.1e-08 | 3.5e-08 | 4.233E-08 | 1.629E-08 | 3.100E-08 | 6.100E-08 |
| 1279 | 64 | I64Y | A | 4e-08 | 1.9e-07 | 5.3e-08 | 9.433E-08 | 8.310E-08 | 4.000E-08 | 1.900E-07 |
| 1280 | 64 | I64V | A | 3e-08 | 1.6e-08 | 3.7e-08 | 2.767E-08 | 1.069E-08 | 1.600E-08 | 3.700E-08 |
| 1281 | 65 | A65A | A | 1.1e-07 | 2.8e-08 | 5.5e-08 | 6.433E-08 | 4.179E-08 | 2.800E-08 | 1.100E-07 |
| 1282 | 65 | A65R | A | 6.5e-08 | 5.4e-08 | 6.9e-08 | 6.267E-08 | 7.767E-09 | 5.400E-08 | 6.900E-08 |
| 1283 | 65 | A65N | A | 4.5e-08 | 1.4e-07 | 3.9e-08 | 7.467E-08 | 5.666E-08 | 3.900E-08 | 1.400E-07 |
| 1284 | 65 | A65D | A | 2.7e-08 | 8e-08 | 2.9e-08 | 4.533E-08 | 3.004E-08 | 2.700E-08 | 8.000E-08 |
| 1285 | 65 | A65C | A | 2.9e-08 | 2.7e-08 | 3.7e-08 | 3.100E-08 | 5.292E-09 | 2.700E-08 | 3.700E-08 |
| 1286 | 65 | A65E | A | 2.3e-08 | 7.1e-08 | 8e-08 | 5.800E-08 | 3.064E-08 | 2.300E-08 | 8.000E-08 |
| 1287 | 65 | A65Q | A | 3.2e-08 | 9.1e-08 | 4.4e-08 | 5.567E-08 | 3.118E-08 | 3.200E-08 | 9.100E-08 |
| 1288 | 65 | A65G | A | 6e-08 | 7.5e-08 | 2.4e-08 | 5.300E-08 | 2.621E-08 | 2.400E-08 | 7.500E-08 |
| 1289 | 65 | A65H | A | 1.2e-07 | 3.8e-08 | 3.3e-08 | 6.367E-08 | 4.885E-08 | 3.300E-08 | 1.200E-07 |
| 1290 | 65 | A65I | A | 5.9e-08 | 4.4e-08 | 8.2e-08 | 6.167E-08 | 1.914E-08 | 4.400E-08 | 8.200E-08 |
| 1291 | 65 | A65L | A | 4.6e-08 | 2.3e-08 | 6e-08 | 4.300E-08 | 1.868E-08 | 2.300E-08 | 6.000E-08 |
| 1292 | 65 | A65K | A | 4e-08 | 1.5e-08 | 4.2e-08 | 3.233E-08 | 1.504E-08 | 1.500E-08 | 4.200E-08 |
| 1293 | 65 | A65M | A | 4.1e-08 | 1.5e-08 | 3.4e-08 | 3.000E-08 | 1.345E-08 | 1.500E-08 | 4.100E-08 |
| 1294 | 65 | A65F | A | 9.8e-08 | 2.9e-08 | 2e-08 | 4.900E-08 | 4.267E-08 | 2.000E-08 | 9.800E-08 |
| 1295 | 65 | A65P | A | 5.2e-08 | 6.8e-08 | 1.8e-08 | 4.600E-08 | 2.553E-08 | 1.800E-08 | 6.800E-08 |
| 1296 | 65 | A65S | A | 8.1e-08 | 5.9e-08 | 5.5e-08 | 6.500E-08 | 1.400E-08 | 5.500E-08 | 8.100E-08 |
| 1297 | 65 | A65T | A | 1.2e-08 | 5.8e-08 | 1.1e-07 | 6.000E-08 | 4.903E-08 | 1.200E-08 | 1.100E-07 |
| 1298 | 65 | A65W | A | 2.6e-08 | 4.4e-08 | 1.5e-07 | 7.333E-08 | 6.700E-08 | 2.600E-08 | 1.500E-07 |
| 1299 | 65 | A65Y | A | 1.1e-07 | 2.4e-08 | 8.5e-08 | 7.300E-08 | 4.424E-08 | 2.400E-08 | 1.100E-07 |
| 1300 | 65 | A65V | A | 6.9e-08 | 4.9e-08 | 8.4e-08 | 6.733E-08 | 1.756E-08 | 4.900E-08 | 8.400E-08 |
| 1301 | 66 | H66A | A | 3.7e-08 | 2.6e-08 | 2.5e-08 | 2.933E-08 | 6.658E-09 | 2.500E-08 | 3.700E-08 |
| 1302 | 66 | H66R | A | 2.7e-08 | 5.7e-08 | 7.5e-08 | 5.300E-08 | 2.425E-08 | 2.700E-08 | 7.500E-08 |
| 1303 | 66 | H66N | A | 3.6e-08 | 3.8e-08 | 6.9e-09 | 2.697E-08 | 1.741E-08 | 6.900E-09 | 3.800E-08 |
| 1304 | 66 | H66D | A | 3e-08 | 6.2e-08 | 4.9e-08 | 4.700E-08 | 1.609E-08 | 3.000E-08 | 6.200E-08 |
| 1305 | 66 | H66C | A | 3.1e-08 | 7.7e-08 | 3.3e-08 | 4.700E-08 | 2.600E-08 | 3.100E-08 | 7.700E-08 |
| 1306 | 66 | H66E | A | 7.1e-08 | 1.1e-07 | 7.3e-08 | 8.467E-08 | 2.196E-08 | 7.100E-08 | 1.100E-07 |
| 1307 | 66 | H66Q | A | 2.8e-08 | 1e-07 | 5.9e-08 | 6.233E-08 | 3.612E-08 | 2.800E-08 | 1.000E-07 |
| 1308 | 66 | H66G | A | 4.8e-08 | 2.5e-08 | 3.2e-08 | 3.500E-08 | 1.179E-08 | 2.500E-08 | 4.800E-08 |
| 1309 | 66 | H66H | A | 2.9e-08 | 1e-07 | 3.8e-08 | 5.567E-08 | 3.866E-08 | 2.900E-08 | 1.000E-07 |
| 1310 | 66 | H66I | A | 2.7e-08 | 2.5e-08 | 3e-08 | 2.733E-08 | 2.517E-09 | 2.500E-08 | 3.000E-08 |
| 1311 | 66 | H66L | A | 7e-08 | 3.4e-08 | 5.5e-08 | 5.300E-08 | 1.808E-08 | 3.400E-08 | 7.000E-08 |
| 1312 | 66 | H66K | A | 6.8e-08 | 6.3e-08 | 8.7e-08 | 7.267E-08 | 1.266E-08 | 6.300E-08 | 8.700E-08 |
| 1313 | 66 | H66M | A | 3.7e-08 | 3.2e-08 | 5.5e-08 | 4.133E-08 | 1.210E-08 | 3.200E-08 | 5.500E-08 |
| 1314 | 66 | H66F | A | 1.7e-08 | 2.7e-08 | 3.4e-08 | 2.600E-08 | 8.544E-09 | 1.700E-08 | 3.400E-08 |
| 1315 | 66 | H66P | A | 4.3e-08 | 1.3e-07 | 5e-08 | 7.433E-08 | 4.834E-08 | 4.300E-08 | 1.300E-07 |
| 1316 | 66 | H66S | A | 2.5e-08 | 4.9e-08 | 3.8e-08 | 3.733E-08 | 1.201E-08 | 2.500E-08 | 4.900E-08 |
| 1317 | 66 | H66T | A | 3.3e-08 | 8.3e-08 | 1.9e-08 | 4.500E-08 | 3.365E-08 | 1.900E-08 | 8.300E-08 |
| 1318 | 66 | H66W | A | 5.9e-08 | 7.9e-08 | 6.5e-08 | 6.767E-08 | 1.026E-08 | 5.900E-08 | 7.900E-08 |
| 1319 | 66 | H66Y | A | 1.5e-08 | 5.9e-08 | 2.4e-08 | 3.267E-08 | 2.325E-08 | 1.500E-08 | 5.900E-08 |
| 1320 | 66 | H66V | A | 4.5e-08 | 4.2e-08 | 6.6e-08 | 5.100E-08 | 1.308E-08 | 4.200E-08 | 6.600E-08 |
| 1321 | 67 | N67A | A | 6e-08 | 8.1e-08 | 1.2e-07 | 8.700E-08 | 3.045E-08 | 6.000E-08 | 1.200E-07 |
| 1322 | 67 | N67R | A | 4.1e-08 | 2.9e-08 | 5.4e-08 | 4.133E-08 | 1.250E-08 | 2.900E-08 | 5.400E-08 |
| 1323 | 67 | N67N | A | 1.6e-08 | 3.3e-08 | 5.8e-08 | 3.567E-08 | 2.113E-08 | 1.600E-08 | 5.800E-08 |
| 1324 | 67 | N67D | A | 1.6e-08 | 5.4e-08 | 5e-08 | 4.000E-08 | 2.088E-08 | 1.600E-08 | 5.400E-08 |
| 1325 | 67 | N67C | A | 4.3e-08 | 9.1e-08 | 3.8e-08 | 5.733E-08 | 2.926E-08 | 3.800E-08 | 9.100E-08 |
| 1326 | 67 | N67E | A | 4.9e-08 | 4.1e-08 | 9.4e-08 | 6.133E-08 | 2.857E-08 | 4.100E-08 | 9.400E-08 |
| 1327 | 67 | N67Q | A | 6.6e-08 | 2.9e-08 | 3e-08 | 4.167E-08 | 2.108E-08 | 2.900E-08 | 6.600E-08 |
| 1328 | 67 | N67G | A | 3.6e-08 | 2.9e-08 | 4e-08 | 3.500E-08 | 5.568E-09 | 2.900E-08 | 4.000E-08 |
| 1329 | 67 | N67H | A | 1.8e-08 | 5.1e-08 | 3e-08 | 3.300E-08 | 1.670E-08 | 1.800E-08 | 5.100E-08 |
| 1330 | 67 | N67I | A | 2.5e-08 | 8.3e-08 | 1.6e-08 | 4.133E-08 | 3.636E-08 | 1.600E-08 | 8.300E-08 |
| 1331 | 67 | N67L | A | 4.8e-08 | 2.1e-08 | 6.1e-08 | 4.333E-08 | 2.040E-08 | 2.100E-08 | 6.100E-08 |
| 1332 | 67 | N67K | A | 5.5e-08 | 1.2e-08 | 3.1e-08 | 3.267E-08 | 2.155E-08 | 1.200E-08 | 5.500E-08 |
| 1333 | 67 | N67M | A | 2.4e-08 | 4.6e-08 | 5.8e-08 | 4.267E-08 | 1.724E-08 | 2.400E-08 | 5.800E-08 |
| 1334 | 67 | N67F | A | 4.2e-08 | 8.2e-08 | 3.5e-08 | 5.300E-08 | 2.536E-08 | 3.500E-08 | 8.200E-08 |
| 1335 | 67 | N67P | A | 1.2e-08 | 3.8e-08 | 1.1e-07 | 5.333E-08 | 5.077E-08 | 1.200E-08 | 1.100E-07 |
| 1336 | 67 | N67S | A | 4.5e-08 | 2.3e-08 | 6.5e-08 | 4.433E-08 | 2.101E-08 | 2.300E-08 | 6.500E-08 |
| 1337 | 67 | N67T | A | 3.4e-08 | 5e-08 | 3.7e-08 | 4.033E-08 | 8.505E-09 | 3.400E-08 | 5.000E-08 |
| 1338 | 67 | N67W | A | 2.2e-08 | 1e-07 | 5e-08 | 5.733E-08 | 3.951E-08 | 2.200E-08 | 1.000E-07 |
| 1339 | 67 | N67Y | A | 1.1e-07 | 4.8e-08 | 2e-08 | 5.933E-08 | 4.606E-08 | 2.000E-08 | 1.100E-07 |
| 1340 | 67 | N67V | A | 2.6e-08 | 1.7e-08 | 1.7e-07 | 7.100E-08 | 8.585E-08 | 1.700E-08 | 1.700E-07 |
| 1341 | 68 | Q68A | A | 5.5e-08 | 1.6e-07 | 7e-08 | 9.500E-08 | 5.679E-08 | 5.500E-08 | 1.600E-07 |
| 1342 | 68 | Q68R | A | 4.6e-08 | 1.2e-08 | 3.7e-08 | 3.167E-08 | 1.762E-08 | 1.200E-08 | 4.600E-08 |
| 1343 | 68 | Q68N | A | 2.8e-08 | 1.5e-08 | 2.4e-08 | 2.233E-08 | 6.658E-09 | 1.500E-08 | 2.800E-08 |
| 1344 | 68 | Q68D | A | 2.3e-08 | 5.1e-08 | 1.6e-07 | 7.800E-08 | 7.238E-08 | 2.300E-08 | 1.600E-07 |
| 1345 | 68 | Q68C | A | 5e-08 | 1.3e-08 | 5e-08 | 3.767E-08 | 2.136E-08 | 1.300E-08 | 5.000E-08 |
| 1346 | 68 | Q68E | A | 5.4e-08 | 4.6e-08 | 7.7e-08 | 5.900E-08 | 1.609E-08 | 4.600E-08 | 7.700E-08 |
| 1347 | 68 | Q68Q | A | 4.9e-08 | 1.6e-08 | 1.8e-08 | 2.767E-08 | 1.850E-08 | 1.600E-08 | 4.900E-08 |
| 1348 | 68 | Q68G | A | 3.9e-08 | 7.7e-08 | 1.1e-08 | 4.233E-08 | 3.313E-08 | 1.100E-08 | 7.700E-08 |
| 1349 | 68 | Q68H | A | 2.8e-08 | 7.3e-08 | 6.8e-08 | 5.633E-08 | 2.466E-08 | 2.800E-08 | 7.300E-08 |
| 1350 | 68 | Q68I | A | 1.2e-07 | 1.8e-08 | 3.3e-08 | 5.700E-08 | 5.507E-08 | 1.800E-08 | 1.200E-07 |
| 1351 | 68 | Q68L | A | 1e-07 | 3.6e-08 | 2.1e-08 | 5.233E-08 | 4.196E-08 | 2.100E-08 | 1.000E-07 |
| 1352 | 68 | Q68K | A | 1.3e-07 | 7e-08 | 1e-07 | 1.000E-07 | 3.000E-08 | 7.000E-08 | 1.300E-07 |
| 1353 | 68 | Q68M | A | 4.5e-08 | 1e-07 | 2.3e-08 | 5.600E-08 | 3.966E-08 | 2.300E-08 | 1.000E-07 |
| 1354 | 68 | Q68F | A | 5.8e-08 | 7.9e-08 | 4.7e-08 | 6.133E-08 | 1.626E-08 | 4.700E-08 | 7.900E-08 |
| 1355 | 68 | Q68P | A | 9e-08 | 6.7e-08 | 6.2e-08 | 7.300E-08 | 1.493E-08 | 6.200E-08 | 9.000E-08 |
| 1356 | 68 | Q68S | A | 8.3e-08 | 5.6e-08 | 8e-08 | 7.300E-08 | 1.480E-08 | 5.600E-08 | 8.300E-08 |
| 1357 | 68 | Q68T | A | 4.5e-08 | 3.6e-08 | 4e-08 | 4.033E-08 | 4.509E-09 | 3.600E-08 | 4.500E-08 |
| 1358 | 68 | Q68W | A | 4.1e-08 | 6e-09 | 1.1e-07 | 5.233E-08 | 5.292E-08 | 6.000E-09 | 1.100E-07 |
| 1359 | 68 | Q68Y | A | 3.3e-07 | 2.9e-08 | 3e-08 | 1.297E-07 | 1.735E-07 | 2.900E-08 | 3.300E-07 |

| | | | | | | | | | | |
|---|---|---|---|---|---|---|---|---|---|---|
| 1360 | 68 | Q68V | A | 2.7e-08 | 5.6e-08 | 4.8e-08 | 4.367E-08 | 1.498E-08 | 2.700E-08 | 5.600E-08 |
| 1361 | 69 | V69A | A | 5.6e-08 | 7.8e-08 | 4.3e-08 | 5.900E-08 | 1.769E-08 | 4.300E-08 | 7.800E-08 |
| 1362 | 69 | V69R | A | 4.5e-08 | 2.6e-08 | 1.5e-08 | 2.867E-08 | 1.518E-08 | 1.500E-08 | 4.500E-08 |
| 1363 | 69 | V69N | A | 6e-08 | 4.2e-08 | 6.1e-08 | 5.433E-08 | 1.069E-08 | 4.200E-08 | 6.100E-08 |
| 1364 | 69 | V69D | A | 2.8e-08 | 3.2e-08 | 6e-08 | 4.000E-08 | 1.744E-08 | 2.800E-08 | 6.000E-08 |
| 1365 | 69 | V69C | A | 5.9e-08 | 5e-08 | 2e-08 | 4.300E-08 | 2.042E-08 | 2.000E-08 | 5.900E-08 |
| 1366 | 69 | V69E | A | 2.8e-08 | 3.3e-08 | 7.5e-08 | 4.533E-08 | 2.581E-08 | 2.800E-08 | 7.500E-08 |
| 1367 | 69 | V69Q | A | 5.7e-08 | 2.4e-08 | 5.3e-08 | 4.467E-08 | 1.801E-08 | 2.400E-08 | 5.700E-08 |
| 1368 | 69 | V69G | A | 2.7e-08 | 1.1e-07 | 6.8e-08 | 6.833E-08 | 4.150E-08 | 2.700E-08 | 1.100E-07 |
| 1369 | 69 | V69H | A | 3.5e-08 | 8.7e-08 | 1.4e-07 | 8.733E-08 | 5.250E-08 | 3.500E-08 | 1.400E-07 |
| 1370 | 69 | V69I | A | 5.3e-08 | 8.3e-08 | 1.1e-07 | 8.200E-08 | 2.851E-08 | 5.300E-08 | 1.100E-07 |
| 1371 | 69 | V69L | A | 2.7e-08 | 5e-08 | 5.1e-08 | 4.267E-08 | 1.358E-08 | 2.700E-08 | 5.100E-08 |
| 1372 | 69 | V69K | A | 1.9e-08 | 4.7e-08 | 3.1e-08 | 3.233E-08 | 1.405E-08 | 1.900E-08 | 4.700E-08 |
| 1373 | 69 | V69M | A | 8.5e-08 | 4.4e-08 | 2.6e-08 | 5.167E-08 | 3.024E-08 | 2.600E-08 | 8.500E-08 |
| 1374 | 69 | V69F | A | 1.9e-07 | 7.3e-08 | 6e-08 | 1.077E-07 | 7.160E-08 | 6.000E-08 | 1.900E-07 |
| 1375 | 69 | V69P | A | 2.6e-08 | 3.1e-08 | 3.6e-08 | 3.100E-08 | 5.000E-09 | 2.600E-08 | 3.600E-08 |
| 1376 | 69 | V69S | A | 3.5e-08 | 9e-08 | 7.2e-08 | 6.567E-08 | 2.804E-08 | 3.500E-08 | 9.000E-08 |
| 1377 | 69 | V69T | A | 4.9e-08 | 9.9e-08 | 2.1e-08 | 5.633E-08 | 3.951E-08 | 2.100E-08 | 9.900E-08 |
| 1378 | 69 | V69W | A | 2.1e-08 | 5.2e-08 | 3.8e-08 | 3.700E-08 | 1.552E-08 | 2.100E-08 | 5.200E-08 |
| 1379 | 69 | V69Y | A | 2.8e-08 | 1.7e-08 | 1.8e-07 | 7.500E-08 | 9.110E-08 | 1.700E-08 | 1.800E-07 |
| 1380 | 69 | V69V | A | 7.7e-08 | 6.3e-08 | 7.3e-08 | 7.100E-08 | 7.211E-09 | 6.300E-08 | 7.700E-08 |
| 1381 | 70 | R70A | A | 1.2e-08 | 1.9e-08 | 1.2e-08 | 1.433E-08 | 4.041E-09 | 1.200E-08 | 1.900E-08 |
| 1382 | 70 | R70R | A | 9.8e-08 | 3.1e-08 | 1.3e-08 | 4.733E-08 | 4.479E-08 | 1.300E-08 | 9.800E-08 |
| 1383 | 70 | R70N | A | 4e-08 | 7.5e-08 | 1.3e-07 | 8.167E-08 | 4.537E-08 | 4.000E-08 | 1.300E-07 |
| 1384 | 70 | R70D | A | 1.5e-08 | 4.5e-08 | 2.3e-08 | 2.767E-08 | 1.553E-08 | 1.500E-08 | 4.500E-08 |
| 1385 | 70 | R70C | A | 5.2e-08 | 4.6e-08 | 8.7e-08 | 6.167E-08 | 2.214E-08 | 4.600E-08 | 8.700E-08 |
| 1386 | 70 | R70E | A | 2.7e-08 | 5.5e-08 | 1.9e-07 | 9.067E-08 | 8.716E-08 | 2.700E-08 | 1.900E-07 |
| 1387 | 70 | R70Q | A | 1.4e-07 | 1.9e-07 | 2.5e-08 | 1.183E-07 | 8.461E-08 | 2.500E-08 | 1.900E-07 |
| 1388 | 70 | R70G | A | 4.5e-08 | 3.9e-08 | 3.3e-08 | 3.900E-08 | 6.000E-09 | 3.300E-08 | 4.500E-08 |
| 1389 | 70 | R70H | A | 8.1e-08 | 8.6e-08 | 8.1e-08 | 8.267E-08 | 2.887E-09 | 8.100E-08 | 8.600E-08 |
| 1390 | 70 | R70I | A | 3.5e-08 | 5e-08 | 2.6e-08 | 3.700E-08 | 1.212E-08 | 2.600E-08 | 5.000E-08 |
| 1391 | 70 | R70L | A | 7.2e-08 | 5.9e-08 | 7e-08 | 6.700E-08 | 7.000E-09 | 5.900E-08 | 7.200E-08 |
| 1392 | 70 | R70K | A | 1.5e-07 | 3e-08 | 8e-08 | 8.667E-08 | 6.028E-08 | 3.000E-08 | 1.500E-07 |
| 1393 | 70 | R70M | A | 1e-08 | 2.2e-08 | 2.9e-08 | 2.033E-08 | 9.609E-09 | 1.000E-08 | 2.900E-08 |
| 1394 | 70 | R70F | A | 2.1e-08 | 8.7e-08 | 4.3e-08 | 5.033E-08 | 3.361E-08 | 2.100E-08 | 8.700E-08 |
| 1395 | 70 | R70P | A | 4.3e-08 | 4.3e-08 | 3.6e-08 | 4.067E-08 | 4.041E-09 | 3.600E-08 | 4.300E-08 |
| 1396 | 70 | R70S | A | 4.3e-08 | 1.6e-08 | 3.1e-08 | 3.000E-08 | 1.353E-08 | 1.600E-08 | 4.300E-08 |
| 1397 | 70 | R70T | A | 7.3e-08 | 8.5e-08 | 1.4e-08 | 5.733E-08 | 3.800E-08 | 1.400E-08 | 8.500E-08 |
| 1398 | 70 | R70W | A | 1.2e-07 | 4.8e-08 | 3.7e-08 | 6.833E-08 | 4.508E-08 | 3.700E-08 | 1.200E-07 |
| 1399 | 70 | R70Y | A | 5.2e-08 | 3.7e-08 | 5.1e-08 | 4.667E-08 | 8.386E-09 | 3.700E-08 | 5.200E-08 |
| 1400 | 70 | R70V | A | 5e-08 | 4.9e-08 | 1.1e-07 | 6.967E-08 | 3.493E-08 | 4.900E-08 | 1.100E-07 |
| 1401 | 71 | Q71A | A | 4.3e-08 | 6.2e-08 | 2.8e-08 | 4.433E-08 | 1.704E-08 | 2.800E-08 | 6.200E-08 |
| 1402 | 71 | Q71R | A | 2.6e-08 | 7.4e-08 | 8.3e-08 | 6.100E-08 | 3.064E-08 | 2.600E-08 | 8.300E-08 |
| 1403 | 71 | Q71N | A | 2.9e-08 | 5.6e-08 | 1.1e-07 | 6.500E-08 | 4.124E-08 | 2.900E-08 | 1.100E-07 |
| 1404 | 71 | Q71D | A | 3.8e-08 | 9.5e-09 | 2.6e-07 | 1.025E-07 | 1.371E-07 | 9.500E-09 | 2.600E-07 |
| 1405 | 71 | Q71C | A | 5.2e-08 | 1.8e-08 | 5.9e-08 | 4.300E-08 | 2.193E-08 | 1.800E-08 | 5.900E-08 |
| 1406 | 71 | Q71E | A | 4.7e-08 | 1.8e-08 | 8.9e-08 | 5.133E-08 | 3.570E-08 | 1.800E-08 | 8.900E-08 |
| 1407 | 71 | Q71Q | A | 2.8e-08 | 3.7e-08 | 6.1e-08 | 4.200E-08 | 1.706E-08 | 2.800E-08 | 6.100E-08 |
| 1408 | 71 | Q71G | A | 3.7e-08 | 8.6e-08 | 2.9e-08 | 5.067E-08 | 3.086E-08 | 2.900E-08 | 8.600E-08 |
| 1409 | 71 | Q71H | A | 1.9e-08 | 2.8e-08 | 3.6e-08 | 2.767E-08 | 8.505E-09 | 1.900E-08 | 3.600E-08 |
| 1410 | 71 | Q71I | A | 2.2e-08 | 5.1e-08 | 4.6e-08 | 3.967E-08 | 1.550E-08 | 2.200E-08 | 5.100E-08 |
| 1411 | 71 | Q71L | A | 3.1e-08 | 3.6e-08 | 7.8e-08 | 4.833E-08 | 2.581E-08 | 3.100E-08 | 7.800E-08 |
| 1412 | 71 | Q71K | A | 4.7e-08 | 4.9e-08 | 6.1e-08 | 5.233E-08 | 7.572E-09 | 4.700E-08 | 6.100E-08 |
| 1413 | 71 | Q71M | A | 1.7e-08 | 5.5e-08 | 2.2e-08 | 3.133E-08 | 2.065E-08 | 1.700E-08 | 5.500E-08 |
| 1414 | 71 | Q71F | A | 4.2e-08 | 5.6e-08 | 2e-08 | 3.933E-08 | 1.815E-08 | 2.000E-08 | 5.600E-08 |
| 1415 | 71 | Q71P | A | 3.2e-08 | 7.6e-08 | 8e-08 | 6.267E-08 | 2.663E-08 | 3.200E-08 | 8.000E-08 |
| 1416 | 71 | Q71S | A | 5.1e-08 | 1.5e-07 | 6.9e-08 | 9.000E-08 | 5.274E-08 | 5.100E-08 | 1.500E-07 |
| 1417 | 71 | Q71T | A | 2.6e-08 | 4.7e-08 | 1.5e-08 | 2.933E-08 | 1.626E-08 | 1.500E-08 | 4.700E-08 |
| 1418 | 71 | Q71W | A | 4.3e-08 | 2.6e-08 | 4.7e-08 | 3.867E-08 | 1.115E-08 | 2.600E-08 | 4.700E-08 |
| 1419 | 71 | Q71Y | A | 5.2e-08 | 3.9e-08 | 5e-08 | 4.700E-08 | 7.000E-09 | 3.900E-08 | 5.200E-08 |
| 1420 | 71 | Q71V | A | 1e-07 | 2.1e-08 | 6.2e-08 | 6.100E-08 | 3.951E-08 | 2.100E-08 | 1.000E-07 |
| 1421 | 72 | V72A | A | 3.8e-08 | 2.2e-07 | 8.6e-08 | 1.147E-07 | 9.433E-08 | 3.800E-08 | 2.200E-07 |
| 1422 | 72 | V72R | A | 3e-08 | 1.2e-08 | 2.8e-08 | 2.333E-08 | 9.866E-09 | 1.200E-08 | 3.000E-08 |
| 1423 | 72 | V72N | A | 3.6e-08 | 3.1e-08 | 5e-08 | 3.900E-08 | 9.849E-09 | 3.100E-08 | 5.000E-08 |
| 1424 | 72 | V72D | A | 6.3e-08 | 8.3e-08 | 2.5e-08 | 5.700E-08 | 2.946E-08 | 2.500E-08 | 8.300E-08 |
| 1425 | 72 | V72C | A | 2.1e-08 | 3.7e-08 | 1.9e-08 | 2.567E-08 | 9.866E-09 | 1.900E-08 | 3.700E-08 |
| 1426 | 72 | V72E | A | 3e-07 | 1.6e-08 | 1.3e-07 | 1.487E-07 | 1.429E-07 | 1.600E-08 | 3.000E-07 |
| 1427 | 72 | V72Q | A | 6e-08 | 1.1e-07 | 5.2e-08 | 7.400E-08 | 3.143E-08 | 5.200E-08 | 1.100E-07 |
| 1428 | 72 | V72G | A | 9e-08 | 8.1e-08 | 1.5e-07 | 1.070E-07 | 3.751E-08 | 8.100E-08 | 1.500E-07 |
| 1429 | 72 | V72H | A | 1e-07 | 5.4e-08 | 6e-08 | 7.133E-08 | 2.501E-08 | 5.400E-08 | 1.000E-07 |
| 1430 | 72 | V72I | A | 6.2e-08 | 2.8e-08 | 5e-08 | 4.667E-08 | 1.724E-08 | 2.800E-08 | 6.200E-08 |
| 1431 | 72 | V72L | A | 3.5e-08 | 2.5e-08 | 5.1e-08 | 3.700E-08 | 1.311E-08 | 2.500E-08 | 5.100E-08 |
| 1432 | 72 | V72K | A | 4.5e-08 | 1.4e-07 | 2e-08 | 6.833E-08 | 6.331E-08 | 2.000E-08 | 1.400E-07 |
| 1433 | 72 | V72M | A | 3.2e-08 | 4.2e-08 | 5e-08 | 4.133E-08 | 9.018E-09 | 3.200E-08 | 5.000E-08 |
| 1434 | 72 | V72F | A | 1.7e-08 | 5.5e-08 | 3.7e-08 | 3.633E-08 | 1.901E-08 | 1.700E-08 | 5.500E-08 |
| 1435 | 72 | V72P | A | 3.4e-08 | 6.9e-08 | 1.7e-08 | 4.000E-08 | 2.651E-08 | 1.700E-08 | 6.900E-08 |
| 1436 | 72 | V72S | A | 1.2e-07 | 2.8e-08 | 4.9e-08 | 6.567E-08 | 4.821E-08 | 2.800E-08 | 1.200E-07 |
| 1437 | 72 | V72T | A | 1.4e-07 | 2.7e-08 | 4.3e-08 | 7.000E-08 | 6.115E-08 | 2.700E-08 | 1.400E-07 |
| 1438 | 72 | V72W | A | 5e-08 | 3.1e-08 | 2.7e-08 | 3.600E-08 | 1.229E-08 | 2.700E-08 | 5.000E-08 |
| 1439 | 72 | V72Y | A | 4.4e-08 | 2.9e-08 | 5.4e-08 | 4.233E-08 | 1.258E-08 | 2.900E-08 | 5.400E-08 |
| 1440 | 72 | V72V | A | 1.3e-08 | 4.9e-08 | 4.5e-08 | 3.567E-08 | 1.973E-08 | 1.300E-08 | 4.900E-08 |
| 1441 | 73 | P73A | A | 8e-08 | 7.4e-08 | 1.9e-08 | 5.767E-08 | 3.362E-08 | 1.900E-08 | 8.000E-08 |
| 1442 | 73 | P73R | A | 1e-08 | 4.9e-08 | 1.3e-08 | 2.400E-08 | 2.170E-08 | 1.000E-08 | 4.900E-08 |
| 1443 | 73 | P73N | A | 3.6e-08 | 4e-08 | 4.4e-08 | 4.000E-08 | 4.000E-09 | 3.600E-08 | 4.400E-08 |
| 1444 | 73 | P73D | A | 4e-08 | 2.5e-08 | 2.6e-08 | 3.033E-08 | 8.386E-09 | 2.500E-08 | 4.000E-08 |

| | | | | | | | | | | |
|---|---|---|---|---|---|---|---|---|---|---|
| 1445 | 73 | P73C | A | 1e-07 | 7.2e-08 | 8.8e-08 | 8.667E-08 | 1.405E-08 | 7.200E-08 | 1.000E-07 |
| 1446 | 73 | P73E | A | 1.9e-08 | 4.8e-08 | 6.3e-08 | 4.333E-08 | 2.237E-08 | 1.900E-08 | 6.300E-08 |
| 1447 | 73 | P73Q | A | 3.3e-08 | 9e-09 | 1.3e-08 | 1.833E-08 | 1.286E-08 | 9.000E-09 | 3.300E-08 |
| 1448 | 73 | P73G | A | 8.9e-09 | 4.9e-08 | 7.6e-09 | 2.183E-08 | 2.354E-08 | 7.600E-09 | 4.900E-08 |
| 1449 | 73 | P73H | A | 3.5e-08 | 3e-08 | 2.1e-08 | 2.867E-08 | 7.095E-09 | 2.100E-08 | 3.500E-08 |
| 1450 | 73 | P73I | A | 2.7e-08 | 8.7e-08 | 5.5e-08 | 5.633E-08 | 3.002E-08 | 2.700E-08 | 8.700E-08 |
| 1451 | 73 | P73L | A | 9.9e-09 | 1.4e-08 | 3.6e-08 | 1.997E-08 | 1.404E-08 | 9.900E-09 | 3.600E-08 |
| 1452 | 73 | P73K | A | 5e-08 | 4.5e-08 | 1.2e-07 | 7.167E-08 | 4.193E-08 | 4.500E-08 | 1.200E-07 |
| 1453 | 73 | P73M | A | 4.4e-08 | 2.5e-08 | 3e-08 | 3.300E-08 | 9.849E-09 | 2.500E-08 | 4.400E-08 |
| 1454 | 73 | P73F | A | 1.6e-08 | 3.4e-08 | 4.3e-08 | 3.100E-08 | 1.375E-08 | 1.600E-08 | 4.300E-08 |
| 1455 | 73 | P73P | A | 7.6e-08 | 8.2e-08 | 5.5e-08 | 7.100E-08 | 1.418E-08 | 5.500E-08 | 8.200E-08 |
| 1456 | 73 | P73S | A | 3.8e-08 | 2e-08 | 2e-08 | 2.600E-08 | 1.039E-08 | 2.000E-08 | 3.800E-08 |
| 1457 | 73 | P73T | A | 3.3e-08 | 5.4e-08 | 3.8e-08 | 4.167E-08 | 1.097E-08 | 3.300E-08 | 5.400E-08 |
| 1458 | 73 | P73W | A | 2.2e-08 | 3.6e-08 | 4.5e-08 | 3.433E-08 | 1.159E-08 | 2.200E-08 | 4.500E-08 |
| 1459 | 73 | P73Y | A | 1.3e-08 | 5.8e-09 | 5.2e-08 | 2.360E-08 | 2.486E-08 | 5.800E-09 | 5.200E-08 |
| 1460 | 73 | P73V | A | 6.5e-08 | 2.3e-08 | 4.6e-08 | 4.467E-08 | 2.103E-08 | 2.300E-08 | 6.500E-08 |
| 1461 | 74 | L74A | A | 2.5e-08 | 8.3e-08 | 2.8e-08 | 4.533E-08 | 3.265E-08 | 2.500E-08 | 8.300E-08 |
| 1462 | 74 | L74R | A | 1.9e-08 | 4.1e-08 | 4.7e-08 | 3.567E-08 | 1.474E-08 | 1.900E-08 | 4.700E-08 |
| 1463 | 74 | L74N | A | 2.3e-08 | 4.5e-08 | 3.1e-08 | 3.300E-08 | 1.114E-08 | 2.300E-08 | 4.500E-08 |
| 1464 | 74 | L74D | A | 3e-08 | 1.1e-08 | 2.3e-08 | 2.133E-08 | 9.609E-09 | 1.100E-08 | 3.000E-08 |
| 1465 | 74 | L74C | A | 3.5e-08 | 2.8e-08 | 1.1e-07 | 5.767E-08 | 4.546E-08 | 2.800E-08 | 1.100E-07 |
| 1466 | 74 | L74E | A | 3.7e-08 | 5.1e-08 | 2.4e-08 | 3.733E-08 | 1.350E-08 | 2.400E-08 | 5.100E-08 |
| 1467 | 74 | L74Q | A | 5.2e-08 | 5e-08 | 4.6e-08 | 4.933E-08 | 3.055E-09 | 4.600E-08 | 5.200E-08 |
| 1468 | 74 | L74G | A | 4.1e-08 | 5.3e-08 | 8e-08 | 5.800E-08 | 1.997E-08 | 4.100E-08 | 8.000E-08 |
| 1469 | 74 | L74H | A | 1.7e-08 | 1.1e-08 | 2.6e-08 | 1.800E-08 | 7.550E-09 | 1.100E-08 | 2.600E-08 |
| 1470 | 74 | L74I | A | 2.4e-08 | 2.6e-08 | 8.4e-08 | 4.467E-08 | 3.408E-08 | 2.400E-08 | 8.400E-08 |
| 1471 | 74 | L74L | A | 1.3e-08 | 1.3e-08 | 5.6e-08 | 2.733E-08 | 2.483E-08 | 1.300E-08 | 5.600E-08 |
| 1472 | 74 | L74K | A | 4.8e-08 | 2.1e-08 | 6.9e-08 | 4.600E-08 | 2.406E-08 | 2.100E-08 | 6.900E-08 |
| 1473 | 74 | L74M | A | 4.9e-08 | 4.1e-08 | 8.6e-09 | 3.287E-08 | 2.139E-08 | 8.600E-09 | 4.900E-08 |
| 1474 | 74 | L74F | A | 2.2e-08 | 3.6e-08 | 5.9e-08 | 3.900E-08 | 1.868E-08 | 2.200E-08 | 5.900E-08 |
| 1475 | 74 | L74P | A | 2.2e-08 | 3.8e-08 | 4.2e-08 | 3.400E-08 | 1.058E-08 | 2.200E-08 | 4.200E-08 |
| 1476 | 74 | L74S | A | 3.1e-08 | 1.5e-08 | 7e-08 | 3.867E-08 | 2.829E-08 | 1.500E-08 | 7.000E-08 |
| 1477 | 74 | L74T | A | 2.2e-08 | 5.7e-08 | 4.6e-08 | 4.167E-08 | 1.790E-08 | 2.200E-08 | 5.700E-08 |
| 1478 | 74 | L74W | A | 4.1e-08 | 3.8e-08 | 3.7e-08 | 3.867E-08 | 2.082E-09 | 3.700E-08 | 4.100E-08 |
| 1479 | 74 | L74Y | A | 5.2e-08 | 9.7e-08 | 3e-08 | 5.967E-08 | 3.415E-08 | 3.000E-08 | 9.700E-08 |
| 1480 | 74 | L74V | A | 6.5e-08 | 4.6e-08 | 5e-08 | 5.367E-08 | 1.002E-08 | 4.600E-08 | 6.500E-08 |
| 1481 | 75 | Q75A | A | 4e-08 | 7.1e-08 | 6.7e-08 | 5.933E-08 | 1.686E-08 | 4.000E-08 | 7.100E-08 |
| 1482 | 75 | Q75R | A | 3e-08 | 4.3e-08 | 4.2e-08 | 3.833E-08 | 7.234E-09 | 3.000E-08 | 4.300E-08 |
| 1483 | 75 | Q75N | A | 9.6e-08 | 1.4e-07 | 2.3e-08 | 8.633E-08 | 5.910E-08 | 2.300E-08 | 1.400E-07 |
| 1484 | 75 | Q75D | A | 2e-08 | 7.3e-08 | 2.1e-08 | 3.800E-08 | 3.032E-08 | 2.000E-08 | 7.300E-08 |
| 1485 | 75 | Q75C | A | 8.7e-08 | 1.3e-07 | 1.6e-08 | 7.767E-08 | 5.757E-08 | 1.600E-08 | 1.300E-07 |
| 1486 | 75 | Q75E | A | 3.5e-08 | 2.4e-08 | 2e-08 | 2.633E-08 | 7.767E-09 | 2.000E-08 | 3.500E-08 |
| 1487 | 75 | Q75Q | A | 2.3e-08 | 4.7e-08 | 1.5e-08 | 2.833E-08 | 1.665E-08 | 1.500E-08 | 4.700E-08 |
| 1488 | 75 | Q75G | A | 4.3e-08 | 1.1e-07 | 7.5e-08 | 7.600E-08 | 3.351E-08 | 4.300E-08 | 1.100E-07 |
| 1489 | 75 | Q75H | A | 1.4e-08 | 1.2e-07 | 7.2e-08 | 6.867E-08 | 5.308E-08 | 1.400E-08 | 1.200E-07 |
| 1490 | 75 | Q75I | A | 4.7e-08 | 2.8e-08 | 6e-08 | 4.500E-08 | 1.609E-08 | 2.800E-08 | 6.000E-08 |
| 1491 | 75 | Q75L | A | 4.1e-08 | 2.2e-08 | 9.6e-08 | 5.300E-08 | 3.843E-08 | 2.200E-08 | 9.600E-08 |
| 1492 | 75 | Q75K | A | 1.3e-07 | 5.5e-08 | 5.9e-08 | 8.133E-08 | 4.219E-08 | 5.500E-08 | 1.300E-07 |
| 1493 | 75 | Q75M | A | 3.1e-08 | 1.8e-08 | 1.2e-07 | 5.633E-08 | 5.552E-08 | 1.800E-08 | 1.200E-07 |
| 1494 | 75 | Q75F | A | 3.6e-08 | 5.4e-08 | 4.2e-08 | 4.400E-08 | 9.165E-09 | 3.600E-08 | 5.400E-08 |
| 1495 | 75 | Q75P | A | 2e-08 | 4.9e-08 | 4.6e-08 | 3.833E-08 | 1.595E-08 | 2.000E-08 | 4.900E-08 |
| 1496 | 75 | Q75S | A | 1.1e-08 | 3.6e-08 | 4.4e-08 | 3.033E-08 | 1.721E-08 | 1.100E-08 | 4.400E-08 |
| 1497 | 75 | Q75T | A | 2.6e-08 | 6.5e-08 | 4.5e-08 | 4.533E-08 | 1.950E-08 | 2.600E-08 | 6.500E-08 |
| 1498 | 75 | Q75W | A | 1.2e-08 | 2.2e-08 | 5e-08 | 2.800E-08 | 1.970E-08 | 1.200E-08 | 5.000E-08 |
| 1499 | 75 | Q75Y | A | 2.9e-08 | 7.1e-08 | 5.7e-08 | 5.233E-08 | 2.139E-08 | 2.900E-08 | 7.100E-08 |
| 1500 | 75 | Q75V | A | 1e-07 | 5.3e-08 | 4.8e-08 | 6.700E-08 | 2.869E-08 | 4.800E-08 | 1.000E-07 |
| 1501 | 76 | R76A | A | 1.1e-07 | 4.5e-08 | 2.7e-08 | 6.067E-08 | 4.366E-08 | 2.700E-08 | 1.100E-07 |
| 1502 | 76 | R76R | A | 2.9e-08 | 3.2e-08 | 2.1e-08 | 2.733E-08 | 5.686E-09 | 2.100E-08 | 3.200E-08 |
| 1503 | 76 | R76N | A | 4.3e-08 | 5.9e-08 | 7.1e-08 | 5.767E-08 | 1.405E-08 | 4.300E-08 | 7.100E-08 |
| 1504 | 76 | R76D | A | 1.7e-08 | 6e-08 | 7.5e-08 | 5.067E-08 | 3.011E-08 | 1.700E-08 | 7.500E-08 |
| 1505 | 76 | R76C | A | 4.7e-08 | 1.9e-08 | 6e-08 | 4.200E-08 | 2.095E-08 | 1.900E-08 | 6.000E-08 |
| 1506 | 76 | R76E | A | 2.5e-08 | 3.6e-08 | 6.9e-08 | 4.333E-08 | 2.290E-08 | 2.500E-08 | 6.900E-08 |
| 1507 | 76 | R76Q | A | 3.4e-08 | 2.3e-08 | 5.2e-08 | 3.633E-08 | 1.464E-08 | 2.300E-08 | 5.200E-08 |
| 1508 | 76 | R76G | A | 6.2e-08 | 5.4e-08 | 3.4e-08 | 5.000E-08 | 1.442E-08 | 3.400E-08 | 6.200E-08 |
| 1509 | 76 | R76H | A | 4.3e-08 | 4.7e-08 | 1.2e-08 | 3.400E-08 | 1.916E-08 | 1.200E-08 | 4.700E-08 |
| 1510 | 76 | R76I | A | 3.7e-08 | 1.2e-08 | 3.5e-08 | 2.800E-08 | 1.389E-08 | 1.200E-08 | 3.700E-08 |
| 1511 | 76 | R76L | A | 4.5e-08 | 4.8e-08 | 2.1e-08 | 3.800E-08 | 1.480E-08 | 2.100E-08 | 4.800E-08 |
| 1512 | 76 | R76K | A | 6.1e-08 | 4.8e-08 | 2.6e-08 | 4.500E-08 | 1.769E-08 | 2.600E-08 | 6.100E-08 |
| 1513 | 76 | R76M | A | 3.4e-08 | 2.9e-08 | 9.4e-08 | 5.233E-08 | 3.617E-08 | 2.900E-08 | 9.400E-08 |
| 1514 | 76 | R76F | A | 3.6e-08 | 4.3e-08 | 6.1e-08 | 4.667E-08 | 1.290E-08 | 3.600E-08 | 6.100E-08 |
| 1515 | 76 | R76P | A | 7.3e-08 | 1e-07 | 7.5e-08 | 8.267E-08 | 1.504E-08 | 7.300E-08 | 1.000E-07 |
| 1516 | 76 | R76S | A | 2.6e-08 | 3e-08 | 6.8e-08 | 4.133E-08 | 2.318E-08 | 2.600E-08 | 6.800E-08 |
| 1517 | 76 | R76T | A | 2.4e-08 | 9.3e-08 | 1.4e-08 | 4.367E-08 | 4.302E-08 | 1.400E-08 | 9.300E-08 |
| 1518 | 76 | R76W | A | 4e-08 | 2.4e-08 | 1.8e-08 | 2.733E-08 | 1.137E-08 | 1.800E-08 | 4.000E-08 |
| 1519 | 76 | R76Y | A | 8.1e-08 | 8.7e-08 | 2.8e-08 | 6.533E-08 | 3.247E-08 | 2.800E-08 | 8.700E-08 |
| 1520 | 76 | R76V | A | 3.8e-08 | 3.4e-08 | 9.4e-08 | 5.533E-08 | 3.355E-08 | 3.400E-08 | 9.400E-08 |
| 1521 | 77 | L77A | A | 1e-07 | 3.8e-08 | 6.1e-08 | 6.633E-08 | 3.134E-08 | 3.800E-08 | 1.000E-07 |
| 1522 | 77 | L77R | A | 9.2e-08 | 5.6e-08 | 9.3e-08 | 8.033E-08 | 2.108E-08 | 5.600E-08 | 9.300E-08 |
| 1523 | 77 | L77N | A | 5.1e-08 | 2e-08 | 6.1e-08 | 4.400E-08 | 2.138E-08 | 2.000E-08 | 6.100E-08 |
| 1524 | 77 | L77D | A | 2e-08 | 1.9e-08 | 5e-08 | 2.967E-08 | 1.762E-08 | 1.900E-08 | 5.000E-08 |
| 1525 | 77 | L77C | A | 4.3e-08 | 4.6e-08 | 4.4e-08 | 4.433E-08 | 1.528E-09 | 4.300E-08 | 4.600E-08 |
| 1526 | 77 | L77E | A | 4.8e-08 | 3.4e-08 | 3.1e-08 | 3.767E-08 | 9.074E-09 | 3.100E-08 | 4.800E-08 |
| 1527 | 77 | L77Q | A | 2e-08 | 2.7e-08 | 9.2e-08 | 4.633E-08 | 3.970E-08 | 2.000E-08 | 9.200E-08 |
| 1528 | 77 | L77G | A | 3.6e-08 | 3.4e-08 | 9.9e-08 | 5.633E-08 | 3.696E-08 | 3.400E-08 | 9.900E-08 |
| 1529 | 77 | L77H | A | 2.5e-08 | 6.2e-08 | 1.8e-08 | 3.500E-08 | 2.364E-08 | 1.800E-08 | 6.200E-08 |

| | | | | | | | | | | |
|---|---|---|---|---|---|---|---|---|---|---|
| 1530 | 77 | L77I | A | 3.9e-08 | 6.1e-08 | 1.8e-07 | 9.333E-08 | 7.586E-08 | 3.900E-08 | 1.800E-07 |
| 1531 | 77 | L77L | A | 3.2e-08 | 3.7e-08 | 3e-08 | 3.300E-08 | 3.606E-09 | 3.000E-08 | 3.700E-08 |
| 1532 | 77 | L77K | A | 9.6e-09 | 2.7e-08 | 1.5e-07 | 6.220E-08 | 7.653E-08 | 9.600E-09 | 1.500E-07 |
| 1533 | 77 | L77M | A | 7.2e-08 | 1.2e-08 | 2.4e-08 | 3.600E-08 | 3.175E-08 | 1.200E-08 | 7.200E-08 |
| 1534 | 77 | L77F | A | 8.8e-08 | 3.3e-08 | 6.5e-08 | 6.200E-08 | 2.762E-08 | 3.300E-08 | 8.800E-08 |
| 1535 | 77 | L77P | A | 9.5e-08 | 4.8e-08 | 6.8e-08 | 7.033E-08 | 2.359E-08 | 4.800E-08 | 9.500E-08 |
| 1536 | 77 | L77S | A | 2.3e-08 | 4.3e-08 | 4e-08 | 3.533E-08 | 1.079E-08 | 2.300E-08 | 4.300E-08 |
| 1537 | 77 | L77T | A | 1.3e-07 | 2.4e-08 | 4.7e-08 | 6.700E-08 | 5.576E-08 | 2.400E-08 | 1.300E-07 |
| 1538 | 77 | L77W | A | 3e-08 | 1.3e-08 | 4.1e-08 | 2.800E-08 | 1.411E-08 | 1.300E-08 | 4.100E-08 |
| 1539 | 77 | L77Y | A | 1.7e-08 | 5.9e-08 | 5.3e-08 | 4.300E-08 | 2.272E-08 | 1.700E-08 | 5.900E-08 |
| 1540 | 77 | L77V | A | 6.5e-08 | 5.8e-08 | 4.6e-08 | 5.633E-08 | 9.609E-09 | 4.600E-08 | 6.500E-08 |
| 1541 | 78 | R78A | A | 5.3e-08 | 3.4e-08 | 4.4e-08 | 4.367E-08 | 9.504E-09 | 3.400E-08 | 5.300E-08 |
| 1542 | 78 | R78R | A | 4.5e-08 | 2.9e-08 | 2.7e-08 | 3.367E-08 | 9.866E-09 | 2.700E-08 | 4.500E-08 |
| 1543 | 78 | R78N | A | 6.3e-08 | 3.1e-08 | 1.1e-08 | 3.500E-08 | 2.623E-08 | 1.100E-08 | 6.300E-08 |
| 1544 | 78 | R78D | A | 8.1e-08 | 1e-08 | 4.2e-08 | 4.433E-08 | 3.556E-08 | 1.000E-08 | 8.100E-08 |
| 1545 | 78 | R78C | A | 9.6e-08 | 2e-08 | 3.4e-08 | 5.000E-08 | 4.045E-08 | 2.000E-08 | 9.600E-08 |
| 1546 | 78 | R78E | A | 6.9e-08 | 4.6e-08 | 9.1e-08 | 6.867E-08 | 2.250E-08 | 4.600E-08 | 9.100E-08 |
| 1547 | 78 | R78Q | A | 4.5e-08 | 7.6e-08 | 2.5e-08 | 4.867E-08 | 2.570E-08 | 2.500E-08 | 7.600E-08 |
| 1548 | 78 | R78G | A | 5.3e-08 | 4.9e-08 | 4.5e-08 | 4.900E-08 | 4.000E-09 | 4.500E-08 | 5.300E-08 |
| 1549 | 78 | R78H | A | 3.2e-08 | 3.1e-08 | 9.4e-08 | 5.233E-08 | 3.609E-08 | 3.100E-08 | 9.400E-08 |
| 1550 | 78 | R78I | A | 1.6e-08 | 3.5e-08 | 2.1e-08 | 2.400E-08 | 9.849E-09 | 1.600E-08 | 3.500E-08 |
| 1551 | 78 | R78L | A | 8.5e-08 | 3.2e-08 | 4e-08 | 5.233E-08 | 2.857E-08 | 3.200E-08 | 8.500E-08 |
| 1552 | 78 | R78K | A | 5.6e-08 | 2.7e-08 | 2.1e-08 | 3.467E-08 | 1.872E-08 | 2.100E-08 | 5.600E-08 |
| 1553 | 78 | R78M | A | 3e-08 | 3e-08 | 4.2e-08 | 3.400E-08 | 6.928E-09 | 3.000E-08 | 4.200E-08 |
| 1554 | 78 | R78F | A | 2.1e-08 | 2.9e-08 | 1e-07 | 5.000E-08 | 4.349E-08 | 2.100E-08 | 1.000E-07 |
| 1555 | 78 | R78P | A | 3e-08 | 1.8e-07 | 2.7e-08 | 7.900E-08 | 8.748E-08 | 2.700E-08 | 1.800E-07 |
| 1556 | 78 | R78S | A | 4e-08 | 4.5e-08 | 9.8e-08 | 6.100E-08 | 3.214E-08 | 4.000E-08 | 9.800E-08 |
| 1557 | 78 | R78T | A | 3.8e-08 | 7.2e-08 | 2.5e-08 | 4.500E-08 | 2.427E-08 | 2.500E-08 | 7.200E-08 |
| 1558 | 78 | R78W | A | 2e-07 | 2.2e-08 | 1.5e-08 | 7.900E-08 | 1.048E-07 | 1.500E-08 | 2.000E-07 |
| 1559 | 78 | R78Y | A | 2.6e-08 | 4.5e-08 | 5.2e-08 | 4.100E-08 | 1.345E-08 | 2.600E-08 | 5.200E-08 |
| 1560 | 78 | R78V | A | 2.2e-08 | 3.1e-08 | 4.5e-08 | 3.267E-08 | 1.159E-08 | 2.200E-08 | 4.500E-08 |
| 1561 | 79 | I79A | A | 6.3e-08 | 4.3e-08 | 2.9e-08 | 4.500E-08 | 1.709E-08 | 2.900E-08 | 6.300E-08 |
| 1562 | 79 | I79R | A | 5.1e-08 | 3.2e-08 | 2e-07 | 9.433E-08 | 9.200E-08 | 3.200E-08 | 2.000E-07 |
| 1563 | 79 | I79N | A | 1.8e-07 | 5.4e-08 | 4e-08 | 9.133E-08 | 7.711E-08 | 4.000E-08 | 1.800E-07 |
| 1564 | 79 | I79D | A | 7.7e-08 | 2.8e-08 | 4.3e-08 | 4.933E-08 | 2.511E-08 | 2.800E-08 | 7.700E-08 |
| 1565 | 79 | I79C | A | 3.1e-08 | 1e-07 | 3.7e-08 | 5.600E-08 | 3.822E-08 | 3.100E-08 | 1.000E-07 |
| 1566 | 79 | I79E | A | 1.2e-07 | 3.7e-08 | 5.2e-08 | 6.967E-08 | 4.423E-08 | 3.700E-08 | 1.200E-07 |
| 1567 | 79 | I79Q | A | 6.9e-08 | 4e-08 | 4.4e-08 | 5.100E-08 | 1.572E-08 | 4.000E-08 | 6.900E-08 |
| 1568 | 79 | I79G | A | 2.2e-08 | 3.3e-08 | 7.2e-08 | 4.233E-08 | 2.627E-08 | 2.200E-08 | 7.200E-08 |
| 1569 | 79 | I79H | A | 1.2e-07 | 4.1e-08 | 4.8e-08 | 6.967E-08 | 4.373E-08 | 4.100E-08 | 1.200E-07 |
| 1570 | 79 | I79I | A | 4.2e-08 | 2.3e-08 | 8.4e-08 | 4.967E-08 | 3.121E-08 | 2.300E-08 | 8.400E-08 |
| 1571 | 79 | I79L | A | 4e-08 | 5.6e-08 | 4.9e-08 | 4.833E-08 | 8.021E-09 | 4.000E-08 | 5.600E-08 |
| 1572 | 79 | I79K | A | 4.1e-08 | 3.9e-08 | 3.2e-08 | 3.733E-08 | 4.726E-09 | 3.200E-08 | 4.100E-08 |
| 1573 | 79 | I79M | A | 6.6e-08 | 3.6e-08 | 5.3e-08 | 5.167E-08 | 1.504E-08 | 3.600E-08 | 6.600E-08 |
| 1574 | 79 | I79F | A | 8.4e-08 | 2.7e-08 | 3.1e-08 | 4.733E-08 | 3.182E-08 | 2.700E-08 | 8.400E-08 |
| 1575 | 79 | I79P | A | 7.5e-08 | 2.4e-08 | 1.9e-07 | 9.633E-08 | 8.503E-08 | 2.400E-08 | 1.900E-07 |
| 1576 | 79 | I79S | A | 4.5e-08 | 6.8e-08 | 7.5e-08 | 6.267E-08 | 1.570E-08 | 4.500E-08 | 7.500E-08 |
| 1577 | 79 | I79T | A | 3.4e-08 | 2.1e-07 | 9.8e-08 | 1.140E-07 | 8.908E-08 | 3.400E-08 | 2.100E-07 |
| 1578 | 79 | I79W | A | 1.9e-07 | 3.9e-08 | 3.2e-08 | 8.700E-08 | 8.927E-08 | 3.200E-08 | 1.900E-07 |
| 1579 | 79 | I79Y | A | 3.5e-08 | 4.2e-08 | 2.5e-08 | 3.400E-08 | 8.544E-09 | 2.500E-08 | 4.200E-08 |
| 1580 | 79 | I79V | A | 6.4e-08 | 6.5e-08 | 5.5e-08 | 6.133E-08 | 5.508E-09 | 5.500E-08 | 6.500E-08 |
| 1581 | 80 | V80A | A | 4.2e-08 | 2.2e-08 | 2.7e-08 | 3.033E-08 | 1.041E-08 | 2.200E-08 | 4.200E-08 |
| 1582 | 80 | V80R | A | 6.7e-08 | 4.1e-08 | 4.6e-08 | 5.133E-08 | 1.380E-08 | 4.100E-08 | 6.700E-08 |
| 1583 | 80 | V80N | A | 2.9e-08 | 4.8e-08 | 4.3e-08 | 4.000E-08 | 9.849E-09 | 2.900E-08 | 4.800E-08 |
| 1584 | 80 | V80D | A | 7.7e-08 | 7.9e-08 | 3.9e-08 | 6.500E-08 | 2.254E-08 | 3.900E-08 | 7.900E-08 |
| 1585 | 80 | V80C | A | 2.1e-08 | 2.4e-08 | 1.7e-08 | 2.067E-08 | 3.512E-09 | 1.700E-08 | 2.400E-08 |
| 1586 | 80 | V80E | A | 3.3e-08 | 7.9e-08 | 4.8e-08 | 5.333E-08 | 2.346E-08 | 3.300E-08 | 7.900E-08 |
| 1587 | 80 | V80Q | A | 4.3e-08 | 6.5e-08 | 4.4e-08 | 5.067E-08 | 1.242E-08 | 4.300E-08 | 6.500E-08 |
| 1588 | 80 | V80G | A | 2.9e-08 | 6e-09 | 4.1e-08 | 2.533E-08 | 1.779E-08 | 6.000E-09 | 4.100E-08 |
| 1589 | 80 | V80H | A | 4e-08 | 1.8e-08 | 3.5e-08 | 3.100E-08 | 1.153E-08 | 1.800E-08 | 4.000E-08 |
| 1590 | 80 | V80I | A | 3.9e-08 | 7.5e-08 | 2.1e-08 | 4.500E-08 | 2.750E-08 | 2.100E-08 | 7.500E-08 |
| 1591 | 80 | V80L | A | 2.3e-08 | 2.2e-08 | 3.8e-08 | 2.767E-08 | 8.963E-09 | 2.200E-08 | 3.800E-08 |
| 1592 | 80 | V80K | A | 4.3e-08 | 6e-08 | 1.2e-08 | 3.833E-08 | 2.434E-08 | 1.200E-08 | 6.000E-08 |
| 1593 | 80 | V80M | A | 6.2e-08 | 6.8e-08 | 4.6e-08 | 5.867E-08 | 1.137E-08 | 4.600E-08 | 6.800E-08 |
| 1594 | 80 | V80F | A | 5.9e-08 | 4e-08 | 6.2e-08 | 5.367E-08 | 1.193E-08 | 4.000E-08 | 6.200E-08 |
| 1595 | 80 | V80P | A | 1.3e-08 | 4.4e-08 | 5.9e-08 | 3.867E-08 | 2.346E-08 | 1.300E-08 | 5.900E-08 |
| 1596 | 80 | V80S | A | 3.6e-08 | 4.1e-08 | 3e-08 | 3.567E-08 | 5.508E-09 | 3.000E-08 | 4.100E-08 |
| 1597 | 80 | V80T | A | 6.4e-08 | 7.4e-08 | 1.9e-08 | 5.233E-08 | 2.930E-08 | 1.900E-08 | 7.400E-08 |
| 1598 | 80 | V80W | A | 2.1e-08 | 4.9e-08 | 8.7e-08 | 5.233E-08 | 3.313E-08 | 2.100E-08 | 8.700E-08 |
| 1599 | 80 | V80Y | A | 5.3e-08 | 5e-08 | 7.7e-09 | 3.690E-08 | 2.533E-08 | 7.700E-09 | 5.300E-08 |
| 1600 | 80 | V80V | A | 5.2e-08 | 5.5e-08 | 2.5e-08 | 4.400E-08 | 1.652E-08 | 2.500E-08 | 5.500E-08 |
| 1601 | 81 | R81A | A | 5.4e-08 | 4.9e-08 | 3.2e-08 | 4.500E-08 | 1.153E-08 | 3.200E-08 | 5.400E-08 |
| 1602 | 81 | R81R | A | 1.6e-08 | 4.6e-08 | 2.3e-08 | 2.833E-08 | 1.570E-08 | 1.600E-08 | 4.600E-08 |
| 1603 | 81 | R81N | A | 5.9e-08 | 2.1e-08 | 3.4e-08 | 3.800E-08 | 1.931E-08 | 2.100E-08 | 5.900E-08 |
| 1604 | 81 | R81D | A | 8.6e-09 | 8.9e-08 | 6.7e-08 | 5.487E-08 | 4.155E-08 | 8.600E-09 | 8.900E-08 |
| 1605 | 81 | R81C | A | 2.2e-08 | 3.2e-08 | 2.9e-08 | 2.767E-08 | 5.132E-09 | 2.200E-08 | 3.200E-08 |
| 1606 | 81 | R81E | A | 1.2e-08 | 1.1e-07 | 7.4e-08 | 6.533E-08 | 4.957E-08 | 1.200E-08 | 1.100E-07 |
| 1607 | 81 | R81Q | A | 5.4e-08 | 6e-08 | 3.6e-08 | 5.000E-08 | 1.249E-08 | 3.600E-08 | 6.000E-08 |
| 1608 | 81 | R81G | A | 2.4e-08 | 5.5e-08 | 4.9e-08 | 4.267E-08 | 1.644E-08 | 2.400E-08 | 5.500E-08 |
| 1609 | 81 | R81H | A | 9.4e-08 | 5.1e-08 | 3.3e-08 | 5.933E-08 | 3.134E-08 | 3.300E-08 | 9.400E-08 |
| 1610 | 81 | R81I | A | 1.7e-08 | 5e-08 | 9e-08 | 5.233E-08 | 3.656E-08 | 1.700E-08 | 9.000E-08 |
| 1611 | 81 | R81L | A | 1.2e-08 | 3.2e-08 | 1.8e-08 | 2.067E-08 | 1.026E-08 | 1.200E-08 | 3.200E-08 |
| 1612 | 81 | R81K | A | 6e-08 | 1.6e-08 | 8.1e-08 | 5.233E-08 | 3.317E-08 | 1.600E-08 | 8.100E-08 |
| 1613 | 81 | R81M | A | 3.3e-08 | 1.4e-08 | 4.3e-08 | 3.000E-08 | 1.473E-08 | 1.400E-08 | 4.300E-08 |
| 1614 | 81 | R81F | A | 4.4e-08 | 5.3e-08 | 1.4e-08 | 3.700E-08 | 2.042E-08 | 1.400E-08 | 5.300E-08 |

| | | | | | | | | | | |
|---|---|---|---|---|---|---|---|---|---|---|
| 1615 | 81 | R81P | A | 2.6e-08 | 5.2e-08 | 3.8e-08 | 3.867E-08 | 1.301E-08 | 2.600E-08 | 5.200E-08 |
| 1616 | 81 | R81S | A | 5.4e-08 | 1.9e-08 | 5e-08 | 4.100E-08 | 1.916E-08 | 1.900E-08 | 5.400E-08 |
| 1617 | 81 | R81T | A | 5.1e-08 | 1.8e-08 | 7.2e-08 | 4.700E-08 | 2.722E-08 | 1.800E-08 | 7.200E-08 |
| 1618 | 81 | R81W | A | 8.6e-08 | 5e-08 | 3.5e-08 | 5.700E-08 | 2.621E-08 | 3.500E-08 | 8.600E-08 |
| 1619 | 81 | R81Y | A | 3.5e-08 | 1.9e-08 | 2.6e-08 | 2.667E-08 | 8.021E-09 | 1.900E-08 | 3.500E-08 |
| 1620 | 81 | R81V | A | 4.9e-08 | 4.5e-08 | 1.4e-07 | 7.800E-08 | 5.373E-08 | 4.500E-08 | 1.400E-07 |
| 1621 | 82 | G82A | A | 1.6e-08 | 3.3e-08 | 6.9e-08 | 3.933E-08 | 2.706E-08 | 1.600E-08 | 6.900E-08 |
| 1622 | 82 | G82R | A | 6.9e-08 | 1.2e-07 | 3.1e-08 | 7.333E-08 | 4.466E-08 | 3.100E-08 | 1.200E-07 |
| 1623 | 82 | G82N | A | 4.2e-08 | 3.4e-08 | 6.2e-08 | 4.600E-08 | 1.442E-08 | 3.400E-08 | 6.200E-08 |
| 1624 | 82 | G82D | A | 4.9e-08 | 9.2e-08 | 3.8e-08 | 5.967E-08 | 2.854E-08 | 3.800E-08 | 9.200E-08 |
| 1625 | 82 | G82C | A | 4.4e-08 | 4.2e-08 | 5.2e-08 | 4.600E-08 | 5.292E-09 | 4.200E-08 | 5.200E-08 |
| 1626 | 82 | G82E | A | 7.7e-08 | 6.6e-08 | 5.3e-08 | 6.533E-08 | 1.201E-08 | 5.300E-08 | 7.700E-08 |
| 1627 | 82 | G82Q | A | 3.7e-08 | 1.1e-07 | 6.1e-08 | 6.933E-08 | 3.721E-08 | 3.700E-08 | 1.100E-07 |
| 1628 | 82 | G82G | A | 1.8e-07 | 5.2e-08 | 8.3e-08 | 1.050E-07 | 6.678E-08 | 5.200E-08 | 1.800E-07 |
| 1629 | 82 | G82H | A | 2e-08 | 4.9e-08 | 4.1e-08 | 3.667E-08 | 1.498E-08 | 2.000E-08 | 4.900E-08 |
| 1630 | 82 | G82I | A | 3.9e-08 | 2.1e-08 | 9.7e-08 | 5.233E-08 | 3.972E-08 | 2.100E-08 | 9.700E-08 |
| 1631 | 82 | G82L | A | 4.4e-08 | 6.2e-08 | 9.1e-08 | 6.567E-08 | 2.371E-08 | 4.400E-08 | 9.100E-08 |
| 1632 | 82 | G82K | A | 6e-09 | 4.1e-08 | 2.6e-08 | 2.433E-08 | 1.756E-08 | 6.000E-09 | 4.100E-08 |
| 1633 | 82 | G82M | A | 6.5e-09 | 2.9e-08 | 3.8e-08 | 2.450E-08 | 1.622E-08 | 6.500E-09 | 3.800E-08 |
| 1634 | 82 | G82F | A | 6.6e-08 | 5.6e-08 | 6.2e-08 | 6.133E-08 | 5.033E-09 | 5.600E-08 | 6.600E-08 |
| 1635 | 82 | G82P | A | 5.9e-08 | 3.5e-07 | 2.3e-08 | 1.440E-07 | 1.793E-07 | 2.300E-08 | 3.500E-07 |
| 1636 | 82 | G82S | A | 9.2e-08 | 6.3e-08 | 3.7e-08 | 6.400E-08 | 2.751E-08 | 3.700E-08 | 9.200E-08 |
| 1637 | 82 | G82T | A | 1.4e-08 | 5.4e-08 | 1.7e-08 | 2.833E-08 | 2.228E-08 | 1.400E-08 | 5.400E-08 |
| 1638 | 82 | G82W | A | 6.6e-08 | 2.2e-08 | 5.3e-08 | 4.700E-08 | 2.261E-08 | 2.200E-08 | 6.600E-08 |
| 1639 | 82 | G82Y | A | 4.7e-08 | 5.3e-08 | 5.9e-08 | 5.300E-08 | 6.000E-09 | 4.700E-08 | 5.900E-08 |
| 1640 | 82 | G82V | A | 5.4e-08 | 6.5e-08 | 3.9e-08 | 5.267E-08 | 1.305E-08 | 3.900E-08 | 6.500E-08 |
| 1641 | 83 | T83A | A | 3.3e-08 | 7.5e-08 | 4.2e-08 | 5.000E-08 | 2.211E-08 | 3.300E-08 | 7.500E-08 |
| 1642 | 83 | T83R | A | 3.1e-08 | 3.9e-08 | 4.1e-08 | 3.700E-08 | 5.292E-09 | 3.100E-08 | 4.100E-08 |
| 1643 | 83 | T83N | A | 7.7e-08 | 5.6e-08 | 1.1e-07 | 8.100E-08 | 2.722E-08 | 5.600E-08 | 1.100E-07 |
| 1644 | 83 | T83D | A | 4.6e-08 | 6.5e-08 | 5.2e-08 | 5.433E-08 | 9.713E-09 | 4.600E-08 | 6.500E-08 |
| 1645 | 83 | T83C | A | 3.4e-08 | 7.2e-08 | 6.4e-08 | 5.667E-08 | 2.003E-08 | 3.400E-08 | 7.200E-08 |
| 1646 | 83 | T83E | A | 6.6e-08 | 6.8e-08 | 2.2e-08 | 5.200E-08 | 2.600E-08 | 2.200E-08 | 6.800E-08 |
| 1647 | 83 | T83Q | A | 5.5e-08 | 1.7e-08 | 4.8e-08 | 4.000E-08 | 2.022E-08 | 1.700E-08 | 5.500E-08 |
| 1648 | 83 | T83G | A | 2.6e-08 | 4.2e-08 | 7.3e-08 | 4.700E-08 | 2.390E-08 | 2.600E-08 | 7.300E-08 |
| 1649 | 83 | T83H | A | 5.7e-08 | 1.3e-07 | 4.6e-08 | 7.767E-08 | 4.565E-08 | 4.600E-08 | 1.300E-07 |
| 1650 | 83 | T83I | A | 7.8e-08 | 2.8e-08 | 3.3e-08 | 4.633E-08 | 2.754E-08 | 2.800E-08 | 7.800E-08 |
| 1651 | 83 | T83L | A | 1.1e-07 | 3.4e-08 | 2.5e-08 | 5.633E-08 | 4.669E-08 | 2.500E-08 | 1.100E-07 |
| 1652 | 83 | T83K | A | 2.6e-08 | 4.7e-08 | 2.9e-08 | 3.400E-08 | 1.136E-08 | 2.600E-08 | 4.700E-08 |
| 1653 | 83 | T83M | A | 7.5e-08 | 2.4e-08 | 4.4e-08 | 4.767E-08 | 2.570E-08 | 2.400E-08 | 7.500E-08 |
| 1654 | 83 | T83F | A | 2.2e-08 | 6.4e-08 | 3e-08 | 3.867E-08 | 2.230E-08 | 2.200E-08 | 6.400E-08 |
| 1655 | 83 | T83P | A | 1.2e-07 | 8.1e-08 | 5.1e-08 | 8.400E-08 | 3.460E-08 | 5.100E-08 | 1.200E-07 |
| 1656 | 83 | T83S | A | 1.2e-07 | 3.1e-08 | 5.1e-08 | 6.733E-08 | 4.669E-08 | 3.100E-08 | 1.200E-07 |
| 1657 | 83 | T83T | A | 1e-07 | 5.1e-08 | 6.7e-08 | 7.267E-08 | 2.499E-08 | 5.100E-08 | 1.000E-07 |
| 1658 | 83 | T83W | A | 3e-08 | 3.2e-08 | 2.8e-08 | 3.000E-08 | 2.000E-09 | 2.800E-08 | 3.200E-08 |
| 1659 | 83 | T83Y | A | 4.2e-08 | 5e-08 | 2.2e-08 | 3.800E-08 | 1.442E-08 | 2.200E-08 | 5.000E-08 |
| 1660 | 83 | T83V | A | 8.1e-08 | 6.8e-08 | 4.8e-08 | 6.567E-08 | 1.662E-08 | 4.800E-08 | 8.100E-08 |
| 1661 | 84 | Q84A | A | 1.8e-08 | 4.1e-08 | 3e-08 | 2.967E-08 | 1.150E-08 | 1.800E-08 | 4.100E-08 |
| 1662 | 84 | Q84R | A | 3.5e-08 | 4.9e-08 | 5.5e-08 | 4.633E-08 | 1.026E-08 | 3.500E-08 | 5.500E-08 |
| 1663 | 84 | Q84N | A | 2.7e-08 | 4.3e-08 | 9.4e-09 | 2.647E-08 | 1.681E-08 | 9.400E-09 | 4.300E-08 |
| 1664 | 84 | Q84D | A | 4.3e-08 | 3.4e-08 | 5.3e-08 | 4.333E-08 | 9.504E-09 | 3.400E-08 | 5.300E-08 |
| 1665 | 84 | Q84C | A | 8.5e-08 | 5.6e-08 | 1.2e-08 | 5.100E-08 | 3.676E-08 | 1.200E-08 | 8.500E-08 |
| 1666 | 84 | Q84E | A | 9e-08 | 2.8e-08 | 6.2e-08 | 6.000E-08 | 3.105E-08 | 2.800E-08 | 9.000E-08 |
| 1667 | 84 | Q84Q | A | 4.3e-08 | 3.5e-08 | 6.8e-08 | 4.867E-08 | 1.721E-08 | 3.500E-08 | 6.800E-08 |
| 1668 | 84 | Q84G | A | 3.7e-08 | 7e-08 | 3.1e-08 | 4.600E-08 | 2.100E-08 | 3.100E-08 | 7.000E-08 |
| 1669 | 84 | Q84H | A | 4.7e-08 | 4.4e-08 | 1.5e-07 | 8.033E-08 | 6.035E-08 | 4.400E-08 | 1.500E-07 |
| 1670 | 84 | Q84I | A | 4e-08 | 6.4e-08 | 5.2e-08 | 5.200E-08 | 1.200E-08 | 4.000E-08 | 6.400E-08 |
| 1671 | 84 | Q84L | A | 1.1e-07 | 4.5e-08 | 2.2e-08 | 5.900E-08 | 4.564E-08 | 2.200E-08 | 1.100E-07 |
| 1672 | 84 | Q84K | A | 5.9e-08 | 5.9e-08 | 8.1e-08 | 6.633E-08 | 1.270E-08 | 5.900E-08 | 8.100E-08 |
| 1673 | 84 | Q84M | A | 4.1e-08 | 3.5e-08 | 3.4e-08 | 3.667E-08 | 3.786E-09 | 3.400E-08 | 4.100E-08 |
| 1674 | 84 | Q84F | A | 9.5e-08 | 2.4e-08 | 3.7e-08 | 5.200E-08 | 3.780E-08 | 2.400E-08 | 9.500E-08 |
| 1675 | 84 | Q84P | A | 2.5e-08 | 1.5e-08 | 2.1e-08 | 2.033E-08 | 5.033E-09 | 1.500E-08 | 2.500E-08 |
| 1676 | 84 | Q84S | A | 3.8e-08 | 4.8e-08 | 2.7e-08 | 3.767E-08 | 1.050E-08 | 2.700E-08 | 4.800E-08 |
| 1677 | 84 | Q84T | A | 9e-08 | 1.1e-08 | 4.5e-08 | 4.867E-08 | 3.963E-08 | 1.100E-08 | 9.000E-08 |
| 1678 | 84 | Q84W | A | 5.2e-08 | 7.2e-08 | 4.8e-08 | 5.733E-08 | 1.286E-08 | 4.800E-08 | 7.200E-08 |
| 1679 | 84 | Q84Y | A | 7e-08 | 1e-07 | 1.8e-08 | 6.267E-08 | 4.149E-08 | 1.800E-08 | 1.000E-07 |
| 1680 | 84 | Q84V | A | 1.2e-07 | 5.3e-08 | 3.4e-08 | 6.900E-08 | 4.518E-08 | 3.400E-08 | 1.200E-07 |
| 1681 | 85 | L85A | A | 2.4e-08 | 1.9e-08 | 2.1e-08 | 2.133E-08 | 2.517E-09 | 1.900E-08 | 2.400E-08 |
| 1682 | 85 | L85R | A | 8.8e-08 | 6.1e-08 | 2.5e-08 | 5.800E-08 | 3.161E-08 | 2.500E-08 | 8.800E-08 |
| 1683 | 85 | L85N | A | 2.1e-08 | 3.4e-08 | 5.4e-08 | 3.633E-08 | 1.662E-08 | 2.100E-08 | 5.400E-08 |
| 1684 | 85 | L85D | A | 2.5e-08 | 7.2e-08 | 4.6e-08 | 4.767E-08 | 2.354E-08 | 2.500E-08 | 7.200E-08 |
| 1685 | 85 | L85C | A | 3.4e-08 | 9.2e-08 | 1e-07 | 7.533E-08 | 3.602E-08 | 3.400E-08 | 1.000E-07 |
| 1686 | 85 | L85E | A | 1.7e-07 | 3.6e-08 | 5.2e-08 | 8.600E-08 | 7.318E-08 | 3.600E-08 | 1.700E-07 |
| 1687 | 85 | L85Q | A | 4.2e-08 | 6.4e-08 | 1.5e-08 | 4.033E-08 | 2.454E-08 | 1.500E-08 | 6.400E-08 |
| 1688 | 85 | L85G | A | 3e-08 | 9.2e-08 | 1.1e-07 | 7.733E-08 | 4.197E-08 | 3.000E-08 | 1.100E-07 |
| 1689 | 85 | L85H | A | 5.1e-08 | 5.6e-08 | 5.8e-08 | 5.500E-08 | 3.606E-09 | 5.100E-08 | 5.800E-08 |
| 1690 | 85 | L85I | A | 3.6e-08 | 5.5e-08 | 4.8e-08 | 4.633E-08 | 9.609E-09 | 3.600E-08 | 5.500E-08 |
| 1691 | 85 | L85L | A | 6.3e-08 | 5.4e-08 | 2e-08 | 4.567E-08 | 2.268E-08 | 2.000E-08 | 6.300E-08 |
| 1692 | 85 | L85K | A | 3.9e-08 | 8.5e-08 | 2.3e-08 | 4.900E-08 | 3.219E-08 | 2.300E-08 | 8.500E-08 |
| 1693 | 85 | L85M | A | 3e-08 | 6e-08 | 2.6e-08 | 3.867E-08 | 1.858E-08 | 2.600E-08 | 6.000E-08 |
| 1694 | 85 | L85F | A | 7.4e-08 | 5.9e-08 | 5.6e-08 | 6.300E-08 | 9.644E-09 | 5.600E-08 | 7.400E-08 |
| 1695 | 85 | L85P | A | 9.4e-08 | 4.6e-08 | 4.5e-08 | 6.167E-08 | 2.801E-08 | 4.500E-08 | 9.400E-08 |
| 1696 | 85 | L85S | A | 1.7e-08 | 4.9e-08 | 2.8e-08 | 3.133E-08 | 1.626E-08 | 1.700E-08 | 4.900E-08 |
| 1697 | 85 | L85T | A | 6.5e-08 | 3.6e-08 | 2.6e-08 | 4.233E-08 | 2.026E-08 | 2.600E-08 | 6.500E-08 |
| 1698 | 85 | L85W | A | 3.6e-08 | 7.8e-08 | 4.3e-08 | 5.233E-08 | 2.250E-08 | 3.600E-08 | 7.800E-08 |
| 1699 | 85 | L85Y | A | 4.5e-08 | 3.2e-08 | 2.4e-08 | 3.367E-08 | 1.060E-08 | 2.400E-08 | 4.500E-08 |

| | | | | | | | | | | |
|---|---|---|---|---|---|---|---|---|---|---|
| 1700 | 85 | L85V | A | 6.5e-08 | 4.9e-08 | 3.8e-08 | 5.067E-08 | 1.358E-08 | 3.800E-08 | 6.500E-08 |
| 1701 | 86 | F86A | A | 2.5e-08 | 3e-08 | 6.9e-08 | 4.133E-08 | 2.409E-08 | 2.500E-08 | 6.900E-08 |
| 1702 | 86 | F86R | A | 1.2e-07 | 8.1e-08 | 1.3e-07 | 1.103E-07 | 2.589E-08 | 8.100E-08 | 1.300E-07 |
| 1703 | 86 | F86N | A | 5.6e-08 | 3.4e-08 | 7.5e-08 | 5.500E-08 | 2.052E-08 | 3.400E-08 | 7.500E-08 |
| 1704 | 86 | F86D | A | 3.8e-08 | 2e-08 | 3.5e-08 | 3.100E-08 | 9.644E-09 | 2.000E-08 | 3.800E-08 |
| 1705 | 86 | F86C | A | 3e-08 | 2.9e-08 | 4.7e-08 | 3.533E-08 | 1.012E-08 | 2.900E-08 | 4.700E-08 |
| 1706 | 86 | F86E | A | 4.4e-08 | 4.9e-08 | 7.5e-08 | 5.600E-08 | 1.664E-08 | 4.400E-08 | 7.500E-08 |
| 1707 | 86 | F86Q | A | 2e-08 | 1.8e-08 | 5.8e-08 | 3.200E-08 | 2.254E-08 | 1.800E-08 | 5.800E-08 |
| 1708 | 86 | F86G | A | 1e-07 | 1.3e-07 | 9.1e-08 | 1.070E-07 | 2.042E-08 | 9.100E-08 | 1.300E-07 |
| 1709 | 86 | F86H | A | 8.5e-08 | 2.9e-08 | 1.5e-08 | 4.300E-08 | 3.704E-08 | 1.500E-08 | 8.500E-08 |
| 1710 | 86 | F86I | A | 7.3e-08 | 5.6e-08 | 2.3e-08 | 5.067E-08 | 2.542E-08 | 2.300E-08 | 7.300E-08 |
| 1711 | 86 | F86L | A | 4.4e-08 | 3.4e-08 | 5.1e-08 | 4.300E-08 | 8.544E-09 | 3.400E-08 | 5.100E-08 |
| 1712 | 86 | F86K | A | 4e-08 | 6.8e-08 | 2.4e-08 | 4.400E-08 | 2.227E-08 | 2.400E-08 | 6.800E-08 |
| 1713 | 86 | F86M | A | 2.6e-08 | 9.6e-08 | 1.1e-08 | 4.433E-08 | 4.537E-08 | 1.100E-08 | 9.600E-08 |
| 1714 | 86 | F86F | A | 3.2e-08 | 5.7e-08 | 3.5e-08 | 4.133E-08 | 1.365E-08 | 3.200E-08 | 5.700E-08 |
| 1715 | 86 | F86P | A | 5.7e-08 | 2.6e-08 | 6.8e-08 | 5.033E-08 | 2.178E-08 | 2.600E-08 | 6.800E-08 |
| 1716 | 86 | F86S | A | 2.4e-08 | 6e-08 | 3e-08 | 3.800E-08 | 1.929E-08 | 2.400E-08 | 6.000E-08 |
| 1717 | 86 | F86T | A | 1.2e-07 | 2.1e-08 | 2.3e-08 | 5.467E-08 | 5.659E-08 | 2.100E-08 | 1.200E-07 |
| 1718 | 86 | F86W | A | 2.3e-08 | 4.8e-08 | 2.2e-08 | 3.100E-08 | 1.473E-08 | 2.200E-08 | 4.800E-08 |
| 1719 | 86 | F86Y | A | 1.6e-08 | 5.1e-08 | 2.8e-08 | 3.167E-08 | 1.779E-08 | 1.600E-08 | 5.100E-08 |
| 1720 | 86 | F86V | A | 3.7e-08 | 8.2e-08 | 4.7e-08 | 5.533E-08 | 2.363E-08 | 3.700E-08 | 8.200E-08 |
| 1721 | 87 | E87A | A | 4.3e-08 | 5.1e-08 | 8.2e-08 | 5.867E-08 | 2.060E-08 | 4.300E-08 | 8.200E-08 |
| 1722 | 87 | E87R | A | 9e-08 | 3.8e-08 | 4.7e-08 | 5.833E-08 | 2.779E-08 | 3.800E-08 | 9.000E-08 |
| 1723 | 87 | E87N | A | 6.2e-08 | 4.1e-08 | 1.6e-07 | 8.767E-08 | 6.352E-08 | 4.100E-08 | 1.600E-07 |
| 1724 | 87 | E87D | A | 5.5e-08 | 4.7e-08 | 5.6e-08 | 5.267E-08 | 4.933E-09 | 4.700E-08 | 5.600E-08 |
| 1725 | 87 | E87C | A | 6.9e-08 | 1.2e-07 | 4.4e-08 | 7.767E-08 | 3.873E-08 | 4.400E-08 | 1.200E-07 |
| 1726 | 87 | E87E | A | 4.9e-08 | 5.4e-08 | 7e-08 | 5.767E-08 | 1.097E-08 | 4.900E-08 | 7.000E-08 |
| 1727 | 87 | E87Q | A | 2.9e-08 | 1.6e-08 | 7.2e-08 | 3.900E-08 | 2.931E-08 | 1.600E-08 | 7.200E-08 |
| 1728 | 87 | E87G | A | 5.2e-08 | 3.1e-08 | 4.1e-08 | 4.133E-08 | 1.050E-08 | 3.100E-08 | 5.200E-08 |
| 1729 | 87 | E87H | A | 1.9e-08 | 2.8e-08 | 5.9e-08 | 3.533E-08 | 2.098E-08 | 1.900E-08 | 5.900E-08 |
| 1730 | 87 | E87I | A | 7.1e-08 | 6.7e-08 | 4.7e-08 | 6.167E-08 | 1.286E-08 | 4.700E-08 | 7.100E-08 |
| 1731 | 87 | E87L | A | 2.9e-08 | 6.2e-08 | 8.5e-08 | 5.867E-08 | 2.815E-08 | 2.900E-08 | 8.500E-08 |
| 1732 | 87 | E87K | A | 9e-08 | 5.6e-08 | 7.7e-08 | 7.433E-08 | 1.716E-08 | 5.600E-08 | 9.000E-08 |
| 1733 | 87 | E87M | A | 3.9e-08 | 2.2e-08 | 2.8e-08 | 2.967E-08 | 8.622E-09 | 2.200E-08 | 3.900E-08 |
| 1734 | 87 | E87F | A | 5.5e-08 | 7.6e-08 | 8.6e-08 | 7.233E-08 | 1.582E-08 | 5.500E-08 | 8.600E-08 |
| 1735 | 87 | E87P | A | 3e-08 | 3.8e-08 | 4.7e-08 | 3.833E-08 | 8.505E-09 | 3.000E-08 | 4.700E-08 |
| 1736 | 87 | E87S | A | 1.7e-08 | 3.6e-08 | 2.8e-08 | 2.700E-08 | 9.539E-09 | 1.700E-08 | 3.600E-08 |
| 1737 | 87 | E87T | A | 5.7e-08 | 1.8e-08 | 4.2e-08 | 3.900E-08 | 1.967E-08 | 1.800E-08 | 5.700E-08 |
| 1738 | 87 | E87W | A | 3.1e-08 | 1.8e-08 | 5e-08 | 3.300E-08 | 1.609E-08 | 1.800E-08 | 5.000E-08 |
| 1739 | 87 | E87Y | A | 6e-08 | 5.9e-08 | 5.3e-08 | 5.733E-08 | 3.786E-09 | 5.300E-08 | 6.000E-08 |
| 1740 | 87 | E87V | A | 3e-08 | 5.3e-08 | 7.1e-08 | 5.133E-08 | 2.055E-08 | 3.000E-08 | 7.100E-08 |
| 1741 | 88 | D88A | A | 6.1e-08 | 8.3e-08 | 3e-08 | 5.800E-08 | 2.663E-08 | 3.000E-08 | 8.300E-08 |
| 1742 | 88 | D88R | A | 8.2e-08 | 3.3e-08 | 2.1e-08 | 4.533E-08 | 3.232E-08 | 2.100E-08 | 8.200E-08 |
| 1743 | 88 | D88N | A | 4.6e-08 | 1.4e-08 | 3.6e-08 | 3.200E-08 | 1.637E-08 | 1.400E-08 | 4.600E-08 |
| 1744 | 88 | D88D | A | 3.7e-08 | 6.4e-08 | 1.4e-08 | 3.833E-08 | 2.503E-08 | 1.400E-08 | 6.400E-08 |
| 1745 | 88 | D88C | A | 3.3e-08 | 3.3e-08 | 1.4e-07 | 6.867E-08 | 6.178E-08 | 3.300E-08 | 1.400E-07 |
| 1746 | 88 | D88E | A | 3.4e-08 | 5.6e-08 | 3.2e-08 | 4.067E-08 | 1.332E-08 | 3.200E-08 | 5.600E-08 |
| 1747 | 88 | D88Q | A | 2.7e-08 | 6.1e-08 | 3e-08 | 3.933E-08 | 1.882E-08 | 2.700E-08 | 6.100E-08 |
| 1748 | 88 | D88G | A | 9.7e-08 | 4.6e-08 | 6.5e-08 | 6.933E-08 | 2.577E-08 | 4.600E-08 | 9.700E-08 |
| 1749 | 88 | D88H | A | 6.4e-08 | 3.9e-08 | 3.2e-08 | 4.500E-08 | 1.682E-08 | 3.200E-08 | 6.400E-08 |
| 1750 | 88 | D88I | A | 1e-07 | 3e-08 | 2.8e-08 | 5.267E-08 | 4.100E-08 | 2.800E-08 | 1.000E-07 |
| 1751 | 88 | D88L | A | 2.8e-08 | 5.5e-08 | 4.3e-08 | 4.200E-08 | 1.353E-08 | 2.800E-08 | 5.500E-08 |
| 1752 | 88 | D88K | A | 4.3e-08 | 4.9e-08 | 2.2e-08 | 3.800E-08 | 1.418E-08 | 2.200E-08 | 4.900E-08 |
| 1753 | 88 | D88M | A | 1.7e-08 | 3.7e-08 | 3.2e-08 | 2.867E-08 | 1.041E-08 | 1.700E-08 | 3.700E-08 |
| 1754 | 88 | D88F | A | 5.8e-08 | 3.3e-08 | 4e-08 | 4.367E-08 | 1.290E-08 | 3.300E-08 | 5.800E-08 |
| 1755 | 88 | D88P | A | 1.6e-07 | 4.6e-08 | 4.1e-08 | 8.233E-08 | 6.731E-08 | 4.100E-08 | 1.600E-07 |
| 1756 | 88 | D88S | A | 2.6e-08 | 9.6e-09 | 4.9e-08 | 2.820E-08 | 1.979E-08 | 9.600E-09 | 4.900E-08 |
| 1757 | 88 | D88T | A | 1.8e-08 | 8.7e-08 | 4.2e-08 | 4.900E-08 | 3.503E-08 | 1.800E-08 | 8.700E-08 |
| 1758 | 88 | D88W | A | 1.3e-08 | 3e-08 | 6e-08 | 3.433E-08 | 2.380E-08 | 1.300E-08 | 6.000E-08 |
| 1759 | 88 | D88Y | A | 3.1e-08 | 4.5e-08 | 3.3e-08 | 3.633E-08 | 7.572E-09 | 3.100E-08 | 4.500E-08 |
| 1760 | 88 | D88V | A | 2.9e-08 | 1.5e-08 | 2.4e-07 | 9.467E-08 | 1.261E-07 | 1.500E-08 | 2.400E-07 |
| 1761 | 89 | N89A | A | 4.3e-08 | 2.6e-08 | 2e-08 | 2.967E-08 | 1.193E-08 | 2.000E-08 | 4.300E-08 |
| 1762 | 89 | N89R | A | 5e-08 | 1.3e-07 | 6e-08 | 8.000E-08 | 4.359E-08 | 5.000E-08 | 1.300E-07 |
| 1763 | 89 | N89N | A | 2.9e-08 | 1e-07 | 7.5e-08 | 6.800E-08 | 3.601E-08 | 2.900E-08 | 1.000E-07 |
| 1764 | 89 | N89D | A | 3.3e-08 | 2.8e-08 | 4e-08 | 3.367E-08 | 6.028E-09 | 2.800E-08 | 4.000E-08 |
| 1765 | 89 | N89C | A | 4.1e-08 | 4.1e-08 | 7.5e-08 | 5.233E-08 | 1.963E-08 | 4.100E-08 | 7.500E-08 |
| 1766 | 89 | N89E | A | 1.4e-08 | 9.3e-08 | 1.8e-08 | 4.167E-08 | 4.450E-08 | 1.400E-08 | 9.300E-08 |
| 1767 | 89 | N89Q | A | 7.7e-08 | 1.1e-07 | 2.4e-08 | 7.033E-08 | 4.339E-08 | 2.400E-08 | 1.100E-07 |
| 1768 | 89 | N89G | A | 3.8e-08 | 8.8e-08 | 3.8e-08 | 5.467E-08 | 2.887E-08 | 3.800E-08 | 8.800E-08 |
| 1769 | 89 | N89H | A | 2.7e-08 | 5.5e-08 | 1.6e-07 | 8.067E-08 | 7.012E-08 | 2.700E-08 | 1.600E-07 |
| 1770 | 89 | N89I | A | 2.3e-08 | 8.6e-08 | 1.1e-07 | 7.300E-08 | 4.493E-08 | 2.300E-08 | 1.100E-07 |
| 1771 | 89 | N89L | A | 7.2e-08 | 4.2e-08 | 3.5e-08 | 4.967E-08 | 1.966E-08 | 3.500E-08 | 7.200E-08 |
| 1772 | 89 | N89K | A | 7e-08 | 2.3e-08 | 6.7e-08 | 5.333E-08 | 2.631E-08 | 2.300E-08 | 7.000E-08 |
| 1773 | 89 | N89M | A | 2e-08 | 8e-08 | 4e-08 | 4.667E-08 | 3.055E-08 | 2.000E-08 | 8.000E-08 |
| 1774 | 89 | N89F | A | 4.1e-08 | 3e-08 | 1.3e-07 | 6.700E-08 | 5.484E-08 | 3.000E-08 | 1.300E-07 |
| 1775 | 89 | N89P | A | 1.2e-07 | 4e-08 | 4.5e-08 | 6.833E-08 | 4.481E-08 | 4.000E-08 | 1.200E-07 |
| 1776 | 89 | N89S | A | 8.8e-08 | 4.9e-08 | 1.9e-08 | 5.200E-08 | 3.460E-08 | 1.900E-08 | 8.800E-08 |
| 1777 | 89 | N89T | A | 3.4e-08 | 2.1e-08 | 3.4e-08 | 2.967E-08 | 7.506E-09 | 2.100E-08 | 3.400E-08 |
| 1778 | 89 | N89W | A | 2.9e-08 | 3.7e-08 | 2.9e-08 | 3.167E-08 | 4.619E-09 | 2.900E-08 | 3.700E-08 |
| 1779 | 89 | N89Y | A | 6.1e-08 | 8.6e-08 | 1.2e-07 | 8.900E-08 | 2.961E-08 | 6.100E-08 | 1.200E-07 |
| 1780 | 89 | N89V | A | 4.8e-08 | 9.6e-08 | 4.6e-08 | 6.333E-08 | 2.831E-08 | 4.600E-08 | 9.600E-08 |
| 1781 | 90 | Y90A | A | 5.2e-08 | 4.8e-08 | 3.1e-08 | 4.367E-08 | 1.115E-08 | 3.100E-08 | 5.200E-08 |
| 1782 | 90 | Y90R | A | 1.1e-07 | 6.8e-08 | 1.9e-08 | 6.567E-08 | 4.554E-08 | 1.900E-08 | 1.100E-07 |
| 1783 | 90 | Y90N | A | 8.4e-08 | 6.5e-08 | 3.7e-08 | 6.200E-08 | 2.364E-08 | 3.700E-08 | 8.400E-08 |
| 1784 | 90 | Y90D | A | 1.2e-07 | 1.6e-07 | 5.3e-08 | 1.110E-07 | 5.406E-08 | 5.300E-08 | 1.600E-07 |

| | | | | | | | | | | |
|---|---|---|---|---|---|---|---|---|---|---|
| 1785 | 90 | Y90C | A | 7.3e-08 | 3.9e-08 | 4.9e-08 | 5.367E-08 | 1.747E-08 | 3.900E-08 | 7.300E-08 |
| 1786 | 90 | Y90E | A | 6e-08 | 2.1e-08 | 8e-08 | 5.367E-08 | 3.001E-08 | 2.100E-08 | 8.000E-08 |
| 1787 | 90 | Y90Q | A | 6.1e-08 | 6.9e-08 | 1e-07 | 7.667E-08 | 2.060E-08 | 6.100E-08 | 1.000E-07 |
| 1788 | 90 | Y90G | A | 6.8e-08 | 4.3e-08 | 2.2e-08 | 4.433E-08 | 2.303E-08 | 2.200E-08 | 6.800E-08 |
| 1789 | 90 | Y90H | A | 3.7e-08 | 5.8e-08 | 2.8e-08 | 4.100E-08 | 1.539E-08 | 2.800E-08 | 5.800E-08 |
| 1790 | 90 | Y90I | A | 9.2e-08 | 3.7e-08 | 7.1e-08 | 6.667E-08 | 2.775E-08 | 3.700E-08 | 9.200E-08 |
| 1791 | 90 | Y90L | A | 4.6e-08 | 7.5e-08 | 3.6e-08 | 5.233E-08 | 2.026E-08 | 3.600E-08 | 7.500E-08 |
| 1792 | 90 | Y90K | A | 7.1e-08 | 5.2e-08 | 2.9e-08 | 5.067E-08 | 2.103E-08 | 2.900E-08 | 7.100E-08 |
| 1793 | 90 | Y90M | A | 1.6e-08 | 1.9e-08 | 2.9e-08 | 2.133E-08 | 6.807E-09 | 1.600E-08 | 2.900E-08 |
| 1794 | 90 | Y90F | A | 2.8e-08 | 2.5e-08 | 2e-08 | 2.433E-08 | 4.041E-09 | 2.000E-08 | 2.800E-08 |
| 1795 | 90 | Y90P | A | 2.2e-08 | 2.6e-08 | 3.5e-08 | 2.767E-08 | 6.658E-09 | 2.200E-08 | 3.500E-08 |
| 1796 | 90 | Y90S | A | 1.7e-08 | 1.3e-08 | 3e-08 | 2.000E-08 | 8.888E-09 | 1.300E-08 | 3.000E-08 |
| 1797 | 90 | Y90T | A | 2e-07 | 3.2e-08 | 3.7e-08 | 8.967E-08 | 9.558E-08 | 3.200E-08 | 2.000E-07 |
| 1798 | 90 | Y90W | A | 8.4e-08 | 2.8e-08 | 2.2e-08 | 4.467E-08 | 3.420E-08 | 2.200E-08 | 8.400E-08 |
| 1799 | 90 | Y90Y | A | 3.4e-08 | 2.6e-08 | 7.8e-08 | 4.600E-08 | 2.800E-08 | 2.600E-08 | 7.800E-08 |
| 1800 | 90 | Y90V | A | 2.1e-08 | 8.1e-08 | 4.1e-08 | 4.767E-08 | 3.055E-08 | 2.100E-08 | 8.100E-08 |
| 1801 | 91 | A91A | A | 1e-07 | 5.2e-08 | 9.1e-08 | 8.100E-08 | 2.551E-08 | 5.200E-08 | 1.000E-07 |
| 1802 | 91 | A91R | A | 4.7e-08 | 6.8e-08 | 1.5e-08 | 4.333E-08 | 2.669E-08 | 1.500E-08 | 6.800E-08 |
| 1803 | 91 | A91N | A | 6.7e-08 | 5.4e-08 | 8.4e-08 | 6.833E-08 | 1.504E-08 | 5.400E-08 | 8.400E-08 |
| 1804 | 91 | A91D | A | 7.6e-08 | 2.9e-08 | 2.9e-08 | 4.467E-08 | 2.714E-08 | 2.900E-08 | 7.600E-08 |
| 1805 | 91 | A91C | A | 1.4e-07 | 6.2e-08 | 2.5e-08 | 7.567E-08 | 5.871E-08 | 2.500E-08 | 1.400E-07 |
| 1806 | 91 | A91E | A | 4.4e-08 | 9.8e-08 | 2.8e-08 | 5.667E-08 | 3.668E-08 | 2.800E-08 | 9.800E-08 |
| 1807 | 91 | A91Q | A | 7e-08 | 7.9e-09 | 4.7e-08 | 4.163E-08 | 3.140E-08 | 7.900E-09 | 7.000E-08 |
| 1808 | 91 | A91G | A | 6.9e-08 | 2.8e-08 | 6.2e-08 | 5.300E-08 | 2.193E-08 | 2.800E-08 | 6.900E-08 |
| 1809 | 91 | A91H | A | 2.4e-07 | 3.8e-08 | 6.6e-08 | 1.147E-07 | 1.094E-07 | 3.800E-08 | 2.400E-07 |
| 1810 | 91 | A91I | A | 2.4e-08 | 4.9e-08 | 6.9e-08 | 4.733E-08 | 2.255E-08 | 2.400E-08 | 6.900E-08 |
| 1811 | 91 | A91L | A | 5.3e-08 | 9.4e-08 | 5.2e-08 | 6.633E-08 | 2.397E-08 | 5.200E-08 | 9.400E-08 |
| 1812 | 91 | A91K | A | 3.7e-08 | 1.2e-07 | 7.7e-08 | 7.800E-08 | 4.151E-08 | 3.700E-08 | 1.200E-07 |
| 1813 | 91 | A91M | A | 2.1e-08 | 8.4e-08 | 3e-08 | 4.500E-08 | 3.407E-08 | 2.100E-08 | 8.400E-08 |
| 1814 | 91 | A91F | A | 1.9e-08 | 3e-08 | 3.3e-08 | 2.733E-08 | 7.371E-09 | 1.900E-08 | 3.300E-08 |
| 1815 | 91 | A91P | A | 9.9e-08 | 3.6e-08 | 2.5e-08 | 5.333E-08 | 3.993E-08 | 2.500E-08 | 9.900E-08 |
| 1816 | 91 | A91S | A | 4.1e-08 | 5e-08 | 5.7e-08 | 4.933E-08 | 8.021E-09 | 4.100E-08 | 5.700E-08 |
| 1817 | 91 | A91T | A | 2.1e-08 | 5.5e-08 | 2.2e-08 | 3.267E-08 | 1.935E-08 | 2.100E-08 | 5.500E-08 |
| 1818 | 91 | A91W | A | 3.1e-08 | 4.8e-08 | 8.6e-08 | 5.500E-08 | 2.816E-08 | 3.100E-08 | 8.600E-08 |
| 1819 | 91 | A91Y | A | 4.5e-08 | 6.7e-08 | 6e-08 | 5.733E-08 | 1.124E-08 | 4.500E-08 | 6.700E-08 |
| 1820 | 91 | A91V | A | 1.3e-07 | 2.7e-08 | 3.9e-08 | 6.533E-08 | 5.632E-08 | 2.700E-08 | 1.300E-07 |
| 1821 | 92 | L92A | A | 5.6e-08 | 1.8e-08 | 3.5e-08 | 3.633E-08 | 1.904E-08 | 1.800E-08 | 5.600E-08 |
| 1822 | 92 | L92R | A | 2.4e-08 | 4.6e-08 | 4.9e-08 | 3.967E-08 | 1.365E-08 | 2.400E-08 | 4.900E-08 |
| 1823 | 92 | L92N | A | 3.9e-08 | 1.4e-08 | 1.2e-08 | 2.167E-08 | 1.504E-08 | 1.200E-08 | 3.900E-08 |
| 1824 | 92 | L92D | A | 2.9e-08 | 2.2e-07 | 3.8e-08 | 9.567E-08 | 1.078E-07 | 2.900E-08 | 2.200E-07 |
| 1825 | 92 | L92C | A | 2.7e-08 | 3.2e-08 | 3.2e-08 | 3.033E-08 | 2.887E-09 | 2.700E-08 | 3.200E-08 |
| 1826 | 92 | L92E | A | 4.1e-08 | 1.6e-08 | 7.6e-08 | 4.433E-08 | 3.014E-08 | 1.600E-08 | 7.600E-08 |
| 1827 | 92 | L92Q | A | 8.7e-08 | 1.1e-07 | 2.4e-08 | 7.367E-08 | 4.452E-08 | 2.400E-08 | 1.100E-07 |
| 1828 | 92 | L92G | A | 6.4e-08 | 8.6e-08 | 2.4e-08 | 5.800E-08 | 3.143E-08 | 2.400E-08 | 8.600E-08 |
| 1829 | 92 | L92H | A | 5e-08 | 6.3e-08 | 5e-08 | 5.433E-08 | 7.506E-09 | 5.000E-08 | 6.300E-08 |
| 1830 | 92 | L92I | A | 3.8e-08 | 2.5e-08 | 5.3e-08 | 3.867E-08 | 1.401E-08 | 2.500E-08 | 5.300E-08 |
| 1831 | 92 | L92L | A | 3.9e-08 | 3.7e-08 | 4.4e-08 | 4.000E-08 | 3.606E-09 | 3.700E-08 | 4.400E-08 |
| 1832 | 92 | L92K | A | 3.7e-08 | 3.5e-08 | 7.4e-08 | 4.867E-08 | 2.196E-08 | 3.500E-08 | 7.400E-08 |
| 1833 | 92 | L92M | A | 4.9e-08 | 6.3e-08 | 5.4e-08 | 5.533E-08 | 7.095E-09 | 4.900E-08 | 6.300E-08 |
| 1834 | 92 | L92F | A | 4.3e-08 | 3e-08 | 3.3e-08 | 3.533E-08 | 6.807E-09 | 3.000E-08 | 4.300E-08 |
| 1835 | 92 | L92P | A | 1.1e-08 | 4.5e-08 | 7.7e-08 | 4.433E-08 | 3.301E-08 | 1.100E-08 | 7.700E-08 |
| 1836 | 92 | L92S | A | 1.3e-08 | 7.2e-08 | 3.3e-08 | 3.933E-08 | 3.001E-08 | 1.300E-08 | 7.200E-08 |
| 1837 | 92 | L92T | A | 2.9e-08 | 7.8e-08 | 2e-08 | 4.233E-08 | 3.121E-08 | 2.000E-08 | 7.800E-08 |
| 1838 | 92 | L92W | A | 3.1e-08 | 2.2e-08 | 4.2e-08 | 3.167E-08 | 1.002E-08 | 2.200E-08 | 4.200E-08 |
| 1839 | 92 | L92Y | A | 5.3e-08 | 2.1e-08 | 4.5e-08 | 3.967E-08 | 1.665E-08 | 2.100E-08 | 5.300E-08 |
| 1840 | 92 | L92V | A | 3.7e-08 | 5.5e-08 | 5.7e-09 | 3.257E-08 | 2.495E-08 | 5.700E-09 | 5.500E-08 |
| 1841 | 93 | A93A | A | 4.4e-08 | 2e-08 | 1.3e-07 | 6.467E-08 | 5.784E-08 | 2.000E-08 | 1.300E-07 |
| 1842 | 93 | A93R | A | 1.7e-08 | 2.6e-08 | 1.1e-07 | 5.100E-08 | 5.129E-08 | 1.700E-08 | 1.100E-07 |
| 1843 | 93 | A93N | A | 2.9e-08 | 5.5e-08 | 3.6e-08 | 4.000E-08 | 1.345E-08 | 2.900E-08 | 5.500E-08 |
| 1844 | 93 | A93D | A | 2.8e-08 | 2.5e-07 | 3.1e-08 | 1.030E-07 | 1.273E-07 | 2.800E-08 | 2.500E-07 |
| 1845 | 93 | A93C | A | 3.2e-08 | 3.4e-08 | 6.1e-08 | 4.233E-08 | 1.620E-08 | 3.200E-08 | 6.100E-08 |
| 1846 | 93 | A93E | A | 1.4e-07 | 3.1e-08 | 5.6e-08 | 7.567E-08 | 5.710E-08 | 3.100E-08 | 1.400E-07 |
| 1847 | 93 | A93Q | A | 4.4e-08 | 3.5e-08 | 4.4e-08 | 4.100E-08 | 5.196E-09 | 3.500E-08 | 4.400E-08 |
| 1848 | 93 | A93G | A | 6.1e-08 | 3.9e-08 | 6.8e-08 | 5.600E-08 | 1.513E-08 | 3.900E-08 | 6.800E-08 |
| 1849 | 93 | A93H | A | 3.7e-08 | 6.4e-08 | 2.3e-08 | 4.133E-08 | 2.084E-08 | 2.300E-08 | 6.400E-08 |
| 1850 | 93 | A93I | A | 4.4e-08 | 2.7e-08 | 1.7e-08 | 2.933E-08 | 1.365E-08 | 1.700E-08 | 4.400E-08 |
| 1851 | 93 | A93L | A | 2.7e-08 | 2.1e-08 | 5.5e-08 | 3.433E-08 | 1.815E-08 | 2.100E-08 | 5.500E-08 |
| 1852 | 93 | A93K | A | 4.6e-08 | 9.3e-09 | 5e-08 | 3.510E-08 | 2.243E-08 | 9.300E-09 | 5.000E-08 |
| 1853 | 93 | A93M | A | 3.3e-08 | 5.2e-08 | 8.4e-08 | 5.633E-08 | 2.577E-08 | 3.300E-08 | 8.400E-08 |
| 1854 | 93 | A93F | A | 5.5e-08 | 5.1e-08 | 2.9e-08 | 4.500E-08 | 1.400E-08 | 2.900E-08 | 5.500E-08 |
| 1855 | 93 | A93P | A | 2.3e-08 | 2.7e-08 | 5.5e-08 | 3.500E-08 | 1.744E-08 | 2.300E-08 | 5.500E-08 |
| 1856 | 93 | A93S | A | 1.6e-08 | 9.2e-08 | 1.5e-08 | 4.100E-08 | 4.417E-08 | 1.500E-08 | 9.200E-08 |
| 1857 | 93 | A93T | A | 3.1e-08 | 2.2e-08 | 4.1e-08 | 3.133E-08 | 9.504E-09 | 2.200E-08 | 4.100E-08 |
| 1858 | 93 | A93W | A | 4.3e-08 | 4.1e-08 | 6.6e-08 | 5.000E-08 | 1.389E-08 | 4.100E-08 | 6.600E-08 |
| 1859 | 93 | A93Y | A | 4.3e-08 | 4e-08 | 4e-08 | 4.100E-08 | 1.732E-09 | 4.000E-08 | 4.300E-08 |
| 1860 | 93 | A93V | A | 7.1e-08 | 3.6e-08 | 6.2e-08 | 5.633E-08 | 1.818E-08 | 3.600E-08 | 7.100E-08 |
| 1861 | 94 | V94A | A | 5.9e-08 | 3.1e-08 | 5.7e-08 | 4.900E-08 | 1.562E-08 | 3.100E-08 | 5.900E-08 |
| 1862 | 94 | V94R | A | 4.1e-08 | 2.1e-08 | 1.5e-07 | 7.067E-08 | 6.943E-08 | 2.100E-08 | 1.500E-07 |
| 1863 | 94 | V94N | A | 3.4e-08 | 5.5e-08 | 5.3e-08 | 4.733E-08 | 1.159E-08 | 3.400E-08 | 5.500E-08 |
| 1864 | 94 | V94D | A | 3.9e-08 | 3.2e-08 | 1.9e-08 | 3.000E-08 | 1.015E-08 | 1.900E-08 | 3.900E-08 |
| 1865 | 94 | V94C | A | 5.3e-08 | 7.4e-08 | 4.1e-08 | 5.600E-08 | 1.670E-08 | 4.100E-08 | 7.400E-08 |
| 1866 | 94 | V94E | A | 2.6e-08 | 3.6e-08 | 5.6e-08 | 3.933E-08 | 1.528E-08 | 2.600E-08 | 5.600E-08 |
| 1867 | 94 | V94Q | A | 1.6e-08 | 3e-08 | 3e-08 | 2.533E-08 | 8.083E-09 | 1.600E-08 | 3.000E-08 |
| 1868 | 94 | V94G | A | 4.2e-08 | 1.7e-08 | 4.9e-08 | 3.600E-08 | 1.682E-08 | 1.700E-08 | 4.900E-08 |
| 1869 | 94 | V94H | A | 2.9e-08 | 2.7e-08 | 6e-08 | 3.867E-08 | 1.850E-08 | 2.700E-08 | 6.000E-08 |

| | | | | | | | | | | |
|---|---|---|---|---|---|---|---|---|---|---|
| 1870 | 94 | V94I | A | 6.1e-08 | 1.7e-08 | 4.2e-08 | 4.000E-08 | 2.207E-08 | 1.700E-08 | 6.100E-08 |
| 1871 | 94 | V94L | A | 1.4e-08 | 8.7e-08 | 2.5e-08 | 4.200E-08 | 3.936E-08 | 1.400E-08 | 8.700E-08 |
| 1872 | 94 | V94K | A | 4.9e-08 | 4.9e-08 | 3.1e-08 | 4.300E-08 | 1.039E-08 | 3.100E-08 | 4.900E-08 |
| 1873 | 94 | V94M | A | 2e-08 | 4.7e-08 | 1.1e-07 | 5.900E-08 | 4.618E-08 | 2.000E-08 | 1.100E-07 |
| 1874 | 94 | V94F | A | 7.5e-08 | 4.6e-08 | 1.3e-07 | 8.367E-08 | 4.267E-08 | 4.600E-08 | 1.300E-07 |
| 1875 | 94 | V94P | A | 3.2e-08 | 1.8e-08 | 7.3e-08 | 4.100E-08 | 2.858E-08 | 1.800E-08 | 7.300E-08 |
| 1876 | 94 | V94S | A | 5.1e-08 | 3.8e-08 | 5.8e-08 | 4.900E-08 | 1.015E-08 | 3.800E-08 | 5.800E-08 |
| 1877 | 94 | V94T | A | 1.9e-08 | 1.3e-08 | 4.3e-08 | 2.500E-08 | 1.587E-08 | 1.300E-08 | 4.300E-08 |
| 1878 | 94 | V94W | A | 1.7e-08 | 5.2e-08 | 2.3e-08 | 3.067E-08 | 1.872E-08 | 1.700E-08 | 5.200E-08 |
| 1879 | 94 | V94Y | A | 6.8e-08 | 7.7e-08 | 5.6e-08 | 6.700E-08 | 1.054E-08 | 5.600E-08 | 7.700E-08 |
| 1880 | 94 | V94V | A | 5.7e-08 | 2.9e-08 | 4.3e-08 | 4.300E-08 | 1.400E-08 | 2.900E-08 | 5.700E-08 |
| 1881 | 95 | L95A | A | 2e-08 | 3e-08 | 7.7e-09 | 1.923E-08 | 1.117E-08 | 7.700E-09 | 3.000E-08 |
| 1882 | 95 | L95R | A | 2.4e-08 | 4.4e-08 | 2.6e-08 | 3.133E-08 | 1.102E-08 | 2.400E-08 | 4.400E-08 |
| 1883 | 95 | L95N | A | 4.7e-08 | 4.4e-08 | 7.3e-08 | 5.467E-08 | 1.595E-08 | 4.400E-08 | 7.300E-08 |
| 1884 | 95 | L95D | A | 3.9e-08 | 1.1e-08 | 3.7e-08 | 2.900E-08 | 1.562E-08 | 1.100E-08 | 3.900E-08 |
| 1885 | 95 | L95C | A | 2.2e-08 | 1.1e-07 | 5.8e-08 | 6.333E-08 | 4.424E-08 | 2.200E-08 | 1.100E-07 |
| 1886 | 95 | L95E | A | 3.1e-08 | 2.4e-08 | 3.3e-08 | 2.933E-08 | 4.726E-09 | 2.400E-08 | 3.300E-08 |
| 1887 | 95 | L95Q | A | 4.4e-08 | 1.4e-08 | 1.7e-08 | 2.500E-08 | 1.652E-08 | 1.400E-08 | 4.400E-08 |
| 1888 | 95 | L95G | A | 9.8e-08 | 4e-08 | 5.8e-08 | 6.533E-08 | 2.969E-08 | 4.000E-08 | 9.800E-08 |
| 1889 | 95 | L95H | A | 3e-08 | 6.8e-08 | 1.2e-08 | 3.667E-08 | 2.859E-08 | 1.200E-08 | 6.800E-08 |
| 1890 | 95 | L95I | A | 3.1e-08 | 2.3e-08 | 5.1e-08 | 3.500E-08 | 1.442E-08 | 2.300E-08 | 5.100E-08 |
| 1891 | 95 | L95L | A | 4.6e-08 | 3e-08 | 5.1e-08 | 4.233E-08 | 1.097E-08 | 3.000E-08 | 5.100E-08 |
| 1892 | 95 | L95K | A | 2.2e-08 | 1.1e-07 | 3.1e-08 | 5.433E-08 | 4.842E-08 | 2.200E-08 | 1.100E-07 |
| 1893 | 95 | L95M | A | 1.1e-07 | 8.6e-08 | 7.4e-08 | 9.000E-08 | 1.833E-08 | 7.400E-08 | 1.100E-07 |
| 1894 | 95 | L95F | A | 3.7e-08 | 8.6e-08 | 6.5e-08 | 6.267E-08 | 2.458E-08 | 3.700E-08 | 8.600E-08 |
| 1895 | 95 | L95P | A | 1e-07 | 3.8e-08 | 7.2e-08 | 7.000E-08 | 3.105E-08 | 3.800E-08 | 1.000E-07 |
| 1896 | 95 | L95S | A | 3.1e-08 | 1.2e-07 | 4.8e-08 | 6.633E-08 | 4.725E-08 | 3.100E-08 | 1.200E-07 |
| 1897 | 95 | L95T | A | 1.9e-08 | 2.8e-08 | 4.5e-08 | 3.067E-08 | 1.320E-08 | 1.900E-08 | 4.500E-08 |
| 1898 | 95 | L95W | A | 2.2e-08 | 5.2e-08 | 1e-07 | 5.800E-08 | 3.934E-08 | 2.200E-08 | 1.000E-07 |
| 1899 | 95 | L95Y | A | 6.1e-08 | 8.3e-08 | 2.7e-08 | 5.700E-08 | 2.821E-08 | 2.700E-08 | 8.300E-08 |
| 1900 | 95 | L95V | A | 9.6e-08 | 7.7e-08 | 1.8e-08 | 6.367E-08 | 4.067E-08 | 1.800E-08 | 9.600E-08 |
| 1901 | 96 | D96A | A | 6.4e-08 | 5.4e-08 | 6.5e-08 | 6.100E-08 | 6.083E-09 | 5.400E-08 | 6.500E-08 |
| 1902 | 96 | D96R | A | 5.5e-08 | 1.6e-08 | 4.9e-08 | 4.000E-08 | 2.100E-08 | 1.600E-08 | 5.500E-08 |
| 1903 | 96 | D96N | A | 5.6e-08 | 2.8e-08 | 4.7e-08 | 4.367E-08 | 1.429E-08 | 2.800E-08 | 5.600E-08 |
| 1904 | 96 | D96D | A | 3e-08 | 5.2e-08 | 2.1e-08 | 3.433E-08 | 1.595E-08 | 2.100E-08 | 5.200E-08 |
| 1905 | 96 | D96C | A | 3.3e-08 | 5.8e-08 | 4.4e-08 | 4.500E-08 | 1.253E-08 | 3.300E-08 | 5.800E-08 |
| 1906 | 96 | D96E | A | 4.1e-08 | 3.6e-08 | 4.6e-08 | 4.100E-08 | 5.000E-09 | 3.600E-08 | 4.600E-08 |
| 1907 | 96 | D96Q | A | 4.3e-08 | 5e-08 | 4.7e-08 | 4.667E-08 | 3.512E-09 | 4.300E-08 | 5.000E-08 |
| 1908 | 96 | D96G | A | 1.9e-08 | 9.1e-08 | 2.3e-08 | 4.433E-08 | 4.046E-08 | 1.900E-08 | 9.100E-08 |
| 1909 | 96 | D96H | A | 2.1e-08 | 4.3e-08 | 3e-08 | 3.133E-08 | 1.106E-08 | 2.100E-08 | 4.300E-08 |
| 1910 | 96 | D96I | A | 1.3e-08 | 3.4e-08 | 7.5e-08 | 4.067E-08 | 3.153E-08 | 1.300E-08 | 7.500E-08 |
| 1911 | 96 | D96L | A | 7.3e-08 | 7.3e-08 | 3.3e-08 | 5.967E-08 | 2.309E-08 | 3.300E-08 | 7.300E-08 |
| 1912 | 96 | D96K | A | 4.7e-08 | 7.9e-08 | 1.4e-08 | 4.667E-08 | 3.250E-08 | 1.400E-08 | 7.900E-08 |
| 1913 | 96 | D96M | A | 1.3e-08 | 4.7e-08 | 3.3e-08 | 3.100E-08 | 1.709E-08 | 1.300E-08 | 4.700E-08 |
| 1914 | 96 | D96F | A | 5.8e-08 | 1.4e-07 | 4.7e-08 | 8.167E-08 | 5.082E-08 | 4.700E-08 | 1.400E-07 |
| 1915 | 96 | D96P | A | 3.6e-08 | 8e-08 | 3.6e-08 | 5.067E-08 | 2.540E-08 | 3.600E-08 | 8.000E-08 |
| 1916 | 96 | D96S | A | 2.9e-08 | 1.2e-08 | 6.3e-08 | 3.467E-08 | 2.597E-08 | 1.200E-08 | 6.300E-08 |
| 1917 | 96 | D96T | A | 1.3e-08 | 1.1e-07 | 3.9e-08 | 5.400E-08 | 5.021E-08 | 1.300E-08 | 1.100E-07 |
| 1918 | 96 | D96W | A | 4.4e-08 | 2e-08 | 5.8e-08 | 4.067E-08 | 1.922E-08 | 2.000E-08 | 5.800E-08 |
| 1919 | 96 | D96Y | A | 2.8e-08 | 4.5e-08 | 3.9e-08 | 3.733E-08 | 8.622E-09 | 2.800E-08 | 4.500E-08 |
| 1920 | 96 | D96V | A | 7.9e-08 | 3.6e-08 | 6.7e-08 | 6.067E-08 | 2.219E-08 | 3.600E-08 | 7.900E-08 |
| 1921 | 97 | N97A | A | 2.3e-07 | 1.6e-08 | 4.7e-08 | 9.767E-08 | 1.156E-07 | 1.600E-08 | 2.300E-07 |
| 1922 | 97 | N97R | A | 1e-07 | 2.8e-08 | 2e-08 | 4.933E-08 | 4.406E-08 | 2.000E-08 | 1.000E-07 |
| 1923 | 97 | N97N | A | 8e-08 | 3.5e-08 | 3.1e-08 | 4.867E-08 | 2.721E-08 | 3.100E-08 | 8.000E-08 |
| 1924 | 97 | N97D | A | 3e-08 | 6e-08 | 1.2e-07 | 7.000E-08 | 4.583E-08 | 3.000E-08 | 1.200E-07 |
| 1925 | 97 | N97C | A | 1.1e-07 | 8.5e-08 | 1.7e-08 | 7.067E-08 | 4.813E-08 | 1.700E-08 | 1.100E-07 |
| 1926 | 97 | N97E | A | 4.8e-08 | 1.8e-08 | 5.2e-08 | 3.933E-08 | 1.858E-08 | 1.800E-08 | 5.200E-08 |
| 1927 | 97 | N97Q | A | 4.6e-08 | 5.2e-08 | 3.8e-08 | 4.533E-08 | 7.024E-09 | 3.800E-08 | 5.200E-08 |
| 1928 | 97 | N97G | A | 1.2e-07 | 5.5e-08 | 2.1e-08 | 6.533E-08 | 5.030E-08 | 2.100E-08 | 1.200E-07 |
| 1929 | 97 | N97H | A | 2.2e-08 | 7.3e-08 | 5.4e-08 | 4.967E-08 | 2.577E-08 | 2.200E-08 | 7.300E-08 |
| 1930 | 97 | N97I | A | 1e-07 | 2.3e-07 | 1.6e-08 | 1.153E-07 | 1.078E-07 | 1.600E-08 | 2.300E-07 |
| 1931 | 97 | N97L | A | 5.1e-08 | 5.1e-08 | 5.5e-08 | 5.233E-08 | 2.309E-09 | 5.100E-08 | 5.500E-08 |
| 1932 | 97 | N97K | A | 4.3e-08 | 5e-08 | 2.9e-08 | 4.067E-08 | 1.069E-08 | 2.900E-08 | 5.000E-08 |
| 1933 | 97 | N97M | A | 8.4e-08 | 4.2e-08 | 1.6e-08 | 4.733E-08 | 3.431E-08 | 1.600E-08 | 8.400E-08 |
| 1934 | 97 | N97F | A | 7.2e-08 | 1.6e-08 | 1.1e-07 | 6.600E-08 | 4.729E-08 | 1.600E-08 | 1.100E-07 |
| 1935 | 97 | N97P | A | 2.6e-08 | 3e-08 | 5.2e-08 | 3.600E-08 | 1.400E-08 | 2.600E-08 | 5.200E-08 |
| 1936 | 97 | N97S | A | 5e-08 | 5.5e-08 | 5.3e-08 | 5.267E-08 | 2.517E-09 | 5.000E-08 | 5.500E-08 |
| 1937 | 97 | N97T | A | 3.2e-08 | 2.3e-08 | 4e-08 | 3.167E-08 | 8.505E-09 | 2.300E-08 | 4.000E-08 |
| 1938 | 97 | N97W | A | 4.4e-08 | 4.9e-08 | 5.8e-08 | 5.033E-08 | 7.095E-09 | 4.400E-08 | 5.800E-08 |
| 1939 | 97 | N97Y | A | 5.3e-08 | 3.9e-08 | 5.3e-08 | 4.833E-08 | 8.083E-09 | 3.900E-08 | 5.300E-08 |
| 1940 | 97 | N97V | A | 2.7e-08 | 5.2e-08 | 4.9e-08 | 4.267E-08 | 1.365E-08 | 2.700E-08 | 5.200E-08 |
| 1941 | 98 | G98A | A | 8.7e-08 | 3.4e-08 | 2.6e-08 | 4.900E-08 | 3.315E-08 | 2.600E-08 | 8.700E-08 |
| 1942 | 98 | G98R | A | 6.1e-08 | 4.5e-08 | 3e-08 | 4.533E-08 | 1.550E-08 | 3.000E-08 | 6.100E-08 |
| 1943 | 98 | G98N | A | 2e-08 | 4.7e-08 | 4e-08 | 3.567E-08 | 1.401E-08 | 2.000E-08 | 4.700E-08 |
| 1944 | 98 | G98D | A | 2.4e-08 | 7.9e-09 | 4e-08 | 2.397E-08 | 1.605E-08 | 7.900E-09 | 4.000E-08 |
| 1945 | 98 | G98C | A | 2.2e-08 | 1.8e-08 | 1.7e-08 | 1.900E-08 | 2.646E-09 | 1.700E-08 | 2.200E-08 |
| 1946 | 98 | G98E | A | 3.9e-08 | 7.6e-08 | 4.4e-08 | 5.300E-08 | 2.007E-08 | 3.900E-08 | 7.600E-08 |
| 1947 | 98 | G98Q | A | 1.3e-08 | 7.3e-08 | 6.5e-08 | 5.033E-08 | 3.258E-08 | 1.300E-08 | 7.300E-08 |
| 1948 | 98 | G98G | A | 6.1e-08 | 1.4e-07 | 1.9e-07 | 1.303E-07 | 6.504E-08 | 6.100E-08 | 1.900E-07 |
| 1949 | 98 | G98H | A | 2e-08 | 2.4e-08 | 8.8e-08 | 4.400E-08 | 3.816E-08 | 2.000E-08 | 8.800E-08 |
| 1950 | 98 | G98I | A | 9.1e-08 | 4e-08 | 1.2e-07 | 8.367E-08 | 4.050E-08 | 4.000E-08 | 1.200E-07 |
| 1951 | 98 | G98L | A | 3.6e-08 | 1.5e-08 | 2.9e-08 | 2.667E-08 | 1.069E-08 | 1.500E-08 | 3.600E-08 |
| 1952 | 98 | G98K | A | 5e-08 | 1.3e-07 | 3.8e-08 | 7.267E-08 | 5.001E-08 | 3.800E-08 | 1.300E-07 |
| 1953 | 98 | G98M | A | 7.9e-08 | 2e-08 | 2.7e-08 | 4.200E-08 | 3.223E-08 | 2.000E-08 | 7.900E-08 |
| 1954 | 98 | G98F | A | 9.8e-08 | 4.4e-08 | 1e-07 | 8.067E-08 | 3.177E-08 | 4.400E-08 | 1.000E-07 |

| | | | | | | | | | | |
|---|---|---|---|---|---|---|---|---|---|---|
| 1955 | 98 | G98P | A | 2.8e-08 | 9e-08 | 4.1e-08 | 5.300E-08 | 3.270E-08 | 2.800E-08 | 9.000E-08 |
| 1956 | 98 | G98S | A | 2.9e-08 | 3.3e-08 | 8.2e-08 | 4.800E-08 | 2.951E-08 | 2.900E-08 | 8.200E-08 |
| 1957 | 98 | G98T | A | 3e-08 | 2e-08 | 4.5e-08 | 3.167E-08 | 1.258E-08 | 2.000E-08 | 4.500E-08 |
| 1958 | 98 | G98W | A | 5.1e-08 | 1.4e-07 | 2.6e-08 | 7.233E-08 | 5.992E-08 | 2.600E-08 | 1.400E-07 |
| 1959 | 98 | G98Y | A | 4e-08 | 1.7e-08 | 5.6e-08 | 3.767E-08 | 1.960E-08 | 1.700E-08 | 5.600E-08 |
| 1960 | 98 | G98V | A | 5.2e-08 | 4e-08 | 1.9e-08 | 3.700E-08 | 1.670E-08 | 1.900E-08 | 5.200E-08 |
| 1961 | 99 | D99A | A | 4.8e-08 | 2.8e-08 | 8.2e-08 | 5.267E-08 | 2.730E-08 | 2.800E-08 | 8.200E-08 |
| 1962 | 99 | D99R | A | 3.6e-07 | 6e-08 | 5.3e-08 | 1.577E-07 | 1.753E-07 | 5.300E-08 | 3.600E-07 |
| 1963 | 99 | D99N | A | 3.6e-08 | 1.1e-07 | 7.4e-09 | 5.113E-08 | 5.295E-08 | 7.400E-09 | 1.100E-07 |
| 1964 | 99 | D99D | A | 2e-08 | 2.4e-08 | 4.3e-08 | 2.900E-08 | 1.229E-08 | 2.000E-08 | 4.300E-08 |
| 1965 | 99 | D99C | A | 4.7e-08 | 3.7e-08 | 4.8e-08 | 4.400E-08 | 6.083E-09 | 3.700E-08 | 4.800E-08 |
| 1966 | 99 | D99E | A | 7.8e-08 | 8e-08 | 4e-08 | 6.600E-08 | 2.254E-08 | 4.000E-08 | 8.000E-08 |
| 1967 | 99 | D99Q | A | 3.7e-08 | 9.8e-08 | 9.4e-08 | 7.633E-08 | 3.412E-08 | 3.700E-08 | 9.800E-08 |
| 1968 | 99 | D99G | A | 1.1e-07 | 1.5e-07 | 4.3e-08 | 1.010E-07 | 5.406E-08 | 4.300E-08 | 1.500E-07 |
| 1969 | 99 | D99H | A | 2.6e-08 | 1.4e-08 | 6.4e-08 | 3.467E-08 | 2.610E-08 | 1.400E-08 | 6.400E-08 |
| 1970 | 99 | D99I | A | 2.3e-08 | 1.2e-07 | 4.2e-08 | 6.167E-08 | 5.140E-08 | 2.300E-08 | 1.200E-07 |
| 1971 | 99 | D99L | A | 2.8e-08 | 1.9e-08 | 6.2e-08 | 3.633E-08 | 2.268E-08 | 1.900E-08 | 6.200E-08 |
| 1972 | 99 | D99K | A | 6.9e-08 | 3.3e-08 | 5.6e-08 | 5.267E-08 | 1.823E-08 | 3.300E-08 | 6.900E-08 |
| 1973 | 99 | D99M | A | 5e-08 | 5.3e-08 | 4.4e-08 | 4.900E-08 | 4.583E-09 | 4.400E-08 | 5.300E-08 |
| 1974 | 99 | D99F | A | 3.5e-08 | 8.3e-08 | 5.9e-08 | 5.900E-08 | 2.400E-08 | 3.500E-08 | 8.300E-08 |
| 1975 | 99 | D99P | A | 8.7e-08 | 5.2e-08 | 5.2e-08 | 6.367E-08 | 2.021E-08 | 5.200E-08 | 8.700E-08 |
| 1976 | 99 | D99S | A | 1.8e-08 | 9.8e-08 | 8.7e-09 | 4.157E-08 | 4.909E-08 | 8.700E-09 | 9.800E-08 |
| 1977 | 99 | D99T | A | 2.2e-08 | 1.1e-07 | 4.1e-08 | 5.767E-08 | 4.631E-08 | 2.200E-08 | 1.100E-07 |
| 1978 | 99 | D99W | A | 2.1e-08 | 4.6e-08 | 3.4e-08 | 3.367E-08 | 1.250E-08 | 2.100E-08 | 4.600E-08 |
| 1979 | 99 | D99Y | A | 7.8e-08 | 8.3e-08 | 6.1e-08 | 7.400E-08 | 1.153E-08 | 6.100E-08 | 8.300E-08 |
| 1980 | 99 | D99V | A | 4.2e-08 | 7.9e-08 | 2.9e-08 | 5.000E-08 | 2.594E-08 | 2.900E-08 | 7.900E-08 |
| 1981 | 100 | P100A | A | 5.4e-08 | 3.8e-08 | 8.3e-08 | 5.833E-08 | 2.281E-08 | 3.800E-08 | 8.300E-08 |
| 1982 | 100 | P100R | A | 3.6e-08 | 4.1e-08 | 2.7e-08 | 3.467E-08 | 7.095E-09 | 2.700E-08 | 4.100E-08 |
| 1983 | 100 | P100N | A | 6.4e-08 | 3.7e-08 | 5.9e-08 | 5.333E-08 | 1.436E-08 | 3.700E-08 | 6.400E-08 |
| 1984 | 100 | P100D | A | 3.8e-08 | 7e-08 | 2.3e-08 | 4.367E-08 | 2.401E-08 | 2.300E-08 | 7.000E-08 |
| 1985 | 100 | P100C | A | 8.2e-08 | 4.7e-08 | 2.9e-08 | 5.267E-08 | 2.695E-08 | 2.900E-08 | 8.200E-08 |
| 1986 | 100 | P100E | A | 3.7e-08 | 3.2e-08 | 1.2e-08 | 2.700E-08 | 1.323E-08 | 1.200E-08 | 3.700E-08 |
| 1987 | 100 | P100Q | A | 3.5e-08 | 5e-08 | 5.9e-08 | 4.800E-08 | 1.212E-08 | 3.500E-08 | 5.900E-08 |
| 1988 | 100 | P100G | A | 7.6e-08 | 1.3e-07 | 6.1e-08 | 8.900E-08 | 3.629E-08 | 6.100E-08 | 1.300E-07 |
| 1989 | 100 | P100H | A | 4.3e-08 | 8.5e-08 | 2.5e-08 | 5.100E-08 | 3.079E-08 | 2.500E-08 | 8.500E-08 |
| 1990 | 100 | P100I | A | 6.6e-08 | 5.2e-08 | 2.3e-08 | 4.700E-08 | 2.193E-08 | 2.300E-08 | 6.600E-08 |
| 1991 | 100 | P100L | A | 5.2e-08 | 8.8e-08 | 1.3e-07 | 9.000E-08 | 3.904E-08 | 5.200E-08 | 1.300E-07 |
| 1992 | 100 | P100K | A | 3.9e-08 | 6.1e-08 | 2.5e-08 | 4.167E-08 | 1.815E-08 | 2.500E-08 | 6.100E-08 |
| 1993 | 100 | P100M | A | 4.3e-08 | 7e-08 | 3.7e-08 | 5.000E-08 | 1.758E-08 | 3.700E-08 | 7.000E-08 |
| 1994 | 100 | P100F | A | 4.4e-08 | 5.5e-08 | 2.7e-08 | 4.200E-08 | 1.411E-08 | 2.700E-08 | 5.500E-08 |
| 1995 | 100 | P100P | A | 7.4e-08 | 2.5e-08 | 1.2e-08 | 3.700E-08 | 3.270E-08 | 1.200E-08 | 7.400E-08 |
| 1996 | 100 | P100S | A | 5.9e-08 | 4e-08 | 5e-08 | 4.967E-08 | 9.504E-09 | 4.000E-08 | 5.900E-08 |
| 1997 | 100 | P100T | A | 6.8e-08 | 3.1e-08 | 1.2e-07 | 7.300E-08 | 4.471E-08 | 3.100E-08 | 1.200E-07 |
| 1998 | 100 | P100W | A | 1.1e-07 | 1.4e-07 | 3.6e-08 | 9.533E-08 | 5.353E-08 | 3.600E-08 | 1.400E-07 |
| 1999 | 100 | P100Y | A | 3e-08 | 6.4e-08 | 2e-08 | 3.800E-08 | 2.307E-08 | 2.000E-08 | 6.400E-08 |
| 2000 | 100 | P100V | A | 9.3e-08 | 4.9e-08 | 4.5e-08 | 6.233E-08 | 2.663E-08 | 4.500E-08 | 9.300E-08 |
| 2001 | 101 | L101A | A | 4.6e-08 | 1e-08 | 2.8e-08 | 2.800E-08 | 1.800E-08 | 1.000E-08 | 4.600E-08 |
| 2002 | 101 | L101R | A | 4.7e-08 | 7.9e-08 | 4.1e-08 | 5.567E-08 | 2.043E-08 | 4.100E-08 | 7.900E-08 |
| 2003 | 101 | L101N | A | 4.4e-08 | 8.7e-08 | 7.7e-08 | 6.933E-08 | 2.250E-08 | 4.400E-08 | 8.700E-08 |
| 2004 | 101 | L101D | A | 3e-08 | 3.5e-08 | 1.4e-08 | 2.633E-08 | 1.097E-08 | 1.400E-08 | 3.500E-08 |
| 2005 | 101 | L101C | A | 6.1e-08 | 8e-08 | 5.4e-08 | 6.500E-08 | 1.345E-08 | 5.400E-08 | 8.000E-08 |
| 2006 | 101 | L101E | A | 2.8e-08 | 3.7e-08 | 4.8e-08 | 3.767E-08 | 1.002E-08 | 2.800E-08 | 4.800E-08 |
| 2007 | 101 | L101Q | A | 4e-08 | 4e-08 | 4e-08 | 4.000E-08 | 8.105E-24 | 4.000E-08 | 4.000E-08 |
| 2008 | 101 | L101G | A | 3.1e-08 | 1.1e-08 | 7.2e-08 | 3.800E-08 | 3.110E-08 | 1.100E-08 | 7.200E-08 |
| 2009 | 101 | L101H | A | 2.5e-08 | 5e-08 | 5.1e-08 | 4.200E-08 | 1.473E-08 | 2.500E-08 | 5.100E-08 |
| 2010 | 101 | L101I | A | 3.8e-08 | 6.3e-08 | 8.8e-08 | 6.300E-08 | 2.500E-08 | 3.800E-08 | 8.800E-08 |
| 2011 | 101 | L101L | A | 7.2e-08 | 1.7e-07 | 5e-08 | 9.733E-08 | 6.389E-08 | 5.000E-08 | 1.700E-07 |
| 2012 | 101 | L101K | A | 8.7e-08 | 1.1e-07 | 2.3e-08 | 7.333E-08 | 4.508E-08 | 2.300E-08 | 1.100E-07 |
| 2013 | 101 | L101M | A | 6e-08 | 3.8e-08 | 8e-08 | 5.933E-08 | 2.101E-08 | 3.800E-08 | 8.000E-08 |
| 2014 | 101 | L101F | A | 1.3e-08 | 5.7e-08 | 9.3e-08 | 5.433E-08 | 4.007E-08 | 1.300E-08 | 9.300E-08 |
| 2015 | 101 | L101P | A | 2.2e-08 | 9.9e-08 | 5.9e-08 | 6.000E-08 | 3.851E-08 | 2.200E-08 | 9.900E-08 |
| 2016 | 101 | L101S | A | 3.3e-08 | 4.1e-08 | 7.4e-08 | 4.933E-08 | 2.173E-08 | 3.300E-08 | 7.400E-08 |
| 2017 | 101 | L101T | A | 6.9e-08 | 7e-08 | 7.8e-08 | 7.233E-08 | 4.933E-09 | 6.900E-08 | 7.800E-08 |
| 2018 | 101 | L101W | A | 9.4e-08 | 1.9e-08 | 4.5e-08 | 5.267E-08 | 3.808E-08 | 1.900E-08 | 9.400E-08 |
| 2019 | 101 | L101Y | A | 5.2e-08 | 6.5e-08 | 4.6e-08 | 5.433E-08 | 9.713E-09 | 4.600E-08 | 6.500E-08 |
| 2020 | 101 | L101V | A | 6.9e-08 | 2.5e-08 | 1.5e-08 | 3.633E-08 | 2.873E-08 | 1.500E-08 | 6.900E-08 |
| 2021 | 102 | N102A | A | 5.9e-08 | 2.7e-08 | 4.5e-08 | 4.367E-08 | 1.604E-08 | 2.700E-08 | 5.900E-08 |
| 2022 | 102 | N102R | A | 2.3e-08 | 3.2e-08 | 6.9e-08 | 4.133E-08 | 2.438E-08 | 2.300E-08 | 6.900E-08 |
| 2023 | 102 | N102N | A | 4.4e-08 | 2.6e-08 | 3.1e-08 | 3.367E-08 | 9.292E-09 | 2.600E-08 | 4.400E-08 |
| 2024 | 102 | N102D | A | 6.5e-08 | 3.3e-08 | 6.2e-08 | 5.333E-08 | 1.767E-08 | 3.300E-08 | 6.500E-08 |
| 2025 | 102 | N102C | A | 7.7e-08 | 6.9e-08 | 1.8e-07 | 1.087E-07 | 6.191E-08 | 6.900E-08 | 1.800E-07 |
| 2026 | 102 | N102E | A | 3e-08 | 7.2e-08 | 1.5e-08 | 3.900E-08 | 2.955E-08 | 1.500E-08 | 7.200E-08 |
| 2027 | 102 | N102Q | A | 1.5e-07 | 4e-08 | 7.8e-08 | 8.933E-08 | 5.587E-08 | 4.000E-08 | 1.500E-07 |
| 2028 | 102 | N102G | A | 8.4e-08 | 4.2e-08 | 4.4e-08 | 5.667E-08 | 2.369E-08 | 4.200E-08 | 8.400E-08 |
| 2029 | 102 | N102H | A | 1.9e-08 | 5.4e-08 | 2.1e-08 | 3.133E-08 | 1.966E-08 | 1.900E-08 | 5.400E-08 |
| 2030 | 102 | N102I | A | 3.1e-08 | 5e-08 | 2.1e-08 | 3.400E-08 | 1.473E-08 | 2.100E-08 | 5.000E-08 |
| 2031 | 102 | N102L | A | 7e-09 | 1.2e-07 | 6.5e-08 | 6.400E-08 | 5.651E-08 | 7.000E-09 | 1.200E-07 |
| 2032 | 102 | N102K | A | 3.8e-08 | 2.8e-08 | 3.8e-08 | 3.467E-08 | 5.774E-09 | 2.800E-08 | 3.800E-08 |
| 2033 | 102 | N102M | A | 1.3e-07 | 1.1e-07 | 3.5e-08 | 9.167E-08 | 5.008E-08 | 3.500E-08 | 1.300E-07 |
| 2034 | 102 | N102F | A | 4.6e-08 | 9.4e-08 | 1.3e-08 | 5.100E-08 | 4.073E-08 | 1.300E-08 | 9.400E-08 |
| 2035 | 102 | N102P | A | 5.8e-08 | 4.4e-08 | 1.1e-07 | 7.067E-08 | 3.478E-08 | 4.400E-08 | 1.100E-07 |
| 2036 | 102 | N102S | A | 7.2e-08 | 3e-08 | 7.9e-08 | 6.033E-08 | 2.650E-08 | 3.000E-08 | 7.900E-08 |
| 2037 | 102 | N102T | A | 3.5e-08 | 4.4e-08 | 1.9e-07 | 8.967E-08 | 8.701E-08 | 3.500E-08 | 1.900E-07 |
| 2038 | 102 | N102W | A | 3.1e-08 | 3.2e-08 | 6.3e-08 | 4.200E-08 | 1.819E-08 | 3.100E-08 | 6.300E-08 |
| 2039 | 102 | N102Y | A | 4.2e-08 | 2.9e-08 | 9.6e-08 | 5.567E-08 | 3.553E-08 | 2.900E-08 | 9.600E-08 |

| | | | | | | | | | | |
|---|---|---|---|---|---|---|---|---|---|---|
| 2040 | 102 | N102V | A | 7.5e-08 | 7e-08 | 3.2e-08 | 5.900E-08 | 2.352E-08 | 3.200E-08 | 7.500E-08 |
| 2041 | 103 | N103A | A | 2.1e-08 | 2.9e-08 | 7e-08 | 4.000E-08 | 2.629E-08 | 2.100E-08 | 7.000E-08 |
| 2042 | 103 | N103R | A | 8.2e-08 | 5.2e-08 | 8.4e-08 | 7.267E-08 | 1.793E-08 | 5.200E-08 | 8.400E-08 |
| 2043 | 103 | N103N | A | 4.1e-08 | 6.6e-08 | 3.3e-08 | 4.667E-08 | 1.721E-08 | 3.300E-08 | 6.600E-08 |
| 2044 | 103 | N103D | A | 7.2e-08 | 2.1e-08 | 4.8e-08 | 4.700E-08 | 2.551E-08 | 2.100E-08 | 7.200E-08 |
| 2045 | 103 | N103C | A | 3.5e-08 | 4.8e-08 | 3.5e-08 | 3.933E-08 | 7.506E-09 | 3.500E-08 | 4.800E-08 |
| 2046 | 103 | N103E | A | 6.5e-08 | 1e-08 | 4.3e-08 | 3.933E-08 | 2.768E-08 | 1.000E-08 | 6.500E-08 |
| 2047 | 103 | N103Q | A | 7.1e-08 | 3.2e-08 | 2.7e-08 | 4.333E-08 | 2.409E-08 | 2.700E-08 | 7.100E-08 |
| 2048 | 103 | N103G | A | 1e-07 | 3.1e-08 | 8.3e-08 | 7.133E-08 | 3.595E-08 | 3.100E-08 | 1.000E-07 |
| 2049 | 103 | N103H | A | 5.5e-08 | 5.1e-08 | 1.4e-07 | 8.200E-08 | 5.027E-08 | 5.100E-08 | 1.400E-07 |
| 2050 | 103 | N103I | A | 3.3e-08 | 3.4e-08 | 3.4e-08 | 3.367E-08 | 5.774E-10 | 3.300E-08 | 3.400E-08 |
| 2051 | 103 | N103L | A | 3.9e-08 | 4.5e-08 | 4e-08 | 4.133E-08 | 3.215E-09 | 3.900E-08 | 4.500E-08 |
| 2052 | 103 | N103K | A | 5.7e-08 | 5.4e-08 | 3.1e-08 | 4.733E-08 | 1.422E-08 | 3.100E-08 | 5.700E-08 |
| 2053 | 103 | N103M | A | 4.3e-08 | 2.6e-08 | 2.2e-08 | 3.033E-08 | 1.115E-08 | 2.200E-08 | 4.300E-08 |
| 2054 | 103 | N103F | A | 4.7e-08 | 2.9e-08 | 6.4e-08 | 4.667E-08 | 1.750E-08 | 2.900E-08 | 6.400E-08 |
| 2055 | 103 | N103P | A | 8.4e-08 | 4.3e-08 | 5.1e-08 | 5.933E-08 | 2.173E-08 | 4.300E-08 | 8.400E-08 |
| 2056 | 103 | N103S | A | 3.4e-08 | 5.7e-08 | 2.9e-08 | 4.000E-08 | 1.493E-08 | 2.900E-08 | 5.700E-08 |
| 2057 | 103 | N103T | A | 1.7e-08 | 4.9e-08 | 2e-07 | 8.867E-08 | 9.774E-08 | 1.700E-08 | 2.000E-07 |
| 2058 | 103 | N103W | A | 6.5e-08 | 8.6e-08 | 3.9e-08 | 6.333E-08 | 2.354E-08 | 3.900E-08 | 8.600E-08 |
| 2059 | 103 | N103Y | A | 1.8e-08 | 1.1e-07 | 2.3e-08 | 5.033E-08 | 5.173E-08 | 1.800E-08 | 1.100E-07 |
| 2060 | 103 | N103V | A | 2.4e-08 | 5.1e-08 | 2.4e-08 | 3.300E-08 | 1.559E-08 | 2.400E-08 | 5.100E-08 |
| 2061 | 104 | T104A | A | 8.9e-08 | 2.7e-08 | 8.6e-08 | 6.733E-08 | 3.496E-08 | 2.700E-08 | 8.900E-08 |
| 2062 | 104 | T104R | A | 7.4e-08 | 3e-08 | 2.4e-08 | 4.267E-08 | 2.730E-08 | 2.400E-08 | 7.400E-08 |
| 2063 | 104 | T104N | A | 2.8e-08 | 4.4e-08 | 9.7e-08 | 5.633E-08 | 3.612E-08 | 2.800E-08 | 9.700E-08 |
| 2064 | 104 | T104D | A | 6.8e-08 | 2.9e-08 | 4.4e-08 | 4.700E-08 | 1.967E-08 | 2.900E-08 | 6.800E-08 |
| 2065 | 104 | T104C | A | 3.3e-08 | 4.8e-08 | 2e-08 | 3.367E-08 | 1.401E-08 | 2.000E-08 | 4.800E-08 |
| 2066 | 104 | T104E | A | 8.4e-08 | 8.2e-08 | 3.6e-08 | 6.733E-08 | 2.715E-08 | 3.600E-08 | 8.400E-08 |
| 2067 | 104 | T104Q | A | 6.9e-08 | 7.5e-08 | 1.2e-08 | 5.200E-08 | 3.477E-08 | 1.200E-08 | 7.500E-08 |
| 2068 | 104 | T104G | A | 5e-08 | 1.1e-07 | 5.3e-08 | 7.100E-08 | 3.381E-08 | 5.000E-08 | 1.100E-07 |
| 2069 | 104 | T104H | A | 2.4e-08 | 9.2e-08 | 5.1e-08 | 5.567E-08 | 3.424E-08 | 2.400E-08 | 9.200E-08 |
| 2070 | 104 | T104I | A | 2.3e-08 | 1.9e-08 | 2.8e-08 | 2.333E-08 | 4.509E-09 | 1.900E-08 | 2.800E-08 |
| 2071 | 104 | T104L | A | 1.8e-08 | 4.7e-08 | 5.7e-08 | 4.067E-08 | 2.026E-08 | 1.800E-08 | 5.700E-08 |
| 2072 | 104 | T104K | A | 4.7e-08 | 8.5e-08 | 4.3e-08 | 5.833E-08 | 2.318E-08 | 4.300E-08 | 8.500E-08 |
| 2073 | 104 | T104M | A | 4.9e-08 | 8.4e-08 | 6.4e-08 | 6.567E-08 | 1.756E-08 | 4.900E-08 | 8.400E-08 |
| 2074 | 104 | T104F | A | 4.7e-08 | 3.4e-08 | 3.8e-08 | 3.967E-08 | 6.658E-09 | 3.400E-08 | 4.700E-08 |
| 2075 | 104 | T104P | A | 5e-08 | 8.1e-08 | 8.2e-08 | 7.100E-08 | 1.819E-08 | 5.000E-08 | 8.200E-08 |
| 2076 | 104 | T104S | A | 6.8e-08 | 2.3e-08 | 9.2e-08 | 6.100E-08 | 3.503E-08 | 2.300E-08 | 9.200E-08 |
| 2077 | 104 | T104T | A | 2.8e-08 | 5.2e-08 | 2.9e-08 | 3.633E-08 | 1.358E-08 | 2.800E-08 | 5.200E-08 |
| 2078 | 104 | T104W | A | 7.3e-08 | 5.2e-08 | 3e-08 | 5.167E-08 | 2.150E-08 | 3.000E-08 | 7.300E-08 |
| 2079 | 104 | T104Y | A | 2.4e-08 | 1.8e-08 | 3.7e-08 | 2.633E-08 | 9.713E-09 | 1.800E-08 | 3.700E-08 |
| 2080 | 104 | T104V | A | 2e-08 | 2.9e-08 | 2.2e-07 | 8.967E-08 | 1.130E-07 | 2.000E-08 | 2.200E-07 |
| 2081 | 105 | T105A | A | 5.4e-08 | 4.7e-08 | 3.9e-08 | 4.667E-08 | 7.506E-09 | 3.900E-08 | 5.400E-08 |
| 2082 | 105 | T105R | A | 1.9e-08 | 3.8e-08 | 4e-08 | 3.233E-08 | 1.159E-08 | 1.900E-08 | 4.000E-08 |
| 2083 | 105 | T105N | A | 3.4e-08 | 3.5e-08 | 3.2e-08 | 3.367E-08 | 1.528E-09 | 3.200E-08 | 3.500E-08 |
| 2084 | 105 | T105D | A | 8.1e-08 | 3.8e-08 | 2.1e-08 | 4.667E-08 | 3.092E-08 | 2.100E-08 | 8.100E-08 |
| 2085 | 105 | T105C | A | 9e-08 | 4.9e-08 | 4.3e-08 | 6.067E-08 | 2.558E-08 | 4.300E-08 | 9.000E-08 |
| 2086 | 105 | T105E | A | 5.6e-08 | 9.9e-08 | 4.6e-08 | 6.700E-08 | 2.816E-08 | 4.600E-08 | 9.900E-08 |
| 2087 | 105 | T105Q | A | 7.3e-08 | 1.1e-08 | 2.6e-08 | 3.667E-08 | 3.235E-08 | 1.100E-08 | 7.300E-08 |
| 2088 | 105 | T105G | A | 1.9e-07 | 8.3e-08 | 1.4e-08 | 9.567E-08 | 8.868E-08 | 1.400E-08 | 1.900E-07 |
| 2089 | 105 | T105H | A | 2.4e-08 | 6.6e-08 | 6.8e-08 | 5.267E-08 | 2.485E-08 | 2.400E-08 | 6.800E-08 |
| 2090 | 105 | T105I | A | 3.6e-08 | 5e-08 | 4.2e-08 | 4.267E-08 | 7.024E-09 | 3.600E-08 | 5.000E-08 |
| 2091 | 105 | T105L | A | 5.6e-08 | 3e-08 | 2.9e-08 | 3.833E-08 | 1.531E-08 | 2.900E-08 | 5.600E-08 |
| 2092 | 105 | T105K | A | 5.8e-08 | 4.5e-08 | 3.3e-08 | 4.533E-08 | 1.250E-08 | 3.300E-08 | 5.800E-08 |
| 2093 | 105 | T105M | A | 8.4e-08 | 2.3e-08 | 8e-08 | 6.233E-08 | 3.412E-08 | 2.300E-08 | 8.400E-08 |
| 2094 | 105 | T105F | A | 1.4e-08 | 3.6e-08 | 1.4e-08 | 2.133E-08 | 1.270E-08 | 1.400E-08 | 3.600E-08 |
| 2095 | 105 | T105P | A | 4.4e-08 | 2.9e-08 | 6.5e-08 | 4.600E-08 | 1.808E-08 | 2.900E-08 | 6.500E-08 |
| 2096 | 105 | T105S | A | 1.6e-08 | 2.8e-08 | 2.9e-08 | 2.433E-08 | 7.234E-09 | 1.600E-08 | 2.900E-08 |
| 2097 | 105 | T105T | A | 1e-08 | 6.1e-08 | 6.1e-08 | 4.400E-08 | 2.944E-08 | 1.000E-08 | 6.100E-08 |
| 2098 | 105 | T105W | A | 4.6e-08 | 4.6e-08 | 6.2e-08 | 5.133E-08 | 9.238E-09 | 4.600E-08 | 6.200E-08 |
| 2099 | 105 | T105Y | A | 9.9e-08 | 2.4e-08 | 4.7e-08 | 5.667E-08 | 3.842E-08 | 2.400E-08 | 9.900E-08 |
| 2100 | 105 | T105V | A | 3.7e-08 | 5.2e-08 | 4.3e-08 | 4.400E-08 | 7.550E-09 | 3.700E-08 | 5.200E-08 |
| 2101 | 106 | P106A | A | 2.4e-08 | 6.6e-08 | 3.7e-08 | 4.233E-08 | 2.150E-08 | 2.400E-08 | 6.600E-08 |
| 2102 | 106 | P106R | A | 8.4e-08 | 2.8e-08 | 4e-08 | 5.067E-08 | 2.948E-08 | 2.800E-08 | 8.400E-08 |
| 2103 | 106 | P106N | A | 5.8e-08 | 3.1e-08 | 3.1e-08 | 4.000E-08 | 1.559E-08 | 3.100E-08 | 5.800E-08 |
| 2104 | 106 | P106D | A | 2.2e-08 | 4.5e-08 | 2.1e-08 | 2.933E-08 | 1.358E-08 | 2.100E-08 | 4.500E-08 |
| 2105 | 106 | P106C | A | 3e-08 | 1.3e-08 | 4.1e-08 | 2.800E-08 | 1.411E-08 | 1.300E-08 | 4.100E-08 |
| 2106 | 106 | P106E | A | 4.9e-08 | 5.5e-08 | 3.5e-08 | 4.633E-08 | 1.026E-08 | 3.500E-08 | 5.500E-08 |
| 2107 | 106 | P106Q | A | 1.4e-08 | 6.4e-08 | 1.5e-08 | 3.100E-08 | 2.858E-08 | 1.400E-08 | 6.400E-08 |
| 2108 | 106 | P106G | A | 3.2e-08 | 1.1e-07 | 6.4e-08 | 6.867E-08 | 3.921E-08 | 3.200E-08 | 1.100E-07 |
| 2109 | 106 | P106H | A | 5.4e-08 | 5.5e-08 | 4.8e-08 | 5.233E-08 | 3.786E-09 | 4.800E-08 | 5.500E-08 |
| 2110 | 106 | P106I | A | 6e-08 | 1e-07 | 5.8e-08 | 7.267E-08 | 2.369E-08 | 5.800E-08 | 1.000E-07 |
| 2111 | 106 | P106L | A | 1.6e-08 | 5.5e-08 | 5.1e-08 | 4.067E-08 | 2.146E-08 | 1.600E-08 | 5.500E-08 |
| 2112 | 106 | P106K | A | 3.2e-08 | 3.3e-08 | 1.3e-08 | 2.600E-08 | 1.127E-08 | 1.300E-08 | 3.300E-08 |
| 2113 | 106 | P106M | A | 8.5e-08 | 2.3e-08 | 5e-08 | 5.267E-08 | 3.109E-08 | 2.300E-08 | 8.500E-08 |
| 2114 | 106 | P106F | A | 9.6e-08 | 7.5e-08 | 9.7e-08 | 8.933E-08 | 1.242E-08 | 7.500E-08 | 9.700E-08 |
| 2115 | 106 | P106P | A | 4.3e-08 | 1.3e-08 | 2.5e-08 | 2.700E-08 | 1.510E-08 | 1.300E-08 | 4.300E-08 |
| 2116 | 106 | P106S | A | 2.8e-08 | 6e-08 | 3.7e-08 | 4.167E-08 | 1.650E-08 | 2.800E-08 | 6.000E-08 |
| 2117 | 106 | P106T | A | 6.2e-08 | 6.7e-08 | 2.9e-08 | 5.267E-08 | 2.065E-08 | 2.900E-08 | 6.700E-08 |
| 2118 | 106 | P106W | A | 2.2e-07 | 6.8e-08 | 2.4e-08 | 1.040E-07 | 1.028E-07 | 2.400E-08 | 2.200E-07 |
| 2119 | 106 | P106Y | A | 9.2e-08 | 6.4e-08 | 4.4e-08 | 6.667E-08 | 2.411E-08 | 4.400E-08 | 9.200E-08 |
| 2120 | 106 | P106V | A | 3.6e-08 | 1.1e-08 | 7.7e-09 | 1.823E-08 | 1.547E-08 | 7.700E-09 | 3.600E-08 |
| 2121 | 107 | V107A | A | 7.4e-08 | 6.9e-08 | 4.6e-08 | 6.300E-08 | 1.493E-08 | 4.600E-08 | 7.400E-08 |
| 2122 | 107 | V107R | A | 3.7e-08 | 3.8e-08 | 1e-07 | 5.833E-08 | 3.609E-08 | 3.700E-08 | 1.000E-07 |
| 2123 | 107 | V107N | A | 3.2e-08 | 3.6e-08 | 1.7e-08 | 2.833E-08 | 1.002E-08 | 1.700E-08 | 3.600E-08 |
| 2124 | 107 | V107D | A | 7.1e-08 | 5.4e-08 | 2.8e-08 | 5.100E-08 | 2.166E-08 | 2.800E-08 | 7.100E-08 |

| | | | | | | | | | | |
|---|---|---|---|---|---|---|---|---|---|---|
| 2125 | 107 | V107C | A | 1.7e-08 | 2.8e-08 | 1.5e-08 | 2.000E-08 | 7.000E-09 | 1.500E-08 | 2.800E-08 |
| 2126 | 107 | V107E | A | 1.2e-07 | 4.4e-08 | 8.1e-08 | 8.167E-08 | 3.800E-08 | 4.400E-08 | 1.200E-07 |
| 2127 | 107 | V107Q | A | 2.6e-08 | 2.4e-08 | 7.4e-08 | 4.133E-08 | 2.831E-08 | 2.400E-08 | 7.400E-08 |
| 2128 | 107 | V107G | A | 1.4e-08 | 4e-08 | 1.8e-08 | 2.400E-08 | 1.400E-08 | 1.400E-08 | 4.000E-08 |
| 2129 | 107 | V107H | A | 2.1e-08 | 3.1e-08 | 6.9e-08 | 4.033E-08 | 2.532E-08 | 2.100E-08 | 6.900E-08 |
| 2130 | 107 | V107I | A | 5.4e-08 | 3.4e-08 | 1.3e-07 | 7.267E-08 | 5.065E-08 | 3.400E-08 | 1.300E-07 |
| 2131 | 107 | V107L | A | 1.3e-07 | 3.4e-08 | 4.2e-08 | 6.867E-08 | 5.327E-08 | 3.400E-08 | 1.300E-07 |
| 2132 | 107 | V107K | A | 1.5e-07 | 1.1e-07 | 5e-08 | 1.033E-07 | 5.033E-08 | 5.000E-08 | 1.500E-07 |
| 2133 | 107 | V107M | A | 2.2e-08 | 2.9e-08 | 7.2e-08 | 4.100E-08 | 2.707E-08 | 2.200E-08 | 7.200E-08 |
| 2134 | 107 | V107F | A | 2.3e-08 | 6.2e-08 | 4.3e-08 | 4.267E-08 | 1.950E-08 | 2.300E-08 | 6.200E-08 |
| 2135 | 107 | V107P | A | 5.1e-08 | 4.4e-08 | 1.3e-07 | 7.500E-08 | 4.776E-08 | 4.400E-08 | 1.300E-07 |
| 2136 | 107 | V107S | A | 3.9e-08 | 4.4e-08 | 8.2e-08 | 5.500E-08 | 2.352E-08 | 3.900E-08 | 8.200E-08 |
| 2137 | 107 | V107T | A | 2.8e-08 | 8.4e-08 | 3.8e-08 | 5.000E-08 | 2.987E-08 | 2.800E-08 | 8.400E-08 |
| 2138 | 107 | V107W | A | 4.4e-08 | 6.1e-08 | 1.5e-08 | 4.000E-08 | 2.326E-08 | 1.500E-08 | 6.100E-08 |
| 2139 | 107 | V107Y | A | 4e-08 | 2.1e-08 | 7.3e-08 | 4.467E-08 | 2.631E-08 | 2.100E-08 | 7.300E-08 |
| 2140 | 107 | V107V | A | 6.9e-08 | 3.1e-08 | 2.4e-08 | 4.133E-08 | 2.421E-08 | 2.400E-08 | 6.900E-08 |
| 2141 | 108 | T108A | A | 1.7e-08 | 7.2e-08 | 6.8e-08 | 5.233E-08 | 3.066E-08 | 1.700E-08 | 7.200E-08 |
| 2142 | 108 | T108R | A | 2.8e-08 | 6.2e-08 | 7.4e-08 | 5.467E-08 | 2.386E-08 | 2.800E-08 | 7.400E-08 |
| 2143 | 108 | T108N | A | 1.4e-07 | 6e-08 | 5.9e-08 | 8.633E-08 | 4.648E-08 | 5.900E-08 | 1.400E-07 |
| 2144 | 108 | T108D | A | 5.1e-08 | 4e-08 | 5.7e-08 | 4.933E-08 | 8.622E-09 | 4.000E-08 | 5.700E-08 |
| 2145 | 108 | T108C | A | 4.9e-08 | 4e-08 | 3.9e-08 | 4.267E-08 | 5.508E-09 | 3.900E-08 | 4.900E-08 |
| 2146 | 108 | T108E | A | 4.2e-08 | 1.2e-07 | 5.3e-08 | 7.167E-08 | 4.222E-08 | 4.200E-08 | 1.200E-07 |
| 2147 | 108 | T108Q | A | 3.6e-08 | 1.1e-08 | 1e-07 | 4.900E-08 | 4.590E-08 | 1.100E-08 | 1.000E-07 |
| 2148 | 108 | T108G | A | 6.1e-08 | 7.8e-08 | 7.7e-08 | 7.200E-08 | 9.539E-09 | 6.100E-08 | 7.800E-08 |
| 2149 | 108 | T108H | A | 1.3e-08 | 5.2e-08 | 5.9e-08 | 4.133E-08 | 2.479E-08 | 1.300E-08 | 5.900E-08 |
| 2150 | 108 | T108I | A | 5.8e-08 | 5.8e-08 | 3.2e-08 | 4.933E-08 | 1.501E-08 | 3.200E-08 | 5.800E-08 |
| 2151 | 108 | T108L | A | 1.2e-08 | 3.3e-08 | 1.9e-08 | 2.133E-08 | 1.069E-08 | 1.200E-08 | 3.300E-08 |
| 2152 | 108 | T108K | A | 1.8e-08 | 4.6e-08 | 4.1e-08 | 3.500E-08 | 1.493E-08 | 1.800E-08 | 4.600E-08 |
| 2153 | 108 | T108M | A | 4.4e-08 | 5.4e-08 | 5.6e-08 | 5.133E-08 | 6.429E-09 | 4.400E-08 | 5.600E-08 |
| 2154 | 108 | T108F | A | 5e-08 | 8.5e-08 | 3.6e-08 | 5.700E-08 | 2.524E-08 | 3.600E-08 | 8.500E-08 |
| 2155 | 108 | T108P | A | 4.6e-08 | 2.6e-08 | 5.8e-08 | 4.333E-08 | 1.617E-08 | 2.600E-08 | 5.800E-08 |
| 2156 | 108 | T108S | A | 3.7e-08 | 3.5e-08 | 5.4e-08 | 4.200E-08 | 1.044E-08 | 3.500E-08 | 5.400E-08 |
| 2157 | 108 | T108T | A | 3.3e-08 | 5.8e-08 | 9.6e-08 | 6.233E-08 | 3.172E-08 | 3.300E-08 | 9.600E-08 |
| 2158 | 108 | T108W | A | 5.4e-08 | 6.2e-08 | 4.9e-08 | 5.500E-08 | 6.557E-09 | 4.900E-08 | 6.200E-08 |
| 2159 | 108 | T108Y | A | 2e-08 | 2.7e-08 | 5.7e-08 | 3.467E-08 | 1.966E-08 | 2.000E-08 | 5.700E-08 |
| 2160 | 108 | T108V | A | 2.2e-08 | 5e-08 | 4.3e-08 | 3.833E-08 | 1.457E-08 | 2.200E-08 | 5.000E-08 |
| 2161 | 109 | G109A | A | 1.8e-08 | 3.4e-08 | 5.1e-08 | 3.433E-08 | 1.650E-08 | 1.800E-08 | 5.100E-08 |
| 2162 | 109 | G109R | A | 4.7e-08 | 4.7e-08 | 2.1e-08 | 3.833E-08 | 1.501E-08 | 2.100E-08 | 4.700E-08 |
| 2163 | 109 | G109N | A | 2.3e-08 | 6.7e-08 | 2.1e-08 | 3.700E-08 | 2.600E-08 | 2.100E-08 | 6.700E-08 |
| 2164 | 109 | G109D | A | 2.5e-08 | 8.7e-08 | 3.4e-08 | 4.867E-08 | 3.350E-08 | 2.500E-08 | 8.700E-08 |
| 2165 | 109 | G109C | A | 5.2e-08 | 6.6e-08 | 2.1e-07 | 1.093E-07 | 8.746E-08 | 5.200E-08 | 2.100E-07 |
| 2166 | 109 | G109E | A | 3.2e-08 | 7.1e-08 | 3.1e-08 | 4.467E-08 | 2.281E-08 | 3.100E-08 | 7.100E-08 |
| 2167 | 109 | G109Q | A | 4.5e-08 | 8e-09 | 3.8e-08 | 3.033E-08 | 1.966E-08 | 8.000E-09 | 4.500E-08 |
| 2168 | 109 | G109G | A | 4.4e-08 | 3.1e-08 | 4.7e-08 | 4.067E-08 | 8.505E-09 | 3.100E-08 | 4.700E-08 |
| 2169 | 109 | G109H | A | 8.6e-08 | 2.5e-08 | 1e-08 | 4.033E-08 | 4.025E-08 | 1.000E-08 | 8.600E-08 |
| 2170 | 109 | G109I | A | 5e-08 | 2e-08 | 3.4e-08 | 3.467E-08 | 1.501E-08 | 2.000E-08 | 5.000E-08 |
| 2171 | 109 | G109L | A | 1.8e-08 | 1.3e-08 | 5.5e-08 | 2.867E-08 | 2.294E-08 | 1.300E-08 | 5.500E-08 |
| 2172 | 109 | G109K | A | 6.2e-08 | 2.4e-08 | 6.5e-09 | 3.083E-08 | 2.837E-08 | 6.500E-09 | 6.200E-08 |
| 2173 | 109 | G109M | A | 7.7e-08 | 2.9e-08 | 4e-08 | 4.867E-08 | 2.515E-08 | 2.900E-08 | 7.700E-08 |
| 2174 | 109 | G109F | A | 4.4e-08 | 7.8e-08 | 8e-08 | 6.733E-08 | 2.023E-08 | 4.400E-08 | 8.000E-08 |
| 2175 | 109 | G109P | A | 1.8e-07 | 3.4e-08 | 3.9e-08 | 8.433E-08 | 8.289E-08 | 3.400E-08 | 1.800E-07 |
| 2176 | 109 | G109S | A | 6.1e-08 | 7.3e-08 | 2.9e-08 | 5.433E-08 | 2.274E-08 | 2.900E-08 | 7.300E-08 |
| 2177 | 109 | G109T | A | 7.2e-08 | 2e-08 | 2.2e-08 | 3.800E-08 | 2.946E-08 | 2.000E-08 | 7.200E-08 |
| 2178 | 109 | G109W | A | 4.6e-08 | 1e-07 | 6.9e-08 | 7.167E-08 | 2.710E-08 | 4.600E-08 | 1.000E-07 |
| 2179 | 109 | G109Y | A | 6.4e-08 | 3.4e-08 | 3.6e-08 | 4.467E-08 | 1.677E-08 | 3.400E-08 | 6.400E-08 |
| 2180 | 109 | G109V | A | 3.8e-08 | 3.3e-08 | 5.4e-08 | 4.167E-08 | 1.097E-08 | 3.300E-08 | 5.400E-08 |
| 2181 | 110 | A110A | A | 7.1e-08 | 2.7e-08 | 1.7e-08 | 3.833E-08 | 2.873E-08 | 1.700E-08 | 7.100E-08 |
| 2182 | 110 | A110R | A | 4.1e-08 | 5.8e-08 | 6e-08 | 5.300E-08 | 1.044E-08 | 4.100E-08 | 6.000E-08 |
| 2183 | 110 | A110N | A | 2.5e-08 | 3.6e-08 | 2.4e-08 | 2.833E-08 | 6.658E-09 | 2.400E-08 | 3.600E-08 |
| 2184 | 110 | A110D | A | 6.6e-08 | 1.8e-07 | 7.7e-08 | 1.077E-07 | 6.288E-08 | 6.600E-08 | 1.800E-07 |
| 2185 | 110 | A110C | A | 1.5e-08 | 8.1e-08 | 4.9e-08 | 4.833E-08 | 3.301E-08 | 1.500E-08 | 8.100E-08 |
| 2186 | 110 | A110E | A | 6.8e-08 | 4.8e-08 | 3.3e-08 | 4.967E-08 | 1.756E-08 | 3.300E-08 | 6.800E-08 |
| 2187 | 110 | A110Q | A | 1.8e-08 | 5.5e-08 | 2.9e-08 | 3.400E-08 | 1.900E-08 | 1.800E-08 | 5.500E-08 |
| 2188 | 110 | A110G | A | 3.8e-08 | 2.5e-08 | 5.2e-08 | 3.833E-08 | 1.350E-08 | 2.500E-08 | 5.200E-08 |
| 2189 | 110 | A110H | A | 6.5e-08 | 3.3e-08 | 4.3e-08 | 4.700E-08 | 1.637E-08 | 3.300E-08 | 6.500E-08 |
| 2190 | 110 | A110I | A | 6.8e-08 | 5.8e-08 | 3.3e-08 | 5.300E-08 | 1.803E-08 | 3.300E-08 | 6.800E-08 |
| 2191 | 110 | A110L | A | 8.3e-08 | 2e-08 | 5.8e-08 | 5.367E-08 | 3.172E-08 | 2.000E-08 | 8.300E-08 |
| 2192 | 110 | A110K | A | 1e-08 | 7.5e-08 | 5.9e-09 | 3.030E-08 | 3.877E-08 | 5.900E-09 | 7.500E-08 |
| 2193 | 110 | A110M | A | 4.5e-08 | 3.1e-08 | 2e-08 | 3.200E-08 | 1.253E-08 | 2.000E-08 | 4.500E-08 |
| 2194 | 110 | A110F | A | 5.2e-08 | 7e-08 | 6.3e-08 | 6.167E-08 | 9.074E-09 | 5.200E-08 | 7.000E-08 |
| 2195 | 110 | A110P | A | 6.1e-08 | 4.7e-08 | 3.2e-08 | 4.667E-08 | 1.450E-08 | 3.200E-08 | 6.100E-08 |
| 2196 | 110 | A110S | A | 9.2e-08 | 8.3e-08 | 4.3e-08 | 7.267E-08 | 2.608E-08 | 4.300E-08 | 9.200E-08 |
| 2197 | 110 | A110T | A | 3.4e-08 | 1.6e-08 | 4.2e-08 | 3.067E-08 | 1.332E-08 | 1.600E-08 | 4.200E-08 |
| 2198 | 110 | A110W | A | 1.9e-08 | 1.2e-08 | 1e-07 | 4.367E-08 | 4.891E-08 | 1.200E-08 | 1.000E-07 |
| 2199 | 110 | A110Y | A | 3.6e-08 | 1e-08 | 5.2e-08 | 3.267E-08 | 2.120E-08 | 1.000E-08 | 5.200E-08 |
| 2200 | 110 | A110V | A | 7.1e-08 | 1.1e-07 | 8.8e-08 | 8.967E-08 | 1.955E-08 | 7.100E-08 | 1.100E-07 |
| 2201 | 111 | S111A | A | 2.6e-08 | 3.2e-08 | 6.3e-08 | 4.033E-08 | 1.986E-08 | 2.600E-08 | 6.300E-08 |
| 2202 | 111 | S111R | A | 7.1e-08 | 1.2e-07 | 2e-07 | 1.303E-07 | 6.512E-08 | 7.100E-08 | 2.000E-07 |
| 2203 | 111 | S111N | A | 3.5e-08 | 8.5e-08 | 2.5e-08 | 4.833E-08 | 3.215E-08 | 2.500E-08 | 8.500E-08 |
| 2204 | 111 | S111D | A | 2.9e-08 | 9.4e-08 | 6.6e-08 | 6.300E-08 | 3.260E-08 | 2.900E-08 | 9.400E-08 |
| 2205 | 111 | S111C | A | 6.1e-08 | 2.6e-08 | 9.4e-08 | 6.033E-08 | 3.400E-08 | 2.600E-08 | 9.400E-08 |
| 2206 | 111 | S111E | A | 4e-08 | 5.1e-08 | 8.6e-08 | 5.900E-08 | 2.402E-08 | 4.000E-08 | 8.600E-08 |
| 2207 | 111 | S111Q | A | 1.7e-08 | 2.1e-08 | 2.3e-08 | 2.033E-08 | 3.055E-09 | 1.700E-08 | 2.300E-08 |
| 2208 | 111 | S111G | A | 5.5e-08 | 5.9e-08 | 1.1e-07 | 7.467E-08 | 3.066E-08 | 5.500E-08 | 1.100E-07 |
| 2209 | 111 | S111H | A | 4.4e-08 | 6.1e-08 | 4.5e-08 | 5.000E-08 | 9.539E-09 | 4.400E-08 | 6.100E-08 |

| | | | | | | | | | | |
|---|---|---|---|---|---|---|---|---|---|---|
| 2210 | 111 | S111I | A | 4.1e-08 | 4.8e-08 | 6.9e-08 | 5.267E-08 | 1.457E-08 | 4.100E-08 | 6.900E-08 |
| 2211 | 111 | S111L | A | 5.9e-08 | 3.7e-08 | 2.8e-08 | 4.133E-08 | 1.595E-08 | 2.800E-08 | 5.900E-08 |
| 2212 | 111 | S111K | A | 2.5e-08 | 4.6e-08 | 3.3e-08 | 3.467E-08 | 1.060E-08 | 2.500E-08 | 4.600E-08 |
| 2213 | 111 | S111M | A | 1.2e-08 | 1.4e-08 | 5.7e-08 | 2.767E-08 | 2.542E-08 | 1.200E-08 | 5.700E-08 |
| 2214 | 111 | S111F | A | 2.4e-08 | 2.5e-08 | 5.6e-08 | 3.500E-08 | 1.819E-08 | 2.400E-08 | 5.600E-08 |
| 2215 | 111 | S111P | A | 6.5e-08 | 4.1e-08 | 5.5e-08 | 5.367E-08 | 1.206E-08 | 4.100E-08 | 6.500E-08 |
| 2216 | 111 | S111S | A | 4.1e-08 | 3.1e-08 | 5.9e-08 | 4.367E-08 | 1.419E-08 | 3.100E-08 | 5.900E-08 |
| 2217 | 111 | S111T | A | 5.5e-08 | 4.3e-08 | 2.4e-08 | 4.067E-08 | 1.563E-08 | 2.400E-08 | 5.500E-08 |
| 2218 | 111 | S111W | A | 5.8e-08 | 2.2e-08 | 8.7e-08 | 5.567E-08 | 3.256E-08 | 2.200E-08 | 8.700E-08 |
| 2219 | 111 | S111Y | A | 6e-08 | 3.6e-08 | 4e-08 | 4.533E-08 | 1.286E-08 | 3.600E-08 | 6.000E-08 |
| 2220 | 111 | S111V | A | 3.8e-08 | 5.8e-08 | 7.5e-08 | 5.700E-08 | 1.852E-08 | 3.800E-08 | 7.500E-08 |
| 2221 | 112 | P112A | A | 5.1e-08 | 6.4e-08 | 2.7e-08 | 4.733E-08 | 1.877E-08 | 2.700E-08 | 6.400E-08 |
| 2222 | 112 | P112R | A | 4.9e-08 | 9.7e-09 | 3.1e-08 | 2.990E-08 | 1.967E-08 | 9.700E-09 | 4.900E-08 |
| 2223 | 112 | P112N | A | 3.9e-08 | 2.6e-08 | 2e-08 | 2.833E-08 | 9.713E-09 | 2.000E-08 | 3.900E-08 |
| 2224 | 112 | P112D | A | 7.6e-08 | 5.6e-08 | 1.2e-07 | 8.400E-08 | 3.274E-08 | 5.600E-08 | 1.200E-07 |
| 2225 | 112 | P112C | A | 4.6e-08 | 2.3e-08 | 5e-08 | 3.967E-08 | 1.457E-08 | 2.300E-08 | 5.000E-08 |
| 2226 | 112 | P112E | A | 9.3e-08 | 2.9e-08 | 1.3e-08 | 4.500E-08 | 4.233E-08 | 1.300E-08 | 9.300E-08 |
| 2227 | 112 | P112Q | A | 2.6e-08 | 3.4e-08 | 3.3e-08 | 3.100E-08 | 4.359E-09 | 2.600E-08 | 3.400E-08 |
| 2228 | 112 | P112G | A | 2.8e-08 | 5.1e-08 | 5.9e-08 | 4.600E-08 | 1.609E-08 | 2.800E-08 | 5.900E-08 |
| 2229 | 112 | P112H | A | 2.9e-08 | 9.6e-08 | 1.5e-07 | 9.167E-08 | 6.062E-08 | 2.900E-08 | 1.500E-07 |
| 2230 | 112 | P112I | A | 7e-08 | 2.9e-08 | 3.6e-08 | 4.500E-08 | 2.193E-08 | 2.900E-08 | 7.000E-08 |
| 2231 | 112 | P112L | A | 7.1e-08 | 1e-07 | 3.8e-08 | 6.967E-08 | 3.102E-08 | 3.800E-08 | 1.000E-07 |
| 2232 | 112 | P112K | A | 7.4e-08 | 5.2e-08 | 9.3e-08 | 7.300E-08 | 2.052E-08 | 5.200E-08 | 9.300E-08 |
| 2233 | 112 | P112M | A | 3.8e-08 | 5.6e-08 | 1.3e-07 | 7.467E-08 | 4.876E-08 | 3.800E-08 | 1.300E-07 |
| 2234 | 112 | P112F | A | 6.5e-08 | 2.5e-08 | 3.4e-08 | 4.133E-08 | 2.098E-08 | 2.500E-08 | 6.500E-08 |
| 2235 | 112 | P112P | A | 4.6e-08 | 8.8e-08 | 6.3e-08 | 6.567E-08 | 2.113E-08 | 4.600E-08 | 8.800E-08 |
| 2236 | 112 | P112S | A | 3e-08 | 7.2e-08 | 2.5e-08 | 4.233E-08 | 2.581E-08 | 2.500E-08 | 7.200E-08 |
| 2237 | 112 | P112T | A | 7.8e-08 | 3.6e-08 | 6.5e-08 | 5.967E-08 | 2.150E-08 | 3.600E-08 | 7.800E-08 |
| 2238 | 112 | P112W | A | 5.7e-08 | 2.9e-08 | 4.9e-08 | 4.500E-08 | 1.442E-08 | 2.900E-08 | 5.700E-08 |
| 2239 | 112 | P112Y | A | 1.1e-07 | 3.1e-08 | 1.8e-08 | 5.300E-08 | 4.979E-08 | 1.800E-08 | 1.100E-07 |
| 2240 | 112 | P112V | A | 1.4e-08 | 5.7e-08 | 4.2e-08 | 3.767E-08 | 2.183E-08 | 1.400E-08 | 5.700E-08 |
| 2241 | 113 | G113A | A | 7.7e-08 | 3.7e-08 | 3.9e-08 | 5.100E-08 | 2.254E-08 | 3.700E-08 | 7.700E-08 |
| 2242 | 113 | G113R | A | 5.2e-08 | 2e-08 | 4.5e-08 | 3.900E-08 | 1.682E-08 | 2.000E-08 | 5.200E-08 |
| 2243 | 113 | G113N | A | 3.5e-08 | 1.7e-08 | 2.9e-08 | 2.700E-08 | 9.165E-09 | 1.700E-08 | 3.500E-08 |
| 2244 | 113 | G113D | A | 2.6e-08 | 2.9e-08 | 4.7e-08 | 3.400E-08 | 1.136E-08 | 2.600E-08 | 4.700E-08 |
| 2245 | 113 | G113C | A | 5.7e-08 | 2.9e-08 | 1.2e-07 | 6.867E-08 | 4.661E-08 | 2.900E-08 | 1.200E-07 |
| 2246 | 113 | G113E | A | 2.5e-08 | 2.2e-08 | 5.6e-08 | 3.433E-08 | 1.882E-08 | 2.200E-08 | 5.600E-08 |
| 2247 | 113 | G113Q | A | 7.2e-08 | 7.9e-08 | 6.7e-08 | 7.267E-08 | 6.028E-09 | 6.700E-08 | 7.900E-08 |
| 2248 | 113 | G113G | A | 3.4e-08 | 1.9e-08 | 8e-08 | 4.433E-08 | 3.179E-08 | 1.900E-08 | 8.000E-08 |
| 2249 | 113 | G113H | A | 4.9e-08 | 5.1e-08 | 5.8e-08 | 5.267E-08 | 4.726E-09 | 4.900E-08 | 5.800E-08 |
| 2250 | 113 | G113I | A | 2.8e-08 | 3.6e-08 | 3.9e-08 | 3.433E-08 | 5.686E-09 | 2.800E-08 | 3.900E-08 |
| 2251 | 113 | G113L | A | 2.5e-08 | 2.7e-08 | 3.5e-08 | 2.900E-08 | 5.292E-09 | 2.500E-08 | 3.500E-08 |
| 2252 | 113 | G113K | A | 1.7e-08 | 2.9e-08 | 2.5e-08 | 2.367E-08 | 6.110E-09 | 1.700E-08 | 2.900E-08 |
| 2253 | 113 | G113M | A | 3.7e-08 | 5.9e-08 | 7e-08 | 5.533E-08 | 1.680E-08 | 3.700E-08 | 7.000E-08 |
| 2254 | 113 | G113F | A | 6.4e-08 | 1.5e-07 | 6.3e-08 | 9.233E-08 | 4.994E-08 | 6.300E-08 | 1.500E-07 |
| 2255 | 113 | G113P | A | 3.7e-08 | 1.5e-08 | 2.5e-08 | 2.567E-08 | 1.102E-08 | 1.500E-08 | 3.700E-08 |
| 2256 | 113 | G113S | A | 1.7e-08 | 5.9e-08 | 3.1e-08 | 3.567E-08 | 2.139E-08 | 1.700E-08 | 5.900E-08 |
| 2257 | 113 | G113T | A | 2.3e-08 | 3.6e-08 | 6.2e-08 | 4.033E-08 | 1.986E-08 | 2.300E-08 | 6.200E-08 |
| 2258 | 113 | G113W | A | 5.8e-08 | 2e-08 | 2.8e-08 | 3.533E-08 | 2.003E-08 | 2.000E-08 | 5.800E-08 |
| 2259 | 113 | G113Y | A | 7e-08 | 5.5e-08 | 1.7e-08 | 4.733E-08 | 2.732E-08 | 1.700E-08 | 7.000E-08 |
| 2260 | 113 | G113V | A | 3.2e-08 | 5.4e-08 | 3.3e-08 | 3.967E-08 | 1.242E-08 | 3.200E-08 | 5.400E-08 |
| 2261 | 114 | G114A | A | 4.4e-08 | 4.9e-08 | 4.4e-08 | 4.567E-08 | 2.887E-09 | 4.400E-08 | 4.900E-08 |
| 2262 | 114 | G114R | A | 3.6e-08 | 4.5e-08 | 1.3e-07 | 7.033E-08 | 5.187E-08 | 3.600E-08 | 1.300E-07 |
| 2263 | 114 | G114N | A | 5.4e-08 | 2.4e-08 | 1e-07 | 5.933E-08 | 3.828E-08 | 2.400E-08 | 1.000E-07 |
| 2264 | 114 | G114D | A | 4.6e-08 | 2.7e-08 | 7.9e-08 | 5.067E-08 | 2.631E-08 | 2.700E-08 | 7.900E-08 |
| 2265 | 114 | G114C | A | 3.4e-08 | 3.3e-08 | 5.2e-08 | 3.967E-08 | 1.069E-08 | 3.300E-08 | 5.200E-08 |
| 2266 | 114 | G114E | A | 6.3e-08 | 2.1e-08 | 3.3e-08 | 3.900E-08 | 2.163E-08 | 2.100E-08 | 6.300E-08 |
| 2267 | 114 | G114Q | A | 7.3e-08 | 7.4e-08 | 1.7e-08 | 5.467E-08 | 3.262E-08 | 1.700E-08 | 7.400E-08 |
| 2268 | 114 | G114G | A | 4.2e-08 | 1.5e-08 | 4.6e-08 | 3.433E-08 | 1.686E-08 | 1.500E-08 | 4.600E-08 |
| 2269 | 114 | G114H | A | 2.1e-08 | 4.1e-08 | 2.4e-08 | 2.867E-08 | 1.079E-08 | 2.100E-08 | 4.100E-08 |
| 2270 | 114 | G114I | A | 7.3e-08 | 5.3e-08 | 3.7e-08 | 5.433E-08 | 1.804E-08 | 3.700E-08 | 7.300E-08 |
| 2271 | 114 | G114L | A | 3.9e-08 | 1e-07 | 3.4e-08 | 5.767E-08 | 3.675E-08 | 3.400E-08 | 1.000E-07 |
| 2272 | 114 | G114K | A | 1.6e-08 | 4.1e-08 | 3.6e-08 | 3.100E-08 | 1.323E-08 | 1.600E-08 | 4.100E-08 |
| 2273 | 114 | G114M | A | 1.9e-08 | 7.5e-08 | 6e-08 | 5.133E-08 | 2.899E-08 | 1.900E-08 | 7.500E-08 |
| 2274 | 114 | G114F | A | 5.7e-08 | 2.4e-08 | 8.7e-08 | 5.600E-08 | 3.151E-08 | 2.400E-08 | 8.700E-08 |
| 2275 | 114 | G114P | A | 4.2e-08 | 6.5e-08 | 6.8e-08 | 5.833E-08 | 1.422E-08 | 4.200E-08 | 6.800E-08 |
| 2276 | 114 | G114S | A | 3.6e-08 | 4.3e-08 | 2e-08 | 3.300E-08 | 1.179E-08 | 2.000E-08 | 4.300E-08 |
| 2277 | 114 | G114T | A | 1.8e-07 | 1.8e-08 | 4.4e-08 | 8.067E-08 | 8.700E-08 | 1.800E-08 | 1.800E-07 |
| 2278 | 114 | G114W | A | 1.3e-08 | 9e-09 | 8.2e-08 | 3.467E-08 | 4.104E-08 | 9.000E-09 | 8.200E-08 |
| 2279 | 114 | G114Y | A | 4.4e-08 | 2.6e-08 | 4.4e-08 | 3.800E-08 | 1.039E-08 | 2.600E-08 | 4.400E-08 |
| 2280 | 114 | G114V | A | 1.2e-08 | 4.9e-08 | 3.4e-08 | 3.167E-08 | 1.861E-08 | 1.200E-08 | 4.900E-08 |
| 2281 | 115 | L115A | A | 8.1e-08 | 3.7e-08 | 4.8e-08 | 5.533E-08 | 2.290E-08 | 3.700E-08 | 8.100E-08 |
| 2282 | 115 | L115R | A | 8.1e-08 | 2.2e-08 | 4.9e-08 | 5.067E-08 | 2.954E-08 | 2.200E-08 | 8.100E-08 |
| 2283 | 115 | L115N | A | 4.9e-08 | 8.3e-08 | 5.2e-08 | 6.133E-08 | 1.882E-08 | 4.900E-08 | 8.300E-08 |
| 2284 | 115 | L115D | A | 8.5e-08 | 1.3e-07 | 3.8e-08 | 8.433E-08 | 4.600E-08 | 3.800E-08 | 1.300E-07 |
| 2285 | 115 | L115C | A | 2.5e-08 | 7e-08 | 3.4e-08 | 4.300E-08 | 2.381E-08 | 2.500E-08 | 7.000E-08 |
| 2286 | 115 | L115E | A | 4.2e-08 | 4.9e-08 | 6.8e-08 | 5.300E-08 | 1.345E-08 | 4.200E-08 | 6.800E-08 |
| 2287 | 115 | L115Q | A | 5.9e-08 | 4.8e-08 | 3e-08 | 4.567E-08 | 1.464E-08 | 3.000E-08 | 5.900E-08 |
| 2288 | 115 | L115G | A | 5.9e-08 | 2.6e-08 | 4.7e-08 | 4.400E-08 | 1.670E-08 | 2.600E-08 | 5.900E-08 |
| 2289 | 115 | L115H | A | 5.5e-08 | 4.9e-08 | 1.2e-08 | 3.867E-08 | 2.329E-08 | 1.200E-08 | 5.500E-08 |
| 2290 | 115 | L115I | A | 3.2e-08 | 5.1e-08 | 3.1e-08 | 3.800E-08 | 1.127E-08 | 3.100E-08 | 5.100E-08 |
| 2291 | 115 | L115L | A | 3.9e-08 | 3.7e-08 | 1.7e-08 | 3.100E-08 | 1.217E-08 | 1.700E-08 | 3.900E-08 |
| 2292 | 115 | L115K | A | 5.8e-08 | 4.1e-08 | 1.8e-08 | 3.900E-08 | 2.007E-08 | 1.800E-08 | 5.800E-08 |
| 2293 | 115 | L115M | A | 2e-08 | 6.1e-08 | 7.5e-08 | 5.200E-08 | 2.858E-08 | 2.000E-08 | 7.500E-08 |
| 2294 | 115 | L115F | A | 4.1e-08 | 4.3e-08 | 3.2e-08 | 3.867E-08 | 5.859E-09 | 3.200E-08 | 4.300E-08 |

| | | | | | | | | | | |
|---|---|---|---|---|---|---|---|---|---|---|
| 2295 | 115 | L115P | A | 3.2e-08 | 3.8e-08 | 2.7e-08 | 3.233E-08 | 5.508E-09 | 2.700E-08 | 3.800E-08 |
| 2296 | 115 | L115S | A | 7.2e-08 | 1.1e-07 | 5.9e-08 | 8.033E-08 | 2.650E-08 | 5.900E-08 | 1.100E-07 |
| 2297 | 115 | L115T | A | 2.4e-08 | 6.9e-08 | 1.4e-07 | 7.767E-08 | 5.848E-08 | 2.400E-08 | 1.400E-07 |
| 2298 | 115 | L115W | A | 2.2e-08 | 2.3e-08 | 3.8e-08 | 2.767E-08 | 8.963E-09 | 2.200E-08 | 3.800E-08 |
| 2299 | 115 | L115Y | A | 3.3e-08 | 3.9e-08 | 7.6e-08 | 4.933E-08 | 2.329E-08 | 3.300E-08 | 7.600E-08 |
| 2300 | 115 | L115V | A | 4.1e-08 | 5.2e-08 | 3.1e-08 | 4.133E-08 | 1.050E-08 | 3.100E-08 | 5.200E-08 |
| 2301 | 116 | R116A | A | 2.9e-08 | 3.2e-08 | 5.5e-08 | 3.867E-08 | 1.422E-08 | 2.900E-08 | 5.500E-08 |
| 2302 | 116 | R116R | A | 2.5e-08 | 1.1e-07 | 4.8e-08 | 6.100E-08 | 4.397E-08 | 2.500E-08 | 1.100E-07 |
| 2303 | 116 | R116N | A | 8.1e-08 | 4.7e-08 | 2.8e-08 | 5.200E-08 | 2.685E-08 | 2.800E-08 | 8.100E-08 |
| 2304 | 116 | R116D | A | 6.7e-08 | 2.6e-08 | 5.5e-08 | 4.933E-08 | 2.108E-08 | 2.600E-08 | 6.700E-08 |
| 2305 | 116 | R116C | A | 1e-07 | 3.8e-08 | 4.7e-08 | 6.167E-08 | 3.350E-08 | 3.800E-08 | 1.000E-07 |
| 2306 | 116 | R116E | A | 3.5e-08 | 6.7e-08 | 3.7e-08 | 4.633E-08 | 1.793E-08 | 3.500E-08 | 6.700E-08 |
| 2307 | 116 | R116Q | A | 7e-08 | 9.5e-08 | 1.2e-07 | 9.500E-08 | 2.500E-08 | 7.000E-08 | 1.200E-07 |
| 2308 | 116 | R116G | A | 1.8e-08 | 7e-09 | 5.1e-08 | 2.533E-08 | 2.290E-08 | 7.000E-09 | 5.100E-08 |
| 2309 | 116 | R116H | A | 3e-08 | 5.9e-08 | 4.6e-08 | 4.500E-08 | 1.453E-08 | 3.000E-08 | 5.900E-08 |
| 2310 | 116 | R116I | A | 1.8e-08 | 7.2e-08 | 2.7e-08 | 3.900E-08 | 2.893E-08 | 1.800E-08 | 7.200E-08 |
| 2311 | 116 | R116L | A | 1e-07 | 4.1e-08 | 3.1e-08 | 5.733E-08 | 3.729E-08 | 3.100E-08 | 1.000E-07 |
| 2312 | 116 | R116K | A | 1.6e-08 | 5.8e-08 | 1.6e-07 | 7.800E-08 | 7.405E-08 | 1.600E-08 | 1.600E-07 |
| 2313 | 116 | R116M | A | 1.4e-08 | 3.1e-08 | 1.2e-07 | 5.500E-08 | 5.693E-08 | 1.400E-08 | 1.200E-07 |
| 2314 | 116 | R116F | A | 1.4e-08 | 1.6e-08 | 4.3e-08 | 2.433E-08 | 1.620E-08 | 1.400E-08 | 4.300E-08 |
| 2315 | 116 | R116P | A | 2.4e-08 | 3.1e-08 | 5.9e-08 | 3.800E-08 | 1.852E-08 | 2.400E-08 | 5.900E-08 |
| 2316 | 116 | R116S | A | 1.2e-07 | 6.5e-08 | 8.7e-08 | 9.067E-08 | 2.768E-08 | 6.500E-08 | 1.200E-07 |
| 2317 | 116 | R116T | A | 2.3e-08 | 2.8e-08 | 7.8e-08 | 4.300E-08 | 3.041E-08 | 2.300E-08 | 7.800E-08 |
| 2318 | 116 | R116W | A | 1.2e-07 | 2e-08 | 6.4e-08 | 6.800E-08 | 5.012E-08 | 2.000E-08 | 1.200E-07 |
| 2319 | 116 | R116Y | A | 2.7e-08 | 9.7e-09 | 4.3e-08 | 2.657E-08 | 1.665E-08 | 9.700E-09 | 4.300E-08 |
| 2320 | 116 | R116V | A | 6e-08 | 2.7e-08 | 6.8e-08 | 5.167E-08 | 2.173E-08 | 2.700E-08 | 6.800E-08 |
| 2321 | 117 | E117A | A | 5.5e-08 | 5.5e-08 | 5.6e-08 | 5.533E-08 | 5.774E-10 | 5.500E-08 | 5.600E-08 |
| 2322 | 117 | E117R | A | 5.1e-08 | 3.9e-08 | 6.1e-08 | 5.033E-08 | 1.102E-08 | 3.900E-08 | 6.100E-08 |
| 2323 | 117 | E117N | A | 9.6e-08 | 8.3e-08 | 6.1e-08 | 8.000E-08 | 1.769E-08 | 6.100E-08 | 9.600E-08 |
| 2324 | 117 | E117D | A | 3.7e-08 | 1.1e-07 | 9.9e-08 | 8.200E-08 | 3.936E-08 | 3.700E-08 | 1.100E-07 |
| 2325 | 117 | E117C | A | 6.8e-08 | 1.2e-08 | 9.2e-08 | 5.733E-08 | 4.105E-08 | 1.200E-08 | 9.200E-08 |
| 2326 | 117 | E117E | A | 7.8e-08 | 9.6e-08 | 5.3e-08 | 7.567E-08 | 2.159E-08 | 5.300E-08 | 9.600E-08 |
| 2327 | 117 | E117Q | A | 9.8e-08 | 2.7e-08 | 7.1e-08 | 6.533E-08 | 3.584E-08 | 2.700E-08 | 9.800E-08 |
| 2328 | 117 | E117G | A | 1.7e-08 | 5.5e-08 | 5.4e-08 | 4.200E-08 | 2.166E-08 | 1.700E-08 | 5.500E-08 |
| 2329 | 117 | E117H | A | 8.3e-08 | 4.1e-08 | 6.2e-08 | 6.200E-08 | 2.100E-08 | 4.100E-08 | 8.300E-08 |
| 2330 | 117 | E117I | A | 4.9e-08 | 4.5e-08 | 5e-08 | 4.800E-08 | 2.646E-09 | 4.500E-08 | 5.000E-08 |
| 2331 | 117 | E117L | A | 2.4e-08 | 3.5e-08 | 3.1e-08 | 3.000E-08 | 5.568E-09 | 2.400E-08 | 3.500E-08 |
| 2332 | 117 | E117K | A | 2.2e-08 | 5.8e-08 | 3.2e-08 | 3.733E-08 | 1.858E-08 | 2.200E-08 | 5.800E-08 |
| 2333 | 117 | E117M | A | 5e-08 | 3.3e-08 | 1.3e-07 | 7.100E-08 | 5.180E-08 | 3.300E-08 | 1.300E-07 |
| 2334 | 117 | E117F | A | 4.4e-08 | 9.4e-08 | 4.4e-08 | 6.067E-08 | 2.887E-08 | 4.400E-08 | 9.400E-08 |
| 2335 | 117 | E117P | A | 2.6e-08 | 2.3e-08 | 7.3e-08 | 4.067E-08 | 2.804E-08 | 2.300E-08 | 7.300E-08 |
| 2336 | 117 | E117S | A | 6e-08 | 8.1e-08 | 9.5e-08 | 7.867E-08 | 1.762E-08 | 6.000E-08 | 9.500E-08 |
| 2337 | 117 | E117T | A | 4.7e-08 | 8e-08 | 1.8e-08 | 4.833E-08 | 3.102E-08 | 1.800E-08 | 8.000E-08 |
| 2338 | 117 | E117W | A | 4.3e-08 | 7.9e-08 | 7.6e-08 | 6.600E-08 | 1.997E-08 | 4.300E-08 | 7.900E-08 |
| 2339 | 117 | E117Y | A | 3.2e-08 | 2.8e-08 | 3.5e-08 | 3.167E-08 | 3.512E-09 | 2.800E-08 | 3.500E-08 |
| 2340 | 117 | E117V | A | 3e-08 | 2.8e-08 | 6.3e-08 | 4.033E-08 | 1.966E-08 | 2.800E-08 | 6.300E-08 |
| 2341 | 118 | L118A | A | 9.8e-08 | 7.8e-08 | 2e-08 | 6.533E-08 | 4.051E-08 | 2.000E-08 | 9.800E-08 |
| 2342 | 118 | L118R | A | 2.7e-08 | 4.4e-08 | 1.4e-07 | 7.033E-08 | 6.093E-08 | 2.700E-08 | 1.400E-07 |
| 2343 | 118 | L118N | A | 1.2e-08 | 8.3e-08 | 1.5e-08 | 3.667E-08 | 4.015E-08 | 1.200E-08 | 8.300E-08 |
| 2344 | 118 | L118D | A | 4.2e-08 | 6e-08 | 4.8e-08 | 5.000E-08 | 9.165E-09 | 4.200E-08 | 6.000E-08 |
| 2345 | 118 | L118C | A | 3.9e-08 | 4e-08 | 5.2e-08 | 4.367E-08 | 7.234E-09 | 3.900E-08 | 5.200E-08 |
| 2346 | 118 | L118E | A | 6.4e-08 | 5.5e-08 | 6.2e-08 | 6.033E-08 | 4.726E-09 | 5.500E-08 | 6.400E-08 |
| 2347 | 118 | L118Q | A | 4.9e-08 | 6.6e-08 | 2e-08 | 4.500E-08 | 2.326E-08 | 2.000E-08 | 6.600E-08 |
| 2348 | 118 | L118G | A | 4.2e-08 | 1.4e-08 | 4.6e-08 | 3.400E-08 | 1.744E-08 | 1.400E-08 | 4.600E-08 |
| 2349 | 118 | L118H | A | 3.1e-08 | 1e-08 | 7.8e-08 | 3.967E-08 | 3.482E-08 | 1.000E-08 | 7.800E-08 |
| 2350 | 118 | L118I | A | 3e-08 | 5e-08 | 9.9e-09 | 2.997E-08 | 2.005E-08 | 9.900E-09 | 5.000E-08 |
| 2351 | 118 | L118L | A | 2.7e-08 | 1e-07 | 2.8e-08 | 5.167E-08 | 4.186E-08 | 2.700E-08 | 1.000E-07 |
| 2352 | 118 | L118K | A | 3.1e-08 | 2.8e-08 | 3.6e-08 | 3.167E-08 | 4.041E-09 | 2.800E-08 | 3.600E-08 |
| 2353 | 118 | L118M | A | 9.1e-08 | 6.4e-08 | 1.2e-08 | 5.567E-08 | 4.015E-08 | 1.200E-08 | 9.100E-08 |
| 2354 | 118 | L118F | A | 1.3e-07 | 1.7e-08 | 4e-08 | 6.233E-08 | 5.972E-08 | 1.700E-08 | 1.300E-07 |
| 2355 | 118 | L118P | A | 1.5e-07 | 5.8e-08 | 2.4e-08 | 7.733E-08 | 6.519E-08 | 2.400E-08 | 1.500E-07 |
| 2356 | 118 | L118S | A | 1.1e-07 | 1.8e-08 | 9e-08 | 7.267E-08 | 4.839E-08 | 1.800E-08 | 1.100E-07 |
| 2357 | 118 | L118T | A | 8.9e-08 | 3.2e-08 | 4.5e-08 | 5.533E-08 | 2.987E-08 | 3.200E-08 | 8.900E-08 |
| 2358 | 118 | L118W | A | 2.5e-08 | 8e-08 | 3.3e-08 | 4.600E-08 | 2.972E-08 | 2.500E-08 | 8.000E-08 |
| 2359 | 118 | L118Y | A | 4.2e-08 | 4.9e-08 | 2.4e-08 | 3.833E-08 | 1.290E-08 | 2.400E-08 | 4.900E-08 |
| 2360 | 118 | L118V | A | 6.9e-08 | 2.5e-08 | 1.3e-07 | 7.467E-08 | 5.273E-08 | 2.500E-08 | 1.300E-07 |
| 2361 | 119 | Q119A | A | 4.5e-08 | 5.5e-08 | 1.7e-08 | 3.900E-08 | 1.970E-08 | 1.700E-08 | 5.500E-08 |
| 2362 | 119 | Q119R | A | 4e-08 | 2.1e-08 | 7.5e-08 | 4.533E-08 | 2.739E-08 | 2.100E-08 | 7.500E-08 |
| 2363 | 119 | Q119N | A | 4.5e-08 | 4.9e-08 | 2.9e-08 | 4.100E-08 | 1.058E-08 | 2.900E-08 | 4.900E-08 |
| 2364 | 119 | Q119D | A | 2.7e-08 | 6.5e-08 | 3e-08 | 4.067E-08 | 2.113E-08 | 2.700E-08 | 6.500E-08 |
| 2365 | 119 | Q119C | A | 3.3e-08 | 6.2e-08 | 4.6e-08 | 4.700E-08 | 1.453E-08 | 3.300E-08 | 6.200E-08 |
| 2366 | 119 | Q119E | A | 2e-08 | 2.4e-08 | 6e-08 | 3.467E-08 | 2.203E-08 | 2.000E-08 | 6.000E-08 |
| 2367 | 119 | Q119Q | A | 3.6e-08 | 3.5e-08 | 5.8e-08 | 4.300E-08 | 1.300E-08 | 3.500E-08 | 5.800E-08 |
| 2368 | 119 | Q119G | A | 3.3e-08 | 9.8e-08 | 8.3e-08 | 7.133E-08 | 3.403E-08 | 3.300E-08 | 9.800E-08 |
| 2369 | 119 | Q119H | A | 1e-08 | 6.6e-08 | 4.7e-08 | 4.100E-08 | 2.848E-08 | 1.000E-08 | 6.600E-08 |
| 2370 | 119 | Q119I | A | 4.6e-08 | 4.3e-08 | 2.7e-08 | 3.867E-08 | 1.021E-08 | 2.700E-08 | 4.600E-08 |
| 2371 | 119 | Q119L | A | 3.7e-08 | 1.9e-08 | 9.1e-08 | 4.900E-08 | 3.747E-08 | 1.900E-08 | 9.100E-08 |
| 2372 | 119 | Q119K | A | 3.5e-08 | 1.8e-08 | 3.5e-08 | 2.933E-08 | 9.815E-09 | 1.800E-08 | 3.500E-08 |
| 2373 | 119 | Q119M | A | 5.4e-08 | 3.2e-08 | 2.8e-08 | 3.800E-08 | 1.400E-08 | 2.800E-08 | 5.400E-08 |
| 2374 | 119 | Q119F | A | 9e-09 | 2e-08 | 5.5e-08 | 2.800E-08 | 2.402E-08 | 9.000E-09 | 5.500E-08 |
| 2375 | 119 | Q119P | A | 3.5e-08 | 2.6e-08 | 1.3e-08 | 2.467E-08 | 1.106E-08 | 1.300E-08 | 3.500E-08 |
| 2376 | 119 | Q119S | A | 1.5e-08 | 3.7e-08 | 2.1e-08 | 2.433E-08 | 1.137E-08 | 1.500E-08 | 3.700E-08 |
| 2377 | 119 | Q119T | A | 2.1e-08 | 6e-08 | 1.5e-08 | 3.200E-08 | 2.443E-08 | 1.500E-08 | 6.000E-08 |
| 2378 | 119 | Q119W | A | 3.9e-08 | 1e-07 | 1.5e-08 | 5.133E-08 | 4.382E-08 | 1.500E-08 | 1.000E-07 |
| 2379 | 119 | Q119Y | A | 3.5e-08 | 7.5e-08 | 7.5e-08 | 6.167E-08 | 2.309E-08 | 3.500E-08 | 7.500E-08 |

| | | | | | | | | | | |
|---|---|---|---|---|---|---|---|---|---|---|
| 2380 | 119 | Q119V | A | 4.8e-08 | 1.5e-07 | 4e-08 | 7.933E-08 | 6.133E-08 | 4.000E-08 | 1.500E-07 |
| 2381 | 120 | L120A | A | 2.5e-08 | 3.3e-08 | 1.7e-08 | 2.500E-08 | 8.000E-09 | 1.700E-08 | 3.300E-08 |
| 2382 | 120 | L120R | A | 3.4e-08 | 4.7e-08 | 2.3e-08 | 3.467E-08 | 1.201E-08 | 2.300E-08 | 4.700E-08 |
| 2383 | 120 | L120N | A | 3.6e-08 | 9.9e-08 | 2.4e-08 | 5.300E-08 | 4.029E-08 | 2.400E-08 | 9.900E-08 |
| 2384 | 120 | L120D | A | 5.8e-08 | 3.8e-08 | 7.3e-08 | 5.633E-08 | 1.756E-08 | 3.800E-08 | 7.300E-08 |
| 2385 | 120 | L120C | A | 9e-08 | 2e-08 | 4.5e-08 | 5.167E-08 | 3.547E-08 | 2.000E-08 | 9.000E-08 |
| 2386 | 120 | L120E | A | 3.7e-08 | 6.8e-08 | 3.5e-08 | 4.667E-08 | 1.850E-08 | 3.500E-08 | 6.800E-08 |
| 2387 | 120 | L120Q | A | 5e-08 | 6.3e-08 | 4.5e-08 | 5.267E-08 | 9.292E-09 | 4.500E-08 | 6.300E-08 |
| 2388 | 120 | L120G | A | 4.3e-08 | 2.9e-08 | 3.2e-08 | 3.467E-08 | 7.371E-09 | 2.900E-08 | 4.300E-08 |
| 2389 | 120 | L120H | A | 5e-08 | 3e-08 | 3.4e-08 | 3.800E-08 | 1.058E-08 | 3.000E-08 | 5.000E-08 |
| 2390 | 120 | L120I | A | 9.6e-08 | 3.8e-08 | 5.6e-08 | 6.333E-08 | 2.969E-08 | 3.800E-08 | 9.600E-08 |
| 2391 | 120 | L120L | A | 6.2e-08 | 4.7e-08 | 2.2e-08 | 4.367E-08 | 2.021E-08 | 2.200E-08 | 6.200E-08 |
| 2392 | 120 | L120K | A | 6.8e-08 | 3.8e-08 | 4.8e-08 | 5.133E-08 | 1.528E-08 | 3.800E-08 | 6.800E-08 |
| 2393 | 120 | L120M | A | 1.5e-08 | 7.9e-08 | 3.6e-08 | 4.333E-08 | 3.262E-08 | 1.500E-08 | 7.900E-08 |
| 2394 | 120 | L120F | A | 2.2e-08 | 5.1e-08 | 6.8e-08 | 4.700E-08 | 2.326E-08 | 2.200E-08 | 6.800E-08 |
| 2395 | 120 | L120P | A | 1.2e-07 | 4.4e-08 | 5.2e-08 | 7.200E-08 | 4.176E-08 | 4.400E-08 | 1.200E-07 |
| 2396 | 120 | L120S | A | 5.7e-08 | 2e-08 | 7.1e-08 | 4.933E-08 | 2.635E-08 | 2.000E-08 | 7.100E-08 |
| 2397 | 120 | L120T | A | 8.9e-08 | 4.2e-08 | 3.6e-08 | 5.567E-08 | 2.902E-08 | 3.600E-08 | 8.900E-08 |
| 2398 | 120 | L120W | A | 4e-08 | 4.8e-08 | 4e-08 | 4.267E-08 | 4.619E-09 | 4.000E-08 | 4.800E-08 |
| 2399 | 120 | L120Y | A | 4e-08 | 1.7e-08 | 1.9e-08 | 2.533E-08 | 1.274E-08 | 1.700E-08 | 4.000E-08 |
| 2400 | 120 | L120V | A | 2.8e-08 | 4.8e-08 | 4.2e-08 | 3.933E-08 | 1.026E-08 | 2.800E-08 | 4.800E-08 |
| 2401 | 121 | R121A | A | 5.8e-08 | 4e-08 | 7.7e-08 | 5.833E-08 | 1.850E-08 | 4.000E-08 | 7.700E-08 |
| 2402 | 121 | R121R | A | 2.2e-08 | 1.1e-07 | 4.6e-08 | 5.933E-08 | 4.549E-08 | 2.200E-08 | 1.100E-07 |
| 2403 | 121 | R121N | A | 2.7e-08 | 6.2e-08 | 1.5e-07 | 7.967E-08 | 6.337E-08 | 2.700E-08 | 1.500E-07 |
| 2404 | 121 | R121D | A | 7e-08 | 2.3e-08 | 3.6e-08 | 4.300E-08 | 2.427E-08 | 2.300E-08 | 7.000E-08 |
| 2405 | 121 | R121C | A | 8.5e-08 | 2.1e-08 | 4e-08 | 4.867E-08 | 3.287E-08 | 2.100E-08 | 8.500E-08 |
| 2406 | 121 | R121E | A | 2.1e-08 | 4.1e-08 | 4e-08 | 3.400E-08 | 1.127E-08 | 2.100E-08 | 4.100E-08 |
| 2407 | 121 | R121Q | A | 1.9e-08 | 8e-08 | 7.3e-08 | 5.733E-08 | 3.338E-08 | 1.900E-08 | 8.000E-08 |
| 2408 | 121 | R121G | A | 1e-08 | 1.9e-08 | 2.8e-08 | 1.900E-08 | 9.000E-09 | 1.000E-08 | 2.800E-08 |
| 2409 | 121 | R121H | A | 5.1e-08 | 1.8e-08 | 5.8e-08 | 4.233E-08 | 2.136E-08 | 1.800E-08 | 5.800E-08 |
| 2410 | 121 | R121I | A | 4.4e-08 | 3.9e-08 | 4.5e-08 | 4.267E-08 | 3.215E-09 | 3.900E-08 | 4.500E-08 |
| 2411 | 121 | R121L | A | 6.8e-08 | 2.1e-08 | 1.7e-08 | 3.533E-08 | 2.836E-08 | 1.700E-08 | 6.800E-08 |
| 2412 | 121 | R121K | A | 3.5e-08 | 4.9e-08 | 8.9e-08 | 5.767E-08 | 2.802E-08 | 3.500E-08 | 8.900E-08 |
| 2413 | 121 | R121M | A | 4.2e-08 | 6.9e-08 | 3.2e-08 | 4.767E-08 | 1.914E-08 | 3.200E-08 | 6.900E-08 |
| 2414 | 121 | R121F | A | 5.5e-08 | 3.2e-08 | 2.9e-08 | 3.867E-08 | 1.422E-08 | 2.900E-08 | 5.500E-08 |
| 2415 | 121 | R121P | A | 1.7e-08 | 8.8e-08 | 4.1e-08 | 4.867E-08 | 3.612E-08 | 1.700E-08 | 8.800E-08 |
| 2416 | 121 | R121S | A | 4.6e-08 | 1.5e-08 | 3.7e-08 | 3.267E-08 | 1.595E-08 | 1.500E-08 | 4.600E-08 |
| 2417 | 121 | R121T | A | 3.6e-08 | 9.5e-09 | 3.2e-08 | 2.583E-08 | 1.429E-08 | 9.500E-09 | 3.600E-08 |
| 2418 | 121 | R121W | A | 7.6e-08 | 2.4e-08 | 2.2e-08 | 4.067E-08 | 3.062E-08 | 2.200E-08 | 7.600E-08 |
| 2419 | 121 | R121Y | A | 3.5e-08 | 7.5e-08 | 3.9e-08 | 4.967E-08 | 2.203E-08 | 3.500E-08 | 7.500E-08 |
| 2420 | 121 | R121V | A | 2.5e-08 | 4.3e-08 | 6.2e-08 | 4.333E-08 | 1.850E-08 | 2.500E-08 | 6.200E-08 |
| 2421 | 122 | S122A | A | 5e-08 | 2.7e-08 | 8.4e-08 | 5.367E-08 | 2.868E-08 | 2.700E-08 | 8.400E-08 |
| 2422 | 122 | S122R | A | 2.2e-08 | 3.2e-08 | 5.7e-08 | 3.700E-08 | 1.803E-08 | 2.200E-08 | 5.700E-08 |
| 2423 | 122 | S122N | A | 3.5e-08 | 3.6e-08 | 2.5e-08 | 3.200E-08 | 6.083E-09 | 2.500E-08 | 3.600E-08 |
| 2424 | 122 | S122D | A | 2.5e-08 | 3e-08 | 9e-09 | 2.133E-08 | 1.097E-08 | 9.000E-09 | 3.000E-08 |
| 2425 | 122 | S122C | A | 4.3e-08 | 4.4e-08 | 4.2e-08 | 4.300E-08 | 1.000E-09 | 4.200E-08 | 4.400E-08 |
| 2426 | 122 | S122E | A | 3.6e-08 | 8.7e-08 | 4.5e-08 | 5.600E-08 | 2.722E-08 | 3.600E-08 | 8.700E-08 |
| 2427 | 122 | S122Q | A | 2e-08 | 3.4e-08 | 5.5e-08 | 3.633E-08 | 1.762E-08 | 2.000E-08 | 5.500E-08 |
| 2428 | 122 | S122G | A | 1.5e-08 | 2.5e-08 | 4.6e-08 | 2.867E-08 | 1.582E-08 | 1.500E-08 | 4.600E-08 |
| 2429 | 122 | S122H | A | 8.8e-08 | 2.9e-08 | 1.2e-07 | 7.900E-08 | 4.616E-08 | 2.900E-08 | 1.200E-07 |
| 2430 | 122 | S122I | A | 5.2e-08 | 9.8e-08 | 2.4e-08 | 5.800E-08 | 3.736E-08 | 2.400E-08 | 9.800E-08 |
| 2431 | 122 | S122L | A | 2.8e-08 | 4.1e-08 | 7.2e-08 | 4.700E-08 | 2.261E-08 | 2.800E-08 | 7.200E-08 |
| 2432 | 122 | S122K | A | 4.1e-08 | 6.7e-08 | 2.4e-08 | 4.400E-08 | 2.166E-08 | 2.400E-08 | 6.700E-08 |
| 2433 | 122 | S122M | A | 2.2e-08 | 3.6e-08 | 4e-08 | 3.267E-08 | 9.452E-09 | 2.200E-08 | 4.000E-08 |
| 2434 | 122 | S122F | A | 8.8e-08 | 1.4e-07 | 6.5e-08 | 9.767E-08 | 3.842E-08 | 6.500E-08 | 1.400E-07 |
| 2435 | 122 | S122P | A | 3e-08 | 3.5e-08 | 3.2e-08 | 3.233E-08 | 2.517E-09 | 3.000E-08 | 3.500E-08 |
| 2436 | 122 | S122S | A | 7.2e-08 | 4e-08 | 4.6e-08 | 5.267E-08 | 1.701E-08 | 4.000E-08 | 7.200E-08 |
| 2437 | 122 | S122T | A | 3.8e-08 | 2.9e-08 | 3.9e-08 | 3.533E-08 | 5.508E-09 | 2.900E-08 | 3.900E-08 |
| 2438 | 122 | S122W | A | 7.3e-08 | 4.4e-08 | 1.2e-07 | 7.900E-08 | 3.835E-08 | 4.400E-08 | 1.200E-07 |
| 2439 | 122 | S122Y | A | 6e-08 | 3.3e-08 | 3e-08 | 4.100E-08 | 1.652E-08 | 3.000E-08 | 6.000E-08 |
| 2440 | 122 | S122V | A | 3.8e-08 | 4.8e-08 | 5.6e-08 | 4.733E-08 | 9.018E-09 | 3.800E-08 | 5.600E-08 |
| 2441 | 123 | L123A | A | 2.2e-08 | 5.8e-08 | 4.4e-08 | 4.133E-08 | 1.815E-08 | 2.200E-08 | 5.800E-08 |
| 2442 | 123 | L123R | A | 1.1e-07 | 6.6e-08 | 4.5e-08 | 7.367E-08 | 3.317E-08 | 4.500E-08 | 1.100E-07 |
| 2443 | 123 | L123N | A | 2.3e-08 | 7.2e-08 | 5.1e-08 | 4.867E-08 | 2.458E-08 | 2.300E-08 | 7.200E-08 |
| 2444 | 123 | L123D | A | 1.9e-08 | 4.6e-08 | 6.2e-08 | 4.233E-08 | 2.173E-08 | 1.900E-08 | 6.200E-08 |
| 2445 | 123 | L123C | A | 5.7e-08 | 3.7e-08 | 2.6e-08 | 4.000E-08 | 1.572E-08 | 2.600E-08 | 5.700E-08 |
| 2446 | 123 | L123E | A | 3.4e-08 | 3.9e-08 | 1.9e-08 | 3.067E-08 | 1.041E-08 | 1.900E-08 | 3.900E-08 |
| 2447 | 123 | L123Q | A | 3.7e-08 | 3.4e-08 | 7.6e-08 | 4.900E-08 | 2.343E-08 | 3.400E-08 | 7.600E-08 |
| 2448 | 123 | L123G | A | 3.5e-08 | 8.1e-08 | 3.1e-08 | 4.900E-08 | 2.778E-08 | 3.100E-08 | 8.100E-08 |
| 2449 | 123 | L123H | A | 2.1e-08 | 5.3e-08 | 3.5e-08 | 3.633E-08 | 1.604E-08 | 2.100E-08 | 5.300E-08 |
| 2450 | 123 | L123I | A | 5.6e-08 | 2.9e-08 | 6.9e-08 | 5.133E-08 | 2.040E-08 | 2.900E-08 | 6.900E-08 |
| 2451 | 123 | L123L | A | 5.2e-08 | 4.7e-08 | 4.8e-08 | 4.900E-08 | 2.646E-09 | 4.700E-08 | 5.200E-08 |
| 2452 | 123 | L123K | A | 1.4e-08 | 4.1e-08 | 2.4e-08 | 2.633E-08 | 1.365E-08 | 1.400E-08 | 4.100E-08 |
| 2453 | 123 | L123M | A | 3e-08 | 4.7e-08 | 4.8e-08 | 4.167E-08 | 1.012E-08 | 3.000E-08 | 4.800E-08 |
| 2454 | 123 | L123F | A | 4.2e-08 | 6.7e-08 | 2.8e-08 | 4.567E-08 | 1.976E-08 | 2.800E-08 | 6.700E-08 |
| 2455 | 123 | L123P | A | 2.1e-08 | 2.9e-08 | 1.7e-08 | 2.233E-08 | 6.110E-09 | 1.700E-08 | 2.900E-08 |
| 2456 | 123 | L123S | A | 3.5e-08 | 1.6e-07 | 5.3e-08 | 8.267E-08 | 6.757E-08 | 3.500E-08 | 1.600E-07 |
| 2457 | 123 | L123T | A | 2.9e-08 | 1.4e-08 | 3.9e-08 | 2.733E-08 | 1.258E-08 | 1.400E-08 | 3.900E-08 |
| 2458 | 123 | L123W | A | 5.6e-08 | 2.7e-08 | 2.6e-08 | 3.633E-08 | 1.704E-08 | 2.600E-08 | 5.600E-08 |
| 2459 | 123 | L123Y | A | 7.6e-08 | 4.8e-08 | 3.9e-08 | 5.433E-08 | 1.930E-08 | 3.900E-08 | 7.600E-08 |
| 2460 | 123 | L123V | A | 2.6e-08 | 5e-08 | 2.9e-08 | 3.500E-08 | 1.308E-08 | 2.600E-08 | 5.000E-08 |
| 2461 | 124 | T124A | A | 1.8e-08 | 3e-08 | 3.1e-08 | 2.633E-08 | 7.234E-09 | 1.800E-08 | 3.100E-08 |
| 2462 | 124 | T124R | A | 2.7e-08 | 2.9e-08 | 4.2e-08 | 3.267E-08 | 8.145E-09 | 2.700E-08 | 4.200E-08 |
| 2463 | 124 | T124N | A | 2.6e-08 | 2.4e-08 | 4.5e-08 | 3.167E-08 | 1.159E-08 | 2.400E-08 | 4.500E-08 |
| 2464 | 124 | T124D | A | 3.7e-08 | 4.2e-08 | 4.3e-08 | 4.067E-08 | 3.215E-09 | 3.700E-08 | 4.300E-08 |

| | | | | | | | | | | |
|---|---|---|---|---|---|---|---|---|---|---|
| 2465 | 124 | T124C | A | 5.2e-08 | 4.9e-08 | 6.7e-08 | 5.600E-08 | 9.644E-09 | 4.900E-08 | 6.700E-08 |
| 2466 | 124 | T124E | A | 2.2e-08 | 3.7e-08 | 4.6e-08 | 3.500E-08 | 1.212E-08 | 2.200E-08 | 4.600E-08 |
| 2467 | 124 | T124Q | A | 5e-08 | 6.1e-08 | 2.7e-08 | 4.600E-08 | 1.735E-08 | 2.700E-08 | 6.100E-08 |
| 2468 | 124 | T124G | A | 3e-08 | 3e-08 | 7.2e-08 | 4.400E-08 | 2.425E-08 | 3.000E-08 | 7.200E-08 |
| 2469 | 124 | T124H | A | 5.6e-08 | 1.7e-08 | 6.6e-08 | 4.633E-08 | 2.589E-08 | 1.700E-08 | 6.600E-08 |
| 2470 | 124 | T124I | A | 4e-08 | 8.6e-08 | 4.7e-08 | 5.767E-08 | 2.479E-08 | 4.000E-08 | 8.600E-08 |
| 2471 | 124 | T124L | A | 1.3e-08 | 2.7e-08 | 1.9e-08 | 1.967E-08 | 7.024E-09 | 1.300E-08 | 2.700E-08 |
| 2472 | 124 | T124K | A | 3.3e-08 | 5.7e-08 | 2e-07 | 9.667E-08 | 9.029E-08 | 3.300E-08 | 2.000E-07 |
| 2473 | 124 | T124M | A | 5.8e-08 | 9.5e-08 | 4.7e-08 | 6.667E-08 | 2.515E-08 | 4.700E-08 | 9.500E-08 |
| 2474 | 124 | T124F | A | 4e-08 | 6.8e-08 | 8.3e-08 | 6.367E-08 | 2.183E-08 | 4.000E-08 | 8.300E-08 |
| 2475 | 124 | T124P | A | 9.6e-08 | 2.1e-08 | 5.6e-08 | 5.767E-08 | 3.753E-08 | 2.100E-08 | 9.600E-08 |
| 2476 | 124 | T124S | A | 3e-08 | 3.4e-08 | 1.2e-08 | 2.533E-08 | 1.172E-08 | 1.200E-08 | 3.400E-08 |
| 2477 | 124 | T124T | A | 5.1e-08 | 2.8e-08 | 3.2e-08 | 3.700E-08 | 1.229E-08 | 2.800E-08 | 5.100E-08 |
| 2478 | 124 | T124W | A | 2.1e-08 | 1.4e-08 | 1.7e-08 | 1.733E-08 | 3.512E-09 | 1.400E-08 | 2.100E-08 |
| 2479 | 124 | T124Y | A | 3.7e-08 | 4.4e-08 | 1.6e-08 | 3.233E-08 | 1.457E-08 | 1.600E-08 | 4.400E-08 |
| 2480 | 124 | T124V | A | 1.6e-07 | 1.3e-07 | 2.4e-08 | 1.047E-07 | 7.145E-08 | 2.400E-08 | 1.600E-07 |
| 2481 | 125 | E125A | A | 3.6e-08 | 7.8e-08 | 7.6e-08 | 6.333E-08 | 2.369E-08 | 3.600E-08 | 7.800E-08 |
| 2482 | 125 | E125R | A | 3.4e-08 | 2.5e-08 | 9.8e-08 | 5.233E-08 | 3.980E-08 | 2.500E-08 | 9.800E-08 |
| 2483 | 125 | E125N | A | 5e-08 | 5.5e-08 | 2e-07 | 1.017E-07 | 8.520E-08 | 5.000E-08 | 2.000E-07 |
| 2484 | 125 | E125D | A | 2.4e-08 | 7.7e-08 | 5.4e-08 | 5.167E-08 | 2.658E-08 | 2.400E-08 | 7.700E-08 |
| 2485 | 125 | E125C | A | 3.5e-08 | 3.7e-08 | 4.6e-08 | 3.933E-08 | 5.859E-09 | 3.500E-08 | 4.600E-08 |
| 2486 | 125 | E125E | A | 3.7e-08 | 9.1e-08 | 3.4e-08 | 5.400E-08 | 3.208E-08 | 3.400E-08 | 9.100E-08 |
| 2487 | 125 | E125Q | A | 6.7e-08 | 3.5e-08 | 4.1e-08 | 4.767E-08 | 1.701E-08 | 3.500E-08 | 6.700E-08 |
| 2488 | 125 | E125G | A | 7.5e-08 | 4.2e-08 | 5.9e-08 | 5.867E-08 | 1.650E-08 | 4.200E-08 | 7.500E-08 |
| 2489 | 125 | E125H | A | 3.1e-08 | 5.5e-08 | 5.9e-08 | 4.833E-08 | 1.514E-08 | 3.100E-08 | 5.900E-08 |
| 2490 | 125 | E125I | A | 1.9e-08 | 4.6e-08 | 6.4e-08 | 4.300E-08 | 2.265E-08 | 1.900E-08 | 6.400E-08 |
| 2491 | 125 | E125L | A | 4.8e-08 | 6.2e-08 | 6.2e-08 | 5.733E-08 | 8.083E-09 | 4.800E-08 | 6.200E-08 |
| 2492 | 125 | E125K | A | 6.3e-08 | 1.6e-07 | 6.7e-08 | 9.667E-08 | 5.488E-08 | 6.300E-08 | 1.600E-07 |
| 2493 | 125 | E125M | A | 1.4e-08 | 4.1e-08 | 2.8e-08 | 2.767E-08 | 1.350E-08 | 1.400E-08 | 4.100E-08 |
| 2494 | 125 | E125F | A | 8.2e-08 | 4.7e-08 | 1.4e-07 | 8.967E-08 | 4.697E-08 | 4.700E-08 | 1.400E-07 |
| 2495 | 125 | E125P | A | 5.3e-08 | 4.5e-08 | 4.4e-08 | 4.733E-08 | 4.933E-09 | 4.400E-08 | 5.300E-08 |
| 2496 | 125 | E125S | A | 5.2e-08 | 3.7e-08 | 4.8e-08 | 4.567E-08 | 7.767E-09 | 3.700E-08 | 5.200E-08 |
| 2497 | 125 | E125T | A | 1.3e-08 | 2.2e-08 | 4.3e-08 | 2.600E-08 | 1.539E-08 | 1.300E-08 | 4.300E-08 |
| 2498 | 125 | E125W | A | 3.1e-08 | 2e-08 | 3.4e-08 | 2.833E-08 | 7.371E-09 | 2.000E-08 | 3.400E-08 |
| 2499 | 125 | E125Y | A | 5.2e-08 | 1.6e-08 | 1.6e-08 | 2.800E-08 | 2.078E-08 | 1.600E-08 | 5.200E-08 |
| 2500 | 125 | E125V | A | 9.9e-08 | 6.8e-08 | 3e-08 | 6.567E-08 | 3.456E-08 | 3.000E-08 | 9.900E-08 |
| 2501 | 126 | I126A | A | 6.5e-08 | 5.4e-08 | 2.2e-08 | 4.700E-08 | 2.234E-08 | 2.200E-08 | 6.500E-08 |
| 2502 | 126 | I126R | A | 1.2e-07 | 2.9e-08 | 4.7e-08 | 6.533E-08 | 4.819E-08 | 2.900E-08 | 1.200E-07 |
| 2503 | 126 | I126N | A | 2.6e-08 | 1.1e-07 | 7.7e-08 | 7.100E-08 | 4.232E-08 | 2.600E-08 | 1.100E-07 |
| 2504 | 126 | I126D | A | 8.4e-08 | 5.2e-08 | 6.9e-08 | 6.833E-08 | 1.601E-08 | 5.200E-08 | 8.400E-08 |
| 2505 | 126 | I126C | A | 1.5e-08 | 2.6e-08 | 7.4e-08 | 3.833E-08 | 3.137E-08 | 1.500E-08 | 7.400E-08 |
| 2506 | 126 | I126E | A | 3.4e-08 | 3.7e-08 | 5.6e-08 | 4.233E-08 | 1.193E-08 | 3.400E-08 | 5.600E-08 |
| 2507 | 126 | I126Q | A | 2.8e-08 | 2.6e-08 | 1.9e-08 | 2.433E-08 | 4.726E-09 | 1.900E-08 | 2.800E-08 |
| 2508 | 126 | I126G | A | 2.7e-08 | 3.7e-08 | 7.7e-08 | 4.700E-08 | 2.646E-08 | 2.700E-08 | 7.700E-08 |
| 2509 | 126 | I126H | A | 7.5e-08 | 5.1e-08 | 5.6e-08 | 6.067E-08 | 1.266E-08 | 5.100E-08 | 7.500E-08 |
| 2510 | 126 | I126I | A | 3.7e-08 | 2.1e-08 | 1.5e-08 | 2.433E-08 | 1.137E-08 | 1.500E-08 | 3.700E-08 |
| 2511 | 126 | I126L | A | 1.1e-07 | 4.2e-08 | 5.7e-08 | 6.967E-08 | 3.573E-08 | 4.200E-08 | 1.100E-07 |
| 2512 | 126 | I126K | A | 6.1e-08 | 2.7e-08 | 6.4e-08 | 5.067E-08 | 2.055E-08 | 2.700E-08 | 6.400E-08 |
| 2513 | 126 | I126M | A | 4.3e-08 | 3.7e-08 | 7.5e-08 | 5.167E-08 | 2.043E-08 | 3.700E-08 | 7.500E-08 |
| 2514 | 126 | I126F | A | 1.5e-08 | 7.5e-08 | 4.6e-08 | 4.533E-08 | 3.001E-08 | 1.500E-08 | 7.500E-08 |
| 2515 | 126 | I126P | A | 5.5e-08 | 4.5e-08 | 5.1e-08 | 5.033E-08 | 5.033E-09 | 4.500E-08 | 5.500E-08 |
| 2516 | 126 | I126S | A | 3.3e-08 | 4.5e-08 | 8.1e-08 | 5.300E-08 | 2.498E-08 | 3.300E-08 | 8.100E-08 |
| 2517 | 126 | I126T | A | 5.6e-08 | 2.4e-08 | 1.3e-07 | 7.000E-08 | 5.437E-08 | 2.400E-08 | 1.300E-07 |
| 2518 | 126 | I126W | A | 2e-08 | 6.5e-08 | 2.8e-08 | 3.767E-08 | 2.401E-08 | 2.000E-08 | 6.500E-08 |
| 2519 | 126 | I126Y | A | 3.3e-08 | 5e-08 | 4.6e-08 | 4.300E-08 | 8.888E-09 | 3.300E-08 | 5.000E-08 |
| 2520 | 126 | I126V | A | 5.1e-08 | 1.9e-08 | 3.1e-08 | 3.367E-08 | 1.617E-08 | 1.900E-08 | 5.100E-08 |
| 2521 | 127 | L127A | A | 2.2e-08 | 3.8e-08 | 5.1e-08 | 3.700E-08 | 1.453E-08 | 2.200E-08 | 5.100E-08 |
| 2522 | 127 | L127R | A | 1.9e-08 | 2.2e-08 | 6.3e-08 | 3.467E-08 | 2.458E-08 | 1.900E-08 | 6.300E-08 |
| 2523 | 127 | L127N | A | 2.6e-08 | 7.6e-08 | 4.8e-08 | 5.000E-08 | 2.506E-08 | 2.600E-08 | 7.600E-08 |
| 2524 | 127 | L127D | A | 8.2e-08 | 3.5e-08 | 3.5e-08 | 5.067E-08 | 2.714E-08 | 3.500E-08 | 8.200E-08 |
| 2525 | 127 | L127C | A | 5e-08 | 1e-07 | 5.1e-08 | 6.700E-08 | 2.858E-08 | 5.000E-08 | 1.000E-07 |
| 2526 | 127 | L127E | A | 8.2e-08 | 1.8e-08 | 2.7e-08 | 4.233E-08 | 3.465E-08 | 1.800E-08 | 8.200E-08 |
| 2527 | 127 | L127Q | A | 3e-08 | 1.1e-07 | 2.9e-08 | 5.633E-08 | 4.648E-08 | 2.900E-08 | 1.100E-07 |
| 2528 | 127 | L127G | A | 5.7e-08 | 6.4e-08 | 5e-08 | 5.700E-08 | 7.000E-09 | 5.000E-08 | 6.400E-08 |
| 2529 | 127 | L127H | A | 5e-08 | 8e-08 | 3.3e-08 | 5.433E-08 | 2.380E-08 | 3.300E-08 | 8.000E-08 |
| 2530 | 127 | L127I | A | 4.4e-08 | 6.2e-08 | 7.6e-08 | 6.067E-08 | 1.604E-08 | 4.400E-08 | 7.600E-08 |
| 2531 | 127 | L127L | A | 8.5e-08 | 1.4e-07 | 3.6e-08 | 8.700E-08 | 5.203E-08 | 3.600E-08 | 1.400E-07 |
| 2532 | 127 | L127K | A | 9.6e-08 | 2.6e-08 | 4.7e-08 | 5.633E-08 | 3.592E-08 | 2.600E-08 | 9.600E-08 |
| 2533 | 127 | L127M | A | 6.4e-08 | 2e-08 | 1.1e-07 | 6.467E-08 | 4.500E-08 | 2.000E-08 | 1.100E-07 |
| 2534 | 127 | L127F | A | 2.3e-08 | 7.3e-08 | 8.3e-08 | 5.967E-08 | 3.215E-08 | 2.300E-08 | 8.300E-08 |
| 2535 | 127 | L127P | A | 2.1e-08 | 2.8e-08 | 4.6e-08 | 3.167E-08 | 1.290E-08 | 2.100E-08 | 4.600E-08 |
| 2536 | 127 | L127S | A | 1.3e-07 | 5.8e-08 | 2.3e-08 | 7.033E-08 | 5.456E-08 | 2.300E-08 | 1.300E-07 |
| 2537 | 127 | L127T | A | 9.8e-08 | 2.5e-08 | 4.2e-08 | 5.500E-08 | 3.820E-08 | 2.500E-08 | 9.800E-08 |
| 2538 | 127 | L127W | A | 1e-08 | 5.5e-08 | 8.9e-08 | 5.133E-08 | 3.963E-08 | 1.000E-08 | 8.900E-08 |
| 2539 | 127 | L127Y | A | 6.8e-08 | 2.3e-08 | 3.4e-08 | 4.167E-08 | 2.346E-08 | 2.300E-08 | 6.800E-08 |
| 2540 | 127 | L127V | A | 6.5e-08 | 2.4e-08 | 8.8e-08 | 5.900E-08 | 3.242E-08 | 2.400E-08 | 8.800E-08 |
| 2541 | 128 | K128A | A | 3.2e-08 | 8e-08 | 8.8e-08 | 6.667E-08 | 3.029E-08 | 3.200E-08 | 8.800E-08 |
| 2542 | 128 | K128R | A | 6.3e-08 | 3.6e-08 | 3e-08 | 4.300E-08 | 1.758E-08 | 3.000E-08 | 6.300E-08 |
| 2543 | 128 | K128N | A | 2.4e-08 | 5.7e-08 | 4.2e-08 | 4.100E-08 | 1.652E-08 | 2.400E-08 | 5.700E-08 |
| 2544 | 128 | K128D | A | 4.9e-08 | 2.9e-08 | 1.8e-08 | 3.200E-08 | 1.572E-08 | 1.800E-08 | 4.900E-08 |
| 2545 | 128 | K128C | A | 2.3e-08 | 8.2e-08 | 8.9e-08 | 6.467E-08 | 3.625E-08 | 2.300E-08 | 8.900E-08 |
| 2546 | 128 | K128E | A | 5e-08 | 3.6e-08 | 6.6e-08 | 5.067E-08 | 1.501E-08 | 3.600E-08 | 6.600E-08 |
| 2547 | 128 | K128Q | A | 2.5e-07 | 4.2e-08 | 1.7e-07 | 1.540E-07 | 1.049E-07 | 4.200E-08 | 2.500E-07 |
| 2548 | 128 | K128G | A | 6.7e-08 | 5e-08 | 3.8e-08 | 5.167E-08 | 1.457E-08 | 3.800E-08 | 6.700E-08 |
| 2549 | 128 | K128H | A | 8.4e-08 | 3.5e-08 | 4.5e-08 | 5.467E-08 | 2.589E-08 | 3.500E-08 | 8.400E-08 |

| | | | | | | | | | | |
|---|---|---|---|---|---|---|---|---|---|---|
| 2550 | 128 | K128I | A | 2.6e-08 | 4.8e-08 | 3.6e-08 | 3.667E-08 | 1.102E-08 | 2.600E-08 | 4.800E-08 |
| 2551 | 128 | K128L | A | 5.8e-08 | 5.2e-08 | 2.8e-08 | 4.600E-08 | 1.587E-08 | 2.800E-08 | 5.800E-08 |
| 2552 | 128 | K128K | A | 2.6e-08 | 4.7e-08 | 2.9e-08 | 3.400E-08 | 1.136E-08 | 2.600E-08 | 4.700E-08 |
| 2553 | 128 | K128M | A | 9.3e-08 | 2.3e-08 | 2.4e-08 | 4.667E-08 | 4.013E-08 | 2.300E-08 | 9.300E-08 |
| 2554 | 128 | K128F | A | 2.3e-08 | 2.5e-08 | 7e-08 | 3.933E-08 | 2.658E-08 | 2.300E-08 | 7.000E-08 |
| 2555 | 128 | K128P | A | 7.9e-08 | 7e-08 | 4.4e-08 | 6.433E-08 | 1.818E-08 | 4.400E-08 | 7.900E-08 |
| 2556 | 128 | K128S | A | 5.3e-08 | 4.1e-08 | 4.2e-08 | 4.533E-08 | 6.658E-09 | 4.100E-08 | 5.300E-08 |
| 2557 | 128 | K128T | A | 5e-08 | 3.5e-08 | 2.8e-08 | 3.767E-08 | 1.124E-08 | 2.800E-08 | 5.000E-08 |
| 2558 | 128 | K128W | A | 7.8e-08 | 6.1e-08 | 1.4e-08 | 5.100E-08 | 3.315E-08 | 1.400E-08 | 7.800E-08 |
| 2559 | 128 | K128Y | A | 8e-08 | 1.9e-08 | 2.4e-08 | 4.100E-08 | 3.387E-08 | 1.900E-08 | 8.000E-08 |
| 2560 | 128 | K128V | A | 2.9e-08 | 1.8e-08 | 1.3e-07 | 5.900E-08 | 6.173E-08 | 1.800E-08 | 1.300E-07 |
| 2561 | 129 | G129A | A | 5.1e-08 | 2.6e-08 | 7.8e-08 | 5.167E-08 | 2.601E-08 | 2.600E-08 | 7.800E-08 |
| 2562 | 129 | G129R | A | 1.3e-07 | 3.2e-08 | 2.2e-08 | 6.133E-08 | 5.968E-08 | 2.200E-08 | 1.300E-07 |
| 2563 | 129 | G129N | A | 1.7e-08 | 2.5e-08 | 1.8e-08 | 2.000E-08 | 4.359E-09 | 1.700E-08 | 2.500E-08 |
| 2564 | 129 | G129D | A | 3.5e-08 | 1.3e-07 | 4.4e-08 | 6.967E-08 | 5.244E-08 | 3.500E-08 | 1.300E-07 |
| 2565 | 129 | G129C | A | 4.5e-08 | 8.4e-08 | 8.6e-08 | 7.167E-08 | 2.312E-08 | 4.500E-08 | 8.600E-08 |
| 2566 | 129 | G129E | A | 3.2e-08 | 7.3e-08 | 6.1e-08 | 5.533E-08 | 2.108E-08 | 3.200E-08 | 7.300E-08 |
| 2567 | 129 | G129Q | A | 7.9e-08 | 5.9e-08 | 1.4e-08 | 5.067E-08 | 3.329E-08 | 1.400E-08 | 7.900E-08 |
| 2568 | 129 | G129G | A | 1.1e-08 | 4.9e-08 | 4.8e-08 | 3.600E-08 | 2.166E-08 | 1.100E-08 | 4.900E-08 |
| 2569 | 129 | G129H | A | 2.5e-08 | 1.8e-08 | 4.9e-08 | 3.067E-08 | 1.626E-08 | 1.800E-08 | 4.900E-08 |
| 2570 | 129 | G129I | A | 8.5e-08 | 4.8e-08 | 2e-08 | 5.100E-08 | 3.260E-08 | 2.000E-08 | 8.500E-08 |
| 2571 | 129 | G129L | A | 1.9e-08 | 3e-08 | 2.2e-08 | 2.367E-08 | 5.686E-09 | 1.900E-08 | 3.000E-08 |
| 2572 | 129 | G129K | A | 2.5e-07 | 1.6e-08 | 1.9e-08 | 9.500E-08 | 1.342E-07 | 1.600E-08 | 2.500E-07 |
| 2573 | 129 | G129M | A | 1.8e-08 | 1.3e-08 | 4.3e-08 | 2.467E-08 | 1.607E-08 | 1.300E-08 | 4.300E-08 |
| 2574 | 129 | G129F | A | 4.8e-08 | 7.5e-08 | 6.7e-08 | 6.333E-08 | 1.387E-08 | 4.800E-08 | 7.500E-08 |
| 2575 | 129 | G129P | A | 1.1e-07 | 9.7e-08 | 3e-08 | 7.900E-08 | 4.293E-08 | 3.000E-08 | 1.100E-07 |
| 2576 | 129 | G129S | A | 2e-08 | 3e-07 | 7.5e-08 | 1.317E-07 | 1.484E-07 | 2.000E-08 | 3.000E-07 |
| 2577 | 129 | G129T | A | 1.1e-07 | 6.8e-08 | 4.1e-08 | 7.300E-08 | 3.477E-08 | 4.100E-08 | 1.100E-07 |
| 2578 | 129 | G129W | A | 3.4e-08 | 8.3e-08 | 4.4e-08 | 5.367E-08 | 2.589E-08 | 3.400E-08 | 8.300E-08 |
| 2579 | 129 | G129Y | A | 5.5e-08 | 3.5e-08 | 6.1e-08 | 5.033E-08 | 1.361E-08 | 3.500E-08 | 6.100E-08 |
| 2580 | 129 | G129V | A | 7.8e-08 | 3.3e-08 | 6.3e-08 | 5.800E-08 | 2.291E-08 | 3.300E-08 | 7.800E-08 |
| 2581 | 130 | G130A | A | 7.8e-08 | 1.2e-07 | 2.4e-08 | 7.400E-08 | 4.812E-08 | 2.400E-08 | 1.200E-07 |
| 2582 | 130 | G130R | A | 5.2e-08 | 2.1e-08 | 2.2e-07 | 9.767E-08 | 1.071E-07 | 2.100E-08 | 2.200E-07 |
| 2583 | 130 | G130N | A | 5.9e-08 | 6.3e-08 | 6.9e-08 | 6.367E-08 | 5.033E-09 | 5.900E-08 | 6.900E-08 |
| 2584 | 130 | G130D | A | 6.7e-08 | 7.1e-08 | 5e-08 | 6.267E-08 | 1.115E-08 | 5.000E-08 | 7.100E-08 |
| 2585 | 130 | G130C | A | 1.8e-08 | 4.9e-08 | 3.2e-08 | 3.300E-08 | 1.552E-08 | 1.800E-08 | 4.900E-08 |
| 2586 | 130 | G130E | A | 9.4e-08 | 2.7e-08 | 5.2e-08 | 5.767E-08 | 3.386E-08 | 2.700E-08 | 9.400E-08 |
| 2587 | 130 | G130Q | A | 5.5e-08 | 8.1e-08 | 3e-08 | 5.533E-08 | 2.550E-08 | 3.000E-08 | 8.100E-08 |
| 2588 | 130 | G130G | A | 9.9e-08 | 1.3e-07 | 9.4e-09 | 7.947E-08 | 6.263E-08 | 9.400E-09 | 1.300E-07 |
| 2589 | 130 | G130H | A | 7.4e-08 | 3.3e-08 | 4.5e-08 | 5.067E-08 | 2.108E-08 | 3.300E-08 | 7.400E-08 |
| 2590 | 130 | G130I | A | 4.6e-08 | 1e-07 | 7.5e-08 | 7.367E-08 | 2.702E-08 | 4.600E-08 | 1.000E-07 |
| 2591 | 130 | G130L | A | 5.6e-08 | 3.1e-08 | 4.1e-08 | 4.267E-08 | 1.258E-08 | 3.100E-08 | 5.600E-08 |
| 2592 | 130 | G130K | A | 3e-08 | 2.5e-08 | 1.2e-07 | 5.833E-08 | 5.346E-08 | 2.500E-08 | 1.200E-07 |
| 2593 | 130 | G130M | A | 3.2e-08 | 8.2e-08 | 4.4e-08 | 5.267E-08 | 2.610E-08 | 3.200E-08 | 8.200E-08 |
| 2594 | 130 | G130F | A | 1.3e-07 | 1.3e-08 | 6.3e-08 | 6.867E-08 | 5.871E-08 | 1.300E-08 | 1.300E-07 |
| 2595 | 130 | G130P | A | 8e-08 | 5.2e-08 | 4.5e-08 | 5.900E-08 | 1.852E-08 | 4.500E-08 | 8.000E-08 |
| 2596 | 130 | G130S | A | 5.6e-08 | 3.2e-08 | 2.7e-08 | 3.833E-08 | 1.550E-08 | 2.700E-08 | 5.600E-08 |
| 2597 | 130 | G130T | A | 5e-08 | 6.8e-08 | 5.8e-08 | 5.867E-08 | 9.018E-09 | 5.000E-08 | 6.800E-08 |
| 2598 | 130 | G130W | A | 6.5e-08 | 3.1e-08 | 3.3e-08 | 4.300E-08 | 1.908E-08 | 3.100E-08 | 6.500E-08 |
| 2599 | 130 | G130Y | A | 2.9e-08 | 5.1e-08 | 1.4e-08 | 3.133E-08 | 1.861E-08 | 1.400E-08 | 5.100E-08 |
| 2600 | 130 | G130V | A | 9.4e-09 | 3.5e-08 | 7.4e-08 | 3.947E-08 | 3.253E-08 | 9.400E-09 | 7.400E-08 |
| 2601 | 131 | V131A | A | 1.6e-08 | 2.6e-08 | 5.7e-08 | 3.300E-08 | 2.138E-08 | 1.600E-08 | 5.700E-08 |
| 2602 | 131 | V131R | A | 5.4e-08 | 3e-08 | 2.8e-08 | 3.733E-08 | 1.447E-08 | 2.800E-08 | 5.400E-08 |
| 2603 | 131 | V131N | A | 5.3e-08 | 4.8e-08 | 7.8e-08 | 5.967E-08 | 1.607E-08 | 4.800E-08 | 7.800E-08 |
| 2604 | 131 | V131D | A | 3.7e-08 | 1.5e-08 | 4.9e-08 | 3.367E-08 | 1.724E-08 | 1.500E-08 | 4.900E-08 |
| 2605 | 131 | V131C | A | 6.6e-08 | 5.4e-08 | 7.6e-08 | 6.533E-08 | 1.102E-08 | 5.400E-08 | 7.600E-08 |
| 2606 | 131 | V131E | A | 2e-08 | 6.9e-08 | 3.5e-08 | 4.133E-08 | 2.511E-08 | 2.000E-08 | 6.900E-08 |
| 2607 | 131 | V131Q | A | 3.1e-08 | 1.2e-07 | 5e-08 | 6.700E-08 | 4.687E-08 | 3.100E-08 | 1.200E-07 |
| 2608 | 131 | V131G | A | 9.8e-08 | 4.2e-08 | 6.3e-08 | 6.767E-08 | 2.829E-08 | 4.200E-08 | 9.800E-08 |
| 2609 | 131 | V131H | A | 1.2e-08 | 2e-08 | 1.3e-08 | 1.500E-08 | 4.359E-09 | 1.200E-08 | 2.000E-08 |
| 2610 | 131 | V131I | A | 4e-08 | 1e-08 | 2.3e-08 | 2.433E-08 | 1.504E-08 | 1.000E-08 | 4.000E-08 |
| 2611 | 131 | V131L | A | 6.4e-08 | 1.6e-07 | 6.1e-08 | 9.500E-08 | 5.631E-08 | 6.100E-08 | 1.600E-07 |
| 2612 | 131 | V131K | A | 5.6e-08 | 5e-08 | 3.6e-08 | 4.733E-08 | 1.026E-08 | 3.600E-08 | 5.600E-08 |
| 2613 | 131 | V131M | A | 3.4e-08 | 4.9e-08 | 4.5e-08 | 4.267E-08 | 7.767E-09 | 3.400E-08 | 4.900E-08 |
| 2614 | 131 | V131F | A | 6.1e-08 | 2.8e-08 | 7.7e-08 | 5.533E-08 | 2.499E-08 | 2.800E-08 | 7.700E-08 |
| 2615 | 131 | V131P | A | 2.9e-08 | 4.8e-08 | 5.1e-08 | 4.267E-08 | 1.193E-08 | 2.900E-08 | 5.100E-08 |
| 2616 | 131 | V131S | A | 4e-08 | 6e-08 | 3.3e-08 | 4.433E-08 | 1.401E-08 | 3.300E-08 | 6.000E-08 |
| 2617 | 131 | V131T | A | 2.1e-08 | 6.1e-08 | 2.8e-08 | 3.667E-08 | 2.136E-08 | 2.100E-08 | 6.100E-08 |
| 2618 | 131 | V131W | A | 1.3e-07 | 4.2e-08 | 7.6e-08 | 8.267E-08 | 4.438E-08 | 4.200E-08 | 1.300E-07 |
| 2619 | 131 | V131Y | A | 3.6e-08 | 5.2e-08 | 3.3e-08 | 4.033E-08 | 1.021E-08 | 3.300E-08 | 5.200E-08 |
| 2620 | 131 | V131V | A | 3.1e-08 | 5.4e-08 | 9e-08 | 5.833E-08 | 2.974E-08 | 3.100E-08 | 9.000E-08 |
| 2621 | 132 | L132A | A | 2.4e-08 | 1.5e-08 | 5.5e-08 | 3.133E-08 | 2.098E-08 | 1.500E-08 | 5.500E-08 |
| 2622 | 132 | L132R | A | 5.9e-08 | 4.3e-08 | 2.7e-08 | 4.300E-08 | 1.600E-08 | 2.700E-08 | 5.900E-08 |
| 2623 | 132 | L132N | A | 2.5e-08 | 2.9e-08 | 1.7e-07 | 7.467E-08 | 8.259E-08 | 2.500E-08 | 1.700E-07 |
| 2624 | 132 | L132D | A | 2.9e-08 | 4.2e-08 | 1.4e-08 | 2.833E-08 | 1.401E-08 | 1.400E-08 | 4.200E-08 |
| 2625 | 132 | L132C | A | 2.4e-08 | 4e-08 | 4.7e-08 | 3.700E-08 | 1.179E-08 | 2.400E-08 | 4.700E-08 |
| 2626 | 132 | L132E | A | 8.7e-08 | 3.9e-08 | 3.9e-08 | 5.500E-08 | 2.771E-08 | 3.900E-08 | 8.700E-08 |
| 2627 | 132 | L132Q | A | 1.8e-08 | 9.3e-08 | 5e-08 | 5.367E-08 | 3.763E-08 | 1.800E-08 | 9.300E-08 |
| 2628 | 132 | L132G | A | 2.7e-08 | 2.6e-08 | 3.5e-08 | 2.933E-08 | 4.933E-09 | 2.600E-08 | 3.500E-08 |
| 2629 | 132 | L132H | A | 6.9e-08 | 2e-08 | 3.8e-08 | 4.233E-08 | 2.479E-08 | 2.000E-08 | 6.900E-08 |
| 2630 | 132 | L132I | A | 1.2e-08 | 4.9e-08 | 5.9e-08 | 4.000E-08 | 2.476E-08 | 1.200E-08 | 5.900E-08 |
| 2631 | 132 | L132L | A | 5.3e-08 | 6.8e-08 | 4.3e-08 | 5.467E-08 | 1.258E-08 | 4.300E-08 | 6.800E-08 |
| 2632 | 132 | L132K | A | 6.6e-08 | 7e-08 | 8.9e-08 | 7.500E-08 | 1.229E-08 | 6.600E-08 | 8.900E-08 |
| 2633 | 132 | L132M | A | 3.3e-08 | 1.6e-08 | 3.6e-08 | 2.833E-08 | 1.079E-08 | 1.600E-08 | 3.600E-08 |
| 2634 | 132 | L132F | A | 1.8e-08 | 1.7e-08 | 3.6e-08 | 2.367E-08 | 1.069E-08 | 1.700E-08 | 3.600E-08 |

| | | | | | | | | | | |
|---|---|---|---|---|---|---|---|---|---|---|
| 2635 | 132 | L132P | A | 4.9e-08 | 1.1e-08 | 4.4e-08 | 3.467E-08 | 2.065E-08 | 1.100E-08 | 4.900E-08 |
| 2636 | 132 | L132S | A | 9e-08 | 2.8e-08 | 1.9e-08 | 4.567E-08 | 3.866E-08 | 1.900E-08 | 9.000E-08 |
| 2637 | 132 | L132T | A | 5e-08 | 8.4e-08 | 3e-08 | 5.467E-08 | 2.730E-08 | 3.000E-08 | 8.400E-08 |
| 2638 | 132 | L132W | A | 5.5e-08 | 4.1e-08 | 3.7e-08 | 4.433E-08 | 9.452E-09 | 3.700E-08 | 5.500E-08 |
| 2639 | 132 | L132Y | A | 3e-08 | 2.4e-08 | 2.8e-08 | 2.733E-08 | 3.055E-09 | 2.400E-08 | 3.000E-08 |
| 2640 | 132 | L132V | A | 7.8e-08 | 3.3e-08 | 8.3e-08 | 6.467E-08 | 2.754E-08 | 3.300E-08 | 8.300E-08 |
| 2641 | 133 | I133A | A | 3.7e-08 | 1.4e-08 | 5.1e-08 | 3.400E-08 | 1.868E-08 | 1.400E-08 | 5.100E-08 |
| 2642 | 133 | I133R | A | 3.3e-08 | 7.6e-09 | 5.8e-08 | 3.287E-08 | 2.520E-08 | 7.600E-09 | 5.800E-08 |
| 2643 | 133 | I133N | A | 2.7e-08 | 5.8e-08 | 3.5e-08 | 4.000E-08 | 1.609E-08 | 2.700E-08 | 5.800E-08 |
| 2644 | 133 | I133D | A | 3.2e-08 | 7.4e-08 | 2e-08 | 4.200E-08 | 2.835E-08 | 2.000E-08 | 7.400E-08 |
| 2645 | 133 | I133C | A | 3.2e-08 | 2e-08 | 9.5e-09 | 2.050E-08 | 1.126E-08 | 9.500E-09 | 3.200E-08 |
| 2646 | 133 | I133E | A | 6.3e-08 | 2.2e-08 | 2.6e-08 | 3.700E-08 | 2.261E-08 | 2.200E-08 | 6.300E-08 |
| 2647 | 133 | I133Q | A | 7e-08 | 1.4e-07 | 5.7e-09 | 7.190E-08 | 6.717E-08 | 5.700E-09 | 1.400E-07 |
| 2648 | 133 | I133G | A | 6.5e-08 | 4.6e-08 | 2.5e-08 | 4.533E-08 | 2.001E-08 | 2.500E-08 | 6.500E-08 |
| 2649 | 133 | I133H | A | 1.9e-08 | 2.7e-08 | 4e-08 | 2.867E-08 | 1.060E-08 | 1.900E-08 | 4.000E-08 |
| 2650 | 133 | I133I | A | 2e-08 | 3.6e-08 | 2e-08 | 2.533E-08 | 9.238E-09 | 2.000E-08 | 3.600E-08 |
| 2651 | 133 | I133L | A | 1.6e-08 | 3.4e-08 | 9.6e-08 | 4.867E-08 | 4.197E-08 | 1.600E-08 | 9.600E-08 |
| 2652 | 133 | I133K | A | 5e-08 | 3.7e-08 | 8.5e-08 | 5.733E-08 | 2.483E-08 | 3.700E-08 | 8.500E-08 |
| 2653 | 133 | I133M | A | 6.6e-08 | 3.7e-08 | 3.3e-08 | 4.533E-08 | 1.801E-08 | 3.300E-08 | 6.600E-08 |
| 2654 | 133 | I133F | A | 5.7e-08 | 2.7e-08 | 5.4e-08 | 4.600E-08 | 1.652E-08 | 2.700E-08 | 5.700E-08 |
| 2655 | 133 | I133P | A | 1.6e-07 | 2.9e-08 | 3.2e-08 | 7.367E-08 | 7.478E-08 | 2.900E-08 | 1.600E-07 |
| 2656 | 133 | I133S | A | 1e-07 | 4.2e-08 | 6.3e-08 | 6.833E-08 | 2.937E-08 | 4.200E-08 | 1.000E-07 |
| 2657 | 133 | I133T | A | 5.8e-08 | 6.5e-08 | 2e-08 | 4.767E-08 | 2.421E-08 | 2.000E-08 | 6.500E-08 |
| 2658 | 133 | I133W | A | 1.7e-07 | 4.5e-08 | 4e-08 | 8.500E-08 | 7.365E-08 | 4.000E-08 | 1.700E-07 |
| 2659 | 133 | I133Y | A | 1.1e-08 | 5.1e-08 | 3.4e-08 | 3.200E-08 | 2.007E-08 | 1.100E-08 | 5.100E-08 |
| 2660 | 133 | I133V | A | 6.5e-08 | 5.6e-08 | 7.7e-08 | 6.600E-08 | 1.054E-08 | 5.600E-08 | 7.700E-08 |
| 2661 | 134 | Q134A | A | 2.8e-08 | 4e-08 | 7.9e-08 | 4.900E-08 | 2.666E-08 | 2.800E-08 | 7.900E-08 |
| 2662 | 134 | Q134R | A | 3.5e-08 | 4.4e-08 | 9.2e-08 | 5.700E-08 | 3.064E-08 | 3.500E-08 | 9.200E-08 |
| 2663 | 134 | Q134N | A | 2.8e-08 | 1.7e-08 | 5.7e-08 | 3.400E-08 | 2.066E-08 | 1.700E-08 | 5.700E-08 |
| 2664 | 134 | Q134D | A | 3.8e-08 | 4.1e-08 | 2.9e-08 | 3.600E-08 | 6.245E-09 | 2.900E-08 | 4.100E-08 |
| 2665 | 134 | Q134C | A | 6.1e-08 | 3.2e-08 | 6.3e-08 | 5.200E-08 | 1.735E-08 | 3.200E-08 | 6.300E-08 |
| 2666 | 134 | Q134E | A | 1.4e-08 | 5.6e-08 | 4.2e-08 | 3.733E-08 | 2.139E-08 | 1.400E-08 | 5.600E-08 |
| 2667 | 134 | Q134Q | A | 1.3e-07 | 1.4e-08 | 8.3e-08 | 7.567E-08 | 5.835E-08 | 1.400E-08 | 1.300E-07 |
| 2668 | 134 | Q134G | A | 4.9e-08 | 6.5e-08 | 6.1e-08 | 5.833E-08 | 8.327E-09 | 4.900E-08 | 6.500E-08 |
| 2669 | 134 | Q134H | A | 8.8e-09 | 3e-08 | 1.4e-08 | 1.760E-08 | 1.105E-08 | 8.800E-09 | 3.000E-08 |
| 2670 | 134 | Q134I | A | 5.4e-08 | 2.5e-08 | 3.2e-08 | 3.700E-08 | 1.513E-08 | 2.500E-08 | 5.400E-08 |
| 2671 | 134 | Q134L | A | 7.2e-08 | 9.1e-08 | 2e-08 | 6.100E-08 | 3.676E-08 | 2.000E-08 | 9.100E-08 |
| 2672 | 134 | Q134K | A | 2.3e-08 | 4.6e-08 | 4.3e-08 | 3.733E-08 | 1.250E-08 | 2.300E-08 | 4.600E-08 |
| 2673 | 134 | Q134M | A | 9.4e-08 | 5.1e-08 | 4.1e-08 | 6.200E-08 | 2.816E-08 | 4.100E-08 | 9.400E-08 |
| 2674 | 134 | Q134F | A | 1.1e-07 | 7.5e-08 | 4.4e-08 | 7.633E-08 | 3.302E-08 | 4.400E-08 | 1.100E-07 |
| 2675 | 134 | Q134P | A | 4.4e-08 | 2.2e-08 | 2.3e-08 | 2.967E-08 | 1.242E-08 | 2.200E-08 | 4.400E-08 |
| 2676 | 134 | Q134S | A | 9.1e-08 | 2.8e-08 | 2.3e-08 | 4.733E-08 | 3.790E-08 | 2.300E-08 | 9.100E-08 |
| 2677 | 134 | Q134T | A | 3.8e-08 | 8.7e-08 | 1.7e-08 | 4.733E-08 | 3.592E-08 | 1.700E-08 | 8.700E-08 |
| 2678 | 134 | Q134W | A | 2.4e-08 | 8.9e-08 | 2.3e-08 | 4.533E-08 | 3.782E-08 | 2.300E-08 | 8.900E-08 |
| 2679 | 134 | Q134Y | A | 6.7e-08 | 1.6e-08 | 4.3e-08 | 4.200E-08 | 2.551E-08 | 1.600E-08 | 6.700E-08 |
| 2680 | 134 | Q134V | A | 7.1e-08 | 2.7e-08 | 5.4e-08 | 5.067E-08 | 2.219E-08 | 2.700E-08 | 7.100E-08 |
| 2681 | 135 | R135A | A | 4.4e-08 | 6.1e-08 | 4.9e-08 | 5.133E-08 | 8.737E-09 | 4.400E-08 | 6.100E-08 |
| 2682 | 135 | R135R | A | 1.3e-07 | 4.3e-08 | 5.2e-08 | 7.500E-08 | 4.784E-08 | 4.300E-08 | 1.300E-07 |
| 2683 | 135 | R135N | A | 4e-08 | 2.4e-08 | 2e-08 | 2.800E-08 | 1.058E-08 | 2.000E-08 | 4.000E-08 |
| 2684 | 135 | R135D | A | 1.1e-07 | 5e-08 | 2.6e-08 | 6.200E-08 | 4.327E-08 | 2.600E-08 | 1.100E-07 |
| 2685 | 135 | R135C | A | 4.1e-08 | 7.2e-08 | 2.8e-08 | 4.700E-08 | 2.261E-08 | 2.800E-08 | 7.200E-08 |
| 2686 | 135 | R135E | A | 7.9e-08 | 3e-08 | 7.9e-08 | 6.267E-08 | 2.829E-08 | 3.000E-08 | 7.900E-08 |
| 2687 | 135 | R135Q | A | 3.2e-08 | 3.9e-08 | 1.5e-08 | 2.867E-08 | 1.234E-08 | 1.500E-08 | 3.900E-08 |
| 2688 | 135 | R135G | A | 5.2e-08 | 1.1e-07 | 6.2e-08 | 7.467E-08 | 3.101E-08 | 5.200E-08 | 1.100E-07 |
| 2689 | 135 | R135H | A | 7e-08 | 1.5e-08 | 3.1e-08 | 3.867E-08 | 2.829E-08 | 1.500E-08 | 7.000E-08 |
| 2690 | 135 | R135I | A | 3.7e-08 | 4.3e-08 | 5.1e-08 | 4.367E-08 | 7.024E-09 | 3.700E-08 | 5.100E-08 |
| 2691 | 135 | R135L | A | 6.6e-08 | 9.1e-08 | 9.6e-08 | 8.433E-08 | 1.607E-08 | 6.600E-08 | 9.600E-08 |
| 2692 | 135 | R135K | A | 9.4e-08 | 1.2e-07 | 1.9e-08 | 7.767E-08 | 5.244E-08 | 1.900E-08 | 1.200E-07 |
| 2693 | 135 | R135M | A | 2.6e-08 | 5e-08 | 4.6e-08 | 4.067E-08 | 1.286E-08 | 2.600E-08 | 5.000E-08 |
| 2694 | 135 | R135F | A | 3.8e-08 | 7.6e-08 | 4.8e-08 | 5.400E-08 | 1.970E-08 | 3.800E-08 | 7.600E-08 |
| 2695 | 135 | R135P | A | 3e-08 | 1.3e-08 | 1.2e-08 | 1.833E-08 | 1.012E-08 | 1.200E-08 | 3.000E-08 |
| 2696 | 135 | R135S | A | 1.1e-07 | 3.1e-08 | 2.1e-08 | 5.400E-08 | 4.875E-08 | 2.100E-08 | 1.100E-07 |
| 2697 | 135 | R135T | A | 4.6e-08 | 1.6e-08 | 2.7e-08 | 2.967E-08 | 1.518E-08 | 1.600E-08 | 4.600E-08 |
| 2698 | 135 | R135W | A | 1.5e-07 | 1.8e-08 | 8.9e-08 | 8.567E-08 | 6.606E-08 | 1.800E-08 | 1.500E-07 |
| 2699 | 135 | R135Y | A | 3.7e-08 | 5.9e-08 | 7.5e-08 | 5.700E-08 | 1.908E-08 | 3.700E-08 | 7.500E-08 |
| 2700 | 135 | R135V | A | 4e-08 | 3.7e-08 | 4.1e-08 | 3.933E-08 | 2.082E-09 | 3.700E-08 | 4.100E-08 |
| 2701 | 136 | N136A | A | 5e-08 | 5e-08 | 3.1e-08 | 4.367E-08 | 1.097E-08 | 3.100E-08 | 5.000E-08 |
| 2702 | 136 | N136R | A | 5.2e-08 | 5.9e-08 | 3.1e-08 | 4.733E-08 | 1.457E-08 | 3.100E-08 | 5.900E-08 |
| 2703 | 136 | N136N | A | 1.5e-07 | 5.4e-08 | 2.9e-08 | 7.767E-08 | 6.388E-08 | 2.900E-08 | 1.500E-07 |
| 2704 | 136 | N136D | A | 2.6e-08 | 2e-07 | 4.8e-08 | 9.133E-08 | 9.475E-08 | 2.600E-08 | 2.000E-07 |
| 2705 | 136 | N136C | A | 4.6e-08 | 5.4e-08 | 2.9e-08 | 4.300E-08 | 1.277E-08 | 2.900E-08 | 5.400E-08 |
| 2706 | 136 | N136E | A | 5.1e-08 | 3.1e-08 | 3.5e-08 | 3.900E-08 | 1.058E-08 | 3.100E-08 | 5.100E-08 |
| 2707 | 136 | N136Q | A | 4e-08 | 3.9e-08 | 3.8e-08 | 3.900E-08 | 1.000E-09 | 3.800E-08 | 4.000E-08 |
| 2708 | 136 | N136G | A | 5.8e-08 | 2.2e-08 | 2e-08 | 3.333E-08 | 2.139E-08 | 2.000E-08 | 5.800E-08 |
| 2709 | 136 | N136H | A | 8.5e-08 | 2.2e-08 | 1.6e-08 | 4.100E-08 | 3.822E-08 | 1.600E-08 | 8.500E-08 |
| 2710 | 136 | N136I | A | 1.5e-08 | 3.4e-08 | 1.7e-07 | 7.300E-08 | 8.454E-08 | 1.500E-08 | 1.700E-07 |
| 2711 | 136 | N136L | A | 4.1e-08 | 4.9e-08 | 2.3e-08 | 3.767E-08 | 1.332E-08 | 2.300E-08 | 4.900E-08 |
| 2712 | 136 | N136K | A | 7.1e-08 | 1.8e-08 | 5.4e-08 | 4.767E-08 | 2.706E-08 | 1.800E-08 | 7.100E-08 |
| 2713 | 136 | N136M | A | 4.7e-08 | 3.5e-08 | 4.6e-08 | 4.267E-08 | 6.658E-09 | 3.500E-08 | 4.700E-08 |
| 2714 | 136 | N136F | A | 9.2e-08 | 2.6e-08 | 2.2e-08 | 4.667E-08 | 3.931E-08 | 2.200E-08 | 9.200E-08 |
| 2715 | 136 | N136P | A | 3.9e-08 | 1.5e-08 | 5e-08 | 3.467E-08 | 1.790E-08 | 1.500E-08 | 5.000E-08 |
| 2716 | 136 | N136S | A | 4.6e-08 | 7.8e-08 | 1e-07 | 7.467E-08 | 2.715E-08 | 4.600E-08 | 1.000E-07 |
| 2717 | 136 | N136T | A | 3.1e-08 | 3.4e-08 | 1.4e-08 | 2.633E-08 | 1.079E-08 | 1.400E-08 | 3.400E-08 |
| 2718 | 136 | N136W | A | 1.8e-08 | 3.9e-08 | 5.1e-08 | 3.600E-08 | 1.670E-08 | 1.800E-08 | 5.100E-08 |
| 2719 | 136 | N136Y | A | 7.2e-08 | 3.3e-08 | 5.2e-08 | 5.233E-08 | 1.950E-08 | 3.300E-08 | 7.200E-08 |

| | | | | | | | | | | |
|---|---|---|---|---|---|---|---|---|---|---|
| 2720 | 136 | N136V | A | 2.8e-08 | 2.2e-08 | 4.2e-08 | 3.067E-08 | 1.026E-08 | 2.200E-08 | 4.200E-08 |
| 2721 | 137 | P137A | A | 1.5e-07 | 2.1e-08 | 9.2e-08 | 8.767E-08 | 6.461E-08 | 2.100E-08 | 1.500E-07 |
| 2722 | 137 | P137R | A | 8e-09 | 4.8e-08 | 3.4e-08 | 3.000E-08 | 2.030E-08 | 8.000E-09 | 4.800E-08 |
| 2723 | 137 | P137N | A | 1.6e-07 | 7e-08 | 9.9e-08 | 1.097E-07 | 4.594E-08 | 7.000E-08 | 1.600E-07 |
| 2724 | 137 | P137D | A | 1.5e-07 | 7.6e-08 | 4.3e-08 | 8.967E-08 | 5.479E-08 | 4.300E-08 | 1.500E-07 |
| 2725 | 137 | P137C | A | 3.2e-08 | 1.4e-08 | 6.3e-08 | 3.633E-08 | 2.479E-08 | 1.400E-08 | 6.300E-08 |
| 2726 | 137 | P137E | A | 3.1e-08 | 4e-08 | 3e-08 | 3.367E-08 | 5.508E-09 | 3.000E-08 | 4.000E-08 |
| 2727 | 137 | P137Q | A | 7.8e-08 | 4e-08 | 8.5e-08 | 6.767E-08 | 2.421E-08 | 4.000E-08 | 8.500E-08 |
| 2728 | 137 | P137G | A | 7.5e-08 | 1.9e-08 | 8.5e-08 | 5.967E-08 | 3.557E-08 | 1.900E-08 | 8.500E-08 |
| 2729 | 137 | P137H | A | 4.7e-08 | 3.7e-08 | 3.7e-08 | 4.033E-08 | 5.774E-09 | 3.700E-08 | 4.700E-08 |
| 2730 | 137 | P137I | A | 3.5e-08 | 6.3e-08 | 7.3e-08 | 5.700E-08 | 1.970E-08 | 3.500E-08 | 7.300E-08 |
| 2731 | 137 | P137L | A | 7e-08 | 3.4e-08 | 3.7e-08 | 4.700E-08 | 1.997E-08 | 3.400E-08 | 7.000E-08 |
| 2732 | 137 | P137K | A | 4.5e-08 | 6e-08 | 2.9e-08 | 4.467E-08 | 1.550E-08 | 2.900E-08 | 6.000E-08 |
| 2733 | 137 | P137M | A | 5.9e-08 | 9.7e-08 | 1.3e-07 | 9.533E-08 | 3.553E-08 | 5.900E-08 | 1.300E-07 |
| 2734 | 137 | P137F | A | 8.2e-08 | 1.3e-07 | 4.1e-08 | 8.433E-08 | 4.455E-08 | 4.100E-08 | 1.300E-07 |
| 2735 | 137 | P137P | A | 4.8e-08 | 6.3e-08 | 1.6e-08 | 4.233E-08 | 2.401E-08 | 1.600E-08 | 6.300E-08 |
| 2736 | 137 | P137S | A | 3.1e-08 | 1.7e-07 | 7.6e-08 | 9.233E-08 | 7.092E-08 | 3.100E-08 | 1.700E-07 |
| 2737 | 137 | P137T | A | 5e-08 | 1.6e-08 | 5.7e-08 | 4.100E-08 | 2.193E-08 | 1.600E-08 | 5.700E-08 |
| 2738 | 137 | P137W | A | 3.6e-08 | 1.1e-07 | 3.3e-08 | 5.967E-08 | 4.362E-08 | 3.300E-08 | 1.100E-07 |
| 2739 | 137 | P137Y | A | 3.5e-08 | 3.3e-08 | 4.2e-08 | 3.667E-08 | 4.726E-09 | 3.300E-08 | 4.200E-08 |
| 2740 | 137 | P137V | A | 6.5e-08 | 1.2e-07 | 5.7e-08 | 8.067E-08 | 3.430E-08 | 5.700E-08 | 1.200E-07 |
| 2741 | 138 | Q138A | A | 4.9e-08 | 4.8e-08 | 2.5e-08 | 4.067E-08 | 1.358E-08 | 2.500E-08 | 4.900E-08 |
| 2742 | 138 | Q138R | A | 3.2e-08 | 1.1e-08 | 2.5e-08 | 2.267E-08 | 1.069E-08 | 1.100E-08 | 3.200E-08 |
| 2743 | 138 | Q138N | A | 2.2e-08 | 4.6e-08 | 1.7e-08 | 2.833E-08 | 1.550E-08 | 1.700E-08 | 4.600E-08 |
| 2744 | 138 | Q138D | A | 2.4e-08 | 4.8e-08 | 3.5e-08 | 3.567E-08 | 1.201E-08 | 2.400E-08 | 4.800E-08 |
| 2745 | 138 | Q138C | A | 5.7e-08 | 4.9e-08 | 2.5e-08 | 4.367E-08 | 1.665E-08 | 2.500E-08 | 5.700E-08 |
| 2746 | 138 | Q138E | A | 3e-08 | 6.4e-08 | 1.1e-07 | 6.800E-08 | 4.015E-08 | 3.000E-08 | 1.100E-07 |
| 2747 | 138 | Q138Q | A | 5.1e-08 | 2.7e-08 | 6.1e-08 | 4.633E-08 | 1.747E-08 | 2.700E-08 | 6.100E-08 |
| 2748 | 138 | Q138G | A | 3.2e-08 | 4.6e-08 | 1.8e-08 | 3.200E-08 | 1.400E-08 | 1.800E-08 | 4.600E-08 |
| 2749 | 138 | Q138H | A | 1.3e-07 | 5.8e-08 | 1.6e-08 | 6.800E-08 | 5.765E-08 | 1.600E-08 | 1.300E-07 |
| 2750 | 138 | Q138I | A | 4.8e-08 | 3.7e-08 | 1.1e-07 | 6.500E-08 | 3.936E-08 | 3.700E-08 | 1.100E-07 |
| 2751 | 138 | Q138L | A | 4.3e-08 | 4e-08 | 1.5e-08 | 3.267E-08 | 1.537E-08 | 1.500E-08 | 4.300E-08 |
| 2752 | 138 | Q138K | A | 1.6e-08 | 2.4e-08 | 2e-08 | 2.000E-08 | 4.000E-09 | 1.600E-08 | 2.400E-08 |
| 2753 | 138 | Q138M | A | 9.5e-09 | 3.3e-08 | 1.3e-08 | 1.850E-08 | 1.268E-08 | 9.500E-09 | 3.300E-08 |
| 2754 | 138 | Q138F | A | 6.8e-08 | 3.2e-08 | 8.5e-08 | 6.167E-08 | 2.706E-08 | 3.200E-08 | 8.500E-08 |
| 2755 | 138 | Q138P | A | 2e-08 | 3.7e-08 | 2.9e-08 | 2.867E-08 | 8.505E-09 | 2.000E-08 | 3.700E-08 |
| 2756 | 138 | Q138S | A | 6.3e-08 | 3.7e-08 | 1.4e-07 | 8.000E-08 | 5.356E-08 | 3.700E-08 | 1.400E-07 |
| 2757 | 138 | Q138T | A | 3.2e-08 | 4.1e-08 | 2.1e-08 | 3.133E-08 | 1.002E-08 | 2.100E-08 | 4.100E-08 |
| 2758 | 138 | Q138W | A | 2.3e-08 | 4.5e-08 | 5.1e-08 | 3.967E-08 | 1.474E-08 | 2.300E-08 | 5.100E-08 |
| 2759 | 138 | Q138Y | A | 7e-08 | 1.6e-08 | 2.7e-08 | 3.767E-08 | 2.854E-08 | 1.600E-08 | 7.000E-08 |
| 2760 | 138 | Q138V | A | 2.6e-08 | 2.8e-08 | 4.8e-08 | 3.400E-08 | 1.217E-08 | 2.600E-08 | 4.800E-08 |
| 2761 | 139 | L139A | A | 2.1e-08 | 7.2e-08 | 4.9e-08 | 4.733E-08 | 2.554E-08 | 2.100E-08 | 7.200E-08 |
| 2762 | 139 | L139R | A | 5.6e-08 | 2.5e-08 | 4.5e-08 | 4.200E-08 | 1.572E-08 | 2.500E-08 | 5.600E-08 |
| 2763 | 139 | L139N | A | 9.9e-08 | 2e-08 | 1.2e-07 | 7.967E-08 | 5.273E-08 | 2.000E-08 | 1.200E-07 |
| 2764 | 139 | L139D | A | 1.3e-08 | 1.7e-08 | 5e-08 | 2.667E-08 | 2.031E-08 | 1.300E-08 | 5.000E-08 |
| 2765 | 139 | L139C | A | 4.3e-08 | 5.8e-08 | 3.2e-08 | 4.433E-08 | 1.305E-08 | 3.200E-08 | 5.800E-08 |
| 2766 | 139 | L139E | A | 6.1e-08 | 2.2e-08 | 4.3e-08 | 4.200E-08 | 1.952E-08 | 2.200E-08 | 6.100E-08 |
| 2767 | 139 | L139Q | A | 5.6e-08 | 6.2e-08 | 1.6e-07 | 9.267E-08 | 5.839E-08 | 5.600E-08 | 1.600E-07 |
| 2768 | 139 | L139G | A | 3.3e-08 | 3.7e-08 | 6.5e-08 | 4.500E-08 | 1.744E-08 | 3.300E-08 | 6.500E-08 |
| 2769 | 139 | L139H | A | 2.4e-08 | 7.4e-08 | 5.2e-08 | 5.000E-08 | 2.506E-08 | 2.400E-08 | 7.400E-08 |
| 2770 | 139 | L139I | A | 4.3e-08 | 8.4e-08 | 2.9e-08 | 5.200E-08 | 2.858E-08 | 2.900E-08 | 8.400E-08 |
| 2771 | 139 | L139L | A | 2.1e-08 | 2.3e-08 | 2.1e-08 | 2.167E-08 | 1.155E-09 | 2.100E-08 | 2.300E-08 |
| 2772 | 139 | L139K | A | 2.5e-08 | 8.7e-08 | 1.8e-08 | 4.333E-08 | 3.798E-08 | 1.800E-08 | 8.700E-08 |
| 2773 | 139 | L139M | A | 5.9e-08 | 2.2e-08 | 3.2e-08 | 3.767E-08 | 1.914E-08 | 2.200E-08 | 5.900E-08 |
| 2774 | 139 | L139F | A | 3e-08 | 5.4e-08 | 7.5e-08 | 5.300E-08 | 2.252E-08 | 3.000E-08 | 7.500E-08 |
| 2775 | 139 | L139P | A | 7.3e-08 | 3.5e-08 | 4.7e-08 | 5.167E-08 | 1.943E-08 | 3.500E-08 | 7.300E-08 |
| 2776 | 139 | L139S | A | 6.2e-08 | 6.2e-08 | 5.8e-08 | 6.067E-08 | 2.309E-09 | 5.800E-08 | 6.200E-08 |
| 2777 | 139 | L139T | A | 6.6e-08 | 1e-07 | 3.2e-08 | 6.600E-08 | 3.400E-08 | 3.200E-08 | 1.000E-07 |
| 2778 | 139 | L139W | A | 5.5e-08 | 4.3e-08 | 4.7e-08 | 4.833E-08 | 6.110E-09 | 4.300E-08 | 5.500E-08 |
| 2779 | 139 | L139Y | A | 6.9e-08 | 1.5e-08 | 4.2e-08 | 4.200E-08 | 2.700E-08 | 1.500E-08 | 6.900E-08 |
| 2780 | 139 | L139V | A | 3.6e-08 | 2.5e-08 | 1.4e-08 | 2.500E-08 | 1.100E-08 | 1.400E-08 | 3.600E-08 |
| 2781 | 140 | C140A | A | 3.3e-08 | 1.8e-08 | 6.6e-08 | 3.900E-08 | 2.456E-08 | 1.800E-08 | 6.600E-08 |
| 2782 | 140 | C140R | A | 3.4e-08 | 3.7e-08 | 1.5e-08 | 2.867E-08 | 1.193E-08 | 1.500E-08 | 3.700E-08 |
| 2783 | 140 | C140N | A | 7.5e-08 | 7.7e-08 | 1.3e-07 | 9.400E-08 | 3.119E-08 | 7.500E-08 | 1.300E-07 |
| 2784 | 140 | C140D | A | 3.7e-08 | 1.8e-08 | 4.2e-08 | 3.233E-08 | 1.266E-08 | 1.800E-08 | 4.200E-08 |
| 2785 | 140 | C140C | A | 5.3e-08 | 1.1e-07 | 2.4e-08 | 6.233E-08 | 4.375E-08 | 2.400E-08 | 1.100E-07 |
| 2786 | 140 | C140E | A | 6.4e-08 | 9.3e-08 | 4.6e-08 | 6.767E-08 | 2.371E-08 | 4.600E-08 | 9.300E-08 |
| 2787 | 140 | C140Q | A | 4.4e-08 | 6e-08 | 4e-08 | 4.800E-08 | 1.058E-08 | 4.000E-08 | 6.000E-08 |
| 2788 | 140 | C140G | A | 4.3e-08 | 2.9e-08 | 5e-08 | 4.067E-08 | 1.069E-08 | 2.900E-08 | 5.000E-08 |
| 2789 | 140 | C140H | A | 5.3e-08 | 4.7e-08 | 6.3e-08 | 5.433E-08 | 8.083E-09 | 4.700E-08 | 6.300E-08 |
| 2790 | 140 | C140I | A | 2.9e-08 | 2.9e-08 | 4.6e-08 | 3.467E-08 | 9.815E-09 | 2.900E-08 | 4.600E-08 |
| 2791 | 140 | C140L | A | 5.9e-08 | 2e-08 | 1.6e-07 | 7.967E-08 | 7.225E-08 | 2.000E-08 | 1.600E-07 |
| 2792 | 140 | C140K | A | 2.9e-08 | 5.5e-08 | 1.1e-07 | 6.467E-08 | 4.136E-08 | 2.900E-08 | 1.100E-07 |
| 2793 | 140 | C140M | A | 7e-08 | 1.6e-08 | 6.9e-08 | 5.167E-08 | 3.089E-08 | 1.600E-08 | 7.000E-08 |
| 2794 | 140 | C140F | A | 7.9e-08 | 5.1e-08 | 1.6e-08 | 4.867E-08 | 3.156E-08 | 1.600E-08 | 7.900E-08 |
| 2795 | 140 | C140P | A | 5.7e-08 | 2.5e-08 | 2.6e-08 | 3.600E-08 | 1.819E-08 | 2.500E-08 | 5.700E-08 |
| 2796 | 140 | C140S | A | 6.3e-08 | 2.5e-08 | 4e-08 | 4.267E-08 | 1.914E-08 | 2.500E-08 | 6.300E-08 |
| 2797 | 140 | C140T | A | 1.9e-08 | 7.1e-08 | 4.8e-08 | 4.600E-08 | 2.606E-08 | 1.900E-08 | 7.100E-08 |
| 2798 | 140 | C140W | A | 4.3e-08 | 1.4e-08 | 1.8e-08 | 2.500E-08 | 1.572E-08 | 1.400E-08 | 4.300E-08 |
| 2799 | 140 | C140Y | A | 5.5e-08 | 4.1e-08 | 4.2e-08 | 4.600E-08 | 7.810E-09 | 4.100E-08 | 5.500E-08 |
| 2800 | 140 | C140V | A | 2.3e-08 | 2.7e-08 | 6.8e-08 | 3.933E-08 | 2.491E-08 | 2.300E-08 | 6.800E-08 |
| 2801 | 141 | Y141A | A | 2.4e-08 | 9.8e-08 | 2.7e-08 | 4.967E-08 | 4.188E-08 | 2.400E-08 | 9.800E-08 |
| 2802 | 141 | Y141R | A | 3.5e-08 | 9.4e-08 | 1.5e-07 | 9.300E-08 | 5.751E-08 | 3.500E-08 | 1.500E-07 |
| 2803 | 141 | Y141N | A | 3e-08 | 3.5e-08 | 4.6e-08 | 3.700E-08 | 8.185E-09 | 3.000E-08 | 4.600E-08 |
| 2804 | 141 | Y141D | A | 3.4e-08 | 3.6e-08 | 2.3e-08 | 3.100E-08 | 7.000E-09 | 2.300E-08 | 3.600E-08 |

| | | | | | | | | | | |
|---|---|---|---|---|---|---|---|---|---|---|
| 2805 | 141 | Y141C | A | 7e-08 | 2.5e-08 | 1e-07 | 6.500E-08 | 3.775E-08 | 2.500E-08 | 1.000E-07 |
| 2806 | 141 | Y141E | A | 4.3e-08 | 2e-08 | 1.2e-07 | 6.100E-08 | 5.237E-08 | 2.000E-08 | 1.200E-07 |
| 2807 | 141 | Y141Q | A | 3.1e-08 | 4.1e-08 | 3e-08 | 3.400E-08 | 6.083E-09 | 3.000E-08 | 4.100E-08 |
| 2808 | 141 | Y141G | A | 4.3e-08 | 7.1e-08 | 2.8e-08 | 4.733E-08 | 2.183E-08 | 2.800E-08 | 7.100E-08 |
| 2809 | 141 | Y141H | A | 4.7e-08 | 3.5e-08 | 2.1e-08 | 3.433E-08 | 1.301E-08 | 2.100E-08 | 4.700E-08 |
| 2810 | 141 | Y141I | A | 1.3e-08 | 6.6e-08 | 3.9e-08 | 3.933E-08 | 2.650E-08 | 1.300E-08 | 6.600E-08 |
| 2811 | 141 | Y141L | A | 6.9e-08 | 5.4e-08 | 4.2e-08 | 5.500E-08 | 1.353E-08 | 4.200E-08 | 6.900E-08 |
| 2812 | 141 | Y141K | A | 3.5e-08 | 2.6e-08 | 6.7e-08 | 4.267E-08 | 2.155E-08 | 2.600E-08 | 6.700E-08 |
| 2813 | 141 | Y141M | A | 4.5e-08 | 2e-08 | 4e-08 | 3.500E-08 | 1.323E-08 | 2.000E-08 | 4.500E-08 |
| 2814 | 141 | Y141F | A | 6.7e-08 | 9.5e-08 | 3.3e-08 | 6.500E-08 | 3.105E-08 | 3.300E-08 | 9.500E-08 |
| 2815 | 141 | Y141P | A | 4.6e-08 | 1.8e-08 | 4.8e-08 | 3.733E-08 | 1.677E-08 | 1.800E-08 | 4.800E-08 |
| 2816 | 141 | Y141S | A | 9.4e-08 | 6.7e-08 | 1e-07 | 8.700E-08 | 1.758E-08 | 6.700E-08 | 1.000E-07 |
| 2817 | 141 | Y141T | A | 2.3e-08 | 8.2e-08 | 6.9e-08 | 5.800E-08 | 3.100E-08 | 2.300E-08 | 8.200E-08 |
| 2818 | 141 | Y141W | A | 5e-08 | 8e-08 | 2.5e-08 | 5.167E-08 | 2.754E-08 | 2.500E-08 | 8.000E-08 |
| 2819 | 141 | Y141Y | A | 4.3e-08 | 6.1e-08 | 7.9e-08 | 6.100E-08 | 1.800E-08 | 4.300E-08 | 7.900E-08 |
| 2820 | 141 | Y141V | A | 8.1e-08 | 5.8e-08 | 4.4e-08 | 6.100E-08 | 1.868E-08 | 4.400E-08 | 8.100E-08 |
| 2821 | 142 | Q142A | A | 7.3e-08 | 5.9e-08 | 7.6e-08 | 6.933E-08 | 9.074E-09 | 5.900E-08 | 7.600E-08 |
| 2822 | 142 | Q142R | A | 7.2e-08 | 3.2e-08 | 6.3e-08 | 5.567E-08 | 2.098E-08 | 3.200E-08 | 7.200E-08 |
| 2823 | 142 | Q142N | A | 2.8e-08 | 8.1e-08 | 5.8e-08 | 5.567E-08 | 2.658E-08 | 2.800E-08 | 8.100E-08 |
| 2824 | 142 | Q142D | A | 3.6e-08 | 7.5e-08 | 7.3e-08 | 6.133E-08 | 2.196E-08 | 3.600E-08 | 7.500E-08 |
| 2825 | 142 | Q142C | A | 6.4e-08 | 7.3e-08 | 1.1e-07 | 8.233E-08 | 2.438E-08 | 6.400E-08 | 1.100E-07 |
| 2826 | 142 | Q142E | A | 4.2e-08 | 6.4e-08 | 1.5e-08 | 4.033E-08 | 2.454E-08 | 1.500E-08 | 6.400E-08 |
| 2827 | 142 | Q142Q | A | 5e-08 | 3.9e-08 | 2.6e-08 | 3.833E-08 | 1.201E-08 | 2.600E-08 | 5.000E-08 |
| 2828 | 142 | Q142G | A | 2.2e-08 | 4.4e-08 | 5.3e-08 | 3.967E-08 | 1.595E-08 | 2.200E-08 | 5.300E-08 |
| 2829 | 142 | Q142H | A | 3.4e-08 | 5.6e-08 | 2.3e-08 | 3.767E-08 | 1.680E-08 | 2.300E-08 | 5.600E-08 |
| 2830 | 142 | Q142I | A | 6.9e-08 | 2.4e-08 | 9.8e-08 | 6.367E-08 | 3.729E-08 | 2.400E-08 | 9.800E-08 |
| 2831 | 142 | Q142L | A | 1e-07 | 2e-08 | 2.1e-08 | 4.700E-08 | 4.590E-08 | 2.000E-08 | 1.000E-07 |
| 2832 | 142 | Q142K | A | 3e-08 | 4.1e-08 | 4.3e-08 | 3.800E-08 | 7.000E-09 | 3.000E-08 | 4.300E-08 |
| 2833 | 142 | Q142M | A | 8e-08 | 3.6e-08 | 7.2e-08 | 6.267E-08 | 2.344E-08 | 3.600E-08 | 8.000E-08 |
| 2834 | 142 | Q142F | A | 4.8e-08 | 3.4e-08 | 2.8e-08 | 3.667E-08 | 1.026E-08 | 2.800E-08 | 4.800E-08 |
| 2835 | 142 | Q142P | A | 7.2e-08 | 3.9e-08 | 1.2e-08 | 4.100E-08 | 3.005E-08 | 1.200E-08 | 7.200E-08 |
| 2836 | 142 | Q142S | A | 5.3e-08 | 1.9e-08 | 8.2e-09 | 2.673E-08 | 2.338E-08 | 8.200E-09 | 5.300E-08 |
| 2837 | 142 | Q142T | A | 7.4e-08 | 9.6e-08 | 4.7e-08 | 7.233E-08 | 2.454E-08 | 4.700E-08 | 9.600E-08 |
| 2838 | 142 | Q142W | A | 3.8e-08 | 3.9e-08 | 4.1e-08 | 3.933E-08 | 1.528E-09 | 3.800E-08 | 4.100E-08 |
| 2839 | 142 | Q142Y | A | 5.4e-08 | 3.1e-08 | 5.3e-08 | 4.600E-08 | 1.300E-08 | 3.100E-08 | 5.400E-08 |
| 2840 | 142 | Q142V | A | 5.1e-08 | 4e-08 | 2e-08 | 3.700E-08 | 1.572E-08 | 2.000E-08 | 5.100E-08 |
| 2841 | 143 | D143A | A | 3.4e-08 | 2.4e-08 | 4.8e-08 | 3.533E-08 | 1.206E-08 | 2.400E-08 | 4.800E-08 |
| 2842 | 143 | D143R | A | 2.4e-08 | 2e-08 | 7.2e-08 | 3.867E-08 | 2.894E-08 | 2.000E-08 | 7.200E-08 |
| 2843 | 143 | D143N | A | 4.4e-08 | 3.2e-08 | 7e-08 | 4.867E-08 | 1.943E-08 | 3.200E-08 | 7.000E-08 |
| 2844 | 143 | D143D | A | 9.4e-08 | 3.1e-08 | 4.8e-08 | 5.767E-08 | 3.259E-08 | 3.100E-08 | 9.400E-08 |
| 2845 | 143 | D143C | A | 5.4e-08 | 2.9e-08 | 1.1e-07 | 6.433E-08 | 4.148E-08 | 2.900E-08 | 1.100E-07 |
| 2846 | 143 | D143E | A | 1.6e-08 | 2.4e-08 | 3.7e-08 | 2.567E-08 | 1.060E-08 | 1.600E-08 | 3.700E-08 |
| 2847 | 143 | D143Q | A | 2.7e-08 | 6.3e-08 | 2.2e-08 | 3.733E-08 | 2.237E-08 | 2.200E-08 | 6.300E-08 |
| 2848 | 143 | D143G | A | 5.2e-08 | 3.1e-08 | 7.5e-08 | 5.267E-08 | 2.201E-08 | 3.100E-08 | 7.500E-08 |
| 2849 | 143 | D143H | A | 6.7e-09 | 6.5e-08 | 4.2e-08 | 3.790E-08 | 2.937E-08 | 6.700E-09 | 6.500E-08 |
| 2850 | 143 | D143I | A | 4.7e-08 | 1.6e-07 | 5.2e-08 | 8.633E-08 | 6.385E-08 | 4.700E-08 | 1.600E-07 |
| 2851 | 143 | D143L | A | 4.2e-08 | 2.7e-08 | 1e-07 | 5.633E-08 | 3.855E-08 | 2.700E-08 | 1.000E-07 |
| 2852 | 143 | D143K | A | 5.1e-08 | 6e-08 | 4.2e-08 | 5.100E-08 | 9.000E-09 | 4.200E-08 | 6.000E-08 |
| 2853 | 143 | D143M | A | 4.1e-08 | 7.7e-08 | 4e-08 | 5.267E-08 | 2.108E-08 | 4.000E-08 | 7.700E-08 |
| 2854 | 143 | D143F | A | 3.6e-08 | 6.2e-08 | 3.8e-08 | 4.533E-08 | 1.447E-08 | 3.600E-08 | 6.200E-08 |
| 2855 | 143 | D143P | A | 7.3e-08 | 3.7e-08 | 9.2e-08 | 6.733E-08 | 2.793E-08 | 3.700E-08 | 9.200E-08 |
| 2856 | 143 | D143S | A | 9e-08 | 3e-08 | 6.4e-08 | 6.133E-08 | 3.009E-08 | 3.000E-08 | 9.000E-08 |
| 2857 | 143 | D143T | A | 1.6e-08 | 5.4e-08 | 3.8e-08 | 3.600E-08 | 1.908E-08 | 1.600E-08 | 5.400E-08 |
| 2858 | 143 | D143W | A | 2.8e-08 | 2.4e-08 | 3.2e-08 | 2.800E-08 | 4.000E-09 | 2.400E-08 | 3.200E-08 |
| 2859 | 143 | D143Y | A | 4.5e-08 | 5.6e-08 | 5.4e-08 | 5.167E-08 | 5.859E-09 | 4.500E-08 | 5.600E-08 |
| 2860 | 143 | D143V | A | 3.3e-08 | 3.8e-08 | 3.4e-08 | 3.500E-08 | 2.646E-09 | 3.300E-08 | 3.800E-08 |
| 2861 | 144 | T144A | A | 8.9e-08 | 1.5e-08 | 5.8e-08 | 5.400E-08 | 3.716E-08 | 1.500E-08 | 8.900E-08 |
| 2862 | 144 | T144R | A | 4e-08 | 2.6e-08 | 7.7e-08 | 4.767E-08 | 2.635E-08 | 2.600E-08 | 7.700E-08 |
| 2863 | 144 | T144N | A | 6.2e-08 | 3.1e-08 | 5.3e-08 | 4.867E-08 | 1.595E-08 | 3.100E-08 | 6.200E-08 |
| 2864 | 144 | T144D | A | 2.6e-08 | 4.4e-08 | 4.4e-08 | 3.800E-08 | 1.039E-08 | 2.600E-08 | 4.400E-08 |
| 2865 | 144 | T144C | A | 1.9e-08 | 2.7e-08 | 1.4e-07 | 6.200E-08 | 6.767E-08 | 1.900E-08 | 1.400E-07 |
| 2866 | 144 | T144E | A | 2.7e-08 | 2.3e-08 | 8.3e-08 | 4.433E-08 | 3.355E-08 | 2.300E-08 | 8.300E-08 |
| 2867 | 144 | T144Q | A | 8.2e-08 | 5.4e-08 | 3.3e-08 | 5.633E-08 | 2.458E-08 | 3.300E-08 | 8.200E-08 |
| 2868 | 144 | T144G | A | 3.6e-08 | 6e-08 | 4e-08 | 4.533E-08 | 1.286E-08 | 3.600E-08 | 6.000E-08 |
| 2869 | 144 | T144H | A | 3.9e-08 | 6.6e-08 | 6.2e-08 | 5.567E-08 | 1.457E-08 | 3.900E-08 | 6.600E-08 |
| 2870 | 144 | T144I | A | 2.9e-08 | 3.3e-08 | 4.2e-08 | 3.467E-08 | 6.658E-09 | 2.900E-08 | 4.200E-08 |
| 2871 | 144 | T144L | A | 4.2e-08 | 7.2e-08 | 7.4e-08 | 6.267E-08 | 1.793E-08 | 4.200E-08 | 7.400E-08 |
| 2872 | 144 | T144K | A | 9.6e-08 | 2.5e-08 | 2e-08 | 4.700E-08 | 4.251E-08 | 2.000E-08 | 9.600E-08 |
| 2873 | 144 | T144M | A | 5.3e-08 | 7.5e-08 | 2.2e-08 | 5.000E-08 | 2.663E-08 | 2.200E-08 | 7.500E-08 |
| 2874 | 144 | T144F | A | 4.1e-08 | 1.3e-08 | 1.9e-08 | 2.433E-08 | 1.474E-08 | 1.300E-08 | 4.100E-08 |
| 2875 | 144 | T144P | A | 2.2e-08 | 6.2e-08 | 1e-07 | 6.133E-08 | 3.900E-08 | 2.200E-08 | 1.000E-07 |
| 2876 | 144 | T144S | A | 8.3e-08 | 3.8e-08 | 4.1e-08 | 5.400E-08 | 2.516E-08 | 3.800E-08 | 8.300E-08 |
| 2877 | 144 | T144T | A | 3.8e-08 | 2.2e-08 | 5.1e-08 | 3.700E-08 | 1.453E-08 | 2.200E-08 | 5.100E-08 |
| 2878 | 144 | T144W | A | 2.5e-08 | 5e-08 | 8.3e-08 | 5.267E-08 | 2.909E-08 | 2.500E-08 | 8.300E-08 |
| 2879 | 144 | T144Y | A | 1.7e-08 | 1e-07 | 4.5e-08 | 5.400E-08 | 4.223E-08 | 1.700E-08 | 1.000E-07 |
| 2880 | 144 | T144V | A | 2.3e-08 | 2.9e-08 | 4.9e-08 | 3.367E-08 | 1.361E-08 | 2.300E-08 | 4.900E-08 |
| 2881 | 145 | I145A | A | 4.4e-08 | 8e-08 | 4.3e-08 | 5.567E-08 | 2.108E-08 | 4.300E-08 | 8.000E-08 |
| 2882 | 145 | I145R | A | 2e-08 | 4.9e-08 | 4.9e-08 | 3.933E-08 | 1.674E-08 | 2.000E-08 | 4.900E-08 |
| 2883 | 145 | I145N | A | 4.4e-08 | 1.1e-07 | 1.2e-08 | 5.533E-08 | 4.997E-08 | 1.200E-08 | 1.100E-07 |
| 2884 | 145 | I145D | A | 5.1e-08 | 1.7e-08 | 9.1e-08 | 5.300E-08 | 3.704E-08 | 1.700E-08 | 9.100E-08 |
| 2885 | 145 | I145C | A | 2.1e-07 | 5.9e-08 | 5.6e-08 | 1.083E-07 | 8.806E-08 | 5.600E-08 | 2.100E-07 |
| 2886 | 145 | I145E | A | 3.2e-08 | 6.9e-08 | 4.4e-08 | 4.833E-08 | 1.888E-08 | 3.200E-08 | 6.900E-08 |
| 2887 | 145 | I145Q | A | 8e-08 | 1.3e-08 | 5.6e-08 | 4.967E-08 | 3.395E-08 | 1.300E-08 | 8.000E-08 |
| 2888 | 145 | I145G | A | 1e-07 | 2.7e-08 | 6.8e-08 | 6.500E-08 | 3.659E-08 | 2.700E-08 | 1.000E-07 |
| 2889 | 145 | I145H | A | 3.9e-08 | 2.5e-08 | 4.6e-08 | 3.667E-08 | 1.069E-08 | 2.500E-08 | 4.600E-08 |

| | | | | | | | | | | |
|---|---|---|---|---|---|---|---|---|---|---|
| 2890 | 145 | I145I | A | 2.3e-08 | 7.7e-08 | 6.2e-08 | 5.400E-08 | 2.787E-08 | 2.300E-08 | 7.700E-08 |
| 2891 | 145 | I145L | A | 7.2e-08 | 6.4e-08 | 7.2e-08 | 6.933E-08 | 4.619E-09 | 6.400E-08 | 7.200E-08 |
| 2892 | 145 | I145K | A | 6.8e-08 | 1e-07 | 5.8e-08 | 7.533E-08 | 2.194E-08 | 5.800E-08 | 1.000E-07 |
| 2893 | 145 | I145M | A | 3.7e-08 | 6.9e-08 | 2.7e-08 | 4.433E-08 | 2.194E-08 | 2.700E-08 | 6.900E-08 |
| 2894 | 145 | I145F | A | 3.2e-08 | 7.8e-08 | 5.5e-08 | 5.500E-08 | 2.300E-08 | 3.200E-08 | 7.800E-08 |
| 2895 | 145 | I145P | A | 2.9e-08 | 1.1e-08 | 4.3e-08 | 2.767E-08 | 1.604E-08 | 1.100E-08 | 4.300E-08 |
| 2896 | 145 | I145S | A | 5.2e-08 | 8.4e-08 | 8.5e-08 | 7.367E-08 | 1.877E-08 | 5.200E-08 | 8.500E-08 |
| 2897 | 145 | I145T | A | 3.1e-08 | 2.2e-08 | 6.3e-08 | 3.867E-08 | 2.155E-08 | 2.200E-08 | 6.300E-08 |
| 2898 | 145 | I145W | A | 2.6e-08 | 3.3e-08 | 8.8e-08 | 4.900E-08 | 3.396E-08 | 2.600E-08 | 8.800E-08 |
| 2899 | 145 | I145Y | A | 5.2e-08 | 4.1e-08 | 6.6e-08 | 5.300E-08 | 1.253E-08 | 4.100E-08 | 6.600E-08 |
| 2900 | 145 | I145V | A | 4.3e-08 | 3.7e-08 | 4.8e-08 | 4.267E-08 | 5.508E-09 | 3.700E-08 | 4.800E-08 |
| 2901 | 146 | L146A | A | 4e-08 | 1.8e-07 | 3.1e-08 | 8.367E-08 | 8.355E-08 | 3.100E-08 | 1.800E-07 |
| 2902 | 146 | L146R | A | 1.7e-08 | 4.6e-08 | 8.7e-08 | 5.000E-08 | 3.517E-08 | 1.700E-08 | 8.700E-08 |
| 2903 | 146 | L146N | A | 4.1e-08 | 2.2e-08 | 6.7e-08 | 4.333E-08 | 2.259E-08 | 2.200E-08 | 6.700E-08 |
| 2904 | 146 | L146D | A | 3.5e-08 | 9.8e-08 | 4.2e-08 | 5.833E-08 | 3.453E-08 | 3.500E-08 | 9.800E-08 |
| 2905 | 146 | L146C | A | 1.5e-08 | 7.4e-08 | 1.7e-08 | 3.533E-08 | 3.350E-08 | 1.500E-08 | 7.400E-08 |
| 2906 | 146 | L146E | A | 1.1e-08 | 7e-08 | 4.6e-08 | 4.233E-08 | 2.967E-08 | 1.100E-08 | 7.000E-08 |
| 2907 | 146 | L146Q | A | 2.2e-08 | 2.5e-08 | 1.2e-07 | 5.567E-08 | 5.573E-08 | 2.200E-08 | 1.200E-07 |
| 2908 | 146 | L146G | A | 3.4e-08 | 5.6e-08 | 5e-08 | 4.667E-08 | 1.137E-08 | 3.400E-08 | 5.600E-08 |
| 2909 | 146 | L146H | A | 5.1e-08 | 4.9e-08 | 4.6e-08 | 4.867E-08 | 2.517E-09 | 4.600E-08 | 5.100E-08 |
| 2910 | 146 | L146I | A | 5.1e-08 | 7e-08 | 1.4e-08 | 4.500E-08 | 2.848E-08 | 1.400E-08 | 7.000E-08 |
| 2911 | 146 | L146L | A | 3.6e-08 | 2e-08 | 8e-08 | 4.533E-08 | 3.107E-08 | 2.000E-08 | 8.000E-08 |
| 2912 | 146 | L146K | A | 6.5e-08 | 8.1e-08 | 3.7e-08 | 6.100E-08 | 2.227E-08 | 3.700E-08 | 8.100E-08 |
| 2913 | 146 | L146M | A | 6.8e-08 | 1.1e-07 | 4.1e-08 | 7.300E-08 | 3.477E-08 | 4.100E-08 | 1.100E-07 |
| 2914 | 146 | L146F | A | 7.2e-08 | 7.7e-08 | 9.2e-08 | 8.033E-08 | 1.041E-08 | 7.200E-08 | 9.200E-08 |
| 2915 | 146 | L146P | A | 3.7e-08 | 2e-07 | 1.1e-07 | 1.157E-07 | 8.165E-08 | 3.700E-08 | 2.000E-07 |
| 2916 | 146 | L146S | A | 9.3e-08 | 2.4e-08 | 2.9e-08 | 4.867E-08 | 3.848E-08 | 2.400E-08 | 9.300E-08 |
| 2917 | 146 | L146T | A | 4.6e-08 | 5.3e-08 | 2e-08 | 3.967E-08 | 1.739E-08 | 2.000E-08 | 5.300E-08 |
| 2918 | 146 | L146W | A | 6.2e-08 | 8.6e-08 | 1.1e-07 | 8.600E-08 | 2.400E-08 | 6.200E-08 | 1.100E-07 |
| 2919 | 146 | L146Y | A | 5.7e-08 | 2.2e-08 | 2.5e-08 | 3.467E-08 | 1.940E-08 | 2.200E-08 | 5.700E-08 |
| 2920 | 146 | L146V | A | 7.7e-08 | 3.2e-08 | 7.2e-08 | 6.033E-08 | 2.466E-08 | 3.200E-08 | 7.700E-08 |
| 2921 | 147 | W147A | A | 5.7e-08 | 2.3e-08 | 6.4e-08 | 4.800E-08 | 2.193E-08 | 2.300E-08 | 6.400E-08 |
| 2922 | 147 | W147R | A | 6.3e-08 | 8.3e-08 | 2.8e-08 | 5.800E-08 | 2.784E-08 | 2.800E-08 | 8.300E-08 |
| 2923 | 147 | W147N | A | 7e-08 | 3.4e-08 | 6.1e-08 | 5.500E-08 | 1.873E-08 | 3.400E-08 | 7.000E-08 |
| 2924 | 147 | W147D | A | 9.3e-09 | 4.9e-08 | 5.6e-08 | 3.810E-08 | 2.519E-08 | 9.300E-09 | 5.600E-08 |
| 2925 | 147 | W147C | A | 7.5e-08 | 1e-07 | 1.7e-07 | 1.150E-07 | 4.924E-08 | 7.500E-08 | 1.700E-07 |
| 2926 | 147 | W147E | A | 3.6e-08 | 2.9e-08 | 3.4e-08 | 3.300E-08 | 3.606E-09 | 2.900E-08 | 3.600E-08 |
| 2927 | 147 | W147Q | A | 6.4e-08 | 4.3e-08 | 2.3e-08 | 4.333E-08 | 2.050E-08 | 2.300E-08 | 6.400E-08 |
| 2928 | 147 | W147G | A | 1.8e-08 | 7.6e-08 | 1.4e-08 | 3.600E-08 | 3.470E-08 | 1.400E-08 | 7.600E-08 |
| 2929 | 147 | W147H | A | 5.4e-08 | 6.3e-08 | 5e-08 | 5.567E-08 | 6.658E-09 | 5.000E-08 | 6.300E-08 |
| 2930 | 147 | W147I | A | 3.2e-08 | 5.5e-08 | 4.7e-08 | 4.467E-08 | 1.168E-08 | 3.200E-08 | 5.500E-08 |
| 2931 | 147 | W147L | A | 9.6e-08 | 4e-08 | 5.3e-08 | 6.300E-08 | 2.931E-08 | 4.000E-08 | 9.600E-08 |
| 2932 | 147 | W147K | A | 3.9e-08 | 4.4e-08 | 2e-08 | 3.433E-08 | 1.266E-08 | 2.000E-08 | 4.400E-08 |
| 2933 | 147 | W147M | A | 1.2e-07 | 4.9e-08 | 1.6e-08 | 6.167E-08 | 5.314E-08 | 1.600E-08 | 1.200E-07 |
| 2934 | 147 | W147F | A | 6e-08 | 2.5e-08 | 5.4e-08 | 4.633E-08 | 1.872E-08 | 2.500E-08 | 6.000E-08 |
| 2935 | 147 | W147P | A | 4e-08 | 5.2e-08 | 1.6e-08 | 3.600E-08 | 1.833E-08 | 1.600E-08 | 5.200E-08 |
| 2936 | 147 | W147S | A | 2.2e-07 | 2.3e-08 | 5.1e-08 | 9.800E-08 | 1.066E-07 | 2.300E-08 | 2.200E-07 |
| 2937 | 147 | W147T | A | 3.3e-08 | 3.5e-08 | 3e-08 | 3.267E-08 | 2.517E-09 | 3.000E-08 | 3.500E-08 |
| 2938 | 147 | W147W | A | 2.9e-08 | 3.9e-08 | 4.6e-08 | 3.800E-08 | 8.544E-09 | 2.900E-08 | 4.600E-08 |
| 2939 | 147 | W147Y | A | 9.4e-08 | 3.9e-08 | 3.2e-08 | 5.500E-08 | 3.396E-08 | 3.200E-08 | 9.400E-08 |
| 2940 | 147 | W147V | A | 1.9e-08 | 2.4e-08 | 3.3e-08 | 2.533E-08 | 7.095E-09 | 1.900E-08 | 3.300E-08 |
| 2941 | 148 | K148A | A | 5.2e-08 | 5.1e-08 | 9.5e-08 | 6.600E-08 | 2.512E-08 | 5.100E-08 | 9.500E-08 |
| 2942 | 148 | K148R | A | 3e-08 | 8.4e-08 | 5.5e-08 | 5.633E-08 | 2.702E-08 | 3.000E-08 | 8.400E-08 |
| 2943 | 148 | K148N | A | 7.9e-08 | 2e-08 | 3.5e-08 | 4.467E-08 | 3.066E-08 | 2.000E-08 | 7.900E-08 |
| 2944 | 148 | K148D | A | 4.8e-08 | 3.1e-08 | 1.7e-08 | 3.200E-08 | 1.552E-08 | 1.700E-08 | 4.800E-08 |
| 2945 | 148 | K148C | A | 5.1e-08 | 1.3e-07 | 4e-08 | 7.367E-08 | 4.910E-08 | 4.000E-08 | 1.300E-07 |
| 2946 | 148 | K148E | A | 2.6e-08 | 9.5e-08 | 6.8e-08 | 6.300E-08 | 3.477E-08 | 2.600E-08 | 9.500E-08 |
| 2947 | 148 | K148Q | A | 2.9e-08 | 5.1e-08 | 3e-08 | 3.667E-08 | 1.242E-08 | 2.900E-08 | 5.100E-08 |
| 2948 | 148 | K148G | A | 9.3e-09 | 5.8e-08 | 7.6e-08 | 4.777E-08 | 3.451E-08 | 9.300E-09 | 7.600E-08 |
| 2949 | 148 | K148H | A | 3.7e-08 | 8.2e-08 | 5.5e-08 | 5.800E-08 | 2.265E-08 | 3.700E-08 | 8.200E-08 |
| 2950 | 148 | K148I | A | 4.4e-08 | 3.7e-08 | 2e-08 | 3.367E-08 | 1.234E-08 | 2.000E-08 | 4.400E-08 |
| 2951 | 148 | K148L | A | 1.8e-08 | 6e-08 | 2.9e-08 | 3.567E-08 | 2.178E-08 | 1.800E-08 | 6.000E-08 |
| 2952 | 148 | K148K | A | 4.2e-08 | 2.9e-08 | 1.6e-08 | 2.900E-08 | 1.300E-08 | 1.600E-08 | 4.200E-08 |
| 2953 | 148 | K148M | A | 1e-07 | 5.2e-08 | 5.7e-08 | 6.967E-08 | 2.639E-08 | 5.200E-08 | 1.000E-07 |
| 2954 | 148 | K148F | A | 8.5e-08 | 1.3e-07 | 9.1e-09 | 7.470E-08 | 6.110E-08 | 9.100E-09 | 1.300E-07 |
| 2955 | 148 | K148P | A | 2.5e-08 | 2.7e-08 | 3e-08 | 2.733E-08 | 2.517E-09 | 2.500E-08 | 3.000E-08 |
| 2956 | 148 | K148S | A | 5.4e-08 | 5.9e-08 | 2.4e-08 | 4.567E-08 | 1.893E-08 | 2.400E-08 | 5.900E-08 |
| 2957 | 148 | K148T | A | 4e-08 | 4.7e-08 | 2.6e-08 | 3.767E-08 | 1.069E-08 | 2.600E-08 | 4.700E-08 |
| 2958 | 148 | K148W | A | 2.5e-08 | 6.5e-08 | 7.2e-08 | 5.400E-08 | 2.536E-08 | 2.500E-08 | 7.200E-08 |
| 2959 | 148 | K148Y | A | 1.2e-08 | 2.2e-08 | 2.5e-08 | 1.967E-08 | 6.807E-09 | 1.200E-08 | 2.500E-08 |
| 2960 | 148 | K148V | A | 3.9e-08 | 7.3e-08 | 1.7e-07 | 9.400E-08 | 6.798E-08 | 3.900E-08 | 1.700E-07 |
| 2961 | 149 | D149A | A | 1.2e-07 | 2.4e-08 | 4.5e-08 | 6.300E-08 | 5.047E-08 | 2.400E-08 | 1.200E-07 |
| 2962 | 149 | D149R | A | 1.6e-08 | 1.3e-08 | 4.6e-08 | 2.500E-08 | 1.825E-08 | 1.300E-08 | 4.600E-08 |
| 2963 | 149 | D149N | A | 3e-08 | 2e-08 | 7.8e-08 | 4.267E-08 | 3.101E-08 | 2.000E-08 | 7.800E-08 |
| 2964 | 149 | D149D | A | 2.3e-08 | 6.1e-09 | 8.5e-08 | 3.803E-08 | 4.154E-08 | 6.100E-09 | 8.500E-08 |
| 2965 | 149 | D149C | A | 3.1e-08 | 5.5e-08 | 2.4e-08 | 3.667E-08 | 1.626E-08 | 2.400E-08 | 5.500E-08 |
| 2966 | 149 | D149E | A | 6.9e-08 | 5e-08 | 7.3e-08 | 6.400E-08 | 1.229E-08 | 5.000E-08 | 7.300E-08 |
| 2967 | 149 | D149Q | A | 3.2e-08 | 5.1e-08 | 6e-08 | 4.767E-08 | 1.429E-08 | 3.200E-08 | 6.000E-08 |
| 2968 | 149 | D149G | A | 3.2e-08 | 5.5e-08 | 7.9e-08 | 5.533E-08 | 2.350E-08 | 3.200E-08 | 7.900E-08 |
| 2969 | 149 | D149H | A | 2.6e-08 | 3.5e-08 | 1.2e-07 | 6.033E-08 | 5.187E-08 | 2.600E-08 | 1.200E-07 |
| 2970 | 149 | D149I | A | 5.1e-08 | 6.6e-08 | 1.7e-07 | 9.567E-08 | 6.481E-08 | 5.100E-08 | 1.700E-07 |
| 2971 | 149 | D149L | A | 2.9e-08 | 9.1e-08 | 5.8e-08 | 5.933E-08 | 3.102E-08 | 2.900E-08 | 9.100E-08 |
| 2972 | 149 | D149K | A | 2.3e-08 | 2.2e-08 | 4.5e-08 | 3.000E-08 | 1.300E-08 | 2.200E-08 | 4.500E-08 |
| 2973 | 149 | D149M | A | 1.6e-08 | 8.3e-09 | 7.5e-08 | 3.310E-08 | 3.649E-08 | 8.300E-09 | 7.500E-08 |
| 2974 | 149 | D149F | A | 6.5e-08 | 4.6e-08 | 2.1e-08 | 4.400E-08 | 2.207E-08 | 2.100E-08 | 6.500E-08 |

| | | | | | | | | | | |
|---|---|---|---|---|---|---|---|---|---|---|
| 2975 | 149 | D149P | A | 2e-08 | 8.6e-08 | 2.9e-08 | 4.500E-08 | 3.579E-08 | 2.000E-08 | 8.600E-08 |
| 2976 | 149 | D149S | A | 1.2e-07 | 1.1e-08 | 4.2e-08 | 5.767E-08 | 5.616E-08 | 1.100E-08 | 1.200E-07 |
| 2977 | 149 | D149T | A | 5.1e-08 | 7.8e-08 | 2e-08 | 4.967E-08 | 2.902E-08 | 2.000E-08 | 7.800E-08 |
| 2978 | 149 | D149W | A | 2.6e-08 | 4e-08 | 9e-08 | 5.200E-08 | 3.365E-08 | 2.600E-08 | 9.000E-08 |
| 2979 | 149 | D149Y | A | 2.5e-08 | 4.9e-08 | 9.3e-08 | 5.567E-08 | 3.449E-08 | 2.500E-08 | 9.300E-08 |
| 2980 | 149 | D149V | A | 5.3e-08 | 2.2e-08 | 5.4e-08 | 4.300E-08 | 1.819E-08 | 2.200E-08 | 5.400E-08 |
| 2981 | 150 | I150A | A | 2.3e-08 | 5.4e-08 | 1.8e-08 | 3.167E-08 | 1.950E-08 | 1.800E-08 | 5.400E-08 |
| 2982 | 150 | I150R | A | 4.2e-08 | 2.2e-08 | 1.7e-07 | 7.800E-08 | 8.030E-08 | 2.200E-08 | 1.700E-07 |
| 2983 | 150 | I150N | A | 8.6e-08 | 3.5e-08 | 5.9e-08 | 6.000E-08 | 2.551E-08 | 3.500E-08 | 8.600E-08 |
| 2984 | 150 | I150D | A | 8e-08 | 3.4e-08 | 2e-08 | 4.467E-08 | 3.139E-08 | 2.000E-08 | 8.000E-08 |
| 2985 | 150 | I150C | A | 2.4e-08 | 7e-08 | 8.4e-08 | 5.933E-08 | 3.139E-08 | 2.400E-08 | 8.400E-08 |
| 2986 | 150 | I150E | A | 3.3e-08 | 2.3e-08 | 4.5e-08 | 3.367E-08 | 1.102E-08 | 2.300E-08 | 4.500E-08 |
| 2987 | 150 | I150Q | A | 3.7e-08 | 6.6e-08 | 2e-08 | 4.100E-08 | 2.326E-08 | 2.000E-08 | 6.600E-08 |
| 2988 | 150 | I150G | A | 8.6e-08 | 5.2e-08 | 2.2e-07 | 1.193E-07 | 8.882E-08 | 5.200E-08 | 2.200E-07 |
| 2989 | 150 | I150H | A | 1.1e-07 | 2.7e-08 | 2.5e-08 | 5.400E-08 | 4.851E-08 | 2.500E-08 | 1.100E-07 |
| 2990 | 150 | I150I | A | 4.3e-08 | 3.2e-08 | 1.2e-08 | 2.900E-08 | 1.572E-08 | 1.200E-08 | 4.300E-08 |
| 2991 | 150 | I150L | A | 4.6e-08 | 2.2e-08 | 5.4e-08 | 4.067E-08 | 1.665E-08 | 2.200E-08 | 5.400E-08 |
| 2992 | 150 | I150K | A | 3e-08 | 2e-08 | 7.5e-08 | 4.167E-08 | 2.930E-08 | 2.000E-08 | 7.500E-08 |
| 2993 | 150 | I150M | A | 2.6e-08 | 7.8e-08 | 9.2e-08 | 6.533E-08 | 3.478E-08 | 2.600E-08 | 9.200E-08 |
| 2994 | 150 | I150F | A | 6.3e-08 | 1.6e-08 | 1.1e-07 | 6.300E-08 | 4.700E-08 | 1.600E-08 | 1.100E-07 |
| 2995 | 150 | I150P | A | 4.2e-08 | 4.4e-08 | 5.1e-08 | 4.567E-08 | 4.726E-09 | 4.200E-08 | 5.100E-08 |
| 2996 | 150 | I150S | A | 3.1e-08 | 4.1e-08 | 4e-08 | 3.733E-08 | 5.508E-09 | 3.100E-08 | 4.100E-08 |
| 2997 | 150 | I150T | A | 5.1e-08 | 4.3e-08 | 5.6e-08 | 5.000E-08 | 6.557E-09 | 4.300E-08 | 5.600E-08 |
| 2998 | 150 | I150W | A | 2.7e-08 | 2e-08 | 3.9e-08 | 2.867E-08 | 9.609E-09 | 2.000E-08 | 3.900E-08 |
| 2999 | 150 | I150Y | A | 2e-08 | 4.3e-08 | 2.9e-08 | 3.067E-08 | 1.159E-08 | 2.000E-08 | 4.300E-08 |
| 3000 | 150 | I150V | A | 8.6e-08 | 4e-08 | 5.9e-08 | 6.167E-08 | 2.312E-08 | 4.000E-08 | 8.600E-08 |
| 3001 | 151 | F151A | A | 5.9e-08 | 3.8e-08 | 3.2e-08 | 4.300E-08 | 1.418E-08 | 3.200E-08 | 5.900E-08 |
| 3002 | 151 | F151R | A | 2.2e-08 | 4.3e-08 | 8.6e-09 | 2.453E-08 | 1.734E-08 | 8.600E-09 | 4.300E-08 |
| 3003 | 151 | F151N | A | 5.3e-08 | 2.9e-08 | 4.2e-08 | 4.133E-08 | 1.201E-08 | 2.900E-08 | 5.300E-08 |
| 3004 | 151 | F151D | A | 3.2e-08 | 6.6e-08 | 5.2e-08 | 5.000E-08 | 1.709E-08 | 3.200E-08 | 6.600E-08 |
| 3005 | 151 | F151C | A | 1.2e-07 | 9.7e-08 | 1.3e-07 | 1.157E-07 | 1.692E-08 | 9.700E-08 | 1.300E-07 |
| 3006 | 151 | F151E | A | 6.1e-08 | 5.2e-08 | 3e-08 | 4.767E-08 | 1.595E-08 | 3.000E-08 | 6.100E-08 |
| 3007 | 151 | F151Q | A | 2.6e-08 | 1.4e-08 | 2.2e-08 | 2.067E-08 | 6.110E-09 | 1.400E-08 | 2.600E-08 |
| 3008 | 151 | F151G | A | 3.3e-08 | 1.6e-08 | 6.5e-08 | 3.800E-08 | 2.488E-08 | 1.600E-08 | 6.500E-08 |
| 3009 | 151 | F151H | A | 1.5e-07 | 5.2e-08 | 4.6e-08 | 8.267E-08 | 5.839E-08 | 4.600E-08 | 1.500E-07 |
| 3010 | 151 | F151I | A | 5e-08 | 6.4e-08 | 1.1e-07 | 7.467E-08 | 3.139E-08 | 5.000E-08 | 1.100E-07 |
| 3011 | 151 | F151L | A | 4.8e-08 | 4.1e-08 | 4.1e-08 | 4.333E-08 | 4.041E-09 | 4.100E-08 | 4.800E-08 |
| 3012 | 151 | F151K | A | 3.6e-08 | 1.1e-07 | 1.9e-08 | 5.500E-08 | 4.838E-08 | 1.900E-08 | 1.100E-07 |
| 3013 | 151 | F151M | A | 8.5e-08 | 1.1e-07 | 5e-08 | 8.167E-08 | 3.014E-08 | 5.000E-08 | 1.100E-07 |
| 3014 | 151 | F151F | A | 2.5e-08 | 2.5e-08 | 6.5e-08 | 3.833E-08 | 2.309E-08 | 2.500E-08 | 6.500E-08 |
| 3015 | 151 | F151P | A | 1.1e-07 | 5.1e-08 | 1.8e-08 | 5.967E-08 | 4.661E-08 | 1.800E-08 | 1.100E-07 |
| 3016 | 151 | F151S | A | 3.6e-08 | 4.5e-08 | 4.3e-08 | 4.133E-08 | 4.726E-09 | 3.600E-08 | 4.500E-08 |
| 3017 | 151 | F151T | A | 2.1e-08 | 3.8e-08 | 5.1e-08 | 3.667E-08 | 1.504E-08 | 2.100E-08 | 5.100E-08 |
| 3018 | 151 | F151W | A | 2.6e-08 | 1e-07 | 1e-07 | 7.533E-08 | 4.272E-08 | 2.600E-08 | 1.000E-07 |
| 3019 | 151 | F151Y | A | 9.2e-08 | 4e-08 | 6.9e-08 | 6.700E-08 | 2.606E-08 | 4.000E-08 | 9.200E-08 |
| 3020 | 151 | F151V | A | 3e-08 | 6.4e-08 | 3.1e-08 | 4.167E-08 | 1.935E-08 | 3.000E-08 | 6.400E-08 |
| 3021 | 152 | H152A | A | 1.1e-07 | 7.3e-08 | 6.1e-08 | 8.133E-08 | 2.554E-08 | 6.100E-08 | 1.100E-07 |
| 3022 | 152 | H152R | A | 2.7e-08 | 3.7e-08 | 2.5e-08 | 2.967E-08 | 6.429E-09 | 2.500E-08 | 3.700E-08 |
| 3023 | 152 | H152N | A | 3.7e-08 | 6.3e-08 | 5.9e-08 | 5.300E-08 | 1.400E-08 | 3.700E-08 | 6.300E-08 |
| 3024 | 152 | H152D | A | 2.1e-08 | 2e-08 | 2.4e-08 | 2.167E-08 | 2.082E-09 | 2.000E-08 | 2.400E-08 |
| 3025 | 152 | H152C | A | 1.7e-08 | 7.4e-08 | 6.4e-08 | 5.167E-08 | 3.044E-08 | 1.700E-08 | 7.400E-08 |
| 3026 | 152 | H152E | A | 1.3e-08 | 3.7e-08 | 5.8e-08 | 3.600E-08 | 2.252E-08 | 1.300E-08 | 5.800E-08 |
| 3027 | 152 | H152Q | A | 2e-08 | 5.1e-08 | 5e-08 | 4.033E-08 | 1.762E-08 | 2.000E-08 | 5.100E-08 |
| 3028 | 152 | H152G | A | 4.4e-08 | 1.1e-07 | 4.6e-08 | 6.667E-08 | 3.754E-08 | 4.400E-08 | 1.100E-07 |
| 3029 | 152 | H152H | A | 3.3e-08 | 9.4e-08 | 8e-09 | 4.500E-08 | 4.424E-08 | 8.000E-09 | 9.400E-08 |
| 3030 | 152 | H152I | A | 2.3e-08 | 6.7e-08 | 5.8e-08 | 4.933E-08 | 2.325E-08 | 2.300E-08 | 6.700E-08 |
| 3031 | 152 | H152L | A | 6.6e-08 | 3.3e-08 | 5.9e-08 | 5.267E-08 | 1.739E-08 | 3.300E-08 | 6.600E-08 |
| 3032 | 152 | H152K | A | 1.1e-07 | 3.9e-08 | 1.1e-07 | 8.633E-08 | 4.099E-08 | 3.900E-08 | 1.100E-07 |
| 3033 | 152 | H152M | A | 1.3e-08 | 4.4e-08 | 7.5e-08 | 4.400E-08 | 3.100E-08 | 1.300E-08 | 7.500E-08 |
| 3034 | 152 | H152F | A | 9.2e-08 | 4.8e-08 | 6.4e-08 | 6.800E-08 | 2.227E-08 | 4.800E-08 | 9.200E-08 |
| 3035 | 152 | H152P | A | 4.5e-08 | 6.2e-08 | 2.4e-08 | 4.367E-08 | 1.904E-08 | 2.400E-08 | 6.200E-08 |
| 3036 | 152 | H152S | A | 2.7e-08 | 7.8e-08 | 8.9e-08 | 6.467E-08 | 3.308E-08 | 2.700E-08 | 8.900E-08 |
| 3037 | 152 | H152T | A | 5.4e-08 | 1.6e-08 | 5.4e-08 | 4.133E-08 | 2.194E-08 | 1.600E-08 | 5.400E-08 |
| 3038 | 152 | H152W | A | 7.2e-08 | 3.4e-08 | 6.5e-08 | 5.700E-08 | 2.022E-08 | 3.400E-08 | 7.200E-08 |
| 3039 | 152 | H152Y | A | 5.1e-08 | 1.3e-07 | 1.1e-07 | 9.700E-08 | 4.107E-08 | 5.100E-08 | 1.300E-07 |
| 3040 | 152 | H152V | A | 2.4e-08 | 3.2e-08 | 2.4e-08 | 2.667E-08 | 4.619E-09 | 2.400E-08 | 3.200E-08 |
| 3041 | 153 | K153A | A | 2.7e-08 | 2.1e-08 | 6.3e-08 | 3.700E-08 | 2.272E-08 | 2.100E-08 | 6.300E-08 |
| 3042 | 153 | K153R | A | 3.4e-08 | 6.7e-09 | 7.8e-08 | 3.957E-08 | 3.597E-08 | 6.700E-09 | 7.800E-08 |
| 3043 | 153 | K153N | A | 9.8e-09 | 4e-08 | 2.3e-08 | 2.427E-08 | 1.514E-08 | 9.800E-09 | 4.000E-08 |
| 3044 | 153 | K153D | A | 3e-08 | 4.7e-08 | 3.2e-08 | 3.633E-08 | 9.292E-09 | 3.000E-08 | 4.700E-08 |
| 3045 | 153 | K153C | A | 2.9e-08 | 3e-08 | 8e-08 | 4.633E-08 | 2.916E-08 | 2.900E-08 | 8.000E-08 |
| 3046 | 153 | K153E | A | 2.3e-08 | 1.1e-08 | 3.8e-08 | 2.400E-08 | 1.353E-08 | 1.100E-08 | 3.800E-08 |
| 3047 | 153 | K153Q | A | 1.8e-08 | 5.7e-08 | 5.2e-08 | 4.233E-08 | 2.122E-08 | 1.800E-08 | 5.700E-08 |
| 3048 | 153 | K153G | A | 7.5e-08 | 4.7e-08 | 1e-07 | 7.400E-08 | 2.651E-08 | 4.700E-08 | 1.000E-07 |
| 3049 | 153 | K153H | A | 2.2e-08 | 7.1e-08 | 3.6e-08 | 4.300E-08 | 2.524E-08 | 2.200E-08 | 7.100E-08 |
| 3050 | 153 | K153I | A | 5e-08 | 7.8e-08 | 4.8e-08 | 5.867E-08 | 1.677E-08 | 4.800E-08 | 7.800E-08 |
| 3051 | 153 | K153L | A | 2.5e-08 | 9.3e-08 | 4.4e-08 | 5.400E-08 | 3.509E-08 | 2.500E-08 | 9.300E-08 |
| 3052 | 153 | K153K | A | 1.1e-07 | 5.2e-08 | 4.7e-08 | 6.967E-08 | 3.502E-08 | 4.700E-08 | 1.100E-07 |
| 3053 | 153 | K153M | A | 8e-08 | 7.4e-08 | 8.4e-08 | 7.933E-08 | 5.033E-09 | 7.400E-08 | 8.400E-08 |
| 3054 | 153 | K153F | A | 3.7e-08 | 2.1e-07 | 7.2e-08 | 1.063E-07 | 9.147E-08 | 3.700E-08 | 2.100E-07 |
| 3055 | 153 | K153P | A | 1.7e-08 | 1.8e-08 | 3.2e-08 | 2.233E-08 | 8.386E-09 | 1.700E-08 | 3.200E-08 |
| 3056 | 153 | K153S | A | 2.5e-08 | 1e-07 | 3.2e-08 | 5.233E-08 | 4.143E-08 | 2.500E-08 | 1.000E-07 |
| 3057 | 153 | K153T | A | 2.8e-08 | 2.3e-08 | 1.1e-07 | 5.367E-08 | 4.885E-08 | 2.300E-08 | 1.100E-07 |
| 3058 | 153 | K153W | A | 4e-08 | 3.9e-08 | 1.9e-08 | 3.267E-08 | 1.185E-08 | 1.900E-08 | 4.000E-08 |
| 3059 | 153 | K153Y | A | 3e-08 | 4.5e-08 | 1.5e-08 | 3.000E-08 | 1.500E-08 | 1.500E-08 | 4.500E-08 |

| | | | | | | | | | | |
|---|---|---|---|---|---|---|---|---|---|---|
| 3060 | 153 | K153V | A | 2e-08 | 4.2e-08 | 4e-08 | 3.400E-08 | 1.217E-08 | 2.000E-08 | 4.200E-08 |
| 3061 | 154 | N154A | A | 7.6e-08 | 3.7e-08 | 1.4e-08 | 4.233E-08 | 3.134E-08 | 1.400E-08 | 7.600E-08 |
| 3062 | 154 | N154R | A | 1.8e-08 | 4.2e-08 | 4.6e-08 | 3.533E-08 | 1.514E-08 | 1.800E-08 | 4.600E-08 |
| 3063 | 154 | N154N | A | 4.4e-08 | 2.3e-08 | 4.7e-08 | 3.800E-08 | 1.308E-08 | 2.300E-08 | 4.700E-08 |
| 3064 | 154 | N154D | A | 7.3e-08 | 2.5e-08 | 1.2e-07 | 7.267E-08 | 4.750E-08 | 2.500E-08 | 1.200E-07 |
| 3065 | 154 | N154C | A | 4.4e-08 | 1.2e-07 | 6.4e-08 | 7.600E-08 | 3.940E-08 | 4.400E-08 | 1.200E-07 |
| 3066 | 154 | N154E | A | 7.9e-08 | 2e-08 | 3.6e-08 | 4.500E-08 | 3.051E-08 | 2.000E-08 | 7.900E-08 |
| 3067 | 154 | N154Q | A | 2.8e-08 | 3e-08 | 6.2e-08 | 4.000E-08 | 1.908E-08 | 2.800E-08 | 6.200E-08 |
| 3068 | 154 | N154G | A | 6.4e-08 | 5.4e-08 | 1e-07 | 7.267E-08 | 2.419E-08 | 5.400E-08 | 1.000E-07 |
| 3069 | 154 | N154H | A | 2.1e-08 | 6.3e-08 | 4.5e-08 | 4.300E-08 | 2.107E-08 | 2.100E-08 | 6.300E-08 |
| 3070 | 154 | N154I | A | 6.1e-08 | 9.6e-09 | 1.2e-08 | 2.753E-08 | 2.901E-08 | 9.600E-09 | 6.100E-08 |
| 3071 | 154 | N154L | A | 2e-08 | 4.3e-08 | 3.5e-08 | 3.267E-08 | 1.168E-08 | 2.000E-08 | 4.300E-08 |
| 3072 | 154 | N154K | A | 5.8e-08 | 8.1e-08 | 1.7e-08 | 5.200E-08 | 3.242E-08 | 1.700E-08 | 8.100E-08 |
| 3073 | 154 | N154M | A | 2.4e-08 | 2.5e-08 | 6e-08 | 3.633E-08 | 2.050E-08 | 2.400E-08 | 6.000E-08 |
| 3074 | 154 | N154F | A | 2e-08 | 1e-07 | 4.8e-08 | 5.600E-08 | 4.060E-08 | 2.000E-08 | 1.000E-07 |
| 3075 | 154 | N154P | A | 6.3e-08 | 7e-08 | 3.7e-08 | 5.667E-08 | 1.739E-08 | 3.700E-08 | 7.000E-08 |
| 3076 | 154 | N154S | A | 3.1e-08 | 2.1e-08 | 3e-08 | 2.733E-08 | 5.508E-09 | 2.100E-08 | 3.100E-08 |
| 3077 | 154 | N154T | A | 6.2e-08 | 2.4e-08 | 7e-08 | 5.200E-08 | 2.458E-08 | 2.400E-08 | 7.000E-08 |
| 3078 | 154 | N154W | A | 2.8e-08 | 4.3e-08 | 1.7e-08 | 2.933E-08 | 1.305E-08 | 1.700E-08 | 4.300E-08 |
| 3079 | 154 | N154Y | A | 2e-08 | 4.5e-08 | 2.2e-08 | 2.900E-08 | 1.389E-08 | 2.000E-08 | 4.500E-08 |
| 3080 | 154 | N154V | A | 1.5e-08 | 4.5e-08 | 1.4e-08 | 2.467E-08 | 1.762E-08 | 1.400E-08 | 4.500E-08 |
| 3081 | 155 | N155A | A | 2.7e-08 | 6.1e-08 | 2.7e-08 | 3.833E-08 | 1.963E-08 | 2.700E-08 | 6.100E-08 |
| 3082 | 155 | N155R | A | 5.3e-08 | 3.3e-08 | 4.4e-08 | 4.333E-08 | 1.002E-08 | 3.300E-08 | 5.300E-08 |
| 3083 | 155 | N155N | A | 2.3e-08 | 3.1e-08 | 4.5e-08 | 3.300E-08 | 1.114E-08 | 2.300E-08 | 4.500E-08 |
| 3084 | 155 | N155D | A | 9.5e-08 | 3.5e-08 | 4.2e-08 | 5.733E-08 | 3.281E-08 | 3.500E-08 | 9.500E-08 |
| 3085 | 155 | N155C | A | 5.5e-08 | 1.6e-08 | 8.7e-08 | 5.267E-08 | 3.556E-08 | 1.600E-08 | 8.700E-08 |
| 3086 | 155 | N155E | A | 5.7e-08 | 4.7e-08 | 7.1e-08 | 5.833E-08 | 1.206E-08 | 4.700E-08 | 7.100E-08 |
| 3087 | 155 | N155Q | A | 1.5e-08 | 5.9e-08 | 1.1e-07 | 6.133E-08 | 4.754E-08 | 1.500E-08 | 1.100E-07 |
| 3088 | 155 | N155G | A | 1.4e-08 | 1.3e-07 | 1.2e-08 | 5.200E-08 | 6.756E-08 | 1.200E-08 | 1.300E-07 |
| 3089 | 155 | N155H | A | 1.2e-07 | 3.7e-08 | 3.8e-08 | 6.500E-08 | 4.763E-08 | 3.700E-08 | 1.200E-07 |
| 3090 | 155 | N155I | A | 1.4e-08 | 1.3e-08 | 2.2e-08 | 1.633E-08 | 4.933E-09 | 1.300E-08 | 2.200E-08 |
| 3091 | 155 | N155L | A | 6.8e-08 | 1e-08 | 2.1e-08 | 3.300E-08 | 3.081E-08 | 1.000E-08 | 6.800E-08 |
| 3092 | 155 | N155K | A | 3.2e-08 | 3.6e-08 | 9.7e-09 | 2.590E-08 | 1.417E-08 | 9.700E-09 | 3.600E-08 |
| 3093 | 155 | N155M | A | 6.3e-08 | 4.8e-08 | 4e-08 | 5.033E-08 | 1.168E-08 | 4.000E-08 | 6.300E-08 |
| 3094 | 155 | N155F | A | 2.9e-08 | 1.2e-08 | 2.5e-08 | 2.200E-08 | 8.888E-09 | 1.200E-08 | 2.900E-08 |
| 3095 | 155 | N155P | A | 3.5e-08 | 4.6e-08 | 5e-08 | 4.367E-08 | 7.767E-09 | 3.500E-08 | 5.000E-08 |
| 3096 | 155 | N155S | A | 5e-08 | 4e-08 | 2.4e-08 | 3.800E-08 | 1.311E-08 | 2.400E-08 | 5.000E-08 |
| 3097 | 155 | N155T | A | 1.8e-08 | 5.8e-08 | 4.5e-08 | 4.033E-08 | 2.040E-08 | 1.800E-08 | 5.800E-08 |
| 3098 | 155 | N155W | A | 8.9e-08 | 6.3e-08 | 2.1e-08 | 5.767E-08 | 3.431E-08 | 2.100E-08 | 8.900E-08 |
| 3099 | 155 | N155Y | A | 1e-07 | 5.3e-08 | 3.4e-08 | 6.233E-08 | 3.398E-08 | 3.400E-08 | 1.000E-07 |
| 3100 | 155 | N155V | A | 4.4e-08 | 3.8e-08 | 1.7e-08 | 3.300E-08 | 1.418E-08 | 1.700E-08 | 4.400E-08 |
| 3101 | 156 | Q156A | A | 1.7e-08 | 2.3e-08 | 2.9e-08 | 2.300E-08 | 6.000E-09 | 1.700E-08 | 2.900E-08 |
| 3102 | 156 | Q156R | A | 6.1e-08 | 6.3e-08 | 2.7e-08 | 5.033E-08 | 2.023E-08 | 2.700E-08 | 6.300E-08 |
| 3103 | 156 | Q156N | A | 3.7e-08 | 2.6e-08 | 6e-08 | 4.100E-08 | 1.735E-08 | 2.600E-08 | 6.000E-08 |
| 3104 | 156 | Q156D | A | 7.3e-08 | 7e-08 | 5.7e-08 | 6.667E-08 | 8.505E-09 | 5.700E-08 | 7.300E-08 |
| 3105 | 156 | Q156C | A | 2.1e-08 | 2.8e-08 | 7.8e-08 | 4.233E-08 | 3.109E-08 | 2.100E-08 | 7.800E-08 |
| 3106 | 156 | Q156E | A | 2.2e-08 | 4.1e-08 | 2.5e-08 | 2.933E-08 | 1.021E-08 | 2.200E-08 | 4.100E-08 |
| 3107 | 156 | Q156Q | A | 3.3e-08 | 1.7e-07 | 7.5e-08 | 9.267E-08 | 7.019E-08 | 3.300E-08 | 1.700E-07 |
| 3108 | 156 | Q156G | A | 6.1e-08 | 3.5e-08 | 4.5e-08 | 4.700E-08 | 1.311E-08 | 3.500E-08 | 6.100E-08 |
| 3109 | 156 | Q156H | A | 3.5e-08 | 4.4e-08 | 1.8e-08 | 3.233E-08 | 1.320E-08 | 1.800E-08 | 4.400E-08 |
| 3110 | 156 | Q156I | A | 5.1e-08 | 4.8e-08 | 3.6e-08 | 4.500E-08 | 7.937E-09 | 3.600E-08 | 5.100E-08 |
| 3111 | 156 | Q156L | A | 3.5e-08 | 5.9e-08 | 4.7e-08 | 4.700E-08 | 1.200E-08 | 3.500E-08 | 5.900E-08 |
| 3112 | 156 | Q156K | A | 6.6e-08 | 2.9e-08 | 1.4e-08 | 3.633E-08 | 2.676E-08 | 1.400E-08 | 6.600E-08 |
| 3113 | 156 | Q156M | A | 2.3e-08 | 2.9e-08 | 2.6e-08 | 2.600E-08 | 3.000E-09 | 2.300E-08 | 2.900E-08 |
| 3114 | 156 | Q156F | A | 4.8e-08 | 9e-08 | 3.7e-08 | 5.833E-08 | 2.797E-08 | 3.700E-08 | 9.000E-08 |
| 3115 | 156 | Q156P | A | 7.2e-08 | 3.9e-08 | 5e-08 | 5.367E-08 | 1.680E-08 | 3.900E-08 | 7.200E-08 |
| 3116 | 156 | Q156S | A | 7.9e-08 | 9.3e-08 | 8.5e-08 | 8.567E-08 | 7.024E-09 | 7.900E-08 | 9.300E-08 |
| 3117 | 156 | Q156T | A | 2.2e-08 | 4e-08 | 3.9e-08 | 3.367E-08 | 1.012E-08 | 2.200E-08 | 4.000E-08 |
| 3118 | 156 | Q156W | A | 2e-08 | 9.5e-08 | 4e-08 | 5.167E-08 | 3.884E-08 | 2.000E-08 | 9.500E-08 |
| 3119 | 156 | Q156Y | A | 3.2e-08 | 2.9e-08 | 6.1e-08 | 4.067E-08 | 1.767E-08 | 2.900E-08 | 6.100E-08 |
| 3120 | 156 | Q156V | A | 1.5e-08 | 6e-08 | 2.1e-08 | 3.200E-08 | 2.443E-08 | 1.500E-08 | 6.000E-08 |
| 3121 | 157 | L157A | A | 4.5e-08 | 8.3e-08 | 5.1e-08 | 5.967E-08 | 2.043E-08 | 4.500E-08 | 8.300E-08 |
| 3122 | 157 | L157R | A | 2.9e-08 | 5.6e-08 | 3.9e-08 | 4.133E-08 | 1.365E-08 | 2.900E-08 | 5.600E-08 |
| 3123 | 157 | L157N | A | 3.4e-08 | 3.8e-08 | 2.7e-08 | 3.300E-08 | 5.568E-09 | 2.700E-08 | 3.800E-08 |
| 3124 | 157 | L157D | A | 4.4e-08 | 2e-08 | 6.3e-08 | 4.233E-08 | 2.155E-08 | 2.000E-08 | 6.300E-08 |
| 3125 | 157 | L157C | A | 4.7e-08 | 4.5e-08 | 3.1e-08 | 4.100E-08 | 8.718E-09 | 3.100E-08 | 4.700E-08 |
| 3126 | 157 | L157E | A | 2e-08 | 3.7e-08 | 2.1e-08 | 2.600E-08 | 9.539E-09 | 2.000E-08 | 3.700E-08 |
| 3127 | 157 | L157Q | A | 2.4e-08 | 5.2e-08 | 3.2e-08 | 3.600E-08 | 1.442E-08 | 2.400E-08 | 5.200E-08 |
| 3128 | 157 | L157G | A | 5.5e-08 | 3.5e-08 | 6.8e-08 | 5.267E-08 | 1.662E-08 | 3.500E-08 | 6.800E-08 |
| 3129 | 157 | L157H | A | 4.9e-08 | 2.1e-08 | 7e-08 | 4.667E-08 | 2.458E-08 | 2.100E-08 | 7.000E-08 |
| 3130 | 157 | L157I | A | 9.5e-08 | 4.2e-08 | 2.2e-08 | 5.300E-08 | 3.772E-08 | 2.200E-08 | 9.500E-08 |
| 3131 | 157 | L157L | A | 4.1e-08 | 3.6e-08 | 1.5e-08 | 3.067E-08 | 1.380E-08 | 1.500E-08 | 4.100E-08 |
| 3132 | 157 | L157K | A | 8.5e-08 | 1.4e-08 | 5.1e-08 | 5.000E-08 | 3.551E-08 | 1.400E-08 | 8.500E-08 |
| 3133 | 157 | L157M | A | 1.6e-08 | 3.1e-08 | 7e-08 | 3.900E-08 | 2.787E-08 | 1.600E-08 | 7.000E-08 |
| 3134 | 157 | L157F | A | 8.2e-08 | 5.1e-08 | 6.5e-09 | 4.650E-08 | 3.795E-08 | 6.500E-09 | 8.200E-08 |
| 3135 | 157 | L157P | A | 6.1e-08 | 3.8e-08 | 2.2e-08 | 4.033E-08 | 1.960E-08 | 2.200E-08 | 6.100E-08 |
| 3136 | 157 | L157S | A | 8.3e-08 | 7.9e-08 | 5.3e-08 | 7.167E-08 | 1.629E-08 | 5.300E-08 | 8.300E-08 |
| 3137 | 157 | L157T | A | 3.8e-08 | 2.2e-08 | 3.8e-08 | 3.267E-08 | 9.238E-09 | 2.200E-08 | 3.800E-08 |
| 3138 | 157 | L157W | A | 5.2e-08 | 3.4e-08 | 1.1e-07 | 6.533E-08 | 3.972E-08 | 3.400E-08 | 1.100E-07 |
| 3139 | 157 | L157Y | A | 1.1e-07 | 1.5e-07 | 2.6e-08 | 9.533E-08 | 6.329E-08 | 2.600E-08 | 1.500E-07 |
| 3140 | 157 | L157V | A | 4.2e-08 | 5.6e-08 | 1.2e-08 | 3.667E-08 | 2.248E-08 | 1.200E-08 | 5.600E-08 |
| 3141 | 158 | A158A | A | 6.9e-08 | 4.3e-08 | 6.7e-08 | 5.967E-08 | 1.447E-08 | 4.300E-08 | 6.900E-08 |
| 3142 | 158 | A158R | A | 8.7e-08 | 4.7e-08 | 6.7e-08 | 6.700E-08 | 2.000E-08 | 4.700E-08 | 8.700E-08 |
| 3143 | 158 | A158N | A | 1.9e-08 | 1.9e-08 | 5.6e-08 | 3.133E-08 | 2.136E-08 | 1.900E-08 | 5.600E-08 |
| 3144 | 158 | A158D | A | 2.3e-08 | 5.3e-08 | 3.4e-08 | 3.667E-08 | 1.518E-08 | 2.300E-08 | 5.300E-08 |

| | | | | | | | | | | |
|---|---|---|---|---|---|---|---|---|---|---|
| 3145 | 158 | A158C | A | 7.1e-08 | 1.2e-08 | 1.1e-07 | 6.433E-08 | 4.934E-08 | 1.200E-08 | 1.100E-07 |
| 3146 | 158 | A158E | A | 7.1e-08 | 4.8e-08 | 7.6e-08 | 6.500E-08 | 1.493E-08 | 4.800E-08 | 7.600E-08 |
| 3147 | 158 | A158Q | A | 2.1e-08 | 3.6e-08 | 3.5e-08 | 3.067E-08 | 8.386E-09 | 2.100E-08 | 3.600E-08 |
| 3148 | 158 | A158G | A | 1.9e-08 | 4.9e-08 | 5.2e-08 | 4.000E-08 | 1.825E-08 | 1.900E-08 | 5.200E-08 |
| 3149 | 158 | A158H | A | 1.1e-08 | 6.8e-08 | 6.1e-08 | 4.667E-08 | 3.109E-08 | 1.100E-08 | 6.800E-08 |
| 3150 | 158 | A158I | A | 8.5e-08 | 4.6e-08 | 7.8e-08 | 6.967E-08 | 2.079E-08 | 4.600E-08 | 8.500E-08 |
| 3151 | 158 | A158L | A | 2.7e-08 | 1.3e-08 | 4e-08 | 2.667E-08 | 1.350E-08 | 1.300E-08 | 4.000E-08 |
| 3152 | 158 | A158K | A | 2.2e-08 | 3.8e-08 | 2.4e-08 | 2.800E-08 | 8.718E-09 | 2.200E-08 | 3.800E-08 |
| 3153 | 158 | A158M | A | 1.8e-08 | 2.5e-08 | 9.1e-08 | 4.467E-08 | 4.028E-08 | 1.800E-08 | 9.100E-08 |
| 3154 | 158 | A158F | A | 3.5e-08 | 9.4e-08 | 5.6e-08 | 6.167E-08 | 2.991E-08 | 3.500E-08 | 9.400E-08 |
| 3155 | 158 | A158P | A | 4.7e-08 | 6.1e-08 | 5.7e-08 | 5.500E-08 | 7.211E-09 | 4.700E-08 | 6.100E-08 |
| 3156 | 158 | A158S | A | 4.7e-09 | 1.1e-07 | 4.5e-08 | 5.323E-08 | 5.313E-08 | 4.700E-09 | 1.100E-07 |
| 3157 | 158 | A158T | A | 6.1e-08 | 9.3e-08 | 4.5e-08 | 6.633E-08 | 2.444E-08 | 4.500E-08 | 9.300E-08 |
| 3158 | 158 | A158W | A | 1.3e-08 | 3.4e-08 | 9.7e-08 | 4.800E-08 | 4.371E-08 | 1.300E-08 | 9.700E-08 |
| 3159 | 158 | A158Y | A | 3.4e-08 | 1.6e-08 | 4.2e-08 | 3.067E-08 | 1.332E-08 | 1.600E-08 | 4.200E-08 |
| 3160 | 158 | A158V | A | 2.4e-08 | 4.7e-08 | 9e-08 | 5.367E-08 | 3.350E-08 | 2.400E-08 | 9.000E-08 |
| 3161 | 159 | L159A | A | 2.4e-08 | 2.6e-08 | 6.8e-08 | 3.933E-08 | 2.485E-08 | 2.400E-08 | 6.800E-08 |
| 3162 | 159 | L159R | A | 5.8e-08 | 7.8e-08 | 9.1e-08 | 7.567E-08 | 1.662E-08 | 5.800E-08 | 9.100E-08 |
| 3163 | 159 | L159N | A | 1.4e-08 | 4.7e-08 | 3.8e-08 | 3.300E-08 | 1.706E-08 | 1.400E-08 | 4.700E-08 |
| 3164 | 159 | L159D | A | 3.8e-08 | 7.5e-08 | 3.5e-08 | 4.933E-08 | 2.228E-08 | 3.500E-08 | 7.500E-08 |
| 3165 | 159 | L159C | A | 8.9e-08 | 4.9e-08 | 4.3e-08 | 6.033E-08 | 2.501E-08 | 4.300E-08 | 8.900E-08 |
| 3166 | 159 | L159E | A | 5.4e-08 | 7.7e-08 | 7e-08 | 6.700E-08 | 1.179E-08 | 5.400E-08 | 7.700E-08 |
| 3167 | 159 | L159Q | A | 1e-07 | 7.2e-08 | 8.6e-08 | 8.600E-08 | 1.400E-08 | 7.200E-08 | 1.000E-07 |
| 3168 | 159 | L159G | A | 1.5e-08 | 5.2e-08 | 4.6e-08 | 3.767E-08 | 1.986E-08 | 1.500E-08 | 5.200E-08 |
| 3169 | 159 | L159H | A | 5.8e-09 | 2.8e-08 | 4.1e-08 | 2.493E-08 | 1.780E-08 | 5.800E-09 | 4.100E-08 |
| 3170 | 159 | L159I | A | 1.8e-08 | 5.6e-08 | 2.8e-08 | 3.400E-08 | 1.970E-08 | 1.800E-08 | 5.600E-08 |
| 3171 | 159 | L159L | A | 6.5e-08 | 4.5e-08 | 1.2e-07 | 7.667E-08 | 3.884E-08 | 4.500E-08 | 1.200E-07 |
| 3172 | 159 | L159K | A | 5.5e-08 | 3.6e-08 | 3.9e-08 | 4.333E-08 | 1.021E-08 | 3.600E-08 | 5.500E-08 |
| 3173 | 159 | L159M | A | 1.9e-08 | 6.3e-08 | 2.2e-07 | 1.007E-07 | 1.057E-07 | 1.900E-08 | 2.200E-07 |
| 3174 | 159 | L159F | A | 1.7e-08 | 4.8e-08 | 1.7e-08 | 2.733E-08 | 1.790E-08 | 1.700E-08 | 4.800E-08 |
| 3175 | 159 | L159P | A | 2.3e-08 | 5.3e-08 | 2.8e-08 | 3.467E-08 | 1.607E-08 | 2.300E-08 | 5.300E-08 |
| 3176 | 159 | L159S | A | 2e-08 | 9.1e-08 | 1.1e-07 | 7.367E-08 | 4.744E-08 | 2.000E-08 | 1.100E-07 |
| 3177 | 159 | L159T | A | 6.4e-08 | 3.6e-08 | 6.5e-08 | 5.500E-08 | 1.646E-08 | 3.600E-08 | 6.500E-08 |
| 3178 | 159 | L159W | A | 5.5e-08 | 8.6e-08 | 7.2e-08 | 7.100E-08 | 1.552E-08 | 5.500E-08 | 8.600E-08 |
| 3179 | 159 | L159Y | A | 8.9e-08 | 2e-08 | 3.6e-08 | 4.833E-08 | 3.612E-08 | 2.000E-08 | 8.900E-08 |
| 3180 | 159 | L159V | A | 1.1e-07 | 1.8e-08 | 7.9e-08 | 6.900E-08 | 4.681E-08 | 1.800E-08 | 1.100E-07 |
| 3181 | 160 | T160A | A | 5.3e-08 | 2.8e-08 | 4.8e-08 | 4.300E-08 | 1.323E-08 | 2.800E-08 | 5.300E-08 |
| 3182 | 160 | T160R | A | 2.3e-08 | 1.8e-07 | 4.2e-08 | 8.167E-08 | 8.569E-08 | 2.300E-08 | 1.800E-07 |
| 3183 | 160 | T160N | A | 5.5e-08 | 6.9e-08 | 2.3e-08 | 4.900E-08 | 2.358E-08 | 2.300E-08 | 6.900E-08 |
| 3184 | 160 | T160D | A | 2e-08 | 2.9e-08 | 1.3e-07 | 5.967E-08 | 6.108E-08 | 2.000E-08 | 1.300E-07 |
| 3185 | 160 | T160C | A | 6.3e-08 | 2.9e-08 | 1.8e-08 | 3.667E-08 | 2.346E-08 | 1.800E-08 | 6.300E-08 |
| 3186 | 160 | T160E | A | 5e-08 | 4.4e-08 | 1e-07 | 6.467E-08 | 3.075E-08 | 4.400E-08 | 1.000E-07 |
| 3187 | 160 | T160Q | A | 1.1e-08 | 5.9e-08 | 3.1e-08 | 3.367E-08 | 2.411E-08 | 1.100E-08 | 5.900E-08 |
| 3188 | 160 | T160G | A | 9.8e-08 | 4.3e-08 | 4e-08 | 6.033E-08 | 3.265E-08 | 4.000E-08 | 9.800E-08 |
| 3189 | 160 | T160H | A | 5.1e-08 | 1.9e-08 | 5.3e-08 | 4.100E-08 | 1.908E-08 | 1.900E-08 | 5.300E-08 |
| 3190 | 160 | T160I | A | 2.2e-08 | 2.8e-08 | 1e-08 | 2.000E-08 | 9.165E-09 | 1.000E-08 | 2.800E-08 |
| 3191 | 160 | T160L | A | 6.9e-08 | 5.6e-08 | 4.6e-08 | 5.700E-08 | 1.153E-08 | 4.600E-08 | 6.900E-08 |
| 3192 | 160 | T160K | A | 2.9e-08 | 1.4e-08 | 6.4e-08 | 3.567E-08 | 2.566E-08 | 1.400E-08 | 6.400E-08 |
| 3193 | 160 | T160M | A | 4.3e-08 | 3.4e-08 | 4e-08 | 3.900E-08 | 4.583E-09 | 3.400E-08 | 4.300E-08 |
| 3194 | 160 | T160F | A | 1.5e-08 | 8.1e-08 | 1.1e-08 | 3.567E-08 | 3.931E-08 | 1.100E-08 | 8.100E-08 |
| 3195 | 160 | T160P | A | 4.5e-08 | 5.4e-09 | 4.8e-08 | 3.280E-08 | 2.378E-08 | 5.400E-09 | 4.800E-08 |
| 3196 | 160 | T160S | A | 1.8e-08 | 4e-08 | 2.2e-08 | 2.667E-08 | 1.172E-08 | 1.800E-08 | 4.000E-08 |
| 3197 | 160 | T160T | A | 6.6e-08 | 1.8e-08 | 3.8e-08 | 4.067E-08 | 2.411E-08 | 1.800E-08 | 6.600E-08 |
| 3198 | 160 | T160W | A | 7.8e-08 | 7.7e-08 | 3.2e-08 | 6.233E-08 | 2.627E-08 | 3.200E-08 | 7.800E-08 |
| 3199 | 160 | T160Y | A | 5.7e-08 | 3.9e-08 | 7.9e-08 | 5.833E-08 | 2.003E-08 | 3.900E-08 | 7.900E-08 |
| 3200 | 160 | T160V | A | 2.7e-08 | 4.5e-08 | 4.5e-08 | 3.900E-08 | 1.039E-08 | 2.700E-08 | 4.500E-08 |
| 3201 | 161 | L161A | A | 3.6e-08 | 1e-07 | 6.1e-08 | 6.567E-08 | 3.225E-08 | 3.600E-08 | 1.000E-07 |
| 3202 | 161 | L161R | A | 4.8e-08 | 6.3e-08 | 6.2e-08 | 5.767E-08 | 8.386E-09 | 4.800E-08 | 6.300E-08 |
| 3203 | 161 | L161N | A | 5.1e-08 | 1.3e-08 | 1.3e-08 | 2.567E-08 | 2.194E-08 | 1.300E-08 | 5.100E-08 |
| 3204 | 161 | L161D | A | 1.5e-08 | 3.7e-08 | 3.6e-08 | 2.933E-08 | 1.242E-08 | 1.500E-08 | 3.700E-08 |
| 3205 | 161 | L161C | A | 4.6e-08 | 3.1e-08 | 3.3e-08 | 3.667E-08 | 8.145E-09 | 3.100E-08 | 4.600E-08 |
| 3206 | 161 | L161E | A | 2.8e-08 | 6.5e-08 | 3.9e-08 | 4.400E-08 | 1.900E-08 | 2.800E-08 | 6.500E-08 |
| 3207 | 161 | L161Q | A | 4.7e-08 | 4.7e-08 | 1.3e-08 | 3.567E-08 | 1.963E-08 | 1.300E-08 | 4.700E-08 |
| 3208 | 161 | L161G | A | 1.4e-08 | 4.5e-08 | 3.8e-08 | 3.233E-08 | 1.626E-08 | 1.400E-08 | 4.500E-08 |
| 3209 | 161 | L161H | A | 1.1e-07 | 3.5e-08 | 1e-08 | 5.167E-08 | 5.204E-08 | 1.000E-08 | 1.100E-07 |
| 3210 | 161 | L161I | A | 3.5e-08 | 3.6e-08 | 4.2e-08 | 3.767E-08 | 3.786E-09 | 3.500E-08 | 4.200E-08 |
| 3211 | 161 | L161L | A | 5.1e-08 | 3.9e-08 | 3.8e-08 | 4.267E-08 | 7.234E-09 | 3.800E-08 | 5.100E-08 |
| 3212 | 161 | L161K | A | 5.8e-08 | 4.3e-08 | 5.2e-08 | 5.100E-08 | 7.550E-09 | 4.300E-08 | 5.800E-08 |
| 3213 | 161 | L161M | A | 2.2e-08 | 2.5e-08 | 4.8e-08 | 3.167E-08 | 1.422E-08 | 2.200E-08 | 4.800E-08 |
| 3214 | 161 | L161F | A | 4.3e-08 | 6.4e-08 | 3e-08 | 4.567E-08 | 1.716E-08 | 3.000E-08 | 6.400E-08 |
| 3215 | 161 | L161P | A | 1.2e-08 | 3.3e-08 | 4.8e-08 | 3.100E-08 | 1.808E-08 | 1.200E-08 | 4.800E-08 |
| 3216 | 161 | L161S | A | 3.8e-08 | 2.9e-08 | 7e-08 | 4.567E-08 | 2.155E-08 | 2.900E-08 | 7.000E-08 |
| 3217 | 161 | L161T | A | 4.3e-08 | 2.4e-08 | 2.8e-08 | 3.167E-08 | 1.002E-08 | 2.400E-08 | 4.300E-08 |
| 3218 | 161 | L161W | A | 4.9e-08 | 5.3e-08 | 5e-08 | 5.067E-08 | 2.082E-09 | 4.900E-08 | 5.300E-08 |
| 3219 | 161 | L161Y | A | 5.7e-08 | 2.3e-08 | 7.5e-08 | 5.167E-08 | 2.641E-08 | 2.300E-08 | 7.500E-08 |
| 3220 | 161 | L161V | A | 8.5e-08 | 8.3e-08 | 4.1e-08 | 6.967E-08 | 2.485E-08 | 4.100E-08 | 8.500E-08 |
| 3221 | 162 | I162A | A | 5.3e-08 | 4.3e-08 | 4.1e-08 | 4.567E-08 | 6.429E-09 | 4.100E-08 | 5.300E-08 |
| 3222 | 162 | I162R | A | 3.8e-08 | 6.5e-08 | 1.9e-08 | 4.067E-08 | 2.312E-08 | 1.900E-08 | 6.500E-08 |
| 3223 | 162 | I162N | A | 2.2e-08 | 4e-08 | 2.7e-08 | 2.967E-08 | 9.292E-09 | 2.200E-08 | 4.000E-08 |
| 3224 | 162 | I162D | A | 3e-08 | 7e-08 | 3.8e-08 | 4.600E-08 | 2.117E-08 | 3.000E-08 | 7.000E-08 |
| 3225 | 162 | I162C | A | 3.6e-08 | 3e-07 | 2.9e-08 | 1.217E-07 | 1.545E-07 | 2.900E-08 | 3.000E-07 |
| 3226 | 162 | I162E | A | 4.3e-08 | 9.1e-08 | 1.8e-08 | 5.067E-08 | 3.710E-08 | 1.800E-08 | 9.100E-08 |
| 3227 | 162 | I162Q | A | 3.9e-08 | 3.6e-08 | 2.6e-08 | 3.367E-08 | 6.807E-09 | 2.600E-08 | 3.900E-08 |
| 3228 | 162 | I162G | A | 1.5e-08 | 5e-08 | 4.7e-08 | 3.733E-08 | 1.940E-08 | 1.500E-08 | 5.000E-08 |
| 3229 | 162 | I162H | A | 2.5e-08 | 2.5e-08 | 1.8e-08 | 2.267E-08 | 4.041E-09 | 1.800E-08 | 2.500E-08 |

| | | | | | | | | | | |
|---|---|---|---|---|---|---|---|---|---|---|
| 3230 | 162 | I162I | A | 5.6e-08 | 3.7e-08 | 3e-08 | 4.100E-08 | 1.345E-08 | 3.000E-08 | 5.600E-08 |
| 3231 | 162 | I162L | A | 1.1e-08 | 3.2e-08 | 1.4e-08 | 1.900E-08 | 1.136E-08 | 1.100E-08 | 3.200E-08 |
| 3232 | 162 | I162K | A | 8.8e-08 | 8.8e-08 | 3.4e-08 | 7.000E-08 | 3.118E-08 | 3.400E-08 | 8.800E-08 |
| 3233 | 162 | I162M | A | 8.3e-08 | 4.1e-08 | 3.4e-08 | 5.267E-08 | 2.650E-08 | 3.400E-08 | 8.300E-08 |
| 3234 | 162 | I162F | A | 7.1e-08 | 2.9e-08 | 6.2e-08 | 5.400E-08 | 2.211E-08 | 2.900E-08 | 7.100E-08 |
| 3235 | 162 | I162P | A | 3.9e-08 | 1.5e-08 | 3.4e-08 | 2.933E-08 | 1.266E-08 | 1.500E-08 | 3.900E-08 |
| 3236 | 162 | I162S | A | 4.3e-08 | 3.6e-08 | 1.8e-08 | 3.233E-08 | 1.290E-08 | 1.800E-08 | 4.300E-08 |
| 3237 | 162 | I162T | A | 7.4e-08 | 1.1e-07 | 7e-08 | 8.467E-08 | 2.203E-08 | 7.000E-08 | 1.100E-07 |
| 3238 | 162 | I162W | A | 1.9e-08 | 3.6e-08 | 2.7e-08 | 2.733E-08 | 8.505E-09 | 1.900E-08 | 3.600E-08 |
| 3239 | 162 | I162Y | A | 1.4e-07 | 8.5e-08 | 5.5e-08 | 9.333E-08 | 4.311E-08 | 5.500E-08 | 1.400E-07 |
| 3240 | 162 | I162V | A | 2e-08 | 6e-08 | 2.1e-08 | 3.367E-08 | 2.281E-08 | 2.000E-08 | 6.000E-08 |
| 3241 | 163 | D163A | A | 1.3e-08 | 7.2e-08 | 4.9e-08 | 4.467E-08 | 2.974E-08 | 1.300E-08 | 7.200E-08 |
| 3242 | 163 | D163R | A | 7e-08 | 9.7e-08 | 1.9e-08 | 6.200E-08 | 3.961E-08 | 1.900E-08 | 9.700E-08 |
| 3243 | 163 | D163N | A | 3.2e-08 | 2.9e-08 | 1.4e-07 | 6.700E-08 | 6.324E-08 | 2.900E-08 | 1.400E-07 |
| 3244 | 163 | D163D | A | 4.5e-08 | 3.8e-08 | 1.2e-07 | 6.767E-08 | 4.546E-08 | 3.800E-08 | 1.200E-07 |
| 3245 | 163 | D163C | A | 4.1e-08 | 2.6e-08 | 4.1e-08 | 3.600E-08 | 8.660E-09 | 2.600E-08 | 4.100E-08 |
| 3246 | 163 | D163E | A | 8.4e-08 | 3.4e-08 | 7.9e-08 | 6.567E-08 | 2.754E-08 | 3.400E-08 | 8.400E-08 |
| 3247 | 163 | D163Q | A | 6.7e-08 | 2.3e-08 | 2.5e-08 | 3.833E-08 | 2.485E-08 | 2.300E-08 | 6.700E-08 |
| 3248 | 163 | D163G | A | 9e-08 | 2e-08 | 9.1e-08 | 6.700E-08 | 4.071E-08 | 2.000E-08 | 9.100E-08 |
| 3249 | 163 | D163H | A | 2.2e-08 | 2e-08 | 2.8e-08 | 2.333E-08 | 4.163E-09 | 2.000E-08 | 2.800E-08 |
| 3250 | 163 | D163I | A | 2.3e-08 | 5e-08 | 8.5e-08 | 5.267E-08 | 3.109E-08 | 2.300E-08 | 8.500E-08 |
| 3251 | 163 | D163L | A | 3.7e-08 | 7e-08 | 7.1e-08 | 5.933E-08 | 1.935E-08 | 3.700E-08 | 7.100E-08 |
| 3252 | 163 | D163K | A | 6.9e-08 | 2.2e-08 | 6.9e-08 | 5.333E-08 | 2.714E-08 | 2.200E-08 | 6.900E-08 |
| 3253 | 163 | D163M | A | 4.3e-08 | 7.7e-08 | 3e-08 | 5.000E-08 | 2.427E-08 | 3.000E-08 | 7.700E-08 |
| 3254 | 163 | D163F | A | 1.1e-07 | 3.6e-08 | 2.8e-08 | 5.800E-08 | 4.521E-08 | 2.800E-08 | 1.100E-07 |
| 3255 | 163 | D163P | A | 2.5e-08 | 6.1e-08 | 7.6e-08 | 5.400E-08 | 2.621E-08 | 2.500E-08 | 7.600E-08 |
| 3256 | 163 | D163S | A | 2.8e-08 | 2.8e-08 | 5.7e-08 | 3.767E-08 | 1.674E-08 | 2.800E-08 | 5.700E-08 |
| 3257 | 163 | D163T | A | 7.6e-08 | 3.1e-08 | 2.8e-08 | 4.500E-08 | 2.689E-08 | 2.800E-08 | 7.600E-08 |
| 3258 | 163 | D163W | A | 2.9e-08 | 1.7e-08 | 6.3e-08 | 3.633E-08 | 2.386E-08 | 1.700E-08 | 6.300E-08 |
| 3259 | 163 | D163Y | A | 2.1e-08 | 2.9e-08 | 2.9e-08 | 2.633E-08 | 4.619E-09 | 2.100E-08 | 2.900E-08 |
| 3260 | 163 | D163V | A | 7.6e-08 | 1.4e-08 | 3e-08 | 4.000E-08 | 3.219E-08 | 1.400E-08 | 7.600E-08 |
| 3261 | 164 | T164A | A | 1.5e-08 | 3.9e-08 | 2.8e-08 | 2.733E-08 | 1.201E-08 | 1.500E-08 | 3.900E-08 |
| 3262 | 164 | T164R | A | 3.4e-08 | 4.3e-08 | 6.2e-08 | 4.633E-08 | 1.429E-08 | 3.400E-08 | 6.200E-08 |
| 3263 | 164 | T164N | A | 6.6e-08 | 1.2e-08 | 3.4e-08 | 3.733E-08 | 2.715E-08 | 1.200E-08 | 6.600E-08 |
| 3264 | 164 | T164D | A | 8e-08 | 1.7e-07 | 1.1e-08 | 8.700E-08 | 7.973E-08 | 1.100E-08 | 1.700E-07 |
| 3265 | 164 | T164C | A | 4.1e-08 | 4.2e-08 | 2.9e-08 | 3.733E-08 | 7.234E-09 | 2.900E-08 | 4.200E-08 |
| 3266 | 164 | T164E | A | 5.7e-08 | 2.5e-08 | 1.6e-08 | 3.267E-08 | 2.155E-08 | 1.600E-08 | 5.700E-08 |
| 3267 | 164 | T164Q | A | 6.9e-08 | 4.8e-08 | 2.3e-08 | 4.667E-08 | 2.303E-08 | 2.300E-08 | 6.900E-08 |
| 3268 | 164 | T164G | A | 3.3e-08 | 5.3e-08 | 7.1e-08 | 5.233E-08 | 1.901E-08 | 3.300E-08 | 7.100E-08 |
| 3269 | 164 | T164H | A | 4.2e-08 | 3.5e-08 | 4.8e-08 | 4.167E-08 | 6.506E-09 | 3.500E-08 | 4.800E-08 |
| 3270 | 164 | T164I | A | 4.3e-08 | 4.6e-08 | 4.3e-08 | 4.400E-08 | 1.732E-09 | 4.300E-08 | 4.600E-08 |
| 3271 | 164 | T164L | A | 1.9e-08 | 4.1e-08 | 4e-08 | 3.333E-08 | 1.242E-08 | 1.900E-08 | 4.100E-08 |
| 3272 | 164 | T164K | A | 5e-08 | 4e-08 | 7.7e-08 | 5.567E-08 | 1.914E-08 | 4.000E-08 | 7.700E-08 |
| 3273 | 164 | T164M | A | 5.1e-08 | 1e-07 | 2.6e-08 | 5.900E-08 | 3.764E-08 | 2.600E-08 | 1.000E-07 |
| 3274 | 164 | T164F | A | 2.2e-08 | 3.8e-08 | 9e-08 | 5.000E-08 | 3.555E-08 | 2.200E-08 | 9.000E-08 |
| 3275 | 164 | T164P | A | 3.2e-08 | 2.1e-08 | 1.3e-08 | 2.200E-08 | 9.539E-09 | 1.300E-08 | 3.200E-08 |
| 3276 | 164 | T164S | A | 1.7e-08 | 2.8e-08 | 2.9e-08 | 2.467E-08 | 6.658E-09 | 1.700E-08 | 2.900E-08 |
| 3277 | 164 | T164T | A | 3.8e-08 | 1.2e-07 | 5e-08 | 6.933E-08 | 4.429E-08 | 3.800E-08 | 1.200E-07 |
| 3278 | 164 | T164W | A | 4.7e-08 | 1.5e-07 | 5.3e-08 | 8.333E-08 | 5.781E-08 | 4.700E-08 | 1.500E-07 |
| 3279 | 164 | T164Y | A | 2.8e-08 | 5.9e-08 | 2.4e-08 | 3.700E-08 | 1.916E-08 | 2.400E-08 | 5.900E-08 |
| 3280 | 164 | T164V | A | 2.8e-08 | 1.2e-07 | 7.3e-08 | 7.367E-08 | 4.600E-08 | 2.800E-08 | 1.200E-07 |
| 3281 | 165 | N165A | A | 2.4e-08 | 4.9e-08 | 1.7e-08 | 3.000E-08 | 1.682E-08 | 1.700E-08 | 4.900E-08 |
| 3282 | 165 | N165R | A | 3.7e-08 | 1.1e-07 | 6.2e-08 | 6.967E-08 | 3.710E-08 | 3.700E-08 | 1.100E-07 |
| 3283 | 165 | N165N | A | 3.5e-08 | 3.2e-08 | 2.5e-08 | 3.067E-08 | 5.132E-09 | 2.500E-08 | 3.500E-08 |
| 3284 | 165 | N165D | A | 3.4e-08 | 2.8e-08 | 2.3e-08 | 2.833E-08 | 5.508E-09 | 2.300E-08 | 3.400E-08 |
| 3285 | 165 | N165C | A | 7e-08 | 4.7e-08 | 4.7e-08 | 5.467E-08 | 1.328E-08 | 4.700E-08 | 7.000E-08 |
| 3286 | 165 | N165E | A | 7.6e-08 | 2.4e-08 | 2.8e-08 | 4.267E-08 | 2.894E-08 | 2.400E-08 | 7.600E-08 |
| 3287 | 165 | N165Q | A | 5.4e-08 | 6.8e-08 | 3.4e-08 | 5.200E-08 | 1.709E-08 | 3.400E-08 | 6.800E-08 |
| 3288 | 165 | N165G | A | 2e-08 | 4e-08 | 2.6e-08 | 2.867E-08 | 1.026E-08 | 2.000E-08 | 4.000E-08 |
| 3289 | 165 | N165H | A | 2.8e-08 | 1.3e-07 | 1.1e-07 | 8.933E-08 | 5.405E-08 | 2.800E-08 | 1.300E-07 |
| 3290 | 165 | N165I | A | 5.1e-08 | 5.4e-08 | 1.7e-07 | 9.167E-08 | 6.786E-08 | 5.100E-08 | 1.700E-07 |
| 3291 | 165 | N165L | A | 5.9e-08 | 2.5e-08 | 2.5e-08 | 3.633E-08 | 1.963E-08 | 2.500E-08 | 5.900E-08 |
| 3292 | 165 | N165K | A | 4.9e-08 | 4.1e-08 | 1.8e-08 | 3.600E-08 | 1.609E-08 | 1.800E-08 | 4.900E-08 |
| 3293 | 165 | N165M | A | 3.1e-08 | 3.7e-08 | 4.1e-08 | 3.633E-08 | 5.033E-09 | 3.100E-08 | 4.100E-08 |
| 3294 | 165 | N165F | A | 2.6e-08 | 5.9e-08 | 1.5e-08 | 3.333E-08 | 2.290E-08 | 1.500E-08 | 5.900E-08 |
| 3295 | 165 | N165P | A | 6.4e-08 | 6.8e-08 | 9.4e-08 | 7.533E-08 | 1.629E-08 | 6.400E-08 | 9.400E-08 |
| 3296 | 165 | N165S | A | 1.1e-07 | 1.4e-08 | 2.1e-08 | 4.833E-08 | 5.352E-08 | 1.400E-08 | 1.100E-07 |
| 3297 | 165 | N165T | A | 3.2e-08 | 1.1e-08 | 1e-07 | 4.767E-08 | 4.652E-08 | 1.100E-08 | 1.000E-07 |
| 3298 | 165 | N165W | A | 6.5e-08 | 4.4e-08 | 3e-08 | 4.633E-08 | 1.762E-08 | 3.000E-08 | 6.500E-08 |
| 3299 | 165 | N165Y | A | 6.2e-08 | 1.3e-08 | 3e-08 | 3.500E-08 | 2.488E-08 | 1.300E-08 | 6.200E-08 |
| 3300 | 165 | N165V | A | 4.2e-08 | 1.4e-08 | 1.8e-07 | 7.867E-08 | 8.887E-08 | 1.400E-08 | 1.800E-07 |
| 3301 | 166 | R166A | A | 3.9e-08 | 5e-08 | 2.4e-08 | 3.767E-08 | 1.305E-08 | 2.400E-08 | 5.000E-08 |
| 3302 | 166 | R166R | A | 2.1e-08 | 4.1e-08 | 1.5e-08 | 2.567E-08 | 1.361E-08 | 1.500E-08 | 4.100E-08 |
| 3303 | 166 | R166N | A | 5e-08 | 4.6e-08 | 3e-08 | 4.200E-08 | 1.058E-08 | 3.000E-08 | 5.000E-08 |
| 3304 | 166 | R166D | A | 8.7e-08 | 8.7e-08 | 5e-08 | 7.467E-08 | 2.136E-08 | 5.000E-08 | 8.700E-08 |
| 3305 | 166 | R166C | A | 3.7e-08 | 3.8e-08 | 1e-08 | 2.833E-08 | 1.589E-08 | 1.000E-08 | 3.800E-08 |
| 3306 | 166 | R166E | A | 3.2e-08 | 7.4e-08 | 2.5e-08 | 4.367E-08 | 2.650E-08 | 2.500E-08 | 7.400E-08 |
| 3307 | 166 | R166Q | A | 5.9e-08 | 3.4e-08 | 5.7e-08 | 5.000E-08 | 1.389E-08 | 3.400E-08 | 5.900E-08 |
| 3308 | 166 | R166G | A | 5.6e-08 | 6e-08 | 6.6e-08 | 6.067E-08 | 5.033E-09 | 5.600E-08 | 6.600E-08 |
| 3309 | 166 | R166H | A | 4e-08 | 3.3e-08 | 1.5e-08 | 2.933E-08 | 1.290E-08 | 1.500E-08 | 4.000E-08 |
| 3310 | 166 | R166I | A | 7.1e-08 | 3.2e-08 | 3.6e-08 | 4.633E-08 | 2.146E-08 | 3.200E-08 | 7.100E-08 |
| 3311 | 166 | R166L | A | 1.2e-08 | 4.8e-08 | 2e-08 | 2.667E-08 | 1.890E-08 | 1.200E-08 | 4.800E-08 |
| 3312 | 166 | R166K | A | 3.7e-08 | 7e-08 | 3.1e-08 | 4.600E-08 | 2.100E-08 | 3.100E-08 | 7.000E-08 |
| 3313 | 166 | R166M | A | 3.3e-08 | 1.1e-08 | 7.9e-08 | 4.100E-08 | 3.470E-08 | 1.100E-08 | 7.900E-08 |
| 3314 | 166 | R166F | A | 2e-08 | 2.8e-08 | 3.6e-08 | 2.800E-08 | 8.000E-09 | 2.000E-08 | 3.600E-08 |

| | | | | | | | | | | |
|---|---|---|---|---|---|---|---|---|---|---|
| 3315 | 166 | R166P | A | 3.5e-08 | 2.6e-08 | 6.2e-08 | 4.100E-08 | 1.873E-08 | 2.600E-08 | 6.200E-08 |
| 3316 | 166 | R166S | A | 2.6e-08 | 1.9e-08 | 2.5e-08 | 2.333E-08 | 3.786E-09 | 1.900E-08 | 2.600E-08 |
| 3317 | 166 | R166T | A | 2.3e-08 | 7.4e-08 | 7.1e-08 | 5.600E-08 | 2.862E-08 | 2.300E-08 | 7.400E-08 |
| 3318 | 166 | R166W | A | 4.5e-08 | 1.3e-08 | 1.5e-08 | 2.433E-08 | 1.793E-08 | 1.300E-08 | 4.500E-08 |
| 3319 | 166 | R166Y | A | 3.5e-08 | 4.4e-08 | 2.4e-08 | 3.433E-08 | 1.002E-08 | 2.400E-08 | 4.400E-08 |
| 3320 | 166 | R166V | A | 7.3e-08 | 2.4e-08 | 2.8e-08 | 4.167E-08 | 2.721E-08 | 2.400E-08 | 7.300E-08 |
| 3321 | 167 | S167A | A | 2e-08 | 2.3e-08 | 1.2e-08 | 1.833E-08 | 5.686E-09 | 1.200E-08 | 2.300E-08 |
| 3322 | 167 | S167R | A | 2.6e-08 | 1.9e-08 | 3.1e-08 | 2.533E-08 | 6.028E-09 | 1.900E-08 | 3.100E-08 |
| 3323 | 167 | S167N | A | 4e-08 | 7.5e-08 | 9.6e-08 | 7.033E-08 | 2.829E-08 | 4.000E-08 | 9.600E-08 |
| 3324 | 167 | S167D | A | 6.3e-08 | 2e-08 | 7.5e-08 | 5.267E-08 | 2.892E-08 | 2.000E-08 | 7.500E-08 |
| 3325 | 167 | S167C | A | 1.7e-07 | 5.2e-08 | 6.5e-08 | 9.567E-08 | 6.470E-08 | 5.200E-08 | 1.700E-07 |
| 3326 | 167 | S167E | A | 2.5e-08 | 2.7e-08 | 4.7e-08 | 3.300E-08 | 1.217E-08 | 2.500E-08 | 4.700E-08 |
| 3327 | 167 | S167Q | A | 1.1e-07 | 3.4e-08 | 2.5e-08 | 5.633E-08 | 4.669E-08 | 2.500E-08 | 1.100E-07 |
| 3328 | 167 | S167G | A | 3.6e-08 | 9.1e-08 | 4.9e-08 | 5.867E-08 | 2.875E-08 | 3.600E-08 | 9.100E-08 |
| 3329 | 167 | S167H | A | 5e-08 | 2.8e-08 | 2.1e-08 | 3.300E-08 | 1.513E-08 | 2.100E-08 | 5.000E-08 |
| 3330 | 167 | S167I | A | 1.1e-08 | 1.5e-07 | 2e-07 | 1.203E-07 | 9.793E-08 | 1.100E-08 | 2.000E-07 |
| 3331 | 167 | S167L | A | 4.4e-08 | 2.8e-08 | 4.2e-08 | 3.800E-08 | 8.718E-09 | 2.800E-08 | 4.400E-08 |
| 3332 | 167 | S167K | A | 2.2e-08 | 4.3e-08 | 1.3e-07 | 6.500E-08 | 5.726E-08 | 2.200E-08 | 1.300E-07 |
| 3333 | 167 | S167M | A | 5.4e-08 | 5.1e-08 | 1.5e-08 | 4.000E-08 | 2.170E-08 | 1.500E-08 | 5.400E-08 |
| 3334 | 167 | S167F | A | 2.3e-08 | 4.7e-08 | 3.7e-08 | 3.567E-08 | 1.206E-08 | 2.300E-08 | 4.700E-08 |
| 3335 | 167 | S167P | A | 4.1e-08 | 4.3e-08 | 9.1e-08 | 5.833E-08 | 2.831E-08 | 4.100E-08 | 9.100E-08 |
| 3336 | 167 | S167S | A | 4.7e-08 | 4.2e-08 | 4.7e-08 | 4.533E-08 | 2.887E-09 | 4.200E-08 | 4.700E-08 |
| 3337 | 167 | S167T | A | 1.7e-08 | 5.9e-08 | 1.3e-07 | 6.867E-08 | 5.712E-08 | 1.700E-08 | 1.300E-07 |
| 3338 | 167 | S167W | A | 8.1e-08 | 4.7e-08 | 6e-08 | 6.267E-08 | 1.716E-08 | 4.700E-08 | 8.100E-08 |
| 3339 | 167 | S167Y | A | 1e-08 | 4e-08 | 5.1e-08 | 3.367E-08 | 2.122E-08 | 1.000E-08 | 5.100E-08 |
| 3340 | 167 | S167V | A | 5.6e-08 | 2.2e-08 | 5e-08 | 4.267E-08 | 1.815E-08 | 2.200E-08 | 5.600E-08 |
| 3341 | 168 | R168A | A | 3.4e-08 | 2.7e-08 | 2.7e-08 | 2.933E-08 | 4.041E-09 | 2.700E-08 | 3.400E-08 |
| 3342 | 168 | R168R | A | 2.5e-08 | 3.9e-08 | 1.1e-07 | 5.800E-08 | 4.557E-08 | 2.500E-08 | 1.100E-07 |
| 3343 | 168 | R168N | A | 8.9e-08 | 6.2e-08 | 4.8e-08 | 6.633E-08 | 2.084E-08 | 4.800E-08 | 8.900E-08 |
| 3344 | 168 | R168D | A | 7.5e-08 | 2.6e-08 | 4.9e-08 | 5.000E-08 | 2.452E-08 | 2.600E-08 | 7.500E-08 |
| 3345 | 168 | R168C | A | 2.1e-08 | 8e-08 | 6.5e-08 | 5.533E-08 | 3.066E-08 | 2.100E-08 | 8.000E-08 |
| 3346 | 168 | R168E | A | 3.4e-08 | 5.1e-08 | 3.1e-08 | 3.867E-08 | 1.079E-08 | 3.100E-08 | 5.100E-08 |
| 3347 | 168 | R168Q | A | 5.3e-08 | 5.5e-08 | 4.2e-08 | 5.000E-08 | 7.000E-09 | 4.200E-08 | 5.500E-08 |
| 3348 | 168 | R168G | A | 5e-08 | 4.4e-08 | 6.9e-08 | 5.433E-08 | 1.305E-08 | 4.400E-08 | 6.900E-08 |
| 3349 | 168 | R168H | A | 2.9e-08 | 2e-08 | 8.2e-08 | 4.367E-08 | 3.350E-08 | 2.000E-08 | 8.200E-08 |
| 3350 | 168 | R168I | A | 5.2e-08 | 5.2e-08 | 7.9e-08 | 6.100E-08 | 1.559E-08 | 5.200E-08 | 7.900E-08 |
| 3351 | 168 | R168L | A | 6.7e-08 | 4.2e-08 | 2.5e-08 | 4.467E-08 | 2.113E-08 | 2.500E-08 | 6.700E-08 |
| 3352 | 168 | R168K | A | 4.4e-08 | 4.1e-08 | 1.4e-08 | 3.300E-08 | 1.652E-08 | 1.400E-08 | 4.400E-08 |
| 3353 | 168 | R168M | A | 2.2e-08 | 2.9e-08 | 1.4e-08 | 2.167E-08 | 7.506E-09 | 1.400E-08 | 2.900E-08 |
| 3354 | 168 | R168F | A | 8.9e-08 | 3.6e-08 | 3.4e-08 | 5.300E-08 | 3.119E-08 | 3.400E-08 | 8.900E-08 |
| 3355 | 168 | R168P | A | 5.9e-08 | 7.3e-08 | 1.4e-08 | 4.867E-08 | 3.083E-08 | 1.400E-08 | 7.300E-08 |
| 3356 | 168 | R168S | A | 7.1e-08 | 8.2e-08 | 2.7e-08 | 6.000E-08 | 2.910E-08 | 2.700E-08 | 8.200E-08 |
| 3357 | 168 | R168T | A | 1.1e-08 | 7.3e-08 | 4.4e-08 | 4.267E-08 | 3.102E-08 | 1.100E-08 | 7.300E-08 |
| 3358 | 168 | R168W | A | 3.5e-08 | 3.6e-08 | 3.3e-08 | 3.467E-08 | 1.528E-09 | 3.300E-08 | 3.600E-08 |
| 3359 | 168 | R168Y | A | 3.4e-08 | 4.9e-08 | 4.6e-08 | 4.300E-08 | 7.937E-09 | 3.400E-08 | 4.900E-08 |
| 3360 | 168 | R168V | A | 2.4e-08 | 5.9e-08 | 4.9e-08 | 4.400E-08 | 1.803E-08 | 2.400E-08 | 5.900E-08 |
| 3361 | 169 | A169A | A | 4.3e-08 | 6e-08 | 1.9e-08 | 4.067E-08 | 2.060E-08 | 1.900E-08 | 6.000E-08 |
| 3362 | 169 | A169R | A | 3.2e-08 | 6.6e-08 | 1.4e-08 | 3.733E-08 | 2.641E-08 | 1.400E-08 | 6.600E-08 |
| 3363 | 169 | A169N | A | 1e-07 | 2.6e-08 | 8.7e-08 | 7.100E-08 | 3.951E-08 | 2.600E-08 | 1.000E-07 |
| 3364 | 169 | A169D | A | 1.5e-08 | 8.1e-08 | 3.8e-08 | 4.467E-08 | 3.350E-08 | 1.500E-08 | 8.100E-08 |
| 3365 | 169 | A169C | A | 5.7e-08 | 4.3e-08 | 3e-08 | 4.333E-08 | 1.350E-08 | 3.000E-08 | 5.700E-08 |
| 3366 | 169 | A169E | A | 5.4e-08 | 8.5e-08 | 2.2e-08 | 5.367E-08 | 3.150E-08 | 2.200E-08 | 8.500E-08 |
| 3367 | 169 | A169Q | A | 2.8e-08 | 2.4e-08 | 7.6e-08 | 4.267E-08 | 2.894E-08 | 2.400E-08 | 7.600E-08 |
| 3368 | 169 | A169G | A | 3.5e-08 | 1.2e-08 | 2e-08 | 2.233E-08 | 1.168E-08 | 1.200E-08 | 3.500E-08 |
| 3369 | 169 | A169H | A | 1.8e-08 | 2.7e-08 | 4.9e-08 | 3.133E-08 | 1.595E-08 | 1.800E-08 | 4.900E-08 |
| 3370 | 169 | A169I | A | 3.5e-08 | 1.4e-08 | 5.2e-08 | 3.367E-08 | 1.904E-08 | 1.400E-08 | 5.200E-08 |
| 3371 | 169 | A169L | A | 4.4e-08 | 1.1e-07 | 2.5e-08 | 5.967E-08 | 4.461E-08 | 2.500E-08 | 1.100E-07 |
| 3372 | 169 | A169K | A | 2e-07 | 6.8e-08 | 1e-07 | 1.227E-07 | 6.886E-08 | 6.800E-08 | 2.000E-07 |
| 3373 | 169 | A169M | A | 5.3e-08 | 6.5e-08 | 8.2e-08 | 6.667E-08 | 1.457E-08 | 5.300E-08 | 8.200E-08 |
| 3374 | 169 | A169F | A | 2.5e-08 | 4.3e-08 | 6.5e-08 | 4.433E-08 | 2.003E-08 | 2.500E-08 | 6.500E-08 |
| 3375 | 169 | A169P | A | 2.8e-08 | 2.2e-08 | 2.1e-08 | 2.367E-08 | 3.786E-09 | 2.100E-08 | 2.800E-08 |
| 3376 | 169 | A169S | A | 3.1e-08 | 2.6e-08 | 1.1e-07 | 5.567E-08 | 4.712E-08 | 2.600E-08 | 1.100E-07 |
| 3377 | 169 | A169T | A | 2.5e-08 | 5.4e-08 | 1.4e-08 | 3.100E-08 | 2.066E-08 | 1.400E-08 | 5.400E-08 |
| 3378 | 169 | A169W | A | 8.1e-08 | 1.1e-07 | 7.5e-08 | 8.867E-08 | 1.872E-08 | 7.500E-08 | 1.100E-07 |
| 3379 | 169 | A169Y | A | 7.7e-08 | 1.1e-08 | 6.6e-08 | 5.133E-08 | 3.536E-08 | 1.100E-08 | 7.700E-08 |
| 3380 | 169 | A169V | A | 1.6e-08 | 7.7e-08 | 3.1e-08 | 4.133E-08 | 3.179E-08 | 1.600E-08 | 7.700E-08 |
| 3381 | 170 | C170A | A | 2.2e-08 | 8.2e-08 | 5e-08 | 5.133E-08 | 3.002E-08 | 2.200E-08 | 8.200E-08 |
| 3382 | 170 | C170R | A | 9.2e-08 | 9.1e-09 | 3.3e-08 | 4.470E-08 | 4.267E-08 | 9.100E-09 | 9.200E-08 |
| 3383 | 170 | C170N | A | 5.7e-08 | 8.5e-08 | 1.4e-08 | 5.200E-08 | 3.576E-08 | 1.400E-08 | 8.500E-08 |
| 3384 | 170 | C170D | A | 3.9e-08 | 1.2e-07 | 7.5e-08 | 7.800E-08 | 4.058E-08 | 3.900E-08 | 1.200E-07 |
| 3385 | 170 | C170C | A | 1.5e-07 | 3.7e-08 | 6.5e-08 | 8.400E-08 | 5.885E-08 | 3.700E-08 | 1.500E-07 |
| 3386 | 170 | C170E | A | 2.8e-08 | 1.7e-08 | 5.9e-08 | 3.467E-08 | 2.178E-08 | 1.700E-08 | 5.900E-08 |
| 3387 | 170 | C170Q | A | 2.3e-08 | 4.3e-08 | 1.2e-07 | 6.200E-08 | 5.122E-08 | 2.300E-08 | 1.200E-07 |
| 3388 | 170 | C170G | A | 1.5e-08 | 4.2e-08 | 1.3e-08 | 2.333E-08 | 1.620E-08 | 1.300E-08 | 4.200E-08 |
| 3389 | 170 | C170H | A | 4e-08 | 3.6e-08 | 7.3e-09 | 2.777E-08 | 1.784E-08 | 7.300E-09 | 4.000E-08 |
| 3390 | 170 | C170I | A | 1.1e-07 | 6.4e-08 | 1.6e-08 | 6.333E-08 | 4.700E-08 | 1.600E-08 | 1.100E-07 |
| 3391 | 170 | C170L | A | 1.2e-08 | 1.3e-08 | 3.5e-08 | 2.000E-08 | 1.300E-08 | 1.200E-08 | 3.500E-08 |
| 3392 | 170 | C170K | A | 4.4e-08 | 4e-08 | 1.2e-07 | 6.800E-08 | 4.508E-08 | 4.000E-08 | 1.200E-07 |
| 3393 | 170 | C170M | A | 3e-08 | 3.1e-08 | 4.6e-08 | 3.567E-08 | 8.963E-09 | 3.000E-08 | 4.600E-08 |
| 3394 | 170 | C170F | A | 4.1e-08 | 4.2e-08 | 3.3e-08 | 3.867E-08 | 4.933E-09 | 3.300E-08 | 4.200E-08 |
| 3395 | 170 | C170P | A | 5.5e-08 | 1.1e-07 | 3.2e-08 | 6.567E-08 | 4.008E-08 | 3.200E-08 | 1.100E-07 |
| 3396 | 170 | C170S | A | 2.8e-08 | 2.6e-08 | 2.4e-08 | 2.600E-08 | 2.000E-09 | 2.400E-08 | 2.800E-08 |
| 3397 | 170 | C170T | A | 3.1e-08 | 5e-08 | 4e-08 | 4.033E-08 | 9.504E-09 | 3.100E-08 | 5.000E-08 |
| 3398 | 170 | C170W | A | 6.4e-08 | 5.2e-08 | 3.3e-08 | 4.967E-08 | 1.563E-08 | 3.300E-08 | 6.400E-08 |
| 3399 | 170 | C170Y | A | 6.1e-08 | 4.8e-08 | 2e-08 | 4.300E-08 | 2.095E-08 | 2.000E-08 | 6.100E-08 |

| | | | | | | | | | | |
|---|---|---|---|---|---|---|---|---|---|---|
| 3400 | 170 | C170V | A | 1.1e-07 | 5.7e-08 | 3.4e-08 | 6.700E-08 | 3.897E-08 | 3.400E-08 | 1.100E-07 |
| 3401 | 171 | H171A | A | 1.6e-08 | 6.6e-08 | 6.6e-08 | 4.933E-08 | 2.887E-08 | 1.600E-08 | 6.600E-08 |
| 3402 | 171 | H171R | A | 5.3e-08 | 4.3e-08 | 5.4e-08 | 5.000E-08 | 6.083E-09 | 4.300E-08 | 5.400E-08 |
| 3403 | 171 | H171N | A | 5.8e-08 | 2e-08 | 3.9e-08 | 3.900E-08 | 1.900E-08 | 2.000E-08 | 5.800E-08 |
| 3404 | 171 | H171D | A | 1e-08 | 2.9e-08 | 1.7e-08 | 1.867E-08 | 9.609E-09 | 1.000E-08 | 2.900E-08 |
| 3405 | 171 | H171C | A | 3.4e-08 | 8.8e-08 | 2.5e-08 | 4.900E-08 | 3.407E-08 | 2.500E-08 | 8.800E-08 |
| 3406 | 171 | H171E | A | 6.5e-08 | 5.5e-08 | 4.2e-08 | 5.400E-08 | 1.153E-08 | 4.200E-08 | 6.500E-08 |
| 3407 | 171 | H171Q | A | 2.2e-08 | 5.6e-08 | 5.1e-08 | 4.300E-08 | 1.836E-08 | 2.200E-08 | 5.600E-08 |
| 3408 | 171 | H171G | A | 2.6e-08 | 1.3e-08 | 1.4e-08 | 1.767E-08 | 7.234E-09 | 1.300E-08 | 2.600E-08 |
| 3409 | 171 | H171H | A | 4.6e-08 | 5.6e-08 | 1.5e-07 | 8.400E-08 | 5.738E-08 | 4.600E-08 | 1.500E-07 |
| 3410 | 171 | H171I | A | 7.8e-08 | 4.2e-08 | 3.1e-08 | 5.033E-08 | 2.458E-08 | 3.100E-08 | 7.800E-08 |
| 3411 | 171 | H171L | A | 6e-08 | 3.8e-08 | 5.2e-08 | 5.000E-08 | 1.114E-08 | 3.800E-08 | 6.000E-08 |
| 3412 | 171 | H171K | A | 4.2e-08 | 5.1e-08 | 6.5e-08 | 5.267E-08 | 1.159E-08 | 4.200E-08 | 6.500E-08 |
| 3413 | 171 | H171M | A | 2.9e-08 | 3e-08 | 3.1e-08 | 3.000E-08 | 1.000E-09 | 2.900E-08 | 3.100E-08 |
| 3414 | 171 | H171F | A | 4.9e-08 | 2e-08 | 6.1e-08 | 4.333E-08 | 2.108E-08 | 2.000E-08 | 6.100E-08 |
| 3415 | 171 | H171P | A | 7e-08 | 3.3e-08 | 3.5e-08 | 4.600E-08 | 2.081E-08 | 3.300E-08 | 7.000E-08 |
| 3416 | 171 | H171S | A | 1.5e-08 | 5.5e-08 | 4.1e-08 | 3.700E-08 | 2.030E-08 | 1.500E-08 | 5.500E-08 |
| 3417 | 171 | H171T | A | 3.7e-08 | 1.3e-07 | 3e-08 | 6.567E-08 | 5.582E-08 | 3.000E-08 | 1.300E-07 |
| 3418 | 171 | H171W | A | 4.9e-08 | 6.4e-08 | 3e-08 | 4.767E-08 | 1.704E-08 | 3.000E-08 | 6.400E-08 |
| 3419 | 171 | H171Y | A | 2.9e-08 | 4.2e-08 | 6.7e-08 | 4.600E-08 | 1.931E-08 | 2.900E-08 | 6.700E-08 |
| 3420 | 171 | H171V | A | 1e-07 | 2.2e-08 | 3.8e-08 | 5.333E-08 | 4.120E-08 | 2.200E-08 | 1.000E-07 |
| 3421 | 172 | P172A | A | 2.5e-08 | 8.5e-08 | 5.6e-08 | 5.533E-08 | 3.001E-08 | 2.500E-08 | 8.500E-08 |
| 3422 | 172 | P172R | A | 1.7e-08 | 8.4e-08 | 1.3e-08 | 3.800E-08 | 3.989E-08 | 1.300E-08 | 8.400E-08 |
| 3423 | 172 | P172N | A | 6.6e-08 | 3e-08 | 5.7e-08 | 5.100E-08 | 1.873E-08 | 3.000E-08 | 6.600E-08 |
| 3424 | 172 | P172D | A | 2.9e-08 | 7.9e-08 | 6.3e-08 | 5.700E-08 | 2.553E-08 | 2.900E-08 | 7.900E-08 |
| 3425 | 172 | P172C | A | 4.4e-08 | 2.8e-08 | 2.9e-08 | 3.367E-08 | 8.963E-09 | 2.800E-08 | 4.400E-08 |
| 3426 | 172 | P172E | A | 9.8e-09 | 7.6e-08 | 1.3e-08 | 3.293E-08 | 3.733E-08 | 9.800E-09 | 7.600E-08 |
| 3427 | 172 | P172Q | A | 1.8e-08 | 3e-08 | 4.5e-08 | 3.100E-08 | 1.353E-08 | 1.800E-08 | 4.500E-08 |
| 3428 | 172 | P172G | A | 9.8e-08 | 5.1e-08 | 1.3e-07 | 9.300E-08 | 3.974E-08 | 5.100E-08 | 1.300E-07 |
| 3429 | 172 | P172H | A | 4.5e-08 | 7.1e-08 | 5.5e-08 | 5.700E-08 | 1.311E-08 | 4.500E-08 | 7.100E-08 |
| 3430 | 172 | P172I | A | 2.5e-08 | 2.1e-08 | 3.5e-08 | 2.700E-08 | 7.211E-09 | 2.100E-08 | 3.500E-08 |
| 3431 | 172 | P172L | A | 4.1e-08 | 6.8e-08 | 3.8e-08 | 4.900E-08 | 1.652E-08 | 3.800E-08 | 6.800E-08 |
| 3432 | 172 | P172K | A | 1.3e-07 | 8.8e-08 | 2.6e-08 | 8.133E-08 | 5.232E-08 | 2.600E-08 | 1.300E-07 |
| 3433 | 172 | P172M | A | 6.6e-08 | 7.5e-08 | 3.9e-08 | 6.000E-08 | 1.873E-08 | 3.900E-08 | 7.500E-08 |
| 3434 | 172 | P172F | A | 2.8e-08 | 1.2e-08 | 2.9e-08 | 2.300E-08 | 9.539E-09 | 1.200E-08 | 2.900E-08 |
| 3435 | 172 | P172P | A | 2.9e-08 | 5.5e-08 | 4.4e-08 | 4.267E-08 | 1.305E-08 | 2.900E-08 | 5.500E-08 |
| 3436 | 172 | P172S | A | 7.2e-08 | 3.3e-08 | 3.1e-08 | 4.533E-08 | 2.312E-08 | 3.100E-08 | 7.200E-08 |
| 3437 | 172 | P172T | A | 3.6e-08 | 1.2e-08 | 3e-08 | 2.600E-08 | 1.249E-08 | 1.200E-08 | 3.600E-08 |
| 3438 | 172 | P172W | A | 2.4e-08 | 2.7e-08 | 3.1e-08 | 2.733E-08 | 3.512E-09 | 2.400E-08 | 3.100E-08 |
| 3439 | 172 | P172Y | A | 3e-08 | 5.2e-08 | 3.5e-08 | 3.900E-08 | 1.153E-08 | 3.000E-08 | 5.200E-08 |
| 3440 | 172 | P172V | A | 2.2e-08 | 3e-08 | 2e-08 | 2.400E-08 | 5.292E-09 | 2.000E-08 | 3.000E-08 |
| 3441 | 173 | C173A | A | 1.4e-08 | 1.4e-08 | 6.6e-08 | 3.133E-08 | 3.002E-08 | 1.400E-08 | 6.600E-08 |
| 3442 | 173 | C173R | A | 6.2e-08 | 3.2e-08 | 1.2e-08 | 3.533E-08 | 2.517E-08 | 1.200E-08 | 6.200E-08 |
| 3443 | 173 | C173N | A | 5.9e-08 | 3.3e-08 | 1.7e-08 | 3.633E-08 | 2.120E-08 | 1.700E-08 | 5.900E-08 |
| 3444 | 173 | C173D | A | 5.2e-08 | 1.7e-07 | 4.1e-08 | 8.767E-08 | 7.151E-08 | 4.100E-08 | 1.700E-07 |
| 3445 | 173 | C173C | A | 3.7e-08 | 7.8e-08 | 5.9e-08 | 5.800E-08 | 2.052E-08 | 3.700E-08 | 7.800E-08 |
| 3446 | 173 | C173E | A | 4.6e-08 | 1.8e-08 | 3.8e-08 | 3.400E-08 | 1.442E-08 | 1.800E-08 | 4.600E-08 |
| 3447 | 173 | C173Q | A | 1.3e-08 | 7.1e-08 | 3e-08 | 3.800E-08 | 2.982E-08 | 1.300E-08 | 7.100E-08 |
| 3448 | 173 | C173G | A | 3.2e-08 | 5.4e-08 | 4.6e-08 | 4.400E-08 | 1.114E-08 | 3.200E-08 | 5.400E-08 |
| 3449 | 173 | C173H | A | 5.4e-08 | 3.8e-08 | 5.5e-08 | 4.900E-08 | 9.539E-09 | 3.800E-08 | 5.500E-08 |
| 3450 | 173 | C173I | A | 3.5e-08 | 1.1e-07 | 1.6e-07 | 1.017E-07 | 6.292E-08 | 3.500E-08 | 1.600E-07 |
| 3451 | 173 | C173L | A | 2.5e-08 | 1.2e-08 | 5.6e-08 | 3.100E-08 | 2.261E-08 | 1.200E-08 | 5.600E-08 |
| 3452 | 173 | C173K | A | 6e-08 | 5.1e-08 | 1.7e-07 | 9.367E-08 | 6.626E-08 | 5.100E-08 | 1.700E-07 |
| 3453 | 173 | C173M | A | 3.9e-08 | 5.9e-08 | 4.9e-08 | 4.900E-08 | 1.000E-08 | 3.900E-08 | 5.900E-08 |
| 3454 | 173 | C173F | A | 1.8e-08 | 2.7e-08 | 1.7e-08 | 2.067E-08 | 5.508E-09 | 1.700E-08 | 2.700E-08 |
| 3455 | 173 | C173P | A | 7.1e-08 | 7.7e-08 | 5.2e-08 | 6.667E-08 | 1.305E-08 | 5.200E-08 | 7.700E-08 |
| 3456 | 173 | C173S | A | 1.1e-08 | 1.2e-07 | 7.8e-08 | 6.967E-08 | 5.498E-08 | 1.100E-08 | 1.200E-07 |
| 3457 | 173 | C173T | A | 1.4e-07 | 3.9e-08 | 8.3e-09 | 6.243E-08 | 6.891E-08 | 8.300E-09 | 1.400E-07 |
| 3458 | 173 | C173W | A | 1.3e-07 | 3.8e-08 | 1.1e-07 | 9.267E-08 | 4.839E-08 | 3.800E-08 | 1.300E-07 |
| 3459 | 173 | C173Y | A | 4.9e-08 | 1.3e-07 | 1.9e-08 | 6.600E-08 | 5.742E-08 | 1.900E-08 | 1.300E-07 |
| 3460 | 173 | C173V | A | 3.7e-08 | 7.4e-08 | 2.8e-08 | 4.633E-08 | 2.438E-08 | 2.800E-08 | 7.400E-08 |
| 3461 | 174 | S174A | A | 5.2e-08 | 9.1e-08 | 3.6e-08 | 5.967E-08 | 2.829E-08 | 3.600E-08 | 9.100E-08 |
| 3462 | 174 | S174R | A | 1.3e-08 | 2.7e-08 | 1.7e-07 | 7.000E-08 | 8.688E-08 | 1.300E-08 | 1.700E-07 |
| 3463 | 174 | S174N | A | 7.6e-08 | 7.4e-08 | 1.5e-08 | 5.500E-08 | 3.466E-08 | 1.500E-08 | 7.600E-08 |
| 3464 | 174 | S174D | A | 2.3e-08 | 3.9e-08 | 6.2e-08 | 4.133E-08 | 1.960E-08 | 2.300E-08 | 6.200E-08 |
| 3465 | 174 | S174C | A | 7.3e-08 | 6e-08 | 2.5e-08 | 5.267E-08 | 2.483E-08 | 2.500E-08 | 7.300E-08 |
| 3466 | 174 | S174E | A | 4.3e-08 | 8.5e-08 | 5.6e-08 | 6.133E-08 | 2.150E-08 | 4.300E-08 | 8.500E-08 |
| 3467 | 174 | S174Q | A | 9.8e-08 | 3.5e-08 | 1.8e-08 | 5.033E-08 | 4.215E-08 | 1.800E-08 | 9.800E-08 |
| 3468 | 174 | S174G | A | 3.3e-08 | 3.4e-08 | 5.8e-08 | 4.167E-08 | 1.415E-08 | 3.300E-08 | 5.800E-08 |
| 3469 | 174 | S174H | A | 1.3e-08 | 5e-08 | 3e-08 | 3.100E-08 | 1.852E-08 | 1.300E-08 | 5.000E-08 |
| 3470 | 174 | S174I | A | 3.9e-08 | 2.6e-08 | 2.1e-08 | 2.867E-08 | 9.292E-09 | 2.100E-08 | 3.900E-08 |
| 3471 | 174 | S174L | A | 2.5e-08 | 8.4e-08 | 2.4e-08 | 4.433E-08 | 3.436E-08 | 2.400E-08 | 8.400E-08 |
| 3472 | 174 | S174K | A | 1e-07 | 7e-08 | 3.5e-08 | 6.833E-08 | 3.253E-08 | 3.500E-08 | 1.000E-07 |
| 3473 | 174 | S174M | A | 7.1e-08 | 6.1e-08 | 7.3e-08 | 6.833E-08 | 6.429E-09 | 6.100E-08 | 7.300E-08 |
| 3474 | 174 | S174F | A | 5.6e-08 | 6.5e-08 | 4.6e-08 | 5.567E-08 | 9.504E-09 | 4.600E-08 | 6.500E-08 |
| 3475 | 174 | S174P | A | 8.3e-08 | 3e-08 | 6.8e-08 | 6.033E-08 | 2.732E-08 | 3.000E-08 | 8.300E-08 |
| 3476 | 174 | S174S | A | 3.8e-08 | 4.6e-08 | 2.1e-08 | 3.500E-08 | 1.277E-08 | 2.100E-08 | 4.600E-08 |
| 3477 | 174 | S174T | A | 2.5e-08 | 2.9e-08 | 8.3e-08 | 4.567E-08 | 3.239E-08 | 2.500E-08 | 8.300E-08 |
| 3478 | 174 | S174W | A | 8.6e-08 | 4.4e-08 | 1.3e-07 | 8.667E-08 | 4.300E-08 | 4.400E-08 | 1.300E-07 |
| 3479 | 174 | S174Y | A | 2.5e-08 | 3.8e-08 | 6.4e-08 | 4.233E-08 | 1.986E-08 | 2.500E-08 | 6.400E-08 |
| 3480 | 174 | S174V | A | 5.4e-08 | 8.8e-08 | 8.2e-08 | 7.467E-08 | 1.815E-08 | 5.400E-08 | 8.800E-08 |
| 3481 | 175 | P175A | A | 4.6e-08 | 3.3e-08 | 6.4e-08 | 4.767E-08 | 1.557E-08 | 3.300E-08 | 6.400E-08 |
| 3482 | 175 | P175R | A | 7.7e-08 | 3.3e-08 | 8.8e-08 | 6.600E-08 | 2.910E-08 | 3.300E-08 | 8.800E-08 |
| 3483 | 175 | P175N | A | 2.9e-08 | 6e-08 | 6.4e-08 | 5.100E-08 | 1.916E-08 | 2.900E-08 | 6.400E-08 |
| 3484 | 175 | P175D | A | 3.9e-08 | 3.2e-08 | 2.8e-08 | 3.300E-08 | 5.568E-09 | 2.800E-08 | 3.900E-08 |

| | | | | | | | | | | |
|---|---|---|---|---|---|---|---|---|---|---|
| 3485 | 175 | P175C | A | 3.6e-08 | 9.5e-08 | 2.2e-08 | 5.100E-08 | 3.874E-08 | 2.200E-08 | 9.500E-08 |
| 3486 | 175 | P175E | A | 7.3e-09 | 4e-08 | 4.7e-08 | 3.143E-08 | 2.119E-08 | 7.300E-09 | 4.700E-08 |
| 3487 | 175 | P175Q | A | 3e-08 | 3.7e-08 | 7.3e-08 | 4.667E-08 | 2.307E-08 | 3.000E-08 | 7.300E-08 |
| 3488 | 175 | P175G | A | 4.7e-08 | 9.4e-08 | 3.9e-08 | 6.000E-08 | 2.972E-08 | 3.900E-08 | 9.400E-08 |
| 3489 | 175 | P175H | A | 7.5e-08 | 2.7e-08 | 4.2e-08 | 4.800E-08 | 2.456E-08 | 2.700E-08 | 7.500E-08 |
| 3490 | 175 | P175I | A | 3e-08 | 4.4e-08 | 2.3e-08 | 3.233E-08 | 1.069E-08 | 2.300E-08 | 4.400E-08 |
| 3491 | 175 | P175L | A | 2.4e-08 | 5.2e-08 | 9.7e-08 | 5.767E-08 | 3.683E-08 | 2.400E-08 | 9.700E-08 |
| 3492 | 175 | P175K | A | 6.4e-08 | 4e-08 | 4.5e-08 | 4.967E-08 | 1.266E-08 | 4.000E-08 | 6.400E-08 |
| 3493 | 175 | P175M | A | 3.7e-08 | 3.4e-08 | 4e-08 | 3.700E-08 | 3.000E-09 | 3.400E-08 | 4.000E-08 |
| 3494 | 175 | P175F | A | 8.2e-08 | 2.7e-08 | 7.5e-08 | 6.133E-08 | 2.994E-08 | 2.700E-08 | 8.200E-08 |
| 3495 | 175 | P175P | A | 1e-07 | 2.1e-07 | 7.7e-08 | 1.290E-07 | 7.108E-08 | 7.700E-08 | 2.100E-07 |
| 3496 | 175 | P175S | A | 1.1e-07 | 3.3e-08 | 7.6e-09 | 5.020E-08 | 5.332E-08 | 7.600E-09 | 1.100E-07 |
| 3497 | 175 | P175T | A | 4.6e-08 | 3e-08 | 3.4e-08 | 3.667E-08 | 8.327E-09 | 3.000E-08 | 4.600E-08 |
| 3498 | 175 | P175W | A | 2.2e-08 | 2.6e-08 | 1.4e-08 | 2.067E-08 | 6.110E-09 | 1.400E-08 | 2.600E-08 |
| 3499 | 175 | P175Y | A | 3.5e-08 | 7.8e-09 | 4.3e-08 | 2.860E-08 | 1.845E-08 | 7.800E-09 | 4.300E-08 |
| 3500 | 175 | P175V | A | 2.4e-08 | 2.3e-07 | 3.1e-08 | 9.500E-08 | 1.170E-07 | 2.400E-08 | 2.300E-07 |
| 3501 | 176 | M176A | A | 1.8e-08 | 5e-08 | 4.5e-08 | 3.767E-08 | 1.721E-08 | 1.800E-08 | 5.000E-08 |
| 3502 | 176 | M176R | A | 1.9e-08 | 6.8e-08 | 3.6e-08 | 4.100E-08 | 2.488E-08 | 1.900E-08 | 6.800E-08 |
| 3503 | 176 | M176N | A | 4.8e-08 | 2.3e-08 | 4.7e-08 | 3.933E-08 | 1.415E-08 | 2.300E-08 | 4.800E-08 |
| 3504 | 176 | M176D | A | 9.2e-08 | 7e-08 | 2.3e-08 | 6.167E-08 | 3.525E-08 | 2.300E-08 | 9.200E-08 |
| 3505 | 176 | M176C | A | 2.2e-08 | 2.5e-08 | 1.8e-08 | 2.167E-08 | 3.512E-09 | 1.800E-08 | 2.500E-08 |
| 3506 | 176 | M176E | A | 2.4e-08 | 9.1e-08 | 4.4e-08 | 5.300E-08 | 3.439E-08 | 2.400E-08 | 9.100E-08 |
| 3507 | 176 | M176Q | A | 5.5e-08 | 3.8e-08 | 1.8e-08 | 3.700E-08 | 1.852E-08 | 1.800E-08 | 5.500E-08 |
| 3508 | 176 | M176G | A | 3.9e-08 | 4.9e-08 | 1e-07 | 6.267E-08 | 3.272E-08 | 3.900E-08 | 1.000E-07 |
| 3509 | 176 | M176H | A | 2.2e-08 | 6.8e-09 | 4.1e-08 | 2.327E-08 | 1.714E-08 | 6.800E-09 | 4.100E-08 |
| 3510 | 176 | M176I | A | 2e-08 | 2.2e-08 | 6.4e-08 | 3.533E-08 | 2.485E-08 | 2.000E-08 | 6.400E-08 |
| 3511 | 176 | M176L | A | 4.6e-08 | 2.1e-08 | 4.2e-08 | 3.633E-08 | 1.343E-08 | 2.100E-08 | 4.600E-08 |
| 3512 | 176 | M176K | A | 3.1e-08 | 7e-08 | 4.3e-08 | 4.800E-08 | 1.997E-08 | 3.100E-08 | 7.000E-08 |
| 3513 | 176 | M176M | A | 5.8e-08 | 2.4e-08 | 2.4e-08 | 3.533E-08 | 1.963E-08 | 2.400E-08 | 5.800E-08 |
| 3514 | 176 | M176F | A | 6.1e-08 | 4.9e-08 | 3.1e-08 | 4.700E-08 | 1.510E-08 | 3.100E-08 | 6.100E-08 |
| 3515 | 176 | M176P | A | 3.6e-08 | 2.3e-08 | 9.1e-08 | 5.000E-08 | 3.610E-08 | 2.300E-08 | 9.100E-08 |
| 3516 | 176 | M176S | A | 3.8e-08 | 5.2e-08 | 1e-08 | 3.333E-08 | 2.139E-08 | 1.000E-08 | 5.200E-08 |
| 3517 | 176 | M176T | A | 3.7e-08 | 6.8e-08 | 5.6e-08 | 5.367E-08 | 1.563E-08 | 3.700E-08 | 6.800E-08 |
| 3518 | 176 | M176W | A | 1.3e-08 | 6.7e-08 | 1.1e-08 | 3.033E-08 | 3.177E-08 | 1.100E-08 | 6.700E-08 |
| 3519 | 176 | M176Y | A | 2.4e-08 | 3.1e-08 | 2.2e-08 | 2.567E-08 | 4.726E-09 | 2.200E-08 | 3.100E-08 |
| 3520 | 176 | M176V | A | 3.2e-08 | 2.9e-08 | 5.2e-08 | 3.767E-08 | 1.250E-08 | 2.900E-08 | 5.200E-08 |
| 3521 | 177 | C177A | A | 1.4e-08 | 2.1e-08 | 2.5e-08 | 2.000E-08 | 5.568E-09 | 1.400E-08 | 2.500E-08 |
| 3522 | 177 | C177R | A | 1.5e-07 | 1.6e-08 | 3e-08 | 6.533E-08 | 7.366E-08 | 1.600E-08 | 1.500E-07 |
| 3523 | 177 | C177N | A | 3.3e-08 | 3.9e-08 | 3.7e-08 | 3.633E-08 | 3.055E-09 | 3.300E-08 | 3.900E-08 |
| 3524 | 177 | C177D | A | 4.8e-08 | 6.1e-08 | 8.5e-08 | 6.467E-08 | 1.877E-08 | 4.800E-08 | 8.500E-08 |
| 3525 | 177 | C177C | A | 5.8e-08 | 5.1e-08 | 3.1e-08 | 4.667E-08 | 1.401E-08 | 3.100E-08 | 5.800E-08 |
| 3526 | 177 | C177E | A | 4.5e-08 | 2.8e-08 | 1.4e-07 | 7.100E-08 | 6.036E-08 | 2.800E-08 | 1.400E-07 |
| 3527 | 177 | C177Q | A | 7.4e-08 | 3.1e-08 | 1.3e-08 | 3.933E-08 | 3.134E-08 | 1.300E-08 | 7.400E-08 |
| 3528 | 177 | C177G | A | 1.8e-08 | 5e-08 | 1.7e-08 | 2.833E-08 | 1.877E-08 | 1.700E-08 | 5.000E-08 |
| 3529 | 177 | C177H | A | 5.2e-08 | 4.6e-08 | 8.7e-08 | 6.167E-08 | 2.214E-08 | 4.600E-08 | 8.700E-08 |
| 3530 | 177 | C177I | A | 3.2e-08 | 1e-07 | 2.4e-08 | 5.200E-08 | 4.176E-08 | 2.400E-08 | 1.000E-07 |
| 3531 | 177 | C177L | A | 2.1e-08 | 4.7e-08 | 4.6e-08 | 3.800E-08 | 1.473E-08 | 2.100E-08 | 4.700E-08 |
| 3532 | 177 | C177K | A | 5.1e-08 | 3.6e-08 | 5.2e-08 | 4.633E-08 | 8.963E-09 | 3.600E-08 | 5.200E-08 |
| 3533 | 177 | C177M | A | 4.2e-08 | 2.7e-08 | 2e-08 | 2.967E-08 | 1.124E-08 | 2.000E-08 | 4.200E-08 |
| 3534 | 177 | C177F | A | 3.5e-08 | 1.3e-08 | 6.5e-08 | 3.767E-08 | 2.610E-08 | 1.300E-08 | 6.500E-08 |
| 3535 | 177 | C177P | A | 7.1e-08 | 1.1e-07 | 5e-08 | 7.700E-08 | 3.045E-08 | 5.000E-08 | 1.100E-07 |
| 3536 | 177 | C177S | A | 6.1e-08 | 3.2e-08 | 4.7e-08 | 4.667E-08 | 1.450E-08 | 3.200E-08 | 6.100E-08 |
| 3537 | 177 | C177T | A | 5.5e-08 | 4.9e-08 | 1.9e-08 | 4.100E-08 | 1.929E-08 | 1.900E-08 | 5.500E-08 |
| 3538 | 177 | C177W | A | 5.5e-08 | 5.1e-08 | 1.5e-08 | 4.033E-08 | 2.203E-08 | 1.500E-08 | 5.500E-08 |
| 3539 | 177 | C177Y | A | 2.9e-08 | 5.6e-08 | 3.5e-08 | 4.000E-08 | 1.418E-08 | 2.900E-08 | 5.600E-08 |
| 3540 | 177 | C177V | A | 2.1e-08 | 4.2e-08 | 4.2e-08 | 3.500E-08 | 1.212E-08 | 2.100E-08 | 4.200E-08 |
| 3541 | 178 | K178A | A | 7.9e-08 | 1.9e-08 | 2.7e-08 | 4.167E-08 | 3.258E-08 | 1.900E-08 | 7.900E-08 |
| 3542 | 178 | K178R | A | 5.1e-08 | 1.1e-08 | 6.2e-08 | 4.133E-08 | 2.684E-08 | 1.100E-08 | 6.200E-08 |
| 3543 | 178 | K178N | A | 6.2e-08 | 4.2e-08 | 4.5e-08 | 4.967E-08 | 1.079E-08 | 4.200E-08 | 6.200E-08 |
| 3544 | 178 | K178D | A | 4.4e-08 | 8.2e-08 | 5.2e-08 | 5.933E-08 | 2.003E-08 | 4.400E-08 | 8.200E-08 |
| 3545 | 178 | K178C | A | 4.2e-08 | 3.5e-08 | 5.5e-08 | 4.400E-08 | 1.015E-08 | 3.500E-08 | 5.500E-08 |
| 3546 | 178 | K178E | A | 2.3e-08 | 4.7e-08 | 6.3e-08 | 4.433E-08 | 2.013E-08 | 2.300E-08 | 6.300E-08 |
| 3547 | 178 | K178Q | A | 2.2e-08 | 2.5e-08 | 1.7e-08 | 2.133E-08 | 4.041E-09 | 1.700E-08 | 2.500E-08 |
| 3548 | 178 | K178G | A | 2.8e-08 | 3.2e-08 | 4.1e-08 | 3.367E-08 | 6.658E-09 | 2.800E-08 | 4.100E-08 |
| 3549 | 178 | K178H | A | 6.1e-09 | 2.5e-08 | 1.2e-07 | 5.037E-08 | 6.104E-08 | 6.100E-09 | 1.200E-07 |
| 3550 | 178 | K178I | A | 6e-08 | 1.8e-08 | 8e-08 | 5.267E-08 | 3.164E-08 | 1.800E-08 | 8.000E-08 |
| 3551 | 178 | K178L | A | 6.6e-08 | 3.3e-08 | 1.5e-08 | 3.800E-08 | 2.587E-08 | 1.500E-08 | 6.600E-08 |
| 3552 | 178 | K178K | A | 4.4e-08 | 5.1e-08 | 1.2e-08 | 3.567E-08 | 2.079E-08 | 1.200E-08 | 5.100E-08 |
| 3553 | 178 | K178M | A | 1.7e-08 | 4.1e-08 | 2.4e-08 | 2.733E-08 | 1.234E-08 | 1.700E-08 | 4.100E-08 |
| 3554 | 178 | K178F | A | 3.1e-08 | 3.8e-08 | 3.5e-08 | 3.467E-08 | 3.512E-09 | 3.100E-08 | 3.800E-08 |
| 3555 | 178 | K178P | A | 1.4e-07 | 5.1e-08 | 9.7e-09 | 6.690E-08 | 6.659E-08 | 9.700E-09 | 1.400E-07 |
| 3556 | 178 | K178S | A | 2.1e-08 | 2.5e-08 | 2.3e-08 | 2.300E-08 | 2.000E-09 | 2.100E-08 | 2.500E-08 |
| 3557 | 178 | K178T | A | 1.9e-08 | 2.2e-08 | 1.6e-08 | 1.900E-08 | 3.000E-09 | 1.600E-08 | 2.200E-08 |
| 3558 | 178 | K178W | A | 1.1e-07 | 3.5e-08 | 7.1e-08 | 7.200E-08 | 3.751E-08 | 3.500E-08 | 1.100E-07 |
| 3559 | 178 | K178Y | A | 3.6e-08 | 3.9e-08 | 7.5e-08 | 5.000E-08 | 2.170E-08 | 3.600E-08 | 7.500E-08 |
| 3560 | 178 | K178V | A | 6.6e-08 | 4.7e-08 | 4.1e-08 | 5.133E-08 | 1.305E-08 | 4.100E-08 | 6.600E-08 |
| 3561 | 179 | G179A | A | 3.7e-08 | 1.1e-07 | 4.7e-08 | 6.467E-08 | 3.958E-08 | 3.700E-08 | 1.100E-07 |
| 3562 | 179 | G179R | A | 2.7e-08 | 6e-08 | 5.4e-08 | 4.700E-08 | 1.758E-08 | 2.700E-08 | 6.000E-08 |
| 3563 | 179 | G179N | A | 4.8e-08 | 7.5e-08 | 8.2e-08 | 6.833E-08 | 1.795E-08 | 4.800E-08 | 8.200E-08 |
| 3564 | 179 | G179D | A | 3.3e-08 | 9.8e-09 | 1.9e-08 | 2.060E-08 | 1.168E-08 | 9.800E-09 | 3.300E-08 |
| 3565 | 179 | G179C | A | 3e-08 | 2.6e-08 | 6.3e-08 | 3.967E-08 | 2.031E-08 | 2.600E-08 | 6.300E-08 |
| 3566 | 179 | G179E | A | 9.4e-09 | 1.8e-08 | 6.3e-08 | 3.013E-08 | 2.879E-08 | 9.400E-09 | 6.300E-08 |
| 3567 | 179 | G179Q | A | 1.2e-07 | 4.5e-08 | 5.2e-08 | 7.233E-08 | 4.143E-08 | 4.500E-08 | 1.200E-07 |
| 3568 | 179 | G179G | A | 5.9e-08 | 5.7e-08 | 6.2e-08 | 5.933E-08 | 2.517E-09 | 5.700E-08 | 6.200E-08 |
| 3569 | 179 | G179H | A | 1e-07 | 3.7e-08 | 9.1e-08 | 7.600E-08 | 3.407E-08 | 3.700E-08 | 1.000E-07 |

| | | | | | | | | | | |
|---|---|---|---|---|---|---|---|---|---|---|
| 3570 | 179 | G179I | A | 4.7e-08 | 5.4e-08 | 1.8e-08 | 3.967E-08 | 1.909E-08 | 1.800E-08 | 5.400E-08 |
| 3571 | 179 | G179L | A | 1e-07 | 3.9e-08 | 3.6e-08 | 5.833E-08 | 3.612E-08 | 3.600E-08 | 1.000E-07 |
| 3572 | 179 | G179K | A | 7.8e-08 | 4.3e-08 | 1.2e-07 | 8.033E-08 | 3.855E-08 | 4.300E-08 | 1.200E-07 |
| 3573 | 179 | G179M | A | 2e-08 | 3.9e-08 | 4.9e-08 | 3.600E-08 | 1.473E-08 | 2.000E-08 | 4.900E-08 |
| 3574 | 179 | G179F | A | 3e-08 | 8.2e-08 | 4.1e-08 | 5.100E-08 | 2.740E-08 | 3.000E-08 | 8.200E-08 |
| 3575 | 179 | G179P | A | 1.5e-08 | 3.5e-08 | 2.8e-08 | 2.600E-08 | 1.015E-08 | 1.500E-08 | 3.500E-08 |
| 3576 | 179 | G179S | A | 6e-08 | 7.4e-08 | 1.3e-08 | 4.900E-08 | 3.195E-08 | 1.300E-08 | 7.400E-08 |
| 3577 | 179 | G179T | A | 2.4e-08 | 5e-08 | 1.7e-08 | 3.033E-08 | 1.739E-08 | 1.700E-08 | 5.000E-08 |
| 3578 | 179 | G179W | A | 2.5e-08 | 6.8e-08 | 1.2e-07 | 7.100E-08 | 4.757E-08 | 2.500E-08 | 1.200E-07 |
| 3579 | 179 | G179Y | A | 1.8e-08 | 3.8e-08 | 6.3e-08 | 3.967E-08 | 2.255E-08 | 1.800E-08 | 6.300E-08 |
| 3580 | 179 | G179V | A | 3.5e-08 | 4.5e-08 | 2.4e-08 | 3.467E-08 | 1.050E-08 | 2.400E-08 | 4.500E-08 |
| 3581 | 180 | S180A | A | 4.7e-08 | 5.4e-08 | 1.8e-08 | 3.967E-08 | 1.909E-08 | 1.800E-08 | 5.400E-08 |
| 3582 | 180 | S180R | A | 7.9e-08 | 6.6e-08 | 1.4e-07 | 9.500E-08 | 3.951E-08 | 6.600E-08 | 1.400E-07 |
| 3583 | 180 | S180N | A | 3.4e-08 | 4.5e-08 | 4.7e-08 | 4.200E-08 | 7.000E-09 | 3.400E-08 | 4.700E-08 |
| 3584 | 180 | S180D | A | 4.9e-08 | 4.7e-08 | 1.6e-07 | 8.533E-08 | 6.467E-08 | 4.700E-08 | 1.600E-07 |
| 3585 | 180 | S180C | A | 3.8e-08 | 2.7e-08 | 2.3e-08 | 2.933E-08 | 7.767E-09 | 2.300E-08 | 3.800E-08 |
| 3586 | 180 | S180E | A | 3e-08 | 6.8e-08 | 4.7e-08 | 4.833E-08 | 1.904E-08 | 3.000E-08 | 6.800E-08 |
| 3587 | 180 | S180Q | A | 3.1e-08 | 3.4e-08 | 4.9e-08 | 3.800E-08 | 9.644E-09 | 3.100E-08 | 4.900E-08 |
| 3588 | 180 | S180G | A | 8.4e-08 | 8.3e-08 | 1.3e-07 | 9.900E-08 | 2.685E-08 | 8.300E-08 | 1.300E-07 |
| 3589 | 180 | S180H | A | 1.1e-07 | 3.4e-08 | 5e-08 | 6.467E-08 | 4.007E-08 | 3.400E-08 | 1.100E-07 |
| 3590 | 180 | S180I | A | 5.1e-08 | 1.9e-08 | 6.9e-08 | 4.633E-08 | 2.532E-08 | 1.900E-08 | 6.900E-08 |
| 3591 | 180 | S180L | A | 8.3e-08 | 3.4e-08 | 1.2e-08 | 4.300E-08 | 3.635E-08 | 1.200E-08 | 8.300E-08 |
| 3592 | 180 | S180K | A | 4.8e-08 | 8e-08 | 8.1e-08 | 6.967E-08 | 1.877E-08 | 4.800E-08 | 8.100E-08 |
| 3593 | 180 | S180M | A | 4.1e-08 | 3.5e-08 | 3.9e-08 | 3.833E-08 | 3.055E-09 | 3.500E-08 | 4.100E-08 |
| 3594 | 180 | S180F | A | 3.3e-08 | 4.2e-08 | 1e-08 | 2.833E-08 | 1.650E-08 | 1.000E-08 | 4.200E-08 |
| 3595 | 180 | S180P | A | 2.9e-08 | 3.3e-08 | 3.4e-08 | 3.200E-08 | 2.646E-09 | 2.900E-08 | 3.400E-08 |
| 3596 | 180 | S180S | A | 4.6e-08 | 4.3e-08 | 4.5e-08 | 4.467E-08 | 1.528E-09 | 4.300E-08 | 4.600E-08 |
| 3597 | 180 | S180T | A | 1.6e-08 | 2e-08 | 1.7e-08 | 1.767E-08 | 2.082E-09 | 1.600E-08 | 2.000E-08 |
| 3598 | 180 | S180W | A | 7.5e-08 | 3e-08 | 5.2e-08 | 5.233E-08 | 2.250E-08 | 3.000E-08 | 7.500E-08 |
| 3599 | 180 | S180Y | A | 2.1e-08 | 4.1e-08 | 3.1e-08 | 3.100E-08 | 1.000E-08 | 2.100E-08 | 4.100E-08 |
| 3600 | 180 | S180V | A | 1.8e-07 | 6.5e-08 | 4.5e-08 | 9.667E-08 | 7.286E-08 | 4.500E-08 | 1.800E-07 |
| 3601 | 181 | R181A | A | 1.2e-07 | 1.1e-07 | 1.7e-08 | 8.233E-08 | 5.680E-08 | 1.700E-08 | 1.200E-07 |
| 3602 | 181 | R181R | A | 1.5e-08 | 8e-08 | 4.7e-08 | 4.733E-08 | 3.250E-08 | 1.500E-08 | 8.000E-08 |
| 3603 | 181 | R181N | A | 6.5e-08 | 7.1e-08 | 7.5e-08 | 7.033E-08 | 5.033E-09 | 6.500E-08 | 7.500E-08 |
| 3604 | 181 | R181D | A | 7.5e-08 | 2.1e-08 | 2.7e-08 | 4.100E-08 | 2.960E-08 | 2.100E-08 | 7.500E-08 |
| 3605 | 181 | R181C | A | 2.3e-08 | 6e-08 | 4.7e-08 | 4.333E-08 | 1.877E-08 | 2.300E-08 | 6.000E-08 |
| 3606 | 181 | R181E | A | 3.2e-08 | 2.5e-08 | 7.6e-08 | 4.433E-08 | 2.765E-08 | 2.500E-08 | 7.600E-08 |
| 3607 | 181 | R181Q | A | 2.9e-08 | 4e-08 | 8.8e-08 | 5.233E-08 | 3.137E-08 | 2.900E-08 | 8.800E-08 |
| 3608 | 181 | R181G | A | 3.3e-08 | 9.5e-08 | 5.6e-08 | 6.133E-08 | 3.134E-08 | 3.300E-08 | 9.500E-08 |
| 3609 | 181 | R181H | A | 6.3e-08 | 4.2e-08 | 3.4e-08 | 4.633E-08 | 1.498E-08 | 3.400E-08 | 6.300E-08 |
| 3610 | 181 | R181I | A | 3.6e-08 | 7.2e-08 | 1.7e-08 | 4.167E-08 | 2.793E-08 | 1.700E-08 | 7.200E-08 |
| 3611 | 181 | R181L | A | 5.7e-08 | 5.2e-08 | 3.4e-08 | 4.767E-08 | 1.210E-08 | 3.400E-08 | 5.700E-08 |
| 3612 | 181 | R181K | A | 8.9e-08 | 2.8e-08 | 1.3e-07 | 8.233E-08 | 5.133E-08 | 2.800E-08 | 1.300E-07 |
| 3613 | 181 | R181M | A | 2e-08 | 8.6e-08 | 9.9e-08 | 6.833E-08 | 4.236E-08 | 2.000E-08 | 9.900E-08 |
| 3614 | 181 | R181F | A | 5.4e-08 | 3.9e-08 | 4.9e-08 | 4.733E-08 | 7.638E-09 | 3.900E-08 | 5.400E-08 |
| 3615 | 181 | R181P | A | 8.4e-08 | 4.2e-08 | 2e-08 | 4.867E-08 | 3.252E-08 | 2.000E-08 | 8.400E-08 |
| 3616 | 181 | R181S | A | 1.9e-08 | 2.5e-08 | 5.5e-08 | 3.300E-08 | 1.929E-08 | 1.900E-08 | 5.500E-08 |
| 3617 | 181 | R181T | A | 3.2e-08 | 3.5e-08 | 3.7e-08 | 3.467E-08 | 2.517E-09 | 3.200E-08 | 3.700E-08 |
| 3618 | 181 | R181W | A | 9.2e-09 | 7.3e-08 | 2.4e-08 | 3.540E-08 | 3.339E-08 | 9.200E-09 | 7.300E-08 |
| 3619 | 181 | R181Y | A | 6e-08 | 2.1e-08 | 7.7e-08 | 5.267E-08 | 2.871E-08 | 2.100E-08 | 7.700E-08 |
| 3620 | 181 | R181V | A | 1.3e-07 | 2e-08 | 4.5e-08 | 6.500E-08 | 5.766E-08 | 2.000E-08 | 1.300E-07 |
| 3621 | 182 | C182A | A | 9e-08 | 4.1e-08 | 3.5e-08 | 5.533E-08 | 3.017E-08 | 3.500E-08 | 9.000E-08 |
| 3622 | 182 | C182R | A | 2.3e-08 | 7.7e-08 | 3.8e-08 | 4.600E-08 | 2.787E-08 | 2.300E-08 | 7.700E-08 |
| 3623 | 182 | C182N | A | 5.3e-08 | 4.1e-08 | 3.7e-08 | 4.367E-08 | 8.327E-09 | 3.700E-08 | 5.300E-08 |
| 3624 | 182 | C182D | A | 3.7e-08 | 3.8e-08 | 5.5e-08 | 4.333E-08 | 1.012E-08 | 3.700E-08 | 5.500E-08 |
| 3625 | 182 | C182C | A | 2.2e-08 | 3.9e-08 | 5.5e-08 | 3.867E-08 | 1.650E-08 | 2.200E-08 | 5.500E-08 |
| 3626 | 182 | C182E | A | 7.7e-08 | 5.6e-08 | 1.3e-08 | 4.867E-08 | 3.262E-08 | 1.300E-08 | 7.700E-08 |
| 3627 | 182 | C182Q | A | 5.8e-08 | 4.2e-08 | 3.7e-08 | 4.567E-08 | 1.097E-08 | 3.700E-08 | 5.800E-08 |
| 3628 | 182 | C182G | A | 7.1e-09 | 2.6e-08 | 5e-08 | 2.770E-08 | 2.150E-08 | 7.100E-09 | 5.000E-08 |
| 3629 | 182 | C182H | A | 4.9e-08 | 3.6e-08 | 2.2e-08 | 3.567E-08 | 1.350E-08 | 2.200E-08 | 4.900E-08 |
| 3630 | 182 | C182I | A | 3.3e-08 | 4.8e-08 | 5.4e-08 | 4.500E-08 | 1.082E-08 | 3.300E-08 | 5.400E-08 |
| 3631 | 182 | C182L | A | 2.5e-08 | 1.5e-08 | 3e-08 | 2.333E-08 | 7.638E-09 | 1.500E-08 | 3.000E-08 |
| 3632 | 182 | C182K | A | 1.3e-08 | 3.2e-08 | 4.2e-08 | 2.900E-08 | 1.473E-08 | 1.300E-08 | 4.200E-08 |
| 3633 | 182 | C182M | A | 5.7e-08 | 4.3e-08 | 4.4e-08 | 4.800E-08 | 7.810E-09 | 4.300E-08 | 5.700E-08 |
| 3634 | 182 | C182F | A | 4e-08 | 1.5e-08 | 7.7e-08 | 4.400E-08 | 3.119E-08 | 1.500E-08 | 7.700E-08 |
| 3635 | 182 | C182P | A | 3.9e-08 | 5.7e-08 | 3.3e-08 | 4.300E-08 | 1.249E-08 | 3.300E-08 | 5.700E-08 |
| 3636 | 182 | C182S | A | 5.8e-08 | 3.9e-08 | 3.6e-08 | 4.433E-08 | 1.193E-08 | 3.600E-08 | 5.800E-08 |
| 3637 | 182 | C182T | A | 4e-08 | 1.8e-08 | 8.5e-08 | 4.767E-08 | 3.415E-08 | 1.800E-08 | 8.500E-08 |
| 3638 | 182 | C182W | A | 4e-08 | 3.7e-08 | 4.1e-08 | 3.933E-08 | 2.082E-09 | 3.700E-08 | 4.100E-08 |
| 3639 | 182 | C182Y | A | 2.8e-08 | 1.2e-08 | 4.4e-08 | 2.800E-08 | 1.600E-08 | 1.200E-08 | 4.400E-08 |
| 3640 | 182 | C182V | A | 5.7e-08 | 9.1e-08 | 7.8e-08 | 7.533E-08 | 1.716E-08 | 5.700E-08 | 9.100E-08 |
| 3641 | 183 | W183A | A | 1.3e-07 | 2.7e-08 | 4.9e-08 | 6.867E-08 | 5.424E-08 | 2.700E-08 | 1.300E-07 |
| 3642 | 183 | W183R | A | 2.5e-08 | 1.9e-08 | 5.4e-08 | 3.267E-08 | 1.872E-08 | 1.900E-08 | 5.400E-08 |
| 3643 | 183 | W183N | A | 5.2e-08 | 2.1e-08 | 1.1e-08 | 2.800E-08 | 2.138E-08 | 1.100E-08 | 5.200E-08 |
| 3644 | 183 | W183D | A | 1.1e-07 | 5.3e-08 | 3.1e-08 | 6.467E-08 | 4.077E-08 | 3.100E-08 | 1.100E-07 |
| 3645 | 183 | W183C | A | 7.2e-08 | 4.3e-08 | 5.2e-08 | 5.567E-08 | 1.484E-08 | 4.300E-08 | 7.200E-08 |
| 3646 | 183 | W183E | A | 3.5e-08 | 1.1e-08 | 4.4e-08 | 3.000E-08 | 1.706E-08 | 1.100E-08 | 4.400E-08 |
| 3647 | 183 | W183Q | A | 5.8e-08 | 7.7e-08 | 5e-08 | 6.167E-08 | 1.387E-08 | 5.000E-08 | 7.700E-08 |
| 3648 | 183 | W183G | A | 3.3e-08 | 8e-08 | 8.2e-08 | 6.500E-08 | 2.773E-08 | 3.300E-08 | 8.200E-08 |
| 3649 | 183 | W183H | A | 3.8e-08 | 5.9e-08 | 3.4e-08 | 4.367E-08 | 1.343E-08 | 3.400E-08 | 5.900E-08 |
| 3650 | 183 | W183I | A | 1.1e-07 | 1e-07 | 2.8e-08 | 7.933E-08 | 4.474E-08 | 2.800E-08 | 1.100E-07 |
| 3651 | 183 | W183L | A | 2e-07 | 1e-08 | 8.4e-08 | 9.800E-08 | 9.577E-08 | 1.000E-08 | 2.000E-07 |
| 3652 | 183 | W183K | A | 6.4e-08 | 4e-08 | 2.9e-08 | 4.433E-08 | 1.790E-08 | 2.900E-08 | 6.400E-08 |
| 3653 | 183 | W183M | A | 4.8e-08 | 1.2e-07 | 8.6e-08 | 8.467E-08 | 3.602E-08 | 4.800E-08 | 1.200E-07 |
| 3654 | 183 | W183F | A | 8.9e-08 | 3.4e-08 | 3.7e-08 | 5.333E-08 | 3.092E-08 | 3.400E-08 | 8.900E-08 |

| | | | | | | | | | | |
|---|---|---|---|---|---|---|---|---|---|---|
| 3655 | 183 | W183P | A | 8.7e-08 | 2.7e-08 | 9e-08 | 6.800E-08 | 3.554E-08 | 2.700E-08 | 9.000E-08 |
| 3656 | 183 | W183S | A | 4.6e-08 | 2.1e-08 | 5.5e-08 | 4.067E-08 | 1.762E-08 | 2.100E-08 | 5.500E-08 |
| 3657 | 183 | W183T | A | 1.3e-07 | 1.6e-07 | 5.3e-08 | 1.143E-07 | 5.519E-08 | 5.300E-08 | 1.600E-07 |
| 3658 | 183 | W183W | A | 5.4e-08 | 3.7e-08 | 6.1e-08 | 5.067E-08 | 1.234E-08 | 3.700E-08 | 6.100E-08 |
| 3659 | 183 | W183Y | A | 6.7e-08 | 6.2e-08 | 5.9e-08 | 6.267E-08 | 4.041E-09 | 5.900E-08 | 6.700E-08 |
| 3660 | 183 | W183V | A | 2.3e-08 | 5.5e-08 | 6.1e-08 | 4.633E-08 | 2.043E-08 | 2.300E-08 | 6.100E-08 |
| 3661 | 184 | G184A | A | 4.6e-08 | 7e-09 | 1.1e-07 | 5.433E-08 | 5.200E-08 | 7.000E-09 | 1.100E-07 |
| 3662 | 184 | G184R | A | 8.5e-08 | 3e-08 | 5.9e-08 | 5.800E-08 | 2.751E-08 | 3.000E-08 | 8.500E-08 |
| 3663 | 184 | G184N | A | 2.2e-08 | 2e-08 | 4.6e-08 | 2.933E-08 | 1.447E-08 | 2.000E-08 | 4.600E-08 |
| 3664 | 184 | G184D | A | 1e-07 | 9.4e-08 | 1.7e-07 | 1.213E-07 | 4.225E-08 | 9.400E-08 | 1.700E-07 |
| 3665 | 184 | G184C | A | 2.4e-08 | 5.6e-08 | 2.5e-08 | 3.500E-08 | 1.819E-08 | 2.400E-08 | 5.600E-08 |
| 3666 | 184 | G184E | A | 3e-08 | 5.7e-08 | 8.5e-08 | 5.733E-08 | 2.750E-08 | 3.000E-08 | 8.500E-08 |
| 3667 | 184 | G184Q | A | 3.2e-08 | 2.1e-08 | 3.2e-08 | 2.833E-08 | 6.351E-09 | 2.100E-08 | 3.200E-08 |
| 3668 | 184 | G184G | A | 5.1e-08 | 4.6e-08 | 1.5e-08 | 3.733E-08 | 1.950E-08 | 1.500E-08 | 5.100E-08 |
| 3669 | 184 | G184H | A | 5e-08 | 4.6e-08 | 5.1e-08 | 4.900E-08 | 2.646E-09 | 4.600E-08 | 5.100E-08 |
| 3670 | 184 | G184I | A | 5.6e-08 | 1.3e-07 | 3.5e-08 | 7.367E-08 | 4.990E-08 | 3.500E-08 | 1.300E-07 |
| 3671 | 184 | G184L | A | 4.9e-08 | 3.3e-08 | 4.4e-08 | 4.200E-08 | 8.185E-09 | 3.300E-08 | 4.900E-08 |
| 3672 | 184 | G184K | A | 8.9e-08 | 1.6e-08 | 2.8e-08 | 4.433E-08 | 3.915E-08 | 1.600E-08 | 8.900E-08 |
| 3673 | 184 | G184M | A | 1.1e-07 | 3.1e-08 | 6.1e-08 | 6.733E-08 | 3.988E-08 | 3.100E-08 | 1.100E-07 |
| 3674 | 184 | G184F | A | 2.7e-08 | 4.5e-08 | 4.7e-08 | 3.967E-08 | 1.102E-08 | 2.700E-08 | 4.700E-08 |
| 3675 | 184 | G184P | A | 2.8e-08 | 5.5e-08 | 2.9e-08 | 3.733E-08 | 1.531E-08 | 2.800E-08 | 5.500E-08 |
| 3676 | 184 | G184S | A | 6.5e-08 | 7.4e-08 | 1.6e-07 | 9.967E-08 | 5.244E-08 | 6.500E-08 | 1.600E-07 |
| 3677 | 184 | G184T | A | 5.1e-08 | 4e-08 | 4.3e-08 | 4.467E-08 | 5.686E-09 | 4.000E-08 | 5.100E-08 |
| 3678 | 184 | G184W | A | 2.1e-08 | 3.1e-08 | 2.3e-08 | 2.500E-08 | 5.292E-09 | 2.100E-08 | 3.100E-08 |
| 3679 | 184 | G184Y | A | 4.2e-08 | 1.5e-08 | 8.2e-08 | 4.633E-08 | 3.371E-08 | 1.500E-08 | 8.200E-08 |
| 3680 | 184 | G184V | A | 7.6e-08 | 8.9e-08 | 4.3e-08 | 6.933E-08 | 2.371E-08 | 4.300E-08 | 8.900E-08 |
| 3681 | 185 | E185A | A | 3.9e-08 | 3.3e-08 | 2.1e-08 | 3.100E-08 | 9.165E-09 | 2.100E-08 | 3.900E-08 |
| 3682 | 185 | E185R | A | 6.8e-08 | 3.4e-08 | 8.2e-08 | 6.133E-08 | 2.468E-08 | 3.400E-08 | 8.200E-08 |
| 3683 | 185 | E185N | A | 1.9e-08 | 7.5e-08 | 5.9e-08 | 5.100E-08 | 2.884E-08 | 1.900E-08 | 7.500E-08 |
| 3684 | 185 | E185D | A | 3.8e-08 | 3.7e-08 | 2.1e-08 | 3.200E-08 | 9.539E-09 | 2.100E-08 | 3.800E-08 |
| 3685 | 185 | E185C | A | 1.3e-08 | 7e-08 | 3.1e-08 | 3.800E-08 | 2.914E-08 | 1.300E-08 | 7.000E-08 |
| 3686 | 185 | E185E | A | 6.1e-08 | 2.1e-08 | 3.9e-08 | 4.033E-08 | 2.003E-08 | 2.100E-08 | 6.100E-08 |
| 3687 | 185 | E185Q | A | 4.9e-08 | 1.8e-07 | 2e-08 | 8.300E-08 | 8.525E-08 | 2.000E-08 | 1.800E-07 |
| 3688 | 185 | E185G | A | 2.6e-08 | 6.5e-08 | 2.2e-08 | 3.767E-08 | 2.376E-08 | 2.200E-08 | 6.500E-08 |
| 3689 | 185 | E185H | A | 3.7e-08 | 1.9e-08 | 3e-08 | 2.867E-08 | 9.074E-09 | 1.900E-08 | 3.700E-08 |
| 3690 | 185 | E185I | A | 5.6e-08 | 3.2e-08 | 4.8e-08 | 4.533E-08 | 1.222E-08 | 3.200E-08 | 5.600E-08 |
| 3691 | 185 | E185L | A | 1.2e-07 | 2e-08 | 6.4e-08 | 6.800E-08 | 5.012E-08 | 2.000E-08 | 1.200E-07 |
| 3692 | 185 | E185K | A | 3.7e-08 | 1.4e-08 | 1.1e-07 | 5.367E-08 | 5.012E-08 | 1.400E-08 | 1.100E-07 |
| 3693 | 185 | E185M | A | 3.1e-08 | 1e-07 | 3.1e-08 | 5.400E-08 | 3.984E-08 | 3.100E-08 | 1.000E-07 |
| 3694 | 185 | E185F | A | 4.2e-08 | 1.1e-07 | 3.8e-08 | 6.333E-08 | 4.046E-08 | 3.800E-08 | 1.100E-07 |
| 3695 | 185 | E185P | A | 5.2e-08 | 8.5e-08 | 7.5e-08 | 7.067E-08 | 1.692E-08 | 5.200E-08 | 8.500E-08 |
| 3696 | 185 | E185S | A | 2.5e-08 | 4.4e-08 | 6.6e-09 | 2.520E-08 | 1.870E-08 | 6.600E-09 | 4.400E-08 |
| 3697 | 185 | E185T | A | 3.3e-08 | 3e-08 | 1.4e-08 | 2.567E-08 | 1.021E-08 | 1.400E-08 | 3.300E-08 |
| 3698 | 185 | E185W | A | 3.5e-08 | 5.4e-08 | 5.7e-08 | 4.867E-08 | 1.193E-08 | 3.500E-08 | 5.700E-08 |
| 3699 | 185 | E185Y | A | 5.2e-08 | 5.2e-08 | 7.6e-08 | 6.000E-08 | 1.386E-08 | 5.200E-08 | 7.600E-08 |
| 3700 | 185 | E185V | A | 8.9e-08 | 9.5e-08 | 3.8e-08 | 7.400E-08 | 3.132E-08 | 3.800E-08 | 9.500E-08 |
| 3701 | 186 | S186A | A | 2.6e-08 | 3.5e-08 | 3.6e-08 | 3.233E-08 | 5.508E-09 | 2.600E-08 | 3.600E-08 |
| 3702 | 186 | S186R | A | 9.3e-09 | 7.4e-08 | 4.3e-08 | 4.210E-08 | 3.236E-08 | 9.300E-09 | 7.400E-08 |
| 3703 | 186 | S186N | A | 2.9e-08 | 1.1e-07 | 4.5e-08 | 6.133E-08 | 4.290E-08 | 2.900E-08 | 1.100E-07 |
| 3704 | 186 | S186D | A | 6.1e-08 | 8.8e-08 | 6.5e-08 | 7.133E-08 | 1.457E-08 | 6.100E-08 | 8.800E-08 |
| 3705 | 186 | S186C | A | 1.5e-08 | 7.9e-08 | 4.2e-08 | 4.533E-08 | 3.213E-08 | 1.500E-08 | 7.900E-08 |
| 3706 | 186 | S186E | A | 5.6e-08 | 8.6e-08 | 8.8e-08 | 7.667E-08 | 1.793E-08 | 5.600E-08 | 8.800E-08 |
| 3707 | 186 | S186Q | A | 1.5e-08 | 4.4e-08 | 2e-08 | 2.633E-08 | 1.550E-08 | 1.500E-08 | 4.400E-08 |
| 3708 | 186 | S186G | A | 1.4e-07 | 2.5e-08 | 8.8e-08 | 8.433E-08 | 5.759E-08 | 2.500E-08 | 1.400E-07 |
| 3709 | 186 | S186H | A | 1.7e-08 | 3.1e-08 | 3.9e-08 | 2.900E-08 | 1.114E-08 | 1.700E-08 | 3.900E-08 |
| 3710 | 186 | S186I | A | 1e-07 | 7.7e-08 | 6.7e-08 | 8.133E-08 | 1.692E-08 | 6.700E-08 | 1.000E-07 |
| 3711 | 186 | S186L | A | 7.7e-08 | 1.9e-08 | 5.4e-08 | 5.000E-08 | 2.921E-08 | 1.900E-08 | 7.700E-08 |
| 3712 | 186 | S186K | A | 4.7e-08 | 5.9e-08 | 9e-08 | 6.533E-08 | 2.219E-08 | 4.700E-08 | 9.000E-08 |
| 3713 | 186 | S186M | A | 6.5e-08 | 3.2e-08 | 1.8e-08 | 3.833E-08 | 2.413E-08 | 1.800E-08 | 6.500E-08 |
| 3714 | 186 | S186F | A | 5.9e-08 | 1.1e-07 | 4.5e-08 | 7.133E-08 | 3.421E-08 | 4.500E-08 | 1.100E-07 |
| 3715 | 186 | S186P | A | 5.8e-08 | 2.4e-08 | 3.6e-08 | 3.933E-08 | 1.724E-08 | 2.400E-08 | 5.800E-08 |
| 3716 | 186 | S186S | A | 7.6e-09 | 4.7e-08 | 9.1e-08 | 4.853E-08 | 4.172E-08 | 7.600E-09 | 9.100E-08 |
| 3717 | 186 | S186T | A | 1.9e-08 | 4.4e-08 | 5.1e-08 | 3.800E-08 | 1.682E-08 | 1.900E-08 | 5.100E-08 |
| 3718 | 186 | S186W | A | 1.7e-07 | 1.4e-08 | 3.4e-08 | 7.267E-08 | 8.488E-08 | 1.400E-08 | 1.700E-07 |
| 3719 | 186 | S186Y | A | 8.4e-08 | 1.1e-08 | 7.7e-08 | 5.733E-08 | 4.028E-08 | 1.100E-08 | 8.400E-08 |
| 3720 | 186 | S186V | A | 4.5e-08 | 4.6e-08 | 1.2e-07 | 7.033E-08 | 4.302E-08 | 4.500E-08 | 1.200E-07 |
| 3721 | 187 | S187A | A | 2.6e-08 | 5.7e-08 | 3.4e-08 | 3.900E-08 | 1.609E-08 | 2.600E-08 | 5.700E-08 |
| 3722 | 187 | S187R | A | 3.3e-08 | 1.4e-08 | 3.4e-08 | 2.700E-08 | 1.127E-08 | 1.400E-08 | 3.400E-08 |
| 3723 | 187 | S187N | A | 4.5e-08 | 7.8e-08 | 5.7e-08 | 6.000E-08 | 1.670E-08 | 4.500E-08 | 7.800E-08 |
| 3724 | 187 | S187D | A | 8.9e-08 | 4.2e-08 | 1.9e-08 | 5.000E-08 | 3.568E-08 | 1.900E-08 | 8.900E-08 |
| 3725 | 187 | S187C | A | 5.7e-08 | 9.3e-08 | 2.1e-08 | 5.700E-08 | 3.600E-08 | 2.100E-08 | 9.300E-08 |
| 3726 | 187 | S187E | A | 1.7e-08 | 3.8e-08 | 1.9e-08 | 2.467E-08 | 1.159E-08 | 1.700E-08 | 3.800E-08 |
| 3727 | 187 | S187Q | A | 6e-08 | 1e-07 | 2.9e-08 | 6.300E-08 | 3.559E-08 | 2.900E-08 | 1.000E-07 |
| 3728 | 187 | S187G | A | 1.7e-07 | 2.7e-08 | 1.3e-07 | 1.090E-07 | 7.378E-08 | 2.700E-08 | 1.700E-07 |
| 3729 | 187 | S187H | A | 3.1e-08 | 9.4e-08 | 4.9e-08 | 5.800E-08 | 3.245E-08 | 3.100E-08 | 9.400E-08 |
| 3730 | 187 | S187I | A | 5.2e-08 | 1e-07 | 6.3e-08 | 7.167E-08 | 2.515E-08 | 5.200E-08 | 1.000E-07 |
| 3731 | 187 | S187L | A | 2.6e-08 | 3.7e-08 | 7.4e-08 | 4.567E-08 | 2.515E-08 | 2.600E-08 | 7.400E-08 |
| 3732 | 187 | S187K | A | 6.9e-08 | 6.1e-08 | 4.3e-08 | 5.767E-08 | 1.332E-08 | 4.300E-08 | 6.900E-08 |
| 3733 | 187 | S187M | A | 8.9e-08 | 1.6e-08 | 3.7e-08 | 4.733E-08 | 3.758E-08 | 1.600E-08 | 8.900E-08 |
| 3734 | 187 | S187F | A | 4e-08 | 3.6e-08 | 5.8e-08 | 4.467E-08 | 1.172E-08 | 3.600E-08 | 5.800E-08 |
| 3735 | 187 | S187P | A | 6.3e-08 | 4e-08 | 2.6e-08 | 4.300E-08 | 1.868E-08 | 2.600E-08 | 6.300E-08 |
| 3736 | 187 | S187S | A | 2.3e-08 | 3.5e-08 | 5.6e-08 | 3.800E-08 | 1.670E-08 | 2.300E-08 | 5.600E-08 |
| 3737 | 187 | S187T | A | 8.8e-08 | 5.6e-08 | 5.5e-08 | 6.633E-08 | 1.877E-08 | 5.500E-08 | 8.800E-08 |
| 3738 | 187 | S187W | A | 1.1e-07 | 3.7e-08 | 3e-08 | 5.900E-08 | 4.431E-08 | 3.000E-08 | 1.100E-07 |
| 3739 | 187 | S187Y | A | 2.8e-08 | 1.9e-08 | 8.8e-08 | 4.500E-08 | 3.751E-08 | 1.900E-08 | 8.800E-08 |

| | | | | | | | | | | |
|---|---|---|---|---|---|---|---|---|---|---|
| 3740 | 187 | S187V | A | 3.6e-08 | 4.3e-08 | 1e-07 | 5.967E-08 | 3.510E-08 | 3.600E-08 | 1.000E-07 |
| 3741 | 188 | E188A | A | 3.1e-08 | 2.7e-08 | 1.5e-07 | 6.933E-08 | 6.989E-08 | 2.700E-08 | 1.500E-07 |
| 3742 | 188 | E188R | A | 3.5e-08 | 3.7e-08 | 7.5e-08 | 4.900E-08 | 2.254E-08 | 3.500E-08 | 7.500E-08 |
| 3743 | 188 | E188N | A | 7.2e-08 | 1.4e-08 | 6.2e-08 | 4.933E-08 | 3.101E-08 | 1.400E-08 | 7.200E-08 |
| 3744 | 188 | E188D | A | 4.1e-08 | 1.3e-07 | 6.5e-08 | 7.867E-08 | 4.605E-08 | 4.100E-08 | 1.300E-07 |
| 3745 | 188 | E188C | A | 3.7e-08 | 5.5e-08 | 5.1e-08 | 4.767E-08 | 9.452E-09 | 3.700E-08 | 5.500E-08 |
| 3746 | 188 | E188E | A | 7e-08 | 3.2e-08 | 2e-08 | 4.067E-08 | 2.610E-08 | 2.000E-08 | 7.000E-08 |
| 3747 | 188 | E188Q | A | 4.1e-08 | 4.3e-08 | 9.1e-08 | 5.833E-08 | 2.831E-08 | 4.100E-08 | 9.100E-08 |
| 3748 | 188 | E188G | A | 6.3e-08 | 5.5e-08 | 2.4e-07 | 1.193E-07 | 1.046E-07 | 5.500E-08 | 2.400E-07 |
| 3749 | 188 | E188H | A | 5.2e-08 | 4.3e-08 | 8.4e-08 | 5.967E-08 | 2.155E-08 | 4.300E-08 | 8.400E-08 |
| 3750 | 188 | E188I | A | 7.7e-08 | 5.5e-08 | 6.6e-08 | 6.600E-08 | 1.100E-08 | 5.500E-08 | 7.700E-08 |
| 3751 | 188 | E188L | A | 5.4e-08 | 5.4e-08 | 1.8e-08 | 4.200E-08 | 2.078E-08 | 1.800E-08 | 5.400E-08 |
| 3752 | 188 | E188K | A | 4.5e-08 | 7.9e-08 | 7.1e-08 | 6.500E-08 | 1.778E-08 | 4.500E-08 | 7.900E-08 |
| 3753 | 188 | E188M | A | 5.1e-08 | 3.2e-08 | 5e-08 | 4.433E-08 | 1.069E-08 | 3.200E-08 | 5.100E-08 |
| 3754 | 188 | E188F | A | 5.7e-08 | 1e-07 | 6.3e-08 | 7.333E-08 | 2.329E-08 | 5.700E-08 | 1.000E-07 |
| 3755 | 188 | E188P | A | 1.7e-08 | 9.1e-08 | 3e-08 | 4.600E-08 | 3.951E-08 | 1.700E-08 | 9.100E-08 |
| 3756 | 188 | E188S | A | 1.5e-07 | 3.2e-08 | 5e-08 | 7.733E-08 | 6.357E-08 | 3.200E-08 | 1.500E-07 |
| 3757 | 188 | E188T | A | 1.8e-07 | 6.3e-08 | 3.1e-08 | 9.133E-08 | 7.844E-08 | 3.100E-08 | 1.800E-07 |
| 3758 | 188 | E188W | A | 4.4e-08 | 3e-08 | 4.1e-08 | 3.833E-08 | 7.371E-09 | 3.000E-08 | 4.400E-08 |
| 3759 | 188 | E188Y | A | 5.9e-08 | 5.1e-08 | 4.1e-08 | 5.033E-08 | 9.018E-09 | 4.100E-08 | 5.900E-08 |
| 3760 | 188 | E188V | A | 6.1e-08 | 1.9e-08 | 3e-08 | 3.667E-08 | 2.178E-08 | 1.900E-08 | 6.100E-08 |
| 3761 | 189 | D189A | A | 7.1e-08 | 2.3e-08 | 5.7e-08 | 5.033E-08 | 2.468E-08 | 2.300E-08 | 7.100E-08 |
| 3762 | 189 | D189R | A | 7.2e-08 | 2.8e-08 | 4.6e-08 | 4.867E-08 | 2.212E-08 | 2.800E-08 | 7.200E-08 |
| 3763 | 189 | D189N | A | 3.3e-08 | 8.6e-08 | 2.4e-08 | 4.767E-08 | 3.350E-08 | 2.400E-08 | 8.600E-08 |
| 3764 | 189 | D189D | A | 4.8e-08 | 2.2e-08 | 6.8e-08 | 4.600E-08 | 2.307E-08 | 2.200E-08 | 6.800E-08 |
| 3765 | 189 | D189C | A | 5.5e-08 | 3.7e-08 | 6e-08 | 5.067E-08 | 1.210E-08 | 3.700E-08 | 6.000E-08 |
| 3766 | 189 | D189E | A | 1.9e-08 | 3.8e-08 | 7.8e-08 | 4.500E-08 | 3.012E-08 | 1.900E-08 | 7.800E-08 |
| 3767 | 189 | D189Q | A | 9.5e-08 | 6.6e-08 | 4.4e-08 | 6.833E-08 | 2.558E-08 | 4.400E-08 | 9.500E-08 |
| 3768 | 189 | D189G | A | 5.2e-08 | 1.3e-07 | 1.1e-08 | 6.433E-08 | 6.045E-08 | 1.100E-08 | 1.300E-07 |
| 3769 | 189 | D189H | A | 9.6e-08 | 1.3e-07 | 9.1e-08 | 1.057E-07 | 2.122E-08 | 9.100E-08 | 1.300E-07 |
| 3770 | 189 | D189I | A | 8.1e-08 | 5.5e-08 | 3.2e-08 | 5.600E-08 | 2.452E-08 | 3.200E-08 | 8.100E-08 |
| 3771 | 189 | D189L | A | 2.7e-08 | 6.3e-08 | 5.6e-08 | 4.867E-08 | 1.909E-08 | 2.700E-08 | 6.300E-08 |
| 3772 | 189 | D189K | A | 1.3e-08 | 2.3e-08 | 3e-08 | 2.200E-08 | 8.544E-09 | 1.300E-08 | 3.000E-08 |
| 3773 | 189 | D189M | A | 9.2e-08 | 6e-08 | 2.5e-08 | 5.900E-08 | 3.351E-08 | 2.500E-08 | 9.200E-08 |
| 3774 | 189 | D189F | A | 5.5e-08 | 5.7e-08 | 3.1e-08 | 4.767E-08 | 1.447E-08 | 3.100E-08 | 5.700E-08 |
| 3775 | 189 | D189P | A | 2.2e-08 | 7.2e-09 | 4.9e-08 | 2.607E-08 | 2.119E-08 | 7.200E-09 | 4.900E-08 |
| 3776 | 189 | D189S | A | 4.6e-08 | 7.3e-08 | 6.1e-08 | 6.000E-08 | 1.353E-08 | 4.600E-08 | 7.300E-08 |
| 3777 | 189 | D189T | A | 1.8e-07 | 9.8e-08 | 4.8e-08 | 1.087E-07 | 6.664E-08 | 4.800E-08 | 1.800E-07 |
| 3778 | 189 | D189W | A | 7.5e-08 | 3.3e-08 | 7.6e-08 | 6.133E-08 | 2.454E-08 | 3.300E-08 | 7.600E-08 |
| 3779 | 189 | D189Y | A | 5.6e-08 | 7.3e-08 | 3.4e-08 | 5.433E-08 | 1.955E-08 | 3.400E-08 | 7.300E-08 |
| 3780 | 189 | D189V | A | 1e-07 | 5e-08 | 6.2e-08 | 7.067E-08 | 2.610E-08 | 5.000E-08 | 1.000E-07 |
| 3781 | 190 | C190A | A | 6.9e-08 | 5.3e-08 | 2.6e-08 | 4.933E-08 | 2.173E-08 | 2.600E-08 | 6.900E-08 |
| 3782 | 190 | C190R | A | 2.7e-08 | 1.8e-08 | 5.2e-08 | 3.233E-08 | 1.762E-08 | 1.800E-08 | 5.200E-08 |
| 3783 | 190 | C190N | A | 2.3e-08 | 6e-08 | 1.3e-08 | 3.200E-08 | 2.476E-08 | 1.300E-08 | 6.000E-08 |
| 3784 | 190 | C190D | A | 4.6e-08 | 2.8e-08 | 2.3e-08 | 3.233E-08 | 1.210E-08 | 2.300E-08 | 4.600E-08 |
| 3785 | 190 | C190C | A | 5e-08 | 3.2e-08 | 3.7e-08 | 3.967E-08 | 9.292E-09 | 3.200E-08 | 5.000E-08 |
| 3786 | 190 | C190E | A | 2.3e-08 | 2.1e-08 | 3.6e-08 | 2.667E-08 | 8.145E-09 | 2.100E-08 | 3.600E-08 |
| 3787 | 190 | C190Q | A | 2.3e-08 | 1.4e-07 | 7.6e-08 | 7.967E-08 | 5.859E-08 | 2.300E-08 | 1.400E-07 |
| 3788 | 190 | C190G | A | 7.9e-08 | 1.4e-08 | 4.9e-08 | 4.733E-08 | 3.253E-08 | 1.400E-08 | 7.900E-08 |
| 3789 | 190 | C190H | A | 4.6e-08 | 1.9e-08 | 2.2e-08 | 2.900E-08 | 1.480E-08 | 1.900E-08 | 4.600E-08 |
| 3790 | 190 | C190I | A | 2.7e-08 | 3.1e-08 | 2.6e-08 | 2.800E-08 | 2.646E-09 | 2.600E-08 | 3.100E-08 |
| 3791 | 190 | C190L | A | 7.3e-08 | 3.3e-08 | 2.4e-08 | 4.333E-08 | 2.608E-08 | 2.400E-08 | 7.300E-08 |
| 3792 | 190 | C190K | A | 7.2e-08 | 2.3e-08 | 1.2e-07 | 7.167E-08 | 4.850E-08 | 2.300E-08 | 1.200E-07 |
| 3793 | 190 | C190M | A | 2.7e-08 | 1.5e-08 | 2.8e-08 | 2.333E-08 | 7.234E-09 | 1.500E-08 | 2.800E-08 |
| 3794 | 190 | C190F | A | 1.5e-08 | 1.8e-08 | 2.2e-08 | 1.833E-08 | 3.512E-09 | 1.500E-08 | 2.200E-08 |
| 3795 | 190 | C190P | A | 1.9e-08 | 3.3e-08 | 6.1e-08 | 3.767E-08 | 2.139E-08 | 1.900E-08 | 6.100E-08 |
| 3796 | 190 | C190S | A | 3.5e-08 | 4.1e-08 | 4.5e-08 | 4.033E-08 | 5.033E-09 | 3.500E-08 | 4.500E-08 |
| 3797 | 190 | C190T | A | 1.6e-08 | 1.9e-08 | 3.7e-08 | 2.400E-08 | 1.136E-08 | 1.600E-08 | 3.700E-08 |
| 3798 | 190 | C190W | A | 3.4e-08 | 1.3e-08 | 6.4e-08 | 3.700E-08 | 2.563E-08 | 1.300E-08 | 6.400E-08 |
| 3799 | 190 | C190Y | A | 6.6e-08 | 3.2e-08 | 4.1e-08 | 4.633E-08 | 1.762E-08 | 3.200E-08 | 6.600E-08 |
| 3800 | 190 | C190V | A | 3.2e-08 | 6.6e-08 | 2.9e-08 | 4.233E-08 | 2.055E-08 | 2.900E-08 | 6.600E-08 |
| 3801 | 191 | Q191A | A | 3.3e-08 | 7.1e-08 | 1.2e-07 | 7.467E-08 | 4.362E-08 | 3.300E-08 | 1.200E-07 |
| 3802 | 191 | Q191R | A | 4.8e-08 | 3.1e-08 | 8.1e-08 | 5.333E-08 | 2.542E-08 | 3.100E-08 | 8.100E-08 |
| 3803 | 191 | Q191N | A | 5.5e-08 | 2.8e-08 | 3.2e-08 | 3.833E-08 | 1.457E-08 | 2.800E-08 | 5.500E-08 |
| 3804 | 191 | Q191D | A | 3.7e-08 | 3.4e-08 | 3.5e-08 | 3.533E-08 | 1.528E-09 | 3.400E-08 | 3.700E-08 |
| 3805 | 191 | Q191C | A | 1.8e-08 | 3e-08 | 2.8e-08 | 2.533E-08 | 6.429E-09 | 1.800E-08 | 3.000E-08 |
| 3806 | 191 | Q191E | A | 2.8e-08 | 4.4e-08 | 4.4e-08 | 3.867E-08 | 9.238E-09 | 2.800E-08 | 4.400E-08 |
| 3807 | 191 | Q191Q | A | 8.1e-08 | 3e-08 | 6.9e-08 | 6.000E-08 | 2.666E-08 | 3.000E-08 | 8.100E-08 |
| 3808 | 191 | Q191G | A | 4.8e-08 | 6e-08 | 2.6e-08 | 4.467E-08 | 1.724E-08 | 2.600E-08 | 6.000E-08 |
| 3809 | 191 | Q191H | A | 2.6e-08 | 5.1e-08 | 6.5e-08 | 4.733E-08 | 1.976E-08 | 2.600E-08 | 6.500E-08 |
| 3810 | 191 | Q191I | A | 1.7e-08 | 1.8e-08 | 5.5e-08 | 3.000E-08 | 2.166E-08 | 1.700E-08 | 5.500E-08 |
| 3811 | 191 | Q191L | A | 4.6e-08 | 9.6e-08 | 2.1e-08 | 5.433E-08 | 3.819E-08 | 2.100E-08 | 9.600E-08 |
| 3812 | 191 | Q191K | A | 4e-08 | 1.4e-08 | 4.6e-08 | 3.333E-08 | 1.701E-08 | 1.400E-08 | 4.600E-08 |
| 3813 | 191 | Q191M | A | 3.9e-08 | 7.7e-08 | 8.9e-08 | 6.833E-08 | 2.610E-08 | 3.900E-08 | 8.900E-08 |
| 3814 | 191 | Q191F | A | 1.6e-08 | 5.4e-08 | 5.8e-08 | 4.267E-08 | 2.318E-08 | 1.600E-08 | 5.800E-08 |
| 3815 | 191 | Q191P | A | 3.1e-08 | 8.2e-08 | 3.3e-08 | 4.867E-08 | 2.888E-08 | 3.100E-08 | 8.200E-08 |
| 3816 | 191 | Q191S | A | 1.5e-07 | 2.6e-08 | 5.4e-08 | 7.667E-08 | 6.503E-08 | 2.600E-08 | 1.500E-07 |
| 3817 | 191 | Q191T | A | 3.1e-08 | 5e-08 | 2.1e-08 | 3.400E-08 | 1.473E-08 | 2.100E-08 | 5.000E-08 |
| 3818 | 191 | Q191W | A | 9.9e-09 | 8.8e-08 | 5e-08 | 4.930E-08 | 3.905E-08 | 9.900E-09 | 8.800E-08 |
| 3819 | 191 | Q191Y | A | 4.4e-08 | 5.7e-08 | 3.2e-08 | 4.433E-08 | 1.250E-08 | 3.200E-08 | 5.700E-08 |
| 3820 | 191 | Q191V | A | 1.1e-08 | 1.8e-07 | 3.4e-08 | 7.500E-08 | 9.166E-08 | 1.100E-08 | 1.800E-07 |
| 3821 | 192 | S192A | A | 3.3e-08 | 1.7e-08 | 4.1e-08 | 3.033E-08 | 1.222E-08 | 1.700E-08 | 4.100E-08 |
| 3822 | 192 | S192R | A | 1.5e-08 | 2.2e-08 | 3.1e-08 | 2.267E-08 | 8.021E-09 | 1.500E-08 | 3.100E-08 |
| 3823 | 192 | S192N | A | 3e-08 | 8.1e-08 | 8.1e-08 | 6.400E-08 | 2.944E-08 | 3.000E-08 | 8.100E-08 |
| 3824 | 192 | S192D | A | 6.3e-08 | 3e-08 | 8.8e-09 | 3.393E-08 | 2.731E-08 | 8.800E-09 | 6.300E-08 |

| | | | | | | | | | | |
|---|---|---|---|---|---|---|---|---|---|---|
| 3825 | 192 | S192C | A | 5.7e-08 | 1.9e-08 | 5.9e-08 | 4.500E-08 | 2.254E-08 | 1.900E-08 | 5.900E-08 |
| 3826 | 192 | S192E | A | 5.8e-08 | 3.2e-08 | 4.2e-08 | 4.400E-08 | 1.311E-08 | 3.200E-08 | 5.800E-08 |
| 3827 | 192 | S192Q | A | 2.5e-08 | 7.3e-08 | 1.9e-08 | 3.900E-08 | 2.960E-08 | 1.900E-08 | 7.300E-08 |
| 3828 | 192 | S192G | A | 2.8e-08 | 4.1e-08 | 7e-08 | 4.633E-08 | 2.150E-08 | 2.800E-08 | 7.000E-08 |
| 3829 | 192 | S192H | A | 7.9e-08 | 3.4e-08 | 5.7e-08 | 5.667E-08 | 2.250E-08 | 3.400E-08 | 7.900E-08 |
| 3830 | 192 | S192I | A | 3e-08 | 8.2e-08 | 4.1e-08 | 5.100E-08 | 2.740E-08 | 3.000E-08 | 8.200E-08 |
| 3831 | 192 | S192L | A | 9e-08 | 3.7e-08 | 4.9e-08 | 5.867E-08 | 2.779E-08 | 3.700E-08 | 9.000E-08 |
| 3832 | 192 | S192K | A | 3.6e-08 | 2.8e-08 | 3.4e-08 | 3.267E-08 | 4.163E-09 | 2.800E-08 | 3.600E-08 |
| 3833 | 192 | S192M | A | 8.9e-08 | 2.7e-08 | 4.2e-08 | 5.267E-08 | 3.235E-08 | 2.700E-08 | 8.900E-08 |
| 3834 | 192 | S192F | A | 6.4e-08 | 4.7e-08 | 2.6e-08 | 4.567E-08 | 1.904E-08 | 2.600E-08 | 6.400E-08 |
| 3835 | 192 | S192P | A | 4.3e-08 | 8.8e-08 | 5.8e-08 | 6.300E-08 | 2.291E-08 | 4.300E-08 | 8.800E-08 |
| 3836 | 192 | S192S | A | 2e-07 | 1e-08 | 4.7e-08 | 8.567E-08 | 1.007E-07 | 1.000E-08 | 2.000E-07 |
| 3837 | 192 | S192T | A | 6.9e-08 | 7e-08 | 3.2e-08 | 5.700E-08 | 2.166E-08 | 3.200E-08 | 7.000E-08 |
| 3838 | 192 | S192W | A | 1.3e-07 | 3.4e-08 | 5.7e-08 | 7.367E-08 | 5.012E-08 | 3.400E-08 | 1.300E-07 |
| 3839 | 192 | S192Y | A | 1.6e-08 | 1e-08 | 3.7e-08 | 2.100E-08 | 1.418E-08 | 1.000E-08 | 3.700E-08 |
| 3840 | 192 | S192V | A | 2.2e-08 | 6e-08 | 3.5e-08 | 3.900E-08 | 1.931E-08 | 2.200E-08 | 6.000E-08 |
| 3841 | 193 | L193A | A | 4.1e-08 | 5.8e-08 | 1.1e-07 | 6.967E-08 | 3.595E-08 | 4.100E-08 | 1.100E-07 |
| 3842 | 193 | L193R | A | 2.8e-08 | 2.1e-08 | 1.3e-08 | 2.067E-08 | 7.506E-09 | 1.300E-08 | 2.800E-08 |
| 3843 | 193 | L193N | A | 5.3e-08 | 4.5e-08 | 4e-08 | 4.600E-08 | 6.557E-09 | 4.000E-08 | 5.300E-08 |
| 3844 | 193 | L193D | A | 5.9e-08 | 2.3e-08 | 2.8e-08 | 3.667E-08 | 1.950E-08 | 2.300E-08 | 5.900E-08 |
| 3845 | 193 | L193C | A | 3.5e-08 | 5.3e-08 | 2.7e-08 | 3.833E-08 | 1.332E-08 | 2.700E-08 | 5.300E-08 |
| 3846 | 193 | L193E | A | 3.1e-08 | 4.2e-08 | 5.7e-08 | 4.333E-08 | 1.305E-08 | 3.100E-08 | 5.700E-08 |
| 3847 | 193 | L193Q | A | 4.2e-08 | 3.2e-08 | 4.4e-08 | 3.933E-08 | 6.429E-09 | 3.200E-08 | 4.400E-08 |
| 3848 | 193 | L193G | A | 2.7e-08 | 7.1e-08 | 2.1e-08 | 3.967E-08 | 2.730E-08 | 2.100E-08 | 7.100E-08 |
| 3849 | 193 | L193H | A | 4.3e-08 | 1.1e-07 | 1.7e-08 | 5.667E-08 | 4.798E-08 | 1.700E-08 | 1.100E-07 |
| 3850 | 193 | L193I | A | 6.2e-08 | 6e-08 | 7.4e-08 | 6.533E-08 | 7.572E-09 | 6.000E-08 | 7.400E-08 |
| 3851 | 193 | L193L | A | 3.6e-08 | 4.9e-08 | 8.1e-08 | 5.533E-08 | 2.316E-08 | 3.600E-08 | 8.100E-08 |
| 3852 | 193 | L193K | A | 6.1e-08 | 5.7e-08 | 6.3e-08 | 6.033E-08 | 3.055E-09 | 5.700E-08 | 6.300E-08 |
| 3853 | 193 | L193M | A | 3.2e-08 | 7.5e-08 | 1.8e-08 | 4.167E-08 | 2.970E-08 | 1.800E-08 | 7.500E-08 |
| 3854 | 193 | L193F | A | 9.2e-08 | 1.9e-08 | 3.5e-08 | 4.867E-08 | 3.837E-08 | 1.900E-08 | 9.200E-08 |
| 3855 | 193 | L193P | A | 4.9e-08 | 2e-08 | 1.3e-08 | 2.733E-08 | 1.909E-08 | 1.300E-08 | 4.900E-08 |
| 3856 | 193 | L193S | A | 3.3e-08 | 2.4e-08 | 6.3e-08 | 4.000E-08 | 2.042E-08 | 2.400E-08 | 6.300E-08 |
| 3857 | 193 | L193T | A | 2.6e-08 | 2.7e-08 | 3.4e-08 | 2.900E-08 | 4.359E-09 | 2.600E-08 | 3.400E-08 |
| 3858 | 193 | L193W | A | 3.9e-08 | 8.4e-08 | 4.1e-08 | 5.467E-08 | 2.542E-08 | 3.900E-08 | 8.400E-08 |
| 3859 | 193 | L193Y | A | 2.7e-08 | 3.8e-08 | 4e-08 | 3.500E-08 | 7.000E-09 | 2.700E-08 | 4.000E-08 |
| 3860 | 193 | L193V | A | 1.9e-08 | 4.1e-08 | 1.6e-08 | 2.533E-08 | 1.365E-08 | 1.600E-08 | 4.100E-08 |
| 3861 | 194 | T194A | A | 1.4e-08 | 3.7e-08 | 1.7e-08 | 2.267E-08 | 1.250E-08 | 1.400E-08 | 3.700E-08 |
| 3862 | 194 | T194R | A | 1.1e-07 | 1.9e-08 | 4.8e-08 | 5.900E-08 | 4.649E-08 | 1.900E-08 | 1.100E-07 |
| 3863 | 194 | T194N | A | 4.3e-08 | 4.7e-08 | 8.1e-08 | 5.700E-08 | 2.088E-08 | 4.300E-08 | 8.100E-08 |
| 3864 | 194 | T194D | A | 5.2e-08 | 8.6e-09 | 7e-08 | 4.353E-08 | 3.156E-08 | 8.600E-09 | 7.000E-08 |
| 3865 | 194 | T194C | A | 1.9e-08 | 2e-08 | 2.3e-08 | 2.067E-08 | 2.082E-09 | 1.900E-08 | 2.300E-08 |
| 3866 | 194 | T194E | A | 1.2e-07 | 2.7e-08 | 4.4e-08 | 6.367E-08 | 4.952E-08 | 2.700E-08 | 1.200E-07 |
| 3867 | 194 | T194Q | A | 6e-08 | 5.3e-08 | 2.1e-08 | 4.467E-08 | 2.079E-08 | 2.100E-08 | 6.000E-08 |
| 3868 | 194 | T194G | A | 2.4e-08 | 6.4e-08 | 2.6e-08 | 3.800E-08 | 2.254E-08 | 2.400E-08 | 6.400E-08 |
| 3869 | 194 | T194H | A | 3.4e-08 | 1.1e-07 | 3.8e-08 | 6.067E-08 | 4.277E-08 | 3.400E-08 | 1.100E-07 |
| 3870 | 194 | T194I | A | 6.1e-08 | 5.2e-08 | 4.9e-08 | 5.400E-08 | 6.245E-09 | 4.900E-08 | 6.100E-08 |
| 3871 | 194 | T194L | A | 2e-08 | 3e-08 | 1e-07 | 5.000E-08 | 4.359E-08 | 2.000E-08 | 1.000E-07 |
| 3872 | 194 | T194K | A | 8.8e-08 | 2.2e-08 | 8.3e-08 | 6.433E-08 | 3.675E-08 | 2.200E-08 | 8.800E-08 |
| 3873 | 194 | T194M | A | 5.4e-08 | 4.9e-08 | 2.2e-08 | 4.167E-08 | 1.721E-08 | 2.200E-08 | 5.400E-08 |
| 3874 | 194 | T194F | A | 3.7e-08 | 5.2e-08 | 5.9e-08 | 4.933E-08 | 1.124E-08 | 3.700E-08 | 5.900E-08 |
| 3875 | 194 | T194P | A | 6.6e-08 | 1.9e-07 | 5.1e-08 | 1.023E-07 | 7.629E-08 | 5.100E-08 | 1.900E-07 |
| 3876 | 194 | T194S | A | 6.2e-08 | 3.3e-08 | 1.2e-07 | 7.167E-08 | 4.430E-08 | 3.300E-08 | 1.200E-07 |
| 3877 | 194 | T194T | A | 2.9e-08 | 5e-08 | 2.7e-08 | 3.533E-08 | 1.274E-08 | 2.700E-08 | 5.000E-08 |
| 3878 | 194 | T194W | A | 4.6e-08 | 6.6e-08 | 9.6e-08 | 6.933E-08 | 2.517E-08 | 4.600E-08 | 9.600E-08 |
| 3879 | 194 | T194Y | A | 1.7e-08 | 5.3e-08 | 2.1e-08 | 3.033E-08 | 1.973E-08 | 1.700E-08 | 5.300E-08 |
| 3880 | 194 | T194V | A | 1.1e-08 | 3.7e-08 | 4.6e-08 | 3.133E-08 | 1.818E-08 | 1.100E-08 | 4.600E-08 |
| 3881 | 195 | R195A | A | 2.4e-08 | 1.3e-07 | 4.5e-08 | 6.633E-08 | 5.613E-08 | 2.400E-08 | 1.300E-07 |
| 3882 | 195 | R195R | A | 1.2e-07 | 4e-08 | 2.4e-08 | 6.133E-08 | 5.143E-08 | 2.400E-08 | 1.200E-07 |
| 3883 | 195 | R195N | A | 2.5e-08 | 1.2e-08 | 2.8e-08 | 2.167E-08 | 8.505E-09 | 1.200E-08 | 2.800E-08 |
| 3884 | 195 | R195D | A | 1.2e-07 | 6.9e-08 | 3.2e-08 | 7.367E-08 | 4.419E-08 | 3.200E-08 | 1.200E-07 |
| 3885 | 195 | R195C | A | 8.1e-08 | 9.9e-08 | 5.6e-08 | 7.867E-08 | 2.159E-08 | 5.600E-08 | 9.900E-08 |
| 3886 | 195 | R195E | A | 6.1e-08 | 5.5e-08 | 1.2e-07 | 7.867E-08 | 3.592E-08 | 5.500E-08 | 1.200E-07 |
| 3887 | 195 | R195Q | A | 4.5e-08 | 2.8e-08 | 3.9e-08 | 3.733E-08 | 8.622E-09 | 2.800E-08 | 4.500E-08 |
| 3888 | 195 | R195G | A | 2.7e-08 | 2.1e-08 | 7.1e-08 | 3.967E-08 | 2.730E-08 | 2.100E-08 | 7.100E-08 |
| 3889 | 195 | R195H | A | 7.7e-08 | 4e-08 | 2e-08 | 4.567E-08 | 2.892E-08 | 2.000E-08 | 7.700E-08 |
| 3890 | 195 | R195I | A | 5.6e-08 | 3.2e-08 | 4.2e-08 | 4.333E-08 | 1.206E-08 | 3.200E-08 | 5.600E-08 |
| 3891 | 195 | R195L | A | 4.4e-08 | 1.5e-08 | 2.2e-08 | 2.700E-08 | 1.513E-08 | 1.500E-08 | 4.400E-08 |
| 3892 | 195 | R195K | A | 6.2e-08 | 2.8e-08 | 4.9e-08 | 4.633E-08 | 1.716E-08 | 2.800E-08 | 6.200E-08 |
| 3893 | 195 | R195M | A | 2.2e-08 | 3.7e-08 | 2.5e-08 | 2.800E-08 | 7.937E-09 | 2.200E-08 | 3.700E-08 |
| 3894 | 195 | R195F | A | 6.8e-08 | 3.3e-08 | 4.2e-08 | 4.767E-08 | 1.818E-08 | 3.300E-08 | 6.800E-08 |
| 3895 | 195 | R195P | A | 4.5e-08 | 4.2e-08 | 9.8e-08 | 6.167E-08 | 3.150E-08 | 4.200E-08 | 9.800E-08 |
| 3896 | 195 | R195S | A | 5.6e-08 | 3.4e-08 | 3.4e-08 | 4.133E-08 | 1.270E-08 | 3.400E-08 | 5.600E-08 |
| 3897 | 195 | R195T | A | 7.2e-08 | 2.2e-08 | 3.1e-08 | 4.167E-08 | 2.665E-08 | 2.200E-08 | 7.200E-08 |
| 3898 | 195 | R195W | A | 1e-07 | 4.2e-08 | 7.5e-08 | 7.233E-08 | 2.909E-08 | 4.200E-08 | 1.000E-07 |
| 3899 | 195 | R195Y | A | 2.9e-08 | 5.7e-08 | 4.4e-08 | 4.333E-08 | 1.401E-08 | 2.900E-08 | 5.700E-08 |
| 3900 | 195 | R195V | A | 4.4e-08 | 2.2e-08 | 5.3e-08 | 3.967E-08 | 1.595E-08 | 2.200E-08 | 5.300E-08 |
| 3901 | 196 | T196A | A | 5.3e-08 | 5.3e-08 | 4e-08 | 4.867E-08 | 7.506E-09 | 4.000E-08 | 5.300E-08 |
| 3902 | 196 | T196R | A | 4.7e-08 | 8e-08 | 1e-08 | 4.567E-08 | 3.502E-08 | 1.000E-08 | 8.000E-08 |
| 3903 | 196 | T196N | A | 4.3e-08 | 3e-08 | 2.9e-08 | 3.400E-08 | 7.810E-09 | 2.900E-08 | 4.300E-08 |
| 3904 | 196 | T196D | A | 7.9e-08 | 3.3e-08 | 5.3e-08 | 5.500E-08 | 2.307E-08 | 3.300E-08 | 7.900E-08 |
| 3905 | 196 | T196C | A | 3.1e-08 | 3.3e-08 | 2.9e-08 | 3.100E-08 | 2.000E-09 | 2.900E-08 | 3.300E-08 |
| 3906 | 196 | T196E | A | 3.9e-08 | 3.7e-08 | 2.7e-08 | 3.433E-08 | 6.429E-09 | 2.700E-08 | 3.900E-08 |
| 3907 | 196 | T196Q | A | 5.3e-08 | 3.9e-08 | 1.2e-08 | 3.467E-08 | 2.084E-08 | 1.200E-08 | 5.300E-08 |
| 3908 | 196 | T196G | A | 4.9e-08 | 2.8e-08 | 3.2e-08 | 3.633E-08 | 1.115E-08 | 2.800E-08 | 4.900E-08 |
| 3909 | 196 | T196H | A | 1.2e-07 | 7.7e-08 | 1.4e-07 | 1.123E-07 | 3.219E-08 | 7.700E-08 | 1.400E-07 |

| | | | | | | | | | | |
|---|---|---|---|---|---|---|---|---|---|---|
| 3910 | 196 | T196I | A | 3.5e-08 | 6.1e-08 | 3.4e-08 | 4.333E-08 | 1.531E-08 | 3.400E-08 | 6.100E-08 |
| 3911 | 196 | T196L | A | 2.2e-08 | 2.2e-08 | 8.1e-08 | 4.167E-08 | 3.406E-08 | 2.200E-08 | 8.100E-08 |
| 3912 | 196 | T196K | A | 3.6e-08 | 7.2e-09 | 2.7e-08 | 2.340E-08 | 1.473E-08 | 7.200E-09 | 3.600E-08 |
| 3913 | 196 | T196M | A | 3.3e-08 | 1.8e-08 | 3.6e-08 | 2.900E-08 | 9.644E-09 | 1.800E-08 | 3.600E-08 |
| 3914 | 196 | T196F | A | 3.1e-08 | 1.1e-07 | 1.3e-08 | 5.133E-08 | 5.160E-08 | 1.300E-08 | 1.100E-07 |
| 3915 | 196 | T196P | A | 2.5e-08 | 4.4e-08 | 6.8e-08 | 4.567E-08 | 2.155E-08 | 2.500E-08 | 6.800E-08 |
| 3916 | 196 | T196S | A | 6e-08 | 7.3e-08 | 2e-08 | 5.100E-08 | 2.762E-08 | 2.000E-08 | 7.300E-08 |
| 3917 | 196 | T196T | A | 6.3e-08 | 4.2e-08 | 5.2e-08 | 5.233E-08 | 1.050E-08 | 4.200E-08 | 6.300E-08 |
| 3918 | 196 | T196W | A | 6.1e-08 | 3.3e-08 | 3e-08 | 4.133E-08 | 1.710E-08 | 3.000E-08 | 6.100E-08 |
| 3919 | 196 | T196Y | A | 6.3e-08 | 3.7e-08 | 1.6e-08 | 3.867E-08 | 2.354E-08 | 1.600E-08 | 6.300E-08 |
| 3920 | 196 | T196V | A | 5.2e-08 | 1.2e-08 | 1.4e-08 | 2.600E-08 | 2.254E-08 | 1.200E-08 | 5.200E-08 |
| 3921 | 197 | V197A | A | 4.5e-08 | 5.1e-08 | 4.7e-08 | 4.767E-08 | 3.055E-09 | 4.500E-08 | 5.100E-08 |
| 3922 | 197 | V197R | A | 2.8e-08 | 6.2e-08 | 3.2e-08 | 4.067E-08 | 1.858E-08 | 2.800E-08 | 6.200E-08 |
| 3923 | 197 | V197N | A | 3e-08 | 2.4e-08 | 1.1e-07 | 5.467E-08 | 4.801E-08 | 2.400E-08 | 1.100E-07 |
| 3924 | 197 | V197D | A | 3.6e-08 | 2.1e-08 | 4.7e-08 | 3.467E-08 | 1.305E-08 | 2.100E-08 | 4.700E-08 |
| 3925 | 197 | V197C | A | 2e-08 | 3.9e-08 | 3.8e-08 | 3.233E-08 | 1.069E-08 | 2.000E-08 | 3.900E-08 |
| 3926 | 197 | V197E | A | 5.5e-08 | 5.7e-08 | 5.2e-08 | 5.467E-08 | 2.517E-09 | 5.200E-08 | 5.700E-08 |
| 3927 | 197 | V197Q | A | 5.3e-08 | 1.8e-08 | 3.2e-08 | 3.433E-08 | 1.762E-08 | 1.800E-08 | 5.300E-08 |
| 3928 | 197 | V197G | A | 5.9e-08 | 1.2e-07 | 2.2e-08 | 6.700E-08 | 4.949E-08 | 2.200E-08 | 1.200E-07 |
| 3929 | 197 | V197H | A | 3.3e-08 | 6.9e-08 | 5.2e-08 | 5.133E-08 | 1.801E-08 | 3.300E-08 | 6.900E-08 |
| 3930 | 197 | V197I | A | 1.5e-08 | 2.1e-08 | 8.3e-09 | 1.477E-08 | 6.353E-09 | 8.300E-09 | 2.100E-08 |
| 3931 | 197 | V197L | A | 3.7e-08 | 6.3e-08 | 5.8e-08 | 5.267E-08 | 1.380E-08 | 3.700E-08 | 6.300E-08 |
| 3932 | 197 | V197K | A | 2.4e-08 | 3e-08 | 6.2e-08 | 3.867E-08 | 2.043E-08 | 2.400E-08 | 6.200E-08 |
| 3933 | 197 | V197M | A | 5.7e-08 | 6e-08 | 1.5e-08 | 4.400E-08 | 2.516E-08 | 1.500E-08 | 6.000E-08 |
| 3934 | 197 | V197F | A | 1.3e-08 | 5.1e-08 | 2.8e-08 | 3.067E-08 | 1.914E-08 | 1.300E-08 | 5.100E-08 |
| 3935 | 197 | V197P | A | 3.8e-08 | 6.1e-08 | 2.5e-08 | 4.133E-08 | 1.823E-08 | 2.500E-08 | 6.100E-08 |
| 3936 | 197 | V197S | A | 3.1e-08 | 3.5e-08 | 3.7e-08 | 3.433E-08 | 3.055E-09 | 3.100E-08 | 3.700E-08 |
| 3937 | 197 | V197T | A | 1.8e-08 | 5e-08 | 1.6e-08 | 2.800E-08 | 1.908E-08 | 1.600E-08 | 5.000E-08 |
| 3938 | 197 | V197W | A | 4.5e-08 | 3.3e-08 | 3.9e-08 | 3.900E-08 | 6.000E-09 | 3.300E-08 | 4.500E-08 |
| 3939 | 197 | V197Y | A | 4.2e-08 | 4.3e-08 | 8.8e-08 | 5.767E-08 | 2.627E-08 | 4.200E-08 | 8.800E-08 |
| 3940 | 197 | V197V | A | 3.1e-08 | 2.8e-08 | 4e-08 | 3.300E-08 | 6.245E-09 | 2.800E-08 | 4.000E-08 |
| 3941 | 198 | C198A | A | 9.5e-08 | 6.3e-08 | 2.1e-08 | 5.967E-08 | 3.711E-08 | 2.100E-08 | 9.500E-08 |
| 3942 | 198 | C198R | A | 3.2e-08 | 9.6e-08 | 2.9e-08 | 5.233E-08 | 3.785E-08 | 2.900E-08 | 9.600E-08 |
| 3943 | 198 | C198N | A | 2.5e-08 | 8.1e-08 | 3.3e-08 | 4.633E-08 | 3.029E-08 | 2.500E-08 | 8.100E-08 |
| 3944 | 198 | C198D | A | 8.7e-08 | 3.8e-08 | 7e-08 | 6.500E-08 | 2.488E-08 | 3.800E-08 | 8.700E-08 |
| 3945 | 198 | C198C | A | 2.1e-08 | 2.7e-08 | 7.4e-08 | 4.067E-08 | 2.902E-08 | 2.100E-08 | 7.400E-08 |
| 3946 | 198 | C198E | A | 4.9e-08 | 9.5e-08 | 4.9e-08 | 6.433E-08 | 2.656E-08 | 4.900E-08 | 9.500E-08 |
| 3947 | 198 | C198Q | A | 6.3e-08 | 9e-08 | 5.5e-08 | 6.933E-08 | 1.834E-08 | 5.500E-08 | 9.000E-08 |
| 3948 | 198 | C198G | A | 3.3e-08 | 3.1e-08 | 3.2e-08 | 3.200E-08 | 1.000E-09 | 3.100E-08 | 3.300E-08 |
| 3949 | 198 | C198H | A | 3e-08 | 4.3e-08 | 5.4e-08 | 4.233E-08 | 1.201E-08 | 3.000E-08 | 5.400E-08 |
| 3950 | 198 | C198I | A | 3.6e-08 | 4.6e-08 | 6.6e-08 | 4.933E-08 | 1.528E-08 | 3.600E-08 | 6.600E-08 |
| 3951 | 198 | C198L | A | 6e-08 | 7.5e-08 | 9.3e-08 | 7.600E-08 | 1.652E-08 | 6.000E-08 | 9.300E-08 |
| 3952 | 198 | C198K | A | 2.9e-08 | 7.7e-08 | 9.2e-08 | 6.600E-08 | 3.291E-08 | 2.900E-08 | 9.200E-08 |
| 3953 | 198 | C198M | A | 3.7e-08 | 4.8e-08 | 2.8e-08 | 3.767E-08 | 1.002E-08 | 2.800E-08 | 4.800E-08 |
| 3954 | 198 | C198F | A | 4.7e-08 | 4e-08 | 1.8e-08 | 3.500E-08 | 1.513E-08 | 1.800E-08 | 4.700E-08 |
| 3955 | 198 | C198P | A | 3.3e-08 | 1.1e-07 | 6.3e-08 | 6.867E-08 | 3.881E-08 | 3.300E-08 | 1.100E-07 |
| 3956 | 198 | C198S | A | 4.9e-08 | 3.4e-08 | 7.8e-09 | 3.027E-08 | 2.085E-08 | 7.800E-09 | 4.900E-08 |
| 3957 | 198 | C198T | A | 5.3e-08 | 7.8e-08 | 3.8e-08 | 5.633E-08 | 2.021E-08 | 3.800E-08 | 7.800E-08 |
| 3958 | 198 | C198W | A | 9.1e-09 | 3.9e-08 | 3.5e-08 | 2.770E-08 | 1.623E-08 | 9.100E-09 | 3.900E-08 |
| 3959 | 198 | C198Y | A | 5.9e-08 | 1.8e-08 | 1.7e-08 | 3.133E-08 | 2.397E-08 | 1.700E-08 | 5.900E-08 |
| 3960 | 198 | C198V | A | 6.1e-08 | 5.1e-08 | 2.1e-08 | 4.433E-08 | 2.082E-08 | 2.100E-08 | 6.100E-08 |
| 3961 | 199 | A199A | A | 1.8e-08 | 1.4e-07 | 2.8e-08 | 6.200E-08 | 6.773E-08 | 1.800E-08 | 1.400E-07 |
| 3962 | 199 | A199R | A | 6.7e-08 | 3.4e-08 | 2.6e-08 | 4.233E-08 | 2.173E-08 | 2.600E-08 | 6.700E-08 |
| 3963 | 199 | A199N | A | 2.3e-08 | 3.3e-08 | 2.6e-08 | 2.733E-08 | 5.132E-09 | 2.300E-08 | 3.300E-08 |
| 3964 | 199 | A199D | A | 1.3e-07 | 2.7e-08 | 1.1e-08 | 5.600E-08 | 6.458E-08 | 1.100E-08 | 1.300E-07 |
| 3965 | 199 | A199C | A | 5.4e-08 | 6.2e-08 | 5.6e-08 | 5.733E-08 | 4.163E-09 | 5.400E-08 | 6.200E-08 |
| 3966 | 199 | A199E | A | 5.3e-08 | 3.1e-08 | 1e-07 | 6.133E-08 | 3.525E-08 | 3.100E-08 | 1.000E-07 |
| 3967 | 199 | A199Q | A | 5.4e-08 | 2.8e-08 | 4.5e-08 | 4.233E-08 | 1.320E-08 | 2.800E-08 | 5.400E-08 |
| 3968 | 199 | A199G | A | 1.8e-07 | 2.9e-08 | 3.4e-08 | 8.100E-08 | 8.577E-08 | 2.900E-08 | 1.800E-07 |
| 3969 | 199 | A199H | A | 9.3e-08 | 1.5e-08 | 7.3e-08 | 6.033E-08 | 4.051E-08 | 1.500E-08 | 9.300E-08 |
| 3970 | 199 | A199I | A | 4.5e-08 | 1.2e-08 | 1e-07 | 5.233E-08 | 4.446E-08 | 1.200E-08 | 1.000E-07 |
| 3971 | 199 | A199L | A | 6.3e-08 | 6.3e-08 | 1.7e-08 | 4.767E-08 | 2.656E-08 | 1.700E-08 | 6.300E-08 |
| 3972 | 199 | A199K | A | 3.9e-08 | 3.8e-08 | 3.9e-08 | 3.867E-08 | 5.774E-10 | 3.800E-08 | 3.900E-08 |
| 3973 | 199 | A199M | A | 4.3e-08 | 2.3e-08 | 5.3e-08 | 3.967E-08 | 1.528E-08 | 2.300E-08 | 5.300E-08 |
| 3974 | 199 | A199F | A | 6.8e-08 | 2.5e-08 | 7.8e-08 | 5.700E-08 | 2.816E-08 | 2.500E-08 | 7.800E-08 |
| 3975 | 199 | A199P | A | 3.5e-08 | 4.6e-08 | 4.5e-08 | 4.200E-08 | 6.083E-09 | 3.500E-08 | 4.600E-08 |
| 3976 | 199 | A199S | A | 5e-08 | 7.5e-08 | 1.8e-08 | 4.767E-08 | 2.857E-08 | 1.800E-08 | 7.500E-08 |
| 3977 | 199 | A199T | A | 1.8e-08 | 6.9e-08 | 1.2e-07 | 6.900E-08 | 5.100E-08 | 1.800E-08 | 1.200E-07 |
| 3978 | 199 | A199W | A | 1.1e-07 | 1.7e-08 | 2.1e-08 | 4.933E-08 | 5.258E-08 | 1.700E-08 | 1.100E-07 |
| 3979 | 199 | A199Y | A | 4.9e-08 | 2.2e-07 | 6.7e-08 | 1.120E-07 | 9.396E-08 | 4.900E-08 | 2.200E-07 |
| 3980 | 199 | A199V | A | 4.2e-08 | 1.1e-08 | 1e-07 | 5.100E-08 | 4.518E-08 | 1.100E-08 | 1.000E-07 |
| 3981 | 200 | G200A | A | 3.5e-08 | 5.8e-08 | 1.4e-07 | 7.767E-08 | 5.519E-08 | 3.500E-08 | 1.400E-07 |
| 3982 | 200 | G200R | A | 2.8e-08 | 3.2e-08 | 1.1e-07 | 5.667E-08 | 4.623E-08 | 2.800E-08 | 1.100E-07 |
| 3983 | 200 | G200N | A | 5.2e-08 | 2.6e-07 | 1.7e-08 | 1.097E-07 | 1.314E-07 | 1.700E-08 | 2.600E-07 |
| 3984 | 200 | G200D | A | 2e-08 | 8.4e-08 | 5.3e-08 | 5.233E-08 | 3.201E-08 | 2.000E-08 | 8.400E-08 |
| 3985 | 200 | G200C | A | 5.5e-08 | 7e-08 | 3.4e-08 | 5.300E-08 | 1.808E-08 | 3.400E-08 | 7.000E-08 |
| 3986 | 200 | G200E | A | 1.8e-08 | 5.5e-08 | 2.3e-07 | 1.010E-07 | 1.132E-07 | 1.800E-08 | 2.300E-07 |
| 3987 | 200 | G200Q | A | 2.8e-08 | 2.9e-08 | 4e-08 | 3.233E-08 | 6.658E-09 | 2.800E-08 | 4.000E-08 |
| 3988 | 200 | G200G | A | 3e-08 | 2.9e-08 | 1.4e-08 | 2.433E-08 | 8.963E-09 | 1.400E-08 | 3.000E-08 |
| 3989 | 200 | G200H | A | 5.7e-08 | 1.8e-08 | 8.4e-08 | 5.300E-08 | 3.318E-08 | 1.800E-08 | 8.400E-08 |
| 3990 | 200 | G200I | A | 6.1e-08 | 2.2e-08 | 3.6e-08 | 3.967E-08 | 1.976E-08 | 2.200E-08 | 6.100E-08 |
| 3991 | 200 | G200L | A | 2.3e-08 | 1.5e-08 | 1.7e-07 | 6.933E-08 | 8.727E-08 | 1.500E-08 | 1.700E-07 |
| 3992 | 200 | G200K | A | 2.1e-08 | 5.8e-08 | 2e-08 | 3.300E-08 | 2.166E-08 | 2.000E-08 | 5.800E-08 |
| 3993 | 200 | G200M | A | 4.1e-08 | 1.8e-08 | 2.7e-08 | 2.867E-08 | 1.159E-08 | 1.800E-08 | 4.100E-08 |
| 3994 | 200 | G200F | A | 2.4e-08 | 3.2e-08 | 3e-08 | 2.867E-08 | 4.163E-09 | 2.400E-08 | 3.200E-08 |

| | | | | | | | | | | |
|---|---|---|---|---|---|---|---|---|---|---|
| 3995 | 200 | G200P | A | 2.7e-08 | 6.5e-08 | 4.1e-08 | 4.433E-08 | 1.922E-08 | 2.700E-08 | 6.500E-08 |
| 3996 | 200 | G200S | A | 3.3e-08 | 8.4e-08 | 3.2e-08 | 4.967E-08 | 2.974E-08 | 3.200E-08 | 8.400E-08 |
| 3997 | 200 | G200T | A | 3.7e-08 | 7e-08 | 5.9e-08 | 5.533E-08 | 1.680E-08 | 3.700E-08 | 7.000E-08 |
| 3998 | 200 | G200W | A | 4.2e-08 | 8.9e-08 | 2.8e-08 | 5.300E-08 | 3.195E-08 | 2.800E-08 | 8.900E-08 |
| 3999 | 200 | G200Y | A | 1.3e-07 | 3.7e-08 | 4.3e-08 | 7.000E-08 | 5.205E-08 | 3.700E-08 | 1.300E-07 |
| 4000 | 200 | G200V | A | 3.9e-08 | 7.4e-08 | 8.4e-08 | 6.567E-08 | 2.363E-08 | 3.900E-08 | 8.400E-08 |
| 4001 | 201 | G201A | A | 8.1e-08 | 3.5e-08 | 3.9e-08 | 5.167E-08 | 2.548E-08 | 3.500E-08 | 8.100E-08 |
| 4002 | 201 | G201R | A | 3.2e-08 | 3.4e-08 | 4.8e-08 | 3.800E-08 | 8.718E-09 | 3.200E-08 | 4.800E-08 |
| 4003 | 201 | G201N | A | 2.5e-08 | 2e-08 | 5.8e-08 | 3.433E-08 | 2.065E-08 | 2.000E-08 | 5.800E-08 |
| 4004 | 201 | G201D | A | 8.1e-08 | 3.8e-08 | 2.3e-08 | 4.733E-08 | 3.011E-08 | 2.300E-08 | 8.100E-08 |
| 4005 | 201 | G201C | A | 6.1e-08 | 7.4e-08 | 6.8e-08 | 6.767E-08 | 6.506E-09 | 6.100E-08 | 7.400E-08 |
| 4006 | 201 | G201E | A | 5.6e-08 | 3.3e-08 | 7.8e-08 | 5.567E-08 | 2.250E-08 | 3.300E-08 | 7.800E-08 |
| 4007 | 201 | G201Q | A | 2.8e-08 | 6.7e-08 | 6.8e-08 | 5.433E-08 | 2.281E-08 | 2.800E-08 | 6.800E-08 |
| 4008 | 201 | G201G | A | 2.9e-08 | 1.4e-07 | 3.1e-08 | 6.667E-08 | 6.352E-08 | 2.900E-08 | 1.400E-07 |
| 4009 | 201 | G201H | A | 5.9e-08 | 6.7e-08 | 4.2e-08 | 5.600E-08 | 1.277E-08 | 4.200E-08 | 6.700E-08 |
| 4010 | 201 | G201I | A | 5e-08 | 2.2e-07 | 3.5e-08 | 1.017E-07 | 1.028E-07 | 3.500E-08 | 2.200E-07 |
| 4011 | 201 | G201L | A | 3.1e-08 | 8.2e-08 | 4.5e-08 | 5.267E-08 | 2.635E-08 | 3.100E-08 | 8.200E-08 |
| 4012 | 201 | G201K | A | 1.1e-07 | 6e-08 | 5.4e-08 | 7.467E-08 | 3.075E-08 | 5.400E-08 | 1.100E-07 |
| 4013 | 201 | G201M | A | 1.6e-08 | 1.4e-08 | 5.1e-08 | 2.700E-08 | 2.081E-08 | 1.400E-08 | 5.100E-08 |
| 4014 | 201 | G201F | A | 6e-08 | 8.9e-08 | 6.7e-08 | 7.200E-08 | 1.513E-08 | 6.000E-08 | 8.900E-08 |
| 4015 | 201 | G201P | A | 5.6e-08 | 2.3e-08 | 3e-08 | 3.633E-08 | 1.739E-08 | 2.300E-08 | 5.600E-08 |
| 4016 | 201 | G201S | A | 6.2e-08 | 9.6e-08 | 2e-08 | 5.933E-08 | 3.807E-08 | 2.000E-08 | 9.600E-08 |
| 4017 | 201 | G201T | A | 2.2e-08 | 9.2e-08 | 6.3e-08 | 5.900E-08 | 3.517E-08 | 2.200E-08 | 9.200E-08 |
| 4018 | 201 | G201W | A | 2.7e-08 | 4.8e-08 | 2e-08 | 3.167E-08 | 1.457E-08 | 2.000E-08 | 4.800E-08 |
| 4019 | 201 | G201Y | A | 4.6e-08 | 1.6e-08 | 9.2e-08 | 5.133E-08 | 3.828E-08 | 1.600E-08 | 9.200E-08 |
| 4020 | 201 | G201V | A | 5.2e-08 | 3.1e-08 | 4e-08 | 4.100E-08 | 1.054E-08 | 3.100E-08 | 5.200E-08 |
| 4021 | 202 | C202A | A | 7.3e-08 | 5.3e-08 | 5.4e-08 | 6.000E-08 | 1.127E-08 | 5.300E-08 | 7.300E-08 |
| 4022 | 202 | C202R | A | 5e-08 | 3.8e-08 | 5.5e-08 | 4.767E-08 | 8.737E-09 | 3.800E-08 | 5.500E-08 |
| 4023 | 202 | C202N | A | 4.4e-08 | 9.7e-08 | 5e-08 | 6.367E-08 | 2.902E-08 | 4.400E-08 | 9.700E-08 |
| 4024 | 202 | C202D | A | 4.3e-08 | 4.9e-08 | 1.3e-07 | 7.400E-08 | 4.859E-08 | 4.300E-08 | 1.300E-07 |
| 4025 | 202 | C202C | A | 6.8e-08 | 2.5e-08 | 5.3e-08 | 4.867E-08 | 2.183E-08 | 2.500E-08 | 6.800E-08 |
| 4026 | 202 | C202E | A | 6.9e-08 | 3.3e-08 | 1e-07 | 6.733E-08 | 3.353E-08 | 3.300E-08 | 1.000E-07 |
| 4027 | 202 | C202Q | A | 1.3e-07 | 8.4e-08 | 1.6e-08 | 7.667E-08 | 5.735E-08 | 1.600E-08 | 1.300E-07 |
| 4028 | 202 | C202G | A | 4.5e-08 | 1.3e-08 | 1.1e-07 | 5.600E-08 | 4.943E-08 | 1.300E-08 | 1.100E-07 |
| 4029 | 202 | C202H | A | 3.7e-08 | 3.1e-08 | 4.7e-08 | 3.833E-08 | 8.083E-09 | 3.100E-08 | 4.700E-08 |
| 4030 | 202 | C202I | A | 5e-08 | 2.7e-08 | 6.3e-08 | 4.667E-08 | 1.823E-08 | 2.700E-08 | 6.300E-08 |
| 4031 | 202 | C202L | A | 4.3e-08 | 3.1e-08 | 1.9e-08 | 3.100E-08 | 1.200E-08 | 1.900E-08 | 4.300E-08 |
| 4032 | 202 | C202K | A | 5.8e-08 | 8.8e-08 | 1.5e-08 | 5.367E-08 | 3.669E-08 | 1.500E-08 | 8.800E-08 |
| 4033 | 202 | C202M | A | 8e-08 | 1.8e-08 | 4.5e-08 | 4.767E-08 | 3.109E-08 | 1.800E-08 | 8.000E-08 |
| 4034 | 202 | C202F | A | 2.5e-08 | 3.9e-08 | 3.3e-08 | 3.233E-08 | 7.024E-09 | 2.500E-08 | 3.900E-08 |
| 4035 | 202 | C202P | A | 1.9e-07 | 7.6e-08 | 2.9e-08 | 9.833E-08 | 8.279E-08 | 2.900E-08 | 1.900E-07 |
| 4036 | 202 | C202S | A | 9.7e-08 | 1.3e-07 | 9.6e-08 | 1.077E-07 | 1.935E-08 | 9.600E-08 | 1.300E-07 |
| 4037 | 202 | C202T | A | 6.9e-08 | 3.3e-08 | 2.7e-08 | 4.300E-08 | 2.272E-08 | 2.700E-08 | 6.900E-08 |
| 4038 | 202 | C202W | A | 3.5e-08 | 1.9e-08 | 2.9e-08 | 2.767E-08 | 8.083E-09 | 1.900E-08 | 3.500E-08 |
| 4039 | 202 | C202Y | A | 4.7e-08 | 1.3e-07 | 3.2e-08 | 6.967E-08 | 5.279E-08 | 3.200E-08 | 1.300E-07 |
| 4040 | 202 | C202V | A | 7.4e-08 | 9.7e-09 | 6.7e-08 | 5.023E-08 | 3.528E-08 | 9.700E-09 | 7.400E-08 |
| 4041 | 203 | A203A | A | 3.6e-08 | 1.1e-07 | 4.2e-08 | 6.267E-08 | 4.110E-08 | 3.600E-08 | 1.100E-07 |
| 4042 | 203 | A203R | A | 2.9e-08 | 8.2e-08 | 3.3e-08 | 4.800E-08 | 2.951E-08 | 2.900E-08 | 8.200E-08 |
| 4043 | 203 | A203N | A | 5.1e-08 | 5.3e-08 | 8.3e-08 | 6.233E-08 | 1.793E-08 | 5.100E-08 | 8.300E-08 |
| 4044 | 203 | A203D | A | 1.2e-07 | 3.2e-08 | 3.9e-08 | 6.367E-08 | 4.891E-08 | 3.200E-08 | 1.200E-07 |
| 4045 | 203 | A203C | A | 2.6e-08 | 9.3e-08 | 1.1e-07 | 7.633E-08 | 4.441E-08 | 2.600E-08 | 1.100E-07 |
| 4046 | 203 | A203E | A | 3.6e-08 | 4e-08 | 2.8e-08 | 3.467E-08 | 6.110E-09 | 2.800E-08 | 4.000E-08 |
| 4047 | 203 | A203Q | A | 2.3e-07 | 6.9e-08 | 4.4e-08 | 1.143E-07 | 1.009E-07 | 4.400E-08 | 2.300E-07 |
| 4048 | 203 | A203G | A | 8e-08 | 3e-08 | 2e-08 | 4.333E-08 | 3.215E-08 | 2.000E-08 | 8.000E-08 |
| 4049 | 203 | A203H | A | 4.4e-08 | 1.5e-07 | 4.1e-08 | 7.833E-08 | 6.208E-08 | 4.100E-08 | 1.500E-07 |
| 4050 | 203 | A203I | A | 4.8e-08 | 3.6e-08 | 1.7e-08 | 3.367E-08 | 1.563E-08 | 1.700E-08 | 4.800E-08 |
| 4051 | 203 | A203L | A | 1.3e-07 | 6.8e-08 | 4.3e-08 | 8.033E-08 | 4.479E-08 | 4.300E-08 | 1.300E-07 |
| 4052 | 203 | A203K | A | 3.1e-08 | 5.6e-08 | 2.3e-07 | 1.057E-07 | 1.084E-07 | 3.100E-08 | 2.300E-07 |
| 4053 | 203 | A203M | A | 6e-08 | 5.7e-08 | 8.6e-08 | 6.767E-08 | 1.595E-08 | 5.700E-08 | 8.600E-08 |
| 4054 | 203 | A203F | A | 3.1e-08 | 2.6e-08 | 3.2e-08 | 2.967E-08 | 3.215E-09 | 2.600E-08 | 3.200E-08 |
| 4055 | 203 | A203P | A | 8.4e-08 | 2e-08 | 1.7e-07 | 9.133E-08 | 7.527E-08 | 2.000E-08 | 1.700E-07 |
| 4056 | 203 | A203S | A | 3.2e-08 | 4.8e-08 | 2.3e-08 | 3.433E-08 | 1.266E-08 | 2.300E-08 | 4.800E-08 |
| 4057 | 203 | A203T | A | 2.9e-08 | 7.2e-08 | 4e-08 | 4.700E-08 | 2.234E-08 | 2.900E-08 | 7.200E-08 |
| 4058 | 203 | A203W | A | 7.8e-08 | 6.9e-08 | 3.1e-08 | 5.933E-08 | 2.495E-08 | 3.100E-08 | 7.800E-08 |
| 4059 | 203 | A203Y | A | 3.3e-08 | 4.1e-08 | 1.1e-08 | 2.833E-08 | 1.553E-08 | 1.100E-08 | 4.100E-08 |
| 4060 | 203 | A203V | A | 6.5e-08 | 6.4e-08 | 2e-08 | 4.967E-08 | 2.570E-08 | 2.000E-08 | 6.500E-08 |
| 4061 | 204 | R204A | A | 2.9e-08 | 1.2e-07 | 4.9e-08 | 6.600E-08 | 4.782E-08 | 2.900E-08 | 1.200E-07 |
| 4062 | 204 | R204R | A | 3.2e-08 | 5.2e-08 | 3.3e-08 | 3.900E-08 | 1.127E-08 | 3.200E-08 | 5.200E-08 |
| 4063 | 204 | R204N | A | 5.2e-08 | 4.4e-08 | 1.5e-08 | 3.700E-08 | 1.947E-08 | 1.500E-08 | 5.200E-08 |
| 4064 | 204 | R204D | A | 4.4e-08 | 6.8e-08 | 1.9e-08 | 4.367E-08 | 2.450E-08 | 1.900E-08 | 6.800E-08 |
| 4065 | 204 | R204C | A | 1.8e-08 | 2e-08 | 8.2e-08 | 4.000E-08 | 3.639E-08 | 1.800E-08 | 8.200E-08 |
| 4066 | 204 | R204E | A | 7.3e-08 | 1.2e-08 | 4.1e-08 | 4.200E-08 | 3.051E-08 | 1.200E-08 | 7.300E-08 |
| 4067 | 204 | R204Q | A | 8e-08 | 4e-08 | 3e-08 | 5.000E-08 | 2.646E-08 | 3.000E-08 | 8.000E-08 |
| 4068 | 204 | R204G | A | 4.2e-08 | 7.2e-08 | 5.2e-08 | 5.533E-08 | 1.528E-08 | 4.200E-08 | 7.200E-08 |
| 4069 | 204 | R204H | A | 5.2e-08 | 1.8e-08 | 1.1e-07 | 6.000E-08 | 4.652E-08 | 1.800E-08 | 1.100E-07 |
| 4070 | 204 | R204I | A | 3e-08 | 3.2e-08 | 2.6e-08 | 2.933E-08 | 3.055E-09 | 2.600E-08 | 3.200E-08 |
| 4071 | 204 | R204L | A | 9.7e-08 | 5.1e-08 | 1.4e-08 | 5.400E-08 | 4.158E-08 | 1.400E-08 | 9.700E-08 |
| 4072 | 204 | R204K | A | 4.1e-08 | 3.5e-08 | 8.4e-08 | 5.333E-08 | 2.673E-08 | 3.500E-08 | 8.400E-08 |
| 4073 | 204 | R204M | A | 9.1e-08 | 2.9e-08 | 2.6e-08 | 4.867E-08 | 3.669E-08 | 2.600E-08 | 9.100E-08 |
| 4074 | 204 | R204F | A | 4.7e-08 | 5e-08 | 7.1e-08 | 5.600E-08 | 1.308E-08 | 4.700E-08 | 7.100E-08 |
| 4075 | 204 | R204P | A | 2.9e-08 | 5.8e-08 | 5.5e-08 | 4.733E-08 | 1.595E-08 | 2.900E-08 | 5.800E-08 |
| 4076 | 204 | R204S | A | 8.8e-08 | 8.6e-08 | 2.8e-08 | 6.733E-08 | 3.408E-08 | 2.800E-08 | 8.800E-08 |
| 4077 | 204 | R204T | A | 4e-08 | 2.1e-08 | 4.7e-08 | 3.600E-08 | 1.345E-08 | 2.100E-08 | 4.700E-08 |
| 4078 | 204 | R204W | A | 3.7e-08 | 2.5e-08 | 1.3e-07 | 6.400E-08 | 5.747E-08 | 2.500E-08 | 1.300E-07 |
| 4079 | 204 | R204Y | A | 4.5e-08 | 6.5e-08 | 2.9e-08 | 4.633E-08 | 1.804E-08 | 2.900E-08 | 6.500E-08 |

| | | | | | | | | | | |
|---|---|---|---|---|---|---|---|---|---|---|
| 4080 | 204 | R204V | A | 9.3e-08 | 1.8e-07 | 5.2e-08 | 1.083E-07 | 6.536E-08 | 5.200E-08 | 1.800E-07 |
| 4081 | 205 | C205A | A | 6.5e-08 | 2.9e-08 | 3e-08 | 4.133E-08 | 2.050E-08 | 2.900E-08 | 6.500E-08 |
| 4082 | 205 | C205R | A | 3.3e-08 | 6.5e-08 | 3.3e-08 | 4.367E-08 | 1.848E-08 | 3.300E-08 | 6.500E-08 |
| 4083 | 205 | C205N | A | 2.8e-08 | 3.7e-08 | 2.9e-08 | 3.133E-08 | 4.933E-09 | 2.800E-08 | 3.700E-08 |
| 4084 | 205 | C205D | A | 1.1e-08 | 1.3e-07 | 7.1e-08 | 7.067E-08 | 5.950E-08 | 1.100E-08 | 1.300E-07 |
| 4085 | 205 | C205C | A | 8.3e-08 | 6.1e-08 | 3.2e-08 | 5.867E-08 | 2.558E-08 | 3.200E-08 | 8.300E-08 |
| 4086 | 205 | C205E | A | 3.9e-08 | 1.6e-07 | 3.1e-08 | 7.667E-08 | 7.228E-08 | 3.100E-08 | 1.600E-07 |
| 4087 | 205 | C205Q | A | 4e-08 | 9.2e-08 | 1.1e-07 | 8.067E-08 | 3.635E-08 | 4.000E-08 | 1.100E-07 |
| 4088 | 205 | C205G | A | 4.2e-08 | 4.4e-08 | 8.6e-08 | 5.733E-08 | 2.485E-08 | 4.200E-08 | 8.600E-08 |
| 4089 | 205 | C205H | A | 6.7e-08 | 5.3e-08 | 5e-08 | 5.667E-08 | 9.074E-09 | 5.000E-08 | 6.700E-08 |
| 4090 | 205 | C205I | A | 1.8e-08 | 1.1e-07 | 3.1e-08 | 5.300E-08 | 4.979E-08 | 1.800E-08 | 1.100E-07 |
| 4091 | 205 | C205L | A | 4.3e-08 | 2.5e-08 | 3.1e-08 | 3.300E-08 | 9.165E-09 | 2.500E-08 | 4.300E-08 |
| 4092 | 205 | C205K | A | 1.2e-07 | 2.1e-08 | 2.2e-08 | 5.433E-08 | 5.687E-08 | 2.100E-08 | 1.200E-07 |
| 4093 | 205 | C205M | A | 1.4e-07 | 6.3e-08 | 3e-08 | 7.767E-08 | 5.645E-08 | 3.000E-08 | 1.400E-07 |
| 4094 | 205 | C205F | A | 3.1e-08 | 2.9e-08 | 3.3e-08 | 3.100E-08 | 2.000E-09 | 2.900E-08 | 3.300E-08 |
| 4095 | 205 | C205P | A | 4.4e-08 | 5.5e-08 | 4.9e-08 | 4.933E-08 | 5.508E-09 | 4.400E-08 | 5.500E-08 |
| 4096 | 205 | C205S | A | 3.2e-08 | 4.2e-08 | 8.2e-08 | 5.200E-08 | 2.646E-08 | 3.200E-08 | 8.200E-08 |
| 4097 | 205 | C205T | A | 6.9e-08 | 4e-08 | 4.4e-08 | 5.100E-08 | 1.572E-08 | 4.000E-08 | 6.900E-08 |
| 4098 | 205 | C205W | A | 4.6e-08 | 2.6e-08 | 4.1e-08 | 3.767E-08 | 1.041E-08 | 2.600E-08 | 4.600E-08 |
| 4099 | 205 | C205Y | A | 2.1e-08 | 3.5e-08 | 7.4e-08 | 4.333E-08 | 2.747E-08 | 2.100E-08 | 7.400E-08 |
| 4100 | 205 | C205V | A | 6.4e-08 | 5.8e-08 | 2.1e-08 | 4.767E-08 | 2.329E-08 | 2.100E-08 | 6.400E-08 |
| 4101 | 206 | K206A | A | 4.3e-08 | 2e-08 | 7.7e-08 | 4.667E-08 | 2.868E-08 | 2.000E-08 | 7.700E-08 |
| 4102 | 206 | K206R | A | 1.9e-08 | 2.5e-08 | 4.5e-08 | 2.967E-08 | 1.361E-08 | 1.900E-08 | 4.500E-08 |
| 4103 | 206 | K206N | A | 7.8e-08 | 2.1e-08 | 3.1e-08 | 4.333E-08 | 3.044E-08 | 2.100E-08 | 7.800E-08 |
| 4104 | 206 | K206D | A | 3.7e-08 | 4e-08 | 4.2e-08 | 3.967E-08 | 2.517E-09 | 3.700E-08 | 4.200E-08 |
| 4105 | 206 | K206C | A | 4.1e-08 | 1.7e-08 | 4e-08 | 3.267E-08 | 1.358E-08 | 1.700E-08 | 4.100E-08 |
| 4106 | 206 | K206E | A | 5.2e-08 | 7.1e-08 | 1e-07 | 7.433E-08 | 2.417E-08 | 5.200E-08 | 1.000E-07 |
| 4107 | 206 | K206Q | A | 4.4e-08 | 5.5e-08 | 3.4e-08 | 4.433E-08 | 1.050E-08 | 3.400E-08 | 5.500E-08 |
| 4108 | 206 | K206G | A | 5.7e-08 | 6.9e-08 | 5e-08 | 5.867E-08 | 9.609E-09 | 5.000E-08 | 6.900E-08 |
| 4109 | 206 | K206H | A | 9.3e-09 | 3.9e-08 | 5.3e-08 | 3.377E-08 | 2.232E-08 | 9.300E-09 | 5.300E-08 |
| 4110 | 206 | K206I | A | 6.1e-08 | 7e-08 | 3.7e-08 | 5.600E-08 | 1.706E-08 | 3.700E-08 | 7.000E-08 |
| 4111 | 206 | K206L | A | 9.5e-08 | 3.8e-08 | 5.3e-08 | 6.200E-08 | 2.955E-08 | 3.800E-08 | 9.500E-08 |
| 4112 | 206 | K206K | A | 5.3e-08 | 5.1e-08 | 2.3e-08 | 4.233E-08 | 1.677E-08 | 2.300E-08 | 5.300E-08 |
| 4113 | 206 | K206M | A | 3.9e-08 | 3.1e-08 | 1.1e-07 | 6.000E-08 | 4.349E-08 | 3.100E-08 | 1.100E-07 |
| 4114 | 206 | K206F | A | 1.8e-08 | 6.9e-08 | 6.2e-08 | 4.967E-08 | 2.765E-08 | 1.800E-08 | 6.900E-08 |
| 4115 | 206 | K206P | A | 5.8e-08 | 9.7e-08 | 6.3e-08 | 7.267E-08 | 2.122E-08 | 5.800E-08 | 9.700E-08 |
| 4116 | 206 | K206S | A | 2.5e-08 | 1.3e-08 | 4.8e-08 | 2.867E-08 | 1.779E-08 | 1.300E-08 | 4.800E-08 |
| 4117 | 206 | K206T | A | 1.2e-07 | 4.7e-08 | 5.6e-08 | 7.433E-08 | 3.980E-08 | 4.700E-08 | 1.200E-07 |
| 4118 | 206 | K206W | A | 1.4e-08 | 4e-08 | 4.2e-08 | 3.200E-08 | 1.562E-08 | 1.400E-08 | 4.200E-08 |
| 4119 | 206 | K206Y | A | 4.6e-08 | 5.8e-08 | 2.7e-08 | 4.367E-08 | 1.563E-08 | 2.700E-08 | 5.800E-08 |
| 4120 | 206 | K206V | A | 4.7e-08 | 1.9e-08 | 2.3e-08 | 2.967E-08 | 1.514E-08 | 1.900E-08 | 4.700E-08 |
| 4121 | 207 | G207A | A | 6.9e-08 | 1.2e-08 | 2.9e-08 | 3.667E-08 | 2.926E-08 | 1.200E-08 | 6.900E-08 |
| 4122 | 207 | G207R | A | 3.8e-08 | 2e-08 | 3e-08 | 2.933E-08 | 9.018E-09 | 2.000E-08 | 3.800E-08 |
| 4123 | 207 | G207N | A | 4.4e-08 | 2.8e-08 | 3.3e-08 | 3.500E-08 | 8.185E-09 | 2.800E-08 | 4.400E-08 |
| 4124 | 207 | G207D | A | 8.5e-08 | 3.1e-08 | 3.9e-08 | 5.167E-08 | 2.914E-08 | 3.100E-08 | 8.500E-08 |
| 4125 | 207 | G207C | A | 2.4e-08 | 6.6e-08 | 4.4e-08 | 4.467E-08 | 2.101E-08 | 2.400E-08 | 6.600E-08 |
| 4126 | 207 | G207E | A | 3e-08 | 1.9e-08 | 3.5e-08 | 2.800E-08 | 8.185E-09 | 1.900E-08 | 3.500E-08 |
| 4127 | 207 | G207Q | A | 3.2e-08 | 4.9e-08 | 6.5e-08 | 4.867E-08 | 1.650E-08 | 3.200E-08 | 6.500E-08 |
| 4128 | 207 | G207G | A | 1.9e-08 | 3.3e-08 | 6.1e-08 | 3.767E-08 | 2.139E-08 | 1.900E-08 | 6.100E-08 |
| 4129 | 207 | G207H | A | 7.2e-08 | 2.7e-08 | 3.7e-08 | 4.533E-08 | 2.363E-08 | 2.700E-08 | 7.200E-08 |
| 4130 | 207 | G207I | A | 3.6e-08 | 1e-07 | 3.8e-08 | 5.800E-08 | 3.639E-08 | 3.600E-08 | 1.000E-07 |
| 4131 | 207 | G207L | A | 9.6e-08 | 3.3e-08 | 1.2e-08 | 4.700E-08 | 4.371E-08 | 1.200E-08 | 9.600E-08 |
| 4132 | 207 | G207K | A | 4.5e-08 | 2.9e-08 | 9.6e-08 | 5.667E-08 | 3.499E-08 | 2.900E-08 | 9.600E-08 |
| 4133 | 207 | G207M | A | 3.8e-08 | 2.6e-08 | 3.9e-08 | 3.433E-08 | 7.234E-09 | 2.600E-08 | 3.900E-08 |
| 4134 | 207 | G207F | A | 9.1e-09 | 3.1e-08 | 7.1e-08 | 3.703E-08 | 3.139E-08 | 9.100E-09 | 7.100E-08 |
| 4135 | 207 | G207P | A | 5.5e-08 | 2.6e-08 | 5.3e-08 | 4.467E-08 | 1.620E-08 | 2.600E-08 | 5.500E-08 |
| 4136 | 207 | G207S | A | 5.3e-08 | 2.2e-08 | 2.7e-08 | 3.400E-08 | 1.664E-08 | 2.200E-08 | 5.300E-08 |
| 4137 | 207 | G207T | A | 8.9e-08 | 6.3e-08 | 4.3e-08 | 6.500E-08 | 2.307E-08 | 4.300E-08 | 8.900E-08 |
| 4138 | 207 | G207W | A | 3.6e-08 | 4.5e-08 | 2.4e-08 | 3.500E-08 | 1.054E-08 | 2.400E-08 | 4.500E-08 |
| 4139 | 207 | G207Y | A | 3.8e-08 | 7.7e-08 | 5.1e-08 | 5.533E-08 | 1.986E-08 | 3.800E-08 | 7.700E-08 |
| 4140 | 207 | G207V | A | 1.4e-08 | 8.7e-08 | 5.7e-08 | 5.267E-08 | 3.669E-08 | 1.400E-08 | 8.700E-08 |
| 4141 | 208 | P208A | A | 9.2e-08 | 6.7e-08 | 2.9e-08 | 6.267E-08 | 3.172E-08 | 2.900E-08 | 9.200E-08 |
| 4142 | 208 | P208R | A | 4e-08 | 9.4e-08 | 3.4e-08 | 5.600E-08 | 3.305E-08 | 3.400E-08 | 9.400E-08 |
| 4143 | 208 | P208N | A | 3.3e-08 | 2.6e-08 | 1.5e-08 | 2.467E-08 | 9.074E-09 | 1.500E-08 | 3.300E-08 |
| 4144 | 208 | P208D | A | 2e-07 | 5.2e-08 | 4.2e-08 | 9.800E-08 | 8.848E-08 | 4.200E-08 | 2.000E-07 |
| 4145 | 208 | P208C | A | 7.9e-08 | 6.1e-08 | 2.4e-08 | 5.467E-08 | 2.804E-08 | 2.400E-08 | 7.900E-08 |
| 4146 | 208 | P208E | A | 3.5e-08 | 3e-08 | 2.5e-08 | 3.000E-08 | 5.000E-09 | 2.500E-08 | 3.500E-08 |
| 4147 | 208 | P208Q | A | 1.4e-07 | 6e-08 | 3.9e-08 | 7.967E-08 | 5.329E-08 | 3.900E-08 | 1.400E-07 |
| 4148 | 208 | P208G | A | 4.4e-08 | 4.1e-08 | 9.4e-08 | 5.967E-08 | 2.977E-08 | 4.100E-08 | 9.400E-08 |
| 4149 | 208 | P208H | A | 3.1e-08 | 5.8e-08 | 1.9e-08 | 3.600E-08 | 1.997E-08 | 1.900E-08 | 5.800E-08 |
| 4150 | 208 | P208I | A | 3.5e-08 | 4.3e-08 | 6.8e-08 | 4.867E-08 | 1.721E-08 | 3.500E-08 | 6.800E-08 |
| 4151 | 208 | P208L | A | 4.7e-08 | 3.9e-08 | 7.2e-08 | 5.267E-08 | 1.721E-08 | 3.900E-08 | 7.200E-08 |
| 4152 | 208 | P208K | A | 2.6e-08 | 4.5e-08 | 3.4e-08 | 3.500E-08 | 9.539E-09 | 2.600E-08 | 4.500E-08 |
| 4153 | 208 | P208M | A | 4e-08 | 2.4e-08 | 3.1e-08 | 3.167E-08 | 8.021E-09 | 2.400E-08 | 4.000E-08 |
| 4154 | 208 | P208F | A | 3.1e-08 | 1.1e-07 | 6.2e-08 | 6.767E-08 | 3.980E-08 | 3.100E-08 | 1.100E-07 |
| 4155 | 208 | P208P | A | 5.3e-08 | 1.9e-08 | 3.7e-08 | 3.633E-08 | 1.701E-08 | 1.900E-08 | 5.300E-08 |
| 4156 | 208 | P208S | A | 2.7e-08 | 5.5e-08 | 4.1e-08 | 4.100E-08 | 1.400E-08 | 2.700E-08 | 5.500E-08 |
| 4157 | 208 | P208T | A | 3.2e-08 | 2.7e-08 | 7.9e-08 | 4.600E-08 | 2.869E-08 | 2.700E-08 | 7.900E-08 |
| 4158 | 208 | P208W | A | 5.5e-08 | 4.7e-08 | 5.1e-08 | 5.100E-08 | 4.000E-09 | 4.700E-08 | 5.500E-08 |
| 4159 | 208 | P208Y | A | 6.1e-08 | 3.3e-08 | 6e-08 | 5.133E-08 | 1.589E-08 | 3.300E-08 | 6.100E-08 |
| 4160 | 208 | P208V | A | 3.8e-08 | 4.6e-08 | 3.6e-08 | 4.000E-08 | 5.292E-09 | 3.600E-08 | 4.600E-08 |
| 4161 | 209 | L209A | A | 7.6e-08 | 1e-07 | 3.7e-08 | 7.100E-08 | 3.180E-08 | 3.700E-08 | 1.000E-07 |
| 4162 | 209 | L209R | A | 3.2e-09 | 5.1e-08 | 7.4e-08 | 4.273E-08 | 3.612E-08 | 3.200E-09 | 7.400E-08 |
| 4163 | 209 | L209N | A | 1.1e-07 | 4.5e-08 | 8e-08 | 7.833E-08 | 3.253E-08 | 4.500E-08 | 1.100E-07 |
| 4164 | 209 | L209D | A | 3e-08 | 2.8e-08 | 1.2e-08 | 2.333E-08 | 9.866E-09 | 1.200E-08 | 3.000E-08 |

| | | | | | | | | | | |
|---|---|---|---|---|---|---|---|---|---|---|
| 4165 | 209 | L209C | A | 2.7e-08 | 3.1e-08 | 9.9e-08 | 5.233E-08 | 4.046E-08 | 2.700E-08 | 9.900E-08 |
| 4166 | 209 | L209E | A | 2.7e-08 | 7.5e-08 | 3.4e-08 | 4.533E-08 | 2.593E-08 | 2.700E-08 | 7.500E-08 |
| 4167 | 209 | L209Q | A | 3.6e-08 | 2.1e-08 | 4.5e-08 | 3.400E-08 | 1.212E-08 | 2.100E-08 | 4.500E-08 |
| 4168 | 209 | L209G | A | 1.8e-08 | 4.8e-08 | 4.4e-08 | 3.667E-08 | 1.629E-08 | 1.800E-08 | 4.800E-08 |
| 4169 | 209 | L209H | A | 1.9e-08 | 4.3e-08 | 3.6e-08 | 3.267E-08 | 1.234E-08 | 1.900E-08 | 4.300E-08 |
| 4170 | 209 | L209I | A | 3e-08 | 2.1e-08 | 5.9e-08 | 3.667E-08 | 1.986E-08 | 2.100E-08 | 5.900E-08 |
| 4171 | 209 | L209L | A | 3.8e-08 | 2.4e-08 | 3e-08 | 3.067E-08 | 7.024E-09 | 2.400E-08 | 3.800E-08 |
| 4172 | 209 | L209K | A | 5.3e-08 | 3.6e-08 | 2.7e-08 | 3.867E-08 | 1.320E-08 | 2.700E-08 | 5.300E-08 |
| 4173 | 209 | L209M | A | 4e-08 | 1.3e-07 | 2.8e-08 | 6.600E-08 | 5.575E-08 | 2.800E-08 | 1.300E-07 |
| 4174 | 209 | L209F | A | 5.2e-08 | 1.6e-07 | 2.3e-08 | 7.833E-08 | 7.220E-08 | 2.300E-08 | 1.600E-07 |
| 4175 | 209 | L209P | A | 2.1e-08 | 3e-08 | 1.2e-07 | 5.700E-08 | 5.474E-08 | 2.100E-08 | 1.200E-07 |
| 4176 | 209 | L209S | A | 4.5e-08 | 8.2e-08 | 3.4e-08 | 5.367E-08 | 2.515E-08 | 3.400E-08 | 8.200E-08 |
| 4177 | 209 | L209T | A | 4.6e-08 | 9e-08 | 5.7e-08 | 6.433E-08 | 2.290E-08 | 4.600E-08 | 9.000E-08 |
| 4178 | 209 | L209W | A | 5.8e-08 | 4.4e-08 | 1.1e-07 | 7.067E-08 | 3.478E-08 | 4.400E-08 | 1.100E-07 |
| 4179 | 209 | L209Y | A | 4.5e-08 | 4.1e-08 | 4e-08 | 4.200E-08 | 2.646E-09 | 4.000E-08 | 4.500E-08 |
| 4180 | 209 | L209V | A | 2.4e-08 | 7.7e-08 | 2.5e-08 | 4.200E-08 | 3.032E-08 | 2.400E-08 | 7.700E-08 |
| 4181 | 210 | P210A | A | 4.5e-08 | 6.9e-08 | 2.3e-08 | 4.567E-08 | 2.301E-08 | 2.300E-08 | 6.900E-08 |
| 4182 | 210 | P210R | A | 2.7e-08 | 2e-08 | 3.8e-08 | 2.833E-08 | 9.074E-09 | 2.000E-08 | 3.800E-08 |
| 4183 | 210 | P210N | A | 2.1e-08 | 7.9e-08 | 1e-08 | 3.667E-08 | 3.707E-08 | 1.000E-08 | 7.900E-08 |
| 4184 | 210 | P210D | A | 5.4e-08 | 1.3e-07 | 5.7e-08 | 8.033E-08 | 4.304E-08 | 5.400E-08 | 1.300E-07 |
| 4185 | 210 | P210C | A | 1.2e-07 | 3.2e-08 | 6.6e-08 | 7.267E-08 | 4.438E-08 | 3.200E-08 | 1.200E-07 |
| 4186 | 210 | P210E | A | 5e-08 | 5.2e-08 | 3.5e-08 | 4.567E-08 | 9.292E-09 | 3.500E-08 | 5.200E-08 |
| 4187 | 210 | P210Q | A | 4.2e-08 | 2.8e-08 | 1.6e-07 | 7.667E-08 | 7.251E-08 | 2.800E-08 | 1.600E-07 |
| 4188 | 210 | P210G | A | 1e-07 | 2.7e-08 | 3.5e-08 | 5.400E-08 | 4.004E-08 | 2.700E-08 | 1.000E-07 |
| 4189 | 210 | P210H | A | 3.4e-08 | 1.1e-08 | 2.7e-08 | 2.400E-08 | 1.179E-08 | 1.100E-08 | 3.400E-08 |
| 4190 | 210 | P210I | A | 2.4e-08 | 5.2e-08 | 1.4e-07 | 7.200E-08 | 6.053E-08 | 2.400E-08 | 1.400E-07 |
| 4191 | 210 | P210L | A | 3.3e-08 | 4.7e-08 | 3.9e-08 | 3.967E-08 | 7.024E-09 | 3.300E-08 | 4.700E-08 |
| 4192 | 210 | P210K | A | 3.9e-08 | 2.6e-08 | 4e-08 | 3.500E-08 | 7.810E-09 | 2.600E-08 | 4.000E-08 |
| 4193 | 210 | P210M | A | 9.9e-09 | 3.9e-08 | 1.6e-08 | 2.163E-08 | 1.535E-08 | 9.900E-09 | 3.900E-08 |
| 4194 | 210 | P210F | A | 5.8e-08 | 3.6e-08 | 1.9e-08 | 3.767E-08 | 1.955E-08 | 1.900E-08 | 5.800E-08 |
| 4195 | 210 | P210P | A | 7.5e-08 | 1.3e-08 | 2.5e-08 | 3.767E-08 | 3.288E-08 | 1.300E-08 | 7.500E-08 |
| 4196 | 210 | P210S | A | 6.9e-08 | 2.3e-08 | 3.4e-08 | 4.200E-08 | 2.402E-08 | 2.300E-08 | 6.900E-08 |
| 4197 | 210 | P210T | A | 6.9e-08 | 1.1e-08 | 1.7e-07 | 8.333E-08 | 8.046E-08 | 1.100E-08 | 1.700E-07 |
| 4198 | 210 | P210W | A | 3.4e-08 | 5.7e-08 | 5e-08 | 4.700E-08 | 1.179E-08 | 3.400E-08 | 5.700E-08 |
| 4199 | 210 | P210Y | A | 1.4e-08 | 2.6e-08 | 3.8e-08 | 2.600E-08 | 1.200E-08 | 1.400E-08 | 3.800E-08 |
| 4200 | 210 | P210V | A | 2.5e-08 | 1.1e-07 | 9.7e-08 | 7.733E-08 | 4.579E-08 | 2.500E-08 | 1.100E-07 |
| 4201 | 211 | T211A | A | 6.2e-08 | 1.8e-08 | 4.8e-08 | 4.267E-08 | 2.248E-08 | 1.800E-08 | 6.200E-08 |
| 4202 | 211 | T211R | A | 8e-08 | 1.6e-08 | 5.2e-08 | 4.933E-08 | 3.208E-08 | 1.600E-08 | 8.000E-08 |
| 4203 | 211 | T211N | A | 6.9e-08 | 4e-08 | 1.9e-08 | 4.267E-08 | 2.511E-08 | 1.900E-08 | 6.900E-08 |
| 4204 | 211 | T211D | A | 1.5e-08 | 5.8e-08 | 2.3e-08 | 3.200E-08 | 2.287E-08 | 1.500E-08 | 5.800E-08 |
| 4205 | 211 | T211C | A | 2.2e-08 | 7e-08 | 7.5e-08 | 5.567E-08 | 2.926E-08 | 2.200E-08 | 7.500E-08 |
| 4206 | 211 | T211E | A | 2.1e-08 | 6.3e-08 | 7.3e-08 | 5.233E-08 | 2.759E-08 | 2.100E-08 | 7.300E-08 |
| 4207 | 211 | T211Q | A | 5.2e-08 | 5.6e-08 | 4e-08 | 4.933E-08 | 8.327E-09 | 4.000E-08 | 5.600E-08 |
| 4208 | 211 | T211G | A | 6.7e-08 | 9.9e-08 | 3.3e-08 | 6.633E-08 | 3.301E-08 | 3.300E-08 | 9.900E-08 |
| 4209 | 211 | T211H | A | 5.6e-08 | 6.4e-08 | 1.8e-08 | 4.600E-08 | 2.458E-08 | 1.800E-08 | 6.400E-08 |
| 4210 | 211 | T211I | A | 8.3e-08 | 2.9e-08 | 3.4e-08 | 4.867E-08 | 2.984E-08 | 2.900E-08 | 8.300E-08 |
| 4211 | 211 | T211L | A | 3.9e-08 | 4.8e-08 | 4.6e-08 | 4.433E-08 | 4.726E-09 | 3.900E-08 | 4.800E-08 |
| 4212 | 211 | T211K | A | 1.4e-08 | 2.5e-08 | 7e-08 | 3.633E-08 | 2.967E-08 | 1.400E-08 | 7.000E-08 |
| 4213 | 211 | T211M | A | 2.3e-08 | 3.8e-08 | 3.3e-08 | 3.133E-08 | 7.638E-09 | 2.300E-08 | 3.800E-08 |
| 4214 | 211 | T211F | A | 8.6e-08 | 8e-08 | 8.6e-08 | 8.400E-08 | 3.464E-09 | 8.000E-08 | 8.600E-08 |
| 4215 | 211 | T211P | A | 3.5e-08 | 4.4e-08 | 3.5e-08 | 3.800E-08 | 5.196E-09 | 3.500E-08 | 4.400E-08 |
| 4216 | 211 | T211S | A | 2.9e-08 | 5.8e-08 | 1.7e-08 | 3.467E-08 | 2.108E-08 | 1.700E-08 | 5.800E-08 |
| 4217 | 211 | T211T | A | 2.1e-08 | 1.2e-07 | 7.1e-08 | 7.067E-08 | 4.950E-08 | 2.100E-08 | 1.200E-07 |
| 4218 | 211 | T211W | A | 8e-08 | 6.4e-08 | 3.2e-08 | 5.867E-08 | 2.444E-08 | 3.200E-08 | 8.000E-08 |
| 4219 | 211 | T211Y | A | 5.8e-08 | 2.3e-08 | 3.9e-08 | 4.000E-08 | 1.752E-08 | 2.300E-08 | 5.800E-08 |
| 4220 | 211 | T211V | A | 3.2e-08 | 2.9e-08 | 6.5e-08 | 4.200E-08 | 1.997E-08 | 2.900E-08 | 6.500E-08 |
| 4221 | 212 | D212A | A | 6.2e-08 | 7.4e-08 | 4.7e-08 | 6.100E-08 | 1.353E-08 | 4.700E-08 | 7.400E-08 |
| 4222 | 212 | D212R | A | 5.9e-08 | 4e-08 | 3.6e-08 | 4.500E-08 | 1.229E-08 | 3.600E-08 | 5.900E-08 |
| 4223 | 212 | D212N | A | 5.6e-08 | 3e-08 | 2.6e-08 | 3.733E-08 | 1.629E-08 | 2.600E-08 | 5.600E-08 |
| 4224 | 212 | D212D | A | 6.2e-08 | 2.5e-07 | 2.5e-08 | 1.123E-07 | 1.206E-07 | 2.500E-08 | 2.500E-07 |
| 4225 | 212 | D212C | A | 4.5e-08 | 3.2e-08 | 3.4e-08 | 3.700E-08 | 7.000E-09 | 3.200E-08 | 4.500E-08 |
| 4226 | 212 | D212E | A | 3.1e-08 | 3.6e-08 | 5.4e-08 | 4.033E-08 | 1.210E-08 | 3.100E-08 | 5.400E-08 |
| 4227 | 212 | D212Q | A | 4.2e-08 | 4e-08 | 5e-08 | 4.400E-08 | 5.292E-09 | 4.000E-08 | 5.000E-08 |
| 4228 | 212 | D212G | A | 3.1e-08 | 4.7e-08 | 4.1e-08 | 3.967E-08 | 8.083E-09 | 3.100E-08 | 4.700E-08 |
| 4229 | 212 | D212H | A | 3.3e-08 | 3.9e-08 | 3.3e-08 | 3.500E-08 | 3.464E-09 | 3.300E-08 | 3.900E-08 |
| 4230 | 212 | D212I | A | 5.4e-08 | 5.2e-08 | 4.2e-08 | 4.933E-08 | 6.429E-09 | 4.200E-08 | 5.400E-08 |
| 4231 | 212 | D212L | A | 1.8e-08 | 3.8e-08 | 6.8e-08 | 4.133E-08 | 2.517E-08 | 1.800E-08 | 6.800E-08 |
| 4232 | 212 | D212K | A | 9e-08 | 6.3e-08 | 5.8e-08 | 7.033E-08 | 1.721E-08 | 5.800E-08 | 9.000E-08 |
| 4233 | 212 | D212M | A | 2.9e-08 | 7.2e-08 | 7.4e-08 | 5.833E-08 | 2.542E-08 | 2.900E-08 | 7.400E-08 |
| 4234 | 212 | D212F | A | 3.4e-08 | 3.8e-08 | 6.3e-08 | 4.500E-08 | 1.572E-08 | 3.400E-08 | 6.300E-08 |
| 4235 | 212 | D212P | A | 2.4e-08 | 2.5e-08 | 1.4e-07 | 6.300E-08 | 6.669E-08 | 2.400E-08 | 1.400E-07 |
| 4236 | 212 | D212S | A | 4e-08 | 4.5e-08 | 2.2e-08 | 3.567E-08 | 1.210E-08 | 2.200E-08 | 4.500E-08 |
| 4237 | 212 | D212T | A | 4e-08 | 2.8e-08 | 3.5e-08 | 3.433E-08 | 6.028E-09 | 2.800E-08 | 4.000E-08 |
| 4238 | 212 | D212W | A | 1.1e-07 | 1.2e-07 | 4e-08 | 9.000E-08 | 4.359E-08 | 4.000E-08 | 1.200E-07 |
| 4239 | 212 | D212Y | A | 1.6e-07 | 1.8e-07 | 3.5e-08 | 1.250E-07 | 7.858E-08 | 3.500E-08 | 1.800E-07 |
| 4240 | 212 | D212V | A | 3.4e-08 | 1.4e-08 | 2.5e-08 | 2.433E-08 | 1.002E-08 | 1.400E-08 | 3.400E-08 |
| 4241 | 213 | C213A | A | 4.9e-08 | 5.4e-08 | 3.6e-08 | 4.633E-08 | 9.292E-09 | 3.600E-08 | 5.400E-08 |
| 4242 | 213 | C213R | A | 3.7e-08 | 3.5e-08 | 3.8e-08 | 3.667E-08 | 1.528E-09 | 3.500E-08 | 3.800E-08 |
| 4243 | 213 | C213N | A | 7e-08 | 2.8e-08 | 1.3e-07 | 7.600E-08 | 5.126E-08 | 2.800E-08 | 1.300E-07 |
| 4244 | 213 | C213D | A | 6.5e-08 | 2.7e-08 | 3.5e-08 | 4.233E-08 | 2.003E-08 | 2.700E-08 | 6.500E-08 |
| 4245 | 213 | C213C | A | 4.3e-08 | 4.4e-08 | 3.5e-08 | 4.067E-08 | 4.933E-09 | 3.500E-08 | 4.400E-08 |
| 4246 | 213 | C213E | A | 3.3e-08 | 8e-08 | 7e-08 | 6.100E-08 | 2.476E-08 | 3.300E-08 | 8.000E-08 |
| 4247 | 213 | C213Q | A | 1.8e-07 | 2e-08 | 7.3e-08 | 9.100E-08 | 8.150E-08 | 2.000E-08 | 1.800E-07 |
| 4248 | 213 | C213G | A | 1.5e-08 | 3e-08 | 4.2e-08 | 2.900E-08 | 1.353E-08 | 1.500E-08 | 4.200E-08 |
| 4249 | 213 | C213H | A | 5.3e-08 | 2.3e-08 | 6e-08 | 4.533E-08 | 1.966E-08 | 2.300E-08 | 6.000E-08 |

| | | | | | | | | | | |
|---|---|---|---|---|---|---|---|---|---|---|
| 4250 | 213 | C213I | A | 3.7e-08 | 3.8e-08 | 1.1e-07 | 6.167E-08 | 4.186E-08 | 3.700E-08 | 1.100E-07 |
| 4251 | 213 | C213L | A | 3.1e-08 | 4.9e-08 | 2.5e-08 | 3.500E-08 | 1.249E-08 | 2.500E-08 | 4.900E-08 |
| 4252 | 213 | C213K | A | 4.1e-08 | 5e-08 | 5.2e-08 | 4.767E-08 | 5.859E-09 | 4.100E-08 | 5.200E-08 |
| 4253 | 213 | C213M | A | 8.3e-08 | 8.5e-08 | 5.7e-08 | 7.500E-08 | 1.562E-08 | 5.700E-08 | 8.500E-08 |
| 4254 | 213 | C213F | A | 7.8e-08 | 3.9e-08 | 4.3e-08 | 5.333E-08 | 2.146E-08 | 3.900E-08 | 7.800E-08 |
| 4255 | 213 | C213P | A | 2.4e-08 | 2.8e-08 | 4.8e-08 | 3.333E-08 | 1.286E-08 | 2.400E-08 | 4.800E-08 |
| 4256 | 213 | C213S | A | 6.1e-08 | 8.1e-08 | 2.1e-08 | 5.433E-08 | 3.055E-08 | 2.100E-08 | 8.100E-08 |
| 4257 | 213 | C213T | A | 5.1e-08 | 1.1e-07 | 2.8e-08 | 6.300E-08 | 4.230E-08 | 2.800E-08 | 1.100E-07 |
| 4258 | 213 | C213W | A | 7.4e-08 | 3.4e-08 | 6.5e-08 | 5.767E-08 | 2.098E-08 | 3.400E-08 | 7.400E-08 |
| 4259 | 213 | C213Y | A | 2.6e-08 | 7.5e-08 | 6.8e-08 | 5.633E-08 | 2.650E-08 | 2.600E-08 | 7.500E-08 |
| 4260 | 213 | C213V | A | 7.3e-08 | 4.3e-08 | 1.8e-08 | 4.467E-08 | 2.754E-08 | 1.800E-08 | 7.300E-08 |
| 4261 | 214 | C214A | A | 1.5e-08 | 3.8e-08 | 5.9e-08 | 3.733E-08 | 2.201E-08 | 1.500E-08 | 5.900E-08 |
| 4262 | 214 | C214R | A | 3.7e-08 | 2.1e-08 | 2.7e-08 | 2.833E-08 | 8.083E-09 | 2.100E-08 | 3.700E-08 |
| 4263 | 214 | C214N | A | 1e-07 | 2.8e-08 | 5e-08 | 5.933E-08 | 3.690E-08 | 2.800E-08 | 1.000E-07 |
| 4264 | 214 | C214D | A | 2.6e-08 | 3.9e-08 | 5.2e-08 | 3.900E-08 | 1.300E-08 | 2.600E-08 | 5.200E-08 |
| 4265 | 214 | C214C | A | 7.8e-08 | 6e-08 | 6.5e-08 | 6.767E-08 | 9.292E-09 | 6.000E-08 | 7.800E-08 |
| 4266 | 214 | C214E | A | 1.6e-08 | 6e-08 | 2.9e-08 | 3.500E-08 | 2.261E-08 | 1.600E-08 | 6.000E-08 |
| 4267 | 214 | C214Q | A | 6.3e-08 | 6.4e-08 | 2.3e-08 | 5.000E-08 | 2.339E-08 | 2.300E-08 | 6.400E-08 |
| 4268 | 214 | C214G | A | 3.7e-08 | 2.2e-08 | 3.5e-08 | 3.133E-08 | 8.145E-09 | 2.200E-08 | 3.700E-08 |
| 4269 | 214 | C214H | A | 6.1e-08 | 4.1e-08 | 6.4e-08 | 5.533E-08 | 1.250E-08 | 4.100E-08 | 6.400E-08 |
| 4270 | 214 | C214I | A | 5.1e-08 | 2.1e-08 | 5.8e-08 | 4.333E-08 | 1.966E-08 | 2.100E-08 | 5.800E-08 |
| 4271 | 214 | C214L | A | 3.6e-08 | 1.9e-08 | 2e-08 | 2.500E-08 | 9.539E-09 | 1.900E-08 | 3.600E-08 |
| 4272 | 214 | C214K | A | 3e-08 | 1.1e-07 | 9.4e-08 | 7.800E-08 | 4.233E-08 | 3.000E-08 | 1.100E-07 |
| 4273 | 214 | C214M | A | 3.5e-08 | 1.3e-07 | 9.9e-08 | 8.800E-08 | 4.845E-08 | 3.500E-08 | 1.300E-07 |
| 4274 | 214 | C214F | A | 6.6e-08 | 9.8e-08 | 3.2e-08 | 6.533E-08 | 3.301E-08 | 3.200E-08 | 9.800E-08 |
| 4275 | 214 | C214P | A | 3.7e-08 | 6.6e-08 | 5.2e-08 | 5.167E-08 | 1.450E-08 | 3.700E-08 | 6.600E-08 |
| 4276 | 214 | C214S | A | 1.1e-07 | 6.9e-08 | 1.4e-07 | 1.063E-07 | 3.564E-08 | 6.900E-08 | 1.400E-07 |
| 4277 | 214 | C214T | A | 3.4e-08 | 4e-08 | 5.7e-08 | 4.367E-08 | 1.193E-08 | 3.400E-08 | 5.700E-08 |
| 4278 | 214 | C214W | A | 2.8e-08 | 1.4e-08 | 4.8e-08 | 3.000E-08 | 1.709E-08 | 1.400E-08 | 4.800E-08 |
| 4279 | 214 | C214Y | A | 3.2e-08 | 9.4e-08 | 1.1e-08 | 4.567E-08 | 4.315E-08 | 1.100E-08 | 9.400E-08 |
| 4280 | 214 | C214V | A | 1.5e-08 | 6.1e-08 | 5.9e-08 | 4.500E-08 | 2.600E-08 | 1.500E-08 | 6.100E-08 |
| 4281 | 215 | H215A | A | 4.7e-08 | 2.6e-08 | 9.6e-08 | 5.633E-08 | 3.592E-08 | 2.600E-08 | 9.600E-08 |
| 4282 | 215 | H215R | A | 3.4e-08 | 3.6e-08 | 2.6e-08 | 3.200E-08 | 5.292E-09 | 2.600E-08 | 3.600E-08 |
| 4283 | 215 | H215N | A | 2e-08 | 7.8e-08 | 2.3e-08 | 4.033E-08 | 3.265E-08 | 2.000E-08 | 7.800E-08 |
| 4284 | 215 | H215D | A | 4.9e-08 | 1.1e-08 | 2.8e-08 | 2.933E-08 | 1.904E-08 | 1.100E-08 | 4.900E-08 |
| 4285 | 215 | H215C | A | 3.4e-08 | 5.6e-08 | 1.3e-07 | 7.333E-08 | 5.029E-08 | 3.400E-08 | 1.300E-07 |
| 4286 | 215 | H215E | A | 9.5e-08 | 4e-08 | 6.6e-08 | 6.700E-08 | 2.751E-08 | 4.000E-08 | 9.500E-08 |
| 4287 | 215 | H215Q | A | 2e-08 | 3.3e-08 | 3.3e-08 | 2.867E-08 | 7.506E-09 | 2.000E-08 | 3.300E-08 |
| 4288 | 215 | H215G | A | 9.8e-08 | 4.3e-08 | 5.7e-08 | 6.600E-08 | 2.858E-08 | 4.300E-08 | 9.800E-08 |
| 4289 | 215 | H215H | A | 1.3e-07 | 1.7e-08 | 4.4e-08 | 6.367E-08 | 5.901E-08 | 1.700E-08 | 1.300E-07 |
| 4290 | 215 | H215I | A | 4.4e-08 | 1e-07 | 5.5e-08 | 6.633E-08 | 2.967E-08 | 4.400E-08 | 1.000E-07 |
| 4291 | 215 | H215L | A | 6.5e-08 | 4.8e-08 | 5.3e-08 | 5.533E-08 | 8.737E-09 | 4.800E-08 | 6.500E-08 |
| 4292 | 215 | H215K | A | 1.1e-07 | 5.1e-08 | 1.3e-08 | 5.800E-08 | 4.888E-08 | 1.300E-08 | 1.100E-07 |
| 4293 | 215 | H215M | A | 6.3e-08 | 2.2e-08 | 1.7e-08 | 3.400E-08 | 2.524E-08 | 1.700E-08 | 6.300E-08 |
| 4294 | 215 | H215F | A | 1.6e-08 | 9.6e-08 | 9.4e-08 | 6.867E-08 | 4.562E-08 | 1.600E-08 | 9.600E-08 |
| 4295 | 215 | H215P | A | 7.4e-08 | 1.2e-07 | 5.9e-08 | 8.433E-08 | 3.179E-08 | 5.900E-08 | 1.200E-07 |
| 4296 | 215 | H215S | A | 3.5e-08 | 6.4e-08 | 6.9e-08 | 5.600E-08 | 1.836E-08 | 3.500E-08 | 6.900E-08 |
| 4297 | 215 | H215T | A | 2.7e-08 | 4e-08 | 4.3e-08 | 3.667E-08 | 8.505E-09 | 2.700E-08 | 4.300E-08 |
| 4298 | 215 | H215W | A | 2.3e-08 | 7.2e-08 | 8.7e-08 | 6.067E-08 | 3.347E-08 | 2.300E-08 | 8.700E-08 |
| 4299 | 215 | H215Y | A | 7.5e-08 | 1.5e-08 | 1.5e-08 | 3.500E-08 | 3.464E-08 | 1.500E-08 | 7.500E-08 |
| 4300 | 215 | H215V | A | 6e-08 | 3.7e-08 | 3.8e-08 | 4.500E-08 | 1.300E-08 | 3.700E-08 | 6.000E-08 |
| 4301 | 216 | E216A | A | 3e-08 | 6.8e-08 | 2.6e-08 | 4.133E-08 | 2.318E-08 | 2.600E-08 | 6.800E-08 |
| 4302 | 216 | E216R | A | 4e-08 | 7.3e-08 | 1.1e-07 | 7.433E-08 | 3.502E-08 | 4.000E-08 | 1.100E-07 |
| 4303 | 216 | E216N | A | 1.7e-08 | 2.7e-08 | 2.2e-08 | 2.200E-08 | 5.000E-09 | 1.700E-08 | 2.700E-08 |
| 4304 | 216 | E216D | A | 1.9e-08 | 7.7e-08 | 3.3e-08 | 4.300E-08 | 3.027E-08 | 1.900E-08 | 7.700E-08 |
| 4305 | 216 | E216C | A | 6e-08 | 4.1e-08 | 1e-07 | 6.700E-08 | 3.012E-08 | 4.100E-08 | 1.000E-07 |
| 4306 | 216 | E216E | A | 3.6e-08 | 3.4e-08 | 3.4e-08 | 3.467E-08 | 1.155E-09 | 3.400E-08 | 3.600E-08 |
| 4307 | 216 | E216Q | A | 3.8e-08 | 1.1e-07 | 5.8e-08 | 6.867E-08 | 3.717E-08 | 3.800E-08 | 1.100E-07 |
| 4308 | 216 | E216G | A | 1.2e-07 | 5e-08 | 2.7e-08 | 6.567E-08 | 4.844E-08 | 2.700E-08 | 1.200E-07 |
| 4309 | 216 | E216H | A | 2.8e-08 | 3.5e-08 | 7e-08 | 4.433E-08 | 2.250E-08 | 2.800E-08 | 7.000E-08 |
| 4310 | 216 | E216I | A | 6.7e-08 | 1.6e-08 | 2.3e-08 | 3.533E-08 | 2.765E-08 | 1.600E-08 | 6.700E-08 |
| 4311 | 216 | E216L | A | 2.5e-08 | 1.2e-07 | 1.2e-07 | 8.833E-08 | 5.485E-08 | 2.500E-08 | 1.200E-07 |
| 4312 | 216 | E216K | A | 2.1e-08 | 9.9e-08 | 6.5e-08 | 6.167E-08 | 3.911E-08 | 2.100E-08 | 9.900E-08 |
| 4313 | 216 | E216M | A | 2.6e-08 | 2.1e-08 | 7.9e-08 | 4.200E-08 | 3.214E-08 | 2.100E-08 | 7.900E-08 |
| 4314 | 216 | E216F | A | 3.2e-08 | 2.5e-08 | 2.1e-08 | 2.600E-08 | 5.568E-09 | 2.100E-08 | 3.200E-08 |
| 4315 | 216 | E216P | A | 7.1e-08 | 2.8e-08 | 8.7e-08 | 6.200E-08 | 3.051E-08 | 2.800E-08 | 8.700E-08 |
| 4316 | 216 | E216S | A | 1.6e-07 | 3.9e-08 | 1.1e-07 | 1.030E-07 | 6.080E-08 | 3.900E-08 | 1.600E-07 |
| 4317 | 216 | E216T | A | 7.1e-08 | 4.3e-08 | 7.1e-08 | 6.167E-08 | 1.617E-08 | 4.300E-08 | 7.100E-08 |
| 4318 | 216 | E216W | A | 2.8e-08 | 5e-08 | 3.6e-08 | 3.800E-08 | 1.114E-08 | 2.800E-08 | 5.000E-08 |
| 4319 | 216 | E216Y | A | 5.9e-08 | 3.3e-08 | 1.8e-08 | 3.667E-08 | 2.074E-08 | 1.800E-08 | 5.900E-08 |
| 4320 | 216 | E216V | A | 6.5e-08 | 6e-08 | 5.2e-08 | 5.900E-08 | 6.557E-09 | 5.200E-08 | 6.500E-08 |
| 4321 | 217 | Q217A | A | 2.8e-08 | 6e-08 | 6.5e-08 | 5.100E-08 | 2.007E-08 | 2.800E-08 | 6.500E-08 |
| 4322 | 217 | Q217R | A | 9.8e-09 | 7.9e-08 | 7.4e-08 | 5.427E-08 | 3.859E-08 | 9.800E-09 | 7.900E-08 |
| 4323 | 217 | Q217N | A | 2e-08 | 5.2e-08 | 1.7e-07 | 8.067E-08 | 7.900E-08 | 2.000E-08 | 1.700E-07 |
| 4324 | 217 | Q217D | A | 7.9e-08 | 4e-08 | 4e-08 | 5.300E-08 | 2.252E-08 | 4.000E-08 | 7.900E-08 |
| 4325 | 217 | Q217C | A | 3.3e-08 | 6.6e-08 | 3.4e-08 | 4.433E-08 | 1.877E-08 | 3.300E-08 | 6.600E-08 |
| 4326 | 217 | Q217E | A | 2e-08 | 4.7e-08 | 5e-08 | 3.900E-08 | 1.652E-08 | 2.000E-08 | 5.000E-08 |
| 4327 | 217 | Q217Q | A | 5.8e-08 | 2.6e-08 | 1.7e-08 | 3.367E-08 | 2.155E-08 | 1.700E-08 | 5.800E-08 |
| 4328 | 217 | Q217G | A | 4.1e-08 | 3.9e-08 | 4.8e-08 | 4.267E-08 | 4.726E-09 | 3.900E-08 | 4.800E-08 |
| 4329 | 217 | Q217H | A | 1.7e-07 | 2.9e-08 | 8e-09 | 6.900E-08 | 8.810E-08 | 8.000E-09 | 1.700E-07 |
| 4330 | 217 | Q217I | A | 2.4e-08 | 5.9e-08 | 2.5e-08 | 3.600E-08 | 1.992E-08 | 2.400E-08 | 5.900E-08 |
| 4331 | 217 | Q217L | A | 7.9e-08 | 4.8e-08 | 3.4e-08 | 5.367E-08 | 2.303E-08 | 3.400E-08 | 7.900E-08 |
| 4332 | 217 | Q217K | A | 2.2e-08 | 3.6e-08 | 4.2e-08 | 3.333E-08 | 1.026E-08 | 2.200E-08 | 4.200E-08 |
| 4333 | 217 | Q217M | A | 3.4e-08 | 4.7e-08 | 3.7e-08 | 3.933E-08 | 6.807E-09 | 3.400E-08 | 4.700E-08 |
| 4334 | 217 | Q217F | A | 3e-08 | 5.4e-08 | 5.5e-08 | 4.633E-08 | 1.415E-08 | 3.000E-08 | 5.500E-08 |

| | | | | | | | | | | |
|---|---|---|---|---|---|---|---|---|---|---|
| 4335 | 217 | Q217P | A | 2.6e-08 | 4.9e-08 | 3e-08 | 3.500E-08 | 1.229E-08 | 2.600E-08 | 4.900E-08 |
| 4336 | 217 | Q217S | A | 9e-08 | 6.9e-08 | 1.1e-07 | 8.967E-08 | 2.050E-08 | 6.900E-08 | 1.100E-07 |
| 4337 | 217 | Q217T | A | 2.3e-08 | 5.7e-08 | 6e-08 | 4.667E-08 | 2.055E-08 | 2.300E-08 | 6.000E-08 |
| 4338 | 217 | Q217W | A | 2.1e-08 | 4.5e-08 | 8.1e-08 | 4.900E-08 | 3.020E-08 | 2.100E-08 | 8.100E-08 |
| 4339 | 217 | Q217Y | A | 3.1e-08 | 3.2e-08 | 9.3e-08 | 5.200E-08 | 3.551E-08 | 3.100E-08 | 9.300E-08 |
| 4340 | 217 | Q217V | A | 1e-08 | 4.1e-08 | 2.8e-08 | 2.633E-08 | 1.557E-08 | 1.000E-08 | 4.100E-08 |
| 4341 | 218 | C218A | A | 5.9e-08 | 1.7e-07 | 7.8e-08 | 1.023E-07 | 5.937E-08 | 5.900E-08 | 1.700E-07 |
| 4342 | 218 | C218R | A | 5.5e-08 | 2.7e-08 | 5.8e-08 | 4.667E-08 | 1.710E-08 | 2.700E-08 | 5.800E-08 |
| 4343 | 218 | C218N | A | 1.4e-07 | 3.3e-08 | 3.7e-08 | 7.000E-08 | 6.065E-08 | 3.300E-08 | 1.400E-07 |
| 4344 | 218 | C218D | A | 7.8e-08 | 1.7e-08 | 8e-08 | 5.833E-08 | 3.581E-08 | 1.700E-08 | 8.000E-08 |
| 4345 | 218 | C218C | A | 2.8e-08 | 2.9e-08 | 4.6e-08 | 3.433E-08 | 1.012E-08 | 2.800E-08 | 4.600E-08 |
| 4346 | 218 | C218E | A | 2.8e-08 | 6.1e-08 | 4.7e-08 | 4.533E-08 | 1.656E-08 | 2.800E-08 | 6.100E-08 |
| 4347 | 218 | C218Q | A | 5.3e-08 | 2.9e-08 | 5.2e-08 | 4.467E-08 | 1.358E-08 | 2.900E-08 | 5.300E-08 |
| 4348 | 218 | C218G | A | 1.4e-08 | 8.9e-08 | 7.1e-08 | 5.800E-08 | 3.915E-08 | 1.400E-08 | 8.900E-08 |
| 4349 | 218 | C218H | A | 3.1e-08 | 1e-07 | 8.2e-08 | 7.100E-08 | 3.579E-08 | 3.100E-08 | 1.000E-07 |
| 4350 | 218 | C218I | A | 3.7e-08 | 1.1e-08 | 3.9e-08 | 2.900E-08 | 1.562E-08 | 1.100E-08 | 3.900E-08 |
| 4351 | 218 | C218L | A | 5e-08 | 1.5e-07 | 1.1e-07 | 1.033E-07 | 5.033E-08 | 5.000E-08 | 1.500E-07 |
| 4352 | 218 | C218K | A | 6.3e-08 | 8.4e-08 | 8.2e-08 | 7.633E-08 | 1.159E-08 | 6.300E-08 | 8.400E-08 |
| 4353 | 218 | C218M | A | 2.3e-08 | 4e-08 | 5.8e-08 | 4.033E-08 | 1.750E-08 | 2.300E-08 | 5.800E-08 |
| 4354 | 218 | C218F | A | 3.1e-08 | 5.4e-08 | 8.8e-08 | 5.767E-08 | 2.868E-08 | 3.100E-08 | 8.800E-08 |
| 4355 | 218 | C218P | A | 4.7e-08 | 9.6e-09 | 6.3e-08 | 3.987E-08 | 2.741E-08 | 9.600E-09 | 6.300E-08 |
| 4356 | 218 | C218S | A | 3.1e-08 | 4.6e-08 | 8.5e-08 | 5.400E-08 | 2.787E-08 | 3.100E-08 | 8.500E-08 |
| 4357 | 218 | C218T | A | 4.1e-08 | 8.2e-08 | 6.9e-08 | 6.400E-08 | 2.095E-08 | 4.100E-08 | 8.200E-08 |
| 4358 | 218 | C218W | A | 5.1e-08 | 4.3e-08 | 2.7e-08 | 4.033E-08 | 1.222E-08 | 2.700E-08 | 5.100E-08 |
| 4359 | 218 | C218Y | A | 7e-08 | 5.3e-08 | 6.4e-08 | 6.233E-08 | 8.622E-09 | 5.300E-08 | 7.000E-08 |
| 4360 | 218 | C218V | A | 2.2e-08 | 7.8e-09 | 2.9e-08 | 1.960E-08 | 1.080E-08 | 7.800E-09 | 2.900E-08 |
| 4361 | 219 | A219A | A | 1.2e-08 | 2.4e-08 | 2.8e-08 | 2.133E-08 | 8.327E-09 | 1.200E-08 | 2.800E-08 |
| 4362 | 219 | A219R | A | 1.1e-07 | 1.5e-08 | 2.8e-08 | 5.100E-08 | 5.151E-08 | 1.500E-08 | 1.100E-07 |
| 4363 | 219 | A219N | A | 1.9e-08 | 5.9e-08 | 2e-08 | 3.267E-08 | 2.281E-08 | 1.900E-08 | 5.900E-08 |
| 4364 | 219 | A219D | A | 4.2e-08 | 3.6e-08 | 2.3e-08 | 3.367E-08 | 9.713E-09 | 2.300E-08 | 4.200E-08 |
| 4365 | 219 | A219C | A | 6.7e-08 | 7.6e-08 | 1.2e-07 | 8.767E-08 | 2.836E-08 | 6.700E-08 | 1.200E-07 |
| 4366 | 219 | A219E | A | 8.3e-08 | 2.3e-08 | 3e-08 | 4.533E-08 | 3.281E-08 | 2.300E-08 | 8.300E-08 |
| 4367 | 219 | A219Q | A | 5.3e-08 | 6.1e-08 | 3.8e-08 | 5.067E-08 | 1.168E-08 | 3.800E-08 | 6.100E-08 |
| 4368 | 219 | A219G | A | 8.3e-08 | 6.1e-08 | 6.9e-08 | 7.100E-08 | 1.114E-08 | 6.100E-08 | 8.300E-08 |
| 4369 | 219 | A219H | A | 1.6e-08 | 5.5e-08 | 3e-08 | 3.367E-08 | 1.976E-08 | 1.600E-08 | 5.500E-08 |
| 4370 | 219 | A219I | A | 9.1e-08 | 2.6e-08 | 3.3e-08 | 5.000E-08 | 3.568E-08 | 2.600E-08 | 9.100E-08 |
| 4371 | 219 | A219L | A | 5.3e-08 | 5.1e-08 | 1.4e-08 | 3.933E-08 | 2.196E-08 | 1.400E-08 | 5.300E-08 |
| 4372 | 219 | A219K | A | 3.3e-08 | 1.8e-08 | 1.8e-08 | 2.300E-08 | 8.660E-09 | 1.800E-08 | 3.300E-08 |
| 4373 | 219 | A219M | A | 1.2e-07 | 3.6e-08 | 6.7e-08 | 7.433E-08 | 4.248E-08 | 3.600E-08 | 1.200E-07 |
| 4374 | 219 | A219F | A | 9.3e-08 | 4.4e-08 | 2.7e-08 | 5.467E-08 | 3.427E-08 | 2.700E-08 | 9.300E-08 |
| 4375 | 219 | A219P | A | 7.1e-08 | 3.4e-08 | 7.4e-08 | 5.967E-08 | 2.228E-08 | 3.400E-08 | 7.400E-08 |
| 4376 | 219 | A219S | A | 5.9e-08 | 5.2e-08 | 6.8e-08 | 5.967E-08 | 8.021E-09 | 5.200E-08 | 6.800E-08 |
| 4377 | 219 | A219T | A | 1e-07 | 9.8e-08 | 6.3e-08 | 8.700E-08 | 2.081E-08 | 6.300E-08 | 1.000E-07 |
| 4378 | 219 | A219W | A | 1.1e-07 | 5.2e-08 | 5e-08 | 7.067E-08 | 3.408E-08 | 5.000E-08 | 1.100E-07 |
| 4379 | 219 | A219Y | A | 8.7e-09 | 6.5e-08 | 1.2e-07 | 6.457E-08 | 5.565E-08 | 8.700E-09 | 1.200E-07 |
| 4380 | 219 | A219V | A | 2.2e-07 | 1.8e-08 | 1.7e-08 | 8.500E-08 | 1.169E-07 | 1.700E-08 | 2.200E-07 |
| 4381 | 220 | A220A | A | 2.2e-08 | 6.6e-09 | 6.2e-08 | 3.020E-08 | 2.860E-08 | 6.600E-09 | 6.200E-08 |
| 4382 | 220 | A220R | A | 6e-08 | 2.3e-08 | 3.1e-08 | 3.800E-08 | 1.947E-08 | 2.300E-08 | 6.000E-08 |
| 4383 | 220 | A220N | A | 2.7e-08 | 2.8e-08 | 7.2e-08 | 4.233E-08 | 2.570E-08 | 2.700E-08 | 7.200E-08 |
| 4384 | 220 | A220D | A | 7.1e-08 | 3e-08 | 2.7e-08 | 4.267E-08 | 2.458E-08 | 2.700E-08 | 7.100E-08 |
| 4385 | 220 | A220C | A | 5.4e-08 | 6e-08 | 6.5e-08 | 5.967E-08 | 5.508E-09 | 5.400E-08 | 6.500E-08 |
| 4386 | 220 | A220E | A | 9.2e-08 | 2.2e-08 | 8.2e-08 | 6.533E-08 | 3.786E-08 | 2.200E-08 | 9.200E-08 |
| 4387 | 220 | A220Q | A | 5.1e-08 | 4.1e-08 | 4.3e-08 | 4.500E-08 | 5.292E-09 | 4.100E-08 | 5.100E-08 |
| 4388 | 220 | A220G | A | 1.8e-08 | 2.3e-08 | 3.9e-08 | 2.667E-08 | 1.097E-08 | 1.800E-08 | 3.900E-08 |
| 4389 | 220 | A220H | A | 3.8e-08 | 2.8e-08 | 4.7e-08 | 3.767E-08 | 9.504E-09 | 2.800E-08 | 4.700E-08 |
| 4390 | 220 | A220I | A | 2.2e-08 | 4e-08 | 3.2e-08 | 3.133E-08 | 9.018E-09 | 2.200E-08 | 4.000E-08 |
| 4391 | 220 | A220L | A | 2.2e-08 | 5.5e-08 | 3e-08 | 3.567E-08 | 1.721E-08 | 2.200E-08 | 5.500E-08 |
| 4392 | 220 | A220K | A | 5.9e-09 | 6.4e-08 | 3.3e-08 | 3.430E-08 | 2.907E-08 | 5.900E-09 | 6.400E-08 |
| 4393 | 220 | A220M | A | 4.1e-08 | 2.7e-08 | 1.3e-07 | 6.600E-08 | 5.587E-08 | 2.700E-08 | 1.300E-07 |
| 4394 | 220 | A220F | A | 1.5e-08 | 4.6e-08 | 2.8e-08 | 2.967E-08 | 1.557E-08 | 1.500E-08 | 4.600E-08 |
| 4395 | 220 | A220P | A | 4e-08 | 4.7e-08 | 2.7e-08 | 3.800E-08 | 1.015E-08 | 2.700E-08 | 4.700E-08 |
| 4396 | 220 | A220S | A | 4e-08 | 1.8e-08 | 3.7e-08 | 3.167E-08 | 1.193E-08 | 1.800E-08 | 4.000E-08 |
| 4397 | 220 | A220T | A | 3.3e-08 | 5.3e-08 | 1.8e-08 | 3.467E-08 | 1.756E-08 | 1.800E-08 | 5.300E-08 |
| 4398 | 220 | A220W | A | 1.5e-08 | 5.7e-08 | 2.2e-08 | 3.133E-08 | 2.250E-08 | 1.500E-08 | 5.700E-08 |
| 4399 | 220 | A220Y | A | 5.6e-08 | 5.3e-08 | 2e-08 | 4.300E-08 | 1.997E-08 | 2.000E-08 | 5.600E-08 |
| 4400 | 220 | A220V | A | 2.3e-08 | 6.8e-08 | 3e-08 | 4.033E-08 | 2.421E-08 | 2.300E-08 | 6.800E-08 |
| 4401 | 221 | G221A | A | 1.1e-07 | 3.9e-08 | 4.6e-08 | 6.500E-08 | 3.913E-08 | 3.900E-08 | 1.100E-07 |
| 4402 | 221 | G221R | A | 6.3e-08 | 1.7e-08 | 2.3e-08 | 3.433E-08 | 2.501E-08 | 1.700E-08 | 6.300E-08 |
| 4403 | 221 | G221N | A | 5.9e-08 | 5.2e-08 | 2.7e-08 | 4.600E-08 | 1.682E-08 | 2.700E-08 | 5.900E-08 |
| 4404 | 221 | G221D | A | 6.8e-08 | 6.4e-08 | 7e-08 | 6.733E-08 | 3.055E-09 | 6.400E-08 | 7.000E-08 |
| 4405 | 221 | G221C | A | 4.2e-08 | 6e-08 | 4.4e-08 | 4.867E-08 | 9.866E-09 | 4.200E-08 | 6.000E-08 |
| 4406 | 221 | G221E | A | 9.6e-08 | 3.1e-08 | 3e-08 | 5.233E-08 | 3.782E-08 | 3.000E-08 | 9.600E-08 |
| 4407 | 221 | G221Q | A | 2.8e-08 | 9e-09 | 4.5e-08 | 2.733E-08 | 1.801E-08 | 9.000E-09 | 4.500E-08 |
| 4408 | 221 | G221G | A | 5.5e-08 | 3.9e-08 | 3.1e-08 | 4.167E-08 | 1.222E-08 | 3.100E-08 | 5.500E-08 |
| 4409 | 221 | G221H | A | 3.8e-08 | 7e-08 | 6.1e-08 | 5.633E-08 | 1.650E-08 | 3.800E-08 | 7.000E-08 |
| 4410 | 221 | G221I | A | 1.1e-07 | 5.6e-08 | 5.6e-08 | 7.400E-08 | 3.118E-08 | 5.600E-08 | 1.100E-07 |
| 4411 | 221 | G221L | A | 3.4e-08 | 7.2e-08 | 5e-08 | 5.200E-08 | 1.908E-08 | 3.400E-08 | 7.200E-08 |
| 4412 | 221 | G221K | A | 4.1e-08 | 3e-08 | 3e-08 | 3.367E-08 | 6.351E-09 | 3.000E-08 | 4.100E-08 |
| 4413 | 221 | G221M | A | 4e-08 | 9.9e-08 | 9.9e-08 | 7.933E-08 | 3.406E-08 | 4.000E-08 | 9.900E-08 |
| 4414 | 221 | G221F | A | 3.7e-08 | 2.6e-08 | 4.5e-08 | 3.600E-08 | 9.539E-09 | 2.600E-08 | 4.500E-08 |
| 4415 | 221 | G221P | A | 1e-07 | 9.7e-08 | 3.7e-08 | 7.800E-08 | 3.554E-08 | 3.700E-08 | 1.000E-07 |
| 4416 | 221 | G221S | A | 1.6e-08 | 3.4e-08 | 5.6e-08 | 3.533E-08 | 2.003E-08 | 1.600E-08 | 5.600E-08 |
| 4417 | 221 | G221T | A | 3.1e-08 | 4.8e-08 | 3.8e-08 | 3.900E-08 | 8.544E-09 | 3.100E-08 | 4.800E-08 |
| 4418 | 221 | G221W | A | 7.4e-08 | 1.4e-08 | 7.2e-08 | 5.333E-08 | 3.408E-08 | 1.400E-08 | 7.400E-08 |
| 4419 | 221 | G221Y | A | 1.1e-08 | 6.3e-08 | 3.6e-08 | 3.667E-08 | 2.601E-08 | 1.100E-08 | 6.300E-08 |

| | | | | | | | | | | |
|---|---|---|---|---|---|---|---|---|---|---|
| 4420 | 221 | G221V | A | 3.3e-08 | 2.4e-08 | 6e-08 | 3.900E-08 | 1.873E-08 | 2.400E-08 | 6.000E-08 |
| 4421 | 222 | C222A | A | 3.1e-08 | 1.2e-07 | 4.2e-08 | 6.433E-08 | 4.852E-08 | 3.100E-08 | 1.200E-07 |
| 4422 | 222 | C222R | A | 2.1e-08 | 2.3e-08 | 3.4e-08 | 2.600E-08 | 7.000E-09 | 2.100E-08 | 3.400E-08 |
| 4423 | 222 | C222N | A | 2.1e-07 | 6.4e-08 | 4.3e-08 | 1.057E-07 | 9.096E-08 | 4.300E-08 | 2.100E-07 |
| 4424 | 222 | C222D | A | 1.7e-08 | 6.1e-08 | 1.3e-07 | 6.933E-08 | 5.696E-08 | 1.700E-08 | 1.300E-07 |
| 4425 | 222 | C222C | A | 1e-07 | 1.9e-08 | 2.7e-08 | 4.867E-08 | 4.464E-08 | 1.900E-08 | 1.000E-07 |
| 4426 | 222 | C222E | A | 1.8e-08 | 4.5e-08 | 2.7e-08 | 3.000E-08 | 1.375E-08 | 1.800E-08 | 4.500E-08 |
| 4427 | 222 | C222Q | A | 4.9e-08 | 9.7e-08 | 2.8e-08 | 5.800E-08 | 3.537E-08 | 2.800E-08 | 9.700E-08 |
| 4428 | 222 | C222G | A | 4.4e-08 | 2.5e-08 | 3.2e-08 | 3.367E-08 | 9.609E-09 | 2.500E-08 | 4.400E-08 |
| 4429 | 222 | C222H | A | 5.3e-08 | 2e-08 | 2.9e-08 | 3.400E-08 | 1.706E-08 | 2.000E-08 | 5.300E-08 |
| 4430 | 222 | C222I | A | 2.5e-08 | 4.1e-08 | 3.1e-08 | 3.233E-08 | 8.083E-09 | 2.500E-08 | 4.100E-08 |
| 4431 | 222 | C222L | A | 3.8e-08 | 1e-07 | 4.5e-08 | 6.100E-08 | 3.396E-08 | 3.800E-08 | 1.000E-07 |
| 4432 | 222 | C222K | A | 1.5e-08 | 9.7e-08 | 1.1e-07 | 7.400E-08 | 5.151E-08 | 1.500E-08 | 1.100E-07 |
| 4433 | 222 | C222M | A | 2.6e-08 | 7.2e-08 | 6.1e-08 | 5.300E-08 | 2.402E-08 | 2.600E-08 | 7.200E-08 |
| 4434 | 222 | C222F | A | 4e-08 | 4.6e-08 | 7e-08 | 5.200E-08 | 1.587E-08 | 4.000E-08 | 7.000E-08 |
| 4435 | 222 | C222P | A | 1e-07 | 2.7e-08 | 3.3e-08 | 5.333E-08 | 4.053E-08 | 2.700E-08 | 1.000E-07 |
| 4436 | 222 | C222S | A | 5.2e-08 | 5.1e-08 | 3.7e-08 | 4.667E-08 | 8.386E-09 | 3.700E-08 | 5.200E-08 |
| 4437 | 222 | C222T | A | 3.5e-08 | 3.2e-08 | 6.2e-08 | 4.300E-08 | 1.652E-08 | 3.200E-08 | 6.200E-08 |
| 4438 | 222 | C222W | A | 5.3e-08 | 1e-07 | 1.7e-08 | 5.667E-08 | 4.162E-08 | 1.700E-08 | 1.000E-07 |
| 4439 | 222 | C222Y | A | 2.2e-08 | 2.5e-08 | 5.3e-08 | 3.333E-08 | 1.710E-08 | 2.200E-08 | 5.300E-08 |
| 4440 | 222 | C222V | A | 2.6e-08 | 4.1e-08 | 5.7e-08 | 4.133E-08 | 1.550E-08 | 2.600E-08 | 5.700E-08 |
| 4441 | 223 | T223A | A | 6.8e-08 | 6.2e-08 | 1e-07 | 7.667E-08 | 2.043E-08 | 6.200E-08 | 1.000E-07 |
| 4442 | 223 | T223R | A | 6.6e-08 | 1e-07 | 1.8e-08 | 6.133E-08 | 4.120E-08 | 1.800E-08 | 1.000E-07 |
| 4443 | 223 | T223N | A | 3e-08 | 2.9e-08 | 1.9e-08 | 2.600E-08 | 6.083E-09 | 1.900E-08 | 3.000E-08 |
| 4444 | 223 | T223D | A | 2e-08 | 6.9e-08 | 3.4e-08 | 4.100E-08 | 2.524E-08 | 2.000E-08 | 6.900E-08 |
| 4445 | 223 | T223C | A | 4.1e-08 | 2.5e-08 | 2.4e-08 | 3.000E-08 | 9.539E-09 | 2.400E-08 | 4.100E-08 |
| 4446 | 223 | T223E | A | 7.4e-08 | 3.3e-08 | 2.5e-08 | 4.400E-08 | 2.629E-08 | 2.500E-08 | 7.400E-08 |
| 4447 | 223 | T223Q | A | 2e-08 | 8.4e-09 | 4.5e-08 | 2.447E-08 | 1.870E-08 | 8.400E-09 | 4.500E-08 |
| 4448 | 223 | T223G | A | 5.4e-08 | 2.1e-07 | 3.7e-08 | 1.003E-07 | 9.535E-08 | 3.700E-08 | 2.100E-07 |
| 4449 | 223 | T223H | A | 1.2e-08 | 3.9e-08 | 4.4e-08 | 3.167E-08 | 1.721E-08 | 1.200E-08 | 4.400E-08 |
| 4450 | 223 | T223I | A | 4.3e-08 | 5e-08 | 2.7e-08 | 4.000E-08 | 1.179E-08 | 2.700E-08 | 5.000E-08 |
| 4451 | 223 | T223L | A | 4.6e-08 | 1.9e-08 | 1.7e-08 | 2.733E-08 | 1.620E-08 | 1.700E-08 | 4.600E-08 |
| 4452 | 223 | T223K | A | 5.1e-08 | 3.2e-08 | 5.4e-08 | 4.567E-08 | 1.193E-08 | 3.200E-08 | 5.400E-08 |
| 4453 | 223 | T223M | A | 4.2e-08 | 4.1e-08 | 1.1e-07 | 6.433E-08 | 3.955E-08 | 4.100E-08 | 1.100E-07 |
| 4454 | 223 | T223F | A | 3.4e-08 | 1.3e-08 | 5.1e-08 | 3.267E-08 | 1.904E-08 | 1.300E-08 | 5.100E-08 |
| 4455 | 223 | T223P | A | 2.4e-08 | 3.1e-08 | 2e-08 | 2.500E-08 | 5.568E-09 | 2.000E-08 | 3.100E-08 |
| 4456 | 223 | T223S | A | 1.7e-08 | 3.4e-08 | 2.4e-08 | 2.500E-08 | 8.544E-09 | 1.700E-08 | 3.400E-08 |
| 4457 | 223 | T223T | A | 7.5e-08 | 2.6e-08 | 5.1e-08 | 5.067E-08 | 2.450E-08 | 2.600E-08 | 7.500E-08 |
| 4458 | 223 | T223W | A | 4.2e-08 | 5.2e-08 | 1.4e-07 | 7.800E-08 | 5.393E-08 | 4.200E-08 | 1.400E-07 |
| 4459 | 223 | T223Y | A | 1.9e-08 | 4.6e-08 | 2.3e-08 | 2.933E-08 | 1.457E-08 | 1.900E-08 | 4.600E-08 |
| 4460 | 223 | T223V | A | 4.9e-08 | 1.1e-08 | 3e-08 | 3.000E-08 | 1.900E-08 | 1.100E-08 | 4.900E-08 |
| 4461 | 224 | G224A | A | 1.7e-08 | 2.6e-08 | 3.5e-08 | 2.600E-08 | 9.000E-09 | 1.700E-08 | 3.500E-08 |
| 4462 | 224 | G224R | A | 3.5e-08 | 1.5e-08 | 1.2e-07 | 5.667E-08 | 5.575E-08 | 1.500E-08 | 1.200E-07 |
| 4463 | 224 | G224N | A | 3.3e-08 | 3.4e-08 | 7.3e-08 | 4.667E-08 | 2.281E-08 | 3.300E-08 | 7.300E-08 |
| 4464 | 224 | G224D | A | 6e-08 | 2.5e-08 | 1.7e-08 | 3.400E-08 | 2.287E-08 | 1.700E-08 | 6.000E-08 |
| 4465 | 224 | G224C | A | 4.3e-08 | 2.9e-08 | 4.5e-08 | 3.900E-08 | 8.718E-09 | 2.900E-08 | 4.500E-08 |
| 4466 | 224 | G224E | A | 4.6e-08 | 6.5e-08 | 3.4e-08 | 4.833E-08 | 1.563E-08 | 3.400E-08 | 6.500E-08 |
| 4467 | 224 | G224Q | A | 1.9e-08 | 4.6e-08 | 1e-07 | 5.500E-08 | 4.124E-08 | 1.900E-08 | 1.000E-07 |
| 4468 | 224 | G224G | A | 9.4e-09 | 2.4e-08 | 3.8e-08 | 2.380E-08 | 1.430E-08 | 9.400E-09 | 3.800E-08 |
| 4469 | 224 | G224H | A | 3.2e-08 | 9.4e-08 | 6.2e-08 | 6.267E-08 | 3.101E-08 | 3.200E-08 | 9.400E-08 |
| 4470 | 224 | G224I | A | 2.9e-08 | 5.6e-08 | 3.5e-08 | 4.000E-08 | 1.418E-08 | 2.900E-08 | 5.600E-08 |
| 4471 | 224 | G224L | A | 3.3e-08 | 3.6e-08 | 1.2e-08 | 2.700E-08 | 1.308E-08 | 1.200E-08 | 3.600E-08 |
| 4472 | 224 | G224K | A | 3.7e-08 | 5.1e-08 | 1.9e-08 | 3.567E-08 | 1.604E-08 | 1.900E-08 | 5.100E-08 |
| 4473 | 224 | G224M | A | 4.3e-08 | 6.1e-08 | 6.8e-08 | 5.733E-08 | 1.290E-08 | 4.300E-08 | 6.800E-08 |
| 4474 | 224 | G224F | A | 2.3e-08 | 6.8e-08 | 4.6e-08 | 4.567E-08 | 2.250E-08 | 2.300E-08 | 6.800E-08 |
| 4475 | 224 | G224P | A | 3.8e-08 | 9.9e-09 | 7.4e-08 | 4.063E-08 | 3.213E-08 | 9.900E-09 | 7.400E-08 |
| 4476 | 224 | G224S | A | 3e-08 | 1.6e-08 | 4.9e-08 | 3.167E-08 | 1.656E-08 | 1.600E-08 | 4.900E-08 |
| 4477 | 224 | G224T | A | 9.4e-08 | 6.6e-08 | 6.4e-08 | 7.467E-08 | 1.677E-08 | 6.400E-08 | 9.400E-08 |
| 4478 | 224 | G224W | A | 1.6e-08 | 2.2e-08 | 1.8e-08 | 1.867E-08 | 3.055E-09 | 1.600E-08 | 2.200E-08 |
| 4479 | 224 | G224Y | A | 2.5e-08 | 1.3e-07 | 3.9e-08 | 6.467E-08 | 5.701E-08 | 2.500E-08 | 1.300E-07 |
| 4480 | 224 | G224V | A | 2.8e-08 | 2.7e-08 | 3.2e-08 | 2.900E-08 | 2.646E-09 | 2.700E-08 | 3.200E-08 |
| 4481 | 225 | P225A | A | 2.5e-08 | 4.7e-08 | 4.2e-08 | 3.800E-08 | 1.153E-08 | 2.500E-08 | 4.700E-08 |
| 4482 | 225 | P225R | A | 2.7e-08 | 4.6e-08 | 2.7e-08 | 3.333E-08 | 1.097E-08 | 2.700E-08 | 4.600E-08 |
| 4483 | 225 | P225N | A | 6.5e-08 | 4.8e-08 | 4.5e-08 | 5.267E-08 | 1.079E-08 | 4.500E-08 | 6.500E-08 |
| 4484 | 225 | P225D | A | 1e-07 | 1.9e-08 | 3e-08 | 4.967E-08 | 4.394E-08 | 1.900E-08 | 1.000E-07 |
| 4485 | 225 | P225C | A | 6.1e-08 | 7.1e-08 | 5.9e-08 | 6.367E-08 | 6.429E-09 | 5.900E-08 | 7.100E-08 |
| 4486 | 225 | P225E | A | 9.5e-08 | 1.5e-08 | 7.9e-08 | 6.300E-08 | 4.233E-08 | 1.500E-08 | 9.500E-08 |
| 4487 | 225 | P225Q | A | 3.6e-08 | 1.1e-08 | 1.8e-08 | 2.167E-08 | 1.290E-08 | 1.100E-08 | 3.600E-08 |
| 4488 | 225 | P225G | A | 1.1e-07 | 4.2e-08 | 3e-08 | 6.067E-08 | 4.314E-08 | 3.000E-08 | 1.100E-07 |
| 4489 | 225 | P225H | A | 6.2e-08 | 3.1e-08 | 1.3e-08 | 3.533E-08 | 2.479E-08 | 1.300E-08 | 6.200E-08 |
| 4490 | 225 | P225I | A | 1.8e-08 | 4e-08 | 2.5e-08 | 2.767E-08 | 1.124E-08 | 1.800E-08 | 4.000E-08 |
| 4491 | 225 | P225L | A | 9.2e-09 | 5.8e-08 | 6.4e-08 | 4.373E-08 | 3.006E-08 | 9.200E-09 | 6.400E-08 |
| 4492 | 225 | P225K | A | 6e-08 | 1.9e-08 | 4.9e-08 | 4.267E-08 | 2.122E-08 | 1.900E-08 | 6.000E-08 |
| 4493 | 225 | P225M | A | 1.2e-08 | 4.3e-08 | 7.3e-08 | 4.267E-08 | 3.050E-08 | 1.200E-08 | 7.300E-08 |
| 4494 | 225 | P225F | A | 6.6e-08 | 3.6e-08 | 4.4e-08 | 4.867E-08 | 1.553E-08 | 3.600E-08 | 6.600E-08 |
| 4495 | 225 | P225P | A | 7.5e-08 | 6.6e-08 | 7.3e-08 | 7.133E-08 | 4.726E-09 | 6.600E-08 | 7.500E-08 |
| 4496 | 225 | P225S | A | 4.9e-08 | 2.6e-08 | 7.3e-08 | 4.933E-08 | 2.350E-08 | 2.600E-08 | 7.300E-08 |
| 4497 | 225 | P225T | A | 4e-08 | 2.6e-08 | 3.8e-08 | 3.467E-08 | 7.572E-09 | 2.600E-08 | 4.000E-08 |
| 4498 | 225 | P225W | A | 4.3e-08 | 4.2e-08 | 2.4e-08 | 3.633E-08 | 1.069E-08 | 2.400E-08 | 4.300E-08 |
| 4499 | 225 | P225Y | A | 2.8e-08 | 8e-08 | 2.5e-08 | 4.433E-08 | 3.092E-08 | 2.500E-08 | 8.000E-08 |
| 4500 | 225 | P225V | A | 1.9e-08 | 5e-08 | 7.1e-08 | 4.667E-08 | 2.616E-08 | 1.900E-08 | 7.100E-08 |
| 4501 | 226 | K226A | A | 3.8e-08 | 4.4e-08 | 3.1e-08 | 3.767E-08 | 6.506E-09 | 3.100E-08 | 4.400E-08 |
| 4502 | 226 | K226R | A | 5.2e-08 | 6.5e-08 | 3.2e-08 | 4.967E-08 | 1.662E-08 | 3.200E-08 | 6.500E-08 |
| 4503 | 226 | K226N | A | 7.4e-08 | 7.9e-08 | 2.1e-08 | 5.800E-08 | 3.214E-08 | 2.100E-08 | 7.900E-08 |
| 4504 | 226 | K226D | A | 1e-07 | 3.7e-08 | 3.5e-08 | 5.733E-08 | 3.696E-08 | 3.500E-08 | 1.000E-07 |

| | | | | | | | | | | |
|---|---|---|---|---|---|---|---|---|---|---|
| 4505 | 226 | K226C | A | 4.1e-08 | 1.1e-07 | 5.8e-08 | 6.967E-08 | 3.595E-08 | 4.100E-08 | 1.100E-07 |
| 4506 | 226 | K226E | A | 3.9e-08 | 4e-08 | 3.6e-08 | 3.833E-08 | 2.082E-09 | 3.600E-08 | 4.000E-08 |
| 4507 | 226 | K226Q | A | 6e-08 | 1.3e-08 | 2.8e-08 | 3.367E-08 | 2.401E-08 | 1.300E-08 | 6.000E-08 |
| 4508 | 226 | K226G | A | 2.1e-08 | 2.9e-08 | 2.4e-08 | 2.467E-08 | 4.041E-09 | 2.100E-08 | 2.900E-08 |
| 4509 | 226 | K226H | A | 7e-08 | 3.2e-08 | 4.3e-08 | 4.833E-08 | 1.955E-08 | 3.200E-08 | 7.000E-08 |
| 4510 | 226 | K226I | A | 3.1e-08 | 3.5e-08 | 6.9e-08 | 4.500E-08 | 2.088E-08 | 3.100E-08 | 6.900E-08 |
| 4511 | 226 | K226L | A | 1.1e-07 | 1e-07 | 2e-08 | 7.667E-08 | 4.933E-08 | 2.000E-08 | 1.100E-07 |
| 4512 | 226 | K226K | A | 6e-08 | 5.8e-08 | 7.5e-08 | 6.433E-08 | 9.292E-09 | 5.800E-08 | 7.500E-08 |
| 4513 | 226 | K226M | A | 4.8e-08 | 2.3e-08 | 1e-08 | 2.700E-08 | 1.931E-08 | 1.000E-08 | 4.800E-08 |
| 4514 | 226 | K226F | A | 2.9e-08 | 9.3e-08 | 2.9e-08 | 5.033E-08 | 3.695E-08 | 2.900E-08 | 9.300E-08 |
| 4515 | 226 | K226P | A | 1.3e-07 | 3.2e-08 | 5.5e-08 | 7.233E-08 | 5.125E-08 | 3.200E-08 | 1.300E-07 |
| 4516 | 226 | K226S | A | 2.9e-08 | 2.2e-08 | 1e-07 | 5.033E-08 | 4.315E-08 | 2.200E-08 | 1.000E-07 |
| 4517 | 226 | K226T | A | 2.3e-08 | 8.6e-08 | 3.1e-08 | 4.667E-08 | 3.430E-08 | 2.300E-08 | 8.600E-08 |
| 4518 | 226 | K226W | A | 5.1e-08 | 5.4e-08 | 2.9e-08 | 4.467E-08 | 1.365E-08 | 2.900E-08 | 5.400E-08 |
| 4519 | 226 | K226Y | A | 3.1e-08 | 7.4e-08 | 1.5e-08 | 4.000E-08 | 3.051E-08 | 1.500E-08 | 7.400E-08 |
| 4520 | 226 | K226V | A | 9.5e-08 | 4.1e-08 | 3.5e-08 | 5.700E-08 | 3.305E-08 | 3.500E-08 | 9.500E-08 |
| 4521 | 227 | H227A | A | 5.7e-08 | 4.3e-08 | 5.7e-08 | 5.233E-08 | 8.083E-09 | 4.300E-08 | 5.700E-08 |
| 4522 | 227 | H227R | A | 4.1e-08 | 6.1e-08 | 2.8e-08 | 4.333E-08 | 1.662E-08 | 2.800E-08 | 6.100E-08 |
| 4523 | 227 | H227N | A | 4.6e-08 | 1.7e-08 | 3.4e-08 | 3.233E-08 | 1.457E-08 | 1.700E-08 | 4.600E-08 |
| 4524 | 227 | H227D | A | 3.1e-08 | 4.2e-08 | 5.1e-08 | 4.133E-08 | 1.002E-08 | 3.100E-08 | 5.100E-08 |
| 4525 | 227 | H227C | A | 2e-08 | 6.3e-08 | 6.4e-08 | 4.900E-08 | 2.512E-08 | 2.000E-08 | 6.400E-08 |
| 4526 | 227 | H227E | A | 6.9e-08 | 6.5e-08 | 4.2e-08 | 5.867E-08 | 1.457E-08 | 4.200E-08 | 6.900E-08 |
| 4527 | 227 | H227Q | A | 5.9e-08 | 8.4e-08 | 8.9e-09 | 5.063E-08 | 3.824E-08 | 8.900E-09 | 8.400E-08 |
| 4528 | 227 | H227G | A | 5.3e-08 | 5.3e-08 | 1.1e-07 | 7.200E-08 | 3.291E-08 | 5.300E-08 | 1.100E-07 |
| 4529 | 227 | H227H | A | 2.9e-08 | 2.4e-08 | 1.2e-07 | 5.767E-08 | 5.404E-08 | 2.400E-08 | 1.200E-07 |
| 4530 | 227 | H227I | A | 3.6e-08 | 1.4e-08 | 4.4e-08 | 3.133E-08 | 1.553E-08 | 1.400E-08 | 4.400E-08 |
| 4531 | 227 | H227L | A | 8.1e-08 | 5.4e-08 | 1.3e-08 | 4.933E-08 | 3.424E-08 | 1.300E-08 | 8.100E-08 |
| 4532 | 227 | H227K | A | 3.4e-08 | 5.1e-08 | 2.9e-08 | 3.800E-08 | 1.153E-08 | 2.900E-08 | 5.100E-08 |
| 4533 | 227 | H227M | A | 4.9e-08 | 9.1e-08 | 4.7e-08 | 6.233E-08 | 2.485E-08 | 4.700E-08 | 9.100E-08 |
| 4534 | 227 | H227F | A | 4.6e-08 | 5.4e-08 | 4.3e-08 | 4.767E-08 | 5.686E-09 | 4.300E-08 | 5.400E-08 |
| 4535 | 227 | H227P | A | 3.6e-08 | 9.3e-08 | 1.4e-07 | 8.967E-08 | 5.208E-08 | 3.600E-08 | 1.400E-07 |
| 4536 | 227 | H227S | A | 7.1e-08 | 4.6e-08 | 2e-08 | 4.567E-08 | 2.550E-08 | 2.000E-08 | 7.100E-08 |
| 4537 | 227 | H227T | A | 3.9e-08 | 7.1e-08 | 4e-08 | 5.000E-08 | 1.819E-08 | 3.900E-08 | 7.100E-08 |
| 4538 | 227 | H227W | A | 5.9e-08 | 4.3e-08 | 2.2e-08 | 4.133E-08 | 1.856E-08 | 2.200E-08 | 5.900E-08 |
| 4539 | 227 | H227Y | A | 3.5e-08 | 6.8e-08 | 7.4e-08 | 5.900E-08 | 2.100E-08 | 3.500E-08 | 7.400E-08 |
| 4540 | 227 | H227V | A | 2.2e-08 | 5.2e-08 | 3e-08 | 3.467E-08 | 1.553E-08 | 2.200E-08 | 5.200E-08 |
| 4541 | 228 | S228A | A | 3.7e-08 | 1.2e-07 | 8.5e-08 | 8.067E-08 | 4.167E-08 | 3.700E-08 | 1.200E-07 |
| 4542 | 228 | S228R | A | 7.4e-08 | 3.2e-08 | 3.5e-08 | 4.700E-08 | 2.343E-08 | 3.200E-08 | 7.400E-08 |
| 4543 | 228 | S228N | A | 3e-08 | 2.3e-08 | 4.6e-08 | 3.300E-08 | 1.179E-08 | 2.300E-08 | 4.600E-08 |
| 4544 | 228 | S228D | A | 9.1e-08 | 2.1e-08 | 8.4e-08 | 6.533E-08 | 3.855E-08 | 2.100E-08 | 9.100E-08 |
| 4545 | 228 | S228C | A | 2.8e-08 | 2.7e-08 | 1.1e-07 | 5.500E-08 | 4.763E-08 | 2.700E-08 | 1.100E-07 |
| 4546 | 228 | S228E | A | 2e-08 | 2.7e-08 | 6.1e-08 | 3.600E-08 | 2.193E-08 | 2.000E-08 | 6.100E-08 |
| 4547 | 228 | S228Q | A | 3.8e-08 | 3e-08 | 8.4e-08 | 5.067E-08 | 2.914E-08 | 3.000E-08 | 8.400E-08 |
| 4548 | 228 | S228G | A | 4e-08 | 1.8e-08 | 2.8e-08 | 2.867E-08 | 1.102E-08 | 1.800E-08 | 4.000E-08 |
| 4549 | 228 | S228H | A | 7.8e-08 | 8.2e-08 | 7.2e-08 | 7.733E-08 | 5.033E-09 | 7.200E-08 | 8.200E-08 |
| 4550 | 228 | S228I | A | 6e-08 | 3.7e-08 | 2.4e-08 | 4.033E-08 | 1.823E-08 | 2.400E-08 | 6.000E-08 |
| 4551 | 228 | S228L | A | 4.1e-08 | 5.3e-08 | 1.2e-08 | 3.533E-08 | 2.108E-08 | 1.200E-08 | 5.300E-08 |
| 4552 | 228 | S228K | A | 3.7e-08 | 5.3e-08 | 8.3e-08 | 5.767E-08 | 2.335E-08 | 3.700E-08 | 8.300E-08 |
| 4553 | 228 | S228M | A | 2.3e-08 | 4.8e-08 | 3e-08 | 3.367E-08 | 1.290E-08 | 2.300E-08 | 4.800E-08 |
| 4554 | 228 | S228F | A | 2.7e-08 | 5.7e-08 | 3.7e-08 | 4.033E-08 | 1.528E-08 | 2.700E-08 | 5.700E-08 |
| 4555 | 228 | S228P | A | 1.4e-08 | 1.1e-07 | 1.5e-08 | 4.633E-08 | 5.514E-08 | 1.400E-08 | 1.100E-07 |
| 4556 | 228 | S228S | A | 4.8e-08 | 5e-08 | 2.9e-08 | 4.233E-08 | 1.159E-08 | 2.900E-08 | 5.000E-08 |
| 4557 | 228 | S228T | A | 8.4e-08 | 8.1e-08 | 2.1e-08 | 6.200E-08 | 3.554E-08 | 2.100E-08 | 8.400E-08 |
| 4558 | 228 | S228W | A | 6.4e-08 | 6.8e-08 | 8.9e-08 | 7.367E-08 | 1.343E-08 | 6.400E-08 | 8.900E-08 |
| 4559 | 228 | S228Y | A | 1.8e-07 | 6.4e-08 | 7.6e-08 | 1.067E-07 | 6.379E-08 | 6.400E-08 | 1.800E-07 |
| 4560 | 228 | S228V | A | 1.9e-08 | 6.7e-08 | 4.5e-08 | 4.367E-08 | 2.403E-08 | 1.900E-08 | 6.700E-08 |
| 4561 | 229 | D229A | A | 3.9e-08 | 4.3e-08 | 2.9e-08 | 3.700E-08 | 7.211E-09 | 2.900E-08 | 4.300E-08 |
| 4562 | 229 | D229R | A | 5e-08 | 5.3e-08 | 7.8e-09 | 3.693E-08 | 2.527E-08 | 7.800E-09 | 5.300E-08 |
| 4563 | 229 | D229N | A | 3.3e-08 | 2e-08 | 2.8e-08 | 2.700E-08 | 6.557E-09 | 2.000E-08 | 3.300E-08 |
| 4564 | 229 | D229D | A | 5.2e-08 | 6.3e-08 | 4.1e-08 | 5.200E-08 | 1.100E-08 | 4.100E-08 | 6.300E-08 |
| 4565 | 229 | D229C | A | 5.3e-08 | 1.3e-07 | 2e-08 | 6.767E-08 | 5.645E-08 | 2.000E-08 | 1.300E-07 |
| 4566 | 229 | D229E | A | 5.2e-08 | 4.5e-08 | 1.8e-08 | 3.833E-08 | 1.795E-08 | 1.800E-08 | 5.200E-08 |
| 4567 | 229 | D229Q | A | 4.5e-08 | 1.6e-08 | 6.6e-08 | 4.233E-08 | 2.511E-08 | 1.600E-08 | 6.600E-08 |
| 4568 | 229 | D229G | A | 3.2e-08 | 4.9e-08 | 1.1e-07 | 6.367E-08 | 4.102E-08 | 3.200E-08 | 1.100E-07 |
| 4569 | 229 | D229H | A | 6e-08 | 1.7e-08 | 2.7e-08 | 3.467E-08 | 2.250E-08 | 1.700E-08 | 6.000E-08 |
| 4570 | 229 | D229I | A | 6.7e-08 | 4.2e-08 | 9.2e-08 | 6.700E-08 | 2.500E-08 | 4.200E-08 | 9.200E-08 |
| 4571 | 229 | D229L | A | 7.6e-08 | 1.2e-07 | 9e-08 | 9.533E-08 | 2.248E-08 | 7.600E-08 | 1.200E-07 |
| 4572 | 229 | D229K | A | 3.5e-08 | 3.3e-08 | 5.6e-08 | 4.133E-08 | 1.274E-08 | 3.300E-08 | 5.600E-08 |
| 4573 | 229 | D229M | A | 6.5e-08 | 1.3e-07 | 4.2e-08 | 7.900E-08 | 4.564E-08 | 4.200E-08 | 1.300E-07 |
| 4574 | 229 | D229F | A | 2.4e-08 | 2.6e-08 | 4.5e-08 | 3.167E-08 | 1.159E-08 | 2.400E-08 | 4.500E-08 |
| 4575 | 229 | D229P | A | 1.5e-07 | 5.3e-08 | 4e-08 | 8.100E-08 | 6.011E-08 | 4.000E-08 | 1.500E-07 |
| 4576 | 229 | D229S | A | 1.8e-07 | 3e-08 | 2.8e-08 | 7.933E-08 | 8.719E-08 | 2.800E-08 | 1.800E-07 |
| 4577 | 229 | D229T | A | 8.7e-09 | 5.2e-08 | 4.4e-08 | 3.490E-08 | 2.304E-08 | 8.700E-09 | 5.200E-08 |
| 4578 | 229 | D229W | A | 4.8e-08 | 5.3e-08 | 1.4e-08 | 3.833E-08 | 2.122E-08 | 1.400E-08 | 5.300E-08 |
| 4579 | 229 | D229Y | A | 7.9e-08 | 8.9e-09 | 2.3e-08 | 3.697E-08 | 3.708E-08 | 8.900E-09 | 7.900E-08 |
| 4580 | 229 | D229V | A | 4.2e-08 | 3.1e-08 | 9.5e-08 | 5.600E-08 | 3.422E-08 | 3.100E-08 | 9.500E-08 |
| 4581 | 230 | C230A | A | 2.9e-08 | 7.4e-08 | 5.6e-08 | 5.300E-08 | 2.265E-08 | 2.900E-08 | 7.400E-08 |
| 4582 | 230 | C230R | A | 6.4e-08 | 3.1e-08 | 3.5e-08 | 4.333E-08 | 1.801E-08 | 3.100E-08 | 6.400E-08 |
| 4583 | 230 | C230N | A | 3.2e-08 | 3.4e-08 | 4.4e-08 | 3.667E-08 | 6.429E-09 | 3.200E-08 | 4.400E-08 |
| 4584 | 230 | C230D | A | 2.6e-08 | 8.2e-08 | 6.3e-08 | 5.700E-08 | 2.848E-08 | 2.600E-08 | 8.200E-08 |
| 4585 | 230 | C230C | A | 6.7e-08 | 5.3e-08 | 1.1e-07 | 7.667E-08 | 2.970E-08 | 5.300E-08 | 1.100E-07 |
| 4586 | 230 | C230E | A | 3.4e-08 | 5.6e-08 | 1e-08 | 3.333E-08 | 2.301E-08 | 1.000E-08 | 5.600E-08 |
| 4587 | 230 | C230Q | A | 7.1e-08 | 2.3e-08 | 9.8e-08 | 6.400E-08 | 3.799E-08 | 2.300E-08 | 9.800E-08 |
| 4588 | 230 | C230G | A | 8.5e-08 | 5.9e-08 | 1e-07 | 8.133E-08 | 2.074E-08 | 5.900E-08 | 1.000E-07 |
| 4589 | 230 | C230H | A | 6.3e-08 | 1.2e-08 | 1.4e-08 | 2.967E-08 | 2.888E-08 | 1.200E-08 | 6.300E-08 |

| | | | | | | | | | | |
|---|---|---|---|---|---|---|---|---|---|---|
| 4590 | 230 | C230I | A | 2.7e-08 | 7.6e-08 | 3.1e-08 | 4.467E-08 | 2.721E-08 | 2.700E-08 | 7.600E-08 |
| 4591 | 230 | C230L | A | 2.1e-08 | 2.1e-08 | 4.6e-08 | 2.933E-08 | 1.443E-08 | 2.100E-08 | 4.600E-08 |
| 4592 | 230 | C230K | A | 7.3e-08 | 1.2e-07 | 6.8e-08 | 8.700E-08 | 2.869E-08 | 6.800E-08 | 1.200E-07 |
| 4593 | 230 | C230M | A | 4.6e-08 | 4.8e-08 | 3.1e-08 | 4.167E-08 | 9.292E-09 | 3.100E-08 | 4.800E-08 |
| 4594 | 230 | C230F | A | 3.1e-08 | 9.9e-08 | 4.6e-08 | 5.867E-08 | 3.573E-08 | 3.100E-08 | 9.900E-08 |
| 4595 | 230 | C230P | A | 4.6e-08 | 1.2e-07 | 2.2e-08 | 6.267E-08 | 5.108E-08 | 2.200E-08 | 1.200E-07 |
| 4596 | 230 | C230S | A | 3.4e-08 | 5.6e-08 | 5.1e-08 | 4.700E-08 | 1.153E-08 | 3.400E-08 | 5.600E-08 |
| 4597 | 230 | C230T | A | 2.1e-08 | 2e-08 | 9.9e-08 | 4.667E-08 | 4.532E-08 | 2.000E-08 | 9.900E-08 |
| 4598 | 230 | C230W | A | 8e-08 | 4.6e-08 | 5.6e-08 | 6.067E-08 | 1.747E-08 | 4.600E-08 | 8.000E-08 |
| 4599 | 230 | C230Y | A | 6.7e-08 | 9.3e-08 | 8.1e-08 | 8.033E-08 | 1.301E-08 | 6.700E-08 | 9.300E-08 |
| 4600 | 230 | C230V | A | 8.6e-08 | 3.1e-08 | 1.9e-07 | 1.023E-07 | 8.075E-08 | 3.100E-08 | 1.900E-07 |
| 4601 | 231 | L231A | A | 4.8e-08 | 6.1e-08 | 1.1e-08 | 4.000E-08 | 2.594E-08 | 1.100E-08 | 6.100E-08 |
| 4602 | 231 | L231R | A | 2.1e-08 | 7.8e-08 | 1.2e-07 | 7.300E-08 | 4.969E-08 | 2.100E-08 | 1.200E-07 |
| 4603 | 231 | L231N | A | 9e-08 | 6e-08 | 1.3e-08 | 5.433E-08 | 3.881E-08 | 1.300E-08 | 9.000E-08 |
| 4604 | 231 | L231D | A | 3e-08 | 8.9e-08 | 8.9e-08 | 6.933E-08 | 3.406E-08 | 3.000E-08 | 8.900E-08 |
| 4605 | 231 | L231C | A | 6.2e-08 | 1.8e-08 | 5.1e-08 | 4.367E-08 | 2.290E-08 | 1.800E-08 | 6.200E-08 |
| 4606 | 231 | L231E | A | 2.2e-08 | 1.7e-08 | 2.9e-08 | 2.267E-08 | 6.028E-09 | 1.700E-08 | 2.900E-08 |
| 4607 | 231 | L231Q | A | 3.5e-08 | 1.5e-08 | 3e-08 | 2.667E-08 | 1.041E-08 | 1.500E-08 | 3.500E-08 |
| 4608 | 231 | L231G | A | 4.3e-08 | 1.7e-08 | 9.8e-08 | 5.267E-08 | 4.136E-08 | 1.700E-08 | 9.800E-08 |
| 4609 | 231 | L231H | A | 4.2e-08 | 4.7e-08 | 6.6e-08 | 5.167E-08 | 1.266E-08 | 4.200E-08 | 6.600E-08 |
| 4610 | 231 | L231I | A | 4.7e-08 | 8.7e-08 | 2.6e-08 | 5.333E-08 | 3.099E-08 | 2.600E-08 | 8.700E-08 |
| 4611 | 231 | L231L | A | 2.1e-08 | 5.3e-08 | 5.6e-08 | 4.333E-08 | 1.940E-08 | 2.100E-08 | 5.600E-08 |
| 4612 | 231 | L231K | A | 3.9e-08 | 9.1e-08 | 4.2e-08 | 5.733E-08 | 2.919E-08 | 3.900E-08 | 9.100E-08 |
| 4613 | 231 | L231M | A | 4.5e-08 | 3.2e-08 | 2.5e-08 | 3.400E-08 | 1.015E-08 | 2.500E-08 | 4.500E-08 |
| 4614 | 231 | L231F | A | 5.7e-08 | 5.8e-08 | 7.2e-08 | 6.233E-08 | 8.386E-09 | 5.700E-08 | 7.200E-08 |
| 4615 | 231 | L231P | A | 3.8e-08 | 2e-08 | 6e-08 | 3.933E-08 | 2.003E-08 | 2.000E-08 | 6.000E-08 |
| 4616 | 231 | L231S | A | 2.7e-08 | 4.9e-08 | 3.5e-08 | 3.700E-08 | 1.114E-08 | 2.700E-08 | 4.900E-08 |
| 4617 | 231 | L231T | A | 3.1e-08 | 1e-07 | 4.1e-08 | 5.733E-08 | 3.729E-08 | 3.100E-08 | 1.000E-07 |
| 4618 | 231 | L231W | A | 3.3e-08 | 4.2e-08 | 4.2e-08 | 3.900E-08 | 5.196E-09 | 3.300E-08 | 4.200E-08 |
| 4619 | 231 | L231Y | A | 4.9e-08 | 7.6e-08 | 3.8e-08 | 5.433E-08 | 1.955E-08 | 3.800E-08 | 7.600E-08 |
| 4620 | 231 | L231V | A | 3.5e-08 | 1.3e-07 | 4.6e-08 | 7.033E-08 | 5.196E-08 | 3.500E-08 | 1.300E-07 |
| 4621 | 232 | A232A | A | 9.1e-08 | 3.5e-08 | 4.8e-08 | 5.800E-08 | 2.931E-08 | 3.500E-08 | 9.100E-08 |
| 4622 | 232 | A232R | A | 5.1e-08 | 4.3e-08 | 2.1e-08 | 3.833E-08 | 1.553E-08 | 2.100E-08 | 5.100E-08 |
| 4623 | 232 | A232N | A | 3.3e-08 | 2.9e-08 | 3e-08 | 3.067E-08 | 2.082E-09 | 2.900E-08 | 3.300E-08 |
| 4624 | 232 | A232D | A | 4.8e-08 | 3.2e-08 | 3.9e-08 | 3.967E-08 | 8.021E-09 | 3.200E-08 | 4.800E-08 |
| 4625 | 232 | A232C | A | 8.6e-09 | 1.7e-08 | 3.3e-08 | 1.953E-08 | 1.240E-08 | 8.600E-09 | 3.300E-08 |
| 4626 | 232 | A232E | A | 8.8e-09 | 1.8e-07 | 4.4e-08 | 7.760E-08 | 9.041E-08 | 8.800E-09 | 1.800E-07 |
| 4627 | 232 | A232Q | A | 2.4e-08 | 8.8e-08 | 2.1e-08 | 4.433E-08 | 3.785E-08 | 2.100E-08 | 8.800E-08 |
| 4628 | 232 | A232G | A | 3.3e-08 | 1.2e-07 | 8e-08 | 7.767E-08 | 4.355E-08 | 3.300E-08 | 1.200E-07 |
| 4629 | 232 | A232H | A | 2.2e-08 | 3.1e-08 | 4.8e-08 | 3.367E-08 | 1.320E-08 | 2.200E-08 | 4.800E-08 |
| 4630 | 232 | A232I | A | 3.3e-08 | 2.6e-08 | 4.5e-08 | 3.467E-08 | 9.609E-09 | 2.600E-08 | 4.500E-08 |
| 4631 | 232 | A232L | A | 8.3e-08 | 3.3e-08 | 6.9e-08 | 6.167E-08 | 2.579E-08 | 3.300E-08 | 8.300E-08 |
| 4632 | 232 | A232K | A | 7.1e-08 | 2.4e-08 | 3.9e-08 | 4.467E-08 | 2.401E-08 | 2.400E-08 | 7.100E-08 |
| 4633 | 232 | A232M | A | 3e-08 | 4e-08 | 1e-07 | 5.667E-08 | 3.786E-08 | 3.000E-08 | 1.000E-07 |
| 4634 | 232 | A232F | A | 8.3e-08 | 8.7e-08 | 7.7e-08 | 8.233E-08 | 5.033E-09 | 7.700E-08 | 8.700E-08 |
| 4635 | 232 | A232P | A | 5.5e-08 | 1.1e-07 | 3e-08 | 6.500E-08 | 4.093E-08 | 3.000E-08 | 1.100E-07 |
| 4636 | 232 | A232S | A | 3.7e-08 | 4.1e-08 | 1.2e-07 | 6.600E-08 | 4.681E-08 | 3.700E-08 | 1.200E-07 |
| 4637 | 232 | A232T | A | 3.2e-08 | 4.9e-08 | 6.1e-08 | 4.733E-08 | 1.457E-08 | 3.200E-08 | 6.100E-08 |
| 4638 | 232 | A232W | A | 6.1e-08 | 3e-08 | 5.1e-08 | 4.733E-08 | 1.582E-08 | 3.000E-08 | 6.100E-08 |
| 4639 | 232 | A232Y | A | 1.1e-07 | 4.2e-08 | 7e-08 | 7.400E-08 | 3.418E-08 | 4.200E-08 | 1.100E-07 |
| 4640 | 232 | A232V | A | 7.9e-08 | 5e-08 | 4e-08 | 5.633E-08 | 2.026E-08 | 4.000E-08 | 7.900E-08 |
| 4641 | 233 | C233A | A | 4.1e-08 | 1.8e-08 | 3e-08 | 2.967E-08 | 1.150E-08 | 1.800E-08 | 4.100E-08 |
| 4642 | 233 | C233R | A | 1.6e-08 | 3.4e-08 | 4e-08 | 3.000E-08 | 1.249E-08 | 1.600E-08 | 4.000E-08 |
| 4643 | 233 | C233N | A | 4.1e-08 | 2.3e-08 | 7e-08 | 4.467E-08 | 2.371E-08 | 2.300E-08 | 7.000E-08 |
| 4644 | 233 | C233D | A | 1.8e-08 | 3.3e-08 | 3.1e-08 | 2.733E-08 | 8.145E-09 | 1.800E-08 | 3.300E-08 |
| 4645 | 233 | C233C | A | 7.5e-08 | 3.6e-08 | 4.4e-08 | 5.167E-08 | 2.060E-08 | 3.600E-08 | 7.500E-08 |
| 4646 | 233 | C233E | A | 3.9e-08 | 4.4e-08 | 6.1e-08 | 4.800E-08 | 1.153E-08 | 3.900E-08 | 6.100E-08 |
| 4647 | 233 | C233Q | A | 4.2e-08 | 7.1e-09 | 4.4e-08 | 3.103E-08 | 2.075E-08 | 7.100E-09 | 4.400E-08 |
| 4648 | 233 | C233G | A | 5e-08 | 9.4e-08 | 2.4e-08 | 5.600E-08 | 3.538E-08 | 2.400E-08 | 9.400E-08 |
| 4649 | 233 | C233H | A | 4.1e-08 | 6.5e-08 | 4.2e-08 | 4.933E-08 | 1.358E-08 | 4.100E-08 | 6.500E-08 |
| 4650 | 233 | C233I | A | 1.1e-08 | 6.1e-08 | 1.9e-08 | 3.033E-08 | 2.686E-08 | 1.100E-08 | 6.100E-08 |
| 4651 | 233 | C233L | A | 3e-08 | 1.1e-07 | 2.7e-08 | 5.567E-08 | 4.708E-08 | 2.700E-08 | 1.100E-07 |
| 4652 | 233 | C233K | A | 3.6e-08 | 3.2e-08 | 3.6e-08 | 3.467E-08 | 2.309E-09 | 3.200E-08 | 3.600E-08 |
| 4653 | 233 | C233M | A | 5.6e-08 | 3.9e-08 | 2.7e-08 | 4.067E-08 | 1.457E-08 | 2.700E-08 | 5.600E-08 |
| 4654 | 233 | C233F | A | 8.7e-08 | 4.9e-08 | 2.7e-08 | 5.433E-08 | 3.035E-08 | 2.700E-08 | 8.700E-08 |
| 4655 | 233 | C233P | A | 2.7e-08 | 5.2e-08 | 5.6e-08 | 4.500E-08 | 1.572E-08 | 2.700E-08 | 5.600E-08 |
| 4656 | 233 | C233S | A | 2e-08 | 1.3e-07 | 7.3e-08 | 7.433E-08 | 5.501E-08 | 2.000E-08 | 1.300E-07 |
| 4657 | 233 | C233T | A | 3.9e-08 | 4.9e-08 | 3.5e-08 | 4.100E-08 | 7.211E-09 | 3.500E-08 | 4.900E-08 |
| 4658 | 233 | C233W | A | 1.9e-08 | 7.9e-08 | 1.1e-07 | 6.933E-08 | 4.626E-08 | 1.900E-08 | 1.100E-07 |
| 4659 | 233 | C233Y | A | 4.7e-08 | 4.6e-08 | 3.4e-08 | 4.233E-08 | 7.234E-09 | 3.400E-08 | 4.700E-08 |
| 4660 | 233 | C233V | A | 2.3e-08 | 1.8e-07 | 3.5e-08 | 7.933E-08 | 8.739E-08 | 2.300E-08 | 1.800E-07 |
| 4661 | 234 | L234A | A | 9e-08 | 4e-08 | 4.8e-08 | 5.933E-08 | 2.686E-08 | 4.000E-08 | 9.000E-08 |
| 4662 | 234 | L234R | A | 8.7e-08 | 1e-07 | 6.3e-08 | 8.333E-08 | 1.877E-08 | 6.300E-08 | 1.000E-07 |
| 4663 | 234 | L234N | A | 1.1e-08 | 3.5e-08 | 3.1e-08 | 2.567E-08 | 1.286E-08 | 1.100E-08 | 3.500E-08 |
| 4664 | 234 | L234D | A | 2e-08 | 3.3e-08 | 2.4e-08 | 2.567E-08 | 6.658E-09 | 2.000E-08 | 3.300E-08 |
| 4665 | 234 | L234C | A | 5.8e-08 | 1.1e-07 | 1.5e-08 | 6.100E-08 | 4.757E-08 | 1.500E-08 | 1.100E-07 |
| 4666 | 234 | L234E | A | 4.3e-08 | 4.6e-08 | 9e-08 | 5.967E-08 | 2.631E-08 | 4.300E-08 | 9.000E-08 |
| 4667 | 234 | L234Q | A | 7.4e-08 | 7.7e-08 | 4.3e-08 | 6.467E-08 | 1.882E-08 | 4.300E-08 | 7.700E-08 |
| 4668 | 234 | L234G | A | 3e-08 | 4e-08 | 2.7e-08 | 3.233E-08 | 6.807E-09 | 2.700E-08 | 4.000E-08 |
| 4669 | 234 | L234H | A | 3.7e-08 | 4.8e-08 | 1.3e-08 | 3.267E-08 | 1.790E-08 | 1.300E-08 | 4.800E-08 |
| 4670 | 234 | L234I | A | 5.9e-08 | 2e-08 | 3.9e-08 | 3.933E-08 | 1.950E-08 | 2.000E-08 | 5.900E-08 |
| 4671 | 234 | L234L | A | 3.5e-08 | 3.4e-08 | 5e-08 | 3.967E-08 | 8.963E-09 | 3.400E-08 | 5.000E-08 |
| 4672 | 234 | L234K | A | 5.1e-08 | 6.3e-08 | 6.2e-08 | 5.867E-08 | 6.658E-09 | 5.100E-08 | 6.300E-08 |
| 4673 | 234 | L234M | A | 4.3e-08 | 5.5e-08 | 4e-08 | 4.600E-08 | 7.937E-09 | 4.000E-08 | 5.500E-08 |
| 4674 | 234 | L234F | A | 2.1e-08 | 2e-08 | 3.7e-08 | 2.600E-08 | 9.539E-09 | 2.000E-08 | 3.700E-08 |

| | | | | | | | | | | |
|---|---|---|---|---|---|---|---|---|---|---|
| 4675 | 234 | L234P | A | 3.3e-08 | 2.7e-08 | 2e-08 | 2.667E-08 | 6.506E-09 | 2.000E-08 | 3.300E-08 |
| 4676 | 234 | L234S | A | 1.6e-08 | 5.1e-08 | 2.2e-08 | 2.967E-08 | 1.872E-08 | 1.600E-08 | 5.100E-08 |
| 4677 | 234 | L234T | A | 3.5e-08 | 2.6e-08 | 4.9e-08 | 3.667E-08 | 1.159E-08 | 2.600E-08 | 4.900E-08 |
| 4678 | 234 | L234W | A | 4.6e-08 | 7.6e-08 | 3e-08 | 5.067E-08 | 2.335E-08 | 3.000E-08 | 7.600E-08 |
| 4679 | 234 | L234Y | A | 1e-07 | 1.5e-08 | 4.5e-08 | 5.333E-08 | 4.311E-08 | 1.500E-08 | 1.000E-07 |
| 4680 | 234 | L234V | A | 3.7e-08 | 4.3e-08 | 3.9e-08 | 3.967E-08 | 3.055E-09 | 3.700E-08 | 4.300E-08 |
| 4681 | 235 | H235A | A | 5.9e-08 | 3.4e-08 | 3.4e-08 | 4.233E-08 | 1.443E-08 | 3.400E-08 | 5.900E-08 |
| 4682 | 235 | H235R | A | 2.2e-08 | 4.8e-08 | 2e-08 | 3.000E-08 | 1.562E-08 | 2.000E-08 | 4.800E-08 |
| 4683 | 235 | H235N | A | 6.8e-08 | 7e-08 | 2.6e-08 | 5.467E-08 | 2.485E-08 | 2.600E-08 | 7.000E-08 |
| 4684 | 235 | H235D | A | 7.1e-08 | 1.9e-08 | 1.2e-08 | 3.400E-08 | 3.223E-08 | 1.200E-08 | 7.100E-08 |
| 4685 | 235 | H235C | A | 4.7e-08 | 2.8e-08 | 5.6e-08 | 4.367E-08 | 1.429E-08 | 2.800E-08 | 5.600E-08 |
| 4686 | 235 | H235E | A | 1.9e-08 | 4.1e-08 | 5.9e-08 | 3.967E-08 | 2.003E-08 | 1.900E-08 | 5.900E-08 |
| 4687 | 235 | H235Q | A | 4e-08 | 2.4e-08 | 1.1e-07 | 5.800E-08 | 4.574E-08 | 2.400E-08 | 1.100E-07 |
| 4688 | 235 | H235G | A | 2.6e-08 | 5.1e-08 | 8.4e-08 | 5.367E-08 | 2.909E-08 | 2.600E-08 | 8.400E-08 |
| 4689 | 235 | H235H | A | 8.9e-08 | 6.1e-08 | 4.6e-08 | 6.533E-08 | 2.183E-08 | 4.600E-08 | 8.900E-08 |
| 4690 | 235 | H235I | A | 7.4e-08 | 1.2e-08 | 4.1e-08 | 4.233E-08 | 3.102E-08 | 1.200E-08 | 7.400E-08 |
| 4691 | 235 | H235L | A | 1.8e-08 | 8.1e-08 | 4.6e-08 | 4.833E-08 | 3.156E-08 | 1.800E-08 | 8.100E-08 |
| 4692 | 235 | H235K | A | 5.6e-08 | 5e-08 | 3e-08 | 4.533E-08 | 1.361E-08 | 3.000E-08 | 5.600E-08 |
| 4693 | 235 | H235M | A | 1.2e-08 | 3.8e-08 | 3.5e-08 | 2.833E-08 | 1.422E-08 | 1.200E-08 | 3.800E-08 |
| 4694 | 235 | H235F | A | 1.8e-08 | 7.3e-08 | 4e-08 | 4.367E-08 | 2.768E-08 | 1.800E-08 | 7.300E-08 |
| 4695 | 235 | H235P | A | 6.7e-08 | 4.4e-08 | 1.8e-08 | 4.300E-08 | 2.452E-08 | 1.800E-08 | 6.700E-08 |
| 4696 | 235 | H235S | A | 3.3e-08 | 2.7e-08 | 3e-08 | 3.000E-08 | 3.000E-09 | 2.700E-08 | 3.300E-08 |
| 4697 | 235 | H235T | A | 2.8e-08 | 6e-08 | 5.7e-08 | 4.833E-08 | 1.767E-08 | 2.800E-08 | 6.000E-08 |
| 4698 | 235 | H235W | A | 1.8e-08 | 2.1e-08 | 1.2e-08 | 1.700E-08 | 4.583E-09 | 1.200E-08 | 2.100E-08 |
| 4699 | 235 | H235Y | A | 5.9e-08 | 2.5e-08 | 4.8e-08 | 4.400E-08 | 1.735E-08 | 2.500E-08 | 5.900E-08 |
| 4700 | 235 | H235V | A | 3.8e-08 | 2.7e-08 | 7.9e-08 | 4.800E-08 | 2.740E-08 | 2.700E-08 | 7.900E-08 |
| 4701 | 236 | F236A | A | 6.5e-08 | 7.6e-08 | 7.2e-08 | 7.100E-08 | 5.568E-09 | 6.500E-08 | 7.600E-08 |
| 4702 | 236 | F236R | A | 3.6e-08 | 4.2e-08 | 7.1e-08 | 4.967E-08 | 1.872E-08 | 3.600E-08 | 7.100E-08 |
| 4703 | 236 | F236N | A | 3.6e-08 | 7.8e-08 | 1e-07 | 7.133E-08 | 3.252E-08 | 3.600E-08 | 1.000E-07 |
| 4704 | 236 | F236D | A | 1.2e-07 | 9.6e-08 | 1e-07 | 1.053E-07 | 1.286E-08 | 9.600E-08 | 1.200E-07 |
| 4705 | 236 | F236C | A | 1.4e-08 | 4.4e-08 | 1.2e-07 | 5.933E-08 | 5.464E-08 | 1.400E-08 | 1.200E-07 |
| 4706 | 236 | F236E | A | 1e-07 | 1.9e-08 | 9.4e-08 | 7.100E-08 | 4.513E-08 | 1.900E-08 | 1.000E-07 |
| 4707 | 236 | F236Q | A | 1.9e-07 | 4.4e-08 | 1.2e-07 | 1.180E-07 | 7.302E-08 | 4.400E-08 | 1.900E-07 |
| 4708 | 236 | F236G | A | 9.6e-08 | 4.2e-08 | 6.5e-08 | 6.767E-08 | 2.710E-08 | 4.200E-08 | 9.600E-08 |
| 4709 | 236 | F236H | A | 3.4e-08 | 3.3e-08 | 1.6e-07 | 7.567E-08 | 7.304E-08 | 3.300E-08 | 1.600E-07 |
| 4710 | 236 | F236I | A | 4.8e-08 | 5.5e-08 | 6.3e-08 | 5.533E-08 | 7.506E-09 | 4.800E-08 | 6.300E-08 |
| 4711 | 236 | F236L | A | 4.7e-08 | 3.2e-08 | 3.4e-08 | 3.767E-08 | 8.145E-09 | 3.200E-08 | 4.700E-08 |
| 4712 | 236 | F236K | A | 2.5e-08 | 4.2e-08 | 2.9e-08 | 3.200E-08 | 8.888E-09 | 2.500E-08 | 4.200E-08 |
| 4713 | 236 | F236M | A | 3.8e-08 | 2.7e-08 | 2e-08 | 2.833E-08 | 9.074E-09 | 2.000E-08 | 3.800E-08 |
| 4714 | 236 | F236F | A | 4.5e-08 | 4e-08 | 1.8e-08 | 3.433E-08 | 1.436E-08 | 1.800E-08 | 4.500E-08 |
| 4715 | 236 | F236P | A | 1.7e-07 | 9.2e-08 | 8.6e-08 | 1.160E-07 | 4.686E-08 | 8.600E-08 | 1.700E-07 |
| 4716 | 236 | F236S | A | 1.1e-07 | 8.1e-08 | 9.4e-08 | 9.500E-08 | 1.453E-08 | 8.100E-08 | 1.100E-07 |
| 4717 | 236 | F236T | A | 2.3e-08 | 4.9e-08 | 6e-08 | 4.400E-08 | 1.900E-08 | 2.300E-08 | 6.000E-08 |
| 4718 | 236 | F236W | A | 4.2e-08 | 1.3e-08 | 4.9e-08 | 3.467E-08 | 1.909E-08 | 1.300E-08 | 4.900E-08 |
| 4719 | 236 | F236Y | A | 3.6e-08 | 1.8e-08 | 4.2e-08 | 3.200E-08 | 1.249E-08 | 1.800E-08 | 4.200E-08 |
| 4720 | 236 | F236V | A | 5.5e-08 | 6.1e-08 | 4.6e-08 | 5.400E-08 | 7.550E-09 | 4.600E-08 | 6.100E-08 |
| 4721 | 237 | N237A | A | 5.2e-08 | 4.7e-08 | 1e-07 | 6.633E-08 | 2.926E-08 | 4.700E-08 | 1.000E-07 |
| 4722 | 237 | N237R | A | 3.2e-08 | 1.1e-07 | 4.3e-08 | 6.167E-08 | 4.222E-08 | 3.200E-08 | 1.100E-07 |
| 4723 | 237 | N237N | A | 5.2e-08 | 3.5e-08 | 2.5e-08 | 3.733E-08 | 1.365E-08 | 2.500E-08 | 5.200E-08 |
| 4724 | 237 | N237D | A | 1.7e-08 | 7.2e-08 | 1.6e-08 | 3.500E-08 | 3.205E-08 | 1.600E-08 | 7.200E-08 |
| 4725 | 237 | N237C | A | 6.3e-08 | 6e-08 | 3.9e-08 | 5.400E-08 | 1.308E-08 | 3.900E-08 | 6.300E-08 |
| 4726 | 237 | N237E | A | 3e-08 | 2.7e-08 | 2.2e-08 | 2.633E-08 | 4.041E-09 | 2.200E-08 | 3.000E-08 |
| 4727 | 237 | N237Q | A | 5.3e-08 | 8.1e-09 | 3e-08 | 3.037E-08 | 2.245E-08 | 8.100E-09 | 5.300E-08 |
| 4728 | 237 | N237G | A | 9.7e-08 | 2.1e-08 | 2.7e-08 | 4.833E-08 | 4.225E-08 | 2.100E-08 | 9.700E-08 |
| 4729 | 237 | N237H | A | 1.8e-08 | 4.9e-08 | 7.9e-08 | 4.867E-08 | 3.050E-08 | 1.800E-08 | 7.900E-08 |
| 4730 | 237 | N237I | A | 4.5e-08 | 5.5e-08 | 5e-08 | 5.000E-08 | 5.000E-09 | 4.500E-08 | 5.500E-08 |
| 4731 | 237 | N237L | A | 3.7e-08 | 9e-08 | 8e-09 | 4.500E-08 | 4.158E-08 | 8.000E-09 | 9.000E-08 |
| 4732 | 237 | N237K | A | 6.8e-08 | 3.1e-08 | 4.8e-08 | 4.900E-08 | 1.852E-08 | 3.100E-08 | 6.800E-08 |
| 4733 | 237 | N237M | A | 1.1e-07 | 3.1e-08 | 2.2e-08 | 5.433E-08 | 4.842E-08 | 2.200E-08 | 1.100E-07 |
| 4734 | 237 | N237F | A | 5.1e-08 | 6e-08 | 4.6e-08 | 5.233E-08 | 7.095E-09 | 4.600E-08 | 6.000E-08 |
| 4735 | 237 | N237P | A | 2.9e-08 | 4.1e-08 | 2.5e-08 | 3.167E-08 | 8.327E-09 | 2.500E-08 | 4.100E-08 |
| 4736 | 237 | N237S | A | 4.5e-08 | 1.6e-08 | 2.8e-08 | 2.967E-08 | 1.457E-08 | 1.600E-08 | 4.500E-08 |
| 4737 | 237 | N237T | A | 6.1e-08 | 5.3e-08 | 4.7e-08 | 5.367E-08 | 7.024E-09 | 4.700E-08 | 6.100E-08 |
| 4738 | 237 | N237W | A | 8.1e-08 | 2.8e-08 | 9.9e-08 | 6.933E-08 | 3.691E-08 | 2.800E-08 | 9.900E-08 |
| 4739 | 237 | N237Y | A | 5.2e-08 | 5.1e-08 | 7.1e-08 | 5.800E-08 | 1.127E-08 | 5.100E-08 | 7.100E-08 |
| 4740 | 237 | N237V | A | 2.1e-08 | 2.4e-08 | 6.7e-08 | 3.733E-08 | 2.574E-08 | 2.100E-08 | 6.700E-08 |
| 4741 | 238 | H238A | A | 7.4e-08 | 8.1e-08 | 7.5e-08 | 7.667E-08 | 3.786E-09 | 7.400E-08 | 8.100E-08 |
| 4742 | 238 | H238R | A | 4.9e-08 | 2.7e-08 | 3.7e-08 | 3.767E-08 | 1.102E-08 | 2.700E-08 | 4.900E-08 |
| 4743 | 238 | H238N | A | 6.8e-08 | 6.7e-08 | 3.5e-08 | 5.667E-08 | 1.877E-08 | 3.500E-08 | 6.800E-08 |
| 4744 | 238 | H238D | A | 3.2e-08 | 3.9e-08 | 5.1e-08 | 4.067E-08 | 9.609E-09 | 3.200E-08 | 5.100E-08 |
| 4745 | 238 | H238C | A | 1.7e-08 | 2.1e-08 | 1.4e-08 | 1.733E-08 | 3.512E-09 | 1.400E-08 | 2.100E-08 |
| 4746 | 238 | H238E | A | 5.6e-08 | 3.4e-08 | 6.3e-08 | 5.100E-08 | 1.513E-08 | 3.400E-08 | 6.300E-08 |
| 4747 | 238 | H238Q | A | 3.3e-08 | 4.3e-08 | 1.1e-07 | 6.200E-08 | 4.187E-08 | 3.300E-08 | 1.100E-07 |
| 4748 | 238 | H238G | A | 2.1e-08 | 8e-08 | 4.9e-08 | 5.000E-08 | 2.951E-08 | 2.100E-08 | 8.000E-08 |
| 4749 | 238 | H238H | A | 3.5e-08 | 4.7e-08 | 6.9e-08 | 5.033E-08 | 1.724E-08 | 3.500E-08 | 6.900E-08 |
| 4750 | 238 | H238I | A | 6.5e-08 | 6.8e-08 | 1.6e-08 | 4.967E-08 | 2.919E-08 | 1.600E-08 | 6.800E-08 |
| 4751 | 238 | H238L | A | 3.9e-08 | 6.8e-08 | 1e-07 | 6.900E-08 | 3.051E-08 | 3.900E-08 | 1.000E-07 |
| 4752 | 238 | H238K | A | 4e-08 | 2.2e-08 | 3.8e-08 | 3.333E-08 | 9.866E-09 | 2.200E-08 | 4.000E-08 |
| 4753 | 238 | H238M | A | 3.3e-08 | 3.9e-08 | 5.2e-08 | 4.133E-08 | 9.713E-09 | 3.300E-08 | 5.200E-08 |
| 4754 | 238 | H238F | A | 4.4e-08 | 1.1e-08 | 3.7e-08 | 3.067E-08 | 1.739E-08 | 1.100E-08 | 4.400E-08 |
| 4755 | 238 | H238P | A | 2.9e-08 | 5.4e-08 | 9.3e-08 | 5.867E-08 | 3.225E-08 | 2.900E-08 | 9.300E-08 |
| 4756 | 238 | H238S | A | 1.2e-07 | 6.4e-08 | 8.8e-08 | 9.067E-08 | 2.810E-08 | 6.400E-08 | 1.200E-07 |
| 4757 | 238 | H238T | A | 3.6e-08 | 5.5e-09 | 6.2e-08 | 3.450E-08 | 2.828E-08 | 5.500E-09 | 6.200E-08 |
| 4758 | 238 | H238W | A | 3.5e-08 | 4e-08 | 8.3e-09 | 2.777E-08 | 1.704E-08 | 8.300E-09 | 4.000E-08 |
| 4759 | 238 | H238Y | A | 4.4e-08 | 8.4e-08 | 7.7e-08 | 6.833E-08 | 2.136E-08 | 4.400E-08 | 8.400E-08 |

| | | | | | | | | | | |
|---|---|---|---|---|---|---|---|---|---|---|
| 4760 | 238 | H238V | A | 8.6e-09 | 3.2e-08 | 3.3e-08 | 2.453E-08 | 1.381E-08 | 8.600E-09 | 3.300E-08 |
| 4761 | 239 | S239A | A | 8e-08 | 1.4e-07 | 3.4e-08 | 8.467E-08 | 5.315E-08 | 3.400E-08 | 1.400E-07 |
| 4762 | 239 | S239R | A | 7e-08 | 4.8e-08 | 1e-07 | 7.267E-08 | 2.610E-08 | 4.800E-08 | 1.000E-07 |
| 4763 | 239 | S239N | A | 2.7e-08 | 9.2e-08 | 2.8e-08 | 4.900E-08 | 3.724E-08 | 2.700E-08 | 9.200E-08 |
| 4764 | 239 | S239D | A | 4.5e-08 | 5e-08 | 1.1e-07 | 6.833E-08 | 3.617E-08 | 4.500E-08 | 1.100E-07 |
| 4765 | 239 | S239C | A | 5.7e-08 | 3e-08 | 3.4e-08 | 4.033E-08 | 1.457E-08 | 3.000E-08 | 5.700E-08 |
| 4766 | 239 | S239E | A | 6.2e-08 | 1.4e-07 | 2.6e-08 | 7.600E-08 | 5.828E-08 | 2.600E-08 | 1.400E-07 |
| 4767 | 239 | S239Q | A | 5.3e-08 | 3.9e-08 | 4e-08 | 4.400E-08 | 7.810E-09 | 3.900E-08 | 5.300E-08 |
| 4768 | 239 | S239G | A | 3.6e-08 | 3.8e-08 | 5e-08 | 4.133E-08 | 7.572E-09 | 3.600E-08 | 5.000E-08 |
| 4769 | 239 | S239H | A | 1e-08 | 1.7e-08 | 3.6e-08 | 2.100E-08 | 1.345E-08 | 1.000E-08 | 3.600E-08 |
| 4770 | 239 | S239I | A | 3e-08 | 4.3e-08 | 4.4e-08 | 3.900E-08 | 7.810E-09 | 3.000E-08 | 4.400E-08 |
| 4771 | 239 | S239L | A | 1.9e-08 | 4.9e-08 | 7.2e-08 | 4.667E-08 | 2.658E-08 | 1.900E-08 | 7.200E-08 |
| 4772 | 239 | S239K | A | 3.8e-08 | 3.2e-08 | 4.1e-08 | 3.700E-08 | 4.583E-09 | 3.200E-08 | 4.100E-08 |
| 4773 | 239 | S239M | A | 1.2e-07 | 7.6e-08 | 3.6e-08 | 7.733E-08 | 4.202E-08 | 3.600E-08 | 1.200E-07 |
| 4774 | 239 | S239F | A | 1.1e-07 | 2.5e-08 | 4e-08 | 5.833E-08 | 4.537E-08 | 2.500E-08 | 1.100E-07 |
| 4775 | 239 | S239P | A | 6e-08 | 6.4e-08 | 5.7e-08 | 6.033E-08 | 3.512E-09 | 5.700E-08 | 6.400E-08 |
| 4776 | 239 | S239S | A | 5.9e-08 | 8.6e-08 | 2.6e-08 | 5.700E-08 | 3.005E-08 | 2.600E-08 | 8.600E-08 |
| 4777 | 239 | S239T | A | 1.6e-08 | 1.3e-07 | 1.6e-08 | 5.400E-08 | 6.582E-08 | 1.600E-08 | 1.300E-07 |
| 4778 | 239 | S239W | A | 2.8e-08 | 1e-07 | 8e-08 | 6.933E-08 | 3.717E-08 | 2.800E-08 | 1.000E-07 |
| 4779 | 239 | S239Y | A | 1.3e-07 | 1.7e-08 | 4.5e-08 | 6.400E-08 | 5.885E-08 | 1.700E-08 | 1.300E-07 |
| 4780 | 239 | S239V | A | 1.7e-08 | 1e-08 | 5.1e-08 | 2.600E-08 | 2.193E-08 | 1.000E-08 | 5.100E-08 |
| 4781 | 240 | G240A | A | 2.2e-08 | 2.8e-08 | 6.7e-08 | 3.900E-08 | 2.443E-08 | 2.200E-08 | 6.700E-08 |
| 4782 | 240 | G240R | A | 2.1e-08 | 1.2e-07 | 8.5e-09 | 4.983E-08 | 6.109E-08 | 8.500E-09 | 1.200E-07 |
| 4783 | 240 | G240N | A | 4.1e-08 | 3.3e-08 | 2.1e-08 | 3.167E-08 | 1.007E-08 | 2.100E-08 | 4.100E-08 |
| 4784 | 240 | G240D | A | 2.2e-08 | 2.3e-08 | 5.8e-08 | 3.433E-08 | 2.050E-08 | 2.200E-08 | 5.800E-08 |
| 4785 | 240 | G240C | A | 3.9e-08 | 5.3e-08 | 2e-08 | 3.733E-08 | 1.656E-08 | 2.000E-08 | 5.300E-08 |
| 4786 | 240 | G240E | A | 1.6e-08 | 5.6e-08 | 2.3e-08 | 3.167E-08 | 2.136E-08 | 1.600E-08 | 5.600E-08 |
| 4787 | 240 | G240Q | A | 1.5e-08 | 1.4e-07 | 1.4e-08 | 5.633E-08 | 7.246E-08 | 1.400E-08 | 1.400E-07 |
| 4788 | 240 | G240G | A | 5.7e-08 | 3.1e-08 | 4.7e-08 | 4.500E-08 | 1.311E-08 | 3.100E-08 | 5.700E-08 |
| 4789 | 240 | G240H | A | 3.1e-08 | 1.4e-07 | 5.2e-08 | 7.433E-08 | 5.783E-08 | 3.100E-08 | 1.400E-07 |
| 4790 | 240 | G240I | A | 3.7e-08 | 5.5e-08 | 1.1e-07 | 6.733E-08 | 3.803E-08 | 3.700E-08 | 1.100E-07 |
| 4791 | 240 | G240L | A | 4.6e-08 | 2.9e-08 | 7.7e-08 | 5.067E-08 | 2.434E-08 | 2.900E-08 | 7.700E-08 |
| 4792 | 240 | G240K | A | 5e-08 | 8.5e-08 | 4.4e-08 | 5.967E-08 | 2.214E-08 | 4.400E-08 | 8.500E-08 |
| 4793 | 240 | G240M | A | 4.6e-08 | 5.2e-08 | 1.1e-08 | 3.633E-08 | 2.214E-08 | 1.100E-08 | 5.200E-08 |
| 4794 | 240 | G240F | A | 4e-08 | 3.2e-08 | 1.3e-08 | 2.833E-08 | 1.387E-08 | 1.300E-08 | 4.000E-08 |
| 4795 | 240 | G240P | A | 4.8e-08 | 2.6e-08 | 9.4e-08 | 5.600E-08 | 3.470E-08 | 2.600E-08 | 9.400E-08 |
| 4796 | 240 | G240S | A | 6.9e-08 | 1e-08 | 5.4e-08 | 4.433E-08 | 3.066E-08 | 1.000E-08 | 6.900E-08 |
| 4797 | 240 | G240T | A | 4e-08 | 3e-08 | 4.6e-08 | 3.867E-08 | 8.083E-09 | 3.000E-08 | 4.600E-08 |
| 4798 | 240 | G240W | A | 2.9e-08 | 1.2e-07 | 2.3e-08 | 5.733E-08 | 5.435E-08 | 2.300E-08 | 1.200E-07 |
| 4799 | 240 | G240Y | A | 2.9e-08 | 5.1e-08 | 1.2e-08 | 3.067E-08 | 1.955E-08 | 1.200E-08 | 5.100E-08 |
| 4800 | 240 | G240V | A | 7.5e-08 | 8.1e-08 | 3.5e-08 | 6.367E-08 | 2.501E-08 | 3.500E-08 | 8.100E-08 |
| 4801 | 241 | I241A | A | 4.5e-08 | 3.2e-08 | 2.6e-08 | 3.433E-08 | 9.713E-09 | 2.600E-08 | 4.500E-08 |
| 4802 | 241 | I241R | A | 3.9e-08 | 6.9e-08 | 4.9e-08 | 5.233E-08 | 1.528E-08 | 3.900E-08 | 6.900E-08 |
| 4803 | 241 | I241N | A | 1.2e-07 | 6.3e-08 | 7.6e-08 | 8.633E-08 | 2.987E-08 | 6.300E-08 | 1.200E-07 |
| 4804 | 241 | I241D | A | 6.1e-08 | 4.9e-08 | 3.5e-08 | 4.833E-08 | 1.301E-08 | 3.500E-08 | 6.100E-08 |
| 4805 | 241 | I241C | A | 2.8e-08 | 2.3e-08 | 1.5e-07 | 6.700E-08 | 7.192E-08 | 2.300E-08 | 1.500E-07 |
| 4806 | 241 | I241E | A | 8.8e-08 | 2.9e-08 | 7.3e-09 | 4.143E-08 | 4.176E-08 | 7.300E-09 | 8.800E-08 |
| 4807 | 241 | I241Q | A | 1.9e-08 | 5.5e-08 | 2.2e-08 | 3.200E-08 | 1.997E-08 | 1.900E-08 | 5.500E-08 |
| 4808 | 241 | I241G | A | 4.1e-08 | 8e-08 | 1.2e-08 | 4.433E-08 | 3.412E-08 | 1.200E-08 | 8.000E-08 |
| 4809 | 241 | I241H | A | 2.9e-08 | 3.5e-08 | 2.4e-08 | 2.933E-08 | 5.508E-09 | 2.400E-08 | 3.500E-08 |
| 4810 | 241 | I241I | A | 6.8e-08 | 2.7e-08 | 5.3e-08 | 4.933E-08 | 2.074E-08 | 2.700E-08 | 6.800E-08 |
| 4811 | 241 | I241L | A | 8e-08 | 1.9e-08 | 3.4e-08 | 4.433E-08 | 3.179E-08 | 1.900E-08 | 8.000E-08 |
| 4812 | 241 | I241K | A | 2.1e-08 | 4.4e-08 | 3.1e-08 | 3.200E-08 | 1.153E-08 | 2.100E-08 | 4.400E-08 |
| 4813 | 241 | I241M | A | 5.2e-08 | 7.1e-08 | 2.1e-08 | 4.800E-08 | 2.524E-08 | 2.100E-08 | 7.100E-08 |
| 4814 | 241 | I241F | A | 1.1e-07 | 1.6e-08 | 3.5e-08 | 5.367E-08 | 4.970E-08 | 1.600E-08 | 1.100E-07 |
| 4815 | 241 | I241P | A | 2.2e-08 | 5.5e-08 | 6.1e-08 | 4.600E-08 | 2.100E-08 | 2.200E-08 | 6.100E-08 |
| 4816 | 241 | I241S | A | 2.3e-08 | 5.3e-08 | 1.4e-07 | 7.200E-08 | 6.077E-08 | 2.300E-08 | 1.400E-07 |
| 4817 | 241 | I241T | A | 3.5e-08 | 3.1e-08 | 9.5e-08 | 5.367E-08 | 3.585E-08 | 3.100E-08 | 9.500E-08 |
| 4818 | 241 | I241W | A | 1.9e-08 | 2.3e-08 | 1.2e-07 | 5.400E-08 | 5.719E-08 | 1.900E-08 | 1.200E-07 |
| 4819 | 241 | I241Y | A | 2.8e-08 | 3.7e-08 | 7.5e-08 | 4.667E-08 | 2.495E-08 | 2.800E-08 | 7.500E-08 |
| 4820 | 241 | I241V | A | 4.4e-08 | 5.3e-08 | 1.8e-08 | 3.833E-08 | 1.818E-08 | 1.800E-08 | 5.300E-08 |
| 4821 | 242 | C242A | A | 7.5e-08 | 2.1e-08 | 4.5e-08 | 4.700E-08 | 2.706E-08 | 2.100E-08 | 7.500E-08 |
| 4822 | 242 | C242R | A | 6e-08 | 1.1e-07 | 5.8e-08 | 7.600E-08 | 2.946E-08 | 5.800E-08 | 1.100E-07 |
| 4823 | 242 | C242N | A | 2.1e-08 | 4e-08 | 2.9e-08 | 3.000E-08 | 9.539E-09 | 2.100E-08 | 4.000E-08 |
| 4824 | 242 | C242D | A | 5.6e-08 | 4.2e-08 | 3e-08 | 4.267E-08 | 1.301E-08 | 3.000E-08 | 5.600E-08 |
| 4825 | 242 | C242C | A | 2.7e-08 | 6.6e-08 | 5.6e-08 | 4.967E-08 | 2.026E-08 | 2.700E-08 | 6.600E-08 |
| 4826 | 242 | C242E | A | 9.6e-08 | 3.6e-08 | 3.9e-08 | 5.700E-08 | 3.381E-08 | 3.600E-08 | 9.600E-08 |
| 4827 | 242 | C242Q | A | 1.6e-08 | 1.7e-08 | 6.6e-08 | 3.300E-08 | 2.858E-08 | 1.600E-08 | 6.600E-08 |
| 4828 | 242 | C242G | A | 6.1e-08 | 1.1e-07 | 8.1e-08 | 8.400E-08 | 2.464E-08 | 6.100E-08 | 1.100E-07 |
| 4829 | 242 | C242H | A | 6.4e-08 | 7.2e-08 | 5.8e-08 | 6.467E-08 | 7.024E-09 | 5.800E-08 | 7.200E-08 |
| 4830 | 242 | C242I | A | 2.7e-08 | 7.6e-08 | 2.7e-08 | 4.333E-08 | 2.829E-08 | 2.700E-08 | 7.600E-08 |
| 4831 | 242 | C242L | A | 2.7e-08 | 2.2e-08 | 3.6e-08 | 2.833E-08 | 7.095E-09 | 2.200E-08 | 3.600E-08 |
| 4832 | 242 | C242K | A | 3.3e-08 | 4.6e-08 | 6.6e-08 | 4.833E-08 | 1.662E-08 | 3.300E-08 | 6.600E-08 |
| 4833 | 242 | C242M | A | 2.9e-08 | 8.8e-08 | 3.2e-08 | 4.967E-08 | 3.323E-08 | 2.900E-08 | 8.800E-08 |
| 4834 | 242 | C242F | A | 5.5e-08 | 6.1e-08 | 3.6e-08 | 5.067E-08 | 1.305E-08 | 3.600E-08 | 6.100E-08 |
| 4835 | 242 | C242P | A | 3.9e-08 | 6.7e-08 | 3.1e-08 | 4.567E-08 | 1.890E-08 | 3.100E-08 | 6.700E-08 |
| 4836 | 242 | C242S | A | 3.6e-08 | 6.5e-08 | 4.8e-08 | 4.967E-08 | 1.457E-08 | 3.600E-08 | 6.500E-08 |
| 4837 | 242 | C242T | A | 3.4e-08 | 2e-08 | 2.1e-08 | 2.500E-08 | 7.810E-09 | 2.000E-08 | 3.400E-08 |
| 4838 | 242 | C242W | A | 9.7e-08 | 4e-08 | 4.5e-08 | 6.067E-08 | 3.156E-08 | 4.000E-08 | 9.700E-08 |
| 4839 | 242 | C242Y | A | 6.8e-08 | 7.8e-08 | 2.5e-08 | 5.700E-08 | 2.816E-08 | 2.500E-08 | 7.800E-08 |
| 4840 | 242 | C242V | A | 5e-08 | 1e-07 | 2.4e-08 | 5.800E-08 | 3.863E-08 | 2.400E-08 | 1.000E-07 |
| 4841 | 243 | E243A | A | 2e-08 | 2.3e-08 | 2.4e-08 | 2.233E-08 | 2.082E-09 | 2.000E-08 | 2.400E-08 |
| 4842 | 243 | E243R | A | 4.1e-08 | 4.3e-08 | 9.4e-08 | 5.933E-08 | 3.004E-08 | 4.100E-08 | 9.400E-08 |
| 4843 | 243 | E243N | A | 3.7e-08 | 3.6e-08 | 2.8e-08 | 3.367E-08 | 4.933E-09 | 2.800E-08 | 3.700E-08 |
| 4844 | 243 | E243D | A | 2.3e-07 | 3.7e-08 | 3.7e-08 | 1.013E-07 | 1.114E-07 | 3.700E-08 | 2.300E-07 |

| | | | | | | | | | | |
|---|---|---|---|---|---|---|---|---|---|---|
| 4845 | 243 | E243C | A | 5.4e-08 | 3.7e-08 | 1.8e-08 | 3.633E-08 | 1.801E-08 | 1.800E-08 | 5.400E-08 |
| 4846 | 243 | E243E | A | 4.7e-08 | 9.8e-08 | 6.2e-08 | 6.900E-08 | 2.621E-08 | 4.700E-08 | 9.800E-08 |
| 4847 | 243 | E243Q | A | 6.6e-08 | 7e-08 | 4.5e-08 | 6.033E-08 | 1.343E-08 | 4.500E-08 | 7.000E-08 |
| 4848 | 243 | E243G | A | 2.5e-08 | 1.8e-08 | 4.9e-08 | 3.067E-08 | 1.626E-08 | 1.800E-08 | 4.900E-08 |
| 4849 | 243 | E243H | A | 2.5e-08 | 2.7e-08 | 6e-08 | 3.733E-08 | 1.966E-08 | 2.500E-08 | 6.000E-08 |
| 4850 | 243 | E243I | A | 3.8e-08 | 2.4e-08 | 4.4e-08 | 3.533E-08 | 1.026E-08 | 2.400E-08 | 4.400E-08 |
| 4851 | 243 | E243L | A | 6.4e-08 | 3e-08 | 3.3e-08 | 4.233E-08 | 1.882E-08 | 3.000E-08 | 6.400E-08 |
| 4852 | 243 | E243K | A | 6.6e-08 | 5.7e-08 | 2.5e-08 | 4.933E-08 | 2.155E-08 | 2.500E-08 | 6.600E-08 |
| 4853 | 243 | E243M | A | 6.6e-08 | 1.1e-07 | 4.4e-08 | 7.333E-08 | 3.361E-08 | 4.400E-08 | 1.100E-07 |
| 4854 | 243 | E243F | A | 1.1e-07 | 3.5e-08 | 4.3e-08 | 6.267E-08 | 4.119E-08 | 3.500E-08 | 1.100E-07 |
| 4855 | 243 | E243P | A | 5.4e-08 | 4.1e-08 | 6.6e-08 | 5.367E-08 | 1.250E-08 | 4.100E-08 | 6.600E-08 |
| 4856 | 243 | E243S | A | 5.1e-08 | 2.1e-08 | 4.3e-08 | 3.833E-08 | 1.553E-08 | 2.100E-08 | 5.100E-08 |
| 4857 | 243 | E243T | A | 1.1e-07 | 2.7e-08 | 4.4e-08 | 6.033E-08 | 4.384E-08 | 2.700E-08 | 1.100E-07 |
| 4858 | 243 | E243W | A | 1.8e-08 | 5.6e-08 | 7.2e-08 | 4.867E-08 | 2.774E-08 | 1.800E-08 | 7.200E-08 |
| 4859 | 243 | E243Y | A | 1.2e-07 | 6.5e-08 | 3.6e-08 | 7.367E-08 | 4.267E-08 | 3.600E-08 | 1.200E-07 |
| 4860 | 243 | E243V | A | 4.2e-08 | 3.6e-08 | 3.4e-08 | 3.733E-08 | 4.163E-09 | 3.400E-08 | 4.200E-08 |
| 4861 | 244 | L244A | A | 2.3e-08 | 1.5e-08 | 3.9e-08 | 2.567E-08 | 1.222E-08 | 1.500E-08 | 3.900E-08 |
| 4862 | 244 | L244R | A | 4.1e-08 | 1.5e-07 | 9.6e-08 | 9.567E-08 | 5.450E-08 | 4.100E-08 | 1.500E-07 |
| 4863 | 244 | L244N | A | 9.3e-08 | 1e-07 | 9.8e-08 | 9.700E-08 | 3.606E-09 | 9.300E-08 | 1.000E-07 |
| 4864 | 244 | L244D | A | 2.1e-08 | 2.7e-08 | 3.8e-08 | 2.867E-08 | 8.622E-09 | 2.100E-08 | 3.800E-08 |
| 4865 | 244 | L244C | A | 6.3e-08 | 2.2e-08 | 3.3e-08 | 3.933E-08 | 2.122E-08 | 2.200E-08 | 6.300E-08 |
| 4866 | 244 | L244E | A | 7.8e-08 | 4.9e-08 | 4.3e-08 | 5.667E-08 | 1.872E-08 | 4.300E-08 | 7.800E-08 |
| 4867 | 244 | L244Q | A | 1.3e-07 | 2.4e-08 | 1.7e-07 | 1.080E-07 | 7.545E-08 | 2.400E-08 | 1.700E-07 |
| 4868 | 244 | L244G | A | 5.6e-08 | 6.4e-08 | 5.1e-08 | 5.700E-08 | 6.557E-09 | 5.100E-08 | 6.400E-08 |
| 4869 | 244 | L244H | A | 5.5e-08 | 7e-08 | 6.7e-08 | 6.400E-08 | 7.937E-09 | 5.500E-08 | 7.000E-08 |
| 4870 | 244 | L244I | A | 1.5e-08 | 8.4e-08 | 9.2e-09 | 3.607E-08 | 4.161E-08 | 9.200E-09 | 8.400E-08 |
| 4871 | 244 | L244L | A | 3.6e-08 | 4.7e-08 | 8.1e-08 | 5.467E-08 | 2.346E-08 | 3.600E-08 | 8.100E-08 |
| 4872 | 244 | L244K | A | 8.8e-08 | 3.5e-08 | 9.5e-08 | 7.267E-08 | 3.281E-08 | 3.500E-08 | 9.500E-08 |
| 4873 | 244 | L244M | A | 6.5e-08 | 3.2e-08 | 1.8e-08 | 3.833E-08 | 2.413E-08 | 1.800E-08 | 6.500E-08 |
| 4874 | 244 | L244F | A | 4e-08 | 1.7e-08 | 9.7e-08 | 5.133E-08 | 4.119E-08 | 1.700E-08 | 9.700E-08 |
| 4875 | 244 | L244P | A | 6.6e-08 | 3.4e-08 | 8.2e-08 | 6.067E-08 | 2.444E-08 | 3.400E-08 | 8.200E-08 |
| 4876 | 244 | L244S | A | 1.2e-07 | 1.2e-07 | 2.4e-07 | 1.600E-07 | 6.928E-08 | 1.200E-07 | 2.400E-07 |
| 4877 | 244 | L244T | A | 9.7e-08 | 9.3e-08 | 1.5e-07 | 1.133E-07 | 3.182E-08 | 9.300E-08 | 1.500E-07 |
| 4878 | 244 | L244W | A | 2.7e-08 | 8.6e-08 | 9.9e-09 | 4.097E-08 | 3.993E-08 | 9.900E-09 | 8.600E-08 |
| 4879 | 244 | L244Y | A | 5e-08 | 2.3e-08 | 5e-08 | 4.100E-08 | 1.559E-08 | 2.300E-08 | 5.000E-08 |
| 4880 | 244 | L244V | A | 5.2e-08 | 1.7e-08 | 8.1e-08 | 5.000E-08 | 3.205E-08 | 1.700E-08 | 8.100E-08 |
| 4881 | 245 | H245A | A | 6.3e-08 | 5e-08 | 1.4e-08 | 4.233E-08 | 2.538E-08 | 1.400E-08 | 6.300E-08 |
| 4882 | 245 | H245R | A | 6.2e-08 | 3.2e-08 | 8.4e-08 | 5.933E-08 | 2.610E-08 | 3.200E-08 | 8.400E-08 |
| 4883 | 245 | H245N | A | 1.1e-07 | 5.8e-08 | 3.9e-08 | 6.900E-08 | 3.676E-08 | 3.900E-08 | 1.100E-07 |
| 4884 | 245 | H245D | A | 3.1e-08 | 7.1e-08 | 8.3e-08 | 6.167E-08 | 2.723E-08 | 3.100E-08 | 8.300E-08 |
| 4885 | 245 | H245C | A | 3.9e-08 | 5.3e-08 | 6.9e-08 | 5.367E-08 | 1.501E-08 | 3.900E-08 | 6.900E-08 |
| 4886 | 245 | H245E | A | 6.9e-08 | 7.8e-08 | 6.3e-08 | 7.000E-08 | 7.550E-09 | 6.300E-08 | 7.800E-08 |
| 4887 | 245 | H245Q | A | 1.2e-07 | 9.8e-08 | 8.5e-08 | 1.010E-07 | 1.769E-08 | 8.500E-08 | 1.200E-07 |
| 4888 | 245 | H245G | A | 5.2e-08 | 1.6e-08 | 1.7e-08 | 2.833E-08 | 2.050E-08 | 1.600E-08 | 5.200E-08 |
| 4889 | 245 | H245H | A | 2.6e-08 | 1e-07 | 8.6e-08 | 7.067E-08 | 3.931E-08 | 2.600E-08 | 1.000E-07 |
| 4890 | 245 | H245I | A | 3.6e-08 | 5.6e-08 | 1.8e-08 | 3.667E-08 | 1.901E-08 | 1.800E-08 | 5.600E-08 |
| 4891 | 245 | H245L | A | 8.8e-09 | 3.2e-08 | 3.2e-08 | 2.427E-08 | 1.339E-08 | 8.800E-09 | 3.200E-08 |
| 4892 | 245 | H245K | A | 1.8e-08 | 3.9e-08 | 4.8e-08 | 3.500E-08 | 1.539E-08 | 1.800E-08 | 4.800E-08 |
| 4893 | 245 | H245M | A | 3.3e-08 | 2.9e-08 | 8.9e-09 | 2.363E-08 | 1.292E-08 | 8.900E-09 | 3.300E-08 |
| 4894 | 245 | H245F | A | 1.7e-08 | 9.5e-09 | 2.4e-08 | 1.683E-08 | 7.251E-09 | 9.500E-09 | 2.400E-08 |
| 4895 | 245 | H245P | A | 7e-08 | 5.4e-08 | 2.2e-08 | 4.867E-08 | 2.444E-08 | 2.200E-08 | 7.000E-08 |
| 4896 | 245 | H245S | A | 4.2e-08 | 6.6e-08 | 2.6e-07 | 1.227E-07 | 1.195E-07 | 4.200E-08 | 2.600E-07 |
| 4897 | 245 | H245T | A | 6.8e-08 | 1.1e-07 | 7.3e-08 | 8.367E-08 | 2.294E-08 | 6.800E-08 | 1.100E-07 |
| 4898 | 245 | H245W | A | 1.7e-08 | 1.4e-08 | 4.3e-08 | 2.467E-08 | 1.595E-08 | 1.400E-08 | 4.300E-08 |
| 4899 | 245 | H245Y | A | 1.9e-08 | 1.4e-08 | 1.9e-08 | 1.733E-08 | 2.887E-09 | 1.400E-08 | 1.900E-08 |
| 4900 | 245 | H245V | A | 5.5e-08 | 5.2e-08 | 1.2e-07 | 7.567E-08 | 3.842E-08 | 5.200E-08 | 1.200E-07 |
| 4901 | 246 | C246A | A | 3.9e-08 | 1.5e-08 | 1.6e-07 | 7.133E-08 | 7.772E-08 | 1.500E-08 | 1.600E-07 |
| 4902 | 246 | C246R | A | 8.8e-08 | 1.4e-07 | 6.7e-08 | 9.833E-08 | 3.758E-08 | 6.700E-08 | 1.400E-07 |
| 4903 | 246 | C246N | A | 1.7e-07 | 1.4e-07 | 2.4e-07 | 1.833E-07 | 5.132E-08 | 1.400E-07 | 2.400E-07 |
| 4904 | 246 | C246D | A | 5.4e-08 | 1.6e-07 | 1.6e-07 | 1.247E-07 | 6.120E-08 | 5.400E-08 | 1.600E-07 |
| 4905 | 246 | C246C | A | 3.5e-08 | 6.9e-08 | 4.7e-08 | 5.033E-08 | 1.724E-08 | 3.500E-08 | 6.900E-08 |
| 4906 | 246 | C246E | A | 3.6e-08 | 2.5e-07 | 1e-08 | 9.867E-08 | 1.317E-07 | 1.000E-08 | 2.500E-07 |
| 4907 | 246 | C246Q | A | 4.9e-08 | 2e-07 | 4.4e-07 | 2.297E-07 | 1.972E-07 | 4.900E-08 | 4.400E-07 |
| 4908 | 246 | C246G | A | 1.1e-07 | 9.2e-08 | 3.1e-08 | 7.767E-08 | 4.140E-08 | 3.100E-08 | 1.100E-07 |
| 4909 | 246 | C246H | A | 1.3e-07 | 1.3e-07 | 3.6e-07 | 2.067E-07 | 1.328E-07 | 1.300E-07 | 3.600E-07 |
| 4910 | 246 | C246I | A | 7.8e-08 | 1.5e-08 | 5.9e-08 | 5.067E-08 | 3.232E-08 | 1.500E-08 | 7.800E-08 |
| 4911 | 246 | C246L | A | 1.2e-08 | 2e-08 | 2.5e-08 | 1.900E-08 | 6.557E-09 | 1.200E-08 | 2.500E-08 |
| 4912 | 246 | C246K | A | 6.6e-08 | 2.5e-07 | 1.5e-07 | 1.553E-07 | 9.212E-08 | 6.600E-08 | 2.500E-07 |
| 4913 | 246 | C246M | A | 9.3e-08 | 7.7e-08 | 1.7e-07 | 1.133E-07 | 4.972E-08 | 7.700E-08 | 1.700E-07 |
| 4914 | 246 | C246F | A | 2.5e-08 | 1.8e-08 | 4.7e-08 | 3.000E-08 | 1.513E-08 | 1.800E-08 | 4.700E-08 |
| 4915 | 246 | C246P | A | 1.2e-07 | 5.9e-08 | 9.4e-08 | 9.100E-08 | 3.061E-08 | 5.900E-08 | 1.200E-07 |
| 4916 | 246 | C246S | A | 1.5e-07 | 4e-07 | 1.7e-07 | 2.400E-07 | 1.389E-07 | 1.500E-07 | 4.000E-07 |
| 4917 | 246 | C246T | A | 3e-07 | 2e-07 | 1.6e-07 | 2.200E-07 | 7.211E-08 | 1.600E-07 | 3.000E-07 |
| 4918 | 246 | C246W | A | 3.8e-08 | 5.2e-08 | 1.5e-07 | 8.000E-08 | 6.102E-08 | 3.800E-08 | 1.500E-07 |
| 4919 | 246 | C246Y | A | 4.3e-08 | 1.7e-07 | 3.6e-08 | 8.300E-08 | 7.543E-08 | 3.600E-08 | 1.700E-07 |
| 4920 | 246 | C246V | A | 9.5e-08 | 1e-07 | 1.4e-08 | 6.967E-08 | 4.827E-08 | 1.400E-08 | 1.000E-07 |
| 4921 | 247 | P247A | A | 7.1e-08 | 5.3e-08 | 7e-08 | 6.467E-08 | 1.012E-08 | 5.300E-08 | 7.100E-08 |
| 4922 | 247 | P247R | A | 6.3e-08 | 5.7e-08 | 2e-07 | 1.067E-07 | 8.088E-08 | 5.700E-08 | 2.000E-07 |
| 4923 | 247 | P247N | A | 1.1e-07 | 1.4e-07 | 8.7e-08 | 1.123E-07 | 2.658E-08 | 8.700E-08 | 1.400E-07 |
| 4924 | 247 | P247D | A | 4.8e-08 | 1.3e-07 | 5.8e-08 | 7.867E-08 | 4.474E-08 | 4.800E-08 | 1.300E-07 |
| 4925 | 247 | P247C | A | 5e-08 | 9.5e-08 | 1.2e-07 | 8.833E-08 | 3.547E-08 | 5.000E-08 | 1.200E-07 |
| 4926 | 247 | P247E | A | 5.3e-08 | 2.5e-08 | 4.6e-08 | 4.133E-08 | 1.457E-08 | 2.500E-08 | 5.300E-08 |
| 4927 | 247 | P247Q | A | 9.6e-08 | 6.6e-08 | 1.6e-07 | 1.073E-07 | 4.801E-08 | 6.600E-08 | 1.600E-07 |
| 4928 | 247 | P247G | A | 8.2e-08 | 3.6e-08 | 3.2e-08 | 5.000E-08 | 2.778E-08 | 3.200E-08 | 8.200E-08 |
| 4929 | 247 | P247H | A | 8.5e-08 | 1.5e-07 | 6.7e-08 | 1.007E-07 | 4.366E-08 | 6.700E-08 | 1.500E-07 |

| | | | | | | | | | | |
|---|---|---|---|---|---|---|---|---|---|---|
| 4930 | 247 | P247I | A | 1.5e-07 | 7.1e-08 | 2.5e-08 | 8.200E-08 | 6.322E-08 | 2.500E-08 | 1.500E-07 |
| 4931 | 247 | P247L | A | 1.7e-08 | 2.4e-08 | 5e-08 | 3.033E-08 | 1.739E-08 | 1.700E-08 | 5.000E-08 |
| 4932 | 247 | P247K | A | 6e-08 | 8.3e-08 | 8.9e-08 | 7.733E-08 | 1.531E-08 | 6.000E-08 | 8.900E-08 |
| 4933 | 247 | P247M | A | 4.4e-08 | 3.6e-08 | 3.8e-08 | 3.933E-08 | 4.163E-09 | 3.600E-08 | 4.400E-08 |
| 4934 | 247 | P247F | A | 7.6e-08 | 1.6e-07 | 3.8e-08 | 9.133E-08 | 6.243E-08 | 3.800E-08 | 1.600E-07 |
| 4935 | 247 | P247P | A | 3.7e-08 | 3.6e-08 | 2.8e-08 | 3.367E-08 | 4.933E-09 | 2.800E-08 | 3.700E-08 |
| 4936 | 247 | P247S | A | 8.7e-08 | 6.4e-08 | 7.3e-08 | 7.467E-08 | 1.159E-08 | 6.400E-08 | 8.700E-08 |
| 4937 | 247 | P247T | A | 9.6e-08 | 1.2e-07 | 7.2e-08 | 9.600E-08 | 2.400E-08 | 7.200E-08 | 1.200E-07 |
| 4938 | 247 | P247W | A | 1.9e-08 | 9.6e-09 | 1.1e-07 | 4.620E-08 | 5.545E-08 | 9.600E-09 | 1.100E-07 |
| 4939 | 247 | P247Y | A | 2.9e-08 | 5.9e-08 | 2.1e-08 | 3.633E-08 | 2.003E-08 | 2.100E-08 | 5.900E-08 |
| 4940 | 247 | P247V | A | 2.1e-08 | 2.8e-08 | 4.3e-08 | 3.067E-08 | 1.124E-08 | 2.100E-08 | 4.300E-08 |
| 4941 | 248 | A248A | A | 3.8e-08 | 6.8e-08 | 4.3e-08 | 4.967E-08 | 1.607E-08 | 3.800E-08 | 6.800E-08 |
| 4942 | 248 | A248R | A | 3.9e-08 | 2.9e-08 | 1.9e-08 | 2.900E-08 | 1.000E-08 | 1.900E-08 | 3.900E-08 |
| 4943 | 248 | A248N | A | 2.9e-07 | 1.2e-07 | 9.8e-08 | 1.693E-07 | 1.051E-07 | 9.800E-08 | 2.900E-07 |
| 4944 | 248 | A248D | A | 7.9e-08 | 5.5e-08 | 1.8e-07 | 1.047E-07 | 6.634E-08 | 5.500E-08 | 1.800E-07 |
| 4945 | 248 | A248C | A | 3e-08 | 5.3e-08 | 5.7e-08 | 4.667E-08 | 1.457E-08 | 3.000E-08 | 5.700E-08 |
| 4946 | 248 | A248E | A | 5.6e-08 | 6.6e-08 | 4.9e-08 | 5.700E-08 | 8.544E-09 | 4.900E-08 | 6.600E-08 |
| 4947 | 248 | A248Q | A | 1.1e-07 | 1.8e-07 | 7.4e-08 | 1.213E-07 | 5.390E-08 | 7.400E-08 | 1.800E-07 |
| 4948 | 248 | A248G | A | 9.7e-08 | 9.6e-08 | 1.1e-07 | 1.010E-07 | 7.810E-09 | 9.600E-08 | 1.100E-07 |
| 4949 | 248 | A248H | A | 3.7e-08 | 4.1e-08 | 1.7e-08 | 3.167E-08 | 1.286E-08 | 1.700E-08 | 4.100E-08 |
| 4950 | 248 | A248I | A | 4.5e-08 | 8.3e-08 | 5e-08 | 5.933E-08 | 2.065E-08 | 4.500E-08 | 8.300E-08 |
| 4951 | 248 | A248L | A | 2.9e-08 | 2.5e-08 | 1.4e-08 | 2.267E-08 | 7.767E-09 | 1.400E-08 | 2.900E-08 |
| 4952 | 248 | A248K | A | 7e-08 | 1.5e-07 | 5.6e-08 | 9.200E-08 | 5.071E-08 | 5.600E-08 | 1.500E-07 |
| 4953 | 248 | A248M | A | 5.2e-08 | 6.6e-08 | 1.1e-07 | 7.600E-08 | 3.027E-08 | 5.200E-08 | 1.100E-07 |
| 4954 | 248 | A248F | A | 6.7e-08 | 6.1e-08 | 1.2e-08 | 4.667E-08 | 3.017E-08 | 1.200E-08 | 6.700E-08 |
| 4955 | 248 | A248P | A | 4.6e-08 | 4.4e-08 | 3.8e-08 | 4.267E-08 | 4.163E-09 | 3.800E-08 | 4.600E-08 |
| 4956 | 248 | A248S | A | 8.3e-08 | 8.1e-08 | 8.3e-08 | 8.233E-08 | 1.155E-09 | 8.100E-08 | 8.300E-08 |
| 4957 | 248 | A248T | A | 2.5e-08 | 6.1e-08 | 1e-07 | 6.200E-08 | 3.751E-08 | 2.500E-08 | 1.000E-07 |
| 4958 | 248 | A248W | A | 1e-07 | 7.3e-08 | 4e-08 | 7.100E-08 | 3.005E-08 | 4.000E-08 | 1.000E-07 |
| 4959 | 248 | A248Y | A | 1.8e-08 | 5.7e-08 | 6.3e-08 | 4.600E-08 | 2.443E-08 | 1.800E-08 | 6.300E-08 |
| 4960 | 248 | A248V | A | 3e-08 | 8.8e-08 | 2.9e-08 | 4.900E-08 | 3.378E-08 | 2.900E-08 | 8.800E-08 |
| 4961 | 249 | L249A | A | 7.9e-08 | 3.9e-08 | 5e-08 | 5.600E-08 | 2.066E-08 | 3.900E-08 | 7.900E-08 |
| 4962 | 249 | L249R | A | 1.4e-07 | 1e-07 | 7.2e-08 | 1.040E-07 | 3.418E-08 | 7.200E-08 | 1.400E-07 |
| 4963 | 249 | L249N | A | 9.9e-08 | 1.6e-07 | 1.7e-08 | 9.200E-08 | 7.176E-08 | 1.700E-08 | 1.600E-07 |
| 4964 | 249 | L249D | A | 1.7e-07 | 2.2e-08 | 1.6e-08 | 6.933E-08 | 8.723E-08 | 1.600E-08 | 1.700E-07 |
| 4965 | 249 | L249C | A | 3.4e-08 | 7.1e-08 | 3.5e-08 | 4.667E-08 | 2.108E-08 | 3.400E-08 | 7.100E-08 |
| 4966 | 249 | L249E | A | 7.8e-08 | 5.3e-08 | 1.2e-07 | 8.367E-08 | 3.386E-08 | 5.300E-08 | 1.200E-07 |
| 4967 | 249 | L249Q | A | 1.5e-07 | 2.7e-07 | 9.7e-08 | 1.723E-07 | 8.864E-08 | 9.700E-08 | 2.700E-07 |
| 4968 | 249 | L249G | A | 8.7e-08 | 9.5e-08 | 4e-08 | 7.400E-08 | 2.972E-08 | 4.000E-08 | 9.500E-08 |
| 4969 | 249 | L249H | A | 1.1e-07 | 9.6e-08 | 6.8e-08 | 9.133E-08 | 2.139E-08 | 6.800E-08 | 1.100E-07 |
| 4970 | 249 | L249I | A | 6e-08 | 2.3e-08 | 6.5e-08 | 4.933E-08 | 2.294E-08 | 2.300E-08 | 6.500E-08 |
| 4971 | 249 | L249L | A | 1.5e-08 | 2.5e-08 | 6.5e-08 | 3.500E-08 | 2.646E-08 | 1.500E-08 | 6.500E-08 |
| 4972 | 249 | L249K | A | 1.9e-07 | 1.2e-07 | 4e-08 | 1.167E-07 | 7.506E-08 | 4.000E-08 | 1.900E-07 |
| 4973 | 249 | L249M | A | 9e-08 | 3.3e-08 | 3.8e-08 | 5.367E-08 | 3.156E-08 | 3.300E-08 | 9.000E-08 |
| 4974 | 249 | L249F | A | 2.2e-08 | 2.3e-07 | 4.8e-08 | 1.000E-07 | 1.133E-07 | 2.200E-08 | 2.300E-07 |
| 4975 | 249 | L249P | A | 1.7e-08 | 1.5e-08 | 2.8e-08 | 2.000E-08 | 7.000E-09 | 1.500E-08 | 2.800E-08 |
| 4976 | 249 | L249S | A | 6e-08 | 1.6e-07 | 2.6e-07 | 1.600E-07 | 1.000E-07 | 6.000E-08 | 2.600E-07 |
| 4977 | 249 | L249T | A | 8.2e-08 | 1.3e-07 | 1.2e-07 | 1.107E-07 | 2.532E-08 | 8.200E-08 | 1.300E-07 |
| 4978 | 249 | L249W | A | 7.8e-08 | 9e-08 | 3.2e-08 | 6.667E-08 | 3.062E-08 | 3.200E-08 | 9.000E-08 |
| 4979 | 249 | L249Y | A | 8.1e-09 | 5.3e-08 | 3.3e-08 | 3.137E-08 | 2.249E-08 | 8.100E-09 | 5.300E-08 |
| 4980 | 249 | L249V | A | 1e-07 | 2.4e-08 | 3.9e-08 | 5.433E-08 | 4.025E-08 | 2.400E-08 | 1.000E-07 |
| 4981 | 250 | V250A | A | 3.6e-08 | 4.9e-08 | 2.6e-08 | 3.700E-08 | 1.153E-08 | 2.600E-08 | 4.900E-08 |
| 4982 | 250 | V250R | A | 3.9e-08 | 4.9e-08 | 4e-08 | 4.267E-08 | 5.508E-09 | 3.900E-08 | 4.900E-08 |
| 4983 | 250 | V250N | A | 6.9e-08 | 2.9e-08 | 3e-08 | 4.267E-08 | 2.281E-08 | 2.900E-08 | 6.900E-08 |
| 4984 | 250 | V250D | A | 1.2e-07 | 3.5e-08 | 1.4e-08 | 5.633E-08 | 5.613E-08 | 1.400E-08 | 1.200E-07 |
| 4985 | 250 | V250C | A | 2.8e-08 | 3.3e-08 | 3.1e-08 | 3.067E-08 | 2.517E-09 | 2.800E-08 | 3.300E-08 |
| 4986 | 250 | V250E | A | 2.5e-07 | 4.6e-08 | 1.6e-08 | 1.040E-07 | 1.273E-07 | 1.600E-08 | 2.500E-07 |
| 4987 | 250 | V250Q | A | 7.5e-08 | 6.4e-08 | 8.9e-08 | 7.600E-08 | 1.253E-08 | 6.400E-08 | 8.900E-08 |
| 4988 | 250 | V250G | A | 5.4e-08 | 3.4e-08 | 4.3e-08 | 4.367E-08 | 1.002E-08 | 3.400E-08 | 5.400E-08 |
| 4989 | 250 | V250H | A | 6.6e-08 | 4.9e-08 | 2.3e-08 | 4.600E-08 | 2.166E-08 | 2.300E-08 | 6.600E-08 |
| 4990 | 250 | V250I | A | 5.7e-08 | 3.1e-08 | 1.4e-07 | 7.600E-08 | 5.693E-08 | 3.100E-08 | 1.400E-07 |
| 4991 | 250 | V250L | A | 2.7e-08 | 1.7e-08 | 1.4e-08 | 1.933E-08 | 6.807E-09 | 1.400E-08 | 2.700E-08 |
| 4992 | 250 | V250K | A | 5.2e-08 | 3.6e-08 | 1.8e-08 | 3.533E-08 | 1.701E-08 | 1.800E-08 | 5.200E-08 |
| 4993 | 250 | V250M | A | 4.2e-08 | 5.9e-08 | 4.9e-08 | 5.000E-08 | 8.544E-09 | 4.200E-08 | 5.900E-08 |
| 4994 | 250 | V250F | A | 8.4e-08 | 8e-08 | 1.5e-08 | 5.967E-08 | 3.873E-08 | 1.500E-08 | 8.400E-08 |
| 4995 | 250 | V250P | A | 4.2e-08 | 2.8e-08 | 3.1e-08 | 3.367E-08 | 7.371E-09 | 2.800E-08 | 4.200E-08 |
| 4996 | 250 | V250S | A | 4.8e-08 | 8.2e-08 | 1.8e-08 | 4.933E-08 | 3.202E-08 | 1.800E-08 | 8.200E-08 |
| 4997 | 250 | V250T | A | 1e-07 | 3.8e-08 | 3e-08 | 5.600E-08 | 3.831E-08 | 3.000E-08 | 1.000E-07 |
| 4998 | 250 | V250W | A | 3.8e-08 | 5.3e-08 | 5e-08 | 4.700E-08 | 7.937E-09 | 3.800E-08 | 5.300E-08 |
| 4999 | 250 | V250Y | A | 1.8e-08 | 3.2e-08 | 9.5e-09 | 1.983E-08 | 1.136E-08 | 9.500E-09 | 3.200E-08 |
| 5000 | 250 | V250V | A | 4e-08 | 3.3e-08 | 5.3e-08 | 4.200E-08 | 1.015E-08 | 3.300E-08 | 5.300E-08 |
| 5001 | 251 | T251A | A | 4.6e-08 | 3.4e-08 | 5e-08 | 4.333E-08 | 8.327E-09 | 3.400E-08 | 5.000E-08 |
| 5002 | 251 | T251R | A | 5.4e-08 | 1.1e-07 | 1.2e-07 | 9.467E-08 | 3.557E-08 | 5.400E-08 | 1.200E-07 |
| 5003 | 251 | T251N | A | 8.7e-09 | 4e-08 | 5.5e-08 | 3.457E-08 | 2.362E-08 | 8.700E-09 | 5.500E-08 |
| 5004 | 251 | T251D | A | 3.5e-08 | 1.7e-08 | 4e-08 | 3.067E-08 | 1.210E-08 | 1.700E-08 | 4.000E-08 |
| 5005 | 251 | T251C | A | 7.7e-08 | 2e-08 | 4.8e-08 | 4.833E-08 | 2.850E-08 | 2.000E-08 | 7.700E-08 |
| 5006 | 251 | T251E | A | 3.9e-08 | 1.8e-08 | 5.9e-08 | 3.867E-08 | 2.050E-08 | 1.800E-08 | 5.900E-08 |
| 5007 | 251 | T251Q | A | 6.4e-08 | 4.3e-08 | 5.9e-08 | 5.533E-08 | 1.097E-08 | 4.300E-08 | 6.400E-08 |
| 5008 | 251 | T251G | A | 4.2e-08 | 7.4e-08 | 6.4e-08 | 6.000E-08 | 1.637E-08 | 4.200E-08 | 7.400E-08 |
| 5009 | 251 | T251H | A | 1.1e-08 | 2.5e-08 | 4e-08 | 2.533E-08 | 1.450E-08 | 1.100E-08 | 4.000E-08 |
| 5010 | 251 | T251I | A | 1e-07 | 1.6e-07 | 5.2e-08 | 1.040E-07 | 5.411E-08 | 5.200E-08 | 1.600E-07 |
| 5011 | 251 | T251L | A | 5.7e-08 | 8.4e-08 | 1.1e-07 | 8.367E-08 | 2.650E-08 | 5.700E-08 | 1.100E-07 |
| 5012 | 251 | T251K | A | 9.8e-08 | 1.6e-07 | 3.7e-08 | 9.833E-08 | 6.150E-08 | 3.700E-08 | 1.600E-07 |
| 5013 | 251 | T251M | A | 5.9e-08 | 7.2e-08 | 8.8e-08 | 7.300E-08 | 1.453E-08 | 5.900E-08 | 8.800E-08 |
| 5014 | 251 | T251F | A | 4e-08 | 2.1e-08 | 3.4e-08 | 3.167E-08 | 9.713E-09 | 2.100E-08 | 4.000E-08 |

| | | | | | | | | | | |
|---|---|---|---|---|---|---|---|---|---|---|
| 5015 | 251 | T251P | A | 5.8e-08 | 2.6e-08 | 5e-08 | 4.467E-08 | 1.665E-08 | 2.600E-08 | 5.800E-08 |
| 5016 | 251 | T251S | A | 4.9e-08 | 3e-08 | 4.6e-08 | 4.167E-08 | 1.021E-08 | 3.000E-08 | 4.900E-08 |
| 5017 | 251 | T251T | A | 2.1e-08 | 7.2e-08 | 1.5e-08 | 3.600E-08 | 3.132E-08 | 1.500E-08 | 7.200E-08 |
| 5018 | 251 | T251W | A | 1e-08 | 1.6e-07 | 1.7e-08 | 6.233E-08 | 8.465E-08 | 1.000E-08 | 1.600E-07 |
| 5019 | 251 | T251Y | A | 1.1e-08 | 4.7e-08 | 2.3e-09 | 2.010E-08 | 2.370E-08 | 2.300E-09 | 4.700E-08 |
| 5020 | 251 | T251V | A | 4.8e-08 | 2.2e-08 | 4.5e-08 | 3.833E-08 | 1.422E-08 | 2.200E-08 | 4.800E-08 |
| 5021 | 252 | Y252A | A | 6.8e-08 | 5.2e-08 | 1e-07 | 7.333E-08 | 2.444E-08 | 5.200E-08 | 1.000E-07 |
| 5022 | 252 | Y252R | A | 2.3e-08 | 2.4e-08 | 2.1e-08 | 2.267E-08 | 1.528E-09 | 2.100E-08 | 2.400E-08 |
| 5023 | 252 | Y252N | A | 2.2e-08 | 1.7e-08 | 2e-08 | 1.967E-08 | 2.517E-09 | 1.700E-08 | 2.200E-08 |
| 5024 | 252 | Y252D | A | 6.1e-08 | 1.6e-08 | 4.1e-08 | 3.933E-08 | 2.255E-08 | 1.600E-08 | 6.100E-08 |
| 5025 | 252 | Y252C | A | 8.9e-08 | 2.7e-08 | 4.8e-09 | 4.027E-08 | 4.364E-08 | 4.800E-09 | 8.900E-08 |
| 5026 | 252 | Y252E | A | 2.5e-08 | 2.5e-08 | 4.4e-08 | 3.133E-08 | 1.097E-08 | 2.500E-08 | 4.400E-08 |
| 5027 | 252 | Y252Q | A | 3.2e-08 | 1.1e-08 | 1.7e-08 | 2.000E-08 | 1.082E-08 | 1.100E-08 | 3.200E-08 |
| 5028 | 252 | Y252G | A | 3e-08 | 3.4e-08 | 2.7e-08 | 3.033E-08 | 3.512E-09 | 2.700E-08 | 3.400E-08 |
| 5029 | 252 | Y252H | A | 5.4e-08 | 3e-08 | 4.5e-08 | 4.300E-08 | 1.212E-08 | 3.000E-08 | 5.400E-08 |
| 5030 | 252 | Y252I | A | 1.3e-08 | 3.7e-08 | 2.6e-08 | 2.533E-08 | 1.201E-08 | 1.300E-08 | 3.700E-08 |
| 5031 | 252 | Y252L | A | 6.6e-08 | 5.5e-08 | 3.2e-08 | 5.100E-08 | 1.735E-08 | 3.200E-08 | 6.600E-08 |
| 5032 | 252 | Y252K | A | 4e-08 | 2.1e-08 | 2.4e-08 | 2.833E-08 | 1.021E-08 | 2.100E-08 | 4.000E-08 |
| 5033 | 252 | Y252M | A | 2.2e-08 | 1.1e-08 | 3.4e-08 | 2.233E-08 | 1.150E-08 | 1.100E-08 | 3.400E-08 |
| 5034 | 252 | Y252F | A | 2.5e-08 | 7.2e-08 | 6.5e-08 | 5.400E-08 | 2.536E-08 | 2.500E-08 | 7.200E-08 |
| 5035 | 252 | Y252P | A | 2.8e-08 | 5.3e-08 | 7.8e-08 | 5.300E-08 | 2.500E-08 | 2.800E-08 | 7.800E-08 |
| 5036 | 252 | Y252S | A | 3e-08 | 6e-08 | 4.8e-08 | 4.600E-08 | 1.510E-08 | 3.000E-08 | 6.000E-08 |
| 5037 | 252 | Y252T | A | 1.3e-08 | 1.9e-08 | 2.9e-08 | 2.033E-08 | 8.083E-09 | 1.300E-08 | 2.900E-08 |
| 5038 | 252 | Y252W | A | 3.5e-08 | 2.1e-08 | 9.3e-08 | 4.967E-08 | 3.818E-08 | 2.100E-08 | 9.300E-08 |
| 5039 | 252 | Y252Y | A | 7.8e-08 | 5.3e-08 | 8.7e-08 | 7.267E-08 | 1.762E-08 | 5.300E-08 | 8.700E-08 |
| 5040 | 252 | Y252V | A | 2.6e-08 | 2.3e-08 | 3.9e-08 | 2.933E-08 | 8.505E-09 | 2.300E-08 | 3.900E-08 |
| 5041 | 253 | N253A | A | 2.3e-08 | 4.5e-08 | 4.3e-08 | 3.700E-08 | 1.217E-08 | 2.300E-08 | 4.500E-08 |
| 5042 | 253 | N253R | A | 5e-08 | 5.2e-08 | 8.5e-08 | 6.233E-08 | 1.966E-08 | 5.000E-08 | 8.500E-08 |
| 5043 | 253 | N253N | A | 5.7e-08 | 5.3e-08 | 6.6e-08 | 5.867E-08 | 6.658E-09 | 5.300E-08 | 6.600E-08 |
| 5044 | 253 | N253D | A | 2.9e-08 | 7.4e-08 | 5.7e-08 | 5.333E-08 | 2.272E-08 | 2.900E-08 | 7.400E-08 |
| 5045 | 253 | N253C | A | 5.9e-09 | 6.7e-08 | 3.1e-08 | 3.463E-08 | 3.071E-08 | 5.900E-09 | 6.700E-08 |
| 5046 | 253 | N253E | A | 1.2e-07 | 2.8e-08 | 6.2e-08 | 7.000E-08 | 4.652E-08 | 2.800E-08 | 1.200E-07 |
| 5047 | 253 | N253Q | A | 4e-08 | 1.5e-07 | 4.4e-08 | 7.800E-08 | 6.239E-08 | 4.000E-08 | 1.500E-07 |
| 5048 | 253 | N253G | A | 1e-07 | 5.1e-08 | 1.5e-08 | 5.533E-08 | 4.267E-08 | 1.500E-08 | 1.000E-07 |
| 5049 | 253 | N253H | A | 6.6e-08 | 1.1e-07 | 3.4e-08 | 7.000E-08 | 3.816E-08 | 3.400E-08 | 1.100E-07 |
| 5050 | 253 | N253I | A | 5.6e-08 | 6.8e-08 | 2.7e-08 | 5.033E-08 | 2.108E-08 | 2.700E-08 | 6.800E-08 |
| 5051 | 253 | N253L | A | 1.7e-08 | 7.6e-08 | 9.5e-08 | 6.267E-08 | 4.067E-08 | 1.700E-08 | 9.500E-08 |
| 5052 | 253 | N253K | A | 4.7e-08 | 2.8e-08 | 6.8e-08 | 4.767E-08 | 2.001E-08 | 2.800E-08 | 6.800E-08 |
| 5053 | 253 | N253M | A | 8.4e-08 | 3.8e-08 | 1.6e-07 | 9.400E-08 | 6.161E-08 | 3.800E-08 | 1.600E-07 |
| 5054 | 253 | N253F | A | 3.6e-08 | 5.5e-08 | 6.1e-08 | 5.067E-08 | 1.305E-08 | 3.600E-08 | 6.100E-08 |
| 5055 | 253 | N253P | A | 1.3e-07 | 3.5e-08 | 2.8e-08 | 6.433E-08 | 5.698E-08 | 2.800E-08 | 1.300E-07 |
| 5056 | 253 | N253S | A | 1.6e-08 | 1.1e-07 | 7.6e-08 | 6.733E-08 | 4.760E-08 | 1.600E-08 | 1.100E-07 |
| 5057 | 253 | N253T | A | 1.8e-08 | 1.6e-08 | 1.3e-07 | 5.467E-08 | 6.525E-08 | 1.600E-08 | 1.300E-07 |
| 5058 | 253 | N253W | A | 3.1e-08 | 4.7e-08 | 3.4e-08 | 3.733E-08 | 8.505E-09 | 3.100E-08 | 4.700E-08 |
| 5059 | 253 | N253Y | A | 7.3e-08 | 3.1e-08 | 4e-08 | 4.800E-08 | 2.211E-08 | 3.100E-08 | 7.300E-08 |
| 5060 | 253 | N253V | A | 9.6e-08 | 6.6e-08 | 2.8e-08 | 6.333E-08 | 3.408E-08 | 2.800E-08 | 9.600E-08 |
| 5061 | 254 | T254A | A | 1.6e-08 | 1.4e-08 | 1.2e-08 | 1.400E-08 | 2.000E-09 | 1.200E-08 | 1.600E-08 |
| 5062 | 254 | T254R | A | 1.3e-07 | 2e-08 | 1.3e-08 | 5.433E-08 | 6.562E-08 | 1.300E-08 | 1.300E-07 |
| 5063 | 254 | T254N | A | 5.7e-08 | 7.1e-08 | 3.6e-08 | 5.467E-08 | 1.762E-08 | 3.600E-08 | 7.100E-08 |
| 5064 | 254 | T254D | A | 3.4e-08 | 1.9e-08 | 3.8e-08 | 3.033E-08 | 1.002E-08 | 1.900E-08 | 3.800E-08 |
| 5065 | 254 | T254C | A | 1.2e-08 | 2.2e-08 | 3e-08 | 2.133E-08 | 9.018E-09 | 1.200E-08 | 3.000E-08 |
| 5066 | 254 | T254E | A | 6.8e-08 | 5.3e-08 | 7.7e-09 | 4.290E-08 | 3.139E-08 | 7.700E-09 | 6.800E-08 |
| 5067 | 254 | T254Q | A | 3.2e-08 | 1.6e-08 | 4.1e-08 | 2.967E-08 | 1.266E-08 | 1.600E-08 | 4.100E-08 |
| 5068 | 254 | T254G | A | 1.7e-08 | 1.5e-08 | 2.7e-08 | 1.967E-08 | 6.429E-09 | 1.500E-08 | 2.700E-08 |
| 5069 | 254 | T254H | A | 1.5e-08 | 2.5e-08 | 1.2e-08 | 1.733E-08 | 6.807E-09 | 1.200E-08 | 2.500E-08 |
| 5070 | 254 | T254I | A | 2.1e-08 | 9.9e-09 | 3e-08 | 2.030E-08 | 1.007E-08 | 9.900E-09 | 3.000E-08 |
| 5071 | 254 | T254L | A | 1.5e-08 | 6e-09 | 1.5e-08 | 1.200E-08 | 5.196E-09 | 6.000E-09 | 1.500E-08 |
| 5072 | 254 | T254K | A | 1.6e-08 | 2e-08 | 1.7e-08 | 1.767E-08 | 2.082E-09 | 1.600E-08 | 2.000E-08 |
| 5073 | 254 | T254M | A | 1e-08 | 2e-08 | 3.3e-08 | 2.100E-08 | 1.153E-08 | 1.000E-08 | 3.300E-08 |
| 5074 | 254 | T254F | A | 5.7e-09 | 1.9e-08 | 4.7e-08 | 2.390E-08 | 2.108E-08 | 5.700E-09 | 4.700E-08 |
| 5075 | 254 | T254P | A | 1.3e-08 | 2.7e-08 | 2.6e-08 | 2.200E-08 | 7.810E-09 | 1.300E-08 | 2.700E-08 |
| 5076 | 254 | T254S | A | 3.3e-08 | 3.8e-08 | 9.9e-08 | 5.667E-08 | 3.675E-08 | 3.300E-08 | 9.900E-08 |
| 5077 | 254 | T254T | A | 8.2e-08 | 2.4e-08 | 4.9e-08 | 5.167E-08 | 2.909E-08 | 2.400E-08 | 8.200E-08 |
| 5078 | 254 | T254W | A | 1.7e-08 | 1.8e-08 | 1.3e-08 | 1.600E-08 | 2.646E-09 | 1.300E-08 | 1.800E-08 |
| 5079 | 254 | T254Y | A | 1.8e-08 | 2.4e-08 | 2e-08 | 2.067E-08 | 3.055E-09 | 1.800E-08 | 2.400E-08 |
| 5080 | 254 | T254V | A | 1.4e-08 | 7.2e-08 | 2.2e-08 | 3.600E-08 | 3.143E-08 | 1.400E-08 | 7.200E-08 |
| 5081 | 255 | D255A | A | 5.1e-08 | 2.4e-08 | 1.3e-08 | 2.933E-08 | 1.955E-08 | 1.300E-08 | 5.100E-08 |
| 5082 | 255 | D255R | A | 4.2e-08 | 4.4e-08 | 2.9e-08 | 3.833E-08 | 8.145E-09 | 2.900E-08 | 4.400E-08 |
| 5083 | 255 | D255N | A | 3e-07 | 3.9e-08 | 2.8e-08 | 1.223E-07 | 1.540E-07 | 2.800E-08 | 3.000E-07 |
| 5084 | 255 | D255D | A | 2.6e-08 | 6.8e-08 | 1.3e-08 | 3.567E-08 | 2.875E-08 | 1.300E-08 | 6.800E-08 |
| 5085 | 255 | D255C | A | 3e-08 | 9.9e-08 | 5e-08 | 5.967E-08 | 3.550E-08 | 3.000E-08 | 9.900E-08 |
| 5086 | 255 | D255E | A | 8.3e-08 | 5.5e-08 | 3.7e-08 | 5.833E-08 | 2.318E-08 | 3.700E-08 | 8.300E-08 |
| 5087 | 255 | D255Q | A | 5.8e-08 | 7.8e-08 | 8.6e-08 | 7.400E-08 | 1.442E-08 | 5.800E-08 | 8.600E-08 |
| 5088 | 255 | D255G | A | 7.6e-09 | 4.6e-08 | 2e-08 | 2.453E-08 | 1.960E-08 | 7.600E-09 | 4.600E-08 |
| 5089 | 255 | D255H | A | 1.2e-08 | 9.1e-08 | 4.8e-08 | 5.033E-08 | 3.955E-08 | 1.200E-08 | 9.100E-08 |
| 5090 | 255 | D255I | A | 3.5e-08 | 3.2e-08 | 1.6e-08 | 2.767E-08 | 1.021E-08 | 1.600E-08 | 3.500E-08 |
| 5091 | 255 | D255L | A | 2.4e-08 | 1e-08 | 6.4e-08 | 3.267E-08 | 2.802E-08 | 1.000E-08 | 6.400E-08 |
| 5092 | 255 | D255K | A | 2.8e-08 | 2.8e-08 | 6.1e-08 | 3.900E-08 | 1.905E-08 | 2.800E-08 | 6.100E-08 |
| 5093 | 255 | D255M | A | 4.1e-08 | 2.9e-08 | 4.8e-08 | 3.933E-08 | 9.609E-09 | 2.900E-08 | 4.800E-08 |
| 5094 | 255 | D255F | A | 2.5e-08 | 5.1e-08 | 2.3e-08 | 3.300E-08 | 1.562E-08 | 2.300E-08 | 5.100E-08 |
| 5095 | 255 | D255P | A | 3.7e-08 | 6.7e-08 | 1.3e-08 | 3.900E-08 | 2.706E-08 | 1.300E-08 | 6.700E-08 |
| 5096 | 255 | D255S | A | 7.2e-08 | 5.6e-08 | 1.2e-07 | 8.267E-08 | 3.331E-08 | 5.600E-08 | 1.200E-07 |
| 5097 | 255 | D255T | A | 1.1e-07 | 7.4e-08 | 2.6e-08 | 7.000E-08 | 4.214E-08 | 2.600E-08 | 1.100E-07 |
| 5098 | 255 | D255W | A | 7e-08 | 4.1e-08 | 3.8e-08 | 4.967E-08 | 1.767E-08 | 3.800E-08 | 7.000E-08 |
| 5099 | 255 | D255Y | A | 4.5e-08 | 1.5e-08 | 2.7e-08 | 2.900E-08 | 1.510E-08 | 1.500E-08 | 4.500E-08 |

| | | | | | | | | | | |
|---|---|---|---|---|---|---|---|---|---|---|
| 5100 | 255 | D255V | A | 1.7e-08 | 6.8e-08 | 1.4e-08 | 3.300E-08 | 3.035E-08 | 1.400E-08 | 6.800E-08 |
| 5101 | 256 | T256A | A | 2.6e-08 | 7.7e-08 | 3.5e-08 | 4.600E-08 | 2.722E-08 | 2.600E-08 | 7.700E-08 |
| 5102 | 256 | T256R | A | 3e-08 | 3.1e-08 | 3.7e-08 | 3.267E-08 | 3.786E-09 | 3.000E-08 | 3.700E-08 |
| 5103 | 256 | T256N | A | 8.6e-08 | 3.2e-08 | 6.3e-08 | 6.033E-08 | 2.710E-08 | 3.200E-08 | 8.600E-08 |
| 5104 | 256 | T256D | A | 1e-08 | 8e-08 | 4.6e-08 | 4.533E-08 | 3.500E-08 | 1.000E-08 | 8.000E-08 |
| 5105 | 256 | T256C | A | 1.8e-08 | 3.1e-08 | 6e-08 | 3.633E-08 | 2.150E-08 | 1.800E-08 | 6.000E-08 |
| 5106 | 256 | T256E | A | 2e-08 | 3.3e-08 | 3.8e-08 | 3.033E-08 | 9.292E-09 | 2.000E-08 | 3.800E-08 |
| 5107 | 256 | T256Q | A | 7e-08 | 9.3e-08 | 3.3e-08 | 6.533E-08 | 3.027E-08 | 3.300E-08 | 9.300E-08 |
| 5108 | 256 | T256G | A | 2.1e-08 | 1.7e-08 | 1.9e-08 | 1.900E-08 | 2.000E-09 | 1.700E-08 | 2.100E-08 |
| 5109 | 256 | T256H | A | 3.5e-08 | 2.7e-08 | 8.3e-08 | 4.833E-08 | 3.029E-08 | 2.700E-08 | 8.300E-08 |
| 5110 | 256 | T256I | A | 3e-08 | 1.2e-08 | 2.3e-08 | 2.167E-08 | 9.074E-09 | 1.200E-08 | 3.000E-08 |
| 5111 | 256 | T256L | A | 2.8e-08 | 3.5e-08 | 3.1e-08 | 3.133E-08 | 3.512E-09 | 2.800E-08 | 3.500E-08 |
| 5112 | 256 | T256K | A | 3.9e-08 | 7.1e-08 | 2.8e-08 | 4.600E-08 | 2.234E-08 | 2.800E-08 | 7.100E-08 |
| 5113 | 256 | T256M | A | 2.4e-08 | 1.1e-08 | 2.1e-08 | 1.867E-08 | 6.807E-09 | 1.100E-08 | 2.400E-08 |
| 5114 | 256 | T256F | A | 4.4e-08 | 6.8e-09 | 2e-08 | 2.360E-08 | 1.886E-08 | 6.800E-09 | 4.400E-08 |
| 5115 | 256 | T256P | A | 1e-08 | 1.5e-08 | 3.3e-08 | 1.933E-08 | 1.210E-08 | 1.000E-08 | 3.300E-08 |
| 5116 | 256 | T256S | A | 3.2e-08 | 6.3e-08 | 2.7e-08 | 4.067E-08 | 1.950E-08 | 2.700E-08 | 6.300E-08 |
| 5117 | 256 | T256T | A | 1.3e-08 | 2.5e-08 | 3.2e-08 | 2.333E-08 | 9.609E-09 | 1.300E-08 | 3.200E-08 |
| 5118 | 256 | T256W | A | 7.2e-08 | 2.3e-08 | 1.5e-07 | 8.167E-08 | 6.405E-08 | 2.300E-08 | 1.500E-07 |
| 5119 | 256 | T256Y | A | 2.5e-08 | 3.4e-08 | 5.2e-09 | 2.140E-08 | 1.473E-08 | 5.200E-09 | 3.400E-08 |
| 5120 | 256 | T256V | A | 1.4e-08 | 3.2e-08 | 4.7e-08 | 3.100E-08 | 1.652E-08 | 1.400E-08 | 4.700E-08 |
| 5121 | 257 | F257A | A | 9.8e-08 | 5.3e-08 | 7e-08 | 7.367E-08 | 2.272E-08 | 5.300E-08 | 9.800E-08 |
| 5122 | 257 | F257R | A | 9.5e-08 | 1.1e-07 | 8.2e-08 | 9.567E-08 | 1.401E-08 | 8.200E-08 | 1.100E-07 |
| 5123 | 257 | F257N | A | 7.5e-08 | 5.3e-08 | 3.8e-08 | 5.533E-08 | 1.861E-08 | 3.800E-08 | 7.500E-08 |
| 5124 | 257 | F257D | A | 7.5e-08 | 2.3e-08 | 1.1e-07 | 6.933E-08 | 4.378E-08 | 2.300E-08 | 1.100E-07 |
| 5125 | 257 | F257C | A | 1.2e-07 | 4e-08 | 9.1e-08 | 8.367E-08 | 4.050E-08 | 4.000E-08 | 1.200E-07 |
| 5126 | 257 | F257E | A | 1.8e-08 | 6.2e-08 | 6e-08 | 4.667E-08 | 2.485E-08 | 1.800E-08 | 6.200E-08 |
| 5127 | 257 | F257Q | A | 2.5e-07 | 1.4e-07 | 4.1e-08 | 1.437E-07 | 1.045E-07 | 4.100E-08 | 2.500E-07 |
| 5128 | 257 | F257G | A | 7.5e-08 | 3.1e-08 | 7.6e-08 | 6.067E-08 | 2.570E-08 | 3.100E-08 | 7.600E-08 |
| 5129 | 257 | F257H | A | 2.2e-08 | 1.8e-07 | 5.8e-08 | 8.667E-08 | 8.281E-08 | 2.200E-08 | 1.800E-07 |
| 5130 | 257 | F257I | A | 9.5e-08 | 1.1e-07 | 1.6e-07 | 1.217E-07 | 3.403E-08 | 9.500E-08 | 1.600E-07 |
| 5131 | 257 | F257L | A | 1.6e-07 | 5.2e-08 | 8e-08 | 9.733E-08 | 5.605E-08 | 5.200E-08 | 1.600E-07 |
| 5132 | 257 | F257K | A | 8.5e-08 | 1.4e-07 | 6.5e-08 | 9.667E-08 | 3.884E-08 | 6.500E-08 | 1.400E-07 |
| 5133 | 257 | F257M | A | 1.6e-07 | 2.5e-08 | 2.9e-08 | 7.133E-08 | 7.681E-08 | 2.500E-08 | 1.600E-07 |
| 5134 | 257 | F257F | A | 4.5e-08 | 4.3e-08 | 6.5e-08 | 5.100E-08 | 1.217E-08 | 4.300E-08 | 6.500E-08 |
| 5135 | 257 | F257P | A | 3.6e-08 | 5e-08 | 6.2e-08 | 4.933E-08 | 1.301E-08 | 3.600E-08 | 6.200E-08 |
| 5136 | 257 | F257S | A | 3.7e-08 | 7.5e-08 | 5.7e-08 | 5.633E-08 | 1.901E-08 | 3.700E-08 | 7.500E-08 |
| 5137 | 257 | F257T | A | 7.8e-08 | 1e-07 | 7.3e-08 | 8.367E-08 | 1.436E-08 | 7.300E-08 | 1.000E-07 |
| 5138 | 257 | F257W | A | 2e-08 | 3.2e-08 | 2.8e-08 | 2.667E-08 | 6.110E-09 | 2.000E-08 | 3.200E-08 |
| 5139 | 257 | F257Y | A | 3.8e-08 | 6.3e-08 | 1.1e-07 | 7.033E-08 | 3.656E-08 | 3.800E-08 | 1.100E-07 |
| 5140 | 257 | F257V | A | 5.3e-08 | 5.9e-08 | 4.6e-08 | 5.267E-08 | 6.506E-09 | 4.600E-08 | 5.900E-08 |
| 5141 | 258 | E258A | A | 6.9e-08 | 4.5e-08 | 5.2e-08 | 5.533E-08 | 1.234E-08 | 4.500E-08 | 6.900E-08 |
| 5142 | 258 | E258R | A | 3.7e-08 | 5e-08 | 2.4e-08 | 3.700E-08 | 1.300E-08 | 2.400E-08 | 5.000E-08 |
| 5143 | 258 | E258N | A | 2.7e-08 | 5.5e-08 | 1.8e-08 | 3.333E-08 | 1.930E-08 | 1.800E-08 | 5.500E-08 |
| 5144 | 258 | E258D | A | 5.4e-08 | 6.3e-08 | 2.5e-08 | 4.733E-08 | 1.986E-08 | 2.500E-08 | 6.300E-08 |
| 5145 | 258 | E258C | A | 5e-08 | 5e-08 | 4.9e-08 | 4.967E-08 | 5.774E-10 | 4.900E-08 | 5.000E-08 |
| 5146 | 258 | E258E | A | 8.4e-08 | 2.2e-08 | 4.6e-08 | 5.067E-08 | 3.126E-08 | 2.200E-08 | 8.400E-08 |
| 5147 | 258 | E258Q | A | 1.1e-07 | 3.3e-08 | 3.8e-08 | 6.033E-08 | 4.309E-08 | 3.300E-08 | 1.100E-07 |
| 5148 | 258 | E258G | A | 3.4e-08 | 2.9e-08 | 3e-08 | 3.100E-08 | 2.646E-09 | 2.900E-08 | 3.400E-08 |
| 5149 | 258 | E258H | A | 7.3e-08 | 2.8e-08 | 4.6e-08 | 4.900E-08 | 2.265E-08 | 2.800E-08 | 7.300E-08 |
| 5150 | 258 | E258I | A | 5.4e-08 | 3.4e-08 | 6.3e-08 | 5.033E-08 | 1.484E-08 | 3.400E-08 | 6.300E-08 |
| 5151 | 258 | E258L | A | 4.5e-08 | 1.1e-08 | 5.9e-08 | 3.833E-08 | 2.468E-08 | 1.100E-08 | 5.900E-08 |
| 5152 | 258 | E258K | A | 6.7e-08 | 1.8e-08 | 5.5e-08 | 4.667E-08 | 2.554E-08 | 1.800E-08 | 6.700E-08 |
| 5153 | 258 | E258M | A | 6.7e-08 | 6.2e-08 | 7.2e-08 | 6.700E-08 | 5.000E-09 | 6.200E-08 | 7.200E-08 |
| 5154 | 258 | E258F | A | 4.7e-08 | 9.9e-08 | 4e-08 | 6.200E-08 | 3.223E-08 | 4.000E-08 | 9.900E-08 |
| 5155 | 258 | E258P | A | 2.3e-08 | 4.3e-08 | 1.4e-07 | 6.867E-08 | 6.258E-08 | 2.300E-08 | 1.400E-07 |
| 5156 | 258 | E258S | A | 4.7e-08 | 2.6e-08 | 5.6e-08 | 4.300E-08 | 1.539E-08 | 2.600E-08 | 5.600E-08 |
| 5157 | 258 | E258T | A | 2.3e-08 | 1.1e-07 | 3.6e-08 | 5.633E-08 | 4.693E-08 | 2.300E-08 | 1.100E-07 |
| 5158 | 258 | E258W | A | 2.6e-08 | 3.8e-08 | 1.1e-08 | 2.500E-08 | 1.353E-08 | 1.100E-08 | 3.800E-08 |
| 5159 | 258 | E258Y | A | 2.3e-08 | 4e-08 | 1.6e-08 | 2.633E-08 | 1.234E-08 | 1.600E-08 | 4.000E-08 |
| 5160 | 258 | E258V | A | 1.3e-08 | 6.1e-08 | 1.9e-08 | 3.100E-08 | 2.615E-08 | 1.300E-08 | 6.100E-08 |
| 5161 | 259 | S259A | A | 8.9e-08 | 3.3e-08 | 5e-08 | 5.733E-08 | 2.871E-08 | 3.300E-08 | 8.900E-08 |
| 5162 | 259 | S259R | A | 1.4e-07 | 5.3e-08 | 1.7e-08 | 7.000E-08 | 6.324E-08 | 1.700E-08 | 1.400E-07 |
| 5163 | 259 | S259N | A | 6.3e-08 | 6.1e-08 | 3.1e-08 | 5.167E-08 | 1.793E-08 | 3.100E-08 | 6.300E-08 |
| 5164 | 259 | S259D | A | 3.9e-08 | 5.4e-08 | 4.5e-08 | 4.600E-08 | 7.550E-09 | 3.900E-08 | 5.400E-08 |
| 5165 | 259 | S259C | A | 7e-08 | 6.6e-08 | 1.1e-07 | 8.200E-08 | 2.433E-08 | 6.600E-08 | 1.100E-07 |
| 5166 | 259 | S259E | A | 1.3e-07 | 3.2e-08 | 4e-08 | 6.733E-08 | 5.442E-08 | 3.200E-08 | 1.300E-07 |
| 5167 | 259 | S259Q | A | 1.8e-08 | 2.2e-08 | 4.7e-08 | 2.900E-08 | 1.572E-08 | 1.800E-08 | 4.700E-08 |
| 5168 | 259 | S259G | A | 4.7e-08 | 5.6e-08 | 3.7e-08 | 4.667E-08 | 9.504E-09 | 3.700E-08 | 5.600E-08 |
| 5169 | 259 | S259H | A | 2.2e-08 | 1.2e-07 | 3.2e-08 | 5.800E-08 | 5.393E-08 | 2.200E-08 | 1.200E-07 |
| 5170 | 259 | S259I | A | 5.4e-08 | 2e-08 | 8.4e-08 | 5.267E-08 | 3.202E-08 | 2.000E-08 | 8.400E-08 |
| 5171 | 259 | S259L | A | 2e-08 | 5.2e-08 | 3.5e-08 | 3.567E-08 | 1.601E-08 | 2.000E-08 | 5.200E-08 |
| 5172 | 259 | S259K | A | 3e-08 | 2.6e-08 | 5.7e-08 | 3.767E-08 | 1.686E-08 | 2.600E-08 | 5.700E-08 |
| 5173 | 259 | S259M | A | 3.5e-08 | 3.4e-08 | 2.5e-08 | 3.133E-08 | 5.508E-09 | 2.500E-08 | 3.500E-08 |
| 5174 | 259 | S259F | A | 3.5e-08 | 1.3e-07 | 1.2e-08 | 5.900E-08 | 6.255E-08 | 1.200E-08 | 1.300E-07 |
| 5175 | 259 | S259P | A | 3.4e-08 | 4.2e-08 | 3.2e-08 | 3.600E-08 | 5.292E-09 | 3.200E-08 | 4.200E-08 |
| 5176 | 259 | S259S | A | 2.9e-08 | 4.5e-08 | 6.1e-08 | 4.500E-08 | 1.600E-08 | 2.900E-08 | 6.100E-08 |
| 5177 | 259 | S259T | A | 2.8e-08 | 7.3e-08 | 3.7e-08 | 4.600E-08 | 2.381E-08 | 2.800E-08 | 7.300E-08 |
| 5178 | 259 | S259W | A | 1.3e-08 | 4.5e-08 | 2.8e-08 | 2.867E-08 | 1.601E-08 | 1.300E-08 | 4.500E-08 |
| 5179 | 259 | S259Y | A | 8.7e-08 | 3.2e-08 | 1.6e-07 | 9.300E-08 | 6.421E-08 | 3.200E-08 | 1.600E-07 |
| 5180 | 259 | S259V | A | 3.5e-08 | 8.5e-08 | 4.6e-08 | 5.533E-08 | 2.627E-08 | 3.500E-08 | 8.500E-08 |
| 5181 | 260 | M260A | A | 3.6e-08 | 2.3e-08 | 7.1e-08 | 4.333E-08 | 2.483E-08 | 2.300E-08 | 7.100E-08 |
| 5182 | 260 | M260R | A | 9.3e-08 | 2.9e-08 | 2.5e-08 | 4.900E-08 | 3.816E-08 | 2.500E-08 | 9.300E-08 |
| 5183 | 260 | M260N | A | 4.6e-08 | 1.8e-08 | 3.2e-08 | 3.200E-08 | 1.400E-08 | 1.800E-08 | 4.600E-08 |
| 5184 | 260 | M260D | A | 3.5e-08 | 2.2e-08 | 9.1e-08 | 4.933E-08 | 3.667E-08 | 2.200E-08 | 9.100E-08 |

| | | | | | | | | | | |
|---|---|---|---|---|---|---|---|---|---|---|
| 5185 | 260 | M260C | A | 7.2e-08 | 2.8e-08 | 1.9e-08 | 3.967E-08 | 2.836E-08 | 1.900E-08 | 7.200E-08 |
| 5186 | 260 | M260E | A | 1.1e-07 | 4.7e-08 | 5e-08 | 6.900E-08 | 3.554E-08 | 4.700E-08 | 1.100E-07 |
| 5187 | 260 | M260Q | A | 3e-08 | 1.8e-08 | 6.3e-08 | 3.700E-08 | 2.330E-08 | 1.800E-08 | 6.300E-08 |
| 5188 | 260 | M260G | A | 4.7e-08 | 1.3e-08 | 6.8e-08 | 4.267E-08 | 2.775E-08 | 1.300E-08 | 6.800E-08 |
| 5189 | 260 | M260H | A | 2.7e-08 | 3.5e-08 | 3.8e-08 | 3.333E-08 | 5.686E-09 | 2.700E-08 | 3.800E-08 |
| 5190 | 260 | M260I | A | 4.2e-08 | 1.5e-08 | 4e-08 | 3.233E-08 | 1.504E-08 | 1.500E-08 | 4.200E-08 |
| 5191 | 260 | M260L | A | 5.5e-08 | 3.6e-08 | 4.7e-08 | 4.600E-08 | 9.539E-09 | 3.600E-08 | 5.500E-08 |
| 5192 | 260 | M260K | A | 5.2e-08 | 4.4e-08 | 2.4e-08 | 4.000E-08 | 1.442E-08 | 2.400E-08 | 5.200E-08 |
| 5193 | 260 | M260M | A | 1.6e-08 | 2.3e-08 | 6.9e-08 | 3.600E-08 | 2.879E-08 | 1.600E-08 | 6.900E-08 |
| 5194 | 260 | M260F | A | 2.6e-08 | 5.8e-08 | 1.8e-07 | 8.800E-08 | 8.126E-08 | 2.600E-08 | 1.800E-07 |
| 5195 | 260 | M260P | A | 3.5e-08 | 4.3e-08 | 3.3e-08 | 3.700E-08 | 5.292E-09 | 3.300E-08 | 4.300E-08 |
| 5196 | 260 | M260S | A | 2.6e-08 | 1.1e-07 | 4.8e-08 | 6.133E-08 | 4.356E-08 | 2.600E-08 | 1.100E-07 |
| 5197 | 260 | M260T | A | 2.7e-08 | 3.3e-08 | 3.4e-08 | 3.133E-08 | 3.786E-09 | 2.700E-08 | 3.400E-08 |
| 5198 | 260 | M260W | A | 1.7e-08 | 6.3e-08 | 2e-08 | 3.333E-08 | 2.574E-08 | 1.700E-08 | 6.300E-08 |
| 5199 | 260 | M260Y | A | 2.9e-08 | 9.2e-08 | 5.9e-08 | 6.000E-08 | 3.151E-08 | 2.900E-08 | 9.200E-08 |
| 5200 | 260 | M260V | A | 5e-08 | 2.4e-08 | 1.6e-07 | 7.800E-08 | 7.219E-08 | 2.400E-08 | 1.600E-07 |
| 5201 | 261 | P261A | A | 1.5e-08 | 2.3e-08 | 2.2e-08 | 2.000E-08 | 4.359E-09 | 1.500E-08 | 2.300E-08 |
| 5202 | 261 | P261R | A | 5.6e-08 | 8.1e-08 | 4.5e-08 | 6.067E-08 | 1.845E-08 | 4.500E-08 | 8.100E-08 |
| 5203 | 261 | P261N | A | 2.4e-08 | 6.3e-08 | 2.8e-08 | 3.833E-08 | 2.146E-08 | 2.400E-08 | 6.300E-08 |
| 5204 | 261 | P261D | A | 5e-08 | 7.3e-08 | 3.2e-08 | 5.167E-08 | 2.055E-08 | 3.200E-08 | 7.300E-08 |
| 5205 | 261 | P261C | A | 2.5e-08 | 1.3e-08 | 3.3e-08 | 2.367E-08 | 1.007E-08 | 1.300E-08 | 3.300E-08 |
| 5206 | 261 | P261E | A | 2.8e-08 | 3.9e-08 | 1.7e-08 | 2.800E-08 | 1.100E-08 | 1.700E-08 | 3.900E-08 |
| 5207 | 261 | P261Q | A | 1.7e-08 | 3.6e-08 | 2.6e-08 | 2.633E-08 | 9.504E-09 | 1.700E-08 | 3.600E-08 |
| 5208 | 261 | P261G | A | 2.8e-08 | 2.6e-08 | 3.4e-08 | 2.933E-08 | 4.163E-09 | 2.600E-08 | 3.400E-08 |
| 5209 | 261 | P261H | A | 8.9e-08 | 6.2e-08 | 3.8e-08 | 6.300E-08 | 2.551E-08 | 3.800E-08 | 8.900E-08 |
| 5210 | 261 | P261I | A | 6.3e-08 | 6.1e-08 | 8.9e-08 | 7.100E-08 | 1.562E-08 | 6.100E-08 | 8.900E-08 |
| 5211 | 261 | P261L | A | 1.2e-08 | 2.9e-08 | 2.3e-08 | 2.133E-08 | 8.622E-09 | 1.200E-08 | 2.900E-08 |
| 5212 | 261 | P261K | A | 4.3e-08 | 5.8e-08 | 5.9e-08 | 5.333E-08 | 8.963E-09 | 4.300E-08 | 5.900E-08 |
| 5213 | 261 | P261M | A | 4.1e-08 | 4.4e-08 | 2.1e-08 | 3.533E-08 | 1.250E-08 | 2.100E-08 | 4.400E-08 |
| 5214 | 261 | P261F | A | 8.2e-08 | 1.2e-07 | 2.1e-08 | 7.433E-08 | 4.994E-08 | 2.100E-08 | 1.200E-07 |
| 5215 | 261 | P261P | A | 1.1e-08 | 9.8e-08 | 3e-08 | 4.633E-08 | 4.574E-08 | 1.100E-08 | 9.800E-08 |
| 5216 | 261 | P261S | A | 4.4e-08 | 5.6e-08 | 3.9e-08 | 4.633E-08 | 8.737E-09 | 3.900E-08 | 5.600E-08 |
| 5217 | 261 | P261T | A | 6.5e-08 | 2.7e-08 | 5.6e-08 | 4.933E-08 | 1.986E-08 | 2.700E-08 | 6.500E-08 |
| 5218 | 261 | P261W | A | 4.1e-08 | 4.3e-08 | 2.2e-08 | 3.533E-08 | 1.159E-08 | 2.200E-08 | 4.300E-08 |
| 5219 | 261 | P261Y | A | 6.9e-08 | 4e-08 | 4.3e-08 | 5.067E-08 | 1.595E-08 | 4.000E-08 | 6.900E-08 |
| 5220 | 261 | P261V | A | 3.5e-08 | 7.1e-08 | 4.9e-08 | 5.167E-08 | 1.815E-08 | 3.500E-08 | 7.100E-08 |
| 5221 | 262 | N262A | A | 1.7e-08 | 3.1e-08 | 3.1e-08 | 2.633E-08 | 8.083E-09 | 1.700E-08 | 3.100E-08 |
| 5222 | 262 | N262R | A | 2.1e-08 | 5.4e-08 | 4.5e-08 | 4.000E-08 | 1.706E-08 | 2.100E-08 | 5.400E-08 |
| 5223 | 262 | N262N | A | 5.6e-08 | 2.5e-08 | 5.1e-08 | 4.400E-08 | 1.664E-08 | 2.500E-08 | 5.600E-08 |
| 5224 | 262 | N262D | A | 1.1e-07 | 4.9e-08 | 7.7e-08 | 7.867E-08 | 3.053E-08 | 4.900E-08 | 1.100E-07 |
| 5225 | 262 | N262C | A | 1.6e-08 | 3.7e-08 | 7.1e-08 | 4.133E-08 | 2.775E-08 | 1.600E-08 | 7.100E-08 |
| 5226 | 262 | N262E | A | 3.7e-08 | 2.7e-08 | 2e-08 | 2.800E-08 | 8.544E-09 | 2.000E-08 | 3.700E-08 |
| 5227 | 262 | N262Q | A | 1.4e-07 | 1.2e-08 | 4.4e-08 | 6.533E-08 | 6.661E-08 | 1.200E-08 | 1.400E-07 |
| 5228 | 262 | N262G | A | 6.2e-08 | 4.9e-08 | 4.8e-08 | 5.300E-08 | 7.810E-09 | 4.800E-08 | 6.200E-08 |
| 5229 | 262 | N262H | A | 3.6e-08 | 2.4e-08 | 4.7e-08 | 3.567E-08 | 1.150E-08 | 2.400E-08 | 4.700E-08 |
| 5230 | 262 | N262I | A | 5.8e-08 | 3e-08 | 9.2e-08 | 6.000E-08 | 3.105E-08 | 3.000E-08 | 9.200E-08 |
| 5231 | 262 | N262L | A | 6.4e-08 | 1.7e-08 | 1.4e-08 | 3.167E-08 | 2.804E-08 | 1.400E-08 | 6.400E-08 |
| 5232 | 262 | N262K | A | 3.3e-08 | 7.5e-08 | 5.7e-08 | 5.500E-08 | 2.107E-08 | 3.300E-08 | 7.500E-08 |
| 5233 | 262 | N262M | A | 1.9e-08 | 3.2e-08 | 1e-07 | 5.033E-08 | 4.350E-08 | 1.900E-08 | 1.000E-07 |
| 5234 | 262 | N262F | A | 6.7e-08 | 2.3e-08 | 2.2e-08 | 3.733E-08 | 2.570E-08 | 2.200E-08 | 6.700E-08 |
| 5235 | 262 | N262P | A | 1.3e-07 | 7.2e-08 | 5.8e-08 | 8.667E-08 | 3.818E-08 | 5.800E-08 | 1.300E-07 |
| 5236 | 262 | N262S | A | 7.8e-08 | 8e-08 | 4.7e-08 | 6.833E-08 | 1.850E-08 | 4.700E-08 | 8.000E-08 |
| 5237 | 262 | N262T | A | 3.1e-08 | 2.5e-08 | 5.6e-08 | 3.733E-08 | 1.644E-08 | 2.500E-08 | 5.600E-08 |
| 5238 | 262 | N262W | A | 3.5e-08 | 1.1e-07 | 1.9e-08 | 5.467E-08 | 4.858E-08 | 1.900E-08 | 1.100E-07 |
| 5239 | 262 | N262Y | A | 3.8e-08 | 5.5e-08 | 5.7e-08 | 5.000E-08 | 1.044E-08 | 3.800E-08 | 5.700E-08 |
| 5240 | 262 | N262V | A | 7.3e-08 | 6.3e-08 | 7e-08 | 6.867E-08 | 5.132E-09 | 6.300E-08 | 7.300E-08 |
| 5241 | 263 | P263A | A | 2.8e-08 | 9.6e-08 | 3.8e-08 | 5.400E-08 | 3.672E-08 | 2.800E-08 | 9.600E-08 |
| 5242 | 263 | P263R | A | 9e-08 | 3.5e-08 | 2e-08 | 4.833E-08 | 3.686E-08 | 2.000E-08 | 9.000E-08 |
| 5243 | 263 | P263N | A | 4e-08 | 2.3e-08 | 2.6e-08 | 2.967E-08 | 9.074E-09 | 2.300E-08 | 4.000E-08 |
| 5244 | 263 | P263D | A | 4.6e-08 | 9.2e-08 | 5.3e-08 | 6.367E-08 | 2.479E-08 | 4.600E-08 | 9.200E-08 |
| 5245 | 263 | P263C | A | 3.8e-08 | 9.3e-08 | 7.2e-08 | 6.767E-08 | 2.775E-08 | 3.800E-08 | 9.300E-08 |
| 5246 | 263 | P263E | A | 1.4e-08 | 5.1e-08 | 1e-07 | 5.500E-08 | 4.314E-08 | 1.400E-08 | 1.000E-07 |
| 5247 | 263 | P263Q | A | 1.2e-07 | 2.4e-08 | 4.5e-08 | 6.300E-08 | 5.047E-08 | 2.400E-08 | 1.200E-07 |
| 5248 | 263 | P263G | A | 3.8e-08 | 6e-08 | 5e-08 | 4.933E-08 | 1.102E-08 | 3.800E-08 | 6.000E-08 |
| 5249 | 263 | P263H | A | 3e-08 | 5.9e-08 | 2.7e-08 | 3.867E-08 | 1.767E-08 | 2.700E-08 | 5.900E-08 |
| 5250 | 263 | P263I | A | 1.8e-08 | 3.6e-08 | 6.5e-09 | 2.017E-08 | 1.487E-08 | 6.500E-09 | 3.600E-08 |
| 5251 | 263 | P263L | A | 4.8e-08 | 4.4e-08 | 7.1e-09 | 3.303E-08 | 2.255E-08 | 7.100E-09 | 4.800E-08 |
| 5252 | 263 | P263K | A | 3.6e-08 | 2.4e-08 | 7.5e-08 | 4.500E-08 | 2.666E-08 | 2.400E-08 | 7.500E-08 |
| 5253 | 263 | P263M | A | 2e-08 | 6.3e-08 | 1.2e-07 | 6.767E-08 | 5.016E-08 | 2.000E-08 | 1.200E-07 |
| 5254 | 263 | P263F | A | 3.8e-08 | 7.3e-08 | 4.9e-08 | 5.333E-08 | 1.790E-08 | 3.800E-08 | 7.300E-08 |
| 5255 | 263 | P263P | A | 4.2e-08 | 4.2e-08 | 7.6e-08 | 5.333E-08 | 1.963E-08 | 4.200E-08 | 7.600E-08 |
| 5256 | 263 | P263S | A | 6.1e-08 | 2.1e-08 | 2.5e-08 | 3.567E-08 | 2.203E-08 | 2.100E-08 | 6.100E-08 |
| 5257 | 263 | P263T | A | 4.1e-08 | 7.6e-08 | 2.4e-08 | 4.700E-08 | 2.651E-08 | 2.400E-08 | 7.600E-08 |
| 5258 | 263 | P263W | A | 9.2e-08 | 4.9e-08 | 7.2e-08 | 7.100E-08 | 2.152E-08 | 4.900E-08 | 9.200E-08 |
| 5259 | 263 | P263Y | A | 1.5e-07 | 7.8e-08 | 6.4e-08 | 9.733E-08 | 4.614E-08 | 6.400E-08 | 1.500E-07 |
| 5260 | 263 | P263V | A | 6.5e-08 | 2.5e-08 | 8.2e-08 | 5.733E-08 | 2.926E-08 | 2.500E-08 | 8.200E-08 |
| 5261 | 264 | E264A | A | 4.4e-08 | 2.4e-08 | 8.8e-08 | 5.200E-08 | 3.274E-08 | 2.400E-08 | 8.800E-08 |
| 5262 | 264 | E264R | A | 6.6e-08 | 2.8e-08 | 5.3e-08 | 4.900E-08 | 1.931E-08 | 2.800E-08 | 6.600E-08 |
| 5263 | 264 | E264N | A | 1.6e-08 | 1.3e-08 | 4.9e-08 | 2.600E-08 | 1.997E-08 | 1.300E-08 | 4.900E-08 |
| 5264 | 264 | E264D | A | 3.8e-08 | 1.9e-08 | 1.2e-07 | 5.900E-08 | 5.367E-08 | 1.900E-08 | 1.200E-07 |
| 5265 | 264 | E264C | A | 7.7e-08 | 4.6e-08 | 2.2e-08 | 4.833E-08 | 2.757E-08 | 2.200E-08 | 7.700E-08 |
| 5266 | 264 | E264E | A | 4.8e-08 | 4.1e-08 | 5.4e-08 | 4.767E-08 | 6.506E-09 | 4.100E-08 | 5.400E-08 |
| 5267 | 264 | E264Q | A | 5.5e-08 | 8.7e-08 | 7.7e-08 | 7.300E-08 | 1.637E-08 | 5.500E-08 | 8.700E-08 |
| 5268 | 264 | E264G | A | 1.8e-08 | 6.6e-08 | 6e-08 | 4.800E-08 | 2.615E-08 | 1.800E-08 | 6.600E-08 |
| 5269 | 264 | E264H | A | 2.3e-08 | 8.2e-08 | 8.5e-08 | 6.333E-08 | 3.496E-08 | 2.300E-08 | 8.500E-08 |

| | | | | | | | | | | |
|---|---|---|---|---|---|---|---|---|---|---|
| 5270 | 264 | E264I | A | 5.9e-08 | 4.4e-08 | 2.8e-08 | 4.367E-08 | 1.550E-08 | 2.800E-08 | 5.900E-08 |
| 5271 | 264 | E264L | A | 8.5e-08 | 3.2e-08 | 7.4e-08 | 6.367E-08 | 2.797E-08 | 3.200E-08 | 8.500E-08 |
| 5272 | 264 | E264K | A | 4.9e-08 | 7.4e-08 | 3.6e-08 | 5.300E-08 | 1.931E-08 | 3.600E-08 | 7.400E-08 |
| 5273 | 264 | E264M | A | 7.7e-08 | 2.1e-08 | 1e-07 | 6.600E-08 | 4.063E-08 | 2.100E-08 | 1.000E-07 |
| 5274 | 264 | E264F | A | 2.5e-08 | 5e-08 | 1.4e-07 | 7.167E-08 | 6.048E-08 | 2.500E-08 | 1.400E-07 |
| 5275 | 264 | E264P | A | 1.8e-08 | 5.5e-08 | 4.5e-08 | 3.933E-08 | 1.914E-08 | 1.800E-08 | 5.500E-08 |
| 5276 | 264 | E264S | A | 2.8e-08 | 1.5e-07 | 4.5e-08 | 7.433E-08 | 6.608E-08 | 2.800E-08 | 1.500E-07 |
| 5277 | 264 | E264T | A | 6.2e-08 | 4.5e-08 | 6.1e-08 | 5.600E-08 | 9.539E-09 | 4.500E-08 | 6.200E-08 |
| 5278 | 264 | E264W | A | 4.5e-08 | 7.7e-08 | 5e-08 | 5.733E-08 | 1.721E-08 | 4.500E-08 | 7.700E-08 |
| 5279 | 264 | E264Y | A | 5.6e-08 | 2.9e-08 | 4e-08 | 4.167E-08 | 1.358E-08 | 2.900E-08 | 5.600E-08 |
| 5280 | 264 | E264V | A | 2.8e-08 | 5.4e-08 | 9.8e-08 | 6.000E-08 | 3.538E-08 | 2.800E-08 | 9.800E-08 |
| 5281 | 265 | G265A | A | 2.6e-08 | 4.7e-08 | 5.6e-08 | 4.300E-08 | 1.539E-08 | 2.600E-08 | 5.600E-08 |
| 5282 | 265 | G265R | A | 4.2e-08 | 1.1e-07 | 5.7e-08 | 6.967E-08 | 3.573E-08 | 4.200E-08 | 1.100E-07 |
| 5283 | 265 | G265N | A | 4.8e-08 | 1.6e-08 | 4.2e-08 | 3.533E-08 | 1.701E-08 | 1.600E-08 | 4.800E-08 |
| 5284 | 265 | G265D | A | 1.1e-07 | 1e-07 | 4.2e-08 | 8.400E-08 | 3.672E-08 | 4.200E-08 | 1.100E-07 |
| 5285 | 265 | G265C | A | 5.7e-08 | 3.3e-08 | 1.1e-08 | 3.367E-08 | 2.301E-08 | 1.100E-08 | 5.700E-08 |
| 5286 | 265 | G265E | A | 3.2e-08 | 7.6e-08 | 2e-08 | 4.267E-08 | 2.948E-08 | 2.000E-08 | 7.600E-08 |
| 5287 | 265 | G265Q | A | 3.3e-08 | 8.2e-08 | 4.2e-08 | 5.233E-08 | 2.608E-08 | 3.300E-08 | 8.200E-08 |
| 5288 | 265 | G265G | A | 6.1e-08 | 1.9e-08 | 2.4e-08 | 3.467E-08 | 2.294E-08 | 1.900E-08 | 6.100E-08 |
| 5289 | 265 | G265H | A | 3.3e-08 | 2.5e-08 | 5.8e-08 | 3.867E-08 | 1.721E-08 | 2.500E-08 | 5.800E-08 |
| 5290 | 265 | G265I | A | 1.9e-08 | 2.5e-08 | 3.1e-08 | 2.500E-08 | 6.000E-09 | 1.900E-08 | 3.100E-08 |
| 5291 | 265 | G265L | A | 2.1e-08 | 5.2e-08 | 4.7e-08 | 4.000E-08 | 1.664E-08 | 2.100E-08 | 5.200E-08 |
| 5292 | 265 | G265K | A | 3e-08 | 3.8e-08 | 8.3e-08 | 5.033E-08 | 2.857E-08 | 3.000E-08 | 8.300E-08 |
| 5293 | 265 | G265M | A | 5.6e-08 | 3.6e-08 | 5.5e-08 | 4.900E-08 | 1.127E-08 | 3.600E-08 | 5.600E-08 |
| 5294 | 265 | G265F | A | 3.6e-08 | 4.4e-08 | 7.4e-09 | 2.913E-08 | 1.924E-08 | 7.400E-09 | 4.400E-08 |
| 5295 | 265 | G265P | A | 2.5e-08 | 2.9e-08 | 5.8e-08 | 3.733E-08 | 1.801E-08 | 2.500E-08 | 5.800E-08 |
| 5296 | 265 | G265S | A | 9.3e-08 | 7.5e-08 | 6.6e-08 | 7.800E-08 | 1.375E-08 | 6.600E-08 | 9.300E-08 |
| 5297 | 265 | G265T | A | 4.2e-08 | 4.6e-08 | 7.5e-08 | 5.433E-08 | 1.801E-08 | 4.200E-08 | 7.500E-08 |
| 5298 | 265 | G265W | A | 1.1e-08 | 3e-08 | 1.3e-08 | 1.800E-08 | 1.044E-08 | 1.100E-08 | 3.000E-08 |
| 5299 | 265 | G265Y | A | 2.1e-08 | 3.1e-08 | 9.3e-08 | 4.833E-08 | 3.900E-08 | 2.100E-08 | 9.300E-08 |
| 5300 | 265 | G265V | A | 4.2e-08 | 2.3e-08 | 7.9e-08 | 4.800E-08 | 2.848E-08 | 2.300E-08 | 7.900E-08 |
| 5301 | 266 | R266A | A | 3.3e-08 | 4.2e-08 | 4.9e-08 | 4.133E-08 | 8.021E-09 | 3.300E-08 | 4.900E-08 |
| 5302 | 266 | R266R | A | 3.7e-08 | 2.6e-08 | 5e-08 | 3.767E-08 | 1.201E-08 | 2.600E-08 | 5.000E-08 |
| 5303 | 266 | R266N | A | 7.2e-08 | 1.2e-08 | 5.3e-08 | 4.567E-08 | 3.066E-08 | 1.200E-08 | 7.200E-08 |
| 5304 | 266 | R266D | A | 4.6e-08 | 2.3e-07 | 1.5e-07 | 1.420E-07 | 9.226E-08 | 4.600E-08 | 2.300E-07 |
| 5305 | 266 | R266C | A | 1.6e-08 | 2.9e-08 | 3.9e-08 | 2.800E-08 | 1.153E-08 | 1.600E-08 | 3.900E-08 |
| 5306 | 266 | R266E | A | 1.5e-08 | 4.6e-08 | 8.2e-08 | 4.767E-08 | 3.353E-08 | 1.500E-08 | 8.200E-08 |
| 5307 | 266 | R266Q | A | 8.4e-08 | 1e-07 | 7.9e-08 | 8.767E-08 | 1.097E-08 | 7.900E-08 | 1.000E-07 |
| 5308 | 266 | R266G | A | 3.6e-08 | 1.8e-08 | 2.9e-08 | 2.767E-08 | 9.074E-09 | 1.800E-08 | 3.600E-08 |
| 5309 | 266 | R266H | A | 5.1e-08 | 2e-08 | 2.6e-08 | 3.233E-08 | 1.644E-08 | 2.000E-08 | 5.100E-08 |
| 5310 | 266 | R266I | A | 1.6e-08 | 4.9e-08 | 1.8e-08 | 2.767E-08 | 1.850E-08 | 1.600E-08 | 4.900E-08 |
| 5311 | 266 | R266L | A | 6e-08 | 2.3e-08 | 2.1e-08 | 3.467E-08 | 2.196E-08 | 2.100E-08 | 6.000E-08 |
| 5312 | 266 | R266K | A | 5.4e-08 | 2.6e-08 | 2.7e-08 | 3.567E-08 | 1.589E-08 | 2.600E-08 | 5.400E-08 |
| 5313 | 266 | R266M | A | 4e-08 | 9.9e-09 | 2.7e-08 | 2.563E-08 | 1.510E-08 | 9.900E-09 | 4.000E-08 |
| 5314 | 266 | R266F | A | 4.9e-08 | 3e-08 | 3.7e-08 | 3.867E-08 | 9.609E-09 | 3.000E-08 | 4.900E-08 |
| 5315 | 266 | R266P | A | 6.3e-08 | 4.6e-08 | 2.8e-08 | 4.567E-08 | 1.750E-08 | 2.800E-08 | 6.300E-08 |
| 5316 | 266 | R266S | A | 7.1e-08 | 3.6e-08 | 2.3e-08 | 4.333E-08 | 2.483E-08 | 2.300E-08 | 7.100E-08 |
| 5317 | 266 | R266T | A | 1.2e-07 | 3.5e-08 | 6.9e-08 | 7.467E-08 | 4.278E-08 | 3.500E-08 | 1.200E-07 |
| 5318 | 266 | R266W | A | 3.9e-08 | 4.7e-08 | 1.8e-08 | 3.467E-08 | 1.498E-08 | 1.800E-08 | 4.700E-08 |
| 5319 | 266 | R266Y | A | 2.5e-08 | 1.5e-08 | 1.8e-08 | 1.933E-08 | 5.132E-09 | 1.500E-08 | 2.500E-08 |
| 5320 | 266 | R266V | A | 3e-08 | 4.1e-08 | 3.4e-08 | 3.500E-08 | 5.568E-09 | 3.000E-08 | 4.100E-08 |
| 5321 | 267 | Y267A | A | 2.6e-08 | 5.2e-08 | 2.8e-08 | 3.533E-08 | 1.447E-08 | 2.600E-08 | 5.200E-08 |
| 5322 | 267 | Y267R | A | 3e-08 | 1.8e-07 | 2.5e-08 | 7.833E-08 | 8.808E-08 | 2.500E-08 | 1.800E-07 |
| 5323 | 267 | Y267N | A | 3.8e-08 | 3.3e-08 | 4e-08 | 3.700E-08 | 3.606E-09 | 3.300E-08 | 4.000E-08 |
| 5324 | 267 | Y267D | A | 9.4e-08 | 7.5e-08 | 1e-07 | 8.967E-08 | 1.305E-08 | 7.500E-08 | 1.000E-07 |
| 5325 | 267 | Y267C | A | 8.9e-08 | 8e-08 | 7.5e-08 | 8.133E-08 | 7.095E-09 | 7.500E-08 | 8.900E-08 |
| 5326 | 267 | Y267E | A | 1.8e-08 | 6.3e-08 | 7.8e-08 | 5.300E-08 | 3.122E-08 | 1.800E-08 | 7.800E-08 |
| 5327 | 267 | Y267Q | A | 9.6e-08 | 4.8e-08 | 9.8e-08 | 8.067E-08 | 2.831E-08 | 4.800E-08 | 9.800E-08 |
| 5328 | 267 | Y267G | A | 4.3e-08 | 1e-07 | 5.8e-08 | 6.700E-08 | 2.955E-08 | 4.300E-08 | 1.000E-07 |
| 5329 | 267 | Y267H | A | 7.3e-08 | 3.1e-08 | 4.4e-08 | 4.933E-08 | 2.150E-08 | 3.100E-08 | 7.300E-08 |
| 5330 | 267 | Y267I | A | 3.8e-08 | 2e-08 | 8.3e-08 | 4.700E-08 | 3.245E-08 | 2.000E-08 | 8.300E-08 |
| 5331 | 267 | Y267L | A | 2.3e-08 | 9.7e-08 | 3.4e-08 | 5.133E-08 | 3.993E-08 | 2.300E-08 | 9.700E-08 |
| 5332 | 267 | Y267K | A | 4.8e-08 | 3.3e-08 | 3.5e-08 | 3.867E-08 | 8.145E-09 | 3.300E-08 | 4.800E-08 |
| 5333 | 267 | Y267M | A | 3e-08 | 1.4e-07 | 5.3e-08 | 7.433E-08 | 5.802E-08 | 3.000E-08 | 1.400E-07 |
| 5334 | 267 | Y267F | A | 3.4e-08 | 4e-08 | 4.7e-08 | 4.033E-08 | 6.506E-09 | 3.400E-08 | 4.700E-08 |
| 5335 | 267 | Y267P | A | 5.9e-08 | 4.4e-08 | 5.4e-08 | 5.233E-08 | 7.638E-09 | 4.400E-08 | 5.900E-08 |
| 5336 | 267 | Y267S | A | 5.6e-08 | 4.5e-08 | 6.3e-08 | 5.467E-08 | 9.074E-09 | 4.500E-08 | 6.300E-08 |
| 5337 | 267 | Y267T | A | 4.8e-08 | 1e-07 | 1.8e-07 | 1.093E-07 | 6.649E-08 | 4.800E-08 | 1.800E-07 |
| 5338 | 267 | Y267W | A | 3e-08 | 5.9e-08 | 1.1e-08 | 3.333E-08 | 2.417E-08 | 1.100E-08 | 5.900E-08 |
| 5339 | 267 | Y267Y | A | 2.2e-08 | 6.8e-08 | 8.8e-09 | 3.293E-08 | 3.108E-08 | 8.800E-09 | 6.800E-08 |
| 5340 | 267 | Y267V | A | 5.6e-08 | 5.9e-08 | 1.7e-08 | 4.400E-08 | 2.343E-08 | 1.700E-08 | 5.900E-08 |
| 5341 | 268 | T268A | A | 1e-08 | 7.9e-09 | 1.4e-08 | 1.063E-08 | 3.099E-09 | 7.900E-09 | 1.400E-08 |
| 5342 | 268 | T268R | A | 3.6e-08 | 1.7e-08 | 2.9e-08 | 2.733E-08 | 9.609E-09 | 1.700E-08 | 3.600E-08 |
| 5343 | 268 | T268N | A | 5.4e-08 | 1.2e-07 | 6.7e-08 | 8.033E-08 | 3.496E-08 | 5.400E-08 | 1.200E-07 |
| 5344 | 268 | T268D | A | 1.5e-08 | 3.6e-08 | 1.7e-08 | 2.267E-08 | 1.159E-08 | 1.500E-08 | 3.600E-08 |
| 5345 | 268 | T268C | A | 8.2e-09 | 6e-09 | 8.5e-09 | 7.567E-09 | 1.365E-09 | 6.000E-09 | 8.500E-09 |
| 5346 | 268 | T268E | A | 1.6e-08 | 2e-08 | 2.1e-08 | 1.900E-08 | 2.646E-09 | 1.600E-08 | 2.100E-08 |
| 5347 | 268 | T268Q | A | 8.7e-09 | 4.2e-08 | 8.5e-08 | 4.523E-08 | 3.825E-08 | 8.700E-09 | 8.500E-08 |
| 5348 | 268 | T268G | A | 8.8e-09 | 1.4e-08 | 1.5e-08 | 1.260E-08 | 3.329E-09 | 8.800E-09 | 1.500E-08 |
| 5349 | 268 | T268H | A | 3.1e-08 | 1.7e-08 | 1.3e-08 | 2.033E-08 | 9.452E-09 | 1.300E-08 | 3.100E-08 |
| 5350 | 268 | T268I | A | 1e-08 | 1.9e-08 | 1.3e-08 | 1.400E-08 | 4.583E-09 | 1.000E-08 | 1.900E-08 |
| 5351 | 268 | T268L | A | 1.3e-08 | 1.7e-08 | 5.7e-09 | 1.190E-08 | 5.730E-09 | 5.700E-09 | 1.700E-08 |
| 5352 | 268 | T268K | A | 5.4e-08 | 3.4e-08 | 4e-09 | 3.067E-08 | 2.517E-08 | 4.000E-09 | 5.400E-08 |
| 5353 | 268 | T268M | A | 3.6e-08 | 3e-08 | 1.3e-08 | 2.633E-08 | 1.193E-08 | 1.300E-08 | 3.600E-08 |
| 5354 | 268 | T268F | A | 1.1e-08 | 1.5e-08 | 6.2e-09 | 1.073E-08 | 4.406E-09 | 6.200E-09 | 1.500E-08 |

| | | | | | | | | | | |
|---|---|---|---|---|---|---|---|---|---|---|
| 5355 | 268 | T268P | A | 3.4e-09 | 5e-09 | 2.5e-08 | 1.113E-08 | 1.204E-08 | 3.400E-09 | 2.500E-08 |
| 5356 | 268 | T268S | A | 1e-08 | 8.4e-09 | 2.1e-08 | 1.313E-08 | 6.860E-09 | 8.400E-09 | 2.100E-08 |
| 5357 | 268 | T268T | A | 6.4e-08 | 3e-08 | 3.7e-08 | 4.367E-08 | 1.795E-08 | 3.000E-08 | 6.400E-08 |
| 5358 | 268 | T268W | A | 1.3e-08 | 1.4e-08 | 1.8e-08 | 1.500E-08 | 2.646E-09 | 1.300E-08 | 1.800E-08 |
| 5359 | 268 | T268Y | A | 6.7e-09 | 2.7e-08 | 5.5e-09 | 1.307E-08 | 1.208E-08 | 5.500E-09 | 2.700E-08 |
| 5360 | 268 | T268V | A | 3.5e-09 | 7.9e-09 | 8.4e-09 | 6.600E-09 | 2.696E-09 | 3.500E-09 | 8.400E-09 |
| 5361 | 269 | F269A | A | 8.8e-08 | 3.1e-08 | 1.9e-08 | 4.600E-08 | 3.686E-08 | 1.900E-08 | 8.800E-08 |
| 5362 | 269 | F269R | A | 1.2e-07 | 5.4e-08 | 3.7e-08 | 7.033E-08 | 4.384E-08 | 3.700E-08 | 1.200E-07 |
| 5363 | 269 | F269N | A | 2.6e-08 | 3.4e-08 | 3.3e-08 | 3.100E-08 | 4.359E-09 | 2.600E-08 | 3.400E-08 |
| 5364 | 269 | F269D | A | 7.1e-08 | 6.1e-08 | 6.1e-08 | 6.433E-08 | 5.774E-09 | 6.100E-08 | 7.100E-08 |
| 5365 | 269 | F269C | A | 6.7e-08 | 6.9e-08 | 6.4e-08 | 6.667E-08 | 2.517E-09 | 6.400E-08 | 6.900E-08 |
| 5366 | 269 | F269E | A | 3.2e-08 | 7.9e-08 | 4.8e-08 | 5.300E-08 | 2.390E-08 | 3.200E-08 | 7.900E-08 |
| 5367 | 269 | F269Q | A | 7.6e-08 | 8.6e-08 | 3.6e-08 | 6.600E-08 | 2.646E-08 | 3.600E-08 | 8.600E-08 |
| 5368 | 269 | F269G | A | 9.6e-09 | 5e-08 | 1.5e-08 | 2.487E-08 | 2.193E-08 | 9.600E-09 | 5.000E-08 |
| 5369 | 269 | F269H | A | 2.6e-08 | 4.1e-08 | 6.6e-08 | 4.433E-08 | 2.021E-08 | 2.600E-08 | 6.600E-08 |
| 5370 | 269 | F269I | A | 1.1e-07 | 6.2e-08 | 3.3e-08 | 6.833E-08 | 3.889E-08 | 3.300E-08 | 1.100E-07 |
| 5371 | 269 | F269L | A | 1.3e-07 | 6e-08 | 2.6e-08 | 7.200E-08 | 5.303E-08 | 2.600E-08 | 1.300E-07 |
| 5372 | 269 | F269K | A | 1.2e-07 | 4.7e-08 | 4.3e-08 | 7.000E-08 | 4.335E-08 | 4.300E-08 | 1.200E-07 |
| 5373 | 269 | F269M | A | 2.1e-08 | 6.5e-08 | 4.6e-08 | 4.400E-08 | 2.207E-08 | 2.100E-08 | 6.500E-08 |
| 5374 | 269 | F269F | A | 3.7e-08 | 2e-08 | 8.6e-08 | 4.767E-08 | 3.427E-08 | 2.000E-08 | 8.600E-08 |
| 5375 | 269 | F269P | A | 3.1e-08 | 4.9e-08 | 5.8e-08 | 4.600E-08 | 1.375E-08 | 3.100E-08 | 5.800E-08 |
| 5376 | 269 | F269S | A | 3.4e-08 | 3.6e-08 | 2.3e-08 | 3.100E-08 | 7.000E-09 | 2.300E-08 | 3.600E-08 |
| 5377 | 269 | F269T | A | 3.4e-08 | 8.2e-08 | 4.4e-08 | 5.333E-08 | 2.532E-08 | 3.400E-08 | 8.200E-08 |
| 5378 | 269 | F269W | A | 4.2e-08 | 5.4e-08 | 2.4e-08 | 4.000E-08 | 1.510E-08 | 2.400E-08 | 5.400E-08 |
| 5379 | 269 | F269Y | A | 2.9e-08 | 8.9e-08 | 4.2e-08 | 5.333E-08 | 3.156E-08 | 2.900E-08 | 8.900E-08 |
| 5380 | 269 | F269V | A | 1.8e-08 | 6.8e-08 | 3.5e-08 | 4.033E-08 | 2.542E-08 | 1.800E-08 | 6.800E-08 |
| 5381 | 270 | G270A | A | 3.5e-08 | 4.5e-08 | 3.3e-08 | 3.767E-08 | 6.429E-09 | 3.300E-08 | 4.500E-08 |
| 5382 | 270 | G270R | A | 5.2e-08 | 6.6e-08 | 7e-08 | 6.267E-08 | 9.452E-09 | 5.200E-08 | 7.000E-08 |
| 5383 | 270 | G270N | A | 7.3e-08 | 1.3e-07 | 6e-08 | 8.767E-08 | 3.723E-08 | 6.000E-08 | 1.300E-07 |
| 5384 | 270 | G270D | A | 7.1e-08 | 4.9e-08 | 7.3e-08 | 6.433E-08 | 1.332E-08 | 4.900E-08 | 7.300E-08 |
| 5385 | 270 | G270C | A | 9.2e-08 | 8.6e-08 | 5e-08 | 7.600E-08 | 2.272E-08 | 5.000E-08 | 9.200E-08 |
| 5386 | 270 | G270E | A | 6.6e-08 | 6.2e-08 | 1.6e-08 | 4.800E-08 | 2.778E-08 | 1.600E-08 | 6.600E-08 |
| 5387 | 270 | G270Q | A | 5.3e-08 | 4.2e-08 | 1.9e-08 | 3.800E-08 | 1.735E-08 | 1.900E-08 | 5.300E-08 |
| 5388 | 270 | G270G | A | 1e-07 | 5.3e-08 | 9.2e-08 | 8.167E-08 | 2.515E-08 | 5.300E-08 | 1.000E-07 |
| 5389 | 270 | G270H | A | 3.3e-08 | 2.4e-08 | 4.9e-08 | 3.533E-08 | 1.266E-08 | 2.400E-08 | 4.900E-08 |
| 5390 | 270 | G270I | A | 1.4e-07 | 2.8e-08 | 1e-07 | 8.933E-08 | 5.676E-08 | 2.800E-08 | 1.400E-07 |
| 5391 | 270 | G270L | A | 4.9e-08 | 5.5e-09 | 3.4e-08 | 2.950E-08 | 2.210E-08 | 5.500E-09 | 4.900E-08 |
| 5392 | 270 | G270K | A | 1.2e-08 | 3.2e-08 | 5e-08 | 3.133E-08 | 1.901E-08 | 1.200E-08 | 5.000E-08 |
| 5393 | 270 | G270M | A | 4.6e-08 | 1.3e-08 | 7.1e-08 | 4.333E-08 | 2.909E-08 | 1.300E-08 | 7.100E-08 |
| 5394 | 270 | G270F | A | 1.1e-07 | 3.2e-08 | 3e-08 | 5.733E-08 | 4.562E-08 | 3.000E-08 | 1.100E-07 |
| 5395 | 270 | G270P | A | 4.3e-08 | 1.7e-08 | 2.6e-08 | 2.867E-08 | 1.320E-08 | 1.700E-08 | 4.300E-08 |
| 5396 | 270 | G270S | A | 5.7e-08 | 8.6e-08 | 4.4e-08 | 6.233E-08 | 2.150E-08 | 4.400E-08 | 8.600E-08 |
| 5397 | 270 | G270T | A | 1e-07 | 7e-08 | 5.5e-08 | 7.500E-08 | 2.291E-08 | 5.500E-08 | 1.000E-07 |
| 5398 | 270 | G270W | A | 2.1e-08 | 5.2e-08 | 6.4e-08 | 4.567E-08 | 2.219E-08 | 2.100E-08 | 6.400E-08 |
| 5399 | 270 | G270Y | A | 7.7e-08 | 2.6e-08 | 1.7e-08 | 4.000E-08 | 3.236E-08 | 1.700E-08 | 7.700E-08 |
| 5400 | 270 | G270V | A | 2.5e-08 | 6e-08 | 3.1e-08 | 3.867E-08 | 1.872E-08 | 2.500E-08 | 6.000E-08 |
| 5401 | 271 | A271A | A | 9e-08 | 5e-08 | 3.6e-08 | 5.867E-08 | 2.802E-08 | 3.600E-08 | 9.000E-08 |
| 5402 | 271 | A271R | A | 6.6e-08 | 2.1e-08 | 2.7e-08 | 3.800E-08 | 2.443E-08 | 2.100E-08 | 6.600E-08 |
| 5403 | 271 | A271N | A | 4.3e-08 | 1.8e-08 | 1.8e-08 | 2.633E-08 | 1.443E-08 | 1.800E-08 | 4.300E-08 |
| 5404 | 271 | A271D | A | 1e-07 | 3.2e-08 | 3.8e-08 | 5.667E-08 | 3.765E-08 | 3.200E-08 | 1.000E-07 |
| 5405 | 271 | A271C | A | 2.9e-08 | 4.6e-08 | 3.4e-08 | 3.633E-08 | 8.737E-09 | 2.900E-08 | 4.600E-08 |
| 5406 | 271 | A271E | A | 4.1e-08 | 2.7e-08 | 8.9e-08 | 5.233E-08 | 3.252E-08 | 2.700E-08 | 8.900E-08 |
| 5407 | 271 | A271Q | A | 1.3e-08 | 2.9e-08 | 1.6e-08 | 1.933E-08 | 8.505E-09 | 1.300E-08 | 2.900E-08 |
| 5408 | 271 | A271G | A | 5.2e-08 | 3e-08 | 3.6e-08 | 3.933E-08 | 1.137E-08 | 3.000E-08 | 5.200E-08 |
| 5409 | 271 | A271H | A | 9.4e-08 | 2.6e-08 | 3.3e-08 | 5.100E-08 | 3.740E-08 | 2.600E-08 | 9.400E-08 |
| 5410 | 271 | A271I | A | 7.5e-08 | 1.8e-08 | 2.7e-08 | 4.000E-08 | 3.064E-08 | 1.800E-08 | 7.500E-08 |
| 5411 | 271 | A271L | A | 9.4e-09 | 5.5e-08 | 3.5e-08 | 3.313E-08 | 2.286E-08 | 9.400E-09 | 5.500E-08 |
| 5412 | 271 | A271K | A | 1.1e-07 | 3.2e-08 | 4.1e-08 | 6.100E-08 | 4.267E-08 | 3.200E-08 | 1.100E-07 |
| 5413 | 271 | A271M | A | 2.1e-08 | 3.6e-08 | 5.4e-08 | 3.700E-08 | 1.652E-08 | 2.100E-08 | 5.400E-08 |
| 5414 | 271 | A271F | A | 7.2e-08 | 3.9e-08 | 2.9e-08 | 4.667E-08 | 2.250E-08 | 2.900E-08 | 7.200E-08 |
| 5415 | 271 | A271P | A | 8.2e-08 | 7.3e-08 | 5.5e-08 | 7.000E-08 | 1.375E-08 | 5.500E-08 | 8.200E-08 |
| 5416 | 271 | A271S | A | 7.1e-08 | 4.5e-08 | 4e-08 | 5.200E-08 | 1.664E-08 | 4.000E-08 | 7.100E-08 |
| 5417 | 271 | A271T | A | 1.3e-08 | 6.9e-08 | 5.7e-08 | 4.633E-08 | 2.948E-08 | 1.300E-08 | 6.900E-08 |
| 5418 | 271 | A271W | A | 3.3e-08 | 2.1e-08 | 1.7e-08 | 2.367E-08 | 8.327E-09 | 1.700E-08 | 3.300E-08 |
| 5419 | 271 | A271Y | A | 1.2e-07 | 3.3e-08 | 8.5e-08 | 7.933E-08 | 4.378E-08 | 3.300E-08 | 1.200E-07 |
| 5420 | 271 | A271V | A | 3.4e-08 | 6.3e-08 | 2.8e-08 | 4.167E-08 | 1.872E-08 | 2.800E-08 | 6.300E-08 |
| 5421 | 272 | S272A | A | 6.8e-08 | 3.8e-08 | 1.5e-08 | 4.033E-08 | 2.658E-08 | 1.500E-08 | 6.800E-08 |
| 5422 | 272 | S272R | A | 4.6e-08 | 4.4e-08 | 2.3e-08 | 3.767E-08 | 1.274E-08 | 2.300E-08 | 4.600E-08 |
| 5423 | 272 | S272N | A | 4.7e-08 | 4.6e-08 | 1.3e-07 | 7.433E-08 | 4.821E-08 | 4.600E-08 | 1.300E-07 |
| 5424 | 272 | S272D | A | 3.2e-08 | 1.2e-07 | 6.4e-08 | 7.200E-08 | 4.454E-08 | 3.200E-08 | 1.200E-07 |
| 5425 | 272 | S272C | A | 7.7e-08 | 2.2e-08 | 1.7e-08 | 3.867E-08 | 3.329E-08 | 1.700E-08 | 7.700E-08 |
| 5426 | 272 | S272E | A | 2.2e-08 | 2.4e-08 | 3.9e-08 | 2.833E-08 | 9.292E-09 | 2.200E-08 | 3.900E-08 |
| 5427 | 272 | S272Q | A | 2.7e-08 | 8.5e-08 | 5.5e-08 | 5.567E-08 | 2.901E-08 | 2.700E-08 | 8.500E-08 |
| 5428 | 272 | S272G | A | 3e-08 | 3.4e-08 | 2.4e-08 | 2.933E-08 | 5.033E-09 | 2.400E-08 | 3.400E-08 |
| 5429 | 272 | S272H | A | 4.7e-08 | 3.7e-08 | 3.8e-08 | 4.067E-08 | 5.508E-09 | 3.700E-08 | 4.700E-08 |
| 5430 | 272 | S272I | A | 7e-08 | 1.3e-07 | 4.4e-08 | 8.133E-08 | 4.411E-08 | 4.400E-08 | 1.300E-07 |
| 5431 | 272 | S272L | A | 7.7e-08 | 1.1e-08 | 4.6e-08 | 4.467E-08 | 3.302E-08 | 1.100E-08 | 7.700E-08 |
| 5432 | 272 | S272K | A | 5.4e-08 | 9e-09 | 8.2e-08 | 4.833E-08 | 3.683E-08 | 9.000E-09 | 8.200E-08 |
| 5433 | 272 | S272M | A | 1.2e-07 | 2.4e-08 | 4e-08 | 6.133E-08 | 5.143E-08 | 2.400E-08 | 1.200E-07 |
| 5434 | 272 | S272F | A | 3e-08 | 6.7e-08 | 5.1e-08 | 4.933E-08 | 1.856E-08 | 3.000E-08 | 6.700E-08 |
| 5435 | 272 | S272P | A | 1.4e-08 | 6.2e-08 | 5.1e-08 | 4.233E-08 | 2.515E-08 | 1.400E-08 | 6.200E-08 |
| 5436 | 272 | S272S | A | 5.9e-08 | 4.3e-08 | 3.5e-08 | 4.567E-08 | 1.222E-08 | 3.500E-08 | 5.900E-08 |
| 5437 | 272 | S272T | A | 8.4e-08 | 2.3e-08 | 3.6e-08 | 4.767E-08 | 3.213E-08 | 2.300E-08 | 8.400E-08 |
| 5438 | 272 | S272W | A | 4.3e-08 | 5e-08 | 6.8e-08 | 5.367E-08 | 1.290E-08 | 4.300E-08 | 6.800E-08 |
| 5439 | 272 | S272Y | A | 3.6e-08 | 5.8e-08 | 2.5e-08 | 3.967E-08 | 1.680E-08 | 2.500E-08 | 5.800E-08 |

| | | | | | | | | | | |
|---|---|---|---|---|---|---|---|---|---|---|
| 5440 | 272 | S272V | A | 3.5e-08 | 2.2e-08 | 3.8e-08 | 3.167E-08 | 8.505E-09 | 2.200E-08 | 3.800E-08 |
| 5441 | 273 | C273A | A | 4.6e-08 | 3.4e-08 | 6.5e-08 | 4.833E-08 | 1.563E-08 | 3.400E-08 | 6.500E-08 |
| 5442 | 273 | C273R | A | 6.6e-08 | 1.3e-07 | 8.5e-08 | 9.367E-08 | 3.287E-08 | 6.600E-08 | 1.300E-07 |
| 5443 | 273 | C273N | A | 3.9e-08 | 5.1e-08 | 1.1e-07 | 6.667E-08 | 3.800E-08 | 3.900E-08 | 1.100E-07 |
| 5444 | 273 | C273D | A | 1.5e-08 | 9.1e-08 | 1.1e-07 | 7.200E-08 | 5.027E-08 | 1.500E-08 | 1.100E-07 |
| 5445 | 273 | C273C | A | 4.9e-08 | 6.2e-08 | 5.6e-08 | 5.567E-08 | 6.506E-09 | 4.900E-08 | 6.200E-08 |
| 5446 | 273 | C273E | A | 1.2e-07 | 5.3e-08 | 5.9e-08 | 7.733E-08 | 3.707E-08 | 5.300E-08 | 1.200E-07 |
| 5447 | 273 | C273Q | A | 1.7e-07 | 6.4e-08 | 4.7e-08 | 9.367E-08 | 6.665E-08 | 4.700E-08 | 1.700E-07 |
| 5448 | 273 | C273G | A | 1.3e-07 | 4.7e-08 | 1.8e-08 | 6.500E-08 | 5.813E-08 | 1.800E-08 | 1.300E-07 |
| 5449 | 273 | C273H | A | 3.6e-08 | 7e-08 | 8.7e-08 | 6.433E-08 | 2.597E-08 | 3.600E-08 | 8.700E-08 |
| 5450 | 273 | C273I | A | 4.6e-08 | 3.9e-08 | 3e-08 | 3.833E-08 | 8.021E-09 | 3.000E-08 | 4.600E-08 |
| 5451 | 273 | C273L | A | 2.6e-08 | 1.4e-07 | 2.8e-08 | 6.467E-08 | 6.525E-08 | 2.600E-08 | 1.400E-07 |
| 5452 | 273 | C273K | A | 1.8e-08 | 3.6e-08 | 4e-08 | 3.133E-08 | 1.172E-08 | 1.800E-08 | 4.000E-08 |
| 5453 | 273 | C273M | A | 3.2e-08 | 2e-08 | 5.3e-08 | 3.500E-08 | 1.670E-08 | 2.000E-08 | 5.300E-08 |
| 5454 | 273 | C273F | A | 6.4e-08 | 5e-08 | 2.9e-08 | 4.767E-08 | 1.762E-08 | 2.900E-08 | 6.400E-08 |
| 5455 | 273 | C273P | A | 4e-08 | 6.5e-08 | 1.1e-07 | 7.167E-08 | 3.547E-08 | 4.000E-08 | 1.100E-07 |
| 5456 | 273 | C273S | A | 5.7e-08 | 5.9e-08 | 5.8e-08 | 5.800E-08 | 1.000E-09 | 5.700E-08 | 5.900E-08 |
| 5457 | 273 | C273T | A | 3.3e-08 | 5.1e-08 | 2.8e-08 | 3.733E-08 | 1.210E-08 | 2.800E-08 | 5.100E-08 |
| 5458 | 273 | C273W | A | 4.5e-08 | 5.4e-08 | 4.9e-08 | 4.933E-08 | 4.509E-09 | 4.500E-08 | 5.400E-08 |
| 5459 | 273 | C273Y | A | 3.1e-08 | 8.8e-08 | 2.4e-08 | 4.767E-08 | 3.510E-08 | 2.400E-08 | 8.800E-08 |
| 5460 | 273 | C273V | A | 2.9e-08 | 6e-08 | 7.2e-08 | 5.367E-08 | 2.219E-08 | 2.900E-08 | 7.200E-08 |
| 5461 | 274 | V274A | A | 6.4e-08 | 5.1e-08 | 3.5e-08 | 5.000E-08 | 1.453E-08 | 3.500E-08 | 6.400E-08 |
| 5462 | 274 | V274R | A | 2.1e-08 | 4.3e-08 | 8.2e-08 | 4.867E-08 | 3.089E-08 | 2.100E-08 | 8.200E-08 |
| 5463 | 274 | V274N | A | 1.1e-07 | 3.6e-08 | 3.1e-08 | 5.900E-08 | 4.424E-08 | 3.100E-08 | 1.100E-07 |
| 5464 | 274 | V274D | A | 4.2e-08 | 2.7e-08 | 8.6e-08 | 5.167E-08 | 3.066E-08 | 2.700E-08 | 8.600E-08 |
| 5465 | 274 | V274C | A | 4.1e-08 | 1.1e-07 | 3.1e-08 | 6.067E-08 | 4.302E-08 | 3.100E-08 | 1.100E-07 |
| 5466 | 274 | V274E | A | 5.3e-08 | 1.2e-07 | 3.1e-08 | 6.800E-08 | 4.636E-08 | 3.100E-08 | 1.200E-07 |
| 5467 | 274 | V274Q | A | 1.2e-07 | 4.1e-08 | 6.2e-08 | 7.433E-08 | 4.092E-08 | 4.100E-08 | 1.200E-07 |
| 5468 | 274 | V274G | A | 4.1e-08 | 7.5e-08 | 2.1e-08 | 4.567E-08 | 2.730E-08 | 2.100E-08 | 7.500E-08 |
| 5469 | 274 | V274H | A | 4.5e-08 | 3.7e-08 | 2.1e-08 | 3.433E-08 | 1.222E-08 | 2.100E-08 | 4.500E-08 |
| 5470 | 274 | V274I | A | 9e-08 | 1.1e-08 | 6.1e-08 | 5.400E-08 | 3.996E-08 | 1.100E-08 | 9.000E-08 |
| 5471 | 274 | V274L | A | 5.4e-08 | 2.7e-08 | 1.8e-08 | 3.300E-08 | 1.873E-08 | 1.800E-08 | 5.400E-08 |
| 5472 | 274 | V274K | A | 4.5e-08 | 2.7e-08 | 1.3e-07 | 6.733E-08 | 5.501E-08 | 2.700E-08 | 1.300E-07 |
| 5473 | 274 | V274M | A | 5.9e-08 | 9.3e-08 | 8.4e-09 | 5.347E-08 | 4.257E-08 | 8.400E-09 | 9.300E-08 |
| 5474 | 274 | V274F | A | 3.4e-08 | 3.9e-08 | 3.4e-08 | 3.567E-08 | 2.887E-09 | 3.400E-08 | 3.900E-08 |
| 5475 | 274 | V274P | A | 1.2e-07 | 5.3e-08 | 3.6e-08 | 6.967E-08 | 4.441E-08 | 3.600E-08 | 1.200E-07 |
| 5476 | 274 | V274S | A | 6.4e-08 | 4.1e-08 | 6.9e-08 | 5.800E-08 | 1.493E-08 | 4.100E-08 | 6.900E-08 |
| 5477 | 274 | V274T | A | 1e-07 | 5.1e-08 | 4.4e-08 | 6.500E-08 | 3.051E-08 | 4.400E-08 | 1.000E-07 |
| 5478 | 274 | V274W | A | 6.9e-08 | 2.6e-08 | 2e-08 | 3.833E-08 | 2.673E-08 | 2.000E-08 | 6.900E-08 |
| 5479 | 274 | V274Y | A | 2.7e-08 | 3.7e-08 | 4.8e-08 | 3.733E-08 | 1.050E-08 | 2.700E-08 | 4.800E-08 |
| 5480 | 274 | V274V | A | 7.6e-08 | 2.2e-08 | 5e-08 | 4.933E-08 | 2.701E-08 | 2.200E-08 | 7.600E-08 |
| 5481 | 275 | T275A | A | 1.5e-07 | 3.7e-08 | 3.5e-08 | 7.400E-08 | 6.583E-08 | 3.500E-08 | 1.500E-07 |
| 5482 | 275 | T275R | A | 3.6e-08 | 6e-08 | 1.1e-07 | 6.867E-08 | 3.775E-08 | 3.600E-08 | 1.100E-07 |
| 5483 | 275 | T275N | A | 4.1e-08 | 3.5e-08 | 3.5e-08 | 3.700E-08 | 3.464E-09 | 3.500E-08 | 4.100E-08 |
| 5484 | 275 | T275D | A | 2e-08 | 3.1e-08 | 2.2e-08 | 2.433E-08 | 5.859E-09 | 2.000E-08 | 3.100E-08 |
| 5485 | 275 | T275C | A | 4.2e-08 | 3.8e-08 | 4.4e-08 | 4.133E-08 | 3.055E-09 | 3.800E-08 | 4.400E-08 |
| 5486 | 275 | T275E | A | 1.2e-07 | 3.5e-08 | 9e-08 | 8.167E-08 | 4.311E-08 | 3.500E-08 | 1.200E-07 |
| 5487 | 275 | T275Q | A | 2.2e-08 | 3e-08 | 1.5e-08 | 2.233E-08 | 7.506E-09 | 1.500E-08 | 3.000E-08 |
| 5488 | 275 | T275G | A | 8.5e-08 | 3.3e-08 | 5.5e-08 | 5.767E-08 | 2.610E-08 | 3.300E-08 | 8.500E-08 |
| 5489 | 275 | T275H | A | 6.9e-08 | 2.2e-08 | 8.2e-08 | 5.767E-08 | 3.156E-08 | 2.200E-08 | 8.200E-08 |
| 5490 | 275 | T275I | A | 1.4e-07 | 2.8e-08 | 6.1e-08 | 7.633E-08 | 5.755E-08 | 2.800E-08 | 1.400E-07 |
| 5491 | 275 | T275L | A | 4.1e-08 | 2.6e-08 | 5.4e-08 | 4.033E-08 | 1.401E-08 | 2.600E-08 | 5.400E-08 |
| 5492 | 275 | T275K | A | 1.7e-08 | 2e-08 | 5.2e-08 | 2.967E-08 | 1.940E-08 | 1.700E-08 | 5.200E-08 |
| 5493 | 275 | T275M | A | 2.5e-08 | 7.7e-08 | 4.6e-08 | 4.933E-08 | 2.616E-08 | 2.500E-08 | 7.700E-08 |
| 5494 | 275 | T275F | A | 3.3e-08 | 3.6e-08 | 2.6e-08 | 3.167E-08 | 5.132E-09 | 2.600E-08 | 3.600E-08 |
| 5495 | 275 | T275P | A | 3.8e-08 | 1.9e-08 | 1.1e-07 | 5.567E-08 | 4.800E-08 | 1.900E-08 | 1.100E-07 |
| 5496 | 275 | T275S | A | 3.6e-08 | 5.2e-08 | 6.3e-08 | 5.033E-08 | 1.358E-08 | 3.600E-08 | 6.300E-08 |
| 5497 | 275 | T275T | A | 3.4e-08 | 3.5e-08 | 4.8e-08 | 3.900E-08 | 7.810E-09 | 3.400E-08 | 4.800E-08 |
| 5498 | 275 | T275W | A | 2.1e-08 | 2.8e-08 | 4.5e-08 | 3.133E-08 | 1.234E-08 | 2.100E-08 | 4.500E-08 |
| 5499 | 275 | T275Y | A | 6.8e-08 | 3e-08 | 1.9e-08 | 3.900E-08 | 2.571E-08 | 1.900E-08 | 6.800E-08 |
| 5500 | 275 | T275V | A | 6.5e-08 | 3.9e-08 | 4.2e-08 | 4.867E-08 | 1.422E-08 | 3.900E-08 | 6.500E-08 |
| 5501 | 276 | A276A | A | 5.8e-08 | 6.9e-08 | 3.7e-08 | 5.467E-08 | 1.626E-08 | 3.700E-08 | 6.900E-08 |
| 5502 | 276 | A276R | A | 1e-08 | 3.9e-08 | 5.4e-08 | 3.433E-08 | 2.237E-08 | 1.000E-08 | 5.400E-08 |
| 5503 | 276 | A276N | A | 1.5e-07 | 7.5e-08 | 1.5e-08 | 8.000E-08 | 6.764E-08 | 1.500E-08 | 1.500E-07 |
| 5504 | 276 | A276D | A | 5e-08 | 7.7e-08 | 5.3e-08 | 6.000E-08 | 1.480E-08 | 5.000E-08 | 7.700E-08 |
| 5505 | 276 | A276C | A | 2.5e-07 | 6e-09 | 1.3e-08 | 8.967E-08 | 1.389E-07 | 6.000E-09 | 2.500E-07 |
| 5506 | 276 | A276E | A | 6.4e-08 | 7.4e-08 | 8.1e-08 | 7.300E-08 | 8.544E-09 | 6.400E-08 | 8.100E-08 |
| 5507 | 276 | A276Q | A | 5.8e-08 | 5e-08 | 5.6e-08 | 5.467E-08 | 4.163E-09 | 5.000E-08 | 5.800E-08 |
| 5508 | 276 | A276G | A | 4.1e-08 | 3.4e-08 | 1.2e-07 | 6.500E-08 | 4.776E-08 | 3.400E-08 | 1.200E-07 |
| 5509 | 276 | A276H | A | 5.8e-08 | 1.7e-07 | 6.1e-08 | 9.633E-08 | 6.381E-08 | 5.800E-08 | 1.700E-07 |
| 5510 | 276 | A276I | A | 6.1e-08 | 3.5e-08 | 5.7e-08 | 5.100E-08 | 1.400E-08 | 3.500E-08 | 6.100E-08 |
| 5511 | 276 | A276L | A | 3e-08 | 5.4e-08 | 3.7e-08 | 4.033E-08 | 1.234E-08 | 3.000E-08 | 5.400E-08 |
| 5512 | 276 | A276K | A | 1.1e-07 | 3.8e-08 | 1.9e-07 | 1.127E-07 | 7.604E-08 | 3.800E-08 | 1.900E-07 |
| 5513 | 276 | A276M | A | 3.9e-08 | 3e-08 | 3.5e-08 | 3.467E-08 | 4.509E-09 | 3.000E-08 | 3.900E-08 |
| 5514 | 276 | A276F | A | 1.7e-08 | 4.9e-08 | 3.6e-08 | 3.400E-08 | 1.609E-08 | 1.700E-08 | 4.900E-08 |
| 5515 | 276 | A276P | A | 4.7e-08 | 3.3e-08 | 4e-08 | 4.000E-08 | 7.000E-09 | 3.300E-08 | 4.700E-08 |
| 5516 | 276 | A276S | A | 2e-08 | 2.3e-08 | 6.9e-08 | 3.733E-08 | 2.747E-08 | 2.000E-08 | 6.900E-08 |
| 5517 | 276 | A276T | A | 6.4e-08 | 9e-08 | 4.5e-08 | 6.633E-08 | 2.259E-08 | 4.500E-08 | 9.000E-08 |
| 5518 | 276 | A276W | A | 5e-08 | 2.7e-08 | 3.5e-08 | 3.733E-08 | 1.168E-08 | 2.700E-08 | 5.000E-08 |
| 5519 | 276 | A276Y | A | 3.2e-08 | 2.2e-08 | 7.3e-08 | 4.233E-08 | 2.702E-08 | 2.200E-08 | 7.300E-08 |
| 5520 | 276 | A276V | A | 1.5e-08 | 4.6e-08 | 2.8e-08 | 2.967E-08 | 1.557E-08 | 1.500E-08 | 4.600E-08 |
| 5521 | 277 | C277A | A | 1.8e-08 | 4.7e-08 | 2e-08 | 2.833E-08 | 1.620E-08 | 1.800E-08 | 4.700E-08 |
| 5522 | 277 | C277R | A | 7e-08 | 1.4e-07 | 1.4e-07 | 1.167E-07 | 4.041E-08 | 7.000E-08 | 1.400E-07 |
| 5523 | 277 | C277N | A | 8.5e-08 | 5.1e-08 | 2.6e-08 | 5.400E-08 | 2.961E-08 | 2.600E-08 | 8.500E-08 |
| 5524 | 277 | C277D | A | 5.8e-08 | 2.8e-08 | 5.4e-08 | 4.667E-08 | 1.629E-08 | 2.800E-08 | 5.800E-08 |

| | | | | | | | | | | |
|---|---|---|---|---|---|---|---|---|---|---|
| 5525 | 277 | C277C | A | 6.5e-08 | 2.8e-08 | 1.5e-08 | 3.600E-08 | 2.594E-08 | 1.500E-08 | 6.500E-08 |
| 5526 | 277 | C277E | A | 2.9e-08 | 3e-08 | 4.1e-08 | 3.333E-08 | 6.658E-09 | 2.900E-08 | 4.100E-08 |
| 5527 | 277 | C277Q | A | 1.9e-08 | 3.6e-08 | 5.6e-08 | 3.700E-08 | 1.852E-08 | 1.900E-08 | 5.600E-08 |
| 5528 | 277 | C277G | A | 6.2e-08 | 5.5e-08 | 5e-08 | 5.567E-08 | 6.028E-09 | 5.000E-08 | 6.200E-08 |
| 5529 | 277 | C277H | A | 8.4e-08 | 1.5e-08 | 3.6e-08 | 4.500E-08 | 3.537E-08 | 1.500E-08 | 8.400E-08 |
| 5530 | 277 | C277I | A | 1.7e-08 | 2.5e-08 | 1.6e-08 | 1.933E-08 | 4.933E-09 | 1.600E-08 | 2.500E-08 |
| 5531 | 277 | C277L | A | 5e-08 | 7.5e-08 | 4.1e-08 | 5.533E-08 | 1.762E-08 | 4.100E-08 | 7.500E-08 |
| 5532 | 277 | C277K | A | 5.5e-08 | 2.8e-08 | 5.7e-08 | 4.667E-08 | 1.620E-08 | 2.800E-08 | 5.700E-08 |
| 5533 | 277 | C277M | A | 2.5e-08 | 1.5e-08 | 1.9e-08 | 1.967E-08 | 5.033E-09 | 1.500E-08 | 2.500E-08 |
| 5534 | 277 | C277F | A | 1.4e-07 | 5.7e-08 | 1.2e-07 | 1.057E-07 | 4.332E-08 | 5.700E-08 | 1.400E-07 |
| 5535 | 277 | C277P | A | 6.6e-08 | 2.1e-08 | 1.2e-07 | 6.900E-08 | 4.957E-08 | 2.100E-08 | 1.200E-07 |
| 5536 | 277 | C277S | A | 4.1e-08 | 4.7e-08 | 7.5e-08 | 5.433E-08 | 1.815E-08 | 4.100E-08 | 7.500E-08 |
| 5537 | 277 | C277T | A | 3.2e-08 | 1.1e-07 | 8.6e-08 | 7.600E-08 | 3.995E-08 | 3.200E-08 | 1.100E-07 |
| 5538 | 277 | C277W | A | 3.5e-08 | 5.1e-08 | 1.8e-08 | 3.467E-08 | 1.650E-08 | 1.800E-08 | 5.100E-08 |
| 5539 | 277 | C277Y | A | 2.9e-08 | 2.8e-08 | 2.3e-08 | 2.667E-08 | 3.215E-09 | 2.300E-08 | 2.900E-08 |
| 5540 | 277 | C277V | A | 3.7e-08 | 7.5e-09 | 4.9e-08 | 3.117E-08 | 2.136E-08 | 7.500E-09 | 4.900E-08 |
| 5541 | 278 | P278A | A | 3.3e-08 | 1.1e-08 | 2.7e-08 | 2.367E-08 | 1.137E-08 | 1.100E-08 | 3.300E-08 |
| 5542 | 278 | P278R | A | 5.5e-08 | 3.4e-08 | 5.1e-08 | 4.667E-08 | 1.115E-08 | 3.400E-08 | 5.500E-08 |
| 5543 | 278 | P278N | A | 4.8e-08 | 4.4e-08 | 6.9e-08 | 5.367E-08 | 1.343E-08 | 4.400E-08 | 6.900E-08 |
| 5544 | 278 | P278D | A | 3.2e-08 | 4.5e-08 | 5.5e-08 | 4.400E-08 | 1.153E-08 | 3.200E-08 | 5.500E-08 |
| 5545 | 278 | P278C | A | 4.4e-08 | 2.1e-08 | 3.4e-08 | 3.300E-08 | 1.153E-08 | 2.100E-08 | 4.400E-08 |
| 5546 | 278 | P278E | A | 6.5e-08 | 8.9e-08 | 8.8e-08 | 8.067E-08 | 1.358E-08 | 6.500E-08 | 8.900E-08 |
| 5547 | 278 | P278Q | A | 6.8e-08 | 9.5e-08 | 2.2e-08 | 6.167E-08 | 3.691E-08 | 2.200E-08 | 9.500E-08 |
| 5548 | 278 | P278G | A | 3.1e-08 | 3.1e-08 | 5e-08 | 3.733E-08 | 1.097E-08 | 3.100E-08 | 5.000E-08 |
| 5549 | 278 | P278H | A | 4.2e-08 | 2.3e-08 | 1.2e-07 | 6.167E-08 | 5.140E-08 | 2.300E-08 | 1.200E-07 |
| 5550 | 278 | P278I | A | 3.7e-08 | 2e-08 | 1.1e-08 | 2.267E-08 | 1.320E-08 | 1.100E-08 | 3.700E-08 |
| 5551 | 278 | P278L | A | 2.8e-08 | 4.7e-08 | 1.9e-08 | 3.133E-08 | 1.429E-08 | 1.900E-08 | 4.700E-08 |
| 5552 | 278 | P278K | A | 2.4e-08 | 5.7e-08 | 5.1e-08 | 4.400E-08 | 1.758E-08 | 2.400E-08 | 5.700E-08 |
| 5553 | 278 | P278M | A | 7.7e-08 | 2.4e-08 | 2.2e-08 | 4.100E-08 | 3.119E-08 | 2.200E-08 | 7.700E-08 |
| 5554 | 278 | P278F | A | 1.9e-08 | 2.5e-08 | 3.8e-08 | 2.733E-08 | 9.713E-09 | 1.900E-08 | 3.800E-08 |
| 5555 | 278 | P278P | A | 3.2e-08 | 4.3e-08 | 1.7e-08 | 3.067E-08 | 1.305E-08 | 1.700E-08 | 4.300E-08 |
| 5556 | 278 | P278S | A | 6.7e-08 | 2.4e-08 | 2.3e-08 | 3.800E-08 | 2.512E-08 | 2.300E-08 | 6.700E-08 |
| 5557 | 278 | P278T | A | 1.1e-08 | 8e-09 | 2.3e-08 | 1.400E-08 | 7.937E-09 | 8.000E-09 | 2.300E-08 |
| 5558 | 278 | P278W | A | 1.9e-08 | 7.9e-08 | 5e-08 | 4.933E-08 | 3.001E-08 | 1.900E-08 | 7.900E-08 |
| 5559 | 278 | P278Y | A | 1.4e-08 | 7.4e-08 | 6e-08 | 4.933E-08 | 3.139E-08 | 1.400E-08 | 7.400E-08 |
| 5560 | 278 | P278V | A | 2.1e-08 | 3.5e-08 | 3.1e-08 | 2.900E-08 | 7.211E-09 | 2.100E-08 | 3.500E-08 |
| 5561 | 279 | Y279A | A | 3.3e-08 | 8.8e-08 | 3.8e-08 | 5.300E-08 | 3.041E-08 | 3.300E-08 | 8.800E-08 |
| 5562 | 279 | Y279R | A | 1.1e-07 | 5.1e-08 | 3.5e-08 | 6.533E-08 | 3.950E-08 | 3.500E-08 | 1.100E-07 |
| 5563 | 279 | Y279N | A | 3.1e-08 | 7.8e-08 | 4.4e-08 | 5.100E-08 | 2.427E-08 | 3.100E-08 | 7.800E-08 |
| 5564 | 279 | Y279D | A | 3.6e-08 | 1.2e-07 | 2.5e-08 | 6.033E-08 | 5.196E-08 | 2.500E-08 | 1.200E-07 |
| 5565 | 279 | Y279C | A | 2.5e-08 | 8.6e-08 | 1.4e-07 | 8.367E-08 | 5.754E-08 | 2.500E-08 | 1.400E-07 |
| 5566 | 279 | Y279E | A | 5.7e-08 | 5.2e-08 | 4.8e-08 | 5.233E-08 | 4.509E-09 | 4.800E-08 | 5.700E-08 |
| 5567 | 279 | Y279Q | A | 2.7e-08 | 2.4e-08 | 1.8e-08 | 2.300E-08 | 4.583E-09 | 1.800E-08 | 2.700E-08 |
| 5568 | 279 | Y279G | A | 7.9e-08 | 6.1e-08 | 6e-08 | 6.667E-08 | 1.069E-08 | 6.000E-08 | 7.900E-08 |
| 5569 | 279 | Y279H | A | 2.1e-08 | 7.9e-08 | 3e-08 | 4.333E-08 | 3.121E-08 | 2.100E-08 | 7.900E-08 |
| 5570 | 279 | Y279I | A | 3.9e-08 | 8.5e-08 | 1.7e-07 | 9.800E-08 | 6.646E-08 | 3.900E-08 | 1.700E-07 |
| 5571 | 279 | Y279L | A | 3.3e-08 | 1.7e-08 | 8.1e-08 | 4.367E-08 | 3.331E-08 | 1.700E-08 | 8.100E-08 |
| 5572 | 279 | Y279K | A | 6.4e-08 | 5.8e-08 | 4.4e-08 | 5.533E-08 | 1.026E-08 | 4.400E-08 | 6.400E-08 |
| 5573 | 279 | Y279M | A | 6.8e-08 | 2.4e-08 | 1.4e-07 | 7.733E-08 | 5.856E-08 | 2.400E-08 | 1.400E-07 |
| 5574 | 279 | Y279F | A | 4.1e-08 | 5.9e-08 | 3e-08 | 4.333E-08 | 1.464E-08 | 3.000E-08 | 5.900E-08 |
| 5575 | 279 | Y279P | A | 4.1e-08 | 5.4e-08 | 7.6e-08 | 5.700E-08 | 1.769E-08 | 4.100E-08 | 7.600E-08 |
| 5576 | 279 | Y279S | A | 1.1e-07 | 7.3e-08 | 4.3e-08 | 7.533E-08 | 3.356E-08 | 4.300E-08 | 1.100E-07 |
| 5577 | 279 | Y279T | A | 3.1e-08 | 1.1e-07 | 6.8e-08 | 6.967E-08 | 3.953E-08 | 3.100E-08 | 1.100E-07 |
| 5578 | 279 | Y279W | A | 3.6e-08 | 4.2e-08 | 6.7e-08 | 4.833E-08 | 1.644E-08 | 3.600E-08 | 6.700E-08 |
| 5579 | 279 | Y279Y | A | 1e-08 | 3.8e-08 | 7e-08 | 3.933E-08 | 3.002E-08 | 1.000E-08 | 7.000E-08 |
| 5580 | 279 | Y279V | A | 7.3e-08 | 4.1e-08 | 3.6e-08 | 5.000E-08 | 2.007E-08 | 3.600E-08 | 7.300E-08 |
| 5581 | 280 | N280A | A | 6.2e-08 | 3.1e-08 | 2.6e-08 | 3.967E-08 | 1.950E-08 | 2.600E-08 | 6.200E-08 |
| 5582 | 280 | N280R | A | 7.1e-08 | 2.5e-08 | 1.9e-08 | 3.833E-08 | 2.845E-08 | 1.900E-08 | 7.100E-08 |
| 5583 | 280 | N280N | A | 4e-08 | 1.4e-07 | 3.1e-08 | 7.033E-08 | 6.050E-08 | 3.100E-08 | 1.400E-07 |
| 5584 | 280 | N280D | A | 2.9e-08 | 3.8e-08 | 1.1e-07 | 5.900E-08 | 4.440E-08 | 2.900E-08 | 1.100E-07 |
| 5585 | 280 | N280C | A | 9.6e-08 | 7e-08 | 1.4e-08 | 6.000E-08 | 4.190E-08 | 1.400E-08 | 9.600E-08 |
| 5586 | 280 | N280E | A | 1.9e-08 | 2.4e-08 | 1.7e-07 | 7.100E-08 | 8.577E-08 | 1.900E-08 | 1.700E-07 |
| 5587 | 280 | N280Q | A | 6.1e-08 | 3.4e-08 | 5.4e-08 | 4.967E-08 | 1.401E-08 | 3.400E-08 | 6.100E-08 |
| 5588 | 280 | N280G | A | 5.6e-08 | 1e-07 | 4.2e-08 | 6.600E-08 | 3.027E-08 | 4.200E-08 | 1.000E-07 |
| 5589 | 280 | N280H | A | 4e-08 | 3.9e-08 | 5.1e-08 | 4.333E-08 | 6.658E-09 | 3.900E-08 | 5.100E-08 |
| 5590 | 280 | N280I | A | 3.5e-08 | 2e-08 | 8.5e-08 | 4.667E-08 | 3.403E-08 | 2.000E-08 | 8.500E-08 |
| 5591 | 280 | N280L | A | 1.1e-07 | 3.4e-08 | 2.2e-07 | 1.213E-07 | 9.352E-08 | 3.400E-08 | 2.200E-07 |
| 5592 | 280 | N280K | A | 7.8e-08 | 4.2e-08 | 2.3e-08 | 4.767E-08 | 2.793E-08 | 2.300E-08 | 7.800E-08 |
| 5593 | 280 | N280M | A | 4.3e-08 | 4.2e-08 | 4.1e-08 | 4.200E-08 | 1.000E-09 | 4.100E-08 | 4.300E-08 |
| 5594 | 280 | N280F | A | 5.1e-08 | 1e-07 | 2.9e-08 | 6.000E-08 | 3.635E-08 | 2.900E-08 | 1.000E-07 |
| 5595 | 280 | N280P | A | 4.8e-08 | 6.2e-08 | 1.3e-08 | 4.100E-08 | 2.524E-08 | 1.300E-08 | 6.200E-08 |
| 5596 | 280 | N280S | A | 2.2e-08 | 8.5e-08 | 1.2e-07 | 7.567E-08 | 4.966E-08 | 2.200E-08 | 1.200E-07 |
| 5597 | 280 | N280T | A | 2.8e-08 | 7.1e-08 | 3.2e-08 | 4.367E-08 | 2.376E-08 | 2.800E-08 | 7.100E-08 |
| 5598 | 280 | N280W | A | 2.4e-08 | 4.1e-08 | 4.8e-08 | 3.767E-08 | 1.234E-08 | 2.400E-08 | 4.800E-08 |
| 5599 | 280 | N280Y | A | 5.3e-08 | 7.9e-08 | 2.7e-08 | 5.300E-08 | 2.600E-08 | 2.700E-08 | 7.900E-08 |
| 5600 | 280 | N280V | A | 3.2e-08 | 2.6e-08 | 5.5e-08 | 3.767E-08 | 1.531E-08 | 2.600E-08 | 5.500E-08 |
| 5601 | 281 | Y281A | A | 1.4e-08 | 2.2e-08 | 3.3e-08 | 2.300E-08 | 9.539E-09 | 1.400E-08 | 3.300E-08 |
| 5602 | 281 | Y281R | A | 4.1e-08 | 4.7e-08 | 3.8e-08 | 4.200E-08 | 4.583E-09 | 3.800E-08 | 4.700E-08 |
| 5603 | 281 | Y281N | A | 4.6e-08 | 2.5e-08 | 2.3e-08 | 3.133E-08 | 1.274E-08 | 2.300E-08 | 4.600E-08 |
| 5604 | 281 | Y281D | A | 1.6e-08 | 5.6e-08 | 1.1e-08 | 2.767E-08 | 2.466E-08 | 1.100E-08 | 5.600E-08 |
| 5605 | 281 | Y281C | A | 3.1e-08 | 3.6e-08 | 1.6e-08 | 2.767E-08 | 1.041E-08 | 1.600E-08 | 3.600E-08 |
| 5606 | 281 | Y281E | A | 1.3e-08 | 4.6e-08 | 7.1e-08 | 4.333E-08 | 2.909E-08 | 1.300E-08 | 7.100E-08 |
| 5607 | 281 | Y281Q | A | 2e-08 | 7.1e-08 | 2.9e-08 | 4.000E-08 | 2.722E-08 | 2.000E-08 | 7.100E-08 |
| 5608 | 281 | Y281G | A | 4.5e-08 | 1.8e-08 | 9.4e-08 | 5.233E-08 | 3.853E-08 | 1.800E-08 | 9.400E-08 |
| 5609 | 281 | Y281H | A | 4.4e-08 | 5.3e-08 | 5.8e-08 | 5.167E-08 | 7.095E-09 | 4.400E-08 | 5.800E-08 |

| | | | | | | | | | | |
|---|---|---|---|---|---|---|---|---|---|---|
| 5610 | 281 | Y281I | A | 2.4e-08 | 4.3e-08 | 8.2e-08 | 4.967E-08 | 2.957E-08 | 2.400E-08 | 8.200E-08 |
| 5611 | 281 | Y281L | A | 8.1e-08 | 2.6e-08 | 5.6e-08 | 5.433E-08 | 2.754E-08 | 2.600E-08 | 8.100E-08 |
| 5612 | 281 | Y281K | A | 5.9e-08 | 3.4e-08 | 4.5e-08 | 4.600E-08 | 1.253E-08 | 3.400E-08 | 5.900E-08 |
| 5613 | 281 | Y281M | A | 2.2e-08 | 3.5e-08 | 2.2e-08 | 2.633E-08 | 7.506E-09 | 2.200E-08 | 3.500E-08 |
| 5614 | 281 | Y281F | A | 5.4e-08 | 1.7e-08 | 2.7e-08 | 3.267E-08 | 1.914E-08 | 1.700E-08 | 5.400E-08 |
| 5615 | 281 | Y281P | A | 5.9e-08 | 5.6e-08 | 5.6e-08 | 5.700E-08 | 1.732E-09 | 5.600E-08 | 5.900E-08 |
| 5616 | 281 | Y281S | A | 7.4e-08 | 2.9e-08 | 6.4e-08 | 5.567E-08 | 2.363E-08 | 2.900E-08 | 7.400E-08 |
| 5617 | 281 | Y281T | A | 2.9e-08 | 2.1e-08 | 4.7e-08 | 3.233E-08 | 1.332E-08 | 2.100E-08 | 4.700E-08 |
| 5618 | 281 | Y281W | A | 2.6e-08 | 1.3e-07 | 2.6e-08 | 6.067E-08 | 6.004E-08 | 2.600E-08 | 1.300E-07 |
| 5619 | 281 | Y281Y | A | 3.3e-08 | 1.6e-08 | 5.7e-08 | 3.533E-08 | 2.060E-08 | 1.600E-08 | 5.700E-08 |
| 5620 | 281 | Y281V | A | 4e-08 | 3.2e-08 | 3.7e-08 | 3.633E-08 | 4.041E-09 | 3.200E-08 | 4.000E-08 |
| 5621 | 282 | L282A | A | 7.2e-08 | 5.4e-08 | 1.9e-08 | 4.833E-08 | 2.695E-08 | 1.900E-08 | 7.200E-08 |
| 5622 | 282 | L282R | A | 4e-08 | 4.7e-08 | 9e-08 | 5.900E-08 | 2.707E-08 | 4.000E-08 | 9.000E-08 |
| 5623 | 282 | L282N | A | 5.9e-08 | 4.4e-08 | 2.7e-08 | 4.333E-08 | 1.601E-08 | 2.700E-08 | 5.900E-08 |
| 5624 | 282 | L282D | A | 5.6e-08 | 6.1e-08 | 1.2e-07 | 7.900E-08 | 3.559E-08 | 5.600E-08 | 1.200E-07 |
| 5625 | 282 | L282C | A | 2.1e-08 | 8.8e-08 | 5.2e-08 | 5.367E-08 | 3.353E-08 | 2.100E-08 | 8.800E-08 |
| 5626 | 282 | L282E | A | 3.9e-08 | 3.8e-08 | 5e-08 | 4.233E-08 | 6.658E-09 | 3.800E-08 | 5.000E-08 |
| 5627 | 282 | L282Q | A | 7.4e-09 | 5.4e-08 | 3.5e-08 | 3.213E-08 | 2.343E-08 | 7.400E-09 | 5.400E-08 |
| 5628 | 282 | L282G | A | 3.2e-08 | 2.6e-08 | 3.2e-08 | 3.000E-08 | 3.464E-09 | 2.600E-08 | 3.200E-08 |
| 5629 | 282 | L282H | A | 1.6e-07 | 2.5e-08 | 5.5e-08 | 8.000E-08 | 7.089E-08 | 2.500E-08 | 1.600E-07 |
| 5630 | 282 | L282I | A | 5.1e-08 | 8.6e-08 | 8.8e-08 | 7.500E-08 | 2.081E-08 | 5.100E-08 | 8.800E-08 |
| 5631 | 282 | L282L | A | 1.9e-08 | 6.9e-08 | 7.1e-08 | 5.300E-08 | 2.946E-08 | 1.900E-08 | 7.100E-08 |
| 5632 | 282 | L282K | A | 4.1e-08 | 1.4e-07 | 2.1e-08 | 6.733E-08 | 6.372E-08 | 2.100E-08 | 1.400E-07 |
| 5633 | 282 | L282M | A | 7e-08 | 2.8e-08 | 6.9e-09 | 3.497E-08 | 3.212E-08 | 6.900E-09 | 7.000E-08 |
| 5634 | 282 | L282F | A | 6.6e-08 | 5.3e-08 | 3.2e-08 | 5.033E-08 | 1.716E-08 | 3.200E-08 | 6.600E-08 |
| 5635 | 282 | L282P | A | 6.1e-08 | 3.2e-08 | 1.1e-07 | 6.767E-08 | 3.943E-08 | 3.200E-08 | 1.100E-07 |
| 5636 | 282 | L282S | A | 5.4e-08 | 5.1e-08 | 5.4e-08 | 5.300E-08 | 1.732E-09 | 5.100E-08 | 5.400E-08 |
| 5637 | 282 | L282T | A | 3.2e-08 | 3.8e-08 | 3.6e-08 | 3.533E-08 | 3.055E-09 | 3.200E-08 | 3.800E-08 |
| 5638 | 282 | L282W | A | 9.5e-08 | 3.6e-08 | 7.3e-08 | 6.800E-08 | 2.982E-08 | 3.600E-08 | 9.500E-08 |
| 5639 | 282 | L282Y | A | 2.6e-08 | 5.9e-08 | 3.5e-08 | 4.000E-08 | 1.706E-08 | 2.600E-08 | 5.900E-08 |
| 5640 | 282 | L282V | A | 6.4e-08 | 5.3e-08 | 7.8e-08 | 6.500E-08 | 1.253E-08 | 5.300E-08 | 7.800E-08 |
| 5641 | 283 | S283A | A | 6.4e-08 | 1.8e-08 | 3.6e-08 | 3.933E-08 | 2.318E-08 | 1.800E-08 | 6.400E-08 |
| 5642 | 283 | S283R | A | 6.4e-08 | 6.7e-08 | 2.1e-07 | 1.137E-07 | 8.344E-08 | 6.400E-08 | 2.100E-07 |
| 5643 | 283 | S283N | A | 4e-08 | 3.4e-08 | 2.2e-08 | 3.200E-08 | 9.165E-09 | 2.200E-08 | 4.000E-08 |
| 5644 | 283 | S283D | A | 5.3e-08 | 1.9e-07 | 3.7e-08 | 9.333E-08 | 8.410E-08 | 3.700E-08 | 1.900E-07 |
| 5645 | 283 | S283C | A | 4.5e-08 | 3.2e-08 | 1.4e-07 | 7.233E-08 | 5.896E-08 | 3.200E-08 | 1.400E-07 |
| 5646 | 283 | S283E | A | 5.7e-08 | 2.4e-08 | 8.1e-08 | 5.400E-08 | 2.862E-08 | 2.400E-08 | 8.100E-08 |
| 5647 | 283 | S283Q | A | 7.8e-08 | 2.7e-08 | 2.2e-08 | 4.233E-08 | 3.099E-08 | 2.200E-08 | 7.800E-08 |
| 5648 | 283 | S283G | A | 3.3e-08 | 1.9e-08 | 4.1e-08 | 3.100E-08 | 1.114E-08 | 1.900E-08 | 4.100E-08 |
| 5649 | 283 | S283H | A | 7e-08 | 1.6e-08 | 3.8e-08 | 4.133E-08 | 2.715E-08 | 1.600E-08 | 7.000E-08 |
| 5650 | 283 | S283I | A | 2.2e-08 | 1e-07 | 1.1e-07 | 7.733E-08 | 4.818E-08 | 2.200E-08 | 1.100E-07 |
| 5651 | 283 | S283L | A | 4.4e-08 | 4.3e-08 | 1.2e-08 | 3.300E-08 | 1.819E-08 | 1.200E-08 | 4.400E-08 |
| 5652 | 283 | S283K | A | 3.6e-08 | 6.2e-08 | 3.5e-08 | 4.433E-08 | 1.531E-08 | 3.500E-08 | 6.200E-08 |
| 5653 | 283 | S283M | A | 5.5e-08 | 1.3e-08 | 3.2e-08 | 3.333E-08 | 2.103E-08 | 1.300E-08 | 5.500E-08 |
| 5654 | 283 | S283F | A | 2.3e-08 | 3.4e-08 | 3.9e-08 | 3.200E-08 | 8.185E-09 | 2.300E-08 | 3.900E-08 |
| 5655 | 283 | S283P | A | 1.1e-07 | 2.5e-08 | 1.1e-07 | 8.167E-08 | 4.907E-08 | 2.500E-08 | 1.100E-07 |
| 5656 | 283 | S283S | A | 2e-08 | 1.5e-08 | 1.1e-07 | 4.833E-08 | 5.346E-08 | 1.500E-08 | 1.100E-07 |
| 5657 | 283 | S283T | A | 9.4e-08 | 2.6e-08 | 1.6e-08 | 4.533E-08 | 4.244E-08 | 1.600E-08 | 9.400E-08 |
| 5658 | 283 | S283W | A | 1.6e-07 | 3.9e-08 | 9.2e-08 | 9.700E-08 | 6.065E-08 | 3.900E-08 | 1.600E-07 |
| 5659 | 283 | S283Y | A | 3.7e-08 | 3.9e-08 | 2.2e-08 | 3.267E-08 | 9.292E-09 | 2.200E-08 | 3.900E-08 |
| 5660 | 283 | S283V | A | 1.7e-08 | 1.1e-07 | 4.8e-08 | 5.833E-08 | 4.735E-08 | 1.700E-08 | 1.100E-07 |
| 5661 | 284 | T284A | A | 5.4e-08 | 7.8e-08 | 2.3e-08 | 5.167E-08 | 2.757E-08 | 2.300E-08 | 7.800E-08 |
| 5662 | 284 | T284R | A | 2.9e-08 | 2.1e-08 | 2.9e-08 | 2.633E-08 | 4.619E-09 | 2.100E-08 | 2.900E-08 |
| 5663 | 284 | T284N | A | 5.8e-08 | 4.3e-08 | 1.2e-07 | 7.367E-08 | 4.082E-08 | 4.300E-08 | 1.200E-07 |
| 5664 | 284 | T284D | A | 2.4e-08 | 1.7e-08 | 2e-08 | 2.033E-08 | 3.512E-09 | 1.700E-08 | 2.400E-08 |
| 5665 | 284 | T284C | A | 2.5e-08 | 5.4e-08 | 3.6e-08 | 3.833E-08 | 1.464E-08 | 2.500E-08 | 5.400E-08 |
| 5666 | 284 | T284E | A | 7.1e-08 | 2.7e-08 | 5.1e-08 | 4.967E-08 | 2.203E-08 | 2.700E-08 | 7.100E-08 |
| 5667 | 284 | T284Q | A | 4.1e-08 | 5e-08 | 3.3e-08 | 4.133E-08 | 8.505E-09 | 3.300E-08 | 5.000E-08 |
| 5668 | 284 | T284G | A | 4.9e-08 | 5.5e-08 | 1.1e-07 | 7.133E-08 | 3.362E-08 | 4.900E-08 | 1.100E-07 |
| 5669 | 284 | T284H | A | 3.7e-08 | 2.7e-08 | 4.6e-08 | 3.667E-08 | 9.504E-09 | 2.700E-08 | 4.600E-08 |
| 5670 | 284 | T284I | A | 5.5e-08 | 5.2e-08 | 1.8e-08 | 4.167E-08 | 2.055E-08 | 1.800E-08 | 5.500E-08 |
| 5671 | 284 | T284L | A | 3.6e-08 | 3.2e-08 | 2.9e-08 | 3.233E-08 | 3.512E-09 | 2.900E-08 | 3.600E-08 |
| 5672 | 284 | T284K | A | 9.8e-08 | 1.4e-08 | 4.7e-08 | 5.300E-08 | 4.232E-08 | 1.400E-08 | 9.800E-08 |
| 5673 | 284 | T284M | A | 2.1e-08 | 6.4e-09 | 3.6e-08 | 2.113E-08 | 1.480E-08 | 6.400E-09 | 3.600E-08 |
| 5674 | 284 | T284F | A | 1.5e-08 | 2e-08 | 5.1e-08 | 2.867E-08 | 1.950E-08 | 1.500E-08 | 5.100E-08 |
| 5675 | 284 | T284P | A | 4.6e-08 | 6.6e-08 | 8.1e-08 | 6.433E-08 | 1.756E-08 | 4.600E-08 | 8.100E-08 |
| 5676 | 284 | T284S | A | 1.3e-08 | 3.1e-08 | 1.2e-07 | 5.467E-08 | 5.729E-08 | 1.300E-08 | 1.200E-07 |
| 5677 | 284 | T284T | A | 1.3e-07 | 6.5e-08 | 6.2e-08 | 8.567E-08 | 3.842E-08 | 6.200E-08 | 1.300E-07 |
| 5678 | 284 | T284W | A | 2.7e-08 | 2e-08 | 2.8e-08 | 2.500E-08 | 4.359E-09 | 2.000E-08 | 2.800E-08 |
| 5679 | 284 | T284Y | A | 2e-08 | 2.9e-08 | 3.9e-08 | 2.933E-08 | 9.504E-09 | 2.000E-08 | 3.900E-08 |
| 5680 | 284 | T284V | A | 2.6e-08 | 3.5e-08 | 4.1e-08 | 3.400E-08 | 7.550E-09 | 2.600E-08 | 4.100E-08 |
| 5681 | 285 | D285A | A | 3.4e-08 | 9.4e-08 | 8.3e-08 | 7.033E-08 | 3.194E-08 | 3.400E-08 | 9.400E-08 |
| 5682 | 285 | D285R | A | 2e-08 | 2.1e-08 | 6.6e-08 | 3.567E-08 | 2.627E-08 | 2.000E-08 | 6.600E-08 |
| 5683 | 285 | D285N | A | 6.2e-08 | 7.7e-08 | 3.1e-08 | 5.667E-08 | 2.346E-08 | 3.100E-08 | 7.700E-08 |
| 5684 | 285 | D285D | A | 8e-08 | 1e-07 | 3.9e-08 | 7.300E-08 | 3.110E-08 | 3.900E-08 | 1.000E-07 |
| 5685 | 285 | D285C | A | 5.2e-08 | 7.6e-08 | 2.5e-08 | 5.100E-08 | 2.551E-08 | 2.500E-08 | 7.600E-08 |
| 5686 | 285 | D285E | A | 1.2e-07 | 1.6e-08 | 8.5e-08 | 7.367E-08 | 5.292E-08 | 1.600E-08 | 1.200E-07 |
| 5687 | 285 | D285Q | A | 1.6e-08 | 3e-08 | 6.2e-08 | 3.600E-08 | 2.358E-08 | 1.600E-08 | 6.200E-08 |
| 5688 | 285 | D285G | A | 5.3e-08 | 5.7e-08 | 4.8e-08 | 5.267E-08 | 4.509E-09 | 4.800E-08 | 5.700E-08 |
| 5689 | 285 | D285H | A | 2.7e-08 | 7.2e-08 | 3.7e-08 | 4.533E-08 | 2.363E-08 | 2.700E-08 | 7.200E-08 |
| 5690 | 285 | D285I | A | 1e-07 | 2.9e-07 | 5.7e-08 | 1.490E-07 | 1.240E-07 | 5.700E-08 | 2.900E-07 |
| 5691 | 285 | D285L | A | 5.9e-08 | 2.3e-08 | 5.4e-09 | 2.913E-08 | 2.732E-08 | 5.400E-09 | 5.900E-08 |
| 5692 | 285 | D285K | A | 1.1e-08 | 7.4e-08 | 1.5e-08 | 3.333E-08 | 3.528E-08 | 1.100E-08 | 7.400E-08 |
| 5693 | 285 | D285M | A | 1.1e-08 | 7.5e-08 | 8.7e-08 | 5.767E-08 | 4.086E-08 | 1.100E-08 | 8.700E-08 |
| 5694 | 285 | D285F | A | 1e-07 | 4.4e-08 | 3.6e-08 | 6.000E-08 | 3.487E-08 | 3.600E-08 | 1.000E-07 |

| | | | | | | | | | | |
|---|---|---|---|---|---|---|---|---|---|---|
| 5695 | 285 | D285P | A | 9.2e-09 | 1.2e-07 | 8.7e-08 | 7.207E-08 | 5.689E-08 | 9.200E-09 | 1.200E-07 |
| 5696 | 285 | D285S | A | 1.8e-08 | 7.6e-08 | 5.4e-08 | 4.933E-08 | 2.928E-08 | 1.800E-08 | 7.600E-08 |
| 5697 | 285 | D285T | A | 6.8e-08 | 3.7e-08 | 1.7e-08 | 4.067E-08 | 2.570E-08 | 1.700E-08 | 6.800E-08 |
| 5698 | 285 | D285W | A | 7.5e-08 | 3.4e-08 | 5.6e-08 | 5.500E-08 | 2.052E-08 | 3.400E-08 | 7.500E-08 |
| 5699 | 285 | D285Y | A | 7.2e-08 | 4.7e-08 | 6.3e-08 | 6.067E-08 | 1.266E-08 | 4.700E-08 | 7.200E-08 |
| 5700 | 285 | D285V | A | 2.1e-08 | 2.1e-08 | 3.2e-08 | 2.467E-08 | 6.351E-09 | 2.100E-08 | 3.200E-08 |
| 5701 | 286 | V286A | A | 9.9e-08 | 7e-08 | 2.1e-08 | 6.333E-08 | 3.943E-08 | 2.100E-08 | 9.900E-08 |
| 5702 | 286 | V286R | A | 9.9e-08 | 1.7e-07 | 5.5e-08 | 1.080E-07 | 5.803E-08 | 5.500E-08 | 1.700E-07 |
| 5703 | 286 | V286N | A | 5.8e-07 | 5.7e-07 | 9.4e-07 | 6.967E-07 | 2.108E-07 | 5.700E-07 | 9.400E-07 |
| 5704 | 286 | V286D | A | 2e-07 | 5.2e-08 | 5.5e-08 | 1.023E-07 | 8.460E-08 | 5.200E-08 | 2.000E-07 |
| 5705 | 286 | V286C | A | 1.6e-08 | 8e-08 | 2.2e-08 | 3.933E-08 | 3.535E-08 | 1.600E-08 | 8.000E-08 |
| 5706 | 286 | V286E | A | 1.4e-07 | 8.5e-08 | 1.1e-07 | 1.117E-07 | 2.754E-08 | 8.500E-08 | 1.400E-07 |
| 5707 | 286 | V286Q | A | 5e-07 | 4.3e-07 | 3.2e-07 | 4.167E-07 | 9.074E-08 | 3.200E-07 | 5.000E-07 |
| 5708 | 286 | V286G | A | 9.5e-08 | 1.6e-07 | 1.1e-07 | 1.217E-07 | 3.403E-08 | 9.500E-08 | 1.600E-07 |
| 5709 | 286 | V286H | A | 5.8e-08 | 8.8e-08 | 1.9e-07 | 1.120E-07 | 6.920E-08 | 5.800E-08 | 1.900E-07 |
| 5710 | 286 | V286I | A | 2.8e-08 | 2.5e-08 | 1.4e-07 | 6.433E-08 | 6.555E-08 | 2.500E-08 | 1.400E-07 |
| 5711 | 286 | V286L | A | 1.9e-08 | 2.1e-08 | 8.5e-08 | 4.167E-08 | 3.754E-08 | 1.900E-08 | 8.500E-08 |
| 5712 | 286 | V286K | A | 1.4e-07 | 1.2e-07 | 1.4e-07 | 1.333E-07 | 1.155E-08 | 1.200E-07 | 1.400E-07 |
| 5713 | 286 | V286M | A | 3.2e-08 | 3e-08 | 4.5e-08 | 3.567E-08 | 8.145E-09 | 3.000E-08 | 4.500E-08 |
| 5714 | 286 | V286F | A | 4e-08 | 1.7e-08 | 1e-07 | 5.233E-08 | 4.285E-08 | 1.700E-08 | 1.000E-07 |
| 5715 | 286 | V286P | A | 2.8e-08 | 1.4e-08 | 1.6e-08 | 1.933E-08 | 7.572E-09 | 1.400E-08 | 2.800E-08 |
| 5716 | 286 | V286S | A | 2.7e-07 | 5.6e-07 | 3.6e-07 | 3.967E-07 | 1.484E-07 | 2.700E-07 | 5.600E-07 |
| 5717 | 286 | V286T | A | 4.8e-07 | 2.8e-07 | 1.4e-06 | 7.200E-07 | 5.973E-07 | 2.800E-07 | 1.400E-06 |
| 5718 | 286 | V286W | A | 1.8e-08 | 6.7e-09 | 4.1e-08 | 2.190E-08 | 1.748E-08 | 6.700E-09 | 4.100E-08 |
| 5719 | 286 | V286Y | A | 3.6e-08 | 3.4e-08 | 1.4e-08 | 2.800E-08 | 1.217E-08 | 1.400E-08 | 3.600E-08 |
| 5720 | 286 | V286V | A | 6.9e-08 | 4e-08 | 5e-08 | 5.300E-08 | 1.473E-08 | 4.000E-08 | 6.900E-08 |
| 5721 | 287 | G287A | A | 9.5e-08 | 2.6e-08 | 6e-08 | 6.033E-08 | 3.450E-08 | 2.600E-08 | 9.500E-08 |
| 5722 | 287 | G287R | A | 4.3e-08 | 5.2e-08 | 9.3e-08 | 6.267E-08 | 2.665E-08 | 4.300E-08 | 9.300E-08 |
| 5723 | 287 | G287N | A | 4.9e-08 | 8.2e-08 | 9.6e-08 | 7.567E-08 | 2.413E-08 | 4.900E-08 | 9.600E-08 |
| 5724 | 287 | G287D | A | 9.1e-08 | 7e-08 | 2.6e-07 | 1.403E-07 | 1.042E-07 | 7.000E-08 | 2.600E-07 |
| 5725 | 287 | G287C | A | 2.2e-08 | 1.9e-08 | 3.3e-08 | 2.467E-08 | 7.371E-09 | 1.900E-08 | 3.300E-08 |
| 5726 | 287 | G287E | A | 1.6e-07 | 1.3e-07 | 1.1e-07 | 1.333E-07 | 2.517E-08 | 1.100E-07 | 1.600E-07 |
| 5727 | 287 | G287Q | A | 3.4e-07 | 8.8e-08 | 2.1e-07 | 2.127E-07 | 1.260E-07 | 8.800E-08 | 3.400E-07 |
| 5728 | 287 | G287G | A | 1.6e-07 | 1.8e-08 | 1.7e-08 | 6.500E-08 | 8.227E-08 | 1.700E-08 | 1.600E-07 |
| 5729 | 287 | G287H | A | 2.9e-08 | 9.8e-08 | 9e-08 | 7.233E-08 | 3.774E-08 | 2.900E-08 | 9.800E-08 |
| 5730 | 287 | G287I | A | 4.5e-08 | 1.3e-08 | 3.3e-08 | 3.033E-08 | 1.617E-08 | 1.300E-08 | 4.500E-08 |
| 5731 | 287 | G287L | A | 4.8e-08 | 1.8e-08 | 4.2e-08 | 3.600E-08 | 1.587E-08 | 1.800E-08 | 4.800E-08 |
| 5732 | 287 | G287K | A | 3.2e-08 | 8.8e-08 | 3e-08 | 5.000E-08 | 3.292E-08 | 3.000E-08 | 8.800E-08 |
| 5733 | 287 | G287M | A | 2.9e-08 | 6.4e-08 | 4.7e-08 | 4.667E-08 | 1.750E-08 | 2.900E-08 | 6.400E-08 |
| 5734 | 287 | G287F | A | 3.2e-08 | 3.3e-08 | 6.2e-08 | 4.233E-08 | 1.704E-08 | 3.200E-08 | 6.200E-08 |
| 5735 | 287 | G287P | A | 9.8e-09 | 2e-08 | 8.9e-09 | 1.290E-08 | 6.165E-09 | 8.900E-09 | 2.000E-08 |
| 5736 | 287 | G287S | A | 4.1e-08 | 4.6e-08 | 1.6e-07 | 8.233E-08 | 6.731E-08 | 4.100E-08 | 1.600E-07 |
| 5737 | 287 | G287T | A | 9.9e-08 | 7.4e-08 | 6.9e-07 | 2.877E-07 | 3.487E-07 | 7.400E-08 | 6.900E-07 |
| 5738 | 287 | G287W | A | 2.5e-08 | 3.5e-08 | 4.3e-08 | 3.433E-08 | 9.018E-09 | 2.500E-08 | 4.300E-08 |
| 5739 | 287 | G287Y | A | 1.4e-08 | 6.4e-08 | 6.7e-08 | 4.833E-08 | 2.977E-08 | 1.400E-08 | 6.700E-08 |
| 5740 | 287 | G287V | A | 7.6e-08 | 8.9e-08 | 4.2e-08 | 6.900E-08 | 2.427E-08 | 4.200E-08 | 8.900E-08 |
| 5741 | 288 | S288A | A | 9.4e-09 | 6e-09 | 7.3e-09 | 7.567E-09 | 1.716E-09 | 6.000E-09 | 9.400E-09 |
| 5742 | 288 | S288R | A | 4.6e-08 | 2e-08 | 3.5e-08 | 3.367E-08 | 1.305E-08 | 2.000E-08 | 4.600E-08 |
| 5743 | 288 | S288N | A | 2.3e-07 | 2e-08 | 6.8e-08 | 1.060E-07 | 1.100E-07 | 2.000E-08 | 2.300E-07 |
| 5744 | 288 | S288D | A | 4.6e-08 | 3.4e-08 | 1.4e-08 | 3.133E-08 | 1.617E-08 | 1.400E-08 | 4.600E-08 |
| 5745 | 288 | S288C | A | 1.1e-08 | 6.8e-09 | 8.6e-09 | 8.800E-09 | 2.107E-09 | 6.800E-09 | 1.100E-08 |
| 5746 | 288 | S288E | A | 1.1e-08 | 2.2e-08 | 3e-08 | 2.100E-08 | 9.539E-09 | 1.100E-08 | 3.000E-08 |
| 5747 | 288 | S288Q | A | 4.1e-08 | 3.8e-08 | 7.2e-08 | 5.033E-08 | 1.882E-08 | 3.800E-08 | 7.200E-08 |
| 5748 | 288 | S288G | A | 2e-08 | 1e-08 | 1.9e-08 | 1.633E-08 | 5.508E-09 | 1.000E-08 | 2.000E-08 |
| 5749 | 288 | S288H | A | 3.3e-08 | 1.5e-08 | 1.1e-08 | 1.967E-08 | 1.172E-08 | 1.100E-08 | 3.300E-08 |
| 5750 | 288 | S288I | A | 1.4e-08 | 2.8e-08 | 1.4e-08 | 1.867E-08 | 8.083E-09 | 1.400E-08 | 2.800E-08 |
| 5751 | 288 | S288L | A | 1.7e-08 | 1.5e-08 | 2.9e-08 | 2.033E-08 | 7.572E-09 | 1.500E-08 | 2.900E-08 |
| 5752 | 288 | S288K | A | 1.7e-08 | 3.6e-08 | 2.7e-09 | 1.857E-08 | 1.671E-08 | 2.700E-09 | 3.600E-08 |
| 5753 | 288 | S288M | A | 6.9e-09 | 2.7e-08 | 7e-09 | 1.363E-08 | 1.158E-08 | 6.900E-09 | 2.700E-08 |
| 5754 | 288 | S288F | A | 1.9e-08 | 3.7e-09 | 6.4e-09 | 9.700E-09 | 8.166E-09 | 3.700E-09 | 1.900E-08 |
| 5755 | 288 | S288P | A | 1.3e-08 | 1.1e-08 | 3.4e-08 | 1.933E-08 | 1.274E-08 | 1.100E-08 | 3.400E-08 |
| 5756 | 288 | S288S | A | 3.4e-08 | 1.5e-08 | 5.9e-08 | 3.600E-08 | 2.207E-08 | 1.500E-08 | 5.900E-08 |
| 5757 | 288 | S288T | A | 3.7e-08 | 2.5e-08 | 5.8e-08 | 4.000E-08 | 1.670E-08 | 2.500E-08 | 5.800E-08 |
| 5758 | 288 | S288W | A | 9.7e-09 | 6.9e-09 | 1.1e-08 | 9.200E-09 | 2.095E-09 | 6.900E-09 | 1.100E-08 |
| 5759 | 288 | S288Y | A | 6.3e-09 | 4.5e-09 | 2.3e-08 | 1.127E-08 | 1.020E-08 | 4.500E-09 | 2.300E-08 |
| 5760 | 288 | S288V | A | 4e-09 | 3.3e-09 | 9.3e-09 | 5.533E-09 | 3.281E-09 | 3.300E-09 | 9.300E-09 |
| 5761 | 289 | C289A | A | 6.3e-08 | 1.6e-07 | 3.5e-08 | 8.600E-08 | 6.560E-08 | 3.500E-08 | 1.600E-07 |
| 5762 | 289 | C289R | A | 3.8e-08 | 4.9e-08 | 8.2e-08 | 5.633E-08 | 2.290E-08 | 3.800E-08 | 8.200E-08 |
| 5763 | 289 | C289N | A | 1.7e-07 | 7.9e-08 | 5.8e-08 | 1.023E-07 | 5.953E-08 | 5.800E-08 | 1.700E-07 |
| 5764 | 289 | C289D | A | 2.2e-08 | 1e-07 | 4.9e-08 | 5.700E-08 | 3.961E-08 | 2.200E-08 | 1.000E-07 |
| 5765 | 289 | C289C | A | 7.7e-08 | 4.6e-08 | 1e-07 | 7.433E-08 | 2.710E-08 | 4.600E-08 | 1.000E-07 |
| 5766 | 289 | C289E | A | 5.9e-08 | 3.6e-08 | 3.2e-08 | 4.233E-08 | 1.457E-08 | 3.200E-08 | 5.900E-08 |
| 5767 | 289 | C289Q | A | 8.7e-09 | 5.7e-08 | 1.3e-07 | 6.523E-08 | 6.107E-08 | 8.700E-09 | 1.300E-07 |
| 5768 | 289 | C289G | A | 2.2e-08 | 4.8e-08 | 5.8e-08 | 4.267E-08 | 1.858E-08 | 2.200E-08 | 5.800E-08 |
| 5769 | 289 | C289H | A | 4.1e-08 | 9.9e-08 | 2.9e-08 | 5.633E-08 | 3.743E-08 | 2.900E-08 | 9.900E-08 |
| 5770 | 289 | C289I | A | 6e-08 | 2e-08 | 6.1e-08 | 4.700E-08 | 2.339E-08 | 2.000E-08 | 6.100E-08 |
| 5771 | 289 | C289L | A | 4.6e-08 | 1.7e-08 | 3.2e-08 | 3.167E-08 | 1.450E-08 | 1.700E-08 | 4.600E-08 |
| 5772 | 289 | C289K | A | 3.2e-08 | 4.7e-08 | 5.5e-08 | 4.467E-08 | 1.168E-08 | 3.200E-08 | 5.500E-08 |
| 5773 | 289 | C289M | A | 5.4e-08 | 2.7e-08 | 3.6e-08 | 3.900E-08 | 1.375E-08 | 2.700E-08 | 5.400E-08 |
| 5774 | 289 | C289F | A | 3.1e-08 | 4e-08 | 3.7e-08 | 3.600E-08 | 4.583E-09 | 3.100E-08 | 4.000E-08 |
| 5775 | 289 | C289P | A | 1.1e-07 | 5.9e-08 | 3.9e-08 | 6.933E-08 | 3.661E-08 | 3.900E-08 | 1.100E-07 |
| 5776 | 289 | C289S | A | 3e-08 | 2.8e-08 | 2.8e-08 | 2.867E-08 | 1.155E-09 | 2.800E-08 | 3.000E-08 |
| 5777 | 289 | C289T | A | 2.4e-08 | 3.6e-08 | 4.6e-08 | 3.533E-08 | 1.102E-08 | 2.400E-08 | 4.600E-08 |
| 5778 | 289 | C289W | A | 6.7e-08 | 2.2e-08 | 1.2e-08 | 3.367E-08 | 2.930E-08 | 1.200E-08 | 6.700E-08 |
| 5779 | 289 | C289Y | A | 7.1e-08 | 1.4e-07 | 3.2e-08 | 8.100E-08 | 5.469E-08 | 3.200E-08 | 1.400E-07 |

| | | | | | | | | | | |
|---|---|---|---|---|---|---|---|---|---|---|
| 5780 | 289 | C289V | A | 7.9e-08 | 1.4e-07 | 1.2e-08 | 7.700E-08 | 6.402E-08 | 1.200E-08 | 1.400E-07 |
| 5781 | 290 | T290A | A | 4.9e-09 | 1.8e-08 | 6.9e-09 | 9.933E-09 | 7.057E-09 | 4.900E-09 | 1.800E-08 |
| 5782 | 290 | T290R | A | 2.1e-08 | 1.1e-08 | 7.5e-09 | 1.317E-08 | 7.006E-09 | 7.500E-09 | 2.100E-08 |
| 5783 | 290 | T290N | A | 4.5e-08 | 1.2e-08 | 7.1e-08 | 4.267E-08 | 2.957E-08 | 1.200E-08 | 7.100E-08 |
| 5784 | 290 | T290D | A | 2.2e-08 | 1.8e-08 | 1.9e-08 | 1.967E-08 | 2.082E-09 | 1.800E-08 | 2.200E-08 |
| 5785 | 290 | T290C | A | 1.6e-08 | 7.5e-09 | 1.7e-09 | 8.400E-09 | 7.192E-09 | 1.700E-09 | 1.600E-08 |
| 5786 | 290 | T290E | A | 2.4e-08 | 2.5e-08 | 4e-08 | 2.967E-08 | 8.963E-09 | 2.400E-08 | 4.000E-08 |
| 5787 | 290 | T290Q | A | 2.9e-08 | 5.1e-08 | 8.5e-08 | 5.500E-08 | 2.821E-08 | 2.900E-08 | 8.500E-08 |
| 5788 | 290 | T290G | A | 7.3e-09 | 2.2e-08 | 7.3e-09 | 1.220E-08 | 8.487E-09 | 7.300E-09 | 2.200E-08 |
| 5789 | 290 | T290H | A | 5.5e-09 | 1e-08 | 5.5e-08 | 2.350E-08 | 2.737E-08 | 5.500E-09 | 5.500E-08 |
| 5790 | 290 | T290I | A | 4.7e-09 | 5.6e-09 | 2e-09 | 4.100E-09 | 1.873E-09 | 2.000E-09 | 5.600E-09 |
| 5791 | 290 | T290L | A | 1.5e-08 | 1.2e-08 | 1.6e-08 | 1.433E-08 | 2.082E-09 | 1.200E-08 | 1.600E-08 |
| 5792 | 290 | T290K | A | 1.7e-08 | 1.5e-08 | 4.7e-08 | 2.633E-08 | 1.793E-08 | 1.500E-08 | 4.700E-08 |
| 5793 | 290 | T290M | A | 9.6e-09 | 1.1e-08 | 1.1e-08 | 1.053E-08 | 8.083E-10 | 9.600E-09 | 1.100E-08 |
| 5794 | 290 | T290F | A | 5.4e-09 | 1.8e-08 | 1.2e-08 | 1.180E-08 | 6.302E-09 | 5.400E-09 | 1.800E-08 |
| 5795 | 290 | T290P | A | 8.8e-09 | 1.4e-08 | 2.1e-08 | 1.460E-08 | 6.122E-09 | 8.800E-09 | 2.100E-08 |
| 5796 | 290 | T290S | A | 3e-08 | 3.2e-08 | 2.5e-08 | 2.900E-08 | 3.606E-09 | 2.500E-08 | 3.200E-08 |
| 5797 | 290 | T290T | A | 3.6e-08 | 3.8e-08 | 6.4e-08 | 4.600E-08 | 1.562E-08 | 3.600E-08 | 6.400E-08 |
| 5798 | 290 | T290W | A | 2.1e-08 | 9e-09 | 9.3e-09 | 1.310E-08 | 6.843E-09 | 9.000E-09 | 2.100E-08 |
| 5799 | 290 | T290Y | A | 2e-08 | 7.4e-09 | 8.6e-09 | 1.200E-08 | 6.954E-09 | 7.400E-09 | 2.000E-08 |
| 5800 | 290 | T290V | A | 6e-08 | 3.1e-08 | 8.1e-09 | 3.303E-08 | 2.601E-08 | 8.100E-09 | 6.000E-08 |
| 5801 | 291 | L291A | A | 2.7e-08 | 4.6e-08 | 1.3e-08 | 2.867E-08 | 1.656E-08 | 1.300E-08 | 4.600E-08 |
| 5802 | 291 | L291R | A | 8.6e-08 | 2.2e-08 | 3e-08 | 4.600E-08 | 3.487E-08 | 2.200E-08 | 8.600E-08 |
| 5803 | 291 | L291N | A | 1e-07 | 2.1e-08 | 1.2e-07 | 8.033E-08 | 5.235E-08 | 2.100E-08 | 1.200E-07 |
| 5804 | 291 | L291D | A | 9e-09 | 1e-07 | 3.6e-08 | 4.833E-08 | 4.674E-08 | 9.000E-09 | 1.000E-07 |
| 5805 | 291 | L291C | A | 2.2e-08 | 4.4e-08 | 9.9e-08 | 5.500E-08 | 3.966E-08 | 2.200E-08 | 9.900E-08 |
| 5806 | 291 | L291E | A | 8.3e-08 | 2.3e-08 | 4.1e-08 | 4.900E-08 | 3.079E-08 | 2.300E-08 | 8.300E-08 |
| 5807 | 291 | L291Q | A | 5.5e-08 | 3.2e-08 | 2.2e-08 | 3.633E-08 | 1.692E-08 | 2.200E-08 | 5.500E-08 |
| 5808 | 291 | L291G | A | 6.9e-08 | 4.7e-08 | 8.3e-08 | 6.633E-08 | 1.815E-08 | 4.700E-08 | 8.300E-08 |
| 5809 | 291 | L291H | A | 8.4e-08 | 6.1e-08 | 4.2e-08 | 6.233E-08 | 2.103E-08 | 4.200E-08 | 8.400E-08 |
| 5810 | 291 | L291I | A | 6.7e-08 | 7.4e-08 | 1.7e-08 | 5.267E-08 | 3.109E-08 | 1.700E-08 | 7.400E-08 |
| 5811 | 291 | L291L | A | 5.1e-08 | 3.9e-08 | 3.9e-08 | 4.300E-08 | 6.928E-09 | 3.900E-08 | 5.100E-08 |
| 5812 | 291 | L291K | A | 2.5e-08 | 4.1e-08 | 6.2e-08 | 4.267E-08 | 1.856E-08 | 2.500E-08 | 6.200E-08 |
| 5813 | 291 | L291M | A | 2.9e-07 | 1.9e-07 | 6e-08 | 1.800E-07 | 1.153E-07 | 6.000E-08 | 2.900E-07 |
| 5814 | 291 | L291F | A | 4.6e-08 | 2.5e-08 | 5.9e-08 | 4.333E-08 | 1.716E-08 | 2.500E-08 | 5.900E-08 |
| 5815 | 291 | L291P | A | 3e-08 | 7.4e-08 | 6.8e-08 | 5.733E-08 | 2.386E-08 | 3.000E-08 | 7.400E-08 |
| 5816 | 291 | L291S | A | 3.4e-08 | 6.5e-08 | 2.1e-08 | 4.000E-08 | 2.261E-08 | 2.100E-08 | 6.500E-08 |
| 5817 | 291 | L291T | A | 3.6e-08 | 2.5e-08 | 3.2e-08 | 3.100E-08 | 5.568E-09 | 2.500E-08 | 3.600E-08 |
| 5818 | 291 | L291W | A | 1.2e-07 | 9.8e-09 | 9.2e-08 | 7.393E-08 | 5.728E-08 | 9.800E-09 | 1.200E-07 |
| 5819 | 291 | L291Y | A | 2.8e-08 | 2.8e-08 | 6.2e-08 | 3.933E-08 | 1.963E-08 | 2.800E-08 | 6.200E-08 |
| 5820 | 291 | L291V | A | 4.7e-08 | 3.5e-08 | 3.2e-08 | 3.800E-08 | 7.937E-09 | 3.200E-08 | 4.700E-08 |
| 5821 | 292 | V292A | A | 3.1e-08 | 2.8e-08 | 3.5e-08 | 3.133E-08 | 3.512E-09 | 2.800E-08 | 3.500E-08 |
| 5822 | 292 | V292R | A | 2.9e-08 | 6.6e-08 | 5.8e-08 | 5.100E-08 | 1.947E-08 | 2.900E-08 | 6.600E-08 |
| 5823 | 292 | V292N | A | 7.1e-08 | 5.2e-08 | 5.7e-08 | 6.000E-08 | 9.849E-09 | 5.200E-08 | 7.100E-08 |
| 5824 | 292 | V292D | A | 6e-08 | 9.1e-08 | 7.3e-08 | 7.467E-08 | 1.557E-08 | 6.000E-08 | 9.100E-08 |
| 5825 | 292 | V292C | A | 5.5e-08 | 9e-08 | 2.3e-08 | 5.600E-08 | 3.351E-08 | 2.300E-08 | 9.000E-08 |
| 5826 | 292 | V292E | A | 3.2e-08 | 2.1e-08 | 2.4e-08 | 2.567E-08 | 5.686E-09 | 2.100E-08 | 3.200E-08 |
| 5827 | 292 | V292Q | A | 3.4e-08 | 8.2e-08 | 2.3e-08 | 4.633E-08 | 3.137E-08 | 2.300E-08 | 8.200E-08 |
| 5828 | 292 | V292G | A | 6.9e-08 | 3.4e-08 | 7.3e-08 | 5.867E-08 | 2.146E-08 | 3.400E-08 | 7.300E-08 |
| 5829 | 292 | V292H | A | 5.7e-08 | 5.9e-08 | 3.9e-08 | 5.167E-08 | 1.102E-08 | 3.900E-08 | 5.900E-08 |
| 5830 | 292 | V292I | A | 6.6e-08 | 5.3e-08 | 2.9e-08 | 4.933E-08 | 1.877E-08 | 2.900E-08 | 6.600E-08 |
| 5831 | 292 | V292L | A | 4.4e-08 | 2.9e-08 | 5e-08 | 4.100E-08 | 1.082E-08 | 2.900E-08 | 5.000E-08 |
| 5832 | 292 | V292K | A | 8.1e-08 | 4.3e-08 | 8.3e-09 | 4.410E-08 | 3.636E-08 | 8.300E-09 | 8.100E-08 |
| 5833 | 292 | V292M | A | 2.7e-08 | 4.1e-08 | 3.8e-08 | 3.533E-08 | 7.371E-09 | 2.700E-08 | 4.100E-08 |
| 5834 | 292 | V292F | A | 7.3e-08 | 2.6e-08 | 7.1e-08 | 5.667E-08 | 2.658E-08 | 2.600E-08 | 7.300E-08 |
| 5835 | 292 | V292P | A | 4.6e-08 | 1.8e-07 | 2.5e-08 | 8.367E-08 | 8.409E-08 | 2.500E-08 | 1.800E-07 |
| 5836 | 292 | V292S | A | 3.1e-08 | 7.6e-08 | 5.9e-08 | 5.533E-08 | 2.272E-08 | 3.100E-08 | 7.600E-08 |
| 5837 | 292 | V292T | A | 1e-07 | 5e-08 | 2.8e-07 | 1.433E-07 | 1.210E-07 | 5.000E-08 | 2.800E-07 |
| 5838 | 292 | V292W | A | 6.3e-08 | 4.9e-08 | 7.3e-08 | 6.167E-08 | 1.206E-08 | 4.900E-08 | 7.300E-08 |
| 5839 | 292 | V292Y | A | 5.7e-08 | 5.4e-08 | 4.8e-08 | 5.300E-08 | 4.583E-09 | 4.800E-08 | 5.700E-08 |
| 5840 | 292 | V292V | A | 4e-08 | 2.1e-08 | 9.3e-08 | 5.133E-08 | 3.731E-08 | 2.100E-08 | 9.300E-08 |
| 5841 | 293 | C293A | A | 2.4e-08 | 1.7e-08 | 3.8e-08 | 2.633E-08 | 1.069E-08 | 1.700E-08 | 3.800E-08 |
| 5842 | 293 | C293R | A | 2.2e-08 | 4e-08 | 2.6e-08 | 2.933E-08 | 9.452E-09 | 2.200E-08 | 4.000E-08 |
| 5843 | 293 | C293N | A | 3.8e-08 | 5.4e-08 | 2.8e-08 | 4.000E-08 | 1.311E-08 | 2.800E-08 | 5.400E-08 |
| 5844 | 293 | C293D | A | 3.5e-08 | 1.1e-07 | 8.6e-08 | 7.700E-08 | 3.830E-08 | 3.500E-08 | 1.100E-07 |
| 5845 | 293 | C293C | A | 5.2e-08 | 1e-07 | 6.5e-08 | 7.233E-08 | 2.483E-08 | 5.200E-08 | 1.000E-07 |
| 5846 | 293 | C293E | A | 4.7e-08 | 2.1e-08 | 5.3e-08 | 4.033E-08 | 1.701E-08 | 2.100E-08 | 5.300E-08 |
| 5847 | 293 | C293Q | A | 3.2e-08 | 3.1e-08 | 3.5e-08 | 3.267E-08 | 2.082E-09 | 3.100E-08 | 3.500E-08 |
| 5848 | 293 | C293G | A | 3.1e-08 | 2.1e-08 | 6.5e-08 | 3.900E-08 | 2.307E-08 | 2.100E-08 | 6.500E-08 |
| 5849 | 293 | C293H | A | 4.1e-08 | 2.6e-08 | 3e-08 | 3.233E-08 | 7.767E-09 | 2.600E-08 | 4.100E-08 |
| 5850 | 293 | C293I | A | 4.7e-08 | 3.6e-08 | 1e-08 | 3.100E-08 | 1.900E-08 | 1.000E-08 | 4.700E-08 |
| 5851 | 293 | C293L | A | 3.9e-08 | 7e-08 | 1.5e-07 | 8.633E-08 | 5.727E-08 | 3.900E-08 | 1.500E-07 |
| 5852 | 293 | C293K | A | 3e-08 | 2.8e-08 | 3e-08 | 2.933E-08 | 1.155E-09 | 2.800E-08 | 3.000E-08 |
| 5853 | 293 | C293M | A | 1.9e-08 | 4.5e-08 | 8.4e-08 | 4.933E-08 | 3.272E-08 | 1.900E-08 | 8.400E-08 |
| 5854 | 293 | C293F | A | 1e-07 | 2e-08 | 4.8e-08 | 5.600E-08 | 4.060E-08 | 2.000E-08 | 1.000E-07 |
| 5855 | 293 | C293P | A | 6e-08 | 3.4e-08 | 3.1e-08 | 4.167E-08 | 1.595E-08 | 3.100E-08 | 6.000E-08 |
| 5856 | 293 | C293S | A | 1e-08 | 1.5e-07 | 4.7e-08 | 6.900E-08 | 7.255E-08 | 1.000E-08 | 1.500E-07 |
| 5857 | 293 | C293T | A | 5.4e-08 | 7.5e-08 | 3.8e-08 | 5.567E-08 | 1.856E-08 | 3.800E-08 | 7.500E-08 |
| 5858 | 293 | C293W | A | 6.9e-08 | 7.8e-08 | 8.9e-08 | 7.867E-08 | 1.002E-08 | 6.900E-08 | 8.900E-08 |
| 5859 | 293 | C293Y | A | 2.4e-08 | 4.6e-08 | 4.4e-08 | 3.800E-08 | 1.217E-08 | 2.400E-08 | 4.600E-08 |
| 5860 | 293 | C293V | A | 5.7e-08 | 1.9e-07 | 5.3e-08 | 1.000E-07 | 7.797E-08 | 5.300E-08 | 1.900E-07 |
| 5861 | 294 | P294A | A | 9.1e-08 | 5.7e-08 | 8.1e-08 | 7.633E-08 | 1.747E-08 | 5.700E-08 | 9.100E-08 |
| 5862 | 294 | P294R | A | 3.2e-08 | 4.8e-08 | 2.8e-08 | 3.600E-08 | 1.058E-08 | 2.800E-08 | 4.800E-08 |
| 5863 | 294 | P294N | A | 1.8e-08 | 7.2e-08 | 2.7e-08 | 3.900E-08 | 2.893E-08 | 1.800E-08 | 7.200E-08 |
| 5864 | 294 | P294D | A | 6.3e-08 | 1.8e-07 | 3.1e-08 | 9.133E-08 | 7.844E-08 | 3.100E-08 | 1.800E-07 |

| | | | | | | | | | | |
|---|---|---|---|---|---|---|---|---|---|---|
| 5865 | 294 | P294C | A | 3.9e-08 | 6.7e-08 | 1.7e-07 | 9.200E-08 | 6.899E-08 | 3.900E-08 | 1.700E-07 |
| 5866 | 294 | P294E | A | 2.2e-08 | 1.5e-08 | 2.8e-08 | 2.167E-08 | 6.506E-09 | 1.500E-08 | 2.800E-08 |
| 5867 | 294 | P294Q | A | 3.8e-08 | 8.9e-08 | 2.1e-08 | 4.933E-08 | 3.539E-08 | 2.100E-08 | 8.900E-08 |
| 5868 | 294 | P294G | A | 2.4e-08 | 4.5e-08 | 6.5e-08 | 4.467E-08 | 2.050E-08 | 2.400E-08 | 6.500E-08 |
| 5869 | 294 | P294H | A | 1.9e-08 | 9.6e-09 | 1.4e-08 | 1.420E-08 | 4.703E-09 | 9.600E-09 | 1.900E-08 |
| 5870 | 294 | P294I | A | 3.8e-08 | 9.7e-08 | 3.8e-08 | 5.767E-08 | 3.406E-08 | 3.800E-08 | 9.700E-08 |
| 5871 | 294 | P294L | A | 2.5e-08 | 3e-08 | 2.3e-08 | 2.600E-08 | 3.606E-09 | 2.300E-08 | 3.000E-08 |
| 5872 | 294 | P294K | A | 6.1e-08 | 6e-08 | 6.2e-08 | 6.100E-08 | 1.000E-09 | 6.000E-08 | 6.200E-08 |
| 5873 | 294 | P294M | A | 1.9e-08 | 3.9e-08 | 1.2e-08 | 2.333E-08 | 1.401E-08 | 1.200E-08 | 3.900E-08 |
| 5874 | 294 | P294F | A | 9.3e-08 | 3.7e-08 | 3.6e-08 | 5.533E-08 | 3.262E-08 | 3.600E-08 | 9.300E-08 |
| 5875 | 294 | P294P | A | 5.2e-08 | 3.6e-08 | 4.3e-08 | 4.367E-08 | 8.021E-09 | 3.600E-08 | 5.200E-08 |
| 5876 | 294 | P294S | A | 9.2e-09 | 2.9e-08 | 2.7e-08 | 2.173E-08 | 1.090E-08 | 9.200E-09 | 2.900E-08 |
| 5877 | 294 | P294T | A | 3.1e-08 | 1.2e-08 | 4e-08 | 2.767E-08 | 1.429E-08 | 1.200E-08 | 4.000E-08 |
| 5878 | 294 | P294W | A | 3.2e-08 | 2e-08 | 1.5e-08 | 2.233E-08 | 8.737E-09 | 1.500E-08 | 3.200E-08 |
| 5879 | 294 | P294Y | A | 5.5e-08 | 4.3e-08 | 8.6e-08 | 6.133E-08 | 2.219E-08 | 4.300E-08 | 8.600E-08 |
| 5880 | 294 | P294V | A | 6.1e-08 | 2.1e-07 | 4.9e-08 | 1.067E-07 | 8.969E-08 | 4.900E-08 | 2.100E-07 |
| 5881 | 295 | L295A | A | 4.7e-08 | 4e-08 | 6.2e-08 | 4.967E-08 | 1.124E-08 | 4.000E-08 | 6.200E-08 |
| 5882 | 295 | L295R | A | 3.6e-08 | 5e-08 | 2.4e-08 | 3.667E-08 | 1.301E-08 | 2.400E-08 | 5.000E-08 |
| 5883 | 295 | L295N | A | 1.1e-08 | 1.7e-08 | 5.7e-09 | 1.123E-08 | 5.654E-09 | 5.700E-09 | 1.700E-08 |
| 5884 | 295 | L295D | A | 4.5e-08 | 1.5e-08 | 3.4e-08 | 3.133E-08 | 1.518E-08 | 1.500E-08 | 4.500E-08 |
| 5885 | 295 | L295C | A | 4.4e-08 | 6.5e-08 | 3.3e-08 | 4.733E-08 | 1.626E-08 | 3.300E-08 | 6.500E-08 |
| 5886 | 295 | L295E | A | 1.3e-08 | 1.4e-08 | 2.9e-08 | 1.867E-08 | 8.963E-09 | 1.300E-08 | 2.900E-08 |
| 5887 | 295 | L295Q | A | 9.1e-09 | 2.5e-08 | 1e-08 | 1.470E-08 | 8.931E-09 | 9.100E-09 | 2.500E-08 |
| 5888 | 295 | L295G | A | 2.2e-08 | 7.4e-08 | 1.1e-07 | 6.867E-08 | 4.424E-08 | 2.200E-08 | 1.100E-07 |
| 5889 | 295 | L295H | A | 3e-08 | 1.3e-07 | 4.7e-08 | 6.900E-08 | 5.351E-08 | 3.000E-08 | 1.300E-07 |
| 5890 | 295 | L295I | A | 2.5e-08 | 4.5e-08 | 2.7e-08 | 3.233E-08 | 1.102E-08 | 2.500E-08 | 4.500E-08 |
| 5891 | 295 | L295L | A | 5.3e-08 | 5.6e-08 | 9.4e-08 | 6.767E-08 | 2.285E-08 | 5.300E-08 | 9.400E-08 |
| 5892 | 295 | L295K | A | 4.5e-08 | 7.9e-09 | 6.1e-08 | 3.797E-08 | 2.724E-08 | 7.900E-09 | 6.100E-08 |
| 5893 | 295 | L295M | A | 5.4e-08 | 3.1e-08 | 2.5e-08 | 3.667E-08 | 1.531E-08 | 2.500E-08 | 5.400E-08 |
| 5894 | 295 | L295F | A | 7.5e-08 | 2.4e-08 | 1.8e-08 | 3.900E-08 | 3.132E-08 | 1.800E-08 | 7.500E-08 |
| 5895 | 295 | L295P | A | 4.5e-08 | 3.6e-08 | 3e-08 | 3.700E-08 | 7.550E-09 | 3.000E-08 | 4.500E-08 |
| 5896 | 295 | L295S | A | 2.9e-08 | 1.8e-08 | 2.3e-08 | 2.333E-08 | 5.508E-09 | 1.800E-08 | 2.900E-08 |
| 5897 | 295 | L295T | A | 1.6e-08 | 2.7e-08 | 1.8e-08 | 2.033E-08 | 5.859E-09 | 1.600E-08 | 2.700E-08 |
| 5898 | 295 | L295W | A | 3.7e-08 | 6.4e-08 | 1.8e-07 | 9.367E-08 | 7.598E-08 | 3.700E-08 | 1.800E-07 |
| 5899 | 295 | L295Y | A | 1e-07 | 2.5e-08 | 8.7e-08 | 7.067E-08 | 4.008E-08 | 2.500E-08 | 1.000E-07 |
| 5900 | 295 | L295V | A | 5.6e-08 | 4.9e-08 | 2.6e-08 | 4.367E-08 | 1.570E-08 | 2.600E-08 | 5.600E-08 |
| 5901 | 296 | H296A | A | 1.1e-07 | 1.5e-07 | 5.8e-08 | 1.060E-07 | 4.613E-08 | 5.800E-08 | 1.500E-07 |
| 5902 | 296 | H296R | A | 8.1e-08 | 4.5e-08 | 6.1e-08 | 6.233E-08 | 1.804E-08 | 4.500E-08 | 8.100E-08 |
| 5903 | 296 | H296N | A | 5.5e-09 | 1.5e-08 | 7.5e-09 | 9.333E-09 | 5.008E-09 | 5.500E-09 | 1.500E-08 |
| 5904 | 296 | H296D | A | 4.9e-08 | 4.4e-08 | 4.5e-08 | 4.600E-08 | 2.646E-09 | 4.400E-08 | 4.900E-08 |
| 5905 | 296 | H296C | A | 6e-08 | 2e-07 | 8.8e-08 | 1.160E-07 | 7.408E-08 | 6.000E-08 | 2.000E-07 |
| 5906 | 296 | H296E | A | 1.8e-08 | 2.5e-08 | 1.5e-08 | 1.933E-08 | 5.132E-09 | 1.500E-08 | 2.500E-08 |
| 5907 | 296 | H296Q | A | 2.3e-08 | 7e-09 | 4e-08 | 2.333E-08 | 1.650E-08 | 7.000E-09 | 4.000E-08 |
| 5908 | 296 | H296G | A | 1.8e-08 | 4.3e-08 | 4.1e-08 | 3.400E-08 | 1.389E-08 | 1.800E-08 | 4.300E-08 |
| 5909 | 296 | H296H | A | 6.1e-08 | 1.8e-08 | 1e-07 | 5.967E-08 | 4.102E-08 | 1.800E-08 | 1.000E-07 |
| 5910 | 296 | H296I | A | 8.2e-08 | 2.2e-08 | 8.2e-08 | 6.200E-08 | 3.464E-08 | 2.200E-08 | 8.200E-08 |
| 5911 | 296 | H296L | A | 5.1e-08 | 1.3e-07 | 6.3e-08 | 8.133E-08 | 4.257E-08 | 5.100E-08 | 1.300E-07 |
| 5912 | 296 | H296K | A | 4.2e-08 | 3.7e-08 | 6.3e-08 | 4.733E-08 | 1.380E-08 | 3.700E-08 | 6.300E-08 |
| 5913 | 296 | H296M | A | 9.6e-08 | 6.3e-08 | 1.4e-07 | 9.967E-08 | 3.863E-08 | 6.300E-08 | 1.400E-07 |
| 5914 | 296 | H296F | A | 1.1e-07 | 6.3e-08 | 8.2e-08 | 8.500E-08 | 2.364E-08 | 6.300E-08 | 1.100E-07 |
| 5915 | 296 | H296P | A | 9.9e-08 | 4.4e-08 | 3.2e-07 | 1.543E-07 | 1.461E-07 | 4.400E-08 | 3.200E-07 |
| 5916 | 296 | H296S | A | 2.6e-08 | 2.6e-08 | 1.2e-08 | 2.133E-08 | 8.083E-09 | 1.200E-08 | 2.600E-08 |
| 5917 | 296 | H296T | A | 4.2e-09 | 3.2e-08 | 1.4e-08 | 1.673E-08 | 1.410E-08 | 4.200E-09 | 3.200E-08 |
| 5918 | 296 | H296W | A | 5.9e-08 | 1.1e-07 | 1.5e-07 | 1.063E-07 | 4.561E-08 | 5.900E-08 | 1.500E-07 |
| 5919 | 296 | H296Y | A | 7.1e-08 | 5e-08 | 8.7e-08 | 6.933E-08 | 1.856E-08 | 5.000E-08 | 8.700E-08 |
| 5920 | 296 | H296V | A | 6.7e-08 | 5.2e-08 | 1.8e-07 | 9.967E-08 | 6.997E-08 | 5.200E-08 | 1.800E-07 |
| 5921 | 297 | N297A | A | 6e-08 | 2.3e-07 | 1.9e-07 | 1.600E-07 | 8.888E-08 | 6.000E-08 | 2.300E-07 |
| 5922 | 297 | N297R | A | 2.6e-07 | 3.6e-08 | 3.1e-08 | 1.090E-07 | 1.308E-07 | 3.100E-08 | 2.600E-07 |
| 5923 | 297 | N297N | A | 1.8e-08 | 2.3e-08 | 3.6e-08 | 2.567E-08 | 9.292E-09 | 1.800E-08 | 3.600E-08 |
| 5924 | 297 | N297D | A | 2.5e-08 | 4.9e-08 | 6.5e-08 | 4.633E-08 | 2.013E-08 | 2.500E-08 | 6.500E-08 |
| 5925 | 297 | N297C | A | 8.6e-08 | 2.9e-07 | 1.6e-07 | 1.787E-07 | 1.033E-07 | 8.600E-08 | 2.900E-07 |
| 5926 | 297 | N297E | A | 7.8e-08 | 1.2e-07 | 4.9e-08 | 8.233E-08 | 3.570E-08 | 4.900E-08 | 1.200E-07 |
| 5927 | 297 | N297Q | A | 4e-08 | 6.8e-08 | 6.5e-08 | 5.767E-08 | 1.537E-08 | 4.000E-08 | 6.800E-08 |
| 5928 | 297 | N297G | A | 9.2e-08 | 2.5e-07 | 1.6e-07 | 1.673E-07 | 7.925E-08 | 9.200E-08 | 2.500E-07 |
| 5929 | 297 | N297H | A | 1.4e-07 | 1.4e-07 | 6.5e-08 | 1.150E-07 | 4.330E-08 | 6.500E-08 | 1.400E-07 |
| 5930 | 297 | N297I | A | 1.2e-07 | 2.1e-07 | 1.4e-07 | 1.567E-07 | 4.726E-08 | 1.200E-07 | 2.100E-07 |
| 5931 | 297 | N297L | A | 1.1e-07 | 8.9e-08 | 1.5e-07 | 1.163E-07 | 3.099E-08 | 8.900E-08 | 1.500E-07 |
| 5932 | 297 | N297K | A | 7.9e-08 | 1.1e-07 | 6.6e-08 | 8.500E-08 | 2.261E-08 | 6.600E-08 | 1.100E-07 |
| 5933 | 297 | N297M | A | 1.1e-07 | 8.8e-08 | 1.7e-07 | 1.227E-07 | 4.244E-08 | 8.800E-08 | 1.700E-07 |
| 5934 | 297 | N297F | A | 5.2e-08 | 5.4e-08 | 2.3e-07 | 1.120E-07 | 1.022E-07 | 5.200E-08 | 2.300E-07 |
| 5935 | 297 | N297P | A | 7.1e-08 | 4.8e-07 | 8.1e-08 | 2.107E-07 | 2.333E-07 | 7.100E-08 | 4.800E-07 |
| 5936 | 297 | N297S | A | 3.7e-08 | 1.5e-07 | 1e-07 | 9.567E-08 | 5.662E-08 | 3.700E-08 | 1.500E-07 |
| 5937 | 297 | N297T | A | 1.2e-07 | 1.6e-07 | 3.4e-08 | 1.047E-07 | 6.438E-08 | 3.400E-08 | 1.600E-07 |
| 5938 | 297 | N297W | A | 5.8e-08 | 9e-08 | 1.3e-07 | 9.267E-08 | 3.607E-08 | 5.800E-08 | 1.300E-07 |
| 5939 | 297 | N297Y | A | 5.8e-08 | 6.4e-08 | 6.4e-08 | 6.200E-08 | 3.464E-09 | 5.800E-08 | 6.400E-08 |
| 5940 | 297 | N297V | A | 9.7e-08 | 9.3e-08 | 1.6e-07 | 1.167E-07 | 3.758E-08 | 9.300E-08 | 1.600E-07 |
| 5941 | 298 | Q298A | A | 4.1e-08 | 1.8e-08 | 6.1e-08 | 4.000E-08 | 2.152E-08 | 1.800E-08 | 6.100E-08 |
| 5942 | 298 | Q298R | A | 3.7e-08 | 5.7e-08 | 2.2e-08 | 3.867E-08 | 1.756E-08 | 2.200E-08 | 5.700E-08 |
| 5943 | 298 | Q298N | A | 5.6e-08 | 1.7e-08 | 3.3e-08 | 3.533E-08 | 1.960E-08 | 1.700E-08 | 5.600E-08 |
| 5944 | 298 | Q298D | A | 7.8e-08 | 2.2e-07 | 6.5e-08 | 1.210E-07 | 8.598E-08 | 6.500E-08 | 2.200E-07 |
| 5945 | 298 | Q298C | A | 2.1e-08 | 3.8e-08 | 2.6e-08 | 2.833E-08 | 8.737E-09 | 2.100E-08 | 3.800E-08 |
| 5946 | 298 | Q298E | A | 2.7e-08 | 5.8e-08 | 5.5e-08 | 4.667E-08 | 1.710E-08 | 2.700E-08 | 5.800E-08 |
| 5947 | 298 | Q298Q | A | 8.1e-08 | 1.3e-08 | 2.7e-08 | 4.033E-08 | 3.591E-08 | 1.300E-08 | 8.100E-08 |
| 5948 | 298 | Q298G | A | 3e-08 | 3.7e-08 | 3.2e-08 | 3.300E-08 | 3.606E-09 | 3.000E-08 | 3.700E-08 |
| 5949 | 298 | Q298H | A | 8.4e-08 | 7.8e-08 | 8.7e-08 | 8.300E-08 | 4.583E-09 | 7.800E-08 | 8.700E-08 |

| | | | | | | | | | | |
|---|---|---|---|---|---|---|---|---|---|---|
| 5950 | 298 | Q298I | A | 3.6e-08 | 1.2e-07 | 1.1e-07 | 8.867E-08 | 4.588E-08 | 3.600E-08 | 1.200E-07 |
| 5951 | 298 | Q298L | A | 9.7e-08 | 5.8e-08 | 4.2e-08 | 6.567E-08 | 2.829E-08 | 4.200E-08 | 9.700E-08 |
| 5952 | 298 | Q298K | A | 4.1e-08 | 2.5e-07 | 7.1e-08 | 1.207E-07 | 1.130E-07 | 4.100E-08 | 2.500E-07 |
| 5953 | 298 | Q298M | A | 7.6e-08 | 3.6e-08 | 1.5e-07 | 8.733E-08 | 5.784E-08 | 3.600E-08 | 1.500E-07 |
| 5954 | 298 | Q298F | A | 1.8e-07 | 7.1e-08 | 3.4e-08 | 9.500E-08 | 7.590E-08 | 3.400E-08 | 1.800E-07 |
| 5955 | 298 | Q298P | A | 5.4e-08 | 3.4e-08 | 9.7e-08 | 6.167E-08 | 3.219E-08 | 3.400E-08 | 9.700E-08 |
| 5956 | 298 | Q298S | A | 8.6e-08 | 2.6e-08 | 2.6e-08 | 4.600E-08 | 3.464E-08 | 2.600E-08 | 8.600E-08 |
| 5957 | 298 | Q298T | A | 1.1e-08 | 9.8e-08 | 1.6e-08 | 4.167E-08 | 4.885E-08 | 1.100E-08 | 9.800E-08 |
| 5958 | 298 | Q298W | A | 4.3e-08 | 4.9e-08 | 4.6e-08 | 4.600E-08 | 3.000E-09 | 4.300E-08 | 4.900E-08 |
| 5959 | 298 | Q298Y | A | 2.5e-08 | 5.5e-08 | 1.6e-08 | 3.200E-08 | 2.042E-08 | 1.600E-08 | 5.500E-08 |
| 5960 | 298 | Q298V | A | 3.8e-08 | 5.1e-08 | 6.3e-08 | 5.067E-08 | 1.250E-08 | 3.800E-08 | 6.300E-08 |
| 5961 | 299 | E299A | A | 7.2e-09 | 3e-08 | 1.3e-08 | 1.673E-08 | 1.185E-08 | 7.200E-09 | 3.000E-08 |
| 5962 | 299 | E299R | A | 5.4e-08 | 7.4e-08 | 3.5e-08 | 5.433E-08 | 1.950E-08 | 3.500E-08 | 7.400E-08 |
| 5963 | 299 | E299N | A | 9.9e-08 | 7e-08 | 1.6e-08 | 6.167E-08 | 4.212E-08 | 1.600E-08 | 9.900E-08 |
| 5964 | 299 | E299D | A | 6.8e-08 | 5.6e-08 | 8e-08 | 6.800E-08 | 1.200E-08 | 5.600E-08 | 8.000E-08 |
| 5965 | 299 | E299C | A | 4.5e-08 | 2.4e-08 | 9.5e-08 | 5.467E-08 | 3.647E-08 | 2.400E-08 | 9.500E-08 |
| 5966 | 299 | E299E | A | 1e-07 | 2.4e-07 | 6.4e-08 | 1.347E-07 | 9.298E-08 | 6.400E-08 | 2.400E-07 |
| 5967 | 299 | E299Q | A | 2.6e-08 | 5.9e-08 | 1.2e-07 | 6.833E-08 | 4.769E-08 | 2.600E-08 | 1.200E-07 |
| 5968 | 299 | E299G | A | 1.8e-08 | 2.1e-08 | 5.1e-08 | 3.000E-08 | 1.825E-08 | 1.800E-08 | 5.100E-08 |
| 5969 | 299 | E299H | A | 2.9e-08 | 3.9e-08 | 4.6e-08 | 3.800E-08 | 8.544E-09 | 2.900E-08 | 4.600E-08 |
| 5970 | 299 | E299I | A | 1.5e-08 | 3.1e-08 | 2.4e-08 | 2.333E-08 | 8.021E-09 | 1.500E-08 | 3.100E-08 |
| 5971 | 299 | E299L | A | 1.9e-08 | 2.4e-08 | 3.9e-08 | 2.733E-08 | 1.041E-08 | 1.900E-08 | 3.900E-08 |
| 5972 | 299 | E299K | A | 3.7e-08 | 1.5e-08 | 2.7e-08 | 2.633E-08 | 1.102E-08 | 1.500E-08 | 3.700E-08 |
| 5973 | 299 | E299M | A | 4e-08 | 1.2e-07 | 2.9e-08 | 6.300E-08 | 4.967E-08 | 2.900E-08 | 1.200E-07 |
| 5974 | 299 | E299F | A | 4.1e-08 | 9.2e-09 | 3.2e-08 | 2.740E-08 | 1.639E-08 | 9.200E-09 | 4.100E-08 |
| 5975 | 299 | E299P | A | 1.7e-08 | 9.4e-08 | 3.2e-08 | 4.767E-08 | 4.082E-08 | 1.700E-08 | 9.400E-08 |
| 5976 | 299 | E299S | A | 4.9e-08 | 7.1e-08 | 3.2e-08 | 5.067E-08 | 1.955E-08 | 3.200E-08 | 7.100E-08 |
| 5977 | 299 | E299T | A | 5.6e-09 | 3.7e-08 | 5.3e-08 | 3.187E-08 | 2.411E-08 | 5.600E-09 | 5.300E-08 |
| 5978 | 299 | E299W | A | 6.3e-08 | 3.5e-08 | 2.3e-08 | 4.033E-08 | 2.053E-08 | 2.300E-08 | 6.300E-08 |
| 5979 | 299 | E299Y | A | 1.9e-08 | 3.5e-08 | 1.2e-07 | 5.800E-08 | 5.429E-08 | 1.900E-08 | 1.200E-07 |
| 5980 | 299 | E299V | A | 2.6e-08 | 4e-08 | 3.2e-08 | 3.267E-08 | 7.024E-09 | 2.600E-08 | 4.000E-08 |
| 5981 | 300 | V300A | A | 2.4e-08 | 3.6e-08 | 1.3e-08 | 2.433E-08 | 1.150E-08 | 1.300E-08 | 3.600E-08 |
| 5982 | 300 | V300R | A | 1.1e-07 | 9.4e-08 | 6.1e-08 | 8.833E-08 | 2.499E-08 | 6.100E-08 | 1.100E-07 |
| 5983 | 300 | V300N | A | 5.8e-08 | 2.2e-08 | 1.4e-07 | 7.333E-08 | 6.048E-08 | 2.200E-08 | 1.400E-07 |
| 5984 | 300 | V300D | A | 1e-07 | 5.9e-08 | 6.4e-08 | 7.433E-08 | 2.237E-08 | 5.900E-08 | 1.000E-07 |
| 5985 | 300 | V300C | A | 6.1e-08 | 4.2e-08 | 4.8e-08 | 5.033E-08 | 9.713E-09 | 4.200E-08 | 6.100E-08 |
| 5986 | 300 | V300E | A | 5.9e-08 | 2.9e-08 | 5.4e-08 | 4.733E-08 | 1.607E-08 | 2.900E-08 | 5.900E-08 |
| 5987 | 300 | V300Q | A | 6.5e-08 | 2.7e-08 | 5.8e-08 | 5.000E-08 | 2.022E-08 | 2.700E-08 | 6.500E-08 |
| 5988 | 300 | V300G | A | 3.4e-08 | 4.1e-08 | 1.2e-07 | 6.500E-08 | 4.776E-08 | 3.400E-08 | 1.200E-07 |
| 5989 | 300 | V300H | A | 3.8e-08 | 4.7e-08 | 2.7e-08 | 3.733E-08 | 1.002E-08 | 2.700E-08 | 4.700E-08 |
| 5990 | 300 | V300I | A | 3e-08 | 5.6e-08 | 4.8e-08 | 4.467E-08 | 1.332E-08 | 3.000E-08 | 5.600E-08 |
| 5991 | 300 | V300L | A | 2e-08 | 8.2e-08 | 5.1e-08 | 5.100E-08 | 3.100E-08 | 2.000E-08 | 8.200E-08 |
| 5992 | 300 | V300K | A | 2e-08 | 3.7e-08 | 1.3e-08 | 2.333E-08 | 1.234E-08 | 1.300E-08 | 3.700E-08 |
| 5993 | 300 | V300M | A | 5.3e-08 | 5.2e-08 | 2.7e-08 | 4.400E-08 | 1.473E-08 | 2.700E-08 | 5.300E-08 |
| 5994 | 300 | V300F | A | 3.7e-08 | 2.1e-08 | 9.5e-08 | 5.100E-08 | 3.894E-08 | 2.100E-08 | 9.500E-08 |
| 5995 | 300 | V300P | A | 8e-08 | 6.2e-08 | 4.2e-08 | 6.133E-08 | 1.901E-08 | 4.200E-08 | 8.000E-08 |
| 5996 | 300 | V300S | A | 3.1e-08 | 4e-08 | 4.2e-08 | 3.767E-08 | 5.859E-09 | 3.100E-08 | 4.200E-08 |
| 5997 | 300 | V300T | A | 3.7e-08 | 2.9e-08 | 3e-08 | 3.200E-08 | 4.359E-09 | 2.900E-08 | 3.700E-08 |
| 5998 | 300 | V300W | A | 4.3e-08 | 6.1e-08 | 1.4e-08 | 3.933E-08 | 2.371E-08 | 1.400E-08 | 6.100E-08 |
| 5999 | 300 | V300Y | A | 2.6e-08 | 4.2e-08 | 4.7e-08 | 3.833E-08 | 1.097E-08 | 2.600E-08 | 4.700E-08 |
| 6000 | 300 | V300V | A | 1.9e-08 | 5.5e-08 | 6.7e-08 | 4.700E-08 | 2.498E-08 | 1.900E-08 | 6.700E-08 |
| 6001 | 301 | T301A | A | 5.1e-08 | 7.4e-08 | 5.2e-08 | 5.900E-08 | 1.300E-08 | 5.100E-08 | 7.400E-08 |
| 6002 | 301 | T301R | A | 2.8e-08 | 9.7e-08 | 3.7e-08 | 5.400E-08 | 3.751E-08 | 2.800E-08 | 9.700E-08 |
| 6003 | 301 | T301N | A | 7.4e-08 | 4.4e-08 | 5.4e-08 | 5.733E-08 | 1.528E-08 | 4.400E-08 | 7.400E-08 |
| 6004 | 301 | T301D | A | 5.1e-08 | 2.7e-08 | 9e-08 | 5.600E-08 | 3.180E-08 | 2.700E-08 | 9.000E-08 |
| 6005 | 301 | T301C | A | 7.6e-08 | 2.4e-08 | 6.6e-08 | 5.533E-08 | 2.759E-08 | 2.400E-08 | 7.600E-08 |
| 6006 | 301 | T301E | A | 6e-08 | 2.4e-08 | 6.7e-08 | 5.033E-08 | 2.307E-08 | 2.400E-08 | 6.700E-08 |
| 6007 | 301 | T301Q | A | 6.2e-08 | 7.9e-08 | 3.3e-08 | 5.800E-08 | 2.326E-08 | 3.300E-08 | 7.900E-08 |
| 6008 | 301 | T301G | A | 4.9e-08 | 4.5e-08 | 3.1e-08 | 4.167E-08 | 9.452E-09 | 3.100E-08 | 4.900E-08 |
| 6009 | 301 | T301H | A | 2.5e-07 | 6.2e-08 | 7.6e-08 | 1.293E-07 | 1.047E-07 | 6.200E-08 | 2.500E-07 |
| 6010 | 301 | T301I | A | 8.2e-08 | 1.8e-08 | 2e-07 | 1.000E-07 | 9.233E-08 | 1.800E-08 | 2.000E-07 |
| 6011 | 301 | T301L | A | 1.1e-07 | 6.3e-08 | 8.3e-08 | 8.533E-08 | 2.359E-08 | 6.300E-08 | 1.100E-07 |
| 6012 | 301 | T301K | A | 3.5e-08 | 9e-08 | 9.3e-08 | 7.267E-08 | 3.265E-08 | 3.500E-08 | 9.300E-08 |
| 6013 | 301 | T301M | A | 1.1e-08 | 2.7e-08 | 2.5e-08 | 2.100E-08 | 8.718E-09 | 1.100E-08 | 2.700E-08 |
| 6014 | 301 | T301F | A | 1.8e-07 | 4.3e-08 | 4.3e-08 | 8.867E-08 | 7.910E-08 | 4.300E-08 | 1.800E-07 |
| 6015 | 301 | T301P | A | 2.9e-08 | 1.4e-08 | 4.2e-08 | 2.833E-08 | 1.401E-08 | 1.400E-08 | 4.200E-08 |
| 6016 | 301 | T301S | A | 4.9e-08 | 8.3e-08 | 2.4e-08 | 5.200E-08 | 2.961E-08 | 2.400E-08 | 8.300E-08 |
| 6017 | 301 | T301T | A | 4.8e-08 | 4.1e-08 | 5.3e-08 | 4.733E-08 | 6.028E-09 | 4.100E-08 | 5.300E-08 |
| 6018 | 301 | T301W | A | 3.6e-08 | 4.3e-08 | 4.8e-08 | 4.233E-08 | 6.028E-09 | 3.600E-08 | 4.800E-08 |
| 6019 | 301 | T301Y | A | 4.2e-08 | 2.1e-08 | 7.6e-08 | 4.633E-08 | 2.775E-08 | 2.100E-08 | 7.600E-08 |
| 6020 | 301 | T301V | A | 4.7e-08 | 3.1e-08 | 2.7e-08 | 3.500E-08 | 1.058E-08 | 2.700E-08 | 4.700E-08 |
| 6021 | 302 | A302A | A | 8.3e-08 | 8.6e-08 | 7.7e-08 | 8.200E-08 | 4.583E-09 | 7.700E-08 | 8.600E-08 |
| 6022 | 302 | A302R | A | 2.1e-08 | 2.7e-08 | 5.1e-08 | 3.300E-08 | 1.587E-08 | 2.100E-08 | 5.100E-08 |
| 6023 | 302 | A302N | A | 5.3e-08 | 2.2e-08 | 4.5e-08 | 4.000E-08 | 1.609E-08 | 2.200E-08 | 5.300E-08 |
| 6024 | 302 | A302D | A | 1.1e-07 | 9.4e-08 | 5.8e-08 | 8.733E-08 | 2.663E-08 | 5.800E-08 | 1.100E-07 |
| 6025 | 302 | A302C | A | 1.1e-07 | 7.6e-08 | 5e-08 | 7.867E-08 | 3.009E-08 | 5.000E-08 | 1.100E-07 |
| 6026 | 302 | A302E | A | 3.3e-08 | 7.3e-08 | 2.6e-08 | 4.400E-08 | 2.536E-08 | 2.600E-08 | 7.300E-08 |
| 6027 | 302 | A302Q | A | 1.3e-07 | 1.5e-08 | 3.5e-08 | 6.000E-08 | 6.144E-08 | 1.500E-08 | 1.300E-07 |
| 6028 | 302 | A302G | A | 1.1e-07 | 6.1e-08 | 5.3e-08 | 7.467E-08 | 3.086E-08 | 5.300E-08 | 1.100E-07 |
| 6029 | 302 | A302H | A | 4e-08 | 6.6e-08 | 5.2e-08 | 5.267E-08 | 1.301E-08 | 4.000E-08 | 6.600E-08 |
| 6030 | 302 | A302I | A | 3.9e-08 | 1.1e-08 | 3.7e-08 | 2.900E-08 | 1.562E-08 | 1.100E-08 | 3.900E-08 |
| 6031 | 302 | A302L | A | 5.8e-08 | 1.3e-08 | 1.9e-08 | 3.000E-08 | 2.443E-08 | 1.300E-08 | 5.800E-08 |
| 6032 | 302 | A302K | A | 1.3e-07 | 3.1e-08 | 6.6e-08 | 7.567E-08 | 5.020E-08 | 3.100E-08 | 1.300E-07 |
| 6033 | 302 | A302M | A | 3.8e-08 | 4.6e-08 | 1.1e-07 | 6.467E-08 | 3.946E-08 | 3.800E-08 | 1.100E-07 |
| 6034 | 302 | A302F | A | 6.4e-08 | 3.1e-08 | 8.4e-08 | 5.967E-08 | 2.676E-08 | 3.100E-08 | 8.400E-08 |

| | | | | | | | | | | |
|---|---|---|---|---|---|---|---|---|---|---|
| 6035 | 302 | A302P | A | 8.2e-08 | 7.7e-08 | 5.7e-08 | 7.200E-08 | 1.323E-08 | 5.700E-08 | 8.200E-08 |
| 6036 | 302 | A302S | A | 6.7e-08 | 4.3e-08 | 1.7e-08 | 4.233E-08 | 2.501E-08 | 1.700E-08 | 6.700E-08 |
| 6037 | 302 | A302T | A | 5.8e-08 | 6.5e-08 | 5.1e-08 | 5.800E-08 | 7.000E-09 | 5.100E-08 | 6.500E-08 |
| 6038 | 302 | A302W | A | 3.3e-08 | 3.7e-08 | 1.4e-08 | 2.800E-08 | 1.229E-08 | 1.400E-08 | 3.700E-08 |
| 6039 | 302 | A302Y | A | 1.8e-08 | 2.2e-08 | 3.4e-08 | 2.467E-08 | 8.327E-09 | 1.800E-08 | 3.400E-08 |
| 6040 | 302 | A302V | A | 9.1e-08 | 6.8e-08 | 2.5e-08 | 6.133E-08 | 3.350E-08 | 2.500E-08 | 9.100E-08 |
| 6041 | 303 | E303A | A | 3.4e-08 | 2.1e-08 | 5e-08 | 3.500E-08 | 1.453E-08 | 2.100E-08 | 5.000E-08 |
| 6042 | 303 | E303R | A | 3.8e-08 | 1.7e-08 | 1.5e-08 | 2.333E-08 | 1.274E-08 | 1.500E-08 | 3.800E-08 |
| 6043 | 303 | E303N | A | 3.4e-08 | 1.9e-08 | 7.9e-08 | 4.400E-08 | 3.122E-08 | 1.900E-08 | 7.900E-08 |
| 6044 | 303 | E303D | A | 6.9e-08 | 2.8e-08 | 5.6e-08 | 5.100E-08 | 2.095E-08 | 2.800E-08 | 6.900E-08 |
| 6045 | 303 | E303C | A | 9.2e-08 | 4.5e-08 | 4.8e-08 | 6.167E-08 | 2.631E-08 | 4.500E-08 | 9.200E-08 |
| 6046 | 303 | E303E | A | 2.2e-07 | 1.5e-08 | 1.8e-08 | 8.433E-08 | 1.175E-07 | 1.500E-08 | 2.200E-07 |
| 6047 | 303 | E303Q | A | 2.4e-08 | 4.9e-08 | 4.4e-08 | 3.900E-08 | 1.323E-08 | 2.400E-08 | 4.900E-08 |
| 6048 | 303 | E303G | A | 2.6e-08 | 4.7e-08 | 1.1e-07 | 6.100E-08 | 4.371E-08 | 2.600E-08 | 1.100E-07 |
| 6049 | 303 | E303H | A | 1.8e-08 | 4.5e-08 | 9.2e-08 | 5.167E-08 | 3.745E-08 | 1.800E-08 | 9.200E-08 |
| 6050 | 303 | E303I | A | 1.8e-08 | 5.4e-08 | 4.2e-08 | 3.800E-08 | 1.833E-08 | 1.800E-08 | 5.400E-08 |
| 6051 | 303 | E303L | A | 2.8e-08 | 3.1e-08 | 2.9e-08 | 2.933E-08 | 1.528E-09 | 2.800E-08 | 3.100E-08 |
| 6052 | 303 | E303K | A | 3.6e-08 | 1.1e-07 | 3.2e-08 | 5.933E-08 | 4.392E-08 | 3.200E-08 | 1.100E-07 |
| 6053 | 303 | E303M | A | 2.4e-08 | 4.3e-08 | 6.9e-08 | 4.533E-08 | 2.259E-08 | 2.400E-08 | 6.900E-08 |
| 6054 | 303 | E303F | A | 1.2e-08 | 1.1e-08 | 2.4e-08 | 1.567E-08 | 7.234E-09 | 1.100E-08 | 2.400E-08 |
| 6055 | 303 | E303P | A | 3.8e-08 | 1.2e-08 | 4.4e-08 | 3.133E-08 | 1.701E-08 | 1.200E-08 | 4.400E-08 |
| 6056 | 303 | E303S | A | 9e-08 | 4.7e-08 | 2.2e-08 | 5.300E-08 | 3.439E-08 | 2.200E-08 | 9.000E-08 |
| 6057 | 303 | E303T | A | 5.2e-08 | 2e-08 | 7.6e-08 | 4.933E-08 | 2.810E-08 | 2.000E-08 | 7.600E-08 |
| 6058 | 303 | E303W | A | 1.6e-08 | 5.5e-08 | 2.9e-08 | 3.333E-08 | 1.986E-08 | 1.600E-08 | 5.500E-08 |
| 6059 | 303 | E303Y | A | 1.4e-08 | 8.6e-09 | 1.2e-07 | 4.753E-08 | 6.282E-08 | 8.600E-09 | 1.200E-07 |
| 6060 | 303 | E303V | A | 2.8e-08 | 1.5e-07 | 9.1e-08 | 8.967E-08 | 6.101E-08 | 2.800E-08 | 1.500E-07 |
| 6061 | 304 | D304A | A | 7.7e-08 | 1.8e-08 | 1.4e-07 | 7.833E-08 | 6.101E-08 | 1.800E-08 | 1.400E-07 |
| 6062 | 304 | D304R | A | 2.5e-08 | 5.9e-08 | 2.6e-08 | 3.667E-08 | 1.935E-08 | 2.500E-08 | 5.900E-08 |
| 6063 | 304 | D304N | A | 5.6e-08 | 9.8e-08 | 3.1e-08 | 6.167E-08 | 3.386E-08 | 3.100E-08 | 9.800E-08 |
| 6064 | 304 | D304D | A | 8.5e-08 | 5.4e-08 | 2.8e-08 | 5.567E-08 | 2.854E-08 | 2.800E-08 | 8.500E-08 |
| 6065 | 304 | D304C | A | 2.3e-08 | 1.9e-07 | 3.6e-08 | 8.300E-08 | 9.289E-08 | 2.300E-08 | 1.900E-07 |
| 6066 | 304 | D304E | A | 1.5e-07 | 3.9e-08 | 6.1e-08 | 8.333E-08 | 5.877E-08 | 3.900E-08 | 1.500E-07 |
| 6067 | 304 | D304Q | A | 2.7e-08 | 1.5e-08 | 1.4e-08 | 1.867E-08 | 7.234E-09 | 1.400E-08 | 2.700E-08 |
| 6068 | 304 | D304G | A | 6e-08 | 5.4e-08 | 3.5e-08 | 4.967E-08 | 1.305E-08 | 3.500E-08 | 6.000E-08 |
| 6069 | 304 | D304H | A | 5.4e-08 | 3.6e-08 | 7.9e-08 | 5.633E-08 | 2.159E-08 | 3.600E-08 | 7.900E-08 |
| 6070 | 304 | D304I | A | 1.8e-08 | 5.1e-08 | 2.6e-08 | 3.167E-08 | 1.721E-08 | 1.800E-08 | 5.100E-08 |
| 6071 | 304 | D304L | A | 2.2e-08 | 6e-08 | 1.9e-08 | 3.367E-08 | 2.285E-08 | 1.900E-08 | 6.000E-08 |
| 6072 | 304 | D304K | A | 1.7e-08 | 4.7e-08 | 4e-08 | 3.467E-08 | 1.570E-08 | 1.700E-08 | 4.700E-08 |
| 6073 | 304 | D304M | A | 4.3e-08 | 8.4e-08 | 2.5e-08 | 5.067E-08 | 3.024E-08 | 2.500E-08 | 8.400E-08 |
| 6074 | 304 | D304F | A | 6.7e-08 | 4.2e-08 | 1e-07 | 6.967E-08 | 2.909E-08 | 4.200E-08 | 1.000E-07 |
| 6075 | 304 | D304P | A | 1.5e-08 | 4.2e-08 | 7.4e-08 | 4.367E-08 | 2.954E-08 | 1.500E-08 | 7.400E-08 |
| 6076 | 304 | D304S | A | 4.2e-08 | 2.5e-08 | 3e-08 | 3.233E-08 | 8.737E-09 | 2.500E-08 | 4.200E-08 |
| 6077 | 304 | D304T | A | 7.3e-08 | 3e-08 | 2.5e-08 | 4.267E-08 | 2.639E-08 | 2.500E-08 | 7.300E-08 |
| 6078 | 304 | D304W | A | 7.9e-08 | 9e-08 | 4e-08 | 6.967E-08 | 2.627E-08 | 4.000E-08 | 9.000E-08 |
| 6079 | 304 | D304Y | A | 5.1e-08 | 3.4e-08 | 4.1e-08 | 4.200E-08 | 8.544E-09 | 3.400E-08 | 5.100E-08 |
| 6080 | 304 | D304V | A | 2.5e-08 | 2.2e-08 | 6.1e-08 | 3.600E-08 | 2.170E-08 | 2.200E-08 | 6.100E-08 |
| 6081 | 305 | G305A | A | 3.9e-08 | 4.9e-08 | 1.2e-08 | 3.333E-08 | 1.914E-08 | 1.200E-08 | 4.900E-08 |
| 6082 | 305 | G305R | A | 6.6e-08 | 5.1e-08 | 2.4e-08 | 4.700E-08 | 2.128E-08 | 2.400E-08 | 6.600E-08 |
| 6083 | 305 | G305N | A | 3.2e-08 | 3.5e-08 | 4.3e-08 | 3.667E-08 | 5.686E-09 | 3.200E-08 | 4.300E-08 |
| 6084 | 305 | G305D | A | 1.3e-07 | 3.3e-08 | 7.8e-08 | 8.033E-08 | 4.854E-08 | 3.300E-08 | 1.300E-07 |
| 6085 | 305 | G305C | A | 7.9e-08 | 5.2e-08 | 8.6e-09 | 4.653E-08 | 3.552E-08 | 8.600E-09 | 7.900E-08 |
| 6086 | 305 | G305E | A | 1.9e-08 | 2.9e-08 | 4.3e-08 | 3.033E-08 | 1.206E-08 | 1.900E-08 | 4.300E-08 |
| 6087 | 305 | G305Q | A | 1.6e-07 | 1.1e-07 | 8.8e-08 | 1.193E-07 | 3.690E-08 | 8.800E-08 | 1.600E-07 |
| 6088 | 305 | G305G | A | 1.1e-08 | 5.5e-08 | 1.3e-07 | 6.533E-08 | 6.017E-08 | 1.100E-08 | 1.300E-07 |
| 6089 | 305 | G305H | A | 3.1e-08 | 6.3e-08 | 1.2e-07 | 7.133E-08 | 4.508E-08 | 3.100E-08 | 1.200E-07 |
| 6090 | 305 | G305I | A | 4.2e-08 | 3.6e-08 | 1.5e-08 | 3.100E-08 | 1.418E-08 | 1.500E-08 | 4.200E-08 |
| 6091 | 305 | G305L | A | 8.6e-08 | 4.3e-08 | 1e-07 | 7.633E-08 | 2.970E-08 | 4.300E-08 | 1.000E-07 |
| 6092 | 305 | G305K | A | 1e-07 | 3.2e-08 | 1.8e-07 | 1.040E-07 | 7.408E-08 | 3.200E-08 | 1.800E-07 |
| 6093 | 305 | G305M | A | 4.1e-08 | 3.2e-08 | 5.2e-08 | 4.167E-08 | 1.002E-08 | 3.200E-08 | 5.200E-08 |
| 6094 | 305 | G305F | A | 3.8e-08 | 1.2e-07 | 4.1e-08 | 6.633E-08 | 4.650E-08 | 3.800E-08 | 1.200E-07 |
| 6095 | 305 | G305P | A | 4.6e-08 | 6.6e-08 | 3.6e-08 | 4.933E-08 | 1.528E-08 | 3.600E-08 | 6.600E-08 |
| 6096 | 305 | G305S | A | 5.8e-08 | 2.9e-08 | 6.3e-08 | 5.000E-08 | 1.836E-08 | 2.900E-08 | 6.300E-08 |
| 6097 | 305 | G305T | A | 1.2e-07 | 1.1e-07 | 2.1e-08 | 8.367E-08 | 5.450E-08 | 2.100E-08 | 1.200E-07 |
| 6098 | 305 | G305W | A | 7.8e-08 | 2.2e-08 | 6.9e-08 | 5.633E-08 | 3.007E-08 | 2.200E-08 | 7.800E-08 |
| 6099 | 305 | G305Y | A | 1.8e-08 | 5.1e-08 | 5.1e-08 | 4.000E-08 | 1.905E-08 | 1.800E-08 | 5.100E-08 |
| 6100 | 305 | G305V | A | 1.7e-08 | 3.3e-08 | 3.8e-08 | 2.933E-08 | 1.097E-08 | 1.700E-08 | 3.800E-08 |
| 6101 | 306 | T306A | A | 2.3e-08 | 2.6e-08 | 3.3e-08 | 2.733E-08 | 5.132E-09 | 2.300E-08 | 3.300E-08 |
| 6102 | 306 | T306R | A | 9.1e-08 | 6.5e-08 | 5.1e-08 | 6.900E-08 | 2.030E-08 | 5.100E-08 | 9.100E-08 |
| 6103 | 306 | T306N | A | 6e-08 | 8.5e-08 | 1.6e-08 | 5.367E-08 | 3.493E-08 | 1.600E-08 | 8.500E-08 |
| 6104 | 306 | T306D | A | 3.5e-08 | 1.8e-08 | 4.7e-08 | 3.333E-08 | 1.457E-08 | 1.800E-08 | 4.700E-08 |
| 6105 | 306 | T306C | A | 2.8e-08 | 3e-08 | 6e-08 | 3.933E-08 | 1.793E-08 | 2.800E-08 | 6.000E-08 |
| 6106 | 306 | T306E | A | 4.6e-08 | 2.5e-08 | 3.8e-08 | 3.633E-08 | 1.060E-08 | 2.500E-08 | 4.600E-08 |
| 6107 | 306 | T306Q | A | 5.8e-08 | 1.7e-08 | 5.1e-08 | 4.200E-08 | 2.193E-08 | 1.700E-08 | 5.800E-08 |
| 6108 | 306 | T306G | A | 3.4e-08 | 1.9e-08 | 3.3e-08 | 2.867E-08 | 8.386E-09 | 1.900E-08 | 3.400E-08 |
| 6109 | 306 | T306H | A | 3.2e-08 | 5.4e-08 | 3e-08 | 3.867E-08 | 1.332E-08 | 3.000E-08 | 5.400E-08 |
| 6110 | 306 | T306I | A | 7.3e-08 | 2e-08 | 4.5e-08 | 4.600E-08 | 2.651E-08 | 2.000E-08 | 7.300E-08 |
| 6111 | 306 | T306L | A | 2.6e-08 | 1.7e-08 | 2.4e-08 | 2.233E-08 | 4.726E-09 | 1.700E-08 | 2.600E-08 |
| 6112 | 306 | T306K | A | 9.3e-08 | 3.9e-08 | 2.1e-08 | 5.100E-08 | 3.747E-08 | 2.100E-08 | 9.300E-08 |
| 6113 | 306 | T306M | A | 4.2e-08 | 4.2e-08 | 6.2e-08 | 4.867E-08 | 1.155E-08 | 4.200E-08 | 6.200E-08 |
| 6114 | 306 | T306F | A | 4.3e-08 | 2e-08 | 3.1e-08 | 3.133E-08 | 1.150E-08 | 2.000E-08 | 4.300E-08 |
| 6115 | 306 | T306P | A | 4.3e-08 | 5.3e-08 | 4.2e-08 | 4.600E-08 | 6.083E-09 | 4.200E-08 | 5.300E-08 |
| 6116 | 306 | T306S | A | 2.5e-08 | 5.2e-08 | 1.3e-07 | 6.900E-08 | 5.453E-08 | 2.500E-08 | 1.300E-07 |
| 6117 | 306 | T306T | A | 2.1e-08 | 9.7e-08 | 6.9e-08 | 6.233E-08 | 3.844E-08 | 2.100E-08 | 9.700E-08 |
| 6118 | 306 | T306W | A | 7e-08 | 5.4e-08 | 2.4e-08 | 4.933E-08 | 2.335E-08 | 2.400E-08 | 7.000E-08 |
| 6119 | 306 | T306Y | A | 3.7e-08 | 2.1e-08 | 6.9e-08 | 4.233E-08 | 2.444E-08 | 2.100E-08 | 6.900E-08 |

| | | | | | | | | | | |
|---|---|---|---|---|---|---|---|---|---|---|
| 6120 | 306 | T306V | A | 6.7e-08 | 6.7e-08 | 1.5e-08 | 4.967E-08 | 3.002E-08 | 1.500E-08 | 6.700E-08 |
| 6121 | 307 | Q307A | A | 1.4e-08 | 1.9e-08 | 5e-08 | 2.767E-08 | 1.950E-08 | 1.400E-08 | 5.000E-08 |
| 6122 | 307 | Q307R | A | 3.5e-08 | 4.6e-08 | 3.8e-08 | 3.967E-08 | 5.686E-09 | 3.500E-08 | 4.600E-08 |
| 6123 | 307 | Q307N | A | 2.6e-08 | 2.5e-08 | 9e-08 | 4.700E-08 | 3.724E-08 | 2.500E-08 | 9.000E-08 |
| 6124 | 307 | Q307D | A | 6.6e-08 | 1.3e-07 | 2.7e-08 | 7.433E-08 | 5.200E-08 | 2.700E-08 | 1.300E-07 |
| 6125 | 307 | Q307C | A | 3.7e-08 | 2.3e-08 | 3.3e-08 | 3.100E-08 | 7.211E-09 | 2.300E-08 | 3.700E-08 |
| 6126 | 307 | Q307E | A | 2.9e-08 | 4.9e-08 | 5.4e-08 | 4.400E-08 | 1.323E-08 | 2.900E-08 | 5.400E-08 |
| 6127 | 307 | Q307Q | A | 2.5e-08 | 8e-08 | 4.4e-08 | 4.967E-08 | 2.793E-08 | 2.500E-08 | 8.000E-08 |
| 6128 | 307 | Q307G | A | 3.5e-08 | 4.2e-08 | 8.9e-08 | 5.533E-08 | 2.937E-08 | 3.500E-08 | 8.900E-08 |
| 6129 | 307 | Q307H | A | 2.4e-08 | 2.7e-08 | 9.9e-08 | 5.000E-08 | 4.246E-08 | 2.400E-08 | 9.900E-08 |
| 6130 | 307 | Q307I | A | 4.6e-08 | 4.6e-08 | 5.3e-08 | 4.833E-08 | 4.041E-09 | 4.600E-08 | 5.300E-08 |
| 6131 | 307 | Q307L | A | 3.5e-08 | 2.4e-08 | 6.4e-08 | 4.100E-08 | 2.066E-08 | 2.400E-08 | 6.400E-08 |
| 6132 | 307 | Q307K | A | 5.7e-08 | 5.1e-08 | 3.3e-08 | 4.700E-08 | 1.249E-08 | 3.300E-08 | 5.700E-08 |
| 6133 | 307 | Q307M | A | 1.4e-08 | 3.9e-08 | 4.9e-08 | 3.400E-08 | 1.803E-08 | 1.400E-08 | 4.900E-08 |
| 6134 | 307 | Q307F | A | 2.5e-08 | 4.4e-08 | 4.2e-08 | 3.700E-08 | 1.044E-08 | 2.500E-08 | 4.400E-08 |
| 6135 | 307 | Q307P | A | 4.9e-08 | 6.4e-08 | 5.9e-09 | 3.963E-08 | 3.016E-08 | 5.900E-09 | 6.400E-08 |
| 6136 | 307 | Q307S | A | 3.2e-08 | 4.5e-08 | 9.2e-08 | 5.633E-08 | 3.156E-08 | 3.200E-08 | 9.200E-08 |
| 6137 | 307 | Q307T | A | 3.9e-08 | 1e-07 | 3.7e-08 | 5.867E-08 | 3.581E-08 | 3.700E-08 | 1.000E-07 |
| 6138 | 307 | Q307W | A | 1.6e-08 | 3e-08 | 2e-08 | 2.200E-08 | 7.211E-09 | 1.600E-08 | 3.000E-08 |
| 6139 | 307 | Q307Y | A | 2.9e-08 | 6.2e-08 | 1.2e-07 | 7.033E-08 | 4.607E-08 | 2.900E-08 | 1.200E-07 |
| 6140 | 307 | Q307V | A | 1.3e-08 | 5.5e-08 | 5.3e-08 | 4.033E-08 | 2.369E-08 | 1.300E-08 | 5.500E-08 |
| 6141 | 308 | R308A | A | 6.4e-08 | 3.9e-08 | 6.6e-08 | 5.633E-08 | 1.504E-08 | 3.900E-08 | 6.600E-08 |
| 6142 | 308 | R308R | A | 1e-07 | 3e-08 | 2.8e-08 | 5.267E-08 | 4.100E-08 | 2.800E-08 | 1.000E-07 |
| 6143 | 308 | R308N | A | 8e-08 | 3.6e-08 | 6.6e-08 | 6.067E-08 | 2.248E-08 | 3.600E-08 | 8.000E-08 |
| 6144 | 308 | R308D | A | 5.9e-08 | 5.2e-08 | 1.2e-08 | 4.100E-08 | 2.536E-08 | 1.200E-08 | 5.900E-08 |
| 6145 | 308 | R308C | A | 6.5e-08 | 1.2e-07 | 2.4e-08 | 6.967E-08 | 4.817E-08 | 2.400E-08 | 1.200E-07 |
| 6146 | 308 | R308E | A | 3.5e-08 | 2.2e-08 | 8.6e-08 | 4.767E-08 | 3.383E-08 | 2.200E-08 | 8.600E-08 |
| 6147 | 308 | R308Q | A | 5.3e-08 | 4.1e-08 | 1.7e-08 | 3.700E-08 | 1.833E-08 | 1.700E-08 | 5.300E-08 |
| 6148 | 308 | R308G | A | 3.1e-08 | 7e-08 | 1.6e-07 | 8.700E-08 | 6.616E-08 | 3.100E-08 | 1.600E-07 |
| 6149 | 308 | R308H | A | 3.6e-08 | 3.7e-08 | 7.2e-08 | 4.833E-08 | 2.050E-08 | 3.600E-08 | 7.200E-08 |
| 6150 | 308 | R308I | A | 7.4e-08 | 4.3e-08 | 7e-08 | 6.233E-08 | 1.686E-08 | 4.300E-08 | 7.400E-08 |
| 6151 | 308 | R308L | A | 5.4e-08 | 1.7e-08 | 4.7e-08 | 3.933E-08 | 1.966E-08 | 1.700E-08 | 5.400E-08 |
| 6152 | 308 | R308K | A | 2e-08 | 6.7e-08 | 2.5e-08 | 3.733E-08 | 2.581E-08 | 2.000E-08 | 6.700E-08 |
| 6153 | 308 | R308M | A | 2.4e-08 | 7.1e-08 | 3.3e-08 | 4.267E-08 | 2.495E-08 | 2.400E-08 | 7.100E-08 |
| 6154 | 308 | R308F | A | 7.4e-08 | 1e-07 | 3.2e-08 | 6.867E-08 | 3.431E-08 | 3.200E-08 | 1.000E-07 |
| 6155 | 308 | R308P | A | 3.5e-08 | 1.8e-08 | 1.9e-07 | 8.100E-08 | 9.478E-08 | 1.800E-08 | 1.900E-07 |
| 6156 | 308 | R308S | A | 2.1e-08 | 4.7e-08 | 3.1e-08 | 3.300E-08 | 1.311E-08 | 2.100E-08 | 4.700E-08 |
| 6157 | 308 | R308T | A | 1.5e-08 | 3.1e-08 | 4.9e-08 | 3.167E-08 | 1.701E-08 | 1.500E-08 | 4.900E-08 |
| 6158 | 308 | R308W | A | 6e-08 | 2.8e-08 | 9.2e-08 | 6.000E-08 | 3.200E-08 | 2.800E-08 | 9.200E-08 |
| 6159 | 308 | R308Y | A | 4.7e-08 | 4e-08 | 1.1e-07 | 6.567E-08 | 3.855E-08 | 4.000E-08 | 1.100E-07 |
| 6160 | 308 | R308V | A | 4.8e-08 | 8.6e-08 | 1.5e-08 | 4.967E-08 | 3.553E-08 | 1.500E-08 | 8.600E-08 |
| 6161 | 309 | C309A | A | 5.2e-08 | 5.8e-08 | 3.4e-08 | 4.800E-08 | 1.249E-08 | 3.400E-08 | 5.800E-08 |
| 6162 | 309 | C309R | A | 3.4e-08 | 4.8e-08 | 4.1e-08 | 4.100E-08 | 7.000E-09 | 3.400E-08 | 4.800E-08 |
| 6163 | 309 | C309N | A | 6.2e-08 | 4.8e-08 | 5.1e-08 | 5.367E-08 | 7.371E-09 | 4.800E-08 | 6.200E-08 |
| 6164 | 309 | C309D | A | 6e-08 | 1.6e-07 | 1.4e-07 | 1.200E-07 | 5.292E-08 | 6.000E-08 | 1.600E-07 |
| 6165 | 309 | C309C | A | 5e-08 | 3.4e-08 | 1.9e-08 | 3.433E-08 | 1.550E-08 | 1.900E-08 | 5.000E-08 |
| 6166 | 309 | C309E | A | 5.9e-08 | 2.4e-08 | 1.1e-08 | 3.133E-08 | 2.483E-08 | 1.100E-08 | 5.900E-08 |
| 6167 | 309 | C309Q | A | 5.1e-08 | 9.2e-08 | 3.2e-08 | 5.833E-08 | 3.066E-08 | 3.200E-08 | 9.200E-08 |
| 6168 | 309 | C309G | A | 1.8e-08 | 3.7e-08 | 3.5e-08 | 3.000E-08 | 1.044E-08 | 1.800E-08 | 3.700E-08 |
| 6169 | 309 | C309H | A | 6.4e-08 | 9.1e-08 | 4.2e-08 | 6.567E-08 | 2.454E-08 | 4.200E-08 | 9.100E-08 |
| 6170 | 309 | C309I | A | 3.5e-08 | 3.4e-08 | 3.5e-08 | 3.467E-08 | 5.774E-10 | 3.400E-08 | 3.500E-08 |
| 6171 | 309 | C309L | A | 4e-08 | 1.6e-08 | 9e-08 | 4.867E-08 | 3.775E-08 | 1.600E-08 | 9.000E-08 |
| 6172 | 309 | C309K | A | 3.6e-08 | 9e-09 | 3.5e-08 | 2.667E-08 | 1.531E-08 | 9.000E-09 | 3.600E-08 |
| 6173 | 309 | C309M | A | 6e-08 | 2.1e-08 | 4.7e-08 | 4.267E-08 | 1.986E-08 | 2.100E-08 | 6.000E-08 |
| 6174 | 309 | C309F | A | 3.5e-08 | 4.3e-08 | 2.7e-08 | 3.500E-08 | 8.000E-09 | 2.700E-08 | 4.300E-08 |
| 6175 | 309 | C309P | A | 4.2e-08 | 9.2e-08 | 2e-08 | 5.133E-08 | 3.690E-08 | 2.000E-08 | 9.200E-08 |
| 6176 | 309 | C309S | A | 1.1e-07 | 7.4e-08 | 1.7e-08 | 6.700E-08 | 4.689E-08 | 1.700E-08 | 1.100E-07 |
| 6177 | 309 | C309T | A | 2.6e-08 | 5.6e-08 | 2e-08 | 3.400E-08 | 1.929E-08 | 2.000E-08 | 5.600E-08 |
| 6178 | 309 | C309W | A | 3.5e-08 | 1.6e-08 | 1e-07 | 5.033E-08 | 4.405E-08 | 1.600E-08 | 1.000E-07 |
| 6179 | 309 | C309Y | A | 2.9e-08 | 1.8e-07 | 4.3e-08 | 8.400E-08 | 8.343E-08 | 2.900E-08 | 1.800E-07 |
| 6180 | 309 | C309V | A | 7.7e-08 | 5.9e-08 | 2e-08 | 5.200E-08 | 2.914E-08 | 2.000E-08 | 7.700E-08 |
| 6181 | 310 | E310A | A | 2.8e-08 | 6e-08 | 4e-08 | 4.267E-08 | 1.617E-08 | 2.800E-08 | 6.000E-08 |
| 6182 | 310 | E310R | A | 5.1e-08 | 3.9e-08 | 8.7e-08 | 5.900E-08 | 2.498E-08 | 3.900E-08 | 8.700E-08 |
| 6183 | 310 | E310N | A | 4.5e-08 | 3.7e-08 | 2.3e-08 | 3.500E-08 | 1.114E-08 | 2.300E-08 | 4.500E-08 |
| 6184 | 310 | E310D | A | 2e-08 | 6.4e-08 | 3.3e-08 | 3.900E-08 | 2.261E-08 | 2.000E-08 | 6.400E-08 |
| 6185 | 310 | E310C | A | 1.1e-07 | 2.5e-08 | 2.5e-08 | 5.333E-08 | 4.907E-08 | 2.500E-08 | 1.100E-07 |
| 6186 | 310 | E310E | A | 6.9e-08 | 1.4e-08 | 4.9e-08 | 4.400E-08 | 2.784E-08 | 1.400E-08 | 6.900E-08 |
| 6187 | 310 | E310Q | A | 1e-07 | 8.5e-08 | 2.6e-08 | 7.033E-08 | 3.912E-08 | 2.600E-08 | 1.000E-07 |
| 6188 | 310 | E310G | A | 2.7e-08 | 5e-08 | 6.1e-08 | 4.600E-08 | 1.735E-08 | 2.700E-08 | 6.100E-08 |
| 6189 | 310 | E310H | A | 3.4e-08 | 7.2e-08 | 5.8e-08 | 5.467E-08 | 1.922E-08 | 3.400E-08 | 7.200E-08 |
| 6190 | 310 | E310I | A | 1.7e-07 | 8.1e-09 | 6.6e-08 | 8.137E-08 | 8.204E-08 | 8.100E-09 | 1.700E-07 |
| 6191 | 310 | E310L | A | 6.4e-08 | 2.3e-08 | 5e-08 | 4.567E-08 | 2.084E-08 | 2.300E-08 | 6.400E-08 |
| 6192 | 310 | E310K | A | 2.4e-08 | 2.7e-08 | 6.2e-08 | 3.767E-08 | 2.113E-08 | 2.400E-08 | 6.200E-08 |
| 6193 | 310 | E310M | A | 7.6e-08 | 2.1e-08 | 5.2e-08 | 4.967E-08 | 2.757E-08 | 2.100E-08 | 7.600E-08 |
| 6194 | 310 | E310F | A | 3.8e-08 | 1.6e-08 | 2.7e-08 | 2.700E-08 | 1.100E-08 | 1.600E-08 | 3.800E-08 |
| 6195 | 310 | E310P | A | 3.2e-08 | 3.2e-08 | 8.4e-08 | 4.933E-08 | 3.002E-08 | 3.200E-08 | 8.400E-08 |
| 6196 | 310 | E310S | A | 5.9e-08 | 1.2e-07 | 1e-07 | 9.300E-08 | 3.110E-08 | 5.900E-08 | 1.200E-07 |
| 6197 | 310 | E310T | A | 4e-08 | 5.8e-08 | 6.2e-08 | 5.333E-08 | 1.172E-08 | 4.000E-08 | 6.200E-08 |
| 6198 | 310 | E310W | A | 3.9e-08 | 5.5e-08 | 3.9e-08 | 4.433E-08 | 9.238E-09 | 3.900E-08 | 5.500E-08 |
| 6199 | 310 | E310Y | A | 3.4e-08 | 2e-08 | 4.7e-08 | 3.367E-08 | 1.350E-08 | 2.000E-08 | 4.700E-08 |
| 6200 | 310 | E310V | A | 3.2e-08 | 6.1e-08 | 1.3e-07 | 7.433E-08 | 5.034E-08 | 3.200E-08 | 1.300E-07 |
| 6201 | 311 | K311A | A | 5.7e-08 | 7.1e-08 | 7.4e-08 | 6.733E-08 | 9.074E-09 | 5.700E-08 | 7.400E-08 |
| 6202 | 311 | K311R | A | 3.8e-08 | 2.1e-08 | 4.9e-08 | 3.600E-08 | 1.411E-08 | 2.100E-08 | 4.900E-08 |
| 6203 | 311 | K311N | A | 2.8e-08 | 2.5e-08 | 5.7e-08 | 3.667E-08 | 1.767E-08 | 2.500E-08 | 5.700E-08 |
| 6204 | 311 | K311D | A | 7.2e-08 | 3.4e-08 | 3.5e-08 | 4.700E-08 | 2.166E-08 | 3.400E-08 | 7.200E-08 |

| | | | | | | | | | | |
|---|---|---|---|---|---|---|---|---|---|---|
| 6205 | 311 | K311C | A | 2.3e-08 | 1.2e-07 | 1e-07 | 8.100E-08 | 5.122E-08 | 2.300E-08 | 1.200E-07 |
| 6206 | 311 | K311E | A | 3.5e-08 | 3.3e-08 | 7.9e-08 | 4.900E-08 | 2.600E-08 | 3.300E-08 | 7.900E-08 |
| 6207 | 311 | K311Q | A | 1.8e-07 | 4.8e-08 | 2.4e-08 | 8.400E-08 | 8.400E-08 | 2.400E-08 | 1.800E-07 |
| 6208 | 311 | K311G | A | 6.1e-08 | 1.1e-07 | 2.1e-08 | 6.400E-08 | 4.458E-08 | 2.100E-08 | 1.100E-07 |
| 6209 | 311 | K311H | A | 4.2e-08 | 5.6e-08 | 4.8e-08 | 4.867E-08 | 7.024E-09 | 4.200E-08 | 5.600E-08 |
| 6210 | 311 | K311I | A | 4.1e-08 | 7.6e-08 | 2.8e-08 | 4.833E-08 | 2.483E-08 | 2.800E-08 | 7.600E-08 |
| 6211 | 311 | K311L | A | 1.5e-07 | 1.9e-08 | 5.5e-08 | 7.467E-08 | 6.768E-08 | 1.900E-08 | 1.500E-07 |
| 6212 | 311 | K311K | A | 1.8e-08 | 4.7e-08 | 5.3e-08 | 3.933E-08 | 1.872E-08 | 1.800E-08 | 5.300E-08 |
| 6213 | 311 | K311M | A | 1.3e-08 | 3.2e-08 | 7.3e-09 | 1.743E-08 | 1.293E-08 | 7.300E-09 | 3.200E-08 |
| 6214 | 311 | K311F | A | 2.1e-07 | 3.7e-08 | 6.6e-08 | 1.043E-07 | 9.265E-08 | 3.700E-08 | 2.100E-07 |
| 6215 | 311 | K311P | A | 7.4e-08 | 2.7e-08 | 5.3e-08 | 5.133E-08 | 2.354E-08 | 2.700E-08 | 7.400E-08 |
| 6216 | 311 | K311S | A | 2.4e-08 | 3.1e-08 | 6.3e-08 | 3.933E-08 | 2.079E-08 | 2.400E-08 | 6.300E-08 |
| 6217 | 311 | K311T | A | 1.8e-08 | 2.9e-08 | 9e-08 | 4.567E-08 | 3.879E-08 | 1.800E-08 | 9.000E-08 |
| 6218 | 311 | K311W | A | 7.8e-08 | 2.2e-08 | 2e-08 | 4.000E-08 | 3.292E-08 | 2.000E-08 | 7.800E-08 |
| 6219 | 311 | K311Y | A | 1.1e-07 | 3e-08 | 3.7e-08 | 5.900E-08 | 4.431E-08 | 3.000E-08 | 1.100E-07 |
| 6220 | 311 | K311V | A | 1.6e-08 | 3.7e-08 | 2.8e-08 | 2.700E-08 | 1.054E-08 | 1.600E-08 | 3.700E-08 |
| 6221 | 312 | C312A | A | 3.1e-08 | 3.4e-08 | 3.5e-08 | 3.333E-08 | 2.082E-09 | 3.100E-08 | 3.500E-08 |
| 6222 | 312 | C312R | A | 9.2e-08 | 4.9e-08 | 3.4e-08 | 5.833E-08 | 3.011E-08 | 3.400E-08 | 9.200E-08 |
| 6223 | 312 | C312N | A | 4.2e-08 | 8.2e-08 | 1.8e-08 | 4.733E-08 | 3.233E-08 | 1.800E-08 | 8.200E-08 |
| 6224 | 312 | C312D | A | 3.1e-08 | 5.2e-08 | 6.1e-08 | 4.800E-08 | 1.539E-08 | 3.100E-08 | 6.100E-08 |
| 6225 | 312 | C312C | A | 5.1e-08 | 4.2e-08 | 8.4e-08 | 5.900E-08 | 2.211E-08 | 4.200E-08 | 8.400E-08 |
| 6226 | 312 | C312E | A | 7e-08 | 2.9e-08 | 3e-08 | 4.300E-08 | 2.339E-08 | 2.900E-08 | 7.000E-08 |
| 6227 | 312 | C312Q | A | 2.5e-08 | 4.3e-08 | 3.3e-08 | 3.367E-08 | 9.018E-09 | 2.500E-08 | 4.300E-08 |
| 6228 | 312 | C312G | A | 1.4e-08 | 8.7e-08 | 4.8e-08 | 4.967E-08 | 3.653E-08 | 1.400E-08 | 8.700E-08 |
| 6229 | 312 | C312H | A | 1.6e-07 | 5.7e-08 | 1.2e-07 | 1.123E-07 | 5.193E-08 | 5.700E-08 | 1.600E-07 |
| 6230 | 312 | C312I | A | 3.8e-08 | 1.4e-07 | 3.3e-08 | 7.033E-08 | 6.038E-08 | 3.300E-08 | 1.400E-07 |
| 6231 | 312 | C312L | A | 4.7e-08 | 4e-08 | 5.3e-08 | 4.667E-08 | 6.506E-09 | 4.000E-08 | 5.300E-08 |
| 6232 | 312 | C312K | A | 7.2e-08 | 1.7e-08 | 4.6e-08 | 4.500E-08 | 2.751E-08 | 1.700E-08 | 7.200E-08 |
| 6233 | 312 | C312M | A | 4e-08 | 8.4e-08 | 2.2e-08 | 4.867E-08 | 3.190E-08 | 2.200E-08 | 8.400E-08 |
| 6234 | 312 | C312F | A | 1.3e-08 | 7.7e-08 | 6.7e-08 | 5.233E-08 | 3.443E-08 | 1.300E-08 | 7.700E-08 |
| 6235 | 312 | C312P | A | 3.8e-08 | 3.1e-08 | 1.7e-08 | 2.867E-08 | 1.069E-08 | 1.700E-08 | 3.800E-08 |
| 6236 | 312 | C312S | A | 5.8e-08 | 5.8e-08 | 5.1e-08 | 5.567E-08 | 4.041E-09 | 5.100E-08 | 5.800E-08 |
| 6237 | 312 | C312T | A | 8.8e-08 | 8.1e-08 | 4.1e-08 | 7.000E-08 | 2.536E-08 | 4.100E-08 | 8.800E-08 |
| 6238 | 312 | C312W | A | 7.6e-08 | 6.5e-08 | 4.4e-08 | 6.167E-08 | 1.626E-08 | 4.400E-08 | 7.600E-08 |
| 6239 | 312 | C312Y | A | 6.2e-08 | 3.3e-08 | 4e-08 | 4.500E-08 | 1.513E-08 | 3.300E-08 | 6.200E-08 |
| 6240 | 312 | C312V | A | 3.3e-08 | 6.3e-08 | 6.9e-08 | 5.500E-08 | 1.929E-08 | 3.300E-08 | 6.900E-08 |
| 6241 | 313 | S313A | A | 2.6e-08 | 4.6e-08 | 3e-07 | 1.240E-07 | 1.527E-07 | 2.600E-08 | 3.000E-07 |
| 6242 | 313 | S313R | A | 3.2e-08 | 3.3e-08 | 5.5e-08 | 4.000E-08 | 1.300E-08 | 3.200E-08 | 5.500E-08 |
| 6243 | 313 | S313N | A | 2.8e-08 | 4.8e-08 | 2.7e-08 | 3.433E-08 | 1.185E-08 | 2.700E-08 | 4.800E-08 |
| 6244 | 313 | S313D | A | 1.1e-07 | 9.3e-08 | 3.2e-08 | 7.833E-08 | 4.102E-08 | 3.200E-08 | 1.100E-07 |
| 6245 | 313 | S313C | A | 3.8e-08 | 1.6e-08 | 1.1e-08 | 2.167E-08 | 1.436E-08 | 1.100E-08 | 3.800E-08 |
| 6246 | 313 | S313E | A | 4.2e-08 | 1.6e-08 | 2.7e-08 | 2.833E-08 | 1.305E-08 | 1.600E-08 | 4.200E-08 |
| 6247 | 313 | S313Q | A | 1.8e-08 | 9e-08 | 2.2e-08 | 4.333E-08 | 4.046E-08 | 1.800E-08 | 9.000E-08 |
| 6248 | 313 | S313G | A | 9.5e-09 | 1.6e-08 | 8.9e-08 | 3.817E-08 | 4.414E-08 | 9.500E-09 | 8.900E-08 |
| 6249 | 313 | S313H | A | 1.6e-08 | 7.1e-08 | 3.5e-08 | 4.067E-08 | 2.793E-08 | 1.600E-08 | 7.100E-08 |
| 6250 | 313 | S313I | A | 2.8e-08 | 5.7e-09 | 5.2e-08 | 2.857E-08 | 2.316E-08 | 5.700E-09 | 5.200E-08 |
| 6251 | 313 | S313L | A | 9.5e-08 | 1.2e-07 | 4.2e-08 | 8.567E-08 | 3.983E-08 | 4.200E-08 | 1.200E-07 |
| 6252 | 313 | S313K | A | 1.9e-08 | 7.2e-08 | 4.7e-08 | 4.600E-08 | 2.651E-08 | 1.900E-08 | 7.200E-08 |
| 6253 | 313 | S313M | A | 5.4e-08 | 1e-07 | 7.6e-08 | 7.667E-08 | 2.301E-08 | 5.400E-08 | 1.000E-07 |
| 6254 | 313 | S313F | A | 3.8e-08 | 3.8e-08 | 4.4e-08 | 4.000E-08 | 3.464E-09 | 3.800E-08 | 4.400E-08 |
| 6255 | 313 | S313P | A | 1.5e-07 | 4.7e-08 | 1.8e-08 | 7.167E-08 | 6.937E-08 | 1.800E-08 | 1.500E-07 |
| 6256 | 313 | S313S | A | 2.8e-08 | 4e-08 | 3.3e-08 | 3.367E-08 | 6.028E-09 | 2.800E-08 | 4.000E-08 |
| 6257 | 313 | S313T | A | 3.6e-08 | 2.1e-07 | 5.3e-08 | 9.967E-08 | 9.593E-08 | 3.600E-08 | 2.100E-07 |
| 6258 | 313 | S313W | A | 6e-08 | 3.9e-08 | 5.9e-08 | 5.267E-08 | 1.185E-08 | 3.900E-08 | 6.000E-08 |
| 6259 | 313 | S313Y | A | 2.5e-08 | 2.2e-08 | 6.6e-08 | 3.767E-08 | 2.458E-08 | 2.200E-08 | 6.600E-08 |
| 6260 | 313 | S313V | A | 2.3e-08 | 4.1e-08 | 7.7e-08 | 4.700E-08 | 2.750E-08 | 2.300E-08 | 7.700E-08 |
| 6261 | 314 | K314A | A | 2.4e-08 | 7e-08 | 7.9e-08 | 5.767E-08 | 2.950E-08 | 2.400E-08 | 7.900E-08 |
| 6262 | 314 | K314R | A | 1.7e-07 | 5.1e-08 | 4.5e-08 | 8.867E-08 | 7.050E-08 | 4.500E-08 | 1.700E-07 |
| 6263 | 314 | K314N | A | 2.2e-08 | 1e-07 | 6.1e-08 | 6.100E-08 | 3.900E-08 | 2.200E-08 | 1.000E-07 |
| 6264 | 314 | K314D | A | 8e-08 | 7.4e-08 | 7.5e-08 | 7.633E-08 | 3.215E-09 | 7.400E-08 | 8.000E-08 |
| 6265 | 314 | K314C | A | 5.7e-08 | 7.7e-08 | 9e-08 | 7.467E-08 | 1.662E-08 | 5.700E-08 | 9.000E-08 |
| 6266 | 314 | K314E | A | 7.1e-08 | 1.3e-07 | 6.1e-08 | 8.733E-08 | 3.729E-08 | 6.100E-08 | 1.300E-07 |
| 6267 | 314 | K314Q | A | 1.6e-07 | 7.9e-08 | 5.9e-08 | 9.933E-08 | 5.348E-08 | 5.900E-08 | 1.600E-07 |
| 6268 | 314 | K314G | A | 5.2e-08 | 1e-08 | 7.2e-08 | 4.467E-08 | 3.164E-08 | 1.000E-08 | 7.200E-08 |
| 6269 | 314 | K314H | A | 3.2e-08 | 6.6e-08 | 4e-08 | 4.600E-08 | 1.778E-08 | 3.200E-08 | 6.600E-08 |
| 6270 | 314 | K314I | A | 1.1e-08 | 3.4e-08 | 2.5e-08 | 2.333E-08 | 1.159E-08 | 1.100E-08 | 3.400E-08 |
| 6271 | 314 | K314L | A | 3.9e-08 | 3.1e-08 | 3.3e-08 | 3.433E-08 | 4.163E-09 | 3.100E-08 | 3.900E-08 |
| 6272 | 314 | K314K | A | 5e-08 | 5.1e-08 | 3e-08 | 4.367E-08 | 1.185E-08 | 3.000E-08 | 5.100E-08 |
| 6273 | 314 | K314M | A | 6.8e-08 | 3.6e-08 | 6.9e-08 | 5.767E-08 | 1.877E-08 | 3.600E-08 | 6.900E-08 |
| 6274 | 314 | K314F | A | 3.9e-08 | 4.1e-08 | 5.2e-08 | 4.400E-08 | 7.000E-09 | 3.900E-08 | 5.200E-08 |
| 6275 | 314 | K314P | A | 1.1e-08 | 9.9e-08 | 1.4e-07 | 8.333E-08 | 6.591E-08 | 1.100E-08 | 1.400E-07 |
| 6276 | 314 | K314S | A | 3e-08 | 2.6e-08 | 1.1e-07 | 5.533E-08 | 4.738E-08 | 2.600E-08 | 1.100E-07 |
| 6277 | 314 | K314T | A | 7.6e-08 | 9.5e-08 | 1.5e-08 | 6.200E-08 | 4.180E-08 | 1.500E-08 | 9.500E-08 |
| 6278 | 314 | K314W | A | 2e-08 | 7.1e-08 | 7.1e-08 | 5.400E-08 | 2.944E-08 | 2.000E-08 | 7.100E-08 |
| 6279 | 314 | K314Y | A | 8e-08 | 6.1e-08 | 5.3e-08 | 6.467E-08 | 1.387E-08 | 5.300E-08 | 8.000E-08 |
| 6280 | 314 | K314V | A | 5.9e-08 | 2.6e-08 | 1e-08 | 3.167E-08 | 2.499E-08 | 1.000E-08 | 5.900E-08 |
| 6281 | 315 | P315A | A | 3.1e-08 | 5.7e-08 | 5.2e-08 | 4.667E-08 | 1.380E-08 | 3.100E-08 | 5.700E-08 |
| 6282 | 315 | P315R | A | 5.3e-08 | 2.6e-08 | 2.9e-08 | 3.600E-08 | 1.480E-08 | 2.600E-08 | 5.300E-08 |
| 6283 | 315 | P315N | A | 3.5e-08 | 7.4e-08 | 3.9e-08 | 4.933E-08 | 2.146E-08 | 3.500E-08 | 7.400E-08 |
| 6284 | 315 | P315D | A | 5.6e-08 | 6.1e-08 | 6.7e-08 | 6.133E-08 | 5.508E-09 | 5.600E-08 | 6.700E-08 |
| 6285 | 315 | P315C | A | 2.5e-08 | 6.6e-08 | 5.5e-08 | 4.867E-08 | 2.122E-08 | 2.500E-08 | 6.600E-08 |
| 6286 | 315 | P315E | A | 3.1e-08 | 7.7e-08 | 2.1e-08 | 4.300E-08 | 2.987E-08 | 2.100E-08 | 7.700E-08 |
| 6287 | 315 | P315Q | A | 5.2e-08 | 3e-08 | 8e-08 | 5.400E-08 | 2.506E-08 | 3.000E-08 | 8.000E-08 |
| 6288 | 315 | P315G | A | 2.4e-08 | 4.7e-08 | 4.3e-08 | 3.800E-08 | 1.229E-08 | 2.400E-08 | 4.700E-08 |
| 6289 | 315 | P315H | A | 2.5e-08 | 5e-08 | 1.7e-08 | 3.067E-08 | 1.721E-08 | 1.700E-08 | 5.000E-08 |

| | | | | | | | | | | |
|---|---|---|---|---|---|---|---|---|---|---|
| 6290 | 315 | P315I | A | 1.6e-07 | 4.2e-08 | 5.5e-08 | 8.567E-08 | 6.470E-08 | 4.200E-08 | 1.600E-07 |
| 6291 | 315 | P315L | A | 3.6e-08 | 3.3e-08 | 3.5e-08 | 3.467E-08 | 1.528E-09 | 3.300E-08 | 3.600E-08 |
| 6292 | 315 | P315K | A | 3.5e-08 | 3.8e-08 | 2.6e-08 | 3.300E-08 | 6.245E-09 | 2.600E-08 | 3.800E-08 |
| 6293 | 315 | P315M | A | 8.7e-08 | 9.9e-08 | 2.5e-08 | 7.033E-08 | 3.972E-08 | 2.500E-08 | 9.900E-08 |
| 6294 | 315 | P315F | A | 4.6e-08 | 5.8e-08 | 2.1e-08 | 4.167E-08 | 1.888E-08 | 2.100E-08 | 5.800E-08 |
| 6295 | 315 | P315P | A | 6.2e-08 | 2.1e-08 | 3.6e-08 | 3.967E-08 | 2.074E-08 | 2.100E-08 | 6.200E-08 |
| 6296 | 315 | P315S | A | 4.6e-08 | 5e-08 | 4.3e-08 | 4.633E-08 | 3.512E-09 | 4.300E-08 | 5.000E-08 |
| 6297 | 315 | P315T | A | 3.4e-08 | 2.5e-08 | 4.4e-08 | 3.433E-08 | 9.504E-09 | 2.500E-08 | 4.400E-08 |
| 6298 | 315 | P315W | A | 6.2e-08 | 2.6e-08 | 3.3e-08 | 4.033E-08 | 1.909E-08 | 2.600E-08 | 6.200E-08 |
| 6299 | 315 | P315Y | A | 7.6e-08 | 7.3e-08 | 8.7e-08 | 7.867E-08 | 7.371E-09 | 7.300E-08 | 8.700E-08 |
| 6300 | 315 | P315V | A | 7.8e-08 | 2.8e-08 | 6.4e-08 | 5.667E-08 | 2.579E-08 | 2.800E-08 | 7.800E-08 |
| 6301 | 316 | C316A | A | 4.5e-08 | 4.9e-08 | 1.1e-07 | 6.800E-08 | 3.643E-08 | 4.500E-08 | 1.100E-07 |
| 6302 | 316 | C316R | A | 6e-08 | 5.9e-08 | 5.3e-08 | 5.733E-08 | 3.786E-09 | 5.300E-08 | 6.000E-08 |
| 6303 | 316 | C316N | A | 1.5e-07 | 6e-08 | 1.2e-08 | 7.400E-08 | 7.006E-08 | 1.200E-08 | 1.500E-07 |
| 6304 | 316 | C316D | A | 3.9e-08 | 2.8e-08 | 1.1e-07 | 5.900E-08 | 4.451E-08 | 2.800E-08 | 1.100E-07 |
| 6305 | 316 | C316C | A | 7.8e-08 | 6.8e-08 | 3.2e-08 | 5.933E-08 | 2.419E-08 | 3.200E-08 | 7.800E-08 |
| 6306 | 316 | C316E | A | 9.4e-08 | 5.4e-08 | 3.3e-08 | 6.033E-08 | 3.099E-08 | 3.300E-08 | 9.400E-08 |
| 6307 | 316 | C316Q | A | 6.2e-08 | 2.7e-08 | 8.7e-08 | 5.867E-08 | 3.014E-08 | 2.700E-08 | 8.700E-08 |
| 6308 | 316 | C316G | A | 2.3e-08 | 4.5e-08 | 4e-08 | 3.600E-08 | 1.153E-08 | 2.300E-08 | 4.500E-08 |
| 6309 | 316 | C316H | A | 3.1e-08 | 8.5e-08 | 4.9e-08 | 5.500E-08 | 2.750E-08 | 3.100E-08 | 8.500E-08 |
| 6310 | 316 | C316I | A | 1.5e-07 | 8.6e-08 | 3.4e-08 | 9.000E-08 | 5.810E-08 | 3.400E-08 | 1.500E-07 |
| 6311 | 316 | C316L | A | 1.1e-08 | 1.3e-08 | 3.9e-08 | 2.100E-08 | 1.562E-08 | 1.100E-08 | 3.900E-08 |
| 6312 | 316 | C316K | A | 2.6e-08 | 1e-07 | 8e-08 | 6.867E-08 | 3.828E-08 | 2.600E-08 | 1.000E-07 |
| 6313 | 316 | C316M | A | 4.4e-08 | 1.3e-07 | 3.5e-08 | 6.967E-08 | 5.244E-08 | 3.500E-08 | 1.300E-07 |
| 6314 | 316 | C316F | A | 2.6e-08 | 3.3e-08 | 6e-08 | 3.967E-08 | 1.795E-08 | 2.600E-08 | 6.000E-08 |
| 6315 | 316 | C316P | A | 6.3e-08 | 3.7e-08 | 1.2e-07 | 7.333E-08 | 4.245E-08 | 3.700E-08 | 1.200E-07 |
| 6316 | 316 | C316S | A | 7e-08 | 3.8e-08 | 4.8e-08 | 5.200E-08 | 1.637E-08 | 3.800E-08 | 7.000E-08 |
| 6317 | 316 | C316T | A | 7.1e-08 | 2.2e-08 | 2.8e-08 | 4.033E-08 | 2.673E-08 | 2.200E-08 | 7.100E-08 |
| 6318 | 316 | C316W | A | 1.8e-08 | 9.3e-08 | 8.7e-08 | 6.600E-08 | 4.168E-08 | 1.800E-08 | 9.300E-08 |
| 6319 | 316 | C316Y | A | 4.9e-08 | 3.6e-08 | 8.2e-08 | 5.567E-08 | 2.371E-08 | 3.600E-08 | 8.200E-08 |
| 6320 | 316 | C316V | A | 2.6e-08 | 4.6e-08 | 3.3e-08 | 3.500E-08 | 1.015E-08 | 2.600E-08 | 4.600E-08 |
| 6321 | 317 | A317A | A | 2.8e-08 | 2.3e-08 | 7.3e-08 | 4.133E-08 | 2.754E-08 | 2.300E-08 | 7.300E-08 |
| 6322 | 317 | A317R | A | 5.7e-08 | 1.1e-07 | 2.1e-08 | 6.267E-08 | 4.477E-08 | 2.100E-08 | 1.100E-07 |
| 6323 | 317 | A317N | A | 4.4e-08 | 6.9e-08 | 1.6e-08 | 4.300E-08 | 2.651E-08 | 1.600E-08 | 6.900E-08 |
| 6324 | 317 | A317D | A | 3.8e-08 | 3.1e-08 | 3.1e-08 | 3.333E-08 | 4.041E-09 | 3.100E-08 | 3.800E-08 |
| 6325 | 317 | A317C | A | 5.6e-08 | 4.7e-08 | 9.1e-08 | 6.467E-08 | 2.325E-08 | 4.700E-08 | 9.100E-08 |
| 6326 | 317 | A317E | A | 1.2e-08 | 4.5e-08 | 4.7e-08 | 3.467E-08 | 1.966E-08 | 1.200E-08 | 4.700E-08 |
| 6327 | 317 | A317Q | A | 3.8e-08 | 1.8e-08 | 5.6e-08 | 3.733E-08 | 1.901E-08 | 1.800E-08 | 5.600E-08 |
| 6328 | 317 | A317G | A | 4.4e-08 | 4.6e-08 | 6.7e-08 | 5.233E-08 | 1.274E-08 | 4.400E-08 | 6.700E-08 |
| 6329 | 317 | A317H | A | 3.7e-08 | 2.8e-08 | 3e-08 | 3.167E-08 | 4.726E-09 | 2.800E-08 | 3.700E-08 |
| 6330 | 317 | A317I | A | 4.8e-08 | 4.3e-08 | 5e-08 | 4.700E-08 | 3.606E-09 | 4.300E-08 | 5.000E-08 |
| 6331 | 317 | A317L | A | 5.6e-08 | 3.5e-08 | 9e-08 | 6.033E-08 | 2.775E-08 | 3.500E-08 | 9.000E-08 |
| 6332 | 317 | A317K | A | 3.6e-08 | 6.2e-08 | 4.2e-08 | 4.667E-08 | 1.361E-08 | 3.600E-08 | 6.200E-08 |
| 6333 | 317 | A317M | A | 3.6e-08 | 3.9e-08 | 2.4e-08 | 3.300E-08 | 7.937E-09 | 2.400E-08 | 3.900E-08 |
| 6334 | 317 | A317F | A | 1.8e-08 | 4.5e-08 | 4.9e-08 | 3.733E-08 | 1.686E-08 | 1.800E-08 | 4.900E-08 |
| 6335 | 317 | A317P | A | 6.5e-08 | 6.2e-08 | 1.7e-08 | 4.800E-08 | 2.689E-08 | 1.700E-08 | 6.500E-08 |
| 6336 | 317 | A317S | A | 2.2e-08 | 1.2e-08 | 1.9e-08 | 1.767E-08 | 5.132E-09 | 1.200E-08 | 2.200E-08 |
| 6337 | 317 | A317T | A | 6.1e-08 | 2.6e-08 | 2.4e-08 | 3.700E-08 | 2.081E-08 | 2.400E-08 | 6.100E-08 |
| 6338 | 317 | A317W | A | 5.9e-08 | 7.7e-08 | 8.4e-08 | 7.333E-08 | 1.290E-08 | 5.900E-08 | 8.400E-08 |
| 6339 | 317 | A317Y | A | 1.5e-07 | 4.9e-08 | 5.9e-08 | 8.600E-08 | 5.565E-08 | 4.900E-08 | 1.500E-07 |
| 6340 | 317 | A317V | A | 3.5e-08 | 5.8e-08 | 1.6e-08 | 3.633E-08 | 2.103E-08 | 1.600E-08 | 5.800E-08 |
| 6341 | 318 | R318A | A | 5.5e-08 | 3.4e-08 | 9.4e-08 | 6.100E-08 | 3.045E-08 | 3.400E-08 | 9.400E-08 |
| 6342 | 318 | R318R | A | 1e-07 | 5e-08 | 5.3e-08 | 6.767E-08 | 2.804E-08 | 5.000E-08 | 1.000E-07 |
| 6343 | 318 | R318N | A | 6.9e-08 | 6.8e-08 | 4.5e-08 | 6.067E-08 | 1.358E-08 | 4.500E-08 | 6.900E-08 |
| 6344 | 318 | R318D | A | 5.9e-08 | 1.5e-07 | 6e-08 | 8.967E-08 | 5.225E-08 | 5.900E-08 | 1.500E-07 |
| 6345 | 318 | R318C | A | 6.9e-08 | 4.8e-08 | 3.8e-08 | 5.167E-08 | 1.582E-08 | 3.800E-08 | 6.900E-08 |
| 6346 | 318 | R318E | A | 1.2e-07 | 6.5e-08 | 8.2e-08 | 8.900E-08 | 2.816E-08 | 6.500E-08 | 1.200E-07 |
| 6347 | 318 | R318Q | A | 7.8e-08 | 4.9e-08 | 6.1e-08 | 6.267E-08 | 1.457E-08 | 4.900E-08 | 7.800E-08 |
| 6348 | 318 | R318G | A | 7.9e-08 | 1.4e-07 | 2.2e-08 | 8.033E-08 | 5.901E-08 | 2.200E-08 | 1.400E-07 |
| 6349 | 318 | R318H | A | 5.8e-08 | 8.2e-08 | 5.6e-08 | 6.533E-08 | 1.447E-08 | 5.600E-08 | 8.200E-08 |
| 6350 | 318 | R318I | A | 3.3e-08 | 3.6e-08 | 3.6e-08 | 3.500E-08 | 1.732E-09 | 3.300E-08 | 3.600E-08 |
| 6351 | 318 | R318L | A | 4.8e-08 | 4.7e-08 | 1.7e-08 | 3.733E-08 | 1.762E-08 | 1.700E-08 | 4.800E-08 |
| 6352 | 318 | R318K | A | 6.1e-08 | 5.3e-08 | 6.8e-08 | 6.067E-08 | 7.506E-09 | 5.300E-08 | 6.800E-08 |
| 6353 | 318 | R318M | A | 2.6e-08 | 9.5e-08 | 1.8e-08 | 4.633E-08 | 4.234E-08 | 1.800E-08 | 9.500E-08 |
| 6354 | 318 | R318F | A | 6.6e-08 | 8e-08 | 4.9e-08 | 6.500E-08 | 1.552E-08 | 4.900E-08 | 8.000E-08 |
| 6355 | 318 | R318P | A | 5.5e-08 | 1.1e-07 | 3.2e-08 | 6.567E-08 | 4.008E-08 | 3.200E-08 | 1.100E-07 |
| 6356 | 318 | R318S | A | 2.3e-08 | 9e-08 | 3.8e-08 | 5.033E-08 | 3.516E-08 | 2.300E-08 | 9.000E-08 |
| 6357 | 318 | R318T | A | 1.3e-08 | 6.2e-08 | 1e-07 | 5.833E-08 | 4.362E-08 | 1.300E-08 | 1.000E-07 |
| 6358 | 318 | R318W | A | 8.9e-08 | 3.1e-08 | 1e-07 | 7.333E-08 | 3.707E-08 | 3.100E-08 | 1.000E-07 |
| 6359 | 318 | R318Y | A | 1.2e-07 | 1.2e-07 | 2e-08 | 8.667E-08 | 5.774E-08 | 2.000E-08 | 1.200E-07 |
| 6360 | 318 | R318V | A | 6.7e-08 | 4.2e-08 | 3.7e-08 | 4.867E-08 | 1.607E-08 | 3.700E-08 | 6.700E-08 |
| 6361 | 319 | V319A | A | 1.8e-08 | 3.4e-08 | 1.5e-08 | 2.233E-08 | 1.021E-08 | 1.500E-08 | 3.400E-08 |
| 6362 | 319 | V319R | A | 5.8e-08 | 4.5e-08 | 4.6e-08 | 4.967E-08 | 7.234E-09 | 4.500E-08 | 5.800E-08 |
| 6363 | 319 | V319N | A | 2.1e-08 | 6.9e-08 | 4.5e-08 | 4.500E-08 | 2.400E-08 | 2.100E-08 | 6.900E-08 |
| 6364 | 319 | V319D | A | 4.2e-08 | 2.2e-08 | 4e-08 | 3.467E-08 | 1.102E-08 | 2.200E-08 | 4.200E-08 |
| 6365 | 319 | V319C | A | 6.8e-08 | 2.1e-08 | 3.1e-08 | 4.000E-08 | 2.476E-08 | 2.100E-08 | 6.800E-08 |
| 6366 | 319 | V319E | A | 2.9e-08 | 2.6e-08 | 3.6e-08 | 3.033E-08 | 5.132E-09 | 2.600E-08 | 3.600E-08 |
| 6367 | 319 | V319Q | A | 4.4e-08 | 4e-08 | 5.6e-08 | 4.667E-08 | 8.327E-09 | 4.000E-08 | 5.600E-08 |
| 6368 | 319 | V319G | A | 4.2e-08 | 4.8e-08 | 1.5e-07 | 8.000E-08 | 6.070E-08 | 4.200E-08 | 1.500E-07 |
| 6369 | 319 | V319H | A | 1.2e-08 | 6.5e-08 | 8.2e-08 | 5.300E-08 | 3.651E-08 | 1.200E-08 | 8.200E-08 |
| 6370 | 319 | V319I | A | 6.8e-08 | 5.6e-08 | 3.5e-08 | 5.300E-08 | 1.670E-08 | 3.500E-08 | 6.800E-08 |
| 6371 | 319 | V319L | A | 3.9e-08 | 1e-07 | 5.4e-08 | 6.433E-08 | 3.179E-08 | 3.900E-08 | 1.000E-07 |
| 6372 | 319 | V319K | A | 2.2e-08 | 4.1e-08 | 8.6e-08 | 4.967E-08 | 3.287E-08 | 2.200E-08 | 8.600E-08 |
| 6373 | 319 | V319M | A | 2.7e-08 | 1.7e-08 | 4e-08 | 2.800E-08 | 1.153E-08 | 1.700E-08 | 4.000E-08 |
| 6374 | 319 | V319F | A | 5.1e-08 | 5.6e-08 | 5.7e-08 | 5.467E-08 | 3.215E-09 | 5.100E-08 | 5.700E-08 |

| | | | | | | | | | | |
|---|---|---|---|---|---|---|---|---|---|---|
| 6375 | 319 | V319P | A | 5.8e-08 | 2.4e-08 | 4e-08 | 4.067E-08 | 1.701E-08 | 2.400E-08 | 5.800E-08 |
| 6376 | 319 | V319S | A | 4.4e-08 | 4.4e-08 | 4.3e-08 | 4.367E-08 | 5.774E-10 | 4.300E-08 | 4.400E-08 |
| 6377 | 319 | V319T | A | 1.1e-07 | 1e-07 | 2.3e-08 | 7.767E-08 | 4.761E-08 | 2.300E-08 | 1.100E-07 |
| 6378 | 319 | V319W | A | 2.4e-08 | 5.3e-08 | 3.7e-08 | 3.800E-08 | 1.453E-08 | 2.400E-08 | 5.300E-08 |
| 6379 | 319 | V319Y | A | 1.9e-08 | 6e-08 | 3.5e-08 | 3.800E-08 | 2.066E-08 | 1.900E-08 | 6.000E-08 |
| 6380 | 319 | V319V | A | 5.9e-08 | 3.7e-08 | 2.1e-08 | 3.900E-08 | 1.908E-08 | 2.100E-08 | 5.900E-08 |
| 6381 | 320 | C320A | A | 2.8e-08 | 3.1e-08 | 1.8e-08 | 2.567E-08 | 6.807E-09 | 1.800E-08 | 3.100E-08 |
| 6382 | 320 | C320R | A | 2e-08 | 4.2e-08 | 5.1e-08 | 3.767E-08 | 1.595E-08 | 2.000E-08 | 5.100E-08 |
| 6383 | 320 | C320N | A | 1.7e-08 | 2.2e-08 | 6.9e-08 | 3.600E-08 | 2.869E-08 | 1.700E-08 | 6.900E-08 |
| 6384 | 320 | C320D | A | 7.2e-08 | 4.6e-08 | 4.3e-08 | 5.367E-08 | 1.595E-08 | 4.300E-08 | 7.200E-08 |
| 6385 | 320 | C320C | A | 8.8e-08 | 5.9e-08 | 5.7e-08 | 6.800E-08 | 1.735E-08 | 5.700E-08 | 8.800E-08 |
| 6386 | 320 | C320E | A | 2.3e-08 | 4.1e-08 | 9e-08 | 5.133E-08 | 3.467E-08 | 2.300E-08 | 9.000E-08 |
| 6387 | 320 | C320Q | A | 3.7e-08 | 9.8e-08 | 7.4e-08 | 6.967E-08 | 3.073E-08 | 3.700E-08 | 9.800E-08 |
| 6388 | 320 | C320G | A | 9.4e-08 | 7.5e-09 | 1.8e-08 | 3.983E-08 | 4.720E-08 | 7.500E-09 | 9.400E-08 |
| 6389 | 320 | C320H | A | 3e-08 | 1.1e-07 | 9e-09 | 4.967E-08 | 5.329E-08 | 9.000E-09 | 1.100E-07 |
| 6390 | 320 | C320I | A | 2.1e-08 | 4.1e-08 | 3.1e-08 | 3.100E-08 | 1.000E-08 | 2.100E-08 | 4.100E-08 |
| 6391 | 320 | C320L | A | 4.7e-08 | 2.9e-08 | 5e-08 | 4.200E-08 | 1.136E-08 | 2.900E-08 | 5.000E-08 |
| 6392 | 320 | C320K | A | 2.7e-08 | 4.1e-08 | 1.7e-08 | 2.833E-08 | 1.206E-08 | 1.700E-08 | 4.100E-08 |
| 6393 | 320 | C320M | A | 5.6e-08 | 5.2e-08 | 5e-08 | 5.267E-08 | 3.055E-09 | 5.000E-08 | 5.600E-08 |
| 6394 | 320 | C320F | A | 1.4e-08 | 3e-08 | 5.9e-08 | 3.433E-08 | 2.281E-08 | 1.400E-08 | 5.900E-08 |
| 6395 | 320 | C320P | A | 1e-07 | 7.2e-08 | 3.4e-08 | 6.867E-08 | 3.313E-08 | 3.400E-08 | 1.000E-07 |
| 6396 | 320 | C320S | A | 1.8e-08 | 5.8e-08 | 4.4e-08 | 4.000E-08 | 2.030E-08 | 1.800E-08 | 5.800E-08 |
| 6397 | 320 | C320T | A | 5.2e-08 | 5.5e-08 | 5.8e-08 | 5.500E-08 | 3.000E-09 | 5.200E-08 | 5.800E-08 |
| 6398 | 320 | C320W | A | 3.2e-08 | 5.6e-08 | 1.5e-08 | 3.433E-08 | 2.060E-08 | 1.500E-08 | 5.600E-08 |
| 6399 | 320 | C320Y | A | 4.6e-08 | 3.7e-08 | 4.1e-08 | 4.133E-08 | 4.509E-09 | 3.700E-08 | 4.600E-08 |
| 6400 | 320 | C320V | A | 5.3e-08 | 4.2e-08 | 5.5e-08 | 5.000E-08 | 7.000E-09 | 4.200E-08 | 5.500E-08 |
| 6401 | 321 | Y321A | A | 4.9e-08 | 5.7e-08 | 1.3e-08 | 3.967E-08 | 2.344E-08 | 1.300E-08 | 5.700E-08 |
| 6402 | 321 | Y321R | A | 2.3e-08 | 2.3e-08 | 4.5e-08 | 3.033E-08 | 1.270E-08 | 2.300E-08 | 4.500E-08 |
| 6403 | 321 | Y321N | A | 2.5e-08 | 2.6e-08 | 1.5e-08 | 2.200E-08 | 6.083E-09 | 1.500E-08 | 2.600E-08 |
| 6404 | 321 | Y321D | A | 1.1e-07 | 2.6e-08 | 3.3e-08 | 5.633E-08 | 4.661E-08 | 2.600E-08 | 1.100E-07 |
| 6405 | 321 | Y321C | A | 1.3e-07 | 4.9e-08 | 4.6e-08 | 7.500E-08 | 4.766E-08 | 4.600E-08 | 1.300E-07 |
| 6406 | 321 | Y321E | A | 2.2e-08 | 1.1e-08 | 3.7e-08 | 2.333E-08 | 1.305E-08 | 1.100E-08 | 3.700E-08 |
| 6407 | 321 | Y321Q | A | 1.2e-08 | 3.2e-08 | 2.2e-08 | 2.200E-08 | 1.000E-08 | 1.200E-08 | 3.200E-08 |
| 6408 | 321 | Y321G | A | 5.8e-08 | 9e-08 | 6.9e-08 | 7.233E-08 | 1.626E-08 | 5.800E-08 | 9.000E-08 |
| 6409 | 321 | Y321H | A | 1.4e-07 | 1.4e-08 | 3.4e-08 | 6.267E-08 | 6.772E-08 | 1.400E-08 | 1.400E-07 |
| 6410 | 321 | Y321I | A | 1.5e-08 | 8.7e-08 | 3.2e-08 | 4.467E-08 | 3.763E-08 | 1.500E-08 | 8.700E-08 |
| 6411 | 321 | Y321L | A | 1.1e-07 | 2.6e-08 | 2.7e-08 | 5.433E-08 | 4.821E-08 | 2.600E-08 | 1.100E-07 |
| 6412 | 321 | Y321K | A | 7.5e-08 | 1.3e-08 | 2.3e-08 | 3.700E-08 | 3.329E-08 | 1.300E-08 | 7.500E-08 |
| 6413 | 321 | Y321M | A | 2e-08 | 6e-08 | 2.1e-08 | 3.367E-08 | 2.281E-08 | 2.000E-08 | 6.000E-08 |
| 6414 | 321 | Y321F | A | 5.7e-08 | 8.8e-08 | 2.9e-08 | 5.800E-08 | 2.951E-08 | 2.900E-08 | 8.800E-08 |
| 6415 | 321 | Y321P | A | 5.7e-08 | 1.2e-07 | 9.1e-08 | 8.933E-08 | 3.153E-08 | 5.700E-08 | 1.200E-07 |
| 6416 | 321 | Y321S | A | 6.5e-08 | 5.4e-08 | 6.6e-08 | 6.167E-08 | 6.658E-09 | 5.400E-08 | 6.600E-08 |
| 6417 | 321 | Y321T | A | 6.3e-08 | 3.4e-08 | 2.3e-08 | 4.000E-08 | 2.066E-08 | 2.300E-08 | 6.300E-08 |
| 6418 | 321 | Y321W | A | 4.7e-08 | 1.2e-07 | 1.5e-08 | 6.067E-08 | 5.382E-08 | 1.500E-08 | 1.200E-07 |
| 6419 | 321 | Y321Y | A | 1.4e-08 | 2.3e-08 | 7.6e-08 | 3.767E-08 | 3.350E-08 | 1.400E-08 | 7.600E-08 |
| 6420 | 321 | Y321V | A | 5.9e-08 | 5.5e-08 | 6.2e-08 | 5.867E-08 | 3.512E-09 | 5.500E-08 | 6.200E-08 |
| 6421 | 322 | G322A | A | 2.7e-08 | 3.1e-08 | 2.5e-08 | 2.767E-08 | 3.055E-09 | 2.500E-08 | 3.100E-08 |
| 6422 | 322 | G322R | A | 1.4e-08 | 2.1e-08 | 1.8e-08 | 1.767E-08 | 3.512E-09 | 1.400E-08 | 2.100E-08 |
| 6423 | 322 | G322N | A | 3.9e-08 | 9.8e-08 | 1.2e-07 | 8.567E-08 | 4.188E-08 | 3.900E-08 | 1.200E-07 |
| 6424 | 322 | G322D | A | 3e-08 | 1.2e-08 | 4.3e-08 | 2.833E-08 | 1.557E-08 | 1.200E-08 | 4.300E-08 |
| 6425 | 322 | G322C | A | 8.2e-08 | 5.3e-08 | 5.9e-08 | 6.467E-08 | 1.531E-08 | 5.300E-08 | 8.200E-08 |
| 6426 | 322 | G322E | A | 3.2e-08 | 3.8e-08 | 6.3e-08 | 4.433E-08 | 1.644E-08 | 3.200E-08 | 6.300E-08 |
| 6427 | 322 | G322Q | A | 6.8e-08 | 3.5e-08 | 2.9e-08 | 4.400E-08 | 2.100E-08 | 2.900E-08 | 6.800E-08 |
| 6428 | 322 | G322G | A | 2.8e-08 | 2.6e-08 | 3.4e-08 | 2.933E-08 | 4.163E-09 | 2.600E-08 | 3.400E-08 |
| 6429 | 322 | G322H | A | 8.7e-08 | 7.5e-08 | 5.1e-08 | 7.100E-08 | 1.833E-08 | 5.100E-08 | 8.700E-08 |
| 6430 | 322 | G322I | A | 6.1e-08 | 6.4e-08 | 8.6e-08 | 7.033E-08 | 1.365E-08 | 6.100E-08 | 8.600E-08 |
| 6431 | 322 | G322L | A | 3.6e-08 | 7.6e-08 | 2.2e-08 | 4.467E-08 | 2.802E-08 | 2.200E-08 | 7.600E-08 |
| 6432 | 322 | G322K | A | 4.6e-08 | 5.7e-08 | 8e-08 | 6.100E-08 | 1.735E-08 | 4.600E-08 | 8.000E-08 |
| 6433 | 322 | G322M | A | 4.8e-08 | 1.6e-08 | 6.9e-08 | 4.433E-08 | 2.669E-08 | 1.600E-08 | 6.900E-08 |
| 6434 | 322 | G322F | A | 7.8e-08 | 5.2e-08 | 3.4e-08 | 5.467E-08 | 2.212E-08 | 3.400E-08 | 7.800E-08 |
| 6435 | 322 | G322P | A | 5.6e-08 | 2.7e-08 | 9.5e-08 | 5.933E-08 | 3.412E-08 | 2.700E-08 | 9.500E-08 |
| 6436 | 322 | G322S | A | 3.8e-08 | 3.7e-08 | 9.7e-08 | 5.733E-08 | 3.436E-08 | 3.700E-08 | 9.700E-08 |
| 6437 | 322 | G322T | A | 4.8e-08 | 5.6e-08 | 6.3e-08 | 5.567E-08 | 7.506E-09 | 4.800E-08 | 6.300E-08 |
| 6438 | 322 | G322W | A | 9.7e-09 | 5.2e-08 | 3.2e-08 | 3.123E-08 | 2.116E-08 | 9.700E-09 | 5.200E-08 |
| 6439 | 322 | G322Y | A | 4.6e-08 | 1.2e-08 | 5.1e-08 | 3.633E-08 | 2.122E-08 | 1.200E-08 | 5.100E-08 |
| 6440 | 322 | G322V | A | 6.3e-08 | 6.7e-08 | 4.2e-08 | 5.733E-08 | 1.343E-08 | 4.200E-08 | 6.700E-08 |
| 6441 | 323 | L323A | A | 1.3e-08 | 7.1e-08 | 7e-08 | 5.133E-08 | 3.320E-08 | 1.300E-08 | 7.100E-08 |
| 6442 | 323 | L323R | A | 5.8e-08 | 6.5e-08 | 3.5e-08 | 5.267E-08 | 1.570E-08 | 3.500E-08 | 6.500E-08 |
| 6443 | 323 | L323N | A | 5.3e-08 | 5e-08 | 1.8e-08 | 4.033E-08 | 1.940E-08 | 1.800E-08 | 5.300E-08 |
| 6444 | 323 | L323D | A | 3.4e-08 | 4e-08 | 1.9e-08 | 3.100E-08 | 1.082E-08 | 1.900E-08 | 4.000E-08 |
| 6445 | 323 | L323C | A | 5.1e-08 | 4.5e-08 | 3.5e-08 | 4.367E-08 | 8.083E-09 | 3.500E-08 | 5.100E-08 |
| 6446 | 323 | L323E | A | 2.6e-08 | 5.4e-08 | 2e-08 | 3.333E-08 | 1.815E-08 | 2.000E-08 | 5.400E-08 |
| 6447 | 323 | L323Q | A | 3.6e-08 | 5.2e-08 | 3e-08 | 3.933E-08 | 1.137E-08 | 3.000E-08 | 5.200E-08 |
| 6448 | 323 | L323G | A | 1.8e-08 | 1.6e-08 | 4.1e-08 | 2.500E-08 | 1.389E-08 | 1.600E-08 | 4.100E-08 |
| 6449 | 323 | L323H | A | 5.9e-08 | 5.6e-08 | 3.7e-08 | 5.067E-08 | 1.193E-08 | 3.700E-08 | 5.900E-08 |
| 6450 | 323 | L323I | A | 1e-07 | 4.9e-08 | 3.2e-08 | 6.033E-08 | 3.539E-08 | 3.200E-08 | 1.000E-07 |
| 6451 | 323 | L323L | A | 4.1e-08 | 3e-08 | 1.2e-08 | 2.767E-08 | 1.464E-08 | 1.200E-08 | 4.100E-08 |
| 6452 | 323 | L323K | A | 2.5e-08 | 6.1e-08 | 1.1e-07 | 6.533E-08 | 4.267E-08 | 2.500E-08 | 1.100E-07 |
| 6453 | 323 | L323M | A | 8.5e-08 | 2e-08 | 3.1e-08 | 4.533E-08 | 3.479E-08 | 2.000E-08 | 8.500E-08 |
| 6454 | 323 | L323F | A | 2.8e-08 | 4.6e-08 | 4.6e-08 | 4.000E-08 | 1.039E-08 | 2.800E-08 | 4.600E-08 |
| 6455 | 323 | L323P | A | 4.8e-08 | 7.2e-08 | 4.1e-08 | 5.367E-08 | 1.626E-08 | 4.100E-08 | 7.200E-08 |
| 6456 | 323 | L323S | A | 7e-08 | 6.1e-08 | 1.5e-07 | 9.367E-08 | 4.899E-08 | 6.100E-08 | 1.500E-07 |
| 6457 | 323 | L323T | A | 7.5e-08 | 8e-08 | 1.2e-08 | 5.567E-08 | 3.790E-08 | 1.200E-08 | 8.000E-08 |
| 6458 | 323 | L323W | A | 1.3e-07 | 3.5e-08 | 1.7e-08 | 6.067E-08 | 6.072E-08 | 1.700E-08 | 1.300E-07 |
| 6459 | 323 | L323Y | A | 4.7e-08 | 3.6e-08 | 5.4e-08 | 4.567E-08 | 9.074E-09 | 3.600E-08 | 5.400E-08 |

| | | | | | | | | | | |
|---|---|---|---|---|---|---|---|---|---|---|
| 6460 | 323 | L323V | A | 3.3e-08 | 1.9e-08 | 6.6e-08 | 3.933E-08 | 2.413E-08 | 1.900E-08 | 6.600E-08 |
| 6461 | 324 | G324A | A | 1.5e-08 | 2.5e-08 | 2.5e-08 | 2.167E-08 | 5.774E-09 | 1.500E-08 | 2.500E-08 |
| 6462 | 324 | G324R | A | 3.2e-08 | 3.6e-08 | 1.7e-08 | 2.833E-08 | 1.002E-08 | 1.700E-08 | 3.600E-08 |
| 6463 | 324 | G324N | A | 7.9e-08 | 2e-08 | 4e-08 | 4.633E-08 | 3.001E-08 | 2.000E-08 | 7.900E-08 |
| 6464 | 324 | G324D | A | 1.5e-08 | 1.1e-07 | 4.2e-08 | 5.567E-08 | 4.895E-08 | 1.500E-08 | 1.100E-07 |
| 6465 | 324 | G324C | A | 7.9e-09 | 1.1e-07 | 6.4e-08 | 6.063E-08 | 5.113E-08 | 7.900E-09 | 1.100E-07 |
| 6466 | 324 | G324E | A | 9.9e-08 | 1.6e-08 | 5.4e-08 | 5.633E-08 | 4.155E-08 | 1.600E-08 | 9.900E-08 |
| 6467 | 324 | G324Q | A | 8.5e-08 | 3.4e-08 | 3.1e-08 | 5.000E-08 | 3.035E-08 | 3.100E-08 | 8.500E-08 |
| 6468 | 324 | G324G | A | 4.1e-08 | 1.6e-08 | 2.8e-08 | 2.833E-08 | 1.250E-08 | 1.600E-08 | 4.100E-08 |
| 6469 | 324 | G324H | A | 5.1e-08 | 1.4e-08 | 1.1e-07 | 5.833E-08 | 4.842E-08 | 1.400E-08 | 1.100E-07 |
| 6470 | 324 | G324I | A | 7.2e-08 | 7.8e-08 | 2.1e-08 | 5.700E-08 | 3.132E-08 | 2.100E-08 | 7.800E-08 |
| 6471 | 324 | G324L | A | 5.2e-08 | 3.7e-08 | 3.1e-08 | 4.000E-08 | 1.082E-08 | 3.100E-08 | 5.200E-08 |
| 6472 | 324 | G324K | A | 3.7e-08 | 6.4e-08 | 6e-08 | 5.367E-08 | 1.457E-08 | 3.700E-08 | 6.400E-08 |
| 6473 | 324 | G324M | A | 9.9e-08 | 4.5e-08 | 3.4e-08 | 5.933E-08 | 3.479E-08 | 3.400E-08 | 9.900E-08 |
| 6474 | 324 | G324F | A | 3.7e-08 | 2.1e-08 | 2.2e-08 | 2.667E-08 | 8.963E-09 | 2.100E-08 | 3.700E-08 |
| 6475 | 324 | G324P | A | 5e-08 | 8.2e-08 | 6e-08 | 6.400E-08 | 1.637E-08 | 5.000E-08 | 8.200E-08 |
| 6476 | 324 | G324S | A | 5.4e-08 | 5.6e-08 | 4.5e-08 | 5.167E-08 | 5.859E-09 | 4.500E-08 | 5.600E-08 |
| 6477 | 324 | G324T | A | 4.4e-08 | 1.2e-08 | 2.2e-08 | 2.600E-08 | 1.637E-08 | 1.200E-08 | 4.400E-08 |
| 6478 | 324 | G324W | A | 2.7e-08 | 8.8e-08 | 6.7e-08 | 6.067E-08 | 3.099E-08 | 2.700E-08 | 8.800E-08 |
| 6479 | 324 | G324Y | A | 2.6e-08 | 5.1e-08 | 6.2e-08 | 4.633E-08 | 1.845E-08 | 2.600E-08 | 6.200E-08 |
| 6480 | 324 | G324V | A | 4.6e-08 | 3e-08 | 2.9e-08 | 3.500E-08 | 9.539E-09 | 2.900E-08 | 4.600E-08 |
| 6481 | 325 | M325A | A | 9.1e-08 | 5.6e-08 | 2.9e-08 | 5.867E-08 | 3.109E-08 | 2.900E-08 | 9.100E-08 |
| 6482 | 325 | M325R | A | 1.5e-07 | 4.9e-08 | 8.3e-08 | 9.400E-08 | 5.139E-08 | 4.900E-08 | 1.500E-07 |
| 6483 | 325 | M325N | A | 4e-08 | 9.2e-08 | 1.9e-08 | 5.033E-08 | 3.758E-08 | 1.900E-08 | 9.200E-08 |
| 6484 | 325 | M325D | A | 6.8e-08 | 3.3e-08 | 1.8e-08 | 3.967E-08 | 2.566E-08 | 1.800E-08 | 6.800E-08 |
| 6485 | 325 | M325C | A | 2.4e-08 | 4.7e-08 | 3.5e-08 | 3.533E-08 | 1.150E-08 | 2.400E-08 | 4.700E-08 |
| 6486 | 325 | M325E | A | 6.1e-08 | 2.6e-08 | 3.8e-08 | 4.167E-08 | 1.779E-08 | 2.600E-08 | 6.100E-08 |
| 6487 | 325 | M325Q | A | 3.1e-08 | 3.5e-08 | 1.6e-08 | 2.733E-08 | 1.002E-08 | 1.600E-08 | 3.500E-08 |
| 6488 | 325 | M325G | A | 2.3e-08 | 3.5e-08 | 4.6e-08 | 3.467E-08 | 1.150E-08 | 2.300E-08 | 4.600E-08 |
| 6489 | 325 | M325H | A | 9.7e-09 | 3.3e-08 | 2.1e-08 | 2.123E-08 | 1.165E-08 | 9.700E-09 | 3.300E-08 |
| 6490 | 325 | M325I | A | 4.8e-08 | 3.6e-08 | 3.8e-08 | 4.067E-08 | 6.429E-09 | 3.600E-08 | 4.800E-08 |
| 6491 | 325 | M325L | A | 5.5e-08 | 2.5e-08 | 4.6e-08 | 4.200E-08 | 1.539E-08 | 2.500E-08 | 5.500E-08 |
| 6492 | 325 | M325K | A | 3.3e-08 | 1.5e-07 | 1.3e-08 | 6.533E-08 | 7.400E-08 | 1.300E-08 | 1.500E-07 |
| 6493 | 325 | M325M | A | 2.7e-08 | 3.8e-08 | 6.2e-08 | 4.233E-08 | 1.790E-08 | 2.700E-08 | 6.200E-08 |
| 6494 | 325 | M325F | A | 3.5e-08 | 2.3e-08 | 6.1e-08 | 3.967E-08 | 1.943E-08 | 2.300E-08 | 6.100E-08 |
| 6495 | 325 | M325P | A | 1.4e-08 | 3.2e-08 | 4.8e-08 | 3.133E-08 | 1.701E-08 | 1.400E-08 | 4.800E-08 |
| 6496 | 325 | M325S | A | 3.5e-08 | 4.4e-08 | 7.8e-08 | 5.233E-08 | 2.268E-08 | 3.500E-08 | 7.800E-08 |
| 6497 | 325 | M325T | A | 2.5e-08 | 6.7e-08 | 5.6e-08 | 4.933E-08 | 2.178E-08 | 2.500E-08 | 6.700E-08 |
| 6498 | 325 | M325W | A | 9.8e-08 | 1.5e-07 | 8.5e-08 | 1.110E-07 | 3.439E-08 | 8.500E-08 | 1.500E-07 |
| 6499 | 325 | M325Y | A | 2.1e-08 | 1.8e-08 | 6.1e-08 | 3.333E-08 | 2.401E-08 | 1.800E-08 | 6.100E-08 |
| 6500 | 325 | M325V | A | 3.4e-08 | 1.1e-07 | 4.8e-08 | 6.400E-08 | 4.045E-08 | 3.400E-08 | 1.100E-07 |
| 6501 | 326 | E326A | A | 1.7e-08 | 2.9e-08 | 4e-08 | 2.867E-08 | 1.150E-08 | 1.700E-08 | 4.000E-08 |
| 6502 | 326 | E326R | A | 7.7e-08 | 4e-08 | 5e-08 | 5.567E-08 | 1.914E-08 | 4.000E-08 | 7.700E-08 |
| 6503 | 326 | E326N | A | 3.9e-08 | 6e-08 | 7.2e-08 | 5.700E-08 | 1.670E-08 | 3.900E-08 | 7.200E-08 |
| 6504 | 326 | E326D | A | 1.2e-08 | 1.1e-07 | 5.6e-08 | 5.933E-08 | 4.908E-08 | 1.200E-08 | 1.100E-07 |
| 6505 | 326 | E326C | A | 3.5e-08 | 3.9e-08 | 5.1e-08 | 4.167E-08 | 8.327E-09 | 3.500E-08 | 5.100E-08 |
| 6506 | 326 | E326E | A | 5.2e-08 | 3.5e-08 | 7.3e-08 | 5.333E-08 | 1.904E-08 | 3.500E-08 | 7.300E-08 |
| 6507 | 326 | E326Q | A | 7.9e-08 | 3.3e-08 | 6.6e-08 | 5.933E-08 | 2.371E-08 | 3.300E-08 | 7.900E-08 |
| 6508 | 326 | E326G | A | 7.6e-09 | 5.5e-08 | 6e-08 | 4.087E-08 | 2.892E-08 | 7.600E-09 | 6.000E-08 |
| 6509 | 326 | E326H | A | 3.3e-08 | 8.3e-08 | 2.1e-08 | 4.567E-08 | 3.288E-08 | 2.100E-08 | 8.300E-08 |
| 6510 | 326 | E326I | A | 2.4e-08 | 1.3e-07 | 6e-08 | 7.133E-08 | 5.390E-08 | 2.400E-08 | 1.300E-07 |
| 6511 | 326 | E326L | A | 5e-08 | 2.1e-08 | 5.5e-08 | 4.200E-08 | 1.836E-08 | 2.100E-08 | 5.500E-08 |
| 6512 | 326 | E326K | A | 5.5e-08 | 9.1e-08 | 2.5e-08 | 5.700E-08 | 3.305E-08 | 2.500E-08 | 9.100E-08 |
| 6513 | 326 | E326M | A | 1e-08 | 1.3e-07 | 1.5e-08 | 5.167E-08 | 6.788E-08 | 1.000E-08 | 1.300E-07 |
| 6514 | 326 | E326F | A | 3.3e-08 | 1.3e-08 | 2.1e-08 | 2.233E-08 | 1.007E-08 | 1.300E-08 | 3.300E-08 |
| 6515 | 326 | E326P | A | 1e-07 | 4.5e-08 | 3.7e-08 | 6.067E-08 | 3.430E-08 | 3.700E-08 | 1.000E-07 |
| 6516 | 326 | E326S | A | 3e-08 | 1.3e-07 | 3.5e-08 | 6.500E-08 | 5.635E-08 | 3.000E-08 | 1.300E-07 |
| 6517 | 326 | E326T | A | 9.7e-08 | 2.5e-08 | 4.6e-08 | 5.600E-08 | 3.703E-08 | 2.500E-08 | 9.700E-08 |
| 6518 | 326 | E326W | A | 7.3e-08 | 8.7e-08 | 3e-08 | 6.333E-08 | 2.970E-08 | 3.000E-08 | 8.700E-08 |
| 6519 | 326 | E326Y | A | 2.4e-08 | 7.5e-08 | 5.1e-08 | 5.000E-08 | 2.551E-08 | 2.400E-08 | 7.500E-08 |
| 6520 | 326 | E326V | A | 2.3e-08 | 5.4e-08 | 2.2e-08 | 3.300E-08 | 1.819E-08 | 2.200E-08 | 5.400E-08 |
| 6521 | 327 | H327A | A | 8e-08 | 5.2e-08 | 5.8e-08 | 6.333E-08 | 1.474E-08 | 5.200E-08 | 8.000E-08 |
| 6522 | 327 | H327R | A | 8.2e-08 | 1.1e-07 | 2.4e-08 | 7.200E-08 | 4.386E-08 | 2.400E-08 | 1.100E-07 |
| 6523 | 327 | H327N | A | 3.9e-08 | 2.6e-08 | 5.5e-08 | 4.000E-08 | 1.453E-08 | 2.600E-08 | 5.500E-08 |
| 6524 | 327 | H327D | A | 2.1e-08 | 1.8e-08 | 6.1e-08 | 3.333E-08 | 2.401E-08 | 1.800E-08 | 6.100E-08 |
| 6525 | 327 | H327C | A | 3.6e-08 | 3.7e-08 | 3.5e-08 | 3.600E-08 | 1.000E-09 | 3.500E-08 | 3.700E-08 |
| 6526 | 327 | H327E | A | 3e-08 | 1.6e-08 | 1.4e-08 | 2.000E-08 | 8.718E-09 | 1.400E-08 | 3.000E-08 |
| 6527 | 327 | H327Q | A | 2.1e-08 | 3e-08 | 7.6e-08 | 4.233E-08 | 2.950E-08 | 2.100E-08 | 7.600E-08 |
| 6528 | 327 | H327G | A | 3.9e-08 | 4.2e-08 | 2.3e-08 | 3.467E-08 | 1.021E-08 | 2.300E-08 | 4.200E-08 |
| 6529 | 327 | H327H | A | 3.6e-08 | 2.1e-08 | 7.7e-08 | 4.467E-08 | 2.899E-08 | 2.100E-08 | 7.700E-08 |
| 6530 | 327 | H327I | A | 3.2e-08 | 8.6e-08 | 1.9e-08 | 4.567E-08 | 3.553E-08 | 1.900E-08 | 8.600E-08 |
| 6531 | 327 | H327L | A | 3.9e-08 | 3.1e-08 | 9.1e-08 | 5.367E-08 | 3.258E-08 | 3.100E-08 | 9.100E-08 |
| 6532 | 327 | H327K | A | 8.2e-08 | 8.7e-08 | 1.3e-08 | 6.067E-08 | 4.136E-08 | 1.300E-08 | 8.700E-08 |
| 6533 | 327 | H327M | A | 3.7e-08 | 5.6e-08 | 3.3e-08 | 4.200E-08 | 1.229E-08 | 3.300E-08 | 5.600E-08 |
| 6534 | 327 | H327F | A | 6.1e-08 | 1e-07 | 7.1e-08 | 7.733E-08 | 2.026E-08 | 6.100E-08 | 1.000E-07 |
| 6535 | 327 | H327P | A | 2.3e-08 | 1e-07 | 1.4e-07 | 8.767E-08 | 5.947E-08 | 2.300E-08 | 1.400E-07 |
| 6536 | 327 | H327S | A | 2.3e-08 | 3.1e-08 | 3.9e-08 | 3.100E-08 | 8.000E-09 | 2.300E-08 | 3.900E-08 |
| 6537 | 327 | H327T | A | 7.9e-08 | 8e-08 | 3.4e-08 | 6.433E-08 | 2.627E-08 | 3.400E-08 | 8.000E-08 |
| 6538 | 327 | H327W | A | 3.9e-08 | 2.4e-08 | 1.8e-07 | 8.100E-08 | 8.606E-08 | 2.400E-08 | 1.800E-07 |
| 6539 | 327 | H327Y | A | 3.8e-08 | 3.6e-08 | 2.3e-08 | 3.233E-08 | 8.145E-09 | 2.300E-08 | 3.800E-08 |
| 6540 | 327 | H327V | A | 2.6e-08 | 2.7e-08 | 6e-08 | 3.767E-08 | 1.935E-08 | 2.600E-08 | 6.000E-08 |
| 6541 | 328 | L328A | A | 9.8e-08 | 6.9e-08 | 3.1e-08 | 6.600E-08 | 3.360E-08 | 3.100E-08 | 9.800E-08 |
| 6542 | 328 | L328R | A | 7e-08 | 7.6e-08 | 1.4e-08 | 5.333E-08 | 3.420E-08 | 1.400E-08 | 7.600E-08 |
| 6543 | 328 | L328N | A | 9.3e-08 | 5.1e-08 | 9.3e-08 | 7.900E-08 | 2.425E-08 | 5.100E-08 | 9.300E-08 |
| 6544 | 328 | L328D | A | 1.5e-08 | 6.5e-08 | 2.3e-08 | 3.433E-08 | 2.686E-08 | 1.500E-08 | 6.500E-08 |

| | | | | | | | | | | |
|---|---|---|---|---|---|---|---|---|---|---|
| 6545 | 328 | L328C | A | 3.3e-08 | 8.1e-08 | 5.5e-08 | 5.633E-08 | 2.403E-08 | 3.300E-08 | 8.100E-08 |
| 6546 | 328 | L328E | A | 2.8e-08 | 6e-08 | 7.3e-08 | 5.367E-08 | 2.316E-08 | 2.800E-08 | 7.300E-08 |
| 6547 | 328 | L328Q | A | 4.5e-08 | 1.2e-08 | 1.9e-08 | 2.533E-08 | 1.739E-08 | 1.200E-08 | 4.500E-08 |
| 6548 | 328 | L328G | A | 3.6e-08 | 2e-07 | 4.9e-08 | 9.500E-08 | 9.116E-08 | 3.600E-08 | 2.000E-07 |
| 6549 | 328 | L328H | A | 3.7e-08 | 6.3e-08 | 1.4e-08 | 3.800E-08 | 2.452E-08 | 1.400E-08 | 6.300E-08 |
| 6550 | 328 | L328I | A | 2.5e-08 | 4.5e-08 | 2.2e-07 | 9.667E-08 | 1.073E-07 | 2.500E-08 | 2.200E-07 |
| 6551 | 328 | L328L | A | 6.3e-08 | 7.6e-08 | 5.3e-08 | 6.400E-08 | 1.153E-08 | 5.300E-08 | 7.600E-08 |
| 6552 | 328 | L328K | A | 6.6e-08 | 2.6e-08 | 2.5e-08 | 3.900E-08 | 2.339E-08 | 2.500E-08 | 6.600E-08 |
| 6553 | 328 | L328M | A | 2.8e-08 | 1.8e-07 | 7.1e-08 | 9.300E-08 | 7.835E-08 | 2.800E-08 | 1.800E-07 |
| 6554 | 328 | L328F | A | 4.5e-08 | 6.1e-08 | 3.2e-08 | 4.600E-08 | 1.453E-08 | 3.200E-08 | 6.100E-08 |
| 6555 | 328 | L328P | A | 2.2e-08 | 3.8e-08 | 2.8e-08 | 2.933E-08 | 8.083E-09 | 2.200E-08 | 3.800E-08 |
| 6556 | 328 | L328S | A | 2.8e-08 | 5e-08 | 5.5e-08 | 4.433E-08 | 1.436E-08 | 2.800E-08 | 5.500E-08 |
| 6557 | 328 | L328T | A | 9.3e-08 | 6.1e-08 | 2.2e-08 | 5.867E-08 | 3.556E-08 | 2.200E-08 | 9.300E-08 |
| 6558 | 328 | L328W | A | 5.3e-08 | 5.5e-08 | 4.3e-08 | 5.033E-08 | 6.429E-09 | 4.300E-08 | 5.500E-08 |
| 6559 | 328 | L328Y | A | 3.7e-08 | 2.7e-08 | 8.4e-08 | 4.933E-08 | 3.044E-08 | 2.700E-08 | 8.400E-08 |
| 6560 | 328 | L328V | A | 3.4e-08 | 5.3e-08 | 3.6e-08 | 4.100E-08 | 1.044E-08 | 3.400E-08 | 5.300E-08 |
| 6561 | 329 | R329A | A | 4e-08 | 6.6e-08 | 5.8e-08 | 5.467E-08 | 1.332E-08 | 4.000E-08 | 6.600E-08 |
| 6562 | 329 | R329R | A | 5.1e-08 | 3.7e-08 | 7.2e-08 | 5.333E-08 | 1.762E-08 | 3.700E-08 | 7.200E-08 |
| 6563 | 329 | R329N | A | 4.7e-08 | 1.2e-07 | 2.6e-08 | 6.433E-08 | 4.934E-08 | 2.600E-08 | 1.200E-07 |
| 6564 | 329 | R329D | A | 4e-08 | 7e-08 | 2.1e-08 | 4.367E-08 | 2.470E-08 | 2.100E-08 | 7.000E-08 |
| 6565 | 329 | R329C | A | 3.8e-08 | 1.5e-08 | 2.3e-08 | 2.533E-08 | 1.168E-08 | 1.500E-08 | 3.800E-08 |
| 6566 | 329 | R329E | A | 3.7e-08 | 3.4e-08 | 8.8e-08 | 5.300E-08 | 3.035E-08 | 3.400E-08 | 8.800E-08 |
| 6567 | 329 | R329Q | A | 3.6e-08 | 4e-08 | 9.1e-08 | 5.567E-08 | 3.066E-08 | 3.600E-08 | 9.100E-08 |
| 6568 | 329 | R329G | A | 3.5e-08 | 1e-07 | 4.4e-08 | 5.967E-08 | 3.522E-08 | 3.500E-08 | 1.000E-07 |
| 6569 | 329 | R329H | A | 8.2e-08 | 4.3e-08 | 5.6e-08 | 6.033E-08 | 1.986E-08 | 4.300E-08 | 8.200E-08 |
| 6570 | 329 | R329I | A | 2.8e-08 | 5.8e-08 | 9.7e-08 | 6.100E-08 | 3.460E-08 | 2.800E-08 | 9.700E-08 |
| 6571 | 329 | R329L | A | 5.8e-08 | 6.8e-08 | 2.3e-08 | 4.967E-08 | 2.363E-08 | 2.300E-08 | 6.800E-08 |
| 6572 | 329 | R329K | A | 1e-08 | 6.7e-08 | 2.8e-08 | 3.500E-08 | 2.914E-08 | 1.000E-08 | 6.700E-08 |
| 6573 | 329 | R329M | A | 3.8e-08 | 6.3e-08 | 3.5e-08 | 4.533E-08 | 1.537E-08 | 3.500E-08 | 6.300E-08 |
| 6574 | 329 | R329F | A | 1.7e-08 | 4.5e-08 | 2.4e-08 | 2.867E-08 | 1.457E-08 | 1.700E-08 | 4.500E-08 |
| 6575 | 329 | R329P | A | 4.4e-08 | 3.6e-08 | 2e-08 | 3.333E-08 | 1.222E-08 | 2.000E-08 | 4.400E-08 |
| 6576 | 329 | R329S | A | 3.3e-08 | 2.7e-08 | 6.8e-08 | 4.267E-08 | 2.214E-08 | 2.700E-08 | 6.800E-08 |
| 6577 | 329 | R329T | A | 6.5e-08 | 3e-08 | 7.1e-08 | 5.533E-08 | 2.214E-08 | 3.000E-08 | 7.100E-08 |
| 6578 | 329 | R329W | A | 4.9e-08 | 4.4e-08 | 1.6e-08 | 3.633E-08 | 1.779E-08 | 1.600E-08 | 4.900E-08 |
| 6579 | 329 | R329Y | A | 9.8e-08 | 2.9e-08 | 4.2e-08 | 5.633E-08 | 3.667E-08 | 2.900E-08 | 9.800E-08 |
| 6580 | 329 | R329V | A | 6.1e-08 | 1.7e-08 | 1.6e-08 | 3.133E-08 | 2.570E-08 | 1.600E-08 | 6.100E-08 |
| 6581 | 330 | E330A | A | 5.5e-08 | 4.9e-08 | 2.5e-08 | 4.300E-08 | 1.587E-08 | 2.500E-08 | 5.500E-08 |
| 6582 | 330 | E330R | A | 2.4e-08 | 2.1e-08 | 3.3e-08 | 2.600E-08 | 6.245E-09 | 2.100E-08 | 3.300E-08 |
| 6583 | 330 | E330N | A | 6.4e-08 | 1.9e-08 | 5.3e-08 | 4.533E-08 | 2.346E-08 | 1.900E-08 | 6.400E-08 |
| 6584 | 330 | E330D | A | 2.5e-08 | 3e-08 | 3.5e-08 | 3.000E-08 | 5.000E-09 | 2.500E-08 | 3.500E-08 |
| 6585 | 330 | E330C | A | 3.6e-08 | 2.2e-08 | 3.6e-08 | 3.133E-08 | 8.083E-09 | 2.200E-08 | 3.600E-08 |
| 6586 | 330 | E330E | A | 1.8e-08 | 1.1e-07 | 1.1e-08 | 4.633E-08 | 5.525E-08 | 1.100E-08 | 1.100E-07 |
| 6587 | 330 | E330Q | A | 3e-08 | 4.2e-08 | 1.4e-07 | 7.067E-08 | 6.034E-08 | 3.000E-08 | 1.400E-07 |
| 6588 | 330 | E330G | A | 3.7e-08 | 4.2e-08 | 4.3e-08 | 4.067E-08 | 3.215E-09 | 3.700E-08 | 4.300E-08 |
| 6589 | 330 | E330H | A | 1.7e-08 | 1.9e-08 | 2.8e-08 | 2.133E-08 | 5.859E-09 | 1.700E-08 | 2.800E-08 |
| 6590 | 330 | E330I | A | 4e-08 | 5.6e-08 | 3.3e-08 | 4.300E-08 | 1.179E-08 | 3.300E-08 | 5.600E-08 |
| 6591 | 330 | E330L | A | 6.1e-08 | 3.4e-08 | 4.1e-08 | 4.533E-08 | 1.401E-08 | 3.400E-08 | 6.100E-08 |
| 6592 | 330 | E330K | A | 5.5e-08 | 3e-08 | 5.3e-08 | 4.600E-08 | 1.389E-08 | 3.000E-08 | 5.500E-08 |
| 6593 | 330 | E330M | A | 4.8e-08 | 9e-08 | 1.4e-08 | 5.067E-08 | 3.807E-08 | 1.400E-08 | 9.000E-08 |
| 6594 | 330 | E330F | A | 4e-08 | 2.6e-08 | 3.4e-08 | 3.333E-08 | 7.024E-09 | 2.600E-08 | 4.000E-08 |
| 6595 | 330 | E330P | A | 4.9e-08 | 1.2e-07 | 6.5e-08 | 7.800E-08 | 3.724E-08 | 4.900E-08 | 1.200E-07 |
| 6596 | 330 | E330S | A | 4.3e-08 | 7.4e-08 | 3.5e-08 | 5.067E-08 | 2.060E-08 | 3.500E-08 | 7.400E-08 |
| 6597 | 330 | E330T | A | 5.3e-08 | 1.8e-08 | 1e-07 | 5.700E-08 | 4.115E-08 | 1.800E-08 | 1.000E-07 |
| 6598 | 330 | E330W | A | 2.3e-08 | 3e-08 | 5.8e-08 | 3.700E-08 | 1.852E-08 | 2.300E-08 | 5.800E-08 |
| 6599 | 330 | E330Y | A | 4.3e-08 | 8.7e-08 | 6.1e-08 | 6.367E-08 | 2.212E-08 | 4.300E-08 | 8.700E-08 |
| 6600 | 330 | E330V | A | 1.7e-08 | 5.9e-08 | 6.7e-08 | 4.767E-08 | 2.686E-08 | 1.700E-08 | 6.700E-08 |
| 6601 | 331 | V331A | A | 4.8e-08 | 3.6e-08 | 6.4e-08 | 4.933E-08 | 1.405E-08 | 3.600E-08 | 6.400E-08 |
| 6602 | 331 | V331R | A | 4e-08 | 5.7e-08 | 2.4e-08 | 4.033E-08 | 1.650E-08 | 2.400E-08 | 5.700E-08 |
| 6603 | 331 | V331N | A | 1.1e-07 | 1.6e-08 | 4e-08 | 5.533E-08 | 4.884E-08 | 1.600E-08 | 1.100E-07 |
| 6604 | 331 | V331D | A | 1.5e-07 | 1.2e-07 | 3.4e-08 | 1.013E-07 | 6.021E-08 | 3.400E-08 | 1.500E-07 |
| 6605 | 331 | V331C | A | 8.5e-08 | 3.9e-08 | 7.6e-08 | 6.667E-08 | 2.438E-08 | 3.900E-08 | 8.500E-08 |
| 6606 | 331 | V331E | A | 1.7e-08 | 3.5e-08 | 2.9e-08 | 2.700E-08 | 9.165E-09 | 1.700E-08 | 3.500E-08 |
| 6607 | 331 | V331Q | A | 5.9e-08 | 3.2e-08 | 5.1e-08 | 4.733E-08 | 1.387E-08 | 3.200E-08 | 5.900E-08 |
| 6608 | 331 | V331G | A | 3.1e-08 | 4.3e-08 | 8.4e-08 | 5.267E-08 | 2.779E-08 | 3.100E-08 | 8.400E-08 |
| 6609 | 331 | V331H | A | 3.5e-08 | 2.8e-08 | 4.1e-08 | 3.467E-08 | 6.506E-09 | 2.800E-08 | 4.100E-08 |
| 6610 | 331 | V331I | A | 4.1e-08 | 9.2e-08 | 1.8e-08 | 5.033E-08 | 3.787E-08 | 1.800E-08 | 9.200E-08 |
| 6611 | 331 | V331L | A | 2.9e-08 | 3e-08 | 3.9e-08 | 3.267E-08 | 5.508E-09 | 2.900E-08 | 3.900E-08 |
| 6612 | 331 | V331K | A | 5.7e-08 | 2.4e-08 | 5e-08 | 4.367E-08 | 1.739E-08 | 2.400E-08 | 5.700E-08 |
| 6613 | 331 | V331M | A | 3e-08 | 3.3e-08 | 7.2e-08 | 4.500E-08 | 2.343E-08 | 3.000E-08 | 7.200E-08 |
| 6614 | 331 | V331F | A | 2.6e-08 | 3.6e-08 | 7.6e-08 | 4.600E-08 | 2.646E-08 | 2.600E-08 | 7.600E-08 |
| 6615 | 331 | V331P | A | 3.9e-08 | 2.5e-08 | 4.9e-08 | 3.767E-08 | 1.206E-08 | 2.500E-08 | 4.900E-08 |
| 6616 | 331 | V331S | A | 3.9e-08 | 3.6e-08 | 3.9e-08 | 3.800E-08 | 1.732E-09 | 3.600E-08 | 3.900E-08 |
| 6617 | 331 | V331T | A | 2.7e-08 | 5.1e-08 | 7.1e-08 | 4.967E-08 | 2.203E-08 | 2.700E-08 | 7.100E-08 |
| 6618 | 331 | V331W | A | 2.5e-08 | 5e-08 | 3.6e-08 | 3.700E-08 | 1.253E-08 | 2.500E-08 | 5.000E-08 |
| 6619 | 331 | V331Y | A | 7.1e-08 | 2.5e-08 | 2.8e-08 | 4.133E-08 | 2.574E-08 | 2.500E-08 | 7.100E-08 |
| 6620 | 331 | V331V | A | 5.6e-08 | 2.2e-08 | 2.4e-08 | 3.400E-08 | 1.908E-08 | 2.200E-08 | 5.600E-08 |
| 6621 | 332 | R332A | A | 6.9e-08 | 3.7e-08 | 5.5e-08 | 5.367E-08 | 1.604E-08 | 3.700E-08 | 6.900E-08 |
| 6622 | 332 | R332R | A | 1.4e-08 | 2.9e-08 | 1.3e-07 | 5.767E-08 | 6.309E-08 | 1.400E-08 | 1.300E-07 |
| 6623 | 332 | R332N | A | 5.4e-08 | 6.2e-08 | 4.9e-08 | 5.500E-08 | 6.557E-09 | 4.900E-08 | 6.200E-08 |
| 6624 | 332 | R332D | A | 1.5e-07 | 1.9e-08 | 1.5e-07 | 1.063E-07 | 7.563E-08 | 1.900E-08 | 1.500E-07 |
| 6625 | 332 | R332C | A | 2.3e-08 | 3.7e-08 | 3.5e-08 | 3.167E-08 | 7.572E-09 | 2.300E-08 | 3.700E-08 |
| 6626 | 332 | R332E | A | 6e-08 | 1.4e-07 | 6.4e-08 | 8.800E-08 | 4.508E-08 | 6.000E-08 | 1.400E-07 |
| 6627 | 332 | R332Q | A | 1.8e-08 | 6.5e-08 | 8.2e-08 | 5.500E-08 | 3.315E-08 | 1.800E-08 | 8.200E-08 |
| 6628 | 332 | R332G | A | 9.1e-08 | 3.8e-08 | 2.2e-08 | 5.033E-08 | 3.612E-08 | 2.200E-08 | 9.100E-08 |
| 6629 | 332 | R332H | A | 3.2e-08 | 4e-08 | 3.4e-08 | 3.533E-08 | 4.163E-09 | 3.200E-08 | 4.000E-08 |

| | | | | | | | | | | |
|---|---|---|---|---|---|---|---|---|---|---|
| 6630 | 332 | R332I | A | 6.1e-08 | 6.4e-08 | 3.3e-08 | 5.267E-08 | 1.710E-08 | 3.300E-08 | 6.400E-08 |
| 6631 | 332 | R332L | A | 5.7e-08 | 6e-08 | 5.6e-08 | 5.767E-08 | 2.082E-09 | 5.600E-08 | 6.000E-08 |
| 6632 | 332 | R332K | A | 2.6e-08 | 2.1e-08 | 3e-08 | 2.567E-08 | 4.509E-09 | 2.100E-08 | 3.000E-08 |
| 6633 | 332 | R332M | A | 1.8e-08 | 9e-08 | 1.2e-07 | 7.600E-08 | 5.242E-08 | 1.800E-08 | 1.200E-07 |
| 6634 | 332 | R332F | A | 5.2e-08 | 7.2e-08 | 2.3e-08 | 4.900E-08 | 2.464E-08 | 2.300E-08 | 7.200E-08 |
| 6635 | 332 | R332P | A | 4.7e-08 | 3.5e-08 | 8e-08 | 5.400E-08 | 2.330E-08 | 3.500E-08 | 8.000E-08 |
| 6636 | 332 | R332S | A | 6.5e-08 | 3.3e-08 | 2.4e-08 | 4.067E-08 | 2.155E-08 | 2.400E-08 | 6.500E-08 |
| 6637 | 332 | R332T | A | 7.8e-08 | 3.2e-08 | 5.3e-08 | 5.433E-08 | 2.303E-08 | 3.200E-08 | 7.800E-08 |
| 6638 | 332 | R332W | A | 9.1e-09 | 2.5e-08 | 3.2e-08 | 2.203E-08 | 1.173E-08 | 9.100E-09 | 3.200E-08 |
| 6639 | 332 | R332Y | A | 5e-08 | 5.3e-08 | 1.2e-07 | 7.433E-08 | 3.958E-08 | 5.000E-08 | 1.200E-07 |
| 6640 | 332 | R332V | A | 6e-08 | 4.3e-08 | 7.1e-08 | 5.800E-08 | 1.411E-08 | 4.300E-08 | 7.100E-08 |
| 6641 | 333 | A333A | A | 4.4e-08 | 5.2e-08 | 3.6e-08 | 4.400E-08 | 8.000E-09 | 3.600E-08 | 5.200E-08 |
| 6642 | 333 | A333R | A | 3.5e-08 | 4e-08 | 3.8e-08 | 3.767E-08 | 2.517E-09 | 3.500E-08 | 4.000E-08 |
| 6643 | 333 | A333N | A | 1.9e-08 | 5.1e-09 | 2e-08 | 1.470E-08 | 8.329E-09 | 5.100E-09 | 2.000E-08 |
| 6644 | 333 | A333D | A | 4.2e-08 | 1.9e-08 | 5.5e-08 | 3.867E-08 | 1.823E-08 | 1.900E-08 | 5.500E-08 |
| 6645 | 333 | A333C | A | 4.8e-08 | 1.1e-08 | 3.8e-08 | 3.233E-08 | 1.914E-08 | 1.100E-08 | 4.800E-08 |
| 6646 | 333 | A333E | A | 1.9e-08 | 1.2e-07 | 1.7e-07 | 1.030E-07 | 7.692E-08 | 1.900E-08 | 1.700E-07 |
| 6647 | 333 | A333Q | A | 4e-08 | 3.3e-08 | 2.8e-08 | 3.367E-08 | 6.028E-09 | 2.800E-08 | 4.000E-08 |
| 6648 | 333 | A333G | A | 3.7e-08 | 1.3e-08 | 1.1e-07 | 5.333E-08 | 5.052E-08 | 1.300E-08 | 1.100E-07 |
| 6649 | 333 | A333H | A | 1.1e-07 | 9.1e-08 | 3.6e-08 | 7.900E-08 | 3.843E-08 | 3.600E-08 | 1.100E-07 |
| 6650 | 333 | A333I | A | 3.7e-08 | 4.2e-08 | 1.1e-07 | 6.300E-08 | 4.078E-08 | 3.700E-08 | 1.100E-07 |
| 6651 | 333 | A333L | A | 4.4e-08 | 8.4e-08 | 4.4e-08 | 5.733E-08 | 2.309E-08 | 4.400E-08 | 8.400E-08 |
| 6652 | 333 | A333K | A | 4.6e-08 | 5.8e-08 | 6.4e-08 | 5.600E-08 | 9.165E-09 | 4.600E-08 | 6.400E-08 |
| 6653 | 333 | A333M | A | 4.3e-08 | 6.1e-08 | 3.6e-08 | 4.667E-08 | 1.290E-08 | 3.600E-08 | 6.100E-08 |
| 6654 | 333 | A333F | A | 6.9e-08 | 2.9e-08 | 1.1e-08 | 3.633E-08 | 2.969E-08 | 1.100E-08 | 6.900E-08 |
| 6655 | 333 | A333P | A | 1.4e-08 | 8.3e-08 | 5.4e-08 | 5.033E-08 | 3.465E-08 | 1.400E-08 | 8.300E-08 |
| 6656 | 333 | A333S | A | 4.3e-08 | 4.4e-08 | 4.3e-08 | 4.333E-08 | 5.774E-10 | 4.300E-08 | 4.400E-08 |
| 6657 | 333 | A333T | A | 7.8e-08 | 4.6e-08 | 2.9e-08 | 5.100E-08 | 2.488E-08 | 2.900E-08 | 7.800E-08 |
| 6658 | 333 | A333W | A | 4.3e-08 | 6.3e-08 | 1e-07 | 6.867E-08 | 2.892E-08 | 4.300E-08 | 1.000E-07 |
| 6659 | 333 | A333Y | A | 6.4e-08 | 4e-08 | 9.4e-08 | 6.600E-08 | 2.706E-08 | 4.000E-08 | 9.400E-08 |
| 6660 | 333 | A333V | A | 4.6e-08 | 3.8e-08 | 1e-07 | 6.133E-08 | 3.372E-08 | 3.800E-08 | 1.000E-07 |
| 6661 | 334 | V334A | A | 5.9e-08 | 4.4e-08 | 6.1e-08 | 5.467E-08 | 9.292E-09 | 4.400E-08 | 6.100E-08 |
| 6662 | 334 | V334R | A | 9.6e-09 | 4e-08 | 5e-08 | 3.320E-08 | 2.104E-08 | 9.600E-09 | 5.000E-08 |
| 6663 | 334 | V334N | A | 2.1e-08 | 3.5e-08 | 5e-08 | 3.533E-08 | 1.450E-08 | 2.100E-08 | 5.000E-08 |
| 6664 | 334 | V334D | A | 5.3e-08 | 2.1e-08 | 4.6e-08 | 4.000E-08 | 1.682E-08 | 2.100E-08 | 5.300E-08 |
| 6665 | 334 | V334C | A | 1.6e-08 | 3.8e-08 | 4e-08 | 3.133E-08 | 1.332E-08 | 1.600E-08 | 4.000E-08 |
| 6666 | 334 | V334E | A | 8.7e-08 | 3.3e-08 | 6.4e-08 | 6.133E-08 | 2.710E-08 | 3.300E-08 | 8.700E-08 |
| 6667 | 334 | V334Q | A | 6.2e-08 | 1.7e-08 | 3.9e-08 | 3.933E-08 | 2.250E-08 | 1.700E-08 | 6.200E-08 |
| 6668 | 334 | V334G | A | 2.5e-08 | 1.1e-07 | 3.1e-08 | 5.533E-08 | 4.744E-08 | 2.500E-08 | 1.100E-07 |
| 6669 | 334 | V334H | A | 3.5e-08 | 1.4e-08 | 9.3e-08 | 4.733E-08 | 4.092E-08 | 1.400E-08 | 9.300E-08 |
| 6670 | 334 | V334I | A | 3.2e-08 | 5.2e-08 | 4.4e-08 | 4.267E-08 | 1.007E-08 | 3.200E-08 | 5.200E-08 |
| 6671 | 334 | V334L | A | 3.5e-08 | 3.8e-08 | 8.6e-08 | 5.300E-08 | 2.862E-08 | 3.500E-08 | 8.600E-08 |
| 6672 | 334 | V334K | A | 5.6e-08 | 9e-08 | 3.9e-08 | 6.167E-08 | 2.597E-08 | 3.900E-08 | 9.000E-08 |
| 6673 | 334 | V334M | A | 3.1e-08 | 2.5e-08 | 2.6e-08 | 2.733E-08 | 3.215E-09 | 2.500E-08 | 3.100E-08 |
| 6674 | 334 | V334F | A | 4.2e-08 | 4e-08 | 2.1e-08 | 3.433E-08 | 1.159E-08 | 2.100E-08 | 4.200E-08 |
| 6675 | 334 | V334P | A | 1.8e-07 | 3.5e-08 | 6e-08 | 9.167E-08 | 7.751E-08 | 3.500E-08 | 1.800E-07 |
| 6676 | 334 | V334S | A | 5.3e-08 | 2.8e-08 | 7e-08 | 5.033E-08 | 2.113E-08 | 2.800E-08 | 7.000E-08 |
| 6677 | 334 | V334T | A | 1.8e-08 | 3.4e-08 | 5.5e-08 | 3.567E-08 | 1.856E-08 | 1.800E-08 | 5.500E-08 |
| 6678 | 334 | V334W | A | 3.7e-08 | 4.2e-08 | 1.1e-07 | 6.300E-08 | 4.078E-08 | 3.700E-08 | 1.100E-07 |
| 6679 | 334 | V334Y | A | 6.2e-08 | 7.2e-08 | 2.2e-08 | 5.200E-08 | 2.646E-08 | 2.200E-08 | 7.200E-08 |
| 6680 | 334 | V334V | A | 4.1e-08 | 2.9e-08 | 3.4e-08 | 3.467E-08 | 6.028E-09 | 2.900E-08 | 4.100E-08 |
| 6681 | 335 | T335A | A | 3.4e-08 | 2.5e-08 | 6.5e-08 | 4.133E-08 | 2.098E-08 | 2.500E-08 | 6.500E-08 |
| 6682 | 335 | T335R | A | 4.1e-08 | 2.2e-08 | 6.2e-08 | 4.167E-08 | 2.001E-08 | 2.200E-08 | 6.200E-08 |
| 6683 | 335 | T335N | A | 3.1e-08 | 5.3e-08 | 6.8e-08 | 5.067E-08 | 1.861E-08 | 3.100E-08 | 6.800E-08 |
| 6684 | 335 | T335D | A | 6.9e-08 | 5.3e-08 | 2.3e-08 | 4.833E-08 | 2.335E-08 | 2.300E-08 | 6.900E-08 |
| 6685 | 335 | T335C | A | 7.2e-08 | 8.7e-08 | 2.5e-08 | 6.133E-08 | 3.235E-08 | 2.500E-08 | 8.700E-08 |
| 6686 | 335 | T335E | A | 1.6e-08 | 2.7e-08 | 3.5e-08 | 2.600E-08 | 9.539E-09 | 1.600E-08 | 3.500E-08 |
| 6687 | 335 | T335Q | A | 4.1e-08 | 4.1e-08 | 6.4e-08 | 4.867E-08 | 1.328E-08 | 4.100E-08 | 6.400E-08 |
| 6688 | 335 | T335G | A | 1.1e-07 | 7.3e-08 | 2.2e-08 | 6.833E-08 | 4.419E-08 | 2.200E-08 | 1.100E-07 |
| 6689 | 335 | T335H | A | 6.1e-08 | 8.3e-08 | 1.1e-07 | 8.467E-08 | 2.454E-08 | 6.100E-08 | 1.100E-07 |
| 6690 | 335 | T335I | A | 3.1e-08 | 1e-07 | 6.1e-08 | 6.400E-08 | 3.460E-08 | 3.100E-08 | 1.000E-07 |
| 6691 | 335 | T335L | A | 5.5e-08 | 7.2e-08 | 7.1e-08 | 6.600E-08 | 9.539E-09 | 5.500E-08 | 7.200E-08 |
| 6692 | 335 | T335K | A | 5.1e-08 | 7.4e-08 | 9.8e-08 | 7.433E-08 | 2.350E-08 | 5.100E-08 | 9.800E-08 |
| 6693 | 335 | T335M | A | 3.4e-08 | 7.4e-08 | 7.3e-08 | 6.033E-08 | 2.281E-08 | 3.400E-08 | 7.400E-08 |
| 6694 | 335 | T335F | A | 7.9e-08 | 6e-08 | 5.4e-08 | 6.433E-08 | 1.305E-08 | 5.400E-08 | 7.900E-08 |
| 6695 | 335 | T335P | A | 2.3e-08 | 8.5e-08 | 7.5e-08 | 6.100E-08 | 3.329E-08 | 2.300E-08 | 8.500E-08 |
| 6696 | 335 | T335S | A | 1.7e-08 | 5.7e-08 | 4.2e-08 | 3.867E-08 | 2.021E-08 | 1.700E-08 | 5.700E-08 |
| 6697 | 335 | T335T | A | 2.3e-08 | 2.8e-08 | 3.5e-08 | 2.867E-08 | 6.028E-09 | 2.300E-08 | 3.500E-08 |
| 6698 | 335 | T335W | A | 3.4e-08 | 4.2e-08 | 3.9e-08 | 3.833E-08 | 4.041E-09 | 3.400E-08 | 4.200E-08 |
| 6699 | 335 | T335Y | A | 4.9e-08 | 2.9e-08 | 1.9e-08 | 3.233E-08 | 1.528E-08 | 1.900E-08 | 4.900E-08 |
| 6700 | 335 | T335V | A | 4.6e-08 | 3.9e-08 | 3.5e-08 | 4.000E-08 | 5.568E-09 | 3.500E-08 | 4.600E-08 |
| 6701 | 336 | S336A | A | 2.5e-08 | 2.4e-08 | 3e-08 | 2.633E-08 | 3.215E-09 | 2.400E-08 | 3.000E-08 |
| 6702 | 336 | S336R | A | 5.6e-08 | 1e-07 | 1e-07 | 8.533E-08 | 2.540E-08 | 5.600E-08 | 1.000E-07 |
| 6703 | 336 | S336N | A | 3.1e-08 | 2.1e-08 | 6e-08 | 3.733E-08 | 2.026E-08 | 2.100E-08 | 6.000E-08 |
| 6704 | 336 | S336D | A | 1.9e-08 | 2.4e-08 | 7.1e-08 | 3.800E-08 | 2.869E-08 | 1.900E-08 | 7.100E-08 |
| 6705 | 336 | S336C | A | 8.4e-08 | 4.5e-08 | 2.3e-07 | 1.197E-07 | 9.752E-08 | 4.500E-08 | 2.300E-07 |
| 6706 | 336 | S336E | A | 4.5e-08 | 2.8e-08 | 4.2e-08 | 3.833E-08 | 9.074E-09 | 2.800E-08 | 4.500E-08 |
| 6707 | 336 | S336Q | A | 5.5e-08 | 4.5e-08 | 6.3e-08 | 5.433E-08 | 9.018E-09 | 4.500E-08 | 6.300E-08 |
| 6708 | 336 | S336G | A | 3.2e-08 | 7.9e-08 | 1.7e-07 | 9.367E-08 | 7.016E-08 | 3.200E-08 | 1.700E-07 |
| 6709 | 336 | S336H | A | 5.2e-08 | 4e-08 | 1.1e-08 | 3.433E-08 | 2.108E-08 | 1.100E-08 | 5.200E-08 |
| 6710 | 336 | S336I | A | 3e-08 | 4.6e-08 | 5.1e-08 | 4.233E-08 | 1.097E-08 | 3.000E-08 | 5.100E-08 |
| 6711 | 336 | S336L | A | 2.4e-08 | 1.8e-08 | 5.6e-08 | 3.267E-08 | 2.043E-08 | 1.800E-08 | 5.600E-08 |
| 6712 | 336 | S336K | A | 9.6e-08 | 8.2e-08 | 6.3e-08 | 8.033E-08 | 1.656E-08 | 6.300E-08 | 9.600E-08 |
| 6713 | 336 | S336M | A | 3e-08 | 2.8e-08 | 1.5e-08 | 2.433E-08 | 8.145E-09 | 1.500E-08 | 3.000E-08 |
| 6714 | 336 | S336F | A | 6.3e-08 | 1.6e-08 | 2.7e-08 | 3.533E-08 | 2.458E-08 | 1.600E-08 | 6.300E-08 |

| | | | | | | | | | | |
|---|---|---|---|---|---|---|---|---|---|---|
| 6715 | 336 | S336P | A | 3.9e-08 | 6.1e-08 | 3.9e-08 | 4.633E-08 | 1.270E-08 | 3.900E-08 | 6.100E-08 |
| 6716 | 336 | S336S | A | 4.5e-08 | 3e-08 | 4.7e-08 | 4.067E-08 | 9.292E-09 | 3.000E-08 | 4.700E-08 |
| 6717 | 336 | S336T | A | 3.4e-08 | 4.7e-08 | 7.4e-08 | 5.167E-08 | 2.040E-08 | 3.400E-08 | 7.400E-08 |
| 6718 | 336 | S336W | A | 1.1e-07 | 4.5e-08 | 1.6e-08 | 5.700E-08 | 4.814E-08 | 1.600E-08 | 1.100E-07 |
| 6719 | 336 | S336Y | A | 1e-07 | 2.5e-08 | 6.1e-08 | 6.200E-08 | 3.751E-08 | 2.500E-08 | 1.000E-07 |
| 6720 | 336 | S336V | A | 6.1e-08 | 4.9e-08 | 3.2e-08 | 4.733E-08 | 1.457E-08 | 3.200E-08 | 6.100E-08 |
| 6721 | 337 | A337A | A | 4.7e-08 | 8.6e-08 | 9.7e-09 | 4.757E-08 | 3.815E-08 | 9.700E-09 | 8.600E-08 |
| 6722 | 337 | A337R | A | 5e-08 | 6.3e-08 | 1e-07 | 7.100E-08 | 2.594E-08 | 5.000E-08 | 1.000E-07 |
| 6723 | 337 | A337N | A | 2.5e-08 | 4e-08 | 2.8e-08 | 3.100E-08 | 7.937E-09 | 2.500E-08 | 4.000E-08 |
| 6724 | 337 | A337D | A | 4.3e-08 | 4.6e-08 | 7.3e-08 | 5.400E-08 | 1.652E-08 | 4.300E-08 | 7.300E-08 |
| 6725 | 337 | A337C | A | 5.3e-08 | 1.9e-08 | 1.9e-08 | 3.033E-08 | 1.963E-08 | 1.900E-08 | 5.300E-08 |
| 6726 | 337 | A337E | A | 3.3e-08 | 8.1e-08 | 6.9e-08 | 6.100E-08 | 2.498E-08 | 3.300E-08 | 8.100E-08 |
| 6727 | 337 | A337Q | A | 7.3e-08 | 2.6e-08 | 3.1e-08 | 4.333E-08 | 2.581E-08 | 2.600E-08 | 7.300E-08 |
| 6728 | 337 | A337G | A | 3.2e-08 | 7.3e-08 | 1.6e-08 | 4.033E-08 | 2.940E-08 | 1.600E-08 | 7.300E-08 |
| 6729 | 337 | A337H | A | 8.7e-08 | 2.1e-08 | 8.5e-08 | 6.433E-08 | 3.754E-08 | 2.100E-08 | 8.700E-08 |
| 6730 | 337 | A337I | A | 4.6e-08 | 4.4e-08 | 2.6e-08 | 3.867E-08 | 1.102E-08 | 2.600E-08 | 4.600E-08 |
| 6731 | 337 | A337L | A | 4.4e-08 | 4.4e-08 | 5.7e-08 | 4.833E-08 | 7.506E-09 | 4.400E-08 | 5.700E-08 |
| 6732 | 337 | A337K | A | 2.4e-08 | 1.8e-08 | 5.9e-08 | 3.367E-08 | 2.214E-08 | 1.800E-08 | 5.900E-08 |
| 6733 | 337 | A337M | A | 5.4e-08 | 6.2e-08 | 6.4e-08 | 6.000E-08 | 5.292E-09 | 5.400E-08 | 6.400E-08 |
| 6734 | 337 | A337F | A | 2.4e-08 | 2.2e-08 | 4.8e-08 | 3.133E-08 | 1.447E-08 | 2.200E-08 | 4.800E-08 |
| 6735 | 337 | A337P | A | 4e-08 | 2.2e-08 | 2.1e-08 | 2.767E-08 | 1.069E-08 | 2.100E-08 | 4.000E-08 |
| 6736 | 337 | A337S | A | 3.2e-08 | 2e-08 | 8.4e-08 | 4.533E-08 | 3.402E-08 | 2.000E-08 | 8.400E-08 |
| 6737 | 337 | A337T | A | 4.4e-08 | 9.9e-08 | 4e-08 | 6.100E-08 | 3.297E-08 | 4.000E-08 | 9.900E-08 |
| 6738 | 337 | A337W | A | 1.3e-07 | 2.1e-08 | 3.2e-08 | 6.100E-08 | 6.001E-08 | 2.100E-08 | 1.300E-07 |
| 6739 | 337 | A337Y | A | 2.6e-08 | 1.4e-08 | 4.6e-08 | 2.867E-08 | 1.617E-08 | 1.400E-08 | 4.600E-08 |
| 6740 | 337 | A337V | A | 7.4e-08 | 2.3e-08 | 3.4e-08 | 4.367E-08 | 2.684E-08 | 2.300E-08 | 7.400E-08 |
| 6741 | 338 | N338A | A | 4.4e-08 | 1.9e-08 | 3.7e-08 | 3.333E-08 | 1.290E-08 | 1.900E-08 | 4.400E-08 |
| 6742 | 338 | N338R | A | 4.3e-08 | 1.7e-08 | 2.9e-08 | 2.967E-08 | 1.301E-08 | 1.700E-08 | 4.300E-08 |
| 6743 | 338 | N338N | A | 6.2e-08 | 6e-08 | 6.3e-08 | 6.167E-08 | 1.528E-09 | 6.000E-08 | 6.300E-08 |
| 6744 | 338 | N338D | A | 5.9e-08 | 3.1e-08 | 2.1e-08 | 3.700E-08 | 1.970E-08 | 2.100E-08 | 5.900E-08 |
| 6745 | 338 | N338C | A | 6e-08 | 5.9e-08 | 3.3e-08 | 5.067E-08 | 1.531E-08 | 3.300E-08 | 6.000E-08 |
| 6746 | 338 | N338E | A | 4.6e-08 | 1e-07 | 2.6e-08 | 5.733E-08 | 3.828E-08 | 2.600E-08 | 1.000E-07 |
| 6747 | 338 | N338Q | A | 4.3e-08 | 2.2e-08 | 5.8e-08 | 4.100E-08 | 1.808E-08 | 2.200E-08 | 5.800E-08 |
| 6748 | 338 | N338G | A | 2.2e-08 | 2.6e-08 | 7e-08 | 3.933E-08 | 2.663E-08 | 2.200E-08 | 7.000E-08 |
| 6749 | 338 | N338H | A | 5.2e-08 | 1e-08 | 1.3e-07 | 6.400E-08 | 6.089E-08 | 1.000E-08 | 1.300E-07 |
| 6750 | 338 | N338I | A | 6e-08 | 5.2e-08 | 3.4e-08 | 4.867E-08 | 1.332E-08 | 3.400E-08 | 6.000E-08 |
| 6751 | 338 | N338L | A | 2.5e-08 | 3.1e-08 | 1.2e-07 | 5.867E-08 | 5.320E-08 | 2.500E-08 | 1.200E-07 |
| 6752 | 338 | N338K | A | 5e-08 | 2.6e-08 | 9.7e-09 | 2.857E-08 | 2.027E-08 | 9.700E-09 | 5.000E-08 |
| 6753 | 338 | N338M | A | 6.4e-08 | 7.7e-09 | 2.1e-08 | 3.090E-08 | 2.943E-08 | 7.700E-09 | 6.400E-08 |
| 6754 | 338 | N338F | A | 3.5e-08 | 4e-08 | 3.8e-08 | 3.767E-08 | 2.517E-09 | 3.500E-08 | 4.000E-08 |
| 6755 | 338 | N338P | A | 1.8e-08 | 1.6e-08 | 1e-07 | 4.467E-08 | 4.793E-08 | 1.600E-08 | 1.000E-07 |
| 6756 | 338 | N338S | A | 9.6e-08 | 1.2e-08 | 1.8e-08 | 4.200E-08 | 4.686E-08 | 1.200E-08 | 9.600E-08 |
| 6757 | 338 | N338T | A | 6.3e-08 | 5.6e-08 | 6.4e-08 | 6.100E-08 | 4.359E-09 | 5.600E-08 | 6.400E-08 |
| 6758 | 338 | N338W | A | 5.8e-08 | 2.5e-08 | 5.5e-08 | 4.600E-08 | 1.825E-08 | 2.500E-08 | 5.800E-08 |
| 6759 | 338 | N338Y | A | 5.7e-08 | 6.5e-08 | 2.6e-08 | 4.933E-08 | 2.060E-08 | 2.600E-08 | 6.500E-08 |
| 6760 | 338 | N338V | A | 8.6e-08 | 5e-08 | 6.3e-08 | 6.633E-08 | 1.823E-08 | 5.000E-08 | 8.600E-08 |
| 6761 | 339 | I339A | A | 7.9e-08 | 1.9e-08 | 8.8e-08 | 6.200E-08 | 3.751E-08 | 1.900E-08 | 8.800E-08 |
| 6762 | 339 | I339R | A | 7.5e-08 | 6.9e-08 | 5.2e-08 | 6.533E-08 | 1.193E-08 | 5.200E-08 | 7.500E-08 |
| 6763 | 339 | I339N | A | 2.3e-08 | 5.1e-08 | 4.5e-08 | 3.967E-08 | 1.474E-08 | 2.300E-08 | 5.100E-08 |
| 6764 | 339 | I339D | A | 5e-08 | 2.9e-08 | 2.9e-08 | 3.600E-08 | 1.212E-08 | 2.900E-08 | 5.000E-08 |
| 6765 | 339 | I339C | A | 5.4e-08 | 3.9e-08 | 6.6e-08 | 5.300E-08 | 1.353E-08 | 3.900E-08 | 6.600E-08 |
| 6766 | 339 | I339E | A | 3.8e-08 | 2.8e-08 | 5.7e-08 | 4.100E-08 | 1.473E-08 | 2.800E-08 | 5.700E-08 |
| 6767 | 339 | I339Q | A | 9.6e-08 | 6.2e-08 | 2.7e-08 | 6.167E-08 | 3.450E-08 | 2.700E-08 | 9.600E-08 |
| 6768 | 339 | I339G | A | 9.4e-08 | 3.5e-08 | 9e-08 | 7.300E-08 | 3.297E-08 | 3.500E-08 | 9.400E-08 |
| 6769 | 339 | I339H | A | 7.8e-08 | 3.8e-08 | 2.1e-08 | 4.567E-08 | 2.926E-08 | 2.100E-08 | 7.800E-08 |
| 6770 | 339 | I339I | A | 4.2e-08 | 4.4e-08 | 6.5e-08 | 5.033E-08 | 1.274E-08 | 4.200E-08 | 6.500E-08 |
| 6771 | 339 | I339L | A | 5.7e-08 | 2.7e-08 | 4.3e-08 | 4.233E-08 | 1.501E-08 | 2.700E-08 | 5.700E-08 |
| 6772 | 339 | I339K | A | 4.8e-08 | 1.1e-07 | 1.5e-08 | 5.767E-08 | 4.823E-08 | 1.500E-08 | 1.100E-07 |
| 6773 | 339 | I339M | A | 2.8e-08 | 4.2e-08 | 3.2e-08 | 3.400E-08 | 7.211E-09 | 2.800E-08 | 4.200E-08 |
| 6774 | 339 | I339F | A | 5.5e-08 | 1.6e-08 | 6.4e-08 | 4.500E-08 | 2.551E-08 | 1.600E-08 | 6.400E-08 |
| 6775 | 339 | I339P | A | 7.3e-08 | 8.1e-08 | 1e-08 | 5.467E-08 | 3.889E-08 | 1.000E-08 | 8.100E-08 |
| 6776 | 339 | I339S | A | 4.3e-08 | 2.1e-08 | 1.5e-08 | 2.633E-08 | 1.474E-08 | 1.500E-08 | 4.300E-08 |
| 6777 | 339 | I339T | A | 6.6e-08 | 1.8e-08 | 4.9e-08 | 4.433E-08 | 2.434E-08 | 1.800E-08 | 6.600E-08 |
| 6778 | 339 | I339W | A | 8.2e-09 | 1.8e-08 | 1.7e-07 | 6.540E-08 | 9.072E-08 | 8.200E-09 | 1.700E-07 |
| 6779 | 339 | I339Y | A | 1.4e-08 | 6.7e-08 | 3.8e-08 | 3.967E-08 | 2.654E-08 | 1.400E-08 | 6.700E-08 |
| 6780 | 339 | I339V | A | 1.9e-08 | 2.9e-08 | 2.4e-08 | 2.400E-08 | 5.000E-09 | 1.900E-08 | 2.900E-08 |
| 6781 | 340 | Q340A | A | 5.2e-08 | 8.9e-09 | 3.9e-08 | 3.330E-08 | 2.211E-08 | 8.900E-09 | 5.200E-08 |
| 6782 | 340 | Q340R | A | 5.2e-08 | 3.3e-08 | 3.9e-08 | 4.133E-08 | 9.713E-09 | 3.300E-08 | 5.200E-08 |
| 6783 | 340 | Q340N | A | 3.6e-07 | 3.3e-08 | 4.8e-08 | 1.470E-07 | 1.846E-07 | 3.300E-08 | 3.600E-07 |
| 6784 | 340 | Q340D | A | 1.2e-08 | 1.7e-08 | 5e-08 | 2.633E-08 | 2.065E-08 | 1.200E-08 | 5.000E-08 |
| 6785 | 340 | Q340C | A | 4.1e-08 | 4.7e-08 | 9.9e-08 | 6.233E-08 | 3.190E-08 | 4.100E-08 | 9.900E-08 |
| 6786 | 340 | Q340E | A | 5.9e-08 | 4.1e-08 | 1.7e-08 | 3.900E-08 | 2.107E-08 | 1.700E-08 | 5.900E-08 |
| 6787 | 340 | Q340Q | A | 4.2e-08 | 4.8e-08 | 2.1e-08 | 3.700E-08 | 1.418E-08 | 2.100E-08 | 4.800E-08 |
| 6788 | 340 | Q340G | A | 3.1e-08 | 7.8e-08 | 5.9e-07 | 2.330E-07 | 3.101E-07 | 3.100E-08 | 5.900E-07 |
| 6789 | 340 | Q340H | A | 4.2e-08 | 1.1e-07 | 5e-08 | 6.733E-08 | 3.717E-08 | 4.200E-08 | 1.100E-07 |
| 6790 | 340 | Q340I | A | 4.6e-08 | 5.1e-08 | 5.4e-08 | 5.033E-08 | 4.041E-09 | 4.600E-08 | 5.400E-08 |
| 6791 | 340 | Q340L | A | 5.8e-08 | 3.2e-08 | 2.8e-08 | 3.933E-08 | 1.629E-08 | 2.800E-08 | 5.800E-08 |
| 6792 | 340 | Q340K | A | 7.5e-08 | 2.2e-08 | 1.2e-08 | 3.633E-08 | 3.386E-08 | 1.200E-08 | 7.500E-08 |
| 6793 | 340 | Q340M | A | 5.8e-08 | 4.9e-08 | 1.2e-08 | 3.967E-08 | 2.438E-08 | 1.200E-08 | 5.800E-08 |
| 6794 | 340 | Q340F | A | 3.3e-08 | 8.7e-08 | 4.9e-08 | 5.633E-08 | 2.774E-08 | 3.300E-08 | 8.700E-08 |
| 6795 | 340 | Q340P | A | 1.8e-08 | 9.9e-08 | 6.7e-09 | 4.123E-08 | 5.035E-08 | 6.700E-09 | 9.900E-08 |
| 6796 | 340 | Q340S | A | 7.3e-08 | 5.6e-08 | 6.9e-08 | 6.600E-08 | 8.888E-09 | 5.600E-08 | 7.300E-08 |
| 6797 | 340 | Q340T | A | 1.3e-08 | 1e-07 | 8.4e-08 | 6.567E-08 | 4.631E-08 | 1.300E-08 | 1.000E-07 |
| 6798 | 340 | Q340W | A | 4.6e-08 | 7.4e-08 | 5e-08 | 5.667E-08 | 1.514E-08 | 4.600E-08 | 7.400E-08 |
| 6799 | 340 | Q340Y | A | 1.7e-08 | 2.8e-08 | 1e-07 | 4.833E-08 | 4.508E-08 | 1.700E-08 | 1.000E-07 |

| | | | | | | | | | | |
|---|---|---|---|---|---|---|---|---|---|---|
| 6800 | 340 | Q340V | A | 2e-08 | 1.2e-08 | 4.1e-08 | 2.433E-08 | 1.498E-08 | 1.200E-08 | 4.100E-08 |
| 6801 | 341 | E341A | A | 1.7e-08 | 5.6e-08 | 4.6e-08 | 3.967E-08 | 2.026E-08 | 1.700E-08 | 5.600E-08 |
| 6802 | 341 | E341R | A | 4.4e-08 | 3.2e-08 | 5.8e-08 | 4.467E-08 | 1.301E-08 | 3.200E-08 | 5.800E-08 |
| 6803 | 341 | E341N | A | 5.4e-08 | 2.2e-08 | 4.4e-08 | 4.000E-08 | 1.637E-08 | 2.200E-08 | 5.400E-08 |
| 6804 | 341 | E341D | A | 2.8e-08 | 2.6e-08 | 1.7e-08 | 2.367E-08 | 5.859E-09 | 1.700E-08 | 2.800E-08 |
| 6805 | 341 | E341C | A | 1.9e-08 | 4.3e-08 | 3.9e-08 | 3.367E-08 | 1.286E-08 | 1.900E-08 | 4.300E-08 |
| 6806 | 341 | E341E | A | 3.6e-08 | 8.9e-08 | 9e-08 | 7.167E-08 | 3.089E-08 | 3.600E-08 | 9.000E-08 |
| 6807 | 341 | E341Q | A | 3.1e-08 | 3.8e-08 | 2.3e-08 | 3.067E-08 | 7.506E-09 | 2.300E-08 | 3.800E-08 |
| 6808 | 341 | E341G | A | 6.7e-08 | 1.2e-08 | 3.9e-08 | 3.933E-08 | 2.750E-08 | 1.200E-08 | 6.700E-08 |
| 6809 | 341 | E341H | A | 2.6e-08 | 2.3e-08 | 3.4e-08 | 2.767E-08 | 5.686E-09 | 2.300E-08 | 3.400E-08 |
| 6810 | 341 | E341I | A | 2.6e-08 | 4.2e-08 | 8.4e-08 | 5.067E-08 | 2.996E-08 | 2.600E-08 | 8.400E-08 |
| 6811 | 341 | E341L | A | 3.6e-08 | 3.6e-08 | 6.3e-08 | 4.500E-08 | 1.559E-08 | 3.600E-08 | 6.300E-08 |
| 6812 | 341 | E341K | A | 1.8e-07 | 5.2e-08 | 6.4e-08 | 9.867E-08 | 7.069E-08 | 5.200E-08 | 1.800E-07 |
| 6813 | 341 | E341M | A | 2.8e-08 | 5.1e-08 | 4.8e-08 | 4.233E-08 | 1.250E-08 | 2.800E-08 | 5.100E-08 |
| 6814 | 341 | E341F | A | 1.7e-08 | 7.5e-08 | 2.2e-08 | 3.800E-08 | 3.214E-08 | 1.700E-08 | 7.500E-08 |
| 6815 | 341 | E341P | A | 1.7e-08 | 8.1e-08 | 2e-08 | 3.933E-08 | 3.612E-08 | 1.700E-08 | 8.100E-08 |
| 6816 | 341 | E341S | A | 4.3e-08 | 8.6e-08 | 1.6e-07 | 9.633E-08 | 5.918E-08 | 4.300E-08 | 1.600E-07 |
| 6817 | 341 | E341T | A | 3.5e-08 | 4.6e-08 | 2.4e-08 | 3.500E-08 | 1.100E-08 | 2.400E-08 | 4.600E-08 |
| 6818 | 341 | E341W | A | 7.7e-09 | 5e-08 | 4.9e-08 | 3.557E-08 | 2.414E-08 | 7.700E-09 | 5.000E-08 |
| 6819 | 341 | E341Y | A | 3.2e-08 | 3.9e-08 | 8.2e-08 | 5.100E-08 | 2.707E-08 | 3.200E-08 | 8.200E-08 |
| 6820 | 341 | E341V | A | 4.4e-08 | 7.2e-08 | 2.2e-08 | 4.600E-08 | 2.506E-08 | 2.200E-08 | 7.200E-08 |
| 6821 | 342 | F342A | A | 7.9e-08 | 5.2e-08 | 3.8e-08 | 5.633E-08 | 2.084E-08 | 3.800E-08 | 7.900E-08 |
| 6822 | 342 | F342R | A | 8.6e-08 | 8.5e-09 | 3.6e-08 | 4.350E-08 | 3.929E-08 | 8.500E-09 | 8.600E-08 |
| 6823 | 342 | F342N | A | 4.1e-08 | 4.6e-08 | 2.4e-08 | 3.700E-08 | 1.153E-08 | 2.400E-08 | 4.600E-08 |
| 6824 | 342 | F342D | A | 6.4e-08 | 7.1e-08 | 5.7e-08 | 6.400E-08 | 7.000E-09 | 5.700E-08 | 7.100E-08 |
| 6825 | 342 | F342C | A | 1.6e-07 | 1.6e-08 | 7e-08 | 8.200E-08 | 7.275E-08 | 1.600E-08 | 1.600E-07 |
| 6826 | 342 | F342E | A | 3e-08 | 3.8e-08 | 3.8e-08 | 3.533E-08 | 4.619E-09 | 3.000E-08 | 3.800E-08 |
| 6827 | 342 | F342Q | A | 1.6e-08 | 4.5e-08 | 1.1e-08 | 2.400E-08 | 1.836E-08 | 1.100E-08 | 4.500E-08 |
| 6828 | 342 | F342G | A | 9.5e-08 | 2.7e-07 | 4.3e-08 | 1.360E-07 | 1.189E-07 | 4.300E-08 | 2.700E-07 |
| 6829 | 342 | F342H | A | 2.5e-08 | 1.2e-08 | 3.7e-08 | 2.467E-08 | 1.250E-08 | 1.200E-08 | 3.700E-08 |
| 6830 | 342 | F342I | A | 7.5e-08 | 3e-08 | 6e-08 | 5.500E-08 | 2.291E-08 | 3.000E-08 | 7.500E-08 |
| 6831 | 342 | F342L | A | 6.6e-08 | 6.4e-08 | 5.4e-08 | 6.133E-08 | 6.429E-09 | 5.400E-08 | 6.600E-08 |
| 6832 | 342 | F342K | A | 7e-08 | 8.6e-08 | 1.6e-08 | 5.733E-08 | 3.668E-08 | 1.600E-08 | 8.600E-08 |
| 6833 | 342 | F342M | A | 3.3e-08 | 8.1e-08 | 6.8e-08 | 6.067E-08 | 2.483E-08 | 3.300E-08 | 8.100E-08 |
| 6834 | 342 | F342F | A | 6.7e-08 | 6.1e-08 | 5.9e-09 | 4.463E-08 | 3.368E-08 | 5.900E-09 | 6.700E-08 |
| 6835 | 342 | F342P | A | 5.5e-08 | 2.1e-08 | 6.2e-08 | 4.600E-08 | 2.193E-08 | 2.100E-08 | 6.200E-08 |
| 6836 | 342 | F342S | A | 5.2e-08 | 8.8e-08 | 1.1e-07 | 8.333E-08 | 2.928E-08 | 5.200E-08 | 1.100E-07 |
| 6837 | 342 | F342T | A | 2.2e-08 | 6.4e-08 | 1.9e-08 | 3.500E-08 | 2.516E-08 | 1.900E-08 | 6.400E-08 |
| 6838 | 342 | F342W | A | 5.6e-08 | 2.1e-08 | 5.6e-08 | 4.433E-08 | 2.021E-08 | 2.100E-08 | 5.600E-08 |
| 6839 | 342 | F342Y | A | 7.9e-08 | 1e-07 | 4.6e-08 | 7.500E-08 | 2.722E-08 | 4.600E-08 | 1.000E-07 |
| 6840 | 342 | F342V | A | 5.5e-08 | 9.6e-08 | 2.2e-08 | 5.767E-08 | 3.707E-08 | 2.200E-08 | 9.600E-08 |
| 6841 | 343 | A343A | A | 2e-08 | 1.4e-08 | 4.5e-08 | 2.633E-08 | 1.644E-08 | 1.400E-08 | 4.500E-08 |
| 6842 | 343 | A343R | A | 1.5e-08 | 4.5e-08 | 4.5e-08 | 3.500E-08 | 1.732E-08 | 1.500E-08 | 4.500E-08 |
| 6843 | 343 | A343N | A | 2.4e-08 | 9.5e-09 | 9.9e-08 | 4.417E-08 | 4.804E-08 | 9.500E-09 | 9.900E-08 |
| 6844 | 343 | A343D | A | 2.7e-08 | 1.5e-08 | 2.5e-08 | 2.233E-08 | 6.429E-09 | 1.500E-08 | 2.700E-08 |
| 6845 | 343 | A343C | A | 6.8e-08 | 2.6e-08 | 3.3e-08 | 4.233E-08 | 2.250E-08 | 2.600E-08 | 6.800E-08 |
| 6846 | 343 | A343E | A | 5.6e-08 | 7.3e-08 | 4e-08 | 5.633E-08 | 1.650E-08 | 4.000E-08 | 7.300E-08 |
| 6847 | 343 | A343Q | A | 5.5e-08 | 2.7e-08 | 4e-08 | 4.067E-08 | 1.401E-08 | 2.700E-08 | 5.500E-08 |
| 6848 | 343 | A343G | A | 3.7e-08 | 4.9e-08 | 4.3e-08 | 4.300E-08 | 6.000E-09 | 3.700E-08 | 4.900E-08 |
| 6849 | 343 | A343H | A | 2.8e-08 | 3.3e-08 | 2.8e-08 | 2.967E-08 | 2.887E-09 | 2.800E-08 | 3.300E-08 |
| 6850 | 343 | A343I | A | 2e-08 | 1.4e-07 | 1.1e-07 | 9.000E-08 | 6.245E-08 | 2.000E-08 | 1.400E-07 |
| 6851 | 343 | A343L | A | 1.5e-08 | 6e-08 | 3.8e-08 | 3.767E-08 | 2.250E-08 | 1.500E-08 | 6.000E-08 |
| 6852 | 343 | A343K | A | 8.2e-08 | 4.7e-08 | 3.1e-08 | 5.333E-08 | 2.608E-08 | 3.100E-08 | 8.200E-08 |
| 6853 | 343 | A343M | A | 2.7e-08 | 2.5e-08 | 2.5e-08 | 2.567E-08 | 1.155E-09 | 2.500E-08 | 2.700E-08 |
| 6854 | 343 | A343F | A | 3.2e-08 | 5.2e-08 | 1.1e-08 | 3.167E-08 | 2.050E-08 | 1.100E-08 | 5.200E-08 |
| 6855 | 343 | A343P | A | 2.5e-08 | 6.8e-08 | 2e-08 | 3.767E-08 | 2.639E-08 | 2.000E-08 | 6.800E-08 |
| 6856 | 343 | A343S | A | 6.3e-08 | 5e-08 | 3.5e-08 | 4.933E-08 | 1.401E-08 | 3.500E-08 | 6.300E-08 |
| 6857 | 343 | A343T | A | 9.1e-08 | 3.8e-08 | 2.5e-08 | 5.133E-08 | 3.496E-08 | 2.500E-08 | 9.100E-08 |
| 6858 | 343 | A343W | A | 8.7e-08 | 4.3e-08 | 2.2e-08 | 5.067E-08 | 3.317E-08 | 2.200E-08 | 8.700E-08 |
| 6859 | 343 | A343Y | A | 6.3e-08 | 2.5e-08 | 5.8e-08 | 4.867E-08 | 2.065E-08 | 2.500E-08 | 6.300E-08 |
| 6860 | 343 | A343V | A | 3.9e-08 | 1.8e-08 | 6.4e-08 | 4.033E-08 | 2.303E-08 | 1.800E-08 | 6.400E-08 |
| 6861 | 344 | G344A | A | 1.7e-07 | 5.9e-08 | 5.9e-08 | 9.600E-08 | 6.409E-08 | 5.900E-08 | 1.700E-07 |
| 6862 | 344 | G344R | A | 4.5e-08 | 9e-08 | 1e-07 | 7.833E-08 | 2.930E-08 | 4.500E-08 | 1.000E-07 |
| 6863 | 344 | G344N | A | 3.4e-08 | 4.1e-08 | 5.7e-08 | 4.400E-08 | 1.179E-08 | 3.400E-08 | 5.700E-08 |
| 6864 | 344 | G344D | A | 1e-07 | 3.5e-08 | 1.3e-08 | 4.933E-08 | 4.524E-08 | 1.300E-08 | 1.000E-07 |
| 6865 | 344 | G344C | A | 5.4e-08 | 4.3e-08 | 5.3e-08 | 5.000E-08 | 6.083E-09 | 4.300E-08 | 5.400E-08 |
| 6866 | 344 | G344E | A | 2.4e-08 | 2.6e-08 | 2e-08 | 2.333E-08 | 3.055E-09 | 2.000E-08 | 2.600E-08 |
| 6867 | 344 | G344Q | A | 2.9e-08 | 7.3e-08 | 6.1e-08 | 5.433E-08 | 2.274E-08 | 2.900E-08 | 7.300E-08 |
| 6868 | 344 | G344G | A | 6.3e-08 | 6.3e-08 | 5.3e-08 | 5.967E-08 | 5.774E-09 | 5.300E-08 | 6.300E-08 |
| 6869 | 344 | G344H | A | 2.5e-08 | 2.4e-08 | 4.4e-08 | 3.100E-08 | 1.127E-08 | 2.400E-08 | 4.400E-08 |
| 6870 | 344 | G344I | A | 5.2e-08 | 4.5e-08 | 9.7e-09 | 3.557E-08 | 2.267E-08 | 9.700E-09 | 5.200E-08 |
| 6871 | 344 | G344L | A | 7.4e-08 | 3.5e-08 | 3.8e-08 | 4.900E-08 | 2.170E-08 | 3.500E-08 | 7.400E-08 |
| 6872 | 344 | G344K | A | 2.9e-08 | 7.4e-09 | 3.5e-08 | 2.380E-08 | 1.452E-08 | 7.400E-09 | 3.500E-08 |
| 6873 | 344 | G344M | A | 6.1e-08 | 1.1e-08 | 2.4e-08 | 3.200E-08 | 2.594E-08 | 1.100E-08 | 6.100E-08 |
| 6874 | 344 | G344F | A | 1.4e-08 | 1e-07 | 3.7e-08 | 5.033E-08 | 4.452E-08 | 1.400E-08 | 1.000E-07 |
| 6875 | 344 | G344P | A | 1.5e-07 | 2.7e-08 | 2.7e-08 | 6.800E-08 | 7.101E-08 | 2.700E-08 | 1.500E-07 |
| 6876 | 344 | G344S | A | 4.9e-08 | 4.6e-08 | 4.6e-08 | 4.700E-08 | 1.732E-09 | 4.600E-08 | 4.900E-08 |
| 6877 | 344 | G344T | A | 3.3e-08 | 3.9e-08 | 2.7e-08 | 3.300E-08 | 6.000E-09 | 2.700E-08 | 3.900E-08 |
| 6878 | 344 | G344W | A | 7.7e-08 | 2.4e-08 | 6.3e-08 | 5.467E-08 | 2.747E-08 | 2.400E-08 | 7.700E-08 |
| 6879 | 344 | G344Y | A | 4.8e-08 | 4e-08 | 9.2e-09 | 3.240E-08 | 2.049E-08 | 9.200E-09 | 4.800E-08 |
| 6880 | 344 | G344V | A | 5.1e-08 | 2.6e-08 | 4.1e-08 | 3.933E-08 | 1.258E-08 | 2.600E-08 | 5.100E-08 |
| 6881 | 345 | C345A | A | 3.2e-08 | 2e-08 | 4.8e-08 | 3.333E-08 | 1.405E-08 | 2.000E-08 | 4.800E-08 |
| 6882 | 345 | C345R | A | 1.7e-08 | 5.7e-08 | 4e-08 | 3.800E-08 | 2.007E-08 | 1.700E-08 | 5.700E-08 |
| 6883 | 345 | C345N | A | 8.5e-08 | 2.1e-07 | 2.8e-08 | 1.077E-07 | 9.309E-08 | 2.800E-08 | 2.100E-07 |
| 6884 | 345 | C345D | A | 5.9e-08 | 7.2e-08 | 1.5e-08 | 4.867E-08 | 2.987E-08 | 1.500E-08 | 7.200E-08 |

| | | | | | | | | | | |
|---|---|---|---|---|---|---|---|---|---|---|
| 6885 | 345 | C345C | A | 3.7e-08 | 1.1e-08 | 3e-08 | 2.600E-08 | 1.345E-08 | 1.100E-08 | 3.700E-08 |
| 6886 | 345 | C345E | A | 2.6e-08 | 5.7e-08 | 2.7e-08 | 3.667E-08 | 1.762E-08 | 2.600E-08 | 5.700E-08 |
| 6887 | 345 | C345Q | A | 7.9e-08 | 4.5e-08 | 1.4e-08 | 4.600E-08 | 3.251E-08 | 1.400E-08 | 7.900E-08 |
| 6888 | 345 | C345G | A | 6.3e-08 | 2.3e-08 | 6.1e-09 | 3.070E-08 | 2.922E-08 | 6.100E-09 | 6.300E-08 |
| 6889 | 345 | C345H | A | 3.7e-08 | 4.9e-08 | 4e-08 | 4.200E-08 | 6.245E-09 | 3.700E-08 | 4.900E-08 |
| 6890 | 345 | C345I | A | 5.8e-08 | 7.8e-08 | 2.4e-08 | 5.333E-08 | 2.730E-08 | 2.400E-08 | 7.800E-08 |
| 6891 | 345 | C345L | A | 6.4e-08 | 1.5e-08 | 4.2e-08 | 4.033E-08 | 2.454E-08 | 1.500E-08 | 6.400E-08 |
| 6892 | 345 | C345K | A | 5.3e-08 | 5.6e-08 | 1.9e-08 | 4.267E-08 | 2.055E-08 | 1.900E-08 | 5.600E-08 |
| 6893 | 345 | C345M | A | 4.3e-08 | 1.9e-08 | 3.3e-08 | 3.167E-08 | 1.206E-08 | 1.900E-08 | 4.300E-08 |
| 6894 | 345 | C345F | A | 3.3e-08 | 1.3e-07 | 3.5e-08 | 6.600E-08 | 5.543E-08 | 3.300E-08 | 1.300E-07 |
| 6895 | 345 | C345P | A | 2e-08 | 4.8e-08 | 4.5e-08 | 3.767E-08 | 1.537E-08 | 2.000E-08 | 4.800E-08 |
| 6896 | 345 | C345S | A | 1.1e-08 | 3e-08 | 1.1e-07 | 5.033E-08 | 5.254E-08 | 1.100E-08 | 1.100E-07 |
| 6897 | 345 | C345T | A | 3.4e-08 | 3e-08 | 6e-08 | 4.133E-08 | 1.629E-08 | 3.000E-08 | 6.000E-08 |
| 6898 | 345 | C345W | A | 5.7e-08 | 3e-08 | 2.1e-08 | 3.600E-08 | 1.873E-08 | 2.100E-08 | 5.700E-08 |
| 6899 | 345 | C345Y | A | 4.1e-08 | 1.1e-07 | 1.5e-08 | 5.533E-08 | 4.910E-08 | 1.500E-08 | 1.100E-07 |
| 6900 | 345 | C345V | A | 1.2e-07 | 2.6e-08 | 3.7e-08 | 6.100E-08 | 5.139E-08 | 2.600E-08 | 1.200E-07 |
| 6901 | 346 | K346A | A | 3.7e-08 | 4.4e-08 | 5.4e-08 | 4.500E-08 | 8.544E-09 | 3.700E-08 | 5.400E-08 |
| 6902 | 346 | K346R | A | 6.2e-08 | 3.2e-08 | 4.5e-08 | 4.633E-08 | 1.504E-08 | 3.200E-08 | 6.200E-08 |
| 6903 | 346 | K346N | A | 5.6e-08 | 4.2e-08 | 1e-07 | 6.600E-08 | 3.027E-08 | 4.200E-08 | 1.000E-07 |
| 6904 | 346 | K346D | A | 2.9e-08 | 7.7e-08 | 9.5e-08 | 6.700E-08 | 3.412E-08 | 2.900E-08 | 9.500E-08 |
| 6905 | 346 | K346C | A | 2.4e-08 | 2e-08 | 1.4e-08 | 1.933E-08 | 5.033E-09 | 1.400E-08 | 2.400E-08 |
| 6906 | 346 | K346E | A | 4.7e-08 | 5.2e-08 | 1.7e-08 | 3.867E-08 | 1.893E-08 | 1.700E-08 | 5.200E-08 |
| 6907 | 346 | K346Q | A | 5.6e-08 | 4.5e-08 | 4.1e-08 | 4.733E-08 | 7.767E-09 | 4.100E-08 | 5.600E-08 |
| 6908 | 346 | K346G | A | 6.7e-08 | 4.6e-08 | 5.8e-08 | 5.700E-08 | 1.054E-08 | 4.600E-08 | 6.700E-08 |
| 6909 | 346 | K346H | A | 3.8e-08 | 9e-08 | 9.8e-08 | 7.533E-08 | 3.258E-08 | 3.800E-08 | 9.800E-08 |
| 6910 | 346 | K346I | A | 3.6e-08 | 1.4e-08 | 3e-08 | 2.667E-08 | 1.137E-08 | 1.400E-08 | 3.600E-08 |
| 6911 | 346 | K346L | A | 4.2e-08 | 6.7e-08 | 1.8e-08 | 4.233E-08 | 2.450E-08 | 1.800E-08 | 6.700E-08 |
| 6912 | 346 | K346K | A | 2.3e-08 | 8.7e-08 | 5.1e-08 | 5.367E-08 | 3.208E-08 | 2.300E-08 | 8.700E-08 |
| 6913 | 346 | K346M | A | 3.4e-08 | 6.1e-08 | 6.9e-08 | 5.467E-08 | 1.834E-08 | 3.400E-08 | 6.900E-08 |
| 6914 | 346 | K346F | A | 6.2e-08 | 2.4e-08 | 5.6e-08 | 4.733E-08 | 2.043E-08 | 2.400E-08 | 6.200E-08 |
| 6915 | 346 | K346P | A | 6.3e-08 | 3.7e-08 | 2.5e-08 | 4.167E-08 | 1.943E-08 | 2.500E-08 | 6.300E-08 |
| 6916 | 346 | K346S | A | 2.3e-08 | 5.1e-08 | 3.5e-08 | 3.633E-08 | 1.405E-08 | 2.300E-08 | 5.100E-08 |
| 6917 | 346 | K346T | A | 6.7e-08 | 5.1e-08 | 2e-08 | 4.600E-08 | 2.390E-08 | 2.000E-08 | 6.700E-08 |
| 6918 | 346 | K346W | A | 8.5e-08 | 3.8e-08 | 3.7e-08 | 5.333E-08 | 2.743E-08 | 3.700E-08 | 8.500E-08 |
| 6919 | 346 | K346Y | A | 7.5e-08 | 3.5e-08 | 7.7e-08 | 6.233E-08 | 2.369E-08 | 3.500E-08 | 7.700E-08 |
| 6920 | 346 | K346V | A | 3.7e-08 | 3.7e-08 | 1.3e-07 | 6.800E-08 | 5.369E-08 | 3.700E-08 | 1.300E-07 |
| 6921 | 347 | K347A | A | 8.9e-08 | 6.3e-08 | 1.5e-07 | 1.007E-07 | 4.466E-08 | 6.300E-08 | 1.500E-07 |
| 6922 | 347 | K347R | A | 3.1e-08 | 1.8e-08 | 4.8e-08 | 3.233E-08 | 1.504E-08 | 1.800E-08 | 4.800E-08 |
| 6923 | 347 | K347N | A | 6.2e-08 | 5.6e-08 | 2e-08 | 4.600E-08 | 2.272E-08 | 2.000E-08 | 6.200E-08 |
| 6924 | 347 | K347D | A | 9.7e-08 | 5.5e-08 | 9.7e-08 | 8.300E-08 | 2.425E-08 | 5.500E-08 | 9.700E-08 |
| 6925 | 347 | K347C | A | 4.5e-08 | 5.7e-08 | 3.5e-08 | 4.567E-08 | 1.102E-08 | 3.500E-08 | 5.700E-08 |
| 6926 | 347 | K347E | A | 9.9e-08 | 8.3e-08 | 3.7e-08 | 7.300E-08 | 3.219E-08 | 3.700E-08 | 9.900E-08 |
| 6927 | 347 | K347Q | A | 8.3e-08 | 3.8e-08 | 3e-08 | 5.033E-08 | 2.857E-08 | 3.000E-08 | 8.300E-08 |
| 6928 | 347 | K347G | A | 1e-07 | 8.1e-08 | 5e-08 | 7.700E-08 | 2.524E-08 | 5.000E-08 | 1.000E-07 |
| 6929 | 347 | K347H | A | 1.5e-08 | 2.6e-08 | 2.2e-08 | 2.100E-08 | 5.568E-09 | 1.500E-08 | 2.600E-08 |
| 6930 | 347 | K347I | A | 4.1e-08 | 4.4e-08 | 4.6e-08 | 4.367E-08 | 2.517E-09 | 4.100E-08 | 4.600E-08 |
| 6931 | 347 | K347L | A | 5e-08 | 2.7e-08 | 3e-08 | 3.567E-08 | 1.250E-08 | 2.700E-08 | 5.000E-08 |
| 6932 | 347 | K347K | A | 4.2e-08 | 3.9e-08 | 4.7e-08 | 4.267E-08 | 4.041E-09 | 3.900E-08 | 4.700E-08 |
| 6933 | 347 | K347M | A | 2.3e-08 | 3e-08 | 4.4e-08 | 3.233E-08 | 1.069E-08 | 2.300E-08 | 4.400E-08 |
| 6934 | 347 | K347F | A | 4.3e-08 | 4.1e-08 | 5.6e-08 | 4.667E-08 | 8.145E-09 | 4.100E-08 | 5.600E-08 |
| 6935 | 347 | K347P | A | 5.4e-08 | 9.3e-08 | 2.1e-08 | 5.600E-08 | 3.604E-08 | 2.100E-08 | 9.300E-08 |
| 6936 | 347 | K347S | A | 2.4e-08 | 5.2e-08 | 6.4e-08 | 4.667E-08 | 2.053E-08 | 2.400E-08 | 6.400E-08 |
| 6937 | 347 | K347T | A | 9e-08 | 2.9e-08 | 3.6e-08 | 5.167E-08 | 3.338E-08 | 2.900E-08 | 9.000E-08 |
| 6938 | 347 | K347W | A | 3.3e-08 | 2.5e-08 | 3.8e-08 | 3.200E-08 | 6.557E-09 | 2.500E-08 | 3.800E-08 |
| 6939 | 347 | K347Y | A | 1e-07 | 5.8e-08 | 4.7e-08 | 6.833E-08 | 2.797E-08 | 4.700E-08 | 1.000E-07 |
| 6940 | 347 | K347V | A | 2.7e-08 | 4.7e-08 | 4.4e-08 | 3.933E-08 | 1.079E-08 | 2.700E-08 | 4.700E-08 |
| 6941 | 348 | I348A | A | 3.2e-08 | 1e-07 | 4.9e-08 | 6.033E-08 | 3.539E-08 | 3.200E-08 | 1.000E-07 |
| 6942 | 348 | I348R | A | 3.7e-08 | 2.7e-08 | 8.3e-08 | 4.900E-08 | 2.987E-08 | 2.700E-08 | 8.300E-08 |
| 6943 | 348 | I348N | A | 4.8e-08 | 4.1e-08 | 1.7e-08 | 3.533E-08 | 1.626E-08 | 1.700E-08 | 4.800E-08 |
| 6944 | 348 | I348D | A | 6.4e-08 | 4.9e-08 | 7.5e-08 | 6.267E-08 | 1.305E-08 | 4.900E-08 | 7.500E-08 |
| 6945 | 348 | I348C | A | 4.5e-08 | 5.7e-08 | 9.8e-08 | 6.667E-08 | 2.779E-08 | 4.500E-08 | 9.800E-08 |
| 6946 | 348 | I348E | A | 3.1e-08 | 4.8e-08 | 3.3e-08 | 3.733E-08 | 9.292E-09 | 3.100E-08 | 4.800E-08 |
| 6947 | 348 | I348Q | A | 1.6e-08 | 4.8e-08 | 6.7e-08 | 4.367E-08 | 2.577E-08 | 1.600E-08 | 6.700E-08 |
| 6948 | 348 | I348G | A | 1.4e-08 | 4.9e-08 | 7.2e-08 | 4.500E-08 | 2.921E-08 | 1.400E-08 | 7.200E-08 |
| 6949 | 348 | I348H | A | 3.4e-08 | 8.6e-08 | 5.5e-08 | 5.833E-08 | 2.616E-08 | 3.400E-08 | 8.600E-08 |
| 6950 | 348 | I348I | A | 1.7e-08 | 8.8e-08 | 7.2e-08 | 5.900E-08 | 3.724E-08 | 1.700E-08 | 8.800E-08 |
| 6951 | 348 | I348L | A | 3.6e-08 | 1.9e-08 | 3.5e-08 | 3.000E-08 | 9.539E-09 | 1.900E-08 | 3.600E-08 |
| 6952 | 348 | I348K | A | 1.7e-08 | 5.4e-08 | 2.3e-08 | 3.133E-08 | 1.986E-08 | 1.700E-08 | 5.400E-08 |
| 6953 | 348 | I348M | A | 1.6e-07 | 9.5e-08 | 2e-08 | 9.167E-08 | 7.006E-08 | 2.000E-08 | 1.600E-07 |
| 6954 | 348 | I348F | A | 3.8e-08 | 5e-08 | 6.1e-08 | 4.967E-08 | 1.150E-08 | 3.800E-08 | 6.100E-08 |
| 6955 | 348 | I348P | A | 3e-08 | 1.9e-08 | 5.2e-08 | 3.367E-08 | 1.680E-08 | 1.900E-08 | 5.200E-08 |
| 6956 | 348 | I348S | A | 2.4e-08 | 1.7e-08 | 8.8e-08 | 4.300E-08 | 3.913E-08 | 1.700E-08 | 8.800E-08 |
| 6957 | 348 | I348T | A | 3.6e-08 | 2.3e-08 | 8.7e-08 | 4.867E-08 | 3.383E-08 | 2.300E-08 | 8.700E-08 |
| 6958 | 348 | I348W | A | 1.6e-08 | 9.4e-08 | 6.5e-08 | 5.833E-08 | 3.943E-08 | 1.600E-08 | 9.400E-08 |
| 6959 | 348 | I348Y | A | 4.8e-08 | 5.3e-08 | 4.1e-08 | 4.733E-08 | 6.028E-09 | 4.100E-08 | 5.300E-08 |
| 6960 | 348 | I348V | A | 3.2e-08 | 5.8e-08 | 4.1e-08 | 4.367E-08 | 1.320E-08 | 3.200E-08 | 5.800E-08 |
| 6961 | 349 | F349A | A | 4.8e-08 | 1.3e-08 | 4.7e-08 | 3.600E-08 | 1.992E-08 | 1.300E-08 | 4.800E-08 |
| 6962 | 349 | F349R | A | 2.3e-08 | 5.1e-08 | 2e-08 | 3.133E-08 | 1.710E-08 | 2.000E-08 | 5.100E-08 |
| 6963 | 349 | F349N | A | 2.4e-08 | 1.9e-08 | 6.6e-08 | 3.633E-08 | 2.581E-08 | 1.900E-08 | 6.600E-08 |
| 6964 | 349 | F349D | A | 2.5e-08 | 5.1e-08 | 1.3e-07 | 6.867E-08 | 5.468E-08 | 2.500E-08 | 1.300E-07 |
| 6965 | 349 | F349C | A | 7.8e-08 | 5.1e-08 | 1.1e-07 | 7.967E-08 | 2.954E-08 | 5.100E-08 | 1.100E-07 |
| 6966 | 349 | F349E | A | 2.9e-08 | 4.5e-08 | 1.4e-08 | 2.933E-08 | 1.550E-08 | 1.400E-08 | 4.500E-08 |
| 6967 | 349 | F349Q | A | 6.7e-08 | 3.2e-08 | 2.8e-08 | 4.233E-08 | 2.146E-08 | 2.800E-08 | 6.700E-08 |
| 6968 | 349 | F349G | A | 4.9e-08 | 2.2e-08 | 4.1e-08 | 3.733E-08 | 1.387E-08 | 2.200E-08 | 4.900E-08 |
| 6969 | 349 | F349H | A | 2.8e-08 | 4.4e-08 | 2.1e-08 | 3.100E-08 | 1.179E-08 | 2.100E-08 | 4.400E-08 |

| | | | | | | | | | | |
|---|---|---|---|---|---|---|---|---|---|---|
| 6970 | 349 | F349I | A | 7.7e-08 | 3.8e-08 | 5.2e-08 | 5.567E-08 | 1.976E-08 | 3.800E-08 | 7.700E-08 |
| 6971 | 349 | F349L | A | 7.8e-08 | 2e-08 | 6.2e-08 | 5.333E-08 | 2.996E-08 | 2.000E-08 | 7.800E-08 |
| 6972 | 349 | F349K | A | 3.6e-08 | 6.2e-08 | 2.8e-08 | 4.200E-08 | 1.778E-08 | 2.800E-08 | 6.200E-08 |
| 6973 | 349 | F349M | A | 6e-08 | 2.6e-08 | 1.8e-08 | 3.467E-08 | 2.230E-08 | 1.800E-08 | 6.000E-08 |
| 6974 | 349 | F349F | A | 5.7e-08 | 3.2e-08 | 8e-08 | 5.633E-08 | 2.401E-08 | 3.200E-08 | 8.000E-08 |
| 6975 | 349 | F349P | A | 5.6e-08 | 1.9e-08 | 5.4e-08 | 4.300E-08 | 2.081E-08 | 1.900E-08 | 5.600E-08 |
| 6976 | 349 | F349S | A | 1.5e-08 | 1.3e-07 | 3.9e-08 | 6.133E-08 | 6.067E-08 | 1.500E-08 | 1.300E-07 |
| 6977 | 349 | F349T | A | 4.5e-08 | 2.2e-08 | 5e-08 | 3.900E-08 | 1.493E-08 | 2.200E-08 | 5.000E-08 |
| 6978 | 349 | F349W | A | 4e-08 | 2.4e-08 | 4e-08 | 3.467E-08 | 9.238E-09 | 2.400E-08 | 4.000E-08 |
| 6979 | 349 | F349Y | A | 4.9e-08 | 5.4e-08 | 8e-08 | 6.100E-08 | 1.664E-08 | 4.900E-08 | 8.000E-08 |
| 6980 | 349 | F349V | A | 2.4e-08 | 4.8e-08 | 4.6e-08 | 3.933E-08 | 1.332E-08 | 2.400E-08 | 4.800E-08 |
| 6981 | 350 | G350A | A | 3.8e-08 | 5.7e-08 | 4.8e-08 | 4.767E-08 | 9.504E-09 | 3.800E-08 | 5.700E-08 |
| 6982 | 350 | G350R | A | 2.5e-08 | 1.4e-08 | 8.8e-08 | 4.233E-08 | 3.993E-08 | 1.400E-08 | 8.800E-08 |
| 6983 | 350 | G350N | A | 7.4e-08 | 6.2e-08 | 8.8e-08 | 7.467E-08 | 1.301E-08 | 6.200E-08 | 8.800E-08 |
| 6984 | 350 | G350D | A | 4.7e-08 | 4.1e-08 | 5.1e-09 | 3.103E-08 | 2.266E-08 | 5.100E-09 | 4.700E-08 |
| 6985 | 350 | G350C | A | 2.8e-08 | 5.5e-08 | 2e-08 | 3.433E-08 | 1.834E-08 | 2.000E-08 | 5.500E-08 |
| 6986 | 350 | G350E | A | 2.6e-08 | 4e-08 | 8.6e-08 | 5.067E-08 | 3.139E-08 | 2.600E-08 | 8.600E-08 |
| 6987 | 350 | G350Q | A | 1.4e-07 | 5.6e-08 | 1e-07 | 9.867E-08 | 4.202E-08 | 5.600E-08 | 1.400E-07 |
| 6988 | 350 | G350G | A | 3.8e-08 | 2.4e-08 | 8e-08 | 4.733E-08 | 2.914E-08 | 2.400E-08 | 8.000E-08 |
| 6989 | 350 | G350H | A | 1.9e-08 | 5.2e-08 | 6.6e-08 | 4.567E-08 | 2.413E-08 | 1.900E-08 | 6.600E-08 |
| 6990 | 350 | G350I | A | 6.1e-08 | 2.8e-08 | 3.8e-08 | 4.233E-08 | 1.692E-08 | 2.800E-08 | 6.100E-08 |
| 6991 | 350 | G350L | A | 2.3e-08 | 9.2e-08 | 3.9e-08 | 5.133E-08 | 3.612E-08 | 2.300E-08 | 9.200E-08 |
| 6992 | 350 | G350K | A | 3.4e-08 | 1.4e-08 | 2.4e-08 | 2.400E-08 | 1.000E-08 | 1.400E-08 | 3.400E-08 |
| 6993 | 350 | G350M | A | 7.1e-08 | 7.4e-08 | 3.7e-08 | 6.067E-08 | 2.055E-08 | 3.700E-08 | 7.400E-08 |
| 6994 | 350 | G350F | A | 4.1e-08 | 3.8e-08 | 4e-08 | 3.967E-08 | 1.528E-09 | 3.800E-08 | 4.100E-08 |
| 6995 | 350 | G350P | A | 5.9e-08 | 1.4e-08 | 4.9e-08 | 4.067E-08 | 2.363E-08 | 1.400E-08 | 5.900E-08 |
| 6996 | 350 | G350S | A | 5.9e-08 | 1.1e-07 | 1.9e-08 | 6.267E-08 | 4.561E-08 | 1.900E-08 | 1.100E-07 |
| 6997 | 350 | G350T | A | 1e-07 | 7.6e-08 | 3.7e-08 | 7.100E-08 | 3.180E-08 | 3.700E-08 | 1.000E-07 |
| 6998 | 350 | G350W | A | 3.2e-08 | 3.1e-08 | 1.2e-07 | 6.100E-08 | 5.110E-08 | 3.100E-08 | 1.200E-07 |
| 6999 | 350 | G350Y | A | 2.4e-08 | 1.4e-07 | 4.3e-08 | 6.900E-08 | 6.222E-08 | 2.400E-08 | 1.400E-07 |
| 7000 | 350 | G350V | A | 8.9e-08 | 1.2e-07 | 4.1e-08 | 8.333E-08 | 3.980E-08 | 4.100E-08 | 1.200E-07 |
| 7001 | 351 | S351A | A | 1.1e-08 | 9.8e-08 | 6.2e-08 | 5.700E-08 | 4.371E-08 | 1.100E-08 | 9.800E-08 |
| 7002 | 351 | S351R | A | 6.8e-08 | 1.9e-08 | 7.3e-08 | 5.333E-08 | 2.984E-08 | 1.900E-08 | 7.300E-08 |
| 7003 | 351 | S351N | A | 4.2e-08 | 1.3e-08 | 5.5e-08 | 3.667E-08 | 2.150E-08 | 1.300E-08 | 5.500E-08 |
| 7004 | 351 | S351D | A | 6.2e-08 | 3.8e-08 | 5.3e-08 | 5.100E-08 | 1.212E-08 | 3.800E-08 | 6.200E-08 |
| 7005 | 351 | S351C | A | 7.7e-08 | 3.3e-08 | 2.9e-08 | 4.633E-08 | 2.663E-08 | 2.900E-08 | 7.700E-08 |
| 7006 | 351 | S351E | A | 5.2e-08 | 4.3e-08 | 7.5e-08 | 5.667E-08 | 1.650E-08 | 4.300E-08 | 7.500E-08 |
| 7007 | 351 | S351Q | A | 4e-08 | 4.1e-08 | 9.3e-08 | 5.800E-08 | 3.032E-08 | 4.000E-08 | 9.300E-08 |
| 7008 | 351 | S351G | A | 4.4e-08 | 8.9e-08 | 4.7e-08 | 6.000E-08 | 2.516E-08 | 4.400E-08 | 8.900E-08 |
| 7009 | 351 | S351H | A | 6.8e-08 | 1.9e-08 | 9.2e-08 | 5.967E-08 | 3.721E-08 | 1.900E-08 | 9.200E-08 |
| 7010 | 351 | S351I | A | 2e-08 | 2e-08 | 2.9e-08 | 2.300E-08 | 5.196E-09 | 2.000E-08 | 2.900E-08 |
| 7011 | 351 | S351L | A | 3.2e-08 | 8.2e-08 | 4.1e-08 | 5.167E-08 | 2.665E-08 | 3.200E-08 | 8.200E-08 |
| 7012 | 351 | S351K | A | 1.5e-07 | 6.1e-08 | 3.5e-08 | 8.200E-08 | 6.031E-08 | 3.500E-08 | 1.500E-07 |
| 7013 | 351 | S351M | A | 9.6e-08 | 5.7e-08 | 2.2e-08 | 5.833E-08 | 3.702E-08 | 2.200E-08 | 9.600E-08 |
| 7014 | 351 | S351F | A | 5.3e-08 | 5.9e-08 | 1.2e-07 | 7.733E-08 | 3.707E-08 | 5.300E-08 | 1.200E-07 |
| 7015 | 351 | S351P | A | 3.5e-08 | 1.1e-07 | 8.3e-08 | 7.600E-08 | 3.799E-08 | 3.500E-08 | 1.100E-07 |
| 7016 | 351 | S351S | A | 3.5e-08 | 9.9e-08 | 2.4e-08 | 5.267E-08 | 4.050E-08 | 2.400E-08 | 9.900E-08 |
| 7017 | 351 | S351T | A | 7.8e-08 | 3.8e-08 | 2.2e-08 | 4.600E-08 | 2.884E-08 | 2.200E-08 | 7.800E-08 |
| 7018 | 351 | S351W | A | 3.3e-08 | 1.5e-07 | 1.2e-07 | 1.010E-07 | 6.077E-08 | 3.300E-08 | 1.500E-07 |
| 7019 | 351 | S351Y | A | 9e-09 | 1.4e-08 | 2.9e-08 | 1.733E-08 | 1.041E-08 | 9.000E-09 | 2.900E-08 |
| 7020 | 351 | S351V | A | 2.7e-08 | 2.5e-08 | 1e-07 | 5.067E-08 | 4.274E-08 | 2.500E-08 | 1.000E-07 |
| 7021 | 352 | L352A | A | 2.4e-08 | 2e-08 | 6.4e-08 | 3.600E-08 | 2.433E-08 | 2.000E-08 | 6.400E-08 |
| 7022 | 352 | L352R | A | 1e-08 | 4.6e-08 | 3.1e-08 | 2.900E-08 | 1.808E-08 | 1.000E-08 | 4.600E-08 |
| 7023 | 352 | L352N | A | 1e-07 | 3.1e-08 | 8.4e-08 | 7.167E-08 | 3.612E-08 | 3.100E-08 | 1.000E-07 |
| 7024 | 352 | L352D | A | 7.1e-08 | 6.7e-08 | 3.8e-08 | 5.867E-08 | 1.801E-08 | 3.800E-08 | 7.100E-08 |
| 7025 | 352 | L352C | A | 5.7e-08 | 4.5e-08 | 3.4e-08 | 4.533E-08 | 1.150E-08 | 3.400E-08 | 5.700E-08 |
| 7026 | 352 | L352E | A | 2.2e-08 | 7.7e-08 | 5.7e-08 | 5.200E-08 | 2.784E-08 | 2.200E-08 | 7.700E-08 |
| 7027 | 352 | L352Q | A | 3.6e-08 | 9.7e-08 | 2.1e-08 | 5.133E-08 | 4.025E-08 | 2.100E-08 | 9.700E-08 |
| 7028 | 352 | L352G | A | 2.9e-08 | 1e-07 | 3.4e-08 | 5.433E-08 | 3.963E-08 | 2.900E-08 | 1.000E-07 |
| 7029 | 352 | L352H | A | 4.4e-08 | 3.1e-08 | 4.9e-08 | 4.133E-08 | 9.292E-09 | 3.100E-08 | 4.900E-08 |
| 7030 | 352 | L352I | A | 2.7e-08 | 2.4e-08 | 4.1e-08 | 3.067E-08 | 9.074E-09 | 2.400E-08 | 4.100E-08 |
| 7031 | 352 | L352L | A | 5e-08 | 3.2e-08 | 5.6e-08 | 4.600E-08 | 1.249E-08 | 3.200E-08 | 5.600E-08 |
| 7032 | 352 | L352K | A | 3e-08 | 1.6e-08 | 5.2e-08 | 3.267E-08 | 1.815E-08 | 1.600E-08 | 5.200E-08 |
| 7033 | 352 | L352M | A | 5.5e-08 | 5.7e-08 | 1.6e-08 | 4.267E-08 | 2.312E-08 | 1.600E-08 | 5.700E-08 |
| 7034 | 352 | L352F | A | 1.7e-08 | 1.5e-08 | 2.2e-08 | 1.800E-08 | 3.606E-09 | 1.500E-08 | 2.200E-08 |
| 7035 | 352 | L352P | A | 2.9e-08 | 5.8e-08 | 1.9e-08 | 3.533E-08 | 2.026E-08 | 1.900E-08 | 5.800E-08 |
| 7036 | 352 | L352S | A | 3e-08 | 3.8e-08 | 3.8e-08 | 3.533E-08 | 4.619E-09 | 3.000E-08 | 3.800E-08 |
| 7037 | 352 | L352T | A | 5.5e-08 | 9.5e-08 | 8.8e-08 | 7.933E-08 | 2.136E-08 | 5.500E-08 | 9.500E-08 |
| 7038 | 352 | L352W | A | 1.5e-08 | 4.1e-08 | 2.2e-08 | 2.600E-08 | 1.345E-08 | 1.500E-08 | 4.100E-08 |
| 7039 | 352 | L352Y | A | 3.1e-08 | 3.7e-08 | 3.7e-08 | 3.500E-08 | 3.464E-09 | 3.100E-08 | 3.700E-08 |
| 7040 | 352 | L352V | A | 3.5e-08 | 7.6e-08 | 5.6e-08 | 5.567E-08 | 2.050E-08 | 3.500E-08 | 7.600E-08 |
| 7041 | 353 | A353A | A | 6.1e-08 | 3.7e-08 | 4.2e-08 | 4.667E-08 | 1.266E-08 | 3.700E-08 | 6.100E-08 |
| 7042 | 353 | A353R | A | 3.8e-08 | 3.7e-08 | 1.6e-08 | 3.033E-08 | 1.242E-08 | 1.600E-08 | 3.800E-08 |
| 7043 | 353 | A353N | A | 3.5e-08 | 4e-08 | 7.2e-08 | 4.900E-08 | 2.007E-08 | 3.500E-08 | 7.200E-08 |
| 7044 | 353 | A353D | A | 5.1e-08 | 2.6e-08 | 3e-08 | 3.567E-08 | 1.343E-08 | 2.600E-08 | 5.100E-08 |
| 7045 | 353 | A353C | A | 1.7e-08 | 7e-08 | 1.8e-08 | 3.500E-08 | 3.032E-08 | 1.700E-08 | 7.000E-08 |
| 7046 | 353 | A353E | A | 6.7e-08 | 2.7e-08 | 3.7e-08 | 4.367E-08 | 2.082E-08 | 2.700E-08 | 6.700E-08 |
| 7047 | 353 | A353Q | A | 2.3e-08 | 3.5e-08 | 1.5e-08 | 2.433E-08 | 1.007E-08 | 1.500E-08 | 3.500E-08 |
| 7048 | 353 | A353G | A | 4.3e-08 | 5.6e-08 | 4e-08 | 4.633E-08 | 8.505E-09 | 4.000E-08 | 5.600E-08 |
| 7049 | 353 | A353H | A | 2.2e-08 | 3.2e-08 | 7e-08 | 4.133E-08 | 2.532E-08 | 2.200E-08 | 7.000E-08 |
| 7050 | 353 | A353I | A | 6.4e-08 | 6e-08 | 6.2e-08 | 6.200E-08 | 2.000E-09 | 6.000E-08 | 6.400E-08 |
| 7051 | 353 | A353L | A | 7.4e-08 | 9.9e-08 | 4.1e-08 | 7.133E-08 | 2.909E-08 | 4.100E-08 | 9.900E-08 |
| 7052 | 353 | A353K | A | 7.8e-08 | 1.1e-07 | 3.4e-08 | 7.400E-08 | 3.816E-08 | 3.400E-08 | 1.100E-07 |
| 7053 | 353 | A353M | A | 7.1e-08 | 1.2e-07 | 5.2e-08 | 8.100E-08 | 3.509E-08 | 5.200E-08 | 1.200E-07 |
| 7054 | 353 | A353F | A | 8.6e-08 | 1.8e-08 | 6.1e-08 | 5.500E-08 | 3.439E-08 | 1.800E-08 | 8.600E-08 |

| | | | | | | | | | | |
|---|---|---|---|---|---|---|---|---|---|---|
| 7055 | 353 | A353P | A | 5.2e-08 | 5.7e-08 | 7.5e-08 | 6.133E-08 | 1.210E-08 | 5.200E-08 | 7.500E-08 |
| 7056 | 353 | A353S | A | 2.9e-08 | 5.1e-08 | 2.8e-08 | 3.600E-08 | 1.300E-08 | 2.800E-08 | 5.100E-08 |
| 7057 | 353 | A353T | A | 7.5e-08 | 2.2e-08 | 1.2e-07 | 7.233E-08 | 4.905E-08 | 2.200E-08 | 1.200E-07 |
| 7058 | 353 | A353W | A | 6.9e-09 | 4e-08 | 1.1e-08 | 1.930E-08 | 1.804E-08 | 6.900E-09 | 4.000E-08 |
| 7059 | 353 | A353Y | A | 4.8e-08 | 5.4e-08 | 5.4e-08 | 5.200E-08 | 3.464E-09 | 4.800E-08 | 5.400E-08 |
| 7060 | 353 | A353V | A | 5.9e-08 | 1.5e-07 | 3.7e-08 | 8.200E-08 | 5.991E-08 | 3.700E-08 | 1.500E-07 |
| 7061 | 354 | F354A | A | 8.4e-08 | 3.2e-08 | 3.5e-08 | 5.033E-08 | 2.919E-08 | 3.200E-08 | 8.400E-08 |
| 7062 | 354 | F354R | A | 4.2e-08 | 9.2e-08 | 3e-08 | 5.467E-08 | 3.288E-08 | 3.000E-08 | 9.200E-08 |
| 7063 | 354 | F354N | A | 7.5e-08 | 3.9e-08 | 2.9e-08 | 4.767E-08 | 2.419E-08 | 2.900E-08 | 7.500E-08 |
| 7064 | 354 | F354D | A | 1.5e-07 | 5.2e-08 | 8.6e-08 | 9.600E-08 | 4.976E-08 | 5.200E-08 | 1.500E-07 |
| 7065 | 354 | F354C | A | 1e-07 | 4.4e-08 | 2.1e-08 | 5.500E-08 | 4.063E-08 | 2.100E-08 | 1.000E-07 |
| 7066 | 354 | F354E | A | 1.5e-07 | 1.4e-07 | 3.8e-08 | 1.093E-07 | 6.198E-08 | 3.800E-08 | 1.500E-07 |
| 7067 | 354 | F354Q | A | 2.5e-08 | 2.6e-08 | 5.9e-08 | 3.667E-08 | 1.935E-08 | 2.500E-08 | 5.900E-08 |
| 7068 | 354 | F354G | A | 3.3e-08 | 1.7e-08 | 5.7e-08 | 3.567E-08 | 2.013E-08 | 1.700E-08 | 5.700E-08 |
| 7069 | 354 | F354H | A | 4.7e-08 | 3.1e-08 | 5.9e-08 | 4.567E-08 | 1.405E-08 | 3.100E-08 | 5.900E-08 |
| 7070 | 354 | F354I | A | 6.3e-08 | 1.7e-08 | 2.3e-08 | 3.433E-08 | 2.501E-08 | 1.700E-08 | 6.300E-08 |
| 7071 | 354 | F354L | A | 4.5e-08 | 3.9e-08 | 3.5e-08 | 3.967E-08 | 5.033E-09 | 3.500E-08 | 4.500E-08 |
| 7072 | 354 | F354K | A | 7.3e-08 | 3.6e-08 | 3.1e-08 | 4.667E-08 | 2.294E-08 | 3.100E-08 | 7.300E-08 |
| 7073 | 354 | F354M | A | 6.3e-08 | 9.7e-08 | 2.7e-08 | 6.233E-08 | 3.500E-08 | 2.700E-08 | 9.700E-08 |
| 7074 | 354 | F354F | A | 4.7e-08 | 2.2e-08 | 9.2e-08 | 5.367E-08 | 3.547E-08 | 2.200E-08 | 9.200E-08 |
| 7075 | 354 | F354P | A | 1.5e-07 | 3.9e-08 | 5.9e-08 | 8.267E-08 | 5.916E-08 | 3.900E-08 | 1.500E-07 |
| 7076 | 354 | F354S | A | 2.3e-08 | 3.7e-08 | 2.4e-08 | 2.800E-08 | 7.810E-09 | 2.300E-08 | 3.700E-08 |
| 7077 | 354 | F354T | A | 2.9e-08 | 8.2e-08 | 7.2e-08 | 6.100E-08 | 2.816E-08 | 2.900E-08 | 8.200E-08 |
| 7078 | 354 | F354W | A | 1.4e-08 | 7.2e-08 | 2.1e-08 | 3.567E-08 | 3.166E-08 | 1.400E-08 | 7.200E-08 |
| 7079 | 354 | F354Y | A | 1.6e-08 | 5.1e-08 | 3.3e-08 | 3.333E-08 | 1.750E-08 | 1.600E-08 | 5.100E-08 |
| 7080 | 354 | F354V | A | 3.4e-08 | 5.4e-08 | 5e-08 | 4.600E-08 | 1.058E-08 | 3.400E-08 | 5.400E-08 |
| 7081 | 355 | L355A | A | 4.3e-08 | 5.2e-08 | 3.6e-08 | 4.367E-08 | 8.021E-09 | 3.600E-08 | 5.200E-08 |
| 7082 | 355 | L355R | A | 1.8e-08 | 5.1e-08 | 4.9e-08 | 3.933E-08 | 1.850E-08 | 1.800E-08 | 5.100E-08 |
| 7083 | 355 | L355N | A | 6.7e-08 | 4e-08 | 2.5e-08 | 4.400E-08 | 2.128E-08 | 2.500E-08 | 6.700E-08 |
| 7084 | 355 | L355D | A | 6e-08 | 2.6e-08 | 6e-08 | 4.867E-08 | 1.963E-08 | 2.600E-08 | 6.000E-08 |
| 7085 | 355 | L355C | A | 6.5e-08 | 4.5e-08 | 1.7e-08 | 4.233E-08 | 2.411E-08 | 1.700E-08 | 6.500E-08 |
| 7086 | 355 | L355E | A | 2.6e-08 | 3e-08 | 3.1e-08 | 2.900E-08 | 2.646E-09 | 2.600E-08 | 3.100E-08 |
| 7087 | 355 | L355Q | A | 2.8e-08 | 7.3e-08 | 4.5e-08 | 4.867E-08 | 2.272E-08 | 2.800E-08 | 7.300E-08 |
| 7088 | 355 | L355G | A | 7.4e-08 | 1.5e-07 | 4.8e-08 | 9.067E-08 | 5.300E-08 | 4.800E-08 | 1.500E-07 |
| 7089 | 355 | L355H | A | 3.3e-08 | 8.3e-08 | 4.6e-08 | 5.400E-08 | 2.594E-08 | 3.300E-08 | 8.300E-08 |
| 7090 | 355 | L355I | A | 9.2e-08 | 5.1e-08 | 4.6e-08 | 6.300E-08 | 2.524E-08 | 4.600E-08 | 9.200E-08 |
| 7091 | 355 | L355L | A | 4e-08 | 8e-08 | 4.1e-08 | 5.367E-08 | 2.281E-08 | 4.000E-08 | 8.000E-08 |
| 7092 | 355 | L355K | A | 7.2e-08 | 2.4e-08 | 4.7e-08 | 4.767E-08 | 2.401E-08 | 2.400E-08 | 7.200E-08 |
| 7093 | 355 | L355M | A | 3.9e-08 | 6.5e-08 | 2.5e-08 | 4.300E-08 | 2.030E-08 | 2.500E-08 | 6.500E-08 |
| 7094 | 355 | L355F | A | 5.8e-08 | 6.9e-08 | 1.2e-08 | 4.633E-08 | 3.024E-08 | 1.200E-08 | 6.900E-08 |
| 7095 | 355 | L355P | A | 6.4e-08 | 6.7e-08 | 6.1e-08 | 6.400E-08 | 3.000E-09 | 6.100E-08 | 6.700E-08 |
| 7096 | 355 | L355S | A | 4.9e-08 | 7.6e-08 | 6e-08 | 6.167E-08 | 1.358E-08 | 4.900E-08 | 7.600E-08 |
| 7097 | 355 | L355T | A | 5.5e-08 | 5e-08 | 9e-08 | 6.500E-08 | 2.179E-08 | 5.000E-08 | 9.000E-08 |
| 7098 | 355 | L355W | A | 3.7e-08 | 5.5e-08 | 2.5e-08 | 3.900E-08 | 1.510E-08 | 2.500E-08 | 5.500E-08 |
| 7099 | 355 | L355Y | A | 3.7e-08 | 2.6e-08 | 4.9e-08 | 3.733E-08 | 1.150E-08 | 2.600E-08 | 4.900E-08 |
| 7100 | 355 | L355V | A | 3.6e-08 | 5e-08 | 2.1e-08 | 3.567E-08 | 1.450E-08 | 2.100E-08 | 5.000E-08 |
| 7101 | 356 | P356A | A | 3e-08 | 3.8e-08 | 2.9e-08 | 3.233E-08 | 4.933E-09 | 2.900E-08 | 3.800E-08 |
| 7102 | 356 | P356R | A | 3.1e-08 | 4.7e-08 | 3.2e-08 | 3.667E-08 | 8.963E-09 | 3.100E-08 | 4.700E-08 |
| 7103 | 356 | P356N | A | 2.1e-08 | 4.5e-08 | 5.4e-08 | 4.000E-08 | 1.706E-08 | 2.100E-08 | 5.400E-08 |
| 7104 | 356 | P356D | A | 5.5e-08 | 9.9e-08 | 4.6e-08 | 6.667E-08 | 2.836E-08 | 4.600E-08 | 9.900E-08 |
| 7105 | 356 | P356C | A | 2.9e-08 | 3e-08 | 6.8e-08 | 4.233E-08 | 2.223E-08 | 2.900E-08 | 6.800E-08 |
| 7106 | 356 | P356E | A | 3.4e-08 | 8.8e-08 | 3.1e-08 | 5.100E-08 | 3.208E-08 | 3.100E-08 | 8.800E-08 |
| 7107 | 356 | P356Q | A | 3.4e-08 | 9.1e-08 | 3.8e-08 | 5.433E-08 | 3.182E-08 | 3.400E-08 | 9.100E-08 |
| 7108 | 356 | P356G | A | 8.3e-08 | 2.8e-08 | 4.5e-08 | 5.200E-08 | 2.816E-08 | 2.800E-08 | 8.300E-08 |
| 7109 | 356 | P356H | A | 7.6e-08 | 4.4e-08 | 4.5e-08 | 5.500E-08 | 1.819E-08 | 4.400E-08 | 7.600E-08 |
| 7110 | 356 | P356I | A | 8.8e-08 | 7e-08 | 7.9e-08 | 7.900E-08 | 9.000E-09 | 7.000E-08 | 8.800E-08 |
| 7111 | 356 | P356L | A | 3.8e-08 | 2.1e-08 | 6.5e-08 | 4.133E-08 | 2.219E-08 | 2.100E-08 | 6.500E-08 |
| 7112 | 356 | P356K | A | 4.4e-08 | 6.5e-08 | 4.2e-08 | 5.033E-08 | 1.274E-08 | 4.200E-08 | 6.500E-08 |
| 7113 | 356 | P356M | A | 4.6e-08 | 5.2e-08 | 1.7e-08 | 3.833E-08 | 1.872E-08 | 1.700E-08 | 5.200E-08 |
| 7114 | 356 | P356F | A | 2.8e-07 | 7.5e-08 | 3.6e-08 | 1.303E-07 | 1.311E-07 | 3.600E-08 | 2.800E-07 |
| 7115 | 356 | P356P | A | 6.7e-08 | 4e-08 | 4.3e-08 | 5.000E-08 | 1.480E-08 | 4.000E-08 | 6.700E-08 |
| 7116 | 356 | P356S | A | 1.2e-07 | 1e-07 | 4e-08 | 8.667E-08 | 4.163E-08 | 4.000E-08 | 1.200E-07 |
| 7117 | 356 | P356T | A | 2.6e-08 | 7.9e-08 | 3.7e-08 | 4.733E-08 | 2.797E-08 | 2.600E-08 | 7.900E-08 |
| 7118 | 356 | P356W | A | 3.5e-08 | 6.7e-08 | 6.2e-08 | 5.467E-08 | 1.721E-08 | 3.500E-08 | 6.700E-08 |
| 7119 | 356 | P356Y | A | 5.2e-08 | 7.4e-08 | 3.1e-08 | 5.233E-08 | 2.150E-08 | 3.100E-08 | 7.400E-08 |
| 7120 | 356 | P356V | A | 2.9e-08 | 3.7e-08 | 8.6e-08 | 5.067E-08 | 3.086E-08 | 2.900E-08 | 8.600E-08 |
| 7121 | 357 | E357A | A | 1.2e-08 | 4.1e-08 | 5.8e-08 | 3.700E-08 | 2.326E-08 | 1.200E-08 | 5.800E-08 |
| 7122 | 357 | E357R | A | 3.8e-08 | 1.9e-08 | 2.7e-08 | 2.800E-08 | 9.539E-09 | 1.900E-08 | 3.800E-08 |
| 7123 | 357 | E357N | A | 5.5e-08 | 6.4e-08 | 2.9e-08 | 4.933E-08 | 1.818E-08 | 2.900E-08 | 6.400E-08 |
| 7124 | 357 | E357D | A | 9.5e-08 | 1.7e-08 | 2.8e-08 | 4.667E-08 | 4.222E-08 | 1.700E-08 | 9.500E-08 |
| 7125 | 357 | E357C | A | 1.7e-08 | 2.8e-08 | 2.8e-08 | 2.433E-08 | 6.351E-09 | 1.700E-08 | 2.800E-08 |
| 7126 | 357 | E357E | A | 5.7e-08 | 9.5e-08 | 3.8e-08 | 6.333E-08 | 2.902E-08 | 3.800E-08 | 9.500E-08 |
| 7127 | 357 | E357Q | A | 4.8e-08 | 7.1e-08 | 4.9e-08 | 5.600E-08 | 1.300E-08 | 4.800E-08 | 7.100E-08 |
| 7128 | 357 | E357G | A | 8.8e-08 | 1.3e-07 | 3.4e-08 | 8.400E-08 | 4.812E-08 | 3.400E-08 | 1.300E-07 |
| 7129 | 357 | E357H | A | 6.4e-08 | 4.4e-08 | 6.7e-08 | 5.833E-08 | 1.250E-08 | 4.400E-08 | 6.700E-08 |
| 7130 | 357 | E357I | A | 5.9e-08 | 8.7e-08 | 1.3e-07 | 9.200E-08 | 3.576E-08 | 5.900E-08 | 1.300E-07 |
| 7131 | 357 | E357L | A | 7.4e-08 | 6.7e-08 | 1.1e-07 | 8.367E-08 | 2.307E-08 | 6.700E-08 | 1.100E-07 |
| 7132 | 357 | E357K | A | 1.8e-08 | 3.1e-08 | 4.9e-08 | 3.267E-08 | 1.557E-08 | 1.800E-08 | 4.900E-08 |
| 7133 | 357 | E357M | A | 7.3e-08 | 3.4e-08 | 7.5e-08 | 6.067E-08 | 2.312E-08 | 3.400E-08 | 7.500E-08 |
| 7134 | 357 | E357F | A | 3.4e-08 | 1.9e-08 | 7.8e-08 | 4.367E-08 | 3.066E-08 | 1.900E-08 | 7.800E-08 |
| 7135 | 357 | E357P | A | 6.1e-08 | 8.1e-08 | 2.6e-08 | 5.600E-08 | 2.784E-08 | 2.600E-08 | 8.100E-08 |
| 7136 | 357 | E357S | A | 1.4e-07 | 5.2e-08 | 4.4e-08 | 7.867E-08 | 5.327E-08 | 4.400E-08 | 1.400E-07 |
| 7137 | 357 | E357T | A | 9.2e-08 | 4e-08 | 3.2e-08 | 5.467E-08 | 3.258E-08 | 3.200E-08 | 9.200E-08 |
| 7138 | 357 | E357W | A | 1.7e-07 | 1.2e-08 | 7.6e-08 | 8.600E-08 | 7.947E-08 | 1.200E-08 | 1.700E-07 |
| 7139 | 357 | E357Y | A | 5.9e-08 | 8.1e-08 | 5.4e-08 | 6.467E-08 | 1.436E-08 | 5.400E-08 | 8.100E-08 |

| | | | | | | | | | | |
|---|---|---|---|---|---|---|---|---|---|---|
| 7140 | 357 | E357V | A | 3.8e-08 | 2.4e-08 | 2.6e-08 | 2.933E-08 | 7.572E-09 | 2.400E-08 | 3.800E-08 |
| 7141 | 358 | S358A | A | 3.3e-08 | 1.6e-08 | 1.3e-07 | 5.967E-08 | 6.150E-08 | 1.600E-08 | 1.300E-07 |
| 7142 | 358 | S358R | A | 2.1e-08 | 8.5e-08 | 2.1e-08 | 4.233E-08 | 3.695E-08 | 2.100E-08 | 8.500E-08 |
| 7143 | 358 | S358N | A | 1.6e-08 | 5.1e-08 | 2e-08 | 2.900E-08 | 1.916E-08 | 1.600E-08 | 5.100E-08 |
| 7144 | 358 | S358D | A | 5.5e-08 | 5.7e-08 | 3.6e-08 | 4.933E-08 | 1.159E-08 | 3.600E-08 | 5.700E-08 |
| 7145 | 358 | S358C | A | 4.9e-08 | 1.9e-08 | 3.2e-08 | 3.333E-08 | 1.504E-08 | 1.900E-08 | 4.900E-08 |
| 7146 | 358 | S358E | A | 5.4e-08 | 1.9e-08 | 3.4e-08 | 3.567E-08 | 1.756E-08 | 1.900E-08 | 5.400E-08 |
| 7147 | 358 | S358Q | A | 6.7e-08 | 2.3e-08 | 7.6e-08 | 5.533E-08 | 2.836E-08 | 2.300E-08 | 7.600E-08 |
| 7148 | 358 | S358G | A | 3.7e-08 | 1.7e-08 | 2e-08 | 2.467E-08 | 1.079E-08 | 1.700E-08 | 3.700E-08 |
| 7149 | 358 | S358H | A | 4.6e-08 | 5.8e-08 | 6.5e-08 | 5.633E-08 | 9.609E-09 | 4.600E-08 | 6.500E-08 |
| 7150 | 358 | S358I | A | 4.6e-08 | 8.8e-08 | 7.7e-08 | 7.033E-08 | 2.178E-08 | 4.600E-08 | 8.800E-08 |
| 7151 | 358 | S358L | A | 4.9e-08 | 4.8e-08 | 4.5e-08 | 4.733E-08 | 2.082E-09 | 4.500E-08 | 4.900E-08 |
| 7152 | 358 | S358K | A | 1.2e-07 | 2.2e-08 | 2.2e-08 | 5.467E-08 | 5.658E-08 | 2.200E-08 | 1.200E-07 |
| 7153 | 358 | S358M | A | 5.2e-08 | 1.7e-07 | 1.4e-08 | 7.867E-08 | 8.135E-08 | 1.400E-08 | 1.700E-07 |
| 7154 | 358 | S358F | A | 5.3e-08 | 3.8e-08 | 2.2e-08 | 3.767E-08 | 1.550E-08 | 2.200E-08 | 5.300E-08 |
| 7155 | 358 | S358P | A | 3.5e-08 | 3.2e-08 | 4.3e-08 | 3.667E-08 | 5.686E-09 | 3.200E-08 | 4.300E-08 |
| 7156 | 358 | S358S | A | 2.8e-08 | 1.4e-07 | 2.5e-08 | 6.433E-08 | 6.555E-08 | 2.500E-08 | 1.400E-07 |
| 7157 | 358 | S358T | A | 2.5e-08 | 1.9e-07 | 4.6e-08 | 8.700E-08 | 8.982E-08 | 2.500E-08 | 1.900E-07 |
| 7158 | 358 | S358W | A | 4.7e-08 | 5.9e-08 | 5.5e-08 | 5.367E-08 | 6.110E-09 | 4.700E-08 | 5.900E-08 |
| 7159 | 358 | S358Y | A | 4.3e-08 | 5.1e-08 | 3.1e-08 | 4.167E-08 | 1.007E-08 | 3.100E-08 | 5.100E-08 |
| 7160 | 358 | S358V | A | 3.5e-08 | 2.2e-08 | 5.9e-08 | 3.867E-08 | 1.877E-08 | 2.200E-08 | 5.900E-08 |
| 7161 | 359 | F359A | A | 3.5e-08 | 5.7e-08 | 9e-08 | 6.067E-08 | 2.768E-08 | 3.500E-08 | 9.000E-08 |
| 7162 | 359 | F359R | A | 3.8e-08 | 2.6e-08 | 3.3e-08 | 3.233E-08 | 6.028E-09 | 2.600E-08 | 3.800E-08 |
| 7163 | 359 | F359N | A | 3.4e-08 | 7.2e-08 | 3.3e-08 | 4.633E-08 | 2.223E-08 | 3.300E-08 | 7.200E-08 |
| 7164 | 359 | F359D | A | 3.1e-08 | 2.3e-08 | 3.1e-08 | 2.833E-08 | 4.619E-09 | 2.300E-08 | 3.100E-08 |
| 7165 | 359 | F359C | A | 1.1e-07 | 2.8e-08 | 7.2e-08 | 7.000E-08 | 4.104E-08 | 2.800E-08 | 1.100E-07 |
| 7166 | 359 | F359E | A | 6.9e-08 | 3.3e-08 | 2.5e-08 | 4.233E-08 | 2.344E-08 | 2.500E-08 | 6.900E-08 |
| 7167 | 359 | F359Q | A | 2.9e-08 | 4.2e-08 | 5.1e-08 | 4.067E-08 | 1.106E-08 | 2.900E-08 | 5.100E-08 |
| 7168 | 359 | F359G | A | 1e-07 | 3.6e-08 | 3.7e-08 | 5.767E-08 | 3.667E-08 | 3.600E-08 | 1.000E-07 |
| 7169 | 359 | F359H | A | 1.1e-08 | 1.5e-08 | 1.6e-07 | 6.200E-08 | 8.489E-08 | 1.100E-08 | 1.600E-07 |
| 7170 | 359 | F359I | A | 3.3e-08 | 5.3e-08 | 6.2e-08 | 4.933E-08 | 1.484E-08 | 3.300E-08 | 6.200E-08 |
| 7171 | 359 | F359L | A | 3e-08 | 3.5e-08 | 2.9e-08 | 3.133E-08 | 3.215E-09 | 2.900E-08 | 3.500E-08 |
| 7172 | 359 | F359K | A | 5.3e-08 | 6.3e-08 | 1e-07 | 7.200E-08 | 2.476E-08 | 5.300E-08 | 1.000E-07 |
| 7173 | 359 | F359M | A | 3.7e-08 | 2.5e-08 | 4.4e-08 | 3.533E-08 | 9.609E-09 | 2.500E-08 | 4.400E-08 |
| 7174 | 359 | F359F | A | 5.5e-08 | 7e-09 | 1.2e-08 | 2.467E-08 | 2.639E-08 | 7.000E-09 | 5.500E-08 |
| 7175 | 359 | F359P | A | 4.6e-08 | 6.8e-08 | 7.5e-08 | 6.300E-08 | 1.513E-08 | 4.600E-08 | 7.500E-08 |
| 7176 | 359 | F359S | A | 2e-08 | 6.6e-08 | 1.3e-08 | 3.300E-08 | 2.879E-08 | 1.300E-08 | 6.600E-08 |
| 7177 | 359 | F359T | A | 7.8e-08 | 3e-08 | 4.1e-08 | 4.967E-08 | 2.515E-08 | 3.000E-08 | 7.800E-08 |
| 7178 | 359 | F359W | A | 7.2e-08 | 6.2e-08 | 4.7e-08 | 6.033E-08 | 1.258E-08 | 4.700E-08 | 7.200E-08 |
| 7179 | 359 | F359Y | A | 6.9e-08 | 4.2e-08 | 3.9e-08 | 5.000E-08 | 1.652E-08 | 3.900E-08 | 6.900E-08 |
| 7180 | 359 | F359V | A | 3.2e-08 | 1.1e-07 | 3.2e-08 | 5.800E-08 | 4.503E-08 | 3.200E-08 | 1.100E-07 |
| 7181 | 360 | D360A | A | 3e-08 | 4.5e-08 | 1.6e-08 | 3.033E-08 | 1.450E-08 | 1.600E-08 | 4.500E-08 |
| 7182 | 360 | D360R | A | 7.4e-08 | 5.1e-08 | 1.1e-07 | 7.833E-08 | 2.974E-08 | 5.100E-08 | 1.100E-07 |
| 7183 | 360 | D360N | A | 2.9e-08 | 2.9e-08 | 4e-08 | 3.267E-08 | 6.351E-09 | 2.900E-08 | 4.000E-08 |
| 7184 | 360 | D360D | A | 8.5e-08 | 6.5e-08 | 9.2e-08 | 8.067E-08 | 1.401E-08 | 6.500E-08 | 9.200E-08 |
| 7185 | 360 | D360C | A | 6.8e-08 | 1.5e-07 | 3.1e-08 | 8.300E-08 | 6.090E-08 | 3.100E-08 | 1.500E-07 |
| 7186 | 360 | D360E | A | 4.2e-08 | 2.3e-08 | 4.6e-08 | 3.700E-08 | 1.229E-08 | 2.300E-08 | 4.600E-08 |
| 7187 | 360 | D360Q | A | 3.1e-08 | 7.7e-08 | 2.4e-08 | 4.400E-08 | 2.879E-08 | 2.400E-08 | 7.700E-08 |
| 7188 | 360 | D360G | A | 7e-08 | 3.6e-08 | 1.2e-08 | 3.933E-08 | 2.914E-08 | 1.200E-08 | 7.000E-08 |
| 7189 | 360 | D360H | A | 8.9e-08 | 3.5e-08 | 5.7e-08 | 6.033E-08 | 2.715E-08 | 3.500E-08 | 8.900E-08 |
| 7190 | 360 | D360I | A | 6.1e-08 | 8e-08 | 3.6e-08 | 5.900E-08 | 2.207E-08 | 3.600E-08 | 8.000E-08 |
| 7191 | 360 | D360L | A | 2e-08 | 3.1e-08 | 5.4e-08 | 3.500E-08 | 1.735E-08 | 2.000E-08 | 5.400E-08 |
| 7192 | 360 | D360K | A | 1.9e-08 | 8.2e-08 | 1.9e-08 | 4.000E-08 | 3.637E-08 | 1.900E-08 | 8.200E-08 |
| 7193 | 360 | D360M | A | 5.7e-08 | 8.2e-08 | 3.4e-08 | 5.767E-08 | 2.401E-08 | 3.400E-08 | 8.200E-08 |
| 7194 | 360 | D360F | A | 2.1e-08 | 2.5e-08 | 3.5e-08 | 2.700E-08 | 7.211E-09 | 2.100E-08 | 3.500E-08 |
| 7195 | 360 | D360P | A | 3.4e-08 | 1.4e-07 | 3e-08 | 6.800E-08 | 6.239E-08 | 3.000E-08 | 1.400E-07 |
| 7196 | 360 | D360S | A | 6.1e-08 | 4.8e-08 | 4.5e-08 | 5.133E-08 | 8.505E-09 | 4.500E-08 | 6.100E-08 |
| 7197 | 360 | D360T | A | 1.3e-08 | 1e-07 | 2.1e-08 | 4.467E-08 | 4.809E-08 | 1.300E-08 | 1.000E-07 |
| 7198 | 360 | D360W | A | 8.3e-08 | 3.5e-08 | 8e-08 | 6.600E-08 | 2.689E-08 | 3.500E-08 | 8.300E-08 |
| 7199 | 360 | D360Y | A | 3.6e-08 | 1.5e-08 | 1.8e-08 | 2.300E-08 | 1.136E-08 | 1.500E-08 | 3.600E-08 |
| 7200 | 360 | D360V | A | 1.6e-08 | 4.1e-08 | 5.8e-08 | 3.833E-08 | 2.113E-08 | 1.600E-08 | 5.800E-08 |
| 7201 | 361 | G361A | A | 6.2e-08 | 2.8e-08 | 4.4e-08 | 4.467E-08 | 1.701E-08 | 2.800E-08 | 6.200E-08 |
| 7202 | 361 | G361R | A | 3.6e-08 | 6.4e-08 | 2.1e-08 | 4.033E-08 | 2.183E-08 | 2.100E-08 | 6.400E-08 |
| 7203 | 361 | G361N | A | 1.9e-08 | 4.3e-08 | 2.3e-08 | 2.833E-08 | 1.286E-08 | 1.900E-08 | 4.300E-08 |
| 7204 | 361 | G361D | A | 1.7e-08 | 3e-08 | 3.6e-08 | 2.767E-08 | 9.713E-09 | 1.700E-08 | 3.600E-08 |
| 7205 | 361 | G361C | A | 6.1e-08 | 9.6e-09 | 5.8e-08 | 4.287E-08 | 2.885E-08 | 9.600E-09 | 6.100E-08 |
| 7206 | 361 | G361E | A | 5.7e-08 | 4.2e-08 | 7.8e-08 | 5.900E-08 | 1.808E-08 | 4.200E-08 | 7.800E-08 |
| 7207 | 361 | G361Q | A | 1.3e-08 | 6.1e-08 | 4.2e-08 | 3.867E-08 | 2.417E-08 | 1.300E-08 | 6.100E-08 |
| 7208 | 361 | G361G | A | 1.1e-07 | 4.8e-08 | 1.4e-08 | 5.733E-08 | 4.868E-08 | 1.400E-08 | 1.100E-07 |
| 7209 | 361 | G361H | A | 7e-08 | 4.8e-08 | 5e-08 | 5.600E-08 | 1.217E-08 | 4.800E-08 | 7.000E-08 |
| 7210 | 361 | G361I | A | 5.8e-08 | 2e-08 | 2.1e-08 | 3.300E-08 | 2.166E-08 | 2.000E-08 | 5.800E-08 |
| 7211 | 361 | G361L | A | 4.3e-08 | 5.1e-08 | 4.2e-08 | 4.533E-08 | 4.933E-09 | 4.200E-08 | 5.100E-08 |
| 7212 | 361 | G361K | A | 3.6e-08 | 4.6e-08 | 2.6e-08 | 3.600E-08 | 1.000E-08 | 2.600E-08 | 4.600E-08 |
| 7213 | 361 | G361M | A | 5.2e-08 | 6e-08 | 2.4e-08 | 4.533E-08 | 1.890E-08 | 2.400E-08 | 6.000E-08 |
| 7214 | 361 | G361F | A | 2.5e-08 | 1.5e-08 | 2.2e-08 | 2.067E-08 | 5.132E-09 | 1.500E-08 | 2.500E-08 |
| 7215 | 361 | G361P | A | 9.1e-08 | 3.7e-08 | 9.6e-08 | 7.467E-08 | 3.272E-08 | 3.700E-08 | 9.600E-08 |
| 7216 | 361 | G361S | A | 1.2e-07 | 3.6e-08 | 4.2e-08 | 6.600E-08 | 4.686E-08 | 3.600E-08 | 1.200E-07 |
| 7217 | 361 | G361T | A | 5.1e-08 | 5.6e-08 | 2.3e-08 | 4.333E-08 | 1.779E-08 | 2.300E-08 | 5.600E-08 |
| 7218 | 361 | G361W | A | 3.6e-08 | 3.7e-08 | 4.4e-08 | 3.900E-08 | 4.359E-09 | 3.600E-08 | 4.400E-08 |
| 7219 | 361 | G361Y | A | 3.8e-08 | 7.7e-08 | 8.3e-08 | 6.600E-08 | 2.443E-08 | 3.800E-08 | 8.300E-08 |
| 7220 | 361 | G361V | A | 4.9e-08 | 1e-07 | 1.6e-08 | 5.500E-08 | 4.232E-08 | 1.600E-08 | 1.000E-07 |
| 7221 | 362 | D362A | A | 6.5e-08 | 6.3e-08 | 3e-08 | 5.267E-08 | 1.966E-08 | 3.000E-08 | 6.500E-08 |
| 7222 | 362 | D362R | A | 4.3e-08 | 8.5e-08 | 2.9e-08 | 5.233E-08 | 2.914E-08 | 2.900E-08 | 8.500E-08 |
| 7223 | 362 | D362N | A | 4.1e-08 | 4.2e-08 | 3.1e-08 | 3.800E-08 | 6.083E-09 | 3.100E-08 | 4.200E-08 |
| 7224 | 362 | D362D | A | 3.7e-08 | 2.2e-08 | 1e-07 | 5.300E-08 | 4.139E-08 | 2.200E-08 | 1.000E-07 |

| | | | | | | | | | | |
|---|---|---|---|---|---|---|---|---|---|---|
| 7225 | 362 | D362C | A | 9.4e-08 | 3e-08 | 4.1e-08 | 5.500E-08 | 3.422E-08 | 3.000E-08 | 9.400E-08 |
| 7226 | 362 | D362E | A | 5.1e-08 | 8.9e-08 | 2.3e-07 | 1.233E-07 | 9.431E-08 | 5.100E-08 | 2.300E-07 |
| 7227 | 362 | D362Q | A | 3.7e-08 | 3.2e-08 | 1.3e-07 | 6.633E-08 | 5.519E-08 | 3.200E-08 | 1.300E-07 |
| 7228 | 362 | D362G | A | 1.6e-08 | 6.9e-08 | 1.2e-07 | 6.833E-08 | 5.200E-08 | 1.600E-08 | 1.200E-07 |
| 7229 | 362 | D362H | A | 5.9e-08 | 1.6e-07 | 2.2e-08 | 8.033E-08 | 7.143E-08 | 2.200E-08 | 1.600E-07 |
| 7230 | 362 | D362I | A | 2e-08 | 2e-08 | 6.2e-08 | 3.400E-08 | 2.425E-08 | 2.000E-08 | 6.200E-08 |
| 7231 | 362 | D362L | A | 1.8e-08 | 3.5e-08 | 9.8e-08 | 5.033E-08 | 4.215E-08 | 1.800E-08 | 9.800E-08 |
| 7232 | 362 | D362K | A | 3.9e-08 | 5.2e-08 | 3.5e-08 | 4.200E-08 | 8.888E-09 | 3.500E-08 | 5.200E-08 |
| 7233 | 362 | D362M | A | 5.5e-08 | 3.4e-08 | 3.7e-08 | 4.200E-08 | 1.136E-08 | 3.400E-08 | 5.500E-08 |
| 7234 | 362 | D362F | A | 5.1e-08 | 6e-08 | 3.2e-08 | 4.767E-08 | 1.429E-08 | 3.200E-08 | 6.000E-08 |
| 7235 | 362 | D362P | A | 5.7e-08 | 6.5e-08 | 6.6e-08 | 6.267E-08 | 4.933E-09 | 5.700E-08 | 6.600E-08 |
| 7236 | 362 | D362S | A | 6.7e-08 | 4.3e-08 | 8.7e-08 | 6.567E-08 | 2.203E-08 | 4.300E-08 | 8.700E-08 |
| 7237 | 362 | D362T | A | 1.5e-07 | 9.6e-08 | 4.5e-08 | 9.700E-08 | 5.251E-08 | 4.500E-08 | 1.500E-07 |
| 7238 | 362 | D362W | A | 4.6e-08 | 2.8e-08 | 7.5e-08 | 4.967E-08 | 2.371E-08 | 2.800E-08 | 7.500E-08 |
| 7239 | 362 | D362Y | A | 4.9e-08 | 2.4e-08 | 8.2e-08 | 5.167E-08 | 2.909E-08 | 2.400E-08 | 8.200E-08 |
| 7240 | 362 | D362V | A | 2.2e-08 | 5.3e-08 | 7.2e-08 | 4.900E-08 | 2.524E-08 | 2.200E-08 | 7.200E-08 |
| 7241 | 363 | P363A | A | 3.4e-08 | 6.8e-08 | 1.1e-08 | 3.767E-08 | 2.868E-08 | 1.100E-08 | 6.800E-08 |
| 7242 | 363 | P363R | A | 4.1e-08 | 4.4e-08 | 8.9e-09 | 3.130E-08 | 1.946E-08 | 8.900E-09 | 4.400E-08 |
| 7243 | 363 | P363N | A | 1.8e-08 | 4.3e-08 | 6.5e-08 | 4.200E-08 | 2.352E-08 | 1.800E-08 | 6.500E-08 |
| 7244 | 363 | P363D | A | 5.1e-08 | 6.7e-08 | 7.1e-08 | 6.300E-08 | 1.058E-08 | 5.100E-08 | 7.100E-08 |
| 7245 | 363 | P363C | A | 7.4e-08 | 7.1e-08 | 4.8e-08 | 6.433E-08 | 1.422E-08 | 4.800E-08 | 7.400E-08 |
| 7246 | 363 | P363E | A | 7.4e-08 | 5.3e-08 | 2.9e-08 | 5.200E-08 | 2.252E-08 | 2.900E-08 | 7.400E-08 |
| 7247 | 363 | P363Q | A | 3.5e-08 | 4.3e-08 | 4.1e-08 | 3.967E-08 | 4.163E-09 | 3.500E-08 | 4.300E-08 |
| 7248 | 363 | P363G | A | 6.9e-08 | 3.7e-08 | 6.9e-08 | 5.833E-08 | 1.848E-08 | 3.700E-08 | 6.900E-08 |
| 7249 | 363 | P363H | A | 9.1e-08 | 3.3e-08 | 1.9e-08 | 4.767E-08 | 3.818E-08 | 1.900E-08 | 9.100E-08 |
| 7250 | 363 | P363I | A | 2.8e-08 | 4.9e-08 | 3.5e-08 | 3.733E-08 | 1.069E-08 | 2.800E-08 | 4.900E-08 |
| 7251 | 363 | P363L | A | 3.3e-08 | 6.3e-08 | 1.6e-07 | 8.533E-08 | 6.638E-08 | 3.300E-08 | 1.600E-07 |
| 7252 | 363 | P363K | A | 2.8e-08 | 3.1e-08 | 4.8e-08 | 3.567E-08 | 1.079E-08 | 2.800E-08 | 4.800E-08 |
| 7253 | 363 | P363M | A | 7.5e-08 | 5.8e-08 | 2.2e-08 | 5.167E-08 | 2.706E-08 | 2.200E-08 | 7.500E-08 |
| 7254 | 363 | P363F | A | 3.6e-08 | 2.2e-08 | 3.7e-08 | 3.167E-08 | 8.386E-09 | 2.200E-08 | 3.700E-08 |
| 7255 | 363 | P363P | A | 9e-08 | 5.8e-08 | 4.9e-08 | 6.567E-08 | 2.155E-08 | 4.900E-08 | 9.000E-08 |
| 7256 | 363 | P363S | A | 7.8e-08 | 9.9e-09 | 2.9e-08 | 3.897E-08 | 3.513E-08 | 9.900E-09 | 7.800E-08 |
| 7257 | 363 | P363T | A | 4e-08 | 5.6e-08 | 5.1e-08 | 4.900E-08 | 8.185E-09 | 4.000E-08 | 5.600E-08 |
| 7258 | 363 | P363W | A | 3.1e-08 | 4.1e-08 | 5.7e-08 | 4.300E-08 | 1.311E-08 | 3.100E-08 | 5.700E-08 |
| 7259 | 363 | P363Y | A | 5.9e-08 | 9.4e-08 | 1.6e-08 | 5.633E-08 | 3.907E-08 | 1.600E-08 | 9.400E-08 |
| 7260 | 363 | P363V | A | 2.8e-08 | 7.4e-08 | 4.4e-08 | 4.867E-08 | 2.335E-08 | 2.800E-08 | 7.400E-08 |
| 7261 | 364 | A364A | A | 8.8e-08 | 4.5e-08 | 3.1e-08 | 5.467E-08 | 2.970E-08 | 3.100E-08 | 8.800E-08 |
| 7262 | 364 | A364R | A | 5.3e-08 | 6.1e-08 | 3.3e-08 | 4.900E-08 | 1.442E-08 | 3.300E-08 | 6.100E-08 |
| 7263 | 364 | A364N | A | 3.6e-08 | 9.4e-08 | 1.4e-07 | 9.000E-08 | 5.212E-08 | 3.600E-08 | 1.400E-07 |
| 7264 | 364 | A364D | A | 8.6e-08 | 2.2e-08 | 3e-08 | 4.600E-08 | 3.487E-08 | 2.200E-08 | 8.600E-08 |
| 7265 | 364 | A364C | A | 5.8e-08 | 7.9e-08 | 1.9e-08 | 5.200E-08 | 3.045E-08 | 1.900E-08 | 7.900E-08 |
| 7266 | 364 | A364E | A | 8.2e-08 | 4.7e-08 | 3e-08 | 5.300E-08 | 2.651E-08 | 3.000E-08 | 8.200E-08 |
| 7267 | 364 | A364Q | A | 8.2e-08 | 3e-08 | 1.2e-07 | 7.733E-08 | 4.518E-08 | 3.000E-08 | 1.200E-07 |
| 7268 | 364 | A364G | A | 3.7e-08 | 4.5e-08 | 5.7e-08 | 4.633E-08 | 1.007E-08 | 3.700E-08 | 5.700E-08 |
| 7269 | 364 | A364H | A | 6e-08 | 2.1e-08 | 6.8e-08 | 4.967E-08 | 2.515E-08 | 2.100E-08 | 6.800E-08 |
| 7270 | 364 | A364I | A | 4e-08 | 5.1e-08 | 5.4e-08 | 4.833E-08 | 7.371E-09 | 4.000E-08 | 5.400E-08 |
| 7271 | 364 | A364L | A | 3.6e-08 | 5.4e-08 | 1.9e-08 | 3.633E-08 | 1.750E-08 | 1.900E-08 | 5.400E-08 |
| 7272 | 364 | A364K | A | 2.7e-08 | 3e-08 | 4.6e-08 | 3.433E-08 | 1.021E-08 | 2.700E-08 | 4.600E-08 |
| 7273 | 364 | A364M | A | 2e-08 | 3.5e-08 | 4.1e-08 | 3.200E-08 | 1.082E-08 | 2.000E-08 | 4.100E-08 |
| 7274 | 364 | A364F | A | 2e-08 | 3e-08 | 8.8e-08 | 4.600E-08 | 3.672E-08 | 2.000E-08 | 8.800E-08 |
| 7275 | 364 | A364P | A | 4.4e-08 | 4.4e-08 | 5e-08 | 4.600E-08 | 3.464E-09 | 4.400E-08 | 5.000E-08 |
| 7276 | 364 | A364S | A | 1.1e-08 | 1.1e-08 | 4.2e-08 | 2.133E-08 | 1.790E-08 | 1.100E-08 | 4.200E-08 |
| 7277 | 364 | A364T | A | 5.6e-08 | 8.3e-08 | 1e-07 | 7.967E-08 | 2.219E-08 | 5.600E-08 | 1.000E-07 |
| 7278 | 364 | A364W | A | 3e-08 | 6.5e-08 | 2.3e-08 | 3.933E-08 | 2.250E-08 | 2.300E-08 | 6.500E-08 |
| 7279 | 364 | A364Y | A | 4.4e-08 | 7.6e-08 | 2.6e-08 | 4.867E-08 | 2.532E-08 | 2.600E-08 | 7.600E-08 |
| 7280 | 364 | A364V | A | 4.6e-08 | 2.7e-08 | 3e-08 | 3.433E-08 | 1.021E-08 | 2.700E-08 | 4.600E-08 |
| 7281 | 365 | S365A | A | 2e-08 | 4.2e-08 | 3.7e-08 | 3.300E-08 | 1.153E-08 | 2.000E-08 | 4.200E-08 |
| 7282 | 365 | S365R | A | 1.8e-08 | 2.7e-08 | 2.3e-08 | 2.267E-08 | 4.509E-09 | 1.800E-08 | 2.700E-08 |
| 7283 | 365 | S365N | A | 4.8e-08 | 3.4e-08 | 5.6e-08 | 4.600E-08 | 1.114E-08 | 3.400E-08 | 5.600E-08 |
| 7284 | 365 | S365D | A | 3.8e-08 | 1e-07 | 8.1e-08 | 7.300E-08 | 3.176E-08 | 3.800E-08 | 1.000E-07 |
| 7285 | 365 | S365C | A | 5e-08 | 5e-09 | 2.4e-08 | 2.633E-08 | 2.259E-08 | 5.000E-09 | 5.000E-08 |
| 7286 | 365 | S365E | A | 2.4e-08 | 6.5e-08 | 4.5e-08 | 4.467E-08 | 2.050E-08 | 2.400E-08 | 6.500E-08 |
| 7287 | 365 | S365Q | A | 2.1e-08 | 1.8e-08 | 2.3e-08 | 2.067E-08 | 2.517E-09 | 1.800E-08 | 2.300E-08 |
| 7288 | 365 | S365G | A | 4.8e-08 | 1.1e-07 | 4.5e-08 | 6.767E-08 | 3.669E-08 | 4.500E-08 | 1.100E-07 |
| 7289 | 365 | S365H | A | 2.7e-08 | 1.7e-08 | 6.5e-08 | 3.633E-08 | 2.532E-08 | 1.700E-08 | 6.500E-08 |
| 7290 | 365 | S365I | A | 2.4e-08 | 9.4e-08 | 4.4e-08 | 5.400E-08 | 3.606E-08 | 2.400E-08 | 9.400E-08 |
| 7291 | 365 | S365L | A | 4.7e-08 | 1.6e-08 | 3e-08 | 3.100E-08 | 1.552E-08 | 1.600E-08 | 4.700E-08 |
| 7292 | 365 | S365K | A | 5.5e-08 | 9.6e-08 | 4.3e-08 | 6.467E-08 | 2.779E-08 | 4.300E-08 | 9.600E-08 |
| 7293 | 365 | S365M | A | 2.2e-08 | 1.7e-07 | 2e-08 | 7.067E-08 | 8.603E-08 | 2.000E-08 | 1.700E-07 |
| 7294 | 365 | S365F | A | 4.8e-08 | 9.5e-08 | 1e-07 | 8.100E-08 | 2.869E-08 | 4.800E-08 | 1.000E-07 |
| 7295 | 365 | S365P | A | 3.7e-08 | 5.2e-08 | 7.9e-08 | 5.600E-08 | 2.128E-08 | 3.700E-08 | 7.900E-08 |
| 7296 | 365 | S365S | A | 4.9e-08 | 4.1e-08 | 3.7e-08 | 4.233E-08 | 6.110E-09 | 3.700E-08 | 4.900E-08 |
| 7297 | 365 | S365T | A | 2.8e-08 | 4.7e-08 | 6.3e-08 | 4.600E-08 | 1.752E-08 | 2.800E-08 | 6.300E-08 |
| 7298 | 365 | S365W | A | 5e-08 | 7.9e-08 | 1.1e-07 | 7.967E-08 | 3.001E-08 | 5.000E-08 | 1.100E-07 |
| 7299 | 365 | S365Y | A | 4.1e-08 | 2.9e-08 | 2.3e-08 | 3.100E-08 | 9.165E-09 | 2.300E-08 | 4.100E-08 |
| 7300 | 365 | S365V | A | 4.6e-08 | 3.7e-08 | 3.7e-08 | 4.000E-08 | 5.196E-09 | 3.700E-08 | 4.600E-08 |
| 7301 | 366 | N366A | A | 8.8e-08 | 3e-08 | 4.3e-08 | 5.367E-08 | 3.044E-08 | 3.000E-08 | 8.800E-08 |
| 7302 | 366 | N366R | A | 5.4e-08 | 4.7e-08 | 2.8e-08 | 4.300E-08 | 1.345E-08 | 2.800E-08 | 5.400E-08 |
| 7303 | 366 | N366N | A | 2.5e-08 | 4e-08 | 2.5e-08 | 3.000E-08 | 8.660E-09 | 2.500E-08 | 4.000E-08 |
| 7304 | 366 | N366D | A | 6.7e-08 | 3.1e-08 | 1.8e-08 | 3.867E-08 | 2.538E-08 | 1.800E-08 | 6.700E-08 |
| 7305 | 366 | N366C | A | 3.6e-08 | 6.2e-08 | 3.3e-08 | 4.367E-08 | 1.595E-08 | 3.300E-08 | 6.200E-08 |
| 7306 | 366 | N366E | A | 4.3e-08 | 4.5e-08 | 6e-08 | 4.933E-08 | 9.292E-09 | 4.300E-08 | 6.000E-08 |
| 7307 | 366 | N366Q | A | 5.4e-08 | 1.2e-08 | 3.2e-08 | 3.267E-08 | 2.101E-08 | 1.200E-08 | 5.400E-08 |
| 7308 | 366 | N366G | A | 2.7e-07 | 9.9e-08 | 3.5e-08 | 1.347E-07 | 1.215E-07 | 3.500E-08 | 2.700E-07 |
| 7309 | 366 | N366H | A | 4.2e-08 | 5.7e-08 | 4e-08 | 4.633E-08 | 9.292E-09 | 4.000E-08 | 5.700E-08 |

| | | | | | | | | | | |
|---|---|---|---|---|---|---|---|---|---|---|
| 7310 | 366 | N366I | A | 2.5e-08 | 4.2e-08 | 2.1e-08 | 2.933E-08 | 1.115E-08 | 2.100E-08 | 4.200E-08 |
| 7311 | 366 | N366L | A | 3e-08 | 2e-08 | 5.8e-08 | 3.600E-08 | 1.970E-08 | 2.000E-08 | 5.800E-08 |
| 7312 | 366 | N366K | A | 6.3e-08 | 2.5e-08 | 2.9e-08 | 3.900E-08 | 2.088E-08 | 2.500E-08 | 6.300E-08 |
| 7313 | 366 | N366M | A | 2.7e-08 | 5.9e-08 | 2e-08 | 3.533E-08 | 2.079E-08 | 2.000E-08 | 5.900E-08 |
| 7314 | 366 | N366F | A | 2.5e-08 | 6.4e-08 | 3.3e-08 | 4.067E-08 | 2.060E-08 | 2.500E-08 | 6.400E-08 |
| 7315 | 366 | N366P | A | 1.5e-08 | 6.8e-08 | 6.4e-08 | 4.900E-08 | 2.951E-08 | 1.500E-08 | 6.800E-08 |
| 7316 | 366 | N366S | A | 1.4e-08 | 9.6e-08 | 5.1e-08 | 5.367E-08 | 4.106E-08 | 1.400E-08 | 9.600E-08 |
| 7317 | 366 | N366T | A | 4.1e-08 | 1.7e-08 | 1.9e-08 | 2.567E-08 | 1.332E-08 | 1.700E-08 | 4.100E-08 |
| 7318 | 366 | N366W | A | 3.1e-08 | 1.6e-08 | 1.5e-08 | 2.067E-08 | 8.963E-09 | 1.500E-08 | 3.100E-08 |
| 7319 | 366 | N366Y | A | 1.6e-08 | 5.8e-08 | 2.6e-08 | 3.333E-08 | 2.194E-08 | 1.600E-08 | 5.800E-08 |
| 7320 | 366 | N366V | A | 6.5e-08 | 1.8e-08 | 6e-08 | 4.767E-08 | 2.581E-08 | 1.800E-08 | 6.500E-08 |
| 7321 | 367 | T367A | A | 3.6e-08 | 4e-08 | 4.1e-08 | 3.900E-08 | 2.646E-09 | 3.600E-08 | 4.100E-08 |
| 7322 | 367 | T367R | A | 3.4e-08 | 6.4e-08 | 6.3e-08 | 5.367E-08 | 1.704E-08 | 3.400E-08 | 6.400E-08 |
| 7323 | 367 | T367N | A | 1.1e-08 | 3.1e-08 | 4.7e-08 | 2.967E-08 | 1.804E-08 | 1.100E-08 | 4.700E-08 |
| 7324 | 367 | T367D | A | 4.9e-08 | 5.5e-08 | 5.9e-08 | 5.433E-08 | 5.033E-09 | 4.900E-08 | 5.900E-08 |
| 7325 | 367 | T367C | A | 3.2e-08 | 2.5e-08 | 3.6e-08 | 3.100E-08 | 5.568E-09 | 2.500E-08 | 3.600E-08 |
| 7326 | 367 | T367E | A | 2.3e-08 | 9e-08 | 2.6e-08 | 4.633E-08 | 3.785E-08 | 2.300E-08 | 9.000E-08 |
| 7327 | 367 | T367Q | A | 3.6e-08 | 3.2e-08 | 6.5e-08 | 4.433E-08 | 1.801E-08 | 3.200E-08 | 6.500E-08 |
| 7328 | 367 | T367G | A | 1e-07 | 1.2e-07 | 8.4e-08 | 1.013E-07 | 1.804E-08 | 8.400E-08 | 1.200E-07 |
| 7329 | 367 | T367H | A | 2.6e-08 | 2.1e-08 | 5e-08 | 3.233E-08 | 1.550E-08 | 2.100E-08 | 5.000E-08 |
| 7330 | 367 | T367I | A | 3.2e-08 | 2.4e-08 | 2.9e-08 | 2.833E-08 | 4.041E-09 | 2.400E-08 | 3.200E-08 |
| 7331 | 367 | T367L | A | 4.3e-08 | 2.5e-08 | 3.1e-08 | 3.300E-08 | 9.165E-09 | 2.500E-08 | 4.300E-08 |
| 7332 | 367 | T367K | A | 5.7e-08 | 7.2e-08 | 6.1e-08 | 6.333E-08 | 7.767E-09 | 5.700E-08 | 7.200E-08 |
| 7333 | 367 | T367M | A | 8.4e-08 | 1.7e-07 | 1.1e-07 | 1.213E-07 | 4.411E-08 | 8.400E-08 | 1.700E-07 |
| 7334 | 367 | T367F | A | 2.2e-08 | 4.4e-08 | 2.5e-08 | 3.033E-08 | 1.193E-08 | 2.200E-08 | 4.400E-08 |
| 7335 | 367 | T367P | A | 6.9e-08 | 3e-08 | 2.5e-08 | 4.133E-08 | 2.409E-08 | 2.500E-08 | 6.900E-08 |
| 7336 | 367 | T367S | A | 2.9e-08 | 5.9e-08 | 2.6e-08 | 3.800E-08 | 1.825E-08 | 2.600E-08 | 5.900E-08 |
| 7337 | 367 | T367T | A | 4.1e-08 | 4.7e-08 | 1.2e-08 | 3.333E-08 | 1.872E-08 | 1.200E-08 | 4.700E-08 |
| 7338 | 367 | T367W | A | 3.8e-08 | 1.1e-07 | 2.9e-08 | 5.900E-08 | 4.440E-08 | 2.900E-08 | 1.100E-07 |
| 7339 | 367 | T367Y | A | 4.4e-08 | 4.1e-08 | 1.3e-08 | 3.267E-08 | 1.710E-08 | 1.300E-08 | 4.400E-08 |
| 7340 | 367 | T367V | A | 1.8e-08 | 2.7e-08 | 9e-08 | 4.500E-08 | 3.923E-08 | 1.800E-08 | 9.000E-08 |
| 7341 | 368 | A368A | A | 6.5e-08 | 1e-07 | 3.2e-08 | 6.567E-08 | 3.400E-08 | 3.200E-08 | 1.000E-07 |
| 7342 | 368 | A368R | A | 5.9e-08 | 4.8e-08 | 1.2e-07 | 7.567E-08 | 3.879E-08 | 4.800E-08 | 1.200E-07 |
| 7343 | 368 | A368N | A | 2.4e-08 | 2.8e-08 | 3.3e-08 | 2.833E-08 | 4.509E-09 | 2.400E-08 | 3.300E-08 |
| 7344 | 368 | A368D | A | 1.9e-08 | 6e-08 | 1.7e-08 | 3.200E-08 | 2.427E-08 | 1.700E-08 | 6.000E-08 |
| 7345 | 368 | A368C | A | 1.4e-08 | 5.6e-08 | 6.1e-08 | 4.367E-08 | 2.581E-08 | 1.400E-08 | 6.100E-08 |
| 7346 | 368 | A368E | A | 6e-08 | 3.3e-08 | 6.4e-08 | 5.233E-08 | 1.686E-08 | 3.300E-08 | 6.400E-08 |
| 7347 | 368 | A368Q | A | 3.9e-08 | 2.9e-08 | 3.1e-08 | 3.300E-08 | 5.292E-09 | 2.900E-08 | 3.900E-08 |
| 7348 | 368 | A368G | A | 5.3e-08 | 1e-07 | 6.7e-08 | 7.333E-08 | 2.413E-08 | 5.300E-08 | 1.000E-07 |
| 7349 | 368 | A368H | A | 2.7e-08 | 3.6e-08 | 2.7e-08 | 3.000E-08 | 5.196E-09 | 2.700E-08 | 3.600E-08 |
| 7350 | 368 | A368I | A | 1.9e-08 | 1.1e-08 | 4.3e-08 | 2.433E-08 | 1.665E-08 | 1.100E-08 | 4.300E-08 |
| 7351 | 368 | A368L | A | 5.8e-08 | 2.4e-08 | 1.4e-08 | 3.200E-08 | 2.307E-08 | 1.400E-08 | 5.800E-08 |
| 7352 | 368 | A368K | A | 3.4e-08 | 4.2e-08 | 1.5e-07 | 7.533E-08 | 6.479E-08 | 3.400E-08 | 1.500E-07 |
| 7353 | 368 | A368M | A | 8.2e-08 | 5.3e-08 | 6.8e-08 | 6.767E-08 | 1.450E-08 | 5.300E-08 | 8.200E-08 |
| 7354 | 368 | A368F | A | 5.1e-08 | 9.1e-08 | 9.7e-09 | 5.057E-08 | 4.065E-08 | 9.700E-09 | 9.100E-08 |
| 7355 | 368 | A368P | A | 4.4e-08 | 2.6e-08 | 4.6e-08 | 3.867E-08 | 1.102E-08 | 2.600E-08 | 4.600E-08 |
| 7356 | 368 | A368S | A | 2.4e-08 | 3.6e-08 | 3.1e-08 | 3.033E-08 | 6.028E-09 | 2.400E-08 | 3.600E-08 |
| 7357 | 368 | A368T | A | 3.1e-08 | 3.2e-08 | 2.5e-08 | 2.933E-08 | 3.786E-09 | 2.500E-08 | 3.200E-08 |
| 7358 | 368 | A368W | A | 1.7e-08 | 1e-07 | 7.3e-08 | 6.333E-08 | 4.234E-08 | 1.700E-08 | 1.000E-07 |
| 7359 | 368 | A368Y | A | 1.6e-08 | 9.6e-09 | 2.4e-08 | 1.653E-08 | 7.215E-09 | 9.600E-09 | 2.400E-08 |
| 7360 | 368 | A368V | A | 6.8e-08 | 4.4e-08 | 8.6e-08 | 6.600E-08 | 2.107E-08 | 4.400E-08 | 8.600E-08 |
| 7361 | 369 | P369A | A | 4.2e-08 | 5.7e-08 | 1.6e-08 | 3.833E-08 | 2.074E-08 | 1.600E-08 | 5.700E-08 |
| 7362 | 369 | P369R | A | 3.2e-08 | 6.6e-08 | 4.7e-08 | 4.833E-08 | 1.704E-08 | 3.200E-08 | 6.600E-08 |
| 7363 | 369 | P369N | A | 5.1e-08 | 3.2e-08 | 6.3e-08 | 4.867E-08 | 1.563E-08 | 3.200E-08 | 6.300E-08 |
| 7364 | 369 | P369D | A | 1.3e-07 | 3.8e-08 | 7e-08 | 7.933E-08 | 4.670E-08 | 3.800E-08 | 1.300E-07 |
| 7365 | 369 | P369C | A | 2.6e-08 | 8e-08 | 4.9e-08 | 5.167E-08 | 2.710E-08 | 2.600E-08 | 8.000E-08 |
| 7366 | 369 | P369E | A | 4.8e-08 | 8.4e-08 | 9.6e-08 | 7.600E-08 | 2.498E-08 | 4.800E-08 | 9.600E-08 |
| 7367 | 369 | P369Q | A | 3.3e-08 | 2.4e-08 | 4.2e-08 | 3.300E-08 | 9.000E-09 | 2.400E-08 | 4.200E-08 |
| 7368 | 369 | P369G | A | 4.9e-08 | 7.1e-08 | 6e-08 | 6.000E-08 | 1.100E-08 | 4.900E-08 | 7.100E-08 |
| 7369 | 369 | P369H | A | 2.9e-08 | 8.1e-08 | 3.1e-08 | 4.700E-08 | 2.946E-08 | 2.900E-08 | 8.100E-08 |
| 7370 | 369 | P369I | A | 8.5e-08 | 6.1e-08 | 5e-08 | 6.533E-08 | 1.790E-08 | 5.000E-08 | 8.500E-08 |
| 7371 | 369 | P369L | A | 3.8e-08 | 3.6e-08 | 4.4e-08 | 3.933E-08 | 4.163E-09 | 3.600E-08 | 4.400E-08 |
| 7372 | 369 | P369K | A | 7.1e-08 | 4.2e-08 | 2.9e-08 | 4.733E-08 | 2.150E-08 | 2.900E-08 | 7.100E-08 |
| 7373 | 369 | P369M | A | 7.9e-08 | 3.4e-08 | 4.2e-08 | 5.167E-08 | 2.401E-08 | 3.400E-08 | 7.900E-08 |
| 7374 | 369 | P369F | A | 3.9e-08 | 7.3e-08 | 8.2e-08 | 6.467E-08 | 2.268E-08 | 3.900E-08 | 8.200E-08 |
| 7375 | 369 | P369P | A | 4.2e-08 | 4.8e-08 | 4e-08 | 4.333E-08 | 4.163E-09 | 4.000E-08 | 4.800E-08 |
| 7376 | 369 | P369S | A | 1.5e-08 | 5.1e-08 | 3.7e-08 | 3.433E-08 | 1.815E-08 | 1.500E-08 | 5.100E-08 |
| 7377 | 369 | P369T | A | 5e-08 | 4.4e-08 | 1.1e-07 | 6.800E-08 | 3.650E-08 | 4.400E-08 | 1.100E-07 |
| 7378 | 369 | P369W | A | 4.6e-08 | 2.1e-08 | 4.8e-08 | 3.833E-08 | 1.504E-08 | 2.100E-08 | 4.800E-08 |
| 7379 | 369 | P369Y | A | 3.8e-08 | 2.9e-08 | 6.1e-08 | 4.267E-08 | 1.650E-08 | 2.900E-08 | 6.100E-08 |
| 7380 | 369 | P369V | A | 1.2e-07 | 4.6e-08 | 5.4e-08 | 7.333E-08 | 4.061E-08 | 4.600E-08 | 1.200E-07 |
| 7381 | 370 | L370A | A | 3.1e-08 | 2.6e-08 | 8.5e-08 | 4.733E-08 | 3.272E-08 | 2.600E-08 | 8.500E-08 |
| 7382 | 370 | L370R | A | 4.4e-08 | 7.3e-08 | 6.1e-08 | 5.933E-08 | 1.457E-08 | 4.400E-08 | 7.300E-08 |
| 7383 | 370 | L370N | A | 4.2e-08 | 2.8e-08 | 3.7e-08 | 3.567E-08 | 7.095E-09 | 2.800E-08 | 4.200E-08 |
| 7384 | 370 | L370D | A | 1.9e-08 | 2.5e-08 | 4.8e-08 | 3.067E-08 | 1.531E-08 | 1.900E-08 | 4.800E-08 |
| 7385 | 370 | L370C | A | 9.4e-08 | 6.3e-08 | 8.1e-08 | 7.933E-08 | 1.557E-08 | 6.300E-08 | 9.400E-08 |
| 7386 | 370 | L370E | A | 9.7e-08 | 2.1e-08 | 3.8e-08 | 5.200E-08 | 3.989E-08 | 2.100E-08 | 9.700E-08 |
| 7387 | 370 | L370Q | A | 4e-08 | 1.7e-08 | 5.6e-08 | 3.767E-08 | 1.960E-08 | 1.700E-08 | 5.600E-08 |
| 7388 | 370 | L370G | A | 4e-08 | 5.1e-08 | 3.5e-08 | 4.200E-08 | 8.185E-09 | 3.500E-08 | 5.100E-08 |
| 7389 | 370 | L370H | A | 2e-08 | 7.8e-08 | 6.4e-08 | 5.400E-08 | 3.027E-08 | 2.000E-08 | 7.800E-08 |
| 7390 | 370 | L370I | A | 2.4e-08 | 3.6e-08 | 3.5e-08 | 3.167E-08 | 6.658E-09 | 2.400E-08 | 3.600E-08 |
| 7391 | 370 | L370L | A | 7.6e-08 | 5.9e-08 | 5.2e-08 | 6.233E-08 | 1.234E-08 | 5.200E-08 | 7.600E-08 |
| 7392 | 370 | L370K | A | 2.2e-08 | 3.8e-08 | 2.7e-08 | 2.900E-08 | 8.185E-09 | 2.200E-08 | 3.800E-08 |
| 7393 | 370 | L370M | A | 5.8e-08 | 5.8e-08 | 1.7e-08 | 4.433E-08 | 2.367E-08 | 1.700E-08 | 5.800E-08 |
| 7394 | 370 | L370F | A | 4.4e-08 | 6.2e-08 | 3.1e-08 | 4.567E-08 | 1.557E-08 | 3.100E-08 | 6.200E-08 |

| | | | | | | | | | | |
|---|---|---|---|---|---|---|---|---|---|---|
| 7395 | 370 | L370P | A | 1.1e-07 | 8.8e-08 | 2.1e-08 | 7.300E-08 | 4.636E-08 | 2.100E-08 | 1.100E-07 |
| 7396 | 370 | L370S | A | 3.1e-08 | 2.9e-08 | 3.7e-08 | 3.233E-08 | 4.163E-09 | 2.900E-08 | 3.700E-08 |
| 7397 | 370 | L370T | A | 3.8e-08 | 9.5e-08 | 2.8e-08 | 5.367E-08 | 3.614E-08 | 2.800E-08 | 9.500E-08 |
| 7398 | 370 | L370W | A | 3.8e-08 | 2.4e-08 | 4.9e-08 | 3.700E-08 | 1.253E-08 | 2.400E-08 | 4.900E-08 |
| 7399 | 370 | L370Y | A | 8.3e-08 | 3.5e-08 | 2e-08 | 4.600E-08 | 3.291E-08 | 2.000E-08 | 8.300E-08 |
| 7400 | 370 | L370V | A | 6e-08 | 6.4e-08 | 8.3e-08 | 6.900E-08 | 1.229E-08 | 6.000E-08 | 8.300E-08 |
| 7401 | 371 | Q371A | A | 1.2e-07 | 5.5e-08 | 1.6e-08 | 6.367E-08 | 5.254E-08 | 1.600E-08 | 1.200E-07 |
| 7402 | 371 | Q371R | A | 3.3e-08 | 3.9e-08 | 6.9e-08 | 4.700E-08 | 1.929E-08 | 3.300E-08 | 6.900E-08 |
| 7403 | 371 | Q371N | A | 4.9e-08 | 4e-08 | 3.7e-08 | 4.200E-08 | 6.245E-09 | 3.700E-08 | 4.900E-08 |
| 7404 | 371 | Q371D | A | 3.2e-08 | 4.1e-08 | 4.1e-08 | 3.800E-08 | 5.196E-09 | 3.200E-08 | 4.100E-08 |
| 7405 | 371 | Q371C | A | 6e-08 | 7.4e-09 | 1.5e-07 | 7.247E-08 | 7.211E-08 | 7.400E-09 | 1.500E-07 |
| 7406 | 371 | Q371E | A | 4.2e-08 | 5.7e-08 | 6.3e-08 | 5.400E-08 | 1.082E-08 | 4.200E-08 | 6.300E-08 |
| 7407 | 371 | Q371Q | A | 9.2e-08 | 7.7e-08 | 1.6e-07 | 1.097E-07 | 4.423E-08 | 7.700E-08 | 1.600E-07 |
| 7408 | 371 | Q371G | A | 6.4e-08 | 1.2e-08 | 4.4e-08 | 4.000E-08 | 2.623E-08 | 1.200E-08 | 6.400E-08 |
| 7409 | 371 | Q371H | A | 5.7e-08 | 1.2e-07 | 1.1e-08 | 6.267E-08 | 5.472E-08 | 1.100E-08 | 1.200E-07 |
| 7410 | 371 | Q371I | A | 3e-08 | 8.4e-08 | 7.4e-08 | 6.267E-08 | 2.873E-08 | 3.000E-08 | 8.400E-08 |
| 7411 | 371 | Q371L | A | 3.1e-08 | 4.5e-08 | 3.7e-08 | 3.767E-08 | 7.024E-09 | 3.100E-08 | 4.500E-08 |
| 7412 | 371 | Q371K | A | 4.3e-08 | 3.6e-08 | 4.6e-08 | 4.167E-08 | 5.132E-09 | 3.600E-08 | 4.600E-08 |
| 7413 | 371 | Q371M | A | 7e-08 | 3.1e-08 | 3.6e-08 | 4.567E-08 | 2.122E-08 | 3.100E-08 | 7.000E-08 |
| 7414 | 371 | Q371F | A | 1.9e-08 | 6.9e-08 | 2.5e-08 | 3.767E-08 | 2.730E-08 | 1.900E-08 | 6.900E-08 |
| 7415 | 371 | Q371P | A | 3.1e-08 | 2.3e-08 | 6.8e-08 | 4.067E-08 | 2.401E-08 | 2.300E-08 | 6.800E-08 |
| 7416 | 371 | Q371S | A | 6.3e-08 | 1.8e-07 | 4.8e-08 | 9.700E-08 | 7.227E-08 | 4.800E-08 | 1.800E-07 |
| 7417 | 371 | Q371T | A | 1.6e-08 | 3.9e-08 | 2.3e-08 | 2.600E-08 | 1.179E-08 | 1.600E-08 | 3.900E-08 |
| 7418 | 371 | Q371W | A | 3.7e-08 | 6.2e-08 | 6.6e-08 | 5.500E-08 | 1.572E-08 | 3.700E-08 | 6.600E-08 |
| 7419 | 371 | Q371Y | A | 5.2e-08 | 3.8e-08 | 3.8e-08 | 4.267E-08 | 8.083E-09 | 3.800E-08 | 5.200E-08 |
| 7420 | 371 | Q371V | A | 8.3e-09 | 3e-08 | 5.6e-08 | 3.143E-08 | 2.388E-08 | 8.300E-09 | 5.600E-08 |
| 7421 | 372 | P372A | A | 1.4e-07 | 7.7e-08 | 3.6e-08 | 8.433E-08 | 5.239E-08 | 3.600E-08 | 1.400E-07 |
| 7422 | 372 | P372R | A | 1.9e-08 | 7.9e-08 | 4.1e-08 | 4.633E-08 | 3.035E-08 | 1.900E-08 | 7.900E-08 |
| 7423 | 372 | P372N | A | 2.6e-08 | 1.8e-08 | 3.1e-08 | 2.500E-08 | 6.557E-09 | 1.800E-08 | 3.100E-08 |
| 7424 | 372 | P372D | A | 4.1e-08 | 2.1e-08 | 4.7e-08 | 3.633E-08 | 1.361E-08 | 2.100E-08 | 4.700E-08 |
| 7425 | 372 | P372C | A | 8.1e-08 | 4.8e-08 | 3.3e-08 | 5.400E-08 | 2.456E-08 | 3.300E-08 | 8.100E-08 |
| 7426 | 372 | P372E | A | 4.6e-08 | 3.6e-08 | 8.3e-08 | 5.500E-08 | 2.476E-08 | 3.600E-08 | 8.300E-08 |
| 7427 | 372 | P372Q | A | 3e-08 | 3.5e-08 | 6.6e-08 | 4.367E-08 | 1.950E-08 | 3.000E-08 | 6.600E-08 |
| 7428 | 372 | P372G | A | 1.8e-08 | 9.8e-08 | 3e-08 | 4.867E-08 | 4.314E-08 | 1.800E-08 | 9.800E-08 |
| 7429 | 372 | P372H | A | 9.6e-09 | 6.7e-08 | 7.3e-08 | 4.987E-08 | 3.500E-08 | 9.600E-09 | 7.300E-08 |
| 7430 | 372 | P372I | A | 1.5e-08 | 3.3e-08 | 3.8e-08 | 2.867E-08 | 1.210E-08 | 1.500E-08 | 3.800E-08 |
| 7431 | 372 | P372L | A | 3.3e-08 | 8.2e-08 | 7.6e-08 | 6.367E-08 | 2.673E-08 | 3.300E-08 | 8.200E-08 |
| 7432 | 372 | P372K | A | 5.4e-08 | 3.4e-08 | 7.3e-08 | 5.367E-08 | 1.950E-08 | 3.400E-08 | 7.300E-08 |
| 7433 | 372 | P372M | A | 1.4e-08 | 6e-08 | 1.9e-08 | 3.100E-08 | 2.524E-08 | 1.400E-08 | 6.000E-08 |
| 7434 | 372 | P372F | A | 4.4e-08 | 4.3e-08 | 2.3e-08 | 3.667E-08 | 1.185E-08 | 2.300E-08 | 4.400E-08 |
| 7435 | 372 | P372P | A | 8.7e-08 | 7.6e-08 | 1.8e-08 | 6.033E-08 | 3.707E-08 | 1.800E-08 | 8.700E-08 |
| 7436 | 372 | P372S | A | 2.6e-08 | 7.1e-08 | 5.7e-08 | 5.133E-08 | 2.303E-08 | 2.600E-08 | 7.100E-08 |
| 7437 | 372 | P372T | A | 3.3e-08 | 1.7e-08 | 3.4e-08 | 2.800E-08 | 9.539E-09 | 1.700E-08 | 3.400E-08 |
| 7438 | 372 | P372W | A | 2.3e-08 | 2.5e-08 | 6e-08 | 3.600E-08 | 2.081E-08 | 2.300E-08 | 6.000E-08 |
| 7439 | 372 | P372Y | A | 1.3e-07 | 5.3e-08 | 3.4e-08 | 7.233E-08 | 5.084E-08 | 3.400E-08 | 1.300E-07 |
| 7440 | 372 | P372V | A | 2.3e-08 | 1.1e-07 | 5.1e-08 | 6.133E-08 | 4.441E-08 | 2.300E-08 | 1.100E-07 |
| 7441 | 373 | E373A | A | 7.2e-08 | 4.5e-08 | 6.4e-08 | 6.033E-08 | 1.387E-08 | 4.500E-08 | 7.200E-08 |
| 7442 | 373 | E373R | A | 2.4e-08 | 3e-08 | 6.1e-08 | 3.833E-08 | 1.986E-08 | 2.400E-08 | 6.100E-08 |
| 7443 | 373 | E373N | A | 4e-08 | 1e-07 | 2.4e-08 | 5.467E-08 | 4.007E-08 | 2.400E-08 | 1.000E-07 |
| 7444 | 373 | E373D | A | 4.1e-08 | 2.1e-08 | 7.4e-08 | 4.533E-08 | 2.676E-08 | 2.100E-08 | 7.400E-08 |
| 7445 | 373 | E373C | A | 1.2e-07 | 6.4e-08 | 4.4e-08 | 7.600E-08 | 3.940E-08 | 4.400E-08 | 1.200E-07 |
| 7446 | 373 | E373E | A | 5.8e-08 | 7.4e-08 | 3.7e-08 | 5.633E-08 | 1.856E-08 | 3.700E-08 | 7.400E-08 |
| 7447 | 373 | E373Q | A | 6e-08 | 3.4e-08 | 5.6e-08 | 5.000E-08 | 1.400E-08 | 3.400E-08 | 6.000E-08 |
| 7448 | 373 | E373G | A | 5.1e-08 | 2.5e-08 | 5.2e-08 | 4.267E-08 | 1.531E-08 | 2.500E-08 | 5.200E-08 |
| 7449 | 373 | E373H | A | 2.8e-08 | 6.8e-08 | 9.8e-08 | 6.467E-08 | 3.512E-08 | 2.800E-08 | 9.800E-08 |
| 7450 | 373 | E373I | A | 2.2e-08 | 1.5e-07 | 2.9e-08 | 6.700E-08 | 7.197E-08 | 2.200E-08 | 1.500E-07 |
| 7451 | 373 | E373L | A | 6.1e-08 | 3.8e-08 | 2.8e-08 | 4.233E-08 | 1.692E-08 | 2.800E-08 | 6.100E-08 |
| 7452 | 373 | E373K | A | 6e-08 | 9.4e-08 | 1.2e-07 | 9.133E-08 | 3.009E-08 | 6.000E-08 | 1.200E-07 |
| 7453 | 373 | E373M | A | 1.5e-08 | 6.4e-08 | 3.2e-08 | 3.700E-08 | 2.488E-08 | 1.500E-08 | 6.400E-08 |
| 7454 | 373 | E373F | A | 1e-07 | 2.8e-08 | 2.4e-08 | 5.067E-08 | 4.277E-08 | 2.400E-08 | 1.000E-07 |
| 7455 | 373 | E373P | A | 1.2e-08 | 8.1e-08 | 3.7e-08 | 4.333E-08 | 3.493E-08 | 1.200E-08 | 8.100E-08 |
| 7456 | 373 | E373S | A | 7.5e-08 | 3.9e-08 | 4.2e-08 | 5.200E-08 | 1.997E-08 | 3.900E-08 | 7.500E-08 |
| 7457 | 373 | E373T | A | 2.9e-08 | 8.4e-08 | 2.3e-08 | 4.533E-08 | 3.362E-08 | 2.300E-08 | 8.400E-08 |
| 7458 | 373 | E373W | A | 8.1e-08 | 4.6e-08 | 6.8e-08 | 6.500E-08 | 1.769E-08 | 4.600E-08 | 8.100E-08 |
| 7459 | 373 | E373Y | A | 6.5e-08 | 2e-08 | 3.2e-08 | 3.900E-08 | 2.330E-08 | 2.000E-08 | 6.500E-08 |
| 7460 | 373 | E373V | A | 4.1e-08 | 1.6e-08 | 6.7e-08 | 4.133E-08 | 2.550E-08 | 1.600E-08 | 6.700E-08 |
| 7461 | 374 | Q374A | A | 2.7e-08 | 4.8e-08 | 9.2e-08 | 5.567E-08 | 3.317E-08 | 2.700E-08 | 9.200E-08 |
| 7462 | 374 | Q374R | A | 3e-08 | 4.9e-08 | 1.4e-08 | 3.100E-08 | 1.752E-08 | 1.400E-08 | 4.900E-08 |
| 7463 | 374 | Q374N | A | 3.7e-08 | 3.4e-08 | 2.4e-08 | 3.167E-08 | 6.807E-09 | 2.400E-08 | 3.700E-08 |
| 7464 | 374 | Q374D | A | 7.5e-08 | 3.9e-08 | 4e-08 | 5.133E-08 | 2.050E-08 | 3.900E-08 | 7.500E-08 |
| 7465 | 374 | Q374C | A | 9.7e-08 | 1.7e-08 | 1.6e-07 | 9.133E-08 | 7.167E-08 | 1.700E-08 | 1.600E-07 |
| 7466 | 374 | Q374E | A | 6e-08 | 5.6e-08 | 5.1e-08 | 5.567E-08 | 4.509E-09 | 5.100E-08 | 6.000E-08 |
| 7467 | 374 | Q374Q | A | 4.4e-08 | 4.3e-08 | 3.5e-08 | 4.067E-08 | 4.933E-09 | 3.500E-08 | 4.400E-08 |
| 7468 | 374 | Q374G | A | 6.8e-08 | 3.7e-08 | 2.5e-08 | 4.333E-08 | 2.219E-08 | 2.500E-08 | 6.800E-08 |
| 7469 | 374 | Q374H | A | 1.1e-07 | 3.9e-08 | 4.8e-08 | 6.567E-08 | 3.866E-08 | 3.900E-08 | 1.100E-07 |
| 7470 | 374 | Q374I | A | 6.6e-08 | 6.2e-08 | 6.6e-08 | 6.467E-08 | 2.309E-09 | 6.200E-08 | 6.600E-08 |
| 7471 | 374 | Q374L | A | 4.2e-08 | 1.3e-08 | 3e-08 | 2.833E-08 | 1.457E-08 | 1.300E-08 | 4.200E-08 |
| 7472 | 374 | Q374K | A | 2.8e-08 | 2.5e-08 | 2.9e-08 | 2.733E-08 | 2.082E-09 | 2.500E-08 | 2.900E-08 |
| 7473 | 374 | Q374M | A | 6.4e-08 | 7.7e-08 | 5.2e-08 | 6.433E-08 | 1.250E-08 | 5.200E-08 | 7.700E-08 |
| 7474 | 374 | Q374F | A | 2.6e-08 | 2.4e-08 | 1.1e-07 | 5.333E-08 | 4.908E-08 | 2.400E-08 | 1.100E-07 |
| 7475 | 374 | Q374P | A | 2.6e-08 | 7.5e-08 | 5.5e-08 | 5.200E-08 | 2.464E-08 | 2.600E-08 | 7.500E-08 |
| 7476 | 374 | Q374S | A | 5.1e-08 | 5.2e-08 | 6.6e-08 | 5.633E-08 | 8.386E-09 | 5.100E-08 | 6.600E-08 |
| 7477 | 374 | Q374T | A | 3.2e-08 | 2.5e-08 | 2.9e-08 | 2.867E-08 | 3.512E-09 | 2.500E-08 | 3.200E-08 |
| 7478 | 374 | Q374W | A | 5.2e-08 | 2.2e-08 | 2e-08 | 3.133E-08 | 1.793E-08 | 2.000E-08 | 5.200E-08 |
| 7479 | 374 | Q374Y | A | 4.2e-08 | 2e-08 | 4.6e-08 | 3.600E-08 | 1.400E-08 | 2.000E-08 | 4.600E-08 |

| | | | | | | | | | | |
|---|---|---|---|---|---|---|---|---|---|---|
| 7480 | 374 | Q374V | A | 3.5e-08 | 6.7e-08 | 2.3e-08 | 4.167E-08 | 2.274E-08 | 2.300E-08 | 6.700E-08 |
| 7481 | 375 | L375A | A | 3.9e-08 | 3.4e-08 | 2.6e-08 | 3.300E-08 | 6.557E-09 | 2.600E-08 | 3.900E-08 |
| 7482 | 375 | L375R | A | 2.9e-08 | 4.8e-08 | 1.8e-08 | 3.167E-08 | 1.518E-08 | 1.800E-08 | 4.800E-08 |
| 7483 | 375 | L375N | A | 4.5e-08 | 3.8e-08 | 4.1e-08 | 4.133E-08 | 3.512E-09 | 3.800E-08 | 4.500E-08 |
| 7484 | 375 | L375D | A | 1.6e-08 | 2.9e-08 | 5.6e-08 | 3.367E-08 | 2.040E-08 | 1.600E-08 | 5.600E-08 |
| 7485 | 375 | L375C | A | 3.3e-08 | 9.9e-08 | 8.9e-08 | 7.367E-08 | 3.557E-08 | 3.300E-08 | 9.900E-08 |
| 7486 | 375 | L375E | A | 2.4e-08 | 3.1e-08 | 1e-07 | 5.167E-08 | 4.200E-08 | 2.400E-08 | 1.000E-07 |
| 7487 | 375 | L375Q | A | 6.8e-08 | 3.1e-08 | 4.4e-08 | 4.767E-08 | 1.877E-08 | 3.100E-08 | 6.800E-08 |
| 7488 | 375 | L375G | A | 1.3e-08 | 7.6e-08 | 7.4e-08 | 5.433E-08 | 3.581E-08 | 1.300E-08 | 7.600E-08 |
| 7489 | 375 | L375H | A | 2.9e-08 | 5.4e-08 | 9.8e-08 | 6.033E-08 | 3.493E-08 | 2.900E-08 | 9.800E-08 |
| 7490 | 375 | L375I | A | 2.8e-08 | 6.2e-08 | 5.1e-08 | 4.700E-08 | 1.735E-08 | 2.800E-08 | 6.200E-08 |
| 7491 | 375 | L375L | A | 3.4e-08 | 1.1e-07 | 3.5e-08 | 5.967E-08 | 4.359E-08 | 3.400E-08 | 1.100E-07 |
| 7492 | 375 | L375K | A | 3.6e-08 | 1.1e-07 | 3.9e-08 | 6.167E-08 | 4.188E-08 | 3.600E-08 | 1.100E-07 |
| 7493 | 375 | L375M | A | 1.4e-08 | 4.5e-08 | 4.8e-08 | 3.567E-08 | 1.882E-08 | 1.400E-08 | 4.800E-08 |
| 7494 | 375 | L375F | A | 5.9e-08 | 2.9e-08 | 4.9e-08 | 4.567E-08 | 1.528E-08 | 2.900E-08 | 5.900E-08 |
| 7495 | 375 | L375P | A | 6.5e-08 | 2.6e-08 | 7.7e-08 | 5.600E-08 | 2.666E-08 | 2.600E-08 | 7.700E-08 |
| 7496 | 375 | L375S | A | 3.9e-08 | 5.5e-08 | 1.5e-07 | 8.133E-08 | 6.000E-08 | 3.900E-08 | 1.500E-07 |
| 7497 | 375 | L375T | A | 4.9e-08 | 4e-08 | 3.8e-08 | 4.233E-08 | 5.859E-09 | 3.800E-08 | 4.900E-08 |
| 7498 | 375 | L375W | A | 2e-08 | 8.2e-08 | 3.2e-08 | 4.467E-08 | 3.288E-08 | 2.000E-08 | 8.200E-08 |
| 7499 | 375 | L375Y | A | 1.6e-08 | 3.3e-08 | 4.7e-08 | 3.200E-08 | 1.552E-08 | 1.600E-08 | 4.700E-08 |
| 7500 | 375 | L375V | A | 5.7e-08 | 4.9e-08 | 6.4e-08 | 5.667E-08 | 7.506E-09 | 4.900E-08 | 6.400E-08 |
| 7501 | 376 | Q376A | A | 2.3e-08 | 6.7e-08 | 2.5e-08 | 3.833E-08 | 2.485E-08 | 2.300E-08 | 6.700E-08 |
| 7502 | 376 | Q376R | A | 7.1e-08 | 6.2e-08 | 5.8e-08 | 6.367E-08 | 6.658E-09 | 5.800E-08 | 7.100E-08 |
| 7503 | 376 | Q376N | A | 1.2e-07 | 5.1e-08 | 2.7e-08 | 6.600E-08 | 4.828E-08 | 2.700E-08 | 1.200E-07 |
| 7504 | 376 | Q376D | A | 2.9e-08 | 4.8e-08 | 5.7e-08 | 4.467E-08 | 1.429E-08 | 2.900E-08 | 5.700E-08 |
| 7505 | 376 | Q376C | A | 4.4e-08 | 5.6e-08 | 5.4e-08 | 5.133E-08 | 6.429E-09 | 4.400E-08 | 5.600E-08 |
| 7506 | 376 | Q376E | A | 2.8e-08 | 1.8e-08 | 1.9e-08 | 2.167E-08 | 5.508E-09 | 1.800E-08 | 2.800E-08 |
| 7507 | 376 | Q376Q | A | 3.5e-08 | 1.6e-08 | 5.6e-08 | 3.567E-08 | 2.001E-08 | 1.600E-08 | 5.600E-08 |
| 7508 | 376 | Q376G | A | 1.1e-07 | 4.3e-08 | 2e-08 | 5.767E-08 | 4.676E-08 | 2.000E-08 | 1.100E-07 |
| 7509 | 376 | Q376H | A | 8.4e-08 | 2.9e-08 | 5.1e-08 | 5.467E-08 | 2.768E-08 | 2.900E-08 | 8.400E-08 |
| 7510 | 376 | Q376I | A | 1.9e-07 | 4.9e-08 | 2.9e-08 | 8.933E-08 | 8.775E-08 | 2.900E-08 | 1.900E-07 |
| 7511 | 376 | Q376L | A | 4.2e-08 | 1.3e-07 | 8.6e-08 | 8.600E-08 | 4.400E-08 | 4.200E-08 | 1.300E-07 |
| 7512 | 376 | Q376K | A | 4.3e-08 | 4.1e-08 | 3.2e-08 | 3.867E-08 | 5.859E-09 | 3.200E-08 | 4.300E-08 |
| 7513 | 376 | Q376M | A | 5.7e-08 | 4.4e-08 | 1.9e-08 | 4.000E-08 | 1.931E-08 | 1.900E-08 | 5.700E-08 |
| 7514 | 376 | Q376F | A | 2.7e-08 | 3.8e-08 | 3.5e-08 | 3.333E-08 | 5.686E-09 | 2.700E-08 | 3.800E-08 |
| 7515 | 376 | Q376P | A | 6.1e-08 | 5.2e-08 | 5e-08 | 5.433E-08 | 5.859E-09 | 5.000E-08 | 6.100E-08 |
| 7516 | 376 | Q376S | A | 4.7e-08 | 8.6e-08 | 2.9e-08 | 5.400E-08 | 2.914E-08 | 2.900E-08 | 8.600E-08 |
| 7517 | 376 | Q376T | A | 6.5e-08 | 2.1e-08 | 1.4e-08 | 3.333E-08 | 2.765E-08 | 1.400E-08 | 6.500E-08 |
| 7518 | 376 | Q376W | A | 3.4e-08 | 2e-08 | 4.4e-08 | 3.267E-08 | 1.206E-08 | 2.000E-08 | 4.400E-08 |
| 7519 | 376 | Q376Y | A | 7.7e-08 | 1.5e-08 | 1.8e-08 | 3.667E-08 | 3.496E-08 | 1.500E-08 | 7.700E-08 |
| 7520 | 376 | Q376V | A | 7.2e-08 | 6.9e-08 | 1.2e-07 | 8.700E-08 | 2.862E-08 | 6.900E-08 | 1.200E-07 |
| 7521 | 377 | V377A | A | 2.2e-08 | 7.1e-08 | 4.7e-08 | 4.667E-08 | 2.450E-08 | 2.200E-08 | 7.100E-08 |
| 7522 | 377 | V377R | A | 2.4e-08 | 4.9e-08 | 3.9e-08 | 3.733E-08 | 1.258E-08 | 2.400E-08 | 4.900E-08 |
| 7523 | 377 | V377N | A | 3.1e-08 | 2.6e-08 | 3e-08 | 2.900E-08 | 2.646E-09 | 2.600E-08 | 3.100E-08 |
| 7524 | 377 | V377D | A | 2.8e-08 | 9.8e-08 | 5.9e-08 | 6.167E-08 | 3.508E-08 | 2.800E-08 | 9.800E-08 |
| 7525 | 377 | V377C | A | 6.2e-08 | 3.6e-08 | 4.2e-08 | 4.667E-08 | 1.361E-08 | 3.600E-08 | 6.200E-08 |
| 7526 | 377 | V377E | A | 1.1e-07 | 3.4e-08 | 5.6e-08 | 6.667E-08 | 3.911E-08 | 3.400E-08 | 1.100E-07 |
| 7527 | 377 | V377Q | A | 2.1e-08 | 1.4e-08 | 4.1e-08 | 2.533E-08 | 1.401E-08 | 1.400E-08 | 4.100E-08 |
| 7528 | 377 | V377G | A | 8.9e-08 | 2.3e-08 | 9.8e-09 | 4.060E-08 | 4.243E-08 | 9.800E-09 | 8.900E-08 |
| 7529 | 377 | V377H | A | 6.1e-08 | 3e-08 | 6.6e-08 | 5.233E-08 | 1.950E-08 | 3.000E-08 | 6.600E-08 |
| 7530 | 377 | V377I | A | 5.8e-08 | 6.2e-08 | 2.3e-08 | 4.767E-08 | 2.146E-08 | 2.300E-08 | 6.200E-08 |
| 7531 | 377 | V377L | A | 5.1e-08 | 9.7e-08 | 1.6e-08 | 5.467E-08 | 4.062E-08 | 1.600E-08 | 9.700E-08 |
| 7532 | 377 | V377K | A | 1.9e-07 | 5.9e-08 | 9.4e-08 | 1.143E-07 | 6.783E-08 | 5.900E-08 | 1.900E-07 |
| 7533 | 377 | V377M | A | 1.3e-08 | 3.8e-08 | 1e-07 | 5.033E-08 | 4.479E-08 | 1.300E-08 | 1.000E-07 |
| 7534 | 377 | V377F | A | 3.6e-08 | 1.7e-08 | 2.3e-08 | 2.533E-08 | 9.713E-09 | 1.700E-08 | 3.600E-08 |
| 7535 | 377 | V377P | A | 5e-08 | 7.2e-08 | 3e-08 | 5.067E-08 | 2.101E-08 | 3.000E-08 | 7.200E-08 |
| 7536 | 377 | V377S | A | 6.6e-08 | 2.7e-08 | 8.6e-08 | 5.967E-08 | 3.001E-08 | 2.700E-08 | 8.600E-08 |
| 7537 | 377 | V377T | A | 2.8e-08 | 1.2e-08 | 6.8e-08 | 3.600E-08 | 2.884E-08 | 1.200E-08 | 6.800E-08 |
| 7538 | 377 | V377W | A | 5.8e-08 | 1.1e-08 | 8.8e-08 | 5.233E-08 | 3.881E-08 | 1.100E-08 | 8.800E-08 |
| 7539 | 377 | V377Y | A | 5.6e-08 | 5e-08 | 3.7e-08 | 4.767E-08 | 9.713E-09 | 3.700E-08 | 5.600E-08 |
| 7540 | 377 | V377V | A | 7.3e-08 | 5.7e-08 | 4e-08 | 5.667E-08 | 1.650E-08 | 4.000E-08 | 7.300E-08 |
| 7541 | 378 | F378A | A | 6.2e-08 | 5.8e-08 | 2e-08 | 4.667E-08 | 2.318E-08 | 2.000E-08 | 6.200E-08 |
| 7542 | 378 | F378R | A | 2.3e-08 | 2.1e-08 | 2.2e-08 | 2.200E-08 | 1.000E-09 | 2.100E-08 | 2.300E-08 |
| 7543 | 378 | F378N | A | 1.2e-07 | 1.9e-08 | 4.9e-08 | 6.267E-08 | 5.187E-08 | 1.900E-08 | 1.200E-07 |
| 7544 | 378 | F378D | A | 3.9e-08 | 9.1e-08 | 5.5e-08 | 6.167E-08 | 2.663E-08 | 3.900E-08 | 9.100E-08 |
| 7545 | 378 | F378C | A | 1.6e-08 | 3.9e-08 | 8.7e-08 | 4.733E-08 | 3.623E-08 | 1.600E-08 | 8.700E-08 |
| 7546 | 378 | F378E | A | 1.5e-08 | 6.9e-08 | 3.1e-08 | 3.833E-08 | 2.774E-08 | 1.500E-08 | 6.900E-08 |
| 7547 | 378 | F378Q | A | 3.1e-08 | 3.6e-08 | 4.8e-08 | 3.833E-08 | 8.737E-09 | 3.100E-08 | 4.800E-08 |
| 7548 | 378 | F378G | A | 7.3e-08 | 5.4e-08 | 6.4e-08 | 6.367E-08 | 9.504E-09 | 5.400E-08 | 7.300E-08 |
| 7549 | 378 | F378H | A | 5.4e-08 | 4.1e-08 | 6.1e-08 | 5.200E-08 | 1.015E-08 | 4.100E-08 | 6.100E-08 |
| 7550 | 378 | F378I | A | 2.4e-08 | 2.2e-08 | 1.4e-07 | 6.200E-08 | 6.756E-08 | 2.200E-08 | 1.400E-07 |
| 7551 | 378 | F378L | A | 9.7e-08 | 4.3e-08 | 6.7e-08 | 6.900E-08 | 2.706E-08 | 4.300E-08 | 9.700E-08 |
| 7552 | 378 | F378K | A | 1.2e-07 | 3.6e-08 | 6.8e-08 | 7.467E-08 | 4.239E-08 | 3.600E-08 | 1.200E-07 |
| 7553 | 378 | F378M | A | 2.1e-08 | 1.8e-08 | 7e-08 | 3.633E-08 | 2.919E-08 | 1.800E-08 | 7.000E-08 |
| 7554 | 378 | F378F | A | 1.1e-08 | 4.8e-08 | 5.3e-08 | 3.733E-08 | 2.294E-08 | 1.100E-08 | 5.300E-08 |
| 7555 | 378 | F378P | A | 1.8e-08 | 6.2e-08 | 5.8e-08 | 4.600E-08 | 2.433E-08 | 1.800E-08 | 6.200E-08 |
| 7556 | 378 | F378S | A | 3.4e-08 | 2.9e-08 | 1.1e-07 | 5.767E-08 | 4.539E-08 | 2.900E-08 | 1.100E-07 |
| 7557 | 378 | F378T | A | 3e-08 | 5.9e-08 | 4.4e-08 | 4.433E-08 | 1.450E-08 | 3.000E-08 | 5.900E-08 |
| 7558 | 378 | F378W | A | 1.5e-08 | 2.9e-08 | 2.2e-08 | 2.200E-08 | 7.000E-09 | 1.500E-08 | 2.900E-08 |
| 7559 | 378 | F378Y | A | 1.1e-08 | 5.5e-08 | 9.6e-08 | 5.400E-08 | 4.251E-08 | 1.100E-08 | 9.600E-08 |
| 7560 | 378 | F378V | A | 8.4e-08 | 5.4e-08 | 3.3e-08 | 5.700E-08 | 2.563E-08 | 3.300E-08 | 8.400E-08 |
| 7561 | 379 | E379A | A | 3.6e-08 | 6.2e-08 | 2e-08 | 3.933E-08 | 2.120E-08 | 2.000E-08 | 6.200E-08 |
| 7562 | 379 | E379R | A | 3.6e-08 | 3.7e-08 | 2.4e-08 | 3.233E-08 | 7.234E-09 | 2.400E-08 | 3.700E-08 |
| 7563 | 379 | E379N | A | 5.5e-09 | 5e-08 | 1.7e-07 | 7.517E-08 | 8.509E-08 | 5.500E-09 | 1.700E-07 |
| 7564 | 379 | E379D | A | 3.7e-08 | 5.4e-08 | 1.5e-07 | 8.033E-08 | 6.093E-08 | 3.700E-08 | 1.500E-07 |

| | | | | | | | | | | |
|---|---|---|---|---|---|---|---|---|---|---|
| 7565 | 379 | E379C | A | 3e-08 | 4.8e-08 | 9.2e-08 | 5.667E-08 | 3.190E-08 | 3.000E-08 | 9.200E-08 |
| 7566 | 379 | E379E | A | 6.3e-08 | 1.8e-08 | 4.4e-08 | 4.167E-08 | 2.259E-08 | 1.800E-08 | 6.300E-08 |
| 7567 | 379 | E379Q | A | 3.2e-08 | 8.6e-08 | 7.7e-08 | 6.500E-08 | 2.893E-08 | 3.200E-08 | 8.600E-08 |
| 7568 | 379 | E379G | A | 5.2e-08 | 3.8e-08 | 4.7e-08 | 4.567E-08 | 7.095E-09 | 3.800E-08 | 5.200E-08 |
| 7569 | 379 | E379H | A | 4.6e-08 | 2.7e-08 | 8.6e-08 | 5.300E-08 | 3.012E-08 | 2.700E-08 | 8.600E-08 |
| 7570 | 379 | E379I | A | 3.5e-08 | 4.9e-08 | 7.1e-08 | 5.167E-08 | 1.815E-08 | 3.500E-08 | 7.100E-08 |
| 7571 | 379 | E379L | A | 7.2e-08 | 2.6e-08 | 2.2e-08 | 4.000E-08 | 2.778E-08 | 2.200E-08 | 7.200E-08 |
| 7572 | 379 | E379K | A | 4.3e-08 | 5e-08 | 3.6e-08 | 4.300E-08 | 7.000E-09 | 3.600E-08 | 5.000E-08 |
| 7573 | 379 | E379M | A | 5.1e-08 | 3.9e-08 | 4.5e-08 | 4.500E-08 | 6.000E-09 | 3.900E-08 | 5.100E-08 |
| 7574 | 379 | E379F | A | 4e-08 | 4.4e-08 | 7.2e-08 | 5.200E-08 | 1.744E-08 | 4.000E-08 | 7.200E-08 |
| 7575 | 379 | E379P | A | 4.1e-08 | 5.3e-08 | 1.1e-07 | 6.800E-08 | 3.686E-08 | 4.100E-08 | 1.100E-07 |
| 7576 | 379 | E379S | A | 5.9e-08 | 6e-08 | 4.5e-08 | 5.467E-08 | 8.386E-09 | 4.500E-08 | 6.000E-08 |
| 7577 | 379 | E379T | A | 5.6e-08 | 8e-08 | 7.9e-08 | 7.167E-08 | 1.358E-08 | 5.600E-08 | 8.000E-08 |
| 7578 | 379 | E379W | A | 5.4e-08 | 1.4e-07 | 3.8e-08 | 7.733E-08 | 5.486E-08 | 3.800E-08 | 1.400E-07 |
| 7579 | 379 | E379Y | A | 1.8e-07 | 5.9e-08 | 2.8e-08 | 8.900E-08 | 8.032E-08 | 2.800E-08 | 1.800E-07 |
| 7580 | 379 | E379V | A | 4.1e-08 | 6.6e-08 | 6.3e-08 | 5.667E-08 | 1.365E-08 | 4.100E-08 | 6.600E-08 |
| 7581 | 380 | T380A | A | 3.1e-07 | 8.3e-08 | 6e-08 | 1.510E-07 | 1.382E-07 | 6.000E-08 | 3.100E-07 |
| 7582 | 380 | T380R | A | 4.2e-08 | 4.4e-08 | 1.5e-07 | 7.867E-08 | 6.178E-08 | 4.200E-08 | 1.500E-07 |
| 7583 | 380 | T380N | A | 3.1e-08 | 3.9e-08 | 4.1e-08 | 3.700E-08 | 5.292E-09 | 3.100E-08 | 4.100E-08 |
| 7584 | 380 | T380D | A | 1.8e-08 | 9.1e-08 | 3.5e-08 | 4.800E-08 | 3.820E-08 | 1.800E-08 | 9.100E-08 |
| 7585 | 380 | T380C | A | 5.4e-08 | 2.2e-08 | 4.7e-08 | 4.100E-08 | 1.682E-08 | 2.200E-08 | 5.400E-08 |
| 7586 | 380 | T380E | A | 3e-08 | 4.6e-08 | 2e-08 | 3.200E-08 | 1.311E-08 | 2.000E-08 | 4.600E-08 |
| 7587 | 380 | T380Q | A | 9.4e-08 | 4.8e-08 | 2.6e-08 | 5.600E-08 | 3.470E-08 | 2.600E-08 | 9.400E-08 |
| 7588 | 380 | T380G | A | 1.6e-08 | 5.1e-08 | 8.5e-08 | 5.067E-08 | 3.450E-08 | 1.600E-08 | 8.500E-08 |
| 7589 | 380 | T380H | A | 3.4e-08 | 7.4e-09 | 2.2e-08 | 2.113E-08 | 1.332E-08 | 7.400E-09 | 3.400E-08 |
| 7590 | 380 | T380I | A | 6e-08 | 6.3e-08 | 3.5e-08 | 5.267E-08 | 1.537E-08 | 3.500E-08 | 6.300E-08 |
| 7591 | 380 | T380L | A | 4.9e-08 | 4.7e-08 | 3.4e-08 | 4.333E-08 | 8.145E-09 | 3.400E-08 | 4.900E-08 |
| 7592 | 380 | T380K | A | 2.7e-08 | 3.4e-08 | 5.1e-09 | 2.203E-08 | 1.508E-08 | 5.100E-09 | 3.400E-08 |
| 7593 | 380 | T380M | A | 3e-08 | 5.2e-08 | 3.5e-08 | 3.900E-08 | 1.153E-08 | 3.000E-08 | 5.200E-08 |
| 7594 | 380 | T380F | A | 3.6e-08 | 5.8e-08 | 8e-08 | 5.800E-08 | 2.200E-08 | 3.600E-08 | 8.000E-08 |
| 7595 | 380 | T380P | A | 5.4e-08 | 3.9e-08 | 1.9e-08 | 3.733E-08 | 1.756E-08 | 1.900E-08 | 5.400E-08 |
| 7596 | 380 | T380S | A | 4e-08 | 6.9e-08 | 3.9e-08 | 4.933E-08 | 1.704E-08 | 3.900E-08 | 6.900E-08 |
| 7597 | 380 | T380T | A | 9e-08 | 5.6e-08 | 6.6e-08 | 7.067E-08 | 1.747E-08 | 5.600E-08 | 9.000E-08 |
| 7598 | 380 | T380W | A | 2.7e-08 | 1e-08 | 4.9e-08 | 2.867E-08 | 1.955E-08 | 1.000E-08 | 4.900E-08 |
| 7599 | 380 | T380Y | A | 7.1e-08 | 8.8e-08 | 5.7e-08 | 7.200E-08 | 1.552E-08 | 5.700E-08 | 8.800E-08 |
| 7600 | 380 | T380V | A | 6e-08 | 2.9e-08 | 3.2e-08 | 4.033E-08 | 1.710E-08 | 2.900E-08 | 6.000E-08 |
| 7601 | 381 | L381A | A | 5e-08 | 6.7e-08 | 4.4e-08 | 5.367E-08 | 1.193E-08 | 4.400E-08 | 6.700E-08 |
| 7602 | 381 | L381R | A | 1.5e-08 | 7.6e-08 | 3.3e-08 | 4.133E-08 | 3.134E-08 | 1.500E-08 | 7.600E-08 |
| 7603 | 381 | L381N | A | 5.6e-08 | 3.4e-08 | 5.1e-08 | 4.700E-08 | 1.153E-08 | 3.400E-08 | 5.600E-08 |
| 7604 | 381 | L381D | A | 3.1e-08 | 2.6e-08 | 3.8e-08 | 3.167E-08 | 6.028E-09 | 2.600E-08 | 3.800E-08 |
| 7605 | 381 | L381C | A | 2.2e-08 | 2.7e-08 | 9e-08 | 4.633E-08 | 3.790E-08 | 2.200E-08 | 9.000E-08 |
| 7606 | 381 | L381E | A | 4.7e-08 | 3e-08 | 3.4e-08 | 3.700E-08 | 8.888E-09 | 3.000E-08 | 4.700E-08 |
| 7607 | 381 | L381Q | A | 2.1e-08 | 3.2e-08 | 8.4e-08 | 4.567E-08 | 3.365E-08 | 2.100E-08 | 8.400E-08 |
| 7608 | 381 | L381G | A | 5.1e-08 | 3.7e-08 | 3.5e-08 | 4.100E-08 | 8.718E-09 | 3.500E-08 | 5.100E-08 |
| 7609 | 381 | L381H | A | 3.6e-08 | 1.5e-07 | 4.8e-08 | 7.800E-08 | 6.264E-08 | 3.600E-08 | 1.500E-07 |
| 7610 | 381 | L381I | A | 6.9e-08 | 5.4e-08 | 4.7e-08 | 5.667E-08 | 1.124E-08 | 4.700E-08 | 6.900E-08 |
| 7611 | 381 | L381L | A | 2.2e-08 | 6.3e-08 | 8.7e-08 | 5.733E-08 | 3.287E-08 | 2.200E-08 | 8.700E-08 |
| 7612 | 381 | L381K | A | 2.3e-08 | 2.4e-08 | 7.4e-08 | 4.033E-08 | 2.916E-08 | 2.300E-08 | 7.400E-08 |
| 7613 | 381 | L381M | A | 4.2e-08 | 1.1e-07 | 1.1e-07 | 8.733E-08 | 3.926E-08 | 4.200E-08 | 1.100E-07 |
| 7614 | 381 | L381F | A | 3.2e-08 | 6.8e-08 | 7e-08 | 5.667E-08 | 2.139E-08 | 3.200E-08 | 7.000E-08 |
| 7615 | 381 | L381P | A | 4.2e-08 | 5.4e-08 | 2.7e-08 | 4.100E-08 | 1.353E-08 | 2.700E-08 | 5.400E-08 |
| 7616 | 381 | L381S | A | 1.1e-07 | 6.4e-08 | 3.2e-08 | 6.867E-08 | 3.921E-08 | 3.200E-08 | 1.100E-07 |
| 7617 | 381 | L381T | A | 6.8e-08 | 5.3e-08 | 8.9e-08 | 7.000E-08 | 1.808E-08 | 5.300E-08 | 8.900E-08 |
| 7618 | 381 | L381W | A | 6.7e-08 | 3e-08 | 4.6e-08 | 4.767E-08 | 1.856E-08 | 3.000E-08 | 6.700E-08 |
| 7619 | 381 | L381Y | A | 5.4e-08 | 6.7e-08 | 1.3e-08 | 4.467E-08 | 2.818E-08 | 1.300E-08 | 6.700E-08 |
| 7620 | 381 | L381V | A | 1.6e-08 | 3e-08 | 6.4e-08 | 3.667E-08 | 2.468E-08 | 1.600E-08 | 6.400E-08 |
| 7621 | 382 | E382A | A | 7.4e-08 | 4.3e-08 | 2.5e-08 | 4.733E-08 | 2.479E-08 | 2.500E-08 | 7.400E-08 |
| 7622 | 382 | E382R | A | 2.3e-08 | 4e-08 | 2.5e-08 | 2.933E-08 | 9.292E-09 | 2.300E-08 | 4.000E-08 |
| 7623 | 382 | E382N | A | 1.9e-08 | 2.7e-08 | 3.4e-08 | 2.667E-08 | 7.506E-09 | 1.900E-08 | 3.400E-08 |
| 7624 | 382 | E382D | A | 1.3e-08 | 3.5e-08 | 3.5e-08 | 2.767E-08 | 1.270E-08 | 1.300E-08 | 3.500E-08 |
| 7625 | 382 | E382C | A | 1.9e-08 | 1.6e-08 | 4.1e-08 | 2.533E-08 | 1.365E-08 | 1.600E-08 | 4.100E-08 |
| 7626 | 382 | E382E | A | 4.6e-08 | 3.3e-08 | 1.1e-07 | 6.300E-08 | 4.122E-08 | 3.300E-08 | 1.100E-07 |
| 7627 | 382 | E382Q | A | 5.2e-08 | 2.4e-08 | 8.1e-08 | 5.233E-08 | 2.850E-08 | 2.400E-08 | 8.100E-08 |
| 7628 | 382 | E382G | A | 2.4e-08 | 3.8e-08 | 8.8e-08 | 5.000E-08 | 3.365E-08 | 2.400E-08 | 8.800E-08 |
| 7629 | 382 | E382H | A | 1.8e-08 | 2.6e-08 | 2.7e-08 | 2.367E-08 | 4.933E-09 | 1.800E-08 | 2.700E-08 |
| 7630 | 382 | E382I | A | 2.5e-08 | 3e-08 | 3.6e-08 | 3.033E-08 | 5.508E-09 | 2.500E-08 | 3.600E-08 |
| 7631 | 382 | E382L | A | 8.5e-08 | 3.7e-08 | 1.1e-08 | 4.433E-08 | 3.754E-08 | 1.100E-08 | 8.500E-08 |
| 7632 | 382 | E382K | A | 4.9e-08 | 1.4e-08 | 2.5e-08 | 2.933E-08 | 1.790E-08 | 1.400E-08 | 4.900E-08 |
| 7633 | 382 | E382M | A | 2.2e-08 | 9.7e-08 | 7.3e-08 | 6.400E-08 | 3.830E-08 | 2.200E-08 | 9.700E-08 |
| 7634 | 382 | E382F | A | 1.4e-07 | 8.2e-08 | 7.9e-08 | 1.003E-07 | 3.439E-08 | 7.900E-08 | 1.400E-07 |
| 7635 | 382 | E382P | A | 6.7e-08 | 4.3e-08 | 5.5e-08 | 5.500E-08 | 1.200E-08 | 4.300E-08 | 6.700E-08 |
| 7636 | 382 | E382S | A | 4.9e-08 | 8.1e-08 | 4.5e-08 | 5.833E-08 | 1.973E-08 | 4.500E-08 | 8.100E-08 |
| 7637 | 382 | E382T | A | 4e-08 | 2e-08 | 5.9e-08 | 3.967E-08 | 1.950E-08 | 2.000E-08 | 5.900E-08 |
| 7638 | 382 | E382W | A | 6.7e-08 | 9.7e-08 | 3.3e-08 | 6.567E-08 | 3.202E-08 | 3.300E-08 | 9.700E-08 |
| 7639 | 382 | E382Y | A | 9e-08 | 8.4e-08 | 1.3e-07 | 1.013E-07 | 2.501E-08 | 8.400E-08 | 1.300E-07 |
| 7640 | 382 | E382V | A | 5.3e-08 | 7.3e-08 | 6.1e-08 | 6.233E-08 | 1.007E-08 | 5.300E-08 | 7.300E-08 |
| 7641 | 383 | E383A | A | 3.7e-08 | 3.2e-08 | 7.8e-08 | 4.900E-08 | 2.524E-08 | 3.200E-08 | 7.800E-08 |
| 7642 | 383 | E383R | A | 8.2e-08 | 7.2e-08 | 1.8e-08 | 5.733E-08 | 3.443E-08 | 1.800E-08 | 8.200E-08 |
| 7643 | 383 | E383N | A | 3e-08 | 3.3e-08 | 2.1e-08 | 2.800E-08 | 6.245E-09 | 2.100E-08 | 3.300E-08 |
| 7644 | 383 | E383D | A | 1.8e-08 | 5.7e-08 | 6.9e-08 | 4.800E-08 | 2.666E-08 | 1.800E-08 | 6.900E-08 |
| 7645 | 383 | E383C | A | 6.1e-08 | 5.5e-08 | 4.4e-08 | 5.333E-08 | 8.622E-09 | 4.400E-08 | 6.100E-08 |
| 7646 | 383 | E383E | A | 3.5e-08 | 4e-08 | 5.7e-08 | 4.400E-08 | 1.153E-08 | 3.500E-08 | 5.700E-08 |
| 7647 | 383 | E383Q | A | 9.7e-08 | 3.2e-08 | 4.8e-08 | 5.900E-08 | 3.387E-08 | 3.200E-08 | 9.700E-08 |
| 7648 | 383 | E383G | A | 3e-07 | 4.1e-08 | 5e-08 | 1.303E-07 | 1.470E-07 | 4.100E-08 | 3.000E-07 |
| 7649 | 383 | E383H | A | 6.4e-08 | 4.8e-08 | 5e-08 | 5.400E-08 | 8.718E-09 | 4.800E-08 | 6.400E-08 |

| | | | | | | | | | | |
|---|---|---|---|---|---|---|---|---|---|---|
| 7650 | 383 | E383I | A | 1.5e-08 | 3.3e-08 | 3.2e-08 | 2.667E-08 | 1.012E-08 | 1.500E-08 | 3.300E-08 |
| 7651 | 383 | E383L | A | 3.5e-08 | 4.5e-08 | 1.1e-07 | 6.333E-08 | 4.072E-08 | 3.500E-08 | 1.100E-07 |
| 7652 | 383 | E383K | A | 1.9e-08 | 5.3e-08 | 6.2e-08 | 4.467E-08 | 2.268E-08 | 1.900E-08 | 6.200E-08 |
| 7653 | 383 | E383M | A | 4.6e-08 | 1.7e-07 | 4.2e-08 | 8.600E-08 | 7.277E-08 | 4.200E-08 | 1.700E-07 |
| 7654 | 383 | E383F | A | 2.5e-08 | 7.1e-08 | 6.3e-08 | 5.300E-08 | 2.458E-08 | 2.500E-08 | 7.100E-08 |
| 7655 | 383 | E383P | A | 2.2e-08 | 5.5e-08 | 3.9e-08 | 3.867E-08 | 1.650E-08 | 2.200E-08 | 5.500E-08 |
| 7656 | 383 | E383S | A | 5.1e-08 | 2.1e-08 | 9.3e-09 | 2.710E-08 | 2.151E-08 | 9.300E-09 | 5.100E-08 |
| 7657 | 383 | E383T | A | 5.5e-08 | 1.9e-08 | 2.9e-08 | 3.433E-08 | 1.858E-08 | 1.900E-08 | 5.500E-08 |
| 7658 | 383 | E383W | A | 1.9e-08 | 1.2e-08 | 2.1e-08 | 1.733E-08 | 4.726E-09 | 1.200E-08 | 2.100E-08 |
| 7659 | 383 | E383Y | A | 2.6e-08 | 3.2e-08 | 4.8e-08 | 3.533E-08 | 1.137E-08 | 2.600E-08 | 4.800E-08 |
| 7660 | 383 | E383V | A | 4.2e-08 | 5.6e-08 | 5.4e-08 | 5.067E-08 | 7.572E-09 | 4.200E-08 | 5.600E-08 |
| 7661 | 384 | I384A | A | 8.6e-08 | 5.6e-08 | 6.9e-08 | 7.033E-08 | 1.504E-08 | 5.600E-08 | 8.600E-08 |
| 7662 | 384 | I384R | A | 8.4e-08 | 7.2e-08 | 1.5e-08 | 5.700E-08 | 3.686E-08 | 1.500E-08 | 8.400E-08 |
| 7663 | 384 | I384N | A | 2.6e-08 | 7.1e-08 | 1.6e-08 | 3.767E-08 | 2.930E-08 | 1.600E-08 | 7.100E-08 |
| 7664 | 384 | I384D | A | 4.9e-08 | 2.8e-08 | 3.8e-08 | 3.833E-08 | 1.050E-08 | 2.800E-08 | 4.900E-08 |
| 7665 | 384 | I384C | A | 5.5e-08 | 5.7e-08 | 8.6e-08 | 6.600E-08 | 1.735E-08 | 5.500E-08 | 8.600E-08 |
| 7666 | 384 | I384E | A | 2e-08 | 4.1e-08 | 1.1e-07 | 5.700E-08 | 4.709E-08 | 2.000E-08 | 1.100E-07 |
| 7667 | 384 | I384Q | A | 3.4e-08 | 8.7e-08 | 3.2e-08 | 5.100E-08 | 3.119E-08 | 3.200E-08 | 8.700E-08 |
| 7668 | 384 | I384G | A | 4.2e-08 | 4.5e-08 | 4.4e-08 | 4.367E-08 | 1.528E-09 | 4.200E-08 | 4.500E-08 |
| 7669 | 384 | I384H | A | 2.3e-08 | 4.6e-08 | 6.3e-08 | 4.400E-08 | 2.007E-08 | 2.300E-08 | 6.300E-08 |
| 7670 | 384 | I384I | A | 3.3e-08 | 4.5e-08 | 7.8e-08 | 5.200E-08 | 2.330E-08 | 3.300E-08 | 7.800E-08 |
| 7671 | 384 | I384L | A | 4.6e-08 | 3.6e-08 | 4.5e-08 | 4.233E-08 | 5.508E-09 | 3.600E-08 | 4.600E-08 |
| 7672 | 384 | I384K | A | 3.3e-08 | 2.8e-08 | 3.2e-08 | 3.100E-08 | 2.646E-09 | 2.800E-08 | 3.300E-08 |
| 7673 | 384 | I384M | A | 3.2e-08 | 5.2e-08 | 2.7e-08 | 3.700E-08 | 1.323E-08 | 2.700E-08 | 5.200E-08 |
| 7674 | 384 | I384F | A | 2.3e-08 | 5.6e-08 | 1e-07 | 5.967E-08 | 3.863E-08 | 2.300E-08 | 1.000E-07 |
| 7675 | 384 | I384P | A | 1.2e-07 | 3.5e-08 | 3.7e-08 | 6.400E-08 | 4.851E-08 | 3.500E-08 | 1.200E-07 |
| 7676 | 384 | I384S | A | 2.4e-08 | 3.6e-08 | 6e-08 | 4.000E-08 | 1.833E-08 | 2.400E-08 | 6.000E-08 |
| 7677 | 384 | I384T | A | 3.7e-08 | 4.1e-08 | 1.2e-07 | 6.600E-08 | 4.681E-08 | 3.700E-08 | 1.200E-07 |
| 7678 | 384 | I384W | A | 5.8e-08 | 1.9e-08 | 1.3e-08 | 3.000E-08 | 2.443E-08 | 1.300E-08 | 5.800E-08 |
| 7679 | 384 | I384Y | A | 2.8e-08 | 5.3e-08 | 1.5e-08 | 3.200E-08 | 1.931E-08 | 1.500E-08 | 5.300E-08 |
| 7680 | 384 | I384V | A | 5.5e-08 | 5.4e-08 | 4.9e-08 | 5.267E-08 | 3.215E-09 | 4.900E-08 | 5.500E-08 |
| 7681 | 385 | T385A | A | 2.5e-08 | 4.5e-08 | 6.9e-08 | 4.633E-08 | 2.203E-08 | 2.500E-08 | 6.900E-08 |
| 7682 | 385 | T385R | A | 3.5e-08 | 5.5e-08 | 4.2e-08 | 4.400E-08 | 1.015E-08 | 3.500E-08 | 5.500E-08 |
| 7683 | 385 | T385N | A | 6.1e-08 | 3.2e-08 | 1.4e-08 | 3.567E-08 | 2.371E-08 | 1.400E-08 | 6.100E-08 |
| 7684 | 385 | T385D | A | 4.2e-08 | 4.5e-08 | 2.6e-08 | 3.767E-08 | 1.021E-08 | 2.600E-08 | 4.500E-08 |
| 7685 | 385 | T385C | A | 9.3e-08 | 4.3e-08 | 7.4e-08 | 7.000E-08 | 2.524E-08 | 4.300E-08 | 9.300E-08 |
| 7686 | 385 | T385E | A | 2.1e-08 | 4.9e-08 | 6e-08 | 4.333E-08 | 2.011E-08 | 2.100E-08 | 6.000E-08 |
| 7687 | 385 | T385Q | A | 6e-08 | 5.2e-08 | 4.3e-08 | 5.167E-08 | 8.505E-09 | 4.300E-08 | 6.000E-08 |
| 7688 | 385 | T385G | A | 6.4e-08 | 5.7e-08 | 2e-08 | 4.700E-08 | 2.364E-08 | 2.000E-08 | 6.400E-08 |
| 7689 | 385 | T385H | A | 7.1e-09 | 4.9e-08 | 2.3e-08 | 2.637E-08 | 2.115E-08 | 7.100E-09 | 4.900E-08 |
| 7690 | 385 | T385I | A | 2.2e-08 | 8.5e-08 | 6.7e-08 | 5.800E-08 | 3.245E-08 | 2.200E-08 | 8.500E-08 |
| 7691 | 385 | T385L | A | 1.2e-07 | 2.7e-08 | 2.6e-08 | 5.767E-08 | 5.398E-08 | 2.600E-08 | 1.200E-07 |
| 7692 | 385 | T385K | A | 3.3e-08 | 1.4e-08 | 4.4e-08 | 3.033E-08 | 1.518E-08 | 1.400E-08 | 4.400E-08 |
| 7693 | 385 | T385M | A | 3.6e-08 | 2.4e-08 | 9.4e-08 | 5.133E-08 | 3.743E-08 | 2.400E-08 | 9.400E-08 |
| 7694 | 385 | T385F | A | 1.4e-08 | 5.7e-08 | 1.1e-08 | 2.733E-08 | 2.574E-08 | 1.100E-08 | 5.700E-08 |
| 7695 | 385 | T385P | A | 7.3e-08 | 7.5e-08 | 1.3e-08 | 5.367E-08 | 3.523E-08 | 1.300E-08 | 7.500E-08 |
| 7696 | 385 | T385S | A | 3.5e-08 | 5.1e-08 | 5e-08 | 4.533E-08 | 8.963E-09 | 3.500E-08 | 5.100E-08 |
| 7697 | 385 | T385T | A | 1.1e-07 | 1.9e-08 | 2.7e-08 | 5.200E-08 | 5.039E-08 | 1.900E-08 | 1.100E-07 |
| 7698 | 385 | T385W | A | 3.2e-08 | 5.9e-08 | 7.6e-08 | 5.567E-08 | 2.219E-08 | 3.200E-08 | 7.600E-08 |
| 7699 | 385 | T385Y | A | 6.7e-08 | 1.2e-08 | 4.4e-08 | 4.100E-08 | 2.762E-08 | 1.200E-08 | 6.700E-08 |
| 7700 | 385 | T385V | A | 2.4e-08 | 7e-08 | 2.5e-08 | 3.967E-08 | 2.627E-08 | 2.400E-08 | 7.000E-08 |
| 7701 | 386 | G386A | A | 9.3e-08 | 9.5e-08 | 9.6e-08 | 9.467E-08 | 1.528E-09 | 9.300E-08 | 9.600E-08 |
| 7702 | 386 | G386R | A | 2.4e-08 | 5.6e-08 | 1.5e-07 | 7.667E-08 | 6.549E-08 | 2.400E-08 | 1.500E-07 |
| 7703 | 386 | G386N | A | 8.3e-08 | 3.3e-08 | 1.5e-08 | 4.367E-08 | 3.523E-08 | 1.500E-08 | 8.300E-08 |
| 7704 | 386 | G386D | A | 5e-08 | 6.7e-08 | 3.1e-08 | 4.933E-08 | 1.801E-08 | 3.100E-08 | 6.700E-08 |
| 7705 | 386 | G386C | A | 5.9e-08 | 1e-07 | 4.6e-08 | 6.833E-08 | 2.818E-08 | 4.600E-08 | 1.000E-07 |
| 7706 | 386 | G386E | A | 3.7e-08 | 8.5e-08 | 5.1e-08 | 5.767E-08 | 2.468E-08 | 3.700E-08 | 8.500E-08 |
| 7707 | 386 | G386Q | A | 2.5e-07 | 4.9e-08 | 3.1e-08 | 1.100E-07 | 1.216E-07 | 3.100E-08 | 2.500E-07 |
| 7708 | 386 | G386G | A | 9.1e-08 | 1.9e-08 | 4e-08 | 5.000E-08 | 3.703E-08 | 1.900E-08 | 9.100E-08 |
| 7709 | 386 | G386H | A | 2.3e-08 | 4.5e-08 | 7.4e-08 | 4.733E-08 | 2.558E-08 | 2.300E-08 | 7.400E-08 |
| 7710 | 386 | G386I | A | 8e-08 | 4.6e-08 | 1.2e-08 | 4.600E-08 | 3.400E-08 | 1.200E-08 | 8.000E-08 |
| 7711 | 386 | G386L | A | 7.3e-08 | 6e-08 | 2.7e-08 | 5.333E-08 | 2.371E-08 | 2.700E-08 | 7.300E-08 |
| 7712 | 386 | G386K | A | 3.3e-08 | 8.4e-08 | 1.7e-07 | 9.567E-08 | 6.924E-08 | 3.300E-08 | 1.700E-07 |
| 7713 | 386 | G386M | A | 9.6e-09 | 4.5e-08 | 2.2e-08 | 2.553E-08 | 1.796E-08 | 9.600E-09 | 4.500E-08 |
| 7714 | 386 | G386F | A | 2.2e-08 | 7.4e-08 | 3.1e-08 | 4.233E-08 | 2.779E-08 | 2.200E-08 | 7.400E-08 |
| 7715 | 386 | G386P | A | 6e-08 | 6.9e-08 | 4.6e-08 | 5.833E-08 | 1.159E-08 | 4.600E-08 | 6.900E-08 |
| 7716 | 386 | G386S | A | 9e-08 | 3e-08 | 4.3e-08 | 5.433E-08 | 3.156E-08 | 3.000E-08 | 9.000E-08 |
| 7717 | 386 | G386T | A | 1.1e-08 | 1e-07 | 2.1e-08 | 4.400E-08 | 4.875E-08 | 1.100E-08 | 1.000E-07 |
| 7718 | 386 | G386W | A | 4.9e-08 | 6.6e-08 | 7.2e-08 | 6.233E-08 | 1.193E-08 | 4.900E-08 | 7.200E-08 |
| 7719 | 386 | G386Y | A | 3.9e-08 | 1.9e-08 | 1.1e-08 | 2.300E-08 | 1.442E-08 | 1.100E-08 | 3.900E-08 |
| 7720 | 386 | G386V | A | 8.2e-08 | 7.4e-08 | 9.3e-08 | 8.300E-08 | 9.539E-09 | 7.400E-08 | 9.300E-08 |
| 7721 | 387 | Y387A | A | 3.1e-08 | 9.5e-08 | 3.1e-08 | 5.233E-08 | 3.695E-08 | 3.100E-08 | 9.500E-08 |
| 7722 | 387 | Y387R | A | 7.6e-08 | 4.4e-08 | 4.4e-08 | 5.467E-08 | 1.848E-08 | 4.400E-08 | 7.600E-08 |
| 7723 | 387 | Y387N | A | 1e-07 | 4.3e-08 | 1e-08 | 5.100E-08 | 4.553E-08 | 1.000E-08 | 1.000E-07 |
| 7724 | 387 | Y387D | A | 2.5e-08 | 4.6e-08 | 2.9e-08 | 3.333E-08 | 1.115E-08 | 2.500E-08 | 4.600E-08 |
| 7725 | 387 | Y387C | A | 6.3e-08 | 1.1e-08 | 4.1e-08 | 3.833E-08 | 2.610E-08 | 1.100E-08 | 6.300E-08 |
| 7726 | 387 | Y387E | A | 1.7e-07 | 7.6e-08 | 5.5e-08 | 1.003E-07 | 6.124E-08 | 5.500E-08 | 1.700E-07 |
| 7727 | 387 | Y387Q | A | 3.3e-08 | 6e-08 | 1.2e-07 | 7.100E-08 | 4.453E-08 | 3.300E-08 | 1.200E-07 |
| 7728 | 387 | Y387G | A | 1.2e-08 | 5.4e-08 | 5e-08 | 3.867E-08 | 2.318E-08 | 1.200E-08 | 5.400E-08 |
| 7729 | 387 | Y387H | A | 2.2e-08 | 5e-08 | 3.4e-08 | 3.533E-08 | 1.405E-08 | 2.200E-08 | 5.000E-08 |
| 7730 | 387 | Y387I | A | 2.4e-08 | 7.6e-08 | 3.4e-08 | 4.467E-08 | 2.759E-08 | 2.400E-08 | 7.600E-08 |
| 7731 | 387 | Y387L | A | 3.3e-08 | 4.1e-08 | 3.5e-08 | 3.633E-08 | 4.163E-09 | 3.300E-08 | 4.100E-08 |
| 7732 | 387 | Y387K | A | 3e-08 | 1e-07 | 8e-08 | 7.000E-08 | 3.606E-08 | 3.000E-08 | 1.000E-07 |
| 7733 | 387 | Y387M | A | 1.4e-08 | 3.7e-08 | 1.1e-08 | 2.067E-08 | 1.422E-08 | 1.100E-08 | 3.700E-08 |
| 7734 | 387 | Y387F | A | 2.1e-08 | 1.6e-08 | 1e-07 | 4.567E-08 | 4.712E-08 | 1.600E-08 | 1.000E-07 |

| | | | | | | | | | | |
|---|---|---|---|---|---|---|---|---|---|---|
| 7735 | 387 | Y387P | A | 2.8e-08 | 7.6e-08 | 2.7e-08 | 4.367E-08 | 2.801E-08 | 2.700E-08 | 7.600E-08 |
| 7736 | 387 | Y387S | A | 1.4e-08 | 1.3e-07 | 9.4e-08 | 7.933E-08 | 5.937E-08 | 1.400E-08 | 1.300E-07 |
| 7737 | 387 | Y387T | A | 4.3e-08 | 5.4e-08 | 3.7e-08 | 4.467E-08 | 8.622E-09 | 3.700E-08 | 5.400E-08 |
| 7738 | 387 | Y387W | A | 4e-08 | 4.7e-08 | 3.8e-08 | 4.167E-08 | 4.726E-09 | 3.800E-08 | 4.700E-08 |
| 7739 | 387 | Y387Y | A | 6.7e-08 | 6e-08 | 9.7e-08 | 7.467E-08 | 1.966E-08 | 6.000E-08 | 9.700E-08 |
| 7740 | 387 | Y387V | A | 3e-08 | 3.2e-08 | 7.6e-08 | 4.600E-08 | 2.600E-08 | 3.000E-08 | 7.600E-08 |
| 7741 | 388 | L388A | A | 1.2e-07 | 6.5e-08 | 7.8e-08 | 8.767E-08 | 2.875E-08 | 6.500E-08 | 1.200E-07 |
| 7742 | 388 | L388R | A | 4.6e-08 | 3.5e-08 | 1.1e-08 | 3.067E-08 | 1.790E-08 | 1.100E-08 | 4.600E-08 |
| 7743 | 388 | L388N | A | 2.9e-08 | 2.5e-08 | 3e-08 | 2.800E-08 | 2.646E-09 | 2.500E-08 | 3.000E-08 |
| 7744 | 388 | L388D | A | 5.3e-08 | 2.6e-08 | 2.6e-08 | 3.500E-08 | 1.559E-08 | 2.600E-08 | 5.300E-08 |
| 7745 | 388 | L388C | A | 2.7e-08 | 4.9e-08 | 9.6e-09 | 2.853E-08 | 1.974E-08 | 9.600E-09 | 4.900E-08 |
| 7746 | 388 | L388E | A | 3.3e-08 | 5e-08 | 7.7e-08 | 5.333E-08 | 2.219E-08 | 3.300E-08 | 7.700E-08 |
| 7747 | 388 | L388Q | A | 5.1e-08 | 1.9e-08 | 2.2e-08 | 3.067E-08 | 1.767E-08 | 1.900E-08 | 5.100E-08 |
| 7748 | 388 | L388G | A | 5.3e-08 | 6.4e-08 | 5.8e-08 | 5.833E-08 | 5.508E-09 | 5.300E-08 | 6.400E-08 |
| 7749 | 388 | L388H | A | 6.4e-08 | 5.8e-08 | 1.5e-08 | 4.567E-08 | 2.673E-08 | 1.500E-08 | 6.400E-08 |
| 7750 | 388 | L388I | A | 1.8e-07 | 5.9e-08 | 3.5e-08 | 9.133E-08 | 7.772E-08 | 3.500E-08 | 1.800E-07 |
| 7751 | 388 | L388L | A | 8.5e-08 | 1.9e-08 | 7.6e-09 | 3.720E-08 | 4.179E-08 | 7.600E-09 | 8.500E-08 |
| 7752 | 388 | L388K | A | 4.5e-08 | 3.6e-08 | 1.7e-08 | 3.267E-08 | 1.429E-08 | 1.700E-08 | 4.500E-08 |
| 7753 | 388 | L388M | A | 4.5e-08 | 4.3e-08 | 1.7e-07 | 8.600E-08 | 7.275E-08 | 4.300E-08 | 1.700E-07 |
| 7754 | 388 | L388F | A | 3.1e-08 | 4.6e-08 | 7.3e-08 | 5.000E-08 | 2.128E-08 | 3.100E-08 | 7.300E-08 |
| 7755 | 388 | L388P | A | 9e-08 | 2.9e-08 | 3.6e-08 | 5.167E-08 | 3.338E-08 | 2.900E-08 | 9.000E-08 |
| 7756 | 388 | L388S | A | 6.5e-08 | 2.9e-08 | 2.5e-08 | 3.967E-08 | 2.203E-08 | 2.500E-08 | 6.500E-08 |
| 7757 | 388 | L388T | A | 2.3e-08 | 1.2e-08 | 6.4e-08 | 3.300E-08 | 2.740E-08 | 1.200E-08 | 6.400E-08 |
| 7758 | 388 | L388W | A | 3.7e-08 | 8.9e-08 | 2.3e-08 | 4.967E-08 | 3.478E-08 | 2.300E-08 | 8.900E-08 |
| 7759 | 388 | L388Y | A | 2.9e-08 | 2.3e-08 | 2.7e-08 | 2.633E-08 | 3.055E-09 | 2.300E-08 | 2.900E-08 |
| 7760 | 388 | L388V | A | 5.8e-08 | 4.2e-08 | 1.3e-08 | 3.767E-08 | 2.281E-08 | 1.300E-08 | 5.800E-08 |
| 7761 | 389 | Y389A | A | 1.7e-08 | 2.7e-08 | 3.6e-08 | 2.667E-08 | 9.504E-09 | 1.700E-08 | 3.600E-08 |
| 7762 | 389 | Y389R | A | 6.4e-08 | 5.9e-08 | 1e-07 | 7.433E-08 | 2.237E-08 | 5.900E-08 | 1.000E-07 |
| 7763 | 389 | Y389N | A | 4.1e-08 | 1.2e-07 | 9e-08 | 8.367E-08 | 3.988E-08 | 4.100E-08 | 1.200E-07 |
| 7764 | 389 | Y389D | A | 3.6e-08 | 4.4e-08 | 4.4e-08 | 4.133E-08 | 4.619E-09 | 3.600E-08 | 4.400E-08 |
| 7765 | 389 | Y389C | A | 6.9e-08 | 2.5e-08 | 2.3e-08 | 3.900E-08 | 2.600E-08 | 2.300E-08 | 6.900E-08 |
| 7766 | 389 | Y389E | A | 1.5e-08 | 3.3e-08 | 5.8e-08 | 3.533E-08 | 2.159E-08 | 1.500E-08 | 5.800E-08 |
| 7767 | 389 | Y389Q | A | 5.3e-08 | 1.9e-08 | 4.5e-08 | 3.900E-08 | 1.778E-08 | 1.900E-08 | 5.300E-08 |
| 7768 | 389 | Y389G | A | 7.1e-08 | 6.5e-08 | 4.6e-08 | 6.067E-08 | 1.305E-08 | 4.600E-08 | 7.100E-08 |
| 7769 | 389 | Y389H | A | 2.2e-08 | 2.1e-08 | 8.3e-08 | 4.200E-08 | 3.551E-08 | 2.100E-08 | 8.300E-08 |
| 7770 | 389 | Y389I | A | 8.9e-08 | 1.6e-07 | 5e-08 | 9.967E-08 | 5.577E-08 | 5.000E-08 | 1.600E-07 |
| 7771 | 389 | Y389L | A | 1e-07 | 7.5e-08 | 5.2e-08 | 7.567E-08 | 2.401E-08 | 5.200E-08 | 1.000E-07 |
| 7772 | 389 | Y389K | A | 3.3e-08 | 5.7e-08 | 4.5e-08 | 4.500E-08 | 1.200E-08 | 3.300E-08 | 5.700E-08 |
| 7773 | 389 | Y389M | A | 5.3e-08 | 2.6e-08 | 6.4e-08 | 4.767E-08 | 1.955E-08 | 2.600E-08 | 6.400E-08 |
| 7774 | 389 | Y389F | A | 8.4e-08 | 5e-08 | 2.3e-08 | 5.233E-08 | 3.057E-08 | 2.300E-08 | 8.400E-08 |
| 7775 | 389 | Y389P | A | 1.6e-07 | 7.1e-08 | 1.1e-07 | 1.137E-07 | 4.461E-08 | 7.100E-08 | 1.600E-07 |
| 7776 | 389 | Y389S | A | 4.4e-08 | 5.7e-08 | 7.9e-08 | 6.000E-08 | 1.769E-08 | 4.400E-08 | 7.900E-08 |
| 7777 | 389 | Y389T | A | 2.6e-08 | 7.6e-08 | 2.1e-08 | 4.100E-08 | 3.041E-08 | 2.100E-08 | 7.600E-08 |
| 7778 | 389 | Y389W | A | 5.7e-08 | 1.7e-08 | 3e-08 | 3.467E-08 | 2.040E-08 | 1.700E-08 | 5.700E-08 |
| 7779 | 389 | Y389Y | A | 5.9e-08 | 5e-08 | 1.3e-08 | 4.067E-08 | 2.438E-08 | 1.300E-08 | 5.900E-08 |
| 7780 | 389 | Y389V | A | 7.2e-08 | 2.1e-08 | 2e-08 | 3.767E-08 | 2.974E-08 | 2.000E-08 | 7.200E-08 |
| 7781 | 390 | I390A | A | 2.2e-08 | 3e-08 | 3.6e-08 | 2.933E-08 | 7.024E-09 | 2.200E-08 | 3.600E-08 |
| 7782 | 390 | I390R | A | 2.5e-08 | 4.3e-08 | 6.6e-08 | 4.467E-08 | 2.055E-08 | 2.500E-08 | 6.600E-08 |
| 7783 | 390 | I390N | A | 1.5e-08 | 6e-08 | 1.5e-08 | 3.000E-08 | 2.598E-08 | 1.500E-08 | 6.000E-08 |
| 7784 | 390 | I390D | A | 1.1e-07 | 2.3e-08 | 1.5e-08 | 4.933E-08 | 5.269E-08 | 1.500E-08 | 1.100E-07 |
| 7785 | 390 | I390C | A | 2e-08 | 3.5e-08 | 7.2e-08 | 4.233E-08 | 2.676E-08 | 2.000E-08 | 7.200E-08 |
| 7786 | 390 | I390E | A | 6.7e-08 | 4e-08 | 3.8e-08 | 4.833E-08 | 1.620E-08 | 3.800E-08 | 6.700E-08 |
| 7787 | 390 | I390Q | A | 5.2e-08 | 7.1e-08 | 4.6e-08 | 5.633E-08 | 1.305E-08 | 4.600E-08 | 7.100E-08 |
| 7788 | 390 | I390G | A | 7.5e-09 | 4.1e-08 | 3.6e-08 | 2.817E-08 | 1.807E-08 | 7.500E-09 | 4.100E-08 |
| 7789 | 390 | I390H | A | 1.2e-08 | 5.7e-08 | 1.1e-08 | 2.667E-08 | 2.627E-08 | 1.100E-08 | 5.700E-08 |
| 7790 | 390 | I390I | A | 5.6e-08 | 2.2e-08 | 7.9e-08 | 5.233E-08 | 2.868E-08 | 2.200E-08 | 7.900E-08 |
| 7791 | 390 | I390L | A | 5.6e-08 | 1.9e-08 | 4.7e-08 | 4.067E-08 | 1.930E-08 | 1.900E-08 | 5.600E-08 |
| 7792 | 390 | I390K | A | 4.2e-08 | 4.3e-08 | 3.3e-08 | 3.933E-08 | 5.508E-09 | 3.300E-08 | 4.300E-08 |
| 7793 | 390 | I390M | A | 2.2e-08 | 5e-08 | 2.6e-08 | 3.267E-08 | 1.514E-08 | 2.200E-08 | 5.000E-08 |
| 7794 | 390 | I390F | A | 4.4e-08 | 6.8e-08 | 5.2e-08 | 5.467E-08 | 1.222E-08 | 4.400E-08 | 6.800E-08 |
| 7795 | 390 | I390P | A | 7.1e-08 | 1.8e-08 | 3.7e-08 | 4.200E-08 | 2.685E-08 | 1.800E-08 | 7.100E-08 |
| 7796 | 390 | I390S | A | 7.7e-08 | 1.1e-08 | 3.4e-08 | 4.067E-08 | 3.350E-08 | 1.100E-08 | 7.700E-08 |
| 7797 | 390 | I390T | A | 3.7e-08 | 2.9e-08 | 5.7e-08 | 4.100E-08 | 1.442E-08 | 2.900E-08 | 5.700E-08 |
| 7798 | 390 | I390W | A | 4.3e-08 | 4.4e-08 | 1.4e-07 | 7.567E-08 | 5.572E-08 | 4.300E-08 | 1.400E-07 |
| 7799 | 390 | I390Y | A | 3.2e-08 | 7.9e-08 | 9.2e-08 | 6.767E-08 | 3.156E-08 | 3.200E-08 | 9.200E-08 |
| 7800 | 390 | I390V | A | 3.3e-08 | 1.4e-07 | 4.1e-08 | 7.133E-08 | 5.960E-08 | 3.300E-08 | 1.400E-07 |
| 7801 | 391 | S391A | A | 4e-08 | 5.9e-08 | 2.8e-08 | 4.233E-08 | 1.563E-08 | 2.800E-08 | 5.900E-08 |
| 7802 | 391 | S391R | A | 1.1e-07 | 2.8e-08 | 5.1e-08 | 6.300E-08 | 4.230E-08 | 2.800E-08 | 1.100E-07 |
| 7803 | 391 | S391N | A | 7.5e-08 | 1.9e-08 | 7.2e-08 | 5.533E-08 | 3.150E-08 | 1.900E-08 | 7.500E-08 |
| 7804 | 391 | S391D | A | 1e-07 | 2.5e-08 | 7.1e-08 | 6.533E-08 | 3.782E-08 | 2.500E-08 | 1.000E-07 |
| 7805 | 391 | S391C | A | 4.4e-08 | 8.8e-08 | 6.7e-08 | 6.633E-08 | 2.201E-08 | 4.400E-08 | 8.800E-08 |
| 7806 | 391 | S391E | A | 5.1e-08 | 1.3e-07 | 3.3e-08 | 7.133E-08 | 5.160E-08 | 3.300E-08 | 1.300E-07 |
| 7807 | 391 | S391Q | A | 4.3e-08 | 8.8e-09 | 3.9e-08 | 3.027E-08 | 1.870E-08 | 8.800E-09 | 4.300E-08 |
| 7808 | 391 | S391G | A | 6.4e-08 | 6.4e-08 | 1.5e-07 | 9.267E-08 | 4.965E-08 | 6.400E-08 | 1.500E-07 |
| 7809 | 391 | S391H | A | 2.4e-08 | 3.2e-08 | 6.9e-08 | 4.167E-08 | 2.401E-08 | 2.400E-08 | 6.900E-08 |
| 7810 | 391 | S391I | A | 4.6e-08 | 5.2e-08 | 3.3e-08 | 4.367E-08 | 9.713E-09 | 3.300E-08 | 5.200E-08 |
| 7811 | 391 | S391L | A | 5.8e-08 | 3e-08 | 1.3e-07 | 7.267E-08 | 5.159E-08 | 3.000E-08 | 1.300E-07 |
| 7812 | 391 | S391K | A | 4.2e-08 | 2.2e-07 | 5e-08 | 1.040E-07 | 1.005E-07 | 4.200E-08 | 2.200E-07 |
| 7813 | 391 | S391M | A | 2.7e-08 | 3.9e-08 | 2.1e-08 | 2.900E-08 | 9.165E-09 | 2.100E-08 | 3.900E-08 |
| 7814 | 391 | S391F | A | 1.1e-07 | 4.4e-08 | 4e-08 | 6.467E-08 | 3.931E-08 | 4.000E-08 | 1.100E-07 |
| 7815 | 391 | S391P | A | 5.2e-08 | 6.2e-08 | 6.3e-08 | 5.900E-08 | 6.083E-09 | 5.200E-08 | 6.300E-08 |
| 7816 | 391 | S391S | A | 4.8e-08 | 2.2e-08 | 7.1e-08 | 4.700E-08 | 2.452E-08 | 2.200E-08 | 7.100E-08 |
| 7817 | 391 | S391T | A | 4.5e-08 | 1.8e-08 | 1.4e-08 | 2.567E-08 | 1.686E-08 | 1.400E-08 | 4.500E-08 |
| 7818 | 391 | S391W | A | 3.3e-08 | 8.8e-08 | 1.6e-08 | 4.567E-08 | 3.763E-08 | 1.600E-08 | 8.800E-08 |
| 7819 | 391 | S391Y | A | 3.9e-08 | 7.8e-08 | 6.8e-08 | 6.167E-08 | 2.026E-08 | 3.900E-08 | 7.800E-08 |

| | | | | | | | | | | |
|---|---|---|---|---|---|---|---|---|---|---|
| 7820 | 391 | S391V | A | 3.6e-08 | 3e-08 | 1.8e-08 | 2.800E-08 | 9.165E-09 | 1.800E-08 | 3.600E-08 |
| 7821 | 392 | A392A | A | 4.4e-08 | 3.6e-08 | 2e-08 | 3.333E-08 | 1.222E-08 | 2.000E-08 | 4.400E-08 |
| 7822 | 392 | A392R | A | 6.1e-08 | 4.7e-08 | 1.2e-08 | 4.000E-08 | 2.524E-08 | 1.200E-08 | 6.100E-08 |
| 7823 | 392 | A392N | A | 3.7e-08 | 6.5e-08 | 3e-08 | 4.400E-08 | 1.852E-08 | 3.000E-08 | 6.500E-08 |
| 7824 | 392 | A392D | A | 2.2e-08 | 2.7e-08 | 2.5e-08 | 2.467E-08 | 2.517E-09 | 2.200E-08 | 2.700E-08 |
| 7825 | 392 | A392C | A | 1.2e-08 | 1.7e-07 | 1.1e-08 | 6.433E-08 | 9.151E-08 | 1.100E-08 | 1.700E-07 |
| 7826 | 392 | A392E | A | 4.1e-08 | 2.8e-08 | 1e-07 | 5.633E-08 | 3.837E-08 | 2.800E-08 | 1.000E-07 |
| 7827 | 392 | A392Q | A | 4.4e-08 | 4.1e-08 | 2.1e-08 | 3.533E-08 | 1.250E-08 | 2.100E-08 | 4.400E-08 |
| 7828 | 392 | A392G | A | 1.6e-08 | 3.5e-08 | 2.8e-08 | 2.633E-08 | 9.609E-09 | 1.600E-08 | 3.500E-08 |
| 7829 | 392 | A392H | A | 1.1e-07 | 4.7e-08 | 2.6e-08 | 6.100E-08 | 4.371E-08 | 2.600E-08 | 1.100E-07 |
| 7830 | 392 | A392I | A | 2.1e-08 | 2.7e-08 | 4.9e-08 | 3.233E-08 | 1.474E-08 | 2.100E-08 | 4.900E-08 |
| 7831 | 392 | A392L | A | 3.9e-08 | 2.2e-08 | 4.4e-08 | 3.500E-08 | 1.153E-08 | 2.200E-08 | 4.400E-08 |
| 7832 | 392 | A392K | A | 1.6e-07 | 4e-08 | 8.1e-08 | 9.367E-08 | 6.099E-08 | 4.000E-08 | 1.600E-07 |
| 7833 | 392 | A392M | A | 4.4e-08 | 3.4e-08 | 2e-08 | 3.267E-08 | 1.206E-08 | 2.000E-08 | 4.400E-08 |
| 7834 | 392 | A392F | A | 4.3e-08 | 7e-08 | 2e-08 | 4.433E-08 | 2.503E-08 | 2.000E-08 | 7.000E-08 |
| 7835 | 392 | A392P | A | 2.7e-08 | 2.8e-08 | 2.1e-08 | 2.533E-08 | 3.786E-09 | 2.100E-08 | 2.800E-08 |
| 7836 | 392 | A392S | A | 4e-08 | 1.5e-08 | 4.7e-09 | 1.990E-08 | 1.815E-08 | 4.700E-09 | 4.000E-08 |
| 7837 | 392 | A392T | A | 1.2e-08 | 6.4e-08 | 1.9e-08 | 3.167E-08 | 2.822E-08 | 1.200E-08 | 6.400E-08 |
| 7838 | 392 | A392W | A | 5.2e-08 | 1.7e-08 | 7.2e-08 | 4.700E-08 | 2.784E-08 | 1.700E-08 | 7.200E-08 |
| 7839 | 392 | A392Y | A | 1.3e-08 | 4.2e-08 | 7.6e-08 | 4.367E-08 | 3.153E-08 | 1.300E-08 | 7.600E-08 |
| 7840 | 392 | A392V | A | 1.6e-08 | 8.1e-08 | 7.6e-09 | 3.487E-08 | 4.017E-08 | 7.600E-09 | 8.100E-08 |
| 7841 | 393 | W393A | A | 5.4e-08 | 1.2e-07 | 3.5e-08 | 6.967E-08 | 4.461E-08 | 3.500E-08 | 1.200E-07 |
| 7842 | 393 | W393R | A | 6.8e-08 | 5.1e-08 | 5.5e-08 | 5.800E-08 | 8.888E-09 | 5.100E-08 | 6.800E-08 |
| 7843 | 393 | W393N | A | 3.3e-08 | 6.3e-08 | 3.3e-08 | 4.300E-08 | 1.732E-08 | 3.300E-08 | 6.300E-08 |
| 7844 | 393 | W393D | A | 4.2e-08 | 7.1e-08 | 4.1e-08 | 5.133E-08 | 1.704E-08 | 4.100E-08 | 7.100E-08 |
| 7845 | 393 | W393C | A | 6e-08 | 5.5e-08 | 9.7e-08 | 7.067E-08 | 2.294E-08 | 5.500E-08 | 9.700E-08 |
| 7846 | 393 | W393E | A | 2.5e-08 | 1.3e-08 | 2.7e-08 | 2.167E-08 | 7.572E-09 | 1.300E-08 | 2.700E-08 |
| 7847 | 393 | W393Q | A | 1.7e-08 | 4.3e-08 | 1.7e-07 | 7.667E-08 | 8.187E-08 | 1.700E-08 | 1.700E-07 |
| 7848 | 393 | W393G | A | 5.3e-08 | 4.6e-08 | 2.1e-08 | 4.000E-08 | 1.682E-08 | 2.100E-08 | 5.300E-08 |
| 7849 | 393 | W393H | A | 4.1e-08 | 8.3e-08 | 1.8e-08 | 4.733E-08 | 3.296E-08 | 1.800E-08 | 8.300E-08 |
| 7850 | 393 | W393I | A | 2.7e-08 | 4.1e-08 | 2.9e-08 | 3.233E-08 | 7.572E-09 | 2.700E-08 | 4.100E-08 |
| 7851 | 393 | W393L | A | 1.3e-08 | 3.6e-08 | 1.1e-07 | 5.300E-08 | 5.069E-08 | 1.300E-08 | 1.100E-07 |
| 7852 | 393 | W393K | A | 3.7e-08 | 3.9e-08 | 2.8e-08 | 3.467E-08 | 5.859E-09 | 2.800E-08 | 3.900E-08 |
| 7853 | 393 | W393M | A | 4.3e-08 | 1.3e-07 | 5.6e-08 | 7.633E-08 | 4.693E-08 | 4.300E-08 | 1.300E-07 |
| 7854 | 393 | W393F | A | 6e-08 | 7.2e-08 | 2.7e-08 | 5.300E-08 | 2.330E-08 | 2.700E-08 | 7.200E-08 |
| 7855 | 393 | W393P | A | 5.4e-08 | 8.4e-08 | 7.5e-08 | 7.100E-08 | 1.539E-08 | 5.400E-08 | 8.400E-08 |
| 7856 | 393 | W393S | A | 3.5e-08 | 3.2e-08 | 6.8e-08 | 4.500E-08 | 1.997E-08 | 3.200E-08 | 6.800E-08 |
| 7857 | 393 | W393T | A | 4.4e-08 | 1.8e-08 | 4.2e-08 | 3.467E-08 | 1.447E-08 | 1.800E-08 | 4.400E-08 |
| 7858 | 393 | W393W | A | 3.1e-08 | 1.6e-07 | 3.7e-08 | 7.600E-08 | 7.281E-08 | 3.100E-08 | 1.600E-07 |
| 7859 | 393 | W393Y | A | 1.1e-08 | 2.9e-08 | 4.5e-08 | 2.833E-08 | 1.701E-08 | 1.100E-08 | 4.500E-08 |
| 7860 | 393 | W393V | A | 7.8e-08 | 7.2e-08 | 1.1e-07 | 8.667E-08 | 2.043E-08 | 7.200E-08 | 1.100E-07 |
| 7861 | 394 | P394A | A | 1.3e-07 | 4e-08 | 3.8e-08 | 6.933E-08 | 5.255E-08 | 3.800E-08 | 1.300E-07 |
| 7862 | 394 | P394R | A | 8.4e-08 | 1.2e-08 | 5.5e-08 | 5.033E-08 | 3.623E-08 | 1.200E-08 | 8.400E-08 |
| 7863 | 394 | P394N | A | 4.1e-08 | 5.9e-08 | 4.6e-08 | 4.867E-08 | 9.292E-09 | 4.100E-08 | 5.900E-08 |
| 7864 | 394 | P394D | A | 4.6e-08 | 3.6e-08 | 4.1e-08 | 4.100E-08 | 5.000E-09 | 3.600E-08 | 4.600E-08 |
| 7865 | 394 | P394C | A | 4.3e-08 | 3.2e-08 | 2.8e-08 | 3.433E-08 | 7.767E-09 | 2.800E-08 | 4.300E-08 |
| 7866 | 394 | P394E | A | 1.2e-08 | 5e-08 | 6e-08 | 4.067E-08 | 2.532E-08 | 1.200E-08 | 6.000E-08 |
| 7867 | 394 | P394Q | A | 4.3e-08 | 1.7e-08 | 2e-08 | 2.667E-08 | 1.422E-08 | 1.700E-08 | 4.300E-08 |
| 7868 | 394 | P394G | A | 4.6e-08 | 2.9e-08 | 3.3e-08 | 3.600E-08 | 8.888E-09 | 2.900E-08 | 4.600E-08 |
| 7869 | 394 | P394H | A | 2.5e-08 | 3.6e-08 | 6.1e-08 | 4.067E-08 | 1.845E-08 | 2.500E-08 | 6.100E-08 |
| 7870 | 394 | P394I | A | 3.4e-08 | 6.9e-08 | 1e-07 | 6.767E-08 | 3.302E-08 | 3.400E-08 | 1.000E-07 |
| 7871 | 394 | P394L | A | 1.6e-07 | 8.9e-08 | 7.5e-08 | 1.080E-07 | 4.557E-08 | 7.500E-08 | 1.600E-07 |
| 7872 | 394 | P394K | A | 3.6e-08 | 3.7e-08 | 4.4e-08 | 3.900E-08 | 4.359E-09 | 3.600E-08 | 4.400E-08 |
| 7873 | 394 | P394M | A | 6.1e-08 | 1.4e-08 | 3.6e-08 | 3.700E-08 | 2.352E-08 | 1.400E-08 | 6.100E-08 |
| 7874 | 394 | P394F | A | 3.4e-08 | 5.6e-08 | 4.2e-08 | 4.400E-08 | 1.114E-08 | 3.400E-08 | 5.600E-08 |
| 7875 | 394 | P394P | A | 6.1e-08 | 3.4e-08 | 1.5e-08 | 3.667E-08 | 2.312E-08 | 1.500E-08 | 6.100E-08 |
| 7876 | 394 | P394S | A | 9.9e-08 | 5.9e-08 | 2.6e-08 | 6.133E-08 | 3.656E-08 | 2.600E-08 | 9.900E-08 |
| 7877 | 394 | P394T | A | 4.3e-08 | 4.4e-08 | 3.6e-08 | 4.100E-08 | 4.359E-09 | 3.600E-08 | 4.400E-08 |
| 7878 | 394 | P394W | A | 6.9e-08 | 7.9e-08 | 1.4e-08 | 5.400E-08 | 3.500E-08 | 1.400E-08 | 7.900E-08 |
| 7879 | 394 | P394Y | A | 4.4e-08 | 3.1e-08 | 2.8e-08 | 3.433E-08 | 8.505E-09 | 2.800E-08 | 4.400E-08 |
| 7880 | 394 | P394V | A | 2.1e-08 | 2.7e-08 | 6.1e-08 | 3.633E-08 | 2.157E-08 | 2.100E-08 | 6.100E-08 |
| 7881 | 395 | D395A | A | 4.5e-08 | 6.8e-08 | 2.2e-08 | 4.500E-08 | 2.300E-08 | 2.200E-08 | 6.800E-08 |
| 7882 | 395 | D395R | A | 3.6e-08 | 2e-08 | 5.6e-08 | 3.733E-08 | 1.804E-08 | 2.000E-08 | 5.600E-08 |
| 7883 | 395 | D395N | A | 5.3e-08 | 3.5e-08 | 2.1e-08 | 3.633E-08 | 1.604E-08 | 2.100E-08 | 5.300E-08 |
| 7884 | 395 | D395D | A | 5.8e-08 | 4.5e-08 | 5.1e-08 | 5.133E-08 | 6.506E-09 | 4.500E-08 | 5.800E-08 |
| 7885 | 395 | D395C | A | 1.9e-08 | 3.3e-08 | 2.8e-08 | 2.667E-08 | 7.095E-09 | 1.900E-08 | 3.300E-08 |
| 7886 | 395 | D395E | A | 2.3e-08 | 2.1e-08 | 2.5e-08 | 2.300E-08 | 2.000E-09 | 2.100E-08 | 2.500E-08 |
| 7887 | 395 | D395Q | A | 1.2e-07 | 2.7e-08 | 1.8e-08 | 5.500E-08 | 5.647E-08 | 1.800E-08 | 1.200E-07 |
| 7888 | 395 | D395G | A | 5.6e-08 | 5.8e-08 | 4.3e-08 | 5.233E-08 | 8.145E-09 | 4.300E-08 | 5.800E-08 |
| 7889 | 395 | D395H | A | 4.4e-08 | 8.1e-08 | 5.6e-08 | 6.033E-08 | 1.888E-08 | 4.400E-08 | 8.100E-08 |
| 7890 | 395 | D395I | A | 1.5e-08 | 5.6e-08 | 5.8e-08 | 4.300E-08 | 2.427E-08 | 1.500E-08 | 5.800E-08 |
| 7891 | 395 | D395L | A | 2e-08 | 7e-08 | 7.8e-08 | 5.600E-08 | 3.143E-08 | 2.000E-08 | 7.800E-08 |
| 7892 | 395 | D395K | A | 8e-08 | 6e-08 | 4.2e-08 | 6.067E-08 | 1.901E-08 | 4.200E-08 | 8.000E-08 |
| 7893 | 395 | D395M | A | 3.6e-08 | 9.4e-08 | 1.3e-07 | 8.667E-08 | 4.743E-08 | 3.600E-08 | 1.300E-07 |
| 7894 | 395 | D395F | A | 4.3e-08 | 5.5e-08 | 4.5e-08 | 4.767E-08 | 6.429E-09 | 4.300E-08 | 5.500E-08 |
| 7895 | 395 | D395P | A | 4.9e-08 | 3.2e-08 | 7.1e-08 | 5.067E-08 | 1.955E-08 | 3.200E-08 | 7.100E-08 |
| 7896 | 395 | D395S | A | 1.1e-07 | 5.2e-08 | 2.3e-08 | 6.167E-08 | 4.430E-08 | 2.300E-08 | 1.100E-07 |
| 7897 | 395 | D395T | A | 3.7e-08 | 2.7e-08 | 6.9e-08 | 4.433E-08 | 2.194E-08 | 2.700E-08 | 6.900E-08 |
| 7898 | 395 | D395W | A | 2.4e-08 | 4e-08 | 3.6e-08 | 3.333E-08 | 8.327E-09 | 2.400E-08 | 4.000E-08 |
| 7899 | 395 | D395Y | A | 3.2e-08 | 2.1e-08 | 3.3e-08 | 2.867E-08 | 6.658E-09 | 2.100E-08 | 3.300E-08 |
| 7900 | 395 | D395V | A | 5.9e-08 | 4.5e-08 | 7.1e-08 | 5.833E-08 | 1.301E-08 | 4.500E-08 | 7.100E-08 |
| 7901 | 396 | S396A | A | 3.3e-08 | 3.6e-08 | 6.6e-08 | 4.500E-08 | 1.825E-08 | 3.300E-08 | 6.600E-08 |
| 7902 | 396 | S396R | A | 3.1e-08 | 5.7e-08 | 5.2e-08 | 4.667E-08 | 1.380E-08 | 3.100E-08 | 5.700E-08 |
| 7903 | 396 | S396N | A | 6e-08 | 1.3e-07 | 6.4e-08 | 8.467E-08 | 3.931E-08 | 6.000E-08 | 1.300E-07 |
| 7904 | 396 | S396D | A | 9.5e-08 | 9.1e-08 | 5.3e-08 | 7.967E-08 | 2.318E-08 | 5.300E-08 | 9.500E-08 |

| | | | | | | | | | | |
|---|---|---|---|---|---|---|---|---|---|---|
| 7905 | 396 | S396C | A | 2e-08 | 3.1e-08 | 4.2e-08 | 3.100E-08 | 1.100E-08 | 2.000E-08 | 4.200E-08 |
| 7906 | 396 | S396E | A | 3e-08 | 7.7e-08 | 4.5e-08 | 5.067E-08 | 2.401E-08 | 3.000E-08 | 7.700E-08 |
| 7907 | 396 | S396Q | A | 3.8e-08 | 3.4e-08 | 1.9e-08 | 3.033E-08 | 1.002E-08 | 1.900E-08 | 3.800E-08 |
| 7908 | 396 | S396G | A | 3.5e-08 | 3.2e-08 | 5.3e-08 | 4.000E-08 | 1.136E-08 | 3.200E-08 | 5.300E-08 |
| 7909 | 396 | S396H | A | 5.5e-08 | 3.4e-08 | 6.8e-08 | 5.233E-08 | 1.716E-08 | 3.400E-08 | 6.800E-08 |
| 7910 | 396 | S396I | A | 9.1e-08 | 2.9e-08 | 3.4e-08 | 5.133E-08 | 3.444E-08 | 2.900E-08 | 9.100E-08 |
| 7911 | 396 | S396L | A | 1.4e-08 | 2.1e-08 | 7.7e-08 | 3.733E-08 | 3.453E-08 | 1.400E-08 | 7.700E-08 |
| 7912 | 396 | S396K | A | 3.8e-08 | 4.8e-08 | 5.9e-08 | 4.833E-08 | 1.050E-08 | 3.800E-08 | 5.900E-08 |
| 7913 | 396 | S396M | A | 2.8e-08 | 1.3e-07 | 9.6e-08 | 8.467E-08 | 5.194E-08 | 2.800E-08 | 1.300E-07 |
| 7914 | 396 | S396F | A | 2.8e-08 | 5e-08 | 5.7e-08 | 4.500E-08 | 1.513E-08 | 2.800E-08 | 5.700E-08 |
| 7915 | 396 | S396P | A | 4.3e-08 | 3.7e-08 | 5.2e-08 | 4.400E-08 | 7.550E-09 | 3.700E-08 | 5.200E-08 |
| 7916 | 396 | S396S | A | 3e-08 | 3.5e-08 | 1.8e-08 | 2.767E-08 | 8.737E-09 | 1.800E-08 | 3.500E-08 |
| 7917 | 396 | S396T | A | 2.5e-08 | 4e-08 | 7.4e-08 | 4.633E-08 | 2.511E-08 | 2.500E-08 | 7.400E-08 |
| 7918 | 396 | S396W | A | 3e-08 | 1.1e-07 | 3.9e-08 | 5.967E-08 | 4.382E-08 | 3.000E-08 | 1.100E-07 |
| 7919 | 396 | S396Y | A | 1.7e-08 | 9.7e-08 | 1.3e-08 | 4.233E-08 | 4.738E-08 | 1.300E-08 | 9.700E-08 |
| 7920 | 396 | S396V | A | 2.1e-08 | 4.3e-08 | 4.6e-08 | 3.667E-08 | 1.365E-08 | 2.100E-08 | 4.600E-08 |
| 7921 | 397 | L397A | A | 5.2e-08 | 8.2e-08 | 3.2e-08 | 5.533E-08 | 2.517E-08 | 3.200E-08 | 8.200E-08 |
| 7922 | 397 | L397R | A | 4.1e-08 | 1.1e-07 | 7.4e-08 | 7.500E-08 | 3.451E-08 | 4.100E-08 | 1.100E-07 |
| 7923 | 397 | L397N | A | 3.1e-08 | 3e-08 | 2.9e-08 | 3.000E-08 | 1.000E-09 | 2.900E-08 | 3.100E-08 |
| 7924 | 397 | L397D | A | 4.1e-08 | 4.4e-08 | 5.9e-08 | 4.800E-08 | 9.644E-09 | 4.100E-08 | 5.900E-08 |
| 7925 | 397 | L397C | A | 1.2e-07 | 2.4e-08 | 3.1e-08 | 5.833E-08 | 5.352E-08 | 2.400E-08 | 1.200E-07 |
| 7926 | 397 | L397E | A | 3.1e-08 | 6.5e-08 | 3e-08 | 4.200E-08 | 1.992E-08 | 3.000E-08 | 6.500E-08 |
| 7927 | 397 | L397Q | A | 1.8e-08 | 3.6e-08 | 2.2e-08 | 2.533E-08 | 9.452E-09 | 1.800E-08 | 3.600E-08 |
| 7928 | 397 | L397G | A | 1.5e-08 | 2.5e-08 | 9.1e-08 | 4.367E-08 | 4.130E-08 | 1.500E-08 | 9.100E-08 |
| 7929 | 397 | L397H | A | 4.1e-08 | 3.8e-08 | 4.3e-08 | 4.067E-08 | 2.517E-09 | 3.800E-08 | 4.300E-08 |
| 7930 | 397 | L397I | A | 4.4e-08 | 7.2e-08 | 4.2e-08 | 5.267E-08 | 1.677E-08 | 4.200E-08 | 7.200E-08 |
| 7931 | 397 | L397L | A | 2.5e-08 | 3.4e-08 | 8.2e-08 | 4.700E-08 | 3.064E-08 | 2.500E-08 | 8.200E-08 |
| 7932 | 397 | L397K | A | 2.3e-08 | 2.2e-08 | 3.6e-08 | 2.700E-08 | 7.810E-09 | 2.200E-08 | 3.600E-08 |
| 7933 | 397 | L397M | A | 1.5e-08 | 6.6e-08 | 6.6e-08 | 4.900E-08 | 2.944E-08 | 1.500E-08 | 6.600E-08 |
| 7934 | 397 | L397F | A | 1.1e-07 | 8.3e-08 | 1.3e-07 | 1.077E-07 | 2.359E-08 | 8.300E-08 | 1.300E-07 |
| 7935 | 397 | L397P | A | 6.5e-08 | 4.2e-08 | 6.5e-08 | 5.733E-08 | 1.328E-08 | 4.200E-08 | 6.500E-08 |
| 7936 | 397 | L397S | A | 7.6e-08 | 1.9e-08 | 2.3e-08 | 3.933E-08 | 3.182E-08 | 1.900E-08 | 7.600E-08 |
| 7937 | 397 | L397T | A | 4e-08 | 5.2e-08 | 3.2e-08 | 4.133E-08 | 1.007E-08 | 3.200E-08 | 5.200E-08 |
| 7938 | 397 | L397W | A | 1.9e-08 | 5e-08 | 4.3e-08 | 3.733E-08 | 1.626E-08 | 1.900E-08 | 5.000E-08 |
| 7939 | 397 | L397Y | A | 7.3e-08 | 8.3e-08 | 5.9e-08 | 7.167E-08 | 1.206E-08 | 5.900E-08 | 8.300E-08 |
| 7940 | 397 | L397V | A | 7.1e-08 | 4e-08 | 9.1e-08 | 6.733E-08 | 2.570E-08 | 4.000E-08 | 9.100E-08 |
| 7941 | 398 | P398A | A | 4.2e-08 | 6e-08 | 4.9e-08 | 5.033E-08 | 9.074E-09 | 4.200E-08 | 6.000E-08 |
| 7942 | 398 | P398R | A | 4.4e-08 | 4.9e-08 | 4e-08 | 4.433E-08 | 4.509E-09 | 4.000E-08 | 4.900E-08 |
| 7943 | 398 | P398N | A | 4.4e-08 | 5.5e-08 | 2.2e-08 | 4.033E-08 | 1.680E-08 | 2.200E-08 | 5.500E-08 |
| 7944 | 398 | P398D | A | 5e-08 | 1.5e-07 | 5.2e-08 | 8.400E-08 | 5.717E-08 | 5.000E-08 | 1.500E-07 |
| 7945 | 398 | P398C | A | 5.1e-08 | 6.9e-08 | 2.8e-08 | 4.933E-08 | 2.055E-08 | 2.800E-08 | 6.900E-08 |
| 7946 | 398 | P398E | A | 1e-08 | 3e-08 | 3.8e-08 | 2.600E-08 | 1.442E-08 | 1.000E-08 | 3.800E-08 |
| 7947 | 398 | P398Q | A | 1.6e-08 | 3.4e-08 | 3.9e-08 | 2.967E-08 | 1.210E-08 | 1.600E-08 | 3.900E-08 |
| 7948 | 398 | P398G | A | 4.4e-08 | 6.3e-08 | 3e-08 | 4.567E-08 | 1.656E-08 | 3.000E-08 | 6.300E-08 |
| 7949 | 398 | P398H | A | 8.8e-08 | 3.6e-08 | 2.5e-08 | 4.967E-08 | 3.365E-08 | 2.500E-08 | 8.800E-08 |
| 7950 | 398 | P398I | A | 7.6e-08 | 2.1e-08 | 3.2e-08 | 4.300E-08 | 2.910E-08 | 2.100E-08 | 7.600E-08 |
| 7951 | 398 | P398L | A | 2.2e-08 | 3.1e-08 | 4.1e-08 | 3.133E-08 | 9.504E-09 | 2.200E-08 | 4.100E-08 |
| 7952 | 398 | P398K | A | 2.5e-08 | 5.8e-08 | 4.3e-08 | 4.200E-08 | 1.652E-08 | 2.500E-08 | 5.800E-08 |
| 7953 | 398 | P398M | A | 2.7e-08 | 4.5e-08 | 3.3e-08 | 3.500E-08 | 9.165E-09 | 2.700E-08 | 4.500E-08 |
| 7954 | 398 | P398F | A | 1.6e-07 | 3.4e-08 | 7.9e-08 | 9.100E-08 | 6.385E-08 | 3.400E-08 | 1.600E-07 |
| 7955 | 398 | P398P | A | 2.4e-08 | 2.4e-08 | 2.9e-08 | 2.567E-08 | 2.887E-09 | 2.400E-08 | 2.900E-08 |
| 7956 | 398 | P398S | A | 3.1e-08 | 6.8e-08 | 4.9e-08 | 4.933E-08 | 1.850E-08 | 3.100E-08 | 6.800E-08 |
| 7957 | 398 | P398T | A | 8.2e-08 | 4.3e-08 | 2.5e-08 | 5.000E-08 | 2.914E-08 | 2.500E-08 | 8.200E-08 |
| 7958 | 398 | P398W | A | 3.1e-08 | 2.4e-08 | 4.8e-08 | 3.433E-08 | 1.234E-08 | 2.400E-08 | 4.800E-08 |
| 7959 | 398 | P398Y | A | 3e-08 | 4.6e-08 | 5.5e-08 | 4.367E-08 | 1.266E-08 | 3.000E-08 | 5.500E-08 |
| 7960 | 398 | P398V | A | 6.4e-08 | 4.1e-08 | 1.7e-08 | 4.067E-08 | 2.350E-08 | 1.700E-08 | 6.400E-08 |
| 7961 | 399 | D399A | A | 2.3e-08 | 9.2e-08 | 5.1e-08 | 5.533E-08 | 3.470E-08 | 2.300E-08 | 9.200E-08 |
| 7962 | 399 | D399R | A | 3.8e-08 | 1.5e-07 | 5.4e-08 | 8.067E-08 | 6.058E-08 | 3.800E-08 | 1.500E-07 |
| 7963 | 399 | D399N | A | 4.4e-08 | 3.1e-08 | 8.2e-08 | 5.233E-08 | 2.650E-08 | 3.100E-08 | 8.200E-08 |
| 7964 | 399 | D399D | A | 6.8e-08 | 5.5e-08 | 8.9e-08 | 7.067E-08 | 1.716E-08 | 5.500E-08 | 8.900E-08 |
| 7965 | 399 | D399C | A | 2.4e-08 | 3.1e-08 | 6e-08 | 3.833E-08 | 1.909E-08 | 2.400E-08 | 6.000E-08 |
| 7966 | 399 | D399E | A | 5.5e-08 | 5e-08 | 8e-08 | 6.167E-08 | 1.607E-08 | 5.000E-08 | 8.000E-08 |
| 7967 | 399 | D399Q | A | 9.5e-08 | 1.2e-07 | 2.6e-08 | 8.033E-08 | 4.869E-08 | 2.600E-08 | 1.200E-07 |
| 7968 | 399 | D399G | A | 7.1e-08 | 2.7e-08 | 3.8e-08 | 4.533E-08 | 2.290E-08 | 2.700E-08 | 7.100E-08 |
| 7969 | 399 | D399H | A | 4e-08 | 3.7e-08 | 4.2e-08 | 3.967E-08 | 2.517E-09 | 3.700E-08 | 4.200E-08 |
| 7970 | 399 | D399I | A | 6.4e-08 | 2.4e-08 | 5.2e-08 | 4.667E-08 | 2.053E-08 | 2.400E-08 | 6.400E-08 |
| 7971 | 399 | D399L | A | 3.2e-08 | 3.6e-08 | 1.2e-08 | 2.667E-08 | 1.286E-08 | 1.200E-08 | 3.600E-08 |
| 7972 | 399 | D399K | A | 1.9e-08 | 8.8e-08 | 3.5e-08 | 4.733E-08 | 3.612E-08 | 1.900E-08 | 8.800E-08 |
| 7973 | 399 | D399M | A | 3.6e-08 | 2.9e-08 | 5.1e-08 | 3.867E-08 | 1.124E-08 | 2.900E-08 | 5.100E-08 |
| 7974 | 399 | D399F | A | 5.1e-08 | 5.1e-08 | 4.2e-08 | 4.800E-08 | 5.196E-09 | 4.200E-08 | 5.100E-08 |
| 7975 | 399 | D399P | A | 5.3e-08 | 4e-08 | 6.6e-08 | 5.300E-08 | 1.300E-08 | 4.000E-08 | 6.600E-08 |
| 7976 | 399 | D399S | A | 6e-08 | 3.5e-08 | 5.7e-08 | 5.067E-08 | 1.365E-08 | 3.500E-08 | 6.000E-08 |
| 7977 | 399 | D399T | A | 6.5e-08 | 6.2e-08 | 4.2e-08 | 5.633E-08 | 1.250E-08 | 4.200E-08 | 6.500E-08 |
| 7978 | 399 | D399W | A | 9.6e-08 | 3.2e-08 | 3.6e-08 | 5.467E-08 | 3.585E-08 | 3.200E-08 | 9.600E-08 |
| 7979 | 399 | D399Y | A | 4.2e-08 | 4e-08 | 3.7e-08 | 3.967E-08 | 2.517E-09 | 3.700E-08 | 4.200E-08 |
| 7980 | 399 | D399V | A | 3.6e-08 | 4.6e-08 | 2.4e-08 | 3.533E-08 | 1.102E-08 | 2.400E-08 | 4.600E-08 |
| 7981 | 400 | L400A | A | 5.6e-08 | 5.7e-08 | 8.7e-08 | 6.667E-08 | 1.762E-08 | 5.600E-08 | 8.700E-08 |
| 7982 | 400 | L400R | A | 1.3e-08 | 2.7e-08 | 2.9e-08 | 2.300E-08 | 8.718E-09 | 1.300E-08 | 2.900E-08 |
| 7983 | 400 | L400N | A | 3.6e-08 | 3.2e-08 | 9.5e-08 | 5.433E-08 | 3.528E-08 | 3.200E-08 | 9.500E-08 |
| 7984 | 400 | L400D | A | 5.9e-08 | 7e-08 | 6e-08 | 6.300E-08 | 6.083E-09 | 5.900E-08 | 7.000E-08 |
| 7985 | 400 | L400C | A | 4.5e-08 | 1.8e-08 | 7.3e-08 | 4.533E-08 | 2.750E-08 | 1.800E-08 | 7.300E-08 |
| 7986 | 400 | L400E | A | 3.4e-08 | 2.4e-08 | 4e-08 | 3.267E-08 | 8.083E-09 | 2.400E-08 | 4.000E-08 |
| 7987 | 400 | L400Q | A | 3.9e-08 | 2.9e-08 | 2.7e-08 | 3.167E-08 | 6.429E-09 | 2.700E-08 | 3.900E-08 |
| 7988 | 400 | L400G | A | 5.1e-08 | 8.5e-08 | 3.3e-08 | 5.633E-08 | 2.641E-08 | 3.300E-08 | 8.500E-08 |
| 7989 | 400 | L400H | A | 5.4e-08 | 7e-08 | 2.9e-08 | 5.100E-08 | 2.066E-08 | 2.900E-08 | 7.000E-08 |

| | | | | | | | | | | |
|---|---|---|---|---|---|---|---|---|---|---|
| 7990 | 400 | L400I | A | 6.9e-08 | 9e-08 | 1.1e-07 | 8.967E-08 | 2.050E-08 | 6.900E-08 | 1.100E-07 |
| 7991 | 400 | L400L | A | 4.3e-08 | 8e-08 | 2.7e-08 | 5.000E-08 | 2.718E-08 | 2.700E-08 | 8.000E-08 |
| 7992 | 400 | L400K | A | 2.1e-08 | 2.4e-08 | 7.2e-08 | 3.900E-08 | 2.862E-08 | 2.100E-08 | 7.200E-08 |
| 7993 | 400 | L400M | A | 4.4e-08 | 3.4e-08 | 8.8e-08 | 5.533E-08 | 2.873E-08 | 3.400E-08 | 8.800E-08 |
| 7994 | 400 | L400F | A | 8.8e-08 | 7.3e-08 | 1.8e-07 | 1.137E-07 | 5.793E-08 | 7.300E-08 | 1.800E-07 |
| 7995 | 400 | L400P | A | 4.9e-08 | 1.8e-08 | 7e-08 | 4.567E-08 | 2.616E-08 | 1.800E-08 | 7.000E-08 |
| 7996 | 400 | L400S | A | 2.4e-08 | 2.8e-08 | 3.8e-08 | 3.000E-08 | 7.211E-09 | 2.400E-08 | 3.800E-08 |
| 7997 | 400 | L400T | A | 7.5e-08 | 4.7e-08 | 4.7e-08 | 5.633E-08 | 1.617E-08 | 4.700E-08 | 7.500E-08 |
| 7998 | 400 | L400W | A | 3.3e-08 | 3.3e-08 | 1.1e-08 | 2.567E-08 | 1.270E-08 | 1.100E-08 | 3.300E-08 |
| 7999 | 400 | L400Y | A | 3.8e-08 | 3.5e-08 | 1.1e-07 | 6.100E-08 | 4.246E-08 | 3.500E-08 | 1.100E-07 |
| 8000 | 400 | L400V | A | 7.7e-09 | 1.1e-07 | 7.4e-08 | 6.390E-08 | 5.189E-08 | 7.700E-09 | 1.100E-07 |
| 8001 | 401 | S401A | A | 3.1e-08 | 6.9e-08 | 4.4e-08 | 4.800E-08 | 1.931E-08 | 3.100E-08 | 6.900E-08 |
| 8002 | 401 | S401R | A | 3.1e-08 | 2.5e-08 | 5.6e-08 | 3.733E-08 | 1.644E-08 | 2.500E-08 | 5.600E-08 |
| 8003 | 401 | S401N | A | 2.3e-08 | 2.4e-08 | 5.1e-08 | 3.267E-08 | 1.589E-08 | 2.300E-08 | 5.100E-08 |
| 8004 | 401 | S401D | A | 3.9e-08 | 4.1e-08 | 6.5e-08 | 4.833E-08 | 1.447E-08 | 3.900E-08 | 6.500E-08 |
| 8005 | 401 | S401C | A | 1.1e-07 | 2.7e-08 | 6.3e-08 | 6.667E-08 | 4.162E-08 | 2.700E-08 | 1.100E-07 |
| 8006 | 401 | S401E | A | 7.6e-08 | 6.4e-08 | 2.1e-08 | 5.367E-08 | 2.892E-08 | 2.100E-08 | 7.600E-08 |
| 8007 | 401 | S401Q | A | 2.7e-08 | 6.6e-08 | 4.8e-08 | 4.700E-08 | 1.952E-08 | 2.700E-08 | 6.600E-08 |
| 8008 | 401 | S401G | A | 3.8e-08 | 3.7e-08 | 1.1e-07 | 6.167E-08 | 4.186E-08 | 3.700E-08 | 1.100E-07 |
| 8009 | 401 | S401H | A | 2.9e-08 | 1.8e-08 | 4e-08 | 2.900E-08 | 1.100E-08 | 1.800E-08 | 4.000E-08 |
| 8010 | 401 | S401I | A | 7.2e-08 | 9.6e-08 | 1.3e-07 | 9.933E-08 | 2.914E-08 | 7.200E-08 | 1.300E-07 |
| 8011 | 401 | S401L | A | 5.3e-08 | 7.9e-08 | 4.4e-08 | 5.867E-08 | 1.818E-08 | 4.400E-08 | 7.900E-08 |
| 8012 | 401 | S401K | A | 5.6e-08 | 6.5e-08 | 6.8e-08 | 6.300E-08 | 6.245E-09 | 5.600E-08 | 6.800E-08 |
| 8013 | 401 | S401M | A | 2.4e-08 | 2.5e-08 | 4.4e-08 | 3.100E-08 | 1.127E-08 | 2.400E-08 | 4.400E-08 |
| 8014 | 401 | S401F | A | 9.2e-08 | 6.6e-08 | 3.1e-08 | 6.300E-08 | 3.061E-08 | 3.100E-08 | 9.200E-08 |
| 8015 | 401 | S401P | A | 7.8e-08 | 2e-08 | 7.7e-08 | 5.833E-08 | 3.320E-08 | 2.000E-08 | 7.800E-08 |
| 8016 | 401 | S401S | A | 2.5e-08 | 1e-07 | 5.1e-08 | 5.867E-08 | 3.808E-08 | 2.500E-08 | 1.000E-07 |
| 8017 | 401 | S401T | A | 3e-08 | 6.3e-08 | 4.3e-08 | 4.533E-08 | 1.662E-08 | 3.000E-08 | 6.300E-08 |
| 8018 | 401 | S401W | A | 3.3e-08 | 8.8e-08 | 2.9e-08 | 5.000E-08 | 3.297E-08 | 2.900E-08 | 8.800E-08 |
| 8019 | 401 | S401Y | A | 2.5e-08 | 1.9e-08 | 7.2e-08 | 3.867E-08 | 2.902E-08 | 1.900E-08 | 7.200E-08 |
| 8020 | 401 | S401V | A | 5e-08 | 3.4e-08 | 5.5e-08 | 4.633E-08 | 1.097E-08 | 3.400E-08 | 5.500E-08 |
| 8021 | 402 | V402A | A | 4.2e-08 | 8.6e-08 | 2.3e-08 | 5.033E-08 | 3.232E-08 | 2.300E-08 | 8.600E-08 |
| 8022 | 402 | V402R | A | 4.7e-08 | 6.3e-08 | 1.9e-08 | 4.300E-08 | 2.227E-08 | 1.900E-08 | 6.300E-08 |
| 8023 | 402 | V402N | A | 8.7e-08 | 4.9e-08 | 9.9e-08 | 7.833E-08 | 2.610E-08 | 4.900E-08 | 9.900E-08 |
| 8024 | 402 | V402D | A | 3.8e-08 | 5.5e-08 | 3.6e-08 | 4.300E-08 | 1.044E-08 | 3.600E-08 | 5.500E-08 |
| 8025 | 402 | V402C | A | 8.9e-08 | 1e-07 | 5.7e-08 | 8.200E-08 | 2.234E-08 | 5.700E-08 | 1.000E-07 |
| 8026 | 402 | V402E | A | 6.5e-08 | 4e-08 | 1.1e-07 | 7.167E-08 | 3.547E-08 | 4.000E-08 | 1.100E-07 |
| 8027 | 402 | V402Q | A | 8.4e-08 | 4.7e-08 | 5.2e-08 | 6.100E-08 | 2.007E-08 | 4.700E-08 | 8.400E-08 |
| 8028 | 402 | V402G | A | 1.1e-07 | 3.4e-08 | 1.8e-08 | 5.400E-08 | 4.915E-08 | 1.800E-08 | 1.100E-07 |
| 8029 | 402 | V402H | A | 2.6e-08 | 4.7e-08 | 4.9e-08 | 4.067E-08 | 1.274E-08 | 2.600E-08 | 4.900E-08 |
| 8030 | 402 | V402I | A | 1e-07 | 9.1e-08 | 3.7e-08 | 7.600E-08 | 3.407E-08 | 3.700E-08 | 1.000E-07 |
| 8031 | 402 | V402L | A | 7e-08 | 9e-08 | 2.9e-08 | 6.300E-08 | 3.110E-08 | 2.900E-08 | 9.000E-08 |
| 8032 | 402 | V402K | A | 7.4e-08 | 1.1e-07 | 5.2e-08 | 7.867E-08 | 2.928E-08 | 5.200E-08 | 1.100E-07 |
| 8033 | 402 | V402M | A | 5.3e-08 | 2.9e-08 | 2e-08 | 3.400E-08 | 1.706E-08 | 2.000E-08 | 5.300E-08 |
| 8034 | 402 | V402F | A | 2.1e-08 | 4.5e-08 | 6e-08 | 4.200E-08 | 1.967E-08 | 2.100E-08 | 6.000E-08 |
| 8035 | 402 | V402P | A | 5.6e-08 | 4.5e-08 | 7.1e-08 | 5.733E-08 | 1.305E-08 | 4.500E-08 | 7.100E-08 |
| 8036 | 402 | V402S | A | 2.2e-08 | 1.3e-07 | 2.6e-08 | 5.933E-08 | 6.123E-08 | 2.200E-08 | 1.300E-07 |
| 8037 | 402 | V402T | A | 3.9e-08 | 6.4e-08 | 2.4e-08 | 4.233E-08 | 2.021E-08 | 2.400E-08 | 6.400E-08 |
| 8038 | 402 | V402W | A | 3e-08 | 5.2e-08 | 3.9e-08 | 4.033E-08 | 1.106E-08 | 3.000E-08 | 5.200E-08 |
| 8039 | 402 | V402Y | A | 3.4e-08 | 2.6e-08 | 2.5e-08 | 2.833E-08 | 4.933E-09 | 2.500E-08 | 3.400E-08 |
| 8040 | 402 | V402V | A | 9.1e-08 | 3e-08 | 3.2e-08 | 5.100E-08 | 3.466E-08 | 3.000E-08 | 9.100E-08 |
| 8041 | 403 | F403A | A | 1e-07 | 2.4e-08 | 3.2e-08 | 5.200E-08 | 4.176E-08 | 2.400E-08 | 1.000E-07 |
| 8042 | 403 | F403R | A | 6e-08 | 2.7e-08 | 2.6e-08 | 3.767E-08 | 1.935E-08 | 2.600E-08 | 6.000E-08 |
| 8043 | 403 | F403N | A | 4.7e-08 | 4.1e-08 | 6e-08 | 4.933E-08 | 9.713E-09 | 4.100E-08 | 6.000E-08 |
| 8044 | 403 | F403D | A | 2.8e-08 | 9.1e-08 | 8.9e-08 | 6.933E-08 | 3.581E-08 | 2.800E-08 | 9.100E-08 |
| 8045 | 403 | F403C | A | 7.4e-08 | 3.4e-08 | 5.4e-08 | 5.400E-08 | 2.000E-08 | 3.400E-08 | 7.400E-08 |
| 8046 | 403 | F403E | A | 8.3e-08 | 3.7e-08 | 1.8e-08 | 4.600E-08 | 3.342E-08 | 1.800E-08 | 8.300E-08 |
| 8047 | 403 | F403Q | A | 2e-07 | 4.5e-08 | 2.1e-07 | 1.517E-07 | 9.251E-08 | 4.500E-08 | 2.100E-07 |
| 8048 | 403 | F403G | A | 3.8e-08 | 5.8e-08 | 1.8e-08 | 3.800E-08 | 2.000E-08 | 1.800E-08 | 5.800E-08 |
| 8049 | 403 | F403H | A | 3.8e-08 | 4.7e-08 | 2.9e-08 | 3.800E-08 | 9.000E-09 | 2.900E-08 | 4.700E-08 |
| 8050 | 403 | F403I | A | 1.2e-07 | 8.3e-08 | 4.8e-08 | 8.367E-08 | 3.600E-08 | 4.800E-08 | 1.200E-07 |
| 8051 | 403 | F403L | A | 3.3e-08 | 8.4e-08 | 4.4e-08 | 5.367E-08 | 2.684E-08 | 3.300E-08 | 8.400E-08 |
| 8052 | 403 | F403K | A | 5.9e-08 | 2.8e-08 | 3.8e-08 | 4.167E-08 | 1.582E-08 | 2.800E-08 | 5.900E-08 |
| 8053 | 403 | F403M | A | 5e-08 | 4e-08 | 5.8e-08 | 4.933E-08 | 9.018E-09 | 4.000E-08 | 5.800E-08 |
| 8054 | 403 | F403F | A | 1.5e-07 | 7.4e-08 | 3.1e-08 | 8.500E-08 | 6.026E-08 | 3.100E-08 | 1.500E-07 |
| 8055 | 403 | F403P | A | 2.1e-08 | 3.9e-08 | 7.8e-08 | 4.600E-08 | 2.914E-08 | 2.100E-08 | 7.800E-08 |
| 8056 | 403 | F403S | A | 3.6e-08 | 2.1e-08 | 3e-08 | 2.900E-08 | 7.550E-09 | 2.100E-08 | 3.600E-08 |
| 8057 | 403 | F403T | A | 1.2e-07 | 6.9e-08 | 2.1e-08 | 7.000E-08 | 4.951E-08 | 2.100E-08 | 1.200E-07 |
| 8058 | 403 | F403W | A | 2.4e-08 | 3.7e-08 | 3.8e-08 | 3.300E-08 | 7.810E-09 | 2.400E-08 | 3.800E-08 |
| 8059 | 403 | F403Y | A | 4.2e-08 | 3e-08 | 4.4e-08 | 3.867E-08 | 7.572E-09 | 3.000E-08 | 4.400E-08 |
| 8060 | 403 | F403V | A | 3.4e-08 | 2.3e-08 | 4.4e-08 | 3.367E-08 | 1.050E-08 | 2.300E-08 | 4.400E-08 |
| 8061 | 404 | Q404A | A | 2.1e-08 | 4.6e-08 | 3.9e-08 | 3.533E-08 | 1.290E-08 | 2.100E-08 | 4.600E-08 |
| 8062 | 404 | Q404R | A | 3.8e-08 | 2.7e-08 | 4.1e-08 | 3.533E-08 | 7.371E-09 | 2.700E-08 | 4.100E-08 |
| 8063 | 404 | Q404N | A | 5.5e-08 | 5.5e-08 | 1.5e-08 | 4.167E-08 | 2.309E-08 | 1.500E-08 | 5.500E-08 |
| 8064 | 404 | Q404D | A | 3.4e-08 | 5e-08 | 9.7e-08 | 6.033E-08 | 3.275E-08 | 3.400E-08 | 9.700E-08 |
| 8065 | 404 | Q404C | A | 3.5e-08 | 7.8e-08 | 4.5e-08 | 5.267E-08 | 2.250E-08 | 3.500E-08 | 7.800E-08 |
| 8066 | 404 | Q404E | A | 7.4e-08 | 2.3e-08 | 5.2e-08 | 4.967E-08 | 2.558E-08 | 2.300E-08 | 7.400E-08 |
| 8067 | 404 | Q404Q | A | 2.4e-08 | 3e-08 | 2.6e-08 | 2.667E-08 | 3.055E-09 | 2.400E-08 | 3.000E-08 |
| 8068 | 404 | Q404G | A | 7.8e-08 | 2e-08 | 2.9e-08 | 4.233E-08 | 3.121E-08 | 2.000E-08 | 7.800E-08 |
| 8069 | 404 | Q404H | A | 6.3e-08 | 5.8e-08 | 2.1e-08 | 4.733E-08 | 2.294E-08 | 2.100E-08 | 6.300E-08 |
| 8070 | 404 | Q404I | A | 2.3e-08 | 6.8e-08 | 7.8e-08 | 5.633E-08 | 2.930E-08 | 2.300E-08 | 7.800E-08 |
| 8071 | 404 | Q404L | A | 3.1e-08 | 5.8e-08 | 1.9e-08 | 3.600E-08 | 1.997E-08 | 1.900E-08 | 5.800E-08 |
| 8072 | 404 | Q404K | A | 6.6e-08 | 7.6e-09 | 3.7e-08 | 3.687E-08 | 2.920E-08 | 7.600E-09 | 6.600E-08 |
| 8073 | 404 | Q404M | A | 6.9e-08 | 9.9e-08 | 3.3e-08 | 6.700E-08 | 3.305E-08 | 3.300E-08 | 9.900E-08 |
| 8074 | 404 | Q404F | A | 8.8e-08 | 1.1e-07 | 5.1e-08 | 8.300E-08 | 2.982E-08 | 5.100E-08 | 1.100E-07 |

| | | | | | | | | | | |
|---|---|---|---|---|---|---|---|---|---|---|
| 8075 | 404 | Q404P | A | 1e-07 | 3.1e-08 | 5.1e-08 | 6.067E-08 | 3.550E-08 | 3.100E-08 | 1.000E-07 |
| 8076 | 404 | Q404S | A | 4.4e-08 | 3e-08 | 9.5e-08 | 5.633E-08 | 3.421E-08 | 3.000E-08 | 9.500E-08 |
| 8077 | 404 | Q404T | A | 8e-08 | 4.1e-08 | 5.4e-08 | 5.833E-08 | 1.986E-08 | 4.100E-08 | 8.000E-08 |
| 8078 | 404 | Q404W | A | 9.5e-08 | 4.9e-08 | 2.5e-08 | 5.633E-08 | 3.557E-08 | 2.500E-08 | 9.500E-08 |
| 8079 | 404 | Q404Y | A | 5.5e-08 | 4.7e-08 | 4.9e-08 | 5.033E-08 | 4.163E-09 | 4.700E-08 | 5.500E-08 |
| 8080 | 404 | Q404V | A | 4.7e-08 | 3.8e-08 | 1.1e-07 | 6.500E-08 | 3.923E-08 | 3.800E-08 | 1.100E-07 |
| 8081 | 405 | N405A | A | 3.5e-08 | 3.7e-08 | 1.6e-08 | 2.933E-08 | 1.159E-08 | 1.600E-08 | 3.700E-08 |
| 8082 | 405 | N405R | A | 8.7e-08 | 5.4e-08 | 6.9e-08 | 7.000E-08 | 1.652E-08 | 5.400E-08 | 8.700E-08 |
| 8083 | 405 | N405N | A | 3.4e-08 | 2.1e-08 | 4.7e-08 | 3.400E-08 | 1.300E-08 | 2.100E-08 | 4.700E-08 |
| 8084 | 405 | N405D | A | 4.3e-08 | 7.3e-08 | 1.6e-07 | 9.200E-08 | 6.077E-08 | 4.300E-08 | 1.600E-07 |
| 8085 | 405 | N405C | A | 7.6e-08 | 3.3e-08 | 2.7e-08 | 4.533E-08 | 2.673E-08 | 2.700E-08 | 7.600E-08 |
| 8086 | 405 | N405E | A | 7.3e-08 | 4.4e-08 | 1.6e-08 | 4.433E-08 | 2.850E-08 | 1.600E-08 | 7.300E-08 |
| 8087 | 405 | N405Q | A | 1.5e-08 | 3.3e-08 | 1.9e-08 | 2.233E-08 | 9.452E-09 | 1.500E-08 | 3.300E-08 |
| 8088 | 405 | N405G | A | 5.7e-08 | 5.6e-08 | 2.6e-08 | 4.633E-08 | 1.762E-08 | 2.600E-08 | 5.700E-08 |
| 8089 | 405 | N405H | A | 4.8e-08 | 3.7e-08 | 2.7e-08 | 3.733E-08 | 1.050E-08 | 2.700E-08 | 4.800E-08 |
| 8090 | 405 | N405I | A | 8.6e-08 | 2.5e-08 | 5.1e-08 | 5.400E-08 | 3.061E-08 | 2.500E-08 | 8.600E-08 |
| 8091 | 405 | N405L | A | 1.1e-07 | 6.4e-08 | 3.6e-08 | 7.000E-08 | 3.736E-08 | 3.600E-08 | 1.100E-07 |
| 8092 | 405 | N405K | A | 6.1e-08 | 7.3e-08 | 1.5e-07 | 9.467E-08 | 4.829E-08 | 6.100E-08 | 1.500E-07 |
| 8093 | 405 | N405M | A | 2.2e-08 | 1.4e-07 | 3.4e-08 | 6.533E-08 | 6.494E-08 | 2.200E-08 | 1.400E-07 |
| 8094 | 405 | N405F | A | 3.4e-08 | 6.4e-08 | 1.5e-07 | 8.267E-08 | 6.021E-08 | 3.400E-08 | 1.500E-07 |
| 8095 | 405 | N405P | A | 3.4e-08 | 4.7e-08 | 6.2e-08 | 4.767E-08 | 1.401E-08 | 3.400E-08 | 6.200E-08 |
| 8096 | 405 | N405S | A | 7.6e-08 | 1.5e-07 | 3.8e-08 | 8.800E-08 | 5.696E-08 | 3.800E-08 | 1.500E-07 |
| 8097 | 405 | N405T | A | 9.5e-08 | 5.5e-08 | 1e-07 | 8.333E-08 | 2.466E-08 | 5.500E-08 | 1.000E-07 |
| 8098 | 405 | N405W | A | 1.2e-08 | 8.5e-08 | 5.7e-08 | 5.133E-08 | 3.683E-08 | 1.200E-08 | 8.500E-08 |
| 8099 | 405 | N405Y | A | 8.3e-08 | 1.7e-08 | 5.8e-08 | 5.267E-08 | 3.332E-08 | 1.700E-08 | 8.300E-08 |
| 8100 | 405 | N405V | A | 5.9e-08 | 5.4e-08 | 1.1e-07 | 7.433E-08 | 3.099E-08 | 5.400E-08 | 1.100E-07 |
| 8101 | 406 | L406A | A | 3.7e-08 | 6.7e-08 | 1.4e-07 | 8.133E-08 | 5.297E-08 | 3.700E-08 | 1.400E-07 |
| 8102 | 406 | L406R | A | 5.1e-08 | 5.2e-08 | 5.2e-08 | 5.167E-08 | 5.774E-10 | 5.100E-08 | 5.200E-08 |
| 8103 | 406 | L406N | A | 1.3e-08 | 9.3e-08 | 3.9e-08 | 4.833E-08 | 4.081E-08 | 1.300E-08 | 9.300E-08 |
| 8104 | 406 | L406D | A | 5.6e-08 | 6.4e-08 | 2.7e-08 | 4.900E-08 | 1.947E-08 | 2.700E-08 | 6.400E-08 |
| 8105 | 406 | L406C | A | 2.3e-08 | 2e-08 | 6.2e-08 | 3.500E-08 | 2.343E-08 | 2.000E-08 | 6.200E-08 |
| 8106 | 406 | L406E | A | 6.4e-08 | 2.8e-08 | 6e-08 | 5.067E-08 | 1.973E-08 | 2.800E-08 | 6.400E-08 |
| 8107 | 406 | L406Q | A | 3.7e-08 | 5.4e-08 | 2.1e-08 | 3.733E-08 | 1.650E-08 | 2.100E-08 | 5.400E-08 |
| 8108 | 406 | L406G | A | 8.1e-08 | 4.3e-08 | 1e-07 | 7.467E-08 | 2.902E-08 | 4.300E-08 | 1.000E-07 |
| 8109 | 406 | L406H | A | 3.2e-08 | 4.7e-08 | 5.4e-08 | 4.433E-08 | 1.124E-08 | 3.200E-08 | 5.400E-08 |
| 8110 | 406 | L406I | A | 7.4e-08 | 3.1e-08 | 3.1e-08 | 4.533E-08 | 2.483E-08 | 3.100E-08 | 7.400E-08 |
| 8111 | 406 | L406L | A | 3e-08 | 4.3e-08 | 9.3e-08 | 5.533E-08 | 3.326E-08 | 3.000E-08 | 9.300E-08 |
| 8112 | 406 | L406K | A | 4.5e-08 | 2.6e-08 | 5.6e-08 | 4.233E-08 | 1.518E-08 | 2.600E-08 | 5.600E-08 |
| 8113 | 406 | L406M | A | 6e-08 | 8.4e-08 | 3.4e-08 | 5.933E-08 | 2.501E-08 | 3.400E-08 | 8.400E-08 |
| 8114 | 406 | L406F | A | 1.1e-07 | 5e-08 | 5.4e-08 | 7.133E-08 | 3.355E-08 | 5.000E-08 | 1.100E-07 |
| 8115 | 406 | L406P | A | 3.8e-08 | 3.8e-08 | 1e-07 | 5.867E-08 | 3.580E-08 | 3.800E-08 | 1.000E-07 |
| 8116 | 406 | L406S | A | 2.6e-08 | 6.9e-08 | 3.5e-08 | 4.333E-08 | 2.268E-08 | 2.600E-08 | 6.900E-08 |
| 8117 | 406 | L406T | A | 3.5e-08 | 3.1e-08 | 2e-08 | 2.867E-08 | 7.767E-09 | 2.000E-08 | 3.500E-08 |
| 8118 | 406 | L406W | A | 6e-08 | 3.5e-08 | 3.7e-08 | 4.400E-08 | 1.389E-08 | 3.500E-08 | 6.000E-08 |
| 8119 | 406 | L406Y | A | 1.2e-07 | 5.2e-08 | 1e-07 | 9.067E-08 | 3.495E-08 | 5.200E-08 | 1.200E-07 |
| 8120 | 406 | L406V | A | 6.3e-08 | 7.5e-08 | 4.7e-08 | 6.167E-08 | 1.405E-08 | 4.700E-08 | 7.500E-08 |
| 8121 | 407 | Q407A | A | 6e-08 | 2e-08 | 4e-08 | 4.000E-08 | 2.000E-08 | 2.000E-08 | 6.000E-08 |
| 8122 | 407 | Q407R | A | 3.6e-08 | 2.7e-08 | 3.9e-08 | 3.400E-08 | 6.245E-09 | 2.700E-08 | 3.900E-08 |
| 8123 | 407 | Q407N | A | 3e-08 | 3.5e-08 | 6.3e-08 | 4.267E-08 | 1.779E-08 | 3.000E-08 | 6.300E-08 |
| 8124 | 407 | Q407D | A | 2.4e-08 | 9.3e-08 | 3.4e-08 | 5.033E-08 | 3.729E-08 | 2.400E-08 | 9.300E-08 |
| 8125 | 407 | Q407C | A | 4e-08 | 4.2e-08 | 6e-08 | 4.733E-08 | 1.102E-08 | 4.000E-08 | 6.000E-08 |
| 8126 | 407 | Q407E | A | 6e-08 | 3.4e-08 | 4.4e-08 | 4.600E-08 | 1.311E-08 | 3.400E-08 | 6.000E-08 |
| 8127 | 407 | Q407Q | A | 8e-08 | 3.6e-08 | 3.7e-08 | 5.100E-08 | 2.512E-08 | 3.600E-08 | 8.000E-08 |
| 8128 | 407 | Q407G | A | 2e-08 | 2.5e-08 | 3e-08 | 2.500E-08 | 5.000E-09 | 2.000E-08 | 3.000E-08 |
| 8129 | 407 | Q407H | A | 6.1e-08 | 8.5e-08 | 1.8e-08 | 5.467E-08 | 3.395E-08 | 1.800E-08 | 8.500E-08 |
| 8130 | 407 | Q407I | A | 3e-08 | 4.8e-08 | 3.6e-08 | 3.800E-08 | 9.165E-09 | 3.000E-08 | 4.800E-08 |
| 8131 | 407 | Q407L | A | 3.9e-08 | 2.4e-08 | 7.8e-08 | 4.700E-08 | 2.787E-08 | 2.400E-08 | 7.800E-08 |
| 8132 | 407 | Q407K | A | 7e-08 | 2.5e-08 | 9.3e-08 | 6.267E-08 | 3.459E-08 | 2.500E-08 | 9.300E-08 |
| 8133 | 407 | Q407M | A | 1.5e-07 | 7.9e-08 | 1.1e-07 | 1.130E-07 | 3.559E-08 | 7.900E-08 | 1.500E-07 |
| 8134 | 407 | Q407F | A | 1.6e-08 | 5.9e-08 | 1.7e-08 | 3.067E-08 | 2.454E-08 | 1.600E-08 | 5.900E-08 |
| 8135 | 407 | Q407P | A | 7e-08 | 7.2e-08 | 4.3e-08 | 6.167E-08 | 1.620E-08 | 4.300E-08 | 7.200E-08 |
| 8136 | 407 | Q407S | A | 6.2e-08 | 9.9e-08 | 3.6e-08 | 6.567E-08 | 3.166E-08 | 3.600E-08 | 9.900E-08 |
| 8137 | 407 | Q407T | A | 3e-08 | 7.2e-08 | 2.3e-08 | 4.167E-08 | 2.650E-08 | 2.300E-08 | 7.200E-08 |
| 8138 | 407 | Q407W | A | 4.1e-08 | 7.1e-08 | 5.7e-08 | 5.633E-08 | 1.501E-08 | 4.100E-08 | 7.100E-08 |
| 8139 | 407 | Q407Y | A | 2.9e-08 | 4.4e-08 | 2.6e-08 | 3.300E-08 | 9.644E-09 | 2.600E-08 | 4.400E-08 |
| 8140 | 407 | Q407V | A | 2.5e-08 | 5.3e-08 | 5.7e-08 | 4.500E-08 | 1.744E-08 | 2.500E-08 | 5.700E-08 |
| 8141 | 408 | V408A | A | 5.2e-08 | 8e-08 | 6.2e-08 | 6.467E-08 | 1.419E-08 | 5.200E-08 | 8.000E-08 |
| 8142 | 408 | V408R | A | 2.5e-08 | 1.6e-08 | 2.8e-08 | 2.300E-08 | 6.245E-09 | 1.600E-08 | 2.800E-08 |
| 8143 | 408 | V408N | A | 3.4e-08 | 6.2e-08 | 2.4e-08 | 4.000E-08 | 1.970E-08 | 2.400E-08 | 6.200E-08 |
| 8144 | 408 | V408D | A | 7.1e-09 | 8.3e-08 | 6.9e-08 | 5.303E-08 | 4.039E-08 | 7.100E-09 | 8.300E-08 |
| 8145 | 408 | V408C | A | 3.9e-08 | 3.7e-08 | 3.2e-08 | 3.600E-08 | 3.606E-09 | 3.200E-08 | 3.900E-08 |
| 8146 | 408 | V408E | A | 6.9e-08 | 2.7e-08 | 7.4e-08 | 5.667E-08 | 2.581E-08 | 2.700E-08 | 7.400E-08 |
| 8147 | 408 | V408Q | A | 2.7e-08 | 4.9e-08 | 3.5e-08 | 3.700E-08 | 1.114E-08 | 2.700E-08 | 4.900E-08 |
| 8148 | 408 | V408G | A | 6.4e-08 | 5.4e-08 | 3e-08 | 4.933E-08 | 1.747E-08 | 3.000E-08 | 6.400E-08 |
| 8149 | 408 | V408H | A | 1.6e-07 | 5.3e-08 | 3e-08 | 8.100E-08 | 6.938E-08 | 3.000E-08 | 1.600E-07 |
| 8150 | 408 | V408I | A | 4.6e-08 | 2.1e-08 | 3.1e-08 | 3.267E-08 | 1.258E-08 | 2.100E-08 | 4.600E-08 |
| 8151 | 408 | V408L | A | 3.2e-08 | 6.3e-08 | 2.4e-08 | 3.967E-08 | 2.060E-08 | 2.400E-08 | 6.300E-08 |
| 8152 | 408 | V408K | A | 1.2e-08 | 1.9e-08 | 1.5e-07 | 6.033E-08 | 7.773E-08 | 1.200E-08 | 1.500E-07 |
| 8153 | 408 | V408M | A | 1.6e-08 | 1.3e-08 | 2.8e-08 | 1.900E-08 | 7.937E-09 | 1.300E-08 | 2.800E-08 |
| 8154 | 408 | V408F | A | 5e-08 | 4.3e-08 | 2.5e-08 | 3.933E-08 | 1.290E-08 | 2.500E-08 | 5.000E-08 |
| 8155 | 408 | V408P | A | 3.9e-08 | 5.6e-08 | 3.1e-08 | 4.200E-08 | 1.277E-08 | 3.100E-08 | 5.600E-08 |
| 8156 | 408 | V408S | A | 1e-08 | 4.3e-08 | 2e-08 | 2.433E-08 | 1.692E-08 | 1.000E-08 | 4.300E-08 |
| 8157 | 408 | V408T | A | 3.2e-08 | 5.9e-08 | 3e-08 | 4.033E-08 | 1.620E-08 | 3.000E-08 | 5.900E-08 |
| 8158 | 408 | V408W | A | 2.9e-08 | 7.6e-08 | 4.2e-08 | 4.900E-08 | 2.427E-08 | 2.900E-08 | 7.600E-08 |
| 8159 | 408 | V408Y | A | 3.3e-08 | 6.4e-08 | 2e-08 | 3.900E-08 | 2.261E-08 | 2.000E-08 | 6.400E-08 |

| | | | | | | | | | | |
|---|---|---|---|---|---|---|---|---|---|---|
| 8160 | 408 | V408V | A | 5.3e-08 | 1.2e-07 | 3.4e-08 | 6.900E-08 | 4.518E-08 | 3.400E-08 | 1.200E-07 |
| 8161 | 409 | I409A | A | 4.6e-08 | 4.2e-08 | 9.5e-08 | 6.100E-08 | 2.951E-08 | 4.200E-08 | 9.500E-08 |
| 8162 | 409 | I409R | A | 2.8e-08 | 3e-08 | 7.2e-08 | 4.333E-08 | 2.485E-08 | 2.800E-08 | 7.200E-08 |
| 8163 | 409 | I409N | A | 1.2e-08 | 7.4e-08 | 2.1e-08 | 3.567E-08 | 3.350E-08 | 1.200E-08 | 7.400E-08 |
| 8164 | 409 | I409D | A | 3.7e-08 | 7.1e-08 | 1.7e-07 | 9.267E-08 | 6.910E-08 | 3.700E-08 | 1.700E-07 |
| 8165 | 409 | I409C | A | 1.3e-07 | 6.2e-08 | 4.4e-08 | 7.867E-08 | 4.536E-08 | 4.400E-08 | 1.300E-07 |
| 8166 | 409 | I409E | A | 4.7e-08 | 2e-08 | 2.6e-08 | 3.100E-08 | 1.418E-08 | 2.000E-08 | 4.700E-08 |
| 8167 | 409 | I409Q | A | 4.1e-08 | 3.6e-08 | 3.7e-08 | 3.800E-08 | 2.646E-09 | 3.600E-08 | 4.100E-08 |
| 8168 | 409 | I409G | A | 3.4e-08 | 5.1e-08 | 5e-08 | 4.500E-08 | 9.539E-09 | 3.400E-08 | 5.100E-08 |
| 8169 | 409 | I409H | A | 5.6e-08 | 6.6e-08 | 1.2e-07 | 8.067E-08 | 3.443E-08 | 5.600E-08 | 1.200E-07 |
| 8170 | 409 | I409I | A | 5.9e-08 | 4.6e-08 | 4.8e-08 | 5.100E-08 | 7.000E-09 | 4.600E-08 | 5.900E-08 |
| 8171 | 409 | I409L | A | 3.6e-08 | 6.7e-08 | 8.4e-08 | 6.233E-08 | 2.434E-08 | 3.600E-08 | 8.400E-08 |
| 8172 | 409 | I409K | A | 2.1e-08 | 6.7e-08 | 4e-08 | 4.267E-08 | 2.312E-08 | 2.100E-08 | 6.700E-08 |
| 8173 | 409 | I409M | A | 1.7e-08 | 5.1e-08 | 3.8e-08 | 3.533E-08 | 1.716E-08 | 1.700E-08 | 5.100E-08 |
| 8174 | 409 | I409F | A | 4.1e-08 | 3.6e-08 | 2.3e-08 | 3.333E-08 | 9.292E-09 | 2.300E-08 | 4.100E-08 |
| 8175 | 409 | I409P | A | 3.9e-08 | 1.4e-07 | 2.9e-08 | 6.933E-08 | 6.140E-08 | 2.900E-08 | 1.400E-07 |
| 8176 | 409 | I409S | A | 4.4e-08 | 1.6e-08 | 5.1e-08 | 3.700E-08 | 1.852E-08 | 1.600E-08 | 5.100E-08 |
| 8177 | 409 | I409T | A | 5.8e-08 | 8e-08 | 1.8e-07 | 1.060E-07 | 6.502E-08 | 5.800E-08 | 1.800E-07 |
| 8178 | 409 | I409W | A | 2.9e-08 | 4.4e-08 | 3.9e-08 | 3.733E-08 | 7.638E-09 | 2.900E-08 | 4.400E-08 |
| 8179 | 409 | I409Y | A | 5.7e-08 | 3e-08 | 2.9e-08 | 3.867E-08 | 1.589E-08 | 2.900E-08 | 5.700E-08 |
| 8180 | 409 | I409V | A | 2.1e-08 | 5.5e-08 | 2.9e-08 | 3.500E-08 | 1.778E-08 | 2.100E-08 | 5.500E-08 |
| 8181 | 410 | R410A | A | 5.4e-08 | 4.7e-08 | 3.6e-08 | 4.567E-08 | 9.074E-09 | 3.600E-08 | 5.400E-08 |
| 8182 | 410 | R410R | A | 2.2e-08 | 3.6e-08 | 1.9e-08 | 2.567E-08 | 9.074E-09 | 1.900E-08 | 3.600E-08 |
| 8183 | 410 | R410N | A | 4.8e-08 | 9.5e-08 | 4.8e-08 | 6.367E-08 | 2.714E-08 | 4.800E-08 | 9.500E-08 |
| 8184 | 410 | R410D | A | 4.6e-08 | 1.6e-08 | 6.4e-08 | 4.200E-08 | 2.425E-08 | 1.600E-08 | 6.400E-08 |
| 8185 | 410 | R410C | A | 3.2e-08 | 6.8e-08 | 2.4e-08 | 4.133E-08 | 2.344E-08 | 2.400E-08 | 6.800E-08 |
| 8186 | 410 | R410E | A | 8.9e-08 | 2.6e-08 | 5.3e-08 | 5.600E-08 | 3.161E-08 | 2.600E-08 | 8.900E-08 |
| 8187 | 410 | R410Q | A | 1.9e-08 | 1.7e-08 | 3e-08 | 2.200E-08 | 7.000E-09 | 1.700E-08 | 3.000E-08 |
| 8188 | 410 | R410G | A | 1.9e-08 | 3.6e-08 | 7.1e-08 | 4.200E-08 | 2.651E-08 | 1.900E-08 | 7.100E-08 |
| 8189 | 410 | R410H | A | 4.1e-08 | 7.7e-08 | 1.6e-08 | 4.467E-08 | 3.066E-08 | 1.600E-08 | 7.700E-08 |
| 8190 | 410 | R410I | A | 6.5e-08 | 2.2e-08 | 4e-08 | 4.233E-08 | 2.159E-08 | 2.200E-08 | 6.500E-08 |
| 8191 | 410 | R410L | A | 1.9e-08 | 4.4e-08 | 2.5e-08 | 2.933E-08 | 1.305E-08 | 1.900E-08 | 4.400E-08 |
| 8192 | 410 | R410K | A | 5.2e-08 | 2.9e-08 | 9.3e-08 | 5.800E-08 | 3.242E-08 | 2.900E-08 | 9.300E-08 |
| 8193 | 410 | R410M | A | 7.6e-08 | 6.1e-08 | 5.2e-08 | 6.300E-08 | 1.212E-08 | 5.200E-08 | 7.600E-08 |
| 8194 | 410 | R410F | A | 3.7e-08 | 2.9e-08 | 5.9e-08 | 4.167E-08 | 1.553E-08 | 2.900E-08 | 5.900E-08 |
| 8195 | 410 | R410P | A | 1.5e-08 | 3.1e-08 | 5e-08 | 3.200E-08 | 1.752E-08 | 1.500E-08 | 5.000E-08 |
| 8196 | 410 | R410S | A | 1.2e-07 | 5.7e-08 | 1.7e-08 | 6.467E-08 | 5.193E-08 | 1.700E-08 | 1.200E-07 |
| 8197 | 410 | R410T | A | 2.2e-07 | 4e-08 | 1.1e-07 | 1.233E-07 | 9.074E-08 | 4.000E-08 | 2.200E-07 |
| 8198 | 410 | R410W | A | 2.6e-08 | 1.2e-07 | 6.1e-08 | 6.900E-08 | 4.751E-08 | 2.600E-08 | 1.200E-07 |
| 8199 | 410 | R410Y | A | 4.7e-08 | 3.5e-08 | 7.6e-08 | 5.267E-08 | 2.108E-08 | 3.500E-08 | 7.600E-08 |
| 8200 | 410 | R410V | A | 1.1e-08 | 4.2e-08 | 4.7e-08 | 3.333E-08 | 1.950E-08 | 1.100E-08 | 4.700E-08 |
| 8201 | 411 | G411A | A | 8.7e-08 | 5.5e-08 | 1.8e-08 | 5.333E-08 | 3.453E-08 | 1.800E-08 | 8.700E-08 |
| 8202 | 411 | G411R | A | 2.3e-08 | 3.6e-08 | 2.2e-08 | 2.700E-08 | 7.810E-09 | 2.200E-08 | 3.600E-08 |
| 8203 | 411 | G411N | A | 1.6e-08 | 3.8e-08 | 2e-08 | 2.467E-08 | 1.172E-08 | 1.600E-08 | 3.800E-08 |
| 8204 | 411 | G411D | A | 2.6e-08 | 5.8e-08 | 3.2e-08 | 3.867E-08 | 1.701E-08 | 2.600E-08 | 5.800E-08 |
| 8205 | 411 | G411C | A | 2.8e-08 | 2.2e-08 | 6.5e-08 | 3.833E-08 | 2.329E-08 | 2.200E-08 | 6.500E-08 |
| 8206 | 411 | G411E | A | 4.3e-08 | 4.2e-08 | 4e-08 | 4.167E-08 | 1.528E-09 | 4.000E-08 | 4.300E-08 |
| 8207 | 411 | G411Q | A | 7.8e-08 | 5.7e-08 | 6.2e-08 | 6.567E-08 | 1.097E-08 | 5.700E-08 | 7.800E-08 |
| 8208 | 411 | G411G | A | 4.2e-08 | 2.6e-08 | 1.4e-08 | 2.733E-08 | 1.405E-08 | 1.400E-08 | 4.200E-08 |
| 8209 | 411 | G411H | A | 3.7e-08 | 8.7e-09 | 4.6e-08 | 3.057E-08 | 1.946E-08 | 8.700E-09 | 4.600E-08 |
| 8210 | 411 | G411I | A | 3.2e-08 | 3.6e-08 | 5.1e-08 | 3.967E-08 | 1.002E-08 | 3.200E-08 | 5.100E-08 |
| 8211 | 411 | G411L | A | 2.7e-08 | 3.6e-08 | 3.8e-08 | 3.367E-08 | 5.859E-09 | 2.700E-08 | 3.800E-08 |
| 8212 | 411 | G411K | A | 1e-07 | 3.9e-08 | 4.2e-08 | 6.033E-08 | 3.439E-08 | 3.900E-08 | 1.000E-07 |
| 8213 | 411 | G411M | A | 2.1e-08 | 8.9e-08 | 6.4e-08 | 5.800E-08 | 3.439E-08 | 2.100E-08 | 8.900E-08 |
| 8214 | 411 | G411F | A | 6.2e-09 | 4e-08 | 8.3e-08 | 4.307E-08 | 3.849E-08 | 6.200E-09 | 8.300E-08 |
| 8215 | 411 | G411P | A | 3.2e-08 | 5.2e-08 | 2.5e-08 | 3.633E-08 | 1.401E-08 | 2.500E-08 | 5.200E-08 |
| 8216 | 411 | G411S | A | 5.3e-08 | 4.3e-08 | 1.5e-08 | 3.700E-08 | 1.970E-08 | 1.500E-08 | 5.300E-08 |
| 8217 | 411 | G411T | A | 5.1e-08 | 6.4e-08 | 4.6e-08 | 5.367E-08 | 9.292E-09 | 4.600E-08 | 6.400E-08 |
| 8218 | 411 | G411W | A | 3.1e-08 | 4e-08 | 8.2e-08 | 5.100E-08 | 2.722E-08 | 3.100E-08 | 8.200E-08 |
| 8219 | 411 | G411Y | A | 1.9e-08 | 8.9e-08 | 6.2e-08 | 5.667E-08 | 3.530E-08 | 1.900E-08 | 8.900E-08 |
| 8220 | 411 | G411V | A | 5.1e-08 | 4.9e-08 | 1e-08 | 3.667E-08 | 2.312E-08 | 1.000E-08 | 5.100E-08 |
| 8221 | 412 | R412A | A | 4.2e-08 | 1.3e-07 | 3.8e-08 | 7.000E-08 | 5.200E-08 | 3.800E-08 | 1.300E-07 |
| 8222 | 412 | R412R | A | 1.8e-08 | 1.9e-08 | 1.8e-08 | 1.833E-08 | 5.774E-10 | 1.800E-08 | 1.900E-08 |
| 8223 | 412 | R412N | A | 6e-08 | 5.8e-08 | 3.7e-08 | 5.167E-08 | 1.274E-08 | 3.700E-08 | 6.000E-08 |
| 8224 | 412 | R412D | A | 6.4e-08 | 4.7e-08 | 2.1e-08 | 4.400E-08 | 2.166E-08 | 2.100E-08 | 6.400E-08 |
| 8225 | 412 | R412C | A | 9.6e-08 | 6.2e-08 | 1.5e-08 | 5.767E-08 | 4.067E-08 | 1.500E-08 | 9.600E-08 |
| 8226 | 412 | R412E | A | 7.8e-08 | 8.5e-08 | 8.4e-08 | 8.233E-08 | 3.786E-09 | 7.800E-08 | 8.500E-08 |
| 8227 | 412 | R412Q | A | 1.8e-08 | 6.9e-08 | 4.5e-08 | 4.400E-08 | 2.551E-08 | 1.800E-08 | 6.900E-08 |
| 8228 | 412 | R412G | A | 8.6e-08 | 3e-08 | 7.2e-08 | 6.267E-08 | 2.914E-08 | 3.000E-08 | 8.600E-08 |
| 8229 | 412 | R412H | A | 4.7e-08 | 2.5e-08 | 4e-08 | 3.733E-08 | 1.124E-08 | 2.500E-08 | 4.700E-08 |
| 8230 | 412 | R412I | A | 5.7e-08 | 1.5e-08 | 4.9e-08 | 4.033E-08 | 2.230E-08 | 1.500E-08 | 5.700E-08 |
| 8231 | 412 | R412L | A | 2.8e-08 | 1.3e-07 | 3e-08 | 6.267E-08 | 5.832E-08 | 2.800E-08 | 1.300E-07 |
| 8232 | 412 | R412K | A | 2.9e-08 | 3.3e-08 | 2.2e-08 | 2.800E-08 | 5.568E-09 | 2.200E-08 | 3.300E-08 |
| 8233 | 412 | R412M | A | 4.2e-08 | 1e-08 | 1.4e-08 | 2.200E-08 | 1.744E-08 | 1.000E-08 | 4.200E-08 |
| 8234 | 412 | R412F | A | 4.7e-08 | 6.9e-09 | 8e-09 | 2.063E-08 | 2.284E-08 | 6.900E-09 | 4.700E-08 |
| 8235 | 412 | R412P | A | 2.3e-08 | 8.6e-08 | 6.6e-08 | 5.833E-08 | 3.219E-08 | 2.300E-08 | 8.600E-08 |
| 8236 | 412 | R412S | A | 7.2e-08 | 3.9e-08 | 6.2e-08 | 5.767E-08 | 1.692E-08 | 3.900E-08 | 7.200E-08 |
| 8237 | 412 | R412T | A | 7.6e-08 | 1.3e-07 | 3.8e-08 | 8.133E-08 | 4.623E-08 | 3.800E-08 | 1.300E-07 |
| 8238 | 412 | R412W | A | 4.7e-08 | 1.1e-07 | 1e-07 | 8.567E-08 | 3.386E-08 | 4.700E-08 | 1.100E-07 |
| 8239 | 412 | R412Y | A | 2.7e-08 | 2.3e-08 | 7.2e-08 | 4.067E-08 | 2.721E-08 | 2.300E-08 | 7.200E-08 |
| 8240 | 412 | R412V | A | 3.7e-08 | 2.8e-08 | 2.7e-08 | 3.067E-08 | 5.508E-09 | 2.700E-08 | 3.700E-08 |
| 8241 | 413 | I413A | A | 8.3e-08 | 8.8e-08 | 3.2e-08 | 6.767E-08 | 3.099E-08 | 3.200E-08 | 8.800E-08 |
| 8242 | 413 | I413R | A | 7.9e-08 | 7.2e-08 | 5e-08 | 6.700E-08 | 1.513E-08 | 5.000E-08 | 7.900E-08 |
| 8243 | 413 | I413N | A | 1.1e-07 | 5.8e-08 | 4.9e-08 | 7.233E-08 | 3.293E-08 | 4.900E-08 | 1.100E-07 |
| 8244 | 413 | I413D | A | 6.6e-08 | 5e-08 | 2e-08 | 4.533E-08 | 2.335E-08 | 2.000E-08 | 6.600E-08 |

| | | | | | | | | | | |
|---|---|---|---|---|---|---|---|---|---|---|
| 8245 | 413 | I413C | A | 4.8e-08 | 5.4e-08 | 1e-08 | 3.733E-08 | 2.386E-08 | 1.000E-08 | 5.400E-08 |
| 8246 | 413 | I413E | A | 1.5e-08 | 1.8e-08 | 1.1e-07 | 4.767E-08 | 5.400E-08 | 1.500E-08 | 1.100E-07 |
| 8247 | 413 | I413Q | A | 5.7e-08 | 6.8e-08 | 5.6e-08 | 6.033E-08 | 6.658E-09 | 5.600E-08 | 6.800E-08 |
| 8248 | 413 | I413G | A | 6.1e-08 | 3.5e-08 | 5.5e-08 | 5.033E-08 | 1.361E-08 | 3.500E-08 | 6.100E-08 |
| 8249 | 413 | I413H | A | 2e-08 | 6.6e-08 | 9.9e-08 | 6.167E-08 | 3.968E-08 | 2.000E-08 | 9.900E-08 |
| 8250 | 413 | I413I | A | 2.9e-08 | 8.7e-09 | 9.1e-08 | 4.290E-08 | 4.287E-08 | 8.700E-09 | 9.100E-08 |
| 8251 | 413 | I413L | A | 5.8e-08 | 8.5e-09 | 6.1e-08 | 4.250E-08 | 2.948E-08 | 8.500E-09 | 6.100E-08 |
| 8252 | 413 | I413K | A | 4e-08 | 9.6e-08 | 2.8e-08 | 5.467E-08 | 3.630E-08 | 2.800E-08 | 9.600E-08 |
| 8253 | 413 | I413M | A | 2e-08 | 1.1e-07 | 5e-08 | 6.000E-08 | 4.583E-08 | 2.000E-08 | 1.100E-07 |
| 8254 | 413 | I413F | A | 6.1e-09 | 5.1e-08 | 4.9e-08 | 3.537E-08 | 2.537E-08 | 6.100E-09 | 5.100E-08 |
| 8255 | 413 | I413P | A | 6.1e-08 | 2.1e-08 | 3.5e-08 | 3.900E-08 | 2.030E-08 | 2.100E-08 | 6.100E-08 |
| 8256 | 413 | I413S | A | 2.1e-08 | 5.8e-08 | 2.1e-08 | 3.333E-08 | 2.136E-08 | 2.100E-08 | 5.800E-08 |
| 8257 | 413 | I413T | A | 2.3e-08 | 5.7e-08 | 4.9e-08 | 4.300E-08 | 1.778E-08 | 2.300E-08 | 5.700E-08 |
| 8258 | 413 | I413W | A | 2e-07 | 7.8e-08 | 6.1e-08 | 1.130E-07 | 7.582E-08 | 6.100E-08 | 2.000E-07 |
| 8259 | 413 | I413Y | A | 1.4e-08 | 1.2e-07 | 1.8e-08 | 5.067E-08 | 6.008E-08 | 1.400E-08 | 1.200E-07 |
| 8260 | 413 | I413V | A | 2.9e-08 | 5.1e-08 | 1.2e-07 | 6.667E-08 | 4.748E-08 | 2.900E-08 | 1.200E-07 |
| 8261 | 414 | L414A | A | 5.8e-08 | 3.4e-08 | 2.6e-08 | 3.933E-08 | 1.665E-08 | 2.600E-08 | 5.800E-08 |
| 8262 | 414 | L414R | A | 2.6e-08 | 1.7e-07 | 9.8e-08 | 9.800E-08 | 7.200E-08 | 2.600E-08 | 1.700E-07 |
| 8263 | 414 | L414N | A | 2.1e-08 | 2.2e-08 | 2.6e-08 | 2.300E-08 | 2.646E-09 | 2.100E-08 | 2.600E-08 |
| 8264 | 414 | L414D | A | 3.4e-08 | 6.3e-08 | 6.3e-08 | 5.333E-08 | 1.674E-08 | 3.400E-08 | 6.300E-08 |
| 8265 | 414 | L414C | A | 6.2e-08 | 1.2e-07 | 4.6e-08 | 7.600E-08 | 3.894E-08 | 4.600E-08 | 1.200E-07 |
| 8266 | 414 | L414E | A | 2.7e-08 | 2.5e-08 | 4.2e-08 | 3.133E-08 | 9.292E-09 | 2.500E-08 | 4.200E-08 |
| 8267 | 414 | L414Q | A | 5.5e-08 | 3.4e-08 | 2.4e-08 | 3.767E-08 | 1.582E-08 | 2.400E-08 | 5.500E-08 |
| 8268 | 414 | L414G | A | 5.8e-08 | 4.4e-08 | 4.5e-08 | 4.900E-08 | 7.810E-09 | 4.400E-08 | 5.800E-08 |
| 8269 | 414 | L414H | A | 3e-08 | 5.9e-08 | 2.2e-08 | 3.700E-08 | 1.947E-08 | 2.200E-08 | 5.900E-08 |
| 8270 | 414 | L414I | A | 3.2e-08 | 2.8e-08 | 1.2e-08 | 2.400E-08 | 1.058E-08 | 1.200E-08 | 3.200E-08 |
| 8271 | 414 | L414L | A | 3.5e-08 | 5e-08 | 4.8e-08 | 4.433E-08 | 8.145E-09 | 3.500E-08 | 5.000E-08 |
| 8272 | 414 | L414K | A | 1.5e-08 | 3e-08 | 7.4e-09 | 1.747E-08 | 1.150E-08 | 7.400E-09 | 3.000E-08 |
| 8273 | 414 | L414M | A | 4.5e-08 | 6.2e-08 | 5.1e-09 | 3.737E-08 | 2.921E-08 | 5.100E-09 | 6.200E-08 |
| 8274 | 414 | L414F | A | 2.6e-08 | 5.4e-08 | 5.1e-08 | 4.367E-08 | 1.537E-08 | 2.600E-08 | 5.400E-08 |
| 8275 | 414 | L414P | A | 4.7e-08 | 3.9e-08 | 5.5e-08 | 4.700E-08 | 8.000E-09 | 3.900E-08 | 5.500E-08 |
| 8276 | 414 | L414S | A | 2.5e-08 | 5.4e-08 | 7.4e-08 | 5.100E-08 | 2.464E-08 | 2.500E-08 | 7.400E-08 |
| 8277 | 414 | L414T | A | 2e-08 | 1.5e-08 | 8.4e-09 | 1.447E-08 | 5.818E-09 | 8.400E-09 | 2.000E-08 |
| 8278 | 414 | L414W | A | 1e-07 | 4.9e-08 | 1.9e-08 | 5.600E-08 | 4.095E-08 | 1.900E-08 | 1.000E-07 |
| 8279 | 414 | L414Y | A | 1.9e-08 | 8.4e-08 | 4.9e-08 | 5.067E-08 | 3.253E-08 | 1.900E-08 | 8.400E-08 |
| 8280 | 414 | L414V | A | 7.1e-08 | 1e-07 | 4.6e-08 | 7.233E-08 | 2.702E-08 | 4.600E-08 | 1.000E-07 |
| 8281 | 415 | H415A | A | 2.1e-08 | 4e-08 | 3e-08 | 3.033E-08 | 9.504E-09 | 2.100E-08 | 4.000E-08 |
| 8282 | 415 | H415R | A | 9e-08 | 5.6e-08 | 1.9e-08 | 5.500E-08 | 3.551E-08 | 1.900E-08 | 9.000E-08 |
| 8283 | 415 | H415N | A | 6.2e-08 | 4.4e-08 | 9.8e-08 | 6.800E-08 | 2.750E-08 | 4.400E-08 | 9.800E-08 |
| 8284 | 415 | H415D | A | 6.7e-08 | 2.6e-08 | 2.8e-08 | 4.033E-08 | 2.312E-08 | 2.600E-08 | 6.700E-08 |
| 8285 | 415 | H415C | A | 1.3e-07 | 4e-08 | 6.4e-08 | 7.800E-08 | 4.660E-08 | 4.000E-08 | 1.300E-07 |
| 8286 | 415 | H415E | A | 1.6e-08 | 4.1e-08 | 2.9e-08 | 2.867E-08 | 1.250E-08 | 1.600E-08 | 4.100E-08 |
| 8287 | 415 | H415Q | A | 8.5e-09 | 3.2e-08 | 3.7e-08 | 2.583E-08 | 1.522E-08 | 8.500E-09 | 3.700E-08 |
| 8288 | 415 | H415G | A | 2.2e-08 | 6.8e-08 | 4.8e-08 | 4.600E-08 | 2.307E-08 | 2.200E-08 | 6.800E-08 |
| 8289 | 415 | H415H | A | 4.1e-08 | 7e-08 | 4.2e-08 | 5.100E-08 | 1.646E-08 | 4.100E-08 | 7.000E-08 |
| 8290 | 415 | H415I | A | 6.9e-08 | 3e-08 | 5.6e-08 | 5.167E-08 | 1.986E-08 | 3.000E-08 | 6.900E-08 |
| 8291 | 415 | H415L | A | 3.9e-08 | 6e-08 | 6.2e-08 | 5.367E-08 | 1.274E-08 | 3.900E-08 | 6.200E-08 |
| 8292 | 415 | H415K | A | 1.9e-08 | 3.5e-08 | 1.7e-08 | 2.367E-08 | 9.866E-09 | 1.700E-08 | 3.500E-08 |
| 8293 | 415 | H415M | A | 8.5e-08 | 4e-08 | 6.5e-08 | 6.333E-08 | 2.255E-08 | 4.000E-08 | 8.500E-08 |
| 8294 | 415 | H415F | A | 4.8e-08 | 4.1e-08 | 8.9e-08 | 5.933E-08 | 2.593E-08 | 4.100E-08 | 8.900E-08 |
| 8295 | 415 | H415P | A | 1.4e-07 | 2.9e-08 | 6.4e-08 | 7.767E-08 | 5.675E-08 | 2.900E-08 | 1.400E-07 |
| 8296 | 415 | H415S | A | 3.9e-08 | 2.1e-08 | 2.1e-08 | 2.700E-08 | 1.039E-08 | 2.100E-08 | 3.900E-08 |
| 8297 | 415 | H415T | A | 5e-08 | 2e-08 | 2.6e-08 | 3.200E-08 | 1.587E-08 | 2.000E-08 | 5.000E-08 |
| 8298 | 415 | H415W | A | 9.4e-08 | 4.7e-08 | 3.5e-08 | 5.867E-08 | 3.118E-08 | 3.500E-08 | 9.400E-08 |
| 8299 | 415 | H415Y | A | 3.1e-08 | 2.4e-08 | 2.4e-08 | 2.633E-08 | 4.041E-09 | 2.400E-08 | 3.100E-08 |
| 8300 | 415 | H415V | A | 3.6e-08 | 2.7e-08 | 9.6e-08 | 5.300E-08 | 3.751E-08 | 2.700E-08 | 9.600E-08 |
| 8301 | 416 | N416A | A | 1.9e-08 | 3.5e-08 | 2.3e-08 | 2.567E-08 | 8.327E-09 | 1.900E-08 | 3.500E-08 |
| 8302 | 416 | N416R | A | 9.1e-08 | 2.7e-08 | 8.3e-08 | 6.700E-08 | 3.487E-08 | 2.700E-08 | 9.100E-08 |
| 8303 | 416 | N416N | A | 3e-08 | 6.3e-08 | 5.9e-08 | 5.067E-08 | 1.801E-08 | 3.000E-08 | 6.300E-08 |
| 8304 | 416 | N416D | A | 4.4e-08 | 2.6e-08 | 6.8e-08 | 4.600E-08 | 2.107E-08 | 2.600E-08 | 6.800E-08 |
| 8305 | 416 | N416C | A | 2.2e-08 | 6.4e-08 | 2.7e-08 | 3.767E-08 | 2.294E-08 | 2.200E-08 | 6.400E-08 |
| 8306 | 416 | N416E | A | 2.5e-08 | 2e-08 | 3.8e-08 | 2.767E-08 | 9.292E-09 | 2.000E-08 | 3.800E-08 |
| 8307 | 416 | N416Q | A | 2.4e-08 | 1.6e-07 | 9.9e-08 | 9.433E-08 | 6.812E-08 | 2.400E-08 | 1.600E-07 |
| 8308 | 416 | N416G | A | 1.4e-07 | 2.7e-08 | 4.5e-08 | 7.067E-08 | 6.072E-08 | 2.700E-08 | 1.400E-07 |
| 8309 | 416 | N416H | A | 1.7e-08 | 1.4e-08 | 3.1e-08 | 2.067E-08 | 9.074E-09 | 1.400E-08 | 3.100E-08 |
| 8310 | 416 | N416I | A | 3.8e-08 | 3.8e-08 | 8.4e-08 | 5.333E-08 | 2.656E-08 | 3.800E-08 | 8.400E-08 |
| 8311 | 416 | N416L | A | 3.1e-08 | 4.8e-08 | 9.7e-08 | 5.867E-08 | 3.427E-08 | 3.100E-08 | 9.700E-08 |
| 8312 | 416 | N416K | A | 9.6e-08 | 2.8e-07 | 4.4e-08 | 1.400E-07 | 1.240E-07 | 4.400E-08 | 2.800E-07 |
| 8313 | 416 | N416M | A | 3.8e-08 | 1e-07 | 7.6e-08 | 7.133E-08 | 3.126E-08 | 3.800E-08 | 1.000E-07 |
| 8314 | 416 | N416F | A | 3.7e-08 | 6.9e-08 | 3.7e-08 | 4.767E-08 | 1.848E-08 | 3.700E-08 | 6.900E-08 |
| 8315 | 416 | N416P | A | 1.7e-08 | 5.2e-08 | 3.7e-08 | 3.533E-08 | 1.756E-08 | 1.700E-08 | 5.200E-08 |
| 8316 | 416 | N416S | A | 5.1e-08 | 5.9e-08 | 4.1e-08 | 5.033E-08 | 9.018E-09 | 4.100E-08 | 5.900E-08 |
| 8317 | 416 | N416T | A | 8.2e-08 | 3e-08 | 2.9e-08 | 4.700E-08 | 3.032E-08 | 2.900E-08 | 8.200E-08 |
| 8318 | 416 | N416W | A | 4.6e-08 | 1.1e-08 | 1.3e-08 | 2.333E-08 | 1.966E-08 | 1.100E-08 | 4.600E-08 |
| 8319 | 416 | N416Y | A | 3.9e-08 | 5.8e-08 | 3.4e-08 | 4.367E-08 | 1.266E-08 | 3.400E-08 | 5.800E-08 |
| 8320 | 416 | N416V | A | 1.4e-08 | 1.6e-08 | 2.3e-08 | 1.767E-08 | 4.726E-09 | 1.400E-08 | 2.300E-08 |
| 8321 | 417 | G417A | A | 7.1e-08 | 7.3e-08 | 7e-08 | 7.133E-08 | 1.528E-09 | 7.000E-08 | 7.300E-08 |
| 8322 | 417 | G417R | A | 2.2e-08 | 3.4e-08 | 3.4e-08 | 3.000E-08 | 6.928E-09 | 2.200E-08 | 3.400E-08 |
| 8323 | 417 | G417N | A | 2.8e-08 | 5.9e-08 | 6.2e-08 | 4.967E-08 | 1.882E-08 | 2.800E-08 | 6.200E-08 |
| 8324 | 417 | G417D | A | 3.9e-08 | 1.2e-07 | 9.6e-08 | 8.500E-08 | 4.161E-08 | 3.900E-08 | 1.200E-07 |
| 8325 | 417 | G417C | A | 4.4e-08 | 1.8e-08 | 2.3e-08 | 2.833E-08 | 1.380E-08 | 1.800E-08 | 4.400E-08 |
| 8326 | 417 | G417E | A | 6.3e-08 | 5e-08 | 7.8e-08 | 6.367E-08 | 1.401E-08 | 5.000E-08 | 7.800E-08 |
| 8327 | 417 | G417Q | A | 5.5e-08 | 4.4e-08 | 2e-08 | 3.967E-08 | 1.790E-08 | 2.000E-08 | 5.500E-08 |
| 8328 | 417 | G417G | A | 4.1e-08 | 1.5e-08 | 5.2e-08 | 3.600E-08 | 1.900E-08 | 1.500E-08 | 5.200E-08 |
| 8329 | 417 | G417H | A | 5.8e-08 | 4.2e-08 | 1.6e-07 | 8.667E-08 | 6.401E-08 | 4.200E-08 | 1.600E-07 |

| | | | | | | | | | | |
|---|---|---|---|---|---|---|---|---|---|---|
| 8330 | 417 | G417I | A | 1.3e-08 | 2.1e-08 | 2.6e-08 | 2.000E-08 | 6.557E-09 | 1.300E-08 | 2.600E-08 |
| 8331 | 417 | G417L | A | 4.6e-08 | 4.2e-08 | 3.7e-08 | 4.167E-08 | 4.509E-09 | 3.700E-08 | 4.600E-08 |
| 8332 | 417 | G417K | A | 7e-08 | 4.5e-08 | 7.3e-08 | 6.267E-08 | 1.537E-08 | 4.500E-08 | 7.300E-08 |
| 8333 | 417 | G417M | A | 4.5e-08 | 6.1e-08 | 1.9e-08 | 4.167E-08 | 2.120E-08 | 1.900E-08 | 6.100E-08 |
| 8334 | 417 | G417F | A | 1.3e-07 | 1e-07 | 1.5e-08 | 8.167E-08 | 5.965E-08 | 1.500E-08 | 1.300E-07 |
| 8335 | 417 | G417P | A | 5.2e-08 | 8.3e-08 | 7.7e-08 | 7.067E-08 | 1.644E-08 | 5.200E-08 | 8.300E-08 |
| 8336 | 417 | G417S | A | 3.2e-08 | 7.8e-08 | 1.9e-08 | 4.300E-08 | 3.100E-08 | 1.900E-08 | 7.800E-08 |
| 8337 | 417 | G417T | A | 2.7e-08 | 3.4e-08 | 4.3e-08 | 3.467E-08 | 8.021E-09 | 2.700E-08 | 4.300E-08 |
| 8338 | 417 | G417W | A | 2e-08 | 2.2e-08 | 5e-08 | 3.067E-08 | 1.677E-08 | 2.000E-08 | 5.000E-08 |
| 8339 | 417 | G417Y | A | 7e-08 | 2.8e-08 | 3.8e-08 | 4.533E-08 | 2.194E-08 | 2.800E-08 | 7.000E-08 |
| 8340 | 417 | G417V | A | 5.3e-08 | 2.9e-08 | 2.5e-08 | 3.567E-08 | 1.514E-08 | 2.500E-08 | 5.300E-08 |
| 8341 | 418 | A418A | A | 5.3e-08 | 4.7e-08 | 8.7e-08 | 6.233E-08 | 2.157E-08 | 4.700E-08 | 8.700E-08 |
| 8342 | 418 | A418R | A | 1.6e-08 | 4.1e-08 | 5.8e-08 | 3.833E-08 | 2.113E-08 | 1.600E-08 | 5.800E-08 |
| 8343 | 418 | A418N | A | 3.6e-08 | 6.5e-08 | 5.6e-08 | 5.233E-08 | 1.484E-08 | 3.600E-08 | 6.500E-08 |
| 8344 | 418 | A418D | A | 2.7e-08 | 3.3e-08 | 2.4e-08 | 2.800E-08 | 4.583E-09 | 2.400E-08 | 3.300E-08 |
| 8345 | 418 | A418C | A | 2.3e-08 | 5.5e-08 | 6e-08 | 4.600E-08 | 2.007E-08 | 2.300E-08 | 6.000E-08 |
| 8346 | 418 | A418E | A | 2.2e-08 | 4.3e-08 | 3e-08 | 3.167E-08 | 1.060E-08 | 2.200E-08 | 4.300E-08 |
| 8347 | 418 | A418Q | A | 4e-08 | 2.1e-08 | 3e-08 | 3.033E-08 | 9.504E-09 | 2.100E-08 | 4.000E-08 |
| 8348 | 418 | A418G | A | 1.3e-08 | 3.7e-08 | 2.4e-08 | 2.467E-08 | 1.201E-08 | 1.300E-08 | 3.700E-08 |
| 8349 | 418 | A418H | A | 7.4e-08 | 3.7e-08 | 3.8e-08 | 4.967E-08 | 2.108E-08 | 3.700E-08 | 7.400E-08 |
| 8350 | 418 | A418I | A | 2.9e-08 | 6e-08 | 2.5e-08 | 3.800E-08 | 1.916E-08 | 2.500E-08 | 6.000E-08 |
| 8351 | 418 | A418L | A | 5.9e-08 | 8.6e-08 | 4.4e-08 | 6.300E-08 | 2.128E-08 | 4.400E-08 | 8.600E-08 |
| 8352 | 418 | A418K | A | 4.4e-08 | 1.2e-07 | 3.7e-08 | 6.700E-08 | 4.603E-08 | 3.700E-08 | 1.200E-07 |
| 8353 | 418 | A418M | A | 5.4e-08 | 3e-08 | 6e-08 | 4.800E-08 | 1.587E-08 | 3.000E-08 | 6.000E-08 |
| 8354 | 418 | A418F | A | 3e-08 | 4.1e-08 | 4.9e-08 | 4.000E-08 | 9.539E-09 | 3.000E-08 | 4.900E-08 |
| 8355 | 418 | A418P | A | 1.4e-08 | 4.7e-08 | 3.3e-08 | 3.133E-08 | 1.656E-08 | 1.400E-08 | 4.700E-08 |
| 8356 | 418 | A418S | A | 3.6e-08 | 5e-08 | 8.5e-08 | 5.700E-08 | 2.524E-08 | 3.600E-08 | 8.500E-08 |
| 8357 | 418 | A418T | A | 2.6e-08 | 2e-08 | 6.6e-08 | 3.733E-08 | 2.501E-08 | 2.000E-08 | 6.600E-08 |
| 8358 | 418 | A418W | A | 4.6e-08 | 4.4e-08 | 2.6e-08 | 3.867E-08 | 1.102E-08 | 2.600E-08 | 4.600E-08 |
| 8359 | 418 | A418Y | A | 3e-08 | 6e-08 | 4.6e-08 | 4.533E-08 | 1.501E-08 | 3.000E-08 | 6.000E-08 |
| 8360 | 418 | A418V | A | 2.3e-08 | 3.2e-08 | 4.9e-08 | 3.467E-08 | 1.320E-08 | 2.300E-08 | 4.900E-08 |
| 8361 | 419 | Y419A | A | 9.2e-09 | 2.3e-08 | 5.4e-08 | 2.873E-08 | 2.294E-08 | 9.200E-09 | 5.400E-08 |
| 8362 | 419 | Y419R | A | 2.6e-08 | 2.1e-08 | 4.4e-08 | 3.033E-08 | 1.210E-08 | 2.100E-08 | 4.400E-08 |
| 8363 | 419 | Y419N | A | 2.9e-08 | 4.4e-08 | 3.9e-08 | 3.733E-08 | 7.638E-09 | 2.900E-08 | 4.400E-08 |
| 8364 | 419 | Y419D | A | 5.4e-08 | 7e-08 | 1.1e-07 | 7.800E-08 | 2.884E-08 | 5.400E-08 | 1.100E-07 |
| 8365 | 419 | Y419C | A | 6.4e-08 | 1.9e-08 | 9.6e-08 | 5.967E-08 | 3.868E-08 | 1.900E-08 | 9.600E-08 |
| 8366 | 419 | Y419E | A | 2.1e-08 | 7e-08 | 6.6e-08 | 5.233E-08 | 2.721E-08 | 2.100E-08 | 7.000E-08 |
| 8367 | 419 | Y419Q | A | 8e-08 | 3.4e-08 | 3.3e-08 | 4.900E-08 | 2.685E-08 | 3.300E-08 | 8.000E-08 |
| 8368 | 419 | Y419G | A | 5.9e-08 | 6.6e-08 | 7.1e-08 | 6.533E-08 | 6.028E-09 | 5.900E-08 | 7.100E-08 |
| 8369 | 419 | Y419H | A | 6.6e-08 | 4.5e-08 | 5.9e-08 | 5.667E-08 | 1.069E-08 | 4.500E-08 | 6.600E-08 |
| 8370 | 419 | Y419I | A | 2.5e-08 | 5.2e-08 | 1.3e-08 | 3.000E-08 | 1.997E-08 | 1.300E-08 | 5.200E-08 |
| 8371 | 419 | Y419L | A | 3.6e-08 | 3.1e-08 | 2.4e-08 | 3.033E-08 | 6.028E-09 | 2.400E-08 | 3.600E-08 |
| 8372 | 419 | Y419K | A | 4.4e-08 | 4.1e-08 | 5.7e-08 | 4.733E-08 | 8.505E-09 | 4.100E-08 | 5.700E-08 |
| 8373 | 419 | Y419M | A | 1.2e-07 | 4.8e-08 | 3.3e-08 | 6.700E-08 | 4.651E-08 | 3.300E-08 | 1.200E-07 |
| 8374 | 419 | Y419F | A | 6.4e-08 | 2.7e-08 | 3e-08 | 4.033E-08 | 2.055E-08 | 2.700E-08 | 6.400E-08 |
| 8375 | 419 | Y419P | A | 2.4e-08 | 1.9e-08 | 4e-08 | 2.767E-08 | 1.097E-08 | 1.900E-08 | 4.000E-08 |
| 8376 | 419 | Y419S | A | 6.6e-08 | 2.4e-08 | 3.3e-08 | 4.100E-08 | 2.211E-08 | 2.400E-08 | 6.600E-08 |
| 8377 | 419 | Y419T | A | 6.6e-08 | 5.7e-08 | 2.8e-08 | 5.033E-08 | 1.986E-08 | 2.800E-08 | 6.600E-08 |
| 8378 | 419 | Y419W | A | 7.3e-08 | 4.3e-08 | 1.7e-08 | 4.433E-08 | 2.802E-08 | 1.700E-08 | 7.300E-08 |
| 8379 | 419 | Y419Y | A | 1.9e-08 | 2.8e-08 | 5.3e-08 | 3.333E-08 | 1.762E-08 | 1.900E-08 | 5.300E-08 |
| 8380 | 419 | Y419V | A | 3.8e-08 | 3.5e-08 | 2.8e-08 | 3.367E-08 | 5.132E-09 | 2.800E-08 | 3.800E-08 |
| 8381 | 420 | S420A | A | 1.1e-08 | 7.2e-08 | 5.9e-08 | 4.733E-08 | 3.213E-08 | 1.100E-08 | 7.200E-08 |
| 8382 | 420 | S420R | A | 2.7e-07 | 7.3e-08 | 2.1e-08 | 1.213E-07 | 1.313E-07 | 2.100E-08 | 2.700E-07 |
| 8383 | 420 | S420N | A | 2.9e-08 | 5.5e-08 | 2.9e-08 | 3.767E-08 | 1.501E-08 | 2.900E-08 | 5.500E-08 |
| 8384 | 420 | S420D | A | 8.1e-08 | 3.9e-08 | 1.3e-08 | 4.433E-08 | 3.431E-08 | 1.300E-08 | 8.100E-08 |
| 8385 | 420 | S420C | A | 5.6e-08 | 2.1e-08 | 3.8e-08 | 3.833E-08 | 1.750E-08 | 2.100E-08 | 5.600E-08 |
| 8386 | 420 | S420E | A | 6.2e-08 | 5.6e-08 | 1.2e-07 | 7.933E-08 | 3.535E-08 | 5.600E-08 | 1.200E-07 |
| 8387 | 420 | S420Q | A | 2.5e-08 | 5.1e-08 | 3.7e-08 | 3.767E-08 | 1.301E-08 | 2.500E-08 | 5.100E-08 |
| 8388 | 420 | S420G | A | 3.2e-08 | 6.5e-08 | 1.1e-07 | 6.900E-08 | 3.915E-08 | 3.200E-08 | 1.100E-07 |
| 8389 | 420 | S420H | A | 5.1e-08 | 2.8e-08 | 6.9e-08 | 4.933E-08 | 2.055E-08 | 2.800E-08 | 6.900E-08 |
| 8390 | 420 | S420I | A | 6.3e-08 | 2.4e-08 | 1.1e-08 | 3.267E-08 | 2.706E-08 | 1.100E-08 | 6.300E-08 |
| 8391 | 420 | S420L | A | 6.2e-08 | 2.8e-08 | 4.6e-08 | 4.533E-08 | 1.701E-08 | 2.800E-08 | 6.200E-08 |
| 8392 | 420 | S420K | A | 4.9e-08 | 4.7e-08 | 1.8e-07 | 9.200E-08 | 7.622E-08 | 4.700E-08 | 1.800E-07 |
| 8393 | 420 | S420M | A | 4.2e-08 | 9e-08 | 1.6e-08 | 4.933E-08 | 3.754E-08 | 1.600E-08 | 9.000E-08 |
| 8394 | 420 | S420F | A | 4.5e-08 | 6.8e-08 | 4.3e-08 | 5.200E-08 | 1.389E-08 | 4.300E-08 | 6.800E-08 |
| 8395 | 420 | S420P | A | 1.2e-08 | 4.2e-08 | 3.2e-08 | 2.867E-08 | 1.528E-08 | 1.200E-08 | 4.200E-08 |
| 8396 | 420 | S420S | A | 4.4e-08 | 1e-07 | 2.8e-08 | 5.733E-08 | 3.781E-08 | 2.800E-08 | 1.000E-07 |
| 8397 | 420 | S420T | A | 8e-08 | 2.9e-08 | 3.8e-08 | 4.900E-08 | 2.722E-08 | 2.900E-08 | 8.000E-08 |
| 8398 | 420 | S420W | A | 1.6e-08 | 1.2e-07 | 1.8e-08 | 5.133E-08 | 5.948E-08 | 1.600E-08 | 1.200E-07 |
| 8399 | 420 | S420Y | A | 9.7e-08 | 2.1e-08 | 3.7e-08 | 5.167E-08 | 4.007E-08 | 2.100E-08 | 9.700E-08 |
| 8400 | 420 | S420V | A | 2.9e-08 | 2.3e-08 | 5e-08 | 3.400E-08 | 1.418E-08 | 2.300E-08 | 5.000E-08 |
| 8401 | 421 | L421A | A | 2.1e-08 | 3.3e-08 | 2.5e-08 | 2.633E-08 | 6.110E-09 | 2.100E-08 | 3.300E-08 |
| 8402 | 421 | L421R | A | 3.4e-08 | 9.6e-08 | 6e-08 | 6.333E-08 | 3.113E-08 | 3.400E-08 | 9.600E-08 |
| 8403 | 421 | L421N | A | 4.4e-08 | 6.2e-08 | 1.1e-07 | 7.200E-08 | 3.412E-08 | 4.400E-08 | 1.100E-07 |
| 8404 | 421 | L421D | A | 6.6e-08 | 3.1e-08 | 2.9e-08 | 4.200E-08 | 2.081E-08 | 2.900E-08 | 6.600E-08 |
| 8405 | 421 | L421C | A | 8.7e-09 | 2.5e-08 | 1.6e-08 | 1.657E-08 | 8.165E-09 | 8.700E-09 | 2.500E-08 |
| 8406 | 421 | L421E | A | 5.1e-08 | 1.9e-08 | 5e-08 | 4.000E-08 | 1.819E-08 | 1.900E-08 | 5.100E-08 |
| 8407 | 421 | L421Q | A | 3e-08 | 5.4e-08 | 2.8e-08 | 3.733E-08 | 1.447E-08 | 2.800E-08 | 5.400E-08 |
| 8408 | 421 | L421G | A | 3.6e-08 | 2.3e-08 | 2e-08 | 2.633E-08 | 8.505E-09 | 2.000E-08 | 3.600E-08 |
| 8409 | 421 | L421H | A | 2.6e-08 | 1e-08 | 4.2e-08 | 2.600E-08 | 1.600E-08 | 1.000E-08 | 4.200E-08 |
| 8410 | 421 | L421I | A | 3.7e-08 | 5.5e-08 | 3.6e-08 | 4.267E-08 | 1.069E-08 | 3.600E-08 | 5.500E-08 |
| 8411 | 421 | L421L | A | 7.3e-08 | 4e-08 | 3.4e-08 | 4.900E-08 | 2.100E-08 | 3.400E-08 | 7.300E-08 |
| 8412 | 421 | L421K | A | 2.7e-08 | 3.6e-08 | 1.8e-08 | 2.700E-08 | 9.000E-09 | 1.800E-08 | 3.600E-08 |
| 8413 | 421 | L421M | A | 2.6e-08 | 3.4e-08 | 5.4e-08 | 3.800E-08 | 1.442E-08 | 2.600E-08 | 5.400E-08 |
| 8414 | 421 | L421F | A | 6.1e-08 | 1e-07 | 1.6e-08 | 5.900E-08 | 4.204E-08 | 1.600E-08 | 1.000E-07 |

| | | | | | | | | | | |
|---|---|---|---|---|---|---|---|---|---|---|
| 8415 | 421 | L421P | A | 3.6e-08 | 4e-08 | 6.4e-08 | 4.667E-08 | 1.514E-08 | 3.600E-08 | 6.400E-08 |
| 8416 | 421 | L421S | A | 3.8e-08 | 1.1e-07 | 5.1e-08 | 6.633E-08 | 3.837E-08 | 3.800E-08 | 1.100E-07 |
| 8417 | 421 | L421T | A | 6.4e-08 | 7.3e-08 | 2.2e-08 | 5.300E-08 | 2.722E-08 | 2.200E-08 | 7.300E-08 |
| 8418 | 421 | L421W | A | 3.8e-08 | 6.5e-08 | 5e-08 | 5.100E-08 | 1.353E-08 | 3.800E-08 | 6.500E-08 |
| 8419 | 421 | L421Y | A | 1.5e-08 | 2.6e-08 | 8.3e-08 | 4.133E-08 | 3.650E-08 | 1.500E-08 | 8.300E-08 |
| 8420 | 421 | L421V | A | 2.4e-08 | 3e-08 | 3.3e-08 | 2.900E-08 | 4.583E-09 | 2.400E-08 | 3.300E-08 |
| 8421 | 422 | T422A | A | 4.2e-08 | 3.3e-08 | 3.5e-08 | 3.667E-08 | 4.726E-09 | 3.300E-08 | 4.200E-08 |
| 8422 | 422 | T422R | A | 3e-08 | 2.9e-08 | 1.1e-08 | 2.333E-08 | 1.069E-08 | 1.100E-08 | 3.000E-08 |
| 8423 | 422 | T422N | A | 1.7e-08 | 4.2e-08 | 5.7e-08 | 3.867E-08 | 2.021E-08 | 1.700E-08 | 5.700E-08 |
| 8424 | 422 | T422D | A | 2.3e-08 | 5.6e-08 | 4.3e-08 | 4.067E-08 | 1.662E-08 | 2.300E-08 | 5.600E-08 |
| 8425 | 422 | T422C | A | 3e-08 | 1.6e-08 | 8.4e-08 | 4.333E-08 | 3.591E-08 | 1.600E-08 | 8.400E-08 |
| 8426 | 422 | T422E | A | 3.4e-08 | 1.9e-08 | 7.3e-08 | 4.200E-08 | 2.787E-08 | 1.900E-08 | 7.300E-08 |
| 8427 | 422 | T422Q | A | 7.9e-08 | 3.9e-08 | 4.1e-08 | 5.300E-08 | 2.254E-08 | 3.900E-08 | 7.900E-08 |
| 8428 | 422 | T422G | A | 3.9e-08 | 2.6e-08 | 2.6e-08 | 3.033E-08 | 7.506E-09 | 2.600E-08 | 3.900E-08 |
| 8429 | 422 | T422H | A | 5.5e-08 | 6.6e-08 | 4e-08 | 5.367E-08 | 1.305E-08 | 4.000E-08 | 6.600E-08 |
| 8430 | 422 | T422I | A | 5.6e-08 | 2.2e-08 | 1e-07 | 5.933E-08 | 3.911E-08 | 2.200E-08 | 1.000E-07 |
| 8431 | 422 | T422L | A | 5.9e-08 | 6.3e-08 | 3.2e-08 | 5.133E-08 | 1.686E-08 | 3.200E-08 | 6.300E-08 |
| 8432 | 422 | T422K | A | 3.3e-08 | 2.9e-08 | 3.4e-08 | 3.200E-08 | 2.646E-09 | 2.900E-08 | 3.400E-08 |
| 8433 | 422 | T422M | A | 2.5e-08 | 3e-08 | 6.9e-08 | 4.133E-08 | 2.409E-08 | 2.500E-08 | 6.900E-08 |
| 8434 | 422 | T422F | A | 4e-08 | 2.2e-08 | 1.1e-08 | 2.433E-08 | 1.464E-08 | 1.100E-08 | 4.000E-08 |
| 8435 | 422 | T422P | A | 1.5e-07 | 7.9e-08 | 3.1e-08 | 8.667E-08 | 5.987E-08 | 3.100E-08 | 1.500E-07 |
| 8436 | 422 | T422S | A | 2.8e-08 | 4.8e-08 | 4e-08 | 3.867E-08 | 1.007E-08 | 2.800E-08 | 4.800E-08 |
| 8437 | 422 | T422T | A | 2.4e-08 | 2e-08 | 3.6e-08 | 2.667E-08 | 8.327E-09 | 2.000E-08 | 3.600E-08 |
| 8438 | 422 | T422W | A | 1.1e-07 | 2.6e-08 | 7.7e-08 | 7.100E-08 | 4.232E-08 | 2.600E-08 | 1.100E-07 |
| 8439 | 422 | T422Y | A | 6.8e-08 | 3.1e-08 | 3.3e-08 | 4.400E-08 | 2.081E-08 | 3.100E-08 | 6.800E-08 |
| 8440 | 422 | T422V | A | 2.3e-08 | 3.2e-08 | 3.3e-08 | 2.933E-08 | 5.508E-09 | 2.300E-08 | 3.300E-08 |
| 8441 | 423 | L423A | A | 6.1e-08 | 4.2e-08 | 3.1e-08 | 4.467E-08 | 1.518E-08 | 3.100E-08 | 6.100E-08 |
| 8442 | 423 | L423R | A | 3.2e-08 | 1.4e-07 | 2.5e-08 | 6.567E-08 | 6.447E-08 | 2.500E-08 | 1.400E-07 |
| 8443 | 423 | L423N | A | 2.9e-08 | 9.5e-08 | 3.5e-08 | 5.300E-08 | 3.650E-08 | 2.900E-08 | 9.500E-08 |
| 8444 | 423 | L423D | A | 2.5e-08 | 6.8e-08 | 3.2e-08 | 4.167E-08 | 2.307E-08 | 2.500E-08 | 6.800E-08 |
| 8445 | 423 | L423C | A | 4.1e-08 | 6e-08 | 4.6e-08 | 4.900E-08 | 9.849E-09 | 4.100E-08 | 6.000E-08 |
| 8446 | 423 | L423E | A | 3.2e-08 | 9e-08 | 2.9e-08 | 5.033E-08 | 3.439E-08 | 2.900E-08 | 9.000E-08 |
| 8447 | 423 | L423Q | A | 7.5e-08 | 1.6e-08 | 4.3e-08 | 4.467E-08 | 2.954E-08 | 1.600E-08 | 7.500E-08 |
| 8448 | 423 | L423G | A | 8.8e-08 | 3.7e-08 | 1e-07 | 7.500E-08 | 3.345E-08 | 3.700E-08 | 1.000E-07 |
| 8449 | 423 | L423H | A | 2.5e-08 | 3.4e-08 | 2.9e-08 | 2.933E-08 | 4.509E-09 | 2.500E-08 | 3.400E-08 |
| 8450 | 423 | L423I | A | 5.5e-08 | 2.2e-08 | 8.7e-08 | 5.467E-08 | 3.250E-08 | 2.200E-08 | 8.700E-08 |
| 8451 | 423 | L423L | A | 6.8e-08 | 8.4e-08 | 5.2e-08 | 6.800E-08 | 1.600E-08 | 5.200E-08 | 8.400E-08 |
| 8452 | 423 | L423K | A | 4.8e-08 | 5.1e-08 | 3.5e-08 | 4.467E-08 | 8.505E-09 | 3.500E-08 | 5.100E-08 |
| 8453 | 423 | L423M | A | 7.1e-08 | 3.2e-08 | 1e-07 | 6.767E-08 | 3.412E-08 | 3.200E-08 | 1.000E-07 |
| 8454 | 423 | L423F | A | 2e-08 | 6.4e-08 | 4.1e-08 | 4.167E-08 | 2.201E-08 | 2.000E-08 | 6.400E-08 |
| 8455 | 423 | L423P | A | 4.9e-08 | 2.1e-08 | 7.9e-08 | 4.967E-08 | 2.901E-08 | 2.100E-08 | 7.900E-08 |
| 8456 | 423 | L423S | A | 2.7e-08 | 3.3e-08 | 3.5e-08 | 3.167E-08 | 4.163E-09 | 2.700E-08 | 3.500E-08 |
| 8457 | 423 | L423T | A | 5.9e-08 | 2.6e-08 | 4.6e-08 | 4.367E-08 | 1.662E-08 | 2.600E-08 | 5.900E-08 |
| 8458 | 423 | L423W | A | 8.3e-08 | 3.6e-08 | 3.5e-08 | 5.133E-08 | 2.743E-08 | 3.500E-08 | 8.300E-08 |
| 8459 | 423 | L423Y | A | 1.6e-08 | 3.1e-08 | 5e-08 | 3.233E-08 | 1.704E-08 | 1.600E-08 | 5.000E-08 |
| 8460 | 423 | L423V | A | 3.4e-08 | 2.6e-08 | 5.7e-08 | 3.900E-08 | 1.609E-08 | 2.600E-08 | 5.700E-08 |
| 8461 | 424 | Q424A | A | 3.7e-07 | 4.7e-08 | 2e-08 | 1.457E-07 | 1.947E-07 | 2.000E-08 | 3.700E-07 |
| 8462 | 424 | Q424R | A | 4e-08 | 5.1e-08 | 2.7e-08 | 3.933E-08 | 1.201E-08 | 2.700E-08 | 5.100E-08 |
| 8463 | 424 | Q424N | A | 3.6e-08 | 4.4e-08 | 1e-07 | 6.000E-08 | 3.487E-08 | 3.600E-08 | 1.000E-07 |
| 8464 | 424 | Q424D | A | 3.7e-08 | 6.1e-08 | 1.8e-08 | 3.867E-08 | 2.155E-08 | 1.800E-08 | 6.100E-08 |
| 8465 | 424 | Q424C | A | 2.6e-08 | 1.1e-08 | 5.6e-08 | 3.100E-08 | 2.291E-08 | 1.100E-08 | 5.600E-08 |
| 8466 | 424 | Q424E | A | 2.2e-08 | 3.2e-08 | 8.6e-08 | 4.667E-08 | 3.443E-08 | 2.200E-08 | 8.600E-08 |
| 8467 | 424 | Q424Q | A | 5.1e-08 | 7.8e-08 | 4.8e-08 | 5.900E-08 | 1.652E-08 | 4.800E-08 | 7.800E-08 |
| 8468 | 424 | Q424G | A | 3.5e-08 | 2.3e-08 | 3.6e-08 | 3.133E-08 | 7.234E-09 | 2.300E-08 | 3.600E-08 |
| 8469 | 424 | Q424H | A | 1.7e-08 | 2.9e-08 | 1.5e-08 | 2.033E-08 | 7.572E-09 | 1.500E-08 | 2.900E-08 |
| 8470 | 424 | Q424I | A | 2.2e-08 | 7.3e-08 | 2.4e-08 | 3.967E-08 | 2.888E-08 | 2.200E-08 | 7.300E-08 |
| 8471 | 424 | Q424L | A | 3.8e-08 | 6.2e-08 | 4.1e-08 | 4.700E-08 | 1.308E-08 | 3.800E-08 | 6.200E-08 |
| 8472 | 424 | Q424K | A | 5.6e-08 | 3.2e-08 | 2.2e-08 | 3.667E-08 | 1.747E-08 | 2.200E-08 | 5.600E-08 |
| 8473 | 424 | Q424M | A | 3.1e-08 | 2.9e-08 | 4.2e-08 | 3.400E-08 | 7.000E-09 | 2.900E-08 | 4.200E-08 |
| 8474 | 424 | Q424F | A | 4.1e-08 | 3e-08 | 6.8e-08 | 4.633E-08 | 1.955E-08 | 3.000E-08 | 6.800E-08 |
| 8475 | 424 | Q424P | A | 3.5e-08 | 2.6e-08 | 4e-08 | 3.367E-08 | 7.095E-09 | 2.600E-08 | 4.000E-08 |
| 8476 | 424 | Q424S | A | 5.7e-08 | 2.2e-08 | 1.9e-08 | 3.267E-08 | 2.113E-08 | 1.900E-08 | 5.700E-08 |
| 8477 | 424 | Q424T | A | 2.3e-08 | 2.1e-08 | 3.4e-08 | 2.600E-08 | 7.000E-09 | 2.100E-08 | 3.400E-08 |
| 8478 | 424 | Q424W | A | 1.5e-08 | 2.8e-08 | 6e-08 | 3.433E-08 | 2.316E-08 | 1.500E-08 | 6.000E-08 |
| 8479 | 424 | Q424Y | A | 5.4e-08 | 1.6e-08 | 2.8e-08 | 3.267E-08 | 1.943E-08 | 1.600E-08 | 5.400E-08 |
| 8480 | 424 | Q424V | A | 2.6e-08 | 2.8e-08 | 6.7e-08 | 4.033E-08 | 2.312E-08 | 2.600E-08 | 6.700E-08 |
| 8481 | 425 | G425A | A | 9.3e-08 | 3.1e-08 | 2.8e-08 | 5.067E-08 | 3.669E-08 | 2.800E-08 | 9.300E-08 |
| 8482 | 425 | G425R | A | 1.9e-08 | 1.1e-07 | 3.9e-08 | 5.600E-08 | 4.782E-08 | 1.900E-08 | 1.100E-07 |
| 8483 | 425 | G425N | A | 2.9e-08 | 2.9e-08 | 7.9e-08 | 4.567E-08 | 2.887E-08 | 2.900E-08 | 7.900E-08 |
| 8484 | 425 | G425D | A | 4.1e-08 | 4.2e-08 | 2.6e-08 | 3.633E-08 | 8.963E-09 | 2.600E-08 | 4.200E-08 |
| 8485 | 425 | G425C | A | 3.3e-08 | 8.4e-08 | 2.8e-08 | 4.833E-08 | 3.099E-08 | 2.800E-08 | 8.400E-08 |
| 8486 | 425 | G425E | A | 3.1e-08 | 5.6e-08 | 5.3e-08 | 4.667E-08 | 1.365E-08 | 3.100E-08 | 5.600E-08 |
| 8487 | 425 | G425Q | A | 3.8e-08 | 8.4e-08 | 6.4e-08 | 6.200E-08 | 2.307E-08 | 3.800E-08 | 8.400E-08 |
| 8488 | 425 | G425G | A | 6.3e-08 | 1.2e-07 | 6.6e-08 | 8.300E-08 | 3.208E-08 | 6.300E-08 | 1.200E-07 |
| 8489 | 425 | G425H | A | 8.3e-09 | 1.1e-07 | 5.2e-08 | 5.677E-08 | 5.102E-08 | 8.300E-09 | 1.100E-07 |
| 8490 | 425 | G425I | A | 8.7e-08 | 3.1e-08 | 2.1e-08 | 4.633E-08 | 3.557E-08 | 2.100E-08 | 8.700E-08 |
| 8491 | 425 | G425L | A | 5.2e-08 | 3.1e-08 | 9.7e-08 | 6.000E-08 | 3.372E-08 | 3.100E-08 | 9.700E-08 |
| 8492 | 425 | G425K | A | 3.4e-08 | 1.8e-08 | 3.3e-08 | 2.833E-08 | 8.963E-09 | 1.800E-08 | 3.400E-08 |
| 8493 | 425 | G425M | A | 3.5e-08 | 5.1e-08 | 4.7e-08 | 4.433E-08 | 8.327E-09 | 3.500E-08 | 5.100E-08 |
| 8494 | 425 | G425F | A | 1.8e-08 | 6.7e-09 | 8e-08 | 3.490E-08 | 3.946E-08 | 6.700E-09 | 8.000E-08 |
| 8495 | 425 | G425P | A | 4.5e-08 | 2.3e-08 | 1.6e-08 | 2.800E-08 | 1.513E-08 | 1.600E-08 | 4.500E-08 |
| 8496 | 425 | G425S | A | 1.6e-08 | 3.1e-08 | 1.5e-08 | 2.067E-08 | 8.963E-09 | 1.500E-08 | 3.100E-08 |
| 8497 | 425 | G425T | A | 3.3e-08 | 3e-08 | 3e-08 | 3.100E-08 | 1.732E-09 | 3.000E-08 | 3.300E-08 |
| 8498 | 425 | G425W | A | 3.5e-08 | 5.7e-08 | 2.9e-08 | 4.033E-08 | 1.474E-08 | 2.900E-08 | 5.700E-08 |
| 8499 | 425 | G425Y | A | 9.1e-08 | 5.3e-08 | 4.8e-08 | 6.400E-08 | 2.352E-08 | 4.800E-08 | 9.100E-08 |

| | | | | | | | | | | |
|---|---|---|---|---|---|---|---|---|---|---|
| 8500 | 425 | G425V | A | 7.1e-08 | 4.3e-08 | 3.9e-08 | 5.100E-08 | 1.744E-08 | 3.900E-08 | 7.100E-08 |
| 8501 | 426 | L426A | A | 4.9e-08 | 2.5e-08 | 8.2e-08 | 5.200E-08 | 2.862E-08 | 2.500E-08 | 8.200E-08 |
| 8502 | 426 | L426R | A | 2.3e-08 | 1.9e-08 | 3.5e-08 | 2.567E-08 | 8.327E-09 | 1.900E-08 | 3.500E-08 |
| 8503 | 426 | L426N | A | 1.2e-08 | 3.3e-08 | 4.2e-08 | 2.900E-08 | 1.539E-08 | 1.200E-08 | 4.200E-08 |
| 8504 | 426 | L426D | A | 6.3e-08 | 8e-08 | 4e-08 | 6.100E-08 | 2.007E-08 | 4.000E-08 | 8.000E-08 |
| 8505 | 426 | L426C | A | 3.8e-08 | 8.1e-08 | 4.1e-08 | 5.333E-08 | 2.401E-08 | 3.800E-08 | 8.100E-08 |
| 8506 | 426 | L426E | A | 6.7e-08 | 5.4e-08 | 4.3e-08 | 5.467E-08 | 1.201E-08 | 4.300E-08 | 6.700E-08 |
| 8507 | 426 | L426Q | A | 1.6e-08 | 2.6e-08 | 1.1e-07 | 5.067E-08 | 5.163E-08 | 1.600E-08 | 1.100E-07 |
| 8508 | 426 | L426G | A | 2.1e-08 | 1.5e-08 | 3.4e-08 | 2.333E-08 | 9.713E-09 | 1.500E-08 | 3.400E-08 |
| 8509 | 426 | L426H | A | 2.9e-08 | 8e-08 | 2.5e-08 | 4.467E-08 | 3.066E-08 | 2.500E-08 | 8.000E-08 |
| 8510 | 426 | L426I | A | 2e-08 | 1.9e-07 | 3.2e-08 | 8.067E-08 | 9.488E-08 | 2.000E-08 | 1.900E-07 |
| 8511 | 426 | L426L | A | 4.4e-08 | 4.8e-08 | 8.6e-08 | 5.933E-08 | 2.318E-08 | 4.400E-08 | 8.600E-08 |
| 8512 | 426 | L426K | A | 2.6e-08 | 6.8e-08 | 3.6e-08 | 4.333E-08 | 2.194E-08 | 2.600E-08 | 6.800E-08 |
| 8513 | 426 | L426M | A | 3.8e-08 | 6.5e-08 | 2.8e-08 | 4.367E-08 | 1.914E-08 | 2.800E-08 | 6.500E-08 |
| 8514 | 426 | L426F | A | 4.7e-08 | 4.3e-08 | 7.6e-08 | 5.533E-08 | 1.801E-08 | 4.300E-08 | 7.600E-08 |
| 8515 | 426 | L426P | A | 2.2e-08 | 7.2e-08 | 4.4e-08 | 4.600E-08 | 2.506E-08 | 2.200E-08 | 7.200E-08 |
| 8516 | 426 | L426S | A | 3.3e-08 | 2.2e-08 | 1.9e-08 | 2.467E-08 | 7.371E-09 | 1.900E-08 | 3.300E-08 |
| 8517 | 426 | L426T | A | 4e-08 | 4.5e-08 | 9.8e-08 | 6.100E-08 | 3.214E-08 | 4.000E-08 | 9.800E-08 |
| 8518 | 426 | L426W | A | 1.3e-07 | 1.9e-08 | 2.4e-08 | 5.767E-08 | 6.269E-08 | 1.900E-08 | 1.300E-07 |
| 8519 | 426 | L426Y | A | 1.3e-07 | 3.4e-08 | 3.9e-08 | 6.767E-08 | 5.404E-08 | 3.400E-08 | 1.300E-07 |
| 8520 | 426 | L426V | A | 7.2e-08 | 6.4e-08 | 2.3e-08 | 5.300E-08 | 2.629E-08 | 2.300E-08 | 7.200E-08 |
| 8521 | 427 | G427A | A | 8.4e-08 | 4.4e-08 | 2.6e-08 | 5.133E-08 | 2.969E-08 | 2.600E-08 | 8.400E-08 |
| 8522 | 427 | G427R | A | 2.9e-08 | 6.6e-08 | 8e-09 | 3.433E-08 | 2.937E-08 | 8.000E-09 | 6.600E-08 |
| 8523 | 427 | G427N | A | 5.5e-08 | 1e-07 | 6.5e-08 | 7.333E-08 | 2.363E-08 | 5.500E-08 | 1.000E-07 |
| 8524 | 427 | G427D | A | 2e-08 | 4.5e-08 | 2.9e-08 | 3.133E-08 | 1.266E-08 | 2.000E-08 | 4.500E-08 |
| 8525 | 427 | G427C | A | 3.2e-08 | 4.2e-08 | 1.6e-08 | 3.000E-08 | 1.311E-08 | 1.600E-08 | 4.200E-08 |
| 8526 | 427 | G427E | A | 4.7e-08 | 5.7e-08 | 2.4e-08 | 4.267E-08 | 1.692E-08 | 2.400E-08 | 5.700E-08 |
| 8527 | 427 | G427Q | A | 4.5e-08 | 1.1e-07 | 2.2e-08 | 5.900E-08 | 4.564E-08 | 2.200E-08 | 1.100E-07 |
| 8528 | 427 | G427G | A | 5.4e-08 | 2e-08 | 6.1e-08 | 4.500E-08 | 2.193E-08 | 2.000E-08 | 6.100E-08 |
| 8529 | 427 | G427H | A | 4e-08 | 2.2e-08 | 7e-08 | 4.400E-08 | 2.425E-08 | 2.200E-08 | 7.000E-08 |
| 8530 | 427 | G427I | A | 7.5e-08 | 4e-08 | 6.5e-08 | 6.000E-08 | 1.803E-08 | 4.000E-08 | 7.500E-08 |
| 8531 | 427 | G427L | A | 4.2e-08 | 2.6e-08 | 7.3e-08 | 4.700E-08 | 2.390E-08 | 2.600E-08 | 7.300E-08 |
| 8532 | 427 | G427K | A | 4.3e-08 | 3.7e-08 | 1.7e-07 | 8.333E-08 | 7.512E-08 | 3.700E-08 | 1.700E-07 |
| 8533 | 427 | G427M | A | 5.4e-08 | 1e-07 | 3.1e-08 | 6.167E-08 | 3.513E-08 | 3.100E-08 | 1.000E-07 |
| 8534 | 427 | G427F | A | 4.3e-08 | 1.4e-07 | 4.4e-08 | 7.567E-08 | 5.572E-08 | 4.300E-08 | 1.400E-07 |
| 8535 | 427 | G427P | A | 1e-07 | 3.6e-08 | 3.9e-08 | 5.833E-08 | 3.612E-08 | 3.600E-08 | 1.000E-07 |
| 8536 | 427 | G427S | A | 1.2e-07 | 6.5e-08 | 1.4e-08 | 6.633E-08 | 5.301E-08 | 1.400E-08 | 1.200E-07 |
| 8537 | 427 | G427T | A | 1.2e-08 | 6.8e-08 | 2.3e-08 | 3.433E-08 | 2.967E-08 | 1.200E-08 | 6.800E-08 |
| 8538 | 427 | G427W | A | 1.1e-07 | 2.6e-08 | 1.2e-08 | 4.933E-08 | 5.300E-08 | 1.200E-08 | 1.100E-07 |
| 8539 | 427 | G427Y | A | 1.1e-07 | 5e-08 | 5.4e-08 | 7.133E-08 | 3.355E-08 | 5.000E-08 | 1.100E-07 |
| 8540 | 427 | G427V | A | 6.2e-08 | 7.6e-08 | 6.4e-08 | 6.733E-08 | 7.572E-09 | 6.200E-08 | 7.600E-08 |
| 8541 | 428 | I428A | A | 1.3e-07 | 4.1e-08 | 2.8e-08 | 6.633E-08 | 5.552E-08 | 2.800E-08 | 1.300E-07 |
| 8542 | 428 | I428R | A | 1.4e-07 | 2.7e-08 | 6e-08 | 7.567E-08 | 5.811E-08 | 2.700E-08 | 1.400E-07 |
| 8543 | 428 | I428N | A | 2.2e-08 | 3.8e-08 | 1.7e-08 | 2.567E-08 | 1.097E-08 | 1.700E-08 | 3.800E-08 |
| 8544 | 428 | I428D | A | 6.9e-08 | 1.3e-08 | 4.2e-08 | 4.133E-08 | 2.801E-08 | 1.300E-08 | 6.900E-08 |
| 8545 | 428 | I428C | A | 5.8e-08 | 2.7e-08 | 3.2e-08 | 3.900E-08 | 1.664E-08 | 2.700E-08 | 5.800E-08 |
| 8546 | 428 | I428E | A | 4.1e-08 | 9.6e-08 | 8.6e-08 | 7.433E-08 | 2.930E-08 | 4.100E-08 | 9.600E-08 |
| 8547 | 428 | I428Q | A | 7.3e-08 | 4.3e-08 | 3.2e-08 | 4.933E-08 | 2.122E-08 | 3.200E-08 | 7.300E-08 |
| 8548 | 428 | I428G | A | 3.7e-08 | 3.2e-08 | 1.2e-07 | 6.300E-08 | 4.943E-08 | 3.200E-08 | 1.200E-07 |
| 8549 | 428 | I428H | A | 5e-08 | 7.6e-08 | 4.6e-08 | 5.733E-08 | 1.629E-08 | 4.600E-08 | 7.600E-08 |
| 8550 | 428 | I428I | A | 2.5e-08 | 6.1e-08 | 6.5e-08 | 5.033E-08 | 2.203E-08 | 2.500E-08 | 6.500E-08 |
| 8551 | 428 | I428L | A | 1.3e-08 | 1.4e-08 | 2.5e-08 | 1.733E-08 | 6.658E-09 | 1.300E-08 | 2.500E-08 |
| 8552 | 428 | I428K | A | 4e-08 | 2.6e-08 | 6.1e-08 | 4.233E-08 | 1.762E-08 | 2.600E-08 | 6.100E-08 |
| 8553 | 428 | I428M | A | 3.5e-08 | 8.1e-08 | 5e-08 | 5.533E-08 | 2.346E-08 | 3.500E-08 | 8.100E-08 |
| 8554 | 428 | I428F | A | 8.2e-08 | 1.2e-07 | 1.5e-07 | 1.173E-07 | 3.408E-08 | 8.200E-08 | 1.500E-07 |
| 8555 | 428 | I428P | A | 5.9e-08 | 6e-08 | 3.5e-08 | 5.133E-08 | 1.415E-08 | 3.500E-08 | 6.000E-08 |
| 8556 | 428 | I428S | A | 1.1e-07 | 2.6e-08 | 2.9e-08 | 5.500E-08 | 4.766E-08 | 2.600E-08 | 1.100E-07 |
| 8557 | 428 | I428T | A | 2.8e-08 | 5.1e-08 | 6.6e-08 | 4.833E-08 | 1.914E-08 | 2.800E-08 | 6.600E-08 |
| 8558 | 428 | I428W | A | 8e-08 | 2.8e-08 | 1.5e-08 | 4.100E-08 | 3.439E-08 | 1.500E-08 | 8.000E-08 |
| 8559 | 428 | I428Y | A | 3.9e-08 | 2.2e-08 | 3.9e-08 | 3.333E-08 | 9.815E-09 | 2.200E-08 | 3.900E-08 |
| 8560 | 428 | I428V | A | 4.3e-08 | 2.6e-08 | 1.5e-08 | 2.800E-08 | 1.411E-08 | 1.500E-08 | 4.300E-08 |
| 8561 | 429 | S429A | A | 8.9e-09 | 4e-08 | 4.8e-08 | 3.230E-08 | 2.066E-08 | 8.900E-09 | 4.800E-08 |
| 8562 | 429 | S429R | A | 2.6e-08 | 5.9e-08 | 5e-08 | 4.500E-08 | 1.706E-08 | 2.600E-08 | 5.900E-08 |
| 8563 | 429 | S429N | A | 5.5e-08 | 3.8e-08 | 2.9e-08 | 4.067E-08 | 1.320E-08 | 2.900E-08 | 5.500E-08 |
| 8564 | 429 | S429D | A | 1.8e-08 | 4.1e-08 | 2.6e-08 | 2.833E-08 | 1.168E-08 | 1.800E-08 | 4.100E-08 |
| 8565 | 429 | S429C | A | 2e-08 | 5.6e-08 | 4.1e-08 | 3.900E-08 | 1.808E-08 | 2.000E-08 | 5.600E-08 |
| 8566 | 429 | S429E | A | 3.3e-08 | 3.3e-08 | 8.1e-08 | 4.900E-08 | 2.771E-08 | 3.300E-08 | 8.100E-08 |
| 8567 | 429 | S429Q | A | 4e-08 | 6.4e-08 | 8.5e-08 | 6.300E-08 | 2.252E-08 | 4.000E-08 | 8.500E-08 |
| 8568 | 429 | S429G | A | 4.2e-08 | 3.5e-08 | 3e-07 | 1.257E-07 | 1.510E-07 | 3.500E-08 | 3.000E-07 |
| 8569 | 429 | S429H | A | 3.7e-08 | 4.6e-08 | 5.6e-08 | 4.633E-08 | 9.504E-09 | 3.700E-08 | 5.600E-08 |
| 8570 | 429 | S429I | A | 4.4e-08 | 6.4e-08 | 5e-08 | 5.267E-08 | 1.026E-08 | 4.400E-08 | 6.400E-08 |
| 8571 | 429 | S429L | A | 8.2e-08 | 3e-08 | 1e-07 | 7.067E-08 | 3.635E-08 | 3.000E-08 | 1.000E-07 |
| 8572 | 429 | S429K | A | 1.1e-08 | 7.9e-08 | 4.6e-08 | 4.533E-08 | 3.400E-08 | 1.100E-08 | 7.900E-08 |
| 8573 | 429 | S429M | A | 1.9e-08 | 5.7e-08 | 7.2e-08 | 4.933E-08 | 2.732E-08 | 1.900E-08 | 7.200E-08 |
| 8574 | 429 | S429F | A | 5.3e-08 | 4.7e-08 | 5.9e-08 | 5.300E-08 | 6.000E-09 | 4.700E-08 | 5.900E-08 |
| 8575 | 429 | S429P | A | 3.3e-08 | 6.2e-08 | 1.6e-08 | 3.700E-08 | 2.326E-08 | 1.600E-08 | 6.200E-08 |
| 8576 | 429 | S429S | A | 8.2e-08 | 3.1e-08 | 3.4e-08 | 4.900E-08 | 2.862E-08 | 3.100E-08 | 8.200E-08 |
| 8577 | 429 | S429T | A | 5.9e-08 | 2.3e-08 | 2.8e-08 | 3.667E-08 | 1.950E-08 | 2.300E-08 | 5.900E-08 |
| 8578 | 429 | S429W | A | 1e-07 | 6.5e-08 | 1.1e-07 | 9.167E-08 | 2.363E-08 | 6.500E-08 | 1.100E-07 |
| 8579 | 429 | S429Y | A | 5.5e-08 | 5.3e-08 | 1.3e-07 | 7.933E-08 | 4.389E-08 | 5.300E-08 | 1.300E-07 |
| 8580 | 429 | S429V | A | 4.3e-08 | 2.5e-08 | 8.5e-08 | 5.100E-08 | 3.079E-08 | 2.500E-08 | 8.500E-08 |
| 8581 | 430 | W430A | A | 6.6e-08 | 1.3e-07 | 4.1e-08 | 7.900E-08 | 4.590E-08 | 4.100E-08 | 1.300E-07 |
| 8582 | 430 | W430R | A | 5e-08 | 3.7e-08 | 4.3e-08 | 4.333E-08 | 6.506E-09 | 3.700E-08 | 5.000E-08 |
| 8583 | 430 | W430N | A | 8.6e-09 | 1.7e-08 | 3e-08 | 1.853E-08 | 1.078E-08 | 8.600E-09 | 3.000E-08 |
| 8584 | 430 | W430D | A | 5.9e-08 | 2.4e-08 | 3.5e-08 | 3.933E-08 | 1.790E-08 | 2.400E-08 | 5.900E-08 |

| | | | | | | | | | | |
|---|---|---|---|---|---|---|---|---|---|---|
| 8585 | 430 | W430C | A | 3.1e-08 | 1.3e-07 | 4.1e-08 | 6.733E-08 | 5.450E-08 | 3.100E-08 | 1.300E-07 |
| 8586 | 430 | W430E | A | 4e-08 | 1.4e-07 | 4e-08 | 7.333E-08 | 5.774E-08 | 4.000E-08 | 1.400E-07 |
| 8587 | 430 | W430Q | A | 5.9e-08 | 4.5e-08 | 3.1e-08 | 4.500E-08 | 1.400E-08 | 3.100E-08 | 5.900E-08 |
| 8588 | 430 | W430G | A | 2.1e-08 | 3.5e-08 | 5e-08 | 3.533E-08 | 1.450E-08 | 2.100E-08 | 5.000E-08 |
| 8589 | 430 | W430H | A | 6.1e-08 | 2.8e-08 | 7.2e-08 | 5.367E-08 | 2.290E-08 | 2.800E-08 | 7.200E-08 |
| 8590 | 430 | W430I | A | 3e-08 | 1e-07 | 3.6e-08 | 5.533E-08 | 3.880E-08 | 3.000E-08 | 1.000E-07 |
| 8591 | 430 | W430L | A | 8.9e-08 | 7.9e-08 | 7.8e-08 | 8.200E-08 | 6.083E-09 | 7.800E-08 | 8.900E-08 |
| 8592 | 430 | W430K | A | 3.9e-08 | 2.2e-08 | 1.9e-08 | 2.667E-08 | 1.079E-08 | 1.900E-08 | 3.900E-08 |
| 8593 | 430 | W430M | A | 3.4e-08 | 7.1e-08 | 2.9e-08 | 4.467E-08 | 2.294E-08 | 2.900E-08 | 7.100E-08 |
| 8594 | 430 | W430F | A | 3.9e-08 | 2e-08 | 3.3e-08 | 3.067E-08 | 9.713E-09 | 2.000E-08 | 3.900E-08 |
| 8595 | 430 | W430P | A | 7.6e-08 | 1.8e-08 | 5.4e-08 | 4.933E-08 | 2.928E-08 | 1.800E-08 | 7.600E-08 |
| 8596 | 430 | W430S | A | 2.2e-08 | 1.2e-07 | 2.7e-08 | 5.633E-08 | 5.519E-08 | 2.200E-08 | 1.200E-07 |
| 8597 | 430 | W430T | A | 6.2e-08 | 3.4e-08 | 2.5e-07 | 1.153E-07 | 1.175E-07 | 3.400E-08 | 2.500E-07 |
| 8598 | 430 | W430W | A | 1.4e-08 | 1.6e-08 | 2.1e-08 | 1.700E-08 | 3.606E-09 | 1.400E-08 | 2.100E-08 |
| 8599 | 430 | W430Y | A | 3.5e-08 | 7.1e-08 | 7.8e-08 | 6.133E-08 | 2.307E-08 | 3.500E-08 | 7.800E-08 |
| 8600 | 430 | W430V | A | 3.2e-08 | 1.2e-07 | 4.5e-08 | 6.567E-08 | 4.750E-08 | 3.200E-08 | 1.200E-07 |
| 8601 | 431 | L431A | A | 9.2e-08 | 2.5e-08 | 5.8e-08 | 5.833E-08 | 3.350E-08 | 2.500E-08 | 9.200E-08 |
| 8602 | 431 | L431R | A | 2.3e-08 | 2.9e-08 | 3.9e-08 | 3.033E-08 | 8.083E-09 | 2.300E-08 | 3.900E-08 |
| 8603 | 431 | L431N | A | 1.6e-08 | 5.4e-08 | 2.3e-08 | 3.100E-08 | 2.022E-08 | 1.600E-08 | 5.400E-08 |
| 8604 | 431 | L431D | A | 5.7e-08 | 3.8e-08 | 2.1e-08 | 3.867E-08 | 1.801E-08 | 2.100E-08 | 5.700E-08 |
| 8605 | 431 | L431C | A | 1.6e-08 | 1e-07 | 3.3e-08 | 4.967E-08 | 4.441E-08 | 1.600E-08 | 1.000E-07 |
| 8606 | 431 | L431E | A | 9.4e-08 | 5.3e-08 | 2.4e-08 | 5.700E-08 | 3.517E-08 | 2.400E-08 | 9.400E-08 |
| 8607 | 431 | L431Q | A | 2.1e-08 | 7.6e-08 | 8.9e-09 | 3.530E-08 | 3.576E-08 | 8.900E-09 | 7.600E-08 |
| 8608 | 431 | L431G | A | 2.5e-08 | 6.6e-08 | 9.1e-08 | 6.067E-08 | 3.332E-08 | 2.500E-08 | 9.100E-08 |
| 8609 | 431 | L431H | A | 2.6e-08 | 1.3e-08 | 1.2e-07 | 5.300E-08 | 5.839E-08 | 1.300E-08 | 1.200E-07 |
| 8610 | 431 | L431I | A | 1e-07 | 8.3e-08 | 1.9e-08 | 6.733E-08 | 4.271E-08 | 1.900E-08 | 1.000E-07 |
| 8611 | 431 | L431L | A | 3.9e-08 | 2.6e-08 | 5.2e-08 | 3.900E-08 | 1.300E-08 | 2.600E-08 | 5.200E-08 |
| 8612 | 431 | L431K | A | 4.5e-08 | 1.4e-08 | 3.5e-08 | 3.133E-08 | 1.582E-08 | 1.400E-08 | 4.500E-08 |
| 8613 | 431 | L431M | A | 3.5e-08 | 4.7e-08 | 3.5e-08 | 3.900E-08 | 6.928E-09 | 3.500E-08 | 4.700E-08 |
| 8614 | 431 | L431F | A | 1.5e-08 | 5.3e-08 | 3.1e-08 | 3.300E-08 | 1.908E-08 | 1.500E-08 | 5.300E-08 |
| 8615 | 431 | L431P | A | 2.9e-08 | 1e-07 | 3.5e-08 | 5.467E-08 | 3.937E-08 | 2.900E-08 | 1.000E-07 |
| 8616 | 431 | L431S | A | 3.6e-08 | 5.1e-08 | 5.6e-08 | 4.767E-08 | 1.041E-08 | 3.600E-08 | 5.600E-08 |
| 8617 | 431 | L431T | A | 4.5e-08 | 4.6e-08 | 3.4e-08 | 4.167E-08 | 6.658E-09 | 3.400E-08 | 4.600E-08 |
| 8618 | 431 | L431W | A | 3.3e-08 | 5.1e-08 | 1e-07 | 6.133E-08 | 3.467E-08 | 3.300E-08 | 1.000E-07 |
| 8619 | 431 | L431Y | A | 6.1e-08 | 3.5e-08 | 4.4e-08 | 4.667E-08 | 1.320E-08 | 3.500E-08 | 6.100E-08 |
| 8620 | 431 | L431V | A | 4.9e-08 | 5.1e-08 | 4.4e-08 | 4.800E-08 | 3.606E-09 | 4.400E-08 | 5.100E-08 |
| 8621 | 432 | G432A | A | 6.9e-08 | 4.3e-08 | 4e-08 | 5.067E-08 | 1.595E-08 | 4.000E-08 | 6.900E-08 |
| 8622 | 432 | G432R | A | 2.3e-08 | 3.4e-08 | 1.9e-08 | 2.533E-08 | 7.767E-09 | 1.900E-08 | 3.400E-08 |
| 8623 | 432 | G432N | A | 4e-08 | 2.7e-08 | 7e-08 | 4.567E-08 | 2.205E-08 | 2.700E-08 | 7.000E-08 |
| 8624 | 432 | G432D | A | 4.6e-08 | 4.2e-08 | 9.5e-09 | 3.250E-08 | 2.002E-08 | 9.500E-09 | 4.600E-08 |
| 8625 | 432 | G432C | A | 2.6e-08 | 6.2e-08 | 4.2e-08 | 4.333E-08 | 1.804E-08 | 2.600E-08 | 6.200E-08 |
| 8626 | 432 | G432E | A | 4.7e-08 | 3.9e-08 | 3.5e-08 | 4.033E-08 | 6.110E-09 | 3.500E-08 | 4.700E-08 |
| 8627 | 432 | G432Q | A | 8.1e-08 | 5.6e-08 | 9e-08 | 7.567E-08 | 1.762E-08 | 5.600E-08 | 9.000E-08 |
| 8628 | 432 | G432G | A | 1.1e-08 | 4.5e-08 | 5.2e-08 | 3.600E-08 | 2.193E-08 | 1.100E-08 | 5.200E-08 |
| 8629 | 432 | G432H | A | 1.9e-08 | 1.5e-08 | 2.6e-08 | 2.000E-08 | 5.568E-09 | 1.500E-08 | 2.600E-08 |
| 8630 | 432 | G432I | A | 1.7e-08 | 5.4e-08 | 2.7e-08 | 3.267E-08 | 1.914E-08 | 1.700E-08 | 5.400E-08 |
| 8631 | 432 | G432L | A | 8.2e-08 | 7.6e-08 | 5e-08 | 6.933E-08 | 1.701E-08 | 5.000E-08 | 8.200E-08 |
| 8632 | 432 | G432K | A | 6e-08 | 3.7e-08 | 7.5e-08 | 5.733E-08 | 1.914E-08 | 3.700E-08 | 7.500E-08 |
| 8633 | 432 | G432M | A | 3.6e-08 | 8.5e-08 | 7.1e-08 | 6.400E-08 | 2.524E-08 | 3.600E-08 | 8.500E-08 |
| 8634 | 432 | G432F | A | 1e-07 | 3.9e-08 | 9.1e-08 | 7.667E-08 | 3.293E-08 | 3.900E-08 | 1.000E-07 |
| 8635 | 432 | G432P | A | 5.3e-08 | 5.7e-08 | 7.2e-08 | 6.067E-08 | 1.002E-08 | 5.300E-08 | 7.200E-08 |
| 8636 | 432 | G432S | A | 7.2e-08 | 6.7e-08 | 4.1e-08 | 6.000E-08 | 1.664E-08 | 4.100E-08 | 7.200E-08 |
| 8637 | 432 | G432T | A | 3.5e-08 | 4e-08 | 3.7e-08 | 3.733E-08 | 2.517E-09 | 3.500E-08 | 4.000E-08 |
| 8638 | 432 | G432W | A | 4.8e-08 | 2.3e-08 | 8.1e-09 | 2.637E-08 | 2.016E-08 | 8.100E-09 | 4.800E-08 |
| 8639 | 432 | G432Y | A | 3.8e-08 | 3.7e-08 | 8.7e-08 | 5.400E-08 | 2.858E-08 | 3.700E-08 | 8.700E-08 |
| 8640 | 432 | G432V | A | 5.4e-08 | 3.8e-08 | 3.6e-08 | 4.267E-08 | 9.866E-09 | 3.600E-08 | 5.400E-08 |
| 8641 | 433 | L433A | A | 5.6e-08 | 2.4e-08 | 2.4e-08 | 3.467E-08 | 1.848E-08 | 2.400E-08 | 5.600E-08 |
| 8642 | 433 | L433R | A | 1.2e-07 | 8.4e-08 | 5.3e-09 | 6.977E-08 | 5.866E-08 | 5.300E-09 | 1.200E-07 |
| 8643 | 433 | L433N | A | 6.4e-08 | 4.4e-08 | 1.4e-08 | 4.067E-08 | 2.517E-08 | 1.400E-08 | 6.400E-08 |
| 8644 | 433 | L433D | A | 3.9e-08 | 4.8e-08 | 6.7e-08 | 5.133E-08 | 1.429E-08 | 3.900E-08 | 6.700E-08 |
| 8645 | 433 | L433C | A | 2.6e-08 | 3.6e-08 | 7.2e-08 | 4.467E-08 | 2.419E-08 | 2.600E-08 | 7.200E-08 |
| 8646 | 433 | L433E | A | 4.2e-08 | 5.7e-08 | 9.4e-08 | 6.433E-08 | 2.676E-08 | 4.200E-08 | 9.400E-08 |
| 8647 | 433 | L433Q | A | 2.8e-08 | 3.1e-08 | 4.6e-08 | 3.500E-08 | 9.644E-09 | 2.800E-08 | 4.600E-08 |
| 8648 | 433 | L433G | A | 7.6e-08 | 4.9e-08 | 5.1e-08 | 5.867E-08 | 1.504E-08 | 4.900E-08 | 7.600E-08 |
| 8649 | 433 | L433H | A | 7.5e-08 | 3.6e-08 | 2.2e-08 | 4.433E-08 | 2.747E-08 | 2.200E-08 | 7.500E-08 |
| 8650 | 433 | L433I | A | 5.3e-08 | 4.4e-08 | 2.1e-08 | 3.933E-08 | 1.650E-08 | 2.100E-08 | 5.300E-08 |
| 8651 | 433 | L433L | A | 8.5e-08 | 1e-07 | 2.3e-08 | 6.933E-08 | 4.082E-08 | 2.300E-08 | 1.000E-07 |
| 8652 | 433 | L433K | A | 1.2e-08 | 2.1e-08 | 3.1e-08 | 2.133E-08 | 9.504E-09 | 1.200E-08 | 3.100E-08 |
| 8653 | 433 | L433M | A | 6.3e-08 | 1.5e-07 | 3.1e-08 | 8.133E-08 | 6.158E-08 | 3.100E-08 | 1.500E-07 |
| 8654 | 433 | L433F | A | 1.8e-08 | 2.3e-08 | 4.8e-08 | 2.967E-08 | 1.607E-08 | 1.800E-08 | 4.800E-08 |
| 8655 | 433 | L433P | A | 3.1e-08 | 8.3e-08 | 2.1e-08 | 4.500E-08 | 3.329E-08 | 2.100E-08 | 8.300E-08 |
| 8656 | 433 | L433S | A | 2e-07 | 2.7e-08 | 2.3e-08 | 8.333E-08 | 1.011E-07 | 2.300E-08 | 2.000E-07 |
| 8657 | 433 | L433T | A | 3e-08 | 9.6e-08 | 8.9e-08 | 7.167E-08 | 3.625E-08 | 3.000E-08 | 9.600E-08 |
| 8658 | 433 | L433W | A | 6e-08 | 2.6e-08 | 8.4e-08 | 5.667E-08 | 2.914E-08 | 2.600E-08 | 8.400E-08 |
| 8659 | 433 | L433Y | A | 3.2e-08 | 7.2e-08 | 4.1e-08 | 4.833E-08 | 2.098E-08 | 3.200E-08 | 7.200E-08 |
| 8660 | 433 | L433V | A | 5e-08 | 1.7e-08 | 9.8e-08 | 5.500E-08 | 4.073E-08 | 1.700E-08 | 9.800E-08 |
| 8661 | 434 | R434A | A | 5.5e-08 | 4.5e-08 | 7.5e-08 | 5.833E-08 | 1.528E-08 | 4.500E-08 | 7.500E-08 |
| 8662 | 434 | R434R | A | 1.6e-08 | 4.9e-08 | 1.2e-08 | 2.567E-08 | 2.031E-08 | 1.200E-08 | 4.900E-08 |
| 8663 | 434 | R434N | A | 7.8e-08 | 4.5e-08 | 7.5e-08 | 6.600E-08 | 1.825E-08 | 4.500E-08 | 7.800E-08 |
| 8664 | 434 | R434D | A | 3.1e-08 | 1.3e-07 | 3.2e-08 | 6.433E-08 | 5.687E-08 | 3.100E-08 | 1.300E-07 |
| 8665 | 434 | R434C | A | 5.6e-08 | 9.3e-08 | 2.6e-08 | 5.833E-08 | 3.356E-08 | 2.600E-08 | 9.300E-08 |
| 8666 | 434 | R434E | A | 4.5e-08 | 6.2e-08 | 6.3e-08 | 5.667E-08 | 1.012E-08 | 4.500E-08 | 6.300E-08 |
| 8667 | 434 | R434Q | A | 1e-08 | 7.7e-08 | 5.8e-08 | 4.833E-08 | 3.453E-08 | 1.000E-08 | 7.700E-08 |
| 8668 | 434 | R434G | A | 1.1e-07 | 2.8e-08 | 3.7e-08 | 5.833E-08 | 4.497E-08 | 2.800E-08 | 1.100E-07 |
| 8669 | 434 | R434H | A | 7.3e-08 | 4.6e-08 | 3.8e-08 | 5.233E-08 | 1.834E-08 | 3.800E-08 | 7.300E-08 |

| | | | | | | | | | | |
|---|---|---|---|---|---|---|---|---|---|---|
| 8670 | 434 | R434I | A | 1.8e-08 | 4.5e-08 | 1.7e-08 | 2.667E-08 | 1.589E-08 | 1.700E-08 | 4.500E-08 |
| 8671 | 434 | R434L | A | 1.1e-07 | 8.7e-08 | 5.9e-08 | 8.533E-08 | 2.554E-08 | 5.900E-08 | 1.100E-07 |
| 8672 | 434 | R434K | A | 3.2e-08 | 3.6e-08 | 2.5e-08 | 3.100E-08 | 5.568E-09 | 2.500E-08 | 3.600E-08 |
| 8673 | 434 | R434M | A | 2.1e-08 | 3.4e-08 | 4.2e-08 | 3.233E-08 | 1.060E-08 | 2.100E-08 | 4.200E-08 |
| 8674 | 434 | R434F | A | 2.5e-08 | 3.5e-08 | 2.8e-08 | 2.933E-08 | 5.132E-09 | 2.500E-08 | 3.500E-08 |
| 8675 | 434 | R434P | A | 5.5e-08 | 2.9e-08 | 1.4e-07 | 7.467E-08 | 5.805E-08 | 2.900E-08 | 1.400E-07 |
| 8676 | 434 | R434S | A | 8.9e-09 | 4.6e-08 | 1.7e-08 | 2.397E-08 | 1.951E-08 | 8.900E-09 | 4.600E-08 |
| 8677 | 434 | R434T | A | 2.9e-08 | 7.3e-08 | 4.2e-08 | 4.800E-08 | 2.261E-08 | 2.900E-08 | 7.300E-08 |
| 8678 | 434 | R434W | A | 6.8e-08 | 2.3e-08 | 5.2e-08 | 4.767E-08 | 2.281E-08 | 2.300E-08 | 6.800E-08 |
| 8679 | 434 | R434Y | A | 2.2e-08 | 9.1e-08 | 3e-08 | 4.767E-08 | 3.774E-08 | 2.200E-08 | 9.100E-08 |
| 8680 | 434 | R434V | A | 6.7e-08 | 3.2e-08 | 5.2e-08 | 5.033E-08 | 1.756E-08 | 3.200E-08 | 6.700E-08 |
| 8681 | 435 | S435A | A | 6.6e-08 | 4.5e-08 | 6.6e-08 | 5.900E-08 | 1.212E-08 | 4.500E-08 | 6.600E-08 |
| 8682 | 435 | S435R | A | 2.2e-08 | 5.7e-08 | 8.3e-08 | 5.400E-08 | 3.061E-08 | 2.200E-08 | 8.300E-08 |
| 8683 | 435 | S435N | A | 1.6e-07 | 2.1e-08 | 5.8e-08 | 7.967E-08 | 7.199E-08 | 2.100E-08 | 1.600E-07 |
| 8684 | 435 | S435D | A | 3.6e-08 | 6.8e-08 | 5.7e-08 | 5.367E-08 | 1.626E-08 | 3.600E-08 | 6.800E-08 |
| 8685 | 435 | S435C | A | 3.5e-08 | 2.3e-08 | 7.7e-08 | 4.500E-08 | 2.835E-08 | 2.300E-08 | 7.700E-08 |
| 8686 | 435 | S435E | A | 3.9e-08 | 8.3e-08 | 2.3e-08 | 4.833E-08 | 3.107E-08 | 2.300E-08 | 8.300E-08 |
| 8687 | 435 | S435Q | A | 2.9e-08 | 1.2e-07 | 8e-08 | 7.633E-08 | 4.561E-08 | 2.900E-08 | 1.200E-07 |
| 8688 | 435 | S435G | A | 4e-08 | 9.9e-08 | 3.7e-08 | 5.867E-08 | 3.496E-08 | 3.700E-08 | 9.900E-08 |
| 8689 | 435 | S435H | A | 9.7e-08 | 5.3e-08 | 3.5e-08 | 6.167E-08 | 3.190E-08 | 3.500E-08 | 9.700E-08 |
| 8690 | 435 | S435I | A | 3.9e-08 | 1.6e-08 | 3e-08 | 2.833E-08 | 1.159E-08 | 1.600E-08 | 3.900E-08 |
| 8691 | 435 | S435L | A | 1.1e-08 | 3.8e-08 | 7.8e-08 | 4.233E-08 | 3.371E-08 | 1.100E-08 | 7.800E-08 |
| 8692 | 435 | S435K | A | 7.1e-08 | 3.7e-08 | 7.4e-08 | 6.067E-08 | 2.055E-08 | 3.700E-08 | 7.400E-08 |
| 8693 | 435 | S435M | A | 8.9e-08 | 3e-08 | 5.2e-08 | 5.700E-08 | 2.982E-08 | 3.000E-08 | 8.900E-08 |
| 8694 | 435 | S435F | A | 2.8e-08 | 3.7e-08 | 2.7e-08 | 3.067E-08 | 5.508E-09 | 2.700E-08 | 3.700E-08 |
| 8695 | 435 | S435P | A | 1e-07 | 3.1e-08 | 5.8e-08 | 6.300E-08 | 3.477E-08 | 3.100E-08 | 1.000E-07 |
| 8696 | 435 | S435S | A | 3.5e-08 | 2.1e-08 | 5.9e-08 | 3.833E-08 | 1.922E-08 | 2.100E-08 | 5.900E-08 |
| 8697 | 435 | S435T | A | 7.3e-08 | 6.2e-08 | 4e-08 | 5.833E-08 | 1.680E-08 | 4.000E-08 | 7.300E-08 |
| 8698 | 435 | S435W | A | 4.1e-08 | 8.8e-08 | 1.7e-08 | 4.867E-08 | 3.612E-08 | 1.700E-08 | 8.800E-08 |
| 8699 | 435 | S435Y | A | 4e-08 | 4.4e-08 | 2.7e-08 | 3.700E-08 | 8.888E-09 | 2.700E-08 | 4.400E-08 |
| 8700 | 435 | S435V | A | 3.8e-08 | 3.9e-08 | 8.4e-08 | 5.367E-08 | 2.627E-08 | 3.800E-08 | 8.400E-08 |
| 8701 | 436 | L436A | A | 1.9e-08 | 6.5e-08 | 2.5e-08 | 3.633E-08 | 2.501E-08 | 1.900E-08 | 6.500E-08 |
| 8702 | 436 | L436R | A | 2.5e-08 | 1.6e-08 | 5e-08 | 3.033E-08 | 1.762E-08 | 1.600E-08 | 5.000E-08 |
| 8703 | 436 | L436N | A | 1.2e-07 | 5.1e-08 | 3.2e-08 | 6.767E-08 | 4.631E-08 | 3.200E-08 | 1.200E-07 |
| 8704 | 436 | L436D | A | 2.8e-08 | 5.1e-08 | 3.9e-08 | 3.933E-08 | 1.150E-08 | 2.800E-08 | 5.100E-08 |
| 8705 | 436 | L436C | A | 3.8e-08 | 6.4e-08 | 6.9e-08 | 5.700E-08 | 1.664E-08 | 3.800E-08 | 6.900E-08 |
| 8706 | 436 | L436E | A | 7.8e-08 | 5.8e-08 | 1.9e-08 | 5.167E-08 | 3.001E-08 | 1.900E-08 | 7.800E-08 |
| 8707 | 436 | L436Q | A | 3.1e-08 | 3.2e-08 | 9.2e-08 | 5.167E-08 | 3.493E-08 | 3.100E-08 | 9.200E-08 |
| 8708 | 436 | L436G | A | 4.9e-08 | 3e-08 | 5.4e-08 | 4.433E-08 | 1.266E-08 | 3.000E-08 | 5.400E-08 |
| 8709 | 436 | L436H | A | 1.4e-08 | 4.7e-08 | 2.2e-08 | 2.767E-08 | 1.721E-08 | 1.400E-08 | 4.700E-08 |
| 8710 | 436 | L436I | A | 1.5e-08 | 5.5e-08 | 1.9e-07 | 8.667E-08 | 9.170E-08 | 1.500E-08 | 1.900E-07 |
| 8711 | 436 | L436L | A | 5.2e-08 | 8.4e-09 | 4.7e-08 | 3.580E-08 | 2.386E-08 | 8.400E-09 | 5.200E-08 |
| 8712 | 436 | L436K | A | 8.9e-08 | 2.1e-08 | 1e-07 | 7.000E-08 | 4.279E-08 | 2.100E-08 | 1.000E-07 |
| 8713 | 436 | L436M | A | 8.8e-08 | 6.1e-08 | 2e-08 | 5.633E-08 | 3.424E-08 | 2.000E-08 | 8.800E-08 |
| 8714 | 436 | L436F | A | 2.8e-08 | 4.2e-08 | 4.8e-08 | 3.933E-08 | 1.026E-08 | 2.800E-08 | 4.800E-08 |
| 8715 | 436 | L436P | A | 2.4e-08 | 1.6e-07 | 2.4e-08 | 6.933E-08 | 7.852E-08 | 2.400E-08 | 1.600E-07 |
| 8716 | 436 | L436S | A | 5e-08 | 2.6e-08 | 4.7e-08 | 4.100E-08 | 1.308E-08 | 2.600E-08 | 5.000E-08 |
| 8717 | 436 | L436T | A | 3.6e-08 | 1.3e-07 | 1.3e-07 | 9.867E-08 | 5.427E-08 | 3.600E-08 | 1.300E-07 |
| 8718 | 436 | L436W | A | 6.1e-08 | 2.5e-08 | 3.4e-08 | 4.000E-08 | 1.873E-08 | 2.500E-08 | 6.100E-08 |
| 8719 | 436 | L436Y | A | 4.7e-08 | 7.7e-08 | 3.6e-08 | 5.333E-08 | 2.122E-08 | 3.600E-08 | 7.700E-08 |
| 8720 | 436 | L436V | A | 6.6e-08 | 7.3e-08 | 6.2e-08 | 6.700E-08 | 5.568E-09 | 6.200E-08 | 7.300E-08 |
| 8721 | 437 | R437A | A | 1e-08 | 3.1e-08 | 2.7e-08 | 2.267E-08 | 1.115E-08 | 1.000E-08 | 3.100E-08 |
| 8722 | 437 | R437R | A | 1.1e-07 | 3.1e-08 | 6.5e-08 | 6.867E-08 | 3.963E-08 | 3.100E-08 | 1.100E-07 |
| 8723 | 437 | R437N | A | 2e-08 | 9.3e-08 | 2e-08 | 4.433E-08 | 4.215E-08 | 2.000E-08 | 9.300E-08 |
| 8724 | 437 | R437D | A | 1.8e-08 | 3.4e-08 | 9.5e-08 | 4.900E-08 | 4.063E-08 | 1.800E-08 | 9.500E-08 |
| 8725 | 437 | R437C | A | 1.8e-07 | 2.5e-08 | 3.3e-08 | 7.933E-08 | 8.727E-08 | 2.500E-08 | 1.800E-07 |
| 8726 | 437 | R437E | A | 1.1e-07 | 1e-07 | 1.5e-07 | 1.200E-07 | 2.646E-08 | 1.000E-07 | 1.500E-07 |
| 8727 | 437 | R437Q | A | 1.2e-08 | 1e-07 | 4e-08 | 5.067E-08 | 4.496E-08 | 1.200E-08 | 1.000E-07 |
| 8728 | 437 | R437G | A | 6.8e-08 | 4.4e-08 | 8.8e-08 | 6.667E-08 | 2.203E-08 | 4.400E-08 | 8.800E-08 |
| 8729 | 437 | R437H | A | 5e-08 | 2e-08 | 7.5e-08 | 4.833E-08 | 2.754E-08 | 2.000E-08 | 7.500E-08 |
| 8730 | 437 | R437I | A | 3.6e-08 | 2.1e-08 | 4.2e-08 | 3.300E-08 | 1.082E-08 | 2.100E-08 | 4.200E-08 |
| 8731 | 437 | R437L | A | 3.9e-08 | 1.7e-08 | 1.2e-08 | 2.267E-08 | 1.436E-08 | 1.200E-08 | 3.900E-08 |
| 8732 | 437 | R437K | A | 4.1e-08 | 5.4e-08 | 4.8e-08 | 4.767E-08 | 6.506E-09 | 4.100E-08 | 5.400E-08 |
| 8733 | 437 | R437M | A | 3.3e-08 | 1.2e-07 | 1.6e-07 | 1.043E-07 | 6.493E-08 | 3.300E-08 | 1.600E-07 |
| 8734 | 437 | R437F | A | 4.5e-08 | 4.1e-08 | 4.9e-08 | 4.500E-08 | 4.000E-09 | 4.100E-08 | 4.900E-08 |
| 8735 | 437 | R437P | A | 3.9e-08 | 3.4e-08 | 3.6e-08 | 3.633E-08 | 2.517E-09 | 3.400E-08 | 3.900E-08 |
| 8736 | 437 | R437S | A | 5.9e-08 | 6.1e-08 | 2.8e-08 | 4.933E-08 | 1.850E-08 | 2.800E-08 | 6.100E-08 |
| 8737 | 437 | R437T | A | 4.2e-08 | 3.6e-08 | 1e-07 | 5.933E-08 | 3.535E-08 | 3.600E-08 | 1.000E-07 |
| 8738 | 437 | R437W | A | 2e-08 | 1.2e-07 | 3.6e-08 | 5.867E-08 | 5.372E-08 | 2.000E-08 | 1.200E-07 |
| 8739 | 437 | R437Y | A | 1.9e-08 | 2.1e-08 | 3.7e-08 | 2.567E-08 | 9.866E-09 | 1.900E-08 | 3.700E-08 |
| 8740 | 437 | R437V | A | 3.6e-08 | 3.1e-08 | 3e-08 | 3.233E-08 | 3.215E-09 | 3.000E-08 | 3.600E-08 |
| 8741 | 438 | E438A | A | 2e-08 | 1e-07 | 6.7e-08 | 6.233E-08 | 4.020E-08 | 2.000E-08 | 1.000E-07 |
| 8742 | 438 | E438R | A | 9e-08 | 5.7e-08 | 3.7e-08 | 6.133E-08 | 2.676E-08 | 3.700E-08 | 9.000E-08 |
| 8743 | 438 | E438N | A | 1.4e-08 | 8.3e-08 | 3.5e-08 | 4.400E-08 | 3.537E-08 | 1.400E-08 | 8.300E-08 |
| 8744 | 438 | E438D | A | 1.8e-08 | 6.2e-08 | 8.9e-08 | 5.633E-08 | 3.584E-08 | 1.800E-08 | 8.900E-08 |
| 8745 | 438 | E438C | A | 4.4e-08 | 4.1e-08 | 9.2e-08 | 5.900E-08 | 2.862E-08 | 4.100E-08 | 9.200E-08 |
| 8746 | 438 | E438E | A | 4.2e-08 | 4.9e-08 | 4.6e-09 | 3.187E-08 | 2.387E-08 | 4.600E-09 | 4.900E-08 |
| 8747 | 438 | E438Q | A | 6.2e-08 | 5.3e-08 | 4.1e-08 | 5.200E-08 | 1.054E-08 | 4.100E-08 | 6.200E-08 |
| 8748 | 438 | E438G | A | 2e-08 | 4.7e-08 | 2.8e-08 | 3.167E-08 | 1.387E-08 | 2.000E-08 | 4.700E-08 |
| 8749 | 438 | E438H | A | 9.3e-09 | 5e-08 | 1.5e-08 | 2.477E-08 | 2.204E-08 | 9.300E-09 | 5.000E-08 |
| 8750 | 438 | E438I | A | 9.8e-08 | 6e-08 | 2.9e-08 | 6.233E-08 | 3.456E-08 | 2.900E-08 | 9.800E-08 |
| 8751 | 438 | E438L | A | 5.1e-08 | 1e-07 | 1e-07 | 8.367E-08 | 2.829E-08 | 5.100E-08 | 1.000E-07 |
| 8752 | 438 | E438K | A | 1.6e-08 | 3.7e-08 | 4.3e-08 | 3.200E-08 | 1.418E-08 | 1.600E-08 | 4.300E-08 |
| 8753 | 438 | E438M | A | 2.9e-08 | 3.2e-08 | 1.5e-08 | 2.533E-08 | 9.074E-09 | 1.500E-08 | 3.200E-08 |
| 8754 | 438 | E438F | A | 2.1e-08 | 2.8e-08 | 1.3e-07 | 5.967E-08 | 6.101E-08 | 2.100E-08 | 1.300E-07 |

| | | | | | | | | | | |
|---|---|---|---|---|---|---|---|---|---|---|
| 8755 | 438 | E438P | A | 4.6e-08 | 5.7e-08 | 5.1e-08 | 5.133E-08 | 5.508E-09 | 4.600E-08 | 5.700E-08 |
| 8756 | 438 | E438S | A | 6.5e-08 | 7.2e-08 | 8.2e-08 | 7.300E-08 | 8.544E-09 | 6.500E-08 | 8.200E-08 |
| 8757 | 438 | E438T | A | 7.9e-08 | 3.9e-08 | 2.8e-08 | 4.867E-08 | 2.684E-08 | 2.800E-08 | 7.900E-08 |
| 8758 | 438 | E438W | A | 6.8e-08 | 3.7e-08 | 4.6e-08 | 5.033E-08 | 1.595E-08 | 3.700E-08 | 6.800E-08 |
| 8759 | 438 | E438Y | A | 5.2e-08 | 1.1e-07 | 4.2e-08 | 6.800E-08 | 3.672E-08 | 4.200E-08 | 1.100E-07 |
| 8760 | 438 | E438V | A | 6.5e-08 | 2.3e-07 | 6.5e-08 | 1.200E-07 | 9.526E-08 | 6.500E-08 | 2.300E-07 |
| 8761 | 439 | L439A | A | 4.6e-08 | 2.8e-08 | 4.9e-08 | 4.100E-08 | 1.136E-08 | 2.800E-08 | 4.900E-08 |
| 8762 | 439 | L439R | A | 4e-08 | 7.9e-08 | 1.8e-08 | 4.567E-08 | 3.089E-08 | 1.800E-08 | 7.900E-08 |
| 8763 | 439 | L439N | A | 6.4e-08 | 5.4e-08 | 5.5e-08 | 5.767E-08 | 5.508E-09 | 5.400E-08 | 6.400E-08 |
| 8764 | 439 | L439D | A | 5e-08 | 1.1e-07 | 3.8e-08 | 6.600E-08 | 3.857E-08 | 3.800E-08 | 1.100E-07 |
| 8765 | 439 | L439C | A | 2.1e-08 | 4.8e-08 | 4.1e-08 | 3.667E-08 | 1.401E-08 | 2.100E-08 | 4.800E-08 |
| 8766 | 439 | L439E | A | 6.9e-08 | 5.5e-08 | 3.8e-08 | 5.400E-08 | 1.552E-08 | 3.800E-08 | 6.900E-08 |
| 8767 | 439 | L439Q | A | 1.7e-08 | 5e-08 | 4.3e-08 | 3.667E-08 | 1.739E-08 | 1.700E-08 | 5.000E-08 |
| 8768 | 439 | L439G | A | 1.2e-07 | 2.2e-08 | 5.3e-08 | 6.500E-08 | 5.009E-08 | 2.200E-08 | 1.200E-07 |
| 8769 | 439 | L439H | A | 6.7e-08 | 3e-08 | 3e-08 | 4.233E-08 | 2.136E-08 | 3.000E-08 | 6.700E-08 |
| 8770 | 439 | L439I | A | 3.2e-08 | 4.8e-08 | 2e-08 | 3.333E-08 | 1.405E-08 | 2.000E-08 | 4.800E-08 |
| 8771 | 439 | L439L | A | 6.5e-08 | 8.2e-08 | 2.1e-08 | 5.600E-08 | 3.148E-08 | 2.100E-08 | 8.200E-08 |
| 8772 | 439 | L439K | A | 3.5e-08 | 2.1e-08 | 3e-08 | 2.867E-08 | 7.095E-09 | 2.100E-08 | 3.500E-08 |
| 8773 | 439 | L439M | A | 1.1e-08 | 4.1e-08 | 6.9e-08 | 4.033E-08 | 2.901E-08 | 1.100E-08 | 6.900E-08 |
| 8774 | 439 | L439F | A | 1.1e-07 | 3e-07 | 3.8e-08 | 1.493E-07 | 1.354E-07 | 3.800E-08 | 3.000E-07 |
| 8775 | 439 | L439P | A | 4e-08 | 7.9e-08 | 1.8e-08 | 4.567E-08 | 3.089E-08 | 1.800E-08 | 7.900E-08 |
| 8776 | 439 | L439S | A | 3.3e-08 | 2.7e-08 | 6.3e-08 | 4.100E-08 | 1.929E-08 | 2.700E-08 | 6.300E-08 |
| 8777 | 439 | L439T | A | 9.4e-08 | 4.1e-08 | 3.8e-08 | 5.767E-08 | 3.150E-08 | 3.800E-08 | 9.400E-08 |
| 8778 | 439 | L439W | A | 8.2e-08 | 1.6e-08 | 1.4e-08 | 3.733E-08 | 3.870E-08 | 1.400E-08 | 8.200E-08 |
| 8779 | 439 | L439Y | A | 6e-08 | 1.3e-07 | 4.2e-08 | 7.733E-08 | 4.649E-08 | 4.200E-08 | 1.300E-07 |
| 8780 | 439 | L439V | A | 2.3e-08 | 5.9e-08 | 2.6e-08 | 3.600E-08 | 1.997E-08 | 2.300E-08 | 5.900E-08 |
| 8781 | 440 | G440A | A | 5.1e-08 | 6.6e-08 | 5.1e-08 | 5.600E-08 | 8.660E-09 | 5.100E-08 | 6.600E-08 |
| 8782 | 440 | G440R | A | 4.5e-08 | 2.5e-08 | 8.3e-08 | 5.100E-08 | 2.946E-08 | 2.500E-08 | 8.300E-08 |
| 8783 | 440 | G440N | A | 1.1e-07 | 2.8e-08 | 5.4e-08 | 6.400E-08 | 4.190E-08 | 2.800E-08 | 1.100E-07 |
| 8784 | 440 | G440D | A | 1.3e-07 | 1.8e-07 | 2.3e-08 | 1.110E-07 | 8.021E-08 | 2.300E-08 | 1.800E-07 |
| 8785 | 440 | G440C | A | 1.4e-07 | 4.1e-08 | 1.7e-08 | 6.600E-08 | 6.520E-08 | 1.700E-08 | 1.400E-07 |
| 8786 | 440 | G440E | A | 4.9e-08 | 8.8e-08 | 3.3e-08 | 5.667E-08 | 2.829E-08 | 3.300E-08 | 8.800E-08 |
| 8787 | 440 | G440Q | A | 5e-08 | 2.1e-08 | 2.8e-08 | 3.300E-08 | 1.513E-08 | 2.100E-08 | 5.000E-08 |
| 8788 | 440 | G440G | A | 2.3e-08 | 6.5e-08 | 3.8e-08 | 4.200E-08 | 2.128E-08 | 2.300E-08 | 6.500E-08 |
| 8789 | 440 | G440H | A | 7.2e-08 | 4.7e-08 | 4e-08 | 5.300E-08 | 1.682E-08 | 4.000E-08 | 7.200E-08 |
| 8790 | 440 | G440I | A | 7.3e-08 | 3.9e-08 | 4.9e-08 | 5.367E-08 | 1.747E-08 | 3.900E-08 | 7.300E-08 |
| 8791 | 440 | G440L | A | 7.3e-08 | 2.7e-08 | 4.1e-08 | 4.700E-08 | 2.358E-08 | 2.700E-08 | 7.300E-08 |
| 8792 | 440 | G440K | A | 6.2e-08 | 2.3e-08 | 4e-08 | 4.167E-08 | 1.955E-08 | 2.300E-08 | 6.200E-08 |
| 8793 | 440 | G440M | A | 5.8e-08 | 8.5e-08 | 3.6e-08 | 5.967E-08 | 2.454E-08 | 3.600E-08 | 8.500E-08 |
| 8794 | 440 | G440F | A | 2.5e-08 | 2.8e-08 | 3.3e-08 | 2.867E-08 | 4.041E-09 | 2.500E-08 | 3.300E-08 |
| 8795 | 440 | G440P | A | 2.5e-08 | 4.1e-08 | 1e-07 | 5.533E-08 | 3.950E-08 | 2.500E-08 | 1.000E-07 |
| 8796 | 440 | G440S | A | 4.9e-08 | 2.5e-08 | 2.7e-08 | 3.367E-08 | 1.332E-08 | 2.500E-08 | 4.900E-08 |
| 8797 | 440 | G440T | A | 3.1e-08 | 3.7e-08 | 9.3e-09 | 2.577E-08 | 1.457E-08 | 9.300E-09 | 3.700E-08 |
| 8798 | 440 | G440W | A | 7e-08 | 2.1e-08 | 9.2e-08 | 6.100E-08 | 3.635E-08 | 2.100E-08 | 9.200E-08 |
| 8799 | 440 | G440Y | A | 2.9e-08 | 5.9e-08 | 4.2e-08 | 4.333E-08 | 1.504E-08 | 2.900E-08 | 5.900E-08 |
| 8800 | 440 | G440V | A | 2.9e-08 | 1.4e-08 | 5e-08 | 3.100E-08 | 1.808E-08 | 1.400E-08 | 5.000E-08 |
| 8801 | 441 | S441A | A | 4e-08 | 4.1e-08 | 4.4e-08 | 4.167E-08 | 2.082E-09 | 4.000E-08 | 4.400E-08 |
| 8802 | 441 | S441R | A | 2.4e-08 | 5.3e-08 | 3.4e-08 | 3.700E-08 | 1.473E-08 | 2.400E-08 | 5.300E-08 |
| 8803 | 441 | S441N | A | 3.5e-08 | 5.2e-08 | 4.6e-08 | 4.433E-08 | 8.622E-09 | 3.500E-08 | 5.200E-08 |
| 8804 | 441 | S441D | A | 3.6e-08 | 5.4e-08 | 1.3e-08 | 3.433E-08 | 2.055E-08 | 1.300E-08 | 5.400E-08 |
| 8805 | 441 | S441C | A | 2.3e-08 | 2.5e-08 | 3.1e-08 | 2.633E-08 | 4.163E-09 | 2.300E-08 | 3.100E-08 |
| 8806 | 441 | S441E | A | 4.3e-08 | 4.7e-08 | 1.8e-08 | 3.600E-08 | 1.572E-08 | 1.800E-08 | 4.700E-08 |
| 8807 | 441 | S441Q | A | 1.3e-08 | 8.7e-08 | 1.5e-08 | 3.833E-08 | 4.216E-08 | 1.300E-08 | 8.700E-08 |
| 8808 | 441 | S441G | A | 2.4e-08 | 1.5e-08 | 8.7e-08 | 4.200E-08 | 3.923E-08 | 1.500E-08 | 8.700E-08 |
| 8809 | 441 | S441H | A | 8.9e-09 | 2.2e-08 | 2.4e-08 | 1.830E-08 | 8.202E-09 | 8.900E-09 | 2.400E-08 |
| 8810 | 441 | S441I | A | 2.1e-08 | 2.5e-08 | 8.5e-08 | 4.367E-08 | 3.585E-08 | 2.100E-08 | 8.500E-08 |
| 8811 | 441 | S441L | A | 2.5e-08 | 6e-08 | 5e-08 | 4.500E-08 | 1.803E-08 | 2.500E-08 | 6.000E-08 |
| 8812 | 441 | S441K | A | 2.9e-08 | 3.2e-08 | 2e-08 | 2.700E-08 | 6.245E-09 | 2.000E-08 | 3.200E-08 |
| 8813 | 441 | S441M | A | 1.1e-07 | 3.4e-08 | 7.2e-08 | 7.200E-08 | 3.800E-08 | 3.400E-08 | 1.100E-07 |
| 8814 | 441 | S441F | A | 3.2e-08 | 2.2e-08 | 3.4e-08 | 2.933E-08 | 6.429E-09 | 2.200E-08 | 3.400E-08 |
| 8815 | 441 | S441P | A | 5.2e-08 | 3.4e-08 | 5.4e-08 | 4.667E-08 | 1.102E-08 | 3.400E-08 | 5.400E-08 |
| 8816 | 441 | S441S | A | 6.2e-08 | 5.9e-08 | 7.1e-08 | 6.400E-08 | 6.245E-09 | 5.900E-08 | 7.100E-08 |
| 8817 | 441 | S441T | A | 6.4e-08 | 1.8e-08 | 1.7e-08 | 3.300E-08 | 2.685E-08 | 1.700E-08 | 6.400E-08 |
| 8818 | 441 | S441W | A | 7.4e-08 | 6e-08 | 4.4e-08 | 5.933E-08 | 1.501E-08 | 4.400E-08 | 7.400E-08 |
| 8819 | 441 | S441Y | A | 8.7e-08 | 5.5e-08 | 4.9e-08 | 6.367E-08 | 2.043E-08 | 4.900E-08 | 8.700E-08 |
| 8820 | 441 | S441V | A | 6.4e-08 | 2e-08 | 8.6e-08 | 5.667E-08 | 3.361E-08 | 2.000E-08 | 8.600E-08 |
| 8821 | 442 | G442A | A | 3.6e-08 | 2.9e-08 | 1.1e-07 | 5.833E-08 | 4.488E-08 | 2.900E-08 | 1.100E-07 |
| 8822 | 442 | G442R | A | 4.4e-08 | 4e-08 | 3.4e-08 | 3.933E-08 | 5.033E-09 | 3.400E-08 | 4.400E-08 |
| 8823 | 442 | G442N | A | 2.4e-08 | 2e-08 | 4.3e-08 | 2.900E-08 | 1.229E-08 | 2.000E-08 | 4.300E-08 |
| 8824 | 442 | G442D | A | 8.6e-08 | 2.4e-08 | 4.3e-08 | 5.100E-08 | 3.176E-08 | 2.400E-08 | 8.600E-08 |
| 8825 | 442 | G442C | A | 6.4e-08 | 3.7e-08 | 1.9e-08 | 4.000E-08 | 2.265E-08 | 1.900E-08 | 6.400E-08 |
| 8826 | 442 | G442E | A | 2.4e-08 | 1.1e-08 | 2.9e-08 | 2.133E-08 | 9.292E-09 | 1.100E-08 | 2.900E-08 |
| 8827 | 442 | G442Q | A | 9.7e-08 | 1.6e-08 | 5.7e-08 | 5.667E-08 | 4.050E-08 | 1.600E-08 | 9.700E-08 |
| 8828 | 442 | G442G | A | 2.1e-08 | 8.1e-08 | 2.8e-08 | 4.333E-08 | 3.281E-08 | 2.100E-08 | 8.100E-08 |
| 8829 | 442 | G442H | A | 2.2e-08 | 7e-08 | 2.4e-08 | 3.867E-08 | 2.715E-08 | 2.200E-08 | 7.000E-08 |
| 8830 | 442 | G442I | A | 5.4e-08 | 4.1e-08 | 1.8e-08 | 3.767E-08 | 1.823E-08 | 1.800E-08 | 5.400E-08 |
| 8831 | 442 | G442L | A | 1.1e-07 | 4.8e-08 | 3.9e-08 | 6.567E-08 | 3.866E-08 | 3.900E-08 | 1.100E-07 |
| 8832 | 442 | G442K | A | 5.5e-08 | 1e-08 | 1.5e-08 | 2.667E-08 | 2.466E-08 | 1.000E-08 | 5.500E-08 |
| 8833 | 442 | G442M | A | 1.3e-07 | 2.5e-08 | 7.2e-08 | 7.567E-08 | 5.260E-08 | 2.500E-08 | 1.300E-07 |
| 8834 | 442 | G442F | A | 5.7e-08 | 9.2e-08 | 4.7e-08 | 6.533E-08 | 2.363E-08 | 4.700E-08 | 9.200E-08 |
| 8835 | 442 | G442P | A | 4.6e-08 | 7.2e-08 | 4.8e-08 | 5.533E-08 | 1.447E-08 | 4.600E-08 | 7.200E-08 |
| 8836 | 442 | G442S | A | 8e-08 | 1e-07 | 2.5e-08 | 6.833E-08 | 3.884E-08 | 2.500E-08 | 1.000E-07 |
| 8837 | 442 | G442T | A | 8.5e-08 | 4e-08 | 2.5e-08 | 5.000E-08 | 3.122E-08 | 2.500E-08 | 8.500E-08 |
| 8838 | 442 | G442W | A | 8.2e-08 | 2.5e-08 | 9e-08 | 6.567E-08 | 3.544E-08 | 2.500E-08 | 9.000E-08 |
| 8839 | 442 | G442Y | A | 5e-08 | 6.3e-08 | 1.2e-07 | 7.767E-08 | 3.723E-08 | 5.000E-08 | 1.200E-07 |

| | | | | | | | | | | |
|---|---|---|---|---|---|---|---|---|---|---|
| 8840 | 442 | G442V | A | 3.4e-08 | 6.6e-08 | 5e-08 | 5.000E-08 | 1.600E-08 | 3.400E-08 | 6.600E-08 |
| 8841 | 443 | L443A | A | 8.9e-08 | 1.2e-07 | 1.3e-07 | 1.130E-07 | 2.138E-08 | 8.900E-08 | 1.300E-07 |
| 8842 | 443 | L443R | A | 6.9e-08 | 1.4e-08 | 5.2e-08 | 4.500E-08 | 2.816E-08 | 1.400E-08 | 6.900E-08 |
| 8843 | 443 | L443N | A | 1.6e-08 | 2.7e-08 | 1.9e-08 | 2.067E-08 | 5.686E-09 | 1.600E-08 | 2.700E-08 |
| 8844 | 443 | L443D | A | 3.6e-08 | 1.2e-08 | 6.6e-08 | 3.800E-08 | 2.706E-08 | 1.200E-08 | 6.600E-08 |
| 8845 | 443 | L443C | A | 8.5e-08 | 4.6e-08 | 5.7e-08 | 6.267E-08 | 2.011E-08 | 4.600E-08 | 8.500E-08 |
| 8846 | 443 | L443E | A | 3e-08 | 6.2e-08 | 8e-08 | 5.733E-08 | 2.532E-08 | 3.000E-08 | 8.000E-08 |
| 8847 | 443 | L443Q | A | 1.3e-07 | 4.6e-08 | 4e-08 | 7.200E-08 | 5.032E-08 | 4.000E-08 | 1.300E-07 |
| 8848 | 443 | L443G | A | 6.3e-08 | 6e-08 | 2e-08 | 4.767E-08 | 2.401E-08 | 2.000E-08 | 6.300E-08 |
| 8849 | 443 | L443H | A | 6e-08 | 2.6e-07 | 4.6e-08 | 1.220E-07 | 1.197E-07 | 4.600E-08 | 2.600E-07 |
| 8850 | 443 | L443I | A | 1.1e-07 | 8.2e-08 | 8.9e-08 | 9.367E-08 | 1.457E-08 | 8.200E-08 | 1.100E-07 |
| 8851 | 443 | L443L | A | 2.5e-08 | 1e-08 | 3.8e-08 | 2.433E-08 | 1.401E-08 | 1.000E-08 | 3.800E-08 |
| 8852 | 443 | L443K | A | 2.4e-08 | 9.9e-08 | 5.3e-08 | 5.867E-08 | 3.782E-08 | 2.400E-08 | 9.900E-08 |
| 8853 | 443 | L443M | A | 3.9e-08 | 5.4e-08 | 2.7e-08 | 4.000E-08 | 1.353E-08 | 2.700E-08 | 5.400E-08 |
| 8854 | 443 | L443F | A | 2.7e-08 | 2.1e-08 | 4.7e-08 | 3.167E-08 | 1.361E-08 | 2.100E-08 | 4.700E-08 |
| 8855 | 443 | L443P | A | 4.5e-08 | 6.8e-08 | 1e-07 | 7.100E-08 | 2.762E-08 | 4.500E-08 | 1.000E-07 |
| 8856 | 443 | L443S | A | 4.7e-08 | 5.4e-08 | 4.7e-08 | 4.933E-08 | 4.041E-09 | 4.700E-08 | 5.400E-08 |
| 8857 | 443 | L443T | A | 1.6e-08 | 4.6e-08 | 5.7e-08 | 3.967E-08 | 2.122E-08 | 1.600E-08 | 5.700E-08 |
| 8858 | 443 | L443W | A | 9.9e-08 | 4.3e-08 | 1.1e-07 | 8.400E-08 | 3.593E-08 | 4.300E-08 | 1.100E-07 |
| 8859 | 443 | L443Y | A | 3.4e-08 | 3.2e-08 | 3.1e-08 | 3.233E-08 | 1.528E-09 | 3.100E-08 | 3.400E-08 |
| 8860 | 443 | L443V | A | 3.2e-08 | 3.8e-08 | 2e-08 | 3.000E-08 | 9.165E-09 | 2.000E-08 | 3.800E-08 |
| 8861 | 444 | A444A | A | 4.1e-08 | 5.8e-08 | 4.2e-08 | 4.700E-08 | 9.539E-09 | 4.100E-08 | 5.800E-08 |
| 8862 | 444 | A444R | A | 4.4e-08 | 4.4e-08 | 2e-08 | 3.600E-08 | 1.386E-08 | 2.000E-08 | 4.400E-08 |
| 8863 | 444 | A444N | A | 2e-08 | 3.3e-08 | 1.1e-08 | 2.133E-08 | 1.106E-08 | 1.100E-08 | 3.300E-08 |
| 8864 | 444 | A444D | A | 5.1e-08 | 5.6e-08 | 4.5e-08 | 5.067E-08 | 5.508E-09 | 4.500E-08 | 5.600E-08 |
| 8865 | 444 | A444C | A | 3.8e-08 | 6.7e-08 | 5.1e-08 | 5.200E-08 | 1.453E-08 | 3.800E-08 | 6.700E-08 |
| 8866 | 444 | A444E | A | 3.1e-08 | 4.2e-08 | 3.6e-08 | 3.633E-08 | 5.508E-09 | 3.100E-08 | 4.200E-08 |
| 8867 | 444 | A444Q | A | 4e-08 | 2.9e-08 | 5e-08 | 3.967E-08 | 1.050E-08 | 2.900E-08 | 5.000E-08 |
| 8868 | 444 | A444G | A | 5.7e-08 | 2.7e-08 | 3.1e-08 | 3.833E-08 | 1.629E-08 | 2.700E-08 | 5.700E-08 |
| 8869 | 444 | A444H | A | 2.5e-08 | 7.8e-08 | 1.3e-08 | 3.867E-08 | 3.459E-08 | 1.300E-08 | 7.800E-08 |
| 8870 | 444 | A444I | A | 5e-08 | 6.3e-08 | 2.7e-08 | 4.667E-08 | 1.823E-08 | 2.700E-08 | 6.300E-08 |
| 8871 | 444 | A444L | A | 2.8e-08 | 3.5e-08 | 2.5e-08 | 2.933E-08 | 5.132E-09 | 2.500E-08 | 3.500E-08 |
| 8872 | 444 | A444K | A | 3.9e-08 | 2.1e-08 | 4.2e-08 | 3.400E-08 | 1.136E-08 | 2.100E-08 | 4.200E-08 |
| 8873 | 444 | A444M | A | 7.1e-08 | 5.8e-08 | 5.9e-08 | 6.267E-08 | 7.234E-09 | 5.800E-08 | 7.100E-08 |
| 8874 | 444 | A444F | A | 1e-08 | 1.1e-07 | 1.5e-08 | 4.500E-08 | 5.635E-08 | 1.000E-08 | 1.100E-07 |
| 8875 | 444 | A444P | A | 8.7e-08 | 4.1e-08 | 2e-08 | 4.933E-08 | 3.427E-08 | 2.000E-08 | 8.700E-08 |
| 8876 | 444 | A444S | A | 3.2e-08 | 3.8e-08 | 1.9e-08 | 2.967E-08 | 9.713E-09 | 1.900E-08 | 3.800E-08 |
| 8877 | 444 | A444T | A | 3.5e-08 | 5.1e-08 | 3.4e-08 | 4.000E-08 | 9.539E-09 | 3.400E-08 | 5.100E-08 |
| 8878 | 444 | A444W | A | 5.1e-08 | 5.8e-08 | 7.3e-08 | 6.067E-08 | 1.124E-08 | 5.100E-08 | 7.300E-08 |
| 8879 | 444 | A444Y | A | 3.7e-08 | 3.4e-08 | 7.3e-08 | 4.800E-08 | 2.170E-08 | 3.400E-08 | 7.300E-08 |
| 8880 | 444 | A444V | A | 2.4e-08 | 7.2e-08 | 6.7e-08 | 5.433E-08 | 2.639E-08 | 2.400E-08 | 7.200E-08 |
| 8881 | 445 | L445A | A | 1.1e-07 | 2.5e-08 | 8.1e-08 | 7.200E-08 | 4.321E-08 | 2.500E-08 | 1.100E-07 |
| 8882 | 445 | L445R | A | 1.2e-08 | 6.5e-08 | 3e-08 | 3.567E-08 | 2.695E-08 | 1.200E-08 | 6.500E-08 |
| 8883 | 445 | L445N | A | 6.9e-08 | 1.8e-08 | 2.7e-08 | 3.800E-08 | 2.722E-08 | 1.800E-08 | 6.900E-08 |
| 8884 | 445 | L445D | A | 1.3e-08 | 2.4e-08 | 8.3e-08 | 4.000E-08 | 3.764E-08 | 1.300E-08 | 8.300E-08 |
| 8885 | 445 | L445C | A | 5.7e-08 | 4.4e-08 | 5.4e-08 | 5.167E-08 | 6.807E-09 | 4.400E-08 | 5.700E-08 |
| 8886 | 445 | L445E | A | 5.1e-08 | 8.4e-08 | 2.2e-08 | 5.233E-08 | 3.102E-08 | 2.200E-08 | 8.400E-08 |
| 8887 | 445 | L445Q | A | 2.6e-08 | 3e-08 | 2.8e-08 | 2.800E-08 | 2.000E-09 | 2.600E-08 | 3.000E-08 |
| 8888 | 445 | L445G | A | 5.2e-08 | 6.4e-09 | 3.7e-08 | 3.180E-08 | 2.324E-08 | 6.400E-09 | 5.200E-08 |
| 8889 | 445 | L445H | A | 4e-08 | 2.8e-08 | 6.9e-08 | 4.567E-08 | 2.108E-08 | 2.800E-08 | 6.900E-08 |
| 8890 | 445 | L445I | A | 8.7e-08 | 5.7e-08 | 3.8e-08 | 6.067E-08 | 2.470E-08 | 3.800E-08 | 8.700E-08 |
| 8891 | 445 | L445L | A | 3.2e-08 | 3.9e-08 | 3.6e-08 | 3.567E-08 | 3.512E-09 | 3.200E-08 | 3.900E-08 |
| 8892 | 445 | L445K | A | 2.2e-08 | 3e-08 | 4.3e-08 | 3.167E-08 | 1.060E-08 | 2.200E-08 | 4.300E-08 |
| 8893 | 445 | L445M | A | 1.3e-07 | 3.6e-08 | 2.2e-08 | 6.267E-08 | 5.873E-08 | 2.200E-08 | 1.300E-07 |
| 8894 | 445 | L445F | A | 1.1e-07 | 7.1e-08 | 4.6e-08 | 7.567E-08 | 3.225E-08 | 4.600E-08 | 1.100E-07 |
| 8895 | 445 | L445P | A | 1.8e-08 | 3.5e-08 | 3.3e-08 | 2.867E-08 | 9.292E-09 | 1.800E-08 | 3.500E-08 |
| 8896 | 445 | L445S | A | 5.9e-08 | 5.7e-08 | 5e-08 | 5.533E-08 | 4.726E-09 | 5.000E-08 | 5.900E-08 |
| 8897 | 445 | L445T | A | 1.4e-08 | 1.2e-07 | 5e-08 | 6.133E-08 | 5.390E-08 | 1.400E-08 | 1.200E-07 |
| 8898 | 445 | L445W | A | 4.4e-08 | 3.8e-08 | 3.6e-08 | 3.933E-08 | 4.163E-09 | 3.600E-08 | 4.400E-08 |
| 8899 | 445 | L445Y | A | 6.9e-08 | 5.7e-08 | 1.7e-08 | 4.767E-08 | 2.723E-08 | 1.700E-08 | 6.900E-08 |
| 8900 | 445 | L445V | A | 8e-08 | 4e-08 | 5.7e-08 | 5.900E-08 | 2.007E-08 | 4.000E-08 | 8.000E-08 |
| 8901 | 446 | I446A | A | 2.4e-08 | 2.1e-08 | 7.8e-08 | 4.100E-08 | 3.208E-08 | 2.100E-08 | 7.800E-08 |
| 8902 | 446 | I446R | A | 4.4e-08 | 4.1e-08 | 4.6e-09 | 2.987E-08 | 2.193E-08 | 4.600E-09 | 4.400E-08 |
| 8903 | 446 | I446N | A | 2.4e-08 | 7.1e-08 | 1.3e-08 | 3.600E-08 | 3.081E-08 | 1.300E-08 | 7.100E-08 |
| 8904 | 446 | I446D | A | 1.1e-07 | 4.2e-08 | 4.4e-08 | 6.533E-08 | 3.870E-08 | 4.200E-08 | 1.100E-07 |
| 8905 | 446 | I446C | A | 7e-08 | 4.7e-08 | 1.4e-07 | 8.567E-08 | 4.844E-08 | 4.700E-08 | 1.400E-07 |
| 8906 | 446 | I446E | A | 7.3e-08 | 1.8e-08 | 1.6e-08 | 3.567E-08 | 3.235E-08 | 1.600E-08 | 7.300E-08 |
| 8907 | 446 | I446Q | A | 2.4e-08 | 1.6e-08 | 5e-08 | 3.000E-08 | 1.778E-08 | 1.600E-08 | 5.000E-08 |
| 8908 | 446 | I446G | A | 2.1e-08 | 5e-08 | 7.7e-08 | 4.933E-08 | 2.801E-08 | 2.100E-08 | 7.700E-08 |
| 8909 | 446 | I446H | A | 5.7e-08 | 5e-08 | 4e-08 | 4.900E-08 | 8.544E-09 | 4.000E-08 | 5.700E-08 |
| 8910 | 446 | I446I | A | 5.6e-08 | 7.7e-08 | 2.5e-08 | 5.267E-08 | 2.616E-08 | 2.500E-08 | 7.700E-08 |
| 8911 | 446 | I446L | A | 4.9e-08 | 6.8e-08 | 9.3e-09 | 4.210E-08 | 2.995E-08 | 9.300E-09 | 6.800E-08 |
| 8912 | 446 | I446K | A | 3.9e-08 | 6.8e-08 | 8.2e-08 | 6.300E-08 | 2.193E-08 | 3.900E-08 | 8.200E-08 |
| 8913 | 446 | I446M | A | 2.5e-08 | 2.8e-08 | 3.4e-08 | 2.900E-08 | 4.583E-09 | 2.500E-08 | 3.400E-08 |
| 8914 | 446 | I446F | A | 6.2e-08 | 4.2e-08 | 1.5e-08 | 3.967E-08 | 2.359E-08 | 1.500E-08 | 6.200E-08 |
| 8915 | 446 | I446P | A | 2.4e-08 | 5.2e-08 | 5e-08 | 4.200E-08 | 1.562E-08 | 2.400E-08 | 5.200E-08 |
| 8916 | 446 | I446S | A | 1.3e-07 | 2.3e-08 | 9.6e-08 | 8.300E-08 | 5.467E-08 | 2.300E-08 | 1.300E-07 |
| 8917 | 446 | I446T | A | 5.1e-08 | 3.2e-08 | 3.8e-08 | 4.033E-08 | 9.713E-09 | 3.200E-08 | 5.100E-08 |
| 8918 | 446 | I446W | A | 6.9e-08 | 4.2e-08 | 6.5e-08 | 5.867E-08 | 1.457E-08 | 4.200E-08 | 6.900E-08 |
| 8919 | 446 | I446Y | A | 5.5e-08 | 3.8e-08 | 4.3e-08 | 4.533E-08 | 8.737E-09 | 3.800E-08 | 5.500E-08 |
| 8920 | 446 | I446V | A | 4.6e-08 | 3.1e-08 | 6e-08 | 4.567E-08 | 1.450E-08 | 3.100E-08 | 6.000E-08 |
| 8921 | 447 | H447A | A | 3.5e-08 | 3.1e-08 | 3.3e-08 | 3.300E-08 | 2.000E-09 | 3.100E-08 | 3.500E-08 |
| 8922 | 447 | H447R | A | 3.8e-08 | 2.7e-08 | 2.7e-08 | 3.067E-08 | 6.351E-09 | 2.700E-08 | 3.800E-08 |
| 8923 | 447 | H447N | A | 6.9e-08 | 6.3e-08 | 3.7e-08 | 5.633E-08 | 1.701E-08 | 3.700E-08 | 6.900E-08 |
| 8924 | 447 | H447D | A | 1.8e-08 | 1.9e-07 | 7.5e-08 | 9.433E-08 | 8.761E-08 | 1.800E-08 | 1.900E-07 |

| | | | | | | | | | | |
|---|---|---|---|---|---|---|---|---|---|---|
| 8925 | 447 | H447C | A | 3.1e-08 | 3.1e-08 | 5.7e-08 | 3.967E-08 | 1.501E-08 | 3.100E-08 | 5.700E-08 |
| 8926 | 447 | H447E | A | 2.1e-08 | 5.7e-08 | 5.9e-08 | 4.567E-08 | 2.139E-08 | 2.100E-08 | 5.900E-08 |
| 8927 | 447 | H447Q | A | 2.5e-08 | 2.4e-08 | 2.7e-08 | 2.533E-08 | 1.528E-09 | 2.400E-08 | 2.700E-08 |
| 8928 | 447 | H447G | A | 5.1e-08 | 3.8e-08 | 3.5e-08 | 4.133E-08 | 8.505E-09 | 3.500E-08 | 5.100E-08 |
| 8929 | 447 | H447H | A | 3.4e-08 | 1.2e-07 | 3.5e-08 | 6.300E-08 | 4.937E-08 | 3.400E-08 | 1.200E-07 |
| 8930 | 447 | H447I | A | 1.7e-08 | 1.8e-08 | 7.2e-08 | 3.567E-08 | 3.147E-08 | 1.700E-08 | 7.200E-08 |
| 8931 | 447 | H447L | A | 1.7e-07 | 2.9e-08 | 2.7e-08 | 7.533E-08 | 8.199E-08 | 2.700E-08 | 1.700E-07 |
| 8932 | 447 | H447K | A | 3.2e-08 | 6.7e-08 | 3.9e-08 | 4.600E-08 | 1.852E-08 | 3.200E-08 | 6.700E-08 |
| 8933 | 447 | H447M | A | 7.5e-08 | 8.1e-08 | 1.3e-08 | 5.633E-08 | 3.765E-08 | 1.300E-08 | 8.100E-08 |
| 8934 | 447 | H447F | A | 2.6e-08 | 7.9e-08 | 3.7e-08 | 4.733E-08 | 2.797E-08 | 2.600E-08 | 7.900E-08 |
| 8935 | 447 | H447P | A | 3.7e-08 | 4.1e-08 | 8.6e-08 | 5.467E-08 | 2.721E-08 | 3.700E-08 | 8.600E-08 |
| 8936 | 447 | H447S | A | 6.4e-08 | 5.5e-08 | 2.8e-08 | 4.900E-08 | 1.873E-08 | 2.800E-08 | 6.400E-08 |
| 8937 | 447 | H447T | A | 7.9e-08 | 8.7e-08 | 5.3e-08 | 7.300E-08 | 1.778E-08 | 5.300E-08 | 8.700E-08 |
| 8938 | 447 | H447W | A | 4.6e-08 | 3.4e-08 | 4.9e-08 | 4.300E-08 | 7.937E-09 | 3.400E-08 | 4.900E-08 |
| 8939 | 447 | H447Y | A | 8.9e-08 | 4.2e-08 | 5e-08 | 6.033E-08 | 2.515E-08 | 4.200E-08 | 8.900E-08 |
| 8940 | 447 | H447V | A | 6.4e-08 | 8.3e-08 | 3.9e-08 | 6.200E-08 | 2.207E-08 | 3.900E-08 | 8.300E-08 |
| 8941 | 448 | H448A | A | 1.2e-07 | 5.2e-08 | 1.7e-08 | 6.300E-08 | 5.237E-08 | 1.700E-08 | 1.200E-07 |
| 8942 | 448 | H448R | A | 4e-08 | 1.1e-07 | 5.7e-08 | 6.900E-08 | 3.651E-08 | 4.000E-08 | 1.100E-07 |
| 8943 | 448 | H448N | A | 3.1e-08 | 6e-08 | 5e-08 | 4.700E-08 | 1.473E-08 | 3.100E-08 | 6.000E-08 |
| 8944 | 448 | H448D | A | 5.1e-08 | 1.2e-07 | 2.7e-08 | 6.600E-08 | 4.828E-08 | 2.700E-08 | 1.200E-07 |
| 8945 | 448 | H448C | A | 2.6e-08 | 6e-08 | 5.6e-08 | 4.733E-08 | 1.858E-08 | 2.600E-08 | 6.000E-08 |
| 8946 | 448 | H448E | A | 5.8e-08 | 1.2e-07 | 4.9e-08 | 7.567E-08 | 3.866E-08 | 4.900E-08 | 1.200E-07 |
| 8947 | 448 | H448Q | A | 2.9e-08 | 1.1e-07 | 8.8e-08 | 7.567E-08 | 4.188E-08 | 2.900E-08 | 1.100E-07 |
| 8948 | 448 | H448G | A | 5.6e-08 | 2.4e-08 | 4.9e-08 | 4.300E-08 | 1.682E-08 | 2.400E-08 | 5.600E-08 |
| 8949 | 448 | H448H | A | 5.3e-08 | 4e-08 | 4.7e-08 | 4.667E-08 | 6.506E-09 | 4.000E-08 | 5.300E-08 |
| 8950 | 448 | H448I | A | 4.8e-08 | 3e-08 | 1.5e-07 | 7.600E-08 | 6.471E-08 | 3.000E-08 | 1.500E-07 |
| 8951 | 448 | H448L | A | 2.7e-08 | 1.2e-07 | 2e-08 | 5.567E-08 | 5.582E-08 | 2.000E-08 | 1.200E-07 |
| 8952 | 448 | H448K | A | 6.3e-08 | 3.8e-08 | 6.3e-08 | 5.467E-08 | 1.443E-08 | 3.800E-08 | 6.300E-08 |
| 8953 | 448 | H448M | A | 4e-08 | 6.5e-08 | 8.5e-08 | 6.333E-08 | 2.255E-08 | 4.000E-08 | 8.500E-08 |
| 8954 | 448 | H448F | A | 4.4e-08 | 5.2e-08 | 6.1e-08 | 5.233E-08 | 8.505E-09 | 4.400E-08 | 6.100E-08 |
| 8955 | 448 | H448P | A | 7.5e-08 | 3.3e-08 | 4e-08 | 4.933E-08 | 2.250E-08 | 3.300E-08 | 7.500E-08 |
| 8956 | 448 | H448S | A | 3.9e-08 | 7.4e-08 | 3.3e-08 | 4.867E-08 | 2.214E-08 | 3.300E-08 | 7.400E-08 |
| 8957 | 448 | H448T | A | 5e-08 | 4.1e-08 | 2.2e-08 | 3.767E-08 | 1.429E-08 | 2.200E-08 | 5.000E-08 |
| 8958 | 448 | H448W | A | 1.8e-08 | 6.7e-08 | 2e-08 | 3.500E-08 | 2.773E-08 | 1.800E-08 | 6.700E-08 |
| 8959 | 448 | H448Y | A | 5.1e-08 | 3.2e-08 | 3.3e-08 | 3.867E-08 | 1.069E-08 | 3.200E-08 | 5.100E-08 |
| 8960 | 448 | H448V | A | 2.2e-08 | 8e-08 | 7.3e-08 | 5.833E-08 | 3.166E-08 | 2.200E-08 | 8.000E-08 |
| 8961 | 449 | N449A | A | 4.1e-08 | 2.2e-08 | 7.6e-08 | 4.633E-08 | 2.739E-08 | 2.200E-08 | 7.600E-08 |
| 8962 | 449 | N449R | A | 2e-08 | 1.3e-08 | 3.6e-08 | 2.300E-08 | 1.179E-08 | 1.300E-08 | 3.600E-08 |
| 8963 | 449 | N449N | A | 3.1e-08 | 1.5e-07 | 6.5e-08 | 8.200E-08 | 6.129E-08 | 3.100E-08 | 1.500E-07 |
| 8964 | 449 | N449D | A | 3e-08 | 9.8e-08 | 1.7e-08 | 4.833E-08 | 4.350E-08 | 1.700E-08 | 9.800E-08 |
| 8965 | 449 | N449C | A | 3.7e-08 | 1.8e-08 | 2.5e-08 | 2.667E-08 | 9.609E-09 | 1.800E-08 | 3.700E-08 |
| 8966 | 449 | N449E | A | 5.4e-08 | 7.7e-08 | 8.1e-08 | 7.067E-08 | 1.457E-08 | 5.400E-08 | 8.100E-08 |
| 8967 | 449 | N449Q | A | 3.1e-08 | 3.9e-08 | 1.5e-07 | 7.333E-08 | 6.652E-08 | 3.100E-08 | 1.500E-07 |
| 8968 | 449 | N449G | A | 4.7e-08 | 5e-08 | 4.8e-08 | 4.833E-08 | 1.528E-09 | 4.700E-08 | 5.000E-08 |
| 8969 | 449 | N449H | A | 3e-08 | 8.7e-08 | 2.8e-08 | 4.833E-08 | 3.350E-08 | 2.800E-08 | 8.700E-08 |
| 8970 | 449 | N449I | A | 7.2e-08 | 8.3e-08 | 4.6e-08 | 6.700E-08 | 1.900E-08 | 4.600E-08 | 8.300E-08 |
| 8971 | 449 | N449L | A | 4.2e-08 | 1.4e-08 | 5e-08 | 3.533E-08 | 1.890E-08 | 1.400E-08 | 5.000E-08 |
| 8972 | 449 | N449K | A | 9.8e-09 | 2.9e-08 | 5.7e-08 | 3.193E-08 | 2.374E-08 | 9.800E-09 | 5.700E-08 |
| 8973 | 449 | N449M | A | 1.3e-08 | 2.7e-08 | 1.3e-07 | 5.667E-08 | 6.389E-08 | 1.300E-08 | 1.300E-07 |
| 8974 | 449 | N449F | A | 3.5e-08 | 2.6e-08 | 2.8e-08 | 2.967E-08 | 4.726E-09 | 2.600E-08 | 3.500E-08 |
| 8975 | 449 | N449P | A | 2.1e-08 | 7.1e-08 | 4.5e-08 | 4.567E-08 | 2.501E-08 | 2.100E-08 | 7.100E-08 |
| 8976 | 449 | N449S | A | 2.4e-08 | 9.8e-08 | 4.9e-08 | 5.700E-08 | 3.764E-08 | 2.400E-08 | 9.800E-08 |
| 8977 | 449 | N449T | A | 1e-07 | 4.6e-08 | 5.8e-08 | 6.800E-08 | 2.835E-08 | 4.600E-08 | 1.000E-07 |
| 8978 | 449 | N449W | A | 1.6e-08 | 4.9e-08 | 2.6e-08 | 3.033E-08 | 1.692E-08 | 1.600E-08 | 4.900E-08 |
| 8979 | 449 | N449Y | A | 3.3e-08 | 3.4e-08 | 5.1e-08 | 3.933E-08 | 1.012E-08 | 3.300E-08 | 5.100E-08 |
| 8980 | 449 | N449V | A | 4.1e-08 | 2.8e-08 | 9.2e-08 | 5.367E-08 | 3.383E-08 | 2.800E-08 | 9.200E-08 |
| 8981 | 450 | T450A | A | 5.1e-08 | 6.7e-08 | 5e-08 | 5.600E-08 | 9.539E-09 | 5.000E-08 | 6.700E-08 |
| 8982 | 450 | T450R | A | 5.3e-08 | 5.7e-08 | 1.8e-08 | 4.267E-08 | 2.146E-08 | 1.800E-08 | 5.700E-08 |
| 8983 | 450 | T450N | A | 5.6e-08 | 3.1e-08 | 4.2e-08 | 4.300E-08 | 1.253E-08 | 3.100E-08 | 5.600E-08 |
| 8984 | 450 | T450D | A | 3.5e-08 | 3.9e-08 | 5e-08 | 4.133E-08 | 7.767E-09 | 3.500E-08 | 5.000E-08 |
| 8985 | 450 | T450C | A | 3.7e-08 | 5e-08 | 4e-08 | 4.233E-08 | 6.807E-09 | 3.700E-08 | 5.000E-08 |
| 8986 | 450 | T450E | A | 8.1e-08 | 8.9e-08 | 5.6e-08 | 7.533E-08 | 1.721E-08 | 5.600E-08 | 8.900E-08 |
| 8987 | 450 | T450Q | A | 5.3e-08 | 2.5e-08 | 9.2e-08 | 5.667E-08 | 3.365E-08 | 2.500E-08 | 9.200E-08 |
| 8988 | 450 | T450G | A | 1.9e-08 | 2.2e-08 | 2.3e-08 | 2.133E-08 | 2.082E-09 | 1.900E-08 | 2.300E-08 |
| 8989 | 450 | T450H | A | 4.6e-08 | 4e-08 | 1.1e-08 | 3.233E-08 | 1.872E-08 | 1.100E-08 | 4.600E-08 |
| 8990 | 450 | T450I | A | 1.3e-08 | 6e-08 | 3.9e-08 | 3.733E-08 | 2.354E-08 | 1.300E-08 | 6.000E-08 |
| 8991 | 450 | T450L | A | 6.9e-08 | 4.1e-08 | 2e-08 | 4.333E-08 | 2.458E-08 | 2.000E-08 | 6.900E-08 |
| 8992 | 450 | T450K | A | 1.7e-07 | 4e-08 | 5e-08 | 8.667E-08 | 7.234E-08 | 4.000E-08 | 1.700E-07 |
| 8993 | 450 | T450M | A | 2.6e-08 | 5.7e-08 | 7.6e-08 | 5.300E-08 | 2.524E-08 | 2.600E-08 | 7.600E-08 |
| 8994 | 450 | T450F | A | 3.8e-08 | 3.1e-08 | 2.9e-08 | 3.267E-08 | 4.726E-09 | 2.900E-08 | 3.800E-08 |
| 8995 | 450 | T450P | A | 3.1e-08 | 3e-08 | 8.8e-08 | 4.967E-08 | 3.320E-08 | 3.000E-08 | 8.800E-08 |
| 8996 | 450 | T450S | A | 1.8e-08 | 4.9e-08 | 2.8e-08 | 3.167E-08 | 1.582E-08 | 1.800E-08 | 4.900E-08 |
| 8997 | 450 | T450T | A | 1.9e-08 | 3.5e-08 | 7.8e-08 | 4.400E-08 | 3.051E-08 | 1.900E-08 | 7.800E-08 |
| 8998 | 450 | T450W | A | 1e-07 | 3.8e-08 | 9.6e-08 | 7.800E-08 | 3.470E-08 | 3.800E-08 | 1.000E-07 |
| 8999 | 450 | T450Y | A | 5.2e-08 | 8.8e-08 | 1.4e-07 | 9.333E-08 | 4.424E-08 | 5.200E-08 | 1.400E-07 |
| 9000 | 450 | T450V | A | 4e-08 | 3.5e-08 | 8.4e-08 | 5.300E-08 | 2.696E-08 | 3.500E-08 | 8.400E-08 |
| 9001 | 451 | H451A | A | 7.2e-08 | 3.7e-08 | 5.5e-08 | 5.467E-08 | 1.750E-08 | 3.700E-08 | 7.200E-08 |
| 9002 | 451 | H451R | A | 5.4e-08 | 7.1e-08 | 4.4e-08 | 5.633E-08 | 1.365E-08 | 4.400E-08 | 7.100E-08 |
| 9003 | 451 | H451N | A | 3.2e-08 | 7.7e-08 | 5.3e-08 | 5.400E-08 | 2.252E-08 | 3.200E-08 | 7.700E-08 |
| 9004 | 451 | H451D | A | 4.4e-08 | 4e-08 | 2.4e-08 | 3.600E-08 | 1.058E-08 | 2.400E-08 | 4.400E-08 |
| 9005 | 451 | H451C | A | 9.4e-08 | 3.7e-08 | 7.3e-08 | 6.800E-08 | 2.883E-08 | 3.700E-08 | 9.400E-08 |
| 9006 | 451 | H451E | A | 1.5e-08 | 3.5e-08 | 5.9e-08 | 3.633E-08 | 2.203E-08 | 1.500E-08 | 5.900E-08 |
| 9007 | 451 | H451Q | A | 3.5e-08 | 6.2e-08 | 4.8e-08 | 4.833E-08 | 1.350E-08 | 3.500E-08 | 6.200E-08 |
| 9008 | 451 | H451G | A | 1.1e-07 | 5.1e-08 | 6.9e-08 | 7.667E-08 | 3.024E-08 | 5.100E-08 | 1.100E-07 |
| 9009 | 451 | H451H | A | 2.5e-08 | 3.9e-08 | 2.6e-08 | 3.000E-08 | 7.810E-09 | 2.500E-08 | 3.900E-08 |

| | | | | | | | | | | |
|---|---|---|---|---|---|---|---|---|---|---|
| 9010 | 451 | H451I | A | 4.6e-08 | 7.9e-08 | 5.8e-08 | 6.100E-08 | 1.670E-08 | 4.600E-08 | 7.900E-08 |
| 9011 | 451 | H451L | A | 2e-08 | 5.6e-08 | 1.2e-07 | 6.533E-08 | 5.065E-08 | 2.000E-08 | 1.200E-07 |
| 9012 | 451 | H451K | A | 7.7e-08 | 4e-08 | 1.8e-08 | 4.500E-08 | 2.982E-08 | 1.800E-08 | 7.700E-08 |
| 9013 | 451 | H451M | A | 2e-08 | 9e-08 | 2.9e-08 | 4.633E-08 | 3.808E-08 | 2.000E-08 | 9.000E-08 |
| 9014 | 451 | H451F | A | 5.6e-08 | 8.1e-08 | 2.1e-08 | 5.267E-08 | 3.014E-08 | 2.100E-08 | 8.100E-08 |
| 9015 | 451 | H451P | A | 4.8e-08 | 5.8e-08 | 8.2e-08 | 6.267E-08 | 1.747E-08 | 4.800E-08 | 8.200E-08 |
| 9016 | 451 | H451S | A | 3.3e-08 | 1.2e-07 | 1.8e-08 | 5.700E-08 | 5.507E-08 | 1.800E-08 | 1.200E-07 |
| 9017 | 451 | H451T | A | 2.1e-08 | 3e-08 | 7.1e-08 | 4.067E-08 | 2.665E-08 | 2.100E-08 | 7.100E-08 |
| 9018 | 451 | H451W | A | 2e-08 | 7.5e-08 | 7.1e-08 | 5.533E-08 | 3.066E-08 | 2.000E-08 | 7.500E-08 |
| 9019 | 451 | H451Y | A | 7.5e-08 | 3.7e-08 | 1.1e-07 | 7.400E-08 | 3.651E-08 | 3.700E-08 | 1.100E-07 |
| 9020 | 451 | H451V | A | 5.2e-08 | 6e-08 | 5.3e-08 | 5.500E-08 | 4.359E-09 | 5.200E-08 | 6.000E-08 |
| 9021 | 452 | L452A | A | 3.3e-08 | 6.6e-08 | 5.4e-08 | 5.100E-08 | 1.670E-08 | 3.300E-08 | 6.600E-08 |
| 9022 | 452 | L452R | A | 4e-08 | 2.2e-08 | 5.7e-08 | 3.967E-08 | 1.750E-08 | 2.200E-08 | 5.700E-08 |
| 9023 | 452 | L452N | A | 3e-08 | 4.4e-08 | 8.1e-08 | 5.167E-08 | 2.635E-08 | 3.000E-08 | 8.100E-08 |
| 9024 | 452 | L452D | A | 8.3e-08 | 5.6e-08 | 4.4e-08 | 6.100E-08 | 1.997E-08 | 4.400E-08 | 8.300E-08 |
| 9025 | 452 | L452C | A | 1.9e-08 | 1.8e-08 | 4.5e-08 | 2.733E-08 | 1.531E-08 | 1.800E-08 | 4.500E-08 |
| 9026 | 452 | L452E | A | 5.9e-08 | 3.8e-08 | 2.5e-08 | 4.067E-08 | 1.716E-08 | 2.500E-08 | 5.900E-08 |
| 9027 | 452 | L452Q | A | 3.2e-08 | 1.8e-07 | 4.5e-08 | 8.567E-08 | 8.195E-08 | 3.200E-08 | 1.800E-07 |
| 9028 | 452 | L452G | A | 3.8e-08 | 3.9e-08 | 3.7e-08 | 3.800E-08 | 1.000E-09 | 3.700E-08 | 3.900E-08 |
| 9029 | 452 | L452H | A | 5.7e-08 | 1.8e-07 | 8.7e-08 | 1.080E-07 | 6.413E-08 | 5.700E-08 | 1.800E-07 |
| 9030 | 452 | L452I | A | 8.5e-09 | 3.4e-08 | 6.5e-08 | 3.583E-08 | 2.829E-08 | 8.500E-09 | 6.500E-08 |
| 9031 | 452 | L452L | A | 9.9e-08 | 2.2e-08 | 2.8e-08 | 4.967E-08 | 4.283E-08 | 2.200E-08 | 9.900E-08 |
| 9032 | 452 | L452K | A | 1.9e-08 | 4e-08 | 9e-08 | 4.967E-08 | 3.647E-08 | 1.900E-08 | 9.000E-08 |
| 9033 | 452 | L452M | A | 6.6e-08 | 4.3e-08 | 2.8e-08 | 4.567E-08 | 1.914E-08 | 2.800E-08 | 6.600E-08 |
| 9034 | 452 | L452F | A | 6.5e-08 | 3.7e-08 | 9.8e-09 | 3.727E-08 | 2.760E-08 | 9.800E-09 | 6.500E-08 |
| 9035 | 452 | L452P | A | 5.3e-08 | 1.3e-08 | 8e-09 | 2.467E-08 | 2.466E-08 | 8.000E-09 | 5.300E-08 |
| 9036 | 452 | L452S | A | 1.8e-08 | 1.5e-08 | 8.1e-08 | 3.800E-08 | 3.727E-08 | 1.500E-08 | 8.100E-08 |
| 9037 | 452 | L452T | A | 4e-08 | 2.1e-08 | 3.9e-08 | 3.333E-08 | 1.069E-08 | 2.100E-08 | 4.000E-08 |
| 9038 | 452 | L452W | A | 1.1e-08 | 6.6e-08 | 6e-08 | 4.567E-08 | 3.017E-08 | 1.100E-08 | 6.600E-08 |
| 9039 | 452 | L452Y | A | 3.3e-08 | 5.1e-08 | 5.9e-08 | 4.767E-08 | 1.332E-08 | 3.300E-08 | 5.900E-08 |
| 9040 | 452 | L452V | A | 5.5e-08 | 2.5e-08 | 6.3e-08 | 4.767E-08 | 2.003E-08 | 2.500E-08 | 6.300E-08 |
| 9041 | 453 | C453A | A | 1.1e-08 | 5.3e-08 | 6e-08 | 4.133E-08 | 2.650E-08 | 1.100E-08 | 6.000E-08 |
| 9042 | 453 | C453R | A | 4.9e-08 | 1e-07 | 2e-08 | 5.633E-08 | 4.050E-08 | 2.000E-08 | 1.000E-07 |
| 9043 | 453 | C453N | A | 1.8e-08 | 4.8e-08 | 1.5e-08 | 2.700E-08 | 1.825E-08 | 1.500E-08 | 4.800E-08 |
| 9044 | 453 | C453D | A | 4.5e-08 | 5.5e-08 | 5.8e-08 | 5.267E-08 | 6.807E-09 | 4.500E-08 | 5.800E-08 |
| 9045 | 453 | C453C | A | 1.7e-08 | 3.3e-08 | 6.5e-08 | 3.833E-08 | 2.444E-08 | 1.700E-08 | 6.500E-08 |
| 9046 | 453 | C453E | A | 3.8e-08 | 6.3e-08 | 4.5e-08 | 4.867E-08 | 1.290E-08 | 3.800E-08 | 6.300E-08 |
| 9047 | 453 | C453Q | A | 5.2e-08 | 1.9e-08 | 2.2e-08 | 3.100E-08 | 1.825E-08 | 1.900E-08 | 5.200E-08 |
| 9048 | 453 | C453G | A | 9.1e-08 | 3.7e-08 | 1.7e-08 | 4.833E-08 | 3.828E-08 | 1.700E-08 | 9.100E-08 |
| 9049 | 453 | C453H | A | 5.1e-08 | 1.3e-08 | 7.3e-08 | 4.567E-08 | 3.035E-08 | 1.300E-08 | 7.300E-08 |
| 9050 | 453 | C453I | A | 3.4e-08 | 1.4e-07 | 2.1e-08 | 6.500E-08 | 6.528E-08 | 2.100E-08 | 1.400E-07 |
| 9051 | 453 | C453L | A | 4.7e-08 | 1.1e-07 | 4.3e-08 | 6.667E-08 | 3.758E-08 | 4.300E-08 | 1.100E-07 |
| 9052 | 453 | C453K | A | 1.8e-08 | 2.6e-08 | 1e-07 | 4.800E-08 | 4.521E-08 | 1.800E-08 | 1.000E-07 |
| 9053 | 453 | C453M | A | 1e-07 | 2.9e-08 | 2.4e-08 | 5.100E-08 | 4.251E-08 | 2.400E-08 | 1.000E-07 |
| 9054 | 453 | C453F | A | 2.2e-08 | 1.2e-07 | 3e-08 | 5.733E-08 | 5.442E-08 | 2.200E-08 | 1.200E-07 |
| 9055 | 453 | C453P | A | 2.7e-08 | 4.5e-08 | 5.4e-08 | 4.200E-08 | 1.375E-08 | 2.700E-08 | 5.400E-08 |
| 9056 | 453 | C453S | A | 4e-08 | 2.9e-08 | 1.8e-08 | 2.900E-08 | 1.100E-08 | 1.800E-08 | 4.000E-08 |
| 9057 | 453 | C453T | A | 1.1e-07 | 5.9e-08 | 1.2e-07 | 9.633E-08 | 3.272E-08 | 5.900E-08 | 1.200E-07 |
| 9058 | 453 | C453W | A | 3e-08 | 9.1e-08 | 3.4e-08 | 5.167E-08 | 3.412E-08 | 3.000E-08 | 9.100E-08 |
| 9059 | 453 | C453Y | A | 3e-08 | 4.8e-08 | 3.4e-08 | 3.733E-08 | 9.452E-09 | 3.000E-08 | 4.800E-08 |
| 9060 | 453 | C453V | A | 3.7e-08 | 2.9e-08 | 6.2e-08 | 4.267E-08 | 1.721E-08 | 2.900E-08 | 6.200E-08 |
| 9061 | 454 | F454A | A | 5.8e-08 | 7e-08 | 1.8e-08 | 4.867E-08 | 2.723E-08 | 1.800E-08 | 7.000E-08 |
| 9062 | 454 | F454R | A | 5.3e-08 | 1.3e-08 | 3.4e-08 | 3.333E-08 | 2.001E-08 | 1.300E-08 | 5.300E-08 |
| 9063 | 454 | F454N | A | 1.2e-07 | 4e-08 | 9.1e-08 | 8.367E-08 | 4.050E-08 | 4.000E-08 | 1.200E-07 |
| 9064 | 454 | F454D | A | 8.9e-09 | 1.8e-08 | 8.5e-08 | 3.730E-08 | 4.156E-08 | 8.900E-09 | 8.500E-08 |
| 9065 | 454 | F454C | A | 2.5e-08 | 1.3e-07 | 2.4e-08 | 5.967E-08 | 6.091E-08 | 2.400E-08 | 1.300E-07 |
| 9066 | 454 | F454E | A | 2.6e-08 | 2.5e-08 | 4e-08 | 3.033E-08 | 8.386E-09 | 2.500E-08 | 4.000E-08 |
| 9067 | 454 | F454Q | A | 1.5e-08 | 2.9e-08 | 1.5e-07 | 6.467E-08 | 7.423E-08 | 1.500E-08 | 1.500E-07 |
| 9068 | 454 | F454G | A | 6.5e-08 | 3.7e-08 | 3.9e-08 | 4.700E-08 | 1.562E-08 | 3.700E-08 | 6.500E-08 |
| 9069 | 454 | F454H | A | 3.8e-08 | 5.3e-08 | 4.3e-08 | 4.467E-08 | 7.638E-09 | 3.800E-08 | 5.300E-08 |
| 9070 | 454 | F454I | A | 1.6e-08 | 3.4e-08 | 4.3e-08 | 3.100E-08 | 1.375E-08 | 1.600E-08 | 4.300E-08 |
| 9071 | 454 | F454L | A | 5.3e-08 | 8.7e-08 | 3.6e-08 | 5.867E-08 | 2.597E-08 | 3.600E-08 | 8.700E-08 |
| 9072 | 454 | F454K | A | 6.6e-08 | 5.2e-08 | 2.6e-08 | 4.800E-08 | 2.030E-08 | 2.600E-08 | 6.600E-08 |
| 9073 | 454 | F454M | A | 5.6e-08 | 2e-08 | 4.3e-08 | 3.967E-08 | 1.823E-08 | 2.000E-08 | 5.600E-08 |
| 9074 | 454 | F454F | A | 4e-08 | 5.3e-08 | 8.2e-08 | 5.833E-08 | 2.150E-08 | 4.000E-08 | 8.200E-08 |
| 9075 | 454 | F454P | A | 5.6e-08 | 9.3e-08 | 1.1e-08 | 5.333E-08 | 4.106E-08 | 1.100E-08 | 9.300E-08 |
| 9076 | 454 | F454S | A | 4.4e-08 | 4.1e-08 | 1.8e-07 | 8.833E-08 | 7.940E-08 | 4.100E-08 | 1.800E-07 |
| 9077 | 454 | F454T | A | 2.1e-07 | 3.1e-08 | 3e-08 | 9.033E-08 | 1.036E-07 | 3.000E-08 | 2.100E-07 |
| 9078 | 454 | F454W | A | 7e-08 | 8.4e-08 | 3.2e-08 | 6.200E-08 | 2.691E-08 | 3.200E-08 | 8.400E-08 |
| 9079 | 454 | F454Y | A | 8.8e-08 | 2.3e-08 | 2e-08 | 4.367E-08 | 3.842E-08 | 2.000E-08 | 8.800E-08 |
| 9080 | 454 | F454V | A | 1.9e-08 | 1.6e-08 | 6.5e-08 | 3.333E-08 | 2.747E-08 | 1.600E-08 | 6.500E-08 |
| 9081 | 455 | V455A | A | 6.8e-08 | 4.1e-08 | 3.7e-08 | 4.867E-08 | 1.686E-08 | 3.700E-08 | 6.800E-08 |
| 9082 | 455 | V455R | A | 4.6e-08 | 4e-08 | 4.2e-08 | 4.267E-08 | 3.055E-09 | 4.000E-08 | 4.600E-08 |
| 9083 | 455 | V455N | A | 3.1e-08 | 8.7e-08 | 1.8e-08 | 4.533E-08 | 3.667E-08 | 1.800E-08 | 8.700E-08 |
| 9084 | 455 | V455D | A | 2.5e-08 | 7.7e-08 | 3.1e-08 | 4.433E-08 | 2.845E-08 | 2.500E-08 | 7.700E-08 |
| 9085 | 455 | V455C | A | 3e-08 | 2.1e-08 | 4.2e-08 | 3.100E-08 | 1.054E-08 | 2.100E-08 | 4.200E-08 |
| 9086 | 455 | V455E | A | 4.6e-08 | 5.6e-08 | 2.7e-08 | 4.300E-08 | 1.473E-08 | 2.700E-08 | 5.600E-08 |
| 9087 | 455 | V455Q | A | 5.4e-08 | 4.5e-08 | 2.9e-08 | 4.267E-08 | 1.266E-08 | 2.900E-08 | 5.400E-08 |
| 9088 | 455 | V455G | A | 3.5e-08 | 3e-08 | 6.6e-08 | 4.367E-08 | 1.950E-08 | 3.000E-08 | 6.600E-08 |
| 9089 | 455 | V455H | A | 2.7e-08 | 3.8e-08 | 6.3e-08 | 4.267E-08 | 1.845E-08 | 2.700E-08 | 6.300E-08 |
| 9090 | 455 | V455I | A | 3.2e-08 | 4.1e-08 | 2.9e-08 | 3.400E-08 | 6.245E-09 | 2.900E-08 | 4.100E-08 |
| 9091 | 455 | V455L | A | 7.5e-08 | 2.1e-07 | 9.9e-09 | 9.830E-08 | 1.021E-07 | 9.900E-09 | 2.100E-07 |
| 9092 | 455 | V455K | A | 1.2e-07 | 4e-08 | 3.4e-08 | 6.467E-08 | 4.801E-08 | 3.400E-08 | 1.200E-07 |
| 9093 | 455 | V455M | A | 2.5e-08 | 7.9e-08 | 4.1e-08 | 4.833E-08 | 2.774E-08 | 2.500E-08 | 7.900E-08 |
| 9094 | 455 | V455F | A | 1.7e-08 | 4.8e-08 | 3.9e-08 | 3.467E-08 | 1.595E-08 | 1.700E-08 | 4.800E-08 |

| | | | | | | | | | | |
|---|---|---|---|---|---|---|---|---|---|---|
| 9095 | 455 | V455P | A | 5.1e-08 | 1.5e-08 | 7.9e-08 | 4.833E-08 | 3.208E-08 | 1.500E-08 | 7.900E-08 |
| 9096 | 455 | V455S | A | 4.9e-08 | 5.9e-08 | 4.4e-08 | 5.067E-08 | 7.638E-09 | 4.400E-08 | 5.900E-08 |
| 9097 | 455 | V455T | A | 5.8e-08 | 2.7e-08 | 1.8e-08 | 3.433E-08 | 2.098E-08 | 1.800E-08 | 5.800E-08 |
| 9098 | 455 | V455W | A | 2.6e-08 | 4.5e-08 | 3.7e-08 | 3.600E-08 | 9.539E-09 | 2.600E-08 | 4.500E-08 |
| 9099 | 455 | V455Y | A | 8.8e-08 | 1.5e-08 | 5.2e-08 | 5.167E-08 | 3.650E-08 | 1.500E-08 | 8.800E-08 |
| 9100 | 455 | V455V | A | 4.7e-08 | 5.7e-08 | 2.4e-08 | 4.267E-08 | 1.692E-08 | 2.400E-08 | 5.700E-08 |
| 9101 | 456 | H456A | A | 7.5e-08 | 1.3e-07 | 1.8e-07 | 1.283E-07 | 5.252E-08 | 7.500E-08 | 1.800E-07 |
| 9102 | 456 | H456R | A | 1.1e-08 | 1.1e-07 | 2.5e-08 | 4.867E-08 | 5.358E-08 | 1.100E-08 | 1.100E-07 |
| 9103 | 456 | H456N | A | 1.4e-08 | 3.2e-08 | 5.7e-08 | 3.433E-08 | 2.159E-08 | 1.400E-08 | 5.700E-08 |
| 9104 | 456 | H456D | A | 7.8e-08 | 3.4e-08 | 1.8e-07 | 9.733E-08 | 7.490E-08 | 3.400E-08 | 1.800E-07 |
| 9105 | 456 | H456C | A | 2.4e-08 | 7.9e-08 | 4.9e-08 | 5.067E-08 | 2.754E-08 | 2.400E-08 | 7.900E-08 |
| 9106 | 456 | H456E | A | 6.8e-08 | 3.7e-08 | 4.8e-08 | 5.100E-08 | 1.572E-08 | 3.700E-08 | 6.800E-08 |
| 9107 | 456 | H456Q | A | 3e-08 | 5.4e-08 | 2.1e-08 | 3.500E-08 | 1.706E-08 | 2.100E-08 | 5.400E-08 |
| 9108 | 456 | H456G | A | 4.8e-08 | 8.3e-08 | 3.4e-08 | 5.500E-08 | 2.524E-08 | 3.400E-08 | 8.300E-08 |
| 9109 | 456 | H456H | A | 3.7e-08 | 4.2e-08 | 5e-08 | 4.300E-08 | 6.557E-09 | 3.700E-08 | 5.000E-08 |
| 9110 | 456 | H456I | A | 4.8e-08 | 6.5e-08 | 7.9e-09 | 4.030E-08 | 2.932E-08 | 7.900E-09 | 6.500E-08 |
| 9111 | 456 | H456L | A | 1.1e-07 | 6.1e-08 | 2.7e-08 | 6.600E-08 | 4.173E-08 | 2.700E-08 | 1.100E-07 |
| 9112 | 456 | H456K | A | 2.7e-08 | 4e-08 | 4.7e-08 | 3.800E-08 | 1.015E-08 | 2.700E-08 | 4.700E-08 |
| 9113 | 456 | H456M | A | 6.9e-08 | 3.7e-08 | 6.7e-08 | 5.767E-08 | 1.793E-08 | 3.700E-08 | 6.900E-08 |
| 9114 | 456 | H456F | A | 8.8e-08 | 2.1e-08 | 3.3e-08 | 4.733E-08 | 3.573E-08 | 2.100E-08 | 8.800E-08 |
| 9115 | 456 | H456P | A | 9.2e-08 | 6.5e-08 | 1.5e-07 | 1.023E-07 | 4.343E-08 | 6.500E-08 | 1.500E-07 |
| 9116 | 456 | H456S | A | 4.1e-08 | 5.7e-08 | 2.4e-08 | 4.067E-08 | 1.650E-08 | 2.400E-08 | 5.700E-08 |
| 9117 | 456 | H456T | A | 1e-08 | 3e-08 | 2.3e-08 | 2.100E-08 | 1.015E-08 | 1.000E-08 | 3.000E-08 |
| 9118 | 456 | H456W | A | 2.2e-08 | 2.6e-08 | 1.5e-08 | 2.100E-08 | 5.568E-09 | 1.500E-08 | 2.600E-08 |
| 9119 | 456 | H456Y | A | 6.9e-08 | 2.5e-08 | 3.2e-08 | 4.200E-08 | 2.364E-08 | 2.500E-08 | 6.900E-08 |
| 9120 | 456 | H456V | A | 2.4e-08 | 1.1e-07 | 8.8e-08 | 7.400E-08 | 4.468E-08 | 2.400E-08 | 1.100E-07 |
| 9121 | 457 | T457A | A | 1.4e-07 | 2.4e-08 | 6.5e-08 | 7.633E-08 | 5.882E-08 | 2.400E-08 | 1.400E-07 |
| 9122 | 457 | T457R | A | 1.8e-08 | 3.1e-08 | 6.4e-08 | 3.767E-08 | 2.371E-08 | 1.800E-08 | 6.400E-08 |
| 9123 | 457 | T457N | A | 4.4e-08 | 7.1e-08 | 5.6e-08 | 5.700E-08 | 1.353E-08 | 4.400E-08 | 7.100E-08 |
| 9124 | 457 | T457D | A | 6.1e-08 | 3.7e-08 | 8.9e-08 | 6.233E-08 | 2.603E-08 | 3.700E-08 | 8.900E-08 |
| 9125 | 457 | T457C | A | 3.4e-08 | 4.9e-08 | 5.8e-08 | 4.700E-08 | 1.212E-08 | 3.400E-08 | 5.800E-08 |
| 9126 | 457 | T457E | A | 3.2e-08 | 2.4e-08 | 2.7e-08 | 2.767E-08 | 4.041E-09 | 2.400E-08 | 3.200E-08 |
| 9127 | 457 | T457Q | A | 2.9e-08 | 5e-08 | 2.1e-08 | 3.333E-08 | 1.498E-08 | 2.100E-08 | 5.000E-08 |
| 9128 | 457 | T457G | A | 3.7e-08 | 5.8e-09 | 3.8e-08 | 2.693E-08 | 1.831E-08 | 5.800E-09 | 3.800E-08 |
| 9129 | 457 | T457H | A | 4.5e-08 | 2.3e-07 | 8.1e-08 | 1.187E-07 | 9.808E-08 | 4.500E-08 | 2.300E-07 |
| 9130 | 457 | T457I | A | 5.5e-08 | 6.9e-08 | 6.5e-08 | 6.300E-08 | 7.211E-09 | 5.500E-08 | 6.900E-08 |
| 9131 | 457 | T457L | A | 5e-08 | 1.3e-07 | 5.8e-08 | 7.933E-08 | 4.406E-08 | 5.000E-08 | 1.300E-07 |
| 9132 | 457 | T457K | A | 5.4e-08 | 4.8e-08 | 2e-08 | 4.067E-08 | 1.815E-08 | 2.000E-08 | 5.400E-08 |
| 9133 | 457 | T457M | A | 5e-08 | 6.5e-08 | 3e-08 | 4.833E-08 | 1.756E-08 | 3.000E-08 | 6.500E-08 |
| 9134 | 457 | T457F | A | 9.4e-09 | 1.2e-07 | 5.5e-08 | 6.147E-08 | 5.558E-08 | 9.400E-09 | 1.200E-07 |
| 9135 | 457 | T457P | A | 8.4e-08 | 2.2e-08 | 3.2e-08 | 4.600E-08 | 3.329E-08 | 2.200E-08 | 8.400E-08 |
| 9136 | 457 | T457S | A | 3.4e-08 | 2.8e-08 | 1.7e-08 | 2.633E-08 | 8.622E-09 | 1.700E-08 | 3.400E-08 |
| 9137 | 457 | T457T | A | 2.9e-08 | 2.2e-08 | 3.6e-08 | 2.900E-08 | 7.000E-09 | 2.200E-08 | 3.600E-08 |
| 9138 | 457 | T457W | A | 4e-08 | 4.1e-08 | 2.1e-08 | 3.400E-08 | 1.127E-08 | 2.100E-08 | 4.100E-08 |
| 9139 | 457 | T457Y | A | 4.9e-08 | 4.1e-08 | 6.6e-08 | 5.200E-08 | 1.277E-08 | 4.100E-08 | 6.600E-08 |
| 9140 | 457 | T457V | A | 5.3e-08 | 3.2e-08 | 3.6e-08 | 4.033E-08 | 1.115E-08 | 3.200E-08 | 5.300E-08 |
| 9141 | 458 | V458A | A | 1.4e-07 | 8.8e-08 | 7.7e-08 | 1.017E-07 | 3.365E-08 | 7.700E-08 | 1.400E-07 |
| 9142 | 458 | V458R | A | 8.6e-08 | 4.6e-08 | 1.9e-08 | 5.033E-08 | 3.371E-08 | 1.900E-08 | 8.600E-08 |
| 9143 | 458 | V458N | A | 2e-08 | 2e-08 | 5.6e-08 | 3.200E-08 | 2.078E-08 | 2.000E-08 | 5.600E-08 |
| 9144 | 458 | V458D | A | 2.2e-08 | 3.6e-08 | 5.1e-08 | 3.633E-08 | 1.450E-08 | 2.200E-08 | 5.100E-08 |
| 9145 | 458 | V458C | A | 6.4e-08 | 6.2e-08 | 2.1e-08 | 4.900E-08 | 2.427E-08 | 2.100E-08 | 6.400E-08 |
| 9146 | 458 | V458E | A | 5.5e-08 | 1.8e-08 | 5.7e-08 | 4.333E-08 | 2.196E-08 | 1.800E-08 | 5.700E-08 |
| 9147 | 458 | V458Q | A | 3e-08 | 2.2e-08 | 4e-08 | 3.067E-08 | 9.018E-09 | 2.200E-08 | 4.000E-08 |
| 9148 | 458 | V458G | A | 5e-08 | 6.4e-08 | 3.6e-08 | 5.000E-08 | 1.400E-08 | 3.600E-08 | 6.400E-08 |
| 9149 | 458 | V458H | A | 2.8e-08 | 8.3e-08 | 6.9e-08 | 6.000E-08 | 2.858E-08 | 2.800E-08 | 8.300E-08 |
| 9150 | 458 | V458I | A | 4e-08 | 2.2e-08 | 9.2e-08 | 5.133E-08 | 3.635E-08 | 2.200E-08 | 9.200E-08 |
| 9151 | 458 | V458L | A | 3.6e-08 | 7.4e-08 | 3.4e-08 | 4.800E-08 | 2.254E-08 | 3.400E-08 | 7.400E-08 |
| 9152 | 458 | V458K | A | 6.6e-08 | 4.2e-08 | 3.8e-08 | 4.867E-08 | 1.514E-08 | 3.800E-08 | 6.600E-08 |
| 9153 | 458 | V458M | A | 1.9e-07 | 3.8e-08 | 2.6e-08 | 8.467E-08 | 9.142E-08 | 2.600E-08 | 1.900E-07 |
| 9154 | 458 | V458F | A | 4.3e-08 | 7.5e-08 | 6.8e-08 | 6.200E-08 | 1.682E-08 | 4.300E-08 | 7.500E-08 |
| 9155 | 458 | V458P | A | 6.9e-08 | 4.6e-08 | 3.9e-08 | 5.133E-08 | 1.570E-08 | 3.900E-08 | 6.900E-08 |
| 9156 | 458 | V458S | A | 2.7e-08 | 1.3e-08 | 2.9e-08 | 2.300E-08 | 8.718E-09 | 1.300E-08 | 2.900E-08 |
| 9157 | 458 | V458T | A | 2.7e-08 | 4.3e-08 | 1.2e-07 | 6.333E-08 | 4.972E-08 | 2.700E-08 | 1.200E-07 |
| 9158 | 458 | V458W | A | 3.1e-08 | 2.9e-08 | 3.6e-08 | 3.200E-08 | 3.606E-09 | 2.900E-08 | 3.600E-08 |
| 9159 | 458 | V458Y | A | 6.7e-08 | 6.7e-08 | 2.1e-08 | 5.167E-08 | 2.656E-08 | 2.100E-08 | 6.700E-08 |
| 9160 | 458 | V458V | A | 9.5e-08 | 6.2e-08 | 2.9e-08 | 6.200E-08 | 3.300E-08 | 2.900E-08 | 9.500E-08 |
| 9161 | 459 | P459A | A | 1.2e-08 | 1.4e-08 | 6.6e-08 | 3.067E-08 | 3.062E-08 | 1.200E-08 | 6.600E-08 |
| 9162 | 459 | P459R | A | 2.2e-08 | 8.1e-08 | 2.5e-08 | 4.267E-08 | 3.323E-08 | 2.200E-08 | 8.100E-08 |
| 9163 | 459 | P459N | A | 3.7e-08 | 2.4e-08 | 3.4e-08 | 3.167E-08 | 6.807E-09 | 2.400E-08 | 3.700E-08 |
| 9164 | 459 | P459D | A | 5.9e-08 | 3.9e-08 | 3.4e-08 | 4.400E-08 | 1.323E-08 | 3.400E-08 | 5.900E-08 |
| 9165 | 459 | P459C | A | 3e-08 | 4e-08 | 4.4e-08 | 3.800E-08 | 7.211E-09 | 3.000E-08 | 4.400E-08 |
| 9166 | 459 | P459E | A | 4.3e-08 | 7.1e-08 | 5.2e-08 | 5.533E-08 | 1.429E-08 | 4.300E-08 | 7.100E-08 |
| 9167 | 459 | P459Q | A | 2.8e-08 | 2.2e-08 | 2.5e-07 | 1.000E-07 | 1.299E-07 | 2.200E-08 | 2.500E-07 |
| 9168 | 459 | P459G | A | 1.4e-08 | 4.2e-08 | 1.1e-07 | 5.533E-08 | 4.937E-08 | 1.400E-08 | 1.100E-07 |
| 9169 | 459 | P459H | A | 6.7e-08 | 4.5e-08 | 5.8e-08 | 5.667E-08 | 1.106E-08 | 4.500E-08 | 6.700E-08 |
| 9170 | 459 | P459I | A | 5.4e-08 | 2.8e-08 | 3.5e-08 | 3.900E-08 | 1.345E-08 | 2.800E-08 | 5.400E-08 |
| 9171 | 459 | P459L | A | 8.7e-08 | 4e-08 | 3.4e-08 | 5.367E-08 | 2.902E-08 | 3.400E-08 | 8.700E-08 |
| 9172 | 459 | P459K | A | 2.2e-08 | 4e-08 | 2.1e-08 | 2.767E-08 | 1.069E-08 | 2.100E-08 | 4.000E-08 |
| 9173 | 459 | P459M | A | 5.8e-08 | 5.8e-08 | 3.9e-08 | 5.167E-08 | 1.097E-08 | 3.900E-08 | 5.800E-08 |
| 9174 | 459 | P459F | A | 9.7e-08 | 7.4e-08 | 3.6e-08 | 6.900E-08 | 3.081E-08 | 3.600E-08 | 9.700E-08 |
| 9175 | 459 | P459P | A | 4.4e-08 | 2.5e-08 | 2.6e-08 | 3.167E-08 | 1.069E-08 | 2.500E-08 | 4.400E-08 |
| 9176 | 459 | P459S | A | 3.4e-08 | 2.6e-08 | 3.3e-08 | 3.100E-08 | 4.359E-09 | 2.600E-08 | 3.400E-08 |
| 9177 | 459 | P459T | A | 9.1e-08 | 6e-08 | 3.1e-08 | 6.067E-08 | 3.001E-08 | 3.100E-08 | 9.100E-08 |
| 9178 | 459 | P459W | A | 1.4e-08 | 6.9e-08 | 3.7e-08 | 4.000E-08 | 2.762E-08 | 1.400E-08 | 6.900E-08 |
| 9179 | 459 | P459Y | A | 2.7e-08 | 1e-07 | 2.9e-08 | 5.200E-08 | 4.158E-08 | 2.700E-08 | 1.000E-07 |

| | | | | | | | | | | |
|---|---|---|---|---|---|---|---|---|---|---|
| 9180 | 459 | P459V | A | 6.9e-08 | 3.4e-08 | 2.4e-08 | 4.233E-08 | 2.363E-08 | 2.400E-08 | 6.900E-08 |
| 9181 | 460 | W460A | A | 2.8e-08 | 5.2e-08 | 6.3e-08 | 4.767E-08 | 1.790E-08 | 2.800E-08 | 6.300E-08 |
| 9182 | 460 | W460R | A | 6.7e-08 | 2.8e-08 | 3.1e-08 | 4.200E-08 | 2.170E-08 | 2.800E-08 | 6.700E-08 |
| 9183 | 460 | W460N | A | 1.7e-08 | 3.2e-08 | 2.2e-08 | 2.367E-08 | 7.638E-09 | 1.700E-08 | 3.200E-08 |
| 9184 | 460 | W460D | A | 5.3e-08 | 2.8e-08 | 2.7e-08 | 3.600E-08 | 1.473E-08 | 2.700E-08 | 5.300E-08 |
| 9185 | 460 | W460C | A | 1.1e-07 | 5.7e-08 | 6.8e-08 | 7.833E-08 | 2.797E-08 | 5.700E-08 | 1.100E-07 |
| 9186 | 460 | W460E | A | 1.9e-07 | 9.7e-08 | 5.6e-08 | 1.143E-07 | 6.866E-08 | 5.600E-08 | 1.900E-07 |
| 9187 | 460 | W460Q | A | 7.1e-08 | 2.6e-08 | 2.3e-08 | 4.000E-08 | 2.689E-08 | 2.300E-08 | 7.100E-08 |
| 9188 | 460 | W460G | A | 2.6e-08 | 1.6e-08 | 8e-08 | 4.067E-08 | 3.443E-08 | 1.600E-08 | 8.000E-08 |
| 9189 | 460 | W460H | A | 7.2e-08 | 4.9e-08 | 4e-08 | 5.367E-08 | 1.650E-08 | 4.000E-08 | 7.200E-08 |
| 9190 | 460 | W460I | A | 3.2e-08 | 2.2e-08 | 3.1e-08 | 2.833E-08 | 5.508E-09 | 2.200E-08 | 3.200E-08 |
| 9191 | 460 | W460L | A | 5.5e-08 | 3.9e-08 | 2.9e-08 | 4.100E-08 | 1.311E-08 | 2.900E-08 | 5.500E-08 |
| 9192 | 460 | W460K | A | 7.5e-08 | 2.8e-08 | 5.5e-08 | 5.267E-08 | 2.359E-08 | 2.800E-08 | 7.500E-08 |
| 9193 | 460 | W460M | A | 3.8e-08 | 1.6e-08 | 3.6e-08 | 3.000E-08 | 1.217E-08 | 1.600E-08 | 3.800E-08 |
| 9194 | 460 | W460F | A | 6.1e-08 | 6e-08 | 3.1e-08 | 5.067E-08 | 1.704E-08 | 3.100E-08 | 6.100E-08 |
| 9195 | 460 | W460P | A | 4.4e-08 | 5.7e-08 | 4e-08 | 4.700E-08 | 8.888E-09 | 4.000E-08 | 5.700E-08 |
| 9196 | 460 | W460S | A | 1.8e-08 | 2.7e-08 | 4.3e-08 | 2.933E-08 | 1.266E-08 | 1.800E-08 | 4.300E-08 |
| 9197 | 460 | W460T | A | 5.7e-08 | 3.5e-08 | 2.5e-08 | 3.900E-08 | 1.637E-08 | 2.500E-08 | 5.700E-08 |
| 9198 | 460 | W460W | A | 2.2e-08 | 2.9e-08 | 2.2e-08 | 2.433E-08 | 4.041E-09 | 2.200E-08 | 2.900E-08 |
| 9199 | 460 | W460Y | A | 7.4e-08 | 2e-08 | 1.5e-08 | 3.633E-08 | 3.272E-08 | 1.500E-08 | 7.400E-08 |
| 9200 | 460 | W460V | A | 4e-08 | 1.7e-08 | 4.1e-08 | 3.267E-08 | 1.358E-08 | 1.700E-08 | 4.100E-08 |
| 9201 | 461 | D461A | A | 1.3e-08 | 3e-08 | 4.1e-08 | 2.800E-08 | 1.411E-08 | 1.300E-08 | 4.100E-08 |
| 9202 | 461 | D461R | A | 3.1e-08 | 3.1e-08 | 3.2e-08 | 3.133E-08 | 5.774E-10 | 3.100E-08 | 3.200E-08 |
| 9203 | 461 | D461N | A | 2.4e-08 | 6.5e-08 | 1.9e-08 | 3.600E-08 | 2.524E-08 | 1.900E-08 | 6.500E-08 |
| 9204 | 461 | D461D | A | 9.1e-08 | 1.6e-08 | 7.6e-08 | 6.100E-08 | 3.969E-08 | 1.600E-08 | 9.100E-08 |
| 9205 | 461 | D461C | A | 4.7e-08 | 4.6e-08 | 9.8e-09 | 3.427E-08 | 2.119E-08 | 9.800E-09 | 4.700E-08 |
| 9206 | 461 | D461E | A | 4.1e-08 | 6.9e-08 | 3.6e-08 | 4.867E-08 | 1.779E-08 | 3.600E-08 | 6.900E-08 |
| 9207 | 461 | D461Q | A | 7e-08 | 3.3e-08 | 2.4e-08 | 4.233E-08 | 2.438E-08 | 2.400E-08 | 7.000E-08 |
| 9208 | 461 | D461G | A | 1e-07 | 1.8e-08 | 3.6e-08 | 5.133E-08 | 4.310E-08 | 1.800E-08 | 1.000E-07 |
| 9209 | 461 | D461H | A | 6.2e-08 | 4.9e-08 | 2.9e-08 | 4.667E-08 | 1.662E-08 | 2.900E-08 | 6.200E-08 |
| 9210 | 461 | D461I | A | 1.3e-08 | 1.3e-07 | 1.8e-08 | 5.367E-08 | 6.615E-08 | 1.300E-08 | 1.300E-07 |
| 9211 | 461 | D461L | A | 1.2e-08 | 4.3e-08 | 2e-08 | 2.500E-08 | 1.609E-08 | 1.200E-08 | 4.300E-08 |
| 9212 | 461 | D461K | A | 2.9e-08 | 2.3e-08 | 3.6e-08 | 2.933E-08 | 6.506E-09 | 2.300E-08 | 3.600E-08 |
| 9213 | 461 | D461M | A | 4.1e-08 | 7.2e-08 | 5.5e-08 | 5.600E-08 | 1.552E-08 | 4.100E-08 | 7.200E-08 |
| 9214 | 461 | D461F | A | 7.5e-08 | 6.1e-08 | 2.6e-08 | 5.400E-08 | 2.524E-08 | 2.600E-08 | 7.500E-08 |
| 9215 | 461 | D461P | A | 1.2e-07 | 3.7e-08 | 2.3e-08 | 6.000E-08 | 5.243E-08 | 2.300E-08 | 1.200E-07 |
| 9216 | 461 | D461S | A | 2.6e-08 | 4.4e-08 | 7.5e-08 | 4.833E-08 | 2.479E-08 | 2.600E-08 | 7.500E-08 |
| 9217 | 461 | D461T | A | 5.6e-08 | 4.3e-08 | 2.6e-08 | 4.167E-08 | 1.504E-08 | 2.600E-08 | 5.600E-08 |
| 9218 | 461 | D461W | A | 1.8e-08 | 3.9e-08 | 1.5e-08 | 2.400E-08 | 1.308E-08 | 1.500E-08 | 3.900E-08 |
| 9219 | 461 | D461Y | A | 3.3e-08 | 3.5e-08 | 3.8e-08 | 3.533E-08 | 2.517E-09 | 3.300E-08 | 3.800E-08 |
| 9220 | 461 | D461V | A | 3.5e-08 | 3.2e-08 | 2.9e-08 | 3.200E-08 | 3.000E-09 | 2.900E-08 | 3.500E-08 |
| 9221 | 462 | Q462A | A | 1.1e-08 | 3.4e-08 | 5.8e-08 | 3.433E-08 | 2.350E-08 | 1.100E-08 | 5.800E-08 |
| 9222 | 462 | Q462R | A | 2.3e-08 | 1.9e-08 | 5e-08 | 3.067E-08 | 1.686E-08 | 1.900E-08 | 5.000E-08 |
| 9223 | 462 | Q462N | A | 7.1e-08 | 1.1e-08 | 1.5e-08 | 3.233E-08 | 3.355E-08 | 1.100E-08 | 7.100E-08 |
| 9224 | 462 | Q462D | A | 1.6e-08 | 9.5e-08 | 1.7e-08 | 4.267E-08 | 4.532E-08 | 1.600E-08 | 9.500E-08 |
| 9225 | 462 | Q462C | A | 8.8e-08 | 8.6e-08 | 1.6e-07 | 1.113E-07 | 4.216E-08 | 8.600E-08 | 1.600E-07 |
| 9226 | 462 | Q462E | A | 1.8e-08 | 8.4e-08 | 4.6e-08 | 4.933E-08 | 3.313E-08 | 1.800E-08 | 8.400E-08 |
| 9227 | 462 | Q462Q | A | 4.5e-08 | 3.9e-08 | 6.8e-08 | 5.067E-08 | 1.531E-08 | 3.900E-08 | 6.800E-08 |
| 9228 | 462 | Q462G | A | 1.5e-08 | 5.6e-08 | 2.9e-08 | 3.333E-08 | 2.084E-08 | 1.500E-08 | 5.600E-08 |
| 9229 | 462 | Q462H | A | 2.5e-08 | 5.9e-08 | 2.7e-08 | 3.700E-08 | 1.908E-08 | 2.500E-08 | 5.900E-08 |
| 9230 | 462 | Q462I | A | 2.4e-08 | 6e-08 | 1.2e-08 | 3.200E-08 | 2.498E-08 | 1.200E-08 | 6.000E-08 |
| 9231 | 462 | Q462L | A | 3e-08 | 6.6e-08 | 8.9e-08 | 6.167E-08 | 2.974E-08 | 3.000E-08 | 8.900E-08 |
| 9232 | 462 | Q462K | A | 5.5e-08 | 2.5e-08 | 3e-08 | 3.667E-08 | 1.607E-08 | 2.500E-08 | 5.500E-08 |
| 9233 | 462 | Q462M | A | 5.4e-08 | 2.8e-08 | 8.3e-08 | 5.500E-08 | 2.751E-08 | 2.800E-08 | 8.300E-08 |
| 9234 | 462 | Q462F | A | 4.2e-08 | 2.2e-08 | 8.4e-08 | 4.933E-08 | 3.164E-08 | 2.200E-08 | 8.400E-08 |
| 9235 | 462 | Q462P | A | 3e-08 | 1.7e-08 | 1.6e-08 | 2.100E-08 | 7.810E-09 | 1.600E-08 | 3.000E-08 |
| 9236 | 462 | Q462S | A | 2.6e-08 | 5.5e-08 | 2.5e-08 | 3.533E-08 | 1.704E-08 | 2.500E-08 | 5.500E-08 |
| 9237 | 462 | Q462T | A | 3e-08 | 4.3e-08 | 5.4e-08 | 4.233E-08 | 1.201E-08 | 3.000E-08 | 5.400E-08 |
| 9238 | 462 | Q462W | A | 4.1e-08 | 3.3e-08 | 5.2e-08 | 4.200E-08 | 9.539E-09 | 3.300E-08 | 5.200E-08 |
| 9239 | 462 | Q462Y | A | 8.5e-08 | 9.3e-08 | 2.5e-08 | 6.767E-08 | 3.717E-08 | 2.500E-08 | 9.300E-08 |
| 9240 | 462 | Q462V | A | 2.1e-08 | 1.3e-07 | 5.5e-08 | 6.867E-08 | 5.577E-08 | 2.100E-08 | 1.300E-07 |
| 9241 | 463 | L463A | A | 2.8e-08 | 6.5e-08 | 6.1e-08 | 5.133E-08 | 2.031E-08 | 2.800E-08 | 6.500E-08 |
| 9242 | 463 | L463R | A | 1.5e-08 | 8.9e-08 | 7e-08 | 5.800E-08 | 3.843E-08 | 1.500E-08 | 8.900E-08 |
| 9243 | 463 | L463N | A | 5.1e-08 | 1.2e-08 | 5.1e-08 | 3.800E-08 | 2.252E-08 | 1.200E-08 | 5.100E-08 |
| 9244 | 463 | L463D | A | 1.4e-08 | 5.3e-08 | 3.5e-08 | 3.400E-08 | 1.952E-08 | 1.400E-08 | 5.300E-08 |
| 9245 | 463 | L463C | A | 2.1e-07 | 4.7e-08 | 6.2e-08 | 1.063E-07 | 9.009E-08 | 4.700E-08 | 2.100E-07 |
| 9246 | 463 | L463E | A | 3.5e-08 | 6.2e-08 | 3.1e-08 | 4.267E-08 | 1.686E-08 | 3.100E-08 | 6.200E-08 |
| 9247 | 463 | L463Q | A | 3.6e-08 | 6.9e-08 | 5e-08 | 5.167E-08 | 1.656E-08 | 3.600E-08 | 6.900E-08 |
| 9248 | 463 | L463G | A | 3e-08 | 4e-08 | 7.5e-08 | 4.833E-08 | 2.363E-08 | 3.000E-08 | 7.500E-08 |
| 9249 | 463 | L463H | A | 5.9e-08 | 1.1e-08 | 2.8e-08 | 3.267E-08 | 2.434E-08 | 1.100E-08 | 5.900E-08 |
| 9250 | 463 | L463I | A | 3e-08 | 4.1e-08 | 1.1e-07 | 6.033E-08 | 4.336E-08 | 3.000E-08 | 1.100E-07 |
| 9251 | 463 | L463L | A | 2.2e-08 | 5.1e-08 | 2.4e-08 | 3.233E-08 | 1.620E-08 | 2.200E-08 | 5.100E-08 |
| 9252 | 463 | L463K | A | 2.5e-08 | 1.1e-07 | 6.1e-08 | 6.533E-08 | 4.267E-08 | 2.500E-08 | 1.100E-07 |
| 9253 | 463 | L463M | A | 2.4e-08 | 7e-08 | 6.3e-08 | 5.233E-08 | 2.479E-08 | 2.400E-08 | 7.000E-08 |
| 9254 | 463 | L463F | A | 2.3e-08 | 7.1e-08 | 9.5e-08 | 6.300E-08 | 3.666E-08 | 2.300E-08 | 9.500E-08 |
| 9255 | 463 | L463P | A | 2.8e-08 | 2.8e-08 | 4.5e-08 | 3.367E-08 | 9.815E-09 | 2.800E-08 | 4.500E-08 |
| 9256 | 463 | L463S | A | 7.1e-08 | 4.5e-08 | 2.7e-07 | 1.287E-07 | 1.231E-07 | 4.500E-08 | 2.700E-07 |
| 9257 | 463 | L463T | A | 4.1e-08 | 3.6e-08 | 3.8e-08 | 3.833E-08 | 2.517E-09 | 3.600E-08 | 4.100E-08 |
| 9258 | 463 | L463W | A | 6.4e-08 | 3.6e-08 | 5.4e-08 | 5.133E-08 | 1.419E-08 | 3.600E-08 | 6.400E-08 |
| 9259 | 463 | L463Y | A | 5.3e-08 | 6.8e-08 | 5e-08 | 5.700E-08 | 9.644E-09 | 5.000E-08 | 6.800E-08 |
| 9260 | 463 | L463V | A | 3.5e-08 | 3.7e-08 | 2.1e-07 | 9.400E-08 | 1.005E-07 | 3.500E-08 | 2.100E-07 |
| 9261 | 464 | F464A | A | 2.9e-08 | 3.4e-08 | 5.5e-08 | 3.933E-08 | 1.380E-08 | 2.900E-08 | 5.500E-08 |
| 9262 | 464 | F464R | A | 6.5e-08 | 4.4e-08 | 3.6e-08 | 4.833E-08 | 1.498E-08 | 3.600E-08 | 6.500E-08 |
| 9263 | 464 | F464N | A | 1.3e-07 | 6e-08 | 9.3e-08 | 9.433E-08 | 3.502E-08 | 6.000E-08 | 1.300E-07 |
| 9264 | 464 | F464D | A | 5e-08 | 4.4e-08 | 4.5e-08 | 4.633E-08 | 3.215E-09 | 4.400E-08 | 5.000E-08 |

| | | | | | | | | | | |
|---|---|---|---|---|---|---|---|---|---|---|
| 9265 | 464 | F464C | A | 3.2e-08 | 6.2e-08 | 6.3e-08 | 5.233E-08 | 1.762E-08 | 3.200E-08 | 6.300E-08 |
| 9266 | 464 | F464E | A | 3.5e-08 | 1.5e-08 | 2.2e-08 | 2.400E-08 | 1.015E-08 | 1.500E-08 | 3.500E-08 |
| 9267 | 464 | F464Q | A | 5.7e-08 | 4.8e-08 | 7.6e-08 | 6.033E-08 | 1.429E-08 | 4.800E-08 | 7.600E-08 |
| 9268 | 464 | F464G | A | 4e-08 | 8.2e-08 | 2.7e-08 | 4.967E-08 | 2.875E-08 | 2.700E-08 | 8.200E-08 |
| 9269 | 464 | F464H | A | 3.8e-08 | 4.8e-08 | 2.2e-08 | 3.600E-08 | 1.311E-08 | 2.200E-08 | 4.800E-08 |
| 9270 | 464 | F464I | A | 3.2e-08 | 7.8e-08 | 6.3e-08 | 5.767E-08 | 2.346E-08 | 3.200E-08 | 7.800E-08 |
| 9271 | 464 | F464L | A | 4.5e-08 | 3.5e-08 | 2.6e-08 | 3.533E-08 | 9.504E-09 | 2.600E-08 | 4.500E-08 |
| 9272 | 464 | F464K | A | 6.9e-08 | 4.5e-08 | 4.4e-08 | 5.267E-08 | 1.415E-08 | 4.400E-08 | 6.900E-08 |
| 9273 | 464 | F464M | A | 2.5e-08 | 5.6e-08 | 1e-07 | 6.033E-08 | 3.769E-08 | 2.500E-08 | 1.000E-07 |
| 9274 | 464 | F464F | A | 3.9e-08 | 1.7e-08 | 3.8e-08 | 3.133E-08 | 1.242E-08 | 1.700E-08 | 3.900E-08 |
| 9275 | 464 | F464P | A | 3.6e-08 | 1.2e-07 | 2.2e-08 | 5.933E-08 | 5.300E-08 | 2.200E-08 | 1.200E-07 |
| 9276 | 464 | F464S | A | 7.3e-08 | 2.1e-08 | 6.2e-08 | 5.200E-08 | 2.740E-08 | 2.100E-08 | 7.300E-08 |
| 9277 | 464 | F464T | A | 2.7e-08 | 3.2e-08 | 5.8e-08 | 3.900E-08 | 1.664E-08 | 2.700E-08 | 5.800E-08 |
| 9278 | 464 | F464W | A | 3.6e-08 | 5.5e-08 | 4.9e-08 | 4.667E-08 | 9.713E-09 | 3.600E-08 | 5.500E-08 |
| 9279 | 464 | F464Y | A | 5.9e-08 | 3.5e-08 | 3.6e-08 | 4.333E-08 | 1.358E-08 | 3.500E-08 | 5.900E-08 |
| 9280 | 464 | F464V | A | 7.5e-08 | 2e-08 | 6.4e-08 | 5.300E-08 | 2.910E-08 | 2.000E-08 | 7.500E-08 |
| 9281 | 465 | R465A | A | 8.1e-08 | 3e-08 | 1.7e-08 | 4.267E-08 | 3.383E-08 | 1.700E-08 | 8.100E-08 |
| 9282 | 465 | R465R | A | 8.3e-08 | 3.1e-08 | 4.6e-08 | 5.333E-08 | 2.676E-08 | 3.100E-08 | 8.300E-08 |
| 9283 | 465 | R465N | A | 6.5e-08 | 3e-08 | 2.6e-08 | 4.033E-08 | 2.146E-08 | 2.600E-08 | 6.500E-08 |
| 9284 | 465 | R465D | A | 9.4e-08 | 2.8e-08 | 9e-08 | 7.067E-08 | 3.700E-08 | 2.800E-08 | 9.400E-08 |
| 9285 | 465 | R465C | A | 4.6e-08 | 1.8e-08 | 3.2e-08 | 3.200E-08 | 1.400E-08 | 1.800E-08 | 4.600E-08 |
| 9286 | 465 | R465E | A | 4.3e-08 | 8e-08 | 3.7e-08 | 5.333E-08 | 2.329E-08 | 3.700E-08 | 8.000E-08 |
| 9287 | 465 | R465Q | A | 1.9e-08 | 1.5e-07 | 6e-08 | 7.633E-08 | 6.701E-08 | 1.900E-08 | 1.500E-07 |
| 9288 | 465 | R465G | A | 2.7e-08 | 3.4e-08 | 2.4e-08 | 2.833E-08 | 5.132E-09 | 2.400E-08 | 3.400E-08 |
| 9289 | 465 | R465H | A | 3.7e-08 | 6.4e-08 | 4.6e-08 | 4.900E-08 | 1.375E-08 | 3.700E-08 | 6.400E-08 |
| 9290 | 465 | R465I | A | 7.4e-08 | 4.1e-08 | 6e-08 | 5.833E-08 | 1.656E-08 | 4.100E-08 | 7.400E-08 |
| 9291 | 465 | R465L | A | 2.5e-08 | 5.8e-08 | 3.6e-08 | 3.967E-08 | 1.680E-08 | 2.500E-08 | 5.800E-08 |
| 9292 | 465 | R465K | A | 3.2e-08 | 2.1e-08 | 1.5e-08 | 2.267E-08 | 8.622E-09 | 1.500E-08 | 3.200E-08 |
| 9293 | 465 | R465M | A | 1.1e-07 | 3.5e-08 | 2.7e-08 | 5.733E-08 | 4.579E-08 | 2.700E-08 | 1.100E-07 |
| 9294 | 465 | R465F | A | 3.7e-08 | 9.7e-08 | 3.3e-08 | 5.567E-08 | 3.585E-08 | 3.300E-08 | 9.700E-08 |
| 9295 | 465 | R465P | A | 5.1e-08 | 1.1e-07 | 1.8e-07 | 1.137E-07 | 6.458E-08 | 5.100E-08 | 1.800E-07 |
| 9296 | 465 | R465S | A | 3.3e-08 | 5e-08 | 2.1e-08 | 3.467E-08 | 1.457E-08 | 2.100E-08 | 5.000E-08 |
| 9297 | 465 | R465T | A | 2.8e-08 | 2.7e-08 | 3.8e-08 | 3.100E-08 | 6.083E-09 | 2.700E-08 | 3.800E-08 |
| 9298 | 465 | R465W | A | 1.6e-08 | 4.8e-08 | 6.1e-08 | 4.167E-08 | 2.316E-08 | 1.600E-08 | 6.100E-08 |
| 9299 | 465 | R465Y | A | 3.7e-08 | 1.8e-08 | 3.8e-08 | 3.100E-08 | 1.127E-08 | 1.800E-08 | 3.800E-08 |
| 9300 | 465 | R465V | A | 1e-07 | 6.1e-08 | 4.6e-08 | 6.900E-08 | 2.787E-08 | 4.600E-08 | 1.000E-07 |
| 9301 | 466 | N466A | A | 5.1e-08 | 6.7e-08 | 3.7e-08 | 5.167E-08 | 1.501E-08 | 3.700E-08 | 6.700E-08 |
| 9302 | 466 | N466R | A | 9.1e-08 | 4.2e-08 | 2e-08 | 5.100E-08 | 3.635E-08 | 2.000E-08 | 9.100E-08 |
| 9303 | 466 | N466N | A | 6.2e-08 | 7.7e-08 | 4.3e-08 | 6.067E-08 | 1.704E-08 | 4.300E-08 | 7.700E-08 |
| 9304 | 466 | N466D | A | 2.2e-08 | 4.4e-08 | 2.2e-08 | 2.933E-08 | 1.270E-08 | 2.200E-08 | 4.400E-08 |
| 9305 | 466 | N466C | A | 2.3e-08 | 1.7e-08 | 3.8e-08 | 2.600E-08 | 1.082E-08 | 1.700E-08 | 3.800E-08 |
| 9306 | 466 | N466E | A | 3.5e-08 | 2.7e-08 | 4.2e-08 | 3.467E-08 | 7.506E-09 | 2.700E-08 | 4.200E-08 |
| 9307 | 466 | N466Q | A | 5.7e-08 | 3.5e-08 | 7.1e-08 | 5.433E-08 | 1.815E-08 | 3.500E-08 | 7.100E-08 |
| 9308 | 466 | N466G | A | 5.1e-08 | 1.5e-08 | 5.3e-08 | 3.967E-08 | 2.139E-08 | 1.500E-08 | 5.300E-08 |
| 9309 | 466 | N466H | A | 3.4e-08 | 9e-08 | 9.9e-09 | 4.463E-08 | 4.110E-08 | 9.900E-09 | 9.000E-08 |
| 9310 | 466 | N466I | A | 5.5e-08 | 5.1e-08 | 4.1e-08 | 4.900E-08 | 7.211E-09 | 4.100E-08 | 5.500E-08 |
| 9311 | 466 | N466L | A | 5.2e-08 | 4.9e-08 | 2.5e-08 | 4.200E-08 | 1.480E-08 | 2.500E-08 | 5.200E-08 |
| 9312 | 466 | N466K | A | 4.9e-08 | 2e-08 | 5.1e-08 | 4.000E-08 | 1.735E-08 | 2.000E-08 | 5.100E-08 |
| 9313 | 466 | N466M | A | 1.5e-07 | 6.4e-08 | 5.6e-08 | 9.000E-08 | 5.212E-08 | 5.600E-08 | 1.500E-07 |
| 9314 | 466 | N466F | A | 1.2e-07 | 7.4e-08 | 4.8e-08 | 8.067E-08 | 3.646E-08 | 4.800E-08 | 1.200E-07 |
| 9315 | 466 | N466P | A | 3.7e-08 | 8.7e-09 | 4.3e-08 | 2.957E-08 | 1.832E-08 | 8.700E-09 | 4.300E-08 |
| 9316 | 466 | N466S | A | 2.4e-08 | 2.4e-08 | 2.2e-08 | 2.333E-08 | 1.155E-09 | 2.200E-08 | 2.400E-08 |
| 9317 | 466 | N466T | A | 6.1e-08 | 1.2e-07 | 5.9e-08 | 8.000E-08 | 3.466E-08 | 5.900E-08 | 1.200E-07 |
| 9318 | 466 | N466W | A | 1.4e-08 | 1.6e-08 | 1.4e-08 | 1.467E-08 | 1.155E-09 | 1.400E-08 | 1.600E-08 |
| 9319 | 466 | N466Y | A | 4.2e-08 | 3.3e-08 | 3.2e-08 | 3.567E-08 | 5.508E-09 | 3.200E-08 | 4.200E-08 |
| 9320 | 466 | N466V | A | 1.3e-07 | 2.7e-08 | 2.8e-08 | 6.167E-08 | 5.918E-08 | 2.700E-08 | 1.300E-07 |
| 9321 | 467 | P467A | A | 7.7e-08 | 2.9e-08 | 8.1e-08 | 6.233E-08 | 2.894E-08 | 2.900E-08 | 8.100E-08 |
| 9322 | 467 | P467R | A | 2.4e-08 | 3.6e-08 | 5.6e-08 | 3.867E-08 | 1.617E-08 | 2.400E-08 | 5.600E-08 |
| 9323 | 467 | P467N | A | 7.2e-08 | 4.9e-08 | 3.4e-08 | 5.167E-08 | 1.914E-08 | 3.400E-08 | 7.200E-08 |
| 9324 | 467 | P467D | A | 9.4e-08 | 2.6e-08 | 9.3e-08 | 7.100E-08 | 3.897E-08 | 2.600E-08 | 9.400E-08 |
| 9325 | 467 | P467C | A | 1.1e-07 | 6.2e-08 | 3.2e-08 | 6.800E-08 | 3.934E-08 | 3.200E-08 | 1.100E-07 |
| 9326 | 467 | P467E | A | 3.2e-08 | 2.9e-08 | 5.5e-08 | 3.867E-08 | 1.422E-08 | 2.900E-08 | 5.500E-08 |
| 9327 | 467 | P467Q | A | 2e-08 | 3.4e-08 | 3.2e-08 | 2.867E-08 | 7.572E-09 | 2.000E-08 | 3.400E-08 |
| 9328 | 467 | P467G | A | 2.9e-08 | 2.2e-08 | 6.2e-08 | 3.767E-08 | 2.136E-08 | 2.200E-08 | 6.200E-08 |
| 9329 | 467 | P467H | A | 2.7e-08 | 5.7e-08 | 6.9e-08 | 5.100E-08 | 2.163E-08 | 2.700E-08 | 6.900E-08 |
| 9330 | 467 | P467I | A | 5.8e-08 | 3e-08 | 4.4e-08 | 4.400E-08 | 1.400E-08 | 3.000E-08 | 5.800E-08 |
| 9331 | 467 | P467L | A | 3.4e-08 | 5.1e-08 | 8.1e-08 | 5.533E-08 | 2.380E-08 | 3.400E-08 | 8.100E-08 |
| 9332 | 467 | P467K | A | 7.4e-08 | 8.4e-08 | 4e-08 | 6.600E-08 | 2.307E-08 | 4.000E-08 | 8.400E-08 |
| 9333 | 467 | P467M | A | 7.4e-08 | 1.3e-07 | 3.8e-08 | 8.067E-08 | 4.636E-08 | 3.800E-08 | 1.300E-07 |
| 9334 | 467 | P467F | A | 1.6e-08 | 5.3e-08 | 6.7e-08 | 4.533E-08 | 2.635E-08 | 1.600E-08 | 6.700E-08 |
| 9335 | 467 | P467P | A | 9.1e-08 | 6.7e-08 | 2.7e-08 | 6.167E-08 | 3.233E-08 | 2.700E-08 | 9.100E-08 |
| 9336 | 467 | P467S | A | 8.6e-08 | 5.1e-08 | 8e-08 | 7.233E-08 | 1.872E-08 | 5.100E-08 | 8.600E-08 |
| 9337 | 467 | P467T | A | 4.2e-08 | 5.3e-08 | 5.5e-08 | 5.000E-08 | 7.000E-09 | 4.200E-08 | 5.500E-08 |
| 9338 | 467 | P467W | A | 4.8e-08 | 8.3e-08 | 3.8e-08 | 5.633E-08 | 2.363E-08 | 3.800E-08 | 8.300E-08 |
| 9339 | 467 | P467Y | A | 4e-08 | 1.2e-07 | 3.5e-08 | 6.500E-08 | 4.770E-08 | 3.500E-08 | 1.200E-07 |
| 9340 | 467 | P467V | A | 6.7e-08 | 7.3e-08 | 3e-08 | 5.667E-08 | 2.329E-08 | 3.000E-08 | 7.300E-08 |
| 9341 | 468 | H468A | A | 4e-08 | 3.6e-08 | 5.3e-08 | 4.300E-08 | 8.888E-09 | 3.600E-08 | 5.300E-08 |
| 9342 | 468 | H468R | A | 9.8e-08 | 3.5e-08 | 7.7e-08 | 7.000E-08 | 3.208E-08 | 3.500E-08 | 9.800E-08 |
| 9343 | 468 | H468N | A | 2.9e-08 | 8.9e-08 | 5e-08 | 5.600E-08 | 3.045E-08 | 2.900E-08 | 8.900E-08 |
| 9344 | 468 | H468D | A | 3.2e-08 | 7.7e-08 | 3e-08 | 4.633E-08 | 2.658E-08 | 3.000E-08 | 7.700E-08 |
| 9345 | 468 | H468C | A | 4.6e-08 | 2.8e-08 | 5.6e-08 | 4.333E-08 | 1.419E-08 | 2.800E-08 | 5.600E-08 |
| 9346 | 468 | H468E | A | 1.1e-08 | 7.9e-08 | 4.9e-08 | 4.633E-08 | 3.408E-08 | 1.100E-08 | 7.900E-08 |
| 9347 | 468 | H468Q | A | 7.6e-09 | 3.7e-08 | 3.4e-08 | 2.620E-08 | 1.618E-08 | 7.600E-09 | 3.700E-08 |
| 9348 | 468 | H468G | A | 4.5e-08 | 2.1e-08 | 9.5e-08 | 5.367E-08 | 3.775E-08 | 2.100E-08 | 9.500E-08 |
| 9349 | 468 | H468H | A | 2.4e-08 | 3.1e-08 | 1.8e-07 | 7.833E-08 | 8.812E-08 | 2.400E-08 | 1.800E-07 |

| | | | | | | | | | | |
|---|---|---|---|---|---|---|---|---|---|---|
| 9350 | 468 | H468I | A | 1.6e-08 | 4.3e-08 | 7.1e-08 | 4.333E-08 | 2.750E-08 | 1.600E-08 | 7.100E-08 |
| 9351 | 468 | H468L | A | 4.5e-08 | 9.3e-08 | 5.3e-08 | 6.367E-08 | 2.572E-08 | 4.500E-08 | 9.300E-08 |
| 9352 | 468 | H468K | A | 3.9e-08 | 2.6e-08 | 1.8e-08 | 2.767E-08 | 1.060E-08 | 1.800E-08 | 3.900E-08 |
| 9353 | 468 | H468M | A | 1.5e-07 | 2.7e-08 | 6.3e-08 | 8.000E-08 | 6.324E-08 | 2.700E-08 | 1.500E-07 |
| 9354 | 468 | H468F | A | 3.5e-08 | 3.9e-08 | 2.8e-08 | 3.400E-08 | 5.568E-09 | 2.800E-08 | 3.900E-08 |
| 9355 | 468 | H468P | A | 4.9e-08 | 3e-08 | 1.8e-08 | 3.233E-08 | 1.563E-08 | 1.800E-08 | 4.900E-08 |
| 9356 | 468 | H468S | A | 3.9e-08 | 5.3e-08 | 3.7e-08 | 4.300E-08 | 8.718E-09 | 3.700E-08 | 5.300E-08 |
| 9357 | 468 | H468T | A | 1.8e-07 | 6.5e-08 | 7.3e-08 | 1.060E-07 | 6.421E-08 | 6.500E-08 | 1.800E-07 |
| 9358 | 468 | H468W | A | 1.9e-08 | 4.9e-08 | 5.5e-08 | 4.100E-08 | 1.929E-08 | 1.900E-08 | 5.500E-08 |
| 9359 | 468 | H468Y | A | 7.5e-08 | 1.5e-07 | 5e-08 | 9.167E-08 | 5.204E-08 | 5.000E-08 | 1.500E-07 |
| 9360 | 468 | H468V | A | 2.9e-08 | 4.8e-08 | 3e-08 | 3.567E-08 | 1.069E-08 | 2.900E-08 | 4.800E-08 |
| 9361 | 469 | Q469A | A | 5.9e-08 | 3.6e-08 | 7.4e-08 | 5.633E-08 | 1.914E-08 | 3.600E-08 | 7.400E-08 |
| 9362 | 469 | Q469R | A | 4.1e-08 | 5e-08 | 4.3e-08 | 4.467E-08 | 4.726E-09 | 4.100E-08 | 5.000E-08 |
| 9363 | 469 | Q469N | A | 6.5e-08 | 4.8e-08 | 1.7e-08 | 4.333E-08 | 2.434E-08 | 1.700E-08 | 6.500E-08 |
| 9364 | 469 | Q469D | A | 6.8e-08 | 1.3e-08 | 2.4e-08 | 3.500E-08 | 2.910E-08 | 1.300E-08 | 6.800E-08 |
| 9365 | 469 | Q469C | A | 9.9e-08 | 1.9e-08 | 5.6e-08 | 5.800E-08 | 4.004E-08 | 1.900E-08 | 9.900E-08 |
| 9366 | 469 | Q469E | A | 6.7e-08 | 2.1e-08 | 6e-08 | 4.933E-08 | 2.479E-08 | 2.100E-08 | 6.700E-08 |
| 9367 | 469 | Q469Q | A | 5.5e-08 | 6.2e-08 | 4.5e-08 | 5.400E-08 | 8.544E-09 | 4.500E-08 | 6.200E-08 |
| 9368 | 469 | Q469G | A | 8.9e-08 | 8.3e-08 | 2.5e-08 | 6.567E-08 | 3.535E-08 | 2.500E-08 | 8.900E-08 |
| 9369 | 469 | Q469H | A | 6.8e-08 | 5.3e-08 | 2.8e-08 | 4.967E-08 | 2.021E-08 | 2.800E-08 | 6.800E-08 |
| 9370 | 469 | Q469I | A | 7.2e-08 | 3e-08 | 6.7e-08 | 5.633E-08 | 2.294E-08 | 3.000E-08 | 7.200E-08 |
| 9371 | 469 | Q469L | A | 2.4e-08 | 7e-08 | 1.1e-07 | 6.800E-08 | 4.303E-08 | 2.400E-08 | 1.100E-07 |
| 9372 | 469 | Q469K | A | 2.3e-08 | 2.7e-08 | 2e-08 | 2.333E-08 | 3.512E-09 | 2.000E-08 | 2.700E-08 |
| 9373 | 469 | Q469M | A | 2.1e-07 | 4.6e-08 | 2.3e-08 | 9.300E-08 | 1.020E-07 | 2.300E-08 | 2.100E-07 |
| 9374 | 469 | Q469F | A | 4.2e-08 | 7.9e-08 | 4.3e-08 | 5.467E-08 | 2.108E-08 | 4.200E-08 | 7.900E-08 |
| 9375 | 469 | Q469P | A | 2.7e-08 | 1.3e-07 | 7.2e-08 | 7.633E-08 | 5.164E-08 | 2.700E-08 | 1.300E-07 |
| 9376 | 469 | Q469S | A | 1.3e-07 | 5.7e-08 | 3.1e-08 | 7.267E-08 | 5.133E-08 | 3.100E-08 | 1.300E-07 |
| 9377 | 469 | Q469T | A | 9.7e-08 | 5.1e-08 | 2.7e-08 | 5.833E-08 | 3.557E-08 | 2.700E-08 | 9.700E-08 |
| 9378 | 469 | Q469W | A | 9.9e-08 | 1.3e-07 | 3.1e-08 | 8.667E-08 | 5.064E-08 | 3.100E-08 | 1.300E-07 |
| 9379 | 469 | Q469Y | A | 1.2e-08 | 6.8e-08 | 8.5e-08 | 5.500E-08 | 3.820E-08 | 1.200E-08 | 8.500E-08 |
| 9380 | 469 | Q469V | A | 7.4e-08 | 1.2e-07 | 2e-07 | 1.313E-07 | 6.376E-08 | 7.400E-08 | 2.000E-07 |
| 9381 | 470 | A470A | A | 4.4e-08 | 1e-07 | 3.5e-08 | 5.967E-08 | 3.522E-08 | 3.500E-08 | 1.000E-07 |
| 9382 | 470 | A470R | A | 7.4e-08 | 3.1e-08 | 2.6e-08 | 4.367E-08 | 2.639E-08 | 2.600E-08 | 7.400E-08 |
| 9383 | 470 | A470N | A | 5.3e-08 | 2.5e-08 | 2.1e-08 | 3.300E-08 | 1.744E-08 | 2.100E-08 | 5.300E-08 |
| 9384 | 470 | A470D | A | 1.8e-08 | 2.2e-08 | 2.5e-08 | 2.167E-08 | 3.512E-09 | 1.800E-08 | 2.500E-08 |
| 9385 | 470 | A470C | A | 4.1e-08 | 1.2e-07 | 6.3e-08 | 7.467E-08 | 4.077E-08 | 4.100E-08 | 1.200E-07 |
| 9386 | 470 | A470E | A | 1e-08 | 3e-08 | 2.9e-08 | 2.300E-08 | 1.127E-08 | 1.000E-08 | 3.000E-08 |
| 9387 | 470 | A470Q | A | 2.8e-08 | 4.2e-08 | 8.6e-08 | 5.200E-08 | 3.027E-08 | 2.800E-08 | 8.600E-08 |
| 9388 | 470 | A470G | A | 2.8e-08 | 4.9e-08 | 7e-08 | 4.900E-08 | 2.100E-08 | 2.800E-08 | 7.000E-08 |
| 9389 | 470 | A470H | A | 6.3e-08 | 4.6e-08 | 3e-08 | 4.633E-08 | 1.650E-08 | 3.000E-08 | 6.300E-08 |
| 9390 | 470 | A470I | A | 4.1e-08 | 5.3e-08 | 2.8e-08 | 4.067E-08 | 1.250E-08 | 2.800E-08 | 5.300E-08 |
| 9391 | 470 | A470L | A | 7.1e-08 | 8.9e-08 | 2.7e-08 | 6.233E-08 | 3.190E-08 | 2.700E-08 | 8.900E-08 |
| 9392 | 470 | A470K | A | 3.2e-08 | 5.3e-08 | 3.1e-08 | 3.867E-08 | 1.242E-08 | 3.100E-08 | 5.300E-08 |
| 9393 | 470 | A470M | A | 5.4e-08 | 1.1e-08 | 1.1e-07 | 5.833E-08 | 4.964E-08 | 1.100E-08 | 1.100E-07 |
| 9394 | 470 | A470F | A | 6e-08 | 4.9e-08 | 4.9e-08 | 5.267E-08 | 6.351E-09 | 4.900E-08 | 6.000E-08 |
| 9395 | 470 | A470P | A | 1.8e-08 | 2.9e-08 | 3.9e-08 | 2.867E-08 | 1.050E-08 | 1.800E-08 | 3.900E-08 |
| 9396 | 470 | A470S | A | 1.1e-07 | 1.8e-08 | 6.7e-08 | 6.500E-08 | 4.603E-08 | 1.800E-08 | 1.100E-07 |
| 9397 | 470 | A470T | A | 8.4e-08 | 9e-08 | 2.4e-08 | 6.600E-08 | 3.650E-08 | 2.400E-08 | 9.000E-08 |
| 9398 | 470 | A470W | A | 4.3e-08 | 5.9e-08 | 3.9e-08 | 4.700E-08 | 1.058E-08 | 3.900E-08 | 5.900E-08 |
| 9399 | 470 | A470Y | A | 3.2e-08 | 3.9e-08 | 4.3e-08 | 3.800E-08 | 5.568E-09 | 3.200E-08 | 4.300E-08 |
| 9400 | 470 | A470V | A | 2.4e-08 | 1.7e-07 | 6.2e-08 | 8.533E-08 | 7.575E-08 | 2.400E-08 | 1.700E-07 |
| 9401 | 471 | L471A | A | 4.7e-08 | 2.4e-08 | 6.2e-08 | 4.433E-08 | 1.914E-08 | 2.400E-08 | 6.200E-08 |
| 9402 | 471 | L471R | A | 8.4e-08 | 5.3e-08 | 2.1e-08 | 5.267E-08 | 3.150E-08 | 2.100E-08 | 8.400E-08 |
| 9403 | 471 | L471N | A | 8.3e-08 | 3e-08 | 3.3e-08 | 4.867E-08 | 2.977E-08 | 3.000E-08 | 8.300E-08 |
| 9404 | 471 | L471D | A | 2e-08 | 3.3e-08 | 2.8e-08 | 2.700E-08 | 6.557E-09 | 2.000E-08 | 3.300E-08 |
| 9405 | 471 | L471C | A | 2.6e-08 | 7.2e-08 | 5.4e-08 | 5.067E-08 | 2.318E-08 | 2.600E-08 | 7.200E-08 |
| 9406 | 471 | L471E | A | 2.3e-08 | 1.2e-07 | 3.6e-08 | 5.967E-08 | 5.265E-08 | 2.300E-08 | 1.200E-07 |
| 9407 | 471 | L471Q | A | 1.2e-08 | 2.3e-08 | 4.3e-08 | 2.600E-08 | 1.572E-08 | 1.200E-08 | 4.300E-08 |
| 9408 | 471 | L471G | A | 1.1e-07 | 4.9e-08 | 3.1e-08 | 6.333E-08 | 4.140E-08 | 3.100E-08 | 1.100E-07 |
| 9409 | 471 | L471H | A | 6.8e-08 | 3.1e-08 | 1.3e-08 | 3.733E-08 | 2.804E-08 | 1.300E-08 | 6.800E-08 |
| 9410 | 471 | L471I | A | 2.6e-08 | 3.6e-08 | 6.2e-08 | 4.133E-08 | 1.858E-08 | 2.600E-08 | 6.200E-08 |
| 9411 | 471 | L471L | A | 5.9e-08 | 2.2e-08 | 6.6e-08 | 4.900E-08 | 2.364E-08 | 2.200E-08 | 6.600E-08 |
| 9412 | 471 | L471K | A | 5.7e-08 | 2.5e-08 | 4.3e-08 | 4.167E-08 | 1.604E-08 | 2.500E-08 | 5.700E-08 |
| 9413 | 471 | L471M | A | 8.4e-08 | 1.4e-08 | 6.8e-08 | 5.533E-08 | 3.668E-08 | 1.400E-08 | 8.400E-08 |
| 9414 | 471 | L471F | A | 1.7e-08 | 7.2e-08 | 5.7e-08 | 4.867E-08 | 2.843E-08 | 1.700E-08 | 7.200E-08 |
| 9415 | 471 | L471P | A | 1.3e-07 | 2.6e-08 | 2.6e-08 | 6.067E-08 | 6.004E-08 | 2.600E-08 | 1.300E-07 |
| 9416 | 471 | L471S | A | 3.2e-08 | 5.6e-08 | 4.6e-08 | 4.467E-08 | 1.206E-08 | 3.200E-08 | 5.600E-08 |
| 9417 | 471 | L471T | A | 5.4e-08 | 2.7e-08 | 2.4e-08 | 3.500E-08 | 1.652E-08 | 2.400E-08 | 5.400E-08 |
| 9418 | 471 | L471W | A | 6.4e-08 | 3.4e-08 | 3.3e-08 | 4.367E-08 | 1.762E-08 | 3.300E-08 | 6.400E-08 |
| 9419 | 471 | L471Y | A | 9.9e-08 | 1.1e-07 | 2.9e-08 | 7.933E-08 | 4.394E-08 | 2.900E-08 | 1.100E-07 |
| 9420 | 471 | L471V | A | 2.5e-08 | 3.2e-08 | 9.2e-08 | 4.967E-08 | 3.683E-08 | 2.500E-08 | 9.200E-08 |
| 9421 | 472 | L472A | A | 4.7e-08 | 3.7e-08 | 3e-08 | 3.800E-08 | 8.544E-09 | 3.000E-08 | 4.700E-08 |
| 9422 | 472 | L472R | A | 3.9e-08 | 6.3e-08 | 3.1e-08 | 4.433E-08 | 1.665E-08 | 3.100E-08 | 6.300E-08 |
| 9423 | 472 | L472N | A | 4.3e-08 | 1.4e-07 | 2.5e-08 | 6.933E-08 | 6.186E-08 | 2.500E-08 | 1.400E-07 |
| 9424 | 472 | L472D | A | 5.5e-08 | 1.6e-08 | 5.5e-08 | 4.200E-08 | 2.252E-08 | 1.600E-08 | 5.500E-08 |
| 9425 | 472 | L472C | A | 5.2e-08 | 8.6e-08 | 5.6e-08 | 6.467E-08 | 1.858E-08 | 5.200E-08 | 8.600E-08 |
| 9426 | 472 | L472E | A | 2.8e-08 | 2.6e-08 | 3.3e-08 | 2.900E-08 | 3.606E-09 | 2.600E-08 | 3.300E-08 |
| 9427 | 472 | L472Q | A | 1.1e-07 | 7e-08 | 7.2e-08 | 8.400E-08 | 2.254E-08 | 7.000E-08 | 1.100E-07 |
| 9428 | 472 | L472G | A | 4.2e-08 | 4.6e-08 | 5.5e-08 | 4.767E-08 | 6.658E-09 | 4.200E-08 | 5.500E-08 |
| 9429 | 472 | L472H | A | 8.6e-09 | 3.5e-08 | 1e-07 | 4.787E-08 | 4.704E-08 | 8.600E-09 | 1.000E-07 |
| 9430 | 472 | L472I | A | 3.6e-08 | 4.3e-08 | 3.2e-08 | 3.700E-08 | 5.568E-09 | 3.200E-08 | 4.300E-08 |
| 9431 | 472 | L472L | A | 7.5e-08 | 6.2e-08 | 4.7e-08 | 6.133E-08 | 1.401E-08 | 4.700E-08 | 7.500E-08 |
| 9432 | 472 | L472K | A | 1.8e-08 | 7.1e-08 | 6.5e-08 | 5.133E-08 | 2.902E-08 | 1.800E-08 | 7.100E-08 |
| 9433 | 472 | L472M | A | 1.7e-08 | 6.4e-08 | 4.2e-08 | 4.100E-08 | 2.352E-08 | 1.700E-08 | 6.400E-08 |
| 9434 | 472 | L472F | A | 2e-08 | 3.6e-08 | 8e-08 | 4.533E-08 | 3.107E-08 | 2.000E-08 | 8.000E-08 |

| | | | | | | | | | | |
|---|---|---|---|---|---|---|---|---|---|---|
| 9435 | 472 | L472P | A | 4.1e-08 | 1.2e-07 | 4.2e-08 | 6.767E-08 | 4.532E-08 | 4.100E-08 | 1.200E-07 |
| 9436 | 472 | L472S | A | 4e-08 | 4.4e-08 | 1.6e-07 | 8.133E-08 | 6.816E-08 | 4.000E-08 | 1.600E-07 |
| 9437 | 472 | L472T | A | 2.4e-08 | 7.7e-08 | 4.4e-08 | 4.833E-08 | 2.676E-08 | 2.400E-08 | 7.700E-08 |
| 9438 | 472 | L472W | A | 2.8e-08 | 5.4e-08 | 1.9e-08 | 3.367E-08 | 1.818E-08 | 1.900E-08 | 5.400E-08 |
| 9439 | 472 | L472Y | A | 1.2e-07 | 1.8e-08 | 1.7e-07 | 1.027E-07 | 7.747E-08 | 1.800E-08 | 1.700E-07 |
| 9440 | 472 | L472V | A | 3.9e-08 | 6.5e-08 | 1.6e-07 | 8.800E-08 | 6.369E-08 | 3.900E-08 | 1.600E-07 |
| 9441 | 473 | H473A | A | 5.4e-08 | 1.2e-08 | 4.5e-08 | 3.700E-08 | 2.211E-08 | 1.200E-08 | 5.400E-08 |
| 9442 | 473 | H473R | A | 5.9e-08 | 2.7e-08 | 4.8e-08 | 4.467E-08 | 1.626E-08 | 2.700E-08 | 5.900E-08 |
| 9443 | 473 | H473N | A | 7e-08 | 1.3e-08 | 5.3e-08 | 4.533E-08 | 2.926E-08 | 1.300E-08 | 7.000E-08 |
| 9444 | 473 | H473D | A | 2.2e-08 | 1.3e-07 | 3.6e-08 | 6.267E-08 | 5.873E-08 | 2.200E-08 | 1.300E-07 |
| 9445 | 473 | H473C | A | 3.6e-08 | 5.5e-08 | 4.9e-08 | 4.667E-08 | 9.713E-09 | 3.600E-08 | 5.500E-08 |
| 9446 | 473 | H473E | A | 4.2e-08 | 1.1e-08 | 3.2e-08 | 2.833E-08 | 1.582E-08 | 1.100E-08 | 4.200E-08 |
| 9447 | 473 | H473Q | A | 2.9e-08 | 5.1e-08 | 2.4e-08 | 3.467E-08 | 1.436E-08 | 2.400E-08 | 5.100E-08 |
| 9448 | 473 | H473G | A | 3e-08 | 2.7e-08 | 5.5e-08 | 3.733E-08 | 1.537E-08 | 2.700E-08 | 5.500E-08 |
| 9449 | 473 | H473H | A | 1.4e-08 | 4.8e-08 | 4.4e-08 | 3.533E-08 | 1.858E-08 | 1.400E-08 | 4.800E-08 |
| 9450 | 473 | H473I | A | 3.1e-08 | 5.8e-08 | 3.3e-08 | 4.067E-08 | 1.504E-08 | 3.100E-08 | 5.800E-08 |
| 9451 | 473 | H473L | A | 4.5e-08 | 9.4e-09 | 4.2e-08 | 3.213E-08 | 1.974E-08 | 9.400E-09 | 4.500E-08 |
| 9452 | 473 | H473K | A | 5.2e-08 | 2.7e-08 | 9.5e-08 | 5.800E-08 | 3.439E-08 | 2.700E-08 | 9.500E-08 |
| 9453 | 473 | H473M | A | 6.5e-08 | 6.6e-08 | 9e-08 | 7.367E-08 | 1.415E-08 | 6.500E-08 | 9.000E-08 |
| 9454 | 473 | H473F | A | 9.9e-08 | 2.9e-08 | 3.9e-08 | 5.567E-08 | 3.786E-08 | 2.900E-08 | 9.900E-08 |
| 9455 | 473 | H473P | A | 3.2e-08 | 1.3e-08 | 1.2e-08 | 1.900E-08 | 1.127E-08 | 1.200E-08 | 3.200E-08 |
| 9456 | 473 | H473S | A | 3.2e-08 | 3.2e-08 | 8.9e-08 | 5.100E-08 | 3.291E-08 | 3.200E-08 | 8.900E-08 |
| 9457 | 473 | H473T | A | 1e-07 | 4.3e-08 | 4e-08 | 6.100E-08 | 3.381E-08 | 4.000E-08 | 1.000E-07 |
| 9458 | 473 | H473W | A | 4.1e-08 | 4.7e-08 | 5.9e-08 | 4.900E-08 | 9.165E-09 | 4.100E-08 | 5.900E-08 |
| 9459 | 473 | H473Y | A | 6.6e-08 | 2.3e-08 | 3.2e-08 | 4.033E-08 | 2.268E-08 | 2.300E-08 | 6.600E-08 |
| 9460 | 473 | H473V | A | 1.1e-07 | 1.4e-08 | 6.2e-08 | 6.200E-08 | 4.800E-08 | 1.400E-08 | 1.100E-07 |
| 9461 | 474 | T474A | A | 5.3e-08 | 3.8e-08 | 2.3e-08 | 3.800E-08 | 1.500E-08 | 2.300E-08 | 5.300E-08 |
| 9462 | 474 | T474R | A | 3.2e-08 | 5.1e-08 | 4e-08 | 4.100E-08 | 9.539E-09 | 3.200E-08 | 5.100E-08 |
| 9463 | 474 | T474N | A | 3e-08 | 5.7e-08 | 9.1e-08 | 5.933E-08 | 3.057E-08 | 3.000E-08 | 9.100E-08 |
| 9464 | 474 | T474D | A | 6.1e-08 | 2.7e-08 | 1.2e-08 | 3.333E-08 | 2.511E-08 | 1.200E-08 | 6.100E-08 |
| 9465 | 474 | T474C | A | 1.3e-07 | 9.6e-08 | 5.9e-08 | 9.500E-08 | 3.551E-08 | 5.900E-08 | 1.300E-07 |
| 9466 | 474 | T474E | A | 1.7e-08 | 3.3e-08 | 6.1e-08 | 3.700E-08 | 2.227E-08 | 1.700E-08 | 6.100E-08 |
| 9467 | 474 | T474Q | A | 4.7e-08 | 3.5e-08 | 3.1e-08 | 3.767E-08 | 8.327E-09 | 3.100E-08 | 4.700E-08 |
| 9468 | 474 | T474G | A | 9.9e-08 | 1e-07 | 3e-08 | 7.633E-08 | 4.013E-08 | 3.000E-08 | 1.000E-07 |
| 9469 | 474 | T474H | A | 6.8e-08 | 4.7e-08 | 9.5e-08 | 7.000E-08 | 2.406E-08 | 4.700E-08 | 9.500E-08 |
| 9470 | 474 | T474I | A | 2.5e-08 | 4e-08 | 6.5e-08 | 4.333E-08 | 2.021E-08 | 2.500E-08 | 6.500E-08 |
| 9471 | 474 | T474L | A | 2.2e-08 | 3e-08 | 2.5e-08 | 2.567E-08 | 4.041E-09 | 2.200E-08 | 3.000E-08 |
| 9472 | 474 | T474K | A | 3.1e-08 | 2.4e-08 | 1.4e-08 | 2.300E-08 | 8.544E-09 | 1.400E-08 | 3.100E-08 |
| 9473 | 474 | T474M | A | 8.1e-08 | 8.8e-08 | 4.1e-08 | 7.000E-08 | 2.536E-08 | 4.100E-08 | 8.800E-08 |
| 9474 | 474 | T474F | A | 4.1e-09 | 3.3e-08 | 3.9e-08 | 2.537E-08 | 1.866E-08 | 4.100E-09 | 3.900E-08 |
| 9475 | 474 | T474P | A | 8.4e-08 | 6.2e-08 | 7.7e-08 | 7.433E-08 | 1.124E-08 | 6.200E-08 | 8.400E-08 |
| 9476 | 474 | T474S | A | 6.8e-08 | 2.1e-08 | 2.5e-08 | 3.800E-08 | 2.606E-08 | 2.100E-08 | 6.800E-08 |
| 9477 | 474 | T474T | A | 3.9e-08 | 5.6e-08 | 4.2e-08 | 4.567E-08 | 9.074E-09 | 3.900E-08 | 5.600E-08 |
| 9478 | 474 | T474W | A | 1.5e-07 | 6.1e-08 | 3.7e-08 | 8.267E-08 | 5.953E-08 | 3.700E-08 | 1.500E-07 |
| 9479 | 474 | T474Y | A | 3.2e-08 | 3e-08 | 1.5e-08 | 2.567E-08 | 9.292E-09 | 1.500E-08 | 3.200E-08 |
| 9480 | 474 | T474V | A | 4.4e-08 | 1.2e-08 | 4.4e-08 | 3.333E-08 | 1.848E-08 | 1.200E-08 | 4.400E-08 |
| 9481 | 475 | A475A | A | 3.7e-08 | 6.1e-08 | 4.8e-08 | 4.867E-08 | 1.201E-08 | 3.700E-08 | 6.100E-08 |
| 9482 | 475 | A475R | A | 2.1e-08 | 9.6e-08 | 4.9e-08 | 5.533E-08 | 3.790E-08 | 2.100E-08 | 9.600E-08 |
| 9483 | 475 | A475N | A | 5.9e-08 | 2.8e-08 | 2.9e-08 | 3.867E-08 | 1.762E-08 | 2.800E-08 | 5.900E-08 |
| 9484 | 475 | A475D | A | 7.1e-08 | 3.9e-08 | 2.6e-08 | 4.533E-08 | 2.316E-08 | 2.600E-08 | 7.100E-08 |
| 9485 | 475 | A475C | A | 4.3e-08 | 8.4e-08 | 4.4e-08 | 5.700E-08 | 2.339E-08 | 4.300E-08 | 8.400E-08 |
| 9486 | 475 | A475E | A | 1.7e-08 | 1.5e-08 | 1.5e-08 | 1.567E-08 | 1.155E-09 | 1.500E-08 | 1.700E-08 |
| 9487 | 475 | A475Q | A | 1.7e-08 | 6.6e-08 | 4.7e-08 | 4.333E-08 | 2.470E-08 | 1.700E-08 | 6.600E-08 |
| 9488 | 475 | A475G | A | 1.1e-07 | 2.3e-08 | 7.1e-08 | 6.800E-08 | 4.358E-08 | 2.300E-08 | 1.100E-07 |
| 9489 | 475 | A475H | A | 4.4e-08 | 5.4e-08 | 3.8e-08 | 4.533E-08 | 8.083E-09 | 3.800E-08 | 5.400E-08 |
| 9490 | 475 | A475I | A | 4.4e-08 | 7.5e-08 | 4.5e-08 | 5.467E-08 | 1.762E-08 | 4.400E-08 | 7.500E-08 |
| 9491 | 475 | A475L | A | 4.1e-08 | 4.2e-08 | 4.7e-08 | 4.333E-08 | 3.215E-09 | 4.100E-08 | 4.700E-08 |
| 9492 | 475 | A475K | A | 7.5e-08 | 8.6e-08 | 3.1e-08 | 6.400E-08 | 2.910E-08 | 3.100E-08 | 8.600E-08 |
| 9493 | 475 | A475M | A | 1e-08 | 9.1e-09 | 1.5e-08 | 1.137E-08 | 3.179E-09 | 9.100E-09 | 1.500E-08 |
| 9494 | 475 | A475F | A | 4e-08 | 1.8e-08 | 1.9e-08 | 2.567E-08 | 1.242E-08 | 1.800E-08 | 4.000E-08 |
| 9495 | 475 | A475P | A | 3.3e-08 | 4.5e-08 | 1.5e-07 | 7.600E-08 | 6.437E-08 | 3.300E-08 | 1.500E-07 |
| 9496 | 475 | A475S | A | 8.9e-08 | 3.5e-08 | 1.5e-08 | 4.633E-08 | 3.828E-08 | 1.500E-08 | 8.900E-08 |
| 9497 | 475 | A475T | A | 3.9e-08 | 5.2e-08 | 1.5e-07 | 8.033E-08 | 6.068E-08 | 3.900E-08 | 1.500E-07 |
| 9498 | 475 | A475W | A | 3.4e-08 | 3.2e-08 | 7.9e-08 | 4.833E-08 | 2.658E-08 | 3.200E-08 | 7.900E-08 |
| 9499 | 475 | A475Y | A | 3.8e-08 | 6.5e-08 | 4.5e-08 | 4.933E-08 | 1.401E-08 | 3.800E-08 | 6.500E-08 |
| 9500 | 475 | A475V | A | 3.1e-08 | 3.1e-08 | 5.2e-08 | 3.800E-08 | 1.212E-08 | 3.100E-08 | 5.200E-08 |
| 9501 | 476 | N476A | A | 9.6e-08 | 4.8e-08 | 2.5e-08 | 5.633E-08 | 3.623E-08 | 2.500E-08 | 9.600E-08 |
| 9502 | 476 | N476R | A | 4e-08 | 3.8e-08 | 1.2e-07 | 6.600E-08 | 4.678E-08 | 3.800E-08 | 1.200E-07 |
| 9503 | 476 | N476N | A | 2.2e-08 | 5.9e-08 | 6.7e-08 | 4.933E-08 | 2.401E-08 | 2.200E-08 | 6.700E-08 |
| 9504 | 476 | N476D | A | 5.9e-08 | 1.5e-07 | 3e-08 | 7.967E-08 | 6.261E-08 | 3.000E-08 | 1.500E-07 |
| 9505 | 476 | N476C | A | 1.2e-08 | 5.6e-08 | 3e-08 | 3.267E-08 | 2.212E-08 | 1.200E-08 | 5.600E-08 |
| 9506 | 476 | N476E | A | 3e-08 | 5.4e-08 | 9.7e-08 | 6.033E-08 | 3.395E-08 | 3.000E-08 | 9.700E-08 |
| 9507 | 476 | N476Q | A | 4e-08 | 2.1e-08 | 5.5e-08 | 3.867E-08 | 1.704E-08 | 2.100E-08 | 5.500E-08 |
| 9508 | 476 | N476G | A | 6.2e-08 | 6.4e-08 | 5.9e-08 | 6.167E-08 | 2.517E-09 | 5.900E-08 | 6.400E-08 |
| 9509 | 476 | N476H | A | 2.8e-08 | 3.4e-08 | 4.2e-08 | 3.467E-08 | 7.024E-09 | 2.800E-08 | 4.200E-08 |
| 9510 | 476 | N476I | A | 1.3e-08 | 4.6e-08 | 4.9e-08 | 3.600E-08 | 1.997E-08 | 1.300E-08 | 4.900E-08 |
| 9511 | 476 | N476L | A | 6.3e-08 | 2.9e-08 | 8.5e-08 | 5.900E-08 | 2.821E-08 | 2.900E-08 | 8.500E-08 |
| 9512 | 476 | N476K | A | 5.7e-08 | 1.9e-08 | 4.7e-08 | 4.100E-08 | 1.970E-08 | 1.900E-08 | 5.700E-08 |
| 9513 | 476 | N476M | A | 3.7e-08 | 4.4e-08 | 5e-08 | 4.367E-08 | 6.506E-09 | 3.700E-08 | 5.000E-08 |
| 9514 | 476 | N476F | A | 5.4e-08 | 5.6e-08 | 1.6e-08 | 4.200E-08 | 2.254E-08 | 1.600E-08 | 5.600E-08 |
| 9515 | 476 | N476P | A | 6.1e-08 | 3.9e-08 | 3.5e-08 | 4.500E-08 | 1.400E-08 | 3.500E-08 | 6.100E-08 |
| 9516 | 476 | N476S | A | 2.4e-08 | 4.5e-08 | 6.2e-08 | 4.367E-08 | 1.904E-08 | 2.400E-08 | 6.200E-08 |
| 9517 | 476 | N476T | A | 4.7e-08 | 4.4e-08 | 6.9e-08 | 5.333E-08 | 1.365E-08 | 4.400E-08 | 6.900E-08 |
| 9518 | 476 | N476W | A | 4.5e-08 | 2.4e-08 | 4.8e-08 | 3.900E-08 | 1.308E-08 | 2.400E-08 | 4.800E-08 |
| 9519 | 476 | N476Y | A | 6.9e-08 | 1.1e-07 | 2e-08 | 6.633E-08 | 4.506E-08 | 2.000E-08 | 1.100E-07 |

| | | | | | | | | | | |
|---|---|---|---|---|---|---|---|---|---|---|
| 9520 | 476 | N476V | A | 2.7e-08 | 2.8e-08 | 1.4e-08 | 2.300E-08 | 7.810E-09 | 1.400E-08 | 2.800E-08 |
| 9521 | 477 | R477A | A | 4.1e-08 | 4.9e-08 | 5.9e-08 | 4.967E-08 | 9.018E-09 | 4.100E-08 | 5.900E-08 |
| 9522 | 477 | R477R | A | 3.8e-08 | 5e-08 | 2.4e-08 | 3.733E-08 | 1.301E-08 | 2.400E-08 | 5.000E-08 |
| 9523 | 477 | R477N | A | 8.4e-08 | 4.2e-08 | 2.9e-08 | 5.167E-08 | 2.875E-08 | 2.900E-08 | 8.400E-08 |
| 9524 | 477 | R477D | A | 2.7e-08 | 2e-08 | 6.2e-08 | 3.633E-08 | 2.250E-08 | 2.000E-08 | 6.200E-08 |
| 9525 | 477 | R477C | A | 4.3e-08 | 1.6e-08 | 1.8e-08 | 2.567E-08 | 1.504E-08 | 1.600E-08 | 4.300E-08 |
| 9526 | 477 | R477E | A | 8.9e-08 | 1.8e-08 | 9.4e-08 | 6.700E-08 | 4.251E-08 | 1.800E-08 | 9.400E-08 |
| 9527 | 477 | R477Q | A | 5.2e-08 | 8.3e-08 | 4.6e-08 | 6.033E-08 | 1.986E-08 | 4.600E-08 | 8.300E-08 |
| 9528 | 477 | R477G | A | 2.3e-08 | 4.1e-08 | 3.9e-08 | 3.433E-08 | 9.866E-09 | 2.300E-08 | 4.100E-08 |
| 9529 | 477 | R477H | A | 7.8e-09 | 6.6e-08 | 1.9e-08 | 3.093E-08 | 3.088E-08 | 7.800E-09 | 6.600E-08 |
| 9530 | 477 | R477I | A | 1.8e-08 | 4.7e-08 | 1e-08 | 2.500E-08 | 1.947E-08 | 1.000E-08 | 4.700E-08 |
| 9531 | 477 | R477L | A | 5.9e-08 | 7.7e-08 | 3.5e-08 | 5.700E-08 | 2.107E-08 | 3.500E-08 | 7.700E-08 |
| 9532 | 477 | R477K | A | 3.5e-08 | 1.4e-08 | 6.1e-08 | 3.667E-08 | 2.354E-08 | 1.400E-08 | 6.100E-08 |
| 9533 | 477 | R477M | A | 2e-08 | 2.9e-08 | 2.6e-08 | 2.500E-08 | 4.583E-09 | 2.000E-08 | 2.900E-08 |
| 9534 | 477 | R477F | A | 1.6e-08 | 1.2e-08 | 4.2e-08 | 2.333E-08 | 1.629E-08 | 1.200E-08 | 4.200E-08 |
| 9535 | 477 | R477P | A | 1e-07 | 6.4e-08 | 2.6e-08 | 6.333E-08 | 3.700E-08 | 2.600E-08 | 1.000E-07 |
| 9536 | 477 | R477S | A | 8.7e-08 | 3.8e-08 | 3.4e-08 | 5.300E-08 | 2.951E-08 | 3.400E-08 | 8.700E-08 |
| 9537 | 477 | R477T | A | 5.8e-08 | 2.9e-08 | 9.2e-08 | 5.967E-08 | 3.153E-08 | 2.900E-08 | 9.200E-08 |
| 9538 | 477 | R477W | A | 8.5e-08 | 5.1e-08 | 1.1e-07 | 8.200E-08 | 2.961E-08 | 5.100E-08 | 1.100E-07 |
| 9539 | 477 | R477Y | A | 4.5e-08 | 4e-08 | 2.2e-07 | 1.017E-07 | 1.025E-07 | 4.000E-08 | 2.200E-07 |
| 9540 | 477 | R477V | A | 1.2e-07 | 3.6e-08 | 2.5e-08 | 6.033E-08 | 5.196E-08 | 2.500E-08 | 1.200E-07 |
| 9541 | 478 | P478A | A | 5.9e-08 | 5.2e-08 | 6.9e-08 | 6.000E-08 | 8.544E-09 | 5.200E-08 | 6.900E-08 |
| 9542 | 478 | P478R | A | 4.3e-08 | 2e-08 | 5.4e-08 | 3.900E-08 | 1.735E-08 | 2.000E-08 | 5.400E-08 |
| 9543 | 478 | P478N | A | 2.5e-08 | 1.9e-08 | 2.8e-08 | 2.400E-08 | 4.583E-09 | 1.900E-08 | 2.800E-08 |
| 9544 | 478 | P478D | A | 4.2e-08 | 2.7e-08 | 3.3e-08 | 3.400E-08 | 7.550E-09 | 2.700E-08 | 4.200E-08 |
| 9545 | 478 | P478C | A | 2.4e-08 | 1.3e-08 | 4.3e-08 | 2.667E-08 | 1.518E-08 | 1.300E-08 | 4.300E-08 |
| 9546 | 478 | P478E | A | 7.4e-08 | 3.2e-08 | 1e-07 | 6.867E-08 | 3.431E-08 | 3.200E-08 | 1.000E-07 |
| 9547 | 478 | P478Q | A | 3.1e-08 | 6.9e-08 | 4.1e-08 | 4.700E-08 | 1.970E-08 | 3.100E-08 | 6.900E-08 |
| 9548 | 478 | P478G | A | 4e-08 | 3.4e-08 | 2.7e-08 | 3.367E-08 | 6.506E-09 | 2.700E-08 | 4.000E-08 |
| 9549 | 478 | P478H | A | 6.2e-08 | 1.3e-08 | 4.7e-08 | 4.067E-08 | 2.511E-08 | 1.300E-08 | 6.200E-08 |
| 9550 | 478 | P478I | A | 2.8e-08 | 1.1e-07 | 4.5e-08 | 6.100E-08 | 4.328E-08 | 2.800E-08 | 1.100E-07 |
| 9551 | 478 | P478L | A | 3e-08 | 6.5e-08 | 1.2e-07 | 7.167E-08 | 4.537E-08 | 3.000E-08 | 1.200E-07 |
| 9552 | 478 | P478K | A | 7.5e-08 | 1.8e-08 | 1.2e-07 | 7.100E-08 | 5.112E-08 | 1.800E-08 | 1.200E-07 |
| 9553 | 478 | P478M | A | 1.1e-07 | 3.4e-08 | 4.9e-08 | 6.433E-08 | 4.025E-08 | 3.400E-08 | 1.100E-07 |
| 9554 | 478 | P478F | A | 9.5e-08 | 2.7e-08 | 5.1e-08 | 5.767E-08 | 3.449E-08 | 2.700E-08 | 9.500E-08 |
| 9555 | 478 | P478P | A | 3.7e-08 | 1.3e-08 | 3.3e-08 | 2.767E-08 | 1.286E-08 | 1.300E-08 | 3.700E-08 |
| 9556 | 478 | P478S | A | 5.9e-08 | 4.1e-08 | 6e-08 | 5.333E-08 | 1.069E-08 | 4.100E-08 | 6.000E-08 |
| 9557 | 478 | P478T | A | 2.9e-08 | 4.3e-08 | 1.7e-08 | 2.967E-08 | 1.301E-08 | 1.700E-08 | 4.300E-08 |
| 9558 | 478 | P478W | A | 4.1e-08 | 1.4e-08 | 5.7e-08 | 3.733E-08 | 2.173E-08 | 1.400E-08 | 5.700E-08 |
| 9559 | 478 | P478Y | A | 7.1e-08 | 1.9e-08 | 3e-08 | 4.000E-08 | 2.740E-08 | 1.900E-08 | 7.100E-08 |
| 9560 | 478 | P478V | A | 5.7e-08 | 4.2e-08 | 6.4e-08 | 5.433E-08 | 1.124E-08 | 4.200E-08 | 6.400E-08 |
| 9561 | 479 | E479A | A | 5.2e-08 | 3.3e-08 | 3.7e-08 | 4.067E-08 | 1.002E-08 | 3.300E-08 | 5.200E-08 |
| 9562 | 479 | E479R | A | 5.6e-08 | 1.9e-08 | 4.8e-08 | 4.100E-08 | 1.947E-08 | 1.900E-08 | 5.600E-08 |
| 9563 | 479 | E479N | A | 5.2e-09 | 2.1e-08 | 1.4e-08 | 1.340E-08 | 7.917E-09 | 5.200E-09 | 2.100E-08 |
| 9564 | 479 | E479D | A | 3.2e-08 | 3.1e-08 | 6.2e-08 | 4.167E-08 | 1.762E-08 | 3.100E-08 | 6.200E-08 |
| 9565 | 479 | E479C | A | 6.9e-08 | 3.2e-08 | 2e-08 | 4.033E-08 | 2.554E-08 | 2.000E-08 | 6.900E-08 |
| 9566 | 479 | E479E | A | 4.5e-08 | 4.6e-08 | 2.8e-08 | 3.967E-08 | 1.012E-08 | 2.800E-08 | 4.600E-08 |
| 9567 | 479 | E479Q | A | 4.7e-08 | 1.7e-07 | 7.1e-08 | 9.600E-08 | 6.520E-08 | 4.700E-08 | 1.700E-07 |
| 9568 | 479 | E479G | A | 2.2e-08 | 1.9e-07 | 5.1e-08 | 8.767E-08 | 8.980E-08 | 2.200E-08 | 1.900E-07 |
| 9569 | 479 | E479H | A | 4.3e-08 | 7.5e-08 | 6.3e-08 | 6.033E-08 | 1.617E-08 | 4.300E-08 | 7.500E-08 |
| 9570 | 479 | E479I | A | 1.8e-08 | 1.4e-08 | 6.1e-08 | 3.100E-08 | 2.606E-08 | 1.400E-08 | 6.100E-08 |
| 9571 | 479 | E479L | A | 6.7e-08 | 2.8e-08 | 6.9e-08 | 5.467E-08 | 2.312E-08 | 2.800E-08 | 6.900E-08 |
| 9572 | 479 | E479K | A | 6.9e-08 | 3.8e-08 | 7.9e-08 | 6.200E-08 | 2.138E-08 | 3.800E-08 | 7.900E-08 |
| 9573 | 479 | E479M | A | 4.2e-08 | 6.3e-08 | 3.9e-08 | 4.800E-08 | 1.308E-08 | 3.900E-08 | 6.300E-08 |
| 9574 | 479 | E479F | A | 4.3e-08 | 6.1e-08 | 6.2e-08 | 5.533E-08 | 1.069E-08 | 4.300E-08 | 6.200E-08 |
| 9575 | 479 | E479P | A | 3.8e-08 | 7.3e-08 | 6.4e-08 | 5.833E-08 | 1.818E-08 | 3.800E-08 | 7.300E-08 |
| 9576 | 479 | E479S | A | 5.5e-08 | 3.4e-08 | 1.7e-08 | 3.533E-08 | 1.904E-08 | 1.700E-08 | 5.500E-08 |
| 9577 | 479 | E479T | A | 2.4e-08 | 8.2e-08 | 8.2e-08 | 6.267E-08 | 3.349E-08 | 2.400E-08 | 8.200E-08 |
| 9578 | 479 | E479W | A | 1.3e-08 | 5.9e-08 | 2.9e-08 | 3.367E-08 | 2.335E-08 | 1.300E-08 | 5.900E-08 |
| 9579 | 479 | E479Y | A | 4.5e-08 | 5e-08 | 5.1e-08 | 4.867E-08 | 3.215E-09 | 4.500E-08 | 5.100E-08 |
| 9580 | 479 | E479V | A | 1.1e-07 | 4.3e-08 | 3.5e-08 | 6.267E-08 | 4.119E-08 | 3.500E-08 | 1.100E-07 |
| 9581 | 480 | D480A | A | 3.3e-08 | 2.5e-08 | 2.9e-08 | 2.900E-08 | 4.000E-09 | 2.500E-08 | 3.300E-08 |
| 9582 | 480 | D480R | A | 2e-08 | 2e-08 | 4.7e-08 | 2.900E-08 | 1.559E-08 | 2.000E-08 | 4.700E-08 |
| 9583 | 480 | D480N | A | 2.8e-08 | 4.3e-08 | 5.6e-08 | 4.233E-08 | 1.401E-08 | 2.800E-08 | 5.600E-08 |
| 9584 | 480 | D480D | A | 5.5e-08 | 5.9e-08 | 5.7e-08 | 5.700E-08 | 2.000E-09 | 5.500E-08 | 5.900E-08 |
| 9585 | 480 | D480C | A | 6.9e-08 | 2.7e-08 | 3.1e-08 | 4.233E-08 | 2.318E-08 | 2.700E-08 | 6.900E-08 |
| 9586 | 480 | D480E | A | 1.9e-08 | 3.9e-08 | 6.4e-08 | 4.067E-08 | 2.255E-08 | 1.900E-08 | 6.400E-08 |
| 9587 | 480 | D480Q | A | 9.3e-08 | 4.4e-08 | 1.9e-08 | 5.200E-08 | 3.764E-08 | 1.900E-08 | 9.300E-08 |
| 9588 | 480 | D480G | A | 7.2e-08 | 2.6e-08 | 4.7e-08 | 4.833E-08 | 2.303E-08 | 2.600E-08 | 7.200E-08 |
| 9589 | 480 | D480H | A | 2.4e-08 | 1.7e-08 | 4.4e-08 | 2.833E-08 | 1.401E-08 | 1.700E-08 | 4.400E-08 |
| 9590 | 480 | D480I | A | 3.2e-08 | 1.2e-07 | 6.9e-08 | 7.367E-08 | 4.419E-08 | 3.200E-08 | 1.200E-07 |
| 9591 | 480 | D480L | A | 1.3e-07 | 1.6e-08 | 3.1e-08 | 5.900E-08 | 6.194E-08 | 1.600E-08 | 1.300E-07 |
| 9592 | 480 | D480K | A | 9.9e-08 | 5.1e-08 | 4.1e-08 | 6.367E-08 | 3.101E-08 | 4.100E-08 | 9.900E-08 |
| 9593 | 480 | D480M | A | 2.1e-08 | 1.1e-07 | 7.2e-08 | 6.767E-08 | 4.466E-08 | 2.100E-08 | 1.100E-07 |
| 9594 | 480 | D480F | A | 4.1e-08 | 2.9e-08 | 2.1e-08 | 3.033E-08 | 1.007E-08 | 2.100E-08 | 4.100E-08 |
| 9595 | 480 | D480P | A | 6.4e-08 | 3.9e-08 | 1.7e-08 | 4.000E-08 | 2.352E-08 | 1.700E-08 | 6.400E-08 |
| 9596 | 480 | D480S | A | 5.3e-08 | 2.3e-08 | 9.3e-09 | 2.843E-08 | 2.235E-08 | 9.300E-09 | 5.300E-08 |
| 9597 | 480 | D480T | A | 5.1e-08 | 3.4e-08 | 3.5e-08 | 4.000E-08 | 9.539E-09 | 3.400E-08 | 5.100E-08 |
| 9598 | 480 | D480W | A | 6.1e-08 | 3.2e-08 | 2.6e-08 | 3.967E-08 | 1.872E-08 | 2.600E-08 | 6.100E-08 |
| 9599 | 480 | D480Y | A | 4.3e-08 | 6.6e-08 | 5.4e-08 | 5.433E-08 | 1.150E-08 | 4.300E-08 | 6.600E-08 |
| 9600 | 480 | D480V | A | 5.8e-08 | 3.8e-08 | 1.9e-08 | 3.833E-08 | 1.950E-08 | 1.900E-08 | 5.800E-08 |
| 9601 | 481 | E481A | A | 8.2e-08 | 4.8e-08 | 8.7e-08 | 7.233E-08 | 2.122E-08 | 4.800E-08 | 8.700E-08 |
| 9602 | 481 | E481R | A | 2.7e-08 | 3.9e-08 | 2.5e-08 | 3.033E-08 | 7.572E-09 | 2.500E-08 | 3.900E-08 |
| 9603 | 481 | E481N | A | 3e-08 | 3.8e-08 | 9.6e-08 | 5.467E-08 | 3.602E-08 | 3.000E-08 | 9.600E-08 |
| 9604 | 481 | E481D | A | 5.4e-08 | 2e-08 | 1.1e-07 | 6.133E-08 | 4.545E-08 | 2.000E-08 | 1.100E-07 |

| | | | | | | | | | | |
|---|---|---|---|---|---|---|---|---|---|---|
| 9605 | 481 | E481C | A | 3.6e-08 | 2e-08 | 7.6e-08 | 4.400E-08 | 2.884E-08 | 2.000E-08 | 7.600E-08 |
| 9606 | 481 | E481E | A | 3.4e-08 | 2e-08 | 2.8e-08 | 2.733E-08 | 7.024E-09 | 2.000E-08 | 3.400E-08 |
| 9607 | 481 | E481Q | A | 3.7e-08 | 3.3e-08 | 7.6e-08 | 4.867E-08 | 2.376E-08 | 3.300E-08 | 7.600E-08 |
| 9608 | 481 | E481G | A | 2.2e-08 | 1.3e-08 | 2.1e-08 | 1.867E-08 | 4.933E-09 | 1.300E-08 | 2.200E-08 |
| 9609 | 481 | E481H | A | 2.3e-08 | 3.4e-08 | 2.3e-08 | 2.667E-08 | 6.351E-09 | 2.300E-08 | 3.400E-08 |
| 9610 | 481 | E481I | A | 2.9e-08 | 2.1e-08 | 1.6e-08 | 2.200E-08 | 6.557E-09 | 1.600E-08 | 2.900E-08 |
| 9611 | 481 | E481L | A | 2.3e-08 | 6.6e-08 | 4.4e-08 | 4.433E-08 | 2.150E-08 | 2.300E-08 | 6.600E-08 |
| 9612 | 481 | E481K | A | 8e-08 | 5e-08 | 3.1e-08 | 5.367E-08 | 2.470E-08 | 3.100E-08 | 8.000E-08 |
| 9613 | 481 | E481M | A | 2.6e-08 | 3.9e-08 | 7.2e-08 | 4.567E-08 | 2.371E-08 | 2.600E-08 | 7.200E-08 |
| 9614 | 481 | E481F | A | 3.2e-08 | 2.9e-08 | 1.5e-08 | 2.533E-08 | 9.074E-09 | 1.500E-08 | 3.200E-08 |
| 9615 | 481 | E481P | A | 1.7e-08 | 3.2e-08 | 1.3e-08 | 2.067E-08 | 1.002E-08 | 1.300E-08 | 3.200E-08 |
| 9616 | 481 | E481S | A | 2.5e-08 | 2.7e-08 | 1.9e-08 | 2.367E-08 | 4.163E-09 | 1.900E-08 | 2.700E-08 |
| 9617 | 481 | E481T | A | 1.8e-08 | 4.6e-08 | 3e-08 | 3.133E-08 | 1.405E-08 | 1.800E-08 | 4.600E-08 |
| 9618 | 481 | E481W | A | 7.9e-08 | 5.2e-08 | 5.2e-08 | 6.100E-08 | 1.559E-08 | 5.200E-08 | 7.900E-08 |
| 9619 | 481 | E481Y | A | 4.9e-08 | 4.6e-08 | 5.9e-08 | 5.133E-08 | 6.807E-09 | 4.600E-08 | 5.900E-08 |
| 9620 | 481 | E481V | A | 2.8e-08 | 2.7e-08 | 1.1e-07 | 5.500E-08 | 4.763E-08 | 2.700E-08 | 1.100E-07 |
| 9621 | 482 | C482A | A | 5e-08 | 3.6e-08 | 3.6e-08 | 4.067E-08 | 8.083E-09 | 3.600E-08 | 5.000E-08 |
| 9622 | 482 | C482R | A | 9.7e-08 | 3.6e-08 | 1.3e-07 | 8.767E-08 | 4.769E-08 | 3.600E-08 | 1.300E-07 |
| 9623 | 482 | C482N | A | 5.8e-08 | 1.6e-08 | 3.6e-08 | 3.667E-08 | 2.101E-08 | 1.600E-08 | 5.800E-08 |
| 9624 | 482 | C482D | A | 4.3e-08 | 4.5e-08 | 1.9e-08 | 3.567E-08 | 1.447E-08 | 1.900E-08 | 4.500E-08 |
| 9625 | 482 | C482C | A | 1.8e-08 | 4.5e-08 | 3.3e-08 | 3.200E-08 | 1.353E-08 | 1.800E-08 | 4.500E-08 |
| 9626 | 482 | C482E | A | 6.8e-08 | 2.6e-08 | 8.2e-08 | 5.867E-08 | 2.914E-08 | 2.600E-08 | 8.200E-08 |
| 9627 | 482 | C482Q | A | 3.1e-08 | 2.7e-08 | 2.6e-08 | 2.800E-08 | 2.646E-09 | 2.600E-08 | 3.100E-08 |
| 9628 | 482 | C482G | A | 4.5e-08 | 4.3e-08 | 3.3e-08 | 4.033E-08 | 6.429E-09 | 3.300E-08 | 4.500E-08 |
| 9629 | 482 | C482H | A | 5.8e-08 | 4.9e-08 | 5.4e-08 | 5.367E-08 | 4.509E-09 | 4.900E-08 | 5.800E-08 |
| 9630 | 482 | C482I | A | 6.1e-08 | 2.5e-08 | 1.2e-07 | 6.867E-08 | 4.796E-08 | 2.500E-08 | 1.200E-07 |
| 9631 | 482 | C482L | A | 3.1e-08 | 1.8e-08 | 5.8e-08 | 3.567E-08 | 2.040E-08 | 1.800E-08 | 5.800E-08 |
| 9632 | 482 | C482K | A | 9.9e-08 | 4e-08 | 1.1e-07 | 8.300E-08 | 3.764E-08 | 4.000E-08 | 1.100E-07 |
| 9633 | 482 | C482M | A | 5.1e-08 | 4.9e-08 | 1.6e-08 | 3.867E-08 | 1.966E-08 | 1.600E-08 | 5.100E-08 |
| 9634 | 482 | C482F | A | 4.3e-08 | 3.6e-08 | 5.4e-08 | 4.433E-08 | 9.074E-09 | 3.600E-08 | 5.400E-08 |
| 9635 | 482 | C482P | A | 9e-08 | 7.5e-08 | 2.1e-08 | 6.200E-08 | 3.629E-08 | 2.100E-08 | 9.000E-08 |
| 9636 | 482 | C482S | A | 6.1e-08 | 2.1e-08 | 2.5e-08 | 3.567E-08 | 2.203E-08 | 2.100E-08 | 6.100E-08 |
| 9637 | 482 | C482T | A | 2.6e-07 | 5.4e-08 | 2.5e-08 | 1.130E-07 | 1.281E-07 | 2.500E-08 | 2.600E-07 |
| 9638 | 482 | C482W | A | 5.6e-08 | 7.2e-08 | 6.3e-08 | 6.367E-08 | 8.021E-09 | 5.600E-08 | 7.200E-08 |
| 9639 | 482 | C482Y | A | 3.5e-08 | 2.5e-08 | 1.4e-08 | 2.467E-08 | 1.050E-08 | 1.400E-08 | 3.500E-08 |
| 9640 | 482 | C482V | A | 4.3e-08 | 3.6e-08 | 1.9e-08 | 3.267E-08 | 1.234E-08 | 1.900E-08 | 4.300E-08 |
| 9641 | 483 | V483A | A | 4.6e-08 | 5.8e-08 | 6.3e-08 | 5.567E-08 | 8.737E-09 | 4.600E-08 | 6.300E-08 |
| 9642 | 483 | V483R | A | 1.4e-07 | 3.2e-08 | 1.9e-08 | 6.367E-08 | 6.643E-08 | 1.900E-08 | 1.400E-07 |
| 9643 | 483 | V483N | A | 2.3e-08 | 5.8e-08 | 2.9e-08 | 3.667E-08 | 1.872E-08 | 2.300E-08 | 5.800E-08 |
| 9644 | 483 | V483D | A | 7.1e-08 | 8.1e-08 | 3.9e-08 | 6.367E-08 | 2.194E-08 | 3.900E-08 | 8.100E-08 |
| 9645 | 483 | V483C | A | 1.6e-07 | 3e-08 | 1.7e-08 | 6.900E-08 | 7.908E-08 | 1.700E-08 | 1.600E-07 |
| 9646 | 483 | V483E | A | 5.1e-08 | 2.9e-08 | 3e-08 | 3.667E-08 | 1.242E-08 | 2.900E-08 | 5.100E-08 |
| 9647 | 483 | V483Q | A | 1.3e-08 | 1.9e-08 | 4.5e-08 | 2.567E-08 | 1.701E-08 | 1.300E-08 | 4.500E-08 |
| 9648 | 483 | V483G | A | 1.8e-08 | 3e-08 | 3.2e-08 | 2.667E-08 | 7.572E-09 | 1.800E-08 | 3.200E-08 |
| 9649 | 483 | V483H | A | 5.5e-08 | 4.9e-08 | 3e-08 | 4.467E-08 | 1.305E-08 | 3.000E-08 | 5.500E-08 |
| 9650 | 483 | V483I | A | 3.8e-08 | 1.3e-07 | 3.4e-08 | 6.733E-08 | 5.431E-08 | 3.400E-08 | 1.300E-07 |
| 9651 | 483 | V483L | A | 5.1e-08 | 4.6e-08 | 9.5e-09 | 3.550E-08 | 2.266E-08 | 9.500E-09 | 5.100E-08 |
| 9652 | 483 | V483K | A | 3.2e-08 | 4.1e-08 | 2.4e-08 | 3.233E-08 | 8.505E-09 | 2.400E-08 | 4.100E-08 |
| 9653 | 483 | V483M | A | 3.4e-08 | 3.7e-08 | 2.2e-08 | 3.100E-08 | 7.937E-09 | 2.200E-08 | 3.700E-08 |
| 9654 | 483 | V483F | A | 6.3e-08 | 1.4e-07 | 3.5e-08 | 7.933E-08 | 5.437E-08 | 3.500E-08 | 1.400E-07 |
| 9655 | 483 | V483P | A | 4.1e-08 | 5.7e-08 | 4.7e-08 | 4.833E-08 | 8.083E-09 | 4.100E-08 | 5.700E-08 |
| 9656 | 483 | V483S | A | 1.8e-08 | 6.1e-08 | 5.8e-08 | 4.567E-08 | 2.401E-08 | 1.800E-08 | 6.100E-08 |
| 9657 | 483 | V483T | A | 6.2e-08 | 2.2e-08 | 4.1e-08 | 4.167E-08 | 2.001E-08 | 2.200E-08 | 6.200E-08 |
| 9658 | 483 | V483W | A | 2.1e-08 | 1.8e-08 | 8.7e-08 | 4.200E-08 | 3.900E-08 | 1.800E-08 | 8.700E-08 |
| 9659 | 483 | V483Y | A | 3.7e-08 | 5.5e-08 | 8.8e-08 | 6.000E-08 | 2.587E-08 | 3.700E-08 | 8.800E-08 |
| 9660 | 483 | V483V | A | 6.6e-08 | 2.5e-08 | 6.4e-08 | 5.167E-08 | 2.312E-08 | 2.500E-08 | 6.600E-08 |
| 9661 | 484 | G484A | A | 4.1e-08 | 2.5e-08 | 2.3e-08 | 2.967E-08 | 9.866E-09 | 2.300E-08 | 4.100E-08 |
| 9662 | 484 | G484R | A | 1.1e-07 | 2e-08 | 8.3e-08 | 7.100E-08 | 4.618E-08 | 2.000E-08 | 1.100E-07 |
| 9663 | 484 | G484N | A | 6.3e-08 | 5.1e-08 | 9.4e-08 | 6.933E-08 | 2.219E-08 | 5.100E-08 | 9.400E-08 |
| 9664 | 484 | G484D | A | 1.4e-07 | 2e-08 | 8.7e-08 | 8.233E-08 | 6.014E-08 | 2.000E-08 | 1.400E-07 |
| 9665 | 484 | G484C | A | 4.2e-08 | 5.5e-08 | 8.2e-08 | 5.967E-08 | 2.040E-08 | 4.200E-08 | 8.200E-08 |
| 9666 | 484 | G484E | A | 5.3e-08 | 3.9e-08 | 2.6e-08 | 3.933E-08 | 1.350E-08 | 2.600E-08 | 5.300E-08 |
| 9667 | 484 | G484Q | A | 1.6e-08 | 3.8e-08 | 7.6e-08 | 4.333E-08 | 3.035E-08 | 1.600E-08 | 7.600E-08 |
| 9668 | 484 | G484G | A | 8.9e-08 | 2.6e-08 | 3e-08 | 4.833E-08 | 3.528E-08 | 2.600E-08 | 8.900E-08 |
| 9669 | 484 | G484H | A | 3.9e-08 | 8.4e-08 | 1.3e-08 | 4.533E-08 | 3.592E-08 | 1.300E-08 | 8.400E-08 |
| 9670 | 484 | G484I | A | 1e-07 | 6e-08 | 8.3e-08 | 8.100E-08 | 2.007E-08 | 6.000E-08 | 1.000E-07 |
| 9671 | 484 | G484L | A | 6.2e-08 | 6e-08 | 4.8e-08 | 5.667E-08 | 7.572E-09 | 4.800E-08 | 6.200E-08 |
| 9672 | 484 | G484K | A | 4.6e-08 | 2.9e-08 | 7.6e-08 | 5.033E-08 | 2.380E-08 | 2.900E-08 | 7.600E-08 |
| 9673 | 484 | G484M | A | 8.8e-08 | 2.7e-08 | 3.6e-08 | 5.033E-08 | 3.293E-08 | 2.700E-08 | 8.800E-08 |
| 9674 | 484 | G484F | A | 3.8e-08 | 2.7e-08 | 3.6e-08 | 3.367E-08 | 5.859E-09 | 2.700E-08 | 3.800E-08 |
| 9675 | 484 | G484P | A | 6.3e-08 | 2.1e-07 | 4.2e-08 | 1.050E-07 | 9.154E-08 | 4.200E-08 | 2.100E-07 |
| 9676 | 484 | G484S | A | 5.1e-08 | 5.1e-08 | 6.8e-08 | 5.667E-08 | 9.815E-09 | 5.100E-08 | 6.800E-08 |
| 9677 | 484 | G484T | A | 5.2e-08 | 2.1e-08 | 1.3e-08 | 2.867E-08 | 2.060E-08 | 1.300E-08 | 5.200E-08 |
| 9678 | 484 | G484W | A | 2.8e-08 | 7.6e-08 | 9.4e-08 | 6.600E-08 | 3.412E-08 | 2.800E-08 | 9.400E-08 |
| 9679 | 484 | G484Y | A | 9.4e-08 | 3.2e-08 | 3.4e-08 | 5.333E-08 | 3.523E-08 | 3.200E-08 | 9.400E-08 |
| 9680 | 484 | G484V | A | 4e-08 | 4.2e-08 | 4.9e-08 | 4.367E-08 | 4.726E-09 | 4.000E-08 | 4.900E-08 |
| 9681 | 485 | E485A | A | 6.1e-08 | 4.8e-08 | 1.3e-07 | 7.967E-08 | 4.407E-08 | 4.800E-08 | 1.300E-07 |
| 9682 | 485 | E485R | A | 6.1e-08 | 8.6e-08 | 2.4e-08 | 5.700E-08 | 3.119E-08 | 2.400E-08 | 8.600E-08 |
| 9683 | 485 | E485N | A | 5.4e-08 | 1.9e-08 | 2.4e-08 | 3.233E-08 | 1.893E-08 | 1.900E-08 | 5.400E-08 |
| 9684 | 485 | E485D | A | 4.6e-08 | 5.9e-08 | 9.5e-08 | 6.667E-08 | 2.538E-08 | 4.600E-08 | 9.500E-08 |
| 9685 | 485 | E485C | A | 1.5e-08 | 5.9e-08 | 5.2e-08 | 4.200E-08 | 2.364E-08 | 1.500E-08 | 5.900E-08 |
| 9686 | 485 | E485E | A | 6.5e-08 | 2.7e-08 | 2.9e-08 | 4.033E-08 | 2.139E-08 | 2.700E-08 | 6.500E-08 |
| 9687 | 485 | E485Q | A | 3.4e-08 | 2.3e-08 | 4.5e-08 | 3.400E-08 | 1.100E-08 | 2.300E-08 | 4.500E-08 |
| 9688 | 485 | E485G | A | 5.2e-08 | 7.5e-08 | 1.8e-08 | 4.833E-08 | 2.868E-08 | 1.800E-08 | 7.500E-08 |
| 9689 | 485 | E485H | A | 2.3e-08 | 2.3e-08 | 1.6e-08 | 2.067E-08 | 4.041E-09 | 1.600E-08 | 2.300E-08 |

| | | | | | | | | | | |
|---|---|---|---|---|---|---|---|---|---|---|
| 9690 | 485 | E485I | A | 7.5e-08 | 4.7e-08 | 2.5e-08 | 4.900E-08 | 2.506E-08 | 2.500E-08 | 7.500E-08 |
| 9691 | 485 | E485L | A | 9.5e-08 | 4.8e-08 | 2.1e-08 | 5.467E-08 | 3.745E-08 | 2.100E-08 | 9.500E-08 |
| 9692 | 485 | E485K | A | 9.3e-08 | 2.5e-08 | 5.4e-08 | 5.733E-08 | 3.412E-08 | 2.500E-08 | 9.300E-08 |
| 9693 | 485 | E485M | A | 5.2e-08 | 2.7e-08 | 1.6e-07 | 7.967E-08 | 7.068E-08 | 2.700E-08 | 1.600E-07 |
| 9694 | 485 | E485F | A | 3.6e-08 | 5.2e-08 | 3.2e-08 | 4.000E-08 | 1.058E-08 | 3.200E-08 | 5.200E-08 |
| 9695 | 485 | E485P | A | 3.2e-08 | 4e-08 | 3.8e-08 | 3.667E-08 | 4.163E-09 | 3.200E-08 | 4.000E-08 |
| 9696 | 485 | E485S | A | 1.1e-08 | 4.5e-08 | 5.4e-08 | 3.667E-08 | 2.268E-08 | 1.100E-08 | 5.400E-08 |
| 9697 | 485 | E485T | A | 2.4e-08 | 7.9e-08 | 7e-08 | 5.767E-08 | 2.950E-08 | 2.400E-08 | 7.900E-08 |
| 9698 | 485 | E485W | A | 8.8e-08 | 7.4e-08 | 1.5e-08 | 5.900E-08 | 3.874E-08 | 1.500E-08 | 8.800E-08 |
| 9699 | 485 | E485Y | A | 8.5e-08 | 3.9e-08 | 3.6e-08 | 5.333E-08 | 2.747E-08 | 3.600E-08 | 8.500E-08 |
| 9700 | 485 | E485V | A | 4.9e-08 | 7.4e-08 | 2.4e-08 | 4.900E-08 | 2.500E-08 | 2.400E-08 | 7.400E-08 |
| 9701 | 486 | G486A | A | 1.9e-08 | 6.4e-08 | 2.1e-08 | 3.467E-08 | 2.542E-08 | 1.900E-08 | 6.400E-08 |
| 9702 | 486 | G486R | A | 3.4e-08 | 1.2e-07 | 1.6e-08 | 5.667E-08 | 5.558E-08 | 1.600E-08 | 1.200E-07 |
| 9703 | 486 | G486N | A | 2.3e-08 | 3.2e-08 | 2.4e-08 | 2.633E-08 | 4.933E-09 | 2.300E-08 | 3.200E-08 |
| 9704 | 486 | G486D | A | 7.5e-08 | 3.8e-08 | 8.7e-08 | 6.667E-08 | 2.554E-08 | 3.800E-08 | 8.700E-08 |
| 9705 | 486 | G486C | A | 4.7e-08 | 6.5e-08 | 1e-07 | 7.067E-08 | 2.695E-08 | 4.700E-08 | 1.000E-07 |
| 9706 | 486 | G486E | A | 3.8e-08 | 1.5e-08 | 9.3e-08 | 4.867E-08 | 4.008E-08 | 1.500E-08 | 9.300E-08 |
| 9707 | 486 | G486Q | A | 5.8e-08 | 1.7e-08 | 5e-08 | 4.167E-08 | 2.173E-08 | 1.700E-08 | 5.800E-08 |
| 9708 | 486 | G486G | A | 3.7e-08 | 5.1e-08 | 7e-08 | 5.267E-08 | 1.656E-08 | 3.700E-08 | 7.000E-08 |
| 9709 | 486 | G486H | A | 8.4e-08 | 6.1e-08 | 1.1e-07 | 8.500E-08 | 2.452E-08 | 6.100E-08 | 1.100E-07 |
| 9710 | 486 | G486I | A | 2.1e-08 | 6.8e-08 | 1.8e-08 | 3.567E-08 | 2.804E-08 | 1.800E-08 | 6.800E-08 |
| 9711 | 486 | G486L | A | 6.3e-08 | 4.1e-08 | 3.5e-08 | 4.633E-08 | 1.474E-08 | 3.500E-08 | 6.300E-08 |
| 9712 | 486 | G486K | A | 7.8e-08 | 5.9e-08 | 4.4e-08 | 6.033E-08 | 1.704E-08 | 4.400E-08 | 7.800E-08 |
| 9713 | 486 | G486M | A | 9e-08 | 9.1e-08 | 4.2e-08 | 7.433E-08 | 2.801E-08 | 4.200E-08 | 9.100E-08 |
| 9714 | 486 | G486F | A | 2.9e-08 | 1.6e-08 | 1.7e-08 | 2.067E-08 | 7.234E-09 | 1.600E-08 | 2.900E-08 |
| 9715 | 486 | G486P | A | 6.2e-08 | 3.8e-08 | 2.5e-08 | 4.167E-08 | 1.877E-08 | 2.500E-08 | 6.200E-08 |
| 9716 | 486 | G486S | A | 2.9e-08 | 9.6e-08 | 3.8e-08 | 5.433E-08 | 3.636E-08 | 2.900E-08 | 9.600E-08 |
| 9717 | 486 | G486T | A | 7.5e-09 | 5.7e-08 | 4.7e-08 | 3.717E-08 | 2.617E-08 | 7.500E-09 | 5.700E-08 |
| 9718 | 486 | G486W | A | 3.4e-08 | 4.2e-08 | 1.4e-08 | 3.000E-08 | 1.442E-08 | 1.400E-08 | 4.200E-08 |
| 9719 | 486 | G486Y | A | 2.9e-08 | 3.2e-08 | 3.1e-08 | 3.067E-08 | 1.528E-09 | 2.900E-08 | 3.200E-08 |
| 9720 | 486 | G486V | A | 1e-08 | 2.3e-08 | 9e-08 | 4.100E-08 | 4.293E-08 | 1.000E-08 | 9.000E-08 |
| 9721 | 487 | L487A | A | 4.2e-08 | 3.3e-08 | 3.5e-08 | 3.667E-08 | 4.726E-09 | 3.300E-08 | 4.200E-08 |
| 9722 | 487 | L487R | A | 6.3e-08 | 2.2e-08 | 3.7e-08 | 4.067E-08 | 2.074E-08 | 2.200E-08 | 6.300E-08 |
| 9723 | 487 | L487N | A | 7.6e-08 | 4.5e-08 | 2.1e-08 | 4.733E-08 | 2.757E-08 | 2.100E-08 | 7.600E-08 |
| 9724 | 487 | L487D | A | 8e-08 | 4.9e-08 | 1.8e-08 | 4.900E-08 | 3.100E-08 | 1.800E-08 | 8.000E-08 |
| 9725 | 487 | L487C | A | 3.3e-08 | 1.4e-08 | 2.8e-08 | 2.500E-08 | 9.849E-09 | 1.400E-08 | 3.300E-08 |
| 9726 | 487 | L487E | A | 3.9e-08 | 2.5e-08 | 1.5e-08 | 2.633E-08 | 1.206E-08 | 1.500E-08 | 3.900E-08 |
| 9727 | 487 | L487Q | A | 7.5e-08 | 7.5e-08 | 2.2e-08 | 5.733E-08 | 3.060E-08 | 2.200E-08 | 7.500E-08 |
| 9728 | 487 | L487G | A | 4.3e-08 | 4e-08 | 2.6e-08 | 3.633E-08 | 9.074E-09 | 2.600E-08 | 4.300E-08 |
| 9729 | 487 | L487H | A | 1e-08 | 4.2e-08 | 4.3e-08 | 3.167E-08 | 1.877E-08 | 1.000E-08 | 4.300E-08 |
| 9730 | 487 | L487I | A | 2.7e-08 | 7.5e-08 | 4.1e-08 | 4.767E-08 | 2.468E-08 | 2.700E-08 | 7.500E-08 |
| 9731 | 487 | L487L | A | 6.9e-08 | 6.4e-08 | 1.6e-07 | 9.767E-08 | 5.404E-08 | 6.400E-08 | 1.600E-07 |
| 9732 | 487 | L487K | A | 3.2e-08 | 2.4e-08 | 6.7e-08 | 4.100E-08 | 2.287E-08 | 2.400E-08 | 6.700E-08 |
| 9733 | 487 | L487M | A | 4e-08 | 6.6e-08 | 3.2e-08 | 4.600E-08 | 1.778E-08 | 3.200E-08 | 6.600E-08 |
| 9734 | 487 | L487F | A | 5.8e-08 | 4.7e-08 | 2.3e-08 | 4.267E-08 | 1.790E-08 | 2.300E-08 | 5.800E-08 |
| 9735 | 487 | L487P | A | 5e-08 | 4.4e-08 | 4.4e-08 | 4.600E-08 | 3.464E-09 | 4.400E-08 | 5.000E-08 |
| 9736 | 487 | L487S | A | 9e-08 | 8.3e-08 | 1.7e-08 | 6.333E-08 | 4.028E-08 | 1.700E-08 | 9.000E-08 |
| 9737 | 487 | L487T | A | 1.3e-08 | 1.4e-07 | 6.8e-08 | 7.367E-08 | 6.369E-08 | 1.300E-08 | 1.400E-07 |
| 9738 | 487 | L487W | A | 6.6e-08 | 3.8e-08 | 1.1e-07 | 7.133E-08 | 3.630E-08 | 3.800E-08 | 1.100E-07 |
| 9739 | 487 | L487Y | A | 2.1e-08 | 4.4e-08 | 2.4e-08 | 2.967E-08 | 1.250E-08 | 2.100E-08 | 4.400E-08 |
| 9740 | 487 | L487V | A | 4.3e-08 | 6.5e-08 | 5.5e-08 | 5.433E-08 | 1.102E-08 | 4.300E-08 | 6.500E-08 |
| 9741 | 488 | A488A | A | 2.1e-08 | 7e-08 | 3.9e-08 | 4.333E-08 | 2.479E-08 | 2.100E-08 | 7.000E-08 |
| 9742 | 488 | A488R | A | 1.6e-08 | 3.1e-08 | 8.6e-08 | 4.433E-08 | 3.686E-08 | 1.600E-08 | 8.600E-08 |
| 9743 | 488 | A488N | A | 2.2e-08 | 7.2e-08 | 5.8e-08 | 5.067E-08 | 2.579E-08 | 2.200E-08 | 7.200E-08 |
| 9744 | 488 | A488D | A | 6e-08 | 7e-08 | 4.5e-08 | 5.833E-08 | 1.258E-08 | 4.500E-08 | 7.000E-08 |
| 9745 | 488 | A488C | A | 5.5e-08 | 6.4e-08 | 1.3e-07 | 8.300E-08 | 4.095E-08 | 5.500E-08 | 1.300E-07 |
| 9746 | 488 | A488E | A | 2.7e-08 | 7.3e-08 | 3.3e-08 | 4.433E-08 | 2.501E-08 | 2.700E-08 | 7.300E-08 |
| 9747 | 488 | A488Q | A | 9.4e-08 | 3.9e-08 | 9e-08 | 7.433E-08 | 3.066E-08 | 3.900E-08 | 9.400E-08 |
| 9748 | 488 | A488G | A | 6.9e-08 | 3.5e-08 | 7.3e-08 | 5.900E-08 | 2.088E-08 | 3.500E-08 | 7.300E-08 |
| 9749 | 488 | A488H | A | 7.3e-08 | 3.7e-08 | 3.7e-08 | 4.900E-08 | 2.078E-08 | 3.700E-08 | 7.300E-08 |
| 9750 | 488 | A488I | A | 1.7e-08 | 2.4e-08 | 1.4e-08 | 1.833E-08 | 5.132E-09 | 1.400E-08 | 2.400E-08 |
| 9751 | 488 | A488L | A | 1e-07 | 3.5e-08 | 2.2e-08 | 5.233E-08 | 4.179E-08 | 2.200E-08 | 1.000E-07 |
| 9752 | 488 | A488K | A | 5e-08 | 5.8e-08 | 4.5e-08 | 5.100E-08 | 6.557E-09 | 4.500E-08 | 5.800E-08 |
| 9753 | 488 | A488M | A | 1.8e-08 | 1e-07 | 1e-07 | 7.267E-08 | 4.734E-08 | 1.800E-08 | 1.000E-07 |
| 9754 | 488 | A488F | A | 1.8e-07 | 2.5e-08 | 4.6e-08 | 8.367E-08 | 8.409E-08 | 2.500E-08 | 1.800E-07 |
| 9755 | 488 | A488P | A | 4.7e-08 | 1.6e-08 | 8.1e-08 | 4.800E-08 | 3.251E-08 | 1.600E-08 | 8.100E-08 |
| 9756 | 488 | A488S | A | 6.3e-08 | 4e-08 | 5.1e-08 | 5.133E-08 | 1.150E-08 | 4.000E-08 | 6.300E-08 |
| 9757 | 488 | A488T | A | 2.2e-08 | 3.1e-08 | 4.1e-08 | 3.133E-08 | 9.504E-09 | 2.200E-08 | 4.100E-08 |
| 9758 | 488 | A488W | A | 1.5e-07 | 2.7e-08 | 4.9e-08 | 7.533E-08 | 6.559E-08 | 2.700E-08 | 1.500E-07 |
| 9759 | 488 | A488Y | A | 6.5e-08 | 5.5e-08 | 1.5e-08 | 4.500E-08 | 2.646E-08 | 1.500E-08 | 6.500E-08 |
| 9760 | 488 | A488V | A | 7.4e-08 | 2.7e-08 | 7.6e-08 | 5.900E-08 | 2.773E-08 | 2.700E-08 | 7.600E-08 |
| 9761 | 489 | C489A | A | 5.2e-08 | 4.7e-08 | 2.4e-08 | 4.100E-08 | 1.493E-08 | 2.400E-08 | 5.200E-08 |
| 9762 | 489 | C489R | A | 4.7e-08 | 3.9e-08 | 1.3e-07 | 7.200E-08 | 5.039E-08 | 3.900E-08 | 1.300E-07 |
| 9763 | 489 | C489N | A | 3.8e-08 | 5.2e-08 | 1.1e-07 | 6.667E-08 | 3.818E-08 | 3.800E-08 | 1.100E-07 |
| 9764 | 489 | C489D | A | 4.3e-08 | 3e-08 | 1.7e-07 | 8.100E-08 | 7.735E-08 | 3.000E-08 | 1.700E-07 |
| 9765 | 489 | C489C | A | 3.6e-08 | 1.3e-07 | 2.5e-08 | 6.367E-08 | 5.771E-08 | 2.500E-08 | 1.300E-07 |
| 9766 | 489 | C489E | A | 2.7e-08 | 3.2e-08 | 6.6e-08 | 4.167E-08 | 2.122E-08 | 2.700E-08 | 6.600E-08 |
| 9767 | 489 | C489Q | A | 9.1e-08 | 3.9e-08 | 5.3e-08 | 6.100E-08 | 2.691E-08 | 3.900E-08 | 9.100E-08 |
| 9768 | 489 | C489G | A | 2.9e-08 | 9.5e-08 | 4.1e-08 | 5.500E-08 | 3.516E-08 | 2.900E-08 | 9.500E-08 |
| 9769 | 489 | C489H | A | 5.7e-08 | 3.2e-08 | 4.9e-08 | 4.600E-08 | 1.277E-08 | 3.200E-08 | 5.700E-08 |
| 9770 | 489 | C489I | A | 4.5e-08 | 4e-08 | 5.7e-08 | 4.733E-08 | 8.737E-09 | 4.000E-08 | 5.700E-08 |
| 9771 | 489 | C489L | A | 3.2e-08 | 6.3e-08 | 2.5e-08 | 4.000E-08 | 2.022E-08 | 2.500E-08 | 6.300E-08 |
| 9772 | 489 | C489K | A | 4.3e-08 | 3.9e-08 | 7.9e-08 | 5.367E-08 | 2.203E-08 | 3.900E-08 | 7.900E-08 |
| 9773 | 489 | C489M | A | 9.8e-08 | 3.2e-08 | 6.4e-08 | 6.467E-08 | 3.301E-08 | 3.200E-08 | 9.800E-08 |
| 9774 | 489 | C489F | A | 4.5e-08 | 2.8e-08 | 2.5e-08 | 3.267E-08 | 1.079E-08 | 2.500E-08 | 4.500E-08 |

| | | | | | | | | | | |
|---|---|---|---|---|---|---|---|---|---|---|
| 9775 | 489 | C489P | A | 7.2e-08 | 2.3e-08 | 5.9e-08 | 5.133E-08 | 2.538E-08 | 2.300E-08 | 7.200E-08 |
| 9776 | 489 | C489S | A | 3e-08 | 4.5e-08 | 1.3e-07 | 6.833E-08 | 5.393E-08 | 3.000E-08 | 1.300E-07 |
| 9777 | 489 | C489T | A | 5.3e-08 | 8.5e-08 | 9.6e-08 | 7.800E-08 | 2.234E-08 | 5.300E-08 | 9.600E-08 |
| 9778 | 489 | C489W | A | 3.1e-08 | 5e-08 | 8.5e-08 | 5.533E-08 | 2.739E-08 | 3.100E-08 | 8.500E-08 |
| 9779 | 489 | C489Y | A | 9.1e-09 | 5.9e-08 | 3.1e-08 | 3.303E-08 | 2.501E-08 | 9.100E-09 | 5.900E-08 |
| 9780 | 489 | C489V | A | 3.4e-08 | 2.8e-08 | 4.4e-08 | 3.533E-08 | 8.083E-09 | 2.800E-08 | 4.400E-08 |
| 9781 | 490 | H490A | A | 2.9e-08 | 2.2e-08 | 1.1e-07 | 5.367E-08 | 4.891E-08 | 2.200E-08 | 1.100E-07 |
| 9782 | 490 | H490R | A | 6.4e-08 | 2.6e-08 | 3.4e-08 | 4.133E-08 | 2.003E-08 | 2.600E-08 | 6.400E-08 |
| 9783 | 490 | H490N | A | 3.9e-08 | 2.9e-08 | 4.3e-08 | 3.700E-08 | 7.211E-09 | 2.900E-08 | 4.300E-08 |
| 9784 | 490 | H490D | A | 6.1e-08 | 2.8e-08 | 2.6e-08 | 3.833E-08 | 1.966E-08 | 2.600E-08 | 6.100E-08 |
| 9785 | 490 | H490C | A | 5.6e-08 | 6.6e-08 | 6.5e-08 | 6.233E-08 | 5.508E-09 | 5.600E-08 | 6.600E-08 |
| 9786 | 490 | H490E | A | 2.5e-07 | 3.2e-08 | 1.2e-08 | 9.800E-08 | 1.320E-07 | 1.200E-08 | 2.500E-07 |
| 9787 | 490 | H490Q | A | 2.4e-08 | 5.6e-08 | 6.3e-08 | 4.767E-08 | 2.079E-08 | 2.400E-08 | 6.300E-08 |
| 9788 | 490 | H490G | A | 5e-08 | 1.3e-07 | 8e-08 | 8.667E-08 | 4.041E-08 | 5.000E-08 | 1.300E-07 |
| 9789 | 490 | H490H | A | 6.4e-08 | 4.9e-08 | 3.3e-08 | 4.867E-08 | 1.550E-08 | 3.300E-08 | 6.400E-08 |
| 9790 | 490 | H490I | A | 5e-08 | 4.6e-08 | 1e-08 | 3.533E-08 | 2.203E-08 | 1.000E-08 | 5.000E-08 |
| 9791 | 490 | H490L | A | 2.6e-08 | 7e-08 | 9e-08 | 6.200E-08 | 3.274E-08 | 2.600E-08 | 9.000E-08 |
| 9792 | 490 | H490K | A | 2.8e-08 | 3.1e-08 | 4.1e-08 | 3.333E-08 | 6.807E-09 | 2.800E-08 | 4.100E-08 |
| 9793 | 490 | H490M | A | 2.3e-08 | 2e-08 | 9.2e-08 | 4.500E-08 | 4.073E-08 | 2.000E-08 | 9.200E-08 |
| 9794 | 490 | H490F | A | 1.4e-07 | 5.2e-08 | 2.6e-08 | 7.267E-08 | 5.974E-08 | 2.600E-08 | 1.400E-07 |
| 9795 | 490 | H490P | A | 4.8e-08 | 4.5e-08 | 1.4e-07 | 7.767E-08 | 5.400E-08 | 4.500E-08 | 1.400E-07 |
| 9796 | 490 | H490S | A | 5e-08 | 3.9e-08 | 2.9e-08 | 3.933E-08 | 1.050E-08 | 2.900E-08 | 5.000E-08 |
| 9797 | 490 | H490T | A | 4.8e-08 | 7.3e-08 | 9.8e-08 | 7.300E-08 | 2.500E-08 | 4.800E-08 | 9.800E-08 |
| 9798 | 490 | H490W | A | 8.3e-08 | 3.7e-08 | 2.5e-08 | 4.833E-08 | 3.062E-08 | 2.500E-08 | 8.300E-08 |
| 9799 | 490 | H490Y | A | 4.9e-08 | 2e-08 | 2.9e-08 | 3.267E-08 | 1.484E-08 | 2.000E-08 | 4.900E-08 |
| 9800 | 490 | H490V | A | 4.9e-08 | 4.4e-08 | 3.8e-08 | 4.367E-08 | 5.508E-09 | 3.800E-08 | 4.900E-08 |
| 9801 | 491 | Q491A | A | 8.8e-09 | 2.5e-08 | 4.2e-08 | 2.527E-08 | 1.660E-08 | 8.800E-09 | 4.200E-08 |
| 9802 | 491 | Q491R | A | 9.6e-08 | 2.1e-07 | 4.7e-08 | 1.177E-07 | 8.363E-08 | 4.700E-08 | 2.100E-07 |
| 9803 | 491 | Q491N | A | 8.7e-08 | 4.1e-08 | 2.5e-08 | 5.100E-08 | 3.219E-08 | 2.500E-08 | 8.700E-08 |
| 9804 | 491 | Q491D | A | 8e-08 | 5.5e-08 | 2.1e-08 | 5.200E-08 | 2.961E-08 | 2.100E-08 | 8.000E-08 |
| 9805 | 491 | Q491C | A | 2.1e-08 | 4.7e-08 | 3e-08 | 3.267E-08 | 1.320E-08 | 2.100E-08 | 4.700E-08 |
| 9806 | 491 | Q491E | A | 6.2e-08 | 5.2e-08 | 5.1e-08 | 5.500E-08 | 6.083E-09 | 5.100E-08 | 6.200E-08 |
| 9807 | 491 | Q491Q | A | 4.4e-08 | 9.9e-09 | 3e-08 | 2.797E-08 | 1.714E-08 | 9.900E-09 | 4.400E-08 |
| 9808 | 491 | Q491G | A | 9.2e-08 | 1.5e-08 | 9.2e-08 | 6.633E-08 | 4.446E-08 | 1.500E-08 | 9.200E-08 |
| 9809 | 491 | Q491H | A | 2.6e-08 | 2.5e-08 | 3.9e-08 | 3.000E-08 | 7.810E-09 | 2.500E-08 | 3.900E-08 |
| 9810 | 491 | Q491I | A | 3.2e-08 | 9.6e-08 | 2.1e-08 | 4.967E-08 | 4.050E-08 | 2.100E-08 | 9.600E-08 |
| 9811 | 491 | Q491L | A | 6.7e-08 | 9.2e-08 | 2.6e-08 | 6.167E-08 | 3.332E-08 | 2.600E-08 | 9.200E-08 |
| 9812 | 491 | Q491K | A | 3.8e-08 | 1.9e-08 | 8.3e-09 | 2.177E-08 | 1.504E-08 | 8.300E-09 | 3.800E-08 |
| 9813 | 491 | Q491M | A | 5.1e-08 | 3e-08 | 8.3e-08 | 5.467E-08 | 2.669E-08 | 3.000E-08 | 8.300E-08 |
| 9814 | 491 | Q491F | A | 4.4e-08 | 4.3e-08 | 4.7e-08 | 4.467E-08 | 2.082E-09 | 4.300E-08 | 4.700E-08 |
| 9815 | 491 | Q491P | A | 6.9e-08 | 2e-08 | 5.1e-08 | 4.667E-08 | 2.479E-08 | 2.000E-08 | 6.900E-08 |
| 9816 | 491 | Q491S | A | 2.8e-08 | 3.9e-08 | 5.3e-08 | 4.000E-08 | 1.253E-08 | 2.800E-08 | 5.300E-08 |
| 9817 | 491 | Q491T | A | 3.6e-08 | 5.1e-08 | 7.4e-08 | 5.367E-08 | 1.914E-08 | 3.600E-08 | 7.400E-08 |
| 9818 | 491 | Q491W | A | 4.2e-08 | 5.5e-08 | 5.9e-08 | 5.200E-08 | 8.888E-09 | 4.200E-08 | 5.900E-08 |
| 9819 | 491 | Q491Y | A | 5.4e-08 | 4.9e-08 | 5.7e-08 | 5.333E-08 | 4.041E-09 | 4.900E-08 | 5.700E-08 |
| 9820 | 491 | Q491V | A | 6.3e-08 | 3.4e-08 | 4e-08 | 4.567E-08 | 1.531E-08 | 3.400E-08 | 6.300E-08 |
| 9821 | 492 | L492A | A | 3e-08 | 8.2e-08 | 3.3e-08 | 4.833E-08 | 2.919E-08 | 3.000E-08 | 8.200E-08 |
| 9822 | 492 | L492R | A | 1.2e-08 | 1.8e-08 | 6.9e-08 | 3.300E-08 | 3.132E-08 | 1.200E-08 | 6.900E-08 |
| 9823 | 492 | L492N | A | 8.9e-08 | 5.5e-08 | 9.1e-08 | 7.833E-08 | 2.023E-08 | 5.500E-08 | 9.100E-08 |
| 9824 | 492 | L492D | A | 3.4e-08 | 2.9e-08 | 2.9e-08 | 3.067E-08 | 2.887E-09 | 2.900E-08 | 3.400E-08 |
| 9825 | 492 | L492C | A | 5.1e-08 | 3.6e-08 | 3.5e-08 | 4.067E-08 | 8.963E-09 | 3.500E-08 | 5.100E-08 |
| 9826 | 492 | L492E | A | 2e-08 | 8.8e-08 | 4.2e-08 | 5.000E-08 | 3.470E-08 | 2.000E-08 | 8.800E-08 |
| 9827 | 492 | L492Q | A | 1.1e-08 | 5.9e-08 | 2.3e-08 | 3.100E-08 | 2.498E-08 | 1.100E-08 | 5.900E-08 |
| 9828 | 492 | L492G | A | 6e-08 | 5.2e-08 | 5.8e-08 | 5.667E-08 | 4.163E-09 | 5.200E-08 | 6.000E-08 |
| 9829 | 492 | L492H | A | 8.6e-08 | 3.5e-08 | 5.5e-08 | 5.867E-08 | 2.570E-08 | 3.500E-08 | 8.600E-08 |
| 9830 | 492 | L492I | A | 6.5e-08 | 3.4e-08 | 2.1e-08 | 4.000E-08 | 2.261E-08 | 2.100E-08 | 6.500E-08 |
| 9831 | 492 | L492L | A | 7.1e-08 | 5.5e-08 | 1.6e-08 | 4.733E-08 | 2.829E-08 | 1.600E-08 | 7.100E-08 |
| 9832 | 492 | L492K | A | 3.4e-08 | 4.7e-08 | 9.5e-08 | 5.867E-08 | 3.213E-08 | 3.400E-08 | 9.500E-08 |
| 9833 | 492 | L492M | A | 5.2e-08 | 3.3e-08 | 6.3e-08 | 4.933E-08 | 1.518E-08 | 3.300E-08 | 6.300E-08 |
| 9834 | 492 | L492F | A | 5.1e-08 | 3e-08 | 4.3e-08 | 4.133E-08 | 1.060E-08 | 3.000E-08 | 5.100E-08 |
| 9835 | 492 | L492P | A | 3.1e-08 | 2.2e-08 | 2e-08 | 2.433E-08 | 5.859E-09 | 2.000E-08 | 3.100E-08 |
| 9836 | 492 | L492S | A | 1.4e-07 | 2.8e-08 | 2.7e-08 | 6.500E-08 | 6.495E-08 | 2.700E-08 | 1.400E-07 |
| 9837 | 492 | L492T | A | 2e-07 | 8.4e-08 | 7.8e-08 | 1.207E-07 | 6.877E-08 | 7.800E-08 | 2.000E-07 |
| 9838 | 492 | L492W | A | 8.5e-08 | 2.3e-08 | 2.5e-08 | 4.433E-08 | 3.523E-08 | 2.300E-08 | 8.500E-08 |
| 9839 | 492 | L492Y | A | 2.3e-08 | 2e-08 | 1.2e-08 | 1.833E-08 | 5.686E-09 | 1.200E-08 | 2.300E-08 |
| 9840 | 492 | L492V | A | 7.1e-08 | 4e-08 | 5.4e-08 | 5.500E-08 | 1.552E-08 | 4.000E-08 | 7.100E-08 |
| 9841 | 493 | C493A | A | 1.3e-07 | 6.9e-08 | 2.4e-08 | 7.433E-08 | 5.320E-08 | 2.400E-08 | 1.300E-07 |
| 9842 | 493 | C493R | A | 1.1e-07 | 2.2e-08 | 4.2e-08 | 5.800E-08 | 4.613E-08 | 2.200E-08 | 1.100E-07 |
| 9843 | 493 | C493N | A | 4.8e-08 | 4.2e-08 | 1.7e-08 | 3.567E-08 | 1.644E-08 | 1.700E-08 | 4.800E-08 |
| 9844 | 493 | C493D | A | 2.6e-08 | 3.7e-08 | 9.3e-08 | 5.200E-08 | 3.593E-08 | 2.600E-08 | 9.300E-08 |
| 9845 | 493 | C493C | A | 3.6e-08 | 1.2e-07 | 6.4e-08 | 7.333E-08 | 4.277E-08 | 3.600E-08 | 1.200E-07 |
| 9846 | 493 | C493E | A | 1.3e-08 | 2.9e-08 | 2.5e-08 | 2.233E-08 | 8.327E-09 | 1.300E-08 | 2.900E-08 |
| 9847 | 493 | C493Q | A | 3.8e-08 | 3.1e-08 | 2.9e-08 | 3.267E-08 | 4.726E-09 | 2.900E-08 | 3.800E-08 |
| 9848 | 493 | C493G | A | 5.2e-08 | 6.2e-08 | 4.2e-08 | 5.200E-08 | 1.000E-08 | 4.200E-08 | 6.200E-08 |
| 9849 | 493 | C493H | A | 1.1e-08 | 3.5e-08 | 5.2e-08 | 3.267E-08 | 2.060E-08 | 1.100E-08 | 5.200E-08 |
| 9850 | 493 | C493I | A | 5.4e-08 | 3.6e-08 | 3.7e-08 | 4.233E-08 | 1.012E-08 | 3.600E-08 | 5.400E-08 |
| 9851 | 493 | C493L | A | 6.7e-08 | 1.2e-07 | 6.3e-08 | 8.333E-08 | 3.182E-08 | 6.300E-08 | 1.200E-07 |
| 9852 | 493 | C493K | A | 6.6e-08 | 3.2e-08 | 5.2e-08 | 5.000E-08 | 1.709E-08 | 3.200E-08 | 6.600E-08 |
| 9853 | 493 | C493M | A | 4.2e-08 | 3.1e-08 | 9e-08 | 5.433E-08 | 3.137E-08 | 3.100E-08 | 9.000E-08 |
| 9854 | 493 | C493F | A | 4.4e-08 | 1.3e-08 | 1.1e-07 | 5.567E-08 | 4.954E-08 | 1.300E-08 | 1.100E-07 |
| 9855 | 493 | C493P | A | 2.6e-08 | 4.6e-08 | 7.3e-08 | 4.833E-08 | 2.359E-08 | 2.600E-08 | 7.300E-08 |
| 9856 | 493 | C493S | A | 7.7e-08 | 3.3e-08 | 5.2e-08 | 5.400E-08 | 2.207E-08 | 3.300E-08 | 7.700E-08 |
| 9857 | 493 | C493T | A | 8.1e-08 | 2e-08 | 2.1e-08 | 4.067E-08 | 3.493E-08 | 2.000E-08 | 8.100E-08 |
| 9858 | 493 | C493W | A | 1.8e-08 | 1.6e-08 | 2.5e-08 | 1.967E-08 | 4.726E-09 | 1.600E-08 | 2.500E-08 |
| 9859 | 493 | C493Y | A | 6.7e-08 | 3.2e-08 | 4.6e-08 | 4.833E-08 | 1.762E-08 | 3.200E-08 | 6.700E-08 |

| | | | | | | | | | | |
|---|---|---|---|---|---|---|---|---|---|---|
| 9860 | 493 | C493V | A | 1.8e-08 | 4.5e-08 | 1.7e-08 | 2.667E-08 | 1.589E-08 | 1.700E-08 | 4.500E-08 |
| 9861 | 494 | A494A | A | 3.9e-08 | 8.7e-08 | 1e-07 | 7.533E-08 | 3.213E-08 | 3.900E-08 | 1.000E-07 |
| 9862 | 494 | A494R | A | 2.4e-08 | 2.1e-08 | 2.9e-08 | 2.467E-08 | 4.041E-09 | 2.100E-08 | 2.900E-08 |
| 9863 | 494 | A494N | A | 1.3e-08 | 3.3e-08 | 3.1e-08 | 2.567E-08 | 1.102E-08 | 1.300E-08 | 3.300E-08 |
| 9864 | 494 | A494D | A | 1.8e-08 | 4e-08 | 4.7e-08 | 3.500E-08 | 1.513E-08 | 1.800E-08 | 4.700E-08 |
| 9865 | 494 | A494C | A | 5.3e-08 | 5.6e-08 | 2.7e-08 | 4.533E-08 | 1.595E-08 | 2.700E-08 | 5.600E-08 |
| 9866 | 494 | A494E | A | 7.5e-08 | 4.7e-08 | 7.5e-08 | 6.567E-08 | 1.617E-08 | 4.700E-08 | 7.500E-08 |
| 9867 | 494 | A494Q | A | 3.3e-08 | 2.1e-08 | 4.2e-08 | 3.200E-08 | 1.054E-08 | 2.100E-08 | 4.200E-08 |
| 9868 | 494 | A494G | A | 4.1e-08 | 3.4e-08 | 6e-08 | 4.500E-08 | 1.345E-08 | 3.400E-08 | 6.000E-08 |
| 9869 | 494 | A494H | A | 4.6e-08 | 3.2e-08 | 3.1e-08 | 3.633E-08 | 8.386E-09 | 3.100E-08 | 4.600E-08 |
| 9870 | 494 | A494I | A | 4.2e-08 | 1.4e-07 | 7.8e-08 | 8.667E-08 | 4.957E-08 | 4.200E-08 | 1.400E-07 |
| 9871 | 494 | A494L | A | 2.6e-08 | 9e-08 | 5.6e-08 | 5.733E-08 | 3.202E-08 | 2.600E-08 | 9.000E-08 |
| 9872 | 494 | A494K | A | 4.3e-08 | 7.3e-08 | 2.2e-08 | 4.600E-08 | 2.563E-08 | 2.200E-08 | 7.300E-08 |
| 9873 | 494 | A494M | A | 3.8e-08 | 2.8e-08 | 6.5e-08 | 4.367E-08 | 1.914E-08 | 2.800E-08 | 6.500E-08 |
| 9874 | 494 | A494F | A | 6.2e-08 | 4.3e-08 | 5.2e-08 | 5.233E-08 | 9.504E-09 | 4.300E-08 | 6.200E-08 |
| 9875 | 494 | A494P | A | 2.7e-08 | 8.5e-08 | 5.6e-08 | 5.600E-08 | 2.900E-08 | 2.700E-08 | 8.500E-08 |
| 9876 | 494 | A494S | A | 8.2e-08 | 4.7e-08 | 6.5e-08 | 6.467E-08 | 1.750E-08 | 4.700E-08 | 8.200E-08 |
| 9877 | 494 | A494T | A | 6e-08 | 6.6e-08 | 4.1e-08 | 5.567E-08 | 1.305E-08 | 4.100E-08 | 6.600E-08 |
| 9878 | 494 | A494W | A | 6.1e-08 | 4.6e-08 | 4.5e-08 | 5.067E-08 | 8.963E-09 | 4.500E-08 | 6.100E-08 |
| 9879 | 494 | A494Y | A | 3.1e-08 | 1.1e-07 | 1.9e-08 | 5.333E-08 | 4.944E-08 | 1.900E-08 | 1.100E-07 |
| 9880 | 494 | A494V | A | 7.5e-08 | 7.1e-08 | 5.4e-08 | 6.667E-08 | 1.115E-08 | 5.400E-08 | 7.500E-08 |
| 9881 | 495 | R495A | A | 4.5e-08 | 2.6e-08 | 3.6e-08 | 3.567E-08 | 9.504E-09 | 2.600E-08 | 4.500E-08 |
| 9882 | 495 | R495R | A | 7e-08 | 2.7e-08 | 4.5e-08 | 4.733E-08 | 2.159E-08 | 2.700E-08 | 7.000E-08 |
| 9883 | 495 | R495N | A | 2.2e-08 | 5.1e-08 | 4.3e-08 | 3.867E-08 | 1.498E-08 | 2.200E-08 | 5.100E-08 |
| 9884 | 495 | R495D | A | 7.4e-08 | 5.1e-08 | 4.1e-08 | 5.533E-08 | 1.692E-08 | 4.100E-08 | 7.400E-08 |
| 9885 | 495 | R495C | A | 9.8e-08 | 4e-08 | 2.5e-08 | 5.433E-08 | 3.855E-08 | 2.500E-08 | 9.800E-08 |
| 9886 | 495 | R495E | A | 1.2e-08 | 7.1e-08 | 2.1e-08 | 3.467E-08 | 3.179E-08 | 1.200E-08 | 7.100E-08 |
| 9887 | 495 | R495Q | A | 2.9e-08 | 7.8e-09 | 1.8e-08 | 1.827E-08 | 1.060E-08 | 7.800E-09 | 2.900E-08 |
| 9888 | 495 | R495G | A | 3.6e-08 | 4.4e-08 | 2.7e-08 | 3.567E-08 | 8.505E-09 | 2.700E-08 | 4.400E-08 |
| 9889 | 495 | R495H | A | 5.2e-08 | 6.1e-08 | 5.8e-08 | 5.700E-08 | 4.583E-09 | 5.200E-08 | 6.100E-08 |
| 9890 | 495 | R495I | A | 4.8e-08 | 9.2e-08 | 5.1e-08 | 6.367E-08 | 2.458E-08 | 4.800E-08 | 9.200E-08 |
| 9891 | 495 | R495L | A | 3.7e-08 | 7.3e-08 | 3.5e-08 | 4.833E-08 | 2.139E-08 | 3.500E-08 | 7.300E-08 |
| 9892 | 495 | R495K | A | 3.3e-08 | 7.7e-08 | 1.6e-08 | 4.200E-08 | 3.148E-08 | 1.600E-08 | 7.700E-08 |
| 9893 | 495 | R495M | A | 4e-08 | 3.2e-08 | 4.3e-08 | 3.833E-08 | 5.686E-09 | 3.200E-08 | 4.300E-08 |
| 9894 | 495 | R495F | A | 1.6e-07 | 5.1e-08 | 4e-08 | 8.367E-08 | 6.634E-08 | 4.000E-08 | 1.600E-07 |
| 9895 | 495 | R495P | A | 2.3e-08 | 2.6e-08 | 3.6e-08 | 2.833E-08 | 6.807E-09 | 2.300E-08 | 3.600E-08 |
| 9896 | 495 | R495S | A | 5.4e-08 | 3.7e-08 | 8.1e-08 | 5.733E-08 | 2.219E-08 | 3.700E-08 | 8.100E-08 |
| 9897 | 495 | R495T | A | 5.2e-08 | 2.5e-08 | 2.2e-08 | 3.300E-08 | 1.652E-08 | 2.200E-08 | 5.200E-08 |
| 9898 | 495 | R495W | A | 4.4e-08 | 4e-08 | 5.5e-08 | 4.633E-08 | 7.767E-09 | 4.000E-08 | 5.500E-08 |
| 9899 | 495 | R495Y | A | 2e-08 | 2.8e-08 | 8.6e-08 | 4.467E-08 | 3.602E-08 | 2.000E-08 | 8.600E-08 |
| 9900 | 495 | R495V | A | 7.6e-08 | 5.2e-08 | 2.3e-07 | 1.193E-07 | 9.659E-08 | 5.200E-08 | 2.300E-07 |
| 9901 | 496 | G496A | A | 4.2e-08 | 6.5e-08 | 6.3e-08 | 5.667E-08 | 1.274E-08 | 4.200E-08 | 6.500E-08 |
| 9902 | 496 | G496R | A | 7.4e-08 | 7.8e-08 | 5.1e-08 | 6.767E-08 | 1.457E-08 | 5.100E-08 | 7.800E-08 |
| 9903 | 496 | G496N | A | 4.4e-08 | 5.8e-08 | 3.7e-08 | 4.633E-08 | 1.069E-08 | 3.700E-08 | 5.800E-08 |
| 9904 | 496 | G496D | A | 7e-08 | 1.3e-07 | 2e-08 | 7.333E-08 | 5.508E-08 | 2.000E-08 | 1.300E-07 |
| 9905 | 496 | G496C | A | 1.7e-08 | 2.8e-08 | 1.1e-07 | 5.167E-08 | 5.082E-08 | 1.700E-08 | 1.100E-07 |
| 9906 | 496 | G496E | A | 4.2e-08 | 6.7e-08 | 9.2e-08 | 6.700E-08 | 2.500E-08 | 4.200E-08 | 9.200E-08 |
| 9907 | 496 | G496Q | A | 1.1e-07 | 5.9e-08 | 4.7e-08 | 7.200E-08 | 3.345E-08 | 4.700E-08 | 1.100E-07 |
| 9908 | 496 | G496G | A | 2.5e-08 | 2.6e-08 | 4.5e-08 | 3.200E-08 | 1.127E-08 | 2.500E-08 | 4.500E-08 |
| 9909 | 496 | G496H | A | 8.7e-08 | 4.2e-08 | 7.5e-08 | 6.800E-08 | 2.330E-08 | 4.200E-08 | 8.700E-08 |
| 9910 | 496 | G496I | A | 7.5e-08 | 2.3e-08 | 2.4e-08 | 4.067E-08 | 2.974E-08 | 2.300E-08 | 7.500E-08 |
| 9911 | 496 | G496L | A | 1e-07 | 5.3e-08 | 1.2e-07 | 9.100E-08 | 3.439E-08 | 5.300E-08 | 1.200E-07 |
| 9912 | 496 | G496K | A | 2.9e-08 | 3.4e-08 | 3.3e-08 | 3.200E-08 | 2.646E-09 | 2.900E-08 | 3.400E-08 |
| 9913 | 496 | G496M | A | 7.6e-08 | 4.5e-08 | 7.5e-08 | 6.533E-08 | 1.762E-08 | 4.500E-08 | 7.600E-08 |
| 9914 | 496 | G496F | A | 6.7e-08 | 3.2e-08 | 5.5e-08 | 5.133E-08 | 1.779E-08 | 3.200E-08 | 6.700E-08 |
| 9915 | 496 | G496P | A | 2e-08 | 9.4e-08 | 1.7e-08 | 4.367E-08 | 4.362E-08 | 1.700E-08 | 9.400E-08 |
| 9916 | 496 | G496S | A | 1.5e-07 | 6.6e-08 | 7.6e-08 | 9.733E-08 | 4.588E-08 | 6.600E-08 | 1.500E-07 |
| 9917 | 496 | G496T | A | 4.8e-08 | 5.2e-08 | 2.5e-08 | 4.167E-08 | 1.457E-08 | 2.500E-08 | 5.200E-08 |
| 9918 | 496 | G496W | A | 5.5e-08 | 3.5e-08 | 3.5e-08 | 4.167E-08 | 1.155E-08 | 3.500E-08 | 5.500E-08 |
| 9919 | 496 | G496Y | A | 1.8e-08 | 5.6e-08 | 3.1e-08 | 3.500E-08 | 1.931E-08 | 1.800E-08 | 5.600E-08 |
| 9920 | 496 | G496V | A | 1.3e-08 | 3.3e-08 | 1.2e-08 | 1.933E-08 | 1.185E-08 | 1.200E-08 | 3.300E-08 |
| 9921 | 497 | H497A | A | 9.3e-08 | 3.2e-08 | 2e-08 | 4.833E-08 | 3.915E-08 | 2.000E-08 | 9.300E-08 |
| 9922 | 497 | H497R | A | 4e-08 | 6.9e-08 | 4.9e-08 | 5.267E-08 | 1.484E-08 | 4.000E-08 | 6.900E-08 |
| 9923 | 497 | H497N | A | 3.6e-08 | 1.4e-08 | 7.6e-09 | 1.920E-08 | 1.490E-08 | 7.600E-09 | 3.600E-08 |
| 9924 | 497 | H497D | A | 1.3e-07 | 6.5e-08 | 8.5e-08 | 9.333E-08 | 3.329E-08 | 6.500E-08 | 1.300E-07 |
| 9925 | 497 | H497C | A | 6.4e-08 | 3.5e-08 | 2.3e-08 | 4.067E-08 | 2.108E-08 | 2.300E-08 | 6.400E-08 |
| 9926 | 497 | H497E | A | 1e-07 | 6.7e-08 | 2.6e-08 | 6.433E-08 | 3.707E-08 | 2.600E-08 | 1.000E-07 |
| 9927 | 497 | H497Q | A | 6.1e-08 | 3.6e-08 | 3e-08 | 4.233E-08 | 1.644E-08 | 3.000E-08 | 6.100E-08 |
| 9928 | 497 | H497G | A | 2.9e-08 | 4.6e-08 | 4.2e-08 | 3.900E-08 | 8.888E-09 | 2.900E-08 | 4.600E-08 |
| 9929 | 497 | H497H | A | 1.6e-08 | 5.9e-08 | 2.3e-08 | 3.267E-08 | 2.307E-08 | 1.600E-08 | 5.900E-08 |
| 9930 | 497 | H497I | A | 5.5e-08 | 1.5e-08 | 2.6e-08 | 3.200E-08 | 2.066E-08 | 1.500E-08 | 5.500E-08 |
| 9931 | 497 | H497L | A | 4.7e-08 | 3.2e-08 | 4.5e-08 | 4.133E-08 | 8.145E-09 | 3.200E-08 | 4.700E-08 |
| 9932 | 497 | H497K | A | 6.8e-08 | 2.5e-08 | 2.2e-08 | 3.833E-08 | 2.574E-08 | 2.200E-08 | 6.800E-08 |
| 9933 | 497 | H497M | A | 3.1e-08 | 6.6e-08 | 2.7e-08 | 4.133E-08 | 2.146E-08 | 2.700E-08 | 6.600E-08 |
| 9934 | 497 | H497F | A | 2.2e-08 | 8.4e-08 | 7.9e-08 | 6.167E-08 | 3.444E-08 | 2.200E-08 | 8.400E-08 |
| 9935 | 497 | H497P | A | 2.9e-08 | 4.7e-08 | 1.1e-07 | 6.200E-08 | 4.253E-08 | 2.900E-08 | 1.100E-07 |
| 9936 | 497 | H497S | A | 4e-08 | 3.9e-08 | 5.5e-08 | 4.467E-08 | 8.963E-09 | 3.900E-08 | 5.500E-08 |
| 9937 | 497 | H497T | A | 3e-08 | 6.3e-08 | 5.5e-08 | 4.933E-08 | 1.721E-08 | 3.000E-08 | 6.300E-08 |
| 9938 | 497 | H497W | A | 8.1e-08 | 5.7e-08 | 5.1e-08 | 6.300E-08 | 1.587E-08 | 5.100E-08 | 8.100E-08 |
| 9939 | 497 | H497Y | A | 4e-08 | 4.1e-08 | 4.3e-08 | 4.133E-08 | 1.528E-09 | 4.000E-08 | 4.300E-08 |
| 9940 | 497 | H497V | A | 2.3e-08 | 3.2e-08 | 5.9e-08 | 3.800E-08 | 1.873E-08 | 2.300E-08 | 5.900E-08 |
| 9941 | 498 | C498A | A | 2.3e-08 | 8.1e-08 | 2.1e-08 | 4.167E-08 | 3.408E-08 | 2.100E-08 | 8.100E-08 |
| 9942 | 498 | C498R | A | 7.2e-08 | 3.5e-08 | 9.9e-08 | 6.867E-08 | 3.213E-08 | 3.500E-08 | 9.900E-08 |
| 9943 | 498 | C498N | A | 1.1e-07 | 9.5e-08 | 1.3e-08 | 7.267E-08 | 5.221E-08 | 1.300E-08 | 1.100E-07 |
| 9944 | 498 | C498D | A | 2.9e-08 | 6.9e-08 | 2.1e-08 | 3.967E-08 | 2.572E-08 | 2.100E-08 | 6.900E-08 |

| | | | | | | | | | | |
|---|---|---|---|---|---|---|---|---|---|---|
| 9945 | 498 | C498C | A | 5.4e-08 | 2.6e-08 | 4.4e-08 | 4.133E-08 | 1.419E-08 | 2.600E-08 | 5.400E-08 |
| 9946 | 498 | C498E | A | 1.1e-07 | 5e-08 | 1.2e-07 | 9.333E-08 | 3.786E-08 | 5.000E-08 | 1.200E-07 |
| 9947 | 498 | C498Q | A | 5.1e-08 | 6.5e-08 | 1.4e-07 | 8.533E-08 | 4.786E-08 | 5.100E-08 | 1.400E-07 |
| 9948 | 498 | C498G | A | 4.9e-08 | 1.2e-07 | 1.6e-08 | 6.167E-08 | 5.314E-08 | 1.600E-08 | 1.200E-07 |
| 9949 | 498 | C498H | A | 5.9e-08 | 2.6e-08 | 8.1e-09 | 3.103E-08 | 2.582E-08 | 8.100E-09 | 5.900E-08 |
| 9950 | 498 | C498I | A | 9e-08 | 5e-08 | 4.2e-08 | 6.067E-08 | 2.572E-08 | 4.200E-08 | 9.000E-08 |
| 9951 | 498 | C498L | A | 2.1e-08 | 3.4e-08 | 4.6e-08 | 3.367E-08 | 1.250E-08 | 2.100E-08 | 4.600E-08 |
| 9952 | 498 | C498K | A | 3.8e-08 | 1.2e-08 | 1.7e-08 | 2.233E-08 | 1.380E-08 | 1.200E-08 | 3.800E-08 |
| 9953 | 498 | C498M | A | 5e-08 | 8.3e-08 | 3.9e-08 | 5.733E-08 | 2.290E-08 | 3.900E-08 | 8.300E-08 |
| 9954 | 498 | C498F | A | 3.2e-08 | 4.5e-08 | 2.2e-08 | 3.300E-08 | 1.153E-08 | 2.200E-08 | 4.500E-08 |
| 9955 | 498 | C498P | A | 1e-07 | 7.4e-08 | 2.8e-08 | 6.733E-08 | 3.646E-08 | 2.800E-08 | 1.000E-07 |
| 9956 | 498 | C498S | A | 4.3e-08 | 5.8e-08 | 2.3e-08 | 4.133E-08 | 1.756E-08 | 2.300E-08 | 5.800E-08 |
| 9957 | 498 | C498T | A | 3.8e-08 | 2.8e-08 | 3.7e-08 | 3.433E-08 | 5.508E-09 | 2.800E-08 | 3.800E-08 |
| 9958 | 498 | C498W | A | 1.1e-08 | 1.3e-08 | 1e-07 | 4.133E-08 | 5.082E-08 | 1.100E-08 | 1.000E-07 |
| 9959 | 498 | C498Y | A | 1.3e-07 | 1e-07 | 9.2e-08 | 1.073E-07 | 2.003E-08 | 9.200E-08 | 1.300E-07 |
| 9960 | 498 | C498V | A | 4.4e-08 | 8e-08 | 6.7e-08 | 6.367E-08 | 1.823E-08 | 4.400E-08 | 8.000E-08 |
| 9961 | 499 | W499A | A | 3.7e-08 | 4.6e-08 | 6.3e-08 | 4.867E-08 | 1.320E-08 | 3.700E-08 | 6.300E-08 |
| 9962 | 499 | W499R | A | 2.4e-08 | 3e-08 | 2.7e-08 | 2.700E-08 | 3.000E-09 | 2.400E-08 | 3.000E-08 |
| 9963 | 499 | W499N | A | 6.2e-08 | 7.7e-08 | 6.1e-08 | 6.667E-08 | 8.963E-09 | 6.100E-08 | 7.700E-08 |
| 9964 | 499 | W499D | A | 2.1e-07 | 2.2e-08 | 5.5e-08 | 9.567E-08 | 1.004E-07 | 2.200E-08 | 2.100E-07 |
| 9965 | 499 | W499C | A | 9e-08 | 1.8e-08 | 2.8e-08 | 4.533E-08 | 3.900E-08 | 1.800E-08 | 9.000E-08 |
| 9966 | 499 | W499E | A | 5.9e-08 | 8e-08 | 3.2e-08 | 5.700E-08 | 2.406E-08 | 3.200E-08 | 8.000E-08 |
| 9967 | 499 | W499Q | A | 4.3e-08 | 4e-08 | 3.1e-08 | 3.800E-08 | 6.245E-09 | 3.100E-08 | 4.300E-08 |
| 9968 | 499 | W499G | A | 5e-08 | 1.8e-08 | 1e-07 | 5.600E-08 | 4.133E-08 | 1.800E-08 | 1.000E-07 |
| 9969 | 499 | W499H | A | 1.5e-07 | 8.8e-08 | 2.4e-08 | 8.733E-08 | 6.300E-08 | 2.400E-08 | 1.500E-07 |
| 9970 | 499 | W499I | A | 1.2e-08 | 7.6e-08 | 3.1e-08 | 3.967E-08 | 3.287E-08 | 1.200E-08 | 7.600E-08 |
| 9971 | 499 | W499L | A | 8.8e-08 | 5.6e-08 | 3.3e-08 | 5.900E-08 | 2.762E-08 | 3.300E-08 | 8.800E-08 |
| 9972 | 499 | W499K | A | 1.7e-08 | 7.4e-08 | 7.5e-08 | 5.533E-08 | 3.320E-08 | 1.700E-08 | 7.500E-08 |
| 9973 | 499 | W499M | A | 3.7e-08 | 3.5e-08 | 1.1e-08 | 2.767E-08 | 1.447E-08 | 1.100E-08 | 3.700E-08 |
| 9974 | 499 | W499F | A | 1.5e-08 | 4.2e-08 | 3e-08 | 2.900E-08 | 1.353E-08 | 1.500E-08 | 4.200E-08 |
| 9975 | 499 | W499P | A | 2.7e-08 | 9.4e-08 | 3.6e-08 | 5.233E-08 | 3.636E-08 | 2.700E-08 | 9.400E-08 |
| 9976 | 499 | W499S | A | 7.9e-08 | 2.9e-08 | 1.1e-07 | 7.267E-08 | 4.087E-08 | 2.900E-08 | 1.100E-07 |
| 9977 | 499 | W499T | A | 7.7e-08 | 2.1e-08 | 4.3e-08 | 4.700E-08 | 2.821E-08 | 2.100E-08 | 7.700E-08 |
| 9978 | 499 | W499W | A | 1.9e-08 | 4.4e-08 | 5.2e-08 | 3.833E-08 | 1.721E-08 | 1.900E-08 | 5.200E-08 |
| 9979 | 499 | W499Y | A | 5.8e-08 | 2e-08 | 3.8e-08 | 3.867E-08 | 1.901E-08 | 2.000E-08 | 5.800E-08 |
| 9980 | 499 | W499V | A | 7.6e-08 | 2.7e-08 | 4.1e-08 | 4.800E-08 | 2.524E-08 | 2.700E-08 | 7.600E-08 |
| 9981 | 500 | G500A | A | 6.8e-08 | 5.8e-08 | 3.6e-08 | 5.400E-08 | 1.637E-08 | 3.600E-08 | 6.800E-08 |
| 9982 | 500 | G500R | A | 7.2e-08 | 2e-08 | 2.7e-08 | 3.967E-08 | 2.822E-08 | 2.000E-08 | 7.200E-08 |
| 9983 | 500 | G500N | A | 4.9e-08 | 3.4e-08 | 4.2e-08 | 4.167E-08 | 7.506E-09 | 3.400E-08 | 4.900E-08 |
| 9984 | 500 | G500D | A | 3.4e-08 | 3.4e-08 | 6.8e-08 | 4.533E-08 | 1.963E-08 | 3.400E-08 | 6.800E-08 |
| 9985 | 500 | G500C | A | 6.4e-08 | 2.8e-08 | 3.7e-08 | 4.300E-08 | 1.873E-08 | 2.800E-08 | 6.400E-08 |
| 9986 | 500 | G500E | A | 1.6e-08 | 1.6e-08 | 2.9e-08 | 2.033E-08 | 7.506E-09 | 1.600E-08 | 2.900E-08 |
| 9987 | 500 | G500Q | A | 3.6e-08 | 4.3e-08 | 2e-08 | 3.300E-08 | 1.179E-08 | 2.000E-08 | 4.300E-08 |
| 9988 | 500 | G500G | A | 6.6e-08 | 5.7e-08 | 7.8e-08 | 6.700E-08 | 1.054E-08 | 5.700E-08 | 7.800E-08 |
| 9989 | 500 | G500H | A | 4.6e-08 | 1.2e-07 | 6e-08 | 7.533E-08 | 3.931E-08 | 4.600E-08 | 1.200E-07 |
| 9990 | 500 | G500I | A | 7.1e-08 | 8.1e-08 | 4.7e-08 | 6.633E-08 | 1.747E-08 | 4.700E-08 | 8.100E-08 |
| 9991 | 500 | G500L | A | 1.5e-08 | 2.5e-08 | 6.8e-08 | 3.600E-08 | 2.816E-08 | 1.500E-08 | 6.800E-08 |
| 9992 | 500 | G500K | A | 5.2e-08 | 6.2e-08 | 7.5e-08 | 6.300E-08 | 1.153E-08 | 5.200E-08 | 7.500E-08 |
| 9993 | 500 | G500M | A | 5.9e-08 | 2.6e-08 | 3.7e-08 | 4.067E-08 | 1.680E-08 | 2.600E-08 | 5.900E-08 |
| 9994 | 500 | G500F | A | 6.4e-08 | 3.2e-08 | 3.5e-08 | 4.367E-08 | 1.767E-08 | 3.200E-08 | 6.400E-08 |
| 9995 | 500 | G500P | A | 2.5e-08 | 4.2e-08 | 9.6e-08 | 5.433E-08 | 3.707E-08 | 2.500E-08 | 9.600E-08 |
| 9996 | 500 | G500S | A | 5.7e-08 | 4.2e-08 | 2.5e-08 | 4.133E-08 | 1.601E-08 | 2.500E-08 | 5.700E-08 |
| 9997 | 500 | G500T | A | 2.1e-08 | 4.5e-08 | 5.7e-08 | 4.100E-08 | 1.833E-08 | 2.100E-08 | 5.700E-08 |
| 9998 | 500 | G500W | A | 5e-08 | 6.6e-08 | 3.3e-08 | 4.967E-08 | 1.650E-08 | 3.300E-08 | 6.600E-08 |
| 9999 | 500 | G500Y | A | 4.9e-08 | 1.8e-08 | 3.3e-08 | 3.333E-08 | 1.550E-08 | 1.800E-08 | 4.900E-08 |
| 10000 | 500 | G500V | A | 5.4e-08 | 5.8e-08 | 3.9e-08 | 5.033E-08 | 1.002E-08 | 3.900E-08 | 5.800E-08 |
| 10001 | 501 | P501A | A | 2.2e-08 | 1.4e-08 | 1.5e-08 | 1.700E-08 | 4.359E-09 | 1.400E-08 | 2.200E-08 |
| 10002 | 501 | P501R | A | 6.4e-08 | 7.3e-08 | 2.8e-08 | 5.500E-08 | 2.381E-08 | 2.800E-08 | 7.300E-08 |
| 10003 | 501 | P501N | A | 2.4e-08 | 3.9e-08 | 5.4e-08 | 3.900E-08 | 1.500E-08 | 2.400E-08 | 5.400E-08 |
| 10004 | 501 | P501D | A | 3.1e-08 | 7.4e-08 | 5.5e-08 | 5.333E-08 | 2.155E-08 | 3.100E-08 | 7.400E-08 |
| 10005 | 501 | P501C | A | 2e-08 | 1.7e-08 | 6.9e-08 | 3.533E-08 | 2.919E-08 | 1.700E-08 | 6.900E-08 |
| 10006 | 501 | P501E | A | 1.5e-07 | 3.9e-08 | 4.6e-08 | 7.833E-08 | 6.216E-08 | 3.900E-08 | 1.500E-07 |
| 10007 | 501 | P501Q | A | 7.6e-08 | 6.2e-08 | 5.3e-08 | 6.367E-08 | 1.159E-08 | 5.300E-08 | 7.600E-08 |
| 10008 | 501 | P501G | A | 4.3e-08 | 9.8e-08 | 3.9e-08 | 6.000E-08 | 3.297E-08 | 3.900E-08 | 9.800E-08 |
| 10009 | 501 | P501H | A | 2.3e-08 | 7.3e-08 | 7.7e-08 | 5.767E-08 | 3.009E-08 | 2.300E-08 | 7.700E-08 |
| 10010 | 501 | P501I | A | 3e-08 | 4.9e-08 | 5.1e-08 | 4.333E-08 | 1.159E-08 | 3.000E-08 | 5.100E-08 |
| 10011 | 501 | P501L | A | 1.4e-08 | 5.7e-08 | 6.3e-08 | 4.467E-08 | 2.673E-08 | 1.400E-08 | 6.300E-08 |
| 10012 | 501 | P501K | A | 3.6e-08 | 3.7e-08 | 1e-08 | 2.767E-08 | 1.531E-08 | 1.000E-08 | 3.700E-08 |
| 10013 | 501 | P501M | A | 7.8e-09 | 1.1e-08 | 5.5e-08 | 2.460E-08 | 2.638E-08 | 7.800E-09 | 5.500E-08 |
| 10014 | 501 | P501F | A | 2.8e-08 | 5.6e-08 | 6.7e-08 | 5.033E-08 | 2.011E-08 | 2.800E-08 | 6.700E-08 |
| 10015 | 501 | P501P | A | 6e-08 | 6.9e-08 | 2.9e-08 | 5.267E-08 | 2.098E-08 | 2.900E-08 | 6.900E-08 |
| 10016 | 501 | P501S | A | 4.9e-08 | 3.2e-08 | 1.1e-07 | 6.367E-08 | 4.102E-08 | 3.200E-08 | 1.100E-07 |
| 10017 | 501 | P501T | A | 4.7e-08 | 5.4e-08 | 5.3e-08 | 5.133E-08 | 3.786E-09 | 4.700E-08 | 5.400E-08 |
| 10018 | 501 | P501W | A | 4.1e-08 | 2.1e-08 | 7.6e-08 | 4.600E-08 | 2.784E-08 | 2.100E-08 | 7.600E-08 |
| 10019 | 501 | P501Y | A | 4.1e-08 | 8.8e-08 | 4.3e-08 | 5.733E-08 | 2.658E-08 | 4.100E-08 | 8.800E-08 |
| 10020 | 501 | P501V | A | 7.2e-08 | 8.6e-08 | 4e-08 | 6.600E-08 | 2.358E-08 | 4.000E-08 | 8.600E-08 |
| 10021 | 502 | G502A | A | 3.6e-08 | 2.3e-08 | 6.2e-08 | 4.033E-08 | 1.986E-08 | 2.300E-08 | 6.200E-08 |
| 10022 | 502 | G502R | A | 3.1e-08 | 4e-08 | 6.7e-08 | 4.600E-08 | 1.873E-08 | 3.100E-08 | 6.700E-08 |
| 10023 | 502 | G502N | A | 9.5e-08 | 6e-08 | 2.3e-08 | 5.933E-08 | 3.600E-08 | 2.300E-08 | 9.500E-08 |
| 10024 | 502 | G502D | A | 2.3e-08 | 6.3e-08 | 6.3e-08 | 4.967E-08 | 2.309E-08 | 2.300E-08 | 6.300E-08 |
| 10025 | 502 | G502C | A | 4.1e-08 | 3.2e-08 | 2.1e-08 | 3.133E-08 | 1.002E-08 | 2.100E-08 | 4.100E-08 |
| 10026 | 502 | G502E | A | 4.5e-08 | 5e-08 | 1.6e-08 | 3.700E-08 | 1.836E-08 | 1.600E-08 | 5.000E-08 |
| 10027 | 502 | G502Q | A | 4.7e-08 | 9.9e-08 | 2.3e-08 | 5.633E-08 | 3.885E-08 | 2.300E-08 | 9.900E-08 |
| 10028 | 502 | G502G | A | 2.2e-08 | 1.3e-07 | 6.3e-09 | 5.277E-08 | 6.735E-08 | 6.300E-09 | 1.300E-07 |
| 10029 | 502 | G502H | A | 1.8e-08 | 1.9e-08 | 1.6e-08 | 1.767E-08 | 1.528E-09 | 1.600E-08 | 1.900E-08 |

| | | | | | | | | | | |
|---|---|---|---|---|---|---|---|---|---|---|
| 10030 | 502 | G502I | A | 5.7e-08 | 4.7e-08 | 7.9e-08 | 6.100E-08 | 1.637E-08 | 4.700E-08 | 7.900E-08 |
| 10031 | 502 | G502L | A | 2.7e-08 | 1.1e-07 | 3e-08 | 5.567E-08 | 4.708E-08 | 2.700E-08 | 1.100E-07 |
| 10032 | 502 | G502K | A | 1.4e-08 | 5.3e-08 | 4e-08 | 3.567E-08 | 1.986E-08 | 1.400E-08 | 5.300E-08 |
| 10033 | 502 | G502M | A | 1.1e-07 | 6e-08 | 7.1e-08 | 8.033E-08 | 2.627E-08 | 6.000E-08 | 1.100E-07 |
| 10034 | 502 | G502F | A | 4.3e-08 | 1.3e-08 | 3.2e-08 | 2.933E-08 | 1.518E-08 | 1.300E-08 | 4.300E-08 |
| 10035 | 502 | G502P | A | 2.5e-08 | 7.8e-08 | 9.5e-08 | 6.600E-08 | 3.651E-08 | 2.500E-08 | 9.500E-08 |
| 10036 | 502 | G502S | A | 4.6e-08 | 2e-08 | 1.1e-08 | 2.567E-08 | 1.818E-08 | 1.100E-08 | 4.600E-08 |
| 10037 | 502 | G502T | A | 8.8e-08 | 5.2e-08 | 1.9e-08 | 5.300E-08 | 3.451E-08 | 1.900E-08 | 8.800E-08 |
| 10038 | 502 | G502W | A | 2.3e-07 | 6.3e-08 | 5.7e-08 | 1.167E-07 | 9.820E-08 | 5.700E-08 | 2.300E-07 |
| 10039 | 502 | G502Y | A | 4.4e-08 | 4.7e-08 | 3.7e-08 | 4.267E-08 | 5.132E-09 | 3.700E-08 | 4.700E-08 |
| 10040 | 502 | G502V | A | 5.2e-08 | 1.8e-08 | 1.9e-08 | 2.967E-08 | 1.935E-08 | 1.800E-08 | 5.200E-08 |
| 10041 | 503 | P503A | A | 6.6e-08 | 3.1e-08 | 4.6e-08 | 4.767E-08 | 1.756E-08 | 3.100E-08 | 6.600E-08 |
| 10042 | 503 | P503R | A | 4.8e-08 | 5.9e-08 | 1.3e-08 | 4.000E-08 | 2.402E-08 | 1.300E-08 | 5.900E-08 |
| 10043 | 503 | P503N | A | 4.3e-08 | 4.7e-08 | 5.2e-08 | 4.733E-08 | 4.509E-09 | 4.300E-08 | 5.200E-08 |
| 10044 | 503 | P503D | A | 2.7e-08 | 8.6e-08 | 4.3e-08 | 5.200E-08 | 3.051E-08 | 2.700E-08 | 8.600E-08 |
| 10045 | 503 | P503C | A | 5.9e-08 | 4.4e-08 | 2.2e-08 | 4.167E-08 | 1.861E-08 | 2.200E-08 | 5.900E-08 |
| 10046 | 503 | P503E | A | 7.5e-08 | 1.8e-08 | 5.9e-08 | 5.067E-08 | 2.940E-08 | 1.800E-08 | 7.500E-08 |
| 10047 | 503 | P503Q | A | 4.3e-08 | 3.7e-08 | 1.2e-08 | 3.067E-08 | 1.644E-08 | 1.200E-08 | 4.300E-08 |
| 10048 | 503 | P503G | A | 4.6e-08 | 3.9e-08 | 2.7e-08 | 3.733E-08 | 9.609E-09 | 2.700E-08 | 4.600E-08 |
| 10049 | 503 | P503H | A | 4.2e-08 | 8e-08 | 5.5e-08 | 5.900E-08 | 1.931E-08 | 4.200E-08 | 8.000E-08 |
| 10050 | 503 | P503I | A | 4.2e-08 | 7.4e-08 | 5.1e-08 | 5.567E-08 | 1.650E-08 | 4.200E-08 | 7.400E-08 |
| 10051 | 503 | P503L | A | 2.9e-08 | 4.4e-08 | 3.9e-08 | 3.733E-08 | 7.638E-09 | 2.900E-08 | 4.400E-08 |
| 10052 | 503 | P503K | A | 4.6e-08 | 3.1e-08 | 5.3e-08 | 4.333E-08 | 1.124E-08 | 3.100E-08 | 5.300E-08 |
| 10053 | 503 | P503M | A | 5.2e-08 | 7.1e-08 | 4.5e-08 | 5.600E-08 | 1.345E-08 | 4.500E-08 | 7.100E-08 |
| 10054 | 503 | P503F | A | 5.6e-08 | 5.1e-08 | 5.5e-08 | 5.400E-08 | 2.646E-09 | 5.100E-08 | 5.600E-08 |
| 10055 | 503 | P503P | A | 1.9e-08 | 1.2e-08 | 7.3e-08 | 3.467E-08 | 3.338E-08 | 1.200E-08 | 7.300E-08 |
| 10056 | 503 | P503S | A | 7.8e-08 | 1.1e-08 | 5.3e-08 | 4.733E-08 | 3.386E-08 | 1.100E-08 | 7.800E-08 |
| 10057 | 503 | P503T | A | 3.7e-08 | 1.3e-07 | 2.2e-08 | 6.300E-08 | 5.851E-08 | 2.200E-08 | 1.300E-07 |
| 10058 | 503 | P503W | A | 2.5e-08 | 3.6e-08 | 6.8e-08 | 4.300E-08 | 2.234E-08 | 2.500E-08 | 6.800E-08 |
| 10059 | 503 | P503Y | A | 2.4e-08 | 5.7e-08 | 3.4e-08 | 3.833E-08 | 1.692E-08 | 2.400E-08 | 5.700E-08 |
| 10060 | 503 | P503V | A | 1.7e-08 | 5.1e-08 | 5.1e-08 | 3.967E-08 | 1.963E-08 | 1.700E-08 | 5.100E-08 |
| 10061 | 504 | T504A | A | 5.5e-08 | 5.9e-08 | 5.7e-08 | 5.700E-08 | 2.000E-09 | 5.500E-08 | 5.900E-08 |
| 10062 | 504 | T504R | A | 1e-08 | 5.7e-08 | 7.9e-08 | 4.867E-08 | 3.525E-08 | 1.000E-08 | 7.900E-08 |
| 10063 | 504 | T504N | A | 1.1e-07 | 1.6e-08 | 3.4e-08 | 5.333E-08 | 4.989E-08 | 1.600E-08 | 1.100E-07 |
| 10064 | 504 | T504D | A | 3.6e-08 | 8.4e-08 | 3.7e-08 | 5.233E-08 | 2.743E-08 | 3.600E-08 | 8.400E-08 |
| 10065 | 504 | T504C | A | 6.5e-08 | 1.8e-08 | 4e-08 | 4.100E-08 | 2.352E-08 | 1.800E-08 | 6.500E-08 |
| 10066 | 504 | T504E | A | 5.1e-08 | 2.8e-08 | 6.6e-08 | 4.833E-08 | 1.914E-08 | 2.800E-08 | 6.600E-08 |
| 10067 | 504 | T504Q | A | 3.9e-08 | 3.8e-08 | 3.5e-08 | 3.733E-08 | 2.082E-09 | 3.500E-08 | 3.900E-08 |
| 10068 | 504 | T504G | A | 4.3e-08 | 3.3e-08 | 5.2e-08 | 4.267E-08 | 9.504E-09 | 3.300E-08 | 5.200E-08 |
| 10069 | 504 | T504H | A | 1.4e-08 | 4.5e-08 | 4.9e-08 | 3.600E-08 | 1.916E-08 | 1.400E-08 | 4.900E-08 |
| 10070 | 504 | T504I | A | 9.1e-08 | 2.1e-08 | 2e-08 | 4.400E-08 | 4.071E-08 | 2.000E-08 | 9.100E-08 |
| 10071 | 504 | T504L | A | 4.3e-08 | 1.6e-07 | 6.3e-08 | 8.867E-08 | 6.258E-08 | 4.300E-08 | 1.600E-07 |
| 10072 | 504 | T504K | A | 9.8e-08 | 5.3e-08 | 2.9e-08 | 6.000E-08 | 3.503E-08 | 2.900E-08 | 9.800E-08 |
| 10073 | 504 | T504M | A | 2.4e-08 | 3.1e-08 | 5.5e-08 | 3.667E-08 | 1.626E-08 | 2.400E-08 | 5.500E-08 |
| 10074 | 504 | T504F | A | 6.1e-08 | 3.5e-08 | 3.8e-08 | 4.467E-08 | 1.422E-08 | 3.500E-08 | 6.100E-08 |
| 10075 | 504 | T504P | A | 3.9e-08 | 3.5e-08 | 7.3e-08 | 4.900E-08 | 2.088E-08 | 3.500E-08 | 7.300E-08 |
| 10076 | 504 | T504S | A | 4.7e-08 | 3e-08 | 1.4e-07 | 7.233E-08 | 5.921E-08 | 3.000E-08 | 1.400E-07 |
| 10077 | 504 | T504T | A | 4.5e-08 | 3.5e-08 | 5.5e-08 | 4.500E-08 | 1.000E-08 | 3.500E-08 | 5.500E-08 |
| 10078 | 504 | T504W | A | 4e-08 | 3.1e-08 | 5.1e-08 | 4.067E-08 | 1.002E-08 | 3.100E-08 | 5.100E-08 |
| 10079 | 504 | T504Y | A | 4.2e-08 | 4e-08 | 6.6e-08 | 4.933E-08 | 1.447E-08 | 4.000E-08 | 6.600E-08 |
| 10080 | 504 | T504V | A | 2.9e-08 | 3.4e-08 | 4.6e-08 | 3.633E-08 | 8.737E-09 | 2.900E-08 | 4.600E-08 |
| 10081 | 505 | Q505A | A | 3.4e-08 | 2e-08 | 7.9e-08 | 4.433E-08 | 3.083E-08 | 2.000E-08 | 7.900E-08 |
| 10082 | 505 | Q505R | A | 6.5e-08 | 8.1e-08 | 7.6e-08 | 7.400E-08 | 8.185E-09 | 6.500E-08 | 8.100E-08 |
| 10083 | 505 | Q505N | A | 2.6e-08 | 5.1e-08 | 4.2e-08 | 3.967E-08 | 1.266E-08 | 2.600E-08 | 5.100E-08 |
| 10084 | 505 | Q505D | A | 1.2e-07 | 6.4e-08 | 1e-07 | 9.467E-08 | 2.838E-08 | 6.400E-08 | 1.200E-07 |
| 10085 | 505 | Q505C | A | 8.2e-08 | 1.1e-07 | 8.8e-08 | 9.333E-08 | 1.474E-08 | 8.200E-08 | 1.100E-07 |
| 10086 | 505 | Q505E | A | 2.8e-08 | 1.5e-08 | 3.6e-08 | 2.633E-08 | 1.060E-08 | 1.500E-08 | 3.600E-08 |
| 10087 | 505 | Q505Q | A | 5.3e-08 | 6e-08 | 2.1e-07 | 1.077E-07 | 8.869E-08 | 5.300E-08 | 2.100E-07 |
| 10088 | 505 | Q505G | A | 4.3e-08 | 7.5e-09 | 4.3e-08 | 3.117E-08 | 2.050E-08 | 7.500E-09 | 4.300E-08 |
| 10089 | 505 | Q505H | A | 1.3e-08 | 3.7e-08 | 4.5e-08 | 3.167E-08 | 1.665E-08 | 1.300E-08 | 4.500E-08 |
| 10090 | 505 | Q505I | A | 5.5e-08 | 6.3e-08 | 8.5e-08 | 6.767E-08 | 1.553E-08 | 5.500E-08 | 8.500E-08 |
| 10091 | 505 | Q505L | A | 9.1e-08 | 1.5e-07 | 1.1e-07 | 1.170E-07 | 3.012E-08 | 9.100E-08 | 1.500E-07 |
| 10092 | 505 | Q505K | A | 4.3e-08 | 4.6e-08 | 7.9e-08 | 5.600E-08 | 1.997E-08 | 4.300E-08 | 7.900E-08 |
| 10093 | 505 | Q505M | A | 4.6e-08 | 7.4e-08 | 2.2e-08 | 4.733E-08 | 2.603E-08 | 2.200E-08 | 7.400E-08 |
| 10094 | 505 | Q505F | A | 5e-08 | 8e-08 | 4.5e-08 | 5.833E-08 | 1.893E-08 | 4.500E-08 | 8.000E-08 |
| 10095 | 505 | Q505P | A | 7.2e-08 | 7.1e-08 | 5.7e-08 | 6.667E-08 | 8.386E-09 | 5.700E-08 | 7.200E-08 |
| 10096 | 505 | Q505S | A | 2.2e-08 | 5e-08 | 3.5e-08 | 3.567E-08 | 1.401E-08 | 2.200E-08 | 5.000E-08 |
| 10097 | 505 | Q505T | A | 5.7e-08 | 1.5e-07 | 1.7e-08 | 7.467E-08 | 6.824E-08 | 1.700E-08 | 1.500E-07 |
| 10098 | 505 | Q505W | A | 9.4e-08 | 6.7e-08 | 1e-07 | 8.700E-08 | 1.758E-08 | 6.700E-08 | 1.000E-07 |
| 10099 | 505 | Q505Y | A | 2.1e-08 | 3.1e-08 | 3.9e-08 | 3.033E-08 | 9.018E-09 | 2.100E-08 | 3.900E-08 |
| 10100 | 505 | Q505V | A | 3.8e-08 | 6e-08 | 6.7e-08 | 5.500E-08 | 1.513E-08 | 3.800E-08 | 6.700E-08 |
| 10101 | 506 | C506A | A | 5.5e-08 | 5.8e-08 | 6.5e-08 | 5.933E-08 | 5.132E-09 | 5.500E-08 | 6.500E-08 |
| 10102 | 506 | C506R | A | 7e-08 | 3.5e-08 | 3.4e-08 | 4.633E-08 | 2.050E-08 | 3.400E-08 | 7.000E-08 |
| 10103 | 506 | C506N | A | 2.7e-08 | 3e-08 | 3.7e-08 | 3.133E-08 | 5.132E-09 | 2.700E-08 | 3.700E-08 |
| 10104 | 506 | C506D | A | 5e-08 | 1.6e-08 | 8.4e-08 | 5.000E-08 | 3.400E-08 | 1.600E-08 | 8.400E-08 |
| 10105 | 506 | C506C | A | 6.7e-08 | 1.2e-07 | 4.5e-08 | 7.733E-08 | 3.855E-08 | 4.500E-08 | 1.200E-07 |
| 10106 | 506 | C506E | A | 5.9e-08 | 2.9e-08 | 1e-07 | 6.267E-08 | 3.564E-08 | 2.900E-08 | 1.000E-07 |
| 10107 | 506 | C506Q | A | 3.7e-08 | 4e-08 | 3.3e-08 | 3.667E-08 | 3.512E-09 | 3.300E-08 | 4.000E-08 |
| 10108 | 506 | C506G | A | 4.3e-08 | 3e-08 | 4.8e-08 | 4.033E-08 | 9.292E-09 | 3.000E-08 | 4.800E-08 |
| 10109 | 506 | C506H | A | 3.8e-08 | 6.7e-08 | 3.6e-08 | 4.700E-08 | 1.735E-08 | 3.600E-08 | 6.700E-08 |
| 10110 | 506 | C506I | A | 4.7e-08 | 9.5e-08 | 3.5e-08 | 5.900E-08 | 3.175E-08 | 3.500E-08 | 9.500E-08 |
| 10111 | 506 | C506L | A | 1.1e-07 | 3.8e-08 | 1.9e-08 | 5.567E-08 | 4.800E-08 | 1.900E-08 | 1.100E-07 |
| 10112 | 506 | C506K | A | 3.2e-08 | 3.2e-08 | 2.6e-08 | 3.000E-08 | 3.464E-09 | 2.600E-08 | 3.200E-08 |
| 10113 | 506 | C506M | A | 2e-08 | 7.8e-08 | 3.1e-08 | 4.300E-08 | 3.081E-08 | 2.000E-08 | 7.800E-08 |
| 10114 | 506 | C506F | A | 4.7e-08 | 4.5e-08 | 2.5e-08 | 3.900E-08 | 1.217E-08 | 2.500E-08 | 4.700E-08 |

| | | | | | | | | | | |
|---|---|---|---|---|---|---|---|---|---|---|
| 10115 | 506 | C506P | A | 4.3e-08 | 2.1e-08 | 3.2e-08 | 3.200E-08 | 1.100E-08 | 2.100E-08 | 4.300E-08 |
| 10116 | 506 | C506S | A | 1.2e-07 | 3.2e-08 | 2.8e-08 | 6.000E-08 | 5.200E-08 | 2.800E-08 | 1.200E-07 |
| 10117 | 506 | C506T | A | 1.5e-08 | 3.2e-07 | 4.3e-08 | 1.260E-07 | 1.686E-07 | 1.500E-08 | 3.200E-07 |
| 10118 | 506 | C506W | A | 5.9e-08 | 6.9e-08 | 5.1e-08 | 5.967E-08 | 9.018E-09 | 5.100E-08 | 6.900E-08 |
| 10119 | 506 | C506Y | A | 1.3e-07 | 3.4e-08 | 4.1e-08 | 6.833E-08 | 5.352E-08 | 3.400E-08 | 1.300E-07 |
| 10120 | 506 | C506V | A | 4.1e-08 | 4.4e-08 | 2.9e-08 | 3.800E-08 | 7.937E-09 | 2.900E-08 | 4.400E-08 |
| 10121 | 507 | V507A | A | 3.6e-08 | 2.6e-08 | 4.3e-08 | 3.500E-08 | 8.544E-09 | 2.600E-08 | 4.300E-08 |
| 10122 | 507 | V507R | A | 3.5e-08 | 7.2e-08 | 3.8e-08 | 4.833E-08 | 2.055E-08 | 3.500E-08 | 7.200E-08 |
| 10123 | 507 | V507N | A | 3.4e-08 | 8.4e-08 | 5.5e-08 | 5.767E-08 | 2.511E-08 | 3.400E-08 | 8.400E-08 |
| 10124 | 507 | V507D | A | 3.7e-08 | 2.1e-07 | 7e-08 | 1.057E-07 | 9.185E-08 | 3.700E-08 | 2.100E-07 |
| 10125 | 507 | V507C | A | 4e-08 | 4.6e-08 | 5.5e-08 | 4.700E-08 | 7.550E-09 | 4.000E-08 | 5.500E-08 |
| 10126 | 507 | V507E | A | 2.7e-08 | 7.4e-08 | 4.5e-08 | 4.867E-08 | 2.371E-08 | 2.700E-08 | 7.400E-08 |
| 10127 | 507 | V507Q | A | 4.9e-08 | 3.5e-08 | 3.1e-08 | 3.833E-08 | 9.452E-09 | 3.100E-08 | 4.900E-08 |
| 10128 | 507 | V507G | A | 5.3e-08 | 1.1e-07 | 1.6e-07 | 1.077E-07 | 5.354E-08 | 5.300E-08 | 1.600E-07 |
| 10129 | 507 | V507H | A | 8.4e-08 | 5e-08 | 3.8e-08 | 5.733E-08 | 2.386E-08 | 3.800E-08 | 8.400E-08 |
| 10130 | 507 | V507I | A | 6.5e-08 | 5.1e-08 | 2e-08 | 4.533E-08 | 2.303E-08 | 2.000E-08 | 6.500E-08 |
| 10131 | 507 | V507L | A | 5.1e-08 | 6.1e-08 | 4.1e-08 | 5.100E-08 | 1.000E-08 | 4.100E-08 | 6.100E-08 |
| 10132 | 507 | V507K | A | 5.9e-08 | 2.8e-08 | 1.5e-08 | 3.400E-08 | 2.261E-08 | 1.500E-08 | 5.900E-08 |
| 10133 | 507 | V507M | A | 1.1e-08 | 7.6e-09 | 1.4e-08 | 1.087E-08 | 3.202E-09 | 7.600E-09 | 1.400E-08 |
| 10134 | 507 | V507F | A | 2.4e-08 | 3.4e-08 | 9.3e-08 | 5.033E-08 | 3.729E-08 | 2.400E-08 | 9.300E-08 |
| 10135 | 507 | V507P | A | 6.4e-08 | 8.9e-08 | 9.1e-08 | 8.133E-08 | 1.504E-08 | 6.400E-08 | 9.100E-08 |
| 10136 | 507 | V507S | A | 3.1e-08 | 6.7e-08 | 3.8e-08 | 4.533E-08 | 1.909E-08 | 3.100E-08 | 6.700E-08 |
| 10137 | 507 | V507T | A | 2.8e-08 | 5.4e-08 | 2.9e-08 | 3.700E-08 | 1.473E-08 | 2.800E-08 | 5.400E-08 |
| 10138 | 507 | V507W | A | 7.5e-08 | 6.3e-08 | 1.3e-08 | 5.033E-08 | 3.288E-08 | 1.300E-08 | 7.500E-08 |
| 10139 | 507 | V507Y | A | 1.2e-07 | 9.8e-08 | 2.2e-08 | 8.000E-08 | 5.142E-08 | 2.200E-08 | 1.200E-07 |
| 10140 | 507 | V507V | A | 1.7e-08 | 2.6e-08 | 5.3e-08 | 3.200E-08 | 1.873E-08 | 1.700E-08 | 5.300E-08 |
| 10141 | 508 | N508A | A | 3.2e-08 | 2.3e-08 | 2.6e-08 | 2.700E-08 | 4.583E-09 | 2.300E-08 | 3.200E-08 |
| 10142 | 508 | N508R | A | 5e-08 | 3.5e-08 | 4.1e-08 | 4.200E-08 | 7.550E-09 | 3.500E-08 | 5.000E-08 |
| 10143 | 508 | N508N | A | 4e-08 | 3e-08 | 3.2e-08 | 3.400E-08 | 5.292E-09 | 3.000E-08 | 4.000E-08 |
| 10144 | 508 | N508D | A | 1.8e-08 | 8.2e-08 | 3.7e-08 | 4.567E-08 | 3.287E-08 | 1.800E-08 | 8.200E-08 |
| 10145 | 508 | N508C | A | 1.4e-08 | 3.3e-08 | 3.5e-08 | 2.733E-08 | 1.159E-08 | 1.400E-08 | 3.500E-08 |
| 10146 | 508 | N508E | A | 2.3e-08 | 6.7e-08 | 5.1e-08 | 4.700E-08 | 2.227E-08 | 2.300E-08 | 6.700E-08 |
| 10147 | 508 | N508Q | A | 5.4e-08 | 2.8e-08 | 6.3e-08 | 4.833E-08 | 1.818E-08 | 2.800E-08 | 6.300E-08 |
| 10148 | 508 | N508G | A | 7e-08 | 5.5e-08 | 4.6e-08 | 5.700E-08 | 1.212E-08 | 4.600E-08 | 7.000E-08 |
| 10149 | 508 | N508H | A | 9.3e-09 | 7.5e-08 | 9.1e-08 | 5.843E-08 | 4.330E-08 | 9.300E-09 | 9.100E-08 |
| 10150 | 508 | N508I | A | 6e-08 | 1.8e-08 | 8e-08 | 5.267E-08 | 3.164E-08 | 1.800E-08 | 8.000E-08 |
| 10151 | 508 | N508L | A | 3.1e-08 | 3.6e-08 | 3.7e-08 | 3.467E-08 | 3.215E-09 | 3.100E-08 | 3.700E-08 |
| 10152 | 508 | N508K | A | 4.3e-08 | 6.9e-08 | 4.6e-08 | 5.267E-08 | 1.422E-08 | 4.300E-08 | 6.900E-08 |
| 10153 | 508 | N508M | A | 3.9e-08 | 3.6e-08 | 4.1e-08 | 3.867E-08 | 2.517E-09 | 3.600E-08 | 4.100E-08 |
| 10154 | 508 | N508F | A | 1.3e-08 | 9.8e-08 | 5.6e-08 | 5.567E-08 | 4.250E-08 | 1.300E-08 | 9.800E-08 |
| 10155 | 508 | N508P | A | 1.8e-08 | 2.9e-08 | 3.5e-08 | 2.733E-08 | 8.622E-09 | 1.800E-08 | 3.500E-08 |
| 10156 | 508 | N508S | A | 5.7e-08 | 3.6e-08 | 2.4e-08 | 3.900E-08 | 1.670E-08 | 2.400E-08 | 5.700E-08 |
| 10157 | 508 | N508T | A | 2.8e-08 | 2.1e-08 | 9.3e-08 | 4.733E-08 | 3.970E-08 | 2.100E-08 | 9.300E-08 |
| 10158 | 508 | N508W | A | 6.9e-08 | 2.1e-08 | 7e-08 | 5.333E-08 | 2.801E-08 | 2.100E-08 | 7.000E-08 |
| 10159 | 508 | N508Y | A | 3.9e-08 | 4.1e-08 | 2.4e-08 | 3.467E-08 | 9.292E-09 | 2.400E-08 | 4.100E-08 |
| 10160 | 508 | N508V | A | 7.5e-08 | 7.9e-08 | 6.3e-08 | 7.233E-08 | 8.327E-09 | 6.300E-08 | 7.900E-08 |
| 10161 | 509 | C509A | A | 6.1e-09 | 3.1e-08 | 5.5e-08 | 3.070E-08 | 2.445E-08 | 6.100E-09 | 5.500E-08 |
| 10162 | 509 | C509R | A | 4.5e-08 | 8.4e-08 | 6.8e-08 | 6.567E-08 | 1.960E-08 | 4.500E-08 | 8.400E-08 |
| 10163 | 509 | C509N | A | 5.9e-08 | 1.2e-07 | 4.8e-08 | 7.567E-08 | 3.879E-08 | 4.800E-08 | 1.200E-07 |
| 10164 | 509 | C509D | A | 6.1e-08 | 8.7e-08 | 1.1e-07 | 8.600E-08 | 2.452E-08 | 6.100E-08 | 1.100E-07 |
| 10165 | 509 | C509C | A | 4.7e-08 | 7e-08 | 2.2e-08 | 4.633E-08 | 2.401E-08 | 2.200E-08 | 7.000E-08 |
| 10166 | 509 | C509E | A | 2.6e-08 | 6.2e-08 | 4.5e-08 | 4.433E-08 | 1.801E-08 | 2.600E-08 | 6.200E-08 |
| 10167 | 509 | C509Q | A | 3.8e-08 | 5.2e-08 | 6.3e-08 | 5.100E-08 | 1.253E-08 | 3.800E-08 | 6.300E-08 |
| 10168 | 509 | C509G | A | 3.4e-08 | 2.3e-08 | 2.6e-08 | 2.767E-08 | 5.686E-09 | 2.300E-08 | 3.400E-08 |
| 10169 | 509 | C509H | A | 4.4e-08 | 6.4e-08 | 4.8e-08 | 5.200E-08 | 1.058E-08 | 4.400E-08 | 6.400E-08 |
| 10170 | 509 | C509I | A | 1.8e-08 | 4.2e-08 | 7.9e-08 | 4.633E-08 | 3.073E-08 | 1.800E-08 | 7.900E-08 |
| 10171 | 509 | C509L | A | 3.5e-08 | 5.6e-08 | 4.5e-08 | 4.533E-08 | 1.050E-08 | 3.500E-08 | 5.600E-08 |
| 10172 | 509 | C509K | A | 1.1e-07 | 6.5e-08 | 6.3e-08 | 7.933E-08 | 2.658E-08 | 6.300E-08 | 1.100E-07 |
| 10173 | 509 | C509M | A | 4.4e-08 | 4.7e-08 | 4.7e-08 | 4.600E-08 | 1.732E-09 | 4.400E-08 | 4.700E-08 |
| 10174 | 509 | C509F | A | 2.5e-08 | 3e-08 | 6.7e-08 | 4.067E-08 | 2.294E-08 | 2.500E-08 | 6.700E-08 |
| 10175 | 509 | C509P | A | 3.2e-08 | 3.6e-08 | 4.5e-08 | 3.767E-08 | 6.658E-09 | 3.200E-08 | 4.500E-08 |
| 10176 | 509 | C509S | A | 3e-08 | 6.5e-08 | 2.3e-08 | 3.933E-08 | 2.250E-08 | 2.300E-08 | 6.500E-08 |
| 10177 | 509 | C509T | A | 3.8e-08 | 1e-07 | 1.3e-07 | 8.933E-08 | 4.692E-08 | 3.800E-08 | 1.300E-07 |
| 10178 | 509 | C509W | A | 5.9e-08 | 1.3e-08 | 5.8e-08 | 4.333E-08 | 2.627E-08 | 1.300E-08 | 5.900E-08 |
| 10179 | 509 | C509Y | A | 5.5e-08 | 5e-08 | 5.7e-08 | 5.400E-08 | 3.606E-09 | 5.000E-08 | 5.700E-08 |
| 10180 | 509 | C509V | A | 3.3e-08 | 1.5e-08 | 6.6e-08 | 3.800E-08 | 2.587E-08 | 1.500E-08 | 6.600E-08 |
| 10181 | 510 | S510A | A | 1.1e-07 | 2.4e-08 | 6.4e-08 | 6.600E-08 | 4.303E-08 | 2.400E-08 | 1.100E-07 |
| 10182 | 510 | S510R | A | 9.4e-08 | 5.4e-08 | 5.4e-08 | 6.733E-08 | 2.309E-08 | 5.400E-08 | 9.400E-08 |
| 10183 | 510 | S510N | A | 9.7e-08 | 2.3e-08 | 3.3e-08 | 5.100E-08 | 4.015E-08 | 2.300E-08 | 9.700E-08 |
| 10184 | 510 | S510D | A | 4.2e-08 | 1.3e-07 | 4.9e-08 | 7.367E-08 | 4.891E-08 | 4.200E-08 | 1.300E-07 |
| 10185 | 510 | S510C | A | 3.1e-08 | 4.7e-08 | 5.1e-08 | 4.300E-08 | 1.058E-08 | 3.100E-08 | 5.100E-08 |
| 10186 | 510 | S510E | A | 2.2e-08 | 2.9e-08 | 1e-07 | 5.033E-08 | 4.315E-08 | 2.200E-08 | 1.000E-07 |
| 10187 | 510 | S510Q | A | 1.1e-07 | 1.5e-08 | 5e-08 | 5.833E-08 | 4.805E-08 | 1.500E-08 | 1.100E-07 |
| 10188 | 510 | S510G | A | 3.4e-08 | 3.2e-08 | 3.7e-08 | 3.433E-08 | 2.517E-09 | 3.200E-08 | 3.700E-08 |
| 10189 | 510 | S510H | A | 6.1e-08 | 6e-08 | 3.8e-08 | 5.300E-08 | 1.300E-08 | 3.800E-08 | 6.100E-08 |
| 10190 | 510 | S510I | A | 6.1e-08 | 2.2e-08 | 2.4e-08 | 3.567E-08 | 2.196E-08 | 2.200E-08 | 6.100E-08 |
| 10191 | 510 | S510L | A | 3.5e-08 | 2e-08 | 1.7e-08 | 2.400E-08 | 9.644E-09 | 1.700E-08 | 3.500E-08 |
| 10192 | 510 | S510K | A | 5.5e-08 | 4.4e-08 | 1.8e-08 | 3.900E-08 | 1.900E-08 | 1.800E-08 | 5.500E-08 |
| 10193 | 510 | S510M | A | 2.1e-08 | 5.5e-08 | 9.4e-08 | 5.667E-08 | 3.653E-08 | 2.100E-08 | 9.400E-08 |
| 10194 | 510 | S510F | A | 7.6e-08 | 2.1e-08 | 4.4e-08 | 4.700E-08 | 2.762E-08 | 2.100E-08 | 7.600E-08 |
| 10195 | 510 | S510P | A | 3.9e-08 | 1.1e-08 | 2.8e-08 | 2.600E-08 | 1.411E-08 | 1.100E-08 | 3.900E-08 |
| 10196 | 510 | S510S | A | 6.7e-08 | 2.8e-08 | 1.6e-08 | 3.700E-08 | 2.666E-08 | 1.600E-08 | 6.700E-08 |
| 10197 | 510 | S510T | A | 4.7e-08 | 5.1e-08 | 3.4e-08 | 4.400E-08 | 8.888E-09 | 3.400E-08 | 5.100E-08 |
| 10198 | 510 | S510W | A | 3.6e-08 | 5e-08 | 2.9e-08 | 3.833E-08 | 1.069E-08 | 2.900E-08 | 5.000E-08 |
| 10199 | 510 | S510Y | A | 6.1e-08 | 2.1e-08 | 5e-08 | 4.400E-08 | 2.066E-08 | 2.100E-08 | 6.100E-08 |

| | | | | | | | | | | |
|---|---|---|---|---|---|---|---|---|---|---|
| 10200 | 510 | S510V | A | 4.3e-08 | 4.1e-08 | 5e-08 | 4.467E-08 | 4.726E-09 | 4.100E-08 | 5.000E-08 |
| 10201 | 511 | Q511A | A | 3.8e-08 | 6.5e-08 | 3.4e-08 | 4.567E-08 | 1.686E-08 | 3.400E-08 | 6.500E-08 |
| 10202 | 511 | Q511R | A | 1.1e-07 | 2.6e-08 | 7.7e-08 | 7.100E-08 | 4.232E-08 | 2.600E-08 | 1.100E-07 |
| 10203 | 511 | Q511N | A | 1.3e-07 | 8.7e-08 | 3.3e-08 | 8.333E-08 | 4.860E-08 | 3.300E-08 | 1.300E-07 |
| 10204 | 511 | Q511D | A | 2.6e-08 | 1.5e-07 | 4.4e-08 | 7.333E-08 | 6.700E-08 | 2.600E-08 | 1.500E-07 |
| 10205 | 511 | Q511C | A | 7.6e-08 | 4.5e-08 | 6.9e-08 | 6.333E-08 | 1.626E-08 | 4.500E-08 | 7.600E-08 |
| 10206 | 511 | Q511E | A | 8.7e-08 | 2.8e-08 | 3.8e-08 | 5.100E-08 | 3.158E-08 | 2.800E-08 | 8.700E-08 |
| 10207 | 511 | Q511Q | A | 2.4e-08 | 3.6e-08 | 6.3e-08 | 4.100E-08 | 1.997E-08 | 2.400E-08 | 6.300E-08 |
| 10208 | 511 | Q511G | A | 4.3e-08 | 2e-08 | 2.5e-08 | 2.933E-08 | 1.210E-08 | 2.000E-08 | 4.300E-08 |
| 10209 | 511 | Q511H | A | 1.1e-08 | 5e-08 | 2.6e-08 | 2.900E-08 | 1.967E-08 | 1.100E-08 | 5.000E-08 |
| 10210 | 511 | Q511I | A | 4.7e-08 | 1e-08 | 1.4e-07 | 6.567E-08 | 6.698E-08 | 1.000E-08 | 1.400E-07 |
| 10211 | 511 | Q511L | A | 5.5e-08 | 9.6e-08 | 7.4e-08 | 7.500E-08 | 2.052E-08 | 5.500E-08 | 9.600E-08 |
| 10212 | 511 | Q511K | A | 3.8e-08 | 5.5e-08 | 3.3e-08 | 4.200E-08 | 1.153E-08 | 3.300E-08 | 5.500E-08 |
| 10213 | 511 | Q511M | A | 7.7e-08 | 4.4e-08 | 8.1e-08 | 6.733E-08 | 2.031E-08 | 4.400E-08 | 8.100E-08 |
| 10214 | 511 | Q511F | A | 5.2e-08 | 5.5e-08 | 9.6e-08 | 6.767E-08 | 2.458E-08 | 5.200E-08 | 9.600E-08 |
| 10215 | 511 | Q511P | A | 3.5e-08 | 2.7e-08 | 1e-07 | 5.400E-08 | 4.004E-08 | 2.700E-08 | 1.000E-07 |
| 10216 | 511 | Q511S | A | 5.1e-08 | 8e-08 | 6.7e-08 | 6.600E-08 | 1.453E-08 | 5.100E-08 | 8.000E-08 |
| 10217 | 511 | Q511T | A | 2.1e-08 | 4.5e-08 | 2.8e-08 | 3.133E-08 | 1.234E-08 | 2.100E-08 | 4.500E-08 |
| 10218 | 511 | Q511W | A | 5.9e-08 | 8.6e-08 | 3.2e-08 | 5.900E-08 | 2.700E-08 | 3.200E-08 | 8.600E-08 |
| 10219 | 511 | Q511Y | A | 4.5e-08 | 8.1e-08 | 4e-08 | 5.533E-08 | 2.237E-08 | 4.000E-08 | 8.100E-08 |
| 10220 | 511 | Q511V | A | 3.3e-08 | 9.4e-08 | 7.3e-08 | 6.667E-08 | 3.099E-08 | 3.300E-08 | 9.400E-08 |
| 10221 | 512 | F512A | A | 5.6e-08 | 1.8e-08 | 3.8e-08 | 3.733E-08 | 1.901E-08 | 1.800E-08 | 5.600E-08 |
| 10222 | 512 | F512R | A | 4.7e-08 | 2.9e-08 | 6e-08 | 4.533E-08 | 1.557E-08 | 2.900E-08 | 6.000E-08 |
| 10223 | 512 | F512N | A | 2.4e-08 | 4.4e-08 | 6e-08 | 4.267E-08 | 1.804E-08 | 2.400E-08 | 6.000E-08 |
| 10224 | 512 | F512D | A | 2.4e-08 | 4.5e-08 | 7.4e-08 | 4.767E-08 | 2.511E-08 | 2.400E-08 | 7.400E-08 |
| 10225 | 512 | F512C | A | 4e-08 | 5.1e-08 | 3.1e-08 | 4.067E-08 | 1.002E-08 | 3.100E-08 | 5.100E-08 |
| 10226 | 512 | F512E | A | 9.7e-08 | 4.6e-08 | 5e-08 | 6.433E-08 | 2.836E-08 | 4.600E-08 | 9.700E-08 |
| 10227 | 512 | F512Q | A | 3.4e-08 | 2e-08 | 3.9e-08 | 3.100E-08 | 9.849E-09 | 2.000E-08 | 3.900E-08 |
| 10228 | 512 | F512G | A | 1.8e-08 | 6.2e-08 | 5.3e-08 | 4.433E-08 | 2.325E-08 | 1.800E-08 | 6.200E-08 |
| 10229 | 512 | F512H | A | 4.9e-08 | 2.6e-08 | 1e-07 | 5.833E-08 | 3.787E-08 | 2.600E-08 | 1.000E-07 |
| 10230 | 512 | F512I | A | 9.9e-08 | 4.1e-08 | 3.6e-08 | 5.867E-08 | 3.502E-08 | 3.600E-08 | 9.900E-08 |
| 10231 | 512 | F512L | A | 2.6e-08 | 3e-08 | 3.7e-08 | 3.100E-08 | 5.568E-09 | 2.600E-08 | 3.700E-08 |
| 10232 | 512 | F512K | A | 9e-08 | 3.3e-08 | 2.6e-08 | 4.967E-08 | 3.510E-08 | 2.600E-08 | 9.000E-08 |
| 10233 | 512 | F512M | A | 1.8e-08 | 5.3e-08 | 6.7e-08 | 4.600E-08 | 2.524E-08 | 1.800E-08 | 6.700E-08 |
| 10234 | 512 | F512F | A | 2.3e-08 | 3.8e-08 | 2.2e-08 | 2.767E-08 | 8.963E-09 | 2.200E-08 | 3.800E-08 |
| 10235 | 512 | F512P | A | 6.3e-08 | 4.6e-08 | 4.9e-08 | 5.267E-08 | 9.074E-09 | 4.600E-08 | 6.300E-08 |
| 10236 | 512 | F512S | A | 6.9e-08 | 3.2e-08 | 1.8e-08 | 3.967E-08 | 2.635E-08 | 1.800E-08 | 6.900E-08 |
| 10237 | 512 | F512T | A | 1.7e-08 | 1e-07 | 6.2e-08 | 5.967E-08 | 4.155E-08 | 1.700E-08 | 1.000E-07 |
| 10238 | 512 | F512W | A | 1.8e-08 | 9.1e-08 | 4.8e-08 | 5.233E-08 | 3.669E-08 | 1.800E-08 | 9.100E-08 |
| 10239 | 512 | F512Y | A | 6.3e-08 | 2.5e-08 | 3.2e-08 | 4.000E-08 | 2.022E-08 | 2.500E-08 | 6.300E-08 |
| 10240 | 512 | F512V | A | 5.5e-08 | 1.7e-08 | 3.5e-08 | 3.567E-08 | 1.901E-08 | 1.700E-08 | 5.500E-08 |
| 10241 | 513 | L513A | A | 2.2e-08 | 3.6e-08 | 5e-08 | 3.600E-08 | 1.400E-08 | 2.200E-08 | 5.000E-08 |
| 10242 | 513 | L513R | A | 4.9e-08 | 3.4e-08 | 5.4e-08 | 4.567E-08 | 1.041E-08 | 3.400E-08 | 5.400E-08 |
| 10243 | 513 | L513N | A | 2.9e-08 | 2.6e-08 | 3.4e-08 | 2.967E-08 | 4.041E-09 | 2.600E-08 | 3.400E-08 |
| 10244 | 513 | L513D | A | 5.2e-08 | 1.2e-07 | 6.8e-08 | 8.000E-08 | 3.555E-08 | 5.200E-08 | 1.200E-07 |
| 10245 | 513 | L513C | A | 2.5e-07 | 1.8e-08 | 3.5e-08 | 1.010E-07 | 1.293E-07 | 1.800E-08 | 2.500E-07 |
| 10246 | 513 | L513E | A | 9.7e-08 | 2.8e-08 | 3.4e-08 | 5.300E-08 | 3.822E-08 | 2.800E-08 | 9.700E-08 |
| 10247 | 513 | L513Q | A | 5.6e-08 | 6.5e-08 | 7.1e-08 | 6.400E-08 | 7.550E-09 | 5.600E-08 | 7.100E-08 |
| 10248 | 513 | L513G | A | 1.7e-08 | 8.3e-09 | 9e-08 | 3.843E-08 | 4.487E-08 | 8.300E-09 | 9.000E-08 |
| 10249 | 513 | L513H | A | 1e-07 | 5.4e-08 | 3.4e-08 | 6.267E-08 | 3.384E-08 | 3.400E-08 | 1.000E-07 |
| 10250 | 513 | L513I | A | 5.9e-08 | 6e-08 | 7.5e-08 | 6.467E-08 | 8.963E-09 | 5.900E-08 | 7.500E-08 |
| 10251 | 513 | L513L | A | 8e-08 | 5.2e-08 | 5.8e-08 | 6.333E-08 | 1.474E-08 | 5.200E-08 | 8.000E-08 |
| 10252 | 513 | L513K | A | 4.6e-08 | 6.2e-08 | 4.5e-08 | 5.100E-08 | 9.539E-09 | 4.500E-08 | 6.200E-08 |
| 10253 | 513 | L513M | A | 6.4e-08 | 1.1e-07 | 4.6e-08 | 7.333E-08 | 3.301E-08 | 4.600E-08 | 1.100E-07 |
| 10254 | 513 | L513F | A | 5.3e-08 | 8.5e-08 | 2.8e-08 | 5.533E-08 | 2.857E-08 | 2.800E-08 | 8.500E-08 |
| 10255 | 513 | L513P | A | 2.9e-08 | 4.9e-08 | 2.7e-08 | 3.500E-08 | 1.217E-08 | 2.700E-08 | 4.900E-08 |
| 10256 | 513 | L513S | A | 3.2e-08 | 1.7e-08 | 4e-08 | 2.967E-08 | 1.168E-08 | 1.700E-08 | 4.000E-08 |
| 10257 | 513 | L513T | A | 9.1e-08 | 6.8e-08 | 4.2e-08 | 6.700E-08 | 2.452E-08 | 4.200E-08 | 9.100E-08 |
| 10258 | 513 | L513W | A | 3.1e-08 | 5.8e-08 | 4.1e-08 | 4.333E-08 | 1.365E-08 | 3.100E-08 | 5.800E-08 |
| 10259 | 513 | L513Y | A | 7.5e-08 | 4.8e-08 | 6.6e-08 | 6.300E-08 | 1.375E-08 | 4.800E-08 | 7.500E-08 |
| 10260 | 513 | L513V | A | 3.9e-08 | 2e-08 | 3.4e-08 | 3.100E-08 | 9.849E-09 | 2.000E-08 | 3.900E-08 |
| 10261 | 514 | R514A | A | 1.7e-08 | 2.1e-08 | 5.3e-08 | 3.033E-08 | 1.973E-08 | 1.700E-08 | 5.300E-08 |
| 10262 | 514 | R514R | A | 1.3e-08 | 2.9e-08 | 6.8e-08 | 3.667E-08 | 2.829E-08 | 1.300E-08 | 6.800E-08 |
| 10263 | 514 | R514N | A | 5e-08 | 4.4e-08 | 4.2e-08 | 4.533E-08 | 4.163E-09 | 4.200E-08 | 5.000E-08 |
| 10264 | 514 | R514D | A | 3.8e-08 | 2.8e-08 | 3.9e-08 | 3.500E-08 | 6.083E-09 | 2.800E-08 | 3.900E-08 |
| 10265 | 514 | R514C | A | 6.1e-08 | 7.6e-08 | 3.3e-08 | 5.667E-08 | 2.183E-08 | 3.300E-08 | 7.600E-08 |
| 10266 | 514 | R514E | A | 3.4e-08 | 3.2e-08 | 1.9e-08 | 2.833E-08 | 8.145E-09 | 1.900E-08 | 3.400E-08 |
| 10267 | 514 | R514Q | A | 6.2e-08 | 2.6e-08 | 2.9e-08 | 3.900E-08 | 1.997E-08 | 2.600E-08 | 6.200E-08 |
| 10268 | 514 | R514G | A | 6.7e-09 | 9.2e-08 | 1.3e-07 | 7.623E-08 | 6.314E-08 | 6.700E-09 | 1.300E-07 |
| 10269 | 514 | R514H | A | 5.9e-08 | 4.6e-08 | 1.1e-08 | 3.867E-08 | 2.483E-08 | 1.100E-08 | 5.900E-08 |
| 10270 | 514 | R514I | A | 2.6e-08 | 2.2e-07 | 3.7e-08 | 9.433E-08 | 1.090E-07 | 2.600E-08 | 2.200E-07 |
| 10271 | 514 | R514L | A | 9.7e-08 | 5.1e-08 | 5.1e-08 | 6.633E-08 | 2.656E-08 | 5.100E-08 | 9.700E-08 |
| 10272 | 514 | R514K | A | 3e-08 | 5.2e-08 | 5.8e-08 | 4.667E-08 | 1.474E-08 | 3.000E-08 | 5.800E-08 |
| 10273 | 514 | R514M | A | 2.9e-08 | 2.4e-08 | 5.9e-08 | 3.733E-08 | 1.893E-08 | 2.400E-08 | 5.900E-08 |
| 10274 | 514 | R514F | A | 5e-08 | 7.6e-08 | 7.2e-08 | 6.600E-08 | 1.400E-08 | 5.000E-08 | 7.600E-08 |
| 10275 | 514 | R514P | A | 4.5e-08 | 5e-08 | 6.5e-08 | 5.333E-08 | 1.041E-08 | 4.500E-08 | 6.500E-08 |
| 10276 | 514 | R514S | A | 3.6e-08 | 1.9e-08 | 4.5e-08 | 3.333E-08 | 1.320E-08 | 1.900E-08 | 4.500E-08 |
| 10277 | 514 | R514T | A | 8e-08 | 6.5e-08 | 3.2e-08 | 5.900E-08 | 2.456E-08 | 3.200E-08 | 8.000E-08 |
| 10278 | 514 | R514W | A | 3.6e-08 | 5.9e-08 | 9.2e-09 | 3.473E-08 | 2.492E-08 | 9.200E-09 | 5.900E-08 |
| 10279 | 514 | R514Y | A | 8e-08 | 4.3e-08 | 4.5e-08 | 5.600E-08 | 2.081E-08 | 4.300E-08 | 8.000E-08 |
| 10280 | 514 | R514V | A | 5.4e-08 | 4.4e-08 | 4.5e-08 | 4.767E-08 | 5.508E-09 | 4.400E-08 | 5.400E-08 |
| 10281 | 515 | G515A | A | 6.3e-08 | 6.3e-08 | 3.6e-08 | 5.400E-08 | 1.559E-08 | 3.600E-08 | 6.300E-08 |
| 10282 | 515 | G515R | A | 5.8e-08 | 2e-08 | 7.9e-08 | 5.233E-08 | 2.991E-08 | 2.000E-08 | 7.900E-08 |
| 10283 | 515 | G515N | A | 3.3e-08 | 5.2e-08 | 2.4e-08 | 3.633E-08 | 1.429E-08 | 2.400E-08 | 5.200E-08 |
| 10284 | 515 | G515D | A | 2.3e-08 | 3.2e-08 | 8.4e-08 | 4.633E-08 | 3.293E-08 | 2.300E-08 | 8.400E-08 |

| | | | | | | | | | | |
|---|---|---|---|---|---|---|---|---|---|---|
| 10285 | 515 | G515C | A | 1e-07 | 7.5e-08 | 2.2e-08 | 6.567E-08 | 3.983E-08 | 2.200E-08 | 1.000E-07 |
| 10286 | 515 | G515E | A | 3.4e-08 | 5.9e-08 | 1.6e-08 | 3.633E-08 | 2.159E-08 | 1.600E-08 | 5.900E-08 |
| 10287 | 515 | G515Q | A | 4.4e-08 | 6.6e-08 | 6.1e-08 | 5.700E-08 | 1.153E-08 | 4.400E-08 | 6.600E-08 |
| 10288 | 515 | G515G | A | 3.7e-08 | 4.6e-08 | 3.4e-08 | 3.900E-08 | 6.245E-09 | 3.400E-08 | 4.600E-08 |
| 10289 | 515 | G515H | A | 4.6e-08 | 3.1e-08 | 4.4e-08 | 4.033E-08 | 8.145E-09 | 3.100E-08 | 4.600E-08 |
| 10290 | 515 | G515I | A | 3.9e-07 | 5.4e-08 | 5.4e-08 | 1.660E-07 | 1.940E-07 | 5.400E-08 | 3.900E-07 |
| 10291 | 515 | G515L | A | 6.6e-08 | 3.6e-08 | 3.8e-08 | 4.667E-08 | 1.677E-08 | 3.600E-08 | 6.600E-08 |
| 10292 | 515 | G515K | A | 1.2e-07 | 4.9e-08 | 2.6e-08 | 6.500E-08 | 4.900E-08 | 2.600E-08 | 1.200E-07 |
| 10293 | 515 | G515M | A | 9.5e-08 | 5.4e-08 | 7.2e-08 | 7.367E-08 | 2.055E-08 | 5.400E-08 | 9.500E-08 |
| 10294 | 515 | G515F | A | 1.5e-08 | 4.7e-08 | 4.9e-08 | 3.700E-08 | 1.908E-08 | 1.500E-08 | 4.900E-08 |
| 10295 | 515 | G515P | A | 5.4e-08 | 5.3e-08 | 5.2e-08 | 5.300E-08 | 1.000E-09 | 5.200E-08 | 5.400E-08 |
| 10296 | 515 | G515S | A | 1.7e-08 | 3e-08 | 2.1e-08 | 2.267E-08 | 6.658E-09 | 1.700E-08 | 3.000E-08 |
| 10297 | 515 | G515T | A | 2.3e-08 | 1e-07 | 5.1e-08 | 5.800E-08 | 3.897E-08 | 2.300E-08 | 1.000E-07 |
| 10298 | 515 | G515W | A | 1.1e-07 | 7.5e-08 | 2.7e-08 | 7.067E-08 | 4.167E-08 | 2.700E-08 | 1.100E-07 |
| 10299 | 515 | G515Y | A | 5.1e-08 | 1.4e-07 | 5.6e-08 | 8.233E-08 | 5.000E-08 | 5.100E-08 | 1.400E-07 |
| 10300 | 515 | G515V | A | 2.2e-08 | 4.8e-08 | 2.1e-07 | 9.333E-08 | 1.019E-07 | 2.200E-08 | 2.100E-07 |
| 10301 | 516 | Q516A | A | 6.3e-08 | 5.9e-08 | 4.5e-08 | 5.567E-08 | 9.452E-09 | 4.500E-08 | 6.300E-08 |
| 10302 | 516 | Q516R | A | 6.9e-08 | 4e-08 | 2.7e-08 | 4.533E-08 | 2.150E-08 | 2.700E-08 | 6.900E-08 |
| 10303 | 516 | Q516N | A | 4.6e-08 | 1.1e-08 | 7.5e-08 | 4.400E-08 | 3.205E-08 | 1.100E-08 | 7.500E-08 |
| 10304 | 516 | Q516D | A | 6.8e-08 | 7.4e-08 | 9.3e-08 | 7.833E-08 | 1.305E-08 | 6.800E-08 | 9.300E-08 |
| 10305 | 516 | Q516C | A | 1.6e-08 | 1.5e-08 | 3.1e-08 | 2.067E-08 | 8.963E-09 | 1.500E-08 | 3.100E-08 |
| 10306 | 516 | Q516E | A | 3.9e-08 | 7.6e-08 | 3.5e-08 | 5.000E-08 | 2.261E-08 | 3.500E-08 | 7.600E-08 |
| 10307 | 516 | Q516Q | A | 4.5e-08 | 1.2e-07 | 3.1e-09 | 5.603E-08 | 5.923E-08 | 3.100E-09 | 1.200E-07 |
| 10308 | 516 | Q516G | A | 5.2e-08 | 1.4e-07 | 4.7e-08 | 7.967E-08 | 5.231E-08 | 4.700E-08 | 1.400E-07 |
| 10309 | 516 | Q516H | A | 1.1e-07 | 6.7e-08 | 5.4e-08 | 7.700E-08 | 2.931E-08 | 5.400E-08 | 1.100E-07 |
| 10310 | 516 | Q516I | A | 3.6e-08 | 2.8e-08 | 9.2e-08 | 5.200E-08 | 3.487E-08 | 2.800E-08 | 9.200E-08 |
| 10311 | 516 | Q516L | A | 2.2e-08 | 4.2e-08 | 1.6e-08 | 2.667E-08 | 1.361E-08 | 1.600E-08 | 4.200E-08 |
| 10312 | 516 | Q516K | A | 1.5e-08 | 3.7e-08 | 3.1e-08 | 2.767E-08 | 1.137E-08 | 1.500E-08 | 3.700E-08 |
| 10313 | 516 | Q516M | A | 6.3e-08 | 4.6e-08 | 6.3e-08 | 5.733E-08 | 9.815E-09 | 4.600E-08 | 6.300E-08 |
| 10314 | 516 | Q516F | A | 4.5e-08 | 2.8e-08 | 4.3e-08 | 3.867E-08 | 9.292E-09 | 2.800E-08 | 4.500E-08 |
| 10315 | 516 | Q516P | A | 7.3e-08 | 3.6e-08 | 2.2e-08 | 4.367E-08 | 2.635E-08 | 2.200E-08 | 7.300E-08 |
| 10316 | 516 | Q516S | A | 5.9e-08 | 1e-07 | 2.4e-08 | 6.100E-08 | 3.804E-08 | 2.400E-08 | 1.000E-07 |
| 10317 | 516 | Q516T | A | 4.1e-08 | 6.4e-08 | 5.8e-08 | 5.433E-08 | 1.193E-08 | 4.100E-08 | 6.400E-08 |
| 10318 | 516 | Q516W | A | 3.2e-08 | 5.4e-08 | 2e-08 | 3.533E-08 | 1.724E-08 | 2.000E-08 | 5.400E-08 |
| 10319 | 516 | Q516Y | A | 4.5e-08 | 2.7e-08 | 8e-08 | 5.067E-08 | 2.695E-08 | 2.700E-08 | 8.000E-08 |
| 10320 | 516 | Q516V | A | 4.5e-08 | 4.9e-08 | 4.2e-08 | 4.533E-08 | 3.512E-09 | 4.200E-08 | 4.900E-08 |
| 10321 | 517 | E517A | A | 4.9e-08 | 7.2e-08 | 6.4e-08 | 6.167E-08 | 1.168E-08 | 4.900E-08 | 7.200E-08 |
| 10322 | 517 | E517R | A | 5.2e-08 | 4.3e-08 | 9e-08 | 6.167E-08 | 2.495E-08 | 4.300E-08 | 9.000E-08 |
| 10323 | 517 | E517N | A | 5.6e-08 | 5.2e-08 | 5.1e-08 | 5.300E-08 | 2.646E-09 | 5.100E-08 | 5.600E-08 |
| 10324 | 517 | E517D | A | 4.2e-08 | 5.4e-08 | 1.2e-08 | 3.600E-08 | 2.163E-08 | 1.200E-08 | 5.400E-08 |
| 10325 | 517 | E517C | A | 1.1e-07 | 4.8e-08 | 2.9e-08 | 6.233E-08 | 4.236E-08 | 2.900E-08 | 1.100E-07 |
| 10326 | 517 | E517E | A | 1.9e-08 | 1e-07 | 5.4e-08 | 5.767E-08 | 4.062E-08 | 1.900E-08 | 1.000E-07 |
| 10327 | 517 | E517Q | A | 6.3e-08 | 4.9e-08 | 4.2e-08 | 5.133E-08 | 1.069E-08 | 4.200E-08 | 6.300E-08 |
| 10328 | 517 | E517G | A | 5.9e-08 | 1.9e-08 | 2.7e-08 | 3.500E-08 | 2.117E-08 | 1.900E-08 | 5.900E-08 |
| 10329 | 517 | E517H | A | 3.5e-08 | 2.9e-08 | 4.8e-08 | 3.733E-08 | 9.713E-09 | 2.900E-08 | 4.800E-08 |
| 10330 | 517 | E517I | A | 1.4e-07 | 4.5e-08 | 1.6e-08 | 6.700E-08 | 6.486E-08 | 1.600E-08 | 1.400E-07 |
| 10331 | 517 | E517L | A | 3.1e-08 | 5.1e-08 | 3.1e-08 | 3.767E-08 | 1.155E-08 | 3.100E-08 | 5.100E-08 |
| 10332 | 517 | E517K | A | 5.5e-08 | 8.2e-08 | 1.3e-08 | 5.000E-08 | 3.477E-08 | 1.300E-08 | 8.200E-08 |
| 10333 | 517 | E517M | A | 3.6e-08 | 3.7e-08 | 8.1e-08 | 5.133E-08 | 2.570E-08 | 3.600E-08 | 8.100E-08 |
| 10334 | 517 | E517F | A | 5e-08 | 2.9e-08 | 1.5e-08 | 3.133E-08 | 1.762E-08 | 1.500E-08 | 5.000E-08 |
| 10335 | 517 | E517P | A | 2.5e-08 | 5.2e-08 | 1.8e-08 | 3.167E-08 | 1.795E-08 | 1.800E-08 | 5.200E-08 |
| 10336 | 517 | E517S | A | 4.3e-08 | 4.2e-08 | 9.6e-09 | 3.153E-08 | 1.900E-08 | 9.600E-09 | 4.300E-08 |
| 10337 | 517 | E517T | A | 4e-08 | 1.4e-08 | 6.2e-08 | 3.867E-08 | 2.403E-08 | 1.400E-08 | 6.200E-08 |
| 10338 | 517 | E517W | A | 3.9e-08 | 2.2e-08 | 3e-08 | 3.033E-08 | 8.505E-09 | 2.200E-08 | 3.900E-08 |
| 10339 | 517 | E517Y | A | 2e-08 | 3.9e-08 | 3.2e-08 | 3.033E-08 | 9.609E-09 | 2.000E-08 | 3.900E-08 |
| 10340 | 517 | E517V | A | 7.9e-08 | 2e-08 | 8.8e-08 | 6.233E-08 | 3.694E-08 | 2.000E-08 | 8.800E-08 |
| 10341 | 518 | C518A | A | 1.5e-07 | 6.3e-08 | 5e-08 | 8.767E-08 | 5.437E-08 | 5.000E-08 | 1.500E-07 |
| 10342 | 518 | C518R | A | 4.2e-08 | 5.6e-08 | 2.9e-08 | 4.233E-08 | 1.350E-08 | 2.900E-08 | 5.600E-08 |
| 10343 | 518 | C518N | A | 3.9e-08 | 1.3e-07 | 1.9e-08 | 6.267E-08 | 5.916E-08 | 1.900E-08 | 1.300E-07 |
| 10344 | 518 | C518D | A | 4.6e-08 | 7.1e-08 | 6.1e-08 | 5.933E-08 | 1.258E-08 | 4.600E-08 | 7.100E-08 |
| 10345 | 518 | C518C | A | 1.2e-07 | 3.9e-08 | 1.4e-07 | 9.967E-08 | 5.348E-08 | 3.900E-08 | 1.400E-07 |
| 10346 | 518 | C518E | A | 5.2e-08 | 3.6e-08 | 3e-08 | 3.933E-08 | 1.137E-08 | 3.000E-08 | 5.200E-08 |
| 10347 | 518 | C518Q | A | 1.2e-07 | 3.5e-08 | 5.8e-08 | 7.100E-08 | 4.397E-08 | 3.500E-08 | 1.200E-07 |
| 10348 | 518 | C518G | A | 8.5e-08 | 4.6e-08 | 6.5e-08 | 6.533E-08 | 1.950E-08 | 4.600E-08 | 8.500E-08 |
| 10349 | 518 | C518H | A | 4.7e-08 | 6.4e-08 | 2.3e-08 | 4.467E-08 | 2.060E-08 | 2.300E-08 | 6.400E-08 |
| 10350 | 518 | C518I | A | 6.4e-08 | 5.6e-08 | 4.8e-08 | 5.600E-08 | 8.000E-09 | 4.800E-08 | 6.400E-08 |
| 10351 | 518 | C518L | A | 1.2e-07 | 2e-08 | 8.5e-08 | 7.500E-08 | 5.074E-08 | 2.000E-08 | 1.200E-07 |
| 10352 | 518 | C518K | A | 4.6e-08 | 1.1e-07 | 3.6e-08 | 6.400E-08 | 4.015E-08 | 3.600E-08 | 1.100E-07 |
| 10353 | 518 | C518M | A | 8.7e-08 | 8.5e-08 | 1e-07 | 9.067E-08 | 8.145E-09 | 8.500E-08 | 1.000E-07 |
| 10354 | 518 | C518F | A | 1.3e-07 | 4.4e-08 | 2.7e-08 | 6.700E-08 | 5.522E-08 | 2.700E-08 | 1.300E-07 |
| 10355 | 518 | C518P | A | 1e-07 | 2.3e-07 | 1.5e-08 | 1.150E-07 | 1.083E-07 | 1.500E-08 | 2.300E-07 |
| 10356 | 518 | C518S | A | 7.4e-08 | 3.5e-08 | 2.2e-08 | 4.367E-08 | 2.706E-08 | 2.200E-08 | 7.400E-08 |
| 10357 | 518 | C518T | A | 7.3e-08 | 4.5e-08 | 4e-08 | 5.267E-08 | 1.779E-08 | 4.000E-08 | 7.300E-08 |
| 10358 | 518 | C518W | A | 4.7e-08 | 4.2e-08 | 1.6e-08 | 3.500E-08 | 1.664E-08 | 1.600E-08 | 4.700E-08 |
| 10359 | 518 | C518Y | A | 2.1e-08 | 1.6e-08 | 3.4e-08 | 2.367E-08 | 9.292E-09 | 1.600E-08 | 3.400E-08 |
| 10360 | 518 | C518V | A | 6.8e-08 | 6.3e-08 | 2.7e-08 | 5.267E-08 | 2.237E-08 | 2.700E-08 | 6.800E-08 |
| 10361 | 519 | V519A | A | 3.7e-08 | 5.1e-08 | 3.6e-08 | 4.133E-08 | 8.386E-09 | 3.600E-08 | 5.100E-08 |
| 10362 | 519 | V519R | A | 3.7e-08 | 3.1e-08 | 5.6e-08 | 4.133E-08 | 1.305E-08 | 3.100E-08 | 5.600E-08 |
| 10363 | 519 | V519N | A | 2.1e-08 | 3.5e-08 | 3.2e-08 | 2.933E-08 | 7.371E-09 | 2.100E-08 | 3.500E-08 |
| 10364 | 519 | V519D | A | 2.7e-08 | 4.2e-08 | 4.3e-08 | 3.733E-08 | 8.963E-09 | 2.700E-08 | 4.300E-08 |
| 10365 | 519 | V519C | A | 4.3e-08 | 1.8e-08 | 2.5e-08 | 2.867E-08 | 1.290E-08 | 1.800E-08 | 4.300E-08 |
| 10366 | 519 | V519E | A | 3.9e-08 | 1.7e-08 | 3.1e-08 | 2.900E-08 | 1.114E-08 | 1.700E-08 | 3.900E-08 |
| 10367 | 519 | V519Q | A | 2.3e-08 | 2.9e-08 | 8.9e-08 | 4.700E-08 | 3.650E-08 | 2.300E-08 | 8.900E-08 |
| 10368 | 519 | V519G | A | 8.1e-08 | 1.9e-08 | 7.1e-08 | 5.700E-08 | 3.329E-08 | 1.900E-08 | 8.100E-08 |
| 10369 | 519 | V519H | A | 1.9e-08 | 9.3e-08 | 3.2e-08 | 4.800E-08 | 3.951E-08 | 1.900E-08 | 9.300E-08 |

| | | | | | | | | | | |
|---|---|---|---|---|---|---|---|---|---|---|
| 10370 | 519 | V519I | A | 8.7e-08 | 3e-08 | 1.1e-08 | 4.267E-08 | 3.955E-08 | 1.100E-08 | 8.700E-08 |
| 10371 | 519 | V519L | A | 4.1e-08 | 3.4e-08 | 4.5e-08 | 4.000E-08 | 5.568E-09 | 3.400E-08 | 4.500E-08 |
| 10372 | 519 | V519K | A | 1.3e-08 | 7e-08 | 4.1e-08 | 4.133E-08 | 2.850E-08 | 1.300E-08 | 7.000E-08 |
| 10373 | 519 | V519M | A | 3.3e-08 | 3e-08 | 4e-08 | 3.433E-08 | 5.132E-09 | 3.000E-08 | 4.000E-08 |
| 10374 | 519 | V519F | A | 3.6e-08 | 2.3e-08 | 1.6e-08 | 2.500E-08 | 1.015E-08 | 1.600E-08 | 3.600E-08 |
| 10375 | 519 | V519P | A | 1.3e-08 | 3.1e-08 | 8.2e-08 | 4.200E-08 | 3.579E-08 | 1.300E-08 | 8.200E-08 |
| 10376 | 519 | V519S | A | 2.4e-08 | 4.4e-08 | 5.6e-08 | 4.133E-08 | 1.617E-08 | 2.400E-08 | 5.600E-08 |
| 10377 | 519 | V519T | A | 1.5e-07 | 4.1e-08 | 3.3e-08 | 7.467E-08 | 6.536E-08 | 3.300E-08 | 1.500E-07 |
| 10378 | 519 | V519W | A | 1.3e-07 | 3e-08 | 6.7e-08 | 7.567E-08 | 5.056E-08 | 3.000E-08 | 1.300E-07 |
| 10379 | 519 | V519Y | A | 2.7e-08 | 3.2e-08 | 2e-08 | 2.633E-08 | 6.028E-09 | 2.000E-08 | 3.200E-08 |
| 10380 | 519 | V519V | A | 1.3e-07 | 7.6e-08 | 2.1e-08 | 7.567E-08 | 5.450E-08 | 2.100E-08 | 1.300E-07 |
| 10381 | 520 | E520A | A | 5.5e-08 | 2.5e-08 | 2.7e-08 | 3.567E-08 | 1.677E-08 | 2.500E-08 | 5.500E-08 |
| 10382 | 520 | E520R | A | 6.8e-08 | 4e-08 | 3.5e-08 | 4.767E-08 | 1.779E-08 | 3.500E-08 | 6.800E-08 |
| 10383 | 520 | E520N | A | 2.3e-08 | 3.9e-08 | 4.3e-08 | 3.500E-08 | 1.058E-08 | 2.300E-08 | 4.300E-08 |
| 10384 | 520 | E520D | A | 1.1e-08 | 3.3e-08 | 8.9e-08 | 4.433E-08 | 4.022E-08 | 1.100E-08 | 8.900E-08 |
| 10385 | 520 | E520C | A | 5.1e-08 | 3.7e-08 | 7.7e-08 | 5.500E-08 | 2.030E-08 | 3.700E-08 | 7.700E-08 |
| 10386 | 520 | E520E | A | 2.9e-08 | 5.5e-08 | 5.3e-08 | 4.567E-08 | 1.447E-08 | 2.900E-08 | 5.500E-08 |
| 10387 | 520 | E520Q | A | 9e-08 | 2.9e-08 | 6.9e-08 | 6.267E-08 | 3.099E-08 | 2.900E-08 | 9.000E-08 |
| 10388 | 520 | E520G | A | 4.8e-08 | 3.7e-08 | 8.6e-08 | 5.700E-08 | 2.571E-08 | 3.700E-08 | 8.600E-08 |
| 10389 | 520 | E520H | A | 4.4e-08 | 3.7e-08 | 4.1e-08 | 4.067E-08 | 3.512E-09 | 3.700E-08 | 4.400E-08 |
| 10390 | 520 | E520I | A | 7.5e-08 | 1.4e-08 | 9.1e-08 | 6.000E-08 | 4.063E-08 | 1.400E-08 | 9.100E-08 |
| 10391 | 520 | E520L | A | 6.6e-08 | 4e-08 | 1e-07 | 6.867E-08 | 3.009E-08 | 4.000E-08 | 1.000E-07 |
| 10392 | 520 | E520K | A | 8.7e-08 | 3.9e-08 | 2.2e-08 | 4.933E-08 | 3.371E-08 | 2.200E-08 | 8.700E-08 |
| 10393 | 520 | E520M | A | 5.5e-08 | 5.8e-08 | 2.8e-08 | 4.700E-08 | 1.652E-08 | 2.800E-08 | 5.800E-08 |
| 10394 | 520 | E520F | A | 9.2e-08 | 1.3e-08 | 2.9e-08 | 4.467E-08 | 4.177E-08 | 1.300E-08 | 9.200E-08 |
| 10395 | 520 | E520P | A | 2.4e-08 | 2.6e-08 | 1.1e-07 | 5.333E-08 | 4.908E-08 | 2.400E-08 | 1.100E-07 |
| 10396 | 520 | E520S | A | 1.7e-08 | 7.6e-08 | 3.5e-08 | 4.267E-08 | 3.024E-08 | 1.700E-08 | 7.600E-08 |
| 10397 | 520 | E520T | A | 3.7e-08 | 2.6e-08 | 8.7e-08 | 5.000E-08 | 3.251E-08 | 2.600E-08 | 8.700E-08 |
| 10398 | 520 | E520W | A | 2.8e-08 | 3.1e-08 | 2e-08 | 2.633E-08 | 5.686E-09 | 2.000E-08 | 3.100E-08 |
| 10399 | 520 | E520Y | A | 8.8e-08 | 6.5e-08 | 5.5e-08 | 6.933E-08 | 1.692E-08 | 5.500E-08 | 8.800E-08 |
| 10400 | 520 | E520V | A | 6.3e-08 | 1.9e-08 | 3.3e-08 | 3.833E-08 | 2.248E-08 | 1.900E-08 | 6.300E-08 |
| 10401 | 521 | E521A | A | 1.4e-07 | 1.1e-08 | 3.3e-08 | 6.133E-08 | 6.901E-08 | 1.100E-08 | 1.400E-07 |
| 10402 | 521 | E521R | A | 5.5e-08 | 6.2e-08 | 6.2e-08 | 5.967E-08 | 4.041E-09 | 5.500E-08 | 6.200E-08 |
| 10403 | 521 | E521N | A | 5.3e-08 | 1.6e-08 | 6.8e-08 | 4.567E-08 | 2.676E-08 | 1.600E-08 | 6.800E-08 |
| 10404 | 521 | E521D | A | 5.9e-08 | 6.4e-08 | 3.4e-08 | 5.233E-08 | 1.607E-08 | 3.400E-08 | 6.400E-08 |
| 10405 | 521 | E521C | A | 5.7e-08 | 6e-08 | 9.8e-08 | 7.167E-08 | 2.285E-08 | 5.700E-08 | 9.800E-08 |
| 10406 | 521 | E521E | A | 4.3e-08 | 1.2e-07 | 4.3e-08 | 6.867E-08 | 4.446E-08 | 4.300E-08 | 1.200E-07 |
| 10407 | 521 | E521Q | A | 5.4e-08 | 2.6e-08 | 7.9e-08 | 5.300E-08 | 2.651E-08 | 2.600E-08 | 7.900E-08 |
| 10408 | 521 | E521G | A | 4e-08 | 1.2e-08 | 4e-08 | 3.067E-08 | 1.617E-08 | 1.200E-08 | 4.000E-08 |
| 10409 | 521 | E521H | A | 4.1e-08 | 2.4e-08 | 6e-08 | 4.167E-08 | 1.801E-08 | 2.400E-08 | 6.000E-08 |
| 10410 | 521 | E521I | A | 5.8e-08 | 7.5e-08 | 1.6e-08 | 4.967E-08 | 3.037E-08 | 1.600E-08 | 7.500E-08 |
| 10411 | 521 | E521L | A | 9.5e-08 | 2.6e-08 | 4.2e-08 | 5.433E-08 | 3.612E-08 | 2.600E-08 | 9.500E-08 |
| 10412 | 521 | E521K | A | 7.1e-08 | 6.6e-08 | 1.4e-08 | 5.033E-08 | 3.156E-08 | 1.400E-08 | 7.100E-08 |
| 10413 | 521 | E521M | A | 3.5e-08 | 1.9e-08 | 2.9e-08 | 2.767E-08 | 8.083E-09 | 1.900E-08 | 3.500E-08 |
| 10414 | 521 | E521F | A | 2.9e-08 | 3.7e-08 | 1.2e-07 | 6.200E-08 | 5.039E-08 | 2.900E-08 | 1.200E-07 |
| 10415 | 521 | E521P | A | 3.1e-08 | 7e-08 | 2.3e-08 | 4.133E-08 | 2.515E-08 | 2.300E-08 | 7.000E-08 |
| 10416 | 521 | E521S | A | 2.4e-08 | 5e-08 | 3.7e-08 | 3.700E-08 | 1.300E-08 | 2.400E-08 | 5.000E-08 |
| 10417 | 521 | E521T | A | 3.7e-08 | 9.5e-08 | 2.2e-08 | 5.133E-08 | 3.855E-08 | 2.200E-08 | 9.500E-08 |
| 10418 | 521 | E521W | A | 2.3e-08 | 3.9e-08 | 1.7e-08 | 2.633E-08 | 1.137E-08 | 1.700E-08 | 3.900E-08 |
| 10419 | 521 | E521Y | A | 9.8e-09 | 4.1e-08 | 2.8e-08 | 2.627E-08 | 1.567E-08 | 9.800E-09 | 4.100E-08 |
| 10420 | 521 | E521V | A | 5.9e-08 | 7.1e-08 | 5.9e-08 | 6.300E-08 | 6.928E-09 | 5.900E-08 | 7.100E-08 |
| 10421 | 522 | C522A | A | 3.8e-08 | 2.7e-08 | 1.3e-08 | 2.600E-08 | 1.253E-08 | 1.300E-08 | 3.800E-08 |
| 10422 | 522 | C522R | A | 1.6e-08 | 1e-07 | 4e-08 | 5.200E-08 | 4.327E-08 | 1.600E-08 | 1.000E-07 |
| 10423 | 522 | C522N | A | 3.8e-08 | 3.7e-08 | 3.4e-08 | 3.633E-08 | 2.082E-09 | 3.400E-08 | 3.800E-08 |
| 10424 | 522 | C522D | A | 2.2e-08 | 4.4e-08 | 3.9e-08 | 3.500E-08 | 1.153E-08 | 2.200E-08 | 4.400E-08 |
| 10425 | 522 | C522C | A | 3.5e-08 | 3e-08 | 1.6e-07 | 7.500E-08 | 7.365E-08 | 3.000E-08 | 1.600E-07 |
| 10426 | 522 | C522E | A | 3.9e-08 | 5.1e-08 | 2.1e-08 | 3.700E-08 | 1.510E-08 | 2.100E-08 | 5.100E-08 |
| 10427 | 522 | C522Q | A | 4.5e-08 | 3.4e-08 | 6.4e-08 | 4.767E-08 | 1.518E-08 | 3.400E-08 | 6.400E-08 |
| 10428 | 522 | C522G | A | 2.3e-08 | 1.9e-08 | 6.2e-08 | 3.467E-08 | 2.376E-08 | 1.900E-08 | 6.200E-08 |
| 10429 | 522 | C522H | A | 4e-08 | 1.4e-07 | 8.4e-08 | 8.800E-08 | 5.012E-08 | 4.000E-08 | 1.400E-07 |
| 10430 | 522 | C522I | A | 5.5e-08 | 4e-08 | 6.2e-08 | 5.233E-08 | 1.124E-08 | 4.000E-08 | 6.200E-08 |
| 10431 | 522 | C522L | A | 6e-08 | 3e-08 | 1e-07 | 6.333E-08 | 3.512E-08 | 3.000E-08 | 1.000E-07 |
| 10432 | 522 | C522K | A | 7.4e-08 | 3.2e-08 | 6.9e-08 | 5.833E-08 | 2.294E-08 | 3.200E-08 | 7.400E-08 |
| 10433 | 522 | C522M | A | 6.3e-08 | 4.5e-08 | 3.5e-08 | 4.767E-08 | 1.419E-08 | 3.500E-08 | 6.300E-08 |
| 10434 | 522 | C522F | A | 3.8e-08 | 2.6e-07 | 4.3e-08 | 1.137E-07 | 1.268E-07 | 3.800E-08 | 2.600E-07 |
| 10435 | 522 | C522P | A | 8.3e-08 | 1e-07 | 7.5e-08 | 8.600E-08 | 1.277E-08 | 7.500E-08 | 1.000E-07 |
| 10436 | 522 | C522S | A | 7.5e-08 | 6.2e-08 | 1.1e-08 | 4.933E-08 | 3.383E-08 | 1.100E-08 | 7.500E-08 |
| 10437 | 522 | C522T | A | 3.9e-08 | 2e-08 | 4.1e-08 | 3.333E-08 | 1.159E-08 | 2.000E-08 | 4.100E-08 |
| 10438 | 522 | C522W | A | 1.3e-08 | 3e-08 | 2.7e-08 | 2.333E-08 | 9.074E-09 | 1.300E-08 | 3.000E-08 |
| 10439 | 522 | C522Y | A | 9.6e-08 | 3.5e-08 | 3.9e-08 | 5.667E-08 | 3.412E-08 | 3.500E-08 | 9.600E-08 |
| 10440 | 522 | C522V | A | 4.1e-08 | 2e-08 | 4.9e-08 | 3.667E-08 | 1.498E-08 | 2.000E-08 | 4.900E-08 |
| 10441 | 523 | R523A | A | 4.9e-08 | 3.1e-08 | 3e-08 | 3.667E-08 | 1.069E-08 | 3.000E-08 | 4.900E-08 |
| 10442 | 523 | R523R | A | 4.1e-08 | 9.1e-08 | 5.8e-08 | 6.333E-08 | 2.542E-08 | 4.100E-08 | 9.100E-08 |
| 10443 | 523 | R523N | A | 2.2e-08 | 3.6e-08 | 5.6e-08 | 3.800E-08 | 1.709E-08 | 2.200E-08 | 5.600E-08 |
| 10444 | 523 | R523D | A | 4.1e-08 | 4.7e-08 | 7.3e-08 | 5.367E-08 | 1.701E-08 | 4.100E-08 | 7.300E-08 |
| 10445 | 523 | R523C | A | 3.2e-08 | 4.7e-08 | 8.6e-08 | 5.500E-08 | 2.787E-08 | 3.200E-08 | 8.600E-08 |
| 10446 | 523 | R523E | A | 5.3e-08 | 3.4e-08 | 5.5e-08 | 4.733E-08 | 1.159E-08 | 3.400E-08 | 5.500E-08 |
| 10447 | 523 | R523Q | A | 1.2e-08 | 3.2e-08 | 4.3e-08 | 2.900E-08 | 1.572E-08 | 1.200E-08 | 4.300E-08 |
| 10448 | 523 | R523G | A | 1.2e-07 | 5.2e-08 | 8.2e-08 | 8.467E-08 | 3.408E-08 | 5.200E-08 | 1.200E-07 |
| 10449 | 523 | R523H | A | 4.2e-08 | 6.2e-08 | 2.6e-08 | 4.333E-08 | 1.804E-08 | 2.600E-08 | 6.200E-08 |
| 10450 | 523 | R523I | A | 1.9e-08 | 4.7e-08 | 4e-08 | 3.533E-08 | 1.457E-08 | 1.900E-08 | 4.700E-08 |
| 10451 | 523 | R523L | A | 2.2e-08 | 4.4e-08 | 2e-08 | 2.867E-08 | 1.332E-08 | 2.000E-08 | 4.400E-08 |
| 10452 | 523 | R523K | A | 5.1e-08 | 7.2e-08 | 4.1e-08 | 5.467E-08 | 1.582E-08 | 4.100E-08 | 7.200E-08 |
| 10453 | 523 | R523M | A | 1e-07 | 3.2e-08 | 3.7e-08 | 5.633E-08 | 3.790E-08 | 3.200E-08 | 1.000E-07 |
| 10454 | 523 | R523F | A | 5e-08 | 7.2e-08 | 8.4e-08 | 6.867E-08 | 1.724E-08 | 5.000E-08 | 8.400E-08 |

| | | | | | | | | | | |
|---|---|---|---|---|---|---|---|---|---|---|
| 10455 | 523 | R523P | A | 2.3e-08 | 7e-08 | 3.7e-08 | 4.333E-08 | 2.413E-08 | 2.300E-08 | 7.000E-08 |
| 10456 | 523 | R523S | A | 3.3e-08 | 5.5e-08 | 3.4e-08 | 4.067E-08 | 1.242E-08 | 3.300E-08 | 5.500E-08 |
| 10457 | 523 | R523T | A | 4.1e-08 | 3.1e-08 | 1e-07 | 5.733E-08 | 3.729E-08 | 3.100E-08 | 1.000E-07 |
| 10458 | 523 | R523W | A | 9.1e-08 | 1.9e-08 | 1e-08 | 4.000E-08 | 4.440E-08 | 1.000E-08 | 9.100E-08 |
| 10459 | 523 | R523Y | A | 5.3e-08 | 6.1e-08 | 7.5e-08 | 6.300E-08 | 1.114E-08 | 5.300E-08 | 7.500E-08 |
| 10460 | 523 | R523V | A | 4.6e-08 | 3.4e-08 | 2.4e-08 | 3.467E-08 | 1.102E-08 | 2.400E-08 | 4.600E-08 |
| 10461 | 524 | V524A | A | 6.1e-08 | 7.2e-08 | 2.3e-08 | 5.200E-08 | 2.571E-08 | 2.300E-08 | 7.200E-08 |
| 10462 | 524 | V524R | A | 1.5e-07 | 3.4e-08 | 1.3e-07 | 1.047E-07 | 6.201E-08 | 3.400E-08 | 1.500E-07 |
| 10463 | 524 | V524N | A | 4.3e-08 | 2.6e-08 | 1.4e-08 | 2.767E-08 | 1.457E-08 | 1.400E-08 | 4.300E-08 |
| 10464 | 524 | V524D | A | 3.6e-08 | 2.5e-08 | 1.6e-08 | 2.567E-08 | 1.002E-08 | 1.600E-08 | 3.600E-08 |
| 10465 | 524 | V524C | A | 7.4e-08 | 6.1e-08 | 2.2e-08 | 5.233E-08 | 2.706E-08 | 2.200E-08 | 7.400E-08 |
| 10466 | 524 | V524E | A | 2.6e-08 | 4.1e-08 | 8.1e-08 | 4.933E-08 | 2.843E-08 | 2.600E-08 | 8.100E-08 |
| 10467 | 524 | V524Q | A | 2.5e-08 | 1.4e-07 | 3.3e-08 | 6.600E-08 | 6.421E-08 | 2.500E-08 | 1.400E-07 |
| 10468 | 524 | V524G | A | 8.6e-08 | 3.7e-08 | 3.8e-08 | 5.367E-08 | 2.801E-08 | 3.700E-08 | 8.600E-08 |
| 10469 | 524 | V524H | A | 2.6e-08 | 5.1e-08 | 4.5e-08 | 4.067E-08 | 1.305E-08 | 2.600E-08 | 5.100E-08 |
| 10470 | 524 | V524I | A | 4.6e-08 | 3.6e-08 | 2.2e-08 | 3.467E-08 | 1.206E-08 | 2.200E-08 | 4.600E-08 |
| 10471 | 524 | V524L | A | 3.9e-08 | 5.3e-08 | 6e-08 | 5.067E-08 | 1.069E-08 | 3.900E-08 | 6.000E-08 |
| 10472 | 524 | V524K | A | 6.9e-08 | 7.2e-08 | 2.1e-08 | 5.400E-08 | 2.862E-08 | 2.100E-08 | 7.200E-08 |
| 10473 | 524 | V524M | A | 7.4e-08 | 9.1e-08 | 2.9e-08 | 6.467E-08 | 3.204E-08 | 2.900E-08 | 9.100E-08 |
| 10474 | 524 | V524F | A | 7.8e-09 | 1.5e-07 | 4.3e-08 | 6.693E-08 | 7.406E-08 | 7.800E-09 | 1.500E-07 |
| 10475 | 524 | V524P | A | 1.1e-07 | 7.1e-08 | 3.4e-08 | 7.167E-08 | 3.800E-08 | 3.400E-08 | 1.100E-07 |
| 10476 | 524 | V524S | A | 2.8e-08 | 1.6e-08 | 6.4e-08 | 3.600E-08 | 2.498E-08 | 1.600E-08 | 6.400E-08 |
| 10477 | 524 | V524T | A | 4.1e-08 | 3.8e-08 | 1.6e-08 | 3.167E-08 | 1.365E-08 | 1.600E-08 | 4.100E-08 |
| 10478 | 524 | V524W | A | 3.1e-08 | 5.4e-08 | 2.3e-08 | 3.600E-08 | 1.609E-08 | 2.300E-08 | 5.400E-08 |
| 10479 | 524 | V524Y | A | 3.8e-08 | 2.3e-08 | 5.7e-08 | 3.933E-08 | 1.704E-08 | 2.300E-08 | 5.700E-08 |
| 10480 | 524 | V524V | A | 2.6e-08 | 1.7e-08 | 6.4e-08 | 3.567E-08 | 2.495E-08 | 1.700E-08 | 6.400E-08 |
| 10481 | 525 | L525A | A | 6.6e-08 | 3e-08 | 3.3e-08 | 4.300E-08 | 1.997E-08 | 3.000E-08 | 6.600E-08 |
| 10482 | 525 | L525R | A | 5e-08 | 9e-08 | 8.1e-08 | 7.367E-08 | 2.098E-08 | 5.000E-08 | 9.000E-08 |
| 10483 | 525 | L525N | A | 3e-08 | 3.3e-08 | 1.9e-08 | 2.733E-08 | 7.371E-09 | 1.900E-08 | 3.300E-08 |
| 10484 | 525 | L525D | A | 6.9e-08 | 5.4e-08 | 4.3e-08 | 5.533E-08 | 1.305E-08 | 4.300E-08 | 6.900E-08 |
| 10485 | 525 | L525C | A | 8.7e-08 | 3.9e-08 | 1.7e-07 | 9.867E-08 | 6.627E-08 | 3.900E-08 | 1.700E-07 |
| 10486 | 525 | L525E | A | 3.7e-08 | 7e-08 | 2.9e-08 | 4.533E-08 | 2.173E-08 | 2.900E-08 | 7.000E-08 |
| 10487 | 525 | L525Q | A | 3.8e-08 | 3.1e-08 | 3e-08 | 3.300E-08 | 4.359E-09 | 3.000E-08 | 3.800E-08 |
| 10488 | 525 | L525G | A | 1.9e-07 | 2.3e-08 | 4.9e-08 | 8.733E-08 | 8.986E-08 | 2.300E-08 | 1.900E-07 |
| 10489 | 525 | L525H | A | 6e-08 | 1.1e-07 | 1.1e-07 | 9.333E-08 | 2.887E-08 | 6.000E-08 | 1.100E-07 |
| 10490 | 525 | L525I | A | 6.1e-08 | 2.6e-08 | 2.7e-08 | 3.800E-08 | 1.992E-08 | 2.600E-08 | 6.100E-08 |
| 10491 | 525 | L525L | A | 8.7e-08 | 2.9e-08 | 3e-08 | 4.867E-08 | 3.320E-08 | 2.900E-08 | 8.700E-08 |
| 10492 | 525 | L525K | A | 1.1e-07 | 1.9e-08 | 4.2e-08 | 5.700E-08 | 4.732E-08 | 1.900E-08 | 1.100E-07 |
| 10493 | 525 | L525M | A | 1.8e-08 | 5.2e-08 | 5.7e-08 | 4.233E-08 | 2.122E-08 | 1.800E-08 | 5.700E-08 |
| 10494 | 525 | L525F | A | 8e-08 | 2.5e-08 | 2.1e-08 | 4.200E-08 | 3.297E-08 | 2.100E-08 | 8.000E-08 |
| 10495 | 525 | L525P | A | 4.1e-08 | 3.8e-08 | 2.8e-08 | 3.567E-08 | 6.807E-09 | 2.800E-08 | 4.100E-08 |
| 10496 | 525 | L525S | A | 2.3e-08 | 1.8e-08 | 2.4e-08 | 2.167E-08 | 3.215E-09 | 1.800E-08 | 2.400E-08 |
| 10497 | 525 | L525T | A | 1.1e-07 | 3.9e-08 | 9.5e-08 | 8.133E-08 | 3.742E-08 | 3.900E-08 | 1.100E-07 |
| 10498 | 525 | L525W | A | 3.4e-08 | 3.9e-08 | 1.6e-08 | 2.967E-08 | 1.210E-08 | 1.600E-08 | 3.900E-08 |
| 10499 | 525 | L525Y | A | 4.5e-08 | 7.7e-08 | 2.6e-08 | 4.933E-08 | 2.577E-08 | 2.600E-08 | 7.700E-08 |
| 10500 | 525 | L525V | A | 9.7e-08 | 3e-08 | 2e-08 | 4.900E-08 | 4.187E-08 | 2.000E-08 | 9.700E-08 |
| 10501 | 526 | Q526A | A | 6.3e-08 | 7.1e-08 | 8.1e-08 | 7.167E-08 | 9.018E-09 | 6.300E-08 | 8.100E-08 |
| 10502 | 526 | Q526R | A | 8.5e-08 | 3.1e-08 | 1.1e-07 | 7.533E-08 | 4.038E-08 | 3.100E-08 | 1.100E-07 |
| 10503 | 526 | Q526N | A | 2.3e-08 | 7e-08 | 6.2e-09 | 3.307E-08 | 3.307E-08 | 6.200E-09 | 7.000E-08 |
| 10504 | 526 | Q526D | A | 1.2e-08 | 7.5e-08 | 6.5e-08 | 5.067E-08 | 3.386E-08 | 1.200E-08 | 7.500E-08 |
| 10505 | 526 | Q526C | A | 6.2e-08 | 4.2e-08 | 4.5e-08 | 4.967E-08 | 1.079E-08 | 4.200E-08 | 6.200E-08 |
| 10506 | 526 | Q526E | A | 1.3e-07 | 5.4e-08 | 2.7e-08 | 7.033E-08 | 5.341E-08 | 2.700E-08 | 1.300E-07 |
| 10507 | 526 | Q526Q | A | 2.2e-08 | 9.6e-08 | 5.1e-08 | 5.633E-08 | 3.729E-08 | 2.200E-08 | 9.600E-08 |
| 10508 | 526 | Q526G | A | 2.6e-08 | 1.3e-07 | 8.9e-08 | 8.167E-08 | 5.239E-08 | 2.600E-08 | 1.300E-07 |
| 10509 | 526 | Q526H | A | 1.1e-07 | 7.6e-08 | 4e-08 | 7.533E-08 | 3.500E-08 | 4.000E-08 | 1.100E-07 |
| 10510 | 526 | Q526I | A | 3.6e-08 | 9.8e-08 | 4.5e-08 | 5.967E-08 | 3.350E-08 | 3.600E-08 | 9.800E-08 |
| 10511 | 526 | Q526L | A | 3.6e-08 | 1.6e-08 | 7.1e-08 | 4.100E-08 | 2.784E-08 | 1.600E-08 | 7.100E-08 |
| 10512 | 526 | Q526K | A | 2.4e-08 | 6.2e-08 | 2e-08 | 3.533E-08 | 2.318E-08 | 2.000E-08 | 6.200E-08 |
| 10513 | 526 | Q526M | A | 8.7e-08 | 3.8e-08 | 2e-08 | 4.833E-08 | 3.467E-08 | 2.000E-08 | 8.700E-08 |
| 10514 | 526 | Q526F | A | 2.8e-08 | 8.7e-08 | 2.3e-08 | 4.600E-08 | 3.559E-08 | 2.300E-08 | 8.700E-08 |
| 10515 | 526 | Q526P | A | 2e-08 | 3.6e-08 | 3.4e-08 | 3.000E-08 | 8.718E-09 | 2.000E-08 | 3.600E-08 |
| 10516 | 526 | Q526S | A | 7e-08 | 5.2e-08 | 3.3e-08 | 5.167E-08 | 1.850E-08 | 3.300E-08 | 7.000E-08 |
| 10517 | 526 | Q526T | A | 1.1e-07 | 5.9e-08 | 6.4e-08 | 7.767E-08 | 2.811E-08 | 5.900E-08 | 1.100E-07 |
| 10518 | 526 | Q526W | A | 6e-08 | 5.8e-08 | 2.8e-08 | 4.867E-08 | 1.793E-08 | 2.800E-08 | 6.000E-08 |
| 10519 | 526 | Q526Y | A | 2.1e-08 | 5.8e-08 | 6e-08 | 4.633E-08 | 2.196E-08 | 2.100E-08 | 6.000E-08 |
| 10520 | 526 | Q526V | A | 1.1e-07 | 2e-08 | 3.3e-08 | 5.433E-08 | 4.864E-08 | 2.000E-08 | 1.100E-07 |
| 10521 | 527 | G527A | A | 2.8e-08 | 1.6e-07 | 4.9e-08 | 7.900E-08 | 7.093E-08 | 2.800E-08 | 1.600E-07 |
| 10522 | 527 | G527R | A | 4.9e-08 | 2.4e-08 | 3.5e-08 | 3.600E-08 | 1.253E-08 | 2.400E-08 | 4.900E-08 |
| 10523 | 527 | G527N | A | 1.7e-08 | 9.9e-08 | 6.9e-08 | 6.167E-08 | 4.149E-08 | 1.700E-08 | 9.900E-08 |
| 10524 | 527 | G527D | A | 7e-08 | 3.7e-08 | 2.9e-08 | 4.533E-08 | 2.173E-08 | 2.900E-08 | 7.000E-08 |
| 10525 | 527 | G527C | A | 3e-08 | 2.8e-08 | 3.5e-08 | 3.100E-08 | 3.606E-09 | 2.800E-08 | 3.500E-08 |
| 10526 | 527 | G527E | A | 7.9e-08 | 1.8e-08 | 3e-08 | 4.233E-08 | 3.232E-08 | 1.800E-08 | 7.900E-08 |
| 10527 | 527 | G527Q | A | 2.9e-08 | 4.3e-08 | 4.6e-08 | 3.933E-08 | 9.074E-09 | 2.900E-08 | 4.600E-08 |
| 10528 | 527 | G527G | A | 1.2e-07 | 4.2e-08 | 6.4e-08 | 7.533E-08 | 4.022E-08 | 4.200E-08 | 1.200E-07 |
| 10529 | 527 | G527H | A | 1.4e-08 | 5.1e-08 | 2.9e-08 | 3.133E-08 | 1.861E-08 | 1.400E-08 | 5.100E-08 |
| 10530 | 527 | G527I | A | 7.2e-08 | 2.9e-08 | 3e-08 | 4.367E-08 | 2.454E-08 | 2.900E-08 | 7.200E-08 |
| 10531 | 527 | G527L | A | 8.3e-08 | 2.3e-08 | 8.9e-08 | 6.500E-08 | 3.650E-08 | 2.300E-08 | 8.900E-08 |
| 10532 | 527 | G527K | A | 2.6e-08 | 1.2e-07 | 8.1e-08 | 7.567E-08 | 4.723E-08 | 2.600E-08 | 1.200E-07 |
| 10533 | 527 | G527M | A | 3.5e-08 | 5.3e-08 | 1.4e-07 | 7.600E-08 | 5.615E-08 | 3.500E-08 | 1.400E-07 |
| 10534 | 527 | G527F | A | 5.9e-08 | 7.8e-08 | 5.2e-08 | 6.300E-08 | 1.345E-08 | 5.200E-08 | 7.800E-08 |
| 10535 | 527 | G527P | A | 7.3e-08 | 2.1e-08 | 4.2e-08 | 4.533E-08 | 2.616E-08 | 2.100E-08 | 7.300E-08 |
| 10536 | 527 | G527S | A | 3.2e-08 | 3.9e-08 | 2.9e-08 | 3.333E-08 | 5.132E-09 | 2.900E-08 | 3.900E-08 |
| 10537 | 527 | G527T | A | 1e-07 | 4.9e-08 | 3.4e-08 | 6.100E-08 | 3.460E-08 | 3.400E-08 | 1.000E-07 |
| 10538 | 527 | G527W | A | 4.9e-08 | 4.4e-08 | 8.1e-08 | 5.800E-08 | 2.007E-08 | 4.400E-08 | 8.100E-08 |
| 10539 | 527 | G527Y | A | 4e-08 | 2.4e-08 | 5.2e-08 | 3.867E-08 | 1.405E-08 | 2.400E-08 | 5.200E-08 |

| | | | | | | | | | | |
|---|---|---|---|---|---|---|---|---|---|---|
| 10540 | 527 | G527V | A | 4.7e-08 | 3.3e-08 | 2.2e-08 | 3.400E-08 | 1.253E-08 | 2.200E-08 | 4.700E-08 |
| 10541 | 528 | L528A | A | 2.3e-08 | 3e-08 | 1.7e-08 | 2.333E-08 | 6.506E-09 | 1.700E-08 | 3.000E-08 |
| 10542 | 528 | L528R | A | 2.6e-08 | 2.4e-08 | 2.1e-08 | 2.367E-08 | 2.517E-09 | 2.100E-08 | 2.600E-08 |
| 10543 | 528 | L528N | A | 1.4e-08 | 5.7e-08 | 1.8e-08 | 2.967E-08 | 2.376E-08 | 1.400E-08 | 5.700E-08 |
| 10544 | 528 | L528D | A | 2.6e-08 | 2.4e-08 | 5.9e-08 | 3.633E-08 | 1.966E-08 | 2.400E-08 | 5.900E-08 |
| 10545 | 528 | L528C | A | 1.8e-07 | 4.5e-08 | 3.8e-08 | 8.767E-08 | 8.004E-08 | 3.800E-08 | 1.800E-07 |
| 10546 | 528 | L528E | A | 1.1e-07 | 4.5e-08 | 3.4e-08 | 6.300E-08 | 4.107E-08 | 3.400E-08 | 1.100E-07 |
| 10547 | 528 | L528Q | A | 3.8e-08 | 2.2e-08 | 4.7e-08 | 3.567E-08 | 1.266E-08 | 2.200E-08 | 4.700E-08 |
| 10548 | 528 | L528G | A | 8.1e-08 | 7.6e-08 | 4e-08 | 6.567E-08 | 2.237E-08 | 4.000E-08 | 8.100E-08 |
| 10549 | 528 | L528H | A | 5.1e-08 | 5.2e-08 | 1.9e-08 | 4.067E-08 | 1.877E-08 | 1.900E-08 | 5.200E-08 |
| 10550 | 528 | L528I | A | 9.7e-08 | 3.4e-08 | 7.3e-08 | 6.800E-08 | 3.180E-08 | 3.400E-08 | 9.700E-08 |
| 10551 | 528 | L528L | A | 3.7e-08 | 6.9e-08 | 3e-08 | 4.533E-08 | 2.079E-08 | 3.000E-08 | 6.900E-08 |
| 10552 | 528 | L528K | A | 3.8e-08 | 4.9e-08 | 4.1e-08 | 4.267E-08 | 5.686E-09 | 3.800E-08 | 4.900E-08 |
| 10553 | 528 | L528M | A | 7.1e-08 | 4.1e-08 | 4.6e-08 | 5.267E-08 | 1.607E-08 | 4.100E-08 | 7.100E-08 |
| 10554 | 528 | L528F | A | 2.1e-08 | 8e-08 | 3.2e-08 | 4.433E-08 | 3.137E-08 | 2.100E-08 | 8.000E-08 |
| 10555 | 528 | L528P | A | 1.8e-08 | 2.3e-08 | 4.6e-08 | 2.900E-08 | 1.493E-08 | 1.800E-08 | 4.600E-08 |
| 10556 | 528 | L528S | A | 3.6e-08 | 6.7e-08 | 4e-08 | 4.767E-08 | 1.686E-08 | 3.600E-08 | 6.700E-08 |
| 10557 | 528 | L528T | A | 1e-07 | 7.2e-08 | 1.3e-07 | 1.007E-07 | 2.901E-08 | 7.200E-08 | 1.300E-07 |
| 10558 | 528 | L528W | A | 2.4e-08 | 4.8e-08 | 2.5e-08 | 3.233E-08 | 1.358E-08 | 2.400E-08 | 4.800E-08 |
| 10559 | 528 | L528Y | A | 4e-08 | 4.1e-08 | 1e-07 | 6.033E-08 | 3.436E-08 | 4.000E-08 | 1.000E-07 |
| 10560 | 528 | L528V | A | 1.1e-07 | 4.4e-08 | 9.7e-08 | 8.367E-08 | 3.496E-08 | 4.400E-08 | 1.100E-07 |
| 10561 | 529 | P529A | A | 8.5e-08 | 2.6e-08 | 8.3e-08 | 6.467E-08 | 3.350E-08 | 2.600E-08 | 8.500E-08 |
| 10562 | 529 | P529R | A | 2.1e-08 | 3.4e-08 | 4.3e-08 | 3.267E-08 | 1.106E-08 | 2.100E-08 | 4.300E-08 |
| 10563 | 529 | P529N | A | 1.1e-07 | 2.9e-08 | 3.9e-08 | 5.933E-08 | 4.416E-08 | 2.900E-08 | 1.100E-07 |
| 10564 | 529 | P529D | A | 3.3e-08 | 3.9e-08 | 6e-08 | 4.400E-08 | 1.418E-08 | 3.300E-08 | 6.000E-08 |
| 10565 | 529 | P529C | A | 2.9e-08 | 2.2e-08 | 3.7e-08 | 2.933E-08 | 7.506E-09 | 2.200E-08 | 3.700E-08 |
| 10566 | 529 | P529E | A | 4.2e-08 | 3.9e-08 | 4.7e-08 | 4.267E-08 | 4.041E-09 | 3.900E-08 | 4.700E-08 |
| 10567 | 529 | P529Q | A | 7.7e-08 | 7.4e-08 | 3.6e-08 | 6.233E-08 | 2.285E-08 | 3.600E-08 | 7.700E-08 |
| 10568 | 529 | P529G | A | 2.5e-08 | 5.2e-08 | 7.4e-08 | 5.033E-08 | 2.454E-08 | 2.500E-08 | 7.400E-08 |
| 10569 | 529 | P529H | A | 5.2e-08 | 6.1e-08 | 5.2e-08 | 5.500E-08 | 5.196E-09 | 5.200E-08 | 6.100E-08 |
| 10570 | 529 | P529I | A | 4e-08 | 8.6e-08 | 5.5e-08 | 6.033E-08 | 2.346E-08 | 4.000E-08 | 8.600E-08 |
| 10571 | 529 | P529L | A | 4.7e-08 | 2.1e-08 | 1.1e-07 | 5.933E-08 | 4.576E-08 | 2.100E-08 | 1.100E-07 |
| 10572 | 529 | P529K | A | 3e-08 | 4.4e-08 | 6.9e-08 | 4.767E-08 | 1.976E-08 | 3.000E-08 | 6.900E-08 |
| 10573 | 529 | P529M | A | 7.7e-08 | 4e-08 | 4.8e-08 | 5.500E-08 | 1.947E-08 | 4.000E-08 | 7.700E-08 |
| 10574 | 529 | P529F | A | 3.2e-08 | 4.5e-08 | 4.2e-08 | 3.967E-08 | 6.807E-09 | 3.200E-08 | 4.500E-08 |
| 10575 | 529 | P529P | A | 4.4e-08 | 4.6e-08 | 1.2e-07 | 7.000E-08 | 4.331E-08 | 4.400E-08 | 1.200E-07 |
| 10576 | 529 | P529S | A | 8.9e-08 | 1.4e-08 | 2.8e-08 | 4.367E-08 | 3.988E-08 | 1.400E-08 | 8.900E-08 |
| 10577 | 529 | P529T | A | 6.1e-08 | 1.4e-08 | 3.3e-08 | 3.600E-08 | 2.364E-08 | 1.400E-08 | 6.100E-08 |
| 10578 | 529 | P529W | A | 7.2e-08 | 5.3e-08 | 1.6e-08 | 4.700E-08 | 2.848E-08 | 1.600E-08 | 7.200E-08 |
| 10579 | 529 | P529Y | A | 5.6e-08 | 4.9e-08 | 2.8e-08 | 4.433E-08 | 1.457E-08 | 2.800E-08 | 5.600E-08 |
| 10580 | 529 | P529V | A | 3e-08 | 6.1e-08 | 2.5e-08 | 3.867E-08 | 1.950E-08 | 2.500E-08 | 6.100E-08 |
| 10581 | 530 | R530A | A | 6.5e-08 | 1.1e-07 | 1.7e-08 | 6.400E-08 | 4.651E-08 | 1.700E-08 | 1.100E-07 |
| 10582 | 530 | R530R | A | 9.3e-08 | 5.1e-08 | 2.6e-08 | 5.667E-08 | 3.386E-08 | 2.600E-08 | 9.300E-08 |
| 10583 | 530 | R530N | A | 4.7e-08 | 4.7e-08 | 3.9e-08 | 4.433E-08 | 4.619E-09 | 3.900E-08 | 4.700E-08 |
| 10584 | 530 | R530D | A | 3.2e-08 | 2.8e-08 | 8.6e-08 | 4.867E-08 | 3.239E-08 | 2.800E-08 | 8.600E-08 |
| 10585 | 530 | R530C | A | 5.5e-08 | 2e-08 | 9.6e-08 | 5.700E-08 | 3.804E-08 | 2.000E-08 | 9.600E-08 |
| 10586 | 530 | R530E | A | 9.1e-08 | 4.8e-08 | 5.3e-08 | 6.400E-08 | 2.352E-08 | 4.800E-08 | 9.100E-08 |
| 10587 | 530 | R530Q | A | 1.3e-08 | 3e-08 | 2.2e-08 | 2.167E-08 | 8.505E-09 | 1.300E-08 | 3.000E-08 |
| 10588 | 530 | R530G | A | 2.8e-08 | 4e-08 | 5e-08 | 3.933E-08 | 1.102E-08 | 2.800E-08 | 5.000E-08 |
| 10589 | 530 | R530H | A | 1.3e-08 | 3.7e-08 | 3.4e-08 | 2.800E-08 | 1.308E-08 | 1.300E-08 | 3.700E-08 |
| 10590 | 530 | R530I | A | 4.7e-08 | 8.2e-08 | 3.3e-08 | 5.400E-08 | 2.524E-08 | 3.300E-08 | 8.200E-08 |
| 10591 | 530 | R530L | A | 5.2e-08 | 2.9e-08 | 3.4e-08 | 3.833E-08 | 1.210E-08 | 2.900E-08 | 5.200E-08 |
| 10592 | 530 | R530K | A | 6.9e-08 | 4.9e-08 | 1.2e-07 | 7.933E-08 | 3.661E-08 | 4.900E-08 | 1.200E-07 |
| 10593 | 530 | R530M | A | 9.4e-08 | 9.1e-09 | 1.5e-08 | 3.937E-08 | 4.741E-08 | 9.100E-09 | 9.400E-08 |
| 10594 | 530 | R530F | A | 2.1e-08 | 8.5e-08 | 5.7e-08 | 5.433E-08 | 3.208E-08 | 2.100E-08 | 8.500E-08 |
| 10595 | 530 | R530P | A | 3.6e-08 | 4.3e-08 | 1.1e-07 | 6.300E-08 | 4.085E-08 | 3.600E-08 | 1.100E-07 |
| 10596 | 530 | R530S | A | 1.9e-08 | 4.9e-08 | 3.5e-08 | 3.433E-08 | 1.501E-08 | 1.900E-08 | 4.900E-08 |
| 10597 | 530 | R530T | A | 5.3e-08 | 6.6e-08 | 2.8e-08 | 4.900E-08 | 1.931E-08 | 2.800E-08 | 6.600E-08 |
| 10598 | 530 | R530W | A | 2.8e-08 | 5.9e-08 | 2.9e-08 | 3.867E-08 | 1.762E-08 | 2.800E-08 | 5.900E-08 |
| 10599 | 530 | R530Y | A | 9.3e-08 | 3.3e-08 | 2.5e-08 | 5.033E-08 | 3.717E-08 | 2.500E-08 | 9.300E-08 |
| 10600 | 530 | R530V | A | 3.1e-08 | 7e-08 | 1.1e-07 | 7.033E-08 | 3.950E-08 | 3.100E-08 | 1.100E-07 |
| 10601 | 531 | E531A | A | 4.8e-08 | 8.8e-08 | 8.3e-08 | 7.300E-08 | 2.179E-08 | 4.800E-08 | 8.800E-08 |
| 10602 | 531 | E531R | A | 2.8e-08 | 2.3e-08 | 2.8e-08 | 2.633E-08 | 2.887E-09 | 2.300E-08 | 2.800E-08 |
| 10603 | 531 | E531N | A | 4e-08 | 1.5e-08 | 7.3e-08 | 4.267E-08 | 2.909E-08 | 1.500E-08 | 7.300E-08 |
| 10604 | 531 | E531D | A | 3.1e-08 | 5.4e-08 | 3.8e-08 | 4.100E-08 | 1.179E-08 | 3.100E-08 | 5.400E-08 |
| 10605 | 531 | E531C | A | 4.2e-08 | 3e-08 | 5.1e-08 | 4.100E-08 | 1.054E-08 | 3.000E-08 | 5.100E-08 |
| 10606 | 531 | E531E | A | 1.8e-08 | 2.3e-08 | 3.5e-08 | 2.533E-08 | 8.737E-09 | 1.800E-08 | 3.500E-08 |
| 10607 | 531 | E531Q | A | 2.8e-08 | 7.1e-08 | 6.7e-08 | 5.533E-08 | 2.376E-08 | 2.800E-08 | 7.100E-08 |
| 10608 | 531 | E531G | A | 4.8e-08 | 9.2e-08 | 8.6e-08 | 7.533E-08 | 2.386E-08 | 4.800E-08 | 9.200E-08 |
| 10609 | 531 | E531H | A | 7.2e-08 | 4.5e-08 | 8.2e-08 | 6.633E-08 | 1.914E-08 | 4.500E-08 | 8.200E-08 |
| 10610 | 531 | E531I | A | 3.2e-08 | 3.9e-08 | 3.4e-08 | 3.500E-08 | 3.606E-09 | 3.200E-08 | 3.900E-08 |
| 10611 | 531 | E531L | A | 5.9e-08 | 1.6e-08 | 2.6e-08 | 3.367E-08 | 2.250E-08 | 1.600E-08 | 5.900E-08 |
| 10612 | 531 | E531K | A | 3.3e-08 | 3.2e-08 | 5.8e-08 | 4.100E-08 | 1.473E-08 | 3.200E-08 | 5.800E-08 |
| 10613 | 531 | E531M | A | 3.3e-08 | 2e-08 | 7.7e-08 | 4.333E-08 | 2.987E-08 | 2.000E-08 | 7.700E-08 |
| 10614 | 531 | E531F | A | 9.5e-08 | 7.3e-08 | 3.9e-08 | 6.900E-08 | 2.821E-08 | 3.900E-08 | 9.500E-08 |
| 10615 | 531 | E531P | A | 4.8e-08 | 2.8e-08 | 3e-08 | 3.533E-08 | 1.102E-08 | 2.800E-08 | 4.800E-08 |
| 10616 | 531 | E531S | A | 1.4e-08 | 6.4e-08 | 5.7e-08 | 4.500E-08 | 2.707E-08 | 1.400E-08 | 6.400E-08 |
| 10617 | 531 | E531T | A | 1.8e-08 | 4.6e-08 | 4.1e-08 | 3.500E-08 | 1.493E-08 | 1.800E-08 | 4.600E-08 |
| 10618 | 531 | E531W | A | 1.6e-08 | 2.1e-08 | 1.6e-08 | 1.767E-08 | 2.887E-09 | 1.600E-08 | 2.100E-08 |
| 10619 | 531 | E531Y | A | 1.9e-08 | 3.1e-08 | 7.8e-08 | 4.267E-08 | 3.118E-08 | 1.900E-08 | 7.800E-08 |
| 10620 | 531 | E531V | A | 9.9e-08 | 6.2e-08 | 1.9e-08 | 6.000E-08 | 4.004E-08 | 1.900E-08 | 9.900E-08 |
| 10621 | 532 | Y532A | A | 2.6e-08 | 4.1e-08 | 3e-08 | 3.233E-08 | 7.767E-09 | 2.600E-08 | 4.100E-08 |
| 10622 | 532 | Y532R | A | 2.1e-08 | 4.3e-08 | 3.4e-08 | 3.267E-08 | 1.106E-08 | 2.100E-08 | 4.300E-08 |
| 10623 | 532 | Y532N | A | 5.4e-08 | 3.1e-08 | 6e-08 | 4.833E-08 | 1.531E-08 | 3.100E-08 | 6.000E-08 |
| 10624 | 532 | Y532D | A | 8.7e-08 | 4.4e-08 | 6.1e-08 | 6.400E-08 | 2.166E-08 | 4.400E-08 | 8.700E-08 |

| | | | | | | | | | | |
|---|---|---|---|---|---|---|---|---|---|---|
| 10625 | 532 | Y532C | A | 4e-08 | 3.7e-08 | 3.6e-08 | 3.767E-08 | 2.082E-09 | 3.600E-08 | 4.000E-08 |
| 10626 | 532 | Y532E | A | 2.2e-08 | 1.2e-07 | 4.9e-08 | 6.367E-08 | 5.062E-08 | 2.200E-08 | 1.200E-07 |
| 10627 | 532 | Y532Q | A | 3.8e-08 | 9.1e-08 | 1.7e-08 | 4.867E-08 | 3.814E-08 | 1.700E-08 | 9.100E-08 |
| 10628 | 532 | Y532G | A | 2.7e-08 | 4.9e-08 | 7.6e-08 | 5.067E-08 | 2.454E-08 | 2.700E-08 | 7.600E-08 |
| 10629 | 532 | Y532H | A | 1.2e-07 | 4.8e-08 | 3.3e-08 | 6.700E-08 | 4.651E-08 | 3.300E-08 | 1.200E-07 |
| 10630 | 532 | Y532I | A | 7.4e-08 | 5.1e-08 | 4.1e-08 | 5.533E-08 | 1.692E-08 | 4.100E-08 | 7.400E-08 |
| 10631 | 532 | Y532L | A | 2.8e-08 | 5.4e-08 | 7.1e-08 | 5.100E-08 | 2.166E-08 | 2.800E-08 | 7.100E-08 |
| 10632 | 532 | Y532K | A | 2.8e-08 | 5.2e-08 | 1.1e-07 | 6.333E-08 | 4.216E-08 | 2.800E-08 | 1.100E-07 |
| 10633 | 532 | Y532M | A | 9.2e-09 | 3.4e-08 | 8.2e-08 | 4.173E-08 | 3.701E-08 | 9.200E-09 | 8.200E-08 |
| 10634 | 532 | Y532F | A | 5.5e-08 | 4.3e-08 | 4.7e-08 | 4.833E-08 | 6.110E-09 | 4.300E-08 | 5.500E-08 |
| 10635 | 532 | Y532P | A | 4.5e-08 | 5.2e-08 | 5.8e-08 | 5.167E-08 | 6.506E-09 | 4.500E-08 | 5.800E-08 |
| 10636 | 532 | Y532S | A | 3.6e-08 | 2.7e-08 | 6.5e-08 | 4.267E-08 | 1.986E-08 | 2.700E-08 | 6.500E-08 |
| 10637 | 532 | Y532T | A | 3.5e-08 | 7e-08 | 6.9e-08 | 5.800E-08 | 1.992E-08 | 3.500E-08 | 7.000E-08 |
| 10638 | 532 | Y532W | A | 1.6e-08 | 1.8e-08 | 8.7e-08 | 4.033E-08 | 4.043E-08 | 1.600E-08 | 8.700E-08 |
| 10639 | 532 | Y532Y | A | 9.9e-08 | 3.4e-08 | 3.8e-08 | 5.700E-08 | 3.643E-08 | 3.400E-08 | 9.900E-08 |
| 10640 | 532 | Y532V | A | 9.8e-08 | 5.3e-08 | 3.2e-08 | 6.100E-08 | 3.372E-08 | 3.200E-08 | 9.800E-08 |
| 10641 | 533 | V533A | A | 5.5e-08 | 5.8e-08 | 3e-08 | 4.767E-08 | 1.537E-08 | 3.000E-08 | 5.800E-08 |
| 10642 | 533 | V533R | A | 4.4e-08 | 3.3e-08 | 9.3e-08 | 5.667E-08 | 3.194E-08 | 3.300E-08 | 9.300E-08 |
| 10643 | 533 | V533N | A | 4.6e-08 | 2e-08 | 2.7e-08 | 3.100E-08 | 1.345E-08 | 2.000E-08 | 4.600E-08 |
| 10644 | 533 | V533D | A | 2.1e-08 | 2.2e-08 | 2.6e-08 | 2.300E-08 | 2.646E-09 | 2.100E-08 | 2.600E-08 |
| 10645 | 533 | V533C | A | 1.6e-07 | 3.1e-08 | 4.4e-08 | 7.833E-08 | 7.102E-08 | 3.100E-08 | 1.600E-07 |
| 10646 | 533 | V533E | A | 2.5e-08 | 3.7e-08 | 1.4e-07 | 6.733E-08 | 6.322E-08 | 2.500E-08 | 1.400E-07 |
| 10647 | 533 | V533Q | A | 7.4e-08 | 1.3e-08 | 3.4e-08 | 4.033E-08 | 3.099E-08 | 1.300E-08 | 7.400E-08 |
| 10648 | 533 | V533G | A | 1.8e-08 | 2.2e-07 | 1.2e-07 | 1.193E-07 | 1.010E-07 | 1.800E-08 | 2.200E-07 |
| 10649 | 533 | V533H | A | 4.9e-08 | 5e-08 | 8.7e-08 | 6.200E-08 | 2.166E-08 | 4.900E-08 | 8.700E-08 |
| 10650 | 533 | V533I | A | 7.1e-08 | 3e-08 | 1.1e-08 | 3.733E-08 | 3.066E-08 | 1.100E-08 | 7.100E-08 |
| 10651 | 533 | V533L | A | 6.7e-08 | 5.4e-08 | 5.4e-08 | 5.833E-08 | 7.506E-09 | 5.400E-08 | 6.700E-08 |
| 10652 | 533 | V533K | A | 2.9e-08 | 4e-08 | 7.3e-08 | 4.733E-08 | 2.290E-08 | 2.900E-08 | 7.300E-08 |
| 10653 | 533 | V533M | A | 4.2e-08 | 2e-08 | 2.9e-08 | 3.033E-08 | 1.106E-08 | 2.000E-08 | 4.200E-08 |
| 10654 | 533 | V533F | A | 7.2e-08 | 2.6e-08 | 2e-08 | 3.933E-08 | 2.845E-08 | 2.000E-08 | 7.200E-08 |
| 10655 | 533 | V533P | A | 6.3e-08 | 7.1e-08 | 3e-08 | 5.467E-08 | 2.173E-08 | 3.000E-08 | 7.100E-08 |
| 10656 | 533 | V533S | A | 4.7e-08 | 4.6e-08 | 4.6e-08 | 4.633E-08 | 5.774E-10 | 4.600E-08 | 4.700E-08 |
| 10657 | 533 | V533T | A | 3.5e-08 | 4.7e-08 | 6.2e-08 | 4.800E-08 | 1.353E-08 | 3.500E-08 | 6.200E-08 |
| 10658 | 533 | V533W | A | 4.9e-08 | 2.5e-08 | 1e-07 | 5.800E-08 | 3.830E-08 | 2.500E-08 | 1.000E-07 |
| 10659 | 533 | V533Y | A | 4.1e-08 | 6.9e-08 | 6.8e-08 | 5.933E-08 | 1.589E-08 | 4.100E-08 | 6.900E-08 |
| 10660 | 533 | V533V | A | 4.9e-08 | 6.4e-08 | 6.7e-08 | 6.000E-08 | 9.644E-09 | 4.900E-08 | 6.700E-08 |
| 10661 | 534 | N534A | A | 1.8e-07 | 1.4e-08 | 4.7e-08 | 8.033E-08 | 8.788E-08 | 1.400E-08 | 1.800E-07 |
| 10662 | 534 | N534R | A | 2e-08 | 6.6e-08 | 7.1e-08 | 5.233E-08 | 2.811E-08 | 2.000E-08 | 7.100E-08 |
| 10663 | 534 | N534N | A | 1.7e-08 | 2.8e-08 | 3.9e-08 | 2.800E-08 | 1.100E-08 | 1.700E-08 | 3.900E-08 |
| 10664 | 534 | N534D | A | 6.7e-08 | 4.2e-08 | 6.4e-08 | 5.767E-08 | 1.365E-08 | 4.200E-08 | 6.700E-08 |
| 10665 | 534 | N534C | A | 1.1e-08 | 9.3e-09 | 2.6e-08 | 1.543E-08 | 9.190E-09 | 9.300E-09 | 2.600E-08 |
| 10666 | 534 | N534E | A | 5.3e-08 | 1.3e-08 | 2.9e-08 | 3.167E-08 | 2.013E-08 | 1.300E-08 | 5.300E-08 |
| 10667 | 534 | N534Q | A | 4e-08 | 3.4e-08 | 3.1e-08 | 3.500E-08 | 4.583E-09 | 3.100E-08 | 4.000E-08 |
| 10668 | 534 | N534G | A | 1.6e-08 | 5.5e-08 | 5.6e-08 | 4.233E-08 | 2.281E-08 | 1.600E-08 | 5.600E-08 |
| 10669 | 534 | N534H | A | 3.7e-08 | 5.9e-08 | 5.7e-08 | 5.100E-08 | 1.217E-08 | 3.700E-08 | 5.900E-08 |
| 10670 | 534 | N534I | A | 3.1e-08 | 3.7e-08 | 1.8e-08 | 2.867E-08 | 9.713E-09 | 1.800E-08 | 3.700E-08 |
| 10671 | 534 | N534L | A | 3.5e-08 | 6.8e-08 | 2.9e-08 | 4.400E-08 | 2.100E-08 | 2.900E-08 | 6.800E-08 |
| 10672 | 534 | N534K | A | 6.6e-08 | 3.4e-08 | 2.9e-08 | 4.300E-08 | 2.007E-08 | 2.900E-08 | 6.600E-08 |
| 10673 | 534 | N534M | A | 4.1e-08 | 1.2e-07 | 5.5e-08 | 7.200E-08 | 4.215E-08 | 4.100E-08 | 1.200E-07 |
| 10674 | 534 | N534F | A | 1.1e-08 | 2.7e-08 | 3.8e-08 | 2.533E-08 | 1.358E-08 | 1.100E-08 | 3.800E-08 |
| 10675 | 534 | N534P | A | 4.2e-08 | 3e-08 | 1.8e-08 | 3.000E-08 | 1.200E-08 | 1.800E-08 | 4.200E-08 |
| 10676 | 534 | N534S | A | 5.7e-08 | 7.8e-08 | 3.8e-08 | 5.767E-08 | 2.001E-08 | 3.800E-08 | 7.800E-08 |
| 10677 | 534 | N534T | A | 7.6e-08 | 1.1e-07 | 5.5e-08 | 8.033E-08 | 2.775E-08 | 5.500E-08 | 1.100E-07 |
| 10678 | 534 | N534W | A | 5.6e-08 | 1.1e-07 | 5.2e-08 | 7.267E-08 | 3.239E-08 | 5.200E-08 | 1.100E-07 |
| 10679 | 534 | N534Y | A | 4e-08 | 3.2e-08 | 5.7e-08 | 4.300E-08 | 1.277E-08 | 3.200E-08 | 5.700E-08 |
| 10680 | 534 | N534V | A | 1.2e-07 | 4.1e-08 | 5.2e-08 | 7.100E-08 | 4.279E-08 | 4.100E-08 | 1.200E-07 |
| 10681 | 535 | A535A | A | 5.3e-08 | 3e-08 | 2.5e-08 | 3.600E-08 | 1.493E-08 | 2.500E-08 | 5.300E-08 |
| 10682 | 535 | A535R | A | 4e-08 | 4.3e-08 | 5.5e-08 | 4.600E-08 | 7.937E-09 | 4.000E-08 | 5.500E-08 |
| 10683 | 535 | A535N | A | 7.6e-08 | 2.6e-08 | 9.3e-08 | 6.500E-08 | 3.483E-08 | 2.600E-08 | 9.300E-08 |
| 10684 | 535 | A535D | A | 3.4e-08 | 9e-09 | 1.3e-07 | 5.767E-08 | 6.388E-08 | 9.000E-09 | 1.300E-07 |
| 10685 | 535 | A535C | A | 1.4e-08 | 6.9e-08 | 4e-08 | 4.100E-08 | 2.751E-08 | 1.400E-08 | 6.900E-08 |
| 10686 | 535 | A535E | A | 4.2e-08 | 6.8e-08 | 1e-07 | 7.000E-08 | 2.905E-08 | 4.200E-08 | 1.000E-07 |
| 10687 | 535 | A535Q | A | 5.8e-08 | 1.9e-07 | 3.1e-08 | 9.300E-08 | 8.508E-08 | 3.100E-08 | 1.900E-07 |
| 10688 | 535 | A535G | A | 3.5e-08 | 2.8e-08 | 3.4e-08 | 3.233E-08 | 3.786E-09 | 2.800E-08 | 3.500E-08 |
| 10689 | 535 | A535H | A | 2.8e-08 | 6.3e-08 | 2.9e-08 | 4.000E-08 | 1.992E-08 | 2.800E-08 | 6.300E-08 |
| 10690 | 535 | A535I | A | 2.3e-08 | 4.1e-08 | 5.3e-08 | 3.900E-08 | 1.510E-08 | 2.300E-08 | 5.300E-08 |
| 10691 | 535 | A535L | A | 2e-08 | 8.7e-08 | 6.8e-08 | 5.833E-08 | 3.453E-08 | 2.000E-08 | 8.700E-08 |
| 10692 | 535 | A535K | A | 1e-07 | 2.5e-08 | 6e-08 | 6.167E-08 | 3.753E-08 | 2.500E-08 | 1.000E-07 |
| 10693 | 535 | A535M | A | 4.5e-08 | 2.7e-08 | 4.1e-08 | 3.767E-08 | 9.452E-09 | 2.700E-08 | 4.500E-08 |
| 10694 | 535 | A535F | A | 4.7e-08 | 4.7e-08 | 6.7e-08 | 5.367E-08 | 1.155E-08 | 4.700E-08 | 6.700E-08 |
| 10695 | 535 | A535P | A | 9.3e-08 | 5.1e-08 | 2.8e-08 | 5.733E-08 | 3.296E-08 | 2.800E-08 | 9.300E-08 |
| 10696 | 535 | A535S | A | 2.5e-08 | 3e-08 | 1.6e-08 | 2.367E-08 | 7.095E-09 | 1.600E-08 | 3.000E-08 |
| 10697 | 535 | A535T | A | 4.2e-08 | 4.9e-08 | 4e-08 | 4.367E-08 | 4.726E-09 | 4.000E-08 | 4.900E-08 |
| 10698 | 535 | A535W | A | 6e-08 | 1.3e-07 | 1.6e-08 | 6.867E-08 | 5.749E-08 | 1.600E-08 | 1.300E-07 |
| 10699 | 535 | A535Y | A | 5.8e-08 | 1.1e-07 | 4.6e-08 | 7.133E-08 | 3.402E-08 | 4.600E-08 | 1.100E-07 |
| 10700 | 535 | A535V | A | 4.6e-08 | 3.6e-08 | 4.7e-08 | 4.300E-08 | 6.083E-09 | 3.600E-08 | 4.700E-08 |
| 10701 | 536 | R536A | A | 3.6e-08 | 3.7e-08 | 1.2e-07 | 6.433E-08 | 4.821E-08 | 3.600E-08 | 1.200E-07 |
| 10702 | 536 | R536R | A | 2.1e-08 | 6.2e-08 | 3.4e-08 | 3.900E-08 | 2.095E-08 | 2.100E-08 | 6.200E-08 |
| 10703 | 536 | R536N | A | 3.8e-08 | 6.1e-08 | 3.5e-08 | 4.467E-08 | 1.422E-08 | 3.500E-08 | 6.100E-08 |
| 10704 | 536 | R536D | A | 7.7e-08 | 6e-08 | 8e-08 | 7.233E-08 | 1.079E-08 | 6.000E-08 | 8.000E-08 |
| 10705 | 536 | R536C | A | 5.3e-08 | 2.6e-08 | 4.4e-08 | 4.100E-08 | 1.375E-08 | 2.600E-08 | 5.300E-08 |
| 10706 | 536 | R536E | A | 3.2e-08 | 3.1e-08 | 4.4e-08 | 3.567E-08 | 7.234E-09 | 3.100E-08 | 4.400E-08 |
| 10707 | 536 | R536Q | A | 2e-08 | 4.4e-08 | 3.9e-08 | 3.433E-08 | 1.266E-08 | 2.000E-08 | 4.400E-08 |
| 10708 | 536 | R536G | A | 1.2e-07 | 7.2e-08 | 4.4e-08 | 7.867E-08 | 3.844E-08 | 4.400E-08 | 1.200E-07 |
| 10709 | 536 | R536H | A | 3.4e-08 | 7.9e-08 | 5.9e-08 | 5.733E-08 | 2.255E-08 | 3.400E-08 | 7.900E-08 |

| | | | | | | | | | | |
|---|---|---|---|---|---|---|---|---|---|---|
| 10710 | 536 | R536I | A | 3.1e-08 | 3.7e-08 | 5.1e-08 | 3.967E-08 | 1.026E-08 | 3.100E-08 | 5.100E-08 |
| 10711 | 536 | R536L | A | 4.9e-08 | 2.8e-08 | 3.6e-08 | 3.767E-08 | 1.060E-08 | 2.800E-08 | 4.900E-08 |
| 10712 | 536 | R536K | A | 9.5e-08 | 6.2e-08 | 3.1e-08 | 6.267E-08 | 3.201E-08 | 3.100E-08 | 9.500E-08 |
| 10713 | 536 | R536M | A | 3.5e-08 | 2.9e-08 | 1.5e-08 | 2.633E-08 | 1.026E-08 | 1.500E-08 | 3.500E-08 |
| 10714 | 536 | R536F | A | 7e-08 | 8.2e-08 | 1.1e-08 | 5.433E-08 | 3.800E-08 | 1.100E-08 | 8.200E-08 |
| 10715 | 536 | R536P | A | 3.2e-08 | 1.3e-08 | 1.1e-08 | 1.867E-08 | 1.159E-08 | 1.100E-08 | 3.200E-08 |
| 10716 | 536 | R536S | A | 7.7e-08 | 3e-08 | 9.5e-08 | 6.733E-08 | 3.356E-08 | 3.000E-08 | 9.500E-08 |
| 10717 | 536 | R536T | A | 3.6e-08 | 2.5e-08 | 5.2e-08 | 3.767E-08 | 1.358E-08 | 2.500E-08 | 5.200E-08 |
| 10718 | 536 | R536W | A | 2.6e-08 | 8.3e-08 | 4.8e-08 | 5.233E-08 | 2.875E-08 | 2.600E-08 | 8.300E-08 |
| 10719 | 536 | R536Y | A | 9.1e-08 | 4.5e-08 | 2.9e-08 | 5.500E-08 | 3.219E-08 | 2.900E-08 | 9.100E-08 |
| 10720 | 536 | R536V | A | 4.5e-08 | 2.6e-08 | 3.7e-08 | 3.600E-08 | 9.539E-09 | 2.600E-08 | 4.500E-08 |
| 10721 | 537 | H537A | A | 4.3e-08 | 7.2e-08 | 3.6e-08 | 5.033E-08 | 1.909E-08 | 3.600E-08 | 7.200E-08 |
| 10722 | 537 | H537R | A | 3.9e-08 | 1.3e-07 | 2.5e-08 | 6.467E-08 | 5.701E-08 | 2.500E-08 | 1.300E-07 |
| 10723 | 537 | H537N | A | 4.1e-08 | 4.4e-08 | 4.3e-08 | 4.267E-08 | 1.528E-09 | 4.100E-08 | 4.400E-08 |
| 10724 | 537 | H537D | A | 9.2e-08 | 3.5e-08 | 6.6e-08 | 6.433E-08 | 2.854E-08 | 3.500E-08 | 9.200E-08 |
| 10725 | 537 | H537C | A | 7.9e-08 | 1.9e-08 | 3e-08 | 4.267E-08 | 3.194E-08 | 1.900E-08 | 7.900E-08 |
| 10726 | 537 | H537E | A | 2.1e-08 | 4.5e-08 | 5e-08 | 3.867E-08 | 1.550E-08 | 2.100E-08 | 5.000E-08 |
| 10727 | 537 | H537Q | A | 2.5e-08 | 2.2e-08 | 4.8e-08 | 3.167E-08 | 1.422E-08 | 2.200E-08 | 4.800E-08 |
| 10728 | 537 | H537G | A | 3.3e-08 | 6.6e-08 | 4.2e-08 | 4.700E-08 | 1.706E-08 | 3.300E-08 | 6.600E-08 |
| 10729 | 537 | H537H | A | 3.1e-08 | 6.4e-08 | 3.6e-08 | 4.367E-08 | 1.779E-08 | 3.100E-08 | 6.400E-08 |
| 10730 | 537 | H537I | A | 4.5e-08 | 7e-08 | 1.1e-07 | 7.500E-08 | 3.279E-08 | 4.500E-08 | 1.100E-07 |
| 10731 | 537 | H537L | A | 3.1e-08 | 5.7e-08 | 2.1e-08 | 3.633E-08 | 1.858E-08 | 2.100E-08 | 5.700E-08 |
| 10732 | 537 | H537K | A | 1.4e-07 | 9.3e-09 | 2.8e-08 | 5.910E-08 | 7.068E-08 | 9.300E-09 | 1.400E-07 |
| 10733 | 537 | H537M | A | 3.6e-08 | 3.9e-08 | 2.3e-07 | 1.017E-07 | 1.112E-07 | 3.600E-08 | 2.300E-07 |
| 10734 | 537 | H537F | A | 2.9e-08 | 2.7e-08 | 6.4e-08 | 4.000E-08 | 2.081E-08 | 2.700E-08 | 6.400E-08 |
| 10735 | 537 | H537P | A | 1.3e-07 | 4.9e-08 | 1.2e-07 | 9.967E-08 | 4.416E-08 | 4.900E-08 | 1.300E-07 |
| 10736 | 537 | H537S | A | 2.3e-08 | 4.7e-08 | 1.6e-08 | 2.867E-08 | 1.626E-08 | 1.600E-08 | 4.700E-08 |
| 10737 | 537 | H537T | A | 3.8e-08 | 3.4e-08 | 8.5e-08 | 5.233E-08 | 2.836E-08 | 3.400E-08 | 8.500E-08 |
| 10738 | 537 | H537W | A | 1.1e-07 | 1.1e-07 | 3.9e-08 | 8.633E-08 | 4.099E-08 | 3.900E-08 | 1.100E-07 |
| 10739 | 537 | H537Y | A | 1.5e-08 | 6.5e-08 | 5.9e-08 | 4.633E-08 | 2.730E-08 | 1.500E-08 | 6.500E-08 |
| 10740 | 537 | H537V | A | 5.5e-08 | 5.4e-08 | 2.1e-08 | 4.333E-08 | 1.935E-08 | 2.100E-08 | 5.500E-08 |
| 10741 | 538 | C538A | A | 4e-08 | 5.2e-08 | 3.5e-08 | 4.233E-08 | 8.737E-09 | 3.500E-08 | 5.200E-08 |
| 10742 | 538 | C538R | A | 4.9e-08 | 1.9e-08 | 2.5e-08 | 3.100E-08 | 1.587E-08 | 1.900E-08 | 4.900E-08 |
| 10743 | 538 | C538N | A | 3.6e-08 | 6.8e-08 | 2.4e-08 | 4.267E-08 | 2.274E-08 | 2.400E-08 | 6.800E-08 |
| 10744 | 538 | C538D | A | 3.2e-08 | 1.9e-08 | 1.7e-08 | 2.267E-08 | 8.145E-09 | 1.700E-08 | 3.200E-08 |
| 10745 | 538 | C538C | A | 2.2e-08 | 6.6e-08 | 5.1e-08 | 4.633E-08 | 2.237E-08 | 2.200E-08 | 6.600E-08 |
| 10746 | 538 | C538E | A | 5.7e-08 | 5.1e-08 | 3.9e-08 | 4.900E-08 | 9.165E-09 | 3.900E-08 | 5.700E-08 |
| 10747 | 538 | C538Q | A | 3.4e-08 | 1e-07 | 3.4e-08 | 5.600E-08 | 3.811E-08 | 3.400E-08 | 1.000E-07 |
| 10748 | 538 | C538G | A | 4.6e-08 | 4e-08 | 1.6e-08 | 3.400E-08 | 1.587E-08 | 1.600E-08 | 4.600E-08 |
| 10749 | 538 | C538H | A | 6e-08 | 7.7e-08 | 4.9e-08 | 6.200E-08 | 1.411E-08 | 4.900E-08 | 7.700E-08 |
| 10750 | 538 | C538I | A | 4.9e-08 | 7.2e-08 | 3.6e-08 | 5.233E-08 | 1.823E-08 | 3.600E-08 | 7.200E-08 |
| 10751 | 538 | C538L | A | 7.8e-08 | 3.2e-08 | 2.5e-08 | 4.500E-08 | 2.879E-08 | 2.500E-08 | 7.800E-08 |
| 10752 | 538 | C538K | A | 3.1e-08 | 3.5e-08 | 8.5e-08 | 5.033E-08 | 3.009E-08 | 3.100E-08 | 8.500E-08 |
| 10753 | 538 | C538M | A | 6.3e-08 | 3e-08 | 2.9e-08 | 4.067E-08 | 1.935E-08 | 2.900E-08 | 6.300E-08 |
| 10754 | 538 | C538F | A | 5.7e-08 | 5.5e-08 | 2.9e-08 | 4.700E-08 | 1.562E-08 | 2.900E-08 | 5.700E-08 |
| 10755 | 538 | C538P | A | 9.5e-08 | 1.9e-08 | 1.7e-08 | 4.367E-08 | 4.447E-08 | 1.700E-08 | 9.500E-08 |
| 10756 | 538 | C538S | A | 5.1e-08 | 1.3e-08 | 5.3e-08 | 3.900E-08 | 2.254E-08 | 1.300E-08 | 5.300E-08 |
| 10757 | 538 | C538T | A | 5.1e-08 | 1.3e-07 | 3.9e-08 | 7.333E-08 | 4.944E-08 | 3.900E-08 | 1.300E-07 |
| 10758 | 538 | C538W | A | 6.2e-08 | 1e-07 | 2.7e-08 | 6.300E-08 | 3.651E-08 | 2.700E-08 | 1.000E-07 |
| 10759 | 538 | C538Y | A | 2.5e-08 | 2.2e-08 | 4.8e-08 | 3.167E-08 | 1.422E-08 | 2.200E-08 | 4.800E-08 |
| 10760 | 538 | C538V | A | 3.8e-08 | 4.8e-08 | 3.4e-08 | 4.000E-08 | 7.211E-09 | 3.400E-08 | 4.800E-08 |
| 10761 | 539 | L539A | A | 1e-07 | 7.9e-08 | 1.3e-08 | 6.400E-08 | 4.540E-08 | 1.300E-08 | 1.000E-07 |
| 10762 | 539 | L539R | A | 2.7e-08 | 4.3e-08 | 2.4e-08 | 3.133E-08 | 1.021E-08 | 2.400E-08 | 4.300E-08 |
| 10763 | 539 | L539N | A | 3.1e-08 | 1e-07 | 7.6e-08 | 6.900E-08 | 3.503E-08 | 3.100E-08 | 1.000E-07 |
| 10764 | 539 | L539D | A | 1.2e-08 | 5.8e-08 | 3.8e-08 | 3.600E-08 | 2.307E-08 | 1.200E-08 | 5.800E-08 |
| 10765 | 539 | L539C | A | 3.1e-08 | 8.7e-08 | 6.2e-08 | 6.000E-08 | 2.805E-08 | 3.100E-08 | 8.700E-08 |
| 10766 | 539 | L539E | A | 2.7e-08 | 4.7e-08 | 5.8e-08 | 4.400E-08 | 1.572E-08 | 2.700E-08 | 5.800E-08 |
| 10767 | 539 | L539Q | A | 1.9e-08 | 2e-08 | 3.1e-08 | 2.333E-08 | 6.658E-09 | 1.900E-08 | 3.100E-08 |
| 10768 | 539 | L539G | A | 5.5e-08 | 1e-07 | 6.3e-08 | 7.267E-08 | 2.401E-08 | 5.500E-08 | 1.000E-07 |
| 10769 | 539 | L539H | A | 1.7e-08 | 4.3e-08 | 4e-08 | 3.333E-08 | 1.422E-08 | 1.700E-08 | 4.300E-08 |
| 10770 | 539 | L539I | A | 3.6e-08 | 3.6e-08 | 1.8e-07 | 8.400E-08 | 8.314E-08 | 3.600E-08 | 1.800E-07 |
| 10771 | 539 | L539L | A | 4.6e-08 | 4.5e-08 | 9.5e-08 | 6.200E-08 | 2.858E-08 | 4.500E-08 | 9.500E-08 |
| 10772 | 539 | L539K | A | 4.6e-08 | 4.4e-08 | 5.6e-08 | 4.867E-08 | 6.429E-09 | 4.400E-08 | 5.600E-08 |
| 10773 | 539 | L539M | A | 3.4e-08 | 1.2e-07 | 6.8e-08 | 7.400E-08 | 4.331E-08 | 3.400E-08 | 1.200E-07 |
| 10774 | 539 | L539F | A | 3.2e-08 | 3.5e-08 | 7.4e-08 | 4.700E-08 | 2.343E-08 | 3.200E-08 | 7.400E-08 |
| 10775 | 539 | L539P | A | 4.8e-08 | 4e-08 | 4.9e-08 | 4.567E-08 | 4.933E-09 | 4.000E-08 | 4.900E-08 |
| 10776 | 539 | L539S | A | 2.8e-08 | 3.6e-08 | 2.8e-08 | 3.067E-08 | 4.619E-09 | 2.800E-08 | 3.600E-08 |
| 10777 | 539 | L539T | A | 9.5e-08 | 3.9e-08 | 9.5e-08 | 7.633E-08 | 3.233E-08 | 3.900E-08 | 9.500E-08 |
| 10778 | 539 | L539W | A | 3.2e-08 | 1.7e-08 | 2.9e-08 | 2.600E-08 | 7.937E-09 | 1.700E-08 | 3.200E-08 |
| 10779 | 539 | L539Y | A | 7.5e-08 | 9.2e-08 | 3.4e-08 | 6.700E-08 | 2.982E-08 | 3.400E-08 | 9.200E-08 |
| 10780 | 539 | L539V | A | 3e-08 | 5.9e-08 | 3e-08 | 3.967E-08 | 1.674E-08 | 3.000E-08 | 5.900E-08 |
| 10781 | 540 | P540A | A | 3.5e-08 | 3.9e-08 | 3.6e-08 | 3.667E-08 | 2.082E-09 | 3.500E-08 | 3.900E-08 |
| 10782 | 540 | P540R | A | 5.5e-08 | 2.1e-08 | 6.5e-08 | 4.700E-08 | 2.307E-08 | 2.100E-08 | 6.500E-08 |
| 10783 | 540 | P540N | A | 3.9e-08 | 2.1e-08 | 9.9e-08 | 5.300E-08 | 4.084E-08 | 2.100E-08 | 9.900E-08 |
| 10784 | 540 | P540D | A | 3.7e-08 | 2.1e-08 | 3.2e-08 | 3.000E-08 | 8.185E-09 | 2.100E-08 | 3.700E-08 |
| 10785 | 540 | P540C | A | 1.2e-08 | 3.4e-08 | 5.5e-08 | 3.367E-08 | 2.150E-08 | 1.200E-08 | 5.500E-08 |
| 10786 | 540 | P540E | A | 6.8e-08 | 5.1e-08 | 8.7e-08 | 6.867E-08 | 1.801E-08 | 5.100E-08 | 8.700E-08 |
| 10787 | 540 | P540Q | A | 9.3e-08 | 5.7e-08 | 6.1e-08 | 7.033E-08 | 1.973E-08 | 5.700E-08 | 9.300E-08 |
| 10788 | 540 | P540G | A | 4.3e-08 | 4.3e-08 | 1.7e-08 | 3.433E-08 | 1.501E-08 | 1.700E-08 | 4.300E-08 |
| 10789 | 540 | P540H | A | 7.3e-08 | 5.1e-08 | 4.8e-08 | 5.733E-08 | 1.365E-08 | 4.800E-08 | 7.300E-08 |
| 10790 | 540 | P540I | A | 5.5e-08 | 7.6e-08 | 1.5e-07 | 9.367E-08 | 4.990E-08 | 5.500E-08 | 1.500E-07 |
| 10791 | 540 | P540L | A | 1.3e-08 | 6.5e-08 | 2.6e-08 | 3.467E-08 | 2.706E-08 | 1.300E-08 | 6.500E-08 |
| 10792 | 540 | P540K | A | 1.2e-07 | 4e-08 | 7.1e-08 | 7.700E-08 | 4.034E-08 | 4.000E-08 | 1.200E-07 |
| 10793 | 540 | P540M | A | 1.4e-08 | 7.8e-08 | 5.4e-08 | 4.867E-08 | 3.233E-08 | 1.400E-08 | 7.800E-08 |
| 10794 | 540 | P540F | A | 4e-08 | 1.1e-07 | 7.8e-08 | 7.600E-08 | 3.504E-08 | 4.000E-08 | 1.100E-07 |

| | | | | | | | | | | |
|---|---|---|---|---|---|---|---|---|---|---|
| 10795 | 540 | P540P | A | 4.6e-08 | 1.5e-07 | 1e-07 | 9.867E-08 | 5.201E-08 | 4.600E-08 | 1.500E-07 |
| 10796 | 540 | P540S | A | 3e-08 | 4.6e-08 | 6.6e-08 | 4.733E-08 | 1.804E-08 | 3.000E-08 | 6.600E-08 |
| 10797 | 540 | P540T | A | 3.4e-08 | 3.2e-08 | 1.1e-07 | 5.867E-08 | 4.447E-08 | 3.200E-08 | 1.100E-07 |
| 10798 | 540 | P540W | A | 4.8e-08 | 4.7e-08 | 6.7e-08 | 5.400E-08 | 1.127E-08 | 4.700E-08 | 6.700E-08 |
| 10799 | 540 | P540Y | A | 2.7e-08 | 5.3e-08 | 2.6e-08 | 3.533E-08 | 1.531E-08 | 2.600E-08 | 5.300E-08 |
| 10800 | 540 | P540V | A | 4e-08 | 1.7e-08 | 4.9e-08 | 3.533E-08 | 1.650E-08 | 1.700E-08 | 4.900E-08 |
| 10801 | 541 | C541A | A | 7.5e-08 | 3e-08 | 5.9e-08 | 5.467E-08 | 2.281E-08 | 3.000E-08 | 7.500E-08 |
| 10802 | 541 | C541R | A | 9e-08 | 2.5e-08 | 2.2e-08 | 4.567E-08 | 3.842E-08 | 2.200E-08 | 9.000E-08 |
| 10803 | 541 | C541N | A | 1.8e-08 | 6.6e-08 | 1.5e-08 | 3.300E-08 | 2.862E-08 | 1.500E-08 | 6.600E-08 |
| 10804 | 541 | C541D | A | 3.5e-08 | 1.2e-07 | 3.9e-08 | 6.467E-08 | 4.796E-08 | 3.500E-08 | 1.200E-07 |
| 10805 | 541 | C541C | A | 2.8e-08 | 3.7e-08 | 3.8e-08 | 3.433E-08 | 5.508E-09 | 2.800E-08 | 3.800E-08 |
| 10806 | 541 | C541E | A | 9e-08 | 2.5e-08 | 1.5e-07 | 8.833E-08 | 6.252E-08 | 2.500E-08 | 1.500E-07 |
| 10807 | 541 | C541Q | A | 5e-08 | 4.5e-08 | 2.5e-08 | 4.000E-08 | 1.323E-08 | 2.500E-08 | 5.000E-08 |
| 10808 | 541 | C541G | A | 6.7e-08 | 1.5e-07 | 1e-07 | 1.057E-07 | 4.179E-08 | 6.700E-08 | 1.500E-07 |
| 10809 | 541 | C541H | A | 1.8e-08 | 1.4e-07 | 6.8e-08 | 7.533E-08 | 6.133E-08 | 1.800E-08 | 1.400E-07 |
| 10810 | 541 | C541I | A | 5.3e-09 | 3.5e-08 | 2.2e-08 | 2.077E-08 | 1.489E-08 | 5.300E-09 | 3.500E-08 |
| 10811 | 541 | C541L | A | 3.8e-08 | 5.9e-08 | 2.4e-08 | 4.033E-08 | 1.762E-08 | 2.400E-08 | 5.900E-08 |
| 10812 | 541 | C541K | A | 3.1e-08 | 7.2e-08 | 5.6e-08 | 5.300E-08 | 2.066E-08 | 3.100E-08 | 7.200E-08 |
| 10813 | 541 | C541M | A | 3.4e-08 | 5.2e-08 | 3.8e-08 | 4.133E-08 | 9.452E-09 | 3.400E-08 | 5.200E-08 |
| 10814 | 541 | C541F | A | 4.8e-08 | 7.8e-08 | 1.5e-07 | 9.200E-08 | 5.242E-08 | 4.800E-08 | 1.500E-07 |
| 10815 | 541 | C541P | A | 2.4e-08 | 3.5e-08 | 4.1e-08 | 3.333E-08 | 8.622E-09 | 2.400E-08 | 4.100E-08 |
| 10816 | 541 | C541S | A | 3.9e-08 | 5.2e-08 | 5.1e-08 | 4.733E-08 | 7.234E-09 | 3.900E-08 | 5.200E-08 |
| 10817 | 541 | C541T | A | 6.3e-08 | 4.8e-08 | 1.7e-08 | 4.267E-08 | 2.346E-08 | 1.700E-08 | 6.300E-08 |
| 10818 | 541 | C541W | A | 4.4e-08 | 6.7e-08 | 6.6e-08 | 5.900E-08 | 1.300E-08 | 4.400E-08 | 6.700E-08 |
| 10819 | 541 | C541Y | A | 3.8e-08 | 3.7e-08 | 8.2e-08 | 5.233E-08 | 2.570E-08 | 3.700E-08 | 8.200E-08 |
| 10820 | 541 | C541V | A | 3.9e-08 | 4.3e-08 | 6.5e-08 | 4.900E-08 | 1.400E-08 | 3.900E-08 | 6.500E-08 |
| 10821 | 542 | H542A | A | 7.4e-08 | 4.5e-08 | 2.1e-08 | 4.667E-08 | 2.654E-08 | 2.100E-08 | 7.400E-08 |
| 10822 | 542 | H542R | A | 9.4e-08 | 6.3e-08 | 1.4e-08 | 5.700E-08 | 4.034E-08 | 1.400E-08 | 9.400E-08 |
| 10823 | 542 | H542N | A | 7.5e-08 | 2.7e-08 | 5.1e-08 | 5.100E-08 | 2.400E-08 | 2.700E-08 | 7.500E-08 |
| 10824 | 542 | H542D | A | 1.9e-08 | 6e-08 | 6.1e-08 | 4.667E-08 | 2.397E-08 | 1.900E-08 | 6.100E-08 |
| 10825 | 542 | H542C | A | 2e-08 | 3.1e-08 | 4.3e-08 | 3.133E-08 | 1.150E-08 | 2.000E-08 | 4.300E-08 |
| 10826 | 542 | H542E | A | 2.9e-08 | 7.5e-08 | 6.4e-08 | 5.600E-08 | 2.402E-08 | 2.900E-08 | 7.500E-08 |
| 10827 | 542 | H542Q | A | 7.5e-08 | 3.2e-08 | 3.9e-08 | 4.867E-08 | 2.307E-08 | 3.200E-08 | 7.500E-08 |
| 10828 | 542 | H542G | A | 2.9e-08 | 1e-07 | 3e-08 | 5.300E-08 | 4.071E-08 | 2.900E-08 | 1.000E-07 |
| 10829 | 542 | H542H | A | 7.2e-08 | 3e-08 | 4.3e-08 | 4.833E-08 | 2.150E-08 | 3.000E-08 | 7.200E-08 |
| 10830 | 542 | H542I | A | 4.2e-08 | 5.3e-08 | 3.4e-08 | 4.300E-08 | 9.539E-09 | 3.400E-08 | 5.300E-08 |
| 10831 | 542 | H542L | A | 6.4e-08 | 2.2e-08 | 2.7e-08 | 3.767E-08 | 2.294E-08 | 2.200E-08 | 6.400E-08 |
| 10832 | 542 | H542K | A | 7.3e-08 | 5.3e-08 | 6e-08 | 6.200E-08 | 1.015E-08 | 5.300E-08 | 7.300E-08 |
| 10833 | 542 | H542M | A | 5.4e-08 | 9.3e-08 | 5e-08 | 6.567E-08 | 2.376E-08 | 5.000E-08 | 9.300E-08 |
| 10834 | 542 | H542F | A | 7.9e-08 | 1.5e-08 | 2.1e-08 | 3.833E-08 | 3.535E-08 | 1.500E-08 | 7.900E-08 |
| 10835 | 542 | H542P | A | 3.7e-08 | 3.8e-08 | 2.8e-08 | 3.433E-08 | 5.508E-09 | 2.800E-08 | 3.800E-08 |
| 10836 | 542 | H542S | A | 3.3e-08 | 1.1e-07 | 7.9e-08 | 7.400E-08 | 3.874E-08 | 3.300E-08 | 1.100E-07 |
| 10837 | 542 | H542T | A | 1e-08 | 6.2e-08 | 3.6e-08 | 3.600E-08 | 2.600E-08 | 1.000E-08 | 6.200E-08 |
| 10838 | 542 | H542W | A | 7.9e-08 | 3.2e-08 | 8.7e-08 | 6.600E-08 | 2.972E-08 | 3.200E-08 | 8.700E-08 |
| 10839 | 542 | H542Y | A | 3.1e-08 | 6e-08 | 4.9e-08 | 4.667E-08 | 1.464E-08 | 3.100E-08 | 6.000E-08 |
| 10840 | 542 | H542V | A | 1.5e-07 | 4e-08 | 3.7e-08 | 7.567E-08 | 6.439E-08 | 3.700E-08 | 1.500E-07 |
| 10841 | 543 | P543A | A | 5.6e-08 | 2.2e-08 | 5.1e-08 | 4.300E-08 | 1.836E-08 | 2.200E-08 | 5.600E-08 |
| 10842 | 543 | P543R | A | 2.3e-08 | 6.3e-08 | 2.6e-08 | 3.733E-08 | 2.228E-08 | 2.300E-08 | 6.300E-08 |
| 10843 | 543 | P543N | A | 3.5e-08 | 5.9e-08 | 4.7e-08 | 4.700E-08 | 1.200E-08 | 3.500E-08 | 5.900E-08 |
| 10844 | 543 | P543D | A | 2.6e-08 | 2.6e-08 | 2.8e-08 | 2.667E-08 | 1.155E-09 | 2.600E-08 | 2.800E-08 |
| 10845 | 543 | P543C | A | 4.8e-08 | 5.6e-08 | 4.3e-08 | 4.900E-08 | 6.557E-09 | 4.300E-08 | 5.600E-08 |
| 10846 | 543 | P543E | A | 1.9e-08 | 3.6e-08 | 4.7e-08 | 3.400E-08 | 1.411E-08 | 1.900E-08 | 4.700E-08 |
| 10847 | 543 | P543Q | A | 3.1e-08 | 5.7e-08 | 3.2e-08 | 4.000E-08 | 1.473E-08 | 3.100E-08 | 5.700E-08 |
| 10848 | 543 | P543G | A | 3.7e-08 | 1.6e-07 | 5.5e-08 | 8.400E-08 | 6.643E-08 | 3.700E-08 | 1.600E-07 |
| 10849 | 543 | P543H | A | 9e-08 | 2.9e-08 | 5.1e-08 | 5.667E-08 | 3.089E-08 | 2.900E-08 | 9.000E-08 |
| 10850 | 543 | P543I | A | 4.7e-08 | 4.7e-08 | 1e-07 | 6.467E-08 | 3.060E-08 | 4.700E-08 | 1.000E-07 |
| 10851 | 543 | P543L | A | 3.8e-08 | 3.5e-08 | 5.6e-08 | 4.300E-08 | 1.136E-08 | 3.500E-08 | 5.600E-08 |
| 10852 | 543 | P543K | A | 4.9e-08 | 2.7e-08 | 3.5e-08 | 3.700E-08 | 1.114E-08 | 2.700E-08 | 4.900E-08 |
| 10853 | 543 | P543M | A | 4.4e-08 | 1.2e-08 | 3.2e-08 | 2.933E-08 | 1.617E-08 | 1.200E-08 | 4.400E-08 |
| 10854 | 543 | P543F | A | 3.6e-08 | 8.4e-08 | 7.1e-08 | 6.367E-08 | 2.483E-08 | 3.600E-08 | 8.400E-08 |
| 10855 | 543 | P543P | A | 1.5e-08 | 2.7e-08 | 2.7e-08 | 2.300E-08 | 6.928E-09 | 1.500E-08 | 2.700E-08 |
| 10856 | 543 | P543S | A | 3.3e-08 | 2e-08 | 3.6e-08 | 2.967E-08 | 8.505E-09 | 2.000E-08 | 3.600E-08 |
| 10857 | 543 | P543T | A | 5.3e-08 | 1.8e-08 | 4e-08 | 3.700E-08 | 1.769E-08 | 1.800E-08 | 5.300E-08 |
| 10858 | 543 | P543W | A | 2.5e-08 | 6.9e-08 | 4.6e-08 | 4.667E-08 | 2.201E-08 | 2.500E-08 | 6.900E-08 |
| 10859 | 543 | P543Y | A | 4.8e-08 | 7.1e-08 | 4.9e-08 | 5.600E-08 | 1.300E-08 | 4.800E-08 | 7.100E-08 |
| 10860 | 543 | P543V | A | 3.6e-08 | 5.8e-08 | 4.4e-08 | 4.600E-08 | 1.114E-08 | 3.600E-08 | 5.800E-08 |
| 10861 | 544 | E544A | A | 1.3e-08 | 6.1e-08 | 1.2e-07 | 6.467E-08 | 5.359E-08 | 1.300E-08 | 1.200E-07 |
| 10862 | 544 | E544R | A | 5.3e-08 | 5.5e-08 | 4e-08 | 4.933E-08 | 8.145E-09 | 4.000E-08 | 5.500E-08 |
| 10863 | 544 | E544N | A | 6.6e-08 | 4.6e-08 | 5.4e-08 | 5.533E-08 | 1.007E-08 | 4.600E-08 | 6.600E-08 |
| 10864 | 544 | E544D | A | 8.3e-08 | 1.6e-08 | 1.2e-07 | 7.300E-08 | 5.272E-08 | 1.600E-08 | 1.200E-07 |
| 10865 | 544 | E544C | A | 5.4e-08 | 7e-08 | 6.1e-08 | 6.167E-08 | 8.021E-09 | 5.400E-08 | 7.000E-08 |
| 10866 | 544 | E544E | A | 4.3e-08 | 5.7e-08 | 4.9e-08 | 4.967E-08 | 7.024E-09 | 4.300E-08 | 5.700E-08 |
| 10867 | 544 | E544Q | A | 8.8e-08 | 5.2e-08 | 4.8e-08 | 6.267E-08 | 2.203E-08 | 4.800E-08 | 8.800E-08 |
| 10868 | 544 | E544G | A | 7.2e-08 | 5.2e-08 | 6.2e-08 | 6.200E-08 | 1.000E-08 | 5.200E-08 | 7.200E-08 |
| 10869 | 544 | E544H | A | 3.7e-08 | 3.7e-08 | 7.5e-08 | 4.967E-08 | 2.194E-08 | 3.700E-08 | 7.500E-08 |
| 10870 | 544 | E544I | A | 2.9e-08 | 3.8e-08 | 3.7e-08 | 3.467E-08 | 4.933E-09 | 2.900E-08 | 3.800E-08 |
| 10871 | 544 | E544L | A | 4e-08 | 8.7e-08 | 4.2e-08 | 5.633E-08 | 2.658E-08 | 4.000E-08 | 8.700E-08 |
| 10872 | 544 | E544K | A | 6.2e-08 | 2.5e-08 | 7.7e-08 | 5.467E-08 | 2.676E-08 | 2.500E-08 | 7.700E-08 |
| 10873 | 544 | E544M | A | 4.8e-08 | 5.3e-08 | 3.5e-08 | 4.533E-08 | 9.292E-09 | 3.500E-08 | 5.300E-08 |
| 10874 | 544 | E544F | A | 2.2e-08 | 5.6e-08 | 1.8e-07 | 8.600E-08 | 8.316E-08 | 2.200E-08 | 1.800E-07 |
| 10875 | 544 | E544P | A | 4.2e-08 | 4.2e-08 | 2.2e-08 | 3.533E-08 | 1.155E-08 | 2.200E-08 | 4.200E-08 |
| 10876 | 544 | E544S | A | 9.1e-08 | 4.7e-08 | 1.1e-07 | 8.267E-08 | 3.232E-08 | 4.700E-08 | 1.100E-07 |
| 10877 | 544 | E544T | A | 3.4e-08 | 5.8e-08 | 6.4e-08 | 5.200E-08 | 1.587E-08 | 3.400E-08 | 6.400E-08 |
| 10878 | 544 | E544W | A | 9.2e-08 | 1.3e-08 | 1.7e-08 | 4.067E-08 | 4.450E-08 | 1.300E-08 | 9.200E-08 |
| 10879 | 544 | E544Y | A | 1.1e-07 | 5.6e-08 | 8.4e-08 | 8.333E-08 | 2.701E-08 | 5.600E-08 | 1.100E-07 |

| | | | | | | | | | | |
|---|---|---|---|---|---|---|---|---|---|---|
| 10880 | 544 | E544V | A | 5.4e-08 | 1.6e-08 | 6.9e-08 | 4.633E-08 | 2.732E-08 | 1.600E-08 | 6.900E-08 |
| 10881 | 545 | C545A | A | 3.6e-08 | 7.7e-08 | 1.2e-08 | 4.167E-08 | 3.287E-08 | 1.200E-08 | 7.700E-08 |
| 10882 | 545 | C545R | A | 2.1e-07 | 1.1e-07 | 3.4e-08 | 1.180E-07 | 8.827E-08 | 3.400E-08 | 2.100E-07 |
| 10883 | 545 | C545N | A | 1.2e-07 | 8.4e-08 | 5.3e-08 | 8.567E-08 | 3.353E-08 | 5.300E-08 | 1.200E-07 |
| 10884 | 545 | C545D | A | 4.6e-08 | 1.5e-07 | 6.4e-08 | 8.667E-08 | 5.558E-08 | 4.600E-08 | 1.500E-07 |
| 10885 | 545 | C545C | A | 4.1e-08 | 2.7e-08 | 3.8e-08 | 3.533E-08 | 7.371E-09 | 2.700E-08 | 4.100E-08 |
| 10886 | 545 | C545E | A | 7.1e-08 | 5.8e-08 | 3.1e-08 | 5.333E-08 | 2.040E-08 | 3.100E-08 | 7.100E-08 |
| 10887 | 545 | C545Q | A | 1.6e-07 | 1.5e-08 | 2.7e-08 | 6.733E-08 | 8.048E-08 | 1.500E-08 | 1.600E-07 |
| 10888 | 545 | C545G | A | 1.5e-08 | 1e-07 | 2e-08 | 4.500E-08 | 4.770E-08 | 1.500E-08 | 1.000E-07 |
| 10889 | 545 | C545H | A | 2.7e-08 | 6e-08 | 4.6e-08 | 4.433E-08 | 1.656E-08 | 2.700E-08 | 6.000E-08 |
| 10890 | 545 | C545I | A | 6e-08 | 3.4e-08 | 3.6e-08 | 4.333E-08 | 1.447E-08 | 3.400E-08 | 6.000E-08 |
| 10891 | 545 | C545L | A | 7.5e-08 | 1.9e-07 | 9.1e-08 | 1.187E-07 | 6.229E-08 | 7.500E-08 | 1.900E-07 |
| 10892 | 545 | C545K | A | 7.2e-08 | 8.4e-08 | 5.8e-08 | 7.133E-08 | 1.301E-08 | 5.800E-08 | 8.400E-08 |
| 10893 | 545 | C545M | A | 6.2e-08 | 1.7e-08 | 3.5e-08 | 3.800E-08 | 2.265E-08 | 1.700E-08 | 6.200E-08 |
| 10894 | 545 | C545F | A | 2.6e-08 | 2.8e-08 | 4e-08 | 3.133E-08 | 7.572E-09 | 2.600E-08 | 4.000E-08 |
| 10895 | 545 | C545P | A | 2.5e-08 | 2e-08 | 4.8e-08 | 3.100E-08 | 1.493E-08 | 2.000E-08 | 4.800E-08 |
| 10896 | 545 | C545S | A | 6.8e-08 | 2e-08 | 2.3e-08 | 3.700E-08 | 2.689E-08 | 2.000E-08 | 6.800E-08 |
| 10897 | 545 | C545T | A | 1.3e-08 | 2.7e-08 | 8.3e-08 | 4.100E-08 | 3.704E-08 | 1.300E-08 | 8.300E-08 |
| 10898 | 545 | C545W | A | 6.3e-08 | 7.1e-08 | 2.4e-08 | 5.267E-08 | 2.515E-08 | 2.400E-08 | 7.100E-08 |
| 10899 | 545 | C545Y | A | 6.4e-08 | 3.2e-08 | 7.2e-08 | 5.600E-08 | 2.117E-08 | 3.200E-08 | 7.200E-08 |
| 10900 | 545 | C545V | A | 4.7e-08 | 4.7e-08 | 4.1e-08 | 4.500E-08 | 3.464E-09 | 4.100E-08 | 4.700E-08 |
| 10901 | 546 | Q546A | A | 3.8e-08 | 2.5e-08 | 2.8e-08 | 3.033E-08 | 6.807E-09 | 2.500E-08 | 3.800E-08 |
| 10902 | 546 | Q546R | A | 5.5e-08 | 3.3e-08 | 4.5e-08 | 4.433E-08 | 1.102E-08 | 3.300E-08 | 5.500E-08 |
| 10903 | 546 | Q546N | A | 3.5e-08 | 5.4e-08 | 8.1e-08 | 5.667E-08 | 2.312E-08 | 3.500E-08 | 8.100E-08 |
| 10904 | 546 | Q546D | A | 3.2e-08 | 3.9e-08 | 3.2e-08 | 3.433E-08 | 4.041E-09 | 3.200E-08 | 3.900E-08 |
| 10905 | 546 | Q546C | A | 1.3e-08 | 7.6e-08 | 3.6e-08 | 4.167E-08 | 3.188E-08 | 1.300E-08 | 7.600E-08 |
| 10906 | 546 | Q546E | A | 9e-08 | 6.1e-08 | 1.4e-08 | 5.500E-08 | 3.835E-08 | 1.400E-08 | 9.000E-08 |
| 10907 | 546 | Q546Q | A | 8.6e-08 | 6.5e-08 | 5.5e-08 | 6.867E-08 | 1.582E-08 | 5.500E-08 | 8.600E-08 |
| 10908 | 546 | Q546G | A | 1.9e-08 | 5.5e-08 | 4.5e-08 | 3.967E-08 | 1.858E-08 | 1.900E-08 | 5.500E-08 |
| 10909 | 546 | Q546H | A | 9.4e-08 | 7.5e-08 | 8.5e-08 | 8.467E-08 | 9.504E-09 | 7.500E-08 | 9.400E-08 |
| 10910 | 546 | Q546I | A | 9.1e-08 | 3.1e-08 | 4.2e-08 | 5.467E-08 | 3.194E-08 | 3.100E-08 | 9.100E-08 |
| 10911 | 546 | Q546L | A | 2.6e-08 | 3.9e-08 | 2.4e-08 | 2.967E-08 | 8.145E-09 | 2.400E-08 | 3.900E-08 |
| 10912 | 546 | Q546K | A | 2.5e-08 | 1.3e-07 | 4.7e-08 | 6.733E-08 | 5.537E-08 | 2.500E-08 | 1.300E-07 |
| 10913 | 546 | Q546M | A | 4.8e-08 | 5.3e-08 | 4.2e-08 | 4.767E-08 | 5.508E-09 | 4.200E-08 | 5.300E-08 |
| 10914 | 546 | Q546F | A | 2.5e-08 | 3.5e-08 | 1.6e-08 | 2.533E-08 | 9.504E-09 | 1.600E-08 | 3.500E-08 |
| 10915 | 546 | Q546P | A | 3.1e-08 | 5.5e-08 | 1.6e-08 | 3.400E-08 | 1.967E-08 | 1.600E-08 | 5.500E-08 |
| 10916 | 546 | Q546S | A | 5.6e-08 | 8.6e-08 | 4.4e-08 | 6.200E-08 | 2.163E-08 | 4.400E-08 | 8.600E-08 |
| 10917 | 546 | Q546T | A | 2.4e-08 | 3.2e-08 | 6.5e-08 | 4.033E-08 | 2.173E-08 | 2.400E-08 | 6.500E-08 |
| 10918 | 546 | Q546W | A | 2.4e-08 | 3.5e-08 | 8.3e-08 | 4.733E-08 | 3.137E-08 | 2.400E-08 | 8.300E-08 |
| 10919 | 546 | Q546Y | A | 4.8e-08 | 1.8e-08 | 7.6e-08 | 4.733E-08 | 2.901E-08 | 1.800E-08 | 7.600E-08 |
| 10920 | 546 | Q546V | A | 3.3e-08 | 2.6e-08 | 3.4e-08 | 3.100E-08 | 4.359E-09 | 2.600E-08 | 3.400E-08 |
| 10921 | 547 | P547A | A | 2.6e-08 | 1.5e-08 | 6.2e-08 | 3.433E-08 | 2.458E-08 | 1.500E-08 | 6.200E-08 |
| 10922 | 547 | P547R | A | 4.3e-08 | 8e-08 | 7.7e-08 | 6.667E-08 | 2.055E-08 | 4.300E-08 | 8.000E-08 |
| 10923 | 547 | P547N | A | 4e-08 | 6.7e-08 | 4.6e-08 | 5.100E-08 | 1.418E-08 | 4.000E-08 | 6.700E-08 |
| 10924 | 547 | P547D | A | 3.5e-08 | 3e-08 | 4.2e-08 | 3.567E-08 | 6.028E-09 | 3.000E-08 | 4.200E-08 |
| 10925 | 547 | P547C | A | 7.6e-09 | 3.1e-08 | 2e-08 | 1.953E-08 | 1.171E-08 | 7.600E-09 | 3.100E-08 |
| 10926 | 547 | P547E | A | 3.4e-08 | 9.5e-09 | 3.6e-08 | 2.650E-08 | 1.476E-08 | 9.500E-09 | 3.600E-08 |
| 10927 | 547 | P547Q | A | 2e-08 | 4.6e-08 | 2.9e-08 | 3.167E-08 | 1.320E-08 | 2.000E-08 | 4.600E-08 |
| 10928 | 547 | P547G | A | 1.4e-08 | 1.1e-07 | 5.4e-08 | 5.933E-08 | 4.822E-08 | 1.400E-08 | 1.100E-07 |
| 10929 | 547 | P547H | A | 2.9e-08 | 8.5e-08 | 1.1e-07 | 7.467E-08 | 4.148E-08 | 2.900E-08 | 1.100E-07 |
| 10930 | 547 | P547I | A | 4e-08 | 3.2e-08 | 4.8e-08 | 4.000E-08 | 8.000E-09 | 3.200E-08 | 4.800E-08 |
| 10931 | 547 | P547L | A | 1.5e-08 | 8e-08 | 4.3e-08 | 4.600E-08 | 3.260E-08 | 1.500E-08 | 8.000E-08 |
| 10932 | 547 | P547K | A | 2e-08 | 3.6e-08 | 2.4e-08 | 2.667E-08 | 8.327E-09 | 2.000E-08 | 3.600E-08 |
| 10933 | 547 | P547M | A | 7.6e-08 | 7.4e-08 | 1.2e-07 | 9.000E-08 | 2.600E-08 | 7.400E-08 | 1.200E-07 |
| 10934 | 547 | P547F | A | 3.1e-08 | 4.2e-08 | 1.1e-07 | 6.100E-08 | 4.279E-08 | 3.100E-08 | 1.100E-07 |
| 10935 | 547 | P547P | A | 2.3e-08 | 3.8e-08 | 3.8e-08 | 3.300E-08 | 8.660E-09 | 2.300E-08 | 3.800E-08 |
| 10936 | 547 | P547S | A | 6.9e-08 | 5.6e-08 | 2.8e-08 | 5.100E-08 | 2.095E-08 | 2.800E-08 | 6.900E-08 |
| 10937 | 547 | P547T | A | 9.8e-08 | 2.3e-08 | 3.7e-08 | 5.267E-08 | 3.988E-08 | 2.300E-08 | 9.800E-08 |
| 10938 | 547 | P547W | A | 2.3e-08 | 5.2e-08 | 1.9e-08 | 3.133E-08 | 1.801E-08 | 1.900E-08 | 5.200E-08 |
| 10939 | 547 | P547Y | A | 2.2e-08 | 5.4e-08 | 2.6e-08 | 3.400E-08 | 1.744E-08 | 2.200E-08 | 5.400E-08 |
| 10940 | 547 | P547V | A | 1.4e-07 | 2.6e-08 | 9.6e-08 | 8.733E-08 | 5.749E-08 | 2.600E-08 | 1.400E-07 |
| 10941 | 548 | Q548A | A | 3.8e-08 | 2.5e-08 | 5.3e-08 | 3.867E-08 | 1.401E-08 | 2.500E-08 | 5.300E-08 |
| 10942 | 548 | Q548R | A | 3.4e-08 | 5.9e-08 | 2.7e-08 | 4.000E-08 | 1.682E-08 | 2.700E-08 | 5.900E-08 |
| 10943 | 548 | Q548N | A | 1.8e-08 | 4.4e-08 | 2.4e-08 | 2.867E-08 | 1.361E-08 | 1.800E-08 | 4.400E-08 |
| 10944 | 548 | Q548D | A | 2.6e-08 | 1.4e-08 | 9.6e-08 | 4.533E-08 | 4.429E-08 | 1.400E-08 | 9.600E-08 |
| 10945 | 548 | Q548C | A | 5.1e-08 | 2.2e-08 | 1.2e-07 | 6.433E-08 | 5.034E-08 | 2.200E-08 | 1.200E-07 |
| 10946 | 548 | Q548E | A | 6.4e-08 | 5.2e-08 | 1.5e-08 | 4.367E-08 | 2.554E-08 | 1.500E-08 | 6.400E-08 |
| 10947 | 548 | Q548Q | A | 2.5e-08 | 1.5e-08 | 6.9e-08 | 3.633E-08 | 2.873E-08 | 1.500E-08 | 6.900E-08 |
| 10948 | 548 | Q548G | A | 2.7e-08 | 7.3e-08 | 2.3e-08 | 4.100E-08 | 2.778E-08 | 2.300E-08 | 7.300E-08 |
| 10949 | 548 | Q548H | A | 1.6e-08 | 3.4e-08 | 5.5e-08 | 3.500E-08 | 1.952E-08 | 1.600E-08 | 5.500E-08 |
| 10950 | 548 | Q548I | A | 3.5e-08 | 4.1e-08 | 4.5e-08 | 4.033E-08 | 5.033E-09 | 3.500E-08 | 4.500E-08 |
| 10951 | 548 | Q548L | A | 3.8e-08 | 2.6e-08 | 7.3e-08 | 4.567E-08 | 2.442E-08 | 2.600E-08 | 7.300E-08 |
| 10952 | 548 | Q548K | A | 5.2e-08 | 3e-08 | 2.1e-08 | 3.433E-08 | 1.595E-08 | 2.100E-08 | 5.200E-08 |
| 10953 | 548 | Q548M | A | 3.3e-08 | 5.1e-08 | 2.4e-08 | 3.600E-08 | 1.375E-08 | 2.400E-08 | 5.100E-08 |
| 10954 | 548 | Q548F | A | 4.2e-08 | 1.1e-08 | 3.5e-08 | 2.933E-08 | 1.626E-08 | 1.100E-08 | 4.200E-08 |
| 10955 | 548 | Q548P | A | 3.1e-08 | 7.8e-08 | 6.2e-08 | 5.700E-08 | 2.390E-08 | 3.100E-08 | 7.800E-08 |
| 10956 | 548 | Q548S | A | 2.5e-08 | 4.2e-08 | 5.1e-08 | 3.933E-08 | 1.320E-08 | 2.500E-08 | 5.100E-08 |
| 10957 | 548 | Q548T | A | 3.4e-08 | 5.6e-08 | 1.6e-08 | 3.533E-08 | 2.003E-08 | 1.600E-08 | 5.600E-08 |
| 10958 | 548 | Q548W | A | 1.6e-08 | 1e-07 | 2.3e-08 | 4.633E-08 | 4.661E-08 | 1.600E-08 | 1.000E-07 |
| 10959 | 548 | Q548Y | A | 4.9e-08 | 2.9e-08 | 4e-08 | 3.933E-08 | 1.002E-08 | 2.900E-08 | 4.900E-08 |
| 10960 | 548 | Q548V | A | 4.1e-08 | 2.4e-08 | 6.8e-08 | 4.433E-08 | 2.219E-08 | 2.400E-08 | 6.800E-08 |
| 10961 | 549 | N549A | A | 6.4e-08 | 8.6e-08 | 5.6e-08 | 6.867E-08 | 1.553E-08 | 5.600E-08 | 8.600E-08 |
| 10962 | 549 | N549R | A | 7.6e-09 | 3.7e-08 | 7.8e-08 | 4.087E-08 | 3.536E-08 | 7.600E-09 | 7.800E-08 |
| 10963 | 549 | N549N | A | 4.4e-08 | 4.3e-08 | 3e-08 | 3.900E-08 | 7.810E-09 | 3.000E-08 | 4.400E-08 |
| 10964 | 549 | N549D | A | 6.1e-08 | 7.5e-08 | 4.9e-08 | 6.167E-08 | 1.301E-08 | 4.900E-08 | 7.500E-08 |

| | | | | | | | | | | |
|---|---|---|---|---|---|---|---|---|---|---|
| 10965 | 549 | N549C | A | 1.9e-08 | 3.7e-08 | 9.5e-08 | 5.033E-08 | 3.972E-08 | 1.900E-08 | 9.500E-08 |
| 10966 | 549 | N549E | A | 4.6e-08 | 3.1e-08 | 1e-07 | 5.900E-08 | 3.629E-08 | 3.100E-08 | 1.000E-07 |
| 10967 | 549 | N549Q | A | 4.1e-08 | 7.6e-08 | 3.8e-08 | 5.167E-08 | 2.113E-08 | 3.800E-08 | 7.600E-08 |
| 10968 | 549 | N549G | A | 5.4e-08 | 3.6e-08 | 6.2e-08 | 5.067E-08 | 1.332E-08 | 3.600E-08 | 6.200E-08 |
| 10969 | 549 | N549H | A | 2.6e-08 | 4.2e-08 | 5.9e-08 | 4.233E-08 | 1.650E-08 | 2.600E-08 | 5.900E-08 |
| 10970 | 549 | N549I | A | 2.3e-08 | 3.6e-08 | 2.7e-08 | 2.867E-08 | 6.658E-09 | 2.300E-08 | 3.600E-08 |
| 10971 | 549 | N549L | A | 2.8e-08 | 8.9e-08 | 6.5e-08 | 6.067E-08 | 3.073E-08 | 2.800E-08 | 8.900E-08 |
| 10972 | 549 | N549K | A | 6e-08 | 1e-08 | 6.1e-08 | 4.367E-08 | 2.916E-08 | 1.000E-08 | 6.100E-08 |
| 10973 | 549 | N549M | A | 6.2e-08 | 1.3e-07 | 2.7e-08 | 7.300E-08 | 5.237E-08 | 2.700E-08 | 1.300E-07 |
| 10974 | 549 | N549F | A | 3.9e-08 | 1.1e-07 | 1.7e-08 | 5.533E-08 | 4.860E-08 | 1.700E-08 | 1.100E-07 |
| 10975 | 549 | N549P | A | 2.2e-08 | 3.3e-08 | 9.2e-08 | 4.900E-08 | 3.764E-08 | 2.200E-08 | 9.200E-08 |
| 10976 | 549 | N549S | A | 2e-08 | 1.1e-07 | 5.6e-08 | 6.200E-08 | 4.530E-08 | 2.000E-08 | 1.100E-07 |
| 10977 | 549 | N549T | A | 4.8e-08 | 2.1e-08 | 5e-08 | 3.967E-08 | 1.620E-08 | 2.100E-08 | 5.000E-08 |
| 10978 | 549 | N549W | A | 1.2e-08 | 4.7e-08 | 4.3e-08 | 3.400E-08 | 1.916E-08 | 1.200E-08 | 4.700E-08 |
| 10979 | 549 | N549Y | A | 1.2e-08 | 3.3e-08 | 3.5e-08 | 2.667E-08 | 1.274E-08 | 1.200E-08 | 3.500E-08 |
| 10980 | 549 | N549V | A | 8.9e-08 | 4.1e-08 | 4.4e-08 | 5.800E-08 | 2.689E-08 | 4.100E-08 | 8.900E-08 |
| 10981 | 550 | G550A | A | 4e-08 | 2.2e-08 | 9.3e-08 | 5.167E-08 | 3.691E-08 | 2.200E-08 | 9.300E-08 |
| 10982 | 550 | G550R | A | 6.6e-08 | 2.6e-08 | 2.9e-08 | 4.033E-08 | 2.228E-08 | 2.600E-08 | 6.600E-08 |
| 10983 | 550 | G550N | A | 3.7e-08 | 8e-08 | 4.9e-08 | 5.533E-08 | 2.219E-08 | 3.700E-08 | 8.000E-08 |
| 10984 | 550 | G550D | A | 4e-08 | 3.3e-08 | 6.2e-08 | 4.500E-08 | 1.513E-08 | 3.300E-08 | 6.200E-08 |
| 10985 | 550 | G550C | A | 3.7e-08 | 2.1e-07 | 1.7e-08 | 8.800E-08 | 1.061E-07 | 1.700E-08 | 2.100E-07 |
| 10986 | 550 | G550E | A | 1.5e-08 | 3.8e-08 | 2e-08 | 2.433E-08 | 1.210E-08 | 1.500E-08 | 3.800E-08 |
| 10987 | 550 | G550Q | A | 8.5e-08 | 1e-08 | 6e-08 | 5.167E-08 | 3.819E-08 | 1.000E-08 | 8.500E-08 |
| 10988 | 550 | G550G | A | 4.3e-08 | 7e-08 | 4.6e-08 | 5.300E-08 | 1.480E-08 | 4.300E-08 | 7.000E-08 |
| 10989 | 550 | G550H | A | 4.7e-08 | 1.6e-08 | 6.8e-08 | 4.367E-08 | 2.616E-08 | 1.600E-08 | 6.800E-08 |
| 10990 | 550 | G550I | A | 1.6e-08 | 8.9e-08 | 1.8e-07 | 9.500E-08 | 8.216E-08 | 1.600E-08 | 1.800E-07 |
| 10991 | 550 | G550L | A | 1.9e-08 | 6.1e-08 | 9.7e-09 | 2.990E-08 | 2.733E-08 | 9.700E-09 | 6.100E-08 |
| 10992 | 550 | G550K | A | 1.4e-07 | 2.2e-08 | 2.8e-08 | 6.333E-08 | 6.646E-08 | 2.200E-08 | 1.400E-07 |
| 10993 | 550 | G550M | A | 6e-08 | 1.5e-07 | 1e-07 | 1.033E-07 | 4.509E-08 | 6.000E-08 | 1.500E-07 |
| 10994 | 550 | G550F | A | 1.8e-08 | 8.2e-08 | 4.3e-08 | 4.767E-08 | 3.225E-08 | 1.800E-08 | 8.200E-08 |
| 10995 | 550 | G550P | A | 5.9e-08 | 4.2e-08 | 5.7e-08 | 5.267E-08 | 9.292E-09 | 4.200E-08 | 5.900E-08 |
| 10996 | 550 | G550S | A | 6.5e-08 | 1e-07 | 2.4e-08 | 6.300E-08 | 3.804E-08 | 2.400E-08 | 1.000E-07 |
| 10997 | 550 | G550T | A | 1e-07 | 6.2e-08 | 4.9e-08 | 7.033E-08 | 2.650E-08 | 4.900E-08 | 1.000E-07 |
| 10998 | 550 | G550W | A | 7.4e-08 | 5.1e-08 | 2.6e-08 | 5.033E-08 | 2.401E-08 | 2.600E-08 | 7.400E-08 |
| 10999 | 550 | G550Y | A | 7.5e-08 | 1.3e-08 | 4.3e-08 | 4.367E-08 | 3.101E-08 | 1.300E-08 | 7.500E-08 |
| 11000 | 550 | G550V | A | 6.4e-08 | 6.1e-08 | 2.9e-08 | 5.133E-08 | 1.940E-08 | 2.900E-08 | 6.400E-08 |
| 11001 | 551 | S551A | A | 7.8e-08 | 4.2e-08 | 4.9e-08 | 5.633E-08 | 1.909E-08 | 4.200E-08 | 7.800E-08 |
| 11002 | 551 | S551R | A | 6.2e-08 | 6.6e-08 | 2.4e-08 | 5.067E-08 | 2.318E-08 | 2.400E-08 | 6.600E-08 |
| 11003 | 551 | S551N | A | 4.9e-08 | 8.7e-08 | 1.5e-08 | 5.033E-08 | 3.602E-08 | 1.500E-08 | 8.700E-08 |
| 11004 | 551 | S551D | A | 6.2e-08 | 1.6e-08 | 3.8e-08 | 3.867E-08 | 2.301E-08 | 1.600E-08 | 6.200E-08 |
| 11005 | 551 | S551C | A | 8.5e-08 | 5.1e-08 | 1.3e-08 | 4.967E-08 | 3.602E-08 | 1.300E-08 | 8.500E-08 |
| 11006 | 551 | S551E | A | 2.5e-08 | 4.5e-08 | 1e-07 | 5.667E-08 | 3.884E-08 | 2.500E-08 | 1.000E-07 |
| 11007 | 551 | S551Q | A | 2e-08 | 5.3e-08 | 3.1e-08 | 3.467E-08 | 1.680E-08 | 2.000E-08 | 5.300E-08 |
| 11008 | 551 | S551G | A | 9.1e-08 | 5.6e-08 | 2.1e-08 | 5.600E-08 | 3.500E-08 | 2.100E-08 | 9.100E-08 |
| 11009 | 551 | S551H | A | 7.9e-08 | 2.9e-08 | 2.3e-08 | 4.367E-08 | 3.075E-08 | 2.300E-08 | 7.900E-08 |
| 11010 | 551 | S551I | A | 2.4e-08 | 1.8e-08 | 3.3e-08 | 2.500E-08 | 7.550E-09 | 1.800E-08 | 3.300E-08 |
| 11011 | 551 | S551L | A | 4.4e-08 | 4e-08 | 2.3e-08 | 3.567E-08 | 1.115E-08 | 2.300E-08 | 4.400E-08 |
| 11012 | 551 | S551K | A | 3.2e-08 | 4.3e-08 | 3.4e-08 | 3.633E-08 | 5.859E-09 | 3.200E-08 | 4.300E-08 |
| 11013 | 551 | S551M | A | 3.4e-08 | 5.3e-08 | 3.9e-08 | 4.200E-08 | 9.849E-09 | 3.400E-08 | 5.300E-08 |
| 11014 | 551 | S551F | A | 2e-08 | 9e-08 | 6.9e-08 | 5.967E-08 | 3.592E-08 | 2.000E-08 | 9.000E-08 |
| 11015 | 551 | S551P | A | 2.4e-08 | 7.1e-08 | 6.3e-08 | 5.267E-08 | 2.515E-08 | 2.400E-08 | 7.100E-08 |
| 11016 | 551 | S551S | A | 3.5e-08 | 4.1e-08 | 2.6e-08 | 3.400E-08 | 7.550E-09 | 2.600E-08 | 4.100E-08 |
| 11017 | 551 | S551T | A | 2.9e-08 | 4.1e-08 | 1.3e-07 | 6.667E-08 | 5.518E-08 | 2.900E-08 | 1.300E-07 |
| 11018 | 551 | S551W | A | 3.3e-08 | 2.3e-08 | 5.6e-08 | 3.733E-08 | 1.692E-08 | 2.300E-08 | 5.600E-08 |
| 11019 | 551 | S551Y | A | 4e-08 | 9.7e-08 | 2.7e-08 | 5.467E-08 | 3.723E-08 | 2.700E-08 | 9.700E-08 |
| 11020 | 551 | S551V | A | 8.9e-08 | 1.4e-07 | 2.5e-08 | 8.467E-08 | 5.762E-08 | 2.500E-08 | 1.400E-07 |
| 11021 | 552 | V552A | A | 4.6e-08 | 3.7e-08 | 1.2e-08 | 3.167E-08 | 1.762E-08 | 1.200E-08 | 4.600E-08 |
| 11022 | 552 | V552R | A | 1e-07 | 1.9e-08 | 2.5e-08 | 4.800E-08 | 4.513E-08 | 1.900E-08 | 1.000E-07 |
| 11023 | 552 | V552N | A | 3.6e-08 | 4.1e-08 | 3e-08 | 3.567E-08 | 5.508E-09 | 3.000E-08 | 4.100E-08 |
| 11024 | 552 | V552D | A | 6.7e-08 | 1e-08 | 5.4e-08 | 4.367E-08 | 2.987E-08 | 1.000E-08 | 6.700E-08 |
| 11025 | 552 | V552C | A | 7.7e-08 | 6.5e-08 | 2.4e-08 | 5.533E-08 | 2.779E-08 | 2.400E-08 | 7.700E-08 |
| 11026 | 552 | V552E | A | 3.6e-08 | 7.5e-08 | 5.3e-08 | 5.467E-08 | 1.955E-08 | 3.600E-08 | 7.500E-08 |
| 11027 | 552 | V552Q | A | 5.9e-08 | 1.9e-08 | 5.7e-08 | 4.500E-08 | 2.254E-08 | 1.900E-08 | 5.900E-08 |
| 11028 | 552 | V552G | A | 3e-08 | 4.1e-08 | 8.2e-08 | 5.100E-08 | 2.740E-08 | 3.000E-08 | 8.200E-08 |
| 11029 | 552 | V552H | A | 4.3e-08 | 2.7e-08 | 3.5e-08 | 3.500E-08 | 8.000E-09 | 2.700E-08 | 4.300E-08 |
| 11030 | 552 | V552I | A | 1e-07 | 2.2e-08 | 4.4e-08 | 5.533E-08 | 4.022E-08 | 2.200E-08 | 1.000E-07 |
| 11031 | 552 | V552L | A | 2.5e-07 | 2.3e-08 | 4.6e-08 | 1.063E-07 | 1.249E-07 | 2.300E-08 | 2.500E-07 |
| 11032 | 552 | V552K | A | 1.9e-07 | 2e-07 | 2.5e-08 | 1.383E-07 | 9.828E-08 | 2.500E-08 | 2.000E-07 |
| 11033 | 552 | V552M | A | 6.8e-08 | 3.4e-08 | 1.7e-08 | 3.967E-08 | 2.597E-08 | 1.700E-08 | 6.800E-08 |
| 11034 | 552 | V552F | A | 3.7e-08 | 5.3e-08 | 5.3e-08 | 4.767E-08 | 9.238E-09 | 3.700E-08 | 5.300E-08 |
| 11035 | 552 | V552P | A | 2.7e-08 | 2.5e-08 | 2.9e-08 | 2.700E-08 | 2.000E-09 | 2.500E-08 | 2.900E-08 |
| 11036 | 552 | V552S | A | 6.8e-08 | 6.7e-08 | 1.9e-08 | 5.133E-08 | 2.801E-08 | 1.900E-08 | 6.800E-08 |
| 11037 | 552 | V552T | A | 8.8e-08 | 2e-08 | 9.5e-08 | 6.767E-08 | 4.143E-08 | 2.000E-08 | 9.500E-08 |
| 11038 | 552 | V552W | A | 4.1e-08 | 3.7e-08 | 5.1e-08 | 4.300E-08 | 7.211E-09 | 3.700E-08 | 5.100E-08 |
| 11039 | 552 | V552Y | A | 3.4e-08 | 5.4e-08 | 2.7e-08 | 3.833E-08 | 1.401E-08 | 2.700E-08 | 5.400E-08 |
| 11040 | 552 | V552V | A | 3.9e-08 | 8.1e-08 | 1.6e-08 | 4.533E-08 | 3.296E-08 | 1.600E-08 | 8.100E-08 |
| 11041 | 553 | T553A | A | 2e-08 | 2.3e-08 | 6.2e-08 | 3.500E-08 | 2.343E-08 | 2.000E-08 | 6.200E-08 |
| 11042 | 553 | T553R | A | 4e-08 | 3.8e-08 | 5.1e-08 | 4.300E-08 | 7.000E-09 | 3.800E-08 | 5.100E-08 |
| 11043 | 553 | T553N | A | 2.2e-08 | 6.9e-08 | 3.5e-08 | 4.200E-08 | 2.427E-08 | 2.200E-08 | 6.900E-08 |
| 11044 | 553 | T553D | A | 1.9e-08 | 6.6e-08 | 1e-07 | 6.167E-08 | 4.067E-08 | 1.900E-08 | 1.000E-07 |
| 11045 | 553 | T553C | A | 9.6e-08 | 7.4e-08 | 5.8e-08 | 7.600E-08 | 1.908E-08 | 5.800E-08 | 9.600E-08 |
| 11046 | 553 | T553E | A | 4.1e-08 | 3.5e-08 | 8e-08 | 5.200E-08 | 2.443E-08 | 3.500E-08 | 8.000E-08 |
| 11047 | 553 | T553Q | A | 1.7e-08 | 6.8e-08 | 5.3e-08 | 4.600E-08 | 2.621E-08 | 1.700E-08 | 6.800E-08 |
| 11048 | 553 | T553G | A | 3.5e-08 | 3.9e-08 | 2.7e-08 | 3.367E-08 | 6.110E-09 | 2.700E-08 | 3.900E-08 |
| 11049 | 553 | T553H | A | 6.4e-08 | 1.5e-08 | 4.9e-08 | 4.267E-08 | 2.511E-08 | 1.500E-08 | 6.400E-08 |

| | | | | | | | | | | |
|---|---|---|---|---|---|---|---|---|---|---|
| 11050 | 553 | T553I | A | 5.7e-08 | 6.9e-08 | 8.3e-08 | 6.967E-08 | 1.301E-08 | 5.700E-08 | 8.300E-08 |
| 11051 | 553 | T553L | A | 1.9e-08 | 6e-08 | 5.6e-08 | 4.500E-08 | 2.261E-08 | 1.900E-08 | 6.000E-08 |
| 11052 | 553 | T553K | A | 5.5e-08 | 4.6e-08 | 1.4e-08 | 3.833E-08 | 2.155E-08 | 1.400E-08 | 5.500E-08 |
| 11053 | 553 | T553M | A | 3.8e-08 | 9.6e-09 | 6.6e-08 | 3.787E-08 | 2.820E-08 | 9.600E-09 | 6.600E-08 |
| 11054 | 553 | T553F | A | 8.2e-08 | 3.8e-08 | 1.5e-08 | 4.500E-08 | 3.404E-08 | 1.500E-08 | 8.200E-08 |
| 11055 | 553 | T553P | A | 4.3e-08 | 2.5e-08 | 9.8e-08 | 5.533E-08 | 3.803E-08 | 2.500E-08 | 9.800E-08 |
| 11056 | 553 | T553S | A | 3.1e-08 | 3.7e-08 | 8.6e-08 | 5.133E-08 | 3.017E-08 | 3.100E-08 | 8.600E-08 |
| 11057 | 553 | T553T | A | 4.2e-08 | 1.5e-08 | 3e-08 | 2.900E-08 | 1.353E-08 | 1.500E-08 | 4.200E-08 |
| 11058 | 553 | T553W | A | 2.1e-08 | 3.8e-08 | 4.6e-08 | 3.500E-08 | 1.277E-08 | 2.100E-08 | 4.600E-08 |
| 11059 | 553 | T553Y | A | 3e-08 | 3.4e-08 | 4.1e-08 | 3.500E-08 | 5.568E-09 | 3.000E-08 | 4.100E-08 |
| 11060 | 553 | T553V | A | 2.8e-08 | 5.8e-08 | 8.3e-08 | 5.633E-08 | 2.754E-08 | 2.800E-08 | 8.300E-08 |
| 11061 | 554 | C554A | A | 6.9e-08 | 2.4e-08 | 2.9e-08 | 4.067E-08 | 2.466E-08 | 2.400E-08 | 6.900E-08 |
| 11062 | 554 | C554R | A | 7.3e-08 | 5.3e-08 | 5.1e-08 | 5.900E-08 | 1.217E-08 | 5.100E-08 | 7.300E-08 |
| 11063 | 554 | C554N | A | 5.1e-08 | 3.4e-08 | 1.6e-08 | 3.367E-08 | 1.750E-08 | 1.600E-08 | 5.100E-08 |
| 11064 | 554 | C554D | A | 3.5e-08 | 3.2e-08 | 4.8e-08 | 3.833E-08 | 8.505E-09 | 3.200E-08 | 4.800E-08 |
| 11065 | 554 | C554C | A | 1.4e-08 | 4.6e-08 | 2.4e-08 | 2.800E-08 | 1.637E-08 | 1.400E-08 | 4.600E-08 |
| 11066 | 554 | C554E | A | 3e-08 | 4.4e-08 | 5.5e-08 | 4.300E-08 | 1.253E-08 | 3.000E-08 | 5.500E-08 |
| 11067 | 554 | C554Q | A | 6.3e-08 | 9.8e-08 | 4e-08 | 6.700E-08 | 2.921E-08 | 4.000E-08 | 9.800E-08 |
| 11068 | 554 | C554G | A | 3.7e-08 | 1.6e-08 | 2.5e-08 | 2.600E-08 | 1.054E-08 | 1.600E-08 | 3.700E-08 |
| 11069 | 554 | C554H | A | 1e-07 | 4.3e-08 | 3e-08 | 5.767E-08 | 3.723E-08 | 3.000E-08 | 1.000E-07 |
| 11070 | 554 | C554I | A | 2e-08 | 7.7e-08 | 8.4e-08 | 6.033E-08 | 3.510E-08 | 2.000E-08 | 8.400E-08 |
| 11071 | 554 | C554L | A | 7.1e-08 | 8.1e-08 | 4e-08 | 6.400E-08 | 2.138E-08 | 4.000E-08 | 8.100E-08 |
| 11072 | 554 | C554K | A | 1.8e-08 | 5.2e-08 | 1e-07 | 5.667E-08 | 4.120E-08 | 1.800E-08 | 1.000E-07 |
| 11073 | 554 | C554M | A | 1.5e-08 | 2.8e-08 | 1.1e-07 | 5.100E-08 | 5.151E-08 | 1.500E-08 | 1.100E-07 |
| 11074 | 554 | C554F | A | 2.6e-08 | 3e-08 | 2.6e-08 | 2.733E-08 | 2.309E-09 | 2.600E-08 | 3.000E-08 |
| 11075 | 554 | C554P | A | 7.4e-08 | 9.3e-08 | 6.1e-08 | 7.600E-08 | 1.609E-08 | 6.100E-08 | 9.300E-08 |
| 11076 | 554 | C554S | A | 5.5e-08 | 6.1e-08 | 3.6e-08 | 5.067E-08 | 1.305E-08 | 3.600E-08 | 6.100E-08 |
| 11077 | 554 | C554T | A | 2.7e-08 | 4.8e-08 | 3.6e-08 | 3.700E-08 | 1.054E-08 | 2.700E-08 | 4.800E-08 |
| 11078 | 554 | C554W | A | 5.3e-08 | 1.3e-07 | 5.3e-08 | 7.867E-08 | 4.446E-08 | 5.300E-08 | 1.300E-07 |
| 11079 | 554 | C554Y | A | 1.4e-08 | 6.5e-08 | 3.4e-08 | 3.767E-08 | 2.570E-08 | 1.400E-08 | 6.500E-08 |
| 11080 | 554 | C554V | A | 4.6e-08 | 2.4e-08 | 5e-08 | 4.000E-08 | 1.400E-08 | 2.400E-08 | 5.000E-08 |
| 11081 | 555 | F555A | A | 3.4e-08 | 6.4e-08 | 4e-08 | 4.600E-08 | 1.587E-08 | 3.400E-08 | 6.400E-08 |
| 11082 | 555 | F555R | A | 7.9e-08 | 3.3e-08 | 8.4e-08 | 6.533E-08 | 2.811E-08 | 3.300E-08 | 8.400E-08 |
| 11083 | 555 | F555N | A | 6.9e-08 | 6.3e-08 | 3.6e-08 | 5.600E-08 | 1.758E-08 | 3.600E-08 | 6.900E-08 |
| 11084 | 555 | F555D | A | 4.4e-08 | 6.5e-08 | 8e-08 | 6.300E-08 | 1.808E-08 | 4.400E-08 | 8.000E-08 |
| 11085 | 555 | F555C | A | 2.1e-08 | 2.4e-08 | 4.2e-08 | 2.900E-08 | 1.136E-08 | 2.100E-08 | 4.200E-08 |
| 11086 | 555 | F555E | A | 3.8e-08 | 8.8e-08 | 4.3e-08 | 5.633E-08 | 2.754E-08 | 3.800E-08 | 8.800E-08 |
| 11087 | 555 | F555Q | A | 1.8e-08 | 2e-08 | 4.2e-08 | 2.667E-08 | 1.332E-08 | 1.800E-08 | 4.200E-08 |
| 11088 | 555 | F555G | A | 9.9e-08 | 2.4e-08 | 2.9e-08 | 5.067E-08 | 4.193E-08 | 2.400E-08 | 9.900E-08 |
| 11089 | 555 | F555H | A | 5.2e-08 | 7.1e-08 | 4.1e-08 | 5.467E-08 | 1.518E-08 | 4.100E-08 | 7.100E-08 |
| 11090 | 555 | F555I | A | 6.6e-08 | 6.8e-08 | 1.5e-07 | 9.467E-08 | 4.793E-08 | 6.600E-08 | 1.500E-07 |
| 11091 | 555 | F555L | A | 4.1e-08 | 3.1e-08 | 1.5e-08 | 2.900E-08 | 1.311E-08 | 1.500E-08 | 4.100E-08 |
| 11092 | 555 | F555K | A | 1e-07 | 2.4e-08 | 9e-08 | 7.133E-08 | 4.130E-08 | 2.400E-08 | 1.000E-07 |
| 11093 | 555 | F555M | A | 3.7e-08 | 2.4e-08 | 8.3e-08 | 4.800E-08 | 3.100E-08 | 2.400E-08 | 8.300E-08 |
| 11094 | 555 | F555F | A | 2.7e-08 | 4.2e-08 | 5e-08 | 3.967E-08 | 1.168E-08 | 2.700E-08 | 5.000E-08 |
| 11095 | 555 | F555P | A | 8.4e-08 | 1.7e-08 | 1.4e-07 | 8.033E-08 | 6.158E-08 | 1.700E-08 | 1.400E-07 |
| 11096 | 555 | F555S | A | 2.8e-08 | 2.9e-08 | 2.8e-08 | 2.833E-08 | 5.774E-10 | 2.800E-08 | 2.900E-08 |
| 11097 | 555 | F555T | A | 4.2e-08 | 3.8e-08 | 7.4e-08 | 5.133E-08 | 1.973E-08 | 3.800E-08 | 7.400E-08 |
| 11098 | 555 | F555W | A | 1.5e-08 | 3.2e-08 | 1e-07 | 4.900E-08 | 4.498E-08 | 1.500E-08 | 1.000E-07 |
| 11099 | 555 | F555Y | A | 4.6e-08 | 3.4e-08 | 4.3e-08 | 4.100E-08 | 6.245E-09 | 3.400E-08 | 4.600E-08 |
| 11100 | 555 | F555V | A | 2.1e-08 | 2.3e-08 | 8.4e-08 | 4.267E-08 | 3.581E-08 | 2.100E-08 | 8.400E-08 |
| 11101 | 556 | G556A | A | 8.5e-08 | 1.8e-08 | 1.7e-08 | 4.000E-08 | 3.897E-08 | 1.700E-08 | 8.500E-08 |
| 11102 | 556 | G556R | A | 1.3e-08 | 4.7e-08 | 5.6e-08 | 3.867E-08 | 2.268E-08 | 1.300E-08 | 5.600E-08 |
| 11103 | 556 | G556N | A | 7.3e-08 | 2.3e-08 | 7.3e-08 | 5.633E-08 | 2.887E-08 | 2.300E-08 | 7.300E-08 |
| 11104 | 556 | G556D | A | 4.4e-08 | 3.1e-08 | 2.1e-07 | 9.500E-08 | 9.980E-08 | 3.100E-08 | 2.100E-07 |
| 11105 | 556 | G556C | A | 1.9e-08 | 4.4e-08 | 6.2e-08 | 4.167E-08 | 2.159E-08 | 1.900E-08 | 6.200E-08 |
| 11106 | 556 | G556E | A | 7.4e-08 | 4.6e-08 | 4.7e-08 | 5.567E-08 | 1.589E-08 | 4.600E-08 | 7.400E-08 |
| 11107 | 556 | G556Q | A | 3.3e-08 | 2e-08 | 3.4e-08 | 2.900E-08 | 7.810E-09 | 2.000E-08 | 3.400E-08 |
| 11108 | 556 | G556G | A | 1.1e-08 | 2.8e-08 | 6.7e-08 | 3.533E-08 | 2.871E-08 | 1.100E-08 | 6.700E-08 |
| 11109 | 556 | G556H | A | 7.1e-08 | 1.7e-08 | 7.1e-08 | 5.300E-08 | 3.118E-08 | 1.700E-08 | 7.100E-08 |
| 11110 | 556 | G556I | A | 4.8e-08 | 1.5e-08 | 1.2e-08 | 2.500E-08 | 1.997E-08 | 1.200E-08 | 4.800E-08 |
| 11111 | 556 | G556L | A | 4.8e-08 | 4.8e-08 | 6.8e-08 | 5.467E-08 | 1.155E-08 | 4.800E-08 | 6.800E-08 |
| 11112 | 556 | G556K | A | 2.6e-08 | 5.9e-08 | 2.4e-08 | 3.633E-08 | 1.966E-08 | 2.400E-08 | 5.900E-08 |
| 11113 | 556 | G556M | A | 8.4e-08 | 5.8e-08 | 7.9e-08 | 7.367E-08 | 1.380E-08 | 5.800E-08 | 8.400E-08 |
| 11114 | 556 | G556F | A | 1.8e-08 | 2.5e-08 | 3.1e-08 | 2.467E-08 | 6.506E-09 | 1.800E-08 | 3.100E-08 |
| 11115 | 556 | G556P | A | 3.9e-08 | 1.9e-08 | 4.6e-08 | 3.467E-08 | 1.401E-08 | 1.900E-08 | 4.600E-08 |
| 11116 | 556 | G556S | A | 2e-08 | 3.6e-08 | 3.4e-08 | 3.000E-08 | 8.718E-09 | 2.000E-08 | 3.600E-08 |
| 11117 | 556 | G556T | A | 2.7e-08 | 3.1e-08 | 4.7e-08 | 3.500E-08 | 1.058E-08 | 2.700E-08 | 4.700E-08 |
| 11118 | 556 | G556W | A | 7.8e-08 | 5.3e-08 | 4e-08 | 5.700E-08 | 1.931E-08 | 4.000E-08 | 7.800E-08 |
| 11119 | 556 | G556Y | A | 5.8e-08 | 5.6e-08 | 4.8e-08 | 5.400E-08 | 5.292E-09 | 4.800E-08 | 5.800E-08 |
| 11120 | 556 | G556V | A | 4e-08 | 3.6e-08 | 5.9e-08 | 4.500E-08 | 1.229E-08 | 3.600E-08 | 5.900E-08 |
| 11121 | 557 | P557A | A | 3.4e-08 | 2.1e-08 | 5.9e-08 | 3.800E-08 | 1.931E-08 | 2.100E-08 | 5.900E-08 |
| 11122 | 557 | P557R | A | 7.6e-08 | 2.2e-08 | 4.5e-08 | 4.767E-08 | 2.710E-08 | 2.200E-08 | 7.600E-08 |
| 11123 | 557 | P557N | A | 2e-08 | 5.6e-08 | 1.4e-07 | 7.200E-08 | 6.158E-08 | 2.000E-08 | 1.400E-07 |
| 11124 | 557 | P557D | A | 1.8e-08 | 3.8e-08 | 2.7e-08 | 2.767E-08 | 1.002E-08 | 1.800E-08 | 3.800E-08 |
| 11125 | 557 | P557C | A | 4.5e-08 | 1.4e-07 | 1.2e-08 | 6.567E-08 | 6.646E-08 | 1.200E-08 | 1.400E-07 |
| 11126 | 557 | P557E | A | 1.5e-08 | 3.3e-08 | 4.6e-08 | 3.133E-08 | 1.557E-08 | 1.500E-08 | 4.600E-08 |
| 11127 | 557 | P557Q | A | 5.3e-08 | 5.6e-08 | 1.2e-07 | 7.633E-08 | 3.785E-08 | 5.300E-08 | 1.200E-07 |
| 11128 | 557 | P557G | A | 6.1e-08 | 8.6e-08 | 1.6e-08 | 5.433E-08 | 3.547E-08 | 1.600E-08 | 8.600E-08 |
| 11129 | 557 | P557H | A | 4.2e-08 | 7.1e-08 | 3.5e-08 | 4.933E-08 | 1.909E-08 | 3.500E-08 | 7.100E-08 |
| 11130 | 557 | P557I | A | 2.7e-08 | 8.7e-08 | 2.5e-08 | 4.633E-08 | 3.523E-08 | 2.500E-08 | 8.700E-08 |
| 11131 | 557 | P557L | A | 4.6e-08 | 2.2e-08 | 3.6e-08 | 3.467E-08 | 1.206E-08 | 2.200E-08 | 4.600E-08 |
| 11132 | 557 | P557K | A | 3.2e-08 | 8.1e-08 | 3e-08 | 4.767E-08 | 2.888E-08 | 3.000E-08 | 8.100E-08 |
| 11133 | 557 | P557M | A | 4.2e-08 | 3.2e-08 | 7.3e-08 | 4.900E-08 | 2.138E-08 | 3.200E-08 | 7.300E-08 |
| 11134 | 557 | P557F | A | 4e-08 | 1.3e-07 | 4e-08 | 7.000E-08 | 5.196E-08 | 4.000E-08 | 1.300E-07 |

| | | | | | | | | | | |
|---|---|---|---|---|---|---|---|---|---|---|
| 11135 | 557 | P557P | A | 6.3e-08 | 3e-08 | 2.6e-08 | 3.967E-08 | 2.031E-08 | 2.600E-08 | 6.300E-08 |
| 11136 | 557 | P557S | A | 6.6e-08 | 7.7e-08 | 5.6e-08 | 6.633E-08 | 1.050E-08 | 5.600E-08 | 7.700E-08 |
| 11137 | 557 | P557T | A | 4.5e-08 | 6.3e-08 | 4e-08 | 4.933E-08 | 1.210E-08 | 4.000E-08 | 6.300E-08 |
| 11138 | 557 | P557W | A | 2e-08 | 9.9e-08 | 3.1e-08 | 5.000E-08 | 4.279E-08 | 2.000E-08 | 9.900E-08 |
| 11139 | 557 | P557Y | A | 8.9e-08 | 6.9e-08 | 7.8e-08 | 7.867E-08 | 1.002E-08 | 6.900E-08 | 8.900E-08 |
| 11140 | 557 | P557V | A | 1.2e-08 | 2.8e-08 | 3e-08 | 2.333E-08 | 9.866E-09 | 1.200E-08 | 3.000E-08 |
| 11141 | 558 | E558A | A | 8.4e-08 | 3.8e-08 | 4.1e-08 | 5.433E-08 | 2.574E-08 | 3.800E-08 | 8.400E-08 |
| 11142 | 558 | E558R | A | 1.5e-08 | 4.4e-08 | 1.6e-08 | 2.500E-08 | 1.646E-08 | 1.500E-08 | 4.400E-08 |
| 11143 | 558 | E558N | A | 4.4e-08 | 1.1e-08 | 5.6e-08 | 3.700E-08 | 2.330E-08 | 1.100E-08 | 5.600E-08 |
| 11144 | 558 | E558D | A | 4.4e-08 | 1.4e-08 | 3.1e-08 | 2.967E-08 | 1.504E-08 | 1.400E-08 | 4.400E-08 |
| 11145 | 558 | E558C | A | 8.9e-08 | 2e-08 | 2.9e-08 | 4.600E-08 | 3.751E-08 | 2.000E-08 | 8.900E-08 |
| 11146 | 558 | E558E | A | 2.6e-08 | 2.3e-08 | 8.5e-08 | 4.467E-08 | 3.496E-08 | 2.300E-08 | 8.500E-08 |
| 11147 | 558 | E558Q | A | 4.9e-09 | 3.9e-08 | 6.1e-08 | 3.497E-08 | 2.827E-08 | 4.900E-09 | 6.100E-08 |
| 11148 | 558 | E558G | A | 3e-08 | 3e-08 | 7.1e-08 | 4.367E-08 | 2.367E-08 | 3.000E-08 | 7.100E-08 |
| 11149 | 558 | E558H | A | 1.9e-08 | 5.8e-08 | 2.5e-08 | 3.400E-08 | 2.100E-08 | 1.900E-08 | 5.800E-08 |
| 11150 | 558 | E558I | A | 2.8e-08 | 2.1e-07 | 1.5e-08 | 8.433E-08 | 1.090E-07 | 1.500E-08 | 2.100E-07 |
| 11151 | 558 | E558L | A | 7.5e-08 | 4.1e-08 | 6.7e-08 | 6.100E-08 | 1.778E-08 | 4.100E-08 | 7.500E-08 |
| 11152 | 558 | E558K | A | 2.5e-08 | 3.9e-08 | 2.6e-08 | 3.000E-08 | 7.810E-09 | 2.500E-08 | 3.900E-08 |
| 11153 | 558 | E558M | A | 3.9e-08 | 4e-08 | 2.6e-08 | 3.500E-08 | 7.810E-09 | 2.600E-08 | 4.000E-08 |
| 11154 | 558 | E558F | A | 4.5e-08 | 4.9e-08 | 3.9e-08 | 4.433E-08 | 5.033E-09 | 3.900E-08 | 4.900E-08 |
| 11155 | 558 | E558P | A | 7.3e-08 | 1.1e-07 | 3e-08 | 7.100E-08 | 4.004E-08 | 3.000E-08 | 1.100E-07 |
| 11156 | 558 | E558S | A | 7.5e-08 | 4e-08 | 5.9e-08 | 5.800E-08 | 1.752E-08 | 4.000E-08 | 7.500E-08 |
| 11157 | 558 | E558T | A | 4.8e-08 | 7.7e-08 | 8.6e-08 | 7.033E-08 | 1.986E-08 | 4.800E-08 | 8.600E-08 |
| 11158 | 558 | E558W | A | 2.2e-08 | 6.5e-08 | 6.7e-08 | 5.133E-08 | 2.542E-08 | 2.200E-08 | 6.700E-08 |
| 11159 | 558 | E558Y | A | 4.5e-08 | 6e-08 | 1.3e-07 | 7.833E-08 | 4.537E-08 | 4.500E-08 | 1.300E-07 |
| 11160 | 558 | E558V | A | 3.1e-08 | 4.8e-08 | 8.7e-08 | 5.533E-08 | 2.871E-08 | 3.100E-08 | 8.700E-08 |
| 11161 | 559 | A559A | A | 5.3e-08 | 8.8e-08 | 3.1e-08 | 5.733E-08 | 2.875E-08 | 3.100E-08 | 8.800E-08 |
| 11162 | 559 | A559R | A | 4.1e-08 | 4.9e-08 | 8.2e-08 | 5.733E-08 | 2.173E-08 | 4.100E-08 | 8.200E-08 |
| 11163 | 559 | A559N | A | 4.3e-08 | 1.7e-08 | 5.6e-08 | 3.867E-08 | 1.986E-08 | 1.700E-08 | 5.600E-08 |
| 11164 | 559 | A559D | A | 6.2e-08 | 5.9e-08 | 4.4e-08 | 5.500E-08 | 9.644E-09 | 4.400E-08 | 6.200E-08 |
| 11165 | 559 | A559C | A | 4.6e-08 | 4.3e-08 | 6e-09 | 3.167E-08 | 2.228E-08 | 6.000E-09 | 4.600E-08 |
| 11166 | 559 | A559E | A | 4e-08 | 5.4e-08 | 2.7e-08 | 4.033E-08 | 1.350E-08 | 2.700E-08 | 5.400E-08 |
| 11167 | 559 | A559Q | A | 9.2e-09 | 4.6e-08 | 4.8e-08 | 3.440E-08 | 2.185E-08 | 9.200E-09 | 4.800E-08 |
| 11168 | 559 | A559G | A | 2.9e-08 | 1.3e-08 | 2.1e-08 | 2.100E-08 | 8.000E-09 | 1.300E-08 | 2.900E-08 |
| 11169 | 559 | A559H | A | 5.5e-08 | 6.4e-08 | 7.1e-08 | 6.333E-08 | 8.021E-09 | 5.500E-08 | 7.100E-08 |
| 11170 | 559 | A559I | A | 4.6e-08 | 5.3e-08 | 8.6e-08 | 6.167E-08 | 2.136E-08 | 4.600E-08 | 8.600E-08 |
| 11171 | 559 | A559L | A | 9.4e-08 | 9.2e-09 | 3.2e-08 | 4.507E-08 | 4.388E-08 | 9.200E-09 | 9.400E-08 |
| 11172 | 559 | A559K | A | 1.1e-07 | 8.5e-08 | 9.3e-08 | 9.600E-08 | 1.277E-08 | 8.500E-08 | 1.100E-07 |
| 11173 | 559 | A559M | A | 3.5e-08 | 1e-07 | 1.6e-08 | 5.033E-08 | 4.405E-08 | 1.600E-08 | 1.000E-07 |
| 11174 | 559 | A559F | A | 4.8e-08 | 5e-08 | 5e-08 | 4.933E-08 | 1.155E-09 | 4.800E-08 | 5.000E-08 |
| 11175 | 559 | A559P | A | 9.1e-08 | 5e-08 | 5.6e-08 | 6.567E-08 | 2.214E-08 | 5.000E-08 | 9.100E-08 |
| 11176 | 559 | A559S | A | 5.3e-08 | 2.6e-08 | 1.1e-08 | 3.000E-08 | 2.128E-08 | 1.100E-08 | 5.300E-08 |
| 11177 | 559 | A559T | A | 6.5e-08 | 4.8e-08 | 4.4e-08 | 5.233E-08 | 1.115E-08 | 4.400E-08 | 6.500E-08 |
| 11178 | 559 | A559W | A | 5.9e-08 | 6.9e-09 | 7.2e-08 | 4.597E-08 | 3.445E-08 | 6.900E-09 | 7.200E-08 |
| 11179 | 559 | A559Y | A | 5.1e-08 | 7.5e-08 | 6.7e-08 | 6.433E-08 | 1.222E-08 | 5.100E-08 | 7.500E-08 |
| 11180 | 559 | A559V | A | 5.2e-08 | 7.4e-08 | 8.1e-08 | 6.900E-08 | 1.513E-08 | 5.200E-08 | 8.100E-08 |
| 11181 | 560 | D560A | A | 2.6e-08 | 3.2e-08 | 7e-08 | 4.267E-08 | 2.386E-08 | 2.600E-08 | 7.000E-08 |
| 11182 | 560 | D560R | A | 4.3e-08 | 3.3e-08 | 5.2e-08 | 4.267E-08 | 9.504E-09 | 3.300E-08 | 5.200E-08 |
| 11183 | 560 | D560N | A | 3.6e-08 | 5.5e-08 | 2.2e-08 | 3.767E-08 | 1.656E-08 | 2.200E-08 | 5.500E-08 |
| 11184 | 560 | D560D | A | 6.7e-08 | 6.1e-08 | 8.3e-08 | 7.033E-08 | 1.137E-08 | 6.100E-08 | 8.300E-08 |
| 11185 | 560 | D560C | A | 4.5e-08 | 8.2e-08 | 6e-08 | 6.233E-08 | 1.861E-08 | 4.500E-08 | 8.200E-08 |
| 11186 | 560 | D560E | A | 2.1e-08 | 1.8e-08 | 5.5e-08 | 3.133E-08 | 2.055E-08 | 1.800E-08 | 5.500E-08 |
| 11187 | 560 | D560Q | A | 8.7e-08 | 7.9e-08 | 3.6e-08 | 6.733E-08 | 2.743E-08 | 3.600E-08 | 8.700E-08 |
| 11188 | 560 | D560G | A | 4.7e-08 | 1.6e-08 | 3.8e-08 | 3.367E-08 | 1.595E-08 | 1.600E-08 | 4.700E-08 |
| 11189 | 560 | D560H | A | 1.4e-07 | 1.8e-08 | 1.2e-08 | 5.667E-08 | 7.223E-08 | 1.200E-08 | 1.400E-07 |
| 11190 | 560 | D560I | A | 8.3e-08 | 6.6e-08 | 1.9e-08 | 5.600E-08 | 3.315E-08 | 1.900E-08 | 8.300E-08 |
| 11191 | 560 | D560L | A | 5.3e-08 | 1.9e-07 | 3.6e-08 | 9.300E-08 | 8.443E-08 | 3.600E-08 | 1.900E-07 |
| 11192 | 560 | D560K | A | 6.1e-08 | 3.1e-08 | 1.7e-08 | 3.633E-08 | 2.248E-08 | 1.700E-08 | 6.100E-08 |
| 11193 | 560 | D560M | A | 7.9e-08 | 2.7e-08 | 7.7e-08 | 6.100E-08 | 2.946E-08 | 2.700E-08 | 7.900E-08 |
| 11194 | 560 | D560F | A | 1.1e-07 | 6.2e-08 | 3.2e-08 | 6.800E-08 | 3.934E-08 | 3.200E-08 | 1.100E-07 |
| 11195 | 560 | D560P | A | 2.1e-08 | 9.1e-08 | 8.8e-08 | 6.667E-08 | 3.958E-08 | 2.100E-08 | 9.100E-08 |
| 11196 | 560 | D560S | A | 3e-08 | 1.2e-07 | 3.9e-08 | 6.300E-08 | 4.957E-08 | 3.000E-08 | 1.200E-07 |
| 11197 | 560 | D560T | A | 4.3e-08 | 8.7e-08 | 4e-08 | 5.667E-08 | 2.631E-08 | 4.000E-08 | 8.700E-08 |
| 11198 | 560 | D560W | A | 6e-08 | 3.3e-08 | 5.7e-08 | 5.000E-08 | 1.480E-08 | 3.300E-08 | 6.000E-08 |
| 11199 | 560 | D560Y | A | 7e-08 | 5.6e-08 | 2.1e-08 | 4.900E-08 | 2.524E-08 | 2.100E-08 | 7.000E-08 |
| 11200 | 560 | D560V | A | 1.7e-08 | 3.5e-08 | 3e-08 | 2.733E-08 | 9.292E-09 | 1.700E-08 | 3.500E-08 |
| 11201 | 561 | Q561A | A | 3.5e-08 | 7.6e-08 | 2.2e-08 | 4.433E-08 | 2.818E-08 | 2.200E-08 | 7.600E-08 |
| 11202 | 561 | Q561R | A | 5.4e-08 | 8.8e-08 | 2.3e-08 | 5.500E-08 | 3.251E-08 | 2.300E-08 | 8.800E-08 |
| 11203 | 561 | Q561N | A | 4.6e-08 | 1.8e-08 | 7.1e-08 | 4.500E-08 | 2.651E-08 | 1.800E-08 | 7.100E-08 |
| 11204 | 561 | Q561D | A | 4.6e-08 | 1e-07 | 3.7e-08 | 6.100E-08 | 3.407E-08 | 3.700E-08 | 1.000E-07 |
| 11205 | 561 | Q561C | A | 2.6e-08 | 6.6e-08 | 5e-08 | 4.733E-08 | 2.013E-08 | 2.600E-08 | 6.600E-08 |
| 11206 | 561 | Q561E | A | 3.2e-08 | 4.4e-08 | 2.1e-08 | 3.233E-08 | 1.150E-08 | 2.100E-08 | 4.400E-08 |
| 11207 | 561 | Q561Q | A | 1e-08 | 4.4e-08 | 7.2e-08 | 4.200E-08 | 3.105E-08 | 1.000E-08 | 7.200E-08 |
| 11208 | 561 | Q561G | A | 2.3e-08 | 3.3e-08 | 8.3e-08 | 4.633E-08 | 3.215E-08 | 2.300E-08 | 8.300E-08 |
| 11209 | 561 | Q561H | A | 4.9e-08 | 1.8e-08 | 4.2e-08 | 3.633E-08 | 1.626E-08 | 1.800E-08 | 4.900E-08 |
| 11210 | 561 | Q561I | A | 5e-08 | 1e-08 | 5e-08 | 3.667E-08 | 2.309E-08 | 1.000E-08 | 5.000E-08 |
| 11211 | 561 | Q561L | A | 9.7e-08 | 8.3e-08 | 4.8e-08 | 7.600E-08 | 2.524E-08 | 4.800E-08 | 9.700E-08 |
| 11212 | 561 | Q561K | A | 4.5e-08 | 7.9e-08 | 2.4e-08 | 4.933E-08 | 2.775E-08 | 2.400E-08 | 7.900E-08 |
| 11213 | 561 | Q561M | A | 2.9e-08 | 4.5e-08 | 3.7e-08 | 3.700E-08 | 8.000E-09 | 2.900E-08 | 4.500E-08 |
| 11214 | 561 | Q561F | A | 4.8e-08 | 4.3e-08 | 7.6e-08 | 5.567E-08 | 1.779E-08 | 4.300E-08 | 7.600E-08 |
| 11215 | 561 | Q561P | A | 5.6e-08 | 4.2e-08 | 2.9e-08 | 4.233E-08 | 1.350E-08 | 2.900E-08 | 5.600E-08 |
| 11216 | 561 | Q561S | A | 3.3e-08 | 4.2e-08 | 1.1e-07 | 6.167E-08 | 4.210E-08 | 3.300E-08 | 1.100E-07 |
| 11217 | 561 | Q561T | A | 1.4e-08 | 4.9e-08 | 1.1e-07 | 5.767E-08 | 4.858E-08 | 1.400E-08 | 1.100E-07 |
| 11218 | 561 | Q561W | A | 3.4e-08 | 4.7e-08 | 2.6e-08 | 3.567E-08 | 1.060E-08 | 2.600E-08 | 4.700E-08 |
| 11219 | 561 | Q561Y | A | 5.7e-08 | 1.5e-08 | 2.4e-08 | 3.200E-08 | 2.211E-08 | 1.500E-08 | 5.700E-08 |

| | | | | | | | | | | |
|---|---|---|---|---|---|---|---|---|---|---|
| 11220 | 561 | Q561V | A | 4.3e-08 | 6.7e-08 | 3.7e-08 | 4.900E-08 | 1.587E-08 | 3.700E-08 | 6.700E-08 |
| 11221 | 562 | C562A | A | 2.9e-08 | 2.5e-08 | 4.8e-08 | 3.400E-08 | 1.229E-08 | 2.500E-08 | 4.800E-08 |
| 11222 | 562 | C562R | A | 7.4e-08 | 1e-07 | 6.9e-08 | 8.100E-08 | 1.664E-08 | 6.900E-08 | 1.000E-07 |
| 11223 | 562 | C562N | A | 1.5e-07 | 8.5e-08 | 5.7e-08 | 9.733E-08 | 4.771E-08 | 5.700E-08 | 1.500E-07 |
| 11224 | 562 | C562D | A | 1.5e-08 | 4.6e-08 | 1.9e-08 | 2.667E-08 | 1.686E-08 | 1.500E-08 | 4.600E-08 |
| 11225 | 562 | C562C | A | 6.9e-08 | 5.5e-08 | 2e-08 | 4.800E-08 | 2.524E-08 | 2.000E-08 | 6.900E-08 |
| 11226 | 562 | C562E | A | 4.1e-08 | 8.7e-08 | 2.6e-08 | 5.133E-08 | 3.179E-08 | 2.600E-08 | 8.700E-08 |
| 11227 | 562 | C562Q | A | 4.9e-08 | 1.3e-07 | 3.2e-08 | 7.033E-08 | 5.237E-08 | 3.200E-08 | 1.300E-07 |
| 11228 | 562 | C562G | A | 5.7e-08 | 1.4e-07 | 5.9e-08 | 8.533E-08 | 4.735E-08 | 5.700E-08 | 1.400E-07 |
| 11229 | 562 | C562H | A | 1.1e-07 | 1.4e-07 | 5.5e-08 | 1.017E-07 | 4.311E-08 | 5.500E-08 | 1.400E-07 |
| 11230 | 562 | C562I | A | 5e-08 | 5.4e-08 | 4.3e-08 | 4.900E-08 | 5.568E-09 | 4.300E-08 | 5.400E-08 |
| 11231 | 562 | C562L | A | 2.5e-08 | 2.5e-07 | 5.9e-08 | 1.113E-07 | 1.213E-07 | 2.500E-08 | 2.500E-07 |
| 11232 | 562 | C562K | A | 6.2e-08 | 3.4e-08 | 1.2e-08 | 3.600E-08 | 2.506E-08 | 1.200E-08 | 6.200E-08 |
| 11233 | 562 | C562M | A | 2.5e-08 | 8.1e-08 | 4.3e-08 | 4.967E-08 | 2.859E-08 | 2.500E-08 | 8.100E-08 |
| 11234 | 562 | C562F | A | 2.2e-08 | 6.6e-08 | 4.4e-08 | 4.400E-08 | 2.200E-08 | 2.200E-08 | 6.600E-08 |
| 11235 | 562 | C562P | A | 4.8e-08 | 6.7e-08 | 9.4e-08 | 6.967E-08 | 2.312E-08 | 4.800E-08 | 9.400E-08 |
| 11236 | 562 | C562S | A | 8.5e-08 | 6.3e-08 | 5.5e-08 | 6.767E-08 | 1.553E-08 | 5.500E-08 | 8.500E-08 |
| 11237 | 562 | C562T | A | 3.5e-08 | 4.3e-08 | 4.6e-08 | 4.133E-08 | 5.686E-09 | 3.500E-08 | 4.600E-08 |
| 11238 | 562 | C562W | A | 2.1e-08 | 4.3e-08 | 2.8e-08 | 3.067E-08 | 1.124E-08 | 2.100E-08 | 4.300E-08 |
| 11239 | 562 | C562Y | A | 5.1e-08 | 6.1e-08 | 4.5e-08 | 5.233E-08 | 8.083E-09 | 4.500E-08 | 6.100E-08 |
| 11240 | 562 | C562V | A | 6.6e-08 | 1.6e-08 | 4.4e-08 | 4.200E-08 | 2.506E-08 | 1.600E-08 | 6.600E-08 |
| 11241 | 563 | V563A | A | 6.2e-08 | 1.2e-08 | 6.9e-08 | 4.767E-08 | 3.109E-08 | 1.200E-08 | 6.900E-08 |
| 11242 | 563 | V563R | A | 9.6e-08 | 4.9e-08 | 3.3e-08 | 5.933E-08 | 3.275E-08 | 3.300E-08 | 9.600E-08 |
| 11243 | 563 | V563N | A | 6.2e-08 | 9.8e-08 | 2.7e-08 | 6.233E-08 | 3.550E-08 | 2.700E-08 | 9.800E-08 |
| 11244 | 563 | V563D | A | 4.5e-08 | 6.2e-08 | 1.5e-07 | 8.567E-08 | 5.636E-08 | 4.500E-08 | 1.500E-07 |
| 11245 | 563 | V563C | A | 7.8e-08 | 2.9e-08 | 1.5e-08 | 4.067E-08 | 3.308E-08 | 1.500E-08 | 7.800E-08 |
| 11246 | 563 | V563E | A | 3.7e-08 | 2.4e-08 | 2.1e-08 | 2.733E-08 | 8.505E-09 | 2.100E-08 | 3.700E-08 |
| 11247 | 563 | V563Q | A | 5e-08 | 6e-08 | 3.4e-08 | 4.800E-08 | 1.311E-08 | 3.400E-08 | 6.000E-08 |
| 11248 | 563 | V563G | A | 2.6e-08 | 5.1e-08 | 8.4e-08 | 5.367E-08 | 2.909E-08 | 2.600E-08 | 8.400E-08 |
| 11249 | 563 | V563H | A | 1.4e-07 | 5.3e-09 | 4.2e-08 | 6.243E-08 | 6.964E-08 | 5.300E-09 | 1.400E-07 |
| 11250 | 563 | V563I | A | 5.2e-08 | 2.7e-08 | 1.8e-08 | 3.233E-08 | 1.762E-08 | 1.800E-08 | 5.200E-08 |
| 11251 | 563 | V563L | A | 3e-08 | 1.5e-08 | 9.3e-08 | 4.600E-08 | 4.139E-08 | 1.500E-08 | 9.300E-08 |
| 11252 | 563 | V563K | A | 1.4e-08 | 1e-07 | 2.6e-08 | 4.667E-08 | 4.658E-08 | 1.400E-08 | 1.000E-07 |
| 11253 | 563 | V563M | A | 6.5e-08 | 3.4e-08 | 1.7e-08 | 3.867E-08 | 2.434E-08 | 1.700E-08 | 6.500E-08 |
| 11254 | 563 | V563F | A | 7e-08 | 7.8e-08 | 3.1e-08 | 5.967E-08 | 2.515E-08 | 3.100E-08 | 7.800E-08 |
| 11255 | 563 | V563P | A | 1.8e-07 | 6.1e-08 | 5.9e-08 | 1.000E-07 | 6.929E-08 | 5.900E-08 | 1.800E-07 |
| 11256 | 563 | V563S | A | 4.3e-08 | 3.9e-08 | 4.4e-08 | 4.200E-08 | 2.646E-09 | 3.900E-08 | 4.400E-08 |
| 11257 | 563 | V563T | A | 2e-08 | 4.3e-08 | 4.4e-08 | 3.567E-08 | 1.358E-08 | 2.000E-08 | 4.400E-08 |
| 11258 | 563 | V563W | A | 3.7e-08 | 4.6e-08 | 6.6e-08 | 4.967E-08 | 1.484E-08 | 3.700E-08 | 6.600E-08 |
| 11259 | 563 | V563Y | A | 5.2e-08 | 2.8e-07 | 2.9e-08 | 1.203E-07 | 1.388E-07 | 2.900E-08 | 2.800E-07 |
| 11260 | 563 | V563V | A | 1.2e-07 | 3.9e-08 | 1.2e-07 | 9.300E-08 | 4.677E-08 | 3.900E-08 | 1.200E-07 |
| 11261 | 564 | A564A | A | 1.5e-08 | 1.3e-08 | 2.7e-08 | 1.833E-08 | 7.572E-09 | 1.300E-08 | 2.700E-08 |
| 11262 | 564 | A564R | A | 5.7e-08 | 6.9e-08 | 1.3e-07 | 8.533E-08 | 3.915E-08 | 5.700E-08 | 1.300E-07 |
| 11263 | 564 | A564N | A | 4.6e-08 | 1.6e-08 | 2.6e-08 | 2.933E-08 | 1.528E-08 | 1.600E-08 | 4.600E-08 |
| 11264 | 564 | A564D | A | 1.4e-08 | 6.5e-08 | 3.4e-08 | 3.767E-08 | 2.570E-08 | 1.400E-08 | 6.500E-08 |
| 11265 | 564 | A564C | A | 1.1e-08 | 1.3e-07 | 4.8e-08 | 6.300E-08 | 6.090E-08 | 1.100E-08 | 1.300E-07 |
| 11266 | 564 | A564E | A | 4.4e-08 | 8.7e-08 | 4.5e-08 | 5.867E-08 | 2.454E-08 | 4.400E-08 | 8.700E-08 |
| 11267 | 564 | A564Q | A | 1e-08 | 5.7e-08 | 1.1e-07 | 5.900E-08 | 5.003E-08 | 1.000E-08 | 1.100E-07 |
| 11268 | 564 | A564G | A | 2.7e-08 | 1.7e-08 | 2.7e-08 | 2.367E-08 | 5.774E-09 | 1.700E-08 | 2.700E-08 |
| 11269 | 564 | A564H | A | 1.3e-07 | 4.2e-08 | 3e-08 | 6.733E-08 | 5.460E-08 | 3.000E-08 | 1.300E-07 |
| 11270 | 564 | A564I | A | 5.9e-08 | 5.9e-08 | 1.1e-07 | 7.600E-08 | 2.944E-08 | 5.900E-08 | 1.100E-07 |
| 11271 | 564 | A564L | A | 2e-08 | 2e-08 | 4.8e-08 | 2.933E-08 | 1.617E-08 | 2.000E-08 | 4.800E-08 |
| 11272 | 564 | A564K | A | 6.2e-08 | 2.4e-08 | 3.5e-08 | 4.033E-08 | 1.955E-08 | 2.400E-08 | 6.200E-08 |
| 11273 | 564 | A564M | A | 8.6e-08 | 5.9e-08 | 1e-08 | 5.167E-08 | 3.853E-08 | 1.000E-08 | 8.600E-08 |
| 11274 | 564 | A564F | A | 2.1e-08 | 7.2e-08 | 8.2e-08 | 5.833E-08 | 3.272E-08 | 2.100E-08 | 8.200E-08 |
| 11275 | 564 | A564P | A | 5.1e-08 | 1e-07 | 4.8e-08 | 6.633E-08 | 2.919E-08 | 4.800E-08 | 1.000E-07 |
| 11276 | 564 | A564S | A | 8.1e-08 | 6.8e-08 | 4.3e-08 | 6.400E-08 | 1.931E-08 | 4.300E-08 | 8.100E-08 |
| 11277 | 564 | A564T | A | 1.8e-08 | 1.7e-08 | 6.6e-08 | 3.367E-08 | 2.801E-08 | 1.700E-08 | 6.600E-08 |
| 11278 | 564 | A564W | A | 4e-08 | 1.7e-08 | 1.1e-08 | 2.267E-08 | 1.531E-08 | 1.100E-08 | 4.000E-08 |
| 11279 | 564 | A564Y | A | 3.6e-08 | 4.1e-08 | 4.9e-08 | 4.200E-08 | 6.557E-09 | 3.600E-08 | 4.900E-08 |
| 11280 | 564 | A564V | A | 1.8e-08 | 4.9e-08 | 2.5e-08 | 3.067E-08 | 1.626E-08 | 1.800E-08 | 4.900E-08 |
| 11281 | 565 | C565A | A | 1.9e-08 | 3.4e-08 | 5.4e-08 | 3.567E-08 | 1.756E-08 | 1.900E-08 | 5.400E-08 |
| 11282 | 565 | C565R | A | 1.2e-08 | 7.3e-08 | 3.8e-08 | 4.100E-08 | 3.061E-08 | 1.200E-08 | 7.300E-08 |
| 11283 | 565 | C565N | A | 3.9e-08 | 4.7e-08 | 1.9e-08 | 3.500E-08 | 1.442E-08 | 1.900E-08 | 4.700E-08 |
| 11284 | 565 | C565D | A | 7e-08 | 5e-08 | 6.7e-08 | 6.233E-08 | 1.079E-08 | 5.000E-08 | 7.000E-08 |
| 11285 | 565 | C565C | A | 3e-08 | 3.2e-08 | 4.8e-08 | 3.667E-08 | 9.866E-09 | 3.000E-08 | 4.800E-08 |
| 11286 | 565 | C565E | A | 2.9e-08 | 4.1e-08 | 2.7e-08 | 3.233E-08 | 7.572E-09 | 2.700E-08 | 4.100E-08 |
| 11287 | 565 | C565Q | A | 4.4e-08 | 5.5e-08 | 2.2e-08 | 4.033E-08 | 1.680E-08 | 2.200E-08 | 5.500E-08 |
| 11288 | 565 | C565G | A | 4.6e-08 | 1.2e-07 | 2.1e-08 | 6.233E-08 | 5.148E-08 | 2.100E-08 | 1.200E-07 |
| 11289 | 565 | C565H | A | 7.9e-08 | 3.4e-08 | 4.4e-08 | 5.233E-08 | 2.363E-08 | 3.400E-08 | 7.900E-08 |
| 11290 | 565 | C565I | A | 9.7e-08 | 5.2e-08 | 2.6e-08 | 5.833E-08 | 3.592E-08 | 2.600E-08 | 9.700E-08 |
| 11291 | 565 | C565L | A | 9.6e-08 | 3.4e-08 | 5.3e-08 | 6.100E-08 | 3.176E-08 | 3.400E-08 | 9.600E-08 |
| 11292 | 565 | C565K | A | 3.9e-08 | 6.1e-08 | 2.2e-08 | 4.067E-08 | 1.955E-08 | 2.200E-08 | 6.100E-08 |
| 11293 | 565 | C565M | A | 3.8e-08 | 6.1e-08 | 4.6e-08 | 4.833E-08 | 1.168E-08 | 3.800E-08 | 6.100E-08 |
| 11294 | 565 | C565F | A | 4.6e-08 | 2.8e-08 | 2.8e-08 | 3.400E-08 | 1.039E-08 | 2.800E-08 | 4.600E-08 |
| 11295 | 565 | C565P | A | 8.4e-08 | 5.5e-08 | 3.3e-08 | 5.733E-08 | 2.558E-08 | 3.300E-08 | 8.400E-08 |
| 11296 | 565 | C565S | A | 2.8e-08 | 2.3e-08 | 1.6e-08 | 2.233E-08 | 6.028E-09 | 1.600E-08 | 2.800E-08 |
| 11297 | 565 | C565T | A | 1.5e-08 | 5.1e-08 | 9.6e-09 | 2.520E-08 | 2.251E-08 | 9.600E-09 | 5.100E-08 |
| 11298 | 565 | C565W | A | 5.3e-08 | 2.2e-08 | 3.8e-08 | 3.767E-08 | 1.550E-08 | 2.200E-08 | 5.300E-08 |
| 11299 | 565 | C565Y | A | 2.3e-08 | 7e-08 | 1.6e-07 | 8.433E-08 | 6.962E-08 | 2.300E-08 | 1.600E-07 |
| 11300 | 565 | C565V | A | 2.8e-08 | 3.1e-08 | 2.1e-08 | 2.667E-08 | 5.132E-09 | 2.100E-08 | 3.100E-08 |
| 11301 | 566 | A566A | A | 5.3e-08 | 6e-08 | 1.9e-08 | 4.400E-08 | 2.193E-08 | 1.900E-08 | 6.000E-08 |
| 11302 | 566 | A566R | A | 3.5e-08 | 5.4e-08 | 7.2e-08 | 5.367E-08 | 1.850E-08 | 3.500E-08 | 7.200E-08 |
| 11303 | 566 | A566N | A | 5.3e-08 | 3.4e-08 | 2.6e-08 | 3.767E-08 | 1.387E-08 | 2.600E-08 | 5.300E-08 |
| 11304 | 566 | A566D | A | 3.4e-08 | 1.1e-07 | 3.6e-08 | 6.000E-08 | 4.331E-08 | 3.400E-08 | 1.100E-07 |

| | | | | | | | | | | |
|---|---|---|---|---|---|---|---|---|---|---|
| 11305 | 566 | A566C | A | 2.8e-08 | 2.7e-08 | 4.4e-08 | 3.300E-08 | 9.539E-09 | 2.700E-08 | 4.400E-08 |
| 11306 | 566 | A566E | A | 9e-08 | 1.2e-08 | 6.1e-08 | 5.433E-08 | 3.943E-08 | 1.200E-08 | 9.000E-08 |
| 11307 | 566 | A566Q | A | 5.2e-08 | 2.6e-08 | 2.1e-08 | 3.300E-08 | 1.664E-08 | 2.100E-08 | 5.200E-08 |
| 11308 | 566 | A566G | A | 1.9e-08 | 5.3e-08 | 3.9e-08 | 3.700E-08 | 1.709E-08 | 1.900E-08 | 5.300E-08 |
| 11309 | 566 | A566H | A | 4.8e-08 | 3.4e-08 | 3.8e-08 | 4.000E-08 | 7.211E-09 | 3.400E-08 | 4.800E-08 |
| 11310 | 566 | A566I | A | 3.8e-08 | 3.8e-08 | 8e-08 | 5.200E-08 | 2.425E-08 | 3.800E-08 | 8.000E-08 |
| 11311 | 566 | A566L | A | 1.3e-08 | 4.9e-08 | 3e-08 | 3.067E-08 | 1.801E-08 | 1.300E-08 | 4.900E-08 |
| 11312 | 566 | A566K | A | 2.9e-08 | 9.8e-08 | 4.3e-08 | 5.667E-08 | 3.647E-08 | 2.900E-08 | 9.800E-08 |
| 11313 | 566 | A566M | A | 5.2e-08 | 8e-08 | 4.7e-08 | 5.967E-08 | 1.779E-08 | 4.700E-08 | 8.000E-08 |
| 11314 | 566 | A566F | A | 4.9e-08 | 4.9e-08 | 2.2e-08 | 4.000E-08 | 1.559E-08 | 2.200E-08 | 4.900E-08 |
| 11315 | 566 | A566P | A | 1.6e-08 | 3.3e-08 | 2.4e-08 | 2.433E-08 | 8.505E-09 | 1.600E-08 | 3.300E-08 |
| 11316 | 566 | A566S | A | 4.4e-08 | 6e-08 | 2.3e-08 | 4.233E-08 | 1.856E-08 | 2.300E-08 | 6.000E-08 |
| 11317 | 566 | A566T | A | 6.6e-08 | 3.8e-08 | 3e-08 | 4.467E-08 | 1.890E-08 | 3.000E-08 | 6.600E-08 |
| 11318 | 566 | A566W | A | 2.9e-08 | 3e-08 | 4.2e-08 | 3.367E-08 | 7.234E-09 | 2.900E-08 | 4.200E-08 |
| 11319 | 566 | A566Y | A | 4e-08 | 1.1e-07 | 2e-08 | 5.667E-08 | 4.726E-08 | 2.000E-08 | 1.100E-07 |
| 11320 | 566 | A566V | A | 4.3e-08 | 2.9e-08 | 1.1e-07 | 6.067E-08 | 4.329E-08 | 2.900E-08 | 1.100E-07 |
| 11321 | 567 | H567A | A | 5.9e-08 | 1.7e-08 | 3e-08 | 3.533E-08 | 2.150E-08 | 1.700E-08 | 5.900E-08 |
| 11322 | 567 | H567R | A | 1.7e-08 | 4.5e-08 | 1.5e-08 | 2.567E-08 | 1.677E-08 | 1.500E-08 | 4.500E-08 |
| 11323 | 567 | H567N | A | 1.1e-07 | 4.7e-08 | 4e-08 | 6.567E-08 | 3.855E-08 | 4.000E-08 | 1.100E-07 |
| 11324 | 567 | H567D | A | 3e-08 | 5.1e-08 | 2.1e-08 | 3.400E-08 | 1.539E-08 | 2.100E-08 | 5.100E-08 |
| 11325 | 567 | H567C | A | 3.5e-08 | 9.9e-08 | 5e-08 | 6.133E-08 | 3.347E-08 | 3.500E-08 | 9.900E-08 |
| 11326 | 567 | H567E | A | 3.7e-08 | 3.4e-08 | 2.2e-08 | 3.100E-08 | 7.937E-09 | 2.200E-08 | 3.700E-08 |
| 11327 | 567 | H567Q | A | 3.5e-08 | 1.1e-07 | 9e-08 | 7.833E-08 | 3.884E-08 | 3.500E-08 | 1.100E-07 |
| 11328 | 567 | H567G | A | 4.1e-08 | 6.6e-08 | 3.4e-08 | 4.700E-08 | 1.682E-08 | 3.400E-08 | 6.600E-08 |
| 11329 | 567 | H567H | A | 3.8e-08 | 4.3e-08 | 4.4e-08 | 4.167E-08 | 3.215E-09 | 3.800E-08 | 4.400E-08 |
| 11330 | 567 | H567I | A | 4.6e-08 | 5e-08 | 5.1e-08 | 4.900E-08 | 2.646E-09 | 4.600E-08 | 5.100E-08 |
| 11331 | 567 | H567L | A | 5.4e-08 | 3.8e-08 | 2.8e-08 | 4.000E-08 | 1.311E-08 | 2.800E-08 | 5.400E-08 |
| 11332 | 567 | H567K | A | 5.1e-08 | 7e-08 | 5.1e-08 | 5.733E-08 | 1.097E-08 | 5.100E-08 | 7.000E-08 |
| 11333 | 567 | H567M | A | 1e-07 | 2.5e-08 | 2.8e-08 | 5.100E-08 | 4.246E-08 | 2.500E-08 | 1.000E-07 |
| 11334 | 567 | H567F | A | 4.8e-08 | 6.5e-08 | 7.8e-08 | 6.367E-08 | 1.504E-08 | 4.800E-08 | 7.800E-08 |
| 11335 | 567 | H567P | A | 7.9e-08 | 3e-08 | 4.9e-08 | 5.267E-08 | 2.470E-08 | 3.000E-08 | 7.900E-08 |
| 11336 | 567 | H567S | A | 5e-08 | 8.7e-08 | 1.3e-07 | 8.900E-08 | 4.004E-08 | 5.000E-08 | 1.300E-07 |
| 11337 | 567 | H567T | A | 6.5e-08 | 1.1e-08 | 4.9e-08 | 4.167E-08 | 2.774E-08 | 1.100E-08 | 6.500E-08 |
| 11338 | 567 | H567W | A | 5.8e-08 | 3.6e-08 | 3.9e-08 | 4.433E-08 | 1.193E-08 | 3.600E-08 | 5.800E-08 |
| 11339 | 567 | H567Y | A | 2.5e-08 | 7.4e-08 | 4.7e-08 | 4.867E-08 | 2.454E-08 | 2.500E-08 | 7.400E-08 |
| 11340 | 567 | H567V | A | 4.3e-08 | 7.7e-08 | 6.6e-08 | 6.200E-08 | 1.735E-08 | 4.300E-08 | 7.700E-08 |
| 11341 | 568 | Y568A | A | 2.9e-08 | 8.2e-08 | 2e-08 | 4.367E-08 | 3.350E-08 | 2.000E-08 | 8.200E-08 |
| 11342 | 568 | Y568R | A | 2.6e-08 | 1.1e-07 | 3e-08 | 5.533E-08 | 4.738E-08 | 2.600E-08 | 1.100E-07 |
| 11343 | 568 | Y568N | A | 5.1e-08 | 7.7e-08 | 6.8e-08 | 6.533E-08 | 1.320E-08 | 5.100E-08 | 7.700E-08 |
| 11344 | 568 | Y568D | A | 1.2e-07 | 3.5e-08 | 2.7e-08 | 6.067E-08 | 5.154E-08 | 2.700E-08 | 1.200E-07 |
| 11345 | 568 | Y568C | A | 1.1e-07 | 4.7e-08 | 1.2e-07 | 9.233E-08 | 3.958E-08 | 4.700E-08 | 1.200E-07 |
| 11346 | 568 | Y568E | A | 7.1e-08 | 7e-08 | 1.4e-07 | 9.367E-08 | 4.013E-08 | 7.000E-08 | 1.400E-07 |
| 11347 | 568 | Y568Q | A | 3.4e-08 | 8.4e-08 | 4.3e-08 | 5.367E-08 | 2.665E-08 | 3.400E-08 | 8.400E-08 |
| 11348 | 568 | Y568G | A | 2.3e-08 | 2.2e-08 | 4e-08 | 2.833E-08 | 1.012E-08 | 2.200E-08 | 4.000E-08 |
| 11349 | 568 | Y568H | A | 2.3e-08 | 5.4e-08 | 4.7e-08 | 4.133E-08 | 1.626E-08 | 2.300E-08 | 5.400E-08 |
| 11350 | 568 | Y568I | A | 2.6e-08 | 2.5e-08 | 6.9e-08 | 4.000E-08 | 2.512E-08 | 2.500E-08 | 6.900E-08 |
| 11351 | 568 | Y568L | A | 1.8e-08 | 2e-08 | 2.4e-08 | 2.067E-08 | 3.055E-09 | 1.800E-08 | 2.400E-08 |
| 11352 | 568 | Y568K | A | 2.6e-08 | 5.1e-08 | 8.2e-08 | 5.300E-08 | 2.805E-08 | 2.600E-08 | 8.200E-08 |
| 11353 | 568 | Y568M | A | 6.3e-08 | 4.2e-08 | 6.6e-08 | 5.700E-08 | 1.308E-08 | 4.200E-08 | 6.600E-08 |
| 11354 | 568 | Y568F | A | 1.5e-08 | 4.1e-08 | 5.5e-08 | 3.700E-08 | 2.030E-08 | 1.500E-08 | 5.500E-08 |
| 11355 | 568 | Y568P | A | 7.6e-08 | 7.4e-09 | 2.1e-08 | 3.480E-08 | 3.632E-08 | 7.400E-09 | 7.600E-08 |
| 11356 | 568 | Y568S | A | 3.4e-08 | 2.8e-08 | 2.6e-08 | 2.933E-08 | 4.163E-09 | 2.600E-08 | 3.400E-08 |
| 11357 | 568 | Y568T | A | 2.3e-08 | 6.2e-08 | 8e-08 | 5.500E-08 | 2.914E-08 | 2.300E-08 | 8.000E-08 |
| 11358 | 568 | Y568W | A | 4.8e-08 | 1.1e-07 | 3.9e-08 | 6.567E-08 | 3.866E-08 | 3.900E-08 | 1.100E-07 |
| 11359 | 568 | Y568Y | A | 1.1e-07 | 3.9e-08 | 4.5e-08 | 6.467E-08 | 3.937E-08 | 3.900E-08 | 1.100E-07 |
| 11360 | 568 | Y568V | A | 6e-08 | 4.7e-08 | 5.4e-08 | 5.367E-08 | 6.506E-09 | 4.700E-08 | 6.000E-08 |
| 11361 | 569 | K569A | A | 7.3e-08 | 3.8e-08 | 1.5e-08 | 4.200E-08 | 2.921E-08 | 1.500E-08 | 7.300E-08 |
| 11362 | 569 | K569R | A | 1.5e-08 | 2.8e-08 | 1.4e-08 | 1.900E-08 | 7.810E-09 | 1.400E-08 | 2.800E-08 |
| 11363 | 569 | K569N | A | 2.5e-08 | 4.2e-08 | 5.1e-08 | 3.933E-08 | 1.320E-08 | 2.500E-08 | 5.100E-08 |
| 11364 | 569 | K569D | A | 3.9e-08 | 8.4e-08 | 4.1e-08 | 5.467E-08 | 2.542E-08 | 3.900E-08 | 8.400E-08 |
| 11365 | 569 | K569C | A | 7.4e-08 | 2e-08 | 2.9e-08 | 4.100E-08 | 2.893E-08 | 2.000E-08 | 7.400E-08 |
| 11366 | 569 | K569E | A | 6.3e-08 | 1.3e-08 | 2.1e-08 | 3.233E-08 | 2.686E-08 | 1.300E-08 | 6.300E-08 |
| 11367 | 569 | K569Q | A | 4.3e-08 | 4.3e-08 | 2.8e-08 | 3.800E-08 | 8.660E-09 | 2.800E-08 | 4.300E-08 |
| 11368 | 569 | K569G | A | 6.2e-08 | 5.6e-08 | 4.7e-08 | 5.500E-08 | 7.550E-09 | 4.700E-08 | 6.200E-08 |
| 11369 | 569 | K569H | A | 2e-08 | 5.5e-08 | 5.7e-08 | 4.400E-08 | 2.081E-08 | 2.000E-08 | 5.700E-08 |
| 11370 | 569 | K569I | A | 4.4e-08 | 1.4e-08 | 2.3e-08 | 2.700E-08 | 1.539E-08 | 1.400E-08 | 4.400E-08 |
| 11371 | 569 | K569L | A | 4.8e-08 | 5.9e-08 | 8e-08 | 6.233E-08 | 1.626E-08 | 4.800E-08 | 8.000E-08 |
| 11372 | 569 | K569K | A | 3.4e-08 | 3.6e-08 | 9e-08 | 5.333E-08 | 3.177E-08 | 3.400E-08 | 9.000E-08 |
| 11373 | 569 | K569M | A | 1.3e-07 | 3.4e-08 | 1.6e-08 | 6.000E-08 | 6.129E-08 | 1.600E-08 | 1.300E-07 |
| 11374 | 569 | K569F | A | 3.3e-08 | 4.2e-08 | 4.1e-08 | 3.867E-08 | 4.933E-09 | 3.300E-08 | 4.200E-08 |
| 11375 | 569 | K569P | A | 1.6e-08 | 3.6e-08 | 5.5e-08 | 3.567E-08 | 1.950E-08 | 1.600E-08 | 5.500E-08 |
| 11376 | 569 | K569S | A | 3.5e-08 | 6.5e-08 | 1.6e-08 | 3.867E-08 | 2.470E-08 | 1.600E-08 | 6.500E-08 |
| 11377 | 569 | K569T | A | 5.3e-08 | 3.6e-08 | 4.1e-08 | 4.333E-08 | 8.737E-09 | 3.600E-08 | 5.300E-08 |
| 11378 | 569 | K569W | A | 6.5e-08 | 4.9e-08 | 3.3e-08 | 4.900E-08 | 1.600E-08 | 3.300E-08 | 6.500E-08 |
| 11379 | 569 | K569Y | A | 7.1e-08 | 2.4e-08 | 3.5e-08 | 4.333E-08 | 2.458E-08 | 2.400E-08 | 7.100E-08 |
| 11380 | 569 | K569V | A | 1.7e-07 | 3.7e-08 | 4.8e-08 | 8.500E-08 | 7.382E-08 | 3.700E-08 | 1.700E-07 |
| 11381 | 570 | D570A | A | 4.1e-08 | 3.7e-08 | 3.9e-08 | 3.900E-08 | 2.000E-09 | 3.700E-08 | 4.100E-08 |
| 11382 | 570 | D570R | A | 5.1e-08 | 4.6e-08 | 7.1e-08 | 5.600E-08 | 1.323E-08 | 4.600E-08 | 7.100E-08 |
| 11383 | 570 | D570N | A | 6.5e-08 | 9e-08 | 3.6e-08 | 6.367E-08 | 2.702E-08 | 3.600E-08 | 9.000E-08 |
| 11384 | 570 | D570D | A | 8.6e-08 | 5e-08 | 4.3e-08 | 5.967E-08 | 2.307E-08 | 4.300E-08 | 8.600E-08 |
| 11385 | 570 | D570C | A | 1.5e-08 | 7.2e-08 | 4.4e-08 | 4.367E-08 | 2.850E-08 | 1.500E-08 | 7.200E-08 |
| 11386 | 570 | D570E | A | 3.5e-08 | 2e-08 | 8e-08 | 4.500E-08 | 3.122E-08 | 2.000E-08 | 8.000E-08 |
| 11387 | 570 | D570Q | A | 4.7e-08 | 5.3e-08 | 4.5e-08 | 4.833E-08 | 4.163E-09 | 4.500E-08 | 5.300E-08 |
| 11388 | 570 | D570G | A | 1.3e-07 | 7.4e-08 | 2.7e-07 | 1.580E-07 | 1.010E-07 | 7.400E-08 | 2.700E-07 |
| 11389 | 570 | D570H | A | 7.5e-08 | 3.5e-08 | 8.9e-08 | 6.633E-08 | 2.802E-08 | 3.500E-08 | 8.900E-08 |

| | | | | | | | | | | |
|---|---|---|---|---|---|---|---|---|---|---|
| 11390 | 570 | D570I | A | 8.5e-08 | 3.6e-08 | 3.9e-08 | 5.333E-08 | 2.747E-08 | 3.600E-08 | 8.500E-08 |
| 11391 | 570 | D570L | A | 2.3e-08 | 5.1e-08 | 8.8e-08 | 5.400E-08 | 3.260E-08 | 2.300E-08 | 8.800E-08 |
| 11392 | 570 | D570K | A | 7.1e-08 | 3.5e-08 | 4.7e-08 | 5.100E-08 | 1.833E-08 | 3.500E-08 | 7.100E-08 |
| 11393 | 570 | D570M | A | 1.5e-07 | 6e-08 | 4.6e-08 | 8.533E-08 | 5.644E-08 | 4.600E-08 | 1.500E-07 |
| 11394 | 570 | D570F | A | 6.2e-08 | 3.2e-08 | 4.2e-08 | 4.533E-08 | 1.528E-08 | 3.200E-08 | 6.200E-08 |
| 11395 | 570 | D570P | A | 2.1e-08 | 5.1e-08 | 3.6e-08 | 3.600E-08 | 1.500E-08 | 2.100E-08 | 5.100E-08 |
| 11396 | 570 | D570S | A | 6.3e-08 | 3.8e-08 | 6.5e-08 | 5.533E-08 | 1.504E-08 | 3.800E-08 | 6.500E-08 |
| 11397 | 570 | D570T | A | 4.9e-08 | 2.8e-08 | 5.5e-08 | 4.400E-08 | 1.418E-08 | 2.800E-08 | 5.500E-08 |
| 11398 | 570 | D570W | A | 1.7e-08 | 3.3e-08 | 8.2e-09 | 1.940E-08 | 1.257E-08 | 8.200E-09 | 3.300E-08 |
| 11399 | 570 | D570Y | A | 1.3e-08 | 2.6e-08 | 5.4e-08 | 3.100E-08 | 2.095E-08 | 1.300E-08 | 5.400E-08 |
| 11400 | 570 | D570V | A | 4.7e-08 | 5.6e-08 | 4.8e-08 | 5.033E-08 | 4.933E-09 | 4.700E-08 | 5.600E-08 |
| 11401 | 571 | P571A | A | 3.6e-08 | 3.2e-08 | 7.2e-08 | 4.667E-08 | 2.203E-08 | 3.200E-08 | 7.200E-08 |
| 11402 | 571 | P571R | A | 4.7e-08 | 7.7e-08 | 2.5e-08 | 4.967E-08 | 2.610E-08 | 2.500E-08 | 7.700E-08 |
| 11403 | 571 | P571N | A | 4.9e-08 | 4.7e-08 | 5.6e-08 | 5.067E-08 | 4.726E-09 | 4.700E-08 | 5.600E-08 |
| 11404 | 571 | P571D | A | 6.9e-08 | 4.4e-08 | 7.7e-08 | 6.333E-08 | 1.721E-08 | 4.400E-08 | 7.700E-08 |
| 11405 | 571 | P571C | A | 5.1e-08 | 4.7e-09 | 2.8e-07 | 1.119E-07 | 1.474E-07 | 4.700E-09 | 2.800E-07 |
| 11406 | 571 | P571E | A | 2.8e-08 | 6.6e-08 | 4.6e-08 | 4.667E-08 | 1.901E-08 | 2.800E-08 | 6.600E-08 |
| 11407 | 571 | P571Q | A | 3.4e-08 | 2.4e-08 | 4.8e-08 | 3.533E-08 | 1.206E-08 | 2.400E-08 | 4.800E-08 |
| 11408 | 571 | P571G | A | 4.4e-08 | 3.8e-08 | 9e-08 | 5.733E-08 | 2.845E-08 | 3.800E-08 | 9.000E-08 |
| 11409 | 571 | P571H | A | 2.8e-08 | 2.1e-08 | 2.3e-08 | 2.400E-08 | 3.606E-09 | 2.100E-08 | 2.800E-08 |
| 11410 | 571 | P571I | A | 8e-08 | 2.9e-08 | 5.6e-08 | 5.500E-08 | 2.551E-08 | 2.900E-08 | 8.000E-08 |
| 11411 | 571 | P571L | A | 6.7e-08 | 2.8e-08 | 4.8e-08 | 4.767E-08 | 1.950E-08 | 2.800E-08 | 6.700E-08 |
| 11412 | 571 | P571K | A | 2.1e-08 | 8.3e-08 | 8.7e-08 | 6.367E-08 | 3.700E-08 | 2.100E-08 | 8.700E-08 |
| 11413 | 571 | P571M | A | 8e-08 | 8.3e-08 | 3.7e-08 | 6.667E-08 | 2.574E-08 | 3.700E-08 | 8.300E-08 |
| 11414 | 571 | P571F | A | 1.4e-07 | 6.3e-09 | 3.3e-08 | 5.977E-08 | 7.075E-08 | 6.300E-09 | 1.400E-07 |
| 11415 | 571 | P571P | A | 2.5e-08 | 8.5e-08 | 3.6e-08 | 4.867E-08 | 3.194E-08 | 2.500E-08 | 8.500E-08 |
| 11416 | 571 | P571S | A | 1.6e-08 | 9.3e-08 | 1.3e-08 | 4.067E-08 | 4.535E-08 | 1.300E-08 | 9.300E-08 |
| 11417 | 571 | P571T | A | 3e-08 | 3e-08 | 3.9e-08 | 3.300E-08 | 5.196E-09 | 3.000E-08 | 3.900E-08 |
| 11418 | 571 | P571W | A | 1.7e-08 | 1.6e-07 | 1.1e-07 | 9.567E-08 | 7.257E-08 | 1.700E-08 | 1.600E-07 |
| 11419 | 571 | P571Y | A | 2.1e-08 | 2e-08 | 5.3e-08 | 3.133E-08 | 1.877E-08 | 2.000E-08 | 5.300E-08 |
| 11420 | 571 | P571V | A | 1.8e-08 | 2.4e-08 | 5.2e-08 | 3.133E-08 | 1.815E-08 | 1.800E-08 | 5.200E-08 |
| 11421 | 572 | P572A | A | 3.5e-08 | 5.9e-08 | 3.6e-08 | 4.333E-08 | 1.358E-08 | 3.500E-08 | 5.900E-08 |
| 11422 | 572 | P572R | A | 6.1e-08 | 3.7e-08 | 4e-08 | 4.600E-08 | 1.308E-08 | 3.700E-08 | 6.100E-08 |
| 11423 | 572 | P572N | A | 1.9e-08 | 2.5e-08 | 4.8e-08 | 3.067E-08 | 1.531E-08 | 1.900E-08 | 4.800E-08 |
| 11424 | 572 | P572D | A | 6.9e-08 | 1.3e-08 | 6.8e-08 | 5.000E-08 | 3.205E-08 | 1.300E-08 | 6.900E-08 |
| 11425 | 572 | P572C | A | 2.1e-08 | 2.9e-08 | 3.4e-08 | 2.800E-08 | 6.557E-09 | 2.100E-08 | 3.400E-08 |
| 11426 | 572 | P572E | A | 3.6e-08 | 5.5e-08 | 5.5e-08 | 4.867E-08 | 1.097E-08 | 3.600E-08 | 5.500E-08 |
| 11427 | 572 | P572Q | A | 3.5e-08 | 1.2e-07 | 2.3e-08 | 5.933E-08 | 5.288E-08 | 2.300E-08 | 1.200E-07 |
| 11428 | 572 | P572G | A | 1.4e-07 | 5e-08 | 8.5e-08 | 9.167E-08 | 4.537E-08 | 5.000E-08 | 1.400E-07 |
| 11429 | 572 | P572H | A | 2.2e-08 | 6e-08 | 5.6e-08 | 4.600E-08 | 2.088E-08 | 2.200E-08 | 6.000E-08 |
| 11430 | 572 | P572I | A | 2.1e-08 | 2.7e-08 | 2.8e-08 | 2.533E-08 | 3.786E-09 | 2.100E-08 | 2.800E-08 |
| 11431 | 572 | P572L | A | 2.4e-08 | 7.4e-08 | 3e-08 | 4.267E-08 | 2.730E-08 | 2.400E-08 | 7.400E-08 |
| 11432 | 572 | P572K | A | 7.7e-08 | 7.7e-08 | 4.6e-08 | 6.667E-08 | 1.790E-08 | 4.600E-08 | 7.700E-08 |
| 11433 | 572 | P572M | A | 3.5e-08 | 2.5e-08 | 3.5e-08 | 3.167E-08 | 5.774E-09 | 2.500E-08 | 3.500E-08 |
| 11434 | 572 | P572F | A | 7.8e-08 | 3.7e-08 | 6.8e-08 | 6.100E-08 | 2.138E-08 | 3.700E-08 | 7.800E-08 |
| 11435 | 572 | P572P | A | 4.3e-08 | 3.4e-08 | 3.7e-08 | 3.800E-08 | 4.583E-09 | 3.400E-08 | 4.300E-08 |
| 11436 | 572 | P572S | A | 6e-08 | 3.3e-08 | 1.6e-08 | 3.633E-08 | 2.219E-08 | 1.600E-08 | 6.000E-08 |
| 11437 | 572 | P572T | A | 5.5e-08 | 1.6e-08 | 3.7e-08 | 3.600E-08 | 1.952E-08 | 1.600E-08 | 5.500E-08 |
| 11438 | 572 | P572W | A | 3.8e-08 | 1.2e-08 | 3.4e-08 | 2.800E-08 | 1.400E-08 | 1.200E-08 | 3.800E-08 |
| 11439 | 572 | P572Y | A | 7.9e-08 | 8e-08 | 5e-08 | 6.967E-08 | 1.704E-08 | 5.000E-08 | 8.000E-08 |
| 11440 | 572 | P572V | A | 1.1e-07 | 6.1e-08 | 8e-08 | 8.367E-08 | 2.470E-08 | 6.100E-08 | 1.100E-07 |
| 11441 | 573 | F573A | A | 1.9e-08 | 7.5e-08 | 4.6e-08 | 4.667E-08 | 2.801E-08 | 1.900E-08 | 7.500E-08 |
| 11442 | 573 | F573R | A | 6.2e-08 | 3.3e-08 | 8.5e-08 | 6.000E-08 | 2.606E-08 | 3.300E-08 | 8.500E-08 |
| 11443 | 573 | F573N | A | 2.1e-08 | 9.1e-08 | 5.8e-08 | 5.667E-08 | 3.502E-08 | 2.100E-08 | 9.100E-08 |
| 11444 | 573 | F573D | A | 3.5e-08 | 5e-08 | 3.7e-08 | 4.067E-08 | 8.145E-09 | 3.500E-08 | 5.000E-08 |
| 11445 | 573 | F573C | A | 2.1e-08 | 2.2e-08 | 5.6e-08 | 3.300E-08 | 1.992E-08 | 2.100E-08 | 5.600E-08 |
| 11446 | 573 | F573E | A | 2.5e-08 | 3e-08 | 4.3e-08 | 3.267E-08 | 9.292E-09 | 2.500E-08 | 4.300E-08 |
| 11447 | 573 | F573Q | A | 4.4e-08 | 5.5e-08 | 5.1e-08 | 5.000E-08 | 5.568E-09 | 4.400E-08 | 5.500E-08 |
| 11448 | 573 | F573G | A | 3.8e-08 | 2e-08 | 1.1e-07 | 5.600E-08 | 4.762E-08 | 2.000E-08 | 1.100E-07 |
| 11449 | 573 | F573H | A | 6e-08 | 6.2e-08 | 6e-08 | 6.067E-08 | 1.155E-09 | 6.000E-08 | 6.200E-08 |
| 11450 | 573 | F573I | A | 4.2e-08 | 4.1e-08 | 2.5e-08 | 3.600E-08 | 9.539E-09 | 2.500E-08 | 4.200E-08 |
| 11451 | 573 | F573L | A | 5.8e-08 | 8.7e-08 | 7.5e-08 | 7.333E-08 | 1.457E-08 | 5.800E-08 | 8.700E-08 |
| 11452 | 573 | F573K | A | 2.3e-08 | 6.3e-08 | 3.7e-08 | 4.100E-08 | 2.030E-08 | 2.300E-08 | 6.300E-08 |
| 11453 | 573 | F573M | A | 1.5e-08 | 1.8e-08 | 5.1e-08 | 2.800E-08 | 1.997E-08 | 1.500E-08 | 5.100E-08 |
| 11454 | 573 | F573F | A | 1.4e-07 | 4.1e-08 | 2.6e-08 | 6.900E-08 | 6.194E-08 | 2.600E-08 | 1.400E-07 |
| 11455 | 573 | F573P | A | 1e-08 | 1e-07 | 8.2e-08 | 6.400E-08 | 4.762E-08 | 1.000E-08 | 1.000E-07 |
| 11456 | 573 | F573S | A | 2.9e-08 | 5.4e-08 | 3.5e-08 | 3.933E-08 | 1.305E-08 | 2.900E-08 | 5.400E-08 |
| 11457 | 573 | F573T | A | 4.5e-08 | 5.9e-08 | 3.6e-08 | 4.667E-08 | 1.159E-08 | 3.600E-08 | 5.900E-08 |
| 11458 | 573 | F573W | A | 6.4e-08 | 2.1e-08 | 3.3e-08 | 3.933E-08 | 2.219E-08 | 2.100E-08 | 6.400E-08 |
| 11459 | 573 | F573Y | A | 4.3e-08 | 4.2e-08 | 1.7e-08 | 3.400E-08 | 1.473E-08 | 1.700E-08 | 4.300E-08 |
| 11460 | 573 | F573V | A | 3.8e-08 | 6.5e-08 | 5.3e-08 | 5.200E-08 | 1.353E-08 | 3.800E-08 | 6.500E-08 |
| 11461 | 574 | C574A | A | 3.7e-08 | 7.6e-08 | 1.1e-07 | 7.433E-08 | 3.653E-08 | 3.700E-08 | 1.100E-07 |
| 11462 | 574 | C574R | A | 1.1e-07 | 6.1e-08 | 4.2e-08 | 7.100E-08 | 3.509E-08 | 4.200E-08 | 1.100E-07 |
| 11463 | 574 | C574N | A | 4.2e-08 | 2.6e-08 | 3.5e-08 | 3.433E-08 | 8.021E-09 | 2.600E-08 | 4.200E-08 |
| 11464 | 574 | C574D | A | 1.2e-08 | 1.1e-08 | 7.6e-08 | 3.300E-08 | 3.724E-08 | 1.100E-08 | 7.600E-08 |
| 11465 | 574 | C574C | A | 2.7e-08 | 7.2e-08 | 2.5e-08 | 4.133E-08 | 2.658E-08 | 2.500E-08 | 7.200E-08 |
| 11466 | 574 | C574E | A | 2.3e-08 | 4.4e-08 | 6.2e-08 | 4.300E-08 | 1.952E-08 | 2.300E-08 | 6.200E-08 |
| 11467 | 574 | C574Q | A | 7.1e-08 | 4.6e-08 | 3.7e-08 | 5.133E-08 | 1.762E-08 | 3.700E-08 | 7.100E-08 |
| 11468 | 574 | C574G | A | 6.1e-08 | 1e-08 | 6e-08 | 4.367E-08 | 2.916E-08 | 1.000E-08 | 6.100E-08 |
| 11469 | 574 | C574H | A | 3.4e-08 | 3.7e-08 | 5.5e-08 | 4.200E-08 | 1.136E-08 | 3.400E-08 | 5.500E-08 |
| 11470 | 574 | C574I | A | 4.3e-08 | 3.2e-08 | 2e-08 | 3.167E-08 | 1.150E-08 | 2.000E-08 | 4.300E-08 |
| 11471 | 574 | C574L | A | 7.1e-08 | 3.2e-08 | 1.4e-08 | 3.900E-08 | 2.914E-08 | 1.400E-08 | 7.100E-08 |
| 11472 | 574 | C574K | A | 4.9e-08 | 1.6e-08 | 4.6e-08 | 3.700E-08 | 1.825E-08 | 1.600E-08 | 4.900E-08 |
| 11473 | 574 | C574M | A | 8.8e-08 | 6.1e-08 | 3e-08 | 5.967E-08 | 2.902E-08 | 3.000E-08 | 8.800E-08 |
| 11474 | 574 | C574F | A | 8.4e-08 | 2.8e-08 | 3.6e-08 | 4.933E-08 | 3.029E-08 | 2.800E-08 | 8.400E-08 |

| | | | | | | | | | | |
|---|---|---|---|---|---|---|---|---|---|---|
| 11475 | 574 | C574P | A | 1.6e-08 | 4.4e-08 | 1.3e-08 | 2.433E-08 | 1.710E-08 | 1.300E-08 | 4.400E-08 |
| 11476 | 574 | C574S | A | 5.4e-08 | 5.4e-08 | 4.5e-08 | 5.100E-08 | 5.196E-09 | 4.500E-08 | 5.400E-08 |
| 11477 | 574 | C574T | A | 7e-08 | 4.5e-08 | 4e-08 | 5.167E-08 | 1.607E-08 | 4.000E-08 | 7.000E-08 |
| 11478 | 574 | C574W | A | 2.4e-08 | 8.3e-08 | 8.2e-08 | 6.300E-08 | 3.378E-08 | 2.400E-08 | 8.300E-08 |
| 11479 | 574 | C574Y | A | 1.5e-07 | 2.4e-08 | 6.2e-08 | 7.867E-08 | 6.463E-08 | 2.400E-08 | 1.500E-07 |
| 11480 | 574 | C574V | A | 1.7e-08 | 7.3e-08 | 6.7e-08 | 5.233E-08 | 3.075E-08 | 1.700E-08 | 7.300E-08 |
| 11481 | 575 | V575A | A | 4.7e-08 | 2.6e-08 | 7.7e-08 | 5.000E-08 | 2.563E-08 | 2.600E-08 | 7.700E-08 |
| 11482 | 575 | V575R | A | 4.1e-08 | 4.8e-08 | 3e-08 | 3.967E-08 | 9.074E-09 | 3.000E-08 | 4.800E-08 |
| 11483 | 575 | V575N | A | 5.4e-08 | 1e-08 | 2.2e-08 | 2.867E-08 | 2.274E-08 | 1.000E-08 | 5.400E-08 |
| 11484 | 575 | V575D | A | 4.2e-08 | 3.9e-08 | 2e-08 | 3.367E-08 | 1.193E-08 | 2.000E-08 | 4.200E-08 |
| 11485 | 575 | V575C | A | 4.6e-08 | 5.4e-08 | 1.8e-08 | 3.933E-08 | 1.890E-08 | 1.800E-08 | 5.400E-08 |
| 11486 | 575 | V575E | A | 2.9e-08 | 4.1e-08 | 3.9e-08 | 3.633E-08 | 6.429E-09 | 2.900E-08 | 4.100E-08 |
| 11487 | 575 | V575Q | A | 2.5e-08 | 6.5e-08 | 6.9e-08 | 5.300E-08 | 2.433E-08 | 2.500E-08 | 6.900E-08 |
| 11488 | 575 | V575G | A | 2.1e-08 | 5.4e-08 | 5.7e-08 | 4.400E-08 | 1.997E-08 | 2.100E-08 | 5.700E-08 |
| 11489 | 575 | V575H | A | 3.6e-08 | 3.5e-08 | 3.6e-08 | 3.567E-08 | 5.774E-10 | 3.500E-08 | 3.600E-08 |
| 11490 | 575 | V575I | A | 4.1e-08 | 3.8e-08 | 2.1e-08 | 3.333E-08 | 1.079E-08 | 2.100E-08 | 4.100E-08 |
| 11491 | 575 | V575L | A | 5.5e-08 | 2.3e-08 | 1.7e-08 | 3.167E-08 | 2.043E-08 | 1.700E-08 | 5.500E-08 |
| 11492 | 575 | V575K | A | 3.1e-08 | 1.7e-08 | 3.6e-08 | 2.800E-08 | 9.849E-09 | 1.700E-08 | 3.600E-08 |
| 11493 | 575 | V575M | A | 1.3e-08 | 4.7e-09 | 3.1e-08 | 1.623E-08 | 1.344E-08 | 4.700E-09 | 3.100E-08 |
| 11494 | 575 | V575F | A | 1e-07 | 4.3e-08 | 5.1e-08 | 6.467E-08 | 3.086E-08 | 4.300E-08 | 1.000E-07 |
| 11495 | 575 | V575P | A | 1.7e-08 | 5.8e-08 | 2.7e-08 | 3.400E-08 | 2.138E-08 | 1.700E-08 | 5.800E-08 |
| 11496 | 575 | V575S | A | 5.4e-08 | 3.9e-08 | 2.5e-08 | 3.933E-08 | 1.450E-08 | 2.500E-08 | 5.400E-08 |
| 11497 | 575 | V575T | A | 2.9e-08 | 2.6e-08 | 1.7e-08 | 2.400E-08 | 6.245E-09 | 1.700E-08 | 2.900E-08 |
| 11498 | 575 | V575W | A | 3.7e-08 | 3.8e-08 | 1.9e-07 | 8.833E-08 | 8.805E-08 | 3.700E-08 | 1.900E-07 |
| 11499 | 575 | V575Y | A | 5.3e-08 | 9.4e-08 | 4.6e-08 | 6.433E-08 | 2.593E-08 | 4.600E-08 | 9.400E-08 |
| 11500 | 575 | V575V | A | 2.4e-08 | 3.2e-08 | 5.8e-08 | 3.800E-08 | 1.778E-08 | 2.400E-08 | 5.800E-08 |
| 11501 | 576 | A576A | A | 5.3e-08 | 7.1e-08 | 4e-08 | 5.467E-08 | 1.557E-08 | 4.000E-08 | 7.100E-08 |
| 11502 | 576 | A576R | A | 8.9e-08 | 2.8e-08 | 7e-08 | 6.233E-08 | 3.121E-08 | 2.800E-08 | 8.900E-08 |
| 11503 | 576 | A576N | A | 9.2e-08 | 1.8e-08 | 2.8e-08 | 4.600E-08 | 4.015E-08 | 1.800E-08 | 9.200E-08 |
| 11504 | 576 | A576D | A | 6.6e-08 | 2.8e-08 | 3e-08 | 4.133E-08 | 2.139E-08 | 2.800E-08 | 6.600E-08 |
| 11505 | 576 | A576C | A | 8.8e-08 | 3.8e-08 | 1.9e-08 | 4.833E-08 | 3.564E-08 | 1.900E-08 | 8.800E-08 |
| 11506 | 576 | A576E | A | 3.5e-08 | 3.1e-08 | 3.5e-08 | 3.367E-08 | 2.309E-09 | 3.100E-08 | 3.500E-08 |
| 11507 | 576 | A576Q | A | 2e-08 | 1.8e-08 | 5.9e-08 | 3.233E-08 | 2.312E-08 | 1.800E-08 | 5.900E-08 |
| 11508 | 576 | A576G | A | 4.1e-08 | 2e-08 | 4.7e-08 | 3.600E-08 | 1.418E-08 | 2.000E-08 | 4.700E-08 |
| 11509 | 576 | A576H | A | 1.9e-08 | 4.9e-08 | 2.1e-08 | 2.967E-08 | 1.677E-08 | 1.900E-08 | 4.900E-08 |
| 11510 | 576 | A576I | A | 8.9e-08 | 5.3e-08 | 4.4e-08 | 6.200E-08 | 2.381E-08 | 4.400E-08 | 8.900E-08 |
| 11511 | 576 | A576L | A | 3.7e-08 | 1.8e-08 | 6.7e-08 | 4.067E-08 | 2.470E-08 | 1.800E-08 | 6.700E-08 |
| 11512 | 576 | A576K | A | 2e-08 | 4.5e-08 | 3.7e-08 | 3.400E-08 | 1.277E-08 | 2.000E-08 | 4.500E-08 |
| 11513 | 576 | A576M | A | 1.6e-08 | 4.6e-08 | 5.1e-08 | 3.767E-08 | 1.893E-08 | 1.600E-08 | 5.100E-08 |
| 11514 | 576 | A576F | A | 5.7e-08 | 2.9e-08 | 7.7e-08 | 5.433E-08 | 2.411E-08 | 2.900E-08 | 7.700E-08 |
| 11515 | 576 | A576P | A | 3.2e-08 | 8e-08 | 2.2e-08 | 4.467E-08 | 3.101E-08 | 2.200E-08 | 8.000E-08 |
| 11516 | 576 | A576S | A | 4.8e-08 | 2.5e-08 | 2.3e-08 | 3.200E-08 | 1.389E-08 | 2.300E-08 | 4.800E-08 |
| 11517 | 576 | A576T | A | 9e-08 | 2.7e-08 | 2.8e-08 | 4.833E-08 | 3.609E-08 | 2.700E-08 | 9.000E-08 |
| 11518 | 576 | A576W | A | 4.2e-08 | 6.4e-08 | 6.8e-08 | 5.800E-08 | 1.400E-08 | 4.200E-08 | 6.800E-08 |
| 11519 | 576 | A576Y | A | 8.4e-08 | 1.6e-08 | 3e-08 | 4.333E-08 | 3.591E-08 | 1.600E-08 | 8.400E-08 |
| 11520 | 576 | A576V | A | 2.9e-08 | 2.5e-08 | 4.9e-08 | 3.433E-08 | 1.286E-08 | 2.500E-08 | 4.900E-08 |
| 11521 | 577 | R577A | A | 1.4e-08 | 1.2e-07 | 4.4e-08 | 5.933E-08 | 5.464E-08 | 1.400E-08 | 1.200E-07 |
| 11522 | 577 | R577R | A | 7.2e-08 | 1.8e-08 | 4.9e-08 | 4.633E-08 | 2.710E-08 | 1.800E-08 | 7.200E-08 |
| 11523 | 577 | R577N | A | 1.8e-08 | 1.4e-08 | 8e-08 | 3.733E-08 | 3.700E-08 | 1.400E-08 | 8.000E-08 |
| 11524 | 577 | R577D | A | 5.8e-08 | 5.8e-08 | 5.5e-08 | 5.700E-08 | 1.732E-09 | 5.500E-08 | 5.800E-08 |
| 11525 | 577 | R577C | A | 9.6e-08 | 8.2e-08 | 4.6e-08 | 7.467E-08 | 2.579E-08 | 4.600E-08 | 9.600E-08 |
| 11526 | 577 | R577E | A | 7.4e-08 | 7.7e-08 | 3.1e-08 | 6.067E-08 | 2.574E-08 | 3.100E-08 | 7.700E-08 |
| 11527 | 577 | R577Q | A | 6.8e-08 | 3.4e-08 | 1.8e-08 | 4.000E-08 | 2.553E-08 | 1.800E-08 | 6.800E-08 |
| 11528 | 577 | R577G | A | 1.1e-07 | 1.1e-08 | 1.1e-07 | 7.700E-08 | 5.716E-08 | 1.100E-08 | 1.100E-07 |
| 11529 | 577 | R577H | A | 5.3e-08 | 4.7e-08 | 7.1e-08 | 5.700E-08 | 1.249E-08 | 4.700E-08 | 7.100E-08 |
| 11530 | 577 | R577I | A | 1.1e-07 | 6.5e-08 | 7.4e-08 | 8.300E-08 | 2.381E-08 | 6.500E-08 | 1.100E-07 |
| 11531 | 577 | R577L | A | 2.7e-08 | 4.6e-08 | 1.9e-08 | 3.067E-08 | 1.387E-08 | 1.900E-08 | 4.600E-08 |
| 11532 | 577 | R577K | A | 1.1e-07 | 4.7e-08 | 1.7e-08 | 5.800E-08 | 4.747E-08 | 1.700E-08 | 1.100E-07 |
| 11533 | 577 | R577M | A | 4.9e-08 | 1.3e-08 | 3e-08 | 3.067E-08 | 1.801E-08 | 1.300E-08 | 4.900E-08 |
| 11534 | 577 | R577F | A | 7e-08 | 5.7e-08 | 6.5e-08 | 6.400E-08 | 6.557E-09 | 5.700E-08 | 7.000E-08 |
| 11535 | 577 | R577P | A | 6.6e-08 | 6.3e-08 | 6.2e-08 | 6.367E-08 | 2.082E-09 | 6.200E-08 | 6.600E-08 |
| 11536 | 577 | R577S | A | 3.3e-08 | 4.6e-08 | 9e-08 | 5.633E-08 | 2.987E-08 | 3.300E-08 | 9.000E-08 |
| 11537 | 577 | R577T | A | 4.4e-08 | 4.4e-08 | 5.7e-08 | 4.833E-08 | 7.506E-09 | 4.400E-08 | 5.700E-08 |
| 11538 | 577 | R577W | A | 1.7e-08 | 1.9e-08 | 9.5e-09 | 1.517E-08 | 5.008E-09 | 9.500E-09 | 1.900E-08 |
| 11539 | 577 | R577Y | A | 2.7e-08 | 6.7e-08 | 2.3e-08 | 3.900E-08 | 2.433E-08 | 2.300E-08 | 6.700E-08 |
| 11540 | 577 | R577V | A | 2.7e-08 | 2e-08 | 2.1e-08 | 2.267E-08 | 3.786E-09 | 2.000E-08 | 2.700E-08 |
| 11541 | 578 | C578A | A | 4.6e-08 | 3.5e-08 | 6.2e-08 | 4.767E-08 | 1.358E-08 | 3.500E-08 | 6.200E-08 |
| 11542 | 578 | C578R | A | 2.3e-08 | 2.4e-08 | 3.4e-08 | 2.700E-08 | 6.083E-09 | 2.300E-08 | 3.400E-08 |
| 11543 | 578 | C578N | A | 2.9e-08 | 2.2e-08 | 1.8e-08 | 2.300E-08 | 5.568E-09 | 1.800E-08 | 2.900E-08 |
| 11544 | 578 | C578D | A | 1.2e-08 | 3.7e-08 | 9.9e-08 | 4.933E-08 | 4.479E-08 | 1.200E-08 | 9.900E-08 |
| 11545 | 578 | C578C | A | 3.2e-08 | 9.1e-08 | 7.1e-08 | 6.467E-08 | 3.001E-08 | 3.200E-08 | 9.100E-08 |
| 11546 | 578 | C578E | A | 3.6e-08 | 4.2e-08 | 4.3e-08 | 4.033E-08 | 3.786E-09 | 3.600E-08 | 4.300E-08 |
| 11547 | 578 | C578Q | A | 1.3e-07 | 5.2e-08 | 7.2e-08 | 8.467E-08 | 4.051E-08 | 5.200E-08 | 1.300E-07 |
| 11548 | 578 | C578G | A | 6.9e-08 | 4.6e-08 | 5.9e-08 | 5.800E-08 | 1.153E-08 | 4.600E-08 | 6.900E-08 |
| 11549 | 578 | C578H | A | 8.8e-08 | 3.6e-08 | 2.1e-08 | 4.833E-08 | 3.516E-08 | 2.100E-08 | 8.800E-08 |
| 11550 | 578 | C578I | A | 4.6e-08 | 5.2e-08 | 3.8e-08 | 4.533E-08 | 7.024E-09 | 3.800E-08 | 5.200E-08 |
| 11551 | 578 | C578L | A | 4.2e-08 | 3.7e-08 | 4.3e-08 | 4.067E-08 | 3.215E-09 | 3.700E-08 | 4.300E-08 |
| 11552 | 578 | C578K | A | 4.4e-08 | 5.7e-08 | 3.8e-08 | 4.633E-08 | 9.713E-09 | 3.800E-08 | 5.700E-08 |
| 11553 | 578 | C578M | A | 4e-08 | 5e-08 | 6.7e-08 | 5.233E-08 | 1.365E-08 | 4.000E-08 | 6.700E-08 |
| 11554 | 578 | C578F | A | 4.1e-08 | 2.4e-08 | 3.7e-08 | 3.400E-08 | 8.888E-09 | 2.400E-08 | 4.100E-08 |
| 11555 | 578 | C578P | A | 9.9e-09 | 7.2e-08 | 1.5e-08 | 3.230E-08 | 3.448E-08 | 9.900E-09 | 7.200E-08 |
| 11556 | 578 | C578S | A | 4e-08 | 2.1e-08 | 1.2e-08 | 2.433E-08 | 1.429E-08 | 1.200E-08 | 4.000E-08 |
| 11557 | 578 | C578T | A | 2.2e-08 | 1.6e-08 | 3e-08 | 2.267E-08 | 7.024E-09 | 1.600E-08 | 3.000E-08 |
| 11558 | 578 | C578W | A | 5.8e-08 | 2.8e-08 | 4.6e-08 | 4.400E-08 | 1.510E-08 | 2.800E-08 | 5.800E-08 |
| 11559 | 578 | C578Y | A | 2.9e-08 | 3.4e-08 | 3.9e-08 | 3.400E-08 | 5.000E-09 | 2.900E-08 | 3.900E-08 |

| | | | | | | | | | | |
|---|---|---|---|---|---|---|---|---|---|---|
| 11560 | 578 | C578V | A | 2.6e-08 | 2.9e-08 | 3.3e-08 | 2.933E-08 | 3.512E-09 | 2.600E-08 | 3.300E-08 |
| 11561 | 579 | P579A | A | 1.4e-08 | 7.2e-08 | 8.8e-08 | 5.800E-08 | 3.894E-08 | 1.400E-08 | 8.800E-08 |
| 11562 | 579 | P579R | A | 7.2e-08 | 3.1e-08 | 5.9e-08 | 5.400E-08 | 2.095E-08 | 3.100E-08 | 7.200E-08 |
| 11563 | 579 | P579N | A | 8e-08 | 1.3e-08 | 3.1e-08 | 4.133E-08 | 3.467E-08 | 1.300E-08 | 8.000E-08 |
| 11564 | 579 | P579D | A | 5.4e-08 | 1.3e-07 | 1.7e-08 | 6.700E-08 | 5.761E-08 | 1.700E-08 | 1.300E-07 |
| 11565 | 579 | P579C | A | 2.1e-08 | 4.9e-08 | 1.4e-08 | 2.800E-08 | 1.852E-08 | 1.400E-08 | 4.900E-08 |
| 11566 | 579 | P579E | A | 4e-08 | 3.1e-08 | 3.3e-08 | 3.467E-08 | 4.726E-09 | 3.100E-08 | 4.000E-08 |
| 11567 | 579 | P579Q | A | 5.9e-08 | 2e-08 | 1e-07 | 5.967E-08 | 4.000E-08 | 2.000E-08 | 1.000E-07 |
| 11568 | 579 | P579G | A | 7.7e-08 | 8.4e-08 | 5.5e-08 | 7.200E-08 | 1.513E-08 | 5.500E-08 | 8.400E-08 |
| 11569 | 579 | P579H | A | 7.3e-08 | 1.2e-08 | 7.1e-08 | 5.200E-08 | 3.466E-08 | 1.200E-08 | 7.300E-08 |
| 11570 | 579 | P579I | A | 2.3e-08 | 5.2e-08 | 4e-08 | 3.833E-08 | 1.457E-08 | 2.300E-08 | 5.200E-08 |
| 11571 | 579 | P579L | A | 9.9e-08 | 4.1e-08 | 2.6e-08 | 5.533E-08 | 3.855E-08 | 2.600E-08 | 9.900E-08 |
| 11572 | 579 | P579K | A | 7.9e-08 | 1.3e-07 | 2.2e-08 | 7.700E-08 | 5.403E-08 | 2.200E-08 | 1.300E-07 |
| 11573 | 579 | P579M | A | 9.1e-08 | 3.4e-08 | 5.6e-08 | 6.033E-08 | 2.875E-08 | 3.400E-08 | 9.100E-08 |
| 11574 | 579 | P579F | A | 3.6e-08 | 5.7e-08 | 1e-07 | 6.433E-08 | 3.262E-08 | 3.600E-08 | 1.000E-07 |
| 11575 | 579 | P579P | A | 1.1e-08 | 8.1e-08 | 1.8e-08 | 3.667E-08 | 3.855E-08 | 1.100E-08 | 8.100E-08 |
| 11576 | 579 | P579S | A | 6.4e-08 | 4.7e-08 | 1.4e-08 | 4.167E-08 | 2.542E-08 | 1.400E-08 | 6.400E-08 |
| 11577 | 579 | P579T | A | 6.7e-08 | 5.9e-08 | 3.3e-08 | 5.300E-08 | 1.778E-08 | 3.300E-08 | 6.700E-08 |
| 11578 | 579 | P579W | A | 4.2e-08 | 9.2e-08 | 2.2e-08 | 5.200E-08 | 3.606E-08 | 2.200E-08 | 9.200E-08 |
| 11579 | 579 | P579Y | A | 3.2e-08 | 4.3e-08 | 3.4e-08 | 3.633E-08 | 5.859E-09 | 3.200E-08 | 4.300E-08 |
| 11580 | 579 | P579V | A | 6.4e-08 | 4.6e-08 | 6.2e-08 | 5.733E-08 | 9.866E-09 | 4.600E-08 | 6.400E-08 |
| 11581 | 580 | S580A | A | 5.7e-08 | 2.6e-08 | 5.5e-08 | 4.600E-08 | 1.735E-08 | 2.600E-08 | 5.700E-08 |
| 11582 | 580 | S580R | A | 9.1e-08 | 1.1e-07 | 3.3e-08 | 7.800E-08 | 4.011E-08 | 3.300E-08 | 1.100E-07 |
| 11583 | 580 | S580N | A | 4.9e-08 | 3.1e-08 | 2.1e-08 | 3.367E-08 | 1.419E-08 | 2.100E-08 | 4.900E-08 |
| 11584 | 580 | S580D | A | 2.9e-08 | 3.7e-08 | 4.6e-08 | 3.733E-08 | 8.505E-09 | 2.900E-08 | 4.600E-08 |
| 11585 | 580 | S580C | A | 1.2e-07 | 2.5e-08 | 3.6e-08 | 6.033E-08 | 5.196E-08 | 2.500E-08 | 1.200E-07 |
| 11586 | 580 | S580E | A | 6.1e-08 | 5.9e-08 | 1.1e-07 | 7.667E-08 | 2.888E-08 | 5.900E-08 | 1.100E-07 |
| 11587 | 580 | S580Q | A | 2.2e-08 | 7.2e-08 | 5.8e-08 | 5.067E-08 | 2.579E-08 | 2.200E-08 | 7.200E-08 |
| 11588 | 580 | S580G | A | 4.1e-08 | 8.5e-08 | 9.1e-08 | 7.233E-08 | 2.730E-08 | 4.100E-08 | 9.100E-08 |
| 11589 | 580 | S580H | A | 1.4e-08 | 2.9e-08 | 1.2e-07 | 5.433E-08 | 5.736E-08 | 1.400E-08 | 1.200E-07 |
| 11590 | 580 | S580I | A | 1.1e-08 | 3.8e-08 | 1.4e-07 | 6.300E-08 | 6.804E-08 | 1.100E-08 | 1.400E-07 |
| 11591 | 580 | S580L | A | 4.1e-08 | 3.3e-08 | 4.4e-08 | 3.933E-08 | 5.686E-09 | 3.300E-08 | 4.400E-08 |
| 11592 | 580 | S580K | A | 4.3e-08 | 2.3e-08 | 2.2e-07 | 9.533E-08 | 1.084E-07 | 2.300E-08 | 2.200E-07 |
| 11593 | 580 | S580M | A | 6.7e-08 | 4.8e-08 | 6.8e-08 | 6.100E-08 | 1.127E-08 | 4.800E-08 | 6.800E-08 |
| 11594 | 580 | S580F | A | 3.2e-08 | 2.2e-08 | 2.1e-08 | 2.500E-08 | 6.083E-09 | 2.100E-08 | 3.200E-08 |
| 11595 | 580 | S580P | A | 1.6e-08 | 2.1e-08 | 5.9e-08 | 3.200E-08 | 2.352E-08 | 1.600E-08 | 5.900E-08 |
| 11596 | 580 | S580S | A | 2.9e-08 | 4.5e-08 | 2.7e-08 | 3.367E-08 | 9.866E-09 | 2.700E-08 | 4.500E-08 |
| 11597 | 580 | S580T | A | 4.2e-08 | 4.2e-08 | 1.8e-08 | 3.400E-08 | 1.386E-08 | 1.800E-08 | 4.200E-08 |
| 11598 | 580 | S580W | A | 1.8e-08 | 4.2e-08 | 2.6e-08 | 2.867E-08 | 1.222E-08 | 1.800E-08 | 4.200E-08 |
| 11599 | 580 | S580Y | A | 3.4e-08 | 3.8e-08 | 5.3e-08 | 4.167E-08 | 1.002E-08 | 3.400E-08 | 5.300E-08 |
| 11600 | 580 | S580V | A | 4.9e-08 | 3.9e-08 | 1.9e-08 | 3.567E-08 | 1.528E-08 | 1.900E-08 | 4.900E-08 |
| 11601 | 581 | G581A | A | 7.4e-08 | 2.8e-08 | 1e-07 | 6.733E-08 | 3.646E-08 | 2.800E-08 | 1.000E-07 |
| 11602 | 581 | G581R | A | 4.3e-08 | 1.1e-07 | 5.1e-08 | 6.800E-08 | 3.659E-08 | 4.300E-08 | 1.100E-07 |
| 11603 | 581 | G581N | A | 3.6e-08 | 6.1e-08 | 3.4e-08 | 4.367E-08 | 1.504E-08 | 3.400E-08 | 6.100E-08 |
| 11604 | 581 | G581D | A | 3.6e-08 | 4.1e-08 | 4.4e-08 | 4.033E-08 | 4.041E-09 | 3.600E-08 | 4.400E-08 |
| 11605 | 581 | G581C | A | 4.7e-08 | 3.9e-08 | 5e-08 | 4.533E-08 | 5.686E-09 | 3.900E-08 | 5.000E-08 |
| 11606 | 581 | G581E | A | 7e-08 | 7.6e-08 | 5.5e-08 | 6.700E-08 | 1.082E-08 | 5.500E-08 | 7.600E-08 |
| 11607 | 581 | G581Q | A | 1.2e-08 | 2.3e-08 | 6.7e-08 | 3.400E-08 | 2.910E-08 | 1.200E-08 | 6.700E-08 |
| 11608 | 581 | G581G | A | 3.7e-08 | 4e-08 | 4.4e-08 | 4.033E-08 | 3.512E-09 | 3.700E-08 | 4.400E-08 |
| 11609 | 581 | G581H | A | 5.8e-08 | 6e-08 | 7.9e-08 | 6.567E-08 | 1.159E-08 | 5.800E-08 | 7.900E-08 |
| 11610 | 581 | G581I | A | 4e-08 | 1.2e-08 | 1.8e-08 | 2.333E-08 | 1.474E-08 | 1.200E-08 | 4.000E-08 |
| 11611 | 581 | G581L | A | 4.1e-08 | 2.2e-08 | 1.2e-08 | 2.500E-08 | 1.473E-08 | 1.200E-08 | 4.100E-08 |
| 11612 | 581 | G581K | A | 2.4e-08 | 4e-08 | 4.8e-08 | 3.733E-08 | 1.222E-08 | 2.400E-08 | 4.800E-08 |
| 11613 | 581 | G581M | A | 1.1e-08 | 7.9e-08 | 1.9e-08 | 3.633E-08 | 3.717E-08 | 1.100E-08 | 7.900E-08 |
| 11614 | 581 | G581F | A | 1e-07 | 4.3e-08 | 8.4e-08 | 7.567E-08 | 2.940E-08 | 4.300E-08 | 1.000E-07 |
| 11615 | 581 | G581P | A | 7.5e-08 | 1e-08 | 9.3e-08 | 5.933E-08 | 4.366E-08 | 1.000E-08 | 9.300E-08 |
| 11616 | 581 | G581S | A | 1e-07 | 4.3e-08 | 2.2e-08 | 5.500E-08 | 4.036E-08 | 2.200E-08 | 1.000E-07 |
| 11617 | 581 | G581T | A | 3.4e-08 | 4.8e-08 | 6.1e-08 | 4.767E-08 | 1.350E-08 | 3.400E-08 | 6.100E-08 |
| 11618 | 581 | G581W | A | 1.6e-07 | 2.2e-08 | 5.6e-08 | 7.933E-08 | 7.190E-08 | 2.200E-08 | 1.600E-07 |
| 11619 | 581 | G581Y | A | 2.9e-08 | 1.9e-08 | 2.2e-08 | 2.333E-08 | 5.132E-09 | 1.900E-08 | 2.900E-08 |
| 11620 | 581 | G581V | A | 2.9e-08 | 2.3e-08 | 4.8e-08 | 3.333E-08 | 1.305E-08 | 2.300E-08 | 4.800E-08 |
| 11621 | 582 | V582A | A | 2.4e-08 | 1.7e-08 | 2.2e-08 | 2.100E-08 | 3.606E-09 | 1.700E-08 | 2.400E-08 |
| 11622 | 582 | V582R | A | 4.3e-08 | 7.1e-08 | 3.2e-08 | 4.867E-08 | 2.011E-08 | 3.200E-08 | 7.100E-08 |
| 11623 | 582 | V582N | A | 1.4e-08 | 1.9e-07 | 9.6e-08 | 1.000E-07 | 8.807E-08 | 1.400E-08 | 1.900E-07 |
| 11624 | 582 | V582D | A | 2.5e-08 | 1.7e-08 | 1.7e-08 | 1.967E-08 | 4.619E-09 | 1.700E-08 | 2.500E-08 |
| 11625 | 582 | V582C | A | 4.7e-08 | 2.9e-08 | 1.5e-08 | 3.033E-08 | 1.604E-08 | 1.500E-08 | 4.700E-08 |
| 11626 | 582 | V582E | A | 1.8e-08 | 1.9e-08 | 2.7e-08 | 2.133E-08 | 4.933E-09 | 1.800E-08 | 2.700E-08 |
| 11627 | 582 | V582Q | A | 4.4e-08 | 4.5e-08 | 3.7e-08 | 4.200E-08 | 4.359E-09 | 3.700E-08 | 4.500E-08 |
| 11628 | 582 | V582G | A | 3.3e-08 | 9e-08 | 3.4e-08 | 5.233E-08 | 3.262E-08 | 3.300E-08 | 9.000E-08 |
| 11629 | 582 | V582H | A | 2.6e-08 | 3.7e-08 | 4.9e-08 | 3.733E-08 | 1.150E-08 | 2.600E-08 | 4.900E-08 |
| 11630 | 582 | V582I | A | 4.5e-08 | 8.1e-08 | 3.5e-08 | 5.367E-08 | 2.419E-08 | 3.500E-08 | 8.100E-08 |
| 11631 | 582 | V582L | A | 6.1e-08 | 2.9e-08 | 6.9e-08 | 5.300E-08 | 2.117E-08 | 2.900E-08 | 6.900E-08 |
| 11632 | 582 | V582K | A | 3.4e-08 | 2.7e-08 | 7.4e-08 | 4.500E-08 | 2.536E-08 | 2.700E-08 | 7.400E-08 |
| 11633 | 582 | V582M | A | 6.7e-08 | 3.2e-08 | 1.1e-08 | 3.667E-08 | 2.829E-08 | 1.100E-08 | 6.700E-08 |
| 11634 | 582 | V582F | A | 3.9e-08 | 4.5e-08 | 2.7e-08 | 3.700E-08 | 9.165E-09 | 2.700E-08 | 4.500E-08 |
| 11635 | 582 | V582P | A | 2.4e-08 | 3.1e-08 | 2.2e-08 | 2.567E-08 | 4.726E-09 | 2.200E-08 | 3.100E-08 |
| 11636 | 582 | V582S | A | 9.6e-08 | 1e-07 | 6.2e-08 | 8.600E-08 | 2.088E-08 | 6.200E-08 | 1.000E-07 |
| 11637 | 582 | V582T | A | 2.9e-08 | 1.4e-07 | 4.4e-08 | 7.100E-08 | 6.022E-08 | 2.900E-08 | 1.400E-07 |
| 11638 | 582 | V582W | A | 7.2e-08 | 3.4e-08 | 6.9e-08 | 5.833E-08 | 2.113E-08 | 3.400E-08 | 7.200E-08 |
| 11639 | 582 | V582Y | A | 9.3e-08 | 4.9e-08 | 2.6e-08 | 5.600E-08 | 3.404E-08 | 2.600E-08 | 9.300E-08 |
| 11640 | 582 | V582V | A | 4.9e-08 | 6.6e-08 | 3.2e-08 | 4.900E-08 | 1.700E-08 | 3.200E-08 | 6.600E-08 |
| 11641 | 583 | K583A | A | 3e-08 | 3e-08 | 1.9e-08 | 2.633E-08 | 6.351E-09 | 1.900E-08 | 3.000E-08 |
| 11642 | 583 | K583R | A | 2.1e-08 | 5.1e-08 | 3.4e-08 | 3.533E-08 | 1.504E-08 | 2.100E-08 | 5.100E-08 |
| 11643 | 583 | K583N | A | 9.9e-08 | 7.4e-08 | 3.4e-08 | 6.900E-08 | 3.279E-08 | 3.400E-08 | 9.900E-08 |
| 11644 | 583 | K583D | A | 3.8e-08 | 5.1e-08 | 6.9e-08 | 5.267E-08 | 1.557E-08 | 3.800E-08 | 6.900E-08 |

| | | | | | | | | | | |
|---|---|---|---|---|---|---|---|---|---|---|
| 11645 | 583 | K583C | A | 2.2e-08 | 8.5e-09 | 3e-08 | 2.017E-08 | 1.087E-08 | 8.500E-09 | 3.000E-08 |
| 11646 | 583 | K583E | A | 8e-08 | 6.4e-08 | 2.9e-08 | 5.767E-08 | 2.608E-08 | 2.900E-08 | 8.000E-08 |
| 11647 | 583 | K583Q | A | 2.3e-08 | 1e-07 | 6.7e-08 | 6.333E-08 | 3.863E-08 | 2.300E-08 | 1.000E-07 |
| 11648 | 583 | K583G | A | 1.5e-08 | 7.4e-08 | 3.7e-08 | 4.200E-08 | 2.982E-08 | 1.500E-08 | 7.400E-08 |
| 11649 | 583 | K583H | A | 7.2e-08 | 1.4e-08 | 1.3e-08 | 3.300E-08 | 3.378E-08 | 1.300E-08 | 7.200E-08 |
| 11650 | 583 | K583I | A | 2.8e-08 | 1.8e-08 | 1.2e-07 | 5.533E-08 | 5.623E-08 | 1.800E-08 | 1.200E-07 |
| 11651 | 583 | K583L | A | 4.9e-08 | 3.8e-08 | 3.8e-08 | 4.167E-08 | 6.351E-09 | 3.800E-08 | 4.900E-08 |
| 11652 | 583 | K583K | A | 2.7e-08 | 1.5e-08 | 4.8e-08 | 3.000E-08 | 1.670E-08 | 1.500E-08 | 4.800E-08 |
| 11653 | 583 | K583M | A | 3.9e-08 | 2.2e-08 | 9e-08 | 5.033E-08 | 3.539E-08 | 2.200E-08 | 9.000E-08 |
| 11654 | 583 | K583F | A | 3.1e-08 | 3.5e-08 | 5.4e-08 | 4.000E-08 | 1.229E-08 | 3.100E-08 | 5.400E-08 |
| 11655 | 583 | K583P | A | 7.5e-08 | 3.1e-08 | 5.2e-08 | 5.267E-08 | 2.201E-08 | 3.100E-08 | 7.500E-08 |
| 11656 | 583 | K583S | A | 2.7e-08 | 5.4e-08 | 3.3e-08 | 3.800E-08 | 1.418E-08 | 2.700E-08 | 5.400E-08 |
| 11657 | 583 | K583T | A | 2.7e-08 | 4.2e-08 | 4.5e-08 | 3.800E-08 | 9.644E-09 | 2.700E-08 | 4.500E-08 |
| 11658 | 583 | K583W | A | 1.3e-08 | 4.1e-08 | 3.6e-08 | 3.000E-08 | 1.493E-08 | 1.300E-08 | 4.100E-08 |
| 11659 | 583 | K583Y | A | 4e-08 | 7.1e-08 | 9.6e-08 | 6.900E-08 | 2.805E-08 | 4.000E-08 | 9.600E-08 |
| 11660 | 583 | K583V | A | 4.7e-08 | 5.5e-08 | 9.3e-08 | 6.500E-08 | 2.458E-08 | 4.700E-08 | 9.300E-08 |
| 11661 | 584 | P584A | A | 5.9e-08 | 6.8e-08 | 3.7e-08 | 5.467E-08 | 1.595E-08 | 3.700E-08 | 6.800E-08 |
| 11662 | 584 | P584R | A | 1.4e-08 | 7.7e-08 | 9.8e-08 | 6.300E-08 | 4.371E-08 | 1.400E-08 | 9.800E-08 |
| 11663 | 584 | P584N | A | 2.9e-08 | 2e-08 | 5.2e-08 | 3.367E-08 | 1.650E-08 | 2.000E-08 | 5.200E-08 |
| 11664 | 584 | P584D | A | 7.3e-08 | 2.6e-08 | 4e-08 | 4.633E-08 | 2.413E-08 | 2.600E-08 | 7.300E-08 |
| 11665 | 584 | P584C | A | 3e-08 | 3.7e-08 | 9.4e-08 | 5.367E-08 | 3.510E-08 | 3.000E-08 | 9.400E-08 |
| 11666 | 584 | P584E | A | 3.5e-08 | 7.5e-08 | 1.9e-08 | 4.300E-08 | 2.884E-08 | 1.900E-08 | 7.500E-08 |
| 11667 | 584 | P584Q | A | 8e-08 | 5.5e-08 | 1e-07 | 7.833E-08 | 2.255E-08 | 5.500E-08 | 1.000E-07 |
| 11668 | 584 | P584G | A | 3e-08 | 2.3e-08 | 3.5e-08 | 2.933E-08 | 6.028E-09 | 2.300E-08 | 3.500E-08 |
| 11669 | 584 | P584H | A | 1.6e-07 | 3.6e-08 | 3.3e-08 | 7.633E-08 | 7.247E-08 | 3.300E-08 | 1.600E-07 |
| 11670 | 584 | P584I | A | 4.9e-08 | 3.3e-08 | 4.2e-08 | 4.133E-08 | 8.021E-09 | 3.300E-08 | 4.900E-08 |
| 11671 | 584 | P584L | A | 6.6e-08 | 1.2e-08 | 5.5e-08 | 4.433E-08 | 2.854E-08 | 1.200E-08 | 6.600E-08 |
| 11672 | 584 | P584K | A | 3.9e-08 | 3.7e-08 | 7.3e-08 | 4.967E-08 | 2.023E-08 | 3.700E-08 | 7.300E-08 |
| 11673 | 584 | P584M | A | 4.7e-08 | 9.7e-08 | 4.6e-08 | 6.333E-08 | 2.916E-08 | 4.600E-08 | 9.700E-08 |
| 11674 | 584 | P584F | A | 4.8e-08 | 4.4e-08 | 4.2e-08 | 4.467E-08 | 3.055E-09 | 4.200E-08 | 4.800E-08 |
| 11675 | 584 | P584P | A | 4.2e-08 | 4.3e-08 | 1.6e-07 | 8.167E-08 | 6.784E-08 | 4.200E-08 | 1.600E-07 |
| 11676 | 584 | P584S | A | 3e-08 | 1.7e-08 | 8.1e-08 | 4.267E-08 | 3.383E-08 | 1.700E-08 | 8.100E-08 |
| 11677 | 584 | P584T | A | 2.5e-08 | 1.5e-08 | 2e-08 | 2.000E-08 | 5.000E-09 | 1.500E-08 | 2.500E-08 |
| 11678 | 584 | P584W | A | 2.5e-08 | 3.2e-08 | 3.4e-08 | 3.033E-08 | 4.726E-09 | 2.500E-08 | 3.400E-08 |
| 11679 | 584 | P584Y | A | 2e-08 | 5.8e-08 | 4e-08 | 3.933E-08 | 1.901E-08 | 2.000E-08 | 5.800E-08 |
| 11680 | 584 | P584V | A | 2.4e-08 | 5.7e-08 | 4.9e-08 | 4.333E-08 | 1.721E-08 | 2.400E-08 | 5.700E-08 |
| 11681 | 585 | D585A | A | 4.9e-08 | 2.1e-08 | 3e-08 | 3.333E-08 | 1.429E-08 | 2.100E-08 | 4.900E-08 |
| 11682 | 585 | D585R | A | 5.2e-08 | 4.9e-08 | 2.2e-08 | 4.100E-08 | 1.652E-08 | 2.200E-08 | 5.200E-08 |
| 11683 | 585 | D585N | A | 3.1e-08 | 3.1e-08 | 3.5e-08 | 3.233E-08 | 2.309E-09 | 3.100E-08 | 3.500E-08 |
| 11684 | 585 | D585D | A | 2.1e-08 | 5.6e-08 | 1e-07 | 5.900E-08 | 3.959E-08 | 2.100E-08 | 1.000E-07 |
| 11685 | 585 | D585C | A | 9e-08 | 6.9e-08 | 7.3e-08 | 7.733E-08 | 1.115E-08 | 6.900E-08 | 9.000E-08 |
| 11686 | 585 | D585E | A | 5.5e-08 | 2.4e-08 | 3e-08 | 3.633E-08 | 1.644E-08 | 2.400E-08 | 5.500E-08 |
| 11687 | 585 | D585Q | A | 9.7e-08 | 4.5e-08 | 2e-08 | 5.400E-08 | 3.928E-08 | 2.000E-08 | 9.700E-08 |
| 11688 | 585 | D585G | A | 6.8e-08 | 5.4e-08 | 2.5e-08 | 4.900E-08 | 2.193E-08 | 2.500E-08 | 6.800E-08 |
| 11689 | 585 | D585H | A | 1.9e-07 | 2.8e-08 | 6e-08 | 9.267E-08 | 8.580E-08 | 2.800E-08 | 1.900E-07 |
| 11690 | 585 | D585I | A | 4.6e-08 | 6e-08 | 3.2e-08 | 4.600E-08 | 1.400E-08 | 3.200E-08 | 6.000E-08 |
| 11691 | 585 | D585L | A | 2.6e-08 | 7e-09 | 2.7e-08 | 2.000E-08 | 1.127E-08 | 7.000E-09 | 2.700E-08 |
| 11692 | 585 | D585K | A | 8.1e-08 | 2.7e-08 | 6.7e-08 | 5.833E-08 | 2.802E-08 | 2.700E-08 | 8.100E-08 |
| 11693 | 585 | D585M | A | 4.6e-08 | 3.2e-08 | 6.3e-08 | 4.700E-08 | 1.552E-08 | 3.200E-08 | 6.300E-08 |
| 11694 | 585 | D585F | A | 2.9e-08 | 3.6e-08 | 4.5e-08 | 3.667E-08 | 8.021E-09 | 2.900E-08 | 4.500E-08 |
| 11695 | 585 | D585P | A | 2.8e-08 | 3.2e-08 | 7.7e-08 | 4.567E-08 | 2.721E-08 | 2.800E-08 | 7.700E-08 |
| 11696 | 585 | D585S | A | 1.7e-08 | 2.6e-08 | 2.3e-08 | 2.200E-08 | 4.583E-09 | 1.700E-08 | 2.600E-08 |
| 11697 | 585 | D585T | A | 7.5e-08 | 3.9e-08 | 6.3e-08 | 5.900E-08 | 1.833E-08 | 3.900E-08 | 7.500E-08 |
| 11698 | 585 | D585W | A | 4.8e-08 | 1.2e-08 | 2.5e-08 | 2.833E-08 | 1.823E-08 | 1.200E-08 | 4.800E-08 |
| 11699 | 585 | D585Y | A | 3.7e-08 | 5.2e-08 | 2.3e-08 | 3.733E-08 | 1.450E-08 | 2.300E-08 | 5.200E-08 |
| 11700 | 585 | D585V | A | 1.3e-07 | 3e-08 | 8.1e-08 | 8.033E-08 | 5.000E-08 | 3.000E-08 | 1.300E-07 |
| 11701 | 586 | L586A | A | 3.3e-08 | 1.1e-08 | 1.6e-08 | 2.000E-08 | 1.153E-08 | 1.100E-08 | 3.300E-08 |
| 11702 | 586 | L586R | A | 4.4e-08 | 6.8e-08 | 4.4e-08 | 5.200E-08 | 1.386E-08 | 4.400E-08 | 6.800E-08 |
| 11703 | 586 | L586N | A | 6.5e-08 | 1.1e-07 | 3.7e-08 | 7.067E-08 | 3.683E-08 | 3.700E-08 | 1.100E-07 |
| 11704 | 586 | L586D | A | 1.8e-07 | 7.1e-08 | 2.8e-08 | 9.300E-08 | 7.835E-08 | 2.800E-08 | 1.800E-07 |
| 11705 | 586 | L586C | A | 2.2e-08 | 3.4e-08 | 5.2e-08 | 3.600E-08 | 1.510E-08 | 2.200E-08 | 5.200E-08 |
| 11706 | 586 | L586E | A | 5.7e-08 | 6.7e-08 | 7.5e-08 | 6.633E-08 | 9.018E-09 | 5.700E-08 | 7.500E-08 |
| 11707 | 586 | L586Q | A | 6.7e-08 | 4.2e-08 | 6.8e-08 | 5.900E-08 | 1.473E-08 | 4.200E-08 | 6.800E-08 |
| 11708 | 586 | L586G | A | 3.6e-08 | 1.1e-07 | 3.5e-08 | 6.033E-08 | 4.302E-08 | 3.500E-08 | 1.100E-07 |
| 11709 | 586 | L586H | A | 8.2e-08 | 1.8e-08 | 5.2e-08 | 5.067E-08 | 3.202E-08 | 1.800E-08 | 8.200E-08 |
| 11710 | 586 | L586I | A | 3.5e-08 | 3.5e-08 | 4.4e-08 | 3.800E-08 | 5.196E-09 | 3.500E-08 | 4.400E-08 |
| 11711 | 586 | L586L | A | 4.3e-08 | 3.3e-09 | 2.4e-08 | 2.343E-08 | 1.986E-08 | 3.300E-09 | 4.300E-08 |
| 11712 | 586 | L586K | A | 4e-08 | 3.2e-08 | 4e-08 | 3.733E-08 | 4.619E-09 | 3.200E-08 | 4.000E-08 |
| 11713 | 586 | L586M | A | 1.2e-07 | 5.3e-08 | 3.2e-08 | 6.833E-08 | 4.596E-08 | 3.200E-08 | 1.200E-07 |
| 11714 | 586 | L586F | A | 2.6e-08 | 5.6e-08 | 5.6e-08 | 4.600E-08 | 1.732E-08 | 2.600E-08 | 5.600E-08 |
| 11715 | 586 | L586P | A | 2.1e-08 | 5.2e-08 | 3.2e-08 | 3.500E-08 | 1.572E-08 | 2.100E-08 | 5.200E-08 |
| 11716 | 586 | L586S | A | 8.5e-08 | 7.7e-08 | 4.9e-08 | 7.033E-08 | 1.890E-08 | 4.900E-08 | 8.500E-08 |
| 11717 | 586 | L586T | A | 6.8e-08 | 5e-08 | 1.3e-07 | 8.267E-08 | 4.197E-08 | 5.000E-08 | 1.300E-07 |
| 11718 | 586 | L586W | A | 6.1e-08 | 2.6e-08 | 2.6e-08 | 3.767E-08 | 2.021E-08 | 2.600E-08 | 6.100E-08 |
| 11719 | 586 | L586Y | A | 7.1e-08 | 2.2e-08 | 2.1e-08 | 3.800E-08 | 2.858E-08 | 2.100E-08 | 7.100E-08 |
| 11720 | 586 | L586V | A | 8.6e-09 | 5.5e-08 | 1.8e-08 | 2.720E-08 | 2.453E-08 | 8.600E-09 | 5.500E-08 |
| 11721 | 587 | S587A | A | 3.6e-08 | 3.4e-08 | 5.4e-08 | 4.133E-08 | 1.102E-08 | 3.400E-08 | 5.400E-08 |
| 11722 | 587 | S587R | A | 6.2e-08 | 7.4e-09 | 8.5e-08 | 5.147E-08 | 3.986E-08 | 7.400E-09 | 8.500E-08 |
| 11723 | 587 | S587N | A | 6.9e-08 | 1.1e-07 | 5.4e-08 | 7.767E-08 | 2.899E-08 | 5.400E-08 | 1.100E-07 |
| 11724 | 587 | S587D | A | 3e-08 | 6.4e-08 | 4.3e-08 | 4.567E-08 | 1.716E-08 | 3.000E-08 | 6.400E-08 |
| 11725 | 587 | S587C | A | 8.5e-09 | 2.1e-08 | 4.5e-08 | 2.483E-08 | 1.855E-08 | 8.500E-09 | 4.500E-08 |
| 11726 | 587 | S587E | A | 4.2e-08 | 5.9e-08 | 1.3e-07 | 7.700E-08 | 4.668E-08 | 4.200E-08 | 1.300E-07 |
| 11727 | 587 | S587Q | A | 5.1e-08 | 5.9e-08 | 1.9e-08 | 4.300E-08 | 2.117E-08 | 1.900E-08 | 5.900E-08 |
| 11728 | 587 | S587G | A | 4.4e-08 | 2.8e-08 | 1.9e-08 | 3.033E-08 | 1.266E-08 | 1.900E-08 | 4.400E-08 |
| 11729 | 587 | S587H | A | 1.3e-08 | 5.3e-08 | 9.8e-08 | 5.467E-08 | 4.252E-08 | 1.300E-08 | 9.800E-08 |

| | | | | | | | | | | |
|---|---|---|---|---|---|---|---|---|---|---|
| 11730 | 587 | S587I | A | 7.9e-08 | 3.7e-08 | 4e-08 | 5.200E-08 | 2.343E-08 | 3.700E-08 | 7.900E-08 |
| 11731 | 587 | S587L | A | 6.9e-08 | 1.1e-07 | 7.3e-08 | 8.400E-08 | 2.261E-08 | 6.900E-08 | 1.100E-07 |
| 11732 | 587 | S587K | A | 8e-08 | 2.1e-08 | 1.1e-07 | 7.033E-08 | 4.528E-08 | 2.100E-08 | 1.100E-07 |
| 11733 | 587 | S587M | A | 7.5e-08 | 4.3e-08 | 2.3e-08 | 4.700E-08 | 2.623E-08 | 2.300E-08 | 7.500E-08 |
| 11734 | 587 | S587F | A | 1.1e-07 | 3.5e-08 | 1.2e-07 | 8.833E-08 | 4.646E-08 | 3.500E-08 | 1.200E-07 |
| 11735 | 587 | S587P | A | 2e-08 | 3.8e-08 | 3.2e-08 | 3.000E-08 | 9.165E-09 | 2.000E-08 | 3.800E-08 |
| 11736 | 587 | S587S | A | 1e-07 | 3.2e-08 | 6.4e-08 | 6.533E-08 | 3.402E-08 | 3.200E-08 | 1.000E-07 |
| 11737 | 587 | S587T | A | 4.4e-08 | 8.3e-08 | 3.9e-08 | 5.533E-08 | 2.409E-08 | 3.900E-08 | 8.300E-08 |
| 11738 | 587 | S587W | A | 2.4e-08 | 7.4e-08 | 1.3e-07 | 7.600E-08 | 5.303E-08 | 2.400E-08 | 1.300E-07 |
| 11739 | 587 | S587Y | A | 4.9e-08 | 2.9e-08 | 3.6e-08 | 3.800E-08 | 1.015E-08 | 2.900E-08 | 4.900E-08 |
| 11740 | 587 | S587V | A | 3.4e-08 | 1.2e-08 | 3.9e-08 | 2.833E-08 | 1.436E-08 | 1.200E-08 | 3.900E-08 |
| 11741 | 588 | Y588A | A | 1.5e-08 | 4.8e-08 | 1.1e-07 | 5.767E-08 | 4.823E-08 | 1.500E-08 | 1.100E-07 |
| 11742 | 588 | Y588R | A | 1.7e-08 | 4.6e-08 | 6e-08 | 4.100E-08 | 2.193E-08 | 1.700E-08 | 6.000E-08 |
| 11743 | 588 | Y588N | A | 1.1e-07 | 1.9e-07 | 2e-08 | 1.067E-07 | 8.505E-08 | 2.000E-08 | 1.900E-07 |
| 11744 | 588 | Y588D | A | 1.2e-07 | 6.9e-08 | 1.3e-07 | 1.063E-07 | 3.272E-08 | 6.900E-08 | 1.300E-07 |
| 11745 | 588 | Y588C | A | 4.1e-08 | 7.9e-08 | 3.5e-08 | 5.167E-08 | 2.386E-08 | 3.500E-08 | 7.900E-08 |
| 11746 | 588 | Y588E | A | 4.9e-08 | 5.4e-08 | 3.7e-08 | 4.667E-08 | 8.737E-09 | 3.700E-08 | 5.400E-08 |
| 11747 | 588 | Y588Q | A | 4.6e-08 | 4.7e-08 | 3.8e-08 | 4.367E-08 | 4.933E-09 | 3.800E-08 | 4.700E-08 |
| 11748 | 588 | Y588G | A | 3.2e-08 | 3.2e-08 | 4.1e-08 | 3.500E-08 | 5.196E-09 | 3.200E-08 | 4.100E-08 |
| 11749 | 588 | Y588H | A | 1.1e-07 | 2.5e-08 | 5e-08 | 6.167E-08 | 4.368E-08 | 2.500E-08 | 1.100E-07 |
| 11750 | 588 | Y588I | A | 1.1e-07 | 3.4e-08 | 8.8e-09 | 5.093E-08 | 5.268E-08 | 8.800E-09 | 1.100E-07 |
| 11751 | 588 | Y588L | A | 5.4e-08 | 1.1e-07 | 3.4e-08 | 6.600E-08 | 3.940E-08 | 3.400E-08 | 1.100E-07 |
| 11752 | 588 | Y588K | A | 8.2e-08 | 5.2e-08 | 5e-08 | 6.133E-08 | 1.793E-08 | 5.000E-08 | 8.200E-08 |
| 11753 | 588 | Y588M | A | 4e-08 | 4.1e-08 | 2.4e-08 | 3.500E-08 | 9.539E-09 | 2.400E-08 | 4.100E-08 |
| 11754 | 588 | Y588F | A | 1.5e-07 | 6.8e-08 | 3.1e-07 | 1.760E-07 | 1.231E-07 | 6.800E-08 | 3.100E-07 |
| 11755 | 588 | Y588P | A | 3.5e-08 | 6.3e-08 | 7.3e-08 | 5.700E-08 | 1.970E-08 | 3.500E-08 | 7.300E-08 |
| 11756 | 588 | Y588S | A | 9e-08 | 2.2e-08 | 4.7e-08 | 5.300E-08 | 3.439E-08 | 2.200E-08 | 9.000E-08 |
| 11757 | 588 | Y588T | A | 2.4e-08 | 5.9e-08 | 4.6e-08 | 4.300E-08 | 1.769E-08 | 2.400E-08 | 5.900E-08 |
| 11758 | 588 | Y588W | A | 1.6e-08 | 3.3e-08 | 8.7e-08 | 4.533E-08 | 3.707E-08 | 1.600E-08 | 8.700E-08 |
| 11759 | 588 | Y588Y | A | 3.6e-08 | 3.8e-08 | 3.5e-08 | 3.633E-08 | 1.528E-09 | 3.500E-08 | 3.800E-08 |
| 11760 | 588 | Y588V | A | 5e-08 | 6.2e-08 | 1.4e-08 | 4.200E-08 | 2.498E-08 | 1.400E-08 | 6.200E-08 |
| 11761 | 589 | M589A | A | 1e-07 | 1.2e-08 | 2e-08 | 4.400E-08 | 4.866E-08 | 1.200E-08 | 1.000E-07 |
| 11762 | 589 | M589R | A | 3.1e-08 | 1.3e-08 | 9e-09 | 1.767E-08 | 1.172E-08 | 9.000E-09 | 3.100E-08 |
| 11763 | 589 | M589N | A | 2.2e-08 | 4.8e-08 | 5.2e-08 | 4.067E-08 | 1.629E-08 | 2.200E-08 | 5.200E-08 |
| 11764 | 589 | M589D | A | 1.3e-08 | 5e-08 | 1.3e-08 | 2.533E-08 | 2.136E-08 | 1.300E-08 | 5.000E-08 |
| 11765 | 589 | M589C | A | 6.1e-08 | 1.5e-07 | 5.4e-08 | 8.833E-08 | 5.352E-08 | 5.400E-08 | 1.500E-07 |
| 11766 | 589 | M589E | A | 2.5e-07 | 2.1e-08 | 1.7e-08 | 9.600E-08 | 1.334E-07 | 1.700E-08 | 2.500E-07 |
| 11767 | 589 | M589Q | A | 3.9e-08 | 5e-08 | 5.8e-08 | 4.900E-08 | 9.539E-09 | 3.900E-08 | 5.800E-08 |
| 11768 | 589 | M589G | A | 5.4e-08 | 5.5e-08 | 2.7e-08 | 4.533E-08 | 1.589E-08 | 2.700E-08 | 5.500E-08 |
| 11769 | 589 | M589H | A | 7.9e-09 | 6.9e-08 | 4.8e-08 | 4.163E-08 | 3.104E-08 | 7.900E-09 | 6.900E-08 |
| 11770 | 589 | M589I | A | 9.7e-08 | 2.4e-08 | 2.4e-08 | 4.833E-08 | 4.215E-08 | 2.400E-08 | 9.700E-08 |
| 11771 | 589 | M589L | A | 4.3e-08 | 9.6e-08 | 1.2e-08 | 5.033E-08 | 4.248E-08 | 1.200E-08 | 9.600E-08 |
| 11772 | 589 | M589K | A | 8.3e-08 | 3.6e-08 | 7.5e-08 | 6.467E-08 | 2.515E-08 | 3.600E-08 | 8.300E-08 |
| 11773 | 589 | M589M | A | 4e-08 | 4.9e-08 | 6.8e-08 | 5.233E-08 | 1.429E-08 | 4.000E-08 | 6.800E-08 |
| 11774 | 589 | M589F | A | 7.5e-08 | 4.5e-08 | 1.2e-08 | 4.400E-08 | 3.151E-08 | 1.200E-08 | 7.500E-08 |
| 11775 | 589 | M589P | A | 6.1e-08 | 3.6e-08 | 4.7e-08 | 4.800E-08 | 1.253E-08 | 3.600E-08 | 6.100E-08 |
| 11776 | 589 | M589S | A | 2.9e-08 | 2.5e-08 | 2e-08 | 2.467E-08 | 4.509E-09 | 2.000E-08 | 2.900E-08 |
| 11777 | 589 | M589T | A | 4.6e-08 | 5.7e-08 | 9.3e-08 | 6.533E-08 | 2.458E-08 | 4.600E-08 | 9.300E-08 |
| 11778 | 589 | M589W | A | 4.5e-08 | 1e-07 | 2.8e-08 | 5.767E-08 | 3.763E-08 | 2.800E-08 | 1.000E-07 |
| 11779 | 589 | M589Y | A | 4.9e-08 | 2.1e-08 | 4.1e-08 | 3.700E-08 | 1.442E-08 | 2.100E-08 | 4.900E-08 |
| 11780 | 589 | M589V | A | 5.9e-08 | 3.4e-08 | 1.1e-07 | 6.767E-08 | 3.873E-08 | 3.400E-08 | 1.100E-07 |
| 11781 | 590 | P590A | A | 1e-07 | 4.7e-09 | 5.2e-08 | 5.223E-08 | 4.765E-08 | 4.700E-09 | 1.000E-07 |
| 11782 | 590 | P590R | A | 8.8e-08 | 4e-08 | 1e-07 | 7.600E-08 | 3.175E-08 | 4.000E-08 | 1.000E-07 |
| 11783 | 590 | P590N | A | 6.3e-08 | 3.3e-08 | 6.3e-08 | 5.300E-08 | 1.732E-08 | 3.300E-08 | 6.300E-08 |
| 11784 | 590 | P590D | A | 7.3e-08 | 4.7e-08 | 9.9e-08 | 7.300E-08 | 2.600E-08 | 4.700E-08 | 9.900E-08 |
| 11785 | 590 | P590C | A | 5.4e-08 | 5.1e-08 | 2.2e-08 | 4.233E-08 | 1.767E-08 | 2.200E-08 | 5.400E-08 |
| 11786 | 590 | P590E | A | 2.1e-08 | 4.1e-09 | 5.5e-08 | 2.670E-08 | 2.592E-08 | 4.100E-09 | 5.500E-08 |
| 11787 | 590 | P590Q | A | 3.8e-08 | 2.9e-08 | 3.9e-08 | 3.533E-08 | 5.508E-09 | 2.900E-08 | 3.900E-08 |
| 11788 | 590 | P590G | A | 2.4e-08 | 4.4e-08 | 1.7e-08 | 2.833E-08 | 1.401E-08 | 1.700E-08 | 4.400E-08 |
| 11789 | 590 | P590H | A | 3.1e-08 | 1e-07 | 3.1e-08 | 5.400E-08 | 3.984E-08 | 3.100E-08 | 1.000E-07 |
| 11790 | 590 | P590I | A | 3.5e-08 | 7.5e-08 | 3.1e-08 | 4.700E-08 | 2.433E-08 | 3.100E-08 | 7.500E-08 |
| 11791 | 590 | P590L | A | 1.2e-07 | 2.7e-08 | 2.5e-08 | 5.733E-08 | 5.428E-08 | 2.500E-08 | 1.200E-07 |
| 11792 | 590 | P590K | A | 5.1e-08 | 3.8e-08 | 5.9e-08 | 4.933E-08 | 1.060E-08 | 3.800E-08 | 5.900E-08 |
| 11793 | 590 | P590M | A | 3.8e-08 | 2.5e-08 | 6.2e-08 | 4.167E-08 | 1.877E-08 | 2.500E-08 | 6.200E-08 |
| 11794 | 590 | P590F | A | 1.5e-08 | 2e-08 | 3.2e-08 | 2.233E-08 | 8.737E-09 | 1.500E-08 | 3.200E-08 |
| 11795 | 590 | P590P | A | 2.5e-08 | 2e-08 | 8.8e-08 | 4.433E-08 | 3.790E-08 | 2.000E-08 | 8.800E-08 |
| 11796 | 590 | P590S | A | 5.9e-08 | 6.4e-08 | 1.9e-08 | 4.733E-08 | 2.466E-08 | 1.900E-08 | 6.400E-08 |
| 11797 | 590 | P590T | A | 5e-08 | 1.7e-08 | 1.3e-07 | 6.567E-08 | 5.811E-08 | 1.700E-08 | 1.300E-07 |
| 11798 | 590 | P590W | A | 8.1e-08 | 2.4e-08 | 1.9e-08 | 4.133E-08 | 3.444E-08 | 1.900E-08 | 8.100E-08 |
| 11799 | 590 | P590Y | A | 3.5e-08 | 7.4e-08 | 2.4e-08 | 4.433E-08 | 2.627E-08 | 2.400E-08 | 7.400E-08 |
| 11800 | 590 | P590V | A | 3.5e-08 | 1.9e-08 | 6.6e-08 | 4.000E-08 | 2.390E-08 | 1.900E-08 | 6.600E-08 |
| 11801 | 591 | I591A | A | 3.3e-08 | 2e-08 | 1.4e-07 | 6.433E-08 | 6.585E-08 | 2.000E-08 | 1.400E-07 |
| 11802 | 591 | I591R | A | 5.4e-08 | 1.9e-08 | 8.2e-08 | 5.167E-08 | 3.156E-08 | 1.900E-08 | 8.200E-08 |
| 11803 | 591 | I591N | A | 3.6e-08 | 2.9e-08 | 8.8e-08 | 5.100E-08 | 3.223E-08 | 2.900E-08 | 8.800E-08 |
| 11804 | 591 | I591D | A | 2.6e-08 | 3.7e-08 | 4.1e-08 | 3.467E-08 | 7.767E-09 | 2.600E-08 | 4.100E-08 |
| 11805 | 591 | I591C | A | 2.5e-08 | 2.9e-08 | 1.8e-08 | 2.400E-08 | 5.568E-09 | 1.800E-08 | 2.900E-08 |
| 11806 | 591 | I591E | A | 4.6e-08 | 5.4e-08 | 6.7e-08 | 5.567E-08 | 1.060E-08 | 4.600E-08 | 6.700E-08 |
| 11807 | 591 | I591Q | A | 1.7e-08 | 5.3e-08 | 6.7e-08 | 4.567E-08 | 2.579E-08 | 1.700E-08 | 6.700E-08 |
| 11808 | 591 | I591G | A | 5.4e-08 | 5.7e-08 | 4e-08 | 5.033E-08 | 9.074E-09 | 4.000E-08 | 5.700E-08 |
| 11809 | 591 | I591H | A | 1.7e-08 | 1.1e-07 | 1.3e-07 | 8.567E-08 | 6.030E-08 | 1.700E-08 | 1.300E-07 |
| 11810 | 591 | I591I | A | 5.8e-08 | 5.2e-08 | 4.3e-08 | 5.100E-08 | 7.550E-09 | 4.300E-08 | 5.800E-08 |
| 11811 | 591 | I591L | A | 1.5e-07 | 3e-08 | 2.5e-08 | 6.833E-08 | 7.077E-08 | 2.500E-08 | 1.500E-07 |
| 11812 | 591 | I591K | A | 7.5e-08 | 9.4e-09 | 2.9e-08 | 3.780E-08 | 3.367E-08 | 9.400E-09 | 7.500E-08 |
| 11813 | 591 | I591M | A | 5.2e-08 | 1.2e-07 | 3.5e-08 | 6.900E-08 | 4.498E-08 | 3.500E-08 | 1.200E-07 |
| 11814 | 591 | I591F | A | 5.6e-08 | 2.1e-08 | 3.9e-08 | 3.867E-08 | 1.750E-08 | 2.100E-08 | 5.600E-08 |

| | | | | | | | | | | |
|---|---|---|---|---|---|---|---|---|---|---|
| 11815 | 591 | I591P | A | 5.7e-08 | 2.2e-07 | 7.7e-08 | 1.180E-07 | 8.890E-08 | 5.700E-08 | 2.200E-07 |
| 11816 | 591 | I591S | A | 4.8e-08 | 5.9e-08 | 5.8e-08 | 5.500E-08 | 6.083E-09 | 4.800E-08 | 5.900E-08 |
| 11817 | 591 | I591T | A | 1e-07 | 1.9e-08 | 5.6e-08 | 5.833E-08 | 4.055E-08 | 1.900E-08 | 1.000E-07 |
| 11818 | 591 | I591W | A | 4e-08 | 7.1e-08 | 2.3e-08 | 4.467E-08 | 2.434E-08 | 2.300E-08 | 7.100E-08 |
| 11819 | 591 | I591Y | A | 4.1e-08 | 2.6e-08 | 3.7e-08 | 3.467E-08 | 7.767E-09 | 2.600E-08 | 4.100E-08 |
| 11820 | 591 | I591V | A | 8.5e-08 | 3.7e-08 | 4.5e-08 | 5.567E-08 | 2.572E-08 | 3.700E-08 | 8.500E-08 |
| 11821 | 592 | W592A | A | 1.5e-08 | 5.4e-08 | 1.5e-08 | 2.800E-08 | 2.252E-08 | 1.500E-08 | 5.400E-08 |
| 11822 | 592 | W592R | A | 1.2e-07 | 4.8e-08 | 5.1e-08 | 7.300E-08 | 4.073E-08 | 4.800E-08 | 1.200E-07 |
| 11823 | 592 | W592N | A | 1.2e-08 | 4.4e-08 | 6e-08 | 3.867E-08 | 2.444E-08 | 1.200E-08 | 6.000E-08 |
| 11824 | 592 | W592D | A | 9.1e-08 | 7.1e-08 | 2.9e-08 | 6.367E-08 | 3.164E-08 | 2.900E-08 | 9.100E-08 |
| 11825 | 592 | W592C | A | 3.6e-08 | 1.9e-08 | 1.2e-08 | 2.233E-08 | 1.234E-08 | 1.200E-08 | 3.600E-08 |
| 11826 | 592 | W592E | A | 2.6e-08 | 4.1e-08 | 3.5e-08 | 3.400E-08 | 7.550E-09 | 2.600E-08 | 4.100E-08 |
| 11827 | 592 | W592Q | A | 2.8e-08 | 3e-08 | 8.2e-08 | 4.667E-08 | 3.062E-08 | 2.800E-08 | 8.200E-08 |
| 11828 | 592 | W592G | A | 2.1e-08 | 1e-07 | 5.6e-08 | 5.900E-08 | 3.959E-08 | 2.100E-08 | 1.000E-07 |
| 11829 | 592 | W592H | A | 1.2e-08 | 3.1e-08 | 3.8e-08 | 2.700E-08 | 1.345E-08 | 1.200E-08 | 3.800E-08 |
| 11830 | 592 | W592I | A | 5.6e-08 | 1.1e-08 | 3.6e-08 | 3.433E-08 | 2.255E-08 | 1.100E-08 | 5.600E-08 |
| 11831 | 592 | W592L | A | 6.5e-08 | 3.1e-08 | 2.5e-08 | 4.033E-08 | 2.157E-08 | 2.500E-08 | 6.500E-08 |
| 11832 | 592 | W592K | A | 4.1e-08 | 8.1e-08 | 4e-08 | 5.400E-08 | 2.339E-08 | 4.000E-08 | 8.100E-08 |
| 11833 | 592 | W592M | A | 1.1e-07 | 4.8e-08 | 3.8e-08 | 6.533E-08 | 3.900E-08 | 3.800E-08 | 1.100E-07 |
| 11834 | 592 | W592F | A | 2.9e-08 | 4.2e-08 | 1.2e-07 | 6.367E-08 | 4.922E-08 | 2.900E-08 | 1.200E-07 |
| 11835 | 592 | W592P | A | 3e-08 | 5.3e-08 | 6.1e-08 | 4.800E-08 | 1.609E-08 | 3.000E-08 | 6.100E-08 |
| 11836 | 592 | W592S | A | 8.4e-08 | 4.5e-08 | 1.7e-08 | 4.867E-08 | 3.365E-08 | 1.700E-08 | 8.400E-08 |
| 11837 | 592 | W592T | A | 2.4e-08 | 6.7e-08 | 4.6e-08 | 4.567E-08 | 2.150E-08 | 2.400E-08 | 6.700E-08 |
| 11838 | 592 | W592W | A | 7.6e-08 | 3.2e-08 | 5.3e-08 | 5.367E-08 | 2.201E-08 | 3.200E-08 | 7.600E-08 |
| 11839 | 592 | W592Y | A | 1.4e-08 | 3.9e-08 | 6.1e-08 | 3.800E-08 | 2.352E-08 | 1.400E-08 | 6.100E-08 |
| 11840 | 592 | W592V | A | 6.2e-08 | 8.1e-08 | 8.3e-08 | 7.533E-08 | 1.159E-08 | 6.200E-08 | 8.300E-08 |
| 11841 | 593 | K593A | A | 5.2e-08 | 4.6e-08 | 2.5e-08 | 4.100E-08 | 1.418E-08 | 2.500E-08 | 5.200E-08 |
| 11842 | 593 | K593R | A | 3.8e-08 | 2.3e-08 | 1e-07 | 5.367E-08 | 4.082E-08 | 2.300E-08 | 1.000E-07 |
| 11843 | 593 | K593N | A | 1.9e-08 | 5.5e-08 | 1.5e-08 | 2.967E-08 | 2.203E-08 | 1.500E-08 | 5.500E-08 |
| 11844 | 593 | K593D | A | 3.1e-08 | 3.1e-08 | 3.7e-08 | 3.300E-08 | 3.464E-09 | 3.100E-08 | 3.700E-08 |
| 11845 | 593 | K593C | A | 5.2e-08 | 3.2e-08 | 4.7e-08 | 4.367E-08 | 1.041E-08 | 3.200E-08 | 5.200E-08 |
| 11846 | 593 | K593E | A | 4.4e-08 | 3.1e-08 | 5.6e-08 | 4.367E-08 | 1.250E-08 | 3.100E-08 | 5.600E-08 |
| 11847 | 593 | K593Q | A | 6.6e-08 | 4.4e-08 | 3.5e-08 | 4.833E-08 | 1.595E-08 | 3.500E-08 | 6.600E-08 |
| 11848 | 593 | K593G | A | 3.4e-08 | 2.3e-08 | 4.7e-08 | 3.467E-08 | 1.201E-08 | 2.300E-08 | 4.700E-08 |
| 11849 | 593 | K593H | A | 2.8e-08 | 2.7e-08 | 6.3e-08 | 3.933E-08 | 2.050E-08 | 2.700E-08 | 6.300E-08 |
| 11850 | 593 | K593I | A | 2.9e-08 | 3.1e-08 | 2.2e-08 | 2.733E-08 | 4.726E-09 | 2.200E-08 | 3.100E-08 |
| 11851 | 593 | K593L | A | 5.6e-08 | 3.8e-08 | 7.1e-08 | 5.500E-08 | 1.652E-08 | 3.800E-08 | 7.100E-08 |
| 11852 | 593 | K593K | A | 4e-08 | 5.5e-08 | 1.8e-08 | 3.767E-08 | 1.861E-08 | 1.800E-08 | 5.500E-08 |
| 11853 | 593 | K593M | A | 4.9e-08 | 1.5e-08 | 6.1e-08 | 4.167E-08 | 2.386E-08 | 1.500E-08 | 6.100E-08 |
| 11854 | 593 | K593F | A | 3.2e-08 | 7.5e-08 | 3.9e-08 | 4.867E-08 | 2.307E-08 | 3.200E-08 | 7.500E-08 |
| 11855 | 593 | K593P | A | 6e-08 | 5.1e-08 | 1.5e-08 | 4.200E-08 | 2.381E-08 | 1.500E-08 | 6.000E-08 |
| 11856 | 593 | K593S | A | 1.2e-07 | 2.4e-08 | 1.4e-07 | 9.467E-08 | 6.201E-08 | 2.400E-08 | 1.400E-07 |
| 11857 | 593 | K593T | A | 7.8e-08 | 2.8e-08 | 1.3e-07 | 7.867E-08 | 5.100E-08 | 2.800E-08 | 1.300E-07 |
| 11858 | 593 | K593W | A | 5.6e-08 | 8.3e-08 | 3.3e-08 | 5.733E-08 | 2.503E-08 | 3.300E-08 | 8.300E-08 |
| 11859 | 593 | K593Y | A | 5.2e-08 | 1.2e-08 | 1.6e-08 | 2.667E-08 | 2.203E-08 | 1.200E-08 | 5.200E-08 |
| 11860 | 593 | K593V | A | 3.7e-08 | 9.6e-08 | 3.3e-08 | 5.533E-08 | 3.528E-08 | 3.300E-08 | 9.600E-08 |
| 11861 | 594 | F594A | A | 4.1e-08 | 4.5e-08 | 2.2e-08 | 3.600E-08 | 1.229E-08 | 2.200E-08 | 4.500E-08 |
| 11862 | 594 | F594R | A | 5.5e-08 | 5.2e-08 | 9.9e-08 | 6.867E-08 | 2.631E-08 | 5.200E-08 | 9.900E-08 |
| 11863 | 594 | F594N | A | 3.2e-08 | 7e-08 | 5.1e-08 | 5.100E-08 | 1.900E-08 | 3.200E-08 | 7.000E-08 |
| 11864 | 594 | F594D | A | 5.9e-08 | 1.1e-07 | 4.3e-08 | 7.067E-08 | 3.499E-08 | 4.300E-08 | 1.100E-07 |
| 11865 | 594 | F594C | A | 5.5e-08 | 7.3e-08 | 5.6e-08 | 6.133E-08 | 1.012E-08 | 5.500E-08 | 7.300E-08 |
| 11866 | 594 | F594E | A | 2.3e-08 | 9.3e-08 | 5.7e-08 | 5.767E-08 | 3.500E-08 | 2.300E-08 | 9.300E-08 |
| 11867 | 594 | F594Q | A | 7.9e-08 | 3.5e-08 | 1.5e-08 | 4.300E-08 | 3.274E-08 | 1.500E-08 | 7.900E-08 |
| 11868 | 594 | F594G | A | 8.1e-08 | 4.5e-08 | 2.7e-08 | 5.100E-08 | 2.750E-08 | 2.700E-08 | 8.100E-08 |
| 11869 | 594 | F594H | A | 2.9e-08 | 6.6e-08 | 2.7e-08 | 4.067E-08 | 2.196E-08 | 2.700E-08 | 6.600E-08 |
| 11870 | 594 | F594I | A | 3e-08 | 2.5e-08 | 5.4e-08 | 3.633E-08 | 1.550E-08 | 2.500E-08 | 5.400E-08 |
| 11871 | 594 | F594L | A | 4.4e-08 | 3.8e-08 | 2.5e-08 | 3.567E-08 | 9.713E-09 | 2.500E-08 | 4.400E-08 |
| 11872 | 594 | F594K | A | 1.7e-08 | 3.4e-08 | 4.5e-08 | 3.200E-08 | 1.411E-08 | 1.700E-08 | 4.500E-08 |
| 11873 | 594 | F594M | A | 7.7e-08 | 1.3e-07 | 8.3e-08 | 9.667E-08 | 2.902E-08 | 7.700E-08 | 1.300E-07 |
| 11874 | 594 | F594F | A | 2.2e-08 | 1.5e-08 | 4.5e-08 | 2.733E-08 | 1.570E-08 | 1.500E-08 | 4.500E-08 |
| 11875 | 594 | F594P | A | 8.2e-08 | 1.8e-08 | 1.8e-07 | 9.333E-08 | 8.159E-08 | 1.800E-08 | 1.800E-07 |
| 11876 | 594 | F594S | A | 3.2e-08 | 3.9e-08 | 1.7e-08 | 2.933E-08 | 1.124E-08 | 1.700E-08 | 3.900E-08 |
| 11877 | 594 | F594T | A | 7.5e-08 | 5.1e-08 | 2.4e-08 | 5.000E-08 | 2.551E-08 | 2.400E-08 | 7.500E-08 |
| 11878 | 594 | F594W | A | 1.1e-07 | 9e-08 | 6.4e-08 | 8.800E-08 | 2.307E-08 | 6.400E-08 | 1.100E-07 |
| 11879 | 594 | F594Y | A | 8.2e-08 | 2.8e-08 | 1.8e-08 | 4.267E-08 | 3.443E-08 | 1.800E-08 | 8.200E-08 |
| 11880 | 594 | F594V | A | 8.2e-08 | 1.9e-08 | 1e-07 | 6.700E-08 | 4.253E-08 | 1.900E-08 | 1.000E-07 |
| 11881 | 595 | P595A | A | 1.8e-08 | 4.9e-08 | 1.9e-08 | 2.867E-08 | 1.762E-08 | 1.800E-08 | 4.900E-08 |
| 11882 | 595 | P595R | A | 3.3e-08 | 2.2e-08 | 3.1e-08 | 2.867E-08 | 5.859E-09 | 2.200E-08 | 3.300E-08 |
| 11883 | 595 | P595N | A | 9e-08 | 3.7e-08 | 2.2e-08 | 4.967E-08 | 3.573E-08 | 2.200E-08 | 9.000E-08 |
| 11884 | 595 | P595D | A | 7.7e-08 | 1e-07 | 4.4e-08 | 7.367E-08 | 2.815E-08 | 4.400E-08 | 1.000E-07 |
| 11885 | 595 | P595C | A | 3.3e-08 | 2.8e-08 | 3.7e-08 | 3.267E-08 | 4.509E-09 | 2.800E-08 | 3.700E-08 |
| 11886 | 595 | P595E | A | 1.4e-07 | 2.8e-08 | 5e-08 | 7.267E-08 | 5.934E-08 | 2.800E-08 | 1.400E-07 |
| 11887 | 595 | P595Q | A | 1.7e-08 | 1.3e-07 | 4.8e-08 | 6.500E-08 | 5.839E-08 | 1.700E-08 | 1.300E-07 |
| 11888 | 595 | P595G | A | 2e-08 | 3.5e-08 | 1.7e-08 | 2.400E-08 | 9.644E-09 | 1.700E-08 | 3.500E-08 |
| 11889 | 595 | P595H | A | 2.3e-08 | 6.5e-08 | 3.1e-08 | 3.967E-08 | 2.230E-08 | 2.300E-08 | 6.500E-08 |
| 11890 | 595 | P595I | A | 1.9e-08 | 1.1e-08 | 5.9e-08 | 2.967E-08 | 2.572E-08 | 1.100E-08 | 5.900E-08 |
| 11891 | 595 | P595L | A | 5.1e-08 | 4e-08 | 4.3e-08 | 4.467E-08 | 5.686E-09 | 4.000E-08 | 5.100E-08 |
| 11892 | 595 | P595K | A | 1e-08 | 2.2e-08 | 4.1e-08 | 2.433E-08 | 1.563E-08 | 1.000E-08 | 4.100E-08 |
| 11893 | 595 | P595M | A | 8e-08 | 1.6e-08 | 2e-08 | 3.867E-08 | 3.585E-08 | 1.600E-08 | 8.000E-08 |
| 11894 | 595 | P595F | A | 5.1e-08 | 8.8e-08 | 2.6e-08 | 5.500E-08 | 3.119E-08 | 2.600E-08 | 8.800E-08 |
| 11895 | 595 | P595P | A | 2.9e-08 | 3.5e-08 | 4e-08 | 3.467E-08 | 5.508E-09 | 2.900E-08 | 4.000E-08 |
| 11896 | 595 | P595S | A | 2.7e-08 | 5.4e-08 | 2.3e-08 | 3.467E-08 | 1.686E-08 | 2.300E-08 | 5.400E-08 |
| 11897 | 595 | P595T | A | 1.3e-08 | 7e-08 | 3.4e-08 | 3.900E-08 | 2.883E-08 | 1.300E-08 | 7.000E-08 |
| 11898 | 595 | P595W | A | 6.3e-08 | 1.4e-07 | 6.6e-08 | 8.967E-08 | 4.362E-08 | 6.300E-08 | 1.400E-07 |
| 11899 | 595 | P595Y | A | 7.6e-08 | 1.1e-07 | 2.1e-08 | 6.900E-08 | 4.491E-08 | 2.100E-08 | 1.100E-07 |

| | | | | | | | | | | |
|---|---|---|---|---|---|---|---|---|---|---|
| 11900 | 595 | P595V | A | 2.2e-08 | 2.6e-08 | 2e-08 | 2.267E-08 | 3.055E-09 | 2.000E-08 | 2.600E-08 |
| 11901 | 596 | D596A | A | 1.4e-07 | 6.4e-08 | 2.2e-08 | 7.533E-08 | 5.981E-08 | 2.200E-08 | 1.400E-07 |
| 11902 | 596 | D596R | A | 1.8e-07 | 3.5e-08 | 7.2e-08 | 9.567E-08 | 7.534E-08 | 3.500E-08 | 1.800E-07 |
| 11903 | 596 | D596N | A | 8e-09 | 3.5e-08 | 7.7e-08 | 4.000E-08 | 3.477E-08 | 8.000E-09 | 7.700E-08 |
| 11904 | 596 | D596D | A | 3.4e-08 | 2.6e-08 | 4.1e-08 | 3.367E-08 | 7.506E-09 | 2.600E-08 | 4.100E-08 |
| 11905 | 596 | D596C | A | 8e-08 | 4.2e-08 | 7.5e-08 | 6.567E-08 | 2.065E-08 | 4.200E-08 | 8.000E-08 |
| 11906 | 596 | D596E | A | 3.3e-08 | 2.4e-08 | 6.5e-08 | 4.067E-08 | 2.155E-08 | 2.400E-08 | 6.500E-08 |
| 11907 | 596 | D596Q | A | 8e-08 | 4.4e-08 | 8.6e-08 | 7.000E-08 | 2.272E-08 | 4.400E-08 | 8.600E-08 |
| 11908 | 596 | D596G | A | 2e-08 | 2e-08 | 4.2e-08 | 2.733E-08 | 1.270E-08 | 2.000E-08 | 4.200E-08 |
| 11909 | 596 | D596H | A | 5e-08 | 3.7e-08 | 3.6e-08 | 4.100E-08 | 7.810E-09 | 3.600E-08 | 5.000E-08 |
| 11910 | 596 | D596I | A | 4.7e-08 | 3.5e-08 | 8.9e-09 | 3.030E-08 | 1.948E-08 | 8.900E-09 | 4.700E-08 |
| 11911 | 596 | D596L | A | 7.5e-08 | 4.3e-08 | 1.7e-08 | 4.500E-08 | 2.905E-08 | 1.700E-08 | 7.500E-08 |
| 11912 | 596 | D596K | A | 4.9e-08 | 4.3e-08 | 4.6e-08 | 4.600E-08 | 3.000E-09 | 4.300E-08 | 4.900E-08 |
| 11913 | 596 | D596M | A | 5.5e-08 | 7e-08 | 6.2e-08 | 6.233E-08 | 7.506E-09 | 5.500E-08 | 7.000E-08 |
| 11914 | 596 | D596F | A | 2.5e-08 | 3.8e-08 | 4.1e-08 | 3.467E-08 | 8.505E-09 | 2.500E-08 | 4.100E-08 |
| 11915 | 596 | D596P | A | 3.6e-08 | 3e-08 | 7.3e-08 | 4.633E-08 | 2.329E-08 | 3.000E-08 | 7.300E-08 |
| 11916 | 596 | D596S | A | 3.7e-08 | 3.1e-08 | 8.6e-09 | 2.553E-08 | 1.497E-08 | 8.600E-09 | 3.700E-08 |
| 11917 | 596 | D596T | A | 3.4e-08 | 7.6e-08 | 2e-08 | 4.333E-08 | 2.914E-08 | 2.000E-08 | 7.600E-08 |
| 11918 | 596 | D596W | A | 6.3e-08 | 1e-08 | 5e-08 | 4.100E-08 | 2.762E-08 | 1.000E-08 | 6.300E-08 |
| 11919 | 596 | D596Y | A | 3.1e-08 | 3.2e-08 | 1.9e-07 | 8.433E-08 | 9.151E-08 | 3.100E-08 | 1.900E-07 |
| 11920 | 596 | D596V | A | 4.6e-08 | 3.4e-08 | 7.1e-08 | 5.033E-08 | 1.888E-08 | 3.400E-08 | 7.100E-08 |
| 11921 | 597 | E597A | A | 3.2e-08 | 4.7e-08 | 1.6e-08 | 3.167E-08 | 1.550E-08 | 1.600E-08 | 4.700E-08 |
| 11922 | 597 | E597R | A | 8.7e-08 | 6.5e-08 | 3e-08 | 6.067E-08 | 2.875E-08 | 3.000E-08 | 8.700E-08 |
| 11923 | 597 | E597N | A | 1.6e-08 | 2.9e-08 | 3.3e-08 | 2.600E-08 | 8.888E-09 | 1.600E-08 | 3.300E-08 |
| 11924 | 597 | E597D | A | 3e-08 | 5e-08 | 1.5e-08 | 3.167E-08 | 1.756E-08 | 1.500E-08 | 5.000E-08 |
| 11925 | 597 | E597C | A | 2.4e-08 | 3.1e-08 | 9.1e-08 | 4.867E-08 | 3.683E-08 | 2.400E-08 | 9.100E-08 |
| 11926 | 597 | E597E | A | 4.6e-08 | 2.4e-08 | 5.3e-08 | 4.100E-08 | 1.513E-08 | 2.400E-08 | 5.300E-08 |
| 11927 | 597 | E597Q | A | 4.9e-08 | 5.9e-08 | 6.8e-08 | 5.867E-08 | 9.504E-09 | 4.900E-08 | 6.800E-08 |
| 11928 | 597 | E597G | A | 5.7e-08 | 2.3e-08 | 1.3e-07 | 7.000E-08 | 5.467E-08 | 2.300E-08 | 1.300E-07 |
| 11929 | 597 | E597H | A | 2.7e-08 | 8.9e-08 | 3.8e-08 | 5.133E-08 | 3.308E-08 | 2.700E-08 | 8.900E-08 |
| 11930 | 597 | E597I | A | 1.2e-07 | 1.6e-08 | 2.7e-08 | 5.433E-08 | 5.713E-08 | 1.600E-08 | 1.200E-07 |
| 11931 | 597 | E597L | A | 3.5e-08 | 3.3e-08 | 3.2e-08 | 3.333E-08 | 1.528E-09 | 3.200E-08 | 3.500E-08 |
| 11932 | 597 | E597K | A | 5.1e-08 | 2.6e-08 | 2.9e-08 | 3.533E-08 | 1.365E-08 | 2.600E-08 | 5.100E-08 |
| 11933 | 597 | E597M | A | 1.6e-08 | 4.1e-08 | 2.6e-08 | 2.767E-08 | 1.258E-08 | 1.600E-08 | 4.100E-08 |
| 11934 | 597 | E597F | A | 4.9e-08 | 4.8e-08 | 1.8e-08 | 3.833E-08 | 1.762E-08 | 1.800E-08 | 4.900E-08 |
| 11935 | 597 | E597P | A | 3.7e-08 | 6e-08 | 3.7e-08 | 4.467E-08 | 1.328E-08 | 3.700E-08 | 6.000E-08 |
| 11936 | 597 | E597S | A | 4.2e-08 | 8.4e-08 | 5.1e-08 | 5.900E-08 | 2.211E-08 | 4.200E-08 | 8.400E-08 |
| 11937 | 597 | E597T | A | 4.5e-08 | 4e-08 | 1.8e-08 | 3.433E-08 | 1.436E-08 | 1.800E-08 | 4.500E-08 |
| 11938 | 597 | E597W | A | 5.6e-08 | 1.2e-07 | 5.2e-08 | 7.600E-08 | 3.816E-08 | 5.200E-08 | 1.200E-07 |
| 11939 | 597 | E597Y | A | 3.2e-08 | 9.8e-09 | 2e-08 | 2.060E-08 | 1.111E-08 | 9.800E-09 | 3.200E-08 |
| 11940 | 597 | E597V | A | 8.4e-08 | 1.2e-07 | 3.5e-08 | 7.967E-08 | 4.267E-08 | 3.500E-08 | 1.200E-07 |
| 11941 | 598 | E598A | A | 3.1e-08 | 4.7e-08 | 2.2e-08 | 3.333E-08 | 1.266E-08 | 2.200E-08 | 4.700E-08 |
| 11942 | 598 | E598R | A | 1.7e-07 | 3.2e-08 | 1e-07 | 1.007E-07 | 6.900E-08 | 3.200E-08 | 1.700E-07 |
| 11943 | 598 | E598N | A | 5.1e-08 | 4.6e-08 | 1.4e-08 | 3.700E-08 | 2.007E-08 | 1.400E-08 | 5.100E-08 |
| 11944 | 598 | E598D | A | 1.1e-08 | 3.6e-08 | 2.8e-08 | 2.500E-08 | 1.277E-08 | 1.100E-08 | 3.600E-08 |
| 11945 | 598 | E598C | A | 4.7e-08 | 2.4e-08 | 6.3e-08 | 4.467E-08 | 1.960E-08 | 2.400E-08 | 6.300E-08 |
| 11946 | 598 | E598E | A | 7.6e-08 | 6.3e-08 | 1.1e-08 | 5.000E-08 | 3.439E-08 | 1.100E-08 | 7.600E-08 |
| 11947 | 598 | E598Q | A | 7e-08 | 6.7e-08 | 1.2e-07 | 8.567E-08 | 2.977E-08 | 6.700E-08 | 1.200E-07 |
| 11948 | 598 | E598G | A | 4e-08 | 1.7e-08 | 7.7e-08 | 4.467E-08 | 3.027E-08 | 1.700E-08 | 7.700E-08 |
| 11949 | 598 | E598H | A | 4.9e-08 | 1.3e-07 | 6.1e-08 | 8.000E-08 | 4.371E-08 | 4.900E-08 | 1.300E-07 |
| 11950 | 598 | E598I | A | 4.5e-08 | 5.2e-08 | 3.1e-08 | 4.267E-08 | 1.069E-08 | 3.100E-08 | 5.200E-08 |
| 11951 | 598 | E598L | A | 5.9e-08 | 2.5e-08 | 2.1e-08 | 3.500E-08 | 2.088E-08 | 2.100E-08 | 5.900E-08 |
| 11952 | 598 | E598K | A | 7.3e-08 | 2.2e-08 | 5.2e-08 | 4.900E-08 | 2.563E-08 | 2.200E-08 | 7.300E-08 |
| 11953 | 598 | E598M | A | 2.6e-08 | 1.2e-08 | 6.4e-08 | 3.400E-08 | 2.691E-08 | 1.200E-08 | 6.400E-08 |
| 11954 | 598 | E598F | A | 3.9e-08 | 3.5e-08 | 4.3e-08 | 3.900E-08 | 4.000E-09 | 3.500E-08 | 4.300E-08 |
| 11955 | 598 | E598P | A | 2.7e-08 | 6.8e-08 | 6.7e-08 | 5.400E-08 | 2.339E-08 | 2.700E-08 | 6.800E-08 |
| 11956 | 598 | E598S | A | 2.2e-08 | 6.6e-08 | 2.8e-08 | 3.867E-08 | 2.386E-08 | 2.200E-08 | 6.600E-08 |
| 11957 | 598 | E598T | A | 3.3e-08 | 4.9e-08 | 4.3e-08 | 4.167E-08 | 8.083E-09 | 3.300E-08 | 4.900E-08 |
| 11958 | 598 | E598W | A | 7.3e-08 | 8.5e-08 | 1.4e-07 | 9.933E-08 | 3.573E-08 | 7.300E-08 | 1.400E-07 |
| 11959 | 598 | E598Y | A | 6e-08 | 5e-08 | 4.1e-08 | 5.033E-08 | 9.504E-09 | 4.100E-08 | 6.000E-08 |
| 11960 | 598 | E598V | A | 1.7e-07 | 3.9e-08 | 1.8e-08 | 7.567E-08 | 8.237E-08 | 1.800E-08 | 1.700E-07 |
| 11961 | 599 | G599A | A | 5.2e-08 | 4.6e-08 | 2.9e-08 | 4.233E-08 | 1.193E-08 | 2.900E-08 | 5.200E-08 |
| 11962 | 599 | G599R | A | 2.5e-08 | 3.6e-08 | 6.4e-08 | 4.167E-08 | 2.011E-08 | 2.500E-08 | 6.400E-08 |
| 11963 | 599 | G599N | A | 2.3e-08 | 4.7e-08 | 2.2e-08 | 3.067E-08 | 1.415E-08 | 2.200E-08 | 4.700E-08 |
| 11964 | 599 | G599D | A | 4.7e-08 | 1.8e-07 | 3.5e-08 | 8.733E-08 | 8.048E-08 | 3.500E-08 | 1.800E-07 |
| 11965 | 599 | G599C | A | 5e-08 | 3.4e-08 | 1.9e-08 | 3.433E-08 | 1.550E-08 | 1.900E-08 | 5.000E-08 |
| 11966 | 599 | G599E | A | 3.7e-08 | 1.2e-07 | 6.2e-08 | 7.300E-08 | 4.258E-08 | 3.700E-08 | 1.200E-07 |
| 11967 | 599 | G599Q | A | 2e-08 | 5.1e-08 | 6.8e-08 | 4.633E-08 | 2.434E-08 | 2.000E-08 | 6.800E-08 |
| 11968 | 599 | G599G | A | 6.4e-08 | 6.9e-08 | 5.7e-08 | 6.333E-08 | 6.028E-09 | 5.700E-08 | 6.900E-08 |
| 11969 | 599 | G599H | A | 3.9e-08 | 7.3e-08 | 3.1e-08 | 4.767E-08 | 2.230E-08 | 3.100E-08 | 7.300E-08 |
| 11970 | 599 | G599I | A | 5.3e-08 | 1.1e-07 | 3.9e-08 | 6.733E-08 | 3.761E-08 | 3.900E-08 | 1.100E-07 |
| 11971 | 599 | G599L | A | 4.1e-08 | 6.5e-08 | 5.1e-08 | 5.233E-08 | 1.206E-08 | 4.100E-08 | 6.500E-08 |
| 11972 | 599 | G599K | A | 5.7e-08 | 7e-08 | 1.1e-08 | 4.600E-08 | 3.100E-08 | 1.100E-08 | 7.000E-08 |
| 11973 | 599 | G599M | A | 1e-07 | 1.9e-08 | 2.9e-08 | 4.933E-08 | 4.416E-08 | 1.900E-08 | 1.000E-07 |
| 11974 | 599 | G599F | A | 4.6e-08 | 3.8e-08 | 3.2e-08 | 3.867E-08 | 7.024E-09 | 3.200E-08 | 4.600E-08 |
| 11975 | 599 | G599P | A | 7.1e-08 | 6.9e-08 | 6.9e-08 | 6.967E-08 | 1.155E-09 | 6.900E-08 | 7.100E-08 |
| 11976 | 599 | G599S | A | 3.3e-08 | 2.9e-08 | 2e-08 | 2.733E-08 | 6.658E-09 | 2.000E-08 | 3.300E-08 |
| 11977 | 599 | G599T | A | 3.7e-08 | 3.8e-08 | 6.4e-08 | 4.633E-08 | 1.531E-08 | 3.700E-08 | 6.400E-08 |
| 11978 | 599 | G599W | A | 5.2e-08 | 2.7e-08 | 9.4e-08 | 5.767E-08 | 3.386E-08 | 2.700E-08 | 9.400E-08 |
| 11979 | 599 | G599Y | A | 7.6e-08 | 5.2e-08 | 3.5e-08 | 5.433E-08 | 2.060E-08 | 3.500E-08 | 7.600E-08 |
| 11980 | 599 | G599V | A | 3.2e-08 | 7e-08 | 6e-08 | 5.400E-08 | 1.970E-08 | 3.200E-08 | 7.000E-08 |
| 11981 | 600 | A600A | A | 3.8e-08 | 1.3e-07 | 3.2e-08 | 6.667E-08 | 5.493E-08 | 3.200E-08 | 1.300E-07 |
| 11982 | 600 | A600R | A | 5.7e-08 | 5.9e-08 | 3.4e-08 | 5.000E-08 | 1.389E-08 | 3.400E-08 | 5.900E-08 |
| 11983 | 600 | A600N | A | 1.9e-08 | 8.8e-08 | 1.5e-07 | 8.567E-08 | 6.553E-08 | 1.900E-08 | 1.500E-07 |
| 11984 | 600 | A600D | A | 9.4e-08 | 1.9e-08 | 3.5e-08 | 4.933E-08 | 3.950E-08 | 1.900E-08 | 9.400E-08 |

| | | | | | | | | | | |
|---|---|---|---|---|---|---|---|---|---|---|
| 11985 | 600 | A600C | A | 3.1e-08 | 8.1e-08 | 4.2e-08 | 5.133E-08 | 2.627E-08 | 3.100E-08 | 8.100E-08 |
| 11986 | 600 | A600E | A | 3e-08 | 1.8e-07 | 3.2e-08 | 8.067E-08 | 8.603E-08 | 3.000E-08 | 1.800E-07 |
| 11987 | 600 | A600Q | A | 6.3e-08 | 2.1e-08 | 4.2e-08 | 4.200E-08 | 2.100E-08 | 2.100E-08 | 6.300E-08 |
| 11988 | 600 | A600G | A | 1.3e-07 | 6.2e-08 | 5.7e-08 | 8.300E-08 | 4.078E-08 | 5.700E-08 | 1.300E-07 |
| 11989 | 600 | A600H | A | 3.3e-08 | 3.7e-08 | 1.9e-08 | 2.967E-08 | 9.452E-09 | 1.900E-08 | 3.700E-08 |
| 11990 | 600 | A600I | A | 1.6e-08 | 5.8e-08 | 1e-07 | 5.800E-08 | 4.200E-08 | 1.600E-08 | 1.000E-07 |
| 11991 | 600 | A600L | A | 2.3e-08 | 1.2e-07 | 2.3e-08 | 5.533E-08 | 5.600E-08 | 2.300E-08 | 1.200E-07 |
| 11992 | 600 | A600K | A | 6.9e-08 | 1.8e-08 | 2.2e-08 | 3.633E-08 | 2.836E-08 | 1.800E-08 | 6.900E-08 |
| 11993 | 600 | A600M | A | 7.3e-08 | 4.5e-08 | 5.5e-08 | 5.767E-08 | 1.419E-08 | 4.500E-08 | 7.300E-08 |
| 11994 | 600 | A600F | A | 4e-08 | 5.9e-08 | 2.2e-08 | 4.033E-08 | 1.850E-08 | 2.200E-08 | 5.900E-08 |
| 11995 | 600 | A600P | A | 2e-08 | 5e-08 | 1.9e-08 | 2.967E-08 | 1.762E-08 | 1.900E-08 | 5.000E-08 |
| 11996 | 600 | A600S | A | 3.3e-08 | 2.5e-08 | 3e-08 | 2.933E-08 | 4.041E-09 | 2.500E-08 | 3.300E-08 |
| 11997 | 600 | A600T | A | 4.6e-08 | 4.2e-08 | 1.3e-07 | 7.267E-08 | 4.969E-08 | 4.200E-08 | 1.300E-07 |
| 11998 | 600 | A600W | A | 2.3e-08 | 2.9e-08 | 7e-08 | 4.067E-08 | 2.558E-08 | 2.300E-08 | 7.000E-08 |
| 11999 | 600 | A600Y | A | 1.8e-08 | 9e-08 | 3.7e-08 | 4.833E-08 | 3.731E-08 | 1.800E-08 | 9.000E-08 |
| 12000 | 600 | A600V | A | 2.1e-08 | 4.4e-08 | 4.6e-08 | 3.700E-08 | 1.389E-08 | 2.100E-08 | 4.600E-08 |
| 12001 | 601 | C601A | A | 2.5e-08 | 7.2e-08 | 1.8e-08 | 3.833E-08 | 2.937E-08 | 1.800E-08 | 7.200E-08 |
| 12002 | 601 | C601R | A | 5.4e-08 | 1.5e-08 | 1.6e-08 | 2.833E-08 | 2.223E-08 | 1.500E-08 | 5.400E-08 |
| 12003 | 601 | C601N | A | 4.1e-08 | 2e-08 | 1.7e-07 | 7.700E-08 | 8.122E-08 | 2.000E-08 | 1.700E-07 |
| 12004 | 601 | C601D | A | 3.5e-08 | 1.2e-08 | 4e-08 | 2.900E-08 | 1.493E-08 | 1.200E-08 | 4.000E-08 |
| 12005 | 601 | C601C | A | 3.8e-08 | 4.6e-08 | 4.9e-08 | 4.433E-08 | 5.686E-09 | 3.800E-08 | 4.900E-08 |
| 12006 | 601 | C601E | A | 2.1e-08 | 3.6e-08 | 4e-08 | 3.233E-08 | 1.002E-08 | 2.100E-08 | 4.000E-08 |
| 12007 | 601 | C601Q | A | 9.7e-08 | 5.5e-08 | 3.5e-08 | 6.233E-08 | 3.164E-08 | 3.500E-08 | 9.700E-08 |
| 12008 | 601 | C601G | A | 3.4e-08 | 2.5e-08 | 8.7e-08 | 4.867E-08 | 3.350E-08 | 2.500E-08 | 8.700E-08 |
| 12009 | 601 | C601H | A | 3.3e-08 | 2.1e-08 | 3.5e-08 | 2.967E-08 | 7.572E-09 | 2.100E-08 | 3.500E-08 |
| 12010 | 601 | C601I | A | 5.8e-08 | 8.3e-08 | 2.6e-08 | 5.567E-08 | 2.857E-08 | 2.600E-08 | 8.300E-08 |
| 12011 | 601 | C601L | A | 5e-08 | 1.8e-08 | 3.7e-08 | 3.500E-08 | 1.609E-08 | 1.800E-08 | 5.000E-08 |
| 12012 | 601 | C601K | A | 8.1e-08 | 1.2e-07 | 3.8e-08 | 7.967E-08 | 4.102E-08 | 3.800E-08 | 1.200E-07 |
| 12013 | 601 | C601M | A | 5.1e-08 | 4.4e-08 | 6.5e-08 | 5.333E-08 | 1.069E-08 | 4.400E-08 | 6.500E-08 |
| 12014 | 601 | C601F | A | 2.1e-08 | 5.9e-08 | 6.2e-08 | 4.733E-08 | 2.285E-08 | 2.100E-08 | 6.200E-08 |
| 12015 | 601 | C601P | A | 8.3e-08 | 4.5e-08 | 3.9e-08 | 5.567E-08 | 2.386E-08 | 3.900E-08 | 8.300E-08 |
| 12016 | 601 | C601S | A | 2.5e-08 | 4.9e-08 | 2.7e-08 | 3.367E-08 | 1.332E-08 | 2.500E-08 | 4.900E-08 |
| 12017 | 601 | C601T | A | 8.4e-08 | 5.4e-08 | 8.2e-08 | 7.333E-08 | 1.677E-08 | 5.400E-08 | 8.400E-08 |
| 12018 | 601 | C601W | A | 2.9e-08 | 5.6e-08 | 3.5e-08 | 4.000E-08 | 1.418E-08 | 2.900E-08 | 5.600E-08 |
| 12019 | 601 | C601Y | A | 2.2e-08 | 8.7e-09 | 1.4e-08 | 1.490E-08 | 6.696E-09 | 8.700E-09 | 2.200E-08 |
| 12020 | 601 | C601V | A | 4.3e-08 | 4.4e-08 | 6e-08 | 4.900E-08 | 9.539E-09 | 4.300E-08 | 6.000E-08 |
| 12021 | 602 | Q602A | A | 1.6e-07 | 5.4e-08 | 2e-08 | 7.800E-08 | 7.302E-08 | 2.000E-08 | 1.600E-07 |
| 12022 | 602 | Q602R | A | 3.3e-08 | 7.2e-08 | 2.9e-08 | 4.467E-08 | 2.376E-08 | 2.900E-08 | 7.200E-08 |
| 12023 | 602 | Q602N | A | 5e-08 | 1.1e-07 | 3.4e-08 | 6.467E-08 | 4.007E-08 | 3.400E-08 | 1.100E-07 |
| 12024 | 602 | Q602D | A | 2.8e-08 | 4.7e-08 | 6.2e-08 | 4.567E-08 | 1.704E-08 | 2.800E-08 | 6.200E-08 |
| 12025 | 602 | Q602C | A | 2.5e-08 | 1.9e-08 | 5.7e-08 | 3.367E-08 | 2.043E-08 | 1.900E-08 | 5.700E-08 |
| 12026 | 602 | Q602E | A | 3.1e-08 | 1.8e-08 | 5.5e-08 | 3.467E-08 | 1.877E-08 | 1.800E-08 | 5.500E-08 |
| 12027 | 602 | Q602Q | A | 6.4e-08 | 5.4e-08 | 2.8e-08 | 4.867E-08 | 1.858E-08 | 2.800E-08 | 6.400E-08 |
| 12028 | 602 | Q602G | A | 4.7e-08 | 2.4e-08 | 3.9e-08 | 3.667E-08 | 1.168E-08 | 2.400E-08 | 4.700E-08 |
| 12029 | 602 | Q602H | A | 1.4e-08 | 9.3e-08 | 1.3e-08 | 4.000E-08 | 4.590E-08 | 1.300E-08 | 9.300E-08 |
| 12030 | 602 | Q602I | A | 2.6e-08 | 6.2e-08 | 1.3e-07 | 7.267E-08 | 5.281E-08 | 2.600E-08 | 1.300E-07 |
| 12031 | 602 | Q602L | A | 8.5e-08 | 4.2e-08 | 1.4e-08 | 4.700E-08 | 3.576E-08 | 1.400E-08 | 8.500E-08 |
| 12032 | 602 | Q602K | A | 3.9e-08 | 5.5e-08 | 4.7e-08 | 4.700E-08 | 8.000E-09 | 3.900E-08 | 5.500E-08 |
| 12033 | 602 | Q602M | A | 7.8e-08 | 2.4e-08 | 8.2e-08 | 6.133E-08 | 3.239E-08 | 2.400E-08 | 8.200E-08 |
| 12034 | 602 | Q602F | A | 1.2e-08 | 7.9e-08 | 2.1e-07 | 1.003E-07 | 1.007E-07 | 1.200E-08 | 2.100E-07 |
| 12035 | 602 | Q602P | A | 9.3e-08 | 3.8e-08 | 4.9e-08 | 6.000E-08 | 2.910E-08 | 3.800E-08 | 9.300E-08 |
| 12036 | 602 | Q602S | A | 6.2e-08 | 4.7e-08 | 1.1e-07 | 7.300E-08 | 3.291E-08 | 4.700E-08 | 1.100E-07 |
| 12037 | 602 | Q602T | A | 2.3e-08 | 8.6e-08 | 3.1e-08 | 4.667E-08 | 3.430E-08 | 2.300E-08 | 8.600E-08 |
| 12038 | 602 | Q602W | A | 8.5e-08 | 1.7e-08 | 3.5e-08 | 4.567E-08 | 3.523E-08 | 1.700E-08 | 8.500E-08 |
| 12039 | 602 | Q602Y | A | 3.5e-08 | 2.8e-08 | 1.2e-07 | 6.100E-08 | 5.122E-08 | 2.800E-08 | 1.200E-07 |
| 12040 | 602 | Q602V | A | 7.4e-08 | 7e-08 | 1.4e-08 | 5.267E-08 | 3.355E-08 | 1.400E-08 | 7.400E-08 |
| 12041 | 603 | P603A | A | 5.3e-08 | 1.8e-08 | 6.4e-08 | 4.500E-08 | 2.402E-08 | 1.800E-08 | 6.400E-08 |
| 12042 | 603 | P603R | A | 8e-08 | 2.8e-08 | 2.9e-08 | 4.567E-08 | 2.974E-08 | 2.800E-08 | 8.000E-08 |
| 12043 | 603 | P603N | A | 5.7e-08 | 2e-08 | 1.2e-07 | 6.567E-08 | 5.056E-08 | 2.000E-08 | 1.200E-07 |
| 12044 | 603 | P603D | A | 9e-08 | 4.9e-08 | 2.5e-08 | 5.467E-08 | 3.287E-08 | 2.500E-08 | 9.000E-08 |
| 12045 | 603 | P603C | A | 1.4e-07 | 2.9e-08 | 6.3e-08 | 7.733E-08 | 5.687E-08 | 2.900E-08 | 1.400E-07 |
| 12046 | 603 | P603E | A | 5.6e-08 | 1.1e-08 | 3.8e-08 | 3.500E-08 | 2.265E-08 | 1.100E-08 | 5.600E-08 |
| 12047 | 603 | P603Q | A | 4.7e-08 | 3.3e-08 | 2e-08 | 3.333E-08 | 1.350E-08 | 2.000E-08 | 4.700E-08 |
| 12048 | 603 | P603G | A | 4e-08 | 4.5e-08 | 6.5e-08 | 5.000E-08 | 1.323E-08 | 4.000E-08 | 6.500E-08 |
| 12049 | 603 | P603H | A | 6.9e-09 | 1.7e-08 | 9.2e-08 | 3.863E-08 | 4.649E-08 | 6.900E-09 | 9.200E-08 |
| 12050 | 603 | P603I | A | 1.6e-08 | 2.8e-08 | 3.8e-08 | 2.733E-08 | 1.102E-08 | 1.600E-08 | 3.800E-08 |
| 12051 | 603 | P603L | A | 7.6e-08 | 6.1e-08 | 1.5e-08 | 5.067E-08 | 3.179E-08 | 1.500E-08 | 7.600E-08 |
| 12052 | 603 | P603K | A | 6.1e-08 | 6.3e-08 | 4.3e-08 | 5.567E-08 | 1.102E-08 | 4.300E-08 | 6.300E-08 |
| 12053 | 603 | P603M | A | 8.8e-08 | 7.3e-08 | 7.2e-08 | 7.767E-08 | 8.963E-09 | 7.200E-08 | 8.800E-08 |
| 12054 | 603 | P603F | A | 3e-08 | 3.3e-08 | 1.7e-08 | 2.667E-08 | 8.505E-09 | 1.700E-08 | 3.300E-08 |
| 12055 | 603 | P603P | A | 1.9e-08 | 5.6e-08 | 2.5e-08 | 3.333E-08 | 1.986E-08 | 1.900E-08 | 5.600E-08 |
| 12056 | 603 | P603S | A | 8.3e-08 | 2.7e-08 | 1.9e-08 | 4.300E-08 | 3.487E-08 | 1.900E-08 | 8.300E-08 |
| 12057 | 603 | P603T | A | 1.7e-07 | 3.5e-08 | 7e-08 | 9.167E-08 | 7.006E-08 | 3.500E-08 | 1.700E-07 |
| 12058 | 603 | P603W | A | 3.4e-08 | 6.8e-08 | 3.8e-08 | 4.667E-08 | 1.858E-08 | 3.400E-08 | 6.800E-08 |
| 12059 | 603 | P603Y | A | 3.8e-08 | 6.4e-08 | 5.6e-08 | 5.267E-08 | 1.332E-08 | 3.800E-08 | 6.400E-08 |
| 12060 | 603 | P603V | A | 6.9e-08 | 3.3e-08 | 7.3e-08 | 5.833E-08 | 2.203E-08 | 3.300E-08 | 7.300E-08 |
| 12061 | 604 | C604A | A | 1.2e-07 | 1.7e-08 | 3.5e-08 | 5.733E-08 | 5.501E-08 | 1.700E-08 | 1.200E-07 |
| 12062 | 604 | C604R | A | 4e-08 | 3.7e-08 | 1.4e-08 | 3.033E-08 | 1.422E-08 | 1.400E-08 | 4.000E-08 |
| 12063 | 604 | C604N | A | 5.3e-08 | 5e-08 | 3e-08 | 4.433E-08 | 1.250E-08 | 3.000E-08 | 5.300E-08 |
| 12064 | 604 | C604D | A | 3.9e-08 | 4.9e-08 | 3.8e-08 | 4.200E-08 | 6.083E-09 | 3.800E-08 | 4.900E-08 |
| 12065 | 604 | C604C | A | 2.8e-08 | 9.1e-08 | 3.3e-08 | 5.067E-08 | 3.502E-08 | 2.800E-08 | 9.100E-08 |
| 12066 | 604 | C604E | A | 4.9e-08 | 7.3e-08 | 1.7e-08 | 4.633E-08 | 2.810E-08 | 1.700E-08 | 7.300E-08 |
| 12067 | 604 | C604Q | A | 1.8e-08 | 9.1e-08 | 9e-08 | 6.633E-08 | 4.186E-08 | 1.800E-08 | 9.100E-08 |
| 12068 | 604 | C604G | A | 4.2e-08 | 2.8e-08 | 3.7e-08 | 3.567E-08 | 7.095E-09 | 2.800E-08 | 4.200E-08 |
| 12069 | 604 | C604H | A | 2.7e-08 | 3.8e-08 | 5.6e-08 | 4.033E-08 | 1.464E-08 | 2.700E-08 | 5.600E-08 |

| | | | | | | | | | | |
|---|---|---|---|---|---|---|---|---|---|---|
| 12070 | 604 | C604I | A | 2e-08 | 1.1e-07 | 2.2e-08 | 5.067E-08 | 5.139E-08 | 2.000E-08 | 1.100E-07 |
| 12071 | 604 | C604L | A | 3.9e-08 | 1.5e-07 | 1.6e-08 | 6.833E-08 | 7.165E-08 | 1.600E-08 | 1.500E-07 |
| 12072 | 604 | C604K | A | 3.2e-08 | 1.7e-08 | 4.1e-08 | 3.000E-08 | 1.212E-08 | 1.700E-08 | 4.100E-08 |
| 12073 | 604 | C604M | A | 5.8e-08 | 1.6e-08 | 2.3e-08 | 3.233E-08 | 2.250E-08 | 1.600E-08 | 5.800E-08 |
| 12074 | 604 | C604F | A | 7.5e-08 | 4e-08 | 3.4e-08 | 4.967E-08 | 2.214E-08 | 3.400E-08 | 7.500E-08 |
| 12075 | 604 | C604P | A | 2.4e-08 | 1.1e-07 | 6.7e-08 | 6.700E-08 | 4.300E-08 | 2.400E-08 | 1.100E-07 |
| 12076 | 604 | C604S | A | 2.5e-08 | 3.6e-08 | 3.9e-08 | 3.333E-08 | 7.371E-09 | 2.500E-08 | 3.900E-08 |
| 12077 | 604 | C604T | A | 6.4e-08 | 6.5e-08 | 4.6e-08 | 5.833E-08 | 1.069E-08 | 4.600E-08 | 6.500E-08 |
| 12078 | 604 | C604W | A | 2.6e-08 | 1.1e-07 | 5.7e-08 | 6.433E-08 | 4.248E-08 | 2.600E-08 | 1.100E-07 |
| 12079 | 604 | C604Y | A | 3e-08 | 9.8e-08 | 2.6e-08 | 5.133E-08 | 4.046E-08 | 2.600E-08 | 9.800E-08 |
| 12080 | 604 | C604V | A | 5.5e-08 | 1.9e-08 | 5.5e-08 | 4.300E-08 | 2.078E-08 | 1.900E-08 | 5.500E-08 |
| 12081 | 605 | P605A | A | 3.1e-08 | 3.1e-08 | 4.6e-08 | 3.600E-08 | 8.660E-09 | 3.100E-08 | 4.600E-08 |
| 12082 | 605 | P605R | A | 5.4e-08 | 5.1e-08 | 2.4e-08 | 4.300E-08 | 1.652E-08 | 2.400E-08 | 5.400E-08 |
| 12083 | 605 | P605N | A | 2.8e-08 | 4.7e-08 | 3.5e-08 | 3.667E-08 | 9.609E-09 | 2.800E-08 | 4.700E-08 |
| 12084 | 605 | P605D | A | 1.2e-07 | 1.3e-08 | 3.3e-08 | 5.533E-08 | 5.689E-08 | 1.300E-08 | 1.200E-07 |
| 12085 | 605 | P605C | A | 1.3e-07 | 4.5e-08 | 1.3e-08 | 6.267E-08 | 6.047E-08 | 1.300E-08 | 1.300E-07 |
| 12086 | 605 | P605E | A | 1.2e-08 | 5.8e-08 | 1.6e-07 | 7.667E-08 | 7.575E-08 | 1.200E-08 | 1.600E-07 |
| 12087 | 605 | P605Q | A | 3.1e-08 | 1.6e-08 | 4.8e-08 | 3.167E-08 | 1.601E-08 | 1.600E-08 | 4.800E-08 |
| 12088 | 605 | P605G | A | 3.1e-08 | 1.8e-07 | 1.7e-08 | 7.600E-08 | 9.034E-08 | 1.700E-08 | 1.800E-07 |
| 12089 | 605 | P605H | A | 5.1e-08 | 6e-08 | 5.8e-08 | 5.633E-08 | 4.726E-09 | 5.100E-08 | 6.000E-08 |
| 12090 | 605 | P605I | A | 3.7e-08 | 2.3e-08 | 7.3e-08 | 4.433E-08 | 2.579E-08 | 2.300E-08 | 7.300E-08 |
| 12091 | 605 | P605L | A | 1.8e-08 | 2.8e-08 | 2.1e-08 | 2.233E-08 | 5.132E-09 | 1.800E-08 | 2.800E-08 |
| 12092 | 605 | P605K | A | 6.1e-08 | 2.1e-08 | 4.8e-08 | 4.333E-08 | 2.040E-08 | 2.100E-08 | 6.100E-08 |
| 12093 | 605 | P605M | A | 8.8e-08 | 6.3e-08 | 7.3e-08 | 7.467E-08 | 1.258E-08 | 6.300E-08 | 8.800E-08 |
| 12094 | 605 | P605F | A | 5.4e-08 | 3.3e-08 | 2.8e-08 | 3.833E-08 | 1.380E-08 | 2.800E-08 | 5.400E-08 |
| 12095 | 605 | P605P | A | 7.3e-08 | 6.5e-08 | 2.7e-07 | 1.360E-07 | 1.161E-07 | 6.500E-08 | 2.700E-07 |
| 12096 | 605 | P605S | A | 1.3e-07 | 6.1e-08 | 4e-08 | 7.700E-08 | 4.709E-08 | 4.000E-08 | 1.300E-07 |
| 12097 | 605 | P605T | A | 9.3e-08 | 1.1e-08 | 2.7e-08 | 4.367E-08 | 4.347E-08 | 1.100E-08 | 9.300E-08 |
| 12098 | 605 | P605W | A | 2e-08 | 3.2e-08 | 1.2e-07 | 5.733E-08 | 5.460E-08 | 2.000E-08 | 1.200E-07 |
| 12099 | 605 | P605Y | A | 1.3e-08 | 2.3e-08 | 4.4e-08 | 2.667E-08 | 1.582E-08 | 1.300E-08 | 4.400E-08 |
| 12100 | 605 | P605V | A | 2.6e-08 | 1e-08 | 6.9e-08 | 3.500E-08 | 3.051E-08 | 1.000E-08 | 6.900E-08 |
| 12101 | 606 | I606A | A | 4.1e-08 | 3.9e-08 | 3.6e-08 | 3.867E-08 | 2.517E-09 | 3.600E-08 | 4.100E-08 |
| 12102 | 606 | I606R | A | 3e-08 | 1.8e-08 | 9.4e-08 | 4.733E-08 | 4.086E-08 | 1.800E-08 | 9.400E-08 |
| 12103 | 606 | I606N | A | 5.4e-08 | 1.9e-08 | 2.8e-08 | 3.367E-08 | 1.818E-08 | 1.900E-08 | 5.400E-08 |
| 12104 | 606 | I606D | A | 6.1e-08 | 7.1e-08 | 6.2e-08 | 6.467E-08 | 5.508E-09 | 6.100E-08 | 7.100E-08 |
| 12105 | 606 | I606C | A | 3.7e-08 | 2.6e-08 | 1.3e-07 | 6.433E-08 | 5.713E-08 | 2.600E-08 | 1.300E-07 |
| 12106 | 606 | I606E | A | 1.8e-08 | 3.9e-08 | 3.9e-08 | 3.200E-08 | 1.212E-08 | 1.800E-08 | 3.900E-08 |
| 12107 | 606 | I606Q | A | 4.8e-08 | 3e-08 | 2.2e-08 | 3.333E-08 | 1.332E-08 | 2.200E-08 | 4.800E-08 |
| 12108 | 606 | I606G | A | 3e-08 | 3.7e-08 | 3e-08 | 3.233E-08 | 4.041E-09 | 3.000E-08 | 3.700E-08 |
| 12109 | 606 | I606H | A | 7e-08 | 3.5e-08 | 3.6e-08 | 4.700E-08 | 1.992E-08 | 3.500E-08 | 7.000E-08 |
| 12110 | 606 | I606I | A | 6e-08 | 5.5e-08 | 2.2e-08 | 4.567E-08 | 2.065E-08 | 2.200E-08 | 6.000E-08 |
| 12111 | 606 | I606L | A | 4.6e-08 | 4e-08 | 5.1e-08 | 4.567E-08 | 5.508E-09 | 4.000E-08 | 5.100E-08 |
| 12112 | 606 | I606K | A | 7.4e-08 | 1.3e-07 | 4.5e-08 | 8.300E-08 | 4.321E-08 | 4.500E-08 | 1.300E-07 |
| 12113 | 606 | I606M | A | 1.9e-08 | 2.4e-08 | 9.3e-08 | 4.533E-08 | 4.136E-08 | 1.900E-08 | 9.300E-08 |
| 12114 | 606 | I606F | A | 4.5e-08 | 2.6e-08 | 1.2e-07 | 6.367E-08 | 4.970E-08 | 2.600E-08 | 1.200E-07 |
| 12115 | 606 | I606P | A | 2.8e-08 | 1.8e-08 | 2.9e-08 | 2.500E-08 | 6.083E-09 | 1.800E-08 | 2.900E-08 |
| 12116 | 606 | I606S | A | 2.3e-08 | 6.9e-08 | 5.1e-08 | 4.767E-08 | 2.318E-08 | 2.300E-08 | 6.900E-08 |
| 12117 | 606 | I606T | A | 6.9e-08 | 7e-08 | 6.3e-08 | 6.733E-08 | 3.786E-09 | 6.300E-08 | 7.000E-08 |
| 12118 | 606 | I606W | A | 1.3e-08 | 3.6e-08 | 3.5e-08 | 2.800E-08 | 1.300E-08 | 1.300E-08 | 3.600E-08 |
| 12119 | 606 | I606Y | A | 1.6e-08 | 4.3e-08 | 8.8e-09 | 2.260E-08 | 1.803E-08 | 8.800E-09 | 4.300E-08 |
| 12120 | 606 | I606V | A | 2e-08 | 6.9e-08 | 2.2e-08 | 3.700E-08 | 2.773E-08 | 2.000E-08 | 6.900E-08 |
| 12121 | 607 | N607A | A | 4.2e-08 | 2e-08 | 2e-08 | 2.733E-08 | 1.270E-08 | 2.000E-08 | 4.200E-08 |
| 12122 | 607 | N607R | A | 3.8e-08 | 6.8e-08 | 4.3e-08 | 4.967E-08 | 1.607E-08 | 3.800E-08 | 6.800E-08 |
| 12123 | 607 | N607N | A | 4.2e-08 | 2.5e-08 | 3.1e-08 | 3.267E-08 | 8.622E-09 | 2.500E-08 | 4.200E-08 |
| 12124 | 607 | N607D | A | 2.5e-08 | 1.6e-08 | 6.3e-08 | 3.467E-08 | 2.495E-08 | 1.600E-08 | 6.300E-08 |
| 12125 | 607 | N607C | A | 4.9e-08 | 3.6e-08 | 5.1e-08 | 4.533E-08 | 8.145E-09 | 3.600E-08 | 5.100E-08 |
| 12126 | 607 | N607E | A | 3.8e-08 | 8.7e-08 | 7.5e-08 | 6.667E-08 | 2.554E-08 | 3.800E-08 | 8.700E-08 |
| 12127 | 607 | N607Q | A | 2e-08 | 1.8e-08 | 1.5e-07 | 6.267E-08 | 7.564E-08 | 1.800E-08 | 1.500E-07 |
| 12128 | 607 | N607G | A | 7.4e-08 | 5.7e-08 | 9.9e-08 | 7.667E-08 | 2.113E-08 | 5.700E-08 | 9.900E-08 |
| 12129 | 607 | N607H | A | 7.3e-08 | 6.8e-08 | 6e-08 | 6.700E-08 | 6.557E-09 | 6.000E-08 | 7.300E-08 |
| 12130 | 607 | N607I | A | 4.8e-08 | 2.5e-08 | 1.1e-07 | 6.100E-08 | 4.397E-08 | 2.500E-08 | 1.100E-07 |
| 12131 | 607 | N607L | A | 7e-08 | 5.1e-08 | 6.5e-08 | 6.200E-08 | 9.849E-09 | 5.100E-08 | 7.000E-08 |
| 12132 | 607 | N607K | A | 4.5e-08 | 9.8e-08 | 2.5e-08 | 5.600E-08 | 3.772E-08 | 2.500E-08 | 9.800E-08 |
| 12133 | 607 | N607M | A | 3.9e-08 | 3.6e-08 | 2.4e-08 | 3.300E-08 | 7.937E-09 | 2.400E-08 | 3.900E-08 |
| 12134 | 607 | N607F | A | 1.3e-07 | 4.5e-08 | 4.9e-08 | 7.467E-08 | 4.796E-08 | 4.500E-08 | 1.300E-07 |
| 12135 | 607 | N607P | A | 9e-08 | 3.9e-08 | 1.1e-07 | 7.967E-08 | 3.661E-08 | 3.900E-08 | 1.100E-07 |
| 12136 | 607 | N607S | A | 6.9e-08 | 4.2e-08 | 9.2e-08 | 6.767E-08 | 2.503E-08 | 4.200E-08 | 9.200E-08 |
| 12137 | 607 | N607T | A | 1.6e-08 | 7.8e-08 | 2.9e-08 | 4.100E-08 | 3.270E-08 | 1.600E-08 | 7.800E-08 |
| 12138 | 607 | N607W | A | 8.1e-09 | 2.1e-08 | 5.3e-08 | 2.737E-08 | 2.312E-08 | 8.100E-09 | 5.300E-08 |
| 12139 | 607 | N607Y | A | 5.4e-08 | 7.8e-08 | 2.9e-08 | 5.367E-08 | 2.450E-08 | 2.900E-08 | 7.800E-08 |
| 12140 | 607 | N607V | A | 7.2e-08 | 2.6e-08 | 1.8e-08 | 3.867E-08 | 2.914E-08 | 1.800E-08 | 7.200E-08 |
| 12141 | 608 | C608A | A | 7.4e-08 | 3.1e-08 | 3.6e-08 | 4.700E-08 | 2.352E-08 | 3.100E-08 | 7.400E-08 |
| 12142 | 608 | C608R | A | 2e-08 | 3.6e-08 | 9e-08 | 4.867E-08 | 3.668E-08 | 2.000E-08 | 9.000E-08 |
| 12143 | 608 | C608N | A | 6.8e-08 | 1.5e-08 | 1.6e-08 | 3.300E-08 | 3.032E-08 | 1.500E-08 | 6.800E-08 |
| 12144 | 608 | C608D | A | 5.6e-08 | 1.3e-07 | 2.4e-08 | 7.000E-08 | 5.437E-08 | 2.400E-08 | 1.300E-07 |
| 12145 | 608 | C608C | A | 6.4e-08 | 1.1e-08 | 6.4e-08 | 4.633E-08 | 3.060E-08 | 1.100E-08 | 6.400E-08 |
| 12146 | 608 | C608E | A | 1.4e-07 | 1.4e-07 | 4.7e-08 | 1.090E-07 | 5.369E-08 | 4.700E-08 | 1.400E-07 |
| 12147 | 608 | C608Q | A | 4.1e-08 | 3.1e-08 | 5.1e-08 | 4.100E-08 | 1.000E-08 | 3.100E-08 | 5.100E-08 |
| 12148 | 608 | C608G | A | 4.4e-08 | 5.2e-08 | 2.2e-08 | 3.933E-08 | 1.553E-08 | 2.200E-08 | 5.200E-08 |
| 12149 | 608 | C608H | A | 3.1e-08 | 3.9e-08 | 1.3e-07 | 6.667E-08 | 5.499E-08 | 3.100E-08 | 1.300E-07 |
| 12150 | 608 | C608I | A | 5e-08 | 5.6e-08 | 5.7e-08 | 5.433E-08 | 3.786E-09 | 5.000E-08 | 5.700E-08 |
| 12151 | 608 | C608L | A | 5.7e-08 | 3.5e-08 | 5.8e-08 | 5.000E-08 | 1.300E-08 | 3.500E-08 | 5.800E-08 |
| 12152 | 608 | C608K | A | 7.6e-08 | 6.4e-08 | 8.6e-08 | 7.533E-08 | 1.102E-08 | 6.400E-08 | 8.600E-08 |
| 12153 | 608 | C608M | A | 1.1e-07 | 3.4e-08 | 2.8e-08 | 5.733E-08 | 4.571E-08 | 2.800E-08 | 1.100E-07 |
| 12154 | 608 | C608F | A | 1.3e-08 | 7.4e-08 | 8.6e-08 | 5.767E-08 | 3.915E-08 | 1.300E-08 | 8.600E-08 |

| | | | | | | | | | | |
|---|---|---|---|---|---|---|---|---|---|---|
| 12155 | 608 | C608P | A | 2.6e-08 | 5e-08 | 2.7e-08 | 3.433E-08 | 1.358E-08 | 2.600E-08 | 5.000E-08 |
| 12156 | 608 | C608S | A | 1e-08 | 6.1e-09 | 3.8e-08 | 1.803E-08 | 1.740E-08 | 6.100E-09 | 3.800E-08 |
| 12157 | 608 | C608T | A | 1.8e-07 | 7.2e-08 | 5.9e-08 | 1.037E-07 | 6.643E-08 | 5.900E-08 | 1.800E-07 |
| 12158 | 608 | C608W | A | 6.7e-08 | 3.5e-08 | 3.2e-08 | 4.467E-08 | 1.940E-08 | 3.200E-08 | 6.700E-08 |
| 12159 | 608 | C608Y | A | 7.4e-08 | 3.2e-08 | 2.3e-08 | 4.300E-08 | 2.722E-08 | 2.300E-08 | 7.400E-08 |
| 12160 | 608 | C608V | A | 2.7e-07 | 3.8e-08 | 7.4e-08 | 1.273E-07 | 1.249E-07 | 3.800E-08 | 2.700E-07 |
| 12161 | 609 | T609A | A | 3.8e-08 | 7.8e-08 | 6e-08 | 5.867E-08 | 2.003E-08 | 3.800E-08 | 7.800E-08 |
| 12162 | 609 | T609R | A | 5.1e-08 | 3.6e-08 | 3.8e-08 | 4.167E-08 | 8.145E-09 | 3.600E-08 | 5.100E-08 |
| 12163 | 609 | T609N | A | 4.2e-08 | 4.2e-08 | 2.6e-08 | 3.667E-08 | 9.238E-09 | 2.600E-08 | 4.200E-08 |
| 12164 | 609 | T609D | A | 4.9e-08 | 3.8e-08 | 3.6e-08 | 4.100E-08 | 7.000E-09 | 3.600E-08 | 4.900E-08 |
| 12165 | 609 | T609C | A | 4.3e-08 | 2.7e-08 | 3.8e-08 | 3.600E-08 | 8.185E-09 | 2.700E-08 | 4.300E-08 |
| 12166 | 609 | T609E | A | 1.7e-07 | 3.8e-08 | 3.5e-08 | 8.100E-08 | 7.709E-08 | 3.500E-08 | 1.700E-07 |
| 12167 | 609 | T609Q | A | 6.3e-08 | 5e-08 | 3.4e-08 | 4.900E-08 | 1.453E-08 | 3.400E-08 | 6.300E-08 |
| 12168 | 609 | T609G | A | 3e-08 | 6.1e-08 | 3.2e-08 | 4.100E-08 | 1.735E-08 | 3.000E-08 | 6.100E-08 |
| 12169 | 609 | T609H | A | 4.1e-08 | 4.3e-08 | 1.2e-07 | 6.800E-08 | 4.504E-08 | 4.100E-08 | 1.200E-07 |
| 12170 | 609 | T609I | A | 7.3e-08 | 3.9e-08 | 9.1e-09 | 4.037E-08 | 3.197E-08 | 9.100E-09 | 7.300E-08 |
| 12171 | 609 | T609L | A | 7.5e-08 | 4.4e-08 | 8.6e-08 | 6.833E-08 | 2.178E-08 | 4.400E-08 | 8.600E-08 |
| 12172 | 609 | T609K | A | 3.6e-08 | 7.5e-08 | 4.7e-08 | 5.267E-08 | 2.011E-08 | 3.600E-08 | 7.500E-08 |
| 12173 | 609 | T609M | A | 2.6e-08 | 7.2e-08 | 1.2e-07 | 7.267E-08 | 4.700E-08 | 2.600E-08 | 1.200E-07 |
| 12174 | 609 | T609F | A | 5.6e-08 | 1.1e-08 | 1.5e-08 | 2.733E-08 | 2.491E-08 | 1.100E-08 | 5.600E-08 |
| 12175 | 609 | T609P | A | 3.6e-08 | 3.1e-08 | 3.8e-07 | 1.490E-07 | 2.001E-07 | 3.100E-08 | 3.800E-07 |
| 12176 | 609 | T609S | A | 6.3e-08 | 3.1e-08 | 2.7e-08 | 4.033E-08 | 1.973E-08 | 2.700E-08 | 6.300E-08 |
| 12177 | 609 | T609T | A | 1.5e-07 | 8.6e-08 | 9.6e-08 | 1.107E-07 | 3.443E-08 | 8.600E-08 | 1.500E-07 |
| 12178 | 609 | T609W | A | 4.4e-08 | 8.8e-08 | 3.7e-08 | 5.633E-08 | 2.765E-08 | 3.700E-08 | 8.800E-08 |
| 12179 | 609 | T609Y | A | 3.3e-08 | 3.1e-08 | 4.6e-08 | 3.667E-08 | 8.145E-09 | 3.100E-08 | 4.600E-08 |
| 12180 | 609 | T609V | A | 3.4e-08 | 4.8e-08 | 4.2e-08 | 4.133E-08 | 7.024E-09 | 3.400E-08 | 4.800E-08 |
| 12181 | 610 | H610A | A | 2.2e-08 | 2.7e-08 | 1.4e-07 | 6.300E-08 | 6.673E-08 | 2.200E-08 | 1.400E-07 |
| 12182 | 610 | H610R | A | 5.8e-08 | 7.5e-08 | 1.6e-08 | 4.967E-08 | 3.037E-08 | 1.600E-08 | 7.500E-08 |
| 12183 | 610 | H610N | A | 4.5e-08 | 4.4e-08 | 9.3e-08 | 6.067E-08 | 2.801E-08 | 4.400E-08 | 9.300E-08 |
| 12184 | 610 | H610D | A | 4.4e-08 | 2.1e-08 | 3.7e-08 | 3.400E-08 | 1.179E-08 | 2.100E-08 | 4.400E-08 |
| 12185 | 610 | H610C | A | 2.3e-08 | 4.7e-08 | 2.1e-07 | 9.333E-08 | 1.017E-07 | 2.300E-08 | 2.100E-07 |
| 12186 | 610 | H610E | A | 9.9e-08 | 4.1e-08 | 5e-08 | 6.333E-08 | 3.121E-08 | 4.100E-08 | 9.900E-08 |
| 12187 | 610 | H610Q | A | 1.8e-07 | 2.4e-08 | 6.1e-08 | 8.833E-08 | 8.151E-08 | 2.400E-08 | 1.800E-07 |
| 12188 | 610 | H610G | A | 3.9e-08 | 5.8e-08 | 3.7e-08 | 4.467E-08 | 1.159E-08 | 3.700E-08 | 5.800E-08 |
| 12189 | 610 | H610H | A | 1.9e-08 | 2.8e-08 | 2.2e-08 | 2.300E-08 | 4.583E-09 | 1.900E-08 | 2.800E-08 |
| 12190 | 610 | H610I | A | 3.6e-08 | 1e-07 | 3.2e-08 | 5.600E-08 | 3.816E-08 | 3.200E-08 | 1.000E-07 |
| 12191 | 610 | H610L | A | 1.2e-07 | 2.1e-08 | 3.5e-08 | 5.867E-08 | 5.358E-08 | 2.100E-08 | 1.200E-07 |
| 12192 | 610 | H610K | A | 4e-08 | 4.1e-08 | 2e-08 | 3.367E-08 | 1.185E-08 | 2.000E-08 | 4.100E-08 |
| 12193 | 610 | H610M | A | 6.7e-08 | 3e-08 | 2.7e-08 | 4.133E-08 | 2.228E-08 | 2.700E-08 | 6.700E-08 |
| 12194 | 610 | H610F | A | 4.8e-08 | 7.3e-08 | 1.4e-08 | 4.500E-08 | 2.961E-08 | 1.400E-08 | 7.300E-08 |
| 12195 | 610 | H610P | A | 5.5e-08 | 2.4e-08 | 1.3e-07 | 6.967E-08 | 5.450E-08 | 2.400E-08 | 1.300E-07 |
| 12196 | 610 | H610S | A | 4.8e-08 | 1.9e-08 | 5.5e-08 | 4.067E-08 | 1.909E-08 | 1.900E-08 | 5.500E-08 |
| 12197 | 610 | H610T | A | 1.2e-08 | 2.5e-08 | 3.5e-08 | 2.400E-08 | 1.153E-08 | 1.200E-08 | 3.500E-08 |
| 12198 | 610 | H610W | A | 8.1e-08 | 7.8e-08 | 3e-08 | 6.300E-08 | 2.862E-08 | 3.000E-08 | 8.100E-08 |
| 12199 | 610 | H610Y | A | 1.7e-08 | 5.7e-08 | 4.8e-08 | 4.067E-08 | 2.098E-08 | 1.700E-08 | 5.700E-08 |
| 12200 | 610 | H610V | A | 6.1e-08 | 3.3e-08 | 6.3e-08 | 5.233E-08 | 1.677E-08 | 3.300E-08 | 6.300E-08 |
| 12201 | 611 | S611A | A | 3.9e-08 | 3.7e-08 | 6.5e-08 | 4.700E-08 | 1.562E-08 | 3.700E-08 | 6.500E-08 |
| 12202 | 611 | S611R | A | 1.5e-08 | 1.8e-08 | 4.5e-08 | 2.600E-08 | 1.652E-08 | 1.500E-08 | 4.500E-08 |
| 12203 | 611 | S611N | A | 1.5e-08 | 3.6e-08 | 1.7e-08 | 2.267E-08 | 1.159E-08 | 1.500E-08 | 3.600E-08 |
| 12204 | 611 | S611D | A | 2.6e-08 | 5.1e-08 | 5.4e-08 | 4.367E-08 | 1.537E-08 | 2.600E-08 | 5.400E-08 |
| 12205 | 611 | S611C | A | 4e-08 | 2.8e-08 | 3.7e-08 | 3.500E-08 | 6.245E-09 | 2.800E-08 | 4.000E-08 |
| 12206 | 611 | S611E | A | 4.8e-08 | 3.2e-08 | 8.5e-08 | 5.500E-08 | 2.718E-08 | 3.200E-08 | 8.500E-08 |
| 12207 | 611 | S611Q | A | 1.2e-08 | 5.6e-08 | 4e-08 | 3.600E-08 | 2.227E-08 | 1.200E-08 | 5.600E-08 |
| 12208 | 611 | S611G | A | 4.2e-08 | 2.6e-08 | 3.2e-08 | 3.333E-08 | 8.083E-09 | 2.600E-08 | 4.200E-08 |
| 12209 | 611 | S611H | A | 5.2e-08 | 6.3e-08 | 1.3e-07 | 8.167E-08 | 4.222E-08 | 5.200E-08 | 1.300E-07 |
| 12210 | 611 | S611I | A | 2.5e-08 | 3.3e-08 | 2.5e-08 | 2.767E-08 | 4.619E-09 | 2.500E-08 | 3.300E-08 |
| 12211 | 611 | S611L | A | 4.1e-08 | 4.4e-08 | 5.6e-08 | 4.700E-08 | 7.937E-09 | 4.100E-08 | 5.600E-08 |
| 12212 | 611 | S611K | A | 2.5e-08 | 5.1e-08 | 3.7e-08 | 3.767E-08 | 1.301E-08 | 2.500E-08 | 5.100E-08 |
| 12213 | 611 | S611M | A | 4.5e-08 | 7.8e-08 | 2.4e-08 | 4.900E-08 | 2.722E-08 | 2.400E-08 | 7.800E-08 |
| 12214 | 611 | S611F | A | 5.2e-08 | 1.3e-08 | 7.8e-08 | 4.767E-08 | 3.272E-08 | 1.300E-08 | 7.800E-08 |
| 12215 | 611 | S611P | A | 5.8e-08 | 3.4e-08 | 8e-08 | 5.733E-08 | 2.301E-08 | 3.400E-08 | 8.000E-08 |
| 12216 | 611 | S611S | A | 6.3e-08 | 1.8e-08 | 3.6e-08 | 3.900E-08 | 2.265E-08 | 1.800E-08 | 6.300E-08 |
| 12217 | 611 | S611T | A | 6.2e-08 | 6.5e-08 | 3.3e-08 | 5.333E-08 | 1.767E-08 | 3.300E-08 | 6.500E-08 |
| 12218 | 611 | S611W | A | 6e-08 | 4.4e-08 | 3.3e-08 | 4.567E-08 | 1.358E-08 | 3.300E-08 | 6.000E-08 |
| 12219 | 611 | S611Y | A | 6.5e-08 | 6.5e-08 | 2.9e-08 | 5.300E-08 | 2.078E-08 | 2.900E-08 | 6.500E-08 |
| 12220 | 611 | S611V | A | 5.8e-08 | 4.2e-08 | 3.2e-08 | 4.400E-08 | 1.311E-08 | 3.200E-08 | 5.800E-08 |
| 12221 | 612 | C612A | A | 6e-08 | 2.7e-08 | 3.8e-08 | 4.167E-08 | 1.680E-08 | 2.700E-08 | 6.000E-08 |
| 12222 | 612 | C612R | A | 2.2e-08 | 6.1e-08 | 7.1e-08 | 5.133E-08 | 2.589E-08 | 2.200E-08 | 7.100E-08 |
| 12223 | 612 | C612N | A | 1.6e-08 | 3.2e-08 | 7.6e-08 | 4.133E-08 | 3.107E-08 | 1.600E-08 | 7.600E-08 |
| 12224 | 612 | C612D | A | 4.9e-08 | 5.6e-08 | 4.6e-09 | 3.653E-08 | 2.788E-08 | 4.600E-09 | 5.600E-08 |
| 12225 | 612 | C612C | A | 2.8e-08 | 3.7e-08 | 1.1e-07 | 5.833E-08 | 4.497E-08 | 2.800E-08 | 1.100E-07 |
| 12226 | 612 | C612E | A | 7.2e-08 | 1.7e-07 | 5e-08 | 9.733E-08 | 6.389E-08 | 5.000E-08 | 1.700E-07 |
| 12227 | 612 | C612Q | A | 7.8e-08 | 5.1e-08 | 5e-08 | 5.967E-08 | 1.589E-08 | 5.000E-08 | 7.800E-08 |
| 12228 | 612 | C612G | A | 7.9e-08 | 8.5e-08 | 1.3e-08 | 5.900E-08 | 3.995E-08 | 1.300E-08 | 8.500E-08 |
| 12229 | 612 | C612H | A | 3.4e-08 | 5.7e-08 | 8.3e-08 | 5.800E-08 | 2.452E-08 | 3.400E-08 | 8.300E-08 |
| 12230 | 612 | C612I | A | 7.6e-08 | 5.1e-08 | 3.3e-08 | 5.333E-08 | 2.159E-08 | 3.300E-08 | 7.600E-08 |
| 12231 | 612 | C612L | A | 4.1e-08 | 2e-08 | 1.4e-08 | 2.500E-08 | 1.418E-08 | 1.400E-08 | 4.100E-08 |
| 12232 | 612 | C612K | A | 4.2e-08 | 2.1e-08 | 2.1e-08 | 2.800E-08 | 1.212E-08 | 2.100E-08 | 4.200E-08 |
| 12233 | 612 | C612M | A | 6.8e-08 | 2.9e-08 | 9.3e-08 | 6.333E-08 | 3.225E-08 | 2.900E-08 | 9.300E-08 |
| 12234 | 612 | C612F | A | 3.7e-08 | 6.8e-08 | 1.6e-08 | 4.033E-08 | 2.616E-08 | 1.600E-08 | 6.800E-08 |
| 12235 | 612 | C612P | A | 8.1e-08 | 1.4e-08 | 1.1e-07 | 6.833E-08 | 4.924E-08 | 1.400E-08 | 1.100E-07 |
| 12236 | 612 | C612S | A | 4.3e-08 | 8.9e-08 | 6.9e-08 | 6.700E-08 | 2.307E-08 | 4.300E-08 | 8.900E-08 |
| 12237 | 612 | C612T | A | 9.4e-08 | 9.5e-08 | 6.9e-08 | 8.600E-08 | 1.473E-08 | 6.900E-08 | 9.500E-08 |
| 12238 | 612 | C612W | A | 1.9e-08 | 2.1e-08 | 9.3e-08 | 4.433E-08 | 4.216E-08 | 1.900E-08 | 9.300E-08 |
| 12239 | 612 | C612Y | A | 9.9e-08 | 2.4e-08 | 4.7e-08 | 5.667E-08 | 3.842E-08 | 2.400E-08 | 9.900E-08 |

| | | | | | | | | | | |
|---|---|---|---|---|---|---|---|---|---|---|
| 12240 | 612 | C612V | A | 5.5e-08 | 1.5e-08 | 2.4e-08 | 3.133E-08 | 2.098E-08 | 1.500E-08 | 5.500E-08 |
| 12241 | 613 | V613A | A | 4.8e-08 | 1.1e-07 | 2.9e-08 | 6.233E-08 | 4.236E-08 | 2.900E-08 | 1.100E-07 |
| 12242 | 613 | V613R | A | 4.9e-08 | 5.7e-08 | 3e-08 | 4.533E-08 | 1.387E-08 | 3.000E-08 | 5.700E-08 |
| 12243 | 613 | V613N | A | 2.1e-08 | 3.6e-08 | 6.1e-08 | 3.933E-08 | 2.021E-08 | 2.100E-08 | 6.100E-08 |
| 12244 | 613 | V613D | A | 1.5e-08 | 3.3e-08 | 3.7e-08 | 2.833E-08 | 1.172E-08 | 1.500E-08 | 3.700E-08 |
| 12245 | 613 | V613C | A | 2.5e-08 | 2.8e-08 | 1.6e-08 | 2.300E-08 | 6.245E-09 | 1.600E-08 | 2.800E-08 |
| 12246 | 613 | V613E | A | 4.8e-08 | 1e-07 | 1.2e-08 | 5.333E-08 | 4.424E-08 | 1.200E-08 | 1.000E-07 |
| 12247 | 613 | V613Q | A | 2e-08 | 5.4e-08 | 7.8e-08 | 5.067E-08 | 2.914E-08 | 2.000E-08 | 7.800E-08 |
| 12248 | 613 | V613G | A | 1.6e-08 | 6.4e-08 | 9.9e-08 | 5.967E-08 | 4.167E-08 | 1.600E-08 | 9.900E-08 |
| 12249 | 613 | V613H | A | 5.3e-08 | 4.7e-08 | 3.1e-08 | 4.367E-08 | 1.137E-08 | 3.100E-08 | 5.300E-08 |
| 12250 | 613 | V613I | A | 6.5e-08 | 2.1e-08 | 4.4e-08 | 4.333E-08 | 2.201E-08 | 2.100E-08 | 6.500E-08 |
| 12251 | 613 | V613L | A | 4.9e-08 | 4.7e-08 | 7.3e-08 | 5.633E-08 | 1.447E-08 | 4.700E-08 | 7.300E-08 |
| 12252 | 613 | V613K | A | 8.2e-09 | 1.2e-07 | 6.9e-08 | 6.573E-08 | 5.597E-08 | 8.200E-09 | 1.200E-07 |
| 12253 | 613 | V613M | A | 1.2e-08 | 2e-08 | 2.1e-08 | 1.767E-08 | 4.933E-09 | 1.200E-08 | 2.100E-08 |
| 12254 | 613 | V613F | A | 7e-08 | 5.2e-08 | 7.2e-08 | 6.467E-08 | 1.102E-08 | 5.200E-08 | 7.200E-08 |
| 12255 | 613 | V613P | A | 2e-08 | 3e-08 | 3.1e-08 | 2.700E-08 | 6.083E-09 | 2.000E-08 | 3.100E-08 |
| 12256 | 613 | V613S | A | 5.4e-08 | 4.3e-08 | 5.3e-08 | 5.000E-08 | 6.083E-09 | 4.300E-08 | 5.400E-08 |
| 12257 | 613 | V613T | A | 7.5e-08 | 1.2e-08 | 5.9e-08 | 4.867E-08 | 3.275E-08 | 1.200E-08 | 7.500E-08 |
| 12258 | 613 | V613W | A | 1.1e-08 | 5e-08 | 2.4e-08 | 2.833E-08 | 1.986E-08 | 1.100E-08 | 5.000E-08 |
| 12259 | 613 | V613Y | A | 1.4e-08 | 3.8e-08 | 1.2e-08 | 2.133E-08 | 1.447E-08 | 1.200E-08 | 3.800E-08 |
| 12260 | 613 | V613V | A | 6.5e-08 | 3.2e-08 | 2.8e-08 | 4.167E-08 | 2.031E-08 | 2.800E-08 | 6.500E-08 |
| 12261 | 614 | D614A | A | 4.8e-08 | 4e-08 | 7.8e-08 | 5.533E-08 | 2.003E-08 | 4.000E-08 | 7.800E-08 |
| 12262 | 614 | D614R | A | 5.2e-08 | 1.2e-08 | 3.3e-08 | 3.233E-08 | 2.001E-08 | 1.200E-08 | 5.200E-08 |
| 12263 | 614 | D614N | A | 3.2e-08 | 2.7e-08 | 4.4e-08 | 3.433E-08 | 8.737E-09 | 2.700E-08 | 4.400E-08 |
| 12264 | 614 | D614D | A | 2e-08 | 1.4e-07 | 3.9e-08 | 6.633E-08 | 6.450E-08 | 2.000E-08 | 1.400E-07 |
| 12265 | 614 | D614C | A | 2.3e-08 | 5.4e-08 | 2.2e-07 | 9.900E-08 | 1.059E-07 | 2.300E-08 | 2.200E-07 |
| 12266 | 614 | D614E | A | 4.5e-08 | 5.3e-08 | 3.3e-08 | 4.367E-08 | 1.007E-08 | 3.300E-08 | 5.300E-08 |
| 12267 | 614 | D614Q | A | 9.2e-08 | 5.1e-08 | 5.8e-08 | 6.700E-08 | 2.193E-08 | 5.100E-08 | 9.200E-08 |
| 12268 | 614 | D614G | A | 3.5e-08 | 2.6e-08 | 4.8e-08 | 3.633E-08 | 1.106E-08 | 2.600E-08 | 4.800E-08 |
| 12269 | 614 | D614H | A | 4.7e-08 | 2.1e-08 | 4.5e-08 | 3.767E-08 | 1.447E-08 | 2.100E-08 | 4.700E-08 |
| 12270 | 614 | D614I | A | 3.2e-08 | 4.4e-08 | 5.6e-08 | 4.400E-08 | 1.200E-08 | 3.200E-08 | 5.600E-08 |
| 12271 | 614 | D614L | A | 6.4e-08 | 5.1e-08 | 4.9e-08 | 5.467E-08 | 8.145E-09 | 4.900E-08 | 6.400E-08 |
| 12272 | 614 | D614K | A | 3.9e-08 | 6.2e-08 | 3.3e-08 | 4.467E-08 | 1.531E-08 | 3.300E-08 | 6.200E-08 |
| 12273 | 614 | D614M | A | 1e-07 | 1.4e-08 | 4.5e-08 | 5.300E-08 | 4.355E-08 | 1.400E-08 | 1.000E-07 |
| 12274 | 614 | D614F | A | 3.2e-08 | 5.4e-08 | 4.9e-08 | 4.500E-08 | 1.153E-08 | 3.200E-08 | 5.400E-08 |
| 12275 | 614 | D614P | A | 6.5e-08 | 5.9e-08 | 2.8e-08 | 5.067E-08 | 1.986E-08 | 2.800E-08 | 6.500E-08 |
| 12276 | 614 | D614S | A | 5.4e-08 | 1.9e-08 | 1.5e-08 | 2.933E-08 | 2.146E-08 | 1.500E-08 | 5.400E-08 |
| 12277 | 614 | D614T | A | 2.9e-08 | 1.7e-08 | 5.8e-08 | 3.467E-08 | 2.108E-08 | 1.700E-08 | 5.800E-08 |
| 12278 | 614 | D614W | A | 5e-08 | 7.1e-08 | 3.7e-08 | 5.267E-08 | 1.716E-08 | 3.700E-08 | 7.100E-08 |
| 12279 | 614 | D614Y | A | 3.3e-08 | 3e-08 | 4.4e-08 | 3.567E-08 | 7.371E-09 | 3.000E-08 | 4.400E-08 |
| 12280 | 614 | D614V | A | 6.6e-08 | 4.2e-08 | 9.7e-08 | 6.833E-08 | 2.757E-08 | 4.200E-08 | 9.700E-08 |
| 12281 | 615 | L615A | A | 1.3e-07 | 6.7e-08 | 6.1e-08 | 8.600E-08 | 3.822E-08 | 6.100E-08 | 1.300E-07 |
| 12282 | 615 | L615R | A | 1.5e-08 | 1.5e-08 | 4.1e-08 | 2.367E-08 | 1.501E-08 | 1.500E-08 | 4.100E-08 |
| 12283 | 615 | L615N | A | 1.4e-08 | 4e-08 | 2.6e-08 | 2.667E-08 | 1.301E-08 | 1.400E-08 | 4.000E-08 |
| 12284 | 615 | L615D | A | 7.7e-08 | 6.6e-08 | 9.9e-08 | 8.067E-08 | 1.680E-08 | 6.600E-08 | 9.900E-08 |
| 12285 | 615 | L615C | A | 6.7e-08 | 2.7e-08 | 5.4e-08 | 4.933E-08 | 2.040E-08 | 2.700E-08 | 6.700E-08 |
| 12286 | 615 | L615E | A | 2.3e-08 | 1.6e-08 | 5.9e-08 | 3.267E-08 | 2.307E-08 | 1.600E-08 | 5.900E-08 |
| 12287 | 615 | L615Q | A | 8.7e-08 | 8e-08 | 4e-08 | 6.900E-08 | 2.536E-08 | 4.000E-08 | 8.700E-08 |
| 12288 | 615 | L615G | A | 6.1e-08 | 2.7e-08 | 1.2e-07 | 6.933E-08 | 4.706E-08 | 2.700E-08 | 1.200E-07 |
| 12289 | 615 | L615H | A | 3.8e-08 | 8.5e-08 | 5.8e-08 | 6.033E-08 | 2.359E-08 | 3.800E-08 | 8.500E-08 |
| 12290 | 615 | L615I | A | 6.3e-08 | 2.7e-08 | 2.5e-08 | 3.833E-08 | 2.139E-08 | 2.500E-08 | 6.300E-08 |
| 12291 | 615 | L615L | A | 2.6e-08 | 4.3e-08 | 3.4e-08 | 3.433E-08 | 8.505E-09 | 2.600E-08 | 4.300E-08 |
| 12292 | 615 | L615K | A | 4.6e-08 | 7.7e-08 | 7.7e-08 | 6.667E-08 | 1.790E-08 | 4.600E-08 | 7.700E-08 |
| 12293 | 615 | L615M | A | 5.3e-08 | 4.3e-08 | 1.4e-08 | 3.667E-08 | 2.026E-08 | 1.400E-08 | 5.300E-08 |
| 12294 | 615 | L615F | A | 5.7e-08 | 5.9e-08 | 2.9e-08 | 4.833E-08 | 1.677E-08 | 2.900E-08 | 5.900E-08 |
| 12295 | 615 | L615P | A | 1.9e-08 | 1.5e-07 | 2.5e-08 | 6.467E-08 | 7.396E-08 | 1.900E-08 | 1.500E-07 |
| 12296 | 615 | L615S | A | 8.6e-08 | 2.4e-08 | 5e-08 | 5.333E-08 | 3.113E-08 | 2.400E-08 | 8.600E-08 |
| 12297 | 615 | L615T | A | 7.3e-08 | 3.4e-08 | 6.2e-08 | 5.633E-08 | 2.011E-08 | 3.400E-08 | 7.300E-08 |
| 12298 | 615 | L615W | A | 3.4e-08 | 2.7e-08 | 2.5e-08 | 2.867E-08 | 4.726E-09 | 2.500E-08 | 3.400E-08 |
| 12299 | 615 | L615Y | A | 4e-08 | 6.7e-08 | 1.4e-08 | 4.033E-08 | 2.650E-08 | 1.400E-08 | 6.700E-08 |
| 12300 | 615 | L615V | A | 1.1e-07 | 3.1e-08 | 4.1e-08 | 6.067E-08 | 4.302E-08 | 3.100E-08 | 1.100E-07 |
| 12301 | 616 | D616A | A | 1.5e-07 | 2.2e-08 | 8.5e-08 | 8.567E-08 | 6.400E-08 | 2.200E-08 | 1.500E-07 |
| 12302 | 616 | D616R | A | 6e-08 | 2.3e-08 | 1.1e-07 | 6.433E-08 | 4.366E-08 | 2.300E-08 | 1.100E-07 |
| 12303 | 616 | D616N | A | 5.4e-08 | 5.8e-08 | 9.4e-08 | 6.867E-08 | 2.203E-08 | 5.400E-08 | 9.400E-08 |
| 12304 | 616 | D616D | A | 8.4e-09 | 5.8e-08 | 1.5e-08 | 2.713E-08 | 2.693E-08 | 8.400E-09 | 5.800E-08 |
| 12305 | 616 | D616C | A | 5.3e-08 | 7.2e-08 | 1.1e-08 | 4.533E-08 | 3.121E-08 | 1.100E-08 | 7.200E-08 |
| 12306 | 616 | D616E | A | 4.8e-08 | 7.9e-08 | 5.9e-08 | 6.200E-08 | 1.572E-08 | 4.800E-08 | 7.900E-08 |
| 12307 | 616 | D616Q | A | 6.4e-08 | 6.8e-08 | 2.5e-08 | 5.233E-08 | 2.376E-08 | 2.500E-08 | 6.800E-08 |
| 12308 | 616 | D616G | A | 3e-08 | 1e-07 | 2e-08 | 5.000E-08 | 4.359E-08 | 2.000E-08 | 1.000E-07 |
| 12309 | 616 | D616H | A | 1e-07 | 8.7e-08 | 7.1e-08 | 8.600E-08 | 1.453E-08 | 7.100E-08 | 1.000E-07 |
| 12310 | 616 | D616I | A | 1e-07 | 4.4e-08 | 1.6e-08 | 5.333E-08 | 4.277E-08 | 1.600E-08 | 1.000E-07 |
| 12311 | 616 | D616L | A | 3.5e-08 | 6.3e-08 | 4e-08 | 4.600E-08 | 1.493E-08 | 3.500E-08 | 6.300E-08 |
| 12312 | 616 | D616K | A | 4.9e-08 | 1.7e-08 | 3.7e-08 | 3.433E-08 | 1.617E-08 | 1.700E-08 | 4.900E-08 |
| 12313 | 616 | D616M | A | 1e-07 | 7.9e-08 | 7.4e-08 | 8.433E-08 | 1.380E-08 | 7.400E-08 | 1.000E-07 |
| 12314 | 616 | D616F | A | 1.1e-08 | 3.8e-08 | 2.9e-08 | 2.600E-08 | 1.375E-08 | 1.100E-08 | 3.800E-08 |
| 12315 | 616 | D616P | A | 1.1e-07 | 2.7e-08 | 1.1e-07 | 8.233E-08 | 4.792E-08 | 2.700E-08 | 1.100E-07 |
| 12316 | 616 | D616S | A | 5.4e-08 | 2.1e-08 | 3.9e-08 | 3.800E-08 | 1.652E-08 | 2.100E-08 | 5.400E-08 |
| 12317 | 616 | D616T | A | 8.3e-09 | 2.9e-08 | 2.6e-08 | 2.110E-08 | 1.119E-08 | 8.300E-09 | 2.900E-08 |
| 12318 | 616 | D616W | A | 5.6e-08 | 7.1e-08 | 1e-07 | 7.567E-08 | 2.237E-08 | 5.600E-08 | 1.000E-07 |
| 12319 | 616 | D616Y | A | 2.7e-08 | 2.2e-08 | 3.8e-08 | 2.900E-08 | 8.185E-09 | 2.200E-08 | 3.800E-08 |
| 12320 | 616 | D616V | A | 7.5e-08 | 3.1e-08 | 2.6e-08 | 4.400E-08 | 2.696E-08 | 2.600E-08 | 7.500E-08 |
| 12321 | 617 | D617A | A | 3.9e-08 | 4.4e-08 | 6.8e-08 | 5.033E-08 | 1.550E-08 | 3.900E-08 | 6.800E-08 |
| 12322 | 617 | D617R | A | 5.2e-08 | 9.7e-09 | 4.3e-08 | 3.490E-08 | 2.228E-08 | 9.700E-09 | 5.200E-08 |
| 12323 | 617 | D617N | A | 3e-08 | 1.2e-07 | 7.6e-08 | 7.533E-08 | 4.500E-08 | 3.000E-08 | 1.200E-07 |
| 12324 | 617 | D617D | A | 3.5e-08 | 7.7e-08 | 2.3e-08 | 4.500E-08 | 2.835E-08 | 2.300E-08 | 7.700E-08 |

| | | | | | | | | | | |
|---|---|---|---|---|---|---|---|---|---|---|
| 12325 | 617 | D617C | A | 5.8e-08 | 4.6e-08 | 7.5e-08 | 5.967E-08 | 1.457E-08 | 4.600E-08 | 7.500E-08 |
| 12326 | 617 | D617E | A | 3.1e-08 | 7.8e-08 | 2.3e-08 | 4.400E-08 | 2.972E-08 | 2.300E-08 | 7.800E-08 |
| 12327 | 617 | D617Q | A | 6.6e-08 | 1.2e-07 | 3.6e-08 | 7.400E-08 | 4.257E-08 | 3.600E-08 | 1.200E-07 |
| 12328 | 617 | D617G | A | 4.4e-08 | 1.8e-08 | 4.7e-08 | 3.633E-08 | 1.595E-08 | 1.800E-08 | 4.700E-08 |
| 12329 | 617 | D617H | A | 6e-08 | 6.3e-08 | 2.5e-08 | 4.933E-08 | 2.113E-08 | 2.500E-08 | 6.300E-08 |
| 12330 | 617 | D617I | A | 2.2e-08 | 6.7e-08 | 4.1e-08 | 4.333E-08 | 2.259E-08 | 2.200E-08 | 6.700E-08 |
| 12331 | 617 | D617L | A | 1.1e-07 | 1.2e-07 | 2.1e-08 | 8.367E-08 | 5.450E-08 | 2.100E-08 | 1.200E-07 |
| 12332 | 617 | D617K | A | 4.7e-08 | 6.2e-08 | 4.8e-08 | 5.233E-08 | 8.386E-09 | 4.700E-08 | 6.200E-08 |
| 12333 | 617 | D617M | A | 2.4e-08 | 2.4e-08 | 2.9e-08 | 2.567E-08 | 2.887E-09 | 2.400E-08 | 2.900E-08 |
| 12334 | 617 | D617F | A | 4.9e-08 | 2.8e-08 | 2e-08 | 3.233E-08 | 1.498E-08 | 2.000E-08 | 4.900E-08 |
| 12335 | 617 | D617P | A | 3.8e-08 | 7.8e-08 | 2.1e-08 | 4.567E-08 | 2.926E-08 | 2.100E-08 | 7.800E-08 |
| 12336 | 617 | D617S | A | 3.4e-08 | 7.1e-08 | 1.4e-08 | 3.967E-08 | 2.892E-08 | 1.400E-08 | 7.100E-08 |
| 12337 | 617 | D617T | A | 3.5e-08 | 2.6e-08 | 1.1e-07 | 5.700E-08 | 4.612E-08 | 2.600E-08 | 1.100E-07 |
| 12338 | 617 | D617W | A | 4.5e-08 | 8.7e-08 | 7.6e-08 | 6.933E-08 | 2.178E-08 | 4.500E-08 | 8.700E-08 |
| 12339 | 617 | D617Y | A | 3.8e-08 | 1.2e-07 | 4.4e-08 | 6.733E-08 | 4.571E-08 | 3.800E-08 | 1.200E-07 |
| 12340 | 617 | D617V | A | 5.7e-08 | 2.1e-08 | 7.1e-08 | 4.967E-08 | 2.579E-08 | 2.100E-08 | 7.100E-08 |
| 12341 | 618 | K618A | A | 5.1e-08 | 1.1e-08 | 6.7e-08 | 4.300E-08 | 2.884E-08 | 1.100E-08 | 6.700E-08 |
| 12342 | 618 | K618R | A | 2.4e-08 | 3.9e-08 | 3.4e-08 | 3.233E-08 | 7.638E-09 | 2.400E-08 | 3.900E-08 |
| 12343 | 618 | K618N | A | 3.8e-08 | 2.6e-08 | 1e-07 | 5.467E-08 | 3.972E-08 | 2.600E-08 | 1.000E-07 |
| 12344 | 618 | K618D | A | 6.1e-08 | 5e-08 | 7.7e-08 | 6.267E-08 | 1.358E-08 | 5.000E-08 | 7.700E-08 |
| 12345 | 618 | K618C | A | 3.4e-08 | 8.7e-08 | 5.2e-08 | 5.767E-08 | 2.695E-08 | 3.400E-08 | 8.700E-08 |
| 12346 | 618 | K618E | A | 8.2e-08 | 5.9e-08 | 4e-08 | 6.033E-08 | 2.103E-08 | 4.000E-08 | 8.200E-08 |
| 12347 | 618 | K618Q | A | 2.7e-08 | 3.3e-08 | 6.6e-08 | 4.200E-08 | 2.100E-08 | 2.700E-08 | 6.600E-08 |
| 12348 | 618 | K618G | A | 3e-08 | 3.7e-08 | 6.6e-08 | 4.433E-08 | 1.909E-08 | 3.000E-08 | 6.600E-08 |
| 12349 | 618 | K618H | A | 2.6e-08 | 7.7e-08 | 1.3e-07 | 7.767E-08 | 5.200E-08 | 2.600E-08 | 1.300E-07 |
| 12350 | 618 | K618I | A | 5.3e-08 | 7.9e-09 | 2.6e-08 | 2.897E-08 | 2.270E-08 | 7.900E-09 | 5.300E-08 |
| 12351 | 618 | K618L | A | 4.3e-08 | 2.3e-08 | 6e-08 | 4.200E-08 | 1.852E-08 | 2.300E-08 | 6.000E-08 |
| 12352 | 618 | K618K | A | 7.2e-08 | 9.5e-08 | 5.5e-08 | 7.400E-08 | 2.007E-08 | 5.500E-08 | 9.500E-08 |
| 12353 | 618 | K618M | A | 8.4e-08 | 9.1e-08 | 5.1e-08 | 7.533E-08 | 2.136E-08 | 5.100E-08 | 9.100E-08 |
| 12354 | 618 | K618F | A | 4.8e-08 | 5.2e-08 | 5.7e-08 | 5.233E-08 | 4.509E-09 | 4.800E-08 | 5.700E-08 |
| 12355 | 618 | K618P | A | 2.8e-08 | 3.4e-08 | 6.3e-08 | 4.167E-08 | 1.872E-08 | 2.800E-08 | 6.300E-08 |
| 12356 | 618 | K618S | A | 2.8e-08 | 5.3e-08 | 4.3e-08 | 4.133E-08 | 1.258E-08 | 2.800E-08 | 5.300E-08 |
| 12357 | 618 | K618T | A | 2.4e-08 | 2.7e-08 | 6.8e-08 | 3.967E-08 | 2.458E-08 | 2.400E-08 | 6.800E-08 |
| 12358 | 618 | K618W | A | 2.8e-08 | 4.4e-08 | 5.4e-08 | 4.200E-08 | 1.311E-08 | 2.800E-08 | 5.400E-08 |
| 12359 | 618 | K618Y | A | 4.7e-08 | 2.6e-08 | 3.4e-08 | 3.567E-08 | 1.060E-08 | 2.600E-08 | 4.700E-08 |
| 12360 | 618 | K618V | A | 6.1e-08 | 6.1e-08 | 1.2e-07 | 8.067E-08 | 3.406E-08 | 6.100E-08 | 1.200E-07 |
| 12361 | 619 | G619A | A | 3.7e-08 | 6.2e-08 | 4.4e-08 | 4.767E-08 | 1.290E-08 | 3.700E-08 | 6.200E-08 |
| 12362 | 619 | G619R | A | 5.2e-08 | 2.3e-08 | 2.8e-08 | 3.433E-08 | 1.550E-08 | 2.300E-08 | 5.200E-08 |
| 12363 | 619 | G619N | A | 5e-08 | 2.3e-08 | 1.5e-08 | 2.933E-08 | 1.834E-08 | 1.500E-08 | 5.000E-08 |
| 12364 | 619 | G619D | A | 8.6e-08 | 1.8e-08 | 4.1e-08 | 4.833E-08 | 3.459E-08 | 1.800E-08 | 8.600E-08 |
| 12365 | 619 | G619C | A | 2.3e-08 | 4.5e-08 | 3.9e-08 | 3.567E-08 | 1.137E-08 | 2.300E-08 | 4.500E-08 |
| 12366 | 619 | G619E | A | 5.9e-08 | 4.4e-08 | 3.4e-08 | 4.567E-08 | 1.258E-08 | 3.400E-08 | 5.900E-08 |
| 12367 | 619 | G619Q | A | 3.9e-08 | 3.7e-08 | 2.3e-08 | 3.300E-08 | 8.718E-09 | 2.300E-08 | 3.900E-08 |
| 12368 | 619 | G619G | A | 2.2e-08 | 8.1e-08 | 1.9e-08 | 4.067E-08 | 3.496E-08 | 1.900E-08 | 8.100E-08 |
| 12369 | 619 | G619H | A | 9.9e-08 | 5.8e-08 | 2.5e-08 | 6.067E-08 | 3.707E-08 | 2.500E-08 | 9.900E-08 |
| 12370 | 619 | G619I | A | 4.5e-08 | 2.7e-08 | 2.3e-08 | 3.167E-08 | 1.172E-08 | 2.300E-08 | 4.500E-08 |
| 12371 | 619 | G619L | A | 7.1e-08 | 2.7e-08 | 1.2e-07 | 7.267E-08 | 4.652E-08 | 2.700E-08 | 1.200E-07 |
| 12372 | 619 | G619K | A | 1.5e-07 | 5.3e-08 | 5e-08 | 8.433E-08 | 5.689E-08 | 5.000E-08 | 1.500E-07 |
| 12373 | 619 | G619M | A | 8.1e-08 | 1.6e-07 | 2.7e-08 | 8.933E-08 | 6.689E-08 | 2.700E-08 | 1.600E-07 |
| 12374 | 619 | G619F | A | 4.8e-08 | 4e-08 | 1.4e-07 | 7.600E-08 | 5.557E-08 | 4.000E-08 | 1.400E-07 |
| 12375 | 619 | G619P | A | 5.2e-08 | 1e-08 | 3.3e-08 | 3.167E-08 | 2.103E-08 | 1.000E-08 | 5.200E-08 |
| 12376 | 619 | G619S | A | 5.9e-08 | 4.9e-08 | 9.3e-09 | 3.910E-08 | 2.629E-08 | 9.300E-09 | 5.900E-08 |
| 12377 | 619 | G619T | A | 1.3e-08 | 1.2e-07 | 1.4e-07 | 9.100E-08 | 6.829E-08 | 1.300E-08 | 1.400E-07 |
| 12378 | 619 | G619W | A | 3.6e-08 | 5.3e-08 | 3.5e-08 | 4.133E-08 | 1.012E-08 | 3.500E-08 | 5.300E-08 |
| 12379 | 619 | G619Y | A | 2.9e-08 | 1.9e-08 | 9.7e-08 | 4.833E-08 | 4.244E-08 | 1.900E-08 | 9.700E-08 |
| 12380 | 619 | G619V | A | 5.8e-08 | 2.5e-08 | 2.9e-08 | 3.733E-08 | 1.801E-08 | 2.500E-08 | 5.800E-08 |
| 12381 | 620 | C620A | A | 1.3e-07 | 4.4e-08 | 6.2e-08 | 7.867E-08 | 4.536E-08 | 4.400E-08 | 1.300E-07 |
| 12382 | 620 | C620R | A | 4e-08 | 1.2e-07 | 1.2e-07 | 9.333E-08 | 4.619E-08 | 4.000E-08 | 1.200E-07 |
| 12383 | 620 | C620N | A | 9.2e-08 | 2.4e-08 | 5e-08 | 5.533E-08 | 3.431E-08 | 2.400E-08 | 9.200E-08 |
| 12384 | 620 | C620D | A | 7.5e-08 | 3.4e-08 | 7.8e-08 | 6.233E-08 | 2.458E-08 | 3.400E-08 | 7.800E-08 |
| 12385 | 620 | C620C | A | 6.6e-08 | 3.5e-08 | 3.7e-08 | 4.600E-08 | 1.735E-08 | 3.500E-08 | 6.600E-08 |
| 12386 | 620 | C620E | A | 4.8e-08 | 3.3e-08 | 4.9e-08 | 4.333E-08 | 8.963E-09 | 3.300E-08 | 4.900E-08 |
| 12387 | 620 | C620Q | A | 4.3e-08 | 2.3e-08 | 3.1e-08 | 3.233E-08 | 1.007E-08 | 2.300E-08 | 4.300E-08 |
| 12388 | 620 | C620G | A | 5.4e-08 | 1.2e-07 | 7.1e-08 | 8.167E-08 | 3.427E-08 | 5.400E-08 | 1.200E-07 |
| 12389 | 620 | C620H | A | 7.4e-08 | 3.8e-08 | 5.2e-08 | 5.467E-08 | 1.815E-08 | 3.800E-08 | 7.400E-08 |
| 12390 | 620 | C620I | A | 5.4e-08 | 4.4e-08 | 2.1e-08 | 3.967E-08 | 1.692E-08 | 2.100E-08 | 5.400E-08 |
| 12391 | 620 | C620L | A | 2.7e-08 | 4.2e-08 | 3e-08 | 3.300E-08 | 7.937E-09 | 2.700E-08 | 4.200E-08 |
| 12392 | 620 | C620K | A | 7.4e-08 | 6.2e-08 | 2.9e-08 | 5.500E-08 | 2.330E-08 | 2.900E-08 | 7.400E-08 |
| 12393 | 620 | C620M | A | 2.7e-08 | 4.3e-08 | 7.7e-08 | 4.900E-08 | 2.553E-08 | 2.700E-08 | 7.700E-08 |
| 12394 | 620 | C620F | A | 5.8e-08 | 3e-08 | 4.2e-08 | 4.333E-08 | 1.405E-08 | 3.000E-08 | 5.800E-08 |
| 12395 | 620 | C620P | A | 3.7e-08 | 1.6e-08 | 2.4e-08 | 2.567E-08 | 1.060E-08 | 1.600E-08 | 3.700E-08 |
| 12396 | 620 | C620S | A | 4.4e-08 | 4.2e-08 | 2.2e-08 | 3.600E-08 | 1.217E-08 | 2.200E-08 | 4.400E-08 |
| 12397 | 620 | C620T | A | 2.7e-08 | 2.2e-08 | 4.9e-08 | 3.267E-08 | 1.436E-08 | 2.200E-08 | 4.900E-08 |
| 12398 | 620 | C620W | A | 1.1e-07 | 4.4e-08 | 1.1e-07 | 8.800E-08 | 3.811E-08 | 4.400E-08 | 1.100E-07 |
| 12399 | 620 | C620Y | A | 5.2e-08 | 4.3e-08 | 6.5e-08 | 5.333E-08 | 1.106E-08 | 4.300E-08 | 6.500E-08 |
| 12400 | 620 | C620V | A | 2.7e-08 | 1.4e-08 | 3.6e-08 | 2.567E-08 | 1.106E-08 | 1.400E-08 | 3.600E-08 |
| 12401 | 621 | P621A | A | 8.1e-08 | 1.8e-08 | 7.4e-08 | 5.767E-08 | 3.453E-08 | 1.800E-08 | 8.100E-08 |
| 12402 | 621 | P621R | A | 3.6e-08 | 8.5e-08 | 2.9e-08 | 5.000E-08 | 3.051E-08 | 2.900E-08 | 8.500E-08 |
| 12403 | 621 | P621N | A | 6e-08 | 3.7e-08 | 6.1e-08 | 5.267E-08 | 1.358E-08 | 3.700E-08 | 6.100E-08 |
| 12404 | 621 | P621D | A | 3.6e-08 | 1.5e-08 | 4.5e-08 | 3.200E-08 | 1.539E-08 | 1.500E-08 | 4.500E-08 |
| 12405 | 621 | P621C | A | 3.1e-08 | 1e-07 | 4.7e-08 | 5.933E-08 | 3.612E-08 | 3.100E-08 | 1.000E-07 |
| 12406 | 621 | P621E | A | 3.4e-08 | 3.8e-08 | 1.5e-08 | 2.900E-08 | 1.229E-08 | 1.500E-08 | 3.800E-08 |
| 12407 | 621 | P621Q | A | 3.4e-08 | 2.8e-08 | 5.5e-08 | 3.900E-08 | 1.418E-08 | 2.800E-08 | 5.500E-08 |
| 12408 | 621 | P621G | A | 4.4e-08 | 2.6e-08 | 3e-08 | 3.333E-08 | 9.452E-09 | 2.600E-08 | 4.400E-08 |
| 12409 | 621 | P621H | A | 6e-08 | 3.8e-08 | 2.9e-08 | 4.233E-08 | 1.595E-08 | 2.900E-08 | 6.000E-08 |

| | | | | | | | | | | |
|---|---|---|---|---|---|---|---|---|---|---|
| 12410 | 621 | P621I | A | 3.2e-08 | 2.4e-08 | 2.1e-08 | 2.567E-08 | 5.686E-09 | 2.100E-08 | 3.200E-08 |
| 12411 | 621 | P621L | A | 1.8e-08 | 3.4e-08 | 1.3e-08 | 2.167E-08 | 1.097E-08 | 1.300E-08 | 3.400E-08 |
| 12412 | 621 | P621K | A | 4.7e-08 | 5.8e-08 | 2.3e-08 | 4.267E-08 | 1.790E-08 | 2.300E-08 | 5.800E-08 |
| 12413 | 621 | P621M | A | 3.6e-08 | 6.3e-08 | 4.1e-08 | 4.667E-08 | 1.436E-08 | 3.600E-08 | 6.300E-08 |
| 12414 | 621 | P621F | A | 3.1e-08 | 4.9e-08 | 1.4e-07 | 7.333E-08 | 5.843E-08 | 3.100E-08 | 1.400E-07 |
| 12415 | 621 | P621P | A | 7.8e-08 | 1.9e-08 | 3.2e-08 | 4.300E-08 | 3.100E-08 | 1.900E-08 | 7.800E-08 |
| 12416 | 621 | P621S | A | 2.5e-08 | 4.6e-08 | 7.5e-08 | 4.867E-08 | 2.511E-08 | 2.500E-08 | 7.500E-08 |
| 12417 | 621 | P621T | A | 1.2e-08 | 8.6e-08 | 4.4e-08 | 4.733E-08 | 3.711E-08 | 1.200E-08 | 8.600E-08 |
| 12418 | 621 | P621W | A | 5.7e-08 | 1.3e-07 | 3.1e-08 | 7.267E-08 | 5.133E-08 | 3.100E-08 | 1.300E-07 |
| 12419 | 621 | P621Y | A | 1.5e-08 | 5.4e-08 | 1.2e-07 | 6.300E-08 | 5.308E-08 | 1.500E-08 | 1.200E-07 |
| 12420 | 621 | P621V | A | 4.8e-08 | 4.4e-08 | 6.1e-08 | 5.100E-08 | 8.888E-09 | 4.400E-08 | 6.100E-08 |
| 12421 | 622 | A622A | A | 2.2e-08 | 2.6e-08 | 1.6e-08 | 2.133E-08 | 5.033E-09 | 1.600E-08 | 2.600E-08 |
| 12422 | 622 | A622R | A | 6.5e-08 | 1.8e-08 | 5.6e-08 | 4.633E-08 | 2.495E-08 | 1.800E-08 | 6.500E-08 |
| 12423 | 622 | A622N | A | 1.6e-08 | 3.5e-08 | 6e-08 | 3.700E-08 | 2.207E-08 | 1.600E-08 | 6.000E-08 |
| 12424 | 622 | A622D | A | 3.1e-08 | 2.5e-08 | 6.5e-08 | 4.033E-08 | 2.157E-08 | 2.500E-08 | 6.500E-08 |
| 12425 | 622 | A622C | A | 2.9e-08 | 2.9e-08 | 4.1e-08 | 3.300E-08 | 6.928E-09 | 2.900E-08 | 4.100E-08 |
| 12426 | 622 | A622E | A | 4e-08 | 4.4e-08 | 5.1e-08 | 4.500E-08 | 5.568E-09 | 4.000E-08 | 5.100E-08 |
| 12427 | 622 | A622Q | A | 2.3e-08 | 1.2e-07 | 7.8e-08 | 7.367E-08 | 4.864E-08 | 2.300E-08 | 1.200E-07 |
| 12428 | 622 | A622G | A | 2.6e-08 | 3.3e-08 | 1.2e-07 | 5.967E-08 | 5.237E-08 | 2.600E-08 | 1.200E-07 |
| 12429 | 622 | A622H | A | 2e-08 | 3.8e-08 | 3e-08 | 2.933E-08 | 9.018E-09 | 2.000E-08 | 3.800E-08 |
| 12430 | 622 | A622I | A | 6.2e-08 | 4.3e-08 | 9.2e-08 | 6.567E-08 | 2.470E-08 | 4.300E-08 | 9.200E-08 |
| 12431 | 622 | A622L | A | 3.4e-08 | 9.1e-08 | 1.4e-08 | 4.633E-08 | 3.995E-08 | 1.400E-08 | 9.100E-08 |
| 12432 | 622 | A622K | A | 5.8e-08 | 5.5e-08 | 7.4e-08 | 6.233E-08 | 1.021E-08 | 5.500E-08 | 7.400E-08 |
| 12433 | 622 | A622M | A | 7.8e-08 | 1.2e-08 | 3.1e-08 | 4.033E-08 | 3.398E-08 | 1.200E-08 | 7.800E-08 |
| 12434 | 622 | A622F | A | 3e-08 | 4.9e-08 | 4.3e-08 | 4.067E-08 | 9.713E-09 | 3.000E-08 | 4.900E-08 |
| 12435 | 622 | A622P | A | 1.6e-08 | 3.5e-08 | 2.5e-08 | 2.533E-08 | 9.504E-09 | 1.600E-08 | 3.500E-08 |
| 12436 | 622 | A622S | A | 6.7e-08 | 1.1e-07 | 2.9e-08 | 6.867E-08 | 4.053E-08 | 2.900E-08 | 1.100E-07 |
| 12437 | 622 | A622T | A | 5.2e-08 | 1.6e-08 | 8.1e-08 | 4.967E-08 | 3.256E-08 | 1.600E-08 | 8.100E-08 |
| 12438 | 622 | A622W | A | 6.3e-08 | 4.4e-08 | 3.2e-08 | 4.633E-08 | 1.563E-08 | 3.200E-08 | 6.300E-08 |
| 12439 | 622 | A622Y | A | 7.9e-08 | 2.9e-08 | 2.7e-08 | 4.500E-08 | 2.946E-08 | 2.700E-08 | 7.900E-08 |
| 12440 | 622 | A622V | A | 3e-08 | 1e-07 | 5.8e-09 | 4.527E-08 | 4.892E-08 | 5.800E-09 | 1.000E-07 |
| 12441 | 623 | D623A | B | 3.2e-08 | 4.6e-08 | 5.8e-08 | 4.533E-08 | 1.301E-08 | 3.200E-08 | 5.800E-08 |
| 12442 | 623 | D623R | B | 5.3e-08 | 5.8e-08 | 6.1e-08 | 5.733E-08 | 4.041E-09 | 5.300E-08 | 6.100E-08 |
| 12443 | 623 | D623N | B | 1e-08 | 3.3e-08 | 2.4e-08 | 2.233E-08 | 1.159E-08 | 1.000E-08 | 3.300E-08 |
| 12444 | 623 | D623D | B | 4.4e-08 | 2.2e-08 | 2.5e-08 | 3.033E-08 | 1.193E-08 | 2.200E-08 | 4.400E-08 |
| 12445 | 623 | D623C | B | 1.2e-08 | 5.1e-08 | 9.2e-08 | 5.167E-08 | 4.000E-08 | 1.200E-08 | 9.200E-08 |
| 12446 | 623 | D623E | B | 7.7e-08 | 2.2e-08 | 5.2e-08 | 5.033E-08 | 2.754E-08 | 2.200E-08 | 7.700E-08 |
| 12447 | 623 | D623Q | B | 1.2e-08 | 2.8e-08 | 2.6e-08 | 2.200E-08 | 8.718E-09 | 1.200E-08 | 2.800E-08 |
| 12448 | 623 | D623G | B | 1.2e-08 | 5.4e-08 | 2.4e-08 | 3.000E-08 | 2.163E-08 | 1.200E-08 | 5.400E-08 |
| 12449 | 623 | D623H | B | 7.6e-08 | 3.3e-08 | 6.9e-08 | 5.933E-08 | 2.307E-08 | 3.300E-08 | 7.600E-08 |
| 12450 | 623 | D623I | B | 5.1e-08 | 8.8e-08 | 8.4e-08 | 7.433E-08 | 2.031E-08 | 5.100E-08 | 8.800E-08 |
| 12451 | 623 | D623L | B | 2.3e-08 | 1.9e-08 | 7.7e-08 | 3.967E-08 | 3.239E-08 | 1.900E-08 | 7.700E-08 |
| 12452 | 623 | D623K | B | 1.3e-08 | 7.5e-08 | 4e-08 | 4.267E-08 | 3.109E-08 | 1.300E-08 | 7.500E-08 |
| 12453 | 623 | D623M | B | 5.3e-08 | 2.6e-08 | 8.1e-08 | 5.333E-08 | 2.750E-08 | 2.600E-08 | 8.100E-08 |
| 12454 | 623 | D623F | B | 4.9e-08 | 1.3e-08 | 4.2e-08 | 3.467E-08 | 1.909E-08 | 1.300E-08 | 4.900E-08 |
| 12455 | 623 | D623P | B | 3.9e-08 | 8.3e-08 | 4.3e-08 | 5.500E-08 | 2.433E-08 | 3.900E-08 | 8.300E-08 |
| 12456 | 623 | D623S | B | 3.4e-08 | 6.9e-08 | 1.2e-07 | 7.433E-08 | 4.325E-08 | 3.400E-08 | 1.200E-07 |
| 12457 | 623 | D623T | B | 1e-07 | 7.1e-08 | 4.8e-08 | 7.300E-08 | 2.606E-08 | 4.800E-08 | 1.000E-07 |
| 12458 | 623 | D623W | B | 6.5e-08 | 5.6e-08 | 6.1e-08 | 6.067E-08 | 4.509E-09 | 5.600E-08 | 6.500E-08 |
| 12459 | 623 | D623Y | B | 1e-07 | 3.6e-08 | 4.6e-08 | 6.067E-08 | 3.443E-08 | 3.600E-08 | 1.000E-07 |
| 12460 | 623 | D623V | B | 2.9e-08 | 3.2e-08 | 9.3e-08 | 5.133E-08 | 3.612E-08 | 2.900E-08 | 9.300E-08 |
| 12461 | 624 | I624A | B | 2.2e-07 | 6.3e-08 | 4.4e-08 | 1.090E-07 | 9.660E-08 | 4.400E-08 | 2.200E-07 |
| 12462 | 624 | I624R | B | 9.3e-09 | 3.7e-08 | 8.6e-08 | 4.410E-08 | 3.884E-08 | 9.300E-09 | 8.600E-08 |
| 12463 | 624 | I624N | B | 2.5e-08 | 3.9e-08 | 2.3e-08 | 2.900E-08 | 8.718E-09 | 2.300E-08 | 3.900E-08 |
| 12464 | 624 | I624D | B | 6.3e-08 | 5.5e-08 | 1.6e-08 | 4.467E-08 | 2.515E-08 | 1.600E-08 | 6.300E-08 |
| 12465 | 624 | I624C | B | 7e-08 | 1.2e-07 | 3.1e-08 | 7.367E-08 | 4.461E-08 | 3.100E-08 | 1.200E-07 |
| 12466 | 624 | I624E | B | 3.3e-08 | 5.3e-08 | 9.9e-08 | 6.167E-08 | 3.384E-08 | 3.300E-08 | 9.900E-08 |
| 12467 | 624 | I624Q | B | 7.5e-08 | 4.3e-08 | 1.3e-07 | 8.267E-08 | 4.400E-08 | 4.300E-08 | 1.300E-07 |
| 12468 | 624 | I624G | B | 6.3e-08 | 3.1e-08 | 1e-07 | 6.467E-08 | 3.453E-08 | 3.100E-08 | 1.000E-07 |
| 12469 | 624 | I624H | B | 4.4e-08 | 3.1e-08 | 4.2e-08 | 3.900E-08 | 7.000E-09 | 3.100E-08 | 4.400E-08 |
| 12470 | 624 | I624I | B | 9.5e-09 | 2.9e-08 | 3.7e-08 | 2.517E-08 | 1.415E-08 | 9.500E-09 | 3.700E-08 |
| 12471 | 624 | I624L | B | 3.9e-08 | 1.9e-08 | 4.7e-08 | 3.500E-08 | 1.442E-08 | 1.900E-08 | 4.700E-08 |
| 12472 | 624 | I624K | B | 2e-08 | 3.5e-08 | 4.1e-08 | 3.200E-08 | 1.082E-08 | 2.000E-08 | 4.100E-08 |
| 12473 | 624 | I624M | B | 5.9e-08 | 2.8e-08 | 3e-08 | 3.900E-08 | 1.735E-08 | 2.800E-08 | 5.900E-08 |
| 12474 | 624 | I624F | B | 5e-08 | 3.6e-08 | 4.6e-08 | 4.400E-08 | 7.211E-09 | 3.600E-08 | 5.000E-08 |
| 12475 | 624 | I624P | B | 2.3e-08 | 6.8e-08 | 6.7e-09 | 3.257E-08 | 3.175E-08 | 6.700E-09 | 6.800E-08 |
| 12476 | 624 | I624S | B | 4.2e-08 | 5.6e-08 | 6.6e-08 | 5.467E-08 | 1.206E-08 | 4.200E-08 | 6.600E-08 |
| 12477 | 624 | I624T | B | 2.2e-08 | 2.3e-08 | 2.2e-08 | 2.233E-08 | 5.774E-10 | 2.200E-08 | 2.300E-08 |
| 12478 | 624 | I624W | B | 3.1e-08 | 2e-08 | 9.7e-08 | 4.933E-08 | 4.165E-08 | 2.000E-08 | 9.700E-08 |
| 12479 | 624 | I624Y | B | 4.6e-08 | 5.9e-08 | 4.3e-08 | 4.933E-08 | 8.505E-09 | 4.300E-08 | 5.900E-08 |
| 12480 | 624 | I624V | B | 8.3e-08 | 1.3e-08 | 5.3e-08 | 4.967E-08 | 3.512E-08 | 1.300E-08 | 8.300E-08 |
| 12481 | 625 | Q625A | B | 3.2e-08 | 4.8e-08 | 6.3e-08 | 4.767E-08 | 1.550E-08 | 3.200E-08 | 6.300E-08 |
| 12482 | 625 | Q625R | B | 3.9e-08 | 2.6e-08 | 4.4e-08 | 3.633E-08 | 9.292E-09 | 2.600E-08 | 4.400E-08 |
| 12483 | 625 | Q625N | B | 2.5e-08 | 1.3e-07 | 8.1e-08 | 7.867E-08 | 5.254E-08 | 2.500E-08 | 1.300E-07 |
| 12484 | 625 | Q625D | B | 2e-08 | 3.9e-08 | 7e-09 | 2.200E-08 | 1.609E-08 | 7.000E-09 | 3.900E-08 |
| 12485 | 625 | Q625C | B | 4.5e-08 | 3.6e-08 | 1.9e-08 | 3.333E-08 | 1.320E-08 | 1.900E-08 | 4.500E-08 |
| 12486 | 625 | Q625E | B | 6.7e-08 | 4.8e-08 | 7e-08 | 6.167E-08 | 1.193E-08 | 4.800E-08 | 7.000E-08 |
| 12487 | 625 | Q625Q | B | 3.2e-08 | 7.6e-08 | 5.3e-08 | 5.367E-08 | 2.201E-08 | 3.200E-08 | 7.600E-08 |
| 12488 | 625 | Q625G | B | 4.6e-08 | 3.7e-08 | 1.3e-07 | 7.100E-08 | 5.129E-08 | 3.700E-08 | 1.300E-07 |
| 12489 | 625 | Q625H | B | 3.8e-08 | 1.8e-08 | 6.5e-08 | 4.033E-08 | 2.359E-08 | 1.800E-08 | 6.500E-08 |
| 12490 | 625 | Q625I | B | 7.1e-08 | 2.5e-08 | 7.5e-08 | 5.700E-08 | 2.778E-08 | 2.500E-08 | 7.500E-08 |
| 12491 | 625 | Q625L | B | 3.7e-08 | 2.8e-08 | 3.9e-08 | 3.467E-08 | 5.859E-09 | 2.800E-08 | 3.900E-08 |
| 12492 | 625 | Q625K | B | 3.7e-08 | 8.1e-08 | 1.6e-07 | 9.267E-08 | 6.232E-08 | 3.700E-08 | 1.600E-07 |
| 12493 | 625 | Q625M | B | 7e-08 | 8e-08 | 3.5e-08 | 6.167E-08 | 2.363E-08 | 3.500E-08 | 8.000E-08 |
| 12494 | 625 | Q625F | B | 5.9e-08 | 2.8e-08 | 9.9e-08 | 6.200E-08 | 3.559E-08 | 2.800E-08 | 9.900E-08 |

| | | | | | | | | | | |
|---|---|---|---|---|---|---|---|---|---|---|
| 12495 | 625 | Q625P | B | 7.5e-08 | 9.3e-08 | 2.2e-08 | 6.333E-08 | 3.691E-08 | 2.200E-08 | 9.300E-08 |
| 12496 | 625 | Q625S | B | 4.7e-08 | 3.7e-08 | 4.2e-08 | 4.200E-08 | 5.000E-09 | 3.700E-08 | 4.700E-08 |
| 12497 | 625 | Q625T | B | 3.6e-08 | 5e-08 | 1.5e-07 | 7.867E-08 | 6.217E-08 | 3.600E-08 | 1.500E-07 |
| 12498 | 625 | Q625W | B | 8.4e-08 | 8e-08 | 2.9e-08 | 6.433E-08 | 3.066E-08 | 2.900E-08 | 8.400E-08 |
| 12499 | 625 | Q625Y | B | 2.2e-08 | 4.3e-08 | 7.8e-08 | 4.767E-08 | 2.829E-08 | 2.200E-08 | 7.800E-08 |
| 12500 | 625 | Q625V | B | 1.2e-08 | 4.3e-08 | 2.4e-08 | 2.633E-08 | 1.563E-08 | 1.200E-08 | 4.300E-08 |
| 12501 | 626 | M626A | B | 3.6e-08 | 3.5e-08 | 2.8e-08 | 3.300E-08 | 4.359E-09 | 2.800E-08 | 3.600E-08 |
| 12502 | 626 | M626R | B | 3.3e-08 | 7.2e-08 | 5.5e-08 | 5.333E-08 | 1.955E-08 | 3.300E-08 | 7.200E-08 |
| 12503 | 626 | M626N | B | 6.6e-08 | 1.5e-08 | 2.9e-08 | 3.667E-08 | 2.635E-08 | 1.500E-08 | 6.600E-08 |
| 12504 | 626 | M626D | B | 9.9e-08 | 3.4e-08 | 5.2e-08 | 6.167E-08 | 3.356E-08 | 3.400E-08 | 9.900E-08 |
| 12505 | 626 | M626C | B | 5.2e-08 | 5.6e-08 | 2.8e-08 | 4.533E-08 | 1.514E-08 | 2.800E-08 | 5.600E-08 |
| 12506 | 626 | M626E | B | 5.4e-08 | 3.6e-08 | 3.3e-08 | 4.100E-08 | 1.136E-08 | 3.300E-08 | 5.400E-08 |
| 12507 | 626 | M626Q | B | 1.6e-08 | 1.8e-08 | 3.7e-08 | 2.367E-08 | 1.159E-08 | 1.600E-08 | 3.700E-08 |
| 12508 | 626 | M626G | B | 5.6e-08 | 4.3e-08 | 2.1e-08 | 4.000E-08 | 1.769E-08 | 2.100E-08 | 5.600E-08 |
| 12509 | 626 | M626H | B | 2.5e-08 | 3.1e-08 | 5.1e-08 | 3.567E-08 | 1.361E-08 | 2.500E-08 | 5.100E-08 |
| 12510 | 626 | M626I | B | 5.5e-08 | 4.4e-08 | 6e-08 | 5.300E-08 | 8.185E-09 | 4.400E-08 | 6.000E-08 |
| 12511 | 626 | M626L | B | 7.4e-08 | 8.4e-08 | 3.6e-08 | 6.467E-08 | 2.532E-08 | 3.600E-08 | 8.400E-08 |
| 12512 | 626 | M626K | B | 9.2e-08 | 6.3e-08 | 2.2e-08 | 5.900E-08 | 3.517E-08 | 2.200E-08 | 9.200E-08 |
| 12513 | 626 | M626M | B | 6.5e-08 | 9.2e-09 | 3.2e-08 | 3.540E-08 | 2.805E-08 | 9.200E-09 | 6.500E-08 |
| 12514 | 626 | M626F | B | 1.1e-08 | 6.7e-08 | 2.2e-08 | 3.333E-08 | 2.967E-08 | 1.100E-08 | 6.700E-08 |
| 12515 | 626 | M626P | B | 6.6e-08 | 3.7e-08 | 2.1e-08 | 4.133E-08 | 2.281E-08 | 2.100E-08 | 6.600E-08 |
| 12516 | 626 | M626S | B | 3.3e-08 | 2.4e-08 | 3.7e-08 | 3.133E-08 | 6.658E-09 | 2.400E-08 | 3.700E-08 |
| 12517 | 626 | M626T | B | 4.5e-08 | 4.5e-08 | 4.8e-08 | 4.600E-08 | 1.732E-09 | 4.500E-08 | 4.800E-08 |
| 12518 | 626 | M626W | B | 1.8e-08 | 2.2e-08 | 4.2e-08 | 2.733E-08 | 1.286E-08 | 1.800E-08 | 4.200E-08 |
| 12519 | 626 | M626Y | B | 9.7e-08 | 5.2e-08 | 5.9e-08 | 6.933E-08 | 2.421E-08 | 5.200E-08 | 9.700E-08 |
| 12520 | 626 | M626V | B | 6.1e-08 | 6.8e-08 | 6.5e-08 | 6.467E-08 | 3.512E-09 | 6.100E-08 | 6.800E-08 |
| 12521 | 627 | T627A | B | 2.4e-08 | 2.6e-08 | 2.8e-08 | 2.600E-08 | 2.000E-09 | 2.400E-08 | 2.800E-08 |
| 12522 | 627 | T627R | B | 5.2e-08 | 3.7e-08 | 5.6e-08 | 4.833E-08 | 1.002E-08 | 3.700E-08 | 5.600E-08 |
| 12523 | 627 | T627N | B | 1.7e-08 | 3.3e-08 | 9.4e-08 | 4.800E-08 | 4.063E-08 | 1.700E-08 | 9.400E-08 |
| 12524 | 627 | T627D | B | 3.3e-08 | 9e-08 | 6.5e-08 | 6.267E-08 | 2.857E-08 | 3.300E-08 | 9.000E-08 |
| 12525 | 627 | T627C | B | 2.7e-08 | 1.3e-07 | 3.4e-08 | 6.367E-08 | 5.755E-08 | 2.700E-08 | 1.300E-07 |
| 12526 | 627 | T627E | B | 2.3e-08 | 3.9e-08 | 2.4e-08 | 2.867E-08 | 8.963E-09 | 2.300E-08 | 3.900E-08 |
| 12527 | 627 | T627Q | B | 1.8e-07 | 3.5e-08 | 7.6e-09 | 7.420E-08 | 9.264E-08 | 7.600E-09 | 1.800E-07 |
| 12528 | 627 | T627G | B | 8.2e-08 | 6.7e-08 | 8e-08 | 7.633E-08 | 8.145E-09 | 6.700E-08 | 8.200E-08 |
| 12529 | 627 | T627H | B | 5e-08 | 1.1e-08 | 2.1e-08 | 2.733E-08 | 2.026E-08 | 1.100E-08 | 5.000E-08 |
| 12530 | 627 | T627I | B | 4.6e-08 | 5.8e-08 | 4.7e-08 | 5.033E-08 | 6.658E-09 | 4.600E-08 | 5.800E-08 |
| 12531 | 627 | T627L | B | 1.4e-08 | 2.4e-08 | 5.1e-08 | 2.967E-08 | 1.914E-08 | 1.400E-08 | 5.100E-08 |
| 12532 | 627 | T627K | B | 4.7e-08 | 7.9e-08 | 8.8e-08 | 7.133E-08 | 2.155E-08 | 4.700E-08 | 8.800E-08 |
| 12533 | 627 | T627M | B | 3.1e-08 | 5.6e-08 | 7e-08 | 5.233E-08 | 1.976E-08 | 3.100E-08 | 7.000E-08 |
| 12534 | 627 | T627F | B | 7.5e-08 | 6.5e-08 | 2.2e-08 | 5.400E-08 | 2.816E-08 | 2.200E-08 | 7.500E-08 |
| 12535 | 627 | T627P | B | 2.2e-08 | 4.6e-08 | 2.7e-08 | 3.167E-08 | 1.266E-08 | 2.200E-08 | 4.600E-08 |
| 12536 | 627 | T627S | B | 5.5e-08 | 7e-08 | 7e-08 | 6.500E-08 | 8.660E-09 | 5.500E-08 | 7.000E-08 |
| 12537 | 627 | T627T | B | 1.9e-08 | 1.7e-08 | 2.9e-08 | 2.167E-08 | 6.429E-09 | 1.700E-08 | 2.900E-08 |
| 12538 | 627 | T627W | B | 5.3e-08 | 6.6e-08 | 4e-08 | 5.300E-08 | 1.300E-08 | 4.000E-08 | 6.600E-08 |
| 12539 | 627 | T627Y | B | 1.7e-08 | 5.4e-08 | 5.3e-08 | 4.133E-08 | 2.108E-08 | 1.700E-08 | 5.400E-08 |
| 12540 | 627 | T627V | B | 5.8e-08 | 6.4e-08 | 4.6e-08 | 5.600E-08 | 9.165E-09 | 4.600E-08 | 6.400E-08 |
| 12541 | 628 | Q628A | B | 2.2e-08 | 6.9e-08 | 1.3e-07 | 7.367E-08 | 5.415E-08 | 2.200E-08 | 1.300E-07 |
| 12542 | 628 | Q628R | B | 5.1e-08 | 1.5e-08 | 3.3e-08 | 3.300E-08 | 1.800E-08 | 1.500E-08 | 5.100E-08 |
| 12543 | 628 | Q628N | B | 3.6e-08 | 1.2e-07 | 4.3e-08 | 6.633E-08 | 4.661E-08 | 3.600E-08 | 1.200E-07 |
| 12544 | 628 | Q628D | B | 4.5e-08 | 6.2e-08 | 3.6e-08 | 4.767E-08 | 1.320E-08 | 3.600E-08 | 6.200E-08 |
| 12545 | 628 | Q628C | B | 4.9e-08 | 1.2e-07 | 2.2e-08 | 6.367E-08 | 5.062E-08 | 2.200E-08 | 1.200E-07 |
| 12546 | 628 | Q628E | B | 6.6e-08 | 4.7e-08 | 6.6e-08 | 5.967E-08 | 1.097E-08 | 4.700E-08 | 6.600E-08 |
| 12547 | 628 | Q628Q | B | 3.8e-08 | 6.6e-09 | 5.8e-08 | 3.420E-08 | 2.591E-08 | 6.600E-09 | 5.800E-08 |
| 12548 | 628 | Q628G | B | 4.8e-08 | 3e-08 | 3.1e-08 | 3.633E-08 | 1.012E-08 | 3.000E-08 | 4.800E-08 |
| 12549 | 628 | Q628H | B | 3.8e-08 | 4.3e-08 | 3.1e-08 | 3.733E-08 | 6.028E-09 | 3.100E-08 | 4.300E-08 |
| 12550 | 628 | Q628I | B | 6.1e-08 | 9.2e-08 | 2.7e-08 | 6.000E-08 | 3.251E-08 | 2.700E-08 | 9.200E-08 |
| 12551 | 628 | Q628L | B | 3.9e-08 | 2.8e-08 | 2.7e-08 | 3.133E-08 | 6.658E-09 | 2.700E-08 | 3.900E-08 |
| 12552 | 628 | Q628K | B | 6e-08 | 4.4e-08 | 7.4e-08 | 5.933E-08 | 1.501E-08 | 4.400E-08 | 7.400E-08 |
| 12553 | 628 | Q628M | B | 6.9e-08 | 3.4e-08 | 3.1e-08 | 4.467E-08 | 2.113E-08 | 3.100E-08 | 6.900E-08 |
| 12554 | 628 | Q628F | B | 6.4e-08 | 6.9e-08 | 1e-07 | 7.767E-08 | 1.950E-08 | 6.400E-08 | 1.000E-07 |
| 12555 | 628 | Q628P | B | 5.1e-08 | 7.2e-08 | 6.1e-08 | 6.133E-08 | 1.050E-08 | 5.100E-08 | 7.200E-08 |
| 12556 | 628 | Q628S | B | 1.1e-08 | 3.8e-08 | 3.4e-08 | 2.767E-08 | 1.457E-08 | 1.100E-08 | 3.800E-08 |
| 12557 | 628 | Q628T | B | 5.1e-08 | 7.3e-08 | 1.3e-07 | 8.467E-08 | 4.077E-08 | 5.100E-08 | 1.300E-07 |
| 12558 | 628 | Q628W | B | 2.3e-08 | 3.3e-08 | 5.1e-08 | 3.567E-08 | 1.419E-08 | 2.300E-08 | 5.100E-08 |
| 12559 | 628 | Q628Y | B | 6.7e-08 | 9.9e-08 | 6.3e-08 | 7.633E-08 | 1.973E-08 | 6.300E-08 | 9.900E-08 |
| 12560 | 628 | Q628V | B | 1e-08 | 4e-08 | 6e-08 | 3.667E-08 | 2.517E-08 | 1.000E-08 | 6.000E-08 |
| 12561 | 629 | S629A | B | 1.8e-08 | 5.4e-08 | 3.8e-08 | 3.667E-08 | 1.804E-08 | 1.800E-08 | 5.400E-08 |
| 12562 | 629 | S629R | B | 1.3e-07 | 4.2e-08 | 3.5e-08 | 6.900E-08 | 5.294E-08 | 3.500E-08 | 1.300E-07 |
| 12563 | 629 | S629N | B | 5.3e-08 | 6.1e-08 | 4e-08 | 5.133E-08 | 1.060E-08 | 4.000E-08 | 6.100E-08 |
| 12564 | 629 | S629D | B | 6.9e-08 | 8e-08 | 3.4e-08 | 6.100E-08 | 2.402E-08 | 3.400E-08 | 8.000E-08 |
| 12565 | 629 | S629C | B | 2.5e-08 | 1.1e-07 | 6.6e-08 | 6.700E-08 | 4.251E-08 | 2.500E-08 | 1.100E-07 |
| 12566 | 629 | S629E | B | 9e-08 | 5.4e-08 | 5.5e-08 | 6.633E-08 | 2.050E-08 | 5.400E-08 | 9.000E-08 |
| 12567 | 629 | S629Q | B | 8.1e-08 | 6.7e-08 | 5e-08 | 6.600E-08 | 1.552E-08 | 5.000E-08 | 8.100E-08 |
| 12568 | 629 | S629G | B | 2.2e-08 | 9.1e-08 | 5e-08 | 5.433E-08 | 3.470E-08 | 2.200E-08 | 9.100E-08 |
| 12569 | 629 | S629H | B | 3.2e-08 | 2.9e-08 | 4.8e-08 | 3.633E-08 | 1.021E-08 | 2.900E-08 | 4.800E-08 |
| 12570 | 629 | S629I | B | 4.4e-08 | 6.7e-08 | 1.6e-08 | 4.233E-08 | 2.554E-08 | 1.600E-08 | 6.700E-08 |
| 12571 | 629 | S629L | B | 9.3e-08 | 7.1e-08 | 3e-08 | 6.467E-08 | 3.197E-08 | 3.000E-08 | 9.300E-08 |
| 12572 | 629 | S629K | B | 6.1e-08 | 4.1e-08 | 6.5e-08 | 5.567E-08 | 1.286E-08 | 4.100E-08 | 6.500E-08 |
| 12573 | 629 | S629M | B | 7.7e-08 | 9.9e-09 | 5.2e-08 | 4.630E-08 | 3.391E-08 | 9.900E-09 | 7.700E-08 |
| 12574 | 629 | S629F | B | 3.1e-08 | 5.9e-08 | 1.9e-08 | 3.633E-08 | 2.053E-08 | 1.900E-08 | 5.900E-08 |
| 12575 | 629 | S629P | B | 9e-08 | 4.4e-08 | 3.1e-08 | 5.500E-08 | 3.100E-08 | 3.100E-08 | 9.000E-08 |
| 12576 | 629 | S629S | B | 3.2e-08 | 4.5e-08 | 7.3e-09 | 2.810E-08 | 1.915E-08 | 7.300E-09 | 4.500E-08 |
| 12577 | 629 | S629T | B | 3.3e-08 | 6.5e-08 | 7.1e-08 | 5.633E-08 | 2.043E-08 | 3.300E-08 | 7.100E-08 |
| 12578 | 629 | S629W | B | 3e-08 | 9e-08 | 4.7e-08 | 5.567E-08 | 3.092E-08 | 3.000E-08 | 9.000E-08 |
| 12579 | 629 | S629Y | B | 6.4e-08 | 3.2e-08 | 5.3e-08 | 4.967E-08 | 1.626E-08 | 3.200E-08 | 6.400E-08 |

| | | | | | | | | | | |
|---|---|---|---|---|---|---|---|---|---|---|
| 12580 | 629 | S629V | B | 6e-08 | 4.7e-08 | 4.8e-08 | 5.167E-08 | 7.234E-09 | 4.700E-08 | 6.000E-08 |
| 12581 | 630 | P630A | B | 3.5e-08 | 5.5e-08 | 3.1e-08 | 4.033E-08 | 1.286E-08 | 3.100E-08 | 5.500E-08 |
| 12582 | 630 | P630R | B | 2.5e-07 | 3.8e-08 | 7.8e-08 | 1.220E-07 | 1.126E-07 | 3.800E-08 | 2.500E-07 |
| 12583 | 630 | P630N | B | 6.1e-08 | 4.9e-08 | 4.6e-08 | 5.200E-08 | 7.937E-09 | 4.600E-08 | 6.100E-08 |
| 12584 | 630 | P630D | B | 6.1e-08 | 7.5e-08 | 5e-08 | 6.200E-08 | 1.253E-08 | 5.000E-08 | 7.500E-08 |
| 12585 | 630 | P630C | B | 1.1e-07 | 6.5e-08 | 6.2e-08 | 7.900E-08 | 2.689E-08 | 6.200E-08 | 1.100E-07 |
| 12586 | 630 | P630E | B | 7.2e-08 | 3.9e-08 | 7.8e-08 | 6.300E-08 | 2.100E-08 | 3.900E-08 | 7.800E-08 |
| 12587 | 630 | P630Q | B | 7.4e-08 | 2.5e-08 | 5.2e-08 | 5.033E-08 | 2.454E-08 | 2.500E-08 | 7.400E-08 |
| 12588 | 630 | P630G | B | 4.1e-08 | 1.4e-07 | 3e-08 | 7.033E-08 | 6.058E-08 | 3.000E-08 | 1.400E-07 |
| 12589 | 630 | P630H | B | 5.7e-08 | 2.4e-08 | 4.2e-08 | 4.100E-08 | 1.652E-08 | 2.400E-08 | 5.700E-08 |
| 12590 | 630 | P630I | B | 7.7e-08 | 3e-08 | 2.7e-07 | 1.257E-07 | 1.272E-07 | 3.000E-08 | 2.700E-07 |
| 12591 | 630 | P630L | B | 3.9e-08 | 2.6e-08 | 8.7e-08 | 5.067E-08 | 3.213E-08 | 2.600E-08 | 8.700E-08 |
| 12592 | 630 | P630K | B | 3.1e-08 | 2.3e-08 | 4.6e-08 | 3.333E-08 | 1.168E-08 | 2.300E-08 | 4.600E-08 |
| 12593 | 630 | P630M | B | 1.3e-08 | 4.2e-08 | 4.7e-08 | 3.400E-08 | 1.836E-08 | 1.300E-08 | 4.700E-08 |
| 12594 | 630 | P630F | B | 3.1e-08 | 1e-07 | 2.3e-08 | 5.133E-08 | 4.234E-08 | 2.300E-08 | 1.000E-07 |
| 12595 | 630 | P630P | B | 7.6e-08 | 1.8e-08 | 4.6e-08 | 4.667E-08 | 2.901E-08 | 1.800E-08 | 7.600E-08 |
| 12596 | 630 | P630S | B | 2.5e-08 | 3.4e-08 | 9.8e-08 | 5.233E-08 | 3.980E-08 | 2.500E-08 | 9.800E-08 |
| 12597 | 630 | P630T | B | 4.3e-08 | 2.9e-08 | 7.2e-08 | 4.800E-08 | 2.193E-08 | 2.900E-08 | 7.200E-08 |
| 12598 | 630 | P630W | B | 7.5e-08 | 8.3e-09 | 3.1e-08 | 3.810E-08 | 3.391E-08 | 8.300E-09 | 7.500E-08 |
| 12599 | 630 | P630Y | B | 3.4e-08 | 7.6e-08 | 6.7e-08 | 5.900E-08 | 2.211E-08 | 3.400E-08 | 7.600E-08 |
| 12600 | 630 | P630V | B | 1.7e-08 | 2.3e-08 | 6.1e-08 | 3.367E-08 | 2.386E-08 | 1.700E-08 | 6.100E-08 |
| 12601 | 631 | S631A | B | 1.1e-08 | 8.9e-08 | 1.3e-07 | 7.667E-08 | 6.045E-08 | 1.100E-08 | 1.300E-07 |
| 12602 | 631 | S631R | B | 2.7e-08 | 9.7e-08 | 9.4e-08 | 7.267E-08 | 3.958E-08 | 2.700E-08 | 9.700E-08 |
| 12603 | 631 | S631N | B | 2e-08 | 2.6e-08 | 3.2e-08 | 2.600E-08 | 6.000E-09 | 2.000E-08 | 3.200E-08 |
| 12604 | 631 | S631D | B | 1.4e-07 | 4.3e-08 | 5e-08 | 7.767E-08 | 5.410E-08 | 4.300E-08 | 1.400E-07 |
| 12605 | 631 | S631C | B | 5.1e-08 | 8.2e-08 | 8.1e-08 | 7.133E-08 | 1.762E-08 | 5.100E-08 | 8.200E-08 |
| 12606 | 631 | S631E | B | 5.1e-08 | 9.4e-08 | 3.6e-08 | 6.033E-08 | 3.011E-08 | 3.600E-08 | 9.400E-08 |
| 12607 | 631 | S631Q | B | 7.2e-08 | 5.9e-08 | 2.3e-08 | 5.133E-08 | 2.538E-08 | 2.300E-08 | 7.200E-08 |
| 12608 | 631 | S631G | B | 5.1e-08 | 1e-07 | 6.4e-08 | 7.167E-08 | 2.538E-08 | 5.100E-08 | 1.000E-07 |
| 12609 | 631 | S631H | B | 8.9e-08 | 7.1e-08 | 6.4e-08 | 7.467E-08 | 1.290E-08 | 6.400E-08 | 8.900E-08 |
| 12610 | 631 | S631I | B | 2.5e-08 | 5.7e-08 | 2.5e-08 | 3.567E-08 | 1.848E-08 | 2.500E-08 | 5.700E-08 |
| 12611 | 631 | S631L | B | 4.1e-08 | 3.1e-08 | 3.7e-08 | 3.633E-08 | 5.033E-09 | 3.100E-08 | 4.100E-08 |
| 12612 | 631 | S631K | B | 3.1e-08 | 2.5e-08 | 4e-08 | 3.200E-08 | 7.550E-09 | 2.500E-08 | 4.000E-08 |
| 12613 | 631 | S631M | B | 8.7e-08 | 1.7e-08 | 7.1e-08 | 5.833E-08 | 3.668E-08 | 1.700E-08 | 8.700E-08 |
| 12614 | 631 | S631F | B | 7.6e-08 | 1.8e-08 | 4.4e-08 | 4.600E-08 | 2.905E-08 | 1.800E-08 | 7.600E-08 |
| 12615 | 631 | S631P | B | 4.4e-08 | 3.4e-08 | 3.6e-08 | 3.800E-08 | 5.292E-09 | 3.400E-08 | 4.400E-08 |
| 12616 | 631 | S631S | B | 5.9e-08 | 3e-08 | 9.9e-09 | 3.297E-08 | 2.468E-08 | 9.900E-09 | 5.900E-08 |
| 12617 | 631 | S631T | B | 4e-08 | 6.5e-08 | 2.9e-08 | 4.467E-08 | 1.845E-08 | 2.900E-08 | 6.500E-08 |
| 12618 | 631 | S631W | B | 4.1e-08 | 2.1e-08 | 4.4e-08 | 3.533E-08 | 1.250E-08 | 2.100E-08 | 4.400E-08 |
| 12619 | 631 | S631Y | B | 6.9e-08 | 1.4e-07 | 6.1e-08 | 9.000E-08 | 4.349E-08 | 6.100E-08 | 1.400E-07 |
| 12620 | 631 | S631V | B | 1.7e-08 | 7.4e-08 | 1.2e-07 | 7.033E-08 | 5.160E-08 | 1.700E-08 | 1.200E-07 |
| 12621 | 632 | S632A | B | 4e-08 | 8.6e-08 | 3.3e-08 | 5.300E-08 | 2.879E-08 | 3.300E-08 | 8.600E-08 |
| 12622 | 632 | S632R | B | 5.2e-08 | 2.6e-08 | 4.7e-08 | 4.167E-08 | 1.380E-08 | 2.600E-08 | 5.200E-08 |
| 12623 | 632 | S632N | B | 4.9e-08 | 1.7e-08 | 8.8e-08 | 5.133E-08 | 3.556E-08 | 1.700E-08 | 8.800E-08 |
| 12624 | 632 | S632D | B | 3.4e-08 | 3.3e-08 | 3.5e-08 | 3.400E-08 | 1.000E-09 | 3.300E-08 | 3.500E-08 |
| 12625 | 632 | S632C | B | 7.2e-08 | 4e-08 | 6.7e-08 | 5.967E-08 | 1.721E-08 | 4.000E-08 | 7.200E-08 |
| 12626 | 632 | S632E | B | 2.5e-08 | 6.5e-08 | 3.1e-08 | 4.033E-08 | 2.157E-08 | 2.500E-08 | 6.500E-08 |
| 12627 | 632 | S632Q | B | 1.9e-08 | 5.1e-08 | 2.9e-08 | 3.300E-08 | 1.637E-08 | 1.900E-08 | 5.100E-08 |
| 12628 | 632 | S632G | B | 4.4e-08 | 4.9e-08 | 8.3e-08 | 5.867E-08 | 2.122E-08 | 4.400E-08 | 8.300E-08 |
| 12629 | 632 | S632H | B | 3.4e-08 | 5.8e-08 | 2.9e-08 | 4.033E-08 | 1.550E-08 | 2.900E-08 | 5.800E-08 |
| 12630 | 632 | S632I | B | 3.1e-08 | 1.6e-08 | 1.7e-08 | 2.133E-08 | 8.386E-09 | 1.600E-08 | 3.100E-08 |
| 12631 | 632 | S632L | B | 4.5e-08 | 8.8e-08 | 4.8e-08 | 6.033E-08 | 2.401E-08 | 4.500E-08 | 8.800E-08 |
| 12632 | 632 | S632K | B | 3.2e-08 | 3.9e-08 | 3.5e-08 | 3.533E-08 | 3.512E-09 | 3.200E-08 | 3.900E-08 |
| 12633 | 632 | S632M | B | 4.5e-08 | 6.3e-08 | 5.5e-08 | 5.433E-08 | 9.018E-09 | 4.500E-08 | 6.300E-08 |
| 12634 | 632 | S632F | B | 9.5e-08 | 2.9e-08 | 6.5e-08 | 6.300E-08 | 3.305E-08 | 2.900E-08 | 9.500E-08 |
| 12635 | 632 | S632P | B | 1.1e-07 | 5.6e-08 | 7.2e-08 | 7.933E-08 | 2.774E-08 | 5.600E-08 | 1.100E-07 |
| 12636 | 632 | S632S | B | 4.4e-08 | 5e-08 | 6.3e-08 | 5.233E-08 | 9.713E-09 | 4.400E-08 | 6.300E-08 |
| 12637 | 632 | S632T | B | 5.8e-08 | 3.4e-08 | 6e-08 | 5.067E-08 | 1.447E-08 | 3.400E-08 | 6.000E-08 |
| 12638 | 632 | S632W | B | 5.1e-08 | 4.7e-08 | 3e-08 | 4.267E-08 | 1.115E-08 | 3.000E-08 | 5.100E-08 |
| 12639 | 632 | S632Y | B | 3.7e-08 | 2.9e-08 | 4.5e-08 | 3.700E-08 | 8.000E-09 | 2.900E-08 | 4.500E-08 |
| 12640 | 632 | S632V | B | 1.9e-08 | 7.6e-08 | 4.4e-08 | 4.633E-08 | 2.857E-08 | 1.900E-08 | 7.600E-08 |
| 12641 | 633 | L633A | B | 2.8e-08 | 1.2e-08 | 2.6e-08 | 2.200E-08 | 8.718E-09 | 1.200E-08 | 2.800E-08 |
| 12642 | 633 | L633R | B | 5.4e-08 | 2.2e-08 | 3.2e-08 | 3.600E-08 | 1.637E-08 | 2.200E-08 | 5.400E-08 |
| 12643 | 633 | L633N | B | 4.2e-08 | 5.1e-08 | 4.5e-08 | 4.600E-08 | 4.583E-09 | 4.200E-08 | 5.100E-08 |
| 12644 | 633 | L633D | B | 4.7e-08 | 2.8e-08 | 4.4e-08 | 3.967E-08 | 1.021E-08 | 2.800E-08 | 4.700E-08 |
| 12645 | 633 | L633C | B | 1.1e-07 | 5.2e-08 | 1.1e-07 | 9.067E-08 | 3.349E-08 | 5.200E-08 | 1.100E-07 |
| 12646 | 633 | L633E | B | 7.4e-08 | 1.4e-08 | 3.6e-08 | 4.133E-08 | 3.035E-08 | 1.400E-08 | 7.400E-08 |
| 12647 | 633 | L633Q | B | 2.9e-08 | 3.1e-08 | 5.1e-08 | 3.700E-08 | 1.217E-08 | 2.900E-08 | 5.100E-08 |
| 12648 | 633 | L633G | B | 2.4e-08 | 2.3e-08 | 8.3e-08 | 4.333E-08 | 3.436E-08 | 2.300E-08 | 8.300E-08 |
| 12649 | 633 | L633H | B | 2.2e-08 | 1.1e-08 | 5e-08 | 2.767E-08 | 2.011E-08 | 1.100E-08 | 5.000E-08 |
| 12650 | 633 | L633I | B | 9e-08 | 8.5e-08 | 9.7e-08 | 9.067E-08 | 6.028E-09 | 8.500E-08 | 9.700E-08 |
| 12651 | 633 | L633L | B | 5.7e-08 | 1.2e-07 | 3.7e-08 | 7.133E-08 | 4.332E-08 | 3.700E-08 | 1.200E-07 |
| 12652 | 633 | L633K | B | 8.7e-08 | 1.7e-08 | 2.1e-08 | 4.167E-08 | 3.931E-08 | 1.700E-08 | 8.700E-08 |
| 12653 | 633 | L633M | B | 6.9e-08 | 5.7e-08 | 7.3e-08 | 6.633E-08 | 8.327E-09 | 5.700E-08 | 7.300E-08 |
| 12654 | 633 | L633F | B | 2.6e-08 | 1.6e-08 | 5.3e-08 | 3.167E-08 | 1.914E-08 | 1.600E-08 | 5.300E-08 |
| 12655 | 633 | L633P | B | 8.6e-08 | 4.4e-08 | 3.5e-08 | 5.500E-08 | 2.722E-08 | 3.500E-08 | 8.600E-08 |
| 12656 | 633 | L633S | B | 1.8e-08 | 5.1e-08 | 3.4e-08 | 3.433E-08 | 1.650E-08 | 1.800E-08 | 5.100E-08 |
| 12657 | 633 | L633T | B | 4.2e-08 | 1.1e-07 | 9.5e-08 | 8.233E-08 | 3.573E-08 | 4.200E-08 | 1.100E-07 |
| 12658 | 633 | L633W | B | 2e-08 | 9.2e-08 | 1.2e-08 | 4.133E-08 | 4.406E-08 | 1.200E-08 | 9.200E-08 |
| 12659 | 633 | L633Y | B | 1.8e-08 | 7.8e-08 | 5.3e-08 | 4.967E-08 | 3.014E-08 | 1.800E-08 | 7.800E-08 |
| 12660 | 633 | L633V | B | 3.5e-08 | 2.7e-08 | 2.9e-08 | 3.033E-08 | 4.163E-09 | 2.700E-08 | 3.500E-08 |
| 12661 | 634 | S634A | B | 3.1e-08 | 5.1e-08 | 2e-08 | 3.400E-08 | 1.572E-08 | 2.000E-08 | 5.100E-08 |
| 12662 | 634 | S634R | B | 8e-08 | 5.3e-08 | 1.3e-07 | 8.767E-08 | 3.907E-08 | 5.300E-08 | 1.300E-07 |
| 12663 | 634 | S634N | B | 2.7e-08 | 5.9e-08 | 9e-08 | 5.867E-08 | 3.150E-08 | 2.700E-08 | 9.000E-08 |
| 12664 | 634 | S634D | B | 3.4e-08 | 1.3e-08 | 2.1e-08 | 2.267E-08 | 1.060E-08 | 1.300E-08 | 3.400E-08 |

| | | | | | | | | | | |
|---|---|---|---|---|---|---|---|---|---|---|
| 12665 | 634 | S634C | B | 2.6e-08 | 1.4e-08 | 4.3e-08 | 2.767E-08 | 1.457E-08 | 1.400E-08 | 4.300E-08 |
| 12666 | 634 | S634E | B | 1.1e-07 | 3e-08 | 5.4e-08 | 6.467E-08 | 4.105E-08 | 3.000E-08 | 1.100E-07 |
| 12667 | 634 | S634Q | B | 5.9e-08 | 6.3e-08 | 1.2e-08 | 4.467E-08 | 2.836E-08 | 1.200E-08 | 6.300E-08 |
| 12668 | 634 | S634G | B | 5.3e-08 | 3.7e-08 | 1.1e-07 | 6.667E-08 | 3.837E-08 | 3.700E-08 | 1.100E-07 |
| 12669 | 634 | S634H | B | 3.4e-08 | 6.5e-08 | 2.7e-08 | 4.200E-08 | 2.022E-08 | 2.700E-08 | 6.500E-08 |
| 12670 | 634 | S634I | B | 5e-08 | 1e-07 | 2.6e-08 | 5.867E-08 | 3.775E-08 | 2.600E-08 | 1.000E-07 |
| 12671 | 634 | S634L | B | 5.7e-08 | 2.5e-08 | 3.9e-08 | 4.033E-08 | 1.604E-08 | 2.500E-08 | 5.700E-08 |
| 12672 | 634 | S634K | B | 1e-08 | 4.7e-08 | 6.3e-08 | 4.000E-08 | 2.718E-08 | 1.000E-08 | 6.300E-08 |
| 12673 | 634 | S634M | B | 2.8e-08 | 2e-08 | 2.3e-08 | 2.367E-08 | 4.041E-09 | 2.000E-08 | 2.800E-08 |
| 12674 | 634 | S634F | B | 2.6e-08 | 3.4e-08 | 9.2e-08 | 5.067E-08 | 3.602E-08 | 2.600E-08 | 9.200E-08 |
| 12675 | 634 | S634P | B | 3.5e-08 | 6.5e-08 | 5.3e-08 | 5.100E-08 | 1.510E-08 | 3.500E-08 | 6.500E-08 |
| 12676 | 634 | S634S | B | 4.6e-08 | 4.5e-08 | 6.8e-08 | 5.300E-08 | 1.300E-08 | 4.500E-08 | 6.800E-08 |
| 12677 | 634 | S634T | B | 6.6e-08 | 5.3e-08 | 1.7e-08 | 4.533E-08 | 2.538E-08 | 1.700E-08 | 6.600E-08 |
| 12678 | 634 | S634W | B | 4.1e-08 | 9.7e-08 | 5.2e-08 | 6.333E-08 | 2.967E-08 | 4.100E-08 | 9.700E-08 |
| 12679 | 634 | S634Y | B | 1.1e-07 | 6.6e-08 | 1.9e-08 | 6.500E-08 | 4.551E-08 | 1.900E-08 | 1.100E-07 |
| 12680 | 634 | S634V | B | 8.9e-08 | 1e-07 | 2.6e-08 | 7.167E-08 | 3.993E-08 | 2.600E-08 | 1.000E-07 |
| 12681 | 635 | A635A | B | 2.3e-08 | 6.7e-08 | 4.2e-08 | 4.400E-08 | 2.207E-08 | 2.300E-08 | 6.700E-08 |
| 12682 | 635 | A635R | B | 1.1e-07 | 1.4e-07 | 9.9e-08 | 1.163E-07 | 2.122E-08 | 9.900E-08 | 1.400E-07 |
| 12683 | 635 | A635N | B | 1.9e-08 | 7.3e-08 | 4.8e-08 | 4.667E-08 | 2.702E-08 | 1.900E-08 | 7.300E-08 |
| 12684 | 635 | A635D | B | 3.4e-08 | 4.4e-08 | 1.3e-08 | 3.033E-08 | 1.582E-08 | 1.300E-08 | 4.400E-08 |
| 12685 | 635 | A635C | B | 2.1e-08 | 2e-08 | 4.9e-08 | 3.000E-08 | 1.646E-08 | 2.000E-08 | 4.900E-08 |
| 12686 | 635 | A635E | B | 5.4e-08 | 1.2e-07 | 3.5e-08 | 6.967E-08 | 4.461E-08 | 3.500E-08 | 1.200E-07 |
| 12687 | 635 | A635Q | B | 3e-08 | 9.8e-08 | 1.1e-08 | 4.633E-08 | 4.574E-08 | 1.100E-08 | 9.800E-08 |
| 12688 | 635 | A635G | B | 3.9e-08 | 4.6e-08 | 3.7e-08 | 4.067E-08 | 4.726E-09 | 3.700E-08 | 4.600E-08 |
| 12689 | 635 | A635H | B | 2.3e-08 | 7.3e-08 | 3.3e-08 | 4.300E-08 | 2.646E-08 | 2.300E-08 | 7.300E-08 |
| 12690 | 635 | A635I | B | 4.8e-08 | 9.3e-08 | 2.9e-08 | 5.667E-08 | 3.287E-08 | 2.900E-08 | 9.300E-08 |
| 12691 | 635 | A635L | B | 1.1e-07 | 2.6e-08 | 1.6e-08 | 5.067E-08 | 5.163E-08 | 1.600E-08 | 1.100E-07 |
| 12692 | 635 | A635K | B | 1e-07 | 1.9e-08 | 3.3e-08 | 5.067E-08 | 4.329E-08 | 1.900E-08 | 1.000E-07 |
| 12693 | 635 | A635M | B | 4.2e-08 | 3.3e-08 | 6.3e-08 | 4.600E-08 | 1.539E-08 | 3.300E-08 | 6.300E-08 |
| 12694 | 635 | A635F | B | 4.7e-08 | 6.3e-08 | 5.4e-08 | 5.467E-08 | 8.021E-09 | 4.700E-08 | 6.300E-08 |
| 12695 | 635 | A635P | B | 5.4e-08 | 4.8e-08 | 3.4e-08 | 4.533E-08 | 1.026E-08 | 3.400E-08 | 5.400E-08 |
| 12696 | 635 | A635S | B | 1.3e-08 | 2.3e-08 | 1.1e-08 | 1.567E-08 | 6.429E-09 | 1.100E-08 | 2.300E-08 |
| 12697 | 635 | A635T | B | 4e-08 | 7e-08 | 5.5e-08 | 5.500E-08 | 1.500E-08 | 4.000E-08 | 7.000E-08 |
| 12698 | 635 | A635W | B | 7.5e-08 | 4e-08 | 2.1e-08 | 4.533E-08 | 2.739E-08 | 2.100E-08 | 7.500E-08 |
| 12699 | 635 | A635Y | B | 9.9e-08 | 2.5e-08 | 2.6e-08 | 5.000E-08 | 4.244E-08 | 2.500E-08 | 9.900E-08 |
| 12700 | 635 | A635V | B | 4.9e-08 | 2.4e-08 | 3.5e-08 | 3.600E-08 | 1.253E-08 | 2.400E-08 | 4.900E-08 |
| 12701 | 636 | S636A | B | 5.7e-08 | 9.7e-08 | 4.9e-08 | 6.767E-08 | 2.572E-08 | 4.900E-08 | 9.700E-08 |
| 12702 | 636 | S636R | B | 1.1e-07 | 4.9e-08 | 8.4e-08 | 8.100E-08 | 3.061E-08 | 4.900E-08 | 1.100E-07 |
| 12703 | 636 | S636N | B | 2.5e-08 | 7.8e-08 | 6.1e-08 | 5.467E-08 | 2.706E-08 | 2.500E-08 | 7.800E-08 |
| 12704 | 636 | S636D | B | 5.5e-08 | 3.8e-08 | 3.1e-08 | 4.133E-08 | 1.234E-08 | 3.100E-08 | 5.500E-08 |
| 12705 | 636 | S636C | B | 2.1e-08 | 4.5e-08 | 4.8e-08 | 3.800E-08 | 1.480E-08 | 2.100E-08 | 4.800E-08 |
| 12706 | 636 | S636E | B | 5.7e-08 | 5e-08 | 6.4e-08 | 5.700E-08 | 7.000E-09 | 5.000E-08 | 6.400E-08 |
| 12707 | 636 | S636Q | B | 2.6e-08 | 5.8e-08 | 2.1e-08 | 3.500E-08 | 2.007E-08 | 2.100E-08 | 5.800E-08 |
| 12708 | 636 | S636G | B | 3e-08 | 9.9e-08 | 4.8e-08 | 5.900E-08 | 3.579E-08 | 3.000E-08 | 9.900E-08 |
| 12709 | 636 | S636H | B | 8.3e-08 | 2.9e-08 | 8.7e-08 | 6.633E-08 | 3.239E-08 | 2.900E-08 | 8.700E-08 |
| 12710 | 636 | S636I | B | 6.1e-08 | 5.8e-08 | 5.6e-08 | 5.833E-08 | 2.517E-09 | 5.600E-08 | 6.100E-08 |
| 12711 | 636 | S636L | B | 3.7e-08 | 5.5e-08 | 6.9e-08 | 5.367E-08 | 1.604E-08 | 3.700E-08 | 6.900E-08 |
| 12712 | 636 | S636K | B | 5.7e-08 | 3.1e-08 | 1.7e-08 | 3.500E-08 | 2.030E-08 | 1.700E-08 | 5.700E-08 |
| 12713 | 636 | S636M | B | 7.6e-08 | 6.4e-08 | 8.4e-09 | 4.947E-08 | 3.607E-08 | 8.400E-09 | 7.600E-08 |
| 12714 | 636 | S636F | B | 5e-08 | 9.5e-08 | 5.6e-08 | 6.700E-08 | 2.443E-08 | 5.000E-08 | 9.500E-08 |
| 12715 | 636 | S636P | B | 3.1e-08 | 1.9e-08 | 5.9e-08 | 3.633E-08 | 2.053E-08 | 1.900E-08 | 5.900E-08 |
| 12716 | 636 | S636S | B | 2.8e-08 | 7.2e-08 | 9.6e-08 | 6.533E-08 | 3.449E-08 | 2.800E-08 | 9.600E-08 |
| 12717 | 636 | S636T | B | 1.1e-07 | 4.3e-08 | 1.9e-08 | 5.733E-08 | 4.716E-08 | 1.900E-08 | 1.100E-07 |
| 12718 | 636 | S636W | B | 4.8e-08 | 2.5e-08 | 4.4e-08 | 3.900E-08 | 1.229E-08 | 2.500E-08 | 4.800E-08 |
| 12719 | 636 | S636Y | B | 6.1e-08 | 1.7e-08 | 1.9e-08 | 3.233E-08 | 2.485E-08 | 1.700E-08 | 6.100E-08 |
| 12720 | 636 | S636V | B | 8e-08 | 3e-08 | 4.5e-08 | 5.167E-08 | 2.566E-08 | 3.000E-08 | 8.000E-08 |
| 12721 | 637 | V637A | B | 3.8e-08 | 5.4e-08 | 7.3e-08 | 5.500E-08 | 1.752E-08 | 3.800E-08 | 7.300E-08 |
| 12722 | 637 | V637R | B | 3.6e-08 | 5.3e-08 | 6.5e-08 | 5.133E-08 | 1.457E-08 | 3.600E-08 | 6.500E-08 |
| 12723 | 637 | V637N | B | 5.9e-08 | 3.6e-08 | 7.1e-08 | 5.533E-08 | 1.779E-08 | 3.600E-08 | 7.100E-08 |
| 12724 | 637 | V637D | B | 1.3e-08 | 2.2e-08 | 2.3e-08 | 1.933E-08 | 5.508E-09 | 1.300E-08 | 2.300E-08 |
| 12725 | 637 | V637C | B | 5.7e-08 | 6e-08 | 4.6e-08 | 5.433E-08 | 7.371E-09 | 4.600E-08 | 6.000E-08 |
| 12726 | 637 | V637E | B | 9.1e-08 | 1.1e-07 | 1.6e-07 | 1.203E-07 | 3.564E-08 | 9.100E-08 | 1.600E-07 |
| 12727 | 637 | V637Q | B | 5.7e-08 | 2e-08 | 7.6e-08 | 5.100E-08 | 2.848E-08 | 2.000E-08 | 7.600E-08 |
| 12728 | 637 | V637G | B | 3.2e-08 | 4e-08 | 1e-08 | 2.733E-08 | 1.553E-08 | 1.000E-08 | 4.000E-08 |
| 12729 | 637 | V637H | B | 7.5e-08 | 3.4e-08 | 3.7e-08 | 4.867E-08 | 2.285E-08 | 3.400E-08 | 7.500E-08 |
| 12730 | 637 | V637I | B | 2.5e-08 | 5.9e-08 | 1.8e-08 | 3.400E-08 | 2.193E-08 | 1.800E-08 | 5.900E-08 |
| 12731 | 637 | V637L | B | 7.8e-08 | 5.6e-08 | 1.6e-08 | 5.000E-08 | 3.143E-08 | 1.600E-08 | 7.800E-08 |
| 12732 | 637 | V637K | B | 1.4e-08 | 6.6e-08 | 7.1e-09 | 2.903E-08 | 3.220E-08 | 7.100E-09 | 6.600E-08 |
| 12733 | 637 | V637M | B | 2.3e-08 | 2.7e-08 | 3.1e-08 | 2.700E-08 | 4.000E-09 | 2.300E-08 | 3.100E-08 |
| 12734 | 637 | V637F | B | 3.8e-08 | 5.1e-08 | 6.1e-08 | 5.000E-08 | 1.153E-08 | 3.800E-08 | 6.100E-08 |
| 12735 | 637 | V637P | B | 3.4e-08 | 3.2e-08 | 4.2e-08 | 3.600E-08 | 5.292E-09 | 3.200E-08 | 4.200E-08 |
| 12736 | 637 | V637S | B | 2.1e-08 | 6.6e-08 | 3.6e-08 | 4.100E-08 | 2.291E-08 | 2.100E-08 | 6.600E-08 |
| 12737 | 637 | V637T | B | 9e-08 | 2.8e-08 | 4e-08 | 5.267E-08 | 3.288E-08 | 2.800E-08 | 9.000E-08 |
| 12738 | 637 | V637W | B | 3e-08 | 2.2e-08 | 7.4e-08 | 4.200E-08 | 2.800E-08 | 2.200E-08 | 7.400E-08 |
| 12739 | 637 | V637Y | B | 1.2e-07 | 1.4e-08 | 4.2e-08 | 5.867E-08 | 5.493E-08 | 1.400E-08 | 1.200E-07 |
| 12740 | 637 | V637V | B | 3.2e-08 | 8.1e-08 | 2.7e-08 | 4.667E-08 | 2.984E-08 | 2.700E-08 | 8.100E-08 |
| 12741 | 638 | G638A | B | 3.8e-08 | 3.2e-08 | 2.6e-08 | 3.200E-08 | 6.000E-09 | 2.600E-08 | 3.800E-08 |
| 12742 | 638 | G638R | B | 7.7e-08 | 6.9e-08 | 1.6e-08 | 5.400E-08 | 3.315E-08 | 1.600E-08 | 7.700E-08 |
| 12743 | 638 | G638N | B | 6.8e-08 | 3.6e-08 | 1.4e-08 | 3.933E-08 | 2.715E-08 | 1.400E-08 | 6.800E-08 |
| 12744 | 638 | G638D | B | 2.2e-08 | 1.4e-07 | 9.4e-08 | 8.533E-08 | 5.948E-08 | 2.200E-08 | 1.400E-07 |
| 12745 | 638 | G638C | B | 1.4e-07 | 5.7e-08 | 2.3e-08 | 7.333E-08 | 6.019E-08 | 2.300E-08 | 1.400E-07 |
| 12746 | 638 | G638E | B | 1.9e-08 | 6.9e-08 | 6.6e-08 | 5.133E-08 | 2.804E-08 | 1.900E-08 | 6.900E-08 |
| 12747 | 638 | G638Q | B | 3.3e-08 | 5.1e-08 | 8.1e-08 | 5.500E-08 | 2.425E-08 | 3.300E-08 | 8.100E-08 |
| 12748 | 638 | G638G | B | 3.8e-08 | 3.3e-08 | 5.7e-08 | 4.267E-08 | 1.266E-08 | 3.300E-08 | 5.700E-08 |
| 12749 | 638 | G638H | B | 6.7e-08 | 3.3e-08 | 3e-08 | 4.333E-08 | 2.055E-08 | 3.000E-08 | 6.700E-08 |

| | | | | | | | | | | |
|---|---|---|---|---|---|---|---|---|---|---|
| 12750 | 638 | G638I | B | 8.4e-09 | 1.6e-08 | 4.2e-08 | 2.213E-08 | 1.762E-08 | 8.400E-09 | 4.200E-08 |
| 12751 | 638 | G638L | B | 7.8e-08 | 6.8e-08 | 3.8e-08 | 6.133E-08 | 2.082E-08 | 3.800E-08 | 7.800E-08 |
| 12752 | 638 | G638K | B | 5.6e-08 | 3.1e-08 | 1.8e-08 | 3.500E-08 | 1.931E-08 | 1.800E-08 | 5.600E-08 |
| 12753 | 638 | G638M | B | 3.2e-08 | 8.5e-08 | 4.1e-08 | 5.267E-08 | 2.836E-08 | 3.200E-08 | 8.500E-08 |
| 12754 | 638 | G638F | B | 2.9e-08 | 1.8e-08 | 3.2e-08 | 2.633E-08 | 7.371E-09 | 1.800E-08 | 3.200E-08 |
| 12755 | 638 | G638P | B | 3.9e-08 | 6.3e-08 | 6.2e-08 | 5.467E-08 | 1.358E-08 | 3.900E-08 | 6.300E-08 |
| 12756 | 638 | G638S | B | 4.6e-08 | 3.5e-08 | 1.6e-08 | 3.233E-08 | 1.518E-08 | 1.600E-08 | 4.600E-08 |
| 12757 | 638 | G638T | B | 9.3e-08 | 2.6e-08 | 2.4e-08 | 4.767E-08 | 3.927E-08 | 2.400E-08 | 9.300E-08 |
| 12758 | 638 | G638W | B | 2.4e-08 | 4.4e-08 | 2.3e-08 | 3.033E-08 | 1.185E-08 | 2.300E-08 | 4.400E-08 |
| 12759 | 638 | G638Y | B | 3.8e-08 | 3.8e-08 | 4.1e-08 | 3.900E-08 | 1.732E-09 | 3.800E-08 | 4.100E-08 |
| 12760 | 638 | G638V | B | 2.4e-08 | 1.1e-08 | 3e-08 | 2.167E-08 | 9.713E-09 | 1.100E-08 | 3.000E-08 |
| 12761 | 639 | D639A | B | 7.1e-08 | 2.5e-08 | 3.8e-08 | 4.467E-08 | 2.371E-08 | 2.500E-08 | 7.100E-08 |
| 12762 | 639 | D639R | B | 2.3e-08 | 2.3e-08 | 5.2e-08 | 3.267E-08 | 1.674E-08 | 2.300E-08 | 5.200E-08 |
| 12763 | 639 | D639N | B | 1.2e-07 | 2.3e-08 | 1.7e-08 | 5.333E-08 | 5.781E-08 | 1.700E-08 | 1.200E-07 |
| 12764 | 639 | D639D | B | 7.9e-08 | 7.7e-08 | 1.9e-08 | 5.833E-08 | 3.408E-08 | 1.900E-08 | 7.900E-08 |
| 12765 | 639 | D639C | B | 2.9e-08 | 2.1e-08 | 3.8e-08 | 2.933E-08 | 8.505E-09 | 2.100E-08 | 3.800E-08 |
| 12766 | 639 | D639E | B | 1.6e-08 | 9.7e-09 | 3.5e-08 | 2.023E-08 | 1.317E-08 | 9.700E-09 | 3.500E-08 |
| 12767 | 639 | D639Q | B | 8.2e-08 | 5.7e-08 | 1.3e-07 | 8.967E-08 | 3.710E-08 | 5.700E-08 | 1.300E-07 |
| 12768 | 639 | D639G | B | 2.3e-08 | 3.1e-08 | 4.3e-08 | 3.233E-08 | 1.007E-08 | 2.300E-08 | 4.300E-08 |
| 12769 | 639 | D639H | B | 1.2e-07 | 8.2e-08 | 1.5e-08 | 7.233E-08 | 5.316E-08 | 1.500E-08 | 1.200E-07 |
| 12770 | 639 | D639I | B | 8.8e-08 | 3.8e-08 | 7.2e-08 | 6.600E-08 | 2.553E-08 | 3.800E-08 | 8.800E-08 |
| 12771 | 639 | D639L | B | 3.9e-08 | 6.5e-08 | 6.5e-08 | 5.633E-08 | 1.501E-08 | 3.900E-08 | 6.500E-08 |
| 12772 | 639 | D639K | B | 7.4e-09 | 6.3e-08 | 1.8e-08 | 2.947E-08 | 2.952E-08 | 7.400E-09 | 6.300E-08 |
| 12773 | 639 | D639M | B | 2.5e-08 | 4.8e-08 | 7.7e-08 | 5.000E-08 | 2.606E-08 | 2.500E-08 | 7.700E-08 |
| 12774 | 639 | D639F | B | 1.3e-08 | 1e-07 | 1.1e-07 | 7.433E-08 | 5.335E-08 | 1.300E-08 | 1.100E-07 |
| 12775 | 639 | D639P | B | 6.2e-08 | 3.5e-08 | 3.8e-08 | 4.500E-08 | 1.480E-08 | 3.500E-08 | 6.200E-08 |
| 12776 | 639 | D639S | B | 3.8e-08 | 4.9e-08 | 5.7e-08 | 4.800E-08 | 9.539E-09 | 3.800E-08 | 5.700E-08 |
| 12777 | 639 | D639T | B | 1.1e-07 | 1.6e-07 | 4e-08 | 1.033E-07 | 6.028E-08 | 4.000E-08 | 1.600E-07 |
| 12778 | 639 | D639W | B | 6.1e-08 | 9.2e-08 | 3.6e-08 | 6.300E-08 | 2.805E-08 | 3.600E-08 | 9.200E-08 |
| 12779 | 639 | D639Y | B | 5.1e-08 | 3.9e-08 | 5.5e-08 | 4.833E-08 | 8.327E-09 | 3.900E-08 | 5.500E-08 |
| 12780 | 639 | D639V | B | 2.4e-08 | 8.3e-08 | 2.7e-08 | 4.467E-08 | 3.323E-08 | 2.400E-08 | 8.300E-08 |
| 12781 | 640 | R640A | B | 3.6e-08 | 3.1e-08 | 7.6e-08 | 4.767E-08 | 2.466E-08 | 3.100E-08 | 7.600E-08 |
| 12782 | 640 | R640R | B | 4.6e-08 | 4.3e-08 | 2.6e-08 | 3.833E-08 | 1.079E-08 | 2.600E-08 | 4.600E-08 |
| 12783 | 640 | R640N | B | 4e-08 | 5.9e-08 | 1.1e-07 | 6.967E-08 | 3.620E-08 | 4.000E-08 | 1.100E-07 |
| 12784 | 640 | R640D | B | 1.3e-07 | 7e-08 | 6.5e-08 | 8.833E-08 | 3.617E-08 | 6.500E-08 | 1.300E-07 |
| 12785 | 640 | R640C | B | 3.6e-08 | 5.4e-08 | 3.5e-08 | 4.167E-08 | 1.069E-08 | 3.500E-08 | 5.400E-08 |
| 12786 | 640 | R640E | B | 3.6e-08 | 2.3e-08 | 2.6e-08 | 2.833E-08 | 6.807E-09 | 2.300E-08 | 3.600E-08 |
| 12787 | 640 | R640Q | B | 5.5e-08 | 1.4e-08 | 2.9e-08 | 3.267E-08 | 2.074E-08 | 1.400E-08 | 5.500E-08 |
| 12788 | 640 | R640G | B | 5.3e-08 | 4.2e-08 | 3.8e-08 | 4.433E-08 | 7.767E-09 | 3.800E-08 | 5.300E-08 |
| 12789 | 640 | R640H | B | 4.1e-08 | 2.2e-08 | 2.4e-08 | 2.900E-08 | 1.044E-08 | 2.200E-08 | 4.100E-08 |
| 12790 | 640 | R640I | B | 7.5e-08 | 2.6e-08 | 5.4e-08 | 5.167E-08 | 2.458E-08 | 2.600E-08 | 7.500E-08 |
| 12791 | 640 | R640L | B | 6.2e-08 | 1.5e-07 | 9e-08 | 1.007E-07 | 4.496E-08 | 6.200E-08 | 1.500E-07 |
| 12792 | 640 | R640K | B | 6.9e-08 | 7.2e-08 | 2.4e-08 | 5.500E-08 | 2.689E-08 | 2.400E-08 | 7.200E-08 |
| 12793 | 640 | R640M | B | 3.8e-08 | 7.3e-08 | 3e-08 | 4.700E-08 | 2.287E-08 | 3.000E-08 | 7.300E-08 |
| 12794 | 640 | R640F | B | 3.8e-08 | 6e-08 | 2e-08 | 3.933E-08 | 2.003E-08 | 2.000E-08 | 6.000E-08 |
| 12795 | 640 | R640P | B | 2.3e-08 | 2.7e-08 | 4.3e-08 | 3.100E-08 | 1.058E-08 | 2.300E-08 | 4.300E-08 |
| 12796 | 640 | R640S | B | 5.8e-08 | 6.8e-08 | 3.7e-08 | 5.433E-08 | 1.582E-08 | 3.700E-08 | 6.800E-08 |
| 12797 | 640 | R640T | B | 1.3e-07 | 2.9e-08 | 7e-08 | 7.633E-08 | 5.080E-08 | 2.900E-08 | 1.300E-07 |
| 12798 | 640 | R640W | B | 1.4e-08 | 2.3e-08 | 5e-08 | 2.900E-08 | 1.873E-08 | 1.400E-08 | 5.000E-08 |
| 12799 | 640 | R640Y | B | 3.3e-08 | 5.7e-08 | 2.8e-08 | 3.933E-08 | 1.550E-08 | 2.800E-08 | 5.700E-08 |
| 12800 | 640 | R640V | B | 9.9e-09 | 6.5e-08 | 5.5e-08 | 4.330E-08 | 2.935E-08 | 9.900E-09 | 6.500E-08 |
| 12801 | 641 | V641A | B | 8.9e-08 | 5.5e-08 | 5.5e-08 | 6.633E-08 | 1.963E-08 | 5.500E-08 | 8.900E-08 |
| 12802 | 641 | V641R | B | 5.7e-08 | 9.9e-08 | 3.3e-08 | 6.300E-08 | 3.341E-08 | 3.300E-08 | 9.900E-08 |
| 12803 | 641 | V641N | B | 5.8e-08 | 3e-07 | 4.1e-08 | 1.330E-07 | 1.449E-07 | 4.100E-08 | 3.000E-07 |
| 12804 | 641 | V641D | B | 2e-08 | 7.4e-08 | 4.9e-08 | 4.767E-08 | 2.702E-08 | 2.000E-08 | 7.400E-08 |
| 12805 | 641 | V641C | B | 4.5e-08 | 3.9e-08 | 5.3e-08 | 4.567E-08 | 7.024E-09 | 3.900E-08 | 5.300E-08 |
| 12806 | 641 | V641E | B | 1.5e-08 | 5.4e-08 | 9.7e-09 | 2.623E-08 | 2.419E-08 | 9.700E-09 | 5.400E-08 |
| 12807 | 641 | V641Q | B | 4e-08 | 3.6e-08 | 2.8e-08 | 3.467E-08 | 6.110E-09 | 2.800E-08 | 4.000E-08 |
| 12808 | 641 | V641G | B | 4.2e-08 | 8.1e-08 | 2e-08 | 4.767E-08 | 3.089E-08 | 2.000E-08 | 8.100E-08 |
| 12809 | 641 | V641H | B | 5.3e-08 | 4.2e-08 | 3.6e-08 | 4.367E-08 | 8.622E-09 | 3.600E-08 | 5.300E-08 |
| 12810 | 641 | V641I | B | 2.8e-08 | 4.3e-08 | 3.1e-08 | 3.400E-08 | 7.937E-09 | 2.800E-08 | 4.300E-08 |
| 12811 | 641 | V641L | B | 2.4e-08 | 1.7e-08 | 2.2e-08 | 2.100E-08 | 3.606E-09 | 1.700E-08 | 2.400E-08 |
| 12812 | 641 | V641K | B | 1.2e-08 | 3.6e-08 | 3e-08 | 2.600E-08 | 1.249E-08 | 1.200E-08 | 3.600E-08 |
| 12813 | 641 | V641M | B | 5.2e-08 | 4.7e-08 | 4.8e-08 | 4.900E-08 | 2.646E-09 | 4.700E-08 | 5.200E-08 |
| 12814 | 641 | V641F | B | 7.6e-08 | 7.2e-08 | 4.5e-08 | 6.433E-08 | 1.686E-08 | 4.500E-08 | 7.600E-08 |
| 12815 | 641 | V641P | B | 3.5e-08 | 1.1e-07 | 1.3e-08 | 5.267E-08 | 5.086E-08 | 1.300E-08 | 1.100E-07 |
| 12816 | 641 | V641S | B | 3.5e-08 | 4.1e-08 | 4.4e-08 | 4.000E-08 | 4.583E-09 | 3.500E-08 | 4.400E-08 |
| 12817 | 641 | V641T | B | 2e-07 | 4.2e-08 | 1.1e-08 | 8.433E-08 | 1.014E-07 | 1.100E-08 | 2.000E-07 |
| 12818 | 641 | V641W | B | 1e-07 | 7.5e-08 | 3.9e-08 | 7.133E-08 | 3.066E-08 | 3.900E-08 | 1.000E-07 |
| 12819 | 641 | V641Y | B | 7.2e-08 | 1.8e-08 | 4.5e-08 | 4.500E-08 | 2.700E-08 | 1.800E-08 | 7.200E-08 |
| 12820 | 641 | V641V | B | 5.4e-08 | 2.1e-08 | 8.1e-08 | 5.200E-08 | 3.005E-08 | 2.100E-08 | 8.100E-08 |
| 12821 | 642 | T642A | B | 4.2e-08 | 7.4e-08 | 4.3e-08 | 5.300E-08 | 1.819E-08 | 4.200E-08 | 7.400E-08 |
| 12822 | 642 | T642R | B | 2.5e-08 | 2.9e-08 | 2.2e-08 | 2.533E-08 | 3.512E-09 | 2.200E-08 | 2.900E-08 |
| 12823 | 642 | T642N | B | 5.5e-08 | 2.4e-08 | 4.7e-08 | 4.200E-08 | 1.609E-08 | 2.400E-08 | 5.500E-08 |
| 12824 | 642 | T642D | B | 1.7e-08 | 2.3e-08 | 6.2e-08 | 3.400E-08 | 2.443E-08 | 1.700E-08 | 6.200E-08 |
| 12825 | 642 | T642C | B | 4.4e-08 | 1e-07 | 2e-08 | 5.467E-08 | 4.105E-08 | 2.000E-08 | 1.000E-07 |
| 12826 | 642 | T642E | B | 2.7e-08 | 4.7e-08 | 3.9e-08 | 3.767E-08 | 1.007E-08 | 2.700E-08 | 4.700E-08 |
| 12827 | 642 | T642Q | B | 2.5e-08 | 4.1e-08 | 2.4e-08 | 3.000E-08 | 9.539E-09 | 2.400E-08 | 4.100E-08 |
| 12828 | 642 | T642G | B | 3.7e-08 | 3.9e-08 | 6.1e-08 | 4.567E-08 | 1.332E-08 | 3.700E-08 | 6.100E-08 |
| 12829 | 642 | T642H | B | 1.4e-07 | 5.9e-08 | 5.7e-08 | 8.533E-08 | 4.735E-08 | 5.700E-08 | 1.400E-07 |
| 12830 | 642 | T642I | B | 5.9e-08 | 2.7e-08 | 8e-08 | 5.533E-08 | 2.669E-08 | 2.700E-08 | 8.000E-08 |
| 12831 | 642 | T642L | B | 4e-08 | 5.1e-08 | 2.5e-08 | 3.867E-08 | 1.305E-08 | 2.500E-08 | 5.100E-08 |
| 12832 | 642 | T642K | B | 3.6e-08 | 8e-08 | 2.6e-08 | 4.733E-08 | 2.873E-08 | 2.600E-08 | 8.000E-08 |
| 12833 | 642 | T642M | B | 4.8e-08 | 6.5e-08 | 2.8e-08 | 4.700E-08 | 1.852E-08 | 2.800E-08 | 6.500E-08 |
| 12834 | 642 | T642F | B | 7.1e-09 | 4.7e-08 | 2.4e-08 | 2.603E-08 | 2.003E-08 | 7.100E-09 | 4.700E-08 |

| | | | | | | | | | | |
|---|---|---|---|---|---|---|---|---|---|---|
| 12835 | 642 | T642P | B | 5e-08 | 1.7e-08 | 5.5e-08 | 4.067E-08 | 2.065E-08 | 1.700E-08 | 5.500E-08 |
| 12836 | 642 | T642S | B | 3.6e-08 | 7.3e-08 | 4.1e-08 | 5.000E-08 | 2.007E-08 | 3.600E-08 | 7.300E-08 |
| 12837 | 642 | T642T | B | 7.5e-09 | 8.4e-08 | 1.7e-08 | 3.617E-08 | 4.170E-08 | 7.500E-09 | 8.400E-08 |
| 12838 | 642 | T642W | B | 1e-08 | 5.1e-08 | 2.5e-08 | 2.867E-08 | 2.074E-08 | 1.000E-08 | 5.100E-08 |
| 12839 | 642 | T642Y | B | 4.2e-08 | 4.4e-08 | 4.1e-08 | 4.233E-08 | 1.528E-09 | 4.100E-08 | 4.400E-08 |
| 12840 | 642 | T642V | B | 7e-08 | 7.5e-08 | 4.1e-08 | 6.200E-08 | 1.836E-08 | 4.100E-08 | 7.500E-08 |
| 12841 | 643 | I643A | B | 1.3e-07 | 2.3e-08 | 1.4e-08 | 5.567E-08 | 6.453E-08 | 1.400E-08 | 1.300E-07 |
| 12842 | 643 | I643R | B | 8.5e-08 | 3.4e-08 | 1.2e-08 | 4.367E-08 | 3.745E-08 | 1.200E-08 | 8.500E-08 |
| 12843 | 643 | I643N | B | 3.6e-08 | 2.2e-08 | 3.2e-08 | 3.000E-08 | 7.211E-09 | 2.200E-08 | 3.600E-08 |
| 12844 | 643 | I643D | B | 2.4e-08 | 4.4e-08 | 4.8e-08 | 3.867E-08 | 1.286E-08 | 2.400E-08 | 4.800E-08 |
| 12845 | 643 | I643C | B | 1.2e-08 | 3.4e-09 | 4.9e-08 | 2.147E-08 | 2.423E-08 | 3.400E-09 | 4.900E-08 |
| 12846 | 643 | I643E | B | 7.9e-08 | 3.4e-08 | 1.1e-07 | 7.433E-08 | 3.821E-08 | 3.400E-08 | 1.100E-07 |
| 12847 | 643 | I643Q | B | 5e-08 | 7.2e-08 | 2.5e-08 | 4.900E-08 | 2.352E-08 | 2.500E-08 | 7.200E-08 |
| 12848 | 643 | I643G | B | 4.9e-08 | 3.2e-08 | 6e-08 | 4.700E-08 | 1.411E-08 | 3.200E-08 | 6.000E-08 |
| 12849 | 643 | I643H | B | 4.1e-08 | 4.6e-08 | 1.6e-08 | 3.433E-08 | 1.607E-08 | 1.600E-08 | 4.600E-08 |
| 12850 | 643 | I643I | B | 1.3e-07 | 1.8e-08 | 8.9e-08 | 7.900E-08 | 5.667E-08 | 1.800E-08 | 1.300E-07 |
| 12851 | 643 | I643L | B | 3.4e-08 | 6.1e-08 | 1.9e-08 | 3.800E-08 | 2.128E-08 | 1.900E-08 | 6.100E-08 |
| 12852 | 643 | I643K | B | 5.3e-08 | 3.2e-08 | 8.4e-08 | 5.633E-08 | 2.616E-08 | 3.200E-08 | 8.400E-08 |
| 12853 | 643 | I643M | B | 4.5e-08 | 3.7e-08 | 9.3e-08 | 5.833E-08 | 3.029E-08 | 3.700E-08 | 9.300E-08 |
| 12854 | 643 | I643F | B | 9.2e-08 | 3.2e-08 | 5.2e-08 | 5.867E-08 | 3.055E-08 | 3.200E-08 | 9.200E-08 |
| 12855 | 643 | I643P | B | 3.7e-08 | 5.9e-08 | 2.8e-08 | 4.133E-08 | 1.595E-08 | 2.800E-08 | 5.900E-08 |
| 12856 | 643 | I643S | B | 1.3e-08 | 2.1e-08 | 6e-08 | 3.133E-08 | 2.515E-08 | 1.300E-08 | 6.000E-08 |
| 12857 | 643 | I643T | B | 8.5e-08 | 4.2e-08 | 1.5e-08 | 4.733E-08 | 3.530E-08 | 1.500E-08 | 8.500E-08 |
| 12858 | 643 | I643W | B | 6.8e-08 | 1.6e-08 | 5e-08 | 4.467E-08 | 2.641E-08 | 1.600E-08 | 6.800E-08 |
| 12859 | 643 | I643Y | B | 2.2e-08 | 5.1e-08 | 6.1e-08 | 4.467E-08 | 2.026E-08 | 2.200E-08 | 6.100E-08 |
| 12860 | 643 | I643V | B | 7.8e-08 | 5.3e-08 | 4.9e-08 | 6.000E-08 | 1.572E-08 | 4.900E-08 | 7.800E-08 |
| 12861 | 644 | T644A | B | 2.9e-08 | 4.2e-08 | 7.6e-08 | 4.900E-08 | 2.427E-08 | 2.900E-08 | 7.600E-08 |
| 12862 | 644 | T644R | B | 3.6e-08 | 4.8e-08 | 2.9e-08 | 3.767E-08 | 9.609E-09 | 2.900E-08 | 4.800E-08 |
| 12863 | 644 | T644N | B | 2.5e-08 | 3.9e-08 | 4.4e-08 | 3.600E-08 | 9.849E-09 | 2.500E-08 | 4.400E-08 |
| 12864 | 644 | T644D | B | 7.6e-08 | 5e-08 | 1.6e-08 | 4.733E-08 | 3.009E-08 | 1.600E-08 | 7.600E-08 |
| 12865 | 644 | T644C | B | 3.8e-08 | 2.1e-08 | 6e-08 | 3.967E-08 | 1.955E-08 | 2.100E-08 | 6.000E-08 |
| 12866 | 644 | T644E | B | 3.7e-08 | 4.4e-08 | 2.6e-08 | 3.567E-08 | 9.074E-09 | 2.600E-08 | 4.400E-08 |
| 12867 | 644 | T644Q | B | 1.8e-08 | 3.4e-08 | 6.4e-08 | 3.867E-08 | 2.335E-08 | 1.800E-08 | 6.400E-08 |
| 12868 | 644 | T644G | B | 6.5e-08 | 2.7e-08 | 6.4e-08 | 5.200E-08 | 2.166E-08 | 2.700E-08 | 6.500E-08 |
| 12869 | 644 | T644H | B | 4.1e-08 | 3.8e-08 | 4.6e-08 | 4.167E-08 | 4.041E-09 | 3.800E-08 | 4.600E-08 |
| 12870 | 644 | T644I | B | 3.6e-08 | 5.2e-08 | 9.5e-09 | 3.250E-08 | 2.147E-08 | 9.500E-09 | 5.200E-08 |
| 12871 | 644 | T644L | B | 3.4e-08 | 4.5e-08 | 2.6e-08 | 3.500E-08 | 9.539E-09 | 2.600E-08 | 4.500E-08 |
| 12872 | 644 | T644K | B | 1.8e-08 | 8e-08 | 6.2e-08 | 5.333E-08 | 3.190E-08 | 1.800E-08 | 8.000E-08 |
| 12873 | 644 | T644M | B | 4.4e-08 | 3.3e-08 | 9.8e-08 | 5.833E-08 | 3.479E-08 | 3.300E-08 | 9.800E-08 |
| 12874 | 644 | T644F | B | 5.4e-08 | 5.2e-08 | 1.6e-07 | 8.867E-08 | 6.178E-08 | 5.200E-08 | 1.600E-07 |
| 12875 | 644 | T644P | B | 6e-08 | 3.1e-08 | 3.5e-08 | 4.200E-08 | 1.572E-08 | 3.100E-08 | 6.000E-08 |
| 12876 | 644 | T644S | B | 4.1e-08 | 5.1e-08 | 3.4e-08 | 4.200E-08 | 8.544E-09 | 3.400E-08 | 5.100E-08 |
| 12877 | 644 | T644T | B | 5.5e-08 | 1.4e-08 | 2.7e-08 | 3.200E-08 | 2.095E-08 | 1.400E-08 | 5.500E-08 |
| 12878 | 644 | T644W | B | 2.5e-08 | 1.4e-08 | 2e-08 | 1.967E-08 | 5.508E-09 | 1.400E-08 | 2.500E-08 |
| 12879 | 644 | T644Y | B | 4.3e-08 | 6.5e-08 | 4.9e-08 | 5.233E-08 | 1.137E-08 | 4.300E-08 | 6.500E-08 |
| 12880 | 644 | T644V | B | 6.9e-08 | 3.9e-08 | 3e-08 | 4.600E-08 | 2.042E-08 | 3.000E-08 | 6.900E-08 |
| 12881 | 645 | C645A | B | 7.1e-08 | 3.6e-08 | 7.5e-08 | 6.067E-08 | 2.146E-08 | 3.600E-08 | 7.500E-08 |
| 12882 | 645 | C645R | B | 2e-08 | 6.2e-08 | 8.1e-08 | 5.433E-08 | 3.121E-08 | 2.000E-08 | 8.100E-08 |
| 12883 | 645 | C645N | B | 2.6e-07 | 2e-08 | 2.6e-08 | 1.020E-07 | 1.369E-07 | 2.000E-08 | 2.600E-07 |
| 12884 | 645 | C645D | B | 5.8e-08 | 7.1e-08 | 5.6e-08 | 6.167E-08 | 8.145E-09 | 5.600E-08 | 7.100E-08 |
| 12885 | 645 | C645C | B | 7.9e-08 | 6.2e-08 | 3.3e-08 | 5.800E-08 | 2.326E-08 | 3.300E-08 | 7.900E-08 |
| 12886 | 645 | C645E | B | 2.1e-08 | 1.6e-08 | 4.5e-08 | 2.733E-08 | 1.550E-08 | 1.600E-08 | 4.500E-08 |
| 12887 | 645 | C645Q | B | 3.6e-08 | 4.7e-08 | 7.7e-08 | 5.333E-08 | 2.122E-08 | 3.600E-08 | 7.700E-08 |
| 12888 | 645 | C645G | B | 1.2e-07 | 5.1e-08 | 6.5e-08 | 7.867E-08 | 3.647E-08 | 5.100E-08 | 1.200E-07 |
| 12889 | 645 | C645H | B | 4.2e-08 | 3.6e-08 | 6e-08 | 4.600E-08 | 1.249E-08 | 3.600E-08 | 6.000E-08 |
| 12890 | 645 | C645I | B | 4.1e-08 | 2.4e-08 | 3.3e-08 | 3.267E-08 | 8.505E-09 | 2.400E-08 | 4.100E-08 |
| 12891 | 645 | C645L | B | 2.9e-08 | 1.4e-08 | 6.2e-08 | 3.500E-08 | 2.456E-08 | 1.400E-08 | 6.200E-08 |
| 12892 | 645 | C645K | B | 4.4e-08 | 2.6e-08 | 5.7e-08 | 4.233E-08 | 1.557E-08 | 2.600E-08 | 5.700E-08 |
| 12893 | 645 | C645M | B | 5e-08 | 2e-08 | 6.3e-08 | 4.433E-08 | 2.205E-08 | 2.000E-08 | 6.300E-08 |
| 12894 | 645 | C645F | B | 3.3e-08 | 3.4e-08 | 7.1e-08 | 4.600E-08 | 2.166E-08 | 3.300E-08 | 7.100E-08 |
| 12895 | 645 | C645P | B | 7.3e-08 | 1.3e-08 | 3.6e-08 | 4.067E-08 | 3.027E-08 | 1.300E-08 | 7.300E-08 |
| 12896 | 645 | C645S | B | 5.4e-08 | 5.4e-08 | 3.6e-08 | 4.800E-08 | 1.039E-08 | 3.600E-08 | 5.400E-08 |
| 12897 | 645 | C645T | B | 1.5e-08 | 1e-07 | 2.5e-08 | 4.667E-08 | 4.646E-08 | 1.500E-08 | 1.000E-07 |
| 12898 | 645 | C645W | B | 1.7e-08 | 1.1e-07 | 1.4e-07 | 8.900E-08 | 6.413E-08 | 1.700E-08 | 1.400E-07 |
| 12899 | 645 | C645Y | B | 7.1e-08 | 3.5e-08 | 6e-08 | 5.533E-08 | 1.845E-08 | 3.500E-08 | 7.100E-08 |
| 12900 | 645 | C645V | B | 9.3e-08 | 4.2e-08 | 6.9e-08 | 6.800E-08 | 2.551E-08 | 4.200E-08 | 9.300E-08 |
| 12901 | 646 | K646A | B | 4.5e-08 | 4.4e-08 | 2.6e-08 | 3.833E-08 | 1.069E-08 | 2.600E-08 | 4.500E-08 |
| 12902 | 646 | K646R | B | 5.1e-08 | 1.6e-08 | 1.1e-07 | 5.900E-08 | 4.751E-08 | 1.600E-08 | 1.100E-07 |
| 12903 | 646 | K646N | B | 1e-07 | 1e-08 | 2e-08 | 4.333E-08 | 4.933E-08 | 1.000E-08 | 1.000E-07 |
| 12904 | 646 | K646D | B | 2.1e-08 | 9.8e-08 | 5.8e-08 | 5.900E-08 | 3.851E-08 | 2.100E-08 | 9.800E-08 |
| 12905 | 646 | K646C | B | 3.9e-08 | 4.8e-08 | 4.9e-08 | 4.533E-08 | 5.508E-09 | 3.900E-08 | 4.900E-08 |
| 12906 | 646 | K646E | B | 2.1e-08 | 6.3e-08 | 1.5e-07 | 7.800E-08 | 6.580E-08 | 2.100E-08 | 1.500E-07 |
| 12907 | 646 | K646Q | B | 1.4e-08 | 6.3e-08 | 7.1e-08 | 4.933E-08 | 3.086E-08 | 1.400E-08 | 7.100E-08 |
| 12908 | 646 | K646G | B | 5.5e-08 | 5e-08 | 6.5e-08 | 5.667E-08 | 7.638E-09 | 5.000E-08 | 6.500E-08 |
| 12909 | 646 | K646H | B | 2.6e-08 | 4.8e-08 | 6.6e-08 | 4.667E-08 | 2.003E-08 | 2.600E-08 | 6.600E-08 |
| 12910 | 646 | K646I | B | 3.8e-08 | 5.3e-08 | 1.2e-07 | 7.033E-08 | 4.366E-08 | 3.800E-08 | 1.200E-07 |
| 12911 | 646 | K646L | B | 4.6e-08 | 2.7e-08 | 8.9e-08 | 5.400E-08 | 3.176E-08 | 2.700E-08 | 8.900E-08 |
| 12912 | 646 | K646K | B | 1.2e-07 | 5e-08 | 3.2e-08 | 6.733E-08 | 4.649E-08 | 3.200E-08 | 1.200E-07 |
| 12913 | 646 | K646M | B | 2.3e-08 | 2.3e-08 | 2e-08 | 2.200E-08 | 1.732E-09 | 2.000E-08 | 2.300E-08 |
| 12914 | 646 | K646F | B | 1.8e-08 | 1.2e-07 | 5.8e-08 | 6.533E-08 | 5.139E-08 | 1.800E-08 | 1.200E-07 |
| 12915 | 646 | K646P | B | 8.8e-08 | 3.9e-08 | 2.6e-08 | 5.100E-08 | 3.270E-08 | 2.600E-08 | 8.800E-08 |
| 12916 | 646 | K646S | B | 1.8e-08 | 5.4e-08 | 2.7e-08 | 3.300E-08 | 1.873E-08 | 1.800E-08 | 5.400E-08 |
| 12917 | 646 | K646T | B | 5.1e-08 | 1.3e-08 | 7.5e-08 | 4.633E-08 | 3.126E-08 | 1.300E-08 | 7.500E-08 |
| 12918 | 646 | K646W | B | 5.2e-08 | 2.4e-08 | 5.6e-08 | 4.400E-08 | 1.744E-08 | 2.400E-08 | 5.600E-08 |
| 12919 | 646 | K646Y | B | 1.9e-08 | 3.9e-08 | 3.1e-08 | 2.967E-08 | 1.007E-08 | 1.900E-08 | 3.900E-08 |

| | | | | | | | | | | |
|---|---|---|---|---|---|---|---|---|---|---|
| 12920 | 646 | K646V | B | 4.2e-08 | 8.5e-08 | 2.5e-08 | 5.067E-08 | 3.092E-08 | 2.500E-08 | 8.500E-08 |
| 12921 | 647 | A647A | B | 3.6e-08 | 7.2e-08 | 5e-08 | 5.267E-08 | 1.815E-08 | 3.600E-08 | 7.200E-08 |
| 12922 | 647 | A647R | B | 3.1e-08 | 6.7e-08 | 5e-08 | 4.933E-08 | 1.801E-08 | 3.100E-08 | 6.700E-08 |
| 12923 | 647 | A647N | B | 2.5e-08 | 6e-08 | 4.1e-08 | 4.200E-08 | 1.752E-08 | 2.500E-08 | 6.000E-08 |
| 12924 | 647 | A647D | B | 1.2e-08 | 2.2e-08 | 4.8e-08 | 2.733E-08 | 1.858E-08 | 1.200E-08 | 4.800E-08 |
| 12925 | 647 | A647C | B | 5e-08 | 2.7e-08 | 7.6e-08 | 5.100E-08 | 2.452E-08 | 2.700E-08 | 7.600E-08 |
| 12926 | 647 | A647E | B | 4.5e-08 | 3.9e-08 | 3.8e-08 | 4.067E-08 | 3.786E-09 | 3.800E-08 | 4.500E-08 |
| 12927 | 647 | A647Q | B | 2.4e-08 | 3.3e-08 | 7.6e-08 | 4.433E-08 | 2.779E-08 | 2.400E-08 | 7.600E-08 |
| 12928 | 647 | A647G | B | 6.5e-08 | 5.7e-08 | 1.4e-08 | 4.533E-08 | 2.743E-08 | 1.400E-08 | 6.500E-08 |
| 12929 | 647 | A647H | B | 4.1e-08 | 5.4e-08 | 2.6e-08 | 4.033E-08 | 1.401E-08 | 2.600E-08 | 5.400E-08 |
| 12930 | 647 | A647I | B | 8.4e-08 | 4e-08 | 3.4e-08 | 5.267E-08 | 2.730E-08 | 3.400E-08 | 8.400E-08 |
| 12931 | 647 | A647L | B | 8.8e-08 | 3.5e-08 | 5.2e-08 | 5.833E-08 | 2.706E-08 | 3.500E-08 | 8.800E-08 |
| 12932 | 647 | A647K | B | 8.5e-08 | 3e-08 | 2.7e-08 | 4.733E-08 | 3.265E-08 | 2.700E-08 | 8.500E-08 |
| 12933 | 647 | A647M | B | 4.9e-08 | 7e-08 | 4.4e-08 | 5.433E-08 | 1.380E-08 | 4.400E-08 | 7.000E-08 |
| 12934 | 647 | A647F | B | 4.4e-08 | 5.2e-08 | 8.8e-08 | 6.133E-08 | 2.344E-08 | 4.400E-08 | 8.800E-08 |
| 12935 | 647 | A647P | B | 1e-07 | 7.5e-08 | 7.4e-08 | 8.300E-08 | 1.473E-08 | 7.400E-08 | 1.000E-07 |
| 12936 | 647 | A647S | B | 7.8e-08 | 4.9e-08 | 7.2e-08 | 6.633E-08 | 1.531E-08 | 4.900E-08 | 7.800E-08 |
| 12937 | 647 | A647T | B | 4e-08 | 3.7e-08 | 8.5e-08 | 5.400E-08 | 2.689E-08 | 3.700E-08 | 8.500E-08 |
| 12938 | 647 | A647W | B | 4.4e-08 | 5e-08 | 4.5e-08 | 4.633E-08 | 3.215E-09 | 4.400E-08 | 5.000E-08 |
| 12939 | 647 | A647Y | B | 3.3e-08 | 4.4e-08 | 8e-08 | 5.233E-08 | 2.458E-08 | 3.300E-08 | 8.000E-08 |
| 12940 | 647 | A647V | B | 7.5e-08 | 3.7e-08 | 6.4e-08 | 5.867E-08 | 1.955E-08 | 3.700E-08 | 7.500E-08 |
| 12941 | 648 | S648A | B | 3.2e-08 | 1.8e-08 | 5.6e-08 | 3.533E-08 | 1.922E-08 | 1.800E-08 | 5.600E-08 |
| 12942 | 648 | S648R | B | 1.6e-08 | 2.2e-08 | 1.9e-08 | 1.900E-08 | 3.000E-09 | 1.600E-08 | 2.200E-08 |
| 12943 | 648 | S648N | B | 3.3e-08 | 3.8e-08 | 1.1e-07 | 6.033E-08 | 4.309E-08 | 3.300E-08 | 1.100E-07 |
| 12944 | 648 | S648D | B | 1e-07 | 3.1e-08 | 2.4e-08 | 5.167E-08 | 4.200E-08 | 2.400E-08 | 1.000E-07 |
| 12945 | 648 | S648C | B | 5.8e-08 | 7.2e-08 | 3.3e-08 | 5.433E-08 | 1.976E-08 | 3.300E-08 | 7.200E-08 |
| 12946 | 648 | S648E | B | 9.7e-08 | 5.2e-08 | 1.1e-07 | 8.633E-08 | 3.044E-08 | 5.200E-08 | 1.100E-07 |
| 12947 | 648 | S648Q | B | 1.2e-08 | 5.3e-08 | 3.6e-08 | 3.367E-08 | 2.060E-08 | 1.200E-08 | 5.300E-08 |
| 12948 | 648 | S648G | B | 2.4e-08 | 6.3e-08 | 1.2e-07 | 6.900E-08 | 4.828E-08 | 2.400E-08 | 1.200E-07 |
| 12949 | 648 | S648H | B | 4.8e-08 | 1.2e-08 | 7.2e-08 | 4.400E-08 | 3.020E-08 | 1.200E-08 | 7.200E-08 |
| 12950 | 648 | S648I | B | 3.1e-08 | 6.5e-08 | 1.2e-07 | 7.200E-08 | 4.491E-08 | 3.100E-08 | 1.200E-07 |
| 12951 | 648 | S648L | B | 4.1e-08 | 5e-08 | 1.3e-07 | 7.367E-08 | 4.899E-08 | 4.100E-08 | 1.300E-07 |
| 12952 | 648 | S648K | B | 5.1e-08 | 1.2e-08 | 2.7e-08 | 3.000E-08 | 1.967E-08 | 1.200E-08 | 5.100E-08 |
| 12953 | 648 | S648M | B | 1.1e-08 | 3e-08 | 3.1e-08 | 2.400E-08 | 1.127E-08 | 1.100E-08 | 3.100E-08 |
| 12954 | 648 | S648F | B | 4e-08 | 6.4e-08 | 6.2e-08 | 5.533E-08 | 1.332E-08 | 4.000E-08 | 6.400E-08 |
| 12955 | 648 | S648P | B | 7.3e-08 | 7.4e-08 | 3.8e-08 | 6.167E-08 | 2.050E-08 | 3.800E-08 | 7.400E-08 |
| 12956 | 648 | S648S | B | 4.3e-08 | 1.6e-07 | 2.3e-08 | 7.533E-08 | 7.400E-08 | 2.300E-08 | 1.600E-07 |
| 12957 | 648 | S648T | B | 2.9e-08 | 2.7e-08 | 4.8e-08 | 3.467E-08 | 1.159E-08 | 2.700E-08 | 4.800E-08 |
| 12958 | 648 | S648W | B | 6e-08 | 2e-08 | 4.7e-08 | 4.233E-08 | 2.040E-08 | 2.000E-08 | 6.000E-08 |
| 12959 | 648 | S648Y | B | 4.4e-08 | 8.3e-08 | 3.1e-08 | 5.267E-08 | 2.706E-08 | 3.100E-08 | 8.300E-08 |
| 12960 | 648 | S648V | B | 1.4e-07 | 4.2e-08 | 6.2e-08 | 8.133E-08 | 5.178E-08 | 4.200E-08 | 1.400E-07 |
| 12961 | 649 | Q649A | B | 5.1e-08 | 5.6e-08 | 3.3e-08 | 4.667E-08 | 1.210E-08 | 3.300E-08 | 5.600E-08 |
| 12962 | 649 | Q649R | B | 5.7e-08 | 3.3e-08 | 2e-08 | 3.667E-08 | 1.877E-08 | 2.000E-08 | 5.700E-08 |
| 12963 | 649 | Q649N | B | 5.7e-08 | 2.5e-08 | 2.4e-08 | 3.533E-08 | 1.877E-08 | 2.400E-08 | 5.700E-08 |
| 12964 | 649 | Q649D | B | 4.7e-08 | 3e-08 | 8.2e-08 | 5.300E-08 | 2.651E-08 | 3.000E-08 | 8.200E-08 |
| 12965 | 649 | Q649C | B | 3.2e-08 | 4.5e-08 | 5.2e-08 | 4.300E-08 | 1.015E-08 | 3.200E-08 | 5.200E-08 |
| 12966 | 649 | Q649E | B | 3.6e-08 | 8.7e-08 | 3.6e-08 | 5.300E-08 | 2.944E-08 | 3.600E-08 | 8.700E-08 |
| 12967 | 649 | Q649Q | B | 1.1e-07 | 7.6e-08 | 5.7e-08 | 8.100E-08 | 2.685E-08 | 5.700E-08 | 1.100E-07 |
| 12968 | 649 | Q649G | B | 9e-08 | 1.1e-07 | 2.6e-08 | 7.533E-08 | 4.388E-08 | 2.600E-08 | 1.100E-07 |
| 12969 | 649 | Q649H | B | 5.4e-08 | 3.9e-08 | 8.7e-08 | 6.000E-08 | 2.456E-08 | 3.900E-08 | 8.700E-08 |
| 12970 | 649 | Q649I | B | 3.1e-08 | 2.3e-08 | 4.3e-08 | 3.233E-08 | 1.007E-08 | 2.300E-08 | 4.300E-08 |
| 12971 | 649 | Q649L | B | 1.7e-08 | 2.7e-08 | 7.9e-08 | 4.100E-08 | 3.329E-08 | 1.700E-08 | 7.900E-08 |
| 12972 | 649 | Q649K | B | 2.3e-08 | 1.2e-07 | 2.1e-08 | 5.467E-08 | 5.659E-08 | 2.100E-08 | 1.200E-07 |
| 12973 | 649 | Q649M | B | 3.8e-08 | 4.5e-08 | 2.8e-08 | 3.700E-08 | 8.544E-09 | 2.800E-08 | 4.500E-08 |
| 12974 | 649 | Q649F | B | 4.3e-08 | 4.4e-08 | 3.7e-08 | 4.133E-08 | 3.786E-09 | 3.700E-08 | 4.400E-08 |
| 12975 | 649 | Q649P | B | 5.1e-08 | 4.3e-08 | 6.4e-08 | 5.267E-08 | 1.060E-08 | 4.300E-08 | 6.400E-08 |
| 12976 | 649 | Q649S | B | 1e-07 | 3.6e-08 | 6.5e-08 | 6.700E-08 | 3.205E-08 | 3.600E-08 | 1.000E-07 |
| 12977 | 649 | Q649T | B | 2.7e-08 | 4.2e-08 | 3.4e-08 | 3.433E-08 | 7.506E-09 | 2.700E-08 | 4.200E-08 |
| 12978 | 649 | Q649W | B | 1.7e-07 | 5.2e-08 | 6.6e-08 | 9.600E-08 | 6.447E-08 | 5.200E-08 | 1.700E-07 |
| 12979 | 649 | Q649Y | B | 7e-08 | 2.1e-08 | 6.9e-08 | 5.333E-08 | 2.801E-08 | 2.100E-08 | 7.000E-08 |
| 12980 | 649 | Q649V | B | 5.5e-08 | 3.7e-08 | 4e-08 | 4.400E-08 | 9.644E-09 | 3.700E-08 | 5.500E-08 |
| 12981 | 650 | D650A | B | 1.4e-08 | 1.9e-08 | 8e-08 | 3.767E-08 | 3.675E-08 | 1.400E-08 | 8.000E-08 |
| 12982 | 650 | D650R | B | 7e-08 | 1.3e-07 | 2.2e-08 | 7.400E-08 | 5.411E-08 | 2.200E-08 | 1.300E-07 |
| 12983 | 650 | D650N | B | 9.2e-08 | 3.5e-08 | 7.6e-08 | 6.767E-08 | 2.940E-08 | 3.500E-08 | 9.200E-08 |
| 12984 | 650 | D650D | B | 5.6e-08 | 2.6e-08 | 3e-08 | 3.733E-08 | 1.629E-08 | 2.600E-08 | 5.600E-08 |
| 12985 | 650 | D650C | B | 2.9e-08 | 5.6e-08 | 6.5e-08 | 5.000E-08 | 1.873E-08 | 2.900E-08 | 6.500E-08 |
| 12986 | 650 | D650E | B | 7.3e-08 | 8.2e-08 | 4.8e-08 | 6.767E-08 | 1.762E-08 | 4.800E-08 | 8.200E-08 |
| 12987 | 650 | D650Q | B | 4.2e-08 | 3.1e-08 | 3.3e-08 | 3.533E-08 | 5.859E-09 | 3.100E-08 | 4.200E-08 |
| 12988 | 650 | D650G | B | 2.5e-08 | 4.2e-08 | 9.4e-09 | 2.547E-08 | 1.631E-08 | 9.400E-09 | 4.200E-08 |
| 12989 | 650 | D650H | B | 6.8e-08 | 6.6e-08 | 5.7e-08 | 6.367E-08 | 5.859E-09 | 5.700E-08 | 6.800E-08 |
| 12990 | 650 | D650I | B | 8.8e-08 | 4.2e-08 | 3.1e-08 | 5.367E-08 | 3.024E-08 | 3.100E-08 | 8.800E-08 |
| 12991 | 650 | D650L | B | 5.6e-08 | 4e-08 | 5e-08 | 4.867E-08 | 8.083E-09 | 4.000E-08 | 5.600E-08 |
| 12992 | 650 | D650K | B | 3.3e-08 | 1.8e-08 | 3.7e-08 | 2.933E-08 | 1.002E-08 | 1.800E-08 | 3.700E-08 |
| 12993 | 650 | D650M | B | 9.2e-08 | 2.5e-08 | 7.2e-08 | 6.300E-08 | 3.439E-08 | 2.500E-08 | 9.200E-08 |
| 12994 | 650 | D650F | B | 3e-08 | 5.3e-08 | 2.1e-08 | 3.467E-08 | 1.650E-08 | 2.100E-08 | 5.300E-08 |
| 12995 | 650 | D650P | B | 5.2e-08 | 1.1e-07 | 5.7e-08 | 7.300E-08 | 3.214E-08 | 5.200E-08 | 1.100E-07 |
| 12996 | 650 | D650S | B | 5.9e-08 | 3e-08 | 5.1e-08 | 4.667E-08 | 1.498E-08 | 3.000E-08 | 5.900E-08 |
| 12997 | 650 | D650T | B | 1.8e-08 | 1.8e-08 | 2.8e-08 | 2.133E-08 | 5.774E-09 | 1.800E-08 | 2.800E-08 |
| 12998 | 650 | D650W | B | 1.3e-08 | 2.1e-08 | 3.8e-08 | 2.400E-08 | 1.277E-08 | 1.300E-08 | 3.800E-08 |
| 12999 | 650 | D650Y | B | 4.6e-08 | 2.2e-08 | 3.5e-08 | 3.433E-08 | 1.201E-08 | 2.200E-08 | 4.600E-08 |
| 13000 | 650 | D650V | B | 8.4e-08 | 2.3e-07 | 3.8e-08 | 1.173E-07 | 1.002E-07 | 3.800E-08 | 2.300E-07 |
| 13001 | 651 | V651A | B | 8e-08 | 4e-08 | 9.2e-08 | 7.067E-08 | 2.723E-08 | 4.000E-08 | 9.200E-08 |
| 13002 | 651 | V651R | B | 5.1e-08 | 2.1e-08 | 2.2e-08 | 3.133E-08 | 1.704E-08 | 2.100E-08 | 5.100E-08 |
| 13003 | 651 | V651N | B | 1.7e-07 | 3e-08 | 2.9e-08 | 7.633E-08 | 8.112E-08 | 2.900E-08 | 1.700E-07 |
| 13004 | 651 | V651D | B | 2.5e-08 | 3.6e-08 | 2.4e-08 | 2.833E-08 | 6.658E-09 | 2.400E-08 | 3.600E-08 |

| | | | | | | | | | | |
|---|---|---|---|---|---|---|---|---|---|---|
| 13005 | 651 | V651C | B | 3.5e-08 | 2.2e-08 | 9.9e-08 | 5.200E-08 | 4.122E-08 | 2.200E-08 | 9.900E-08 |
| 13006 | 651 | V651E | B | 4.1e-08 | 7.6e-08 | 2.2e-08 | 4.633E-08 | 2.739E-08 | 2.200E-08 | 7.600E-08 |
| 13007 | 651 | V651Q | B | 5.7e-08 | 9.1e-08 | 4.9e-08 | 6.567E-08 | 2.230E-08 | 4.900E-08 | 9.100E-08 |
| 13008 | 651 | V651G | B | 2.6e-08 | 4e-08 | 6.9e-08 | 4.500E-08 | 2.193E-08 | 2.600E-08 | 6.900E-08 |
| 13009 | 651 | V651H | B | 2e-08 | 2.8e-08 | 8.1e-08 | 4.300E-08 | 3.315E-08 | 2.000E-08 | 8.100E-08 |
| 13010 | 651 | V651I | B | 4.8e-08 | 6.7e-08 | 2.9e-08 | 4.800E-08 | 1.900E-08 | 2.900E-08 | 6.700E-08 |
| 13011 | 651 | V651L | B | 7.2e-08 | 5.2e-08 | 2.1e-08 | 4.833E-08 | 2.570E-08 | 2.100E-08 | 7.200E-08 |
| 13012 | 651 | V651K | B | 4.6e-08 | 7e-08 | 2.6e-08 | 4.733E-08 | 2.203E-08 | 2.600E-08 | 7.000E-08 |
| 13013 | 651 | V651M | B | 4.1e-08 | 4.9e-08 | 5.7e-08 | 4.900E-08 | 8.000E-09 | 4.100E-08 | 5.700E-08 |
| 13014 | 651 | V651F | B | 3.3e-08 | 5.3e-08 | 4.3e-08 | 4.300E-08 | 1.000E-08 | 3.300E-08 | 5.300E-08 |
| 13015 | 651 | V651P | B | 4.9e-08 | 2.2e-08 | 6.2e-08 | 4.433E-08 | 2.040E-08 | 2.200E-08 | 6.200E-08 |
| 13016 | 651 | V651S | B | 1.5e-08 | 2e-08 | 6.7e-08 | 3.400E-08 | 2.869E-08 | 1.500E-08 | 6.700E-08 |
| 13017 | 651 | V651T | B | 2.5e-08 | 7.6e-08 | 8.4e-08 | 6.167E-08 | 3.201E-08 | 2.500E-08 | 8.400E-08 |
| 13018 | 651 | V651W | B | 3.1e-08 | 7.5e-08 | 4.5e-08 | 5.033E-08 | 2.248E-08 | 3.100E-08 | 7.500E-08 |
| 13019 | 651 | V651Y | B | 2e-08 | 2e-08 | 4.3e-08 | 2.767E-08 | 1.328E-08 | 2.000E-08 | 4.300E-08 |
| 13020 | 651 | V651V | B | 5.8e-08 | 2.9e-08 | 4.3e-08 | 4.333E-08 | 1.450E-08 | 2.900E-08 | 5.800E-08 |
| 13021 | 652 | S652A | B | 8.8e-08 | 5.1e-08 | 3.7e-08 | 5.867E-08 | 2.635E-08 | 3.700E-08 | 8.800E-08 |
| 13022 | 652 | S652R | B | 4.5e-08 | 3.3e-08 | 1.2e-08 | 3.000E-08 | 1.670E-08 | 1.200E-08 | 4.500E-08 |
| 13023 | 652 | S652N | B | 3.2e-08 | 5.1e-08 | 3.2e-08 | 3.833E-08 | 1.097E-08 | 3.200E-08 | 5.100E-08 |
| 13024 | 652 | S652D | B | 3.6e-08 | 3.5e-08 | 4e-08 | 3.700E-08 | 2.646E-09 | 3.500E-08 | 4.000E-08 |
| 13025 | 652 | S652C | B | 5.1e-08 | 1.8e-08 | 1.9e-08 | 2.933E-08 | 1.877E-08 | 1.800E-08 | 5.100E-08 |
| 13026 | 652 | S652E | B | 3.4e-08 | 1.9e-08 | 3.2e-08 | 2.833E-08 | 8.145E-09 | 1.900E-08 | 3.400E-08 |
| 13027 | 652 | S652Q | B | 3.9e-08 | 3.4e-08 | 3.8e-08 | 3.700E-08 | 2.646E-09 | 3.400E-08 | 3.900E-08 |
| 13028 | 652 | S652G | B | 1.2e-08 | 7.9e-08 | 4.6e-08 | 4.567E-08 | 3.350E-08 | 1.200E-08 | 7.900E-08 |
| 13029 | 652 | S652H | B | 7.5e-08 | 5.9e-08 | 7.9e-08 | 7.100E-08 | 1.058E-08 | 5.900E-08 | 7.900E-08 |
| 13030 | 652 | S652I | B | 3.6e-08 | 3.7e-08 | 2.6e-08 | 3.300E-08 | 6.083E-09 | 2.600E-08 | 3.700E-08 |
| 13031 | 652 | S652L | B | 1.2e-07 | 5.5e-08 | 3e-08 | 6.833E-08 | 4.646E-08 | 3.000E-08 | 1.200E-07 |
| 13032 | 652 | S652K | B | 5.2e-08 | 3.1e-08 | 1e-07 | 6.100E-08 | 3.537E-08 | 3.100E-08 | 1.000E-07 |
| 13033 | 652 | S652M | B | 3.3e-08 | 5.7e-08 | 1.6e-08 | 3.533E-08 | 2.060E-08 | 1.600E-08 | 5.700E-08 |
| 13034 | 652 | S652F | B | 7.4e-08 | 8.1e-08 | 4.6e-08 | 6.700E-08 | 1.852E-08 | 4.600E-08 | 8.100E-08 |
| 13035 | 652 | S652P | B | 2.1e-08 | 6e-08 | 4.8e-08 | 4.300E-08 | 1.997E-08 | 2.100E-08 | 6.000E-08 |
| 13036 | 652 | S652S | B | 4.7e-08 | 1.1e-07 | 3e-08 | 6.233E-08 | 4.215E-08 | 3.000E-08 | 1.100E-07 |
| 13037 | 652 | S652T | B | 1.4e-08 | 4.1e-08 | 3.9e-08 | 3.133E-08 | 1.504E-08 | 1.400E-08 | 4.100E-08 |
| 13038 | 652 | S652W | B | 5.3e-08 | 8.2e-09 | 1.3e-07 | 6.373E-08 | 6.161E-08 | 8.200E-09 | 1.300E-07 |
| 13039 | 652 | S652Y | B | 5.2e-08 | 6.6e-08 | 1e-07 | 7.267E-08 | 2.468E-08 | 5.200E-08 | 1.000E-07 |
| 13040 | 652 | S652V | B | 3.4e-08 | 7.1e-08 | 4.3e-08 | 4.933E-08 | 1.930E-08 | 3.400E-08 | 7.100E-08 |
| 13041 | 653 | I653A | B | 5.3e-08 | 4.9e-08 | 3.9e-08 | 4.700E-08 | 7.211E-09 | 3.900E-08 | 5.300E-08 |
| 13042 | 653 | I653R | B | 3.4e-08 | 1.4e-07 | 4.4e-08 | 7.267E-08 | 5.853E-08 | 3.400E-08 | 1.400E-07 |
| 13043 | 653 | I653N | B | 4.8e-08 | 8.8e-08 | 3.7e-08 | 5.767E-08 | 2.684E-08 | 3.700E-08 | 8.800E-08 |
| 13044 | 653 | I653D | B | 1.4e-08 | 3.5e-08 | 4.6e-08 | 3.167E-08 | 1.626E-08 | 1.400E-08 | 4.600E-08 |
| 13045 | 653 | I653C | B | 4.6e-08 | 6.1e-08 | 7.2e-08 | 5.967E-08 | 1.305E-08 | 4.600E-08 | 7.200E-08 |
| 13046 | 653 | I653E | B | 6.2e-08 | 8.6e-08 | 3.2e-08 | 6.000E-08 | 2.706E-08 | 3.200E-08 | 8.600E-08 |
| 13047 | 653 | I653Q | B | 6.6e-08 | 4e-08 | 3.8e-08 | 4.800E-08 | 1.562E-08 | 3.800E-08 | 6.600E-08 |
| 13048 | 653 | I653G | B | 3.8e-08 | 3e-08 | 9.9e-08 | 5.567E-08 | 3.774E-08 | 3.000E-08 | 9.900E-08 |
| 13049 | 653 | I653H | B | 8.2e-08 | 4.8e-08 | 1.7e-07 | 1.000E-07 | 6.296E-08 | 4.800E-08 | 1.700E-07 |
| 13050 | 653 | I653I | B | 4.6e-08 | 5.7e-08 | 2e-08 | 4.100E-08 | 1.900E-08 | 2.000E-08 | 5.700E-08 |
| 13051 | 653 | I653L | B | 7.7e-08 | 4e-08 | 1.8e-08 | 4.500E-08 | 2.982E-08 | 1.800E-08 | 7.700E-08 |
| 13052 | 653 | I653K | B | 5.2e-08 | 1.8e-08 | 3.8e-08 | 3.600E-08 | 1.709E-08 | 1.800E-08 | 5.200E-08 |
| 13053 | 653 | I653M | B | 7.4e-08 | 3.5e-08 | 3.2e-08 | 4.700E-08 | 2.343E-08 | 3.200E-08 | 7.400E-08 |
| 13054 | 653 | I653F | B | 5.9e-08 | 4.2e-08 | 4.7e-08 | 4.933E-08 | 8.737E-09 | 4.200E-08 | 5.900E-08 |
| 13055 | 653 | I653P | B | 5.6e-08 | 2.1e-08 | 3.6e-08 | 3.767E-08 | 1.756E-08 | 2.100E-08 | 5.600E-08 |
| 13056 | 653 | I653S | B | 3.1e-08 | 2.8e-08 | 2.5e-08 | 2.800E-08 | 3.000E-09 | 2.500E-08 | 3.100E-08 |
| 13057 | 653 | I653T | B | 6.7e-08 | 1.3e-07 | 7.6e-08 | 9.100E-08 | 3.407E-08 | 6.700E-08 | 1.300E-07 |
| 13058 | 653 | I653W | B | 5.2e-08 | 4.6e-08 | 3.3e-08 | 4.367E-08 | 9.713E-09 | 3.300E-08 | 5.200E-08 |
| 13059 | 653 | I653Y | B | 6.7e-08 | 4.2e-08 | 1.2e-08 | 4.033E-08 | 2.754E-08 | 1.200E-08 | 6.700E-08 |
| 13060 | 653 | I653V | B | 3.3e-08 | 5.3e-08 | 3.4e-08 | 4.000E-08 | 1.127E-08 | 3.300E-08 | 5.300E-08 |
| 13061 | 654 | G654A | B | 5.5e-08 | 5e-08 | 7.1e-08 | 5.867E-08 | 1.097E-08 | 5.000E-08 | 7.100E-08 |
| 13062 | 654 | G654R | B | 3.4e-08 | 3.9e-08 | 4.6e-08 | 3.967E-08 | 6.028E-09 | 3.400E-08 | 4.600E-08 |
| 13063 | 654 | G654N | B | 3.4e-08 | 2.7e-08 | 3.6e-08 | 3.233E-08 | 4.726E-09 | 2.700E-08 | 3.600E-08 |
| 13064 | 654 | G654D | B | 5.4e-08 | 2.4e-08 | 2.4e-08 | 3.400E-08 | 1.732E-08 | 2.400E-08 | 5.400E-08 |
| 13065 | 654 | G654C | B | 7e-08 | 3.6e-08 | 1.8e-08 | 4.133E-08 | 2.641E-08 | 1.800E-08 | 7.000E-08 |
| 13066 | 654 | G654E | B | 6.4e-08 | 7.6e-08 | 5.8e-08 | 6.600E-08 | 9.165E-09 | 5.800E-08 | 7.600E-08 |
| 13067 | 654 | G654Q | B | 6.1e-08 | 6.1e-08 | 3e-08 | 5.067E-08 | 1.790E-08 | 3.000E-08 | 6.100E-08 |
| 13068 | 654 | G654G | B | 3.1e-08 | 4.2e-08 | 7.3e-08 | 4.867E-08 | 2.178E-08 | 3.100E-08 | 7.300E-08 |
| 13069 | 654 | G654H | B | 5.6e-08 | 3.1e-08 | 1.1e-07 | 6.567E-08 | 4.038E-08 | 3.100E-08 | 1.100E-07 |
| 13070 | 654 | G654I | B | 1.2e-07 | 6.8e-08 | 4.1e-08 | 7.633E-08 | 4.015E-08 | 4.100E-08 | 1.200E-07 |
| 13071 | 654 | G654L | B | 4.7e-08 | 3.9e-08 | 3.2e-08 | 3.933E-08 | 7.506E-09 | 3.200E-08 | 4.700E-08 |
| 13072 | 654 | G654K | B | 4.7e-08 | 1.9e-08 | 2.8e-08 | 3.133E-08 | 1.429E-08 | 1.900E-08 | 4.700E-08 |
| 13073 | 654 | G654M | B | 2.8e-08 | 2.6e-08 | 2.5e-08 | 2.633E-08 | 1.528E-09 | 2.500E-08 | 2.800E-08 |
| 13074 | 654 | G654F | B | 2.3e-08 | 4e-08 | 6.5e-08 | 4.267E-08 | 2.113E-08 | 2.300E-08 | 6.500E-08 |
| 13075 | 654 | G654P | B | 9e-08 | 1.9e-08 | 3e-08 | 4.633E-08 | 3.821E-08 | 1.900E-08 | 9.000E-08 |
| 13076 | 654 | G654S | B | 4.2e-08 | 3.1e-08 | 1.5e-08 | 2.933E-08 | 1.358E-08 | 1.500E-08 | 4.200E-08 |
| 13077 | 654 | G654T | B | 7.5e-09 | 2.5e-08 | 1.9e-08 | 1.717E-08 | 8.893E-09 | 7.500E-09 | 2.500E-08 |
| 13078 | 654 | G654W | B | 3.3e-08 | 9.5e-08 | 5.5e-08 | 6.100E-08 | 3.143E-08 | 3.300E-08 | 9.500E-08 |
| 13079 | 654 | G654Y | B | 3.2e-08 | 4.7e-08 | 1.1e-08 | 3.000E-08 | 1.808E-08 | 1.100E-08 | 4.700E-08 |
| 13080 | 654 | G654V | B | 5.3e-08 | 6.6e-08 | 3e-08 | 4.967E-08 | 1.823E-08 | 3.000E-08 | 6.600E-08 |
| 13081 | 655 | V655A | B | 6.1e-08 | 8.7e-08 | 5.7e-08 | 6.833E-08 | 1.629E-08 | 5.700E-08 | 8.700E-08 |
| 13082 | 655 | V655R | B | 2.5e-08 | 2.7e-08 | 1.8e-07 | 7.733E-08 | 8.892E-08 | 2.500E-08 | 1.800E-07 |
| 13083 | 655 | V655N | B | 1.3e-08 | 3.4e-08 | 3.3e-08 | 2.667E-08 | 1.185E-08 | 1.300E-08 | 3.400E-08 |
| 13084 | 655 | V655D | B | 4e-08 | 1.9e-08 | 5.9e-08 | 3.933E-08 | 2.001E-08 | 1.900E-08 | 5.900E-08 |
| 13085 | 655 | V655C | B | 1.2e-08 | 8.1e-09 | 1.2e-07 | 4.670E-08 | 6.351E-08 | 8.100E-09 | 1.200E-07 |
| 13086 | 655 | V655E | B | 8.4e-08 | 1.8e-07 | 4.5e-08 | 1.030E-07 | 6.948E-08 | 4.500E-08 | 1.800E-07 |
| 13087 | 655 | V655Q | B | 7.8e-08 | 5.4e-08 | 8.1e-08 | 7.100E-08 | 1.480E-08 | 5.400E-08 | 8.100E-08 |
| 13088 | 655 | V655G | B | 3.4e-08 | 2.5e-08 | 4.7e-08 | 3.533E-08 | 1.106E-08 | 2.500E-08 | 4.700E-08 |
| 13089 | 655 | V655H | B | 4e-08 | 5.4e-08 | 4.7e-08 | 4.700E-08 | 7.000E-09 | 4.000E-08 | 5.400E-08 |

| | | | | | | | | | | |
|---|---|---|---|---|---|---|---|---|---|---|
| 13090 | 655 | V655I | B | 3.2e-08 | 4.5e-08 | 5.2e-08 | 4.300E-08 | 1.015E-08 | 3.200E-08 | 5.200E-08 |
| 13091 | 655 | V655L | B | 7.8e-08 | 4.3e-08 | 1e-08 | 4.367E-08 | 3.400E-08 | 1.000E-08 | 7.800E-08 |
| 13092 | 655 | V655K | B | 7.5e-09 | 3.1e-08 | 4.6e-08 | 2.817E-08 | 1.941E-08 | 7.500E-09 | 4.600E-08 |
| 13093 | 655 | V655M | B | 4.5e-08 | 4.9e-08 | 2.6e-08 | 4.000E-08 | 1.229E-08 | 2.600E-08 | 4.900E-08 |
| 13094 | 655 | V655F | B | 2.4e-08 | 4e-08 | 4.1e-08 | 3.500E-08 | 9.539E-09 | 2.400E-08 | 4.100E-08 |
| 13095 | 655 | V655P | B | 5.7e-08 | 5.6e-08 | 4.8e-08 | 5.367E-08 | 4.933E-09 | 4.800E-08 | 5.700E-08 |
| 13096 | 655 | V655S | B | 2.7e-08 | 1.8e-08 | 4e-08 | 2.833E-08 | 1.106E-08 | 1.800E-08 | 4.000E-08 |
| 13097 | 655 | V655T | B | 7.3e-08 | 3.5e-08 | 3.7e-08 | 4.833E-08 | 2.139E-08 | 3.500E-08 | 7.300E-08 |
| 13098 | 655 | V655W | B | 1e-07 | 1.3e-07 | 8.4e-08 | 1.047E-07 | 2.335E-08 | 8.400E-08 | 1.300E-07 |
| 13099 | 655 | V655Y | B | 2e-08 | 5.7e-08 | 2.9e-08 | 3.533E-08 | 1.930E-08 | 2.000E-08 | 5.700E-08 |
| 13100 | 655 | V655V | B | 9.9e-08 | 3.9e-08 | 4.1e-08 | 5.967E-08 | 3.408E-08 | 3.900E-08 | 9.900E-08 |
| 13101 | 656 | A656A | B | 3.8e-08 | 6.9e-08 | 8.5e-08 | 6.400E-08 | 2.390E-08 | 3.800E-08 | 8.500E-08 |
| 13102 | 656 | A656R | B | 3.2e-08 | 7.6e-08 | 7.7e-08 | 6.167E-08 | 2.570E-08 | 3.200E-08 | 7.700E-08 |
| 13103 | 656 | A656N | B | 1.4e-07 | 7.1e-08 | 2.6e-08 | 7.900E-08 | 5.742E-08 | 2.600E-08 | 1.400E-07 |
| 13104 | 656 | A656D | B | 1e-08 | 3.5e-08 | 5.4e-08 | 3.300E-08 | 2.207E-08 | 1.000E-08 | 5.400E-08 |
| 13105 | 656 | A656C | B | 3.5e-08 | 2.6e-07 | 3e-08 | 1.083E-07 | 1.314E-07 | 3.000E-08 | 2.600E-07 |
| 13106 | 656 | A656E | B | 1.8e-07 | 3e-08 | 3.4e-08 | 8.133E-08 | 8.547E-08 | 3.000E-08 | 1.800E-07 |
| 13107 | 656 | A656Q | B | 4.4e-08 | 2.1e-07 | 2.4e-08 | 9.267E-08 | 1.021E-07 | 2.400E-08 | 2.100E-07 |
| 13108 | 656 | A656G | B | 1.8e-08 | 3.7e-08 | 8.3e-08 | 4.600E-08 | 3.342E-08 | 1.800E-08 | 8.300E-08 |
| 13109 | 656 | A656H | B | 4.6e-08 | 7.9e-08 | 2.3e-08 | 4.933E-08 | 2.815E-08 | 2.300E-08 | 7.900E-08 |
| 13110 | 656 | A656I | B | 5.2e-08 | 3.9e-08 | 4.2e-08 | 4.433E-08 | 6.807E-09 | 3.900E-08 | 5.200E-08 |
| 13111 | 656 | A656L | B | 9.6e-08 | 4.5e-08 | 4.4e-08 | 6.167E-08 | 2.974E-08 | 4.400E-08 | 9.600E-08 |
| 13112 | 656 | A656K | B | 4.8e-08 | 4e-08 | 3.8e-08 | 4.200E-08 | 5.292E-09 | 3.800E-08 | 4.800E-08 |
| 13113 | 656 | A656M | B | 1.1e-07 | 4.3e-08 | 1.1e-08 | 5.467E-08 | 5.052E-08 | 1.100E-08 | 1.100E-07 |
| 13114 | 656 | A656F | B | 9.7e-08 | 8.9e-08 | 1.2e-07 | 1.020E-07 | 1.609E-08 | 8.900E-08 | 1.200E-07 |
| 13115 | 656 | A656P | B | 4.9e-08 | 1.1e-07 | 8.8e-08 | 8.233E-08 | 3.089E-08 | 4.900E-08 | 1.100E-07 |
| 13116 | 656 | A656S | B | 4.2e-08 | 1.7e-08 | 3.7e-08 | 3.200E-08 | 1.323E-08 | 1.700E-08 | 4.200E-08 |
| 13117 | 656 | A656T | B | 4e-08 | 4.7e-08 | 2.2e-08 | 3.633E-08 | 1.290E-08 | 2.200E-08 | 4.700E-08 |
| 13118 | 656 | A656W | B | 4.4e-08 | 4.7e-08 | 1.8e-08 | 3.633E-08 | 1.595E-08 | 1.800E-08 | 4.700E-08 |
| 13119 | 656 | A656Y | B | 2.7e-08 | 2.1e-08 | 2.4e-08 | 2.400E-08 | 3.000E-09 | 2.100E-08 | 2.700E-08 |
| 13120 | 656 | A656V | B | 1e-07 | 5.8e-08 | 5.5e-08 | 7.100E-08 | 2.516E-08 | 5.500E-08 | 1.000E-07 |
| 13121 | 657 | W657A | B | 4e-08 | 7.3e-08 | 9.3e-08 | 6.867E-08 | 2.676E-08 | 4.000E-08 | 9.300E-08 |
| 13122 | 657 | W657R | B | 5.2e-08 | 2.9e-08 | 2.6e-08 | 3.567E-08 | 1.422E-08 | 2.600E-08 | 5.200E-08 |
| 13123 | 657 | W657N | B | 2.7e-08 | 6.3e-08 | 7.3e-08 | 5.433E-08 | 2.419E-08 | 2.700E-08 | 7.300E-08 |
| 13124 | 657 | W657D | B | 3e-08 | 7e-08 | 8.5e-08 | 6.167E-08 | 2.843E-08 | 3.000E-08 | 8.500E-08 |
| 13125 | 657 | W657C | B | 5.5e-08 | 1.3e-07 | 6.4e-08 | 8.300E-08 | 4.095E-08 | 5.500E-08 | 1.300E-07 |
| 13126 | 657 | W657E | B | 7.5e-08 | 7.4e-08 | 5.7e-08 | 6.867E-08 | 1.012E-08 | 5.700E-08 | 7.500E-08 |
| 13127 | 657 | W657Q | B | 3.3e-08 | 3e-08 | 5.6e-08 | 3.967E-08 | 1.422E-08 | 3.000E-08 | 5.600E-08 |
| 13128 | 657 | W657G | B | 3.8e-08 | 4.6e-08 | 3.8e-08 | 4.067E-08 | 4.619E-09 | 3.800E-08 | 4.600E-08 |
| 13129 | 657 | W657H | B | 1.1e-07 | 3e-08 | 3.1e-08 | 5.700E-08 | 4.590E-08 | 3.000E-08 | 1.100E-07 |
| 13130 | 657 | W657I | B | 3.8e-08 | 4.6e-08 | 6.9e-08 | 5.100E-08 | 1.609E-08 | 3.800E-08 | 6.900E-08 |
| 13131 | 657 | W657L | B | 3.3e-08 | 6.2e-08 | 8.7e-08 | 6.067E-08 | 2.702E-08 | 3.300E-08 | 8.700E-08 |
| 13132 | 657 | W657K | B | 4.6e-08 | 3e-08 | 2.8e-08 | 3.467E-08 | 9.866E-09 | 2.800E-08 | 4.600E-08 |
| 13133 | 657 | W657M | B | 8.5e-08 | 3.2e-08 | 4.6e-08 | 5.433E-08 | 2.747E-08 | 3.200E-08 | 8.500E-08 |
| 13134 | 657 | W657F | B | 3.1e-08 | 3.3e-08 | 9.9e-08 | 5.433E-08 | 3.870E-08 | 3.100E-08 | 9.900E-08 |
| 13135 | 657 | W657P | B | 9.5e-08 | 7.3e-08 | 6.8e-08 | 7.867E-08 | 1.436E-08 | 6.800E-08 | 9.500E-08 |
| 13136 | 657 | W657S | B | 1.4e-07 | 5e-08 | 1.1e-08 | 6.700E-08 | 6.616E-08 | 1.100E-08 | 1.400E-07 |
| 13137 | 657 | W657T | B | 1.5e-08 | 9.1e-08 | 6.1e-08 | 5.567E-08 | 3.828E-08 | 1.500E-08 | 9.100E-08 |
| 13138 | 657 | W657W | B | 4.6e-08 | 5.1e-08 | 2.8e-08 | 4.167E-08 | 1.210E-08 | 2.800E-08 | 5.100E-08 |
| 13139 | 657 | W657Y | B | 6.3e-08 | 1e-07 | 2e-08 | 6.100E-08 | 4.004E-08 | 2.000E-08 | 1.000E-07 |
| 13140 | 657 | W657V | B | 7.5e-08 | 1.3e-07 | 6.3e-08 | 8.933E-08 | 3.573E-08 | 6.300E-08 | 1.300E-07 |
| 13141 | 658 | Y658A | B | 2.8e-08 | 5.1e-08 | 1.3e-08 | 3.067E-08 | 1.914E-08 | 1.300E-08 | 5.100E-08 |
| 13142 | 658 | Y658R | B | 7e-08 | 8e-08 | 7.7e-08 | 7.567E-08 | 5.132E-09 | 7.000E-08 | 8.000E-08 |
| 13143 | 658 | Y658N | B | 1.5e-08 | 4.3e-08 | 3e-08 | 2.933E-08 | 1.401E-08 | 1.500E-08 | 4.300E-08 |
| 13144 | 658 | Y658D | B | 1.1e-07 | 4.2e-08 | 5.6e-08 | 6.933E-08 | 3.591E-08 | 4.200E-08 | 1.100E-07 |
| 13145 | 658 | Y658C | B | 2.4e-08 | 8.8e-08 | 8.6e-08 | 6.600E-08 | 3.639E-08 | 2.400E-08 | 8.800E-08 |
| 13146 | 658 | Y658E | B | 2e-08 | 8.2e-08 | 6.9e-08 | 5.700E-08 | 3.270E-08 | 2.000E-08 | 8.200E-08 |
| 13147 | 658 | Y658Q | B | 1.6e-08 | 4.6e-08 | 3.7e-08 | 3.300E-08 | 1.539E-08 | 1.600E-08 | 4.600E-08 |
| 13148 | 658 | Y658G | B | 6.3e-08 | 1.9e-07 | 4.4e-08 | 9.900E-08 | 7.938E-08 | 4.400E-08 | 1.900E-07 |
| 13149 | 658 | Y658H | B | 1.3e-07 | 3.4e-08 | 2.5e-08 | 6.300E-08 | 5.820E-08 | 2.500E-08 | 1.300E-07 |
| 13150 | 658 | Y658I | B | 5.3e-08 | 3.7e-08 | 3.5e-08 | 4.167E-08 | 9.866E-09 | 3.500E-08 | 5.300E-08 |
| 13151 | 658 | Y658L | B | 2.7e-08 | 8.3e-08 | 5.5e-08 | 5.500E-08 | 2.800E-08 | 2.700E-08 | 8.300E-08 |
| 13152 | 658 | Y658K | B | 4.4e-08 | 9.4e-08 | 3.9e-08 | 5.900E-08 | 3.041E-08 | 3.900E-08 | 9.400E-08 |
| 13153 | 658 | Y658M | B | 4.5e-08 | 2.6e-08 | 1.6e-07 | 7.700E-08 | 7.251E-08 | 2.600E-08 | 1.600E-07 |
| 13154 | 658 | Y658F | B | 2.8e-08 | 5e-08 | 8e-08 | 5.267E-08 | 2.610E-08 | 2.800E-08 | 8.000E-08 |
| 13155 | 658 | Y658P | B | 1.3e-08 | 7.5e-08 | 6.3e-08 | 5.033E-08 | 3.288E-08 | 1.300E-08 | 7.500E-08 |
| 13156 | 658 | Y658S | B | 1.2e-08 | 4.3e-08 | 3.2e-08 | 2.900E-08 | 1.572E-08 | 1.200E-08 | 4.300E-08 |
| 13157 | 658 | Y658T | B | 3.1e-08 | 2.3e-08 | 3.3e-08 | 2.900E-08 | 5.292E-09 | 2.300E-08 | 3.300E-08 |
| 13158 | 658 | Y658W | B | 9e-08 | 6.3e-08 | 4.4e-08 | 6.567E-08 | 2.312E-08 | 4.400E-08 | 9.000E-08 |
| 13159 | 658 | Y658Y | B | 3e-08 | 4.9e-08 | 2.7e-08 | 3.533E-08 | 1.193E-08 | 2.700E-08 | 4.900E-08 |
| 13160 | 658 | Y658V | B | 3e-08 | 6.4e-08 | 4.7e-08 | 4.700E-08 | 1.700E-08 | 3.000E-08 | 6.400E-08 |
| 13161 | 659 | Q659A | B | 1.7e-08 | 5.9e-08 | 1.1e-08 | 2.900E-08 | 2.615E-08 | 1.100E-08 | 5.900E-08 |
| 13162 | 659 | Q659R | B | 4.3e-08 | 5.8e-08 | 3.9e-08 | 4.667E-08 | 1.002E-08 | 3.900E-08 | 5.800E-08 |
| 13163 | 659 | Q659N | B | 6.9e-08 | 5.9e-08 | 1.7e-08 | 4.833E-08 | 2.759E-08 | 1.700E-08 | 6.900E-08 |
| 13164 | 659 | Q659D | B | 5.1e-08 | 5.5e-08 | 7.4e-08 | 6.000E-08 | 1.229E-08 | 5.100E-08 | 7.400E-08 |
| 13165 | 659 | Q659C | B | 1.5e-08 | 3.3e-08 | 6.9e-08 | 3.900E-08 | 2.750E-08 | 1.500E-08 | 6.900E-08 |
| 13166 | 659 | Q659E | B | 2.9e-08 | 3e-08 | 1e-07 | 5.300E-08 | 4.071E-08 | 2.900E-08 | 1.000E-07 |
| 13167 | 659 | Q659Q | B | 1.6e-08 | 9.2e-08 | 4.9e-08 | 5.233E-08 | 3.811E-08 | 1.600E-08 | 9.200E-08 |
| 13168 | 659 | Q659G | B | 1.6e-08 | 4.8e-08 | 6.7e-08 | 4.367E-08 | 2.577E-08 | 1.600E-08 | 6.700E-08 |
| 13169 | 659 | Q659H | B | 2.4e-08 | 5.5e-08 | 3.3e-08 | 3.733E-08 | 1.595E-08 | 2.400E-08 | 5.500E-08 |
| 13170 | 659 | Q659I | B | 6.9e-08 | 4.6e-08 | 3.2e-08 | 4.900E-08 | 1.868E-08 | 3.200E-08 | 6.900E-08 |
| 13171 | 659 | Q659L | B | 4.9e-08 | 2.9e-08 | 1.3e-07 | 6.933E-08 | 5.348E-08 | 2.900E-08 | 1.300E-07 |
| 13172 | 659 | Q659K | B | 2.8e-08 | 3.3e-08 | 3.7e-08 | 3.267E-08 | 4.509E-09 | 2.800E-08 | 3.700E-08 |
| 13173 | 659 | Q659M | B | 5.1e-08 | 6.8e-08 | 2.4e-08 | 4.767E-08 | 2.219E-08 | 2.400E-08 | 6.800E-08 |
| 13174 | 659 | Q659F | B | 5.5e-08 | 5.2e-08 | 3.3e-08 | 4.667E-08 | 1.193E-08 | 3.300E-08 | 5.500E-08 |

| | | | | | | | | | | |
|---|---|---|---|---|---|---|---|---|---|---|
| 13175 | 659 | Q659P | B | 1.1e-07 | 2.1e-08 | 1.1e-08 | 4.733E-08 | 5.450E-08 | 1.100E-08 | 1.100E-07 |
| 13176 | 659 | Q659S | B | 9.1e-09 | 2.1e-08 | 1.7e-08 | 1.570E-08 | 6.056E-09 | 9.100E-09 | 2.100E-08 |
| 13177 | 659 | Q659T | B | 5e-08 | 1.3e-08 | 1.2e-07 | 6.100E-08 | 5.434E-08 | 1.300E-08 | 1.200E-07 |
| 13178 | 659 | Q659W | B | 6.8e-08 | 2.9e-08 | 6.1e-08 | 5.267E-08 | 2.079E-08 | 2.900E-08 | 6.800E-08 |
| 13179 | 659 | Q659Y | B | 3.1e-08 | 3.5e-08 | 3.4e-08 | 3.333E-08 | 2.082E-09 | 3.100E-08 | 3.500E-08 |
| 13180 | 659 | Q659V | B | 4e-08 | 4.1e-08 | 1.5e-08 | 3.200E-08 | 1.473E-08 | 1.500E-08 | 4.100E-08 |
| 13181 | 660 | Q660A | B | 3.2e-08 | 3.9e-08 | 2.9e-08 | 3.333E-08 | 5.132E-09 | 2.900E-08 | 3.900E-08 |
| 13182 | 660 | Q660R | B | 5.6e-08 | 7e-08 | 2.1e-08 | 4.900E-08 | 2.524E-08 | 2.100E-08 | 7.000E-08 |
| 13183 | 660 | Q660N | B | 2.5e-08 | 6.7e-08 | 6.6e-08 | 5.267E-08 | 2.397E-08 | 2.500E-08 | 6.700E-08 |
| 13184 | 660 | Q660D | B | 4.3e-08 | 3.5e-08 | 9.9e-08 | 5.900E-08 | 3.487E-08 | 3.500E-08 | 9.900E-08 |
| 13185 | 660 | Q660C | B | 4.4e-08 | 6.8e-08 | 6.5e-08 | 5.900E-08 | 1.308E-08 | 4.400E-08 | 6.800E-08 |
| 13186 | 660 | Q660E | B | 4.1e-08 | 7.1e-08 | 4.7e-08 | 5.300E-08 | 1.587E-08 | 4.100E-08 | 7.100E-08 |
| 13187 | 660 | Q660Q | B | 5.8e-08 | 4e-08 | 2.4e-08 | 4.067E-08 | 1.701E-08 | 2.400E-08 | 5.800E-08 |
| 13188 | 660 | Q660G | B | 3.8e-08 | 1.5e-08 | 2.7e-08 | 2.667E-08 | 1.150E-08 | 1.500E-08 | 3.800E-08 |
| 13189 | 660 | Q660H | B | 4.7e-08 | 3.6e-08 | 7e-08 | 5.100E-08 | 1.735E-08 | 3.600E-08 | 7.000E-08 |
| 13190 | 660 | Q660I | B | 5e-08 | 4.1e-08 | 5e-08 | 4.700E-08 | 5.196E-09 | 4.100E-08 | 5.000E-08 |
| 13191 | 660 | Q660L | B | 3.6e-08 | 1.8e-08 | 4.6e-08 | 3.333E-08 | 1.419E-08 | 1.800E-08 | 4.600E-08 |
| 13192 | 660 | Q660K | B | 6e-08 | 7.3e-08 | 5.8e-08 | 6.367E-08 | 8.145E-09 | 5.800E-08 | 7.300E-08 |
| 13193 | 660 | Q660M | B | 7.2e-08 | 1.3e-08 | 3.2e-08 | 3.900E-08 | 3.012E-08 | 1.300E-08 | 7.200E-08 |
| 13194 | 660 | Q660F | B | 3.4e-08 | 5.8e-08 | 4.3e-08 | 4.500E-08 | 1.212E-08 | 3.400E-08 | 5.800E-08 |
| 13195 | 660 | Q660P | B | 9.1e-08 | 1.3e-07 | 5.4e-08 | 9.167E-08 | 3.800E-08 | 5.400E-08 | 1.300E-07 |
| 13196 | 660 | Q660S | B | 1.2e-07 | 2.8e-08 | 1.8e-08 | 5.533E-08 | 5.623E-08 | 1.800E-08 | 1.200E-07 |
| 13197 | 660 | Q660T | B | 1.4e-08 | 1.2e-07 | 2.3e-08 | 5.233E-08 | 5.877E-08 | 1.400E-08 | 1.200E-07 |
| 13198 | 660 | Q660W | B | 3.9e-08 | 4.3e-08 | 3.8e-08 | 4.000E-08 | 2.646E-09 | 3.800E-08 | 4.300E-08 |
| 13199 | 660 | Q660Y | B | 5.7e-08 | 3.1e-08 | 4.9e-08 | 4.567E-08 | 1.332E-08 | 3.100E-08 | 5.700E-08 |
| 13200 | 660 | Q660V | B | 1.5e-08 | 4.2e-08 | 2.8e-08 | 2.833E-08 | 1.350E-08 | 1.500E-08 | 4.200E-08 |
| 13201 | 661 | K661A | B | 7.3e-08 | 8.5e-08 | 4.8e-08 | 6.867E-08 | 1.888E-08 | 4.800E-08 | 8.500E-08 |
| 13202 | 661 | K661R | B | 3.1e-08 | 5.7e-08 | 4.3e-08 | 4.367E-08 | 1.301E-08 | 3.100E-08 | 5.700E-08 |
| 13203 | 661 | K661N | B | 4e-08 | 1.9e-08 | 3.4e-08 | 3.100E-08 | 1.082E-08 | 1.900E-08 | 4.000E-08 |
| 13204 | 661 | K661D | B | 2.5e-08 | 2.6e-08 | 6.2e-09 | 1.907E-08 | 1.115E-08 | 6.200E-09 | 2.600E-08 |
| 13205 | 661 | K661C | B | 2.8e-08 | 5.1e-08 | 3.1e-07 | 1.297E-07 | 1.566E-07 | 2.800E-08 | 3.100E-07 |
| 13206 | 661 | K661E | B | 3.7e-08 | 2.5e-08 | 6.2e-08 | 4.133E-08 | 1.888E-08 | 2.500E-08 | 6.200E-08 |
| 13207 | 661 | K661Q | B | 7.9e-08 | 6.8e-08 | 1.1e-07 | 8.567E-08 | 2.178E-08 | 6.800E-08 | 1.100E-07 |
| 13208 | 661 | K661G | B | 3.5e-08 | 2e-08 | 8.3e-08 | 4.600E-08 | 3.291E-08 | 2.000E-08 | 8.300E-08 |
| 13209 | 661 | K661H | B | 5.1e-08 | 3.4e-08 | 6.2e-08 | 4.900E-08 | 1.411E-08 | 3.400E-08 | 6.200E-08 |
| 13210 | 661 | K661I | B | 4.8e-08 | 1.9e-08 | 5.5e-09 | 2.417E-08 | 2.172E-08 | 5.500E-09 | 4.800E-08 |
| 13211 | 661 | K661L | B | 1.3e-08 | 1.2e-07 | 1.9e-08 | 5.067E-08 | 6.012E-08 | 1.300E-08 | 1.200E-07 |
| 13212 | 661 | K661K | B | 1e-08 | 3.7e-08 | 7.5e-08 | 4.067E-08 | 3.265E-08 | 1.000E-08 | 7.500E-08 |
| 13213 | 661 | K661M | B | 3.5e-08 | 3e-08 | 1e-07 | 5.500E-08 | 3.905E-08 | 3.000E-08 | 1.000E-07 |
| 13214 | 661 | K661F | B | 1.3e-08 | 3.7e-08 | 5.7e-08 | 3.567E-08 | 2.203E-08 | 1.300E-08 | 5.700E-08 |
| 13215 | 661 | K661P | B | 4.5e-08 | 1.4e-07 | 4.5e-08 | 7.667E-08 | 5.485E-08 | 4.500E-08 | 1.400E-07 |
| 13216 | 661 | K661S | B | 7.2e-08 | 4.2e-08 | 1.4e-07 | 8.467E-08 | 5.021E-08 | 4.200E-08 | 1.400E-07 |
| 13217 | 661 | K661T | B | 7.2e-08 | 1.3e-07 | 2.7e-08 | 7.633E-08 | 5.164E-08 | 2.700E-08 | 1.300E-07 |
| 13218 | 661 | K661W | B | 5.9e-08 | 5.6e-08 | 5e-08 | 5.500E-08 | 4.583E-09 | 5.000E-08 | 5.900E-08 |
| 13219 | 661 | K661Y | B | 7.1e-08 | 1.6e-08 | 6.9e-08 | 5.200E-08 | 3.119E-08 | 1.600E-08 | 7.100E-08 |
| 13220 | 661 | K661V | B | 4.8e-08 | 4.2e-08 | 3.5e-08 | 4.167E-08 | 6.506E-09 | 3.500E-08 | 4.800E-08 |
| 13221 | 662 | P662A | B | 2.4e-08 | 2.4e-08 | 2.4e-08 | 2.400E-08 | 0.000E+00 | 2.400E-08 | 2.400E-08 |
| 13222 | 662 | P662R | B | 4.9e-08 | 2.2e-08 | 5.2e-08 | 4.100E-08 | 1.652E-08 | 2.200E-08 | 5.200E-08 |
| 13223 | 662 | P662N | B | 1.1e-08 | 4.1e-08 | 1.3e-07 | 6.067E-08 | 6.189E-08 | 1.100E-08 | 1.300E-07 |
| 13224 | 662 | P662D | B | 9.3e-08 | 2.1e-08 | 3.5e-08 | 4.967E-08 | 3.818E-08 | 2.100E-08 | 9.300E-08 |
| 13225 | 662 | P662C | B | 4.4e-08 | 3.3e-08 | 3e-08 | 3.567E-08 | 7.371E-09 | 3.000E-08 | 4.400E-08 |
| 13226 | 662 | P662E | B | 1.2e-07 | 5.6e-08 | 2.5e-08 | 6.700E-08 | 4.845E-08 | 2.500E-08 | 1.200E-07 |
| 13227 | 662 | P662Q | B | 2e-08 | 3.7e-08 | 5.8e-08 | 3.833E-08 | 1.904E-08 | 2.000E-08 | 5.800E-08 |
| 13228 | 662 | P662G | B | 3.7e-08 | 1.4e-08 | 2.5e-08 | 2.533E-08 | 1.150E-08 | 1.400E-08 | 3.700E-08 |
| 13229 | 662 | P662H | B | 5.3e-08 | 3.7e-08 | 7e-08 | 5.333E-08 | 1.650E-08 | 3.700E-08 | 7.000E-08 |
| 13230 | 662 | P662I | B | 2.8e-08 | 9.3e-08 | 4e-08 | 5.367E-08 | 3.459E-08 | 2.800E-08 | 9.300E-08 |
| 13231 | 662 | P662L | B | 3.1e-08 | 2.3e-08 | 1.8e-08 | 2.400E-08 | 6.557E-09 | 1.800E-08 | 3.100E-08 |
| 13232 | 662 | P662K | B | 2.4e-08 | 1.8e-08 | 3.8e-08 | 2.667E-08 | 1.026E-08 | 1.800E-08 | 3.800E-08 |
| 13233 | 662 | P662M | B | 4.9e-08 | 4.4e-08 | 7.6e-09 | 3.353E-08 | 2.260E-08 | 7.600E-09 | 4.900E-08 |
| 13234 | 662 | P662F | B | 5.1e-08 | 1.5e-08 | 2.1e-08 | 2.900E-08 | 1.929E-08 | 1.500E-08 | 5.100E-08 |
| 13235 | 662 | P662P | B | 2.7e-08 | 2.5e-08 | 3.9e-08 | 3.033E-08 | 7.572E-09 | 2.500E-08 | 3.900E-08 |
| 13236 | 662 | P662S | B | 4.8e-08 | 4.6e-08 | 3.5e-08 | 4.300E-08 | 7.000E-09 | 3.500E-08 | 4.800E-08 |
| 13237 | 662 | P662T | B | 7.5e-08 | 2.4e-08 | 3e-08 | 4.300E-08 | 2.787E-08 | 2.400E-08 | 7.500E-08 |
| 13238 | 662 | P662W | B | 1.9e-08 | 5.9e-08 | 1.1e-07 | 6.267E-08 | 4.561E-08 | 1.900E-08 | 1.100E-07 |
| 13239 | 662 | P662Y | B | 2e-08 | 3.2e-08 | 3.5e-08 | 2.900E-08 | 7.937E-09 | 2.000E-08 | 3.500E-08 |
| 13240 | 662 | P662V | B | 1.2e-07 | 4e-08 | 6.8e-08 | 7.600E-08 | 4.060E-08 | 4.000E-08 | 1.200E-07 |
| 13241 | 663 | G663A | B | 6.7e-08 | 1.5e-08 | 8.4e-08 | 5.533E-08 | 3.595E-08 | 1.500E-08 | 8.400E-08 |
| 13242 | 663 | G663R | B | 4.8e-08 | 5.1e-08 | 4.7e-08 | 4.867E-08 | 2.082E-09 | 4.700E-08 | 5.100E-08 |
| 13243 | 663 | G663N | B | 7.9e-08 | 1.1e-08 | 9.9e-08 | 6.300E-08 | 4.613E-08 | 1.100E-08 | 9.900E-08 |
| 13244 | 663 | G663D | B | 5.7e-08 | 6e-08 | 3.1e-08 | 4.933E-08 | 1.595E-08 | 3.100E-08 | 6.000E-08 |
| 13245 | 663 | G663C | B | 2.9e-08 | 7.7e-08 | 7e-08 | 5.867E-08 | 2.593E-08 | 2.900E-08 | 7.700E-08 |
| 13246 | 663 | G663E | B | 3.6e-08 | 2.9e-08 | 4.3e-08 | 3.600E-08 | 7.000E-09 | 2.900E-08 | 4.300E-08 |
| 13247 | 663 | G663Q | B | 8.6e-08 | 7.8e-08 | 1.1e-07 | 9.133E-08 | 1.665E-08 | 7.800E-08 | 1.100E-07 |
| 13248 | 663 | G663G | B | 2.5e-08 | 3.6e-08 | 3.7e-08 | 3.267E-08 | 6.658E-09 | 2.500E-08 | 3.700E-08 |
| 13249 | 663 | G663H | B | 3e-08 | 6.2e-08 | 5e-08 | 4.733E-08 | 1.617E-08 | 3.000E-08 | 6.200E-08 |
| 13250 | 663 | G663I | B | 5.6e-08 | 4e-08 | 4.7e-08 | 4.767E-08 | 8.021E-09 | 4.000E-08 | 5.600E-08 |
| 13251 | 663 | G663L | B | 2.3e-08 | 1e-07 | 1.6e-08 | 4.633E-08 | 4.661E-08 | 1.600E-08 | 1.000E-07 |
| 13252 | 663 | G663K | B | 2.7e-08 | 2.3e-08 | 5.6e-08 | 3.533E-08 | 1.801E-08 | 2.300E-08 | 5.600E-08 |
| 13253 | 663 | G663M | B | 1.4e-08 | 3.4e-08 | 2.6e-08 | 2.467E-08 | 1.007E-08 | 1.400E-08 | 3.400E-08 |
| 13254 | 663 | G663F | B | 3.4e-08 | 6.6e-08 | 3e-08 | 4.333E-08 | 1.973E-08 | 3.000E-08 | 6.600E-08 |
| 13255 | 663 | G663P | B | 1.2e-08 | 3.1e-08 | 4.3e-08 | 2.867E-08 | 1.563E-08 | 1.200E-08 | 4.300E-08 |
| 13256 | 663 | G663S | B | 1.6e-08 | 6.9e-08 | 2.5e-08 | 3.667E-08 | 2.836E-08 | 1.600E-08 | 6.900E-08 |
| 13257 | 663 | G663T | B | 7.4e-08 | 3.8e-08 | 4.2e-08 | 5.133E-08 | 1.973E-08 | 3.800E-08 | 7.400E-08 |
| 13258 | 663 | G663W | B | 5.9e-08 | 5.2e-08 | 5.6e-08 | 5.567E-08 | 3.512E-09 | 5.200E-08 | 5.900E-08 |
| 13259 | 663 | G663Y | B | 2e-08 | 3.3e-08 | 7.1e-08 | 4.133E-08 | 2.650E-08 | 2.000E-08 | 7.100E-08 |

| | | | | | | | | | | |
|---|---|---|---|---|---|---|---|---|---|---|
| 13260 | 663 | G663V | B | 6e-08 | 2e-08 | 5.3e-08 | 4.433E-08 | 2.136E-08 | 2.000E-08 | 6.000E-08 |
| 13261 | 664 | K664A | B | 2.2e-08 | 5.5e-08 | 5.5e-08 | 4.400E-08 | 1.905E-08 | 2.200E-08 | 5.500E-08 |
| 13262 | 664 | K664R | B | 6.8e-08 | 3.4e-08 | 3.4e-08 | 4.533E-08 | 1.963E-08 | 3.400E-08 | 6.800E-08 |
| 13263 | 664 | K664N | B | 1.9e-08 | 1.1e-07 | 6.9e-08 | 6.600E-08 | 4.557E-08 | 1.900E-08 | 1.100E-07 |
| 13264 | 664 | K664D | B | 2.8e-08 | 6.8e-08 | 4.7e-08 | 4.767E-08 | 2.001E-08 | 2.800E-08 | 6.800E-08 |
| 13265 | 664 | K664C | B | 7.5e-08 | 2.2e-08 | 4e-08 | 4.567E-08 | 2.695E-08 | 2.200E-08 | 7.500E-08 |
| 13266 | 664 | K664E | B | 2.4e-08 | 9.3e-08 | 5.8e-08 | 5.833E-08 | 3.450E-08 | 2.400E-08 | 9.300E-08 |
| 13267 | 664 | K664Q | B | 5e-08 | 2.5e-08 | 9e-08 | 5.500E-08 | 3.279E-08 | 2.500E-08 | 9.000E-08 |
| 13268 | 664 | K664G | B | 2.4e-08 | 2.1e-08 | 6.4e-08 | 3.633E-08 | 2.401E-08 | 2.100E-08 | 6.400E-08 |
| 13269 | 664 | K664H | B | 2.5e-08 | 3.8e-08 | 2.1e-08 | 2.800E-08 | 8.888E-09 | 2.100E-08 | 3.800E-08 |
| 13270 | 664 | K664I | B | 7.5e-08 | 6.4e-08 | 5.1e-08 | 6.333E-08 | 1.201E-08 | 5.100E-08 | 7.500E-08 |
| 13271 | 664 | K664L | B | 2e-08 | 1.8e-08 | 3e-08 | 2.267E-08 | 6.429E-09 | 1.800E-08 | 3.000E-08 |
| 13272 | 664 | K664K | B | 7.5e-08 | 1.9e-08 | 5.6e-08 | 5.000E-08 | 2.848E-08 | 1.900E-08 | 7.500E-08 |
| 13273 | 664 | K664M | B | 3.7e-08 | 2.8e-08 | 3.7e-08 | 3.400E-08 | 5.196E-09 | 2.800E-08 | 3.700E-08 |
| 13274 | 664 | K664F | B | 3.5e-08 | 4.6e-08 | 3.2e-08 | 3.767E-08 | 7.371E-09 | 3.200E-08 | 4.600E-08 |
| 13275 | 664 | K664P | B | 5.1e-08 | 4.1e-08 | 2.6e-08 | 3.933E-08 | 1.258E-08 | 2.600E-08 | 5.100E-08 |
| 13276 | 664 | K664S | B | 3.4e-08 | 5.6e-08 | 2.7e-08 | 3.900E-08 | 1.513E-08 | 2.700E-08 | 5.600E-08 |
| 13277 | 664 | K664T | B | 2.1e-08 | 3e-08 | 1.4e-07 | 6.367E-08 | 6.626E-08 | 2.100E-08 | 1.400E-07 |
| 13278 | 664 | K664W | B | 2.7e-08 | 6.1e-08 | 7.4e-08 | 5.400E-08 | 2.427E-08 | 2.700E-08 | 7.400E-08 |
| 13279 | 664 | K664Y | B | 1.1e-07 | 3.1e-08 | 3.3e-08 | 5.800E-08 | 4.504E-08 | 3.100E-08 | 1.100E-07 |
| 13280 | 664 | K664V | B | 8.6e-08 | 2.9e-08 | 2e-08 | 4.500E-08 | 3.579E-08 | 2.000E-08 | 8.600E-08 |
| 13281 | 665 | A665A | B | 5.6e-08 | 1.4e-07 | 1.1e-07 | 1.020E-07 | 4.257E-08 | 5.600E-08 | 1.400E-07 |
| 13282 | 665 | A665R | B | 1.2e-07 | 1.6e-07 | 2.6e-08 | 1.020E-07 | 6.879E-08 | 2.600E-08 | 1.600E-07 |
| 13283 | 665 | A665N | B | 3.8e-08 | 5.2e-08 | 4.9e-08 | 4.633E-08 | 7.371E-09 | 3.800E-08 | 5.200E-08 |
| 13284 | 665 | A665D | B | 5.1e-08 | 9.7e-08 | 7.1e-08 | 7.300E-08 | 2.307E-08 | 5.100E-08 | 9.700E-08 |
| 13285 | 665 | A665C | B | 3e-08 | 3.7e-08 | 4.9e-08 | 3.867E-08 | 9.609E-09 | 3.000E-08 | 4.900E-08 |
| 13286 | 665 | A665E | B | 2.8e-08 | 4.4e-08 | 8.5e-08 | 5.233E-08 | 2.940E-08 | 2.800E-08 | 8.500E-08 |
| 13287 | 665 | A665Q | B | 3e-08 | 1.2e-07 | 6.9e-08 | 7.300E-08 | 4.513E-08 | 3.000E-08 | 1.200E-07 |
| 13288 | 665 | A665G | B | 8.3e-08 | 4.4e-08 | 3.4e-08 | 5.367E-08 | 2.589E-08 | 3.400E-08 | 8.300E-08 |
| 13289 | 665 | A665H | B | 3.1e-08 | 1.8e-08 | 3.9e-08 | 2.933E-08 | 1.060E-08 | 1.800E-08 | 3.900E-08 |
| 13290 | 665 | A665I | B | 2.6e-08 | 2e-08 | 4e-08 | 2.867E-08 | 1.026E-08 | 2.000E-08 | 4.000E-08 |
| 13291 | 665 | A665L | B | 5.2e-08 | 1e-08 | 1.9e-08 | 2.700E-08 | 2.211E-08 | 1.000E-08 | 5.200E-08 |
| 13292 | 665 | A665K | B | 3.6e-08 | 3.7e-08 | 5.4e-08 | 4.233E-08 | 1.012E-08 | 3.600E-08 | 5.400E-08 |
| 13293 | 665 | A665M | B | 6.6e-08 | 8.9e-08 | 7.2e-08 | 7.567E-08 | 1.193E-08 | 6.600E-08 | 8.900E-08 |
| 13294 | 665 | A665F | B | 8.1e-08 | 3.1e-08 | 3e-08 | 4.733E-08 | 2.916E-08 | 3.000E-08 | 8.100E-08 |
| 13295 | 665 | A665P | B | 1.3e-07 | 5.7e-08 | 6e-08 | 8.233E-08 | 4.131E-08 | 5.700E-08 | 1.300E-07 |
| 13296 | 665 | A665S | B | 1.3e-08 | 6.4e-08 | 5.8e-08 | 4.500E-08 | 2.787E-08 | 1.300E-08 | 6.400E-08 |
| 13297 | 665 | A665T | B | 5.2e-08 | 6.6e-08 | 4.1e-08 | 5.300E-08 | 1.253E-08 | 4.100E-08 | 6.600E-08 |
| 13298 | 665 | A665W | B | 4.1e-08 | 1.9e-08 | 3.4e-08 | 3.133E-08 | 1.124E-08 | 1.900E-08 | 4.100E-08 |
| 13299 | 665 | A665Y | B | 3.1e-08 | 5.4e-08 | 8.4e-08 | 5.633E-08 | 2.658E-08 | 3.100E-08 | 8.400E-08 |
| 13300 | 665 | A665V | B | 3.9e-08 | 6.9e-08 | 7.6e-08 | 6.133E-08 | 1.966E-08 | 3.900E-08 | 7.600E-08 |
| 13301 | 666 | P666A | B | 2.1e-08 | 3.8e-08 | 7.5e-08 | 4.467E-08 | 2.761E-08 | 2.100E-08 | 7.500E-08 |
| 13302 | 666 | P666R | B | 1.2e-08 | 7.6e-08 | 3.9e-08 | 4.233E-08 | 3.213E-08 | 1.200E-08 | 7.600E-08 |
| 13303 | 666 | P666N | B | 5.3e-08 | 9.8e-08 | 2.8e-08 | 5.967E-08 | 3.547E-08 | 2.800E-08 | 9.800E-08 |
| 13304 | 666 | P666D | B | 4.6e-08 | 7.5e-08 | 7e-08 | 6.367E-08 | 1.550E-08 | 4.600E-08 | 7.500E-08 |
| 13305 | 666 | P666C | B | 1.1e-07 | 4.4e-08 | 4.3e-08 | 6.567E-08 | 3.840E-08 | 4.300E-08 | 1.100E-07 |
| 13306 | 666 | P666E | B | 1e-07 | 3.3e-08 | 4.5e-08 | 5.933E-08 | 3.573E-08 | 3.300E-08 | 1.000E-07 |
| 13307 | 666 | P666Q | B | 4.7e-08 | 3.3e-08 | 8.2e-08 | 5.400E-08 | 2.524E-08 | 3.300E-08 | 8.200E-08 |
| 13308 | 666 | P666G | B | 3.6e-08 | 2.5e-08 | 5.9e-08 | 4.000E-08 | 1.735E-08 | 2.500E-08 | 5.900E-08 |
| 13309 | 666 | P666H | B | 8.1e-08 | 3.9e-08 | 8.1e-09 | 4.270E-08 | 3.659E-08 | 8.100E-09 | 8.100E-08 |
| 13310 | 666 | P666I | B | 6.7e-08 | 2.1e-08 | 3.3e-08 | 4.033E-08 | 2.386E-08 | 2.100E-08 | 6.700E-08 |
| 13311 | 666 | P666L | B | 5e-08 | 3.5e-08 | 6.3e-08 | 4.933E-08 | 1.401E-08 | 3.500E-08 | 6.300E-08 |
| 13312 | 666 | P666K | B | 7.9e-08 | 1.2e-08 | 3.5e-08 | 4.200E-08 | 3.404E-08 | 1.200E-08 | 7.900E-08 |
| 13313 | 666 | P666M | B | 7.1e-08 | 1.1e-07 | 2.6e-08 | 6.900E-08 | 4.204E-08 | 2.600E-08 | 1.100E-07 |
| 13314 | 666 | P666F | B | 7.7e-08 | 1.3e-07 | 9.7e-08 | 1.013E-07 | 2.676E-08 | 7.700E-08 | 1.300E-07 |
| 13315 | 666 | P666P | B | 4.4e-08 | 2.7e-08 | 1.2e-08 | 2.767E-08 | 1.601E-08 | 1.200E-08 | 4.400E-08 |
| 13316 | 666 | P666S | B | 2.7e-08 | 1e-07 | 1.3e-07 | 8.567E-08 | 5.297E-08 | 2.700E-08 | 1.300E-07 |
| 13317 | 666 | P666T | B | 6.8e-08 | 8.4e-08 | 6.3e-08 | 7.167E-08 | 1.097E-08 | 6.300E-08 | 8.400E-08 |
| 13318 | 666 | P666W | B | 3.3e-08 | 2.9e-08 | 6.5e-08 | 4.233E-08 | 1.973E-08 | 2.900E-08 | 6.500E-08 |
| 13319 | 666 | P666Y | B | 2.5e-08 | 1e-07 | 8.6e-09 | 4.453E-08 | 4.873E-08 | 8.600E-09 | 1.000E-07 |
| 13320 | 666 | P666V | B | 3.8e-08 | 3.1e-08 | 3.4e-08 | 3.433E-08 | 3.512E-09 | 3.100E-08 | 3.800E-08 |
| 13321 | 667 | K667A | B | 2.2e-08 | 4.7e-08 | 3.7e-08 | 3.533E-08 | 1.258E-08 | 2.200E-08 | 4.700E-08 |
| 13322 | 667 | K667R | B | 5.5e-08 | 2.8e-08 | 2.5e-08 | 3.600E-08 | 1.652E-08 | 2.500E-08 | 5.500E-08 |
| 13323 | 667 | K667N | B | 6e-08 | 1.5e-08 | 4e-08 | 3.833E-08 | 2.255E-08 | 1.500E-08 | 6.000E-08 |
| 13324 | 667 | K667D | B | 1.6e-08 | 9.4e-09 | 8.2e-08 | 3.580E-08 | 4.015E-08 | 9.400E-09 | 8.200E-08 |
| 13325 | 667 | K667C | B | 3.7e-08 | 7.6e-08 | 1.3e-08 | 4.200E-08 | 3.180E-08 | 1.300E-08 | 7.600E-08 |
| 13326 | 667 | K667E | B | 1.9e-08 | 4.6e-08 | 6.2e-08 | 4.233E-08 | 2.173E-08 | 1.900E-08 | 6.200E-08 |
| 13327 | 667 | K667Q | B | 9.7e-08 | 6.1e-08 | 5.1e-08 | 6.967E-08 | 2.419E-08 | 5.100E-08 | 9.700E-08 |
| 13328 | 667 | K667G | B | 2.9e-08 | 2.3e-08 | 6e-08 | 3.733E-08 | 1.986E-08 | 2.300E-08 | 6.000E-08 |
| 13329 | 667 | K667H | B | 2e-08 | 1.8e-08 | 3.2e-08 | 2.333E-08 | 7.572E-09 | 1.800E-08 | 3.200E-08 |
| 13330 | 667 | K667I | B | 1.7e-08 | 7.1e-08 | 4.7e-08 | 4.500E-08 | 2.706E-08 | 1.700E-08 | 7.100E-08 |
| 13331 | 667 | K667L | B | 4.5e-08 | 2.6e-08 | 6.6e-08 | 4.567E-08 | 2.001E-08 | 2.600E-08 | 6.600E-08 |
| 13332 | 667 | K667K | B | 4.5e-08 | 4.7e-08 | 2.4e-08 | 3.867E-08 | 1.274E-08 | 2.400E-08 | 4.700E-08 |
| 13333 | 667 | K667M | B | 3.3e-08 | 1.3e-07 | 2.1e-08 | 6.133E-08 | 5.977E-08 | 2.100E-08 | 1.300E-07 |
| 13334 | 667 | K667F | B | 3.6e-08 | 8.4e-08 | 1.5e-08 | 4.500E-08 | 3.537E-08 | 1.500E-08 | 8.400E-08 |
| 13335 | 667 | K667P | B | 5.1e-08 | 6.1e-08 | 7.6e-08 | 6.267E-08 | 1.258E-08 | 5.100E-08 | 7.600E-08 |
| 13336 | 667 | K667S | B | 2.2e-08 | 4.4e-08 | 4.6e-08 | 3.733E-08 | 1.332E-08 | 2.200E-08 | 4.600E-08 |
| 13337 | 667 | K667T | B | 7.1e-08 | 2.1e-08 | 2.5e-08 | 3.900E-08 | 2.778E-08 | 2.100E-08 | 7.100E-08 |
| 13338 | 667 | K667W | B | 3.6e-08 | 9.4e-08 | 2.2e-08 | 5.067E-08 | 3.818E-08 | 2.200E-08 | 9.400E-08 |
| 13339 | 667 | K667Y | B | 2.8e-08 | 7.2e-08 | 5.4e-08 | 5.133E-08 | 2.212E-08 | 2.800E-08 | 7.200E-08 |
| 13340 | 667 | K667V | B | 4.4e-08 | 4.9e-08 | 4.3e-08 | 4.533E-08 | 3.215E-09 | 4.300E-08 | 4.900E-08 |
| 13341 | 668 | L668A | B | 3e-08 | 1.3e-08 | 5.3e-08 | 3.200E-08 | 2.007E-08 | 1.300E-08 | 5.300E-08 |
| 13342 | 668 | L668R | B | 2.9e-08 | 7.9e-08 | 1.7e-08 | 4.167E-08 | 3.288E-08 | 1.700E-08 | 7.900E-08 |
| 13343 | 668 | L668N | B | 2.8e-08 | 2.4e-08 | 3.5e-08 | 2.900E-08 | 5.568E-09 | 2.400E-08 | 3.500E-08 |
| 13344 | 668 | L668D | B | 1.4e-08 | 7e-08 | 2.7e-08 | 3.700E-08 | 2.931E-08 | 1.400E-08 | 7.000E-08 |

| | | | | | | | | | | |
|---|---|---|---|---|---|---|---|---|---|---|
| 13345 | 668 | L668C | B | 7.2e-08 | 6e-08 | 3.2e-08 | 5.467E-08 | 2.053E-08 | 3.200E-08 | 7.200E-08 |
| 13346 | 668 | L668E | B | 6.2e-08 | 3.3e-08 | 3.4e-08 | 4.300E-08 | 1.646E-08 | 3.300E-08 | 6.200E-08 |
| 13347 | 668 | L668Q | B | 6e-08 | 4.1e-08 | 3.4e-08 | 4.500E-08 | 1.345E-08 | 3.400E-08 | 6.000E-08 |
| 13348 | 668 | L668G | B | 2.7e-08 | 4.6e-08 | 3.9e-08 | 3.733E-08 | 9.609E-09 | 2.700E-08 | 4.600E-08 |
| 13349 | 668 | L668H | B | 5.9e-08 | 1.7e-07 | 8.3e-08 | 1.040E-07 | 5.840E-08 | 5.900E-08 | 1.700E-07 |
| 13350 | 668 | L668I | B | 3e-08 | 4.2e-08 | 8e-09 | 2.667E-08 | 1.724E-08 | 8.000E-09 | 4.200E-08 |
| 13351 | 668 | L668L | B | 3.6e-08 | 4.4e-08 | 4.3e-08 | 4.100E-08 | 4.359E-09 | 3.600E-08 | 4.400E-08 |
| 13352 | 668 | L668K | B | 1.5e-08 | 1e-07 | 4.5e-08 | 5.333E-08 | 4.311E-08 | 1.500E-08 | 1.000E-07 |
| 13353 | 668 | L668M | B | 1.4e-07 | 8.8e-08 | 1.4e-08 | 8.067E-08 | 6.332E-08 | 1.400E-08 | 1.400E-07 |
| 13354 | 668 | L668F | B | 4e-08 | 1.8e-08 | 3.7e-08 | 3.167E-08 | 1.193E-08 | 1.800E-08 | 4.000E-08 |
| 13355 | 668 | L668P | B | 1.2e-07 | 3.2e-08 | 2.9e-08 | 6.033E-08 | 5.169E-08 | 2.900E-08 | 1.200E-07 |
| 13356 | 668 | L668S | B | 4.7e-08 | 2.9e-08 | 3.5e-08 | 3.700E-08 | 9.165E-09 | 2.900E-08 | 4.700E-08 |
| 13357 | 668 | L668T | B | 8.8e-08 | 1.4e-07 | 1.1e-07 | 1.127E-07 | 2.610E-08 | 8.800E-08 | 1.400E-07 |
| 13358 | 668 | L668W | B | 4.1e-08 | 2.6e-08 | 1.6e-07 | 7.567E-08 | 7.342E-08 | 2.600E-08 | 1.600E-07 |
| 13359 | 668 | L668Y | B | 8.5e-08 | 1.4e-08 | 3.5e-08 | 4.467E-08 | 3.647E-08 | 1.400E-08 | 8.500E-08 |
| 13360 | 668 | L668V | B | 3.1e-08 | 6e-08 | 6.1e-08 | 5.067E-08 | 1.704E-08 | 3.100E-08 | 6.100E-08 |
| 13361 | 669 | L669A | B | 6.3e-08 | 4.2e-08 | 4.7e-08 | 5.067E-08 | 1.097E-08 | 4.200E-08 | 6.300E-08 |
| 13362 | 669 | L669R | B | 2.3e-08 | 4.8e-08 | 1.3e-07 | 6.700E-08 | 5.597E-08 | 2.300E-08 | 1.300E-07 |
| 13363 | 669 | L669N | B | 5e-09 | 1.7e-08 | 2.1e-08 | 1.433E-08 | 8.327E-09 | 5.000E-09 | 2.100E-08 |
| 13364 | 669 | L669D | B | 5.9e-08 | 4.6e-08 | 4.7e-08 | 5.067E-08 | 7.234E-09 | 4.600E-08 | 5.900E-08 |
| 13365 | 669 | L669C | B | 8.8e-08 | 9.7e-08 | 5.1e-08 | 7.867E-08 | 2.438E-08 | 5.100E-08 | 9.700E-08 |
| 13366 | 669 | L669E | B | 8.3e-08 | 3.6e-08 | 4.4e-08 | 5.433E-08 | 2.515E-08 | 3.600E-08 | 8.300E-08 |
| 13367 | 669 | L669Q | B | 2.8e-08 | 8.2e-08 | 2.9e-08 | 4.633E-08 | 3.089E-08 | 2.800E-08 | 8.200E-08 |
| 13368 | 669 | L669G | B | 4.2e-08 | 7.1e-08 | 3.9e-08 | 5.067E-08 | 1.767E-08 | 3.900E-08 | 7.100E-08 |
| 13369 | 669 | L669H | B | 6.2e-08 | 4e-08 | 9.7e-08 | 6.633E-08 | 2.875E-08 | 4.000E-08 | 9.700E-08 |
| 13370 | 669 | L669I | B | 6.1e-08 | 8.8e-08 | 2.9e-08 | 5.933E-08 | 2.954E-08 | 2.900E-08 | 8.800E-08 |
| 13371 | 669 | L669L | B | 3.7e-08 | 4.8e-08 | 1.5e-07 | 7.833E-08 | 6.231E-08 | 3.700E-08 | 1.500E-07 |
| 13372 | 669 | L669K | B | 1e-07 | 3.8e-08 | 2.9e-08 | 5.567E-08 | 3.866E-08 | 2.900E-08 | 1.000E-07 |
| 13373 | 669 | L669M | B | 6.5e-08 | 4e-08 | 3.8e-08 | 4.767E-08 | 1.504E-08 | 3.800E-08 | 6.500E-08 |
| 13374 | 669 | L669F | B | 4.5e-08 | 2.4e-08 | 2e-08 | 2.967E-08 | 1.343E-08 | 2.000E-08 | 4.500E-08 |
| 13375 | 669 | L669P | B | 7.5e-08 | 2.1e-08 | 1.1e-08 | 3.567E-08 | 3.443E-08 | 1.100E-08 | 7.500E-08 |
| 13376 | 669 | L669S | B | 6e-08 | 5.5e-08 | 7.6e-08 | 6.367E-08 | 1.097E-08 | 5.500E-08 | 7.600E-08 |
| 13377 | 669 | L669T | B | 1e-08 | 5.3e-08 | 5.4e-08 | 3.900E-08 | 2.512E-08 | 1.000E-08 | 5.400E-08 |
| 13378 | 669 | L669W | B | 3.2e-08 | 3e-08 | 3.2e-08 | 3.133E-08 | 1.155E-09 | 3.000E-08 | 3.200E-08 |
| 13379 | 669 | L669Y | B | 4.5e-08 | 2.6e-08 | 5.8e-08 | 4.300E-08 | 1.609E-08 | 2.600E-08 | 5.800E-08 |
| 13380 | 669 | L669V | B | 1.1e-07 | 3.5e-08 | 3.3e-08 | 5.933E-08 | 4.389E-08 | 3.300E-08 | 1.100E-07 |
| 13381 | 670 | I670A | B | 2.4e-08 | 1.8e-08 | 8.1e-08 | 4.100E-08 | 3.477E-08 | 1.800E-08 | 8.100E-08 |
| 13382 | 670 | I670R | B | 1.8e-07 | 6.3e-08 | 9.8e-08 | 1.137E-07 | 6.005E-08 | 6.300E-08 | 1.800E-07 |
| 13383 | 670 | I670N | B | 8.4e-09 | 4.2e-08 | 2.9e-08 | 2.647E-08 | 1.694E-08 | 8.400E-09 | 4.200E-08 |
| 13384 | 670 | I670D | B | 2.8e-08 | 7.5e-08 | 3.2e-08 | 4.500E-08 | 2.606E-08 | 2.800E-08 | 7.500E-08 |
| 13385 | 670 | I670C | B | 1.5e-08 | 6e-08 | 4.5e-08 | 4.000E-08 | 2.291E-08 | 1.500E-08 | 6.000E-08 |
| 13386 | 670 | I670E | B | 4.2e-08 | 2.8e-08 | 5e-08 | 4.000E-08 | 1.114E-08 | 2.800E-08 | 5.000E-08 |
| 13387 | 670 | I670Q | B | 4.8e-08 | 7.3e-08 | 5.8e-08 | 5.967E-08 | 1.258E-08 | 4.800E-08 | 7.300E-08 |
| 13388 | 670 | I670G | B | 7.1e-08 | 5.6e-08 | 3.8e-08 | 5.500E-08 | 1.652E-08 | 3.800E-08 | 7.100E-08 |
| 13389 | 670 | I670H | B | 1.8e-08 | 2.8e-08 | 7.4e-08 | 4.000E-08 | 2.987E-08 | 1.800E-08 | 7.400E-08 |
| 13390 | 670 | I670I | B | 2.6e-08 | 4.4e-08 | 3.5e-08 | 3.500E-08 | 9.000E-09 | 2.600E-08 | 4.400E-08 |
| 13391 | 670 | I670L | B | 3.4e-08 | 1.5e-08 | 2.9e-08 | 2.600E-08 | 9.849E-09 | 1.500E-08 | 3.400E-08 |
| 13392 | 670 | I670K | B | 2.9e-08 | 2.3e-08 | 3.9e-08 | 3.033E-08 | 8.083E-09 | 2.300E-08 | 3.900E-08 |
| 13393 | 670 | I670M | B | 2.9e-08 | 5.1e-08 | 4.2e-08 | 4.067E-08 | 1.106E-08 | 2.900E-08 | 5.100E-08 |
| 13394 | 670 | I670F | B | 4.8e-08 | 3.9e-08 | 4.3e-08 | 4.333E-08 | 4.509E-09 | 3.900E-08 | 4.800E-08 |
| 13395 | 670 | I670P | B | 1.2e-07 | 7.8e-08 | 1.6e-08 | 7.133E-08 | 5.232E-08 | 1.600E-08 | 1.200E-07 |
| 13396 | 670 | I670S | B | 5e-08 | 6.3e-08 | 4.9e-08 | 5.400E-08 | 7.810E-09 | 4.900E-08 | 6.300E-08 |
| 13397 | 670 | I670T | B | 4.2e-08 | 5.6e-08 | 8.7e-08 | 6.167E-08 | 2.303E-08 | 4.200E-08 | 8.700E-08 |
| 13398 | 670 | I670W | B | 3.1e-08 | 2.4e-08 | 3.4e-08 | 2.967E-08 | 5.132E-09 | 2.400E-08 | 3.400E-08 |
| 13399 | 670 | I670Y | B | 5e-08 | 4.9e-08 | 2.6e-08 | 4.167E-08 | 1.358E-08 | 2.600E-08 | 5.000E-08 |
| 13400 | 670 | I670V | B | 6.1e-08 | 9.5e-08 | 3.6e-08 | 6.400E-08 | 2.961E-08 | 3.600E-08 | 9.500E-08 |
| 13401 | 671 | Y671A | B | 4.4e-08 | 6.3e-08 | 5.1e-08 | 5.267E-08 | 9.609E-09 | 4.400E-08 | 6.300E-08 |
| 13402 | 671 | Y671R | B | 3.3e-08 | 7.1e-08 | 4.2e-08 | 4.867E-08 | 1.986E-08 | 3.300E-08 | 7.100E-08 |
| 13403 | 671 | Y671N | B | 3.5e-08 | 4.1e-08 | 4.2e-08 | 3.933E-08 | 3.786E-09 | 3.500E-08 | 4.200E-08 |
| 13404 | 671 | Y671D | B | 3.3e-08 | 2.7e-08 | 5e-08 | 3.667E-08 | 1.193E-08 | 2.700E-08 | 5.000E-08 |
| 13405 | 671 | Y671C | B | 1.6e-08 | 6.2e-08 | 4.1e-08 | 3.967E-08 | 2.303E-08 | 1.600E-08 | 6.200E-08 |
| 13406 | 671 | Y671E | B | 2.3e-08 | 2.9e-08 | 4e-08 | 3.067E-08 | 8.622E-09 | 2.300E-08 | 4.000E-08 |
| 13407 | 671 | Y671Q | B | 3.4e-08 | 3.2e-08 | 3.5e-08 | 3.367E-08 | 1.528E-09 | 3.200E-08 | 3.500E-08 |
| 13408 | 671 | Y671G | B | 3e-08 | 6.1e-08 | 5.3e-08 | 4.800E-08 | 1.609E-08 | 3.000E-08 | 6.100E-08 |
| 13409 | 671 | Y671H | B | 7.7e-08 | 4.3e-08 | 2.3e-08 | 4.767E-08 | 2.730E-08 | 2.300E-08 | 7.700E-08 |
| 13410 | 671 | Y671I | B | 3.9e-08 | 6.6e-08 | 2.8e-08 | 4.433E-08 | 1.955E-08 | 2.800E-08 | 6.600E-08 |
| 13411 | 671 | Y671L | B | 5.8e-08 | 4.9e-08 | 4.3e-08 | 5.000E-08 | 7.550E-09 | 4.300E-08 | 5.800E-08 |
| 13412 | 671 | Y671K | B | 6.5e-08 | 8.7e-08 | 9.6e-08 | 8.267E-08 | 1.595E-08 | 6.500E-08 | 9.600E-08 |
| 13413 | 671 | Y671M | B | 6.3e-08 | 4.1e-08 | 5.2e-08 | 5.200E-08 | 1.100E-08 | 4.100E-08 | 6.300E-08 |
| 13414 | 671 | Y671F | B | 6.3e-08 | 5.9e-08 | 2.6e-08 | 4.933E-08 | 2.031E-08 | 2.600E-08 | 6.300E-08 |
| 13415 | 671 | Y671P | B | 1.8e-07 | 9e-08 | 2.3e-08 | 9.767E-08 | 7.878E-08 | 2.300E-08 | 1.800E-07 |
| 13416 | 671 | Y671S | B | 4.4e-08 | 3.9e-08 | 6e-08 | 4.767E-08 | 1.097E-08 | 3.900E-08 | 6.000E-08 |
| 13417 | 671 | Y671T | B | 3.1e-08 | 2.7e-08 | 4.7e-08 | 3.500E-08 | 1.058E-08 | 2.700E-08 | 4.700E-08 |
| 13418 | 671 | Y671W | B | 9.3e-08 | 4.5e-08 | 3.9e-08 | 5.900E-08 | 2.960E-08 | 3.900E-08 | 9.300E-08 |
| 13419 | 671 | Y671Y | B | 2.7e-08 | 1e-07 | 2.6e-08 | 5.100E-08 | 4.244E-08 | 2.600E-08 | 1.000E-07 |
| 13420 | 671 | Y671V | B | 2.1e-08 | 1.5e-07 | 5.7e-08 | 7.600E-08 | 6.657E-08 | 2.100E-08 | 1.500E-07 |
| 13421 | 672 | S672A | B | 1.7e-08 | 2.7e-08 | 6.1e-08 | 3.500E-08 | 2.307E-08 | 1.700E-08 | 6.100E-08 |
| 13422 | 672 | S672R | B | 6e-08 | 3.8e-08 | 5.6e-08 | 5.133E-08 | 1.172E-08 | 3.800E-08 | 6.000E-08 |
| 13423 | 672 | S672N | B | 3.2e-08 | 5.2e-08 | 4.2e-08 | 4.200E-08 | 1.000E-08 | 3.200E-08 | 5.200E-08 |
| 13424 | 672 | S672D | B | 2.1e-08 | 3.6e-08 | 7.8e-08 | 4.500E-08 | 2.955E-08 | 2.100E-08 | 7.800E-08 |
| 13425 | 672 | S672C | B | 9.9e-08 | 2e-08 | 4e-08 | 5.300E-08 | 4.107E-08 | 2.000E-08 | 9.900E-08 |
| 13426 | 672 | S672E | B | 1.3e-07 | 7.4e-08 | 4e-08 | 8.133E-08 | 4.545E-08 | 4.000E-08 | 1.300E-07 |
| 13427 | 672 | S672Q | B | 3.7e-08 | 3.6e-08 | 1.1e-07 | 6.100E-08 | 4.244E-08 | 3.600E-08 | 1.100E-07 |
| 13428 | 672 | S672G | B | 4.4e-08 | 3.5e-08 | 2e-08 | 3.300E-08 | 1.212E-08 | 2.000E-08 | 4.400E-08 |
| 13429 | 672 | S672H | B | 5.9e-08 | 1.3e-08 | 1.2e-07 | 6.400E-08 | 5.367E-08 | 1.300E-08 | 1.200E-07 |

| | | | | | | | | | | |
|---|---|---|---|---|---|---|---|---|---|---|
| 13430 | 672 | S672I | B | 3.3e-08 | 7.6e-08 | 5.5e-08 | 5.467E-08 | 2.150E-08 | 3.300E-08 | 7.600E-08 |
| 13431 | 672 | S672L | B | 2.9e-08 | 1.6e-08 | 5.6e-08 | 3.367E-08 | 2.040E-08 | 1.600E-08 | 5.600E-08 |
| 13432 | 672 | S672K | B | 3.2e-08 | 9.6e-08 | 2.2e-08 | 5.000E-08 | 4.015E-08 | 2.200E-08 | 9.600E-08 |
| 13433 | 672 | S672M | B | 4.5e-08 | 4.1e-08 | 2.6e-08 | 3.733E-08 | 1.002E-08 | 2.600E-08 | 4.500E-08 |
| 13434 | 672 | S672F | B | 3.1e-08 | 4e-08 | 3.7e-08 | 3.600E-08 | 4.583E-09 | 3.100E-08 | 4.000E-08 |
| 13435 | 672 | S672P | B | 3.8e-08 | 4.5e-08 | 5e-08 | 4.433E-08 | 6.028E-09 | 3.800E-08 | 5.000E-08 |
| 13436 | 672 | S672S | B | 7e-08 | 5.4e-08 | 5.3e-08 | 5.900E-08 | 9.539E-09 | 5.300E-08 | 7.000E-08 |
| 13437 | 672 | S672T | B | 4.1e-08 | 3.2e-08 | 4e-08 | 3.767E-08 | 4.933E-09 | 3.200E-08 | 4.100E-08 |
| 13438 | 672 | S672W | B | 4.7e-08 | 1.5e-08 | 5.6e-08 | 3.933E-08 | 2.155E-08 | 1.500E-08 | 5.600E-08 |
| 13439 | 672 | S672Y | B | 6.5e-08 | 4.2e-08 | 3.8e-08 | 4.833E-08 | 1.457E-08 | 3.800E-08 | 6.500E-08 |
| 13440 | 672 | S672V | B | 7.4e-08 | 5.5e-08 | 6.3e-08 | 6.400E-08 | 9.539E-09 | 5.500E-08 | 7.400E-08 |
| 13441 | 673 | A673A | B | 4.3e-08 | 1.3e-07 | 3.4e-08 | 6.900E-08 | 5.302E-08 | 3.400E-08 | 1.300E-07 |
| 13442 | 673 | A673R | B | 7.6e-08 | 4.1e-08 | 6.4e-08 | 6.033E-08 | 1.779E-08 | 4.100E-08 | 7.600E-08 |
| 13443 | 673 | A673N | B | 5.1e-08 | 3.5e-08 | 7.8e-08 | 5.467E-08 | 2.173E-08 | 3.500E-08 | 7.800E-08 |
| 13444 | 673 | A673D | B | 4.3e-08 | 3.8e-08 | 1.2e-07 | 6.700E-08 | 4.597E-08 | 3.800E-08 | 1.200E-07 |
| 13445 | 673 | A673C | B | 4.5e-08 | 6.9e-08 | 5.6e-08 | 5.667E-08 | 1.201E-08 | 4.500E-08 | 6.900E-08 |
| 13446 | 673 | A673E | B | 7.2e-08 | 2.5e-08 | 1.8e-08 | 3.833E-08 | 2.937E-08 | 1.800E-08 | 7.200E-08 |
| 13447 | 673 | A673Q | B | 3.7e-08 | 6e-08 | 3.5e-08 | 4.400E-08 | 1.389E-08 | 3.500E-08 | 6.000E-08 |
| 13448 | 673 | A673G | B | 5.9e-08 | 5.1e-08 | 4.1e-08 | 5.033E-08 | 9.018E-09 | 4.100E-08 | 5.900E-08 |
| 13449 | 673 | A673H | B | 3.5e-08 | 2e-08 | 3.1e-08 | 2.867E-08 | 7.767E-09 | 2.000E-08 | 3.500E-08 |
| 13450 | 673 | A673I | B | 2.2e-08 | 2.5e-08 | 8.7e-08 | 4.467E-08 | 3.669E-08 | 2.200E-08 | 8.700E-08 |
| 13451 | 673 | A673L | B | 3.8e-08 | 2.6e-08 | 4.2e-08 | 3.533E-08 | 8.327E-09 | 2.600E-08 | 4.200E-08 |
| 13452 | 673 | A673K | B | 6.9e-08 | 6.1e-08 | 7.2e-08 | 6.733E-08 | 5.686E-09 | 6.100E-08 | 7.200E-08 |
| 13453 | 673 | A673M | B | 2.8e-08 | 4.4e-08 | 4.6e-08 | 3.933E-08 | 9.866E-09 | 2.800E-08 | 4.600E-08 |
| 13454 | 673 | A673F | B | 5.3e-08 | 2e-08 | 1.3e-07 | 6.767E-08 | 5.645E-08 | 2.000E-08 | 1.300E-07 |
| 13455 | 673 | A673P | B | 4.7e-08 | 8e-08 | 4.1e-08 | 5.600E-08 | 2.100E-08 | 4.100E-08 | 8.000E-08 |
| 13456 | 673 | A673S | B | 7.6e-08 | 4.7e-08 | 6.2e-08 | 6.167E-08 | 1.450E-08 | 4.700E-08 | 7.600E-08 |
| 13457 | 673 | A673T | B | 5e-08 | 1.1e-07 | 3.4e-08 | 6.467E-08 | 4.007E-08 | 3.400E-08 | 1.100E-07 |
| 13458 | 673 | A673W | B | 5e-08 | 1.5e-08 | 5.5e-08 | 4.000E-08 | 2.179E-08 | 1.500E-08 | 5.500E-08 |
| 13459 | 673 | A673Y | B | 1.7e-08 | 5.5e-08 | 2.4e-08 | 3.200E-08 | 2.022E-08 | 1.700E-08 | 5.500E-08 |
| 13460 | 673 | A673V | B | 6.9e-08 | 4.2e-08 | 1e-07 | 7.033E-08 | 2.902E-08 | 4.200E-08 | 1.000E-07 |
| 13461 | 674 | S674A | B | 7.3e-08 | 9.3e-08 | 8.4e-09 | 5.813E-08 | 4.422E-08 | 8.400E-09 | 9.300E-08 |
| 13462 | 674 | S674R | B | 6.4e-08 | 1.9e-08 | 2.5e-08 | 3.600E-08 | 2.443E-08 | 1.900E-08 | 6.400E-08 |
| 13463 | 674 | S674N | B | 1e-07 | 4e-08 | 3.4e-08 | 5.800E-08 | 3.650E-08 | 3.400E-08 | 1.000E-07 |
| 13464 | 674 | S674D | B | 2.1e-08 | 5.5e-08 | 1e-08 | 2.867E-08 | 2.346E-08 | 1.000E-08 | 5.500E-08 |
| 13465 | 674 | S674C | B | 2.1e-08 | 2.9e-08 | 1.7e-08 | 2.233E-08 | 6.110E-09 | 1.700E-08 | 2.900E-08 |
| 13466 | 674 | S674E | B | 6.7e-08 | 4.4e-08 | 4.8e-08 | 5.300E-08 | 1.229E-08 | 4.400E-08 | 6.700E-08 |
| 13467 | 674 | S674Q | B | 2.9e-08 | 4.9e-08 | 5.6e-08 | 4.467E-08 | 1.401E-08 | 2.900E-08 | 5.600E-08 |
| 13468 | 674 | S674G | B | 3.6e-08 | 1.8e-08 | 8.1e-08 | 4.500E-08 | 3.245E-08 | 1.800E-08 | 8.100E-08 |
| 13469 | 674 | S674H | B | 8.2e-08 | 7.3e-08 | 4.5e-08 | 6.667E-08 | 1.930E-08 | 4.500E-08 | 8.200E-08 |
| 13470 | 674 | S674I | B | 2.8e-08 | 6.5e-08 | 5.8e-08 | 5.033E-08 | 1.966E-08 | 2.800E-08 | 6.500E-08 |
| 13471 | 674 | S674L | B | 8.3e-08 | 6.2e-08 | 2e-08 | 5.500E-08 | 3.208E-08 | 2.000E-08 | 8.300E-08 |
| 13472 | 674 | S674K | B | 1.1e-08 | 9.1e-09 | 7.7e-08 | 3.237E-08 | 3.867E-08 | 9.100E-09 | 7.700E-08 |
| 13473 | 674 | S674M | B | 4.1e-08 | 3.4e-08 | 2.2e-08 | 3.233E-08 | 9.609E-09 | 2.200E-08 | 4.100E-08 |
| 13474 | 674 | S674F | B | 1e-07 | 4.4e-08 | 6.1e-08 | 6.833E-08 | 2.871E-08 | 4.400E-08 | 1.000E-07 |
| 13475 | 674 | S674P | B | 4.7e-08 | 1.4e-07 | 3.9e-08 | 7.533E-08 | 5.615E-08 | 3.900E-08 | 1.400E-07 |
| 13476 | 674 | S674S | B | 6e-08 | 5.5e-08 | 3.1e-08 | 4.867E-08 | 1.550E-08 | 3.100E-08 | 6.000E-08 |
| 13477 | 674 | S674T | B | 3.4e-08 | 2.2e-08 | 4.7e-08 | 3.433E-08 | 1.250E-08 | 2.200E-08 | 4.700E-08 |
| 13478 | 674 | S674W | B | 9.3e-08 | 7.1e-08 | 3.1e-08 | 6.500E-08 | 3.143E-08 | 3.100E-08 | 9.300E-08 |
| 13479 | 674 | S674Y | B | 1.8e-08 | 4.5e-08 | 6.5e-08 | 4.267E-08 | 2.359E-08 | 1.800E-08 | 6.500E-08 |
| 13480 | 674 | S674V | B | 7.7e-08 | 7.2e-08 | 5.5e-08 | 6.800E-08 | 1.153E-08 | 5.500E-08 | 7.700E-08 |
| 13481 | 675 | Y675A | B | 2.6e-08 | 9.5e-08 | 4.1e-08 | 5.400E-08 | 3.629E-08 | 2.600E-08 | 9.500E-08 |
| 13482 | 675 | Y675R | B | 2.8e-08 | 3.3e-08 | 2.7e-08 | 2.933E-08 | 3.215E-09 | 2.700E-08 | 3.300E-08 |
| 13483 | 675 | Y675N | B | 4e-08 | 4.5e-08 | 5.3e-08 | 4.600E-08 | 6.557E-09 | 4.000E-08 | 5.300E-08 |
| 13484 | 675 | Y675D | B | 6.2e-08 | 5.3e-08 | 5.4e-08 | 5.633E-08 | 4.933E-09 | 5.300E-08 | 6.200E-08 |
| 13485 | 675 | Y675C | B | 5.7e-08 | 3.7e-08 | 4.3e-08 | 4.567E-08 | 1.026E-08 | 3.700E-08 | 5.700E-08 |
| 13486 | 675 | Y675E | B | 8.3e-08 | 6.3e-08 | 2.9e-08 | 5.833E-08 | 2.730E-08 | 2.900E-08 | 8.300E-08 |
| 13487 | 675 | Y675Q | B | 7e-08 | 7.4e-08 | 7.2e-08 | 7.200E-08 | 2.000E-09 | 7.000E-08 | 7.400E-08 |
| 13488 | 675 | Y675G | B | 4.4e-08 | 1.8e-08 | 4.9e-08 | 3.700E-08 | 1.664E-08 | 1.800E-08 | 4.900E-08 |
| 13489 | 675 | Y675H | B | 1.8e-08 | 5.3e-08 | 4.6e-08 | 3.900E-08 | 1.852E-08 | 1.800E-08 | 5.300E-08 |
| 13490 | 675 | Y675I | B | 2.1e-08 | 4.5e-08 | 8.8e-08 | 5.133E-08 | 3.395E-08 | 2.100E-08 | 8.800E-08 |
| 13491 | 675 | Y675L | B | 1.1e-07 | 4.2e-08 | 4e-08 | 6.400E-08 | 3.985E-08 | 4.000E-08 | 1.100E-07 |
| 13492 | 675 | Y675K | B | 1.7e-08 | 3.6e-08 | 2.5e-08 | 2.600E-08 | 9.539E-09 | 1.700E-08 | 3.600E-08 |
| 13493 | 675 | Y675M | B | 2.2e-08 | 1.3e-08 | 5e-08 | 2.833E-08 | 1.930E-08 | 1.300E-08 | 5.000E-08 |
| 13494 | 675 | Y675F | B | 1.5e-08 | 9e-08 | 2.2e-08 | 4.233E-08 | 4.143E-08 | 1.500E-08 | 9.000E-08 |
| 13495 | 675 | Y675P | B | 2e-08 | 2.4e-08 | 1.7e-08 | 2.033E-08 | 3.512E-09 | 1.700E-08 | 2.400E-08 |
| 13496 | 675 | Y675S | B | 6.9e-08 | 3e-08 | 6.1e-08 | 5.333E-08 | 2.060E-08 | 3.000E-08 | 6.900E-08 |
| 13497 | 675 | Y675T | B | 9.5e-08 | 2.7e-08 | 9.1e-08 | 7.100E-08 | 3.816E-08 | 2.700E-08 | 9.500E-08 |
| 13498 | 675 | Y675W | B | 9e-08 | 3.2e-08 | 5.2e-09 | 4.240E-08 | 4.335E-08 | 5.200E-09 | 9.000E-08 |
| 13499 | 675 | Y675Y | B | 7.4e-08 | 3.3e-08 | 2.1e-08 | 4.267E-08 | 2.779E-08 | 2.100E-08 | 7.400E-08 |
| 13500 | 675 | Y675V | B | 1.6e-08 | 3.4e-08 | 4.1e-08 | 3.033E-08 | 1.290E-08 | 1.600E-08 | 4.100E-08 |
| 13501 | 676 | R676A | B | 3.6e-08 | 3.6e-08 | 1.7e-07 | 8.067E-08 | 7.736E-08 | 3.600E-08 | 1.700E-07 |
| 13502 | 676 | R676R | B | 7.2e-08 | 2e-08 | 2.8e-08 | 4.000E-08 | 2.800E-08 | 2.000E-08 | 7.200E-08 |
| 13503 | 676 | R676N | B | 8.6e-08 | 2.4e-08 | 5.1e-08 | 5.367E-08 | 3.109E-08 | 2.400E-08 | 8.600E-08 |
| 13504 | 676 | R676D | B | 5.6e-08 | 3.2e-08 | 2.2e-08 | 3.667E-08 | 1.747E-08 | 2.200E-08 | 5.600E-08 |
| 13505 | 676 | R676C | B | 1.5e-07 | 1.6e-08 | 6.4e-08 | 7.667E-08 | 6.789E-08 | 1.600E-08 | 1.500E-07 |
| 13506 | 676 | R676E | B | 4.5e-08 | 6.9e-08 | 3.4e-08 | 4.933E-08 | 1.790E-08 | 3.400E-08 | 6.900E-08 |
| 13507 | 676 | R676Q | B | 3.4e-08 | 4.7e-08 | 6.3e-08 | 4.800E-08 | 1.453E-08 | 3.400E-08 | 6.300E-08 |
| 13508 | 676 | R676G | B | 1.1e-07 | 6e-08 | 2.5e-08 | 6.500E-08 | 4.272E-08 | 2.500E-08 | 1.100E-07 |
| 13509 | 676 | R676H | B | 3e-08 | 1.3e-07 | 2.4e-08 | 6.133E-08 | 5.954E-08 | 2.400E-08 | 1.300E-07 |
| 13510 | 676 | R676I | B | 3.7e-08 | 6e-08 | 2.1e-08 | 3.933E-08 | 1.960E-08 | 2.100E-08 | 6.000E-08 |
| 13511 | 676 | R676L | B | 2.4e-08 | 3.6e-08 | 3.2e-08 | 3.067E-08 | 6.110E-09 | 2.400E-08 | 3.600E-08 |
| 13512 | 676 | R676K | B | 4.7e-08 | 6.4e-08 | 2.2e-08 | 4.433E-08 | 2.113E-08 | 2.200E-08 | 6.400E-08 |
| 13513 | 676 | R676M | B | 3.5e-08 | 6.2e-08 | 2.4e-08 | 4.033E-08 | 1.955E-08 | 2.400E-08 | 6.200E-08 |
| 13514 | 676 | R676F | B | 5.2e-08 | 6.8e-08 | 2.2e-08 | 4.733E-08 | 2.335E-08 | 2.200E-08 | 6.800E-08 |

| | | | | | | | | | | |
|---|---|---|---|---|---|---|---|---|---|---|
| 13515 | 676 | R676P | B | 8.5e-08 | 6.8e-08 | 3.5e-08 | 6.267E-08 | 2.542E-08 | 3.500E-08 | 8.500E-08 |
| 13516 | 676 | R676S | B | 5.1e-08 | 4.2e-08 | 5e-08 | 4.767E-08 | 4.933E-09 | 4.200E-08 | 5.100E-08 |
| 13517 | 676 | R676T | B | 4e-08 | 1.5e-07 | 2.2e-08 | 7.067E-08 | 6.929E-08 | 2.200E-08 | 1.500E-07 |
| 13518 | 676 | R676W | B | 4.2e-08 | 8.1e-08 | 2.1e-08 | 4.800E-08 | 3.045E-08 | 2.100E-08 | 8.100E-08 |
| 13519 | 676 | R676Y | B | 5.4e-08 | 6e-08 | 7.5e-08 | 6.300E-08 | 1.082E-08 | 5.400E-08 | 7.500E-08 |
| 13520 | 676 | R676V | B | 3.3e-08 | 1.9e-08 | 8.1e-08 | 4.433E-08 | 3.252E-08 | 1.900E-08 | 8.100E-08 |
| 13521 | 677 | Y677A | B | 1.3e-07 | 2.2e-08 | 5.8e-08 | 7.000E-08 | 5.499E-08 | 2.200E-08 | 1.300E-07 |
| 13522 | 677 | Y677R | B | 2.6e-08 | 9.1e-08 | 6.9e-08 | 6.200E-08 | 3.306E-08 | 2.600E-08 | 9.100E-08 |
| 13523 | 677 | Y677N | B | 2.7e-08 | 1.1e-07 | 3.8e-08 | 5.833E-08 | 4.508E-08 | 2.700E-08 | 1.100E-07 |
| 13524 | 677 | Y677D | B | 3.3e-08 | 3e-08 | 6.9e-08 | 4.400E-08 | 2.170E-08 | 3.000E-08 | 6.900E-08 |
| 13525 | 677 | Y677C | B | 4e-08 | 3e-08 | 4.4e-08 | 3.800E-08 | 7.211E-09 | 3.000E-08 | 4.400E-08 |
| 13526 | 677 | Y677E | B | 2.4e-08 | 6.5e-08 | 1.8e-08 | 3.567E-08 | 2.558E-08 | 1.800E-08 | 6.500E-08 |
| 13527 | 677 | Y677Q | B | 9.6e-08 | 3.7e-08 | 4.6e-08 | 5.967E-08 | 3.179E-08 | 3.700E-08 | 9.600E-08 |
| 13528 | 677 | Y677G | B | 5.5e-08 | 8.2e-08 | 4.7e-08 | 6.133E-08 | 1.834E-08 | 4.700E-08 | 8.200E-08 |
| 13529 | 677 | Y677H | B | 5.3e-08 | 7.8e-08 | 2.6e-08 | 5.233E-08 | 2.601E-08 | 2.600E-08 | 7.800E-08 |
| 13530 | 677 | Y677I | B | 3.7e-08 | 1.4e-08 | 5.8e-08 | 3.633E-08 | 2.201E-08 | 1.400E-08 | 5.800E-08 |
| 13531 | 677 | Y677L | B | 3.8e-08 | 1.1e-07 | 4.9e-08 | 6.567E-08 | 3.879E-08 | 3.800E-08 | 1.100E-07 |
| 13532 | 677 | Y677K | B | 3.8e-08 | 7e-08 | 5.2e-08 | 5.333E-08 | 1.604E-08 | 3.800E-08 | 7.000E-08 |
| 13533 | 677 | Y677M | B | 4.7e-08 | 3.4e-08 | 2.4e-08 | 3.500E-08 | 1.153E-08 | 2.400E-08 | 4.700E-08 |
| 13534 | 677 | Y677F | B | 5.5e-08 | 2.7e-08 | 2.4e-08 | 3.533E-08 | 1.710E-08 | 2.400E-08 | 5.500E-08 |
| 13535 | 677 | Y677P | B | 7.3e-08 | 2e-08 | 6e-08 | 5.100E-08 | 2.762E-08 | 2.000E-08 | 7.300E-08 |
| 13536 | 677 | Y677S | B | 2e-08 | 5e-08 | 2.4e-08 | 3.133E-08 | 1.629E-08 | 2.000E-08 | 5.000E-08 |
| 13537 | 677 | Y677T | B | 3.4e-08 | 7e-08 | 3.8e-08 | 4.733E-08 | 1.973E-08 | 3.400E-08 | 7.000E-08 |
| 13538 | 677 | Y677W | B | 6.4e-08 | 3.8e-08 | 7.1e-08 | 5.767E-08 | 1.739E-08 | 3.800E-08 | 7.100E-08 |
| 13539 | 677 | Y677Y | B | 4e-08 | 2.4e-08 | 4.7e-08 | 3.700E-08 | 1.179E-08 | 2.400E-08 | 4.700E-08 |
| 13540 | 677 | Y677V | B | 4.8e-08 | 2.1e-08 | 3.1e-08 | 3.333E-08 | 1.365E-08 | 2.100E-08 | 4.800E-08 |
| 13541 | 678 | T678A | B | 1.4e-08 | 1e-07 | 2.5e-08 | 4.633E-08 | 4.680E-08 | 1.400E-08 | 1.000E-07 |
| 13542 | 678 | T678R | B | 7.8e-08 | 2.9e-08 | 1.2e-07 | 7.567E-08 | 4.554E-08 | 2.900E-08 | 1.200E-07 |
| 13543 | 678 | T678N | B | 2.6e-08 | 2.9e-08 | 8.6e-08 | 4.700E-08 | 3.381E-08 | 2.600E-08 | 8.600E-08 |
| 13544 | 678 | T678D | B | 2.1e-08 | 2.1e-08 | 1.2e-08 | 1.800E-08 | 5.196E-09 | 1.200E-08 | 2.100E-08 |
| 13545 | 678 | T678C | B | 2.9e-08 | 1e-07 | 7.3e-08 | 6.733E-08 | 3.584E-08 | 2.900E-08 | 1.000E-07 |
| 13546 | 678 | T678E | B | 1.2e-07 | 3.5e-08 | 2e-08 | 5.833E-08 | 5.393E-08 | 2.000E-08 | 1.200E-07 |
| 13547 | 678 | T678Q | B | 2.6e-08 | 2.8e-08 | 4e-08 | 3.133E-08 | 7.572E-09 | 2.600E-08 | 4.000E-08 |
| 13548 | 678 | T678G | B | 7.7e-08 | 3.3e-08 | 1.4e-07 | 8.333E-08 | 5.378E-08 | 3.300E-08 | 1.400E-07 |
| 13549 | 678 | T678H | B | 3.6e-08 | 3.1e-08 | 9.6e-08 | 5.433E-08 | 3.617E-08 | 3.100E-08 | 9.600E-08 |
| 13550 | 678 | T678I | B | 3.9e-08 | 5.3e-08 | 4.7e-08 | 4.633E-08 | 7.024E-09 | 3.900E-08 | 5.300E-08 |
| 13551 | 678 | T678L | B | 3.7e-08 | 4.5e-08 | 7.8e-08 | 5.333E-08 | 2.173E-08 | 3.700E-08 | 7.800E-08 |
| 13552 | 678 | T678K | B | 4.3e-08 | 9.2e-08 | 3.4e-08 | 5.633E-08 | 3.121E-08 | 3.400E-08 | 9.200E-08 |
| 13553 | 678 | T678M | B | 6.7e-08 | 2.7e-08 | 3.1e-08 | 4.167E-08 | 2.203E-08 | 2.700E-08 | 6.700E-08 |
| 13554 | 678 | T678F | B | 2e-08 | 3.2e-07 | 5.1e-08 | 1.303E-07 | 1.650E-07 | 2.000E-08 | 3.200E-07 |
| 13555 | 678 | T678P | B | 3.3e-08 | 4.3e-08 | 6.4e-08 | 4.667E-08 | 1.582E-08 | 3.300E-08 | 6.400E-08 |
| 13556 | 678 | T678S | B | 4.6e-08 | 2.6e-08 | 3.2e-08 | 3.467E-08 | 1.026E-08 | 2.600E-08 | 4.600E-08 |
| 13557 | 678 | T678T | B | 5.1e-08 | 5.5e-08 | 3.8e-08 | 4.800E-08 | 8.888E-09 | 3.800E-08 | 5.500E-08 |
| 13558 | 678 | T678W | B | 3.3e-08 | 5.8e-08 | 3.5e-08 | 4.200E-08 | 1.389E-08 | 3.300E-08 | 5.800E-08 |
| 13559 | 678 | T678Y | B | 4.1e-08 | 3.6e-08 | 8.4e-08 | 5.367E-08 | 2.639E-08 | 3.600E-08 | 8.400E-08 |
| 13560 | 678 | T678V | B | 1.2e-08 | 2.7e-08 | 3.1e-08 | 2.333E-08 | 1.002E-08 | 1.200E-08 | 3.100E-08 |
| 13561 | 679 | G679A | B | 1.5e-08 | 4.8e-08 | 1.4e-08 | 2.567E-08 | 1.935E-08 | 1.400E-08 | 4.800E-08 |
| 13562 | 679 | G679R | B | 3.5e-08 | 2.1e-08 | 4.3e-08 | 3.300E-08 | 1.114E-08 | 2.100E-08 | 4.300E-08 |
| 13563 | 679 | G679N | B | 6.7e-08 | 3e-08 | 2e-08 | 3.900E-08 | 2.476E-08 | 2.000E-08 | 6.700E-08 |
| 13564 | 679 | G679D | B | 4.4e-08 | 2.3e-08 | 4.2e-08 | 3.633E-08 | 1.159E-08 | 2.300E-08 | 4.400E-08 |
| 13565 | 679 | G679C | B | 8.2e-08 | 4.2e-08 | 5.6e-08 | 6.000E-08 | 2.030E-08 | 4.200E-08 | 8.200E-08 |
| 13566 | 679 | G679E | B | 3.6e-08 | 9.4e-08 | 4.9e-08 | 5.967E-08 | 3.044E-08 | 3.600E-08 | 9.400E-08 |
| 13567 | 679 | G679Q | B | 2.9e-08 | 2.1e-08 | 4.7e-08 | 3.233E-08 | 1.332E-08 | 2.100E-08 | 4.700E-08 |
| 13568 | 679 | G679G | B | 2.6e-08 | 4.9e-08 | 4.2e-08 | 3.900E-08 | 1.179E-08 | 2.600E-08 | 4.900E-08 |
| 13569 | 679 | G679H | B | 3.4e-08 | 8.1e-08 | 1.7e-08 | 4.400E-08 | 3.315E-08 | 1.700E-08 | 8.100E-08 |
| 13570 | 679 | G679I | B | 8.6e-08 | 4.9e-08 | 6.1e-08 | 6.533E-08 | 1.888E-08 | 4.900E-08 | 8.600E-08 |
| 13571 | 679 | G679L | B | 4.5e-08 | 5.2e-08 | 1.7e-08 | 3.800E-08 | 1.852E-08 | 1.700E-08 | 5.200E-08 |
| 13572 | 679 | G679K | B | 2.1e-08 | 3.3e-08 | 5.9e-08 | 3.767E-08 | 1.943E-08 | 2.100E-08 | 5.900E-08 |
| 13573 | 679 | G679M | B | 1.7e-08 | 3.4e-08 | 7.4e-08 | 4.167E-08 | 2.926E-08 | 1.700E-08 | 7.400E-08 |
| 13574 | 679 | G679F | B | 5.9e-08 | 3.8e-08 | 4.3e-08 | 4.667E-08 | 1.097E-08 | 3.800E-08 | 5.900E-08 |
| 13575 | 679 | G679P | B | 4.4e-08 | 5.3e-08 | 3.3e-08 | 4.333E-08 | 1.002E-08 | 3.300E-08 | 5.300E-08 |
| 13576 | 679 | G679S | B | 3.2e-08 | 2.6e-08 | 2.6e-08 | 2.800E-08 | 3.464E-09 | 2.600E-08 | 3.200E-08 |
| 13577 | 679 | G679T | B | 1.1e-07 | 4.3e-08 | 2.7e-08 | 6.000E-08 | 4.403E-08 | 2.700E-08 | 1.100E-07 |
| 13578 | 679 | G679W | B | 4.8e-08 | 5.7e-08 | 3e-08 | 4.500E-08 | 1.375E-08 | 3.000E-08 | 5.700E-08 |
| 13579 | 679 | G679Y | B | 3.6e-08 | 3.7e-08 | 4.5e-08 | 3.933E-08 | 4.933E-09 | 3.600E-08 | 4.500E-08 |
| 13580 | 679 | G679V | B | 9.3e-08 | 2.8e-08 | 6.4e-08 | 6.167E-08 | 3.256E-08 | 2.800E-08 | 9.300E-08 |
| 13581 | 680 | V680A | B | 5.6e-08 | 6.1e-08 | 2.1e-08 | 4.600E-08 | 2.179E-08 | 2.100E-08 | 6.100E-08 |
| 13582 | 680 | V680R | B | 2.3e-08 | 4.7e-08 | 6.3e-08 | 4.433E-08 | 2.013E-08 | 2.300E-08 | 6.300E-08 |
| 13583 | 680 | V680N | B | 1.3e-08 | 1.4e-07 | 3e-08 | 6.100E-08 | 6.894E-08 | 1.300E-08 | 1.400E-07 |
| 13584 | 680 | V680D | B | 4.9e-08 | 5.1e-08 | 5.6e-08 | 5.200E-08 | 3.606E-09 | 4.900E-08 | 5.600E-08 |
| 13585 | 680 | V680C | B | 6.6e-08 | 7.6e-08 | 7.3e-08 | 7.167E-08 | 5.132E-09 | 6.600E-08 | 7.600E-08 |
| 13586 | 680 | V680E | B | 5.3e-08 | 3.7e-08 | 4.8e-08 | 4.600E-08 | 8.185E-09 | 3.700E-08 | 5.300E-08 |
| 13587 | 680 | V680Q | B | 5.3e-08 | 7.8e-08 | 2.2e-08 | 5.100E-08 | 2.805E-08 | 2.200E-08 | 7.800E-08 |
| 13588 | 680 | V680G | B | 7.2e-08 | 1e-07 | 9e-08 | 8.733E-08 | 1.419E-08 | 7.200E-08 | 1.000E-07 |
| 13589 | 680 | V680H | B | 3e-08 | 3.5e-08 | 8.4e-09 | 2.447E-08 | 1.414E-08 | 8.400E-09 | 3.500E-08 |
| 13590 | 680 | V680I | B | 4.5e-08 | 1.8e-08 | 4e-08 | 3.433E-08 | 1.436E-08 | 1.800E-08 | 4.500E-08 |
| 13591 | 680 | V680L | B | 1.7e-08 | 3e-08 | 9.4e-08 | 4.700E-08 | 4.122E-08 | 1.700E-08 | 9.400E-08 |
| 13592 | 680 | V680K | B | 9.8e-08 | 1.1e-07 | 4.1e-08 | 8.300E-08 | 3.686E-08 | 4.100E-08 | 1.100E-07 |
| 13593 | 680 | V680M | B | 9.1e-08 | 3.3e-08 | 2e-08 | 4.800E-08 | 3.780E-08 | 2.000E-08 | 9.100E-08 |
| 13594 | 680 | V680F | B | 5.2e-08 | 5.1e-08 | 2.6e-08 | 4.300E-08 | 1.473E-08 | 2.600E-08 | 5.200E-08 |
| 13595 | 680 | V680P | B | 3.4e-08 | 2.8e-08 | 2.7e-08 | 2.967E-08 | 3.786E-09 | 2.700E-08 | 3.400E-08 |
| 13596 | 680 | V680S | B | 6.8e-08 | 2.1e-08 | 5.1e-08 | 4.667E-08 | 2.380E-08 | 2.100E-08 | 6.800E-08 |
| 13597 | 680 | V680T | B | 8.6e-08 | 6.8e-08 | 2.1e-08 | 5.833E-08 | 3.356E-08 | 2.100E-08 | 8.600E-08 |
| 13598 | 680 | V680W | B | 1.9e-08 | 3.3e-08 | 4.9e-08 | 3.367E-08 | 1.501E-08 | 1.900E-08 | 4.900E-08 |
| 13599 | 680 | V680Y | B | 6.3e-08 | 4.7e-08 | 2.4e-08 | 4.467E-08 | 1.960E-08 | 2.400E-08 | 6.300E-08 |

| | | | | | | | | | | |
|---|---|---|---|---|---|---|---|---|---|---|
| 13600 | 680 | V680V | B | 1.5e-08 | 7.9e-08 | 5.2e-08 | 4.867E-08 | 3.213E-08 | 1.500E-08 | 7.900E-08 |
| 13601 | 681 | P681A | B | 4.1e-08 | 3.2e-08 | 3.4e-08 | 3.567E-08 | 4.726E-09 | 3.200E-08 | 4.100E-08 |
| 13602 | 681 | P681R | B | 1.1e-07 | 1.9e-08 | 5.3e-08 | 6.067E-08 | 4.598E-08 | 1.900E-08 | 1.100E-07 |
| 13603 | 681 | P681N | B | 2.9e-08 | 6.8e-08 | 1.7e-08 | 3.800E-08 | 2.666E-08 | 1.700E-08 | 6.800E-08 |
| 13604 | 681 | P681D | B | 2.8e-08 | 3.3e-08 | 5.1e-08 | 3.733E-08 | 1.210E-08 | 2.800E-08 | 5.100E-08 |
| 13605 | 681 | P681C | B | 2.9e-08 | 3.6e-08 | 4.1e-08 | 3.533E-08 | 6.028E-09 | 2.900E-08 | 4.100E-08 |
| 13606 | 681 | P681E | B | 5.3e-08 | 2e-08 | 5.1e-08 | 4.133E-08 | 1.850E-08 | 2.000E-08 | 5.300E-08 |
| 13607 | 681 | P681Q | B | 3.4e-08 | 6.3e-08 | 3.6e-08 | 4.433E-08 | 1.620E-08 | 3.400E-08 | 6.300E-08 |
| 13608 | 681 | P681G | B | 7.2e-08 | 2.4e-08 | 3.5e-08 | 4.367E-08 | 2.515E-08 | 2.400E-08 | 7.200E-08 |
| 13609 | 681 | P681H | B | 3.6e-08 | 3.9e-08 | 2.4e-08 | 3.300E-08 | 7.937E-09 | 2.400E-08 | 3.900E-08 |
| 13610 | 681 | P681I | B | 4.8e-08 | 2.9e-08 | 2.9e-08 | 3.533E-08 | 1.097E-08 | 2.900E-08 | 4.800E-08 |
| 13611 | 681 | P681L | B | 3.1e-08 | 4.9e-08 | 4.7e-08 | 4.233E-08 | 9.866E-09 | 3.100E-08 | 4.900E-08 |
| 13612 | 681 | P681K | B | 4.6e-08 | 2.8e-08 | 4.9e-08 | 4.100E-08 | 1.136E-08 | 2.800E-08 | 4.900E-08 |
| 13613 | 681 | P681M | B | 3.5e-08 | 2.4e-08 | 2.8e-08 | 2.900E-08 | 5.568E-09 | 2.400E-08 | 3.500E-08 |
| 13614 | 681 | P681F | B | 3e-08 | 1.2e-07 | 4e-08 | 6.333E-08 | 4.933E-08 | 3.000E-08 | 1.200E-07 |
| 13615 | 681 | P681P | B | 3.5e-08 | 6.8e-08 | 1.9e-08 | 4.067E-08 | 2.499E-08 | 1.900E-08 | 6.800E-08 |
| 13616 | 681 | P681S | B | 8.4e-08 | 5.4e-08 | 7.8e-08 | 7.200E-08 | 1.587E-08 | 5.400E-08 | 8.400E-08 |
| 13617 | 681 | P681T | B | 5e-08 | 7e-08 | 4e-08 | 5.333E-08 | 1.528E-08 | 4.000E-08 | 7.000E-08 |
| 13618 | 681 | P681W | B | 4.5e-08 | 2.9e-08 | 1.7e-08 | 3.033E-08 | 1.405E-08 | 1.700E-08 | 4.500E-08 |
| 13619 | 681 | P681Y | B | 3.4e-08 | 3.4e-08 | 3.8e-08 | 3.533E-08 | 2.309E-09 | 3.400E-08 | 3.800E-08 |
| 13620 | 681 | P681V | B | 3.9e-08 | 6.2e-08 | 1.6e-08 | 3.900E-08 | 2.300E-08 | 1.600E-08 | 6.200E-08 |
| 13621 | 682 | S682A | B | 5.6e-08 | 3.3e-08 | 4.4e-08 | 4.433E-08 | 1.150E-08 | 3.300E-08 | 5.600E-08 |
| 13622 | 682 | S682R | B | 4.1e-08 | 7.3e-08 | 8.6e-08 | 6.667E-08 | 2.316E-08 | 4.100E-08 | 8.600E-08 |
| 13623 | 682 | S682N | B | 5.5e-08 | 6.6e-08 | 7e-08 | 6.367E-08 | 7.767E-09 | 5.500E-08 | 7.000E-08 |
| 13624 | 682 | S682D | B | 7.3e-08 | 6.9e-08 | 6.2e-08 | 6.800E-08 | 5.568E-09 | 6.200E-08 | 7.300E-08 |
| 13625 | 682 | S682C | B | 5.6e-08 | 3.5e-08 | 4.5e-08 | 4.533E-08 | 1.050E-08 | 3.500E-08 | 5.600E-08 |
| 13626 | 682 | S682E | B | 1.4e-07 | 2.7e-08 | 1.5e-08 | 6.067E-08 | 6.897E-08 | 1.500E-08 | 1.400E-07 |
| 13627 | 682 | S682Q | B | 3.4e-08 | 2.5e-08 | 4e-08 | 3.300E-08 | 7.550E-09 | 2.500E-08 | 4.000E-08 |
| 13628 | 682 | S682G | B | 2.7e-08 | 7.5e-08 | 7.8e-08 | 6.000E-08 | 2.862E-08 | 2.700E-08 | 7.800E-08 |
| 13629 | 682 | S682H | B | 4.7e-08 | 9.7e-08 | 3.3e-08 | 5.900E-08 | 3.365E-08 | 3.300E-08 | 9.700E-08 |
| 13630 | 682 | S682I | B | 1.4e-07 | 5e-08 | 4.5e-08 | 7.833E-08 | 5.346E-08 | 4.500E-08 | 1.400E-07 |
| 13631 | 682 | S682L | B | 3.2e-08 | 3.5e-08 | 3.1e-08 | 3.267E-08 | 2.082E-09 | 3.100E-08 | 3.500E-08 |
| 13632 | 682 | S682K | B | 7e-08 | 4.2e-08 | 3.3e-08 | 4.833E-08 | 1.930E-08 | 3.300E-08 | 7.000E-08 |
| 13633 | 682 | S682M | B | 2.6e-08 | 6.5e-08 | 8.3e-08 | 5.800E-08 | 2.914E-08 | 2.600E-08 | 8.300E-08 |
| 13634 | 682 | S682F | B | 6.4e-08 | 3.7e-08 | 3.3e-08 | 4.467E-08 | 1.686E-08 | 3.300E-08 | 6.400E-08 |
| 13635 | 682 | S682P | B | 4.5e-08 | 5.6e-08 | 2.4e-08 | 4.167E-08 | 1.626E-08 | 2.400E-08 | 5.600E-08 |
| 13636 | 682 | S682S | B | 9.4e-08 | 7.7e-08 | 2e-08 | 6.367E-08 | 3.876E-08 | 2.000E-08 | 9.400E-08 |
| 13637 | 682 | S682T | B | 6.6e-08 | 1e-07 | 3.5e-08 | 6.700E-08 | 3.251E-08 | 3.500E-08 | 1.000E-07 |
| 13638 | 682 | S682W | B | 7e-08 | 1.3e-07 | 4.6e-08 | 8.200E-08 | 4.327E-08 | 4.600E-08 | 1.300E-07 |
| 13639 | 682 | S682Y | B | 8.4e-09 | 3.4e-08 | 4.3e-08 | 2.847E-08 | 1.795E-08 | 8.400E-09 | 4.300E-08 |
| 13640 | 682 | S682V | B | 2e-08 | 8e-08 | 2.1e-08 | 4.033E-08 | 3.436E-08 | 2.000E-08 | 8.000E-08 |
| 13641 | 683 | R683A | B | 8.8e-08 | 3.2e-08 | 2.8e-08 | 4.933E-08 | 3.355E-08 | 2.800E-08 | 8.800E-08 |
| 13642 | 683 | R683R | B | 1.1e-07 | 7.7e-08 | 5e-08 | 7.900E-08 | 3.005E-08 | 5.000E-08 | 1.100E-07 |
| 13643 | 683 | R683N | B | 3.1e-08 | 2.5e-08 | 1.3e-07 | 6.200E-08 | 5.897E-08 | 2.500E-08 | 1.300E-07 |
| 13644 | 683 | R683D | B | 2.1e-08 | 7.9e-08 | 2.9e-08 | 4.300E-08 | 3.143E-08 | 2.100E-08 | 7.900E-08 |
| 13645 | 683 | R683C | B | 5.6e-08 | 3.2e-08 | 1.2e-07 | 6.933E-08 | 4.549E-08 | 3.200E-08 | 1.200E-07 |
| 13646 | 683 | R683E | B | 3.4e-08 | 2.3e-08 | 4.4e-08 | 3.367E-08 | 1.050E-08 | 2.300E-08 | 4.400E-08 |
| 13647 | 683 | R683Q | B | 2.6e-08 | 1.4e-08 | 2.8e-08 | 2.267E-08 | 7.572E-09 | 1.400E-08 | 2.800E-08 |
| 13648 | 683 | R683G | B | 6.8e-08 | 2.6e-08 | 5.5e-08 | 4.967E-08 | 2.150E-08 | 2.600E-08 | 6.800E-08 |
| 13649 | 683 | R683H | B | 5.1e-08 | 6.4e-08 | 3.9e-08 | 5.133E-08 | 1.250E-08 | 3.900E-08 | 6.400E-08 |
| 13650 | 683 | R683I | B | 1.4e-08 | 2.5e-08 | 2e-08 | 1.967E-08 | 5.508E-09 | 1.400E-08 | 2.500E-08 |
| 13651 | 683 | R683L | B | 5.5e-08 | 1.1e-07 | 7.7e-08 | 8.067E-08 | 2.768E-08 | 5.500E-08 | 1.100E-07 |
| 13652 | 683 | R683K | B | 2.4e-08 | 4.9e-08 | 3.3e-08 | 3.533E-08 | 1.266E-08 | 2.400E-08 | 4.900E-08 |
| 13653 | 683 | R683M | B | 3.3e-08 | 5.7e-08 | 6.7e-08 | 5.233E-08 | 1.747E-08 | 3.300E-08 | 6.700E-08 |
| 13654 | 683 | R683F | B | 5.1e-08 | 1.1e-07 | 1.6e-08 | 5.900E-08 | 4.751E-08 | 1.600E-08 | 1.100E-07 |
| 13655 | 683 | R683P | B | 4.4e-08 | 4.9e-08 | 8.1e-08 | 5.800E-08 | 2.007E-08 | 4.400E-08 | 8.100E-08 |
| 13656 | 683 | R683S | B | 1.2e-07 | 5e-08 | 3.6e-08 | 6.867E-08 | 4.500E-08 | 3.600E-08 | 1.200E-07 |
| 13657 | 683 | R683T | B | 3.3e-08 | 3.4e-08 | 3.8e-08 | 3.500E-08 | 2.646E-09 | 3.300E-08 | 3.800E-08 |
| 13658 | 683 | R683W | B | 5.4e-08 | 2.7e-08 | 3e-08 | 3.700E-08 | 1.480E-08 | 2.700E-08 | 5.400E-08 |
| 13659 | 683 | R683Y | B | 5.2e-08 | 4.1e-08 | 1.8e-08 | 3.700E-08 | 1.735E-08 | 1.800E-08 | 5.200E-08 |
| 13660 | 683 | R683V | B | 4.3e-08 | 6.1e-08 | 3.5e-08 | 4.633E-08 | 1.332E-08 | 3.500E-08 | 6.100E-08 |
| 13661 | 684 | F684A | B | 6.2e-08 | 7.7e-08 | 3.5e-08 | 5.800E-08 | 2.128E-08 | 3.500E-08 | 7.700E-08 |
| 13662 | 684 | F684R | B | 5e-08 | 1.7e-08 | 1.3e-08 | 2.667E-08 | 2.031E-08 | 1.300E-08 | 5.000E-08 |
| 13663 | 684 | F684N | B | 7.2e-08 | 2.8e-08 | 4.4e-08 | 4.800E-08 | 2.227E-08 | 2.800E-08 | 7.200E-08 |
| 13664 | 684 | F684D | B | 7.8e-08 | 1.2e-08 | 4.8e-08 | 4.600E-08 | 3.305E-08 | 1.200E-08 | 7.800E-08 |
| 13665 | 684 | F684C | B | 2.2e-08 | 5.2e-08 | 9e-08 | 5.467E-08 | 3.408E-08 | 2.200E-08 | 9.000E-08 |
| 13666 | 684 | F684E | B | 5.9e-08 | 2.2e-08 | 3.3e-08 | 3.800E-08 | 1.900E-08 | 2.200E-08 | 5.900E-08 |
| 13667 | 684 | F684Q | B | 2.3e-08 | 8.2e-08 | 4.4e-08 | 4.967E-08 | 2.991E-08 | 2.300E-08 | 8.200E-08 |
| 13668 | 684 | F684G | B | 4.6e-08 | 1.9e-08 | 3.7e-08 | 3.400E-08 | 1.375E-08 | 1.900E-08 | 4.600E-08 |
| 13669 | 684 | F684H | B | 2.8e-08 | 4.1e-08 | 3.1e-08 | 3.333E-08 | 6.807E-09 | 2.800E-08 | 4.100E-08 |
| 13670 | 684 | F684I | B | 2.7e-08 | 4.1e-08 | 6.5e-08 | 4.433E-08 | 1.922E-08 | 2.700E-08 | 6.500E-08 |
| 13671 | 684 | F684L | B | 4.9e-08 | 3e-08 | 3.6e-08 | 3.833E-08 | 9.713E-09 | 3.000E-08 | 4.900E-08 |
| 13672 | 684 | F684K | B | 3.9e-08 | 2.6e-08 | 2.2e-08 | 2.900E-08 | 8.888E-09 | 2.200E-08 | 3.900E-08 |
| 13673 | 684 | F684M | B | 4.5e-08 | 2.7e-08 | 2.7e-08 | 3.300E-08 | 1.039E-08 | 2.700E-08 | 4.500E-08 |
| 13674 | 684 | F684F | B | 8.9e-08 | 3.5e-08 | 6e-08 | 6.133E-08 | 2.702E-08 | 3.500E-08 | 8.900E-08 |
| 13675 | 684 | F684P | B | 8.7e-08 | 4.9e-08 | 3e-08 | 5.533E-08 | 2.902E-08 | 3.000E-08 | 8.700E-08 |
| 13676 | 684 | F684S | B | 1.9e-08 | 5e-08 | 1.6e-08 | 2.833E-08 | 1.882E-08 | 1.600E-08 | 5.000E-08 |
| 13677 | 684 | F684T | B | 3.6e-08 | 6.8e-08 | 5e-08 | 5.133E-08 | 1.604E-08 | 3.600E-08 | 6.800E-08 |
| 13678 | 684 | F684W | B | 1.7e-07 | 3.3e-08 | 8.9e-08 | 9.733E-08 | 6.888E-08 | 3.300E-08 | 1.700E-07 |
| 13679 | 684 | F684Y | B | 8.8e-08 | 5.6e-08 | 1.6e-08 | 5.333E-08 | 3.607E-08 | 1.600E-08 | 8.800E-08 |
| 13680 | 684 | F684V | B | 3e-08 | 5.4e-08 | 2.5e-08 | 3.633E-08 | 1.550E-08 | 2.500E-08 | 5.400E-08 |
| 13681 | 685 | S685A | B | 3.7e-08 | 5.6e-08 | 4.3e-08 | 4.533E-08 | 9.713E-09 | 3.700E-08 | 5.600E-08 |
| 13682 | 685 | S685R | B | 2.1e-08 | 2.8e-08 | 2.4e-08 | 2.433E-08 | 3.512E-09 | 2.100E-08 | 2.800E-08 |
| 13683 | 685 | S685N | B | 5.3e-08 | 8e-08 | 6.4e-08 | 6.567E-08 | 1.358E-08 | 5.300E-08 | 8.000E-08 |
| 13684 | 685 | S685D | B | 4.5e-08 | 1.6e-07 | 2.1e-08 | 7.533E-08 | 7.430E-08 | 2.100E-08 | 1.600E-07 |

| | | | | | | | | | | |
|---|---|---|---|---|---|---|---|---|---|---|
| 13685 | 685 | S685C | B | 2.7e-08 | 3.1e-08 | 3.9e-08 | 3.233E-08 | 6.110E-09 | 2.700E-08 | 3.900E-08 |
| 13686 | 685 | S685E | B | 5.8e-08 | 8.5e-08 | 1e-07 | 8.100E-08 | 2.128E-08 | 5.800E-08 | 1.000E-07 |
| 13687 | 685 | S685Q | B | 1.6e-08 | 2.3e-08 | 4e-08 | 2.633E-08 | 1.234E-08 | 1.600E-08 | 4.000E-08 |
| 13688 | 685 | S685G | B | 5.7e-08 | 2.7e-08 | 5.7e-08 | 4.700E-08 | 1.732E-08 | 2.700E-08 | 5.700E-08 |
| 13689 | 685 | S685H | B | 7.9e-08 | 2e-08 | 3.2e-08 | 4.367E-08 | 3.118E-08 | 2.000E-08 | 7.900E-08 |
| 13690 | 685 | S685I | B | 5.1e-08 | 8.5e-08 | 6.9e-08 | 6.833E-08 | 1.701E-08 | 5.100E-08 | 8.500E-08 |
| 13691 | 685 | S685L | B | 3.8e-08 | 8.5e-08 | 7e-08 | 6.433E-08 | 2.401E-08 | 3.800E-08 | 8.500E-08 |
| 13692 | 685 | S685K | B | 5.8e-08 | 1.2e-07 | 8.6e-08 | 8.800E-08 | 3.105E-08 | 5.800E-08 | 1.200E-07 |
| 13693 | 685 | S685M | B | 1.6e-08 | 9.4e-08 | 1.4e-08 | 4.133E-08 | 4.562E-08 | 1.400E-08 | 9.400E-08 |
| 13694 | 685 | S685F | B | 4.6e-08 | 3.7e-08 | 3.1e-08 | 3.800E-08 | 7.550E-09 | 3.100E-08 | 4.600E-08 |
| 13695 | 685 | S685P | B | 2.3e-08 | 2.1e-08 | 7.4e-08 | 3.933E-08 | 3.004E-08 | 2.100E-08 | 7.400E-08 |
| 13696 | 685 | S685S | B | 4.4e-08 | 3.6e-08 | 3.8e-08 | 3.933E-08 | 4.163E-09 | 3.600E-08 | 4.400E-08 |
| 13697 | 685 | S685T | B | 3.4e-08 | 3.5e-08 | 3.3e-08 | 3.400E-08 | 1.000E-09 | 3.300E-08 | 3.500E-08 |
| 13698 | 685 | S685W | B | 4.4e-08 | 1.4e-08 | 2.4e-08 | 2.733E-08 | 1.528E-08 | 1.400E-08 | 4.400E-08 |
| 13699 | 685 | S685Y | B | 3.4e-08 | 5.2e-08 | 3.2e-08 | 3.933E-08 | 1.102E-08 | 3.200E-08 | 5.200E-08 |
| 13700 | 685 | S685V | B | 1.1e-07 | 1.5e-08 | 2.2e-08 | 4.900E-08 | 5.294E-08 | 1.500E-08 | 1.100E-07 |
| 13701 | 686 | G686A | B | 1.1e-07 | 9.2e-08 | 7.9e-08 | 9.367E-08 | 1.557E-08 | 7.900E-08 | 1.100E-07 |
| 13702 | 686 | G686R | B | 6.1e-08 | 4.3e-08 | 1.8e-08 | 4.067E-08 | 2.159E-08 | 1.800E-08 | 6.100E-08 |
| 13703 | 686 | G686N | B | 3.3e-08 | 1e-07 | 4.3e-08 | 5.867E-08 | 3.614E-08 | 3.300E-08 | 1.000E-07 |
| 13704 | 686 | G686D | B | 8e-09 | 6.1e-08 | 4.7e-08 | 3.867E-08 | 2.747E-08 | 8.000E-09 | 6.100E-08 |
| 13705 | 686 | G686C | B | 1.9e-08 | 4.2e-08 | 2.2e-08 | 2.767E-08 | 1.250E-08 | 1.900E-08 | 4.200E-08 |
| 13706 | 686 | G686E | B | 3.6e-08 | 4e-08 | 6.7e-08 | 4.767E-08 | 1.686E-08 | 3.600E-08 | 6.700E-08 |
| 13707 | 686 | G686Q | B | 1.9e-08 | 4.5e-08 | 2.2e-08 | 2.867E-08 | 1.422E-08 | 1.900E-08 | 4.500E-08 |
| 13708 | 686 | G686G | B | 2.1e-08 | 5.7e-08 | 9.3e-08 | 5.700E-08 | 3.600E-08 | 2.100E-08 | 9.300E-08 |
| 13709 | 686 | G686H | B | 2.1e-08 | 3.8e-08 | 2e-08 | 2.633E-08 | 1.012E-08 | 2.000E-08 | 3.800E-08 |
| 13710 | 686 | G686I | B | 2.7e-08 | 1.3e-07 | 2.1e-08 | 5.933E-08 | 6.127E-08 | 2.100E-08 | 1.300E-07 |
| 13711 | 686 | G686L | B | 4.8e-08 | 4.5e-08 | 1.1e-08 | 3.467E-08 | 2.055E-08 | 1.100E-08 | 4.800E-08 |
| 13712 | 686 | G686K | B | 6.3e-08 | 4e-08 | 5.9e-08 | 5.400E-08 | 1.229E-08 | 4.000E-08 | 6.300E-08 |
| 13713 | 686 | G686M | B | 2.7e-08 | 1.3e-08 | 1.9e-08 | 1.967E-08 | 7.024E-09 | 1.300E-08 | 2.700E-08 |
| 13714 | 686 | G686F | B | 2.9e-08 | 4.2e-08 | 4.5e-08 | 3.867E-08 | 8.505E-09 | 2.900E-08 | 4.500E-08 |
| 13715 | 686 | G686P | B | 6.8e-08 | 4.4e-08 | 4.4e-08 | 5.200E-08 | 1.386E-08 | 4.400E-08 | 6.800E-08 |
| 13716 | 686 | G686S | B | 5.8e-08 | 4e-08 | 8.1e-08 | 5.967E-08 | 2.055E-08 | 4.000E-08 | 8.100E-08 |
| 13717 | 686 | G686T | B | 7.2e-08 | 5.5e-08 | 3.5e-08 | 5.400E-08 | 1.852E-08 | 3.500E-08 | 7.200E-08 |
| 13718 | 686 | G686W | B | 1.7e-07 | 3.6e-08 | 1.3e-08 | 7.300E-08 | 8.479E-08 | 1.300E-08 | 1.700E-07 |
| 13719 | 686 | G686Y | B | 4.9e-08 | 1.8e-08 | 4.9e-08 | 3.867E-08 | 1.790E-08 | 1.800E-08 | 4.900E-08 |
| 13720 | 686 | G686V | B | 1.7e-08 | 4.7e-08 | 1.8e-08 | 2.733E-08 | 1.704E-08 | 1.700E-08 | 4.700E-08 |
| 13721 | 687 | S687A | B | 2.4e-08 | 6.3e-08 | 8e-08 | 5.567E-08 | 2.871E-08 | 2.400E-08 | 8.000E-08 |
| 13722 | 687 | S687R | B | 7.3e-08 | 4.1e-08 | 1.6e-08 | 4.333E-08 | 2.857E-08 | 1.600E-08 | 7.300E-08 |
| 13723 | 687 | S687N | B | 7.5e-08 | 1.1e-07 | 2.1e-08 | 6.867E-08 | 4.484E-08 | 2.100E-08 | 1.100E-07 |
| 13724 | 687 | S687D | B | 2.1e-08 | 7.3e-08 | 8.1e-08 | 5.833E-08 | 3.258E-08 | 2.100E-08 | 8.100E-08 |
| 13725 | 687 | S687C | B | 2.9e-08 | 5e-08 | 3.8e-08 | 3.900E-08 | 1.054E-08 | 2.900E-08 | 5.000E-08 |
| 13726 | 687 | S687E | B | 3.8e-08 | 3.6e-08 | 1.8e-08 | 3.067E-08 | 1.102E-08 | 1.800E-08 | 3.800E-08 |
| 13727 | 687 | S687Q | B | 4.2e-08 | 4.2e-08 | 3.4e-08 | 3.933E-08 | 4.619E-09 | 3.400E-08 | 4.200E-08 |
| 13728 | 687 | S687G | B | 2.8e-08 | 4.6e-08 | 4.9e-08 | 4.100E-08 | 1.136E-08 | 2.800E-08 | 4.900E-08 |
| 13729 | 687 | S687H | B | 1.7e-07 | 8.2e-08 | 7.6e-08 | 1.093E-07 | 5.262E-08 | 7.600E-08 | 1.700E-07 |
| 13730 | 687 | S687I | B | 5.7e-08 | 4.1e-08 | 4.6e-08 | 4.800E-08 | 8.185E-09 | 4.100E-08 | 5.700E-08 |
| 13731 | 687 | S687L | B | 7.6e-08 | 8.5e-08 | 3.8e-08 | 6.633E-08 | 2.495E-08 | 3.800E-08 | 8.500E-08 |
| 13732 | 687 | S687K | B | 5.9e-08 | 8.1e-08 | 1.2e-07 | 8.667E-08 | 3.089E-08 | 5.900E-08 | 1.200E-07 |
| 13733 | 687 | S687M | B | 6.7e-08 | 2.5e-08 | 1.5e-08 | 3.567E-08 | 2.759E-08 | 1.500E-08 | 6.700E-08 |
| 13734 | 687 | S687F | B | 2.4e-08 | 8.9e-08 | 3.2e-08 | 4.833E-08 | 3.544E-08 | 2.400E-08 | 8.900E-08 |
| 13735 | 687 | S687P | B | 3.5e-08 | 3.9e-08 | 1.9e-08 | 3.100E-08 | 1.058E-08 | 1.900E-08 | 3.900E-08 |
| 13736 | 687 | S687S | B | 2.6e-07 | 2.4e-08 | 9.7e-08 | 1.270E-07 | 1.208E-07 | 2.400E-08 | 2.600E-07 |
| 13737 | 687 | S687T | B | 7.1e-08 | 7.9e-08 | 1.8e-08 | 5.600E-08 | 3.315E-08 | 1.800E-08 | 7.900E-08 |
| 13738 | 687 | S687W | B | 1.2e-07 | 1.4e-08 | 1.9e-08 | 5.100E-08 | 5.981E-08 | 1.400E-08 | 1.200E-07 |
| 13739 | 687 | S687Y | B | 6.6e-08 | 1.1e-07 | 4.8e-08 | 7.467E-08 | 3.190E-08 | 4.800E-08 | 1.100E-07 |
| 13740 | 687 | S687V | B | 7.6e-08 | 5.5e-08 | 2.1e-08 | 5.067E-08 | 2.775E-08 | 2.100E-08 | 7.600E-08 |
| 13741 | 688 | G688A | B | 5.8e-08 | 3.6e-08 | 9.6e-08 | 6.333E-08 | 3.035E-08 | 3.600E-08 | 9.600E-08 |
| 13742 | 688 | G688R | B | 1e-08 | 1.4e-08 | 2.5e-08 | 1.633E-08 | 7.767E-09 | 1.000E-08 | 2.500E-08 |
| 13743 | 688 | G688N | B | 1.3e-08 | 1.4e-07 | 3.8e-08 | 6.367E-08 | 6.728E-08 | 1.300E-08 | 1.400E-07 |
| 13744 | 688 | G688D | B | 1e-07 | 7.2e-08 | 4.7e-08 | 7.300E-08 | 2.651E-08 | 4.700E-08 | 1.000E-07 |
| 13745 | 688 | G688C | B | 6e-08 | 5.5e-08 | 8.4e-08 | 6.633E-08 | 1.550E-08 | 5.500E-08 | 8.400E-08 |
| 13746 | 688 | G688E | B | 6.8e-08 | 5.1e-08 | 2e-08 | 4.633E-08 | 2.434E-08 | 2.000E-08 | 6.800E-08 |
| 13747 | 688 | G688Q | B | 8.3e-08 | 3e-08 | 3e-08 | 4.767E-08 | 3.060E-08 | 3.000E-08 | 8.300E-08 |
| 13748 | 688 | G688G | B | 2.5e-08 | 2.5e-08 | 6.3e-08 | 3.767E-08 | 2.194E-08 | 2.500E-08 | 6.300E-08 |
| 13749 | 688 | G688H | B | 2.3e-08 | 1.6e-08 | 1e-07 | 4.633E-08 | 4.661E-08 | 1.600E-08 | 1.000E-07 |
| 13750 | 688 | G688I | B | 6.2e-08 | 1.3e-07 | 4.6e-08 | 7.933E-08 | 4.460E-08 | 4.600E-08 | 1.300E-07 |
| 13751 | 688 | G688L | B | 3.3e-08 | 3e-08 | 4.4e-08 | 3.567E-08 | 7.371E-09 | 3.000E-08 | 4.400E-08 |
| 13752 | 688 | G688K | B | 1.3e-07 | 2.6e-08 | 3.5e-08 | 6.367E-08 | 5.762E-08 | 2.600E-08 | 1.300E-07 |
| 13753 | 688 | G688M | B | 4.5e-08 | 2.5e-08 | 9e-08 | 5.333E-08 | 3.329E-08 | 2.500E-08 | 9.000E-08 |
| 13754 | 688 | G688F | B | 5.6e-08 | 3.1e-08 | 2.8e-08 | 3.833E-08 | 1.537E-08 | 2.800E-08 | 5.600E-08 |
| 13755 | 688 | G688P | B | 2.6e-08 | 3.4e-08 | 3.6e-08 | 3.200E-08 | 5.292E-09 | 2.600E-08 | 3.600E-08 |
| 13756 | 688 | G688S | B | 4.7e-08 | 5.9e-08 | 2.4e-08 | 4.333E-08 | 1.779E-08 | 2.400E-08 | 5.900E-08 |
| 13757 | 688 | G688T | B | 5.8e-08 | 2.6e-08 | 4.1e-08 | 4.167E-08 | 1.601E-08 | 2.600E-08 | 5.800E-08 |
| 13758 | 688 | G688W | B | 5.6e-08 | 4.6e-08 | 1.5e-07 | 8.400E-08 | 5.738E-08 | 4.600E-08 | 1.500E-07 |
| 13759 | 688 | G688Y | B | 5.8e-08 | 9.6e-08 | 6.6e-08 | 7.333E-08 | 2.003E-08 | 5.800E-08 | 9.600E-08 |
| 13760 | 688 | G688V | B | 1.8e-08 | 5.9e-08 | 5e-08 | 4.233E-08 | 2.155E-08 | 1.800E-08 | 5.900E-08 |
| 13761 | 689 | S689A | B | 3.4e-08 | 3.7e-08 | 3.2e-08 | 3.433E-08 | 2.517E-09 | 3.200E-08 | 3.700E-08 |
| 13762 | 689 | S689R | B | 5e-08 | 5.5e-08 | 1.8e-08 | 4.100E-08 | 2.007E-08 | 1.800E-08 | 5.500E-08 |
| 13763 | 689 | S689N | B | 5.4e-08 | 3e-08 | 4.2e-08 | 4.200E-08 | 1.200E-08 | 3.000E-08 | 5.400E-08 |
| 13764 | 689 | S689D | B | 2e-08 | 2.4e-08 | 2.4e-08 | 2.267E-08 | 2.309E-09 | 2.000E-08 | 2.400E-08 |
| 13765 | 689 | S689C | B | 3.2e-08 | 7.5e-08 | 5.8e-08 | 5.500E-08 | 2.166E-08 | 3.200E-08 | 7.500E-08 |
| 13766 | 689 | S689E | B | 2.4e-08 | 7.7e-08 | 3.4e-08 | 4.500E-08 | 2.816E-08 | 2.400E-08 | 7.700E-08 |
| 13767 | 689 | S689Q | B | 8.6e-08 | 1.3e-07 | 5.4e-08 | 9.000E-08 | 3.816E-08 | 5.400E-08 | 1.300E-07 |
| 13768 | 689 | S689G | B | 3.4e-08 | 6.9e-08 | 9.7e-08 | 6.667E-08 | 3.156E-08 | 3.400E-08 | 9.700E-08 |
| 13769 | 689 | S689H | B | 5.2e-08 | 3.9e-08 | 4.1e-08 | 4.400E-08 | 7.000E-09 | 3.900E-08 | 5.200E-08 |

| | | | | | | | | | | |
|---|---|---|---|---|---|---|---|---|---|---|
| 13770 | 689 | S689I | B | 4.6e-08 | 2.8e-08 | 2.9e-08 | 3.433E-08 | 1.012E-08 | 2.800E-08 | 4.600E-08 |
| 13771 | 689 | S689L | B | 2.7e-08 | 3.7e-08 | 1.7e-08 | 2.700E-08 | 1.000E-08 | 1.700E-08 | 3.700E-08 |
| 13772 | 689 | S689K | B | 9.1e-08 | 4e-08 | 4.3e-08 | 5.800E-08 | 2.862E-08 | 4.000E-08 | 9.100E-08 |
| 13773 | 689 | S689M | B | 6.9e-08 | 2.8e-08 | 2e-07 | 9.900E-08 | 8.984E-08 | 2.800E-08 | 2.000E-07 |
| 13774 | 689 | S689F | B | 7e-08 | 2.9e-08 | 5e-08 | 4.967E-08 | 2.050E-08 | 2.900E-08 | 7.000E-08 |
| 13775 | 689 | S689P | B | 1.2e-08 | 6.3e-08 | 1.7e-08 | 3.067E-08 | 2.811E-08 | 1.200E-08 | 6.300E-08 |
| 13776 | 689 | S689S | B | 4.7e-08 | 2e-08 | 4.5e-08 | 3.733E-08 | 1.504E-08 | 2.000E-08 | 4.700E-08 |
| 13777 | 689 | S689T | B | 2.9e-08 | 3.8e-08 | 3.4e-08 | 3.367E-08 | 4.509E-09 | 2.900E-08 | 3.800E-08 |
| 13778 | 689 | S689W | B | 3.7e-08 | 2.9e-08 | 5.8e-08 | 4.133E-08 | 1.498E-08 | 2.900E-08 | 5.800E-08 |
| 13779 | 689 | S689Y | B | 4.5e-08 | 2.2e-08 | 4.7e-08 | 3.800E-08 | 1.389E-08 | 2.200E-08 | 4.700E-08 |
| 13780 | 689 | S689V | B | 1e-07 | 6.8e-08 | 3.7e-08 | 6.833E-08 | 3.150E-08 | 3.700E-08 | 1.000E-07 |
| 13781 | 690 | G690A | B | 4.7e-08 | 2.1e-08 | 2.5e-08 | 3.100E-08 | 1.400E-08 | 2.100E-08 | 4.700E-08 |
| 13782 | 690 | G690R | B | 7.8e-08 | 1.2e-08 | 9e-08 | 6.000E-08 | 4.200E-08 | 1.200E-08 | 9.000E-08 |
| 13783 | 690 | G690N | B | 7.7e-08 | 5.8e-08 | 3.8e-08 | 5.767E-08 | 1.950E-08 | 3.800E-08 | 7.700E-08 |
| 13784 | 690 | G690D | B | 7.9e-08 | 3.4e-08 | 1.4e-08 | 4.233E-08 | 3.329E-08 | 1.400E-08 | 7.900E-08 |
| 13785 | 690 | G690C | B | 6.9e-08 | 6.6e-08 | 3.4e-08 | 5.633E-08 | 1.940E-08 | 3.400E-08 | 6.900E-08 |
| 13786 | 690 | G690E | B | 3.4e-08 | 2e-08 | 1.6e-08 | 2.333E-08 | 9.452E-09 | 1.600E-08 | 3.400E-08 |
| 13787 | 690 | G690Q | B | 3.3e-08 | 2.4e-08 | 4e-08 | 3.233E-08 | 8.021E-09 | 2.400E-08 | 4.000E-08 |
| 13788 | 690 | G690G | B | 4.4e-08 | 3.4e-08 | 3.1e-08 | 3.633E-08 | 6.807E-09 | 3.100E-08 | 4.400E-08 |
| 13789 | 690 | G690H | B | 1.4e-07 | 8.3e-08 | 2.9e-08 | 8.400E-08 | 5.551E-08 | 2.900E-08 | 1.400E-07 |
| 13790 | 690 | G690I | B | 4.3e-08 | 2.4e-08 | 1.5e-07 | 7.233E-08 | 6.793E-08 | 2.400E-08 | 1.500E-07 |
| 13791 | 690 | G690L | B | 2.6e-08 | 2.9e-08 | 1.4e-08 | 2.300E-08 | 7.937E-09 | 1.400E-08 | 2.900E-08 |
| 13792 | 690 | G690K | B | 2.9e-08 | 3.1e-08 | 1.2e-08 | 2.400E-08 | 1.044E-08 | 1.200E-08 | 3.100E-08 |
| 13793 | 690 | G690M | B | 2.6e-08 | 1.1e-07 | 2.9e-08 | 5.500E-08 | 4.766E-08 | 2.600E-08 | 1.100E-07 |
| 13794 | 690 | G690F | B | 9e-08 | 4e-08 | 3.3e-08 | 5.433E-08 | 3.109E-08 | 3.300E-08 | 9.000E-08 |
| 13795 | 690 | G690P | B | 8.7e-09 | 4.9e-08 | 2e-08 | 2.590E-08 | 2.079E-08 | 8.700E-09 | 4.900E-08 |
| 13796 | 690 | G690S | B | 5.8e-08 | 4.8e-08 | 2.2e-07 | 1.087E-07 | 9.655E-08 | 4.800E-08 | 2.200E-07 |
| 13797 | 690 | G690T | B | 1.9e-08 | 4e-08 | 5.9e-08 | 3.933E-08 | 2.001E-08 | 1.900E-08 | 5.900E-08 |
| 13798 | 690 | G690W | B | 1.1e-07 | 3.1e-08 | 7.2e-08 | 7.100E-08 | 3.951E-08 | 3.100E-08 | 1.100E-07 |
| 13799 | 690 | G690Y | B | 3.7e-08 | 3.3e-08 | 2.4e-08 | 3.133E-08 | 6.658E-09 | 2.400E-08 | 3.700E-08 |
| 13800 | 690 | G690V | B | 5.4e-08 | 2.1e-08 | 1.1e-08 | 2.867E-08 | 2.250E-08 | 1.100E-08 | 5.400E-08 |
| 13801 | 691 | T691A | B | 5.4e-08 | 3.8e-08 | 1.2e-07 | 7.067E-08 | 4.347E-08 | 3.800E-08 | 1.200E-07 |
| 13802 | 691 | T691R | B | 1.2e-08 | 6.4e-08 | 3.7e-08 | 3.767E-08 | 2.601E-08 | 1.200E-08 | 6.400E-08 |
| 13803 | 691 | T691N | B | 1.7e-08 | 8.8e-08 | 5.2e-08 | 5.233E-08 | 3.550E-08 | 1.700E-08 | 8.800E-08 |
| 13804 | 691 | T691D | B | 1.2e-07 | 8.5e-08 | 6.7e-08 | 9.067E-08 | 2.695E-08 | 6.700E-08 | 1.200E-07 |
| 13805 | 691 | T691C | B | 4.7e-08 | 3.9e-08 | 2.7e-08 | 3.767E-08 | 1.007E-08 | 2.700E-08 | 4.700E-08 |
| 13806 | 691 | T691E | B | 3.6e-08 | 4.6e-08 | 4.6e-08 | 4.267E-08 | 5.774E-09 | 3.600E-08 | 4.600E-08 |
| 13807 | 691 | T691Q | B | 2.9e-08 | 6.8e-08 | 3.8e-08 | 4.500E-08 | 2.042E-08 | 2.900E-08 | 6.800E-08 |
| 13808 | 691 | T691G | B | 6.6e-08 | 7.8e-08 | 3.7e-08 | 6.033E-08 | 2.108E-08 | 3.700E-08 | 7.800E-08 |
| 13809 | 691 | T691H | B | 4.4e-08 | 3e-08 | 5e-08 | 4.133E-08 | 1.026E-08 | 3.000E-08 | 5.000E-08 |
| 13810 | 691 | T691I | B | 1.1e-08 | 3.5e-08 | 3.2e-08 | 2.600E-08 | 1.308E-08 | 1.100E-08 | 3.500E-08 |
| 13811 | 691 | T691L | B | 2.8e-08 | 5.4e-08 | 9.4e-08 | 5.867E-08 | 3.325E-08 | 2.800E-08 | 9.400E-08 |
| 13812 | 691 | T691K | B | 5.9e-08 | 2.6e-08 | 2.5e-08 | 3.667E-08 | 1.935E-08 | 2.500E-08 | 5.900E-08 |
| 13813 | 691 | T691M | B | 1.7e-08 | 3e-08 | 3.9e-08 | 2.867E-08 | 1.106E-08 | 1.700E-08 | 3.900E-08 |
| 13814 | 691 | T691F | B | 7.1e-08 | 9.1e-08 | 7e-09 | 5.633E-08 | 4.388E-08 | 7.000E-09 | 9.100E-08 |
| 13815 | 691 | T691P | B | 5.8e-08 | 2.5e-08 | 4.9e-08 | 4.400E-08 | 1.706E-08 | 2.500E-08 | 5.800E-08 |
| 13816 | 691 | T691S | B | 3.5e-08 | 1.1e-07 | 8.1e-08 | 7.533E-08 | 3.782E-08 | 3.500E-08 | 1.100E-07 |
| 13817 | 691 | T691T | B | 4.8e-08 | 4.6e-08 | 2.3e-08 | 3.900E-08 | 1.389E-08 | 2.300E-08 | 4.800E-08 |
| 13818 | 691 | T691W | B | 5.4e-08 | 4.5e-08 | 1.5e-08 | 3.800E-08 | 2.042E-08 | 1.500E-08 | 5.400E-08 |
| 13819 | 691 | T691Y | B | 3.4e-08 | 4.6e-08 | 3.3e-08 | 3.767E-08 | 7.234E-09 | 3.300E-08 | 4.600E-08 |
| 13820 | 691 | T691V | B | 2.3e-08 | 1.3e-07 | 3.7e-08 | 6.333E-08 | 5.816E-08 | 2.300E-08 | 1.300E-07 |
| 13821 | 692 | D692A | B | 2.1e-08 | 1.6e-07 | 5.9e-08 | 8.000E-08 | 7.184E-08 | 2.100E-08 | 1.600E-07 |
| 13822 | 692 | D692R | B | 1.2e-07 | 2.5e-08 | 3.1e-08 | 5.867E-08 | 5.320E-08 | 2.500E-08 | 1.200E-07 |
| 13823 | 692 | D692N | B | 5.6e-08 | 5.8e-08 | 2.1e-08 | 4.500E-08 | 2.081E-08 | 2.100E-08 | 5.800E-08 |
| 13824 | 692 | D692D | B | 1.9e-08 | 8.9e-08 | 2.4e-08 | 4.400E-08 | 3.905E-08 | 1.900E-08 | 8.900E-08 |
| 13825 | 692 | D692C | B | 1.1e-07 | 3.3e-08 | 8e-08 | 7.433E-08 | 3.881E-08 | 3.300E-08 | 1.100E-07 |
| 13826 | 692 | D692E | B | 3e-08 | 6.7e-08 | 7.8e-08 | 5.833E-08 | 2.515E-08 | 3.000E-08 | 7.800E-08 |
| 13827 | 692 | D692Q | B | 1.6e-08 | 3.2e-08 | 2.5e-08 | 2.433E-08 | 8.021E-09 | 1.600E-08 | 3.200E-08 |
| 13828 | 692 | D692G | B | 2.9e-08 | 9e-08 | 8.1e-08 | 6.667E-08 | 3.293E-08 | 2.900E-08 | 9.000E-08 |
| 13829 | 692 | D692H | B | 5.2e-08 | 7.8e-08 | 3.3e-08 | 5.433E-08 | 2.259E-08 | 3.300E-08 | 7.800E-08 |
| 13830 | 692 | D692I | B | 6.1e-08 | 6.7e-08 | 5.4e-08 | 6.067E-08 | 6.506E-09 | 5.400E-08 | 6.700E-08 |
| 13831 | 692 | D692L | B | 1.9e-08 | 2e-08 | 2.5e-08 | 2.133E-08 | 3.215E-09 | 1.900E-08 | 2.500E-08 |
| 13832 | 692 | D692K | B | 6.6e-08 | 9.6e-08 | 9.3e-08 | 8.500E-08 | 1.652E-08 | 6.600E-08 | 9.600E-08 |
| 13833 | 692 | D692M | B | 3.1e-08 | 4e-08 | 3.1e-08 | 3.400E-08 | 5.196E-09 | 3.100E-08 | 4.000E-08 |
| 13834 | 692 | D692F | B | 3.2e-08 | 6.3e-08 | 3.5e-08 | 4.333E-08 | 1.710E-08 | 3.200E-08 | 6.300E-08 |
| 13835 | 692 | D692P | B | 1.3e-08 | 3.7e-08 | 1.5e-08 | 2.167E-08 | 1.332E-08 | 1.300E-08 | 3.700E-08 |
| 13836 | 692 | D692S | B | 2e-08 | 1.9e-08 | 2.7e-08 | 2.200E-08 | 4.359E-09 | 1.900E-08 | 2.700E-08 |
| 13837 | 692 | D692T | B | 1.9e-08 | 2.2e-08 | 1.9e-08 | 2.000E-08 | 1.732E-09 | 1.900E-08 | 2.200E-08 |
| 13838 | 692 | D692W | B | 1.8e-08 | 6.5e-08 | 6.2e-08 | 4.833E-08 | 2.631E-08 | 1.800E-08 | 6.500E-08 |
| 13839 | 692 | D692Y | B | 9.1e-08 | 5e-08 | 1.5e-08 | 5.200E-08 | 3.804E-08 | 1.500E-08 | 9.100E-08 |
| 13840 | 692 | D692V | B | 4.2e-08 | 9.4e-08 | 5.6e-08 | 6.400E-08 | 2.691E-08 | 4.200E-08 | 9.400E-08 |
| 13841 | 693 | F693A | B | 3.2e-08 | 1.7e-08 | 5.1e-08 | 3.333E-08 | 1.704E-08 | 1.700E-08 | 5.100E-08 |
| 13842 | 693 | F693R | B | 4.6e-08 | 2.6e-08 | 6.2e-08 | 4.467E-08 | 1.804E-08 | 2.600E-08 | 6.200E-08 |
| 13843 | 693 | F693N | B | 5e-08 | 3.8e-08 | 3.9e-08 | 4.233E-08 | 6.658E-09 | 3.800E-08 | 5.000E-08 |
| 13844 | 693 | F693D | B | 1.9e-08 | 5e-08 | 4.5e-08 | 3.800E-08 | 1.664E-08 | 1.900E-08 | 5.000E-08 |
| 13845 | 693 | F693C | B | 8.9e-08 | 3.5e-08 | 2.5e-08 | 4.967E-08 | 3.443E-08 | 2.500E-08 | 8.900E-08 |
| 13846 | 693 | F693E | B | 5.3e-08 | 6.1e-08 | 8.7e-08 | 6.700E-08 | 1.778E-08 | 5.300E-08 | 8.700E-08 |
| 13847 | 693 | F693Q | B | 3.1e-08 | 5.1e-08 | 2.8e-08 | 3.667E-08 | 1.250E-08 | 2.800E-08 | 5.100E-08 |
| 13848 | 693 | F693G | B | 1.5e-08 | 4.9e-08 | 3.9e-08 | 3.433E-08 | 1.747E-08 | 1.500E-08 | 4.900E-08 |
| 13849 | 693 | F693H | B | 2.8e-08 | 4.2e-08 | 6.4e-08 | 4.467E-08 | 1.815E-08 | 2.800E-08 | 6.400E-08 |
| 13850 | 693 | F693I | B | 3.9e-08 | 3e-08 | 6.5e-08 | 4.467E-08 | 1.818E-08 | 3.000E-08 | 6.500E-08 |
| 13851 | 693 | F693L | B | 2.6e-08 | 3.5e-08 | 3.3e-08 | 3.133E-08 | 4.726E-09 | 2.600E-08 | 3.500E-08 |
| 13852 | 693 | F693K | B | 2.1e-08 | 3.6e-08 | 7.1e-09 | 2.137E-08 | 1.445E-08 | 7.100E-09 | 3.600E-08 |
| 13853 | 693 | F693M | B | 3e-08 | 5e-08 | 3e-08 | 3.667E-08 | 1.155E-08 | 3.000E-08 | 5.000E-08 |
| 13854 | 693 | F693F | B | 6e-08 | 7.5e-08 | 3e-08 | 5.500E-08 | 2.291E-08 | 3.000E-08 | 7.500E-08 |

| | | | | | | | | | | |
|---|---|---|---|---|---|---|---|---|---|---|
| 13855 | 693 | F693P | B | 1.9e-08 | 3e-08 | 4.3e-08 | 3.067E-08 | 1.201E-08 | 1.900E-08 | 4.300E-08 |
| 13856 | 693 | F693S | B | 9.7e-08 | 1.6e-08 | 2.5e-08 | 4.600E-08 | 4.440E-08 | 1.600E-08 | 9.700E-08 |
| 13857 | 693 | F693T | B | 1.2e-08 | 4.7e-08 | 9.1e-08 | 5.000E-08 | 3.959E-08 | 1.200E-08 | 9.100E-08 |
| 13858 | 693 | F693W | B | 7.9e-08 | 6.4e-08 | 1.1e-07 | 8.433E-08 | 2.346E-08 | 6.400E-08 | 1.100E-07 |
| 13859 | 693 | F693Y | B | 6.2e-08 | 5.1e-08 | 5.3e-08 | 5.533E-08 | 5.859E-09 | 5.100E-08 | 6.200E-08 |
| 13860 | 693 | F693V | B | 2.3e-08 | 2.5e-08 | 2.5e-08 | 2.433E-08 | 1.155E-09 | 2.300E-08 | 2.500E-08 |
| 13861 | 694 | T694A | B | 9.5e-08 | 2.3e-08 | 6.9e-08 | 6.233E-08 | 3.646E-08 | 2.300E-08 | 9.500E-08 |
| 13862 | 694 | T694R | B | 8.2e-08 | 3.7e-08 | 5e-08 | 5.633E-08 | 2.316E-08 | 3.700E-08 | 8.200E-08 |
| 13863 | 694 | T694N | B | 4e-08 | 6.2e-08 | 6.8e-08 | 5.667E-08 | 1.474E-08 | 4.000E-08 | 6.800E-08 |
| 13864 | 694 | T694D | B | 1.6e-08 | 1e-07 | 3.3e-08 | 4.967E-08 | 4.441E-08 | 1.600E-08 | 1.000E-07 |
| 13865 | 694 | T694C | B | 3.4e-08 | 4e-08 | 1.4e-07 | 7.133E-08 | 5.954E-08 | 3.400E-08 | 1.400E-07 |
| 13866 | 694 | T694E | B | 9.1e-08 | 4.8e-08 | 4.7e-08 | 6.200E-08 | 2.512E-08 | 4.700E-08 | 9.100E-08 |
| 13867 | 694 | T694Q | B | 3.1e-08 | 3.9e-08 | 1.2e-07 | 6.333E-08 | 4.924E-08 | 3.100E-08 | 1.200E-07 |
| 13868 | 694 | T694G | B | 7.9e-08 | 8.9e-08 | 3.8e-08 | 6.867E-08 | 2.702E-08 | 3.800E-08 | 8.900E-08 |
| 13869 | 694 | T694H | B | 2.9e-08 | 3.2e-08 | 4e-08 | 3.367E-08 | 5.686E-09 | 2.900E-08 | 4.000E-08 |
| 13870 | 694 | T694I | B | 2.8e-08 | 2.4e-08 | 1.4e-08 | 2.200E-08 | 7.211E-09 | 1.400E-08 | 2.800E-08 |
| 13871 | 694 | T694L | B | 3.1e-08 | 4.4e-08 | 5.8e-08 | 4.433E-08 | 1.350E-08 | 3.100E-08 | 5.800E-08 |
| 13872 | 694 | T694K | B | 2.7e-08 | 1.1e-08 | 6.9e-08 | 3.567E-08 | 2.996E-08 | 1.100E-08 | 6.900E-08 |
| 13873 | 694 | T694M | B | 4.5e-08 | 4.7e-08 | 6.9e-08 | 5.367E-08 | 1.332E-08 | 4.500E-08 | 6.900E-08 |
| 13874 | 694 | T694F | B | 6.2e-08 | 3.4e-08 | 7.1e-08 | 5.567E-08 | 1.930E-08 | 3.400E-08 | 7.100E-08 |
| 13875 | 694 | T694P | B | 6.3e-08 | 4.1e-08 | 4.8e-08 | 5.067E-08 | 1.124E-08 | 4.100E-08 | 6.300E-08 |
| 13876 | 694 | T694S | B | 5.7e-08 | 1.7e-08 | 3.6e-08 | 3.667E-08 | 2.001E-08 | 1.700E-08 | 5.700E-08 |
| 13877 | 694 | T694T | B | 1.3e-08 | 3.8e-08 | 4.9e-08 | 3.333E-08 | 1.845E-08 | 1.300E-08 | 4.900E-08 |
| 13878 | 694 | T694W | B | 8.8e-08 | 1.2e-07 | 1.5e-08 | 7.433E-08 | 5.382E-08 | 1.500E-08 | 1.200E-07 |
| 13879 | 694 | T694Y | B | 5.9e-08 | 5.7e-08 | 5.8e-08 | 5.800E-08 | 1.000E-09 | 5.700E-08 | 5.900E-08 |
| 13880 | 694 | T694V | B | 6.3e-08 | 6.5e-08 | 8.7e-08 | 7.167E-08 | 1.332E-08 | 6.300E-08 | 8.700E-08 |
| 13881 | 695 | L695A | B | 5.9e-08 | 3.5e-08 | 7.2e-08 | 5.533E-08 | 1.877E-08 | 3.500E-08 | 7.200E-08 |
| 13882 | 695 | L695R | B | 3.8e-08 | 2.1e-08 | 1.2e-08 | 2.367E-08 | 1.320E-08 | 1.200E-08 | 3.800E-08 |
| 13883 | 695 | L695N | B | 2.5e-08 | 2.7e-08 | 2.5e-08 | 2.567E-08 | 1.155E-09 | 2.500E-08 | 2.700E-08 |
| 13884 | 695 | L695D | B | 4.6e-08 | 3e-08 | 9.1e-08 | 5.567E-08 | 3.163E-08 | 3.000E-08 | 9.100E-08 |
| 13885 | 695 | L695C | B | 4e-08 | 5e-08 | 7.4e-08 | 5.467E-08 | 1.747E-08 | 4.000E-08 | 7.400E-08 |
| 13886 | 695 | L695E | B | 4e-08 | 3.3e-08 | 2e-07 | 9.100E-08 | 9.446E-08 | 3.300E-08 | 2.000E-07 |
| 13887 | 695 | L695Q | B | 7.3e-08 | 2e-08 | 4.7e-08 | 4.667E-08 | 2.650E-08 | 2.000E-08 | 7.300E-08 |
| 13888 | 695 | L695G | B | 6.5e-08 | 1.9e-08 | 8.6e-08 | 5.667E-08 | 3.427E-08 | 1.900E-08 | 8.600E-08 |
| 13889 | 695 | L695H | B | 2.2e-07 | 1.3e-08 | 6.7e-08 | 1.000E-07 | 1.074E-07 | 1.300E-08 | 2.200E-07 |
| 13890 | 695 | L695I | B | 1.5e-07 | 5.2e-08 | 2.1e-08 | 7.433E-08 | 6.734E-08 | 2.100E-08 | 1.500E-07 |
| 13891 | 695 | L695L | B | 3.6e-08 | 6.7e-08 | 5.3e-08 | 5.200E-08 | 1.552E-08 | 3.600E-08 | 6.700E-08 |
| 13892 | 695 | L695K | B | 3.9e-08 | 6.9e-08 | 6.6e-08 | 5.800E-08 | 1.652E-08 | 3.900E-08 | 6.900E-08 |
| 13893 | 695 | L695M | B | 6e-08 | 6.3e-08 | 3.9e-08 | 5.400E-08 | 1.308E-08 | 3.900E-08 | 6.300E-08 |
| 13894 | 695 | L695F | B | 2.8e-08 | 2.3e-08 | 4.4e-08 | 3.167E-08 | 1.097E-08 | 2.300E-08 | 4.400E-08 |
| 13895 | 695 | L695P | B | 9.5e-08 | 5.1e-08 | 1.7e-08 | 5.433E-08 | 3.911E-08 | 1.700E-08 | 9.500E-08 |
| 13896 | 695 | L695S | B | 2.3e-08 | 2.8e-08 | 3.7e-08 | 2.933E-08 | 7.095E-09 | 2.300E-08 | 3.700E-08 |
| 13897 | 695 | L695T | B | 2.6e-08 | 4.5e-08 | 3e-08 | 3.367E-08 | 1.002E-08 | 2.600E-08 | 4.500E-08 |
| 13898 | 695 | L695W | B | 4.3e-08 | 1.1e-07 | 1.4e-08 | 5.567E-08 | 4.924E-08 | 1.400E-08 | 1.100E-07 |
| 13899 | 695 | L695Y | B | 5.3e-08 | 4.8e-08 | 1e-07 | 6.700E-08 | 2.869E-08 | 4.800E-08 | 1.000E-07 |
| 13900 | 695 | L695V | B | 6.3e-08 | 1e-07 | 5.3e-08 | 7.200E-08 | 2.476E-08 | 5.300E-08 | 1.000E-07 |
| 13901 | 696 | T696A | B | 3.3e-08 | 6.2e-08 | 1.2e-07 | 7.167E-08 | 4.430E-08 | 3.300E-08 | 1.200E-07 |
| 13902 | 696 | T696R | B | 3.6e-08 | 4.5e-08 | 2.1e-08 | 3.400E-08 | 1.212E-08 | 2.100E-08 | 4.500E-08 |
| 13903 | 696 | T696N | B | 2.4e-08 | 1.7e-08 | 5.8e-08 | 3.300E-08 | 2.193E-08 | 1.700E-08 | 5.800E-08 |
| 13904 | 696 | T696D | B | 2.7e-08 | 5.4e-08 | 5.5e-08 | 4.533E-08 | 1.589E-08 | 2.700E-08 | 5.500E-08 |
| 13905 | 696 | T696C | B | 3.4e-08 | 2.7e-08 | 4.1e-08 | 3.400E-08 | 7.000E-09 | 2.700E-08 | 4.100E-08 |
| 13906 | 696 | T696E | B | 1.8e-08 | 4.7e-08 | 2.6e-08 | 3.033E-08 | 1.498E-08 | 1.800E-08 | 4.700E-08 |
| 13907 | 696 | T696Q | B | 2e-08 | 6.6e-08 | 4.6e-08 | 4.400E-08 | 2.307E-08 | 2.000E-08 | 6.600E-08 |
| 13908 | 696 | T696G | B | 5.3e-08 | 5.5e-08 | 1.3e-07 | 7.933E-08 | 4.389E-08 | 5.300E-08 | 1.300E-07 |
| 13909 | 696 | T696H | B | 3.8e-08 | 5.3e-08 | 2.4e-08 | 3.833E-08 | 1.450E-08 | 2.400E-08 | 5.300E-08 |
| 13910 | 696 | T696I | B | 3.7e-08 | 3.5e-08 | 2.4e-08 | 3.200E-08 | 7.000E-09 | 2.400E-08 | 3.700E-08 |
| 13911 | 696 | T696L | B | 9.3e-08 | 4.6e-08 | 2.1e-08 | 5.333E-08 | 3.656E-08 | 2.100E-08 | 9.300E-08 |
| 13912 | 696 | T696K | B | 5.1e-08 | 6.1e-08 | 2e-08 | 4.400E-08 | 2.138E-08 | 2.000E-08 | 6.100E-08 |
| 13913 | 696 | T696M | B | 1.5e-08 | 3.6e-08 | 5.6e-08 | 3.567E-08 | 2.050E-08 | 1.500E-08 | 5.600E-08 |
| 13914 | 696 | T696F | B | 1e-07 | 3.3e-08 | 1e-07 | 7.767E-08 | 3.868E-08 | 3.300E-08 | 1.000E-07 |
| 13915 | 696 | T696P | B | 3.2e-08 | 2.2e-08 | 4.9e-08 | 3.433E-08 | 1.365E-08 | 2.200E-08 | 4.900E-08 |
| 13916 | 696 | T696S | B | 5.6e-08 | 5e-08 | 5.4e-09 | 3.713E-08 | 2.765E-08 | 5.400E-09 | 5.600E-08 |
| 13917 | 696 | T696T | B | 2.7e-08 | 4.7e-08 | 2.2e-08 | 3.200E-08 | 1.323E-08 | 2.200E-08 | 4.700E-08 |
| 13918 | 696 | T696W | B | 7.5e-08 | 4.8e-08 | 3.5e-08 | 5.267E-08 | 2.040E-08 | 3.500E-08 | 7.500E-08 |
| 13919 | 696 | T696Y | B | 7.7e-08 | 9.8e-08 | 8.4e-08 | 8.633E-08 | 1.069E-08 | 7.700E-08 | 9.800E-08 |
| 13920 | 696 | T696V | B | 6.8e-08 | 4.6e-08 | 8.2e-08 | 6.533E-08 | 1.815E-08 | 4.600E-08 | 8.200E-08 |
| 13921 | 697 | I697A | B | 5.5e-08 | 3.3e-08 | 6.9e-08 | 5.233E-08 | 1.815E-08 | 3.300E-08 | 6.900E-08 |
| 13922 | 697 | I697R | B | 2.7e-08 | 3e-08 | 7.7e-08 | 4.467E-08 | 2.804E-08 | 2.700E-08 | 7.700E-08 |
| 13923 | 697 | I697N | B | 5.4e-08 | 1.1e-07 | 4.3e-08 | 6.900E-08 | 3.593E-08 | 4.300E-08 | 1.100E-07 |
| 13924 | 697 | I697D | B | 9.9e-08 | 4.5e-08 | 5.8e-08 | 6.733E-08 | 2.818E-08 | 4.500E-08 | 9.900E-08 |
| 13925 | 697 | I697C | B | 4.8e-08 | 3.8e-08 | 8.1e-08 | 5.567E-08 | 2.250E-08 | 3.800E-08 | 8.100E-08 |
| 13926 | 697 | I697E | B | 3.1e-08 | 5e-08 | 1.5e-08 | 3.200E-08 | 1.752E-08 | 1.500E-08 | 5.000E-08 |
| 13927 | 697 | I697Q | B | 6.6e-08 | 1.6e-07 | 2.3e-08 | 8.300E-08 | 7.006E-08 | 2.300E-08 | 1.600E-07 |
| 13928 | 697 | I697G | B | 2.9e-08 | 9.1e-08 | 3.7e-08 | 5.233E-08 | 3.372E-08 | 2.900E-08 | 9.100E-08 |
| 13929 | 697 | I697H | B | 5.5e-08 | 1.7e-08 | 2e-08 | 3.067E-08 | 2.113E-08 | 1.700E-08 | 5.500E-08 |
| 13930 | 697 | I697I | B | 8.6e-08 | 5.8e-08 | 5.3e-08 | 6.567E-08 | 1.779E-08 | 5.300E-08 | 8.600E-08 |
| 13931 | 697 | I697L | B | 1.7e-08 | 3.9e-08 | 1.3e-07 | 6.200E-08 | 5.991E-08 | 1.700E-08 | 1.300E-07 |
| 13932 | 697 | I697K | B | 1.7e-08 | 3.2e-08 | 3.3e-08 | 2.733E-08 | 8.963E-09 | 1.700E-08 | 3.300E-08 |
| 13933 | 697 | I697M | B | 8e-08 | 1.8e-08 | 1.8e-07 | 9.267E-08 | 8.174E-08 | 1.800E-08 | 1.800E-07 |
| 13934 | 697 | I697F | B | 7.3e-08 | 4.8e-08 | 4.3e-08 | 5.467E-08 | 1.607E-08 | 4.300E-08 | 7.300E-08 |
| 13935 | 697 | I697P | B | 1.1e-07 | 4.6e-08 | 3.6e-08 | 6.400E-08 | 4.015E-08 | 3.600E-08 | 1.100E-07 |
| 13936 | 697 | I697S | B | 3.7e-08 | 1.2e-07 | 3.9e-08 | 6.533E-08 | 4.735E-08 | 3.700E-08 | 1.200E-07 |
| 13937 | 697 | I697T | B | 5.3e-08 | 1.8e-08 | 6.5e-08 | 4.533E-08 | 2.442E-08 | 1.800E-08 | 6.500E-08 |
| 13938 | 697 | I697W | B | 4.6e-08 | 5.9e-08 | 5.9e-08 | 5.467E-08 | 7.506E-09 | 4.600E-08 | 5.900E-08 |
| 13939 | 697 | I697Y | B | 5e-08 | 3.6e-08 | 7.5e-08 | 5.367E-08 | 1.976E-08 | 3.600E-08 | 7.500E-08 |

| | | | | | | | | | | |
|---|---|---|---|---|---|---|---|---|---|---|
| 13940 | 697 | I697V | B | 2.2e-08 | 2.2e-08 | 6.9e-08 | 3.767E-08 | 2.714E-08 | 2.200E-08 | 6.900E-08 |
| 13941 | 698 | S698A | B | 3.1e-08 | 1.8e-08 | 2.8e-08 | 2.567E-08 | 6.807E-09 | 1.800E-08 | 3.100E-08 |
| 13942 | 698 | S698R | B | 2.1e-07 | 2.4e-08 | 2.2e-08 | 8.533E-08 | 1.080E-07 | 2.200E-08 | 2.100E-07 |
| 13943 | 698 | S698N | B | 5.5e-08 | 1.7e-08 | 6e-08 | 4.400E-08 | 2.352E-08 | 1.700E-08 | 6.000E-08 |
| 13944 | 698 | S698D | B | 2.2e-08 | 9.2e-08 | 4.2e-08 | 5.200E-08 | 3.606E-08 | 2.200E-08 | 9.200E-08 |
| 13945 | 698 | S698C | B | 4.4e-08 | 1.7e-08 | 1.3e-08 | 2.467E-08 | 1.686E-08 | 1.300E-08 | 4.400E-08 |
| 13946 | 698 | S698E | B | 2.9e-08 | 5e-08 | 6.1e-08 | 4.667E-08 | 1.626E-08 | 2.900E-08 | 6.100E-08 |
| 13947 | 698 | S698Q | B | 2.5e-08 | 3.6e-08 | 3.2e-08 | 3.100E-08 | 5.568E-09 | 2.500E-08 | 3.600E-08 |
| 13948 | 698 | S698G | B | 2.8e-08 | 5.7e-08 | 9.8e-08 | 6.100E-08 | 3.517E-08 | 2.800E-08 | 9.800E-08 |
| 13949 | 698 | S698H | B | 3.8e-08 | 6e-08 | 1.1e-07 | 6.933E-08 | 3.690E-08 | 3.800E-08 | 1.100E-07 |
| 13950 | 698 | S698I | B | 3.8e-08 | 7.2e-08 | 3.6e-08 | 4.867E-08 | 2.023E-08 | 3.600E-08 | 7.200E-08 |
| 13951 | 698 | S698L | B | 1.7e-08 | 3e-08 | 1.2e-07 | 5.567E-08 | 5.609E-08 | 1.700E-08 | 1.200E-07 |
| 13952 | 698 | S698K | B | 1.1e-08 | 4.9e-08 | 9.3e-09 | 2.310E-08 | 2.245E-08 | 9.300E-09 | 4.900E-08 |
| 13953 | 698 | S698M | B | 5e-08 | 5.4e-08 | 1e-07 | 6.800E-08 | 2.778E-08 | 5.000E-08 | 1.000E-07 |
| 13954 | 698 | S698F | B | 1.8e-08 | 8.2e-08 | 3.5e-08 | 4.500E-08 | 3.315E-08 | 1.800E-08 | 8.200E-08 |
| 13955 | 698 | S698P | B | 8.5e-08 | 2e-08 | 5.1e-08 | 5.200E-08 | 3.251E-08 | 2.000E-08 | 8.500E-08 |
| 13956 | 698 | S698S | B | 9e-08 | 2.8e-08 | 5.8e-08 | 5.867E-08 | 3.101E-08 | 2.800E-08 | 9.000E-08 |
| 13957 | 698 | S698T | B | 1.4e-07 | 7.7e-08 | 4.2e-08 | 8.633E-08 | 4.966E-08 | 4.200E-08 | 1.400E-07 |
| 13958 | 698 | S698W | B | 5.9e-08 | 9.3e-08 | 8.1e-08 | 7.767E-08 | 1.724E-08 | 5.900E-08 | 9.300E-08 |
| 13959 | 698 | S698Y | B | 2.8e-08 | 2.3e-08 | 4.4e-08 | 3.167E-08 | 1.097E-08 | 2.300E-08 | 4.400E-08 |
| 13960 | 698 | S698V | B | 2.1e-08 | 1.4e-08 | 2.2e-08 | 1.900E-08 | 4.359E-09 | 1.400E-08 | 2.200E-08 |
| 13961 | 699 | S699A | B | 3.4e-08 | 7.6e-08 | 8.1e-08 | 6.367E-08 | 2.581E-08 | 3.400E-08 | 8.100E-08 |
| 13962 | 699 | S699R | B | 5.4e-08 | 3.7e-08 | 6.5e-08 | 5.200E-08 | 1.411E-08 | 3.700E-08 | 6.500E-08 |
| 13963 | 699 | S699N | B | 3.3e-08 | 3.5e-08 | 4e-08 | 3.600E-08 | 3.606E-09 | 3.300E-08 | 4.000E-08 |
| 13964 | 699 | S699D | B | 3.2e-08 | 4.9e-08 | 1.2e-07 | 6.700E-08 | 4.668E-08 | 3.200E-08 | 1.200E-07 |
| 13965 | 699 | S699C | B | 2.5e-08 | 5.5e-08 | 1.1e-07 | 6.333E-08 | 4.311E-08 | 2.500E-08 | 1.100E-07 |
| 13966 | 699 | S699E | B | 7.4e-08 | 4.1e-08 | 6.8e-08 | 6.100E-08 | 1.758E-08 | 4.100E-08 | 7.400E-08 |
| 13967 | 699 | S699Q | B | 1.4e-08 | 4.5e-08 | 1.9e-08 | 2.600E-08 | 1.664E-08 | 1.400E-08 | 4.500E-08 |
| 13968 | 699 | S699G | B | 8e-09 | 4e-08 | 2.9e-08 | 2.567E-08 | 1.626E-08 | 8.000E-09 | 4.000E-08 |
| 13969 | 699 | S699H | B | 3e-08 | 3.9e-08 | 4e-08 | 3.633E-08 | 5.508E-09 | 3.000E-08 | 4.000E-08 |
| 13970 | 699 | S699I | B | 3.3e-08 | 1.3e-08 | 4.9e-08 | 3.167E-08 | 1.804E-08 | 1.300E-08 | 4.900E-08 |
| 13971 | 699 | S699L | B | 7.6e-08 | 9.1e-08 | 3.1e-08 | 6.600E-08 | 3.122E-08 | 3.100E-08 | 9.100E-08 |
| 13972 | 699 | S699K | B | 2.6e-08 | 4.4e-08 | 5.3e-08 | 4.100E-08 | 1.375E-08 | 2.600E-08 | 5.300E-08 |
| 13973 | 699 | S699M | B | 1.4e-07 | 2.4e-08 | 8.4e-08 | 8.267E-08 | 5.801E-08 | 2.400E-08 | 1.400E-07 |
| 13974 | 699 | S699F | B | 5.4e-08 | 8.6e-08 | 9.2e-09 | 4.973E-08 | 3.858E-08 | 9.200E-09 | 8.600E-08 |
| 13975 | 699 | S699P | B | 8.7e-08 | 5.2e-08 | 3.3e-08 | 5.733E-08 | 2.739E-08 | 3.300E-08 | 8.700E-08 |
| 13976 | 699 | S699S | B | 2.6e-08 | 3e-08 | 1.6e-08 | 2.400E-08 | 7.211E-09 | 1.600E-08 | 3.000E-08 |
| 13977 | 699 | S699T | B | 1.6e-08 | 3.6e-08 | 5.4e-08 | 3.533E-08 | 1.901E-08 | 1.600E-08 | 5.400E-08 |
| 13978 | 699 | S699W | B | 1.4e-08 | 4.5e-08 | 1.2e-08 | 2.367E-08 | 1.850E-08 | 1.200E-08 | 4.500E-08 |
| 13979 | 699 | S699Y | B | 6.4e-08 | 1.7e-08 | 4.1e-08 | 4.067E-08 | 2.350E-08 | 1.700E-08 | 6.400E-08 |
| 13980 | 699 | S699V | B | 4.4e-08 | 2.7e-08 | 1.8e-08 | 2.967E-08 | 1.320E-08 | 1.800E-08 | 4.400E-08 |
| 13981 | 700 | L700A | B | 5.6e-08 | 5.9e-08 | 3.2e-08 | 4.900E-08 | 1.480E-08 | 3.200E-08 | 5.900E-08 |
| 13982 | 700 | L700R | B | 7.8e-08 | 5.3e-08 | 2.8e-08 | 5.300E-08 | 2.500E-08 | 2.800E-08 | 7.800E-08 |
| 13983 | 700 | L700N | B | 1.4e-08 | 7.7e-08 | 1.3e-07 | 7.367E-08 | 5.807E-08 | 1.400E-08 | 1.300E-07 |
| 13984 | 700 | L700D | B | 8.3e-08 | 7.1e-08 | 2.4e-08 | 5.933E-08 | 3.118E-08 | 2.400E-08 | 8.300E-08 |
| 13985 | 700 | L700C | B | 6.3e-08 | 3.1e-08 | 1.3e-08 | 3.567E-08 | 2.532E-08 | 1.300E-08 | 6.300E-08 |
| 13986 | 700 | L700E | B | 6.8e-08 | 8.6e-08 | 1.3e-07 | 9.467E-08 | 3.190E-08 | 6.800E-08 | 1.300E-07 |
| 13987 | 700 | L700Q | B | 2.9e-08 | 8.9e-08 | 7.8e-08 | 6.533E-08 | 3.194E-08 | 2.900E-08 | 8.900E-08 |
| 13988 | 700 | L700G | B | 2.6e-08 | 4.5e-08 | 9e-08 | 5.367E-08 | 3.287E-08 | 2.600E-08 | 9.000E-08 |
| 13989 | 700 | L700H | B | 1.5e-08 | 2.7e-08 | 1.9e-08 | 2.033E-08 | 6.110E-09 | 1.500E-08 | 2.700E-08 |
| 13990 | 700 | L700I | B | 2.8e-08 | 5.3e-08 | 5.1e-08 | 4.400E-08 | 1.389E-08 | 2.800E-08 | 5.300E-08 |
| 13991 | 700 | L700L | B | 1.5e-08 | 4e-08 | 6e-08 | 3.833E-08 | 2.255E-08 | 1.500E-08 | 6.000E-08 |
| 13992 | 700 | L700K | B | 4.2e-08 | 2.4e-08 | 6.8e-08 | 4.467E-08 | 2.212E-08 | 2.400E-08 | 6.800E-08 |
| 13993 | 700 | L700M | B | 9.8e-08 | 2.2e-07 | 3.8e-08 | 1.187E-07 | 9.274E-08 | 3.800E-08 | 2.200E-07 |
| 13994 | 700 | L700F | B | 2.1e-08 | 8.7e-08 | 2.1e-08 | 4.300E-08 | 3.811E-08 | 2.100E-08 | 8.700E-08 |
| 13995 | 700 | L700P | B | 6.9e-08 | 5.8e-08 | 5.4e-08 | 6.033E-08 | 7.767E-09 | 5.400E-08 | 6.900E-08 |
| 13996 | 700 | L700S | B | 5.3e-08 | 4.7e-08 | 4.4e-08 | 4.800E-08 | 4.583E-09 | 4.400E-08 | 5.300E-08 |
| 13997 | 700 | L700T | B | 5.4e-08 | 2.6e-08 | 3.2e-08 | 3.733E-08 | 1.474E-08 | 2.600E-08 | 5.400E-08 |
| 13998 | 700 | L700W | B | 9.7e-08 | 4.6e-08 | 2.4e-08 | 5.567E-08 | 3.745E-08 | 2.400E-08 | 9.700E-08 |
| 13999 | 700 | L700Y | B | 2e-08 | 5.4e-08 | 4.1e-08 | 3.833E-08 | 1.716E-08 | 2.000E-08 | 5.400E-08 |
| 14000 | 700 | L700V | B | 8.5e-08 | 7.2e-08 | 1.2e-07 | 9.233E-08 | 2.483E-08 | 7.200E-08 | 1.200E-07 |
| 14001 | 701 | Q701A | B | 1.8e-08 | 2.5e-08 | 5.4e-08 | 3.233E-08 | 1.909E-08 | 1.800E-08 | 5.400E-08 |
| 14002 | 701 | Q701R | B | 1.4e-08 | 5.9e-08 | 7.2e-08 | 4.833E-08 | 3.044E-08 | 1.400E-08 | 7.200E-08 |
| 14003 | 701 | Q701N | B | 1.6e-08 | 2.8e-08 | 6.1e-08 | 3.500E-08 | 2.330E-08 | 1.600E-08 | 6.100E-08 |
| 14004 | 701 | Q701D | B | 4.2e-08 | 3.8e-08 | 5.8e-08 | 4.600E-08 | 1.058E-08 | 3.800E-08 | 5.800E-08 |
| 14005 | 701 | Q701C | B | 1.1e-07 | 6.5e-08 | 6.1e-08 | 7.867E-08 | 2.721E-08 | 6.100E-08 | 1.100E-07 |
| 14006 | 701 | Q701E | B | 2.6e-08 | 6.3e-08 | 1.1e-07 | 6.633E-08 | 4.210E-08 | 2.600E-08 | 1.100E-07 |
| 14007 | 701 | Q701Q | B | 6.7e-08 | 2.4e-08 | 8.5e-08 | 5.867E-08 | 3.134E-08 | 2.400E-08 | 8.500E-08 |
| 14008 | 701 | Q701G | B | 8.5e-08 | 9.9e-08 | 1.4e-08 | 6.600E-08 | 4.557E-08 | 1.400E-08 | 9.900E-08 |
| 14009 | 701 | Q701H | B | 2.2e-08 | 1.5e-08 | 4.2e-08 | 2.633E-08 | 1.401E-08 | 1.500E-08 | 4.200E-08 |
| 14010 | 701 | Q701I | B | 2e-07 | 3.9e-08 | 4.1e-08 | 9.333E-08 | 9.238E-08 | 3.900E-08 | 2.000E-07 |
| 14011 | 701 | Q701L | B | 1.3e-07 | 2.9e-08 | 3.2e-08 | 6.367E-08 | 5.747E-08 | 2.900E-08 | 1.300E-07 |
| 14012 | 701 | Q701K | B | 2.7e-08 | 5e-08 | 3.4e-08 | 3.700E-08 | 1.179E-08 | 2.700E-08 | 5.000E-08 |
| 14013 | 701 | Q701M | B | 3.7e-08 | 4.6e-08 | 5.8e-08 | 4.700E-08 | 1.054E-08 | 3.700E-08 | 5.800E-08 |
| 14014 | 701 | Q701F | B | 2.2e-08 | 1.1e-07 | 3.9e-08 | 5.700E-08 | 4.668E-08 | 2.200E-08 | 1.100E-07 |
| 14015 | 701 | Q701P | B | 3.1e-08 | 3.9e-08 | 3.5e-08 | 3.500E-08 | 4.000E-09 | 3.100E-08 | 3.900E-08 |
| 14016 | 701 | Q701S | B | 1.6e-07 | 4.8e-08 | 3e-08 | 7.933E-08 | 7.044E-08 | 3.000E-08 | 1.600E-07 |
| 14017 | 701 | Q701T | B | 4.1e-08 | 4.8e-08 | 2.3e-08 | 3.733E-08 | 1.290E-08 | 2.300E-08 | 4.800E-08 |
| 14018 | 701 | Q701W | B | 8.6e-09 | 4.3e-08 | 4.8e-08 | 3.320E-08 | 2.145E-08 | 8.600E-09 | 4.800E-08 |
| 14019 | 701 | Q701Y | B | 4e-08 | 3.2e-08 | 6.3e-08 | 4.500E-08 | 1.609E-08 | 3.200E-08 | 6.300E-08 |
| 14020 | 701 | Q701V | B | 3e-08 | 3.8e-08 | 9.6e-08 | 5.467E-08 | 3.602E-08 | 3.000E-08 | 9.600E-08 |
| 14021 | 702 | P702A | B | 4e-08 | 2.8e-08 | 3.4e-08 | 3.400E-08 | 6.000E-09 | 2.800E-08 | 4.000E-08 |
| 14022 | 702 | P702R | B | 1.9e-08 | 3e-08 | 2.3e-08 | 2.400E-08 | 5.568E-09 | 1.900E-08 | 3.000E-08 |
| 14023 | 702 | P702N | B | 8.3e-08 | 3.4e-08 | 2.5e-08 | 4.733E-08 | 3.121E-08 | 2.500E-08 | 8.300E-08 |
| 14024 | 702 | P702D | B | 5.3e-08 | 1.4e-08 | 6e-08 | 4.233E-08 | 2.479E-08 | 1.400E-08 | 6.000E-08 |

| | | | | | | | | | | |
|---|---|---|---|---|---|---|---|---|---|---|
| 14025 | 702 | P702C | B | 2.1e-08 | 2.5e-08 | 5.1e-08 | 3.233E-08 | 1.629E-08 | 2.100E-08 | 5.100E-08 |
| 14026 | 702 | P702E | B | 4.2e-08 | 6.3e-08 | 3.5e-08 | 4.667E-08 | 1.457E-08 | 3.500E-08 | 6.300E-08 |
| 14027 | 702 | P702Q | B | 1.1e-07 | 2.6e-08 | 2.5e-08 | 5.367E-08 | 4.879E-08 | 2.500E-08 | 1.100E-07 |
| 14028 | 702 | P702G | B | 9.8e-08 | 6.9e-08 | 5.2e-08 | 7.300E-08 | 2.326E-08 | 5.200E-08 | 9.800E-08 |
| 14029 | 702 | P702H | B | 1.3e-07 | 3.5e-08 | 3.1e-08 | 6.533E-08 | 5.604E-08 | 3.100E-08 | 1.300E-07 |
| 14030 | 702 | P702I | B | 2.3e-08 | 5e-08 | 5.1e-08 | 4.133E-08 | 1.589E-08 | 2.300E-08 | 5.100E-08 |
| 14031 | 702 | P702L | B | 2.2e-08 | 6.1e-08 | 3.7e-08 | 4.000E-08 | 1.967E-08 | 2.200E-08 | 6.100E-08 |
| 14032 | 702 | P702K | B | 5.6e-08 | 3.2e-08 | 4.2e-08 | 4.333E-08 | 1.206E-08 | 3.200E-08 | 5.600E-08 |
| 14033 | 702 | P702M | B | 2.9e-07 | 4.9e-08 | 1.9e-08 | 1.193E-07 | 1.486E-07 | 1.900E-08 | 2.900E-07 |
| 14034 | 702 | P702F | B | 1.6e-08 | 7.2e-08 | 2e-08 | 3.600E-08 | 3.124E-08 | 1.600E-08 | 7.200E-08 |
| 14035 | 702 | P702P | B | 2.6e-08 | 4.9e-08 | 2.8e-08 | 3.433E-08 | 1.274E-08 | 2.600E-08 | 4.900E-08 |
| 14036 | 702 | P702S | B | 5.3e-08 | 1.4e-08 | 4.7e-08 | 3.800E-08 | 2.100E-08 | 1.400E-08 | 5.300E-08 |
| 14037 | 702 | P702T | B | 4.8e-08 | 2.9e-08 | 4.1e-08 | 3.933E-08 | 9.609E-09 | 2.900E-08 | 4.800E-08 |
| 14038 | 702 | P702W | B | 3.2e-08 | 3.2e-08 | 3.4e-08 | 3.267E-08 | 1.155E-09 | 3.200E-08 | 3.400E-08 |
| 14039 | 702 | P702Y | B | 2.5e-08 | 2.2e-08 | 2.9e-08 | 2.533E-08 | 3.512E-09 | 2.200E-08 | 2.900E-08 |
| 14040 | 702 | P702V | B | 3.1e-08 | 2.5e-08 | 2.4e-08 | 2.667E-08 | 3.786E-09 | 2.400E-08 | 3.100E-08 |
| 14041 | 703 | E703A | B | 4.2e-08 | 2.4e-08 | 5.5e-08 | 4.033E-08 | 1.557E-08 | 2.400E-08 | 5.500E-08 |
| 14042 | 703 | E703R | B | 2.9e-08 | 5.4e-08 | 7.1e-08 | 5.133E-08 | 2.113E-08 | 2.900E-08 | 7.100E-08 |
| 14043 | 703 | E703N | B | 3.9e-08 | 3.4e-08 | 3.1e-08 | 3.467E-08 | 4.041E-09 | 3.100E-08 | 3.900E-08 |
| 14044 | 703 | E703D | B | 2.3e-08 | 1.7e-08 | 1.4e-08 | 1.800E-08 | 4.583E-09 | 1.400E-08 | 2.300E-08 |
| 14045 | 703 | E703C | B | 4.1e-08 | 7.7e-08 | 4.6e-08 | 5.467E-08 | 1.950E-08 | 4.100E-08 | 7.700E-08 |
| 14046 | 703 | E703E | B | 4.7e-08 | 7e-08 | 7.9e-08 | 6.533E-08 | 1.650E-08 | 4.700E-08 | 7.900E-08 |
| 14047 | 703 | E703Q | B | 4.8e-08 | 6.2e-08 | 3.4e-08 | 4.800E-08 | 1.400E-08 | 3.400E-08 | 6.200E-08 |
| 14048 | 703 | E703G | B | 2.9e-08 | 8e-08 | 2.1e-08 | 4.333E-08 | 3.201E-08 | 2.100E-08 | 8.000E-08 |
| 14049 | 703 | E703H | B | 1.5e-08 | 4.7e-08 | 1.1e-07 | 5.733E-08 | 4.834E-08 | 1.500E-08 | 1.100E-07 |
| 14050 | 703 | E703I | B | 3.5e-08 | 3.8e-08 | 1.4e-07 | 7.100E-08 | 5.977E-08 | 3.500E-08 | 1.400E-07 |
| 14051 | 703 | E703L | B | 7.8e-08 | 8.3e-08 | 1.2e-07 | 9.367E-08 | 2.294E-08 | 7.800E-08 | 1.200E-07 |
| 14052 | 703 | E703K | B | 7e-08 | 4.1e-08 | 3.8e-08 | 4.967E-08 | 1.767E-08 | 3.800E-08 | 7.000E-08 |
| 14053 | 703 | E703M | B | 9.1e-08 | 7.8e-08 | 3.6e-08 | 6.833E-08 | 2.875E-08 | 3.600E-08 | 9.100E-08 |
| 14054 | 703 | E703F | B | 3.8e-08 | 3.1e-08 | 4.7e-08 | 3.867E-08 | 8.021E-09 | 3.100E-08 | 4.700E-08 |
| 14055 | 703 | E703P | B | 8e-08 | 5.1e-08 | 1.6e-07 | 9.700E-08 | 5.645E-08 | 5.100E-08 | 1.600E-07 |
| 14056 | 703 | E703S | B | 1e-08 | 6.3e-08 | 4.4e-08 | 3.900E-08 | 2.685E-08 | 1.000E-08 | 6.300E-08 |
| 14057 | 703 | E703T | B | 6e-08 | 4.4e-08 | 4.6e-08 | 5.000E-08 | 8.718E-09 | 4.400E-08 | 6.000E-08 |
| 14058 | 703 | E703W | B | 2.2e-08 | 5.3e-08 | 1e-08 | 2.833E-08 | 2.219E-08 | 1.000E-08 | 5.300E-08 |
| 14059 | 703 | E703Y | B | 4.8e-08 | 3.4e-08 | 4.7e-08 | 4.300E-08 | 7.810E-09 | 3.400E-08 | 4.800E-08 |
| 14060 | 703 | E703V | B | 9.2e-08 | 2e-08 | 3.7e-08 | 4.967E-08 | 3.763E-08 | 2.000E-08 | 9.200E-08 |
| 14061 | 704 | D704A | B | 3.6e-08 | 6.2e-08 | 2.8e-08 | 4.200E-08 | 1.778E-08 | 2.800E-08 | 6.200E-08 |
| 14062 | 704 | D704R | B | 8e-08 | 3.4e-08 | 3.7e-08 | 5.033E-08 | 2.574E-08 | 3.400E-08 | 8.000E-08 |
| 14063 | 704 | D704N | B | 1.1e-07 | 2.2e-08 | 6.9e-08 | 6.700E-08 | 4.403E-08 | 2.200E-08 | 1.100E-07 |
| 14064 | 704 | D704D | B | 1.8e-08 | 2.5e-08 | 9.8e-09 | 1.760E-08 | 7.608E-09 | 9.800E-09 | 2.500E-08 |
| 14065 | 704 | D704C | B | 2.6e-08 | 3.7e-08 | 7e-08 | 4.433E-08 | 2.290E-08 | 2.600E-08 | 7.000E-08 |
| 14066 | 704 | D704E | B | 9.5e-08 | 6.6e-08 | 2.9e-08 | 6.333E-08 | 3.308E-08 | 2.900E-08 | 9.500E-08 |
| 14067 | 704 | D704Q | B | 3e-08 | 3.8e-08 | 3.8e-08 | 3.533E-08 | 4.619E-09 | 3.000E-08 | 3.800E-08 |
| 14068 | 704 | D704G | B | 2.7e-08 | 6.8e-08 | 3.1e-08 | 4.200E-08 | 2.261E-08 | 2.700E-08 | 6.800E-08 |
| 14069 | 704 | D704H | B | 1.7e-08 | 1.2e-07 | 3.9e-08 | 5.867E-08 | 5.424E-08 | 1.700E-08 | 1.200E-07 |
| 14070 | 704 | D704I | B | 6.5e-08 | 1.3e-07 | 4.5e-08 | 8.000E-08 | 4.444E-08 | 4.500E-08 | 1.300E-07 |
| 14071 | 704 | D704L | B | 4.4e-08 | 7e-08 | 1.4e-07 | 8.467E-08 | 4.965E-08 | 4.400E-08 | 1.400E-07 |
| 14072 | 704 | D704K | B | 6e-08 | 5.8e-08 | 4.9e-08 | 5.567E-08 | 5.859E-09 | 4.900E-08 | 6.000E-08 |
| 14073 | 704 | D704M | B | 2.5e-08 | 6.6e-08 | 2e-07 | 9.700E-08 | 9.153E-08 | 2.500E-08 | 2.000E-07 |
| 14074 | 704 | D704F | B | 6.3e-08 | 1.4e-08 | 2.1e-08 | 3.267E-08 | 2.650E-08 | 1.400E-08 | 6.300E-08 |
| 14075 | 704 | D704P | B | 3.1e-08 | 3.2e-08 | 5.7e-08 | 4.000E-08 | 1.473E-08 | 3.100E-08 | 5.700E-08 |
| 14076 | 704 | D704S | B | 1.9e-07 | 8.3e-08 | 4.8e-08 | 1.070E-07 | 7.398E-08 | 4.800E-08 | 1.900E-07 |
| 14077 | 704 | D704T | B | 2.1e-08 | 3.4e-08 | 4.8e-08 | 3.433E-08 | 1.350E-08 | 2.100E-08 | 4.800E-08 |
| 14078 | 704 | D704W | B | 6.7e-08 | 1.9e-08 | 7.2e-08 | 5.267E-08 | 2.926E-08 | 1.900E-08 | 7.200E-08 |
| 14079 | 704 | D704Y | B | 5.6e-08 | 1e-07 | 5.3e-08 | 6.967E-08 | 2.631E-08 | 5.300E-08 | 1.000E-07 |
| 14080 | 704 | D704V | B | 2.8e-08 | 3.9e-08 | 5.9e-08 | 4.200E-08 | 1.572E-08 | 2.800E-08 | 5.900E-08 |
| 14081 | 705 | F705A | B | 6.5e-08 | 9.2e-08 | 4.8e-08 | 6.833E-08 | 2.219E-08 | 4.800E-08 | 9.200E-08 |
| 14082 | 705 | F705R | B | 4.2e-08 | 8.2e-08 | 2.8e-08 | 5.067E-08 | 2.802E-08 | 2.800E-08 | 8.200E-08 |
| 14083 | 705 | F705N | B | 2.9e-08 | 1e-07 | 3e-08 | 5.300E-08 | 4.071E-08 | 2.900E-08 | 1.000E-07 |
| 14084 | 705 | F705D | B | 4.4e-08 | 5.4e-08 | 5.8e-08 | 5.200E-08 | 7.211E-09 | 4.400E-08 | 5.800E-08 |
| 14085 | 705 | F705C | B | 3.7e-08 | 6.2e-08 | 5.5e-08 | 5.133E-08 | 1.290E-08 | 3.700E-08 | 6.200E-08 |
| 14086 | 705 | F705E | B | 3.9e-08 | 4.7e-08 | 7.6e-08 | 5.400E-08 | 1.947E-08 | 3.900E-08 | 7.600E-08 |
| 14087 | 705 | F705Q | B | 4.8e-08 | 4.9e-08 | 9.5e-08 | 6.400E-08 | 2.685E-08 | 4.800E-08 | 9.500E-08 |
| 14088 | 705 | F705G | B | 5.5e-08 | 4.9e-08 | 2.2e-08 | 4.200E-08 | 1.758E-08 | 2.200E-08 | 5.500E-08 |
| 14089 | 705 | F705H | B | 4.2e-08 | 5.1e-08 | 2.5e-08 | 3.933E-08 | 1.320E-08 | 2.500E-08 | 5.100E-08 |
| 14090 | 705 | F705I | B | 5.3e-08 | 3.5e-08 | 2.5e-08 | 3.767E-08 | 1.419E-08 | 2.500E-08 | 5.300E-08 |
| 14091 | 705 | F705L | B | 2.6e-08 | 7.2e-08 | 5.1e-08 | 4.967E-08 | 2.303E-08 | 2.600E-08 | 7.200E-08 |
| 14092 | 705 | F705K | B | 4.1e-08 | 5e-08 | 2.4e-08 | 3.833E-08 | 1.320E-08 | 2.400E-08 | 5.000E-08 |
| 14093 | 705 | F705M | B | 6.5e-08 | 8.9e-08 | 5.8e-08 | 7.067E-08 | 1.626E-08 | 5.800E-08 | 8.900E-08 |
| 14094 | 705 | F705F | B | 6.2e-08 | 4.4e-08 | 2.9e-08 | 4.500E-08 | 1.652E-08 | 2.900E-08 | 6.200E-08 |
| 14095 | 705 | F705P | B | 1.1e-07 | 7e-08 | 6.1e-08 | 8.033E-08 | 2.608E-08 | 6.100E-08 | 1.100E-07 |
| 14096 | 705 | F705S | B | 2.2e-08 | 7.7e-08 | 6.1e-08 | 5.333E-08 | 2.829E-08 | 2.200E-08 | 7.700E-08 |
| 14097 | 705 | F705T | B | 2.1e-08 | 8.6e-08 | 1.2e-07 | 7.567E-08 | 5.030E-08 | 2.100E-08 | 1.200E-07 |
| 14098 | 705 | F705W | B | 4.7e-08 | 7e-08 | 2e-08 | 4.567E-08 | 2.503E-08 | 2.000E-08 | 7.000E-08 |
| 14099 | 705 | F705Y | B | 6.8e-08 | 5.4e-08 | 2.4e-08 | 4.867E-08 | 2.248E-08 | 2.400E-08 | 6.800E-08 |
| 14100 | 705 | F705V | B | 9.5e-08 | 2.6e-08 | 2.7e-08 | 4.933E-08 | 3.955E-08 | 2.600E-08 | 9.500E-08 |
| 14101 | 706 | A706A | B | 4.1e-08 | 7.8e-08 | 2.6e-08 | 4.833E-08 | 2.676E-08 | 2.600E-08 | 7.800E-08 |
| 14102 | 706 | A706R | B | 1.3e-08 | 9.3e-08 | 4.2e-08 | 4.933E-08 | 4.050E-08 | 1.300E-08 | 9.300E-08 |
| 14103 | 706 | A706N | B | 1.4e-07 | 4.4e-08 | 2.2e-08 | 6.867E-08 | 6.275E-08 | 2.200E-08 | 1.400E-07 |
| 14104 | 706 | A706D | B | 3.6e-08 | 3.2e-08 | 2.5e-08 | 3.100E-08 | 5.568E-09 | 2.500E-08 | 3.600E-08 |
| 14105 | 706 | A706C | B | 5.2e-08 | 3.8e-08 | 1.4e-08 | 3.467E-08 | 1.922E-08 | 1.400E-08 | 5.200E-08 |
| 14106 | 706 | A706E | B | 3.3e-08 | 3.6e-08 | 8e-08 | 4.967E-08 | 2.631E-08 | 3.300E-08 | 8.000E-08 |
| 14107 | 706 | A706Q | B | 2.5e-08 | 4.4e-08 | 5.1e-08 | 4.000E-08 | 1.345E-08 | 2.500E-08 | 5.100E-08 |
| 14108 | 706 | A706G | B | 2.2e-08 | 4.2e-08 | 1.1e-07 | 5.800E-08 | 4.613E-08 | 2.200E-08 | 1.100E-07 |
| 14109 | 706 | A706H | B | 2.1e-08 | 3.6e-08 | 2.3e-08 | 2.667E-08 | 8.145E-09 | 2.100E-08 | 3.600E-08 |

| | | | | | | | | | | |
|---|---|---|---|---|---|---|---|---|---|---|
| 14110 | 706 | A706I | B | 2.2e-08 | 1.2e-07 | 8.7e-08 | 7.633E-08 | 4.986E-08 | 2.200E-08 | 1.200E-07 |
| 14111 | 706 | A706L | B | 5.7e-08 | 3.3e-08 | 3.1e-08 | 4.033E-08 | 1.447E-08 | 3.100E-08 | 5.700E-08 |
| 14112 | 706 | A706K | B | 3.2e-08 | 4.1e-08 | 7.8e-08 | 5.033E-08 | 2.438E-08 | 3.200E-08 | 7.800E-08 |
| 14113 | 706 | A706M | B | 9e-09 | 1.8e-08 | 2.8e-08 | 1.833E-08 | 9.504E-09 | 9.000E-09 | 2.800E-08 |
| 14114 | 706 | A706F | B | 1.4e-08 | 7.8e-08 | 2.6e-08 | 3.933E-08 | 3.402E-08 | 1.400E-08 | 7.800E-08 |
| 14115 | 706 | A706P | B | 3.9e-08 | 5.1e-08 | 2.8e-08 | 3.933E-08 | 1.150E-08 | 2.800E-08 | 5.100E-08 |
| 14116 | 706 | A706S | B | 3e-08 | 6.4e-08 | 9.5e-08 | 6.300E-08 | 3.251E-08 | 3.000E-08 | 9.500E-08 |
| 14117 | 706 | A706T | B | 3.6e-08 | 2.4e-08 | 2.8e-08 | 2.933E-08 | 6.110E-09 | 2.400E-08 | 3.600E-08 |
| 14118 | 706 | A706W | B | 3.4e-08 | 4.6e-08 | 3.6e-08 | 3.867E-08 | 6.429E-09 | 3.400E-08 | 4.600E-08 |
| 14119 | 706 | A706Y | B | 3.6e-08 | 3.2e-08 | 3e-08 | 3.267E-08 | 3.055E-09 | 3.000E-08 | 3.600E-08 |
| 14120 | 706 | A706V | B | 1.1e-07 | 3.1e-08 | 6.8e-08 | 6.967E-08 | 3.953E-08 | 3.100E-08 | 1.100E-07 |
| 14121 | 707 | T707A | B | 3.8e-08 | 4.7e-08 | 1.1e-07 | 6.500E-08 | 3.923E-08 | 3.800E-08 | 1.100E-07 |
| 14122 | 707 | T707R | B | 4.5e-09 | 7.1e-08 | 6.1e-08 | 4.550E-08 | 3.586E-08 | 4.500E-09 | 7.100E-08 |
| 14123 | 707 | T707N | B | 4.1e-08 | 2.7e-08 | 4.4e-08 | 3.733E-08 | 9.074E-09 | 2.700E-08 | 4.400E-08 |
| 14124 | 707 | T707D | B | 6.3e-08 | 8.7e-08 | 1.5e-08 | 5.500E-08 | 3.666E-08 | 1.500E-08 | 8.700E-08 |
| 14125 | 707 | T707C | B | 6.1e-08 | 8.7e-08 | 1.9e-08 | 5.567E-08 | 3.431E-08 | 1.900E-08 | 8.700E-08 |
| 14126 | 707 | T707E | B | 3.5e-08 | 4.8e-08 | 5e-08 | 4.433E-08 | 8.145E-09 | 3.500E-08 | 5.000E-08 |
| 14127 | 707 | T707Q | B | 7.1e-08 | 2.6e-08 | 2.6e-08 | 4.100E-08 | 2.598E-08 | 2.600E-08 | 7.100E-08 |
| 14128 | 707 | T707G | B | 3.1e-08 | 2.9e-08 | 5.5e-08 | 3.833E-08 | 1.447E-08 | 2.900E-08 | 5.500E-08 |
| 14129 | 707 | T707H | B | 2.7e-08 | 9e-08 | 5.1e-08 | 5.600E-08 | 3.180E-08 | 2.700E-08 | 9.000E-08 |
| 14130 | 707 | T707I | B | 2.6e-08 | 6.9e-08 | 7.9e-08 | 5.800E-08 | 2.816E-08 | 2.600E-08 | 7.900E-08 |
| 14131 | 707 | T707L | B | 3.1e-08 | 3.4e-08 | 2.9e-08 | 3.133E-08 | 2.517E-09 | 2.900E-08 | 3.400E-08 |
| 14132 | 707 | T707K | B | 3.8e-08 | 2.9e-08 | 8.1e-08 | 4.933E-08 | 2.779E-08 | 2.900E-08 | 8.100E-08 |
| 14133 | 707 | T707M | B | 5.1e-08 | 1.4e-07 | 1.1e-07 | 1.003E-07 | 4.528E-08 | 5.100E-08 | 1.400E-07 |
| 14134 | 707 | T707F | B | 1.7e-08 | 4e-08 | 5.4e-08 | 3.700E-08 | 1.868E-08 | 1.700E-08 | 5.400E-08 |
| 14135 | 707 | T707P | B | 6.1e-08 | 7.2e-08 | 2.5e-08 | 5.267E-08 | 2.458E-08 | 2.500E-08 | 7.200E-08 |
| 14136 | 707 | T707S | B | 4.3e-08 | 2.5e-08 | 1.9e-08 | 2.900E-08 | 1.249E-08 | 1.900E-08 | 4.300E-08 |
| 14137 | 707 | T707T | B | 1.7e-08 | 3.9e-08 | 2.5e-08 | 2.700E-08 | 1.114E-08 | 1.700E-08 | 3.900E-08 |
| 14138 | 707 | T707W | B | 1.8e-08 | 3.5e-08 | 3.7e-08 | 3.000E-08 | 1.044E-08 | 1.800E-08 | 3.700E-08 |
| 14139 | 707 | T707Y | B | 2.8e-08 | 4.4e-08 | 3.7e-08 | 3.633E-08 | 8.021E-09 | 2.800E-08 | 4.400E-08 |
| 14140 | 707 | T707V | B | 7.6e-08 | 2.8e-08 | 3.7e-08 | 4.700E-08 | 2.551E-08 | 2.800E-08 | 7.600E-08 |
| 14141 | 708 | Y708A | B | 3.3e-08 | 6.9e-08 | 2.3e-08 | 4.167E-08 | 2.419E-08 | 2.300E-08 | 6.900E-08 |
| 14142 | 708 | Y708R | B | 4.9e-08 | 5.9e-08 | 9.6e-08 | 6.800E-08 | 2.476E-08 | 4.900E-08 | 9.600E-08 |
| 14143 | 708 | Y708N | B | 1.1e-08 | 3.3e-08 | 1.7e-08 | 2.033E-08 | 1.137E-08 | 1.100E-08 | 3.300E-08 |
| 14144 | 708 | Y708D | B | 1.2e-08 | 4.4e-08 | 3e-08 | 2.867E-08 | 1.604E-08 | 1.200E-08 | 4.400E-08 |
| 14145 | 708 | Y708C | B | 6.2e-08 | 4.4e-08 | 3.6e-08 | 4.733E-08 | 1.332E-08 | 3.600E-08 | 6.200E-08 |
| 14146 | 708 | Y708E | B | 4.8e-08 | 4.7e-08 | 6.1e-08 | 5.200E-08 | 7.810E-09 | 4.700E-08 | 6.100E-08 |
| 14147 | 708 | Y708Q | B | 6.8e-08 | 1.6e-08 | 8.3e-08 | 5.567E-08 | 3.516E-08 | 1.600E-08 | 8.300E-08 |
| 14148 | 708 | Y708G | B | 5.2e-08 | 6.3e-08 | 9e-08 | 6.833E-08 | 1.955E-08 | 5.200E-08 | 9.000E-08 |
| 14149 | 708 | Y708H | B | 4.2e-08 | 5e-08 | 6.5e-08 | 5.233E-08 | 1.168E-08 | 4.200E-08 | 6.500E-08 |
| 14150 | 708 | Y708I | B | 4.2e-08 | 4.6e-08 | 2.9e-08 | 3.900E-08 | 8.888E-09 | 2.900E-08 | 4.600E-08 |
| 14151 | 708 | Y708L | B | 7e-08 | 1.6e-08 | 6.6e-08 | 5.067E-08 | 3.009E-08 | 1.600E-08 | 7.000E-08 |
| 14152 | 708 | Y708K | B | 7.1e-08 | 1.1e-07 | 1.1e-07 | 9.700E-08 | 2.252E-08 | 7.100E-08 | 1.100E-07 |
| 14153 | 708 | Y708M | B | 3e-08 | 5.1e-08 | 4.6e-08 | 4.233E-08 | 1.097E-08 | 3.000E-08 | 5.100E-08 |
| 14154 | 708 | Y708F | B | 2.7e-08 | 7.7e-08 | 4.9e-08 | 5.100E-08 | 2.506E-08 | 2.700E-08 | 7.700E-08 |
| 14155 | 708 | Y708P | B | 5.8e-08 | 1.6e-08 | 2.7e-08 | 3.367E-08 | 2.178E-08 | 1.600E-08 | 5.800E-08 |
| 14156 | 708 | Y708S | B | 5.3e-08 | 4.1e-08 | 3.2e-08 | 4.200E-08 | 1.054E-08 | 3.200E-08 | 5.300E-08 |
| 14157 | 708 | Y708T | B | 1.4e-07 | 2.2e-08 | 4.7e-08 | 6.967E-08 | 6.218E-08 | 2.200E-08 | 1.400E-07 |
| 14158 | 708 | Y708W | B | 2.8e-08 | 3.1e-08 | 2.7e-08 | 2.867E-08 | 2.082E-09 | 2.700E-08 | 3.100E-08 |
| 14159 | 708 | Y708Y | B | 3.3e-08 | 1.7e-08 | 1.9e-08 | 2.300E-08 | 8.718E-09 | 1.700E-08 | 3.300E-08 |
| 14160 | 708 | Y708V | B | 5.6e-08 | 1.8e-08 | 3.5e-08 | 3.633E-08 | 1.904E-08 | 1.800E-08 | 5.600E-08 |
| 14161 | 709 | Y709A | B | 4.8e-08 | 1.9e-08 | 3.4e-08 | 3.367E-08 | 1.450E-08 | 1.900E-08 | 4.800E-08 |
| 14162 | 709 | Y709R | B | 4.5e-08 | 2.7e-08 | 5.9e-08 | 4.367E-08 | 1.604E-08 | 2.700E-08 | 5.900E-08 |
| 14163 | 709 | Y709N | B | 4.1e-08 | 1.1e-07 | 3.9e-08 | 6.333E-08 | 4.043E-08 | 3.900E-08 | 1.100E-07 |
| 14164 | 709 | Y709D | B | 2e-08 | 6.8e-08 | 1.7e-08 | 3.500E-08 | 2.862E-08 | 1.700E-08 | 6.800E-08 |
| 14165 | 709 | Y709C | B | 3e-08 | 5.5e-08 | 7.5e-08 | 5.333E-08 | 2.255E-08 | 3.000E-08 | 7.500E-08 |
| 14166 | 709 | Y709E | B | 6.1e-08 | 3.8e-08 | 5.1e-08 | 5.000E-08 | 1.153E-08 | 3.800E-08 | 6.100E-08 |
| 14167 | 709 | Y709Q | B | 3e-08 | 5e-08 | 5.2e-08 | 4.400E-08 | 1.217E-08 | 3.000E-08 | 5.200E-08 |
| 14168 | 709 | Y709G | B | 7.9e-08 | 6.3e-08 | 2.5e-08 | 5.567E-08 | 2.774E-08 | 2.500E-08 | 7.900E-08 |
| 14169 | 709 | Y709H | B | 2.3e-08 | 1.1e-07 | 2.9e-08 | 5.400E-08 | 4.859E-08 | 2.300E-08 | 1.100E-07 |
| 14170 | 709 | Y709I | B | 6e-08 | 1.8e-07 | 2.8e-08 | 8.933E-08 | 8.013E-08 | 2.800E-08 | 1.800E-07 |
| 14171 | 709 | Y709L | B | 3.2e-08 | 6.5e-08 | 5.1e-08 | 4.933E-08 | 1.656E-08 | 3.200E-08 | 6.500E-08 |
| 14172 | 709 | Y709K | B | 6.9e-08 | 3.6e-08 | 3e-08 | 4.500E-08 | 2.100E-08 | 3.000E-08 | 6.900E-08 |
| 14173 | 709 | Y709M | B | 5.2e-08 | 9.2e-08 | 1.2e-07 | 8.800E-08 | 3.418E-08 | 5.200E-08 | 1.200E-07 |
| 14174 | 709 | Y709F | B | 1.8e-07 | 3.2e-08 | 1.4e-08 | 7.533E-08 | 9.109E-08 | 1.400E-08 | 1.800E-07 |
| 14175 | 709 | Y709P | B | 3.6e-08 | 2.8e-08 | 2.6e-08 | 3.000E-08 | 5.292E-09 | 2.600E-08 | 3.600E-08 |
| 14176 | 709 | Y709S | B | 3.3e-08 | 4.5e-08 | 9e-09 | 2.900E-08 | 1.833E-08 | 9.000E-09 | 4.500E-08 |
| 14177 | 709 | Y709T | B | 3.6e-08 | 4.4e-08 | 1.9e-08 | 3.300E-08 | 1.277E-08 | 1.900E-08 | 4.400E-08 |
| 14178 | 709 | Y709W | B | 8.6e-08 | 2.2e-08 | 3.4e-08 | 4.733E-08 | 3.402E-08 | 2.200E-08 | 8.600E-08 |
| 14179 | 709 | Y709Y | B | 5.8e-08 | 9.4e-08 | 1.9e-08 | 5.700E-08 | 3.751E-08 | 1.900E-08 | 9.400E-08 |
| 14180 | 709 | Y709V | B | 3.4e-08 | 3.1e-08 | 2.8e-08 | 3.100E-08 | 3.000E-09 | 2.800E-08 | 3.400E-08 |
| 14181 | 710 | C710A | B | 3e-08 | 4.2e-08 | 8.5e-08 | 5.233E-08 | 2.892E-08 | 3.000E-08 | 8.500E-08 |
| 14182 | 710 | C710R | B | 3.6e-08 | 1.8e-08 | 6.2e-08 | 3.867E-08 | 2.212E-08 | 1.800E-08 | 6.200E-08 |
| 14183 | 710 | C710N | B | 1.4e-07 | 2.5e-08 | 9.2e-08 | 8.567E-08 | 5.776E-08 | 2.500E-08 | 1.400E-07 |
| 14184 | 710 | C710D | B | 1.3e-08 | 1.9e-08 | 2.5e-08 | 1.900E-08 | 6.000E-09 | 1.300E-08 | 2.500E-08 |
| 14185 | 710 | C710C | B | 6.4e-08 | 1e-07 | 7.1e-08 | 7.833E-08 | 1.909E-08 | 6.400E-08 | 1.000E-07 |
| 14186 | 710 | C710E | B | 3.6e-08 | 6e-08 | 8.5e-08 | 6.033E-08 | 2.450E-08 | 3.600E-08 | 8.500E-08 |
| 14187 | 710 | C710Q | B | 3.5e-08 | 9.6e-09 | 8.6e-08 | 4.353E-08 | 3.891E-08 | 9.600E-09 | 8.600E-08 |
| 14188 | 710 | C710G | B | 5.2e-08 | 3.3e-08 | 1.1e-08 | 3.200E-08 | 2.052E-08 | 1.100E-08 | 5.200E-08 |
| 14189 | 710 | C710H | B | 3e-08 | 2.6e-08 | 6.4e-08 | 4.000E-08 | 2.088E-08 | 2.600E-08 | 6.400E-08 |
| 14190 | 710 | C710I | B | 1.7e-08 | 1.3e-08 | 3.1e-08 | 2.033E-08 | 9.452E-09 | 1.300E-08 | 3.100E-08 |
| 14191 | 710 | C710L | B | 4.4e-08 | 1.5e-08 | 6.9e-08 | 4.267E-08 | 2.702E-08 | 1.500E-08 | 6.900E-08 |
| 14192 | 710 | C710K | B | 1.3e-07 | 1.1e-08 | 3.4e-08 | 5.833E-08 | 6.312E-08 | 1.100E-08 | 1.300E-07 |
| 14193 | 710 | C710M | B | 6.6e-08 | 3.1e-08 | 6.5e-09 | 3.450E-08 | 2.990E-08 | 6.500E-09 | 6.600E-08 |
| 14194 | 710 | C710F | B | 2.1e-08 | 4.8e-08 | 9.3e-08 | 5.400E-08 | 3.637E-08 | 2.100E-08 | 9.300E-08 |

| | | | | | | | | | | |
|---|---|---|---|---|---|---|---|---|---|---|
| 14195 | 710 | C710P | B | 1.1e-07 | 1e-07 | 4.3e-08 | 8.433E-08 | 3.614E-08 | 4.300E-08 | 1.100E-07 |
| 14196 | 710 | C710S | B | 5e-08 | 3e-08 | 3.4e-08 | 3.800E-08 | 1.058E-08 | 3.000E-08 | 5.000E-08 |
| 14197 | 710 | C710T | B | 3.2e-08 | 8.7e-08 | 2e-08 | 4.633E-08 | 3.573E-08 | 2.000E-08 | 8.700E-08 |
| 14198 | 710 | C710W | B | 2.5e-08 | 5e-08 | 8.9e-08 | 5.467E-08 | 3.225E-08 | 2.500E-08 | 8.900E-08 |
| 14199 | 710 | C710Y | B | 8.9e-08 | 5e-08 | 2e-08 | 5.300E-08 | 3.460E-08 | 2.000E-08 | 8.900E-08 |
| 14200 | 710 | C710V | B | 3e-08 | 5.7e-08 | 3.4e-08 | 4.033E-08 | 1.457E-08 | 3.000E-08 | 5.700E-08 |
| 14201 | 711 | Q711A | B | 2.9e-08 | 2.3e-08 | 1.9e-08 | 2.367E-08 | 5.033E-09 | 1.900E-08 | 2.900E-08 |
| 14202 | 711 | Q711R | B | 1e-07 | 4.2e-08 | 5.5e-08 | 6.567E-08 | 3.044E-08 | 4.200E-08 | 1.000E-07 |
| 14203 | 711 | Q711N | B | 4.7e-08 | 4e-08 | 6.3e-08 | 5.000E-08 | 1.179E-08 | 4.000E-08 | 6.300E-08 |
| 14204 | 711 | Q711D | B | 3.7e-08 | 2.7e-08 | 3.3e-08 | 3.233E-08 | 5.033E-09 | 2.700E-08 | 3.700E-08 |
| 14205 | 711 | Q711C | B | 8.1e-08 | 1.1e-07 | 3.3e-08 | 7.467E-08 | 3.889E-08 | 3.300E-08 | 1.100E-07 |
| 14206 | 711 | Q711E | B | 2.7e-08 | 2.8e-08 | 1.1e-07 | 5.500E-08 | 4.763E-08 | 2.700E-08 | 1.100E-07 |
| 14207 | 711 | Q711Q | B | 4.1e-08 | 1.3e-07 | 5.5e-08 | 7.533E-08 | 4.786E-08 | 4.100E-08 | 1.300E-07 |
| 14208 | 711 | Q711G | B | 6.6e-08 | 1.7e-08 | 6.2e-08 | 4.833E-08 | 2.721E-08 | 1.700E-08 | 6.600E-08 |
| 14209 | 711 | Q711H | B | 4.2e-08 | 3.4e-08 | 4.1e-08 | 3.900E-08 | 4.359E-09 | 3.400E-08 | 4.200E-08 |
| 14210 | 711 | Q711I | B | 3.4e-08 | 9.3e-09 | 2.2e-08 | 2.177E-08 | 1.235E-08 | 9.300E-09 | 3.400E-08 |
| 14211 | 711 | Q711L | B | 2.4e-08 | 8.1e-08 | 4.4e-08 | 4.967E-08 | 2.892E-08 | 2.400E-08 | 8.100E-08 |
| 14212 | 711 | Q711K | B | 2.8e-08 | 1.9e-08 | 1.7e-08 | 2.133E-08 | 5.859E-09 | 1.700E-08 | 2.800E-08 |
| 14213 | 711 | Q711M | B | 1.7e-08 | 1.3e-08 | 8.2e-08 | 3.733E-08 | 3.873E-08 | 1.300E-08 | 8.200E-08 |
| 14214 | 711 | Q711F | B | 3.4e-08 | 1e-07 | 6.1e-08 | 6.500E-08 | 3.318E-08 | 3.400E-08 | 1.000E-07 |
| 14215 | 711 | Q711P | B | 8.9e-08 | 3e-08 | 5.9e-08 | 5.933E-08 | 2.950E-08 | 3.000E-08 | 8.900E-08 |
| 14216 | 711 | Q711S | B | 4.6e-08 | 1.3e-08 | 3.1e-08 | 3.000E-08 | 1.652E-08 | 1.300E-08 | 4.600E-08 |
| 14217 | 711 | Q711T | B | 8.9e-08 | 5.7e-08 | 2.9e-08 | 5.833E-08 | 3.002E-08 | 2.900E-08 | 8.900E-08 |
| 14218 | 711 | Q711W | B | 4.3e-08 | 3.5e-08 | 4e-08 | 3.933E-08 | 4.041E-09 | 3.500E-08 | 4.300E-08 |
| 14219 | 711 | Q711Y | B | 3e-08 | 4.9e-08 | 2.5e-08 | 3.467E-08 | 1.266E-08 | 2.500E-08 | 4.900E-08 |
| 14220 | 711 | Q711V | B | 6.6e-08 | 4.5e-08 | 2.3e-08 | 4.467E-08 | 2.150E-08 | 2.300E-08 | 6.600E-08 |
| 14221 | 712 | Q712A | B | 1.7e-08 | 2.5e-08 | 2.6e-08 | 2.267E-08 | 4.933E-09 | 1.700E-08 | 2.600E-08 |
| 14222 | 712 | Q712R | B | 5e-08 | 2.2e-08 | 2.9e-08 | 3.367E-08 | 1.457E-08 | 2.200E-08 | 5.000E-08 |
| 14223 | 712 | Q712N | B | 3.6e-08 | 4.8e-08 | 5.3e-08 | 4.567E-08 | 8.737E-09 | 3.600E-08 | 5.300E-08 |
| 14224 | 712 | Q712D | B | 4.4e-08 | 3.8e-08 | 1.7e-07 | 8.400E-08 | 7.454E-08 | 3.800E-08 | 1.700E-07 |
| 14225 | 712 | Q712C | B | 1.6e-07 | 1.6e-08 | 4.3e-08 | 7.300E-08 | 7.654E-08 | 1.600E-08 | 1.600E-07 |
| 14226 | 712 | Q712E | B | 3.2e-08 | 2.8e-08 | 5.3e-08 | 3.767E-08 | 1.343E-08 | 2.800E-08 | 5.300E-08 |
| 14227 | 712 | Q712Q | B | 8.5e-09 | 3e-08 | 7.3e-08 | 3.717E-08 | 3.284E-08 | 8.500E-09 | 7.300E-08 |
| 14228 | 712 | Q712G | B | 1e-07 | 2.4e-08 | 2.7e-08 | 5.033E-08 | 4.304E-08 | 2.400E-08 | 1.000E-07 |
| 14229 | 712 | Q712H | B | 1.9e-08 | 3.7e-08 | 1.5e-07 | 6.867E-08 | 7.101E-08 | 1.900E-08 | 1.500E-07 |
| 14230 | 712 | Q712I | B | 7.7e-08 | 9e-08 | 2.7e-08 | 6.467E-08 | 3.326E-08 | 2.700E-08 | 9.000E-08 |
| 14231 | 712 | Q712L | B | 5e-08 | 3.1e-08 | 9.9e-08 | 6.000E-08 | 3.509E-08 | 3.100E-08 | 9.900E-08 |
| 14232 | 712 | Q712K | B | 3.5e-08 | 4.6e-08 | 1e-07 | 6.033E-08 | 3.479E-08 | 3.500E-08 | 1.000E-07 |
| 14233 | 712 | Q712M | B | 7.7e-08 | 3.9e-08 | 3.6e-08 | 5.067E-08 | 2.285E-08 | 3.600E-08 | 7.700E-08 |
| 14234 | 712 | Q712F | B | 3.2e-08 | 7.2e-08 | 3.9e-08 | 4.767E-08 | 2.136E-08 | 3.200E-08 | 7.200E-08 |
| 14235 | 712 | Q712P | B | 2.8e-08 | 4e-08 | 2.1e-08 | 2.967E-08 | 9.609E-09 | 2.100E-08 | 4.000E-08 |
| 14236 | 712 | Q712S | B | 9.4e-08 | 5.4e-08 | 3.2e-08 | 6.000E-08 | 3.143E-08 | 3.200E-08 | 9.400E-08 |
| 14237 | 712 | Q712T | B | 6e-08 | 3.9e-08 | 8.4e-08 | 6.100E-08 | 2.252E-08 | 3.900E-08 | 8.400E-08 |
| 14238 | 712 | Q712W | B | 8.6e-08 | 2.8e-08 | 6.8e-08 | 6.067E-08 | 2.969E-08 | 2.800E-08 | 8.600E-08 |
| 14239 | 712 | Q712Y | B | 1e-07 | 2.1e-08 | 1.1e-07 | 7.700E-08 | 4.875E-08 | 2.100E-08 | 1.100E-07 |
| 14240 | 712 | Q712V | B | 1.2e-07 | 6.6e-08 | 8.4e-08 | 9.000E-08 | 2.750E-08 | 6.600E-08 | 1.200E-07 |
| 14241 | 713 | Y713A | B | 2.5e-07 | 3.1e-08 | 2.5e-08 | 1.020E-07 | 1.282E-07 | 2.500E-08 | 2.500E-07 |
| 14242 | 713 | Y713R | B | 3.4e-08 | 2.5e-08 | 1.4e-08 | 2.433E-08 | 1.002E-08 | 1.400E-08 | 3.400E-08 |
| 14243 | 713 | Y713N | B | 6.5e-08 | 5.1e-08 | 6e-08 | 5.867E-08 | 7.095E-09 | 5.100E-08 | 6.500E-08 |
| 14244 | 713 | Y713D | B | 7.2e-08 | 4.2e-08 | 4.8e-08 | 5.400E-08 | 1.587E-08 | 4.200E-08 | 7.200E-08 |
| 14245 | 713 | Y713C | B | 1.3e-07 | 4.3e-08 | 2.5e-08 | 6.600E-08 | 5.615E-08 | 2.500E-08 | 1.300E-07 |
| 14246 | 713 | Y713E | B | 4.9e-08 | 4.7e-08 | 3.4e-08 | 4.333E-08 | 8.145E-09 | 3.400E-08 | 4.900E-08 |
| 14247 | 713 | Y713Q | B | 2e-08 | 2.6e-08 | 3.3e-08 | 2.633E-08 | 6.506E-09 | 2.000E-08 | 3.300E-08 |
| 14248 | 713 | Y713G | B | 3.8e-08 | 3.7e-08 | 3.5e-08 | 3.667E-08 | 1.528E-09 | 3.500E-08 | 3.800E-08 |
| 14249 | 713 | Y713H | B | 5.4e-08 | 3.1e-08 | 6.1e-08 | 4.867E-08 | 1.570E-08 | 3.100E-08 | 6.100E-08 |
| 14250 | 713 | Y713I | B | 6.5e-08 | 5.6e-08 | 4.1e-08 | 5.400E-08 | 1.212E-08 | 4.100E-08 | 6.500E-08 |
| 14251 | 713 | Y713L | B | 8.9e-08 | 2.2e-08 | 1.3e-08 | 4.133E-08 | 4.153E-08 | 1.300E-08 | 8.900E-08 |
| 14252 | 713 | Y713K | B | 5.5e-08 | 1.2e-07 | 9.8e-08 | 9.100E-08 | 3.306E-08 | 5.500E-08 | 1.200E-07 |
| 14253 | 713 | Y713M | B | 7.8e-08 | 5.2e-08 | 3.7e-08 | 5.567E-08 | 2.074E-08 | 3.700E-08 | 7.800E-08 |
| 14254 | 713 | Y713F | B | 4.1e-08 | 4.4e-08 | 1.1e-07 | 6.500E-08 | 3.900E-08 | 4.100E-08 | 1.100E-07 |
| 14255 | 713 | Y713P | B | 4.9e-08 | 6.2e-08 | 5.5e-08 | 5.533E-08 | 6.506E-09 | 4.900E-08 | 6.200E-08 |
| 14256 | 713 | Y713S | B | 3.7e-08 | 1.4e-08 | 2.1e-08 | 2.400E-08 | 1.179E-08 | 1.400E-08 | 3.700E-08 |
| 14257 | 713 | Y713T | B | 5e-08 | 5.6e-08 | 1e-07 | 6.867E-08 | 2.730E-08 | 5.000E-08 | 1.000E-07 |
| 14258 | 713 | Y713W | B | 4.5e-08 | 6.1e-08 | 3.8e-08 | 4.800E-08 | 1.179E-08 | 3.800E-08 | 6.100E-08 |
| 14259 | 713 | Y713Y | B | 1.9e-08 | 5e-08 | 4.8e-08 | 3.900E-08 | 1.735E-08 | 1.900E-08 | 5.000E-08 |
| 14260 | 713 | Y713V | B | 3.7e-08 | 3.9e-08 | 3.1e-08 | 3.567E-08 | 4.163E-09 | 3.100E-08 | 3.900E-08 |
| 14261 | 714 | Y714A | B | 8.7e-08 | 2.8e-08 | 3.2e-08 | 4.900E-08 | 3.297E-08 | 2.800E-08 | 8.700E-08 |
| 14262 | 714 | Y714R | B | 6e-08 | 1.6e-08 | 6.9e-08 | 4.833E-08 | 2.836E-08 | 1.600E-08 | 6.900E-08 |
| 14263 | 714 | Y714N | B | 8.5e-08 | 1e-07 | 1.4e-08 | 6.633E-08 | 4.594E-08 | 1.400E-08 | 1.000E-07 |
| 14264 | 714 | Y714D | B | 3.3e-08 | 2.4e-08 | 5.8e-08 | 3.833E-08 | 1.762E-08 | 2.400E-08 | 5.800E-08 |
| 14265 | 714 | Y714C | B | 6.7e-08 | 7.7e-08 | 2.2e-08 | 5.533E-08 | 2.930E-08 | 2.200E-08 | 7.700E-08 |
| 14266 | 714 | Y714E | B | 8.4e-09 | 1.1e-08 | 6.4e-08 | 2.780E-08 | 3.138E-08 | 8.400E-09 | 6.400E-08 |
| 14267 | 714 | Y714Q | B | 5e-08 | 6.9e-08 | 4.1e-08 | 5.333E-08 | 1.429E-08 | 4.100E-08 | 6.900E-08 |
| 14268 | 714 | Y714G | B | 5.4e-08 | 5.4e-08 | 1.9e-08 | 4.233E-08 | 2.021E-08 | 1.900E-08 | 5.400E-08 |
| 14269 | 714 | Y714H | B | 5.7e-08 | 4e-08 | 5e-08 | 4.900E-08 | 8.544E-09 | 4.000E-08 | 5.700E-08 |
| 14270 | 714 | Y714I | B | 1e-07 | 4.9e-08 | 7.6e-08 | 7.500E-08 | 2.551E-08 | 4.900E-08 | 1.000E-07 |
| 14271 | 714 | Y714L | B | 3e-08 | 3.1e-08 | 3.7e-08 | 3.267E-08 | 3.786E-09 | 3.000E-08 | 3.700E-08 |
| 14272 | 714 | Y714K | B | 5e-08 | 1.3e-07 | 5e-08 | 7.667E-08 | 4.619E-08 | 5.000E-08 | 1.300E-07 |
| 14273 | 714 | Y714M | B | 6.3e-08 | 8.7e-08 | 4.1e-08 | 6.367E-08 | 2.301E-08 | 4.100E-08 | 8.700E-08 |
| 14274 | 714 | Y714F | B | 7.8e-08 | 6.9e-08 | 1.7e-08 | 5.467E-08 | 3.293E-08 | 1.700E-08 | 7.800E-08 |
| 14275 | 714 | Y714P | B | 5.4e-08 | 3.1e-08 | 1e-07 | 6.167E-08 | 3.513E-08 | 3.100E-08 | 1.000E-07 |
| 14276 | 714 | Y714S | B | 2.1e-08 | 3.2e-08 | 3.5e-08 | 2.933E-08 | 7.371E-09 | 2.100E-08 | 3.500E-08 |
| 14277 | 714 | Y714T | B | 8.2e-08 | 2.2e-08 | 7.2e-08 | 5.867E-08 | 3.215E-08 | 2.200E-08 | 8.200E-08 |
| 14278 | 714 | Y714W | B | 6e-08 | 6.4e-08 | 3.4e-08 | 5.267E-08 | 1.629E-08 | 3.400E-08 | 6.400E-08 |
| 14279 | 714 | Y714Y | B | 7.3e-08 | 2.8e-08 | 8.6e-08 | 6.233E-08 | 3.044E-08 | 2.800E-08 | 8.600E-08 |

| | | | | | | | | | | |
|---|---|---|---|---|---|---|---|---|---|---|
| 14280 | 714 | Y714V | B | 4.1e-08 | 2e-08 | 3.4e-08 | 3.167E-08 | 1.069E-08 | 2.000E-08 | 4.100E-08 |
| 14281 | 715 | I715A | B | 7.9e-08 | 4.3e-08 | 2.2e-08 | 4.800E-08 | 2.883E-08 | 2.200E-08 | 7.900E-08 |
| 14282 | 715 | I715R | B | 4.9e-08 | 7.5e-08 | 2.6e-08 | 5.000E-08 | 2.452E-08 | 2.600E-08 | 7.500E-08 |
| 14283 | 715 | I715N | B | 4.5e-08 | 1.8e-08 | 4.4e-08 | 3.567E-08 | 1.531E-08 | 1.800E-08 | 4.500E-08 |
| 14284 | 715 | I715D | B | 1.1e-07 | 3.1e-08 | 1.3e-08 | 5.133E-08 | 5.160E-08 | 1.300E-08 | 1.100E-07 |
| 14285 | 715 | I715C | B | 2.7e-08 | 8.6e-08 | 5e-08 | 5.433E-08 | 2.974E-08 | 2.700E-08 | 8.600E-08 |
| 14286 | 715 | I715E | B | 1.6e-08 | 2.9e-08 | 6.9e-08 | 3.800E-08 | 2.762E-08 | 1.600E-08 | 6.900E-08 |
| 14287 | 715 | I715Q | B | 4.8e-08 | 6e-08 | 7.5e-08 | 6.100E-08 | 1.353E-08 | 4.800E-08 | 7.500E-08 |
| 14288 | 715 | I715G | B | 9.5e-08 | 2.8e-08 | 6.3e-08 | 6.200E-08 | 3.351E-08 | 2.800E-08 | 9.500E-08 |
| 14289 | 715 | I715H | B | 1.5e-08 | 5.5e-08 | 5.6e-08 | 4.200E-08 | 2.339E-08 | 1.500E-08 | 5.600E-08 |
| 14290 | 715 | I715I | B | 1.7e-07 | 1.2e-08 | 1.4e-08 | 6.533E-08 | 9.065E-08 | 1.200E-08 | 1.700E-07 |
| 14291 | 715 | I715L | B | 4.1e-08 | 1.1e-07 | 1.9e-08 | 5.667E-08 | 4.748E-08 | 1.900E-08 | 1.100E-07 |
| 14292 | 715 | I715K | B | 1.9e-08 | 2.7e-08 | 2.7e-08 | 2.433E-08 | 4.619E-09 | 1.900E-08 | 2.700E-08 |
| 14293 | 715 | I715M | B | 6.3e-08 | 7.2e-08 | 6.1e-08 | 6.533E-08 | 5.859E-09 | 6.100E-08 | 7.200E-08 |
| 14294 | 715 | I715F | B | 2.4e-08 | 4.9e-08 | 6.4e-08 | 4.567E-08 | 2.021E-08 | 2.400E-08 | 6.400E-08 |
| 14295 | 715 | I715P | B | 7e-08 | 2.3e-08 | 5.3e-08 | 4.867E-08 | 2.380E-08 | 2.300E-08 | 7.000E-08 |
| 14296 | 715 | I715S | B | 7e-08 | 3.9e-08 | 6.2e-08 | 5.700E-08 | 1.609E-08 | 3.900E-08 | 7.000E-08 |
| 14297 | 715 | I715T | B | 4.3e-08 | 2.8e-08 | 1.7e-08 | 2.933E-08 | 1.305E-08 | 1.700E-08 | 4.300E-08 |
| 14298 | 715 | I715W | B | 1.4e-08 | 2.9e-08 | 1.2e-07 | 5.433E-08 | 5.736E-08 | 1.400E-08 | 1.200E-07 |
| 14299 | 715 | I715Y | B | 4.4e-08 | 1.5e-08 | 5.7e-08 | 3.867E-08 | 2.150E-08 | 1.500E-08 | 5.700E-08 |
| 14300 | 715 | I715V | B | 5.1e-08 | 5.6e-08 | 8.9e-08 | 6.533E-08 | 2.065E-08 | 5.100E-08 | 8.900E-08 |
| 14301 | 716 | Y716A | B | 3.3e-08 | 2.8e-08 | 2.8e-08 | 2.967E-08 | 2.887E-09 | 2.800E-08 | 3.300E-08 |
| 14302 | 716 | Y716R | B | 3e-08 | 2.8e-08 | 1.8e-08 | 2.533E-08 | 6.429E-09 | 1.800E-08 | 3.000E-08 |
| 14303 | 716 | Y716N | B | 4.2e-08 | 5.4e-08 | 1.2e-07 | 7.200E-08 | 4.200E-08 | 4.200E-08 | 1.200E-07 |
| 14304 | 716 | Y716D | B | 3.6e-08 | 4.4e-08 | 3.3e-08 | 3.767E-08 | 5.686E-09 | 3.300E-08 | 4.400E-08 |
| 14305 | 716 | Y716C | B | 3.9e-08 | 7.2e-08 | 4.6e-08 | 5.233E-08 | 1.739E-08 | 3.900E-08 | 7.200E-08 |
| 14306 | 716 | Y716E | B | 4.2e-08 | 9e-09 | 5.1e-08 | 3.400E-08 | 2.211E-08 | 9.000E-09 | 5.100E-08 |
| 14307 | 716 | Y716Q | B | 4.9e-08 | 3e-08 | 3.1e-08 | 3.667E-08 | 1.069E-08 | 3.000E-08 | 4.900E-08 |
| 14308 | 716 | Y716G | B | 6.2e-08 | 1.4e-08 | 1.1e-07 | 6.200E-08 | 4.800E-08 | 1.400E-08 | 1.100E-07 |
| 14309 | 716 | Y716H | B | 3.9e-08 | 4.1e-08 | 4.8e-08 | 4.267E-08 | 4.726E-09 | 3.900E-08 | 4.800E-08 |
| 14310 | 716 | Y716I | B | 7.1e-08 | 1.8e-08 | 4.3e-08 | 4.400E-08 | 2.651E-08 | 1.800E-08 | 7.100E-08 |
| 14311 | 716 | Y716L | B | 3.5e-08 | 8.3e-08 | 1.8e-07 | 9.933E-08 | 7.387E-08 | 3.500E-08 | 1.800E-07 |
| 14312 | 716 | Y716K | B | 2.8e-08 | 2.5e-08 | 6.5e-08 | 3.933E-08 | 2.228E-08 | 2.500E-08 | 6.500E-08 |
| 14313 | 716 | Y716M | B | 2.1e-08 | 8e-08 | 2e-08 | 4.033E-08 | 3.436E-08 | 2.000E-08 | 8.000E-08 |
| 14314 | 716 | Y716F | B | 2.2e-08 | 5e-08 | 7.5e-08 | 4.900E-08 | 2.651E-08 | 2.200E-08 | 7.500E-08 |
| 14315 | 716 | Y716P | B | 1.6e-08 | 3.8e-08 | 4.4e-08 | 3.267E-08 | 1.474E-08 | 1.600E-08 | 4.400E-08 |
| 14316 | 716 | Y716S | B | 5.6e-08 | 1.5e-08 | 1.9e-08 | 3.000E-08 | 2.261E-08 | 1.500E-08 | 5.600E-08 |
| 14317 | 716 | Y716T | B | 5.4e-08 | 1.1e-07 | 2.9e-08 | 6.433E-08 | 4.148E-08 | 2.900E-08 | 1.100E-07 |
| 14318 | 716 | Y716W | B | 5.3e-08 | 1.3e-08 | 6.4e-08 | 4.333E-08 | 2.684E-08 | 1.300E-08 | 6.400E-08 |
| 14319 | 716 | Y716Y | B | 2.9e-08 | 8.1e-08 | 5.4e-08 | 5.467E-08 | 2.601E-08 | 2.900E-08 | 8.100E-08 |
| 14320 | 716 | Y716V | B | 5.2e-08 | 6e-08 | 2.1e-08 | 4.433E-08 | 2.060E-08 | 2.100E-08 | 6.000E-08 |
| 14321 | 717 | P717A | B | 8.4e-09 | 5.7e-08 | 4.5e-08 | 3.680E-08 | 2.532E-08 | 8.400E-09 | 5.700E-08 |
| 14322 | 717 | P717R | B | 3.4e-08 | 4.8e-08 | 4.8e-08 | 4.333E-08 | 8.083E-09 | 3.400E-08 | 4.800E-08 |
| 14323 | 717 | P717N | B | 3.7e-08 | 3.1e-08 | 4.4e-08 | 3.733E-08 | 6.506E-09 | 3.100E-08 | 4.400E-08 |
| 14324 | 717 | P717D | B | 2.4e-08 | 5.4e-08 | 4.5e-08 | 4.100E-08 | 1.539E-08 | 2.400E-08 | 5.400E-08 |
| 14325 | 717 | P717C | B | 3.3e-08 | 6.3e-08 | 2.1e-08 | 3.900E-08 | 2.163E-08 | 2.100E-08 | 6.300E-08 |
| 14326 | 717 | P717E | B | 7.1e-08 | 9e-08 | 1.7e-07 | 1.103E-07 | 5.254E-08 | 7.100E-08 | 1.700E-07 |
| 14327 | 717 | P717Q | B | 4.1e-08 | 1.6e-08 | 1.6e-07 | 7.233E-08 | 7.694E-08 | 1.600E-08 | 1.600E-07 |
| 14328 | 717 | P717G | B | 5.1e-08 | 4.8e-08 | 4.1e-08 | 4.667E-08 | 5.132E-09 | 4.100E-08 | 5.100E-08 |
| 14329 | 717 | P717H | B | 2.7e-08 | 2.2e-08 | 5.1e-08 | 3.333E-08 | 1.550E-08 | 2.200E-08 | 5.100E-08 |
| 14330 | 717 | P717I | B | 5.2e-08 | 4.2e-08 | 2.5e-08 | 3.967E-08 | 1.365E-08 | 2.500E-08 | 5.200E-08 |
| 14331 | 717 | P717L | B | 3.4e-08 | 6.7e-08 | 4.9e-08 | 5.000E-08 | 1.652E-08 | 3.400E-08 | 6.700E-08 |
| 14332 | 717 | P717K | B | 1.9e-08 | 3e-08 | 1.2e-07 | 5.633E-08 | 5.541E-08 | 1.900E-08 | 1.200E-07 |
| 14333 | 717 | P717M | B | 7.8e-09 | 4.1e-08 | 2.2e-08 | 2.360E-08 | 1.666E-08 | 7.800E-09 | 4.100E-08 |
| 14334 | 717 | P717F | B | 3e-08 | 8.5e-08 | 1.4e-08 | 4.300E-08 | 3.724E-08 | 1.400E-08 | 8.500E-08 |
| 14335 | 717 | P717P | B | 3.2e-08 | 3.1e-08 | 2.4e-08 | 2.900E-08 | 4.359E-09 | 2.400E-08 | 3.200E-08 |
| 14336 | 717 | P717S | B | 3.6e-08 | 9.1e-08 | 2.5e-08 | 5.067E-08 | 3.536E-08 | 2.500E-08 | 9.100E-08 |
| 14337 | 717 | P717T | B | 4.4e-08 | 1.6e-07 | 9.9e-08 | 1.010E-07 | 5.803E-08 | 4.400E-08 | 1.600E-07 |
| 14338 | 717 | P717W | B | 2.1e-08 | 4.2e-08 | 1.2e-07 | 6.100E-08 | 5.216E-08 | 2.100E-08 | 1.200E-07 |
| 14339 | 717 | P717Y | B | 4.5e-08 | 3.7e-08 | 2.4e-08 | 3.533E-08 | 1.060E-08 | 2.400E-08 | 4.500E-08 |
| 14340 | 717 | P717V | B | 2.2e-08 | 5.3e-08 | 3.6e-08 | 3.700E-08 | 1.552E-08 | 2.200E-08 | 5.300E-08 |
| 14341 | 718 | Y718A | B | 3.2e-08 | 3.2e-08 | 3.2e-08 | 3.200E-08 | 0.000E+00 | 3.200E-08 | 3.200E-08 |
| 14342 | 718 | Y718R | B | 1.5e-08 | 2.3e-08 | 4e-08 | 2.600E-08 | 1.277E-08 | 1.500E-08 | 4.000E-08 |
| 14343 | 718 | Y718N | B | 4.2e-08 | 5e-08 | 5.6e-08 | 4.933E-08 | 7.024E-09 | 4.200E-08 | 5.600E-08 |
| 14344 | 718 | Y718D | B | 2e-08 | 3.1e-08 | 3.9e-08 | 3.000E-08 | 9.539E-09 | 2.000E-08 | 3.900E-08 |
| 14345 | 718 | Y718C | B | 4.3e-08 | 2.4e-08 | 6.1e-08 | 4.267E-08 | 1.850E-08 | 2.400E-08 | 6.100E-08 |
| 14346 | 718 | Y718E | B | 3.9e-08 | 4.8e-08 | 3e-08 | 3.900E-08 | 9.000E-09 | 3.000E-08 | 4.800E-08 |
| 14347 | 718 | Y718Q | B | 3.5e-08 | 4.1e-08 | 3e-08 | 3.533E-08 | 5.508E-09 | 3.000E-08 | 4.100E-08 |
| 14348 | 718 | Y718G | B | 1.7e-08 | 5.1e-08 | 4.8e-08 | 3.867E-08 | 1.882E-08 | 1.700E-08 | 5.100E-08 |
| 14349 | 718 | Y718H | B | 2.1e-08 | 5.6e-08 | 8.7e-09 | 2.857E-08 | 2.454E-08 | 8.700E-09 | 5.600E-08 |
| 14350 | 718 | Y718I | B | 3.2e-08 | 3.5e-08 | 2.6e-08 | 3.100E-08 | 4.583E-09 | 2.600E-08 | 3.500E-08 |
| 14351 | 718 | Y718L | B | 5e-08 | 5.1e-08 | 3.3e-08 | 4.467E-08 | 1.012E-08 | 3.300E-08 | 5.100E-08 |
| 14352 | 718 | Y718K | B | 2.6e-08 | 5e-08 | 1.6e-08 | 3.067E-08 | 1.747E-08 | 1.600E-08 | 5.000E-08 |
| 14353 | 718 | Y718M | B | 1.7e-08 | 3.2e-08 | 2.3e-08 | 2.400E-08 | 7.550E-09 | 1.700E-08 | 3.200E-08 |
| 14354 | 718 | Y718F | B | 5.4e-08 | 2.4e-08 | 4.1e-08 | 3.967E-08 | 1.504E-08 | 2.400E-08 | 5.400E-08 |
| 14355 | 718 | Y718P | B | 4.5e-08 | 2.4e-08 | 5.6e-08 | 4.167E-08 | 1.626E-08 | 2.400E-08 | 5.600E-08 |
| 14356 | 718 | Y718S | B | 3.2e-08 | 1.8e-08 | 7.3e-08 | 4.100E-08 | 2.858E-08 | 1.800E-08 | 7.300E-08 |
| 14357 | 718 | Y718T | B | 3.2e-08 | 2.4e-08 | 6.5e-08 | 4.033E-08 | 2.173E-08 | 2.400E-08 | 6.500E-08 |
| 14358 | 718 | Y718W | B | 3.6e-08 | 1.6e-07 | 2e-08 | 7.200E-08 | 7.663E-08 | 2.000E-08 | 1.600E-07 |
| 14359 | 718 | Y718Y | B | 4.2e-08 | 2.4e-08 | 9.9e-08 | 5.500E-08 | 3.915E-08 | 2.400E-08 | 9.900E-08 |
| 14360 | 718 | Y718V | B | 6e-08 | 1.8e-08 | 3.4e-08 | 3.733E-08 | 2.120E-08 | 1.800E-08 | 6.000E-08 |
| 14361 | 719 | T719A | B | 3.2e-08 | 1.8e-08 | 5e-08 | 3.333E-08 | 1.604E-08 | 1.800E-08 | 5.000E-08 |
| 14362 | 719 | T719R | B | 5.6e-08 | 7e-08 | 2.4e-08 | 5.000E-08 | 2.358E-08 | 2.400E-08 | 7.000E-08 |
| 14363 | 719 | T719N | B | 2.3e-08 | 2.2e-08 | 4e-08 | 2.833E-08 | 1.012E-08 | 2.200E-08 | 4.000E-08 |
| 14364 | 719 | T719D | B | 4.1e-08 | 2.9e-08 | 6.6e-08 | 4.533E-08 | 1.888E-08 | 2.900E-08 | 6.600E-08 |

| | | | | | | | | | | |
|---|---|---|---|---|---|---|---|---|---|---|
| 14365 | 719 | T719C | B | 3.1e-08 | 1.5e-08 | 4.1e-08 | 2.900E-08 | 1.311E-08 | 1.500E-08 | 4.100E-08 |
| 14366 | 719 | T719E | B | 2.5e-08 | 2.5e-08 | 3.2e-08 | 2.733E-08 | 4.041E-09 | 2.500E-08 | 3.200E-08 |
| 14367 | 719 | T719Q | B | 2.6e-08 | 3.8e-08 | 5.1e-08 | 3.833E-08 | 1.250E-08 | 2.600E-08 | 5.100E-08 |
| 14368 | 719 | T719G | B | 3.3e-08 | 3.1e-08 | 1.1e-07 | 5.800E-08 | 4.504E-08 | 3.100E-08 | 1.100E-07 |
| 14369 | 719 | T719H | B | 9.8e-08 | 3.3e-08 | 9e-08 | 7.367E-08 | 3.544E-08 | 3.300E-08 | 9.800E-08 |
| 14370 | 719 | T719I | B | 1.5e-08 | 1.7e-08 | 1.2e-07 | 5.067E-08 | 6.005E-08 | 1.500E-08 | 1.200E-07 |
| 14371 | 719 | T719L | B | 1.7e-08 | 1.3e-08 | 5.4e-08 | 2.800E-08 | 2.261E-08 | 1.300E-08 | 5.400E-08 |
| 14372 | 719 | T719K | B | 9.7e-08 | 8.1e-08 | 3.8e-08 | 7.200E-08 | 3.051E-08 | 3.800E-08 | 9.700E-08 |
| 14373 | 719 | T719M | B | 3.6e-08 | 4.5e-08 | 3.9e-08 | 4.000E-08 | 4.583E-09 | 3.600E-08 | 4.500E-08 |
| 14374 | 719 | T719F | B | 5.7e-08 | 4.6e-08 | 2.8e-08 | 4.367E-08 | 1.464E-08 | 2.800E-08 | 5.700E-08 |
| 14375 | 719 | T719P | B | 3.2e-08 | 3.3e-08 | 3.8e-08 | 3.433E-08 | 3.215E-09 | 3.200E-08 | 3.800E-08 |
| 14376 | 719 | T719S | B | 2.4e-08 | 6.8e-08 | 1.7e-08 | 3.633E-08 | 2.765E-08 | 1.700E-08 | 6.800E-08 |
| 14377 | 719 | T719T | B | 1.5e-08 | 1.1e-07 | 2.1e-08 | 4.867E-08 | 5.320E-08 | 1.500E-08 | 1.100E-07 |
| 14378 | 719 | T719W | B | 3.6e-08 | 1.2e-07 | 4.3e-08 | 6.633E-08 | 4.661E-08 | 3.600E-08 | 1.200E-07 |
| 14379 | 719 | T719Y | B | 6.4e-08 | 7.8e-08 | 2.2e-08 | 5.467E-08 | 2.914E-08 | 2.200E-08 | 7.800E-08 |
| 14380 | 719 | T719V | B | 6.4e-08 | 5.1e-08 | 9.9e-08 | 7.133E-08 | 2.483E-08 | 5.100E-08 | 9.900E-08 |
| 14381 | 720 | F720A | B | 9.7e-08 | 5.3e-08 | 5.2e-08 | 6.733E-08 | 2.570E-08 | 5.200E-08 | 9.700E-08 |
| 14382 | 720 | F720R | B | 4.3e-08 | 2.5e-08 | 6.4e-08 | 4.400E-08 | 1.952E-08 | 2.500E-08 | 6.400E-08 |
| 14383 | 720 | F720N | B | 3.9e-08 | 3.1e-08 | 7.8e-08 | 4.933E-08 | 2.515E-08 | 3.100E-08 | 7.800E-08 |
| 14384 | 720 | F720D | B | 5.1e-08 | 4.2e-08 | 6e-08 | 5.100E-08 | 9.000E-09 | 4.200E-08 | 6.000E-08 |
| 14385 | 720 | F720C | B | 5.5e-08 | 2.6e-08 | 4.8e-08 | 4.300E-08 | 1.513E-08 | 2.600E-08 | 5.500E-08 |
| 14386 | 720 | F720E | B | 3.8e-08 | 3e-08 | 5.5e-08 | 4.100E-08 | 1.277E-08 | 3.000E-08 | 5.500E-08 |
| 14387 | 720 | F720Q | B | 5.1e-08 | 1.1e-07 | 1.1e-07 | 9.033E-08 | 3.406E-08 | 5.100E-08 | 1.100E-07 |
| 14388 | 720 | F720G | B | 3.1e-08 | 9.6e-08 | 6.5e-08 | 6.400E-08 | 3.251E-08 | 3.100E-08 | 9.600E-08 |
| 14389 | 720 | F720H | B | 3.7e-08 | 7.4e-08 | 1.9e-08 | 4.333E-08 | 2.804E-08 | 1.900E-08 | 7.400E-08 |
| 14390 | 720 | F720I | B | 3.1e-08 | 5.9e-08 | 3.3e-08 | 4.100E-08 | 1.562E-08 | 3.100E-08 | 5.900E-08 |
| 14391 | 720 | F720L | B | 3.6e-08 | 1.1e-07 | 2.4e-08 | 5.667E-08 | 4.658E-08 | 2.400E-08 | 1.100E-07 |
| 14392 | 720 | F720K | B | 3e-08 | 3e-08 | 3.9e-08 | 3.300E-08 | 5.196E-09 | 3.000E-08 | 3.900E-08 |
| 14393 | 720 | F720M | B | 9.4e-08 | 6.1e-08 | 1e-07 | 8.500E-08 | 2.100E-08 | 6.100E-08 | 1.000E-07 |
| 14394 | 720 | F720F | B | 3.9e-08 | 5.2e-08 | 3.2e-08 | 4.100E-08 | 1.015E-08 | 3.200E-08 | 5.200E-08 |
| 14395 | 720 | F720P | B | 2e-08 | 4.5e-08 | 5.8e-08 | 4.100E-08 | 1.931E-08 | 2.000E-08 | 5.800E-08 |
| 14396 | 720 | F720S | B | 3.7e-08 | 5.8e-08 | 4.8e-08 | 4.767E-08 | 1.050E-08 | 3.700E-08 | 5.800E-08 |
| 14397 | 720 | F720T | B | 3.5e-08 | 7.1e-08 | 4e-08 | 4.867E-08 | 1.950E-08 | 3.500E-08 | 7.100E-08 |
| 14398 | 720 | F720W | B | 4.8e-08 | 4.1e-08 | 4.4e-08 | 4.433E-08 | 3.512E-09 | 4.100E-08 | 4.800E-08 |
| 14399 | 720 | F720Y | B | 3.1e-08 | 3e-08 | 4.8e-08 | 3.633E-08 | 1.012E-08 | 3.000E-08 | 4.800E-08 |
| 14400 | 720 | F720V | B | 7.7e-08 | 2.3e-08 | 2.7e-08 | 4.233E-08 | 3.009E-08 | 2.300E-08 | 7.700E-08 |
| 14401 | 721 | G721A | B | 3.3e-08 | 4.5e-08 | 2.6e-08 | 3.467E-08 | 9.609E-09 | 2.600E-08 | 4.500E-08 |
| 14402 | 721 | G721R | B | 5.3e-08 | 3.1e-08 | 1.8e-08 | 3.400E-08 | 1.769E-08 | 1.800E-08 | 5.300E-08 |
| 14403 | 721 | G721N | B | 2.1e-08 | 3.3e-08 | 1.9e-08 | 2.433E-08 | 7.572E-09 | 1.900E-08 | 3.300E-08 |
| 14404 | 721 | G721D | B | 1.2e-08 | 6.8e-08 | 4.4e-08 | 4.133E-08 | 2.810E-08 | 1.200E-08 | 6.800E-08 |
| 14405 | 721 | G721C | B | 2.9e-08 | 3.5e-08 | 5.3e-08 | 3.900E-08 | 1.249E-08 | 2.900E-08 | 5.300E-08 |
| 14406 | 721 | G721E | B | 2.2e-08 | 5.9e-08 | 7.1e-08 | 5.067E-08 | 2.554E-08 | 2.200E-08 | 7.100E-08 |
| 14407 | 721 | G721Q | B | 5.5e-08 | 3.5e-08 | 4.8e-08 | 4.600E-08 | 1.015E-08 | 3.500E-08 | 5.500E-08 |
| 14408 | 721 | G721G | B | 1.1e-07 | 2.2e-08 | 2.5e-08 | 5.233E-08 | 4.996E-08 | 2.200E-08 | 1.100E-07 |
| 14409 | 721 | G721H | B | 3.6e-08 | 9.6e-08 | 1.6e-08 | 4.933E-08 | 4.163E-08 | 1.600E-08 | 9.600E-08 |
| 14410 | 721 | G721I | B | 7.5e-08 | 4.2e-08 | 2.7e-08 | 4.800E-08 | 2.456E-08 | 2.700E-08 | 7.500E-08 |
| 14411 | 721 | G721L | B | 9.2e-08 | 2.1e-08 | 1.3e-07 | 8.100E-08 | 5.533E-08 | 2.100E-08 | 1.300E-07 |
| 14412 | 721 | G721K | B | 2.1e-08 | 5.2e-08 | 6.1e-08 | 4.467E-08 | 2.098E-08 | 2.100E-08 | 6.100E-08 |
| 14413 | 721 | G721M | B | 3.4e-08 | 3.8e-08 | 1.5e-08 | 2.900E-08 | 1.229E-08 | 1.500E-08 | 3.800E-08 |
| 14414 | 721 | G721F | B | 1.8e-08 | 5.4e-08 | 8.4e-08 | 5.200E-08 | 3.305E-08 | 1.800E-08 | 8.400E-08 |
| 14415 | 721 | G721P | B | 3.1e-08 | 3.9e-08 | 2.5e-08 | 3.167E-08 | 7.024E-09 | 2.500E-08 | 3.900E-08 |
| 14416 | 721 | G721S | B | 7.4e-08 | 2.8e-08 | 1.4e-08 | 3.867E-08 | 3.139E-08 | 1.400E-08 | 7.400E-08 |
| 14417 | 721 | G721T | B | 5.1e-08 | 3.5e-08 | 4.2e-08 | 4.267E-08 | 8.021E-09 | 3.500E-08 | 5.100E-08 |
| 14418 | 721 | G721W | B | 9.4e-08 | 2.9e-08 | 3.4e-08 | 5.233E-08 | 3.617E-08 | 2.900E-08 | 9.400E-08 |
| 14419 | 721 | G721Y | B | 2.4e-08 | 3.3e-08 | 3.2e-08 | 2.967E-08 | 4.933E-09 | 2.400E-08 | 3.300E-08 |
| 14420 | 721 | G721V | B | 5.9e-08 | 2.9e-08 | 4.6e-08 | 4.467E-08 | 1.504E-08 | 2.900E-08 | 5.900E-08 |
| 14421 | 722 | Q722A | B | 7.5e-08 | 3.3e-08 | 4.5e-08 | 5.100E-08 | 2.163E-08 | 3.300E-08 | 7.500E-08 |
| 14422 | 722 | Q722R | B | 4e-08 | 1.5e-08 | 2.6e-08 | 2.700E-08 | 1.253E-08 | 1.500E-08 | 4.000E-08 |
| 14423 | 722 | Q722N | B | 5.8e-08 | 9.8e-08 | 2.2e-08 | 5.933E-08 | 3.802E-08 | 2.200E-08 | 9.800E-08 |
| 14424 | 722 | Q722D | B | 2.7e-08 | 1.2e-07 | 1.1e-07 | 8.567E-08 | 5.105E-08 | 2.700E-08 | 1.200E-07 |
| 14425 | 722 | Q722C | B | 4.6e-08 | 2e-08 | 1.7e-08 | 2.767E-08 | 1.595E-08 | 1.700E-08 | 4.600E-08 |
| 14426 | 722 | Q722E | B | 4.6e-08 | 6.7e-08 | 2.9e-08 | 4.733E-08 | 1.904E-08 | 2.900E-08 | 6.700E-08 |
| 14427 | 722 | Q722Q | B | 4.8e-08 | 1.1e-07 | 1.9e-08 | 5.900E-08 | 4.649E-08 | 1.900E-08 | 1.100E-07 |
| 14428 | 722 | Q722G | B | 8.8e-08 | 5.5e-08 | 2.2e-08 | 5.500E-08 | 3.300E-08 | 2.200E-08 | 8.800E-08 |
| 14429 | 722 | Q722H | B | 5.2e-08 | 8.6e-09 | 1.8e-08 | 2.620E-08 | 2.283E-08 | 8.600E-09 | 5.200E-08 |
| 14430 | 722 | Q722I | B | 1e-07 | 2.2e-08 | 6.7e-08 | 6.300E-08 | 3.915E-08 | 2.200E-08 | 1.000E-07 |
| 14431 | 722 | Q722L | B | 2.7e-08 | 3.2e-08 | 6e-08 | 3.967E-08 | 1.779E-08 | 2.700E-08 | 6.000E-08 |
| 14432 | 722 | Q722K | B | 4e-08 | 6.3e-08 | 4.4e-08 | 4.900E-08 | 1.229E-08 | 4.000E-08 | 6.300E-08 |
| 14433 | 722 | Q722M | B | 2.9e-08 | 5.8e-08 | 4.2e-08 | 4.300E-08 | 1.453E-08 | 2.900E-08 | 5.800E-08 |
| 14434 | 722 | Q722F | B | 9.4e-08 | 6.8e-08 | 3.4e-08 | 6.533E-08 | 3.009E-08 | 3.400E-08 | 9.400E-08 |
| 14435 | 722 | Q722P | B | 2.1e-08 | 1.7e-08 | 1.1e-08 | 1.633E-08 | 5.033E-09 | 1.100E-08 | 2.100E-08 |
| 14436 | 722 | Q722S | B | 2.9e-08 | 2.6e-08 | 3.4e-08 | 2.967E-08 | 4.041E-09 | 2.600E-08 | 3.400E-08 |
| 14437 | 722 | Q722T | B | 9.6e-08 | 7.5e-08 | 4.7e-08 | 7.267E-08 | 2.458E-08 | 4.700E-08 | 9.600E-08 |
| 14438 | 722 | Q722W | B | 4.9e-08 | 1.9e-08 | 4e-08 | 3.600E-08 | 1.539E-08 | 1.900E-08 | 4.900E-08 |
| 14439 | 722 | Q722Y | B | 2e-08 | 8.5e-08 | 4.3e-08 | 4.933E-08 | 3.296E-08 | 2.000E-08 | 8.500E-08 |
| 14440 | 722 | Q722V | B | 4.5e-08 | 2.4e-08 | 3.3e-08 | 3.400E-08 | 1.054E-08 | 2.400E-08 | 4.500E-08 |
| 14441 | 723 | G723A | B | 1.4e-08 | 4.5e-08 | 1.6e-08 | 2.500E-08 | 1.735E-08 | 1.400E-08 | 4.500E-08 |
| 14442 | 723 | G723R | B | 9.9e-09 | 1e-07 | 2.4e-08 | 4.463E-08 | 4.846E-08 | 9.900E-09 | 1.000E-07 |
| 14443 | 723 | G723N | B | 3.1e-08 | 2.7e-08 | 4.6e-08 | 3.467E-08 | 1.002E-08 | 2.700E-08 | 4.600E-08 |
| 14444 | 723 | G723D | B | 2.8e-08 | 1.7e-08 | 3.6e-08 | 2.700E-08 | 9.539E-09 | 1.700E-08 | 3.600E-08 |
| 14445 | 723 | G723C | B | 7.5e-08 | 4e-08 | 9.4e-08 | 6.967E-08 | 2.739E-08 | 4.000E-08 | 9.400E-08 |
| 14446 | 723 | G723E | B | 1.1e-08 | 2.2e-08 | 7.6e-08 | 3.633E-08 | 3.479E-08 | 1.100E-08 | 7.600E-08 |
| 14447 | 723 | G723Q | B | 6e-08 | 3.9e-08 | 5.4e-08 | 5.100E-08 | 1.082E-08 | 3.900E-08 | 6.000E-08 |
| 14448 | 723 | G723G | B | 1.5e-08 | 3.5e-08 | 4.9e-08 | 3.300E-08 | 1.709E-08 | 1.500E-08 | 4.900E-08 |
| 14449 | 723 | G723H | B | 4.1e-08 | 5e-08 | 8.5e-08 | 5.867E-08 | 2.325E-08 | 4.100E-08 | 8.500E-08 |

| | | | | | | | | | | |
|---|---|---|---|---|---|---|---|---|---|---|
| 14450 | 723 | G723I | B | 1.1e-07 | 6.2e-08 | 1.2e-07 | 9.733E-08 | 3.101E-08 | 6.200E-08 | 1.200E-07 |
| 14451 | 723 | G723L | B | 2.8e-08 | 8.3e-08 | 6.8e-08 | 5.967E-08 | 2.843E-08 | 2.800E-08 | 8.300E-08 |
| 14452 | 723 | G723K | B | 3.9e-08 | 7.6e-08 | 6.1e-08 | 5.867E-08 | 1.861E-08 | 3.900E-08 | 7.600E-08 |
| 14453 | 723 | G723M | B | 2e-08 | 9.3e-08 | 4.5e-08 | 5.267E-08 | 3.710E-08 | 2.000E-08 | 9.300E-08 |
| 14454 | 723 | G723F | B | 3.9e-08 | 3.2e-08 | 7.6e-08 | 4.900E-08 | 2.364E-08 | 3.200E-08 | 7.600E-08 |
| 14455 | 723 | G723P | B | 3e-08 | 3.4e-08 | 2.8e-08 | 3.067E-08 | 3.055E-09 | 2.800E-08 | 3.400E-08 |
| 14456 | 723 | G723S | B | 1.2e-07 | 8.8e-08 | 2.7e-08 | 7.833E-08 | 4.725E-08 | 2.700E-08 | 1.200E-07 |
| 14457 | 723 | G723T | B | 6.7e-08 | 4.1e-08 | 4.3e-08 | 5.033E-08 | 1.447E-08 | 4.100E-08 | 6.700E-08 |
| 14458 | 723 | G723W | B | 6.1e-08 | 3.2e-08 | 3.6e-08 | 4.300E-08 | 1.572E-08 | 3.200E-08 | 6.100E-08 |
| 14459 | 723 | G723Y | B | 2.4e-08 | 2.4e-08 | 6.5e-08 | 3.767E-08 | 2.367E-08 | 2.400E-08 | 6.500E-08 |
| 14460 | 723 | G723V | B | 7.5e-08 | 1.3e-08 | 4.5e-08 | 4.433E-08 | 3.101E-08 | 1.300E-08 | 7.500E-08 |
| 14461 | 724 | T724A | B | 3.9e-08 | 4.9e-08 | 2e-08 | 3.600E-08 | 1.473E-08 | 2.000E-08 | 4.900E-08 |
| 14462 | 724 | T724R | B | 6e-08 | 2.9e-08 | 4.6e-08 | 4.500E-08 | 1.552E-08 | 2.900E-08 | 6.000E-08 |
| 14463 | 724 | T724N | B | 2e-08 | 1.7e-08 | 8.8e-08 | 4.167E-08 | 4.015E-08 | 1.700E-08 | 8.800E-08 |
| 14464 | 724 | T724D | B | 1.4e-08 | 3.9e-08 | 1.2e-08 | 2.167E-08 | 1.504E-08 | 1.200E-08 | 3.900E-08 |
| 14465 | 724 | T724C | B | 2.5e-08 | 3.2e-08 | 3.9e-08 | 3.200E-08 | 7.000E-09 | 2.500E-08 | 3.900E-08 |
| 14466 | 724 | T724E | B | 3.3e-08 | 4.2e-08 | 4.9e-08 | 4.133E-08 | 8.021E-09 | 3.300E-08 | 4.900E-08 |
| 14467 | 724 | T724Q | B | 6.8e-08 | 5.5e-08 | 3.7e-08 | 5.333E-08 | 1.557E-08 | 3.700E-08 | 6.800E-08 |
| 14468 | 724 | T724G | B | 3.1e-08 | 3.7e-08 | 2e-08 | 2.933E-08 | 8.622E-09 | 2.000E-08 | 3.700E-08 |
| 14469 | 724 | T724H | B | 6.3e-08 | 6.7e-08 | 7.9e-08 | 6.967E-08 | 8.327E-09 | 6.300E-08 | 7.900E-08 |
| 14470 | 724 | T724I | B | 3e-08 | 1.2e-07 | 3.2e-08 | 6.067E-08 | 5.139E-08 | 3.000E-08 | 1.200E-07 |
| 14471 | 724 | T724L | B | 2.8e-08 | 5.1e-08 | 5.9e-08 | 4.600E-08 | 1.609E-08 | 2.800E-08 | 5.900E-08 |
| 14472 | 724 | T724K | B | 5.4e-09 | 4.9e-08 | 1e-07 | 5.147E-08 | 4.735E-08 | 5.400E-09 | 1.000E-07 |
| 14473 | 724 | T724M | B | 2.9e-08 | 3.9e-08 | 1.3e-08 | 2.700E-08 | 1.311E-08 | 1.300E-08 | 3.900E-08 |
| 14474 | 724 | T724F | B | 1.3e-07 | 7e-08 | 3.7e-08 | 7.900E-08 | 4.715E-08 | 3.700E-08 | 1.300E-07 |
| 14475 | 724 | T724P | B | 3e-08 | 4.6e-08 | 5.2e-08 | 4.267E-08 | 1.137E-08 | 3.000E-08 | 5.200E-08 |
| 14476 | 724 | T724S | B | 1.6e-08 | 3.3e-08 | 3.2e-08 | 2.700E-08 | 9.539E-09 | 1.600E-08 | 3.300E-08 |
| 14477 | 724 | T724T | B | 4.6e-08 | 5.7e-08 | 7.4e-08 | 5.900E-08 | 1.411E-08 | 4.600E-08 | 7.400E-08 |
| 14478 | 724 | T724W | B | 5.8e-08 | 9.9e-08 | 7.6e-08 | 7.767E-08 | 2.055E-08 | 5.800E-08 | 9.900E-08 |
| 14479 | 724 | T724Y | B | 4.6e-08 | 4e-08 | 4.8e-08 | 4.467E-08 | 4.163E-09 | 4.000E-08 | 4.800E-08 |
| 14480 | 724 | T724V | B | 4.1e-08 | 7.5e-08 | 2.2e-08 | 4.600E-08 | 2.685E-08 | 2.200E-08 | 7.500E-08 |
| 14481 | 725 | K725A | B | 4.2e-08 | 2.4e-08 | 8.7e-08 | 5.100E-08 | 3.245E-08 | 2.400E-08 | 8.700E-08 |
| 14482 | 725 | K725R | B | 6.4e-08 | 1.1e-07 | 4.3e-08 | 7.233E-08 | 3.427E-08 | 4.300E-08 | 1.100E-07 |
| 14483 | 725 | K725N | B | 3.2e-08 | 5.9e-08 | 1.6e-08 | 3.567E-08 | 2.173E-08 | 1.600E-08 | 5.900E-08 |
| 14484 | 725 | K725D | B | 1.1e-07 | 3.5e-08 | 4.1e-08 | 6.200E-08 | 4.168E-08 | 3.500E-08 | 1.100E-07 |
| 14485 | 725 | K725C | B | 5.2e-08 | 2.8e-08 | 4.3e-08 | 4.100E-08 | 1.212E-08 | 2.800E-08 | 5.200E-08 |
| 14486 | 725 | K725E | B | 3.2e-08 | 2.9e-08 | 4.3e-08 | 3.467E-08 | 7.371E-09 | 2.900E-08 | 4.300E-08 |
| 14487 | 725 | K725Q | B | 7.4e-08 | 2.3e-08 | 2.8e-08 | 4.167E-08 | 2.811E-08 | 2.300E-08 | 7.400E-08 |
| 14488 | 725 | K725G | B | 1.5e-07 | 7.7e-08 | 3.6e-08 | 8.767E-08 | 5.774E-08 | 3.600E-08 | 1.500E-07 |
| 14489 | 725 | K725H | B | 4.8e-08 | 3.4e-08 | 3.9e-08 | 4.033E-08 | 7.095E-09 | 3.400E-08 | 4.800E-08 |
| 14490 | 725 | K725I | B | 4.3e-08 | 6.8e-08 | 1.9e-08 | 4.333E-08 | 2.450E-08 | 1.900E-08 | 6.800E-08 |
| 14491 | 725 | K725L | B | 3.6e-08 | 1.8e-08 | 8.6e-08 | 4.667E-08 | 3.523E-08 | 1.800E-08 | 8.600E-08 |
| 14492 | 725 | K725K | B | 4.7e-08 | 3.4e-08 | 1e-07 | 6.033E-08 | 3.496E-08 | 3.400E-08 | 1.000E-07 |
| 14493 | 725 | K725M | B | 2.8e-08 | 4.7e-08 | 6.7e-08 | 4.733E-08 | 1.950E-08 | 2.800E-08 | 6.700E-08 |
| 14494 | 725 | K725F | B | 7.2e-08 | 9.1e-08 | 5.3e-08 | 7.200E-08 | 1.900E-08 | 5.300E-08 | 9.100E-08 |
| 14495 | 725 | K725P | B | 6.5e-08 | 6.2e-08 | 1.1e-07 | 7.900E-08 | 2.689E-08 | 6.200E-08 | 1.100E-07 |
| 14496 | 725 | K725S | B | 3.2e-08 | 5.7e-08 | 3.6e-08 | 4.167E-08 | 1.343E-08 | 3.200E-08 | 5.700E-08 |
| 14497 | 725 | K725T | B | 7.1e-08 | 2.2e-08 | 6e-08 | 5.100E-08 | 2.571E-08 | 2.200E-08 | 7.100E-08 |
| 14498 | 725 | K725W | B | 5.5e-08 | 9.1e-09 | 3.2e-08 | 3.203E-08 | 2.295E-08 | 9.100E-09 | 5.500E-08 |
| 14499 | 725 | K725Y | B | 3.8e-08 | 5.7e-08 | 9e-08 | 6.167E-08 | 2.631E-08 | 3.800E-08 | 9.000E-08 |
| 14500 | 725 | K725V | B | 4.6e-08 | 5.7e-08 | 7.4e-08 | 5.900E-08 | 1.411E-08 | 4.600E-08 | 7.400E-08 |
| 14501 | 726 | V726A | B | 4.9e-08 | 2.7e-08 | 1.2e-08 | 2.933E-08 | 1.861E-08 | 1.200E-08 | 4.900E-08 |
| 14502 | 726 | V726R | B | 5.2e-08 | 4.3e-08 | 6.7e-08 | 5.400E-08 | 1.212E-08 | 4.300E-08 | 6.700E-08 |
| 14503 | 726 | V726N | B | 5.7e-08 | 5.4e-08 | 3.6e-08 | 4.900E-08 | 1.136E-08 | 3.600E-08 | 5.700E-08 |
| 14504 | 726 | V726D | B | 5.1e-08 | 2.9e-08 | 4.2e-08 | 4.067E-08 | 1.106E-08 | 2.900E-08 | 5.100E-08 |
| 14505 | 726 | V726C | B | 1.6e-08 | 2.3e-08 | 1.9e-08 | 1.933E-08 | 3.512E-09 | 1.600E-08 | 2.300E-08 |
| 14506 | 726 | V726E | B | 2.8e-08 | 2.2e-08 | 1.1e-08 | 2.033E-08 | 8.622E-09 | 1.100E-08 | 2.800E-08 |
| 14507 | 726 | V726Q | B | 1.7e-08 | 2e-08 | 5.3e-08 | 3.000E-08 | 1.997E-08 | 1.700E-08 | 5.300E-08 |
| 14508 | 726 | V726G | B | 3.7e-08 | 7.2e-08 | 6.4e-08 | 5.767E-08 | 1.834E-08 | 3.700E-08 | 7.200E-08 |
| 14509 | 726 | V726H | B | 1.1e-08 | 6.3e-08 | 5.4e-08 | 4.267E-08 | 2.779E-08 | 1.100E-08 | 6.300E-08 |
| 14510 | 726 | V726I | B | 3.4e-08 | 9.1e-08 | 4e-08 | 5.500E-08 | 3.132E-08 | 3.400E-08 | 9.100E-08 |
| 14511 | 726 | V726L | B | 9e-08 | 2.5e-08 | 3.3e-08 | 4.933E-08 | 3.544E-08 | 2.500E-08 | 9.000E-08 |
| 14512 | 726 | V726K | B | 4.9e-08 | 3.6e-08 | 1.1e-08 | 3.200E-08 | 1.931E-08 | 1.100E-08 | 4.900E-08 |
| 14513 | 726 | V726M | B | 3.3e-08 | 2.7e-08 | 4.5e-08 | 3.500E-08 | 9.165E-09 | 2.700E-08 | 4.500E-08 |
| 14514 | 726 | V726F | B | 1.1e-07 | 6.5e-08 | 2.9e-08 | 6.800E-08 | 4.058E-08 | 2.900E-08 | 1.100E-07 |
| 14515 | 726 | V726P | B | 1.2e-07 | 2.3e-08 | 4.5e-08 | 6.267E-08 | 5.086E-08 | 2.300E-08 | 1.200E-07 |
| 14516 | 726 | V726S | B | 6.2e-08 | 3.1e-08 | 4.5e-08 | 4.600E-08 | 1.552E-08 | 3.100E-08 | 6.200E-08 |
| 14517 | 726 | V726T | B | 5.1e-08 | 3.5e-08 | 2.9e-08 | 3.833E-08 | 1.137E-08 | 2.900E-08 | 5.100E-08 |
| 14518 | 726 | V726W | B | 3.6e-08 | 5.1e-08 | 3.1e-08 | 3.933E-08 | 1.041E-08 | 3.100E-08 | 5.100E-08 |
| 14519 | 726 | V726Y | B | 1.1e-07 | 1.1e-07 | 2.4e-08 | 8.133E-08 | 4.965E-08 | 2.400E-08 | 1.100E-07 |
| 14520 | 726 | V726V | B | 2.3e-08 | 3.3e-08 | 2.4e-08 | 2.667E-08 | 5.508E-09 | 2.300E-08 | 3.300E-08 |
| 14521 | 727 | E727A | B | 5.1e-08 | 7.2e-08 | 2.9e-08 | 5.067E-08 | 2.150E-08 | 2.900E-08 | 7.200E-08 |
| 14522 | 727 | E727R | B | 5.1e-08 | 5.4e-08 | 8.1e-08 | 6.200E-08 | 1.652E-08 | 5.100E-08 | 8.100E-08 |
| 14523 | 727 | E727N | B | 5.4e-08 | 7.2e-08 | 6.2e-08 | 6.267E-08 | 9.018E-09 | 5.400E-08 | 7.200E-08 |
| 14524 | 727 | E727D | B | 8.8e-08 | 5.4e-08 | 6.1e-08 | 6.767E-08 | 1.795E-08 | 5.400E-08 | 8.800E-08 |
| 14525 | 727 | E727C | B | 5.2e-08 | 4.2e-08 | 5.7e-08 | 5.033E-08 | 7.638E-09 | 4.200E-08 | 5.700E-08 |
| 14526 | 727 | E727E | B | 3.8e-08 | 2.1e-08 | 2.6e-08 | 2.833E-08 | 8.737E-09 | 2.100E-08 | 3.800E-08 |
| 14527 | 727 | E727Q | B | 4.1e-08 | 3.3e-08 | 3.7e-08 | 3.700E-08 | 4.000E-09 | 3.300E-08 | 4.100E-08 |
| 14528 | 727 | E727G | B | 3.2e-08 | 2.4e-08 | 3.4e-08 | 3.000E-08 | 5.292E-09 | 2.400E-08 | 3.400E-08 |
| 14529 | 727 | E727H | B | 8.5e-08 | 7.8e-08 | 4.1e-08 | 6.800E-08 | 2.364E-08 | 4.100E-08 | 8.500E-08 |
| 14530 | 727 | E727I | B | 4.5e-08 | 1.1e-07 | 1.6e-08 | 5.700E-08 | 4.814E-08 | 1.600E-08 | 1.100E-07 |
| 14531 | 727 | E727L | B | 3.6e-08 | 4e-08 | 5.2e-08 | 4.267E-08 | 8.327E-09 | 3.600E-08 | 5.200E-08 |
| 14532 | 727 | E727K | B | 5.7e-08 | 4.2e-08 | 7.7e-08 | 5.867E-08 | 1.756E-08 | 4.200E-08 | 7.700E-08 |
| 14533 | 727 | E727M | B | 3.5e-08 | 5.9e-08 | 6.4e-08 | 5.267E-08 | 1.550E-08 | 3.500E-08 | 6.400E-08 |
| 14534 | 727 | E727F | B | 1.5e-08 | 4.1e-08 | 4.5e-08 | 3.367E-08 | 1.629E-08 | 1.500E-08 | 4.500E-08 |

| | | | | | | | | | | |
|---|---|---|---|---|---|---|---|---|---|---|
| 14535 | 727 | E727P | B | 2.6e-08 | 4.5e-08 | 4.2e-08 | 3.767E-08 | 1.021E-08 | 2.600E-08 | 4.500E-08 |
| 14536 | 727 | E727S | B | 7.1e-08 | 3.6e-08 | 3.1e-08 | 4.600E-08 | 2.179E-08 | 3.100E-08 | 7.100E-08 |
| 14537 | 727 | E727T | B | 8.1e-08 | 7.5e-08 | 2.9e-08 | 6.167E-08 | 2.845E-08 | 2.900E-08 | 8.100E-08 |
| 14538 | 727 | E727W | B | 4.4e-08 | 5.2e-08 | 1.2e-07 | 7.200E-08 | 4.176E-08 | 4.400E-08 | 1.200E-07 |
| 14539 | 727 | E727Y | B | 2e-08 | 5.8e-08 | 4.4e-08 | 4.067E-08 | 1.922E-08 | 2.000E-08 | 5.800E-08 |
| 14540 | 727 | E727V | B | 4.4e-08 | 2.5e-08 | 1.4e-08 | 2.767E-08 | 1.518E-08 | 1.400E-08 | 4.400E-08 |
| 14541 | 728 | I728A | B | 5.8e-08 | 2.2e-08 | 5.6e-08 | 4.533E-08 | 2.023E-08 | 2.200E-08 | 5.800E-08 |
| 14542 | 728 | I728R | B | 5.2e-08 | 4.1e-08 | 1.4e-07 | 7.767E-08 | 5.426E-08 | 4.100E-08 | 1.400E-07 |
| 14543 | 728 | I728N | B | 1.5e-08 | 3.7e-08 | 4.7e-08 | 3.300E-08 | 1.637E-08 | 1.500E-08 | 4.700E-08 |
| 14544 | 728 | I728D | B | 3e-08 | 5.6e-08 | 4.4e-08 | 4.333E-08 | 1.301E-08 | 3.000E-08 | 5.600E-08 |
| 14545 | 728 | I728C | B | 4.2e-08 | 2e-08 | 4.6e-08 | 3.600E-08 | 1.400E-08 | 2.000E-08 | 4.600E-08 |
| 14546 | 728 | I728E | B | 8.4e-08 | 2.2e-08 | 5.6e-08 | 5.400E-08 | 3.105E-08 | 2.200E-08 | 8.400E-08 |
| 14547 | 728 | I728Q | B | 5.1e-08 | 4.4e-08 | 4.6e-08 | 4.700E-08 | 3.606E-09 | 4.400E-08 | 5.100E-08 |
| 14548 | 728 | I728G | B | 4.4e-08 | 2.3e-08 | 2.3e-08 | 3.000E-08 | 1.212E-08 | 2.300E-08 | 4.400E-08 |
| 14549 | 728 | I728H | B | 1e-08 | 2.2e-08 | 2.3e-08 | 1.833E-08 | 7.234E-09 | 1.000E-08 | 2.300E-08 |
| 14550 | 728 | I728I | B | 3.8e-08 | 4.1e-08 | 2.6e-08 | 3.500E-08 | 7.937E-09 | 2.600E-08 | 4.100E-08 |
| 14551 | 728 | I728L | B | 4.5e-08 | 2e-08 | 4e-08 | 3.500E-08 | 1.323E-08 | 2.000E-08 | 4.500E-08 |
| 14552 | 728 | I728K | B | 4.8e-08 | 4.8e-08 | 8.4e-08 | 6.000E-08 | 2.078E-08 | 4.800E-08 | 8.400E-08 |
| 14553 | 728 | I728M | B | 3.4e-08 | 1.1e-07 | 4.4e-08 | 6.267E-08 | 4.130E-08 | 3.400E-08 | 1.100E-07 |
| 14554 | 728 | I728F | B | 1.5e-08 | 4.1e-08 | 6.1e-08 | 3.900E-08 | 2.307E-08 | 1.500E-08 | 6.100E-08 |
| 14555 | 728 | I728P | B | 5.7e-08 | 4.2e-08 | 1.1e-07 | 6.967E-08 | 3.573E-08 | 4.200E-08 | 1.100E-07 |
| 14556 | 728 | I728S | B | 9.4e-08 | 4.6e-08 | 5.1e-08 | 6.367E-08 | 2.639E-08 | 4.600E-08 | 9.400E-08 |
| 14557 | 728 | I728T | B | 3e-08 | 6e-08 | 9.4e-08 | 6.133E-08 | 3.202E-08 | 3.000E-08 | 9.400E-08 |
| 14558 | 728 | I728W | B | 4.8e-08 | 4.9e-08 | 9.9e-08 | 6.533E-08 | 2.916E-08 | 4.800E-08 | 9.900E-08 |
| 14559 | 728 | I728Y | B | 4.9e-08 | 3.2e-08 | 4.5e-08 | 4.200E-08 | 8.888E-09 | 3.200E-08 | 4.900E-08 |
| 14560 | 728 | I728V | B | 2.8e-08 | 2.6e-08 | 8.1e-08 | 4.500E-08 | 3.119E-08 | 2.600E-08 | 8.100E-08 |
| 14561 | 729 | K729A | B | 7.1e-08 | 3.1e-08 | 1.7e-08 | 3.967E-08 | 2.802E-08 | 1.700E-08 | 7.100E-08 |
| 14562 | 729 | K729R | B | 1.9e-07 | 2.5e-08 | 6e-08 | 9.167E-08 | 8.694E-08 | 2.500E-08 | 1.900E-07 |
| 14563 | 729 | K729N | B | 4.8e-08 | 1.8e-08 | 6.1e-08 | 4.233E-08 | 2.205E-08 | 1.800E-08 | 6.100E-08 |
| 14564 | 729 | K729D | B | 3e-08 | 6e-08 | 2.4e-08 | 3.800E-08 | 1.929E-08 | 2.400E-08 | 6.000E-08 |
| 14565 | 729 | K729C | B | 5.4e-08 | 7.1e-08 | 4.8e-08 | 5.767E-08 | 1.193E-08 | 4.800E-08 | 7.100E-08 |
| 14566 | 729 | K729E | B | 5.1e-08 | 4.6e-08 | 2.6e-08 | 4.100E-08 | 1.323E-08 | 2.600E-08 | 5.100E-08 |
| 14567 | 729 | K729Q | B | 3.2e-08 | 2.8e-08 | 2.4e-08 | 2.800E-08 | 4.000E-09 | 2.400E-08 | 3.200E-08 |
| 14568 | 729 | K729G | B | 7.1e-08 | 6.1e-08 | 4.3e-08 | 5.833E-08 | 1.419E-08 | 4.300E-08 | 7.100E-08 |
| 14569 | 729 | K729H | B | 5.5e-08 | 6.4e-08 | 3.3e-08 | 5.067E-08 | 1.595E-08 | 3.300E-08 | 6.400E-08 |
| 14570 | 729 | K729I | B | 6e-08 | 5.1e-08 | 3.7e-08 | 4.933E-08 | 1.159E-08 | 3.700E-08 | 6.000E-08 |
| 14571 | 729 | K729L | B | 2e-08 | 5.8e-08 | 3e-08 | 3.600E-08 | 1.970E-08 | 2.000E-08 | 5.800E-08 |
| 14572 | 729 | K729K | B | 3.3e-08 | 1.2e-08 | 5.6e-08 | 3.367E-08 | 2.201E-08 | 1.200E-08 | 5.600E-08 |
| 14573 | 729 | K729M | B | 3.4e-08 | 3.4e-08 | 1.4e-07 | 6.933E-08 | 6.120E-08 | 3.400E-08 | 1.400E-07 |
| 14574 | 729 | K729F | B | 6.1e-08 | 5e-08 | 9.6e-08 | 6.900E-08 | 2.402E-08 | 5.000E-08 | 9.600E-08 |
| 14575 | 729 | K729P | B | 1.8e-08 | 2.6e-08 | 3.6e-08 | 2.667E-08 | 9.018E-09 | 1.800E-08 | 3.600E-08 |
| 14576 | 729 | K729S | B | 7.9e-08 | 3.8e-08 | 9.2e-08 | 6.967E-08 | 2.818E-08 | 3.800E-08 | 9.200E-08 |
| 14577 | 729 | K729T | B | 3.3e-08 | 1.4e-07 | 2.6e-08 | 6.633E-08 | 6.389E-08 | 2.600E-08 | 1.400E-07 |
| 14578 | 729 | K729W | B | 4e-08 | 3.4e-08 | 6.1e-08 | 4.500E-08 | 1.418E-08 | 3.400E-08 | 6.100E-08 |
| 14579 | 729 | K729Y | B | 7.1e-08 | 3.1e-08 | 2.8e-08 | 4.333E-08 | 2.401E-08 | 2.800E-08 | 7.100E-08 |
| 14580 | 729 | K729V | B | 2.1e-08 | 2.4e-08 | 5.1e-08 | 3.200E-08 | 1.652E-08 | 2.100E-08 | 5.100E-08 |
| 14581 | 730 | R730A | B | 4e-08 | 3.8e-08 | 4.6e-08 | 4.133E-08 | 4.163E-09 | 3.800E-08 | 4.600E-08 |
| 14582 | 730 | R730R | B | 2.2e-08 | 1.8e-08 | 3.7e-08 | 2.567E-08 | 1.002E-08 | 1.800E-08 | 3.700E-08 |
| 14583 | 730 | R730N | B | 3.7e-08 | 1.1e-07 | 3.2e-08 | 5.967E-08 | 4.366E-08 | 3.200E-08 | 1.100E-07 |
| 14584 | 730 | R730D | B | 6.3e-08 | 3.3e-08 | 7.3e-08 | 5.633E-08 | 2.082E-08 | 3.300E-08 | 7.300E-08 |
| 14585 | 730 | R730C | B | 6.8e-08 | 4.9e-08 | 5.7e-08 | 5.800E-08 | 9.539E-09 | 4.900E-08 | 6.800E-08 |
| 14586 | 730 | R730E | B | 4.3e-08 | 2.9e-08 | 4.9e-08 | 4.033E-08 | 1.026E-08 | 2.900E-08 | 4.900E-08 |
| 14587 | 730 | R730Q | B | 4.9e-08 | 2.4e-08 | 4.3e-08 | 3.867E-08 | 1.305E-08 | 2.400E-08 | 4.900E-08 |
| 14588 | 730 | R730G | B | 3.4e-08 | 4.1e-08 | 9.9e-08 | 5.800E-08 | 3.568E-08 | 3.400E-08 | 9.900E-08 |
| 14589 | 730 | R730H | B | 3.4e-08 | 2.2e-08 | 4.1e-08 | 3.233E-08 | 9.609E-09 | 2.200E-08 | 4.100E-08 |
| 14590 | 730 | R730I | B | 1.9e-08 | 9.5e-08 | 4.2e-08 | 5.200E-08 | 3.897E-08 | 1.900E-08 | 9.500E-08 |
| 14591 | 730 | R730L | B | 8.7e-08 | 1.7e-08 | 6e-08 | 5.467E-08 | 3.530E-08 | 1.700E-08 | 8.700E-08 |
| 14592 | 730 | R730K | B | 3e-08 | 4e-08 | 4.8e-08 | 3.933E-08 | 9.018E-09 | 3.000E-08 | 4.800E-08 |
| 14593 | 730 | R730M | B | 1.1e-07 | 3.6e-08 | 5.7e-08 | 6.767E-08 | 3.814E-08 | 3.600E-08 | 1.100E-07 |
| 14594 | 730 | R730F | B | 6e-08 | 5e-08 | 2.3e-08 | 4.433E-08 | 1.914E-08 | 2.300E-08 | 6.000E-08 |
| 14595 | 730 | R730P | B | 4.9e-08 | 5.4e-08 | 3.9e-08 | 4.733E-08 | 7.638E-09 | 3.900E-08 | 5.400E-08 |
| 14596 | 730 | R730S | B | 2.8e-08 | 3.1e-08 | 2.6e-08 | 2.833E-08 | 2.517E-09 | 2.600E-08 | 3.100E-08 |
| 14597 | 730 | R730T | B | 5.5e-08 | 1.6e-08 | 2.5e-08 | 3.200E-08 | 2.042E-08 | 1.600E-08 | 5.500E-08 |
| 14598 | 730 | R730W | B | 4.6e-08 | 2.2e-08 | 5.3e-08 | 4.033E-08 | 1.626E-08 | 2.200E-08 | 5.300E-08 |
| 14599 | 730 | R730Y | B | 6.4e-08 | 2.1e-08 | 2.9e-08 | 3.800E-08 | 2.287E-08 | 2.100E-08 | 6.400E-08 |
| 14600 | 730 | R730V | B | 6.3e-08 | 3.5e-08 | 1.8e-08 | 3.867E-08 | 2.272E-08 | 1.800E-08 | 6.300E-08 |
| 14601 | 731 | T731A | B | 1.8e-08 | 5.1e-08 | 1.8e-08 | 2.900E-08 | 1.905E-08 | 1.800E-08 | 5.100E-08 |
| 14602 | 731 | T731R | B | 6.1e-08 | 2.1e-08 | 1.4e-07 | 7.400E-08 | 6.056E-08 | 2.100E-08 | 1.400E-07 |
| 14603 | 731 | T731N | B | 9.7e-09 | 6.2e-08 | 9.2e-08 | 5.457E-08 | 4.165E-08 | 9.700E-09 | 9.200E-08 |
| 14604 | 731 | T731D | B | 3e-08 | 9.4e-08 | 2.1e-08 | 4.833E-08 | 3.980E-08 | 2.100E-08 | 9.400E-08 |
| 14605 | 731 | T731C | B | 9.6e-08 | 2e-08 | 3.8e-08 | 5.133E-08 | 3.972E-08 | 2.000E-08 | 9.600E-08 |
| 14606 | 731 | T731E | B | 6.4e-08 | 3.8e-08 | 4.8e-08 | 5.000E-08 | 1.311E-08 | 3.800E-08 | 6.400E-08 |
| 14607 | 731 | T731Q | B | 1.8e-07 | 1.6e-07 | 5.1e-08 | 1.303E-07 | 6.943E-08 | 5.100E-08 | 1.800E-07 |
| 14608 | 731 | T731G | B | 8e-08 | 6e-08 | 3.9e-08 | 5.967E-08 | 2.050E-08 | 3.900E-08 | 8.000E-08 |
| 14609 | 731 | T731H | B | 1.3e-08 | 2.2e-07 | 5.8e-08 | 9.700E-08 | 1.089E-07 | 1.300E-08 | 2.200E-07 |
| 14610 | 731 | T731I | B | 7.8e-08 | 9.1e-08 | 4.3e-08 | 7.067E-08 | 2.483E-08 | 4.300E-08 | 9.100E-08 |
| 14611 | 731 | T731L | B | 4.9e-08 | 7.1e-08 | 1.4e-08 | 4.467E-08 | 2.875E-08 | 1.400E-08 | 7.100E-08 |
| 14612 | 731 | T731K | B | 4e-08 | 5.2e-08 | 3.1e-08 | 4.100E-08 | 1.054E-08 | 3.100E-08 | 5.200E-08 |
| 14613 | 731 | T731M | B | 5.5e-08 | 5.5e-08 | 4.9e-08 | 5.300E-08 | 3.464E-09 | 4.900E-08 | 5.500E-08 |
| 14614 | 731 | T731F | B | 3.1e-08 | 6.6e-08 | 8e-08 | 5.900E-08 | 2.524E-08 | 3.100E-08 | 8.000E-08 |
| 14615 | 731 | T731P | B | 1.5e-08 | 5.1e-08 | 7.2e-08 | 4.600E-08 | 2.883E-08 | 1.500E-08 | 7.200E-08 |
| 14616 | 731 | T731S | B | 1.1e-07 | 9e-08 | 1.5e-08 | 7.167E-08 | 5.008E-08 | 1.500E-08 | 1.100E-07 |
| 14617 | 731 | T731T | B | 4.3e-08 | 8.7e-08 | 6.3e-08 | 6.433E-08 | 2.203E-08 | 4.300E-08 | 8.700E-08 |
| 14618 | 731 | T731W | B | 4.7e-08 | 8.3e-08 | 1.9e-08 | 4.967E-08 | 3.208E-08 | 1.900E-08 | 8.300E-08 |
| 14619 | 731 | T731Y | B | 1.8e-08 | 1.8e-08 | 4e-08 | 2.533E-08 | 1.270E-08 | 1.800E-08 | 4.000E-08 |

| | | | | | | | | | | |
|---|---|---|---|---|---|---|---|---|---|---|
| 14620 | 731 | T731V | B | 3.7e-08 | 3.6e-08 | 5.2e-08 | 4.167E-08 | 8.963E-09 | 3.600E-08 | 5.200E-08 |
| 14621 | 732 | V732A | B | 2.1e-08 | 4e-08 | 7.3e-09 | 2.277E-08 | 1.642E-08 | 7.300E-09 | 4.000E-08 |
| 14622 | 732 | V732R | B | 5.6e-08 | 5.9e-08 | 3.8e-08 | 5.100E-08 | 1.136E-08 | 3.800E-08 | 5.900E-08 |
| 14623 | 732 | V732N | B | 1.7e-08 | 5.6e-08 | 2.2e-08 | 3.167E-08 | 2.122E-08 | 1.700E-08 | 5.600E-08 |
| 14624 | 732 | V732D | B | 1.9e-08 | 3.2e-08 | 5.8e-08 | 3.633E-08 | 1.986E-08 | 1.900E-08 | 5.800E-08 |
| 14625 | 732 | V732C | B | 3.5e-08 | 2.6e-08 | 6.2e-08 | 4.100E-08 | 1.873E-08 | 2.600E-08 | 6.200E-08 |
| 14626 | 732 | V732E | B | 1.7e-07 | 5.9e-08 | 5.5e-08 | 9.467E-08 | 6.527E-08 | 5.500E-08 | 1.700E-07 |
| 14627 | 732 | V732Q | B | 5.3e-08 | 2.8e-08 | 9.8e-08 | 5.967E-08 | 3.547E-08 | 2.800E-08 | 9.800E-08 |
| 14628 | 732 | V732G | B | 3.4e-08 | 5e-08 | 2.6e-08 | 3.667E-08 | 1.222E-08 | 2.600E-08 | 5.000E-08 |
| 14629 | 732 | V732H | B | 8.7e-08 | 6.9e-08 | 5.1e-08 | 6.900E-08 | 1.800E-08 | 5.100E-08 | 8.700E-08 |
| 14630 | 732 | V732I | B | 1.8e-08 | 3.8e-08 | 7.7e-08 | 4.433E-08 | 3.001E-08 | 1.800E-08 | 7.700E-08 |
| 14631 | 732 | V732L | B | 4.7e-08 | 1.8e-07 | 3.5e-08 | 8.733E-08 | 8.048E-08 | 3.500E-08 | 1.800E-07 |
| 14632 | 732 | V732K | B | 3.7e-08 | 4.7e-08 | 2.8e-08 | 3.733E-08 | 9.504E-09 | 2.800E-08 | 4.700E-08 |
| 14633 | 732 | V732M | B | 5.7e-08 | 3.8e-08 | 2.2e-08 | 3.900E-08 | 1.752E-08 | 2.200E-08 | 5.700E-08 |
| 14634 | 732 | V732F | B | 4.3e-08 | 3.6e-08 | 9e-08 | 5.633E-08 | 2.937E-08 | 3.600E-08 | 9.000E-08 |
| 14635 | 732 | V732P | B | 3.2e-08 | 8e-08 | 3.8e-08 | 5.000E-08 | 2.615E-08 | 3.200E-08 | 8.000E-08 |
| 14636 | 732 | V732S | B | 5e-08 | 6.3e-08 | 1.7e-08 | 4.333E-08 | 2.371E-08 | 1.700E-08 | 6.300E-08 |
| 14637 | 732 | V732T | B | 1.5e-07 | 2.9e-08 | 3.7e-08 | 7.200E-08 | 6.767E-08 | 2.900E-08 | 1.500E-07 |
| 14638 | 732 | V732W | B | 5.9e-08 | 4.3e-08 | 3.6e-08 | 4.600E-08 | 1.179E-08 | 3.600E-08 | 5.900E-08 |
| 14639 | 732 | V732Y | B | 1.3e-08 | 2.9e-08 | 1e-07 | 4.733E-08 | 4.631E-08 | 1.300E-08 | 1.000E-07 |
| 14640 | 732 | V732V | B | 4.9e-08 | 5.6e-08 | 4.6e-08 | 5.033E-08 | 5.132E-09 | 4.600E-08 | 5.600E-08 |
| 14641 | 733 | A733A | B | 2.3e-08 | 4.7e-08 | 3e-08 | 3.333E-08 | 1.234E-08 | 2.300E-08 | 4.700E-08 |
| 14642 | 733 | A733R | B | 5.5e-08 | 6.1e-08 | 4.4e-08 | 5.333E-08 | 8.622E-09 | 4.400E-08 | 6.100E-08 |
| 14643 | 733 | A733N | B | 5.8e-08 | 1.9e-08 | 2.6e-08 | 3.433E-08 | 2.079E-08 | 1.900E-08 | 5.800E-08 |
| 14644 | 733 | A733D | B | 6.3e-08 | 2.2e-08 | 4.3e-08 | 4.267E-08 | 2.050E-08 | 2.200E-08 | 6.300E-08 |
| 14645 | 733 | A733C | B | 7.1e-08 | 4.8e-08 | 6.9e-08 | 6.267E-08 | 1.274E-08 | 4.800E-08 | 7.100E-08 |
| 14646 | 733 | A733E | B | 2e-07 | 3.3e-08 | 1.7e-07 | 1.343E-07 | 8.903E-08 | 3.300E-08 | 2.000E-07 |
| 14647 | 733 | A733Q | B | 4.4e-08 | 6.7e-08 | 1e-07 | 7.033E-08 | 2.815E-08 | 4.400E-08 | 1.000E-07 |
| 14648 | 733 | A733G | B | 7e-08 | 4.9e-08 | 1e-07 | 7.300E-08 | 2.563E-08 | 4.900E-08 | 1.000E-07 |
| 14649 | 733 | A733H | B | 3.4e-08 | 6.1e-08 | 7.9e-08 | 5.800E-08 | 2.265E-08 | 3.400E-08 | 7.900E-08 |
| 14650 | 733 | A733I | B | 1.6e-08 | 1.8e-08 | 1.3e-07 | 5.467E-08 | 6.525E-08 | 1.600E-08 | 1.300E-07 |
| 14651 | 733 | A733L | B | 7.8e-08 | 5.9e-08 | 2.3e-08 | 5.333E-08 | 2.793E-08 | 2.300E-08 | 7.800E-08 |
| 14652 | 733 | A733K | B | 5.8e-08 | 3.7e-08 | 4.9e-08 | 4.800E-08 | 1.054E-08 | 3.700E-08 | 5.800E-08 |
| 14653 | 733 | A733M | B | 2.2e-08 | 5.9e-08 | 6.6e-08 | 4.900E-08 | 2.364E-08 | 2.200E-08 | 6.600E-08 |
| 14654 | 733 | A733F | B | 3.6e-08 | 5.1e-08 | 3.7e-08 | 4.133E-08 | 8.386E-09 | 3.600E-08 | 5.100E-08 |
| 14655 | 733 | A733P | B | 3.1e-08 | 8.6e-08 | 2.7e-08 | 4.800E-08 | 3.297E-08 | 2.700E-08 | 8.600E-08 |
| 14656 | 733 | A733S | B | 2.1e-08 | 1e-07 | 6.5e-08 | 6.200E-08 | 3.959E-08 | 2.100E-08 | 1.000E-07 |
| 14657 | 733 | A733T | B | 2.7e-08 | 8.4e-09 | 6.2e-08 | 3.247E-08 | 2.721E-08 | 8.400E-09 | 6.200E-08 |
| 14658 | 733 | A733W | B | 4.9e-08 | 1.9e-08 | 5e-08 | 3.933E-08 | 1.762E-08 | 1.900E-08 | 5.000E-08 |
| 14659 | 733 | A733Y | B | 5.2e-08 | 1.8e-08 | 3.4e-08 | 3.467E-08 | 1.701E-08 | 1.800E-08 | 5.200E-08 |
| 14660 | 733 | A733V | B | 4.5e-08 | 1.8e-08 | 5e-08 | 3.767E-08 | 1.721E-08 | 1.800E-08 | 5.000E-08 |
| 14661 | 734 | A734A | B | 4.2e-08 | 2.6e-08 | 2.8e-08 | 3.200E-08 | 8.718E-09 | 2.600E-08 | 4.200E-08 |
| 14662 | 734 | A734R | B | 5.9e-08 | 4.5e-08 | 3.8e-08 | 4.733E-08 | 1.069E-08 | 3.800E-08 | 5.900E-08 |
| 14663 | 734 | A734N | B | 2.3e-08 | 2.7e-08 | 2.8e-08 | 2.600E-08 | 2.646E-09 | 2.300E-08 | 2.800E-08 |
| 14664 | 734 | A734D | B | 2e-08 | 2.4e-08 | 8.2e-08 | 4.200E-08 | 3.470E-08 | 2.000E-08 | 8.200E-08 |
| 14665 | 734 | A734C | B | 6.3e-08 | 4.5e-08 | 7.8e-08 | 6.200E-08 | 1.652E-08 | 4.500E-08 | 7.800E-08 |
| 14666 | 734 | A734E | B | 1.3e-07 | 2.2e-08 | 6.6e-08 | 7.267E-08 | 5.431E-08 | 2.200E-08 | 1.300E-07 |
| 14667 | 734 | A734Q | B | 3.5e-08 | 6.7e-08 | 7e-08 | 5.733E-08 | 1.940E-08 | 3.500E-08 | 7.000E-08 |
| 14668 | 734 | A734G | B | 2.7e-08 | 1.7e-08 | 6.3e-08 | 3.567E-08 | 2.419E-08 | 1.700E-08 | 6.300E-08 |
| 14669 | 734 | A734H | B | 6e-08 | 2.8e-08 | 1.3e-08 | 3.367E-08 | 2.401E-08 | 1.300E-08 | 6.000E-08 |
| 14670 | 734 | A734I | B | 3.8e-08 | 3.9e-08 | 2.7e-08 | 3.467E-08 | 6.658E-09 | 2.700E-08 | 3.900E-08 |
| 14671 | 734 | A734L | B | 9e-08 | 6.2e-08 | 2.4e-08 | 5.867E-08 | 3.313E-08 | 2.400E-08 | 9.000E-08 |
| 14672 | 734 | A734K | B | 9.8e-08 | 5e-08 | 5.1e-08 | 6.633E-08 | 2.743E-08 | 5.000E-08 | 9.800E-08 |
| 14673 | 734 | A734M | B | 5.6e-08 | 5.4e-08 | 3.2e-08 | 4.733E-08 | 1.332E-08 | 3.200E-08 | 5.600E-08 |
| 14674 | 734 | A734F | B | 1e-07 | 2.6e-08 | 2.3e-08 | 4.967E-08 | 4.362E-08 | 2.300E-08 | 1.000E-07 |
| 14675 | 734 | A734P | B | 3.5e-08 | 6.5e-08 | 1.5e-08 | 3.833E-08 | 2.517E-08 | 1.500E-08 | 6.500E-08 |
| 14676 | 734 | A734S | B | 3.7e-08 | 6e-08 | 2.9e-08 | 4.200E-08 | 1.609E-08 | 2.900E-08 | 6.000E-08 |
| 14677 | 734 | A734T | B | 1e-08 | 2.4e-08 | 3.3e-08 | 2.233E-08 | 1.159E-08 | 1.000E-08 | 3.300E-08 |
| 14678 | 734 | A734W | B | 2.4e-08 | 5.5e-08 | 1.3e-08 | 3.067E-08 | 2.178E-08 | 1.300E-08 | 5.500E-08 |
| 14679 | 734 | A734Y | B | 5.3e-08 | 5.8e-08 | 7.6e-08 | 6.233E-08 | 1.210E-08 | 5.300E-08 | 7.600E-08 |
| 14680 | 734 | A734V | B | 2.1e-08 | 4.9e-08 | 4.8e-08 | 3.933E-08 | 1.589E-08 | 2.100E-08 | 4.900E-08 |
| 14681 | 735 | P735A | B | 2.3e-08 | 7.6e-08 | 5.1e-08 | 5.000E-08 | 2.651E-08 | 2.300E-08 | 7.600E-08 |
| 14682 | 735 | P735R | B | 1.7e-08 | 3.9e-08 | 4.6e-08 | 3.400E-08 | 1.513E-08 | 1.700E-08 | 4.600E-08 |
| 14683 | 735 | P735N | B | 5.1e-08 | 4.4e-08 | 8.7e-08 | 6.067E-08 | 2.307E-08 | 4.400E-08 | 8.700E-08 |
| 14684 | 735 | P735D | B | 1.3e-07 | 7e-08 | 6.2e-08 | 8.733E-08 | 3.717E-08 | 6.200E-08 | 1.300E-07 |
| 14685 | 735 | P735C | B | 5e-08 | 2.2e-08 | 2.5e-08 | 3.233E-08 | 1.537E-08 | 2.200E-08 | 5.000E-08 |
| 14686 | 735 | P735E | B | 3.7e-08 | 2.8e-08 | 2.1e-08 | 2.867E-08 | 8.021E-09 | 2.100E-08 | 3.700E-08 |
| 14687 | 735 | P735Q | B | 7.4e-08 | 9.8e-08 | 8.8e-08 | 8.667E-08 | 1.206E-08 | 7.400E-08 | 9.800E-08 |
| 14688 | 735 | P735G | B | 3.8e-08 | 5.2e-08 | 2.8e-08 | 3.933E-08 | 1.206E-08 | 2.800E-08 | 5.200E-08 |
| 14689 | 735 | P735H | B | 4.6e-08 | 3.1e-08 | 3.4e-08 | 3.700E-08 | 7.937E-09 | 3.100E-08 | 4.600E-08 |
| 14690 | 735 | P735I | B | 3.1e-08 | 7.6e-08 | 3.4e-08 | 4.700E-08 | 2.516E-08 | 3.100E-08 | 7.600E-08 |
| 14691 | 735 | P735L | B | 1.9e-08 | 5e-08 | 3.9e-08 | 3.600E-08 | 1.572E-08 | 1.900E-08 | 5.000E-08 |
| 14692 | 735 | P735K | B | 4.2e-08 | 2.5e-08 | 3.9e-08 | 3.533E-08 | 9.074E-09 | 2.500E-08 | 4.200E-08 |
| 14693 | 735 | P735M | B | 7.8e-08 | 3.3e-08 | 5.5e-08 | 5.533E-08 | 2.250E-08 | 3.300E-08 | 7.800E-08 |
| 14694 | 735 | P735F | B | 7.7e-08 | 9.3e-08 | 2.7e-08 | 6.567E-08 | 3.443E-08 | 2.700E-08 | 9.300E-08 |
| 14695 | 735 | P735P | B | 5.1e-08 | 5.7e-08 | 6.1e-08 | 5.633E-08 | 5.033E-09 | 5.100E-08 | 6.100E-08 |
| 14696 | 735 | P735S | B | 2.6e-08 | 3.7e-08 | 4.9e-08 | 3.733E-08 | 1.150E-08 | 2.600E-08 | 4.900E-08 |
| 14697 | 735 | P735T | B | 1.4e-07 | 6.4e-08 | 5.5e-08 | 8.633E-08 | 4.669E-08 | 5.500E-08 | 1.400E-07 |
| 14698 | 735 | P735W | B | 3.3e-08 | 1.7e-08 | 2.7e-08 | 2.567E-08 | 8.083E-09 | 1.700E-08 | 3.300E-08 |
| 14699 | 735 | P735Y | B | 5e-08 | 4.8e-08 | 3.3e-08 | 4.367E-08 | 9.292E-09 | 3.300E-08 | 5.000E-08 |
| 14700 | 735 | P735V | B | 5.3e-08 | 6e-08 | 5.5e-08 | 5.600E-08 | 3.606E-09 | 5.300E-08 | 6.000E-08 |
| 14701 | 736 | S736A | B | 8.3e-09 | 2.7e-08 | 3.6e-08 | 2.377E-08 | 1.413E-08 | 8.300E-09 | 3.600E-08 |
| 14702 | 736 | S736R | B | 4.6e-08 | 4.9e-08 | 3.5e-08 | 4.333E-08 | 7.371E-09 | 3.500E-08 | 4.900E-08 |
| 14703 | 736 | S736N | B | 4e-08 | 3e-08 | 5.2e-08 | 4.067E-08 | 1.102E-08 | 3.000E-08 | 5.200E-08 |
| 14704 | 736 | S736D | B | 2.4e-08 | 3.3e-08 | 4.3e-08 | 3.333E-08 | 9.504E-09 | 2.400E-08 | 4.300E-08 |

| | | | | | | | | | | |
|---|---|---|---|---|---|---|---|---|---|---|
| 14705 | 736 | S736C | B | 7.8e-08 | 2e-08 | 2.5e-08 | 4.100E-08 | 3.214E-08 | 2.000E-08 | 7.800E-08 |
| 14706 | 736 | S736E | B | 5.1e-08 | 3.2e-08 | 4.5e-08 | 4.267E-08 | 9.713E-09 | 3.200E-08 | 5.100E-08 |
| 14707 | 736 | S736Q | B | 5e-08 | 5.2e-08 | 1.9e-08 | 4.033E-08 | 1.850E-08 | 1.900E-08 | 5.200E-08 |
| 14708 | 736 | S736G | B | 9.2e-08 | 3e-08 | 9.6e-08 | 7.267E-08 | 3.700E-08 | 3.000E-08 | 9.600E-08 |
| 14709 | 736 | S736H | B | 3.4e-08 | 4.5e-08 | 4.8e-08 | 4.233E-08 | 7.371E-09 | 3.400E-08 | 4.800E-08 |
| 14710 | 736 | S736I | B | 2.9e-08 | 7.7e-08 | 5e-08 | 5.200E-08 | 2.406E-08 | 2.900E-08 | 7.700E-08 |
| 14711 | 736 | S736L | B | 7.4e-08 | 1.6e-08 | 1.2e-07 | 7.000E-08 | 5.212E-08 | 1.600E-08 | 1.200E-07 |
| 14712 | 736 | S736K | B | 6.3e-08 | 3.2e-08 | 4.9e-08 | 4.800E-08 | 1.552E-08 | 3.200E-08 | 6.300E-08 |
| 14713 | 736 | S736M | B | 8.9e-08 | 4.4e-08 | 3.4e-08 | 5.567E-08 | 2.930E-08 | 3.400E-08 | 8.900E-08 |
| 14714 | 736 | S736F | B | 1.3e-07 | 3.8e-08 | 2.7e-08 | 6.500E-08 | 5.656E-08 | 2.700E-08 | 1.300E-07 |
| 14715 | 736 | S736P | B | 2.8e-08 | 7.3e-08 | 2.9e-08 | 4.333E-08 | 2.570E-08 | 2.800E-08 | 7.300E-08 |
| 14716 | 736 | S736S | B | 2.7e-08 | 4.5e-08 | 5.4e-08 | 4.200E-08 | 1.375E-08 | 2.700E-08 | 5.400E-08 |
| 14717 | 736 | S736T | B | 4.6e-08 | 3.2e-08 | 4.5e-08 | 4.100E-08 | 7.810E-09 | 3.200E-08 | 4.600E-08 |
| 14718 | 736 | S736W | B | 3.5e-08 | 1e-08 | 2.3e-08 | 2.267E-08 | 1.250E-08 | 1.000E-08 | 3.500E-08 |
| 14719 | 736 | S736Y | B | 1.6e-08 | 4.4e-08 | 3.6e-08 | 3.200E-08 | 1.442E-08 | 1.600E-08 | 4.400E-08 |
| 14720 | 736 | S736V | B | 7.1e-08 | 4.9e-08 | 3.1e-08 | 5.033E-08 | 2.003E-08 | 3.100E-08 | 7.100E-08 |
| 14721 | 737 | V737A | B | 7.5e-08 | 1.1e-07 | 3e-08 | 7.167E-08 | 4.010E-08 | 3.000E-08 | 1.100E-07 |
| 14722 | 737 | V737R | B | 5e-08 | 1.3e-08 | 4.5e-08 | 3.600E-08 | 2.007E-08 | 1.300E-08 | 5.000E-08 |
| 14723 | 737 | V737N | B | 4.9e-08 | 7.1e-08 | 4.2e-08 | 5.400E-08 | 1.513E-08 | 4.200E-08 | 7.100E-08 |
| 14724 | 737 | V737D | B | 9.6e-08 | 4.6e-08 | 5.6e-08 | 6.600E-08 | 2.646E-08 | 4.600E-08 | 9.600E-08 |
| 14725 | 737 | V737C | B | 3.8e-08 | 9e-08 | 4.2e-08 | 5.667E-08 | 2.894E-08 | 3.800E-08 | 9.000E-08 |
| 14726 | 737 | V737E | B | 3.4e-08 | 7e-08 | 1.1e-07 | 7.133E-08 | 3.802E-08 | 3.400E-08 | 1.100E-07 |
| 14727 | 737 | V737Q | B | 3.6e-08 | 3.6e-08 | 4e-08 | 3.733E-08 | 2.309E-09 | 3.600E-08 | 4.000E-08 |
| 14728 | 737 | V737G | B | 8.3e-08 | 4.5e-08 | 4.3e-08 | 5.700E-08 | 2.254E-08 | 4.300E-08 | 8.300E-08 |
| 14729 | 737 | V737H | B | 3.4e-08 | 1.2e-08 | 3.2e-08 | 2.600E-08 | 1.217E-08 | 1.200E-08 | 3.400E-08 |
| 14730 | 737 | V737I | B | 3.8e-08 | 2e-08 | 9.3e-08 | 5.033E-08 | 3.803E-08 | 2.000E-08 | 9.300E-08 |
| 14731 | 737 | V737L | B | 3.6e-08 | 4.9e-08 | 4.1e-08 | 4.200E-08 | 6.557E-09 | 3.600E-08 | 4.900E-08 |
| 14732 | 737 | V737K | B | 5.1e-08 | 2.5e-08 | 4e-08 | 3.867E-08 | 1.305E-08 | 2.500E-08 | 5.100E-08 |
| 14733 | 737 | V737M | B | 8.4e-08 | 6.8e-08 | 5.8e-08 | 7.000E-08 | 1.311E-08 | 5.800E-08 | 8.400E-08 |
| 14734 | 737 | V737F | B | 7.9e-08 | 6.8e-08 | 1.2e-07 | 8.900E-08 | 2.740E-08 | 6.800E-08 | 1.200E-07 |
| 14735 | 737 | V737P | B | 1.2e-07 | 6.7e-08 | 3.5e-08 | 7.400E-08 | 4.293E-08 | 3.500E-08 | 1.200E-07 |
| 14736 | 737 | V737S | B | 4.6e-08 | 3.1e-08 | 5e-08 | 4.233E-08 | 1.002E-08 | 3.100E-08 | 5.000E-08 |
| 14737 | 737 | V737T | B | 2.5e-08 | 2.5e-08 | 3.7e-08 | 2.900E-08 | 6.928E-09 | 2.500E-08 | 3.700E-08 |
| 14738 | 737 | V737W | B | 7.2e-08 | 5.7e-08 | 1e-07 | 7.633E-08 | 2.183E-08 | 5.700E-08 | 1.000E-07 |
| 14739 | 737 | V737Y | B | 9.9e-08 | 8.3e-08 | 2.3e-08 | 6.833E-08 | 4.007E-08 | 2.300E-08 | 9.900E-08 |
| 14740 | 737 | V737V | B | 4.9e-08 | 2.6e-08 | 1.8e-08 | 3.100E-08 | 1.609E-08 | 1.800E-08 | 4.900E-08 |
| 14741 | 738 | F738A | B | 2.8e-08 | 6.5e-08 | 6.7e-08 | 5.333E-08 | 2.196E-08 | 2.800E-08 | 6.700E-08 |
| 14742 | 738 | F738R | B | 1.9e-08 | 3e-08 | 7.8e-08 | 4.233E-08 | 3.137E-08 | 1.900E-08 | 7.800E-08 |
| 14743 | 738 | F738N | B | 4.8e-08 | 5.7e-08 | 2.3e-08 | 4.267E-08 | 1.762E-08 | 2.300E-08 | 5.700E-08 |
| 14744 | 738 | F738D | B | 2.7e-08 | 1.1e-07 | 3.3e-08 | 5.667E-08 | 4.629E-08 | 2.700E-08 | 1.100E-07 |
| 14745 | 738 | F738C | B | 7.7e-08 | 4.6e-08 | 3e-08 | 5.100E-08 | 2.390E-08 | 3.000E-08 | 7.700E-08 |
| 14746 | 738 | F738E | B | 2.1e-08 | 3.4e-08 | 3.2e-08 | 2.900E-08 | 7.000E-09 | 2.100E-08 | 3.400E-08 |
| 14747 | 738 | F738Q | B | 9.5e-09 | 7.1e-08 | 2.1e-08 | 3.383E-08 | 3.270E-08 | 9.500E-09 | 7.100E-08 |
| 14748 | 738 | F738G | B | 5e-08 | 6.5e-08 | 4.3e-08 | 5.267E-08 | 1.124E-08 | 4.300E-08 | 6.500E-08 |
| 14749 | 738 | F738H | B | 3.6e-08 | 1.9e-08 | 1.2e-07 | 5.833E-08 | 5.408E-08 | 1.900E-08 | 1.200E-07 |
| 14750 | 738 | F738I | B | 4.4e-08 | 6e-08 | 2.4e-08 | 4.267E-08 | 1.804E-08 | 2.400E-08 | 6.000E-08 |
| 14751 | 738 | F738L | B | 4.3e-08 | 2.8e-08 | 2.8e-08 | 3.300E-08 | 8.660E-09 | 2.800E-08 | 4.300E-08 |
| 14752 | 738 | F738K | B | 4.3e-08 | 3.2e-08 | 3.3e-08 | 3.600E-08 | 6.083E-09 | 3.200E-08 | 4.300E-08 |
| 14753 | 738 | F738M | B | 4.7e-08 | 4.5e-08 | 1.6e-07 | 8.400E-08 | 6.583E-08 | 4.500E-08 | 1.600E-07 |
| 14754 | 738 | F738F | B | 2.1e-08 | 1.3e-07 | 5.7e-08 | 6.933E-08 | 5.554E-08 | 2.100E-08 | 1.300E-07 |
| 14755 | 738 | F738P | B | 6.1e-08 | 1.4e-08 | 1.5e-08 | 3.000E-08 | 2.685E-08 | 1.400E-08 | 6.100E-08 |
| 14756 | 738 | F738S | B | 4.3e-08 | 2.3e-08 | 1.5e-08 | 2.700E-08 | 1.442E-08 | 1.500E-08 | 4.300E-08 |
| 14757 | 738 | F738T | B | 7.2e-08 | 3.6e-08 | 1.7e-08 | 4.167E-08 | 2.793E-08 | 1.700E-08 | 7.200E-08 |
| 14758 | 738 | F738W | B | 3.8e-08 | 5.9e-08 | 3.8e-08 | 4.500E-08 | 1.212E-08 | 3.800E-08 | 5.900E-08 |
| 14759 | 738 | F738Y | B | 4.5e-08 | 1.6e-08 | 2.5e-08 | 2.867E-08 | 1.484E-08 | 1.600E-08 | 4.500E-08 |
| 14760 | 738 | F738V | B | 4.7e-08 | 6.4e-08 | 7.6e-08 | 6.233E-08 | 1.457E-08 | 4.700E-08 | 7.600E-08 |
| 14761 | 739 | I739A | B | 7e-08 | 3.1e-08 | 6.3e-08 | 5.467E-08 | 2.079E-08 | 3.100E-08 | 7.000E-08 |
| 14762 | 739 | I739R | B | 3.4e-08 | 4.1e-08 | 1e-08 | 2.833E-08 | 1.626E-08 | 1.000E-08 | 4.100E-08 |
| 14763 | 739 | I739N | B | 7.3e-08 | 7e-08 | 4.4e-08 | 6.233E-08 | 1.595E-08 | 4.400E-08 | 7.300E-08 |
| 14764 | 739 | I739D | B | 5e-08 | 7.8e-08 | 1.3e-08 | 4.700E-08 | 3.260E-08 | 1.300E-08 | 7.800E-08 |
| 14765 | 739 | I739C | B | 5.7e-09 | 5.9e-08 | 1e-08 | 2.490E-08 | 2.961E-08 | 5.700E-09 | 5.900E-08 |
| 14766 | 739 | I739E | B | 3e-08 | 1.4e-07 | 7.6e-08 | 8.200E-08 | 5.524E-08 | 3.000E-08 | 1.400E-07 |
| 14767 | 739 | I739Q | B | 2.6e-08 | 2.9e-08 | 5e-08 | 3.500E-08 | 1.308E-08 | 2.600E-08 | 5.000E-08 |
| 14768 | 739 | I739G | B | 5.5e-08 | 1.1e-08 | 2.6e-08 | 3.067E-08 | 2.237E-08 | 1.100E-08 | 5.500E-08 |
| 14769 | 739 | I739H | B | 2.6e-08 | 4.8e-08 | 4.1e-08 | 3.833E-08 | 1.124E-08 | 2.600E-08 | 4.800E-08 |
| 14770 | 739 | I739I | B | 6.8e-08 | 3.9e-08 | 4e-08 | 4.900E-08 | 1.646E-08 | 3.900E-08 | 6.800E-08 |
| 14771 | 739 | I739L | B | 6.2e-08 | 9.4e-08 | 3.4e-08 | 6.333E-08 | 3.002E-08 | 3.400E-08 | 9.400E-08 |
| 14772 | 739 | I739K | B | 2.8e-08 | 3.2e-08 | 4.3e-08 | 3.433E-08 | 7.767E-09 | 2.800E-08 | 4.300E-08 |
| 14773 | 739 | I739M | B | 4e-08 | 1.8e-08 | 5.4e-08 | 3.733E-08 | 1.815E-08 | 1.800E-08 | 5.400E-08 |
| 14774 | 739 | I739F | B | 6.2e-08 | 5.5e-08 | 6.1e-08 | 5.933E-08 | 3.786E-09 | 5.500E-08 | 6.200E-08 |
| 14775 | 739 | I739P | B | 1.1e-07 | 3.7e-08 | 3.6e-08 | 6.100E-08 | 4.244E-08 | 3.600E-08 | 1.100E-07 |
| 14776 | 739 | I739S | B | 8.8e-08 | 6.1e-09 | 2.6e-08 | 4.003E-08 | 4.272E-08 | 6.100E-09 | 8.800E-08 |
| 14777 | 739 | I739T | B | 1.8e-08 | 3.2e-08 | 2.6e-08 | 2.533E-08 | 7.024E-09 | 1.800E-08 | 3.200E-08 |
| 14778 | 739 | I739W | B | 2.7e-08 | 6.2e-08 | 1.6e-08 | 3.500E-08 | 2.402E-08 | 1.600E-08 | 6.200E-08 |
| 14779 | 739 | I739Y | B | 8.3e-08 | 4e-08 | 2.9e-08 | 5.067E-08 | 2.854E-08 | 2.900E-08 | 8.300E-08 |
| 14780 | 739 | I739V | B | 5.3e-08 | 2.3e-08 | 2.2e-08 | 3.267E-08 | 1.762E-08 | 2.200E-08 | 5.300E-08 |
| 14781 | 740 | F740A | B | 4.5e-08 | 4.3e-08 | 1.1e-07 | 6.600E-08 | 3.812E-08 | 4.300E-08 | 1.100E-07 |
| 14782 | 740 | F740R | B | 3.9e-08 | 3.4e-08 | 1.9e-08 | 3.067E-08 | 1.041E-08 | 1.900E-08 | 3.900E-08 |
| 14783 | 740 | F740N | B | 4.5e-08 | 2.8e-08 | 3.6e-08 | 3.633E-08 | 8.505E-09 | 2.800E-08 | 4.500E-08 |
| 14784 | 740 | F740D | B | 2.9e-08 | 2.5e-08 | 6.7e-08 | 4.033E-08 | 2.318E-08 | 2.500E-08 | 6.700E-08 |
| 14785 | 740 | F740C | B | 7.8e-08 | 1.8e-08 | 3.3e-08 | 4.300E-08 | 3.122E-08 | 1.800E-08 | 7.800E-08 |
| 14786 | 740 | F740E | B | 2.1e-08 | 5.3e-08 | 4.7e-08 | 4.033E-08 | 1.701E-08 | 2.100E-08 | 5.300E-08 |
| 14787 | 740 | F740Q | B | 5.3e-08 | 1.2e-07 | 3.8e-08 | 7.033E-08 | 4.366E-08 | 3.800E-08 | 1.200E-07 |
| 14788 | 740 | F740G | B | 5.6e-08 | 7.5e-08 | 4.4e-08 | 5.833E-08 | 1.563E-08 | 4.400E-08 | 7.500E-08 |
| 14789 | 740 | F740H | B | 1.4e-07 | 1.1e-07 | 2.8e-08 | 9.267E-08 | 5.798E-08 | 2.800E-08 | 1.400E-07 |

| | | | | | | | | | | |
|---|---|---|---|---|---|---|---|---|---|---|
| 14790 | 740 | F740I | B | 4.4e-08 | 5.2e-08 | 2.5e-08 | 4.033E-08 | 1.387E-08 | 2.500E-08 | 5.200E-08 |
| 14791 | 740 | F740L | B | 6.3e-08 | 5.9e-08 | 4.8e-08 | 5.667E-08 | 7.767E-09 | 4.800E-08 | 6.300E-08 |
| 14792 | 740 | F740K | B | 4.7e-08 | 3.6e-08 | 6.7e-08 | 5.000E-08 | 1.572E-08 | 3.600E-08 | 6.700E-08 |
| 14793 | 740 | F740M | B | 4.3e-08 | 6.2e-08 | 5.8e-08 | 5.433E-08 | 1.002E-08 | 4.300E-08 | 6.200E-08 |
| 14794 | 740 | F740F | B | 7.3e-08 | 4.8e-08 | 4.4e-08 | 5.500E-08 | 1.572E-08 | 4.400E-08 | 7.300E-08 |
| 14795 | 740 | F740P | B | 7.3e-08 | 7.4e-08 | 9.9e-08 | 8.200E-08 | 1.473E-08 | 7.300E-08 | 9.900E-08 |
| 14796 | 740 | F740S | B | 1.8e-08 | 1.4e-07 | 9.5e-08 | 8.433E-08 | 6.170E-08 | 1.800E-08 | 1.400E-07 |
| 14797 | 740 | F740T | B | 1.8e-08 | 1.1e-07 | 6e-08 | 6.267E-08 | 4.606E-08 | 1.800E-08 | 1.100E-07 |
| 14798 | 740 | F740W | B | 1.4e-07 | 3.9e-08 | 6.2e-08 | 8.033E-08 | 5.294E-08 | 3.900E-08 | 1.400E-07 |
| 14799 | 740 | F740Y | B | 3.3e-08 | 1.2e-07 | 1.1e-07 | 8.767E-08 | 4.761E-08 | 3.300E-08 | 1.200E-07 |
| 14800 | 740 | F740V | B | 3.1e-08 | 4e-08 | 3.5e-08 | 3.533E-08 | 4.509E-09 | 3.100E-08 | 4.000E-08 |
| 14801 | 741 | P741A | B | 7.6e-08 | 5.4e-08 | 1.3e-07 | 8.667E-08 | 3.911E-08 | 5.400E-08 | 1.300E-07 |
| 14802 | 741 | P741R | B | 2.4e-08 | 3e-08 | 3.7e-08 | 3.033E-08 | 6.506E-09 | 2.400E-08 | 3.700E-08 |
| 14803 | 741 | P741N | B | 6.4e-08 | 4.6e-08 | 4.6e-08 | 5.200E-08 | 1.039E-08 | 4.600E-08 | 6.400E-08 |
| 14804 | 741 | P741D | B | 1.2e-07 | 1.2e-07 | 4.8e-08 | 9.600E-08 | 4.157E-08 | 4.800E-08 | 1.200E-07 |
| 14805 | 741 | P741C | B | 5.8e-08 | 3.9e-08 | 5.9e-08 | 5.200E-08 | 1.127E-08 | 3.900E-08 | 5.900E-08 |
| 14806 | 741 | P741E | B | 4.5e-08 | 7.9e-08 | 6.4e-08 | 6.267E-08 | 1.704E-08 | 4.500E-08 | 7.900E-08 |
| 14807 | 741 | P741Q | B | 2.3e-08 | 1.4e-07 | 4e-08 | 6.767E-08 | 6.322E-08 | 2.300E-08 | 1.400E-07 |
| 14808 | 741 | P741G | B | 1.2e-07 | 3.8e-08 | 6.3e-08 | 7.367E-08 | 4.203E-08 | 3.800E-08 | 1.200E-07 |
| 14809 | 741 | P741H | B | 3.2e-08 | 4.8e-08 | 2.3e-08 | 3.433E-08 | 1.266E-08 | 2.300E-08 | 4.800E-08 |
| 14810 | 741 | P741I | B | 7.4e-09 | 3.4e-08 | 8.4e-08 | 4.180E-08 | 3.889E-08 | 7.400E-09 | 8.400E-08 |
| 14811 | 741 | P741L | B | 1.8e-08 | 2.3e-08 | 7.7e-08 | 3.933E-08 | 3.272E-08 | 1.800E-08 | 7.700E-08 |
| 14812 | 741 | P741K | B | 7.9e-08 | 1.3e-08 | 4.4e-08 | 4.533E-08 | 3.302E-08 | 1.300E-08 | 7.900E-08 |
| 14813 | 741 | P741M | B | 2.8e-08 | 2e-08 | 2.3e-08 | 2.367E-08 | 4.041E-09 | 2.000E-08 | 2.800E-08 |
| 14814 | 741 | P741F | B | 2.5e-08 | 2.5e-08 | 5.7e-08 | 3.567E-08 | 1.848E-08 | 2.500E-08 | 5.700E-08 |
| 14815 | 741 | P741P | B | 4.1e-08 | 5.2e-08 | 3.8e-08 | 4.367E-08 | 7.371E-09 | 3.800E-08 | 5.200E-08 |
| 14816 | 741 | P741S | B | 4.4e-08 | 1.6e-08 | 6.1e-08 | 4.033E-08 | 2.272E-08 | 1.600E-08 | 6.100E-08 |
| 14817 | 741 | P741T | B | 2.8e-08 | 3.2e-08 | 2.5e-08 | 2.833E-08 | 3.512E-09 | 2.500E-08 | 3.200E-08 |
| 14818 | 741 | P741W | B | 4.3e-08 | 6.1e-08 | 1.7e-08 | 4.033E-08 | 2.212E-08 | 1.700E-08 | 6.100E-08 |
| 14819 | 741 | P741Y | B | 6.1e-08 | 3.3e-08 | 2.9e-08 | 4.100E-08 | 1.744E-08 | 2.900E-08 | 6.100E-08 |
| 14820 | 741 | P741V | B | 5.5e-08 | 8.3e-08 | 2.4e-08 | 5.400E-08 | 2.951E-08 | 2.400E-08 | 8.300E-08 |
| 14821 | 742 | P742A | B | 3.5e-08 | 7e-08 | 1.9e-08 | 4.133E-08 | 2.608E-08 | 1.900E-08 | 7.000E-08 |
| 14822 | 742 | P742R | B | 7.5e-08 | 8.9e-08 | 2.9e-08 | 6.433E-08 | 3.139E-08 | 2.900E-08 | 8.900E-08 |
| 14823 | 742 | P742N | B | 1.2e-07 | 4.2e-08 | 3.5e-08 | 6.567E-08 | 4.718E-08 | 3.500E-08 | 1.200E-07 |
| 14824 | 742 | P742D | B | 1.1e-07 | 8.7e-08 | 4e-08 | 7.900E-08 | 3.568E-08 | 4.000E-08 | 1.100E-07 |
| 14825 | 742 | P742C | B | 1.6e-08 | 5e-08 | 5.8e-08 | 4.133E-08 | 2.230E-08 | 1.600E-08 | 5.800E-08 |
| 14826 | 742 | P742E | B | 1e-07 | 6e-08 | 1e-07 | 8.667E-08 | 2.309E-08 | 6.000E-08 | 1.000E-07 |
| 14827 | 742 | P742Q | B | 4.3e-08 | 2.4e-08 | 5.5e-08 | 4.067E-08 | 1.563E-08 | 2.400E-08 | 5.500E-08 |
| 14828 | 742 | P742G | B | 7.6e-08 | 5.4e-08 | 2.5e-08 | 5.167E-08 | 2.558E-08 | 2.500E-08 | 7.600E-08 |
| 14829 | 742 | P742H | B | 1.4e-08 | 4.6e-08 | 4.2e-08 | 3.400E-08 | 1.744E-08 | 1.400E-08 | 4.600E-08 |
| 14830 | 742 | P742I | B | 1.7e-08 | 2.2e-08 | 5.3e-08 | 3.067E-08 | 1.950E-08 | 1.700E-08 | 5.300E-08 |
| 14831 | 742 | P742L | B | 2.4e-08 | 1.7e-08 | 7.9e-08 | 4.000E-08 | 3.396E-08 | 1.700E-08 | 7.900E-08 |
| 14832 | 742 | P742K | B | 4.7e-08 | 3.4e-08 | 2.4e-08 | 3.500E-08 | 1.153E-08 | 2.400E-08 | 4.700E-08 |
| 14833 | 742 | P742M | B | 5.8e-08 | 5.7e-08 | 7.4e-08 | 6.300E-08 | 9.539E-09 | 5.700E-08 | 7.400E-08 |
| 14834 | 742 | P742F | B | 2.9e-08 | 1.8e-08 | 4.8e-08 | 3.167E-08 | 1.518E-08 | 1.800E-08 | 4.800E-08 |
| 14835 | 742 | P742P | B | 1.1e-07 | 7.4e-08 | 3.3e-08 | 7.233E-08 | 3.853E-08 | 3.300E-08 | 1.100E-07 |
| 14836 | 742 | P742S | B | 1.9e-08 | 2.5e-08 | 6.9e-08 | 3.767E-08 | 2.730E-08 | 1.900E-08 | 6.900E-08 |
| 14837 | 742 | P742T | B | 1.6e-08 | 3.6e-08 | 2.7e-08 | 2.633E-08 | 1.002E-08 | 1.600E-08 | 3.600E-08 |
| 14838 | 742 | P742W | B | 3.2e-08 | 8.2e-08 | 7.3e-08 | 6.233E-08 | 2.665E-08 | 3.200E-08 | 8.200E-08 |
| 14839 | 742 | P742Y | B | 6e-08 | 3.1e-08 | 4e-08 | 4.367E-08 | 1.484E-08 | 3.100E-08 | 6.000E-08 |
| 14840 | 742 | P742V | B | 5e-08 | 2.8e-08 | 2e-08 | 3.267E-08 | 1.553E-08 | 2.000E-08 | 5.000E-08 |
| 14841 | 743 | S743A | B | 3.7e-08 | 6.7e-08 | 6.3e-08 | 5.567E-08 | 1.629E-08 | 3.700E-08 | 6.700E-08 |
| 14842 | 743 | S743R | B | 2.3e-08 | 5.3e-08 | 1.2e-07 | 6.533E-08 | 4.966E-08 | 2.300E-08 | 1.200E-07 |
| 14843 | 743 | S743N | B | 6.7e-08 | 8.9e-08 | 1.7e-08 | 5.767E-08 | 3.690E-08 | 1.700E-08 | 8.900E-08 |
| 14844 | 743 | S743D | B | 9.5e-08 | 3e-08 | 3.6e-08 | 5.367E-08 | 3.592E-08 | 3.000E-08 | 9.500E-08 |
| 14845 | 743 | S743C | B | 1.6e-08 | 5.1e-08 | 2e-08 | 2.900E-08 | 1.916E-08 | 1.600E-08 | 5.100E-08 |
| 14846 | 743 | S743E | B | 2.8e-08 | 4e-08 | 5.4e-08 | 4.067E-08 | 1.301E-08 | 2.800E-08 | 5.400E-08 |
| 14847 | 743 | S743Q | B | 3.1e-08 | 3.5e-08 | 4.4e-08 | 3.667E-08 | 6.658E-09 | 3.100E-08 | 4.400E-08 |
| 14848 | 743 | S743G | B | 3.7e-08 | 1.1e-07 | 7.5e-08 | 7.400E-08 | 3.651E-08 | 3.700E-08 | 1.100E-07 |
| 14849 | 743 | S743H | B | 1.6e-08 | 2.2e-08 | 3.9e-08 | 2.567E-08 | 1.193E-08 | 1.600E-08 | 3.900E-08 |
| 14850 | 743 | S743I | B | 6.2e-08 | 3.2e-08 | 1.1e-07 | 6.800E-08 | 3.934E-08 | 3.200E-08 | 1.100E-07 |
| 14851 | 743 | S743L | B | 1e-07 | 2.8e-08 | 2.8e-08 | 5.200E-08 | 4.157E-08 | 2.800E-08 | 1.000E-07 |
| 14852 | 743 | S743K | B | 3.4e-08 | 1.4e-07 | 2e-08 | 6.467E-08 | 6.562E-08 | 2.000E-08 | 1.400E-07 |
| 14853 | 743 | S743M | B | 5.5e-08 | 4.4e-08 | 6.2e-08 | 5.367E-08 | 9.074E-09 | 4.400E-08 | 6.200E-08 |
| 14854 | 743 | S743F | B | 3.8e-08 | 5.3e-08 | 2.9e-08 | 4.000E-08 | 1.212E-08 | 2.900E-08 | 5.300E-08 |
| 14855 | 743 | S743P | B | 2.2e-08 | 3.9e-08 | 6.3e-08 | 4.133E-08 | 2.060E-08 | 2.200E-08 | 6.300E-08 |
| 14856 | 743 | S743S | B | 4.2e-08 | 4.7e-08 | 3.7e-08 | 4.200E-08 | 5.000E-09 | 3.700E-08 | 4.700E-08 |
| 14857 | 743 | S743T | B | 4.8e-08 | 2.1e-08 | 3.3e-08 | 3.400E-08 | 1.353E-08 | 2.100E-08 | 4.800E-08 |
| 14858 | 743 | S743W | B | 1.3e-07 | 3.4e-08 | 3.1e-08 | 6.500E-08 | 5.631E-08 | 3.100E-08 | 1.300E-07 |
| 14859 | 743 | S743Y | B | 5.2e-08 | 1.1e-07 | 1e-07 | 8.733E-08 | 3.101E-08 | 5.200E-08 | 1.100E-07 |
| 14860 | 743 | S743V | B | 2.8e-08 | 6.3e-08 | 5.8e-08 | 4.967E-08 | 1.893E-08 | 2.800E-08 | 6.300E-08 |
| 14861 | 744 | D744A | B | 4.9e-08 | 1.2e-07 | 3.2e-08 | 6.700E-08 | 4.668E-08 | 3.200E-08 | 1.200E-07 |
| 14862 | 744 | D744R | B | 4.8e-08 | 6e-08 | 1.1e-07 | 7.267E-08 | 3.288E-08 | 4.800E-08 | 1.100E-07 |
| 14863 | 744 | D744N | B | 3.1e-08 | 4.1e-08 | 5.1e-08 | 4.100E-08 | 1.000E-08 | 3.100E-08 | 5.100E-08 |
| 14864 | 744 | D744D | B | 2.3e-08 | 3.5e-08 | 3.4e-08 | 3.067E-08 | 6.658E-09 | 2.300E-08 | 3.500E-08 |
| 14865 | 744 | D744C | B | 7.5e-08 | 2.6e-08 | 7.7e-08 | 5.933E-08 | 2.888E-08 | 2.600E-08 | 7.700E-08 |
| 14866 | 744 | D744E | B | 4.7e-08 | 4.3e-08 | 6.5e-08 | 5.167E-08 | 1.172E-08 | 4.300E-08 | 6.500E-08 |
| 14867 | 744 | D744Q | B | 4.5e-08 | 3.4e-08 | 4.5e-08 | 4.133E-08 | 6.351E-09 | 3.400E-08 | 4.500E-08 |
| 14868 | 744 | D744G | B | 2.1e-08 | 2.2e-08 | 5.6e-08 | 3.300E-08 | 1.992E-08 | 2.100E-08 | 5.600E-08 |
| 14869 | 744 | D744H | B | 6.2e-08 | 4.8e-08 | 3.7e-08 | 4.900E-08 | 1.253E-08 | 3.700E-08 | 6.200E-08 |
| 14870 | 744 | D744I | B | 9.2e-08 | 7.4e-08 | 3.8e-08 | 6.800E-08 | 2.750E-08 | 3.800E-08 | 9.200E-08 |
| 14871 | 744 | D744L | B | 6.7e-08 | 1.1e-07 | 5.2e-08 | 7.633E-08 | 3.011E-08 | 5.200E-08 | 1.100E-07 |
| 14872 | 744 | D744K | B | 4e-08 | 3.5e-08 | 6.6e-08 | 4.700E-08 | 1.664E-08 | 3.500E-08 | 6.600E-08 |
| 14873 | 744 | D744M | B | 1.7e-08 | 2.3e-08 | 3e-08 | 2.333E-08 | 6.506E-09 | 1.700E-08 | 3.000E-08 |
| 14874 | 744 | D744F | B | 2.7e-08 | 4.1e-08 | 2.7e-08 | 3.167E-08 | 8.083E-09 | 2.700E-08 | 4.100E-08 |

| | | | | | | | | | | |
|---|---|---|---|---|---|---|---|---|---|---|
| 14875 | 744 | D744P | B | 2.3e-08 | 2.7e-07 | 8.8e-08 | 1.270E-07 | 1.280E-07 | 2.300E-08 | 2.700E-07 |
| 14876 | 744 | D744S | B | 2.6e-08 | 6.3e-08 | 6.3e-08 | 5.067E-08 | 2.136E-08 | 2.600E-08 | 6.300E-08 |
| 14877 | 744 | D744T | B | 8.4e-09 | 4.5e-08 | 9.3e-08 | 4.880E-08 | 4.243E-08 | 8.400E-09 | 9.300E-08 |
| 14878 | 744 | D744W | B | 3.3e-08 | 3.8e-08 | 1.6e-08 | 2.900E-08 | 1.153E-08 | 1.600E-08 | 3.800E-08 |
| 14879 | 744 | D744Y | B | 6.1e-08 | 3.3e-08 | 5.3e-08 | 4.900E-08 | 1.442E-08 | 3.300E-08 | 6.100E-08 |
| 14880 | 744 | D744V | B | 5.1e-08 | 4.4e-08 | 5.6e-08 | 5.033E-08 | 6.028E-09 | 4.400E-08 | 5.600E-08 |
| 14881 | 745 | E745A | B | 3.3e-08 | 1.1e-08 | 3.5e-08 | 2.633E-08 | 1.332E-08 | 1.100E-08 | 3.500E-08 |
| 14882 | 745 | E745R | B | 2.2e-08 | 5.9e-08 | 1.5e-08 | 3.200E-08 | 2.364E-08 | 1.500E-08 | 5.900E-08 |
| 14883 | 745 | E745N | B | 5.2e-08 | 3.7e-08 | 1.3e-07 | 7.300E-08 | 4.993E-08 | 3.700E-08 | 1.300E-07 |
| 14884 | 745 | E745D | B | 1.8e-08 | 4.6e-08 | 2.2e-08 | 2.867E-08 | 1.514E-08 | 1.800E-08 | 4.600E-08 |
| 14885 | 745 | E745C | B | 2.5e-08 | 3.2e-08 | 2.4e-08 | 2.700E-08 | 4.359E-09 | 2.400E-08 | 3.200E-08 |
| 14886 | 745 | E745E | B | 2.5e-08 | 2.3e-08 | 1.2e-08 | 2.000E-08 | 7.000E-09 | 1.200E-08 | 2.500E-08 |
| 14887 | 745 | E745Q | B | 5e-08 | 2.6e-08 | 9.2e-08 | 5.600E-08 | 3.341E-08 | 2.600E-08 | 9.200E-08 |
| 14888 | 745 | E745G | B | 3.1e-08 | 3e-08 | 1.9e-08 | 2.667E-08 | 6.658E-09 | 1.900E-08 | 3.100E-08 |
| 14889 | 745 | E745H | B | 4.1e-08 | 5.9e-08 | 9e-08 | 6.333E-08 | 2.479E-08 | 4.100E-08 | 9.000E-08 |
| 14890 | 745 | E745I | B | 2.6e-08 | 3.6e-08 | 5.3e-08 | 3.833E-08 | 1.365E-08 | 2.600E-08 | 5.300E-08 |
| 14891 | 745 | E745L | B | 2.3e-08 | 9.4e-08 | 3.1e-08 | 4.933E-08 | 3.889E-08 | 2.300E-08 | 9.400E-08 |
| 14892 | 745 | E745K | B | 4.2e-08 | 4.9e-08 | 2.2e-08 | 3.767E-08 | 1.401E-08 | 2.200E-08 | 4.900E-08 |
| 14893 | 745 | E745M | B | 1.3e-08 | 4e-08 | 3.6e-08 | 2.967E-08 | 1.457E-08 | 1.300E-08 | 4.000E-08 |
| 14894 | 745 | E745F | B | 2.8e-08 | 5e-08 | 3.2e-08 | 3.667E-08 | 1.172E-08 | 2.800E-08 | 5.000E-08 |
| 14895 | 745 | E745P | B | 2.1e-08 | 1.6e-08 | 2.5e-08 | 2.067E-08 | 4.509E-09 | 1.600E-08 | 2.500E-08 |
| 14896 | 745 | E745S | B | 9.2e-08 | 9e-08 | 2.3e-08 | 6.833E-08 | 3.927E-08 | 2.300E-08 | 9.200E-08 |
| 14897 | 745 | E745T | B | 5.6e-08 | 9.3e-09 | 7.5e-08 | 4.677E-08 | 3.381E-08 | 9.300E-09 | 7.500E-08 |
| 14898 | 745 | E745W | B | 6.5e-08 | 1.8e-08 | 1.7e-07 | 8.433E-08 | 7.782E-08 | 1.800E-08 | 1.700E-07 |
| 14899 | 745 | E745Y | B | 3.1e-08 | 3.1e-08 | 4.2e-08 | 3.467E-08 | 6.351E-09 | 3.100E-08 | 4.200E-08 |
| 14900 | 745 | E745V | B | 7.2e-08 | 2.6e-08 | 4.2e-08 | 4.667E-08 | 2.335E-08 | 2.600E-08 | 7.200E-08 |
| 14901 | 746 | Q746A | B | 1.3e-07 | 2.6e-08 | 5.9e-08 | 7.167E-08 | 5.314E-08 | 2.600E-08 | 1.300E-07 |
| 14902 | 746 | Q746R | B | 1.2e-07 | 1.8e-08 | 5.4e-08 | 6.400E-08 | 5.173E-08 | 1.800E-08 | 1.200E-07 |
| 14903 | 746 | Q746N | B | 1.6e-08 | 7e-08 | 4.6e-08 | 4.400E-08 | 2.706E-08 | 1.600E-08 | 7.000E-08 |
| 14904 | 746 | Q746D | B | 3e-08 | 5.3e-08 | 2.4e-08 | 3.567E-08 | 1.531E-08 | 2.400E-08 | 5.300E-08 |
| 14905 | 746 | Q746C | B | 5.8e-08 | 3.5e-08 | 3.4e-08 | 4.233E-08 | 1.358E-08 | 3.400E-08 | 5.800E-08 |
| 14906 | 746 | Q746E | B | 4.5e-08 | 2.2e-08 | 4e-08 | 3.567E-08 | 1.210E-08 | 2.200E-08 | 4.500E-08 |
| 14907 | 746 | Q746Q | B | 5.4e-08 | 1.7e-08 | 2.6e-08 | 3.233E-08 | 1.930E-08 | 1.700E-08 | 5.400E-08 |
| 14908 | 746 | Q746G | B | 6.1e-08 | 1.6e-08 | 8.1e-08 | 5.267E-08 | 3.329E-08 | 1.600E-08 | 8.100E-08 |
| 14909 | 746 | Q746H | B | 3.4e-08 | 1.3e-07 | 1.9e-08 | 6.100E-08 | 6.022E-08 | 1.900E-08 | 1.300E-07 |
| 14910 | 746 | Q746I | B | 4.7e-08 | 2.9e-08 | 1e-07 | 5.867E-08 | 3.691E-08 | 2.900E-08 | 1.000E-07 |
| 14911 | 746 | Q746L | B | 9.6e-08 | 4.6e-08 | 2.4e-08 | 5.533E-08 | 3.690E-08 | 2.400E-08 | 9.600E-08 |
| 14912 | 746 | Q746K | B | 4.1e-08 | 1.4e-08 | 8.9e-08 | 4.800E-08 | 3.799E-08 | 1.400E-08 | 8.900E-08 |
| 14913 | 746 | Q746M | B | 7.3e-09 | 6.3e-08 | 3.4e-08 | 3.477E-08 | 2.786E-08 | 7.300E-09 | 6.300E-08 |
| 14914 | 746 | Q746F | B | 3.5e-08 | 2e-07 | 4.3e-08 | 9.267E-08 | 9.304E-08 | 3.500E-08 | 2.000E-07 |
| 14915 | 746 | Q746P | B | 4.1e-08 | 1.1e-07 | 3.6e-08 | 6.233E-08 | 4.136E-08 | 3.600E-08 | 1.100E-07 |
| 14916 | 746 | Q746S | B | 2.6e-08 | 3.9e-08 | 5.1e-08 | 3.867E-08 | 1.250E-08 | 2.600E-08 | 5.100E-08 |
| 14917 | 746 | Q746T | B | 6.7e-08 | 4.8e-08 | 4.2e-08 | 5.233E-08 | 1.305E-08 | 4.200E-08 | 6.700E-08 |
| 14918 | 746 | Q746W | B | 9.8e-08 | 4.6e-08 | 7.6e-08 | 7.333E-08 | 2.610E-08 | 4.600E-08 | 9.800E-08 |
| 14919 | 746 | Q746Y | B | 2e-08 | 1e-07 | 1.6e-07 | 9.333E-08 | 7.024E-08 | 2.000E-08 | 1.600E-07 |
| 14920 | 746 | Q746V | B | 3.2e-08 | 4e-08 | 5.7e-08 | 4.300E-08 | 1.277E-08 | 3.200E-08 | 5.700E-08 |
| 14921 | 747 | L747A | B | 7.1e-08 | 9.9e-08 | 3.5e-08 | 6.833E-08 | 3.208E-08 | 3.500E-08 | 9.900E-08 |
| 14922 | 747 | L747R | B | 9.2e-08 | 6.8e-08 | 7e-08 | 7.667E-08 | 1.332E-08 | 6.800E-08 | 9.200E-08 |
| 14923 | 747 | L747N | B | 6.6e-08 | 6e-08 | 6.1e-08 | 6.233E-08 | 3.215E-09 | 6.000E-08 | 6.600E-08 |
| 14924 | 747 | L747D | B | 2e-08 | 3.9e-08 | 4.8e-08 | 3.567E-08 | 1.429E-08 | 2.000E-08 | 4.800E-08 |
| 14925 | 747 | L747C | B | 6.8e-08 | 3.9e-08 | 6.8e-08 | 5.833E-08 | 1.674E-08 | 3.900E-08 | 6.800E-08 |
| 14926 | 747 | L747E | B | 1.9e-08 | 3.2e-08 | 9.4e-08 | 4.833E-08 | 4.008E-08 | 1.900E-08 | 9.400E-08 |
| 14927 | 747 | L747Q | B | 2.8e-08 | 8.9e-08 | 9.4e-08 | 7.033E-08 | 3.675E-08 | 2.800E-08 | 9.400E-08 |
| 14928 | 747 | L747G | B | 1.6e-08 | 2.5e-08 | 4.3e-08 | 2.800E-08 | 1.375E-08 | 1.600E-08 | 4.300E-08 |
| 14929 | 747 | L747H | B | 2.5e-08 | 4.2e-08 | 3.7e-08 | 3.467E-08 | 8.737E-09 | 2.500E-08 | 4.200E-08 |
| 14930 | 747 | L747I | B | 3.5e-08 | 6.5e-08 | 8.3e-08 | 6.100E-08 | 2.425E-08 | 3.500E-08 | 8.300E-08 |
| 14931 | 747 | L747L | B | 4.3e-08 | 2.5e-08 | 5.6e-08 | 4.133E-08 | 1.557E-08 | 2.500E-08 | 5.600E-08 |
| 14932 | 747 | L747K | B | 1.7e-08 | 4.3e-08 | 1.3e-07 | 6.333E-08 | 5.918E-08 | 1.700E-08 | 1.300E-07 |
| 14933 | 747 | L747M | B | 4.7e-08 | 5.5e-08 | 2.4e-08 | 4.200E-08 | 1.609E-08 | 2.400E-08 | 5.500E-08 |
| 14934 | 747 | L747F | B | 4.2e-08 | 6.9e-08 | 7.8e-08 | 6.300E-08 | 1.873E-08 | 4.200E-08 | 7.800E-08 |
| 14935 | 747 | L747P | B | 1.7e-08 | 2.8e-08 | 1.2e-07 | 5.500E-08 | 5.656E-08 | 1.700E-08 | 1.200E-07 |
| 14936 | 747 | L747S | B | 1.2e-08 | 3.3e-08 | 1e-07 | 4.833E-08 | 4.596E-08 | 1.200E-08 | 1.000E-07 |
| 14937 | 747 | L747T | B | 2e-08 | 6e-08 | 7.6e-08 | 5.200E-08 | 2.884E-08 | 2.000E-08 | 7.600E-08 |
| 14938 | 747 | L747W | B | 8.3e-08 | 3.5e-08 | 5.8e-08 | 5.867E-08 | 2.401E-08 | 3.500E-08 | 8.300E-08 |
| 14939 | 747 | L747Y | B | 2.5e-08 | 3.5e-08 | 3.8e-08 | 3.267E-08 | 6.807E-09 | 2.500E-08 | 3.800E-08 |
| 14940 | 747 | L747V | B | 6.8e-08 | 1e-07 | 1.6e-08 | 6.133E-08 | 4.239E-08 | 1.600E-08 | 1.000E-07 |
| 14941 | 748 | K748A | B | 2.2e-08 | 2.3e-08 | 4.8e-08 | 3.100E-08 | 1.473E-08 | 2.200E-08 | 4.800E-08 |
| 14942 | 748 | K748R | B | 3.6e-08 | 2.7e-08 | 7e-08 | 4.433E-08 | 2.268E-08 | 2.700E-08 | 7.000E-08 |
| 14943 | 748 | K748N | B | 5.2e-08 | 1.1e-07 | 2.4e-08 | 6.200E-08 | 4.386E-08 | 2.400E-08 | 1.100E-07 |
| 14944 | 748 | K748D | B | 7.4e-08 | 7e-08 | 2.7e-08 | 5.700E-08 | 2.606E-08 | 2.700E-08 | 7.400E-08 |
| 14945 | 748 | K748C | B | 1.8e-08 | 5.5e-08 | 2.7e-08 | 3.333E-08 | 1.930E-08 | 1.800E-08 | 5.500E-08 |
| 14946 | 748 | K748E | B | 1.4e-08 | 1.1e-08 | 1.4e-08 | 1.300E-08 | 1.732E-09 | 1.100E-08 | 1.400E-08 |
| 14947 | 748 | K748Q | B | 6.4e-08 | 4.6e-08 | 5.7e-08 | 5.567E-08 | 9.074E-09 | 4.600E-08 | 6.400E-08 |
| 14948 | 748 | K748G | B | 5e-08 | 8.7e-08 | 2.2e-08 | 5.300E-08 | 3.260E-08 | 2.200E-08 | 8.700E-08 |
| 14949 | 748 | K748H | B | 3e-08 | 2.8e-08 | 2.7e-08 | 2.833E-08 | 1.528E-09 | 2.700E-08 | 3.000E-08 |
| 14950 | 748 | K748I | B | 1.1e-07 | 4e-08 | 4.8e-08 | 6.600E-08 | 3.831E-08 | 4.000E-08 | 1.100E-07 |
| 14951 | 748 | K748L | B | 2.7e-08 | 8.1e-08 | 3e-08 | 4.600E-08 | 3.035E-08 | 2.700E-08 | 8.100E-08 |
| 14952 | 748 | K748K | B | 5.3e-08 | 1.8e-08 | 2.9e-08 | 3.333E-08 | 1.790E-08 | 1.800E-08 | 5.300E-08 |
| 14953 | 748 | K748M | B | 3e-08 | 6.9e-08 | 3e-08 | 4.300E-08 | 2.252E-08 | 3.000E-08 | 6.900E-08 |
| 14954 | 748 | K748F | B | 6.6e-08 | 1.1e-07 | 7.3e-08 | 8.300E-08 | 2.364E-08 | 6.600E-08 | 1.100E-07 |
| 14955 | 748 | K748P | B | 5e-08 | 1.8e-08 | 6.5e-08 | 4.433E-08 | 2.401E-08 | 1.800E-08 | 6.500E-08 |
| 14956 | 748 | K748S | B | 2.8e-08 | 2.6e-08 | 7.1e-08 | 4.167E-08 | 2.542E-08 | 2.600E-08 | 7.100E-08 |
| 14957 | 748 | K748T | B | 5.7e-08 | 9.4e-08 | 3.5e-08 | 6.200E-08 | 2.982E-08 | 3.500E-08 | 9.400E-08 |
| 14958 | 748 | K748W | B | 6.8e-08 | 1.3e-07 | 2.7e-08 | 7.500E-08 | 5.186E-08 | 2.700E-08 | 1.300E-07 |
| 14959 | 748 | K748Y | B | 1.3e-08 | 6.2e-08 | 1.1e-07 | 6.167E-08 | 4.850E-08 | 1.300E-08 | 1.100E-07 |

| | | | | | | | | | | |
|---|---|---|---|---|---|---|---|---|---|---|
| 14960 | 748 | K748V | B | 5.8e-08 | 1.8e-08 | 3.9e-08 | 3.833E-08 | 2.001E-08 | 1.800E-08 | 5.800E-08 |
| 14961 | 749 | S749A | B | 6.3e-09 | 3.6e-08 | 4.6e-08 | 2.943E-08 | 2.065E-08 | 6.300E-09 | 4.600E-08 |
| 14962 | 749 | S749R | B | 7.7e-08 | 7.9e-08 | 5.2e-08 | 6.933E-08 | 1.504E-08 | 5.200E-08 | 7.900E-08 |
| 14963 | 749 | S749N | B | 7.3e-08 | 6.3e-08 | 9.8e-09 | 4.860E-08 | 3.397E-08 | 9.800E-09 | 7.300E-08 |
| 14964 | 749 | S749D | B | 2.2e-08 | 5.8e-08 | 4e-08 | 4.000E-08 | 1.800E-08 | 2.200E-08 | 5.800E-08 |
| 14965 | 749 | S749C | B | 1.8e-08 | 6.8e-08 | 3.1e-08 | 3.900E-08 | 2.594E-08 | 1.800E-08 | 6.800E-08 |
| 14966 | 749 | S749E | B | 7.1e-08 | 3.2e-08 | 3.5e-08 | 4.600E-08 | 2.170E-08 | 3.200E-08 | 7.100E-08 |
| 14967 | 749 | S749Q | B | 4.9e-08 | 5.6e-08 | 3.6e-08 | 4.700E-08 | 1.015E-08 | 3.600E-08 | 5.600E-08 |
| 14968 | 749 | S749G | B | 3.8e-08 | 2e-08 | 4.3e-08 | 3.367E-08 | 1.210E-08 | 2.000E-08 | 4.300E-08 |
| 14969 | 749 | S749H | B | 3.8e-08 | 6.6e-08 | 1.3e-08 | 3.900E-08 | 2.651E-08 | 1.300E-08 | 6.600E-08 |
| 14970 | 749 | S749I | B | 3.8e-08 | 9e-09 | 1.1e-07 | 5.233E-08 | 5.200E-08 | 9.000E-09 | 1.100E-07 |
| 14971 | 749 | S749L | B | 4.3e-08 | 3.9e-08 | 4.4e-08 | 4.200E-08 | 2.646E-09 | 3.900E-08 | 4.400E-08 |
| 14972 | 749 | S749K | B | 4.4e-08 | 5e-08 | 6.4e-08 | 5.267E-08 | 1.026E-08 | 4.400E-08 | 6.400E-08 |
| 14973 | 749 | S749M | B | 1.9e-08 | 8.3e-09 | 1.8e-08 | 1.510E-08 | 5.910E-09 | 8.300E-09 | 1.900E-08 |
| 14974 | 749 | S749F | B | 1.6e-08 | 6.3e-09 | 6.6e-08 | 2.943E-08 | 3.204E-08 | 6.300E-09 | 6.600E-08 |
| 14975 | 749 | S749P | B | 3.4e-08 | 2.7e-07 | 3.8e-08 | 1.140E-07 | 1.351E-07 | 3.400E-08 | 2.700E-07 |
| 14976 | 749 | S749S | B | 1.6e-08 | 3.9e-08 | 3.9e-08 | 3.133E-08 | 1.328E-08 | 1.600E-08 | 3.900E-08 |
| 14977 | 749 | S749T | B | 5.7e-08 | 4.3e-08 | 2.2e-08 | 4.067E-08 | 1.762E-08 | 2.200E-08 | 5.700E-08 |
| 14978 | 749 | S749W | B | 8.5e-08 | 7.2e-08 | 5.4e-08 | 7.033E-08 | 1.557E-08 | 5.400E-08 | 8.500E-08 |
| 14979 | 749 | S749Y | B | 2.2e-07 | 6e-08 | 1.1e-08 | 9.700E-08 | 1.093E-07 | 1.100E-08 | 2.200E-07 |
| 14980 | 749 | S749V | B | 2.7e-08 | 3.4e-08 | 9.3e-08 | 5.133E-08 | 3.625E-08 | 2.700E-08 | 9.300E-08 |
| 14981 | 750 | G750A | B | 3e-08 | 3.5e-08 | 7.5e-08 | 4.667E-08 | 2.466E-08 | 3.000E-08 | 7.500E-08 |
| 14982 | 750 | G750R | B | 3.2e-08 | 2.4e-08 | 1.8e-08 | 2.467E-08 | 7.024E-09 | 1.800E-08 | 3.200E-08 |
| 14983 | 750 | G750N | B | 1.8e-08 | 5.2e-08 | 2.2e-08 | 3.067E-08 | 1.858E-08 | 1.800E-08 | 5.200E-08 |
| 14984 | 750 | G750D | B | 3.4e-08 | 9.8e-09 | 4.6e-08 | 2.993E-08 | 1.844E-08 | 9.800E-09 | 4.600E-08 |
| 14985 | 750 | G750C | B | 3.1e-08 | 5.1e-08 | 2.6e-08 | 3.600E-08 | 1.323E-08 | 2.600E-08 | 5.100E-08 |
| 14986 | 750 | G750E | B | 7.8e-08 | 3.5e-08 | 3.6e-08 | 4.967E-08 | 2.454E-08 | 3.500E-08 | 7.800E-08 |
| 14987 | 750 | G750Q | B | 2e-08 | 1e-07 | 2.2e-08 | 4.733E-08 | 4.562E-08 | 2.000E-08 | 1.000E-07 |
| 14988 | 750 | G750G | B | 4.9e-08 | 3.6e-08 | 9.3e-08 | 5.933E-08 | 2.987E-08 | 3.600E-08 | 9.300E-08 |
| 14989 | 750 | G750H | B | 1.6e-07 | 3e-08 | 8.8e-08 | 9.267E-08 | 6.513E-08 | 3.000E-08 | 1.600E-07 |
| 14990 | 750 | G750I | B | 2.8e-08 | 6.2e-08 | 3.2e-08 | 4.067E-08 | 1.858E-08 | 2.800E-08 | 6.200E-08 |
| 14991 | 750 | G750L | B | 1.7e-07 | 5.4e-08 | 7.8e-08 | 1.007E-07 | 6.123E-08 | 5.400E-08 | 1.700E-07 |
| 14992 | 750 | G750K | B | 2.1e-08 | 6.9e-08 | 1.1e-07 | 6.667E-08 | 4.455E-08 | 2.100E-08 | 1.100E-07 |
| 14993 | 750 | G750M | B | 3.1e-08 | 4.1e-08 | 5e-08 | 4.067E-08 | 9.504E-09 | 3.100E-08 | 5.000E-08 |
| 14994 | 750 | G750F | B | 6.3e-08 | 3.5e-08 | 1e-07 | 6.600E-08 | 3.260E-08 | 3.500E-08 | 1.000E-07 |
| 14995 | 750 | G750P | B | 2.5e-08 | 4.3e-08 | 1.3e-07 | 6.600E-08 | 5.615E-08 | 2.500E-08 | 1.300E-07 |
| 14996 | 750 | G750S | B | 5e-08 | 7.2e-08 | 2.2e-08 | 4.800E-08 | 2.506E-08 | 2.200E-08 | 7.200E-08 |
| 14997 | 750 | G750T | B | 2.2e-08 | 6.5e-08 | 5.6e-08 | 4.767E-08 | 2.268E-08 | 2.200E-08 | 6.500E-08 |
| 14998 | 750 | G750W | B | 6.1e-08 | 6.3e-08 | 2.5e-08 | 4.967E-08 | 2.139E-08 | 2.500E-08 | 6.300E-08 |
| 14999 | 750 | G750Y | B | 2.2e-08 | 1.3e-08 | 4e-08 | 2.500E-08 | 1.375E-08 | 1.300E-08 | 4.000E-08 |
| 15000 | 750 | G750V | B | 3.6e-08 | 3.5e-08 | 1.1e-08 | 2.733E-08 | 1.415E-08 | 1.100E-08 | 3.600E-08 |
| 15001 | 751 | T751A | B | 3.5e-08 | 5.7e-08 | 8.4e-08 | 5.867E-08 | 2.454E-08 | 3.500E-08 | 8.400E-08 |
| 15002 | 751 | T751R | B | 3.4e-08 | 4.5e-08 | 4.2e-08 | 4.033E-08 | 5.686E-09 | 3.400E-08 | 4.500E-08 |
| 15003 | 751 | T751N | B | 4.3e-08 | 2.2e-08 | 4.3e-08 | 3.600E-08 | 1.212E-08 | 2.200E-08 | 4.300E-08 |
| 15004 | 751 | T751D | B | 6.3e-08 | 1.9e-07 | 3.3e-08 | 9.533E-08 | 8.334E-08 | 3.300E-08 | 1.900E-07 |
| 15005 | 751 | T751C | B | 5.5e-08 | 2e-08 | 4.3e-08 | 3.933E-08 | 1.779E-08 | 2.000E-08 | 5.500E-08 |
| 15006 | 751 | T751E | B | 4.1e-08 | 2.1e-08 | 2.3e-08 | 2.833E-08 | 1.102E-08 | 2.100E-08 | 4.100E-08 |
| 15007 | 751 | T751Q | B | 4.1e-08 | 5.9e-08 | 5e-08 | 5.000E-08 | 9.000E-09 | 4.100E-08 | 5.900E-08 |
| 15008 | 751 | T751G | B | 4.7e-08 | 1.6e-08 | 3.7e-08 | 3.333E-08 | 1.582E-08 | 1.600E-08 | 4.700E-08 |
| 15009 | 751 | T751H | B | 4.5e-08 | 5.7e-08 | 2.8e-08 | 4.333E-08 | 1.457E-08 | 2.800E-08 | 5.700E-08 |
| 15010 | 751 | T751I | B | 4e-08 | 3.6e-08 | 5.3e-08 | 4.300E-08 | 8.888E-09 | 3.600E-08 | 5.300E-08 |
| 15011 | 751 | T751L | B | 1.3e-07 | 1.6e-08 | 2.9e-08 | 5.833E-08 | 6.240E-08 | 1.600E-08 | 1.300E-07 |
| 15012 | 751 | T751K | B | 2.2e-08 | 3.7e-08 | 7.1e-08 | 4.333E-08 | 2.511E-08 | 2.200E-08 | 7.100E-08 |
| 15013 | 751 | T751M | B | 3.7e-08 | 3.8e-08 | 5.4e-08 | 4.300E-08 | 9.539E-09 | 3.700E-08 | 5.400E-08 |
| 15014 | 751 | T751F | B | 1.4e-08 | 5.7e-08 | 2.6e-08 | 3.233E-08 | 2.219E-08 | 1.400E-08 | 5.700E-08 |
| 15015 | 751 | T751P | B | 3.7e-08 | 7.2e-09 | 7.3e-08 | 3.907E-08 | 3.295E-08 | 7.200E-09 | 7.300E-08 |
| 15016 | 751 | T751S | B | 9.9e-08 | 6.9e-08 | 9.7e-08 | 8.833E-08 | 1.677E-08 | 6.900E-08 | 9.900E-08 |
| 15017 | 751 | T751T | B | 7.5e-08 | 2.3e-08 | 2.5e-08 | 4.100E-08 | 2.946E-08 | 2.300E-08 | 7.500E-08 |
| 15018 | 751 | T751W | B | 7.8e-08 | 4.6e-08 | 3.6e-08 | 5.333E-08 | 2.194E-08 | 3.600E-08 | 7.800E-08 |
| 15019 | 751 | T751Y | B | 4.4e-08 | 1.9e-08 | 5.5e-08 | 3.933E-08 | 1.845E-08 | 1.900E-08 | 5.500E-08 |
| 15020 | 751 | T751V | B | 4.3e-08 | 3.8e-08 | 3.1e-08 | 3.733E-08 | 6.028E-09 | 3.100E-08 | 4.300E-08 |
| 15021 | 752 | A752A | B | 1.1e-07 | 1.1e-07 | 9.1e-08 | 1.037E-07 | 1.097E-08 | 9.100E-08 | 1.100E-07 |
| 15022 | 752 | A752R | B | 4.7e-08 | 4.7e-08 | 6.8e-08 | 5.400E-08 | 1.212E-08 | 4.700E-08 | 6.800E-08 |
| 15023 | 752 | A752N | B | 3.5e-08 | 8.4e-08 | 1.2e-08 | 4.367E-08 | 3.677E-08 | 1.200E-08 | 8.400E-08 |
| 15024 | 752 | A752D | B | 4.3e-08 | 1.1e-08 | 8.4e-08 | 4.600E-08 | 3.659E-08 | 1.100E-08 | 8.400E-08 |
| 15025 | 752 | A752C | B | 6.8e-08 | 9.6e-08 | 2.5e-08 | 6.300E-08 | 3.576E-08 | 2.500E-08 | 9.600E-08 |
| 15026 | 752 | A752E | B | 4e-08 | 3.8e-08 | 5.3e-08 | 4.367E-08 | 8.145E-09 | 3.800E-08 | 5.300E-08 |
| 15027 | 752 | A752Q | B | 5e-08 | 5.4e-08 | 6.5e-08 | 5.633E-08 | 7.767E-09 | 5.000E-08 | 6.500E-08 |
| 15028 | 752 | A752G | B | 4.4e-08 | 8.2e-08 | 2.7e-08 | 5.100E-08 | 2.816E-08 | 2.700E-08 | 8.200E-08 |
| 15029 | 752 | A752H | B | 6.1e-08 | 1.2e-07 | 9.6e-08 | 9.233E-08 | 2.967E-08 | 6.100E-08 | 1.200E-07 |
| 15030 | 752 | A752I | B | 7.8e-08 | 6e-08 | 6e-08 | 6.600E-08 | 1.039E-08 | 6.000E-08 | 7.800E-08 |
| 15031 | 752 | A752L | B | 5.1e-08 | 1.1e-07 | 9.2e-08 | 8.433E-08 | 3.024E-08 | 5.100E-08 | 1.100E-07 |
| 15032 | 752 | A752K | B | 4.5e-08 | 3.4e-08 | 8.5e-08 | 5.467E-08 | 2.684E-08 | 3.400E-08 | 8.500E-08 |
| 15033 | 752 | A752M | B | 2.9e-08 | 2.1e-08 | 2e-08 | 2.333E-08 | 4.933E-09 | 2.000E-08 | 2.900E-08 |
| 15034 | 752 | A752F | B | 6.8e-08 | 2.2e-08 | 1.6e-07 | 8.333E-08 | 7.027E-08 | 2.200E-08 | 1.600E-07 |
| 15035 | 752 | A752P | B | 5.1e-08 | 6.1e-08 | 7.1e-08 | 6.100E-08 | 1.000E-08 | 5.100E-08 | 7.100E-08 |
| 15036 | 752 | A752S | B | 7.5e-08 | 3e-08 | 3.8e-08 | 4.767E-08 | 2.401E-08 | 3.000E-08 | 7.500E-08 |
| 15037 | 752 | A752T | B | 4.9e-08 | 4.4e-08 | 2.7e-08 | 4.000E-08 | 1.153E-08 | 2.700E-08 | 4.900E-08 |
| 15038 | 752 | A752W | B | 3e-08 | 4.8e-08 | 4.6e-08 | 4.133E-08 | 9.866E-09 | 3.000E-08 | 4.800E-08 |
| 15039 | 752 | A752Y | B | 1.1e-07 | 2.7e-08 | 2.5e-08 | 5.400E-08 | 4.851E-08 | 2.500E-08 | 1.100E-07 |
| 15040 | 752 | A752V | B | 1.2e-08 | 2.7e-08 | 4.5e-08 | 2.800E-08 | 1.652E-08 | 1.200E-08 | 4.500E-08 |
| 15041 | 753 | S753A | B | 1e-07 | 3.3e-08 | 6.7e-08 | 6.667E-08 | 3.350E-08 | 3.300E-08 | 1.000E-07 |
| 15042 | 753 | S753R | B | 1.3e-07 | 3.3e-08 | 1.8e-08 | 6.033E-08 | 6.080E-08 | 1.800E-08 | 1.300E-07 |
| 15043 | 753 | S753N | B | 7.1e-08 | 5.1e-08 | 1.1e-07 | 7.733E-08 | 3.001E-08 | 5.100E-08 | 1.100E-07 |
| 15044 | 753 | S753D | B | 2.1e-08 | 1.1e-07 | 5.3e-08 | 6.133E-08 | 4.508E-08 | 2.100E-08 | 1.100E-07 |

| | | | | | | | | | | |
|---|---|---|---|---|---|---|---|---|---|---|
| 15045 | 753 | S753C | B | 1.6e-07 | 4.1e-08 | 2.2e-08 | 7.433E-08 | 7.480E-08 | 2.200E-08 | 1.600E-07 |
| 15046 | 753 | S753E | B | 4.1e-08 | 3.3e-08 | 2.3e-08 | 3.233E-08 | 9.018E-09 | 2.300E-08 | 4.100E-08 |
| 15047 | 753 | S753Q | B | 2.6e-08 | 7.5e-08 | 7.7e-08 | 5.933E-08 | 2.888E-08 | 2.600E-08 | 7.700E-08 |
| 15048 | 753 | S753G | B | 5.3e-08 | 4.9e-08 | 3.6e-08 | 4.600E-08 | 8.888E-09 | 3.600E-08 | 5.300E-08 |
| 15049 | 753 | S753H | B | 3.5e-08 | 3.3e-08 | 7.7e-08 | 4.833E-08 | 2.485E-08 | 3.300E-08 | 7.700E-08 |
| 15050 | 753 | S753I | B | 3.1e-08 | 5.7e-08 | 1.5e-07 | 7.933E-08 | 6.256E-08 | 3.100E-08 | 1.500E-07 |
| 15051 | 753 | S753L | B | 9.8e-08 | 3.5e-08 | 1.7e-08 | 5.000E-08 | 4.253E-08 | 1.700E-08 | 9.800E-08 |
| 15052 | 753 | S753K | B | 4.9e-08 | 1.6e-07 | 4e-08 | 8.300E-08 | 6.684E-08 | 4.000E-08 | 1.600E-07 |
| 15053 | 753 | S753M | B | 4.2e-08 | 5.3e-08 | 4.9e-08 | 4.800E-08 | 5.568E-09 | 4.200E-08 | 5.300E-08 |
| 15054 | 753 | S753F | B | 2.6e-08 | 3.3e-08 | 3e-08 | 2.967E-08 | 3.512E-09 | 2.600E-08 | 3.300E-08 |
| 15055 | 753 | S753P | B | 7.4e-08 | 1e-07 | 2e-08 | 6.467E-08 | 4.081E-08 | 2.000E-08 | 1.000E-07 |
| 15056 | 753 | S753S | B | 1.5e-08 | 7.6e-08 | 3.7e-08 | 4.267E-08 | 3.089E-08 | 1.500E-08 | 7.600E-08 |
| 15057 | 753 | S753T | B | 3.3e-08 | 3e-08 | 6e-08 | 4.100E-08 | 1.652E-08 | 3.000E-08 | 6.000E-08 |
| 15058 | 753 | S753W | B | 3.2e-08 | 1.8e-08 | 1.6e-07 | 7.000E-08 | 7.826E-08 | 1.800E-08 | 1.600E-07 |
| 15059 | 753 | S753Y | B | 6.7e-08 | 1.5e-07 | 2.2e-08 | 7.967E-08 | 6.493E-08 | 2.200E-08 | 1.500E-07 |
| 15060 | 753 | S753V | B | 3.9e-08 | 2.6e-08 | 3e-08 | 3.167E-08 | 6.658E-09 | 2.600E-08 | 3.900E-08 |
| 15061 | 754 | V754A | B | 6.9e-08 | 2.9e-08 | 3.8e-08 | 4.533E-08 | 2.098E-08 | 2.900E-08 | 6.900E-08 |
| 15062 | 754 | V754R | B | 7.2e-08 | 5.5e-08 | 4.9e-08 | 5.867E-08 | 1.193E-08 | 4.900E-08 | 7.200E-08 |
| 15063 | 754 | V754N | B | 1.4e-07 | 7.2e-08 | 5.2e-08 | 8.800E-08 | 4.613E-08 | 5.200E-08 | 1.400E-07 |
| 15064 | 754 | V754D | B | 1.4e-08 | 4.2e-08 | 2.2e-08 | 2.600E-08 | 1.442E-08 | 1.400E-08 | 4.200E-08 |
| 15065 | 754 | V754C | B | 2.7e-08 | 3.1e-08 | 3.6e-08 | 3.133E-08 | 4.509E-09 | 2.700E-08 | 3.600E-08 |
| 15066 | 754 | V754E | B | 9e-08 | 2.6e-08 | 7.8e-08 | 6.467E-08 | 3.402E-08 | 2.600E-08 | 9.000E-08 |
| 15067 | 754 | V754Q | B | 4.4e-08 | 2.5e-08 | 3.9e-08 | 3.600E-08 | 9.849E-09 | 2.500E-08 | 4.400E-08 |
| 15068 | 754 | V754G | B | 9.4e-09 | 6.9e-08 | 3.5e-08 | 3.780E-08 | 2.990E-08 | 9.400E-09 | 6.900E-08 |
| 15069 | 754 | V754H | B | 4.3e-08 | 4.3e-08 | 5.3e-08 | 4.633E-08 | 5.774E-09 | 4.300E-08 | 5.300E-08 |
| 15070 | 754 | V754I | B | 5.4e-08 | 6.7e-08 | 3.1e-08 | 5.067E-08 | 1.823E-08 | 3.100E-08 | 6.700E-08 |
| 15071 | 754 | V754L | B | 2.9e-08 | 1.2e-07 | 4.8e-08 | 6.567E-08 | 4.800E-08 | 2.900E-08 | 1.200E-07 |
| 15072 | 754 | V754K | B | 3.4e-08 | 2.6e-08 | 2.7e-08 | 2.900E-08 | 4.359E-09 | 2.600E-08 | 3.400E-08 |
| 15073 | 754 | V754M | B | 5.7e-08 | 1.6e-08 | 6.6e-08 | 4.633E-08 | 2.665E-08 | 1.600E-08 | 6.600E-08 |
| 15074 | 754 | V754F | B | 7.4e-08 | 3.5e-08 | 6.3e-08 | 5.733E-08 | 2.011E-08 | 3.500E-08 | 7.400E-08 |
| 15075 | 754 | V754P | B | 2.6e-08 | 3.6e-08 | 3.5e-08 | 3.233E-08 | 5.508E-09 | 2.600E-08 | 3.600E-08 |
| 15076 | 754 | V754S | B | 4.8e-08 | 5.3e-08 | 1.4e-08 | 3.833E-08 | 2.122E-08 | 1.400E-08 | 5.300E-08 |
| 15077 | 754 | V754T | B | 5e-08 | 1.1e-07 | 2.4e-08 | 6.133E-08 | 4.411E-08 | 2.400E-08 | 1.100E-07 |
| 15078 | 754 | V754W | B | 7e-08 | 3.7e-08 | 6.1e-08 | 5.600E-08 | 1.706E-08 | 3.700E-08 | 7.000E-08 |
| 15079 | 754 | V754Y | B | 7.9e-08 | 1.3e-07 | 2.8e-08 | 7.900E-08 | 5.100E-08 | 2.800E-08 | 1.300E-07 |
| 15080 | 754 | V754V | B | 1.2e-07 | 5e-08 | 2.9e-08 | 6.633E-08 | 4.765E-08 | 2.900E-08 | 1.200E-07 |
| 15081 | 755 | V755A | B | 2.9e-08 | 3e-08 | 4e-08 | 3.300E-08 | 6.083E-09 | 2.900E-08 | 4.000E-08 |
| 15082 | 755 | V755R | B | 5.7e-08 | 2.9e-08 | 9.6e-08 | 6.067E-08 | 3.365E-08 | 2.900E-08 | 9.600E-08 |
| 15083 | 755 | V755N | B | 6.2e-08 | 1.6e-07 | 6.7e-08 | 9.633E-08 | 5.519E-08 | 6.200E-08 | 1.600E-07 |
| 15084 | 755 | V755D | B | 1.1e-07 | 8.7e-08 | 1.5e-08 | 7.067E-08 | 4.956E-08 | 1.500E-08 | 1.100E-07 |
| 15085 | 755 | V755C | B | 9.9e-09 | 2.1e-08 | 8.7e-08 | 3.930E-08 | 4.168E-08 | 9.900E-09 | 8.700E-08 |
| 15086 | 755 | V755E | B | 2e-08 | 4.5e-08 | 7.4e-08 | 4.633E-08 | 2.702E-08 | 2.000E-08 | 7.400E-08 |
| 15087 | 755 | V755Q | B | 2e-08 | 2.5e-08 | 4.2e-08 | 2.900E-08 | 1.153E-08 | 2.000E-08 | 4.200E-08 |
| 15088 | 755 | V755G | B | 5.2e-08 | 8e-08 | 1.4e-07 | 9.067E-08 | 4.496E-08 | 5.200E-08 | 1.400E-07 |
| 15089 | 755 | V755H | B | 1.5e-08 | 1.5e-08 | 2.9e-08 | 1.967E-08 | 8.083E-09 | 1.500E-08 | 2.900E-08 |
| 15090 | 755 | V755I | B | 5.1e-08 | 1e-07 | 3.6e-08 | 6.233E-08 | 3.347E-08 | 3.600E-08 | 1.000E-07 |
| 15091 | 755 | V755L | B | 2.5e-08 | 3.7e-08 | 4.6e-08 | 3.600E-08 | 1.054E-08 | 2.500E-08 | 4.600E-08 |
| 15092 | 755 | V755K | B | 3.7e-08 | 3.6e-08 | 4.6e-08 | 3.967E-08 | 5.508E-09 | 3.600E-08 | 4.600E-08 |
| 15093 | 755 | V755M | B | 8.4e-09 | 1.9e-08 | 7.2e-08 | 3.313E-08 | 3.407E-08 | 8.400E-09 | 7.200E-08 |
| 15094 | 755 | V755F | B | 5.8e-08 | 4.2e-08 | 4.3e-08 | 4.767E-08 | 8.963E-09 | 4.200E-08 | 5.800E-08 |
| 15095 | 755 | V755P | B | 4.5e-08 | 7.3e-08 | 8.9e-08 | 6.900E-08 | 2.227E-08 | 4.500E-08 | 8.900E-08 |
| 15096 | 755 | V755S | B | 4e-08 | 4.1e-08 | 1.9e-08 | 3.333E-08 | 1.242E-08 | 1.900E-08 | 4.100E-08 |
| 15097 | 755 | V755T | B | 2.6e-08 | 5.9e-08 | 3e-08 | 3.833E-08 | 1.801E-08 | 2.600E-08 | 5.900E-08 |
| 15098 | 755 | V755W | B | 6.3e-08 | 9.1e-08 | 1.2e-07 | 9.133E-08 | 2.850E-08 | 6.300E-08 | 1.200E-07 |
| 15099 | 755 | V755Y | B | 5.2e-08 | 9.7e-08 | 1.9e-08 | 5.600E-08 | 3.915E-08 | 1.900E-08 | 9.700E-08 |
| 15100 | 755 | V755V | B | 5.1e-08 | 5.1e-08 | 7.7e-08 | 5.967E-08 | 1.501E-08 | 5.100E-08 | 7.700E-08 |
| 15101 | 756 | C756A | B | 4.2e-08 | 6e-08 | 1.3e-07 | 7.733E-08 | 4.649E-08 | 4.200E-08 | 1.300E-07 |
| 15102 | 756 | C756R | B | 5.6e-08 | 5.1e-08 | 7.5e-08 | 6.067E-08 | 1.266E-08 | 5.100E-08 | 7.500E-08 |
| 15103 | 756 | C756N | B | 3e-08 | 4.7e-08 | 7.6e-08 | 5.100E-08 | 2.326E-08 | 3.000E-08 | 7.600E-08 |
| 15104 | 756 | C756D | B | 3.2e-08 | 3.2e-08 | 1.8e-08 | 2.733E-08 | 8.083E-09 | 1.800E-08 | 3.200E-08 |
| 15105 | 756 | C756C | B | 2.6e-08 | 5.1e-08 | 4e-08 | 3.900E-08 | 1.253E-08 | 2.600E-08 | 5.100E-08 |
| 15106 | 756 | C756E | B | 7.6e-08 | 8.6e-08 | 2.3e-08 | 6.167E-08 | 3.386E-08 | 2.300E-08 | 8.600E-08 |
| 15107 | 756 | C756Q | B | 5.4e-08 | 6.3e-08 | 2.4e-08 | 4.700E-08 | 2.042E-08 | 2.400E-08 | 6.300E-08 |
| 15108 | 756 | C756G | B | 3.4e-08 | 3.4e-08 | 1.5e-07 | 7.267E-08 | 6.697E-08 | 3.400E-08 | 1.500E-07 |
| 15109 | 756 | C756H | B | 3.1e-08 | 2.2e-08 | 4.3e-08 | 3.200E-08 | 1.054E-08 | 2.200E-08 | 4.300E-08 |
| 15110 | 756 | C756I | B | 1e-07 | 2.5e-08 | 7.4e-08 | 6.633E-08 | 3.808E-08 | 2.500E-08 | 1.000E-07 |
| 15111 | 756 | C756L | B | 6e-08 | 2.1e-08 | 4.2e-08 | 4.100E-08 | 1.952E-08 | 2.100E-08 | 6.000E-08 |
| 15112 | 756 | C756K | B | 4.2e-08 | 2.4e-08 | 4.7e-08 | 3.767E-08 | 1.210E-08 | 2.400E-08 | 4.700E-08 |
| 15113 | 756 | C756M | B | 2.6e-08 | 5.6e-08 | 6.7e-09 | 2.957E-08 | 2.484E-08 | 6.700E-09 | 5.600E-08 |
| 15114 | 756 | C756F | B | 7.8e-08 | 8.2e-08 | 4.6e-08 | 6.867E-08 | 1.973E-08 | 4.600E-08 | 8.200E-08 |
| 15115 | 756 | C756P | B | 4.9e-08 | 4.1e-08 | 2.2e-08 | 3.733E-08 | 1.387E-08 | 2.200E-08 | 4.900E-08 |
| 15116 | 756 | C756S | B | 9.3e-08 | 5.2e-08 | 2.5e-08 | 5.667E-08 | 3.424E-08 | 2.500E-08 | 9.300E-08 |
| 15117 | 756 | C756T | B | 1.1e-07 | 6.2e-08 | 4.3e-08 | 7.167E-08 | 3.453E-08 | 4.300E-08 | 1.100E-07 |
| 15118 | 756 | C756W | B | 6.2e-08 | 2.4e-08 | 3.4e-08 | 4.000E-08 | 1.970E-08 | 2.400E-08 | 6.200E-08 |
| 15119 | 756 | C756Y | B | 5.4e-08 | 4.7e-08 | 2.2e-08 | 4.100E-08 | 1.682E-08 | 2.200E-08 | 5.400E-08 |
| 15120 | 756 | C756V | B | 2e-08 | 3.1e-08 | 6.6e-08 | 3.900E-08 | 2.402E-08 | 2.000E-08 | 6.600E-08 |
| 15121 | 757 | L757A | B | 6.8e-08 | 6.7e-08 | 2e-08 | 5.167E-08 | 2.743E-08 | 2.000E-08 | 6.800E-08 |
| 15122 | 757 | L757R | B | 3.1e-08 | 5.3e-08 | 2.4e-08 | 3.600E-08 | 1.513E-08 | 2.400E-08 | 5.300E-08 |
| 15123 | 757 | L757N | B | 4.2e-08 | 7.9e-08 | 3.6e-08 | 5.233E-08 | 2.329E-08 | 3.600E-08 | 7.900E-08 |
| 15124 | 757 | L757D | B | 5e-08 | 6.8e-08 | 4.4e-08 | 5.400E-08 | 1.249E-08 | 4.400E-08 | 6.800E-08 |
| 15125 | 757 | L757C | B | 2.1e-08 | 6.7e-08 | 2.1e-08 | 3.633E-08 | 2.656E-08 | 2.100E-08 | 6.700E-08 |
| 15126 | 757 | L757E | B | 1.3e-08 | 4.8e-08 | 9.3e-09 | 2.343E-08 | 2.136E-08 | 9.300E-09 | 4.800E-08 |
| 15127 | 757 | L757Q | B | 2.1e-08 | 5.3e-08 | 7.9e-08 | 5.100E-08 | 2.905E-08 | 2.100E-08 | 7.900E-08 |
| 15128 | 757 | L757G | B | 4.1e-08 | 2.7e-08 | 8.6e-08 | 5.133E-08 | 3.083E-08 | 2.700E-08 | 8.600E-08 |
| 15129 | 757 | L757H | B | 4.6e-08 | 3.3e-08 | 5.1e-08 | 4.333E-08 | 9.292E-09 | 3.300E-08 | 5.100E-08 |

| | | | | | | | | | | |
|---|---|---|---|---|---|---|---|---|---|---|
| 15130 | 757 | L757I | B | 4.3e-08 | 1e-07 | 6.5e-08 | 6.933E-08 | 2.875E-08 | 4.300E-08 | 1.000E-07 |
| 15131 | 757 | L757L | B | 4.7e-08 | 5.7e-08 | 5.1e-08 | 5.167E-08 | 5.033E-09 | 4.700E-08 | 5.700E-08 |
| 15132 | 757 | L757K | B | 3.1e-08 | 3.4e-08 | 3.1e-08 | 3.200E-08 | 1.732E-09 | 3.100E-08 | 3.400E-08 |
| 15133 | 757 | L757M | B | 7.9e-08 | 1e-07 | 6.4e-08 | 8.100E-08 | 1.808E-08 | 6.400E-08 | 1.000E-07 |
| 15134 | 757 | L757F | B | 5.5e-08 | 1.6e-08 | 5.7e-08 | 4.267E-08 | 2.312E-08 | 1.600E-08 | 5.700E-08 |
| 15135 | 757 | L757P | B | 8.8e-08 | 1e-08 | 2.5e-08 | 4.100E-08 | 4.139E-08 | 1.000E-08 | 8.800E-08 |
| 15136 | 757 | L757S | B | 3.9e-08 | 1.6e-08 | 2.6e-08 | 2.700E-08 | 1.153E-08 | 1.600E-08 | 3.900E-08 |
| 15137 | 757 | L757T | B | 5.4e-08 | 2.7e-08 | 2.4e-08 | 3.500E-08 | 1.652E-08 | 2.400E-08 | 5.400E-08 |
| 15138 | 757 | L757W | B | 2.1e-08 | 7e-08 | 5.7e-08 | 4.933E-08 | 2.538E-08 | 2.100E-08 | 7.000E-08 |
| 15139 | 757 | L757Y | B | 6.5e-08 | 5e-08 | 4.3e-08 | 5.267E-08 | 1.124E-08 | 4.300E-08 | 6.500E-08 |
| 15140 | 757 | L757V | B | 1.1e-07 | 6.7e-08 | 2.7e-08 | 6.800E-08 | 4.151E-08 | 2.700E-08 | 1.100E-07 |
| 15141 | 758 | L758A | B | 1.3e-08 | 1.3e-08 | 3.9e-08 | 2.167E-08 | 1.501E-08 | 1.300E-08 | 3.900E-08 |
| 15142 | 758 | L758R | B | 6.5e-08 | 9.6e-08 | 5.8e-08 | 7.300E-08 | 2.022E-08 | 5.800E-08 | 9.600E-08 |
| 15143 | 758 | L758N | B | 4.1e-08 | 2.4e-08 | 1.3e-08 | 2.600E-08 | 1.411E-08 | 1.300E-08 | 4.100E-08 |
| 15144 | 758 | L758D | B | 1.8e-07 | 4.5e-08 | 7.8e-08 | 1.010E-07 | 7.038E-08 | 4.500E-08 | 1.800E-07 |
| 15145 | 758 | L758C | B | 2.2e-08 | 3.5e-08 | 5e-08 | 3.567E-08 | 1.401E-08 | 2.200E-08 | 5.000E-08 |
| 15146 | 758 | L758E | B | 3.5e-08 | 4e-08 | 2.9e-08 | 3.467E-08 | 5.508E-09 | 2.900E-08 | 4.000E-08 |
| 15147 | 758 | L758Q | B | 5e-08 | 3e-08 | 4.7e-08 | 4.233E-08 | 1.079E-08 | 3.000E-08 | 5.000E-08 |
| 15148 | 758 | L758G | B | 7.1e-08 | 3.5e-08 | 3.3e-08 | 4.633E-08 | 2.139E-08 | 3.300E-08 | 7.100E-08 |
| 15149 | 758 | L758H | B | 2.1e-08 | 2.4e-08 | 2.3e-08 | 2.267E-08 | 1.528E-09 | 2.100E-08 | 2.400E-08 |
| 15150 | 758 | L758I | B | 7.6e-08 | 5.8e-08 | 4.7e-08 | 6.033E-08 | 1.464E-08 | 4.700E-08 | 7.600E-08 |
| 15151 | 758 | L758L | B | 3.1e-08 | 6.5e-08 | 5.9e-08 | 5.167E-08 | 1.815E-08 | 3.100E-08 | 6.500E-08 |
| 15152 | 758 | L758K | B | 3e-08 | 6.4e-08 | 6.9e-08 | 5.433E-08 | 2.122E-08 | 3.000E-08 | 6.900E-08 |
| 15153 | 758 | L758M | B | 2e-08 | 7.3e-08 | 5.6e-08 | 4.967E-08 | 2.706E-08 | 2.000E-08 | 7.300E-08 |
| 15154 | 758 | L758F | B | 7.2e-08 | 3.6e-08 | 6.7e-08 | 5.833E-08 | 1.950E-08 | 3.600E-08 | 7.200E-08 |
| 15155 | 758 | L758P | B | 2.8e-08 | 9.8e-08 | 2e-08 | 4.867E-08 | 4.291E-08 | 2.000E-08 | 9.800E-08 |
| 15156 | 758 | L758S | B | 8.3e-08 | 4.9e-08 | 5.2e-08 | 6.133E-08 | 1.882E-08 | 4.900E-08 | 8.300E-08 |
| 15157 | 758 | L758T | B | 1.3e-08 | 3.2e-08 | 1e-07 | 4.833E-08 | 4.574E-08 | 1.300E-08 | 1.000E-07 |
| 15158 | 758 | L758W | B | 6.3e-08 | 3.9e-08 | 6.8e-08 | 5.667E-08 | 1.550E-08 | 3.900E-08 | 6.800E-08 |
| 15159 | 758 | L758Y | B | 5.9e-08 | 2e-08 | 4.3e-08 | 4.067E-08 | 1.960E-08 | 2.000E-08 | 5.900E-08 |
| 15160 | 758 | L758V | B | 5.6e-08 | 1.9e-07 | 6.4e-08 | 1.033E-07 | 7.516E-08 | 5.600E-08 | 1.900E-07 |
| 15161 | 759 | N759A | B | 3.4e-08 | 6.5e-08 | 2.4e-08 | 4.100E-08 | 2.138E-08 | 2.400E-08 | 6.500E-08 |
| 15162 | 759 | N759R | B | 3.1e-08 | 1.6e-08 | 1.2e-07 | 5.567E-08 | 5.622E-08 | 1.600E-08 | 1.200E-07 |
| 15163 | 759 | N759N | B | 4.1e-08 | 3.7e-08 | 4.7e-08 | 4.167E-08 | 5.033E-09 | 3.700E-08 | 4.700E-08 |
| 15164 | 759 | N759D | B | 9.3e-08 | 1.7e-08 | 5.4e-08 | 5.467E-08 | 3.800E-08 | 1.700E-08 | 9.300E-08 |
| 15165 | 759 | N759C | B | 5e-08 | 2.1e-08 | 4e-08 | 3.700E-08 | 1.473E-08 | 2.100E-08 | 5.000E-08 |
| 15166 | 759 | N759E | B | 6.6e-08 | 7.7e-08 | 6.4e-08 | 6.900E-08 | 7.000E-09 | 6.400E-08 | 7.700E-08 |
| 15167 | 759 | N759Q | B | 6.5e-08 | 1.4e-07 | 1.3e-08 | 7.267E-08 | 6.385E-08 | 1.300E-08 | 1.400E-07 |
| 15168 | 759 | N759G | B | 1.5e-08 | 4e-08 | 6.3e-08 | 3.933E-08 | 2.401E-08 | 1.500E-08 | 6.300E-08 |
| 15169 | 759 | N759H | B | 2.7e-08 | 3.8e-08 | 3.4e-08 | 3.300E-08 | 5.568E-09 | 2.700E-08 | 3.800E-08 |
| 15170 | 759 | N759I | B | 3.5e-08 | 4.6e-08 | 1.8e-08 | 3.300E-08 | 1.411E-08 | 1.800E-08 | 4.600E-08 |
| 15171 | 759 | N759L | B | 2.2e-08 | 7.5e-08 | 2.1e-07 | 1.023E-07 | 9.693E-08 | 2.200E-08 | 2.100E-07 |
| 15172 | 759 | N759K | B | 1.2e-07 | 8.9e-08 | 4.2e-08 | 8.367E-08 | 3.927E-08 | 4.200E-08 | 1.200E-07 |
| 15173 | 759 | N759M | B | 2.5e-08 | 5.2e-08 | 2.2e-08 | 3.300E-08 | 1.652E-08 | 2.200E-08 | 5.200E-08 |
| 15174 | 759 | N759F | B | 6.2e-08 | 4.2e-08 | 3.3e-08 | 4.567E-08 | 1.484E-08 | 3.300E-08 | 6.200E-08 |
| 15175 | 759 | N759P | B | 3e-08 | 2.9e-08 | 6.8e-08 | 4.233E-08 | 2.223E-08 | 2.900E-08 | 6.800E-08 |
| 15176 | 759 | N759S | B | 6.3e-08 | 1.9e-08 | 3.1e-08 | 3.767E-08 | 2.274E-08 | 1.900E-08 | 6.300E-08 |
| 15177 | 759 | N759T | B | 3.2e-08 | 7.6e-08 | 4.1e-08 | 4.967E-08 | 2.325E-08 | 3.200E-08 | 7.600E-08 |
| 15178 | 759 | N759W | B | 3.6e-08 | 7.6e-08 | 2.2e-08 | 4.467E-08 | 2.802E-08 | 2.200E-08 | 7.600E-08 |
| 15179 | 759 | N759Y | B | 5.6e-08 | 4.7e-08 | 7.5e-09 | 3.683E-08 | 2.580E-08 | 7.500E-09 | 5.600E-08 |
| 15180 | 759 | N759V | B | 2.8e-08 | 5.4e-08 | 3.5e-08 | 3.900E-08 | 1.345E-08 | 2.800E-08 | 5.400E-08 |
| 15181 | 760 | N760A | B | 2.9e-08 | 4.5e-08 | 1.1e-07 | 6.133E-08 | 4.290E-08 | 2.900E-08 | 1.100E-07 |
| 15182 | 760 | N760R | B | 2.5e-08 | 4.3e-08 | 2.8e-08 | 3.200E-08 | 9.644E-09 | 2.500E-08 | 4.300E-08 |
| 15183 | 760 | N760N | B | 3.4e-08 | 3.3e-08 | 1.7e-07 | 7.900E-08 | 7.881E-08 | 3.300E-08 | 1.700E-07 |
| 15184 | 760 | N760D | B | 5.1e-08 | 7.9e-08 | 5.7e-08 | 6.233E-08 | 1.474E-08 | 5.100E-08 | 7.900E-08 |
| 15185 | 760 | N760C | B | 3.6e-08 | 3.1e-08 | 4.2e-08 | 3.633E-08 | 5.508E-09 | 3.100E-08 | 4.200E-08 |
| 15186 | 760 | N760E | B | 2.9e-08 | 4e-08 | 2.1e-08 | 3.000E-08 | 9.539E-09 | 2.100E-08 | 4.000E-08 |
| 15187 | 760 | N760Q | B | 4e-08 | 4.5e-08 | 2.1e-08 | 3.533E-08 | 1.266E-08 | 2.100E-08 | 4.500E-08 |
| 15188 | 760 | N760G | B | 1.1e-08 | 5.6e-08 | 5.5e-08 | 4.067E-08 | 2.570E-08 | 1.100E-08 | 5.600E-08 |
| 15189 | 760 | N760H | B | 1.6e-08 | 5e-08 | 6.3e-08 | 4.300E-08 | 2.427E-08 | 1.600E-08 | 6.300E-08 |
| 15190 | 760 | N760I | B | 5.1e-08 | 4.3e-08 | 6.2e-08 | 5.200E-08 | 9.539E-09 | 4.300E-08 | 6.200E-08 |
| 15191 | 760 | N760L | B | 2.7e-08 | 9.2e-08 | 5.4e-08 | 5.767E-08 | 3.265E-08 | 2.700E-08 | 9.200E-08 |
| 15192 | 760 | N760K | B | 4.3e-08 | 5.1e-08 | 2.3e-08 | 3.900E-08 | 1.442E-08 | 2.300E-08 | 5.100E-08 |
| 15193 | 760 | N760M | B | 4.3e-08 | 1.2e-07 | 4.8e-08 | 7.033E-08 | 4.309E-08 | 4.300E-08 | 1.200E-07 |
| 15194 | 760 | N760F | B | 2.9e-08 | 7.1e-08 | 7.7e-08 | 5.900E-08 | 2.615E-08 | 2.900E-08 | 7.700E-08 |
| 15195 | 760 | N760P | B | 9.3e-08 | 4.7e-08 | 1.6e-08 | 5.200E-08 | 3.874E-08 | 1.600E-08 | 9.300E-08 |
| 15196 | 760 | N760S | B | 1.2e-07 | 1.7e-08 | 1.5e-08 | 5.067E-08 | 6.005E-08 | 1.500E-08 | 1.200E-07 |
| 15197 | 760 | N760T | B | 2.9e-08 | 3.3e-08 | 4.9e-08 | 3.700E-08 | 1.058E-08 | 2.900E-08 | 4.900E-08 |
| 15198 | 760 | N760W | B | 7.1e-08 | 4e-08 | 3e-08 | 4.700E-08 | 2.138E-08 | 3.000E-08 | 7.100E-08 |
| 15199 | 760 | N760Y | B | 2.1e-08 | 2.9e-08 | 1.1e-07 | 5.333E-08 | 4.924E-08 | 2.100E-08 | 1.100E-07 |
| 15200 | 760 | N760V | B | 5.4e-08 | 5.1e-08 | 4.4e-08 | 4.967E-08 | 5.132E-09 | 4.400E-08 | 5.400E-08 |
| 15201 | 761 | F761A | B | 4e-08 | 3.6e-08 | 9.1e-08 | 5.567E-08 | 3.066E-08 | 3.600E-08 | 9.100E-08 |
| 15202 | 761 | F761R | B | 2.4e-08 | 2.5e-08 | 6.2e-08 | 3.700E-08 | 2.166E-08 | 2.400E-08 | 6.200E-08 |
| 15203 | 761 | F761N | B | 5e-08 | 4.7e-08 | 1.3e-08 | 3.667E-08 | 2.055E-08 | 1.300E-08 | 5.000E-08 |
| 15204 | 761 | F761D | B | 4.2e-08 | 3.8e-08 | 4.3e-08 | 4.100E-08 | 2.646E-09 | 3.800E-08 | 4.300E-08 |
| 15205 | 761 | F761C | B | 1.3e-08 | 2.4e-08 | 4.2e-08 | 2.633E-08 | 1.464E-08 | 1.300E-08 | 4.200E-08 |
| 15206 | 761 | F761E | B | 5.1e-08 | 2e-08 | 4.8e-08 | 3.967E-08 | 1.710E-08 | 2.000E-08 | 5.100E-08 |
| 15207 | 761 | F761Q | B | 2.9e-08 | 5.4e-08 | 3.9e-08 | 4.067E-08 | 1.258E-08 | 2.900E-08 | 5.400E-08 |
| 15208 | 761 | F761G | B | 1.1e-07 | 2.1e-08 | 6.7e-08 | 6.600E-08 | 4.451E-08 | 2.100E-08 | 1.100E-07 |
| 15209 | 761 | F761H | B | 3.2e-08 | 5.6e-08 | 7.5e-08 | 5.433E-08 | 2.155E-08 | 3.200E-08 | 7.500E-08 |
| 15210 | 761 | F761I | B | 8.1e-08 | 3.1e-08 | 1.1e-07 | 7.400E-08 | 3.996E-08 | 3.100E-08 | 1.100E-07 |
| 15211 | 761 | F761L | B | 4.8e-08 | 3.4e-08 | 4.8e-08 | 4.333E-08 | 8.083E-09 | 3.400E-08 | 4.800E-08 |
| 15212 | 761 | F761K | B | 8.2e-08 | 1.9e-08 | 5.4e-08 | 5.167E-08 | 3.156E-08 | 1.900E-08 | 8.200E-08 |
| 15213 | 761 | F761M | B | 2e-08 | 5.3e-08 | 6e-08 | 4.433E-08 | 2.136E-08 | 2.000E-08 | 6.000E-08 |
| 15214 | 761 | F761F | B | 3.5e-08 | 2.1e-08 | 2.2e-08 | 2.600E-08 | 7.810E-09 | 2.100E-08 | 3.500E-08 |

| | | | | | | | | | | |
|---|---|---|---|---|---|---|---|---|---|---|
| 15215 | 761 | F761P | B | 4.5e-08 | 9.4e-09 | 4.2e-08 | 3.213E-08 | 1.974E-08 | 9.400E-09 | 4.500E-08 |
| 15216 | 761 | F761S | B | 5e-08 | 3.3e-08 | 4.9e-08 | 4.400E-08 | 9.539E-09 | 3.300E-08 | 5.000E-08 |
| 15217 | 761 | F761T | B | 2.7e-08 | 9e-09 | 4e-08 | 2.533E-08 | 1.557E-08 | 9.000E-09 | 4.000E-08 |
| 15218 | 761 | F761W | B | 1.1e-07 | 7.4e-08 | 2.9e-08 | 7.100E-08 | 4.058E-08 | 2.900E-08 | 1.100E-07 |
| 15219 | 761 | F761Y | B | 2.8e-08 | 4.5e-08 | 3.6e-08 | 3.633E-08 | 8.505E-09 | 2.800E-08 | 4.500E-08 |
| 15220 | 761 | F761V | B | 1.7e-08 | 6.5e-08 | 2.7e-08 | 3.633E-08 | 2.532E-08 | 1.700E-08 | 6.500E-08 |
| 15221 | 762 | Y762A | B | 4.5e-08 | 7.7e-08 | 5.9e-08 | 6.033E-08 | 1.604E-08 | 4.500E-08 | 7.700E-08 |
| 15222 | 762 | Y762R | B | 7.4e-08 | 5.1e-08 | 6.1e-08 | 6.200E-08 | 1.153E-08 | 5.100E-08 | 7.400E-08 |
| 15223 | 762 | Y762N | B | 3e-08 | 1.1e-07 | 1.2e-07 | 8.667E-08 | 4.933E-08 | 3.000E-08 | 1.200E-07 |
| 15224 | 762 | Y762D | B | 3.5e-08 | 3e-08 | 5e-08 | 3.833E-08 | 1.041E-08 | 3.000E-08 | 5.000E-08 |
| 15225 | 762 | Y762C | B | 3.6e-08 | 8.3e-09 | 3.1e-08 | 2.510E-08 | 1.476E-08 | 8.300E-09 | 3.600E-08 |
| 15226 | 762 | Y762E | B | 5.7e-08 | 7.7e-08 | 3.2e-08 | 5.533E-08 | 2.255E-08 | 3.200E-08 | 7.700E-08 |
| 15227 | 762 | Y762Q | B | 6.8e-09 | 2e-08 | 7.1e-08 | 3.260E-08 | 3.390E-08 | 6.800E-09 | 7.100E-08 |
| 15228 | 762 | Y762G | B | 2.9e-08 | 2.9e-08 | 5.8e-08 | 3.867E-08 | 1.674E-08 | 2.900E-08 | 5.800E-08 |
| 15229 | 762 | Y762H | B | 9e-08 | 1.7e-07 | 1.7e-08 | 9.233E-08 | 7.653E-08 | 1.700E-08 | 1.700E-07 |
| 15230 | 762 | Y762I | B | 1.2e-07 | 7.8e-08 | 5.1e-08 | 8.300E-08 | 3.477E-08 | 5.100E-08 | 1.200E-07 |
| 15231 | 762 | Y762L | B | 3.1e-08 | 6.6e-08 | 2.8e-07 | 1.257E-07 | 1.348E-07 | 3.100E-08 | 2.800E-07 |
| 15232 | 762 | Y762K | B | 7.4e-08 | 3.4e-08 | 6.4e-08 | 5.733E-08 | 2.082E-08 | 3.400E-08 | 7.400E-08 |
| 15233 | 762 | Y762M | B | 4.9e-08 | 1.9e-08 | 9.1e-08 | 5.300E-08 | 3.617E-08 | 1.900E-08 | 9.100E-08 |
| 15234 | 762 | Y762F | B | 4.7e-08 | 8.1e-08 | 2.6e-08 | 5.133E-08 | 2.775E-08 | 2.600E-08 | 8.100E-08 |
| 15235 | 762 | Y762P | B | 3.7e-08 | 3.6e-08 | 3.7e-08 | 3.667E-08 | 5.774E-10 | 3.600E-08 | 3.700E-08 |
| 15236 | 762 | Y762S | B | 4.3e-08 | 5e-08 | 4.3e-08 | 4.533E-08 | 4.041E-09 | 4.300E-08 | 5.000E-08 |
| 15237 | 762 | Y762T | B | 5.2e-08 | 3.2e-08 | 1.8e-08 | 3.400E-08 | 1.709E-08 | 1.800E-08 | 5.200E-08 |
| 15238 | 762 | Y762W | B | 6e-08 | 8.6e-08 | 6.7e-08 | 7.100E-08 | 1.345E-08 | 6.000E-08 | 8.600E-08 |
| 15239 | 762 | Y762Y | B | 4.3e-08 | 7.6e-08 | 2.6e-08 | 4.833E-08 | 2.542E-08 | 2.600E-08 | 7.600E-08 |
| 15240 | 762 | Y762V | B | 8.6e-08 | 4.5e-08 | 1.2e-07 | 8.367E-08 | 3.755E-08 | 4.500E-08 | 1.200E-07 |
| 15241 | 763 | P763A | B | 4.7e-08 | 5.3e-08 | 5.8e-08 | 5.267E-08 | 5.508E-09 | 4.700E-08 | 5.800E-08 |
| 15242 | 763 | P763R | B | 3.7e-08 | 2.7e-08 | 6.3e-08 | 4.233E-08 | 1.858E-08 | 2.700E-08 | 6.300E-08 |
| 15243 | 763 | P763N | B | 6.6e-08 | 3.7e-08 | 2.7e-08 | 4.333E-08 | 2.026E-08 | 2.700E-08 | 6.600E-08 |
| 15244 | 763 | P763D | B | 1.9e-08 | 6.4e-08 | 6.1e-08 | 4.800E-08 | 2.516E-08 | 1.900E-08 | 6.400E-08 |
| 15245 | 763 | P763C | B | 3.6e-08 | 3.4e-08 | 8.8e-08 | 5.267E-08 | 3.062E-08 | 3.400E-08 | 8.800E-08 |
| 15246 | 763 | P763E | B | 3.5e-08 | 3.7e-08 | 6.6e-08 | 4.600E-08 | 1.735E-08 | 3.500E-08 | 6.600E-08 |
| 15247 | 763 | P763Q | B | 2.2e-08 | 3.5e-08 | 5.1e-08 | 3.600E-08 | 1.453E-08 | 2.200E-08 | 5.100E-08 |
| 15248 | 763 | P763G | B | 4.1e-08 | 3.9e-08 | 5.9e-08 | 4.633E-08 | 1.102E-08 | 3.900E-08 | 5.900E-08 |
| 15249 | 763 | P763H | B | 3.2e-08 | 6e-08 | 3.8e-08 | 4.333E-08 | 1.474E-08 | 3.200E-08 | 6.000E-08 |
| 15250 | 763 | P763I | B | 1e-07 | 1.3e-08 | 6.6e-08 | 5.967E-08 | 4.384E-08 | 1.300E-08 | 1.000E-07 |
| 15251 | 763 | P763L | B | 2.5e-08 | 1.2e-07 | 1e-07 | 8.167E-08 | 5.008E-08 | 2.500E-08 | 1.200E-07 |
| 15252 | 763 | P763K | B | 9.8e-08 | 8e-08 | 1.7e-08 | 6.500E-08 | 4.253E-08 | 1.700E-08 | 9.800E-08 |
| 15253 | 763 | P763M | B | 5e-08 | 4.8e-08 | 6.1e-08 | 5.300E-08 | 7.000E-09 | 4.800E-08 | 6.100E-08 |
| 15254 | 763 | P763F | B | 5e-08 | 2.3e-08 | 2.6e-08 | 3.300E-08 | 1.480E-08 | 2.300E-08 | 5.000E-08 |
| 15255 | 763 | P763P | B | 3.7e-08 | 1.8e-07 | 3.2e-08 | 8.300E-08 | 8.404E-08 | 3.200E-08 | 1.800E-07 |
| 15256 | 763 | P763S | B | 3.9e-08 | 6.9e-08 | 6.6e-08 | 5.800E-08 | 1.652E-08 | 3.900E-08 | 6.900E-08 |
| 15257 | 763 | P763T | B | 8e-08 | 4.9e-08 | 1.2e-07 | 8.300E-08 | 3.559E-08 | 4.900E-08 | 1.200E-07 |
| 15258 | 763 | P763W | B | 4.2e-08 | 2.4e-08 | 7e-08 | 4.533E-08 | 2.318E-08 | 2.400E-08 | 7.000E-08 |
| 15259 | 763 | P763Y | B | 2.5e-08 | 3.8e-08 | 1.5e-08 | 2.600E-08 | 1.153E-08 | 1.500E-08 | 3.800E-08 |
| 15260 | 763 | P763V | B | 6.6e-08 | 1.3e-07 | 1.5e-08 | 7.033E-08 | 5.762E-08 | 1.500E-08 | 1.300E-07 |
| 15261 | 764 | R764A | B | 8.1e-08 | 4.1e-08 | 2.5e-08 | 4.900E-08 | 2.884E-08 | 2.500E-08 | 8.100E-08 |
| 15262 | 764 | R764R | B | 4.1e-08 | 5.2e-08 | 4.1e-08 | 4.467E-08 | 6.351E-09 | 4.100E-08 | 5.200E-08 |
| 15263 | 764 | R764N | B | 1.6e-08 | 5.8e-08 | 1.7e-07 | 8.133E-08 | 7.961E-08 | 1.600E-08 | 1.700E-07 |
| 15264 | 764 | R764D | B | 8.3e-08 | 5.7e-08 | 4.6e-08 | 6.200E-08 | 1.900E-08 | 4.600E-08 | 8.300E-08 |
| 15265 | 764 | R764C | B | 2.7e-08 | 7.1e-08 | 3.3e-08 | 4.367E-08 | 2.386E-08 | 2.700E-08 | 7.100E-08 |
| 15266 | 764 | R764E | B | 6.8e-08 | 1.8e-08 | 8.8e-08 | 5.800E-08 | 3.606E-08 | 1.800E-08 | 8.800E-08 |
| 15267 | 764 | R764Q | B | 5.6e-08 | 3.9e-08 | 4.7e-08 | 4.733E-08 | 8.505E-09 | 3.900E-08 | 5.600E-08 |
| 15268 | 764 | R764G | B | 6.3e-09 | 2.5e-08 | 4e-08 | 2.377E-08 | 1.688E-08 | 6.300E-09 | 4.000E-08 |
| 15269 | 764 | R764H | B | 1.6e-08 | 7.4e-08 | 3.2e-08 | 4.067E-08 | 2.996E-08 | 1.600E-08 | 7.400E-08 |
| 15270 | 764 | R764I | B | 4.4e-08 | 8e-08 | 4.8e-08 | 5.733E-08 | 1.973E-08 | 4.400E-08 | 8.000E-08 |
| 15271 | 764 | R764L | B | 7.2e-08 | 2.1e-08 | 4.3e-08 | 4.533E-08 | 2.558E-08 | 2.100E-08 | 7.200E-08 |
| 15272 | 764 | R764K | B | 2.8e-08 | 2.3e-08 | 2.4e-08 | 2.500E-08 | 2.646E-09 | 2.300E-08 | 2.800E-08 |
| 15273 | 764 | R764M | B | 1.4e-08 | 6.5e-08 | 4.6e-08 | 4.167E-08 | 2.577E-08 | 1.400E-08 | 6.500E-08 |
| 15274 | 764 | R764F | B | 3.3e-08 | 3.5e-08 | 6.4e-08 | 4.400E-08 | 1.735E-08 | 3.300E-08 | 6.400E-08 |
| 15275 | 764 | R764P | B | 8.8e-08 | 5.3e-08 | 6.9e-08 | 7.000E-08 | 1.752E-08 | 5.300E-08 | 8.800E-08 |
| 15276 | 764 | R764S | B | 2.2e-08 | 1.8e-08 | 4.6e-08 | 2.867E-08 | 1.514E-08 | 1.800E-08 | 4.600E-08 |
| 15277 | 764 | R764T | B | 6.9e-08 | 2e-08 | 5.2e-08 | 4.700E-08 | 2.488E-08 | 2.000E-08 | 6.900E-08 |
| 15278 | 764 | R764W | B | 1.4e-08 | 7.7e-08 | 7.3e-08 | 5.467E-08 | 3.528E-08 | 1.400E-08 | 7.700E-08 |
| 15279 | 764 | R764Y | B | 1.1e-07 | 6.5e-08 | 1.3e-08 | 6.267E-08 | 4.854E-08 | 1.300E-08 | 1.100E-07 |
| 15280 | 764 | R764V | B | 4.4e-08 | 1.9e-08 | 7.5e-08 | 4.600E-08 | 2.805E-08 | 1.900E-08 | 7.500E-08 |
| 15281 | 765 | E765A | B | 5.6e-08 | 7.6e-08 | 1.2e-08 | 4.800E-08 | 3.274E-08 | 1.200E-08 | 7.600E-08 |
| 15282 | 765 | E765R | B | 7.1e-08 | 3.4e-08 | 3.1e-08 | 4.533E-08 | 2.228E-08 | 3.100E-08 | 7.100E-08 |
| 15283 | 765 | E765N | B | 4e-08 | 6.3e-08 | 1.8e-08 | 4.033E-08 | 2.250E-08 | 1.800E-08 | 6.300E-08 |
| 15284 | 765 | E765D | B | 1.1e-08 | 5.2e-08 | 3.9e-08 | 3.400E-08 | 2.095E-08 | 1.100E-08 | 5.200E-08 |
| 15285 | 765 | E765C | B | 2.9e-08 | 2.6e-08 | 2.8e-08 | 2.767E-08 | 1.528E-09 | 2.600E-08 | 2.900E-08 |
| 15286 | 765 | E765E | B | 2.1e-08 | 3.2e-08 | 5.2e-08 | 3.500E-08 | 1.572E-08 | 2.100E-08 | 5.200E-08 |
| 15287 | 765 | E765Q | B | 3e-08 | 4.6e-08 | 5e-08 | 4.200E-08 | 1.058E-08 | 3.000E-08 | 5.000E-08 |
| 15288 | 765 | E765G | B | 9.5e-08 | 1.8e-08 | 4e-08 | 5.100E-08 | 3.966E-08 | 1.800E-08 | 9.500E-08 |
| 15289 | 765 | E765H | B | 6.9e-08 | 6.3e-08 | 4.8e-08 | 6.000E-08 | 1.082E-08 | 4.800E-08 | 6.900E-08 |
| 15290 | 765 | E765I | B | 2.7e-08 | 3.2e-08 | 6.6e-08 | 4.167E-08 | 2.122E-08 | 2.700E-08 | 6.600E-08 |
| 15291 | 765 | E765L | B | 3e-08 | 6e-08 | 2.4e-08 | 3.800E-08 | 1.929E-08 | 2.400E-08 | 6.000E-08 |
| 15292 | 765 | E765K | B | 4.8e-08 | 3.7e-08 | 4e-08 | 4.167E-08 | 5.686E-09 | 3.700E-08 | 4.800E-08 |
| 15293 | 765 | E765M | B | 2.3e-08 | 1.2e-08 | 3.1e-08 | 2.200E-08 | 9.539E-09 | 1.200E-08 | 3.100E-08 |
| 15294 | 765 | E765F | B | 6.6e-08 | 2.7e-08 | 5.1e-08 | 4.800E-08 | 1.967E-08 | 2.700E-08 | 6.600E-08 |
| 15295 | 765 | E765P | B | 3.7e-08 | 3.7e-08 | 3.3e-08 | 3.567E-08 | 2.309E-09 | 3.300E-08 | 3.700E-08 |
| 15296 | 765 | E765S | B | 3.9e-08 | 7.8e-08 | 2.2e-08 | 4.633E-08 | 2.871E-08 | 2.200E-08 | 7.800E-08 |
| 15297 | 765 | E765T | B | 4.8e-08 | 3.9e-08 | 3.7e-08 | 4.133E-08 | 5.859E-09 | 3.700E-08 | 4.800E-08 |
| 15298 | 765 | E765W | B | 2.6e-08 | 3.2e-08 | 4.3e-08 | 3.367E-08 | 8.622E-09 | 2.600E-08 | 4.300E-08 |
| 15299 | 765 | E765Y | B | 5.7e-08 | 7.3e-08 | 2.5e-08 | 5.167E-08 | 2.444E-08 | 2.500E-08 | 7.300E-08 |

| | | | | | | | | | | |
|---|---|---|---|---|---|---|---|---|---|---|
| 15300 | 765 | E765V | B | 6e-08 | 4.7e-08 | 4.4e-08 | 5.033E-08 | 8.505E-09 | 4.400E-08 | 6.000E-08 |
| 15301 | 766 | A766A | B | 5.1e-08 | 4.4e-08 | 5.4e-08 | 4.967E-08 | 5.132E-09 | 4.400E-08 | 5.400E-08 |
| 15302 | 766 | A766R | B | 3.5e-08 | 7.8e-08 | 4.4e-08 | 5.233E-08 | 2.268E-08 | 3.500E-08 | 7.800E-08 |
| 15303 | 766 | A766N | B | 5.4e-08 | 8.5e-08 | 7.2e-08 | 7.033E-08 | 1.557E-08 | 5.400E-08 | 8.500E-08 |
| 15304 | 766 | A766D | B | 4.1e-08 | 9.6e-08 | 3.4e-08 | 5.700E-08 | 3.396E-08 | 3.400E-08 | 9.600E-08 |
| 15305 | 766 | A766C | B | 3.4e-08 | 5e-08 | 6e-08 | 4.800E-08 | 1.311E-08 | 3.400E-08 | 6.000E-08 |
| 15306 | 766 | A766E | B | 3.5e-08 | 1e-07 | 9.2e-08 | 7.567E-08 | 3.544E-08 | 3.500E-08 | 1.000E-07 |
| 15307 | 766 | A766Q | B | 8.1e-08 | 6.9e-08 | 6e-08 | 7.000E-08 | 1.054E-08 | 6.000E-08 | 8.100E-08 |
| 15308 | 766 | A766G | B | 2e-07 | 4.9e-08 | 5.5e-08 | 1.013E-07 | 8.550E-08 | 4.900E-08 | 2.000E-07 |
| 15309 | 766 | A766H | B | 8.6e-08 | 5.9e-08 | 2.1e-08 | 5.533E-08 | 3.265E-08 | 2.100E-08 | 8.600E-08 |
| 15310 | 766 | A766I | B | 4e-08 | 2.1e-08 | 4e-08 | 3.367E-08 | 1.097E-08 | 2.100E-08 | 4.000E-08 |
| 15311 | 766 | A766L | B | 5.6e-08 | 2.8e-08 | 4.3e-08 | 4.233E-08 | 1.401E-08 | 2.800E-08 | 5.600E-08 |
| 15312 | 766 | A766K | B | 3.9e-08 | 2.5e-08 | 1.3e-08 | 2.567E-08 | 1.301E-08 | 1.300E-08 | 3.900E-08 |
| 15313 | 766 | A766M | B | 2.9e-08 | 7.7e-08 | 3.6e-08 | 4.733E-08 | 2.593E-08 | 2.900E-08 | 7.700E-08 |
| 15314 | 766 | A766F | B | 5.3e-08 | 8.1e-08 | 2.7e-08 | 5.367E-08 | 2.701E-08 | 2.700E-08 | 8.100E-08 |
| 15315 | 766 | A766P | B | 7e-08 | 4.9e-08 | 1.3e-07 | 8.300E-08 | 4.204E-08 | 4.900E-08 | 1.300E-07 |
| 15316 | 766 | A766S | B | 1.6e-07 | 4.7e-08 | 1.4e-08 | 7.367E-08 | 7.657E-08 | 1.400E-08 | 1.600E-07 |
| 15317 | 766 | A766T | B | 3.3e-08 | 2.8e-08 | 4e-08 | 3.367E-08 | 6.028E-09 | 2.800E-08 | 4.000E-08 |
| 15318 | 766 | A766W | B | 2.2e-08 | 1.2e-07 | 3.3e-08 | 5.833E-08 | 5.369E-08 | 2.200E-08 | 1.200E-07 |
| 15319 | 766 | A766Y | B | 4.1e-08 | 3.7e-08 | 1.9e-08 | 3.233E-08 | 1.172E-08 | 1.900E-08 | 4.100E-08 |
| 15320 | 766 | A766V | B | 6.4e-08 | 2.9e-08 | 3.9e-08 | 4.400E-08 | 1.803E-08 | 2.900E-08 | 6.400E-08 |
| 15321 | 767 | K767A | B | 3.4e-08 | 1.1e-07 | 6.9e-08 | 7.100E-08 | 3.804E-08 | 3.400E-08 | 1.100E-07 |
| 15322 | 767 | K767R | B | 5.7e-08 | 7.4e-08 | 6e-08 | 6.367E-08 | 9.074E-09 | 5.700E-08 | 7.400E-08 |
| 15323 | 767 | K767N | B | 2.6e-08 | 4.9e-08 | 1e-07 | 5.833E-08 | 3.787E-08 | 2.600E-08 | 1.000E-07 |
| 15324 | 767 | K767D | B | 7.2e-08 | 1.7e-08 | 5.9e-08 | 4.933E-08 | 2.875E-08 | 1.700E-08 | 7.200E-08 |
| 15325 | 767 | K767C | B | 7.7e-08 | 3.6e-08 | 4e-08 | 5.100E-08 | 2.261E-08 | 3.600E-08 | 7.700E-08 |
| 15326 | 767 | K767E | B | 3.4e-08 | 7.6e-08 | 8e-08 | 6.333E-08 | 2.548E-08 | 3.400E-08 | 8.000E-08 |
| 15327 | 767 | K767Q | B | 2.2e-08 | 1.6e-08 | 2.8e-08 | 2.200E-08 | 6.000E-09 | 1.600E-08 | 2.800E-08 |
| 15328 | 767 | K767G | B | 3.9e-08 | 3.9e-08 | 1.6e-08 | 3.133E-08 | 1.328E-08 | 1.600E-08 | 3.900E-08 |
| 15329 | 767 | K767H | B | 1.2e-08 | 1.7e-08 | 8.3e-08 | 3.733E-08 | 3.963E-08 | 1.200E-08 | 8.300E-08 |
| 15330 | 767 | K767I | B | 2.3e-08 | 7.1e-08 | 4e-08 | 4.467E-08 | 2.434E-08 | 2.300E-08 | 7.100E-08 |
| 15331 | 767 | K767L | B | 2.5e-08 | 4.6e-08 | 3.1e-08 | 3.400E-08 | 1.082E-08 | 2.500E-08 | 4.600E-08 |
| 15332 | 767 | K767K | B | 1.2e-07 | 6.6e-08 | 5.2e-08 | 7.933E-08 | 3.591E-08 | 5.200E-08 | 1.200E-07 |
| 15333 | 767 | K767M | B | 1.7e-08 | 3.5e-08 | 4.4e-08 | 3.200E-08 | 1.375E-08 | 1.700E-08 | 4.400E-08 |
| 15334 | 767 | K767F | B | 2.1e-08 | 6.6e-08 | 2.2e-08 | 3.633E-08 | 2.570E-08 | 2.100E-08 | 6.600E-08 |
| 15335 | 767 | K767P | B | 4.2e-08 | 4.2e-08 | 4.1e-08 | 4.167E-08 | 5.774E-10 | 4.100E-08 | 4.200E-08 |
| 15336 | 767 | K767S | B | 5.7e-08 | 1.4e-07 | 8.3e-08 | 9.333E-08 | 4.245E-08 | 5.700E-08 | 1.400E-07 |
| 15337 | 767 | K767T | B | 3.8e-08 | 4.9e-08 | 5.5e-08 | 4.733E-08 | 8.622E-09 | 3.800E-08 | 5.500E-08 |
| 15338 | 767 | K767W | B | 3.5e-08 | 5.1e-08 | 5.5e-08 | 4.700E-08 | 1.058E-08 | 3.500E-08 | 5.500E-08 |
| 15339 | 767 | K767Y | B | 1e-07 | 5.3e-08 | 1e-07 | 8.433E-08 | 2.714E-08 | 5.300E-08 | 1.000E-07 |
| 15340 | 767 | K767V | B | 6.2e-08 | 7.9e-08 | 3.3e-08 | 5.800E-08 | 2.326E-08 | 3.300E-08 | 7.900E-08 |
| 15341 | 768 | V768A | B | 1.3e-07 | 4.8e-08 | 3.5e-08 | 7.100E-08 | 5.151E-08 | 3.500E-08 | 1.300E-07 |
| 15342 | 768 | V768R | B | 5.8e-08 | 3e-08 | 2.4e-08 | 3.733E-08 | 1.815E-08 | 2.400E-08 | 5.800E-08 |
| 15343 | 768 | V768N | B | 5.5e-08 | 6e-08 | 4.7e-08 | 5.400E-08 | 6.557E-09 | 4.700E-08 | 6.000E-08 |
| 15344 | 768 | V768D | B | 8.2e-08 | 3.2e-08 | 1.4e-07 | 8.467E-08 | 5.405E-08 | 3.200E-08 | 1.400E-07 |
| 15345 | 768 | V768C | B | 6.1e-08 | 6.1e-08 | 1.6e-08 | 4.600E-08 | 2.598E-08 | 1.600E-08 | 6.100E-08 |
| 15346 | 768 | V768E | B | 5.4e-08 | 4e-08 | 2.3e-08 | 3.900E-08 | 1.552E-08 | 2.300E-08 | 5.400E-08 |
| 15347 | 768 | V768Q | B | 7.5e-08 | 4.7e-08 | 3.7e-08 | 5.300E-08 | 1.970E-08 | 3.700E-08 | 7.500E-08 |
| 15348 | 768 | V768G | B | 1.5e-08 | 1.2e-07 | 8.1e-08 | 7.200E-08 | 5.308E-08 | 1.500E-08 | 1.200E-07 |
| 15349 | 768 | V768H | B | 1.1e-07 | 5.5e-08 | 6.3e-08 | 7.600E-08 | 2.972E-08 | 5.500E-08 | 1.100E-07 |
| 15350 | 768 | V768I | B | 8.8e-08 | 1.2e-07 | 1.2e-07 | 1.093E-07 | 1.848E-08 | 8.800E-08 | 1.200E-07 |
| 15351 | 768 | V768L | B | 5.8e-08 | 4.8e-08 | 4.9e-08 | 5.167E-08 | 5.508E-09 | 4.800E-08 | 5.800E-08 |
| 15352 | 768 | V768K | B | 2.7e-08 | 1.3e-07 | 3.2e-08 | 6.300E-08 | 5.808E-08 | 2.700E-08 | 1.300E-07 |
| 15353 | 768 | V768M | B | 2.3e-08 | 2.7e-08 | 4.2e-08 | 3.067E-08 | 1.002E-08 | 2.300E-08 | 4.200E-08 |
| 15354 | 768 | V768F | B | 4.1e-08 | 6.2e-08 | 1.4e-08 | 3.900E-08 | 2.406E-08 | 1.400E-08 | 6.200E-08 |
| 15355 | 768 | V768P | B | 3.6e-08 | 2.3e-08 | 1.5e-08 | 2.467E-08 | 1.060E-08 | 1.500E-08 | 3.600E-08 |
| 15356 | 768 | V768S | B | 2.8e-08 | 4.7e-08 | 4.4e-08 | 3.967E-08 | 1.021E-08 | 2.800E-08 | 4.700E-08 |
| 15357 | 768 | V768T | B | 3.9e-08 | 5.4e-08 | 6.2e-08 | 5.167E-08 | 1.168E-08 | 3.900E-08 | 6.200E-08 |
| 15358 | 768 | V768W | B | 3.2e-08 | 4.7e-08 | 4.9e-08 | 4.267E-08 | 9.292E-09 | 3.200E-08 | 4.900E-08 |
| 15359 | 768 | V768Y | B | 5.2e-08 | 4.5e-08 | 3.5e-08 | 4.400E-08 | 8.544E-09 | 3.500E-08 | 5.200E-08 |
| 15360 | 768 | V768V | B | 5.4e-08 | 3.1e-08 | 3.2e-08 | 3.900E-08 | 1.300E-08 | 3.100E-08 | 5.400E-08 |
| 15361 | 769 | Q769A | B | 1.1e-07 | 5.4e-08 | 4.5e-08 | 6.967E-08 | 3.522E-08 | 4.500E-08 | 1.100E-07 |
| 15362 | 769 | Q769R | B | 2.7e-08 | 3.8e-08 | 2.8e-08 | 3.100E-08 | 6.083E-09 | 2.700E-08 | 3.800E-08 |
| 15363 | 769 | Q769N | B | 3.8e-08 | 5.2e-08 | 4e-08 | 4.333E-08 | 7.572E-09 | 3.800E-08 | 5.200E-08 |
| 15364 | 769 | Q769D | B | 1.2e-07 | 1.7e-08 | 4.4e-08 | 6.033E-08 | 5.341E-08 | 1.700E-08 | 1.200E-07 |
| 15365 | 769 | Q769C | B | 4.8e-08 | 5.3e-08 | 7.6e-08 | 5.900E-08 | 1.493E-08 | 4.800E-08 | 7.600E-08 |
| 15366 | 769 | Q769E | B | 4.2e-08 | 1.8e-08 | 6.6e-08 | 4.200E-08 | 2.400E-08 | 1.800E-08 | 6.600E-08 |
| 15367 | 769 | Q769Q | B | 4.8e-08 | 1.1e-08 | 7.1e-08 | 4.333E-08 | 3.027E-08 | 1.100E-08 | 7.100E-08 |
| 15368 | 769 | Q769G | B | 5.5e-08 | 4.7e-08 | 3.8e-08 | 4.667E-08 | 8.505E-09 | 3.800E-08 | 5.500E-08 |
| 15369 | 769 | Q769H | B | 1.4e-08 | 3.3e-08 | 3.7e-08 | 2.800E-08 | 1.229E-08 | 1.400E-08 | 3.700E-08 |
| 15370 | 769 | Q769I | B | 5e-08 | 4.9e-08 | 1.1e-08 | 3.667E-08 | 2.223E-08 | 1.100E-08 | 5.000E-08 |
| 15371 | 769 | Q769L | B | 3.6e-08 | 1e-07 | 3.6e-08 | 5.733E-08 | 3.695E-08 | 3.600E-08 | 1.000E-07 |
| 15372 | 769 | Q769K | B | 7e-08 | 2.7e-08 | 2.9e-08 | 4.200E-08 | 2.427E-08 | 2.700E-08 | 7.000E-08 |
| 15373 | 769 | Q769M | B | 3.7e-08 | 4.1e-08 | 1.5e-08 | 3.100E-08 | 1.400E-08 | 1.500E-08 | 4.100E-08 |
| 15374 | 769 | Q769F | B | 7.7e-08 | 3.1e-08 | 4.8e-08 | 5.200E-08 | 2.326E-08 | 3.100E-08 | 7.700E-08 |
| 15375 | 769 | Q769P | B | 1.5e-08 | 4.9e-08 | 2.1e-08 | 2.833E-08 | 1.815E-08 | 1.500E-08 | 4.900E-08 |
| 15376 | 769 | Q769S | B | 5.7e-08 | 1.1e-07 | 5.2e-08 | 7.300E-08 | 3.214E-08 | 5.200E-08 | 1.100E-07 |
| 15377 | 769 | Q769T | B | 4.2e-08 | 5.1e-08 | 1.8e-08 | 3.700E-08 | 1.706E-08 | 1.800E-08 | 5.100E-08 |
| 15378 | 769 | Q769W | B | 3.2e-08 | 3.9e-08 | 3.2e-08 | 3.433E-08 | 4.041E-09 | 3.200E-08 | 3.900E-08 |
| 15379 | 769 | Q769Y | B | 4.2e-08 | 1.9e-08 | 5.4e-08 | 3.833E-08 | 1.779E-08 | 1.900E-08 | 5.400E-08 |
| 15380 | 769 | Q769V | B | 3.5e-08 | 7e-08 | 1.2e-07 | 7.500E-08 | 4.272E-08 | 3.500E-08 | 1.200E-07 |
| 15381 | 770 | W770A | B | 4.2e-08 | 4.1e-08 | 2e-08 | 3.433E-08 | 1.242E-08 | 2.000E-08 | 4.200E-08 |
| 15382 | 770 | W770R | B | 1.2e-07 | 4.3e-08 | 8e-08 | 8.100E-08 | 3.851E-08 | 4.300E-08 | 1.200E-07 |
| 15383 | 770 | W770N | B | 5.3e-08 | 3.3e-08 | 2.3e-08 | 3.633E-08 | 1.528E-08 | 2.300E-08 | 5.300E-08 |
| 15384 | 770 | W770D | B | 2.8e-08 | 3.3e-08 | 3.4e-08 | 3.167E-08 | 3.215E-09 | 2.800E-08 | 3.400E-08 |

| | | | | | | | | | | |
|---|---|---|---|---|---|---|---|---|---|---|
| 15385 | 770 | W770C | B | 3e-08 | 4.8e-08 | 5.9e-08 | 4.567E-08 | 1.464E-08 | 3.000E-08 | 5.900E-08 |
| 15386 | 770 | W770E | B | 3.4e-08 | 5.2e-08 | 3e-08 | 3.867E-08 | 1.172E-08 | 3.000E-08 | 5.200E-08 |
| 15387 | 770 | W770Q | B | 1.5e-07 | 2.1e-08 | 8.8e-08 | 8.633E-08 | 6.452E-08 | 2.100E-08 | 1.500E-07 |
| 15388 | 770 | W770G | B | 2.1e-08 | 5.4e-08 | 8.3e-08 | 5.267E-08 | 3.102E-08 | 2.100E-08 | 8.300E-08 |
| 15389 | 770 | W770H | B | 2.4e-07 | 6.4e-08 | 1e-07 | 1.347E-07 | 9.298E-08 | 6.400E-08 | 2.400E-07 |
| 15390 | 770 | W770I | B | 2.1e-08 | 1.3e-08 | 9e-09 | 1.433E-08 | 6.110E-09 | 9.000E-09 | 2.100E-08 |
| 15391 | 770 | W770L | B | 4.4e-08 | 3.9e-08 | 1.6e-08 | 3.300E-08 | 1.493E-08 | 1.600E-08 | 4.400E-08 |
| 15392 | 770 | W770K | B | 2.9e-08 | 3.5e-08 | 1.6e-07 | 7.467E-08 | 7.396E-08 | 2.900E-08 | 1.600E-07 |
| 15393 | 770 | W770M | B | 5.5e-08 | 4.2e-08 | 9.3e-08 | 6.333E-08 | 2.650E-08 | 4.200E-08 | 9.300E-08 |
| 15394 | 770 | W770F | B | 4.3e-08 | 3.4e-08 | 5.2e-08 | 4.300E-08 | 9.000E-09 | 3.400E-08 | 5.200E-08 |
| 15395 | 770 | W770P | B | 2.6e-08 | 3e-08 | 2.3e-08 | 2.633E-08 | 3.512E-09 | 2.300E-08 | 3.000E-08 |
| 15396 | 770 | W770S | B | 4.5e-08 | 3.6e-08 | 2.5e-08 | 3.533E-08 | 1.002E-08 | 2.500E-08 | 4.500E-08 |
| 15397 | 770 | W770T | B | 4.7e-08 | 2.5e-08 | 1.8e-08 | 3.000E-08 | 1.513E-08 | 1.800E-08 | 4.700E-08 |
| 15398 | 770 | W770W | B | 5.9e-08 | 4.2e-08 | 1.7e-08 | 3.933E-08 | 2.113E-08 | 1.700E-08 | 5.900E-08 |
| 15399 | 770 | W770Y | B | 9.5e-08 | 5.5e-08 | 4.1e-08 | 6.367E-08 | 2.802E-08 | 4.100E-08 | 9.500E-08 |
| 15400 | 770 | W770V | B | 5.7e-08 | 1e-07 | 3.4e-08 | 6.367E-08 | 3.350E-08 | 3.400E-08 | 1.000E-07 |
| 15401 | 771 | K771A | B | 5.5e-08 | 3.9e-08 | 6.8e-09 | 3.360E-08 | 2.455E-08 | 6.800E-09 | 5.500E-08 |
| 15402 | 771 | K771R | B | 2.3e-08 | 4.8e-08 | 9.3e-08 | 5.467E-08 | 3.547E-08 | 2.300E-08 | 9.300E-08 |
| 15403 | 771 | K771N | B | 2.3e-08 | 1.9e-08 | 7.5e-08 | 3.900E-08 | 3.124E-08 | 1.900E-08 | 7.500E-08 |
| 15404 | 771 | K771D | B | 8.2e-08 | 4.4e-08 | 2.4e-08 | 5.000E-08 | 2.946E-08 | 2.400E-08 | 8.200E-08 |
| 15405 | 771 | K771C | B | 6e-08 | 7.1e-08 | 8.6e-08 | 7.233E-08 | 1.305E-08 | 6.000E-08 | 8.600E-08 |
| 15406 | 771 | K771E | B | 1.8e-08 | 2.2e-08 | 9.3e-09 | 1.643E-08 | 6.493E-09 | 9.300E-09 | 2.200E-08 |
| 15407 | 771 | K771Q | B | 5.6e-08 | 2.4e-08 | 1.3e-07 | 7.000E-08 | 5.437E-08 | 2.400E-08 | 1.300E-07 |
| 15408 | 771 | K771G | B | 5.4e-08 | 2.7e-08 | 1e-07 | 6.033E-08 | 3.691E-08 | 2.700E-08 | 1.000E-07 |
| 15409 | 771 | K771H | B | 2.3e-08 | 2.4e-08 | 1.4e-07 | 6.233E-08 | 6.726E-08 | 2.300E-08 | 1.400E-07 |
| 15410 | 771 | K771I | B | 8e-08 | 6.8e-08 | 7.7e-08 | 7.500E-08 | 6.245E-09 | 6.800E-08 | 8.000E-08 |
| 15411 | 771 | K771L | B | 3.4e-08 | 4.3e-08 | 5.3e-08 | 4.333E-08 | 9.504E-09 | 3.400E-08 | 5.300E-08 |
| 15412 | 771 | K771K | B | 1.1e-07 | 4.2e-08 | 4.3e-08 | 6.500E-08 | 3.897E-08 | 4.200E-08 | 1.100E-07 |
| 15413 | 771 | K771M | B | 3.9e-08 | 4.8e-08 | 7.4e-08 | 5.367E-08 | 1.818E-08 | 3.900E-08 | 7.400E-08 |
| 15414 | 771 | K771F | B | 4e-08 | 1.3e-07 | 1.2e-08 | 6.067E-08 | 6.165E-08 | 1.200E-08 | 1.300E-07 |
| 15415 | 771 | K771P | B | 6.6e-08 | 7.8e-08 | 2.1e-08 | 5.500E-08 | 3.005E-08 | 2.100E-08 | 7.800E-08 |
| 15416 | 771 | K771S | B | 5.3e-08 | 2.8e-08 | 6.9e-08 | 5.000E-08 | 2.066E-08 | 2.800E-08 | 6.900E-08 |
| 15417 | 771 | K771T | B | 4e-08 | 4.1e-08 | 6.6e-08 | 4.900E-08 | 1.473E-08 | 4.000E-08 | 6.600E-08 |
| 15418 | 771 | K771W | B | 3.4e-08 | 3.5e-08 | 4.5e-08 | 3.800E-08 | 6.083E-09 | 3.400E-08 | 4.500E-08 |
| 15419 | 771 | K771Y | B | 8.9e-08 | 4.8e-08 | 2.1e-08 | 5.267E-08 | 3.424E-08 | 2.100E-08 | 8.900E-08 |
| 15420 | 771 | K771V | B | 9e-08 | 5.3e-08 | 8.1e-08 | 7.467E-08 | 1.930E-08 | 5.300E-08 | 9.000E-08 |
| 15421 | 772 | V772A | B | 3e-08 | 3.5e-08 | 7.4e-08 | 4.633E-08 | 2.409E-08 | 3.000E-08 | 7.400E-08 |
| 15422 | 772 | V772R | B | 1e-07 | 4.3e-08 | 2.5e-08 | 5.600E-08 | 3.915E-08 | 2.500E-08 | 1.000E-07 |
| 15423 | 772 | V772N | B | 1.6e-08 | 2.9e-08 | 1.1e-08 | 1.867E-08 | 9.292E-09 | 1.100E-08 | 2.900E-08 |
| 15424 | 772 | V772D | B | 2.5e-08 | 4.5e-08 | 1.1e-08 | 2.700E-08 | 1.709E-08 | 1.100E-08 | 4.500E-08 |
| 15425 | 772 | V772C | B | 9.5e-08 | 1.8e-08 | 2.2e-08 | 4.500E-08 | 4.335E-08 | 1.800E-08 | 9.500E-08 |
| 15426 | 772 | V772E | B | 7.1e-08 | 8.4e-08 | 3.8e-08 | 6.433E-08 | 2.371E-08 | 3.800E-08 | 8.400E-08 |
| 15427 | 772 | V772Q | B | 4.9e-09 | 3.5e-08 | 5.1e-08 | 3.030E-08 | 2.341E-08 | 4.900E-09 | 5.100E-08 |
| 15428 | 772 | V772G | B | 3.9e-08 | 6.9e-08 | 4.7e-08 | 5.167E-08 | 1.553E-08 | 3.900E-08 | 6.900E-08 |
| 15429 | 772 | V772H | B | 1.8e-08 | 3.6e-08 | 4e-08 | 3.133E-08 | 1.172E-08 | 1.800E-08 | 4.000E-08 |
| 15430 | 772 | V772I | B | 3.5e-08 | 1.2e-07 | 3.9e-08 | 6.467E-08 | 4.796E-08 | 3.500E-08 | 1.200E-07 |
| 15431 | 772 | V772L | B | 7.7e-08 | 4.8e-08 | 3.3e-08 | 5.267E-08 | 2.237E-08 | 3.300E-08 | 7.700E-08 |
| 15432 | 772 | V772K | B | 8.4e-08 | 3.6e-08 | 5.2e-08 | 5.733E-08 | 2.444E-08 | 3.600E-08 | 8.400E-08 |
| 15433 | 772 | V772M | B | 5.6e-08 | 4.7e-08 | 1.1e-08 | 3.800E-08 | 2.381E-08 | 1.100E-08 | 5.600E-08 |
| 15434 | 772 | V772F | B | 5.4e-08 | 5.8e-08 | 2.8e-08 | 4.667E-08 | 1.629E-08 | 2.800E-08 | 5.800E-08 |
| 15435 | 772 | V772P | B | 6.2e-08 | 4.5e-08 | 6e-08 | 5.567E-08 | 9.292E-09 | 4.500E-08 | 6.200E-08 |
| 15436 | 772 | V772S | B | 5.9e-08 | 1.1e-07 | 1.7e-08 | 6.200E-08 | 4.657E-08 | 1.700E-08 | 1.100E-07 |
| 15437 | 772 | V772T | B | 4.5e-08 | 4.8e-08 | 4.3e-08 | 4.533E-08 | 2.517E-09 | 4.300E-08 | 4.800E-08 |
| 15438 | 772 | V772W | B | 1e-07 | 1.1e-07 | 4.9e-08 | 8.633E-08 | 3.272E-08 | 4.900E-08 | 1.100E-07 |
| 15439 | 772 | V772Y | B | 4.7e-08 | 1.4e-08 | 3.4e-08 | 3.167E-08 | 1.662E-08 | 1.400E-08 | 4.700E-08 |
| 15440 | 772 | V772V | B | 4.7e-08 | 1.2e-08 | 3e-08 | 2.967E-08 | 1.750E-08 | 1.200E-08 | 4.700E-08 |
| 15441 | 773 | D773A | B | 4.4e-08 | 2.6e-08 | 9.2e-08 | 5.400E-08 | 3.412E-08 | 2.600E-08 | 9.200E-08 |
| 15442 | 773 | D773R | B | 1.1e-08 | 4.6e-08 | 4.7e-08 | 3.467E-08 | 2.050E-08 | 1.100E-08 | 4.700E-08 |
| 15443 | 773 | D773N | B | 3.7e-08 | 1.3e-07 | 5.5e-08 | 7.400E-08 | 4.933E-08 | 3.700E-08 | 1.300E-07 |
| 15444 | 773 | D773D | B | 3e-08 | 2e-08 | 3.8e-08 | 2.933E-08 | 9.018E-09 | 2.000E-08 | 3.800E-08 |
| 15445 | 773 | D773C | B | 2e-08 | 4.8e-08 | 3.5e-08 | 3.433E-08 | 1.401E-08 | 2.000E-08 | 4.800E-08 |
| 15446 | 773 | D773E | B | 6.8e-08 | 1.9e-08 | 6.5e-08 | 5.067E-08 | 2.747E-08 | 1.900E-08 | 6.800E-08 |
| 15447 | 773 | D773Q | B | 1.2e-07 | 7.4e-08 | 5.2e-08 | 8.200E-08 | 3.470E-08 | 5.200E-08 | 1.200E-07 |
| 15448 | 773 | D773G | B | 6.5e-08 | 6.7e-08 | 5.4e-08 | 6.200E-08 | 7.000E-09 | 5.400E-08 | 6.700E-08 |
| 15449 | 773 | D773H | B | 4.9e-08 | 8.4e-08 | 5.5e-08 | 6.267E-08 | 1.872E-08 | 4.900E-08 | 8.400E-08 |
| 15450 | 773 | D773I | B | 3.2e-08 | 3.6e-08 | 4.9e-08 | 3.900E-08 | 8.888E-09 | 3.200E-08 | 4.900E-08 |
| 15451 | 773 | D773L | B | 3.5e-08 | 4e-08 | 1.3e-08 | 2.933E-08 | 1.436E-08 | 1.300E-08 | 4.000E-08 |
| 15452 | 773 | D773K | B | 7.3e-08 | 2.1e-08 | 2.7e-08 | 4.033E-08 | 2.845E-08 | 2.100E-08 | 7.300E-08 |
| 15453 | 773 | D773M | B | 5.4e-08 | 4.2e-08 | 5.9e-08 | 5.167E-08 | 8.737E-09 | 4.200E-08 | 5.900E-08 |
| 15454 | 773 | D773F | B | 6.8e-09 | 3.9e-08 | 6.7e-08 | 3.760E-08 | 3.012E-08 | 6.800E-09 | 6.700E-08 |
| 15455 | 773 | D773P | B | 2.7e-08 | 5e-08 | 1.2e-07 | 6.567E-08 | 4.844E-08 | 2.700E-08 | 1.200E-07 |
| 15456 | 773 | D773S | B | 5.8e-08 | 7.6e-08 | 7.8e-08 | 7.067E-08 | 1.102E-08 | 5.800E-08 | 7.800E-08 |
| 15457 | 773 | D773T | B | 1.7e-08 | 1.1e-07 | 1.1e-07 | 7.900E-08 | 5.369E-08 | 1.700E-08 | 1.100E-07 |
| 15458 | 773 | D773W | B | 4.5e-08 | 1e-07 | 5.2e-08 | 6.567E-08 | 2.994E-08 | 4.500E-08 | 1.000E-07 |
| 15459 | 773 | D773Y | B | 5.4e-08 | 2.7e-08 | 2.7e-08 | 3.600E-08 | 1.559E-08 | 2.700E-08 | 5.400E-08 |
| 15460 | 773 | D773V | B | 5.3e-08 | 5.2e-08 | 3.1e-08 | 4.533E-08 | 1.242E-08 | 3.100E-08 | 5.300E-08 |
| 15461 | 774 | N774A | B | 3.6e-08 | 2.3e-08 | 7.2e-08 | 4.367E-08 | 2.538E-08 | 2.300E-08 | 7.200E-08 |
| 15462 | 774 | N774R | B | 5.8e-08 | 3.3e-08 | 4.4e-08 | 4.500E-08 | 1.253E-08 | 3.300E-08 | 5.800E-08 |
| 15463 | 774 | N774N | B | 2.8e-08 | 1.9e-08 | 2.3e-08 | 2.333E-08 | 4.509E-09 | 1.900E-08 | 2.800E-08 |
| 15464 | 774 | N774D | B | 5.2e-08 | 3.4e-08 | 5.5e-08 | 4.700E-08 | 1.136E-08 | 3.400E-08 | 5.500E-08 |
| 15465 | 774 | N774C | B | 3.4e-08 | 1.2e-08 | 2.7e-08 | 2.433E-08 | 1.124E-08 | 1.200E-08 | 3.400E-08 |
| 15466 | 774 | N774E | B | 3.6e-08 | 2.9e-08 | 6.4e-08 | 4.300E-08 | 1.852E-08 | 2.900E-08 | 6.400E-08 |
| 15467 | 774 | N774Q | B | 1.4e-08 | 6.1e-08 | 1.2e-07 | 6.500E-08 | 5.311E-08 | 1.400E-08 | 1.200E-07 |
| 15468 | 774 | N774G | B | 3.3e-08 | 2e-07 | 2.3e-08 | 8.533E-08 | 9.943E-08 | 2.300E-08 | 2.000E-07 |
| 15469 | 774 | N774H | B | 3.6e-08 | 6.2e-08 | 4.6e-08 | 4.800E-08 | 1.311E-08 | 3.600E-08 | 6.200E-08 |

| | | | | | | | | | | |
|---|---|---|---|---|---|---|---|---|---|---|
| 15470 | 774 | N774I | B | 4.5e-08 | 6.7e-08 | 5.1e-08 | 5.433E-08 | 1.137E-08 | 4.500E-08 | 6.700E-08 |
| 15471 | 774 | N774L | B | 1.6e-08 | 2.8e-08 | 3.2e-08 | 2.533E-08 | 8.327E-09 | 1.600E-08 | 3.200E-08 |
| 15472 | 774 | N774K | B | 4.8e-08 | 6.2e-08 | 5.9e-08 | 5.633E-08 | 7.371E-09 | 4.800E-08 | 6.200E-08 |
| 15473 | 774 | N774M | B | 2.3e-08 | 5e-08 | 1.2e-08 | 2.833E-08 | 1.955E-08 | 1.200E-08 | 5.000E-08 |
| 15474 | 774 | N774F | B | 5.6e-08 | 3.9e-08 | 6.3e-08 | 5.267E-08 | 1.234E-08 | 3.900E-08 | 6.300E-08 |
| 15475 | 774 | N774P | B | 6.3e-08 | 7.9e-08 | 3.7e-08 | 5.967E-08 | 2.120E-08 | 3.700E-08 | 7.900E-08 |
| 15476 | 774 | N774S | B | 1.5e-07 | 1.3e-07 | 7.1e-08 | 1.170E-07 | 4.107E-08 | 7.100E-08 | 1.500E-07 |
| 15477 | 774 | N774T | B | 2.9e-08 | 2e-08 | 1.1e-07 | 5.300E-08 | 4.957E-08 | 2.000E-08 | 1.100E-07 |
| 15478 | 774 | N774W | B | 3e-08 | 7.2e-08 | 6.7e-08 | 5.633E-08 | 2.294E-08 | 3.000E-08 | 7.200E-08 |
| 15479 | 774 | N774Y | B | 6e-08 | 1.1e-08 | 2.8e-08 | 3.300E-08 | 2.488E-08 | 1.100E-08 | 6.000E-08 |
| 15480 | 774 | N774V | B | 2.1e-08 | 5.6e-08 | 1.1e-07 | 6.233E-08 | 4.484E-08 | 2.100E-08 | 1.100E-07 |
| 15481 | 775 | A775A | B | 2.4e-08 | 3.7e-08 | 1.2e-07 | 6.033E-08 | 5.208E-08 | 2.400E-08 | 1.200E-07 |
| 15482 | 775 | A775R | B | 5.6e-08 | 3.1e-08 | 4.4e-08 | 4.367E-08 | 1.250E-08 | 3.100E-08 | 5.600E-08 |
| 15483 | 775 | A775N | B | 1.1e-07 | 2.7e-08 | 1.8e-08 | 5.167E-08 | 5.072E-08 | 1.800E-08 | 1.100E-07 |
| 15484 | 775 | A775D | B | 3.5e-08 | 1.3e-08 | 3.7e-08 | 2.833E-08 | 1.332E-08 | 1.300E-08 | 3.700E-08 |
| 15485 | 775 | A775C | B | 3.8e-08 | 4.8e-08 | 2.2e-08 | 3.600E-08 | 1.311E-08 | 2.200E-08 | 4.800E-08 |
| 15486 | 775 | A775E | B | 2.8e-08 | 5.7e-08 | 3.5e-08 | 4.000E-08 | 1.513E-08 | 2.800E-08 | 5.700E-08 |
| 15487 | 775 | A775Q | B | 1e-08 | 1.3e-07 | 1.5e-08 | 5.167E-08 | 6.788E-08 | 1.000E-08 | 1.300E-07 |
| 15488 | 775 | A775G | B | 2.4e-08 | 4.7e-08 | 3.7e-08 | 3.600E-08 | 1.153E-08 | 2.400E-08 | 4.700E-08 |
| 15489 | 775 | A775H | B | 3e-08 | 1.4e-08 | 7.1e-08 | 3.833E-08 | 2.940E-08 | 1.400E-08 | 7.100E-08 |
| 15490 | 775 | A775I | B | 9.6e-08 | 1e-07 | 3.2e-08 | 7.600E-08 | 3.816E-08 | 3.200E-08 | 1.000E-07 |
| 15491 | 775 | A775L | B | 6.9e-08 | 4.1e-08 | 4.3e-08 | 5.100E-08 | 1.562E-08 | 4.100E-08 | 6.900E-08 |
| 15492 | 775 | A775K | B | 1.1e-07 | 5.8e-08 | 1.1e-07 | 9.267E-08 | 3.002E-08 | 5.800E-08 | 1.100E-07 |
| 15493 | 775 | A775M | B | 4.3e-08 | 2.3e-08 | 8.3e-08 | 4.967E-08 | 3.055E-08 | 2.300E-08 | 8.300E-08 |
| 15494 | 775 | A775F | B | 5.1e-08 | 3.8e-08 | 4e-08 | 4.300E-08 | 7.000E-09 | 3.800E-08 | 5.100E-08 |
| 15495 | 775 | A775P | B | 3.7e-08 | 3.2e-08 | 5.2e-08 | 4.033E-08 | 1.041E-08 | 3.200E-08 | 5.200E-08 |
| 15496 | 775 | A775S | B | 4.4e-08 | 3.6e-08 | 3.6e-08 | 3.867E-08 | 4.619E-09 | 3.600E-08 | 4.400E-08 |
| 15497 | 775 | A775T | B | 7.2e-08 | 1.9e-08 | 3.2e-08 | 4.100E-08 | 2.762E-08 | 1.900E-08 | 7.200E-08 |
| 15498 | 775 | A775W | B | 2.3e-08 | 8.2e-09 | 1.7e-08 | 1.607E-08 | 7.444E-09 | 8.200E-09 | 2.300E-08 |
| 15499 | 775 | A775Y | B | 1.8e-08 | 5e-08 | 1.3e-07 | 6.600E-08 | 5.769E-08 | 1.800E-08 | 1.300E-07 |
| 15500 | 775 | A775V | B | 4.9e-08 | 1.3e-07 | 5.6e-08 | 7.833E-08 | 4.488E-08 | 4.900E-08 | 1.300E-07 |
| 15501 | 776 | L776A | B | 2.4e-08 | 3.3e-08 | 2.4e-08 | 2.700E-08 | 5.196E-09 | 2.400E-08 | 3.300E-08 |
| 15502 | 776 | L776R | B | 2.4e-08 | 2.9e-08 | 2.5e-08 | 2.600E-08 | 2.646E-09 | 2.400E-08 | 2.900E-08 |
| 15503 | 776 | L776N | B | 7.3e-09 | 2.7e-08 | 5e-08 | 2.810E-08 | 2.137E-08 | 7.300E-09 | 5.000E-08 |
| 15504 | 776 | L776D | B | 2e-08 | 7.9e-08 | 7.8e-08 | 5.900E-08 | 3.378E-08 | 2.000E-08 | 7.900E-08 |
| 15505 | 776 | L776C | B | 4.1e-08 | 2e-08 | 6.1e-08 | 4.067E-08 | 2.050E-08 | 2.000E-08 | 6.100E-08 |
| 15506 | 776 | L776E | B | 6.7e-08 | 7.9e-08 | 4.3e-08 | 6.300E-08 | 1.833E-08 | 4.300E-08 | 7.900E-08 |
| 15507 | 776 | L776Q | B | 7.4e-08 | 3.6e-08 | 3e-08 | 4.667E-08 | 2.386E-08 | 3.000E-08 | 7.400E-08 |
| 15508 | 776 | L776G | B | 4.3e-08 | 4e-08 | 2e-08 | 3.433E-08 | 1.250E-08 | 2.000E-08 | 4.300E-08 |
| 15509 | 776 | L776H | B | 9.1e-08 | 4.1e-08 | 1.8e-08 | 5.000E-08 | 3.732E-08 | 1.800E-08 | 9.100E-08 |
| 15510 | 776 | L776I | B | 1e-07 | 2.1e-08 | 2.8e-08 | 4.967E-08 | 4.373E-08 | 2.100E-08 | 1.000E-07 |
| 15511 | 776 | L776L | B | 6.2e-08 | 2.4e-08 | 6.3e-08 | 4.967E-08 | 2.223E-08 | 2.400E-08 | 6.300E-08 |
| 15512 | 776 | L776K | B | 7.5e-08 | 2.5e-08 | 2.4e-08 | 4.133E-08 | 2.916E-08 | 2.400E-08 | 7.500E-08 |
| 15513 | 776 | L776M | B | 1.7e-08 | 2.8e-08 | 4.9e-08 | 3.133E-08 | 1.626E-08 | 1.700E-08 | 4.900E-08 |
| 15514 | 776 | L776F | B | 3.9e-08 | 8.3e-08 | 2.5e-08 | 4.900E-08 | 3.027E-08 | 2.500E-08 | 8.300E-08 |
| 15515 | 776 | L776P | B | 7e-08 | 1.4e-08 | 9.3e-08 | 5.900E-08 | 4.063E-08 | 1.400E-08 | 9.300E-08 |
| 15516 | 776 | L776S | B | 7.6e-08 | 3.7e-08 | 3e-08 | 4.767E-08 | 2.479E-08 | 3.000E-08 | 7.600E-08 |
| 15517 | 776 | L776T | B | 5.6e-08 | 2.7e-08 | 2.7e-08 | 3.667E-08 | 1.674E-08 | 2.700E-08 | 5.600E-08 |
| 15518 | 776 | L776W | B | 5.5e-08 | 7e-08 | 4.5e-08 | 5.667E-08 | 1.258E-08 | 4.500E-08 | 7.000E-08 |
| 15519 | 776 | L776Y | B | 5e-08 | 4.1e-08 | 6.3e-08 | 5.133E-08 | 1.106E-08 | 4.100E-08 | 6.300E-08 |
| 15520 | 776 | L776V | B | 4.6e-08 | 8.8e-08 | 3e-08 | 5.467E-08 | 2.996E-08 | 3.000E-08 | 8.800E-08 |
| 15521 | 777 | Q777A | B | 4.9e-08 | 9.1e-08 | 8.6e-08 | 7.533E-08 | 2.294E-08 | 4.900E-08 | 9.100E-08 |
| 15522 | 777 | Q777R | B | 3.2e-08 | 4.6e-08 | 4.7e-08 | 4.167E-08 | 8.386E-09 | 3.200E-08 | 4.700E-08 |
| 15523 | 777 | Q777N | B | 1.5e-08 | 3.7e-08 | 4.8e-08 | 3.333E-08 | 1.680E-08 | 1.500E-08 | 4.800E-08 |
| 15524 | 777 | Q777D | B | 7.2e-08 | 3.7e-08 | 3.6e-08 | 4.833E-08 | 2.050E-08 | 3.600E-08 | 7.200E-08 |
| 15525 | 777 | Q777C | B | 2.6e-08 | 3.9e-08 | 6e-08 | 4.167E-08 | 1.716E-08 | 2.600E-08 | 6.000E-08 |
| 15526 | 777 | Q777E | B | 1e-07 | 1e-07 | 6.5e-08 | 8.833E-08 | 2.021E-08 | 6.500E-08 | 1.000E-07 |
| 15527 | 777 | Q777Q | B | 8.5e-08 | 4.5e-08 | 5e-08 | 6.000E-08 | 2.179E-08 | 4.500E-08 | 8.500E-08 |
| 15528 | 777 | Q777G | B | 1.8e-08 | 5.3e-08 | 2.4e-08 | 3.167E-08 | 1.872E-08 | 1.800E-08 | 5.300E-08 |
| 15529 | 777 | Q777H | B | 6.2e-08 | 3.2e-08 | 1.1e-08 | 3.500E-08 | 2.563E-08 | 1.100E-08 | 6.200E-08 |
| 15530 | 777 | Q777I | B | 4.6e-08 | 3.1e-08 | 4.9e-08 | 4.200E-08 | 9.644E-09 | 3.100E-08 | 4.900E-08 |
| 15531 | 777 | Q777L | B | 1e-07 | 7.2e-08 | 4.3e-08 | 7.167E-08 | 2.850E-08 | 4.300E-08 | 1.000E-07 |
| 15532 | 777 | Q777K | B | 1.2e-07 | 5e-08 | 5.4e-08 | 7.467E-08 | 3.931E-08 | 5.000E-08 | 1.200E-07 |
| 15533 | 777 | Q777M | B | 5e-08 | 8.1e-08 | 2.2e-08 | 5.100E-08 | 2.951E-08 | 2.200E-08 | 8.100E-08 |
| 15534 | 777 | Q777F | B | 7e-08 | 4.3e-08 | 6.2e-08 | 5.833E-08 | 1.387E-08 | 4.300E-08 | 7.000E-08 |
| 15535 | 777 | Q777P | B | 4.1e-08 | 5.9e-08 | 1.1e-08 | 3.700E-08 | 2.425E-08 | 1.100E-08 | 5.900E-08 |
| 15536 | 777 | Q777S | B | 7.2e-09 | 8.4e-08 | 2e-08 | 3.707E-08 | 4.115E-08 | 7.200E-09 | 8.400E-08 |
| 15537 | 777 | Q777T | B | 6.5e-08 | 1.8e-08 | 2.3e-08 | 3.533E-08 | 2.581E-08 | 1.800E-08 | 6.500E-08 |
| 15538 | 777 | Q777W | B | 2.4e-08 | 2.1e-08 | 9.1e-09 | 1.803E-08 | 7.881E-09 | 9.100E-09 | 2.400E-08 |
| 15539 | 777 | Q777Y | B | 1.5e-08 | 4.6e-08 | 4.4e-08 | 3.500E-08 | 1.735E-08 | 1.500E-08 | 4.600E-08 |
| 15540 | 777 | Q777V | B | 4.6e-08 | 4.6e-08 | 2.9e-08 | 4.033E-08 | 9.815E-09 | 2.900E-08 | 4.600E-08 |
| 15541 | 778 | S778A | B | 3.5e-08 | 6.5e-08 | 5.3e-08 | 5.100E-08 | 1.510E-08 | 3.500E-08 | 6.500E-08 |
| 15542 | 778 | S778R | B | 7.2e-08 | 3.2e-08 | 3e-08 | 4.467E-08 | 2.369E-08 | 3.000E-08 | 7.200E-08 |
| 15543 | 778 | S778N | B | 6e-08 | 2e-08 | 4.5e-08 | 4.167E-08 | 2.021E-08 | 2.000E-08 | 6.000E-08 |
| 15544 | 778 | S778D | B | 2.7e-08 | 4.8e-08 | 3.3e-08 | 3.600E-08 | 1.082E-08 | 2.700E-08 | 4.800E-08 |
| 15545 | 778 | S778C | B | 3.1e-08 | 8.8e-08 | 7.5e-08 | 6.467E-08 | 2.987E-08 | 3.100E-08 | 8.800E-08 |
| 15546 | 778 | S778E | B | 4.9e-08 | 9.3e-08 | 8e-08 | 7.400E-08 | 2.261E-08 | 4.900E-08 | 9.300E-08 |
| 15547 | 778 | S778Q | B | 6.3e-08 | 5.6e-08 | 5.8e-08 | 5.900E-08 | 3.606E-09 | 5.600E-08 | 6.300E-08 |
| 15548 | 778 | S778G | B | 6.5e-08 | 1.1e-07 | 2e-08 | 6.500E-08 | 4.500E-08 | 2.000E-08 | 1.100E-07 |
| 15549 | 778 | S778H | B | 4.2e-08 | 8.3e-08 | 2.9e-08 | 5.133E-08 | 2.818E-08 | 2.900E-08 | 8.300E-08 |
| 15550 | 778 | S778I | B | 1.2e-07 | 7.3e-08 | 1.2e-07 | 1.043E-07 | 2.714E-08 | 7.300E-08 | 1.200E-07 |
| 15551 | 778 | S778L | B | 2.8e-08 | 4.1e-08 | 2.5e-08 | 3.133E-08 | 8.505E-09 | 2.500E-08 | 4.100E-08 |
| 15552 | 778 | S778K | B | 4.4e-08 | 4.5e-08 | 4.2e-08 | 4.367E-08 | 1.528E-09 | 4.200E-08 | 4.500E-08 |
| 15553 | 778 | S778M | B | 3.6e-08 | 1e-07 | 1.9e-08 | 5.167E-08 | 4.271E-08 | 1.900E-08 | 1.000E-07 |
| 15554 | 778 | S778F | B | 5.2e-08 | 8.1e-08 | 3.5e-08 | 5.600E-08 | 2.326E-08 | 3.500E-08 | 8.100E-08 |

| | | | | | | | | | | |
|---|---|---|---|---|---|---|---|---|---|---|
| 15555 | 778 | S778P | B | 2.5e-08 | 6.5e-08 | 3.3e-08 | 4.100E-08 | 2.117E-08 | 2.500E-08 | 6.500E-08 |
| 15556 | 778 | S778S | B | 2e-08 | 5.2e-08 | 5.1e-08 | 4.100E-08 | 1.819E-08 | 2.000E-08 | 5.200E-08 |
| 15557 | 778 | S778T | B | 7.8e-09 | 2.7e-08 | 2.4e-08 | 1.960E-08 | 1.033E-08 | 7.800E-09 | 2.700E-08 |
| 15558 | 778 | S778W | B | 1e-07 | 4.2e-08 | 4.4e-08 | 6.200E-08 | 3.292E-08 | 4.200E-08 | 1.000E-07 |
| 15559 | 778 | S778Y | B | 3.6e-08 | 7e-08 | 5.6e-08 | 5.400E-08 | 1.709E-08 | 3.600E-08 | 7.000E-08 |
| 15560 | 778 | S778V | B | 3.5e-08 | 3.8e-08 | 3.3e-08 | 3.533E-08 | 2.517E-09 | 3.300E-08 | 3.800E-08 |
| 15561 | 779 | G779A | B | 2.9e-08 | 1.1e-07 | 5.4e-08 | 6.433E-08 | 4.148E-08 | 2.900E-08 | 1.100E-07 |
| 15562 | 779 | G779R | B | 1.3e-07 | 8.2e-08 | 2.4e-08 | 7.867E-08 | 5.308E-08 | 2.400E-08 | 1.300E-07 |
| 15563 | 779 | G779N | B | 2.5e-08 | 3.7e-08 | 5.4e-08 | 3.867E-08 | 1.457E-08 | 2.500E-08 | 5.400E-08 |
| 15564 | 779 | G779D | B | 8.9e-08 | 2.4e-08 | 4.2e-08 | 5.167E-08 | 3.356E-08 | 2.400E-08 | 8.900E-08 |
| 15565 | 779 | G779C | B | 3.6e-08 | 3.4e-08 | 2.2e-08 | 3.067E-08 | 7.572E-09 | 2.200E-08 | 3.600E-08 |
| 15566 | 779 | G779E | B | 4.2e-08 | 2.3e-08 | 3.5e-08 | 3.333E-08 | 9.609E-09 | 2.300E-08 | 4.200E-08 |
| 15567 | 779 | G779Q | B | 3.1e-08 | 4.8e-08 | 4.1e-08 | 4.000E-08 | 8.544E-09 | 3.100E-08 | 4.800E-08 |
| 15568 | 779 | G779G | B | 1.2e-07 | 3.9e-08 | 5.4e-08 | 7.100E-08 | 4.309E-08 | 3.900E-08 | 1.200E-07 |
| 15569 | 779 | G779H | B | 2.2e-08 | 7.8e-08 | 6.9e-08 | 5.633E-08 | 3.007E-08 | 2.200E-08 | 7.800E-08 |
| 15570 | 779 | G779I | B | 4.5e-08 | 4.6e-08 | 1.5e-08 | 3.533E-08 | 1.762E-08 | 1.500E-08 | 4.600E-08 |
| 15571 | 779 | G779L | B | 8.5e-08 | 2.4e-08 | 5.2e-08 | 5.367E-08 | 3.053E-08 | 2.400E-08 | 8.500E-08 |
| 15572 | 779 | G779K | B | 8.1e-08 | 4.2e-08 | 2.9e-08 | 5.067E-08 | 2.706E-08 | 2.900E-08 | 8.100E-08 |
| 15573 | 779 | G779M | B | 7.2e-08 | 1.4e-08 | 5.5e-08 | 4.700E-08 | 2.982E-08 | 1.400E-08 | 7.200E-08 |
| 15574 | 779 | G779F | B | 4.7e-08 | 3.5e-08 | 1.9e-08 | 3.367E-08 | 1.405E-08 | 1.900E-08 | 4.700E-08 |
| 15575 | 779 | G779P | B | 5.7e-08 | 6.8e-08 | 4.4e-08 | 5.633E-08 | 1.201E-08 | 4.400E-08 | 6.800E-08 |
| 15576 | 779 | G779S | B | 1.7e-08 | 9.6e-08 | 2.2e-08 | 4.500E-08 | 4.424E-08 | 1.700E-08 | 9.600E-08 |
| 15577 | 779 | G779T | B | 1.8e-08 | 3.3e-08 | 2.2e-08 | 2.433E-08 | 7.767E-09 | 1.800E-08 | 3.300E-08 |
| 15578 | 779 | G779W | B | 5.7e-08 | 1.4e-08 | 4e-08 | 3.700E-08 | 2.166E-08 | 1.400E-08 | 5.700E-08 |
| 15579 | 779 | G779Y | B | 1.7e-07 | 3e-08 | 3.9e-08 | 7.967E-08 | 7.836E-08 | 3.000E-08 | 1.700E-07 |
| 15580 | 779 | G779V | B | 8.3e-08 | 7e-08 | 2e-08 | 5.767E-08 | 3.326E-08 | 2.000E-08 | 8.300E-08 |
| 15581 | 780 | N780A | B | 7.6e-08 | 2.3e-08 | 5.6e-08 | 5.167E-08 | 2.676E-08 | 2.300E-08 | 7.600E-08 |
| 15582 | 780 | N780R | B | 3.1e-08 | 6.8e-08 | 4.9e-08 | 4.933E-08 | 1.850E-08 | 3.100E-08 | 6.800E-08 |
| 15583 | 780 | N780N | B | 3.5e-08 | 3e-08 | 5.4e-08 | 3.967E-08 | 1.266E-08 | 3.000E-08 | 5.400E-08 |
| 15584 | 780 | N780D | B | 3.1e-08 | 4.6e-08 | 3.5e-08 | 3.733E-08 | 7.767E-09 | 3.100E-08 | 4.600E-08 |
| 15585 | 780 | N780C | B | 3.8e-08 | 3.2e-08 | 1.9e-08 | 2.967E-08 | 9.713E-09 | 1.900E-08 | 3.800E-08 |
| 15586 | 780 | N780E | B | 7.1e-08 | 6.1e-08 | 3.2e-08 | 5.467E-08 | 2.026E-08 | 3.200E-08 | 7.100E-08 |
| 15587 | 780 | N780Q | B | 7.1e-08 | 4.6e-08 | 2.7e-08 | 4.800E-08 | 2.207E-08 | 2.700E-08 | 7.100E-08 |
| 15588 | 780 | N780G | B | 9.9e-08 | 6.9e-08 | 3.7e-08 | 6.833E-08 | 3.101E-08 | 3.700E-08 | 9.900E-08 |
| 15589 | 780 | N780H | B | 1.4e-07 | 4.9e-08 | 2.9e-08 | 7.267E-08 | 5.916E-08 | 2.900E-08 | 1.400E-07 |
| 15590 | 780 | N780I | B | 7.7e-08 | 2.7e-08 | 4.1e-08 | 4.833E-08 | 2.579E-08 | 2.700E-08 | 7.700E-08 |
| 15591 | 780 | N780L | B | 4.2e-08 | 1.2e-07 | 2.5e-08 | 6.233E-08 | 5.066E-08 | 2.500E-08 | 1.200E-07 |
| 15592 | 780 | N780K | B | 1.7e-08 | 3.3e-08 | 3.6e-08 | 2.867E-08 | 1.021E-08 | 1.700E-08 | 3.600E-08 |
| 15593 | 780 | N780M | B | 1.2e-07 | 8.9e-08 | 2e-08 | 7.633E-08 | 5.119E-08 | 2.000E-08 | 1.200E-07 |
| 15594 | 780 | N780F | B | 5.5e-08 | 1.5e-08 | 9.2e-08 | 5.400E-08 | 3.851E-08 | 1.500E-08 | 9.200E-08 |
| 15595 | 780 | N780P | B | 1.9e-08 | 2.5e-08 | 2.3e-08 | 2.233E-08 | 3.055E-09 | 1.900E-08 | 2.500E-08 |
| 15596 | 780 | N780S | B | 5.9e-08 | 8.4e-08 | 2e-08 | 5.433E-08 | 3.225E-08 | 2.000E-08 | 8.400E-08 |
| 15597 | 780 | N780T | B | 4.1e-08 | 2.9e-08 | 4.3e-08 | 3.767E-08 | 7.572E-09 | 2.900E-08 | 4.300E-08 |
| 15598 | 780 | N780W | B | 2.1e-08 | 5.3e-08 | 1.2e-07 | 6.467E-08 | 5.052E-08 | 2.100E-08 | 1.200E-07 |
| 15599 | 780 | N780Y | B | 4.7e-08 | 5.7e-08 | 5.9e-08 | 5.433E-08 | 6.429E-09 | 4.700E-08 | 5.900E-08 |
| 15600 | 780 | N780V | B | 3.4e-08 | 7.2e-08 | 1.4e-07 | 8.200E-08 | 5.370E-08 | 3.400E-08 | 1.400E-07 |
| 15601 | 781 | S781A | B | 2.6e-08 | 5.5e-08 | 2.6e-08 | 3.567E-08 | 1.674E-08 | 2.600E-08 | 5.500E-08 |
| 15602 | 781 | S781R | B | 3.1e-08 | 9.4e-08 | 8.4e-08 | 6.967E-08 | 3.386E-08 | 3.100E-08 | 9.400E-08 |
| 15603 | 781 | S781N | B | 2.8e-08 | 4e-08 | 7.6e-08 | 4.800E-08 | 2.498E-08 | 2.800E-08 | 7.600E-08 |
| 15604 | 781 | S781D | B | 6.5e-08 | 1.3e-08 | 5.4e-08 | 4.400E-08 | 2.740E-08 | 1.300E-08 | 6.500E-08 |
| 15605 | 781 | S781C | B | 4.7e-08 | 1.2e-07 | 2.3e-08 | 6.333E-08 | 5.052E-08 | 2.300E-08 | 1.200E-07 |
| 15606 | 781 | S781E | B | 4.6e-08 | 3.2e-08 | 2.9e-08 | 3.567E-08 | 9.074E-09 | 2.900E-08 | 4.600E-08 |
| 15607 | 781 | S781Q | B | 4.7e-08 | 6.3e-08 | 5.2e-08 | 5.400E-08 | 8.185E-09 | 4.700E-08 | 6.300E-08 |
| 15608 | 781 | S781G | B | 6.2e-08 | 4.7e-08 | 1.5e-08 | 4.133E-08 | 2.401E-08 | 1.500E-08 | 6.200E-08 |
| 15609 | 781 | S781H | B | 3.9e-08 | 1.4e-07 | 1.4e-08 | 6.433E-08 | 6.671E-08 | 1.400E-08 | 1.400E-07 |
| 15610 | 781 | S781I | B | 2.6e-08 | 1.1e-07 | 3.4e-08 | 5.667E-08 | 4.636E-08 | 2.600E-08 | 1.100E-07 |
| 15611 | 781 | S781L | B | 5e-08 | 5.5e-08 | 7e-08 | 5.833E-08 | 1.041E-08 | 5.000E-08 | 7.000E-08 |
| 15612 | 781 | S781K | B | 3.6e-08 | 2e-08 | 1.3e-07 | 6.200E-08 | 5.943E-08 | 2.000E-08 | 1.300E-07 |
| 15613 | 781 | S781M | B | 4.3e-08 | 1e-08 | 1.2e-07 | 5.767E-08 | 5.645E-08 | 1.000E-08 | 1.200E-07 |
| 15614 | 781 | S781F | B | 4.1e-08 | 6.9e-08 | 1.2e-08 | 4.067E-08 | 2.850E-08 | 1.200E-08 | 6.900E-08 |
| 15615 | 781 | S781P | B | 3.5e-08 | 4.4e-08 | 8.9e-08 | 5.600E-08 | 2.893E-08 | 3.500E-08 | 8.900E-08 |
| 15616 | 781 | S781S | B | 1.7e-08 | 4.7e-08 | 3.6e-08 | 3.333E-08 | 1.518E-08 | 1.700E-08 | 4.700E-08 |
| 15617 | 781 | S781T | B | 2.1e-08 | 5.3e-08 | 6.7e-08 | 4.700E-08 | 2.358E-08 | 2.100E-08 | 6.700E-08 |
| 15618 | 781 | S781W | B | 4.5e-08 | 2.6e-07 | 1.3e-08 | 1.060E-07 | 1.343E-07 | 1.300E-08 | 2.600E-07 |
| 15619 | 781 | S781Y | B | 4e-08 | 1.4e-07 | 2.6e-08 | 6.867E-08 | 6.217E-08 | 2.600E-08 | 1.400E-07 |
| 15620 | 781 | S781V | B | 3.4e-08 | 5e-08 | 4e-08 | 4.133E-08 | 8.083E-09 | 3.400E-08 | 5.000E-08 |
| 15621 | 782 | Q782A | B | 6.2e-08 | 5.7e-08 | 2.3e-08 | 4.733E-08 | 2.122E-08 | 2.300E-08 | 6.200E-08 |
| 15622 | 782 | Q782R | B | 3.7e-08 | 4.2e-08 | 3e-08 | 3.633E-08 | 6.028E-09 | 3.000E-08 | 4.200E-08 |
| 15623 | 782 | Q782N | B | 3.2e-08 | 3.2e-08 | 4e-08 | 3.467E-08 | 4.619E-09 | 3.200E-08 | 4.000E-08 |
| 15624 | 782 | Q782D | B | 2.5e-08 | 1.1e-07 | 8.3e-09 | 4.777E-08 | 5.454E-08 | 8.300E-09 | 1.100E-07 |
| 15625 | 782 | Q782C | B | 8e-08 | 6.3e-08 | 6.4e-08 | 6.900E-08 | 9.539E-09 | 6.300E-08 | 8.000E-08 |
| 15626 | 782 | Q782E | B | 1.1e-07 | 1.5e-07 | 6.8e-08 | 1.093E-07 | 4.100E-08 | 6.800E-08 | 1.500E-07 |
| 15627 | 782 | Q782Q | B | 3e-08 | 8.6e-08 | 1.6e-08 | 4.400E-08 | 3.704E-08 | 1.600E-08 | 8.600E-08 |
| 15628 | 782 | Q782G | B | 4.8e-08 | 4.6e-08 | 5.2e-08 | 4.867E-08 | 3.055E-09 | 4.600E-08 | 5.200E-08 |
| 15629 | 782 | Q782H | B | 1.1e-07 | 2.6e-08 | 1.1e-07 | 8.200E-08 | 4.850E-08 | 2.600E-08 | 1.100E-07 |
| 15630 | 782 | Q782I | B | 5.6e-08 | 5.5e-08 | 6.5e-08 | 5.867E-08 | 5.508E-09 | 5.500E-08 | 6.500E-08 |
| 15631 | 782 | Q782L | B | 5.9e-08 | 2.3e-08 | 4e-08 | 4.067E-08 | 1.801E-08 | 2.300E-08 | 5.900E-08 |
| 15632 | 782 | Q782K | B | 7.7e-08 | 4.3e-08 | 1.5e-08 | 4.500E-08 | 3.105E-08 | 1.500E-08 | 7.700E-08 |
| 15633 | 782 | Q782M | B | 9.9e-08 | 2.5e-08 | 6.8e-08 | 6.400E-08 | 3.716E-08 | 2.500E-08 | 9.900E-08 |
| 15634 | 782 | Q782F | B | 1.8e-08 | 2.7e-08 | 3.4e-08 | 2.633E-08 | 8.021E-09 | 1.800E-08 | 3.400E-08 |
| 15635 | 782 | Q782P | B | 8.3e-08 | 5e-08 | 1.8e-08 | 5.033E-08 | 3.250E-08 | 1.800E-08 | 8.300E-08 |
| 15636 | 782 | Q782S | B | 2.2e-08 | 2.8e-08 | 7.4e-08 | 4.133E-08 | 2.845E-08 | 2.200E-08 | 7.400E-08 |
| 15637 | 782 | Q782T | B | 2.9e-08 | 5.6e-08 | 1.7e-07 | 8.500E-08 | 7.484E-08 | 2.900E-08 | 1.700E-07 |
| 15638 | 782 | Q782W | B | 1.3e-08 | 4.2e-08 | 3.3e-08 | 2.933E-08 | 1.484E-08 | 1.300E-08 | 4.200E-08 |
| 15639 | 782 | Q782Y | B | 1.9e-08 | 1.6e-08 | 3e-08 | 2.167E-08 | 7.371E-09 | 1.600E-08 | 3.000E-08 |

| | | | | | | | | | | |
|---|---|---|---|---|---|---|---|---|---|---|
| 15640 | 782 | Q782V | B | 3.6e-08 | 5.4e-08 | 2.2e-08 | 3.733E-08 | 1.604E-08 | 2.200E-08 | 5.400E-08 |
| 15641 | 783 | E783A | B | 7e-08 | 1e-07 | 6e-08 | 7.667E-08 | 2.082E-08 | 6.000E-08 | 1.000E-07 |
| 15642 | 783 | E783R | B | 5.4e-08 | 8.6e-09 | 2.9e-08 | 3.053E-08 | 2.274E-08 | 8.600E-09 | 5.400E-08 |
| 15643 | 783 | E783N | B | 5e-08 | 2.7e-08 | 5.6e-08 | 4.433E-08 | 1.531E-08 | 2.700E-08 | 5.600E-08 |
| 15644 | 783 | E783D | B | 7.8e-08 | 3e-08 | 8.6e-08 | 6.467E-08 | 3.029E-08 | 3.000E-08 | 8.600E-08 |
| 15645 | 783 | E783C | B | 3.1e-08 | 5.1e-08 | 8.8e-08 | 5.667E-08 | 2.892E-08 | 3.100E-08 | 8.800E-08 |
| 15646 | 783 | E783E | B | 3.2e-08 | 8e-08 | 3.5e-08 | 4.900E-08 | 2.689E-08 | 3.200E-08 | 8.000E-08 |
| 15647 | 783 | E783Q | B | 4.3e-08 | 6.1e-08 | 3.1e-08 | 4.500E-08 | 1.510E-08 | 3.100E-08 | 6.100E-08 |
| 15648 | 783 | E783G | B | 6.5e-08 | 4.1e-08 | 5.6e-08 | 5.400E-08 | 1.212E-08 | 4.100E-08 | 6.500E-08 |
| 15649 | 783 | E783H | B | 1.3e-08 | 8.8e-08 | 3.8e-08 | 4.633E-08 | 3.819E-08 | 1.300E-08 | 8.800E-08 |
| 15650 | 783 | E783I | B | 6.1e-08 | 2.4e-07 | 1.9e-08 | 1.067E-07 | 1.174E-07 | 1.900E-08 | 2.400E-07 |
| 15651 | 783 | E783L | B | 8.9e-08 | 9.5e-08 | 5.6e-08 | 8.000E-08 | 2.100E-08 | 5.600E-08 | 9.500E-08 |
| 15652 | 783 | E783K | B | 4.4e-08 | 2.5e-08 | 5e-08 | 3.967E-08 | 1.305E-08 | 2.500E-08 | 5.000E-08 |
| 15653 | 783 | E783M | B | 5e-08 | 2.5e-08 | 7e-08 | 4.833E-08 | 2.255E-08 | 2.500E-08 | 7.000E-08 |
| 15654 | 783 | E783F | B | 3.8e-08 | 4.5e-08 | 2.3e-08 | 3.533E-08 | 1.124E-08 | 2.300E-08 | 4.500E-08 |
| 15655 | 783 | E783P | B | 9.4e-08 | 4.3e-08 | 7e-08 | 6.900E-08 | 2.551E-08 | 4.300E-08 | 9.400E-08 |
| 15656 | 783 | E783S | B | 1.2e-07 | 3.6e-08 | 5.1e-08 | 6.900E-08 | 4.480E-08 | 3.600E-08 | 1.200E-07 |
| 15657 | 783 | E783T | B | 3.4e-08 | 8.4e-08 | 2.2e-08 | 4.667E-08 | 3.288E-08 | 2.200E-08 | 8.400E-08 |
| 15658 | 783 | E783W | B | 6.9e-08 | 5.8e-08 | 2.5e-08 | 5.067E-08 | 2.290E-08 | 2.500E-08 | 6.900E-08 |
| 15659 | 783 | E783Y | B | 2.9e-08 | 4.1e-08 | 3e-08 | 3.333E-08 | 6.658E-09 | 2.900E-08 | 4.100E-08 |
| 15660 | 783 | E783V | B | 3.5e-08 | 6.4e-08 | 4.3e-08 | 4.733E-08 | 1.498E-08 | 3.500E-08 | 6.400E-08 |
| 15661 | 784 | S784A | B | 1.2e-07 | 5.5e-08 | 7.7e-08 | 8.400E-08 | 3.306E-08 | 5.500E-08 | 1.200E-07 |
| 15662 | 784 | S784R | B | 2.5e-08 | 5.4e-08 | 6.5e-08 | 4.800E-08 | 2.066E-08 | 2.500E-08 | 6.500E-08 |
| 15663 | 784 | S784N | B | 6.6e-08 | 6.2e-08 | 6.7e-08 | 6.500E-08 | 2.646E-09 | 6.200E-08 | 6.700E-08 |
| 15664 | 784 | S784D | B | 6.2e-08 | 2.2e-08 | 2.1e-08 | 3.500E-08 | 2.339E-08 | 2.100E-08 | 6.200E-08 |
| 15665 | 784 | S784C | B | 2.7e-08 | 6.8e-08 | 4.2e-08 | 4.567E-08 | 2.074E-08 | 2.700E-08 | 6.800E-08 |
| 15666 | 784 | S784E | B | 7.9e-08 | 5.9e-08 | 1.9e-08 | 5.233E-08 | 3.055E-08 | 1.900E-08 | 7.900E-08 |
| 15667 | 784 | S784Q | B | 3.3e-08 | 1e-07 | 3e-08 | 5.433E-08 | 3.958E-08 | 3.000E-08 | 1.000E-07 |
| 15668 | 784 | S784G | B | 2.1e-08 | 7.3e-08 | 2.7e-08 | 4.033E-08 | 2.845E-08 | 2.100E-08 | 7.300E-08 |
| 15669 | 784 | S784H | B | 4.9e-08 | 6.1e-08 | 6.8e-08 | 5.933E-08 | 9.609E-09 | 4.900E-08 | 6.800E-08 |
| 15670 | 784 | S784I | B | 1e-07 | 5.3e-08 | 7.4e-08 | 7.567E-08 | 2.354E-08 | 5.300E-08 | 1.000E-07 |
| 15671 | 784 | S784L | B | 2.8e-08 | 3.6e-08 | 1.6e-08 | 2.667E-08 | 1.007E-08 | 1.600E-08 | 3.600E-08 |
| 15672 | 784 | S784K | B | 4.8e-08 | 1.6e-07 | 8.6e-08 | 9.800E-08 | 5.696E-08 | 4.800E-08 | 1.600E-07 |
| 15673 | 784 | S784M | B | 4.9e-08 | 3.9e-08 | 4.6e-08 | 4.467E-08 | 5.132E-09 | 3.900E-08 | 4.900E-08 |
| 15674 | 784 | S784F | B | 2.7e-08 | 7.9e-08 | 4.4e-08 | 5.000E-08 | 2.651E-08 | 2.700E-08 | 7.900E-08 |
| 15675 | 784 | S784P | B | 6.1e-08 | 2.9e-08 | 8.5e-08 | 5.833E-08 | 2.810E-08 | 2.900E-08 | 8.500E-08 |
| 15676 | 784 | S784S | B | 7e-08 | 2.7e-08 | 3e-08 | 4.233E-08 | 2.401E-08 | 2.700E-08 | 7.000E-08 |
| 15677 | 784 | S784T | B | 9e-08 | 2.8e-08 | 5.2e-08 | 5.667E-08 | 3.126E-08 | 2.800E-08 | 9.000E-08 |
| 15678 | 784 | S784W | B | 2.9e-08 | 5.4e-08 | 2e-08 | 3.433E-08 | 1.762E-08 | 2.000E-08 | 5.400E-08 |
| 15679 | 784 | S784Y | B | 4e-08 | 1.8e-08 | 4.1e-08 | 3.300E-08 | 1.300E-08 | 1.800E-08 | 4.100E-08 |
| 15680 | 784 | S784V | B | 2e-08 | 5.1e-08 | 7.6e-08 | 4.900E-08 | 2.805E-08 | 2.000E-08 | 7.600E-08 |
| 15681 | 785 | V785A | B | 1.3e-08 | 3.8e-08 | 8.6e-08 | 4.567E-08 | 3.710E-08 | 1.300E-08 | 8.600E-08 |
| 15682 | 785 | V785R | B | 5.3e-08 | 2.3e-08 | 3.7e-08 | 3.767E-08 | 1.501E-08 | 2.300E-08 | 5.300E-08 |
| 15683 | 785 | V785N | B | 3.2e-08 | 6.1e-08 | 1.3e-07 | 7.433E-08 | 5.034E-08 | 3.200E-08 | 1.300E-07 |
| 15684 | 785 | V785D | B | 3.6e-08 | 5e-08 | 5e-08 | 4.533E-08 | 8.083E-09 | 3.600E-08 | 5.000E-08 |
| 15685 | 785 | V785C | B | 5.5e-08 | 9e-08 | 4.1e-08 | 6.200E-08 | 2.524E-08 | 4.100E-08 | 9.000E-08 |
| 15686 | 785 | V785E | B | 3.1e-08 | 8.1e-08 | 5.1e-08 | 5.433E-08 | 2.517E-08 | 3.100E-08 | 8.100E-08 |
| 15687 | 785 | V785Q | B | 4.6e-08 | 5.6e-08 | 2.7e-08 | 4.300E-08 | 1.473E-08 | 2.700E-08 | 5.600E-08 |
| 15688 | 785 | V785G | B | 3.4e-08 | 4.6e-08 | 4.2e-08 | 4.067E-08 | 6.110E-09 | 3.400E-08 | 4.600E-08 |
| 15689 | 785 | V785H | B | 2.8e-08 | 3.4e-08 | 4.6e-08 | 3.600E-08 | 9.165E-09 | 2.800E-08 | 4.600E-08 |
| 15690 | 785 | V785I | B | 1e-08 | 5.8e-08 | 5.9e-08 | 4.233E-08 | 2.801E-08 | 1.000E-08 | 5.900E-08 |
| 15691 | 785 | V785L | B | 5.9e-08 | 1.4e-08 | 4.9e-08 | 4.067E-08 | 2.363E-08 | 1.400E-08 | 5.900E-08 |
| 15692 | 785 | V785K | B | 1.1e-07 | 9.7e-08 | 5.1e-08 | 8.600E-08 | 3.100E-08 | 5.100E-08 | 1.100E-07 |
| 15693 | 785 | V785M | B | 2.9e-08 | 4.7e-08 | 9.1e-08 | 5.567E-08 | 3.190E-08 | 2.900E-08 | 9.100E-08 |
| 15694 | 785 | V785F | B | 3.6e-08 | 4.3e-08 | 2.3e-08 | 3.400E-08 | 1.015E-08 | 2.300E-08 | 4.300E-08 |
| 15695 | 785 | V785P | B | 3.3e-08 | 2.5e-08 | 3.1e-08 | 2.967E-08 | 4.163E-09 | 2.500E-08 | 3.300E-08 |
| 15696 | 785 | V785S | B | 2.4e-08 | 7.1e-08 | 3.3e-08 | 4.267E-08 | 2.495E-08 | 2.400E-08 | 7.100E-08 |
| 15697 | 785 | V785T | B | 1.1e-07 | 3.1e-08 | 2.3e-08 | 5.467E-08 | 4.809E-08 | 2.300E-08 | 1.100E-07 |
| 15698 | 785 | V785W | B | 1.5e-08 | 6.6e-08 | 6e-08 | 4.700E-08 | 2.787E-08 | 1.500E-08 | 6.600E-08 |
| 15699 | 785 | V785Y | B | 3.7e-08 | 3.2e-08 | 9.2e-08 | 5.367E-08 | 3.329E-08 | 3.200E-08 | 9.200E-08 |
| 15700 | 785 | V785V | B | 2.3e-08 | 1.4e-08 | 2.9e-08 | 2.200E-08 | 7.550E-09 | 1.400E-08 | 2.900E-08 |
| 15701 | 786 | T786A | B | 3.4e-08 | 3.5e-08 | 1.1e-07 | 5.967E-08 | 4.359E-08 | 3.400E-08 | 1.100E-07 |
| 15702 | 786 | T786R | B | 2.2e-08 | 8.3e-08 | 1.8e-08 | 4.100E-08 | 3.643E-08 | 1.800E-08 | 8.300E-08 |
| 15703 | 786 | T786N | B | 8.6e-09 | 3.5e-08 | 4.4e-08 | 2.920E-08 | 1.840E-08 | 8.600E-09 | 4.400E-08 |
| 15704 | 786 | T786D | B | 1.1e-07 | 4.9e-08 | 8.6e-09 | 5.587E-08 | 5.105E-08 | 8.600E-09 | 1.100E-07 |
| 15705 | 786 | T786C | B | 1.2e-07 | 2.1e-08 | 1.2e-08 | 5.100E-08 | 5.992E-08 | 1.200E-08 | 1.200E-07 |
| 15706 | 786 | T786E | B | 1.6e-08 | 3.3e-08 | 2e-08 | 2.300E-08 | 8.888E-09 | 1.600E-08 | 3.300E-08 |
| 15707 | 786 | T786Q | B | 4.6e-08 | 4.1e-08 | 3.1e-08 | 3.933E-08 | 7.638E-09 | 3.100E-08 | 4.600E-08 |
| 15708 | 786 | T786G | B | 9.5e-08 | 2.6e-08 | 7.9e-08 | 6.667E-08 | 3.612E-08 | 2.600E-08 | 9.500E-08 |
| 15709 | 786 | T786H | B | 8.7e-08 | 2.1e-08 | 1.9e-08 | 4.233E-08 | 3.870E-08 | 1.900E-08 | 8.700E-08 |
| 15710 | 786 | T786I | B | 5.1e-08 | 7.9e-08 | 7e-08 | 6.667E-08 | 1.429E-08 | 5.100E-08 | 7.900E-08 |
| 15711 | 786 | T786L | B | 2.3e-08 | 5.4e-08 | 5e-08 | 4.233E-08 | 1.686E-08 | 2.300E-08 | 5.400E-08 |
| 15712 | 786 | T786K | B | 4.2e-08 | 4.2e-08 | 2.6e-08 | 3.667E-08 | 9.238E-09 | 2.600E-08 | 4.200E-08 |
| 15713 | 786 | T786M | B | 5.9e-08 | 2.4e-08 | 8.1e-08 | 5.467E-08 | 2.875E-08 | 2.400E-08 | 8.100E-08 |
| 15714 | 786 | T786F | B | 1.9e-08 | 9.6e-08 | 5.1e-08 | 5.533E-08 | 3.868E-08 | 1.900E-08 | 9.600E-08 |
| 15715 | 786 | T786P | B | 8.6e-08 | 5.9e-08 | 5.4e-08 | 6.633E-08 | 1.721E-08 | 5.400E-08 | 8.600E-08 |
| 15716 | 786 | T786S | B | 4.5e-08 | 8.1e-09 | 2.2e-08 | 2.503E-08 | 1.864E-08 | 8.100E-09 | 4.500E-08 |
| 15717 | 786 | T786T | B | 8.1e-08 | 5.7e-08 | 5.1e-08 | 6.300E-08 | 1.587E-08 | 5.100E-08 | 8.100E-08 |
| 15718 | 786 | T786W | B | 2.3e-08 | 6.3e-08 | 7.1e-08 | 5.233E-08 | 2.572E-08 | 2.300E-08 | 7.100E-08 |
| 15719 | 786 | T786Y | B | 4.7e-08 | 4.1e-08 | 6.9e-08 | 5.233E-08 | 1.474E-08 | 4.100E-08 | 6.900E-08 |
| 15720 | 786 | T786V | B | 2.1e-08 | 6.9e-08 | 2.7e-08 | 3.900E-08 | 2.615E-08 | 2.100E-08 | 6.900E-08 |
| 15721 | 787 | E787A | B | 3.5e-08 | 1.1e-07 | 6.7e-08 | 7.067E-08 | 3.763E-08 | 3.500E-08 | 1.100E-07 |
| 15722 | 787 | E787R | B | 1e-07 | 4.3e-08 | 2.6e-08 | 5.633E-08 | 3.876E-08 | 2.600E-08 | 1.000E-07 |
| 15723 | 787 | E787N | B | 2.5e-08 | 6e-08 | 1.9e-08 | 3.467E-08 | 2.214E-08 | 1.900E-08 | 6.000E-08 |
| 15724 | 787 | E787D | B | 1.6e-08 | 5.5e-08 | 6.9e-08 | 4.667E-08 | 2.747E-08 | 1.600E-08 | 6.900E-08 |

| | | | | | | | | | | |
|---|---|---|---|---|---|---|---|---|---|---|
| 15725 | 787 | E787C | B | 4.5e-08 | 2.3e-07 | 6.6e-08 | 1.137E-07 | 1.013E-07 | 4.500E-08 | 2.300E-07 |
| 15726 | 787 | E787E | B | 1.9e-08 | 6.1e-08 | 3.7e-08 | 3.900E-08 | 2.107E-08 | 1.900E-08 | 6.100E-08 |
| 15727 | 787 | E787Q | B | 3.5e-08 | 4.2e-08 | 2.9e-08 | 3.533E-08 | 6.506E-09 | 2.900E-08 | 4.200E-08 |
| 15728 | 787 | E787G | B | 5.1e-08 | 1.1e-07 | 2.5e-08 | 6.200E-08 | 4.355E-08 | 2.500E-08 | 1.100E-07 |
| 15729 | 787 | E787H | B | 2.2e-08 | 1.6e-08 | 1.5e-08 | 1.767E-08 | 3.786E-09 | 1.500E-08 | 2.200E-08 |
| 15730 | 787 | E787I | B | 3.1e-08 | 3.4e-08 | 4.3e-08 | 3.600E-08 | 6.245E-09 | 3.100E-08 | 4.300E-08 |
| 15731 | 787 | E787L | B | 7.1e-08 | 4.5e-08 | 3e-08 | 4.867E-08 | 2.074E-08 | 3.000E-08 | 7.100E-08 |
| 15732 | 787 | E787K | B | 5.2e-08 | 2.5e-08 | 4.2e-08 | 3.967E-08 | 1.365E-08 | 2.500E-08 | 5.200E-08 |
| 15733 | 787 | E787M | B | 5.1e-08 | 3e-08 | 3.4e-08 | 3.833E-08 | 1.115E-08 | 3.000E-08 | 5.100E-08 |
| 15734 | 787 | E787F | B | 8.9e-08 | 6.1e-08 | 2.6e-08 | 5.867E-08 | 3.156E-08 | 2.600E-08 | 8.900E-08 |
| 15735 | 787 | E787P | B | 2.3e-08 | 6.2e-08 | 5.9e-08 | 4.800E-08 | 2.170E-08 | 2.300E-08 | 6.200E-08 |
| 15736 | 787 | E787S | B | 2.4e-08 | 2.3e-08 | 3.1e-08 | 2.600E-08 | 4.359E-09 | 2.300E-08 | 3.100E-08 |
| 15737 | 787 | E787T | B | 1.2e-07 | 7.2e-08 | 2e-08 | 7.067E-08 | 5.001E-08 | 2.000E-08 | 1.200E-07 |
| 15738 | 787 | E787W | B | 1.8e-07 | 6e-08 | 2.2e-08 | 8.733E-08 | 8.247E-08 | 2.200E-08 | 1.800E-07 |
| 15739 | 787 | E787Y | B | 6e-08 | 3.5e-08 | 1.8e-08 | 3.767E-08 | 2.113E-08 | 1.800E-08 | 6.000E-08 |
| 15740 | 787 | E787V | B | 5.3e-08 | 1.8e-08 | 4.2e-08 | 3.767E-08 | 1.790E-08 | 1.800E-08 | 5.300E-08 |
| 15741 | 788 | Q788A | B | 2.4e-08 | 3.1e-08 | 1.1e-07 | 5.500E-08 | 4.776E-08 | 2.400E-08 | 1.100E-07 |
| 15742 | 788 | Q788R | B | 3.3e-08 | 4e-08 | 3.5e-08 | 3.600E-08 | 3.606E-09 | 3.300E-08 | 4.000E-08 |
| 15743 | 788 | Q788N | B | 4.3e-08 | 1.1e-07 | 7e-08 | 7.433E-08 | 3.371E-08 | 4.300E-08 | 1.100E-07 |
| 15744 | 788 | Q788D | B | 3.8e-08 | 7.2e-08 | 3.3e-08 | 4.767E-08 | 2.122E-08 | 3.300E-08 | 7.200E-08 |
| 15745 | 788 | Q788C | B | 5.2e-08 | 3.5e-08 | 1.1e-08 | 3.267E-08 | 2.060E-08 | 1.100E-08 | 5.200E-08 |
| 15746 | 788 | Q788E | B | 2.4e-08 | 4.5e-08 | 1.1e-07 | 5.967E-08 | 4.484E-08 | 2.400E-08 | 1.100E-07 |
| 15747 | 788 | Q788Q | B | 2.8e-08 | 3.1e-08 | 2.5e-08 | 2.800E-08 | 3.000E-09 | 2.500E-08 | 3.100E-08 |
| 15748 | 788 | Q788G | B | 3.3e-08 | 4.7e-08 | 8.8e-08 | 5.600E-08 | 2.858E-08 | 3.300E-08 | 8.800E-08 |
| 15749 | 788 | Q788H | B | 3.9e-08 | 5.6e-08 | 8e-08 | 5.833E-08 | 2.060E-08 | 3.900E-08 | 8.000E-08 |
| 15750 | 788 | Q788I | B | 7.6e-08 | 5.2e-08 | 3.8e-08 | 5.533E-08 | 1.922E-08 | 3.800E-08 | 7.600E-08 |
| 15751 | 788 | Q788L | B | 8.2e-08 | 2.1e-08 | 6.9e-08 | 5.733E-08 | 3.213E-08 | 2.100E-08 | 8.200E-08 |
| 15752 | 788 | Q788K | B | 3.1e-08 | 3.8e-08 | 2.1e-08 | 3.000E-08 | 8.544E-09 | 2.100E-08 | 3.800E-08 |
| 15753 | 788 | Q788M | B | 2.8e-08 | 5.1e-08 | 2.5e-08 | 3.467E-08 | 1.422E-08 | 2.500E-08 | 5.100E-08 |
| 15754 | 788 | Q788F | B | 5.2e-08 | 4.5e-08 | 3.8e-08 | 4.500E-08 | 7.000E-09 | 3.800E-08 | 5.200E-08 |
| 15755 | 788 | Q788P | B | 8.7e-08 | 1.6e-08 | 4.5e-08 | 4.933E-08 | 3.570E-08 | 1.600E-08 | 8.700E-08 |
| 15756 | 788 | Q788S | B | 4.4e-08 | 4.1e-08 | 1.4e-08 | 3.300E-08 | 1.652E-08 | 1.400E-08 | 4.400E-08 |
| 15757 | 788 | Q788T | B | 6e-08 | 9.1e-08 | 5.1e-08 | 6.733E-08 | 2.098E-08 | 5.100E-08 | 9.100E-08 |
| 15758 | 788 | Q788W | B | 4.2e-08 | 1.6e-08 | 6.2e-08 | 4.000E-08 | 2.307E-08 | 1.600E-08 | 6.200E-08 |
| 15759 | 788 | Q788Y | B | 8.3e-08 | 3.3e-08 | 3e-08 | 4.867E-08 | 2.977E-08 | 3.000E-08 | 8.300E-08 |
| 15760 | 788 | Q788V | B | 2.2e-08 | 4.3e-08 | 7.9e-08 | 4.800E-08 | 2.883E-08 | 2.200E-08 | 7.900E-08 |
| 15761 | 789 | D789A | B | 9e-08 | 4.9e-08 | 1.3e-08 | 5.067E-08 | 3.853E-08 | 1.300E-08 | 9.000E-08 |
| 15762 | 789 | D789R | B | 1.7e-08 | 6.6e-08 | 6.1e-08 | 4.800E-08 | 2.696E-08 | 1.700E-08 | 6.600E-08 |
| 15763 | 789 | D789N | B | 5e-08 | 4.1e-08 | 2.8e-08 | 3.967E-08 | 1.106E-08 | 2.800E-08 | 5.000E-08 |
| 15764 | 789 | D789D | B | 5e-08 | 1.8e-08 | 4.1e-08 | 3.633E-08 | 1.650E-08 | 1.800E-08 | 5.000E-08 |
| 15765 | 789 | D789C | B | 1.6e-07 | 3.8e-08 | 3.1e-08 | 7.633E-08 | 7.254E-08 | 3.100E-08 | 1.600E-07 |
| 15766 | 789 | D789E | B | 6.5e-08 | 3.1e-08 | 6e-08 | 5.200E-08 | 1.836E-08 | 3.100E-08 | 6.500E-08 |
| 15767 | 789 | D789Q | B | 5.9e-08 | 5.5e-08 | 9.8e-09 | 4.127E-08 | 2.732E-08 | 9.800E-09 | 5.900E-08 |
| 15768 | 789 | D789G | B | 7.6e-08 | 2e-08 | 1.1e-07 | 6.867E-08 | 4.545E-08 | 2.000E-08 | 1.100E-07 |
| 15769 | 789 | D789H | B | 5.8e-08 | 1e-07 | 3.6e-08 | 6.467E-08 | 3.252E-08 | 3.600E-08 | 1.000E-07 |
| 15770 | 789 | D789I | B | 9.7e-08 | 4e-08 | 1.1e-07 | 8.233E-08 | 3.723E-08 | 4.000E-08 | 1.100E-07 |
| 15771 | 789 | D789L | B | 2.4e-08 | 2.7e-08 | 1.9e-08 | 2.333E-08 | 4.041E-09 | 1.900E-08 | 2.700E-08 |
| 15772 | 789 | D789K | B | 7.8e-08 | 5.7e-08 | 2.5e-08 | 5.333E-08 | 2.669E-08 | 2.500E-08 | 7.800E-08 |
| 15773 | 789 | D789M | B | 2.8e-08 | 1.2e-08 | 1.7e-08 | 1.900E-08 | 8.185E-09 | 1.200E-08 | 2.800E-08 |
| 15774 | 789 | D789F | B | 6.1e-08 | 2.5e-08 | 2.3e-08 | 3.633E-08 | 2.139E-08 | 2.300E-08 | 6.100E-08 |
| 15775 | 789 | D789P | B | 5.4e-08 | 9.8e-08 | 3.2e-08 | 6.133E-08 | 3.361E-08 | 3.200E-08 | 9.800E-08 |
| 15776 | 789 | D789S | B | 4.8e-08 | 6.1e-08 | 8.3e-08 | 6.400E-08 | 1.769E-08 | 4.800E-08 | 8.300E-08 |
| 15777 | 789 | D789T | B | 9.1e-09 | 2.2e-08 | 1.6e-08 | 1.570E-08 | 6.455E-09 | 9.100E-09 | 2.200E-08 |
| 15778 | 789 | D789W | B | 2.4e-08 | 2.1e-08 | 2e-08 | 2.167E-08 | 2.082E-09 | 2.000E-08 | 2.400E-08 |
| 15779 | 789 | D789Y | B | 4.1e-08 | 7.5e-08 | 3.3e-08 | 4.967E-08 | 2.230E-08 | 3.300E-08 | 7.500E-08 |
| 15780 | 789 | D789V | B | 1.9e-08 | 1.9e-08 | 4.1e-08 | 2.633E-08 | 1.270E-08 | 1.900E-08 | 4.100E-08 |
| 15781 | 790 | S790A | B | 2.5e-08 | 6.7e-08 | 4.1e-08 | 4.433E-08 | 2.120E-08 | 2.500E-08 | 6.700E-08 |
| 15782 | 790 | S790R | B | 2.5e-08 | 5.3e-08 | 2.3e-08 | 3.367E-08 | 1.677E-08 | 2.300E-08 | 5.300E-08 |
| 15783 | 790 | S790N | B | 2.2e-08 | 3.5e-08 | 2.6e-08 | 2.767E-08 | 6.658E-09 | 2.200E-08 | 3.500E-08 |
| 15784 | 790 | S790D | B | 1.6e-08 | 1.5e-07 | 3.7e-08 | 6.767E-08 | 7.207E-08 | 1.600E-08 | 1.500E-07 |
| 15785 | 790 | S790C | B | 5.4e-08 | 1.7e-07 | 2.8e-08 | 8.400E-08 | 7.560E-08 | 2.800E-08 | 1.700E-07 |
| 15786 | 790 | S790E | B | 7.7e-08 | 1.1e-07 | 7.5e-08 | 8.733E-08 | 1.966E-08 | 7.500E-08 | 1.100E-07 |
| 15787 | 790 | S790Q | B | 6.4e-08 | 4.7e-08 | 5.2e-08 | 5.433E-08 | 8.737E-09 | 4.700E-08 | 6.400E-08 |
| 15788 | 790 | S790G | B | 4.2e-08 | 5.3e-08 | 2e-08 | 3.833E-08 | 1.680E-08 | 2.000E-08 | 5.300E-08 |
| 15789 | 790 | S790H | B | 3.7e-08 | 6.1e-08 | 4.1e-08 | 4.633E-08 | 1.286E-08 | 3.700E-08 | 6.100E-08 |
| 15790 | 790 | S790I | B | 2.2e-08 | 3.9e-08 | 3.1e-08 | 3.067E-08 | 8.505E-09 | 2.200E-08 | 3.900E-08 |
| 15791 | 790 | S790L | B | 6.3e-08 | 6.7e-08 | 2.8e-08 | 5.267E-08 | 2.146E-08 | 2.800E-08 | 6.700E-08 |
| 15792 | 790 | S790K | B | 4.8e-08 | 1.1e-07 | 2.3e-08 | 6.033E-08 | 4.479E-08 | 2.300E-08 | 1.100E-07 |
| 15793 | 790 | S790M | B | 3.7e-08 | 7.5e-08 | 2.8e-08 | 4.667E-08 | 2.495E-08 | 2.800E-08 | 7.500E-08 |
| 15794 | 790 | S790F | B | 3.9e-08 | 7.3e-08 | 2.1e-08 | 4.433E-08 | 2.641E-08 | 2.100E-08 | 7.300E-08 |
| 15795 | 790 | S790P | B | 5.1e-08 | 5.9e-08 | 2.4e-08 | 4.467E-08 | 1.834E-08 | 2.400E-08 | 5.900E-08 |
| 15796 | 790 | S790S | B | 3.3e-08 | 6.6e-08 | 6.2e-08 | 5.367E-08 | 1.801E-08 | 3.300E-08 | 6.600E-08 |
| 15797 | 790 | S790T | B | 1.5e-08 | 2.5e-08 | 2.7e-08 | 2.233E-08 | 6.429E-09 | 1.500E-08 | 2.700E-08 |
| 15798 | 790 | S790W | B | 2.9e-08 | 2.5e-08 | 6.7e-08 | 4.033E-08 | 2.318E-08 | 2.500E-08 | 6.700E-08 |
| 15799 | 790 | S790Y | B | 1.4e-07 | 6e-08 | 3e-08 | 7.667E-08 | 5.686E-08 | 3.000E-08 | 1.400E-07 |
| 15800 | 790 | S790V | B | 3.1e-08 | 3.1e-08 | 4e-08 | 3.400E-08 | 5.196E-09 | 3.100E-08 | 4.000E-08 |
| 15801 | 791 | K791A | B | 4e-08 | 3.3e-08 | 5.4e-08 | 4.233E-08 | 1.069E-08 | 3.300E-08 | 5.400E-08 |
| 15802 | 791 | K791R | B | 2.5e-08 | 9.1e-08 | 3e-08 | 4.867E-08 | 3.675E-08 | 2.500E-08 | 9.100E-08 |
| 15803 | 791 | K791N | B | 2.6e-08 | 8.6e-08 | 3.2e-08 | 4.800E-08 | 3.305E-08 | 2.600E-08 | 8.600E-08 |
| 15804 | 791 | K791D | B | 8.8e-08 | 7.7e-08 | 7.2e-08 | 7.900E-08 | 8.185E-09 | 7.200E-08 | 8.800E-08 |
| 15805 | 791 | K791C | B | 5.2e-08 | 5.9e-08 | 6.7e-08 | 5.933E-08 | 7.506E-09 | 5.200E-08 | 6.700E-08 |
| 15806 | 791 | K791E | B | 4.3e-08 | 1.7e-08 | 2.2e-08 | 2.733E-08 | 1.380E-08 | 1.700E-08 | 4.300E-08 |
| 15807 | 791 | K791Q | B | 5.5e-08 | 6.9e-08 | 4.3e-08 | 5.567E-08 | 1.301E-08 | 4.300E-08 | 6.900E-08 |
| 15808 | 791 | K791G | B | 4.8e-08 | 1e-07 | 5.1e-08 | 6.633E-08 | 2.919E-08 | 4.800E-08 | 1.000E-07 |
| 15809 | 791 | K791H | B | 1.1e-07 | 3.3e-08 | 6.8e-08 | 7.033E-08 | 3.855E-08 | 3.300E-08 | 1.100E-07 |

| | | | | | | | | | | |
|---|---|---|---|---|---|---|---|---|---|---|
| 15810 | 791 | K791I | B | 3.3e-08 | 2.7e-08 | 3.2e-08 | 3.067E-08 | 3.215E-09 | 2.700E-08 | 3.300E-08 |
| 15811 | 791 | K791L | B | 2.1e-08 | 4.1e-08 | 4.9e-08 | 3.700E-08 | 1.442E-08 | 2.100E-08 | 4.900E-08 |
| 15812 | 791 | K791K | B | 3.4e-08 | 1.3e-07 | 1.8e-08 | 6.067E-08 | 6.058E-08 | 1.800E-08 | 1.300E-07 |
| 15813 | 791 | K791M | B | 1.1e-07 | 9e-09 | 5e-08 | 5.633E-08 | 5.080E-08 | 9.000E-09 | 1.100E-07 |
| 15814 | 791 | K791F | B | 2.8e-08 | 5.3e-08 | 2.7e-08 | 3.600E-08 | 1.473E-08 | 2.700E-08 | 5.300E-08 |
| 15815 | 791 | K791P | B | 2.9e-08 | 1.3e-07 | 2.8e-08 | 6.233E-08 | 5.860E-08 | 2.800E-08 | 1.300E-07 |
| 15816 | 791 | K791S | B | 4.1e-08 | 2.3e-08 | 2.8e-08 | 3.067E-08 | 9.292E-09 | 2.300E-08 | 4.100E-08 |
| 15817 | 791 | K791T | B | 4e-08 | 2.2e-08 | 3.9e-08 | 3.367E-08 | 1.012E-08 | 2.200E-08 | 4.000E-08 |
| 15818 | 791 | K791W | B | 3.3e-08 | 3.1e-08 | 6.1e-08 | 4.167E-08 | 1.677E-08 | 3.100E-08 | 6.100E-08 |
| 15819 | 791 | K791Y | B | 1e-08 | 2.7e-08 | 9.5e-08 | 4.400E-08 | 4.498E-08 | 1.000E-08 | 9.500E-08 |
| 15820 | 791 | K791V | B | 5.9e-08 | 4.8e-08 | 6.7e-08 | 5.800E-08 | 9.539E-09 | 4.800E-08 | 6.700E-08 |
| 15821 | 792 | D792A | B | 4.3e-08 | 4e-08 | 5.7e-08 | 4.667E-08 | 9.074E-09 | 4.000E-08 | 5.700E-08 |
| 15822 | 792 | D792R | B | 3.2e-07 | 4e-08 | 4.1e-08 | 1.337E-07 | 1.614E-07 | 4.000E-08 | 3.200E-07 |
| 15823 | 792 | D792N | B | 5e-08 | 9.8e-08 | 6.1e-08 | 6.967E-08 | 2.515E-08 | 5.000E-08 | 9.800E-08 |
| 15824 | 792 | D792D | B | 9.5e-08 | 1.8e-08 | 1.1e-07 | 7.433E-08 | 4.936E-08 | 1.800E-08 | 1.100E-07 |
| 15825 | 792 | D792C | B | 3.8e-08 | 5.6e-08 | 7.5e-08 | 5.633E-08 | 1.850E-08 | 3.800E-08 | 7.500E-08 |
| 15826 | 792 | D792E | B | 4e-08 | 4e-08 | 2.3e-08 | 3.433E-08 | 9.815E-09 | 2.300E-08 | 4.000E-08 |
| 15827 | 792 | D792Q | B | 8.3e-08 | 3.5e-08 | 3.9e-08 | 5.233E-08 | 2.663E-08 | 3.500E-08 | 8.300E-08 |
| 15828 | 792 | D792G | B | 6e-08 | 3.9e-08 | 3.1e-08 | 4.333E-08 | 1.498E-08 | 3.100E-08 | 6.000E-08 |
| 15829 | 792 | D792H | B | 2.2e-08 | 5.4e-08 | 5.3e-08 | 4.300E-08 | 1.819E-08 | 2.200E-08 | 5.400E-08 |
| 15830 | 792 | D792I | B | 5.1e-08 | 7.1e-08 | 6e-08 | 6.067E-08 | 1.002E-08 | 5.100E-08 | 7.100E-08 |
| 15831 | 792 | D792L | B | 1.2e-07 | 5.3e-08 | 4.8e-08 | 7.367E-08 | 4.020E-08 | 4.800E-08 | 1.200E-07 |
| 15832 | 792 | D792K | B | 3.1e-08 | 9.2e-08 | 4.3e-08 | 5.533E-08 | 3.232E-08 | 3.100E-08 | 9.200E-08 |
| 15833 | 792 | D792M | B | 3e-08 | 1.7e-08 | 8.1e-08 | 4.267E-08 | 3.383E-08 | 1.700E-08 | 8.100E-08 |
| 15834 | 792 | D792F | B | 3.4e-08 | 4.1e-08 | 2e-08 | 3.167E-08 | 1.069E-08 | 2.000E-08 | 4.100E-08 |
| 15835 | 792 | D792P | B | 3.7e-08 | 4.7e-08 | 3.8e-08 | 4.067E-08 | 5.508E-09 | 3.700E-08 | 4.700E-08 |
| 15836 | 792 | D792S | B | 2e-08 | 4e-08 | 1.9e-07 | 8.333E-08 | 9.292E-08 | 2.000E-08 | 1.900E-07 |
| 15837 | 792 | D792T | B | 1.5e-07 | 9.6e-08 | 4.8e-08 | 9.800E-08 | 5.103E-08 | 4.800E-08 | 1.500E-07 |
| 15838 | 792 | D792W | B | 4.5e-08 | 5.9e-08 | 4.7e-08 | 5.033E-08 | 7.572E-09 | 4.500E-08 | 5.900E-08 |
| 15839 | 792 | D792Y | B | 4.9e-08 | 1e-08 | 5.1e-08 | 3.667E-08 | 2.312E-08 | 1.000E-08 | 5.100E-08 |
| 15840 | 792 | D792V | B | 8e-08 | 1.3e-07 | 3.1e-08 | 8.033E-08 | 4.950E-08 | 3.100E-08 | 1.300E-07 |
| 15841 | 793 | S793A | B | 3.6e-08 | 3.7e-08 | 5.8e-08 | 4.367E-08 | 1.242E-08 | 3.600E-08 | 5.800E-08 |
| 15842 | 793 | S793R | B | 3.4e-08 | 6.7e-08 | 6.9e-08 | 5.667E-08 | 1.966E-08 | 3.400E-08 | 6.900E-08 |
| 15843 | 793 | S793N | B | 2.7e-08 | 1.8e-08 | 3.3e-08 | 2.600E-08 | 7.550E-09 | 1.800E-08 | 3.300E-08 |
| 15844 | 793 | S793D | B | 3.9e-08 | 7.7e-08 | 4.8e-08 | 5.467E-08 | 1.986E-08 | 3.900E-08 | 7.700E-08 |
| 15845 | 793 | S793C | B | 2.1e-08 | 2.5e-08 | 5.2e-08 | 3.267E-08 | 1.686E-08 | 2.100E-08 | 5.200E-08 |
| 15846 | 793 | S793E | B | 5.4e-08 | 4.1e-08 | 3.3e-08 | 4.267E-08 | 1.060E-08 | 3.300E-08 | 5.400E-08 |
| 15847 | 793 | S793Q | B | 3.9e-08 | 6.6e-08 | 3.3e-08 | 4.600E-08 | 1.758E-08 | 3.300E-08 | 6.600E-08 |
| 15848 | 793 | S793G | B | 5.4e-08 | 6.4e-08 | 4.9e-08 | 5.567E-08 | 7.638E-09 | 4.900E-08 | 6.400E-08 |
| 15849 | 793 | S793H | B | 6.8e-08 | 1.1e-08 | 8.3e-08 | 5.400E-08 | 3.799E-08 | 1.100E-08 | 8.300E-08 |
| 15850 | 793 | S793I | B | 5.2e-08 | 2.9e-08 | 4.7e-08 | 4.267E-08 | 1.210E-08 | 2.900E-08 | 5.200E-08 |
| 15851 | 793 | S793L | B | 2.9e-08 | 8.1e-08 | 1.5e-07 | 8.667E-08 | 6.070E-08 | 2.900E-08 | 1.500E-07 |
| 15852 | 793 | S793K | B | 2.7e-08 | 8.2e-08 | 1.2e-07 | 7.633E-08 | 4.676E-08 | 2.700E-08 | 1.200E-07 |
| 15853 | 793 | S793M | B | 6.7e-08 | 1e-07 | 4.6e-08 | 7.100E-08 | 2.722E-08 | 4.600E-08 | 1.000E-07 |
| 15854 | 793 | S793F | B | 3.4e-08 | 3.5e-08 | 3.2e-08 | 3.367E-08 | 1.528E-09 | 3.200E-08 | 3.500E-08 |
| 15855 | 793 | S793P | B | 9.6e-08 | 9.9e-08 | 1.2e-08 | 6.900E-08 | 4.939E-08 | 1.200E-08 | 9.900E-08 |
| 15856 | 793 | S793S | B | 1.8e-07 | 3.2e-08 | 8.8e-08 | 1.000E-07 | 7.473E-08 | 3.200E-08 | 1.800E-07 |
| 15857 | 793 | S793T | B | 7.3e-08 | 2.1e-08 | 2.2e-08 | 3.867E-08 | 2.974E-08 | 2.100E-08 | 7.300E-08 |
| 15858 | 793 | S793W | B | 3.3e-08 | 1.7e-08 | 6.2e-08 | 3.733E-08 | 2.281E-08 | 1.700E-08 | 6.200E-08 |
| 15859 | 793 | S793Y | B | 5.1e-08 | 4.9e-08 | 5.1e-08 | 5.033E-08 | 1.155E-09 | 4.900E-08 | 5.100E-08 |
| 15860 | 793 | S793V | B | 5.9e-08 | 2.8e-08 | 2.2e-08 | 3.633E-08 | 1.986E-08 | 2.200E-08 | 5.900E-08 |
| 15861 | 794 | T794A | B | 5.1e-08 | 2e-08 | 1.2e-07 | 6.367E-08 | 5.119E-08 | 2.000E-08 | 1.200E-07 |
| 15862 | 794 | T794R | B | 1.9e-08 | 3.9e-08 | 1.8e-08 | 2.533E-08 | 1.185E-08 | 1.800E-08 | 3.900E-08 |
| 15863 | 794 | T794N | B | 8.2e-08 | 3.4e-08 | 2.1e-08 | 4.567E-08 | 3.213E-08 | 2.100E-08 | 8.200E-08 |
| 15864 | 794 | T794D | B | 5.4e-08 | 2.9e-08 | 1.8e-08 | 3.367E-08 | 1.845E-08 | 1.800E-08 | 5.400E-08 |
| 15865 | 794 | T794C | B | 3.9e-08 | 3.7e-08 | 1.4e-08 | 3.000E-08 | 1.389E-08 | 1.400E-08 | 3.900E-08 |
| 15866 | 794 | T794E | B | 4e-08 | 3.6e-08 | 1.2e-07 | 6.533E-08 | 4.738E-08 | 3.600E-08 | 1.200E-07 |
| 15867 | 794 | T794Q | B | 2.1e-08 | 1.1e-07 | 2.8e-08 | 5.300E-08 | 4.949E-08 | 2.100E-08 | 1.100E-07 |
| 15868 | 794 | T794G | B | 2.9e-08 | 7.1e-08 | 1.3e-08 | 3.767E-08 | 2.996E-08 | 1.300E-08 | 7.100E-08 |
| 15869 | 794 | T794H | B | 2.6e-08 | 5.8e-08 | 5.6e-08 | 4.667E-08 | 1.793E-08 | 2.600E-08 | 5.800E-08 |
| 15870 | 794 | T794I | B | 4.9e-08 | 3.7e-08 | 6.7e-08 | 5.100E-08 | 1.510E-08 | 3.700E-08 | 6.700E-08 |
| 15871 | 794 | T794L | B | 3.9e-08 | 1.6e-08 | 2.5e-08 | 2.667E-08 | 1.159E-08 | 1.600E-08 | 3.900E-08 |
| 15872 | 794 | T794K | B | 1.3e-08 | 1.7e-08 | 4.7e-08 | 2.567E-08 | 1.858E-08 | 1.300E-08 | 4.700E-08 |
| 15873 | 794 | T794M | B | 6.4e-08 | 2.1e-08 | 6e-08 | 4.833E-08 | 2.376E-08 | 2.100E-08 | 6.400E-08 |
| 15874 | 794 | T794F | B | 2.8e-08 | 1.2e-07 | 9.8e-08 | 8.200E-08 | 4.804E-08 | 2.800E-08 | 1.200E-07 |
| 15875 | 794 | T794P | B | 2.5e-08 | 2.4e-08 | 3.8e-08 | 2.900E-08 | 7.810E-09 | 2.400E-08 | 3.800E-08 |
| 15876 | 794 | T794S | B | 6e-08 | 3e-08 | 4.5e-08 | 4.500E-08 | 1.500E-08 | 3.000E-08 | 6.000E-08 |
| 15877 | 794 | T794T | B | 1.7e-07 | 2.8e-08 | 5.2e-08 | 8.333E-08 | 7.601E-08 | 2.800E-08 | 1.700E-07 |
| 15878 | 794 | T794W | B | 3.1e-08 | 4.4e-08 | 8.6e-08 | 5.367E-08 | 2.875E-08 | 3.100E-08 | 8.600E-08 |
| 15879 | 794 | T794Y | B | 5.8e-08 | 1.1e-08 | 8.1e-08 | 5.000E-08 | 3.568E-08 | 1.100E-08 | 8.100E-08 |
| 15880 | 794 | T794V | B | 1.7e-07 | 4.1e-08 | 9.1e-08 | 1.007E-07 | 6.504E-08 | 4.100E-08 | 1.700E-07 |
| 15881 | 795 | Y795A | B | 8e-08 | 3.3e-08 | 5e-08 | 5.433E-08 | 2.380E-08 | 3.300E-08 | 8.000E-08 |
| 15882 | 795 | Y795R | B | 2.8e-08 | 3.3e-08 | 3.9e-08 | 3.333E-08 | 5.508E-09 | 2.800E-08 | 3.900E-08 |
| 15883 | 795 | Y795N | B | 2.9e-08 | 2.7e-08 | 8.7e-08 | 4.767E-08 | 3.408E-08 | 2.700E-08 | 8.700E-08 |
| 15884 | 795 | Y795D | B | 6.1e-08 | 9e-08 | 5.4e-08 | 6.833E-08 | 1.909E-08 | 5.400E-08 | 9.000E-08 |
| 15885 | 795 | Y795C | B | 4.6e-08 | 2.8e-08 | 7.9e-08 | 5.100E-08 | 2.587E-08 | 2.800E-08 | 7.900E-08 |
| 15886 | 795 | Y795E | B | 2.1e-08 | 4.4e-08 | 2.8e-08 | 3.100E-08 | 1.179E-08 | 2.100E-08 | 4.400E-08 |
| 15887 | 795 | Y795Q | B | 5.6e-08 | 8e-08 | 2.6e-08 | 5.400E-08 | 2.706E-08 | 2.600E-08 | 8.000E-08 |
| 15888 | 795 | Y795G | B | 2e-08 | 4.8e-08 | 3.4e-08 | 3.400E-08 | 1.400E-08 | 2.000E-08 | 4.800E-08 |
| 15889 | 795 | Y795H | B | 9.9e-09 | 5.2e-08 | 4.9e-08 | 3.697E-08 | 2.349E-08 | 9.900E-09 | 5.200E-08 |
| 15890 | 795 | Y795I | B | 4.5e-08 | 4.7e-08 | 3.7e-08 | 4.300E-08 | 5.292E-09 | 3.700E-08 | 4.700E-08 |
| 15891 | 795 | Y795L | B | 2.5e-08 | 1.9e-08 | 3.7e-08 | 2.700E-08 | 9.165E-09 | 1.900E-08 | 3.700E-08 |
| 15892 | 795 | Y795K | B | 1.6e-08 | 3.7e-08 | 4.6e-08 | 3.300E-08 | 1.539E-08 | 1.600E-08 | 4.600E-08 |
| 15893 | 795 | Y795M | B | 3.3e-08 | 2.1e-08 | 6.5e-08 | 3.967E-08 | 2.274E-08 | 2.100E-08 | 6.500E-08 |
| 15894 | 795 | Y795F | B | 4e-08 | 1.1e-07 | 7.5e-08 | 7.500E-08 | 3.500E-08 | 4.000E-08 | 1.100E-07 |

| | | | | | | | | | | |
|---|---|---|---|---|---|---|---|---|---|---|
| 15895 | 795 | Y795P | B | 3.3e-08 | 3.5e-08 | 3.5e-08 | 3.433E-08 | 1.155E-09 | 3.300E-08 | 3.500E-08 |
| 15896 | 795 | Y795S | B | 3.7e-08 | 5.4e-08 | 4.3e-08 | 4.467E-08 | 8.622E-09 | 3.700E-08 | 5.400E-08 |
| 15897 | 795 | Y795T | B | 1.8e-08 | 1.6e-07 | 1.1e-07 | 9.600E-08 | 7.203E-08 | 1.800E-08 | 1.600E-07 |
| 15898 | 795 | Y795W | B | 4.9e-08 | 5.8e-08 | 8.9e-08 | 6.533E-08 | 2.098E-08 | 4.900E-08 | 8.900E-08 |
| 15899 | 795 | Y795Y | B | 5.4e-08 | 2.7e-08 | 3.8e-08 | 3.967E-08 | 1.358E-08 | 2.700E-08 | 5.400E-08 |
| 15900 | 795 | Y795V | B | 7.1e-08 | 5.1e-08 | 9.7e-08 | 7.300E-08 | 2.307E-08 | 5.100E-08 | 9.700E-08 |
| 15901 | 796 | S796A | B | 3.6e-08 | 3.4e-08 | 3e-08 | 3.333E-08 | 3.055E-09 | 3.000E-08 | 3.600E-08 |
| 15902 | 796 | S796R | B | 1.7e-08 | 3e-08 | 2.5e-08 | 2.400E-08 | 6.557E-09 | 1.700E-08 | 3.000E-08 |
| 15903 | 796 | S796N | B | 2.4e-08 | 6e-08 | 2.2e-08 | 3.533E-08 | 2.139E-08 | 2.200E-08 | 6.000E-08 |
| 15904 | 796 | S796D | B | 2e-08 | 2.9e-08 | 4.5e-08 | 3.133E-08 | 1.266E-08 | 2.000E-08 | 4.500E-08 |
| 15905 | 796 | S796C | B | 9.5e-09 | 2.6e-08 | 1.1e-07 | 4.850E-08 | 5.390E-08 | 9.500E-09 | 1.100E-07 |
| 15906 | 796 | S796E | B | 6.7e-08 | 4.8e-08 | 5.8e-08 | 5.767E-08 | 9.504E-09 | 4.800E-08 | 6.700E-08 |
| 15907 | 796 | S796Q | B | 3.3e-08 | 1.1e-08 | 2e-08 | 2.133E-08 | 1.106E-08 | 1.100E-08 | 3.300E-08 |
| 15908 | 796 | S796G | B | 4.5e-08 | 7.1e-08 | 1.4e-08 | 4.333E-08 | 2.854E-08 | 1.400E-08 | 7.100E-08 |
| 15909 | 796 | S796H | B | 1.4e-07 | 6.7e-08 | 2.9e-08 | 7.867E-08 | 5.641E-08 | 2.900E-08 | 1.400E-07 |
| 15910 | 796 | S796I | B | 1.8e-08 | 3.2e-08 | 1e-07 | 5.000E-08 | 4.386E-08 | 1.800E-08 | 1.000E-07 |
| 15911 | 796 | S796L | B | 6.2e-08 | 9.4e-08 | 2.1e-08 | 5.900E-08 | 3.659E-08 | 2.100E-08 | 9.400E-08 |
| 15912 | 796 | S796K | B | 4.7e-08 | 4.8e-08 | 3.1e-08 | 4.200E-08 | 9.539E-09 | 3.100E-08 | 4.800E-08 |
| 15913 | 796 | S796M | B | 4.2e-08 | 5.2e-08 | 4.1e-08 | 4.500E-08 | 6.083E-09 | 4.100E-08 | 5.200E-08 |
| 15914 | 796 | S796F | B | 6.8e-08 | 2.6e-08 | 2.8e-08 | 4.067E-08 | 2.369E-08 | 2.600E-08 | 6.800E-08 |
| 15915 | 796 | S796P | B | 5.9e-08 | 2.5e-08 | 4.8e-08 | 4.400E-08 | 1.735E-08 | 2.500E-08 | 5.900E-08 |
| 15916 | 796 | S796S | B | 5e-08 | 1.1e-07 | 1.9e-08 | 5.967E-08 | 4.626E-08 | 1.900E-08 | 1.100E-07 |
| 15917 | 796 | S796T | B | 4.3e-08 | 1.8e-08 | 5.9e-08 | 4.000E-08 | 2.066E-08 | 1.800E-08 | 5.900E-08 |
| 15918 | 796 | S796W | B | 2.4e-08 | 3.9e-08 | 3.8e-08 | 3.367E-08 | 8.386E-09 | 2.400E-08 | 3.900E-08 |
| 15919 | 796 | S796Y | B | 6.6e-08 | 9.2e-08 | 4.3e-08 | 6.700E-08 | 2.452E-08 | 4.300E-08 | 9.200E-08 |
| 15920 | 796 | S796V | B | 5e-08 | 1.9e-08 | 4.2e-08 | 3.700E-08 | 1.609E-08 | 1.900E-08 | 5.000E-08 |
| 15921 | 797 | L797A | B | 6.5e-08 | 6.6e-08 | 7e-08 | 6.700E-08 | 2.646E-09 | 6.500E-08 | 7.000E-08 |
| 15922 | 797 | L797R | B | 3.8e-08 | 9.6e-09 | 2.4e-08 | 2.387E-08 | 1.420E-08 | 9.600E-09 | 3.800E-08 |
| 15923 | 797 | L797N | B | 1.8e-08 | 4.2e-08 | 4.9e-08 | 3.633E-08 | 1.626E-08 | 1.800E-08 | 4.900E-08 |
| 15924 | 797 | L797D | B | 2.9e-08 | 4.1e-08 | 3.1e-08 | 3.367E-08 | 6.429E-09 | 2.900E-08 | 4.100E-08 |
| 15925 | 797 | L797C | B | 5.1e-08 | 7.1e-08 | 5.5e-08 | 5.900E-08 | 1.058E-08 | 5.100E-08 | 7.100E-08 |
| 15926 | 797 | L797E | B | 3.6e-08 | 4.8e-08 | 2.7e-08 | 3.700E-08 | 1.054E-08 | 2.700E-08 | 4.800E-08 |
| 15927 | 797 | L797Q | B | 3.9e-08 | 2.5e-08 | 3.3e-08 | 3.233E-08 | 7.024E-09 | 2.500E-08 | 3.900E-08 |
| 15928 | 797 | L797G | B | 4.9e-08 | 3.4e-08 | 1.8e-07 | 8.767E-08 | 8.031E-08 | 3.400E-08 | 1.800E-07 |
| 15929 | 797 | L797H | B | 1.2e-07 | 3.7e-08 | 4.1e-08 | 6.600E-08 | 4.681E-08 | 3.700E-08 | 1.200E-07 |
| 15930 | 797 | L797I | B | 4.8e-08 | 5.4e-08 | 7.2e-08 | 5.800E-08 | 1.249E-08 | 4.800E-08 | 7.200E-08 |
| 15931 | 797 | L797L | B | 1.9e-08 | 4e-08 | 4.9e-08 | 3.600E-08 | 1.539E-08 | 1.900E-08 | 4.900E-08 |
| 15932 | 797 | L797K | B | 7.9e-08 | 5e-08 | 2.6e-08 | 5.167E-08 | 2.654E-08 | 2.600E-08 | 7.900E-08 |
| 15933 | 797 | L797M | B | 1.1e-08 | 9.4e-08 | 7.8e-08 | 6.100E-08 | 4.403E-08 | 1.100E-08 | 9.400E-08 |
| 15934 | 797 | L797F | B | 4.7e-08 | 4.3e-08 | 6.8e-08 | 5.267E-08 | 1.343E-08 | 4.300E-08 | 6.800E-08 |
| 15935 | 797 | L797P | B | 8.3e-08 | 2.8e-08 | 4e-08 | 5.033E-08 | 2.892E-08 | 2.800E-08 | 8.300E-08 |
| 15936 | 797 | L797S | B | 9.8e-09 | 7.9e-08 | 1.2e-07 | 6.960E-08 | 5.570E-08 | 9.800E-09 | 1.200E-07 |
| 15937 | 797 | L797T | B | 5.8e-08 | 1.9e-08 | 8.9e-08 | 5.533E-08 | 3.508E-08 | 1.900E-08 | 8.900E-08 |
| 15938 | 797 | L797W | B | 7.2e-08 | 2.4e-08 | 3.5e-08 | 4.367E-08 | 2.515E-08 | 2.400E-08 | 7.200E-08 |
| 15939 | 797 | L797Y | B | 2.4e-08 | 7.1e-08 | 3.7e-08 | 4.400E-08 | 2.427E-08 | 2.400E-08 | 7.100E-08 |
| 15940 | 797 | L797V | B | 3.2e-08 | 1e-07 | 2.7e-08 | 5.300E-08 | 4.078E-08 | 2.700E-08 | 1.000E-07 |
| 15941 | 798 | S798A | B | 2e-08 | 3.3e-08 | 6.1e-08 | 3.800E-08 | 2.095E-08 | 2.000E-08 | 6.100E-08 |
| 15942 | 798 | S798R | B | 1.1e-08 | 4.4e-08 | 4.4e-08 | 3.300E-08 | 1.905E-08 | 1.100E-08 | 4.400E-08 |
| 15943 | 798 | S798N | B | 5.9e-08 | 3.7e-08 | 2.6e-08 | 4.067E-08 | 1.680E-08 | 2.600E-08 | 5.900E-08 |
| 15944 | 798 | S798D | B | 2.2e-08 | 4.1e-08 | 3.8e-08 | 3.367E-08 | 1.021E-08 | 2.200E-08 | 4.100E-08 |
| 15945 | 798 | S798C | B | 3.5e-08 | 1.9e-08 | 1.6e-07 | 7.133E-08 | 7.720E-08 | 1.900E-08 | 1.600E-07 |
| 15946 | 798 | S798E | B | 7.4e-08 | 1.9e-08 | 4.1e-08 | 4.467E-08 | 2.768E-08 | 1.900E-08 | 7.400E-08 |
| 15947 | 798 | S798Q | B | 7.7e-08 | 4.6e-08 | 1.6e-07 | 9.433E-08 | 5.894E-08 | 4.600E-08 | 1.600E-07 |
| 15948 | 798 | S798G | B | 1.7e-08 | 5.4e-08 | 2.1e-08 | 3.067E-08 | 2.031E-08 | 1.700E-08 | 5.400E-08 |
| 15949 | 798 | S798H | B | 1.5e-08 | 5.2e-08 | 4.1e-08 | 3.600E-08 | 1.900E-08 | 1.500E-08 | 5.200E-08 |
| 15950 | 798 | S798I | B | 5.2e-08 | 1e-07 | 5.4e-08 | 6.867E-08 | 2.715E-08 | 5.200E-08 | 1.000E-07 |
| 15951 | 798 | S798L | B | 7.6e-08 | 2.7e-08 | 2.7e-08 | 4.333E-08 | 2.829E-08 | 2.700E-08 | 7.600E-08 |
| 15952 | 798 | S798K | B | 1.9e-08 | 3.7e-08 | 3.9e-08 | 3.167E-08 | 1.102E-08 | 1.900E-08 | 3.900E-08 |
| 15953 | 798 | S798M | B | 6.9e-08 | 6.6e-08 | 2.9e-08 | 5.467E-08 | 2.228E-08 | 2.900E-08 | 6.900E-08 |
| 15954 | 798 | S798F | B | 4.4e-08 | 1.3e-08 | 8.8e-08 | 4.833E-08 | 3.769E-08 | 1.300E-08 | 8.800E-08 |
| 15955 | 798 | S798P | B | 6.1e-08 | 4.8e-08 | 4.4e-08 | 5.100E-08 | 8.888E-09 | 4.400E-08 | 6.100E-08 |
| 15956 | 798 | S798S | B | 1e-07 | 9.4e-08 | 9.2e-08 | 9.533E-08 | 4.163E-09 | 9.200E-08 | 1.000E-07 |
| 15957 | 798 | S798T | B | 2.3e-08 | 7.1e-08 | 2.9e-08 | 4.100E-08 | 2.615E-08 | 2.300E-08 | 7.100E-08 |
| 15958 | 798 | S798W | B | 5.2e-08 | 3.1e-08 | 4.8e-08 | 4.367E-08 | 1.115E-08 | 3.100E-08 | 5.200E-08 |
| 15959 | 798 | S798Y | B | 1.6e-08 | 5.7e-08 | 6.2e-08 | 4.500E-08 | 2.524E-08 | 1.600E-08 | 6.200E-08 |
| 15960 | 798 | S798V | B | 4.7e-08 | 2.6e-08 | 6.2e-08 | 4.500E-08 | 1.808E-08 | 2.600E-08 | 6.200E-08 |
| 15961 | 799 | S799A | B | 2.4e-08 | 3.4e-08 | 6e-08 | 3.933E-08 | 1.858E-08 | 2.400E-08 | 6.000E-08 |
| 15962 | 799 | S799R | B | 8.5e-09 | 3.1e-08 | 4.8e-08 | 2.917E-08 | 1.981E-08 | 8.500E-09 | 4.800E-08 |
| 15963 | 799 | S799N | B | 2.1e-08 | 6.6e-09 | 2.4e-07 | 8.920E-08 | 1.308E-07 | 6.600E-09 | 2.400E-07 |
| 15964 | 799 | S799D | B | 2.2e-08 | 3.3e-08 | 4.2e-08 | 3.233E-08 | 1.002E-08 | 2.200E-08 | 4.200E-08 |
| 15965 | 799 | S799C | B | 7e-08 | 7e-08 | 1.6e-07 | 1.000E-07 | 5.196E-08 | 7.000E-08 | 1.600E-07 |
| 15966 | 799 | S799E | B | 4.9e-08 | 1.9e-08 | 6.8e-08 | 4.533E-08 | 2.470E-08 | 1.900E-08 | 6.800E-08 |
| 15967 | 799 | S799Q | B | 5.8e-08 | 1.4e-07 | 4e-08 | 7.933E-08 | 5.330E-08 | 4.000E-08 | 1.400E-07 |
| 15968 | 799 | S799G | B | 1.9e-08 | 5.4e-08 | 5.3e-08 | 4.200E-08 | 1.992E-08 | 1.900E-08 | 5.400E-08 |
| 15969 | 799 | S799H | B | 4.1e-08 | 1.5e-08 | 3.6e-08 | 3.067E-08 | 1.380E-08 | 1.500E-08 | 4.100E-08 |
| 15970 | 799 | S799I | B | 8.4e-08 | 2.6e-08 | 5.9e-08 | 5.633E-08 | 2.909E-08 | 2.600E-08 | 8.400E-08 |
| 15971 | 799 | S799L | B | 5.5e-08 | 1e-08 | 4.8e-08 | 3.767E-08 | 2.421E-08 | 1.000E-08 | 5.500E-08 |
| 15972 | 799 | S799K | B | 2.2e-08 | 4.6e-08 | 3.5e-08 | 3.433E-08 | 1.201E-08 | 2.200E-08 | 4.600E-08 |
| 15973 | 799 | S799M | B | 2e-08 | 3.4e-08 | 3.7e-08 | 3.033E-08 | 9.074E-09 | 2.000E-08 | 3.700E-08 |
| 15974 | 799 | S799F | B | 8.3e-08 | 1.4e-07 | 2.9e-08 | 8.400E-08 | 5.551E-08 | 2.900E-08 | 1.400E-07 |
| 15975 | 799 | S799P | B | 1.3e-07 | 5.7e-08 | 4.5e-08 | 7.733E-08 | 4.600E-08 | 4.500E-08 | 1.300E-07 |
| 15976 | 799 | S799S | B | 1.8e-08 | 2.2e-08 | 5.1e-08 | 3.033E-08 | 1.801E-08 | 1.800E-08 | 5.100E-08 |
| 15977 | 799 | S799T | B | 8.2e-08 | 4.2e-08 | 1.6e-08 | 4.667E-08 | 3.325E-08 | 1.600E-08 | 8.200E-08 |
| 15978 | 799 | S799W | B | 2.8e-08 | 1.5e-08 | 4.5e-08 | 2.933E-08 | 1.504E-08 | 1.500E-08 | 4.500E-08 |
| 15979 | 799 | S799Y | B | 2.7e-08 | 3.9e-08 | 8.2e-08 | 4.933E-08 | 2.892E-08 | 2.700E-08 | 8.200E-08 |

| | | | | | | | | | | |
|---|---|---|---|---|---|---|---|---|---|---|
| 15980 | 799 | S799V | B | 2.4e-08 | 1e-08 | 6.4e-08 | 3.267E-08 | 2.802E-08 | 1.000E-08 | 6.400E-08 |
| 15981 | 800 | T800A | B | 4.7e-08 | 6.6e-08 | 4.2e-08 | 5.167E-08 | 1.266E-08 | 4.200E-08 | 6.600E-08 |
| 15982 | 800 | T800R | B | 7.2e-08 | 2.6e-08 | 4.9e-08 | 4.900E-08 | 2.300E-08 | 2.600E-08 | 7.200E-08 |
| 15983 | 800 | T800N | B | 4.2e-08 | 7.6e-08 | 8.7e-08 | 6.833E-08 | 2.346E-08 | 4.200E-08 | 8.700E-08 |
| 15984 | 800 | T800D | B | 2.6e-08 | 1.9e-08 | 5.6e-08 | 3.367E-08 | 1.966E-08 | 1.900E-08 | 5.600E-08 |
| 15985 | 800 | T800C | B | 6e-08 | 1.4e-07 | 2.8e-08 | 7.600E-08 | 5.769E-08 | 2.800E-08 | 1.400E-07 |
| 15986 | 800 | T800E | B | 7e-08 | 3.6e-08 | 1.5e-08 | 4.033E-08 | 2.775E-08 | 1.500E-08 | 7.000E-08 |
| 15987 | 800 | T800Q | B | 4.5e-08 | 8.2e-08 | 5e-08 | 5.900E-08 | 2.007E-08 | 4.500E-08 | 8.200E-08 |
| 15988 | 800 | T800G | B | 7.7e-08 | 4.7e-08 | 5.9e-08 | 6.100E-08 | 1.510E-08 | 4.700E-08 | 7.700E-08 |
| 15989 | 800 | T800H | B | 1.1e-07 | 5.4e-09 | 3.5e-08 | 5.013E-08 | 5.392E-08 | 5.400E-09 | 1.100E-07 |
| 15990 | 800 | T800I | B | 4.4e-08 | 2.7e-08 | 5.8e-08 | 4.300E-08 | 1.552E-08 | 2.700E-08 | 5.800E-08 |
| 15991 | 800 | T800L | B | 6.7e-08 | 1.5e-08 | 4.6e-08 | 4.267E-08 | 2.616E-08 | 1.500E-08 | 6.700E-08 |
| 15992 | 800 | T800K | B | 1e-07 | 7.8e-08 | 3.1e-08 | 6.967E-08 | 3.525E-08 | 3.100E-08 | 1.000E-07 |
| 15993 | 800 | T800M | B | 6.5e-08 | 5.2e-08 | 3e-08 | 4.900E-08 | 1.769E-08 | 3.000E-08 | 6.500E-08 |
| 15994 | 800 | T800F | B | 4.9e-08 | 2.4e-08 | 6.3e-08 | 4.533E-08 | 1.976E-08 | 2.400E-08 | 6.300E-08 |
| 15995 | 800 | T800P | B | 7.4e-08 | 1e-07 | 4.3e-08 | 7.233E-08 | 2.854E-08 | 4.300E-08 | 1.000E-07 |
| 15996 | 800 | T800S | B | 3.9e-08 | 4.3e-08 | 4e-08 | 4.067E-08 | 2.082E-09 | 3.900E-08 | 4.300E-08 |
| 15997 | 800 | T800T | B | 6.4e-08 | 4.1e-08 | 8.2e-08 | 6.233E-08 | 2.055E-08 | 4.100E-08 | 8.200E-08 |
| 15998 | 800 | T800W | B | 2.6e-08 | 4.1e-08 | 4.1e-08 | 3.600E-08 | 8.660E-09 | 2.600E-08 | 4.100E-08 |
| 15999 | 800 | T800Y | B | 4.6e-08 | 4.4e-08 | 2.6e-08 | 3.867E-08 | 1.102E-08 | 2.600E-08 | 4.600E-08 |
| 16000 | 800 | T800V | B | 1.8e-08 | 2.3e-08 | 3.9e-08 | 2.667E-08 | 1.097E-08 | 1.800E-08 | 3.900E-08 |
| 16001 | 801 | L801A | B | 9.2e-09 | 3.6e-08 | 1.8e-08 | 2.107E-08 | 1.366E-08 | 9.200E-09 | 3.600E-08 |
| 16002 | 801 | L801R | B | 9.1e-08 | 6.6e-08 | 4.3e-08 | 6.667E-08 | 2.401E-08 | 4.300E-08 | 9.100E-08 |
| 16003 | 801 | L801N | B | 1.5e-08 | 3.8e-08 | 2e-08 | 2.433E-08 | 1.210E-08 | 1.500E-08 | 3.800E-08 |
| 16004 | 801 | L801D | B | 5.8e-08 | 1.3e-08 | 7.9e-08 | 5.000E-08 | 3.372E-08 | 1.300E-08 | 7.900E-08 |
| 16005 | 801 | L801C | B | 2.2e-08 | 5e-08 | 1.8e-08 | 3.000E-08 | 1.744E-08 | 1.800E-08 | 5.000E-08 |
| 16006 | 801 | L801E | B | 4.3e-08 | 1.1e-08 | 2.9e-08 | 2.767E-08 | 1.604E-08 | 1.100E-08 | 4.300E-08 |
| 16007 | 801 | L801Q | B | 1e-07 | 1.1e-08 | 4.7e-08 | 5.267E-08 | 4.477E-08 | 1.100E-08 | 1.000E-07 |
| 16008 | 801 | L801G | B | 1.3e-08 | 4.2e-08 | 6.3e-08 | 3.933E-08 | 2.511E-08 | 1.300E-08 | 6.300E-08 |
| 16009 | 801 | L801H | B | 3.7e-08 | 4e-08 | 3.3e-08 | 3.667E-08 | 3.512E-09 | 3.300E-08 | 4.000E-08 |
| 16010 | 801 | L801I | B | 2.5e-08 | 4.3e-08 | 6.5e-08 | 4.433E-08 | 2.003E-08 | 2.500E-08 | 6.500E-08 |
| 16011 | 801 | L801L | B | 3.7e-08 | 2.7e-08 | 2.7e-08 | 3.033E-08 | 5.774E-09 | 2.700E-08 | 3.700E-08 |
| 16012 | 801 | L801K | B | 2.1e-08 | 1e-07 | 9.3e-08 | 7.133E-08 | 4.373E-08 | 2.100E-08 | 1.000E-07 |
| 16013 | 801 | L801M | B | 2e-08 | 4.4e-08 | 3.4e-08 | 3.267E-08 | 1.206E-08 | 2.000E-08 | 4.400E-08 |
| 16014 | 801 | L801F | B | 1.1e-07 | 1.3e-08 | 4.6e-08 | 5.633E-08 | 4.932E-08 | 1.300E-08 | 1.100E-07 |
| 16015 | 801 | L801P | B | 2.7e-08 | 8.7e-08 | 8.6e-08 | 6.667E-08 | 3.436E-08 | 2.700E-08 | 8.700E-08 |
| 16016 | 801 | L801S | B | 6.3e-08 | 2.8e-08 | 8.3e-08 | 5.800E-08 | 2.784E-08 | 2.800E-08 | 8.300E-08 |
| 16017 | 801 | L801T | B | 9.3e-08 | 1.7e-07 | 5.4e-08 | 1.057E-07 | 5.903E-08 | 5.400E-08 | 1.700E-07 |
| 16018 | 801 | L801W | B | 6.9e-08 | 2.2e-08 | 3.4e-08 | 4.167E-08 | 2.442E-08 | 2.200E-08 | 6.900E-08 |
| 16019 | 801 | L801Y | B | 3e-08 | 2.1e-08 | 6.1e-08 | 3.733E-08 | 2.098E-08 | 2.100E-08 | 6.100E-08 |
| 16020 | 801 | L801V | B | 2.1e-08 | 9.4e-09 | 6.2e-08 | 3.080E-08 | 2.764E-08 | 9.400E-09 | 6.200E-08 |
| 16021 | 802 | T802A | B | 4.4e-08 | 8.8e-08 | 7.6e-08 | 6.933E-08 | 2.274E-08 | 4.400E-08 | 8.800E-08 |
| 16022 | 802 | T802R | B | 4e-08 | 1.2e-08 | 9.8e-09 | 2.060E-08 | 1.684E-08 | 9.800E-09 | 4.000E-08 |
| 16023 | 802 | T802N | B | 2e-08 | 3.1e-08 | 2.7e-08 | 2.600E-08 | 5.568E-09 | 2.000E-08 | 3.100E-08 |
| 16024 | 802 | T802D | B | 5.3e-08 | 2.6e-08 | 3.1e-08 | 3.667E-08 | 1.436E-08 | 2.600E-08 | 5.300E-08 |
| 16025 | 802 | T802C | B | 4.3e-08 | 4.7e-08 | 8.8e-08 | 5.933E-08 | 2.491E-08 | 4.300E-08 | 8.800E-08 |
| 16026 | 802 | T802E | B | 7.2e-08 | 4.2e-08 | 3.9e-08 | 5.100E-08 | 1.825E-08 | 3.900E-08 | 7.200E-08 |
| 16027 | 802 | T802Q | B | 5.9e-08 | 3.4e-08 | 2.9e-08 | 4.067E-08 | 1.607E-08 | 2.900E-08 | 5.900E-08 |
| 16028 | 802 | T802G | B | 1.2e-07 | 2.3e-08 | 8.4e-08 | 7.567E-08 | 4.903E-08 | 2.300E-08 | 1.200E-07 |
| 16029 | 802 | T802H | B | 3.2e-08 | 4.7e-08 | 9.5e-08 | 5.800E-08 | 3.291E-08 | 3.200E-08 | 9.500E-08 |
| 16030 | 802 | T802I | B | 4.4e-08 | 2.8e-08 | 4.8e-08 | 4.000E-08 | 1.058E-08 | 2.800E-08 | 4.800E-08 |
| 16031 | 802 | T802L | B | 5.6e-08 | 2.2e-08 | 5.2e-08 | 4.333E-08 | 1.858E-08 | 2.200E-08 | 5.600E-08 |
| 16032 | 802 | T802K | B | 5.2e-08 | 6.1e-08 | 6.8e-08 | 6.033E-08 | 8.021E-09 | 5.200E-08 | 6.800E-08 |
| 16033 | 802 | T802M | B | 2.3e-08 | 3e-08 | 6.9e-09 | 1.997E-08 | 1.184E-08 | 6.900E-09 | 3.000E-08 |
| 16034 | 802 | T802F | B | 3e-08 | 4.4e-08 | 1.4e-07 | 7.133E-08 | 5.988E-08 | 3.000E-08 | 1.400E-07 |
| 16035 | 802 | T802P | B | 2.8e-08 | 2.7e-07 | 5.1e-09 | 1.010E-07 | 1.468E-07 | 5.100E-09 | 2.700E-07 |
| 16036 | 802 | T802S | B | 4e-08 | 6.2e-08 | 2.9e-08 | 4.367E-08 | 1.680E-08 | 2.900E-08 | 6.200E-08 |
| 16037 | 802 | T802T | B | 6.2e-08 | 5.6e-08 | 1.5e-08 | 4.433E-08 | 2.558E-08 | 1.500E-08 | 6.200E-08 |
| 16038 | 802 | T802W | B | 3.2e-08 | 4.2e-08 | 5e-08 | 4.133E-08 | 9.018E-09 | 3.200E-08 | 5.000E-08 |
| 16039 | 802 | T802Y | B | 1.8e-08 | 1.7e-08 | 3.9e-08 | 2.467E-08 | 1.242E-08 | 1.700E-08 | 3.900E-08 |
| 16040 | 802 | T802V | B | 4.9e-08 | 2e-08 | 2.4e-08 | 3.100E-08 | 1.572E-08 | 2.000E-08 | 4.900E-08 |
| 16041 | 803 | L803A | B | 7.4e-08 | 6.2e-08 | 6e-08 | 6.533E-08 | 7.572E-09 | 6.000E-08 | 7.400E-08 |
| 16042 | 803 | L803R | B | 7.5e-08 | 9.9e-08 | 4.5e-08 | 7.300E-08 | 2.706E-08 | 4.500E-08 | 9.900E-08 |
| 16043 | 803 | L803N | B | 3.6e-08 | 1.9e-08 | 6.1e-08 | 3.867E-08 | 2.113E-08 | 1.900E-08 | 6.100E-08 |
| 16044 | 803 | L803D | B | 2.8e-08 | 3.8e-08 | 4.5e-08 | 3.700E-08 | 8.544E-09 | 2.800E-08 | 4.500E-08 |
| 16045 | 803 | L803C | B | 6.2e-08 | 1.2e-07 | 6.1e-08 | 8.100E-08 | 3.378E-08 | 6.100E-08 | 1.200E-07 |
| 16046 | 803 | L803E | B | 4.6e-08 | 7e-08 | 3.9e-08 | 5.167E-08 | 1.626E-08 | 3.900E-08 | 7.000E-08 |
| 16047 | 803 | L803Q | B | 2.9e-08 | 3.6e-08 | 3.4e-08 | 3.300E-08 | 3.606E-09 | 2.900E-08 | 3.600E-08 |
| 16048 | 803 | L803G | B | 2e-08 | 3e-08 | 5.3e-08 | 3.433E-08 | 1.692E-08 | 2.000E-08 | 5.300E-08 |
| 16049 | 803 | L803H | B | 2.4e-08 | 5.5e-08 | 3.2e-08 | 3.700E-08 | 1.609E-08 | 2.400E-08 | 5.500E-08 |
| 16050 | 803 | L803I | B | 7.1e-08 | 2.9e-08 | 2.8e-08 | 4.267E-08 | 2.454E-08 | 2.800E-08 | 7.100E-08 |
| 16051 | 803 | L803L | B | 3e-08 | 2.1e-08 | 3.2e-08 | 2.767E-08 | 5.859E-09 | 2.100E-08 | 3.200E-08 |
| 16052 | 803 | L803K | B | 2.3e-08 | 2.3e-08 | 2e-08 | 2.200E-08 | 1.732E-09 | 2.000E-08 | 2.300E-08 |
| 16053 | 803 | L803M | B | 6.1e-08 | 1.1e-07 | 7.6e-08 | 8.233E-08 | 2.511E-08 | 6.100E-08 | 1.100E-07 |
| 16054 | 803 | L803F | B | 2.8e-08 | 4.9e-08 | 1e-07 | 5.900E-08 | 3.703E-08 | 2.800E-08 | 1.000E-07 |
| 16055 | 803 | L803P | B | 5.7e-08 | 4.9e-08 | 5.2e-08 | 5.267E-08 | 4.041E-09 | 4.900E-08 | 5.700E-08 |
| 16056 | 803 | L803S | B | 6e-08 | 2.9e-08 | 5.3e-08 | 4.733E-08 | 1.626E-08 | 2.900E-08 | 6.000E-08 |
| 16057 | 803 | L803T | B | 4.5e-08 | 5.3e-08 | 8.1e-08 | 5.967E-08 | 1.890E-08 | 4.500E-08 | 8.100E-08 |
| 16058 | 803 | L803W | B | 1e-08 | 4.4e-08 | 4.6e-08 | 3.333E-08 | 2.023E-08 | 1.000E-08 | 4.600E-08 |
| 16059 | 803 | L803Y | B | 1.3e-08 | 1.5e-08 | 1.4e-07 | 5.600E-08 | 7.275E-08 | 1.300E-08 | 1.400E-07 |
| 16060 | 803 | L803V | B | 5.3e-08 | 6.3e-08 | 4.2e-08 | 5.267E-08 | 1.050E-08 | 4.200E-08 | 6.300E-08 |
| 16061 | 804 | S804A | B | 1.1e-07 | 4e-08 | 4.1e-08 | 6.367E-08 | 4.013E-08 | 4.000E-08 | 1.100E-07 |
| 16062 | 804 | S804R | B | 5.9e-08 | 5.2e-08 | 1.6e-08 | 4.233E-08 | 2.307E-08 | 1.600E-08 | 5.900E-08 |
| 16063 | 804 | S804N | B | 2.2e-08 | 3.7e-08 | 7.2e-08 | 4.367E-08 | 2.566E-08 | 2.200E-08 | 7.200E-08 |
| 16064 | 804 | S804D | B | 8.2e-08 | 2.4e-08 | 2.1e-08 | 4.233E-08 | 3.439E-08 | 2.100E-08 | 8.200E-08 |

| | | | | | | | | | | |
|---|---|---|---|---|---|---|---|---|---|---|
| 16065 | 804 | S804C | B | 6.7e-08 | 7.5e-08 | 3.3e-08 | 5.833E-08 | 2.230E-08 | 3.300E-08 | 7.500E-08 |
| 16066 | 804 | S804E | B | 1.9e-07 | 6.3e-08 | 7.5e-08 | 1.093E-07 | 7.012E-08 | 6.300E-08 | 1.900E-07 |
| 16067 | 804 | S804Q | B | 7.8e-08 | 6.5e-08 | 6.4e-08 | 6.900E-08 | 7.810E-09 | 6.400E-08 | 7.800E-08 |
| 16068 | 804 | S804G | B | 9.1e-08 | 2.4e-08 | 1.2e-07 | 7.833E-08 | 4.924E-08 | 2.400E-08 | 1.200E-07 |
| 16069 | 804 | S804H | B | 1.7e-08 | 2.7e-08 | 2.1e-08 | 2.167E-08 | 5.033E-09 | 1.700E-08 | 2.700E-08 |
| 16070 | 804 | S804I | B | 7e-08 | 4.7e-08 | 2.5e-08 | 4.733E-08 | 2.250E-08 | 2.500E-08 | 7.000E-08 |
| 16071 | 804 | S804L | B | 4.3e-08 | 1.1e-08 | 3.3e-08 | 2.900E-08 | 1.637E-08 | 1.100E-08 | 4.300E-08 |
| 16072 | 804 | S804K | B | 3.5e-08 | 5.4e-08 | 3e-08 | 3.967E-08 | 1.266E-08 | 3.000E-08 | 5.400E-08 |
| 16073 | 804 | S804M | B | 4.1e-08 | 1.8e-07 | 3.4e-08 | 8.500E-08 | 8.235E-08 | 3.400E-08 | 1.800E-07 |
| 16074 | 804 | S804F | B | 4.3e-08 | 2.8e-08 | 3.7e-08 | 3.600E-08 | 7.550E-09 | 2.800E-08 | 4.300E-08 |
| 16075 | 804 | S804P | B | 2.2e-08 | 2.2e-08 | 3.5e-08 | 2.633E-08 | 7.506E-09 | 2.200E-08 | 3.500E-08 |
| 16076 | 804 | S804S | B | 1e-07 | 8.1e-08 | 2e-08 | 6.700E-08 | 4.180E-08 | 2.000E-08 | 1.000E-07 |
| 16077 | 804 | S804T | B | 5.3e-08 | 9.3e-08 | 7.6e-08 | 7.400E-08 | 2.007E-08 | 5.300E-08 | 9.300E-08 |
| 16078 | 804 | S804W | B | 6.7e-08 | 5.7e-08 | 4.6e-08 | 5.667E-08 | 1.050E-08 | 4.600E-08 | 6.700E-08 |
| 16079 | 804 | S804Y | B | 6.2e-08 | 7.9e-08 | 5.4e-08 | 6.500E-08 | 1.277E-08 | 5.400E-08 | 7.900E-08 |
| 16080 | 804 | S804V | B | 9e-08 | 2.9e-08 | 7.8e-08 | 6.567E-08 | 3.232E-08 | 2.900E-08 | 9.000E-08 |
| 16081 | 805 | K805A | B | 6e-08 | 8e-09 | 3.7e-08 | 3.500E-08 | 2.606E-08 | 8.000E-09 | 6.000E-08 |
| 16082 | 805 | K805R | B | 3e-08 | 7.4e-08 | 3.4e-08 | 4.600E-08 | 2.433E-08 | 3.000E-08 | 7.400E-08 |
| 16083 | 805 | K805N | B | 5.6e-08 | 4.5e-08 | 4.3e-08 | 4.800E-08 | 7.000E-09 | 4.300E-08 | 5.600E-08 |
| 16084 | 805 | K805D | B | 2.5e-08 | 2.7e-08 | 3.3e-08 | 2.833E-08 | 4.163E-09 | 2.500E-08 | 3.300E-08 |
| 16085 | 805 | K805C | B | 6e-08 | 2.1e-08 | 4e-08 | 4.033E-08 | 1.950E-08 | 2.100E-08 | 6.000E-08 |
| 16086 | 805 | K805E | B | 1.7e-08 | 7e-08 | 2.5e-08 | 3.733E-08 | 2.857E-08 | 1.700E-08 | 7.000E-08 |
| 16087 | 805 | K805Q | B | 2.8e-08 | 2.1e-08 | 4e-08 | 2.967E-08 | 9.609E-09 | 2.100E-08 | 4.000E-08 |
| 16088 | 805 | K805G | B | 6.2e-08 | 5.5e-08 | 4.3e-08 | 5.333E-08 | 9.609E-09 | 4.300E-08 | 6.200E-08 |
| 16089 | 805 | K805H | B | 1.5e-08 | 5.9e-08 | 6.2e-08 | 4.533E-08 | 2.631E-08 | 1.500E-08 | 6.200E-08 |
| 16090 | 805 | K805I | B | 7.3e-08 | 1.8e-08 | 7.7e-08 | 5.600E-08 | 3.297E-08 | 1.800E-08 | 7.700E-08 |
| 16091 | 805 | K805L | B | 1.2e-07 | 2.9e-08 | 5.3e-08 | 6.733E-08 | 4.716E-08 | 2.900E-08 | 1.200E-07 |
| 16092 | 805 | K805K | B | 3.5e-08 | 3e-08 | 6.4e-08 | 4.300E-08 | 1.836E-08 | 3.000E-08 | 6.400E-08 |
| 16093 | 805 | K805M | B | 3.8e-08 | 2.1e-08 | 5e-08 | 3.633E-08 | 1.457E-08 | 2.100E-08 | 5.000E-08 |
| 16094 | 805 | K805F | B | 3.9e-08 | 2.5e-08 | 7.8e-08 | 4.733E-08 | 2.747E-08 | 2.500E-08 | 7.800E-08 |
| 16095 | 805 | K805P | B | 4.3e-08 | 1.7e-08 | 2.9e-08 | 2.967E-08 | 1.301E-08 | 1.700E-08 | 4.300E-08 |
| 16096 | 805 | K805S | B | 2.7e-08 | 1.8e-08 | 2.6e-08 | 2.367E-08 | 4.933E-09 | 1.800E-08 | 2.700E-08 |
| 16097 | 805 | K805T | B | 2.4e-08 | 3.3e-08 | 4.8e-08 | 3.500E-08 | 1.212E-08 | 2.400E-08 | 4.800E-08 |
| 16098 | 805 | K805W | B | 5.3e-08 | 1e-08 | 5.2e-08 | 3.833E-08 | 2.454E-08 | 1.000E-08 | 5.300E-08 |
| 16099 | 805 | K805Y | B | 3.2e-08 | 5.8e-08 | 4.3e-08 | 4.433E-08 | 1.305E-08 | 3.200E-08 | 5.800E-08 |
| 16100 | 805 | K805V | B | 3.5e-08 | 3.9e-08 | 3.4e-08 | 3.600E-08 | 2.646E-09 | 3.400E-08 | 3.900E-08 |
| 16101 | 806 | A806A | B | 3.1e-08 | 3.4e-08 | 5.4e-08 | 3.967E-08 | 1.250E-08 | 3.100E-08 | 5.400E-08 |
| 16102 | 806 | A806R | B | 4.8e-08 | 4e-08 | 5.7e-08 | 4.833E-08 | 8.505E-09 | 4.000E-08 | 5.700E-08 |
| 16103 | 806 | A806N | B | 5.5e-08 | 3.4e-08 | 4.1e-08 | 4.333E-08 | 1.069E-08 | 3.400E-08 | 5.500E-08 |
| 16104 | 806 | A806D | B | 6.7e-08 | 4.1e-08 | 1.7e-07 | 9.267E-08 | 6.822E-08 | 4.100E-08 | 1.700E-07 |
| 16105 | 806 | A806C | B | 1.3e-08 | 2.1e-08 | 1.1e-07 | 4.800E-08 | 5.384E-08 | 1.300E-08 | 1.100E-07 |
| 16106 | 806 | A806E | B | 4e-08 | 2e-08 | 4.2e-08 | 3.400E-08 | 1.217E-08 | 2.000E-08 | 4.200E-08 |
| 16107 | 806 | A806Q | B | 4.8e-08 | 3.5e-08 | 2.1e-08 | 3.467E-08 | 1.350E-08 | 2.100E-08 | 4.800E-08 |
| 16108 | 806 | A806G | B | 2.8e-08 | 8.9e-08 | 4e-08 | 5.233E-08 | 3.232E-08 | 2.800E-08 | 8.900E-08 |
| 16109 | 806 | A806H | B | 7.6e-08 | 3.9e-08 | 3.4e-08 | 4.967E-08 | 2.294E-08 | 3.400E-08 | 7.600E-08 |
| 16110 | 806 | A806I | B | 1.6e-08 | 3e-08 | 1.3e-07 | 5.867E-08 | 6.217E-08 | 1.600E-08 | 1.300E-07 |
| 16111 | 806 | A806L | B | 3.1e-08 | 1.1e-08 | 4.1e-08 | 2.767E-08 | 1.528E-08 | 1.100E-08 | 4.100E-08 |
| 16112 | 806 | A806K | B | 4.3e-08 | 1.9e-08 | 3.2e-08 | 3.133E-08 | 1.201E-08 | 1.900E-08 | 4.300E-08 |
| 16113 | 806 | A806M | B | 5.9e-08 | 1.2e-07 | 2.5e-08 | 6.800E-08 | 4.814E-08 | 2.500E-08 | 1.200E-07 |
| 16114 | 806 | A806F | B | 4.3e-08 | 4.2e-08 | 2.9e-08 | 3.800E-08 | 7.810E-09 | 2.900E-08 | 4.300E-08 |
| 16115 | 806 | A806P | B | 3.1e-08 | 2.4e-08 | 2.6e-08 | 2.700E-08 | 3.606E-09 | 2.400E-08 | 3.100E-08 |
| 16116 | 806 | A806S | B | 4.8e-08 | 3.3e-08 | 6.1e-08 | 4.733E-08 | 1.401E-08 | 3.300E-08 | 6.100E-08 |
| 16117 | 806 | A806T | B | 6.7e-08 | 3.3e-08 | 3.4e-08 | 4.467E-08 | 1.935E-08 | 3.300E-08 | 6.700E-08 |
| 16118 | 806 | A806W | B | 4.1e-08 | 4.1e-08 | 2.9e-08 | 3.700E-08 | 6.928E-09 | 2.900E-08 | 4.100E-08 |
| 16119 | 806 | A806Y | B | 2.3e-08 | 5.9e-08 | 1.7e-08 | 3.300E-08 | 2.272E-08 | 1.700E-08 | 5.900E-08 |
| 16120 | 806 | A806V | B | 2.9e-08 | 6.1e-08 | 2.7e-08 | 3.900E-08 | 1.908E-08 | 2.700E-08 | 6.100E-08 |
| 16121 | 807 | D807A | B | 6.6e-08 | 8.8e-08 | 8.6e-08 | 8.000E-08 | 1.217E-08 | 6.600E-08 | 8.800E-08 |
| 16122 | 807 | D807R | B | 2.7e-08 | 3.2e-08 | 5.7e-08 | 3.867E-08 | 1.607E-08 | 2.700E-08 | 5.700E-08 |
| 16123 | 807 | D807N | B | 4.3e-08 | 1.2e-08 | 1.1e-07 | 5.500E-08 | 5.009E-08 | 1.200E-08 | 1.100E-07 |
| 16124 | 807 | D807D | B | 5.5e-08 | 1.2e-08 | 6.1e-08 | 4.267E-08 | 2.673E-08 | 1.200E-08 | 6.100E-08 |
| 16125 | 807 | D807C | B | 2.7e-08 | 2.5e-08 | 1.8e-08 | 2.333E-08 | 4.726E-09 | 1.800E-08 | 2.700E-08 |
| 16126 | 807 | D807E | B | 1.3e-07 | 4.7e-08 | 3.1e-08 | 6.933E-08 | 5.314E-08 | 3.100E-08 | 1.300E-07 |
| 16127 | 807 | D807Q | B | 2.2e-08 | 5.4e-08 | 1.6e-07 | 7.867E-08 | 7.223E-08 | 2.200E-08 | 1.600E-07 |
| 16128 | 807 | D807G | B | 2e-08 | 3.8e-08 | 6.1e-08 | 3.967E-08 | 2.055E-08 | 2.000E-08 | 6.100E-08 |
| 16129 | 807 | D807H | B | 1.7e-08 | 2.7e-08 | 6.9e-08 | 3.767E-08 | 2.759E-08 | 1.700E-08 | 6.900E-08 |
| 16130 | 807 | D807I | B | 1.6e-08 | 7.5e-08 | 8.4e-08 | 5.833E-08 | 3.694E-08 | 1.600E-08 | 8.400E-08 |
| 16131 | 807 | D807L | B | 3.3e-08 | 3.5e-08 | 2.4e-08 | 3.067E-08 | 5.859E-09 | 2.400E-08 | 3.500E-08 |
| 16132 | 807 | D807K | B | 5e-08 | 2.1e-08 | 5e-08 | 4.033E-08 | 1.674E-08 | 2.100E-08 | 5.000E-08 |
| 16133 | 807 | D807M | B | 8.9e-09 | 6.9e-08 | 3.8e-08 | 3.863E-08 | 3.006E-08 | 8.900E-09 | 6.900E-08 |
| 16134 | 807 | D807F | B | 6.2e-08 | 3.4e-08 | 2.9e-08 | 4.167E-08 | 1.779E-08 | 2.900E-08 | 6.200E-08 |
| 16135 | 807 | D807P | B | 3e-08 | 9.7e-08 | 1.8e-08 | 4.833E-08 | 4.257E-08 | 1.800E-08 | 9.700E-08 |
| 16136 | 807 | D807S | B | 4.2e-08 | 5.4e-08 | 7.3e-08 | 5.633E-08 | 1.563E-08 | 4.200E-08 | 7.300E-08 |
| 16137 | 807 | D807T | B | 4.4e-08 | 4.2e-08 | 1.2e-07 | 6.867E-08 | 4.447E-08 | 4.200E-08 | 1.200E-07 |
| 16138 | 807 | D807W | B | 6.9e-08 | 3.4e-08 | 2.7e-08 | 4.333E-08 | 2.250E-08 | 2.700E-08 | 6.900E-08 |
| 16139 | 807 | D807Y | B | 4.2e-08 | 2e-08 | 5.8e-08 | 4.000E-08 | 1.908E-08 | 2.000E-08 | 5.800E-08 |
| 16140 | 807 | D807V | B | 4.8e-08 | 2.6e-08 | 3.7e-08 | 3.700E-08 | 1.100E-08 | 2.600E-08 | 4.800E-08 |
| 16141 | 808 | Y808A | B | 6.6e-08 | 9.4e-08 | 9.4e-08 | 8.467E-08 | 1.617E-08 | 6.600E-08 | 9.400E-08 |
| 16142 | 808 | Y808R | B | 7.6e-08 | 8.8e-08 | 7.4e-08 | 7.933E-08 | 7.572E-09 | 7.400E-08 | 8.800E-08 |
| 16143 | 808 | Y808N | B | 4.7e-08 | 4.9e-08 | 9.4e-09 | 3.513E-08 | 2.231E-08 | 9.400E-09 | 4.900E-08 |
| 16144 | 808 | Y808D | B | 1.5e-07 | 4.8e-08 | 3.4e-08 | 7.733E-08 | 6.332E-08 | 3.400E-08 | 1.500E-07 |
| 16145 | 808 | Y808C | B | 1.3e-07 | 4.7e-08 | 2.8e-08 | 6.833E-08 | 5.424E-08 | 2.800E-08 | 1.300E-07 |
| 16146 | 808 | Y808E | B | 8.2e-08 | 8.4e-08 | 6.6e-08 | 7.733E-08 | 9.866E-09 | 6.600E-08 | 8.400E-08 |
| 16147 | 808 | Y808Q | B | 1.9e-08 | 3.3e-08 | 3.2e-08 | 2.800E-08 | 7.810E-09 | 1.900E-08 | 3.300E-08 |
| 16148 | 808 | Y808G | B | 1.8e-08 | 4.9e-08 | 4.5e-08 | 3.733E-08 | 1.686E-08 | 1.800E-08 | 4.900E-08 |
| 16149 | 808 | Y808H | B | 4.3e-08 | 3.9e-08 | 2.1e-08 | 3.433E-08 | 1.172E-08 | 2.100E-08 | 4.300E-08 |

| | | | | | | | | | | |
|---|---|---|---|---|---|---|---|---|---|---|
| 16150 | 808 | Y808I | B | 6.3e-08 | 5.6e-08 | 6.8e-08 | 6.233E-08 | 6.028E-09 | 5.600E-08 | 6.800E-08 |
| 16151 | 808 | Y808L | B | 5.4e-08 | 3.6e-08 | 3e-08 | 4.000E-08 | 1.249E-08 | 3.000E-08 | 5.400E-08 |
| 16152 | 808 | Y808K | B | 4.5e-08 | 4.9e-08 | 3.6e-08 | 4.333E-08 | 6.658E-09 | 3.600E-08 | 4.900E-08 |
| 16153 | 808 | Y808M | B | 6.1e-08 | 6.6e-08 | 6e-08 | 6.233E-08 | 3.215E-09 | 6.000E-08 | 6.600E-08 |
| 16154 | 808 | Y808F | B | 3.6e-08 | 3e-08 | 7.4e-08 | 4.667E-08 | 2.386E-08 | 3.000E-08 | 7.400E-08 |
| 16155 | 808 | Y808P | B | 6.5e-08 | 8.3e-08 | 5.2e-08 | 6.667E-08 | 1.557E-08 | 5.200E-08 | 8.300E-08 |
| 16156 | 808 | Y808S | B | 6.5e-08 | 6.8e-08 | 3.8e-08 | 5.700E-08 | 1.652E-08 | 3.800E-08 | 6.800E-08 |
| 16157 | 808 | Y808T | B | 1.9e-08 | 6.9e-08 | 4e-08 | 4.267E-08 | 2.511E-08 | 1.900E-08 | 6.900E-08 |
| 16158 | 808 | Y808W | B | 8.1e-08 | 4.4e-08 | 1e-07 | 7.500E-08 | 2.848E-08 | 4.400E-08 | 1.000E-07 |
| 16159 | 808 | Y808Y | B | 5.3e-08 | 2.3e-08 | 3.1e-08 | 3.567E-08 | 1.553E-08 | 2.300E-08 | 5.300E-08 |
| 16160 | 808 | Y808V | B | 3e-08 | 1.8e-08 | 5e-08 | 3.267E-08 | 1.617E-08 | 1.800E-08 | 5.000E-08 |
| 16161 | 809 | E809A | B | 3.3e-08 | 5.2e-08 | 4.4e-08 | 4.300E-08 | 9.539E-09 | 3.300E-08 | 5.200E-08 |
| 16162 | 809 | E809R | B | 2.4e-08 | 4e-08 | 4.6e-08 | 3.667E-08 | 1.137E-08 | 2.400E-08 | 4.600E-08 |
| 16163 | 809 | E809N | B | 7.6e-08 | 2.8e-08 | 3.2e-08 | 4.533E-08 | 2.663E-08 | 2.800E-08 | 7.600E-08 |
| 16164 | 809 | E809D | B | 6.5e-08 | 2.4e-08 | 1.1e-07 | 6.633E-08 | 4.302E-08 | 2.400E-08 | 1.100E-07 |
| 16165 | 809 | E809C | B | 3.9e-08 | 1.3e-08 | 1.6e-08 | 2.267E-08 | 1.422E-08 | 1.300E-08 | 3.900E-08 |
| 16166 | 809 | E809E | B | 4.9e-08 | 1e-07 | 4.4e-08 | 6.433E-08 | 3.099E-08 | 4.400E-08 | 1.000E-07 |
| 16167 | 809 | E809Q | B | 4.3e-08 | 1.9e-08 | 6.2e-08 | 4.133E-08 | 2.155E-08 | 1.900E-08 | 6.200E-08 |
| 16168 | 809 | E809G | B | 1.1e-08 | 5.7e-08 | 2.6e-08 | 3.133E-08 | 2.346E-08 | 1.100E-08 | 5.700E-08 |
| 16169 | 809 | E809H | B | 2.8e-08 | 4.6e-08 | 2.1e-08 | 3.167E-08 | 1.290E-08 | 2.100E-08 | 4.600E-08 |
| 16170 | 809 | E809I | B | 5.2e-08 | 3.4e-08 | 3.7e-08 | 4.100E-08 | 9.644E-09 | 3.400E-08 | 5.200E-08 |
| 16171 | 809 | E809L | B | 4.4e-08 | 1.8e-08 | 2.5e-08 | 2.900E-08 | 1.345E-08 | 1.800E-08 | 4.400E-08 |
| 16172 | 809 | E809K | B | 1.6e-08 | 2.9e-08 | 2e-08 | 2.167E-08 | 6.658E-09 | 1.600E-08 | 2.900E-08 |
| 16173 | 809 | E809M | B | 2e-07 | 7.9e-08 | 2.6e-08 | 1.017E-07 | 8.919E-08 | 2.600E-08 | 2.000E-07 |
| 16174 | 809 | E809F | B | 8.5e-08 | 8.2e-08 | 4.5e-08 | 7.067E-08 | 2.228E-08 | 4.500E-08 | 8.500E-08 |
| 16175 | 809 | E809P | B | 3.2e-08 | 1.1e-07 | 6.4e-08 | 6.867E-08 | 3.921E-08 | 3.200E-08 | 1.100E-07 |
| 16176 | 809 | E809S | B | 9.1e-08 | 4e-08 | 6.3e-08 | 6.467E-08 | 2.554E-08 | 4.000E-08 | 9.100E-08 |
| 16177 | 809 | E809T | B | 4.4e-08 | 2.4e-08 | 2e-08 | 2.933E-08 | 1.286E-08 | 2.000E-08 | 4.400E-08 |
| 16178 | 809 | E809W | B | 5.9e-08 | 5.3e-08 | 4e-08 | 5.067E-08 | 9.713E-09 | 4.000E-08 | 5.900E-08 |
| 16179 | 809 | E809Y | B | 6.3e-08 | 6.6e-08 | 4.4e-08 | 5.767E-08 | 1.193E-08 | 4.400E-08 | 6.600E-08 |
| 16180 | 809 | E809V | B | 3.1e-08 | 6.8e-08 | 4.3e-08 | 4.733E-08 | 1.888E-08 | 3.100E-08 | 6.800E-08 |
| 16181 | 810 | K810A | B | 8.5e-08 | 5.7e-08 | 4e-08 | 6.067E-08 | 2.272E-08 | 4.000E-08 | 8.500E-08 |
| 16182 | 810 | K810R | B | 4e-08 | 1.1e-07 | 2.4e-08 | 5.800E-08 | 4.574E-08 | 2.400E-08 | 1.100E-07 |
| 16183 | 810 | K810N | B | 5.6e-08 | 9e-08 | 4.6e-08 | 6.400E-08 | 2.307E-08 | 4.600E-08 | 9.000E-08 |
| 16184 | 810 | K810D | B | 4.6e-08 | 7.1e-08 | 9.8e-08 | 7.167E-08 | 2.601E-08 | 4.600E-08 | 9.800E-08 |
| 16185 | 810 | K810C | B | 4.3e-08 | 6.3e-08 | 5.8e-08 | 5.467E-08 | 1.041E-08 | 4.300E-08 | 6.300E-08 |
| 16186 | 810 | K810E | B | 1.9e-08 | 3.5e-08 | 1.7e-08 | 2.367E-08 | 9.866E-09 | 1.700E-08 | 3.500E-08 |
| 16187 | 810 | K810Q | B | 3.6e-08 | 2.1e-08 | 5.8e-08 | 3.833E-08 | 1.861E-08 | 2.100E-08 | 5.800E-08 |
| 16188 | 810 | K810G | B | 3.1e-08 | 9.2e-08 | 4.2e-08 | 5.500E-08 | 3.251E-08 | 3.100E-08 | 9.200E-08 |
| 16189 | 810 | K810H | B | 3.7e-08 | 1.9e-07 | 3e-08 | 8.567E-08 | 9.042E-08 | 3.000E-08 | 1.900E-07 |
| 16190 | 810 | K810I | B | 1.9e-08 | 5.9e-08 | 3.1e-08 | 3.633E-08 | 2.053E-08 | 1.900E-08 | 5.900E-08 |
| 16191 | 810 | K810L | B | 4.6e-08 | 6e-08 | 1.5e-07 | 8.533E-08 | 5.644E-08 | 4.600E-08 | 1.500E-07 |
| 16192 | 810 | K810K | B | 4.3e-08 | 1e-07 | 9.9e-08 | 8.067E-08 | 3.262E-08 | 4.300E-08 | 1.000E-07 |
| 16193 | 810 | K810M | B | 7.5e-08 | 4.2e-08 | 3.2e-07 | 1.457E-07 | 1.519E-07 | 4.200E-08 | 3.200E-07 |
| 16194 | 810 | K810F | B | 4.8e-08 | 1.1e-07 | 5.7e-08 | 7.167E-08 | 3.350E-08 | 4.800E-08 | 1.100E-07 |
| 16195 | 810 | K810P | B | 1.8e-08 | 7.3e-08 | 5.4e-08 | 4.833E-08 | 2.793E-08 | 1.800E-08 | 7.300E-08 |
| 16196 | 810 | K810S | B | 3.7e-08 | 7.3e-08 | 2e-08 | 4.333E-08 | 2.706E-08 | 2.000E-08 | 7.300E-08 |
| 16197 | 810 | K810T | B | 6.6e-08 | 5.2e-08 | 1.1e-07 | 7.600E-08 | 3.027E-08 | 5.200E-08 | 1.100E-07 |
| 16198 | 810 | K810W | B | 6.4e-08 | 2.3e-08 | 4.7e-08 | 4.467E-08 | 2.060E-08 | 2.300E-08 | 6.400E-08 |
| 16199 | 810 | K810Y | B | 4.5e-08 | 3.8e-08 | 1.7e-08 | 3.333E-08 | 1.457E-08 | 1.700E-08 | 4.500E-08 |
| 16200 | 810 | K810V | B | 5.3e-08 | 5.8e-08 | 6.8e-08 | 5.967E-08 | 7.638E-09 | 5.300E-08 | 6.800E-08 |
| 16201 | 811 | H811A | B | 4.9e-08 | 2.9e-08 | 5.2e-08 | 4.333E-08 | 1.250E-08 | 2.900E-08 | 5.200E-08 |
| 16202 | 811 | H811R | B | 3.4e-08 | 3.6e-08 | 3.6e-08 | 3.533E-08 | 1.155E-09 | 3.400E-08 | 3.600E-08 |
| 16203 | 811 | H811N | B | 2.6e-08 | 2.3e-08 | 5.5e-08 | 3.467E-08 | 1.767E-08 | 2.300E-08 | 5.500E-08 |
| 16204 | 811 | H811D | B | 2.3e-08 | 4.5e-08 | 7.7e-08 | 4.833E-08 | 2.715E-08 | 2.300E-08 | 7.700E-08 |
| 16205 | 811 | H811C | B | 3.1e-08 | 1.3e-08 | 1.2e-08 | 1.867E-08 | 1.069E-08 | 1.200E-08 | 3.100E-08 |
| 16206 | 811 | H811E | B | 5e-08 | 9.2e-08 | 4.7e-08 | 6.300E-08 | 2.516E-08 | 4.700E-08 | 9.200E-08 |
| 16207 | 811 | H811Q | B | 1.3e-08 | 8.1e-08 | 2.7e-08 | 4.033E-08 | 3.591E-08 | 1.300E-08 | 8.100E-08 |
| 16208 | 811 | H811G | B | 3.5e-08 | 1.6e-07 | 7.4e-08 | 8.967E-08 | 6.396E-08 | 3.500E-08 | 1.600E-07 |
| 16209 | 811 | H811H | B | 4.1e-08 | 2.6e-08 | 4.1e-08 | 3.600E-08 | 8.660E-09 | 2.600E-08 | 4.100E-08 |
| 16210 | 811 | H811I | B | 1.9e-08 | 4.9e-08 | 3.3e-08 | 3.367E-08 | 1.501E-08 | 1.900E-08 | 4.900E-08 |
| 16211 | 811 | H811L | B | 2.2e-08 | 2.7e-08 | 6.7e-08 | 3.867E-08 | 2.466E-08 | 2.200E-08 | 6.700E-08 |
| 16212 | 811 | H811K | B | 5.8e-08 | 1.1e-07 | 7.2e-08 | 8.000E-08 | 2.691E-08 | 5.800E-08 | 1.100E-07 |
| 16213 | 811 | H811M | B | 4.3e-08 | 1.3e-08 | 1.2e-08 | 2.267E-08 | 1.762E-08 | 1.200E-08 | 4.300E-08 |
| 16214 | 811 | H811F | B | 1.2e-07 | 2.7e-08 | 9.4e-08 | 8.033E-08 | 4.798E-08 | 2.700E-08 | 1.200E-07 |
| 16215 | 811 | H811P | B | 8.2e-08 | 1.7e-08 | 5.7e-08 | 5.200E-08 | 3.279E-08 | 1.700E-08 | 8.200E-08 |
| 16216 | 811 | H811S | B | 3.4e-08 | 6.7e-08 | 2.3e-08 | 4.133E-08 | 2.290E-08 | 2.300E-08 | 6.700E-08 |
| 16217 | 811 | H811T | B | 5e-08 | 1.4e-08 | 5.2e-08 | 3.867E-08 | 2.139E-08 | 1.400E-08 | 5.200E-08 |
| 16218 | 811 | H811W | B | 6.1e-08 | 3.1e-08 | 8.5e-08 | 5.900E-08 | 2.706E-08 | 3.100E-08 | 8.500E-08 |
| 16219 | 811 | H811Y | B | 2.6e-08 | 1.7e-08 | 5.1e-08 | 3.133E-08 | 1.762E-08 | 1.700E-08 | 5.100E-08 |
| 16220 | 811 | H811V | B | 1.2e-08 | 3.1e-08 | 7.8e-08 | 4.033E-08 | 3.398E-08 | 1.200E-08 | 7.800E-08 |
| 16221 | 812 | K812A | B | 4.7e-08 | 3.1e-08 | 6.5e-08 | 4.767E-08 | 1.701E-08 | 3.100E-08 | 6.500E-08 |
| 16222 | 812 | K812R | B | 8.1e-08 | 7.2e-08 | 2.8e-08 | 6.033E-08 | 2.836E-08 | 2.800E-08 | 8.100E-08 |
| 16223 | 812 | K812N | B | 2.5e-08 | 2.5e-08 | 6.6e-08 | 3.867E-08 | 2.367E-08 | 2.500E-08 | 6.600E-08 |
| 16224 | 812 | K812D | B | 5e-08 | 1.1e-07 | 4.9e-08 | 6.967E-08 | 3.493E-08 | 4.900E-08 | 1.100E-07 |
| 16225 | 812 | K812C | B | 1.5e-07 | 6.7e-08 | 4e-08 | 8.567E-08 | 5.733E-08 | 4.000E-08 | 1.500E-07 |
| 16226 | 812 | K812E | B | 4.3e-08 | 3.9e-08 | 2.7e-08 | 3.633E-08 | 8.327E-09 | 2.700E-08 | 4.300E-08 |
| 16227 | 812 | K812Q | B | 4.9e-08 | 2.5e-08 | 5.4e-08 | 4.267E-08 | 1.550E-08 | 2.500E-08 | 5.400E-08 |
| 16228 | 812 | K812G | B | 2.3e-07 | 4.4e-08 | 2.5e-07 | 1.747E-07 | 1.136E-07 | 4.400E-08 | 2.500E-07 |
| 16229 | 812 | K812H | B | 1.7e-07 | 9.6e-08 | 7.4e-08 | 1.133E-07 | 5.029E-08 | 7.400E-08 | 1.700E-07 |
| 16230 | 812 | K812I | B | 2.6e-08 | 1.4e-07 | 2.7e-08 | 6.433E-08 | 6.553E-08 | 2.600E-08 | 1.400E-07 |
| 16231 | 812 | K812L | B | 9.5e-08 | 2.9e-08 | 2.1e-08 | 4.833E-08 | 4.061E-08 | 2.100E-08 | 9.500E-08 |
| 16232 | 812 | K812K | B | 2.9e-08 | 6.5e-08 | 1.2e-07 | 7.133E-08 | 4.583E-08 | 2.900E-08 | 1.200E-07 |
| 16233 | 812 | K812M | B | 4.4e-08 | 4.2e-08 | 2.8e-08 | 3.800E-08 | 8.718E-09 | 2.800E-08 | 4.400E-08 |
| 16234 | 812 | K812F | B | 9.4e-08 | 1e-07 | 1.2e-08 | 6.867E-08 | 4.917E-08 | 1.200E-08 | 1.000E-07 |

| | | | | | | | | | | |
|---|---|---|---|---|---|---|---|---|---|---|
| 16235 | 812 | K812P | B | 4.6e-08 | 6.3e-08 | 5.8e-09 | 3.827E-08 | 2.937E-08 | 5.800E-09 | 6.300E-08 |
| 16236 | 812 | K812S | B | 2.5e-08 | 1e-07 | 1.8e-08 | 4.767E-08 | 4.546E-08 | 1.800E-08 | 1.000E-07 |
| 16237 | 812 | K812T | B | 3e-08 | 1.1e-07 | 1e-07 | 8.000E-08 | 4.359E-08 | 3.000E-08 | 1.100E-07 |
| 16238 | 812 | K812W | B | 3.8e-08 | 2.1e-08 | 7.1e-09 | 2.203E-08 | 1.548E-08 | 7.100E-09 | 3.800E-08 |
| 16239 | 812 | K812Y | B | 3e-08 | 4.9e-08 | 5e-08 | 4.300E-08 | 1.127E-08 | 3.000E-08 | 5.000E-08 |
| 16240 | 812 | K812V | B | 9.1e-08 | 3.6e-08 | 1.3e-08 | 4.667E-08 | 4.008E-08 | 1.300E-08 | 9.100E-08 |
| 16241 | 813 | V813A | B | 2.5e-08 | 2.2e-08 | 2.3e-08 | 2.333E-08 | 1.528E-09 | 2.200E-08 | 2.500E-08 |
| 16242 | 813 | V813R | B | 5.3e-08 | 3.4e-08 | 8.1e-08 | 5.600E-08 | 2.364E-08 | 3.400E-08 | 8.100E-08 |
| 16243 | 813 | V813N | B | 7.1e-08 | 2.5e-08 | 9.9e-08 | 6.500E-08 | 3.736E-08 | 2.500E-08 | 9.900E-08 |
| 16244 | 813 | V813D | B | 2.6e-08 | 7.1e-08 | 2e-08 | 3.900E-08 | 2.787E-08 | 2.000E-08 | 7.100E-08 |
| 16245 | 813 | V813C | B | 1.2e-07 | 2.3e-08 | 2e-08 | 5.433E-08 | 5.689E-08 | 2.000E-08 | 1.200E-07 |
| 16246 | 813 | V813E | B | 2.2e-08 | 2.7e-08 | 3e-08 | 2.633E-08 | 4.041E-09 | 2.200E-08 | 3.000E-08 |
| 16247 | 813 | V813Q | B | 5.8e-08 | 1.8e-08 | 4.5e-08 | 4.033E-08 | 2.040E-08 | 1.800E-08 | 5.800E-08 |
| 16248 | 813 | V813G | B | 2.9e-08 | 7.7e-08 | 3.4e-08 | 4.667E-08 | 2.639E-08 | 2.900E-08 | 7.700E-08 |
| 16249 | 813 | V813H | B | 3.8e-08 | 6.1e-08 | 2.5e-08 | 4.133E-08 | 1.823E-08 | 2.500E-08 | 6.100E-08 |
| 16250 | 813 | V813I | B | 3.8e-08 | 3.8e-08 | 7.4e-08 | 5.000E-08 | 2.078E-08 | 3.800E-08 | 7.400E-08 |
| 16251 | 813 | V813L | B | 2.5e-08 | 1e-07 | 6.4e-08 | 6.300E-08 | 3.751E-08 | 2.500E-08 | 1.000E-07 |
| 16252 | 813 | V813K | B | 4.7e-08 | 2.3e-08 | 3.6e-08 | 3.533E-08 | 1.201E-08 | 2.300E-08 | 4.700E-08 |
| 16253 | 813 | V813M | B | 5.3e-08 | 9.4e-08 | 2.3e-08 | 5.667E-08 | 3.564E-08 | 2.300E-08 | 9.400E-08 |
| 16254 | 813 | V813F | B | 1.5e-08 | 4.1e-08 | 7.8e-08 | 4.467E-08 | 3.166E-08 | 1.500E-08 | 7.800E-08 |
| 16255 | 813 | V813P | B | 2e-08 | 2.9e-08 | 6.3e-08 | 3.733E-08 | 2.268E-08 | 2.000E-08 | 6.300E-08 |
| 16256 | 813 | V813S | B | 4.7e-08 | 2.4e-08 | 7.2e-08 | 4.767E-08 | 2.401E-08 | 2.400E-08 | 7.200E-08 |
| 16257 | 813 | V813T | B | 2.5e-08 | 3e-08 | 4.4e-08 | 3.300E-08 | 9.849E-09 | 2.500E-08 | 4.400E-08 |
| 16258 | 813 | V813W | B | 3.4e-08 | 6.9e-08 | 1.1e-07 | 7.100E-08 | 3.804E-08 | 3.400E-08 | 1.100E-07 |
| 16259 | 813 | V813Y | B | 4.8e-08 | 4e-08 | 2.4e-08 | 3.733E-08 | 1.222E-08 | 2.400E-08 | 4.800E-08 |
| 16260 | 813 | V813V | B | 7.6e-08 | 3.7e-08 | 2.8e-08 | 4.700E-08 | 2.551E-08 | 2.800E-08 | 7.600E-08 |
| 16261 | 814 | Y814A | B | 3.1e-08 | 2e-08 | 4.8e-08 | 3.300E-08 | 1.411E-08 | 2.000E-08 | 4.800E-08 |
| 16262 | 814 | Y814R | B | 2.8e-08 | 9.2e-08 | 3.7e-08 | 5.233E-08 | 3.465E-08 | 2.800E-08 | 9.200E-08 |
| 16263 | 814 | Y814N | B | 1.5e-08 | 1.8e-08 | 9.1e-08 | 4.133E-08 | 4.304E-08 | 1.500E-08 | 9.100E-08 |
| 16264 | 814 | Y814D | B | 6e-08 | 2.6e-08 | 1.1e-07 | 6.533E-08 | 4.225E-08 | 2.600E-08 | 1.100E-07 |
| 16265 | 814 | Y814C | B | 3.8e-08 | 2.3e-08 | 4.3e-08 | 3.467E-08 | 1.041E-08 | 2.300E-08 | 4.300E-08 |
| 16266 | 814 | Y814E | B | 4.2e-08 | 5.9e-08 | 4.1e-08 | 4.733E-08 | 1.012E-08 | 4.100E-08 | 5.900E-08 |
| 16267 | 814 | Y814Q | B | 6.8e-08 | 2.7e-08 | 5.6e-08 | 5.033E-08 | 2.108E-08 | 2.700E-08 | 6.800E-08 |
| 16268 | 814 | Y814G | B | 6.2e-08 | 2.7e-08 | 2.9e-08 | 3.933E-08 | 1.966E-08 | 2.700E-08 | 6.200E-08 |
| 16269 | 814 | Y814H | B | 6.7e-08 | 1.3e-07 | 6.4e-08 | 8.700E-08 | 3.727E-08 | 6.400E-08 | 1.300E-07 |
| 16270 | 814 | Y814I | B | 7.5e-08 | 2e-08 | 4.8e-08 | 4.767E-08 | 2.750E-08 | 2.000E-08 | 7.500E-08 |
| 16271 | 814 | Y814L | B | 5.2e-08 | 1.9e-08 | 8.8e-08 | 5.300E-08 | 3.451E-08 | 1.900E-08 | 8.800E-08 |
| 16272 | 814 | Y814K | B | 3.5e-08 | 7.1e-08 | 3.4e-08 | 4.667E-08 | 2.108E-08 | 3.400E-08 | 7.100E-08 |
| 16273 | 814 | Y814M | B | 1.9e-08 | 7.7e-08 | 3.7e-08 | 4.433E-08 | 2.969E-08 | 1.900E-08 | 7.700E-08 |
| 16274 | 814 | Y814F | B | 7.5e-08 | 1.6e-08 | 3.6e-08 | 4.233E-08 | 3.001E-08 | 1.600E-08 | 7.500E-08 |
| 16275 | 814 | Y814P | B | 4.8e-08 | 6.4e-08 | 2.3e-08 | 4.500E-08 | 2.066E-08 | 2.300E-08 | 6.400E-08 |
| 16276 | 814 | Y814S | B | 3e-08 | 4.6e-08 | 1.9e-08 | 3.167E-08 | 1.358E-08 | 1.900E-08 | 4.600E-08 |
| 16277 | 814 | Y814T | B | 9e-08 | 6.6e-08 | 5.6e-08 | 7.067E-08 | 1.747E-08 | 5.600E-08 | 9.000E-08 |
| 16278 | 814 | Y814W | B | 1.2e-07 | 3e-08 | 3.1e-08 | 6.033E-08 | 5.168E-08 | 3.000E-08 | 1.200E-07 |
| 16279 | 814 | Y814Y | B | 9.8e-08 | 3.2e-08 | 1.1e-08 | 4.700E-08 | 4.540E-08 | 1.100E-08 | 9.800E-08 |
| 16280 | 814 | Y814V | B | 2.3e-08 | 6.5e-08 | 1.1e-08 | 3.300E-08 | 2.835E-08 | 1.100E-08 | 6.500E-08 |
| 16281 | 815 | A815A | B | 2.2e-08 | 4.2e-08 | 7.4e-08 | 4.600E-08 | 2.623E-08 | 2.200E-08 | 7.400E-08 |
| 16282 | 815 | A815R | B | 7.1e-08 | 2.6e-08 | 5.7e-08 | 5.133E-08 | 2.303E-08 | 2.600E-08 | 7.100E-08 |
| 16283 | 815 | A815N | B | 3.6e-08 | 3.6e-08 | 6.6e-08 | 4.600E-08 | 1.732E-08 | 3.600E-08 | 6.600E-08 |
| 16284 | 815 | A815D | B | 7.3e-08 | 4.2e-08 | 1.3e-07 | 8.167E-08 | 4.464E-08 | 4.200E-08 | 1.300E-07 |
| 16285 | 815 | A815C | B | 8e-08 | 2.7e-08 | 9.2e-08 | 6.633E-08 | 3.459E-08 | 2.700E-08 | 9.200E-08 |
| 16286 | 815 | A815E | B | 4.6e-08 | 8.5e-09 | 1.4e-08 | 2.283E-08 | 2.025E-08 | 8.500E-09 | 4.600E-08 |
| 16287 | 815 | A815Q | B | 3.5e-08 | 3.6e-08 | 2.6e-08 | 3.233E-08 | 5.508E-09 | 2.600E-08 | 3.600E-08 |
| 16288 | 815 | A815G | B | 3.3e-08 | 3.3e-08 | 5.7e-08 | 4.100E-08 | 1.386E-08 | 3.300E-08 | 5.700E-08 |
| 16289 | 815 | A815H | B | 1.4e-08 | 3.1e-08 | 6.1e-08 | 3.533E-08 | 2.380E-08 | 1.400E-08 | 6.100E-08 |
| 16290 | 815 | A815I | B | 4.1e-08 | 6.6e-08 | 2.6e-08 | 4.433E-08 | 2.021E-08 | 2.600E-08 | 6.600E-08 |
| 16291 | 815 | A815L | B | 9e-08 | 5e-08 | 1.8e-08 | 5.267E-08 | 3.607E-08 | 1.800E-08 | 9.000E-08 |
| 16292 | 815 | A815K | B | 3.1e-08 | 9.5e-08 | 1.6e-07 | 9.533E-08 | 6.450E-08 | 3.100E-08 | 1.600E-07 |
| 16293 | 815 | A815M | B | 5.2e-08 | 2.6e-08 | 4.2e-08 | 4.000E-08 | 1.311E-08 | 2.600E-08 | 5.200E-08 |
| 16294 | 815 | A815F | B | 7.7e-08 | 9e-08 | 6.6e-08 | 7.767E-08 | 1.201E-08 | 6.600E-08 | 9.000E-08 |
| 16295 | 815 | A815P | B | 2.9e-08 | 5.4e-08 | 3e-08 | 3.767E-08 | 1.415E-08 | 2.900E-08 | 5.400E-08 |
| 16296 | 815 | A815S | B | 2.9e-07 | 2.4e-08 | 3.3e-08 | 1.157E-07 | 1.510E-07 | 2.400E-08 | 2.900E-07 |
| 16297 | 815 | A815T | B | 5.3e-08 | 3e-08 | 3.8e-08 | 4.033E-08 | 1.168E-08 | 3.000E-08 | 5.300E-08 |
| 16298 | 815 | A815W | B | 9.7e-08 | 7.3e-08 | 1.4e-07 | 1.033E-07 | 3.395E-08 | 7.300E-08 | 1.400E-07 |
| 16299 | 815 | A815Y | B | 2.4e-08 | 8.5e-08 | 9.4e-09 | 3.947E-08 | 4.010E-08 | 9.400E-09 | 8.500E-08 |
| 16300 | 815 | A815V | B | 5.7e-08 | 4.2e-08 | 7.7e-08 | 5.867E-08 | 1.756E-08 | 4.200E-08 | 7.700E-08 |
| 16301 | 816 | C816A | B | 1.3e-08 | 4.4e-08 | 9.8e-08 | 5.167E-08 | 4.302E-08 | 1.300E-08 | 9.800E-08 |
| 16302 | 816 | C816R | B | 4.5e-08 | 4.4e-08 | 1.2e-07 | 6.967E-08 | 4.359E-08 | 4.400E-08 | 1.200E-07 |
| 16303 | 816 | C816N | B | 2.5e-08 | 9e-08 | 3.4e-08 | 4.967E-08 | 3.522E-08 | 2.500E-08 | 9.000E-08 |
| 16304 | 816 | C816D | B | 3.9e-08 | 9.8e-08 | 3.8e-08 | 5.833E-08 | 3.436E-08 | 3.800E-08 | 9.800E-08 |
| 16305 | 816 | C816C | B | 9.9e-08 | 5e-08 | 3.9e-08 | 6.267E-08 | 3.194E-08 | 3.900E-08 | 9.900E-08 |
| 16306 | 816 | C816E | B | 3.8e-08 | 4.6e-08 | 2e-08 | 3.467E-08 | 1.332E-08 | 2.000E-08 | 4.600E-08 |
| 16307 | 816 | C816Q | B | 3.8e-08 | 2.1e-08 | 2.3e-08 | 2.733E-08 | 9.292E-09 | 2.100E-08 | 3.800E-08 |
| 16308 | 816 | C816G | B | 6.9e-08 | 3e-08 | 6.2e-08 | 5.367E-08 | 2.079E-08 | 3.000E-08 | 6.900E-08 |
| 16309 | 816 | C816H | B | 9.3e-08 | 3.2e-08 | 1.3e-08 | 4.600E-08 | 4.180E-08 | 1.300E-08 | 9.300E-08 |
| 16310 | 816 | C816I | B | 5.2e-08 | 2.2e-08 | 5.2e-08 | 4.200E-08 | 1.732E-08 | 2.200E-08 | 5.200E-08 |
| 16311 | 816 | C816L | B | 3.6e-08 | 9.6e-09 | 5.7e-08 | 3.420E-08 | 2.375E-08 | 9.600E-09 | 5.700E-08 |
| 16312 | 816 | C816K | B | 3.5e-08 | 5.2e-08 | 6.9e-08 | 5.200E-08 | 1.700E-08 | 3.500E-08 | 6.900E-08 |
| 16313 | 816 | C816M | B | 8.7e-08 | 3e-08 | 5.9e-08 | 5.867E-08 | 2.850E-08 | 3.000E-08 | 8.700E-08 |
| 16314 | 816 | C816F | B | 2.7e-08 | 3e-08 | 5.4e-08 | 3.700E-08 | 1.480E-08 | 2.700E-08 | 5.400E-08 |
| 16315 | 816 | C816P | B | 5.5e-08 | 3.4e-08 | 3.2e-08 | 4.033E-08 | 1.274E-08 | 3.200E-08 | 5.500E-08 |
| 16316 | 816 | C816S | B | 3.5e-08 | 3.6e-08 | 1.2e-07 | 6.367E-08 | 4.879E-08 | 3.500E-08 | 1.200E-07 |
| 16317 | 816 | C816T | B | 2.5e-08 | 5.9e-08 | 3.7e-08 | 4.033E-08 | 1.724E-08 | 2.500E-08 | 5.900E-08 |
| 16318 | 816 | C816W | B | 1.9e-08 | 5.8e-08 | 3.5e-08 | 3.733E-08 | 1.960E-08 | 1.900E-08 | 5.800E-08 |
| 16319 | 816 | C816Y | B | 4.6e-08 | 5.1e-08 | 3.6e-08 | 4.433E-08 | 7.638E-09 | 3.600E-08 | 5.100E-08 |

| | | | | | | | | | | |
|---|---|---|---|---|---|---|---|---|---|---|
| 16320 | 816 | C816V | B | 3.1e-08 | 1.9e-08 | 3.8e-08 | 2.933E-08 | 9.609E-09 | 1.900E-08 | 3.800E-08 |
| 16321 | 817 | E817A | B | 4.2e-08 | 4.7e-08 | 4.7e-08 | 4.533E-08 | 2.887E-09 | 4.200E-08 | 4.700E-08 |
| 16322 | 817 | E817R | B | 1.6e-08 | 2.1e-08 | 2.3e-08 | 2.000E-08 | 3.606E-09 | 1.600E-08 | 2.300E-08 |
| 16323 | 817 | E817N | B | 3.6e-08 | 4.3e-08 | 1.5e-07 | 7.633E-08 | 6.389E-08 | 3.600E-08 | 1.500E-07 |
| 16324 | 817 | E817D | B | 1.5e-08 | 1.1e-07 | 1.8e-08 | 4.767E-08 | 5.400E-08 | 1.500E-08 | 1.100E-07 |
| 16325 | 817 | E817C | B | 1.9e-08 | 7e-08 | 1.4e-07 | 7.633E-08 | 6.075E-08 | 1.900E-08 | 1.400E-07 |
| 16326 | 817 | E817E | B | 1.4e-07 | 2e-08 | 6.1e-08 | 7.367E-08 | 6.099E-08 | 2.000E-08 | 1.400E-07 |
| 16327 | 817 | E817Q | B | 2.5e-08 | 4.8e-08 | 1.5e-07 | 7.433E-08 | 6.653E-08 | 2.500E-08 | 1.500E-07 |
| 16328 | 817 | E817G | B | 7.2e-08 | 4.4e-08 | 5.4e-08 | 5.667E-08 | 1.419E-08 | 4.400E-08 | 7.200E-08 |
| 16329 | 817 | E817H | B | 7.9e-08 | 2.7e-08 | 2.8e-08 | 4.467E-08 | 2.974E-08 | 2.700E-08 | 7.900E-08 |
| 16330 | 817 | E817I | B | 6.9e-08 | 5.8e-08 | 1.9e-08 | 4.867E-08 | 2.627E-08 | 1.900E-08 | 6.900E-08 |
| 16331 | 817 | E817L | B | 5.1e-08 | 1.4e-08 | 3.7e-08 | 3.400E-08 | 1.868E-08 | 1.400E-08 | 5.100E-08 |
| 16332 | 817 | E817K | B | 4.6e-08 | 5.7e-08 | 3.5e-08 | 4.600E-08 | 1.100E-08 | 3.500E-08 | 5.700E-08 |
| 16333 | 817 | E817M | B | 3.1e-08 | 4.1e-08 | 3.1e-08 | 3.433E-08 | 5.774E-09 | 3.100E-08 | 4.100E-08 |
| 16334 | 817 | E817F | B | 6.1e-08 | 1e-08 | 4.2e-08 | 3.767E-08 | 2.577E-08 | 1.000E-08 | 6.100E-08 |
| 16335 | 817 | E817P | B | 2.5e-08 | 2.2e-08 | 3.7e-08 | 2.800E-08 | 7.937E-09 | 2.200E-08 | 3.700E-08 |
| 16336 | 817 | E817S | B | 7e-08 | 4e-08 | 4.8e-08 | 5.267E-08 | 1.553E-08 | 4.000E-08 | 7.000E-08 |
| 16337 | 817 | E817T | B | 5.5e-08 | 4.4e-08 | 4.4e-08 | 4.767E-08 | 6.351E-09 | 4.400E-08 | 5.500E-08 |
| 16338 | 817 | E817W | B | 6.7e-08 | 9.2e-08 | 2.2e-08 | 6.033E-08 | 3.547E-08 | 2.200E-08 | 9.200E-08 |
| 16339 | 817 | E817Y | B | 3.3e-08 | 4.2e-08 | 7.8e-08 | 5.100E-08 | 2.381E-08 | 3.300E-08 | 7.800E-08 |
| 16340 | 817 | E817V | B | 6.4e-08 | 8.2e-08 | 2.6e-08 | 5.733E-08 | 2.859E-08 | 2.600E-08 | 8.200E-08 |
| 16341 | 818 | V818A | B | 5.4e-08 | 6e-08 | 2.9e-08 | 4.767E-08 | 1.644E-08 | 2.900E-08 | 6.000E-08 |
| 16342 | 818 | V818R | B | 4.2e-08 | 7.1e-08 | 6.2e-08 | 5.833E-08 | 1.484E-08 | 4.200E-08 | 7.100E-08 |
| 16343 | 818 | V818N | B | 3.8e-08 | 2e-08 | 4.4e-08 | 3.400E-08 | 1.249E-08 | 2.000E-08 | 4.400E-08 |
| 16344 | 818 | V818D | B | 1.8e-08 | 2.8e-08 | 4.5e-08 | 3.033E-08 | 1.365E-08 | 1.800E-08 | 4.500E-08 |
| 16345 | 818 | V818C | B | 9.1e-08 | 6.4e-08 | 1.3e-08 | 5.600E-08 | 3.961E-08 | 1.300E-08 | 9.100E-08 |
| 16346 | 818 | V818E | B | 4.4e-08 | 2.8e-08 | 6.1e-08 | 4.433E-08 | 1.650E-08 | 2.800E-08 | 6.100E-08 |
| 16347 | 818 | V818Q | B | 5.1e-08 | 8.1e-08 | 4.5e-08 | 5.900E-08 | 1.929E-08 | 4.500E-08 | 8.100E-08 |
| 16348 | 818 | V818G | B | 6.2e-08 | 3.3e-08 | 4e-08 | 4.500E-08 | 1.513E-08 | 3.300E-08 | 6.200E-08 |
| 16349 | 818 | V818H | B | 1.9e-08 | 2.7e-08 | 9e-08 | 4.533E-08 | 3.889E-08 | 1.900E-08 | 9.000E-08 |
| 16350 | 818 | V818I | B | 2.1e-08 | 7e-08 | 2.4e-08 | 3.833E-08 | 2.747E-08 | 2.100E-08 | 7.000E-08 |
| 16351 | 818 | V818L | B | 3.1e-08 | 7.6e-08 | 3.6e-08 | 4.767E-08 | 2.466E-08 | 3.100E-08 | 7.600E-08 |
| 16352 | 818 | V818K | B | 4.8e-08 | 7.4e-08 | 3.5e-08 | 5.233E-08 | 1.986E-08 | 3.500E-08 | 7.400E-08 |
| 16353 | 818 | V818M | B | 1.2e-07 | 4e-08 | 1.4e-08 | 5.800E-08 | 5.524E-08 | 1.400E-08 | 1.200E-07 |
| 16354 | 818 | V818F | B | 1.7e-08 | 7.7e-08 | 3.6e-08 | 4.333E-08 | 3.066E-08 | 1.700E-08 | 7.700E-08 |
| 16355 | 818 | V818P | B | 1.2e-08 | 2.5e-08 | 1.2e-07 | 5.233E-08 | 5.896E-08 | 1.200E-08 | 1.200E-07 |
| 16356 | 818 | V818S | B | 6e-08 | 1.9e-08 | 1.2e-07 | 6.633E-08 | 5.080E-08 | 1.900E-08 | 1.200E-07 |
| 16357 | 818 | V818T | B | 9.1e-08 | 3.7e-08 | 1.4e-08 | 4.733E-08 | 3.953E-08 | 1.400E-08 | 9.100E-08 |
| 16358 | 818 | V818W | B | 4.9e-08 | 4.1e-08 | 5.4e-08 | 4.800E-08 | 6.557E-09 | 4.100E-08 | 5.400E-08 |
| 16359 | 818 | V818Y | B | 2.8e-08 | 6.4e-08 | 4.2e-08 | 4.467E-08 | 1.815E-08 | 2.800E-08 | 6.400E-08 |
| 16360 | 818 | V818V | B | 4.9e-08 | 2.8e-08 | 4.6e-08 | 4.100E-08 | 1.136E-08 | 2.800E-08 | 4.900E-08 |
| 16361 | 819 | T819A | B | 3.3e-08 | 2.4e-08 | 1.6e-08 | 2.433E-08 | 8.505E-09 | 1.600E-08 | 3.300E-08 |
| 16362 | 819 | T819R | B | 4e-08 | 3.6e-08 | 5.3e-08 | 4.300E-08 | 8.888E-09 | 3.600E-08 | 5.300E-08 |
| 16363 | 819 | T819N | B | 5e-08 | 4.9e-08 | 6.6e-08 | 5.500E-08 | 9.539E-09 | 4.900E-08 | 6.600E-08 |
| 16364 | 819 | T819D | B | 3.9e-08 | 7.9e-08 | 1.8e-08 | 4.533E-08 | 3.099E-08 | 1.800E-08 | 7.900E-08 |
| 16365 | 819 | T819C | B | 4.6e-08 | 3.8e-08 | 6.5e-08 | 4.967E-08 | 1.387E-08 | 3.800E-08 | 6.500E-08 |
| 16366 | 819 | T819E | B | 2.5e-08 | 2.7e-08 | 8e-08 | 4.400E-08 | 3.119E-08 | 2.500E-08 | 8.000E-08 |
| 16367 | 819 | T819Q | B | 2.1e-08 | 8e-08 | 4.8e-08 | 4.967E-08 | 2.954E-08 | 2.100E-08 | 8.000E-08 |
| 16368 | 819 | T819G | B | 1.8e-08 | 2.1e-08 | 5.6e-08 | 3.167E-08 | 2.113E-08 | 1.800E-08 | 5.600E-08 |
| 16369 | 819 | T819H | B | 3.6e-08 | 5.9e-08 | 1.9e-07 | 9.500E-08 | 8.307E-08 | 3.600E-08 | 1.900E-07 |
| 16370 | 819 | T819I | B | 3.8e-08 | 1.4e-08 | 5.9e-08 | 3.700E-08 | 2.252E-08 | 1.400E-08 | 5.900E-08 |
| 16371 | 819 | T819L | B | 3.7e-08 | 2.8e-08 | 4.6e-08 | 3.700E-08 | 9.000E-09 | 2.800E-08 | 4.600E-08 |
| 16372 | 819 | T819K | B | 3.1e-08 | 3.3e-08 | 5e-08 | 3.800E-08 | 1.044E-08 | 3.100E-08 | 5.000E-08 |
| 16373 | 819 | T819M | B | 2.6e-08 | 3.2e-08 | 4.4e-08 | 3.400E-08 | 9.165E-09 | 2.600E-08 | 4.400E-08 |
| 16374 | 819 | T819F | B | 5e-08 | 1.7e-08 | 3.7e-08 | 3.467E-08 | 1.662E-08 | 1.700E-08 | 5.000E-08 |
| 16375 | 819 | T819P | B | 2.3e-08 | 6.1e-08 | 7.3e-08 | 5.233E-08 | 2.610E-08 | 2.300E-08 | 7.300E-08 |
| 16376 | 819 | T819S | B | 4.7e-08 | 6.6e-08 | 2.6e-08 | 4.633E-08 | 2.001E-08 | 2.600E-08 | 6.600E-08 |
| 16377 | 819 | T819T | B | 3.5e-08 | 1.9e-08 | 5.6e-08 | 3.667E-08 | 1.856E-08 | 1.900E-08 | 5.600E-08 |
| 16378 | 819 | T819W | B | 4.3e-08 | 1.7e-08 | 4.3e-08 | 3.433E-08 | 1.501E-08 | 1.700E-08 | 4.300E-08 |
| 16379 | 819 | T819Y | B | 4.9e-08 | 3.2e-08 | 5.1e-08 | 4.400E-08 | 1.044E-08 | 3.200E-08 | 5.100E-08 |
| 16380 | 819 | T819V | B | 3.2e-08 | 3.2e-08 | 2.3e-08 | 2.900E-08 | 5.196E-09 | 2.300E-08 | 3.200E-08 |
| 16381 | 820 | H820A | B | 3.6e-08 | 4.8e-08 | 3.8e-08 | 4.067E-08 | 6.429E-09 | 3.600E-08 | 4.800E-08 |
| 16382 | 820 | H820R | B | 5.9e-08 | 5.6e-08 | 4.6e-08 | 5.367E-08 | 6.807E-09 | 4.600E-08 | 5.900E-08 |
| 16383 | 820 | H820N | B | 5.8e-08 | 2.1e-08 | 2e-08 | 3.300E-08 | 2.166E-08 | 2.000E-08 | 5.800E-08 |
| 16384 | 820 | H820D | B | 8.4e-08 | 6.5e-08 | 4.4e-08 | 6.433E-08 | 2.001E-08 | 4.400E-08 | 8.400E-08 |
| 16385 | 820 | H820C | B | 2.4e-08 | 3.6e-08 | 3.8e-08 | 3.267E-08 | 7.572E-09 | 2.400E-08 | 3.800E-08 |
| 16386 | 820 | H820E | B | 2.4e-08 | 6.6e-08 | 3.7e-08 | 4.233E-08 | 2.150E-08 | 2.400E-08 | 6.600E-08 |
| 16387 | 820 | H820Q | B | 2e-08 | 7.1e-08 | 5.8e-08 | 4.967E-08 | 2.650E-08 | 2.000E-08 | 7.100E-08 |
| 16388 | 820 | H820G | B | 8.4e-08 | 3.3e-08 | 8e-08 | 6.567E-08 | 2.836E-08 | 3.300E-08 | 8.400E-08 |
| 16389 | 820 | H820H | B | 1.6e-08 | 3.8e-08 | 1.2e-08 | 2.200E-08 | 1.400E-08 | 1.200E-08 | 3.800E-08 |
| 16390 | 820 | H820I | B | 4e-08 | 1.8e-08 | 1.4e-08 | 2.400E-08 | 1.400E-08 | 1.400E-08 | 4.000E-08 |
| 16391 | 820 | H820L | B | 5.6e-08 | 1.4e-08 | 6.4e-08 | 4.467E-08 | 2.686E-08 | 1.400E-08 | 6.400E-08 |
| 16392 | 820 | H820K | B | 2.2e-08 | 7.3e-08 | 2.1e-08 | 3.867E-08 | 2.974E-08 | 2.100E-08 | 7.300E-08 |
| 16393 | 820 | H820M | B | 9.8e-08 | 2.4e-08 | 5.9e-08 | 6.033E-08 | 3.702E-08 | 2.400E-08 | 9.800E-08 |
| 16394 | 820 | H820F | B | 5.5e-08 | 5.2e-08 | 4.4e-08 | 5.033E-08 | 5.686E-09 | 4.400E-08 | 5.500E-08 |
| 16395 | 820 | H820P | B | 4.9e-08 | 1.2e-08 | 3.3e-08 | 3.133E-08 | 1.856E-08 | 1.200E-08 | 4.900E-08 |
| 16396 | 820 | H820S | B | 2.8e-08 | 1.6e-08 | 5e-08 | 3.133E-08 | 1.724E-08 | 1.600E-08 | 5.000E-08 |
| 16397 | 820 | H820T | B | 1.9e-07 | 2.1e-08 | 4.2e-08 | 8.433E-08 | 9.211E-08 | 2.100E-08 | 1.900E-07 |
| 16398 | 820 | H820W | B | 6.4e-08 | 1.3e-07 | 7.2e-08 | 8.867E-08 | 3.602E-08 | 6.400E-08 | 1.300E-07 |
| 16399 | 820 | H820Y | B | 5.2e-08 | 3.7e-08 | 5.2e-08 | 4.700E-08 | 8.660E-09 | 3.700E-08 | 5.200E-08 |
| 16400 | 820 | H820V | B | 2.1e-08 | 7.2e-08 | 1.8e-08 | 3.700E-08 | 3.035E-08 | 1.800E-08 | 7.200E-08 |
| 16401 | 821 | Q821A | B | 5.9e-08 | 6.4e-08 | 5.3e-08 | 5.867E-08 | 5.508E-09 | 5.300E-08 | 6.400E-08 |
| 16402 | 821 | Q821R | B | 4.2e-08 | 4.7e-08 | 4.9e-08 | 4.600E-08 | 3.606E-09 | 4.200E-08 | 4.900E-08 |
| 16403 | 821 | Q821N | B | 2.9e-08 | 5e-08 | 4.9e-08 | 4.267E-08 | 1.185E-08 | 2.900E-08 | 5.000E-08 |
| 16404 | 821 | Q821D | B | 4.8e-08 | 5.6e-08 | 3.6e-08 | 4.667E-08 | 1.007E-08 | 3.600E-08 | 5.600E-08 |

| | | | | | | | | | | |
|---|---|---|---|---|---|---|---|---|---|---|
| 16405 | 821 | Q821C | B | 4e-08 | 4.4e-08 | 3.6e-08 | 4.000E-08 | 4.000E-09 | 3.600E-08 | 4.400E-08 |
| 16406 | 821 | Q821E | B | 4.6e-08 | 6.2e-08 | 2.4e-08 | 4.400E-08 | 1.908E-08 | 2.400E-08 | 6.200E-08 |
| 16407 | 821 | Q821Q | B | 4.7e-08 | 5.8e-08 | 3.3e-08 | 4.600E-08 | 1.253E-08 | 3.300E-08 | 5.800E-08 |
| 16408 | 821 | Q821G | B | 2.6e-08 | 8.3e-08 | 1.3e-07 | 7.967E-08 | 5.208E-08 | 2.600E-08 | 1.300E-07 |
| 16409 | 821 | Q821H | B | 3e-08 | 7.6e-08 | 5e-08 | 5.200E-08 | 2.307E-08 | 3.000E-08 | 7.600E-08 |
| 16410 | 821 | Q821I | B | 1.1e-07 | 7.1e-08 | 2.6e-08 | 6.900E-08 | 4.204E-08 | 2.600E-08 | 1.100E-07 |
| 16411 | 821 | Q821L | B | 8e-08 | 6e-08 | 4.6e-08 | 6.200E-08 | 1.709E-08 | 4.600E-08 | 8.000E-08 |
| 16412 | 821 | Q821K | B | 1.7e-08 | 4.7e-08 | 2.8e-08 | 3.067E-08 | 1.518E-08 | 1.700E-08 | 4.700E-08 |
| 16413 | 821 | Q821M | B | 4.9e-08 | 1.7e-08 | 5.9e-08 | 4.167E-08 | 2.194E-08 | 1.700E-08 | 5.900E-08 |
| 16414 | 821 | Q821F | B | 5.2e-09 | 1.2e-07 | 3.6e-08 | 5.373E-08 | 5.942E-08 | 5.200E-09 | 1.200E-07 |
| 16415 | 821 | Q821P | B | 1.9e-08 | 4.4e-08 | 3.9e-08 | 3.400E-08 | 1.323E-08 | 1.900E-08 | 4.400E-08 |
| 16416 | 821 | Q821S | B | 6.7e-08 | 6.8e-08 | 3.3e-08 | 5.600E-08 | 1.992E-08 | 3.300E-08 | 6.800E-08 |
| 16417 | 821 | Q821T | B | 6.8e-08 | 4.2e-08 | 4.2e-08 | 5.067E-08 | 1.501E-08 | 4.200E-08 | 6.800E-08 |
| 16418 | 821 | Q821W | B | 3.1e-08 | 4.8e-08 | 8.8e-08 | 5.567E-08 | 2.926E-08 | 3.100E-08 | 8.800E-08 |
| 16419 | 821 | Q821Y | B | 1.5e-08 | 2.3e-08 | 8.7e-08 | 4.167E-08 | 3.946E-08 | 1.500E-08 | 8.700E-08 |
| 16420 | 821 | Q821V | B | 2.4e-08 | 1.4e-07 | 6.5e-08 | 7.633E-08 | 5.882E-08 | 2.400E-08 | 1.400E-07 |
| 16421 | 822 | G822A | B | 5.1e-08 | 3.9e-08 | 3.3e-08 | 4.100E-08 | 9.165E-09 | 3.300E-08 | 5.100E-08 |
| 16422 | 822 | G822R | B | 2.7e-08 | 2.7e-08 | 6.6e-08 | 4.000E-08 | 2.252E-08 | 2.700E-08 | 6.600E-08 |
| 16423 | 822 | G822N | B | 5.9e-08 | 8.7e-08 | 2.8e-08 | 5.800E-08 | 2.951E-08 | 2.800E-08 | 8.700E-08 |
| 16424 | 822 | G822D | B | 3.5e-08 | 2.1e-08 | 2.8e-08 | 2.800E-08 | 7.000E-09 | 2.100E-08 | 3.500E-08 |
| 16425 | 822 | G822C | B | 9.9e-08 | 4.3e-08 | 4.4e-08 | 6.200E-08 | 3.205E-08 | 4.300E-08 | 9.900E-08 |
| 16426 | 822 | G822E | B | 7.1e-08 | 5e-08 | 2.6e-08 | 4.900E-08 | 2.252E-08 | 2.600E-08 | 7.100E-08 |
| 16427 | 822 | G822Q | B | 4.5e-08 | 5e-08 | 6.7e-08 | 5.400E-08 | 1.153E-08 | 4.500E-08 | 6.700E-08 |
| 16428 | 822 | G822G | B | 1.4e-07 | 4.5e-08 | 5.1e-08 | 7.867E-08 | 5.320E-08 | 4.500E-08 | 1.400E-07 |
| 16429 | 822 | G822H | B | 5e-08 | 1.4e-08 | 9.1e-08 | 5.167E-08 | 3.853E-08 | 1.400E-08 | 9.100E-08 |
| 16430 | 822 | G822I | B | 1.9e-08 | 2.5e-08 | 3.2e-08 | 2.533E-08 | 6.506E-09 | 1.900E-08 | 3.200E-08 |
| 16431 | 822 | G822L | B | 5.7e-08 | 2.1e-08 | 1.7e-08 | 3.167E-08 | 2.203E-08 | 1.700E-08 | 5.700E-08 |
| 16432 | 822 | G822K | B | 3.6e-08 | 5.3e-08 | 2.4e-08 | 3.767E-08 | 1.457E-08 | 2.400E-08 | 5.300E-08 |
| 16433 | 822 | G822M | B | 1.9e-08 | 1.8e-08 | 5.4e-08 | 3.033E-08 | 2.050E-08 | 1.800E-08 | 5.400E-08 |
| 16434 | 822 | G822F | B | 2.1e-08 | 2e-08 | 4.4e-08 | 2.833E-08 | 1.358E-08 | 2.000E-08 | 4.400E-08 |
| 16435 | 822 | G822P | B | 1.7e-08 | 7.5e-08 | 2.2e-08 | 3.800E-08 | 3.214E-08 | 1.700E-08 | 7.500E-08 |
| 16436 | 822 | G822S | B | 5.9e-08 | 6.3e-08 | 4.3e-08 | 5.500E-08 | 1.058E-08 | 4.300E-08 | 6.300E-08 |
| 16437 | 822 | G822T | B | 5.1e-08 | 2e-08 | 3.2e-08 | 3.433E-08 | 1.563E-08 | 2.000E-08 | 5.100E-08 |
| 16438 | 822 | G822W | B | 3.5e-08 | 8.9e-08 | 1.8e-08 | 4.733E-08 | 3.707E-08 | 1.800E-08 | 8.900E-08 |
| 16439 | 822 | G822Y | B | 7.1e-08 | 4.6e-08 | 2.7e-08 | 4.800E-08 | 2.207E-08 | 2.700E-08 | 7.100E-08 |
| 16440 | 822 | G822V | B | 6.7e-08 | 7.6e-08 | 5.6e-08 | 6.633E-08 | 1.002E-08 | 5.600E-08 | 7.600E-08 |
| 16441 | 823 | L823A | B | 3.3e-08 | 7e-08 | 5.3e-08 | 5.200E-08 | 1.852E-08 | 3.300E-08 | 7.000E-08 |
| 16442 | 823 | L823R | B | 5.7e-08 | 7.4e-08 | 2.9e-08 | 5.333E-08 | 2.272E-08 | 2.900E-08 | 7.400E-08 |
| 16443 | 823 | L823N | B | 6.7e-08 | 3.7e-08 | 5.6e-08 | 5.333E-08 | 1.518E-08 | 3.700E-08 | 6.700E-08 |
| 16444 | 823 | L823D | B | 6.1e-08 | 2.2e-08 | 5.6e-08 | 4.633E-08 | 2.122E-08 | 2.200E-08 | 6.100E-08 |
| 16445 | 823 | L823C | B | 5.4e-08 | 4.3e-08 | 4.3e-08 | 4.667E-08 | 6.351E-09 | 4.300E-08 | 5.400E-08 |
| 16446 | 823 | L823E | B | 2.7e-08 | 4.3e-08 | 2.7e-08 | 3.233E-08 | 9.238E-09 | 2.700E-08 | 4.300E-08 |
| 16447 | 823 | L823Q | B | 2.2e-08 | 2.1e-08 | 5.6e-08 | 3.300E-08 | 1.992E-08 | 2.100E-08 | 5.600E-08 |
| 16448 | 823 | L823G | B | 6.5e-09 | 2.1e-08 | 2.5e-08 | 1.750E-08 | 9.734E-09 | 6.500E-09 | 2.500E-08 |
| 16449 | 823 | L823H | B | 2.5e-08 | 2.6e-08 | 2.1e-08 | 2.400E-08 | 2.646E-09 | 2.100E-08 | 2.600E-08 |
| 16450 | 823 | L823I | B | 4.5e-08 | 5.6e-08 | 1.2e-08 | 3.767E-08 | 2.290E-08 | 1.200E-08 | 5.600E-08 |
| 16451 | 823 | L823L | B | 9e-08 | 5e-08 | 3.4e-08 | 5.800E-08 | 2.884E-08 | 3.400E-08 | 9.000E-08 |
| 16452 | 823 | L823K | B | 2.7e-08 | 1.5e-08 | 5.2e-08 | 3.133E-08 | 1.888E-08 | 1.500E-08 | 5.200E-08 |
| 16453 | 823 | L823M | B | 3.3e-08 | 6.7e-08 | 3.3e-08 | 4.433E-08 | 1.963E-08 | 3.300E-08 | 6.700E-08 |
| 16454 | 823 | L823F | B | 1.7e-08 | 5.5e-08 | 1.9e-08 | 3.033E-08 | 2.139E-08 | 1.700E-08 | 5.500E-08 |
| 16455 | 823 | L823P | B | 8.3e-08 | 3.4e-08 | 4.4e-08 | 5.367E-08 | 2.589E-08 | 3.400E-08 | 8.300E-08 |
| 16456 | 823 | L823S | B | 7e-08 | 2e-07 | 6.8e-08 | 1.127E-07 | 7.564E-08 | 6.800E-08 | 2.000E-07 |
| 16457 | 823 | L823T | B | 4.2e-08 | 4e-08 | 4.6e-08 | 4.267E-08 | 3.055E-09 | 4.000E-08 | 4.600E-08 |
| 16458 | 823 | L823W | B | 5.4e-08 | 9.3e-08 | 5.9e-08 | 6.867E-08 | 2.122E-08 | 5.400E-08 | 9.300E-08 |
| 16459 | 823 | L823Y | B | 6.4e-08 | 2.5e-08 | 3.4e-08 | 4.100E-08 | 2.042E-08 | 2.500E-08 | 6.400E-08 |
| 16460 | 823 | L823V | B | 1.2e-07 | 2.6e-08 | 2e-08 | 5.533E-08 | 5.608E-08 | 2.000E-08 | 1.200E-07 |
| 16461 | 824 | S824A | B | 4.8e-08 | 3.8e-08 | 3.3e-08 | 3.967E-08 | 7.638E-09 | 3.300E-08 | 4.800E-08 |
| 16462 | 824 | S824R | B | 6.7e-08 | 3e-08 | 6.9e-08 | 5.533E-08 | 2.196E-08 | 3.000E-08 | 6.900E-08 |
| 16463 | 824 | S824N | B | 4.4e-08 | 6.3e-08 | 1.2e-07 | 7.567E-08 | 3.955E-08 | 4.400E-08 | 1.200E-07 |
| 16464 | 824 | S824D | B | 6.5e-08 | 4.5e-08 | 5.9e-08 | 5.633E-08 | 1.026E-08 | 4.500E-08 | 6.500E-08 |
| 16465 | 824 | S824C | B | 1.2e-07 | 2.6e-08 | 4e-08 | 6.200E-08 | 5.071E-08 | 2.600E-08 | 1.200E-07 |
| 16466 | 824 | S824E | B | 1.5e-08 | 6.7e-08 | 2.9e-08 | 3.700E-08 | 2.691E-08 | 1.500E-08 | 6.700E-08 |
| 16467 | 824 | S824Q | B | 5.4e-08 | 2.7e-08 | 1.6e-07 | 8.033E-08 | 7.030E-08 | 2.700E-08 | 1.600E-07 |
| 16468 | 824 | S824G | B | 1.2e-08 | 2.2e-08 | 6.2e-08 | 3.200E-08 | 2.646E-08 | 1.200E-08 | 6.200E-08 |
| 16469 | 824 | S824H | B | 4.6e-08 | 3.3e-08 | 3.6e-08 | 3.833E-08 | 6.807E-09 | 3.300E-08 | 4.600E-08 |
| 16470 | 824 | S824I | B | 5.1e-08 | 9.4e-08 | 5.5e-08 | 6.667E-08 | 2.376E-08 | 5.100E-08 | 9.400E-08 |
| 16471 | 824 | S824L | B | 4.6e-08 | 5.3e-08 | 3.5e-08 | 4.467E-08 | 9.074E-09 | 3.500E-08 | 5.300E-08 |
| 16472 | 824 | S824K | B | 3.4e-08 | 3.7e-08 | 3e-08 | 3.367E-08 | 3.512E-09 | 3.000E-08 | 3.700E-08 |
| 16473 | 824 | S824M | B | 5.7e-08 | 5.2e-08 | 7.7e-08 | 6.200E-08 | 1.323E-08 | 5.200E-08 | 7.700E-08 |
| 16474 | 824 | S824F | B | 5.5e-08 | 2.1e-08 | 8.4e-08 | 5.333E-08 | 3.153E-08 | 2.100E-08 | 8.400E-08 |
| 16475 | 824 | S824P | B | 3.7e-08 | 4e-08 | 8e-08 | 5.233E-08 | 2.401E-08 | 3.700E-08 | 8.000E-08 |
| 16476 | 824 | S824S | B | 5e-08 | 4e-08 | 5.6e-08 | 4.867E-08 | 8.083E-09 | 4.000E-08 | 5.600E-08 |
| 16477 | 824 | S824T | B | 3.2e-08 | 2.1e-08 | 2e-08 | 2.433E-08 | 6.658E-09 | 2.000E-08 | 3.200E-08 |
| 16478 | 824 | S824W | B | 1.3e-07 | 9.7e-08 | 4.7e-08 | 9.133E-08 | 4.179E-08 | 4.700E-08 | 1.300E-07 |
| 16479 | 824 | S824Y | B | 5.6e-08 | 9.4e-09 | 1.4e-08 | 2.647E-08 | 2.568E-08 | 9.400E-09 | 5.600E-08 |
| 16480 | 824 | S824V | B | 4.6e-08 | 8.2e-08 | 9.1e-08 | 7.300E-08 | 2.381E-08 | 4.600E-08 | 9.100E-08 |
| 16481 | 825 | S825A | B | 4.2e-08 | 1.6e-08 | 6.3e-08 | 4.033E-08 | 2.354E-08 | 1.600E-08 | 6.300E-08 |
| 16482 | 825 | S825R | B | 4e-08 | 1.4e-07 | 3.1e-08 | 7.033E-08 | 6.050E-08 | 3.100E-08 | 1.400E-07 |
| 16483 | 825 | S825N | B | 4.8e-08 | 7.3e-08 | 6.9e-08 | 6.333E-08 | 1.343E-08 | 4.800E-08 | 7.300E-08 |
| 16484 | 825 | S825D | B | 5.8e-08 | 1.4e-08 | 1e-07 | 5.733E-08 | 4.300E-08 | 1.400E-08 | 1.000E-07 |
| 16485 | 825 | S825C | B | 6.1e-08 | 4.4e-08 | 2e-08 | 4.167E-08 | 2.060E-08 | 2.000E-08 | 6.100E-08 |
| 16486 | 825 | S825E | B | 3e-08 | 1.8e-08 | 2.7e-08 | 2.500E-08 | 6.245E-09 | 1.800E-08 | 3.000E-08 |
| 16487 | 825 | S825Q | B | 3e-08 | 3.9e-08 | 9.5e-08 | 5.467E-08 | 3.522E-08 | 3.000E-08 | 9.500E-08 |
| 16488 | 825 | S825G | B | 4.3e-08 | 2.6e-08 | 8.9e-08 | 5.267E-08 | 3.259E-08 | 2.600E-08 | 8.900E-08 |
| 16489 | 825 | S825H | B | 3.1e-08 | 6.6e-08 | 4.7e-08 | 4.800E-08 | 1.752E-08 | 3.100E-08 | 6.600E-08 |

| | | | | | | | | | | |
|---|---|---|---|---|---|---|---|---|---|---|
| 16490 | 825 | S825I | B | 1.1e-07 | 1.8e-08 | 4.7e-08 | 5.833E-08 | 4.704E-08 | 1.800E-08 | 1.100E-07 |
| 16491 | 825 | S825L | B | 1.3e-08 | 3.3e-08 | 2.6e-08 | 2.400E-08 | 1.015E-08 | 1.300E-08 | 3.300E-08 |
| 16492 | 825 | S825K | B | 6.2e-08 | 7.7e-08 | 4.2e-08 | 6.033E-08 | 1.756E-08 | 4.200E-08 | 7.700E-08 |
| 16493 | 825 | S825M | B | 4e-08 | 8.4e-08 | 6.4e-08 | 6.267E-08 | 2.203E-08 | 4.000E-08 | 8.400E-08 |
| 16494 | 825 | S825F | B | 4e-08 | 6.1e-08 | 5.4e-08 | 5.167E-08 | 1.069E-08 | 4.000E-08 | 6.100E-08 |
| 16495 | 825 | S825P | B | 4.5e-08 | 6e-08 | 8.8e-08 | 6.433E-08 | 2.183E-08 | 4.500E-08 | 8.800E-08 |
| 16496 | 825 | S825S | B | 8.7e-09 | 8.2e-09 | 1.8e-07 | 6.563E-08 | 9.904E-08 | 8.200E-09 | 1.800E-07 |
| 16497 | 825 | S825T | B | 4.2e-08 | 3e-08 | 4.5e-08 | 3.900E-08 | 7.937E-09 | 3.000E-08 | 4.500E-08 |
| 16498 | 825 | S825W | B | 5.9e-08 | 4.7e-08 | 3.9e-08 | 4.833E-08 | 1.007E-08 | 3.900E-08 | 5.900E-08 |
| 16499 | 825 | S825Y | B | 2.2e-08 | 1e-08 | 3.2e-08 | 2.133E-08 | 1.102E-08 | 1.000E-08 | 3.200E-08 |
| 16500 | 825 | S825V | B | 8.4e-08 | 6.1e-08 | 5.6e-08 | 6.700E-08 | 1.493E-08 | 5.600E-08 | 8.400E-08 |
| 16501 | 826 | P826A | B | 7.6e-08 | 1.8e-07 | 3.3e-08 | 9.633E-08 | 7.558E-08 | 3.300E-08 | 1.800E-07 |
| 16502 | 826 | P826R | B | 2.8e-08 | 7.1e-08 | 2.1e-08 | 4.000E-08 | 2.707E-08 | 2.100E-08 | 7.100E-08 |
| 16503 | 826 | P826N | B | 7.7e-08 | 6.9e-08 | 2.2e-08 | 5.600E-08 | 2.972E-08 | 2.200E-08 | 7.700E-08 |
| 16504 | 826 | P826D | B | 1.1e-07 | 3.7e-08 | 7.2e-08 | 7.300E-08 | 3.651E-08 | 3.700E-08 | 1.100E-07 |
| 16505 | 826 | P826C | B | 4.3e-08 | 5.6e-08 | 5.9e-08 | 5.267E-08 | 8.505E-09 | 4.300E-08 | 5.900E-08 |
| 16506 | 826 | P826E | B | 1.2e-08 | 5.6e-08 | 6.7e-08 | 4.500E-08 | 2.910E-08 | 1.200E-08 | 6.700E-08 |
| 16507 | 826 | P826Q | B | 2.2e-08 | 1.1e-07 | 3.3e-08 | 5.500E-08 | 4.795E-08 | 2.200E-08 | 1.100E-07 |
| 16508 | 826 | P826G | B | 2.4e-08 | 5.6e-08 | 1.4e-07 | 7.333E-08 | 5.991E-08 | 2.400E-08 | 1.400E-07 |
| 16509 | 826 | P826H | B | 7.9e-08 | 3.2e-08 | 5.6e-08 | 5.567E-08 | 2.350E-08 | 3.200E-08 | 7.900E-08 |
| 16510 | 826 | P826I | B | 1.4e-08 | 7.6e-09 | 3.9e-08 | 2.020E-08 | 1.659E-08 | 7.600E-09 | 3.900E-08 |
| 16511 | 826 | P826L | B | 1.4e-07 | 5.6e-08 | 2.6e-08 | 7.400E-08 | 5.909E-08 | 2.600E-08 | 1.400E-07 |
| 16512 | 826 | P826K | B | 8.6e-08 | 5.5e-08 | 3.1e-08 | 5.733E-08 | 2.757E-08 | 3.100E-08 | 8.600E-08 |
| 16513 | 826 | P826M | B | 5.4e-08 | 3.5e-08 | 6.6e-08 | 5.167E-08 | 1.563E-08 | 3.500E-08 | 6.600E-08 |
| 16514 | 826 | P826F | B | 3.9e-08 | 6.4e-08 | 3.7e-08 | 4.667E-08 | 1.504E-08 | 3.700E-08 | 6.400E-08 |
| 16515 | 826 | P826P | B | 5.9e-08 | 8.4e-08 | 4.5e-08 | 6.267E-08 | 1.976E-08 | 4.500E-08 | 8.400E-08 |
| 16516 | 826 | P826S | B | 1.8e-08 | 8.8e-08 | 2.3e-08 | 4.300E-08 | 3.905E-08 | 1.800E-08 | 8.800E-08 |
| 16517 | 826 | P826T | B | 4.2e-08 | 1e-08 | 5.7e-08 | 3.633E-08 | 2.401E-08 | 1.000E-08 | 5.700E-08 |
| 16518 | 826 | P826W | B | 2.3e-08 | 2.7e-08 | 4.4e-08 | 3.133E-08 | 1.115E-08 | 2.300E-08 | 4.400E-08 |
| 16519 | 826 | P826Y | B | 8.3e-08 | 1e-07 | 5.2e-08 | 7.833E-08 | 2.434E-08 | 5.200E-08 | 1.000E-07 |
| 16520 | 826 | P826V | B | 2.2e-08 | 7.7e-08 | 1.1e-07 | 6.967E-08 | 4.446E-08 | 2.200E-08 | 1.100E-07 |
| 16521 | 827 | V827A | B | 3.7e-08 | 2.7e-08 | 1.5e-08 | 2.633E-08 | 1.102E-08 | 1.500E-08 | 3.700E-08 |
| 16522 | 827 | V827R | B | 5.8e-08 | 2.4e-08 | 3.8e-08 | 4.000E-08 | 1.709E-08 | 2.400E-08 | 5.800E-08 |
| 16523 | 827 | V827N | B | 6.1e-08 | 9.4e-08 | 3.9e-08 | 6.467E-08 | 2.768E-08 | 3.900E-08 | 9.400E-08 |
| 16524 | 827 | V827D | B | 6.9e-08 | 5.6e-09 | 3.8e-08 | 3.753E-08 | 3.170E-08 | 5.600E-09 | 6.900E-08 |
| 16525 | 827 | V827C | B | 2.6e-08 | 2.7e-08 | 1.7e-08 | 2.333E-08 | 5.508E-09 | 1.700E-08 | 2.700E-08 |
| 16526 | 827 | V827E | B | 2.9e-08 | 4.5e-08 | 2.7e-08 | 3.367E-08 | 9.866E-09 | 2.700E-08 | 4.500E-08 |
| 16527 | 827 | V827Q | B | 3.6e-08 | 6.3e-08 | 5.1e-08 | 5.000E-08 | 1.353E-08 | 3.600E-08 | 6.300E-08 |
| 16528 | 827 | V827G | B | 1.1e-07 | 1.3e-07 | 2.1e-08 | 8.700E-08 | 5.803E-08 | 2.100E-08 | 1.300E-07 |
| 16529 | 827 | V827H | B | 1.5e-07 | 5.7e-08 | 6.1e-08 | 8.933E-08 | 5.258E-08 | 5.700E-08 | 1.500E-07 |
| 16530 | 827 | V827I | B | 5.7e-08 | 5.2e-08 | 1.1e-07 | 7.300E-08 | 3.214E-08 | 5.200E-08 | 1.100E-07 |
| 16531 | 827 | V827L | B | 2.8e-08 | 6.2e-08 | 1.7e-08 | 3.567E-08 | 2.346E-08 | 1.700E-08 | 6.200E-08 |
| 16532 | 827 | V827K | B | 4.6e-08 | 7.3e-08 | 5.2e-08 | 5.700E-08 | 1.418E-08 | 4.600E-08 | 7.300E-08 |
| 16533 | 827 | V827M | B | 5.1e-08 | 3.5e-08 | 2.6e-08 | 3.733E-08 | 1.266E-08 | 2.600E-08 | 5.100E-08 |
| 16534 | 827 | V827F | B | 5.5e-08 | 5.7e-08 | 8.8e-08 | 6.667E-08 | 1.850E-08 | 5.500E-08 | 8.800E-08 |
| 16535 | 827 | V827P | B | 8e-08 | 1.1e-07 | 4.8e-08 | 7.933E-08 | 3.101E-08 | 4.800E-08 | 1.100E-07 |
| 16536 | 827 | V827S | B | 3.1e-08 | 8.6e-08 | 6.7e-08 | 6.133E-08 | 2.793E-08 | 3.100E-08 | 8.600E-08 |
| 16537 | 827 | V827T | B | 5.3e-08 | 1.3e-08 | 1.2e-07 | 6.200E-08 | 5.406E-08 | 1.300E-08 | 1.200E-07 |
| 16538 | 827 | V827W | B | 3.3e-08 | 6e-08 | 4.3e-08 | 4.533E-08 | 1.365E-08 | 3.300E-08 | 6.000E-08 |
| 16539 | 827 | V827Y | B | 5.5e-08 | 3.3e-08 | 1.3e-08 | 3.367E-08 | 2.101E-08 | 1.300E-08 | 5.500E-08 |
| 16540 | 827 | V827V | B | 3.5e-08 | 3.6e-08 | 4.7e-08 | 3.933E-08 | 6.658E-09 | 3.500E-08 | 4.700E-08 |
| 16541 | 828 | T828A | B | 8.8e-08 | 1.1e-07 | 1.7e-07 | 1.227E-07 | 4.244E-08 | 8.800E-08 | 1.700E-07 |
| 16542 | 828 | T828R | B | 5.2e-08 | 2.7e-08 | 2.1e-08 | 3.333E-08 | 1.644E-08 | 2.100E-08 | 5.200E-08 |
| 16543 | 828 | T828N | B | 6.9e-08 | 2.7e-08 | 5.9e-08 | 5.167E-08 | 2.194E-08 | 2.700E-08 | 6.900E-08 |
| 16544 | 828 | T828D | B | 1.6e-07 | 1.4e-07 | 2.7e-08 | 1.090E-07 | 7.171E-08 | 2.700E-08 | 1.600E-07 |
| 16545 | 828 | T828C | B | 4.3e-08 | 6.8e-08 | 2e-08 | 4.367E-08 | 2.401E-08 | 2.000E-08 | 6.800E-08 |
| 16546 | 828 | T828E | B | 2.7e-08 | 3.9e-08 | 1.9e-08 | 2.833E-08 | 1.007E-08 | 1.900E-08 | 3.900E-08 |
| 16547 | 828 | T828Q | B | 1.7e-08 | 4.7e-08 | 2.1e-08 | 2.833E-08 | 1.629E-08 | 1.700E-08 | 4.700E-08 |
| 16548 | 828 | T828G | B | 1.4e-07 | 3.5e-08 | 3.2e-08 | 6.900E-08 | 6.151E-08 | 3.200E-08 | 1.400E-07 |
| 16549 | 828 | T828H | B | 4e-08 | 5.6e-08 | 1e-08 | 3.533E-08 | 2.335E-08 | 1.000E-08 | 5.600E-08 |
| 16550 | 828 | T828I | B | 2.3e-07 | 1.2e-08 | 8.8e-08 | 1.100E-07 | 1.107E-07 | 1.200E-08 | 2.300E-07 |
| 16551 | 828 | T828L | B | 2e-08 | 3.6e-08 | 9.8e-08 | 5.133E-08 | 4.120E-08 | 2.000E-08 | 9.800E-08 |
| 16552 | 828 | T828K | B | 4.9e-08 | 7.4e-08 | 1.7e-08 | 4.667E-08 | 2.857E-08 | 1.700E-08 | 7.400E-08 |
| 16553 | 828 | T828M | B | 6.3e-08 | 5.2e-08 | 4.2e-08 | 5.233E-08 | 1.050E-08 | 4.200E-08 | 6.300E-08 |
| 16554 | 828 | T828F | B | 3e-08 | 4.5e-08 | 1.2e-08 | 2.900E-08 | 1.652E-08 | 1.200E-08 | 4.500E-08 |
| 16555 | 828 | T828P | B | 2e-08 | 1e-07 | 5.7e-08 | 5.900E-08 | 4.004E-08 | 2.000E-08 | 1.000E-07 |
| 16556 | 828 | T828S | B | 6.5e-08 | 5.2e-08 | 1.1e-07 | 7.567E-08 | 3.044E-08 | 5.200E-08 | 1.100E-07 |
| 16557 | 828 | T828T | B | 2.7e-08 | 1.8e-08 | 1.4e-08 | 1.967E-08 | 6.658E-09 | 1.400E-08 | 2.700E-08 |
| 16558 | 828 | T828W | B | 4e-08 | 1.7e-08 | 4.1e-08 | 3.267E-08 | 1.358E-08 | 1.700E-08 | 4.100E-08 |
| 16559 | 828 | T828Y | B | 7.2e-08 | 2.5e-08 | 7.1e-08 | 5.600E-08 | 2.685E-08 | 2.500E-08 | 7.200E-08 |
| 16560 | 828 | T828V | B | 5e-08 | 3.1e-08 | 7.5e-09 | 2.950E-08 | 2.129E-08 | 7.500E-09 | 5.000E-08 |
| 16561 | 829 | K829A | B | 1.5e-08 | 2.6e-08 | 1.3e-08 | 1.800E-08 | 7.000E-09 | 1.300E-08 | 2.600E-08 |
| 16562 | 829 | K829R | B | 2.4e-08 | 8.1e-08 | 5.1e-08 | 5.200E-08 | 2.851E-08 | 2.400E-08 | 8.100E-08 |
| 16563 | 829 | K829N | B | 1e-07 | 1.2e-07 | 6.9e-08 | 9.633E-08 | 2.570E-08 | 6.900E-08 | 1.200E-07 |
| 16564 | 829 | K829D | B | 5.5e-08 | 1.4e-07 | 1.1e-08 | 6.867E-08 | 6.558E-08 | 1.100E-08 | 1.400E-07 |
| 16565 | 829 | K829C | B | 7.3e-08 | 3e-08 | 4.5e-08 | 4.933E-08 | 2.183E-08 | 3.000E-08 | 7.300E-08 |
| 16566 | 829 | K829E | B | 3.2e-08 | 4.6e-08 | 6.7e-08 | 4.833E-08 | 1.762E-08 | 3.200E-08 | 6.700E-08 |
| 16567 | 829 | K829Q | B | 6.5e-08 | 1.3e-08 | 4.1e-08 | 3.967E-08 | 2.603E-08 | 1.300E-08 | 6.500E-08 |
| 16568 | 829 | K829G | B | 2.8e-08 | 2.7e-08 | 5.3e-08 | 3.600E-08 | 1.473E-08 | 2.700E-08 | 5.300E-08 |
| 16569 | 829 | K829H | B | 5e-08 | 2.7e-08 | 2.7e-08 | 3.467E-08 | 1.328E-08 | 2.700E-08 | 5.000E-08 |
| 16570 | 829 | K829I | B | 4e-08 | 3e-08 | 2.1e-08 | 3.033E-08 | 9.504E-09 | 2.100E-08 | 4.000E-08 |
| 16571 | 829 | K829L | B | 3.8e-08 | 9.4e-08 | 8.9e-08 | 7.367E-08 | 3.099E-08 | 3.800E-08 | 9.400E-08 |
| 16572 | 829 | K829K | B | 3.5e-08 | 4.1e-08 | 5.6e-08 | 4.400E-08 | 1.082E-08 | 3.500E-08 | 5.600E-08 |
| 16573 | 829 | K829M | B | 2.6e-08 | 1.8e-08 | 1.6e-08 | 2.000E-08 | 5.292E-09 | 1.600E-08 | 2.600E-08 |
| 16574 | 829 | K829F | B | 9.3e-08 | 2.7e-08 | 2.4e-08 | 4.800E-08 | 3.900E-08 | 2.400E-08 | 9.300E-08 |

| | | | | | | | | | | |
|---|---|---|---|---|---|---|---|---|---|---|
| 16575 | 829 | K829P | B | 8e-08 | 1.9e-08 | 6.4e-08 | 5.433E-08 | 3.163E-08 | 1.900E-08 | 8.000E-08 |
| 16576 | 829 | K829S | B | 4.2e-08 | 6.2e-08 | 4.7e-08 | 5.033E-08 | 1.041E-08 | 4.200E-08 | 6.200E-08 |
| 16577 | 829 | K829T | B | 9.3e-08 | 3.5e-08 | 4.4e-08 | 5.733E-08 | 3.121E-08 | 3.500E-08 | 9.300E-08 |
| 16578 | 829 | K829W | B | 6e-08 | 2.9e-08 | 3.7e-08 | 4.200E-08 | 1.609E-08 | 2.900E-08 | 6.000E-08 |
| 16579 | 829 | K829Y | B | 6.7e-08 | 3.8e-08 | 5.2e-08 | 5.233E-08 | 1.450E-08 | 3.800E-08 | 6.700E-08 |
| 16580 | 829 | K829V | B | 1.4e-08 | 6.7e-08 | 4.5e-08 | 4.200E-08 | 2.663E-08 | 1.400E-08 | 6.700E-08 |
| 16581 | 830 | S830A | B | 1.2e-07 | 4.4e-08 | 8.9e-08 | 8.433E-08 | 3.821E-08 | 4.400E-08 | 1.200E-07 |
| 16582 | 830 | S830R | B | 1.3e-07 | 4.6e-08 | 3.4e-08 | 7.000E-08 | 5.231E-08 | 3.400E-08 | 1.300E-07 |
| 16583 | 830 | S830N | B | 4e-08 | 3.8e-08 | 2.7e-08 | 3.500E-08 | 7.000E-09 | 2.700E-08 | 4.000E-08 |
| 16584 | 830 | S830D | B | 2.5e-08 | 5.2e-08 | 3e-08 | 3.567E-08 | 1.436E-08 | 2.500E-08 | 5.200E-08 |
| 16585 | 830 | S830C | B | 3.9e-08 | 3.3e-08 | 6.6e-08 | 4.600E-08 | 1.758E-08 | 3.300E-08 | 6.600E-08 |
| 16586 | 830 | S830E | B | 2.4e-08 | 3.2e-08 | 1.5e-08 | 2.367E-08 | 8.505E-09 | 1.500E-08 | 3.200E-08 |
| 16587 | 830 | S830Q | B | 3.3e-08 | 2.7e-08 | 1.2e-07 | 6.000E-08 | 5.205E-08 | 2.700E-08 | 1.200E-07 |
| 16588 | 830 | S830G | B | 1.4e-08 | 1.6e-08 | 3.2e-08 | 2.067E-08 | 9.866E-09 | 1.400E-08 | 3.200E-08 |
| 16589 | 830 | S830H | B | 2.1e-08 | 7.9e-08 | 1.6e-08 | 3.867E-08 | 3.502E-08 | 1.600E-08 | 7.900E-08 |
| 16590 | 830 | S830I | B | 4.1e-08 | 8.2e-08 | 3e-08 | 5.100E-08 | 2.740E-08 | 3.000E-08 | 8.200E-08 |
| 16591 | 830 | S830L | B | 7.9e-08 | 8.2e-08 | 3.9e-08 | 6.667E-08 | 2.401E-08 | 3.900E-08 | 8.200E-08 |
| 16592 | 830 | S830K | B | 1.8e-08 | 3.7e-08 | 3.2e-08 | 2.900E-08 | 9.849E-09 | 1.800E-08 | 3.700E-08 |
| 16593 | 830 | S830M | B | 1.2e-08 | 2.5e-08 | 4e-08 | 2.567E-08 | 1.401E-08 | 1.200E-08 | 4.000E-08 |
| 16594 | 830 | S830F | B | 2.8e-08 | 6.3e-08 | 5.3e-08 | 4.800E-08 | 1.803E-08 | 2.800E-08 | 6.300E-08 |
| 16595 | 830 | S830P | B | 5.5e-08 | 2e-08 | 2.3e-08 | 3.267E-08 | 1.940E-08 | 2.000E-08 | 5.500E-08 |
| 16596 | 830 | S830S | B | 9e-08 | 2.3e-08 | 4.9e-08 | 5.400E-08 | 3.378E-08 | 2.300E-08 | 9.000E-08 |
| 16597 | 830 | S830T | B | 5.4e-08 | 2.6e-08 | 5e-08 | 4.333E-08 | 1.514E-08 | 2.600E-08 | 5.400E-08 |
| 16598 | 830 | S830W | B | 6e-08 | 4.9e-08 | 4.7e-08 | 5.200E-08 | 7.000E-09 | 4.700E-08 | 6.000E-08 |
| 16599 | 830 | S830Y | B | 3.4e-08 | 9.6e-08 | 3.9e-08 | 5.633E-08 | 3.444E-08 | 3.400E-08 | 9.600E-08 |
| 16600 | 830 | S830V | B | 1.3e-07 | 5.2e-08 | 6.4e-08 | 8.200E-08 | 4.200E-08 | 5.200E-08 | 1.300E-07 |
| 16601 | 831 | F831A | B | 3.7e-08 | 4.9e-08 | 3.4e-08 | 4.000E-08 | 7.937E-09 | 3.400E-08 | 4.900E-08 |
| 16602 | 831 | F831R | B | 5.1e-08 | 5e-08 | 3.6e-08 | 4.567E-08 | 8.386E-09 | 3.600E-08 | 5.100E-08 |
| 16603 | 831 | F831N | B | 3.3e-08 | 7.3e-08 | 1.8e-08 | 4.133E-08 | 2.843E-08 | 1.800E-08 | 7.300E-08 |
| 16604 | 831 | F831D | B | 4e-08 | 1.3e-07 | 2.1e-08 | 6.367E-08 | 5.823E-08 | 2.100E-08 | 1.300E-07 |
| 16605 | 831 | F831C | B | 6e-08 | 2.9e-08 | 6.6e-08 | 5.167E-08 | 1.986E-08 | 2.900E-08 | 6.600E-08 |
| 16606 | 831 | F831E | B | 1.4e-08 | 4.2e-08 | 2.9e-08 | 2.833E-08 | 1.401E-08 | 1.400E-08 | 4.200E-08 |
| 16607 | 831 | F831Q | B | 4.1e-08 | 8.5e-08 | 7.1e-08 | 6.567E-08 | 2.248E-08 | 4.100E-08 | 8.500E-08 |
| 16608 | 831 | F831G | B | 9.4e-08 | 2.8e-08 | 2.8e-08 | 5.000E-08 | 3.811E-08 | 2.800E-08 | 9.400E-08 |
| 16609 | 831 | F831H | B | 2.2e-08 | 6.4e-08 | 2.5e-08 | 3.700E-08 | 2.343E-08 | 2.200E-08 | 6.400E-08 |
| 16610 | 831 | F831I | B | 1.5e-07 | 5.1e-08 | 4.2e-08 | 8.100E-08 | 5.992E-08 | 4.200E-08 | 1.500E-07 |
| 16611 | 831 | F831L | B | 3.4e-08 | 3.5e-08 | 6.2e-08 | 4.367E-08 | 1.589E-08 | 3.400E-08 | 6.200E-08 |
| 16612 | 831 | F831K | B | 2.1e-08 | 5.7e-08 | 3.4e-08 | 3.733E-08 | 1.823E-08 | 2.100E-08 | 5.700E-08 |
| 16613 | 831 | F831M | B | 4.3e-08 | 1.1e-07 | 5.2e-08 | 6.833E-08 | 3.636E-08 | 4.300E-08 | 1.100E-07 |
| 16614 | 831 | F831F | B | 5.6e-08 | 8.1e-08 | 2.8e-08 | 5.500E-08 | 2.651E-08 | 2.800E-08 | 8.100E-08 |
| 16615 | 831 | F831P | B | 2.2e-08 | 8.5e-08 | 7.7e-08 | 6.133E-08 | 3.430E-08 | 2.200E-08 | 8.500E-08 |
| 16616 | 831 | F831S | B | 5e-08 | 1.7e-08 | 9.6e-08 | 5.433E-08 | 3.968E-08 | 1.700E-08 | 9.600E-08 |
| 16617 | 831 | F831T | B | 2.6e-08 | 9.5e-08 | 1.7e-08 | 4.600E-08 | 4.267E-08 | 1.700E-08 | 9.500E-08 |
| 16618 | 831 | F831W | B | 5.8e-08 | 5.3e-08 | 5.6e-08 | 5.567E-08 | 2.517E-09 | 5.300E-08 | 5.800E-08 |
| 16619 | 831 | F831Y | B | 5.6e-08 | 4.7e-08 | 2.1e-08 | 4.133E-08 | 1.818E-08 | 2.100E-08 | 5.600E-08 |
| 16620 | 831 | F831V | B | 4.2e-08 | 1.7e-08 | 2.8e-08 | 2.900E-08 | 1.253E-08 | 1.700E-08 | 4.200E-08 |
| 16621 | 832 | N832A | B | 5.7e-08 | 8.3e-08 | 3e-08 | 5.667E-08 | 2.650E-08 | 3.000E-08 | 8.300E-08 |
| 16622 | 832 | N832R | B | 4.3e-08 | 5.5e-08 | 3.3e-08 | 4.367E-08 | 1.102E-08 | 3.300E-08 | 5.500E-08 |
| 16623 | 832 | N832N | B | 6.2e-08 | 2.1e-08 | 7.4e-08 | 5.233E-08 | 2.779E-08 | 2.100E-08 | 7.400E-08 |
| 16624 | 832 | N832D | B | 5.9e-08 | 2.7e-08 | 1.4e-07 | 7.533E-08 | 5.824E-08 | 2.700E-08 | 1.400E-07 |
| 16625 | 832 | N832C | B | 6.8e-08 | 1.8e-07 | 6.8e-08 | 1.053E-07 | 6.466E-08 | 6.800E-08 | 1.800E-07 |
| 16626 | 832 | N832E | B | 1.2e-07 | 2e-08 | 9.6e-08 | 7.867E-08 | 5.220E-08 | 2.000E-08 | 1.200E-07 |
| 16627 | 832 | N832Q | B | 1.4e-07 | 4.7e-08 | 2.6e-08 | 7.100E-08 | 6.067E-08 | 2.600E-08 | 1.400E-07 |
| 16628 | 832 | N832G | B | 3.2e-08 | 4e-08 | 7.6e-08 | 4.933E-08 | 2.344E-08 | 3.200E-08 | 7.600E-08 |
| 16629 | 832 | N832H | B | 5.7e-08 | 4.3e-08 | 3.8e-08 | 4.600E-08 | 9.849E-09 | 3.800E-08 | 5.700E-08 |
| 16630 | 832 | N832I | B | 6.1e-08 | 2.3e-08 | 7e-08 | 5.133E-08 | 2.495E-08 | 2.300E-08 | 7.000E-08 |
| 16631 | 832 | N832L | B | 6e-08 | 2.4e-08 | 4.7e-08 | 4.367E-08 | 1.823E-08 | 2.400E-08 | 6.000E-08 |
| 16632 | 832 | N832K | B | 4e-08 | 4.5e-08 | 1.2e-07 | 6.833E-08 | 4.481E-08 | 4.000E-08 | 1.200E-07 |
| 16633 | 832 | N832M | B | 3.5e-08 | 2.9e-08 | 4.1e-08 | 3.500E-08 | 6.000E-09 | 2.900E-08 | 4.100E-08 |
| 16634 | 832 | N832F | B | 4.2e-08 | 9.2e-08 | 1.5e-07 | 9.467E-08 | 5.405E-08 | 4.200E-08 | 1.500E-07 |
| 16635 | 832 | N832P | B | 5.1e-08 | 4.4e-08 | 6.5e-08 | 5.333E-08 | 1.069E-08 | 4.400E-08 | 6.500E-08 |
| 16636 | 832 | N832S | B | 6.4e-08 | 6.5e-08 | 1.7e-08 | 4.867E-08 | 2.743E-08 | 1.700E-08 | 6.500E-08 |
| 16637 | 832 | N832T | B | 4.7e-08 | 9.8e-08 | 1.2e-07 | 8.833E-08 | 3.745E-08 | 4.700E-08 | 1.200E-07 |
| 16638 | 832 | N832W | B | 1e-07 | 1.8e-08 | 2.5e-08 | 4.767E-08 | 4.546E-08 | 1.800E-08 | 1.000E-07 |
| 16639 | 832 | N832Y | B | 6.3e-08 | 7.1e-08 | 1e-07 | 7.800E-08 | 1.947E-08 | 6.300E-08 | 1.000E-07 |
| 16640 | 832 | N832V | B | 2.1e-08 | 4.3e-08 | 6.4e-08 | 4.267E-08 | 2.150E-08 | 2.100E-08 | 6.400E-08 |
| 16641 | 833 | R833A | B | 6.6e-08 | 4.2e-08 | 5.3e-08 | 5.367E-08 | 1.201E-08 | 4.200E-08 | 6.600E-08 |
| 16642 | 833 | R833R | B | 6.6e-08 | 5.7e-08 | 3.9e-08 | 5.400E-08 | 1.375E-08 | 3.900E-08 | 6.600E-08 |
| 16643 | 833 | R833N | B | 8.6e-08 | 7.7e-08 | 8.4e-08 | 8.233E-08 | 4.726E-09 | 7.700E-08 | 8.600E-08 |
| 16644 | 833 | R833D | B | 4.6e-08 | 3.2e-08 | 1.7e-08 | 3.167E-08 | 1.450E-08 | 1.700E-08 | 4.600E-08 |
| 16645 | 833 | R833C | B | 2.7e-08 | 4.1e-08 | 3.6e-08 | 3.467E-08 | 7.095E-09 | 2.700E-08 | 4.100E-08 |
| 16646 | 833 | R833E | B | 4.1e-08 | 9.5e-08 | 2.9e-08 | 5.500E-08 | 3.516E-08 | 2.900E-08 | 9.500E-08 |
| 16647 | 833 | R833Q | B | 9.4e-08 | 3e-08 | 4.3e-08 | 5.567E-08 | 3.383E-08 | 3.000E-08 | 9.400E-08 |
| 16648 | 833 | R833G | B | 4.4e-08 | 7.6e-08 | 4.2e-08 | 5.400E-08 | 1.908E-08 | 4.200E-08 | 7.600E-08 |
| 16649 | 833 | R833H | B | 9.1e-08 | 1.6e-07 | 9.4e-09 | 8.680E-08 | 7.539E-08 | 9.400E-09 | 1.600E-07 |
| 16650 | 833 | R833I | B | 2.1e-08 | 5.6e-08 | 4.9e-08 | 4.200E-08 | 1.852E-08 | 2.100E-08 | 5.600E-08 |
| 16651 | 833 | R833L | B | 6.7e-08 | 2.1e-08 | 1.1e-07 | 6.600E-08 | 4.451E-08 | 2.100E-08 | 1.100E-07 |
| 16652 | 833 | R833K | B | 6.7e-08 | 3.3e-08 | 6.8e-09 | 3.560E-08 | 3.018E-08 | 6.800E-09 | 6.700E-08 |
| 16653 | 833 | R833M | B | 1.6e-07 | 4.9e-08 | 3e-08 | 7.967E-08 | 7.022E-08 | 3.000E-08 | 1.600E-07 |
| 16654 | 833 | R833F | B | 3.8e-08 | 5.2e-08 | 5.8e-08 | 4.933E-08 | 1.026E-08 | 3.800E-08 | 5.800E-08 |
| 16655 | 833 | R833P | B | 3.3e-08 | 5.2e-08 | 1.5e-08 | 3.333E-08 | 1.850E-08 | 1.500E-08 | 5.200E-08 |
| 16656 | 833 | R833S | B | 3.3e-08 | 2.5e-08 | 3.9e-08 | 3.233E-08 | 7.024E-09 | 2.500E-08 | 3.900E-08 |
| 16657 | 833 | R833T | B | 1.3e-08 | 2.6e-08 | 4.1e-08 | 2.667E-08 | 1.401E-08 | 1.300E-08 | 4.100E-08 |
| 16658 | 833 | R833W | B | 2.3e-08 | 1.1e-08 | 8.6e-08 | 4.000E-08 | 4.029E-08 | 1.100E-08 | 8.600E-08 |
| 16659 | 833 | R833Y | B | 3.2e-08 | 1.7e-08 | 3.2e-08 | 2.700E-08 | 8.660E-09 | 1.700E-08 | 3.200E-08 |

| | | | | | | | | | | |
|---|---|---|---|---|---|---|---|---|---|---|
| 16660 | 833 | R833V | B | 2.2e-08 | 3.9e-08 | 5.3e-08 | 3.800E-08 | 1.552E-08 | 2.200E-08 | 5.300E-08 |
| 16661 | 834 | G834A | B | 2.9e-08 | 8.7e-08 | 3.7e-08 | 5.100E-08 | 3.143E-08 | 2.900E-08 | 8.700E-08 |
| 16662 | 834 | G834R | B | 2.2e-08 | 5.4e-08 | 5.4e-08 | 4.333E-08 | 1.848E-08 | 2.200E-08 | 5.400E-08 |
| 16663 | 834 | G834N | B | 6e-08 | 4.6e-08 | 7.3e-08 | 5.967E-08 | 1.350E-08 | 4.600E-08 | 7.300E-08 |
| 16664 | 834 | G834D | B | 6.6e-08 | 6.1e-08 | 2.8e-08 | 5.167E-08 | 2.065E-08 | 2.800E-08 | 6.600E-08 |
| 16665 | 834 | G834C | B | 1.5e-08 | 6.9e-08 | 4.1e-08 | 4.167E-08 | 2.701E-08 | 1.500E-08 | 6.900E-08 |
| 16666 | 834 | G834E | B | 3.2e-08 | 3.2e-08 | 3.8e-08 | 3.400E-08 | 3.464E-09 | 3.200E-08 | 3.800E-08 |
| 16667 | 834 | G834Q | B | 2.1e-08 | 3.5e-08 | 2.3e-08 | 2.633E-08 | 7.572E-09 | 2.100E-08 | 3.500E-08 |
| 16668 | 834 | G834G | B | 3.7e-08 | 2e-08 | 6.9e-08 | 4.200E-08 | 2.488E-08 | 2.000E-08 | 6.900E-08 |
| 16669 | 834 | G834H | B | 6.2e-08 | 4.2e-08 | 4.2e-08 | 4.867E-08 | 1.155E-08 | 4.200E-08 | 6.200E-08 |
| 16670 | 834 | G834I | B | 9.9e-09 | 7.1e-08 | 3.6e-08 | 3.897E-08 | 3.066E-08 | 9.900E-09 | 7.100E-08 |
| 16671 | 834 | G834L | B | 3.5e-08 | 3.7e-08 | 8.5e-08 | 5.233E-08 | 2.831E-08 | 3.500E-08 | 8.500E-08 |
| 16672 | 834 | G834K | B | 1.7e-08 | 1.5e-08 | 3.6e-08 | 2.267E-08 | 1.159E-08 | 1.500E-08 | 3.600E-08 |
| 16673 | 834 | G834M | B | 5.9e-08 | 4.5e-08 | 6.1e-08 | 5.500E-08 | 8.718E-09 | 4.500E-08 | 6.100E-08 |
| 16674 | 834 | G834F | B | 2.9e-08 | 9.3e-08 | 2.1e-08 | 4.767E-08 | 3.946E-08 | 2.100E-08 | 9.300E-08 |
| 16675 | 834 | G834P | B | 7e-08 | 8.7e-08 | 5.5e-08 | 7.067E-08 | 1.601E-08 | 5.500E-08 | 8.700E-08 |
| 16676 | 834 | G834S | B | 4.6e-08 | 6.1e-08 | 5.3e-08 | 5.333E-08 | 7.506E-09 | 4.600E-08 | 6.100E-08 |
| 16677 | 834 | G834T | B | 8.1e-08 | 3.6e-08 | 3.4e-08 | 5.033E-08 | 2.658E-08 | 3.400E-08 | 8.100E-08 |
| 16678 | 834 | G834W | B | 5.1e-08 | 7.4e-08 | 6.3e-08 | 6.267E-08 | 1.150E-08 | 5.100E-08 | 7.400E-08 |
| 16679 | 834 | G834Y | B | 1.5e-08 | 4.2e-08 | 3.9e-08 | 3.200E-08 | 1.480E-08 | 1.500E-08 | 4.200E-08 |
| 16680 | 834 | G834V | B | 5.9e-08 | 4.4e-08 | 2.4e-08 | 4.233E-08 | 1.756E-08 | 2.400E-08 | 5.900E-08 |
| 16681 | 835 | E835A | B | 5.8e-08 | 3e-08 | 6e-08 | 4.933E-08 | 1.677E-08 | 3.000E-08 | 6.000E-08 |
| 16682 | 835 | E835R | B | 1.3e-07 | 5.2e-08 | 3.9e-08 | 7.367E-08 | 4.922E-08 | 3.900E-08 | 1.300E-07 |
| 16683 | 835 | E835N | B | 4.6e-08 | 2.4e-08 | 2.4e-08 | 3.133E-08 | 1.270E-08 | 2.400E-08 | 4.600E-08 |
| 16684 | 835 | E835D | B | 3.4e-08 | 3.2e-08 | 7.3e-08 | 4.633E-08 | 2.312E-08 | 3.200E-08 | 7.300E-08 |
| 16685 | 835 | E835C | B | 2.3e-08 | 4.6e-08 | 3.6e-08 | 3.500E-08 | 1.153E-08 | 2.300E-08 | 4.600E-08 |
| 16686 | 835 | E835E | B | 2.6e-08 | 3.6e-08 | 4.4e-08 | 3.533E-08 | 9.018E-09 | 2.600E-08 | 4.400E-08 |
| 16687 | 835 | E835Q | B | 3.7e-08 | 7.6e-08 | 6.3e-08 | 5.867E-08 | 1.986E-08 | 3.700E-08 | 7.600E-08 |
| 16688 | 835 | E835G | B | 3.7e-08 | 6.2e-08 | 6.6e-08 | 5.500E-08 | 1.572E-08 | 3.700E-08 | 6.600E-08 |
| 16689 | 835 | E835H | B | 2.7e-08 | 2.6e-08 | 6e-08 | 3.767E-08 | 1.935E-08 | 2.600E-08 | 6.000E-08 |
| 16690 | 835 | E835I | B | 1.1e-07 | 5.5e-08 | 2.9e-08 | 6.467E-08 | 4.136E-08 | 2.900E-08 | 1.100E-07 |
| 16691 | 835 | E835L | B | 4.3e-08 | 2.9e-08 | 3.8e-08 | 3.667E-08 | 7.095E-09 | 2.900E-08 | 4.300E-08 |
| 16692 | 835 | E835K | B | 4.9e-08 | 4.3e-08 | 3e-08 | 4.067E-08 | 9.713E-09 | 3.000E-08 | 4.900E-08 |
| 16693 | 835 | E835M | B | 9.2e-08 | 9.9e-08 | 1.6e-08 | 6.900E-08 | 4.603E-08 | 1.600E-08 | 9.900E-08 |
| 16694 | 835 | E835F | B | 7.1e-08 | 3.9e-08 | 4.5e-08 | 5.167E-08 | 1.701E-08 | 3.900E-08 | 7.100E-08 |
| 16695 | 835 | E835P | B | 5.3e-08 | 6.2e-08 | 5.2e-08 | 5.567E-08 | 5.508E-09 | 5.200E-08 | 6.200E-08 |
| 16696 | 835 | E835S | B | 4.3e-08 | 6.1e-08 | 7.8e-08 | 6.067E-08 | 1.750E-08 | 4.300E-08 | 7.800E-08 |
| 16697 | 835 | E835T | B | 2.5e-08 | 2.2e-08 | 5.5e-08 | 3.400E-08 | 1.825E-08 | 2.200E-08 | 5.500E-08 |
| 16698 | 835 | E835W | B | 2.4e-08 | 3.3e-08 | 3.3e-08 | 3.000E-08 | 5.196E-09 | 2.400E-08 | 3.300E-08 |
| 16699 | 835 | E835Y | B | 2.4e-08 | 1.7e-08 | 2e-08 | 2.033E-08 | 3.512E-09 | 1.700E-08 | 2.400E-08 |
| 16700 | 835 | E835V | B | 1.3e-07 | 9.1e-09 | 4.5e-08 | 6.137E-08 | 6.209E-08 | 9.100E-09 | 1.300E-07 |
| 16701 | 836 | C836A | B | 5e-08 | 8.8e-08 | 3e-08 | 5.600E-08 | 2.946E-08 | 3.000E-08 | 8.800E-08 |
| 16702 | 836 | C836R | B | 4.6e-08 | 1.5e-08 | 1.6e-07 | 7.367E-08 | 7.636E-08 | 1.500E-08 | 1.600E-07 |
| 16703 | 836 | C836N | B | 2.6e-08 | 1.5e-08 | 3.2e-08 | 2.433E-08 | 8.622E-09 | 1.500E-08 | 3.200E-08 |
| 16704 | 836 | C836D | B | 4.8e-08 | 2e-08 | 2.6e-08 | 3.133E-08 | 1.474E-08 | 2.000E-08 | 4.800E-08 |
| 16705 | 836 | C836C | B | 4.1e-08 | 2.1e-08 | 5.1e-08 | 3.767E-08 | 1.528E-08 | 2.100E-08 | 5.100E-08 |
| 16706 | 836 | C836E | B | 7.4e-08 | 4.1e-08 | 1.3e-08 | 4.267E-08 | 3.053E-08 | 1.300E-08 | 7.400E-08 |
| 16707 | 836 | C836Q | B | 6.7e-08 | 6.3e-08 | 2.9e-08 | 5.300E-08 | 2.088E-08 | 2.900E-08 | 6.700E-08 |
| 16708 | 836 | C836G | B | 4.9e-08 | 8.9e-08 | 2.3e-08 | 5.367E-08 | 3.325E-08 | 2.300E-08 | 8.900E-08 |
| 16709 | 836 | C836H | B | 4.4e-08 | 1.1e-07 | 8.3e-08 | 7.900E-08 | 3.318E-08 | 4.400E-08 | 1.100E-07 |
| 16710 | 836 | C836I | B | 1.1e-07 | 3.7e-08 | 5.6e-08 | 6.767E-08 | 3.787E-08 | 3.700E-08 | 1.100E-07 |
| 16711 | 836 | C836L | B | 2.1e-08 | 8.1e-08 | 5.5e-08 | 5.233E-08 | 3.009E-08 | 2.100E-08 | 8.100E-08 |
| 16712 | 836 | C836K | B | 2.9e-08 | 5.2e-08 | 3e-08 | 3.700E-08 | 1.300E-08 | 2.900E-08 | 5.200E-08 |
| 16713 | 836 | C836M | B | 3.4e-08 | 5.1e-08 | 1.8e-08 | 3.433E-08 | 1.650E-08 | 1.800E-08 | 5.100E-08 |
| 16714 | 836 | C836F | B | 3.2e-08 | 2.1e-08 | 2.5e-08 | 2.600E-08 | 5.568E-09 | 2.100E-08 | 3.200E-08 |
| 16715 | 836 | C836P | B | 7.9e-08 | 5.4e-08 | 2.1e-08 | 5.133E-08 | 2.909E-08 | 2.100E-08 | 7.900E-08 |
| 16716 | 836 | C836S | B | 5.4e-08 | 3.9e-08 | 4.2e-08 | 4.500E-08 | 7.937E-09 | 3.900E-08 | 5.400E-08 |
| 16717 | 836 | C836T | B | 2.5e-08 | 3.6e-08 | 3.1e-08 | 3.067E-08 | 5.508E-09 | 2.500E-08 | 3.600E-08 |
| 16718 | 836 | C836W | B | 1.5e-07 | 1.7e-08 | 1.3e-07 | 9.900E-08 | 7.171E-08 | 1.700E-08 | 1.500E-07 |
| 16719 | 836 | C836Y | B | 3.3e-08 | 1.7e-08 | 2.8e-08 | 2.600E-08 | 8.185E-09 | 1.700E-08 | 3.300E-08 |
| 16720 | 836 | C836V | B | 3.2e-08 | 4.2e-08 | 2.5e-08 | 3.300E-08 | 8.544E-09 | 2.500E-08 | 4.200E-08 |
| 16721 | 837 | E837A | C | 1.7e-08 | 1.9e-08 | 1e-08 | 1.533E-08 | 4.726E-09 | 1.000E-08 | 1.900E-08 |
| 16722 | 837 | E837R | C | 3e-08 | 1.7e-08 | 1.1e-07 | 5.233E-08 | 5.036E-08 | 1.700E-08 | 1.100E-07 |
| 16723 | 837 | E837N | C | 7.3e-08 | 2.7e-08 | 3.6e-08 | 4.533E-08 | 2.438E-08 | 2.700E-08 | 7.300E-08 |
| 16724 | 837 | E837D | C | 4.4e-08 | 5.2e-08 | 3.6e-08 | 4.400E-08 | 8.000E-09 | 3.600E-08 | 5.200E-08 |
| 16725 | 837 | E837C | C | 9e-08 | 4.1e-08 | 4.5e-08 | 5.867E-08 | 2.721E-08 | 4.100E-08 | 9.000E-08 |
| 16726 | 837 | E837E | C | 2.6e-08 | 2.7e-08 | 5.3e-08 | 3.533E-08 | 1.531E-08 | 2.600E-08 | 5.300E-08 |
| 16727 | 837 | E837Q | C | 2.9e-08 | 1e-07 | 7.2e-08 | 6.700E-08 | 3.576E-08 | 2.900E-08 | 1.000E-07 |
| 16728 | 837 | E837G | C | 5.2e-08 | 7.6e-08 | 3e-08 | 5.267E-08 | 2.301E-08 | 3.000E-08 | 7.600E-08 |
| 16729 | 837 | E837H | C | 8e-08 | 2.8e-08 | 7.1e-08 | 5.967E-08 | 2.779E-08 | 2.800E-08 | 8.000E-08 |
| 16730 | 837 | E837I | C | 5.1e-08 | 2.6e-08 | 4.9e-08 | 4.200E-08 | 1.389E-08 | 2.600E-08 | 5.100E-08 |
| 16731 | 837 | E837L | C | 6.6e-08 | 9.6e-08 | 4.7e-08 | 6.967E-08 | 2.470E-08 | 4.700E-08 | 9.600E-08 |
| 16732 | 837 | E837K | C | 1.4e-08 | 3.9e-08 | 3.9e-08 | 3.067E-08 | 1.443E-08 | 1.400E-08 | 3.900E-08 |
| 16733 | 837 | E837M | C | 2.2e-08 | 4.8e-08 | 7.6e-08 | 4.867E-08 | 2.701E-08 | 2.200E-08 | 7.600E-08 |
| 16734 | 837 | E837F | C | 5.5e-08 | 2.4e-08 | 7.1e-08 | 5.000E-08 | 2.390E-08 | 2.400E-08 | 7.100E-08 |
| 16735 | 837 | E837P | C | 4e-08 | 4.5e-08 | 4.1e-08 | 4.200E-08 | 2.646E-09 | 4.000E-08 | 4.500E-08 |
| 16736 | 837 | E837S | C | 4.1e-08 | 5.2e-08 | 4.3e-08 | 4.533E-08 | 5.859E-09 | 4.100E-08 | 5.200E-08 |
| 16737 | 837 | E837T | C | 8.1e-08 | 1.6e-08 | 2.7e-08 | 4.133E-08 | 3.479E-08 | 1.600E-08 | 8.100E-08 |
| 16738 | 837 | E837W | C | 1.9e-08 | 2.5e-08 | 5.2e-08 | 3.200E-08 | 1.758E-08 | 1.900E-08 | 5.200E-08 |
| 16739 | 837 | E837Y | C | 4.9e-08 | 4.4e-08 | 4.3e-08 | 4.533E-08 | 3.215E-09 | 4.300E-08 | 4.900E-08 |
| 16740 | 837 | E837V | C | 1.4e-08 | 2.6e-08 | 8.1e-08 | 4.033E-08 | 3.573E-08 | 1.400E-08 | 8.100E-08 |
| 16741 | 838 | V838A | C | 6e-08 | 4.7e-08 | 3.9e-08 | 4.867E-08 | 1.060E-08 | 3.900E-08 | 6.000E-08 |
| 16742 | 838 | V838R | C | 4.1e-08 | 1.8e-08 | 8.3e-08 | 4.733E-08 | 3.296E-08 | 1.800E-08 | 8.300E-08 |
| 16743 | 838 | V838N | C | 4.3e-08 | 4.2e-08 | 7.4e-08 | 5.300E-08 | 1.819E-08 | 4.200E-08 | 7.400E-08 |
| 16744 | 838 | V838D | C | 2.6e-08 | 3.4e-08 | 7.7e-08 | 4.567E-08 | 2.743E-08 | 2.600E-08 | 7.700E-08 |

| | | | | | | | | | | |
|---|---|---|---|---|---|---|---|---|---|---|
| 16745 | 838 | V838C | C | 1.1e-07 | 4.3e-08 | 6.1e-08 | 7.133E-08 | 3.467E-08 | 4.300E-08 | 1.100E-07 |
| 16746 | 838 | V838E | C | 3e-08 | 7.9e-08 | 2.8e-08 | 4.567E-08 | 2.888E-08 | 2.800E-08 | 7.900E-08 |
| 16747 | 838 | V838Q | C | 4.9e-08 | 5e-08 | 3.7e-08 | 4.533E-08 | 7.234E-09 | 3.700E-08 | 5.000E-08 |
| 16748 | 838 | V838G | C | 1.5e-07 | 3.3e-08 | 1.7e-08 | 6.667E-08 | 7.261E-08 | 1.700E-08 | 1.500E-07 |
| 16749 | 838 | V838H | C | 7.4e-08 | 8.7e-08 | 3.3e-08 | 6.467E-08 | 2.818E-08 | 3.300E-08 | 8.700E-08 |
| 16750 | 838 | V838I | C | 1.6e-08 | 2.2e-08 | 2.2e-08 | 2.000E-08 | 3.464E-09 | 1.600E-08 | 2.200E-08 |
| 16751 | 838 | V838L | C | 2.3e-08 | 9.6e-09 | 5.8e-08 | 3.020E-08 | 2.499E-08 | 9.600E-09 | 5.800E-08 |
| 16752 | 838 | V838K | C | 6.3e-08 | 4.6e-08 | 2.7e-08 | 4.533E-08 | 1.801E-08 | 2.700E-08 | 6.300E-08 |
| 16753 | 838 | V838M | C | 3.7e-08 | 6.7e-08 | 7.7e-08 | 6.033E-08 | 2.082E-08 | 3.700E-08 | 7.700E-08 |
| 16754 | 838 | V838F | C | 8.1e-08 | 3.6e-08 | 3.4e-08 | 5.033E-08 | 2.658E-08 | 3.400E-08 | 8.100E-08 |
| 16755 | 838 | V838P | C | 4.7e-08 | 4.4e-08 | 2.5e-08 | 3.867E-08 | 1.193E-08 | 2.500E-08 | 4.700E-08 |
| 16756 | 838 | V838S | C | 3.7e-08 | 4.2e-08 | 5.8e-08 | 4.567E-08 | 1.097E-08 | 3.700E-08 | 5.800E-08 |
| 16757 | 838 | V838T | C | 4.4e-08 | 1.8e-08 | 9.5e-08 | 5.233E-08 | 3.917E-08 | 1.800E-08 | 9.500E-08 |
| 16758 | 838 | V838W | C | 3.6e-08 | 3.5e-08 | 5.6e-08 | 4.233E-08 | 1.185E-08 | 3.500E-08 | 5.600E-08 |
| 16759 | 838 | V838Y | C | 4.5e-08 | 3e-08 | 4.5e-08 | 4.000E-08 | 8.660E-09 | 3.000E-08 | 4.500E-08 |
| 16760 | 838 | V838V | C | 5e-08 | 1.7e-08 | 6.9e-08 | 4.533E-08 | 2.631E-08 | 1.700E-08 | 6.900E-08 |
| 16761 | 839 | Q839A | C | 5.6e-08 | 4.3e-08 | 1.5e-07 | 8.300E-08 | 5.839E-08 | 4.300E-08 | 1.500E-07 |
| 16762 | 839 | Q839R | C | 3.6e-08 | 5.1e-08 | 1.3e-08 | 3.333E-08 | 1.914E-08 | 1.300E-08 | 5.100E-08 |
| 16763 | 839 | Q839N | C | 1.1e-07 | 7.5e-08 | 2.1e-08 | 6.867E-08 | 4.484E-08 | 2.100E-08 | 1.100E-07 |
| 16764 | 839 | Q839D | C | 8.2e-08 | 3.2e-08 | 4.1e-08 | 5.167E-08 | 2.665E-08 | 3.200E-08 | 8.200E-08 |
| 16765 | 839 | Q839C | C | 3.4e-08 | 1.2e-08 | 3.5e-08 | 2.700E-08 | 1.300E-08 | 1.200E-08 | 3.500E-08 |
| 16766 | 839 | Q839E | C | 2.5e-08 | 1.1e-07 | 2e-08 | 5.167E-08 | 5.058E-08 | 2.000E-08 | 1.100E-07 |
| 16767 | 839 | Q839Q | C | 5.7e-08 | 6.3e-08 | 3.9e-08 | 5.300E-08 | 1.249E-08 | 3.900E-08 | 6.300E-08 |
| 16768 | 839 | Q839G | C | 6.6e-08 | 6.8e-08 | 2.2e-08 | 5.200E-08 | 2.600E-08 | 2.200E-08 | 6.800E-08 |
| 16769 | 839 | Q839H | C | 8.4e-08 | 4e-08 | 4.5e-08 | 5.633E-08 | 2.409E-08 | 4.000E-08 | 8.400E-08 |
| 16770 | 839 | Q839I | C | 7.4e-08 | 9.4e-08 | 7.3e-08 | 8.033E-08 | 1.185E-08 | 7.300E-08 | 9.400E-08 |
| 16771 | 839 | Q839L | C | 1.6e-08 | 6.5e-08 | 1.2e-08 | 3.100E-08 | 2.951E-08 | 1.200E-08 | 6.500E-08 |
| 16772 | 839 | Q839K | C | 7.6e-08 | 4.5e-08 | 1.3e-08 | 4.467E-08 | 3.150E-08 | 1.300E-08 | 7.600E-08 |
| 16773 | 839 | Q839M | C | 1.5e-08 | 5.1e-08 | 5.9e-08 | 4.167E-08 | 2.344E-08 | 1.500E-08 | 5.900E-08 |
| 16774 | 839 | Q839F | C | 3.6e-08 | 4e-08 | 7.5e-08 | 5.033E-08 | 2.146E-08 | 3.600E-08 | 7.500E-08 |
| 16775 | 839 | Q839P | C | 6.2e-08 | 7e-08 | 8.2e-08 | 7.133E-08 | 1.007E-08 | 6.200E-08 | 8.200E-08 |
| 16776 | 839 | Q839S | C | 2e-08 | 3.2e-08 | 7.5e-08 | 4.233E-08 | 2.892E-08 | 2.000E-08 | 7.500E-08 |
| 16777 | 839 | Q839T | C | 6.4e-08 | 4.7e-08 | 3.4e-08 | 4.833E-08 | 1.504E-08 | 3.400E-08 | 6.400E-08 |
| 16778 | 839 | Q839W | C | 3.3e-08 | 3.5e-08 | 1.7e-08 | 2.833E-08 | 9.866E-09 | 1.700E-08 | 3.500E-08 |
| 16779 | 839 | Q839Y | C | 7.1e-08 | 1.7e-08 | 4.7e-08 | 4.500E-08 | 2.706E-08 | 1.700E-08 | 7.100E-08 |
| 16780 | 839 | Q839V | C | 7e-08 | 9.4e-08 | 3.2e-08 | 6.533E-08 | 3.126E-08 | 3.200E-08 | 9.400E-08 |
| 16781 | 840 | L840A | C | 1.1e-07 | 4.5e-08 | 3.9e-08 | 6.467E-08 | 3.937E-08 | 3.900E-08 | 1.100E-07 |
| 16782 | 840 | L840R | C | 8.1e-08 | 6.9e-08 | 6.9e-08 | 7.300E-08 | 6.928E-09 | 6.900E-08 | 8.100E-08 |
| 16783 | 840 | L840N | C | 4.3e-08 | 7e-08 | 2.3e-08 | 4.533E-08 | 2.359E-08 | 2.300E-08 | 7.000E-08 |
| 16784 | 840 | L840D | C | 4.6e-08 | 1.9e-08 | 3.6e-08 | 3.367E-08 | 1.365E-08 | 1.900E-08 | 4.600E-08 |
| 16785 | 840 | L840C | C | 4.1e-08 | 6.9e-08 | 8.1e-08 | 6.367E-08 | 2.053E-08 | 4.100E-08 | 8.100E-08 |
| 16786 | 840 | L840E | C | 4e-08 | 6e-08 | 3.4e-08 | 4.467E-08 | 1.361E-08 | 3.400E-08 | 6.000E-08 |
| 16787 | 840 | L840Q | C | 7.8e-08 | 3.4e-08 | 6.9e-08 | 6.033E-08 | 2.325E-08 | 3.400E-08 | 7.800E-08 |
| 16788 | 840 | L840G | C | 4.2e-08 | 2e-08 | 1.5e-08 | 2.567E-08 | 1.436E-08 | 1.500E-08 | 4.200E-08 |
| 16789 | 840 | L840H | C | 5.8e-08 | 2.4e-08 | 6.1e-08 | 4.767E-08 | 2.055E-08 | 2.400E-08 | 6.100E-08 |
| 16790 | 840 | L840I | C | 1.8e-08 | 2.2e-08 | 3.5e-08 | 2.500E-08 | 8.888E-09 | 1.800E-08 | 3.500E-08 |
| 16791 | 840 | L840L | C | 3.3e-08 | 4.9e-08 | 5.8e-08 | 4.667E-08 | 1.266E-08 | 3.300E-08 | 5.800E-08 |
| 16792 | 840 | L840K | C | 4.2e-08 | 2.6e-08 | 3.5e-08 | 3.433E-08 | 8.021E-09 | 2.600E-08 | 4.200E-08 |
| 16793 | 840 | L840M | C | 4.6e-08 | 1.5e-08 | 1.3e-08 | 2.467E-08 | 1.850E-08 | 1.300E-08 | 4.600E-08 |
| 16794 | 840 | L840F | C | 2.8e-08 | 7.8e-08 | 3.8e-08 | 4.800E-08 | 2.646E-08 | 2.800E-08 | 7.800E-08 |
| 16795 | 840 | L840P | C | 3.7e-08 | 3.3e-08 | 6.1e-08 | 4.367E-08 | 1.514E-08 | 3.300E-08 | 6.100E-08 |
| 16796 | 840 | L840S | C | 3.7e-08 | 2.6e-08 | 3.4e-08 | 3.233E-08 | 5.686E-09 | 2.600E-08 | 3.700E-08 |
| 16797 | 840 | L840T | C | 3.4e-08 | 4e-08 | 4.3e-08 | 3.900E-08 | 4.583E-09 | 3.400E-08 | 4.300E-08 |
| 16798 | 840 | L840W | C | 1e-07 | 5.4e-08 | 5.1e-08 | 6.833E-08 | 2.747E-08 | 5.100E-08 | 1.000E-07 |
| 16799 | 840 | L840Y | C | 4.3e-08 | 5.6e-08 | 7e-08 | 5.633E-08 | 1.350E-08 | 4.300E-08 | 7.000E-08 |
| 16800 | 840 | L840V | C | 3.9e-08 | 2.1e-08 | 7.3e-08 | 4.433E-08 | 2.641E-08 | 2.100E-08 | 7.300E-08 |
| 16801 | 841 | V841A | C | 4.7e-08 | 2.7e-08 | 1.1e-07 | 6.133E-08 | 4.332E-08 | 2.700E-08 | 1.100E-07 |
| 16802 | 841 | V841R | C | 5.4e-08 | 9e-08 | 3.1e-08 | 5.833E-08 | 2.974E-08 | 3.100E-08 | 9.000E-08 |
| 16803 | 841 | V841N | C | 3.1e-08 | 1.5e-08 | 3.9e-08 | 2.833E-08 | 1.222E-08 | 1.500E-08 | 3.900E-08 |
| 16804 | 841 | V841D | C | 3.8e-08 | 5.1e-08 | 3.2e-08 | 4.033E-08 | 9.713E-09 | 3.200E-08 | 5.100E-08 |
| 16805 | 841 | V841C | C | 3.3e-08 | 8.9e-08 | 5.6e-08 | 5.933E-08 | 2.815E-08 | 3.300E-08 | 8.900E-08 |
| 16806 | 841 | V841E | C | 2.5e-08 | 2.7e-08 | 4.2e-08 | 3.133E-08 | 9.292E-09 | 2.500E-08 | 4.200E-08 |
| 16807 | 841 | V841Q | C | 4.6e-08 | 6e-08 | 8.2e-08 | 6.267E-08 | 1.815E-08 | 4.600E-08 | 8.200E-08 |
| 16808 | 841 | V841G | C | 2.1e-08 | 1.9e-07 | 3.7e-08 | 8.267E-08 | 9.330E-08 | 2.100E-08 | 1.900E-07 |
| 16809 | 841 | V841H | C | 4.8e-08 | 6.1e-08 | 4.3e-08 | 5.067E-08 | 9.292E-09 | 4.300E-08 | 6.100E-08 |
| 16810 | 841 | V841I | C | 3.7e-08 | 3.5e-08 | 7.1e-08 | 4.767E-08 | 2.023E-08 | 3.500E-08 | 7.100E-08 |
| 16811 | 841 | V841L | C | 3.1e-08 | 2.6e-08 | 3.2e-08 | 2.967E-08 | 3.215E-09 | 2.600E-08 | 3.200E-08 |
| 16812 | 841 | V841K | C | 2.8e-08 | 9.2e-08 | 3.1e-08 | 5.033E-08 | 3.612E-08 | 2.800E-08 | 9.200E-08 |
| 16813 | 841 | V841M | C | 2.8e-08 | 3.2e-08 | 2.3e-08 | 2.767E-08 | 4.509E-09 | 2.300E-08 | 3.200E-08 |
| 16814 | 841 | V841F | C | 8.7e-08 | 4.5e-08 | 4.7e-08 | 5.967E-08 | 2.369E-08 | 4.500E-08 | 8.700E-08 |
| 16815 | 841 | V841P | C | 3.9e-08 | 6.4e-08 | 3.6e-08 | 4.633E-08 | 1.537E-08 | 3.600E-08 | 6.400E-08 |
| 16816 | 841 | V841S | C | 7.3e-08 | 3.2e-08 | 9.1e-08 | 6.533E-08 | 3.024E-08 | 3.200E-08 | 9.100E-08 |
| 16817 | 841 | V841T | C | 4.6e-08 | 4.1e-08 | 9.8e-09 | 3.227E-08 | 1.962E-08 | 9.800E-09 | 4.600E-08 |
| 16818 | 841 | V841W | C | 4.4e-08 | 4.8e-08 | 2.7e-08 | 3.967E-08 | 1.115E-08 | 2.700E-08 | 4.800E-08 |
| 16819 | 841 | V841Y | C | 2.2e-08 | 2.8e-08 | 4.1e-08 | 3.033E-08 | 9.713E-09 | 2.200E-08 | 4.100E-08 |
| 16820 | 841 | V841V | C | 4.4e-08 | 4.6e-08 | 3.4e-08 | 4.133E-08 | 6.429E-09 | 3.400E-08 | 4.600E-08 |
| 16821 | 842 | E842A | C | 4.7e-08 | 3.7e-08 | 2.7e-08 | 3.700E-08 | 1.000E-08 | 2.700E-08 | 4.700E-08 |
| 16822 | 842 | E842R | C | 1.1e-07 | 1.4e-08 | 6.1e-08 | 6.167E-08 | 4.800E-08 | 1.400E-08 | 1.100E-07 |
| 16823 | 842 | E842N | C | 8.4e-08 | 3.5e-08 | 6.9e-08 | 6.267E-08 | 2.511E-08 | 3.500E-08 | 8.400E-08 |
| 16824 | 842 | E842D | C | 4e-08 | 4.3e-08 | 6.3e-08 | 4.867E-08 | 1.250E-08 | 4.000E-08 | 6.300E-08 |
| 16825 | 842 | E842C | C | 3.5e-08 | 1.6e-08 | 5.2e-08 | 3.433E-08 | 1.801E-08 | 1.600E-08 | 5.200E-08 |
| 16826 | 842 | E842E | C | 7.7e-08 | 2.5e-08 | 5.5e-08 | 5.233E-08 | 2.610E-08 | 2.500E-08 | 7.700E-08 |
| 16827 | 842 | E842Q | C | 4.6e-08 | 1.6e-08 | 7.8e-08 | 4.667E-08 | 3.101E-08 | 1.600E-08 | 7.800E-08 |
| 16828 | 842 | E842G | C | 6e-08 | 3.7e-08 | 6.5e-08 | 5.400E-08 | 1.493E-08 | 3.700E-08 | 6.500E-08 |
| 16829 | 842 | E842H | C | 4.9e-08 | 1.9e-08 | 1.7e-08 | 2.833E-08 | 1.793E-08 | 1.700E-08 | 4.900E-08 |

| | | | | | | | | | | |
|---|---|---|---|---|---|---|---|---|---|---|
| 16830 | 842 | E842I | C | 4.4e-08 | 2.7e-08 | 2.4e-08 | 3.167E-08 | 1.079E-08 | 2.400E-08 | 4.400E-08 |
| 16831 | 842 | E842L | C | 3.7e-08 | 2e-07 | 1.5e-07 | 1.290E-07 | 8.350E-08 | 3.700E-08 | 2.000E-07 |
| 16832 | 842 | E842K | C | 6.6e-08 | 5.4e-08 | 8.8e-09 | 4.293E-08 | 3.016E-08 | 8.800E-09 | 6.600E-08 |
| 16833 | 842 | E842M | C | 6.4e-08 | 2.2e-08 | 2.5e-08 | 3.700E-08 | 2.343E-08 | 2.200E-08 | 6.400E-08 |
| 16834 | 842 | E842F | C | 6.4e-08 | 7.5e-08 | 1e-08 | 4.967E-08 | 3.479E-08 | 1.000E-08 | 7.500E-08 |
| 16835 | 842 | E842P | C | 1.2e-07 | 3.3e-08 | 4.6e-08 | 6.633E-08 | 4.693E-08 | 3.300E-08 | 1.200E-07 |
| 16836 | 842 | E842S | C | 3.1e-08 | 8.3e-08 | 3.8e-08 | 5.067E-08 | 2.822E-08 | 3.100E-08 | 8.300E-08 |
| 16837 | 842 | E842T | C | 3.4e-08 | 4.7e-08 | 3.2e-08 | 3.767E-08 | 8.145E-09 | 3.200E-08 | 4.700E-08 |
| 16838 | 842 | E842W | C | 3.2e-08 | 2e-08 | 3.1e-08 | 2.767E-08 | 6.658E-09 | 2.000E-08 | 3.200E-08 |
| 16839 | 842 | E842Y | C | 2.9e-08 | 4.5e-08 | 2e-08 | 3.133E-08 | 1.266E-08 | 2.000E-08 | 4.500E-08 |
| 16840 | 842 | E842V | C | 1e-07 | 5.2e-08 | 1.3e-07 | 9.400E-08 | 3.934E-08 | 5.200E-08 | 1.300E-07 |
| 16841 | 843 | S843A | C | 3.6e-08 | 2.1e-08 | 1.1e-07 | 5.567E-08 | 4.765E-08 | 2.100E-08 | 1.100E-07 |
| 16842 | 843 | S843R | C | 4.2e-08 | 1.2e-07 | 1.7e-08 | 5.967E-08 | 5.372E-08 | 1.700E-08 | 1.200E-07 |
| 16843 | 843 | S843N | C | 4.9e-08 | 3.3e-08 | 1.2e-07 | 6.733E-08 | 4.631E-08 | 3.300E-08 | 1.200E-07 |
| 16844 | 843 | S843D | C | 2.3e-08 | 3.2e-08 | 8.4e-08 | 4.633E-08 | 3.293E-08 | 2.300E-08 | 8.400E-08 |
| 16845 | 843 | S843C | C | 2e-08 | 6.3e-08 | 3.2e-08 | 3.833E-08 | 2.219E-08 | 2.000E-08 | 6.300E-08 |
| 16846 | 843 | S843E | C | 6e-08 | 5.5e-08 | 5e-08 | 5.500E-08 | 5.000E-09 | 5.000E-08 | 6.000E-08 |
| 16847 | 843 | S843Q | C | 7.4e-08 | 3.6e-08 | 8.4e-08 | 6.467E-08 | 2.532E-08 | 3.600E-08 | 8.400E-08 |
| 16848 | 843 | S843G | C | 2.8e-08 | 1.6e-08 | 5.2e-08 | 3.200E-08 | 1.833E-08 | 1.600E-08 | 5.200E-08 |
| 16849 | 843 | S843H | C | 1.2e-07 | 5.7e-08 | 1.9e-08 | 6.533E-08 | 5.101E-08 | 1.900E-08 | 1.200E-07 |
| 16850 | 843 | S843I | C | 4.1e-08 | 1.8e-08 | 4.2e-08 | 3.367E-08 | 1.358E-08 | 1.800E-08 | 4.200E-08 |
| 16851 | 843 | S843L | C | 3.2e-08 | 5.4e-08 | 3.9e-08 | 4.167E-08 | 1.124E-08 | 3.200E-08 | 5.400E-08 |
| 16852 | 843 | S843K | C | 3.4e-08 | 3.1e-08 | 8.8e-08 | 5.100E-08 | 3.208E-08 | 3.100E-08 | 8.800E-08 |
| 16853 | 843 | S843M | C | 2.2e-08 | 4.1e-08 | 4.3e-08 | 3.533E-08 | 1.159E-08 | 2.200E-08 | 4.300E-08 |
| 16854 | 843 | S843F | C | 3.8e-08 | 1.7e-08 | 2.8e-08 | 2.767E-08 | 1.050E-08 | 1.700E-08 | 3.800E-08 |
| 16855 | 843 | S843P | C | 3.6e-08 | 2.6e-08 | 2.6e-08 | 2.933E-08 | 5.774E-09 | 2.600E-08 | 3.600E-08 |
| 16856 | 843 | S843S | C | 3.1e-08 | 2.4e-08 | 2.6e-08 | 2.700E-08 | 3.606E-09 | 2.400E-08 | 3.100E-08 |
| 16857 | 843 | S843T | C | 5.2e-08 | 1.5e-08 | 5.3e-08 | 4.000E-08 | 2.166E-08 | 1.500E-08 | 5.300E-08 |
| 16858 | 843 | S843W | C | 2.6e-08 | 6.9e-08 | 2.8e-08 | 4.100E-08 | 2.427E-08 | 2.600E-08 | 6.900E-08 |
| 16859 | 843 | S843Y | C | 4.3e-08 | 2.5e-08 | 1.8e-08 | 2.867E-08 | 1.290E-08 | 1.800E-08 | 4.300E-08 |
| 16860 | 843 | S843V | C | 5.1e-08 | 4.2e-08 | 5e-08 | 4.767E-08 | 4.933E-09 | 4.200E-08 | 5.100E-08 |
| 16861 | 844 | G844A | C | 2e-08 | 5.6e-08 | 2.7e-08 | 3.433E-08 | 1.909E-08 | 2.000E-08 | 5.600E-08 |
| 16862 | 844 | G844R | C | 9.4e-08 | 4.1e-08 | 1.3e-07 | 8.833E-08 | 4.477E-08 | 4.100E-08 | 1.300E-07 |
| 16863 | 844 | G844N | C | 5.2e-08 | 2.1e-08 | 8e-08 | 5.100E-08 | 2.951E-08 | 2.100E-08 | 8.000E-08 |
| 16864 | 844 | G844D | C | 2.7e-08 | 3e-08 | 2.4e-08 | 2.700E-08 | 3.000E-09 | 2.400E-08 | 3.000E-08 |
| 16865 | 844 | G844C | C | 3e-08 | 3.7e-08 | 5.7e-08 | 4.133E-08 | 1.401E-08 | 3.000E-08 | 5.700E-08 |
| 16866 | 844 | G844E | C | 3.5e-08 | 3e-08 | 2.2e-08 | 2.900E-08 | 6.557E-09 | 2.200E-08 | 3.500E-08 |
| 16867 | 844 | G844Q | C | 5e-08 | 4.5e-08 | 7e-08 | 5.500E-08 | 1.323E-08 | 4.500E-08 | 7.000E-08 |
| 16868 | 844 | G844G | C | 7.1e-08 | 4.7e-08 | 6.9e-08 | 6.233E-08 | 1.332E-08 | 4.700E-08 | 7.100E-08 |
| 16869 | 844 | G844H | C | 3.5e-08 | 4.4e-08 | 2.5e-08 | 3.467E-08 | 9.504E-09 | 2.500E-08 | 4.400E-08 |
| 16870 | 844 | G844I | C | 1.8e-08 | 3.5e-08 | 5.3e-08 | 3.533E-08 | 1.750E-08 | 1.800E-08 | 5.300E-08 |
| 16871 | 844 | G844L | C | 1.2e-08 | 3e-08 | 3.2e-08 | 2.467E-08 | 1.102E-08 | 1.200E-08 | 3.200E-08 |
| 16872 | 844 | G844K | C | 3.9e-08 | 7.2e-08 | 2.6e-08 | 4.567E-08 | 2.371E-08 | 2.600E-08 | 7.200E-08 |
| 16873 | 844 | G844M | C | 4.5e-08 | 4.2e-08 | 4.5e-08 | 4.400E-08 | 1.732E-09 | 4.200E-08 | 4.500E-08 |
| 16874 | 844 | G844F | C | 6.5e-08 | 2.7e-08 | 6.5e-08 | 5.233E-08 | 2.194E-08 | 2.700E-08 | 6.500E-08 |
| 16875 | 844 | G844P | C | 5.9e-08 | 8.2e-08 | 2e-08 | 5.367E-08 | 3.134E-08 | 2.000E-08 | 8.200E-08 |
| 16876 | 844 | G844S | C | 4.1e-08 | 7.7e-08 | 9.1e-08 | 6.967E-08 | 2.579E-08 | 4.100E-08 | 9.100E-08 |
| 16877 | 844 | G844T | C | 5.1e-08 | 5.6e-08 | 2.7e-08 | 4.467E-08 | 1.550E-08 | 2.700E-08 | 5.600E-08 |
| 16878 | 844 | G844W | C | 2.6e-08 | 6e-08 | 3.8e-08 | 4.133E-08 | 1.724E-08 | 2.600E-08 | 6.000E-08 |
| 16879 | 844 | G844Y | C | 8.3e-08 | 7.7e-08 | 1.4e-07 | 1.000E-07 | 3.477E-08 | 7.700E-08 | 1.400E-07 |
| 16880 | 844 | G844V | C | 8.1e-08 | 2.6e-08 | 3.5e-08 | 4.733E-08 | 2.950E-08 | 2.600E-08 | 8.100E-08 |
| 16881 | 845 | G845A | C | 4.7e-08 | 2.6e-08 | 4.2e-08 | 3.833E-08 | 1.097E-08 | 2.600E-08 | 4.700E-08 |
| 16882 | 845 | G845R | C | 2e-08 | 7.8e-08 | 1e-07 | 6.600E-08 | 4.133E-08 | 2.000E-08 | 1.000E-07 |
| 16883 | 845 | G845N | C | 3.4e-08 | 3.2e-08 | 2.5e-08 | 3.033E-08 | 4.726E-09 | 2.500E-08 | 3.400E-08 |
| 16884 | 845 | G845D | C | 1.7e-08 | 2.4e-08 | 1.6e-08 | 1.900E-08 | 4.359E-09 | 1.600E-08 | 2.400E-08 |
| 16885 | 845 | G845C | C | 6.3e-08 | 1.3e-08 | 2.2e-08 | 3.267E-08 | 2.665E-08 | 1.300E-08 | 6.300E-08 |
| 16886 | 845 | G845E | C | 4.1e-08 | 7.5e-08 | 9.8e-08 | 7.133E-08 | 2.868E-08 | 4.100E-08 | 9.800E-08 |
| 16887 | 845 | G845Q | C | 3.6e-08 | 3.5e-08 | 8.9e-08 | 5.333E-08 | 3.089E-08 | 3.500E-08 | 8.900E-08 |
| 16888 | 845 | G845G | C | 7.2e-08 | 6.5e-08 | 2.2e-08 | 5.300E-08 | 2.707E-08 | 2.200E-08 | 7.200E-08 |
| 16889 | 845 | G845H | C | 3.1e-08 | 3.7e-08 | 4.8e-08 | 3.867E-08 | 8.622E-09 | 3.100E-08 | 4.800E-08 |
| 16890 | 845 | G845I | C | 3.2e-08 | 6.1e-08 | 5.5e-08 | 4.933E-08 | 1.531E-08 | 3.200E-08 | 6.100E-08 |
| 16891 | 845 | G845L | C | 6.8e-08 | 5.3e-08 | 6.7e-08 | 6.267E-08 | 8.386E-09 | 5.300E-08 | 6.800E-08 |
| 16892 | 845 | G845K | C | 6.5e-08 | 4.2e-08 | 3.6e-08 | 4.767E-08 | 1.531E-08 | 3.600E-08 | 6.500E-08 |
| 16893 | 845 | G845M | C | 4e-08 | 4.7e-08 | 2e-08 | 3.567E-08 | 1.401E-08 | 2.000E-08 | 4.700E-08 |
| 16894 | 845 | G845F | C | 3e-07 | 5.5e-08 | 1.1e-07 | 1.550E-07 | 1.285E-07 | 5.500E-08 | 3.000E-07 |
| 16895 | 845 | G845P | C | 5.1e-08 | 4.5e-08 | 7.3e-08 | 5.633E-08 | 1.474E-08 | 4.500E-08 | 7.300E-08 |
| 16896 | 845 | G845S | C | 4.8e-08 | 9.9e-09 | 5.9e-08 | 3.897E-08 | 2.577E-08 | 9.900E-09 | 5.900E-08 |
| 16897 | 845 | G845T | C | 2.1e-08 | 4.2e-08 | 5.3e-08 | 3.867E-08 | 1.626E-08 | 2.100E-08 | 5.300E-08 |
| 16898 | 845 | G845W | C | 1.3e-07 | 6.6e-08 | 1.4e-08 | 7.000E-08 | 5.810E-08 | 1.400E-08 | 1.300E-07 |
| 16899 | 845 | G845Y | C | 3.2e-08 | 5.4e-08 | 2.4e-08 | 3.667E-08 | 1.553E-08 | 2.400E-08 | 5.400E-08 |
| 16900 | 845 | G845V | C | 5.1e-08 | 5.2e-08 | 4.7e-08 | 5.000E-08 | 2.646E-09 | 4.700E-08 | 5.200E-08 |
| 16901 | 846 | G846A | C | 1.6e-08 | 2.6e-08 | 5.5e-08 | 3.233E-08 | 2.026E-08 | 1.600E-08 | 5.500E-08 |
| 16902 | 846 | G846R | C | 2.6e-08 | 4e-08 | 3.5e-08 | 3.367E-08 | 7.095E-09 | 2.600E-08 | 4.000E-08 |
| 16903 | 846 | G846N | C | 7.8e-08 | 4.1e-08 | 5.4e-08 | 5.767E-08 | 1.877E-08 | 4.100E-08 | 7.800E-08 |
| 16904 | 846 | G846D | C | 6.3e-08 | 7e-08 | 4e-08 | 5.767E-08 | 1.570E-08 | 4.000E-08 | 7.000E-08 |
| 16905 | 846 | G846C | C | 1.7e-08 | 2.7e-08 | 3.9e-08 | 2.767E-08 | 1.102E-08 | 1.700E-08 | 3.900E-08 |
| 16906 | 846 | G846E | C | 5.7e-08 | 4.5e-08 | 4.2e-08 | 4.800E-08 | 7.937E-09 | 4.200E-08 | 5.700E-08 |
| 16907 | 846 | G846Q | C | 3.3e-08 | 5.9e-08 | 5.2e-08 | 4.800E-08 | 1.345E-08 | 3.300E-08 | 5.900E-08 |
| 16908 | 846 | G846G | C | 1.4e-07 | 3.8e-08 | 2.1e-08 | 6.633E-08 | 6.436E-08 | 2.100E-08 | 1.400E-07 |
| 16909 | 846 | G846H | C | 1.1e-07 | 8.5e-08 | 1.3e-07 | 1.083E-07 | 2.255E-08 | 8.500E-08 | 1.300E-07 |
| 16910 | 846 | G846I | C | 2.4e-08 | 5.5e-08 | 3.4e-08 | 3.767E-08 | 1.582E-08 | 2.400E-08 | 5.500E-08 |
| 16911 | 846 | G846L | C | 2e-08 | 4.1e-08 | 4e-08 | 3.367E-08 | 1.185E-08 | 2.000E-08 | 4.100E-08 |
| 16912 | 846 | G846K | C | 8.2e-08 | 7.3e-08 | 5.8e-08 | 7.100E-08 | 1.212E-08 | 5.800E-08 | 8.200E-08 |
| 16913 | 846 | G846M | C | 5e-08 | 3.2e-08 | 5.5e-08 | 4.567E-08 | 1.210E-08 | 3.200E-08 | 5.500E-08 |
| 16914 | 846 | G846F | C | 4.4e-08 | 3.3e-08 | 2.3e-08 | 3.333E-08 | 1.050E-08 | 2.300E-08 | 4.400E-08 |

| | | | | | | | | | | |
|---|---|---|---|---|---|---|---|---|---|---|
| 16915 | 846 | G846P | C | 1.4e-08 | 1.5e-08 | 1.9e-08 | 1.600E-08 | 2.646E-09 | 1.400E-08 | 1.900E-08 |
| 16916 | 846 | G846S | C | 5.1e-08 | 3.2e-08 | 1.4e-08 | 3.233E-08 | 1.850E-08 | 1.400E-08 | 5.100E-08 |
| 16917 | 846 | G846T | C | 2.7e-08 | 9.7e-08 | 1.7e-08 | 4.700E-08 | 4.359E-08 | 1.700E-08 | 9.700E-08 |
| 16918 | 846 | G846W | C | 9.1e-08 | 3.6e-08 | 8.6e-08 | 7.100E-08 | 3.041E-08 | 3.600E-08 | 9.100E-08 |
| 16919 | 846 | G846Y | C | 3.9e-08 | 5.3e-08 | 4.4e-08 | 4.533E-08 | 7.095E-09 | 3.900E-08 | 5.300E-08 |
| 16920 | 846 | G846V | C | 1.6e-08 | 6.7e-08 | 3.6e-08 | 3.967E-08 | 2.570E-08 | 1.600E-08 | 6.700E-08 |
| 16921 | 847 | L847A | C | 6.4e-08 | 3.2e-08 | 2.8e-08 | 4.133E-08 | 1.973E-08 | 2.800E-08 | 6.400E-08 |
| 16922 | 847 | L847R | C | 2e-08 | 2.4e-08 | 3.2e-08 | 2.533E-08 | 6.110E-09 | 2.000E-08 | 3.200E-08 |
| 16923 | 847 | L847N | C | 3.5e-08 | 1.8e-08 | 4.7e-08 | 3.333E-08 | 1.457E-08 | 1.800E-08 | 4.700E-08 |
| 16924 | 847 | L847D | C | 6.1e-08 | 8.5e-08 | 4.1e-08 | 6.233E-08 | 2.203E-08 | 4.100E-08 | 8.500E-08 |
| 16925 | 847 | L847C | C | 4.1e-07 | 2.9e-08 | 5.8e-08 | 1.657E-07 | 2.121E-07 | 2.900E-08 | 4.100E-07 |
| 16926 | 847 | L847E | C | 8.5e-08 | 9e-09 | 6.8e-08 | 5.400E-08 | 3.989E-08 | 9.000E-09 | 8.500E-08 |
| 16927 | 847 | L847Q | C | 5.3e-08 | 5.5e-08 | 6.6e-08 | 5.800E-08 | 7.000E-09 | 5.300E-08 | 6.600E-08 |
| 16928 | 847 | L847G | C | 3.4e-08 | 6.2e-08 | 3.3e-08 | 4.300E-08 | 1.646E-08 | 3.300E-08 | 6.200E-08 |
| 16929 | 847 | L847H | C | 2.7e-08 | 2.9e-08 | 2.7e-08 | 2.767E-08 | 1.155E-09 | 2.700E-08 | 2.900E-08 |
| 16930 | 847 | L847I | C | 3.6e-08 | 5.5e-08 | 1.3e-07 | 7.367E-08 | 4.970E-08 | 3.600E-08 | 1.300E-07 |
| 16931 | 847 | L847L | C | 3.2e-08 | 2.9e-08 | 9.9e-08 | 5.333E-08 | 3.958E-08 | 2.900E-08 | 9.900E-08 |
| 16932 | 847 | L847K | C | 4.3e-08 | 3.2e-08 | 9.5e-08 | 5.667E-08 | 3.365E-08 | 3.200E-08 | 9.500E-08 |
| 16933 | 847 | L847M | C | 3.9e-08 | 3.1e-08 | 3.2e-08 | 3.400E-08 | 4.359E-09 | 3.100E-08 | 3.900E-08 |
| 16934 | 847 | L847F | C | 2.6e-08 | 5.2e-09 | 2.9e-08 | 2.007E-08 | 1.296E-08 | 5.200E-09 | 2.900E-08 |
| 16935 | 847 | L847P | C | 9.8e-08 | 4e-08 | 5.9e-08 | 6.567E-08 | 2.957E-08 | 4.000E-08 | 9.800E-08 |
| 16936 | 847 | L847S | C | 2.5e-08 | 3.5e-08 | 6.5e-08 | 4.167E-08 | 2.082E-08 | 2.500E-08 | 6.500E-08 |
| 16937 | 847 | L847T | C | 6.7e-08 | 4.2e-08 | 2.1e-08 | 4.333E-08 | 2.303E-08 | 2.100E-08 | 6.700E-08 |
| 16938 | 847 | L847W | C | 2.2e-08 | 5.7e-08 | 3.3e-08 | 3.733E-08 | 1.790E-08 | 2.200E-08 | 5.700E-08 |
| 16939 | 847 | L847Y | C | 2.4e-08 | 7.3e-08 | 6.2e-08 | 5.300E-08 | 2.571E-08 | 2.400E-08 | 7.300E-08 |
| 16940 | 847 | L847V | C | 2e-08 | 6e-08 | 2.6e-08 | 3.533E-08 | 2.157E-08 | 2.000E-08 | 6.000E-08 |
| 16941 | 848 | V848A | C | 9.1e-08 | 7.6e-08 | 4.6e-08 | 7.100E-08 | 2.291E-08 | 4.600E-08 | 9.100E-08 |
| 16942 | 848 | V848R | C | 6.8e-09 | 4.2e-08 | 7.7e-08 | 4.193E-08 | 3.510E-08 | 6.800E-09 | 7.700E-08 |
| 16943 | 848 | V848N | C | 5.5e-08 | 6.6e-08 | 5e-08 | 5.700E-08 | 8.185E-09 | 5.000E-08 | 6.600E-08 |
| 16944 | 848 | V848D | C | 7.6e-08 | 4.9e-08 | 7.9e-09 | 4.430E-08 | 3.429E-08 | 7.900E-09 | 7.600E-08 |
| 16945 | 848 | V848C | C | 9e-08 | 6.5e-08 | 8.8e-08 | 8.100E-08 | 1.389E-08 | 6.500E-08 | 9.000E-08 |
| 16946 | 848 | V848E | C | 3.4e-08 | 3.6e-08 | 6.7e-08 | 4.567E-08 | 1.850E-08 | 3.400E-08 | 6.700E-08 |
| 16947 | 848 | V848Q | C | 2.7e-08 | 1.6e-07 | 2.4e-08 | 7.033E-08 | 7.767E-08 | 2.400E-08 | 1.600E-07 |
| 16948 | 848 | V848G | C | 5.2e-08 | 4.1e-08 | 1.8e-08 | 3.700E-08 | 1.735E-08 | 1.800E-08 | 5.200E-08 |
| 16949 | 848 | V848H | C | 6.9e-08 | 2e-08 | 3.3e-08 | 4.067E-08 | 2.538E-08 | 2.000E-08 | 6.900E-08 |
| 16950 | 848 | V848I | C | 6.5e-08 | 7.5e-08 | 6.5e-08 | 6.833E-08 | 5.774E-09 | 6.500E-08 | 7.500E-08 |
| 16951 | 848 | V848L | C | 2.5e-07 | 8.7e-08 | 6.1e-08 | 1.327E-07 | 1.024E-07 | 6.100E-08 | 2.500E-07 |
| 16952 | 848 | V848K | C | 4.9e-08 | 5.1e-08 | 4.7e-08 | 4.900E-08 | 2.000E-09 | 4.700E-08 | 5.100E-08 |
| 16953 | 848 | V848M | C | 5.2e-08 | 1e-07 | 3.7e-08 | 6.300E-08 | 3.291E-08 | 3.700E-08 | 1.000E-07 |
| 16954 | 848 | V848F | C | 1.1e-08 | 4e-08 | 3.6e-08 | 2.900E-08 | 1.572E-08 | 1.100E-08 | 4.000E-08 |
| 16955 | 848 | V848P | C | 4.4e-08 | 5.6e-08 | 2.3e-08 | 4.100E-08 | 1.670E-08 | 2.300E-08 | 5.600E-08 |
| 16956 | 848 | V848S | C | 9.6e-08 | 2.4e-08 | 4.6e-08 | 5.533E-08 | 3.690E-08 | 2.400E-08 | 9.600E-08 |
| 16957 | 848 | V848T | C | 3.1e-08 | 3.1e-08 | 3.4e-08 | 3.200E-08 | 1.732E-09 | 3.100E-08 | 3.400E-08 |
| 16958 | 848 | V848W | C | 4.3e-08 | 5.3e-08 | 7.3e-08 | 5.633E-08 | 1.528E-08 | 4.300E-08 | 7.300E-08 |
| 16959 | 848 | V848Y | C | 2.2e-08 | 2.1e-08 | 5.7e-08 | 3.333E-08 | 2.050E-08 | 2.100E-08 | 5.700E-08 |
| 16960 | 848 | V848V | C | 1.4e-07 | 1.1e-08 | 7.5e-08 | 7.533E-08 | 6.450E-08 | 1.100E-08 | 1.400E-07 |
| 16961 | 849 | Q849A | C | 9.4e-08 | 1.1e-08 | 3.7e-08 | 4.733E-08 | 4.245E-08 | 1.100E-08 | 9.400E-08 |
| 16962 | 849 | Q849R | C | 3e-08 | 5.2e-08 | 5.7e-08 | 4.633E-08 | 1.436E-08 | 3.000E-08 | 5.700E-08 |
| 16963 | 849 | Q849N | C | 2.6e-08 | 2.5e-08 | 4.5e-08 | 3.200E-08 | 1.127E-08 | 2.500E-08 | 4.500E-08 |
| 16964 | 849 | Q849D | C | 1e-08 | 6.6e-08 | 7.6e-08 | 5.067E-08 | 3.557E-08 | 1.000E-08 | 7.600E-08 |
| 16965 | 849 | Q849C | C | 2.3e-08 | 5.1e-08 | 2e-08 | 3.133E-08 | 1.710E-08 | 2.000E-08 | 5.100E-08 |
| 16966 | 849 | Q849E | C | 4.5e-08 | 3.3e-08 | 7e-08 | 4.933E-08 | 1.888E-08 | 3.300E-08 | 7.000E-08 |
| 16967 | 849 | Q849Q | C | 3.8e-08 | 3.8e-08 | 3.2e-08 | 3.600E-08 | 3.464E-09 | 3.200E-08 | 3.800E-08 |
| 16968 | 849 | Q849G | C | 7.8e-08 | 4.5e-08 | 5.6e-08 | 5.967E-08 | 1.680E-08 | 4.500E-08 | 7.800E-08 |
| 16969 | 849 | Q849H | C | 6.7e-08 | 4.8e-08 | 5.4e-08 | 5.633E-08 | 9.713E-09 | 4.800E-08 | 6.700E-08 |
| 16970 | 849 | Q849I | C | 4e-08 | 3.1e-08 | 3.9e-08 | 3.667E-08 | 4.933E-09 | 3.100E-08 | 4.000E-08 |
| 16971 | 849 | Q849L | C | 8.3e-09 | 2.5e-08 | 3.5e-08 | 2.277E-08 | 1.349E-08 | 8.300E-09 | 3.500E-08 |
| 16972 | 849 | Q849K | C | 4.7e-08 | 1e-07 | 3.2e-08 | 5.967E-08 | 3.573E-08 | 3.200E-08 | 1.000E-07 |
| 16973 | 849 | Q849M | C | 2e-08 | 5.1e-08 | 1.5e-07 | 7.367E-08 | 6.790E-08 | 2.000E-08 | 1.500E-07 |
| 16974 | 849 | Q849F | C | 5e-08 | 2.9e-08 | 4.3e-08 | 4.067E-08 | 1.069E-08 | 2.900E-08 | 5.000E-08 |
| 16975 | 849 | Q849P | C | 3.5e-08 | 4.2e-08 | 1.1e-07 | 6.233E-08 | 4.143E-08 | 3.500E-08 | 1.100E-07 |
| 16976 | 849 | Q849S | C | 2.7e-08 | 6e-08 | 4.3e-08 | 4.333E-08 | 1.650E-08 | 2.700E-08 | 6.000E-08 |
| 16977 | 849 | Q849T | C | 3.4e-08 | 2.3e-08 | 9.8e-08 | 5.167E-08 | 4.050E-08 | 2.300E-08 | 9.800E-08 |
| 16978 | 849 | Q849W | C | 3e-08 | 1.6e-08 | 3.3e-08 | 2.633E-08 | 9.074E-09 | 1.600E-08 | 3.300E-08 |
| 16979 | 849 | Q849Y | C | 2.6e-08 | 3.6e-08 | 4.2e-08 | 3.467E-08 | 8.083E-09 | 2.600E-08 | 4.200E-08 |
| 16980 | 849 | Q849V | C | 6.3e-08 | 3.2e-08 | 4.9e-08 | 4.800E-08 | 1.552E-08 | 3.200E-08 | 6.300E-08 |
| 16981 | 850 | P850A | C | 3.4e-08 | 3.3e-08 | 8.9e-08 | 5.200E-08 | 3.205E-08 | 3.300E-08 | 8.900E-08 |
| 16982 | 850 | P850R | C | 6e-08 | 5.4e-08 | 4.6e-08 | 5.333E-08 | 7.024E-09 | 4.600E-08 | 6.000E-08 |
| 16983 | 850 | P850N | C | 3.9e-08 | 5.7e-08 | 1.2e-08 | 3.600E-08 | 2.265E-08 | 1.200E-08 | 5.700E-08 |
| 16984 | 850 | P850D | C | 1e-08 | 6.1e-08 | 5.7e-08 | 4.267E-08 | 2.836E-08 | 1.000E-08 | 6.100E-08 |
| 16985 | 850 | P850C | C | 1.2e-08 | 7.4e-08 | 6e-08 | 4.867E-08 | 3.252E-08 | 1.200E-08 | 7.400E-08 |
| 16986 | 850 | P850E | C | 1e-07 | 4.5e-08 | 3.3e-08 | 5.933E-08 | 3.573E-08 | 3.300E-08 | 1.000E-07 |
| 16987 | 850 | P850Q | C | 3.5e-08 | 3.2e-08 | 4.3e-08 | 3.667E-08 | 5.686E-09 | 3.200E-08 | 4.300E-08 |
| 16988 | 850 | P850G | C | 4.3e-08 | 3.4e-08 | 1.5e-07 | 7.567E-08 | 6.453E-08 | 3.400E-08 | 1.500E-07 |
| 16989 | 850 | P850H | C | 2.1e-08 | 2.7e-08 | 7.6e-08 | 4.133E-08 | 3.017E-08 | 2.100E-08 | 7.600E-08 |
| 16990 | 850 | P850I | C | 3.3e-08 | 1.3e-07 | 3.1e-08 | 6.467E-08 | 5.659E-08 | 3.100E-08 | 1.300E-07 |
| 16991 | 850 | P850L | C | 3.5e-08 | 2.2e-08 | 2.2e-08 | 2.633E-08 | 7.506E-09 | 2.200E-08 | 3.500E-08 |
| 16992 | 850 | P850K | C | 6.9e-08 | 4.4e-08 | 6.2e-08 | 5.833E-08 | 1.290E-08 | 4.400E-08 | 6.900E-08 |
| 16993 | 850 | P850M | C | 6.3e-08 | 5.5e-08 | 4.1e-08 | 5.300E-08 | 1.114E-08 | 4.100E-08 | 6.300E-08 |
| 16994 | 850 | P850F | C | 1.9e-08 | 3.5e-08 | 2.2e-08 | 2.533E-08 | 8.505E-09 | 1.900E-08 | 3.500E-08 |
| 16995 | 850 | P850P | C | 5.1e-08 | 1.6e-07 | 6.6e-08 | 9.233E-08 | 5.908E-08 | 5.100E-08 | 1.600E-07 |
| 16996 | 850 | P850S | C | 1.2e-08 | 7.1e-08 | 6.5e-08 | 4.933E-08 | 3.247E-08 | 1.200E-08 | 7.100E-08 |
| 16997 | 850 | P850T | C | 9.1e-08 | 3.4e-08 | 3.5e-08 | 5.333E-08 | 3.262E-08 | 3.400E-08 | 9.100E-08 |
| 16998 | 850 | P850W | C | 2.1e-08 | 2.3e-08 | 6.1e-08 | 3.500E-08 | 2.254E-08 | 2.100E-08 | 6.100E-08 |
| 16999 | 850 | P850Y | C | 1.8e-08 | 1.1e-07 | 6.5e-08 | 6.433E-08 | 4.600E-08 | 1.800E-08 | 1.100E-07 |

| | | | | | | | | | | |
|---|---|---|---|---|---|---|---|---|---|---|
| 17000 | 850 | P850V | C | 8.1e-08 | 1.4e-07 | 9.7e-09 | 7.690E-08 | 6.525E-08 | 9.700E-09 | 1.400E-07 |
| 17001 | 851 | G851A | C | 4.4e-08 | 8.1e-08 | 5.6e-08 | 6.033E-08 | 1.888E-08 | 4.400E-08 | 8.100E-08 |
| 17002 | 851 | G851R | C | 7.6e-08 | 3.2e-08 | 2.9e-08 | 4.567E-08 | 2.631E-08 | 2.900E-08 | 7.600E-08 |
| 17003 | 851 | G851N | C | 2.1e-08 | 5.6e-08 | 5.7e-08 | 4.467E-08 | 2.050E-08 | 2.100E-08 | 5.700E-08 |
| 17004 | 851 | G851D | C | 2.8e-08 | 5.2e-08 | 5e-08 | 4.333E-08 | 1.332E-08 | 2.800E-08 | 5.200E-08 |
| 17005 | 851 | G851C | C | 4.5e-08 | 4.1e-08 | 2e-08 | 3.533E-08 | 1.343E-08 | 2.000E-08 | 4.500E-08 |
| 17006 | 851 | G851E | C | 1.1e-07 | 8.7e-08 | 6.5e-08 | 8.733E-08 | 2.250E-08 | 6.500E-08 | 1.100E-07 |
| 17007 | 851 | G851Q | C | 3.7e-08 | 3.2e-08 | 1.3e-07 | 6.633E-08 | 5.519E-08 | 3.200E-08 | 1.300E-07 |
| 17008 | 851 | G851G | C | 8.2e-08 | 4.9e-08 | 4.9e-08 | 6.000E-08 | 1.905E-08 | 4.900E-08 | 8.200E-08 |
| 17009 | 851 | G851H | C | 4.5e-08 | 1.4e-08 | 6.8e-08 | 4.233E-08 | 2.710E-08 | 1.400E-08 | 6.800E-08 |
| 17010 | 851 | G851I | C | 3.3e-08 | 4.3e-08 | 3.6e-08 | 3.733E-08 | 5.132E-09 | 3.300E-08 | 4.300E-08 |
| 17011 | 851 | G851L | C | 2e-08 | 3.8e-08 | 3.4e-08 | 3.067E-08 | 9.452E-09 | 2.000E-08 | 3.800E-08 |
| 17012 | 851 | G851K | C | 6.4e-08 | 9.5e-08 | 4.8e-08 | 6.900E-08 | 2.390E-08 | 4.800E-08 | 9.500E-08 |
| 17013 | 851 | G851M | C | 3.6e-08 | 1.9e-08 | 4.6e-08 | 3.367E-08 | 1.365E-08 | 1.900E-08 | 4.600E-08 |
| 17014 | 851 | G851F | C | 1.9e-08 | 2.5e-08 | 3.7e-08 | 2.700E-08 | 9.165E-09 | 1.900E-08 | 3.700E-08 |
| 17015 | 851 | G851P | C | 5.9e-08 | 4.4e-08 | 4.8e-08 | 5.033E-08 | 7.767E-09 | 4.400E-08 | 5.900E-08 |
| 17016 | 851 | G851S | C | 3.7e-08 | 5.2e-08 | 3.3e-08 | 4.067E-08 | 1.002E-08 | 3.300E-08 | 5.200E-08 |
| 17017 | 851 | G851T | C | 4.3e-08 | 3.8e-08 | 4.2e-08 | 4.100E-08 | 2.646E-09 | 3.800E-08 | 4.300E-08 |
| 17018 | 851 | G851W | C | 6.4e-08 | 1.1e-07 | 1.9e-08 | 6.433E-08 | 4.550E-08 | 1.900E-08 | 1.100E-07 |
| 17019 | 851 | G851Y | C | 7e-08 | 5.1e-08 | 7.6e-08 | 6.567E-08 | 1.305E-08 | 5.100E-08 | 7.600E-08 |
| 17020 | 851 | G851V | C | 2e-08 | 4.6e-08 | 2.5e-08 | 3.033E-08 | 1.380E-08 | 2.000E-08 | 4.600E-08 |
| 17021 | 852 | G852A | C | 2.2e-08 | 1.8e-08 | 8.8e-08 | 4.267E-08 | 3.931E-08 | 1.800E-08 | 8.800E-08 |
| 17022 | 852 | G852R | C | 4.4e-08 | 8.7e-08 | 4e-08 | 5.700E-08 | 2.606E-08 | 4.000E-08 | 8.700E-08 |
| 17023 | 852 | G852N | C | 5.7e-08 | 4.6e-08 | 3.9e-08 | 4.733E-08 | 9.074E-09 | 3.900E-08 | 5.700E-08 |
| 17024 | 852 | G852D | C | 2.5e-08 | 2.5e-08 | 5.3e-08 | 3.433E-08 | 1.617E-08 | 2.500E-08 | 5.300E-08 |
| 17025 | 852 | G852C | C | 5.5e-08 | 1.4e-07 | 7.9e-08 | 9.133E-08 | 4.382E-08 | 5.500E-08 | 1.400E-07 |
| 17026 | 852 | G852E | C | 2.2e-08 | 4.1e-08 | 5.7e-08 | 4.000E-08 | 1.752E-08 | 2.200E-08 | 5.700E-08 |
| 17027 | 852 | G852Q | C | 1.9e-08 | 2.8e-08 | 1.4e-08 | 2.033E-08 | 7.095E-09 | 1.400E-08 | 2.800E-08 |
| 17028 | 852 | G852G | C | 1.4e-08 | 5.6e-08 | 2.2e-08 | 3.067E-08 | 2.230E-08 | 1.400E-08 | 5.600E-08 |
| 17029 | 852 | G852H | C | 1.9e-08 | 5.6e-08 | 5.1e-08 | 4.200E-08 | 2.007E-08 | 1.900E-08 | 5.600E-08 |
| 17030 | 852 | G852I | C | 3.5e-08 | 7.1e-08 | 4.2e-08 | 4.933E-08 | 1.909E-08 | 3.500E-08 | 7.100E-08 |
| 17031 | 852 | G852L | C | 4.5e-08 | 8.6e-08 | 8.3e-08 | 7.133E-08 | 2.285E-08 | 4.500E-08 | 8.600E-08 |
| 17032 | 852 | G852K | C | 4.5e-08 | 1.7e-08 | 5.3e-08 | 3.833E-08 | 1.890E-08 | 1.700E-08 | 5.300E-08 |
| 17033 | 852 | G852M | C | 3.7e-08 | 2.8e-08 | 4e-08 | 3.500E-08 | 6.245E-09 | 2.800E-08 | 4.000E-08 |
| 17034 | 852 | G852F | C | 2.1e-08 | 1.7e-07 | 2.3e-08 | 7.133E-08 | 8.545E-08 | 2.100E-08 | 1.700E-07 |
| 17035 | 852 | G852P | C | 7e-08 | 3.9e-08 | 1e-07 | 6.967E-08 | 3.050E-08 | 3.900E-08 | 1.000E-07 |
| 17036 | 852 | G852S | C | 8.7e-08 | 1.6e-08 | 8.8e-08 | 6.367E-08 | 4.128E-08 | 1.600E-08 | 8.800E-08 |
| 17037 | 852 | G852T | C | 2.1e-08 | 6.2e-08 | 6.9e-08 | 5.067E-08 | 2.593E-08 | 2.100E-08 | 6.900E-08 |
| 17038 | 852 | G852W | C | 5.9e-08 | 3.9e-08 | 4e-08 | 4.600E-08 | 1.127E-08 | 3.900E-08 | 5.900E-08 |
| 17039 | 852 | G852Y | C | 2.7e-08 | 2.6e-08 | 5.8e-08 | 3.700E-08 | 1.819E-08 | 2.600E-08 | 5.800E-08 |
| 17040 | 852 | G852V | C | 2.3e-08 | 1.6e-08 | 3.3e-08 | 2.400E-08 | 8.544E-09 | 1.600E-08 | 3.300E-08 |
| 17041 | 853 | S853A | C | 2.3e-08 | 3.3e-08 | 1.5e-07 | 6.867E-08 | 7.061E-08 | 2.300E-08 | 1.500E-07 |
| 17042 | 853 | S853R | C | 8.9e-08 | 1.5e-08 | 8.1e-08 | 6.167E-08 | 4.061E-08 | 1.500E-08 | 8.900E-08 |
| 17043 | 853 | S853N | C | 8.3e-08 | 5.2e-08 | 6.5e-08 | 6.667E-08 | 1.557E-08 | 5.200E-08 | 8.300E-08 |
| 17044 | 853 | S853D | C | 8.9e-08 | 8.1e-09 | 6.1e-08 | 5.270E-08 | 4.108E-08 | 8.100E-09 | 8.900E-08 |
| 17045 | 853 | S853C | C | 5.6e-08 | 2.6e-08 | 3e-08 | 3.733E-08 | 1.629E-08 | 2.600E-08 | 5.600E-08 |
| 17046 | 853 | S853E | C | 5.7e-08 | 3.2e-08 | 3.7e-08 | 4.200E-08 | 1.323E-08 | 3.200E-08 | 5.700E-08 |
| 17047 | 853 | S853Q | C | 6.6e-08 | 1.7e-08 | 2.4e-08 | 3.567E-08 | 2.650E-08 | 1.700E-08 | 6.600E-08 |
| 17048 | 853 | S853G | C | 8.2e-08 | 3.1e-08 | 3.5e-08 | 4.933E-08 | 2.836E-08 | 3.100E-08 | 8.200E-08 |
| 17049 | 853 | S853H | C | 5.7e-08 | 4.5e-08 | 2.5e-08 | 4.233E-08 | 1.617E-08 | 2.500E-08 | 5.700E-08 |
| 17050 | 853 | S853I | C | 4.3e-08 | 3.3e-08 | 6.9e-08 | 4.833E-08 | 1.858E-08 | 3.300E-08 | 6.900E-08 |
| 17051 | 853 | S853L | C | 2.7e-08 | 4e-08 | 7.1e-08 | 4.600E-08 | 2.261E-08 | 2.700E-08 | 7.100E-08 |
| 17052 | 853 | S853K | C | 8e-08 | 6.5e-08 | 4.1e-08 | 6.200E-08 | 1.967E-08 | 4.100E-08 | 8.000E-08 |
| 17053 | 853 | S853M | C | 3.5e-08 | 1.6e-08 | 3.8e-08 | 2.967E-08 | 1.193E-08 | 1.600E-08 | 3.800E-08 |
| 17054 | 853 | S853F | C | 4e-08 | 9.4e-08 | 2.6e-08 | 5.333E-08 | 3.591E-08 | 2.600E-08 | 9.400E-08 |
| 17055 | 853 | S853P | C | 3.4e-08 | 6.1e-08 | 5.5e-08 | 5.000E-08 | 1.418E-08 | 3.400E-08 | 6.100E-08 |
| 17056 | 853 | S853S | C | 3.8e-08 | 5.6e-08 | 5e-08 | 4.800E-08 | 9.165E-09 | 3.800E-08 | 5.600E-08 |
| 17057 | 853 | S853T | C | 2.9e-08 | 4.8e-08 | 1.7e-08 | 3.133E-08 | 1.563E-08 | 1.700E-08 | 4.800E-08 |
| 17058 | 853 | S853W | C | 2.6e-08 | 9.3e-08 | 4.9e-08 | 5.600E-08 | 3.404E-08 | 2.600E-08 | 9.300E-08 |
| 17059 | 853 | S853Y | C | 1.1e-07 | 1.7e-08 | 3.3e-08 | 5.333E-08 | 4.972E-08 | 1.700E-08 | 1.100E-07 |
| 17060 | 853 | S853V | C | 5.9e-08 | 5.4e-08 | 1.1e-08 | 4.133E-08 | 2.639E-08 | 1.100E-08 | 5.900E-08 |
| 17061 | 854 | L854A | C | 2.9e-08 | 3.7e-08 | 3.3e-08 | 3.300E-08 | 4.000E-09 | 2.900E-08 | 3.700E-08 |
| 17062 | 854 | L854R | C | 1.1e-07 | 6.4e-08 | 4.3e-08 | 7.233E-08 | 3.427E-08 | 4.300E-08 | 1.100E-07 |
| 17063 | 854 | L854N | C | 5.7e-08 | 2.4e-08 | 2.8e-08 | 3.633E-08 | 1.801E-08 | 2.400E-08 | 5.700E-08 |
| 17064 | 854 | L854D | C | 3.1e-08 | 7e-08 | 4e-08 | 4.700E-08 | 2.042E-08 | 3.100E-08 | 7.000E-08 |
| 17065 | 854 | L854C | C | 1.6e-08 | 4.1e-08 | 3.5e-08 | 3.067E-08 | 1.305E-08 | 1.600E-08 | 4.100E-08 |
| 17066 | 854 | L854E | C | 1.3e-07 | 4.6e-08 | 4.7e-08 | 7.433E-08 | 4.821E-08 | 4.600E-08 | 1.300E-07 |
| 17067 | 854 | L854Q | C | 1.8e-08 | 2.1e-08 | 3.6e-08 | 2.500E-08 | 9.644E-09 | 1.800E-08 | 3.600E-08 |
| 17068 | 854 | L854G | C | 1.8e-07 | 2e-07 | 5.9e-08 | 1.463E-07 | 7.629E-08 | 5.900E-08 | 2.000E-07 |
| 17069 | 854 | L854H | C | 2.3e-08 | 2.9e-08 | 9e-08 | 4.733E-08 | 3.707E-08 | 2.300E-08 | 9.000E-08 |
| 17070 | 854 | L854I | C | 6.7e-08 | 3.8e-08 | 2.7e-08 | 4.400E-08 | 2.066E-08 | 2.700E-08 | 6.700E-08 |
| 17071 | 854 | L854L | C | 9.6e-08 | 5.3e-08 | 1e-07 | 8.300E-08 | 2.606E-08 | 5.300E-08 | 1.000E-07 |
| 17072 | 854 | L854K | C | 8.5e-08 | 8.8e-08 | 5.1e-08 | 7.467E-08 | 2.055E-08 | 5.100E-08 | 8.800E-08 |
| 17073 | 854 | L854M | C | 2.1e-08 | 4.3e-08 | 1.2e-07 | 6.133E-08 | 5.198E-08 | 2.100E-08 | 1.200E-07 |
| 17074 | 854 | L854F | C | 1.8e-08 | 1.4e-07 | 4.4e-08 | 6.733E-08 | 6.426E-08 | 1.800E-08 | 1.400E-07 |
| 17075 | 854 | L854P | C | 6.2e-08 | 5.1e-08 | 7e-08 | 6.100E-08 | 9.539E-09 | 5.100E-08 | 7.000E-08 |
| 17076 | 854 | L854S | C | 4e-08 | 1.6e-08 | 6.4e-08 | 4.000E-08 | 2.400E-08 | 1.600E-08 | 6.400E-08 |
| 17077 | 854 | L854T | C | 7.4e-08 | 4.4e-08 | 3.5e-08 | 5.100E-08 | 2.042E-08 | 3.500E-08 | 7.400E-08 |
| 17078 | 854 | L854W | C | 2.3e-08 | 4e-08 | 3.6e-08 | 3.300E-08 | 8.888E-09 | 2.300E-08 | 4.000E-08 |
| 17079 | 854 | L854Y | C | 7.5e-08 | 2.7e-08 | 3e-08 | 4.400E-08 | 2.689E-08 | 2.700E-08 | 7.500E-08 |
| 17080 | 854 | L854V | C | 5.1e-08 | 5e-08 | 2.2e-08 | 4.100E-08 | 1.646E-08 | 2.200E-08 | 5.100E-08 |
| 17081 | 855 | R855A | C | 7.2e-08 | 1.8e-07 | 4e-08 | 9.733E-08 | 7.336E-08 | 4.000E-08 | 1.800E-07 |
| 17082 | 855 | R855R | C | 4.1e-08 | 9.3e-08 | 7.9e-08 | 7.100E-08 | 2.691E-08 | 4.100E-08 | 9.300E-08 |
| 17083 | 855 | R855N | C | 5.3e-08 | 1.7e-08 | 9.7e-08 | 5.567E-08 | 4.007E-08 | 1.700E-08 | 9.700E-08 |
| 17084 | 855 | R855D | C | 5.2e-08 | 4.6e-08 | 1.9e-08 | 3.900E-08 | 1.758E-08 | 1.900E-08 | 5.200E-08 |

| | | | | | | | | | | |
|---|---|---|---|---|---|---|---|---|---|---|
| 17085 | 855 | R855C | C | 2.6e-08 | 4.4e-08 | 2.4e-08 | 3.133E-08 | 1.102E-08 | 2.400E-08 | 4.400E-08 |
| 17086 | 855 | R855E | C | 2.6e-08 | 1.8e-08 | 5e-08 | 3.133E-08 | 1.665E-08 | 1.800E-08 | 5.000E-08 |
| 17087 | 855 | R855Q | C | 2.2e-08 | 6.5e-08 | 8e-08 | 5.567E-08 | 3.011E-08 | 2.200E-08 | 8.000E-08 |
| 17088 | 855 | R855G | C | 5.7e-08 | 2.3e-08 | 1.4e-08 | 3.133E-08 | 2.268E-08 | 1.400E-08 | 5.700E-08 |
| 17089 | 855 | R855H | C | 3.9e-08 | 4.7e-08 | 1.7e-07 | 8.533E-08 | 7.343E-08 | 3.900E-08 | 1.700E-07 |
| 17090 | 855 | R855I | C | 9.2e-08 | 4.1e-08 | 1.1e-07 | 8.100E-08 | 3.579E-08 | 4.100E-08 | 1.100E-07 |
| 17091 | 855 | R855L | C | 1.3e-08 | 7.9e-08 | 2.8e-08 | 4.000E-08 | 3.460E-08 | 1.300E-08 | 7.900E-08 |
| 17092 | 855 | R855K | C | 2.5e-08 | 3.5e-08 | 3.9e-08 | 3.300E-08 | 7.211E-09 | 2.500E-08 | 3.900E-08 |
| 17093 | 855 | R855M | C | 6e-08 | 3.2e-08 | 7.1e-08 | 5.433E-08 | 2.011E-08 | 3.200E-08 | 7.100E-08 |
| 17094 | 855 | R855F | C | 4.6e-08 | 1.2e-08 | 6.1e-08 | 3.967E-08 | 2.511E-08 | 1.200E-08 | 6.100E-08 |
| 17095 | 855 | R855P | C | 7e-08 | 2e-08 | 2.8e-08 | 3.933E-08 | 2.686E-08 | 2.000E-08 | 7.000E-08 |
| 17096 | 855 | R855S | C | 4.1e-08 | 2.2e-08 | 8e-08 | 4.767E-08 | 2.957E-08 | 2.200E-08 | 8.000E-08 |
| 17097 | 855 | R855T | C | 7.3e-08 | 5.4e-08 | 3.3e-08 | 5.333E-08 | 2.001E-08 | 3.300E-08 | 7.300E-08 |
| 17098 | 855 | R855W | C | 6e-08 | 5e-08 | 8.2e-08 | 6.400E-08 | 1.637E-08 | 5.000E-08 | 8.200E-08 |
| 17099 | 855 | R855Y | C | 1.7e-08 | 6.6e-08 | 2.9e-08 | 3.733E-08 | 2.554E-08 | 1.700E-08 | 6.600E-08 |
| 17100 | 855 | R855V | C | 1.1e-07 | 4.2e-08 | 3.6e-08 | 6.267E-08 | 4.110E-08 | 3.600E-08 | 1.100E-07 |
| 17101 | 856 | L856A | C | 3e-08 | 3.6e-08 | 3.3e-08 | 3.300E-08 | 3.000E-09 | 3.000E-08 | 3.600E-08 |
| 17102 | 856 | L856R | C | 5.8e-08 | 6.2e-08 | 4.7e-08 | 5.567E-08 | 7.767E-09 | 4.700E-08 | 6.200E-08 |
| 17103 | 856 | L856N | C | 7.8e-08 | 4e-08 | 5.1e-08 | 5.633E-08 | 1.955E-08 | 4.000E-08 | 7.800E-08 |
| 17104 | 856 | L856D | C | 2.8e-08 | 1.1e-07 | 7.2e-08 | 7.000E-08 | 4.104E-08 | 2.800E-08 | 1.100E-07 |
| 17105 | 856 | L856C | C | 4.6e-08 | 4.5e-08 | 2.2e-08 | 3.767E-08 | 1.358E-08 | 2.200E-08 | 4.600E-08 |
| 17106 | 856 | L856E | C | 4.4e-08 | 2.4e-08 | 2.5e-08 | 3.100E-08 | 1.127E-08 | 2.400E-08 | 4.400E-08 |
| 17107 | 856 | L856Q | C | 4.7e-08 | 1.9e-08 | 5.8e-08 | 4.133E-08 | 2.011E-08 | 1.900E-08 | 5.800E-08 |
| 17108 | 856 | L856G | C | 4.2e-08 | 2.7e-08 | 3.2e-08 | 3.367E-08 | 7.638E-09 | 2.700E-08 | 4.200E-08 |
| 17109 | 856 | L856H | C | 1.2e-07 | 4.4e-08 | 3.1e-08 | 6.500E-08 | 4.807E-08 | 3.100E-08 | 1.200E-07 |
| 17110 | 856 | L856I | C | 2.6e-08 | 8.6e-08 | 6.4e-08 | 5.867E-08 | 3.035E-08 | 2.600E-08 | 8.600E-08 |
| 17111 | 856 | L856L | C | 2.2e-08 | 5.8e-08 | 3.4e-08 | 3.800E-08 | 1.833E-08 | 2.200E-08 | 5.800E-08 |
| 17112 | 856 | L856K | C | 8.7e-08 | 2.3e-08 | 4.4e-08 | 5.133E-08 | 3.262E-08 | 2.300E-08 | 8.700E-08 |
| 17113 | 856 | L856M | C | 8.6e-08 | 1.5e-08 | 7e-08 | 5.700E-08 | 3.724E-08 | 1.500E-08 | 8.600E-08 |
| 17114 | 856 | L856F | C | 5.5e-08 | 2.9e-08 | 2.6e-08 | 3.667E-08 | 1.595E-08 | 2.600E-08 | 5.500E-08 |
| 17115 | 856 | L856P | C | 4e-08 | 2.5e-08 | 6.9e-08 | 4.467E-08 | 2.237E-08 | 2.500E-08 | 6.900E-08 |
| 17116 | 856 | L856S | C | 3.5e-08 | 9.3e-08 | 1.4e-07 | 8.933E-08 | 5.260E-08 | 3.500E-08 | 1.400E-07 |
| 17117 | 856 | L856T | C | 5.3e-08 | 1.4e-07 | 4.4e-08 | 7.900E-08 | 5.302E-08 | 4.400E-08 | 1.400E-07 |
| 17118 | 856 | L856W | C | 6.6e-08 | 7e-08 | 2.1e-08 | 5.233E-08 | 2.721E-08 | 2.100E-08 | 7.000E-08 |
| 17119 | 856 | L856Y | C | 7.2e-08 | 7.5e-08 | 3.3e-08 | 6.000E-08 | 2.343E-08 | 3.300E-08 | 7.500E-08 |
| 17120 | 856 | L856V | C | 1e-07 | 2.4e-08 | 2.4e-08 | 4.933E-08 | 4.388E-08 | 2.400E-08 | 1.000E-07 |
| 17121 | 857 | S857A | C | 3.5e-08 | 1.8e-08 | 1.8e-08 | 2.367E-08 | 9.815E-09 | 1.800E-08 | 3.500E-08 |
| 17122 | 857 | S857R | C | 4.4e-08 | 9.1e-08 | 4.3e-08 | 5.933E-08 | 2.743E-08 | 4.300E-08 | 9.100E-08 |
| 17123 | 857 | S857N | C | 1.3e-08 | 3e-08 | 3.5e-08 | 2.600E-08 | 1.153E-08 | 1.300E-08 | 3.500E-08 |
| 17124 | 857 | S857D | C | 1.3e-08 | 9e-08 | 2.9e-08 | 4.400E-08 | 4.063E-08 | 1.300E-08 | 9.000E-08 |
| 17125 | 857 | S857C | C | 5.3e-08 | 3.8e-08 | 4.4e-08 | 4.500E-08 | 7.550E-09 | 3.800E-08 | 5.300E-08 |
| 17126 | 857 | S857E | C | 1.6e-08 | 6.4e-08 | 2.8e-08 | 3.600E-08 | 2.498E-08 | 1.600E-08 | 6.400E-08 |
| 17127 | 857 | S857Q | C | 9.6e-08 | 4.6e-08 | 2.7e-08 | 5.633E-08 | 3.564E-08 | 2.700E-08 | 9.600E-08 |
| 17128 | 857 | S857G | C | 8.2e-08 | 2e-08 | 3.4e-08 | 4.533E-08 | 3.252E-08 | 2.000E-08 | 8.200E-08 |
| 17129 | 857 | S857H | C | 1e-07 | 7.3e-08 | 4.4e-08 | 7.233E-08 | 2.801E-08 | 4.400E-08 | 1.000E-07 |
| 17130 | 857 | S857I | C | 1.3e-08 | 5.3e-08 | 2.3e-08 | 2.967E-08 | 2.082E-08 | 1.300E-08 | 5.300E-08 |
| 17131 | 857 | S857L | C | 2.8e-08 | 3.1e-08 | 9.5e-08 | 5.133E-08 | 3.785E-08 | 2.800E-08 | 9.500E-08 |
| 17132 | 857 | S857K | C | 6e-08 | 5.4e-08 | 8.7e-08 | 6.700E-08 | 1.758E-08 | 5.400E-08 | 8.700E-08 |
| 17133 | 857 | S857M | C | 5.1e-08 | 1.1e-07 | 8.2e-08 | 8.100E-08 | 2.951E-08 | 5.100E-08 | 1.100E-07 |
| 17134 | 857 | S857F | C | 1.9e-08 | 2e-08 | 3.3e-08 | 2.400E-08 | 7.810E-09 | 1.900E-08 | 3.300E-08 |
| 17135 | 857 | S857P | C | 1.1e-07 | 3.8e-08 | 8.7e-08 | 7.833E-08 | 3.677E-08 | 3.800E-08 | 1.100E-07 |
| 17136 | 857 | S857S | C | 5.6e-08 | 3.2e-08 | 7.5e-08 | 5.433E-08 | 2.155E-08 | 3.200E-08 | 7.500E-08 |
| 17137 | 857 | S857T | C | 4.2e-08 | 3.5e-08 | 5.4e-08 | 4.367E-08 | 9.609E-09 | 3.500E-08 | 5.400E-08 |
| 17138 | 857 | S857W | C | 6.5e-08 | 3.2e-08 | 5.6e-08 | 5.100E-08 | 1.706E-08 | 3.200E-08 | 6.500E-08 |
| 17139 | 857 | S857Y | C | 7.5e-08 | 2e-08 | 1.4e-08 | 3.633E-08 | 3.362E-08 | 1.400E-08 | 7.500E-08 |
| 17140 | 857 | S857V | C | 3.3e-08 | 1.2e-07 | 6.4e-08 | 7.233E-08 | 4.409E-08 | 3.300E-08 | 1.200E-07 |
| 17141 | 858 | C858A | C | 4.8e-08 | 2e-08 | 2.1e-08 | 2.967E-08 | 1.589E-08 | 2.000E-08 | 4.800E-08 |
| 17142 | 858 | C858R | C | 4.8e-08 | 4.6e-08 | 3.5e-08 | 4.300E-08 | 7.000E-09 | 3.500E-08 | 4.800E-08 |
| 17143 | 858 | C858N | C | 2.8e-08 | 8.5e-09 | 1.1e-07 | 4.883E-08 | 5.386E-08 | 8.500E-09 | 1.100E-07 |
| 17144 | 858 | C858D | C | 3.5e-08 | 4.7e-08 | 3.4e-08 | 3.867E-08 | 7.234E-09 | 3.400E-08 | 4.700E-08 |
| 17145 | 858 | C858C | C | 3.7e-08 | 4.9e-08 | 2.9e-08 | 3.833E-08 | 1.007E-08 | 2.900E-08 | 4.900E-08 |
| 17146 | 858 | C858E | C | 3.1e-08 | 6.4e-08 | 4e-08 | 4.500E-08 | 1.706E-08 | 3.100E-08 | 6.400E-08 |
| 17147 | 858 | C858Q | C | 6.6e-08 | 2e-08 | 7.1e-08 | 5.233E-08 | 2.811E-08 | 2.000E-08 | 7.100E-08 |
| 17148 | 858 | C858G | C | 4.5e-08 | 1e-08 | 2.2e-08 | 2.567E-08 | 1.779E-08 | 1.000E-08 | 4.500E-08 |
| 17149 | 858 | C858H | C | 7.3e-08 | 6.2e-08 | 5.1e-08 | 6.200E-08 | 1.100E-08 | 5.100E-08 | 7.300E-08 |
| 17150 | 858 | C858I | C | 9e-08 | 5.9e-08 | 3.6e-08 | 6.167E-08 | 2.710E-08 | 3.600E-08 | 9.000E-08 |
| 17151 | 858 | C858L | C | 1.2e-07 | 2.9e-08 | 6.6e-08 | 7.167E-08 | 4.576E-08 | 2.900E-08 | 1.200E-07 |
| 17152 | 858 | C858K | C | 4.8e-08 | 9e-08 | 8.2e-08 | 7.333E-08 | 2.230E-08 | 4.800E-08 | 9.000E-08 |
| 17153 | 858 | C858M | C | 3.1e-08 | 2.1e-08 | 1.3e-07 | 6.067E-08 | 6.025E-08 | 2.100E-08 | 1.300E-07 |
| 17154 | 858 | C858F | C | 6.7e-08 | 3.6e-08 | 3.7e-08 | 4.667E-08 | 1.762E-08 | 3.600E-08 | 6.700E-08 |
| 17155 | 858 | C858P | C | 2.6e-08 | 9.8e-08 | 3e-08 | 5.133E-08 | 4.046E-08 | 2.600E-08 | 9.800E-08 |
| 17156 | 858 | C858S | C | 3.4e-08 | 4.5e-08 | 3e-08 | 3.633E-08 | 7.767E-09 | 3.000E-08 | 4.500E-08 |
| 17157 | 858 | C858T | C | 3e-08 | 1.8e-08 | 4.9e-08 | 3.233E-08 | 1.563E-08 | 1.800E-08 | 4.900E-08 |
| 17158 | 858 | C858W | C | 2.2e-08 | 1e-07 | 2.6e-08 | 4.933E-08 | 4.392E-08 | 2.200E-08 | 1.000E-07 |
| 17159 | 858 | C858Y | C | 3.2e-08 | 1.8e-08 | 2.8e-08 | 2.600E-08 | 7.211E-09 | 1.800E-08 | 3.200E-08 |
| 17160 | 858 | C858V | C | 2.7e-08 | 4.7e-08 | 2.7e-08 | 3.367E-08 | 1.155E-08 | 2.700E-08 | 4.700E-08 |
| 17161 | 859 | A859A | C | 7.2e-08 | 3.4e-08 | 1e-07 | 6.867E-08 | 3.313E-08 | 3.400E-08 | 1.000E-07 |
| 17162 | 859 | A859R | C | 1.9e-08 | 4.8e-08 | 7.1e-08 | 4.600E-08 | 2.606E-08 | 1.900E-08 | 7.100E-08 |
| 17163 | 859 | A859N | C | 1.7e-08 | 3e-08 | 4.2e-08 | 2.967E-08 | 1.250E-08 | 1.700E-08 | 4.200E-08 |
| 17164 | 859 | A859D | C | 6.6e-08 | 7e-08 | 3.5e-08 | 5.700E-08 | 1.916E-08 | 3.500E-08 | 7.000E-08 |
| 17165 | 859 | A859C | C | 5e-08 | 7.7e-08 | 6.3e-08 | 6.333E-08 | 1.350E-08 | 5.000E-08 | 7.700E-08 |
| 17166 | 859 | A859E | C | 2.5e-08 | 2.6e-07 | 3.8e-08 | 1.077E-07 | 1.321E-07 | 2.500E-08 | 2.600E-07 |
| 17167 | 859 | A859Q | C | 1.9e-08 | 1.1e-08 | 4.4e-08 | 2.467E-08 | 1.721E-08 | 1.100E-08 | 4.400E-08 |
| 17168 | 859 | A859G | C | 6e-08 | 6.5e-08 | 1.5e-08 | 4.667E-08 | 2.754E-08 | 1.500E-08 | 6.500E-08 |
| 17169 | 859 | A859H | C | 8.3e-08 | 3.8e-08 | 4.1e-08 | 5.400E-08 | 2.516E-08 | 3.800E-08 | 8.300E-08 |

| | | | | | | | | | | |
|---|---|---|---|---|---|---|---|---|---|---|
| 17170 | 859 | A859I | C | 6.3e-08 | 7.1e-08 | 6e-08 | 6.467E-08 | 5.686E-09 | 6.000E-08 | 7.100E-08 |
| 17171 | 859 | A859L | C | 4e-08 | 5e-08 | 2.6e-08 | 3.867E-08 | 1.206E-08 | 2.600E-08 | 5.000E-08 |
| 17172 | 859 | A859K | C | 8.2e-09 | 2.3e-08 | 4.5e-08 | 2.540E-08 | 1.852E-08 | 8.200E-09 | 4.500E-08 |
| 17173 | 859 | A859M | C | 1.7e-08 | 8.2e-09 | 4.7e-08 | 2.407E-08 | 2.034E-08 | 8.200E-09 | 4.700E-08 |
| 17174 | 859 | A859F | C | 5.2e-08 | 3.2e-08 | 2.6e-08 | 3.667E-08 | 1.361E-08 | 2.600E-08 | 5.200E-08 |
| 17175 | 859 | A859P | C | 3.6e-08 | 2.6e-08 | 1.3e-07 | 6.400E-08 | 5.738E-08 | 2.600E-08 | 1.300E-07 |
| 17176 | 859 | A859S | C | 5.8e-08 | 6.7e-08 | 3e-08 | 5.167E-08 | 1.930E-08 | 3.000E-08 | 6.700E-08 |
| 17177 | 859 | A859T | C | 5.7e-08 | 1.5e-08 | 2.5e-08 | 3.233E-08 | 2.194E-08 | 1.500E-08 | 5.700E-08 |
| 17178 | 859 | A859W | C | 6.8e-08 | 4.7e-08 | 1.4e-08 | 4.300E-08 | 2.722E-08 | 1.400E-08 | 6.800E-08 |
| 17179 | 859 | A859Y | C | 7.4e-08 | 9e-08 | 3.6e-08 | 6.667E-08 | 2.774E-08 | 3.600E-08 | 9.000E-08 |
| 17180 | 859 | A859V | C | 7.1e-08 | 1.5e-08 | 5.8e-08 | 4.800E-08 | 2.931E-08 | 1.500E-08 | 7.100E-08 |
| 17181 | 860 | A860A | C | 3.5e-08 | 5.7e-08 | 4.8e-08 | 4.667E-08 | 1.106E-08 | 3.500E-08 | 5.700E-08 |
| 17182 | 860 | A860R | C | 8.9e-08 | 3.2e-08 | 7.9e-08 | 6.667E-08 | 3.044E-08 | 3.200E-08 | 8.900E-08 |
| 17183 | 860 | A860N | C | 8.2e-08 | 4.8e-08 | 5.8e-08 | 6.267E-08 | 1.747E-08 | 4.800E-08 | 8.200E-08 |
| 17184 | 860 | A860D | C | 3.8e-08 | 2e-08 | 5e-08 | 3.600E-08 | 1.510E-08 | 2.000E-08 | 5.000E-08 |
| 17185 | 860 | A860C | C | 8.9e-08 | 4.2e-08 | 4.4e-08 | 5.833E-08 | 2.658E-08 | 4.200E-08 | 8.900E-08 |
| 17186 | 860 | A860E | C | 2e-08 | 9.5e-08 | 2.4e-08 | 4.633E-08 | 4.219E-08 | 2.000E-08 | 9.500E-08 |
| 17187 | 860 | A860Q | C | 7.6e-08 | 3.8e-08 | 7.7e-08 | 6.367E-08 | 2.223E-08 | 3.800E-08 | 7.700E-08 |
| 17188 | 860 | A860G | C | 4.2e-08 | 8.6e-08 | 3.6e-08 | 5.467E-08 | 2.730E-08 | 3.600E-08 | 8.600E-08 |
| 17189 | 860 | A860H | C | 4.1e-08 | 5.3e-08 | 3.1e-08 | 4.167E-08 | 1.102E-08 | 3.100E-08 | 5.300E-08 |
| 17190 | 860 | A860I | C | 3.2e-08 | 5.8e-08 | 2.2e-08 | 3.733E-08 | 1.858E-08 | 2.200E-08 | 5.800E-08 |
| 17191 | 860 | A860L | C | 2.9e-08 | 7.4e-08 | 4.3e-08 | 4.867E-08 | 2.303E-08 | 2.900E-08 | 7.400E-08 |
| 17192 | 860 | A860K | C | 8.6e-08 | 8.8e-08 | 7.3e-08 | 8.233E-08 | 8.145E-09 | 7.300E-08 | 8.800E-08 |
| 17193 | 860 | A860M | C | 2.5e-08 | 5e-08 | 1.5e-08 | 3.000E-08 | 1.803E-08 | 1.500E-08 | 5.000E-08 |
| 17194 | 860 | A860F | C | 2.9e-08 | 5.3e-08 | 4.1e-08 | 4.100E-08 | 1.200E-08 | 2.900E-08 | 5.300E-08 |
| 17195 | 860 | A860P | C | 3.1e-08 | 2e-08 | 3e-08 | 2.700E-08 | 6.083E-09 | 2.000E-08 | 3.100E-08 |
| 17196 | 860 | A860S | C | 4.3e-08 | 2.6e-08 | 2.3e-08 | 3.067E-08 | 1.079E-08 | 2.300E-08 | 4.300E-08 |
| 17197 | 860 | A860T | C | 3.7e-08 | 2.6e-08 | 4.5e-08 | 3.600E-08 | 9.539E-09 | 2.600E-08 | 4.500E-08 |
| 17198 | 860 | A860W | C | 9.3e-09 | 1.1e-07 | 3.3e-08 | 5.077E-08 | 5.265E-08 | 9.300E-09 | 1.100E-07 |
| 17199 | 860 | A860Y | C | 4.5e-08 | 3.5e-08 | 5.3e-08 | 4.433E-08 | 9.018E-09 | 3.500E-08 | 5.300E-08 |
| 17200 | 860 | A860V | C | 4.6e-08 | 3.4e-08 | 4.7e-08 | 4.233E-08 | 7.234E-09 | 3.400E-08 | 4.700E-08 |
| 17201 | 861 | S861A | C | 4.6e-08 | 1.8e-08 | 8.1e-08 | 4.833E-08 | 3.156E-08 | 1.800E-08 | 8.100E-08 |
| 17202 | 861 | S861R | C | 5.3e-08 | 3.3e-08 | 2.3e-08 | 3.633E-08 | 1.528E-08 | 2.300E-08 | 5.300E-08 |
| 17203 | 861 | S861N | C | 3.3e-08 | 8.3e-08 | 3.4e-08 | 5.000E-08 | 2.858E-08 | 3.300E-08 | 8.300E-08 |
| 17204 | 861 | S861D | C | 2.7e-08 | 1.5e-08 | 3.4e-08 | 2.533E-08 | 9.609E-09 | 1.500E-08 | 3.400E-08 |
| 17205 | 861 | S861C | C | 4.2e-08 | 2e-08 | 4.8e-08 | 3.667E-08 | 1.474E-08 | 2.000E-08 | 4.800E-08 |
| 17206 | 861 | S861E | C | 1.8e-08 | 4.5e-08 | 9.3e-08 | 5.200E-08 | 3.799E-08 | 1.800E-08 | 9.300E-08 |
| 17207 | 861 | S861Q | C | 4.2e-08 | 7.3e-09 | 2.4e-08 | 2.443E-08 | 1.735E-08 | 7.300E-09 | 4.200E-08 |
| 17208 | 861 | S861G | C | 1.8e-07 | 2.1e-08 | 5.7e-08 | 8.600E-08 | 8.337E-08 | 2.100E-08 | 1.800E-07 |
| 17209 | 861 | S861H | C | 4e-08 | 6.4e-08 | 8.3e-08 | 6.233E-08 | 2.155E-08 | 4.000E-08 | 8.300E-08 |
| 17210 | 861 | S861I | C | 5.6e-08 | 3.6e-08 | 6.3e-08 | 5.167E-08 | 1.401E-08 | 3.600E-08 | 6.300E-08 |
| 17211 | 861 | S861L | C | 4.1e-08 | 6.1e-08 | 3.9e-08 | 4.700E-08 | 1.217E-08 | 3.900E-08 | 6.100E-08 |
| 17212 | 861 | S861K | C | 7.3e-08 | 9e-08 | 2.7e-08 | 6.333E-08 | 3.259E-08 | 2.700E-08 | 9.000E-08 |
| 17213 | 861 | S861M | C | 1.7e-08 | 5e-08 | 3.5e-08 | 3.400E-08 | 1.652E-08 | 1.700E-08 | 5.000E-08 |
| 17214 | 861 | S861F | C | 3.5e-08 | 7.1e-08 | 4.4e-08 | 5.000E-08 | 1.873E-08 | 3.500E-08 | 7.100E-08 |
| 17215 | 861 | S861P | C | 2.8e-08 | 1e-07 | 4.5e-08 | 5.767E-08 | 3.763E-08 | 2.800E-08 | 1.000E-07 |
| 17216 | 861 | S861S | C | 1.4e-07 | 4e-08 | 1.4e-08 | 6.467E-08 | 6.652E-08 | 1.400E-08 | 1.400E-07 |
| 17217 | 861 | S861T | C | 6.6e-08 | 1.6e-08 | 1.7e-08 | 3.300E-08 | 2.858E-08 | 1.600E-08 | 6.600E-08 |
| 17218 | 861 | S861W | C | 3.7e-08 | 3.2e-08 | 1.8e-08 | 2.900E-08 | 9.849E-09 | 1.800E-08 | 3.700E-08 |
| 17219 | 861 | S861Y | C | 2.4e-08 | 8.8e-08 | 1.1e-07 | 7.400E-08 | 4.468E-08 | 2.400E-08 | 1.100E-07 |
| 17220 | 861 | S861V | C | 1.1e-07 | 1.1e-07 | 1.1e-07 | 1.100E-07 | 0.000E+00 | 1.100E-07 | 1.100E-07 |
| 17221 | 862 | G862A | C | 4.1e-08 | 6e-08 | 3.5e-08 | 4.533E-08 | 1.305E-08 | 3.500E-08 | 6.000E-08 |
| 17222 | 862 | G862R | C | 1.1e-07 | 8.8e-08 | 1e-07 | 9.933E-08 | 1.102E-08 | 8.800E-08 | 1.100E-07 |
| 17223 | 862 | G862N | C | 5e-08 | 8e-08 | 3e-08 | 5.333E-08 | 2.517E-08 | 3.000E-08 | 8.000E-08 |
| 17224 | 862 | G862D | C | 1e-07 | 1.7e-08 | 4.5e-08 | 5.400E-08 | 4.223E-08 | 1.700E-08 | 1.000E-07 |
| 17225 | 862 | G862C | C | 3e-08 | 7e-08 | 2e-08 | 4.000E-08 | 2.646E-08 | 2.000E-08 | 7.000E-08 |
| 17226 | 862 | G862E | C | 3.5e-08 | 4.9e-08 | 2.2e-08 | 3.533E-08 | 1.350E-08 | 2.200E-08 | 4.900E-08 |
| 17227 | 862 | G862Q | C | 1.3e-07 | 5.2e-08 | 5e-08 | 7.733E-08 | 4.562E-08 | 5.000E-08 | 1.300E-07 |
| 17228 | 862 | G862G | C | 3.9e-08 | 3.6e-08 | 5.3e-08 | 4.267E-08 | 9.074E-09 | 3.600E-08 | 5.300E-08 |
| 17229 | 862 | G862H | C | 1.5e-08 | 3.6e-08 | 7.2e-08 | 4.100E-08 | 2.883E-08 | 1.500E-08 | 7.200E-08 |
| 17230 | 862 | G862I | C | 4.6e-08 | 2.7e-08 | 3.8e-08 | 3.700E-08 | 9.539E-09 | 2.700E-08 | 4.600E-08 |
| 17231 | 862 | G862L | C | 2.3e-08 | 1.1e-07 | 2.5e-08 | 5.267E-08 | 4.966E-08 | 2.300E-08 | 1.100E-07 |
| 17232 | 862 | G862K | C | 1.3e-08 | 1.7e-08 | 3.2e-08 | 2.067E-08 | 1.002E-08 | 1.300E-08 | 3.200E-08 |
| 17233 | 862 | G862M | C | 1.1e-07 | 2.1e-08 | 4.7e-08 | 5.933E-08 | 4.576E-08 | 2.100E-08 | 1.100E-07 |
| 17234 | 862 | G862F | C | 3.9e-08 | 3.3e-08 | 6e-08 | 4.400E-08 | 1.418E-08 | 3.300E-08 | 6.000E-08 |
| 17235 | 862 | G862P | C | 6.7e-08 | 3.8e-08 | 2.8e-08 | 4.433E-08 | 2.026E-08 | 2.800E-08 | 6.700E-08 |
| 17236 | 862 | G862S | C | 3.5e-08 | 3.2e-08 | 6e-08 | 4.233E-08 | 1.537E-08 | 3.200E-08 | 6.000E-08 |
| 17237 | 862 | G862T | C | 5.6e-08 | 7.6e-08 | 4.2e-08 | 5.800E-08 | 1.709E-08 | 4.200E-08 | 7.600E-08 |
| 17238 | 862 | G862W | C | 5.3e-08 | 1.8e-08 | 3.7e-08 | 3.600E-08 | 1.752E-08 | 1.800E-08 | 5.300E-08 |
| 17239 | 862 | G862Y | C | 8.4e-08 | 3.6e-08 | 6.2e-08 | 6.067E-08 | 2.403E-08 | 3.600E-08 | 8.400E-08 |
| 17240 | 862 | G862V | C | 1.3e-08 | 3.8e-08 | 5.3e-08 | 3.467E-08 | 2.021E-08 | 1.300E-08 | 5.300E-08 |
| 17241 | 863 | F863A | C | 7.5e-08 | 7e-08 | 7.8e-08 | 7.433E-08 | 4.041E-09 | 7.000E-08 | 7.800E-08 |
| 17242 | 863 | F863R | C | 3.4e-08 | 6.5e-08 | 2e-08 | 3.967E-08 | 2.303E-08 | 2.000E-08 | 6.500E-08 |
| 17243 | 863 | F863N | C | 5.3e-08 | 5.6e-08 | 9.1e-08 | 6.667E-08 | 2.113E-08 | 5.300E-08 | 9.100E-08 |
| 17244 | 863 | F863D | C | 5.1e-08 | 2.9e-08 | 1.3e-07 | 7.000E-08 | 5.311E-08 | 2.900E-08 | 1.300E-07 |
| 17245 | 863 | F863C | C | 9.6e-08 | 3.5e-08 | 3.6e-08 | 5.567E-08 | 3.493E-08 | 3.500E-08 | 9.600E-08 |
| 17246 | 863 | F863E | C | 2.9e-08 | 4.4e-08 | 5.2e-08 | 4.167E-08 | 1.168E-08 | 2.900E-08 | 5.200E-08 |
| 17247 | 863 | F863Q | C | 1.6e-08 | 2.6e-08 | 1.3e-08 | 1.833E-08 | 6.807E-09 | 1.300E-08 | 2.600E-08 |
| 17248 | 863 | F863G | C | 3.4e-08 | 2.6e-08 | 1.3e-08 | 2.433E-08 | 1.060E-08 | 1.300E-08 | 3.400E-08 |
| 17249 | 863 | F863H | C | 8.6e-08 | 4e-08 | 4.1e-08 | 5.567E-08 | 2.627E-08 | 4.000E-08 | 8.600E-08 |
| 17250 | 863 | F863I | C | 4.9e-08 | 5.1e-08 | 3.2e-08 | 4.400E-08 | 1.044E-08 | 3.200E-08 | 5.100E-08 |
| 17251 | 863 | F863L | C | 3.9e-08 | 3.4e-08 | 4.1e-08 | 3.800E-08 | 3.606E-09 | 3.400E-08 | 4.100E-08 |
| 17252 | 863 | F863K | C | 2.3e-08 | 2.5e-08 | 8.6e-08 | 4.467E-08 | 3.581E-08 | 2.300E-08 | 8.600E-08 |
| 17253 | 863 | F863M | C | 3.7e-08 | 2.8e-08 | 2.5e-08 | 3.000E-08 | 6.245E-09 | 2.500E-08 | 3.700E-08 |
| 17254 | 863 | F863F | C | 2.9e-08 | 2.2e-08 | 4e-08 | 3.033E-08 | 9.074E-09 | 2.200E-08 | 4.000E-08 |

| | | | | | | | | | | |
|---|---|---|---|---|---|---|---|---|---|---|
| 17255 | 863 | F863P | C | 1.1e-07 | 4.4e-08 | 3.7e-08 | 6.367E-08 | 4.028E-08 | 3.700E-08 | 1.100E-07 |
| 17256 | 863 | F863S | C | 8.3e-08 | 6.1e-08 | 7.7e-09 | 5.057E-08 | 3.872E-08 | 7.700E-09 | 8.300E-08 |
| 17257 | 863 | F863T | C | 1.3e-08 | 1.7e-08 | 3.7e-08 | 2.233E-08 | 1.286E-08 | 1.300E-08 | 3.700E-08 |
| 17258 | 863 | F863W | C | 9.2e-09 | 1.5e-08 | 3.4e-08 | 1.940E-08 | 1.297E-08 | 9.200E-09 | 3.400E-08 |
| 17259 | 863 | F863Y | C | 4.4e-08 | 3.3e-08 | 4.9e-08 | 4.200E-08 | 8.185E-09 | 3.300E-08 | 4.900E-08 |
| 17260 | 863 | F863V | C | 3e-08 | 2.9e-08 | 5.5e-08 | 3.800E-08 | 1.473E-08 | 2.900E-08 | 5.500E-08 |
| 17261 | 864 | T864A | C | 7.4e-08 | 2e-08 | 1.6e-08 | 3.667E-08 | 3.239E-08 | 1.600E-08 | 7.400E-08 |
| 17262 | 864 | T864R | C | 2.6e-08 | 2.2e-08 | 2.4e-08 | 2.400E-08 | 2.000E-09 | 2.200E-08 | 2.600E-08 |
| 17263 | 864 | T864N | C | 1e-07 | 1e-07 | 1.6e-08 | 7.200E-08 | 4.850E-08 | 1.600E-08 | 1.000E-07 |
| 17264 | 864 | T864D | C | 2.8e-08 | 3.4e-08 | 5.1e-08 | 3.767E-08 | 1.193E-08 | 2.800E-08 | 5.100E-08 |
| 17265 | 864 | T864C | C | 1.7e-08 | 1.5e-08 | 2.8e-08 | 2.000E-08 | 7.000E-09 | 1.500E-08 | 2.800E-08 |
| 17266 | 864 | T864E | C | 4.8e-08 | 2.8e-08 | 1.3e-08 | 2.967E-08 | 1.756E-08 | 1.300E-08 | 4.800E-08 |
| 17267 | 864 | T864Q | C | 4.8e-08 | 6.3e-08 | 7.3e-08 | 6.133E-08 | 1.258E-08 | 4.800E-08 | 7.300E-08 |
| 17268 | 864 | T864G | C | 4.9e-08 | 4.6e-08 | 1.8e-07 | 9.167E-08 | 7.651E-08 | 4.600E-08 | 1.800E-07 |
| 17269 | 864 | T864H | C | 1.1e-07 | 4.1e-08 | 3.9e-08 | 6.333E-08 | 4.043E-08 | 3.900E-08 | 1.100E-07 |
| 17270 | 864 | T864I | C | 4.7e-08 | 7e-09 | 1.8e-08 | 2.400E-08 | 2.066E-08 | 7.000E-09 | 4.700E-08 |
| 17271 | 864 | T864L | C | 1.5e-08 | 3.6e-08 | 4.5e-08 | 3.200E-08 | 1.539E-08 | 1.500E-08 | 4.500E-08 |
| 17272 | 864 | T864K | C | 4.8e-08 | 3.6e-08 | 6e-08 | 4.800E-08 | 1.200E-08 | 3.600E-08 | 6.000E-08 |
| 17273 | 864 | T864M | C | 1.6e-08 | 5.4e-08 | 1.4e-08 | 2.800E-08 | 2.254E-08 | 1.400E-08 | 5.400E-08 |
| 17274 | 864 | T864F | C | 8.1e-09 | 1.9e-08 | 1.7e-08 | 1.470E-08 | 5.803E-09 | 8.100E-09 | 1.900E-08 |
| 17275 | 864 | T864P | C | 4.4e-08 | 1.3e-07 | 2.4e-08 | 6.600E-08 | 5.632E-08 | 2.400E-08 | 1.300E-07 |
| 17276 | 864 | T864S | C | 3e-08 | 4.5e-08 | 4.1e-08 | 3.867E-08 | 7.767E-09 | 3.000E-08 | 4.500E-08 |
| 17277 | 864 | T864T | C | 2.7e-08 | 7.3e-08 | 4.7e-08 | 4.900E-08 | 2.307E-08 | 2.700E-08 | 7.300E-08 |
| 17278 | 864 | T864W | C | 5.7e-08 | 4.7e-09 | 3e-08 | 3.057E-08 | 2.615E-08 | 4.700E-09 | 5.700E-08 |
| 17279 | 864 | T864Y | C | 2e-08 | 1.6e-08 | 1.2e-08 | 1.600E-08 | 4.000E-09 | 1.200E-08 | 2.000E-08 |
| 17280 | 864 | T864V | C | 6.9e-08 | 2.9e-08 | 2.2e-08 | 4.000E-08 | 2.536E-08 | 2.200E-08 | 6.900E-08 |
| 17281 | 865 | F865A | C | 2.2e-08 | 1.1e-07 | 4e-08 | 5.733E-08 | 4.649E-08 | 2.200E-08 | 1.100E-07 |
| 17282 | 865 | F865R | C | 5.4e-08 | 5.3e-08 | 4e-08 | 4.900E-08 | 7.810E-09 | 4.000E-08 | 5.400E-08 |
| 17283 | 865 | F865N | C | 6.5e-08 | 5.9e-08 | 3.2e-08 | 5.200E-08 | 1.758E-08 | 3.200E-08 | 6.500E-08 |
| 17284 | 865 | F865D | C | 2.8e-08 | 4.6e-08 | 6.4e-08 | 4.600E-08 | 1.800E-08 | 2.800E-08 | 6.400E-08 |
| 17285 | 865 | F865C | C | 9.4e-08 | 1.1e-07 | 1.4e-07 | 1.147E-07 | 2.335E-08 | 9.400E-08 | 1.400E-07 |
| 17286 | 865 | F865E | C | 5.6e-08 | 5.2e-08 | 5.1e-08 | 5.300E-08 | 2.646E-09 | 5.100E-08 | 5.600E-08 |
| 17287 | 865 | F865Q | C | 2.5e-08 | 6.2e-08 | 3.2e-08 | 3.967E-08 | 1.966E-08 | 2.500E-08 | 6.200E-08 |
| 17288 | 865 | F865G | C | 4.2e-08 | 3.7e-08 | 5e-08 | 4.300E-08 | 6.557E-09 | 3.700E-08 | 5.000E-08 |
| 17289 | 865 | F865H | C | 1.6e-08 | 6.2e-08 | 3.9e-08 | 3.900E-08 | 2.300E-08 | 1.600E-08 | 6.200E-08 |
| 17290 | 865 | F865I | C | 3.9e-08 | 3.5e-08 | 4.9e-08 | 4.100E-08 | 7.211E-09 | 3.500E-08 | 4.900E-08 |
| 17291 | 865 | F865L | C | 7.1e-08 | 2.5e-08 | 1.2e-07 | 7.200E-08 | 4.751E-08 | 2.500E-08 | 1.200E-07 |
| 17292 | 865 | F865K | C | 5.2e-08 | 1.8e-08 | 1.1e-07 | 6.000E-08 | 4.652E-08 | 1.800E-08 | 1.100E-07 |
| 17293 | 865 | F865M | C | 2.9e-08 | 2.9e-08 | 4.3e-08 | 3.367E-08 | 8.083E-09 | 2.900E-08 | 4.300E-08 |
| 17294 | 865 | F865F | C | 6.6e-09 | 1e-07 | 4.7e-08 | 5.120E-08 | 4.684E-08 | 6.600E-09 | 1.000E-07 |
| 17295 | 865 | F865P | C | 1.1e-07 | 1.1e-07 | 3.6e-08 | 8.533E-08 | 4.272E-08 | 3.600E-08 | 1.100E-07 |
| 17296 | 865 | F865S | C | 4.4e-08 | 2.7e-08 | 3e-08 | 3.367E-08 | 9.074E-09 | 2.700E-08 | 4.400E-08 |
| 17297 | 865 | F865T | C | 4.3e-08 | 6.9e-08 | 1.7e-08 | 4.300E-08 | 2.600E-08 | 1.700E-08 | 6.900E-08 |
| 17298 | 865 | F865W | C | 3.3e-08 | 6.3e-08 | 5.4e-08 | 5.000E-08 | 1.539E-08 | 3.300E-08 | 6.300E-08 |
| 17299 | 865 | F865Y | C | 1.1e-08 | 6.6e-08 | 1.5e-08 | 3.067E-08 | 3.066E-08 | 1.100E-08 | 6.600E-08 |
| 17300 | 865 | F865V | C | 3.2e-08 | 5.2e-08 | 4.9e-08 | 4.433E-08 | 1.079E-08 | 3.200E-08 | 5.200E-08 |
| 17301 | 866 | T866A | C | 2.4e-08 | 1.4e-08 | 3.4e-08 | 2.400E-08 | 1.000E-08 | 1.400E-08 | 3.400E-08 |
| 17302 | 866 | T866R | C | 2.1e-08 | 1e-08 | 3e-08 | 2.033E-08 | 1.002E-08 | 1.000E-08 | 3.000E-08 |
| 17303 | 866 | T866N | C | 4.4e-08 | 1.8e-08 | 3e-08 | 3.067E-08 | 1.301E-08 | 1.800E-08 | 4.400E-08 |
| 17304 | 866 | T866D | C | 8e-09 | 3.3e-08 | 2.7e-08 | 2.267E-08 | 1.305E-08 | 8.000E-09 | 3.300E-08 |
| 17305 | 866 | T866C | C | 1.4e-08 | 1.4e-08 | 2.7e-09 | 1.023E-08 | 6.524E-09 | 2.700E-09 | 1.400E-08 |
| 17306 | 866 | T866E | C | 2.8e-09 | 1.5e-08 | 2.2e-08 | 1.327E-08 | 9.717E-09 | 2.800E-09 | 2.200E-08 |
| 17307 | 866 | T866Q | C | 1.7e-08 | 2.2e-08 | 1.9e-08 | 1.933E-08 | 2.517E-09 | 1.700E-08 | 2.200E-08 |
| 17308 | 866 | T866G | C | 1.1e-08 | 1.8e-08 | 1e-08 | 1.300E-08 | 4.359E-09 | 1.000E-08 | 1.800E-08 |
| 17309 | 866 | T866H | C | 7.4e-09 | 1.1e-08 | 2.9e-08 | 1.580E-08 | 1.157E-08 | 7.400E-09 | 2.900E-08 |
| 17310 | 866 | T866I | C | 9.5e-09 | 1.3e-08 | 1e-08 | 1.083E-08 | 1.893E-09 | 9.500E-09 | 1.300E-08 |
| 17311 | 866 | T866L | C | 1.7e-08 | 1.3e-08 | 1.5e-08 | 1.500E-08 | 2.000E-09 | 1.300E-08 | 1.700E-08 |
| 17312 | 866 | T866K | C | 1.3e-08 | 1.1e-08 | 1.1e-08 | 1.167E-08 | 1.155E-09 | 1.100E-08 | 1.300E-08 |
| 17313 | 866 | T866M | C | 6.3e-09 | 9.1e-09 | 6e-09 | 7.133E-09 | 1.710E-09 | 6.000E-09 | 9.100E-09 |
| 17314 | 866 | T866F | C | 1.2e-08 | 4.5e-09 | 8.2e-09 | 8.233E-09 | 3.750E-09 | 4.500E-09 | 1.200E-08 |
| 17315 | 866 | T866P | C | 1.7e-08 | 7.7e-09 | 6e-09 | 1.023E-08 | 5.921E-09 | 6.000E-09 | 1.700E-08 |
| 17316 | 866 | T866S | C | 1.9e-08 | 4.7e-08 | 5.6e-08 | 4.067E-08 | 1.930E-08 | 1.900E-08 | 5.600E-08 |
| 17317 | 866 | T866T | C | 8.1e-08 | 1.9e-08 | 1.3e-08 | 3.767E-08 | 3.765E-08 | 1.300E-08 | 8.100E-08 |
| 17318 | 866 | T866W | C | 3.4e-09 | 2.6e-09 | 5.7e-09 | 3.900E-09 | 1.609E-09 | 2.600E-09 | 5.700E-09 |
| 17319 | 866 | T866Y | C | 9.8e-09 | 5.2e-09 | 9.7e-09 | 8.233E-09 | 2.627E-09 | 5.200E-09 | 9.800E-09 |
| 17320 | 866 | T866V | C | 1.2e-08 | 7.3e-09 | 3.9e-09 | 7.733E-09 | 4.067E-09 | 3.900E-09 | 1.200E-08 |
| 17321 | 867 | D867A | C | 1.3e-07 | 4.2e-08 | 5.7e-08 | 7.633E-08 | 4.708E-08 | 4.200E-08 | 1.300E-07 |
| 17322 | 867 | D867R | C | 4.7e-08 | 4.3e-08 | 6.7e-08 | 5.233E-08 | 1.286E-08 | 4.300E-08 | 6.700E-08 |
| 17323 | 867 | D867N | C | 3.4e-08 | 9.6e-08 | 1.6e-08 | 4.867E-08 | 4.197E-08 | 1.600E-08 | 9.600E-08 |
| 17324 | 867 | D867D | C | 2.7e-08 | 4.8e-08 | 3.2e-08 | 3.567E-08 | 1.097E-08 | 2.700E-08 | 4.800E-08 |
| 17325 | 867 | D867C | C | 4.8e-08 | 4e-08 | 7e-08 | 5.267E-08 | 1.553E-08 | 4.000E-08 | 7.000E-08 |
| 17326 | 867 | D867E | C | 1.3e-08 | 4.3e-08 | 1.9e-08 | 2.500E-08 | 1.587E-08 | 1.300E-08 | 4.300E-08 |
| 17327 | 867 | D867Q | C | 5.1e-09 | 5.2e-08 | 8.4e-09 | 2.183E-08 | 2.618E-08 | 5.100E-09 | 5.200E-08 |
| 17328 | 867 | D867G | C | 1.4e-07 | 5.1e-08 | 4e-08 | 7.700E-08 | 5.484E-08 | 4.000E-08 | 1.400E-07 |
| 17329 | 867 | D867H | C | 2.2e-08 | 2e-08 | 8.3e-08 | 4.167E-08 | 3.581E-08 | 2.000E-08 | 8.300E-08 |
| 17330 | 867 | D867I | C | 9.6e-08 | 2e-08 | 2.6e-08 | 4.733E-08 | 4.225E-08 | 2.000E-08 | 9.600E-08 |
| 17331 | 867 | D867L | C | 1.1e-07 | 3.3e-08 | 3.8e-08 | 6.033E-08 | 4.309E-08 | 3.300E-08 | 1.100E-07 |
| 17332 | 867 | D867K | C | 2.6e-08 | 1.9e-08 | 1.1e-07 | 5.167E-08 | 5.064E-08 | 1.900E-08 | 1.100E-07 |
| 17333 | 867 | D867M | C | 3e-08 | 9.3e-08 | 2.8e-08 | 5.033E-08 | 3.696E-08 | 2.800E-08 | 9.300E-08 |
| 17334 | 867 | D867F | C | 4.9e-08 | 5.9e-08 | 4.9e-08 | 5.233E-08 | 5.774E-09 | 4.900E-08 | 5.900E-08 |
| 17335 | 867 | D867P | C | 1.1e-07 | 8.9e-09 | 1.1e-08 | 4.330E-08 | 5.777E-08 | 8.900E-09 | 1.100E-07 |
| 17336 | 867 | D867S | C | 5.4e-08 | 1.5e-07 | 3e-08 | 7.800E-08 | 6.350E-08 | 3.000E-08 | 1.500E-07 |
| 17337 | 867 | D867T | C | 1.8e-08 | 8.3e-08 | 5.8e-08 | 5.300E-08 | 3.279E-08 | 1.800E-08 | 8.300E-08 |
| 17338 | 867 | D867W | C | 6.2e-08 | 6.9e-08 | 2.3e-08 | 5.133E-08 | 2.479E-08 | 2.300E-08 | 6.900E-08 |
| 17339 | 867 | D867Y | C | 4.6e-08 | 2.1e-08 | 2.4e-08 | 3.033E-08 | 1.365E-08 | 2.100E-08 | 4.600E-08 |

| | | | | | | | | | | |
|---|---|---|---|---|---|---|---|---|---|---|
| 17340 | 867 | D867V | C | 5.6e-08 | 3.6e-08 | 1.9e-08 | 3.700E-08 | 1.852E-08 | 1.900E-08 | 5.600E-08 |
| 17341 | 868 | Y868A | C | 1.3e-07 | 4.1e-08 | 3.3e-08 | 6.800E-08 | 5.384E-08 | 3.300E-08 | 1.300E-07 |
| 17342 | 868 | Y868R | C | 2.1e-08 | 4.4e-08 | 2.2e-08 | 2.900E-08 | 1.300E-08 | 2.100E-08 | 4.400E-08 |
| 17343 | 868 | Y868N | C | 4.9e-08 | 2.7e-08 | 2.2e-08 | 3.267E-08 | 1.436E-08 | 2.200E-08 | 4.900E-08 |
| 17344 | 868 | Y868D | C | 4.3e-08 | 7.9e-08 | 1e-07 | 7.400E-08 | 2.883E-08 | 4.300E-08 | 1.000E-07 |
| 17345 | 868 | Y868C | C | 9.6e-08 | 1.1e-07 | 9.9e-08 | 1.017E-07 | 7.371E-09 | 9.600E-08 | 1.100E-07 |
| 17346 | 868 | Y868E | C | 4.6e-08 | 3.5e-08 | 4.6e-08 | 4.233E-08 | 6.351E-09 | 3.500E-08 | 4.600E-08 |
| 17347 | 868 | Y868Q | C | 4e-08 | 5.7e-08 | 2.2e-08 | 3.967E-08 | 1.750E-08 | 2.200E-08 | 5.700E-08 |
| 17348 | 868 | Y868G | C | 2.2e-08 | 7.4e-08 | 4.6e-08 | 4.733E-08 | 2.603E-08 | 2.200E-08 | 7.400E-08 |
| 17349 | 868 | Y868H | C | 3e-08 | 2.5e-08 | 4.7e-08 | 3.400E-08 | 1.153E-08 | 2.500E-08 | 4.700E-08 |
| 17350 | 868 | Y868I | C | 3.6e-08 | 6.4e-08 | 1.8e-07 | 9.333E-08 | 7.635E-08 | 3.600E-08 | 1.800E-07 |
| 17351 | 868 | Y868L | C | 4.6e-08 | 9.9e-08 | 7e-08 | 7.167E-08 | 2.654E-08 | 4.600E-08 | 9.900E-08 |
| 17352 | 868 | Y868K | C | 2.7e-08 | 3.4e-08 | 4.7e-08 | 3.600E-08 | 1.015E-08 | 2.700E-08 | 4.700E-08 |
| 17353 | 868 | Y868M | C | 4.6e-08 | 5e-08 | 3.6e-08 | 4.400E-08 | 7.211E-09 | 3.600E-08 | 5.000E-08 |
| 17354 | 868 | Y868F | C | 3.7e-08 | 6.3e-08 | 1.9e-08 | 3.967E-08 | 2.212E-08 | 1.900E-08 | 6.300E-08 |
| 17355 | 868 | Y868P | C | 8.1e-08 | 5.9e-08 | 8.4e-08 | 7.467E-08 | 1.365E-08 | 5.900E-08 | 8.400E-08 |
| 17356 | 868 | Y868S | C | 2e-08 | 2.6e-08 | 4.4e-08 | 3.000E-08 | 1.249E-08 | 2.000E-08 | 4.400E-08 |
| 17357 | 868 | Y868T | C | 7.6e-08 | 4.3e-08 | 4.4e-08 | 5.433E-08 | 1.877E-08 | 4.300E-08 | 7.600E-08 |
| 17358 | 868 | Y868W | C | 2.6e-08 | 3.8e-08 | 3.8e-08 | 3.400E-08 | 6.928E-09 | 2.600E-08 | 3.800E-08 |
| 17359 | 868 | Y868Y | C | 7e-08 | 5.4e-08 | 3.8e-08 | 5.400E-08 | 1.600E-08 | 3.800E-08 | 7.000E-08 |
| 17360 | 868 | Y868V | C | 4.6e-08 | 8.7e-08 | 5e-08 | 6.100E-08 | 2.261E-08 | 4.600E-08 | 8.700E-08 |
| 17361 | 869 | T869A | C | 3.6e-08 | 5.8e-08 | 2.4e-08 | 3.933E-08 | 1.724E-08 | 2.400E-08 | 5.800E-08 |
| 17362 | 869 | T869R | C | 1.9e-08 | 6.4e-08 | 3.4e-08 | 3.900E-08 | 2.291E-08 | 1.900E-08 | 6.400E-08 |
| 17363 | 869 | T869N | C | 5.3e-08 | 5.6e-08 | 1.7e-08 | 4.200E-08 | 2.170E-08 | 1.700E-08 | 5.600E-08 |
| 17364 | 869 | T869D | C | 7.8e-08 | 1.1e-07 | 1.8e-08 | 6.867E-08 | 4.670E-08 | 1.800E-08 | 1.100E-07 |
| 17365 | 869 | T869C | C | 6.8e-08 | 8.3e-08 | 6.2e-08 | 7.100E-08 | 1.082E-08 | 6.200E-08 | 8.300E-08 |
| 17366 | 869 | T869E | C | 7.4e-08 | 4.1e-08 | 4.9e-08 | 5.467E-08 | 1.721E-08 | 4.100E-08 | 7.400E-08 |
| 17367 | 869 | T869Q | C | 2.3e-08 | 4.6e-08 | 3.9e-08 | 3.600E-08 | 1.179E-08 | 2.300E-08 | 4.600E-08 |
| 17368 | 869 | T869G | C | 2.8e-08 | 1.1e-08 | 2.7e-08 | 2.200E-08 | 9.539E-09 | 1.100E-08 | 2.800E-08 |
| 17369 | 869 | T869H | C | 4.7e-08 | 6e-08 | 4.6e-08 | 5.100E-08 | 7.810E-09 | 4.600E-08 | 6.000E-08 |
| 17370 | 869 | T869I | C | 8.6e-08 | 5.5e-08 | 7.9e-08 | 7.333E-08 | 1.626E-08 | 5.500E-08 | 8.600E-08 |
| 17371 | 869 | T869L | C | 2.1e-08 | 4e-08 | 7.3e-08 | 4.467E-08 | 2.631E-08 | 2.100E-08 | 7.300E-08 |
| 17372 | 869 | T869K | C | 6.4e-08 | 6.3e-08 | 8.8e-08 | 7.167E-08 | 1.415E-08 | 6.300E-08 | 8.800E-08 |
| 17373 | 869 | T869M | C | 6.6e-08 | 1.2e-08 | 1.2e-07 | 6.600E-08 | 5.400E-08 | 1.200E-08 | 1.200E-07 |
| 17374 | 869 | T869F | C | 3.6e-08 | 2.5e-08 | 6.9e-08 | 4.333E-08 | 2.290E-08 | 2.500E-08 | 6.900E-08 |
| 17375 | 869 | T869P | C | 5.2e-08 | 7.4e-08 | 2.8e-07 | 1.353E-07 | 1.258E-07 | 5.200E-08 | 2.800E-07 |
| 17376 | 869 | T869S | C | 6.6e-08 | 5.7e-08 | 1e-07 | 7.433E-08 | 2.268E-08 | 5.700E-08 | 1.000E-07 |
| 17377 | 869 | T869T | C | 1.4e-07 | 2.5e-08 | 5.6e-08 | 7.367E-08 | 5.950E-08 | 2.500E-08 | 1.400E-07 |
| 17378 | 869 | T869W | C | 3.3e-08 | 4.6e-08 | 5.3e-08 | 4.400E-08 | 1.015E-08 | 3.300E-08 | 5.300E-08 |
| 17379 | 869 | T869Y | C | 5.8e-08 | 3.1e-08 | 2.9e-08 | 3.933E-08 | 1.620E-08 | 2.900E-08 | 5.800E-08 |
| 17380 | 869 | T869V | C | 1.6e-07 | 1.6e-08 | 2.2e-08 | 6.600E-08 | 8.146E-08 | 1.600E-08 | 1.600E-07 |
| 17381 | 870 | M870A | C | 3.5e-08 | 3.3e-08 | 5e-08 | 3.933E-08 | 9.292E-09 | 3.300E-08 | 5.000E-08 |
| 17382 | 870 | M870R | C | 1.6e-08 | 3.3e-08 | 8.9e-08 | 4.600E-08 | 3.820E-08 | 1.600E-08 | 8.900E-08 |
| 17383 | 870 | M870N | C | 3.9e-08 | 4.6e-08 | 4.6e-08 | 4.367E-08 | 4.041E-09 | 3.900E-08 | 4.600E-08 |
| 17384 | 870 | M870D | C | 2.7e-08 | 5.6e-08 | 4.4e-08 | 4.233E-08 | 1.457E-08 | 2.700E-08 | 5.600E-08 |
| 17385 | 870 | M870C | C | 2.9e-08 | 4.2e-08 | 5.8e-08 | 4.300E-08 | 1.453E-08 | 2.900E-08 | 5.800E-08 |
| 17386 | 870 | M870E | C | 1.1e-07 | 6.2e-08 | 4.1e-08 | 7.100E-08 | 3.537E-08 | 4.100E-08 | 1.100E-07 |
| 17387 | 870 | M870Q | C | 6.5e-08 | 8.9e-08 | 5.5e-08 | 6.967E-08 | 1.747E-08 | 5.500E-08 | 8.900E-08 |
| 17388 | 870 | M870G | C | 7.1e-08 | 4.4e-08 | 5.3e-08 | 5.600E-08 | 1.375E-08 | 4.400E-08 | 7.100E-08 |
| 17389 | 870 | M870H | C | 6e-08 | 5e-08 | 4.5e-08 | 5.167E-08 | 7.638E-09 | 4.500E-08 | 6.000E-08 |
| 17390 | 870 | M870I | C | 5.5e-08 | 1.5e-08 | 3.5e-08 | 3.500E-08 | 2.000E-08 | 1.500E-08 | 5.500E-08 |
| 17391 | 870 | M870L | C | 4.2e-08 | 2.1e-08 | 2.5e-07 | 1.043E-07 | 1.266E-07 | 2.100E-08 | 2.500E-07 |
| 17392 | 870 | M870K | C | 4.2e-08 | 2.4e-08 | 4.5e-08 | 3.700E-08 | 1.136E-08 | 2.400E-08 | 4.500E-08 |
| 17393 | 870 | M870M | C | 1.8e-08 | 6.5e-08 | 7.1e-08 | 5.133E-08 | 2.902E-08 | 1.800E-08 | 7.100E-08 |
| 17394 | 870 | M870F | C | 2.7e-08 | 2.4e-08 | 3.6e-08 | 2.900E-08 | 6.245E-09 | 2.400E-08 | 3.600E-08 |
| 17395 | 870 | M870P | C | 3.9e-08 | 2.5e-08 | 5.5e-08 | 3.967E-08 | 1.501E-08 | 2.500E-08 | 5.500E-08 |
| 17396 | 870 | M870S | C | 3.1e-08 | 2.4e-08 | 2.6e-08 | 2.700E-08 | 3.606E-09 | 2.400E-08 | 3.100E-08 |
| 17397 | 870 | M870T | C | 9.8e-08 | 6.2e-08 | 8.4e-08 | 8.133E-08 | 1.815E-08 | 6.200E-08 | 9.800E-08 |
| 17398 | 870 | M870W | C | 8.4e-08 | 6.3e-08 | 5.9e-08 | 6.867E-08 | 1.343E-08 | 5.900E-08 | 8.400E-08 |
| 17399 | 870 | M870Y | C | 4.3e-08 | 1.1e-07 | 6.7e-08 | 7.333E-08 | 3.395E-08 | 4.300E-08 | 1.100E-07 |
| 17400 | 870 | M870V | C | 4.5e-08 | 1.4e-08 | 1.7e-08 | 2.533E-08 | 1.710E-08 | 1.400E-08 | 4.500E-08 |
| 17401 | 871 | D871A | C | 9e-08 | 2.6e-08 | 3.6e-08 | 5.067E-08 | 3.443E-08 | 2.600E-08 | 9.000E-08 |
| 17402 | 871 | D871R | C | 2.6e-08 | 3.6e-08 | 4.5e-08 | 3.567E-08 | 9.504E-09 | 2.600E-08 | 4.500E-08 |
| 17403 | 871 | D871N | C | 3e-08 | 3.2e-08 | 7.4e-08 | 4.533E-08 | 2.485E-08 | 3.000E-08 | 7.400E-08 |
| 17404 | 871 | D871D | C | 2.2e-08 | 9.9e-08 | 4.6e-08 | 5.567E-08 | 3.940E-08 | 2.200E-08 | 9.900E-08 |
| 17405 | 871 | D871C | C | 1.1e-07 | 4.1e-08 | 7.5e-08 | 7.533E-08 | 3.450E-08 | 4.100E-08 | 1.100E-07 |
| 17406 | 871 | D871E | C | 5.5e-08 | 7.6e-08 | 7e-08 | 6.700E-08 | 1.082E-08 | 5.500E-08 | 7.600E-08 |
| 17407 | 871 | D871Q | C | 2.2e-08 | 4.6e-08 | 2.6e-08 | 3.133E-08 | 1.286E-08 | 2.200E-08 | 4.600E-08 |
| 17408 | 871 | D871G | C | 5.3e-08 | 3.5e-08 | 1.6e-08 | 3.467E-08 | 1.850E-08 | 1.600E-08 | 5.300E-08 |
| 17409 | 871 | D871H | C | 4.5e-08 | 8.3e-08 | 4.8e-08 | 5.867E-08 | 2.113E-08 | 4.500E-08 | 8.300E-08 |
| 17410 | 871 | D871I | C | 6.2e-08 | 3.7e-08 | 4.1e-08 | 4.667E-08 | 1.343E-08 | 3.700E-08 | 6.200E-08 |
| 17411 | 871 | D871L | C | 8.2e-08 | 1.9e-08 | 5e-08 | 5.033E-08 | 3.150E-08 | 1.900E-08 | 8.200E-08 |
| 17412 | 871 | D871K | C | 5e-08 | 2e-08 | 1.4e-07 | 7.000E-08 | 6.245E-08 | 2.000E-08 | 1.400E-07 |
| 17413 | 871 | D871M | C | 5.6e-08 | 1.5e-07 | 1.4e-07 | 1.153E-07 | 5.163E-08 | 5.600E-08 | 1.500E-07 |
| 17414 | 871 | D871F | C | 6.3e-08 | 4.8e-08 | 1.2e-07 | 7.700E-08 | 3.799E-08 | 4.800E-08 | 1.200E-07 |
| 17415 | 871 | D871P | C | 2.1e-08 | 2.2e-08 | 2.1e-08 | 2.133E-08 | 5.774E-10 | 2.100E-08 | 2.200E-08 |
| 17416 | 871 | D871S | C | 1.7e-08 | 2.3e-08 | 1.1e-07 | 5.000E-08 | 5.205E-08 | 1.700E-08 | 1.100E-07 |
| 17417 | 871 | D871T | C | 5.2e-08 | 1.9e-08 | 8.3e-08 | 5.133E-08 | 3.201E-08 | 1.900E-08 | 8.300E-08 |
| 17418 | 871 | D871W | C | 1.1e-08 | 5.5e-08 | 5e-08 | 3.867E-08 | 2.409E-08 | 1.100E-08 | 5.500E-08 |
| 17419 | 871 | D871Y | C | 8.3e-08 | 8.3e-09 | 7e-08 | 5.377E-08 | 3.991E-08 | 8.300E-09 | 8.300E-08 |
| 17420 | 871 | D871V | C | 6e-08 | 4.1e-08 | 2e-08 | 4.033E-08 | 2.001E-08 | 2.000E-08 | 6.000E-08 |
| 17421 | 872 | W872A | C | 2.9e-08 | 7.9e-08 | 2.8e-08 | 4.533E-08 | 2.916E-08 | 2.800E-08 | 7.900E-08 |
| 17422 | 872 | W872R | C | 4.1e-08 | 2.5e-08 | 4.7e-08 | 3.767E-08 | 1.137E-08 | 2.500E-08 | 4.700E-08 |
| 17423 | 872 | W872N | C | 2.4e-08 | 1.9e-08 | 4.2e-08 | 2.833E-08 | 1.210E-08 | 1.900E-08 | 4.200E-08 |
| 17424 | 872 | W872D | C | 2.7e-08 | 2.7e-08 | 4.5e-08 | 3.300E-08 | 1.039E-08 | 2.700E-08 | 4.500E-08 |

| | | | | | | | | | | |
|---|---|---|---|---|---|---|---|---|---|---|
| 17425 | 872 | W872C | C | 6.1e-08 | 2.1e-08 | 1.1e-08 | 3.100E-08 | 2.646E-08 | 1.100E-08 | 6.100E-08 |
| 17426 | 872 | W872E | C | 5.3e-08 | 7.1e-08 | 5.9e-08 | 6.100E-08 | 9.165E-09 | 5.300E-08 | 7.100E-08 |
| 17427 | 872 | W872Q | C | 5.1e-08 | 4.8e-08 | 7.5e-08 | 5.800E-08 | 1.480E-08 | 4.800E-08 | 7.500E-08 |
| 17428 | 872 | W872G | C | 4.2e-08 | 3.2e-08 | 2.6e-08 | 3.333E-08 | 8.083E-09 | 2.600E-08 | 4.200E-08 |
| 17429 | 872 | W872H | C | 3.3e-08 | 5.2e-08 | 4.2e-08 | 4.233E-08 | 9.504E-09 | 3.300E-08 | 5.200E-08 |
| 17430 | 872 | W872I | C | 3.2e-08 | 8.1e-08 | 3.7e-08 | 5.000E-08 | 2.696E-08 | 3.200E-08 | 8.100E-08 |
| 17431 | 872 | W872L | C | 6.2e-08 | 8.7e-08 | 6e-08 | 6.967E-08 | 1.504E-08 | 6.000E-08 | 8.700E-08 |
| 17432 | 872 | W872K | C | 6.6e-08 | 9.7e-08 | 1.8e-08 | 6.033E-08 | 3.980E-08 | 1.800E-08 | 9.700E-08 |
| 17433 | 872 | W872M | C | 1.4e-08 | 1.1e-07 | 5.7e-08 | 6.033E-08 | 4.809E-08 | 1.400E-08 | 1.100E-07 |
| 17434 | 872 | W872F | C | 9.2e-08 | 1.1e-07 | 4.9e-08 | 8.367E-08 | 3.134E-08 | 4.900E-08 | 1.100E-07 |
| 17435 | 872 | W872P | C | 1.1e-07 | 2.8e-08 | 3.9e-08 | 5.900E-08 | 4.451E-08 | 2.800E-08 | 1.100E-07 |
| 17436 | 872 | W872S | C | 3.2e-08 | 6.2e-08 | 2.7e-08 | 4.033E-08 | 1.893E-08 | 2.700E-08 | 6.200E-08 |
| 17437 | 872 | W872T | C | 2.5e-08 | 6.2e-08 | 5.4e-08 | 4.700E-08 | 1.947E-08 | 2.500E-08 | 6.200E-08 |
| 17438 | 872 | W872W | C | 2e-07 | 8.1e-08 | 1.9e-08 | 1.000E-07 | 9.198E-08 | 1.900E-08 | 2.000E-07 |
| 17439 | 872 | W872Y | C | 1.3e-07 | 6.8e-08 | 4.2e-08 | 8.000E-08 | 4.521E-08 | 4.200E-08 | 1.300E-07 |
| 17440 | 872 | W872V | C | 6.4e-08 | 4.1e-08 | 5.2e-08 | 5.233E-08 | 1.150E-08 | 4.100E-08 | 6.400E-08 |
| 17441 | 873 | V873A | C | 6.6e-08 | 6e-08 | 2.6e-08 | 5.067E-08 | 2.157E-08 | 2.600E-08 | 6.600E-08 |
| 17442 | 873 | V873R | C | 4.1e-08 | 2.6e-08 | 9.7e-09 | 2.557E-08 | 1.565E-08 | 9.700E-09 | 4.100E-08 |
| 17443 | 873 | V873N | C | 3.8e-08 | 8.7e-08 | 6e-08 | 6.167E-08 | 2.454E-08 | 3.800E-08 | 8.700E-08 |
| 17444 | 873 | V873D | C | 1.3e-07 | 3.9e-08 | 4.6e-08 | 7.167E-08 | 5.064E-08 | 3.900E-08 | 1.300E-07 |
| 17445 | 873 | V873C | C | 1.7e-08 | 6.4e-08 | 4.4e-08 | 4.167E-08 | 2.359E-08 | 1.700E-08 | 6.400E-08 |
| 17446 | 873 | V873E | C | 4.6e-08 | 2.1e-08 | 1.4e-07 | 6.900E-08 | 6.275E-08 | 2.100E-08 | 1.400E-07 |
| 17447 | 873 | V873Q | C | 4.7e-08 | 6e-08 | 2.9e-08 | 4.533E-08 | 1.557E-08 | 2.900E-08 | 6.000E-08 |
| 17448 | 873 | V873G | C | 3.1e-08 | 3.3e-08 | 2.2e-08 | 2.867E-08 | 5.859E-09 | 2.200E-08 | 3.300E-08 |
| 17449 | 873 | V873H | C | 2.1e-08 | 4.3e-08 | 3.4e-08 | 3.267E-08 | 1.106E-08 | 2.100E-08 | 4.300E-08 |
| 17450 | 873 | V873I | C | 2.2e-08 | 1e-07 | 3.1e-08 | 5.100E-08 | 4.267E-08 | 2.200E-08 | 1.000E-07 |
| 17451 | 873 | V873L | C | 4.4e-08 | 1.1e-07 | 6.8e-08 | 7.400E-08 | 3.341E-08 | 4.400E-08 | 1.100E-07 |
| 17452 | 873 | V873K | C | 8.4e-08 | 8.3e-08 | 5.1e-08 | 7.267E-08 | 1.877E-08 | 5.100E-08 | 8.400E-08 |
| 17453 | 873 | V873M | C | 7.1e-08 | 5.8e-08 | 3.9e-08 | 5.600E-08 | 1.609E-08 | 3.900E-08 | 7.100E-08 |
| 17454 | 873 | V873F | C | 1.2e-08 | 1.2e-08 | 2.7e-08 | 1.700E-08 | 8.660E-09 | 1.200E-08 | 2.700E-08 |
| 17455 | 873 | V873P | C | 6.2e-08 | 8.8e-08 | 9e-08 | 8.000E-08 | 1.562E-08 | 6.200E-08 | 9.000E-08 |
| 17456 | 873 | V873S | C | 6e-08 | 2.6e-08 | 1.8e-08 | 3.467E-08 | 2.230E-08 | 1.800E-08 | 6.000E-08 |
| 17457 | 873 | V873T | C | 5.8e-08 | 2.2e-08 | 1.5e-08 | 3.167E-08 | 2.307E-08 | 1.500E-08 | 5.800E-08 |
| 17458 | 873 | V873W | C | 6.8e-08 | 8.8e-08 | 3.7e-08 | 6.433E-08 | 2.570E-08 | 3.700E-08 | 8.800E-08 |
| 17459 | 873 | V873Y | C | 4.7e-08 | 4.5e-08 | 9.3e-08 | 6.167E-08 | 2.715E-08 | 4.500E-08 | 9.300E-08 |
| 17460 | 873 | V873V | C | 7.8e-08 | 2.5e-08 | 2.9e-08 | 4.400E-08 | 2.951E-08 | 2.500E-08 | 7.800E-08 |
| 17461 | 874 | R874A | C | 1.9e-08 | 3.3e-08 | 6.2e-08 | 3.800E-08 | 2.193E-08 | 1.900E-08 | 6.200E-08 |
| 17462 | 874 | R874R | C | 3.4e-08 | 5.1e-08 | 3.9e-08 | 4.133E-08 | 8.737E-09 | 3.400E-08 | 5.100E-08 |
| 17463 | 874 | R874N | C | 8e-08 | 9.7e-08 | 7e-08 | 8.233E-08 | 1.365E-08 | 7.000E-08 | 9.700E-08 |
| 17464 | 874 | R874D | C | 8e-08 | 3.6e-08 | 3.5e-08 | 5.033E-08 | 2.570E-08 | 3.500E-08 | 8.000E-08 |
| 17465 | 874 | R874C | C | 1.6e-08 | 3.9e-08 | 2.1e-08 | 2.533E-08 | 1.210E-08 | 1.600E-08 | 3.900E-08 |
| 17466 | 874 | R874E | C | 8.6e-08 | 1.9e-07 | 2.7e-08 | 1.010E-07 | 8.253E-08 | 2.700E-08 | 1.900E-07 |
| 17467 | 874 | R874Q | C | 6.5e-08 | 4.7e-08 | 6.7e-08 | 5.967E-08 | 1.102E-08 | 4.700E-08 | 6.700E-08 |
| 17468 | 874 | R874G | C | 4e-08 | 2.9e-08 | 4.1e-08 | 3.667E-08 | 6.658E-09 | 2.900E-08 | 4.100E-08 |
| 17469 | 874 | R874H | C | 4.3e-08 | 4.9e-08 | 4.6e-08 | 4.600E-08 | 3.000E-09 | 4.300E-08 | 4.900E-08 |
| 17470 | 874 | R874I | C | 4.3e-08 | 1.8e-08 | 3.3e-08 | 3.133E-08 | 1.258E-08 | 1.800E-08 | 4.300E-08 |
| 17471 | 874 | R874L | C | 3.6e-08 | 3.8e-08 | 6.8e-08 | 4.733E-08 | 1.793E-08 | 3.600E-08 | 6.800E-08 |
| 17472 | 874 | R874K | C | 6.9e-08 | 5e-08 | 3e-08 | 4.967E-08 | 1.950E-08 | 3.000E-08 | 6.900E-08 |
| 17473 | 874 | R874M | C | 3.1e-08 | 1.4e-08 | 4.8e-08 | 3.100E-08 | 1.700E-08 | 1.400E-08 | 4.800E-08 |
| 17474 | 874 | R874F | C | 2.6e-08 | 9.5e-08 | 4.8e-08 | 5.633E-08 | 3.525E-08 | 2.600E-08 | 9.500E-08 |
| 17475 | 874 | R874P | C | 1.2e-07 | 2.2e-08 | 5.9e-08 | 6.700E-08 | 4.949E-08 | 2.200E-08 | 1.200E-07 |
| 17476 | 874 | R874S | C | 6.9e-08 | 1.7e-08 | 3.4e-08 | 4.000E-08 | 2.651E-08 | 1.700E-08 | 6.900E-08 |
| 17477 | 874 | R874T | C | 6e-08 | 5.9e-08 | 2.9e-08 | 4.933E-08 | 1.762E-08 | 2.900E-08 | 6.000E-08 |
| 17478 | 874 | R874W | C | 2e-08 | 4.8e-08 | 1.7e-08 | 2.833E-08 | 1.710E-08 | 1.700E-08 | 4.800E-08 |
| 17479 | 874 | R874Y | C | 3.7e-08 | 1.4e-07 | 7.9e-08 | 8.533E-08 | 5.179E-08 | 3.700E-08 | 1.400E-07 |
| 17480 | 874 | R874V | C | 1.6e-08 | 5.4e-08 | 3.3e-08 | 3.433E-08 | 1.904E-08 | 1.600E-08 | 5.400E-08 |
| 17481 | 875 | Q875A | C | 3.6e-08 | 2.8e-08 | 1.2e-07 | 6.133E-08 | 5.096E-08 | 2.800E-08 | 1.200E-07 |
| 17482 | 875 | Q875R | C | 6.3e-08 | 6.2e-08 | 4.6e-08 | 5.700E-08 | 9.539E-09 | 4.600E-08 | 6.300E-08 |
| 17483 | 875 | Q875N | C | 3e-08 | 1.5e-08 | 5.9e-08 | 3.467E-08 | 2.237E-08 | 1.500E-08 | 5.900E-08 |
| 17484 | 875 | Q875D | C | 5e-08 | 8.6e-09 | 4.1e-08 | 3.320E-08 | 2.177E-08 | 8.600E-09 | 5.000E-08 |
| 17485 | 875 | Q875C | C | 2.8e-08 | 3.9e-08 | 5.6e-08 | 4.100E-08 | 1.411E-08 | 2.800E-08 | 5.600E-08 |
| 17486 | 875 | Q875E | C | 1.8e-07 | 6.4e-08 | 3.3e-08 | 9.233E-08 | 7.749E-08 | 3.300E-08 | 1.800E-07 |
| 17487 | 875 | Q875Q | C | 6.4e-08 | 2.3e-08 | 5e-08 | 4.567E-08 | 2.084E-08 | 2.300E-08 | 6.400E-08 |
| 17488 | 875 | Q875G | C | 5.8e-08 | 4e-08 | 6.3e-08 | 5.367E-08 | 1.210E-08 | 4.000E-08 | 6.300E-08 |
| 17489 | 875 | Q875H | C | 8.9e-08 | 4.4e-08 | 7.6e-08 | 6.967E-08 | 2.316E-08 | 4.400E-08 | 8.900E-08 |
| 17490 | 875 | Q875I | C | 2.8e-08 | 5.9e-08 | 7.1e-08 | 5.267E-08 | 2.219E-08 | 2.800E-08 | 7.100E-08 |
| 17491 | 875 | Q875L | C | 5.1e-08 | 4.7e-08 | 2.2e-08 | 4.000E-08 | 1.572E-08 | 2.200E-08 | 5.100E-08 |
| 17492 | 875 | Q875K | C | 4.2e-08 | 5.7e-08 | 4.7e-08 | 4.867E-08 | 7.638E-09 | 4.200E-08 | 5.700E-08 |
| 17493 | 875 | Q875M | C | 2.4e-08 | 5.1e-08 | 5.8e-08 | 4.433E-08 | 1.795E-08 | 2.400E-08 | 5.800E-08 |
| 17494 | 875 | Q875F | C | 5.9e-08 | 6.1e-08 | 5.7e-08 | 5.900E-08 | 2.000E-09 | 5.700E-08 | 6.100E-08 |
| 17495 | 875 | Q875P | C | 5.4e-08 | 1.5e-08 | 2.3e-08 | 3.067E-08 | 2.060E-08 | 1.500E-08 | 5.400E-08 |
| 17496 | 875 | Q875S | C | 4.2e-08 | 1.8e-08 | 2.9e-07 | 1.167E-07 | 1.506E-07 | 1.800E-08 | 2.900E-07 |
| 17497 | 875 | Q875T | C | 3.4e-08 | 2.5e-08 | 3.2e-08 | 3.033E-08 | 4.726E-09 | 2.500E-08 | 3.400E-08 |
| 17498 | 875 | Q875W | C | 5.3e-08 | 3.1e-08 | 4.6e-08 | 4.333E-08 | 1.124E-08 | 3.100E-08 | 5.300E-08 |
| 17499 | 875 | Q875Y | C | 5.2e-08 | 1.5e-08 | 7.7e-08 | 4.800E-08 | 3.119E-08 | 1.500E-08 | 7.700E-08 |
| 17500 | 875 | Q875V | C | 3.3e-08 | 7.9e-08 | 2.7e-08 | 4.633E-08 | 2.845E-08 | 2.700E-08 | 7.900E-08 |
| 17501 | 876 | A876A | C | 3.6e-08 | 2e-08 | 7.1e-08 | 4.233E-08 | 2.608E-08 | 2.000E-08 | 7.100E-08 |
| 17502 | 876 | A876R | C | 5.7e-08 | 2.4e-08 | 7.6e-08 | 5.233E-08 | 2.631E-08 | 2.400E-08 | 7.600E-08 |
| 17503 | 876 | A876N | C | 1e-08 | 5.5e-08 | 2.2e-08 | 2.900E-08 | 2.330E-08 | 1.000E-08 | 5.500E-08 |
| 17504 | 876 | A876D | C | 1.5e-08 | 8.2e-08 | 1.3e-08 | 3.667E-08 | 3.927E-08 | 1.300E-08 | 8.200E-08 |
| 17505 | 876 | A876C | C | 5.1e-08 | 2.9e-08 | 2e-08 | 3.333E-08 | 1.595E-08 | 2.000E-08 | 5.100E-08 |
| 17506 | 876 | A876E | C | 3.8e-08 | 1.9e-08 | 2.1e-08 | 2.600E-08 | 1.044E-08 | 1.900E-08 | 3.800E-08 |
| 17507 | 876 | A876Q | C | 2.1e-08 | 3.4e-08 | 7.7e-08 | 4.400E-08 | 2.931E-08 | 2.100E-08 | 7.700E-08 |
| 17508 | 876 | A876G | C | 7.1e-08 | 6.2e-08 | 2.8e-08 | 5.367E-08 | 2.268E-08 | 2.800E-08 | 7.100E-08 |
| 17509 | 876 | A876H | C | 1.3e-07 | 8.7e-08 | 5e-08 | 8.900E-08 | 4.004E-08 | 5.000E-08 | 1.300E-07 |

| | | | | | | | | | | |
|---|---|---|---|---|---|---|---|---|---|---|
| 17510 | 876 | A876I | C | 7.3e-08 | 4.6e-08 | 4.2e-08 | 5.367E-08 | 1.686E-08 | 4.200E-08 | 7.300E-08 |
| 17511 | 876 | A876L | C | 1.8e-08 | 3.8e-08 | 3.1e-08 | 2.900E-08 | 1.015E-08 | 1.800E-08 | 3.800E-08 |
| 17512 | 876 | A876K | C | 3.1e-08 | 5.6e-08 | 4.5e-08 | 4.400E-08 | 1.253E-08 | 3.100E-08 | 5.600E-08 |
| 17513 | 876 | A876M | C | 5.7e-08 | 1.5e-08 | 1e-07 | 5.733E-08 | 4.250E-08 | 1.500E-08 | 1.000E-07 |
| 17514 | 876 | A876F | C | 7.9e-08 | 4e-08 | 3e-08 | 4.967E-08 | 2.589E-08 | 3.000E-08 | 7.900E-08 |
| 17515 | 876 | A876P | C | 4.4e-08 | 2.1e-08 | 1.8e-08 | 2.767E-08 | 1.422E-08 | 1.800E-08 | 4.400E-08 |
| 17516 | 876 | A876S | C | 4e-08 | 8e-08 | 3.1e-08 | 5.033E-08 | 2.608E-08 | 3.100E-08 | 8.000E-08 |
| 17517 | 876 | A876T | C | 1.2e-07 | 6.2e-08 | 5.8e-08 | 8.000E-08 | 3.470E-08 | 5.800E-08 | 1.200E-07 |
| 17518 | 876 | A876W | C | 2e-08 | 5.7e-08 | 5.2e-08 | 4.300E-08 | 2.007E-08 | 2.000E-08 | 5.700E-08 |
| 17519 | 876 | A876Y | C | 4.1e-08 | 6.8e-08 | 5.1e-08 | 5.333E-08 | 1.365E-08 | 4.100E-08 | 6.800E-08 |
| 17520 | 876 | A876V | C | 6.2e-08 | 7.5e-08 | 7.1e-08 | 6.933E-08 | 6.658E-09 | 6.200E-08 | 7.500E-08 |
| 17521 | 877 | P877A | C | 5.2e-08 | 1.2e-07 | 8.9e-08 | 8.700E-08 | 3.404E-08 | 5.200E-08 | 1.200E-07 |
| 17522 | 877 | P877R | C | 4.6e-08 | 3e-08 | 7e-08 | 4.867E-08 | 2.013E-08 | 3.000E-08 | 7.000E-08 |
| 17523 | 877 | P877N | C | 5.7e-08 | 1.7e-07 | 2.5e-08 | 8.400E-08 | 7.618E-08 | 2.500E-08 | 1.700E-07 |
| 17524 | 877 | P877D | C | 2.4e-08 | 6.3e-08 | 3.9e-08 | 4.200E-08 | 1.967E-08 | 2.400E-08 | 6.300E-08 |
| 17525 | 877 | P877C | C | 8.4e-08 | 9.2e-09 | 2.8e-08 | 4.040E-08 | 3.891E-08 | 9.200E-09 | 8.400E-08 |
| 17526 | 877 | P877E | C | 2.7e-08 | 1.9e-08 | 5.9e-08 | 3.500E-08 | 2.117E-08 | 1.900E-08 | 5.900E-08 |
| 17527 | 877 | P877Q | C | 2.5e-08 | 3.1e-08 | 6e-08 | 3.867E-08 | 1.872E-08 | 2.500E-08 | 6.000E-08 |
| 17528 | 877 | P877G | C | 8.1e-08 | 5.5e-08 | 4.5e-08 | 6.033E-08 | 1.858E-08 | 4.500E-08 | 8.100E-08 |
| 17529 | 877 | P877H | C | 6.4e-08 | 4.8e-08 | 2.5e-08 | 4.567E-08 | 1.960E-08 | 2.500E-08 | 6.400E-08 |
| 17530 | 877 | P877I | C | 3.8e-08 | 2e-08 | 2e-08 | 2.600E-08 | 1.039E-08 | 2.000E-08 | 3.800E-08 |
| 17531 | 877 | P877L | C | 1.5e-07 | 9.5e-08 | 2.8e-08 | 9.100E-08 | 6.110E-08 | 2.800E-08 | 1.500E-07 |
| 17532 | 877 | P877K | C | 2.1e-08 | 4.6e-08 | 2.8e-08 | 3.167E-08 | 1.290E-08 | 2.100E-08 | 4.600E-08 |
| 17533 | 877 | P877M | C | 6.6e-08 | 1.1e-07 | 7.5e-08 | 8.367E-08 | 2.325E-08 | 6.600E-08 | 1.100E-07 |
| 17534 | 877 | P877F | C | 3.2e-08 | 7.1e-08 | 4e-08 | 4.767E-08 | 2.060E-08 | 3.200E-08 | 7.100E-08 |
| 17535 | 877 | P877P | C | 8.9e-08 | 6.2e-08 | 7.1e-08 | 7.400E-08 | 1.375E-08 | 6.200E-08 | 8.900E-08 |
| 17536 | 877 | P877S | C | 4.8e-08 | 6.5e-08 | 3.1e-08 | 4.800E-08 | 1.700E-08 | 3.100E-08 | 6.500E-08 |
| 17537 | 877 | P877T | C | 5.7e-08 | 4.6e-08 | 3.9e-08 | 4.733E-08 | 9.074E-09 | 3.900E-08 | 5.700E-08 |
| 17538 | 877 | P877W | C | 2e-08 | 1.2e-08 | 4.2e-08 | 2.467E-08 | 1.553E-08 | 1.200E-08 | 4.200E-08 |
| 17539 | 877 | P877Y | C | 6.4e-08 | 1.2e-07 | 1.1e-07 | 9.800E-08 | 2.987E-08 | 6.400E-08 | 1.200E-07 |
| 17540 | 877 | P877V | C | 5e-08 | 4.3e-08 | 4.1e-08 | 4.467E-08 | 4.726E-09 | 4.100E-08 | 5.000E-08 |
| 17541 | 878 | G878A | C | 6.6e-08 | 5.2e-08 | 2.9e-08 | 4.900E-08 | 1.868E-08 | 2.900E-08 | 6.600E-08 |
| 17542 | 878 | G878R | C | 5e-08 | 5.3e-08 | 2.6e-08 | 4.300E-08 | 1.480E-08 | 2.600E-08 | 5.300E-08 |
| 17543 | 878 | G878N | C | 5.4e-08 | 3.1e-08 | 5.8e-08 | 4.767E-08 | 1.457E-08 | 3.100E-08 | 5.800E-08 |
| 17544 | 878 | G878D | C | 1.8e-08 | 6.7e-08 | 2e-08 | 3.500E-08 | 2.773E-08 | 1.800E-08 | 6.700E-08 |
| 17545 | 878 | G878C | C | 3.5e-08 | 5.4e-08 | 1e-07 | 6.300E-08 | 3.342E-08 | 3.500E-08 | 1.000E-07 |
| 17546 | 878 | G878E | C | 2.3e-08 | 9.2e-08 | 8.7e-08 | 6.733E-08 | 3.848E-08 | 2.300E-08 | 9.200E-08 |
| 17547 | 878 | G878Q | C | 7.3e-08 | 6.4e-08 | 5.9e-08 | 6.533E-08 | 7.095E-09 | 5.900E-08 | 7.300E-08 |
| 17548 | 878 | G878G | C | 4.2e-08 | 5.8e-08 | 6.6e-08 | 5.533E-08 | 1.222E-08 | 4.200E-08 | 6.600E-08 |
| 17549 | 878 | G878H | C | 2.8e-08 | 4.3e-08 | 5.8e-08 | 4.300E-08 | 1.500E-08 | 2.800E-08 | 5.800E-08 |
| 17550 | 878 | G878I | C | 2.2e-08 | 2.8e-08 | 3e-08 | 2.667E-08 | 4.163E-09 | 2.200E-08 | 3.000E-08 |
| 17551 | 878 | G878L | C | 4.6e-08 | 5.3e-08 | 8.1e-08 | 6.000E-08 | 1.852E-08 | 4.600E-08 | 8.100E-08 |
| 17552 | 878 | G878K | C | 5.6e-08 | 1.1e-07 | 3e-08 | 6.533E-08 | 4.081E-08 | 3.000E-08 | 1.100E-07 |
| 17553 | 878 | G878M | C | 2e-08 | 5.6e-08 | 8.9e-08 | 5.500E-08 | 3.451E-08 | 2.000E-08 | 8.900E-08 |
| 17554 | 878 | G878F | C | 1.8e-08 | 2.1e-08 | 5.9e-08 | 3.267E-08 | 2.285E-08 | 1.800E-08 | 5.900E-08 |
| 17555 | 878 | G878P | C | 5.5e-08 | 8.1e-08 | 8.6e-08 | 7.400E-08 | 1.664E-08 | 5.500E-08 | 8.600E-08 |
| 17556 | 878 | G878S | C | 3.6e-08 | 1.3e-08 | 3.2e-08 | 2.700E-08 | 1.229E-08 | 1.300E-08 | 3.600E-08 |
| 17557 | 878 | G878T | C | 7.1e-08 | 7.4e-08 | 8.8e-08 | 7.767E-08 | 9.074E-09 | 7.100E-08 | 8.800E-08 |
| 17558 | 878 | G878W | C | 1.9e-08 | 1.2e-07 | 4.2e-08 | 6.033E-08 | 5.294E-08 | 1.900E-08 | 1.200E-07 |
| 17559 | 878 | G878Y | C | 4.7e-08 | 3.9e-08 | 5.8e-08 | 4.800E-08 | 9.539E-09 | 3.900E-08 | 5.800E-08 |
| 17560 | 878 | G878V | C | 7.3e-08 | 2.2e-08 | 5.2e-08 | 4.900E-08 | 2.563E-08 | 2.200E-08 | 7.300E-08 |
| 17561 | 879 | K879A | C | 3.5e-08 | 2.6e-08 | 3.2e-08 | 3.100E-08 | 4.583E-09 | 2.600E-08 | 3.500E-08 |
| 17562 | 879 | K879R | C | 1.7e-08 | 1.4e-07 | 7.4e-08 | 7.700E-08 | 6.155E-08 | 1.700E-08 | 1.400E-07 |
| 17563 | 879 | K879N | C | 1.8e-08 | 7.6e-08 | 3e-08 | 4.133E-08 | 3.062E-08 | 1.800E-08 | 7.600E-08 |
| 17564 | 879 | K879D | C | 1.8e-08 | 5.5e-08 | 4.1e-08 | 3.800E-08 | 1.868E-08 | 1.800E-08 | 5.500E-08 |
| 17565 | 879 | K879C | C | 3e-07 | 2.9e-08 | 1e-07 | 1.430E-07 | 1.405E-07 | 2.900E-08 | 3.000E-07 |
| 17566 | 879 | K879E | C | 3.9e-08 | 1.6e-08 | 4.2e-08 | 3.233E-08 | 1.422E-08 | 1.600E-08 | 4.200E-08 |
| 17567 | 879 | K879Q | C | 3.4e-08 | 7.7e-08 | 3e-08 | 4.700E-08 | 2.606E-08 | 3.000E-08 | 7.700E-08 |
| 17568 | 879 | K879G | C | 1.1e-07 | 3.3e-08 | 2.9e-08 | 5.733E-08 | 4.565E-08 | 2.900E-08 | 1.100E-07 |
| 17569 | 879 | K879H | C | 4.2e-08 | 5.9e-08 | 3.5e-08 | 4.533E-08 | 1.234E-08 | 3.500E-08 | 5.900E-08 |
| 17570 | 879 | K879I | C | 3e-08 | 4e-08 | 2.2e-07 | 9.667E-08 | 1.069E-07 | 3.000E-08 | 2.200E-07 |
| 17571 | 879 | K879L | C | 8.9e-09 | 8.9e-08 | 3e-08 | 4.263E-08 | 4.152E-08 | 8.900E-09 | 8.900E-08 |
| 17572 | 879 | K879K | C | 7.4e-08 | 3.4e-08 | 4.2e-08 | 5.000E-08 | 2.117E-08 | 3.400E-08 | 7.400E-08 |
| 17573 | 879 | K879M | C | 2.4e-08 | 7.6e-08 | 1.1e-08 | 3.700E-08 | 3.439E-08 | 1.100E-08 | 7.600E-08 |
| 17574 | 879 | K879F | C | 3.7e-08 | 2.4e-08 | 3.8e-08 | 3.300E-08 | 7.810E-09 | 2.400E-08 | 3.800E-08 |
| 17575 | 879 | K879P | C | 2.6e-08 | 2.1e-08 | 1.6e-08 | 2.100E-08 | 5.000E-09 | 1.600E-08 | 2.600E-08 |
| 17576 | 879 | K879S | C | 3.1e-08 | 7.6e-08 | 2.2e-08 | 4.300E-08 | 2.893E-08 | 2.200E-08 | 7.600E-08 |
| 17577 | 879 | K879T | C | 5.6e-08 | 1.6e-08 | 5.2e-08 | 4.133E-08 | 2.203E-08 | 1.600E-08 | 5.600E-08 |
| 17578 | 879 | K879W | C | 4e-08 | 4.3e-08 | 3.7e-08 | 4.000E-08 | 3.000E-09 | 3.700E-08 | 4.300E-08 |
| 17579 | 879 | K879Y | C | 4.2e-08 | 6.7e-08 | 1.1e-08 | 4.000E-08 | 2.805E-08 | 1.100E-08 | 6.700E-08 |
| 17580 | 879 | K879V | C | 6.6e-08 | 3e-08 | 3.2e-08 | 4.267E-08 | 2.023E-08 | 3.000E-08 | 6.600E-08 |
| 17581 | 880 | G880A | C | 2.9e-08 | 2.6e-08 | 9.3e-08 | 4.933E-08 | 3.785E-08 | 2.600E-08 | 9.300E-08 |
| 17582 | 880 | G880R | C | 2.7e-08 | 4e-08 | 2.6e-08 | 3.100E-08 | 7.810E-09 | 2.600E-08 | 4.000E-08 |
| 17583 | 880 | G880N | C | 4.5e-08 | 6.3e-08 | 2.7e-08 | 4.500E-08 | 1.800E-08 | 2.700E-08 | 6.300E-08 |
| 17584 | 880 | G880D | C | 2.9e-08 | 3.6e-08 | 3.9e-08 | 3.467E-08 | 5.132E-09 | 2.900E-08 | 3.900E-08 |
| 17585 | 880 | G880C | C | 7.5e-08 | 6e-08 | 3.5e-08 | 5.667E-08 | 2.021E-08 | 3.500E-08 | 7.500E-08 |
| 17586 | 880 | G880E | C | 3.5e-08 | 7e-08 | 1.1e-07 | 7.167E-08 | 3.753E-08 | 3.500E-08 | 1.100E-07 |
| 17587 | 880 | G880Q | C | 1.3e-07 | 5.8e-08 | 4.4e-08 | 7.733E-08 | 4.614E-08 | 4.400E-08 | 1.300E-07 |
| 17588 | 880 | G880G | C | 2.9e-08 | 5.3e-08 | 5.8e-08 | 4.667E-08 | 1.550E-08 | 2.900E-08 | 5.800E-08 |
| 17589 | 880 | G880H | C | 6.3e-08 | 2.3e-08 | 1.6e-08 | 3.400E-08 | 2.536E-08 | 1.600E-08 | 6.300E-08 |
| 17590 | 880 | G880I | C | 4e-08 | 7e-08 | 5.6e-08 | 5.533E-08 | 1.501E-08 | 4.000E-08 | 7.000E-08 |
| 17591 | 880 | G880L | C | 3.9e-08 | 1.7e-08 | 8.6e-09 | 2.153E-08 | 1.570E-08 | 8.600E-09 | 3.900E-08 |
| 17592 | 880 | G880K | C | 4.5e-08 | 2.8e-08 | 8.2e-08 | 5.167E-08 | 2.761E-08 | 2.800E-08 | 8.200E-08 |
| 17593 | 880 | G880M | C | 1e-08 | 2.3e-08 | 5.1e-08 | 2.800E-08 | 2.095E-08 | 1.000E-08 | 5.100E-08 |
| 17594 | 880 | G880F | C | 6.6e-08 | 3.2e-08 | 3.1e-08 | 4.300E-08 | 1.992E-08 | 3.100E-08 | 6.600E-08 |

| | | | | | | | | | | |
|---|---|---|---|---|---|---|---|---|---|---|
| 17595 | 880 | G880P | C | 4e-08 | 5e-08 | 4.7e-08 | 4.567E-08 | 5.132E-09 | 4.000E-08 | 5.000E-08 |
| 17596 | 880 | G880S | C | 5.3e-08 | 4.7e-08 | 2.5e-08 | 4.167E-08 | 1.474E-08 | 2.500E-08 | 5.300E-08 |
| 17597 | 880 | G880T | C | 5.2e-08 | 3e-08 | 7.4e-08 | 5.200E-08 | 2.200E-08 | 3.000E-08 | 7.400E-08 |
| 17598 | 880 | G880W | C | 6.7e-08 | 1.5e-08 | 1.9e-08 | 3.367E-08 | 2.894E-08 | 1.500E-08 | 6.700E-08 |
| 17599 | 880 | G880Y | C | 5.1e-08 | 2.2e-07 | 4.6e-08 | 1.057E-07 | 9.905E-08 | 4.600E-08 | 2.200E-07 |
| 17600 | 880 | G880V | C | 4.4e-08 | 3.3e-08 | 2.7e-08 | 3.467E-08 | 8.622E-09 | 2.700E-08 | 4.400E-08 |
| 17601 | 881 | L881A | C | 3.5e-08 | 9.1e-08 | 4.6e-08 | 5.733E-08 | 2.967E-08 | 3.500E-08 | 9.100E-08 |
| 17602 | 881 | L881R | C | 4.4e-08 | 3e-08 | 8.6e-08 | 5.333E-08 | 2.914E-08 | 3.000E-08 | 8.600E-08 |
| 17603 | 881 | L881N | C | 7.4e-08 | 3.5e-08 | 1.2e-07 | 7.633E-08 | 4.255E-08 | 3.500E-08 | 1.200E-07 |
| 17604 | 881 | L881D | C | 9.2e-08 | 2.3e-08 | 7.7e-08 | 6.400E-08 | 3.629E-08 | 2.300E-08 | 9.200E-08 |
| 17605 | 881 | L881C | C | 7.5e-08 | 8.6e-08 | 5.9e-08 | 7.333E-08 | 1.358E-08 | 5.900E-08 | 8.600E-08 |
| 17606 | 881 | L881E | C | 3.7e-08 | 5.5e-08 | 4.1e-08 | 4.433E-08 | 9.452E-09 | 3.700E-08 | 5.500E-08 |
| 17607 | 881 | L881Q | C | 5.1e-08 | 1.3e-07 | 3.5e-08 | 7.200E-08 | 5.086E-08 | 3.500E-08 | 1.300E-07 |
| 17608 | 881 | L881G | C | 3.7e-08 | 4.8e-08 | 5.2e-08 | 4.567E-08 | 7.767E-09 | 3.700E-08 | 5.200E-08 |
| 17609 | 881 | L881H | C | 1.8e-08 | 9.5e-08 | 1.2e-07 | 7.767E-08 | 5.316E-08 | 1.800E-08 | 1.200E-07 |
| 17610 | 881 | L881I | C | 1.4e-07 | 3.5e-08 | 4.9e-08 | 7.467E-08 | 5.701E-08 | 3.500E-08 | 1.400E-07 |
| 17611 | 881 | L881L | C | 4.5e-08 | 9.6e-08 | 2.4e-08 | 5.500E-08 | 3.703E-08 | 2.400E-08 | 9.600E-08 |
| 17612 | 881 | L881K | C | 1.4e-08 | 2.9e-08 | 3.7e-08 | 2.667E-08 | 1.168E-08 | 1.400E-08 | 3.700E-08 |
| 17613 | 881 | L881M | C | 1.3e-08 | 5.2e-08 | 7.8e-08 | 4.767E-08 | 3.272E-08 | 1.300E-08 | 7.800E-08 |
| 17614 | 881 | L881F | C | 7e-08 | 4e-08 | 6.2e-08 | 5.733E-08 | 1.553E-08 | 4.000E-08 | 7.000E-08 |
| 17615 | 881 | L881P | C | 5.6e-08 | 6.7e-08 | 3.3e-08 | 5.200E-08 | 1.735E-08 | 3.300E-08 | 6.700E-08 |
| 17616 | 881 | L881S | C | 1.2e-07 | 1.8e-08 | 4e-08 | 5.933E-08 | 5.368E-08 | 1.800E-08 | 1.200E-07 |
| 17617 | 881 | L881T | C | 3.1e-08 | 4.2e-08 | 1e-07 | 5.767E-08 | 3.707E-08 | 3.100E-08 | 1.000E-07 |
| 17618 | 881 | L881W | C | 1.4e-07 | 2.4e-08 | 3.6e-08 | 6.667E-08 | 6.379E-08 | 2.400E-08 | 1.400E-07 |
| 17619 | 881 | L881Y | C | 7.2e-08 | 8.8e-08 | 3.4e-08 | 6.467E-08 | 2.774E-08 | 3.400E-08 | 8.800E-08 |
| 17620 | 881 | L881V | C | 2.7e-08 | 3.7e-08 | 2.2e-08 | 2.867E-08 | 7.638E-09 | 2.200E-08 | 3.700E-08 |
| 17621 | 882 | E882A | C | 1e-07 | 4.6e-08 | 1.9e-08 | 5.500E-08 | 4.124E-08 | 1.900E-08 | 1.000E-07 |
| 17622 | 882 | E882R | C | 4.4e-08 | 2e-07 | 9.9e-08 | 1.143E-07 | 7.912E-08 | 4.400E-08 | 2.000E-07 |
| 17623 | 882 | E882N | C | 7.1e-08 | 2.9e-08 | 6.3e-08 | 5.433E-08 | 2.230E-08 | 2.900E-08 | 7.100E-08 |
| 17624 | 882 | E882D | C | 4.3e-08 | 5.7e-08 | 5e-08 | 5.000E-08 | 7.000E-09 | 4.300E-08 | 5.700E-08 |
| 17625 | 882 | E882C | C | 6.7e-08 | 2.5e-08 | 8.8e-08 | 6.000E-08 | 3.208E-08 | 2.500E-08 | 8.800E-08 |
| 17626 | 882 | E882E | C | 7.3e-08 | 2.2e-08 | 1.8e-08 | 3.767E-08 | 3.066E-08 | 1.800E-08 | 7.300E-08 |
| 17627 | 882 | E882Q | C | 4e-08 | 4.1e-08 | 4.9e-08 | 4.333E-08 | 4.933E-09 | 4.000E-08 | 4.900E-08 |
| 17628 | 882 | E882G | C | 5.4e-08 | 5.3e-08 | 3.9e-08 | 4.867E-08 | 8.386E-09 | 3.900E-08 | 5.400E-08 |
| 17629 | 882 | E882H | C | 6.1e-08 | 2.6e-08 | 3.5e-08 | 4.067E-08 | 1.818E-08 | 2.600E-08 | 6.100E-08 |
| 17630 | 882 | E882I | C | 9e-08 | 4.3e-08 | 6e-08 | 6.433E-08 | 2.380E-08 | 4.300E-08 | 9.000E-08 |
| 17631 | 882 | E882L | C | 4.6e-08 | 1.9e-08 | 2.5e-08 | 3.000E-08 | 1.418E-08 | 1.900E-08 | 4.600E-08 |
| 17632 | 882 | E882K | C | 4e-08 | 2.6e-08 | 5.2e-08 | 3.933E-08 | 1.301E-08 | 2.600E-08 | 5.200E-08 |
| 17633 | 882 | E882M | C | 1.8e-08 | 9e-08 | 9.1e-08 | 6.633E-08 | 4.186E-08 | 1.800E-08 | 9.100E-08 |
| 17634 | 882 | E882F | C | 1.5e-07 | 2.4e-08 | 1.4e-08 | 6.267E-08 | 7.580E-08 | 1.400E-08 | 1.500E-07 |
| 17635 | 882 | E882P | C | 4.5e-08 | 3.1e-08 | 5.5e-08 | 4.367E-08 | 1.206E-08 | 3.100E-08 | 5.500E-08 |
| 17636 | 882 | E882S | C | 3.1e-08 | 4.4e-08 | 7e-08 | 4.833E-08 | 1.986E-08 | 3.100E-08 | 7.000E-08 |
| 17637 | 882 | E882T | C | 3.7e-08 | 3.6e-08 | 7.1e-08 | 4.800E-08 | 1.992E-08 | 3.600E-08 | 7.100E-08 |
| 17638 | 882 | E882W | C | 7.3e-08 | 4.2e-08 | 1.5e-07 | 8.833E-08 | 5.561E-08 | 4.200E-08 | 1.500E-07 |
| 17639 | 882 | E882Y | C | 5e-08 | 8.4e-09 | 8.4e-08 | 4.747E-08 | 3.786E-08 | 8.400E-09 | 8.400E-08 |
| 17640 | 882 | E882V | C | 5e-08 | 4.1e-08 | 2.5e-08 | 3.867E-08 | 1.266E-08 | 2.500E-08 | 5.000E-08 |
| 17641 | 883 | W883A | C | 3.8e-08 | 5.8e-08 | 5.3e-08 | 4.967E-08 | 1.041E-08 | 3.800E-08 | 5.800E-08 |
| 17642 | 883 | W883R | C | 1.1e-08 | 3.7e-08 | 6.1e-08 | 3.633E-08 | 2.501E-08 | 1.100E-08 | 6.100E-08 |
| 17643 | 883 | W883N | C | 2.3e-08 | 3.9e-08 | 2.4e-08 | 2.867E-08 | 8.963E-09 | 2.300E-08 | 3.900E-08 |
| 17644 | 883 | W883D | C | 8.3e-08 | 3.2e-08 | 1.9e-08 | 4.467E-08 | 3.383E-08 | 1.900E-08 | 8.300E-08 |
| 17645 | 883 | W883C | C | 3.8e-08 | 4.3e-08 | 4e-08 | 4.033E-08 | 2.517E-09 | 3.800E-08 | 4.300E-08 |
| 17646 | 883 | W883E | C | 5.1e-08 | 1.9e-08 | 6.4e-08 | 4.467E-08 | 2.316E-08 | 1.900E-08 | 6.400E-08 |
| 17647 | 883 | W883Q | C | 6.2e-08 | 4.3e-08 | 3.3e-08 | 4.600E-08 | 1.473E-08 | 3.300E-08 | 6.200E-08 |
| 17648 | 883 | W883G | C | 7.1e-08 | 6.1e-08 | 4.5e-08 | 5.900E-08 | 1.311E-08 | 4.500E-08 | 7.100E-08 |
| 17649 | 883 | W883H | C | 6.8e-08 | 1.6e-08 | 3e-08 | 3.800E-08 | 2.691E-08 | 1.600E-08 | 6.800E-08 |
| 17650 | 883 | W883I | C | 4.5e-08 | 8.6e-08 | 1.5e-08 | 4.867E-08 | 3.564E-08 | 1.500E-08 | 8.600E-08 |
| 17651 | 883 | W883L | C | 2.1e-08 | 9.5e-08 | 8.3e-08 | 6.633E-08 | 3.972E-08 | 2.100E-08 | 9.500E-08 |
| 17652 | 883 | W883K | C | 1.7e-08 | 3.3e-08 | 6.6e-08 | 3.867E-08 | 2.499E-08 | 1.700E-08 | 6.600E-08 |
| 17653 | 883 | W883M | C | 3.5e-08 | 5.6e-08 | 2.7e-08 | 3.933E-08 | 1.498E-08 | 2.700E-08 | 5.600E-08 |
| 17654 | 883 | W883F | C | 2e-08 | 4.7e-08 | 8.6e-08 | 5.100E-08 | 3.318E-08 | 2.000E-08 | 8.600E-08 |
| 17655 | 883 | W883P | C | 3.7e-08 | 1.3e-08 | 6.6e-08 | 3.867E-08 | 2.654E-08 | 1.300E-08 | 6.600E-08 |
| 17656 | 883 | W883S | C | 5.7e-08 | 3.2e-08 | 2.2e-08 | 3.700E-08 | 1.803E-08 | 2.200E-08 | 5.700E-08 |
| 17657 | 883 | W883T | C | 2.8e-08 | 2.4e-08 | 2.6e-08 | 2.600E-08 | 2.000E-09 | 2.400E-08 | 2.800E-08 |
| 17658 | 883 | W883W | C | 8e-08 | 3.6e-08 | 4.2e-08 | 5.267E-08 | 2.386E-08 | 3.600E-08 | 8.000E-08 |
| 17659 | 883 | W883Y | C | 7.3e-08 | 2.4e-08 | 4.5e-08 | 4.733E-08 | 2.458E-08 | 2.400E-08 | 7.300E-08 |
| 17660 | 883 | W883V | C | 4.8e-08 | 6.3e-08 | 1.7e-08 | 4.267E-08 | 2.346E-08 | 1.700E-08 | 6.300E-08 |
| 17661 | 884 | V884A | C | 2.8e-08 | 6.5e-08 | 3.1e-08 | 4.133E-08 | 2.055E-08 | 2.800E-08 | 6.500E-08 |
| 17662 | 884 | V884R | C | 6.4e-08 | 6.4e-08 | 6e-08 | 6.267E-08 | 2.309E-09 | 6.000E-08 | 6.400E-08 |
| 17663 | 884 | V884N | C | 6.8e-08 | 3e-08 | 4.5e-08 | 4.767E-08 | 1.914E-08 | 3.000E-08 | 6.800E-08 |
| 17664 | 884 | V884D | C | 5.9e-08 | 2.8e-08 | 1.2e-08 | 3.300E-08 | 2.390E-08 | 1.200E-08 | 5.900E-08 |
| 17665 | 884 | V884C | C | 7.3e-08 | 5.8e-08 | 9e-08 | 7.367E-08 | 1.601E-08 | 5.800E-08 | 9.000E-08 |
| 17666 | 884 | V884E | C | 9e-08 | 2.5e-08 | 2.4e-08 | 4.633E-08 | 3.782E-08 | 2.400E-08 | 9.000E-08 |
| 17667 | 884 | V884Q | C | 3.4e-08 | 8.4e-08 | 4.2e-08 | 5.333E-08 | 2.686E-08 | 3.400E-08 | 8.400E-08 |
| 17668 | 884 | V884G | C | 4.9e-08 | 4.5e-08 | 4.4e-08 | 4.600E-08 | 2.646E-09 | 4.400E-08 | 4.900E-08 |
| 17669 | 884 | V884H | C | 8.4e-08 | 2.5e-08 | 4.9e-08 | 5.267E-08 | 2.967E-08 | 2.500E-08 | 8.400E-08 |
| 17670 | 884 | V884I | C | 3.7e-08 | 2.9e-08 | 5.3e-08 | 3.967E-08 | 1.222E-08 | 2.900E-08 | 5.300E-08 |
| 17671 | 884 | V884L | C | 8.4e-08 | 3.2e-08 | 8.3e-08 | 6.633E-08 | 2.974E-08 | 3.200E-08 | 8.400E-08 |
| 17672 | 884 | V884K | C | 4.1e-08 | 3.9e-08 | 1.6e-08 | 3.200E-08 | 1.389E-08 | 1.600E-08 | 4.100E-08 |
| 17673 | 884 | V884M | C | 6.4e-08 | 4e-08 | 3.3e-08 | 4.567E-08 | 1.626E-08 | 3.300E-08 | 6.400E-08 |
| 17674 | 884 | V884F | C | 5.7e-08 | 3.5e-08 | 5.3e-08 | 4.833E-08 | 1.172E-08 | 3.500E-08 | 5.700E-08 |
| 17675 | 884 | V884P | C | 8e-08 | 4.4e-08 | 5e-08 | 5.800E-08 | 1.929E-08 | 4.400E-08 | 8.000E-08 |
| 17676 | 884 | V884S | C | 5.2e-08 | 1.7e-07 | 4.8e-08 | 9.000E-08 | 6.931E-08 | 4.800E-08 | 1.700E-07 |
| 17677 | 884 | V884T | C | 4.1e-08 | 4.1e-08 | 3.5e-08 | 3.900E-08 | 3.464E-09 | 3.500E-08 | 4.100E-08 |
| 17678 | 884 | V884W | C | 7.3e-08 | 3e-08 | 1.2e-08 | 3.833E-08 | 3.134E-08 | 1.200E-08 | 7.300E-08 |
| 17679 | 884 | V884Y | C | 3.8e-08 | 1.1e-08 | 2.3e-08 | 2.400E-08 | 1.353E-08 | 1.100E-08 | 3.800E-08 |

| | | | | | | | | | | |
|---|---|---|---|---|---|---|---|---|---|---|
| 17680 | 884 | V884V | C | 2.7e-08 | 5.6e-08 | 9.7e-08 | 6.000E-08 | 3.517E-08 | 2.700E-08 | 9.700E-08 |
| 17681 | 885 | A885A | C | 1.3e-07 | 3.7e-08 | 5.5e-08 | 7.400E-08 | 4.933E-08 | 3.700E-08 | 1.300E-07 |
| 17682 | 885 | A885R | C | 5.1e-08 | 2.8e-08 | 4.9e-08 | 4.267E-08 | 1.274E-08 | 2.800E-08 | 5.100E-08 |
| 17683 | 885 | A885N | C | 1.8e-08 | 7.8e-08 | 2e-08 | 3.867E-08 | 3.408E-08 | 1.800E-08 | 7.800E-08 |
| 17684 | 885 | A885D | C | 3e-08 | 5e-08 | 2.1e-07 | 9.667E-08 | 9.866E-08 | 3.000E-08 | 2.100E-07 |
| 17685 | 885 | A885C | C | 5.3e-08 | 5e-08 | 3.3e-08 | 4.533E-08 | 1.079E-08 | 3.300E-08 | 5.300E-08 |
| 17686 | 885 | A885E | C | 8.7e-08 | 2.3e-08 | 1.4e-08 | 4.133E-08 | 3.980E-08 | 1.400E-08 | 8.700E-08 |
| 17687 | 885 | A885Q | C | 8.4e-08 | 2.9e-08 | 4.9e-08 | 5.400E-08 | 2.784E-08 | 2.900E-08 | 8.400E-08 |
| 17688 | 885 | A885G | C | 3.9e-08 | 2.7e-08 | 1.8e-08 | 2.800E-08 | 1.054E-08 | 1.800E-08 | 3.900E-08 |
| 17689 | 885 | A885H | C | 2.7e-08 | 2.2e-07 | 9.6e-08 | 1.143E-07 | 9.780E-08 | 2.700E-08 | 2.200E-07 |
| 17690 | 885 | A885I | C | 1.1e-08 | 4.1e-08 | 2.2e-08 | 2.467E-08 | 1.518E-08 | 1.100E-08 | 4.100E-08 |
| 17691 | 885 | A885L | C | 2.1e-08 | 5.3e-08 | 6.4e-08 | 4.600E-08 | 2.234E-08 | 2.100E-08 | 6.400E-08 |
| 17692 | 885 | A885K | C | 4.8e-08 | 2.3e-07 | 3.1e-08 | 1.030E-07 | 1.103E-07 | 3.100E-08 | 2.300E-07 |
| 17693 | 885 | A885M | C | 6.7e-08 | 7.5e-08 | 2.5e-08 | 5.567E-08 | 2.686E-08 | 2.500E-08 | 7.500E-08 |
| 17694 | 885 | A885F | C | 3.8e-08 | 8.6e-08 | 7.2e-08 | 6.533E-08 | 2.468E-08 | 3.800E-08 | 8.600E-08 |
| 17695 | 885 | A885P | C | 7.1e-08 | 3.3e-08 | 2.2e-08 | 4.200E-08 | 2.571E-08 | 2.200E-08 | 7.100E-08 |
| 17696 | 885 | A885S | C | 3.9e-08 | 6.9e-08 | 8.9e-08 | 6.567E-08 | 2.517E-08 | 3.900E-08 | 8.900E-08 |
| 17697 | 885 | A885T | C | 5.1e-08 | 7.2e-08 | 1.2e-08 | 4.500E-08 | 3.045E-08 | 1.200E-08 | 7.200E-08 |
| 17698 | 885 | A885W | C | 4e-08 | 2.1e-08 | 1.1e-07 | 5.700E-08 | 4.687E-08 | 2.100E-08 | 1.100E-07 |
| 17699 | 885 | A885Y | C | 5.1e-08 | 6.7e-08 | 2.5e-08 | 4.767E-08 | 2.120E-08 | 2.500E-08 | 6.700E-08 |
| 17700 | 885 | A885V | C | 9e-08 | 7.5e-08 | 5.7e-08 | 7.400E-08 | 1.652E-08 | 5.700E-08 | 9.000E-08 |
| 17701 | 886 | D886A | C | 2.5e-08 | 3.3e-08 | 4e-08 | 3.267E-08 | 7.506E-09 | 2.500E-08 | 4.000E-08 |
| 17702 | 886 | D886R | C | 4.9e-08 | 7.7e-08 | 2.1e-08 | 4.900E-08 | 2.800E-08 | 2.100E-08 | 7.700E-08 |
| 17703 | 886 | D886N | C | 2.7e-08 | 3.2e-08 | 4.5e-08 | 3.467E-08 | 9.292E-09 | 2.700E-08 | 4.500E-08 |
| 17704 | 886 | D886D | C | 2.5e-08 | 1.2e-07 | 3.3e-08 | 5.933E-08 | 5.269E-08 | 2.500E-08 | 1.200E-07 |
| 17705 | 886 | D886C | C | 3.2e-08 | 1.9e-08 | 3.7e-08 | 2.933E-08 | 9.292E-09 | 1.900E-08 | 3.700E-08 |
| 17706 | 886 | D886E | C | 4.6e-08 | 7.7e-08 | 2.1e-08 | 4.800E-08 | 2.805E-08 | 2.100E-08 | 7.700E-08 |
| 17707 | 886 | D886Q | C | 1e-07 | 4e-08 | 4.4e-08 | 6.133E-08 | 3.355E-08 | 4.000E-08 | 1.000E-07 |
| 17708 | 886 | D886G | C | 4e-08 | 2.2e-08 | 9e-08 | 5.067E-08 | 3.523E-08 | 2.200E-08 | 9.000E-08 |
| 17709 | 886 | D886H | C | 6.8e-08 | 3e-08 | 5.3e-08 | 5.033E-08 | 1.914E-08 | 3.000E-08 | 6.800E-08 |
| 17710 | 886 | D886I | C | 3.4e-08 | 2.9e-08 | 1.9e-08 | 2.733E-08 | 7.638E-09 | 1.900E-08 | 3.400E-08 |
| 17711 | 886 | D886L | C | 1e-07 | 2.3e-08 | 6.2e-08 | 6.167E-08 | 3.850E-08 | 2.300E-08 | 1.000E-07 |
| 17712 | 886 | D886K | C | 5.9e-08 | 3.1e-08 | 2.6e-08 | 3.867E-08 | 1.779E-08 | 2.600E-08 | 5.900E-08 |
| 17713 | 886 | D886M | C | 6.5e-08 | 5.1e-08 | 4.8e-08 | 5.467E-08 | 9.074E-09 | 4.800E-08 | 6.500E-08 |
| 17714 | 886 | D886F | C | 4.5e-08 | 9.6e-08 | 1.8e-08 | 5.300E-08 | 3.961E-08 | 1.800E-08 | 9.600E-08 |
| 17715 | 886 | D886P | C | 6.6e-08 | 4.4e-08 | 2.1e-08 | 4.367E-08 | 2.250E-08 | 2.100E-08 | 6.600E-08 |
| 17716 | 886 | D886S | C | 2.4e-08 | 5.3e-08 | 5.3e-08 | 4.333E-08 | 1.674E-08 | 2.400E-08 | 5.300E-08 |
| 17717 | 886 | D886T | C | 2.1e-08 | 5.5e-08 | 2.2e-08 | 3.267E-08 | 1.935E-08 | 2.100E-08 | 5.500E-08 |
| 17718 | 886 | D886W | C | 6.6e-08 | 2.3e-08 | 4.4e-08 | 4.433E-08 | 2.150E-08 | 2.300E-08 | 6.600E-08 |
| 17719 | 886 | D886Y | C | 5.3e-08 | 9.8e-08 | 5e-08 | 6.700E-08 | 2.689E-08 | 5.000E-08 | 9.800E-08 |
| 17720 | 886 | D886V | C | 1.6e-08 | 4.2e-08 | 5.2e-08 | 3.667E-08 | 1.858E-08 | 1.600E-08 | 5.200E-08 |
| 17721 | 887 | V887A | C | 1e-07 | 6.6e-08 | 2.6e-08 | 6.400E-08 | 3.704E-08 | 2.600E-08 | 1.000E-07 |
| 17722 | 887 | V887R | C | 3.3e-08 | 4e-08 | 2.9e-08 | 3.400E-08 | 5.568E-09 | 2.900E-08 | 4.000E-08 |
| 17723 | 887 | V887N | C | 6.6e-08 | 1e-07 | 2.1e-08 | 6.233E-08 | 3.963E-08 | 2.100E-08 | 1.000E-07 |
| 17724 | 887 | V887D | C | 3.9e-08 | 2.2e-08 | 2.6e-08 | 2.900E-08 | 8.888E-09 | 2.200E-08 | 3.900E-08 |
| 17725 | 887 | V887C | C | 4.5e-08 | 4.3e-08 | 6.1e-08 | 4.967E-08 | 9.866E-09 | 4.300E-08 | 6.100E-08 |
| 17726 | 887 | V887E | C | 4.3e-08 | 4.4e-08 | 1e-07 | 6.233E-08 | 3.262E-08 | 4.300E-08 | 1.000E-07 |
| 17727 | 887 | V887Q | C | 2.8e-08 | 3.7e-08 | 3.5e-08 | 3.333E-08 | 4.726E-09 | 2.800E-08 | 3.700E-08 |
| 17728 | 887 | V887G | C | 2.2e-08 | 4.1e-08 | 7.6e-08 | 4.633E-08 | 2.739E-08 | 2.200E-08 | 7.600E-08 |
| 17729 | 887 | V887H | C | 1.7e-07 | 3.8e-08 | 1.1e-08 | 7.300E-08 | 8.508E-08 | 1.100E-08 | 1.700E-07 |
| 17730 | 887 | V887I | C | 5.7e-08 | 2.8e-08 | 5.8e-08 | 4.767E-08 | 1.704E-08 | 2.800E-08 | 5.800E-08 |
| 17731 | 887 | V887L | C | 3.4e-08 | 4.9e-08 | 3.3e-08 | 3.867E-08 | 8.963E-09 | 3.300E-08 | 4.900E-08 |
| 17732 | 887 | V887K | C | 5.5e-08 | 5.1e-08 | 2.1e-08 | 4.233E-08 | 1.858E-08 | 2.100E-08 | 5.500E-08 |
| 17733 | 887 | V887M | C | 8.7e-08 | 6.2e-08 | 2.3e-08 | 5.733E-08 | 3.225E-08 | 2.300E-08 | 8.700E-08 |
| 17734 | 887 | V887F | C | 3.7e-08 | 2.8e-08 | 2.2e-08 | 2.900E-08 | 7.550E-09 | 2.200E-08 | 3.700E-08 |
| 17735 | 887 | V887P | C | 4.6e-08 | 6.2e-08 | 5.7e-08 | 5.500E-08 | 8.185E-09 | 4.600E-08 | 6.200E-08 |
| 17736 | 887 | V887S | C | 7.9e-08 | 4.5e-08 | 7.9e-08 | 6.767E-08 | 1.963E-08 | 4.500E-08 | 7.900E-08 |
| 17737 | 887 | V887T | C | 1.8e-08 | 4.1e-08 | 3.6e-08 | 3.167E-08 | 1.210E-08 | 1.800E-08 | 4.100E-08 |
| 17738 | 887 | V887W | C | 4e-08 | 6.2e-08 | 1.4e-08 | 3.867E-08 | 2.403E-08 | 1.400E-08 | 6.200E-08 |
| 17739 | 887 | V887Y | C | 4.4e-08 | 3.9e-08 | 1.2e-08 | 3.167E-08 | 1.721E-08 | 1.200E-08 | 4.400E-08 |
| 17740 | 887 | V887V | C | 1e-07 | 2e-07 | 6e-08 | 1.200E-07 | 7.211E-08 | 6.000E-08 | 2.000E-07 |
| 17741 | 888 | N888A | C | 5.9e-08 | 4.6e-08 | 6e-08 | 5.500E-08 | 7.810E-09 | 4.600E-08 | 6.000E-08 |
| 17742 | 888 | N888R | C | 2.8e-08 | 2.4e-08 | 5.3e-08 | 3.500E-08 | 1.572E-08 | 2.400E-08 | 5.300E-08 |
| 17743 | 888 | N888N | C | 6.7e-08 | 3.6e-08 | 2e-08 | 4.100E-08 | 2.390E-08 | 2.000E-08 | 6.700E-08 |
| 17744 | 888 | N888D | C | 5e-08 | 1.7e-08 | 7.5e-08 | 4.733E-08 | 2.909E-08 | 1.700E-08 | 7.500E-08 |
| 17745 | 888 | N888C | C | 4.6e-08 | 7.7e-08 | 1.2e-07 | 8.100E-08 | 3.716E-08 | 4.600E-08 | 1.200E-07 |
| 17746 | 888 | N888E | C | 1.5e-08 | 3.3e-08 | 4.9e-08 | 3.233E-08 | 1.701E-08 | 1.500E-08 | 4.900E-08 |
| 17747 | 888 | N888Q | C | 1.5e-08 | 1.2e-08 | 5.4e-08 | 2.700E-08 | 2.343E-08 | 1.200E-08 | 5.400E-08 |
| 17748 | 888 | N888G | C | 8.1e-08 | 8.3e-08 | 2.5e-08 | 6.300E-08 | 3.292E-08 | 2.500E-08 | 8.300E-08 |
| 17749 | 888 | N888H | C | 1.5e-08 | 3.5e-08 | 2.6e-08 | 2.533E-08 | 1.002E-08 | 1.500E-08 | 3.500E-08 |
| 17750 | 888 | N888I | C | 6.7e-08 | 9.8e-08 | 4.6e-08 | 7.033E-08 | 2.616E-08 | 4.600E-08 | 9.800E-08 |
| 17751 | 888 | N888L | C | 8.7e-08 | 6.8e-08 | 3.5e-08 | 6.333E-08 | 2.631E-08 | 3.500E-08 | 8.700E-08 |
| 17752 | 888 | N888K | C | 6.9e-08 | 1.2e-08 | 2.6e-08 | 3.567E-08 | 2.970E-08 | 1.200E-08 | 6.900E-08 |
| 17753 | 888 | N888M | C | 1.1e-07 | 1.6e-07 | 4.3e-08 | 1.043E-07 | 5.871E-08 | 4.300E-08 | 1.600E-07 |
| 17754 | 888 | N888F | C | 4.1e-08 | 1.2e-07 | 6.4e-08 | 7.500E-08 | 4.063E-08 | 4.100E-08 | 1.200E-07 |
| 17755 | 888 | N888P | C | 4.8e-08 | 8.2e-08 | 5.1e-08 | 6.033E-08 | 1.882E-08 | 4.800E-08 | 8.200E-08 |
| 17756 | 888 | N888S | C | 7.2e-08 | 4.7e-08 | 6.2e-08 | 6.033E-08 | 1.258E-08 | 4.700E-08 | 7.200E-08 |
| 17757 | 888 | N888T | C | 1.8e-08 | 7.6e-08 | 1.1e-07 | 6.800E-08 | 4.652E-08 | 1.800E-08 | 1.100E-07 |
| 17758 | 888 | N888W | C | 1.9e-08 | 1.2e-07 | 1e-07 | 7.967E-08 | 5.348E-08 | 1.900E-08 | 1.200E-07 |
| 17759 | 888 | N888Y | C | 5.6e-08 | 2.5e-07 | 3e-08 | 1.120E-07 | 1.202E-07 | 3.000E-08 | 2.500E-07 |
| 17760 | 888 | N888V | C | 3.1e-08 | 3e-08 | 7.1e-08 | 4.400E-08 | 2.339E-08 | 3.000E-08 | 7.100E-08 |
| 17761 | 889 | P889A | C | 2.9e-08 | 4.6e-08 | 2.1e-08 | 3.200E-08 | 1.277E-08 | 2.100E-08 | 4.600E-08 |
| 17762 | 889 | P889R | C | 6.4e-08 | 3.2e-08 | 3.2e-08 | 4.267E-08 | 1.848E-08 | 3.200E-08 | 6.400E-08 |
| 17763 | 889 | P889N | C | 4.5e-08 | 9.5e-09 | 6.7e-08 | 4.050E-08 | 2.901E-08 | 9.500E-09 | 6.700E-08 |
| 17764 | 889 | P889D | C | 1.2e-07 | 1.1e-07 | 6.4e-08 | 9.800E-08 | 2.987E-08 | 6.400E-08 | 1.200E-07 |

| | | | | | | | | | | |
|---|---|---|---|---|---|---|---|---|---|---|
| 17765 | 889 | P889C | C | 5.7e-08 | 5e-08 | 8.4e-08 | 6.367E-08 | 1.795E-08 | 5.000E-08 | 8.400E-08 |
| 17766 | 889 | P889E | C | 5.9e-08 | 1.2e-08 | 1.1e-07 | 6.033E-08 | 4.901E-08 | 1.200E-08 | 1.100E-07 |
| 17767 | 889 | P889Q | C | 1.1e-08 | 2.9e-08 | 2.5e-08 | 2.167E-08 | 9.452E-09 | 1.100E-08 | 2.900E-08 |
| 17768 | 889 | P889G | C | 6.7e-08 | 8.9e-08 | 1.4e-07 | 9.867E-08 | 3.745E-08 | 6.700E-08 | 1.400E-07 |
| 17769 | 889 | P889H | C | 6e-08 | 4.3e-08 | 3e-08 | 4.433E-08 | 1.504E-08 | 3.000E-08 | 6.000E-08 |
| 17770 | 889 | P889I | C | 4.2e-08 | 3.4e-08 | 1.4e-08 | 3.000E-08 | 1.442E-08 | 1.400E-08 | 4.200E-08 |
| 17771 | 889 | P889L | C | 6.1e-08 | 4.5e-08 | 6e-08 | 5.533E-08 | 8.963E-09 | 4.500E-08 | 6.100E-08 |
| 17772 | 889 | P889K | C | 8e-08 | 7.3e-08 | 5.7e-08 | 7.000E-08 | 1.179E-08 | 5.700E-08 | 8.000E-08 |
| 17773 | 889 | P889M | C | 6.2e-08 | 2.1e-08 | 2.5e-08 | 3.600E-08 | 2.261E-08 | 2.100E-08 | 6.200E-08 |
| 17774 | 889 | P889F | C | 3.9e-08 | 2.9e-08 | 5.3e-08 | 4.033E-08 | 1.206E-08 | 2.900E-08 | 5.300E-08 |
| 17775 | 889 | P889P | C | 1.8e-08 | 3.4e-08 | 2.4e-08 | 2.533E-08 | 8.083E-09 | 1.800E-08 | 3.400E-08 |
| 17776 | 889 | P889S | C | 6.9e-08 | 1.4e-07 | 3.3e-08 | 8.067E-08 | 5.445E-08 | 3.300E-08 | 1.400E-07 |
| 17777 | 889 | P889T | C | 3.7e-08 | 3.7e-08 | 7.3e-08 | 4.900E-08 | 2.078E-08 | 3.700E-08 | 7.300E-08 |
| 17778 | 889 | P889W | C | 2.4e-08 | 1.2e-08 | 3.1e-08 | 2.233E-08 | 9.609E-09 | 1.200E-08 | 3.100E-08 |
| 17779 | 889 | P889Y | C | 8.2e-09 | 2.5e-08 | 1.2e-08 | 1.507E-08 | 8.810E-09 | 8.200E-09 | 2.500E-08 |
| 17780 | 889 | P889V | C | 6.1e-08 | 1.5e-07 | 1.5e-08 | 7.533E-08 | 6.863E-08 | 1.500E-08 | 1.500E-07 |
| 17781 | 890 | N890A | C | 2.1e-07 | 1.2e-07 | 2.5e-07 | 1.933E-07 | 6.658E-08 | 1.200E-07 | 2.500E-07 |
| 17782 | 890 | N890R | C | 3.1e-08 | 7.2e-08 | 5.3e-08 | 5.200E-08 | 2.052E-08 | 3.100E-08 | 7.200E-08 |
| 17783 | 890 | N890N | C | 1.9e-08 | 4e-08 | 2.7e-08 | 2.867E-08 | 1.060E-08 | 1.900E-08 | 4.000E-08 |
| 17784 | 890 | N890D | C | 8.5e-08 | 8.9e-08 | 9e-08 | 8.800E-08 | 2.646E-09 | 8.500E-08 | 9.000E-08 |
| 17785 | 890 | N890C | C | 5.9e-08 | 1.1e-07 | 7.3e-08 | 8.067E-08 | 2.635E-08 | 5.900E-08 | 1.100E-07 |
| 17786 | 890 | N890E | C | 7.3e-08 | 6e-08 | 1e-07 | 7.767E-08 | 2.040E-08 | 6.000E-08 | 1.000E-07 |
| 17787 | 890 | N890Q | C | 3.5e-08 | 9.1e-08 | 1.6e-07 | 9.533E-08 | 6.261E-08 | 3.500E-08 | 1.600E-07 |
| 17788 | 890 | N890G | C | 2.3e-07 | 1e-07 | 9e-08 | 1.400E-07 | 7.810E-08 | 9.000E-08 | 2.300E-07 |
| 17789 | 890 | N890H | C | 4.4e-08 | 5e-08 | 5.7e-08 | 5.033E-08 | 6.506E-09 | 4.400E-08 | 5.700E-08 |
| 17790 | 890 | N890I | C | 4e-08 | 8e-08 | 1.2e-07 | 8.000E-08 | 4.000E-08 | 4.000E-08 | 1.200E-07 |
| 17791 | 890 | N890L | C | 1.3e-07 | 5.6e-08 | 2e-07 | 1.287E-07 | 7.201E-08 | 5.600E-08 | 2.000E-07 |
| 17792 | 890 | N890K | C | 6.8e-08 | 6.5e-08 | 9.8e-08 | 7.700E-08 | 1.825E-08 | 6.500E-08 | 9.800E-08 |
| 17793 | 890 | N890M | C | 2.1e-07 | 1.2e-07 | 4.3e-07 | 2.533E-07 | 1.595E-07 | 1.200E-07 | 4.300E-07 |
| 17794 | 890 | N890F | C | 8.2e-08 | 3.1e-07 | 1.6e-07 | 1.840E-07 | 1.159E-07 | 8.200E-08 | 3.100E-07 |
| 17795 | 890 | N890P | C | 4e-07 | 1.7e-07 | 2.9e-07 | 2.867E-07 | 1.150E-07 | 1.700E-07 | 4.000E-07 |
| 17796 | 890 | N890S | C | 1.7e-08 | 1.8e-08 | 9.9e-08 | 4.467E-08 | 4.706E-08 | 1.700E-08 | 9.900E-08 |
| 17797 | 890 | N890T | C | 8.8e-08 | 5e-08 | 1.6e-08 | 5.133E-08 | 3.602E-08 | 1.600E-08 | 8.800E-08 |
| 17798 | 890 | N890W | C | 2.1e-07 | 1.1e-07 | 9.5e-08 | 1.383E-07 | 6.252E-08 | 9.500E-08 | 2.100E-07 |
| 17799 | 890 | N890Y | C | 3.1e-07 | 2.2e-07 | 2.2e-07 | 2.500E-07 | 5.196E-08 | 2.200E-07 | 3.100E-07 |
| 17800 | 890 | N890V | C | 1.8e-07 | 1.3e-07 | 7.9e-08 | 1.297E-07 | 5.050E-08 | 7.900E-08 | 1.800E-07 |
| 17801 | 891 | S891A | C | 5.5e-08 | 1.1e-07 | 5.1e-08 | 7.200E-08 | 3.297E-08 | 5.100E-08 | 1.100E-07 |
| 17802 | 891 | S891R | C | 5.8e-08 | 7.4e-08 | 4.3e-08 | 5.833E-08 | 1.550E-08 | 4.300E-08 | 7.400E-08 |
| 17803 | 891 | S891N | C | 3.6e-08 | 1.2e-08 | 2.6e-08 | 2.467E-08 | 1.206E-08 | 1.200E-08 | 3.600E-08 |
| 17804 | 891 | S891D | C | 4.6e-08 | 4e-08 | 3.3e-08 | 3.967E-08 | 6.506E-09 | 3.300E-08 | 4.600E-08 |
| 17805 | 891 | S891C | C | 2.2e-07 | 8e-08 | 3.8e-08 | 1.127E-07 | 9.530E-08 | 3.800E-08 | 2.200E-07 |
| 17806 | 891 | S891E | C | 2.6e-08 | 5.2e-08 | 1.5e-08 | 3.100E-08 | 1.900E-08 | 1.500E-08 | 5.200E-08 |
| 17807 | 891 | S891Q | C | 2.8e-08 | 5.7e-08 | 5.9e-08 | 4.800E-08 | 1.735E-08 | 2.800E-08 | 5.900E-08 |
| 17808 | 891 | S891G | C | 1.1e-07 | 1.6e-07 | 1.4e-07 | 1.367E-07 | 2.517E-08 | 1.100E-07 | 1.600E-07 |
| 17809 | 891 | S891H | C | 3.5e-08 | 5e-08 | 1.4e-08 | 3.300E-08 | 1.808E-08 | 1.400E-08 | 5.000E-08 |
| 17810 | 891 | S891I | C | 9.9e-08 | 6e-08 | 2.1e-08 | 6.000E-08 | 3.900E-08 | 2.100E-08 | 9.900E-08 |
| 17811 | 891 | S891L | C | 5.1e-08 | 1.1e-07 | 1.1e-07 | 9.033E-08 | 3.406E-08 | 5.100E-08 | 1.100E-07 |
| 17812 | 891 | S891K | C | 5.1e-08 | 3.1e-08 | 8e-08 | 5.400E-08 | 2.464E-08 | 3.100E-08 | 8.000E-08 |
| 17813 | 891 | S891M | C | 3.7e-08 | 4.9e-08 | 8.5e-08 | 5.700E-08 | 2.498E-08 | 3.700E-08 | 8.500E-08 |
| 17814 | 891 | S891F | C | 6.6e-08 | 1.9e-07 | 8.3e-08 | 1.130E-07 | 6.722E-08 | 6.600E-08 | 1.900E-07 |
| 17815 | 891 | S891P | C | 1.2e-07 | 6e-08 | 5.1e-08 | 7.700E-08 | 3.751E-08 | 5.100E-08 | 1.200E-07 |
| 17816 | 891 | S891S | C | 3.6e-08 | 6.6e-08 | 6.1e-08 | 5.433E-08 | 1.607E-08 | 3.600E-08 | 6.600E-08 |
| 17817 | 891 | S891T | C | 8.9e-09 | 2.8e-08 | 7.3e-08 | 3.663E-08 | 3.291E-08 | 8.900E-09 | 7.300E-08 |
| 17818 | 891 | S891W | C | 6.4e-08 | 4.6e-08 | 7.2e-08 | 6.067E-08 | 1.332E-08 | 4.600E-08 | 7.200E-08 |
| 17819 | 891 | S891Y | C | 3.5e-08 | 7.6e-08 | 9.2e-08 | 6.767E-08 | 2.940E-08 | 3.500E-08 | 9.200E-08 |
| 17820 | 891 | S891V | C | 3.6e-08 | 1e-07 | 1.1e-07 | 8.200E-08 | 4.015E-08 | 3.600E-08 | 1.100E-07 |
| 17821 | 892 | G892A | C | 5.3e-08 | 3e-08 | 2.2e-08 | 3.500E-08 | 1.609E-08 | 2.200E-08 | 5.300E-08 |
| 17822 | 892 | G892R | C | 8.1e-09 | 4.6e-08 | 2.4e-08 | 2.603E-08 | 1.903E-08 | 8.100E-09 | 4.600E-08 |
| 17823 | 892 | G892N | C | 4.9e-08 | 1.2e-08 | 2.1e-08 | 2.733E-08 | 1.930E-08 | 1.200E-08 | 4.900E-08 |
| 17824 | 892 | G892D | C | 3.7e-08 | 2.6e-08 | 2.1e-08 | 2.800E-08 | 8.185E-09 | 2.100E-08 | 3.700E-08 |
| 17825 | 892 | G892C | C | 9.2e-08 | 5.1e-08 | 4.4e-08 | 6.233E-08 | 2.593E-08 | 4.400E-08 | 9.200E-08 |
| 17826 | 892 | G892E | C | 2.8e-08 | 1.1e-08 | 7.5e-08 | 3.800E-08 | 3.315E-08 | 1.100E-08 | 7.500E-08 |
| 17827 | 892 | G892Q | C | 1.6e-07 | 1.6e-08 | 9e-08 | 8.867E-08 | 7.201E-08 | 1.600E-08 | 1.600E-07 |
| 17828 | 892 | G892G | C | 4.6e-08 | 2.1e-08 | 2.1e-08 | 2.933E-08 | 1.443E-08 | 2.100E-08 | 4.600E-08 |
| 17829 | 892 | G892H | C | 4.9e-08 | 3.8e-08 | 4.6e-08 | 4.433E-08 | 5.686E-09 | 3.800E-08 | 4.900E-08 |
| 17830 | 892 | G892I | C | 4.8e-08 | 3.5e-08 | 1.4e-08 | 3.233E-08 | 1.716E-08 | 1.400E-08 | 4.800E-08 |
| 17831 | 892 | G892L | C | 7.7e-08 | 2.9e-08 | 5.3e-08 | 5.300E-08 | 2.400E-08 | 2.900E-08 | 7.700E-08 |
| 17832 | 892 | G892K | C | 4.2e-08 | 3e-08 | 2.7e-08 | 3.300E-08 | 7.937E-09 | 2.700E-08 | 4.200E-08 |
| 17833 | 892 | G892M | C | 4e-08 | 5.2e-08 | 5.3e-08 | 4.833E-08 | 7.234E-09 | 4.000E-08 | 5.300E-08 |
| 17834 | 892 | G892F | C | 5.6e-08 | 1.8e-08 | 6.6e-09 | 2.687E-08 | 2.587E-08 | 6.600E-09 | 5.600E-08 |
| 17835 | 892 | G892P | C | 8.7e-08 | 1.5e-07 | 5e-08 | 9.567E-08 | 5.056E-08 | 5.000E-08 | 1.500E-07 |
| 17836 | 892 | G892S | C | 6.4e-08 | 4.5e-08 | 2.8e-08 | 4.567E-08 | 1.801E-08 | 2.800E-08 | 6.400E-08 |
| 17837 | 892 | G892T | C | 3.2e-08 | 8.2e-08 | 3.2e-08 | 4.867E-08 | 2.887E-08 | 3.200E-08 | 8.200E-08 |
| 17838 | 892 | G892W | C | 2.1e-08 | 2.1e-08 | 2.3e-08 | 2.167E-08 | 1.155E-09 | 2.100E-08 | 2.300E-08 |
| 17839 | 892 | G892Y | C | 1.7e-08 | 3.5e-08 | 1.2e-08 | 2.133E-08 | 1.210E-08 | 1.200E-08 | 3.500E-08 |
| 17840 | 892 | G892V | C | 2.4e-08 | 2.9e-08 | 6.3e-08 | 3.867E-08 | 2.122E-08 | 2.400E-08 | 6.300E-08 |
| 17841 | 893 | G893A | C | 9.7e-08 | 2e-08 | 8e-08 | 6.567E-08 | 4.045E-08 | 2.000E-08 | 9.700E-08 |
| 17842 | 893 | G893R | C | 4.3e-08 | 4.5e-08 | 2.4e-08 | 3.733E-08 | 1.159E-08 | 2.400E-08 | 4.500E-08 |
| 17843 | 893 | G893N | C | 3.8e-08 | 3.8e-08 | 4.4e-08 | 4.000E-08 | 3.464E-09 | 3.800E-08 | 4.400E-08 |
| 17844 | 893 | G893D | C | 1.1e-07 | 2.1e-08 | 5.8e-08 | 6.300E-08 | 4.471E-08 | 2.100E-08 | 1.100E-07 |
| 17845 | 893 | G893C | C | 3.1e-08 | 3.4e-08 | 2.4e-07 | 1.017E-07 | 1.198E-07 | 3.100E-08 | 2.400E-07 |
| 17846 | 893 | G893E | C | 2.5e-08 | 1.8e-08 | 3.4e-08 | 2.567E-08 | 8.021E-09 | 1.800E-08 | 3.400E-08 |
| 17847 | 893 | G893Q | C | 9e-09 | 2.6e-08 | 3e-08 | 2.167E-08 | 1.115E-08 | 9.000E-09 | 3.000E-08 |
| 17848 | 893 | G893G | C | 2.2e-08 | 6.6e-08 | 7.9e-08 | 5.567E-08 | 2.987E-08 | 2.200E-08 | 7.900E-08 |
| 17849 | 893 | G893H | C | 2.8e-08 | 2.4e-08 | 4.7e-08 | 3.300E-08 | 1.229E-08 | 2.400E-08 | 4.700E-08 |

| | | | | | | | | | | |
|---|---|---|---|---|---|---|---|---|---|---|
| 17850 | 893 | G893I | C | 3.8e-08 | 4.4e-08 | 3e-08 | 3.733E-08 | 7.024E-09 | 3.000E-08 | 4.400E-08 |
| 17851 | 893 | G893L | C | 1.4e-08 | 6.3e-08 | 7.1e-08 | 4.933E-08 | 3.086E-08 | 1.400E-08 | 7.100E-08 |
| 17852 | 893 | G893K | C | 7.6e-09 | 5.3e-08 | 1.5e-08 | 2.520E-08 | 2.436E-08 | 7.600E-09 | 5.300E-08 |
| 17853 | 893 | G893M | C | 7.1e-08 | 1.8e-07 | 4.1e-08 | 9.733E-08 | 7.315E-08 | 4.100E-08 | 1.800E-07 |
| 17854 | 893 | G893F | C | 4.3e-08 | 2e-08 | 9.2e-08 | 5.167E-08 | 3.677E-08 | 2.000E-08 | 9.200E-08 |
| 17855 | 893 | G893P | C | 4.3e-08 | 2.7e-08 | 3.7e-08 | 3.567E-08 | 8.083E-09 | 2.700E-08 | 4.300E-08 |
| 17856 | 893 | G893S | C | 3.1e-08 | 3e-08 | 2.8e-08 | 2.967E-08 | 1.528E-09 | 2.800E-08 | 3.100E-08 |
| 17857 | 893 | G893T | C | 9.5e-08 | 1.7e-08 | 1.6e-08 | 4.267E-08 | 4.532E-08 | 1.600E-08 | 9.500E-08 |
| 17858 | 893 | G893W | C | 4e-08 | 5.4e-08 | 6.7e-08 | 5.367E-08 | 1.350E-08 | 4.000E-08 | 6.700E-08 |
| 17859 | 893 | G893Y | C | 2.3e-08 | 3.9e-08 | 7e-08 | 4.400E-08 | 2.390E-08 | 2.300E-08 | 7.000E-08 |
| 17860 | 893 | G893V | C | 5.7e-08 | 2.1e-08 | 4.8e-08 | 4.200E-08 | 1.873E-08 | 2.100E-08 | 5.700E-08 |
| 17861 | 894 | S894A | C | 5.5e-08 | 1e-08 | 5.6e-08 | 4.033E-08 | 2.627E-08 | 1.000E-08 | 5.600E-08 |
| 17862 | 894 | S894R | C | 4.8e-08 | 3.4e-08 | 8.5e-08 | 5.567E-08 | 2.635E-08 | 3.400E-08 | 8.500E-08 |
| 17863 | 894 | S894N | C | 8.3e-08 | 8.1e-09 | 1.2e-07 | 7.037E-08 | 5.701E-08 | 8.100E-09 | 1.200E-07 |
| 17864 | 894 | S894D | C | 2.1e-08 | 5.6e-08 | 7.2e-08 | 4.967E-08 | 2.608E-08 | 2.100E-08 | 7.200E-08 |
| 17865 | 894 | S894C | C | 6.7e-08 | 7.3e-08 | 3.6e-08 | 5.867E-08 | 1.986E-08 | 3.600E-08 | 7.300E-08 |
| 17866 | 894 | S894E | C | 5.3e-08 | 8.8e-08 | 5e-08 | 6.367E-08 | 2.113E-08 | 5.000E-08 | 8.800E-08 |
| 17867 | 894 | S894Q | C | 4.3e-08 | 7.1e-08 | 4.1e-08 | 5.167E-08 | 1.677E-08 | 4.100E-08 | 7.100E-08 |
| 17868 | 894 | S894G | C | 4.6e-08 | 3.5e-08 | 5.5e-08 | 4.533E-08 | 1.002E-08 | 3.500E-08 | 5.500E-08 |
| 17869 | 894 | S894H | C | 5e-08 | 4.6e-08 | 6.5e-08 | 5.367E-08 | 1.002E-08 | 4.600E-08 | 6.500E-08 |
| 17870 | 894 | S894I | C | 4.5e-08 | 7.3e-08 | 5.8e-08 | 5.867E-08 | 1.401E-08 | 4.500E-08 | 7.300E-08 |
| 17871 | 894 | S894L | C | 1.3e-07 | 3.3e-08 | 1.3e-07 | 9.767E-08 | 5.600E-08 | 3.300E-08 | 1.300E-07 |
| 17872 | 894 | S894K | C | 3.4e-08 | 4.4e-08 | 4.3e-08 | 4.033E-08 | 5.508E-09 | 3.400E-08 | 4.400E-08 |
| 17873 | 894 | S894M | C | 2.5e-07 | 1.2e-07 | 1.7e-07 | 1.800E-07 | 6.557E-08 | 1.200E-07 | 2.500E-07 |
| 17874 | 894 | S894F | C | 7.8e-08 | 1.6e-07 | 1e-08 | 8.267E-08 | 7.511E-08 | 1.000E-08 | 1.600E-07 |
| 17875 | 894 | S894P | C | 1.5e-07 | 1.2e-07 | 7.4e-08 | 1.147E-07 | 3.828E-08 | 7.400E-08 | 1.500E-07 |
| 17876 | 894 | S894S | C | 3.7e-08 | 1.5e-08 | 3.6e-08 | 2.933E-08 | 1.242E-08 | 1.500E-08 | 3.700E-08 |
| 17877 | 894 | S894T | C | 5e-08 | 7.3e-08 | 8.7e-08 | 7.000E-08 | 1.868E-08 | 5.000E-08 | 8.700E-08 |
| 17878 | 894 | S894W | C | 5.4e-08 | 5.2e-08 | 4e-08 | 4.867E-08 | 7.572E-09 | 4.000E-08 | 5.400E-08 |
| 17879 | 894 | S894Y | C | 4.5e-08 | 6.8e-08 | 1.5e-07 | 8.767E-08 | 5.519E-08 | 4.500E-08 | 1.500E-07 |
| 17880 | 894 | S894V | C | 7.8e-08 | 4.2e-08 | 4e-08 | 5.333E-08 | 2.139E-08 | 4.000E-08 | 7.800E-08 |
| 17881 | 895 | I895A | C | 6.2e-09 | 2.5e-08 | 1.7e-08 | 1.607E-08 | 9.435E-09 | 6.200E-09 | 2.500E-08 |
| 17882 | 895 | I895R | C | 2.6e-08 | 2.2e-08 | 1e-08 | 1.933E-08 | 8.327E-09 | 1.000E-08 | 2.600E-08 |
| 17883 | 895 | I895N | C | 5.1e-08 | 1.6e-08 | 4.5e-08 | 3.733E-08 | 1.872E-08 | 1.600E-08 | 5.100E-08 |
| 17884 | 895 | I895D | C | 2.3e-08 | 4.6e-08 | 5.8e-08 | 4.233E-08 | 1.779E-08 | 2.300E-08 | 5.800E-08 |
| 17885 | 895 | I895C | C | 1.8e-08 | 1.2e-07 | 3.3e-08 | 5.700E-08 | 5.507E-08 | 1.800E-08 | 1.200E-07 |
| 17886 | 895 | I895E | C | 1.7e-08 | 2.5e-08 | 5.1e-08 | 3.100E-08 | 1.778E-08 | 1.700E-08 | 5.100E-08 |
| 17887 | 895 | I895Q | C | 1.4e-08 | 1.1e-07 | 4.3e-08 | 5.567E-08 | 4.924E-08 | 1.400E-08 | 1.100E-07 |
| 17888 | 895 | I895G | C | 3.7e-08 | 4e-08 | 4.6e-08 | 4.100E-08 | 4.583E-09 | 3.700E-08 | 4.600E-08 |
| 17889 | 895 | I895H | C | 2.4e-08 | 2.7e-08 | 7.1e-08 | 4.067E-08 | 2.631E-08 | 2.400E-08 | 7.100E-08 |
| 17890 | 895 | I895I | C | 5.7e-08 | 1.8e-08 | 4.4e-08 | 3.967E-08 | 1.986E-08 | 1.800E-08 | 5.700E-08 |
| 17891 | 895 | I895L | C | 3.7e-08 | 3.9e-08 | 3.7e-08 | 3.767E-08 | 1.155E-09 | 3.700E-08 | 3.900E-08 |
| 17892 | 895 | I895K | C | 2.6e-08 | 2.9e-08 | 2.5e-08 | 2.667E-08 | 2.082E-09 | 2.500E-08 | 2.900E-08 |
| 17893 | 895 | I895M | C | 8.4e-08 | 8e-08 | 5.3e-08 | 7.233E-08 | 1.686E-08 | 5.300E-08 | 8.400E-08 |
| 17894 | 895 | I895F | C | 3.4e-08 | 4.8e-08 | 5.4e-08 | 4.533E-08 | 1.026E-08 | 3.400E-08 | 5.400E-08 |
| 17895 | 895 | I895P | C | 3.9e-08 | 3.2e-08 | 4.9e-08 | 4.000E-08 | 8.544E-09 | 3.200E-08 | 4.900E-08 |
| 17896 | 895 | I895S | C | 2.8e-08 | 1e-08 | 1.2e-08 | 1.667E-08 | 9.866E-09 | 1.000E-08 | 2.800E-08 |
| 17897 | 895 | I895T | C | 3.4e-08 | 1.8e-08 | 3.9e-08 | 3.033E-08 | 1.097E-08 | 1.800E-08 | 3.900E-08 |
| 17898 | 895 | I895W | C | 1.3e-07 | 3.9e-08 | 2.8e-08 | 6.567E-08 | 5.599E-08 | 2.800E-08 | 1.300E-07 |
| 17899 | 895 | I895Y | C | 1.7e-08 | 3.2e-08 | 7.6e-08 | 4.167E-08 | 3.066E-08 | 1.700E-08 | 7.600E-08 |
| 17900 | 895 | I895V | C | 9.9e-08 | 5.5e-08 | 2.1e-08 | 5.833E-08 | 3.911E-08 | 2.100E-08 | 9.900E-08 |
| 17901 | 896 | Y896A | C | 7.1e-08 | 8.7e-08 | 1.1e-07 | 8.933E-08 | 1.960E-08 | 7.100E-08 | 1.100E-07 |
| 17902 | 896 | Y896R | C | 3.3e-08 | 3.3e-08 | 1.7e-08 | 2.767E-08 | 9.238E-09 | 1.700E-08 | 3.300E-08 |
| 17903 | 896 | Y896N | C | 3.2e-08 | 3.5e-08 | 5e-08 | 3.900E-08 | 9.644E-09 | 3.200E-08 | 5.000E-08 |
| 17904 | 896 | Y896D | C | 4.7e-08 | 6.6e-08 | 9.9e-09 | 4.097E-08 | 2.853E-08 | 9.900E-09 | 6.600E-08 |
| 17905 | 896 | Y896C | C | 3.4e-08 | 3.1e-08 | 2e-08 | 2.833E-08 | 7.371E-09 | 2.000E-08 | 3.400E-08 |
| 17906 | 896 | Y896E | C | 2.9e-08 | 1.1e-07 | 3.9e-08 | 5.933E-08 | 4.416E-08 | 2.900E-08 | 1.100E-07 |
| 17907 | 896 | Y896Q | C | 4.6e-08 | 9.4e-09 | 2.8e-08 | 2.780E-08 | 1.830E-08 | 9.400E-09 | 4.600E-08 |
| 17908 | 896 | Y896G | C | 4.6e-08 | 4.2e-08 | 3.4e-08 | 4.067E-08 | 6.110E-09 | 3.400E-08 | 4.600E-08 |
| 17909 | 896 | Y896H | C | 4.4e-08 | 6.2e-08 | 9.2e-08 | 6.600E-08 | 2.425E-08 | 4.400E-08 | 9.200E-08 |
| 17910 | 896 | Y896I | C | 1.7e-08 | 6.2e-08 | 1.3e-08 | 3.067E-08 | 2.721E-08 | 1.300E-08 | 6.200E-08 |
| 17911 | 896 | Y896L | C | 5.8e-08 | 7.2e-08 | 2.1e-08 | 5.033E-08 | 2.635E-08 | 2.100E-08 | 7.200E-08 |
| 17912 | 896 | Y896K | C | 5.7e-08 | 1.6e-08 | 3.2e-08 | 3.500E-08 | 2.066E-08 | 1.600E-08 | 5.700E-08 |
| 17913 | 896 | Y896M | C | 1.9e-08 | 5.1e-08 | 3.9e-08 | 3.633E-08 | 1.617E-08 | 1.900E-08 | 5.100E-08 |
| 17914 | 896 | Y896F | C | 1.2e-07 | 7.3e-08 | 4.6e-08 | 7.967E-08 | 3.745E-08 | 4.600E-08 | 1.200E-07 |
| 17915 | 896 | Y896P | C | 9.7e-08 | 6.7e-08 | 7.3e-08 | 7.900E-08 | 1.587E-08 | 6.700E-08 | 9.700E-08 |
| 17916 | 896 | Y896S | C | 5.1e-08 | 5.3e-08 | 4.2e-08 | 4.867E-08 | 5.859E-09 | 4.200E-08 | 5.300E-08 |
| 17917 | 896 | Y896T | C | 2.4e-08 | 3.3e-08 | 6.5e-08 | 4.067E-08 | 2.155E-08 | 2.400E-08 | 6.500E-08 |
| 17918 | 896 | Y896W | C | 6.4e-08 | 4.8e-08 | 3.9e-08 | 5.033E-08 | 1.266E-08 | 3.900E-08 | 6.400E-08 |
| 17919 | 896 | Y896Y | C | 1.3e-08 | 5.6e-08 | 1.9e-08 | 2.933E-08 | 2.329E-08 | 1.300E-08 | 5.600E-08 |
| 17920 | 896 | Y896V | C | 5.8e-08 | 4.2e-08 | 6e-08 | 5.333E-08 | 9.866E-09 | 4.200E-08 | 6.000E-08 |
| 17921 | 897 | N897A | C | 3.1e-08 | 5e-08 | 3.3e-08 | 3.800E-08 | 1.044E-08 | 3.100E-08 | 5.000E-08 |
| 17922 | 897 | N897R | C | 3e-08 | 4e-08 | 8.1e-08 | 5.033E-08 | 2.702E-08 | 3.000E-08 | 8.100E-08 |
| 17923 | 897 | N897N | C | 2.4e-08 | 1.3e-08 | 4.7e-08 | 2.800E-08 | 1.735E-08 | 1.300E-08 | 4.700E-08 |
| 17924 | 897 | N897D | C | 9.4e-08 | 8.4e-08 | 7e-08 | 8.267E-08 | 1.206E-08 | 7.000E-08 | 9.400E-08 |
| 17925 | 897 | N897C | C | 3.1e-08 | 5.3e-08 | 4.1e-08 | 4.167E-08 | 1.102E-08 | 3.100E-08 | 5.300E-08 |
| 17926 | 897 | N897E | C | 5.7e-08 | 6.1e-08 | 2.6e-08 | 4.800E-08 | 1.916E-08 | 2.600E-08 | 6.100E-08 |
| 17927 | 897 | N897Q | C | 4.2e-08 | 8.7e-08 | 3.3e-08 | 5.400E-08 | 2.893E-08 | 3.300E-08 | 8.700E-08 |
| 17928 | 897 | N897G | C | 4.3e-08 | 3.4e-08 | 2.2e-08 | 3.300E-08 | 1.054E-08 | 2.200E-08 | 4.300E-08 |
| 17929 | 897 | N897H | C | 3e-08 | 3.4e-08 | 4.5e-08 | 3.633E-08 | 7.767E-09 | 3.000E-08 | 4.500E-08 |
| 17930 | 897 | N897I | C | 3.2e-08 | 4.3e-08 | 8.7e-08 | 5.400E-08 | 2.910E-08 | 3.200E-08 | 8.700E-08 |
| 17931 | 897 | N897L | C | 5.9e-08 | 3.5e-08 | 3.4e-08 | 4.267E-08 | 1.415E-08 | 3.400E-08 | 5.900E-08 |
| 17932 | 897 | N897K | C | 1e-07 | 3.2e-08 | 4.6e-08 | 5.933E-08 | 3.591E-08 | 3.200E-08 | 1.000E-07 |
| 17933 | 897 | N897M | C | 5.3e-08 | 8.3e-08 | 3.7e-08 | 5.767E-08 | 2.335E-08 | 3.700E-08 | 8.300E-08 |
| 17934 | 897 | N897F | C | 7.1e-08 | 9.1e-08 | 4.6e-08 | 6.933E-08 | 2.255E-08 | 4.600E-08 | 9.100E-08 |

| | | | | | | | | | | |
|---|---|---|---|---|---|---|---|---|---|---|
| 17935 | 897 | N897P | C | 7.7e-08 | 4.5e-08 | 8e-08 | 6.733E-08 | 1.940E-08 | 4.500E-08 | 8.000E-08 |
| 17936 | 897 | N897S | C | 6.3e-08 | 3e-08 | 3.1e-08 | 4.133E-08 | 1.877E-08 | 3.000E-08 | 6.300E-08 |
| 17937 | 897 | N897T | C | 2.6e-08 | 6.6e-08 | 5.1e-08 | 4.767E-08 | 2.021E-08 | 2.600E-08 | 6.600E-08 |
| 17938 | 897 | N897W | C | 5.2e-08 | 4e-08 | 1.2e-07 | 7.067E-08 | 4.314E-08 | 4.000E-08 | 1.200E-07 |
| 17939 | 897 | N897Y | C | 6e-08 | 7.5e-08 | 7e-08 | 6.833E-08 | 7.638E-09 | 6.000E-08 | 7.500E-08 |
| 17940 | 897 | N897V | C | 8.1e-08 | 9.9e-08 | 9.8e-08 | 9.267E-08 | 1.012E-08 | 8.100E-08 | 9.900E-08 |
| 17941 | 898 | Q898A | C | 4.7e-08 | 1.3e-08 | 1.8e-08 | 2.600E-08 | 1.836E-08 | 1.300E-08 | 4.700E-08 |
| 17942 | 898 | Q898R | C | 2.6e-08 | 7e-08 | 1e-08 | 3.533E-08 | 3.107E-08 | 1.000E-08 | 7.000E-08 |
| 17943 | 898 | Q898N | C | 1.7e-08 | 6.4e-08 | 5.5e-08 | 4.533E-08 | 2.495E-08 | 1.700E-08 | 6.400E-08 |
| 17944 | 898 | Q898D | C | 3.1e-08 | 6.9e-08 | 3.4e-08 | 4.467E-08 | 2.113E-08 | 3.100E-08 | 6.900E-08 |
| 17945 | 898 | Q898C | C | 2.5e-08 | 3.8e-08 | 2.9e-08 | 3.067E-08 | 6.658E-09 | 2.500E-08 | 3.800E-08 |
| 17946 | 898 | Q898E | C | 6e-08 | 1e-08 | 1.4e-08 | 2.800E-08 | 2.778E-08 | 1.000E-08 | 6.000E-08 |
| 17947 | 898 | Q898Q | C | 3.1e-08 | 3.4e-08 | 7.6e-08 | 4.700E-08 | 2.516E-08 | 3.100E-08 | 7.600E-08 |
| 17948 | 898 | Q898G | C | 8.2e-09 | 4.5e-08 | 2.6e-08 | 2.640E-08 | 1.840E-08 | 8.200E-09 | 4.500E-08 |
| 17949 | 898 | Q898H | C | 6.4e-08 | 2.3e-08 | 4.4e-08 | 4.367E-08 | 2.050E-08 | 2.300E-08 | 6.400E-08 |
| 17950 | 898 | Q898I | C | 4.8e-08 | 1.5e-08 | 3.1e-08 | 3.133E-08 | 1.650E-08 | 1.500E-08 | 4.800E-08 |
| 17951 | 898 | Q898L | C | 4e-08 | 1.3e-08 | 1.3e-08 | 2.200E-08 | 1.559E-08 | 1.300E-08 | 4.000E-08 |
| 17952 | 898 | Q898K | C | 2.6e-08 | 1.2e-08 | 3e-08 | 2.267E-08 | 9.452E-09 | 1.200E-08 | 3.000E-08 |
| 17953 | 898 | Q898M | C | 1.9e-08 | 3.8e-08 | 6.2e-09 | 2.107E-08 | 1.600E-08 | 6.200E-09 | 3.800E-08 |
| 17954 | 898 | Q898F | C | 1.8e-08 | 5.6e-09 | 5.5e-09 | 9.700E-09 | 7.188E-09 | 5.500E-09 | 1.800E-08 |
| 17955 | 898 | Q898P | C | 3e-08 | 1.6e-08 | 2.6e-08 | 2.400E-08 | 7.211E-09 | 1.600E-08 | 3.000E-08 |
| 17956 | 898 | Q898S | C | 2.7e-08 | 7.4e-08 | 5.5e-08 | 5.200E-08 | 2.364E-08 | 2.700E-08 | 7.400E-08 |
| 17957 | 898 | Q898T | C | 3.5e-08 | 6.5e-08 | 2.1e-08 | 4.033E-08 | 2.248E-08 | 2.100E-08 | 6.500E-08 |
| 17958 | 898 | Q898W | C | 2.2e-08 | 2.1e-08 | 1.9e-08 | 2.067E-08 | 1.528E-09 | 1.900E-08 | 2.200E-08 |
| 17959 | 898 | Q898Y | C | 1.7e-08 | 1.8e-08 | 1.9e-08 | 1.800E-08 | 1.000E-09 | 1.700E-08 | 1.900E-08 |
| 17960 | 898 | Q898V | C | 1.4e-08 | 1.4e-08 | 2.7e-08 | 1.833E-08 | 7.506E-09 | 1.400E-08 | 2.700E-08 |
| 17961 | 899 | R899A | C | 5.3e-08 | 7.1e-08 | 4e-08 | 5.467E-08 | 1.557E-08 | 4.000E-08 | 7.100E-08 |
| 17962 | 899 | R899R | C | 2.7e-08 | 3e-08 | 4e-08 | 3.233E-08 | 6.807E-09 | 2.700E-08 | 4.000E-08 |
| 17963 | 899 | R899N | C | 1.4e-08 | 7.3e-08 | 4.2e-08 | 4.300E-08 | 2.951E-08 | 1.400E-08 | 7.300E-08 |
| 17964 | 899 | R899D | C | 4.4e-08 | 1.5e-08 | 3.1e-08 | 3.000E-08 | 1.453E-08 | 1.500E-08 | 4.400E-08 |
| 17965 | 899 | R899C | C | 9.3e-08 | 9.8e-09 | 6.2e-09 | 3.633E-08 | 4.911E-08 | 6.200E-09 | 9.300E-08 |
| 17966 | 899 | R899E | C | 2.9e-08 | 2.2e-08 | 7.5e-08 | 4.200E-08 | 2.879E-08 | 2.200E-08 | 7.500E-08 |
| 17967 | 899 | R899Q | C | 5.4e-08 | 2.1e-08 | 4.4e-08 | 3.967E-08 | 1.692E-08 | 2.100E-08 | 5.400E-08 |
| 17968 | 899 | R899G | C | 3.7e-08 | 4.6e-08 | 6.5e-08 | 4.933E-08 | 1.429E-08 | 3.700E-08 | 6.500E-08 |
| 17969 | 899 | R899H | C | 3.8e-08 | 8.2e-08 | 3.1e-08 | 5.033E-08 | 2.765E-08 | 3.100E-08 | 8.200E-08 |
| 17970 | 899 | R899I | C | 5.1e-08 | 1.2e-07 | 3.8e-08 | 6.967E-08 | 4.407E-08 | 3.800E-08 | 1.200E-07 |
| 17971 | 899 | R899L | C | 1.7e-08 | 8.2e-08 | 6e-08 | 5.300E-08 | 3.306E-08 | 1.700E-08 | 8.200E-08 |
| 17972 | 899 | R899K | C | 4.1e-08 | 3e-08 | 2e-08 | 3.033E-08 | 1.050E-08 | 2.000E-08 | 4.100E-08 |
| 17973 | 899 | R899M | C | 3.7e-08 | 3e-08 | 1.2e-08 | 2.633E-08 | 1.290E-08 | 1.200E-08 | 3.700E-08 |
| 17974 | 899 | R899F | C | 8.2e-08 | 2e-08 | 7.4e-09 | 3.647E-08 | 3.993E-08 | 7.400E-09 | 8.200E-08 |
| 17975 | 899 | R899P | C | 1.2e-08 | 6.9e-08 | 5.4e-08 | 4.500E-08 | 2.955E-08 | 1.200E-08 | 6.900E-08 |
| 17976 | 899 | R899S | C | 8.5e-08 | 7.9e-08 | 1e-07 | 8.800E-08 | 1.082E-08 | 7.900E-08 | 1.000E-07 |
| 17977 | 899 | R899T | C | 3.7e-08 | 2.2e-08 | 2.8e-08 | 2.900E-08 | 7.550E-09 | 2.200E-08 | 3.700E-08 |
| 17978 | 899 | R899W | C | 2.9e-08 | 7.9e-08 | 4.2e-08 | 5.000E-08 | 2.594E-08 | 2.900E-08 | 7.900E-08 |
| 17979 | 899 | R899Y | C | 6.4e-08 | 4.3e-08 | 4.9e-08 | 5.200E-08 | 1.082E-08 | 4.300E-08 | 6.400E-08 |
| 17980 | 899 | R899V | C | 5.8e-08 | 4.2e-08 | 4.4e-08 | 4.800E-08 | 8.718E-09 | 4.200E-08 | 5.800E-08 |
| 17981 | 900 | F900A | C | 4e-08 | 9.7e-08 | 1e-07 | 7.900E-08 | 3.381E-08 | 4.000E-08 | 1.000E-07 |
| 17982 | 900 | F900R | C | 3.9e-08 | 4.6e-08 | 7.4e-08 | 5.300E-08 | 1.852E-08 | 3.900E-08 | 7.400E-08 |
| 17983 | 900 | F900N | C | 9.7e-08 | 8.2e-08 | 8.6e-08 | 8.833E-08 | 7.767E-09 | 8.200E-08 | 9.700E-08 |
| 17984 | 900 | F900D | C | 6.4e-08 | 5.3e-08 | 5.3e-08 | 5.667E-08 | 6.351E-09 | 5.300E-08 | 6.400E-08 |
| 17985 | 900 | F900C | C | 4.9e-08 | 7.7e-08 | 1.3e-08 | 4.633E-08 | 3.208E-08 | 1.300E-08 | 7.700E-08 |
| 17986 | 900 | F900E | C | 6.5e-08 | 6.5e-08 | 2e-08 | 5.000E-08 | 2.598E-08 | 2.000E-08 | 6.500E-08 |
| 17987 | 900 | F900Q | C | 1e-07 | 3.1e-08 | 5e-08 | 6.033E-08 | 3.564E-08 | 3.100E-08 | 1.000E-07 |
| 17988 | 900 | F900G | C | 1.7e-08 | 7.5e-09 | 2.4e-08 | 1.617E-08 | 8.282E-09 | 7.500E-09 | 2.400E-08 |
| 17989 | 900 | F900H | C | 5.1e-08 | 1.3e-07 | 8.3e-08 | 8.800E-08 | 3.974E-08 | 5.100E-08 | 1.300E-07 |
| 17990 | 900 | F900I | C | 5.9e-08 | 4.1e-08 | 2.6e-08 | 4.200E-08 | 1.652E-08 | 2.600E-08 | 5.900E-08 |
| 17991 | 900 | F900L | C | 5.5e-08 | 2e-08 | 2.3e-08 | 3.267E-08 | 1.940E-08 | 2.000E-08 | 5.500E-08 |
| 17992 | 900 | F900K | C | 3.6e-08 | 3.4e-08 | 4.9e-08 | 3.967E-08 | 8.145E-09 | 3.400E-08 | 4.900E-08 |
| 17993 | 900 | F900M | C | 1.1e-08 | 1.4e-07 | 5.7e-08 | 6.933E-08 | 6.538E-08 | 1.100E-08 | 1.400E-07 |
| 17994 | 900 | F900F | C | 5.6e-08 | 7.2e-08 | 9.2e-08 | 7.333E-08 | 1.804E-08 | 5.600E-08 | 9.200E-08 |
| 17995 | 900 | F900P | C | 3.9e-08 | 6.8e-08 | 2.2e-08 | 4.300E-08 | 2.326E-08 | 2.200E-08 | 6.800E-08 |
| 17996 | 900 | F900S | C | 3.8e-08 | 4.2e-08 | 4.7e-08 | 4.233E-08 | 4.509E-09 | 3.800E-08 | 4.700E-08 |
| 17997 | 900 | F900T | C | 2.4e-08 | 4.5e-08 | 1.6e-07 | 7.633E-08 | 7.321E-08 | 2.400E-08 | 1.600E-07 |
| 17998 | 900 | F900W | C | 8.7e-08 | 3e-08 | 2.4e-08 | 4.700E-08 | 3.477E-08 | 2.400E-08 | 8.700E-08 |
| 17999 | 900 | F900Y | C | 4.4e-08 | 5.9e-08 | 6.1e-08 | 5.467E-08 | 9.292E-09 | 4.400E-08 | 6.100E-08 |
| 18000 | 900 | F900V | C | 1.8e-08 | 1.1e-07 | 5e-08 | 5.933E-08 | 4.670E-08 | 1.800E-08 | 1.100E-07 |
| 18001 | 901 | K901A | C | 4.9e-08 | 5.9e-08 | 3e-08 | 4.600E-08 | 1.473E-08 | 3.000E-08 | 5.900E-08 |
| 18002 | 901 | K901R | C | 5e-08 | 6.5e-08 | 2.7e-08 | 4.733E-08 | 1.914E-08 | 2.700E-08 | 6.500E-08 |
| 18003 | 901 | K901N | C | 2.7e-08 | 5.7e-08 | 4.1e-08 | 4.167E-08 | 1.501E-08 | 2.700E-08 | 5.700E-08 |
| 18004 | 901 | K901D | C | 1.7e-07 | 8e-08 | 4.7e-08 | 9.900E-08 | 6.366E-08 | 4.700E-08 | 1.700E-07 |
| 18005 | 901 | K901C | C | 5.5e-08 | 4.3e-08 | 1.9e-08 | 3.900E-08 | 1.833E-08 | 1.900E-08 | 5.500E-08 |
| 18006 | 901 | K901E | C | 5.5e-08 | 4.4e-08 | 6.3e-08 | 5.400E-08 | 9.539E-09 | 4.400E-08 | 6.300E-08 |
| 18007 | 901 | K901Q | C | 4.4e-08 | 7e-08 | 4.6e-08 | 5.333E-08 | 1.447E-08 | 4.400E-08 | 7.000E-08 |
| 18008 | 901 | K901G | C | 4.3e-08 | 2.5e-08 | 1.7e-08 | 2.833E-08 | 1.332E-08 | 1.700E-08 | 4.300E-08 |
| 18009 | 901 | K901H | C | 4.8e-08 | 1.7e-08 | 7.9e-08 | 4.800E-08 | 3.100E-08 | 1.700E-08 | 7.900E-08 |
| 18010 | 901 | K901I | C | 2.5e-08 | 6.1e-08 | 1.8e-08 | 3.467E-08 | 2.307E-08 | 1.800E-08 | 6.100E-08 |
| 18011 | 901 | K901L | C | 4.3e-08 | 2.1e-08 | 4.7e-08 | 3.700E-08 | 1.400E-08 | 2.100E-08 | 4.700E-08 |
| 18012 | 901 | K901K | C | 6e-08 | 4.1e-08 | 1.3e-07 | 7.700E-08 | 4.687E-08 | 4.100E-08 | 1.300E-07 |
| 18013 | 901 | K901M | C | 1.5e-08 | 2.4e-08 | 3.8e-08 | 2.567E-08 | 1.159E-08 | 1.500E-08 | 3.800E-08 |
| 18014 | 901 | K901F | C | 2.6e-08 | 4.3e-08 | 1.4e-08 | 2.767E-08 | 1.457E-08 | 1.400E-08 | 4.300E-08 |
| 18015 | 901 | K901P | C | 6.5e-08 | 7.1e-08 | 5.7e-08 | 6.433E-08 | 7.024E-09 | 5.700E-08 | 7.100E-08 |
| 18016 | 901 | K901S | C | 4.1e-08 | 8.5e-08 | 9.9e-08 | 7.500E-08 | 3.027E-08 | 4.100E-08 | 9.900E-08 |
| 18017 | 901 | K901T | C | 4.3e-08 | 2.5e-08 | 5.7e-08 | 4.167E-08 | 1.604E-08 | 2.500E-08 | 5.700E-08 |
| 18018 | 901 | K901W | C | 3.7e-08 | 2.6e-08 | 3e-08 | 3.100E-08 | 5.568E-09 | 2.600E-08 | 3.700E-08 |
| 18019 | 901 | K901Y | C | 4.4e-08 | 2.8e-08 | 1.5e-08 | 2.900E-08 | 1.453E-08 | 1.500E-08 | 4.400E-08 |

| | | | | | | | | | | |
|---|---|---|---|---|---|---|---|---|---|---|
| 18020 | 901 | K901V | C | 8e-08 | 2.8e-08 | 2e-08 | 4.267E-08 | 3.258E-08 | 2.000E-08 | 8.000E-08 |
| 18021 | 902 | G902A | C | 2.3e-08 | 3.9e-08 | 2.7e-08 | 2.967E-08 | 8.327E-09 | 2.300E-08 | 3.900E-08 |
| 18022 | 902 | G902R | C | 9.1e-08 | 3.6e-08 | 3.2e-08 | 5.300E-08 | 3.297E-08 | 3.200E-08 | 9.100E-08 |
| 18023 | 902 | G902N | C | 4.1e-08 | 3.3e-08 | 4.7e-08 | 4.033E-08 | 7.024E-09 | 3.300E-08 | 4.700E-08 |
| 18024 | 902 | G902D | C | 6.2e-08 | 4.8e-08 | 3.9e-08 | 4.967E-08 | 1.159E-08 | 3.900E-08 | 6.200E-08 |
| 18025 | 902 | G902C | C | 9.4e-08 | 4.6e-08 | 3.2e-08 | 5.733E-08 | 3.252E-08 | 3.200E-08 | 9.400E-08 |
| 18026 | 902 | G902E | C | 6.4e-08 | 2.4e-08 | 6.5e-08 | 5.100E-08 | 2.339E-08 | 2.400E-08 | 6.500E-08 |
| 18027 | 902 | G902Q | C | 2.4e-08 | 5.4e-08 | 3.5e-08 | 3.767E-08 | 1.518E-08 | 2.400E-08 | 5.400E-08 |
| 18028 | 902 | G902G | C | 3.1e-08 | 6.9e-08 | 4.9e-08 | 4.967E-08 | 1.901E-08 | 3.100E-08 | 6.900E-08 |
| 18029 | 902 | G902H | C | 1.5e-07 | 4.7e-08 | 1.3e-07 | 1.090E-07 | 5.462E-08 | 4.700E-08 | 1.500E-07 |
| 18030 | 902 | G902I | C | 2.3e-08 | 6.1e-08 | 4.9e-08 | 4.433E-08 | 1.943E-08 | 2.300E-08 | 6.100E-08 |
| 18031 | 902 | G902L | C | 3.9e-08 | 5.3e-08 | 1.3e-07 | 7.400E-08 | 4.900E-08 | 3.900E-08 | 1.300E-07 |
| 18032 | 902 | G902K | C | 3.4e-08 | 3.5e-08 | 7.1e-08 | 4.667E-08 | 2.108E-08 | 3.400E-08 | 7.100E-08 |
| 18033 | 902 | G902M | C | 1.4e-07 | 1.5e-08 | 3.5e-08 | 6.333E-08 | 6.714E-08 | 1.500E-08 | 1.400E-07 |
| 18034 | 902 | G902F | C | 6.9e-08 | 5.5e-08 | 1.8e-08 | 4.733E-08 | 2.635E-08 | 1.800E-08 | 6.900E-08 |
| 18035 | 902 | G902P | C | 4.2e-08 | 1.6e-07 | 4.8e-08 | 8.333E-08 | 6.646E-08 | 4.200E-08 | 1.600E-07 |
| 18036 | 902 | G902S | C | 8.6e-08 | 3.8e-08 | 6.3e-08 | 6.233E-08 | 2.401E-08 | 3.800E-08 | 8.600E-08 |
| 18037 | 902 | G902T | C | 7.5e-08 | 5.6e-08 | 4.7e-08 | 5.933E-08 | 1.429E-08 | 4.700E-08 | 7.500E-08 |
| 18038 | 902 | G902W | C | 6.2e-08 | 3.2e-08 | 3e-08 | 4.133E-08 | 1.793E-08 | 3.000E-08 | 6.200E-08 |
| 18039 | 902 | G902Y | C | 9.1e-08 | 3.5e-08 | 1.9e-08 | 4.833E-08 | 3.781E-08 | 1.900E-08 | 9.100E-08 |
| 18040 | 902 | G902V | C | 3.7e-08 | 3.6e-08 | 3.2e-08 | 3.500E-08 | 2.646E-09 | 3.200E-08 | 3.700E-08 |
| 18041 | 903 | R903A | C | 6.1e-08 | 4.4e-08 | 2.9e-08 | 4.467E-08 | 1.601E-08 | 2.900E-08 | 6.100E-08 |
| 18042 | 903 | R903R | C | 7.6e-08 | 9.7e-08 | 6.4e-08 | 7.900E-08 | 1.670E-08 | 6.400E-08 | 9.700E-08 |
| 18043 | 903 | R903N | C | 3.7e-08 | 1.3e-08 | 2.9e-08 | 2.633E-08 | 1.222E-08 | 1.300E-08 | 3.700E-08 |
| 18044 | 903 | R903D | C | 7e-08 | 2.7e-08 | 6.6e-08 | 5.433E-08 | 2.376E-08 | 2.700E-08 | 7.000E-08 |
| 18045 | 903 | R903C | C | 2.9e-08 | 4.1e-08 | 1.7e-08 | 2.900E-08 | 1.200E-08 | 1.700E-08 | 4.100E-08 |
| 18046 | 903 | R903E | C | 1.5e-07 | 1.6e-08 | 3.2e-07 | 1.620E-07 | 1.524E-07 | 1.600E-08 | 3.200E-07 |
| 18047 | 903 | R903Q | C | 2e-08 | 2.9e-08 | 5e-08 | 3.300E-08 | 1.539E-08 | 2.000E-08 | 5.000E-08 |
| 18048 | 903 | R903G | C | 5.6e-08 | 3.8e-08 | 2e-08 | 3.800E-08 | 1.800E-08 | 2.000E-08 | 5.600E-08 |
| 18049 | 903 | R903H | C | 5.9e-08 | 1.1e-07 | 1.9e-08 | 6.267E-08 | 4.561E-08 | 1.900E-08 | 1.100E-07 |
| 18050 | 903 | R903I | C | 7.5e-08 | 2.5e-08 | 7.8e-08 | 5.933E-08 | 2.977E-08 | 2.500E-08 | 7.800E-08 |
| 18051 | 903 | R903L | C | 1.1e-07 | 6.6e-08 | 2.7e-08 | 6.767E-08 | 4.153E-08 | 2.700E-08 | 1.100E-07 |
| 18052 | 903 | R903K | C | 2.3e-08 | 4.8e-08 | 1.5e-07 | 7.367E-08 | 6.728E-08 | 2.300E-08 | 1.500E-07 |
| 18053 | 903 | R903M | C | 1.9e-08 | 2.7e-08 | 2.2e-08 | 2.267E-08 | 4.041E-09 | 1.900E-08 | 2.700E-08 |
| 18054 | 903 | R903F | C | 2.3e-08 | 3.6e-08 | 3.5e-08 | 3.133E-08 | 7.234E-09 | 2.300E-08 | 3.600E-08 |
| 18055 | 903 | R903P | C | 3.7e-08 | 1.8e-08 | 5.8e-08 | 3.767E-08 | 2.001E-08 | 1.800E-08 | 5.800E-08 |
| 18056 | 903 | R903S | C | 1.1e-07 | 2.3e-08 | 1.5e-08 | 4.933E-08 | 5.269E-08 | 1.500E-08 | 1.100E-07 |
| 18057 | 903 | R903T | C | 5.7e-08 | 2.4e-08 | 5.8e-08 | 4.633E-08 | 1.935E-08 | 2.400E-08 | 5.800E-08 |
| 18058 | 903 | R903W | C | 8.2e-08 | 8.9e-08 | 6e-08 | 7.700E-08 | 1.513E-08 | 6.000E-08 | 8.900E-08 |
| 18059 | 903 | R903Y | C | 5.1e-08 | 2.5e-08 | 6.1e-08 | 4.567E-08 | 1.858E-08 | 2.500E-08 | 6.100E-08 |
| 18060 | 903 | R903V | C | 4.6e-08 | 5.8e-08 | 3.7e-08 | 4.700E-08 | 1.054E-08 | 3.700E-08 | 5.800E-08 |
| 18061 | 904 | F904A | C | 4.4e-08 | 3.1e-08 | 2.2e-08 | 3.233E-08 | 1.106E-08 | 2.200E-08 | 4.400E-08 |
| 18062 | 904 | F904R | C | 5.3e-08 | 6.4e-08 | 4.3e-08 | 5.333E-08 | 1.050E-08 | 4.300E-08 | 6.400E-08 |
| 18063 | 904 | F904N | C | 3.4e-08 | 2.6e-08 | 2.5e-08 | 2.833E-08 | 4.933E-09 | 2.500E-08 | 3.400E-08 |
| 18064 | 904 | F904D | C | 6.4e-08 | 3.1e-08 | 2.1e-08 | 3.867E-08 | 2.250E-08 | 2.100E-08 | 6.400E-08 |
| 18065 | 904 | F904C | C | 9.3e-08 | 2.3e-08 | 3.2e-08 | 4.933E-08 | 3.808E-08 | 2.300E-08 | 9.300E-08 |
| 18066 | 904 | F904E | C | 3.4e-08 | 3.8e-08 | 3e-08 | 3.400E-08 | 4.000E-09 | 3.000E-08 | 3.800E-08 |
| 18067 | 904 | F904Q | C | 6.5e-08 | 1.6e-08 | 4.4e-08 | 4.167E-08 | 2.458E-08 | 1.600E-08 | 6.500E-08 |
| 18068 | 904 | F904G | C | 3.5e-08 | 1.5e-08 | 5e-08 | 3.333E-08 | 1.756E-08 | 1.500E-08 | 5.000E-08 |
| 18069 | 904 | F904H | C | 6.2e-08 | 1.6e-08 | 4e-08 | 3.933E-08 | 2.301E-08 | 1.600E-08 | 6.200E-08 |
| 18070 | 904 | F904I | C | 3.7e-08 | 1.5e-08 | 1.6e-08 | 2.267E-08 | 1.242E-08 | 1.500E-08 | 3.700E-08 |
| 18071 | 904 | F904L | C | 5.6e-08 | 3.3e-08 | 3.5e-08 | 4.133E-08 | 1.274E-08 | 3.300E-08 | 5.600E-08 |
| 18072 | 904 | F904K | C | 2e-08 | 2.5e-08 | 4.5e-08 | 3.000E-08 | 1.323E-08 | 2.000E-08 | 4.500E-08 |
| 18073 | 904 | F904M | C | 2.4e-08 | 3.2e-08 | 4.4e-08 | 3.333E-08 | 1.007E-08 | 2.400E-08 | 4.400E-08 |
| 18074 | 904 | F904F | C | 3e-08 | 4.1e-08 | 4.4e-08 | 3.833E-08 | 7.371E-09 | 3.000E-08 | 4.400E-08 |
| 18075 | 904 | F904P | C | 1.5e-08 | 2.1e-08 | 5.7e-08 | 3.100E-08 | 2.272E-08 | 1.500E-08 | 5.700E-08 |
| 18076 | 904 | F904S | C | 3.1e-08 | 2.4e-08 | 5.4e-08 | 3.633E-08 | 1.570E-08 | 2.400E-08 | 5.400E-08 |
| 18077 | 904 | F904T | C | 4.7e-08 | 5.1e-08 | 3.7e-08 | 4.500E-08 | 7.211E-09 | 3.700E-08 | 5.100E-08 |
| 18078 | 904 | F904W | C | 2.4e-08 | 4.6e-08 | 4.9e-08 | 3.967E-08 | 1.365E-08 | 2.400E-08 | 4.900E-08 |
| 18079 | 904 | F904Y | C | 5.3e-08 | 3.2e-08 | 8e-08 | 5.500E-08 | 2.406E-08 | 3.200E-08 | 8.000E-08 |
| 18080 | 904 | F904V | C | 3.4e-08 | 3.1e-08 | 5.6e-08 | 4.033E-08 | 1.365E-08 | 3.100E-08 | 5.600E-08 |
| 18081 | 905 | T905A | C | 6.7e-08 | 3.1e-08 | 3.7e-08 | 4.500E-08 | 1.929E-08 | 3.100E-08 | 6.700E-08 |
| 18082 | 905 | T905R | C | 1.7e-08 | 7.6e-08 | 2.7e-08 | 4.000E-08 | 3.158E-08 | 1.700E-08 | 7.600E-08 |
| 18083 | 905 | T905N | C | 8.9e-09 | 2.6e-08 | 2.1e-08 | 1.863E-08 | 8.792E-09 | 8.900E-09 | 2.600E-08 |
| 18084 | 905 | T905D | C | 5.6e-08 | 4.4e-08 | 9.5e-08 | 6.500E-08 | 2.666E-08 | 4.400E-08 | 9.500E-08 |
| 18085 | 905 | T905C | C | 2.4e-08 | 2.7e-08 | 1.1e-07 | 5.367E-08 | 4.881E-08 | 2.400E-08 | 1.100E-07 |
| 18086 | 905 | T905E | C | 3.6e-08 | 3.5e-08 | 3.2e-08 | 3.433E-08 | 2.082E-09 | 3.200E-08 | 3.600E-08 |
| 18087 | 905 | T905Q | C | 5.6e-08 | 4.3e-08 | 9.5e-08 | 6.467E-08 | 2.706E-08 | 4.300E-08 | 9.500E-08 |
| 18088 | 905 | T905G | C | 5.6e-08 | 3.7e-08 | 1.7e-08 | 3.667E-08 | 1.950E-08 | 1.700E-08 | 5.600E-08 |
| 18089 | 905 | T905H | C | 3.1e-08 | 4.2e-08 | 2.9e-08 | 3.400E-08 | 7.000E-09 | 2.900E-08 | 4.200E-08 |
| 18090 | 905 | T905I | C | 3.3e-08 | 3.6e-08 | 1.7e-08 | 2.867E-08 | 1.021E-08 | 1.700E-08 | 3.600E-08 |
| 18091 | 905 | T905L | C | 4e-08 | 7.6e-08 | 1e-08 | 4.200E-08 | 3.305E-08 | 1.000E-08 | 7.600E-08 |
| 18092 | 905 | T905K | C | 2.4e-08 | 2.2e-08 | 2.3e-08 | 2.300E-08 | 1.000E-09 | 2.200E-08 | 2.400E-08 |
| 18093 | 905 | T905M | C | 9.7e-09 | 1.8e-08 | 3.4e-08 | 2.057E-08 | 1.235E-08 | 9.700E-09 | 3.400E-08 |
| 18094 | 905 | T905F | C | 1.1e-07 | 8.9e-08 | 2.2e-08 | 7.367E-08 | 4.596E-08 | 2.200E-08 | 1.100E-07 |
| 18095 | 905 | T905P | C | 7.3e-08 | 3.6e-08 | 2.6e-08 | 4.500E-08 | 2.476E-08 | 2.600E-08 | 7.300E-08 |
| 18096 | 905 | T905S | C | 4.5e-08 | 2.6e-08 | 7e-08 | 4.700E-08 | 2.207E-08 | 2.600E-08 | 7.000E-08 |
| 18097 | 905 | T905T | C | 5.2e-08 | 2.1e-08 | 3.8e-08 | 3.700E-08 | 1.552E-08 | 2.100E-08 | 5.200E-08 |
| 18098 | 905 | T905W | C | 1.9e-08 | 5.2e-08 | 2.5e-08 | 3.200E-08 | 1.758E-08 | 1.900E-08 | 5.200E-08 |
| 18099 | 905 | T905Y | C | 4.9e-08 | 2.6e-08 | 5.5e-08 | 4.333E-08 | 1.531E-08 | 2.600E-08 | 5.500E-08 |
| 18100 | 905 | T905V | C | 4.7e-08 | 5.3e-08 | 4.7e-08 | 4.900E-08 | 3.464E-09 | 4.700E-08 | 5.300E-08 |
| 18101 | 906 | L906A | C | 4.2e-08 | 2.1e-08 | 2e-08 | 2.767E-08 | 1.242E-08 | 2.000E-08 | 4.200E-08 |
| 18102 | 906 | L906R | C | 1.6e-08 | 4.1e-08 | 1.8e-07 | 7.900E-08 | 8.836E-08 | 1.600E-08 | 1.800E-07 |
| 18103 | 906 | L906N | C | 9.2e-08 | 4.2e-08 | 1.6e-08 | 5.000E-08 | 3.863E-08 | 1.600E-08 | 9.200E-08 |
| 18104 | 906 | L906D | C | 1.2e-08 | 5.9e-08 | 2.4e-08 | 3.167E-08 | 2.442E-08 | 1.200E-08 | 5.900E-08 |

| | | | | | | | | | | |
|---|---|---|---|---|---|---|---|---|---|---|
| 18105 | 906 | L906C | C | 1.8e-08 | 7.2e-08 | 3.8e-08 | 4.267E-08 | 2.730E-08 | 1.800E-08 | 7.200E-08 |
| 18106 | 906 | L906E | C | 3.8e-08 | 2.1e-08 | 1.7e-08 | 2.533E-08 | 1.115E-08 | 1.700E-08 | 3.800E-08 |
| 18107 | 906 | L906Q | C | 3.8e-08 | 5e-08 | 1.2e-08 | 3.333E-08 | 1.943E-08 | 1.200E-08 | 5.000E-08 |
| 18108 | 906 | L906G | C | 1.7e-08 | 2e-08 | 3.9e-08 | 2.533E-08 | 1.193E-08 | 1.700E-08 | 3.900E-08 |
| 18109 | 906 | L906H | C | 1.6e-07 | 4.3e-08 | 3e-08 | 7.767E-08 | 7.160E-08 | 3.000E-08 | 1.600E-07 |
| 18110 | 906 | L906I | C | 4.7e-08 | 9.1e-08 | 3e-08 | 5.600E-08 | 3.148E-08 | 3.000E-08 | 9.100E-08 |
| 18111 | 906 | L906L | C | 1.5e-08 | 1.7e-08 | 3.5e-08 | 2.233E-08 | 1.102E-08 | 1.500E-08 | 3.500E-08 |
| 18112 | 906 | L906K | C | 5.7e-08 | 6.6e-08 | 2e-08 | 4.767E-08 | 2.438E-08 | 2.000E-08 | 6.600E-08 |
| 18113 | 906 | L906M | C | 4.6e-08 | 3.6e-08 | 4.7e-08 | 4.300E-08 | 6.083E-09 | 3.600E-08 | 4.700E-08 |
| 18114 | 906 | L906F | C | 3.1e-08 | 5.5e-08 | 6.4e-08 | 5.000E-08 | 1.706E-08 | 3.100E-08 | 6.400E-08 |
| 18115 | 906 | L906P | C | 7.4e-08 | 2.3e-08 | 4.5e-08 | 4.733E-08 | 2.558E-08 | 2.300E-08 | 7.400E-08 |
| 18116 | 906 | L906S | C | 1.4e-07 | 3.2e-08 | 3e-08 | 6.733E-08 | 6.294E-08 | 3.000E-08 | 1.400E-07 |
| 18117 | 906 | L906T | C | 3.4e-08 | 2.4e-08 | 6.3e-08 | 4.033E-08 | 2.026E-08 | 2.400E-08 | 6.300E-08 |
| 18118 | 906 | L906W | C | 3.7e-08 | 3.2e-08 | 5.9e-08 | 4.267E-08 | 1.436E-08 | 3.200E-08 | 5.900E-08 |
| 18119 | 906 | L906Y | C | 7.4e-08 | 3.1e-08 | 1.2e-07 | 7.500E-08 | 4.451E-08 | 3.100E-08 | 1.200E-07 |
| 18120 | 906 | L906V | C | 1.3e-07 | 8.2e-08 | 1e-07 | 1.040E-07 | 2.425E-08 | 8.200E-08 | 1.300E-07 |
| 18121 | 907 | S907A | C | 5.8e-08 | 4.6e-08 | 3.4e-08 | 4.600E-08 | 1.200E-08 | 3.400E-08 | 5.800E-08 |
| 18122 | 907 | S907R | C | 3.6e-08 | 1.4e-08 | 1.9e-08 | 2.300E-08 | 1.153E-08 | 1.400E-08 | 3.600E-08 |
| 18123 | 907 | S907N | C | 1.7e-08 | 2.1e-08 | 2.1e-08 | 1.967E-08 | 2.309E-09 | 1.700E-08 | 2.100E-08 |
| 18124 | 907 | S907D | C | 6.2e-08 | 2.3e-08 | 4.1e-08 | 4.200E-08 | 1.952E-08 | 2.300E-08 | 6.200E-08 |
| 18125 | 907 | S907C | C | 5e-08 | 4.5e-08 | 5.8e-09 | 3.360E-08 | 2.420E-08 | 5.800E-09 | 5.000E-08 |
| 18126 | 907 | S907E | C | 1.9e-08 | 3.8e-08 | 8.5e-08 | 4.733E-08 | 3.398E-08 | 1.900E-08 | 8.500E-08 |
| 18127 | 907 | S907Q | C | 6.7e-08 | 1.8e-08 | 4.1e-08 | 4.200E-08 | 2.452E-08 | 1.800E-08 | 6.700E-08 |
| 18128 | 907 | S907G | C | 2.7e-08 | 2e-08 | 7.9e-08 | 4.200E-08 | 3.223E-08 | 2.000E-08 | 7.900E-08 |
| 18129 | 907 | S907H | C | 3e-08 | 2.8e-08 | 5.1e-08 | 3.633E-08 | 1.274E-08 | 2.800E-08 | 5.100E-08 |
| 18130 | 907 | S907I | C | 4.2e-08 | 1.5e-08 | 9.4e-08 | 5.033E-08 | 4.015E-08 | 1.500E-08 | 9.400E-08 |
| 18131 | 907 | S907L | C | 6.1e-08 | 3.2e-08 | 3.1e-08 | 4.133E-08 | 1.704E-08 | 3.100E-08 | 6.100E-08 |
| 18132 | 907 | S907K | C | 2e-08 | 8.7e-09 | 3.1e-08 | 1.990E-08 | 1.115E-08 | 8.700E-09 | 3.100E-08 |
| 18133 | 907 | S907M | C | 4.5e-08 | 8.2e-08 | 2.6e-08 | 5.100E-08 | 2.848E-08 | 2.600E-08 | 8.200E-08 |
| 18134 | 907 | S907F | C | 2.9e-08 | 4.4e-08 | 1.8e-08 | 3.033E-08 | 1.305E-08 | 1.800E-08 | 4.400E-08 |
| 18135 | 907 | S907P | C | 6e-08 | 3e-08 | 4.2e-08 | 4.400E-08 | 1.510E-08 | 3.000E-08 | 6.000E-08 |
| 18136 | 907 | S907S | C | 5.6e-08 | 6.8e-08 | 4.9e-08 | 5.767E-08 | 9.609E-09 | 4.900E-08 | 6.800E-08 |
| 18137 | 907 | S907T | C | 5.1e-08 | 3.6e-08 | 3e-08 | 3.900E-08 | 1.082E-08 | 3.000E-08 | 5.100E-08 |
| 18138 | 907 | S907W | C | 4e-08 | 5e-08 | 1.1e-07 | 6.667E-08 | 3.786E-08 | 4.000E-08 | 1.100E-07 |
| 18139 | 907 | S907Y | C | 5.9e-08 | 2.9e-08 | 2.7e-08 | 3.833E-08 | 1.793E-08 | 2.700E-08 | 5.900E-08 |
| 18140 | 907 | S907V | C | 2e-08 | 3.6e-08 | 4.9e-08 | 3.500E-08 | 1.453E-08 | 2.000E-08 | 4.900E-08 |
| 18141 | 908 | V908A | C | 4e-08 | 4.4e-08 | 5.5e-08 | 4.633E-08 | 7.767E-09 | 4.000E-08 | 5.500E-08 |
| 18142 | 908 | V908R | C | 6.9e-08 | 5.1e-08 | 2.9e-08 | 4.967E-08 | 2.003E-08 | 2.900E-08 | 6.900E-08 |
| 18143 | 908 | V908N | C | 3.3e-08 | 9.7e-08 | 8.5e-08 | 7.167E-08 | 3.402E-08 | 3.300E-08 | 9.700E-08 |
| 18144 | 908 | V908D | C | 3.2e-08 | 5.9e-08 | 2.6e-08 | 3.900E-08 | 1.758E-08 | 2.600E-08 | 5.900E-08 |
| 18145 | 908 | V908C | C | 4.1e-08 | 2.6e-08 | 2.3e-08 | 3.000E-08 | 9.644E-09 | 2.300E-08 | 4.100E-08 |
| 18146 | 908 | V908E | C | 2.6e-08 | 4.5e-08 | 2.2e-08 | 3.100E-08 | 1.229E-08 | 2.200E-08 | 4.500E-08 |
| 18147 | 908 | V908Q | C | 1.5e-08 | 1.8e-08 | 3.7e-08 | 2.333E-08 | 1.193E-08 | 1.500E-08 | 3.700E-08 |
| 18148 | 908 | V908G | C | 4.5e-08 | 4.2e-08 | 3.4e-08 | 4.033E-08 | 5.686E-09 | 3.400E-08 | 4.500E-08 |
| 18149 | 908 | V908H | C | 1e-07 | 1.5e-08 | 2.1e-08 | 4.533E-08 | 4.744E-08 | 1.500E-08 | 1.000E-07 |
| 18150 | 908 | V908I | C | 8.7e-08 | 1.5e-07 | 3.7e-08 | 9.133E-08 | 5.662E-08 | 3.700E-08 | 1.500E-07 |
| 18151 | 908 | V908L | C | 4.2e-08 | 3.2e-08 | 3.4e-08 | 3.600E-08 | 5.292E-09 | 3.200E-08 | 4.200E-08 |
| 18152 | 908 | V908K | C | 4.2e-08 | 2.5e-08 | 4.6e-08 | 3.767E-08 | 1.115E-08 | 2.500E-08 | 4.600E-08 |
| 18153 | 908 | V908M | C | 8.3e-08 | 3.2e-08 | 4.8e-08 | 5.433E-08 | 2.608E-08 | 3.200E-08 | 8.300E-08 |
| 18154 | 908 | V908F | C | 4.9e-08 | 6.2e-08 | 2.1e-07 | 1.070E-07 | 8.944E-08 | 4.900E-08 | 2.100E-07 |
| 18155 | 908 | V908P | C | 4.8e-08 | 2.9e-08 | 4.3e-08 | 4.000E-08 | 9.849E-09 | 2.900E-08 | 4.800E-08 |
| 18156 | 908 | V908S | C | 5.9e-08 | 1.8e-08 | 7.3e-08 | 5.000E-08 | 2.858E-08 | 1.800E-08 | 7.300E-08 |
| 18157 | 908 | V908T | C | 7.7e-08 | 9e-08 | 2.1e-08 | 6.267E-08 | 3.667E-08 | 2.100E-08 | 9.000E-08 |
| 18158 | 908 | V908W | C | 3.4e-08 | 7.9e-08 | 3.1e-08 | 4.800E-08 | 2.689E-08 | 3.100E-08 | 7.900E-08 |
| 18159 | 908 | V908Y | C | 2.8e-08 | 3e-08 | 3.1e-08 | 2.967E-08 | 1.528E-09 | 2.800E-08 | 3.100E-08 |
| 18160 | 908 | V908V | C | 3.5e-08 | 2.9e-08 | 1.8e-08 | 2.733E-08 | 8.622E-09 | 1.800E-08 | 3.500E-08 |
| 18161 | 909 | D909A | C | 2.8e-08 | 2.9e-08 | 2.9e-08 | 2.867E-08 | 5.774E-10 | 2.800E-08 | 2.900E-08 |
| 18162 | 909 | D909R | C | 3.2e-08 | 1.7e-08 | 7.3e-08 | 4.067E-08 | 2.899E-08 | 1.700E-08 | 7.300E-08 |
| 18163 | 909 | D909N | C | 4.1e-08 | 5.9e-08 | 1e-07 | 6.667E-08 | 3.024E-08 | 4.100E-08 | 1.000E-07 |
| 18164 | 909 | D909D | C | 2.7e-08 | 1.8e-08 | 3.9e-08 | 2.800E-08 | 1.054E-08 | 1.800E-08 | 3.900E-08 |
| 18165 | 909 | D909C | C | 4.5e-08 | 1.6e-07 | 2.9e-08 | 7.800E-08 | 7.146E-08 | 2.900E-08 | 1.600E-07 |
| 18166 | 909 | D909E | C | 1.6e-08 | 4.2e-08 | 2.1e-08 | 2.633E-08 | 1.380E-08 | 1.600E-08 | 4.200E-08 |
| 18167 | 909 | D909Q | C | 2.4e-08 | 5.7e-08 | 5.5e-08 | 4.533E-08 | 1.850E-08 | 2.400E-08 | 5.700E-08 |
| 18168 | 909 | D909G | C | 4.6e-08 | 9.7e-08 | 1.8e-08 | 5.367E-08 | 4.005E-08 | 1.800E-08 | 9.700E-08 |
| 18169 | 909 | D909H | C | 3e-08 | 1e-07 | 4.9e-08 | 5.967E-08 | 3.620E-08 | 3.000E-08 | 1.000E-07 |
| 18170 | 909 | D909I | C | 5.7e-08 | 6.6e-08 | 1.5e-08 | 4.600E-08 | 2.722E-08 | 1.500E-08 | 6.600E-08 |
| 18171 | 909 | D909L | C | 1.5e-08 | 5.2e-08 | 8.8e-08 | 5.167E-08 | 3.650E-08 | 1.500E-08 | 8.800E-08 |
| 18172 | 909 | D909K | C | 2e-08 | 3.1e-08 | 2e-08 | 2.367E-08 | 6.351E-09 | 2.000E-08 | 3.100E-08 |
| 18173 | 909 | D909M | C | 1.2e-07 | 1.1e-07 | 4.9e-08 | 9.300E-08 | 3.843E-08 | 4.900E-08 | 1.200E-07 |
| 18174 | 909 | D909F | C | 4.4e-08 | 1.8e-07 | 9.4e-09 | 7.780E-08 | 9.018E-08 | 9.400E-09 | 1.800E-07 |
| 18175 | 909 | D909P | C | 7.6e-08 | 1.9e-08 | 1.8e-08 | 3.767E-08 | 3.320E-08 | 1.800E-08 | 7.600E-08 |
| 18176 | 909 | D909S | C | 4e-08 | 4.1e-08 | 4e-08 | 4.033E-08 | 5.774E-10 | 4.000E-08 | 4.100E-08 |
| 18177 | 909 | D909T | C | 3.2e-08 | 4.9e-08 | 4.4e-08 | 4.167E-08 | 8.737E-09 | 3.200E-08 | 4.900E-08 |
| 18178 | 909 | D909W | C | 6.4e-08 | 4.3e-08 | 2.9e-08 | 4.533E-08 | 1.762E-08 | 2.900E-08 | 6.400E-08 |
| 18179 | 909 | D909Y | C | 2.2e-08 | 9e-08 | 4.3e-08 | 5.167E-08 | 3.482E-08 | 2.200E-08 | 9.000E-08 |
| 18180 | 909 | D909V | C | 7.9e-08 | 4.3e-08 | 2.9e-08 | 5.033E-08 | 2.579E-08 | 2.900E-08 | 7.900E-08 |
| 18181 | 910 | R910A | C | 8.2e-08 | 6.1e-08 | 3.1e-08 | 5.800E-08 | 2.563E-08 | 3.100E-08 | 8.200E-08 |
| 18182 | 910 | R910R | C | 1.1e-08 | 3.7e-08 | 5.3e-08 | 3.367E-08 | 2.120E-08 | 1.100E-08 | 5.300E-08 |
| 18183 | 910 | R910N | C | 4e-08 | 7.2e-08 | 4.2e-08 | 5.133E-08 | 1.793E-08 | 4.000E-08 | 7.200E-08 |
| 18184 | 910 | R910D | C | 7e-08 | 4e-08 | 4e-08 | 5.000E-08 | 1.732E-08 | 4.000E-08 | 7.000E-08 |
| 18185 | 910 | R910C | C | 1e-07 | 3.3e-08 | 6.1e-08 | 6.467E-08 | 3.365E-08 | 3.300E-08 | 1.000E-07 |
| 18186 | 910 | R910E | C | 2e-08 | 5.6e-08 | 7.9e-08 | 5.167E-08 | 2.974E-08 | 2.000E-08 | 7.900E-08 |
| 18187 | 910 | R910Q | C | 5.4e-08 | 1.9e-08 | 2.4e-08 | 3.233E-08 | 1.893E-08 | 1.900E-08 | 5.400E-08 |
| 18188 | 910 | R910G | C | 4.2e-08 | 3.7e-08 | 7e-08 | 4.967E-08 | 1.779E-08 | 3.700E-08 | 7.000E-08 |
| 18189 | 910 | R910H | C | 5.3e-08 | 3.6e-08 | 4.9e-08 | 4.600E-08 | 8.888E-09 | 3.600E-08 | 5.300E-08 |

| | | | | | | | | | | |
|---|---|---|---|---|---|---|---|---|---|---|
| 18190 | 910 | R910I | C | 5.3e-08 | 4.9e-08 | 1.1e-07 | 7.067E-08 | 3.412E-08 | 4.900E-08 | 1.100E-07 |
| 18191 | 910 | R910L | C | 4.1e-08 | 3.8e-08 | 2.1e-08 | 3.333E-08 | 1.079E-08 | 2.100E-08 | 4.100E-08 |
| 18192 | 910 | R910K | C | 5.5e-08 | 4.6e-08 | 6.7e-08 | 5.600E-08 | 1.054E-08 | 4.600E-08 | 6.700E-08 |
| 18193 | 910 | R910M | C | 5.4e-08 | 1.2e-07 | 6.9e-08 | 8.100E-08 | 3.460E-08 | 5.400E-08 | 1.200E-07 |
| 18194 | 910 | R910F | C | 2.3e-08 | 4.5e-08 | 3.8e-08 | 3.533E-08 | 1.124E-08 | 2.300E-08 | 4.500E-08 |
| 18195 | 910 | R910P | C | 9.8e-08 | 2.1e-07 | 4.9e-08 | 1.190E-07 | 8.253E-08 | 4.900E-08 | 2.100E-07 |
| 18196 | 910 | R910S | C | 4.1e-08 | 2.3e-08 | 1.1e-07 | 5.800E-08 | 4.592E-08 | 2.300E-08 | 1.100E-07 |
| 18197 | 910 | R910T | C | 5.6e-08 | 4.8e-08 | 1e-07 | 6.800E-08 | 2.800E-08 | 4.800E-08 | 1.000E-07 |
| 18198 | 910 | R910W | C | 8.4e-08 | 7.3e-08 | 2.1e-08 | 5.933E-08 | 3.365E-08 | 2.100E-08 | 8.400E-08 |
| 18199 | 910 | R910Y | C | 7.7e-08 | 7e-08 | 3.4e-08 | 6.033E-08 | 2.307E-08 | 3.400E-08 | 7.700E-08 |
| 18200 | 910 | R910V | C | 3.9e-08 | 7.2e-08 | 1.3e-07 | 8.033E-08 | 4.607E-08 | 3.900E-08 | 1.300E-07 |
| 18201 | 911 | S911A | C | 9.1e-09 | 4.9e-08 | 1.2e-07 | 5.937E-08 | 5.617E-08 | 9.100E-09 | 1.200E-07 |
| 18202 | 911 | S911R | C | 1.5e-08 | 3.1e-08 | 8.3e-09 | 1.810E-08 | 1.166E-08 | 8.300E-09 | 3.100E-08 |
| 18203 | 911 | S911N | C | 3.3e-08 | 1.9e-08 | 1.4e-08 | 2.200E-08 | 9.849E-09 | 1.400E-08 | 3.300E-08 |
| 18204 | 911 | S911D | C | 1e-08 | 5.2e-08 | 8.2e-08 | 4.800E-08 | 3.617E-08 | 1.000E-08 | 8.200E-08 |
| 18205 | 911 | S911C | C | 1.2e-08 | 5.4e-08 | 1.9e-08 | 2.833E-08 | 2.250E-08 | 1.200E-08 | 5.400E-08 |
| 18206 | 911 | S911E | C | 3.3e-08 | 6.7e-08 | 2.3e-08 | 4.100E-08 | 2.307E-08 | 2.300E-08 | 6.700E-08 |
| 18207 | 911 | S911Q | C | 3.7e-08 | 1.1e-08 | 3e-08 | 2.600E-08 | 1.345E-08 | 1.100E-08 | 3.700E-08 |
| 18208 | 911 | S911G | C | 1.2e-08 | 1.6e-08 | 4.2e-08 | 2.333E-08 | 1.629E-08 | 1.200E-08 | 4.200E-08 |
| 18209 | 911 | S911H | C | 2.5e-08 | 4.8e-08 | 1.8e-08 | 3.033E-08 | 1.570E-08 | 1.800E-08 | 4.800E-08 |
| 18210 | 911 | S911I | C | 1.3e-07 | 8.2e-09 | 5.5e-08 | 6.440E-08 | 6.144E-08 | 8.200E-09 | 1.300E-07 |
| 18211 | 911 | S911L | C | 2e-08 | 1.9e-08 | 2.2e-08 | 2.033E-08 | 1.528E-09 | 1.900E-08 | 2.200E-08 |
| 18212 | 911 | S911K | C | 2.2e-08 | 1.7e-08 | 1.8e-08 | 1.900E-08 | 2.646E-09 | 1.700E-08 | 2.200E-08 |
| 18213 | 911 | S911M | C | 1.5e-08 | 1.2e-08 | 3.2e-08 | 1.967E-08 | 1.079E-08 | 1.200E-08 | 3.200E-08 |
| 18214 | 911 | S911F | C | 1.7e-08 | 2.1e-09 | 1.9e-08 | 1.270E-08 | 9.234E-09 | 2.100E-09 | 1.900E-08 |
| 18215 | 911 | S911P | C | 5.1e-08 | 2.6e-08 | 2.7e-08 | 3.467E-08 | 1.415E-08 | 2.600E-08 | 5.100E-08 |
| 18216 | 911 | S911S | C | 3.8e-08 | 1.5e-08 | 2.3e-08 | 2.533E-08 | 1.168E-08 | 1.500E-08 | 3.800E-08 |
| 18217 | 911 | S911T | C | 3.7e-08 | 1.8e-08 | 4.3e-08 | 3.267E-08 | 1.305E-08 | 1.800E-08 | 4.300E-08 |
| 18218 | 911 | S911W | C | 8.6e-09 | 8.3e-09 | 1.2e-08 | 9.633E-09 | 2.055E-09 | 8.300E-09 | 1.200E-08 |
| 18219 | 911 | S911Y | C | 4.9e-08 | 6e-09 | 8.4e-09 | 2.113E-08 | 2.416E-08 | 6.000E-09 | 4.900E-08 |
| 18220 | 911 | S911V | C | 2e-08 | 9.2e-09 | 2.3e-08 | 1.740E-08 | 7.258E-09 | 9.200E-09 | 2.300E-08 |
| 18221 | 912 | K912A | C | 1.4e-07 | 3.1e-08 | 3.6e-08 | 6.900E-08 | 6.154E-08 | 3.100E-08 | 1.400E-07 |
| 18222 | 912 | K912R | C | 1.1e-08 | 5.2e-08 | 3.5e-08 | 3.267E-08 | 2.060E-08 | 1.100E-08 | 5.200E-08 |
| 18223 | 912 | K912N | C | 5e-08 | 4.1e-08 | 5.9e-08 | 5.000E-08 | 9.000E-09 | 4.100E-08 | 5.900E-08 |
| 18224 | 912 | K912D | C | 5.8e-08 | 1.6e-08 | 3.9e-08 | 3.767E-08 | 2.103E-08 | 1.600E-08 | 5.800E-08 |
| 18225 | 912 | K912C | C | 6.8e-08 | 9.3e-08 | 2.9e-08 | 6.333E-08 | 3.225E-08 | 2.900E-08 | 9.300E-08 |
| 18226 | 912 | K912E | C | 1.5e-07 | 6.5e-08 | 7.3e-08 | 9.600E-08 | 4.694E-08 | 6.500E-08 | 1.500E-07 |
| 18227 | 912 | K912Q | C | 5.3e-08 | 5.1e-08 | 1.5e-07 | 8.467E-08 | 5.659E-08 | 5.100E-08 | 1.500E-07 |
| 18228 | 912 | K912G | C | 9.9e-09 | 3.4e-08 | 3.3e-08 | 2.563E-08 | 1.363E-08 | 9.900E-09 | 3.400E-08 |
| 18229 | 912 | K912H | C | 1.2e-07 | 5.3e-08 | 3.2e-08 | 6.833E-08 | 4.596E-08 | 3.200E-08 | 1.200E-07 |
| 18230 | 912 | K912I | C | 4e-08 | 1.5e-07 | 8.1e-08 | 9.033E-08 | 5.559E-08 | 4.000E-08 | 1.500E-07 |
| 18231 | 912 | K912L | C | 3.8e-08 | 5.1e-08 | 1.1e-08 | 3.333E-08 | 2.040E-08 | 1.100E-08 | 5.100E-08 |
| 18232 | 912 | K912K | C | 2.9e-08 | 1.4e-08 | 7.7e-09 | 1.690E-08 | 1.094E-08 | 7.700E-09 | 2.900E-08 |
| 18233 | 912 | K912M | C | 6.8e-08 | 5.9e-08 | 1.7e-08 | 4.800E-08 | 2.722E-08 | 1.700E-08 | 6.800E-08 |
| 18234 | 912 | K912F | C | 5.9e-08 | 7.5e-08 | 4.3e-08 | 5.900E-08 | 1.600E-08 | 4.300E-08 | 7.500E-08 |
| 18235 | 912 | K912P | C | 2.8e-08 | 5.9e-08 | 4.9e-08 | 4.533E-08 | 1.582E-08 | 2.800E-08 | 5.900E-08 |
| 18236 | 912 | K912S | C | 1.3e-07 | 3.9e-08 | 5.6e-08 | 7.500E-08 | 4.838E-08 | 3.900E-08 | 1.300E-07 |
| 18237 | 912 | K912T | C | 1.2e-07 | 9.7e-08 | 6.2e-08 | 9.300E-08 | 2.921E-08 | 6.200E-08 | 1.200E-07 |
| 18238 | 912 | K912W | C | 8.2e-08 | 1.2e-08 | 8.3e-08 | 5.900E-08 | 4.071E-08 | 1.200E-08 | 8.300E-08 |
| 18239 | 912 | K912Y | C | 2.2e-08 | 2e-08 | 6.3e-09 | 1.610E-08 | 8.546E-09 | 6.300E-09 | 2.200E-08 |
| 18240 | 912 | K912V | C | 1.5e-08 | 6.3e-08 | 5e-08 | 4.267E-08 | 2.483E-08 | 1.500E-08 | 6.300E-08 |
| 18241 | 913 | N913A | C | 2.2e-08 | 4.6e-08 | 4.8e-08 | 3.867E-08 | 1.447E-08 | 2.200E-08 | 4.800E-08 |
| 18242 | 913 | N913R | C | 4e-08 | 1.4e-08 | 9.1e-08 | 4.833E-08 | 3.917E-08 | 1.400E-08 | 9.100E-08 |
| 18243 | 913 | N913N | C | 3.3e-08 | 3.1e-08 | 5.9e-08 | 4.100E-08 | 1.562E-08 | 3.100E-08 | 5.900E-08 |
| 18244 | 913 | N913D | C | 5e-08 | 4.7e-08 | 1.1e-08 | 3.600E-08 | 2.170E-08 | 1.100E-08 | 5.000E-08 |
| 18245 | 913 | N913C | C | 5.2e-08 | 6e-08 | 5.3e-08 | 5.500E-08 | 4.359E-09 | 5.200E-08 | 6.000E-08 |
| 18246 | 913 | N913E | C | 3.4e-08 | 3.6e-08 | 3.2e-08 | 3.400E-08 | 2.000E-09 | 3.200E-08 | 3.600E-08 |
| 18247 | 913 | N913Q | C | 6.5e-08 | 1.8e-08 | 7.9e-08 | 5.400E-08 | 3.195E-08 | 1.800E-08 | 7.900E-08 |
| 18248 | 913 | N913G | C | 4.1e-08 | 6.1e-08 | 5.4e-08 | 5.200E-08 | 1.015E-08 | 4.100E-08 | 6.100E-08 |
| 18249 | 913 | N913H | C | 3.5e-08 | 4.1e-08 | 1.6e-08 | 3.067E-08 | 1.305E-08 | 1.600E-08 | 4.100E-08 |
| 18250 | 913 | N913I | C | 3.2e-08 | 2.9e-08 | 7.9e-08 | 4.667E-08 | 2.804E-08 | 2.900E-08 | 7.900E-08 |
| 18251 | 913 | N913L | C | 3.6e-08 | 8.9e-08 | 4.8e-08 | 5.767E-08 | 2.779E-08 | 3.600E-08 | 8.900E-08 |
| 18252 | 913 | N913K | C | 2.9e-08 | 7.5e-08 | 4.4e-08 | 4.933E-08 | 2.346E-08 | 2.900E-08 | 7.500E-08 |
| 18253 | 913 | N913M | C | 2.5e-08 | 1e-08 | 8.6e-08 | 4.033E-08 | 4.025E-08 | 1.000E-08 | 8.600E-08 |
| 18254 | 913 | N913F | C | 1.3e-08 | 1.9e-08 | 2.7e-08 | 1.967E-08 | 7.024E-09 | 1.300E-08 | 2.700E-08 |
| 18255 | 913 | N913P | C | 8.9e-08 | 3.4e-08 | 4.6e-08 | 5.633E-08 | 2.892E-08 | 3.400E-08 | 8.900E-08 |
| 18256 | 913 | N913S | C | 4.4e-08 | 5.8e-08 | 3.9e-08 | 4.700E-08 | 9.849E-09 | 3.900E-08 | 5.800E-08 |
| 18257 | 913 | N913T | C | 4.2e-08 | 7.8e-08 | 6.5e-08 | 6.167E-08 | 1.823E-08 | 4.200E-08 | 7.800E-08 |
| 18258 | 913 | N913W | C | 3.9e-08 | 8.2e-09 | 2.2e-08 | 2.307E-08 | 1.543E-08 | 8.200E-09 | 3.900E-08 |
| 18259 | 913 | N913Y | C | 2.3e-08 | 4.6e-08 | 6e-08 | 4.300E-08 | 1.868E-08 | 2.300E-08 | 6.000E-08 |
| 18260 | 913 | N913V | C | 9.6e-08 | 5.3e-08 | 1.9e-08 | 5.600E-08 | 3.859E-08 | 1.900E-08 | 9.600E-08 |
| 18261 | 914 | T914A | C | 3.3e-08 | 3.8e-08 | 1.8e-07 | 8.367E-08 | 8.346E-08 | 3.300E-08 | 1.800E-07 |
| 18262 | 914 | T914R | C | 3.7e-08 | 5.9e-08 | 2.7e-08 | 4.100E-08 | 1.637E-08 | 2.700E-08 | 5.900E-08 |
| 18263 | 914 | T914N | C | 2e-08 | 2.4e-08 | 2.1e-08 | 2.167E-08 | 2.082E-09 | 2.000E-08 | 2.400E-08 |
| 18264 | 914 | T914D | C | 4e-08 | 8.2e-08 | 5e-08 | 5.733E-08 | 2.194E-08 | 4.000E-08 | 8.200E-08 |
| 18265 | 914 | T914C | C | 4.3e-08 | 2.9e-08 | 1e-07 | 5.733E-08 | 3.761E-08 | 2.900E-08 | 1.000E-07 |
| 18266 | 914 | T914E | C | 9.2e-08 | 6e-08 | 9.7e-08 | 8.300E-08 | 2.007E-08 | 6.000E-08 | 9.700E-08 |
| 18267 | 914 | T914Q | C | 2.7e-08 | 2.7e-08 | 3.5e-08 | 2.967E-08 | 4.619E-09 | 2.700E-08 | 3.500E-08 |
| 18268 | 914 | T914G | C | 1e-07 | 5.9e-08 | 1.8e-08 | 5.900E-08 | 4.100E-08 | 1.800E-08 | 1.000E-07 |
| 18269 | 914 | T914H | C | 3.2e-08 | 2.6e-08 | 5.5e-08 | 3.767E-08 | 1.531E-08 | 2.600E-08 | 5.500E-08 |
| 18270 | 914 | T914I | C | 5.6e-08 | 3.1e-08 | 2.3e-08 | 3.667E-08 | 1.721E-08 | 2.300E-08 | 5.600E-08 |
| 18271 | 914 | T914L | C | 3.9e-08 | 5e-08 | 2.7e-08 | 3.867E-08 | 1.150E-08 | 2.700E-08 | 5.000E-08 |
| 18272 | 914 | T914K | C | 7.3e-08 | 4.1e-08 | 2.5e-08 | 4.633E-08 | 2.444E-08 | 2.500E-08 | 7.300E-08 |
| 18273 | 914 | T914M | C | 7.9e-08 | 1.8e-08 | 6.1e-08 | 5.267E-08 | 3.134E-08 | 1.800E-08 | 7.900E-08 |
| 18274 | 914 | T914F | C | 5e-08 | 3.6e-08 | 4.3e-08 | 4.300E-08 | 7.000E-09 | 3.600E-08 | 5.000E-08 |

| | | | | | | | | | | |
|---|---|---|---|---|---|---|---|---|---|---|
| 18275 | 914 | T914P | C | 6.5e-08 | 4.9e-08 | 4.2e-08 | 5.200E-08 | 1.179E-08 | 4.200E-08 | 6.500E-08 |
| 18276 | 914 | T914S | C | 5.5e-08 | 2e-08 | 6.4e-08 | 4.633E-08 | 2.325E-08 | 2.000E-08 | 6.400E-08 |
| 18277 | 914 | T914T | C | 1.5e-07 | 6.6e-08 | 9.7e-08 | 1.043E-07 | 4.248E-08 | 6.600E-08 | 1.500E-07 |
| 18278 | 914 | T914W | C | 2.4e-08 | 1.4e-08 | 4.9e-08 | 2.900E-08 | 1.803E-08 | 1.400E-08 | 4.900E-08 |
| 18279 | 914 | T914Y | C | 9.1e-08 | 2.6e-08 | 5.6e-08 | 5.767E-08 | 3.253E-08 | 2.600E-08 | 9.100E-08 |
| 18280 | 914 | T914V | C | 8.3e-08 | 5.2e-08 | 6e-08 | 6.500E-08 | 1.609E-08 | 5.200E-08 | 8.300E-08 |
| 18281 | 915 | L915A | C | 2.7e-08 | 9.8e-08 | 6.7e-08 | 6.400E-08 | 3.559E-08 | 2.700E-08 | 9.800E-08 |
| 18282 | 915 | L915R | C | 4.2e-08 | 5.8e-08 | 4.1e-08 | 4.700E-08 | 9.539E-09 | 4.100E-08 | 5.800E-08 |
| 18283 | 915 | L915N | C | 5e-08 | 8.1e-08 | 5.4e-08 | 6.167E-08 | 1.686E-08 | 5.000E-08 | 8.100E-08 |
| 18284 | 915 | L915D | C | 1.1e-07 | 6.9e-08 | 5.5e-08 | 7.800E-08 | 2.858E-08 | 5.500E-08 | 1.100E-07 |
| 18285 | 915 | L915C | C | 4.9e-08 | 5.5e-08 | 5.1e-08 | 5.167E-08 | 3.055E-09 | 4.900E-08 | 5.500E-08 |
| 18286 | 915 | L915E | C | 4.3e-08 | 1.4e-08 | 6.6e-08 | 4.100E-08 | 2.606E-08 | 1.400E-08 | 6.600E-08 |
| 18287 | 915 | L915Q | C | 7.5e-08 | 1.8e-08 | 5.3e-08 | 4.867E-08 | 2.875E-08 | 1.800E-08 | 7.500E-08 |
| 18288 | 915 | L915G | C | 9.2e-09 | 3.1e-08 | 2.7e-08 | 2.240E-08 | 1.161E-08 | 9.200E-09 | 3.100E-08 |
| 18289 | 915 | L915H | C | 3.1e-08 | 2.7e-08 | 6.4e-08 | 4.067E-08 | 2.031E-08 | 2.700E-08 | 6.400E-08 |
| 18290 | 915 | L915I | C | 6.7e-08 | 1.7e-08 | 2.7e-08 | 3.700E-08 | 2.646E-08 | 1.700E-08 | 6.700E-08 |
| 18291 | 915 | L915L | C | 9.8e-08 | 2.3e-08 | 4.1e-08 | 5.400E-08 | 3.915E-08 | 2.300E-08 | 9.800E-08 |
| 18292 | 915 | L915K | C | 4.4e-08 | 2.7e-08 | 5.5e-08 | 4.200E-08 | 1.411E-08 | 2.700E-08 | 5.500E-08 |
| 18293 | 915 | L915M | C | 2e-07 | 1.7e-08 | 6e-08 | 9.233E-08 | 9.569E-08 | 1.700E-08 | 2.000E-07 |
| 18294 | 915 | L915F | C | 3.9e-08 | 1.3e-08 | 6.4e-08 | 3.867E-08 | 2.550E-08 | 1.300E-08 | 6.400E-08 |
| 18295 | 915 | L915P | C | 7.5e-08 | 1.5e-08 | 3.7e-08 | 4.233E-08 | 3.035E-08 | 1.500E-08 | 7.500E-08 |
| 18296 | 915 | L915S | C | 7.4e-08 | 4e-08 | 2.9e-08 | 4.767E-08 | 2.346E-08 | 2.900E-08 | 7.400E-08 |
| 18297 | 915 | L915T | C | 7.9e-08 | 2.1e-08 | 2.7e-08 | 4.233E-08 | 3.190E-08 | 2.100E-08 | 7.900E-08 |
| 18298 | 915 | L915W | C | 3.5e-08 | 4.7e-08 | 5.7e-08 | 4.633E-08 | 1.102E-08 | 3.500E-08 | 5.700E-08 |
| 18299 | 915 | L915Y | C | 8.5e-08 | 1.3e-07 | 3.3e-08 | 8.267E-08 | 4.854E-08 | 3.300E-08 | 1.300E-07 |
| 18300 | 915 | L915V | C | 2.4e-08 | 4.8e-08 | 9.8e-09 | 2.727E-08 | 1.931E-08 | 9.800E-09 | 4.800E-08 |
| 18301 | 916 | Y916A | C | 2.3e-08 | 9.2e-08 | 8.1e-08 | 6.533E-08 | 3.707E-08 | 2.300E-08 | 9.200E-08 |
| 18302 | 916 | Y916R | C | 2.9e-08 | 9.4e-08 | 4.6e-08 | 5.633E-08 | 3.371E-08 | 2.900E-08 | 9.400E-08 |
| 18303 | 916 | Y916N | C | 3.2e-08 | 5e-08 | 2.9e-08 | 3.700E-08 | 1.136E-08 | 2.900E-08 | 5.000E-08 |
| 18304 | 916 | Y916D | C | 3.2e-08 | 8.5e-08 | 1.7e-08 | 4.467E-08 | 3.573E-08 | 1.700E-08 | 8.500E-08 |
| 18305 | 916 | Y916C | C | 4.8e-08 | 1.1e-07 | 9.4e-08 | 8.400E-08 | 3.219E-08 | 4.800E-08 | 1.100E-07 |
| 18306 | 916 | Y916E | C | 3.7e-08 | 3.9e-08 | 5.1e-08 | 4.233E-08 | 7.572E-09 | 3.700E-08 | 5.100E-08 |
| 18307 | 916 | Y916Q | C | 5e-08 | 5.2e-08 | 4.8e-08 | 5.000E-08 | 2.000E-09 | 4.800E-08 | 5.200E-08 |
| 18308 | 916 | Y916G | C | 1.3e-07 | 6.3e-08 | 4.5e-08 | 7.933E-08 | 4.479E-08 | 4.500E-08 | 1.300E-07 |
| 18309 | 916 | Y916H | C | 1.1e-08 | 6.8e-08 | 4.3e-08 | 4.067E-08 | 2.857E-08 | 1.100E-08 | 6.800E-08 |
| 18310 | 916 | Y916I | C | 1.1e-07 | 7.9e-08 | 4.1e-08 | 7.667E-08 | 3.456E-08 | 4.100E-08 | 1.100E-07 |
| 18311 | 916 | Y916L | C | 2.2e-08 | 4.8e-08 | 4.7e-08 | 3.900E-08 | 1.473E-08 | 2.200E-08 | 4.800E-08 |
| 18312 | 916 | Y916K | C | 3.3e-08 | 1.3e-07 | 5.3e-08 | 7.200E-08 | 5.122E-08 | 3.300E-08 | 1.300E-07 |
| 18313 | 916 | Y916M | C | 2e-08 | 2.9e-08 | 6.6e-08 | 3.833E-08 | 2.438E-08 | 2.000E-08 | 6.600E-08 |
| 18314 | 916 | Y916F | C | 8.9e-08 | 1e-07 | 4.6e-08 | 7.833E-08 | 2.854E-08 | 4.600E-08 | 1.000E-07 |
| 18315 | 916 | Y916P | C | 8.2e-08 | 1.7e-08 | 7.8e-08 | 5.900E-08 | 3.643E-08 | 1.700E-08 | 8.200E-08 |
| 18316 | 916 | Y916S | C | 5.6e-08 | 2.7e-08 | 2.1e-08 | 3.467E-08 | 1.872E-08 | 2.100E-08 | 5.600E-08 |
| 18317 | 916 | Y916T | C | 2.5e-08 | 1.2e-07 | 4.8e-08 | 6.433E-08 | 4.956E-08 | 2.500E-08 | 1.200E-07 |
| 18318 | 916 | Y916W | C | 8e-08 | 7.3e-08 | 3.8e-08 | 6.367E-08 | 2.250E-08 | 3.800E-08 | 8.000E-08 |
| 18319 | 916 | Y916Y | C | 2.5e-08 | 5.9e-08 | 3.8e-08 | 4.067E-08 | 1.716E-08 | 2.500E-08 | 5.900E-08 |
| 18320 | 916 | Y916V | C | 7.9e-08 | 1.8e-08 | 6.2e-08 | 5.300E-08 | 3.148E-08 | 1.800E-08 | 7.900E-08 |
| 18321 | 917 | L917A | C | 4.6e-08 | 5.2e-08 | 9.2e-08 | 6.333E-08 | 2.501E-08 | 4.600E-08 | 9.200E-08 |
| 18322 | 917 | L917R | C | 4.2e-08 | 3.3e-08 | 1.6e-08 | 3.033E-08 | 1.320E-08 | 1.600E-08 | 4.200E-08 |
| 18323 | 917 | L917N | C | 5e-08 | 7.6e-08 | 4.3e-08 | 5.633E-08 | 1.739E-08 | 4.300E-08 | 7.600E-08 |
| 18324 | 917 | L917D | C | 3.3e-08 | 2.7e-08 | 4.4e-08 | 3.467E-08 | 8.622E-09 | 2.700E-08 | 4.400E-08 |
| 18325 | 917 | L917C | C | 7.4e-08 | 3.5e-08 | 2.1e-08 | 4.333E-08 | 2.747E-08 | 2.100E-08 | 7.400E-08 |
| 18326 | 917 | L917E | C | 8.5e-08 | 1.8e-08 | 3.6e-08 | 4.633E-08 | 3.467E-08 | 1.800E-08 | 8.500E-08 |
| 18327 | 917 | L917Q | C | 4e-08 | 2.8e-08 | 7.3e-08 | 4.700E-08 | 2.330E-08 | 2.800E-08 | 7.300E-08 |
| 18328 | 917 | L917G | C | 2.9e-08 | 7.2e-08 | 1.7e-07 | 9.033E-08 | 7.227E-08 | 2.900E-08 | 1.700E-07 |
| 18329 | 917 | L917H | C | 6.7e-08 | 7.9e-08 | 7.7e-08 | 7.433E-08 | 6.429E-09 | 6.700E-08 | 7.900E-08 |
| 18330 | 917 | L917I | C | 7.4e-08 | 9.2e-08 | 2e-08 | 6.200E-08 | 3.747E-08 | 2.000E-08 | 9.200E-08 |
| 18331 | 917 | L917L | C | 4.3e-08 | 3.7e-08 | 4.8e-08 | 4.267E-08 | 5.508E-09 | 3.700E-08 | 4.800E-08 |
| 18332 | 917 | L917K | C | 6.6e-08 | 4.4e-08 | 5.6e-08 | 5.533E-08 | 1.102E-08 | 4.400E-08 | 6.600E-08 |
| 18333 | 917 | L917M | C | 3e-08 | 4.2e-08 | 1.4e-08 | 2.867E-08 | 1.405E-08 | 1.400E-08 | 4.200E-08 |
| 18334 | 917 | L917F | C | 1.8e-07 | 2.7e-08 | 7.4e-08 | 9.367E-08 | 7.837E-08 | 2.700E-08 | 1.800E-07 |
| 18335 | 917 | L917P | C | 8.1e-08 | 3.5e-08 | 3.3e-08 | 4.967E-08 | 2.715E-08 | 3.300E-08 | 8.100E-08 |
| 18336 | 917 | L917S | C | 1.6e-07 | 6.6e-08 | 5.9e-08 | 9.500E-08 | 5.640E-08 | 5.900E-08 | 1.600E-07 |
| 18337 | 917 | L917T | C | 3e-08 | 1.6e-08 | 6.3e-08 | 3.633E-08 | 2.413E-08 | 1.600E-08 | 6.300E-08 |
| 18338 | 917 | L917W | C | 3.2e-08 | 1.6e-07 | 2.3e-08 | 7.167E-08 | 7.663E-08 | 2.300E-08 | 1.600E-07 |
| 18339 | 917 | L917Y | C | 2.5e-08 | 1.9e-08 | 2.2e-08 | 2.200E-08 | 3.000E-09 | 1.900E-08 | 2.500E-08 |
| 18340 | 917 | L917V | C | 3.8e-08 | 2.7e-08 | 4.3e-08 | 3.600E-08 | 8.185E-09 | 2.700E-08 | 4.300E-08 |
| 18341 | 918 | Q918A | C | 4.8e-08 | 2.8e-08 | 3e-08 | 3.533E-08 | 1.102E-08 | 2.800E-08 | 4.800E-08 |
| 18342 | 918 | Q918R | C | 6e-08 | 2.1e-08 | 3.3e-08 | 3.800E-08 | 1.997E-08 | 2.100E-08 | 6.000E-08 |
| 18343 | 918 | Q918N | C | 3.6e-08 | 1.9e-07 | 5.1e-08 | 9.233E-08 | 8.491E-08 | 3.600E-08 | 1.900E-07 |
| 18344 | 918 | Q918D | C | 3.8e-08 | 3.5e-08 | 1.4e-07 | 7.100E-08 | 5.977E-08 | 3.500E-08 | 1.400E-07 |
| 18345 | 918 | Q918C | C | 3.6e-08 | 5.7e-08 | 3.5e-08 | 4.267E-08 | 1.242E-08 | 3.500E-08 | 5.700E-08 |
| 18346 | 918 | Q918E | C | 6.9e-09 | 6.2e-08 | 2e-08 | 2.963E-08 | 2.879E-08 | 6.900E-09 | 6.200E-08 |
| 18347 | 918 | Q918Q | C | 5.1e-08 | 1.7e-08 | 5.1e-08 | 3.967E-08 | 1.963E-08 | 1.700E-08 | 5.100E-08 |
| 18348 | 918 | Q918G | C | 5.6e-08 | 1e-07 | 5.2e-08 | 6.933E-08 | 2.663E-08 | 5.200E-08 | 1.000E-07 |
| 18349 | 918 | Q918H | C | 5e-08 | 5.4e-08 | 3.4e-08 | 4.600E-08 | 1.058E-08 | 3.400E-08 | 5.400E-08 |
| 18350 | 918 | Q918I | C | 4.7e-08 | 5.1e-08 | 4.1e-08 | 4.633E-08 | 5.033E-09 | 4.100E-08 | 5.100E-08 |
| 18351 | 918 | Q918L | C | 8.9e-08 | 3.4e-08 | 3.5e-08 | 5.267E-08 | 3.147E-08 | 3.400E-08 | 8.900E-08 |
| 18352 | 918 | Q918K | C | 1.7e-08 | 3.4e-08 | 3.1e-08 | 2.733E-08 | 9.074E-09 | 1.700E-08 | 3.400E-08 |
| 18353 | 918 | Q918M | C | 1.6e-08 | 2e-08 | 6.5e-08 | 3.367E-08 | 2.721E-08 | 1.600E-08 | 6.500E-08 |
| 18354 | 918 | Q918F | C | 8.3e-08 | 1.5e-07 | 4.8e-08 | 9.367E-08 | 5.183E-08 | 4.800E-08 | 1.500E-07 |
| 18355 | 918 | Q918P | C | 3.3e-08 | 7.5e-08 | 1.3e-07 | 7.933E-08 | 4.864E-08 | 3.300E-08 | 1.300E-07 |
| 18356 | 918 | Q918S | C | 3.9e-08 | 4.8e-08 | 4.8e-08 | 4.500E-08 | 5.196E-09 | 3.900E-08 | 4.800E-08 |
| 18357 | 918 | Q918T | C | 2.3e-08 | 7.8e-09 | 5.9e-08 | 2.993E-08 | 2.629E-08 | 7.800E-09 | 5.900E-08 |
| 18358 | 918 | Q918W | C | 1e-08 | 5.6e-08 | 7.7e-08 | 4.767E-08 | 3.427E-08 | 1.000E-08 | 7.700E-08 |
| 18359 | 918 | Q918Y | C | 3.6e-08 | 2.1e-08 | 5.1e-08 | 3.600E-08 | 1.500E-08 | 2.100E-08 | 5.100E-08 |

| | | | | | | | | | | |
|---|---|---|---|---|---|---|---|---|---|---|
| 18360 | 918 | Q918V | C | 4.3e-08 | 5.4e-08 | 1.2e-07 | 7.233E-08 | 4.165E-08 | 4.300E-08 | 1.200E-07 |
| 18361 | 919 | M919A | C | 9.1e-08 | 1.6e-08 | 4.4e-08 | 5.033E-08 | 3.790E-08 | 1.600E-08 | 9.100E-08 |
| 18362 | 919 | M919R | C | 6.4e-08 | 3.5e-08 | 2.3e-08 | 4.067E-08 | 2.108E-08 | 2.300E-08 | 6.400E-08 |
| 18363 | 919 | M919N | C | 1.3e-07 | 2.3e-08 | 4.3e-08 | 6.533E-08 | 5.689E-08 | 2.300E-08 | 1.300E-07 |
| 18364 | 919 | M919D | C | 4.1e-08 | 6.2e-08 | 6.2e-08 | 5.500E-08 | 1.212E-08 | 4.100E-08 | 6.200E-08 |
| 18365 | 919 | M919C | C | 5.6e-08 | 3.4e-08 | 6.7e-08 | 5.233E-08 | 1.680E-08 | 3.400E-08 | 6.700E-08 |
| 18366 | 919 | M919E | C | 4.1e-08 | 4.8e-08 | 1.9e-08 | 3.600E-08 | 1.513E-08 | 1.900E-08 | 4.800E-08 |
| 18367 | 919 | M919Q | C | 7.6e-08 | 2.5e-08 | 3.8e-08 | 4.633E-08 | 2.650E-08 | 2.500E-08 | 7.600E-08 |
| 18368 | 919 | M919G | C | 2.3e-08 | 2.3e-08 | 2e-08 | 2.200E-08 | 1.732E-09 | 2.000E-08 | 2.300E-08 |
| 18369 | 919 | M919H | C | 9.4e-08 | 2.6e-08 | 3.8e-08 | 5.267E-08 | 3.630E-08 | 2.600E-08 | 9.400E-08 |
| 18370 | 919 | M919I | C | 5.4e-08 | 5.6e-08 | 1.3e-07 | 8.000E-08 | 4.331E-08 | 5.400E-08 | 1.300E-07 |
| 18371 | 919 | M919L | C | 3.8e-08 | 2.3e-08 | 4.4e-08 | 3.500E-08 | 1.082E-08 | 2.300E-08 | 4.400E-08 |
| 18372 | 919 | M919K | C | 4.3e-08 | 2.3e-08 | 6.8e-08 | 4.467E-08 | 2.255E-08 | 2.300E-08 | 6.800E-08 |
| 18373 | 919 | M919M | C | 1.7e-08 | 5.5e-08 | 1.5e-08 | 2.900E-08 | 2.254E-08 | 1.500E-08 | 5.500E-08 |
| 18374 | 919 | M919F | C | 5.5e-08 | 1.5e-08 | 5.3e-08 | 4.100E-08 | 2.254E-08 | 1.500E-08 | 5.500E-08 |
| 18375 | 919 | M919P | C | 4e-08 | 3.5e-08 | 8e-08 | 5.167E-08 | 2.466E-08 | 3.500E-08 | 8.000E-08 |
| 18376 | 919 | M919S | C | 6.6e-08 | 5.9e-08 | 7.1e-08 | 6.533E-08 | 6.028E-09 | 5.900E-08 | 7.100E-08 |
| 18377 | 919 | M919T | C | 4.3e-08 | 6.2e-08 | 3.9e-08 | 4.800E-08 | 1.229E-08 | 3.900E-08 | 6.200E-08 |
| 18378 | 919 | M919W | C | 3.9e-08 | 3.6e-08 | 1.8e-08 | 3.100E-08 | 1.136E-08 | 1.800E-08 | 3.900E-08 |
| 18379 | 919 | M919Y | C | 5.5e-08 | 9.8e-08 | 5.5e-08 | 6.933E-08 | 2.483E-08 | 5.500E-08 | 9.800E-08 |
| 18380 | 919 | M919V | C | 1.2e-07 | 5.7e-08 | 2.6e-08 | 6.767E-08 | 4.790E-08 | 2.600E-08 | 1.200E-07 |
| 18381 | 920 | N920A | C | 2.5e-08 | 1.8e-08 | 4.3e-08 | 2.867E-08 | 1.290E-08 | 1.800E-08 | 4.300E-08 |
| 18382 | 920 | N920R | C | 9.4e-08 | 1.2e-07 | 1.2e-07 | 1.113E-07 | 1.501E-08 | 9.400E-08 | 1.200E-07 |
| 18383 | 920 | N920N | C | 2.2e-08 | 8.3e-08 | 5.7e-08 | 5.400E-08 | 3.061E-08 | 2.200E-08 | 8.300E-08 |
| 18384 | 920 | N920D | C | 5.3e-08 | 1.8e-08 | 8.8e-08 | 5.300E-08 | 3.500E-08 | 1.800E-08 | 8.800E-08 |
| 18385 | 920 | N920C | C | 4e-08 | 3.6e-08 | 5e-08 | 4.200E-08 | 7.211E-09 | 3.600E-08 | 5.000E-08 |
| 18386 | 920 | N920E | C | 9.4e-08 | 4.6e-08 | 5.8e-08 | 6.600E-08 | 2.498E-08 | 4.600E-08 | 9.400E-08 |
| 18387 | 920 | N920Q | C | 2.9e-08 | 2.6e-08 | 2.6e-08 | 2.700E-08 | 1.732E-09 | 2.600E-08 | 2.900E-08 |
| 18388 | 920 | N920G | C | 2.5e-08 | 1.1e-07 | 8.2e-08 | 7.233E-08 | 4.332E-08 | 2.500E-08 | 1.100E-07 |
| 18389 | 920 | N920H | C | 8.9e-08 | 4.4e-08 | 2.6e-08 | 5.300E-08 | 3.245E-08 | 2.600E-08 | 8.900E-08 |
| 18390 | 920 | N920I | C | 2.3e-08 | 2.1e-08 | 2.1e-08 | 2.167E-08 | 1.155E-09 | 2.100E-08 | 2.300E-08 |
| 18391 | 920 | N920L | C | 5.7e-08 | 4.1e-08 | 6.5e-08 | 5.433E-08 | 1.222E-08 | 4.100E-08 | 6.500E-08 |
| 18392 | 920 | N920K | C | 2.9e-08 | 5.9e-08 | 3.1e-08 | 3.967E-08 | 1.677E-08 | 2.900E-08 | 5.900E-08 |
| 18393 | 920 | N920M | C | 1.4e-08 | 4.7e-08 | 5.8e-08 | 3.967E-08 | 2.290E-08 | 1.400E-08 | 5.800E-08 |
| 18394 | 920 | N920F | C | 6.1e-08 | 2.4e-08 | 2.1e-08 | 3.533E-08 | 2.228E-08 | 2.100E-08 | 6.100E-08 |
| 18395 | 920 | N920P | C | 2.3e-08 | 2.2e-07 | 2.8e-08 | 9.033E-08 | 1.123E-07 | 2.300E-08 | 2.200E-07 |
| 18396 | 920 | N920S | C | 9.4e-08 | 7.1e-08 | 5.3e-08 | 7.267E-08 | 2.055E-08 | 5.300E-08 | 9.400E-08 |
| 18397 | 920 | N920T | C | 5e-08 | 2.8e-08 | 4.5e-08 | 4.100E-08 | 1.153E-08 | 2.800E-08 | 5.000E-08 |
| 18398 | 920 | N920W | C | 6e-08 | 4.9e-08 | 3.9e-08 | 4.933E-08 | 1.050E-08 | 3.900E-08 | 6.000E-08 |
| 18399 | 920 | N920Y | C | 2.9e-08 | 1.6e-08 | 2.7e-08 | 2.400E-08 | 7.000E-09 | 1.600E-08 | 2.900E-08 |
| 18400 | 920 | N920V | C | 3.8e-08 | 2.2e-07 | 5.1e-08 | 1.030E-07 | 1.015E-07 | 3.800E-08 | 2.200E-07 |
| 18401 | 921 | S921A | C | 4e-08 | 5.3e-08 | 4.8e-08 | 4.700E-08 | 6.557E-09 | 4.000E-08 | 5.300E-08 |
| 18402 | 921 | S921R | C | 3.1e-08 | 4.2e-08 | 5.7e-08 | 4.333E-08 | 1.305E-08 | 3.100E-08 | 5.700E-08 |
| 18403 | 921 | S921N | C | 4.4e-08 | 1.8e-08 | 1.8e-08 | 2.667E-08 | 1.501E-08 | 1.800E-08 | 4.400E-08 |
| 18404 | 921 | S921D | C | 6e-08 | 7.7e-08 | 7.1e-08 | 6.933E-08 | 8.622E-09 | 6.000E-08 | 7.700E-08 |
| 18405 | 921 | S921C | C | 6.1e-08 | 8.3e-08 | 1e-07 | 8.133E-08 | 1.955E-08 | 6.100E-08 | 1.000E-07 |
| 18406 | 921 | S921E | C | 1.4e-08 | 5.4e-08 | 7.7e-08 | 4.833E-08 | 3.188E-08 | 1.400E-08 | 7.700E-08 |
| 18407 | 921 | S921Q | C | 4.3e-08 | 5e-08 | 6.8e-08 | 5.367E-08 | 1.290E-08 | 4.300E-08 | 6.800E-08 |
| 18408 | 921 | S921G | C | 8.6e-08 | 8.6e-08 | 7.2e-08 | 8.133E-08 | 8.083E-09 | 7.200E-08 | 8.600E-08 |
| 18409 | 921 | S921H | C | 2.2e-08 | 3.7e-08 | 8e-08 | 4.633E-08 | 3.011E-08 | 2.200E-08 | 8.000E-08 |
| 18410 | 921 | S921I | C | 2e-07 | 2.5e-08 | 3.3e-08 | 8.600E-08 | 9.881E-08 | 2.500E-08 | 2.000E-07 |
| 18411 | 921 | S921L | C | 3.3e-08 | 8.1e-08 | 9.7e-08 | 7.033E-08 | 3.331E-08 | 3.300E-08 | 9.700E-08 |
| 18412 | 921 | S921K | C | 2.4e-08 | 2.6e-08 | 4.8e-08 | 3.267E-08 | 1.332E-08 | 2.400E-08 | 4.800E-08 |
| 18413 | 921 | S921M | C | 6.9e-08 | 3.9e-08 | 1.9e-08 | 4.233E-08 | 2.517E-08 | 1.900E-08 | 6.900E-08 |
| 18414 | 921 | S921F | C | 2.5e-08 | 6.3e-08 | 2.3e-08 | 3.700E-08 | 2.254E-08 | 2.300E-08 | 6.300E-08 |
| 18415 | 921 | S921P | C | 5.6e-08 | 6.6e-08 | 2.3e-08 | 4.833E-08 | 2.250E-08 | 2.300E-08 | 6.600E-08 |
| 18416 | 921 | S921S | C | 3e-08 | 6.2e-08 | 1.3e-07 | 7.400E-08 | 5.107E-08 | 3.000E-08 | 1.300E-07 |
| 18417 | 921 | S921T | C | 1.8e-08 | 8.1e-09 | 7.4e-08 | 3.337E-08 | 3.554E-08 | 8.100E-09 | 7.400E-08 |
| 18418 | 921 | S921W | C | 3.4e-08 | 7.7e-08 | 3.7e-08 | 4.933E-08 | 2.401E-08 | 3.400E-08 | 7.700E-08 |
| 18419 | 921 | S921Y | C | 8.3e-08 | 2.8e-08 | 8e-08 | 6.367E-08 | 3.092E-08 | 2.800E-08 | 8.300E-08 |
| 18420 | 921 | S921V | C | 1.9e-08 | 5.8e-08 | 4e-08 | 3.900E-08 | 1.952E-08 | 1.900E-08 | 5.800E-08 |
| 18421 | 922 | L922A | C | 5.6e-08 | 3.5e-08 | 3.8e-08 | 4.300E-08 | 1.136E-08 | 3.500E-08 | 5.600E-08 |
| 18422 | 922 | L922R | C | 4.5e-08 | 1.2e-07 | 8e-08 | 8.167E-08 | 3.753E-08 | 4.500E-08 | 1.200E-07 |
| 18423 | 922 | L922N | C | 3e-08 | 3.4e-08 | 3.1e-08 | 3.167E-08 | 2.082E-09 | 3.000E-08 | 3.400E-08 |
| 18424 | 922 | L922D | C | 3e-08 | 8.9e-08 | 8.3e-08 | 6.733E-08 | 3.247E-08 | 3.000E-08 | 8.900E-08 |
| 18425 | 922 | L922C | C | 3.1e-08 | 7.6e-08 | 7.3e-08 | 6.000E-08 | 2.516E-08 | 3.100E-08 | 7.600E-08 |
| 18426 | 922 | L922E | C | 6.5e-08 | 4.4e-08 | 1.5e-07 | 8.633E-08 | 5.613E-08 | 4.400E-08 | 1.500E-07 |
| 18427 | 922 | L922Q | C | 9.4e-08 | 4.4e-08 | 2.7e-08 | 5.500E-08 | 3.483E-08 | 2.700E-08 | 9.400E-08 |
| 18428 | 922 | L922G | C | 8.7e-08 | 2.3e-08 | 5.6e-08 | 5.533E-08 | 3.201E-08 | 2.300E-08 | 8.700E-08 |
| 18429 | 922 | L922H | C | 2.3e-08 | 3.3e-08 | 7.4e-08 | 4.333E-08 | 2.702E-08 | 2.300E-08 | 7.400E-08 |
| 18430 | 922 | L922I | C | 4.4e-08 | 4.7e-08 | 5.2e-08 | 4.767E-08 | 4.041E-09 | 4.400E-08 | 5.200E-08 |
| 18431 | 922 | L922L | C | 2e-08 | 1.1e-07 | 5.1e-08 | 6.033E-08 | 4.572E-08 | 2.000E-08 | 1.100E-07 |
| 18432 | 922 | L922K | C | 1.3e-07 | 3e-08 | 5.3e-08 | 7.100E-08 | 5.237E-08 | 3.000E-08 | 1.300E-07 |
| 18433 | 922 | L922M | C | 3.4e-08 | 4.1e-08 | 5.4e-08 | 4.300E-08 | 1.015E-08 | 3.400E-08 | 5.400E-08 |
| 18434 | 922 | L922F | C | 1.1e-07 | 2e-08 | 9.9e-08 | 7.633E-08 | 4.910E-08 | 2.000E-08 | 1.100E-07 |
| 18435 | 922 | L922P | C | 2.6e-08 | 7e-08 | 4.5e-08 | 4.700E-08 | 2.207E-08 | 2.600E-08 | 7.000E-08 |
| 18436 | 922 | L922S | C | 6.4e-08 | 3.1e-08 | 2.1e-08 | 3.867E-08 | 2.250E-08 | 2.100E-08 | 6.400E-08 |
| 18437 | 922 | L922T | C | 3.9e-08 | 1.9e-08 | 8.9e-08 | 4.900E-08 | 3.606E-08 | 1.900E-08 | 8.900E-08 |
| 18438 | 922 | L922W | C | 3.4e-08 | 1.4e-08 | 6.6e-08 | 3.800E-08 | 2.623E-08 | 1.400E-08 | 6.600E-08 |
| 18439 | 922 | L922Y | C | 8.5e-08 | 2.8e-08 | 3.4e-08 | 4.900E-08 | 3.132E-08 | 2.800E-08 | 8.500E-08 |
| 18440 | 922 | L922V | C | 7.8e-08 | 3e-08 | 6.6e-08 | 5.800E-08 | 2.498E-08 | 3.000E-08 | 7.800E-08 |
| 18441 | 923 | R923A | C | 4.9e-08 | 1.7e-08 | 4.3e-08 | 3.633E-08 | 1.701E-08 | 1.700E-08 | 4.900E-08 |
| 18442 | 923 | R923R | C | 4.9e-08 | 3.3e-08 | 5.1e-08 | 4.433E-08 | 9.866E-09 | 3.300E-08 | 5.100E-08 |
| 18443 | 923 | R923N | C | 3e-08 | 2.9e-08 | 1.1e-08 | 2.333E-08 | 1.069E-08 | 1.100E-08 | 3.000E-08 |
| 18444 | 923 | R923D | C | 3.6e-08 | 2.1e-08 | 4.8e-08 | 3.500E-08 | 1.353E-08 | 2.100E-08 | 4.800E-08 |

| | | | | | | | | | | |
|---|---|---|---|---|---|---|---|---|---|---|
| 18445 | 923 | R923C | C | 4.8e-08 | 2.6e-08 | 6.5e-08 | 4.633E-08 | 1.955E-08 | 2.600E-08 | 6.500E-08 |
| 18446 | 923 | R923E | C | 6.5e-08 | 3.2e-08 | 1.9e-08 | 3.867E-08 | 2.371E-08 | 1.900E-08 | 6.500E-08 |
| 18447 | 923 | R923Q | C | 6e-08 | 5.9e-08 | 9.3e-08 | 7.067E-08 | 1.935E-08 | 5.900E-08 | 9.300E-08 |
| 18448 | 923 | R923G | C | 5.4e-08 | 2.4e-08 | 7.4e-08 | 5.067E-08 | 2.517E-08 | 2.400E-08 | 7.400E-08 |
| 18449 | 923 | R923H | C | 4.1e-08 | 3.4e-08 | 4.3e-08 | 3.933E-08 | 4.726E-09 | 3.400E-08 | 4.300E-08 |
| 18450 | 923 | R923I | C | 2.9e-08 | 3.3e-08 | 8.3e-08 | 4.833E-08 | 3.009E-08 | 2.900E-08 | 8.300E-08 |
| 18451 | 923 | R923L | C | 4.8e-08 | 6.1e-08 | 4.2e-08 | 5.033E-08 | 9.713E-09 | 4.200E-08 | 6.100E-08 |
| 18452 | 923 | R923K | C | 2.8e-08 | 2.1e-08 | 4.4e-08 | 3.100E-08 | 1.179E-08 | 2.100E-08 | 4.400E-08 |
| 18453 | 923 | R923M | C | 4.9e-08 | 6.4e-08 | 1.2e-07 | 7.767E-08 | 3.742E-08 | 4.900E-08 | 1.200E-07 |
| 18454 | 923 | R923F | C | 5e-08 | 4.6e-08 | 4.2e-08 | 4.600E-08 | 4.000E-09 | 4.200E-08 | 5.000E-08 |
| 18455 | 923 | R923P | C | 4.5e-08 | 4.9e-08 | 2.8e-08 | 4.067E-08 | 1.115E-08 | 2.800E-08 | 4.900E-08 |
| 18456 | 923 | R923S | C | 7.3e-09 | 5.9e-08 | 7.2e-08 | 4.610E-08 | 3.422E-08 | 7.300E-09 | 7.200E-08 |
| 18457 | 923 | R923T | C | 6.4e-08 | 4.1e-08 | 3.9e-08 | 4.800E-08 | 1.389E-08 | 3.900E-08 | 6.400E-08 |
| 18458 | 923 | R923W | C | 3.4e-08 | 2.8e-08 | 4.4e-08 | 3.533E-08 | 8.083E-09 | 2.800E-08 | 4.400E-08 |
| 18459 | 923 | R923Y | C | 3e-08 | 1.4e-07 | 9.8e-08 | 8.933E-08 | 5.551E-08 | 3.000E-08 | 1.400E-07 |
| 18460 | 923 | R923V | C | 2.6e-08 | 4e-08 | 1.1e-07 | 5.867E-08 | 4.500E-08 | 2.600E-08 | 1.100E-07 |
| 18461 | 924 | A924A | C | 3.6e-08 | 8.8e-08 | 3.2e-08 | 5.200E-08 | 3.124E-08 | 3.200E-08 | 8.800E-08 |
| 18462 | 924 | A924R | C | 7.8e-08 | 2.8e-08 | 1.3e-07 | 7.867E-08 | 5.100E-08 | 2.800E-08 | 1.300E-07 |
| 18463 | 924 | A924N | C | 1.4e-08 | 5e-08 | 3.3e-08 | 3.233E-08 | 1.801E-08 | 1.400E-08 | 5.000E-08 |
| 18464 | 924 | A924D | C | 5e-08 | 2.3e-08 | 4.1e-08 | 3.800E-08 | 1.375E-08 | 2.300E-08 | 5.000E-08 |
| 18465 | 924 | A924C | C | 1.3e-08 | 5.5e-08 | 1.5e-07 | 7.267E-08 | 7.019E-08 | 1.300E-08 | 1.500E-07 |
| 18466 | 924 | A924E | C | 3.9e-08 | 5.6e-08 | 5.5e-08 | 5.000E-08 | 9.539E-09 | 3.900E-08 | 5.600E-08 |
| 18467 | 924 | A924Q | C | 5e-08 | 4.7e-08 | 1e-07 | 6.567E-08 | 2.977E-08 | 4.700E-08 | 1.000E-07 |
| 18468 | 924 | A924G | C | 2.5e-08 | 1.6e-08 | 8.8e-08 | 4.300E-08 | 3.923E-08 | 1.600E-08 | 8.800E-08 |
| 18469 | 924 | A924H | C | 5.7e-08 | 1.9e-08 | 2.4e-08 | 3.333E-08 | 2.065E-08 | 1.900E-08 | 5.700E-08 |
| 18470 | 924 | A924I | C | 1.8e-08 | 3.5e-08 | 4.8e-08 | 3.367E-08 | 1.504E-08 | 1.800E-08 | 4.800E-08 |
| 18471 | 924 | A924L | C | 1.5e-07 | 7.8e-08 | 3e-08 | 8.600E-08 | 6.040E-08 | 3.000E-08 | 1.500E-07 |
| 18472 | 924 | A924K | C | 3.4e-08 | 2.5e-08 | 6.1e-08 | 4.000E-08 | 1.873E-08 | 2.500E-08 | 6.100E-08 |
| 18473 | 924 | A924M | C | 4.5e-08 | 2.9e-08 | 2.2e-08 | 3.200E-08 | 1.179E-08 | 2.200E-08 | 4.500E-08 |
| 18474 | 924 | A924F | C | 1.6e-08 | 5.3e-08 | 7.6e-08 | 4.833E-08 | 3.027E-08 | 1.600E-08 | 7.600E-08 |
| 18475 | 924 | A924P | C | 7e-08 | 4.4e-08 | 1.7e-08 | 4.367E-08 | 2.650E-08 | 1.700E-08 | 7.000E-08 |
| 18476 | 924 | A924S | C | 3.9e-08 | 3.7e-08 | 3.8e-08 | 3.800E-08 | 1.000E-09 | 3.700E-08 | 3.900E-08 |
| 18477 | 924 | A924T | C | 2.6e-08 | 1e-08 | 2.5e-08 | 2.033E-08 | 8.963E-09 | 1.000E-08 | 2.600E-08 |
| 18478 | 924 | A924W | C | 6e-08 | 2.2e-08 | 4.1e-08 | 4.100E-08 | 1.900E-08 | 2.200E-08 | 6.000E-08 |
| 18479 | 924 | A924Y | C | 1.9e-08 | 4.1e-08 | 8.4e-08 | 4.800E-08 | 3.306E-08 | 1.900E-08 | 8.400E-08 |
| 18480 | 924 | A924V | C | 7.9e-08 | 4.1e-08 | 2.5e-08 | 4.833E-08 | 2.774E-08 | 2.500E-08 | 7.900E-08 |
| 18481 | 925 | E925A | C | 6.2e-08 | 5.1e-08 | 4.7e-08 | 5.333E-08 | 7.767E-09 | 4.700E-08 | 6.200E-08 |
| 18482 | 925 | E925R | C | 2.8e-08 | 1.6e-08 | 2.6e-08 | 2.333E-08 | 6.429E-09 | 1.600E-08 | 2.800E-08 |
| 18483 | 925 | E925N | C | 4.7e-08 | 2.8e-08 | 1.9e-08 | 3.133E-08 | 1.429E-08 | 1.900E-08 | 4.700E-08 |
| 18484 | 925 | E925D | C | 4.4e-08 | 4.7e-08 | 2.5e-08 | 3.867E-08 | 1.193E-08 | 2.500E-08 | 4.700E-08 |
| 18485 | 925 | E925C | C | 7.8e-08 | 8.4e-08 | 4.3e-08 | 6.833E-08 | 2.214E-08 | 4.300E-08 | 8.400E-08 |
| 18486 | 925 | E925E | C | 1.4e-08 | 3.7e-08 | 6.1e-08 | 3.733E-08 | 2.350E-08 | 1.400E-08 | 6.100E-08 |
| 18487 | 925 | E925Q | C | 4.5e-08 | 1.6e-07 | 3.5e-08 | 8.000E-08 | 6.946E-08 | 3.500E-08 | 1.600E-07 |
| 18488 | 925 | E925G | C | 7.3e-08 | 4.9e-08 | 2.7e-08 | 4.967E-08 | 2.301E-08 | 2.700E-08 | 7.300E-08 |
| 18489 | 925 | E925H | C | 5.7e-08 | 7.2e-08 | 2.6e-08 | 5.167E-08 | 2.346E-08 | 2.600E-08 | 7.200E-08 |
| 18490 | 925 | E925I | C | 7e-08 | 4.9e-08 | 3.6e-08 | 5.167E-08 | 1.716E-08 | 3.600E-08 | 7.000E-08 |
| 18491 | 925 | E925L | C | 3.7e-08 | 2.2e-08 | 1.4e-07 | 6.633E-08 | 6.424E-08 | 2.200E-08 | 1.400E-07 |
| 18492 | 925 | E925K | C | 4.6e-08 | 6.9e-08 | 2.4e-08 | 4.633E-08 | 2.250E-08 | 2.400E-08 | 6.900E-08 |
| 18493 | 925 | E925M | C | 5.8e-08 | 2.6e-08 | 3.8e-08 | 4.067E-08 | 1.617E-08 | 2.600E-08 | 5.800E-08 |
| 18494 | 925 | E925F | C | 2.6e-08 | 6e-08 | 2.6e-08 | 3.733E-08 | 1.963E-08 | 2.600E-08 | 6.000E-08 |
| 18495 | 925 | E925P | C | 1e-07 | 6.9e-09 | 4.4e-08 | 5.030E-08 | 4.687E-08 | 6.900E-09 | 1.000E-07 |
| 18496 | 925 | E925S | C | 6.1e-08 | 2.8e-08 | 4.6e-08 | 4.500E-08 | 1.652E-08 | 2.800E-08 | 6.100E-08 |
| 18497 | 925 | E925T | C | 8.7e-09 | 5e-08 | 5.4e-08 | 3.757E-08 | 2.508E-08 | 8.700E-09 | 5.400E-08 |
| 18498 | 925 | E925W | C | 3.1e-08 | 5.8e-08 | 2.6e-08 | 3.833E-08 | 1.721E-08 | 2.600E-08 | 5.800E-08 |
| 18499 | 925 | E925Y | C | 3.6e-08 | 5.2e-08 | 4.2e-08 | 4.333E-08 | 8.083E-09 | 3.600E-08 | 5.200E-08 |
| 18500 | 925 | E925V | C | 4.4e-08 | 1.6e-08 | 3.4e-08 | 3.133E-08 | 1.419E-08 | 1.600E-08 | 4.400E-08 |
| 18501 | 926 | D926A | C | 2.2e-08 | 1.9e-08 | 2.8e-08 | 2.300E-08 | 4.583E-09 | 1.900E-08 | 2.800E-08 |
| 18502 | 926 | D926R | C | 3e-08 | 3.7e-08 | 4.1e-08 | 3.600E-08 | 5.568E-09 | 3.000E-08 | 4.100E-08 |
| 18503 | 926 | D926N | C | 2.2e-08 | 8.8e-09 | 5.4e-08 | 2.827E-08 | 2.324E-08 | 8.800E-09 | 5.400E-08 |
| 18504 | 926 | D926D | C | 3.3e-08 | 3.1e-08 | 3.3e-08 | 3.233E-08 | 1.155E-09 | 3.100E-08 | 3.300E-08 |
| 18505 | 926 | D926C | C | 5.4e-08 | 1.7e-08 | 4.1e-08 | 3.733E-08 | 1.877E-08 | 1.700E-08 | 5.400E-08 |
| 18506 | 926 | D926E | C | 2.9e-08 | 8.5e-08 | 2.8e-08 | 4.733E-08 | 3.262E-08 | 2.800E-08 | 8.500E-08 |
| 18507 | 926 | D926Q | C | 4.3e-08 | 4.9e-08 | 4.6e-08 | 4.600E-08 | 3.000E-09 | 4.300E-08 | 4.900E-08 |
| 18508 | 926 | D926G | C | 3e-08 | 1e-08 | 6.9e-08 | 3.633E-08 | 3.001E-08 | 1.000E-08 | 6.900E-08 |
| 18509 | 926 | D926H | C | 4.3e-08 | 5.6e-08 | 3.2e-08 | 4.367E-08 | 1.201E-08 | 3.200E-08 | 5.600E-08 |
| 18510 | 926 | D926I | C | 4.1e-08 | 8.1e-08 | 2.8e-08 | 5.000E-08 | 2.762E-08 | 2.800E-08 | 8.100E-08 |
| 18511 | 926 | D926L | C | 1.4e-07 | 7.3e-08 | 7.2e-08 | 9.500E-08 | 3.897E-08 | 7.200E-08 | 1.400E-07 |
| 18512 | 926 | D926K | C | 7.1e-08 | 3.1e-08 | 5.8e-08 | 5.333E-08 | 2.040E-08 | 3.100E-08 | 7.100E-08 |
| 18513 | 926 | D926M | C | 2.8e-08 | 2.1e-08 | 3.4e-08 | 2.767E-08 | 6.506E-09 | 2.100E-08 | 3.400E-08 |
| 18514 | 926 | D926F | C | 3.1e-08 | 2.9e-08 | 2.3e-08 | 2.767E-08 | 4.163E-09 | 2.300E-08 | 3.100E-08 |
| 18515 | 926 | D926P | C | 2.6e-08 | 4.4e-08 | 3.4e-08 | 3.467E-08 | 9.018E-09 | 2.600E-08 | 4.400E-08 |
| 18516 | 926 | D926S | C | 6.4e-08 | 3.6e-08 | 1.6e-08 | 3.867E-08 | 2.411E-08 | 1.600E-08 | 6.400E-08 |
| 18517 | 926 | D926T | C | 1.3e-08 | 2.5e-08 | 4.6e-08 | 2.800E-08 | 1.670E-08 | 1.300E-08 | 4.600E-08 |
| 18518 | 926 | D926W | C | 1e-08 | 2.1e-08 | 1.5e-08 | 1.533E-08 | 5.508E-09 | 1.000E-08 | 2.100E-08 |
| 18519 | 926 | D926Y | C | 8.4e-08 | 4.9e-08 | 1.4e-08 | 4.900E-08 | 3.500E-08 | 1.400E-08 | 8.400E-08 |
| 18520 | 926 | D926V | C | 3.2e-08 | 8.1e-08 | 1.1e-07 | 7.433E-08 | 3.943E-08 | 3.200E-08 | 1.100E-07 |
| 18521 | 927 | T927A | C | 7.7e-08 | 7.5e-08 | 8.8e-08 | 8.000E-08 | 7.000E-09 | 7.500E-08 | 8.800E-08 |
| 18522 | 927 | T927R | C | 7.1e-08 | 4.7e-08 | 7e-08 | 6.267E-08 | 1.358E-08 | 4.700E-08 | 7.100E-08 |
| 18523 | 927 | T927N | C | 1.8e-08 | 3.7e-08 | 2e-08 | 2.500E-08 | 1.044E-08 | 1.800E-08 | 3.700E-08 |
| 18524 | 927 | T927D | C | 2.6e-08 | 4.9e-08 | 2.5e-08 | 3.333E-08 | 1.358E-08 | 2.500E-08 | 4.900E-08 |
| 18525 | 927 | T927C | C | 4.1e-08 | 8e-08 | 1.7e-08 | 4.600E-08 | 3.180E-08 | 1.700E-08 | 8.000E-08 |
| 18526 | 927 | T927E | C | 4.9e-08 | 3.3e-08 | 4.6e-08 | 4.267E-08 | 8.505E-09 | 3.300E-08 | 4.900E-08 |
| 18527 | 927 | T927Q | C | 2.3e-08 | 4e-08 | 9.9e-08 | 5.400E-08 | 3.989E-08 | 2.300E-08 | 9.900E-08 |
| 18528 | 927 | T927G | C | 3.7e-08 | 2.5e-08 | 5.7e-08 | 3.967E-08 | 1.617E-08 | 2.500E-08 | 5.700E-08 |
| 18529 | 927 | T927H | C | 1.7e-08 | 3.9e-08 | 4.8e-08 | 3.467E-08 | 1.595E-08 | 1.700E-08 | 4.800E-08 |

| | | | | | | | | | | |
|---|---|---|---|---|---|---|---|---|---|---|
| 18530 | 927 | T927I | C | 6.7e-08 | 2.2e-08 | 3.2e-08 | 4.033E-08 | 2.363E-08 | 2.200E-08 | 6.700E-08 |
| 18531 | 927 | T927L | C | 1.5e-08 | 4.1e-08 | 1.8e-08 | 2.467E-08 | 1.422E-08 | 1.500E-08 | 4.100E-08 |
| 18532 | 927 | T927K | C | 8.4e-08 | 4.8e-08 | 4.2e-08 | 5.800E-08 | 2.272E-08 | 4.200E-08 | 8.400E-08 |
| 18533 | 927 | T927M | C | 2.8e-08 | 4.5e-08 | 1.3e-07 | 6.767E-08 | 5.465E-08 | 2.800E-08 | 1.300E-07 |
| 18534 | 927 | T927F | C | 7.2e-08 | 4.9e-08 | 8.3e-08 | 6.800E-08 | 1.735E-08 | 4.900E-08 | 8.300E-08 |
| 18535 | 927 | T927P | C | 2.4e-08 | 8.4e-08 | 3e-08 | 4.600E-08 | 3.305E-08 | 2.400E-08 | 8.400E-08 |
| 18536 | 927 | T927S | C | 5.2e-08 | 2.5e-08 | 9.8e-08 | 5.833E-08 | 3.691E-08 | 2.500E-08 | 9.800E-08 |
| 18537 | 927 | T927T | C | 4.7e-08 | 4.2e-08 | 1.7e-08 | 3.533E-08 | 1.607E-08 | 1.700E-08 | 4.700E-08 |
| 18538 | 927 | T927W | C | 4.6e-08 | 6.8e-08 | 2.8e-08 | 4.733E-08 | 2.003E-08 | 2.800E-08 | 6.800E-08 |
| 18539 | 927 | T927Y | C | 6.5e-08 | 3.5e-08 | 1.6e-08 | 3.867E-08 | 2.470E-08 | 1.600E-08 | 6.500E-08 |
| 18540 | 927 | T927V | C | 3.4e-08 | 4.4e-08 | 3e-08 | 3.600E-08 | 7.211E-09 | 3.000E-08 | 4.400E-08 |
| 18541 | 928 | A928A | C | 4.7e-08 | 5.8e-08 | 5.2e-08 | 5.233E-08 | 5.508E-09 | 4.700E-08 | 5.800E-08 |
| 18542 | 928 | A928R | C | 8.7e-08 | 3.1e-08 | 2.1e-08 | 4.633E-08 | 3.557E-08 | 2.100E-08 | 8.700E-08 |
| 18543 | 928 | A928N | C | 1.6e-08 | 3.6e-08 | 3.7e-08 | 2.967E-08 | 1.185E-08 | 1.600E-08 | 3.700E-08 |
| 18544 | 928 | A928D | C | 2.7e-08 | 2.6e-08 | 8.1e-09 | 2.037E-08 | 1.064E-08 | 8.100E-09 | 2.700E-08 |
| 18545 | 928 | A928C | C | 3e-08 | 5.1e-08 | 5.7e-08 | 4.600E-08 | 1.418E-08 | 3.000E-08 | 5.700E-08 |
| 18546 | 928 | A928E | C | 4.4e-08 | 1.4e-08 | 4.9e-08 | 3.567E-08 | 1.893E-08 | 1.400E-08 | 4.900E-08 |
| 18547 | 928 | A928Q | C | 2.4e-08 | 3e-08 | 2e-08 | 2.467E-08 | 5.033E-09 | 2.000E-08 | 3.000E-08 |
| 18548 | 928 | A928G | C | 4.5e-08 | 4.8e-08 | 2.7e-08 | 4.000E-08 | 1.136E-08 | 2.700E-08 | 4.800E-08 |
| 18549 | 928 | A928H | C | 9.7e-09 | 1.5e-08 | 1.7e-08 | 1.390E-08 | 3.772E-09 | 9.700E-09 | 1.700E-08 |
| 18550 | 928 | A928I | C | 6.8e-08 | 2.7e-08 | 6.2e-08 | 5.233E-08 | 2.214E-08 | 2.700E-08 | 6.800E-08 |
| 18551 | 928 | A928L | C | 5.6e-08 | 2.6e-08 | 8e-08 | 5.400E-08 | 2.706E-08 | 2.600E-08 | 8.000E-08 |
| 18552 | 928 | A928K | C | 2.7e-08 | 2.3e-08 | 2.9e-08 | 2.633E-08 | 3.055E-09 | 2.300E-08 | 2.900E-08 |
| 18553 | 928 | A928M | C | 3.1e-08 | 5.8e-08 | 1.8e-08 | 3.567E-08 | 2.040E-08 | 1.800E-08 | 5.800E-08 |
| 18554 | 928 | A928F | C | 7.1e-08 | 1.2e-08 | 3.9e-08 | 4.067E-08 | 2.954E-08 | 1.200E-08 | 7.100E-08 |
| 18555 | 928 | A928P | C | 7e-08 | 3.9e-08 | 1.1e-07 | 7.300E-08 | 3.559E-08 | 3.900E-08 | 1.100E-07 |
| 18556 | 928 | A928S | C | 1.9e-07 | 2.4e-08 | 2.2e-08 | 7.867E-08 | 9.642E-08 | 2.200E-08 | 1.900E-07 |
| 18557 | 928 | A928T | C | 3.6e-08 | 2.9e-08 | 2e-08 | 2.833E-08 | 8.021E-09 | 2.000E-08 | 3.600E-08 |
| 18558 | 928 | A928W | C | 2.2e-08 | 5.2e-08 | 1.1e-08 | 2.833E-08 | 2.122E-08 | 1.100E-08 | 5.200E-08 |
| 18559 | 928 | A928Y | C | 8.6e-08 | 1.6e-08 | 3.9e-08 | 4.700E-08 | 3.568E-08 | 1.600E-08 | 8.600E-08 |
| 18560 | 928 | A928V | C | 6.9e-08 | 1.1e-07 | 3.5e-08 | 7.133E-08 | 3.755E-08 | 3.500E-08 | 1.100E-07 |
| 18561 | 929 | V929A | C | 2.1e-08 | 3.8e-08 | 3.9e-08 | 3.267E-08 | 1.012E-08 | 2.100E-08 | 3.900E-08 |
| 18562 | 929 | V929R | C | 3.2e-08 | 5.4e-08 | 2.8e-08 | 3.800E-08 | 1.400E-08 | 2.800E-08 | 5.400E-08 |
| 18563 | 929 | V929N | C | 3e-08 | 3.6e-08 | 2.1e-08 | 2.900E-08 | 7.550E-09 | 2.100E-08 | 3.600E-08 |
| 18564 | 929 | V929D | C | 3e-08 | 7.1e-08 | 5.4e-08 | 5.167E-08 | 2.060E-08 | 3.000E-08 | 7.100E-08 |
| 18565 | 929 | V929C | C | 3.7e-08 | 1.1e-07 | 3.8e-08 | 6.167E-08 | 4.186E-08 | 3.700E-08 | 1.100E-07 |
| 18566 | 929 | V929E | C | 1.1e-07 | 5e-08 | 5.6e-08 | 7.200E-08 | 3.305E-08 | 5.000E-08 | 1.100E-07 |
| 18567 | 929 | V929Q | C | 3.2e-08 | 5.3e-08 | 7.3e-08 | 5.267E-08 | 2.050E-08 | 3.200E-08 | 7.300E-08 |
| 18568 | 929 | V929G | C | 8.4e-08 | 2.5e-08 | 2.4e-08 | 4.433E-08 | 3.436E-08 | 2.400E-08 | 8.400E-08 |
| 18569 | 929 | V929H | C | 2.9e-08 | 2.6e-08 | 4.7e-08 | 3.400E-08 | 1.136E-08 | 2.600E-08 | 4.700E-08 |
| 18570 | 929 | V929I | C | 2.3e-08 | 7.8e-08 | 3.5e-08 | 4.533E-08 | 2.892E-08 | 2.300E-08 | 7.800E-08 |
| 18571 | 929 | V929L | C | 3.7e-08 | 3.5e-08 | 6.4e-08 | 4.533E-08 | 1.620E-08 | 3.500E-08 | 6.400E-08 |
| 18572 | 929 | V929K | C | 2e-08 | 1e-07 | 3.7e-08 | 5.233E-08 | 4.215E-08 | 2.000E-08 | 1.000E-07 |
| 18573 | 929 | V929M | C | 3.2e-08 | 6.8e-08 | 5.6e-08 | 5.200E-08 | 1.833E-08 | 3.200E-08 | 6.800E-08 |
| 18574 | 929 | V929F | C | 6.3e-08 | 8.7e-08 | 2.1e-08 | 5.700E-08 | 3.341E-08 | 2.100E-08 | 8.700E-08 |
| 18575 | 929 | V929P | C | 2.6e-08 | 6.2e-08 | 1.7e-08 | 3.500E-08 | 2.381E-08 | 1.700E-08 | 6.200E-08 |
| 18576 | 929 | V929S | C | 5.4e-08 | 6.1e-08 | 1e-07 | 7.167E-08 | 2.479E-08 | 5.400E-08 | 1.000E-07 |
| 18577 | 929 | V929T | C | 1.5e-08 | 1.1e-07 | 6.3e-08 | 6.267E-08 | 4.750E-08 | 1.500E-08 | 1.100E-07 |
| 18578 | 929 | V929W | C | 5.5e-08 | 3.2e-08 | 8.5e-09 | 3.183E-08 | 2.325E-08 | 8.500E-09 | 5.500E-08 |
| 18579 | 929 | V929Y | C | 3.3e-08 | 9.3e-09 | 4.6e-08 | 2.943E-08 | 1.861E-08 | 9.300E-09 | 4.600E-08 |
| 18580 | 929 | V929V | C | 7e-08 | 4.9e-08 | 1.6e-08 | 4.500E-08 | 2.722E-08 | 1.600E-08 | 7.000E-08 |
| 18581 | 930 | Y930A | C | 1.8e-08 | 1.7e-07 | 9.2e-08 | 9.333E-08 | 7.601E-08 | 1.800E-08 | 1.700E-07 |
| 18582 | 930 | Y930R | C | 1e-07 | 6e-08 | 2.7e-08 | 6.233E-08 | 3.656E-08 | 2.700E-08 | 1.000E-07 |
| 18583 | 930 | Y930N | C | 4.8e-08 | 6.4e-08 | 3.5e-08 | 4.900E-08 | 1.453E-08 | 3.500E-08 | 6.400E-08 |
| 18584 | 930 | Y930D | C | 4.5e-08 | 4.1e-08 | 1.8e-08 | 3.467E-08 | 1.457E-08 | 1.800E-08 | 4.500E-08 |
| 18585 | 930 | Y930C | C | 6.7e-08 | 2.5e-08 | 3.8e-08 | 4.333E-08 | 2.150E-08 | 2.500E-08 | 6.700E-08 |
| 18586 | 930 | Y930E | C | 3e-08 | 4.3e-08 | 1.6e-07 | 7.767E-08 | 7.160E-08 | 3.000E-08 | 1.600E-07 |
| 18587 | 930 | Y930Q | C | 8.3e-08 | 4.1e-08 | 4.7e-08 | 5.700E-08 | 2.272E-08 | 4.100E-08 | 8.300E-08 |
| 18588 | 930 | Y930G | C | 2.8e-08 | 3e-08 | 2.2e-08 | 2.667E-08 | 4.163E-09 | 2.200E-08 | 3.000E-08 |
| 18589 | 930 | Y930H | C | 1.8e-08 | 2e-08 | 8.7e-08 | 4.167E-08 | 3.927E-08 | 1.800E-08 | 8.700E-08 |
| 18590 | 930 | Y930I | C | 1.9e-08 | 5.2e-08 | 3.6e-08 | 3.567E-08 | 1.650E-08 | 1.900E-08 | 5.200E-08 |
| 18591 | 930 | Y930L | C | 3.1e-08 | 9.2e-08 | 7.3e-08 | 6.533E-08 | 3.121E-08 | 3.100E-08 | 9.200E-08 |
| 18592 | 930 | Y930K | C | 3.9e-08 | 6.7e-08 | 6e-08 | 5.533E-08 | 1.457E-08 | 3.900E-08 | 6.700E-08 |
| 18593 | 930 | Y930M | C | 6.5e-08 | 3.4e-08 | 4.1e-08 | 4.667E-08 | 1.626E-08 | 3.400E-08 | 6.500E-08 |
| 18594 | 930 | Y930F | C | 3.1e-08 | 4.2e-08 | 1.4e-08 | 2.900E-08 | 1.411E-08 | 1.400E-08 | 4.200E-08 |
| 18595 | 930 | Y930P | C | 2.3e-08 | 5.8e-08 | 9.9e-08 | 6.000E-08 | 3.804E-08 | 2.300E-08 | 9.900E-08 |
| 18596 | 930 | Y930S | C | 8.6e-08 | 3.8e-08 | 2.7e-08 | 5.033E-08 | 3.137E-08 | 2.700E-08 | 8.600E-08 |
| 18597 | 930 | Y930T | C | 7.5e-08 | 4.2e-08 | 1e-07 | 7.233E-08 | 2.909E-08 | 4.200E-08 | 1.000E-07 |
| 18598 | 930 | Y930W | C | 5.6e-08 | 4e-08 | 9.2e-08 | 6.267E-08 | 2.663E-08 | 4.000E-08 | 9.200E-08 |
| 18599 | 930 | Y930Y | C | 5.4e-08 | 1.5e-08 | 1.4e-08 | 2.767E-08 | 2.281E-08 | 1.400E-08 | 5.400E-08 |
| 18600 | 930 | Y930V | C | 8.7e-08 | 2.7e-08 | 2.3e-08 | 4.567E-08 | 3.585E-08 | 2.300E-08 | 8.700E-08 |
| 18601 | 931 | Y931A | C | 1.3e-07 | 8.5e-08 | 1.4e-08 | 7.633E-08 | 5.848E-08 | 1.400E-08 | 1.300E-07 |
| 18602 | 931 | Y931R | C | 7.1e-08 | 7.5e-08 | 9.6e-09 | 5.187E-08 | 3.666E-08 | 9.600E-09 | 7.500E-08 |
| 18603 | 931 | Y931N | C | 3.2e-08 | 2.6e-08 | 9.2e-08 | 5.000E-08 | 3.650E-08 | 2.600E-08 | 9.200E-08 |
| 18604 | 931 | Y931D | C | 5.7e-09 | 2.5e-08 | 1.5e-08 | 1.523E-08 | 9.652E-09 | 5.700E-09 | 2.500E-08 |
| 18605 | 931 | Y931C | C | 8.6e-08 | 7.3e-08 | 4.9e-08 | 6.933E-08 | 1.877E-08 | 4.900E-08 | 8.600E-08 |
| 18606 | 931 | Y931E | C | 6.5e-08 | 5.4e-08 | 6.7e-08 | 6.200E-08 | 7.000E-09 | 5.400E-08 | 6.700E-08 |
| 18607 | 931 | Y931Q | C | 4.5e-08 | 3.1e-08 | 5.4e-08 | 4.333E-08 | 1.159E-08 | 3.100E-08 | 5.400E-08 |
| 18608 | 931 | Y931G | C | 4.1e-08 | 1e-07 | 3.1e-08 | 5.733E-08 | 3.729E-08 | 3.100E-08 | 1.000E-07 |
| 18609 | 931 | Y931H | C | 1.4e-08 | 8e-08 | 1.5e-07 | 8.133E-08 | 6.801E-08 | 1.400E-08 | 1.500E-07 |
| 18610 | 931 | Y931I | C | 4.5e-08 | 4e-08 | 7.4e-08 | 5.300E-08 | 1.836E-08 | 4.000E-08 | 7.400E-08 |
| 18611 | 931 | Y931L | C | 3.3e-08 | 7.5e-08 | 1.7e-08 | 4.167E-08 | 2.996E-08 | 1.700E-08 | 7.500E-08 |
| 18612 | 931 | Y931K | C | 2.9e-08 | 2.7e-08 | 2.5e-08 | 2.700E-08 | 2.000E-09 | 2.500E-08 | 2.900E-08 |
| 18613 | 931 | Y931M | C | 5.8e-08 | 5.8e-08 | 9.8e-08 | 7.133E-08 | 2.309E-08 | 5.800E-08 | 9.800E-08 |
| 18614 | 931 | Y931F | C | 1.8e-08 | 5.9e-08 | 1.9e-08 | 3.200E-08 | 2.339E-08 | 1.800E-08 | 5.900E-08 |

| | | | | | | | | | | |
|---|---|---|---|---|---|---|---|---|---|---|
| 18615 | 931 | Y931P | C | 1.4e-08 | 6.6e-08 | 5.1e-08 | 4.367E-08 | 2.676E-08 | 1.400E-08 | 6.600E-08 |
| 18616 | 931 | Y931S | C | 3.8e-08 | 5.2e-08 | 1e-07 | 6.333E-08 | 3.252E-08 | 3.800E-08 | 1.000E-07 |
| 18617 | 931 | Y931T | C | 9.5e-08 | 6e-08 | 3.6e-08 | 6.367E-08 | 2.967E-08 | 3.600E-08 | 9.500E-08 |
| 18618 | 931 | Y931W | C | 4.3e-08 | 7.7e-08 | 3.9e-08 | 5.300E-08 | 2.088E-08 | 3.900E-08 | 7.700E-08 |
| 18619 | 931 | Y931Y | C | 6.7e-08 | 8.5e-08 | 8.1e-08 | 7.767E-08 | 9.452E-09 | 6.700E-08 | 8.500E-08 |
| 18620 | 931 | Y931V | C | 5.1e-08 | 3.7e-08 | 4.3e-08 | 4.367E-08 | 7.024E-09 | 3.700E-08 | 5.100E-08 |
| 18621 | 932 | C932A | C | 3.6e-08 | 4.8e-08 | 2.3e-08 | 3.567E-08 | 1.250E-08 | 2.300E-08 | 4.800E-08 |
| 18622 | 932 | C932R | C | 1e-07 | 6.6e-08 | 2.7e-08 | 6.433E-08 | 3.653E-08 | 2.700E-08 | 1.000E-07 |
| 18623 | 932 | C932N | C | 2.4e-08 | 3.3e-08 | 2.3e-08 | 2.667E-08 | 5.508E-09 | 2.300E-08 | 3.300E-08 |
| 18624 | 932 | C932D | C | 2.3e-08 | 7.9e-09 | 7.6e-08 | 3.563E-08 | 3.576E-08 | 7.900E-09 | 7.600E-08 |
| 18625 | 932 | C932C | C | 2.9e-08 | 2.2e-08 | 3.1e-08 | 2.733E-08 | 4.726E-09 | 2.200E-08 | 3.100E-08 |
| 18626 | 932 | C932E | C | 6.6e-08 | 4.9e-08 | 5.9e-09 | 4.030E-08 | 3.098E-08 | 5.900E-09 | 6.600E-08 |
| 18627 | 932 | C932Q | C | 2.3e-08 | 7.6e-08 | 5.5e-08 | 5.133E-08 | 2.669E-08 | 2.300E-08 | 7.600E-08 |
| 18628 | 932 | C932G | C | 1.1e-08 | 1.4e-07 | 1e-08 | 5.367E-08 | 7.477E-08 | 1.000E-08 | 1.400E-07 |
| 18629 | 932 | C932H | C | 7.7e-08 | 7.1e-08 | 2.6e-08 | 5.800E-08 | 2.787E-08 | 2.600E-08 | 7.700E-08 |
| 18630 | 932 | C932I | C | 2.7e-08 | 1.2e-07 | 4.5e-08 | 6.400E-08 | 4.933E-08 | 2.700E-08 | 1.200E-07 |
| 18631 | 932 | C932L | C | 9.6e-08 | 5.1e-08 | 8.9e-08 | 7.867E-08 | 2.421E-08 | 5.100E-08 | 9.600E-08 |
| 18632 | 932 | C932K | C | 3.7e-08 | 5.9e-08 | 1.1e-08 | 3.567E-08 | 2.403E-08 | 1.100E-08 | 5.900E-08 |
| 18633 | 932 | C932M | C | 7.2e-08 | 3.4e-08 | 1.1e-07 | 7.200E-08 | 3.800E-08 | 3.400E-08 | 1.100E-07 |
| 18634 | 932 | C932F | C | 2.3e-08 | 5.2e-08 | 4.1e-08 | 3.867E-08 | 1.464E-08 | 2.300E-08 | 5.200E-08 |
| 18635 | 932 | C932P | C | 4.2e-08 | 6.1e-08 | 9.5e-09 | 3.750E-08 | 2.604E-08 | 9.500E-09 | 6.100E-08 |
| 18636 | 932 | C932S | C | 8.6e-08 | 3.1e-08 | 5.9e-08 | 5.867E-08 | 2.750E-08 | 3.100E-08 | 8.600E-08 |
| 18637 | 932 | C932T | C | 3.8e-08 | 7.1e-08 | 4.9e-08 | 5.267E-08 | 1.680E-08 | 3.800E-08 | 7.100E-08 |
| 18638 | 932 | C932W | C | 2.7e-08 | 1.3e-08 | 1.9e-08 | 1.967E-08 | 7.024E-09 | 1.300E-08 | 2.700E-08 |
| 18639 | 932 | C932Y | C | 8.8e-08 | 4.2e-08 | 3.4e-08 | 5.467E-08 | 2.914E-08 | 3.400E-08 | 8.800E-08 |
| 18640 | 932 | C932V | C | 2.8e-08 | 5e-08 | 4.3e-08 | 4.033E-08 | 1.124E-08 | 2.800E-08 | 5.000E-08 |
| 18641 | 933 | A933A | C | 2.9e-08 | 8.6e-08 | 2e-08 | 4.500E-08 | 3.579E-08 | 2.000E-08 | 8.600E-08 |
| 18642 | 933 | A933R | C | 4.7e-08 | 1e-07 | 4.6e-08 | 6.433E-08 | 3.089E-08 | 4.600E-08 | 1.000E-07 |
| 18643 | 933 | A933N | C | 5.5e-08 | 3.9e-08 | 4.3e-08 | 4.567E-08 | 8.327E-09 | 3.900E-08 | 5.500E-08 |
| 18644 | 933 | A933D | C | 3.7e-08 | 3.7e-08 | 7.1e-08 | 4.833E-08 | 1.963E-08 | 3.700E-08 | 7.100E-08 |
| 18645 | 933 | A933C | C | 7.6e-08 | 3.5e-08 | 4e-08 | 5.033E-08 | 2.237E-08 | 3.500E-08 | 7.600E-08 |
| 18646 | 933 | A933E | C | 8.7e-08 | 5.1e-08 | 6.5e-08 | 6.767E-08 | 1.815E-08 | 5.100E-08 | 8.700E-08 |
| 18647 | 933 | A933Q | C | 2.4e-08 | 4.2e-08 | 2.6e-08 | 3.067E-08 | 9.866E-09 | 2.400E-08 | 4.200E-08 |
| 18648 | 933 | A933G | C | 5.6e-08 | 4.9e-08 | 2.8e-08 | 4.433E-08 | 1.457E-08 | 2.800E-08 | 5.600E-08 |
| 18649 | 933 | A933H | C | 1.4e-07 | 3.4e-08 | 6.4e-08 | 7.933E-08 | 5.464E-08 | 3.400E-08 | 1.400E-07 |
| 18650 | 933 | A933I | C | 5.3e-08 | 3.5e-08 | 7.5e-08 | 5.433E-08 | 2.003E-08 | 3.500E-08 | 7.500E-08 |
| 18651 | 933 | A933L | C | 5e-08 | 2.1e-08 | 3.5e-08 | 3.533E-08 | 1.450E-08 | 2.100E-08 | 5.000E-08 |
| 18652 | 933 | A933K | C | 2.1e-08 | 2.5e-08 | 2.9e-08 | 2.500E-08 | 4.000E-09 | 2.100E-08 | 2.900E-08 |
| 18653 | 933 | A933M | C | 6.7e-08 | 4.2e-08 | 4.1e-08 | 5.000E-08 | 1.473E-08 | 4.100E-08 | 6.700E-08 |
| 18654 | 933 | A933F | C | 4.5e-08 | 5.8e-08 | 3e-08 | 4.433E-08 | 1.401E-08 | 3.000E-08 | 5.800E-08 |
| 18655 | 933 | A933P | C | 5.2e-08 | 3.4e-08 | 4.1e-08 | 4.233E-08 | 9.074E-09 | 3.400E-08 | 5.200E-08 |
| 18656 | 933 | A933S | C | 2.4e-08 | 2.4e-08 | 4.5e-08 | 3.100E-08 | 1.212E-08 | 2.400E-08 | 4.500E-08 |
| 18657 | 933 | A933T | C | 6.5e-08 | 5.8e-08 | 7.5e-08 | 6.600E-08 | 8.544E-09 | 5.800E-08 | 7.500E-08 |
| 18658 | 933 | A933W | C | 3.2e-08 | 2.6e-08 | 3.3e-08 | 3.033E-08 | 3.786E-09 | 2.600E-08 | 3.300E-08 |
| 18659 | 933 | A933Y | C | 6.4e-08 | 4.1e-08 | 4.6e-08 | 5.033E-08 | 1.210E-08 | 4.100E-08 | 6.400E-08 |
| 18660 | 933 | A933V | C | 4.1e-08 | 4.3e-08 | 4.6e-08 | 4.333E-08 | 2.517E-09 | 4.100E-08 | 4.600E-08 |
| 18661 | 934 | R934A | C | 1.2e-08 | 6.2e-09 | 6.1e-08 | 2.640E-08 | 3.010E-08 | 6.200E-09 | 6.100E-08 |
| 18662 | 934 | R934R | C | 6.1e-08 | 4.4e-08 | 4.8e-08 | 5.100E-08 | 8.888E-09 | 4.400E-08 | 6.100E-08 |
| 18663 | 934 | R934N | C | 7.5e-08 | 4.6e-08 | 4.9e-08 | 5.667E-08 | 1.595E-08 | 4.600E-08 | 7.500E-08 |
| 18664 | 934 | R934D | C | 4.5e-08 | 2.3e-08 | 2.3e-08 | 3.033E-08 | 1.270E-08 | 2.300E-08 | 4.500E-08 |
| 18665 | 934 | R934C | C | 3.7e-08 | 3.5e-08 | 3.6e-08 | 3.600E-08 | 1.000E-09 | 3.500E-08 | 3.700E-08 |
| 18666 | 934 | R934E | C | 9.6e-08 | 6.1e-08 | 4.2e-08 | 6.633E-08 | 2.739E-08 | 4.200E-08 | 9.600E-08 |
| 18667 | 934 | R934Q | C | 9.8e-08 | 1e-07 | 3.3e-08 | 7.700E-08 | 3.812E-08 | 3.300E-08 | 1.000E-07 |
| 18668 | 934 | R934G | C | 2.8e-08 | 7.8e-08 | 4.8e-08 | 5.133E-08 | 2.517E-08 | 2.800E-08 | 7.800E-08 |
| 18669 | 934 | R934H | C | 4.1e-08 | 3.7e-08 | 7.3e-08 | 5.033E-08 | 1.973E-08 | 3.700E-08 | 7.300E-08 |
| 18670 | 934 | R934I | C | 2.8e-08 | 8.2e-08 | 1.6e-07 | 9.000E-08 | 6.636E-08 | 2.800E-08 | 1.600E-07 |
| 18671 | 934 | R934L | C | 3.1e-08 | 7.5e-08 | 4.3e-08 | 4.967E-08 | 2.274E-08 | 3.100E-08 | 7.500E-08 |
| 18672 | 934 | R934K | C | 1.2e-07 | 3.1e-08 | 3.4e-08 | 6.167E-08 | 5.054E-08 | 3.100E-08 | 1.200E-07 |
| 18673 | 934 | R934M | C | 2.4e-08 | 7.5e-08 | 4.4e-08 | 4.767E-08 | 2.570E-08 | 2.400E-08 | 7.500E-08 |
| 18674 | 934 | R934F | C | 2e-08 | 5.9e-08 | 5.2e-08 | 4.367E-08 | 2.079E-08 | 2.000E-08 | 5.900E-08 |
| 18675 | 934 | R934P | C | 8.7e-08 | 4.4e-08 | 1.1e-07 | 8.033E-08 | 3.350E-08 | 4.400E-08 | 1.100E-07 |
| 18676 | 934 | R934S | C | 4.1e-08 | 4.6e-08 | 7.3e-08 | 5.333E-08 | 1.721E-08 | 4.100E-08 | 7.300E-08 |
| 18677 | 934 | R934T | C | 8e-08 | 1.2e-07 | 3.9e-08 | 7.967E-08 | 4.050E-08 | 3.900E-08 | 1.200E-07 |
| 18678 | 934 | R934W | C | 3.3e-08 | 2.3e-08 | 4.8e-08 | 3.467E-08 | 1.258E-08 | 2.300E-08 | 4.800E-08 |
| 18679 | 934 | R934Y | C | 9.6e-08 | 5.3e-08 | 1.1e-07 | 8.633E-08 | 2.970E-08 | 5.300E-08 | 1.100E-07 |
| 18680 | 934 | R934V | C | 5e-08 | 1e-07 | 3.5e-08 | 6.167E-08 | 3.403E-08 | 3.500E-08 | 1.000E-07 |
| 18681 | 935 | N935A | C | 5e-08 | 2.2e-08 | 3.6e-08 | 3.600E-08 | 1.400E-08 | 2.200E-08 | 5.000E-08 |
| 18682 | 935 | N935R | C | 6e-08 | 9.6e-09 | 3.3e-08 | 3.420E-08 | 2.522E-08 | 9.600E-09 | 6.000E-08 |
| 18683 | 935 | N935N | C | 2.9e-08 | 4.8e-08 | 3.5e-08 | 3.733E-08 | 9.713E-09 | 2.900E-08 | 4.800E-08 |
| 18684 | 935 | N935D | C | 1.2e-08 | 3.9e-08 | 6e-08 | 3.700E-08 | 2.406E-08 | 1.200E-08 | 6.000E-08 |
| 18685 | 935 | N935C | C | 1.9e-08 | 2.7e-08 | 1.7e-08 | 2.100E-08 | 5.292E-09 | 1.700E-08 | 2.700E-08 |
| 18686 | 935 | N935E | C | 9.7e-09 | 1.6e-08 | 5.4e-08 | 2.657E-08 | 2.397E-08 | 9.700E-09 | 5.400E-08 |
| 18687 | 935 | N935Q | C | 4.8e-08 | 5.2e-08 | 4.7e-08 | 4.900E-08 | 2.646E-09 | 4.700E-08 | 5.200E-08 |
| 18688 | 935 | N935G | C | 2.8e-08 | 6e-08 | 5.7e-08 | 4.833E-08 | 1.767E-08 | 2.800E-08 | 6.000E-08 |
| 18689 | 935 | N935H | C | 2.9e-08 | 1.8e-08 | 2e-08 | 2.233E-08 | 5.859E-09 | 1.800E-08 | 2.900E-08 |
| 18690 | 935 | N935I | C | 4.2e-08 | 5.8e-08 | 4e-08 | 4.667E-08 | 9.866E-09 | 4.000E-08 | 5.800E-08 |
| 18691 | 935 | N935L | C | 2.5e-08 | 5e-08 | 1.6e-08 | 3.033E-08 | 1.762E-08 | 1.600E-08 | 5.000E-08 |
| 18692 | 935 | N935K | C | 7.6e-08 | 2.7e-08 | 2.5e-08 | 4.267E-08 | 2.888E-08 | 2.500E-08 | 7.600E-08 |
| 18693 | 935 | N935M | C | 1.1e-07 | 7.1e-08 | 5.6e-08 | 7.900E-08 | 2.787E-08 | 5.600E-08 | 1.100E-07 |
| 18694 | 935 | N935F | C | 3.3e-08 | 1.9e-08 | 4e-08 | 3.067E-08 | 1.069E-08 | 1.900E-08 | 4.000E-08 |
| 18695 | 935 | N935P | C | 2.1e-08 | 1.1e-07 | 4.3e-08 | 5.800E-08 | 4.636E-08 | 2.100E-08 | 1.100E-07 |
| 18696 | 935 | N935S | C | 2.5e-08 | 5.8e-08 | 1.4e-07 | 7.433E-08 | 5.921E-08 | 2.500E-08 | 1.400E-07 |
| 18697 | 935 | N935T | C | 1.1e-07 | 1.9e-08 | 4.2e-08 | 5.700E-08 | 4.732E-08 | 1.900E-08 | 1.100E-07 |
| 18698 | 935 | N935W | C | 2.5e-08 | 1.4e-08 | 4.4e-08 | 2.767E-08 | 1.518E-08 | 1.400E-08 | 4.400E-08 |
| 18699 | 935 | N935Y | C | 2.5e-08 | 3e-08 | 4.7e-08 | 3.400E-08 | 1.153E-08 | 2.500E-08 | 4.700E-08 |

| | | | | | | | | | | |
|---|---|---|---|---|---|---|---|---|---|---|
| 18700 | 935 | N935V | C | 2.6e-08 | 4.2e-08 | 4.5e-08 | 3.767E-08 | 1.021E-08 | 2.600E-08 | 4.500E-08 |
| 18701 | 936 | L936A | C | 5.1e-08 | 1.8e-08 | 2.4e-08 | 3.100E-08 | 1.758E-08 | 1.800E-08 | 5.100E-08 |
| 18702 | 936 | L936R | C | 5.9e-08 | 5.1e-08 | 2.9e-08 | 4.633E-08 | 1.553E-08 | 2.900E-08 | 5.900E-08 |
| 18703 | 936 | L936N | C | 2.3e-07 | 5.5e-08 | 9.6e-08 | 1.270E-07 | 9.153E-08 | 5.500E-08 | 2.300E-07 |
| 18704 | 936 | L936D | C | 4.9e-08 | 3.9e-08 | 1e-07 | 6.267E-08 | 3.272E-08 | 3.900E-08 | 1.000E-07 |
| 18705 | 936 | L936C | C | 5.1e-08 | 5.5e-08 | 5.6e-08 | 5.400E-08 | 2.646E-09 | 5.100E-08 | 5.600E-08 |
| 18706 | 936 | L936E | C | 6e-08 | 7.7e-08 | 1.4e-07 | 9.233E-08 | 4.215E-08 | 6.000E-08 | 1.400E-07 |
| 18707 | 936 | L936Q | C | 6e-08 | 1.4e-07 | 9.5e-08 | 9.833E-08 | 4.010E-08 | 6.000E-08 | 1.400E-07 |
| 18708 | 936 | L936G | C | 5.3e-08 | 7.3e-08 | 4e-08 | 5.533E-08 | 1.662E-08 | 4.000E-08 | 7.300E-08 |
| 18709 | 936 | L936H | C | 2e-08 | 3.6e-08 | 1.1e-07 | 5.533E-08 | 4.801E-08 | 2.000E-08 | 1.100E-07 |
| 18710 | 936 | L936I | C | 2.8e-08 | 4.2e-08 | 6.1e-08 | 4.367E-08 | 1.656E-08 | 2.800E-08 | 6.100E-08 |
| 18711 | 936 | L936L | C | 6.6e-08 | 1.1e-07 | 3.9e-08 | 7.167E-08 | 3.584E-08 | 3.900E-08 | 1.100E-07 |
| 18712 | 936 | L936K | C | 1.8e-08 | 2.9e-08 | 8e-08 | 4.233E-08 | 3.308E-08 | 1.800E-08 | 8.000E-08 |
| 18713 | 936 | L936M | C | 7.4e-08 | 4.2e-08 | 4.6e-08 | 5.400E-08 | 1.744E-08 | 4.200E-08 | 7.400E-08 |
| 18714 | 936 | L936F | C | 2e-08 | 1.9e-08 | 1.4e-07 | 5.967E-08 | 6.957E-08 | 1.900E-08 | 1.400E-07 |
| 18715 | 936 | L936P | C | 7.4e-08 | 1.8e-08 | 3.2e-08 | 4.133E-08 | 2.914E-08 | 1.800E-08 | 7.400E-08 |
| 18716 | 936 | L936S | C | 4.4e-08 | 3.6e-08 | 5.2e-08 | 4.400E-08 | 8.000E-09 | 3.600E-08 | 5.200E-08 |
| 18717 | 936 | L936T | C | 8.7e-08 | 6.6e-08 | 8.8e-08 | 8.033E-08 | 1.242E-08 | 6.600E-08 | 8.800E-08 |
| 18718 | 936 | L936W | C | 1.8e-07 | 8.5e-08 | 6.3e-08 | 1.093E-07 | 6.218E-08 | 6.300E-08 | 1.800E-07 |
| 18719 | 936 | L936Y | C | 3.7e-08 | 1.8e-08 | 9.1e-08 | 4.867E-08 | 3.787E-08 | 1.800E-08 | 9.100E-08 |
| 18720 | 936 | L936V | C | 2e-08 | 8.4e-08 | 4e-08 | 4.800E-08 | 3.274E-08 | 2.000E-08 | 8.400E-08 |
| 18721 | 937 | G937A | C | 4e-08 | 2.5e-08 | 3.4e-08 | 3.300E-08 | 7.550E-09 | 2.500E-08 | 4.000E-08 |
| 18722 | 937 | G937R | C | 2.7e-08 | 2.3e-08 | 5.5e-08 | 3.500E-08 | 1.744E-08 | 2.300E-08 | 5.500E-08 |
| 18723 | 937 | G937N | C | 2.3e-08 | 2.7e-08 | 4e-08 | 3.000E-08 | 8.888E-09 | 2.300E-08 | 4.000E-08 |
| 18724 | 937 | G937D | C | 5.2e-08 | 3.9e-08 | 1.2e-08 | 3.433E-08 | 2.040E-08 | 1.200E-08 | 5.200E-08 |
| 18725 | 937 | G937C | C | 5.3e-08 | 8.3e-08 | 3e-08 | 5.533E-08 | 2.658E-08 | 3.000E-08 | 8.300E-08 |
| 18726 | 937 | G937E | C | 2.8e-08 | 6.8e-08 | 3.3e-08 | 4.300E-08 | 2.179E-08 | 2.800E-08 | 6.800E-08 |
| 18727 | 937 | G937Q | C | 8.2e-08 | 7e-08 | 3e-08 | 6.067E-08 | 2.723E-08 | 3.000E-08 | 8.200E-08 |
| 18728 | 937 | G937G | C | 2.1e-08 | 4.8e-08 | 9.2e-08 | 5.367E-08 | 3.584E-08 | 2.100E-08 | 9.200E-08 |
| 18729 | 937 | G937H | C | 1.4e-08 | 2.2e-08 | 2.7e-08 | 2.100E-08 | 6.557E-09 | 1.400E-08 | 2.700E-08 |
| 18730 | 937 | G937I | C | 3e-08 | 3.5e-08 | 2.3e-08 | 2.933E-08 | 6.028E-09 | 2.300E-08 | 3.500E-08 |
| 18731 | 937 | G937L | C | 3.8e-08 | 2.7e-08 | 3e-08 | 3.167E-08 | 5.686E-09 | 2.700E-08 | 3.800E-08 |
| 18732 | 937 | G937K | C | 1.4e-08 | 7.8e-08 | 2.2e-08 | 3.800E-08 | 3.487E-08 | 1.400E-08 | 7.800E-08 |
| 18733 | 937 | G937M | C | 6.5e-08 | 5.6e-08 | 1e-08 | 4.367E-08 | 2.950E-08 | 1.000E-08 | 6.500E-08 |
| 18734 | 937 | G937F | C | 2.1e-08 | 4.5e-08 | 4.3e-08 | 3.633E-08 | 1.332E-08 | 2.100E-08 | 4.500E-08 |
| 18735 | 937 | G937P | C | 4.4e-08 | 6.5e-08 | 6.3e-08 | 5.733E-08 | 1.159E-08 | 4.400E-08 | 6.500E-08 |
| 18736 | 937 | G937S | C | 2.1e-08 | 3.3e-08 | 2.9e-08 | 2.767E-08 | 6.110E-09 | 2.100E-08 | 3.300E-08 |
| 18737 | 937 | G937T | C | 2.7e-08 | 4.9e-08 | 5.8e-08 | 4.467E-08 | 1.595E-08 | 2.700E-08 | 5.800E-08 |
| 18738 | 937 | G937W | C | 1.2e-08 | 2.6e-08 | 1.6e-08 | 1.800E-08 | 7.211E-09 | 1.200E-08 | 2.600E-08 |
| 18739 | 937 | G937Y | C | 7.6e-08 | 1.1e-07 | 2.1e-08 | 6.900E-08 | 4.491E-08 | 2.100E-08 | 1.100E-07 |
| 18740 | 937 | G937V | C | 6.6e-08 | 2.1e-08 | 3.8e-08 | 4.167E-08 | 2.272E-08 | 2.100E-08 | 6.600E-08 |
| 18741 | 938 | P938A | C | 1e-07 | 4.2e-08 | 9.6e-08 | 7.933E-08 | 3.239E-08 | 4.200E-08 | 1.000E-07 |
| 18742 | 938 | P938R | C | 1.8e-08 | 4.6e-08 | 4.4e-08 | 3.600E-08 | 1.562E-08 | 1.800E-08 | 4.600E-08 |
| 18743 | 938 | P938N | C | 1e-07 | 5.2e-08 | 1.6e-07 | 1.040E-07 | 5.411E-08 | 5.200E-08 | 1.600E-07 |
| 18744 | 938 | P938D | C | 2.6e-08 | 6.3e-08 | 6.8e-08 | 5.233E-08 | 2.294E-08 | 2.600E-08 | 6.800E-08 |
| 18745 | 938 | P938C | C | 3.5e-08 | 7.9e-08 | 4e-08 | 5.133E-08 | 2.409E-08 | 3.500E-08 | 7.900E-08 |
| 18746 | 938 | P938E | C | 1.3e-07 | 3.7e-08 | 6.1e-08 | 7.600E-08 | 4.828E-08 | 3.700E-08 | 1.300E-07 |
| 18747 | 938 | P938Q | C | 8.6e-08 | 3e-08 | 1.7e-07 | 9.533E-08 | 7.047E-08 | 3.000E-08 | 1.700E-07 |
| 18748 | 938 | P938G | C | 6.5e-08 | 4.8e-08 | 6.1e-08 | 5.800E-08 | 8.888E-09 | 4.800E-08 | 6.500E-08 |
| 18749 | 938 | P938H | C | 4.5e-08 | 3.7e-08 | 3.7e-08 | 3.967E-08 | 4.619E-09 | 3.700E-08 | 4.500E-08 |
| 18750 | 938 | P938I | C | 5.2e-08 | 7.6e-08 | 3.3e-08 | 5.367E-08 | 2.155E-08 | 3.300E-08 | 7.600E-08 |
| 18751 | 938 | P938L | C | 4.5e-08 | 6.3e-08 | 4.8e-08 | 5.200E-08 | 9.644E-09 | 4.500E-08 | 6.300E-08 |
| 18752 | 938 | P938K | C | 7.6e-08 | 4.5e-08 | 1.7e-07 | 9.700E-08 | 6.509E-08 | 4.500E-08 | 1.700E-07 |
| 18753 | 938 | P938M | C | 1.6e-08 | 2.3e-08 | 3.5e-08 | 2.467E-08 | 9.609E-09 | 1.600E-08 | 3.500E-08 |
| 18754 | 938 | P938F | C | 1.6e-08 | 4.1e-08 | 7.5e-08 | 4.400E-08 | 2.961E-08 | 1.600E-08 | 7.500E-08 |
| 18755 | 938 | P938P | C | 6.4e-08 | 3.1e-08 | 5.2e-08 | 4.900E-08 | 1.670E-08 | 3.100E-08 | 6.400E-08 |
| 18756 | 938 | P938S | C | 1.3e-07 | 1.4e-07 | 1.2e-07 | 1.300E-07 | 1.000E-08 | 1.200E-07 | 1.400E-07 |
| 18757 | 938 | P938T | C | 5.2e-08 | 1.1e-07 | 1.6e-07 | 1.073E-07 | 5.405E-08 | 5.200E-08 | 1.600E-07 |
| 18758 | 938 | P938W | C | 5.7e-08 | 7.2e-08 | 3e-08 | 5.300E-08 | 2.128E-08 | 3.000E-08 | 7.200E-08 |
| 18759 | 938 | P938Y | C | 2.4e-08 | 6.9e-08 | 2.6e-08 | 3.967E-08 | 2.542E-08 | 2.400E-08 | 6.900E-08 |
| 18760 | 938 | P938V | C | 6.2e-08 | 1.5e-08 | 3.6e-08 | 3.767E-08 | 2.354E-08 | 1.500E-08 | 6.200E-08 |
| 18761 | 939 | S939A | C | 2.1e-08 | 7.5e-08 | 4.5e-08 | 4.700E-08 | 2.706E-08 | 2.100E-08 | 7.500E-08 |
| 18762 | 939 | S939R | C | 1e-07 | 5e-08 | 4.7e-08 | 6.567E-08 | 2.977E-08 | 4.700E-08 | 1.000E-07 |
| 18763 | 939 | S939N | C | 2.6e-08 | 2.6e-08 | 5.3e-08 | 3.500E-08 | 1.559E-08 | 2.600E-08 | 5.300E-08 |
| 18764 | 939 | S939D | C | 7.9e-08 | 7.5e-08 | 5.4e-08 | 6.933E-08 | 1.343E-08 | 5.400E-08 | 7.900E-08 |
| 18765 | 939 | S939C | C | 1.1e-08 | 4e-08 | 5.6e-08 | 3.567E-08 | 2.281E-08 | 1.100E-08 | 5.600E-08 |
| 18766 | 939 | S939E | C | 6.3e-08 | 2.4e-08 | 2.1e-08 | 3.600E-08 | 2.343E-08 | 2.100E-08 | 6.300E-08 |
| 18767 | 939 | S939Q | C | 5.6e-08 | 4.3e-08 | 5e-08 | 4.967E-08 | 6.506E-09 | 4.300E-08 | 5.600E-08 |
| 18768 | 939 | S939G | C | 1.8e-08 | 4.4e-08 | 3.7e-08 | 3.300E-08 | 1.345E-08 | 1.800E-08 | 4.400E-08 |
| 18769 | 939 | S939H | C | 9.5e-08 | 5.8e-08 | 2.7e-08 | 6.000E-08 | 3.404E-08 | 2.700E-08 | 9.500E-08 |
| 18770 | 939 | S939I | C | 2.8e-08 | 2.7e-08 | 1.8e-08 | 2.433E-08 | 5.508E-09 | 1.800E-08 | 2.800E-08 |
| 18771 | 939 | S939L | C | 3.9e-08 | 1.4e-08 | 5e-08 | 3.433E-08 | 1.845E-08 | 1.400E-08 | 5.000E-08 |
| 18772 | 939 | S939K | C | 4.7e-08 | 6e-08 | 5.1e-08 | 5.267E-08 | 6.658E-09 | 4.700E-08 | 6.000E-08 |
| 18773 | 939 | S939M | C | 8.7e-08 | 1.7e-08 | 3.5e-08 | 4.633E-08 | 3.635E-08 | 1.700E-08 | 8.700E-08 |
| 18774 | 939 | S939F | C | 2.2e-08 | 4.4e-08 | 2.4e-08 | 3.000E-08 | 1.217E-08 | 2.200E-08 | 4.400E-08 |
| 18775 | 939 | S939P | C | 2.4e-08 | 1.2e-07 | 1e-08 | 5.133E-08 | 5.988E-08 | 1.000E-08 | 1.200E-07 |
| 18776 | 939 | S939S | C | 5.5e-08 | 1.4e-08 | 2.5e-08 | 3.133E-08 | 2.122E-08 | 1.400E-08 | 5.500E-08 |
| 18777 | 939 | S939T | C | 3.4e-08 | 1.1e-07 | 3.5e-08 | 5.967E-08 | 4.359E-08 | 3.400E-08 | 1.100E-07 |
| 18778 | 939 | S939W | C | 4.3e-08 | 3.6e-08 | 2e-08 | 3.300E-08 | 1.179E-08 | 2.000E-08 | 4.300E-08 |
| 18779 | 939 | S939Y | C | 6.8e-08 | 9e-09 | 4.1e-08 | 3.933E-08 | 2.954E-08 | 9.000E-09 | 6.800E-08 |
| 18780 | 939 | S939V | C | 1.5e-07 | 5.9e-08 | 1.7e-08 | 7.533E-08 | 6.799E-08 | 1.700E-08 | 1.500E-07 |
| 18781 | 940 | F940A | C | 4.8e-08 | 3.4e-08 | 2.4e-08 | 3.533E-08 | 1.206E-08 | 2.400E-08 | 4.800E-08 |
| 18782 | 940 | F940R | C | 5.8e-08 | 8e-08 | 6e-08 | 6.600E-08 | 1.217E-08 | 5.800E-08 | 8.000E-08 |
| 18783 | 940 | F940N | C | 3.3e-08 | 2.2e-08 | 3.4e-08 | 2.967E-08 | 6.658E-09 | 2.200E-08 | 3.400E-08 |
| 18784 | 940 | F940D | C | 5.4e-08 | 4.2e-08 | 3.2e-08 | 4.267E-08 | 1.102E-08 | 3.200E-08 | 5.400E-08 |

| | | | | | | | | | | |
|---|---|---|---|---|---|---|---|---|---|---|
| 18785 | 940 | F940C | C | 6.8e-08 | 3e-08 | 5.2e-08 | 5.000E-08 | 1.908E-08 | 3.000E-08 | 6.800E-08 |
| 18786 | 940 | F940E | C | 3e-08 | 1e-08 | 3.7e-08 | 2.567E-08 | 1.401E-08 | 1.000E-08 | 3.700E-08 |
| 18787 | 940 | F940Q | C | 5.7e-08 | 1.5e-08 | 3.3e-08 | 3.500E-08 | 2.107E-08 | 1.500E-08 | 5.700E-08 |
| 18788 | 940 | F940G | C | 2.4e-08 | 4.2e-08 | 4.3e-08 | 3.633E-08 | 1.069E-08 | 2.400E-08 | 4.300E-08 |
| 18789 | 940 | F940H | C | 3.3e-08 | 6.5e-08 | 3.7e-08 | 4.500E-08 | 1.744E-08 | 3.300E-08 | 6.500E-08 |
| 18790 | 940 | F940I | C | 5.9e-08 | 3.4e-08 | 5.7e-08 | 5.000E-08 | 1.389E-08 | 3.400E-08 | 5.900E-08 |
| 18791 | 940 | F940L | C | 5e-08 | 2.2e-08 | 3.7e-08 | 3.633E-08 | 1.401E-08 | 2.200E-08 | 5.000E-08 |
| 18792 | 940 | F940K | C | 1.8e-08 | 1.3e-07 | 3.5e-08 | 6.100E-08 | 6.036E-08 | 1.800E-08 | 1.300E-07 |
| 18793 | 940 | F940M | C | 9.1e-08 | 4.4e-08 | 2.8e-08 | 5.433E-08 | 3.275E-08 | 2.800E-08 | 9.100E-08 |
| 18794 | 940 | F940F | C | 3.8e-08 | 2.7e-08 | 1.5e-08 | 2.667E-08 | 1.150E-08 | 1.500E-08 | 3.800E-08 |
| 18795 | 940 | F940P | C | 4.9e-08 | 3.1e-08 | 1.9e-08 | 3.300E-08 | 1.510E-08 | 1.900E-08 | 4.900E-08 |
| 18796 | 940 | F940S | C | 3.1e-08 | 3.5e-08 | 4.6e-08 | 3.733E-08 | 7.767E-09 | 3.100E-08 | 4.600E-08 |
| 18797 | 940 | F940T | C | 2.2e-08 | 7.9e-08 | 9.1e-08 | 6.400E-08 | 3.686E-08 | 2.200E-08 | 9.100E-08 |
| 18798 | 940 | F940W | C | 2e-08 | 4.6e-08 | 5.1e-08 | 3.900E-08 | 1.664E-08 | 2.000E-08 | 5.100E-08 |
| 18799 | 940 | F940Y | C | 1.3e-07 | 2.6e-08 | 3.3e-08 | 6.300E-08 | 5.813E-08 | 2.600E-08 | 1.300E-07 |
| 18800 | 940 | F940V | C | 4.9e-08 | 4.4e-08 | 7.8e-08 | 5.700E-08 | 1.836E-08 | 4.400E-08 | 7.800E-08 |
| 18801 | 941 | Y941A | C | 1.4e-07 | 1.8e-08 | 3.9e-08 | 6.567E-08 | 6.523E-08 | 1.800E-08 | 1.400E-07 |
| 18802 | 941 | Y941R | C | 1.3e-07 | 7.3e-08 | 3.3e-08 | 7.867E-08 | 4.875E-08 | 3.300E-08 | 1.300E-07 |
| 18803 | 941 | Y941N | C | 5.1e-08 | 8.8e-08 | 3.6e-08 | 5.833E-08 | 2.676E-08 | 3.600E-08 | 8.800E-08 |
| 18804 | 941 | Y941D | C | 6.8e-08 | 3e-08 | 5.8e-08 | 5.200E-08 | 1.970E-08 | 3.000E-08 | 6.800E-08 |
| 18805 | 941 | Y941C | C | 3.6e-08 | 1.6e-08 | 8.7e-08 | 4.633E-08 | 3.661E-08 | 1.600E-08 | 8.700E-08 |
| 18806 | 941 | Y941E | C | 1.9e-08 | 3.8e-08 | 8.9e-08 | 4.867E-08 | 3.620E-08 | 1.900E-08 | 8.900E-08 |
| 18807 | 941 | Y941Q | C | 5.2e-08 | 4.1e-08 | 1.8e-08 | 3.700E-08 | 1.735E-08 | 1.800E-08 | 5.200E-08 |
| 18808 | 941 | Y941G | C | 1.8e-08 | 4.5e-08 | 4.2e-08 | 3.500E-08 | 1.480E-08 | 1.800E-08 | 4.500E-08 |
| 18809 | 941 | Y941H | C | 5.7e-08 | 3e-08 | 6.2e-08 | 4.967E-08 | 1.721E-08 | 3.000E-08 | 6.200E-08 |
| 18810 | 941 | Y941I | C | 6.5e-08 | 7.2e-08 | 1.3e-08 | 5.000E-08 | 3.223E-08 | 1.300E-08 | 7.200E-08 |
| 18811 | 941 | Y941L | C | 3.7e-08 | 8.5e-08 | 4.9e-08 | 5.700E-08 | 2.498E-08 | 3.700E-08 | 8.500E-08 |
| 18812 | 941 | Y941K | C | 2.7e-08 | 3e-08 | 1.6e-08 | 2.433E-08 | 7.371E-09 | 1.600E-08 | 3.000E-08 |
| 18813 | 941 | Y941M | C | 4.3e-08 | 6.2e-08 | 2.7e-08 | 4.400E-08 | 1.752E-08 | 2.700E-08 | 6.200E-08 |
| 18814 | 941 | Y941F | C | 1.1e-07 | 3.5e-08 | 8.2e-08 | 7.567E-08 | 3.790E-08 | 3.500E-08 | 1.100E-07 |
| 18815 | 941 | Y941P | C | 6.4e-08 | 2.9e-08 | 2.5e-08 | 3.933E-08 | 2.146E-08 | 2.500E-08 | 6.400E-08 |
| 18816 | 941 | Y941S | C | 5.9e-08 | 9.5e-08 | 3.2e-08 | 6.200E-08 | 3.161E-08 | 3.200E-08 | 9.500E-08 |
| 18817 | 941 | Y941T | C | 7.4e-08 | 9e-08 | 4.8e-08 | 7.067E-08 | 2.120E-08 | 4.800E-08 | 9.000E-08 |
| 18818 | 941 | Y941W | C | 2.6e-08 | 8e-08 | 3.8e-08 | 4.800E-08 | 2.835E-08 | 2.600E-08 | 8.000E-08 |
| 18819 | 941 | Y941Y | C | 2.5e-08 | 5.4e-08 | 4.7e-08 | 4.200E-08 | 1.513E-08 | 2.500E-08 | 5.400E-08 |
| 18820 | 941 | Y941V | C | 5.6e-08 | 8.6e-08 | 1.5e-08 | 5.233E-08 | 3.564E-08 | 1.500E-08 | 8.600E-08 |
| 18821 | 942 | F942A | C | 3.5e-08 | 3.6e-08 | 3.2e-08 | 3.433E-08 | 2.082E-09 | 3.200E-08 | 3.600E-08 |
| 18822 | 942 | F942R | C | 1.7e-08 | 2.1e-08 | 1.4e-07 | 5.933E-08 | 6.989E-08 | 1.700E-08 | 1.400E-07 |
| 18823 | 942 | F942N | C | 3e-08 | 4.6e-08 | 3.5e-08 | 3.700E-08 | 8.185E-09 | 3.000E-08 | 4.600E-08 |
| 18824 | 942 | F942D | C | 1e-07 | 5.5e-08 | 1.9e-08 | 5.800E-08 | 4.058E-08 | 1.900E-08 | 1.000E-07 |
| 18825 | 942 | F942C | C | 5.1e-08 | 2.4e-08 | 5.6e-08 | 4.367E-08 | 1.721E-08 | 2.400E-08 | 5.600E-08 |
| 18826 | 942 | F942E | C | 4.8e-08 | 5.8e-08 | 2e-08 | 4.200E-08 | 1.970E-08 | 2.000E-08 | 5.800E-08 |
| 18827 | 942 | F942Q | C | 1.8e-08 | 4.5e-08 | 1.4e-07 | 6.767E-08 | 6.408E-08 | 1.800E-08 | 1.400E-07 |
| 18828 | 942 | F942G | C | 3.2e-08 | 5.8e-08 | 2.5e-08 | 3.833E-08 | 1.739E-08 | 2.500E-08 | 5.800E-08 |
| 18829 | 942 | F942H | C | 3.4e-08 | 6.8e-08 | 3.7e-08 | 4.633E-08 | 1.882E-08 | 3.400E-08 | 6.800E-08 |
| 18830 | 942 | F942I | C | 3e-08 | 2.1e-08 | 4.7e-08 | 3.267E-08 | 1.320E-08 | 2.100E-08 | 4.700E-08 |
| 18831 | 942 | F942L | C | 3.6e-08 | 3.5e-08 | 3.1e-08 | 3.400E-08 | 2.646E-09 | 3.100E-08 | 3.600E-08 |
| 18832 | 942 | F942K | C | 5.8e-08 | 5.3e-08 | 3.6e-08 | 4.900E-08 | 1.153E-08 | 3.600E-08 | 5.800E-08 |
| 18833 | 942 | F942M | C | 3.2e-08 | 2.4e-08 | 5.1e-08 | 3.567E-08 | 1.387E-08 | 2.400E-08 | 5.100E-08 |
| 18834 | 942 | F942F | C | 6.1e-08 | 3e-08 | 3.6e-08 | 4.233E-08 | 1.644E-08 | 3.000E-08 | 6.100E-08 |
| 18835 | 942 | F942P | C | 1.1e-07 | 4.2e-08 | 2.1e-08 | 5.767E-08 | 4.652E-08 | 2.100E-08 | 1.100E-07 |
| 18836 | 942 | F942S | C | 6.9e-08 | 8.6e-08 | 3.6e-08 | 6.367E-08 | 2.542E-08 | 3.600E-08 | 8.600E-08 |
| 18837 | 942 | F942T | C | 5.4e-08 | 3e-08 | 4.7e-08 | 4.367E-08 | 1.234E-08 | 3.000E-08 | 5.400E-08 |
| 18838 | 942 | F942W | C | 2.1e-08 | 2.2e-08 | 3.8e-08 | 2.700E-08 | 9.539E-09 | 2.100E-08 | 3.800E-08 |
| 18839 | 942 | F942Y | C | 3.1e-08 | 2.4e-08 | 2.1e-08 | 2.533E-08 | 5.132E-09 | 2.100E-08 | 3.100E-08 |
| 18840 | 942 | F942V | C | 8e-08 | 2.2e-08 | 4.9e-08 | 5.033E-08 | 2.902E-08 | 2.200E-08 | 8.000E-08 |
| 18841 | 943 | D943A | C | 1.7e-07 | 2.9e-08 | 5.8e-08 | 8.567E-08 | 7.446E-08 | 2.900E-08 | 1.700E-07 |
| 18842 | 943 | D943R | C | 6.1e-08 | 4.4e-08 | 2.1e-08 | 4.200E-08 | 2.007E-08 | 2.100E-08 | 6.100E-08 |
| 18843 | 943 | D943N | C | 7.1e-08 | 4.8e-08 | 3.7e-08 | 5.200E-08 | 1.735E-08 | 3.700E-08 | 7.100E-08 |
| 18844 | 943 | D943D | C | 5.3e-08 | 9.8e-08 | 8.9e-09 | 5.330E-08 | 4.455E-08 | 8.900E-09 | 9.800E-08 |
| 18845 | 943 | D943C | C | 6.1e-08 | 2.5e-08 | 2.2e-08 | 3.600E-08 | 2.170E-08 | 2.200E-08 | 6.100E-08 |
| 18846 | 943 | D943E | C | 5.9e-08 | 1e-07 | 1.1e-08 | 5.667E-08 | 4.455E-08 | 1.100E-08 | 1.000E-07 |
| 18847 | 943 | D943Q | C | 2.6e-08 | 6.4e-08 | 4.8e-08 | 4.600E-08 | 1.908E-08 | 2.600E-08 | 6.400E-08 |
| 18848 | 943 | D943G | C | 5e-08 | 2.8e-08 | 1.6e-07 | 7.933E-08 | 7.072E-08 | 2.800E-08 | 1.600E-07 |
| 18849 | 943 | D943H | C | 5.7e-08 | 1.2e-07 | 4e-08 | 7.233E-08 | 4.215E-08 | 4.000E-08 | 1.200E-07 |
| 18850 | 943 | D943I | C | 1e-07 | 1e-07 | 1.1e-07 | 1.033E-07 | 5.774E-09 | 1.000E-07 | 1.100E-07 |
| 18851 | 943 | D943L | C | 4.3e-08 | 2.7e-08 | 3.2e-08 | 3.400E-08 | 8.185E-09 | 2.700E-08 | 4.300E-08 |
| 18852 | 943 | D943K | C | 8.1e-08 | 7.4e-08 | 5.9e-08 | 7.133E-08 | 1.124E-08 | 5.900E-08 | 8.100E-08 |
| 18853 | 943 | D943M | C | 7.2e-08 | 4.9e-08 | 2.6e-08 | 4.900E-08 | 2.300E-08 | 2.600E-08 | 7.200E-08 |
| 18854 | 943 | D943F | C | 7.9e-08 | 3.4e-08 | 6.2e-08 | 5.833E-08 | 2.272E-08 | 3.400E-08 | 7.900E-08 |
| 18855 | 943 | D943P | C | 7.3e-08 | 6.4e-08 | 9.3e-08 | 7.667E-08 | 1.484E-08 | 6.400E-08 | 9.300E-08 |
| 18856 | 943 | D943S | C | 6e-08 | 6.8e-08 | 7.9e-08 | 6.900E-08 | 9.539E-09 | 6.000E-08 | 7.900E-08 |
| 18857 | 943 | D943T | C | 4.7e-08 | 6.4e-08 | 3.9e-08 | 5.000E-08 | 1.277E-08 | 3.900E-08 | 6.400E-08 |
| 18858 | 943 | D943W | C | 2.5e-08 | 3.2e-08 | 3.8e-08 | 3.167E-08 | 6.506E-09 | 2.500E-08 | 3.800E-08 |
| 18859 | 943 | D943Y | C | 5.4e-08 | 1.1e-07 | 5.5e-08 | 7.300E-08 | 3.205E-08 | 5.400E-08 | 1.100E-07 |
| 18860 | 943 | D943V | C | 2e-08 | 4.1e-08 | 4.6e-08 | 3.567E-08 | 1.380E-08 | 2.000E-08 | 4.600E-08 |
| 18861 | 944 | Y944A | C | 8.9e-08 | 1.7e-08 | 2.2e-08 | 4.267E-08 | 4.020E-08 | 1.700E-08 | 8.900E-08 |
| 18862 | 944 | Y944R | C | 4.1e-08 | 1.2e-07 | 6.8e-08 | 7.633E-08 | 4.015E-08 | 4.100E-08 | 1.200E-07 |
| 18863 | 944 | Y944N | C | 4.8e-08 | 5.3e-08 | 2.9e-08 | 4.333E-08 | 1.266E-08 | 2.900E-08 | 5.300E-08 |
| 18864 | 944 | Y944D | C | 2.7e-08 | 3.4e-08 | 1.7e-07 | 7.700E-08 | 8.062E-08 | 2.700E-08 | 1.700E-07 |
| 18865 | 944 | Y944C | C | 4.4e-08 | 1.3e-08 | 1.4e-07 | 6.567E-08 | 6.621E-08 | 1.300E-08 | 1.400E-07 |
| 18866 | 944 | Y944E | C | 2e-08 | 9.1e-09 | 5.1e-08 | 2.670E-08 | 2.174E-08 | 9.100E-09 | 5.100E-08 |
| 18867 | 944 | Y944Q | C | 2.7e-08 | 4.9e-08 | 1.1e-08 | 2.900E-08 | 1.908E-08 | 1.100E-08 | 4.900E-08 |
| 18868 | 944 | Y944G | C | 2.1e-07 | 6.5e-08 | 1e-07 | 1.250E-07 | 7.566E-08 | 6.500E-08 | 2.100E-07 |
| 18869 | 944 | Y944H | C | 4.2e-08 | 7.6e-08 | 4e-08 | 5.267E-08 | 2.023E-08 | 4.000E-08 | 7.600E-08 |

| | | | | | | | | | | |
|---|---|---|---|---|---|---|---|---|---|---|
| 18870 | 944 | Y944I | C | 1.5e-08 | 6.6e-08 | 3.4e-08 | 3.833E-08 | 2.577E-08 | 1.500E-08 | 6.600E-08 |
| 18871 | 944 | Y944L | C | 6.4e-08 | 2.5e-08 | 7.1e-08 | 5.333E-08 | 2.479E-08 | 2.500E-08 | 7.100E-08 |
| 18872 | 944 | Y944K | C | 1.3e-08 | 3.4e-08 | 3.2e-08 | 2.633E-08 | 1.159E-08 | 1.300E-08 | 3.400E-08 |
| 18873 | 944 | Y944M | C | 1.7e-08 | 4.7e-08 | 1.6e-07 | 7.467E-08 | 7.541E-08 | 1.700E-08 | 1.600E-07 |
| 18874 | 944 | Y944F | C | 2.3e-08 | 5.4e-08 | 4.6e-09 | 2.720E-08 | 2.497E-08 | 4.600E-09 | 5.400E-08 |
| 18875 | 944 | Y944P | C | 2.3e-08 | 2.8e-08 | 7.4e-08 | 4.167E-08 | 2.811E-08 | 2.300E-08 | 7.400E-08 |
| 18876 | 944 | Y944S | C | 7.7e-08 | 6.1e-08 | 3.7e-08 | 5.833E-08 | 2.013E-08 | 3.700E-08 | 7.700E-08 |
| 18877 | 944 | Y944T | C | 3.4e-08 | 1.3e-07 | 5.8e-08 | 7.400E-08 | 4.996E-08 | 3.400E-08 | 1.300E-07 |
| 18878 | 944 | Y944W | C | 3e-08 | 2.1e-08 | 3.7e-08 | 2.933E-08 | 8.021E-09 | 2.100E-08 | 3.700E-08 |
| 18879 | 944 | Y944Y | C | 4.6e-08 | 6.9e-08 | 3.8e-08 | 5.100E-08 | 1.609E-08 | 3.800E-08 | 6.900E-08 |
| 18880 | 944 | Y944V | C | 1e-07 | 1.7e-08 | 4.4e-08 | 5.367E-08 | 4.234E-08 | 1.700E-08 | 1.000E-07 |
| 18881 | 945 | W945A | C | 3.5e-08 | 3.9e-08 | 5e-08 | 4.133E-08 | 7.767E-09 | 3.500E-08 | 5.000E-08 |
| 18882 | 945 | W945R | C | 7.4e-08 | 5.6e-08 | 1.9e-08 | 4.967E-08 | 2.804E-08 | 1.900E-08 | 7.400E-08 |
| 18883 | 945 | W945N | C | 1.2e-07 | 1.1e-08 | 1.1e-07 | 8.033E-08 | 6.025E-08 | 1.100E-08 | 1.200E-07 |
| 18884 | 945 | W945D | C | 4.3e-08 | 2.1e-08 | 3e-08 | 3.133E-08 | 1.106E-08 | 2.100E-08 | 4.300E-08 |
| 18885 | 945 | W945C | C | 2.4e-08 | 3.9e-08 | 7.2e-08 | 4.500E-08 | 2.456E-08 | 2.400E-08 | 7.200E-08 |
| 18886 | 945 | W945E | C | 5.9e-08 | 3.5e-08 | 7.9e-08 | 5.767E-08 | 2.203E-08 | 3.500E-08 | 7.900E-08 |
| 18887 | 945 | W945Q | C | 2.3e-08 | 4.1e-08 | 4.5e-08 | 3.633E-08 | 1.172E-08 | 2.300E-08 | 4.500E-08 |
| 18888 | 945 | W945G | C | 3.6e-08 | 4.8e-08 | 6.7e-08 | 5.033E-08 | 1.563E-08 | 3.600E-08 | 6.700E-08 |
| 18889 | 945 | W945H | C | 6.4e-08 | 8.7e-09 | 1.1e-07 | 6.090E-08 | 5.072E-08 | 8.700E-09 | 1.100E-07 |
| 18890 | 945 | W945I | C | 4.6e-08 | 5.4e-08 | 3.6e-08 | 4.533E-08 | 9.018E-09 | 3.600E-08 | 5.400E-08 |
| 18891 | 945 | W945L | C | 2.4e-08 | 5.3e-08 | 4.5e-08 | 4.067E-08 | 1.498E-08 | 2.400E-08 | 5.300E-08 |
| 18892 | 945 | W945K | C | 3.7e-08 | 7.4e-08 | 4.2e-08 | 5.100E-08 | 2.007E-08 | 3.700E-08 | 7.400E-08 |
| 18893 | 945 | W945M | C | 6.7e-08 | 2e-08 | 1e-07 | 6.233E-08 | 4.020E-08 | 2.000E-08 | 1.000E-07 |
| 18894 | 945 | W945F | C | 1e-07 | 2.2e-08 | 5.9e-08 | 6.033E-08 | 3.902E-08 | 2.200E-08 | 1.000E-07 |
| 18895 | 945 | W945P | C | 1.8e-08 | 4.1e-08 | 5.2e-08 | 3.700E-08 | 1.735E-08 | 1.800E-08 | 5.200E-08 |
| 18896 | 945 | W945S | C | 1.8e-08 | 2.2e-08 | 1.6e-08 | 1.867E-08 | 3.055E-09 | 1.600E-08 | 2.200E-08 |
| 18897 | 945 | W945T | C | 1.4e-08 | 2.8e-08 | 4e-08 | 2.733E-08 | 1.301E-08 | 1.400E-08 | 4.000E-08 |
| 18898 | 945 | W945W | C | 3.1e-08 | 4.5e-08 | 3.4e-08 | 3.667E-08 | 7.371E-09 | 3.100E-08 | 4.500E-08 |
| 18899 | 945 | W945Y | C | 4e-08 | 5.1e-08 | 3.9e-08 | 4.333E-08 | 6.658E-09 | 3.900E-08 | 5.100E-08 |
| 18900 | 945 | W945V | C | 4.3e-08 | 2.7e-08 | 1.2e-07 | 6.333E-08 | 4.972E-08 | 2.700E-08 | 1.200E-07 |
| 18901 | 946 | G946A | C | 4.4e-08 | 9.7e-08 | 3.2e-08 | 5.767E-08 | 3.459E-08 | 3.200E-08 | 9.700E-08 |
| 18902 | 946 | G946R | C | 1.2e-07 | 1.8e-07 | 3.2e-08 | 1.107E-07 | 7.444E-08 | 3.200E-08 | 1.800E-07 |
| 18903 | 946 | G946N | C | 5e-08 | 2.1e-08 | 2.4e-07 | 1.037E-07 | 1.190E-07 | 2.100E-08 | 2.400E-07 |
| 18904 | 946 | G946D | C | 5.2e-08 | 2.6e-08 | 5.5e-08 | 4.433E-08 | 1.595E-08 | 2.600E-08 | 5.500E-08 |
| 18905 | 946 | G946C | C | 4e-08 | 1.5e-07 | 2e-08 | 7.000E-08 | 7.000E-08 | 2.000E-08 | 1.500E-07 |
| 18906 | 946 | G946E | C | 4.1e-08 | 5.9e-08 | 1e-07 | 6.667E-08 | 3.024E-08 | 4.100E-08 | 1.000E-07 |
| 18907 | 946 | G946Q | C | 4.2e-08 | 4e-08 | 4.5e-08 | 4.233E-08 | 2.517E-09 | 4.000E-08 | 4.500E-08 |
| 18908 | 946 | G946G | C | 3.1e-08 | 1.3e-07 | 2.2e-08 | 6.100E-08 | 5.992E-08 | 2.200E-08 | 1.300E-07 |
| 18909 | 946 | G946H | C | 3.2e-08 | 9.3e-08 | 6.9e-08 | 6.467E-08 | 3.073E-08 | 3.200E-08 | 9.300E-08 |
| 18910 | 946 | G946I | C | 3.7e-08 | 1.3e-08 | 6.6e-08 | 3.867E-08 | 2.654E-08 | 1.300E-08 | 6.600E-08 |
| 18911 | 946 | G946L | C | 2.7e-08 | 4.5e-08 | 6.6e-08 | 4.600E-08 | 1.952E-08 | 2.700E-08 | 6.600E-08 |
| 18912 | 946 | G946K | C | 7.1e-08 | 1.2e-08 | 6.8e-08 | 5.033E-08 | 3.323E-08 | 1.200E-08 | 7.100E-08 |
| 18913 | 946 | G946M | C | 9.3e-09 | 9.4e-08 | 4.1e-08 | 4.810E-08 | 4.279E-08 | 9.300E-09 | 9.400E-08 |
| 18914 | 946 | G946F | C | 4.3e-08 | 4.2e-08 | 1.9e-08 | 3.467E-08 | 1.358E-08 | 1.900E-08 | 4.300E-08 |
| 18915 | 946 | G946P | C | 4.3e-08 | 5.4e-08 | 6.2e-08 | 5.300E-08 | 9.539E-09 | 4.300E-08 | 6.200E-08 |
| 18916 | 946 | G946S | C | 2.5e-08 | 5.5e-08 | 2.8e-08 | 3.600E-08 | 1.652E-08 | 2.500E-08 | 5.500E-08 |
| 18917 | 946 | G946T | C | 5.1e-08 | 6.7e-08 | 3.8e-08 | 5.200E-08 | 1.453E-08 | 3.800E-08 | 6.700E-08 |
| 18918 | 946 | G946W | C | 1.3e-07 | 2.1e-08 | 2.4e-08 | 5.833E-08 | 6.208E-08 | 2.100E-08 | 1.300E-07 |
| 18919 | 946 | G946Y | C | 2.9e-08 | 9.5e-08 | 3.8e-08 | 5.400E-08 | 3.579E-08 | 2.900E-08 | 9.500E-08 |
| 18920 | 946 | G946V | C | 8.1e-08 | 6.5e-08 | 3.4e-08 | 6.000E-08 | 2.390E-08 | 3.400E-08 | 8.100E-08 |
| 18921 | 947 | Q947A | C | 7.6e-08 | 4.1e-08 | 7.2e-08 | 6.300E-08 | 1.916E-08 | 4.100E-08 | 7.600E-08 |
| 18922 | 947 | Q947R | C | 4.6e-08 | 4.7e-08 | 2.9e-08 | 4.067E-08 | 1.012E-08 | 2.900E-08 | 4.700E-08 |
| 18923 | 947 | Q947N | C | 7.3e-08 | 1.8e-08 | 5.5e-08 | 4.867E-08 | 2.804E-08 | 1.800E-08 | 7.300E-08 |
| 18924 | 947 | Q947D | C | 2.9e-08 | 2.7e-08 | 4.4e-08 | 3.333E-08 | 9.292E-09 | 2.700E-08 | 4.400E-08 |
| 18925 | 947 | Q947C | C | 4.4e-08 | 2.2e-08 | 3.3e-08 | 3.300E-08 | 1.100E-08 | 2.200E-08 | 4.400E-08 |
| 18926 | 947 | Q947E | C | 4.5e-08 | 3.7e-08 | 3.7e-08 | 3.967E-08 | 4.619E-09 | 3.700E-08 | 4.500E-08 |
| 18927 | 947 | Q947Q | C | 3.8e-08 | 3.7e-08 | 2.2e-08 | 3.233E-08 | 8.963E-09 | 2.200E-08 | 3.800E-08 |
| 18928 | 947 | Q947G | C | 7.3e-08 | 2.7e-08 | 3.3e-08 | 4.433E-08 | 2.501E-08 | 2.700E-08 | 7.300E-08 |
| 18929 | 947 | Q947H | C | 4.1e-08 | 1.2e-07 | 5.6e-08 | 7.233E-08 | 4.196E-08 | 4.100E-08 | 1.200E-07 |
| 18930 | 947 | Q947I | C | 7.9e-08 | 5.4e-08 | 2.5e-08 | 5.267E-08 | 2.702E-08 | 2.500E-08 | 7.900E-08 |
| 18931 | 947 | Q947L | C | 2.7e-08 | 4.5e-08 | 2.8e-08 | 3.333E-08 | 1.012E-08 | 2.700E-08 | 4.500E-08 |
| 18932 | 947 | Q947K | C | 3.1e-07 | 4.5e-08 | 2.1e-08 | 1.253E-07 | 1.604E-07 | 2.100E-08 | 3.100E-07 |
| 18933 | 947 | Q947M | C | 1.7e-08 | 4.4e-08 | 8.3e-08 | 4.800E-08 | 3.318E-08 | 1.700E-08 | 8.300E-08 |
| 18934 | 947 | Q947F | C | 6.8e-08 | 9.6e-08 | 8.9e-08 | 8.433E-08 | 1.457E-08 | 6.800E-08 | 9.600E-08 |
| 18935 | 947 | Q947P | C | 3.9e-08 | 6.2e-08 | 2.1e-08 | 4.067E-08 | 2.055E-08 | 2.100E-08 | 6.200E-08 |
| 18936 | 947 | Q947S | C | 3.8e-08 | 2.3e-08 | 7.6e-08 | 4.567E-08 | 2.732E-08 | 2.300E-08 | 7.600E-08 |
| 18937 | 947 | Q947T | C | 3.6e-08 | 5.6e-08 | 5.6e-08 | 4.933E-08 | 1.155E-08 | 3.600E-08 | 5.600E-08 |
| 18938 | 947 | Q947W | C | 7.3e-08 | 6.3e-08 | 3.1e-08 | 5.567E-08 | 2.194E-08 | 3.100E-08 | 7.300E-08 |
| 18939 | 947 | Q947Y | C | 2.8e-08 | 9.2e-08 | 3e-08 | 5.000E-08 | 3.639E-08 | 2.800E-08 | 9.200E-08 |
| 18940 | 947 | Q947V | C | 2.9e-08 | 1.2e-08 | 5.1e-08 | 3.067E-08 | 1.955E-08 | 1.200E-08 | 5.100E-08 |
| 18941 | 948 | G948A | C | 1.1e-08 | 8.6e-08 | 8.5e-08 | 6.067E-08 | 4.302E-08 | 1.100E-08 | 8.600E-08 |
| 18942 | 948 | G948R | C | 1.9e-08 | 4e-08 | 1.8e-08 | 2.567E-08 | 1.242E-08 | 1.800E-08 | 4.000E-08 |
| 18943 | 948 | G948N | C | 5.1e-08 | 3.7e-08 | 3.3e-08 | 4.033E-08 | 9.452E-09 | 3.300E-08 | 5.100E-08 |
| 18944 | 948 | G948D | C | 2.2e-08 | 6e-08 | 6.1e-08 | 4.767E-08 | 2.223E-08 | 2.200E-08 | 6.100E-08 |
| 18945 | 948 | G948C | C | 5.8e-08 | 3.2e-08 | 5.7e-08 | 4.900E-08 | 1.473E-08 | 3.200E-08 | 5.800E-08 |
| 18946 | 948 | G948E | C | 4.1e-08 | 5e-08 | 5.4e-08 | 4.833E-08 | 6.658E-09 | 4.100E-08 | 5.400E-08 |
| 18947 | 948 | G948Q | C | 1.2e-07 | 8.1e-08 | 8.6e-08 | 9.567E-08 | 2.122E-08 | 8.100E-08 | 1.200E-07 |
| 18948 | 948 | G948G | C | 7.1e-08 | 3.9e-08 | 6.2e-08 | 5.733E-08 | 1.650E-08 | 3.900E-08 | 7.100E-08 |
| 18949 | 948 | G948H | C | 8.7e-08 | 1.3e-08 | 5.8e-08 | 5.267E-08 | 3.729E-08 | 1.300E-08 | 8.700E-08 |
| 18950 | 948 | G948I | C | 4.4e-08 | 1e-08 | 5.7e-08 | 3.700E-08 | 2.427E-08 | 1.000E-08 | 5.700E-08 |
| 18951 | 948 | G948L | C | 1.2e-08 | 1.7e-08 | 1.5e-08 | 1.467E-08 | 2.517E-09 | 1.200E-08 | 1.700E-08 |
| 18952 | 948 | G948K | C | 7.2e-08 | 5.4e-08 | 4.7e-08 | 5.767E-08 | 1.290E-08 | 4.700E-08 | 7.200E-08 |
| 18953 | 948 | G948M | C | 1.1e-08 | 1.7e-08 | 4e-08 | 2.267E-08 | 1.531E-08 | 1.100E-08 | 4.000E-08 |
| 18954 | 948 | G948F | C | 4e-08 | 6.5e-08 | 6.2e-08 | 5.567E-08 | 1.365E-08 | 4.000E-08 | 6.500E-08 |

| | | | | | | | | | | |
|---|---|---|---|---|---|---|---|---|---|---|
| 18955 | 948 | G948P | C | 6.7e-08 | 7.5e-08 | 1.7e-08 | 5.300E-08 | 3.143E-08 | 1.700E-08 | 7.500E-08 |
| 18956 | 948 | G948S | C | 4.1e-08 | 1.1e-07 | 8.2e-08 | 7.767E-08 | 3.470E-08 | 4.100E-08 | 1.100E-07 |
| 18957 | 948 | G948T | C | 5.3e-08 | 2.9e-08 | 5.3e-08 | 4.500E-08 | 1.386E-08 | 2.900E-08 | 5.300E-08 |
| 18958 | 948 | G948W | C | 1.1e-08 | 2.8e-08 | 2.1e-08 | 2.000E-08 | 8.544E-09 | 1.100E-08 | 2.800E-08 |
| 18959 | 948 | G948Y | C | 3.5e-08 | 1.2e-07 | 6.3e-08 | 7.267E-08 | 4.332E-08 | 3.500E-08 | 1.200E-07 |
| 18960 | 948 | G948V | C | 8.3e-08 | 1.5e-08 | 2.3e-08 | 4.033E-08 | 3.717E-08 | 1.500E-08 | 8.300E-08 |
| 18961 | 949 | T949A | C | 3.1e-08 | 1.8e-08 | 2.4e-08 | 2.433E-08 | 6.506E-09 | 1.800E-08 | 3.100E-08 |
| 18962 | 949 | T949R | C | 8e-08 | 2.1e-08 | 3.2e-08 | 4.433E-08 | 3.137E-08 | 2.100E-08 | 8.000E-08 |
| 18963 | 949 | T949N | C | 2.7e-08 | 3.7e-08 | 7.1e-08 | 4.500E-08 | 2.307E-08 | 2.700E-08 | 7.100E-08 |
| 18964 | 949 | T949D | C | 3.1e-08 | 9.6e-08 | 5.4e-08 | 6.033E-08 | 3.296E-08 | 3.100E-08 | 9.600E-08 |
| 18965 | 949 | T949C | C | 3e-08 | 5.7e-08 | 3.8e-08 | 4.167E-08 | 1.387E-08 | 3.000E-08 | 5.700E-08 |
| 18966 | 949 | T949E | C | 9.7e-08 | 4.8e-08 | 5.3e-08 | 6.600E-08 | 2.696E-08 | 4.800E-08 | 9.700E-08 |
| 18967 | 949 | T949Q | C | 4.4e-08 | 1e-07 | 4.6e-08 | 6.333E-08 | 3.177E-08 | 4.400E-08 | 1.000E-07 |
| 18968 | 949 | T949G | C | 5.7e-08 | 3.4e-08 | 3.7e-08 | 4.267E-08 | 1.250E-08 | 3.400E-08 | 5.700E-08 |
| 18969 | 949 | T949H | C | 7.5e-08 | 6.8e-08 | 3.7e-08 | 6.000E-08 | 2.022E-08 | 3.700E-08 | 7.500E-08 |
| 18970 | 949 | T949I | C | 7.8e-08 | 1.1e-07 | 5.3e-08 | 8.033E-08 | 2.857E-08 | 5.300E-08 | 1.100E-07 |
| 18971 | 949 | T949L | C | 4e-08 | 4.6e-08 | 3.3e-08 | 3.967E-08 | 6.506E-09 | 3.300E-08 | 4.600E-08 |
| 18972 | 949 | T949K | C | 6.7e-08 | 8.3e-08 | 1.9e-08 | 5.633E-08 | 3.331E-08 | 1.900E-08 | 8.300E-08 |
| 18973 | 949 | T949M | C | 3.9e-08 | 9.2e-08 | 4.4e-08 | 5.833E-08 | 2.926E-08 | 3.900E-08 | 9.200E-08 |
| 18974 | 949 | T949F | C | 3e-08 | 3.6e-08 | 6e-08 | 4.200E-08 | 1.587E-08 | 3.000E-08 | 6.000E-08 |
| 18975 | 949 | T949P | C | 4.5e-08 | 9.3e-08 | 3.3e-08 | 5.700E-08 | 3.175E-08 | 3.300E-08 | 9.300E-08 |
| 18976 | 949 | T949S | C | 1.1e-08 | 1.7e-08 | 1.1e-07 | 4.600E-08 | 5.551E-08 | 1.100E-08 | 1.100E-07 |
| 18977 | 949 | T949T | C | 3.8e-08 | 3.3e-08 | 1.3e-07 | 6.700E-08 | 5.462E-08 | 3.300E-08 | 1.300E-07 |
| 18978 | 949 | T949W | C | 4.7e-08 | 5e-08 | 2.9e-08 | 4.200E-08 | 1.136E-08 | 2.900E-08 | 5.000E-08 |
| 18979 | 949 | T949Y | C | 4.4e-08 | 5.3e-08 | 2.2e-08 | 3.967E-08 | 1.595E-08 | 2.200E-08 | 5.300E-08 |
| 18980 | 949 | T949V | C | 4.7e-08 | 3.4e-08 | 3.1e-08 | 3.733E-08 | 8.505E-09 | 3.100E-08 | 4.700E-08 |
| 18981 | 950 | L950A | C | 5.2e-08 | 8.2e-08 | 3.7e-08 | 5.700E-08 | 2.291E-08 | 3.700E-08 | 8.200E-08 |
| 18982 | 950 | L950R | C | 5.1e-08 | 4.8e-08 | 1.8e-08 | 3.900E-08 | 1.825E-08 | 1.800E-08 | 5.100E-08 |
| 18983 | 950 | L950N | C | 2.8e-08 | 1.7e-08 | 2e-08 | 2.167E-08 | 5.686E-09 | 1.700E-08 | 2.800E-08 |
| 18984 | 950 | L950D | C | 1.9e-08 | 1.3e-08 | 5.7e-08 | 2.967E-08 | 2.386E-08 | 1.300E-08 | 5.700E-08 |
| 18985 | 950 | L950C | C | 1.7e-08 | 6.2e-08 | 2.4e-08 | 3.433E-08 | 2.421E-08 | 1.700E-08 | 6.200E-08 |
| 18986 | 950 | L950E | C | 2e-08 | 3.8e-08 | 8e-08 | 4.600E-08 | 3.079E-08 | 2.000E-08 | 8.000E-08 |
| 18987 | 950 | L950Q | C | 6.7e-08 | 3.1e-08 | 1.1e-08 | 3.633E-08 | 2.838E-08 | 1.100E-08 | 6.700E-08 |
| 18988 | 950 | L950G | C | 6e-08 | 2e-08 | 3.6e-08 | 3.867E-08 | 2.013E-08 | 2.000E-08 | 6.000E-08 |
| 18989 | 950 | L950H | C | 2.1e-08 | 4.5e-08 | 7.8e-08 | 4.800E-08 | 2.862E-08 | 2.100E-08 | 7.800E-08 |
| 18990 | 950 | L950I | C | 7.3e-08 | 5.2e-08 | 8.2e-08 | 6.900E-08 | 1.539E-08 | 5.200E-08 | 8.200E-08 |
| 18991 | 950 | L950L | C | 4.5e-08 | 3.9e-08 | 5.5e-08 | 4.633E-08 | 8.083E-09 | 3.900E-08 | 5.500E-08 |
| 18992 | 950 | L950K | C | 3.3e-08 | 7e-08 | 3.8e-08 | 4.700E-08 | 2.007E-08 | 3.300E-08 | 7.000E-08 |
| 18993 | 950 | L950M | C | 1.4e-07 | 4e-08 | 3.4e-08 | 7.133E-08 | 5.954E-08 | 3.400E-08 | 1.400E-07 |
| 18994 | 950 | L950F | C | 9.4e-09 | 9.5e-08 | 6.4e-08 | 5.613E-08 | 4.334E-08 | 9.400E-09 | 9.500E-08 |
| 18995 | 950 | L950P | C | 4e-08 | 1.2e-07 | 6.1e-08 | 7.367E-08 | 4.148E-08 | 4.000E-08 | 1.200E-07 |
| 18996 | 950 | L950S | C | 8.2e-08 | 6.1e-08 | 4.8e-08 | 6.367E-08 | 1.716E-08 | 4.800E-08 | 8.200E-08 |
| 18997 | 950 | L950T | C | 2.4e-08 | 2.5e-08 | 3.7e-08 | 2.867E-08 | 7.234E-09 | 2.400E-08 | 3.700E-08 |
| 18998 | 950 | L950W | C | 4.7e-08 | 3.1e-08 | 5.8e-08 | 4.533E-08 | 1.358E-08 | 3.100E-08 | 5.800E-08 |
| 18999 | 950 | L950Y | C | 2.9e-08 | 1.8e-08 | 3.7e-08 | 2.800E-08 | 9.539E-09 | 1.800E-08 | 3.700E-08 |
| 19000 | 950 | L950V | C | 5.1e-08 | 1.5e-08 | 8.1e-08 | 4.900E-08 | 3.305E-08 | 1.500E-08 | 8.100E-08 |
| 19001 | 951 | V951A | C | 3.7e-08 | 2.8e-08 | 1.7e-08 | 2.733E-08 | 1.002E-08 | 1.700E-08 | 3.700E-08 |
| 19002 | 951 | V951R | C | 3.1e-08 | 7.8e-08 | 9.5e-08 | 6.800E-08 | 3.315E-08 | 3.100E-08 | 9.500E-08 |
| 19003 | 951 | V951N | C | 2.5e-08 | 4.3e-08 | 4e-08 | 3.600E-08 | 9.644E-09 | 2.500E-08 | 4.300E-08 |
| 19004 | 951 | V951D | C | 3.2e-08 | 2.9e-08 | 2.6e-08 | 2.900E-08 | 3.000E-09 | 2.600E-08 | 3.200E-08 |
| 19005 | 951 | V951C | C | 3.5e-08 | 1.5e-08 | 6.4e-08 | 3.800E-08 | 2.464E-08 | 1.500E-08 | 6.400E-08 |
| 19006 | 951 | V951E | C | 1.1e-08 | 5.7e-08 | 3.3e-08 | 3.367E-08 | 2.301E-08 | 1.100E-08 | 5.700E-08 |
| 19007 | 951 | V951Q | C | 5.5e-08 | 5e-08 | 2.1e-08 | 4.200E-08 | 1.836E-08 | 2.100E-08 | 5.500E-08 |
| 19008 | 951 | V951G | C | 1.6e-08 | 3.3e-08 | 6.5e-08 | 3.800E-08 | 2.488E-08 | 1.600E-08 | 6.500E-08 |
| 19009 | 951 | V951H | C | 5.3e-08 | 6.6e-08 | 6.7e-08 | 6.200E-08 | 7.810E-09 | 5.300E-08 | 6.700E-08 |
| 19010 | 951 | V951I | C | 1.9e-08 | 4.4e-08 | 1.7e-08 | 2.667E-08 | 1.504E-08 | 1.700E-08 | 4.400E-08 |
| 19011 | 951 | V951L | C | 2.3e-08 | 2.3e-08 | 6.2e-08 | 3.600E-08 | 2.252E-08 | 2.300E-08 | 6.200E-08 |
| 19012 | 951 | V951K | C | 4.8e-08 | 7.1e-08 | 5.1e-08 | 5.667E-08 | 1.250E-08 | 4.800E-08 | 7.100E-08 |
| 19013 | 951 | V951M | C | 2.9e-08 | 3.2e-08 | 6.5e-08 | 4.200E-08 | 1.997E-08 | 2.900E-08 | 6.500E-08 |
| 19014 | 951 | V951F | C | 5.4e-08 | 1.1e-07 | 6.3e-08 | 7.567E-08 | 3.007E-08 | 5.400E-08 | 1.100E-07 |
| 19015 | 951 | V951P | C | 2.5e-08 | 1.3e-08 | 7.4e-08 | 3.733E-08 | 3.232E-08 | 1.300E-08 | 7.400E-08 |
| 19016 | 951 | V951S | C | 6.6e-08 | 4.2e-08 | 6e-08 | 5.600E-08 | 1.249E-08 | 4.200E-08 | 6.600E-08 |
| 19017 | 951 | V951T | C | 4e-08 | 2.7e-08 | 4.5e-08 | 3.733E-08 | 9.292E-09 | 2.700E-08 | 4.500E-08 |
| 19018 | 951 | V951W | C | 7.1e-08 | 3.9e-08 | 4.8e-08 | 5.267E-08 | 1.650E-08 | 3.900E-08 | 7.100E-08 |
| 19019 | 951 | V951Y | C | 6.7e-08 | 3.2e-08 | 4.5e-08 | 4.800E-08 | 1.769E-08 | 3.200E-08 | 6.700E-08 |
| 19020 | 951 | V951V | C | 4.3e-08 | 3.7e-08 | 1.6e-08 | 3.200E-08 | 1.418E-08 | 1.600E-08 | 4.300E-08 |
| 19021 | 952 | T952A | C | 4.1e-08 | 2.6e-08 | 4e-08 | 3.567E-08 | 8.386E-09 | 2.600E-08 | 4.100E-08 |
| 19022 | 952 | T952R | C | 2.2e-08 | 3.6e-08 | 1.2e-07 | 5.933E-08 | 5.300E-08 | 2.200E-08 | 1.200E-07 |
| 19023 | 952 | T952N | C | 3.6e-08 | 5.8e-08 | 8.9e-08 | 6.100E-08 | 2.663E-08 | 3.600E-08 | 8.900E-08 |
| 19024 | 952 | T952D | C | 5.7e-08 | 2.8e-08 | 7.3e-09 | 3.077E-08 | 2.497E-08 | 7.300E-09 | 5.700E-08 |
| 19025 | 952 | T952C | C | 9.4e-08 | 7.3e-08 | 1.2e-07 | 9.567E-08 | 2.354E-08 | 7.300E-08 | 1.200E-07 |
| 19026 | 952 | T952E | C | 3.5e-08 | 3.7e-08 | 1.7e-08 | 2.967E-08 | 1.102E-08 | 1.700E-08 | 3.700E-08 |
| 19027 | 952 | T952Q | C | 2.2e-08 | 2.2e-08 | 3.2e-08 | 2.533E-08 | 5.774E-09 | 2.200E-08 | 3.200E-08 |
| 19028 | 952 | T952G | C | 8.5e-08 | 6.9e-08 | 6.2e-08 | 7.200E-08 | 1.179E-08 | 6.200E-08 | 8.500E-08 |
| 19029 | 952 | T952H | C | 6.5e-08 | 1.2e-07 | 5.2e-08 | 7.900E-08 | 3.610E-08 | 5.200E-08 | 1.200E-07 |
| 19030 | 952 | T952I | C | 5.5e-08 | 5.6e-08 | 2.1e-08 | 4.400E-08 | 1.992E-08 | 2.100E-08 | 5.600E-08 |
| 19031 | 952 | T952L | C | 9.2e-08 | 5.9e-08 | 6.3e-08 | 7.133E-08 | 1.801E-08 | 5.900E-08 | 9.200E-08 |
| 19032 | 952 | T952K | C | 7.2e-08 | 1.6e-08 | 1.1e-07 | 6.600E-08 | 4.729E-08 | 1.600E-08 | 1.100E-07 |
| 19033 | 952 | T952M | C | 5.6e-08 | 7.2e-08 | 5e-08 | 5.933E-08 | 1.137E-08 | 5.000E-08 | 7.200E-08 |
| 19034 | 952 | T952F | C | 3.8e-08 | 4.8e-08 | 4.3e-08 | 4.300E-08 | 5.000E-09 | 3.800E-08 | 4.800E-08 |
| 19035 | 952 | T952P | C | 3e-08 | 8.6e-08 | 1.6e-07 | 9.200E-08 | 6.521E-08 | 3.000E-08 | 1.600E-07 |
| 19036 | 952 | T952S | C | 6.6e-08 | 3.5e-08 | 7e-08 | 5.700E-08 | 1.916E-08 | 3.500E-08 | 7.000E-08 |
| 19037 | 952 | T952T | C | 4.3e-08 | 1.8e-08 | 1.1e-07 | 5.700E-08 | 4.757E-08 | 1.800E-08 | 1.100E-07 |
| 19038 | 952 | T952W | C | 3.2e-08 | 2.6e-08 | 1.8e-08 | 2.533E-08 | 7.024E-09 | 1.800E-08 | 3.200E-08 |
| 19039 | 952 | T952Y | C | 1.8e-08 | 3.7e-08 | 1.8e-08 | 2.433E-08 | 1.097E-08 | 1.800E-08 | 3.700E-08 |

| | | | | | | | | | | |
|---|---|---|---|---|---|---|---|---|---|---|
| 19040 | 952 | T952V | C | 4e-08 | 1.6e-08 | 2e-08 | 2.533E-08 | 1.286E-08 | 1.600E-08 | 4.000E-08 |
| 19041 | 953 | V953A | C | 2.5e-08 | 2.9e-08 | 2.1e-08 | 2.500E-08 | 4.000E-09 | 2.100E-08 | 2.900E-08 |
| 19042 | 953 | V953R | C | 3.5e-08 | 2.2e-08 | 8.2e-08 | 4.633E-08 | 3.156E-08 | 2.200E-08 | 8.200E-08 |
| 19043 | 953 | V953N | C | 4.2e-08 | 8.5e-08 | 2e-08 | 4.900E-08 | 3.306E-08 | 2.000E-08 | 8.500E-08 |
| 19044 | 953 | V953D | C | 2.8e-08 | 1.9e-08 | 2.1e-08 | 2.267E-08 | 4.726E-09 | 1.900E-08 | 2.800E-08 |
| 19045 | 953 | V953C | C | 2.1e-08 | 1.9e-08 | 3.1e-08 | 2.367E-08 | 6.429E-09 | 1.900E-08 | 3.100E-08 |
| 19046 | 953 | V953E | C | 6.4e-08 | 5.4e-08 | 4.7e-08 | 5.500E-08 | 8.544E-09 | 4.700E-08 | 6.400E-08 |
| 19047 | 953 | V953Q | C | 4e-08 | 1.9e-08 | 3.2e-08 | 3.033E-08 | 1.060E-08 | 1.900E-08 | 4.000E-08 |
| 19048 | 953 | V953G | C | 1.8e-08 | 9.5e-08 | 1.3e-08 | 4.200E-08 | 4.597E-08 | 1.300E-08 | 9.500E-08 |
| 19049 | 953 | V953H | C | 3.3e-08 | 2.5e-08 | 2e-08 | 2.600E-08 | 6.557E-09 | 2.000E-08 | 3.300E-08 |
| 19050 | 953 | V953I | C | 5.6e-08 | 7.7e-08 | 1.8e-08 | 5.033E-08 | 2.991E-08 | 1.800E-08 | 7.700E-08 |
| 19051 | 953 | V953L | C | 2.9e-08 | 1.7e-08 | 6.5e-08 | 3.700E-08 | 2.498E-08 | 1.700E-08 | 6.500E-08 |
| 19052 | 953 | V953K | C | 2e-08 | 1.5e-08 | 2e-08 | 1.833E-08 | 2.887E-09 | 1.500E-08 | 2.000E-08 |
| 19053 | 953 | V953M | C | 7.2e-08 | 1.2e-07 | 8.2e-08 | 9.133E-08 | 2.532E-08 | 7.200E-08 | 1.200E-07 |
| 19054 | 953 | V953F | C | 3.2e-08 | 1.4e-07 | 4.6e-08 | 7.267E-08 | 5.873E-08 | 3.200E-08 | 1.400E-07 |
| 19055 | 953 | V953P | C | 4.4e-08 | 4.5e-08 | 6.1e-08 | 5.000E-08 | 9.539E-09 | 4.400E-08 | 6.100E-08 |
| 19056 | 953 | V953S | C | 4.4e-08 | 7.9e-08 | 7.1e-08 | 6.467E-08 | 1.834E-08 | 4.400E-08 | 7.900E-08 |
| 19057 | 953 | V953T | C | 3.8e-08 | 1.6e-08 | 5.4e-08 | 3.600E-08 | 1.908E-08 | 1.600E-08 | 5.400E-08 |
| 19058 | 953 | V953W | C | 2.9e-08 | 6e-08 | 3.3e-08 | 4.067E-08 | 1.686E-08 | 2.900E-08 | 6.000E-08 |
| 19059 | 953 | V953Y | C | 2.6e-08 | 2e-08 | 4e-08 | 2.867E-08 | 1.026E-08 | 2.000E-08 | 4.000E-08 |
| 19060 | 953 | V953V | C | 2.1e-08 | 2.7e-08 | 4.5e-08 | 3.100E-08 | 1.249E-08 | 2.100E-08 | 4.500E-08 |
| 19061 | 954 | S954A | C | 1.6e-08 | 2.5e-08 | 5.3e-08 | 3.133E-08 | 1.930E-08 | 1.600E-08 | 5.300E-08 |
| 19062 | 954 | S954R | C | 2.6e-08 | 8.4e-08 | 2.6e-08 | 4.533E-08 | 3.349E-08 | 2.600E-08 | 8.400E-08 |
| 19063 | 954 | S954N | C | 3.9e-08 | 3.6e-08 | 4.4e-08 | 3.967E-08 | 4.041E-09 | 3.600E-08 | 4.400E-08 |
| 19064 | 954 | S954D | C | 3.2e-08 | 3.1e-08 | 4.1e-08 | 3.467E-08 | 5.508E-09 | 3.100E-08 | 4.100E-08 |
| 19065 | 954 | S954C | C | 2.8e-08 | 4.1e-08 | 3.4e-08 | 3.433E-08 | 6.506E-09 | 2.800E-08 | 4.100E-08 |
| 19066 | 954 | S954E | C | 1.7e-07 | 3.9e-08 | 1.7e-08 | 7.533E-08 | 8.272E-08 | 1.700E-08 | 1.700E-07 |
| 19067 | 954 | S954Q | C | 2.7e-08 | 4.1e-08 | 9.8e-08 | 5.533E-08 | 3.761E-08 | 2.700E-08 | 9.800E-08 |
| 19068 | 954 | S954G | C | 6.8e-08 | 1.2e-07 | 6.6e-08 | 8.467E-08 | 3.062E-08 | 6.600E-08 | 1.200E-07 |
| 19069 | 954 | S954H | C | 3.9e-08 | 4.8e-08 | 6.8e-08 | 5.167E-08 | 1.484E-08 | 3.900E-08 | 6.800E-08 |
| 19070 | 954 | S954I | C | 1.6e-08 | 2.1e-08 | 3.3e-08 | 2.333E-08 | 8.737E-09 | 1.600E-08 | 3.300E-08 |
| 19071 | 954 | S954L | C | 2.7e-08 | 3.3e-08 | 2.2e-08 | 2.733E-08 | 5.508E-09 | 2.200E-08 | 3.300E-08 |
| 19072 | 954 | S954K | C | 8.3e-08 | 3.4e-08 | 1.7e-08 | 4.467E-08 | 3.427E-08 | 1.700E-08 | 8.300E-08 |
| 19073 | 954 | S954M | C | 1.6e-07 | 8.9e-08 | 2e-08 | 8.967E-08 | 7.000E-08 | 2.000E-08 | 1.600E-07 |
| 19074 | 954 | S954F | C | 4.2e-08 | 6.1e-08 | 4.7e-08 | 5.000E-08 | 9.849E-09 | 4.200E-08 | 6.100E-08 |
| 19075 | 954 | S954P | C | 4.5e-08 | 4e-08 | 7.1e-08 | 5.200E-08 | 1.664E-08 | 4.000E-08 | 7.100E-08 |
| 19076 | 954 | S954S | C | 5.5e-08 | 2.6e-08 | 3.9e-08 | 4.000E-08 | 1.453E-08 | 2.600E-08 | 5.500E-08 |
| 19077 | 954 | S954T | C | 4.9e-08 | 4.8e-08 | 5.2e-08 | 4.967E-08 | 2.082E-09 | 4.800E-08 | 5.200E-08 |
| 19078 | 954 | S954W | C | 4.7e-08 | 3.6e-08 | 5.2e-08 | 4.500E-08 | 8.185E-09 | 3.600E-08 | 5.200E-08 |
| 19079 | 954 | S954Y | C | 2.8e-08 | 6.4e-08 | 5.3e-08 | 4.833E-08 | 1.845E-08 | 2.800E-08 | 6.400E-08 |
| 19080 | 954 | S954V | C | 9.6e-08 | 4.6e-08 | 4.7e-08 | 6.300E-08 | 2.858E-08 | 4.600E-08 | 9.600E-08 |
| 19081 | 955 | S955A | C | 2.1e-08 | 1.1e-07 | 3.8e-08 | 5.633E-08 | 4.725E-08 | 2.100E-08 | 1.100E-07 |
| 19082 | 955 | S955R | C | 2.6e-08 | 1e-07 | 8.6e-08 | 7.067E-08 | 3.931E-08 | 2.600E-08 | 1.000E-07 |
| 19083 | 955 | S955N | C | 2.1e-08 | 5.9e-08 | 6.1e-08 | 4.700E-08 | 2.254E-08 | 2.100E-08 | 6.100E-08 |
| 19084 | 955 | S955D | C | 4.1e-08 | 6.4e-08 | 9.6e-09 | 3.820E-08 | 2.731E-08 | 9.600E-09 | 6.400E-08 |
| 19085 | 955 | S955C | C | 1.4e-08 | 1.4e-08 | 6.6e-08 | 3.133E-08 | 3.002E-08 | 1.400E-08 | 6.600E-08 |
| 19086 | 955 | S955E | C | 3.8e-08 | 4.5e-08 | 5.9e-08 | 4.733E-08 | 1.069E-08 | 3.800E-08 | 5.900E-08 |
| 19087 | 955 | S955Q | C | 8.5e-08 | 6.6e-09 | 1.3e-08 | 3.487E-08 | 4.353E-08 | 6.600E-09 | 8.500E-08 |
| 19088 | 955 | S955G | C | 3.2e-08 | 3.1e-08 | 4.4e-08 | 3.567E-08 | 7.234E-09 | 3.100E-08 | 4.400E-08 |
| 19089 | 955 | S955H | C | 1.6e-08 | 8.5e-08 | 5.2e-08 | 5.100E-08 | 3.451E-08 | 1.600E-08 | 8.500E-08 |
| 19090 | 955 | S955I | C | 2.6e-08 | 6.7e-08 | 7.1e-08 | 5.467E-08 | 2.491E-08 | 2.600E-08 | 7.100E-08 |
| 19091 | 955 | S955L | C | 2.6e-08 | 2.3e-08 | 1.1e-07 | 5.300E-08 | 4.939E-08 | 2.300E-08 | 1.100E-07 |
| 19092 | 955 | S955K | C | 3.7e-08 | 6.6e-08 | 5.9e-08 | 5.400E-08 | 1.513E-08 | 3.700E-08 | 6.600E-08 |
| 19093 | 955 | S955M | C | 2.1e-08 | 7e-08 | 3.7e-08 | 4.267E-08 | 2.499E-08 | 2.100E-08 | 7.000E-08 |
| 19094 | 955 | S955F | C | 7.3e-08 | 2.4e-08 | 7.3e-08 | 5.667E-08 | 2.829E-08 | 2.400E-08 | 7.300E-08 |
| 19095 | 955 | S955P | C | 2.9e-08 | 3.7e-08 | 1.2e-07 | 6.200E-08 | 5.039E-08 | 2.900E-08 | 1.200E-07 |
| 19096 | 955 | S955S | C | 9.1e-08 | 6.5e-08 | 2.4e-08 | 6.000E-08 | 3.378E-08 | 2.400E-08 | 9.100E-08 |
| 19097 | 955 | S955T | C | 3e-08 | 2.5e-08 | 2.8e-08 | 2.767E-08 | 2.517E-09 | 2.500E-08 | 3.000E-08 |
| 19098 | 955 | S955W | C | 5.5e-08 | 8.5e-08 | 2.3e-08 | 5.433E-08 | 3.101E-08 | 2.300E-08 | 8.500E-08 |
| 19099 | 955 | S955Y | C | 4.8e-08 | 4.4e-08 | 3.3e-08 | 4.167E-08 | 7.767E-09 | 3.300E-08 | 4.800E-08 |
| 19100 | 955 | S955V | C | 1.3e-08 | 7.5e-08 | 5.1e-08 | 4.633E-08 | 3.126E-08 | 1.300E-08 | 7.500E-08 |
| 19101 | 956 | A956A | C | 7.1e-08 | 3.5e-08 | 8.5e-08 | 6.367E-08 | 2.579E-08 | 3.500E-08 | 8.500E-08 |
| 19102 | 956 | A956R | C | 4.6e-08 | 2.3e-08 | 8.6e-08 | 5.167E-08 | 3.188E-08 | 2.300E-08 | 8.600E-08 |
| 19103 | 956 | A956N | C | 2.4e-08 | 4.5e-08 | 1.6e-08 | 2.833E-08 | 1.498E-08 | 1.600E-08 | 4.500E-08 |
| 19104 | 956 | A956D | C | 1.2e-08 | 5.5e-08 | 2.5e-08 | 3.067E-08 | 2.205E-08 | 1.200E-08 | 5.500E-08 |
| 19105 | 956 | A956C | C | 2.3e-08 | 4.5e-08 | 4.4e-08 | 3.733E-08 | 1.242E-08 | 2.300E-08 | 4.500E-08 |
| 19106 | 956 | A956E | C | 1.3e-08 | 3.4e-08 | 7.4e-08 | 4.033E-08 | 3.099E-08 | 1.300E-08 | 7.400E-08 |
| 19107 | 956 | A956Q | C | 5.8e-08 | 3.8e-08 | 3.1e-08 | 4.233E-08 | 1.401E-08 | 3.100E-08 | 5.800E-08 |
| 19108 | 956 | A956G | C | 2.8e-08 | 1.2e-07 | 3.9e-08 | 6.233E-08 | 5.024E-08 | 2.800E-08 | 1.200E-07 |
| 19109 | 956 | A956H | C | 3.9e-08 | 6.2e-08 | 4.8e-08 | 4.967E-08 | 1.159E-08 | 3.900E-08 | 6.200E-08 |
| 19110 | 956 | A956I | C | 3.1e-08 | 3.4e-08 | 5.4e-08 | 3.967E-08 | 1.250E-08 | 3.100E-08 | 5.400E-08 |
| 19111 | 956 | A956L | C | 2.1e-08 | 2.2e-08 | 5.6e-08 | 3.300E-08 | 1.992E-08 | 2.100E-08 | 5.600E-08 |
| 19112 | 956 | A956K | C | 6.4e-08 | 2.2e-08 | 9.3e-08 | 5.967E-08 | 3.570E-08 | 2.200E-08 | 9.300E-08 |
| 19113 | 956 | A956M | C | 9.9e-08 | 1.7e-07 | 2.8e-08 | 9.900E-08 | 7.100E-08 | 2.800E-08 | 1.700E-07 |
| 19114 | 956 | A956F | C | 8.6e-08 | 5.7e-08 | 2.3e-08 | 5.533E-08 | 3.153E-08 | 2.300E-08 | 8.600E-08 |
| 19115 | 956 | A956P | C | 3.1e-08 | 9.8e-09 | 5.7e-08 | 3.260E-08 | 2.364E-08 | 9.800E-09 | 5.700E-08 |
| 19116 | 956 | A956S | C | 4e-08 | 4.1e-08 | 2.5e-08 | 3.533E-08 | 8.963E-09 | 2.500E-08 | 4.100E-08 |
| 19117 | 956 | A956T | C | 2.6e-08 | 6.1e-08 | 4e-08 | 4.233E-08 | 1.762E-08 | 2.600E-08 | 6.100E-08 |
| 19118 | 956 | A956W | C | 6.8e-09 | 1.4e-08 | 3.3e-08 | 1.793E-08 | 1.354E-08 | 6.800E-09 | 3.300E-08 |
| 19119 | 956 | A956Y | C | 3.5e-08 | 3.9e-08 | 5.4e-08 | 4.267E-08 | 1.002E-08 | 3.500E-08 | 5.400E-08 |
| 19120 | 956 | A956V | C | 4.1e-08 | 3.8e-08 | 1.7e-08 | 3.200E-08 | 1.308E-08 | 1.700E-08 | 4.100E-08 |
| 19121 | 957 | S957A | C | 5.7e-08 | 3e-08 | 6.6e-08 | 5.100E-08 | 1.873E-08 | 3.000E-08 | 6.600E-08 |
| 19122 | 957 | S957R | C | 3.6e-08 | 1.2e-07 | 3.7e-08 | 6.433E-08 | 4.821E-08 | 3.600E-08 | 1.200E-07 |
| 19123 | 957 | S957N | C | 1.7e-08 | 7.6e-08 | 1.3e-08 | 3.533E-08 | 3.528E-08 | 1.300E-08 | 7.600E-08 |
| 19124 | 957 | S957D | C | 5.1e-08 | 2.3e-08 | 1.8e-08 | 3.067E-08 | 1.779E-08 | 1.800E-08 | 5.100E-08 |

| | | | | | | | | | | |
|---|---|---|---|---|---|---|---|---|---|---|
| 19125 | 957 | S957C | C | 6e-08 | 5.8e-08 | 3.3e-08 | 5.033E-08 | 1.504E-08 | 3.300E-08 | 6.000E-08 |
| 19126 | 957 | S957E | C | 6.2e-08 | 4.9e-08 | 4.2e-08 | 5.100E-08 | 1.015E-08 | 4.200E-08 | 6.200E-08 |
| 19127 | 957 | S957Q | C | 6.9e-08 | 4.5e-08 | 3.3e-08 | 4.900E-08 | 1.833E-08 | 3.300E-08 | 6.900E-08 |
| 19128 | 957 | S957G | C | 1.1e-08 | 1.8e-08 | 3.8e-08 | 2.233E-08 | 1.401E-08 | 1.100E-08 | 3.800E-08 |
| 19129 | 957 | S957H | C | 4.5e-08 | 1e-08 | 3.5e-08 | 3.000E-08 | 1.803E-08 | 1.000E-08 | 4.500E-08 |
| 19130 | 957 | S957I | C | 3.5e-08 | 7.7e-08 | 5.3e-08 | 5.500E-08 | 2.107E-08 | 3.500E-08 | 7.700E-08 |
| 19131 | 957 | S957L | C | 4.9e-08 | 8.2e-08 | 5.3e-08 | 6.133E-08 | 1.801E-08 | 4.900E-08 | 8.200E-08 |
| 19132 | 957 | S957K | C | 2.5e-08 | 9.4e-08 | 1.2e-08 | 4.367E-08 | 4.407E-08 | 1.200E-08 | 9.400E-08 |
| 19133 | 957 | S957M | C | 4e-08 | 1.7e-08 | 2.6e-08 | 2.767E-08 | 1.159E-08 | 1.700E-08 | 4.000E-08 |
| 19134 | 957 | S957F | C | 1.7e-07 | 6.9e-08 | 1.8e-08 | 8.567E-08 | 7.736E-08 | 1.800E-08 | 1.700E-07 |
| 19135 | 957 | S957P | C | 4.2e-08 | 4.6e-08 | 3.1e-08 | 3.967E-08 | 7.767E-09 | 3.100E-08 | 4.600E-08 |
| 19136 | 957 | S957S | C | 4.8e-08 | 3.1e-08 | 4.8e-08 | 4.233E-08 | 9.815E-09 | 3.100E-08 | 4.800E-08 |
| 19137 | 957 | S957T | C | 4.8e-08 | 3.2e-08 | 1.2e-08 | 3.067E-08 | 1.804E-08 | 1.200E-08 | 4.800E-08 |
| 19138 | 957 | S957W | C | 7.7e-08 | 6.6e-08 | 3e-08 | 5.767E-08 | 2.458E-08 | 3.000E-08 | 7.700E-08 |
| 19139 | 957 | S957Y | C | 3.5e-08 | 4.2e-08 | 4.1e-08 | 3.933E-08 | 3.786E-09 | 3.500E-08 | 4.200E-08 |
| 19140 | 957 | S957V | C | 2.6e-08 | 5.4e-08 | 1.1e-07 | 6.333E-08 | 4.277E-08 | 2.600E-08 | 1.100E-07 |
| 19141 | 958 | T958A | C | 2.6e-08 | 1.1e-07 | 1.4e-08 | 5.000E-08 | 5.231E-08 | 1.400E-08 | 1.100E-07 |
| 19142 | 958 | T958R | C | 1.4e-07 | 7.6e-08 | 1.1e-07 | 1.087E-07 | 3.202E-08 | 7.600E-08 | 1.400E-07 |
| 19143 | 958 | T958N | C | 1.1e-07 | 6.5e-08 | 5.6e-08 | 7.700E-08 | 2.893E-08 | 5.600E-08 | 1.100E-07 |
| 19144 | 958 | T958D | C | 4.9e-08 | 8.4e-08 | 6.2e-08 | 6.500E-08 | 1.769E-08 | 4.900E-08 | 8.400E-08 |
| 19145 | 958 | T958C | C | 7e-08 | 3e-08 | 3.5e-08 | 4.500E-08 | 2.179E-08 | 3.000E-08 | 7.000E-08 |
| 19146 | 958 | T958E | C | 7.6e-08 | 3.1e-08 | 1.4e-08 | 4.033E-08 | 3.204E-08 | 1.400E-08 | 7.600E-08 |
| 19147 | 958 | T958Q | C | 4.1e-08 | 7.7e-08 | 4.5e-08 | 5.433E-08 | 1.973E-08 | 4.100E-08 | 7.700E-08 |
| 19148 | 958 | T958G | C | 6.9e-08 | 9.9e-08 | 1.4e-08 | 6.067E-08 | 4.311E-08 | 1.400E-08 | 9.900E-08 |
| 19149 | 958 | T958H | C | 4.5e-08 | 4.7e-08 | 3.4e-08 | 4.200E-08 | 7.000E-09 | 3.400E-08 | 4.700E-08 |
| 19150 | 958 | T958I | C | 7.7e-08 | 5.6e-08 | 6.4e-08 | 6.567E-08 | 1.060E-08 | 5.600E-08 | 7.700E-08 |
| 19151 | 958 | T958L | C | 3.6e-08 | 1.9e-08 | 4.3e-08 | 3.267E-08 | 1.234E-08 | 1.900E-08 | 4.300E-08 |
| 19152 | 958 | T958K | C | 1.1e-07 | 4.4e-08 | 8e-08 | 7.800E-08 | 3.305E-08 | 4.400E-08 | 1.100E-07 |
| 19153 | 958 | T958M | C | 6.1e-08 | 2.1e-08 | 1.9e-08 | 3.367E-08 | 2.369E-08 | 1.900E-08 | 6.100E-08 |
| 19154 | 958 | T958F | C | 3.6e-08 | 5.8e-08 | 1.2e-07 | 7.133E-08 | 4.356E-08 | 3.600E-08 | 1.200E-07 |
| 19155 | 958 | T958P | C | 7.6e-08 | 4.1e-08 | 6.1e-08 | 5.933E-08 | 1.756E-08 | 4.100E-08 | 7.600E-08 |
| 19156 | 958 | T958S | C | 2.1e-08 | 2.7e-08 | 2.5e-08 | 2.433E-08 | 3.055E-09 | 2.100E-08 | 2.700E-08 |
| 19157 | 958 | T958T | C | 1.2e-08 | 5.3e-08 | 1.5e-08 | 2.667E-08 | 2.285E-08 | 1.200E-08 | 5.300E-08 |
| 19158 | 958 | T958W | C | 2.8e-08 | 1e-07 | 3.5e-08 | 5.433E-08 | 3.970E-08 | 2.800E-08 | 1.000E-07 |
| 19159 | 958 | T958Y | C | 3.2e-08 | 4.9e-08 | 5e-08 | 4.367E-08 | 1.012E-08 | 3.200E-08 | 5.000E-08 |
| 19160 | 958 | T958V | C | 2.5e-08 | 3.7e-08 | 3.7e-08 | 3.300E-08 | 6.928E-09 | 2.500E-08 | 3.700E-08 |
| 19161 | 959 | K959A | C | 2e-08 | 7.3e-08 | 5.5e-08 | 4.933E-08 | 2.695E-08 | 2.000E-08 | 7.300E-08 |
| 19162 | 959 | K959R | C | 5.9e-08 | 2.7e-08 | 2e-08 | 3.533E-08 | 2.079E-08 | 2.000E-08 | 5.900E-08 |
| 19163 | 959 | K959N | C | 1.8e-08 | 3.3e-08 | 4e-08 | 3.033E-08 | 1.124E-08 | 1.800E-08 | 4.000E-08 |
| 19164 | 959 | K959D | C | 1.2e-07 | 3.2e-08 | 9.2e-08 | 8.133E-08 | 4.496E-08 | 3.200E-08 | 1.200E-07 |
| 19165 | 959 | K959C | C | 5.4e-08 | 4.3e-08 | 6.2e-08 | 5.300E-08 | 9.539E-09 | 4.300E-08 | 6.200E-08 |
| 19166 | 959 | K959E | C | 3e-08 | 5.4e-08 | 2.6e-08 | 3.667E-08 | 1.514E-08 | 2.600E-08 | 5.400E-08 |
| 19167 | 959 | K959Q | C | 8.4e-09 | 3.2e-08 | 2.9e-08 | 2.313E-08 | 1.285E-08 | 8.400E-09 | 3.200E-08 |
| 19168 | 959 | K959G | C | 4e-08 | 3.5e-08 | 3.1e-08 | 3.533E-08 | 4.509E-09 | 3.100E-08 | 4.000E-08 |
| 19169 | 959 | K959H | C | 2.7e-08 | 5e-08 | 2.8e-08 | 3.500E-08 | 1.300E-08 | 2.700E-08 | 5.000E-08 |
| 19170 | 959 | K959I | C | 6e-08 | 2.2e-08 | 3.7e-08 | 3.967E-08 | 1.914E-08 | 2.200E-08 | 6.000E-08 |
| 19171 | 959 | K959L | C | 3e-08 | 4.1e-08 | 7.6e-08 | 4.900E-08 | 2.402E-08 | 3.000E-08 | 7.600E-08 |
| 19172 | 959 | K959K | C | 1.5e-08 | 5.6e-08 | 5.6e-08 | 4.233E-08 | 2.367E-08 | 1.500E-08 | 5.600E-08 |
| 19173 | 959 | K959M | C | 2.8e-08 | 2.5e-08 | 3e-08 | 2.767E-08 | 2.517E-09 | 2.500E-08 | 3.000E-08 |
| 19174 | 959 | K959F | C | 2.5e-07 | 1.9e-08 | 5.1e-08 | 1.067E-07 | 1.252E-07 | 1.900E-08 | 2.500E-07 |
| 19175 | 959 | K959P | C | 1.9e-08 | 3.1e-08 | 2.4e-08 | 2.467E-08 | 6.028E-09 | 1.900E-08 | 3.100E-08 |
| 19176 | 959 | K959S | C | 4.5e-08 | 6.9e-08 | 3e-08 | 4.800E-08 | 1.967E-08 | 3.000E-08 | 6.900E-08 |
| 19177 | 959 | K959T | C | 7.2e-08 | 4.4e-08 | 3.3e-08 | 4.967E-08 | 2.011E-08 | 3.300E-08 | 7.200E-08 |
| 19178 | 959 | K959W | C | 5.6e-08 | 4.1e-08 | 3.5e-08 | 4.400E-08 | 1.082E-08 | 3.500E-08 | 5.600E-08 |
| 19179 | 959 | K959Y | C | 6.1e-08 | 2.9e-08 | 3.3e-08 | 4.100E-08 | 1.744E-08 | 2.900E-08 | 6.100E-08 |
| 19180 | 959 | K959V | C | 2e-08 | 9.9e-09 | 6.9e-08 | 3.297E-08 | 3.161E-08 | 9.900E-09 | 6.900E-08 |
| 19181 | 960 | G960A | C | 4.1e-08 | 4.5e-08 | 6.4e-08 | 5.000E-08 | 1.229E-08 | 4.100E-08 | 6.400E-08 |
| 19182 | 960 | G960R | C | 3.5e-08 | 1.7e-08 | 5e-08 | 3.400E-08 | 1.652E-08 | 1.700E-08 | 5.000E-08 |
| 19183 | 960 | G960N | C | 2.8e-08 | 3.3e-08 | 4.8e-08 | 3.633E-08 | 1.041E-08 | 2.800E-08 | 4.800E-08 |
| 19184 | 960 | G960D | C | 8.4e-08 | 4.8e-08 | 3.9e-08 | 5.700E-08 | 2.381E-08 | 3.900E-08 | 8.400E-08 |
| 19185 | 960 | G960C | C | 4.7e-08 | 1.8e-08 | 2.6e-08 | 3.033E-08 | 1.498E-08 | 1.800E-08 | 4.700E-08 |
| 19186 | 960 | G960E | C | 1.5e-08 | 1.7e-08 | 7.8e-08 | 3.667E-08 | 3.581E-08 | 1.500E-08 | 7.800E-08 |
| 19187 | 960 | G960Q | C | 3.9e-08 | 3.3e-08 | 3.5e-08 | 3.567E-08 | 3.055E-09 | 3.300E-08 | 3.900E-08 |
| 19188 | 960 | G960G | C | 6.3e-08 | 4.4e-08 | 3.4e-08 | 4.700E-08 | 1.473E-08 | 3.400E-08 | 6.300E-08 |
| 19189 | 960 | G960H | C | 1.3e-08 | 2.7e-08 | 8.3e-08 | 4.100E-08 | 3.704E-08 | 1.300E-08 | 8.300E-08 |
| 19190 | 960 | G960I | C | 2.7e-08 | 1.7e-08 | 7.3e-08 | 3.900E-08 | 2.987E-08 | 1.700E-08 | 7.300E-08 |
| 19191 | 960 | G960L | C | 1.1e-08 | 2.5e-08 | 3.6e-08 | 2.400E-08 | 1.253E-08 | 1.100E-08 | 3.600E-08 |
| 19192 | 960 | G960K | C | 6.1e-08 | 3.1e-08 | 1.8e-08 | 3.667E-08 | 2.205E-08 | 1.800E-08 | 6.100E-08 |
| 19193 | 960 | G960M | C | 4.8e-08 | 3.8e-08 | 2.3e-08 | 3.633E-08 | 1.258E-08 | 2.300E-08 | 4.800E-08 |
| 19194 | 960 | G960F | C | 2.1e-08 | 1.9e-08 | 5.7e-08 | 3.233E-08 | 2.139E-08 | 1.900E-08 | 5.700E-08 |
| 19195 | 960 | G960P | C | 1.7e-08 | 3.4e-08 | 3.2e-08 | 2.767E-08 | 9.292E-09 | 1.700E-08 | 3.400E-08 |
| 19196 | 960 | G960S | C | 4.4e-08 | 4.8e-08 | 1.9e-08 | 3.700E-08 | 1.572E-08 | 1.900E-08 | 4.800E-08 |
| 19197 | 960 | G960T | C | 5.9e-08 | 2.8e-08 | 5e-08 | 4.567E-08 | 1.595E-08 | 2.800E-08 | 5.900E-08 |
| 19198 | 960 | G960W | C | 5.3e-08 | 5.7e-08 | 2e-08 | 4.333E-08 | 2.031E-08 | 2.000E-08 | 5.700E-08 |
| 19199 | 960 | G960Y | C | 7e-08 | 3.8e-08 | 2.3e-08 | 4.367E-08 | 2.401E-08 | 2.300E-08 | 7.000E-08 |
| 19200 | 960 | G960V | C | 2.9e-08 | 1.7e-08 | 7e-08 | 3.867E-08 | 2.779E-08 | 1.700E-08 | 7.000E-08 |
| 19201 | 961 | P961A | C | 3.9e-08 | 1.1e-07 | 4.8e-08 | 6.567E-08 | 3.866E-08 | 3.900E-08 | 1.100E-07 |
| 19202 | 961 | P961R | C | 6.4e-08 | 1e-07 | 4.9e-08 | 7.100E-08 | 2.621E-08 | 4.900E-08 | 1.000E-07 |
| 19203 | 961 | P961N | C | 4.4e-08 | 4e-08 | 4.5e-08 | 4.300E-08 | 2.646E-09 | 4.000E-08 | 4.500E-08 |
| 19204 | 961 | P961D | C | 7.4e-08 | 1.1e-07 | 2.7e-08 | 7.033E-08 | 4.162E-08 | 2.700E-08 | 1.100E-07 |
| 19205 | 961 | P961C | C | 1.5e-08 | 2.4e-08 | 6.6e-08 | 3.500E-08 | 2.722E-08 | 1.500E-08 | 6.600E-08 |
| 19206 | 961 | P961E | C | 4.2e-08 | 3.4e-08 | 3e-08 | 3.533E-08 | 6.110E-09 | 3.000E-08 | 4.200E-08 |
| 19207 | 961 | P961Q | C | 3.9e-08 | 7.3e-08 | 3.5e-08 | 4.900E-08 | 2.088E-08 | 3.500E-08 | 7.300E-08 |
| 19208 | 961 | P961G | C | 1.4e-08 | 3.9e-08 | 4.8e-08 | 3.367E-08 | 1.762E-08 | 1.400E-08 | 4.800E-08 |
| 19209 | 961 | P961H | C | 7.9e-08 | 9.7e-08 | 6.6e-08 | 8.067E-08 | 1.557E-08 | 6.600E-08 | 9.700E-08 |

| | | | | | | | | | | |
|---|---|---|---|---|---|---|---|---|---|---|
| 19210 | 961 | P961I | C | 5.5e-08 | 2.6e-08 | 3.9e-08 | 4.000E-08 | 1.453E-08 | 2.600E-08 | 5.500E-08 |
| 19211 | 961 | P961L | C | 3.6e-08 | 1.5e-08 | 4.7e-08 | 3.267E-08 | 1.626E-08 | 1.500E-08 | 4.700E-08 |
| 19212 | 961 | P961K | C | 1e-07 | 3.2e-08 | 5.5e-08 | 6.233E-08 | 3.459E-08 | 3.200E-08 | 1.000E-07 |
| 19213 | 961 | P961M | C | 3.4e-08 | 6.4e-08 | 7.2e-08 | 5.667E-08 | 2.003E-08 | 3.400E-08 | 7.200E-08 |
| 19214 | 961 | P961F | C | 3.3e-08 | 1e-07 | 1.8e-08 | 5.033E-08 | 4.366E-08 | 1.800E-08 | 1.000E-07 |
| 19215 | 961 | P961P | C | 4.7e-08 | 1.9e-08 | 3.8e-08 | 3.467E-08 | 1.429E-08 | 1.900E-08 | 4.700E-08 |
| 19216 | 961 | P961S | C | 3.5e-08 | 4.2e-08 | 3e-08 | 3.567E-08 | 6.028E-09 | 3.000E-08 | 4.200E-08 |
| 19217 | 961 | P961T | C | 4.3e-08 | 5.3e-08 | 3e-08 | 4.200E-08 | 1.153E-08 | 3.000E-08 | 5.300E-08 |
| 19218 | 961 | P961W | C | 7e-08 | 4.1e-08 | 6e-08 | 5.700E-08 | 1.473E-08 | 4.100E-08 | 7.000E-08 |
| 19219 | 961 | P961Y | C | 4.7e-08 | 3.1e-08 | 5.2e-08 | 4.333E-08 | 1.097E-08 | 3.100E-08 | 5.200E-08 |
| 19220 | 961 | P961V | C | 2.7e-08 | 1.1e-07 | 1.7e-08 | 5.133E-08 | 5.105E-08 | 1.700E-08 | 1.100E-07 |
| 19221 | 962 | S962A | C | 4.5e-09 | 4.8e-08 | 1.1e-07 | 5.417E-08 | 5.302E-08 | 4.500E-09 | 1.100E-07 |
| 19222 | 962 | S962R | C | 2.6e-08 | 3.1e-08 | 3.8e-08 | 3.167E-08 | 6.028E-09 | 2.600E-08 | 3.800E-08 |
| 19223 | 962 | S962N | C | 5.5e-08 | 3.6e-08 | 5.4e-08 | 4.833E-08 | 1.069E-08 | 3.600E-08 | 5.500E-08 |
| 19224 | 962 | S962D | C | 4.4e-08 | 2.2e-08 | 1.4e-08 | 2.667E-08 | 1.553E-08 | 1.400E-08 | 4.400E-08 |
| 19225 | 962 | S962C | C | 4.7e-08 | 1.1e-07 | 2.7e-08 | 6.133E-08 | 4.332E-08 | 2.700E-08 | 1.100E-07 |
| 19226 | 962 | S962E | C | 1.1e-08 | 6.4e-08 | 1e-07 | 5.833E-08 | 4.477E-08 | 1.100E-08 | 1.000E-07 |
| 19227 | 962 | S962Q | C | 6.7e-08 | 2.6e-08 | 1.6e-07 | 8.433E-08 | 6.866E-08 | 2.600E-08 | 1.600E-07 |
| 19228 | 962 | S962G | C | 1.1e-07 | 1.1e-08 | 5.7e-08 | 5.933E-08 | 4.954E-08 | 1.100E-08 | 1.100E-07 |
| 19229 | 962 | S962H | C | 7.9e-08 | 1.5e-07 | 6.1e-08 | 9.667E-08 | 4.706E-08 | 6.100E-08 | 1.500E-07 |
| 19230 | 962 | S962I | C | 8.8e-08 | 3.5e-08 | 4.6e-08 | 5.633E-08 | 2.797E-08 | 3.500E-08 | 8.800E-08 |
| 19231 | 962 | S962L | C | 7.9e-08 | 1.7e-08 | 4.5e-08 | 4.700E-08 | 3.105E-08 | 1.700E-08 | 7.900E-08 |
| 19232 | 962 | S962K | C | 1.8e-08 | 4.1e-08 | 4.1e-08 | 3.333E-08 | 1.328E-08 | 1.800E-08 | 4.100E-08 |
| 19233 | 962 | S962M | C | 6.1e-08 | 1.7e-08 | 6.2e-08 | 4.667E-08 | 2.570E-08 | 1.700E-08 | 6.200E-08 |
| 19234 | 962 | S962F | C | 6.8e-08 | 3.8e-08 | 2.8e-08 | 4.467E-08 | 2.082E-08 | 2.800E-08 | 6.800E-08 |
| 19235 | 962 | S962P | C | 6.4e-08 | 2.9e-08 | 4.2e-08 | 4.500E-08 | 1.769E-08 | 2.900E-08 | 6.400E-08 |
| 19236 | 962 | S962S | C | 9.1e-08 | 4.1e-08 | 7.7e-08 | 6.967E-08 | 2.579E-08 | 4.100E-08 | 9.100E-08 |
| 19237 | 962 | S962T | C | 3.3e-08 | 7.6e-08 | 3.8e-08 | 4.900E-08 | 2.352E-08 | 3.300E-08 | 7.600E-08 |
| 19238 | 962 | S962W | C | 1.3e-07 | 3.8e-08 | 8.7e-08 | 8.500E-08 | 4.603E-08 | 3.800E-08 | 1.300E-07 |
| 19239 | 962 | S962Y | C | 5.2e-08 | 4.1e-08 | 3.6e-08 | 4.300E-08 | 8.185E-09 | 3.600E-08 | 5.200E-08 |
| 19240 | 962 | S962V | C | 7.3e-08 | 1.7e-08 | 2.9e-08 | 3.967E-08 | 2.948E-08 | 1.700E-08 | 7.300E-08 |
| 19241 | 963 | V963A | C | 9.4e-08 | 4.7e-08 | 3e-08 | 5.700E-08 | 3.315E-08 | 3.000E-08 | 9.400E-08 |
| 19242 | 963 | V963R | C | 3.2e-08 | 1.1e-07 | 8.8e-08 | 7.667E-08 | 4.022E-08 | 3.200E-08 | 1.100E-07 |
| 19243 | 963 | V963N | C | 6.8e-08 | 4.5e-08 | 6.2e-08 | 5.833E-08 | 1.193E-08 | 4.500E-08 | 6.800E-08 |
| 19244 | 963 | V963D | C | 5.3e-08 | 5.6e-08 | 6.3e-08 | 5.733E-08 | 5.132E-09 | 5.300E-08 | 6.300E-08 |
| 19245 | 963 | V963C | C | 5.8e-08 | 1.1e-07 | 5.1e-08 | 7.300E-08 | 3.223E-08 | 5.100E-08 | 1.100E-07 |
| 19246 | 963 | V963E | C | 2.7e-08 | 9e-08 | 9.1e-09 | 4.203E-08 | 4.249E-08 | 9.100E-09 | 9.000E-08 |
| 19247 | 963 | V963Q | C | 1.2e-07 | 5.9e-08 | 4.1e-08 | 7.333E-08 | 4.140E-08 | 4.100E-08 | 1.200E-07 |
| 19248 | 963 | V963G | C | 5.3e-08 | 4.3e-08 | 2.1e-08 | 3.900E-08 | 1.637E-08 | 2.100E-08 | 5.300E-08 |
| 19249 | 963 | V963H | C | 7.7e-08 | 7.1e-08 | 1e-07 | 8.267E-08 | 1.531E-08 | 7.100E-08 | 1.000E-07 |
| 19250 | 963 | V963I | C | 5.8e-09 | 5.5e-08 | 2e-08 | 2.693E-08 | 2.532E-08 | 5.800E-09 | 5.500E-08 |
| 19251 | 963 | V963L | C | 6.3e-08 | 1.6e-08 | 8.9e-08 | 5.600E-08 | 3.700E-08 | 1.600E-08 | 8.900E-08 |
| 19252 | 963 | V963K | C | 3e-08 | 6.2e-08 | 1.8e-08 | 3.667E-08 | 2.274E-08 | 1.800E-08 | 6.200E-08 |
| 19253 | 963 | V963M | C | 8.4e-08 | 3.1e-08 | 2.5e-08 | 4.667E-08 | 3.247E-08 | 2.500E-08 | 8.400E-08 |
| 19254 | 963 | V963F | C | 2.7e-08 | 2.1e-08 | 1.1e-07 | 5.267E-08 | 4.974E-08 | 2.100E-08 | 1.100E-07 |
| 19255 | 963 | V963P | C | 6.1e-08 | 2e-08 | 8e-08 | 5.367E-08 | 3.066E-08 | 2.000E-08 | 8.000E-08 |
| 19256 | 963 | V963S | C | 6.9e-08 | 9.6e-08 | 1.8e-08 | 6.100E-08 | 3.961E-08 | 1.800E-08 | 9.600E-08 |
| 19257 | 963 | V963T | C | 5.9e-08 | 7.5e-08 | 5.2e-08 | 6.200E-08 | 1.179E-08 | 5.200E-08 | 7.500E-08 |
| 19258 | 963 | V963W | C | 1.5e-07 | 2.7e-08 | 2e-08 | 6.567E-08 | 7.312E-08 | 2.000E-08 | 1.500E-07 |
| 19259 | 963 | V963Y | C | 2.8e-08 | 5.4e-08 | 5.4e-08 | 4.533E-08 | 1.501E-08 | 2.800E-08 | 5.400E-08 |
| 19260 | 963 | V963V | C | 1.7e-08 | 3.8e-08 | 3.1e-08 | 2.867E-08 | 1.069E-08 | 1.700E-08 | 3.800E-08 |
| 19261 | 964 | F964A | C | 3.1e-08 | 3.5e-08 | 5.2e-08 | 3.933E-08 | 1.115E-08 | 3.100E-08 | 5.200E-08 |
| 19262 | 964 | F964R | C | 1.5e-08 | 6.5e-08 | 6.8e-08 | 4.933E-08 | 2.977E-08 | 1.500E-08 | 6.800E-08 |
| 19263 | 964 | F964N | C | 4.4e-08 | 4.1e-08 | 1.4e-08 | 3.300E-08 | 1.652E-08 | 1.400E-08 | 4.400E-08 |
| 19264 | 964 | F964D | C | 4e-08 | 6.7e-08 | 5e-08 | 5.233E-08 | 1.365E-08 | 4.000E-08 | 6.700E-08 |
| 19265 | 964 | F964C | C | 5.9e-08 | 2.9e-08 | 4.9e-08 | 4.567E-08 | 1.528E-08 | 2.900E-08 | 5.900E-08 |
| 19266 | 964 | F964E | C | 7.2e-08 | 2.2e-08 | 6.7e-08 | 5.367E-08 | 2.754E-08 | 2.200E-08 | 7.200E-08 |
| 19267 | 964 | F964Q | C | 4.8e-08 | 7.6e-08 | 7.2e-08 | 6.533E-08 | 1.514E-08 | 4.800E-08 | 7.600E-08 |
| 19268 | 964 | F964G | C | 3.4e-08 | 2.1e-08 | 1.6e-08 | 2.367E-08 | 9.292E-09 | 1.600E-08 | 3.400E-08 |
| 19269 | 964 | F964H | C | 3.6e-08 | 2.1e-08 | 2.3e-08 | 2.667E-08 | 8.145E-09 | 2.100E-08 | 3.600E-08 |
| 19270 | 964 | F964I | C | 3.2e-08 | 2.4e-08 | 3.8e-08 | 3.133E-08 | 7.024E-09 | 2.400E-08 | 3.800E-08 |
| 19271 | 964 | F964L | C | 1.2e-07 | 5.4e-08 | 1.7e-08 | 6.367E-08 | 5.218E-08 | 1.700E-08 | 1.200E-07 |
| 19272 | 964 | F964K | C | 2.7e-08 | 1.5e-07 | 5.9e-08 | 7.867E-08 | 6.381E-08 | 2.700E-08 | 1.500E-07 |
| 19273 | 964 | F964M | C | 1.8e-08 | 7.1e-08 | 5.6e-08 | 4.833E-08 | 2.732E-08 | 1.800E-08 | 7.100E-08 |
| 19274 | 964 | F964F | C | 6e-08 | 2.2e-07 | 1.5e-08 | 9.833E-08 | 1.077E-07 | 1.500E-08 | 2.200E-07 |
| 19275 | 964 | F964P | C | 2.4e-08 | 6e-08 | 4.6e-08 | 4.333E-08 | 1.815E-08 | 2.400E-08 | 6.000E-08 |
| 19276 | 964 | F964S | C | 3.4e-08 | 3.5e-08 | 6.2e-08 | 4.367E-08 | 1.589E-08 | 3.400E-08 | 6.200E-08 |
| 19277 | 964 | F964T | C | 4e-08 | 7.4e-08 | 2.3e-08 | 4.567E-08 | 2.597E-08 | 2.300E-08 | 7.400E-08 |
| 19278 | 964 | F964W | C | 3.5e-08 | 6.8e-08 | 4e-08 | 4.767E-08 | 1.779E-08 | 3.500E-08 | 6.800E-08 |
| 19279 | 964 | F964Y | C | 7.1e-08 | 6.7e-08 | 4.7e-08 | 6.167E-08 | 1.286E-08 | 4.700E-08 | 7.100E-08 |
| 19280 | 964 | F964V | C | 2.6e-08 | 8.3e-08 | 2.4e-08 | 4.433E-08 | 3.350E-08 | 2.400E-08 | 8.300E-08 |
| 19281 | 965 | P965A | C | 8.2e-08 | 5.7e-08 | 3.9e-08 | 5.933E-08 | 2.159E-08 | 3.900E-08 | 8.200E-08 |
| 19282 | 965 | P965R | C | 4.8e-08 | 4.1e-08 | 8.7e-08 | 5.867E-08 | 2.479E-08 | 4.100E-08 | 8.700E-08 |
| 19283 | 965 | P965N | C | 3.6e-08 | 4.8e-08 | 3.6e-08 | 4.000E-08 | 6.928E-09 | 3.600E-08 | 4.800E-08 |
| 19284 | 965 | P965D | C | 1.4e-08 | 4.3e-08 | 2.6e-08 | 2.767E-08 | 1.457E-08 | 1.400E-08 | 4.300E-08 |
| 19285 | 965 | P965C | C | 4.4e-08 | 4.1e-08 | 1.2e-08 | 3.233E-08 | 1.767E-08 | 1.200E-08 | 4.400E-08 |
| 19286 | 965 | P965E | C | 4.1e-08 | 1.8e-08 | 5.2e-08 | 3.700E-08 | 1.735E-08 | 1.800E-08 | 5.200E-08 |
| 19287 | 965 | P965Q | C | 9.4e-08 | 3.2e-08 | 1.8e-07 | 1.020E-07 | 7.432E-08 | 3.200E-08 | 1.800E-07 |
| 19288 | 965 | P965G | C | 3.1e-08 | 5.5e-08 | 6.9e-08 | 5.167E-08 | 1.922E-08 | 3.100E-08 | 6.900E-08 |
| 19289 | 965 | P965H | C | 6.6e-08 | 8.3e-09 | 4.9e-08 | 4.110E-08 | 2.965E-08 | 8.300E-09 | 6.600E-08 |
| 19290 | 965 | P965I | C | 4.4e-08 | 8.4e-08 | 6.5e-08 | 6.433E-08 | 2.001E-08 | 4.400E-08 | 8.400E-08 |
| 19291 | 965 | P965L | C | 3.3e-08 | 4.7e-08 | 4.6e-08 | 4.200E-08 | 7.810E-09 | 3.300E-08 | 4.700E-08 |
| 19292 | 965 | P965K | C | 3.3e-08 | 1.6e-08 | 3.4e-08 | 2.767E-08 | 1.012E-08 | 1.600E-08 | 3.400E-08 |
| 19293 | 965 | P965M | C | 6e-08 | 4.8e-08 | 4.6e-08 | 5.133E-08 | 7.572E-09 | 4.600E-08 | 6.000E-08 |
| 19294 | 965 | P965F | C | 5.2e-08 | 2.4e-08 | 1.8e-08 | 3.133E-08 | 1.815E-08 | 1.800E-08 | 5.200E-08 |

| | | | | | | | | | | |
|---|---|---|---|---|---|---|---|---|---|---|
| 19295 | 965 | P965P | C | 1.7e-08 | 4.7e-08 | 4.8e-08 | 3.733E-08 | 1.762E-08 | 1.700E-08 | 4.800E-08 |
| 19296 | 965 | P965S | C | 1.3e-07 | 6e-08 | 3e-08 | 7.333E-08 | 5.132E-08 | 3.000E-08 | 1.300E-07 |
| 19297 | 965 | P965T | C | 6.6e-08 | 1.3e-08 | 5.1e-08 | 4.333E-08 | 2.732E-08 | 1.300E-08 | 6.600E-08 |
| 19298 | 965 | P965W | C | 4.9e-08 | 8.2e-08 | 7.7e-08 | 6.933E-08 | 1.779E-08 | 4.900E-08 | 8.200E-08 |
| 19299 | 965 | P965Y | C | 7.3e-08 | 5e-08 | 3e-08 | 5.100E-08 | 2.152E-08 | 3.000E-08 | 7.300E-08 |
| 19300 | 965 | P965V | C | 3.7e-08 | 8.3e-08 | 4.4e-08 | 5.467E-08 | 2.479E-08 | 3.700E-08 | 8.300E-08 |
| 19301 | 966 | L966A | C | 2.9e-08 | 1.7e-08 | 7.3e-08 | 3.967E-08 | 2.948E-08 | 1.700E-08 | 7.300E-08 |
| 19302 | 966 | L966R | C | 1.9e-08 | 2.3e-08 | 1.3e-08 | 1.833E-08 | 5.033E-09 | 1.300E-08 | 2.300E-08 |
| 19303 | 966 | L966N | C | 1.8e-08 | 2.7e-08 | 4.7e-08 | 3.067E-08 | 1.484E-08 | 1.800E-08 | 4.700E-08 |
| 19304 | 966 | L966D | C | 2.1e-07 | 1.4e-08 | 4.6e-08 | 9.000E-08 | 1.051E-07 | 1.400E-08 | 2.100E-07 |
| 19305 | 966 | L966C | C | 4.4e-08 | 5.5e-08 | 4.2e-08 | 4.700E-08 | 7.000E-09 | 4.200E-08 | 5.500E-08 |
| 19306 | 966 | L966E | C | 4.5e-08 | 6.4e-08 | 3.8e-08 | 4.900E-08 | 1.345E-08 | 3.800E-08 | 6.400E-08 |
| 19307 | 966 | L966Q | C | 5.1e-08 | 1.8e-08 | 6.3e-08 | 4.400E-08 | 2.330E-08 | 1.800E-08 | 6.300E-08 |
| 19308 | 966 | L966G | C | 4.3e-08 | 9e-08 | 1.5e-08 | 4.933E-08 | 3.790E-08 | 1.500E-08 | 9.000E-08 |
| 19309 | 966 | L966H | C | 3.4e-08 | 5.8e-08 | 3.1e-08 | 4.100E-08 | 1.480E-08 | 3.100E-08 | 5.800E-08 |
| 19310 | 966 | L966I | C | 5.4e-08 | 2.7e-08 | 4.9e-08 | 4.333E-08 | 1.436E-08 | 2.700E-08 | 5.400E-08 |
| 19311 | 966 | L966L | C | 2.9e-08 | 5.2e-08 | 9.2e-08 | 5.767E-08 | 3.188E-08 | 2.900E-08 | 9.200E-08 |
| 19312 | 966 | L966K | C | 2.5e-08 | 2.1e-08 | 5.7e-08 | 3.433E-08 | 1.973E-08 | 2.100E-08 | 5.700E-08 |
| 19313 | 966 | L966M | C | 7.3e-08 | 3.2e-08 | 7.1e-08 | 5.867E-08 | 2.312E-08 | 3.200E-08 | 7.300E-08 |
| 19314 | 966 | L966F | C | 5.1e-08 | 9.4e-08 | 4.8e-08 | 6.433E-08 | 2.574E-08 | 4.800E-08 | 9.400E-08 |
| 19315 | 966 | L966P | C | 1.4e-08 | 3.3e-08 | 3.3e-08 | 2.667E-08 | 1.097E-08 | 1.400E-08 | 3.300E-08 |
| 19316 | 966 | L966S | C | 4.9e-08 | 6.4e-08 | 2.7e-08 | 4.667E-08 | 1.861E-08 | 2.700E-08 | 6.400E-08 |
| 19317 | 966 | L966T | C | 7.3e-08 | 6.4e-08 | 5.7e-08 | 6.467E-08 | 8.021E-09 | 5.700E-08 | 7.300E-08 |
| 19318 | 966 | L966W | C | 3.7e-08 | 9.5e-08 | 1.8e-08 | 5.000E-08 | 4.011E-08 | 1.800E-08 | 9.500E-08 |
| 19319 | 966 | L966Y | C | 2.5e-08 | 1.9e-08 | 3.9e-08 | 2.767E-08 | 1.026E-08 | 1.900E-08 | 3.900E-08 |
| 19320 | 966 | L966V | C | 3.8e-08 | 6e-08 | 1.9e-08 | 3.900E-08 | 2.052E-08 | 1.900E-08 | 6.000E-08 |
| 19321 | 967 | A967A | C | 5.8e-08 | 6e-08 | 4e-08 | 5.267E-08 | 1.102E-08 | 4.000E-08 | 6.000E-08 |
| 19322 | 967 | A967R | C | 5.9e-08 | 5.2e-08 | 4.1e-08 | 5.067E-08 | 9.074E-09 | 4.100E-08 | 5.900E-08 |
| 19323 | 967 | A967N | C | 5.2e-08 | 2.3e-08 | 1.2e-07 | 6.500E-08 | 4.979E-08 | 2.300E-08 | 1.200E-07 |
| 19324 | 967 | A967D | C | 5.8e-08 | 3.5e-08 | 8.5e-08 | 5.933E-08 | 2.503E-08 | 3.500E-08 | 8.500E-08 |
| 19325 | 967 | A967C | C | 1.2e-08 | 2.6e-08 | 3.8e-08 | 2.533E-08 | 1.301E-08 | 1.200E-08 | 3.800E-08 |
| 19326 | 967 | A967E | C | 6.7e-08 | 5.1e-08 | 8.7e-08 | 6.833E-08 | 1.804E-08 | 5.100E-08 | 8.700E-08 |
| 19327 | 967 | A967Q | C | 2.2e-08 | 5.5e-08 | 3.6e-08 | 3.767E-08 | 1.656E-08 | 2.200E-08 | 5.500E-08 |
| 19328 | 967 | A967G | C | 4.2e-08 | 2.3e-08 | 9.2e-08 | 5.233E-08 | 3.564E-08 | 2.300E-08 | 9.200E-08 |
| 19329 | 967 | A967H | C | 6.1e-08 | 5.1e-08 | 2.4e-08 | 4.533E-08 | 1.914E-08 | 2.400E-08 | 6.100E-08 |
| 19330 | 967 | A967I | C | 6.5e-08 | 4.1e-08 | 4.2e-08 | 4.933E-08 | 1.358E-08 | 4.100E-08 | 6.500E-08 |
| 19331 | 967 | A967L | C | 4.5e-08 | 5.8e-08 | 2.9e-08 | 4.400E-08 | 1.453E-08 | 2.900E-08 | 5.800E-08 |
| 19332 | 967 | A967K | C | 5.6e-08 | 1.4e-08 | 2.2e-08 | 3.067E-08 | 2.230E-08 | 1.400E-08 | 5.600E-08 |
| 19333 | 967 | A967M | C | 4.1e-08 | 3.5e-08 | 3e-08 | 3.533E-08 | 5.508E-09 | 3.000E-08 | 4.100E-08 |
| 19334 | 967 | A967F | C | 1.5e-08 | 5.7e-08 | 4.7e-08 | 3.967E-08 | 2.194E-08 | 1.500E-08 | 5.700E-08 |
| 19335 | 967 | A967P | C | 1.6e-08 | 4.3e-08 | 3.4e-08 | 3.100E-08 | 1.375E-08 | 1.600E-08 | 4.300E-08 |
| 19336 | 967 | A967S | C | 4.8e-08 | 1.7e-08 | 5e-08 | 3.833E-08 | 1.850E-08 | 1.700E-08 | 5.000E-08 |
| 19337 | 967 | A967T | C | 4.3e-08 | 3.5e-08 | 7.1e-08 | 4.967E-08 | 1.890E-08 | 3.500E-08 | 7.100E-08 |
| 19338 | 967 | A967W | C | 4.7e-08 | 1.1e-08 | 5.4e-08 | 3.733E-08 | 2.307E-08 | 1.100E-08 | 5.400E-08 |
| 19339 | 967 | A967Y | C | 6.7e-08 | 2.4e-08 | 1.4e-07 | 7.700E-08 | 5.864E-08 | 2.400E-08 | 1.400E-07 |
| 19340 | 967 | A967V | C | 1.6e-08 | 4e-08 | 1.3e-08 | 2.300E-08 | 1.480E-08 | 1.300E-08 | 4.000E-08 |
| 19341 | 968 | P968A | C | 3e-08 | 3.3e-08 | 2.8e-08 | 3.033E-08 | 2.517E-09 | 2.800E-08 | 3.300E-08 |
| 19342 | 968 | P968R | C | 5.7e-08 | 7.1e-08 | 3.2e-08 | 5.333E-08 | 1.976E-08 | 3.200E-08 | 7.100E-08 |
| 19343 | 968 | P968N | C | 2.5e-08 | 9.8e-08 | 1.7e-08 | 4.667E-08 | 4.464E-08 | 1.700E-08 | 9.800E-08 |
| 19344 | 968 | P968D | C | 4.9e-08 | 3.5e-08 | 1.3e-07 | 7.133E-08 | 5.129E-08 | 3.500E-08 | 1.300E-07 |
| 19345 | 968 | P968C | C | 9.7e-08 | 3.7e-08 | 6.9e-08 | 6.767E-08 | 3.002E-08 | 3.700E-08 | 9.700E-08 |
| 19346 | 968 | P968E | C | 5.5e-08 | 3.5e-08 | 3.3e-08 | 4.100E-08 | 1.217E-08 | 3.300E-08 | 5.500E-08 |
| 19347 | 968 | P968Q | C | 4.1e-08 | 4.2e-08 | 3.5e-08 | 3.933E-08 | 3.786E-09 | 3.500E-08 | 4.200E-08 |
| 19348 | 968 | P968G | C | 3.6e-08 | 1e-07 | 1.2e-07 | 8.533E-08 | 4.388E-08 | 3.600E-08 | 1.200E-07 |
| 19349 | 968 | P968H | C | 1.8e-08 | 2.2e-08 | 3e-08 | 2.333E-08 | 6.110E-09 | 1.800E-08 | 3.000E-08 |
| 19350 | 968 | P968I | C | 3.9e-08 | 3e-08 | 1.1e-08 | 2.667E-08 | 1.429E-08 | 1.100E-08 | 3.900E-08 |
| 19351 | 968 | P968L | C | 2.9e-08 | 6.1e-08 | 4.9e-08 | 4.633E-08 | 1.617E-08 | 2.900E-08 | 6.100E-08 |
| 19352 | 968 | P968K | C | 6.1e-08 | 3.4e-08 | 8.6e-08 | 6.033E-08 | 2.601E-08 | 3.400E-08 | 8.600E-08 |
| 19353 | 968 | P968M | C | 2.8e-08 | 1.1e-08 | 2.9e-08 | 2.267E-08 | 1.012E-08 | 1.100E-08 | 2.900E-08 |
| 19354 | 968 | P968F | C | 3.5e-08 | 3.2e-08 | 4.8e-08 | 3.833E-08 | 8.505E-09 | 3.200E-08 | 4.800E-08 |
| 19355 | 968 | P968P | C | 4.8e-08 | 5.6e-08 | 5.9e-08 | 5.433E-08 | 5.686E-09 | 4.800E-08 | 5.900E-08 |
| 19356 | 968 | P968S | C | 6.2e-08 | 2.3e-08 | 1.7e-08 | 3.400E-08 | 2.443E-08 | 1.700E-08 | 6.200E-08 |
| 19357 | 968 | P968T | C | 1.3e-08 | 1.2e-08 | 2.8e-08 | 1.767E-08 | 8.963E-09 | 1.200E-08 | 2.800E-08 |
| 19358 | 968 | P968W | C | 3.9e-08 | 4e-08 | 4.3e-08 | 4.067E-08 | 2.082E-09 | 3.900E-08 | 4.300E-08 |
| 19359 | 968 | P968Y | C | 8.5e-08 | 3.9e-08 | 1.6e-08 | 4.667E-08 | 3.513E-08 | 1.600E-08 | 8.500E-08 |
| 19360 | 968 | P968V | C | 3.9e-08 | 4.6e-08 | 1.9e-08 | 3.467E-08 | 1.401E-08 | 1.900E-08 | 4.600E-08 |
| 19361 | 969 | S969A | C | 5.5e-08 | 6.6e-08 | 4.8e-08 | 5.633E-08 | 9.074E-09 | 4.800E-08 | 6.600E-08 |
| 19362 | 969 | S969R | C | 2.8e-08 | 1.1e-07 | 3.4e-08 | 5.733E-08 | 4.571E-08 | 2.800E-08 | 1.100E-07 |
| 19363 | 969 | S969N | C | 5.3e-08 | 3.2e-08 | 3.7e-08 | 4.067E-08 | 1.097E-08 | 3.200E-08 | 5.300E-08 |
| 19364 | 969 | S969D | C | 3.6e-08 | 7.4e-08 | 1.3e-07 | 8.000E-08 | 4.729E-08 | 3.600E-08 | 1.300E-07 |
| 19365 | 969 | S969C | C | 3.7e-08 | 1.7e-08 | 2.4e-08 | 2.600E-08 | 1.015E-08 | 1.700E-08 | 3.700E-08 |
| 19366 | 969 | S969E | C | 3e-08 | 4.7e-08 | 4.7e-08 | 4.133E-08 | 9.815E-09 | 3.000E-08 | 4.700E-08 |
| 19367 | 969 | S969Q | C | 4.8e-08 | 4.1e-08 | 8.5e-08 | 5.800E-08 | 2.364E-08 | 4.100E-08 | 8.500E-08 |
| 19368 | 969 | S969G | C | 1.1e-07 | 5.4e-08 | 6.3e-08 | 7.567E-08 | 3.007E-08 | 5.400E-08 | 1.100E-07 |
| 19369 | 969 | S969H | C | 8.9e-08 | 1.3e-07 | 4.6e-08 | 8.833E-08 | 4.200E-08 | 4.600E-08 | 1.300E-07 |
| 19370 | 969 | S969I | C | 1e-07 | 5.7e-08 | 6.5e-08 | 7.400E-08 | 2.287E-08 | 5.700E-08 | 1.000E-07 |
| 19371 | 969 | S969L | C | 4.8e-08 | 3.6e-08 | 2.9e-08 | 3.767E-08 | 9.609E-09 | 2.900E-08 | 4.800E-08 |
| 19372 | 969 | S969K | C | 8.3e-08 | 7.8e-08 | 2.1e-08 | 6.067E-08 | 3.444E-08 | 2.100E-08 | 8.300E-08 |
| 19373 | 969 | S969M | C | 8.2e-08 | 5.2e-08 | 5.4e-08 | 6.267E-08 | 1.677E-08 | 5.200E-08 | 8.200E-08 |
| 19374 | 969 | S969F | C | 1.6e-08 | 9.2e-08 | 2.9e-08 | 4.567E-08 | 4.065E-08 | 1.600E-08 | 9.200E-08 |
| 19375 | 969 | S969P | C | 8.9e-08 | 2.3e-08 | 8.4e-08 | 6.533E-08 | 3.675E-08 | 2.300E-08 | 8.900E-08 |
| 19376 | 969 | S969S | C | 2.4e-08 | 4.4e-08 | 9.1e-08 | 5.300E-08 | 3.439E-08 | 2.400E-08 | 9.100E-08 |
| 19377 | 969 | S969T | C | 6.5e-08 | 1.7e-08 | 7.5e-08 | 5.233E-08 | 3.101E-08 | 1.700E-08 | 7.500E-08 |
| 19378 | 969 | S969W | C | 7.3e-08 | 5.8e-08 | 3e-08 | 5.367E-08 | 2.183E-08 | 3.000E-08 | 7.300E-08 |
| 19379 | 969 | S969Y | C | 3.2e-08 | 7e-08 | 2.6e-08 | 4.267E-08 | 2.386E-08 | 2.600E-08 | 7.000E-08 |

| | | | | | | | | | | |
|---|---|---|---|---|---|---|---|---|---|---|
| 19380 | 969 | S969V | C | 3.8e-08 | 6.7e-08 | 3.9e-08 | 4.800E-08 | 1.646E-08 | 3.800E-08 | 6.700E-08 |
| 19381 | 970 | S970A | C | 4.9e-08 | 1.5e-08 | 9e-08 | 5.133E-08 | 3.755E-08 | 1.500E-08 | 9.000E-08 |
| 19382 | 970 | S970R | C | 3.2e-08 | 8.9e-08 | 8.8e-08 | 6.967E-08 | 3.262E-08 | 3.200E-08 | 8.900E-08 |
| 19383 | 970 | S970N | C | 3.1e-08 | 3e-08 | 6.2e-08 | 4.100E-08 | 1.819E-08 | 3.000E-08 | 6.200E-08 |
| 19384 | 970 | S970D | C | 4.5e-08 | 6.1e-08 | 5.8e-08 | 5.467E-08 | 8.505E-09 | 4.500E-08 | 6.100E-08 |
| 19385 | 970 | S970C | C | 9.5e-08 | 2.7e-08 | 1.7e-08 | 4.633E-08 | 4.244E-08 | 1.700E-08 | 9.500E-08 |
| 19386 | 970 | S970E | C | 1.3e-07 | 5.9e-08 | 9.8e-09 | 6.627E-08 | 6.043E-08 | 9.800E-09 | 1.300E-07 |
| 19387 | 970 | S970Q | C | 4e-08 | 3.9e-08 | 1.1e-07 | 6.300E-08 | 4.071E-08 | 3.900E-08 | 1.100E-07 |
| 19388 | 970 | S970G | C | 7.1e-08 | 9.9e-08 | 3.8e-08 | 6.933E-08 | 3.053E-08 | 3.800E-08 | 9.900E-08 |
| 19389 | 970 | S970H | C | 3.7e-08 | 2.8e-08 | 8.8e-08 | 5.100E-08 | 3.236E-08 | 2.800E-08 | 8.800E-08 |
| 19390 | 970 | S970I | C | 2.9e-08 | 2.2e-08 | 5.8e-08 | 3.633E-08 | 1.909E-08 | 2.200E-08 | 5.800E-08 |
| 19391 | 970 | S970L | C | 3.8e-08 | 4e-08 | 6.5e-08 | 4.767E-08 | 1.504E-08 | 3.800E-08 | 6.500E-08 |
| 19392 | 970 | S970K | C | 4.5e-08 | 2.4e-08 | 2.5e-08 | 3.133E-08 | 1.185E-08 | 2.400E-08 | 4.500E-08 |
| 19393 | 970 | S970M | C | 4.7e-08 | 3.2e-08 | 4.1e-08 | 4.000E-08 | 7.550E-09 | 3.200E-08 | 4.700E-08 |
| 19394 | 970 | S970F | C | 6.1e-08 | 4.7e-08 | 2.1e-08 | 4.300E-08 | 2.030E-08 | 2.100E-08 | 6.100E-08 |
| 19395 | 970 | S970P | C | 7e-08 | 7.9e-08 | 3.3e-08 | 6.067E-08 | 2.438E-08 | 3.300E-08 | 7.900E-08 |
| 19396 | 970 | S970S | C | 3.7e-08 | 6.5e-08 | 1.6e-07 | 8.733E-08 | 6.447E-08 | 3.700E-08 | 1.600E-07 |
| 19397 | 970 | S970T | C | 1.1e-07 | 3.3e-08 | 7.1e-08 | 7.133E-08 | 3.850E-08 | 3.300E-08 | 1.100E-07 |
| 19398 | 970 | S970W | C | 2.6e-08 | 4.6e-08 | 3.9e-08 | 3.700E-08 | 1.015E-08 | 2.600E-08 | 4.600E-08 |
| 19399 | 970 | S970Y | C | 2.8e-08 | 3.2e-08 | 2.3e-08 | 2.767E-08 | 4.509E-09 | 2.300E-08 | 3.200E-08 |
| 19400 | 970 | S970V | C | 4.2e-08 | 1.6e-08 | 7.8e-08 | 4.533E-08 | 3.113E-08 | 1.600E-08 | 7.800E-08 |
| 19401 | 971 | K971A | C | 6.1e-08 | 5.4e-08 | 2e-08 | 4.500E-08 | 2.193E-08 | 2.000E-08 | 6.100E-08 |
| 19402 | 971 | K971R | C | 4.4e-08 | 5.3e-08 | 2.3e-08 | 4.000E-08 | 1.539E-08 | 2.300E-08 | 5.300E-08 |
| 19403 | 971 | K971N | C | 2.8e-08 | 2.7e-08 | 2.3e-08 | 2.600E-08 | 2.646E-09 | 2.300E-08 | 2.800E-08 |
| 19404 | 971 | K971D | C | 1.6e-08 | 3.5e-08 | 3.6e-08 | 2.900E-08 | 1.127E-08 | 1.600E-08 | 3.600E-08 |
| 19405 | 971 | K971C | C | 2.8e-08 | 5.9e-08 | 2.5e-08 | 3.733E-08 | 1.882E-08 | 2.500E-08 | 5.900E-08 |
| 19406 | 971 | K971E | C | 2.8e-08 | 3.2e-08 | 6.4e-08 | 4.133E-08 | 1.973E-08 | 2.800E-08 | 6.400E-08 |
| 19407 | 971 | K971Q | C | 3.8e-08 | 2.3e-08 | 2.1e-08 | 2.733E-08 | 9.292E-09 | 2.100E-08 | 3.800E-08 |
| 19408 | 971 | K971G | C | 6.8e-08 | 3.1e-08 | 4e-08 | 4.633E-08 | 1.930E-08 | 3.100E-08 | 6.800E-08 |
| 19409 | 971 | K971H | C | 1.6e-08 | 4.3e-08 | 6.4e-08 | 4.100E-08 | 2.406E-08 | 1.600E-08 | 6.400E-08 |
| 19410 | 971 | K971I | C | 9.6e-08 | 5.8e-08 | 1.5e-08 | 5.633E-08 | 4.053E-08 | 1.500E-08 | 9.600E-08 |
| 19411 | 971 | K971L | C | 6.1e-08 | 9.3e-08 | 5.1e-08 | 6.833E-08 | 2.194E-08 | 5.100E-08 | 9.300E-08 |
| 19412 | 971 | K971K | C | 2.2e-08 | 9e-08 | 4.2e-08 | 5.133E-08 | 3.495E-08 | 2.200E-08 | 9.000E-08 |
| 19413 | 971 | K971M | C | 1.5e-07 | 7.3e-08 | 7.4e-08 | 9.900E-08 | 4.417E-08 | 7.300E-08 | 1.500E-07 |
| 19414 | 971 | K971F | C | 4.6e-08 | 3.3e-08 | 2.8e-08 | 3.567E-08 | 9.292E-09 | 2.800E-08 | 4.600E-08 |
| 19415 | 971 | K971P | C | 2e-08 | 2.3e-08 | 2.9e-08 | 2.400E-08 | 4.583E-09 | 2.000E-08 | 2.900E-08 |
| 19416 | 971 | K971S | C | 9.9e-08 | 2.7e-08 | 9.3e-08 | 7.300E-08 | 3.995E-08 | 2.700E-08 | 9.900E-08 |
| 19417 | 971 | K971T | C | 8e-08 | 4.5e-08 | 7.6e-08 | 6.700E-08 | 1.916E-08 | 4.500E-08 | 8.000E-08 |
| 19418 | 971 | K971W | C | 6.9e-08 | 8.8e-08 | 7.3e-08 | 7.667E-08 | 1.002E-08 | 6.900E-08 | 8.800E-08 |
| 19419 | 971 | K971Y | C | 6.8e-08 | 8.4e-08 | 3.1e-08 | 6.100E-08 | 2.718E-08 | 3.100E-08 | 8.400E-08 |
| 19420 | 971 | K971V | C | 7.3e-08 | 7.2e-08 | 3.4e-08 | 5.967E-08 | 2.223E-08 | 3.400E-08 | 7.300E-08 |
| 19421 | 972 | S972A | C | 2.2e-08 | 4.3e-08 | 1.1e-07 | 5.833E-08 | 4.596E-08 | 2.200E-08 | 1.100E-07 |
| 19422 | 972 | S972R | C | 1.9e-08 | 2.8e-08 | 5.7e-08 | 3.467E-08 | 1.986E-08 | 1.900E-08 | 5.700E-08 |
| 19423 | 972 | S972N | C | 5.7e-08 | 2.3e-08 | 3.5e-08 | 3.833E-08 | 1.724E-08 | 2.300E-08 | 5.700E-08 |
| 19424 | 972 | S972D | C | 4.1e-08 | 4.7e-08 | 7.2e-08 | 5.333E-08 | 1.644E-08 | 4.100E-08 | 7.200E-08 |
| 19425 | 972 | S972C | C | 8.8e-08 | 4.3e-08 | 1.7e-07 | 1.003E-07 | 6.439E-08 | 4.300E-08 | 1.700E-07 |
| 19426 | 972 | S972E | C | 1.1e-07 | 6.8e-08 | 2.2e-08 | 6.667E-08 | 4.402E-08 | 2.200E-08 | 1.100E-07 |
| 19427 | 972 | S972Q | C | 3.7e-08 | 1.1e-07 | 3.4e-08 | 6.033E-08 | 4.304E-08 | 3.400E-08 | 1.100E-07 |
| 19428 | 972 | S972G | C | 2.2e-08 | 2.6e-08 | 3.3e-08 | 2.700E-08 | 5.568E-09 | 2.200E-08 | 3.300E-08 |
| 19429 | 972 | S972H | C | 4.5e-08 | 2.1e-08 | 4e-08 | 3.533E-08 | 1.266E-08 | 2.100E-08 | 4.500E-08 |
| 19430 | 972 | S972I | C | 3.2e-08 | 5.3e-08 | 3.7e-08 | 4.067E-08 | 1.097E-08 | 3.200E-08 | 5.300E-08 |
| 19431 | 972 | S972L | C | 2.5e-08 | 7.4e-08 | 1.1e-07 | 6.967E-08 | 4.267E-08 | 2.500E-08 | 1.100E-07 |
| 19432 | 972 | S972K | C | 4.1e-08 | 3.9e-08 | 4.8e-08 | 4.267E-08 | 4.726E-09 | 3.900E-08 | 4.800E-08 |
| 19433 | 972 | S972M | C | 1.2e-08 | 5.3e-08 | 1.4e-07 | 6.833E-08 | 6.536E-08 | 1.200E-08 | 1.400E-07 |
| 19434 | 972 | S972F | C | 1.5e-08 | 2.7e-08 | 4e-08 | 2.733E-08 | 1.250E-08 | 1.500E-08 | 4.000E-08 |
| 19435 | 972 | S972P | C | 5.2e-08 | 4e-08 | 3.5e-08 | 4.233E-08 | 8.737E-09 | 3.500E-08 | 5.200E-08 |
| 19436 | 972 | S972S | C | 1.8e-08 | 2.9e-08 | 7.6e-08 | 4.100E-08 | 3.081E-08 | 1.800E-08 | 7.600E-08 |
| 19437 | 972 | S972T | C | 1.6e-08 | 6.8e-08 | 3.4e-08 | 3.933E-08 | 2.641E-08 | 1.600E-08 | 6.800E-08 |
| 19438 | 972 | S972W | C | 1.4e-07 | 4.5e-08 | 7.3e-08 | 8.600E-08 | 4.882E-08 | 4.500E-08 | 1.400E-07 |
| 19439 | 972 | S972Y | C | 3e-08 | 1.4e-08 | 3.6e-08 | 2.667E-08 | 1.137E-08 | 1.400E-08 | 3.600E-08 |
| 19440 | 972 | S972V | C | 4.4e-08 | 3.9e-08 | 5.9e-08 | 4.733E-08 | 1.041E-08 | 3.900E-08 | 5.900E-08 |
| 19441 | 973 | T973A | C | 4.4e-08 | 3e-08 | 5.6e-08 | 4.333E-08 | 1.301E-08 | 3.000E-08 | 5.600E-08 |
| 19442 | 973 | T973R | C | 4.4e-08 | 4.5e-08 | 7.7e-08 | 5.533E-08 | 1.877E-08 | 4.400E-08 | 7.700E-08 |
| 19443 | 973 | T973N | C | 2e-07 | 8.7e-08 | 1e-07 | 1.290E-07 | 6.183E-08 | 8.700E-08 | 2.000E-07 |
| 19444 | 973 | T973D | C | 5.8e-08 | 1.9e-07 | 3.2e-08 | 9.333E-08 | 8.472E-08 | 3.200E-08 | 1.900E-07 |
| 19445 | 973 | T973C | C | 1.8e-08 | 3.3e-08 | 5.2e-08 | 3.433E-08 | 1.704E-08 | 1.800E-08 | 5.200E-08 |
| 19446 | 973 | T973E | C | 6.1e-08 | 5.3e-08 | 5.6e-08 | 5.667E-08 | 4.041E-09 | 5.300E-08 | 6.100E-08 |
| 19447 | 973 | T973Q | C | 3.6e-08 | 4.9e-08 | 1.9e-08 | 3.467E-08 | 1.504E-08 | 1.900E-08 | 4.900E-08 |
| 19448 | 973 | T973G | C | 5.1e-08 | 9.4e-08 | 7.2e-08 | 7.233E-08 | 2.150E-08 | 5.100E-08 | 9.400E-08 |
| 19449 | 973 | T973H | C | 6.9e-08 | 3.8e-08 | 1.9e-08 | 4.200E-08 | 2.524E-08 | 1.900E-08 | 6.900E-08 |
| 19450 | 973 | T973I | C | 1e-07 | 3.9e-08 | 6e-08 | 6.633E-08 | 3.099E-08 | 3.900E-08 | 1.000E-07 |
| 19451 | 973 | T973L | C | 1.5e-08 | 1.8e-08 | 7.5e-08 | 3.600E-08 | 3.381E-08 | 1.500E-08 | 7.500E-08 |
| 19452 | 973 | T973K | C | 1.5e-07 | 4.8e-08 | 4.7e-08 | 8.167E-08 | 5.918E-08 | 4.700E-08 | 1.500E-07 |
| 19453 | 973 | T973M | C | 1.8e-08 | 5e-08 | 2e-08 | 2.933E-08 | 1.793E-08 | 1.800E-08 | 5.000E-08 |
| 19454 | 973 | T973F | C | 2.1e-08 | 4.4e-08 | 9.9e-09 | 2.497E-08 | 1.739E-08 | 9.900E-09 | 4.400E-08 |
| 19455 | 973 | T973P | C | 2.1e-08 | 5.3e-08 | 7.6e-08 | 5.000E-08 | 2.762E-08 | 2.100E-08 | 7.600E-08 |
| 19456 | 973 | T973S | C | 1.8e-08 | 2.3e-08 | 6.3e-08 | 3.467E-08 | 2.466E-08 | 1.800E-08 | 6.300E-08 |
| 19457 | 973 | T973T | C | 1.6e-08 | 3.6e-08 | 5.6e-08 | 3.600E-08 | 2.000E-08 | 1.600E-08 | 5.600E-08 |
| 19458 | 973 | T973W | C | 6e-08 | 3.6e-08 | 7.2e-08 | 5.600E-08 | 1.833E-08 | 3.600E-08 | 7.200E-08 |
| 19459 | 973 | T973Y | C | 1e-07 | 3.6e-08 | 3.9e-08 | 5.833E-08 | 3.612E-08 | 3.600E-08 | 1.000E-07 |
| 19460 | 973 | T973V | C | 3.5e-08 | 5.3e-08 | 7.4e-08 | 5.400E-08 | 1.952E-08 | 3.500E-08 | 7.400E-08 |
| 19461 | 974 | S974A | C | 3.1e-08 | 8.3e-08 | 6.5e-08 | 5.967E-08 | 2.641E-08 | 3.100E-08 | 8.300E-08 |
| 19462 | 974 | S974R | C | 4.1e-08 | 3.7e-08 | 1.6e-07 | 7.933E-08 | 6.989E-08 | 3.700E-08 | 1.600E-07 |
| 19463 | 974 | S974N | C | 4.5e-08 | 2.8e-08 | 1.6e-08 | 2.967E-08 | 1.457E-08 | 1.600E-08 | 4.500E-08 |
| 19464 | 974 | S974D | C | 1.1e-07 | 4.5e-08 | 3.7e-08 | 6.400E-08 | 4.004E-08 | 3.700E-08 | 1.100E-07 |

| | | | | | | | | | | |
|---|---|---|---|---|---|---|---|---|---|---|
| 19465 | 974 | S974C | C | 8.5e-08 | 6.1e-08 | 3.9e-08 | 6.167E-08 | 2.301E-08 | 3.900E-08 | 8.500E-08 |
| 19466 | 974 | S974E | C | 5e-08 | 8.3e-08 | 8.6e-08 | 7.300E-08 | 1.997E-08 | 5.000E-08 | 8.600E-08 |
| 19467 | 974 | S974Q | C | 2.2e-08 | 4.9e-08 | 2.8e-08 | 3.300E-08 | 1.418E-08 | 2.200E-08 | 4.900E-08 |
| 19468 | 974 | S974G | C | 3e-08 | 2.5e-08 | 9.9e-08 | 5.133E-08 | 4.136E-08 | 2.500E-08 | 9.900E-08 |
| 19469 | 974 | S974H | C | 1.7e-08 | 8.9e-08 | 4.3e-08 | 4.967E-08 | 3.646E-08 | 1.700E-08 | 8.900E-08 |
| 19470 | 974 | S974I | C | 3.2e-08 | 1e-07 | 2.1e-08 | 5.100E-08 | 4.279E-08 | 2.100E-08 | 1.000E-07 |
| 19471 | 974 | S974L | C | 7.8e-08 | 4.3e-08 | 1.1e-07 | 7.700E-08 | 3.351E-08 | 4.300E-08 | 1.100E-07 |
| 19472 | 974 | S974K | C | 4.4e-08 | 2.6e-08 | 5.7e-08 | 4.233E-08 | 1.557E-08 | 2.600E-08 | 5.700E-08 |
| 19473 | 974 | S974M | C | 8e-08 | 2.7e-08 | 7.1e-08 | 5.933E-08 | 2.836E-08 | 2.700E-08 | 8.000E-08 |
| 19474 | 974 | S974F | C | 2.1e-08 | 1.2e-07 | 5.7e-08 | 6.600E-08 | 5.011E-08 | 2.100E-08 | 1.200E-07 |
| 19475 | 974 | S974P | C | 6.3e-08 | 7.7e-09 | 4e-08 | 3.690E-08 | 2.778E-08 | 7.700E-09 | 6.300E-08 |
| 19476 | 974 | S974S | C | 3.7e-08 | 5.2e-08 | 6.6e-08 | 5.167E-08 | 1.450E-08 | 3.700E-08 | 6.600E-08 |
| 19477 | 974 | S974T | C | 3.5e-08 | 3.2e-08 | 1.7e-08 | 2.800E-08 | 9.644E-09 | 1.700E-08 | 3.500E-08 |
| 19478 | 974 | S974W | C | 5.1e-08 | 2.8e-08 | 6e-08 | 4.633E-08 | 1.650E-08 | 2.800E-08 | 6.000E-08 |
| 19479 | 974 | S974Y | C | 4.3e-08 | 1.4e-07 | 1.8e-08 | 6.700E-08 | 6.444E-08 | 1.800E-08 | 1.400E-07 |
| 19480 | 974 | S974V | C | 1.2e-08 | 1.7e-08 | 2.8e-08 | 1.900E-08 | 8.185E-09 | 1.200E-08 | 2.800E-08 |
| 19481 | 975 | G975A | C | 1e-08 | 4.3e-08 | 1.9e-08 | 2.400E-08 | 1.706E-08 | 1.000E-08 | 4.300E-08 |
| 19482 | 975 | G975R | C | 9.1e-08 | 6.5e-08 | 3.9e-08 | 6.500E-08 | 2.600E-08 | 3.900E-08 | 9.100E-08 |
| 19483 | 975 | G975N | C | 4.2e-08 | 3.4e-08 | 5.2e-08 | 4.267E-08 | 9.018E-09 | 3.400E-08 | 5.200E-08 |
| 19484 | 975 | G975D | C | 2.7e-08 | 9.4e-08 | 6e-08 | 6.033E-08 | 3.350E-08 | 2.700E-08 | 9.400E-08 |
| 19485 | 975 | G975C | C | 2.4e-08 | 3.2e-08 | 6.4e-08 | 4.000E-08 | 2.117E-08 | 2.400E-08 | 6.400E-08 |
| 19486 | 975 | G975E | C | 6.8e-08 | 4.3e-08 | 2.8e-08 | 4.633E-08 | 2.021E-08 | 2.800E-08 | 6.800E-08 |
| 19487 | 975 | G975Q | C | 3.2e-08 | 2.2e-08 | 6.2e-08 | 3.867E-08 | 2.082E-08 | 2.200E-08 | 6.200E-08 |
| 19488 | 975 | G975G | C | 9.9e-08 | 4.5e-08 | 1.1e-07 | 8.467E-08 | 3.479E-08 | 4.500E-08 | 1.100E-07 |
| 19489 | 975 | G975H | C | 4.1e-08 | 1.7e-08 | 5.2e-08 | 3.667E-08 | 1.790E-08 | 1.700E-08 | 5.200E-08 |
| 19490 | 975 | G975I | C | 2.4e-08 | 8.5e-08 | 2.2e-08 | 4.367E-08 | 3.581E-08 | 2.200E-08 | 8.500E-08 |
| 19491 | 975 | G975L | C | 1.5e-07 | 3.3e-08 | 2.1e-08 | 6.800E-08 | 7.127E-08 | 2.100E-08 | 1.500E-07 |
| 19492 | 975 | G975K | C | 6.8e-08 | 1.7e-08 | 5e-08 | 4.500E-08 | 2.587E-08 | 1.700E-08 | 6.800E-08 |
| 19493 | 975 | G975M | C | 2.9e-08 | 4.1e-08 | 2.5e-08 | 3.167E-08 | 8.327E-09 | 2.500E-08 | 4.100E-08 |
| 19494 | 975 | G975F | C | 3.5e-08 | 5.2e-08 | 2.3e-08 | 3.667E-08 | 1.457E-08 | 2.300E-08 | 5.200E-08 |
| 19495 | 975 | G975P | C | 3e-08 | 3.1e-08 | 1.4e-08 | 2.500E-08 | 9.539E-09 | 1.400E-08 | 3.100E-08 |
| 19496 | 975 | G975S | C | 1.4e-07 | 5.9e-08 | 5e-08 | 8.300E-08 | 4.957E-08 | 5.000E-08 | 1.400E-07 |
| 19497 | 975 | G975T | C | 6.3e-08 | 3.4e-08 | 4.2e-08 | 4.633E-08 | 1.498E-08 | 3.400E-08 | 6.300E-08 |
| 19498 | 975 | G975W | C | 3.2e-08 | 5e-08 | 3.7e-08 | 3.967E-08 | 9.292E-09 | 3.200E-08 | 5.000E-08 |
| 19499 | 975 | G975Y | C | 5.1e-08 | 6e-08 | 3.1e-08 | 4.733E-08 | 1.484E-08 | 3.100E-08 | 6.000E-08 |
| 19500 | 975 | G975V | C | 3.5e-08 | 3.2e-08 | 2.1e-08 | 2.933E-08 | 7.371E-09 | 2.100E-08 | 3.500E-08 |
| 19501 | 976 | G976A | C | 3.8e-08 | 2.4e-08 | 2.2e-08 | 2.800E-08 | 8.718E-09 | 2.200E-08 | 3.800E-08 |
| 19502 | 976 | G976R | C | 3.2e-08 | 2.4e-08 | 2.6e-08 | 2.733E-08 | 4.163E-09 | 2.400E-08 | 3.200E-08 |
| 19503 | 976 | G976N | C | 2.3e-08 | 3.9e-08 | 4.1e-08 | 3.433E-08 | 9.866E-09 | 2.300E-08 | 4.100E-08 |
| 19504 | 976 | G976D | C | 4.4e-08 | 1.1e-07 | 7.1e-08 | 7.500E-08 | 3.318E-08 | 4.400E-08 | 1.100E-07 |
| 19505 | 976 | G976C | C | 1.7e-08 | 9.2e-08 | 8.1e-08 | 6.333E-08 | 4.050E-08 | 1.700E-08 | 9.200E-08 |
| 19506 | 976 | G976E | C | 2.6e-08 | 3.5e-08 | 2e-08 | 2.700E-08 | 7.550E-09 | 2.000E-08 | 3.500E-08 |
| 19507 | 976 | G976Q | C | 1.1e-08 | 3.6e-08 | 4.5e-08 | 3.067E-08 | 1.762E-08 | 1.100E-08 | 4.500E-08 |
| 19508 | 976 | G976G | C | 2.9e-08 | 2.7e-08 | 1.8e-07 | 7.867E-08 | 8.776E-08 | 2.700E-08 | 1.800E-07 |
| 19509 | 976 | G976H | C | 2.6e-08 | 7.6e-08 | 5.6e-08 | 5.267E-08 | 2.517E-08 | 2.600E-08 | 7.600E-08 |
| 19510 | 976 | G976I | C | 3.1e-08 | 2e-08 | 4.2e-08 | 3.100E-08 | 1.100E-08 | 2.000E-08 | 4.200E-08 |
| 19511 | 976 | G976L | C | 3.4e-08 | 5.6e-08 | 3.9e-08 | 4.300E-08 | 1.153E-08 | 3.400E-08 | 5.600E-08 |
| 19512 | 976 | G976K | C | 3e-08 | 7.5e-08 | 8.7e-08 | 6.400E-08 | 3.005E-08 | 3.000E-08 | 8.700E-08 |
| 19513 | 976 | G976M | C | 4.1e-08 | 3.5e-08 | 8.1e-08 | 5.233E-08 | 2.501E-08 | 3.500E-08 | 8.100E-08 |
| 19514 | 976 | G976F | C | 3.3e-08 | 1.3e-07 | 2.8e-08 | 6.367E-08 | 5.750E-08 | 2.800E-08 | 1.300E-07 |
| 19515 | 976 | G976P | C | 3.4e-08 | 3.5e-08 | 2e-08 | 2.967E-08 | 8.386E-09 | 2.000E-08 | 3.500E-08 |
| 19516 | 976 | G976S | C | 3.2e-08 | 5e-08 | 1e-07 | 6.067E-08 | 3.523E-08 | 3.200E-08 | 1.000E-07 |
| 19517 | 976 | G976T | C | 3.9e-08 | 5.4e-08 | 3.1e-08 | 4.133E-08 | 1.168E-08 | 3.100E-08 | 5.400E-08 |
| 19518 | 976 | G976W | C | 9.5e-08 | 3.9e-08 | 6.4e-08 | 6.600E-08 | 2.805E-08 | 3.900E-08 | 9.500E-08 |
| 19519 | 976 | G976Y | C | 2.4e-08 | 2.2e-08 | 3.3e-08 | 2.633E-08 | 5.859E-09 | 2.200E-08 | 3.300E-08 |
| 19520 | 976 | G976V | C | 2.9e-08 | 2.3e-08 | 2.1e-08 | 2.433E-08 | 4.163E-09 | 2.100E-08 | 2.900E-08 |
| 19521 | 977 | T977A | C | 2.5e-08 | 2.4e-08 | 9.7e-08 | 4.867E-08 | 4.186E-08 | 2.400E-08 | 9.700E-08 |
| 19522 | 977 | T977R | C | 3.5e-08 | 2.5e-08 | 6.2e-08 | 4.067E-08 | 1.914E-08 | 2.500E-08 | 6.200E-08 |
| 19523 | 977 | T977N | C | 3.5e-08 | 3.2e-08 | 4.3e-08 | 3.667E-08 | 5.686E-09 | 3.200E-08 | 4.300E-08 |
| 19524 | 977 | T977D | C | 4.5e-08 | 6.7e-08 | 6.1e-08 | 5.767E-08 | 1.137E-08 | 4.500E-08 | 6.700E-08 |
| 19525 | 977 | T977C | C | 4.2e-08 | 8.4e-08 | 4.1e-08 | 5.567E-08 | 2.454E-08 | 4.100E-08 | 8.400E-08 |
| 19526 | 977 | T977E | C | 1.6e-08 | 4.5e-08 | 3.4e-08 | 3.167E-08 | 1.464E-08 | 1.600E-08 | 4.500E-08 |
| 19527 | 977 | T977Q | C | 4.6e-08 | 2.3e-08 | 4.9e-08 | 3.933E-08 | 1.422E-08 | 2.300E-08 | 4.900E-08 |
| 19528 | 977 | T977G | C | 6.2e-08 | 1e-07 | 4.3e-08 | 6.833E-08 | 2.902E-08 | 4.300E-08 | 1.000E-07 |
| 19529 | 977 | T977H | C | 4.8e-08 | 3.4e-08 | 5.5e-08 | 4.567E-08 | 1.069E-08 | 3.400E-08 | 5.500E-08 |
| 19530 | 977 | T977I | C | 1.3e-07 | 2.5e-08 | 4.5e-08 | 6.667E-08 | 5.575E-08 | 2.500E-08 | 1.300E-07 |
| 19531 | 977 | T977L | C | 4.6e-08 | 3.9e-08 | 3.6e-08 | 4.033E-08 | 5.132E-09 | 3.600E-08 | 4.600E-08 |
| 19532 | 977 | T977K | C | 5.8e-08 | 3.9e-08 | 1.6e-08 | 3.767E-08 | 2.103E-08 | 1.600E-08 | 5.800E-08 |
| 19533 | 977 | T977M | C | 5.5e-08 | 2.6e-08 | 6.1e-08 | 4.733E-08 | 1.872E-08 | 2.600E-08 | 6.100E-08 |
| 19534 | 977 | T977F | C | 9e-08 | 3.3e-08 | 9.1e-08 | 7.133E-08 | 3.320E-08 | 3.300E-08 | 9.100E-08 |
| 19535 | 977 | T977P | C | 6.8e-08 | 7.7e-08 | 4.7e-08 | 6.400E-08 | 1.539E-08 | 4.700E-08 | 7.700E-08 |
| 19536 | 977 | T977S | C | 1.6e-08 | 2.8e-08 | 8.1e-08 | 4.167E-08 | 3.459E-08 | 1.600E-08 | 8.100E-08 |
| 19537 | 977 | T977T | C | 5.7e-08 | 2.7e-08 | 2.6e-08 | 3.667E-08 | 1.762E-08 | 2.600E-08 | 5.700E-08 |
| 19538 | 977 | T977W | C | 5.7e-08 | 3.3e-08 | 5e-08 | 4.667E-08 | 1.234E-08 | 3.300E-08 | 5.700E-08 |
| 19539 | 977 | T977Y | C | 2.4e-08 | 6.6e-08 | 5.9e-08 | 4.967E-08 | 2.250E-08 | 2.400E-08 | 6.600E-08 |
| 19540 | 977 | T977V | C | 2.5e-08 | 3.2e-08 | 3.8e-08 | 3.167E-08 | 6.506E-09 | 2.500E-08 | 3.800E-08 |
| 19541 | 978 | A978A | C | 4.2e-08 | 3.1e-08 | 6.7e-08 | 4.667E-08 | 1.845E-08 | 3.100E-08 | 6.700E-08 |
| 19542 | 978 | A978R | C | 4.6e-08 | 4.7e-08 | 2e-08 | 3.767E-08 | 1.531E-08 | 2.000E-08 | 4.700E-08 |
| 19543 | 978 | A978N | C | 7.2e-08 | 1.4e-08 | 3.6e-08 | 4.067E-08 | 2.928E-08 | 1.400E-08 | 7.200E-08 |
| 19544 | 978 | A978D | C | 5.4e-08 | 3.8e-08 | 7.9e-08 | 5.700E-08 | 2.066E-08 | 3.800E-08 | 7.900E-08 |
| 19545 | 978 | A978C | C | 3.2e-08 | 2.1e-08 | 5.8e-08 | 3.700E-08 | 1.900E-08 | 2.100E-08 | 5.800E-08 |
| 19546 | 978 | A978E | C | 1.1e-07 | 6.7e-08 | 4.7e-08 | 7.467E-08 | 3.219E-08 | 4.700E-08 | 1.100E-07 |
| 19547 | 978 | A978Q | C | 9.3e-08 | 3e-08 | 7.5e-08 | 6.600E-08 | 3.245E-08 | 3.000E-08 | 9.300E-08 |
| 19548 | 978 | A978G | C | 4.1e-08 | 3.3e-08 | 2.7e-08 | 3.367E-08 | 7.024E-09 | 2.700E-08 | 4.100E-08 |
| 19549 | 978 | A978H | C | 4.7e-08 | 9.7e-08 | 2.9e-08 | 5.767E-08 | 3.523E-08 | 2.900E-08 | 9.700E-08 |

| | | | | | | | | | | |
|---|---|---|---|---|---|---|---|---|---|---|
| 19550 | 978 | A978I | C | 8.1e-08 | 3.3e-08 | 4.1e-08 | 5.167E-08 | 2.572E-08 | 3.300E-08 | 8.100E-08 |
| 19551 | 978 | A978L | C | 2.5e-08 | 4.5e-08 | 9.9e-09 | 2.663E-08 | 1.761E-08 | 9.900E-09 | 4.500E-08 |
| 19552 | 978 | A978K | C | 6e-08 | 7.9e-08 | 1e-07 | 7.967E-08 | 2.001E-08 | 6.000E-08 | 1.000E-07 |
| 19553 | 978 | A978M | C | 2.3e-08 | 4.4e-08 | 2.5e-08 | 3.067E-08 | 1.159E-08 | 2.300E-08 | 4.400E-08 |
| 19554 | 978 | A978F | C | 9.8e-08 | 1.3e-08 | 9.3e-08 | 6.800E-08 | 4.770E-08 | 1.300E-08 | 9.800E-08 |
| 19555 | 978 | A978P | C | 2.7e-08 | 4.2e-08 | 1.6e-08 | 2.833E-08 | 1.305E-08 | 1.600E-08 | 4.200E-08 |
| 19556 | 978 | A978S | C | 3.5e-08 | 2.3e-08 | 9.6e-09 | 2.253E-08 | 1.271E-08 | 9.600E-09 | 3.500E-08 |
| 19557 | 978 | A978T | C | 5.6e-08 | 4.6e-08 | 5.2e-08 | 5.133E-08 | 5.033E-09 | 4.600E-08 | 5.600E-08 |
| 19558 | 978 | A978W | C | 4.5e-08 | 4.3e-08 | 2.9e-08 | 3.900E-08 | 8.718E-09 | 2.900E-08 | 4.500E-08 |
| 19559 | 978 | A978Y | C | 2.3e-08 | 5.6e-08 | 4.6e-08 | 4.167E-08 | 1.692E-08 | 2.300E-08 | 5.600E-08 |
| 19560 | 978 | A978V | C | 3.1e-08 | 6.6e-08 | 5e-08 | 4.900E-08 | 1.752E-08 | 3.100E-08 | 6.600E-08 |
| 19561 | 979 | A979A | C | 3.8e-08 | 1.3e-07 | 6.2e-08 | 7.667E-08 | 4.772E-08 | 3.800E-08 | 1.300E-07 |
| 19562 | 979 | A979R | C | 5.2e-08 | 4.6e-08 | 1.6e-08 | 3.800E-08 | 1.929E-08 | 1.600E-08 | 5.200E-08 |
| 19563 | 979 | A979N | C | 5.2e-08 | 1.7e-08 | 2.7e-08 | 3.200E-08 | 1.803E-08 | 1.700E-08 | 5.200E-08 |
| 19564 | 979 | A979D | C | 4.9e-08 | 4e-08 | 3.1e-08 | 4.000E-08 | 9.000E-09 | 3.100E-08 | 4.900E-08 |
| 19565 | 979 | A979C | C | 1.2e-07 | 3.4e-08 | 5.1e-08 | 6.833E-08 | 4.554E-08 | 3.400E-08 | 1.200E-07 |
| 19566 | 979 | A979E | C | 3.6e-08 | 3.4e-08 | 9.5e-08 | 5.500E-08 | 3.466E-08 | 3.400E-08 | 9.500E-08 |
| 19567 | 979 | A979Q | C | 1.2e-07 | 1e-08 | 7.7e-08 | 6.900E-08 | 5.543E-08 | 1.000E-08 | 1.200E-07 |
| 19568 | 979 | A979G | C | 4.8e-08 | 2.5e-08 | 4e-08 | 3.767E-08 | 1.168E-08 | 2.500E-08 | 4.800E-08 |
| 19569 | 979 | A979H | C | 1.5e-08 | 5.2e-08 | 4.9e-08 | 3.867E-08 | 2.055E-08 | 1.500E-08 | 5.200E-08 |
| 19570 | 979 | A979I | C | 3.7e-08 | 4.4e-08 | 2.3e-08 | 3.467E-08 | 1.069E-08 | 2.300E-08 | 4.400E-08 |
| 19571 | 979 | A979L | C | 2e-08 | 4.2e-08 | 5.7e-08 | 3.967E-08 | 1.861E-08 | 2.000E-08 | 5.700E-08 |
| 19572 | 979 | A979K | C | 4.8e-08 | 3.3e-08 | 3.3e-08 | 3.800E-08 | 8.660E-09 | 3.300E-08 | 4.800E-08 |
| 19573 | 979 | A979M | C | 7e-08 | 3.3e-08 | 2.9e-08 | 4.400E-08 | 2.261E-08 | 2.900E-08 | 7.000E-08 |
| 19574 | 979 | A979F | C | 5.7e-08 | 1.4e-08 | 5.2e-08 | 4.100E-08 | 2.352E-08 | 1.400E-08 | 5.700E-08 |
| 19575 | 979 | A979P | C | 6.3e-08 | 8.3e-08 | 6.3e-08 | 6.967E-08 | 1.155E-08 | 6.300E-08 | 8.300E-08 |
| 19576 | 979 | A979S | C | 2.7e-08 | 5.5e-08 | 2.1e-08 | 3.433E-08 | 1.815E-08 | 2.100E-08 | 5.500E-08 |
| 19577 | 979 | A979T | C | 1.9e-08 | 3.6e-08 | 6.8e-08 | 4.100E-08 | 2.488E-08 | 1.900E-08 | 6.800E-08 |
| 19578 | 979 | A979W | C | 3.3e-08 | 2.1e-08 | 2.9e-08 | 2.767E-08 | 6.110E-09 | 2.100E-08 | 3.300E-08 |
| 19579 | 979 | A979Y | C | 6.4e-08 | 3.9e-08 | 2.8e-08 | 4.367E-08 | 1.845E-08 | 2.800E-08 | 6.400E-08 |
| 19580 | 979 | A979V | C | 3.3e-08 | 1.6e-08 | 1.1e-08 | 2.000E-08 | 1.153E-08 | 1.100E-08 | 3.300E-08 |
| 19581 | 980 | L980A | C | 1.8e-08 | 6.3e-08 | 8.7e-08 | 5.600E-08 | 3.503E-08 | 1.800E-08 | 8.700E-08 |
| 19582 | 980 | L980R | C | 1.9e-08 | 8.1e-08 | 2.9e-08 | 4.300E-08 | 3.329E-08 | 1.900E-08 | 8.100E-08 |
| 19583 | 980 | L980N | C | 3.1e-08 | 4.7e-08 | 5.6e-08 | 4.467E-08 | 1.266E-08 | 3.100E-08 | 5.600E-08 |
| 19584 | 980 | L980D | C | 4.3e-08 | 6.1e-08 | 8.3e-08 | 6.233E-08 | 2.003E-08 | 4.300E-08 | 8.300E-08 |
| 19585 | 980 | L980C | C | 4e-08 | 6.6e-08 | 2.3e-08 | 4.300E-08 | 2.166E-08 | 2.300E-08 | 6.600E-08 |
| 19586 | 980 | L980E | C | 7.8e-08 | 2.1e-08 | 3.9e-08 | 4.600E-08 | 2.914E-08 | 2.100E-08 | 7.800E-08 |
| 19587 | 980 | L980Q | C | 3.2e-08 | 4.3e-08 | 3.7e-08 | 3.733E-08 | 5.508E-09 | 3.200E-08 | 4.300E-08 |
| 19588 | 980 | L980G | C | 6.1e-08 | 6.8e-08 | 4.2e-08 | 5.700E-08 | 1.345E-08 | 4.200E-08 | 6.800E-08 |
| 19589 | 980 | L980H | C | 5e-08 | 4.3e-08 | 2.9e-08 | 4.067E-08 | 1.069E-08 | 2.900E-08 | 5.000E-08 |
| 19590 | 980 | L980I | C | 3.8e-08 | 8.5e-08 | 9.2e-08 | 7.167E-08 | 2.937E-08 | 3.800E-08 | 9.200E-08 |
| 19591 | 980 | L980L | C | 4.1e-08 | 7.2e-08 | 6e-08 | 5.767E-08 | 1.563E-08 | 4.100E-08 | 7.200E-08 |
| 19592 | 980 | L980K | C | 3e-08 | 4e-08 | 3.1e-08 | 3.367E-08 | 5.508E-09 | 3.000E-08 | 4.000E-08 |
| 19593 | 980 | L980M | C | 2.4e-08 | 4.1e-08 | 7.7e-08 | 4.733E-08 | 2.706E-08 | 2.400E-08 | 7.700E-08 |
| 19594 | 980 | L980F | C | 8.5e-08 | 8.4e-08 | 3.3e-08 | 6.733E-08 | 2.974E-08 | 3.300E-08 | 8.500E-08 |
| 19595 | 980 | L980P | C | 5.4e-08 | 3.7e-08 | 6.6e-08 | 5.233E-08 | 1.457E-08 | 3.700E-08 | 6.600E-08 |
| 19596 | 980 | L980S | C | 2.4e-08 | 1.4e-08 | 8.9e-08 | 4.233E-08 | 4.072E-08 | 1.400E-08 | 8.900E-08 |
| 19597 | 980 | L980T | C | 3.7e-08 | 1.6e-08 | 2.6e-08 | 2.633E-08 | 1.050E-08 | 1.600E-08 | 3.700E-08 |
| 19598 | 980 | L980W | C | 3.1e-08 | 5.1e-08 | 9.6e-08 | 5.933E-08 | 3.329E-08 | 3.100E-08 | 9.600E-08 |
| 19599 | 980 | L980Y | C | 5.5e-08 | 1.9e-08 | 2.4e-08 | 3.267E-08 | 1.950E-08 | 1.900E-08 | 5.500E-08 |
| 19600 | 980 | L980V | C | 4.6e-08 | 2.8e-08 | 2e-08 | 3.133E-08 | 1.332E-08 | 2.000E-08 | 4.600E-08 |
| 19601 | 981 | G981A | C | 5.1e-08 | 2.1e-07 | 7.3e-08 | 1.113E-07 | 8.615E-08 | 5.100E-08 | 2.100E-07 |
| 19602 | 981 | G981R | C | 2.1e-08 | 2.1e-08 | 7.7e-09 | 1.657E-08 | 7.679E-09 | 7.700E-09 | 2.100E-08 |
| 19603 | 981 | G981N | C | 2.9e-08 | 4.9e-08 | 7.9e-09 | 2.863E-08 | 2.055E-08 | 7.900E-09 | 4.900E-08 |
| 19604 | 981 | G981D | C | 5e-08 | 7.9e-08 | 2.2e-08 | 5.033E-08 | 2.850E-08 | 2.200E-08 | 7.900E-08 |
| 19605 | 981 | G981C | C | 9.9e-08 | 1.1e-07 | 2.9e-08 | 7.933E-08 | 4.394E-08 | 2.900E-08 | 1.100E-07 |
| 19606 | 981 | G981E | C | 2.4e-08 | 7.3e-08 | 3.6e-08 | 4.433E-08 | 2.554E-08 | 2.400E-08 | 7.300E-08 |
| 19607 | 981 | G981Q | C | 5.2e-08 | 2.5e-08 | 2.7e-08 | 3.467E-08 | 1.504E-08 | 2.500E-08 | 5.200E-08 |
| 19608 | 981 | G981G | C | 3.8e-08 | 7e-08 | 2.2e-08 | 4.333E-08 | 2.444E-08 | 2.200E-08 | 7.000E-08 |
| 19609 | 981 | G981H | C | 3e-08 | 3.5e-08 | 5.5e-08 | 4.000E-08 | 1.323E-08 | 3.000E-08 | 5.500E-08 |
| 19610 | 981 | G981I | C | 4.1e-08 | 3.2e-08 | 4e-08 | 3.767E-08 | 4.933E-09 | 3.200E-08 | 4.100E-08 |
| 19611 | 981 | G981L | C | 4.6e-08 | 4.1e-08 | 2.1e-08 | 3.600E-08 | 1.323E-08 | 2.100E-08 | 4.600E-08 |
| 19612 | 981 | G981K | C | 4e-08 | 5.8e-08 | 3.2e-08 | 4.333E-08 | 1.332E-08 | 3.200E-08 | 5.800E-08 |
| 19613 | 981 | G981M | C | 2.3e-08 | 4e-08 | 1.1e-07 | 5.767E-08 | 4.611E-08 | 2.300E-08 | 1.100E-07 |
| 19614 | 981 | G981F | C | 1.3e-07 | 5.3e-08 | 8.1e-08 | 8.800E-08 | 3.897E-08 | 5.300E-08 | 1.300E-07 |
| 19615 | 981 | G981P | C | 5.2e-08 | 9.4e-08 | 1.1e-08 | 5.233E-08 | 4.150E-08 | 1.100E-08 | 9.400E-08 |
| 19616 | 981 | G981S | C | 3.4e-08 | 3.5e-08 | 9.5e-08 | 5.467E-08 | 3.493E-08 | 3.400E-08 | 9.500E-08 |
| 19617 | 981 | G981T | C | 2.5e-08 | 1e-07 | 4e-08 | 5.500E-08 | 3.969E-08 | 2.500E-08 | 1.000E-07 |
| 19618 | 981 | G981W | C | 6.9e-08 | 1.8e-08 | 2.1e-08 | 3.600E-08 | 2.862E-08 | 1.800E-08 | 6.900E-08 |
| 19619 | 981 | G981Y | C | 2.7e-08 | 5.4e-08 | 5.1e-08 | 4.400E-08 | 1.480E-08 | 2.700E-08 | 5.400E-08 |
| 19620 | 981 | G981V | C | 3.5e-08 | 4.5e-08 | 5e-08 | 4.333E-08 | 7.638E-09 | 3.500E-08 | 5.000E-08 |
| 19621 | 982 | C982A | C | 2e-08 | 2.7e-08 | 9.3e-08 | 4.667E-08 | 4.028E-08 | 2.000E-08 | 9.300E-08 |
| 19622 | 982 | C982R | C | 5.1e-08 | 3.3e-08 | 2.7e-08 | 3.700E-08 | 1.249E-08 | 2.700E-08 | 5.100E-08 |
| 19623 | 982 | C982N | C | 2.6e-08 | 3.3e-08 | 3.9e-08 | 3.267E-08 | 6.506E-09 | 2.600E-08 | 3.900E-08 |
| 19624 | 982 | C982D | C | 6.7e-08 | 2.2e-08 | 1.5e-07 | 7.967E-08 | 6.493E-08 | 2.200E-08 | 1.500E-07 |
| 19625 | 982 | C982C | C | 5.9e-08 | 4.5e-08 | 6.6e-08 | 5.667E-08 | 1.069E-08 | 4.500E-08 | 6.600E-08 |
| 19626 | 982 | C982E | C | 8.8e-08 | 3.1e-08 | 4.2e-08 | 5.367E-08 | 3.024E-08 | 3.100E-08 | 8.800E-08 |
| 19627 | 982 | C982Q | C | 1.3e-08 | 2.6e-08 | 1e-07 | 4.633E-08 | 4.693E-08 | 1.300E-08 | 1.000E-07 |
| 19628 | 982 | C982G | C | 2.5e-08 | 3.7e-08 | 3.4e-08 | 3.200E-08 | 6.245E-09 | 2.500E-08 | 3.700E-08 |
| 19629 | 982 | C982H | C | 1.6e-08 | 6e-08 | 3.9e-08 | 3.833E-08 | 2.201E-08 | 1.600E-08 | 6.000E-08 |
| 19630 | 982 | C982I | C | 1.4e-07 | 3.7e-08 | 5.4e-08 | 7.700E-08 | 5.522E-08 | 3.700E-08 | 1.400E-07 |
| 19631 | 982 | C982L | C | 6.6e-08 | 8.2e-08 | 5.5e-08 | 6.767E-08 | 1.358E-08 | 5.500E-08 | 8.200E-08 |
| 19632 | 982 | C982K | C | 4.4e-08 | 2.1e-08 | 6.6e-08 | 4.367E-08 | 2.250E-08 | 2.100E-08 | 6.600E-08 |
| 19633 | 982 | C982M | C | 5e-08 | 5.3e-08 | 3.8e-08 | 4.700E-08 | 7.937E-09 | 3.800E-08 | 5.300E-08 |
| 19634 | 982 | C982F | C | 9.5e-08 | 7.1e-08 | 1.8e-08 | 6.133E-08 | 3.940E-08 | 1.800E-08 | 9.500E-08 |

| | | | | | | | | | | |
|---|---|---|---|---|---|---|---|---|---|---|
| 19635 | 982 | C982P | C | 2.1e-08 | 5e-08 | 4.9e-08 | 4.000E-08 | 1.646E-08 | 2.100E-08 | 5.000E-08 |
| 19636 | 982 | C982S | C | 3.7e-08 | 3.1e-08 | 3.5e-08 | 3.433E-08 | 3.055E-09 | 3.100E-08 | 3.700E-08 |
| 19637 | 982 | C982T | C | 2.8e-07 | 1.9e-08 | 1.3e-07 | 1.430E-07 | 1.310E-07 | 1.900E-08 | 2.800E-07 |
| 19638 | 982 | C982W | C | 7.1e-08 | 5.6e-08 | 6.7e-08 | 6.467E-08 | 7.767E-09 | 5.600E-08 | 7.100E-08 |
| 19639 | 982 | C982Y | C | 3.3e-08 | 3e-08 | 4.2e-08 | 3.500E-08 | 6.245E-09 | 3.000E-08 | 4.200E-08 |
| 19640 | 982 | C982V | C | 1.5e-07 | 7.9e-08 | 1.2e-07 | 1.163E-07 | 3.564E-08 | 7.900E-08 | 1.500E-07 |
| 19641 | 983 | L983A | C | 3.5e-08 | 2.6e-08 | 5.1e-08 | 3.733E-08 | 1.266E-08 | 2.600E-08 | 5.100E-08 |
| 19642 | 983 | L983R | C | 4.7e-08 | 5.6e-08 | 5.7e-08 | 5.333E-08 | 5.508E-09 | 4.700E-08 | 5.700E-08 |
| 19643 | 983 | L983N | C | 6.1e-08 | 7.3e-08 | 1.2e-07 | 8.467E-08 | 3.118E-08 | 6.100E-08 | 1.200E-07 |
| 19644 | 983 | L983D | C | 2e-08 | 5.4e-08 | 2.8e-08 | 3.400E-08 | 1.778E-08 | 2.000E-08 | 5.400E-08 |
| 19645 | 983 | L983C | C | 2.7e-08 | 6.3e-08 | 2.5e-08 | 3.833E-08 | 2.139E-08 | 2.500E-08 | 6.300E-08 |
| 19646 | 983 | L983E | C | 2.3e-08 | 4.2e-08 | 6.2e-08 | 4.233E-08 | 1.950E-08 | 2.300E-08 | 6.200E-08 |
| 19647 | 983 | L983Q | C | 8.5e-08 | 2.9e-08 | 2.8e-08 | 4.733E-08 | 3.262E-08 | 2.800E-08 | 8.500E-08 |
| 19648 | 983 | L983G | C | 3.1e-08 | 7.6e-08 | 2.3e-08 | 4.333E-08 | 2.857E-08 | 2.300E-08 | 7.600E-08 |
| 19649 | 983 | L983H | C | 5.3e-08 | 3.7e-08 | 1.4e-08 | 3.467E-08 | 1.960E-08 | 1.400E-08 | 5.300E-08 |
| 19650 | 983 | L983I | C | 2.6e-08 | 2.3e-08 | 3.6e-08 | 2.833E-08 | 6.807E-09 | 2.300E-08 | 3.600E-08 |
| 19651 | 983 | L983L | C | 4.2e-08 | 7.4e-08 | 1.2e-08 | 4.267E-08 | 3.101E-08 | 1.200E-08 | 7.400E-08 |
| 19652 | 983 | L983K | C | 2.9e-08 | 3.3e-08 | 3.5e-08 | 3.233E-08 | 3.055E-09 | 2.900E-08 | 3.500E-08 |
| 19653 | 983 | L983M | C | 1.7e-08 | 4.7e-08 | 4.1e-08 | 3.500E-08 | 1.587E-08 | 1.700E-08 | 4.700E-08 |
| 19654 | 983 | L983F | C | 1e-07 | 1.7e-08 | 1.4e-08 | 4.367E-08 | 4.881E-08 | 1.400E-08 | 1.000E-07 |
| 19655 | 983 | L983P | C | 2.1e-08 | 3.3e-08 | 6.5e-08 | 3.967E-08 | 2.274E-08 | 2.100E-08 | 6.500E-08 |
| 19656 | 983 | L983S | C | 4.6e-08 | 8.1e-08 | 1.4e-07 | 8.900E-08 | 4.751E-08 | 4.600E-08 | 1.400E-07 |
| 19657 | 983 | L983T | C | 7.2e-08 | 1e-07 | 8.2e-08 | 8.467E-08 | 1.419E-08 | 7.200E-08 | 1.000E-07 |
| 19658 | 983 | L983W | C | 1.9e-08 | 1.3e-07 | 1.9e-08 | 5.600E-08 | 6.409E-08 | 1.900E-08 | 1.300E-07 |
| 19659 | 983 | L983Y | C | 4.3e-08 | 2e-08 | 4.7e-08 | 3.667E-08 | 1.457E-08 | 2.000E-08 | 4.700E-08 |
| 19660 | 983 | L983V | C | 3.1e-08 | 5.2e-08 | 2.9e-08 | 3.733E-08 | 1.274E-08 | 2.900E-08 | 5.200E-08 |
| 19661 | 984 | V984A | C | 9.3e-08 | 3.9e-08 | 1.2e-07 | 8.400E-08 | 4.124E-08 | 3.900E-08 | 1.200E-07 |
| 19662 | 984 | V984R | C | 3.1e-08 | 5.7e-08 | 2.5e-08 | 3.767E-08 | 1.701E-08 | 2.500E-08 | 5.700E-08 |
| 19663 | 984 | V984N | C | 5.7e-08 | 2.9e-08 | 5.3e-08 | 4.633E-08 | 1.514E-08 | 2.900E-08 | 5.700E-08 |
| 19664 | 984 | V984D | C | 5.1e-08 | 3.8e-08 | 4.9e-08 | 4.600E-08 | 7.000E-09 | 3.800E-08 | 5.100E-08 |
| 19665 | 984 | V984C | C | 9.9e-08 | 2.3e-08 | 2.6e-08 | 4.933E-08 | 4.304E-08 | 2.300E-08 | 9.900E-08 |
| 19666 | 984 | V984E | C | 4.6e-08 | 4.3e-08 | 3.8e-08 | 4.233E-08 | 4.041E-09 | 3.800E-08 | 4.600E-08 |
| 19667 | 984 | V984Q | C | 7.2e-08 | 6.2e-08 | 1.8e-08 | 5.067E-08 | 2.873E-08 | 1.800E-08 | 7.200E-08 |
| 19668 | 984 | V984G | C | 4e-08 | 3.9e-08 | 6.5e-08 | 4.800E-08 | 1.473E-08 | 3.900E-08 | 6.500E-08 |
| 19669 | 984 | V984H | C | 5.2e-08 | 4.9e-08 | 2.2e-08 | 4.100E-08 | 1.652E-08 | 2.200E-08 | 5.200E-08 |
| 19670 | 984 | V984I | C | 3.3e-08 | 3.9e-08 | 8.3e-08 | 5.167E-08 | 2.730E-08 | 3.300E-08 | 8.300E-08 |
| 19671 | 984 | V984L | C | 8.2e-08 | 4.9e-08 | 8.2e-08 | 7.100E-08 | 1.905E-08 | 4.900E-08 | 8.200E-08 |
| 19672 | 984 | V984K | C | 6.1e-08 | 3.3e-08 | 2.1e-08 | 3.833E-08 | 2.053E-08 | 2.100E-08 | 6.100E-08 |
| 19673 | 984 | V984M | C | 6.1e-08 | 2.5e-08 | 1.2e-07 | 6.867E-08 | 4.796E-08 | 2.500E-08 | 1.200E-07 |
| 19674 | 984 | V984F | C | 7.1e-08 | 3.2e-08 | 5e-08 | 5.100E-08 | 1.952E-08 | 3.200E-08 | 7.100E-08 |
| 19675 | 984 | V984P | C | 2.4e-08 | 3.3e-08 | 5.8e-08 | 3.833E-08 | 1.762E-08 | 2.400E-08 | 5.800E-08 |
| 19676 | 984 | V984S | C | 3.1e-08 | 3.2e-08 | 3.1e-08 | 3.133E-08 | 5.774E-10 | 3.100E-08 | 3.200E-08 |
| 19677 | 984 | V984T | C | 3.4e-08 | 3.4e-08 | 3.2e-08 | 3.333E-08 | 1.155E-09 | 3.200E-08 | 3.400E-08 |
| 19678 | 984 | V984W | C | 1.5e-07 | 1.1e-07 | 4.9e-08 | 1.030E-07 | 5.086E-08 | 4.900E-08 | 1.500E-07 |
| 19679 | 984 | V984Y | C | 8.8e-08 | 3.4e-08 | 4.3e-08 | 5.500E-08 | 2.893E-08 | 3.400E-08 | 8.800E-08 |
| 19680 | 984 | V984V | C | 8.1e-08 | 3.1e-08 | 2.6e-08 | 4.600E-08 | 3.041E-08 | 2.600E-08 | 8.100E-08 |
| 19681 | 985 | K985A | C | 4.3e-08 | 5e-08 | 3e-08 | 4.100E-08 | 1.015E-08 | 3.000E-08 | 5.000E-08 |
| 19682 | 985 | K985R | C | 5.5e-08 | 3.9e-08 | 6e-08 | 5.133E-08 | 1.097E-08 | 3.900E-08 | 6.000E-08 |
| 19683 | 985 | K985N | C | 6.8e-08 | 3.4e-08 | 4.2e-08 | 4.800E-08 | 1.778E-08 | 3.400E-08 | 6.800E-08 |
| 19684 | 985 | K985D | C | 3.7e-08 | 5.5e-08 | 3e-08 | 4.067E-08 | 1.290E-08 | 3.000E-08 | 5.500E-08 |
| 19685 | 985 | K985C | C | 4e-08 | 3.4e-08 | 1.6e-08 | 3.000E-08 | 1.249E-08 | 1.600E-08 | 4.000E-08 |
| 19686 | 985 | K985E | C | 5.6e-08 | 4.7e-08 | 4.6e-08 | 4.967E-08 | 5.508E-09 | 4.600E-08 | 5.600E-08 |
| 19687 | 985 | K985Q | C | 2.9e-08 | 2.7e-08 | 3.6e-08 | 3.067E-08 | 4.726E-09 | 2.700E-08 | 3.600E-08 |
| 19688 | 985 | K985G | C | 1.1e-07 | 9.3e-08 | 3.3e-08 | 7.867E-08 | 4.045E-08 | 3.300E-08 | 1.100E-07 |
| 19689 | 985 | K985H | C | 7.5e-08 | 6.6e-08 | 3.8e-08 | 5.967E-08 | 1.930E-08 | 3.800E-08 | 7.500E-08 |
| 19690 | 985 | K985I | C | 3.7e-08 | 3.8e-08 | 2.2e-08 | 3.233E-08 | 8.963E-09 | 2.200E-08 | 3.800E-08 |
| 19691 | 985 | K985L | C | 1.1e-07 | 2.7e-08 | 3.2e-08 | 5.633E-08 | 4.654E-08 | 2.700E-08 | 1.100E-07 |
| 19692 | 985 | K985K | C | 5.2e-08 | 4.6e-08 | 1.1e-07 | 6.933E-08 | 3.535E-08 | 4.600E-08 | 1.100E-07 |
| 19693 | 985 | K985M | C | 2.4e-08 | 2.6e-08 | 9e-08 | 4.667E-08 | 3.754E-08 | 2.400E-08 | 9.000E-08 |
| 19694 | 985 | K985F | C | 5.2e-08 | 6.4e-08 | 4.8e-08 | 5.467E-08 | 8.327E-09 | 4.800E-08 | 6.400E-08 |
| 19695 | 985 | K985P | C | 2.3e-08 | 4.3e-08 | 2.3e-08 | 2.967E-08 | 1.155E-08 | 2.300E-08 | 4.300E-08 |
| 19696 | 985 | K985S | C | 3.1e-08 | 5.3e-08 | 2.9e-08 | 3.767E-08 | 1.332E-08 | 2.900E-08 | 5.300E-08 |
| 19697 | 985 | K985T | C | 6e-08 | 4.4e-08 | 1.8e-08 | 4.067E-08 | 2.120E-08 | 1.800E-08 | 6.000E-08 |
| 19698 | 985 | K985W | C | 1.4e-08 | 2.1e-08 | 1.1e-08 | 1.533E-08 | 5.132E-09 | 1.100E-08 | 2.100E-08 |
| 19699 | 985 | K985Y | C | 7.3e-08 | 2.4e-08 | 5.2e-08 | 4.967E-08 | 2.458E-08 | 2.400E-08 | 7.300E-08 |
| 19700 | 985 | K985V | C | 3.4e-08 | 3.5e-08 | 8.4e-08 | 5.100E-08 | 2.858E-08 | 3.400E-08 | 8.400E-08 |
| 19701 | 986 | D986A | C | 4.3e-08 | 8e-08 | 5.6e-08 | 5.967E-08 | 1.877E-08 | 4.300E-08 | 8.000E-08 |
| 19702 | 986 | D986R | C | 3.5e-08 | 2.5e-08 | 8.8e-08 | 4.933E-08 | 3.386E-08 | 2.500E-08 | 8.800E-08 |
| 19703 | 986 | D986N | C | 7.9e-08 | 1.2e-08 | 9.5e-08 | 6.200E-08 | 4.403E-08 | 1.200E-08 | 9.500E-08 |
| 19704 | 986 | D986D | C | 3.1e-08 | 9.4e-08 | 6.1e-08 | 6.200E-08 | 3.151E-08 | 3.100E-08 | 9.400E-08 |
| 19705 | 986 | D986C | C | 1.2e-08 | 3.4e-08 | 4.4e-08 | 3.000E-08 | 1.637E-08 | 1.200E-08 | 4.400E-08 |
| 19706 | 986 | D986E | C | 8.5e-08 | 6.4e-08 | 6.7e-08 | 7.200E-08 | 1.136E-08 | 6.400E-08 | 8.500E-08 |
| 19707 | 986 | D986Q | C | 4.5e-08 | 3.7e-08 | 7.8e-08 | 5.333E-08 | 2.173E-08 | 3.700E-08 | 7.800E-08 |
| 19708 | 986 | D986G | C | 5.6e-08 | 1.2e-08 | 2.3e-08 | 3.033E-08 | 2.290E-08 | 1.200E-08 | 5.600E-08 |
| 19709 | 986 | D986H | C | 1.2e-07 | 8.9e-08 | 2.8e-08 | 7.900E-08 | 4.681E-08 | 2.800E-08 | 1.200E-07 |
| 19710 | 986 | D986I | C | 5.6e-08 | 3.1e-08 | 1.4e-08 | 3.367E-08 | 2.113E-08 | 1.400E-08 | 5.600E-08 |
| 19711 | 986 | D986L | C | 3.4e-08 | 1.1e-07 | 6.1e-08 | 6.833E-08 | 3.853E-08 | 3.400E-08 | 1.100E-07 |
| 19712 | 986 | D986K | C | 2.5e-08 | 3.7e-08 | 4.9e-08 | 3.700E-08 | 1.200E-08 | 2.500E-08 | 4.900E-08 |
| 19713 | 986 | D986M | C | 2.1e-08 | 2.2e-08 | 3.7e-08 | 2.667E-08 | 8.963E-09 | 2.100E-08 | 3.700E-08 |
| 19714 | 986 | D986F | C | 3.7e-08 | 2.8e-08 | 4.3e-08 | 3.600E-08 | 7.550E-09 | 2.800E-08 | 4.300E-08 |
| 19715 | 986 | D986P | C | 5.3e-08 | 8.2e-08 | 4.3e-08 | 5.933E-08 | 2.026E-08 | 4.300E-08 | 8.200E-08 |
| 19716 | 986 | D986S | C | 7e-08 | 1.8e-08 | 5.5e-08 | 4.767E-08 | 2.676E-08 | 1.800E-08 | 7.000E-08 |
| 19717 | 986 | D986T | C | 3.2e-08 | 6.3e-08 | 3e-08 | 4.167E-08 | 1.850E-08 | 3.000E-08 | 6.300E-08 |
| 19718 | 986 | D986W | C | 4.5e-08 | 3.5e-08 | 3.9e-08 | 3.967E-08 | 5.033E-09 | 3.500E-08 | 4.500E-08 |
| 19719 | 986 | D986Y | C | 8.5e-08 | 1.2e-07 | 6.5e-08 | 9.000E-08 | 2.784E-08 | 6.500E-08 | 1.200E-07 |

| | | | | | | | | | | |
|---|---|---|---|---|---|---|---|---|---|---|
| 19720 | 986 | D986V | C | 3.6e-08 | 3.2e-08 | 4.7e-08 | 3.833E-08 | 7.767E-09 | 3.200E-08 | 4.700E-08 |
| 19721 | 987 | Y987A | C | 1.9e-08 | 4.4e-08 | 4.5e-08 | 3.600E-08 | 1.473E-08 | 1.900E-08 | 4.500E-08 |
| 19722 | 987 | Y987R | C | 4.7e-08 | 1.1e-07 | 3.6e-08 | 6.433E-08 | 3.993E-08 | 3.600E-08 | 1.100E-07 |
| 19723 | 987 | Y987N | C | 2.5e-08 | 4.3e-08 | 6e-08 | 4.267E-08 | 1.750E-08 | 2.500E-08 | 6.000E-08 |
| 19724 | 987 | Y987D | C | 3.7e-08 | 2.4e-08 | 8.6e-09 | 2.320E-08 | 1.422E-08 | 8.600E-09 | 3.700E-08 |
| 19725 | 987 | Y987C | C | 2.9e-08 | 1.6e-08 | 8.8e-08 | 4.433E-08 | 3.837E-08 | 1.600E-08 | 8.800E-08 |
| 19726 | 987 | Y987E | C | 1.2e-08 | 2.5e-08 | 2.4e-08 | 2.033E-08 | 7.234E-09 | 1.200E-08 | 2.500E-08 |
| 19727 | 987 | Y987Q | C | 1.2e-07 | 5.3e-08 | 3.7e-08 | 7.000E-08 | 4.403E-08 | 3.700E-08 | 1.200E-07 |
| 19728 | 987 | Y987G | C | 7.6e-08 | 5.8e-08 | 5e-08 | 6.133E-08 | 1.332E-08 | 5.000E-08 | 7.600E-08 |
| 19729 | 987 | Y987H | C | 3.5e-08 | 1e-07 | 1.1e-07 | 8.167E-08 | 4.072E-08 | 3.500E-08 | 1.100E-07 |
| 19730 | 987 | Y987I | C | 8.8e-08 | 1.5e-08 | 4.8e-08 | 5.033E-08 | 3.656E-08 | 1.500E-08 | 8.800E-08 |
| 19731 | 987 | Y987L | C | 1.7e-08 | 8.6e-08 | 3.5e-08 | 4.600E-08 | 3.579E-08 | 1.700E-08 | 8.600E-08 |
| 19732 | 987 | Y987K | C | 2.2e-08 | 3e-08 | 5.6e-08 | 3.600E-08 | 1.778E-08 | 2.200E-08 | 5.600E-08 |
| 19733 | 987 | Y987M | C | 4.1e-08 | 1.3e-08 | 1.3e-07 | 6.133E-08 | 6.109E-08 | 1.300E-08 | 1.300E-07 |
| 19734 | 987 | Y987F | C | 3.9e-08 | 5.7e-08 | 5.5e-08 | 5.033E-08 | 9.866E-09 | 3.900E-08 | 5.700E-08 |
| 19735 | 987 | Y987P | C | 5.3e-08 | 9.8e-08 | 3.2e-08 | 6.100E-08 | 3.372E-08 | 3.200E-08 | 9.800E-08 |
| 19736 | 987 | Y987S | C | 6.1e-08 | 1.4e-08 | 1.4e-08 | 2.967E-08 | 2.714E-08 | 1.400E-08 | 6.100E-08 |
| 19737 | 987 | Y987T | C | 4.8e-08 | 3.6e-08 | 1.1e-07 | 6.467E-08 | 3.972E-08 | 3.600E-08 | 1.100E-07 |
| 19738 | 987 | Y987W | C | 6.7e-08 | 2.2e-08 | 4.1e-08 | 4.333E-08 | 2.259E-08 | 2.200E-08 | 6.700E-08 |
| 19739 | 987 | Y987Y | C | 1.8e-08 | 1e-07 | 5.2e-08 | 5.667E-08 | 4.120E-08 | 1.800E-08 | 1.000E-07 |
| 19740 | 987 | Y987V | C | 2e-08 | 2.9e-08 | 5e-08 | 3.300E-08 | 1.539E-08 | 2.000E-08 | 5.000E-08 |
| 19741 | 988 | F988A | C | 2.7e-08 | 6.4e-08 | 2.3e-08 | 3.800E-08 | 2.261E-08 | 2.300E-08 | 6.400E-08 |
| 19742 | 988 | F988R | C | 5.9e-08 | 4.5e-08 | 4.6e-08 | 5.000E-08 | 7.810E-09 | 4.500E-08 | 5.900E-08 |
| 19743 | 988 | F988N | C | 5.1e-08 | 2e-08 | 2e-08 | 3.033E-08 | 1.790E-08 | 2.000E-08 | 5.100E-08 |
| 19744 | 988 | F988D | C | 3.4e-08 | 5.9e-08 | 5.4e-08 | 4.900E-08 | 1.323E-08 | 3.400E-08 | 5.900E-08 |
| 19745 | 988 | F988C | C | 2.5e-08 | 4.7e-08 | 3.1e-08 | 3.433E-08 | 1.137E-08 | 2.500E-08 | 4.700E-08 |
| 19746 | 988 | F988E | C | 2.2e-08 | 3.3e-08 | 2.8e-08 | 2.767E-08 | 5.508E-09 | 2.200E-08 | 3.300E-08 |
| 19747 | 988 | F988Q | C | 8.2e-08 | 2.9e-08 | 8.2e-08 | 6.433E-08 | 3.060E-08 | 2.900E-08 | 8.200E-08 |
| 19748 | 988 | F988G | C | 2.2e-08 | 3e-08 | 1e-07 | 5.067E-08 | 4.291E-08 | 2.200E-08 | 1.000E-07 |
| 19749 | 988 | F988H | C | 2.1e-08 | 8.4e-08 | 4e-08 | 4.833E-08 | 3.232E-08 | 2.100E-08 | 8.400E-08 |
| 19750 | 988 | F988I | C | 1e-07 | 8.1e-08 | 4.6e-08 | 7.567E-08 | 2.739E-08 | 4.600E-08 | 1.000E-07 |
| 19751 | 988 | F988L | C | 5.7e-08 | 3.2e-08 | 7.2e-08 | 5.367E-08 | 2.021E-08 | 3.200E-08 | 7.200E-08 |
| 19752 | 988 | F988K | C | 1.3e-07 | 1.5e-08 | 6.9e-08 | 7.133E-08 | 5.754E-08 | 1.500E-08 | 1.300E-07 |
| 19753 | 988 | F988M | C | 7.7e-08 | 7e-08 | 2.9e-08 | 5.867E-08 | 2.593E-08 | 2.900E-08 | 7.700E-08 |
| 19754 | 988 | F988F | C | 4.9e-08 | 4e-08 | 6.9e-08 | 5.267E-08 | 1.484E-08 | 4.000E-08 | 6.900E-08 |
| 19755 | 988 | F988P | C | 4.6e-08 | 3.5e-08 | 3.6e-08 | 3.900E-08 | 6.083E-09 | 3.500E-08 | 4.600E-08 |
| 19756 | 988 | F988S | C | 6.5e-08 | 4.9e-08 | 2.5e-08 | 4.633E-08 | 2.013E-08 | 2.500E-08 | 6.500E-08 |
| 19757 | 988 | F988T | C | 1.6e-08 | 1.5e-07 | 2.3e-08 | 6.300E-08 | 7.543E-08 | 1.600E-08 | 1.500E-07 |
| 19758 | 988 | F988W | C | 3.6e-08 | 9.8e-08 | 2.7e-08 | 5.367E-08 | 3.866E-08 | 2.700E-08 | 9.800E-08 |
| 19759 | 988 | F988Y | C | 3e-08 | 4.2e-08 | 1.2e-07 | 6.400E-08 | 4.887E-08 | 3.000E-08 | 1.200E-07 |
| 19760 | 988 | F988V | C | 2.8e-08 | 4.1e-08 | 3.8e-08 | 3.567E-08 | 6.807E-09 | 2.800E-08 | 4.100E-08 |
| 19761 | 989 | P989A | C | 7.1e-08 | 3.5e-08 | 1.9e-08 | 4.167E-08 | 2.663E-08 | 1.900E-08 | 7.100E-08 |
| 19762 | 989 | P989R | C | 3.2e-08 | 2.9e-08 | 5.3e-08 | 3.800E-08 | 1.308E-08 | 2.900E-08 | 5.300E-08 |
| 19763 | 989 | P989N | C | 6e-09 | 1.8e-08 | 3.4e-08 | 1.933E-08 | 1.405E-08 | 6.000E-09 | 3.400E-08 |
| 19764 | 989 | P989D | C | 3.6e-08 | 2e-08 | 1.1e-07 | 5.533E-08 | 4.801E-08 | 2.000E-08 | 1.100E-07 |
| 19765 | 989 | P989C | C | 2.9e-08 | 9.3e-08 | 7.2e-08 | 6.467E-08 | 3.262E-08 | 2.900E-08 | 9.300E-08 |
| 19766 | 989 | P989E | C | 7.1e-08 | 1e-07 | 5e-08 | 7.367E-08 | 2.511E-08 | 5.000E-08 | 1.000E-07 |
| 19767 | 989 | P989Q | C | 1.3e-08 | 7.3e-08 | 4.2e-08 | 4.267E-08 | 3.001E-08 | 1.300E-08 | 7.300E-08 |
| 19768 | 989 | P989G | C | 5.3e-08 | 5.3e-08 | 1.2e-08 | 3.933E-08 | 2.367E-08 | 1.200E-08 | 5.300E-08 |
| 19769 | 989 | P989H | C | 5.2e-08 | 1.3e-08 | 5.1e-08 | 3.867E-08 | 2.223E-08 | 1.300E-08 | 5.200E-08 |
| 19770 | 989 | P989I | C | 2.7e-08 | 2.5e-08 | 5.6e-08 | 3.600E-08 | 1.735E-08 | 2.500E-08 | 5.600E-08 |
| 19771 | 989 | P989L | C | 3e-08 | 3.3e-08 | 2.4e-08 | 2.900E-08 | 4.583E-09 | 2.400E-08 | 3.300E-08 |
| 19772 | 989 | P989K | C | 1e-07 | 2.5e-08 | 8.7e-08 | 7.067E-08 | 4.008E-08 | 2.500E-08 | 1.000E-07 |
| 19773 | 989 | P989M | C | 4.4e-08 | 2e-07 | 2.6e-08 | 9.000E-08 | 9.569E-08 | 2.600E-08 | 2.000E-07 |
| 19774 | 989 | P989F | C | 5.7e-08 | 3.5e-08 | 1e-07 | 6.400E-08 | 3.306E-08 | 3.500E-08 | 1.000E-07 |
| 19775 | 989 | P989P | C | 2.7e-08 | 2.5e-08 | 6.3e-08 | 3.833E-08 | 2.139E-08 | 2.500E-08 | 6.300E-08 |
| 19776 | 989 | P989S | C | 7.1e-08 | 2.5e-08 | 6.8e-08 | 5.467E-08 | 2.574E-08 | 2.500E-08 | 7.100E-08 |
| 19777 | 989 | P989T | C | 2.2e-08 | 5.9e-08 | 8.5e-08 | 5.533E-08 | 3.166E-08 | 2.200E-08 | 8.500E-08 |
| 19778 | 989 | P989W | C | 1.8e-08 | 1.9e-07 | 1.7e-07 | 1.260E-07 | 9.406E-08 | 1.800E-08 | 1.900E-07 |
| 19779 | 989 | P989Y | C | 4.3e-08 | 2.7e-08 | 2.2e-08 | 3.067E-08 | 1.097E-08 | 2.200E-08 | 4.300E-08 |
| 19780 | 989 | P989V | C | 1.2e-07 | 4.1e-08 | 4.1e-08 | 6.733E-08 | 4.561E-08 | 4.100E-08 | 1.200E-07 |
| 19781 | 990 | E990A | C | 2.7e-08 | 1.4e-08 | 2.3e-08 | 2.133E-08 | 6.658E-09 | 1.400E-08 | 2.700E-08 |
| 19782 | 990 | E990R | C | 5.6e-08 | 3.2e-08 | 3.1e-08 | 3.967E-08 | 1.415E-08 | 3.100E-08 | 5.600E-08 |
| 19783 | 990 | E990N | C | 4.9e-08 | 5.4e-08 | 2.3e-08 | 4.200E-08 | 1.664E-08 | 2.300E-08 | 5.400E-08 |
| 19784 | 990 | E990D | C | 2.2e-08 | 6.1e-08 | 3.3e-08 | 3.867E-08 | 2.011E-08 | 2.200E-08 | 6.100E-08 |
| 19785 | 990 | E990C | C | 2.2e-08 | 3e-08 | 2.8e-08 | 2.667E-08 | 4.163E-09 | 2.200E-08 | 3.000E-08 |
| 19786 | 990 | E990E | C | 5.8e-08 | 6.7e-08 | 1.5e-08 | 4.667E-08 | 2.779E-08 | 1.500E-08 | 6.700E-08 |
| 19787 | 990 | E990Q | C | 7.4e-08 | 3.2e-08 | 6.9e-08 | 5.833E-08 | 2.294E-08 | 3.200E-08 | 7.400E-08 |
| 19788 | 990 | E990G | C | 3.3e-08 | 4.4e-08 | 4.2e-08 | 3.967E-08 | 5.859E-09 | 3.300E-08 | 4.400E-08 |
| 19789 | 990 | E990H | C | 7.7e-08 | 2.9e-08 | 3e-08 | 4.533E-08 | 2.743E-08 | 2.900E-08 | 7.700E-08 |
| 19790 | 990 | E990I | C | 5.6e-08 | 2.3e-08 | 7.4e-08 | 5.100E-08 | 2.587E-08 | 2.300E-08 | 7.400E-08 |
| 19791 | 990 | E990L | C | 6.7e-08 | 4e-08 | 6.2e-08 | 5.633E-08 | 1.436E-08 | 4.000E-08 | 6.700E-08 |
| 19792 | 990 | E990K | C | 3.5e-08 | 4.4e-08 | 1.6e-07 | 7.967E-08 | 6.972E-08 | 3.500E-08 | 1.600E-07 |
| 19793 | 990 | E990M | C | 3.7e-08 | 1.2e-08 | 6.5e-08 | 3.800E-08 | 2.651E-08 | 1.200E-08 | 6.500E-08 |
| 19794 | 990 | E990F | C | 3.5e-08 | 3.3e-08 | 4.4e-08 | 3.733E-08 | 5.859E-09 | 3.300E-08 | 4.400E-08 |
| 19795 | 990 | E990P | C | 2.8e-08 | 2.8e-08 | 2e-08 | 2.533E-08 | 4.619E-09 | 2.000E-08 | 2.800E-08 |
| 19796 | 990 | E990S | C | 1.7e-08 | 3.8e-08 | 1.3e-08 | 2.267E-08 | 1.343E-08 | 1.300E-08 | 3.800E-08 |
| 19797 | 990 | E990T | C | 1.8e-08 | 6.2e-08 | 6.6e-08 | 4.867E-08 | 2.663E-08 | 1.800E-08 | 6.600E-08 |
| 19798 | 990 | E990W | C | 2.5e-08 | 5.6e-08 | 8.5e-08 | 5.533E-08 | 3.001E-08 | 2.500E-08 | 8.500E-08 |
| 19799 | 990 | E990Y | C | 4.6e-08 | 7.6e-08 | 1.8e-08 | 4.667E-08 | 2.901E-08 | 1.800E-08 | 7.600E-08 |
| 19800 | 990 | E990V | C | 2.9e-08 | 1.6e-08 | 5.6e-08 | 3.367E-08 | 2.040E-08 | 1.600E-08 | 5.600E-08 |
| 19801 | 991 | P991A | C | 3.3e-08 | 1.9e-08 | 2.5e-08 | 2.567E-08 | 7.024E-09 | 1.900E-08 | 3.300E-08 |
| 19802 | 991 | P991R | C | 8.7e-08 | 6.7e-08 | 5.5e-08 | 6.967E-08 | 1.617E-08 | 5.500E-08 | 8.700E-08 |
| 19803 | 991 | P991N | C | 3.9e-08 | 7.7e-08 | 2.7e-08 | 4.767E-08 | 2.610E-08 | 2.700E-08 | 7.700E-08 |
| 19804 | 991 | P991D | C | 3.5e-08 | 2.1e-08 | 3.1e-08 | 2.900E-08 | 7.211E-09 | 2.100E-08 | 3.500E-08 |

| | | | | | | | | | | |
|---|---|---|---|---|---|---|---|---|---|---|
| 19805 | 991 | P991C | C | 5.6e-08 | 3.5e-08 | 3.1e-08 | 4.067E-08 | 1.343E-08 | 3.100E-08 | 5.600E-08 |
| 19806 | 991 | P991E | C | 3.2e-08 | 2.4e-08 | 3e-08 | 2.867E-08 | 4.163E-09 | 2.400E-08 | 3.200E-08 |
| 19807 | 991 | P991Q | C | 5.6e-08 | 9e-08 | 3.2e-08 | 5.933E-08 | 2.914E-08 | 3.200E-08 | 9.000E-08 |
| 19808 | 991 | P991G | C | 6.7e-08 | 3.2e-08 | 1.5e-07 | 8.300E-08 | 6.061E-08 | 3.200E-08 | 1.500E-07 |
| 19809 | 991 | P991H | C | 9.3e-08 | 4e-08 | 2.7e-08 | 5.333E-08 | 3.496E-08 | 2.700E-08 | 9.300E-08 |
| 19810 | 991 | P991I | C | 3.4e-08 | 3.2e-08 | 5.8e-08 | 4.133E-08 | 1.447E-08 | 3.200E-08 | 5.800E-08 |
| 19811 | 991 | P991L | C | 2.4e-08 | 6.7e-08 | 4.8e-08 | 4.633E-08 | 2.155E-08 | 2.400E-08 | 6.700E-08 |
| 19812 | 991 | P991K | C | 2.7e-08 | 3.6e-08 | 6.8e-08 | 4.367E-08 | 2.155E-08 | 2.700E-08 | 6.800E-08 |
| 19813 | 991 | P991M | C | 2.4e-08 | 1e-07 | 1.1e-07 | 7.800E-08 | 4.703E-08 | 2.400E-08 | 1.100E-07 |
| 19814 | 991 | P991F | C | 5.5e-08 | 3.6e-08 | 4.1e-08 | 4.400E-08 | 9.849E-09 | 3.600E-08 | 5.500E-08 |
| 19815 | 991 | P991P | C | 6.7e-08 | 5.7e-08 | 2.7e-08 | 5.033E-08 | 2.082E-08 | 2.700E-08 | 6.700E-08 |
| 19816 | 991 | P991S | C | 9.3e-08 | 4.3e-08 | 1.6e-08 | 5.067E-08 | 3.907E-08 | 1.600E-08 | 9.300E-08 |
| 19817 | 991 | P991T | C | 2.8e-08 | 7.1e-08 | 6.5e-08 | 5.467E-08 | 2.329E-08 | 2.800E-08 | 7.100E-08 |
| 19818 | 991 | P991W | C | 4.2e-08 | 4.9e-08 | 7e-08 | 5.367E-08 | 1.457E-08 | 4.200E-08 | 7.000E-08 |
| 19819 | 991 | P991Y | C | 2.5e-08 | 1.5e-08 | 3.9e-08 | 2.633E-08 | 1.206E-08 | 1.500E-08 | 3.900E-08 |
| 19820 | 991 | P991V | C | 5.9e-08 | 5.5e-08 | 9.9e-08 | 7.100E-08 | 2.433E-08 | 5.500E-08 | 9.900E-08 |
| 19821 | 992 | V992A | C | 2e-08 | 1.7e-08 | 2e-08 | 1.900E-08 | 1.732E-09 | 1.700E-08 | 2.000E-08 |
| 19822 | 992 | V992R | C | 1.7e-08 | 2e-08 | 1.8e-08 | 1.833E-08 | 1.528E-09 | 1.700E-08 | 2.000E-08 |
| 19823 | 992 | V992N | C | 2.3e-08 | 2.8e-08 | 6.7e-08 | 3.933E-08 | 2.409E-08 | 2.300E-08 | 6.700E-08 |
| 19824 | 992 | V992D | C | 2.5e-08 | 3.5e-08 | 3.9e-08 | 3.300E-08 | 7.211E-09 | 2.500E-08 | 3.900E-08 |
| 19825 | 992 | V992C | C | 6.9e-08 | 7.2e-08 | 4.5e-08 | 6.200E-08 | 1.480E-08 | 4.500E-08 | 7.200E-08 |
| 19826 | 992 | V992E | C | 3.1e-08 | 3e-08 | 3.3e-08 | 3.133E-08 | 1.528E-09 | 3.000E-08 | 3.300E-08 |
| 19827 | 992 | V992Q | C | 1.6e-08 | 1.1e-08 | 4.8e-08 | 2.500E-08 | 2.007E-08 | 1.100E-08 | 4.800E-08 |
| 19828 | 992 | V992G | C | 2.1e-08 | 2.1e-08 | 5.8e-08 | 3.333E-08 | 2.136E-08 | 2.100E-08 | 5.800E-08 |
| 19829 | 992 | V992H | C | 4e-08 | 2.1e-08 | 5.4e-08 | 3.833E-08 | 1.656E-08 | 2.100E-08 | 5.400E-08 |
| 19830 | 992 | V992I | C | 3.2e-08 | 3e-08 | 3.6e-08 | 3.267E-08 | 3.055E-09 | 3.000E-08 | 3.600E-08 |
| 19831 | 992 | V992L | C | 7.6e-08 | 2.7e-08 | 7.7e-08 | 6.000E-08 | 2.858E-08 | 2.700E-08 | 7.700E-08 |
| 19832 | 992 | V992K | C | 2.1e-08 | 2.6e-08 | 2.9e-08 | 2.533E-08 | 4.041E-09 | 2.100E-08 | 2.900E-08 |
| 19833 | 992 | V992M | C | 4.3e-08 | 8.6e-08 | 3.4e-08 | 5.433E-08 | 2.779E-08 | 3.400E-08 | 8.600E-08 |
| 19834 | 992 | V992F | C | 2.9e-08 | 2.1e-08 | 1.1e-07 | 5.333E-08 | 4.924E-08 | 2.100E-08 | 1.100E-07 |
| 19835 | 992 | V992P | C | 6.9e-08 | 2.5e-08 | 5.8e-08 | 5.067E-08 | 2.290E-08 | 2.500E-08 | 6.900E-08 |
| 19836 | 992 | V992S | C | 2.3e-08 | 1.9e-08 | 6.1e-08 | 3.433E-08 | 2.318E-08 | 1.900E-08 | 6.100E-08 |
| 19837 | 992 | V992T | C | 4.6e-08 | 6.2e-08 | 2.6e-08 | 4.467E-08 | 1.804E-08 | 2.600E-08 | 6.200E-08 |
| 19838 | 992 | V992W | C | 5.5e-08 | 3.2e-08 | 2.7e-08 | 3.800E-08 | 1.493E-08 | 2.700E-08 | 5.500E-08 |
| 19839 | 992 | V992Y | C | 4.5e-08 | 2.8e-08 | 1.7e-08 | 3.000E-08 | 1.411E-08 | 1.700E-08 | 4.500E-08 |
| 19840 | 992 | V992V | C | 4.1e-08 | 3e-08 | 1.2e-07 | 6.367E-08 | 4.910E-08 | 3.000E-08 | 1.200E-07 |
| 19841 | 993 | T993A | C | 2.3e-08 | 7.5e-08 | 6.2e-08 | 5.333E-08 | 2.706E-08 | 2.300E-08 | 7.500E-08 |
| 19842 | 993 | T993R | C | 4.1e-08 | 4.3e-08 | 9.3e-08 | 5.900E-08 | 2.946E-08 | 4.100E-08 | 9.300E-08 |
| 19843 | 993 | T993N | C | 7.2e-08 | 5.8e-08 | 7.9e-08 | 6.967E-08 | 1.069E-08 | 5.800E-08 | 7.900E-08 |
| 19844 | 993 | T993D | C | 3.2e-08 | 6.5e-08 | 4.8e-08 | 4.833E-08 | 1.650E-08 | 3.200E-08 | 6.500E-08 |
| 19845 | 993 | T993C | C | 2e-08 | 4.5e-08 | 3.7e-08 | 3.400E-08 | 1.277E-08 | 2.000E-08 | 4.500E-08 |
| 19846 | 993 | T993E | C | 3.8e-08 | 7.4e-08 | 3.9e-08 | 5.033E-08 | 2.050E-08 | 3.800E-08 | 7.400E-08 |
| 19847 | 993 | T993Q | C | 4.2e-08 | 6.9e-08 | 7.1e-08 | 6.067E-08 | 1.620E-08 | 4.200E-08 | 7.100E-08 |
| 19848 | 993 | T993G | C | 2.3e-08 | 7.8e-08 | 1.2e-07 | 7.367E-08 | 4.864E-08 | 2.300E-08 | 1.200E-07 |
| 19849 | 993 | T993H | C | 3e-08 | 1.9e-08 | 5.4e-08 | 3.433E-08 | 1.790E-08 | 1.900E-08 | 5.400E-08 |
| 19850 | 993 | T993I | C | 5e-08 | 1.2e-08 | 1.4e-07 | 6.733E-08 | 6.574E-08 | 1.200E-08 | 1.400E-07 |
| 19851 | 993 | T993L | C | 9.5e-08 | 5.8e-08 | 4.3e-08 | 6.533E-08 | 2.676E-08 | 4.300E-08 | 9.500E-08 |
| 19852 | 993 | T993K | C | 2.3e-08 | 5.1e-08 | 1.6e-08 | 3.000E-08 | 1.852E-08 | 1.600E-08 | 5.100E-08 |
| 19853 | 993 | T993M | C | 2.9e-08 | 4.5e-08 | 4.9e-08 | 4.100E-08 | 1.058E-08 | 2.900E-08 | 4.900E-08 |
| 19854 | 993 | T993F | C | 1.4e-08 | 3.8e-08 | 3.2e-08 | 2.800E-08 | 1.249E-08 | 1.400E-08 | 3.800E-08 |
| 19855 | 993 | T993P | C | 4.8e-08 | 3.2e-08 | 9.9e-08 | 5.967E-08 | 3.499E-08 | 3.200E-08 | 9.900E-08 |
| 19856 | 993 | T993S | C | 3.4e-08 | 2.8e-08 | 3.7e-08 | 3.300E-08 | 4.583E-09 | 2.800E-08 | 3.700E-08 |
| 19857 | 993 | T993T | C | 4e-08 | 2.2e-08 | 6.9e-08 | 4.367E-08 | 2.371E-08 | 2.200E-08 | 6.900E-08 |
| 19858 | 993 | T993W | C | 4.9e-08 | 8.3e-08 | 4.1e-08 | 5.767E-08 | 2.230E-08 | 4.100E-08 | 8.300E-08 |
| 19859 | 993 | T993Y | C | 6.4e-08 | 9.7e-08 | 1e-07 | 8.700E-08 | 1.997E-08 | 6.400E-08 | 1.000E-07 |
| 19860 | 993 | T993V | C | 2.6e-08 | 9.2e-08 | 5.7e-08 | 5.833E-08 | 3.302E-08 | 2.600E-08 | 9.200E-08 |
| 19861 | 994 | V994A | C | 2.9e-08 | 5.4e-08 | 4.2e-08 | 4.167E-08 | 1.250E-08 | 2.900E-08 | 5.400E-08 |
| 19862 | 994 | V994R | C | 7.6e-08 | 5e-08 | 1.6e-08 | 4.733E-08 | 3.009E-08 | 1.600E-08 | 7.600E-08 |
| 19863 | 994 | V994N | C | 2.4e-08 | 6.7e-08 | 5.4e-08 | 4.833E-08 | 2.205E-08 | 2.400E-08 | 6.700E-08 |
| 19864 | 994 | V994D | C | 3.1e-08 | 6.8e-08 | 3.1e-08 | 4.333E-08 | 2.136E-08 | 3.100E-08 | 6.800E-08 |
| 19865 | 994 | V994C | C | 2.8e-08 | 4.2e-08 | 7e-08 | 4.667E-08 | 2.139E-08 | 2.800E-08 | 7.000E-08 |
| 19866 | 994 | V994E | C | 1.9e-08 | 2.7e-08 | 5.3e-08 | 3.300E-08 | 1.778E-08 | 1.900E-08 | 5.300E-08 |
| 19867 | 994 | V994Q | C | 4.9e-08 | 3.4e-08 | 2.8e-08 | 3.700E-08 | 1.082E-08 | 2.800E-08 | 4.900E-08 |
| 19868 | 994 | V994G | C | 5.4e-08 | 5.5e-08 | 3.9e-08 | 4.933E-08 | 8.963E-09 | 3.900E-08 | 5.500E-08 |
| 19869 | 994 | V994H | C | 4.3e-08 | 6.3e-08 | 4.4e-08 | 5.000E-08 | 1.127E-08 | 4.300E-08 | 6.300E-08 |
| 19870 | 994 | V994I | C | 2.3e-08 | 8.8e-09 | 2.7e-08 | 1.960E-08 | 9.565E-09 | 8.800E-09 | 2.700E-08 |
| 19871 | 994 | V994L | C | 4.6e-08 | 1.2e-08 | 1.4e-08 | 2.400E-08 | 1.908E-08 | 1.200E-08 | 4.600E-08 |
| 19872 | 994 | V994K | C | 3.7e-08 | 3.7e-08 | 2.8e-08 | 3.400E-08 | 5.196E-09 | 2.800E-08 | 3.700E-08 |
| 19873 | 994 | V994M | C | 3.9e-08 | 1.3e-07 | 3.7e-08 | 6.867E-08 | 5.313E-08 | 3.700E-08 | 1.300E-07 |
| 19874 | 994 | V994F | C | 6.3e-08 | 5.2e-08 | 7.1e-09 | 4.070E-08 | 2.961E-08 | 7.100E-09 | 6.300E-08 |
| 19875 | 994 | V994P | C | 2.8e-08 | 4.9e-08 | 1.8e-08 | 3.167E-08 | 1.582E-08 | 1.800E-08 | 4.900E-08 |
| 19876 | 994 | V994S | C | 2.8e-08 | 1.2e-07 | 3.1e-08 | 5.967E-08 | 5.227E-08 | 2.800E-08 | 1.200E-07 |
| 19877 | 994 | V994T | C | 2.9e-08 | 4.3e-08 | 3.1e-08 | 3.433E-08 | 7.572E-09 | 2.900E-08 | 4.300E-08 |
| 19878 | 994 | V994W | C | 3.3e-08 | 5.6e-08 | 1.5e-07 | 7.967E-08 | 6.199E-08 | 3.300E-08 | 1.500E-07 |
| 19879 | 994 | V994Y | C | 2.6e-08 | 8.4e-08 | 1.6e-08 | 4.200E-08 | 3.672E-08 | 1.600E-08 | 8.400E-08 |
| 19880 | 994 | V994V | C | 1.4e-08 | 4.1e-08 | 4.5e-08 | 3.333E-08 | 1.686E-08 | 1.400E-08 | 4.500E-08 |
| 19881 | 995 | S995A | C | 4.7e-08 | 5e-08 | 3e-08 | 4.233E-08 | 1.079E-08 | 3.000E-08 | 5.000E-08 |
| 19882 | 995 | S995R | C | 4.9e-08 | 7.3e-08 | 2.8e-08 | 5.000E-08 | 2.252E-08 | 2.800E-08 | 7.300E-08 |
| 19883 | 995 | S995N | C | 2.3e-08 | 6.9e-08 | 6.2e-08 | 5.133E-08 | 2.479E-08 | 2.300E-08 | 6.900E-08 |
| 19884 | 995 | S995D | C | 3.5e-08 | 7.5e-08 | 3.5e-08 | 4.833E-08 | 2.309E-08 | 3.500E-08 | 7.500E-08 |
| 19885 | 995 | S995C | C | 4.4e-08 | 2.6e-08 | 3.7e-08 | 3.567E-08 | 9.074E-09 | 2.600E-08 | 4.400E-08 |
| 19886 | 995 | S995E | C | 2.1e-08 | 5.7e-08 | 2e-08 | 3.267E-08 | 2.108E-08 | 2.000E-08 | 5.700E-08 |
| 19887 | 995 | S995Q | C | 2.7e-08 | 5.4e-08 | 2.9e-08 | 3.667E-08 | 1.504E-08 | 2.700E-08 | 5.400E-08 |
| 19888 | 995 | S995G | C | 1.2e-08 | 4.7e-08 | 2.2e-08 | 2.700E-08 | 1.803E-08 | 1.200E-08 | 4.700E-08 |
| 19889 | 995 | S995H | C | 2.4e-08 | 8e-08 | 5e-08 | 5.133E-08 | 2.802E-08 | 2.400E-08 | 8.000E-08 |

| | | | | | | | | | | |
|---|---|---|---|---|---|---|---|---|---|---|
| 19890 | 995 | S995I | C | 5.5e-08 | 4.2e-08 | 8.3e-09 | 3.510E-08 | 2.410E-08 | 8.300E-09 | 5.500E-08 |
| 19891 | 995 | S995L | C | 3.1e-08 | 3.6e-08 | 2.9e-08 | 3.200E-08 | 3.606E-09 | 2.900E-08 | 3.600E-08 |
| 19892 | 995 | S995K | C | 6e-08 | 2.3e-08 | 9.6e-09 | 3.087E-08 | 2.610E-08 | 9.600E-09 | 6.000E-08 |
| 19893 | 995 | S995M | C | 7.2e-08 | 8.9e-08 | 3.1e-08 | 6.400E-08 | 2.982E-08 | 3.100E-08 | 8.900E-08 |
| 19894 | 995 | S995F | C | 2.2e-08 | 2.3e-08 | 4.3e-08 | 2.933E-08 | 1.185E-08 | 2.200E-08 | 4.300E-08 |
| 19895 | 995 | S995P | C | 9.4e-08 | 4.9e-08 | 2e-08 | 5.433E-08 | 3.729E-08 | 2.000E-08 | 9.400E-08 |
| 19896 | 995 | S995S | C | 2e-08 | 3.7e-08 | 5.4e-08 | 3.700E-08 | 1.700E-08 | 2.000E-08 | 5.400E-08 |
| 19897 | 995 | S995T | C | 2.2e-08 | 9e-08 | 5.4e-08 | 5.533E-08 | 3.402E-08 | 2.200E-08 | 9.000E-08 |
| 19898 | 995 | S995W | C | 4.6e-08 | 6.4e-08 | 3.4e-08 | 4.800E-08 | 1.510E-08 | 3.400E-08 | 6.400E-08 |
| 19899 | 995 | S995Y | C | 2.5e-08 | 2.8e-08 | 6.5e-08 | 3.933E-08 | 2.228E-08 | 2.500E-08 | 6.500E-08 |
| 19900 | 995 | S995V | C | 2.8e-08 | 2.7e-08 | 5.4e-08 | 3.633E-08 | 1.531E-08 | 2.700E-08 | 5.400E-08 |
| 19901 | 996 | W996A | C | 3.6e-08 | 1.3e-07 | 2.9e-08 | 6.500E-08 | 5.640E-08 | 2.900E-08 | 1.300E-07 |
| 19902 | 996 | W996R | C | 3.8e-08 | 5e-08 | 5.6e-08 | 4.800E-08 | 9.165E-09 | 3.800E-08 | 5.600E-08 |
| 19903 | 996 | W996N | C | 5.3e-08 | 6.8e-08 | 4.7e-08 | 5.600E-08 | 1.082E-08 | 4.700E-08 | 6.800E-08 |
| 19904 | 996 | W996D | C | 2.1e-08 | 5.1e-08 | 3.7e-08 | 3.633E-08 | 1.501E-08 | 2.100E-08 | 5.100E-08 |
| 19905 | 996 | W996C | C | 6e-08 | 5.2e-08 | 4.3e-08 | 5.167E-08 | 8.505E-09 | 4.300E-08 | 6.000E-08 |
| 19906 | 996 | W996E | C | 8.9e-08 | 4.3e-08 | 4.7e-08 | 5.967E-08 | 2.548E-08 | 4.300E-08 | 8.900E-08 |
| 19907 | 996 | W996Q | C | 6e-08 | 9.8e-08 | 4.8e-08 | 6.867E-08 | 2.610E-08 | 4.800E-08 | 9.800E-08 |
| 19908 | 996 | W996G | C | 1e-07 | 2.7e-08 | 8.3e-08 | 7.000E-08 | 3.820E-08 | 2.700E-08 | 1.000E-07 |
| 19909 | 996 | W996H | C | 2.1e-08 | 3e-08 | 8.9e-08 | 4.667E-08 | 3.694E-08 | 2.100E-08 | 8.900E-08 |
| 19910 | 996 | W996I | C | 6.6e-08 | 1.7e-08 | 2.6e-08 | 3.633E-08 | 2.608E-08 | 1.700E-08 | 6.600E-08 |
| 19911 | 996 | W996L | C | 4.5e-08 | 5.5e-08 | 5.7e-08 | 5.233E-08 | 6.429E-09 | 4.500E-08 | 5.700E-08 |
| 19912 | 996 | W996K | C | 1.5e-08 | 1.1e-08 | 4.2e-08 | 2.267E-08 | 1.686E-08 | 1.100E-08 | 4.200E-08 |
| 19913 | 996 | W996M | C | 1.7e-07 | 7.5e-08 | 3e-08 | 9.167E-08 | 7.147E-08 | 3.000E-08 | 1.700E-07 |
| 19914 | 996 | W996F | C | 2.1e-08 | 5.7e-08 | 4.6e-08 | 4.133E-08 | 1.845E-08 | 2.100E-08 | 5.700E-08 |
| 19915 | 996 | W996P | C | 4.1e-08 | 4.3e-08 | 6.1e-08 | 4.833E-08 | 1.102E-08 | 4.100E-08 | 6.100E-08 |
| 19916 | 996 | W996S | C | 4.2e-08 | 2.7e-08 | 3.8e-08 | 3.567E-08 | 7.767E-09 | 2.700E-08 | 4.200E-08 |
| 19917 | 996 | W996T | C | 5.9e-08 | 2.4e-08 | 1e-07 | 6.100E-08 | 3.804E-08 | 2.400E-08 | 1.000E-07 |
| 19918 | 996 | W996W | C | 6.4e-08 | 2.5e-08 | 1.3e-07 | 7.300E-08 | 5.308E-08 | 2.500E-08 | 1.300E-07 |
| 19919 | 996 | W996Y | C | 7.1e-08 | 9.4e-08 | 4.7e-08 | 7.067E-08 | 2.350E-08 | 4.700E-08 | 9.400E-08 |
| 19920 | 996 | W996V | C | 1.6e-08 | 3e-08 | 2.5e-08 | 2.367E-08 | 7.095E-09 | 1.600E-08 | 3.000E-08 |
| 19921 | 997 | N997A | C | 5.5e-08 | 3.1e-08 | 7.1e-08 | 5.233E-08 | 2.013E-08 | 3.100E-08 | 7.100E-08 |
| 19922 | 997 | N997R | C | 6.6e-08 | 3.8e-08 | 2.2e-07 | 1.080E-07 | 9.800E-08 | 3.800E-08 | 2.200E-07 |
| 19923 | 997 | N997N | C | 2e-08 | 4.2e-08 | 5.3e-08 | 3.833E-08 | 1.680E-08 | 2.000E-08 | 5.300E-08 |
| 19924 | 997 | N997D | C | 2.7e-08 | 5.6e-08 | 3.7e-08 | 4.000E-08 | 1.473E-08 | 2.700E-08 | 5.600E-08 |
| 19925 | 997 | N997C | C | 1.3e-07 | 8.9e-08 | 5.5e-08 | 9.133E-08 | 3.755E-08 | 5.500E-08 | 1.300E-07 |
| 19926 | 997 | N997E | C | 3e-08 | 4e-08 | 6.7e-08 | 4.567E-08 | 1.914E-08 | 3.000E-08 | 6.700E-08 |
| 19927 | 997 | N997Q | C | 3.3e-08 | 2.9e-08 | 1.2e-07 | 6.067E-08 | 5.142E-08 | 2.900E-08 | 1.200E-07 |
| 19928 | 997 | N997G | C | 1.7e-08 | 5.1e-08 | 8.5e-08 | 5.100E-08 | 3.400E-08 | 1.700E-08 | 8.500E-08 |
| 19929 | 997 | N997H | C | 2.4e-08 | 2.4e-08 | 2.2e-08 | 2.333E-08 | 1.155E-09 | 2.200E-08 | 2.400E-08 |
| 19930 | 997 | N997I | C | 1.2e-07 | 6.1e-08 | 3.1e-08 | 7.067E-08 | 4.528E-08 | 3.100E-08 | 1.200E-07 |
| 19931 | 997 | N997L | C | 6e-08 | 1.4e-07 | 1.5e-07 | 1.167E-07 | 4.933E-08 | 6.000E-08 | 1.500E-07 |
| 19932 | 997 | N997K | C | 4.5e-08 | 6.8e-08 | 5e-08 | 5.433E-08 | 1.210E-08 | 4.500E-08 | 6.800E-08 |
| 19933 | 997 | N997M | C | 4.7e-08 | 5.5e-08 | 7.3e-08 | 5.833E-08 | 1.332E-08 | 4.700E-08 | 7.300E-08 |
| 19934 | 997 | N997F | C | 4.7e-08 | 1e-07 | 8.8e-09 | 5.193E-08 | 4.580E-08 | 8.800E-09 | 1.000E-07 |
| 19935 | 997 | N997P | C | 3.2e-08 | 3.2e-08 | 5.5e-08 | 3.967E-08 | 1.328E-08 | 3.200E-08 | 5.500E-08 |
| 19936 | 997 | N997S | C | 5.1e-08 | 3.3e-08 | 1.1e-08 | 3.167E-08 | 2.003E-08 | 1.100E-08 | 5.100E-08 |
| 19937 | 997 | N997T | C | 1e-07 | 2.4e-08 | 5.1e-08 | 5.833E-08 | 3.853E-08 | 2.400E-08 | 1.000E-07 |
| 19938 | 997 | N997W | C | 2.9e-08 | 2.4e-08 | 2.4e-08 | 2.567E-08 | 2.887E-09 | 2.400E-08 | 2.900E-08 |
| 19939 | 997 | N997Y | C | 2.3e-07 | 2.7e-08 | 4.7e-08 | 1.013E-07 | 1.119E-07 | 2.700E-08 | 2.300E-07 |
| 19940 | 997 | N997V | C | 6.5e-08 | 3.7e-08 | 2.1e-08 | 4.100E-08 | 2.227E-08 | 2.100E-08 | 6.500E-08 |
| 19941 | 998 | S998A | C | 8.6e-08 | 6e-08 | 1.4e-08 | 5.333E-08 | 3.646E-08 | 1.400E-08 | 8.600E-08 |
| 19942 | 998 | S998R | C | 4e-08 | 3.9e-08 | 5.7e-08 | 4.533E-08 | 1.012E-08 | 3.900E-08 | 5.700E-08 |
| 19943 | 998 | S998N | C | 2.9e-08 | 2.8e-08 | 6.6e-08 | 4.100E-08 | 2.166E-08 | 2.800E-08 | 6.600E-08 |
| 19944 | 998 | S998D | C | 6.3e-08 | 3.8e-08 | 6.6e-09 | 3.587E-08 | 2.826E-08 | 6.600E-09 | 6.300E-08 |
| 19945 | 998 | S998C | C | 1.8e-07 | 7.9e-08 | 4.2e-08 | 1.003E-07 | 7.143E-08 | 4.200E-08 | 1.800E-07 |
| 19946 | 998 | S998E | C | 4.8e-08 | 4.4e-08 | 4.7e-08 | 4.633E-08 | 2.082E-09 | 4.400E-08 | 4.800E-08 |
| 19947 | 998 | S998Q | C | 1.7e-08 | 3.4e-08 | 1.5e-08 | 2.200E-08 | 1.044E-08 | 1.500E-08 | 3.400E-08 |
| 19948 | 998 | S998G | C | 6.5e-08 | 4.9e-08 | 1.4e-07 | 8.467E-08 | 4.858E-08 | 4.900E-08 | 1.400E-07 |
| 19949 | 998 | S998H | C | 3.3e-08 | 3.4e-08 | 1.4e-08 | 2.700E-08 | 1.127E-08 | 1.400E-08 | 3.400E-08 |
| 19950 | 998 | S998I | C | 3.1e-08 | 1.6e-07 | 5.3e-08 | 8.133E-08 | 6.901E-08 | 3.100E-08 | 1.600E-07 |
| 19951 | 998 | S998L | C | 9.8e-08 | 4e-08 | 1.5e-07 | 9.600E-08 | 5.503E-08 | 4.000E-08 | 1.500E-07 |
| 19952 | 998 | S998K | C | 5.2e-08 | 2.9e-08 | 2.9e-08 | 3.667E-08 | 1.328E-08 | 2.900E-08 | 5.200E-08 |
| 19953 | 998 | S998M | C | 1.9e-08 | 5.5e-08 | 3.3e-08 | 3.567E-08 | 1.815E-08 | 1.900E-08 | 5.500E-08 |
| 19954 | 998 | S998F | C | 1.8e-08 | 1e-07 | 4.3e-08 | 5.367E-08 | 4.203E-08 | 1.800E-08 | 1.000E-07 |
| 19955 | 998 | S998P | C | 5.5e-08 | 2.3e-08 | 3.9e-08 | 3.900E-08 | 1.600E-08 | 2.300E-08 | 5.500E-08 |
| 19956 | 998 | S998S | C | 2.3e-08 | 1.3e-07 | 6.2e-08 | 7.167E-08 | 5.415E-08 | 2.300E-08 | 1.300E-07 |
| 19957 | 998 | S998T | C | 4.7e-08 | 4e-08 | 2.1e-08 | 3.600E-08 | 1.345E-08 | 2.100E-08 | 4.700E-08 |
| 19958 | 998 | S998W | C | 2e-08 | 4e-08 | 1.2e-07 | 6.000E-08 | 5.292E-08 | 2.000E-08 | 1.200E-07 |
| 19959 | 998 | S998Y | C | 7.3e-08 | 8.6e-08 | 6.7e-08 | 7.533E-08 | 9.713E-09 | 6.700E-08 | 8.600E-08 |
| 19960 | 998 | S998V | C | 1.2e-07 | 6.1e-08 | 4.1e-08 | 7.400E-08 | 4.107E-08 | 4.100E-08 | 1.200E-07 |
| 19961 | 999 | G999A | C | 4.5e-08 | 5.2e-08 | 4.5e-08 | 4.733E-08 | 4.041E-09 | 4.500E-08 | 5.200E-08 |
| 19962 | 999 | G999R | C | 3.2e-08 | 1.9e-08 | 2.2e-08 | 2.433E-08 | 6.807E-09 | 1.900E-08 | 3.200E-08 |
| 19963 | 999 | G999N | C | 3.2e-08 | 7.6e-08 | 2.8e-08 | 4.533E-08 | 2.663E-08 | 2.800E-08 | 7.600E-08 |
| 19964 | 999 | G999D | C | 3.8e-08 | 4.3e-08 | 2.1e-08 | 3.400E-08 | 1.153E-08 | 2.100E-08 | 4.300E-08 |
| 19965 | 999 | G999C | C | 1.9e-08 | 2.7e-08 | 5.5e-08 | 3.367E-08 | 1.890E-08 | 1.900E-08 | 5.500E-08 |
| 19966 | 999 | G999E | C | 6.6e-08 | 8.4e-08 | 3.4e-08 | 6.133E-08 | 2.532E-08 | 3.400E-08 | 8.400E-08 |
| 19967 | 999 | G999Q | C | 1.7e-08 | 5.4e-08 | 7.7e-08 | 4.933E-08 | 3.027E-08 | 1.700E-08 | 7.700E-08 |
| 19968 | 999 | G999G | C | 1e-07 | 6.2e-08 | 4.5e-08 | 6.900E-08 | 2.816E-08 | 4.500E-08 | 1.000E-07 |
| 19969 | 999 | G999H | C | 6.4e-08 | 6.4e-08 | 3.8e-08 | 5.533E-08 | 1.501E-08 | 3.800E-08 | 6.400E-08 |
| 19970 | 999 | G999I | C | 3.7e-08 | 5e-08 | 3.2e-08 | 3.967E-08 | 9.292E-09 | 3.200E-08 | 5.000E-08 |
| 19971 | 999 | G999L | C | 3.6e-08 | 6.8e-08 | 4.5e-08 | 4.967E-08 | 1.650E-08 | 3.600E-08 | 6.800E-08 |
| 19972 | 999 | G999K | C | 3e-08 | 8.6e-08 | 3e-08 | 4.867E-08 | 3.233E-08 | 3.000E-08 | 8.600E-08 |
| 19973 | 999 | G999M | C | 1.4e-07 | 3.7e-08 | 4.2e-08 | 7.300E-08 | 5.808E-08 | 3.700E-08 | 1.400E-07 |
| 19974 | 999 | G999F | C | 1.1e-07 | 5.2e-08 | 4e-08 | 6.733E-08 | 3.743E-08 | 4.000E-08 | 1.100E-07 |

| | | | | | | | | | | |
|---|---|---|---|---|---|---|---|---|---|---|
| 19975 | 999 | G999P | C | 3.1e-08 | 3e-08 | 3.1e-08 | 3.067E-08 | 5.774E-10 | 3.000E-08 | 3.100E-08 |
| 19976 | 999 | G999S | C | 3.5e-08 | 2.4e-08 | 1.3e-07 | 6.300E-08 | 5.828E-08 | 2.400E-08 | 1.300E-07 |
| 19977 | 999 | G999T | C | 1.7e-08 | 2.7e-08 | 4.9e-08 | 3.100E-08 | 1.637E-08 | 1.700E-08 | 4.900E-08 |
| 19978 | 999 | G999W | C | 5.7e-08 | 2.6e-08 | 4.3e-08 | 4.200E-08 | 1.552E-08 | 2.600E-08 | 5.700E-08 |
| 19979 | 999 | G999Y | C | 2.9e-08 | 1.8e-08 | 4.2e-08 | 2.967E-08 | 1.201E-08 | 1.800E-08 | 4.200E-08 |
| 19980 | 999 | G999V | C | 6.2e-08 | 4.8e-08 | 1.6e-08 | 4.200E-08 | 2.358E-08 | 1.600E-08 | 6.200E-08 |
| 19981 | 1000 | A1000A | C | 6.2e-08 | 1.2e-08 | 2.8e-08 | 3.400E-08 | 2.553E-08 | 1.200E-08 | 6.200E-08 |
| 19982 | 1000 | A1000R | C | 4.5e-08 | 8.7e-08 | 3.8e-08 | 5.667E-08 | 2.650E-08 | 3.800E-08 | 8.700E-08 |
| 19983 | 1000 | A1000N | C | 4.4e-08 | 4.6e-08 | 3.7e-08 | 4.233E-08 | 4.726E-09 | 3.700E-08 | 4.600E-08 |
| 19984 | 1000 | A1000D | C | 2.1e-08 | 2.9e-08 | 3.8e-08 | 2.933E-08 | 8.505E-09 | 2.100E-08 | 3.800E-08 |
| 19985 | 1000 | A1000C | C | 4.7e-08 | 1.1e-07 | 2.4e-08 | 6.033E-08 | 4.452E-08 | 2.400E-08 | 1.100E-07 |
| 19986 | 1000 | A1000E | C | 3.7e-08 | 1.7e-08 | 3.2e-08 | 2.867E-08 | 1.041E-08 | 1.700E-08 | 3.700E-08 |
| 19987 | 1000 | A1000Q | C | 4.7e-08 | 7e-08 | 2.2e-08 | 4.633E-08 | 2.401E-08 | 2.200E-08 | 7.000E-08 |
| 19988 | 1000 | A1000G | C | 2.6e-08 | 8.7e-08 | 3e-08 | 4.767E-08 | 3.412E-08 | 2.600E-08 | 8.700E-08 |
| 19989 | 1000 | A1000H | C | 1.8e-08 | 4.8e-08 | 9.2e-08 | 5.267E-08 | 3.722E-08 | 1.800E-08 | 9.200E-08 |
| 19990 | 1000 | A1000I | C | 5.5e-08 | 7.2e-08 | 1.8e-08 | 4.833E-08 | 2.761E-08 | 1.800E-08 | 7.200E-08 |
| 19991 | 1000 | A1000L | C | 5.2e-08 | 1.4e-08 | 1.5e-08 | 2.700E-08 | 2.166E-08 | 1.400E-08 | 5.200E-08 |
| 19992 | 1000 | A1000K | C | 3.3e-08 | 3.2e-08 | 8.3e-08 | 4.933E-08 | 2.916E-08 | 3.200E-08 | 8.300E-08 |
| 19993 | 1000 | A1000M | C | 4.3e-08 | 6.1e-08 | 9.4e-08 | 6.600E-08 | 2.587E-08 | 4.300E-08 | 9.400E-08 |
| 19994 | 1000 | A1000F | C | 3.5e-08 | 9e-08 | 5e-08 | 5.833E-08 | 2.843E-08 | 3.500E-08 | 9.000E-08 |
| 19995 | 1000 | A1000P | C | 5.6e-08 | 2.6e-08 | 8.7e-09 | 3.023E-08 | 2.393E-08 | 8.700E-09 | 5.600E-08 |
| 19996 | 1000 | A1000S | C | 1.9e-08 | 7.2e-09 | 7.4e-08 | 3.340E-08 | 3.565E-08 | 7.200E-09 | 7.400E-08 |
| 19997 | 1000 | A1000T | C | 5.9e-08 | 1.8e-08 | 1.8e-08 | 3.167E-08 | 2.367E-08 | 1.800E-08 | 5.900E-08 |
| 19998 | 1000 | A1000W | C | 3.4e-08 | 3.3e-08 | 2.4e-08 | 3.033E-08 | 5.508E-09 | 2.400E-08 | 3.400E-08 |
| 19999 | 1000 | A1000Y | C | 2.4e-08 | 9.1e-09 | 2e-08 | 1.770E-08 | 7.712E-09 | 9.100E-09 | 2.400E-08 |
| 20000 | 1000 | A1000V | C | 6e-08 | 4.7e-08 | 3.3e-08 | 4.667E-08 | 1.350E-08 | 3.300E-08 | 6.000E-08 |
| 20001 | 1001 | L1001A | C | 8.7e-08 | 3.9e-08 | 3.3e-08 | 5.300E-08 | 2.960E-08 | 3.300E-08 | 8.700E-08 |
| 20002 | 1001 | L1001R | C | 3.3e-08 | 5.7e-08 | 1.7e-08 | 3.567E-08 | 2.013E-08 | 1.700E-08 | 5.700E-08 |
| 20003 | 1001 | L1001N | C | 1.7e-08 | 3.4e-08 | 2.6e-08 | 2.567E-08 | 8.505E-09 | 1.700E-08 | 3.400E-08 |
| 20004 | 1001 | L1001D | C | 4.4e-08 | 3.2e-08 | 4.5e-08 | 4.033E-08 | 7.234E-09 | 3.200E-08 | 4.500E-08 |
| 20005 | 1001 | L1001C | C | 1.5e-08 | 1.6e-08 | 5e-08 | 2.700E-08 | 1.992E-08 | 1.500E-08 | 5.000E-08 |
| 20006 | 1001 | L1001E | C | 2.4e-08 | 1.5e-08 | 3.7e-08 | 2.533E-08 | 1.106E-08 | 1.500E-08 | 3.700E-08 |
| 20007 | 1001 | L1001Q | C | 3.7e-08 | 5.4e-08 | 8.2e-08 | 5.767E-08 | 2.272E-08 | 3.700E-08 | 8.200E-08 |
| 20008 | 1001 | L1001G | C | 3.3e-08 | 1.3e-07 | 1.4e-07 | 1.010E-07 | 5.910E-08 | 3.300E-08 | 1.400E-07 |
| 20009 | 1001 | L1001H | C | 3.3e-08 | 6.6e-08 | 3.4e-08 | 4.433E-08 | 1.877E-08 | 3.300E-08 | 6.600E-08 |
| 20010 | 1001 | L1001I | C | 3.4e-08 | 4.8e-08 | 1.9e-08 | 3.367E-08 | 1.450E-08 | 1.900E-08 | 4.800E-08 |
| 20011 | 1001 | L1001L | C | 6.4e-08 | 9e-08 | 6.5e-08 | 7.300E-08 | 1.473E-08 | 6.400E-08 | 9.000E-08 |
| 20012 | 1001 | L1001K | C | 5.6e-08 | 3e-08 | 3.6e-08 | 4.067E-08 | 1.361E-08 | 3.000E-08 | 5.600E-08 |
| 20013 | 1001 | L1001M | C | 1.3e-07 | 3.5e-08 | 2.6e-08 | 6.367E-08 | 5.762E-08 | 2.600E-08 | 1.300E-07 |
| 20014 | 1001 | L1001F | C | 8.7e-08 | 4.4e-08 | 3.8e-08 | 5.633E-08 | 2.673E-08 | 3.800E-08 | 8.700E-08 |
| 20015 | 1001 | L1001P | C | 6.4e-08 | 3.7e-08 | 2.2e-08 | 4.100E-08 | 2.128E-08 | 2.200E-08 | 6.400E-08 |
| 20016 | 1001 | L1001S | C | 2.4e-08 | 2.8e-08 | 7.7e-08 | 4.300E-08 | 2.951E-08 | 2.400E-08 | 7.700E-08 |
| 20017 | 1001 | L1001T | C | 1.7e-08 | 5.4e-08 | 6.8e-08 | 4.633E-08 | 2.635E-08 | 1.700E-08 | 6.800E-08 |
| 20018 | 1001 | L1001W | C | 4.8e-08 | 3.7e-08 | 1.3e-07 | 7.167E-08 | 5.082E-08 | 3.700E-08 | 1.300E-07 |
| 20019 | 1001 | L1001Y | C | 3.5e-08 | 5.7e-08 | 1.2e-07 | 7.067E-08 | 4.412E-08 | 3.500E-08 | 1.200E-07 |
| 20020 | 1001 | L1001V | C | 8.7e-08 | 1.7e-08 | 4.1e-08 | 4.833E-08 | 3.557E-08 | 1.700E-08 | 8.700E-08 |
| 20021 | 1002 | T1002A | C | 5.1e-08 | 4.6e-08 | 3.6e-08 | 4.433E-08 | 7.638E-09 | 3.600E-08 | 5.100E-08 |
| 20022 | 1002 | T1002R | C | 3.4e-08 | 3.9e-08 | 3.6e-08 | 3.633E-08 | 2.517E-09 | 3.400E-08 | 3.900E-08 |
| 20023 | 1002 | T1002N | C | 5.5e-08 | 3.6e-08 | 4.6e-08 | 4.567E-08 | 9.504E-09 | 3.600E-08 | 5.500E-08 |
| 20024 | 1002 | T1002D | C | 7.4e-08 | 3.3e-08 | 1.1e-07 | 7.233E-08 | 3.853E-08 | 3.300E-08 | 1.100E-07 |
| 20025 | 1002 | T1002C | C | 4.3e-08 | 1.8e-07 | 5.4e-08 | 9.233E-08 | 7.612E-08 | 4.300E-08 | 1.800E-07 |
| 20026 | 1002 | T1002E | C | 3.8e-08 | 1.7e-08 | 1.2e-08 | 2.233E-08 | 1.380E-08 | 1.200E-08 | 3.800E-08 |
| 20027 | 1002 | T1002Q | C | 3.6e-08 | 5.8e-08 | 6.2e-08 | 5.200E-08 | 1.400E-08 | 3.600E-08 | 6.200E-08 |
| 20028 | 1002 | T1002G | C | 2.3e-08 | 3.6e-08 | 3.2e-08 | 3.033E-08 | 6.658E-09 | 2.300E-08 | 3.600E-08 |
| 20029 | 1002 | T1002H | C | 2.2e-08 | 2.6e-08 | 5.9e-08 | 3.567E-08 | 2.031E-08 | 2.200E-08 | 5.900E-08 |
| 20030 | 1002 | T1002I | C | 1.5e-08 | 2.4e-08 | 5.3e-08 | 3.067E-08 | 1.986E-08 | 1.500E-08 | 5.300E-08 |
| 20031 | 1002 | T1002L | C | 4.3e-08 | 3.4e-08 | 2.4e-08 | 3.367E-08 | 9.504E-09 | 2.400E-08 | 4.300E-08 |
| 20032 | 1002 | T1002K | C | 3.2e-08 | 4.4e-08 | 4.1e-08 | 3.900E-08 | 6.245E-09 | 3.200E-08 | 4.400E-08 |
| 20033 | 1002 | T1002M | C | 7.3e-08 | 2.9e-08 | 4.5e-08 | 4.900E-08 | 2.227E-08 | 2.900E-08 | 7.300E-08 |
| 20034 | 1002 | T1002F | C | 2e-08 | 2.6e-08 | 2.9e-08 | 2.500E-08 | 4.583E-09 | 2.000E-08 | 2.900E-08 |
| 20035 | 1002 | T1002P | C | 3e-08 | 3.9e-08 | 3.7e-08 | 3.533E-08 | 4.726E-09 | 3.000E-08 | 3.900E-08 |
| 20036 | 1002 | T1002S | C | 1.5e-08 | 6.8e-08 | 8e-08 | 5.433E-08 | 3.459E-08 | 1.500E-08 | 8.000E-08 |
| 20037 | 1002 | T1002T | C | 4.7e-08 | 6.9e-08 | 2.5e-08 | 4.700E-08 | 2.200E-08 | 2.500E-08 | 6.900E-08 |
| 20038 | 1002 | T1002W | C | 6.3e-08 | 7.9e-08 | 3.6e-08 | 5.933E-08 | 2.173E-08 | 3.600E-08 | 7.900E-08 |
| 20039 | 1002 | T1002Y | C | 7.7e-08 | 3.3e-08 | 4.5e-08 | 5.167E-08 | 2.274E-08 | 3.300E-08 | 7.700E-08 |
| 20040 | 1002 | T1002V | C | 3.1e-08 | 4.9e-08 | 2.7e-08 | 3.567E-08 | 1.172E-08 | 2.700E-08 | 4.900E-08 |
| 20041 | 1003 | S1003A | C | 2.3e-08 | 3e-08 | 4.1e-08 | 3.133E-08 | 9.074E-09 | 2.300E-08 | 4.100E-08 |
| 20042 | 1003 | S1003R | C | 1.5e-08 | 1.3e-07 | 3.3e-08 | 5.933E-08 | 6.186E-08 | 1.500E-08 | 1.300E-07 |
| 20043 | 1003 | S1003N | C | 7.5e-08 | 4.6e-08 | 3.2e-08 | 5.100E-08 | 2.193E-08 | 3.200E-08 | 7.500E-08 |
| 20044 | 1003 | S1003D | C | 6.8e-08 | 3.6e-08 | 4.9e-08 | 5.100E-08 | 1.609E-08 | 3.600E-08 | 6.800E-08 |
| 20045 | 1003 | S1003C | C | 9.2e-08 | 1e-07 | 6.1e-08 | 8.433E-08 | 2.060E-08 | 6.100E-08 | 1.000E-07 |
| 20046 | 1003 | S1003E | C | 2.3e-08 | 4.9e-08 | 9.8e-08 | 5.667E-08 | 3.808E-08 | 2.300E-08 | 9.800E-08 |
| 20047 | 1003 | S1003Q | C | 6.3e-08 | 4.1e-08 | 8.5e-08 | 6.300E-08 | 2.200E-08 | 4.100E-08 | 8.500E-08 |
| 20048 | 1003 | S1003G | C | 6.3e-08 | 1.9e-08 | 3.9e-08 | 4.033E-08 | 2.203E-08 | 1.900E-08 | 6.300E-08 |
| 20049 | 1003 | S1003H | C | 2.6e-08 | 1.8e-08 | 6.8e-08 | 3.733E-08 | 2.686E-08 | 1.800E-08 | 6.800E-08 |
| 20050 | 1003 | S1003I | C | 2.7e-08 | 3.1e-08 | 8e-08 | 4.600E-08 | 2.951E-08 | 2.700E-08 | 8.000E-08 |
| 20051 | 1003 | S1003L | C | 1.1e-07 | 5.6e-08 | 2.9e-08 | 6.500E-08 | 4.124E-08 | 2.900E-08 | 1.100E-07 |
| 20052 | 1003 | S1003K | C | 1.1e-07 | 3.8e-08 | 5.4e-08 | 6.733E-08 | 3.781E-08 | 3.800E-08 | 1.100E-07 |
| 20053 | 1003 | S1003M | C | 4.3e-08 | 5.3e-08 | 5e-08 | 4.867E-08 | 5.132E-09 | 4.300E-08 | 5.300E-08 |
| 20054 | 1003 | S1003F | C | 7.2e-08 | 1.2e-07 | 4.3e-08 | 7.833E-08 | 3.889E-08 | 4.300E-08 | 1.200E-07 |
| 20055 | 1003 | S1003P | C | 5.5e-08 | 4.7e-08 | 2.1e-08 | 4.100E-08 | 1.778E-08 | 2.100E-08 | 5.500E-08 |
| 20056 | 1003 | S1003S | C | 4.8e-08 | 2.2e-08 | 1.2e-07 | 6.333E-08 | 5.077E-08 | 2.200E-08 | 1.200E-07 |
| 20057 | 1003 | S1003T | C | 4.9e-08 | 6.5e-08 | 1.6e-07 | 9.133E-08 | 6.000E-08 | 4.900E-08 | 1.600E-07 |
| 20058 | 1003 | S1003W | C | 8.1e-08 | 4.4e-08 | 5e-09 | 4.333E-08 | 3.800E-08 | 5.000E-09 | 8.100E-08 |
| 20059 | 1003 | S1003Y | C | 3.8e-08 | 5.5e-08 | 7.2e-08 | 5.500E-08 | 1.700E-08 | 3.800E-08 | 7.200E-08 |

| | | | | | | | | | | |
|---|---|---|---|---|---|---|---|---|---|---|
| 20060 | 1003 | S1003V | C | 2.4e-08 | 4.5e-08 | 5.3e-08 | 4.067E-08 | 1.498E-08 | 2.400E-08 | 5.300E-08 |
| 20061 | 1004 | G1004A | C | 6.1e-08 | 7.7e-08 | 2.1e-07 | 1.160E-07 | 8.180E-08 | 6.100E-08 | 2.100E-07 |
| 20062 | 1004 | G1004R | C | 1.5e-08 | 3.7e-08 | 9.1e-08 | 4.767E-08 | 3.911E-08 | 1.500E-08 | 9.100E-08 |
| 20063 | 1004 | G1004N | C | 3.9e-08 | 1.4e-08 | 7.2e-08 | 4.167E-08 | 2.909E-08 | 1.400E-08 | 7.200E-08 |
| 20064 | 1004 | G1004D | C | 8.2e-08 | 2.9e-08 | 6.7e-08 | 5.933E-08 | 2.732E-08 | 2.900E-08 | 8.200E-08 |
| 20065 | 1004 | G1004C | C | 8.7e-09 | 7.3e-08 | 4.3e-08 | 4.157E-08 | 3.217E-08 | 8.700E-09 | 7.300E-08 |
| 20066 | 1004 | G1004E | C | 1.5e-07 | 2.3e-08 | 1.2e-07 | 9.767E-08 | 6.638E-08 | 2.300E-08 | 1.500E-07 |
| 20067 | 1004 | G1004Q | C | 2.1e-08 | 5.9e-08 | 2.9e-08 | 3.633E-08 | 2.003E-08 | 2.100E-08 | 5.900E-08 |
| 20068 | 1004 | G1004G | C | 1.9e-08 | 9.5e-08 | 3.4e-08 | 4.933E-08 | 4.025E-08 | 1.900E-08 | 9.500E-08 |
| 20069 | 1004 | G1004H | C | 1.4e-08 | 7e-08 | 5.1e-08 | 4.500E-08 | 2.848E-08 | 1.400E-08 | 7.000E-08 |
| 20070 | 1004 | G1004I | C | 3.4e-08 | 3.8e-08 | 1.6e-08 | 2.933E-08 | 1.172E-08 | 1.600E-08 | 3.800E-08 |
| 20071 | 1004 | G1004L | C | 7.4e-08 | 8.5e-08 | 2.7e-08 | 6.200E-08 | 3.081E-08 | 2.700E-08 | 8.500E-08 |
| 20072 | 1004 | G1004K | C | 3.9e-08 | 5e-08 | 2.9e-08 | 3.933E-08 | 1.050E-08 | 2.900E-08 | 5.000E-08 |
| 20073 | 1004 | G1004M | C | 7e-08 | 4.9e-08 | 4.3e-08 | 5.400E-08 | 1.418E-08 | 4.300E-08 | 7.000E-08 |
| 20074 | 1004 | G1004F | C | 3.6e-08 | 1.3e-08 | 1.1e-07 | 5.300E-08 | 5.069E-08 | 1.300E-08 | 1.100E-07 |
| 20075 | 1004 | G1004P | C | 4.9e-08 | 3.8e-08 | 2.8e-08 | 3.833E-08 | 1.050E-08 | 2.800E-08 | 4.900E-08 |
| 20076 | 1004 | G1004S | C | 5.1e-08 | 3.1e-08 | 1.5e-08 | 3.233E-08 | 1.804E-08 | 1.500E-08 | 5.100E-08 |
| 20077 | 1004 | G1004T | C | 5.2e-08 | 9.8e-08 | 6e-08 | 7.000E-08 | 2.458E-08 | 5.200E-08 | 9.800E-08 |
| 20078 | 1004 | G1004W | C | 6.4e-08 | 2.9e-08 | 4e-08 | 4.433E-08 | 1.790E-08 | 2.900E-08 | 6.400E-08 |
| 20079 | 1004 | G1004Y | C | 3.7e-08 | 5e-08 | 3.7e-08 | 4.133E-08 | 7.506E-09 | 3.700E-08 | 5.000E-08 |
| 20080 | 1004 | G1004V | C | 6.6e-08 | 6.6e-08 | 1.4e-08 | 4.867E-08 | 3.002E-08 | 1.400E-08 | 6.600E-08 |
| 20081 | 1005 | V1005A | C | 6.4e-08 | 6.6e-08 | 5.4e-08 | 6.133E-08 | 6.429E-09 | 5.400E-08 | 6.600E-08 |
| 20082 | 1005 | V1005R | C | 4.2e-08 | 1.3e-07 | 7e-08 | 8.067E-08 | 4.496E-08 | 4.200E-08 | 1.300E-07 |
| 20083 | 1005 | V1005N | C | 1.6e-08 | 1.1e-07 | 2.4e-08 | 5.000E-08 | 5.212E-08 | 1.600E-08 | 1.100E-07 |
| 20084 | 1005 | V1005D | C | 3.1e-08 | 5e-08 | 6.3e-08 | 4.800E-08 | 1.609E-08 | 3.100E-08 | 6.300E-08 |
| 20085 | 1005 | V1005C | C | 2.2e-08 | 8.2e-08 | 4.4e-08 | 4.933E-08 | 3.035E-08 | 2.200E-08 | 8.200E-08 |
| 20086 | 1005 | V1005E | C | 4.9e-08 | 3.9e-08 | 4.9e-08 | 4.567E-08 | 5.774E-09 | 3.900E-08 | 4.900E-08 |
| 20087 | 1005 | V1005Q | C | 4.9e-08 | 3.7e-08 | 1.7e-08 | 3.433E-08 | 1.617E-08 | 1.700E-08 | 4.900E-08 |
| 20088 | 1005 | V1005G | C | 1e-07 | 5.2e-08 | 7.4e-08 | 7.533E-08 | 2.403E-08 | 5.200E-08 | 1.000E-07 |
| 20089 | 1005 | V1005H | C | 3.8e-08 | 1.6e-08 | 2.2e-08 | 2.533E-08 | 1.137E-08 | 1.600E-08 | 3.800E-08 |
| 20090 | 1005 | V1005I | C | 2.7e-08 | 2.8e-08 | 8.1e-08 | 4.533E-08 | 3.089E-08 | 2.700E-08 | 8.100E-08 |
| 20091 | 1005 | V1005L | C | 3e-08 | 6.6e-08 | 7.1e-08 | 5.567E-08 | 2.237E-08 | 3.000E-08 | 7.100E-08 |
| 20092 | 1005 | V1005K | C | 3.7e-08 | 5.3e-08 | 7.5e-08 | 5.500E-08 | 1.908E-08 | 3.700E-08 | 7.500E-08 |
| 20093 | 1005 | V1005M | C | 3.7e-08 | 1.4e-08 | 3.5e-08 | 2.867E-08 | 1.274E-08 | 1.400E-08 | 3.700E-08 |
| 20094 | 1005 | V1005F | C | 3.6e-08 | 2.6e-08 | 8.2e-08 | 4.800E-08 | 2.987E-08 | 2.600E-08 | 8.200E-08 |
| 20095 | 1005 | V1005P | C | 6.1e-08 | 7.8e-08 | 7.6e-08 | 7.167E-08 | 9.292E-09 | 6.100E-08 | 7.800E-08 |
| 20096 | 1005 | V1005S | C | 1.1e-07 | 5.3e-08 | 5.4e-08 | 7.233E-08 | 3.262E-08 | 5.300E-08 | 1.100E-07 |
| 20097 | 1005 | V1005T | C | 6.9e-08 | 7.2e-08 | 1.3e-07 | 9.033E-08 | 3.439E-08 | 6.900E-08 | 1.300E-07 |
| 20098 | 1005 | V1005W | C | 2.2e-08 | 5.8e-08 | 4.3e-08 | 4.100E-08 | 1.808E-08 | 2.200E-08 | 5.800E-08 |
| 20099 | 1005 | V1005Y | C | 9.1e-08 | 2.1e-08 | 4.8e-08 | 5.333E-08 | 3.530E-08 | 2.100E-08 | 9.100E-08 |
| 20100 | 1005 | V1005V | C | 3.9e-08 | 2.4e-08 | 1.1e-07 | 5.767E-08 | 4.594E-08 | 2.400E-08 | 1.100E-07 |
| 20101 | 1006 | H1006A | C | 2.7e-08 | 3.1e-08 | 9.1e-08 | 4.967E-08 | 3.585E-08 | 2.700E-08 | 9.100E-08 |
| 20102 | 1006 | H1006R | C | 4e-08 | 4.9e-08 | 7.8e-08 | 5.567E-08 | 1.986E-08 | 4.000E-08 | 7.800E-08 |
| 20103 | 1006 | H1006N | C | 3.3e-08 | 2.3e-08 | 4.3e-08 | 3.300E-08 | 1.000E-08 | 2.300E-08 | 4.300E-08 |
| 20104 | 1006 | H1006D | C | 9.5e-08 | 6.2e-08 | 7.4e-08 | 7.700E-08 | 1.670E-08 | 6.200E-08 | 9.500E-08 |
| 20105 | 1006 | H1006C | C | 3.1e-08 | 9.8e-08 | 4.4e-08 | 5.767E-08 | 3.553E-08 | 3.100E-08 | 9.800E-08 |
| 20106 | 1006 | H1006E | C | 5e-08 | 5.5e-08 | 4e-08 | 4.833E-08 | 7.638E-09 | 4.000E-08 | 5.500E-08 |
| 20107 | 1006 | H1006Q | C | 1.5e-07 | 8.9e-08 | 4.4e-08 | 9.433E-08 | 5.320E-08 | 4.400E-08 | 1.500E-07 |
| 20108 | 1006 | H1006G | C | 8.3e-08 | 1e-07 | 7.1e-08 | 8.467E-08 | 1.457E-08 | 7.100E-08 | 1.000E-07 |
| 20109 | 1006 | H1006H | C | 6.7e-08 | 2.7e-08 | 2.2e-08 | 3.867E-08 | 2.466E-08 | 2.200E-08 | 6.700E-08 |
| 20110 | 1006 | H1006I | C | 3.2e-08 | 1.4e-08 | 3.8e-08 | 2.800E-08 | 1.249E-08 | 1.400E-08 | 3.800E-08 |
| 20111 | 1006 | H1006L | C | 5.3e-08 | 7.7e-08 | 5.2e-09 | 4.507E-08 | 3.655E-08 | 5.200E-09 | 7.700E-08 |
| 20112 | 1006 | H1006K | C | 6.3e-08 | 3e-08 | 4.7e-08 | 4.667E-08 | 1.650E-08 | 3.000E-08 | 6.300E-08 |
| 20113 | 1006 | H1006M | C | 3e-08 | 2e-08 | 6.3e-08 | 3.767E-08 | 2.250E-08 | 2.000E-08 | 6.300E-08 |
| 20114 | 1006 | H1006F | C | 3.6e-08 | 2.4e-08 | 1e-07 | 5.333E-08 | 4.086E-08 | 2.400E-08 | 1.000E-07 |
| 20115 | 1006 | H1006P | C | 3.2e-08 | 8.5e-08 | 3.8e-08 | 5.167E-08 | 2.902E-08 | 3.200E-08 | 8.500E-08 |
| 20116 | 1006 | H1006S | C | 2.1e-08 | 1.4e-08 | 4.6e-08 | 2.700E-08 | 1.682E-08 | 1.400E-08 | 4.600E-08 |
| 20117 | 1006 | H1006T | C | 2.8e-08 | 4.7e-08 | 5e-08 | 4.167E-08 | 1.193E-08 | 2.800E-08 | 5.000E-08 |
| 20118 | 1006 | H1006W | C | 3.8e-08 | 3e-08 | 2.2e-08 | 3.000E-08 | 8.000E-09 | 2.200E-08 | 3.800E-08 |
| 20119 | 1006 | H1006Y | C | 2.9e-08 | 4.3e-08 | 3.5e-08 | 3.567E-08 | 7.024E-09 | 2.900E-08 | 4.300E-08 |
| 20120 | 1006 | H1006V | C | 2.4e-08 | 1.5e-07 | 3.5e-08 | 6.967E-08 | 6.979E-08 | 2.400E-08 | 1.500E-07 |
| 20121 | 1007 | T1007A | C | 3.5e-08 | 2.5e-08 | 1.6e-08 | 2.533E-08 | 9.504E-09 | 1.600E-08 | 3.500E-08 |
| 20122 | 1007 | T1007R | C | 7.1e-08 | 3.3e-08 | 6.8e-08 | 5.733E-08 | 2.113E-08 | 3.300E-08 | 7.100E-08 |
| 20123 | 1007 | T1007N | C | 9.2e-08 | 2.6e-08 | 6.8e-08 | 6.200E-08 | 3.341E-08 | 2.600E-08 | 9.200E-08 |
| 20124 | 1007 | T1007D | C | 7.7e-08 | 2.3e-08 | 7.8e-08 | 5.933E-08 | 3.147E-08 | 2.300E-08 | 7.800E-08 |
| 20125 | 1007 | T1007C | C | 2.1e-08 | 6e-08 | 8.9e-08 | 5.667E-08 | 3.412E-08 | 2.100E-08 | 8.900E-08 |
| 20126 | 1007 | T1007E | C | 4.6e-08 | 3.5e-08 | 3.3e-08 | 3.800E-08 | 7.000E-09 | 3.300E-08 | 4.600E-08 |
| 20127 | 1007 | T1007Q | C | 2.3e-08 | 2.7e-08 | 2.7e-08 | 2.567E-08 | 2.309E-09 | 2.300E-08 | 2.700E-08 |
| 20128 | 1007 | T1007G | C | 2.3e-08 | 1e-07 | 1.4e-08 | 4.567E-08 | 4.727E-08 | 1.400E-08 | 1.000E-07 |
| 20129 | 1007 | T1007H | C | 3.4e-08 | 5.6e-08 | 6.6e-08 | 5.200E-08 | 1.637E-08 | 3.400E-08 | 6.600E-08 |
| 20130 | 1007 | T1007I | C | 6.1e-08 | 5.6e-08 | 5.1e-08 | 5.600E-08 | 5.000E-09 | 5.100E-08 | 6.100E-08 |
| 20131 | 1007 | T1007L | C | 2.6e-08 | 3.5e-08 | 2.4e-08 | 2.833E-08 | 5.859E-09 | 2.400E-08 | 3.500E-08 |
| 20132 | 1007 | T1007K | C | 7.7e-08 | 2.1e-08 | 1.7e-07 | 8.933E-08 | 7.526E-08 | 2.100E-08 | 1.700E-07 |
| 20133 | 1007 | T1007M | C | 7.4e-08 | 1.9e-08 | 5.5e-08 | 4.933E-08 | 2.793E-08 | 1.900E-08 | 7.400E-08 |
| 20134 | 1007 | T1007F | C | 5.7e-08 | 5.4e-08 | 1.9e-08 | 4.333E-08 | 2.113E-08 | 1.900E-08 | 5.700E-08 |
| 20135 | 1007 | T1007P | C | 3.7e-08 | 4e-08 | 2.2e-08 | 3.300E-08 | 9.644E-09 | 2.200E-08 | 4.000E-08 |
| 20136 | 1007 | T1007S | C | 2.2e-08 | 3.5e-08 | 5.1e-08 | 3.600E-08 | 1.453E-08 | 2.200E-08 | 5.100E-08 |
| 20137 | 1007 | T1007T | C | 3.7e-08 | 5.4e-08 | 8.1e-08 | 5.733E-08 | 2.219E-08 | 3.700E-08 | 8.100E-08 |
| 20138 | 1007 | T1007W | C | 2.7e-08 | 7.8e-08 | 5e-08 | 5.167E-08 | 2.554E-08 | 2.700E-08 | 7.800E-08 |
| 20139 | 1007 | T1007Y | C | 3.4e-08 | 7.1e-08 | 1.8e-08 | 4.100E-08 | 2.718E-08 | 1.800E-08 | 7.100E-08 |
| 20140 | 1007 | T1007V | C | 2.8e-08 | 5.2e-08 | 9e-08 | 5.667E-08 | 3.126E-08 | 2.800E-08 | 9.000E-08 |
| 20141 | 1008 | F1008A | C | 4.9e-08 | 9.5e-08 | 3.3e-08 | 5.900E-08 | 3.219E-08 | 3.300E-08 | 9.500E-08 |
| 20142 | 1008 | F1008R | C | 2.2e-08 | 2.2e-08 | 2.2e-08 | 2.200E-08 | 0.000E+00 | 2.200E-08 | 2.200E-08 |
| 20143 | 1008 | F1008N | C | 3e-08 | 2.6e-08 | 3.2e-08 | 2.933E-08 | 3.055E-09 | 2.600E-08 | 3.200E-08 |
| 20144 | 1008 | F1008D | C | 1.2e-07 | 3.2e-08 | 6.2e-08 | 7.133E-08 | 4.474E-08 | 3.200E-08 | 1.200E-07 |

| | | | | | | | | | | |
|---|---|---|---|---|---|---|---|---|---|---|
| 20145 | 1008 | F1008C | C | 5.2e-08 | 2.6e-08 | 4e-08 | 3.933E-08 | 1.301E-08 | 2.600E-08 | 5.200E-08 |
| 20146 | 1008 | F1008E | C | 3.2e-08 | 6.3e-08 | 1.8e-08 | 3.767E-08 | 2.303E-08 | 1.800E-08 | 6.300E-08 |
| 20147 | 1008 | F1008Q | C | 5.6e-08 | 1.6e-08 | 3.2e-08 | 3.467E-08 | 2.013E-08 | 1.600E-08 | 5.600E-08 |
| 20148 | 1008 | F1008G | C | 4.9e-08 | 1.6e-08 | 1.7e-08 | 2.733E-08 | 1.877E-08 | 1.600E-08 | 4.900E-08 |
| 20149 | 1008 | F1008H | C | 1.3e-08 | 1e-08 | 8.3e-08 | 3.533E-08 | 4.131E-08 | 1.000E-08 | 8.300E-08 |
| 20150 | 1008 | F1008I | C | 3e-08 | 3.7e-08 | 5.1e-08 | 3.933E-08 | 1.069E-08 | 3.000E-08 | 5.100E-08 |
| 20151 | 1008 | F1008L | C | 4.8e-08 | 1e-07 | 8.4e-08 | 7.733E-08 | 2.663E-08 | 4.800E-08 | 1.000E-07 |
| 20152 | 1008 | F1008K | C | 6e-08 | 1.8e-08 | 3.6e-08 | 3.800E-08 | 2.107E-08 | 1.800E-08 | 6.000E-08 |
| 20153 | 1008 | F1008M | C | 3.1e-08 | 2.3e-08 | 3e-08 | 2.800E-08 | 4.359E-09 | 2.300E-08 | 3.100E-08 |
| 20154 | 1008 | F1008F | C | 4.9e-08 | 5.1e-08 | 4.1e-08 | 4.700E-08 | 5.292E-09 | 4.100E-08 | 5.100E-08 |
| 20155 | 1008 | F1008P | C | 8.3e-08 | 2e-08 | 7.8e-08 | 6.033E-08 | 3.502E-08 | 2.000E-08 | 8.300E-08 |
| 20156 | 1008 | F1008S | C | 7.1e-08 | 4.7e-08 | 1.9e-08 | 4.567E-08 | 2.603E-08 | 1.900E-08 | 7.100E-08 |
| 20157 | 1008 | F1008T | C | 5e-08 | 1.3e-07 | 1.1e-07 | 9.667E-08 | 4.163E-08 | 5.000E-08 | 1.300E-07 |
| 20158 | 1008 | F1008W | C | 1.3e-07 | 3.5e-08 | 4.9e-08 | 7.133E-08 | 5.129E-08 | 3.500E-08 | 1.300E-07 |
| 20159 | 1008 | F1008Y | C | 1.6e-08 | 2.9e-08 | 2.6e-08 | 2.367E-08 | 6.807E-09 | 1.600E-08 | 2.900E-08 |
| 20160 | 1008 | F1008V | C | 1e-07 | 2.8e-08 | 3.4e-08 | 5.400E-08 | 3.995E-08 | 2.800E-08 | 1.000E-07 |
| 20161 | 1009 | P1009A | C | 5.6e-08 | 3.9e-08 | 3.6e-08 | 4.367E-08 | 1.079E-08 | 3.600E-08 | 5.600E-08 |
| 20162 | 1009 | P1009R | C | 1e-07 | 2.3e-08 | 4.8e-08 | 5.700E-08 | 3.928E-08 | 2.300E-08 | 1.000E-07 |
| 20163 | 1009 | P1009N | C | 1.8e-08 | 2.8e-08 | 3.7e-08 | 2.767E-08 | 9.504E-09 | 1.800E-08 | 3.700E-08 |
| 20164 | 1009 | P1009D | C | 1.9e-08 | 2.6e-08 | 3.1e-08 | 2.533E-08 | 6.028E-09 | 1.900E-08 | 3.100E-08 |
| 20165 | 1009 | P1009C | C | 7.9e-08 | 2.5e-08 | 4.4e-08 | 4.933E-08 | 2.739E-08 | 2.500E-08 | 7.900E-08 |
| 20166 | 1009 | P1009E | C | 2.2e-08 | 4.4e-08 | 2.8e-08 | 3.133E-08 | 1.137E-08 | 2.200E-08 | 4.400E-08 |
| 20167 | 1009 | P1009Q | C | 1.5e-08 | 5.6e-08 | 3.4e-08 | 3.500E-08 | 2.052E-08 | 1.500E-08 | 5.600E-08 |
| 20168 | 1009 | P1009G | C | 5.3e-08 | 3.8e-08 | 5.8e-08 | 4.967E-08 | 1.041E-08 | 3.800E-08 | 5.800E-08 |
| 20169 | 1009 | P1009H | C | 1.8e-08 | 7.1e-08 | 4.4e-08 | 4.433E-08 | 2.650E-08 | 1.800E-08 | 7.100E-08 |
| 20170 | 1009 | P1009I | C | 3e-08 | 8.1e-08 | 3.2e-08 | 4.767E-08 | 2.888E-08 | 3.000E-08 | 8.100E-08 |
| 20171 | 1009 | P1009L | C | 4.5e-08 | 3.7e-08 | 4.3e-08 | 4.167E-08 | 4.163E-09 | 3.700E-08 | 4.500E-08 |
| 20172 | 1009 | P1009K | C | 4.9e-08 | 2.6e-08 | 3.7e-08 | 3.733E-08 | 1.150E-08 | 2.600E-08 | 4.900E-08 |
| 20173 | 1009 | P1009M | C | 3.8e-08 | 5.9e-08 | 6e-08 | 5.233E-08 | 1.242E-08 | 3.800E-08 | 6.000E-08 |
| 20174 | 1009 | P1009F | C | 1.8e-08 | 3.3e-08 | 9e-08 | 4.700E-08 | 3.799E-08 | 1.800E-08 | 9.000E-08 |
| 20175 | 1009 | P1009P | C | 1.5e-07 | 3.3e-08 | 6.4e-08 | 8.233E-08 | 6.062E-08 | 3.300E-08 | 1.500E-07 |
| 20176 | 1009 | P1009S | C | 5.1e-08 | 1.9e-08 | 1.9e-08 | 2.967E-08 | 1.848E-08 | 1.900E-08 | 5.100E-08 |
| 20177 | 1009 | P1009T | C | 4e-08 | 6.3e-08 | 5.8e-08 | 5.367E-08 | 1.210E-08 | 4.000E-08 | 6.300E-08 |
| 20178 | 1009 | P1009W | C | 3.2e-08 | 6.4e-08 | 6.8e-08 | 5.467E-08 | 1.973E-08 | 3.200E-08 | 6.800E-08 |
| 20179 | 1009 | P1009Y | C | 2.5e-08 | 8.7e-08 | 2e-08 | 4.400E-08 | 3.732E-08 | 2.000E-08 | 8.700E-08 |
| 20180 | 1009 | P1009V | C | 1.8e-08 | 4.1e-08 | 6.5e-08 | 4.133E-08 | 2.350E-08 | 1.800E-08 | 6.500E-08 |
| 20181 | 1010 | A1010A | C | 5e-09 | 9.4e-08 | 1.9e-08 | 3.933E-08 | 4.786E-08 | 5.000E-09 | 9.400E-08 |
| 20182 | 1010 | A1010R | C | 4.4e-08 | 2.1e-08 | 3.2e-08 | 3.233E-08 | 1.150E-08 | 2.100E-08 | 4.400E-08 |
| 20183 | 1010 | A1010N | C | 5.6e-08 | 2.2e-08 | 3.2e-08 | 3.667E-08 | 1.747E-08 | 2.200E-08 | 5.600E-08 |
| 20184 | 1010 | A1010D | C | 5.6e-08 | 4e-08 | 6.8e-08 | 5.467E-08 | 1.405E-08 | 4.000E-08 | 6.800E-08 |
| 20185 | 1010 | A1010C | C | 6.4e-08 | 1.8e-08 | 5.7e-08 | 4.633E-08 | 2.479E-08 | 1.800E-08 | 6.400E-08 |
| 20186 | 1010 | A1010E | C | 4.5e-08 | 5.8e-08 | 5.1e-08 | 5.133E-08 | 6.506E-09 | 4.500E-08 | 5.800E-08 |
| 20187 | 1010 | A1010Q | C | 3.4e-08 | 9.7e-08 | 4e-08 | 5.700E-08 | 3.477E-08 | 3.400E-08 | 9.700E-08 |
| 20188 | 1010 | A1010G | C | 2.5e-08 | 4.3e-08 | 1e-07 | 5.600E-08 | 3.915E-08 | 2.500E-08 | 1.000E-07 |
| 20189 | 1010 | A1010H | C | 2e-08 | 4.4e-08 | 7.6e-08 | 4.667E-08 | 2.810E-08 | 2.000E-08 | 7.600E-08 |
| 20190 | 1010 | A1010I | C | 2e-08 | 2.1e-08 | 6.2e-08 | 3.433E-08 | 2.397E-08 | 2.000E-08 | 6.200E-08 |
| 20191 | 1010 | A1010L | C | 1.9e-08 | 4.2e-08 | 7.1e-08 | 4.400E-08 | 2.606E-08 | 1.900E-08 | 7.100E-08 |
| 20192 | 1010 | A1010K | C | 1.7e-08 | 5.6e-08 | 9.7e-08 | 5.667E-08 | 4.000E-08 | 1.700E-08 | 9.700E-08 |
| 20193 | 1010 | A1010M | C | 4.4e-08 | 1e-07 | 1.6e-08 | 5.333E-08 | 4.277E-08 | 1.600E-08 | 1.000E-07 |
| 20194 | 1010 | A1010F | C | 6.9e-08 | 4.4e-08 | 2.6e-08 | 4.633E-08 | 2.159E-08 | 2.600E-08 | 6.900E-08 |
| 20195 | 1010 | A1010P | C | 6.8e-08 | 6.7e-08 | 9e-08 | 7.500E-08 | 1.300E-08 | 6.700E-08 | 9.000E-08 |
| 20196 | 1010 | A1010S | C | 7.8e-08 | 8.8e-08 | 2.8e-08 | 6.467E-08 | 3.215E-08 | 2.800E-08 | 8.800E-08 |
| 20197 | 1010 | A1010T | C | 2.6e-08 | 5.8e-08 | 1.9e-08 | 3.433E-08 | 2.079E-08 | 1.900E-08 | 5.800E-08 |
| 20198 | 1010 | A1010W | C | 4.3e-08 | 3.8e-08 | 3.6e-08 | 3.900E-08 | 3.606E-09 | 3.600E-08 | 4.300E-08 |
| 20199 | 1010 | A1010Y | C | 3.3e-08 | 4e-08 | 9.3e-08 | 5.533E-08 | 3.281E-08 | 3.300E-08 | 9.300E-08 |
| 20200 | 1010 | A1010V | C | 3.2e-08 | 6.6e-08 | 2.4e-08 | 4.067E-08 | 2.230E-08 | 2.400E-08 | 6.600E-08 |
| 20201 | 1011 | V1011A | C | 2.9e-08 | 2e-08 | 2e-07 | 8.300E-08 | 1.014E-07 | 2.000E-08 | 2.000E-07 |
| 20202 | 1011 | V1011R | C | 2.9e-08 | 3.8e-08 | 9.4e-08 | 5.367E-08 | 3.522E-08 | 2.900E-08 | 9.400E-08 |
| 20203 | 1011 | V1011N | C | 5.8e-08 | 5.1e-08 | 7.5e-08 | 6.133E-08 | 1.234E-08 | 5.100E-08 | 7.500E-08 |
| 20204 | 1011 | V1011D | C | 5.2e-08 | 5.5e-08 | 1.8e-08 | 4.167E-08 | 2.055E-08 | 1.800E-08 | 5.500E-08 |
| 20205 | 1011 | V1011C | C | 3.3e-08 | 1.4e-08 | 5.7e-08 | 3.467E-08 | 2.155E-08 | 1.400E-08 | 5.700E-08 |
| 20206 | 1011 | V1011E | C | 2.9e-08 | 3.1e-08 | 5.3e-08 | 3.767E-08 | 1.332E-08 | 2.900E-08 | 5.300E-08 |
| 20207 | 1011 | V1011Q | C | 8e-08 | 3.4e-08 | 8.2e-08 | 6.533E-08 | 2.715E-08 | 3.400E-08 | 8.200E-08 |
| 20208 | 1011 | V1011G | C | 3e-08 | 6.7e-08 | 4.7e-08 | 4.800E-08 | 1.852E-08 | 3.000E-08 | 6.700E-08 |
| 20209 | 1011 | V1011H | C | 3.5e-08 | 5e-08 | 7.8e-08 | 5.433E-08 | 2.183E-08 | 3.500E-08 | 7.800E-08 |
| 20210 | 1011 | V1011I | C | 1.3e-08 | 6.1e-08 | 1.8e-08 | 3.067E-08 | 2.639E-08 | 1.300E-08 | 6.100E-08 |
| 20211 | 1011 | V1011L | C | 1.5e-08 | 8.7e-08 | 8.7e-08 | 6.300E-08 | 4.157E-08 | 1.500E-08 | 8.700E-08 |
| 20212 | 1011 | V1011K | C | 2.1e-08 | 1.6e-08 | 6.7e-08 | 3.467E-08 | 2.811E-08 | 1.600E-08 | 6.700E-08 |
| 20213 | 1011 | V1011M | C | 6.1e-08 | 5.2e-08 | 4.2e-08 | 5.167E-08 | 9.504E-09 | 4.200E-08 | 6.100E-08 |
| 20214 | 1011 | V1011F | C | 4.5e-08 | 1.4e-08 | 1.1e-07 | 5.633E-08 | 4.899E-08 | 1.400E-08 | 1.100E-07 |
| 20215 | 1011 | V1011P | C | 2.2e-08 | 2e-08 | 2e-08 | 2.067E-08 | 1.155E-09 | 2.000E-08 | 2.200E-08 |
| 20216 | 1011 | V1011S | C | 2.3e-08 | 1.1e-07 | 1e-07 | 7.767E-08 | 4.761E-08 | 2.300E-08 | 1.100E-07 |
| 20217 | 1011 | V1011T | C | 4.8e-08 | 3.3e-08 | 2.4e-08 | 3.500E-08 | 1.212E-08 | 2.400E-08 | 4.800E-08 |
| 20218 | 1011 | V1011W | C | 2.6e-08 | 8e-08 | 9.2e-08 | 6.600E-08 | 3.516E-08 | 2.600E-08 | 9.200E-08 |
| 20219 | 1011 | V1011Y | C | 5.2e-08 | 2.6e-08 | 2.2e-08 | 3.333E-08 | 1.629E-08 | 2.200E-08 | 5.200E-08 |
| 20220 | 1011 | V1011V | C | 5.1e-08 | 4.4e-08 | 2.4e-08 | 3.967E-08 | 1.401E-08 | 2.400E-08 | 5.100E-08 |
| 20221 | 1012 | L1012A | C | 3.4e-08 | 4.6e-08 | 2.2e-08 | 3.400E-08 | 1.200E-08 | 2.200E-08 | 4.600E-08 |
| 20222 | 1012 | L1012R | C | 5.3e-08 | 8.8e-08 | 2.1e-08 | 5.400E-08 | 3.351E-08 | 2.100E-08 | 8.800E-08 |
| 20223 | 1012 | L1012N | C | 2.7e-08 | 2.6e-08 | 5.5e-08 | 3.600E-08 | 1.646E-08 | 2.600E-08 | 5.500E-08 |
| 20224 | 1012 | L1012D | C | 1.7e-08 | 6.9e-08 | 4.1e-08 | 4.233E-08 | 2.603E-08 | 1.700E-08 | 6.900E-08 |
| 20225 | 1012 | L1012C | C | 1.1e-07 | 2e-08 | 4.7e-08 | 5.900E-08 | 4.618E-08 | 2.000E-08 | 1.100E-07 |
| 20226 | 1012 | L1012E | C | 9.4e-08 | 6.3e-08 | 5.6e-08 | 7.100E-08 | 2.022E-08 | 5.600E-08 | 9.400E-08 |
| 20227 | 1012 | L1012Q | C | 1.5e-07 | 6.1e-08 | 3.1e-08 | 8.067E-08 | 6.189E-08 | 3.100E-08 | 1.500E-07 |
| 20228 | 1012 | L1012G | C | 1.6e-08 | 2.9e-08 | 7.6e-08 | 4.033E-08 | 3.156E-08 | 1.600E-08 | 7.600E-08 |
| 20229 | 1012 | L1012H | C | 4.5e-08 | 7.4e-08 | 7.1e-08 | 6.333E-08 | 1.595E-08 | 4.500E-08 | 7.400E-08 |

| | | | | | | | | | | |
|---|---|---|---|---|---|---|---|---|---|---|
| 20230 | 1012 | L1012I | C | 8.9e-08 | 1.3e-08 | 3.6e-08 | 4.600E-08 | 3.897E-08 | 1.300E-08 | 8.900E-08 |
| 20231 | 1012 | L1012L | C | 2.1e-08 | 3.4e-08 | 7.3e-08 | 4.267E-08 | 2.706E-08 | 2.100E-08 | 7.300E-08 |
| 20232 | 1012 | L1012K | C | 1.4e-07 | 6.5e-08 | 5.5e-08 | 8.667E-08 | 4.646E-08 | 5.500E-08 | 1.400E-07 |
| 20233 | 1012 | L1012M | C | 7.5e-08 | 5.2e-08 | 1.2e-08 | 4.633E-08 | 3.188E-08 | 1.200E-08 | 7.500E-08 |
| 20234 | 1012 | L1012F | C | 2.9e-08 | 5.3e-08 | 7.5e-09 | 2.983E-08 | 2.276E-08 | 7.500E-09 | 5.300E-08 |
| 20235 | 1012 | L1012P | C | 1.2e-08 | 6.9e-08 | 4.5e-08 | 4.200E-08 | 2.862E-08 | 1.200E-08 | 6.900E-08 |
| 20236 | 1012 | L1012S | C | 5.3e-08 | 1e-07 | 1.9e-08 | 5.733E-08 | 4.067E-08 | 1.900E-08 | 1.000E-07 |
| 20237 | 1012 | L1012T | C | 1.2e-07 | 4.8e-08 | 5.1e-08 | 7.300E-08 | 4.073E-08 | 4.800E-08 | 1.200E-07 |
| 20238 | 1012 | L1012W | C | 6.5e-08 | 2.6e-08 | 1.5e-08 | 3.533E-08 | 2.627E-08 | 1.500E-08 | 6.500E-08 |
| 20239 | 1012 | L1012Y | C | 5.9e-08 | 8.8e-08 | 1.2e-08 | 5.300E-08 | 3.835E-08 | 1.200E-08 | 8.800E-08 |
| 20240 | 1012 | L1012V | C | 3e-08 | 6.9e-08 | 3.4e-08 | 4.433E-08 | 2.146E-08 | 3.000E-08 | 6.900E-08 |
| 20241 | 1013 | Q1013A | C | 3.4e-08 | 3.7e-08 | 3.8e-08 | 3.633E-08 | 2.082E-09 | 3.400E-08 | 3.800E-08 |
| 20242 | 1013 | Q1013R | C | 1.7e-08 | 2.6e-08 | 4.2e-08 | 2.833E-08 | 1.266E-08 | 1.700E-08 | 4.200E-08 |
| 20243 | 1013 | Q1013N | C | 1.5e-08 | 2.2e-08 | 3.3e-08 | 2.333E-08 | 9.074E-09 | 1.500E-08 | 3.300E-08 |
| 20244 | 1013 | Q1013D | C | 2e-08 | 3.6e-08 | 5.7e-08 | 3.767E-08 | 1.856E-08 | 2.000E-08 | 5.700E-08 |
| 20245 | 1013 | Q1013C | C | 6.5e-08 | 3.9e-08 | 3.1e-08 | 4.500E-08 | 1.778E-08 | 3.100E-08 | 6.500E-08 |
| 20246 | 1013 | Q1013E | C | 4.3e-08 | 3.7e-08 | 5.9e-08 | 4.633E-08 | 1.137E-08 | 3.700E-08 | 5.900E-08 |
| 20247 | 1013 | Q1013Q | C | 1.8e-07 | 7.7e-08 | 2e-08 | 9.233E-08 | 8.109E-08 | 2.000E-08 | 1.800E-07 |
| 20248 | 1013 | Q1013G | C | 5.9e-08 | 4.7e-08 | 8.7e-08 | 6.433E-08 | 2.053E-08 | 4.700E-08 | 8.700E-08 |
| 20249 | 1013 | Q1013H | C | 4e-08 | 2.3e-08 | 2.9e-08 | 3.067E-08 | 8.622E-09 | 2.300E-08 | 4.000E-08 |
| 20250 | 1013 | Q1013I | C | 1.9e-08 | 2.6e-08 | 4.2e-08 | 2.900E-08 | 1.179E-08 | 1.900E-08 | 4.200E-08 |
| 20251 | 1013 | Q1013L | C | 8.3e-08 | 4.9e-08 | 4e-08 | 5.733E-08 | 2.268E-08 | 4.000E-08 | 8.300E-08 |
| 20252 | 1013 | Q1013K | C | 6.9e-08 | 2.4e-08 | 3e-08 | 4.100E-08 | 2.443E-08 | 2.400E-08 | 6.900E-08 |
| 20253 | 1013 | Q1013M | C | 1.3e-08 | 3.5e-08 | 4.2e-08 | 3.000E-08 | 1.513E-08 | 1.300E-08 | 4.200E-08 |
| 20254 | 1013 | Q1013F | C | 4.8e-08 | 3.6e-08 | 1.1e-07 | 6.467E-08 | 3.972E-08 | 3.600E-08 | 1.100E-07 |
| 20255 | 1013 | Q1013P | C | 3.7e-08 | 2.3e-08 | 7.6e-08 | 4.533E-08 | 2.747E-08 | 2.300E-08 | 7.600E-08 |
| 20256 | 1013 | Q1013S | C | 4e-08 | 2.5e-08 | 1.5e-07 | 7.167E-08 | 6.825E-08 | 2.500E-08 | 1.500E-07 |
| 20257 | 1013 | Q1013T | C | 3.3e-08 | 3.9e-08 | 4.1e-08 | 3.767E-08 | 4.163E-09 | 3.300E-08 | 4.100E-08 |
| 20258 | 1013 | Q1013W | C | 7.9e-08 | 2.8e-08 | 4e-08 | 4.900E-08 | 2.666E-08 | 2.800E-08 | 7.900E-08 |
| 20259 | 1013 | Q1013Y | C | 2.6e-08 | 5.4e-08 | 8.1e-08 | 5.367E-08 | 2.750E-08 | 2.600E-08 | 8.100E-08 |
| 20260 | 1013 | Q1013V | C | 5.7e-08 | 2e-08 | 3.2e-08 | 3.633E-08 | 1.888E-08 | 2.000E-08 | 5.700E-08 |
| 20261 | 1014 | S1014A | C | 2.8e-08 | 2.1e-08 | 4.1e-08 | 3.000E-08 | 1.015E-08 | 2.100E-08 | 4.100E-08 |
| 20262 | 1014 | S1014R | C | 3.1e-08 | 4.4e-08 | 3e-08 | 3.500E-08 | 7.810E-09 | 3.000E-08 | 4.400E-08 |
| 20263 | 1014 | S1014N | C | 4.6e-08 | 5.4e-08 | 5.6e-08 | 5.200E-08 | 5.292E-09 | 4.600E-08 | 5.600E-08 |
| 20264 | 1014 | S1014D | C | 3.9e-08 | 7.4e-08 | 4e-08 | 5.100E-08 | 1.992E-08 | 3.900E-08 | 7.400E-08 |
| 20265 | 1014 | S1014C | C | 8e-08 | 7.4e-08 | 4.5e-08 | 6.633E-08 | 1.872E-08 | 4.500E-08 | 8.000E-08 |
| 20266 | 1014 | S1014E | C | 3.1e-08 | 9.4e-08 | 2.5e-08 | 5.000E-08 | 3.822E-08 | 2.500E-08 | 9.400E-08 |
| 20267 | 1014 | S1014Q | C | 2.3e-08 | 3.8e-08 | 2.7e-08 | 2.933E-08 | 7.767E-09 | 2.300E-08 | 3.800E-08 |
| 20268 | 1014 | S1014G | C | 4.1e-08 | 5.2e-08 | 5.2e-08 | 4.833E-08 | 6.351E-09 | 4.100E-08 | 5.200E-08 |
| 20269 | 1014 | S1014H | C | 5e-08 | 4.2e-08 | 7.7e-08 | 5.633E-08 | 1.834E-08 | 4.200E-08 | 7.700E-08 |
| 20270 | 1014 | S1014I | C | 6.4e-08 | 5.3e-08 | 1e-07 | 7.233E-08 | 2.458E-08 | 5.300E-08 | 1.000E-07 |
| 20271 | 1014 | S1014L | C | 5.5e-08 | 2.2e-08 | 5.5e-08 | 4.400E-08 | 1.905E-08 | 2.200E-08 | 5.500E-08 |
| 20272 | 1014 | S1014K | C | 1.2e-07 | 9.9e-08 | 3.9e-08 | 8.600E-08 | 4.204E-08 | 3.900E-08 | 1.200E-07 |
| 20273 | 1014 | S1014M | C | 2.9e-08 | 1.2e-08 | 3.4e-08 | 2.500E-08 | 1.153E-08 | 1.200E-08 | 3.400E-08 |
| 20274 | 1014 | S1014F | C | 8.2e-08 | 4.5e-08 | 3.4e-08 | 5.367E-08 | 2.515E-08 | 3.400E-08 | 8.200E-08 |
| 20275 | 1014 | S1014P | C | 5.4e-08 | 5.7e-08 | 5.4e-08 | 5.500E-08 | 1.732E-09 | 5.400E-08 | 5.700E-08 |
| 20276 | 1014 | S1014S | C | 2.7e-08 | 9e-08 | 3.2e-08 | 4.967E-08 | 3.502E-08 | 2.700E-08 | 9.000E-08 |
| 20277 | 1014 | S1014T | C | 3.8e-08 | 4.4e-08 | 2.3e-08 | 3.500E-08 | 1.082E-08 | 2.300E-08 | 4.400E-08 |
| 20278 | 1014 | S1014W | C | 1e-07 | 2.6e-08 | 6.3e-08 | 6.300E-08 | 3.700E-08 | 2.600E-08 | 1.000E-07 |
| 20279 | 1014 | S1014Y | C | 4.2e-08 | 3.9e-08 | 2e-08 | 3.367E-08 | 1.193E-08 | 2.000E-08 | 4.200E-08 |
| 20280 | 1014 | S1014V | C | 1.8e-08 | 2.1e-08 | 6.5e-09 | 1.517E-08 | 7.654E-09 | 6.500E-09 | 2.100E-08 |
| 20281 | 1015 | S1015A | C | 4.7e-08 | 4.7e-08 | 2.5e-08 | 3.967E-08 | 1.270E-08 | 2.500E-08 | 4.700E-08 |
| 20282 | 1015 | S1015R | C | 1.3e-07 | 4.8e-08 | 1.1e-07 | 9.600E-08 | 4.276E-08 | 4.800E-08 | 1.300E-07 |
| 20283 | 1015 | S1015N | C | 6.3e-08 | 7.3e-08 | 7.2e-08 | 6.933E-08 | 5.508E-09 | 6.300E-08 | 7.300E-08 |
| 20284 | 1015 | S1015D | C | 3.9e-08 | 3.1e-08 | 7.8e-08 | 4.933E-08 | 2.515E-08 | 3.100E-08 | 7.800E-08 |
| 20285 | 1015 | S1015C | C | 9.1e-08 | 8.5e-08 | 5.4e-08 | 7.667E-08 | 1.986E-08 | 5.400E-08 | 9.100E-08 |
| 20286 | 1015 | S1015E | C | 4e-08 | 2.2e-08 | 5.4e-08 | 3.867E-08 | 1.604E-08 | 2.200E-08 | 5.400E-08 |
| 20287 | 1015 | S1015Q | C | 3.4e-08 | 1.5e-07 | 6.9e-08 | 8.433E-08 | 5.950E-08 | 3.400E-08 | 1.500E-07 |
| 20288 | 1015 | S1015G | C | 6e-08 | 4.4e-08 | 9.7e-08 | 6.700E-08 | 2.718E-08 | 4.400E-08 | 9.700E-08 |
| 20289 | 1015 | S1015H | C | 1.9e-08 | 1.4e-08 | 1.1e-07 | 4.767E-08 | 5.404E-08 | 1.400E-08 | 1.100E-07 |
| 20290 | 1015 | S1015I | C | 4.5e-08 | 6.1e-08 | 6.1e-08 | 5.567E-08 | 9.238E-09 | 4.500E-08 | 6.100E-08 |
| 20291 | 1015 | S1015L | C | 3e-08 | 2.5e-08 | 4.1e-08 | 3.200E-08 | 8.185E-09 | 2.500E-08 | 4.100E-08 |
| 20292 | 1015 | S1015K | C | 8.3e-08 | 6.6e-08 | 7.5e-08 | 7.467E-08 | 8.505E-09 | 6.600E-08 | 8.300E-08 |
| 20293 | 1015 | S1015M | C | 2.5e-08 | 3.2e-08 | 4.3e-08 | 3.333E-08 | 9.074E-09 | 2.500E-08 | 4.300E-08 |
| 20294 | 1015 | S1015F | C | 1e-07 | 4.4e-08 | 2.1e-08 | 5.500E-08 | 4.063E-08 | 2.100E-08 | 1.000E-07 |
| 20295 | 1015 | S1015P | C | 4.8e-08 | 1.2e-08 | 5.9e-08 | 3.967E-08 | 2.458E-08 | 1.200E-08 | 5.900E-08 |
| 20296 | 1015 | S1015S | C | 3.1e-08 | 5e-08 | 1.5e-08 | 3.200E-08 | 1.752E-08 | 1.500E-08 | 5.000E-08 |
| 20297 | 1015 | S1015T | C | 2.5e-08 | 5e-08 | 5.7e-08 | 4.400E-08 | 1.682E-08 | 2.500E-08 | 5.700E-08 |
| 20298 | 1015 | S1015W | C | 3.6e-08 | 8.3e-08 | 7.2e-08 | 6.367E-08 | 2.458E-08 | 3.600E-08 | 8.300E-08 |
| 20299 | 1015 | S1015Y | C | 2.9e-08 | 3.5e-08 | 6.3e-08 | 4.233E-08 | 1.815E-08 | 2.900E-08 | 6.300E-08 |
| 20300 | 1015 | S1015V | C | 5.6e-08 | 3.8e-08 | 1.5e-07 | 8.133E-08 | 6.014E-08 | 3.800E-08 | 1.500E-07 |
| 20301 | 1016 | G1016A | C | 1.1e-07 | 3.8e-08 | 6.3e-08 | 7.033E-08 | 3.656E-08 | 3.800E-08 | 1.100E-07 |
| 20302 | 1016 | G1016R | C | 4.2e-08 | 5.4e-08 | 5.2e-08 | 4.933E-08 | 6.429E-09 | 4.200E-08 | 5.400E-08 |
| 20303 | 1016 | G1016N | C | 5.7e-08 | 3.1e-08 | 4.3e-08 | 4.367E-08 | 1.301E-08 | 3.100E-08 | 5.700E-08 |
| 20304 | 1016 | G1016D | C | 4.5e-08 | 9.6e-08 | 4.8e-08 | 6.300E-08 | 2.862E-08 | 4.500E-08 | 9.600E-08 |
| 20305 | 1016 | G1016C | C | 1.5e-08 | 7.4e-08 | 3.4e-08 | 4.100E-08 | 3.012E-08 | 1.500E-08 | 7.400E-08 |
| 20306 | 1016 | G1016E | C | 4.7e-08 | 1.5e-08 | 4.6e-08 | 3.600E-08 | 1.819E-08 | 1.500E-08 | 4.700E-08 |
| 20307 | 1016 | G1016Q | C | 2.2e-08 | 4.7e-08 | 3.6e-08 | 3.500E-08 | 1.253E-08 | 2.200E-08 | 4.700E-08 |
| 20308 | 1016 | G1016G | C | 4.1e-08 | 4.9e-08 | 1.3e-08 | 3.433E-08 | 1.890E-08 | 1.300E-08 | 4.900E-08 |
| 20309 | 1016 | G1016H | C | 5.6e-08 | 3e-08 | 3.6e-08 | 4.067E-08 | 1.361E-08 | 3.000E-08 | 5.600E-08 |
| 20310 | 1016 | G1016I | C | 1.4e-08 | 3.2e-08 | 3.2e-08 | 2.600E-08 | 1.039E-08 | 1.400E-08 | 3.200E-08 |
| 20311 | 1016 | G1016L | C | 1.8e-08 | 3.3e-08 | 1.9e-08 | 2.333E-08 | 8.386E-09 | 1.800E-08 | 3.300E-08 |
| 20312 | 1016 | G1016K | C | 4.3e-08 | 4.7e-08 | 1.5e-08 | 3.500E-08 | 1.744E-08 | 1.500E-08 | 4.700E-08 |
| 20313 | 1016 | G1016M | C | 2.7e-08 | 1.7e-07 | 4.7e-08 | 8.133E-08 | 7.744E-08 | 2.700E-08 | 1.700E-07 |
| 20314 | 1016 | G1016F | C | 1.2e-07 | 5.6e-08 | 1.3e-07 | 1.020E-07 | 4.015E-08 | 5.600E-08 | 1.300E-07 |

| | | | | | | | | | | |
|---|---|---|---|---|---|---|---|---|---|---|
| 20315 | 1016 | G1016P | C | 2.9e-08 | 2.9e-08 | 8.4e-08 | 4.733E-08 | 3.175E-08 | 2.900E-08 | 8.400E-08 |
| 20316 | 1016 | G1016S | C | 6.5e-09 | 4e-08 | 1.2e-08 | 1.950E-08 | 1.797E-08 | 6.500E-09 | 4.000E-08 |
| 20317 | 1016 | G1016T | C | 1.2e-08 | 4.7e-08 | 5.3e-08 | 3.733E-08 | 2.214E-08 | 1.200E-08 | 5.300E-08 |
| 20318 | 1016 | G1016W | C | 4.3e-08 | 5.1e-08 | 2.3e-08 | 3.900E-08 | 1.442E-08 | 2.300E-08 | 5.100E-08 |
| 20319 | 1016 | G1016Y | C | 3.9e-08 | 1e-07 | 4.6e-08 | 6.167E-08 | 3.338E-08 | 3.900E-08 | 1.000E-07 |
| 20320 | 1016 | G1016V | C | 2.5e-08 | 6.7e-08 | 5.4e-08 | 4.867E-08 | 2.150E-08 | 2.500E-08 | 6.700E-08 |
| 20321 | 1017 | L1017A | C | 2e-08 | 3.2e-07 | 7.5e-08 | 1.383E-07 | 1.597E-07 | 2.000E-08 | 3.200E-07 |
| 20322 | 1017 | L1017R | C | 2.7e-08 | 1e-07 | 3.9e-08 | 5.533E-08 | 3.915E-08 | 2.700E-08 | 1.000E-07 |
| 20323 | 1017 | L1017N | C | 1.4e-08 | 3.3e-08 | 6.8e-08 | 3.833E-08 | 2.739E-08 | 1.400E-08 | 6.800E-08 |
| 20324 | 1017 | L1017D | C | 1.1e-07 | 1.3e-07 | 1.2e-08 | 8.400E-08 | 6.315E-08 | 1.200E-08 | 1.300E-07 |
| 20325 | 1017 | L1017C | C | 9.9e-08 | 2.5e-08 | 3.5e-08 | 5.300E-08 | 4.015E-08 | 2.500E-08 | 9.900E-08 |
| 20326 | 1017 | L1017E | C | 3.9e-08 | 1.6e-08 | 3.7e-08 | 3.067E-08 | 1.274E-08 | 1.600E-08 | 3.900E-08 |
| 20327 | 1017 | L1017Q | C | 4.6e-08 | 6.7e-08 | 1e-08 | 4.100E-08 | 2.883E-08 | 1.000E-08 | 6.700E-08 |
| 20328 | 1017 | L1017G | C | 3.1e-08 | 4.9e-08 | 3.4e-08 | 3.800E-08 | 9.644E-09 | 3.100E-08 | 4.900E-08 |
| 20329 | 1017 | L1017H | C | 4.1e-08 | 4.1e-08 | 3.6e-08 | 3.933E-08 | 2.887E-09 | 3.600E-08 | 4.100E-08 |
| 20330 | 1017 | L1017I | C | 3.5e-08 | 4.9e-08 | 3.2e-08 | 3.867E-08 | 9.074E-09 | 3.200E-08 | 4.900E-08 |
| 20331 | 1017 | L1017L | C | 1.7e-08 | 6.5e-08 | 3.9e-08 | 4.033E-08 | 2.403E-08 | 1.700E-08 | 6.500E-08 |
| 20332 | 1017 | L1017K | C | 2.5e-08 | 5.6e-08 | 3.9e-08 | 4.000E-08 | 1.552E-08 | 2.500E-08 | 5.600E-08 |
| 20333 | 1017 | L1017M | C | 7.2e-08 | 7e-08 | 4.1e-08 | 6.100E-08 | 1.735E-08 | 4.100E-08 | 7.200E-08 |
| 20334 | 1017 | L1017F | C | 4.9e-08 | 8.7e-08 | 7.7e-08 | 7.100E-08 | 1.970E-08 | 4.900E-08 | 8.700E-08 |
| 20335 | 1017 | L1017P | C | 2.7e-08 | 7.9e-08 | 9.5e-08 | 6.700E-08 | 3.555E-08 | 2.700E-08 | 9.500E-08 |
| 20336 | 1017 | L1017S | C | 6e-08 | 3.7e-08 | 4e-08 | 4.567E-08 | 1.250E-08 | 3.700E-08 | 6.000E-08 |
| 20337 | 1017 | L1017T | C | 6.3e-08 | 6.8e-08 | 3.5e-08 | 5.533E-08 | 1.779E-08 | 3.500E-08 | 6.800E-08 |
| 20338 | 1017 | L1017W | C | 6e-08 | 1.3e-07 | 2.9e-08 | 7.300E-08 | 5.174E-08 | 2.900E-08 | 1.300E-07 |
| 20339 | 1017 | L1017Y | C | 1.3e-08 | 9.1e-08 | 5.7e-08 | 5.367E-08 | 3.911E-08 | 1.300E-08 | 9.100E-08 |
| 20340 | 1017 | L1017V | C | 2e-08 | 8.5e-08 | 7.1e-08 | 5.867E-08 | 3.421E-08 | 2.000E-08 | 8.500E-08 |
| 20341 | 1018 | Y1018A | C | 2.5e-08 | 2.8e-08 | 3.1e-08 | 2.800E-08 | 3.000E-09 | 2.500E-08 | 3.100E-08 |
| 20342 | 1018 | Y1018R | C | 2.3e-08 | 4.9e-08 | 3.6e-08 | 3.600E-08 | 1.300E-08 | 2.300E-08 | 4.900E-08 |
| 20343 | 1018 | Y1018N | C | 3.1e-08 | 2.5e-08 | 2.1e-08 | 2.567E-08 | 5.033E-09 | 2.100E-08 | 3.100E-08 |
| 20344 | 1018 | Y1018D | C | 1.6e-08 | 8.6e-08 | 1.9e-08 | 4.033E-08 | 3.958E-08 | 1.600E-08 | 8.600E-08 |
| 20345 | 1018 | Y1018C | C | 2.4e-08 | 5.5e-08 | 2.5e-08 | 3.467E-08 | 1.762E-08 | 2.400E-08 | 5.500E-08 |
| 20346 | 1018 | Y1018E | C | 3.1e-08 | 9.1e-08 | 3e-08 | 5.067E-08 | 3.493E-08 | 3.000E-08 | 9.100E-08 |
| 20347 | 1018 | Y1018Q | C | 2.7e-08 | 5e-08 | 9.6e-08 | 5.767E-08 | 3.513E-08 | 2.700E-08 | 9.600E-08 |
| 20348 | 1018 | Y1018G | C | 5.7e-08 | 5.8e-08 | 4.5e-08 | 5.333E-08 | 7.234E-09 | 4.500E-08 | 5.800E-08 |
| 20349 | 1018 | Y1018H | C | 4.7e-08 | 3.8e-08 | 7.5e-09 | 3.083E-08 | 2.070E-08 | 7.500E-09 | 4.700E-08 |
| 20350 | 1018 | Y1018I | C | 4.8e-08 | 2.5e-08 | 3.7e-08 | 3.667E-08 | 1.150E-08 | 2.500E-08 | 4.800E-08 |
| 20351 | 1018 | Y1018L | C | 3e-08 | 2.5e-07 | 3e-08 | 1.033E-07 | 1.270E-07 | 3.000E-08 | 2.500E-07 |
| 20352 | 1018 | Y1018K | C | 3.9e-08 | 1.1e-07 | 7e-08 | 7.300E-08 | 3.559E-08 | 3.900E-08 | 1.100E-07 |
| 20353 | 1018 | Y1018M | C | 7.1e-08 | 4.5e-08 | 3.2e-08 | 4.933E-08 | 1.986E-08 | 3.200E-08 | 7.100E-08 |
| 20354 | 1018 | Y1018F | C | 2e-08 | 3.9e-08 | 8.8e-09 | 2.260E-08 | 1.527E-08 | 8.800E-09 | 3.900E-08 |
| 20355 | 1018 | Y1018P | C | 5e-08 | 5.8e-08 | 7.5e-08 | 6.100E-08 | 1.277E-08 | 5.000E-08 | 7.500E-08 |
| 20356 | 1018 | Y1018S | C | 4.2e-08 | 4.8e-08 | 4.9e-08 | 4.633E-08 | 3.786E-09 | 4.200E-08 | 4.900E-08 |
| 20357 | 1018 | Y1018T | C | 6.2e-08 | 2.6e-08 | 7.1e-08 | 5.300E-08 | 2.381E-08 | 2.600E-08 | 7.100E-08 |
| 20358 | 1018 | Y1018W | C | 4e-08 | 1.3e-08 | 2.5e-08 | 2.600E-08 | 1.353E-08 | 1.300E-08 | 4.000E-08 |
| 20359 | 1018 | Y1018Y | C | 4.3e-08 | 2e-08 | 5e-08 | 3.767E-08 | 1.570E-08 | 2.000E-08 | 5.000E-08 |
| 20360 | 1018 | Y1018V | C | 2.9e-08 | 2.5e-08 | 7.6e-08 | 4.333E-08 | 2.836E-08 | 2.500E-08 | 7.600E-08 |
| 20361 | 1019 | S1019A | C | 2.4e-08 | 5.9e-08 | 5.9e-08 | 4.733E-08 | 2.021E-08 | 2.400E-08 | 5.900E-08 |
| 20362 | 1019 | S1019R | C | 3.8e-08 | 6.1e-08 | 1.8e-08 | 3.900E-08 | 2.152E-08 | 1.800E-08 | 6.100E-08 |
| 20363 | 1019 | S1019N | C | 1.7e-08 | 2.6e-08 | 4.5e-08 | 2.933E-08 | 1.429E-08 | 1.700E-08 | 4.500E-08 |
| 20364 | 1019 | S1019D | C | 4.2e-08 | 5e-08 | 2.7e-08 | 3.967E-08 | 1.168E-08 | 2.700E-08 | 5.000E-08 |
| 20365 | 1019 | S1019C | C | 4.6e-08 | 1e-07 | 5.5e-08 | 6.700E-08 | 2.893E-08 | 4.600E-08 | 1.000E-07 |
| 20366 | 1019 | S1019E | C | 1.5e-07 | 3.7e-08 | 9.8e-08 | 9.500E-08 | 5.656E-08 | 3.700E-08 | 1.500E-07 |
| 20367 | 1019 | S1019Q | C | 4.3e-08 | 7.4e-08 | 1.4e-08 | 4.367E-08 | 3.001E-08 | 1.400E-08 | 7.400E-08 |
| 20368 | 1019 | S1019G | C | 2.1e-08 | 1e-07 | 2.1e-08 | 4.733E-08 | 4.561E-08 | 2.100E-08 | 1.000E-07 |
| 20369 | 1019 | S1019H | C | 8.8e-08 | 8.5e-08 | 6.9e-08 | 8.067E-08 | 1.021E-08 | 6.900E-08 | 8.800E-08 |
| 20370 | 1019 | S1019I | C | 6e-08 | 7.7e-08 | 3.3e-08 | 5.667E-08 | 2.219E-08 | 3.300E-08 | 7.700E-08 |
| 20371 | 1019 | S1019L | C | 3.8e-08 | 2.2e-08 | 1.4e-08 | 2.467E-08 | 1.222E-08 | 1.400E-08 | 3.800E-08 |
| 20372 | 1019 | S1019K | C | 1.4e-07 | 8.3e-08 | 7.7e-09 | 7.690E-08 | 6.636E-08 | 7.700E-09 | 1.400E-07 |
| 20373 | 1019 | S1019M | C | 4.8e-08 | 3.1e-08 | 3.9e-08 | 3.933E-08 | 8.505E-09 | 3.100E-08 | 4.800E-08 |
| 20374 | 1019 | S1019F | C | 6.8e-08 | 4.5e-08 | 6.7e-08 | 6.000E-08 | 1.300E-08 | 4.500E-08 | 6.800E-08 |
| 20375 | 1019 | S1019P | C | 4.9e-08 | 1.6e-08 | 7.3e-08 | 4.600E-08 | 2.862E-08 | 1.600E-08 | 7.300E-08 |
| 20376 | 1019 | S1019S | C | 5.6e-08 | 7.7e-08 | 9.8e-08 | 7.700E-08 | 2.100E-08 | 5.600E-08 | 9.800E-08 |
| 20377 | 1019 | S1019T | C | 2.7e-08 | 9.6e-08 | 9.8e-08 | 7.367E-08 | 4.043E-08 | 2.700E-08 | 9.800E-08 |
| 20378 | 1019 | S1019W | C | 3.3e-08 | 4.5e-08 | 3e-08 | 3.600E-08 | 7.937E-09 | 3.000E-08 | 4.500E-08 |
| 20379 | 1019 | S1019Y | C | 5.9e-08 | 2.4e-08 | 5.6e-08 | 4.633E-08 | 1.940E-08 | 2.400E-08 | 5.900E-08 |
| 20380 | 1019 | S1019V | C | 4.1e-08 | 5.9e-08 | 2.6e-08 | 4.200E-08 | 1.652E-08 | 2.600E-08 | 5.900E-08 |
| 20381 | 1020 | L1020A | C | 2.8e-08 | 3.1e-08 | 2.2e-08 | 2.700E-08 | 4.583E-09 | 2.200E-08 | 3.100E-08 |
| 20382 | 1020 | L1020R | C | 1e-08 | 3.9e-08 | 2.6e-08 | 2.500E-08 | 1.453E-08 | 1.000E-08 | 3.900E-08 |
| 20383 | 1020 | L1020N | C | 4.1e-08 | 6.5e-08 | 3.3e-08 | 4.633E-08 | 1.665E-08 | 3.300E-08 | 6.500E-08 |
| 20384 | 1020 | L1020D | C | 1e-08 | 3.5e-08 | 4e-08 | 2.833E-08 | 1.607E-08 | 1.000E-08 | 4.000E-08 |
| 20385 | 1020 | L1020C | C | 3.3e-08 | 4.1e-08 | 8.2e-08 | 5.200E-08 | 2.629E-08 | 3.300E-08 | 8.200E-08 |
| 20386 | 1020 | L1020E | C | 4.5e-08 | 3.1e-08 | 3.7e-08 | 3.767E-08 | 7.024E-09 | 3.100E-08 | 4.500E-08 |
| 20387 | 1020 | L1020Q | C | 5.3e-08 | 4.7e-08 | 1e-07 | 6.667E-08 | 2.902E-08 | 4.700E-08 | 1.000E-07 |
| 20388 | 1020 | L1020G | C | 5e-08 | 3.1e-08 | 4.5e-08 | 4.200E-08 | 9.849E-09 | 3.100E-08 | 5.000E-08 |
| 20389 | 1020 | L1020H | C | 9.1e-08 | 2.7e-08 | 5e-08 | 5.600E-08 | 3.242E-08 | 2.700E-08 | 9.100E-08 |
| 20390 | 1020 | L1020I | C | 3.8e-08 | 3.8e-08 | 6.7e-08 | 4.767E-08 | 1.674E-08 | 3.800E-08 | 6.700E-08 |
| 20391 | 1020 | L1020L | C | 4.1e-08 | 1.2e-07 | 4.6e-08 | 6.900E-08 | 4.424E-08 | 4.100E-08 | 1.200E-07 |
| 20392 | 1020 | L1020K | C | 8.4e-08 | 1.9e-08 | 1.5e-07 | 8.433E-08 | 6.550E-08 | 1.900E-08 | 1.500E-07 |
| 20393 | 1020 | L1020M | C | 3.5e-08 | 5.1e-08 | 6.6e-08 | 5.067E-08 | 1.550E-08 | 3.500E-08 | 6.600E-08 |
| 20394 | 1020 | L1020F | C | 5.2e-08 | 5e-08 | 6.3e-08 | 5.500E-08 | 7.000E-09 | 5.000E-08 | 6.300E-08 |
| 20395 | 1020 | L1020P | C | 5.3e-08 | 8.3e-09 | 3.8e-08 | 3.310E-08 | 2.275E-08 | 8.300E-09 | 5.300E-08 |
| 20396 | 1020 | L1020S | C | 1.3e-08 | 1.7e-08 | 8.4e-08 | 3.800E-08 | 3.989E-08 | 1.300E-08 | 8.400E-08 |
| 20397 | 1020 | L1020T | C | 1.3e-08 | 4.7e-08 | 2.7e-08 | 2.900E-08 | 1.709E-08 | 1.300E-08 | 4.700E-08 |
| 20398 | 1020 | L1020W | C | 4.1e-08 | 4.9e-08 | 5.6e-08 | 4.867E-08 | 7.506E-09 | 4.100E-08 | 5.600E-08 |
| 20399 | 1020 | L1020Y | C | 6.2e-08 | 4.5e-08 | 7.1e-08 | 5.933E-08 | 1.320E-08 | 4.500E-08 | 7.100E-08 |

| | | | | | | | | | | |
|---|---|---|---|---|---|---|---|---|---|---|
| 20400 | 1020 | L1020V | C | 2.2e-08 | 2.8e-08 | 1.9e-08 | 2.300E-08 | 4.583E-09 | 1.900E-08 | 2.800E-08 |
| 20401 | 1021 | S1021A | C | 3.6e-08 | 3.5e-08 | 7.7e-08 | 4.933E-08 | 2.397E-08 | 3.500E-08 | 7.700E-08 |
| 20402 | 1021 | S1021R | C | 6.3e-08 | 2.6e-08 | 2.4e-08 | 3.767E-08 | 2.196E-08 | 2.400E-08 | 6.300E-08 |
| 20403 | 1021 | S1021N | C | 2.9e-08 | 6.1e-08 | 1.2e-07 | 7.000E-08 | 4.616E-08 | 2.900E-08 | 1.200E-07 |
| 20404 | 1021 | S1021D | C | 4.2e-08 | 4.1e-08 | 2.5e-08 | 3.600E-08 | 9.539E-09 | 2.500E-08 | 4.200E-08 |
| 20405 | 1021 | S1021C | C | 2.6e-08 | 4.1e-08 | 4e-08 | 3.567E-08 | 8.386E-09 | 2.600E-08 | 4.100E-08 |
| 20406 | 1021 | S1021E | C | 3.9e-08 | 4.2e-08 | 3.5e-08 | 3.867E-08 | 3.512E-09 | 3.500E-08 | 4.200E-08 |
| 20407 | 1021 | S1021Q | C | 1.7e-08 | 5.8e-08 | 5.4e-08 | 4.300E-08 | 2.261E-08 | 1.700E-08 | 5.800E-08 |
| 20408 | 1021 | S1021G | C | 2.1e-08 | 4e-08 | 3.1e-08 | 3.067E-08 | 9.504E-09 | 2.100E-08 | 4.000E-08 |
| 20409 | 1021 | S1021H | C | 5.7e-08 | 5.7e-08 | 1.1e-07 | 7.467E-08 | 3.060E-08 | 5.700E-08 | 1.100E-07 |
| 20410 | 1021 | S1021I | C | 5.6e-08 | 2.3e-08 | 2.2e-08 | 3.367E-08 | 1.935E-08 | 2.200E-08 | 5.600E-08 |
| 20411 | 1021 | S1021L | C | 4.4e-08 | 5.4e-08 | 6.7e-08 | 5.500E-08 | 1.153E-08 | 4.400E-08 | 6.700E-08 |
| 20412 | 1021 | S1021K | C | 4.5e-08 | 4e-08 | 7.3e-08 | 5.267E-08 | 1.779E-08 | 4.000E-08 | 7.300E-08 |
| 20413 | 1021 | S1021M | C | 5.6e-08 | 2.4e-08 | 3.6e-08 | 3.867E-08 | 1.617E-08 | 2.400E-08 | 5.600E-08 |
| 20414 | 1021 | S1021F | C | 4.2e-08 | 4.2e-08 | 2.6e-08 | 3.667E-08 | 9.238E-09 | 2.600E-08 | 4.200E-08 |
| 20415 | 1021 | S1021P | C | 3.7e-08 | 2.5e-08 | 3.5e-08 | 3.233E-08 | 6.429E-09 | 2.500E-08 | 3.700E-08 |
| 20416 | 1021 | S1021S | C | 8.8e-08 | 5.7e-08 | 2.4e-08 | 5.633E-08 | 3.201E-08 | 2.400E-08 | 8.800E-08 |
| 20417 | 1021 | S1021T | C | 4.7e-08 | 1.6e-08 | 3.6e-08 | 3.300E-08 | 1.572E-08 | 1.600E-08 | 4.700E-08 |
| 20418 | 1021 | S1021W | C | 3.2e-08 | 9.2e-08 | 3.8e-08 | 5.400E-08 | 3.305E-08 | 3.200E-08 | 9.200E-08 |
| 20419 | 1021 | S1021Y | C | 6.2e-08 | 4.8e-08 | 2.4e-08 | 4.467E-08 | 1.922E-08 | 2.400E-08 | 6.200E-08 |
| 20420 | 1021 | S1021V | C | 2.4e-08 | 3.1e-08 | 8.9e-08 | 4.800E-08 | 3.568E-08 | 2.400E-08 | 8.900E-08 |
| 20421 | 1022 | S1022A | C | 3e-08 | 6.2e-08 | 1.3e-08 | 3.500E-08 | 2.488E-08 | 1.300E-08 | 6.200E-08 |
| 20422 | 1022 | S1022R | C | 5.8e-08 | 1.2e-07 | 2.8e-08 | 6.867E-08 | 4.692E-08 | 2.800E-08 | 1.200E-07 |
| 20423 | 1022 | S1022N | C | 2.3e-08 | 5.3e-08 | 1.1e-07 | 6.200E-08 | 4.419E-08 | 2.300E-08 | 1.100E-07 |
| 20424 | 1022 | S1022D | C | 2.8e-08 | 2.9e-08 | 1.8e-08 | 2.500E-08 | 6.083E-09 | 1.800E-08 | 2.900E-08 |
| 20425 | 1022 | S1022C | C | 3.2e-08 | 5.6e-08 | 3.3e-08 | 4.033E-08 | 1.358E-08 | 3.200E-08 | 5.600E-08 |
| 20426 | 1022 | S1022E | C | 2.9e-08 | 8.1e-08 | 9.8e-08 | 6.933E-08 | 3.595E-08 | 2.900E-08 | 9.800E-08 |
| 20427 | 1022 | S1022Q | C | 6e-08 | 2.6e-08 | 2.5e-08 | 3.700E-08 | 1.992E-08 | 2.500E-08 | 6.000E-08 |
| 20428 | 1022 | S1022G | C | 7.7e-08 | 5.4e-08 | 1.8e-08 | 4.967E-08 | 2.974E-08 | 1.800E-08 | 7.700E-08 |
| 20429 | 1022 | S1022H | C | 2.2e-08 | 4.1e-08 | 1.5e-08 | 2.600E-08 | 1.345E-08 | 1.500E-08 | 4.100E-08 |
| 20430 | 1022 | S1022I | C | 4.3e-08 | 2.5e-08 | 2.6e-08 | 3.133E-08 | 1.012E-08 | 2.500E-08 | 4.300E-08 |
| 20431 | 1022 | S1022L | C | 5.1e-08 | 6.9e-08 | 2.2e-08 | 4.733E-08 | 2.371E-08 | 2.200E-08 | 6.900E-08 |
| 20432 | 1022 | S1022K | C | 1.7e-08 | 3.6e-08 | 3e-08 | 2.767E-08 | 9.713E-09 | 1.700E-08 | 3.600E-08 |
| 20433 | 1022 | S1022M | C | 7.2e-08 | 3.8e-08 | 2.5e-07 | 1.200E-07 | 1.139E-07 | 3.800E-08 | 2.500E-07 |
| 20434 | 1022 | S1022F | C | 7.8e-08 | 4.5e-08 | 3.6e-08 | 5.300E-08 | 2.211E-08 | 3.600E-08 | 7.800E-08 |
| 20435 | 1022 | S1022P | C | 3.4e-08 | 5e-08 | 1.3e-08 | 3.233E-08 | 1.856E-08 | 1.300E-08 | 5.000E-08 |
| 20436 | 1022 | S1022S | C | 5.4e-08 | 2.5e-08 | 8.9e-08 | 5.600E-08 | 3.205E-08 | 2.500E-08 | 8.900E-08 |
| 20437 | 1022 | S1022T | C | 4.6e-08 | 3.2e-08 | 4.4e-08 | 4.067E-08 | 7.572E-09 | 3.200E-08 | 4.600E-08 |
| 20438 | 1022 | S1022W | C | 1e-07 | 6.3e-08 | 3e-08 | 6.433E-08 | 3.502E-08 | 3.000E-08 | 1.000E-07 |
| 20439 | 1022 | S1022Y | C | 1.3e-08 | 6.4e-08 | 1.2e-08 | 2.967E-08 | 2.974E-08 | 1.200E-08 | 6.400E-08 |
| 20440 | 1022 | S1022V | C | 6.1e-08 | 3.1e-08 | 6.2e-08 | 5.133E-08 | 1.762E-08 | 3.100E-08 | 6.200E-08 |
| 20441 | 1023 | V1023A | C | 5.3e-08 | 1.6e-07 | 3.2e-08 | 8.167E-08 | 6.865E-08 | 3.200E-08 | 1.600E-07 |
| 20442 | 1023 | V1023R | C | 4.6e-08 | 4.4e-08 | 5.4e-08 | 4.800E-08 | 5.292E-09 | 4.400E-08 | 5.400E-08 |
| 20443 | 1023 | V1023N | C | 3.7e-08 | 4e-08 | 2.6e-08 | 3.433E-08 | 7.371E-09 | 2.600E-08 | 4.000E-08 |
| 20444 | 1023 | V1023D | C | 6e-08 | 6.7e-08 | 6e-08 | 6.233E-08 | 4.041E-09 | 6.000E-08 | 6.700E-08 |
| 20445 | 1023 | V1023C | C | 2.9e-08 | 7.2e-08 | 3.2e-08 | 4.433E-08 | 2.401E-08 | 2.900E-08 | 7.200E-08 |
| 20446 | 1023 | V1023E | C | 7.8e-09 | 1.3e-08 | 2.5e-08 | 1.527E-08 | 8.821E-09 | 7.800E-09 | 2.500E-08 |
| 20447 | 1023 | V1023Q | C | 8.6e-08 | 4.1e-08 | 4.4e-08 | 5.700E-08 | 2.516E-08 | 4.100E-08 | 8.600E-08 |
| 20448 | 1023 | V1023G | C | 5.5e-08 | 3.4e-08 | 3.8e-08 | 4.233E-08 | 1.115E-08 | 3.400E-08 | 5.500E-08 |
| 20449 | 1023 | V1023H | C | 2.1e-08 | 3e-08 | 3.7e-08 | 2.933E-08 | 8.021E-09 | 2.100E-08 | 3.700E-08 |
| 20450 | 1023 | V1023I | C | 7.1e-08 | 3.2e-08 | 7e-08 | 5.767E-08 | 2.223E-08 | 3.200E-08 | 7.100E-08 |
| 20451 | 1023 | V1023L | C | 9.3e-09 | 1.8e-08 | 6.6e-08 | 3.110E-08 | 3.054E-08 | 9.300E-09 | 6.600E-08 |
| 20452 | 1023 | V1023K | C | 3.2e-08 | 1.2e-08 | 6.4e-09 | 1.680E-08 | 1.346E-08 | 6.400E-09 | 3.200E-08 |
| 20453 | 1023 | V1023M | C | 2.8e-08 | 4.7e-08 | 9.3e-08 | 5.600E-08 | 3.342E-08 | 2.800E-08 | 9.300E-08 |
| 20454 | 1023 | V1023F | C | 6.1e-08 | 2.7e-08 | 5e-08 | 4.600E-08 | 1.735E-08 | 2.700E-08 | 6.100E-08 |
| 20455 | 1023 | V1023P | C | 8e-08 | 2.3e-08 | 3.7e-08 | 4.667E-08 | 2.970E-08 | 2.300E-08 | 8.000E-08 |
| 20456 | 1023 | V1023S | C | 4.6e-08 | 5.2e-08 | 4.5e-08 | 4.767E-08 | 3.786E-09 | 4.500E-08 | 5.200E-08 |
| 20457 | 1023 | V1023T | C | 3.4e-08 | 5.2e-08 | 5.9e-08 | 4.833E-08 | 1.290E-08 | 3.400E-08 | 5.900E-08 |
| 20458 | 1023 | V1023W | C | 8.7e-08 | 3.1e-08 | 3.5e-08 | 5.100E-08 | 3.124E-08 | 3.100E-08 | 8.700E-08 |
| 20459 | 1023 | V1023Y | C | 3.2e-08 | 1.3e-07 | 4.7e-08 | 6.967E-08 | 5.279E-08 | 3.200E-08 | 1.300E-07 |
| 20460 | 1023 | V1023V | C | 7.8e-08 | 1.3e-07 | 3.9e-08 | 8.233E-08 | 4.565E-08 | 3.900E-08 | 1.300E-07 |
| 20461 | 1024 | V1024A | C | 2.3e-08 | 5e-08 | 7.2e-08 | 4.833E-08 | 2.454E-08 | 2.300E-08 | 7.200E-08 |
| 20462 | 1024 | V1024R | C | 3.7e-08 | 2.1e-08 | 6.6e-08 | 4.133E-08 | 2.281E-08 | 2.100E-08 | 6.600E-08 |
| 20463 | 1024 | V1024N | C | 8.7e-08 | 2.3e-08 | 2.4e-08 | 4.467E-08 | 3.667E-08 | 2.300E-08 | 8.700E-08 |
| 20464 | 1024 | V1024D | C | 5e-08 | 7.3e-08 | 4e-08 | 5.433E-08 | 1.692E-08 | 4.000E-08 | 7.300E-08 |
| 20465 | 1024 | V1024C | C | 3.5e-08 | 1.8e-08 | 3.1e-08 | 2.800E-08 | 8.888E-09 | 1.800E-08 | 3.500E-08 |
| 20466 | 1024 | V1024E | C | 5e-08 | 9.1e-08 | 9.1e-08 | 7.733E-08 | 2.367E-08 | 5.000E-08 | 9.100E-08 |
| 20467 | 1024 | V1024Q | C | 6.5e-08 | 6.9e-08 | 3.2e-08 | 5.533E-08 | 2.031E-08 | 3.200E-08 | 6.900E-08 |
| 20468 | 1024 | V1024G | C | 4.6e-08 | 7.7e-08 | 1e-07 | 7.433E-08 | 2.710E-08 | 4.600E-08 | 1.000E-07 |
| 20469 | 1024 | V1024H | C | 6.1e-08 | 3.2e-08 | 4.7e-08 | 4.667E-08 | 1.450E-08 | 3.200E-08 | 6.100E-08 |
| 20470 | 1024 | V1024I | C | 6.4e-08 | 4.8e-08 | 3.7e-08 | 4.967E-08 | 1.358E-08 | 3.700E-08 | 6.400E-08 |
| 20471 | 1024 | V1024L | C | 2.3e-08 | 8.2e-08 | 7.5e-08 | 6.000E-08 | 3.223E-08 | 2.300E-08 | 8.200E-08 |
| 20472 | 1024 | V1024K | C | 7.4e-09 | 4.7e-08 | 2e-08 | 2.480E-08 | 2.023E-08 | 7.400E-09 | 4.700E-08 |
| 20473 | 1024 | V1024M | C | 4.2e-08 | 2.7e-08 | 8.1e-08 | 5.000E-08 | 2.787E-08 | 2.700E-08 | 8.100E-08 |
| 20474 | 1024 | V1024F | C | 1.2e-07 | 3.9e-08 | 3e-08 | 6.300E-08 | 4.957E-08 | 3.000E-08 | 1.200E-07 |
| 20475 | 1024 | V1024P | C | 2.8e-08 | 1e-07 | 1.6e-08 | 4.800E-08 | 4.543E-08 | 1.600E-08 | 1.000E-07 |
| 20476 | 1024 | V1024S | C | 4.5e-08 | 2.1e-08 | 9.9e-08 | 5.500E-08 | 3.995E-08 | 2.100E-08 | 9.900E-08 |
| 20477 | 1024 | V1024T | C | 6.3e-08 | 7.2e-08 | 1.6e-07 | 9.833E-08 | 5.359E-08 | 6.300E-08 | 1.600E-07 |
| 20478 | 1024 | V1024W | C | 4.9e-08 | 2.4e-08 | 5.1e-08 | 4.133E-08 | 1.504E-08 | 2.400E-08 | 5.100E-08 |
| 20479 | 1024 | V1024Y | C | 7.2e-08 | 2.1e-08 | 3.1e-08 | 4.133E-08 | 2.702E-08 | 2.100E-08 | 7.200E-08 |
| 20480 | 1024 | V1024V | C | 1.2e-07 | 2e-08 | 6.1e-08 | 6.700E-08 | 5.027E-08 | 2.000E-08 | 1.200E-07 |
| 20481 | 1025 | T1025A | C | 1.2e-07 | 5.4e-08 | 5.1e-08 | 7.500E-08 | 3.900E-08 | 5.100E-08 | 1.200E-07 |
| 20482 | 1025 | T1025R | C | 4.3e-08 | 2.2e-08 | 2.8e-08 | 3.100E-08 | 1.082E-08 | 2.200E-08 | 4.300E-08 |
| 20483 | 1025 | T1025N | C | 4.7e-08 | 5.3e-08 | 2.6e-08 | 4.200E-08 | 1.418E-08 | 2.600E-08 | 5.300E-08 |
| 20484 | 1025 | T1025D | C | 1.6e-07 | 4.4e-08 | 4.6e-08 | 8.333E-08 | 6.640E-08 | 4.400E-08 | 1.600E-07 |

| | | | | | | | | | | |
|---|---|---|---|---|---|---|---|---|---|---|
| 20485 | 1025 | T1025C | C | 5.1e-08 | 1e-07 | 2.3e-08 | 5.800E-08 | 3.897E-08 | 2.300E-08 | 1.000E-07 |
| 20486 | 1025 | T1025E | C | 3.4e-08 | 6.8e-08 | 1.9e-07 | 9.733E-08 | 8.203E-08 | 3.400E-08 | 1.900E-07 |
| 20487 | 1025 | T1025Q | C | 6.5e-08 | 6.4e-08 | 3e-08 | 5.300E-08 | 1.992E-08 | 3.000E-08 | 6.500E-08 |
| 20488 | 1025 | T1025G | C | 2.1e-08 | 9.7e-08 | 6.8e-08 | 6.200E-08 | 3.835E-08 | 2.100E-08 | 9.700E-08 |
| 20489 | 1025 | T1025H | C | 1.5e-08 | 3.7e-08 | 6.9e-08 | 4.033E-08 | 2.715E-08 | 1.500E-08 | 6.900E-08 |
| 20490 | 1025 | T1025I | C | 5.2e-08 | 4.5e-08 | 1.1e-07 | 6.900E-08 | 3.568E-08 | 4.500E-08 | 1.100E-07 |
| 20491 | 1025 | T1025L | C | 7.8e-08 | 3e-08 | 8.6e-08 | 6.467E-08 | 3.029E-08 | 3.000E-08 | 8.600E-08 |
| 20492 | 1025 | T1025K | C | 1e-07 | 2.5e-08 | 8.8e-08 | 7.100E-08 | 4.029E-08 | 2.500E-08 | 1.000E-07 |
| 20493 | 1025 | T1025M | C | 5.3e-08 | 3.5e-08 | 3.6e-08 | 4.133E-08 | 1.012E-08 | 3.500E-08 | 5.300E-08 |
| 20494 | 1025 | T1025F | C | 2.3e-08 | 3.7e-08 | 5.2e-08 | 3.733E-08 | 1.450E-08 | 2.300E-08 | 5.200E-08 |
| 20495 | 1025 | T1025P | C | 4.6e-08 | 2.6e-08 | 5.3e-08 | 4.167E-08 | 1.401E-08 | 2.600E-08 | 5.300E-08 |
| 20496 | 1025 | T1025S | C | 5.5e-08 | 1.8e-08 | 3e-08 | 3.433E-08 | 1.888E-08 | 1.800E-08 | 5.500E-08 |
| 20497 | 1025 | T1025T | C | 7.3e-08 | 1.7e-07 | 1.9e-08 | 8.733E-08 | 7.651E-08 | 1.900E-08 | 1.700E-07 |
| 20498 | 1025 | T1025W | C | 2.4e-08 | 9.6e-08 | 7.2e-08 | 6.400E-08 | 3.666E-08 | 2.400E-08 | 9.600E-08 |
| 20499 | 1025 | T1025Y | C | 7e-08 | 2.1e-08 | 4.2e-08 | 4.433E-08 | 2.458E-08 | 2.100E-08 | 7.000E-08 |
| 20500 | 1025 | T1025V | C | 4.8e-08 | 2.8e-08 | 8.4e-08 | 5.333E-08 | 2.838E-08 | 2.800E-08 | 8.400E-08 |
| 20501 | 1026 | V1026A | C | 1.1e-07 | 5.9e-08 | 5.5e-08 | 7.467E-08 | 3.066E-08 | 5.500E-08 | 1.100E-07 |
| 20502 | 1026 | V1026R | C | 5.4e-08 | 1.8e-08 | 5.2e-08 | 4.133E-08 | 2.023E-08 | 1.800E-08 | 5.400E-08 |
| 20503 | 1026 | V1026N | C | 7.3e-09 | 5.1e-08 | 1.2e-08 | 2.343E-08 | 2.399E-08 | 7.300E-09 | 5.100E-08 |
| 20504 | 1026 | V1026D | C | 3e-08 | 4.1e-08 | 6.8e-08 | 4.633E-08 | 1.955E-08 | 3.000E-08 | 6.800E-08 |
| 20505 | 1026 | V1026C | C | 9e-09 | 1.5e-08 | 6.6e-08 | 3.000E-08 | 3.132E-08 | 9.000E-09 | 6.600E-08 |
| 20506 | 1026 | V1026E | C | 2.8e-08 | 1.7e-07 | 4.7e-08 | 8.167E-08 | 7.709E-08 | 2.800E-08 | 1.700E-07 |
| 20507 | 1026 | V1026Q | C | 1.7e-08 | 3e-08 | 3.4e-08 | 2.700E-08 | 8.888E-09 | 1.700E-08 | 3.400E-08 |
| 20508 | 1026 | V1026G | C | 2.5e-08 | 5.3e-08 | 6e-08 | 4.600E-08 | 1.852E-08 | 2.500E-08 | 6.000E-08 |
| 20509 | 1026 | V1026H | C | 5.9e-08 | 4.4e-08 | 4.7e-08 | 5.000E-08 | 7.937E-09 | 4.400E-08 | 5.900E-08 |
| 20510 | 1026 | V1026I | C | 3.2e-08 | 2.3e-08 | 4.7e-08 | 3.400E-08 | 1.212E-08 | 2.300E-08 | 4.700E-08 |
| 20511 | 1026 | V1026L | C | 2.4e-08 | 3.6e-08 | 2.3e-08 | 2.767E-08 | 7.234E-09 | 2.300E-08 | 3.600E-08 |
| 20512 | 1026 | V1026K | C | 3e-08 | 5.1e-08 | 4.5e-08 | 4.200E-08 | 1.082E-08 | 3.000E-08 | 5.100E-08 |
| 20513 | 1026 | V1026M | C | 9.2e-08 | 8.4e-08 | 1.4e-08 | 6.333E-08 | 4.291E-08 | 1.400E-08 | 9.200E-08 |
| 20514 | 1026 | V1026F | C | 3.1e-08 | 1e-07 | 6.7e-08 | 6.600E-08 | 3.451E-08 | 3.100E-08 | 1.000E-07 |
| 20515 | 1026 | V1026P | C | 1.2e-07 | 6.6e-08 | 2.2e-08 | 6.933E-08 | 4.908E-08 | 2.200E-08 | 1.200E-07 |
| 20516 | 1026 | V1026S | C | 5.6e-08 | 4.3e-08 | 1.6e-08 | 3.833E-08 | 2.040E-08 | 1.600E-08 | 5.600E-08 |
| 20517 | 1026 | V1026T | C | 3.5e-08 | 4.3e-08 | 2e-08 | 3.267E-08 | 1.168E-08 | 2.000E-08 | 4.300E-08 |
| 20518 | 1026 | V1026W | C | 9.1e-08 | 1.5e-08 | 6.4e-08 | 5.667E-08 | 3.853E-08 | 1.500E-08 | 9.100E-08 |
| 20519 | 1026 | V1026Y | C | 2.4e-08 | 2.9e-08 | 1.3e-08 | 2.200E-08 | 8.185E-09 | 1.300E-08 | 2.900E-08 |
| 20520 | 1026 | V1026V | C | 9.4e-08 | 7.3e-08 | 2.3e-08 | 6.333E-08 | 3.647E-08 | 2.300E-08 | 9.400E-08 |
| 20521 | 1027 | P1027A | C | 4.9e-08 | 1.1e-07 | 4.6e-08 | 6.833E-08 | 3.612E-08 | 4.600E-08 | 1.100E-07 |
| 20522 | 1027 | P1027R | C | 4.2e-08 | 1.4e-08 | 1.6e-07 | 7.200E-08 | 7.749E-08 | 1.400E-08 | 1.600E-07 |
| 20523 | 1027 | P1027N | C | 8e-08 | 2.2e-08 | 3.1e-08 | 4.433E-08 | 3.121E-08 | 2.200E-08 | 8.000E-08 |
| 20524 | 1027 | P1027D | C | 2.8e-08 | 5.3e-08 | 4.2e-08 | 4.100E-08 | 1.253E-08 | 2.800E-08 | 5.300E-08 |
| 20525 | 1027 | P1027C | C | 6.1e-08 | 3.1e-08 | 2.9e-08 | 4.033E-08 | 1.793E-08 | 2.900E-08 | 6.100E-08 |
| 20526 | 1027 | P1027E | C | 5.6e-08 | 2.4e-08 | 2.6e-08 | 3.533E-08 | 1.793E-08 | 2.400E-08 | 5.600E-08 |
| 20527 | 1027 | P1027Q | C | 3.9e-08 | 3e-08 | 5.5e-08 | 4.133E-08 | 1.266E-08 | 3.000E-08 | 5.500E-08 |
| 20528 | 1027 | P1027G | C | 3.9e-08 | 1.2e-08 | 3.4e-08 | 2.833E-08 | 1.436E-08 | 1.200E-08 | 3.900E-08 |
| 20529 | 1027 | P1027H | C | 2.5e-08 | 3.1e-08 | 8.8e-08 | 4.800E-08 | 3.477E-08 | 2.500E-08 | 8.800E-08 |
| 20530 | 1027 | P1027I | C | 3.7e-08 | 3.5e-08 | 2.4e-08 | 3.200E-08 | 7.000E-09 | 2.400E-08 | 3.700E-08 |
| 20531 | 1027 | P1027L | C | 4e-08 | 5.4e-08 | 3.7e-08 | 4.367E-08 | 9.074E-09 | 3.700E-08 | 5.400E-08 |
| 20532 | 1027 | P1027K | C | 1.5e-07 | 3.9e-08 | 1.4e-08 | 6.767E-08 | 7.239E-08 | 1.400E-08 | 1.500E-07 |
| 20533 | 1027 | P1027M | C | 4.8e-08 | 3.5e-08 | 3.2e-08 | 3.833E-08 | 8.505E-09 | 3.200E-08 | 4.800E-08 |
| 20534 | 1027 | P1027F | C | 2.9e-08 | 2.7e-08 | 2.9e-08 | 2.833E-08 | 1.155E-09 | 2.700E-08 | 2.900E-08 |
| 20535 | 1027 | P1027P | C | 6.6e-08 | 7.2e-09 | 5.5e-08 | 4.273E-08 | 3.126E-08 | 7.200E-09 | 6.600E-08 |
| 20536 | 1027 | P1027S | C | 1.9e-08 | 3.1e-08 | 2.4e-08 | 2.467E-08 | 6.028E-09 | 1.900E-08 | 3.100E-08 |
| 20537 | 1027 | P1027T | C | 3.8e-08 | 8.2e-08 | 2.1e-08 | 4.700E-08 | 3.148E-08 | 2.100E-08 | 8.200E-08 |
| 20538 | 1027 | P1027W | C | 4.5e-08 | 1.2e-07 | 3.9e-08 | 6.800E-08 | 4.513E-08 | 3.900E-08 | 1.200E-07 |
| 20539 | 1027 | P1027Y | C | 2.6e-08 | 2.3e-08 | 7.1e-08 | 4.000E-08 | 2.689E-08 | 2.300E-08 | 7.100E-08 |
| 20540 | 1027 | P1027V | C | 3.7e-08 | 6.3e-08 | 4.1e-08 | 4.700E-08 | 1.400E-08 | 3.700E-08 | 6.300E-08 |
| 20541 | 1028 | S1028A | C | 1.6e-08 | 4.2e-08 | 7.1e-08 | 4.300E-08 | 2.751E-08 | 1.600E-08 | 7.100E-08 |
| 20542 | 1028 | S1028R | C | 4.7e-08 | 2.5e-08 | 4.6e-08 | 3.933E-08 | 1.242E-08 | 2.500E-08 | 4.700E-08 |
| 20543 | 1028 | S1028N | C | 6.5e-08 | 4.7e-08 | 8.4e-08 | 6.533E-08 | 1.850E-08 | 4.700E-08 | 8.400E-08 |
| 20544 | 1028 | S1028D | C | 1.9e-08 | 9.3e-08 | 6.2e-08 | 5.800E-08 | 3.716E-08 | 1.900E-08 | 9.300E-08 |
| 20545 | 1028 | S1028C | C | 5.5e-08 | 1.1e-07 | 2e-08 | 6.167E-08 | 4.537E-08 | 2.000E-08 | 1.100E-07 |
| 20546 | 1028 | S1028E | C | 3.6e-08 | 5.1e-08 | 6.9e-08 | 5.200E-08 | 1.652E-08 | 3.600E-08 | 6.900E-08 |
| 20547 | 1028 | S1028Q | C | 4.8e-08 | 7.6e-08 | 1.8e-07 | 1.013E-07 | 6.955E-08 | 4.800E-08 | 1.800E-07 |
| 20548 | 1028 | S1028G | C | 8.2e-08 | 4e-08 | 9.3e-08 | 7.167E-08 | 2.797E-08 | 4.000E-08 | 9.300E-08 |
| 20549 | 1028 | S1028H | C | 1e-07 | 3.1e-08 | 5.2e-08 | 6.100E-08 | 3.537E-08 | 3.100E-08 | 1.000E-07 |
| 20550 | 1028 | S1028I | C | 4.4e-08 | 2.5e-08 | 3e-08 | 3.300E-08 | 9.849E-09 | 2.500E-08 | 4.400E-08 |
| 20551 | 1028 | S1028L | C | 3.8e-08 | 1.1e-07 | 3.2e-08 | 6.000E-08 | 4.341E-08 | 3.200E-08 | 1.100E-07 |
| 20552 | 1028 | S1028K | C | 5.3e-08 | 3.3e-08 | 3.1e-08 | 3.900E-08 | 1.217E-08 | 3.100E-08 | 5.300E-08 |
| 20553 | 1028 | S1028M | C | 2.6e-08 | 1.4e-08 | 1.8e-08 | 1.933E-08 | 6.110E-09 | 1.400E-08 | 2.600E-08 |
| 20554 | 1028 | S1028F | C | 4.2e-08 | 1.7e-08 | 5.7e-08 | 3.867E-08 | 2.021E-08 | 1.700E-08 | 5.700E-08 |
| 20555 | 1028 | S1028P | C | 2.7e-07 | 1.5e-08 | 1.1e-07 | 1.317E-07 | 1.289E-07 | 1.500E-08 | 2.700E-07 |
| 20556 | 1028 | S1028S | C | 1.9e-08 | 1.9e-08 | 5.2e-08 | 3.000E-08 | 1.905E-08 | 1.900E-08 | 5.200E-08 |
| 20557 | 1028 | S1028T | C | 2e-08 | 4.7e-08 | 7.6e-08 | 4.767E-08 | 2.801E-08 | 2.000E-08 | 7.600E-08 |
| 20558 | 1028 | S1028W | C | 4.4e-08 | 7.1e-08 | 6.2e-08 | 5.900E-08 | 1.375E-08 | 4.400E-08 | 7.100E-08 |
| 20559 | 1028 | S1028Y | C | 6.7e-08 | 3.3e-08 | 3.3e-08 | 4.433E-08 | 1.963E-08 | 3.300E-08 | 6.700E-08 |
| 20560 | 1028 | S1028V | C | 7.2e-08 | 5.2e-08 | 2.3e-08 | 4.900E-08 | 2.464E-08 | 2.300E-08 | 7.200E-08 |
| 20561 | 1029 | S1029A | C | 5.3e-08 | 6.2e-08 | 5.5e-08 | 5.667E-08 | 4.726E-09 | 5.300E-08 | 6.200E-08 |
| 20562 | 1029 | S1029R | C | 7.3e-08 | 2.7e-08 | 2e-08 | 4.000E-08 | 2.879E-08 | 2.000E-08 | 7.300E-08 |
| 20563 | 1029 | S1029N | C | 2e-08 | 1.6e-08 | 2.1e-08 | 1.900E-08 | 2.646E-09 | 1.600E-08 | 2.100E-08 |
| 20564 | 1029 | S1029D | C | 6.2e-08 | 8e-08 | 6.4e-08 | 6.867E-08 | 9.866E-09 | 6.200E-08 | 8.000E-08 |
| 20565 | 1029 | S1029C | C | 1.5e-08 | 1.9e-08 | 3.7e-08 | 2.367E-08 | 1.172E-08 | 1.500E-08 | 3.700E-08 |
| 20566 | 1029 | S1029E | C | 2.7e-08 | 2.4e-08 | 4e-08 | 3.033E-08 | 8.505E-09 | 2.400E-08 | 4.000E-08 |
| 20567 | 1029 | S1029Q | C | 1e-07 | 4.6e-08 | 6.2e-08 | 6.933E-08 | 2.774E-08 | 4.600E-08 | 1.000E-07 |
| 20568 | 1029 | S1029G | C | 4.6e-08 | 8.6e-08 | 2.7e-08 | 5.300E-08 | 3.012E-08 | 2.700E-08 | 8.600E-08 |
| 20569 | 1029 | S1029H | C | 5.1e-08 | 2.1e-08 | 6.6e-08 | 4.600E-08 | 2.291E-08 | 2.100E-08 | 6.600E-08 |

| | | | | | | | | | | |
|---|---|---|---|---|---|---|---|---|---|---|
| 20570 | 1029 | S1029I | C | 5.5e-08 | 5.1e-08 | 3.9e-08 | 4.833E-08 | 8.327E-09 | 3.900E-08 | 5.500E-08 |
| 20571 | 1029 | S1029L | C | 3.7e-08 | 3e-08 | 6.7e-08 | 4.467E-08 | 1.966E-08 | 3.000E-08 | 6.700E-08 |
| 20572 | 1029 | S1029K | C | 3.4e-08 | 3.5e-08 | 1.3e-08 | 2.733E-08 | 1.242E-08 | 1.300E-08 | 3.500E-08 |
| 20573 | 1029 | S1029M | C | 3e-08 | 7.7e-08 | 5.5e-08 | 5.400E-08 | 2.352E-08 | 3.000E-08 | 7.700E-08 |
| 20574 | 1029 | S1029F | C | 3.9e-08 | 5.1e-09 | 6.8e-08 | 3.737E-08 | 3.148E-08 | 5.100E-09 | 6.800E-08 |
| 20575 | 1029 | S1029P | C | 2.3e-08 | 2.7e-08 | 4.7e-08 | 3.233E-08 | 1.286E-08 | 2.300E-08 | 4.700E-08 |
| 20576 | 1029 | S1029S | C | 1.1e-08 | 2.9e-08 | 6.4e-08 | 3.467E-08 | 2.695E-08 | 1.100E-08 | 6.400E-08 |
| 20577 | 1029 | S1029T | C | 2.6e-08 | 5.5e-08 | 3.1e-08 | 3.733E-08 | 1.550E-08 | 2.600E-08 | 5.500E-08 |
| 20578 | 1029 | S1029W | C | 1.9e-08 | 8.7e-08 | 7e-08 | 5.867E-08 | 3.539E-08 | 1.900E-08 | 8.700E-08 |
| 20579 | 1029 | S1029Y | C | 2.5e-08 | 3.6e-08 | 3.8e-08 | 3.300E-08 | 7.000E-09 | 2.500E-08 | 3.800E-08 |
| 20580 | 1029 | S1029V | C | 5.6e-08 | 6.7e-08 | 1.6e-07 | 9.433E-08 | 5.713E-08 | 5.600E-08 | 1.600E-07 |
| 20581 | 1030 | S1030A | C | 4.8e-08 | 3.4e-08 | 3.8e-08 | 4.000E-08 | 7.211E-09 | 3.400E-08 | 4.800E-08 |
| 20582 | 1030 | S1030R | C | 4.4e-08 | 3.7e-08 | 5.1e-08 | 4.400E-08 | 7.000E-09 | 3.700E-08 | 5.100E-08 |
| 20583 | 1030 | S1030N | C | 4.5e-08 | 3.2e-08 | 3.6e-08 | 3.767E-08 | 6.658E-09 | 3.200E-08 | 4.500E-08 |
| 20584 | 1030 | S1030D | C | 7.1e-08 | 3.4e-08 | 1e-07 | 6.833E-08 | 3.308E-08 | 3.400E-08 | 1.000E-07 |
| 20585 | 1030 | S1030C | C | 2.8e-08 | 2.7e-08 | 8.4e-08 | 4.633E-08 | 3.262E-08 | 2.700E-08 | 8.400E-08 |
| 20586 | 1030 | S1030E | C | 9.2e-08 | 6.5e-08 | 3.7e-08 | 6.467E-08 | 2.750E-08 | 3.700E-08 | 9.200E-08 |
| 20587 | 1030 | S1030Q | C | 3.7e-08 | 2.2e-08 | 6.4e-08 | 4.100E-08 | 2.128E-08 | 2.200E-08 | 6.400E-08 |
| 20588 | 1030 | S1030G | C | 2.8e-08 | 5.4e-08 | 5.7e-08 | 4.633E-08 | 1.595E-08 | 2.800E-08 | 5.700E-08 |
| 20589 | 1030 | S1030H | C | 6.1e-08 | 4.7e-08 | 3.6e-08 | 4.800E-08 | 1.253E-08 | 3.600E-08 | 6.100E-08 |
| 20590 | 1030 | S1030I | C | 2.2e-08 | 5.3e-08 | 6e-08 | 4.500E-08 | 2.022E-08 | 2.200E-08 | 6.000E-08 |
| 20591 | 1030 | S1030L | C | 6.9e-08 | 2.6e-08 | 3.6e-08 | 4.367E-08 | 2.250E-08 | 2.600E-08 | 6.900E-08 |
| 20592 | 1030 | S1030K | C | 6.6e-08 | 1.5e-08 | 3.3e-08 | 3.800E-08 | 2.587E-08 | 1.500E-08 | 6.600E-08 |
| 20593 | 1030 | S1030M | C | 4.8e-08 | 2.1e-08 | 1.1e-08 | 2.667E-08 | 1.914E-08 | 1.100E-08 | 4.800E-08 |
| 20594 | 1030 | S1030F | C | 5.9e-08 | 5.1e-08 | 3.7e-08 | 4.900E-08 | 1.114E-08 | 3.700E-08 | 5.900E-08 |
| 20595 | 1030 | S1030P | C | 3.6e-08 | 5.2e-08 | 3.9e-08 | 4.233E-08 | 8.505E-09 | 3.600E-08 | 5.200E-08 |
| 20596 | 1030 | S1030S | C | 3.1e-08 | 8.3e-08 | 3.4e-08 | 4.933E-08 | 2.919E-08 | 3.100E-08 | 8.300E-08 |
| 20597 | 1030 | S1030T | C | 5.1e-08 | 7.8e-08 | 4.1e-08 | 5.667E-08 | 1.914E-08 | 4.100E-08 | 7.800E-08 |
| 20598 | 1030 | S1030W | C | 2.5e-08 | 2.1e-08 | 3.2e-08 | 2.600E-08 | 5.568E-09 | 2.100E-08 | 3.200E-08 |
| 20599 | 1030 | S1030Y | C | 5.1e-08 | 8.5e-08 | 1.1e-07 | 8.200E-08 | 2.961E-08 | 5.100E-08 | 1.100E-07 |
| 20600 | 1030 | S1030V | C | 5.3e-08 | 1e-07 | 9.8e-08 | 8.367E-08 | 2.658E-08 | 5.300E-08 | 1.000E-07 |
| 20601 | 1031 | L1031A | C | 5e-08 | 3.6e-08 | 4.1e-08 | 4.233E-08 | 7.095E-09 | 3.600E-08 | 5.000E-08 |
| 20602 | 1031 | L1031R | C | 2.2e-08 | 6.3e-08 | 3.1e-08 | 3.867E-08 | 2.155E-08 | 2.200E-08 | 6.300E-08 |
| 20603 | 1031 | L1031N | C | 3.6e-08 | 2.9e-08 | 1.8e-08 | 2.767E-08 | 9.074E-09 | 1.800E-08 | 3.600E-08 |
| 20604 | 1031 | L1031D | C | 4e-08 | 1.8e-07 | 1.8e-08 | 7.933E-08 | 8.787E-08 | 1.800E-08 | 1.800E-07 |
| 20605 | 1031 | L1031C | C | 4.1e-08 | 1.1e-07 | 4e-08 | 6.367E-08 | 4.013E-08 | 4.000E-08 | 1.100E-07 |
| 20606 | 1031 | L1031E | C | 3.6e-08 | 1.7e-08 | 3.4e-08 | 2.900E-08 | 1.044E-08 | 1.700E-08 | 3.600E-08 |
| 20607 | 1031 | L1031Q | C | 4.9e-08 | 7.3e-08 | 2.9e-08 | 5.033E-08 | 2.203E-08 | 2.900E-08 | 7.300E-08 |
| 20608 | 1031 | L1031G | C | 2.5e-08 | 1.5e-08 | 1.8e-08 | 1.933E-08 | 5.132E-09 | 1.500E-08 | 2.500E-08 |
| 20609 | 1031 | L1031H | C | 1.5e-07 | 7.4e-08 | 4.9e-08 | 9.100E-08 | 5.260E-08 | 4.900E-08 | 1.500E-07 |
| 20610 | 1031 | L1031I | C | 3.5e-08 | 3.1e-08 | 2.7e-08 | 3.100E-08 | 4.000E-09 | 2.700E-08 | 3.500E-08 |
| 20611 | 1031 | L1031L | C | 4.9e-08 | 7.5e-08 | 4.1e-08 | 5.500E-08 | 1.778E-08 | 4.100E-08 | 7.500E-08 |
| 20612 | 1031 | L1031K | C | 2.5e-08 | 6.2e-08 | 4e-08 | 4.233E-08 | 1.861E-08 | 2.500E-08 | 6.200E-08 |
| 20613 | 1031 | L1031M | C | 5.5e-08 | 3e-08 | 6e-08 | 4.833E-08 | 1.607E-08 | 3.000E-08 | 6.000E-08 |
| 20614 | 1031 | L1031F | C | 1.9e-08 | 3.5e-08 | 2.8e-08 | 2.733E-08 | 8.021E-09 | 1.900E-08 | 3.500E-08 |
| 20615 | 1031 | L1031P | C | 2.9e-08 | 1.6e-08 | 5.8e-08 | 3.433E-08 | 2.150E-08 | 1.600E-08 | 5.800E-08 |
| 20616 | 1031 | L1031S | C | 7.5e-08 | 6e-08 | 3.7e-08 | 5.733E-08 | 1.914E-08 | 3.700E-08 | 7.500E-08 |
| 20617 | 1031 | L1031T | C | 8.2e-08 | 1e-08 | 1.1e-07 | 6.733E-08 | 5.159E-08 | 1.000E-08 | 1.100E-07 |
| 20618 | 1031 | L1031W | C | 7.7e-08 | 5.2e-08 | 1.8e-08 | 4.900E-08 | 2.961E-08 | 1.800E-08 | 7.700E-08 |
| 20619 | 1031 | L1031Y | C | 3.4e-08 | 2.1e-08 | 8e-08 | 4.500E-08 | 3.100E-08 | 2.100E-08 | 8.000E-08 |
| 20620 | 1031 | L1031V | C | 7.3e-08 | 2.6e-08 | 5.8e-09 | 3.493E-08 | 3.448E-08 | 5.800E-09 | 7.300E-08 |
| 20621 | 1032 | G1032A | C | 1.9e-08 | 4.1e-08 | 3.7e-08 | 3.233E-08 | 1.172E-08 | 1.900E-08 | 4.100E-08 |
| 20622 | 1032 | G1032R | C | 1.3e-07 | 4.4e-08 | 2.9e-08 | 6.767E-08 | 5.450E-08 | 2.900E-08 | 1.300E-07 |
| 20623 | 1032 | G1032N | C | 3.7e-08 | 5.6e-08 | 5e-08 | 4.767E-08 | 9.713E-09 | 3.700E-08 | 5.600E-08 |
| 20624 | 1032 | G1032D | C | 1.5e-07 | 4e-08 | 4.3e-08 | 7.767E-08 | 6.266E-08 | 4.000E-08 | 1.500E-07 |
| 20625 | 1032 | G1032C | C | 1.1e-08 | 3.4e-08 | 5.7e-08 | 3.400E-08 | 2.300E-08 | 1.100E-08 | 5.700E-08 |
| 20626 | 1032 | G1032E | C | 5.8e-08 | 2.5e-08 | 3.6e-08 | 3.967E-08 | 1.680E-08 | 2.500E-08 | 5.800E-08 |
| 20627 | 1032 | G1032Q | C | 5.8e-08 | 3.5e-08 | 4.8e-08 | 4.700E-08 | 1.153E-08 | 3.500E-08 | 5.800E-08 |
| 20628 | 1032 | G1032G | C | 3.8e-08 | 1.5e-08 | 4.9e-08 | 3.400E-08 | 1.735E-08 | 1.500E-08 | 4.900E-08 |
| 20629 | 1032 | G1032H | C | 6e-08 | 4.2e-08 | 1.1e-08 | 3.767E-08 | 2.479E-08 | 1.100E-08 | 6.000E-08 |
| 20630 | 1032 | G1032I | C | 3.8e-08 | 3.8e-08 | 5.8e-08 | 4.467E-08 | 1.155E-08 | 3.800E-08 | 5.800E-08 |
| 20631 | 1032 | G1032L | C | 3e-08 | 8.2e-08 | 4.2e-08 | 5.133E-08 | 2.723E-08 | 3.000E-08 | 8.200E-08 |
| 20632 | 1032 | G1032K | C | 7.5e-08 | 3.2e-08 | 4.1e-08 | 4.933E-08 | 2.268E-08 | 3.200E-08 | 7.500E-08 |
| 20633 | 1032 | G1032M | C | 3.6e-08 | 4.4e-08 | 6.3e-08 | 4.767E-08 | 1.387E-08 | 3.600E-08 | 6.300E-08 |
| 20634 | 1032 | G1032F | C | 1e-07 | 1.1e-07 | 2.1e-08 | 7.700E-08 | 4.875E-08 | 2.100E-08 | 1.100E-07 |
| 20635 | 1032 | G1032P | C | 2.6e-08 | 7.4e-08 | 4.3e-08 | 4.767E-08 | 2.434E-08 | 2.600E-08 | 7.400E-08 |
| 20636 | 1032 | G1032S | C | 8e-08 | 4.5e-08 | 8e-08 | 6.833E-08 | 2.021E-08 | 4.500E-08 | 8.000E-08 |
| 20637 | 1032 | G1032T | C | 6.4e-08 | 4.5e-08 | 3.5e-08 | 4.800E-08 | 1.473E-08 | 3.500E-08 | 6.400E-08 |
| 20638 | 1032 | G1032W | C | 2.1e-08 | 2.2e-08 | 5.9e-08 | 3.400E-08 | 2.166E-08 | 2.100E-08 | 5.900E-08 |
| 20639 | 1032 | G1032Y | C | 2.3e-08 | 1.1e-07 | 1.3e-08 | 4.867E-08 | 5.335E-08 | 1.300E-08 | 1.100E-07 |
| 20640 | 1032 | G1032V | C | 7.4e-08 | 5.5e-08 | 3.7e-08 | 5.533E-08 | 1.850E-08 | 3.700E-08 | 7.400E-08 |
| 20641 | 1033 | T1033A | C | 2e-08 | 1.5e-08 | 7.1e-08 | 3.533E-08 | 3.099E-08 | 1.500E-08 | 7.100E-08 |
| 20642 | 1033 | T1033R | C | 2.2e-08 | 6e-08 | 4.4e-08 | 4.200E-08 | 1.908E-08 | 2.200E-08 | 6.000E-08 |
| 20643 | 1033 | T1033N | C | 2.1e-08 | 3.7e-08 | 6.6e-08 | 4.133E-08 | 2.281E-08 | 2.100E-08 | 6.600E-08 |
| 20644 | 1033 | T1033D | C | 2.8e-08 | 3.5e-08 | 6.5e-08 | 4.267E-08 | 1.966E-08 | 2.800E-08 | 6.500E-08 |
| 20645 | 1033 | T1033C | C | 2.7e-08 | 7.4e-08 | 4.9e-08 | 5.000E-08 | 2.352E-08 | 2.700E-08 | 7.400E-08 |
| 20646 | 1033 | T1033E | C | 5.2e-08 | 5.3e-08 | 1.2e-07 | 7.500E-08 | 3.897E-08 | 5.200E-08 | 1.200E-07 |
| 20647 | 1033 | T1033Q | C | 3.7e-08 | 1.3e-07 | 4.2e-08 | 6.967E-08 | 5.231E-08 | 3.700E-08 | 1.300E-07 |
| 20648 | 1033 | T1033G | C | 8.1e-08 | 5.8e-08 | 1.6e-08 | 5.167E-08 | 3.296E-08 | 1.600E-08 | 8.100E-08 |
| 20649 | 1033 | T1033H | C | 3e-08 | 2.8e-08 | 4.5e-08 | 3.433E-08 | 9.292E-09 | 2.800E-08 | 4.500E-08 |
| 20650 | 1033 | T1033I | C | 2.8e-08 | 8.3e-08 | 6.4e-08 | 5.833E-08 | 2.793E-08 | 2.800E-08 | 8.300E-08 |
| 20651 | 1033 | T1033L | C | 4.6e-08 | 1.8e-08 | 5.5e-08 | 3.967E-08 | 1.930E-08 | 1.800E-08 | 5.500E-08 |
| 20652 | 1033 | T1033K | C | 1.7e-08 | 3.5e-08 | 3.8e-08 | 3.000E-08 | 1.136E-08 | 1.700E-08 | 3.800E-08 |
| 20653 | 1033 | T1033M | C | 2e-08 | 4e-08 | 3.8e-08 | 3.267E-08 | 1.102E-08 | 2.000E-08 | 4.000E-08 |
| 20654 | 1033 | T1033F | C | 4.2e-08 | 9.4e-08 | 2.1e-08 | 5.233E-08 | 3.758E-08 | 2.100E-08 | 9.400E-08 |

| | | | | | | | | | | |
|---|---|---|---|---|---|---|---|---|---|---|
| 20655 | 1033 | T1033P | C | 8.5e-08 | 5.9e-08 | 6.6e-08 | 7.000E-08 | 1.345E-08 | 5.900E-08 | 8.500E-08 |
| 20656 | 1033 | T1033S | C | 2.8e-08 | 3.8e-08 | 3.5e-08 | 3.367E-08 | 5.132E-09 | 2.800E-08 | 3.800E-08 |
| 20657 | 1033 | T1033T | C | 1.8e-08 | 4.1e-08 | 5.8e-08 | 3.900E-08 | 2.007E-08 | 1.800E-08 | 5.800E-08 |
| 20658 | 1033 | T1033W | C | 7.4e-08 | 5.2e-08 | 4.7e-08 | 5.767E-08 | 1.436E-08 | 4.700E-08 | 7.400E-08 |
| 20659 | 1033 | T1033Y | C | 7.7e-08 | 1e-07 | 3.6e-08 | 7.100E-08 | 3.242E-08 | 3.600E-08 | 1.000E-07 |
| 20660 | 1033 | T1033V | C | 3.1e-08 | 1.9e-08 | 2.6e-08 | 2.533E-08 | 6.028E-09 | 1.900E-08 | 3.100E-08 |
| 20661 | 1034 | Q1034A | C | 5.6e-08 | 8.2e-08 | 1.1e-07 | 8.267E-08 | 2.701E-08 | 5.600E-08 | 1.100E-07 |
| 20662 | 1034 | Q1034R | C | 2.8e-08 | 1.3e-08 | 5.9e-08 | 3.333E-08 | 2.346E-08 | 1.300E-08 | 5.900E-08 |
| 20663 | 1034 | Q1034N | C | 1.4e-08 | 5.2e-08 | 3.1e-08 | 3.233E-08 | 1.904E-08 | 1.400E-08 | 5.200E-08 |
| 20664 | 1034 | Q1034D | C | 6.1e-08 | 5.9e-08 | 9.9e-08 | 7.300E-08 | 2.254E-08 | 5.900E-08 | 9.900E-08 |
| 20665 | 1034 | Q1034C | C | 5e-08 | 2.7e-08 | 5e-08 | 4.233E-08 | 1.328E-08 | 2.700E-08 | 5.000E-08 |
| 20666 | 1034 | Q1034E | C | 5.3e-08 | 4.6e-08 | 2.6e-08 | 4.167E-08 | 1.401E-08 | 2.600E-08 | 5.300E-08 |
| 20667 | 1034 | Q1034Q | C | 9.6e-08 | 3.4e-08 | 5.1e-08 | 6.033E-08 | 3.204E-08 | 3.400E-08 | 9.600E-08 |
| 20668 | 1034 | Q1034G | C | 2.7e-08 | 4.9e-08 | 2.7e-08 | 3.433E-08 | 1.270E-08 | 2.700E-08 | 4.900E-08 |
| 20669 | 1034 | Q1034H | C | 4.1e-08 | 2.7e-08 | 2e-08 | 2.933E-08 | 1.069E-08 | 2.000E-08 | 4.100E-08 |
| 20670 | 1034 | Q1034I | C | 1.6e-08 | 8.4e-08 | 1.6e-08 | 3.867E-08 | 3.926E-08 | 1.600E-08 | 8.400E-08 |
| 20671 | 1034 | Q1034L | C | 6.9e-08 | 5.5e-08 | 7.4e-08 | 6.600E-08 | 9.849E-09 | 5.500E-08 | 7.400E-08 |
| 20672 | 1034 | Q1034K | C | 3e-08 | 2.7e-08 | 3.5e-08 | 3.067E-08 | 4.041E-09 | 2.700E-08 | 3.500E-08 |
| 20673 | 1034 | Q1034M | C | 6.7e-08 | 1e-07 | 4.1e-08 | 6.933E-08 | 2.957E-08 | 4.100E-08 | 1.000E-07 |
| 20674 | 1034 | Q1034F | C | 5.4e-08 | 1.9e-08 | 2.8e-08 | 3.367E-08 | 1.818E-08 | 1.900E-08 | 5.400E-08 |
| 20675 | 1034 | Q1034P | C | 7.4e-08 | 7e-08 | 6.1e-08 | 6.833E-08 | 6.658E-09 | 6.100E-08 | 7.400E-08 |
| 20676 | 1034 | Q1034S | C | 5.6e-08 | 2.3e-08 | 1.1e-07 | 6.300E-08 | 4.392E-08 | 2.300E-08 | 1.100E-07 |
| 20677 | 1034 | Q1034T | C | 6.1e-08 | 7.8e-08 | 3.8e-08 | 5.900E-08 | 2.007E-08 | 3.800E-08 | 7.800E-08 |
| 20678 | 1034 | Q1034W | C | 2.3e-08 | 5.5e-08 | 1.3e-08 | 3.033E-08 | 2.194E-08 | 1.300E-08 | 5.500E-08 |
| 20679 | 1034 | Q1034Y | C | 2.5e-07 | 5.5e-08 | 1.3e-08 | 1.060E-07 | 1.265E-07 | 1.300E-08 | 2.500E-07 |
| 20680 | 1034 | Q1034V | C | 4.9e-08 | 7.7e-08 | 1.6e-08 | 4.733E-08 | 3.053E-08 | 1.600E-08 | 7.700E-08 |
| 20681 | 1035 | T1035A | C | 8.9e-08 | 3.8e-08 | 5.4e-08 | 6.033E-08 | 2.608E-08 | 3.800E-08 | 8.900E-08 |
| 20682 | 1035 | T1035R | C | 1.7e-08 | 4.5e-08 | 2.6e-08 | 2.933E-08 | 1.429E-08 | 1.700E-08 | 4.500E-08 |
| 20683 | 1035 | T1035N | C | 6.5e-08 | 3.3e-08 | 4.4e-08 | 4.733E-08 | 1.626E-08 | 3.300E-08 | 6.500E-08 |
| 20684 | 1035 | T1035D | C | 4.2e-08 | 4.4e-08 | 9.2e-08 | 5.933E-08 | 2.831E-08 | 4.200E-08 | 9.200E-08 |
| 20685 | 1035 | T1035C | C | 2.3e-08 | 7.5e-08 | 6.5e-08 | 5.433E-08 | 2.759E-08 | 2.300E-08 | 7.500E-08 |
| 20686 | 1035 | T1035E | C | 3.3e-08 | 8.1e-08 | 4.8e-08 | 5.400E-08 | 2.456E-08 | 3.300E-08 | 8.100E-08 |
| 20687 | 1035 | T1035Q | C | 3.7e-08 | 1.2e-07 | 7.9e-08 | 7.867E-08 | 4.150E-08 | 3.700E-08 | 1.200E-07 |
| 20688 | 1035 | T1035G | C | 6.3e-08 | 3.9e-08 | 8.9e-08 | 6.367E-08 | 2.501E-08 | 3.900E-08 | 8.900E-08 |
| 20689 | 1035 | T1035H | C | 2.1e-08 | 4.2e-08 | 4.1e-08 | 3.467E-08 | 1.185E-08 | 2.100E-08 | 4.200E-08 |
| 20690 | 1035 | T1035I | C | 3.9e-08 | 5.1e-08 | 3.3e-08 | 4.100E-08 | 9.165E-09 | 3.300E-08 | 5.100E-08 |
| 20691 | 1035 | T1035L | C | 6.4e-08 | 3.9e-08 | 3.3e-08 | 4.533E-08 | 1.644E-08 | 3.300E-08 | 6.400E-08 |
| 20692 | 1035 | T1035K | C | 5e-08 | 7.1e-08 | 9.9e-08 | 7.333E-08 | 2.458E-08 | 5.000E-08 | 9.900E-08 |
| 20693 | 1035 | T1035M | C | 5.8e-08 | 5e-08 | 6.1e-08 | 5.633E-08 | 5.686E-09 | 5.000E-08 | 6.100E-08 |
| 20694 | 1035 | T1035F | C | 3.4e-08 | 1.1e-07 | 4.9e-08 | 6.433E-08 | 4.025E-08 | 3.400E-08 | 1.100E-07 |
| 20695 | 1035 | T1035P | C | 1.1e-07 | 6e-08 | 3.2e-08 | 6.733E-08 | 3.951E-08 | 3.200E-08 | 1.100E-07 |
| 20696 | 1035 | T1035S | C | 6.1e-08 | 5e-08 | 4.1e-08 | 5.067E-08 | 1.002E-08 | 4.100E-08 | 6.100E-08 |
| 20697 | 1035 | T1035T | C | 3e-07 | 3.2e-08 | 2e-08 | 1.173E-07 | 1.583E-07 | 2.000E-08 | 3.000E-07 |
| 20698 | 1035 | T1035W | C | 1.6e-08 | 3.6e-08 | 2.8e-08 | 2.667E-08 | 1.007E-08 | 1.600E-08 | 3.600E-08 |
| 20699 | 1035 | T1035Y | C | 2.8e-08 | 1.5e-07 | 2e-07 | 1.260E-07 | 8.848E-08 | 2.800E-08 | 2.000E-07 |
| 20700 | 1035 | T1035V | C | 4.5e-08 | 6.6e-08 | 6.7e-08 | 5.933E-08 | 1.242E-08 | 4.500E-08 | 6.700E-08 |
| 20701 | 1036 | Y1036A | C | 3.5e-08 | 6.4e-08 | 7.3e-08 | 5.733E-08 | 1.986E-08 | 3.500E-08 | 7.300E-08 |
| 20702 | 1036 | Y1036R | C | 1.4e-07 | 4.5e-09 | 7e-08 | 7.150E-08 | 6.776E-08 | 4.500E-09 | 1.400E-07 |
| 20703 | 1036 | Y1036N | C | 3.2e-08 | 8.9e-08 | 5.6e-08 | 5.900E-08 | 2.862E-08 | 3.200E-08 | 8.900E-08 |
| 20704 | 1036 | Y1036D | C | 8.1e-08 | 2.5e-08 | 4.5e-08 | 5.033E-08 | 2.838E-08 | 2.500E-08 | 8.100E-08 |
| 20705 | 1036 | Y1036C | C | 2e-08 | 4.7e-08 | 4e-08 | 3.567E-08 | 1.401E-08 | 2.000E-08 | 4.700E-08 |
| 20706 | 1036 | Y1036E | C | 4.2e-08 | 4.2e-08 | 3.9e-08 | 4.100E-08 | 1.732E-09 | 3.900E-08 | 4.200E-08 |
| 20707 | 1036 | Y1036Q | C | 4.8e-08 | 2.9e-08 | 2.3e-08 | 3.333E-08 | 1.305E-08 | 2.300E-08 | 4.800E-08 |
| 20708 | 1036 | Y1036G | C | 6e-08 | 4e-08 | 1.7e-07 | 9.000E-08 | 7.000E-08 | 4.000E-08 | 1.700E-07 |
| 20709 | 1036 | Y1036H | C | 1.4e-07 | 3.3e-08 | 5.2e-08 | 7.500E-08 | 5.709E-08 | 3.300E-08 | 1.400E-07 |
| 20710 | 1036 | Y1036I | C | 1.7e-08 | 2.6e-08 | 4.2e-08 | 2.833E-08 | 1.266E-08 | 1.700E-08 | 4.200E-08 |
| 20711 | 1036 | Y1036L | C | 5.4e-08 | 3.4e-08 | 3.1e-08 | 3.967E-08 | 1.250E-08 | 3.100E-08 | 5.400E-08 |
| 20712 | 1036 | Y1036K | C | 4.7e-08 | 4.9e-08 | 3.2e-08 | 4.267E-08 | 9.292E-09 | 3.200E-08 | 4.900E-08 |
| 20713 | 1036 | Y1036M | C | 9e-08 | 2.9e-08 | 3.4e-08 | 5.100E-08 | 3.387E-08 | 2.900E-08 | 9.000E-08 |
| 20714 | 1036 | Y1036F | C | 6.1e-08 | 5.3e-08 | 6.4e-08 | 5.933E-08 | 5.686E-09 | 5.300E-08 | 6.400E-08 |
| 20715 | 1036 | Y1036P | C | 1.1e-07 | 6.4e-08 | 6.1e-08 | 7.833E-08 | 2.747E-08 | 6.100E-08 | 1.100E-07 |
| 20716 | 1036 | Y1036S | C | 3.1e-08 | 2.6e-08 | 2.3e-08 | 2.667E-08 | 4.041E-09 | 2.300E-08 | 3.100E-08 |
| 20717 | 1036 | Y1036T | C | 1.5e-08 | 4.2e-08 | 1.9e-07 | 8.233E-08 | 9.421E-08 | 1.500E-08 | 1.900E-07 |
| 20718 | 1036 | Y1036W | C | 1.5e-07 | 6.1e-08 | 8e-08 | 9.700E-08 | 4.687E-08 | 6.100E-08 | 1.500E-07 |
| 20719 | 1036 | Y1036Y | C | 4.1e-08 | 2.7e-08 | 6.2e-08 | 4.333E-08 | 1.762E-08 | 2.700E-08 | 6.200E-08 |
| 20720 | 1036 | Y1036V | C | 4.1e-08 | 2.9e-08 | 6.1e-08 | 4.367E-08 | 1.617E-08 | 2.900E-08 | 6.100E-08 |
| 20721 | 1037 | I1037A | C | 8.3e-08 | 4.3e-08 | 5e-08 | 5.867E-08 | 2.136E-08 | 4.300E-08 | 8.300E-08 |
| 20722 | 1037 | I1037R | C | 5.6e-08 | 8.3e-08 | 5.3e-08 | 6.400E-08 | 1.652E-08 | 5.300E-08 | 8.300E-08 |
| 20723 | 1037 | I1037N | C | 4.2e-08 | 6.8e-08 | 2.2e-08 | 4.400E-08 | 2.307E-08 | 2.200E-08 | 6.800E-08 |
| 20724 | 1037 | I1037D | C | 4.5e-08 | 5.8e-08 | 1.9e-08 | 4.067E-08 | 1.986E-08 | 1.900E-08 | 5.800E-08 |
| 20725 | 1037 | I1037C | C | 3.9e-08 | 4.8e-08 | 3.5e-08 | 4.067E-08 | 6.658E-09 | 3.500E-08 | 4.800E-08 |
| 20726 | 1037 | I1037E | C | 8e-08 | 3e-08 | 3.3e-08 | 4.767E-08 | 2.804E-08 | 3.000E-08 | 8.000E-08 |
| 20727 | 1037 | I1037Q | C | 5.3e-08 | 3.3e-08 | 3.9e-08 | 4.167E-08 | 1.026E-08 | 3.300E-08 | 5.300E-08 |
| 20728 | 1037 | I1037G | C | 4.1e-08 | 4.6e-08 | 3.7e-08 | 4.133E-08 | 4.509E-09 | 3.700E-08 | 4.600E-08 |
| 20729 | 1037 | I1037H | C | 2.6e-08 | 6.6e-08 | 5.5e-08 | 4.900E-08 | 2.066E-08 | 2.600E-08 | 6.600E-08 |
| 20730 | 1037 | I1037I | C | 6.3e-08 | 6.4e-08 | 9.3e-08 | 7.333E-08 | 1.704E-08 | 6.300E-08 | 9.300E-08 |
| 20731 | 1037 | I1037L | C | 3.8e-08 | 1.4e-08 | 2e-08 | 2.400E-08 | 1.249E-08 | 1.400E-08 | 3.800E-08 |
| 20732 | 1037 | I1037K | C | 4.7e-08 | 2.5e-08 | 2.5e-08 | 3.233E-08 | 1.270E-08 | 2.500E-08 | 4.700E-08 |
| 20733 | 1037 | I1037M | C | 2.9e-08 | 1e-07 | 2.2e-08 | 5.033E-08 | 4.315E-08 | 2.200E-08 | 1.000E-07 |
| 20734 | 1037 | I1037F | C | 3.8e-08 | 2.7e-08 | 2.5e-08 | 3.000E-08 | 7.000E-09 | 2.500E-08 | 3.800E-08 |
| 20735 | 1037 | I1037P | C | 2.3e-08 | 4.7e-08 | 2.4e-08 | 3.133E-08 | 1.358E-08 | 2.300E-08 | 4.700E-08 |
| 20736 | 1037 | I1037S | C | 3.8e-08 | 2.8e-08 | 4.6e-08 | 3.733E-08 | 9.018E-09 | 2.800E-08 | 4.600E-08 |
| 20737 | 1037 | I1037T | C | 4e-08 | 4.6e-08 | 3e-08 | 3.867E-08 | 8.083E-09 | 3.000E-08 | 4.600E-08 |
| 20738 | 1037 | I1037W | C | 2.8e-08 | 1.5e-07 | 7.4e-08 | 8.400E-08 | 6.161E-08 | 2.800E-08 | 1.500E-07 |
| 20739 | 1037 | I1037Y | C | 3.8e-08 | 2.6e-08 | 5.3e-08 | 3.900E-08 | 1.353E-08 | 2.600E-08 | 5.300E-08 |

| | | | | | | | | | | |
|---|---|---|---|---|---|---|---|---|---|---|
| 20740 | 1037 | I1037V | C | 3.3e-08 | 5.5e-08 | 9.7e-08 | 6.167E-08 | 3.252E-08 | 3.300E-08 | 9.700E-08 |
| 20741 | 1038 | C1038A | C | 2.6e-08 | 2.7e-08 | 3.4e-08 | 2.900E-08 | 4.359E-09 | 2.600E-08 | 3.400E-08 |
| 20742 | 1038 | C1038R | C | 7.3e-08 | 4.6e-08 | 5.7e-08 | 5.867E-08 | 1.358E-08 | 4.600E-08 | 7.300E-08 |
| 20743 | 1038 | C1038N | C | 1e-07 | 1.9e-08 | 5.1e-08 | 5.667E-08 | 4.080E-08 | 1.900E-08 | 1.000E-07 |
| 20744 | 1038 | C1038D | C | 2.4e-08 | 7.2e-08 | 2.1e-08 | 3.900E-08 | 2.862E-08 | 2.100E-08 | 7.200E-08 |
| 20745 | 1038 | C1038C | C | 6.1e-08 | 8.4e-08 | 3.4e-08 | 5.967E-08 | 2.503E-08 | 3.400E-08 | 8.400E-08 |
| 20746 | 1038 | C1038E | C | 9.1e-09 | 6.2e-08 | 2.5e-08 | 3.203E-08 | 2.714E-08 | 9.100E-09 | 6.200E-08 |
| 20747 | 1038 | C1038Q | C | 9.6e-08 | 5.2e-08 | 4.7e-08 | 6.500E-08 | 2.696E-08 | 4.700E-08 | 9.600E-08 |
| 20748 | 1038 | C1038G | C | 4.4e-08 | 3.8e-08 | 7.1e-08 | 5.100E-08 | 1.758E-08 | 3.800E-08 | 7.100E-08 |
| 20749 | 1038 | C1038H | C | 7e-08 | 2.2e-08 | 2.7e-08 | 3.967E-08 | 2.639E-08 | 2.200E-08 | 7.000E-08 |
| 20750 | 1038 | C1038I | C | 7.4e-08 | 5.3e-08 | 1.4e-08 | 4.700E-08 | 3.045E-08 | 1.400E-08 | 7.400E-08 |
| 20751 | 1038 | C1038L | C | 7.7e-08 | 4.7e-08 | 7.9e-08 | 6.767E-08 | 1.793E-08 | 4.700E-08 | 7.900E-08 |
| 20752 | 1038 | C1038K | C | 4.8e-08 | 1.2e-07 | 7.5e-08 | 8.100E-08 | 3.637E-08 | 4.800E-08 | 1.200E-07 |
| 20753 | 1038 | C1038M | C | 6.2e-08 | 1.1e-08 | 5.8e-08 | 4.367E-08 | 2.836E-08 | 1.100E-08 | 6.200E-08 |
| 20754 | 1038 | C1038F | C | 4.4e-08 | 1.3e-07 | 6.1e-08 | 7.833E-08 | 4.554E-08 | 4.400E-08 | 1.300E-07 |
| 20755 | 1038 | C1038P | C | 8.2e-08 | 2.9e-08 | 4.5e-08 | 5.200E-08 | 2.718E-08 | 2.900E-08 | 8.200E-08 |
| 20756 | 1038 | C1038S | C | 6e-08 | 9.8e-08 | 5.3e-08 | 7.033E-08 | 2.421E-08 | 5.300E-08 | 9.800E-08 |
| 20757 | 1038 | C1038T | C | 7.5e-08 | 8.5e-08 | 3.2e-08 | 6.400E-08 | 2.816E-08 | 3.200E-08 | 8.500E-08 |
| 20758 | 1038 | C1038W | C | 4e-08 | 1.5e-08 | 2.4e-08 | 2.633E-08 | 1.266E-08 | 1.500E-08 | 4.000E-08 |
| 20759 | 1038 | C1038Y | C | 4.6e-08 | 2.4e-08 | 3.6e-08 | 3.533E-08 | 1.102E-08 | 2.400E-08 | 4.600E-08 |
| 20760 | 1038 | C1038V | C | 4.9e-08 | 4.8e-08 | 3.6e-08 | 4.433E-08 | 7.234E-09 | 3.600E-08 | 4.900E-08 |
| 20761 | 1039 | N1039A | C | 1.8e-07 | 3.9e-08 | 4.3e-08 | 8.733E-08 | 8.028E-08 | 3.900E-08 | 1.800E-07 |
| 20762 | 1039 | N1039R | C | 3e-08 | 5.5e-08 | 3.2e-08 | 3.900E-08 | 1.389E-08 | 3.000E-08 | 5.500E-08 |
| 20763 | 1039 | N1039N | C | 5.7e-08 | 5.9e-08 | 1.1e-08 | 4.233E-08 | 2.715E-08 | 1.100E-08 | 5.900E-08 |
| 20764 | 1039 | N1039D | C | 3e-08 | 5.2e-08 | 3.5e-08 | 3.900E-08 | 1.153E-08 | 3.000E-08 | 5.200E-08 |
| 20765 | 1039 | N1039C | C | 2.7e-08 | 5.8e-08 | 2.1e-08 | 3.533E-08 | 1.986E-08 | 2.100E-08 | 5.800E-08 |
| 20766 | 1039 | N1039E | C | 4.1e-08 | 5e-08 | 2.7e-08 | 3.933E-08 | 1.159E-08 | 2.700E-08 | 5.000E-08 |
| 20767 | 1039 | N1039Q | C | 5.4e-08 | 9.8e-08 | 5.2e-08 | 6.800E-08 | 2.600E-08 | 5.200E-08 | 9.800E-08 |
| 20768 | 1039 | N1039G | C | 2.7e-08 | 1e-07 | 7.7e-08 | 6.800E-08 | 3.732E-08 | 2.700E-08 | 1.000E-07 |
| 20769 | 1039 | N1039H | C | 2.5e-08 | 5.3e-08 | 3.1e-08 | 3.633E-08 | 1.474E-08 | 2.500E-08 | 5.300E-08 |
| 20770 | 1039 | N1039I | C | 2.4e-08 | 4.8e-08 | 5.3e-08 | 4.167E-08 | 1.550E-08 | 2.400E-08 | 5.300E-08 |
| 20771 | 1039 | N1039L | C | 1.2e-07 | 3.4e-08 | 3.3e-08 | 6.233E-08 | 4.994E-08 | 3.300E-08 | 1.200E-07 |
| 20772 | 1039 | N1039K | C | 1.6e-07 | 1.8e-08 | 6.4e-08 | 8.067E-08 | 7.245E-08 | 1.800E-08 | 1.600E-07 |
| 20773 | 1039 | N1039M | C | 2.9e-08 | 2.5e-08 | 4.6e-08 | 3.333E-08 | 1.115E-08 | 2.500E-08 | 4.600E-08 |
| 20774 | 1039 | N1039F | C | 1.5e-08 | 2.3e-08 | 1.1e-07 | 4.933E-08 | 5.269E-08 | 1.500E-08 | 1.100E-07 |
| 20775 | 1039 | N1039P | C | 7.6e-08 | 2.6e-08 | 3.2e-08 | 4.467E-08 | 2.730E-08 | 2.600E-08 | 7.600E-08 |
| 20776 | 1039 | N1039S | C | 2.4e-08 | 2.9e-08 | 4.8e-08 | 3.367E-08 | 1.266E-08 | 2.400E-08 | 4.800E-08 |
| 20777 | 1039 | N1039T | C | 9.7e-08 | 4.2e-08 | 3.5e-08 | 5.800E-08 | 3.396E-08 | 3.500E-08 | 9.700E-08 |
| 20778 | 1039 | N1039W | C | 2.2e-08 | 3.8e-08 | 4.9e-08 | 3.633E-08 | 1.358E-08 | 2.200E-08 | 4.900E-08 |
| 20779 | 1039 | N1039Y | C | 4e-08 | 3.7e-08 | 4.6e-08 | 4.100E-08 | 4.583E-09 | 3.700E-08 | 4.600E-08 |
| 20780 | 1039 | N1039V | C | 7.4e-08 | 7.7e-08 | 2.1e-08 | 5.733E-08 | 3.150E-08 | 2.100E-08 | 7.700E-08 |
| 20781 | 1040 | V1040A | C | 8.3e-08 | 2.8e-08 | 2.4e-08 | 4.500E-08 | 3.297E-08 | 2.400E-08 | 8.300E-08 |
| 20782 | 1040 | V1040R | C | 3.5e-08 | 4.4e-08 | 8.1e-08 | 5.333E-08 | 2.438E-08 | 3.500E-08 | 8.100E-08 |
| 20783 | 1040 | V1040N | C | 3.2e-08 | 3.6e-08 | 6.2e-08 | 4.333E-08 | 1.629E-08 | 3.200E-08 | 6.200E-08 |
| 20784 | 1040 | V1040D | C | 4.4e-08 | 4.5e-08 | 2.4e-08 | 3.767E-08 | 1.185E-08 | 2.400E-08 | 4.500E-08 |
| 20785 | 1040 | V1040C | C | 8.5e-08 | 7.4e-08 | 2.9e-08 | 6.267E-08 | 2.967E-08 | 2.900E-08 | 8.500E-08 |
| 20786 | 1040 | V1040E | C | 6.4e-08 | 1.4e-07 | 3.4e-08 | 7.933E-08 | 5.464E-08 | 3.400E-08 | 1.400E-07 |
| 20787 | 1040 | V1040Q | C | 3.3e-08 | 4.4e-08 | 6.4e-08 | 4.700E-08 | 1.572E-08 | 3.300E-08 | 6.400E-08 |
| 20788 | 1040 | V1040G | C | 4.7e-08 | 7.2e-08 | 3.5e-08 | 5.133E-08 | 1.888E-08 | 3.500E-08 | 7.200E-08 |
| 20789 | 1040 | V1040H | C | 4.8e-08 | 7e-08 | 2.4e-08 | 4.733E-08 | 2.301E-08 | 2.400E-08 | 7.000E-08 |
| 20790 | 1040 | V1040I | C | 8.3e-08 | 3.4e-08 | 9.2e-08 | 6.967E-08 | 3.121E-08 | 3.400E-08 | 9.200E-08 |
| 20791 | 1040 | V1040L | C | 1.1e-07 | 1e-07 | 3.9e-08 | 8.300E-08 | 3.843E-08 | 3.900E-08 | 1.100E-07 |
| 20792 | 1040 | V1040K | C | 7.1e-08 | 3.2e-08 | 5.1e-08 | 5.133E-08 | 1.950E-08 | 3.200E-08 | 7.100E-08 |
| 20793 | 1040 | V1040M | C | 1e-07 | 3.4e-08 | 3.9e-08 | 5.767E-08 | 3.675E-08 | 3.400E-08 | 1.000E-07 |
| 20794 | 1040 | V1040F | C | 3.7e-08 | 1.6e-08 | 2e-08 | 2.433E-08 | 1.115E-08 | 1.600E-08 | 3.700E-08 |
| 20795 | 1040 | V1040P | C | 4.2e-08 | 6.5e-08 | 1.8e-08 | 4.167E-08 | 2.350E-08 | 1.800E-08 | 6.500E-08 |
| 20796 | 1040 | V1040S | C | 3.9e-08 | 1.1e-07 | 1.3e-07 | 9.300E-08 | 4.782E-08 | 3.900E-08 | 1.300E-07 |
| 20797 | 1040 | V1040T | C | 6.5e-08 | 3.5e-08 | 2.6e-08 | 4.200E-08 | 2.042E-08 | 2.600E-08 | 6.500E-08 |
| 20798 | 1040 | V1040W | C | 6e-08 | 3.7e-08 | 3.7e-08 | 4.467E-08 | 1.328E-08 | 3.700E-08 | 6.000E-08 |
| 20799 | 1040 | V1040Y | C | 1.5e-08 | 7.9e-08 | 4.1e-08 | 4.500E-08 | 3.219E-08 | 1.500E-08 | 7.900E-08 |
| 20800 | 1040 | V1040V | C | 1.9e-08 | 3.2e-08 | 1e-07 | 5.033E-08 | 4.350E-08 | 1.900E-08 | 1.000E-07 |
| 20801 | 1041 | N1041A | C | 2.9e-08 | 3.8e-08 | 5.4e-08 | 4.033E-08 | 1.266E-08 | 2.900E-08 | 5.400E-08 |
| 20802 | 1041 | N1041R | C | 3.5e-08 | 6.3e-08 | 2.3e-08 | 4.033E-08 | 2.053E-08 | 2.300E-08 | 6.300E-08 |
| 20803 | 1041 | N1041N | C | 7e-08 | 2.5e-08 | 3.8e-08 | 4.433E-08 | 2.316E-08 | 2.500E-08 | 7.000E-08 |
| 20804 | 1041 | N1041D | C | 4.3e-08 | 1.7e-08 | 1.8e-08 | 2.600E-08 | 1.473E-08 | 1.700E-08 | 4.300E-08 |
| 20805 | 1041 | N1041C | C | 9.4e-08 | 5.9e-08 | 4.8e-08 | 6.700E-08 | 2.402E-08 | 4.800E-08 | 9.400E-08 |
| 20806 | 1041 | N1041E | C | 3.2e-08 | 1.6e-08 | 4.7e-08 | 3.167E-08 | 1.550E-08 | 1.600E-08 | 4.700E-08 |
| 20807 | 1041 | N1041Q | C | 3.9e-08 | 3.8e-08 | 2.4e-08 | 3.367E-08 | 8.386E-09 | 2.400E-08 | 3.900E-08 |
| 20808 | 1041 | N1041G | C | 4.7e-08 | 2.1e-08 | 3e-08 | 3.267E-08 | 1.320E-08 | 2.100E-08 | 4.700E-08 |
| 20809 | 1041 | N1041H | C | 3.2e-08 | 9e-08 | 9.1e-08 | 7.100E-08 | 3.378E-08 | 3.200E-08 | 9.100E-08 |
| 20810 | 1041 | N1041I | C | 3.2e-08 | 6.7e-08 | 2.2e-08 | 4.033E-08 | 2.363E-08 | 2.200E-08 | 6.700E-08 |
| 20811 | 1041 | N1041L | C | 4e-08 | 3.2e-08 | 3.8e-08 | 3.667E-08 | 4.163E-09 | 3.200E-08 | 4.000E-08 |
| 20812 | 1041 | N1041K | C | 1e-07 | 6.7e-08 | 6.5e-08 | 7.733E-08 | 1.966E-08 | 6.500E-08 | 1.000E-07 |
| 20813 | 1041 | N1041M | C | 9.4e-08 | 5.1e-08 | 6.4e-08 | 6.967E-08 | 2.205E-08 | 5.100E-08 | 9.400E-08 |
| 20814 | 1041 | N1041F | C | 5.2e-08 | 6e-08 | 6.1e-08 | 5.767E-08 | 4.933E-09 | 5.200E-08 | 6.100E-08 |
| 20815 | 1041 | N1041P | C | 3.2e-08 | 4.4e-08 | 6.4e-08 | 4.667E-08 | 1.617E-08 | 3.200E-08 | 6.400E-08 |
| 20816 | 1041 | N1041S | C | 6e-08 | 3.2e-08 | 4.2e-08 | 4.467E-08 | 1.419E-08 | 3.200E-08 | 6.000E-08 |
| 20817 | 1041 | N1041T | C | 2.8e-08 | 1.4e-08 | 1e-07 | 4.733E-08 | 4.614E-08 | 1.400E-08 | 1.000E-07 |
| 20818 | 1041 | N1041W | C | 5e-08 | 3.2e-08 | 5.7e-08 | 4.633E-08 | 1.290E-08 | 3.200E-08 | 5.700E-08 |
| 20819 | 1041 | N1041Y | C | 4.6e-08 | 2.1e-08 | 1.6e-08 | 2.767E-08 | 1.607E-08 | 1.600E-08 | 4.600E-08 |
| 20820 | 1041 | N1041V | C | 2.2e-08 | 2.6e-08 | 8.9e-08 | 4.567E-08 | 3.758E-08 | 2.200E-08 | 8.900E-08 |
| 20821 | 1042 | H1042A | C | 5.7e-08 | 6.7e-08 | 3.1e-08 | 5.167E-08 | 1.858E-08 | 3.100E-08 | 6.700E-08 |
| 20822 | 1042 | H1042R | C | 3.6e-08 | 3.8e-08 | 9.3e-08 | 5.567E-08 | 3.235E-08 | 3.600E-08 | 9.300E-08 |
| 20823 | 1042 | H1042N | C | 3e-08 | 2.1e-08 | 4e-08 | 3.033E-08 | 9.504E-09 | 2.100E-08 | 4.000E-08 |
| 20824 | 1042 | H1042D | C | 7.7e-08 | 2.8e-08 | 4e-08 | 4.833E-08 | 2.554E-08 | 2.800E-08 | 7.700E-08 |

| | | | | | | | | | | |
|---|---|---|---|---|---|---|---|---|---|---|
| 20825 | 1042 | H1042C | C | 7.6e-08 | 3.9e-08 | 5.4e-08 | 5.633E-08 | 1.861E-08 | 3.900E-08 | 7.600E-08 |
| 20826 | 1042 | H1042E | C | 5.6e-08 | 5e-08 | 5.5e-08 | 5.367E-08 | 3.215E-09 | 5.000E-08 | 5.600E-08 |
| 20827 | 1042 | H1042Q | C | 6.1e-08 | 5.2e-08 | 4.9e-08 | 5.400E-08 | 6.245E-09 | 4.900E-08 | 6.100E-08 |
| 20828 | 1042 | H1042G | C | 3e-08 | 5.5e-08 | 9.1e-08 | 5.867E-08 | 3.066E-08 | 3.000E-08 | 9.100E-08 |
| 20829 | 1042 | H1042H | C | 5.4e-08 | 9.4e-08 | 1.9e-08 | 5.567E-08 | 3.753E-08 | 1.900E-08 | 9.400E-08 |
| 20830 | 1042 | H1042I | C | 2.7e-08 | 5e-08 | 1.1e-07 | 6.233E-08 | 4.285E-08 | 2.700E-08 | 1.100E-07 |
| 20831 | 1042 | H1042L | C | 4.7e-08 | 4.8e-08 | 9.6e-08 | 6.367E-08 | 2.801E-08 | 4.700E-08 | 9.600E-08 |
| 20832 | 1042 | H1042K | C | 1.2e-08 | 2.4e-08 | 6.6e-08 | 3.400E-08 | 2.835E-08 | 1.200E-08 | 6.600E-08 |
| 20833 | 1042 | H1042M | C | 5.9e-08 | 3.3e-08 | 3.7e-08 | 4.300E-08 | 1.400E-08 | 3.300E-08 | 5.900E-08 |
| 20834 | 1042 | H1042F | C | 2.9e-08 | 3.5e-08 | 2.9e-08 | 3.100E-08 | 3.464E-09 | 2.900E-08 | 3.500E-08 |
| 20835 | 1042 | H1042P | C | 2e-08 | 7.2e-08 | 2.4e-08 | 3.867E-08 | 2.894E-08 | 2.000E-08 | 7.200E-08 |
| 20836 | 1042 | H1042S | C | 2.5e-08 | 7.4e-08 | 2e-08 | 3.967E-08 | 2.984E-08 | 2.000E-08 | 7.400E-08 |
| 20837 | 1042 | H1042T | C | 4.3e-08 | 1.3e-07 | 4.3e-08 | 7.200E-08 | 5.023E-08 | 4.300E-08 | 1.300E-07 |
| 20838 | 1042 | H1042W | C | 7.7e-09 | 1.6e-08 | 4e-08 | 2.123E-08 | 1.677E-08 | 7.700E-09 | 4.000E-08 |
| 20839 | 1042 | H1042Y | C | 5.1e-08 | 2.3e-08 | 5.6e-08 | 4.333E-08 | 1.779E-08 | 2.300E-08 | 5.600E-08 |
| 20840 | 1042 | H1042V | C | 6.9e-08 | 3e-08 | 5.4e-08 | 5.100E-08 | 1.967E-08 | 3.000E-08 | 6.900E-08 |
| 20841 | 1043 | K1043A | C | 3e-08 | 6.1e-08 | 2.9e-08 | 4.000E-08 | 1.819E-08 | 2.900E-08 | 6.100E-08 |
| 20842 | 1043 | K1043R | C | 2.2e-08 | 5.4e-08 | 6.3e-09 | 2.743E-08 | 2.431E-08 | 6.300E-09 | 5.400E-08 |
| 20843 | 1043 | K1043N | C | 2.7e-08 | 1.9e-08 | 1e-07 | 4.867E-08 | 4.464E-08 | 1.900E-08 | 1.000E-07 |
| 20844 | 1043 | K1043D | C | 9.5e-08 | 4e-08 | 8.1e-08 | 7.200E-08 | 2.858E-08 | 4.000E-08 | 9.500E-08 |
| 20845 | 1043 | K1043C | C | 5.5e-08 | 5.2e-08 | 6.8e-08 | 5.833E-08 | 8.505E-09 | 5.200E-08 | 6.800E-08 |
| 20846 | 1043 | K1043E | C | 1.1e-08 | 3.6e-08 | 6.2e-08 | 3.633E-08 | 2.550E-08 | 1.100E-08 | 6.200E-08 |
| 20847 | 1043 | K1043Q | C | 6.3e-08 | 3.1e-08 | 5.5e-08 | 4.967E-08 | 1.665E-08 | 3.100E-08 | 6.300E-08 |
| 20848 | 1043 | K1043G | C | 1.5e-07 | 4.5e-08 | 4.1e-08 | 7.867E-08 | 6.181E-08 | 4.100E-08 | 1.500E-07 |
| 20849 | 1043 | K1043H | C | 7.1e-08 | 8.5e-08 | 1.9e-08 | 5.833E-08 | 3.478E-08 | 1.900E-08 | 8.500E-08 |
| 20850 | 1043 | K1043I | C | 7.2e-08 | 4.3e-08 | 5.4e-08 | 5.633E-08 | 1.464E-08 | 4.300E-08 | 7.200E-08 |
| 20851 | 1043 | K1043L | C | 3.2e-08 | 3.6e-08 | 3.3e-08 | 3.367E-08 | 2.082E-09 | 3.200E-08 | 3.600E-08 |
| 20852 | 1043 | K1043K | C | 6.9e-08 | 4.3e-08 | 2.8e-08 | 4.667E-08 | 2.074E-08 | 2.800E-08 | 6.900E-08 |
| 20853 | 1043 | K1043M | C | 8.2e-08 | 2.2e-08 | 1.6e-08 | 4.000E-08 | 3.650E-08 | 1.600E-08 | 8.200E-08 |
| 20854 | 1043 | K1043F | C | 4e-08 | 7.7e-08 | 4.9e-08 | 5.533E-08 | 1.930E-08 | 4.000E-08 | 7.700E-08 |
| 20855 | 1043 | K1043P | C | 2.6e-08 | 5.7e-08 | 1.6e-08 | 3.300E-08 | 2.138E-08 | 1.600E-08 | 5.700E-08 |
| 20856 | 1043 | K1043S | C | 1.3e-08 | 2.4e-08 | 5.8e-08 | 3.167E-08 | 2.346E-08 | 1.300E-08 | 5.800E-08 |
| 20857 | 1043 | K1043T | C | 5.3e-08 | 5.5e-08 | 1.1e-07 | 7.267E-08 | 3.235E-08 | 5.300E-08 | 1.100E-07 |
| 20858 | 1043 | K1043W | C | 5.5e-08 | 1.7e-08 | 9.1e-08 | 5.433E-08 | 3.700E-08 | 1.700E-08 | 9.100E-08 |
| 20859 | 1043 | K1043Y | C | 3.9e-08 | 4.5e-08 | 2.9e-08 | 3.767E-08 | 8.083E-09 | 2.900E-08 | 4.500E-08 |
| 20860 | 1043 | K1043V | C | 3.1e-08 | 7e-08 | 6.3e-08 | 5.467E-08 | 2.079E-08 | 3.100E-08 | 7.000E-08 |
| 20861 | 1044 | P1044A | C | 1.3e-07 | 5.6e-08 | 8.5e-08 | 9.033E-08 | 3.729E-08 | 5.600E-08 | 1.300E-07 |
| 20862 | 1044 | P1044R | C | 2.3e-08 | 2.7e-08 | 2.2e-08 | 2.400E-08 | 2.646E-09 | 2.200E-08 | 2.700E-08 |
| 20863 | 1044 | P1044N | C | 9e-08 | 8.1e-08 | 7.7e-08 | 8.267E-08 | 6.658E-09 | 7.700E-08 | 9.000E-08 |
| 20864 | 1044 | P1044D | C | 4e-08 | 4.3e-08 | 2.6e-08 | 3.633E-08 | 9.074E-09 | 2.600E-08 | 4.300E-08 |
| 20865 | 1044 | P1044C | C | 5.1e-08 | 2.8e-08 | 2.4e-08 | 3.433E-08 | 1.457E-08 | 2.400E-08 | 5.100E-08 |
| 20866 | 1044 | P1044E | C | 1.1e-08 | 5.3e-08 | 2.6e-08 | 3.000E-08 | 2.128E-08 | 1.100E-08 | 5.300E-08 |
| 20867 | 1044 | P1044Q | C | 4.2e-08 | 5.6e-08 | 2.5e-08 | 4.100E-08 | 1.552E-08 | 2.500E-08 | 5.600E-08 |
| 20868 | 1044 | P1044G | C | 1.4e-08 | 4.3e-08 | 2.1e-08 | 2.600E-08 | 1.513E-08 | 1.400E-08 | 4.300E-08 |
| 20869 | 1044 | P1044H | C | 1.6e-08 | 2.4e-08 | 4.2e-08 | 2.733E-08 | 1.332E-08 | 1.600E-08 | 4.200E-08 |
| 20870 | 1044 | P1044I | C | 4.3e-08 | 5.2e-08 | 2.1e-08 | 3.867E-08 | 1.595E-08 | 2.100E-08 | 5.200E-08 |
| 20871 | 1044 | P1044L | C | 3.5e-08 | 2.8e-08 | 3.7e-08 | 3.333E-08 | 4.726E-09 | 2.800E-08 | 3.700E-08 |
| 20872 | 1044 | P1044K | C | 3.3e-08 | 4.9e-08 | 2.5e-08 | 3.567E-08 | 1.222E-08 | 2.500E-08 | 4.900E-08 |
| 20873 | 1044 | P1044M | C | 4.4e-08 | 8.5e-08 | 5.1e-08 | 6.000E-08 | 2.193E-08 | 4.400E-08 | 8.500E-08 |
| 20874 | 1044 | P1044F | C | 4.5e-08 | 2.3e-08 | 5.3e-08 | 4.033E-08 | 1.553E-08 | 2.300E-08 | 5.300E-08 |
| 20875 | 1044 | P1044P | C | 2.1e-08 | 5.6e-08 | 7.2e-08 | 4.967E-08 | 2.608E-08 | 2.100E-08 | 7.200E-08 |
| 20876 | 1044 | P1044S | C | 3.1e-08 | 5.8e-08 | 1e-07 | 6.300E-08 | 3.477E-08 | 3.100E-08 | 1.000E-07 |
| 20877 | 1044 | P1044T | C | 4.8e-08 | 5.2e-08 | 5.1e-08 | 5.033E-08 | 2.082E-09 | 4.800E-08 | 5.200E-08 |
| 20878 | 1044 | P1044W | C | 4.4e-08 | 1.4e-08 | 7.2e-08 | 4.333E-08 | 2.901E-08 | 1.400E-08 | 7.200E-08 |
| 20879 | 1044 | P1044Y | C | 1.4e-08 | 6.2e-08 | 3.1e-08 | 3.567E-08 | 2.434E-08 | 1.400E-08 | 6.200E-08 |
| 20880 | 1044 | P1044V | C | 3.4e-08 | 2.4e-08 | 3.7e-08 | 3.167E-08 | 6.807E-09 | 2.400E-08 | 3.700E-08 |
| 20881 | 1045 | S1045A | C | 5.2e-08 | 3.7e-08 | 7.9e-08 | 5.600E-08 | 2.128E-08 | 3.700E-08 | 7.900E-08 |
| 20882 | 1045 | S1045R | C | 5.2e-08 | 1.7e-08 | 3.5e-08 | 3.467E-08 | 1.750E-08 | 1.700E-08 | 5.200E-08 |
| 20883 | 1045 | S1045N | C | 6e-08 | 4.9e-08 | 2.4e-08 | 4.433E-08 | 1.845E-08 | 2.400E-08 | 6.000E-08 |
| 20884 | 1045 | S1045D | C | 3.5e-08 | 2.2e-08 | 2.8e-08 | 2.833E-08 | 6.506E-09 | 2.200E-08 | 3.500E-08 |
| 20885 | 1045 | S1045C | C | 6e-08 | 4.5e-08 | 3.4e-08 | 4.633E-08 | 1.305E-08 | 3.400E-08 | 6.000E-08 |
| 20886 | 1045 | S1045E | C | 7.2e-08 | 5.5e-08 | 4.3e-08 | 5.667E-08 | 1.457E-08 | 4.300E-08 | 7.200E-08 |
| 20887 | 1045 | S1045Q | C | 4.3e-08 | 4.4e-08 | 4.4e-08 | 4.367E-08 | 5.774E-10 | 4.300E-08 | 4.400E-08 |
| 20888 | 1045 | S1045G | C | 4.9e-08 | 3.8e-08 | 5.1e-08 | 4.600E-08 | 7.000E-09 | 3.800E-08 | 5.100E-08 |
| 20889 | 1045 | S1045H | C | 4.4e-08 | 5.5e-08 | 6.6e-08 | 5.500E-08 | 1.100E-08 | 4.400E-08 | 6.600E-08 |
| 20890 | 1045 | S1045I | C | 7.7e-08 | 2.6e-08 | 3.7e-08 | 4.667E-08 | 2.684E-08 | 2.600E-08 | 7.700E-08 |
| 20891 | 1045 | S1045L | C | 7.2e-08 | 5.7e-08 | 1.7e-08 | 4.867E-08 | 2.843E-08 | 1.700E-08 | 7.200E-08 |
| 20892 | 1045 | S1045K | C | 6.7e-08 | 4.7e-08 | 4.1e-08 | 5.167E-08 | 1.361E-08 | 4.100E-08 | 6.700E-08 |
| 20893 | 1045 | S1045M | C | 4.9e-08 | 5.8e-08 | 6.8e-08 | 5.833E-08 | 9.504E-09 | 4.900E-08 | 6.800E-08 |
| 20894 | 1045 | S1045F | C | 6.1e-08 | 5.2e-08 | 2.7e-08 | 4.667E-08 | 1.762E-08 | 2.700E-08 | 6.100E-08 |
| 20895 | 1045 | S1045P | C | 4.5e-08 | 3.6e-08 | 2.5e-08 | 3.533E-08 | 1.002E-08 | 2.500E-08 | 4.500E-08 |
| 20896 | 1045 | S1045S | C | 2.7e-08 | 4.3e-08 | 3.2e-08 | 3.400E-08 | 8.185E-09 | 2.700E-08 | 4.300E-08 |
| 20897 | 1045 | S1045T | C | 3.2e-08 | 6.9e-08 | 1.6e-07 | 8.700E-08 | 6.587E-08 | 3.200E-08 | 1.600E-07 |
| 20898 | 1045 | S1045W | C | 6.7e-08 | 2.2e-08 | 2.8e-08 | 3.900E-08 | 2.443E-08 | 2.200E-08 | 6.700E-08 |
| 20899 | 1045 | S1045Y | C | 6.3e-08 | 1.6e-08 | 2.2e-08 | 3.367E-08 | 2.558E-08 | 1.600E-08 | 6.300E-08 |
| 20900 | 1045 | S1045V | C | 4.1e-08 | 5.5e-08 | 6.4e-08 | 5.333E-08 | 1.159E-08 | 4.100E-08 | 6.400E-08 |
| 20901 | 1046 | N1046A | C | 2.8e-08 | 2.9e-08 | 7.7e-08 | 4.467E-08 | 2.801E-08 | 2.800E-08 | 7.700E-08 |
| 20902 | 1046 | N1046R | C | 4.1e-08 | 5.5e-08 | 3.7e-08 | 4.433E-08 | 9.452E-09 | 3.700E-08 | 5.500E-08 |
| 20903 | 1046 | N1046N | C | 4.2e-08 | 3.3e-08 | 1.3e-08 | 2.933E-08 | 1.484E-08 | 1.300E-08 | 4.200E-08 |
| 20904 | 1046 | N1046D | C | 4.4e-08 | 1e-07 | 2.7e-08 | 5.700E-08 | 3.820E-08 | 2.700E-08 | 1.000E-07 |
| 20905 | 1046 | N1046C | C | 1.4e-08 | 1.1e-07 | 6.5e-08 | 6.300E-08 | 4.803E-08 | 1.400E-08 | 1.100E-07 |
| 20906 | 1046 | N1046E | C | 6.3e-08 | 2.5e-08 | 5.6e-08 | 4.800E-08 | 2.022E-08 | 2.500E-08 | 6.300E-08 |
| 20907 | 1046 | N1046Q | C | 1.3e-07 | 2.4e-08 | 4.5e-08 | 6.633E-08 | 5.613E-08 | 2.400E-08 | 1.300E-07 |
| 20908 | 1046 | N1046G | C | 1e-07 | 4.7e-08 | 1.9e-08 | 5.533E-08 | 4.114E-08 | 1.900E-08 | 1.000E-07 |
| 20909 | 1046 | N1046H | C | 2e-08 | 3.7e-08 | 5.1e-08 | 3.600E-08 | 1.552E-08 | 2.000E-08 | 5.100E-08 |

| | | | | | | | | | | |
|---|---|---|---|---|---|---|---|---|---|---|
| 20910 | 1046 | N1046I | C | 9.7e-08 | 2.9e-08 | 4.4e-08 | 5.667E-08 | 3.573E-08 | 2.900E-08 | 9.700E-08 |
| 20911 | 1046 | N1046L | C | 2.5e-08 | 3.9e-08 | 4.6e-08 | 3.667E-08 | 1.069E-08 | 2.500E-08 | 4.600E-08 |
| 20912 | 1046 | N1046K | C | 4.3e-08 | 1.8e-07 | 5.5e-08 | 9.267E-08 | 7.587E-08 | 4.300E-08 | 1.800E-07 |
| 20913 | 1046 | N1046M | C | 7.7e-08 | 2e-08 | 7.1e-08 | 5.600E-08 | 3.132E-08 | 2.000E-08 | 7.700E-08 |
| 20914 | 1046 | N1046F | C | 3.4e-08 | 3.4e-08 | 4.6e-08 | 3.800E-08 | 6.928E-09 | 3.400E-08 | 4.600E-08 |
| 20915 | 1046 | N1046P | C | 2.1e-08 | 2.4e-08 | 4.8e-08 | 3.100E-08 | 1.480E-08 | 2.100E-08 | 4.800E-08 |
| 20916 | 1046 | N1046S | C | 5.8e-09 | 3.2e-08 | 4.5e-08 | 2.760E-08 | 1.997E-08 | 5.800E-09 | 4.500E-08 |
| 20917 | 1046 | N1046T | C | 3e-08 | 5.6e-08 | 4.7e-08 | 4.433E-08 | 1.320E-08 | 3.000E-08 | 5.600E-08 |
| 20918 | 1046 | N1046W | C | 5.6e-08 | 4.9e-08 | 7.9e-08 | 6.133E-08 | 1.570E-08 | 4.900E-08 | 7.900E-08 |
| 20919 | 1046 | N1046Y | C | 2.9e-08 | 1.1e-07 | 9.9e-08 | 7.933E-08 | 4.394E-08 | 2.900E-08 | 1.100E-07 |
| 20920 | 1046 | N1046V | C | 4.3e-08 | 2.3e-08 | 5.4e-08 | 4.000E-08 | 1.572E-08 | 2.300E-08 | 5.400E-08 |
| 20921 | 1047 | T1047A | C | 2.8e-08 | 5.6e-08 | 1.8e-08 | 3.400E-08 | 1.970E-08 | 1.800E-08 | 5.600E-08 |
| 20922 | 1047 | T1047R | C | 5e-08 | 1e-07 | 4.3e-08 | 6.433E-08 | 3.109E-08 | 4.300E-08 | 1.000E-07 |
| 20923 | 1047 | T1047N | C | 3.5e-08 | 6.3e-08 | 4.2e-08 | 4.667E-08 | 1.457E-08 | 3.500E-08 | 6.300E-08 |
| 20924 | 1047 | T1047D | C | 2.1e-08 | 4e-08 | 2.3e-08 | 2.800E-08 | 1.044E-08 | 2.100E-08 | 4.000E-08 |
| 20925 | 1047 | T1047C | C | 7e-08 | 1e-08 | 1.3e-07 | 7.000E-08 | 6.000E-08 | 1.000E-08 | 1.300E-07 |
| 20926 | 1047 | T1047E | C | 4.9e-08 | 4.4e-08 | 3.9e-08 | 4.400E-08 | 5.000E-09 | 3.900E-08 | 4.900E-08 |
| 20927 | 1047 | T1047Q | C | 7.6e-08 | 4.8e-08 | 2.8e-08 | 5.067E-08 | 2.411E-08 | 2.800E-08 | 7.600E-08 |
| 20928 | 1047 | T1047G | C | 3.9e-08 | 9.3e-08 | 4e-08 | 5.733E-08 | 3.089E-08 | 3.900E-08 | 9.300E-08 |
| 20929 | 1047 | T1047H | C | 6.8e-08 | 4.4e-08 | 2.5e-08 | 4.567E-08 | 2.155E-08 | 2.500E-08 | 6.800E-08 |
| 20930 | 1047 | T1047I | C | 3.1e-08 | 1e-07 | 2.9e-08 | 5.333E-08 | 4.043E-08 | 2.900E-08 | 1.000E-07 |
| 20931 | 1047 | T1047L | C | 5.1e-08 | 4e-08 | 1e-07 | 6.367E-08 | 3.194E-08 | 4.000E-08 | 1.000E-07 |
| 20932 | 1047 | T1047K | C | 2.7e-08 | 2.8e-08 | 5.2e-08 | 3.567E-08 | 1.415E-08 | 2.700E-08 | 5.200E-08 |
| 20933 | 1047 | T1047M | C | 1.1e-07 | 8.2e-08 | 7.9e-08 | 9.033E-08 | 1.710E-08 | 7.900E-08 | 1.100E-07 |
| 20934 | 1047 | T1047F | C | 3.6e-08 | 3.1e-08 | 1.9e-08 | 2.867E-08 | 8.737E-09 | 1.900E-08 | 3.600E-08 |
| 20935 | 1047 | T1047P | C | 3.1e-08 | 2e-08 | 4.6e-08 | 3.233E-08 | 1.305E-08 | 2.000E-08 | 4.600E-08 |
| 20936 | 1047 | T1047S | C | 4.3e-08 | 5.9e-08 | 6.3e-08 | 5.500E-08 | 1.058E-08 | 4.300E-08 | 6.300E-08 |
| 20937 | 1047 | T1047T | C | 3.7e-08 | 4.9e-08 | 1.2e-07 | 6.867E-08 | 4.486E-08 | 3.700E-08 | 1.200E-07 |
| 20938 | 1047 | T1047W | C | 3.6e-08 | 4.2e-08 | 2.2e-08 | 3.333E-08 | 1.026E-08 | 2.200E-08 | 4.200E-08 |
| 20939 | 1047 | T1047Y | C | 1.5e-07 | 2.9e-08 | 7.5e-08 | 8.467E-08 | 6.108E-08 | 2.900E-08 | 1.500E-07 |
| 20940 | 1047 | T1047V | C | 2.1e-08 | 5.7e-08 | 2.3e-08 | 3.367E-08 | 2.023E-08 | 2.100E-08 | 5.700E-08 |
| 20941 | 1048 | K1048A | C | 4.3e-08 | 5.3e-08 | 1.5e-07 | 8.200E-08 | 5.910E-08 | 4.300E-08 | 1.500E-07 |
| 20942 | 1048 | K1048R | C | 1.4e-07 | 1e-07 | 4.8e-08 | 9.600E-08 | 4.613E-08 | 4.800E-08 | 1.400E-07 |
| 20943 | 1048 | K1048N | C | 2.6e-08 | 1.5e-08 | 2.3e-08 | 2.133E-08 | 5.686E-09 | 1.500E-08 | 2.600E-08 |
| 20944 | 1048 | K1048D | C | 4e-08 | 1.8e-08 | 8.6e-08 | 4.800E-08 | 3.470E-08 | 1.800E-08 | 8.600E-08 |
| 20945 | 1048 | K1048C | C | 6.6e-08 | 2.2e-08 | 5.3e-08 | 4.700E-08 | 2.261E-08 | 2.200E-08 | 6.600E-08 |
| 20946 | 1048 | K1048E | C | 3.7e-08 | 2.7e-08 | 6.1e-08 | 4.167E-08 | 1.747E-08 | 2.700E-08 | 6.100E-08 |
| 20947 | 1048 | K1048Q | C | 6.8e-08 | 3.5e-08 | 4.1e-08 | 4.800E-08 | 1.758E-08 | 3.500E-08 | 6.800E-08 |
| 20948 | 1048 | K1048G | C | 4.7e-08 | 8.9e-08 | 3.6e-08 | 5.733E-08 | 2.797E-08 | 3.600E-08 | 8.900E-08 |
| 20949 | 1048 | K1048H | C | 4.6e-08 | 3.7e-08 | 5e-08 | 4.433E-08 | 6.658E-09 | 3.700E-08 | 5.000E-08 |
| 20950 | 1048 | K1048I | C | 7.7e-08 | 1.7e-08 | 5.5e-08 | 4.967E-08 | 3.035E-08 | 1.700E-08 | 7.700E-08 |
| 20951 | 1048 | K1048L | C | 8.1e-09 | 3.5e-08 | 6.6e-08 | 3.637E-08 | 2.897E-08 | 8.100E-09 | 6.600E-08 |
| 20952 | 1048 | K1048K | C | 4e-08 | 3.5e-08 | 2.7e-08 | 3.400E-08 | 6.557E-09 | 2.700E-08 | 4.000E-08 |
| 20953 | 1048 | K1048M | C | 4.7e-08 | 2.2e-08 | 1.9e-08 | 2.933E-08 | 1.537E-08 | 1.900E-08 | 4.700E-08 |
| 20954 | 1048 | K1048F | C | 2.7e-08 | 2.2e-08 | 5.5e-08 | 3.467E-08 | 1.779E-08 | 2.200E-08 | 5.500E-08 |
| 20955 | 1048 | K1048P | C | 5.2e-08 | 6.2e-09 | 4.2e-08 | 3.340E-08 | 2.408E-08 | 6.200E-09 | 5.200E-08 |
| 20956 | 1048 | K1048S | C | 9.5e-08 | 5.5e-08 | 4e-08 | 6.333E-08 | 2.843E-08 | 4.000E-08 | 9.500E-08 |
| 20957 | 1048 | K1048T | C | 6.1e-08 | 5.4e-08 | 5.3e-08 | 5.600E-08 | 4.359E-09 | 5.300E-08 | 6.100E-08 |
| 20958 | 1048 | K1048W | C | 2.6e-08 | 1.5e-08 | 3.4e-08 | 2.500E-08 | 9.539E-09 | 1.500E-08 | 3.400E-08 |
| 20959 | 1048 | K1048Y | C | 5e-08 | 2.9e-08 | 1.1e-08 | 3.000E-08 | 1.952E-08 | 1.100E-08 | 5.000E-08 |
| 20960 | 1048 | K1048V | C | 2e-08 | 3.7e-08 | 3.6e-08 | 3.100E-08 | 9.539E-09 | 2.000E-08 | 3.700E-08 |
| 20961 | 1049 | V1049A | C | 8.6e-08 | 6.9e-08 | 7.7e-08 | 7.733E-08 | 8.505E-09 | 6.900E-08 | 8.600E-08 |
| 20962 | 1049 | V1049R | C | 7.5e-08 | 4e-08 | 3.9e-08 | 5.133E-08 | 2.050E-08 | 3.900E-08 | 7.500E-08 |
| 20963 | 1049 | V1049N | C | 7.1e-08 | 6.6e-08 | 1.9e-08 | 5.200E-08 | 2.869E-08 | 1.900E-08 | 7.100E-08 |
| 20964 | 1049 | V1049D | C | 3.6e-08 | 5.4e-08 | 3.1e-08 | 4.033E-08 | 1.210E-08 | 3.100E-08 | 5.400E-08 |
| 20965 | 1049 | V1049C | C | 8.5e-08 | 1.9e-08 | 2.8e-08 | 4.400E-08 | 3.579E-08 | 1.900E-08 | 8.500E-08 |
| 20966 | 1049 | V1049E | C | 3e-08 | 7e-08 | 4.6e-08 | 4.867E-08 | 2.013E-08 | 3.000E-08 | 7.000E-08 |
| 20967 | 1049 | V1049Q | C | 5e-08 | 1.8e-08 | 5.6e-08 | 4.133E-08 | 2.043E-08 | 1.800E-08 | 5.600E-08 |
| 20968 | 1049 | V1049G | C | 4.1e-08 | 2.3e-08 | 1.9e-08 | 2.767E-08 | 1.172E-08 | 1.900E-08 | 4.100E-08 |
| 20969 | 1049 | V1049H | C | 3.6e-08 | 4.2e-08 | 2.5e-08 | 3.433E-08 | 8.622E-09 | 2.500E-08 | 4.200E-08 |
| 20970 | 1049 | V1049I | C | 3.1e-08 | 6.5e-08 | 3.3e-08 | 4.300E-08 | 1.908E-08 | 3.100E-08 | 6.500E-08 |
| 20971 | 1049 | V1049L | C | 3.6e-08 | 9.1e-08 | 2.6e-08 | 5.100E-08 | 3.500E-08 | 2.600E-08 | 9.100E-08 |
| 20972 | 1049 | V1049K | C | 4.3e-08 | 6.1e-08 | 1.3e-08 | 3.900E-08 | 2.425E-08 | 1.300E-08 | 6.100E-08 |
| 20973 | 1049 | V1049M | C | 4e-08 | 4.9e-08 | 1.1e-07 | 6.633E-08 | 3.808E-08 | 4.000E-08 | 1.100E-07 |
| 20974 | 1049 | V1049F | C | 7e-08 | 1.6e-07 | 1.6e-07 | 1.300E-07 | 5.196E-08 | 7.000E-08 | 1.600E-07 |
| 20975 | 1049 | V1049P | C | 8.6e-08 | 2.4e-08 | 2.4e-08 | 4.467E-08 | 3.580E-08 | 2.400E-08 | 8.600E-08 |
| 20976 | 1049 | V1049S | C | 6.8e-08 | 1.9e-07 | 2e-08 | 9.267E-08 | 8.764E-08 | 2.000E-08 | 1.900E-07 |
| 20977 | 1049 | V1049T | C | 3.4e-08 | 4.7e-08 | 5.1e-08 | 4.400E-08 | 8.888E-09 | 3.400E-08 | 5.100E-08 |
| 20978 | 1049 | V1049W | C | 3e-08 | 3e-08 | 5.2e-08 | 3.733E-08 | 1.270E-08 | 3.000E-08 | 5.200E-08 |
| 20979 | 1049 | V1049Y | C | 2.3e-08 | 2e-08 | 5.2e-08 | 3.167E-08 | 1.767E-08 | 2.000E-08 | 5.200E-08 |
| 20980 | 1049 | V1049V | C | 1.3e-08 | 1.7e-08 | 3.7e-08 | 2.233E-08 | 1.286E-08 | 1.300E-08 | 3.700E-08 |
| 20981 | 1050 | D1050A | C | 4.6e-08 | 5.9e-08 | 4.1e-08 | 4.867E-08 | 9.292E-09 | 4.100E-08 | 5.900E-08 |
| 20982 | 1050 | D1050R | C | 6.4e-08 | 6.4e-08 | 2.2e-08 | 5.000E-08 | 2.425E-08 | 2.200E-08 | 6.400E-08 |
| 20983 | 1050 | D1050N | C | 8.4e-08 | 2.8e-08 | 8.8e-08 | 6.667E-08 | 3.355E-08 | 2.800E-08 | 8.800E-08 |
| 20984 | 1050 | D1050D | C | 3e-08 | 3.2e-08 | 3.7e-08 | 3.300E-08 | 3.606E-09 | 3.000E-08 | 3.700E-08 |
| 20985 | 1050 | D1050C | C | 7.6e-08 | 1.4e-07 | 3.4e-08 | 8.333E-08 | 5.338E-08 | 3.400E-08 | 1.400E-07 |
| 20986 | 1050 | D1050E | C | 5.8e-08 | 5.4e-08 | 2.7e-08 | 4.633E-08 | 1.686E-08 | 2.700E-08 | 5.800E-08 |
| 20987 | 1050 | D1050Q | C | 6.9e-08 | 1.9e-08 | 6.9e-08 | 5.233E-08 | 2.887E-08 | 1.900E-08 | 6.900E-08 |
| 20988 | 1050 | D1050G | C | 9.1e-08 | 1.3e-07 | 2.1e-07 | 1.437E-07 | 6.067E-08 | 9.100E-08 | 2.100E-07 |
| 20989 | 1050 | D1050H | C | 2.7e-08 | 2.1e-08 | 8e-08 | 4.267E-08 | 3.247E-08 | 2.100E-08 | 8.000E-08 |
| 20990 | 1050 | D1050I | C | 4.1e-08 | 3e-08 | 5.6e-08 | 4.233E-08 | 1.305E-08 | 3.000E-08 | 5.600E-08 |
| 20991 | 1050 | D1050L | C | 1.6e-08 | 1.8e-08 | 2.8e-08 | 2.067E-08 | 6.429E-09 | 1.600E-08 | 2.800E-08 |
| 20992 | 1050 | D1050K | C | 1.4e-08 | 3.6e-08 | 1e-07 | 5.000E-08 | 4.468E-08 | 1.400E-08 | 1.000E-07 |
| 20993 | 1050 | D1050M | C | 3.5e-08 | 1e-07 | 6.4e-08 | 6.633E-08 | 3.256E-08 | 3.500E-08 | 1.000E-07 |
| 20994 | 1050 | D1050F | C | 4.3e-08 | 2.3e-08 | 1.6e-08 | 2.733E-08 | 1.401E-08 | 1.600E-08 | 4.300E-08 |

| | | | | | | | | | | |
|---|---|---|---|---|---|---|---|---|---|---|
| 20995 | 1050 | D1050P | C | 5.5e-08 | 4e-08 | 4.5e-08 | 4.667E-08 | 7.638E-09 | 4.000E-08 | 5.500E-08 |
| 20996 | 1050 | D1050S | C | 2.8e-08 | 5.4e-08 | 9e-08 | 5.733E-08 | 3.113E-08 | 2.800E-08 | 9.000E-08 |
| 20997 | 1050 | D1050T | C | 6.7e-08 | 1e-07 | 3.6e-08 | 6.767E-08 | 3.201E-08 | 3.600E-08 | 1.000E-07 |
| 20998 | 1050 | D1050W | C | 4e-08 | 4.9e-08 | 1.6e-08 | 3.500E-08 | 1.706E-08 | 1.600E-08 | 4.900E-08 |
| 20999 | 1050 | D1050Y | C | 5.3e-08 | 5e-08 | 2.7e-08 | 4.333E-08 | 1.422E-08 | 2.700E-08 | 5.300E-08 |
| 21000 | 1050 | D1050V | C | 1.4e-07 | 3.8e-08 | 2.8e-08 | 6.867E-08 | 6.198E-08 | 2.800E-08 | 1.400E-07 |
| 21001 | 1051 | K1051A | C | 9.4e-08 | 8.9e-08 | 7.3e-08 | 8.533E-08 | 1.097E-08 | 7.300E-08 | 9.400E-08 |
| 21002 | 1051 | K1051R | C | 5.4e-08 | 1.7e-08 | 3.8e-08 | 3.633E-08 | 1.856E-08 | 1.700E-08 | 5.400E-08 |
| 21003 | 1051 | K1051N | C | 2.3e-08 | 2.2e-08 | 5.1e-08 | 3.200E-08 | 1.646E-08 | 2.200E-08 | 5.100E-08 |
| 21004 | 1051 | K1051D | C | 1.7e-08 | 4.7e-08 | 1.5e-08 | 2.633E-08 | 1.793E-08 | 1.500E-08 | 4.700E-08 |
| 21005 | 1051 | K1051C | C | 1.4e-07 | 2.5e-08 | 7.3e-08 | 7.933E-08 | 5.776E-08 | 2.500E-08 | 1.400E-07 |
| 21006 | 1051 | K1051E | C | 4.8e-08 | 9.9e-09 | 8.3e-08 | 4.697E-08 | 3.656E-08 | 9.900E-09 | 8.300E-08 |
| 21007 | 1051 | K1051Q | C | 3.4e-08 | 5.4e-08 | 7e-08 | 5.267E-08 | 1.804E-08 | 3.400E-08 | 7.000E-08 |
| 21008 | 1051 | K1051G | C | 1.7e-08 | 4.6e-08 | 2.8e-08 | 3.033E-08 | 1.464E-08 | 1.700E-08 | 4.600E-08 |
| 21009 | 1051 | K1051H | C | 6.4e-08 | 8.3e-08 | 1.5e-08 | 5.400E-08 | 3.509E-08 | 1.500E-08 | 8.300E-08 |
| 21010 | 1051 | K1051I | C | 2.6e-08 | 9e-08 | 9.3e-08 | 6.967E-08 | 3.785E-08 | 2.600E-08 | 9.300E-08 |
| 21011 | 1051 | K1051L | C | 1.8e-08 | 3e-08 | 6.5e-08 | 3.767E-08 | 2.442E-08 | 1.800E-08 | 6.500E-08 |
| 21012 | 1051 | K1051K | C | 2.3e-08 | 3.8e-08 | 2e-08 | 2.700E-08 | 9.644E-09 | 2.000E-08 | 3.800E-08 |
| 21013 | 1051 | K1051M | C | 6.4e-08 | 4.8e-08 | 1.7e-08 | 4.300E-08 | 2.390E-08 | 1.700E-08 | 6.400E-08 |
| 21014 | 1051 | K1051F | C | 4.9e-08 | 7e-08 | 4e-08 | 5.300E-08 | 1.539E-08 | 4.000E-08 | 7.000E-08 |
| 21015 | 1051 | K1051P | C | 5e-08 | 3.1e-08 | 1e-07 | 6.033E-08 | 3.564E-08 | 3.100E-08 | 1.000E-07 |
| 21016 | 1051 | K1051S | C | 8.1e-08 | 2.1e-08 | 2.6e-08 | 4.267E-08 | 3.329E-08 | 2.100E-08 | 8.100E-08 |
| 21017 | 1051 | K1051T | C | 6.6e-08 | 2.5e-08 | 6.3e-08 | 5.133E-08 | 2.285E-08 | 2.500E-08 | 6.600E-08 |
| 21018 | 1051 | K1051W | C | 9.1e-08 | 4e-08 | 3.6e-08 | 5.567E-08 | 3.066E-08 | 3.600E-08 | 9.100E-08 |
| 21019 | 1051 | K1051Y | C | 4e-08 | 2.3e-08 | 4.5e-08 | 3.600E-08 | 1.153E-08 | 2.300E-08 | 4.500E-08 |
| 21020 | 1051 | K1051V | C | 2.7e-08 | 6.9e-08 | 2e-08 | 3.867E-08 | 2.650E-08 | 2.000E-08 | 6.900E-08 |
| 21021 | 1052 | K1052A | C | 6.1e-08 | 2.2e-08 | 4.6e-08 | 4.300E-08 | 1.967E-08 | 2.200E-08 | 6.100E-08 |
| 21022 | 1052 | K1052R | C | 4.8e-08 | 4.9e-08 | 5.9e-08 | 5.200E-08 | 6.083E-09 | 4.800E-08 | 5.900E-08 |
| 21023 | 1052 | K1052N | C | 2.6e-08 | 5.8e-08 | 5.9e-08 | 4.767E-08 | 1.877E-08 | 2.600E-08 | 5.900E-08 |
| 21024 | 1052 | K1052D | C | 1.2e-08 | 2.4e-08 | 7.1e-08 | 3.567E-08 | 3.118E-08 | 1.200E-08 | 7.100E-08 |
| 21025 | 1052 | K1052C | C | 3.3e-08 | 6.3e-08 | 2.1e-08 | 3.900E-08 | 2.163E-08 | 2.100E-08 | 6.300E-08 |
| 21026 | 1052 | K1052E | C | 2.1e-08 | 9.1e-08 | 2.9e-08 | 4.700E-08 | 3.831E-08 | 2.100E-08 | 9.100E-08 |
| 21027 | 1052 | K1052Q | C | 8.4e-08 | 7.2e-09 | 4.9e-08 | 4.673E-08 | 3.845E-08 | 7.200E-09 | 8.400E-08 |
| 21028 | 1052 | K1052G | C | 1.2e-08 | 2e-08 | 1.7e-07 | 6.733E-08 | 8.900E-08 | 1.200E-08 | 1.700E-07 |
| 21029 | 1052 | K1052H | C | 5.2e-08 | 5.4e-08 | 2.9e-08 | 4.500E-08 | 1.389E-08 | 2.900E-08 | 5.400E-08 |
| 21030 | 1052 | K1052I | C | 5.3e-08 | 3.4e-08 | 5.2e-08 | 4.633E-08 | 1.069E-08 | 3.400E-08 | 5.300E-08 |
| 21031 | 1052 | K1052L | C | 9.1e-08 | 3.1e-08 | 5.3e-08 | 5.833E-08 | 3.035E-08 | 3.100E-08 | 9.100E-08 |
| 21032 | 1052 | K1052K | C | 3.2e-08 | 1.7e-08 | 9e-08 | 4.633E-08 | 3.855E-08 | 1.700E-08 | 9.000E-08 |
| 21033 | 1052 | K1052M | C | 2e-08 | 2.3e-08 | 2.7e-08 | 2.333E-08 | 3.512E-09 | 2.000E-08 | 2.700E-08 |
| 21034 | 1052 | K1052F | C | 6e-08 | 4.6e-08 | 8.9e-08 | 6.500E-08 | 2.193E-08 | 4.600E-08 | 8.900E-08 |
| 21035 | 1052 | K1052P | C | 8e-08 | 2.8e-08 | 4.1e-08 | 4.967E-08 | 2.706E-08 | 2.800E-08 | 8.000E-08 |
| 21036 | 1052 | K1052S | C | 4e-08 | 4.3e-08 | 3.7e-08 | 4.000E-08 | 3.000E-09 | 3.700E-08 | 4.300E-08 |
| 21037 | 1052 | K1052T | C | 5.2e-08 | 8.2e-08 | 3.7e-08 | 5.700E-08 | 2.291E-08 | 3.700E-08 | 8.200E-08 |
| 21038 | 1052 | K1052W | C | 2.9e-08 | 7.8e-08 | 7e-08 | 5.900E-08 | 2.629E-08 | 2.900E-08 | 7.800E-08 |
| 21039 | 1052 | K1052Y | C | 3.1e-08 | 3.2e-08 | 4.6e-08 | 3.633E-08 | 8.386E-09 | 3.100E-08 | 4.600E-08 |
| 21040 | 1052 | K1052V | C | 1.5e-08 | 2.8e-08 | 2.1e-08 | 2.133E-08 | 6.506E-09 | 1.500E-08 | 2.800E-08 |
| 21041 | 1053 | V1053A | C | 7.9e-08 | 3.8e-08 | 3.2e-08 | 4.967E-08 | 2.558E-08 | 3.200E-08 | 7.900E-08 |
| 21042 | 1053 | V1053R | C | 3.5e-08 | 7.1e-08 | 2.8e-08 | 4.467E-08 | 2.307E-08 | 2.800E-08 | 7.100E-08 |
| 21043 | 1053 | V1053N | C | 3.2e-08 | 9.4e-08 | 5.3e-08 | 5.967E-08 | 3.153E-08 | 3.200E-08 | 9.400E-08 |
| 21044 | 1053 | V1053D | C | 7.3e-08 | 1.7e-08 | 2.3e-08 | 3.767E-08 | 3.075E-08 | 1.700E-08 | 7.300E-08 |
| 21045 | 1053 | V1053C | C | 2.3e-08 | 3.1e-08 | 7.1e-08 | 4.167E-08 | 2.572E-08 | 2.300E-08 | 7.100E-08 |
| 21046 | 1053 | V1053E | C | 3.1e-08 | 4.6e-08 | 2.8e-08 | 3.500E-08 | 9.644E-09 | 2.800E-08 | 4.600E-08 |
| 21047 | 1053 | V1053Q | C | 4e-08 | 3.1e-08 | 5.2e-08 | 4.100E-08 | 1.054E-08 | 3.100E-08 | 5.200E-08 |
| 21048 | 1053 | V1053G | C | 1.4e-08 | 8.9e-08 | 5.1e-08 | 5.133E-08 | 3.750E-08 | 1.400E-08 | 8.900E-08 |
| 21049 | 1053 | V1053H | C | 4.2e-08 | 2.7e-08 | 7.2e-08 | 4.700E-08 | 2.291E-08 | 2.700E-08 | 7.200E-08 |
| 21050 | 1053 | V1053I | C | 5.5e-08 | 5e-08 | 5.1e-08 | 5.200E-08 | 2.646E-09 | 5.000E-08 | 5.500E-08 |
| 21051 | 1053 | V1053L | C | 1.2e-08 | 1.9e-08 | 1.3e-08 | 1.467E-08 | 3.786E-09 | 1.200E-08 | 1.900E-08 |
| 21052 | 1053 | V1053K | C | 3.9e-08 | 1.6e-08 | 9.3e-08 | 4.933E-08 | 3.953E-08 | 1.600E-08 | 9.300E-08 |
| 21053 | 1053 | V1053M | C | 2.6e-08 | 6e-08 | 7.6e-08 | 5.400E-08 | 2.553E-08 | 2.600E-08 | 7.600E-08 |
| 21054 | 1053 | V1053F | C | 3.2e-08 | 1.3e-07 | 3.8e-08 | 6.667E-08 | 5.493E-08 | 3.200E-08 | 1.300E-07 |
| 21055 | 1053 | V1053P | C | 4e-08 | 2.6e-08 | 3.2e-08 | 3.267E-08 | 7.024E-09 | 2.600E-08 | 4.000E-08 |
| 21056 | 1053 | V1053S | C | 3e-08 | 1.1e-08 | 1.6e-08 | 1.900E-08 | 9.849E-09 | 1.100E-08 | 3.000E-08 |
| 21057 | 1053 | V1053T | C | 3.4e-08 | 6e-08 | 2.4e-08 | 3.933E-08 | 1.858E-08 | 2.400E-08 | 6.000E-08 |
| 21058 | 1053 | V1053W | C | 6.3e-08 | 4.2e-08 | 4.3e-08 | 4.933E-08 | 1.185E-08 | 4.200E-08 | 6.300E-08 |
| 21059 | 1053 | V1053Y | C | 2.6e-08 | 4.8e-08 | 2e-08 | 3.133E-08 | 1.474E-08 | 2.000E-08 | 4.800E-08 |
| 21060 | 1053 | V1053V | C | 7e-08 | 4.6e-08 | 7.7e-08 | 6.433E-08 | 1.626E-08 | 4.600E-08 | 7.700E-08 |
| 21061 | 1054 | E1054A | C | 7.3e-08 | 8.6e-08 | 6.6e-08 | 7.500E-08 | 1.015E-08 | 6.600E-08 | 8.600E-08 |
| 21062 | 1054 | E1054R | C | 2.2e-08 | 3.6e-08 | 3.2e-08 | 3.000E-08 | 7.211E-09 | 2.200E-08 | 3.600E-08 |
| 21063 | 1054 | E1054N | C | 3.9e-08 | 4.5e-08 | 1.3e-07 | 7.133E-08 | 5.090E-08 | 3.900E-08 | 1.300E-07 |
| 21064 | 1054 | E1054D | C | 6.7e-08 | 3.2e-08 | 1.6e-08 | 3.833E-08 | 2.608E-08 | 1.600E-08 | 6.700E-08 |
| 21065 | 1054 | E1054C | C | 1.2e-07 | 8.5e-08 | 3.9e-08 | 8.133E-08 | 4.062E-08 | 3.900E-08 | 1.200E-07 |
| 21066 | 1054 | E1054E | C | 2.6e-08 | 4e-08 | 4e-08 | 3.533E-08 | 8.083E-09 | 2.600E-08 | 4.000E-08 |
| 21067 | 1054 | E1054Q | C | 4e-08 | 2e-08 | 5e-08 | 3.667E-08 | 1.528E-08 | 2.000E-08 | 5.000E-08 |
| 21068 | 1054 | E1054G | C | 4.3e-08 | 5.1e-08 | 8.5e-08 | 5.967E-08 | 2.230E-08 | 4.300E-08 | 8.500E-08 |
| 21069 | 1054 | E1054H | C | 1.8e-08 | 4.9e-08 | 6.2e-08 | 4.300E-08 | 2.261E-08 | 1.800E-08 | 6.200E-08 |
| 21070 | 1054 | E1054I | C | 5.3e-08 | 5e-08 | 4.1e-08 | 4.800E-08 | 6.245E-09 | 4.100E-08 | 5.300E-08 |
| 21071 | 1054 | E1054L | C | 6.1e-08 | 3.1e-08 | 3.5e-08 | 4.233E-08 | 1.629E-08 | 3.100E-08 | 6.100E-08 |
| 21072 | 1054 | E1054K | C | 5.4e-08 | 5.6e-08 | 5.8e-08 | 5.600E-08 | 2.000E-09 | 5.400E-08 | 5.800E-08 |
| 21073 | 1054 | E1054M | C | 8.5e-09 | 7.9e-08 | 1.5e-08 | 3.417E-08 | 3.896E-08 | 8.500E-09 | 7.900E-08 |
| 21074 | 1054 | E1054F | C | 1.3e-08 | 3.8e-08 | 4.3e-08 | 3.133E-08 | 1.607E-08 | 1.300E-08 | 4.300E-08 |
| 21075 | 1054 | E1054P | C | 1.2e-08 | 5.1e-08 | 2.1e-08 | 2.800E-08 | 2.042E-08 | 1.200E-08 | 5.100E-08 |
| 21076 | 1054 | E1054S | C | 1.6e-08 | 2.7e-08 | 1.4e-07 | 6.100E-08 | 6.864E-08 | 1.600E-08 | 1.400E-07 |
| 21077 | 1054 | E1054T | C | 1.2e-07 | 6.3e-08 | 3.3e-08 | 7.200E-08 | 4.419E-08 | 3.300E-08 | 1.200E-07 |
| 21078 | 1054 | E1054W | C | 6.6e-08 | 4.8e-08 | 4e-08 | 5.133E-08 | 1.332E-08 | 4.000E-08 | 6.600E-08 |
| 21079 | 1054 | E1054Y | C | 9.1e-08 | 1e-08 | 5.3e-08 | 5.133E-08 | 4.053E-08 | 1.000E-08 | 9.100E-08 |

| | | | | | | | | | | |
|---|---|---|---|---|---|---|---|---|---|---|
| 21080 | 1054 | E1054V | C | 9e-08 | 7.5e-08 | 6e-08 | 7.500E-08 | 1.500E-08 | 6.000E-08 | 9.000E-08 |
| 21081 | 1055 | P1055A | C | 3.4e-08 | 4.6e-08 | 5.5e-08 | 4.500E-08 | 1.054E-08 | 3.400E-08 | 5.500E-08 |
| 21082 | 1055 | P1055R | C | 2.4e-08 | 3.3e-08 | 4.4e-08 | 3.367E-08 | 1.002E-08 | 2.400E-08 | 4.400E-08 |
| 21083 | 1055 | P1055N | C | 6.1e-08 | 2.6e-08 | 2.9e-08 | 3.867E-08 | 1.940E-08 | 2.600E-08 | 6.100E-08 |
| 21084 | 1055 | P1055D | C | 2.2e-08 | 5.2e-08 | 4.9e-08 | 4.100E-08 | 1.652E-08 | 2.200E-08 | 5.200E-08 |
| 21085 | 1055 | P1055C | C | 6.9e-08 | 7.5e-08 | 5.4e-08 | 6.600E-08 | 1.082E-08 | 5.400E-08 | 7.500E-08 |
| 21086 | 1055 | P1055E | C | 2.9e-08 | 1.6e-08 | 7.4e-08 | 3.967E-08 | 3.044E-08 | 1.600E-08 | 7.400E-08 |
| 21087 | 1055 | P1055Q | C | 2.1e-08 | 7.5e-08 | 8.2e-09 | 3.473E-08 | 3.545E-08 | 8.200E-09 | 7.500E-08 |
| 21088 | 1055 | P1055G | C | 5.9e-08 | 2.2e-08 | 5.2e-08 | 4.433E-08 | 1.966E-08 | 2.200E-08 | 5.900E-08 |
| 21089 | 1055 | P1055H | C | 9.3e-08 | 4e-08 | 4.1e-08 | 5.800E-08 | 3.032E-08 | 4.000E-08 | 9.300E-08 |
| 21090 | 1055 | P1055I | C | 3.4e-08 | 1e-07 | 4.5e-08 | 5.967E-08 | 3.536E-08 | 3.400E-08 | 1.000E-07 |
| 21091 | 1055 | P1055L | C | 1.8e-08 | 3.5e-08 | 2.1e-08 | 2.467E-08 | 9.074E-09 | 1.800E-08 | 3.500E-08 |
| 21092 | 1055 | P1055K | C | 4.8e-08 | 1.6e-08 | 9.1e-08 | 5.167E-08 | 3.763E-08 | 1.600E-08 | 9.100E-08 |
| 21093 | 1055 | P1055M | C | 2.9e-08 | 6e-08 | 7.2e-08 | 5.367E-08 | 2.219E-08 | 2.900E-08 | 7.200E-08 |
| 21094 | 1055 | P1055F | C | 3.8e-08 | 1.1e-07 | 1.1e-07 | 8.600E-08 | 4.157E-08 | 3.800E-08 | 1.100E-07 |
| 21095 | 1055 | P1055P | C | 3.8e-08 | 4.8e-08 | 6.5e-08 | 5.033E-08 | 1.365E-08 | 3.800E-08 | 6.500E-08 |
| 21096 | 1055 | P1055S | C | 6.2e-08 | 4.9e-08 | 3.4e-08 | 4.833E-08 | 1.401E-08 | 3.400E-08 | 6.200E-08 |
| 21097 | 1055 | P1055T | C | 1e-07 | 5.2e-08 | 4.8e-08 | 6.667E-08 | 2.894E-08 | 4.800E-08 | 1.000E-07 |
| 21098 | 1055 | P1055W | C | 2.6e-08 | 3e-08 | 3.4e-08 | 3.000E-08 | 4.000E-09 | 2.600E-08 | 3.400E-08 |
| 21099 | 1055 | P1055Y | C | 2.6e-08 | 4.9e-08 | 5.9e-08 | 4.467E-08 | 1.692E-08 | 2.600E-08 | 5.900E-08 |
| 21100 | 1055 | P1055V | C | 1.6e-08 | 6.4e-08 | 8.2e-08 | 5.400E-08 | 3.412E-08 | 1.600E-08 | 8.200E-08 |
| 21101 | 1056 | K1056A | C | 8e-08 | 1.8e-08 | 7.6e-08 | 5.800E-08 | 3.470E-08 | 1.800E-08 | 8.000E-08 |
| 21102 | 1056 | K1056R | C | 3.8e-08 | 3.9e-08 | 3e-08 | 3.567E-08 | 4.933E-09 | 3.000E-08 | 3.900E-08 |
| 21103 | 1056 | K1056N | C | 3.8e-08 | 3.5e-08 | 3.2e-08 | 3.500E-08 | 3.000E-09 | 3.200E-08 | 3.800E-08 |
| 21104 | 1056 | K1056D | C | 4.6e-08 | 9.4e-08 | 4.6e-08 | 6.200E-08 | 2.771E-08 | 4.600E-08 | 9.400E-08 |
| 21105 | 1056 | K1056C | C | 3e-08 | 7.3e-08 | 2.6e-08 | 4.300E-08 | 2.606E-08 | 2.600E-08 | 7.300E-08 |
| 21106 | 1056 | K1056E | C | 2.9e-08 | 1.3e-07 | 5.8e-08 | 7.233E-08 | 5.200E-08 | 2.900E-08 | 1.300E-07 |
| 21107 | 1056 | K1056Q | C | 7.5e-08 | 2.6e-08 | 2.7e-08 | 4.267E-08 | 2.801E-08 | 2.600E-08 | 7.500E-08 |
| 21108 | 1056 | K1056G | C | 3e-08 | 4.8e-08 | 7e-08 | 4.933E-08 | 2.003E-08 | 3.000E-08 | 7.000E-08 |
| 21109 | 1056 | K1056H | C | 2.5e-08 | 1.1e-07 | 1e-07 | 7.833E-08 | 4.646E-08 | 2.500E-08 | 1.100E-07 |
| 21110 | 1056 | K1056I | C | 2.1e-08 | 6.1e-08 | 5.7e-08 | 4.633E-08 | 2.203E-08 | 2.100E-08 | 6.100E-08 |
| 21111 | 1056 | K1056L | C | 3.1e-08 | 4.8e-08 | 1.6e-08 | 3.167E-08 | 1.601E-08 | 1.600E-08 | 4.800E-08 |
| 21112 | 1056 | K1056K | C | 4.4e-08 | 6.4e-08 | 5e-08 | 5.267E-08 | 1.026E-08 | 4.400E-08 | 6.400E-08 |
| 21113 | 1056 | K1056M | C | 3.3e-08 | 3.9e-08 | 6.4e-08 | 4.533E-08 | 1.644E-08 | 3.300E-08 | 6.400E-08 |
| 21114 | 1056 | K1056F | C | 8.2e-08 | 7.2e-08 | 2.4e-08 | 5.933E-08 | 3.101E-08 | 2.400E-08 | 8.200E-08 |
| 21115 | 1056 | K1056P | C | 5.5e-08 | 2.5e-08 | 3.8e-08 | 3.933E-08 | 1.504E-08 | 2.500E-08 | 5.500E-08 |
| 21116 | 1056 | K1056S | C | 4.7e-08 | 5.3e-08 | 7.9e-08 | 5.967E-08 | 1.701E-08 | 4.700E-08 | 7.900E-08 |
| 21117 | 1056 | K1056T | C | 3.8e-08 | 4.6e-08 | 2.6e-08 | 3.667E-08 | 1.007E-08 | 2.600E-08 | 4.600E-08 |
| 21118 | 1056 | K1056W | C | 4.4e-08 | 6.7e-08 | 5e-08 | 5.367E-08 | 1.193E-08 | 4.400E-08 | 6.700E-08 |
| 21119 | 1056 | K1056Y | C | 4.1e-08 | 3.1e-08 | 7.2e-08 | 4.800E-08 | 2.138E-08 | 3.100E-08 | 7.200E-08 |
| 21120 | 1056 | K1056V | C | 5.1e-08 | 1.1e-08 | 4.3e-08 | 3.500E-08 | 2.117E-08 | 1.100E-08 | 5.100E-08 |
| 21121 | 1057 | S1057A | C | 8.9e-09 | 6.2e-08 | 7.8e-08 | 4.963E-08 | 3.617E-08 | 8.900E-09 | 7.800E-08 |
| 21122 | 1057 | S1057R | C | 2e-07 | 4e-08 | 4.1e-08 | 9.367E-08 | 9.209E-08 | 4.000E-08 | 2.000E-07 |
| 21123 | 1057 | S1057N | C | 3.7e-08 | 1.6e-07 | 5.2e-08 | 8.300E-08 | 6.710E-08 | 3.700E-08 | 1.600E-07 |
| 21124 | 1057 | S1057D | C | 4.9e-08 | 9.4e-08 | 1.3e-08 | 5.200E-08 | 4.058E-08 | 1.300E-08 | 9.400E-08 |
| 21125 | 1057 | S1057C | C | 1.5e-07 | 1.7e-08 | 4.9e-08 | 7.200E-08 | 6.942E-08 | 1.700E-08 | 1.500E-07 |
| 21126 | 1057 | S1057E | C | 2.7e-08 | 2.3e-08 | 3.1e-08 | 2.700E-08 | 4.000E-09 | 2.300E-08 | 3.100E-08 |
| 21127 | 1057 | S1057Q | C | 5.5e-08 | 9.9e-09 | 3.6e-08 | 3.363E-08 | 2.264E-08 | 9.900E-09 | 5.500E-08 |
| 21128 | 1057 | S1057G | C | 4.1e-08 | 7.2e-08 | 5.1e-08 | 5.467E-08 | 1.582E-08 | 4.100E-08 | 7.200E-08 |
| 21129 | 1057 | S1057H | C | 3.7e-08 | 4.7e-08 | 3.6e-08 | 4.000E-08 | 6.083E-09 | 3.600E-08 | 4.700E-08 |
| 21130 | 1057 | S1057I | C | 2e-08 | 1e-07 | 1.6e-08 | 4.533E-08 | 4.738E-08 | 1.600E-08 | 1.000E-07 |
| 21131 | 1057 | S1057L | C | 6.9e-08 | 3.4e-08 | 3.6e-08 | 4.633E-08 | 1.966E-08 | 3.400E-08 | 6.900E-08 |
| 21132 | 1057 | S1057K | C | 3.3e-08 | 9.3e-08 | 2.4e-08 | 5.000E-08 | 3.751E-08 | 2.400E-08 | 9.300E-08 |
| 21133 | 1057 | S1057M | C | 2.4e-07 | 9.6e-08 | 3.5e-08 | 1.237E-07 | 1.053E-07 | 3.500E-08 | 2.400E-07 |
| 21134 | 1057 | S1057F | C | 3e-08 | 6.3e-08 | 2.2e-08 | 3.833E-08 | 2.173E-08 | 2.200E-08 | 6.300E-08 |
| 21135 | 1057 | S1057P | C | 2.5e-08 | 2.3e-08 | 2.3e-08 | 2.367E-08 | 1.155E-09 | 2.300E-08 | 2.500E-08 |
| 21136 | 1057 | S1057S | C | 7.2e-08 | 5.3e-08 | 4.7e-08 | 5.733E-08 | 1.305E-08 | 4.700E-08 | 7.200E-08 |
| 21137 | 1057 | S1057T | C | 5.6e-08 | 4.1e-08 | 4.7e-08 | 4.800E-08 | 7.550E-09 | 4.100E-08 | 5.600E-08 |
| 21138 | 1057 | S1057W | C | 3e-08 | 1.1e-07 | 6.8e-08 | 6.933E-08 | 4.002E-08 | 3.000E-08 | 1.100E-07 |
| 21139 | 1057 | S1057Y | C | 7.3e-08 | 2e-07 | 8.7e-08 | 1.200E-07 | 6.963E-08 | 7.300E-08 | 2.000E-07 |
| 21140 | 1057 | S1057V | C | 5.4e-08 | 2.9e-08 | 3.6e-08 | 3.967E-08 | 1.290E-08 | 2.900E-08 | 5.400E-08 |
| 21141 | 1058 | C1058A | C | 2.6e-08 | 3.1e-08 | 6.7e-08 | 4.133E-08 | 2.237E-08 | 2.600E-08 | 6.700E-08 |
| 21142 | 1058 | C1058R | C | 7.7e-08 | 1.1e-08 | 9.4e-08 | 6.067E-08 | 4.384E-08 | 1.100E-08 | 9.400E-08 |
| 21143 | 1058 | C1058N | C | 2.1e-08 | 1.4e-08 | 6.6e-08 | 3.367E-08 | 2.822E-08 | 1.400E-08 | 6.600E-08 |
| 21144 | 1058 | C1058D | C | 9.6e-08 | 6.2e-08 | 8e-09 | 5.533E-08 | 4.438E-08 | 8.000E-09 | 9.600E-08 |
| 21145 | 1058 | C1058C | C | 4.3e-08 | 3.4e-08 | 4e-08 | 3.900E-08 | 4.583E-09 | 3.400E-08 | 4.300E-08 |
| 21146 | 1058 | C1058E | C | 8.7e-08 | 3.1e-08 | 8.6e-08 | 6.800E-08 | 3.205E-08 | 3.100E-08 | 8.700E-08 |
| 21147 | 1058 | C1058Q | C | 1.9e-08 | 3.5e-08 | 5.3e-08 | 3.567E-08 | 1.701E-08 | 1.900E-08 | 5.300E-08 |
| 21148 | 1058 | C1058G | C | 2.6e-08 | 2.5e-08 | 3.6e-08 | 2.900E-08 | 6.083E-09 | 2.500E-08 | 3.600E-08 |
| 21149 | 1058 | C1058H | C | 8.5e-08 | 4.5e-08 | 1.5e-08 | 4.833E-08 | 3.512E-08 | 1.500E-08 | 8.500E-08 |
| 21150 | 1058 | C1058I | C | 6.7e-08 | 5.9e-08 | 6e-08 | 6.200E-08 | 4.359E-09 | 5.900E-08 | 6.700E-08 |
| 21151 | 1058 | C1058L | C | 3e-08 | 4e-08 | 2.5e-08 | 3.167E-08 | 7.638E-09 | 2.500E-08 | 4.000E-08 |
| 21152 | 1058 | C1058K | C | 3.9e-08 | 2.3e-08 | 4.4e-08 | 3.533E-08 | 1.097E-08 | 2.300E-08 | 4.400E-08 |
| 21153 | 1058 | C1058M | C | 1.5e-07 | 5.9e-08 | 6.9e-08 | 9.267E-08 | 4.990E-08 | 5.900E-08 | 1.500E-07 |
| 21154 | 1058 | C1058F | C | 1.9e-08 | 4.3e-08 | 3.3e-08 | 3.167E-08 | 1.206E-08 | 1.900E-08 | 4.300E-08 |
| 21155 | 1058 | C1058P | C | 7e-08 | 3.7e-08 | 5.1e-08 | 5.267E-08 | 1.656E-08 | 3.700E-08 | 7.000E-08 |
| 21156 | 1058 | C1058S | C | 8.3e-08 | 2.6e-08 | 1.9e-08 | 4.267E-08 | 3.510E-08 | 1.900E-08 | 8.300E-08 |
| 21157 | 1058 | C1058T | C | 6.6e-09 | 3.4e-08 | 4.4e-08 | 2.820E-08 | 1.936E-08 | 6.600E-09 | 4.400E-08 |
| 21158 | 1058 | C1058W | C | 3.2e-08 | 2.3e-08 | 4.2e-08 | 3.233E-08 | 9.504E-09 | 2.300E-08 | 4.200E-08 |
| 21159 | 1058 | C1058Y | C | 3e-08 | 5.7e-08 | 1e-07 | 6.233E-08 | 3.530E-08 | 3.000E-08 | 1.000E-07 |
| 21160 | 1058 | C1058V | C | 3.6e-08 | 2.5e-08 | 4.8e-08 | 3.633E-08 | 1.150E-08 | 2.500E-08 | 4.800E-08 |
| 21161 | 1059 | D1059A | D | 6.9e-08 | 5e-08 | 2.4e-08 | 4.767E-08 | 2.259E-08 | 2.400E-08 | 6.900E-08 |
| 21162 | 1059 | D1059R | D | 5.4e-08 | 2.2e-08 | 5e-08 | 4.200E-08 | 1.744E-08 | 2.200E-08 | 5.400E-08 |
| 21163 | 1059 | D1059N | D | 5.7e-08 | 7.4e-08 | 3.6e-08 | 5.567E-08 | 1.904E-08 | 3.600E-08 | 7.400E-08 |
| 21164 | 1059 | D1059D | D | 5.6e-08 | 2.6e-08 | 3.3e-08 | 3.833E-08 | 1.570E-08 | 2.600E-08 | 5.600E-08 |

| | | | | | | | | | | |
|---|---|---|---|---|---|---|---|---|---|---|
| 21165 | 1059 | D1059C | D | 5.5e-08 | 1.3e-08 | 2.9e-08 | 3.233E-08 | 2.120E-08 | 1.300E-08 | 5.500E-08 |
| 21166 | 1059 | D1059E | D | 3.3e-08 | 1.1e-08 | 3e-08 | 2.467E-08 | 1.193E-08 | 1.100E-08 | 3.300E-08 |
| 21167 | 1059 | D1059Q | D | 9e-08 | 8.1e-08 | 2.8e-08 | 6.633E-08 | 3.350E-08 | 2.800E-08 | 9.000E-08 |
| 21168 | 1059 | D1059G | D | 1.3e-08 | 5.1e-08 | 2.9e-08 | 3.100E-08 | 1.908E-08 | 1.300E-08 | 5.100E-08 |
| 21169 | 1059 | D1059H | D | 1.2e-07 | 4.4e-08 | 8.1e-08 | 8.167E-08 | 3.800E-08 | 4.400E-08 | 1.200E-07 |
| 21170 | 1059 | D1059I | D | 6.8e-08 | 6.1e-08 | 8.1e-08 | 7.000E-08 | 1.015E-08 | 6.100E-08 | 8.100E-08 |
| 21171 | 1059 | D1059L | D | 3.7e-08 | 2.2e-08 | 5.8e-08 | 3.900E-08 | 1.808E-08 | 2.200E-08 | 5.800E-08 |
| 21172 | 1059 | D1059K | D | 2.7e-08 | 5.4e-08 | 5.2e-08 | 4.433E-08 | 1.504E-08 | 2.700E-08 | 5.400E-08 |
| 21173 | 1059 | D1059M | D | 7.3e-09 | 1.2e-07 | 9.4e-08 | 7.377E-08 | 5.901E-08 | 7.300E-09 | 1.200E-07 |
| 21174 | 1059 | D1059F | D | 5.1e-08 | 1.1e-07 | 1e-07 | 8.700E-08 | 3.158E-08 | 5.100E-08 | 1.100E-07 |
| 21175 | 1059 | D1059P | D | 4.5e-08 | 4.1e-08 | 3.4e-08 | 4.000E-08 | 5.568E-09 | 3.400E-08 | 4.500E-08 |
| 21176 | 1059 | D1059S | D | 4.8e-08 | 2.7e-08 | 5.9e-08 | 4.467E-08 | 1.626E-08 | 2.700E-08 | 5.900E-08 |
| 21177 | 1059 | D1059T | D | 5.5e-08 | 4.7e-08 | 1e-07 | 6.733E-08 | 2.857E-08 | 4.700E-08 | 1.000E-07 |
| 21178 | 1059 | D1059W | D | 1e-07 | 3.9e-08 | 8e-08 | 7.300E-08 | 3.110E-08 | 3.900E-08 | 1.000E-07 |
| 21179 | 1059 | D1059Y | D | 6e-08 | 1.2e-07 | 5e-08 | 7.667E-08 | 3.786E-08 | 5.000E-08 | 1.200E-07 |
| 21180 | 1059 | D1059V | D | 1.6e-08 | 2e-08 | 5.2e-08 | 2.933E-08 | 1.973E-08 | 1.600E-08 | 5.200E-08 |
| 21181 | 1060 | I1060A | D | 6.3e-08 | 1.1e-07 | 2.1e-08 | 6.467E-08 | 4.452E-08 | 2.100E-08 | 1.100E-07 |
| 21182 | 1060 | I1060R | D | 6.2e-08 | 1.6e-08 | 3.9e-08 | 3.900E-08 | 2.300E-08 | 1.600E-08 | 6.200E-08 |
| 21183 | 1060 | I1060N | D | 1.1e-07 | 2.7e-08 | 4.5e-08 | 6.067E-08 | 4.366E-08 | 2.700E-08 | 1.100E-07 |
| 21184 | 1060 | I1060D | D | 5.9e-08 | 8e-08 | 3.6e-08 | 5.833E-08 | 2.201E-08 | 3.600E-08 | 8.000E-08 |
| 21185 | 1060 | I1060C | D | 5.4e-08 | 8.3e-09 | 5e-08 | 3.743E-08 | 2.531E-08 | 8.300E-09 | 5.400E-08 |
| 21186 | 1060 | I1060E | D | 5.8e-08 | 6.4e-08 | 4.2e-08 | 5.467E-08 | 1.137E-08 | 4.200E-08 | 6.400E-08 |
| 21187 | 1060 | I1060Q | D | 9.1e-09 | 4.9e-08 | 5e-08 | 3.603E-08 | 2.333E-08 | 9.100E-09 | 5.000E-08 |
| 21188 | 1060 | I1060G | D | 3.2e-08 | 1.5e-07 | 4.9e-08 | 7.700E-08 | 6.379E-08 | 3.200E-08 | 1.500E-07 |
| 21189 | 1060 | I1060H | D | 2.5e-08 | 4.4e-08 | 3.5e-08 | 3.467E-08 | 9.504E-09 | 2.500E-08 | 4.400E-08 |
| 21190 | 1060 | I1060I | D | 2.5e-08 | 1.2e-08 | 3.8e-08 | 2.500E-08 | 1.300E-08 | 1.200E-08 | 3.800E-08 |
| 21191 | 1060 | I1060L | D | 4e-08 | 4.3e-08 | 6.1e-08 | 4.800E-08 | 1.136E-08 | 4.000E-08 | 6.100E-08 |
| 21192 | 1060 | I1060K | D | 3.4e-08 | 3.7e-08 | 7.7e-08 | 4.933E-08 | 2.401E-08 | 3.400E-08 | 7.700E-08 |
| 21193 | 1060 | I1060M | D | 2e-08 | 2.4e-08 | 4.4e-08 | 2.933E-08 | 1.286E-08 | 2.000E-08 | 4.400E-08 |
| 21194 | 1060 | I1060F | D | 4.2e-08 | 4e-08 | 3.3e-08 | 3.833E-08 | 4.726E-09 | 3.300E-08 | 4.200E-08 |
| 21195 | 1060 | I1060P | D | 2.2e-07 | 1.6e-07 | 3.3e-08 | 1.377E-07 | 9.548E-08 | 3.300E-08 | 2.200E-07 |
| 21196 | 1060 | I1060S | D | 8.7e-08 | 1.1e-07 | 4.8e-08 | 8.167E-08 | 3.134E-08 | 4.800E-08 | 1.100E-07 |
| 21197 | 1060 | I1060T | D | 7.4e-08 | 4.8e-08 | 3.3e-08 | 5.167E-08 | 2.074E-08 | 3.300E-08 | 7.400E-08 |
| 21198 | 1060 | I1060W | D | 1.9e-08 | 4.2e-08 | 4.6e-08 | 3.567E-08 | 1.457E-08 | 1.900E-08 | 4.600E-08 |
| 21199 | 1060 | I1060Y | D | 3.7e-08 | 8.1e-08 | 6.1e-08 | 5.967E-08 | 2.203E-08 | 3.700E-08 | 8.100E-08 |
| 21200 | 1060 | I1060V | D | 1.5e-08 | 2.8e-08 | 2e-08 | 2.100E-08 | 6.557E-09 | 1.500E-08 | 2.800E-08 |
| 21201 | 1061 | Q1061A | D | 6.7e-08 | 2.4e-08 | 4.2e-08 | 4.433E-08 | 2.159E-08 | 2.400E-08 | 6.700E-08 |
| 21202 | 1061 | Q1061R | D | 6.2e-08 | 4.8e-08 | 9.5e-08 | 6.833E-08 | 2.413E-08 | 4.800E-08 | 9.500E-08 |
| 21203 | 1061 | Q1061N | D | 2.1e-08 | 3.7e-08 | 7.4e-08 | 4.400E-08 | 2.718E-08 | 2.100E-08 | 7.400E-08 |
| 21204 | 1061 | Q1061D | D | 8.2e-09 | 2.9e-08 | 2e-08 | 1.907E-08 | 1.043E-08 | 8.200E-09 | 2.900E-08 |
| 21205 | 1061 | Q1061C | D | 5e-08 | 1.6e-08 | 2.6e-08 | 3.067E-08 | 1.747E-08 | 1.600E-08 | 5.000E-08 |
| 21206 | 1061 | Q1061E | D | 3.2e-08 | 2.9e-08 | 4.3e-08 | 3.467E-08 | 7.371E-09 | 2.900E-08 | 4.300E-08 |
| 21207 | 1061 | Q1061Q | D | 5.5e-08 | 3.5e-08 | 4.3e-08 | 4.433E-08 | 1.007E-08 | 3.500E-08 | 5.500E-08 |
| 21208 | 1061 | Q1061G | D | 6.3e-08 | 4e-08 | 1.5e-07 | 8.433E-08 | 5.802E-08 | 4.000E-08 | 1.500E-07 |
| 21209 | 1061 | Q1061H | D | 4.7e-08 | 4.2e-08 | 5.7e-08 | 4.867E-08 | 7.638E-09 | 4.200E-08 | 5.700E-08 |
| 21210 | 1061 | Q1061I | D | 4.6e-08 | 2.8e-08 | 9.2e-08 | 5.533E-08 | 3.301E-08 | 2.800E-08 | 9.200E-08 |
| 21211 | 1061 | Q1061L | D | 2.5e-08 | 2.3e-08 | 1.5e-08 | 2.100E-08 | 5.292E-09 | 1.500E-08 | 2.500E-08 |
| 21212 | 1061 | Q1061K | D | 4.5e-08 | 4.2e-08 | 2.8e-08 | 3.833E-08 | 9.074E-09 | 2.800E-08 | 4.500E-08 |
| 21213 | 1061 | Q1061M | D | 3.4e-08 | 1e-08 | 2.4e-08 | 2.267E-08 | 1.206E-08 | 1.000E-08 | 3.400E-08 |
| 21214 | 1061 | Q1061F | D | 4.4e-08 | 7e-08 | 7.5e-08 | 6.300E-08 | 1.664E-08 | 4.400E-08 | 7.500E-08 |
| 21215 | 1061 | Q1061P | D | 2e-08 | 2.9e-08 | 8.2e-08 | 4.367E-08 | 3.350E-08 | 2.000E-08 | 8.200E-08 |
| 21216 | 1061 | Q1061S | D | 4e-08 | 9.2e-08 | 7.7e-08 | 6.967E-08 | 2.676E-08 | 4.000E-08 | 9.200E-08 |
| 21217 | 1061 | Q1061T | D | 1.1e-07 | 2.2e-08 | 4e-08 | 5.733E-08 | 4.649E-08 | 2.200E-08 | 1.100E-07 |
| 21218 | 1061 | Q1061W | D | 4.1e-08 | 3.2e-08 | 3.2e-08 | 3.500E-08 | 5.196E-09 | 3.200E-08 | 4.100E-08 |
| 21219 | 1061 | Q1061Y | D | 7e-08 | 2.5e-08 | 2.7e-08 | 4.067E-08 | 2.542E-08 | 2.500E-08 | 7.000E-08 |
| 21220 | 1061 | Q1061V | D | 1.1e-07 | 4.4e-08 | 6.5e-08 | 7.300E-08 | 3.372E-08 | 4.400E-08 | 1.100E-07 |
| 21221 | 1062 | M1062A | D | 3.1e-08 | 2e-08 | 8e-08 | 4.367E-08 | 3.194E-08 | 2.000E-08 | 8.000E-08 |
| 21222 | 1062 | M1062R | D | 4.2e-08 | 4.7e-08 | 1.5e-08 | 3.467E-08 | 1.721E-08 | 1.500E-08 | 4.700E-08 |
| 21223 | 1062 | M1062N | D | 1.3e-07 | 6e-08 | 4.2e-08 | 7.733E-08 | 4.649E-08 | 4.200E-08 | 1.300E-07 |
| 21224 | 1062 | M1062D | D | 3.6e-08 | 2.6e-08 | 6.4e-08 | 4.200E-08 | 1.970E-08 | 2.600E-08 | 6.400E-08 |
| 21225 | 1062 | M1062C | D | 2.5e-08 | 6.3e-08 | 1.8e-08 | 3.533E-08 | 2.421E-08 | 1.800E-08 | 6.300E-08 |
| 21226 | 1062 | M1062E | D | 3.5e-08 | 2.1e-08 | 3.8e-08 | 3.133E-08 | 9.074E-09 | 2.100E-08 | 3.800E-08 |
| 21227 | 1062 | M1062Q | D | 2.3e-08 | 5.2e-08 | 9.5e-08 | 5.667E-08 | 3.623E-08 | 2.300E-08 | 9.500E-08 |
| 21228 | 1062 | M1062G | D | 2e-08 | 4.3e-08 | 4.3e-08 | 3.533E-08 | 1.328E-08 | 2.000E-08 | 4.300E-08 |
| 21229 | 1062 | M1062H | D | 4.9e-08 | 4.1e-08 | 2.9e-08 | 3.967E-08 | 1.007E-08 | 2.900E-08 | 4.900E-08 |
| 21230 | 1062 | M1062I | D | 3.4e-08 | 4.2e-08 | 4.7e-08 | 4.100E-08 | 6.557E-09 | 3.400E-08 | 4.700E-08 |
| 21231 | 1062 | M1062L | D | 2.6e-08 | 2.2e-08 | 5.5e-08 | 3.433E-08 | 1.801E-08 | 2.200E-08 | 5.500E-08 |
| 21232 | 1062 | M1062K | D | 6.2e-08 | 4.6e-08 | 2.1e-07 | 1.060E-07 | 9.042E-08 | 4.600E-08 | 2.100E-07 |
| 21233 | 1062 | M1062M | D | 2.9e-08 | 8.2e-08 | 1.3e-07 | 8.033E-08 | 5.052E-08 | 2.900E-08 | 1.300E-07 |
| 21234 | 1062 | M1062F | D | 2e-08 | 1.1e-07 | 3.1e-08 | 5.367E-08 | 4.910E-08 | 2.000E-08 | 1.100E-07 |
| 21235 | 1062 | M1062P | D | 1.8e-08 | 5.6e-08 | 3.7e-08 | 3.700E-08 | 1.900E-08 | 1.800E-08 | 5.600E-08 |
| 21236 | 1062 | M1062S | D | 4.4e-08 | 1.9e-08 | 2.9e-08 | 3.067E-08 | 1.258E-08 | 1.900E-08 | 4.400E-08 |
| 21237 | 1062 | M1062T | D | 4.3e-08 | 2.4e-08 | 3.2e-08 | 3.300E-08 | 9.539E-09 | 2.400E-08 | 4.300E-08 |
| 21238 | 1062 | M1062W | D | 2.5e-08 | 3.9e-08 | 4e-08 | 3.467E-08 | 8.386E-09 | 2.500E-08 | 4.000E-08 |
| 21239 | 1062 | M1062Y | D | 6.5e-08 | 9.8e-08 | 3.9e-08 | 6.733E-08 | 2.957E-08 | 3.900E-08 | 9.800E-08 |
| 21240 | 1062 | M1062V | D | 3.8e-08 | 5.2e-08 | 9.3e-08 | 6.100E-08 | 2.858E-08 | 3.800E-08 | 9.300E-08 |
| 21241 | 1063 | T1063A | D | 6e-08 | 1.1e-08 | 4.8e-08 | 3.967E-08 | 2.554E-08 | 1.100E-08 | 6.000E-08 |
| 21242 | 1063 | T1063R | D | 2.1e-08 | 1.1e-07 | 1.8e-08 | 4.967E-08 | 5.227E-08 | 1.800E-08 | 1.100E-07 |
| 21243 | 1063 | T1063N | D | 1.1e-07 | 3.7e-08 | 5.7e-08 | 6.800E-08 | 3.772E-08 | 3.700E-08 | 1.100E-07 |
| 21244 | 1063 | T1063D | D | 4.5e-08 | 4.2e-08 | 5.1e-08 | 4.600E-08 | 4.583E-09 | 4.200E-08 | 5.100E-08 |
| 21245 | 1063 | T1063C | D | 5.4e-08 | 1.1e-08 | 4.4e-08 | 3.633E-08 | 2.250E-08 | 1.100E-08 | 5.400E-08 |
| 21246 | 1063 | T1063E | D | 3.4e-08 | 4.7e-08 | 2.3e-08 | 3.467E-08 | 1.201E-08 | 2.300E-08 | 4.700E-08 |
| 21247 | 1063 | T1063Q | D | 3.1e-08 | 4.8e-08 | 4.4e-08 | 4.100E-08 | 8.888E-09 | 3.100E-08 | 4.800E-08 |
| 21248 | 1063 | T1063G | D | 1.2e-08 | 4.9e-08 | 3.3e-08 | 3.133E-08 | 1.856E-08 | 1.200E-08 | 4.900E-08 |
| 21249 | 1063 | T1063H | D | 5.7e-08 | 9.7e-08 | 5.2e-08 | 6.867E-08 | 2.466E-08 | 5.200E-08 | 9.700E-08 |

| | | | | | | | | | | |
|---|---|---|---|---|---|---|---|---|---|---|
| 21250 | 1063 | T1063I | D | 1.3e-07 | 4.5e-08 | 1e-07 | 9.167E-08 | 4.311E-08 | 4.500E-08 | 1.300E-07 |
| 21251 | 1063 | T1063L | D | 2.8e-08 | 7e-08 | 8.8e-08 | 6.200E-08 | 3.079E-08 | 2.800E-08 | 8.800E-08 |
| 21252 | 1063 | T1063K | D | 1.8e-08 | 1.9e-08 | 6e-08 | 3.233E-08 | 2.397E-08 | 1.800E-08 | 6.000E-08 |
| 21253 | 1063 | T1063M | D | 3.5e-08 | 3.1e-08 | 7.1e-08 | 4.567E-08 | 2.203E-08 | 3.100E-08 | 7.100E-08 |
| 21254 | 1063 | T1063F | D | 1.6e-08 | 5.8e-08 | 6.6e-08 | 4.667E-08 | 2.686E-08 | 1.600E-08 | 6.600E-08 |
| 21255 | 1063 | T1063P | D | 2.5e-08 | 9.9e-08 | 1.9e-08 | 4.767E-08 | 4.456E-08 | 1.900E-08 | 9.900E-08 |
| 21256 | 1063 | T1063S | D | 4.7e-08 | 1.1e-08 | 7.2e-08 | 4.333E-08 | 3.066E-08 | 1.100E-08 | 7.200E-08 |
| 21257 | 1063 | T1063T | D | 3e-08 | 4.3e-08 | 1.1e-07 | 6.100E-08 | 4.293E-08 | 3.000E-08 | 1.100E-07 |
| 21258 | 1063 | T1063W | D | 2.7e-08 | 4.3e-08 | 4.9e-08 | 3.967E-08 | 1.137E-08 | 2.700E-08 | 4.900E-08 |
| 21259 | 1063 | T1063Y | D | 4e-08 | 6e-08 | 2.7e-08 | 4.233E-08 | 1.662E-08 | 2.700E-08 | 6.000E-08 |
| 21260 | 1063 | T1063V | D | 2e-08 | 7.5e-09 | 5.9e-08 | 2.883E-08 | 2.686E-08 | 7.500E-09 | 5.900E-08 |
| 21261 | 1064 | Q1064A | D | 1.1e-07 | 1.2e-08 | 3.1e-08 | 5.100E-08 | 5.197E-08 | 1.200E-08 | 1.100E-07 |
| 21262 | 1064 | Q1064R | D | 4.9e-08 | 1.1e-07 | 1.5e-08 | 5.800E-08 | 4.814E-08 | 1.500E-08 | 1.100E-07 |
| 21263 | 1064 | Q1064N | D | 5.1e-08 | 1.3e-08 | 2.8e-08 | 3.067E-08 | 1.914E-08 | 1.300E-08 | 5.100E-08 |
| 21264 | 1064 | Q1064D | D | 3.6e-08 | 7.7e-08 | 6.5e-08 | 5.933E-08 | 2.108E-08 | 3.600E-08 | 7.700E-08 |
| 21265 | 1064 | Q1064C | D | 2.5e-08 | 4.2e-08 | 7.6e-08 | 4.767E-08 | 2.597E-08 | 2.500E-08 | 7.600E-08 |
| 21266 | 1064 | Q1064E | D | 1.9e-08 | 1.2e-07 | 5.9e-08 | 6.600E-08 | 5.086E-08 | 1.900E-08 | 1.200E-07 |
| 21267 | 1064 | Q1064Q | D | 1.8e-08 | 3.5e-08 | 3.1e-08 | 2.800E-08 | 8.888E-09 | 1.800E-08 | 3.500E-08 |
| 21268 | 1064 | Q1064G | D | 2.3e-08 | 3.8e-08 | 2.7e-08 | 2.933E-08 | 7.767E-09 | 2.300E-08 | 3.800E-08 |
| 21269 | 1064 | Q1064H | D | 2.7e-08 | 9.9e-09 | 1e-07 | 4.563E-08 | 4.785E-08 | 9.900E-09 | 1.000E-07 |
| 21270 | 1064 | Q1064I | D | 2.4e-08 | 3.5e-08 | 6.3e-08 | 4.067E-08 | 2.011E-08 | 2.400E-08 | 6.300E-08 |
| 21271 | 1064 | Q1064L | D | 2e-08 | 6.7e-08 | 2.1e-08 | 3.600E-08 | 2.685E-08 | 2.000E-08 | 6.700E-08 |
| 21272 | 1064 | Q1064K | D | 7.2e-08 | 2.9e-08 | 3.3e-08 | 4.467E-08 | 2.376E-08 | 2.900E-08 | 7.200E-08 |
| 21273 | 1064 | Q1064M | D | 2.3e-08 | 4.3e-08 | 2.6e-08 | 3.067E-08 | 1.079E-08 | 2.300E-08 | 4.300E-08 |
| 21274 | 1064 | Q1064F | D | 2.9e-08 | 6.7e-08 | 2.1e-08 | 3.900E-08 | 2.458E-08 | 2.100E-08 | 6.700E-08 |
| 21275 | 1064 | Q1064P | D | 6.8e-08 | 2.4e-08 | 2.5e-08 | 3.900E-08 | 2.512E-08 | 2.400E-08 | 6.800E-08 |
| 21276 | 1064 | Q1064S | D | 2.9e-08 | 5.3e-08 | 1.2e-08 | 3.133E-08 | 2.060E-08 | 1.200E-08 | 5.300E-08 |
| 21277 | 1064 | Q1064T | D | 7.1e-08 | 3.6e-08 | 7.5e-08 | 6.067E-08 | 2.146E-08 | 3.600E-08 | 7.500E-08 |
| 21278 | 1064 | Q1064W | D | 9.3e-08 | 2.9e-07 | 5.4e-08 | 1.457E-07 | 1.265E-07 | 5.400E-08 | 2.900E-07 |
| 21279 | 1064 | Q1064Y | D | 6e-08 | 3e-08 | 7.3e-08 | 5.433E-08 | 2.205E-08 | 3.000E-08 | 7.300E-08 |
| 21280 | 1064 | Q1064V | D | 3e-08 | 5.2e-08 | 8.1e-08 | 5.433E-08 | 2.558E-08 | 3.000E-08 | 8.100E-08 |
| 21281 | 1065 | S1065A | D | 1.8e-08 | 5.7e-08 | 4.6e-08 | 4.033E-08 | 2.011E-08 | 1.800E-08 | 5.700E-08 |
| 21282 | 1065 | S1065R | D | 4e-08 | 1.4e-08 | 1.1e-08 | 2.167E-08 | 1.595E-08 | 1.100E-08 | 4.000E-08 |
| 21283 | 1065 | S1065N | D | 1.5e-08 | 5.7e-08 | 8.1e-08 | 5.100E-08 | 3.341E-08 | 1.500E-08 | 8.100E-08 |
| 21284 | 1065 | S1065D | D | 5e-08 | 4.3e-08 | 2.3e-08 | 3.867E-08 | 1.401E-08 | 2.300E-08 | 5.000E-08 |
| 21285 | 1065 | S1065C | D | 3.9e-08 | 6.4e-08 | 9.2e-08 | 6.500E-08 | 2.651E-08 | 3.900E-08 | 9.200E-08 |
| 21286 | 1065 | S1065E | D | 3.4e-08 | 1e-08 | 8.9e-08 | 4.433E-08 | 4.050E-08 | 1.000E-08 | 8.900E-08 |
| 21287 | 1065 | S1065Q | D | 5.6e-08 | 2.4e-08 | 4.5e-08 | 4.167E-08 | 1.626E-08 | 2.400E-08 | 5.600E-08 |
| 21288 | 1065 | S1065G | D | 3.2e-08 | 1.4e-08 | 5.3e-08 | 3.300E-08 | 1.952E-08 | 1.400E-08 | 5.300E-08 |
| 21289 | 1065 | S1065H | D | 1.1e-08 | 3e-08 | 1.9e-07 | 7.700E-08 | 9.832E-08 | 1.100E-08 | 1.900E-07 |
| 21290 | 1065 | S1065I | D | 4e-08 | 5.4e-08 | 1.4e-07 | 7.800E-08 | 5.415E-08 | 4.000E-08 | 1.400E-07 |
| 21291 | 1065 | S1065L | D | 8.7e-08 | 4.4e-08 | 4e-08 | 5.700E-08 | 2.606E-08 | 4.000E-08 | 8.700E-08 |
| 21292 | 1065 | S1065K | D | 2.5e-08 | 6.1e-08 | 3.3e-08 | 3.967E-08 | 1.890E-08 | 2.500E-08 | 6.100E-08 |
| 21293 | 1065 | S1065M | D | 4.9e-08 | 3.1e-08 | 2.8e-08 | 3.600E-08 | 1.136E-08 | 2.800E-08 | 4.900E-08 |
| 21294 | 1065 | S1065F | D | 5.9e-08 | 5.4e-08 | 8.8e-08 | 6.700E-08 | 1.836E-08 | 5.400E-08 | 8.800E-08 |
| 21295 | 1065 | S1065P | D | 1.8e-08 | 4.6e-08 | 1.8e-08 | 2.733E-08 | 1.617E-08 | 1.800E-08 | 4.600E-08 |
| 21296 | 1065 | S1065S | D | 5.2e-08 | 2e-08 | 5.8e-08 | 4.333E-08 | 2.043E-08 | 2.000E-08 | 5.800E-08 |
| 21297 | 1065 | S1065T | D | 2.9e-08 | 6.4e-08 | 3e-08 | 4.100E-08 | 1.992E-08 | 2.900E-08 | 6.400E-08 |
| 21298 | 1065 | S1065W | D | 3.6e-08 | 3.1e-08 | 5.1e-08 | 3.933E-08 | 1.041E-08 | 3.100E-08 | 5.100E-08 |
| 21299 | 1065 | S1065Y | D | 4.5e-08 | 5.8e-08 | 3.7e-08 | 4.667E-08 | 1.060E-08 | 3.700E-08 | 5.800E-08 |
| 21300 | 1065 | S1065V | D | 9.4e-08 | 9.7e-08 | 8.3e-08 | 9.133E-08 | 7.371E-09 | 8.300E-08 | 9.700E-08 |
| 21301 | 1066 | P1066A | D | 1.8e-08 | 3.3e-08 | 3.6e-08 | 2.900E-08 | 9.644E-09 | 1.800E-08 | 3.600E-08 |
| 21302 | 1066 | P1066R | D | 7e-08 | 4.4e-08 | 5.4e-08 | 5.600E-08 | 1.311E-08 | 4.400E-08 | 7.000E-08 |
| 21303 | 1066 | P1066N | D | 8.1e-08 | 7.6e-08 | 1.4e-07 | 9.900E-08 | 3.559E-08 | 7.600E-08 | 1.400E-07 |
| 21304 | 1066 | P1066D | D | 3.3e-08 | 2.1e-08 | 1e-08 | 2.133E-08 | 1.150E-08 | 1.000E-08 | 3.300E-08 |
| 21305 | 1066 | P1066C | D | 1e-08 | 7.4e-08 | 4.2e-08 | 4.200E-08 | 3.200E-08 | 1.000E-08 | 7.400E-08 |
| 21306 | 1066 | P1066E | D | 4.1e-08 | 2.6e-08 | 4.8e-08 | 3.833E-08 | 1.124E-08 | 2.600E-08 | 4.800E-08 |
| 21307 | 1066 | P1066Q | D | 2.8e-08 | 1.6e-07 | 5.3e-08 | 8.033E-08 | 7.012E-08 | 2.800E-08 | 1.600E-07 |
| 21308 | 1066 | P1066G | D | 6.8e-08 | 5.3e-08 | 5.8e-08 | 5.967E-08 | 7.638E-09 | 5.300E-08 | 6.800E-08 |
| 21309 | 1066 | P1066H | D | 1.9e-08 | 5.8e-08 | 8.4e-08 | 5.367E-08 | 3.272E-08 | 1.900E-08 | 8.400E-08 |
| 21310 | 1066 | P1066I | D | 4.8e-08 | 5.1e-08 | 4.2e-08 | 4.700E-08 | 4.583E-09 | 4.200E-08 | 5.100E-08 |
| 21311 | 1066 | P1066L | D | 2e-08 | 7.4e-08 | 7.9e-08 | 5.767E-08 | 3.272E-08 | 2.000E-08 | 7.900E-08 |
| 21312 | 1066 | P1066K | D | 1.7e-08 | 2.2e-08 | 3e-08 | 2.300E-08 | 6.557E-09 | 1.700E-08 | 3.000E-08 |
| 21313 | 1066 | P1066M | D | 1.4e-08 | 1.3e-07 | 3.1e-08 | 5.833E-08 | 6.264E-08 | 1.400E-08 | 1.300E-07 |
| 21314 | 1066 | P1066F | D | 1.6e-07 | 6.6e-08 | 2e-08 | 8.200E-08 | 7.136E-08 | 2.000E-08 | 1.600E-07 |
| 21315 | 1066 | P1066P | D | 2.5e-08 | 4.8e-08 | 3.3e-08 | 3.533E-08 | 1.168E-08 | 2.500E-08 | 4.800E-08 |
| 21316 | 1066 | P1066S | D | 1.5e-08 | 5.1e-08 | 8.4e-08 | 5.000E-08 | 3.451E-08 | 1.500E-08 | 8.400E-08 |
| 21317 | 1066 | P1066T | D | 4.1e-08 | 8.3e-08 | 2.9e-08 | 5.100E-08 | 2.835E-08 | 2.900E-08 | 8.300E-08 |
| 21318 | 1066 | P1066W | D | 4.3e-08 | 1.1e-08 | 1.7e-08 | 2.367E-08 | 1.701E-08 | 1.100E-08 | 4.300E-08 |
| 21319 | 1066 | P1066Y | D | 1.1e-07 | 4.4e-08 | 3e-08 | 6.133E-08 | 4.272E-08 | 3.000E-08 | 1.100E-07 |
| 21320 | 1066 | P1066V | D | 5.6e-08 | 8.1e-08 | 3.6e-08 | 5.767E-08 | 2.255E-08 | 3.600E-08 | 8.100E-08 |
| 21321 | 1067 | S1067A | D | 2.1e-08 | 6.5e-08 | 2.4e-08 | 3.667E-08 | 2.458E-08 | 2.100E-08 | 6.500E-08 |
| 21322 | 1067 | S1067R | D | 1.2e-07 | 6e-08 | 2.2e-08 | 6.733E-08 | 4.941E-08 | 2.200E-08 | 1.200E-07 |
| 21323 | 1067 | S1067N | D | 4.1e-08 | 2.8e-08 | 5.7e-08 | 4.200E-08 | 1.453E-08 | 2.800E-08 | 5.700E-08 |
| 21324 | 1067 | S1067D | D | 3.5e-08 | 4.9e-08 | 1.1e-07 | 6.467E-08 | 3.988E-08 | 3.500E-08 | 1.100E-07 |
| 21325 | 1067 | S1067C | D | 4.4e-08 | 7e-08 | 6.1e-08 | 5.833E-08 | 1.320E-08 | 4.400E-08 | 7.000E-08 |
| 21326 | 1067 | S1067E | D | 5.8e-08 | 3.5e-08 | 3.8e-08 | 4.367E-08 | 1.250E-08 | 3.500E-08 | 5.800E-08 |
| 21327 | 1067 | S1067Q | D | 2.3e-08 | 2.3e-08 | 5.5e-08 | 3.367E-08 | 1.848E-08 | 2.300E-08 | 5.500E-08 |
| 21328 | 1067 | S1067G | D | 7.5e-08 | 7.8e-08 | 7.9e-08 | 7.733E-08 | 2.082E-09 | 7.500E-08 | 7.900E-08 |
| 21329 | 1067 | S1067H | D | 5.4e-08 | 1.6e-08 | 1.8e-08 | 2.933E-08 | 2.139E-08 | 1.600E-08 | 5.400E-08 |
| 21330 | 1067 | S1067I | D | 2.2e-07 | 1.3e-08 | 3.5e-08 | 8.933E-08 | 1.137E-07 | 1.300E-08 | 2.200E-07 |
| 21331 | 1067 | S1067L | D | 3.9e-08 | 4.1e-08 | 4.2e-08 | 4.067E-08 | 1.528E-09 | 3.900E-08 | 4.200E-08 |
| 21332 | 1067 | S1067K | D | 6.6e-08 | 5.6e-08 | 2.1e-08 | 4.767E-08 | 2.363E-08 | 2.100E-08 | 6.600E-08 |
| 21333 | 1067 | S1067M | D | 5.9e-08 | 5.8e-08 | 4.9e-08 | 5.533E-08 | 5.508E-09 | 4.900E-08 | 5.900E-08 |
| 21334 | 1067 | S1067F | D | 3.1e-08 | 7.3e-08 | 9e-08 | 6.467E-08 | 3.037E-08 | 3.100E-08 | 9.000E-08 |

| | | | | | | | | | | |
|---|---|---|---|---|---|---|---|---|---|---|
| 21335 | 1067 | S1067P | D | 5.5e-08 | 7.6e-08 | 7.8e-08 | 6.967E-08 | 1.274E-08 | 5.500E-08 | 7.800E-08 |
| 21336 | 1067 | S1067S | D | 5.3e-08 | 2e-08 | 3.9e-08 | 3.733E-08 | 1.656E-08 | 2.000E-08 | 5.300E-08 |
| 21337 | 1067 | S1067T | D | 1.8e-08 | 7.3e-08 | 7.4e-08 | 5.500E-08 | 3.205E-08 | 1.800E-08 | 7.400E-08 |
| 21338 | 1067 | S1067W | D | 1.8e-08 | 4.5e-08 | 1.1e-07 | 5.767E-08 | 4.729E-08 | 1.800E-08 | 1.100E-07 |
| 21339 | 1067 | S1067Y | D | 4e-08 | 7.1e-08 | 1.6e-08 | 4.233E-08 | 2.757E-08 | 1.600E-08 | 7.100E-08 |
| 21340 | 1067 | S1067V | D | 4.1e-08 | 6e-08 | 8.7e-09 | 3.657E-08 | 2.594E-08 | 8.700E-09 | 6.000E-08 |
| 21341 | 1068 | S1068A | D | 7.1e-08 | 4.7e-08 | 3.3e-08 | 5.033E-08 | 1.922E-08 | 3.300E-08 | 7.100E-08 |
| 21342 | 1068 | S1068R | D | 5.4e-08 | 5.2e-08 | 1.9e-08 | 4.167E-08 | 1.966E-08 | 1.900E-08 | 5.400E-08 |
| 21343 | 1068 | S1068N | D | 1.5e-07 | 3.3e-08 | 1.2e-07 | 1.010E-07 | 6.077E-08 | 3.300E-08 | 1.500E-07 |
| 21344 | 1068 | S1068D | D | 2.8e-08 | 5.2e-08 | 2e-08 | 3.333E-08 | 1.665E-08 | 2.000E-08 | 5.200E-08 |
| 21345 | 1068 | S1068C | D | 3e-08 | 5.7e-08 | 9.8e-08 | 6.167E-08 | 3.424E-08 | 3.000E-08 | 9.800E-08 |
| 21346 | 1068 | S1068E | D | 5e-08 | 1.8e-08 | 4.8e-08 | 3.867E-08 | 1.793E-08 | 1.800E-08 | 5.000E-08 |
| 21347 | 1068 | S1068Q | D | 4.6e-08 | 3.5e-08 | 5e-08 | 4.367E-08 | 7.767E-09 | 3.500E-08 | 5.000E-08 |
| 21348 | 1068 | S1068G | D | 6.7e-08 | 9.2e-08 | 5.2e-08 | 7.033E-08 | 2.021E-08 | 5.200E-08 | 9.200E-08 |
| 21349 | 1068 | S1068H | D | 1.4e-07 | 3.8e-08 | 3.7e-08 | 7.167E-08 | 5.918E-08 | 3.700E-08 | 1.400E-07 |
| 21350 | 1068 | S1068I | D | 5.4e-08 | 6.4e-08 | 1.6e-07 | 9.267E-08 | 5.853E-08 | 5.400E-08 | 1.600E-07 |
| 21351 | 1068 | S1068L | D | 4.7e-08 | 5.3e-08 | 9.1e-08 | 6.367E-08 | 2.386E-08 | 4.700E-08 | 9.100E-08 |
| 21352 | 1068 | S1068K | D | 5e-08 | 4.6e-08 | 4.1e-08 | 4.567E-08 | 4.509E-09 | 4.100E-08 | 5.000E-08 |
| 21353 | 1068 | S1068M | D | 2.6e-08 | 3.2e-08 | 5.7e-08 | 3.833E-08 | 1.644E-08 | 2.600E-08 | 5.700E-08 |
| 21354 | 1068 | S1068F | D | 1e-07 | 5.1e-08 | 9e-09 | 5.333E-08 | 4.554E-08 | 9.000E-09 | 1.000E-07 |
| 21355 | 1068 | S1068P | D | 8.1e-08 | 3.8e-08 | 4.3e-08 | 5.400E-08 | 2.352E-08 | 3.800E-08 | 8.100E-08 |
| 21356 | 1068 | S1068S | D | 4.9e-08 | 6.6e-08 | 2.3e-08 | 4.600E-08 | 2.166E-08 | 2.300E-08 | 6.600E-08 |
| 21357 | 1068 | S1068T | D | 8.8e-08 | 6.3e-08 | 9.2e-08 | 8.100E-08 | 1.572E-08 | 6.300E-08 | 9.200E-08 |
| 21358 | 1068 | S1068W | D | 1.3e-07 | 4.4e-08 | 1.3e-07 | 1.013E-07 | 4.965E-08 | 4.400E-08 | 1.300E-07 |
| 21359 | 1068 | S1068Y | D | 2e-08 | 3.4e-08 | 3.1e-08 | 2.833E-08 | 7.371E-09 | 2.000E-08 | 3.400E-08 |
| 21360 | 1068 | S1068V | D | 1.1e-07 | 6.2e-08 | 3.4e-08 | 6.867E-08 | 3.844E-08 | 3.400E-08 | 1.100E-07 |
| 21361 | 1069 | L1069A | D | 6.1e-08 | 6.5e-08 | 6.5e-08 | 6.367E-08 | 2.309E-09 | 6.100E-08 | 6.500E-08 |
| 21362 | 1069 | L1069R | D | 1.4e-08 | 6.3e-08 | 8.5e-08 | 5.400E-08 | 3.635E-08 | 1.400E-08 | 8.500E-08 |
| 21363 | 1069 | L1069N | D | 4.3e-08 | 8.8e-08 | 1.5e-08 | 4.867E-08 | 3.683E-08 | 1.500E-08 | 8.800E-08 |
| 21364 | 1069 | L1069D | D | 3.8e-08 | 3.6e-08 | 2.7e-08 | 3.367E-08 | 5.859E-09 | 2.700E-08 | 3.800E-08 |
| 21365 | 1069 | L1069C | D | 2.1e-08 | 4e-08 | 1.7e-08 | 2.600E-08 | 1.229E-08 | 1.700E-08 | 4.000E-08 |
| 21366 | 1069 | L1069E | D | 3.7e-08 | 1.2e-08 | 2.6e-08 | 2.500E-08 | 1.253E-08 | 1.200E-08 | 3.700E-08 |
| 21367 | 1069 | L1069Q | D | 6.1e-08 | 1.2e-08 | 3.6e-08 | 3.633E-08 | 2.450E-08 | 1.200E-08 | 6.100E-08 |
| 21368 | 1069 | L1069G | D | 6e-08 | 7.6e-08 | 2.7e-08 | 5.433E-08 | 2.499E-08 | 2.700E-08 | 7.600E-08 |
| 21369 | 1069 | L1069H | D | 5e-08 | 2.9e-08 | 2.1e-08 | 3.333E-08 | 1.498E-08 | 2.100E-08 | 5.000E-08 |
| 21370 | 1069 | L1069I | D | 2.8e-08 | 1.2e-07 | 3.1e-08 | 5.967E-08 | 5.227E-08 | 2.800E-08 | 1.200E-07 |
| 21371 | 1069 | L1069L | D | 3.1e-08 | 5.2e-08 | 3.9e-08 | 4.067E-08 | 1.060E-08 | 3.100E-08 | 5.200E-08 |
| 21372 | 1069 | L1069K | D | 3.8e-08 | 3.6e-08 | 2.4e-08 | 3.267E-08 | 7.572E-09 | 2.400E-08 | 3.800E-08 |
| 21373 | 1069 | L1069M | D | 4.9e-08 | 4.1e-08 | 3.4e-08 | 4.133E-08 | 7.506E-09 | 3.400E-08 | 4.900E-08 |
| 21374 | 1069 | L1069F | D | 9.4e-08 | 5.6e-08 | 1.8e-08 | 5.600E-08 | 3.800E-08 | 1.800E-08 | 9.400E-08 |
| 21375 | 1069 | L1069P | D | 6.3e-08 | 2.4e-08 | 3.4e-08 | 4.033E-08 | 2.026E-08 | 2.400E-08 | 6.300E-08 |
| 21376 | 1069 | L1069S | D | 7e-08 | 2.6e-08 | 2.7e-08 | 4.100E-08 | 2.512E-08 | 2.600E-08 | 7.000E-08 |
| 21377 | 1069 | L1069T | D | 5.6e-08 | 4.5e-08 | 1.3e-08 | 3.800E-08 | 2.234E-08 | 1.300E-08 | 5.600E-08 |
| 21378 | 1069 | L1069W | D | 4.2e-08 | 1.6e-08 | 2.2e-08 | 2.667E-08 | 1.361E-08 | 1.600E-08 | 4.200E-08 |
| 21379 | 1069 | L1069Y | D | 4.9e-08 | 3.3e-08 | 7.2e-08 | 5.133E-08 | 1.960E-08 | 3.300E-08 | 7.200E-08 |
| 21380 | 1069 | L1069V | D | 5.8e-08 | 2.4e-08 | 1.2e-07 | 6.733E-08 | 4.868E-08 | 2.400E-08 | 1.200E-07 |
| 21381 | 1070 | S1070A | D | 2.5e-08 | 7.3e-08 | 1.2e-07 | 7.267E-08 | 4.750E-08 | 2.500E-08 | 1.200E-07 |
| 21382 | 1070 | S1070R | D | 6e-08 | 7.9e-08 | 4.7e-08 | 6.200E-08 | 1.609E-08 | 4.700E-08 | 7.900E-08 |
| 21383 | 1070 | S1070N | D | 7.3e-08 | 7.7e-08 | 7.9e-08 | 7.633E-08 | 3.055E-09 | 7.300E-08 | 7.900E-08 |
| 21384 | 1070 | S1070D | D | 4.1e-08 | 3.4e-08 | 4.4e-08 | 3.967E-08 | 5.132E-09 | 3.400E-08 | 4.400E-08 |
| 21385 | 1070 | S1070C | D | 3.8e-08 | 4e-08 | 2.4e-08 | 3.400E-08 | 8.718E-09 | 2.400E-08 | 4.000E-08 |
| 21386 | 1070 | S1070E | D | 3.4e-08 | 5.3e-08 | 5.7e-08 | 4.800E-08 | 1.229E-08 | 3.400E-08 | 5.700E-08 |
| 21387 | 1070 | S1070Q | D | 9.5e-08 | 5.6e-08 | 4.9e-08 | 6.667E-08 | 2.479E-08 | 4.900E-08 | 9.500E-08 |
| 21388 | 1070 | S1070G | D | 4.2e-08 | 9e-08 | 3.2e-08 | 5.467E-08 | 3.101E-08 | 3.200E-08 | 9.000E-08 |
| 21389 | 1070 | S1070H | D | 4.9e-08 | 3.6e-08 | 3.4e-08 | 3.967E-08 | 8.145E-09 | 3.400E-08 | 4.900E-08 |
| 21390 | 1070 | S1070I | D | 3.7e-08 | 6.2e-08 | 1.6e-08 | 3.833E-08 | 2.303E-08 | 1.600E-08 | 6.200E-08 |
| 21391 | 1070 | S1070L | D | 3.4e-08 | 2.5e-07 | 1.6e-07 | 1.480E-07 | 1.085E-07 | 3.400E-08 | 2.500E-07 |
| 21392 | 1070 | S1070K | D | 5.6e-08 | 3.5e-08 | 4e-08 | 4.367E-08 | 1.097E-08 | 3.500E-08 | 5.600E-08 |
| 21393 | 1070 | S1070M | D | 1.4e-07 | 4.9e-08 | 3.2e-08 | 7.367E-08 | 5.807E-08 | 3.200E-08 | 1.400E-07 |
| 21394 | 1070 | S1070F | D | 2.7e-08 | 4.8e-08 | 2.7e-08 | 3.400E-08 | 1.212E-08 | 2.700E-08 | 4.800E-08 |
| 21395 | 1070 | S1070P | D | 8.6e-08 | 2.9e-08 | 3.9e-08 | 5.133E-08 | 3.044E-08 | 2.900E-08 | 8.600E-08 |
| 21396 | 1070 | S1070S | D | 1.6e-08 | 1.8e-08 | 1.2e-08 | 1.533E-08 | 3.055E-09 | 1.200E-08 | 1.800E-08 |
| 21397 | 1070 | S1070T | D | 3.3e-08 | 5.9e-08 | 3.1e-08 | 4.100E-08 | 1.562E-08 | 3.100E-08 | 5.900E-08 |
| 21398 | 1070 | S1070W | D | 3.1e-08 | 7.6e-08 | 7.5e-08 | 6.067E-08 | 2.570E-08 | 3.100E-08 | 7.600E-08 |
| 21399 | 1070 | S1070Y | D | 2.5e-08 | 3.5e-08 | 3.5e-08 | 3.167E-08 | 5.774E-09 | 2.500E-08 | 3.500E-08 |
| 21400 | 1070 | S1070V | D | 2.3e-08 | 4.7e-08 | 7.9e-08 | 4.967E-08 | 2.810E-08 | 2.300E-08 | 7.900E-08 |
| 21401 | 1071 | A1071A | D | 1.8e-08 | 2.7e-08 | 4.7e-08 | 3.067E-08 | 1.484E-08 | 1.800E-08 | 4.700E-08 |
| 21402 | 1071 | A1071R | D | 8.9e-08 | 9.7e-09 | 3.9e-08 | 4.590E-08 | 4.010E-08 | 9.700E-09 | 8.900E-08 |
| 21403 | 1071 | A1071N | D | 2.9e-08 | 2.2e-08 | 2.3e-08 | 2.467E-08 | 3.786E-09 | 2.200E-08 | 2.900E-08 |
| 21404 | 1071 | A1071D | D | 6.7e-08 | 8.9e-08 | 6.1e-08 | 7.233E-08 | 1.474E-08 | 6.100E-08 | 8.900E-08 |
| 21405 | 1071 | A1071C | D | 4.8e-08 | 7.1e-08 | 2.6e-08 | 4.833E-08 | 2.250E-08 | 2.600E-08 | 7.100E-08 |
| 21406 | 1071 | A1071E | D | 5.2e-08 | 1.4e-08 | 7.5e-08 | 4.700E-08 | 3.081E-08 | 1.400E-08 | 7.500E-08 |
| 21407 | 1071 | A1071Q | D | 1.3e-08 | 2e-08 | 2.3e-08 | 1.867E-08 | 5.132E-09 | 1.300E-08 | 2.300E-08 |
| 21408 | 1071 | A1071G | D | 1e-07 | 6.3e-08 | 7.5e-08 | 7.933E-08 | 1.888E-08 | 6.300E-08 | 1.000E-07 |
| 21409 | 1071 | A1071H | D | 2.6e-08 | 2.9e-08 | 3.1e-08 | 2.867E-08 | 2.517E-09 | 2.600E-08 | 3.100E-08 |
| 21410 | 1071 | A1071I | D | 2.4e-08 | 4.6e-08 | 9.8e-08 | 5.600E-08 | 3.800E-08 | 2.400E-08 | 9.800E-08 |
| 21411 | 1071 | A1071L | D | 3.2e-08 | 5.2e-08 | 3.8e-08 | 4.067E-08 | 1.026E-08 | 3.200E-08 | 5.200E-08 |
| 21412 | 1071 | A1071K | D | 9e-08 | 7e-08 | 4.7e-08 | 6.900E-08 | 2.152E-08 | 4.700E-08 | 9.000E-08 |
| 21413 | 1071 | A1071M | D | 4.4e-08 | 4.3e-08 | 3.8e-08 | 4.167E-08 | 3.215E-09 | 3.800E-08 | 4.400E-08 |
| 21414 | 1071 | A1071F | D | 1.5e-08 | 2.8e-08 | 2.5e-08 | 2.267E-08 | 6.807E-09 | 1.500E-08 | 2.800E-08 |
| 21415 | 1071 | A1071P | D | 5.5e-08 | 4.8e-08 | 4.5e-08 | 4.933E-08 | 5.132E-09 | 4.500E-08 | 5.500E-08 |
| 21416 | 1071 | A1071S | D | 7.8e-08 | 5e-08 | 1e-07 | 7.600E-08 | 2.506E-08 | 5.000E-08 | 1.000E-07 |
| 21417 | 1071 | A1071T | D | 4.4e-08 | 2e-08 | 6.3e-08 | 4.233E-08 | 2.155E-08 | 2.000E-08 | 6.300E-08 |
| 21418 | 1071 | A1071W | D | 7.2e-08 | 2.5e-08 | 4.6e-08 | 4.767E-08 | 2.354E-08 | 2.500E-08 | 7.200E-08 |
| 21419 | 1071 | A1071Y | D | 2.4e-08 | 2.5e-08 | 1.8e-08 | 2.233E-08 | 3.786E-09 | 1.800E-08 | 2.500E-08 |

| | | | | | | | | | | |
|---|---|---|---|---|---|---|---|---|---|---|
| 21420 | 1071 | A1071V | D | 3.6e-08 | 3.4e-08 | 1.4e-07 | 7.000E-08 | 6.063E-08 | 3.400E-08 | 1.400E-07 |
| 21421 | 1072 | S1072A | D | 3.2e-08 | 6.8e-08 | 3.5e-08 | 4.500E-08 | 1.997E-08 | 3.200E-08 | 6.800E-08 |
| 21422 | 1072 | S1072R | D | 8.2e-08 | 2.6e-08 | 3.4e-08 | 4.733E-08 | 3.029E-08 | 2.600E-08 | 8.200E-08 |
| 21423 | 1072 | S1072N | D | 1.7e-07 | 7.6e-08 | 2.2e-08 | 8.933E-08 | 7.490E-08 | 2.200E-08 | 1.700E-07 |
| 21424 | 1072 | S1072D | D | 6.3e-08 | 3.3e-08 | 4.5e-08 | 4.700E-08 | 1.510E-08 | 3.300E-08 | 6.300E-08 |
| 21425 | 1072 | S1072C | D | 3.5e-08 | 7.3e-08 | 1.8e-08 | 4.200E-08 | 2.816E-08 | 1.800E-08 | 7.300E-08 |
| 21426 | 1072 | S1072E | D | 5.6e-08 | 3.2e-08 | 8.7e-08 | 5.833E-08 | 2.757E-08 | 3.200E-08 | 8.700E-08 |
| 21427 | 1072 | S1072Q | D | 4e-08 | 7.8e-08 | 5.9e-08 | 5.900E-08 | 1.900E-08 | 4.000E-08 | 7.800E-08 |
| 21428 | 1072 | S1072G | D | 5.2e-08 | 1.1e-07 | 3.4e-08 | 6.533E-08 | 3.972E-08 | 3.400E-08 | 1.100E-07 |
| 21429 | 1072 | S1072H | D | 5.1e-08 | 3e-08 | 9.6e-08 | 5.900E-08 | 3.372E-08 | 3.000E-08 | 9.600E-08 |
| 21430 | 1072 | S1072I | D | 6.8e-08 | 3.6e-08 | 6.6e-08 | 5.667E-08 | 1.793E-08 | 3.600E-08 | 6.800E-08 |
| 21431 | 1072 | S1072L | D | 9.1e-08 | 7.5e-08 | 1.6e-08 | 6.067E-08 | 3.950E-08 | 1.600E-08 | 9.100E-08 |
| 21432 | 1072 | S1072K | D | 4.4e-08 | 4.7e-08 | 4.3e-08 | 4.467E-08 | 2.082E-09 | 4.300E-08 | 4.700E-08 |
| 21433 | 1072 | S1072M | D | 3.8e-08 | 1.1e-07 | 3.5e-08 | 6.100E-08 | 4.246E-08 | 3.500E-08 | 1.100E-07 |
| 21434 | 1072 | S1072F | D | 4.8e-08 | 1.1e-07 | 8.3e-08 | 8.033E-08 | 3.109E-08 | 4.800E-08 | 1.100E-07 |
| 21435 | 1072 | S1072P | D | 8.6e-08 | 9.3e-08 | 3.5e-08 | 7.133E-08 | 3.166E-08 | 3.500E-08 | 9.300E-08 |
| 21436 | 1072 | S1072S | D | 9.6e-08 | 3.6e-08 | 6.5e-08 | 6.567E-08 | 3.001E-08 | 3.600E-08 | 9.600E-08 |
| 21437 | 1072 | S1072T | D | 1.2e-08 | 4e-08 | 2.8e-08 | 2.667E-08 | 1.405E-08 | 1.200E-08 | 4.000E-08 |
| 21438 | 1072 | S1072W | D | 9.3e-08 | 5.1e-08 | 3.4e-08 | 5.933E-08 | 3.037E-08 | 3.400E-08 | 9.300E-08 |
| 21439 | 1072 | S1072Y | D | 1.1e-08 | 5.8e-08 | 1.5e-08 | 2.800E-08 | 2.606E-08 | 1.100E-08 | 5.800E-08 |
| 21440 | 1072 | S1072V | D | 8.5e-08 | 3.7e-08 | 2.5e-08 | 4.900E-08 | 3.175E-08 | 2.500E-08 | 8.500E-08 |
| 21441 | 1073 | V1073A | D | 1.5e-08 | 1.5e-07 | 8.5e-09 | 5.783E-08 | 7.988E-08 | 8.500E-09 | 1.500E-07 |
| 21442 | 1073 | V1073R | D | 9.5e-08 | 7.7e-08 | 1.5e-08 | 6.233E-08 | 4.197E-08 | 1.500E-08 | 9.500E-08 |
| 21443 | 1073 | V1073N | D | 1.8e-08 | 3.5e-08 | 3.5e-08 | 2.933E-08 | 9.815E-09 | 1.800E-08 | 3.500E-08 |
| 21444 | 1073 | V1073D | D | 2.4e-08 | 6.9e-08 | 1.4e-07 | 7.767E-08 | 5.848E-08 | 2.400E-08 | 1.400E-07 |
| 21445 | 1073 | V1073C | D | 1.1e-07 | 2e-08 | 4.5e-08 | 5.833E-08 | 4.646E-08 | 2.000E-08 | 1.100E-07 |
| 21446 | 1073 | V1073E | D | 7.5e-08 | 4.2e-08 | 9.5e-08 | 7.067E-08 | 2.676E-08 | 4.200E-08 | 9.500E-08 |
| 21447 | 1073 | V1073Q | D | 1.9e-08 | 5.2e-08 | 1.8e-08 | 2.967E-08 | 1.935E-08 | 1.800E-08 | 5.200E-08 |
| 21448 | 1073 | V1073G | D | 8.3e-08 | 2.8e-08 | 2.3e-08 | 4.467E-08 | 3.329E-08 | 2.300E-08 | 8.300E-08 |
| 21449 | 1073 | V1073H | D | 7.9e-08 | 2.6e-08 | 4.8e-08 | 5.100E-08 | 2.663E-08 | 2.600E-08 | 7.900E-08 |
| 21450 | 1073 | V1073I | D | 3.4e-08 | 6e-08 | 2.7e-08 | 4.033E-08 | 1.739E-08 | 2.700E-08 | 6.000E-08 |
| 21451 | 1073 | V1073L | D | 5.9e-08 | 5.2e-08 | 3.1e-08 | 4.733E-08 | 1.457E-08 | 3.100E-08 | 5.900E-08 |
| 21452 | 1073 | V1073K | D | 4.6e-08 | 1.9e-08 | 6.2e-08 | 4.233E-08 | 2.173E-08 | 1.900E-08 | 6.200E-08 |
| 21453 | 1073 | V1073M | D | 4.6e-08 | 6.9e-08 | 8.6e-08 | 6.700E-08 | 2.007E-08 | 4.600E-08 | 8.600E-08 |
| 21454 | 1073 | V1073F | D | 3.2e-08 | 7.2e-08 | 1.8e-08 | 4.067E-08 | 2.802E-08 | 1.800E-08 | 7.200E-08 |
| 21455 | 1073 | V1073P | D | 5.2e-08 | 7.1e-08 | 5.6e-08 | 5.967E-08 | 1.002E-08 | 5.200E-08 | 7.100E-08 |
| 21456 | 1073 | V1073S | D | 3.1e-08 | 2.4e-08 | 3.3e-08 | 2.933E-08 | 4.726E-09 | 2.400E-08 | 3.300E-08 |
| 21457 | 1073 | V1073T | D | 3.1e-08 | 1.5e-08 | 4.6e-08 | 3.067E-08 | 1.550E-08 | 1.500E-08 | 4.600E-08 |
| 21458 | 1073 | V1073W | D | 2.3e-08 | 3.8e-08 | 4.3e-08 | 3.467E-08 | 1.041E-08 | 2.300E-08 | 4.300E-08 |
| 21459 | 1073 | V1073Y | D | 4.8e-08 | 3.5e-08 | 4.5e-08 | 4.267E-08 | 6.807E-09 | 3.500E-08 | 4.800E-08 |
| 21460 | 1073 | V1073V | D | 2.5e-08 | 3.5e-08 | 1.2e-07 | 6.000E-08 | 5.220E-08 | 2.500E-08 | 1.200E-07 |
| 21461 | 1074 | G1074A | D | 7.3e-08 | 3.4e-08 | 4.3e-08 | 5.000E-08 | 2.042E-08 | 3.400E-08 | 7.300E-08 |
| 21462 | 1074 | G1074R | D | 4.7e-08 | 2e-08 | 2.5e-08 | 3.067E-08 | 1.436E-08 | 2.000E-08 | 4.700E-08 |
| 21463 | 1074 | G1074N | D | 3.2e-08 | 4.1e-08 | 1.9e-08 | 3.067E-08 | 1.106E-08 | 1.900E-08 | 4.100E-08 |
| 21464 | 1074 | G1074D | D | 4.1e-08 | 2.6e-08 | 4.9e-08 | 3.867E-08 | 1.168E-08 | 2.600E-08 | 4.900E-08 |
| 21465 | 1074 | G1074C | D | 2e-08 | 5.7e-08 | 4.8e-08 | 4.167E-08 | 1.930E-08 | 2.000E-08 | 5.700E-08 |
| 21466 | 1074 | G1074E | D | 8.9e-08 | 1.1e-08 | 2.3e-08 | 4.100E-08 | 4.200E-08 | 1.100E-08 | 8.900E-08 |
| 21467 | 1074 | G1074Q | D | 4.7e-08 | 1.1e-07 | 5.2e-08 | 6.967E-08 | 3.502E-08 | 4.700E-08 | 1.100E-07 |
| 21468 | 1074 | G1074G | D | 1.2e-08 | 2e-08 | 2.2e-08 | 1.800E-08 | 5.292E-09 | 1.200E-08 | 2.200E-08 |
| 21469 | 1074 | G1074H | D | 6.4e-08 | 4.7e-08 | 1.3e-08 | 4.133E-08 | 2.597E-08 | 1.300E-08 | 6.400E-08 |
| 21470 | 1074 | G1074I | D | 7.5e-08 | 5.6e-08 | 5.4e-08 | 6.167E-08 | 1.159E-08 | 5.400E-08 | 7.500E-08 |
| 21471 | 1074 | G1074L | D | 3e-08 | 2.9e-08 | 3.5e-08 | 3.133E-08 | 3.215E-09 | 2.900E-08 | 3.500E-08 |
| 21472 | 1074 | G1074K | D | 3.7e-08 | 4.8e-08 | 7.4e-08 | 5.300E-08 | 1.900E-08 | 3.700E-08 | 7.400E-08 |
| 21473 | 1074 | G1074M | D | 4e-08 | 2.6e-08 | 5e-08 | 3.867E-08 | 1.206E-08 | 2.600E-08 | 5.000E-08 |
| 21474 | 1074 | G1074F | D | 1.5e-08 | 2.3e-08 | 3.4e-08 | 2.400E-08 | 9.539E-09 | 1.500E-08 | 3.400E-08 |
| 21475 | 1074 | G1074P | D | 3e-08 | 1e-07 | 8.9e-08 | 7.300E-08 | 3.764E-08 | 3.000E-08 | 1.000E-07 |
| 21476 | 1074 | G1074S | D | 3.6e-08 | 5.5e-08 | 3.5e-08 | 4.200E-08 | 1.127E-08 | 3.500E-08 | 5.500E-08 |
| 21477 | 1074 | G1074T | D | 4.8e-08 | 1.1e-08 | 7.4e-08 | 4.433E-08 | 3.166E-08 | 1.100E-08 | 7.400E-08 |
| 21478 | 1074 | G1074W | D | 9.1e-09 | 3.3e-08 | 6.4e-08 | 3.537E-08 | 2.753E-08 | 9.100E-09 | 6.400E-08 |
| 21479 | 1074 | G1074Y | D | 4.9e-08 | 3.6e-08 | 3.8e-08 | 4.100E-08 | 7.000E-09 | 3.600E-08 | 4.900E-08 |
| 21480 | 1074 | G1074V | D | 2.5e-08 | 3e-08 | 3.9e-08 | 3.133E-08 | 7.095E-09 | 2.500E-08 | 3.900E-08 |
| 21481 | 1075 | D1075A | D | 6.4e-08 | 2.7e-08 | 2.4e-08 | 3.833E-08 | 2.228E-08 | 2.400E-08 | 6.400E-08 |
| 21482 | 1075 | D1075R | D | 8.6e-08 | 4.1e-08 | 7.7e-08 | 6.800E-08 | 2.381E-08 | 4.100E-08 | 8.600E-08 |
| 21483 | 1075 | D1075N | D | 2.5e-08 | 1.2e-07 | 4.1e-08 | 6.200E-08 | 5.086E-08 | 2.500E-08 | 1.200E-07 |
| 21484 | 1075 | D1075D | D | 2.8e-08 | 1.9e-07 | 5.5e-08 | 9.100E-08 | 8.679E-08 | 2.800E-08 | 1.900E-07 |
| 21485 | 1075 | D1075C | D | 5.9e-08 | 2.6e-08 | 4.2e-08 | 4.233E-08 | 1.650E-08 | 2.600E-08 | 5.900E-08 |
| 21486 | 1075 | D1075E | D | 4.1e-08 | 3.2e-08 | 9e-08 | 5.433E-08 | 3.121E-08 | 3.200E-08 | 9.000E-08 |
| 21487 | 1075 | D1075Q | D | 4.5e-08 | 7.4e-08 | 2.4e-08 | 4.767E-08 | 2.511E-08 | 2.400E-08 | 7.400E-08 |
| 21488 | 1075 | D1075G | D | 3.5e-08 | 1.3e-07 | 1.9e-08 | 6.133E-08 | 6.000E-08 | 1.900E-08 | 1.300E-07 |
| 21489 | 1075 | D1075H | D | 1.5e-07 | 5.6e-08 | 4.7e-08 | 8.433E-08 | 5.705E-08 | 4.700E-08 | 1.500E-07 |
| 21490 | 1075 | D1075I | D | 6.1e-08 | 3.8e-08 | 3.7e-08 | 4.533E-08 | 1.358E-08 | 3.700E-08 | 6.100E-08 |
| 21491 | 1075 | D1075L | D | 3.3e-08 | 4.4e-08 | 7.7e-08 | 5.133E-08 | 2.290E-08 | 3.300E-08 | 7.700E-08 |
| 21492 | 1075 | D1075K | D | 7.2e-08 | 7.2e-08 | 2.7e-08 | 5.700E-08 | 2.598E-08 | 2.700E-08 | 7.200E-08 |
| 21493 | 1075 | D1075M | D | 3.5e-08 | 4.1e-08 | 5.3e-08 | 4.300E-08 | 9.165E-09 | 3.500E-08 | 5.300E-08 |
| 21494 | 1075 | D1075F | D | 9.6e-08 | 4.1e-08 | 2.3e-08 | 5.333E-08 | 3.803E-08 | 2.300E-08 | 9.600E-08 |
| 21495 | 1075 | D1075P | D | 2.9e-08 | 4.9e-08 | 7e-08 | 4.933E-08 | 2.050E-08 | 2.900E-08 | 7.000E-08 |
| 21496 | 1075 | D1075S | D | 1.1e-08 | 5.5e-08 | 3.3e-08 | 3.300E-08 | 2.200E-08 | 1.100E-08 | 5.500E-08 |
| 21497 | 1075 | D1075T | D | 2.1e-07 | 2.1e-08 | 4.2e-08 | 9.100E-08 | 1.036E-07 | 2.100E-08 | 2.100E-07 |
| 21498 | 1075 | D1075W | D | 4.3e-08 | 2.4e-08 | 1.8e-08 | 2.833E-08 | 1.305E-08 | 1.800E-08 | 4.300E-08 |
| 21499 | 1075 | D1075Y | D | 2e-08 | 2.4e-08 | 2.6e-08 | 2.333E-08 | 3.055E-09 | 2.000E-08 | 2.600E-08 |
| 21500 | 1075 | D1075V | D | 5.8e-08 | 3.8e-08 | 1.5e-08 | 3.700E-08 | 2.152E-08 | 1.500E-08 | 5.800E-08 |
| 21501 | 1076 | R1076A | D | 1.8e-08 | 3.9e-08 | 3.8e-08 | 3.167E-08 | 1.185E-08 | 1.800E-08 | 3.900E-08 |
| 21502 | 1076 | R1076R | D | 8e-08 | 3e-08 | 8.6e-09 | 3.953E-08 | 3.664E-08 | 8.600E-09 | 8.000E-08 |
| 21503 | 1076 | R1076N | D | 3.2e-08 | 6.7e-08 | 1.9e-07 | 9.633E-08 | 8.298E-08 | 3.200E-08 | 1.900E-07 |
| 21504 | 1076 | R1076D | D | 3.9e-08 | 4.4e-08 | 2.6e-08 | 3.633E-08 | 9.292E-09 | 2.600E-08 | 4.400E-08 |

| | | | | | | | | | | |
|---|---|---|---|---|---|---|---|---|---|---|
| 21505 | 1076 | R1076C | D | 3.4e-08 | 3.4e-08 | 2.5e-08 | 3.100E-08 | 5.196E-09 | 2.500E-08 | 3.400E-08 |
| 21506 | 1076 | R1076E | D | 1.8e-08 | 1.3e-08 | 1.1e-08 | 1.400E-08 | 3.606E-09 | 1.100E-08 | 1.800E-08 |
| 21507 | 1076 | R1076Q | D | 3.3e-08 | 5.9e-08 | 9.3e-09 | 3.377E-08 | 2.486E-08 | 9.300E-09 | 5.900E-08 |
| 21508 | 1076 | R1076G | D | 3.5e-08 | 5.1e-08 | 1.2e-08 | 3.267E-08 | 1.960E-08 | 1.200E-08 | 5.100E-08 |
| 21509 | 1076 | R1076H | D | 4.1e-08 | 4.6e-08 | 1.9e-08 | 3.533E-08 | 1.436E-08 | 1.900E-08 | 4.600E-08 |
| 21510 | 1076 | R1076I | D | 2.6e-08 | 1.9e-08 | 2.6e-08 | 2.367E-08 | 4.041E-09 | 1.900E-08 | 2.600E-08 |
| 21511 | 1076 | R1076L | D | 1.1e-08 | 1.5e-08 | 4.2e-08 | 2.267E-08 | 1.686E-08 | 1.100E-08 | 4.200E-08 |
| 21512 | 1076 | R1076K | D | 1.2e-08 | 1e-07 | 8.1e-08 | 6.433E-08 | 4.631E-08 | 1.200E-08 | 1.000E-07 |
| 21513 | 1076 | R1076M | D | 2.9e-08 | 4.7e-08 | 1.2e-08 | 2.933E-08 | 1.750E-08 | 1.200E-08 | 4.700E-08 |
| 21514 | 1076 | R1076F | D | 1.5e-08 | 1.1e-07 | 3e-08 | 5.167E-08 | 5.107E-08 | 1.500E-08 | 1.100E-07 |
| 21515 | 1076 | R1076P | D | 3.8e-08 | 3.8e-08 | 9.8e-08 | 5.800E-08 | 3.464E-08 | 3.800E-08 | 9.800E-08 |
| 21516 | 1076 | R1076S | D | 5.1e-08 | 1.1e-07 | 5.9e-08 | 7.333E-08 | 3.201E-08 | 5.100E-08 | 1.100E-07 |
| 21517 | 1076 | R1076T | D | 2.2e-08 | 4e-08 | 8.7e-08 | 4.967E-08 | 3.356E-08 | 2.200E-08 | 8.700E-08 |
| 21518 | 1076 | R1076W | D | 1.9e-08 | 7.5e-08 | 2.5e-08 | 3.967E-08 | 3.075E-08 | 1.900E-08 | 7.500E-08 |
| 21519 | 1076 | R1076Y | D | 1.2e-07 | 2.6e-08 | 6.2e-08 | 6.933E-08 | 4.743E-08 | 2.600E-08 | 1.200E-07 |
| 21520 | 1076 | R1076V | D | 3.1e-08 | 6.8e-08 | 3.9e-08 | 4.600E-08 | 1.947E-08 | 3.100E-08 | 6.800E-08 |
| 21521 | 1077 | V1077A | D | 3.6e-08 | 5.9e-08 | 3.7e-08 | 4.400E-08 | 1.300E-08 | 3.600E-08 | 5.900E-08 |
| 21522 | 1077 | V1077R | D | 3e-08 | 2.5e-08 | 9.3e-08 | 4.933E-08 | 3.790E-08 | 2.500E-08 | 9.300E-08 |
| 21523 | 1077 | V1077N | D | 3.1e-08 | 3.1e-08 | 1.8e-07 | 8.067E-08 | 8.603E-08 | 3.100E-08 | 1.800E-07 |
| 21524 | 1077 | V1077D | D | 2.7e-08 | 1.8e-07 | 2.1e-08 | 7.600E-08 | 9.012E-08 | 2.100E-08 | 1.800E-07 |
| 21525 | 1077 | V1077C | D | 3.2e-08 | 2.9e-08 | 5.5e-08 | 3.867E-08 | 1.422E-08 | 2.900E-08 | 5.500E-08 |
| 21526 | 1077 | V1077E | D | 2.4e-08 | 3.9e-08 | 4.2e-08 | 3.500E-08 | 9.644E-09 | 2.400E-08 | 4.200E-08 |
| 21527 | 1077 | V1077Q | D | 3.3e-08 | 2.9e-08 | 1.8e-08 | 2.667E-08 | 7.767E-09 | 1.800E-08 | 3.300E-08 |
| 21528 | 1077 | V1077G | D | 4.8e-08 | 4.8e-08 | 7.1e-08 | 5.567E-08 | 1.328E-08 | 4.800E-08 | 7.100E-08 |
| 21529 | 1077 | V1077H | D | 4.4e-07 | 3e-08 | 4.2e-08 | 1.707E-07 | 2.333E-07 | 3.000E-08 | 4.400E-07 |
| 21530 | 1077 | V1077I | D | 5.6e-08 | 1.7e-08 | 3.7e-08 | 3.667E-08 | 1.950E-08 | 1.700E-08 | 5.600E-08 |
| 21531 | 1077 | V1077L | D | 3.3e-08 | 3.1e-08 | 5e-08 | 3.800E-08 | 1.044E-08 | 3.100E-08 | 5.000E-08 |
| 21532 | 1077 | V1077K | D | 3.3e-08 | 5.6e-08 | 2.9e-08 | 3.933E-08 | 1.457E-08 | 2.900E-08 | 5.600E-08 |
| 21533 | 1077 | V1077M | D | 1.4e-08 | 4.7e-08 | 3.2e-08 | 3.100E-08 | 1.652E-08 | 1.400E-08 | 4.700E-08 |
| 21534 | 1077 | V1077F | D | 3.3e-08 | 4.2e-08 | 4.5e-08 | 4.000E-08 | 6.245E-09 | 3.300E-08 | 4.500E-08 |
| 21535 | 1077 | V1077P | D | 4.4e-08 | 2.1e-08 | 4.3e-08 | 3.600E-08 | 1.300E-08 | 2.100E-08 | 4.400E-08 |
| 21536 | 1077 | V1077S | D | 9.3e-08 | 5.6e-08 | 3.5e-08 | 6.133E-08 | 2.937E-08 | 3.500E-08 | 9.300E-08 |
| 21537 | 1077 | V1077T | D | 6.6e-08 | 4.3e-08 | 3.2e-08 | 4.700E-08 | 1.735E-08 | 3.200E-08 | 6.600E-08 |
| 21538 | 1077 | V1077W | D | 3.8e-08 | 5.7e-08 | 2.3e-08 | 3.933E-08 | 1.704E-08 | 2.300E-08 | 5.700E-08 |
| 21539 | 1077 | V1077Y | D | 2.8e-08 | 1.2e-08 | 3e-08 | 2.333E-08 | 9.866E-09 | 1.200E-08 | 3.000E-08 |
| 21540 | 1077 | V1077V | D | 2.5e-08 | 9.9e-09 | 3.6e-08 | 2.363E-08 | 1.310E-08 | 9.900E-09 | 3.600E-08 |
| 21541 | 1078 | T1078A | D | 5.4e-08 | 6.3e-08 | 2.1e-08 | 4.600E-08 | 2.211E-08 | 2.100E-08 | 6.300E-08 |
| 21542 | 1078 | T1078R | D | 1.5e-07 | 2.6e-08 | 2.2e-08 | 6.600E-08 | 7.277E-08 | 2.200E-08 | 1.500E-07 |
| 21543 | 1078 | T1078N | D | 2.2e-08 | 5.1e-08 | 3.8e-08 | 3.700E-08 | 1.453E-08 | 2.200E-08 | 5.100E-08 |
| 21544 | 1078 | T1078D | D | 2.8e-08 | 4e-08 | 4.1e-08 | 3.633E-08 | 7.234E-09 | 2.800E-08 | 4.100E-08 |
| 21545 | 1078 | T1078C | D | 7.5e-08 | 3.2e-08 | 3.3e-08 | 4.667E-08 | 2.454E-08 | 3.200E-08 | 7.500E-08 |
| 21546 | 1078 | T1078E | D | 7.3e-08 | 9.1e-08 | 2.4e-08 | 6.267E-08 | 3.467E-08 | 2.400E-08 | 9.100E-08 |
| 21547 | 1078 | T1078Q | D | 4e-08 | 2.7e-08 | 3.7e-08 | 3.467E-08 | 6.807E-09 | 2.700E-08 | 4.000E-08 |
| 21548 | 1078 | T1078G | D | 3.6e-08 | 1.2e-08 | 3.1e-08 | 2.633E-08 | 1.266E-08 | 1.200E-08 | 3.600E-08 |
| 21549 | 1078 | T1078H | D | 1.5e-08 | 3e-08 | 9.2e-08 | 4.567E-08 | 4.082E-08 | 1.500E-08 | 9.200E-08 |
| 21550 | 1078 | T1078I | D | 2e-08 | 1.5e-08 | 5.6e-08 | 3.033E-08 | 2.237E-08 | 1.500E-08 | 5.600E-08 |
| 21551 | 1078 | T1078L | D | 3.8e-08 | 6.5e-08 | 5.8e-08 | 5.367E-08 | 1.401E-08 | 3.800E-08 | 6.500E-08 |
| 21552 | 1078 | T1078K | D | 4.5e-08 | 8.5e-08 | 4.2e-08 | 5.733E-08 | 2.401E-08 | 4.200E-08 | 8.500E-08 |
| 21553 | 1078 | T1078M | D | 3.6e-08 | 9.5e-08 | 4.6e-08 | 5.900E-08 | 3.158E-08 | 3.600E-08 | 9.500E-08 |
| 21554 | 1078 | T1078F | D | 3.4e-08 | 5e-08 | 6.2e-08 | 4.867E-08 | 1.405E-08 | 3.400E-08 | 6.200E-08 |
| 21555 | 1078 | T1078P | D | 4.1e-08 | 3.2e-08 | 6e-08 | 4.433E-08 | 1.429E-08 | 3.200E-08 | 6.000E-08 |
| 21556 | 1078 | T1078S | D | 6.4e-08 | 6.6e-08 | 2.8e-08 | 5.267E-08 | 2.139E-08 | 2.800E-08 | 6.600E-08 |
| 21557 | 1078 | T1078T | D | 2.6e-08 | 4.2e-08 | 4.3e-08 | 3.700E-08 | 9.539E-09 | 2.600E-08 | 4.300E-08 |
| 21558 | 1078 | T1078W | D | 5.2e-08 | 5.9e-08 | 5e-08 | 5.367E-08 | 4.726E-09 | 5.000E-08 | 5.900E-08 |
| 21559 | 1078 | T1078Y | D | 4.5e-08 | 5.2e-08 | 2.4e-08 | 4.033E-08 | 1.457E-08 | 2.400E-08 | 5.200E-08 |
| 21560 | 1078 | T1078V | D | 7.7e-08 | 3.4e-08 | 5.4e-08 | 5.500E-08 | 2.152E-08 | 3.400E-08 | 7.700E-08 |
| 21561 | 1079 | I1079A | D | 4.3e-08 | 6.8e-08 | 2.7e-08 | 4.600E-08 | 2.066E-08 | 2.700E-08 | 6.800E-08 |
| 21562 | 1079 | I1079R | D | 4.7e-08 | 2e-08 | 9.4e-08 | 5.367E-08 | 3.745E-08 | 2.000E-08 | 9.400E-08 |
| 21563 | 1079 | I1079N | D | 2.4e-08 | 6.4e-08 | 4.4e-08 | 4.400E-08 | 2.000E-08 | 2.400E-08 | 6.400E-08 |
| 21564 | 1079 | I1079D | D | 2.8e-08 | 3.6e-08 | 2.8e-08 | 3.067E-08 | 4.619E-09 | 2.800E-08 | 3.600E-08 |
| 21565 | 1079 | I1079C | D | 6e-08 | 3e-08 | 2e-08 | 3.667E-08 | 2.082E-08 | 2.000E-08 | 6.000E-08 |
| 21566 | 1079 | I1079E | D | 1.3e-08 | 3.1e-08 | 4.5e-08 | 2.967E-08 | 1.604E-08 | 1.300E-08 | 4.500E-08 |
| 21567 | 1079 | I1079Q | D | 1.3e-08 | 3.6e-08 | 1.5e-08 | 2.133E-08 | 1.274E-08 | 1.300E-08 | 3.600E-08 |
| 21568 | 1079 | I1079G | D | 4e-08 | 4.4e-08 | 5.3e-08 | 4.567E-08 | 6.658E-09 | 4.000E-08 | 5.300E-08 |
| 21569 | 1079 | I1079H | D | 2.2e-08 | 4e-08 | 3e-08 | 3.067E-08 | 9.018E-09 | 2.200E-08 | 4.000E-08 |
| 21570 | 1079 | I1079I | D | 8.2e-08 | 2.6e-08 | 7e-08 | 5.933E-08 | 2.948E-08 | 2.600E-08 | 8.200E-08 |
| 21571 | 1079 | I1079L | D | 7.9e-08 | 7.7e-08 | 7.6e-08 | 7.733E-08 | 1.528E-09 | 7.600E-08 | 7.900E-08 |
| 21572 | 1079 | I1079K | D | 4.5e-08 | 9.9e-08 | 7.9e-08 | 7.433E-08 | 2.730E-08 | 4.500E-08 | 9.900E-08 |
| 21573 | 1079 | I1079M | D | 4.2e-08 | 2e-08 | 3.9e-08 | 3.367E-08 | 1.193E-08 | 2.000E-08 | 4.200E-08 |
| 21574 | 1079 | I1079F | D | 3.8e-08 | 4.2e-08 | 2e-08 | 3.333E-08 | 1.172E-08 | 2.000E-08 | 4.200E-08 |
| 21575 | 1079 | I1079P | D | 9.8e-08 | 3.9e-08 | 4.6e-08 | 6.100E-08 | 3.223E-08 | 3.900E-08 | 9.800E-08 |
| 21576 | 1079 | I1079S | D | 8.7e-08 | 4.7e-08 | 1.6e-08 | 5.000E-08 | 3.559E-08 | 1.600E-08 | 8.700E-08 |
| 21577 | 1079 | I1079T | D | 3.5e-08 | 1.4e-08 | 2.8e-08 | 2.567E-08 | 1.069E-08 | 1.400E-08 | 3.500E-08 |
| 21578 | 1079 | I1079W | D | 4.3e-08 | 3.6e-08 | 7.6e-08 | 5.167E-08 | 2.136E-08 | 3.600E-08 | 7.600E-08 |
| 21579 | 1079 | I1079Y | D | 3.4e-08 | 6.5e-08 | 2.2e-08 | 4.033E-08 | 2.219E-08 | 2.200E-08 | 6.500E-08 |
| 21580 | 1079 | I1079V | D | 3.6e-08 | 6.8e-08 | 6.2e-08 | 5.533E-08 | 1.701E-08 | 3.600E-08 | 6.800E-08 |
| 21581 | 1080 | T1080A | D | 5e-08 | 1.2e-07 | 3e-08 | 6.667E-08 | 4.726E-08 | 3.000E-08 | 1.200E-07 |
| 21582 | 1080 | T1080R | D | 5.1e-08 | 2.5e-08 | 3.6e-08 | 3.733E-08 | 1.305E-08 | 2.500E-08 | 5.100E-08 |
| 21583 | 1080 | T1080N | D | 1.5e-07 | 2.6e-08 | 2.7e-07 | 1.487E-07 | 1.220E-07 | 2.600E-08 | 2.700E-07 |
| 21584 | 1080 | T1080D | D | 3.3e-08 | 2.3e-08 | 6.3e-08 | 3.967E-08 | 2.082E-08 | 2.300E-08 | 6.300E-08 |
| 21585 | 1080 | T1080C | D | 1.9e-08 | 1.9e-08 | 2.9e-08 | 2.233E-08 | 5.774E-09 | 1.900E-08 | 2.900E-08 |
| 21586 | 1080 | T1080E | D | 6.9e-08 | 5.2e-08 | 4.2e-08 | 5.433E-08 | 1.365E-08 | 4.200E-08 | 6.900E-08 |
| 21587 | 1080 | T1080Q | D | 5.4e-08 | 2.9e-08 | 9.2e-08 | 5.833E-08 | 3.172E-08 | 2.900E-08 | 9.200E-08 |
| 21588 | 1080 | T1080G | D | 3.3e-08 | 9.7e-08 | 2.4e-08 | 5.133E-08 | 3.980E-08 | 2.400E-08 | 9.700E-08 |
| 21589 | 1080 | T1080H | D | 9.1e-08 | 7.1e-08 | 4.1e-08 | 6.767E-08 | 2.517E-08 | 4.100E-08 | 9.100E-08 |

| | | | | | | | | | | |
|---|---|---|---|---|---|---|---|---|---|---|
| 21590 | 1080 | T1080I | D | 2.9e-08 | 5.5e-08 | 5.6e-08 | 4.667E-08 | 1.531E-08 | 2.900E-08 | 5.600E-08 |
| 21591 | 1080 | T1080L | D | 5.5e-08 | 6.3e-08 | 1e-07 | 7.267E-08 | 2.401E-08 | 5.500E-08 | 1.000E-07 |
| 21592 | 1080 | T1080K | D | 5.6e-08 | 7.6e-08 | 6.4e-08 | 6.533E-08 | 1.007E-08 | 5.600E-08 | 7.600E-08 |
| 21593 | 1080 | T1080M | D | 2.8e-08 | 3.8e-08 | 3.4e-08 | 3.333E-08 | 5.033E-09 | 2.800E-08 | 3.800E-08 |
| 21594 | 1080 | T1080F | D | 5.4e-08 | 3.7e-08 | 1.3e-07 | 7.367E-08 | 4.952E-08 | 3.700E-08 | 1.300E-07 |
| 21595 | 1080 | T1080P | D | 2.9e-08 | 1.6e-08 | 6.1e-08 | 3.533E-08 | 2.316E-08 | 1.600E-08 | 6.100E-08 |
| 21596 | 1080 | T1080S | D | 2.3e-08 | 3.1e-08 | 1.8e-08 | 2.400E-08 | 6.557E-09 | 1.800E-08 | 3.100E-08 |
| 21597 | 1080 | T1080T | D | 1.1e-08 | 2.9e-08 | 6.5e-08 | 3.500E-08 | 2.750E-08 | 1.100E-08 | 6.500E-08 |
| 21598 | 1080 | T1080W | D | 3.5e-08 | 4.3e-08 | 5.7e-08 | 4.500E-08 | 1.114E-08 | 3.500E-08 | 5.700E-08 |
| 21599 | 1080 | T1080Y | D | 4e-08 | 4.8e-08 | 2.4e-08 | 3.733E-08 | 1.222E-08 | 2.400E-08 | 4.800E-08 |
| 21600 | 1080 | T1080V | D | 4.2e-08 | 6.4e-08 | 8.4e-08 | 6.333E-08 | 2.101E-08 | 4.200E-08 | 8.400E-08 |
| 21601 | 1081 | C1081A | D | 9.5e-08 | 7.4e-08 | 3.6e-08 | 6.833E-08 | 2.991E-08 | 3.600E-08 | 9.500E-08 |
| 21602 | 1081 | C1081R | D | 2.9e-08 | 5.1e-08 | 3e-08 | 3.667E-08 | 1.242E-08 | 2.900E-08 | 5.100E-08 |
| 21603 | 1081 | C1081N | D | 6.7e-08 | 2.9e-08 | 2.3e-08 | 3.967E-08 | 2.386E-08 | 2.300E-08 | 6.700E-08 |
| 21604 | 1081 | C1081D | D | 1.1e-07 | 6.5e-08 | 2.5e-08 | 6.667E-08 | 4.252E-08 | 2.500E-08 | 1.100E-07 |
| 21605 | 1081 | C1081C | D | 2.9e-08 | 5e-08 | 3.1e-08 | 3.667E-08 | 1.159E-08 | 2.900E-08 | 5.000E-08 |
| 21606 | 1081 | C1081E | D | 4.3e-08 | 2.9e-08 | 2e-07 | 9.067E-08 | 9.494E-08 | 2.900E-08 | 2.000E-07 |
| 21607 | 1081 | C1081Q | D | 1.9e-08 | 5.3e-08 | 1.3e-07 | 6.733E-08 | 5.687E-08 | 1.900E-08 | 1.300E-07 |
| 21608 | 1081 | C1081G | D | 1.8e-08 | 3.2e-08 | 1.8e-08 | 2.267E-08 | 8.083E-09 | 1.800E-08 | 3.200E-08 |
| 21609 | 1081 | C1081H | D | 6.7e-08 | 3.2e-08 | 4.4e-08 | 4.767E-08 | 1.779E-08 | 3.200E-08 | 6.700E-08 |
| 21610 | 1081 | C1081I | D | 3.4e-08 | 9.4e-08 | 6.7e-08 | 6.500E-08 | 3.005E-08 | 3.400E-08 | 9.400E-08 |
| 21611 | 1081 | C1081L | D | 5.3e-08 | 8.9e-08 | 4.6e-08 | 6.267E-08 | 2.307E-08 | 4.600E-08 | 8.900E-08 |
| 21612 | 1081 | C1081K | D | 6.1e-08 | 9.3e-08 | 5.9e-08 | 7.100E-08 | 1.908E-08 | 5.900E-08 | 9.300E-08 |
| 21613 | 1081 | C1081M | D | 7.4e-08 | 4.4e-08 | 5.1e-08 | 5.633E-08 | 1.570E-08 | 4.400E-08 | 7.400E-08 |
| 21614 | 1081 | C1081F | D | 1.3e-07 | 3.2e-08 | 2.2e-08 | 6.133E-08 | 5.968E-08 | 2.200E-08 | 1.300E-07 |
| 21615 | 1081 | C1081P | D | 2.6e-08 | 1.5e-07 | 2.9e-08 | 6.833E-08 | 7.074E-08 | 2.600E-08 | 1.500E-07 |
| 21616 | 1081 | C1081S | D | 1.2e-07 | 4.9e-08 | 3.5e-08 | 6.800E-08 | 4.557E-08 | 3.500E-08 | 1.200E-07 |
| 21617 | 1081 | C1081T | D | 1.1e-07 | 8.4e-08 | 6.2e-08 | 8.533E-08 | 2.403E-08 | 6.200E-08 | 1.100E-07 |
| 21618 | 1081 | C1081W | D | 4.1e-08 | 8e-08 | 4.8e-08 | 5.633E-08 | 2.079E-08 | 4.100E-08 | 8.000E-08 |
| 21619 | 1081 | C1081Y | D | 3.2e-08 | 1.3e-08 | 3.9e-08 | 2.800E-08 | 1.345E-08 | 1.300E-08 | 3.900E-08 |
| 21620 | 1081 | C1081V | D | 1e-07 | 4.4e-08 | 1.2e-08 | 5.200E-08 | 4.454E-08 | 1.200E-08 | 1.000E-07 |
| 21621 | 1082 | R1082A | D | 1.5e-08 | 4.8e-08 | 3.4e-08 | 3.233E-08 | 1.656E-08 | 1.500E-08 | 4.800E-08 |
| 21622 | 1082 | R1082R | D | 9.6e-08 | 2e-08 | 7.2e-08 | 6.267E-08 | 3.885E-08 | 2.000E-08 | 9.600E-08 |
| 21623 | 1082 | R1082N | D | 1e-08 | 3.7e-08 | 7.1e-08 | 3.933E-08 | 3.057E-08 | 1.000E-08 | 7.100E-08 |
| 21624 | 1082 | R1082D | D | 6.2e-08 | 4.5e-08 | 4.2e-08 | 4.967E-08 | 1.079E-08 | 4.200E-08 | 6.200E-08 |
| 21625 | 1082 | R1082C | D | 4.2e-08 | 4.8e-08 | 3.9e-08 | 4.300E-08 | 4.583E-09 | 3.900E-08 | 4.800E-08 |
| 21626 | 1082 | R1082E | D | 3.8e-08 | 4.7e-08 | 4.9e-08 | 4.467E-08 | 5.859E-09 | 3.800E-08 | 4.900E-08 |
| 21627 | 1082 | R1082Q | D | 1.7e-08 | 7.8e-08 | 4.9e-08 | 4.800E-08 | 3.051E-08 | 1.700E-08 | 7.800E-08 |
| 21628 | 1082 | R1082G | D | 1.4e-08 | 2.4e-08 | 4.5e-08 | 2.767E-08 | 1.582E-08 | 1.400E-08 | 4.500E-08 |
| 21629 | 1082 | R1082H | D | 2.4e-08 | 6.3e-08 | 9e-08 | 5.900E-08 | 3.318E-08 | 2.400E-08 | 9.000E-08 |
| 21630 | 1082 | R1082I | D | 9.8e-08 | 2.9e-08 | 2.1e-08 | 4.933E-08 | 4.234E-08 | 2.100E-08 | 9.800E-08 |
| 21631 | 1082 | R1082L | D | 6.3e-08 | 2.8e-08 | 9.1e-08 | 6.067E-08 | 3.156E-08 | 2.800E-08 | 9.100E-08 |
| 21632 | 1082 | R1082K | D | 3.5e-08 | 6.7e-08 | 3.3e-08 | 4.500E-08 | 1.908E-08 | 3.300E-08 | 6.700E-08 |
| 21633 | 1082 | R1082M | D | 1e-07 | 5.5e-08 | 3.1e-08 | 6.200E-08 | 3.503E-08 | 3.100E-08 | 1.000E-07 |
| 21634 | 1082 | R1082F | D | 3.8e-08 | 2.9e-08 | 4.2e-08 | 3.633E-08 | 6.658E-09 | 2.900E-08 | 4.200E-08 |
| 21635 | 1082 | R1082P | D | 4e-08 | 2.3e-08 | 3.5e-08 | 3.267E-08 | 8.737E-09 | 2.300E-08 | 4.000E-08 |
| 21636 | 1082 | R1082S | D | 6.6e-08 | 4e-08 | 3.7e-08 | 4.767E-08 | 1.595E-08 | 3.700E-08 | 6.600E-08 |
| 21637 | 1082 | R1082T | D | 4.2e-08 | 1.5e-08 | 5.2e-08 | 3.633E-08 | 1.914E-08 | 1.500E-08 | 5.200E-08 |
| 21638 | 1082 | R1082W | D | 2.1e-08 | 1.1e-07 | 4.2e-08 | 5.767E-08 | 4.652E-08 | 2.100E-08 | 1.100E-07 |
| 21639 | 1082 | R1082Y | D | 3.8e-08 | 7.1e-08 | 2.6e-08 | 4.500E-08 | 2.330E-08 | 2.600E-08 | 7.100E-08 |
| 21640 | 1082 | R1082V | D | 6.3e-08 | 7.7e-09 | 3.4e-08 | 3.490E-08 | 2.766E-08 | 7.700E-09 | 6.300E-08 |
| 21641 | 1083 | A1083A | D | 2.8e-08 | 2.8e-08 | 2.7e-08 | 2.767E-08 | 5.774E-10 | 2.700E-08 | 2.800E-08 |
| 21642 | 1083 | A1083R | D | 4.8e-08 | 3.1e-08 | 6.6e-08 | 4.833E-08 | 1.750E-08 | 3.100E-08 | 6.600E-08 |
| 21643 | 1083 | A1083N | D | 9.4e-08 | 5e-08 | 3.8e-08 | 6.067E-08 | 2.948E-08 | 3.800E-08 | 9.400E-08 |
| 21644 | 1083 | A1083D | D | 3.8e-08 | 3.2e-08 | 1e-07 | 5.667E-08 | 3.765E-08 | 3.200E-08 | 1.000E-07 |
| 21645 | 1083 | A1083C | D | 4.5e-08 | 5.6e-08 | 7.3e-08 | 5.800E-08 | 1.411E-08 | 4.500E-08 | 7.300E-08 |
| 21646 | 1083 | A1083E | D | 6e-08 | 2.1e-08 | 4.6e-08 | 4.233E-08 | 1.976E-08 | 2.100E-08 | 6.000E-08 |
| 21647 | 1083 | A1083Q | D | 2.9e-08 | 1.1e-07 | 5.2e-08 | 6.367E-08 | 4.174E-08 | 2.900E-08 | 1.100E-07 |
| 21648 | 1083 | A1083G | D | 6.8e-08 | 3.3e-08 | 4.6e-08 | 4.900E-08 | 1.769E-08 | 3.300E-08 | 6.800E-08 |
| 21649 | 1083 | A1083H | D | 5e-08 | 2.2e-08 | 2.7e-08 | 3.300E-08 | 1.493E-08 | 2.200E-08 | 5.000E-08 |
| 21650 | 1083 | A1083I | D | 2.4e-08 | 4.9e-08 | 8.2e-08 | 5.167E-08 | 2.909E-08 | 2.400E-08 | 8.200E-08 |
| 21651 | 1083 | A1083L | D | 5.8e-08 | 3e-08 | 8.9e-08 | 5.900E-08 | 2.951E-08 | 3.000E-08 | 8.900E-08 |
| 21652 | 1083 | A1083K | D | 8e-08 | 7.7e-08 | 4.4e-08 | 6.700E-08 | 1.997E-08 | 4.400E-08 | 8.000E-08 |
| 21653 | 1083 | A1083M | D | 4.8e-08 | 1.7e-08 | 9.3e-08 | 5.267E-08 | 3.821E-08 | 1.700E-08 | 9.300E-08 |
| 21654 | 1083 | A1083F | D | 4.2e-08 | 3.4e-08 | 1e-07 | 5.867E-08 | 3.602E-08 | 3.400E-08 | 1.000E-07 |
| 21655 | 1083 | A1083P | D | 5e-08 | 3.3e-08 | 4.4e-08 | 4.233E-08 | 8.622E-09 | 3.300E-08 | 5.000E-08 |
| 21656 | 1083 | A1083S | D | 4.9e-08 | 6.5e-08 | 3.5e-08 | 4.967E-08 | 1.501E-08 | 3.500E-08 | 6.500E-08 |
| 21657 | 1083 | A1083T | D | 3.4e-08 | 1.5e-07 | 6.5e-08 | 8.300E-08 | 6.006E-08 | 3.400E-08 | 1.500E-07 |
| 21658 | 1083 | A1083W | D | 1.1e-07 | 9.6e-08 | 4.1e-08 | 8.233E-08 | 3.647E-08 | 4.100E-08 | 1.100E-07 |
| 21659 | 1083 | A1083Y | D | 4.4e-08 | 3.8e-08 | 2.5e-08 | 3.567E-08 | 9.713E-09 | 2.500E-08 | 4.400E-08 |
| 21660 | 1083 | A1083V | D | 9.3e-08 | 2e-08 | 5.4e-08 | 5.567E-08 | 3.653E-08 | 2.000E-08 | 9.300E-08 |
| 21661 | 1084 | S1084A | D | 6.3e-08 | 1.7e-07 | 5.3e-08 | 9.533E-08 | 6.486E-08 | 5.300E-08 | 1.700E-07 |
| 21662 | 1084 | S1084R | D | 4.9e-08 | 4.3e-08 | 7.8e-08 | 5.667E-08 | 1.872E-08 | 4.300E-08 | 7.800E-08 |
| 21663 | 1084 | S1084N | D | 3.5e-08 | 7.6e-08 | 4.8e-08 | 5.300E-08 | 2.095E-08 | 3.500E-08 | 7.600E-08 |
| 21664 | 1084 | S1084D | D | 3.4e-08 | 4e-08 | 3.7e-08 | 3.700E-08 | 3.000E-09 | 3.400E-08 | 4.000E-08 |
| 21665 | 1084 | S1084C | D | 7.8e-08 | 1.4e-07 | 2.6e-08 | 8.133E-08 | 5.707E-08 | 2.600E-08 | 1.400E-07 |
| 21666 | 1084 | S1084E | D | 5.2e-08 | 4e-08 | 3.8e-08 | 4.333E-08 | 7.572E-09 | 3.800E-08 | 5.200E-08 |
| 21667 | 1084 | S1084Q | D | 3.1e-08 | 2.3e-08 | 7.5e-08 | 4.300E-08 | 2.800E-08 | 2.300E-08 | 7.500E-08 |
| 21668 | 1084 | S1084G | D | 5.9e-08 | 1.9e-08 | 4.4e-08 | 4.067E-08 | 2.021E-08 | 1.900E-08 | 5.900E-08 |
| 21669 | 1084 | S1084H | D | 5.8e-08 | 4.4e-08 | 3.9e-08 | 4.700E-08 | 9.849E-09 | 3.900E-08 | 5.800E-08 |
| 21670 | 1084 | S1084I | D | 3.3e-08 | 1.3e-07 | 8.6e-08 | 8.300E-08 | 4.857E-08 | 3.300E-08 | 1.300E-07 |
| 21671 | 1084 | S1084L | D | 7.2e-08 | 1.2e-08 | 5.4e-08 | 4.600E-08 | 3.079E-08 | 1.200E-08 | 7.200E-08 |
| 21672 | 1084 | S1084K | D | 7.4e-08 | 2e-08 | 3.5e-08 | 4.300E-08 | 2.787E-08 | 2.000E-08 | 7.400E-08 |
| 21673 | 1084 | S1084M | D | 5.8e-08 | 2.2e-08 | 7.6e-08 | 5.200E-08 | 2.750E-08 | 2.200E-08 | 7.600E-08 |
| 21674 | 1084 | S1084F | D | 1.8e-08 | 6.9e-08 | 1.2e-08 | 3.300E-08 | 3.132E-08 | 1.200E-08 | 6.900E-08 |

| | | | | | | | | | | |
|---|---|---|---|---|---|---|---|---|---|---|
| 21675 | 1084 | S1084P | D | 1.9e-08 | 7.9e-08 | 4.4e-08 | 4.733E-08 | 3.014E-08 | 1.900E-08 | 7.900E-08 |
| 21676 | 1084 | S1084S | D | 9.4e-08 | 7.6e-08 | 1.1e-07 | 9.333E-08 | 1.701E-08 | 7.600E-08 | 1.100E-07 |
| 21677 | 1084 | S1084T | D | 1.2e-08 | 1.2e-08 | 5.8e-08 | 2.733E-08 | 2.656E-08 | 1.200E-08 | 5.800E-08 |
| 21678 | 1084 | S1084W | D | 2.6e-08 | 9.6e-08 | 4e-08 | 5.400E-08 | 3.704E-08 | 2.600E-08 | 9.600E-08 |
| 21679 | 1084 | S1084Y | D | 7.3e-08 | 7.5e-08 | 7.6e-08 | 7.467E-08 | 1.528E-09 | 7.300E-08 | 7.600E-08 |
| 21680 | 1084 | S1084V | D | 1.1e-07 | 8.9e-08 | 1.9e-07 | 1.297E-07 | 5.329E-08 | 8.900E-08 | 1.900E-07 |
| 21681 | 1085 | Q1085A | D | 1.2e-07 | 3.5e-08 | 4.4e-08 | 6.633E-08 | 4.669E-08 | 3.500E-08 | 1.200E-07 |
| 21682 | 1085 | Q1085R | D | 8.4e-08 | 5.2e-08 | 3.2e-08 | 5.600E-08 | 2.623E-08 | 3.200E-08 | 8.400E-08 |
| 21683 | 1085 | Q1085N | D | 3.8e-08 | 3.2e-08 | 7.4e-08 | 4.800E-08 | 2.272E-08 | 3.200E-08 | 7.400E-08 |
| 21684 | 1085 | Q1085D | D | 4e-08 | 6.4e-08 | 2.4e-08 | 4.267E-08 | 2.013E-08 | 2.400E-08 | 6.400E-08 |
| 21685 | 1085 | Q1085C | D | 2.3e-08 | 5.5e-08 | 5.8e-08 | 4.533E-08 | 1.940E-08 | 2.300E-08 | 5.800E-08 |
| 21686 | 1085 | Q1085E | D | 4.4e-08 | 4.1e-08 | 3e-08 | 3.833E-08 | 7.371E-09 | 3.000E-08 | 4.400E-08 |
| 21687 | 1085 | Q1085Q | D | 8.7e-08 | 2.9e-08 | 1.5e-07 | 8.867E-08 | 6.052E-08 | 2.900E-08 | 1.500E-07 |
| 21688 | 1085 | Q1085G | D | 1.3e-07 | 3.8e-08 | 3.1e-08 | 6.633E-08 | 5.525E-08 | 3.100E-08 | 1.300E-07 |
| 21689 | 1085 | Q1085H | D | 7.6e-08 | 1.5e-08 | 1.5e-08 | 3.533E-08 | 3.522E-08 | 1.500E-08 | 7.600E-08 |
| 21690 | 1085 | Q1085I | D | 2.2e-08 | 5.5e-08 | 7.2e-08 | 4.967E-08 | 2.542E-08 | 2.200E-08 | 7.200E-08 |
| 21691 | 1085 | Q1085L | D | 1.7e-08 | 4.9e-08 | 2.8e-08 | 3.133E-08 | 1.626E-08 | 1.700E-08 | 4.900E-08 |
| 21692 | 1085 | Q1085K | D | 2.6e-08 | 5.5e-08 | 1.7e-08 | 3.267E-08 | 1.986E-08 | 1.700E-08 | 5.500E-08 |
| 21693 | 1085 | Q1085M | D | 9.4e-08 | 9.1e-09 | 4.1e-08 | 4.803E-08 | 4.288E-08 | 9.100E-09 | 9.400E-08 |
| 21694 | 1085 | Q1085F | D | 2.1e-08 | 7.7e-08 | 1.1e-08 | 3.633E-08 | 3.557E-08 | 1.100E-08 | 7.700E-08 |
| 21695 | 1085 | Q1085P | D | 1e-07 | 2.2e-07 | 5.8e-08 | 1.260E-07 | 8.407E-08 | 5.800E-08 | 2.200E-07 |
| 21696 | 1085 | Q1085S | D | 2.1e-08 | 7.9e-08 | 1.6e-08 | 3.867E-08 | 3.502E-08 | 1.600E-08 | 7.900E-08 |
| 21697 | 1085 | Q1085T | D | 7.5e-08 | 4.8e-08 | 3.7e-08 | 5.333E-08 | 1.955E-08 | 3.700E-08 | 7.500E-08 |
| 21698 | 1085 | Q1085W | D | 4.8e-08 | 9.5e-08 | 4.1e-08 | 6.133E-08 | 2.937E-08 | 4.100E-08 | 9.500E-08 |
| 21699 | 1085 | Q1085Y | D | 8.5e-08 | 4.6e-08 | 2.6e-08 | 5.233E-08 | 3.001E-08 | 2.600E-08 | 8.500E-08 |
| 21700 | 1085 | Q1085V | D | 9.4e-08 | 2.3e-08 | 2.6e-08 | 4.767E-08 | 4.015E-08 | 2.300E-08 | 9.400E-08 |
| 21701 | 1086 | D1086A | D | 7.7e-08 | 8e-08 | 4.3e-08 | 6.667E-08 | 2.055E-08 | 4.300E-08 | 8.000E-08 |
| 21702 | 1086 | D1086R | D | 6.3e-08 | 1.7e-08 | 3.3e-08 | 3.767E-08 | 2.335E-08 | 1.700E-08 | 6.300E-08 |
| 21703 | 1086 | D1086N | D | 8.6e-08 | 7e-08 | 6.3e-08 | 7.300E-08 | 1.179E-08 | 6.300E-08 | 8.600E-08 |
| 21704 | 1086 | D1086D | D | 3.6e-08 | 9.9e-08 | 3.1e-08 | 5.533E-08 | 3.790E-08 | 3.100E-08 | 9.900E-08 |
| 21705 | 1086 | D1086C | D | 1.2e-07 | 4.6e-08 | 5.4e-08 | 7.333E-08 | 4.061E-08 | 4.600E-08 | 1.200E-07 |
| 21706 | 1086 | D1086E | D | 5.5e-08 | 9e-08 | 4.8e-08 | 6.433E-08 | 2.250E-08 | 4.800E-08 | 9.000E-08 |
| 21707 | 1086 | D1086Q | D | 6.5e-08 | 1.7e-08 | 3.1e-08 | 3.767E-08 | 2.468E-08 | 1.700E-08 | 6.500E-08 |
| 21708 | 1086 | D1086G | D | 3.5e-08 | 7.5e-08 | 1.5e-08 | 4.167E-08 | 3.055E-08 | 1.500E-08 | 7.500E-08 |
| 21709 | 1086 | D1086H | D | 5.4e-08 | 3.9e-08 | 3.4e-08 | 4.233E-08 | 1.041E-08 | 3.400E-08 | 5.400E-08 |
| 21710 | 1086 | D1086I | D | 4.4e-08 | 8.2e-08 | 2.1e-08 | 4.900E-08 | 3.081E-08 | 2.100E-08 | 8.200E-08 |
| 21711 | 1086 | D1086L | D | 3.4e-08 | 3.2e-08 | 4.2e-08 | 3.600E-08 | 5.292E-09 | 3.200E-08 | 4.200E-08 |
| 21712 | 1086 | D1086K | D | 2.6e-08 | 4.3e-08 | 2.5e-08 | 3.133E-08 | 1.012E-08 | 2.500E-08 | 4.300E-08 |
| 21713 | 1086 | D1086M | D | 8.4e-08 | 6.1e-08 | 6.8e-08 | 7.100E-08 | 1.179E-08 | 6.100E-08 | 8.400E-08 |
| 21714 | 1086 | D1086F | D | 3.2e-08 | 5e-08 | 5.5e-08 | 4.567E-08 | 1.210E-08 | 3.200E-08 | 5.500E-08 |
| 21715 | 1086 | D1086P | D | 5.6e-08 | 6e-08 | 6.4e-08 | 6.000E-08 | 4.000E-09 | 5.600E-08 | 6.400E-08 |
| 21716 | 1086 | D1086S | D | 3.7e-08 | 4.3e-08 | 3.1e-08 | 3.700E-08 | 6.000E-09 | 3.100E-08 | 4.300E-08 |
| 21717 | 1086 | D1086T | D | 2.5e-08 | 7.7e-08 | 1.5e-07 | 8.400E-08 | 6.279E-08 | 2.500E-08 | 1.500E-07 |
| 21718 | 1086 | D1086W | D | 5.1e-08 | 3e-08 | 5e-08 | 4.367E-08 | 1.185E-08 | 3.000E-08 | 5.100E-08 |
| 21719 | 1086 | D1086Y | D | 1.2e-07 | 2.7e-08 | 4.8e-08 | 6.500E-08 | 4.877E-08 | 2.700E-08 | 1.200E-07 |
| 21720 | 1086 | D1086V | D | 8.9e-08 | 1.6e-07 | 4.1e-08 | 9.667E-08 | 5.987E-08 | 4.100E-08 | 1.600E-07 |
| 21721 | 1087 | V1087A | D | 7.7e-08 | 5.5e-08 | 2.8e-08 | 5.333E-08 | 2.454E-08 | 2.800E-08 | 7.700E-08 |
| 21722 | 1087 | V1087R | D | 4.8e-08 | 2.9e-08 | 5.2e-08 | 4.300E-08 | 1.229E-08 | 2.900E-08 | 5.200E-08 |
| 21723 | 1087 | V1087N | D | 5e-08 | 3.6e-08 | 7e-08 | 5.200E-08 | 1.709E-08 | 3.600E-08 | 7.000E-08 |
| 21724 | 1087 | V1087D | D | 2.9e-08 | 4.9e-08 | 3.7e-08 | 3.833E-08 | 1.007E-08 | 2.900E-08 | 4.900E-08 |
| 21725 | 1087 | V1087C | D | 5.8e-08 | 2.6e-08 | 2.5e-08 | 3.633E-08 | 1.877E-08 | 2.500E-08 | 5.800E-08 |
| 21726 | 1087 | V1087E | D | 2e-08 | 5.1e-08 | 2.3e-08 | 3.133E-08 | 1.710E-08 | 2.000E-08 | 5.100E-08 |
| 21727 | 1087 | V1087Q | D | 1.4e-07 | 1.3e-08 | 4.1e-09 | 5.237E-08 | 7.602E-08 | 4.100E-09 | 1.400E-07 |
| 21728 | 1087 | V1087G | D | 3.4e-08 | 5.5e-08 | 5.1e-08 | 4.667E-08 | 1.115E-08 | 3.400E-08 | 5.500E-08 |
| 21729 | 1087 | V1087H | D | 5.9e-08 | 3.3e-08 | 5.9e-08 | 5.033E-08 | 1.501E-08 | 3.300E-08 | 5.900E-08 |
| 21730 | 1087 | V1087I | D | 2.8e-08 | 8.7e-08 | 1.8e-08 | 4.433E-08 | 3.729E-08 | 1.800E-08 | 8.700E-08 |
| 21731 | 1087 | V1087L | D | 8.2e-08 | 1.5e-08 | 4.2e-08 | 4.633E-08 | 3.371E-08 | 1.500E-08 | 8.200E-08 |
| 21732 | 1087 | V1087K | D | 3.3e-08 | 4.7e-08 | 4e-08 | 4.000E-08 | 7.000E-09 | 3.300E-08 | 4.700E-08 |
| 21733 | 1087 | V1087M | D | 2.9e-08 | 5.9e-08 | 2.8e-08 | 3.867E-08 | 1.762E-08 | 2.800E-08 | 5.900E-08 |
| 21734 | 1087 | V1087F | D | 2e-08 | 5.9e-08 | 2.2e-08 | 3.367E-08 | 2.196E-08 | 2.000E-08 | 5.900E-08 |
| 21735 | 1087 | V1087P | D | 3.4e-08 | 3e-08 | 4.2e-08 | 3.533E-08 | 6.110E-09 | 3.000E-08 | 4.200E-08 |
| 21736 | 1087 | V1087S | D | 1.9e-08 | 5.2e-08 | 6e-08 | 4.367E-08 | 2.173E-08 | 1.900E-08 | 6.000E-08 |
| 21737 | 1087 | V1087T | D | 1.7e-08 | 5.2e-08 | 2.8e-08 | 3.233E-08 | 1.790E-08 | 1.700E-08 | 5.200E-08 |
| 21738 | 1087 | V1087W | D | 4.8e-08 | 2.1e-08 | 5.7e-08 | 4.200E-08 | 1.873E-08 | 2.100E-08 | 5.700E-08 |
| 21739 | 1087 | V1087Y | D | 3.1e-08 | 7.4e-08 | 5.8e-08 | 5.433E-08 | 2.173E-08 | 3.100E-08 | 7.400E-08 |
| 21740 | 1087 | V1087V | D | 3.5e-08 | 1.8e-07 | 1.9e-08 | 7.800E-08 | 8.870E-08 | 1.900E-08 | 1.800E-07 |
| 21741 | 1088 | N1088A | D | 7.3e-08 | 4.5e-08 | 2.6e-08 | 4.800E-08 | 2.364E-08 | 2.600E-08 | 7.300E-08 |
| 21742 | 1088 | N1088R | D | 7.3e-08 | 7e-08 | 6.1e-08 | 6.800E-08 | 6.245E-09 | 6.100E-08 | 7.300E-08 |
| 21743 | 1088 | N1088N | D | 4.5e-08 | 4.5e-08 | 5e-08 | 4.667E-08 | 2.887E-09 | 4.500E-08 | 5.000E-08 |
| 21744 | 1088 | N1088D | D | 6.7e-08 | 7.2e-08 | 4.5e-08 | 6.133E-08 | 1.436E-08 | 4.500E-08 | 7.200E-08 |
| 21745 | 1088 | N1088C | D | 2e-08 | 9.1e-08 | 6.7e-08 | 5.933E-08 | 3.612E-08 | 2.000E-08 | 9.100E-08 |
| 21746 | 1088 | N1088E | D | 3.1e-08 | 8.9e-08 | 4.9e-08 | 5.633E-08 | 2.969E-08 | 3.100E-08 | 8.900E-08 |
| 21747 | 1088 | N1088Q | D | 6.5e-08 | 6.7e-08 | 1.3e-07 | 8.733E-08 | 3.696E-08 | 6.500E-08 | 1.300E-07 |
| 21748 | 1088 | N1088G | D | 5.2e-08 | 2.9e-08 | 8e-08 | 5.367E-08 | 2.554E-08 | 2.900E-08 | 8.000E-08 |
| 21749 | 1088 | N1088H | D | 1.7e-08 | 3.3e-08 | 5.4e-08 | 3.467E-08 | 1.856E-08 | 1.700E-08 | 5.400E-08 |
| 21750 | 1088 | N1088I | D | 4.2e-08 | 5.8e-08 | 2.3e-08 | 4.100E-08 | 1.752E-08 | 2.300E-08 | 5.800E-08 |
| 21751 | 1088 | N1088L | D | 1.8e-08 | 2.3e-08 | 1.1e-07 | 5.033E-08 | 5.173E-08 | 1.800E-08 | 1.100E-07 |
| 21752 | 1088 | N1088K | D | 8.9e-09 | 6e-08 | 3.8e-08 | 3.563E-08 | 2.563E-08 | 8.900E-09 | 6.000E-08 |
| 21753 | 1088 | N1088M | D | 3.5e-08 | 3.4e-08 | 9.4e-08 | 5.433E-08 | 3.436E-08 | 3.400E-08 | 9.400E-08 |
| 21754 | 1088 | N1088F | D | 3.5e-08 | 2.1e-08 | 4.2e-08 | 3.267E-08 | 1.069E-08 | 2.100E-08 | 4.200E-08 |
| 21755 | 1088 | N1088P | D | 1.8e-08 | 6.7e-08 | 7.5e-08 | 5.333E-08 | 3.086E-08 | 1.800E-08 | 7.500E-08 |
| 21756 | 1088 | N1088S | D | 3.6e-08 | 5.6e-08 | 3.4e-08 | 4.200E-08 | 1.217E-08 | 3.400E-08 | 5.600E-08 |
| 21757 | 1088 | N1088T | D | 4.3e-08 | 6.6e-08 | 7.2e-08 | 6.033E-08 | 1.531E-08 | 4.300E-08 | 7.200E-08 |
| 21758 | 1088 | N1088W | D | 5e-08 | 1e-07 | 6.2e-08 | 7.067E-08 | 2.610E-08 | 5.000E-08 | 1.000E-07 |
| 21759 | 1088 | N1088Y | D | 5.6e-08 | 1e-07 | 5.3e-08 | 6.967E-08 | 2.631E-08 | 5.300E-08 | 1.000E-07 |

| | | | | | | | | | | |
|---|---|---|---|---|---|---|---|---|---|---|
| 21760 | 1088 | N1088V | D | 8e-08 | 6.4e-08 | 4e-08 | 6.133E-08 | 2.013E-08 | 4.000E-08 | 8.000E-08 |
| 21761 | 1089 | T1089A | D | 3.9e-08 | 6.2e-08 | 4.2e-08 | 4.767E-08 | 1.250E-08 | 3.900E-08 | 6.200E-08 |
| 21762 | 1089 | T1089R | D | 5.2e-08 | 4.5e-08 | 1.5e-08 | 3.733E-08 | 1.966E-08 | 1.500E-08 | 5.200E-08 |
| 21763 | 1089 | T1089N | D | 4e-08 | 2.5e-08 | 4.9e-08 | 3.800E-08 | 1.212E-08 | 2.500E-08 | 4.900E-08 |
| 21764 | 1089 | T1089D | D | 7.3e-08 | 7.9e-08 | 4.5e-08 | 6.567E-08 | 1.815E-08 | 4.500E-08 | 7.900E-08 |
| 21765 | 1089 | T1089C | D | 2.4e-08 | 7e-08 | 2.1e-08 | 3.833E-08 | 2.747E-08 | 2.100E-08 | 7.000E-08 |
| 21766 | 1089 | T1089E | D | 3.9e-08 | 7.2e-08 | 3e-08 | 4.700E-08 | 2.211E-08 | 3.000E-08 | 7.200E-08 |
| 21767 | 1089 | T1089Q | D | 7.8e-08 | 1.8e-08 | 2.8e-08 | 4.133E-08 | 3.215E-08 | 1.800E-08 | 7.800E-08 |
| 21768 | 1089 | T1089G | D | 5.4e-08 | 1.6e-08 | 4.1e-08 | 3.700E-08 | 1.931E-08 | 1.600E-08 | 5.400E-08 |
| 21769 | 1089 | T1089H | D | 8.1e-08 | 1.6e-08 | 1.9e-08 | 3.867E-08 | 3.669E-08 | 1.600E-08 | 8.100E-08 |
| 21770 | 1089 | T1089I | D | 2.4e-08 | 2.9e-08 | 4.7e-08 | 3.333E-08 | 1.210E-08 | 2.400E-08 | 4.700E-08 |
| 21771 | 1089 | T1089L | D | 2.4e-08 | 5.6e-08 | 4.4e-08 | 4.133E-08 | 1.617E-08 | 2.400E-08 | 5.600E-08 |
| 21772 | 1089 | T1089K | D | 3.5e-08 | 1e-07 | 2.4e-08 | 5.300E-08 | 4.107E-08 | 2.400E-08 | 1.000E-07 |
| 21773 | 1089 | T1089M | D | 3.1e-08 | 1.1e-07 | 4.4e-08 | 6.167E-08 | 4.236E-08 | 3.100E-08 | 1.100E-07 |
| 21774 | 1089 | T1089F | D | 4.6e-08 | 5e-08 | 2.4e-08 | 4.000E-08 | 1.400E-08 | 2.400E-08 | 5.000E-08 |
| 21775 | 1089 | T1089P | D | 2.8e-08 | 2.9e-08 | 1.2e-07 | 5.900E-08 | 5.283E-08 | 2.800E-08 | 1.200E-07 |
| 21776 | 1089 | T1089S | D | 1.4e-08 | 3.4e-08 | 6.2e-08 | 3.667E-08 | 2.411E-08 | 1.400E-08 | 6.200E-08 |
| 21777 | 1089 | T1089T | D | 1.2e-07 | 3.3e-08 | 1.3e-07 | 9.433E-08 | 5.335E-08 | 3.300E-08 | 1.300E-07 |
| 21778 | 1089 | T1089W | D | 7.5e-08 | 5.8e-08 | 4.4e-08 | 5.900E-08 | 1.552E-08 | 4.400E-08 | 7.500E-08 |
| 21779 | 1089 | T1089Y | D | 4.2e-08 | 1.1e-07 | 3.2e-08 | 6.133E-08 | 4.244E-08 | 3.200E-08 | 1.100E-07 |
| 21780 | 1089 | T1089V | D | 4.6e-08 | 4.3e-08 | 2.3e-08 | 3.733E-08 | 1.250E-08 | 2.300E-08 | 4.600E-08 |
| 21781 | 1090 | A1090A | D | 3.2e-08 | 3.5e-08 | 7.7e-08 | 4.800E-08 | 2.516E-08 | 3.200E-08 | 7.700E-08 |
| 21782 | 1090 | A1090R | D | 6.8e-08 | 9.4e-09 | 9.8e-08 | 5.847E-08 | 4.506E-08 | 9.400E-09 | 9.800E-08 |
| 21783 | 1090 | A1090N | D | 1.7e-08 | 8.8e-08 | 4.5e-08 | 5.000E-08 | 3.576E-08 | 1.700E-08 | 8.800E-08 |
| 21784 | 1090 | A1090D | D | 3.4e-08 | 2.8e-08 | 3.3e-08 | 3.167E-08 | 3.215E-09 | 2.800E-08 | 3.400E-08 |
| 21785 | 1090 | A1090C | D | 3.2e-08 | 7.4e-08 | 2.8e-08 | 4.467E-08 | 2.548E-08 | 2.800E-08 | 7.400E-08 |
| 21786 | 1090 | A1090E | D | 7.8e-08 | 1.4e-07 | 2.5e-08 | 8.100E-08 | 5.756E-08 | 2.500E-08 | 1.400E-07 |
| 21787 | 1090 | A1090Q | D | 7.5e-08 | 3.2e-08 | 1.2e-07 | 7.567E-08 | 4.400E-08 | 3.200E-08 | 1.200E-07 |
| 21788 | 1090 | A1090G | D | 5.3e-08 | 2.6e-08 | 1.1e-07 | 6.300E-08 | 4.288E-08 | 2.600E-08 | 1.100E-07 |
| 21789 | 1090 | A1090H | D | 3.9e-08 | 1.7e-08 | 2.3e-08 | 2.633E-08 | 1.137E-08 | 1.700E-08 | 3.900E-08 |
| 21790 | 1090 | A1090I | D | 2.5e-08 | 3.1e-08 | 7e-08 | 4.200E-08 | 2.443E-08 | 2.500E-08 | 7.000E-08 |
| 21791 | 1090 | A1090L | D | 6.9e-08 | 6.1e-08 | 3.2e-08 | 5.400E-08 | 1.947E-08 | 3.200E-08 | 6.900E-08 |
| 21792 | 1090 | A1090K | D | 5.1e-08 | 5.2e-08 | 2e-08 | 4.100E-08 | 1.819E-08 | 2.000E-08 | 5.200E-08 |
| 21793 | 1090 | A1090M | D | 1.5e-08 | 5.5e-08 | 8.1e-08 | 5.033E-08 | 3.325E-08 | 1.500E-08 | 8.100E-08 |
| 21794 | 1090 | A1090F | D | 6.4e-08 | 2.6e-08 | 1.4e-08 | 3.467E-08 | 2.610E-08 | 1.400E-08 | 6.400E-08 |
| 21795 | 1090 | A1090P | D | 2.8e-08 | 4.3e-08 | 6.5e-08 | 4.533E-08 | 1.861E-08 | 2.800E-08 | 6.500E-08 |
| 21796 | 1090 | A1090S | D | 2.2e-08 | 3.6e-08 | 8.4e-08 | 4.733E-08 | 3.252E-08 | 2.200E-08 | 8.400E-08 |
| 21797 | 1090 | A1090T | D | 3.7e-08 | 9.2e-08 | 4.7e-08 | 5.867E-08 | 2.930E-08 | 3.700E-08 | 9.200E-08 |
| 21798 | 1090 | A1090W | D | 3.8e-08 | 6.4e-08 | 3.5e-08 | 4.567E-08 | 1.595E-08 | 3.500E-08 | 6.400E-08 |
| 21799 | 1090 | A1090Y | D | 5.2e-08 | 6.5e-08 | 5.5e-08 | 5.733E-08 | 6.807E-09 | 5.200E-08 | 6.500E-08 |
| 21800 | 1090 | A1090V | D | 5.2e-08 | 7.6e-08 | 1.9e-08 | 4.900E-08 | 2.862E-08 | 1.900E-08 | 7.600E-08 |
| 21801 | 1091 | V1091A | D | 6.3e-08 | 6.6e-08 | 3e-08 | 5.300E-08 | 1.997E-08 | 3.000E-08 | 6.600E-08 |
| 21802 | 1091 | V1091R | D | 1.7e-08 | 1.1e-07 | 1.1e-07 | 7.900E-08 | 5.369E-08 | 1.700E-08 | 1.100E-07 |
| 21803 | 1091 | V1091N | D | 2.9e-08 | 4e-08 | 6.1e-08 | 4.333E-08 | 1.626E-08 | 2.900E-08 | 6.100E-08 |
| 21804 | 1091 | V1091D | D | 2.5e-08 | 1.4e-08 | 2.2e-08 | 2.033E-08 | 5.686E-09 | 1.400E-08 | 2.500E-08 |
| 21805 | 1091 | V1091C | D | 3.7e-08 | 3.5e-08 | 5.7e-08 | 4.300E-08 | 1.217E-08 | 3.500E-08 | 5.700E-08 |
| 21806 | 1091 | V1091E | D | 3.2e-08 | 3.9e-08 | 5.2e-08 | 4.100E-08 | 1.015E-08 | 3.200E-08 | 5.200E-08 |
| 21807 | 1091 | V1091Q | D | 5e-08 | 1.1e-07 | 5.8e-08 | 7.267E-08 | 3.258E-08 | 5.000E-08 | 1.100E-07 |
| 21808 | 1091 | V1091G | D | 3.9e-08 | 1.8e-08 | 2.6e-08 | 2.767E-08 | 1.060E-08 | 1.800E-08 | 3.900E-08 |
| 21809 | 1091 | V1091H | D | 6.7e-08 | 3.9e-08 | 5.2e-08 | 5.267E-08 | 1.401E-08 | 3.900E-08 | 6.700E-08 |
| 21810 | 1091 | V1091I | D | 1e-07 | 3.2e-08 | 1.1e-07 | 8.067E-08 | 4.244E-08 | 3.200E-08 | 1.100E-07 |
| 21811 | 1091 | V1091L | D | 3.7e-08 | 1.1e-07 | 6.8e-08 | 7.167E-08 | 3.664E-08 | 3.700E-08 | 1.100E-07 |
| 21812 | 1091 | V1091K | D | 9.9e-08 | 7.3e-08 | 5.2e-08 | 7.467E-08 | 2.354E-08 | 5.200E-08 | 9.900E-08 |
| 21813 | 1091 | V1091M | D | 2.1e-08 | 4.4e-08 | 2.5e-08 | 3.000E-08 | 1.229E-08 | 2.100E-08 | 4.400E-08 |
| 21814 | 1091 | V1091F | D | 8e-08 | 1.1e-07 | 3e-08 | 7.333E-08 | 4.041E-08 | 3.000E-08 | 1.100E-07 |
| 21815 | 1091 | V1091P | D | 8.1e-08 | 7.9e-08 | 2.8e-08 | 6.267E-08 | 3.004E-08 | 2.800E-08 | 8.100E-08 |
| 21816 | 1091 | V1091S | D | 9e-08 | 7.9e-08 | 2.8e-08 | 6.567E-08 | 3.308E-08 | 2.800E-08 | 9.000E-08 |
| 21817 | 1091 | V1091T | D | 5.3e-08 | 1.6e-08 | 2.7e-08 | 3.200E-08 | 1.900E-08 | 1.600E-08 | 5.300E-08 |
| 21818 | 1091 | V1091W | D | 5.5e-08 | 2.5e-08 | 5.6e-08 | 4.533E-08 | 1.762E-08 | 2.500E-08 | 5.600E-08 |
| 21819 | 1091 | V1091Y | D | 5.2e-08 | 4e-08 | 4.6e-08 | 4.600E-08 | 6.000E-09 | 4.000E-08 | 5.200E-08 |
| 21820 | 1091 | V1091V | D | 3.4e-08 | 2.8e-08 | 2.2e-08 | 2.800E-08 | 6.000E-09 | 2.200E-08 | 3.400E-08 |
| 21821 | 1092 | A1092A | D | 4e-08 | 3.8e-08 | 4.8e-08 | 4.200E-08 | 5.292E-09 | 3.800E-08 | 4.800E-08 |
| 21822 | 1092 | A1092R | D | 3.7e-08 | 5.2e-08 | 7.5e-08 | 5.467E-08 | 1.914E-08 | 3.700E-08 | 7.500E-08 |
| 21823 | 1092 | A1092N | D | 1.6e-08 | 1e-08 | 7.8e-08 | 3.467E-08 | 3.765E-08 | 1.000E-08 | 7.800E-08 |
| 21824 | 1092 | A1092D | D | 2.5e-08 | 4.6e-08 | 2e-08 | 3.033E-08 | 1.380E-08 | 2.000E-08 | 4.600E-08 |
| 21825 | 1092 | A1092C | D | 6.7e-08 | 6.1e-08 | 3.6e-08 | 5.467E-08 | 1.644E-08 | 3.600E-08 | 6.700E-08 |
| 21826 | 1092 | A1092E | D | 5e-08 | 9.6e-08 | 2e-07 | 1.153E-07 | 7.685E-08 | 5.000E-08 | 2.000E-07 |
| 21827 | 1092 | A1092Q | D | 2.1e-08 | 1.5e-08 | 1.9e-08 | 1.833E-08 | 3.055E-09 | 1.500E-08 | 2.100E-08 |
| 21828 | 1092 | A1092G | D | 3.3e-08 | 4.1e-08 | 6.7e-08 | 4.700E-08 | 1.778E-08 | 3.300E-08 | 6.700E-08 |
| 21829 | 1092 | A1092H | D | 1.4e-08 | 2.6e-08 | 8.8e-08 | 4.267E-08 | 3.972E-08 | 1.400E-08 | 8.800E-08 |
| 21830 | 1092 | A1092I | D | 2.7e-08 | 1.9e-08 | 3.3e-08 | 2.633E-08 | 7.024E-09 | 1.900E-08 | 3.300E-08 |
| 21831 | 1092 | A1092L | D | 7.6e-08 | 3.3e-08 | 1.9e-08 | 4.267E-08 | 2.970E-08 | 1.900E-08 | 7.600E-08 |
| 21832 | 1092 | A1092K | D | 7.6e-08 | 1.2e-07 | 6.6e-08 | 8.733E-08 | 2.873E-08 | 6.600E-08 | 1.200E-07 |
| 21833 | 1092 | A1092M | D | 8.2e-08 | 6.9e-08 | 3.8e-08 | 6.300E-08 | 2.261E-08 | 3.800E-08 | 8.200E-08 |
| 21834 | 1092 | A1092F | D | 1.9e-08 | 4.5e-08 | 1.1e-07 | 5.800E-08 | 4.687E-08 | 1.900E-08 | 1.100E-07 |
| 21835 | 1092 | A1092P | D | 2.9e-08 | 3.2e-08 | 3.4e-08 | 3.167E-08 | 2.517E-09 | 2.900E-08 | 3.400E-08 |
| 21836 | 1092 | A1092S | D | 3.9e-08 | 1.9e-08 | 7.3e-08 | 4.367E-08 | 2.730E-08 | 1.900E-08 | 7.300E-08 |
| 21837 | 1092 | A1092T | D | 5.1e-08 | 3.6e-08 | 3.5e-08 | 4.067E-08 | 8.963E-09 | 3.500E-08 | 5.100E-08 |
| 21838 | 1092 | A1092W | D | 5.5e-08 | 4.4e-08 | 3.4e-08 | 4.433E-08 | 1.050E-08 | 3.400E-08 | 5.500E-08 |
| 21839 | 1092 | A1092Y | D | 1.4e-07 | 4.9e-08 | 2.3e-08 | 7.067E-08 | 6.144E-08 | 2.300E-08 | 1.400E-07 |
| 21840 | 1092 | A1092V | D | 4.4e-08 | 8.6e-08 | 6.1e-08 | 6.367E-08 | 2.113E-08 | 4.400E-08 | 8.600E-08 |
| 21841 | 1093 | W1093A | D | 1.1e-07 | 4.6e-08 | 6.6e-08 | 7.400E-08 | 3.274E-08 | 4.600E-08 | 1.100E-07 |
| 21842 | 1093 | W1093R | D | 5.4e-08 | 3.3e-08 | 7.9e-08 | 5.533E-08 | 2.303E-08 | 3.300E-08 | 7.900E-08 |
| 21843 | 1093 | W1093N | D | 2.3e-08 | 1.9e-08 | 4.4e-08 | 2.867E-08 | 1.343E-08 | 1.900E-08 | 4.400E-08 |
| 21844 | 1093 | W1093D | D | 2.4e-08 | 2.5e-08 | 2.8e-08 | 2.567E-08 | 2.082E-09 | 2.400E-08 | 2.800E-08 |

| | | | | | | | | | | |
|---|---|---|---|---|---|---|---|---|---|---|
| 21845 | 1093 | W1093C | D | 4e-08 | 1.8e-08 | 2.9e-08 | 2.900E-08 | 1.100E-08 | 1.800E-08 | 4.000E-08 |
| 21846 | 1093 | W1093E | D | 1.6e-08 | 7.6e-08 | 6.7e-08 | 5.300E-08 | 3.236E-08 | 1.600E-08 | 7.600E-08 |
| 21847 | 1093 | W1093Q | D | 6.1e-08 | 5.2e-08 | 5.7e-08 | 5.667E-08 | 4.509E-09 | 5.200E-08 | 6.100E-08 |
| 21848 | 1093 | W1093G | D | 4.4e-08 | 2.6e-08 | 1e-07 | 5.667E-08 | 3.859E-08 | 2.600E-08 | 1.000E-07 |
| 21849 | 1093 | W1093H | D | 6.9e-08 | 4.6e-08 | 4.9e-08 | 5.467E-08 | 1.250E-08 | 4.600E-08 | 6.900E-08 |
| 21850 | 1093 | W1093I | D | 5.2e-08 | 2.9e-08 | 1.6e-08 | 3.233E-08 | 1.823E-08 | 1.600E-08 | 5.200E-08 |
| 21851 | 1093 | W1093L | D | 3.4e-08 | 5.5e-08 | 5.3e-08 | 4.733E-08 | 1.159E-08 | 3.400E-08 | 5.500E-08 |
| 21852 | 1093 | W1093K | D | 2.2e-08 | 5.2e-08 | 3.9e-08 | 3.767E-08 | 1.504E-08 | 2.200E-08 | 5.200E-08 |
| 21853 | 1093 | W1093M | D | 3.1e-08 | 2.4e-08 | 4.6e-08 | 3.367E-08 | 1.124E-08 | 2.400E-08 | 4.600E-08 |
| 21854 | 1093 | W1093F | D | 3.3e-08 | 3.5e-08 | 4.8e-08 | 3.867E-08 | 8.145E-09 | 3.300E-08 | 4.800E-08 |
| 21855 | 1093 | W1093P | D | 3.6e-08 | 7.4e-08 | 2e-08 | 4.333E-08 | 2.774E-08 | 2.000E-08 | 7.400E-08 |
| 21856 | 1093 | W1093S | D | 2.1e-08 | 6.9e-08 | 5.1e-08 | 4.700E-08 | 2.425E-08 | 2.100E-08 | 6.900E-08 |
| 21857 | 1093 | W1093T | D | 7.1e-08 | 3.9e-08 | 3.6e-08 | 4.867E-08 | 1.940E-08 | 3.600E-08 | 7.100E-08 |
| 21858 | 1093 | W1093W | D | 8e-08 | 8.8e-08 | 5.2e-08 | 7.333E-08 | 1.890E-08 | 5.200E-08 | 8.800E-08 |
| 21859 | 1093 | W1093Y | D | 3.7e-08 | 4.2e-08 | 5.2e-08 | 4.367E-08 | 7.638E-09 | 3.700E-08 | 5.200E-08 |
| 21860 | 1093 | W1093V | D | 8.9e-08 | 7.4e-08 | 4.2e-08 | 6.833E-08 | 2.401E-08 | 4.200E-08 | 8.900E-08 |
| 21861 | 1094 | Y1094A | D | 4.4e-08 | 3.8e-08 | 1.1e-08 | 3.100E-08 | 1.758E-08 | 1.100E-08 | 4.400E-08 |
| 21862 | 1094 | Y1094R | D | 5.9e-08 | 6e-08 | 1.9e-08 | 4.600E-08 | 2.339E-08 | 1.900E-08 | 6.000E-08 |
| 21863 | 1094 | Y1094N | D | 4.2e-08 | 7.3e-08 | 2.2e-08 | 4.567E-08 | 2.570E-08 | 2.200E-08 | 7.300E-08 |
| 21864 | 1094 | Y1094D | D | 1e-07 | 4.1e-08 | 2.4e-08 | 5.500E-08 | 3.989E-08 | 2.400E-08 | 1.000E-07 |
| 21865 | 1094 | Y1094C | D | 3.5e-08 | 3.3e-08 | 3.7e-08 | 3.500E-08 | 2.000E-09 | 3.300E-08 | 3.700E-08 |
| 21866 | 1094 | Y1094E | D | 4.1e-08 | 3.7e-08 | 6.7e-08 | 4.833E-08 | 1.629E-08 | 3.700E-08 | 6.700E-08 |
| 21867 | 1094 | Y1094Q | D | 9.2e-08 | 7.7e-08 | 2.6e-08 | 6.500E-08 | 3.460E-08 | 2.600E-08 | 9.200E-08 |
| 21868 | 1094 | Y1094G | D | 2.7e-08 | 5.3e-08 | 4.3e-08 | 4.100E-08 | 1.311E-08 | 2.700E-08 | 5.300E-08 |
| 21869 | 1094 | Y1094H | D | 4.8e-08 | 3.9e-08 | 4.1e-08 | 4.267E-08 | 4.726E-09 | 3.900E-08 | 4.800E-08 |
| 21870 | 1094 | Y1094I | D | 3.6e-08 | 1.4e-07 | 7.5e-08 | 8.367E-08 | 5.254E-08 | 3.600E-08 | 1.400E-07 |
| 21871 | 1094 | Y1094L | D | 6.6e-08 | 5.5e-08 | 8.8e-09 | 4.327E-08 | 3.035E-08 | 8.800E-09 | 6.600E-08 |
| 21872 | 1094 | Y1094K | D | 6.6e-08 | 4.7e-08 | 3.3e-08 | 4.867E-08 | 1.656E-08 | 3.300E-08 | 6.600E-08 |
| 21873 | 1094 | Y1094M | D | 2e-08 | 6.5e-08 | 4.6e-08 | 4.367E-08 | 2.259E-08 | 2.000E-08 | 6.500E-08 |
| 21874 | 1094 | Y1094F | D | 2e-08 | 1.2e-08 | 3.5e-08 | 2.233E-08 | 1.168E-08 | 1.200E-08 | 3.500E-08 |
| 21875 | 1094 | Y1094P | D | 5.9e-08 | 6.5e-08 | 5.3e-08 | 5.900E-08 | 6.000E-09 | 5.300E-08 | 6.500E-08 |
| 21876 | 1094 | Y1094S | D | 9.7e-09 | 6.3e-08 | 5.7e-08 | 4.323E-08 | 2.920E-08 | 9.700E-09 | 6.300E-08 |
| 21877 | 1094 | Y1094T | D | 2.6e-08 | 2.1e-08 | 1.7e-08 | 2.133E-08 | 4.509E-09 | 1.700E-08 | 2.600E-08 |
| 21878 | 1094 | Y1094W | D | 1e-07 | 5.1e-08 | 6.1e-08 | 7.067E-08 | 2.589E-08 | 5.100E-08 | 1.000E-07 |
| 21879 | 1094 | Y1094Y | D | 7.3e-09 | 4.6e-08 | 8.8e-08 | 4.710E-08 | 4.036E-08 | 7.300E-09 | 8.800E-08 |
| 21880 | 1094 | Y1094V | D | 4e-08 | 7.6e-08 | 3.3e-08 | 4.967E-08 | 2.307E-08 | 3.300E-08 | 7.600E-08 |
| 21881 | 1095 | Q1095A | D | 6.5e-08 | 7.3e-09 | 4.3e-08 | 3.843E-08 | 2.912E-08 | 7.300E-09 | 6.500E-08 |
| 21882 | 1095 | Q1095R | D | 5.4e-08 | 5.2e-08 | 3e-08 | 4.533E-08 | 1.332E-08 | 3.000E-08 | 5.400E-08 |
| 21883 | 1095 | Q1095N | D | 6.8e-08 | 4.3e-08 | 4.5e-08 | 5.200E-08 | 1.389E-08 | 4.300E-08 | 6.800E-08 |
| 21884 | 1095 | Q1095D | D | 3.1e-08 | 1.3e-08 | 3.4e-08 | 2.600E-08 | 1.136E-08 | 1.300E-08 | 3.400E-08 |
| 21885 | 1095 | Q1095C | D | 6e-08 | 4.5e-08 | 8.6e-08 | 6.367E-08 | 2.074E-08 | 4.500E-08 | 8.600E-08 |
| 21886 | 1095 | Q1095E | D | 5.8e-08 | 3.3e-08 | 6e-08 | 5.033E-08 | 1.504E-08 | 3.300E-08 | 6.000E-08 |
| 21887 | 1095 | Q1095Q | D | 2.8e-08 | 3.1e-08 | 4.3e-08 | 3.400E-08 | 7.937E-09 | 2.800E-08 | 4.300E-08 |
| 21888 | 1095 | Q1095G | D | 5e-08 | 5.9e-08 | 1.1e-07 | 7.300E-08 | 3.236E-08 | 5.000E-08 | 1.100E-07 |
| 21889 | 1095 | Q1095H | D | 4e-08 | 5.1e-08 | 3.3e-08 | 4.133E-08 | 9.074E-09 | 3.300E-08 | 5.100E-08 |
| 21890 | 1095 | Q1095I | D | 3.2e-08 | 8.3e-08 | 5.5e-08 | 5.667E-08 | 2.554E-08 | 3.200E-08 | 8.300E-08 |
| 21891 | 1095 | Q1095L | D | 4.5e-08 | 5.9e-08 | 2.6e-08 | 4.333E-08 | 1.656E-08 | 2.600E-08 | 5.900E-08 |
| 21892 | 1095 | Q1095K | D | 4e-08 | 2.3e-08 | 1e-07 | 5.433E-08 | 4.045E-08 | 2.300E-08 | 1.000E-07 |
| 21893 | 1095 | Q1095M | D | 4e-08 | 3.1e-08 | 3e-08 | 3.367E-08 | 5.508E-09 | 3.000E-08 | 4.000E-08 |
| 21894 | 1095 | Q1095F | D | 2.3e-08 | 5.4e-08 | 4.8e-08 | 4.167E-08 | 1.644E-08 | 2.300E-08 | 5.400E-08 |
| 21895 | 1095 | Q1095P | D | 7.4e-08 | 9.9e-08 | 1.6e-08 | 6.300E-08 | 4.258E-08 | 1.600E-08 | 9.900E-08 |
| 21896 | 1095 | Q1095S | D | 7.8e-08 | 2.2e-08 | 6.2e-08 | 5.400E-08 | 2.884E-08 | 2.200E-08 | 7.800E-08 |
| 21897 | 1095 | Q1095T | D | 3.1e-08 | 2.2e-08 | 3.7e-08 | 3.000E-08 | 7.550E-09 | 2.200E-08 | 3.700E-08 |
| 21898 | 1095 | Q1095W | D | 8.8e-08 | 7.2e-08 | 3.4e-08 | 6.467E-08 | 2.774E-08 | 3.400E-08 | 8.800E-08 |
| 21899 | 1095 | Q1095Y | D | 6.2e-08 | 5.8e-08 | 4.9e-08 | 5.633E-08 | 6.658E-09 | 4.900E-08 | 6.200E-08 |
| 21900 | 1095 | Q1095V | D | 2.8e-08 | 8e-08 | 1.7e-07 | 9.267E-08 | 7.184E-08 | 2.800E-08 | 1.700E-07 |
| 21901 | 1096 | Q1096A | D | 7.8e-08 | 4.3e-08 | 4.9e-08 | 5.667E-08 | 1.872E-08 | 4.300E-08 | 7.800E-08 |
| 21902 | 1096 | Q1096R | D | 7.8e-08 | 7.3e-08 | 6.5e-08 | 7.200E-08 | 6.557E-09 | 6.500E-08 | 7.800E-08 |
| 21903 | 1096 | Q1096N | D | 3.8e-08 | 3.4e-08 | 7.7e-08 | 4.967E-08 | 2.376E-08 | 3.400E-08 | 7.700E-08 |
| 21904 | 1096 | Q1096D | D | 5.2e-08 | 5e-08 | 4.3e-08 | 4.833E-08 | 4.726E-09 | 4.300E-08 | 5.200E-08 |
| 21905 | 1096 | Q1096C | D | 4.6e-08 | 7e-08 | 3.8e-08 | 5.133E-08 | 1.665E-08 | 3.800E-08 | 7.000E-08 |
| 21906 | 1096 | Q1096E | D | 3.6e-08 | 5.6e-08 | 1.1e-07 | 6.733E-08 | 3.828E-08 | 3.600E-08 | 1.100E-07 |
| 21907 | 1096 | Q1096Q | D | 2.8e-08 | 5.7e-08 | 5.5e-08 | 4.667E-08 | 1.620E-08 | 2.800E-08 | 5.700E-08 |
| 21908 | 1096 | Q1096G | D | 6.6e-08 | 4e-08 | 2.5e-08 | 4.367E-08 | 2.074E-08 | 2.500E-08 | 6.600E-08 |
| 21909 | 1096 | Q1096H | D | 5.5e-08 | 1.2e-08 | 1.9e-07 | 8.567E-08 | 9.288E-08 | 1.200E-08 | 1.900E-07 |
| 21910 | 1096 | Q1096I | D | 9.1e-08 | 6.2e-08 | 5.3e-08 | 6.867E-08 | 1.986E-08 | 5.300E-08 | 9.100E-08 |
| 21911 | 1096 | Q1096L | D | 2.8e-08 | 1.2e-07 | 7.9e-08 | 7.567E-08 | 4.609E-08 | 2.800E-08 | 1.200E-07 |
| 21912 | 1096 | Q1096K | D | 1.6e-07 | 1.9e-08 | 1.2e-08 | 6.367E-08 | 8.350E-08 | 1.200E-08 | 1.600E-07 |
| 21913 | 1096 | Q1096M | D | 3.6e-08 | 7e-08 | 9.4e-08 | 6.667E-08 | 2.914E-08 | 3.600E-08 | 9.400E-08 |
| 21914 | 1096 | Q1096F | D | 5.4e-08 | 4.5e-08 | 3.3e-08 | 4.400E-08 | 1.054E-08 | 3.300E-08 | 5.400E-08 |
| 21915 | 1096 | Q1096P | D | 4.3e-08 | 6.3e-08 | 5.7e-08 | 5.433E-08 | 1.026E-08 | 4.300E-08 | 6.300E-08 |
| 21916 | 1096 | Q1096S | D | 2.6e-08 | 6.9e-08 | 6.3e-08 | 5.267E-08 | 2.329E-08 | 2.600E-08 | 6.900E-08 |
| 21917 | 1096 | Q1096T | D | 7.5e-08 | 8.7e-08 | 3.9e-08 | 6.700E-08 | 2.498E-08 | 3.900E-08 | 8.700E-08 |
| 21918 | 1096 | Q1096W | D | 2.3e-08 | 6e-08 | 5.4e-08 | 4.567E-08 | 1.986E-08 | 2.300E-08 | 6.000E-08 |
| 21919 | 1096 | Q1096Y | D | 8.3e-08 | 3.9e-08 | 5.6e-08 | 5.933E-08 | 2.219E-08 | 3.900E-08 | 8.300E-08 |
| 21920 | 1096 | Q1096V | D | 1.2e-07 | 4.2e-08 | 8.9e-09 | 5.697E-08 | 5.704E-08 | 8.900E-09 | 1.200E-07 |
| 21921 | 1097 | K1097A | D | 2.1e-08 | 4.9e-08 | 1.5e-08 | 2.833E-08 | 1.815E-08 | 1.500E-08 | 4.900E-08 |
| 21922 | 1097 | K1097R | D | 6.9e-08 | 2.7e-08 | 3.1e-08 | 4.233E-08 | 2.318E-08 | 2.700E-08 | 6.900E-08 |
| 21923 | 1097 | K1097N | D | 2.8e-08 | 3.8e-08 | 3.4e-08 | 3.333E-08 | 5.033E-09 | 2.800E-08 | 3.800E-08 |
| 21924 | 1097 | K1097D | D | 1.2e-08 | 2.1e-08 | 4.1e-08 | 2.467E-08 | 1.484E-08 | 1.200E-08 | 4.100E-08 |
| 21925 | 1097 | K1097C | D | 2.4e-08 | 7.6e-08 | 1.3e-08 | 3.767E-08 | 3.365E-08 | 1.300E-08 | 7.600E-08 |
| 21926 | 1097 | K1097E | D | 1.7e-08 | 3.1e-08 | 2.7e-08 | 2.500E-08 | 7.211E-09 | 1.700E-08 | 3.100E-08 |
| 21927 | 1097 | K1097Q | D | 2.8e-08 | 5.3e-08 | 3.2e-08 | 3.767E-08 | 1.343E-08 | 2.800E-08 | 5.300E-08 |
| 21928 | 1097 | K1097G | D | 1.8e-07 | 4.5e-08 | 4.6e-08 | 9.033E-08 | 7.766E-08 | 4.500E-08 | 1.800E-07 |
| 21929 | 1097 | K1097H | D | 2.9e-08 | 6.5e-08 | 1e-08 | 3.467E-08 | 2.793E-08 | 1.000E-08 | 6.500E-08 |

| | | | | | | | | | | |
|---|---|---|---|---|---|---|---|---|---|---|
| 21930 | 1097 | K1097I | D | 3.1e-08 | 5.7e-08 | 3.4e-08 | 4.067E-08 | 1.422E-08 | 3.100E-08 | 5.700E-08 |
| 21931 | 1097 | K1097L | D | 7.8e-08 | 8.4e-08 | 6.5e-08 | 7.567E-08 | 9.713E-09 | 6.500E-08 | 8.400E-08 |
| 21932 | 1097 | K1097K | D | 7e-08 | 2.8e-08 | 3.4e-08 | 4.400E-08 | 2.272E-08 | 2.800E-08 | 7.000E-08 |
| 21933 | 1097 | K1097M | D | 7.4e-08 | 3.4e-08 | 3.6e-08 | 4.800E-08 | 2.254E-08 | 3.400E-08 | 7.400E-08 |
| 21934 | 1097 | K1097F | D | 3.5e-08 | 2.1e-08 | 3.6e-08 | 3.067E-08 | 8.386E-09 | 2.100E-08 | 3.600E-08 |
| 21935 | 1097 | K1097P | D | 3e-08 | 2.7e-08 | 3.7e-08 | 3.133E-08 | 5.132E-09 | 2.700E-08 | 3.700E-08 |
| 21936 | 1097 | K1097S | D | 6.2e-08 | 5.7e-08 | 7.3e-08 | 6.400E-08 | 8.185E-09 | 5.700E-08 | 7.300E-08 |
| 21937 | 1097 | K1097T | D | 4.1e-08 | 3.8e-08 | 3.7e-08 | 3.867E-08 | 2.082E-09 | 3.700E-08 | 4.100E-08 |
| 21938 | 1097 | K1097W | D | 1.1e-07 | 1.1e-07 | 2.4e-08 | 8.133E-08 | 4.965E-08 | 2.400E-08 | 1.100E-07 |
| 21939 | 1097 | K1097Y | D | 5.6e-08 | 9.4e-08 | 3.5e-08 | 6.167E-08 | 2.991E-08 | 3.500E-08 | 9.400E-08 |
| 21940 | 1097 | K1097V | D | 3.4e-08 | 4.5e-08 | 8.3e-08 | 5.400E-08 | 2.571E-08 | 3.400E-08 | 8.300E-08 |
| 21941 | 1098 | P1098A | D | 3.2e-08 | 9.4e-08 | 3.9e-08 | 5.500E-08 | 3.396E-08 | 3.200E-08 | 9.400E-08 |
| 21942 | 1098 | P1098R | D | 8.1e-08 | 5.1e-08 | 1.9e-08 | 5.033E-08 | 3.101E-08 | 1.900E-08 | 8.100E-08 |
| 21943 | 1098 | P1098N | D | 4.2e-08 | 2.4e-08 | 1.9e-08 | 2.833E-08 | 1.210E-08 | 1.900E-08 | 4.200E-08 |
| 21944 | 1098 | P1098D | D | 3.9e-08 | 2.2e-08 | 7.4e-08 | 4.500E-08 | 2.651E-08 | 2.200E-08 | 7.400E-08 |
| 21945 | 1098 | P1098C | D | 4.5e-08 | 5.4e-08 | 2.9e-08 | 4.267E-08 | 1.266E-08 | 2.900E-08 | 5.400E-08 |
| 21946 | 1098 | P1098E | D | 1.4e-08 | 5.2e-08 | 4.7e-08 | 3.767E-08 | 2.065E-08 | 1.400E-08 | 5.200E-08 |
| 21947 | 1098 | P1098Q | D | 4.3e-08 | 2.8e-08 | 1.8e-08 | 2.967E-08 | 1.258E-08 | 1.800E-08 | 4.300E-08 |
| 21948 | 1098 | P1098G | D | 4.9e-08 | 1e-07 | 4.4e-08 | 6.433E-08 | 3.099E-08 | 4.400E-08 | 1.000E-07 |
| 21949 | 1098 | P1098H | D | 3.3e-08 | 3.5e-08 | 3.1e-08 | 3.300E-08 | 2.000E-09 | 3.100E-08 | 3.500E-08 |
| 21950 | 1098 | P1098I | D | 1.7e-07 | 9.2e-08 | 3.7e-08 | 9.967E-08 | 6.683E-08 | 3.700E-08 | 1.700E-07 |
| 21951 | 1098 | P1098L | D | 1.2e-08 | 4.4e-08 | 5.9e-08 | 3.833E-08 | 2.401E-08 | 1.200E-08 | 5.900E-08 |
| 21952 | 1098 | P1098K | D | 3.1e-08 | 4.5e-08 | 1.1e-07 | 6.200E-08 | 4.215E-08 | 3.100E-08 | 1.100E-07 |
| 21953 | 1098 | P1098M | D | 2.9e-08 | 4.9e-08 | 9.2e-08 | 5.667E-08 | 3.219E-08 | 2.900E-08 | 9.200E-08 |
| 21954 | 1098 | P1098F | D | 1.7e-08 | 2.9e-08 | 6.5e-08 | 3.700E-08 | 2.498E-08 | 1.700E-08 | 6.500E-08 |
| 21955 | 1098 | P1098P | D | 1e-07 | 5.4e-08 | 2.1e-08 | 5.833E-08 | 3.968E-08 | 2.100E-08 | 1.000E-07 |
| 21956 | 1098 | P1098S | D | 8.3e-08 | 9e-08 | 7.2e-08 | 8.167E-08 | 9.074E-09 | 7.200E-08 | 9.000E-08 |
| 21957 | 1098 | P1098T | D | 2.3e-08 | 3.4e-08 | 3.3e-08 | 3.000E-08 | 6.083E-09 | 2.300E-08 | 3.400E-08 |
| 21958 | 1098 | P1098W | D | 3.1e-08 | 8e-08 | 4.7e-08 | 5.267E-08 | 2.499E-08 | 3.100E-08 | 8.000E-08 |
| 21959 | 1098 | P1098Y | D | 3.5e-08 | 7.4e-08 | 3.9e-08 | 4.933E-08 | 2.146E-08 | 3.500E-08 | 7.400E-08 |
| 21960 | 1098 | P1098V | D | 1.3e-08 | 9.5e-09 | 3.6e-08 | 1.950E-08 | 1.440E-08 | 9.500E-09 | 3.600E-08 |
| 21961 | 1099 | G1099A | D | 4.4e-08 | 3.1e-08 | 3.4e-08 | 3.633E-08 | 6.807E-09 | 3.100E-08 | 4.400E-08 |
| 21962 | 1099 | G1099R | D | 1.9e-08 | 3.4e-08 | 4.4e-08 | 3.233E-08 | 1.258E-08 | 1.900E-08 | 4.400E-08 |
| 21963 | 1099 | G1099N | D | 2.9e-08 | 3.5e-08 | 6.2e-08 | 4.200E-08 | 1.758E-08 | 2.900E-08 | 6.200E-08 |
| 21964 | 1099 | G1099D | D | 2.1e-08 | 4.5e-08 | 1.8e-08 | 2.800E-08 | 1.480E-08 | 1.800E-08 | 4.500E-08 |
| 21965 | 1099 | G1099C | D | 1.3e-08 | 5.4e-08 | 1.5e-07 | 7.233E-08 | 7.032E-08 | 1.300E-08 | 1.500E-07 |
| 21966 | 1099 | G1099E | D | 3.1e-08 | 5e-08 | 3.4e-08 | 3.833E-08 | 1.021E-08 | 3.100E-08 | 5.000E-08 |
| 21967 | 1099 | G1099Q | D | 6e-08 | 2.6e-08 | 7.2e-08 | 5.267E-08 | 2.386E-08 | 2.600E-08 | 7.200E-08 |
| 21968 | 1099 | G1099G | D | 7.5e-08 | 4.7e-08 | 3.4e-08 | 5.200E-08 | 2.095E-08 | 3.400E-08 | 7.500E-08 |
| 21969 | 1099 | G1099H | D | 4.7e-08 | 4.5e-08 | 2.3e-08 | 3.833E-08 | 1.332E-08 | 2.300E-08 | 4.700E-08 |
| 21970 | 1099 | G1099I | D | 5.2e-08 | 4.8e-08 | 4.1e-08 | 4.700E-08 | 5.568E-09 | 4.100E-08 | 5.200E-08 |
| 21971 | 1099 | G1099L | D | 6.8e-08 | 4.1e-08 | 2.6e-08 | 4.500E-08 | 2.128E-08 | 2.600E-08 | 6.800E-08 |
| 21972 | 1099 | G1099K | D | 1.2e-07 | 9.3e-08 | 2.6e-08 | 7.967E-08 | 4.840E-08 | 2.600E-08 | 1.200E-07 |
| 21973 | 1099 | G1099M | D | 3.5e-08 | 1.5e-07 | 4.9e-08 | 7.800E-08 | 6.275E-08 | 3.500E-08 | 1.500E-07 |
| 21974 | 1099 | G1099F | D | 4e-08 | 2.4e-08 | 3e-08 | 3.133E-08 | 8.083E-09 | 2.400E-08 | 4.000E-08 |
| 21975 | 1099 | G1099P | D | 8.3e-08 | 1.4e-08 | 4.5e-08 | 4.733E-08 | 3.456E-08 | 1.400E-08 | 8.300E-08 |
| 21976 | 1099 | G1099S | D | 3.4e-08 | 1.9e-08 | 5.1e-08 | 3.467E-08 | 1.601E-08 | 1.900E-08 | 5.100E-08 |
| 21977 | 1099 | G1099T | D | 9.1e-08 | 1.9e-08 | 1.9e-08 | 4.300E-08 | 4.157E-08 | 1.900E-08 | 9.100E-08 |
| 21978 | 1099 | G1099W | D | 3e-08 | 2.9e-08 | 3.9e-08 | 3.267E-08 | 5.508E-09 | 2.900E-08 | 3.900E-08 |
| 21979 | 1099 | G1099Y | D | 1.9e-08 | 2.1e-08 | 1.6e-08 | 1.867E-08 | 2.517E-09 | 1.600E-08 | 2.100E-08 |
| 21980 | 1099 | G1099V | D | 9e-08 | 3.9e-08 | 3.4e-08 | 5.433E-08 | 3.099E-08 | 3.400E-08 | 9.000E-08 |
| 21981 | 1100 | K1100A | D | 3.9e-08 | 3.4e-08 | 3.4e-08 | 3.567E-08 | 2.887E-09 | 3.400E-08 | 3.900E-08 |
| 21982 | 1100 | K1100R | D | 6.7e-08 | 3.2e-08 | 2.1e-08 | 4.000E-08 | 2.402E-08 | 2.100E-08 | 6.700E-08 |
| 21983 | 1100 | K1100N | D | 1.3e-07 | 8.2e-08 | 1.1e-07 | 1.073E-07 | 2.411E-08 | 8.200E-08 | 1.300E-07 |
| 21984 | 1100 | K1100D | D | 2.3e-08 | 4e-08 | 2.3e-08 | 2.867E-08 | 9.815E-09 | 2.300E-08 | 4.000E-08 |
| 21985 | 1100 | K1100C | D | 6.4e-08 | 9.2e-08 | 5.3e-08 | 6.967E-08 | 2.011E-08 | 5.300E-08 | 9.200E-08 |
| 21986 | 1100 | K1100E | D | 2.8e-08 | 5.9e-08 | 1.4e-07 | 7.567E-08 | 5.783E-08 | 2.800E-08 | 1.400E-07 |
| 21987 | 1100 | K1100Q | D | 3.8e-08 | 2.3e-08 | 1.9e-08 | 2.667E-08 | 1.002E-08 | 1.900E-08 | 3.800E-08 |
| 21988 | 1100 | K1100G | D | 1.7e-08 | 7.5e-08 | 1.4e-07 | 7.733E-08 | 6.153E-08 | 1.700E-08 | 1.400E-07 |
| 21989 | 1100 | K1100H | D | 4.1e-08 | 7.5e-09 | 1.7e-08 | 2.183E-08 | 1.727E-08 | 7.500E-09 | 4.100E-08 |
| 21990 | 1100 | K1100I | D | 4.3e-08 | 3.6e-08 | 1.9e-08 | 3.267E-08 | 1.234E-08 | 1.900E-08 | 4.300E-08 |
| 21991 | 1100 | K1100L | D | 2.4e-08 | 1.6e-08 | 1e-07 | 4.667E-08 | 4.636E-08 | 1.600E-08 | 1.000E-07 |
| 21992 | 1100 | K1100K | D | 2.7e-08 | 2.1e-08 | 4.5e-08 | 3.100E-08 | 1.249E-08 | 2.100E-08 | 4.500E-08 |
| 21993 | 1100 | K1100M | D | 3.7e-08 | 8.2e-08 | 1.2e-07 | 7.967E-08 | 4.155E-08 | 3.700E-08 | 1.200E-07 |
| 21994 | 1100 | K1100F | D | 7e-08 | 3.7e-08 | 6.3e-09 | 3.777E-08 | 3.186E-08 | 6.300E-09 | 7.000E-08 |
| 21995 | 1100 | K1100P | D | 7.5e-09 | 4.7e-08 | 5.2e-08 | 3.550E-08 | 2.438E-08 | 7.500E-09 | 5.200E-08 |
| 21996 | 1100 | K1100S | D | 7.6e-08 | 4.2e-08 | 5.2e-08 | 5.667E-08 | 1.747E-08 | 4.200E-08 | 7.600E-08 |
| 21997 | 1100 | K1100T | D | 4e-08 | 2.5e-08 | 1.9e-08 | 2.800E-08 | 1.082E-08 | 1.900E-08 | 4.000E-08 |
| 21998 | 1100 | K1100W | D | 3e-08 | 5.9e-08 | 6.9e-08 | 5.267E-08 | 2.026E-08 | 3.000E-08 | 6.900E-08 |
| 21999 | 1100 | K1100Y | D | 1.7e-07 | 3.6e-08 | 1.8e-08 | 7.467E-08 | 8.305E-08 | 1.800E-08 | 1.700E-07 |
| 22000 | 1100 | K1100V | D | 2.6e-08 | 3.1e-08 | 4.6e-08 | 3.433E-08 | 1.041E-08 | 2.600E-08 | 4.600E-08 |
| 22001 | 1101 | A1101A | D | 5.8e-08 | 1.5e-08 | 5.3e-08 | 4.200E-08 | 2.352E-08 | 1.500E-08 | 5.800E-08 |
| 22002 | 1101 | A1101R | D | 2.9e-08 | 6e-08 | 4.4e-08 | 4.433E-08 | 1.550E-08 | 2.900E-08 | 6.000E-08 |
| 22003 | 1101 | A1101N | D | 8.1e-08 | 7.8e-08 | 5.3e-08 | 7.067E-08 | 1.537E-08 | 5.300E-08 | 8.100E-08 |
| 22004 | 1101 | A1101D | D | 7.4e-08 | 7.1e-08 | 2.9e-08 | 5.800E-08 | 2.516E-08 | 2.900E-08 | 7.400E-08 |
| 22005 | 1101 | A1101C | D | 2.8e-08 | 2.8e-08 | 1.2e-08 | 2.267E-08 | 9.238E-09 | 1.200E-08 | 2.800E-08 |
| 22006 | 1101 | A1101E | D | 3.2e-08 | 4.9e-08 | 6.9e-08 | 5.000E-08 | 1.852E-08 | 3.200E-08 | 6.900E-08 |
| 22007 | 1101 | A1101Q | D | 4.4e-08 | 5.9e-08 | 1.9e-08 | 4.067E-08 | 2.021E-08 | 1.900E-08 | 5.900E-08 |
| 22008 | 1101 | A1101G | D | 5e-08 | 6e-08 | 5.9e-08 | 5.633E-08 | 5.508E-09 | 5.000E-08 | 6.000E-08 |
| 22009 | 1101 | A1101H | D | 7e-08 | 3.1e-08 | 7.4e-08 | 5.833E-08 | 2.376E-08 | 3.100E-08 | 7.400E-08 |
| 22010 | 1101 | A1101I | D | 8.9e-08 | 4.2e-08 | 4.8e-08 | 5.967E-08 | 2.558E-08 | 4.200E-08 | 8.900E-08 |
| 22011 | 1101 | A1101L | D | 2.7e-08 | 1.1e-07 | 5.3e-08 | 6.333E-08 | 4.245E-08 | 2.700E-08 | 1.100E-07 |
| 22012 | 1101 | A1101K | D | 7.7e-08 | 1.7e-08 | 4.2e-08 | 4.533E-08 | 3.014E-08 | 1.700E-08 | 7.700E-08 |
| 22013 | 1101 | A1101M | D | 5.2e-08 | 5.8e-08 | 2.7e-08 | 4.567E-08 | 1.644E-08 | 2.700E-08 | 5.800E-08 |
| 22014 | 1101 | A1101F | D | 4.1e-08 | 3.9e-08 | 2.2e-08 | 3.400E-08 | 1.044E-08 | 2.200E-08 | 4.100E-08 |

| | | | | | | | | | | |
|---|---|---|---|---|---|---|---|---|---|---|
| 22015 | 1101 | A1101P | D | 1.1e-07 | 3.9e-08 | 5.2e-08 | 6.700E-08 | 3.780E-08 | 3.900E-08 | 1.100E-07 |
| 22016 | 1101 | A1101S | D | 1.1e-08 | 9.9e-08 | 7.6e-08 | 6.200E-08 | 4.564E-08 | 1.100E-08 | 9.900E-08 |
| 22017 | 1101 | A1101T | D | 6e-08 | 2.3e-08 | 6e-08 | 4.767E-08 | 2.136E-08 | 2.300E-08 | 6.000E-08 |
| 22018 | 1101 | A1101W | D | 3e-08 | 4.2e-08 | 4.7e-08 | 3.967E-08 | 8.737E-09 | 3.000E-08 | 4.700E-08 |
| 22019 | 1101 | A1101Y | D | 6.1e-08 | 2.4e-08 | 2.3e-08 | 3.600E-08 | 2.166E-08 | 2.300E-08 | 6.100E-08 |
| 22020 | 1101 | A1101V | D | 2.6e-07 | 4.6e-08 | 4.9e-08 | 1.183E-07 | 1.227E-07 | 4.600E-08 | 2.600E-07 |
| 22021 | 1102 | P1102A | D | 6.1e-08 | 2.1e-07 | 4.9e-08 | 1.067E-07 | 8.969E-08 | 4.900E-08 | 2.100E-07 |
| 22022 | 1102 | P1102R | D | 2.9e-08 | 2.5e-08 | 2.7e-08 | 2.700E-08 | 2.000E-09 | 2.500E-08 | 2.900E-08 |
| 22023 | 1102 | P1102N | D | 1.7e-08 | 6.6e-08 | 3.7e-08 | 4.000E-08 | 2.464E-08 | 1.700E-08 | 6.600E-08 |
| 22024 | 1102 | P1102D | D | 5.7e-08 | 3.6e-08 | 4.3e-08 | 4.533E-08 | 1.069E-08 | 3.600E-08 | 5.700E-08 |
| 22025 | 1102 | P1102C | D | 2.6e-08 | 6.6e-08 | 6.5e-08 | 5.233E-08 | 2.281E-08 | 2.600E-08 | 6.600E-08 |
| 22026 | 1102 | P1102E | D | 5.5e-08 | 3.8e-08 | 6.1e-08 | 5.133E-08 | 1.193E-08 | 3.800E-08 | 6.100E-08 |
| 22027 | 1102 | P1102Q | D | 4.5e-08 | 5.6e-08 | 9.3e-09 | 3.677E-08 | 2.441E-08 | 9.300E-09 | 5.600E-08 |
| 22028 | 1102 | P1102G | D | 1.8e-08 | 1.7e-08 | 3e-08 | 2.167E-08 | 7.234E-09 | 1.700E-08 | 3.000E-08 |
| 22029 | 1102 | P1102H | D | 9.8e-08 | 5.5e-08 | 1.1e-07 | 8.767E-08 | 2.892E-08 | 5.500E-08 | 1.100E-07 |
| 22030 | 1102 | P1102I | D | 3.4e-08 | 8e-08 | 2.2e-08 | 4.533E-08 | 3.062E-08 | 2.200E-08 | 8.000E-08 |
| 22031 | 1102 | P1102L | D | 8.9e-08 | 2.9e-08 | 4.7e-08 | 5.500E-08 | 3.079E-08 | 2.900E-08 | 8.900E-08 |
| 22032 | 1102 | P1102K | D | 1.7e-08 | 8.2e-09 | 6.6e-08 | 3.040E-08 | 3.114E-08 | 8.200E-09 | 6.600E-08 |
| 22033 | 1102 | P1102M | D | 5e-08 | 3.5e-08 | 1.9e-08 | 3.467E-08 | 1.550E-08 | 1.900E-08 | 5.000E-08 |
| 22034 | 1102 | P1102F | D | 8.3e-08 | 1.3e-07 | 3.9e-08 | 8.400E-08 | 4.551E-08 | 3.900E-08 | 1.300E-07 |
| 22035 | 1102 | P1102P | D | 4.9e-08 | 4.4e-08 | 3.8e-08 | 4.367E-08 | 5.508E-09 | 3.800E-08 | 4.900E-08 |
| 22036 | 1102 | P1102S | D | 3.1e-08 | 7e-08 | 2.9e-08 | 4.333E-08 | 2.312E-08 | 2.900E-08 | 7.000E-08 |
| 22037 | 1102 | P1102T | D | 8.8e-08 | 6.2e-08 | 1.6e-08 | 5.533E-08 | 3.646E-08 | 1.600E-08 | 8.800E-08 |
| 22038 | 1102 | P1102W | D | 2.7e-08 | 4.5e-08 | 2.2e-08 | 3.133E-08 | 1.210E-08 | 2.200E-08 | 4.500E-08 |
| 22039 | 1102 | P1102Y | D | 3.1e-08 | 7.9e-08 | 2.7e-08 | 4.567E-08 | 2.894E-08 | 2.700E-08 | 7.900E-08 |
| 22040 | 1102 | P1102V | D | 5.8e-08 | 1.1e-07 | 1.5e-08 | 6.100E-08 | 4.757E-08 | 1.500E-08 | 1.100E-07 |
| 22041 | 1103 | K1103A | D | 8.5e-08 | 3e-08 | 3.3e-08 | 4.933E-08 | 3.092E-08 | 3.000E-08 | 8.500E-08 |
| 22042 | 1103 | K1103R | D | 3.4e-08 | 3.6e-08 | 1.7e-07 | 8.000E-08 | 7.795E-08 | 3.400E-08 | 1.700E-07 |
| 22043 | 1103 | K1103N | D | 4e-08 | 3.5e-08 | 3.8e-08 | 3.767E-08 | 2.517E-09 | 3.500E-08 | 4.000E-08 |
| 22044 | 1103 | K1103D | D | 1.5e-08 | 5.8e-08 | 3.5e-08 | 3.600E-08 | 2.152E-08 | 1.500E-08 | 5.800E-08 |
| 22045 | 1103 | K1103C | D | 3e-08 | 5e-08 | 2.9e-08 | 3.633E-08 | 1.185E-08 | 2.900E-08 | 5.000E-08 |
| 22046 | 1103 | K1103E | D | 2.8e-08 | 3.1e-08 | 1e-07 | 5.300E-08 | 4.073E-08 | 2.800E-08 | 1.000E-07 |
| 22047 | 1103 | K1103Q | D | 3.5e-08 | 1.9e-08 | 2.9e-08 | 2.767E-08 | 8.083E-09 | 1.900E-08 | 3.500E-08 |
| 22048 | 1103 | K1103G | D | 6.3e-08 | 5.4e-08 | 8.4e-08 | 6.700E-08 | 1.539E-08 | 5.400E-08 | 8.400E-08 |
| 22049 | 1103 | K1103H | D | 1e-07 | 3.6e-08 | 6.3e-08 | 6.633E-08 | 3.213E-08 | 3.600E-08 | 1.000E-07 |
| 22050 | 1103 | K1103I | D | 1.3e-08 | 8.7e-08 | 1.4e-08 | 3.800E-08 | 4.244E-08 | 1.300E-08 | 8.700E-08 |
| 22051 | 1103 | K1103L | D | 6.7e-08 | 1.1e-07 | 1.2e-07 | 9.900E-08 | 2.816E-08 | 6.700E-08 | 1.200E-07 |
| 22052 | 1103 | K1103K | D | 7.6e-08 | 2.2e-08 | 4e-08 | 4.600E-08 | 2.750E-08 | 2.200E-08 | 7.600E-08 |
| 22053 | 1103 | K1103M | D | 9e-08 | 4.3e-08 | 1.3e-08 | 4.867E-08 | 3.881E-08 | 1.300E-08 | 9.000E-08 |
| 22054 | 1103 | K1103F | D | 9.8e-08 | 6.6e-08 | 2.7e-08 | 6.367E-08 | 3.556E-08 | 2.700E-08 | 9.800E-08 |
| 22055 | 1103 | K1103P | D | 5.3e-08 | 5.4e-08 | 2.5e-08 | 4.400E-08 | 1.646E-08 | 2.500E-08 | 5.400E-08 |
| 22056 | 1103 | K1103S | D | 7.4e-08 | 3.1e-08 | 4.3e-08 | 4.933E-08 | 2.219E-08 | 3.100E-08 | 7.400E-08 |
| 22057 | 1103 | K1103T | D | 3.9e-08 | 4.2e-08 | 7.2e-08 | 5.100E-08 | 1.825E-08 | 3.900E-08 | 7.200E-08 |
| 22058 | 1103 | K1103W | D | 1.2e-08 | 9.6e-08 | 3.3e-08 | 4.700E-08 | 4.371E-08 | 1.200E-08 | 9.600E-08 |
| 22059 | 1103 | K1103Y | D | 8.5e-08 | 1.8e-08 | 1.8e-07 | 9.433E-08 | 8.140E-08 | 1.800E-08 | 1.800E-07 |
| 22060 | 1103 | K1103V | D | 5.2e-08 | 1.8e-08 | 1.4e-07 | 7.000E-08 | 6.296E-08 | 1.800E-08 | 1.400E-07 |
| 22061 | 1104 | L1104A | D | 3.7e-08 | 2e-08 | 4.6e-08 | 3.433E-08 | 1.320E-08 | 2.000E-08 | 4.600E-08 |
| 22062 | 1104 | L1104R | D | 6.3e-08 | 8.2e-08 | 6.2e-08 | 6.900E-08 | 1.127E-08 | 6.200E-08 | 8.200E-08 |
| 22063 | 1104 | L1104N | D | 1.2e-07 | 4.7e-08 | 1.1e-07 | 9.233E-08 | 3.958E-08 | 4.700E-08 | 1.200E-07 |
| 22064 | 1104 | L1104D | D | 1e-07 | 7.3e-08 | 5.9e-08 | 7.733E-08 | 2.084E-08 | 5.900E-08 | 1.000E-07 |
| 22065 | 1104 | L1104C | D | 3e-08 | 3.2e-08 | 5.4e-08 | 3.867E-08 | 1.332E-08 | 3.000E-08 | 5.400E-08 |
| 22066 | 1104 | L1104E | D | 4.2e-08 | 4.3e-08 | 4.4e-08 | 4.300E-08 | 1.000E-09 | 4.200E-08 | 4.400E-08 |
| 22067 | 1104 | L1104Q | D | 1.2e-08 | 4.5e-08 | 4.1e-08 | 3.267E-08 | 1.801E-08 | 1.200E-08 | 4.500E-08 |
| 22068 | 1104 | L1104G | D | 2.4e-08 | 4.6e-08 | 4.6e-08 | 3.867E-08 | 1.270E-08 | 2.400E-08 | 4.600E-08 |
| 22069 | 1104 | L1104H | D | 2.8e-08 | 8.3e-08 | 5.5e-08 | 5.533E-08 | 2.750E-08 | 2.800E-08 | 8.300E-08 |
| 22070 | 1104 | L1104I | D | 3.9e-08 | 1.2e-07 | 1.7e-08 | 5.867E-08 | 5.424E-08 | 1.700E-08 | 1.200E-07 |
| 22071 | 1104 | L1104L | D | 2.6e-08 | 4.4e-08 | 2.1e-08 | 3.033E-08 | 1.210E-08 | 2.100E-08 | 4.400E-08 |
| 22072 | 1104 | L1104K | D | 3.2e-08 | 2.7e-08 | 6.4e-08 | 4.100E-08 | 2.007E-08 | 2.700E-08 | 6.400E-08 |
| 22073 | 1104 | L1104M | D | 4.3e-08 | 1.8e-07 | 3.1e-08 | 8.467E-08 | 8.278E-08 | 3.100E-08 | 1.800E-07 |
| 22074 | 1104 | L1104F | D | 7.6e-08 | 4.4e-08 | 7.2e-08 | 6.400E-08 | 1.744E-08 | 4.400E-08 | 7.600E-08 |
| 22075 | 1104 | L1104P | D | 2.8e-08 | 5.1e-08 | 2e-08 | 3.300E-08 | 1.609E-08 | 2.000E-08 | 5.100E-08 |
| 22076 | 1104 | L1104S | D | 7.6e-08 | 2.4e-08 | 9e-08 | 6.333E-08 | 3.478E-08 | 2.400E-08 | 9.000E-08 |
| 22077 | 1104 | L1104T | D | 2.9e-08 | 7.5e-08 | 4.9e-08 | 5.100E-08 | 2.307E-08 | 2.900E-08 | 7.500E-08 |
| 22078 | 1104 | L1104W | D | 2e-08 | 4.1e-08 | 5.1e-08 | 3.733E-08 | 1.582E-08 | 2.000E-08 | 5.100E-08 |
| 22079 | 1104 | L1104Y | D | 2.9e-08 | 1.9e-08 | 5.3e-08 | 3.367E-08 | 1.747E-08 | 1.900E-08 | 5.300E-08 |
| 22080 | 1104 | L1104V | D | 1.9e-08 | 3.4e-08 | 5.7e-08 | 3.667E-08 | 1.914E-08 | 1.900E-08 | 5.700E-08 |
| 22081 | 1105 | L1105A | D | 2.9e-08 | 4e-08 | 4.8e-08 | 3.900E-08 | 9.539E-09 | 2.900E-08 | 4.800E-08 |
| 22082 | 1105 | L1105R | D | 7.4e-08 | 9.4e-08 | 5.3e-08 | 7.367E-08 | 2.050E-08 | 5.300E-08 | 9.400E-08 |
| 22083 | 1105 | L1105N | D | 3.3e-08 | 2.1e-08 | 1.5e-07 | 6.800E-08 | 7.127E-08 | 2.100E-08 | 1.500E-07 |
| 22084 | 1105 | L1105D | D | 2.9e-08 | 4.4e-08 | 2.2e-08 | 3.167E-08 | 1.124E-08 | 2.200E-08 | 4.400E-08 |
| 22085 | 1105 | L1105C | D | 6.8e-08 | 2.9e-08 | 3.9e-08 | 4.533E-08 | 2.026E-08 | 2.900E-08 | 6.800E-08 |
| 22086 | 1105 | L1105E | D | 8.5e-08 | 4.8e-08 | 1.8e-08 | 5.033E-08 | 3.356E-08 | 1.800E-08 | 8.500E-08 |
| 22087 | 1105 | L1105Q | D | 4e-08 | 1.4e-07 | 3.5e-08 | 7.167E-08 | 5.923E-08 | 3.500E-08 | 1.400E-07 |
| 22088 | 1105 | L1105G | D | 7.8e-08 | 9.1e-08 | 4.2e-08 | 7.033E-08 | 2.538E-08 | 4.200E-08 | 9.100E-08 |
| 22089 | 1105 | L1105H | D | 4.4e-08 | 5.9e-08 | 3.1e-08 | 4.467E-08 | 1.401E-08 | 3.100E-08 | 5.900E-08 |
| 22090 | 1105 | L1105I | D | 8.6e-08 | 1.2e-08 | 1.8e-08 | 3.867E-08 | 4.110E-08 | 1.200E-08 | 8.600E-08 |
| 22091 | 1105 | L1105L | D | 3.3e-08 | 1.9e-08 | 3.7e-08 | 2.967E-08 | 9.452E-09 | 1.900E-08 | 3.700E-08 |
| 22092 | 1105 | L1105K | D | 3.6e-08 | 3.6e-08 | 9.5e-09 | 2.717E-08 | 1.530E-08 | 9.500E-09 | 3.600E-08 |
| 22093 | 1105 | L1105M | D | 2.7e-08 | 6.7e-08 | 2.3e-08 | 3.900E-08 | 2.433E-08 | 2.300E-08 | 6.700E-08 |
| 22094 | 1105 | L1105F | D | 3.3e-08 | 2.8e-08 | 4.6e-08 | 3.567E-08 | 9.292E-09 | 2.800E-08 | 4.600E-08 |
| 22095 | 1105 | L1105P | D | 3.6e-08 | 5.6e-08 | 4.7e-08 | 4.633E-08 | 1.002E-08 | 3.600E-08 | 5.600E-08 |
| 22096 | 1105 | L1105S | D | 9.1e-08 | 3.3e-08 | 1.4e-08 | 4.600E-08 | 4.011E-08 | 1.400E-08 | 9.100E-08 |
| 22097 | 1105 | L1105T | D | 1.9e-08 | 4.4e-08 | 6.1e-08 | 4.133E-08 | 2.113E-08 | 1.900E-08 | 6.100E-08 |
| 22098 | 1105 | L1105W | D | 4.7e-08 | 2e-08 | 1.1e-07 | 5.900E-08 | 4.618E-08 | 2.000E-08 | 1.100E-07 |
| 22099 | 1105 | L1105Y | D | 1.5e-08 | 2.9e-08 | 1.8e-07 | 7.467E-08 | 9.149E-08 | 1.500E-08 | 1.800E-07 |

| | | | | | | | | | | |
|---|---|---|---|---|---|---|---|---|---|---|
| 22100 | 1105 | L1105V | D | 5.4e-08 | 7.7e-09 | 3.3e-08 | 3.157E-08 | 2.318E-08 | 7.700E-09 | 5.400E-08 |
| 22101 | 1106 | I1106A | D | 3.4e-08 | 3.6e-08 | 6.2e-08 | 4.400E-08 | 1.562E-08 | 3.400E-08 | 6.200E-08 |
| 22102 | 1106 | I1106R | D | 6.2e-08 | 7.6e-08 | 2.3e-08 | 5.367E-08 | 2.747E-08 | 2.300E-08 | 7.600E-08 |
| 22103 | 1106 | I1106N | D | 4.6e-08 | 5.5e-08 | 3e-08 | 4.367E-08 | 1.266E-08 | 3.000E-08 | 5.500E-08 |
| 22104 | 1106 | I1106D | D | 1e-07 | 3.3e-08 | 1.7e-08 | 5.000E-08 | 4.403E-08 | 1.700E-08 | 1.000E-07 |
| 22105 | 1106 | I1106C | D | 2.6e-08 | 8.6e-09 | 4.1e-08 | 2.520E-08 | 1.621E-08 | 8.600E-09 | 4.100E-08 |
| 22106 | 1106 | I1106E | D | 6.8e-08 | 2.5e-08 | 4.3e-08 | 4.533E-08 | 2.159E-08 | 2.500E-08 | 6.800E-08 |
| 22107 | 1106 | I1106Q | D | 2.9e-08 | 7.6e-08 | 3.2e-08 | 4.567E-08 | 2.631E-08 | 2.900E-08 | 7.600E-08 |
| 22108 | 1106 | I1106G | D | 8.6e-08 | 2.4e-08 | 3.9e-08 | 4.967E-08 | 3.235E-08 | 2.400E-08 | 8.600E-08 |
| 22109 | 1106 | I1106H | D | 1.2e-08 | 5.5e-08 | 2.5e-08 | 3.067E-08 | 2.205E-08 | 1.200E-08 | 5.500E-08 |
| 22110 | 1106 | I1106I | D | 5e-08 | 3.3e-08 | 4e-08 | 4.100E-08 | 8.544E-09 | 3.300E-08 | 5.000E-08 |
| 22111 | 1106 | I1106L | D | 1.3e-07 | 6.4e-08 | 3.6e-08 | 7.667E-08 | 4.826E-08 | 3.600E-08 | 1.300E-07 |
| 22112 | 1106 | I1106K | D | 3.4e-08 | 1.5e-07 | 3.9e-08 | 7.433E-08 | 6.558E-08 | 3.400E-08 | 1.500E-07 |
| 22113 | 1106 | I1106M | D | 3.1e-08 | 1e-07 | 2.4e-08 | 5.167E-08 | 4.200E-08 | 2.400E-08 | 1.000E-07 |
| 22114 | 1106 | I1106F | D | 4.2e-08 | 5e-08 | 2.5e-07 | 1.140E-07 | 1.178E-07 | 4.200E-08 | 2.500E-07 |
| 22115 | 1106 | I1106P | D | 5e-08 | 6.8e-08 | 3.3e-08 | 5.033E-08 | 1.750E-08 | 3.300E-08 | 6.800E-08 |
| 22116 | 1106 | I1106S | D | 3.5e-08 | 5.7e-08 | 2.7e-08 | 3.967E-08 | 1.553E-08 | 2.700E-08 | 5.700E-08 |
| 22117 | 1106 | I1106T | D | 4e-08 | 1.1e-07 | 1.6e-08 | 5.533E-08 | 4.884E-08 | 1.600E-08 | 1.100E-07 |
| 22118 | 1106 | I1106W | D | 4.1e-08 | 1e-07 | 3.1e-08 | 5.733E-08 | 3.729E-08 | 3.100E-08 | 1.000E-07 |
| 22119 | 1106 | I1106Y | D | 2.5e-08 | 4.7e-08 | 1.5e-07 | 7.400E-08 | 6.673E-08 | 2.500E-08 | 1.500E-07 |
| 22120 | 1106 | I1106V | D | 5.6e-08 | 3.3e-08 | 1.1e-07 | 6.633E-08 | 3.953E-08 | 3.300E-08 | 1.100E-07 |
| 22121 | 1107 | Y1107A | D | 7.4e-08 | 1.6e-08 | 1.7e-08 | 3.567E-08 | 3.320E-08 | 1.600E-08 | 7.400E-08 |
| 22122 | 1107 | Y1107R | D | 3.4e-08 | 2.2e-08 | 5.6e-08 | 3.733E-08 | 1.724E-08 | 2.200E-08 | 5.600E-08 |
| 22123 | 1107 | Y1107N | D | 3e-08 | 7.5e-08 | 6e-08 | 5.500E-08 | 2.291E-08 | 3.000E-08 | 7.500E-08 |
| 22124 | 1107 | Y1107D | D | 1.9e-08 | 6.5e-08 | 6.5e-08 | 4.967E-08 | 2.656E-08 | 1.900E-08 | 6.500E-08 |
| 22125 | 1107 | Y1107C | D | 2.6e-08 | 5.3e-08 | 6.4e-08 | 4.767E-08 | 1.955E-08 | 2.600E-08 | 6.400E-08 |
| 22126 | 1107 | Y1107E | D | 2.7e-08 | 2.5e-08 | 2.6e-08 | 2.600E-08 | 1.000E-09 | 2.500E-08 | 2.700E-08 |
| 22127 | 1107 | Y1107Q | D | 6.8e-08 | 3e-08 | 2.4e-08 | 4.067E-08 | 2.386E-08 | 2.400E-08 | 6.800E-08 |
| 22128 | 1107 | Y1107G | D | 2.3e-08 | 2.8e-08 | 1.4e-07 | 6.367E-08 | 6.615E-08 | 2.300E-08 | 1.400E-07 |
| 22129 | 1107 | Y1107H | D | 2.8e-08 | 3.5e-08 | 1.9e-08 | 2.733E-08 | 8.021E-09 | 1.900E-08 | 3.500E-08 |
| 22130 | 1107 | Y1107I | D | 5.6e-08 | 2e-07 | 3.6e-08 | 9.733E-08 | 8.947E-08 | 3.600E-08 | 2.000E-07 |
| 22131 | 1107 | Y1107L | D | 3.2e-08 | 2.1e-07 | 9.3e-08 | 1.117E-07 | 9.046E-08 | 3.200E-08 | 2.100E-07 |
| 22132 | 1107 | Y1107K | D | 7.5e-08 | 3e-08 | 7.1e-08 | 5.867E-08 | 2.491E-08 | 3.000E-08 | 7.500E-08 |
| 22133 | 1107 | Y1107M | D | 9.2e-08 | 3.4e-08 | 3.8e-08 | 5.467E-08 | 3.239E-08 | 3.400E-08 | 9.200E-08 |
| 22134 | 1107 | Y1107F | D | 2.1e-08 | 1.5e-08 | 7.3e-08 | 3.633E-08 | 3.190E-08 | 1.500E-08 | 7.300E-08 |
| 22135 | 1107 | Y1107P | D | 5e-08 | 9.5e-08 | 6.7e-08 | 7.067E-08 | 2.272E-08 | 5.000E-08 | 9.500E-08 |
| 22136 | 1107 | Y1107S | D | 8.3e-09 | 1.4e-08 | 4.1e-08 | 2.110E-08 | 1.747E-08 | 8.300E-09 | 4.100E-08 |
| 22137 | 1107 | Y1107T | D | 3.3e-08 | 5.5e-08 | 7.3e-08 | 5.367E-08 | 2.003E-08 | 3.300E-08 | 7.300E-08 |
| 22138 | 1107 | Y1107W | D | 4.6e-08 | 6.6e-08 | 3.1e-08 | 4.767E-08 | 1.756E-08 | 3.100E-08 | 6.600E-08 |
| 22139 | 1107 | Y1107Y | D | 3.2e-08 | 6e-09 | 5.1e-08 | 2.967E-08 | 2.259E-08 | 6.000E-09 | 5.100E-08 |
| 22140 | 1107 | Y1107V | D | 4.4e-08 | 1.1e-08 | 2.1e-08 | 2.533E-08 | 1.692E-08 | 1.100E-08 | 4.400E-08 |
| 22141 | 1108 | S1108A | D | 6.7e-08 | 2.2e-08 | 2.9e-08 | 3.933E-08 | 2.421E-08 | 2.200E-08 | 6.700E-08 |
| 22142 | 1108 | S1108R | D | 9.9e-09 | 2e-08 | 3.6e-08 | 2.197E-08 | 1.316E-08 | 9.900E-09 | 3.600E-08 |
| 22143 | 1108 | S1108N | D | 3.6e-08 | 2.9e-08 | 3.5e-08 | 3.333E-08 | 3.786E-09 | 2.900E-08 | 3.600E-08 |
| 22144 | 1108 | S1108D | D | 3.8e-08 | 5.4e-08 | 3.8e-08 | 4.333E-08 | 9.238E-09 | 3.800E-08 | 5.400E-08 |
| 22145 | 1108 | S1108C | D | 2e-08 | 2e-08 | 7.4e-08 | 3.800E-08 | 3.118E-08 | 2.000E-08 | 7.400E-08 |
| 22146 | 1108 | S1108E | D | 6.9e-08 | 6.3e-08 | 2.8e-08 | 5.333E-08 | 2.214E-08 | 2.800E-08 | 6.900E-08 |
| 22147 | 1108 | S1108Q | D | 9.8e-08 | 7e-08 | 5.6e-08 | 7.467E-08 | 2.139E-08 | 5.600E-08 | 9.800E-08 |
| 22148 | 1108 | S1108G | D | 3.4e-08 | 5.4e-08 | 2.8e-08 | 3.867E-08 | 1.361E-08 | 2.800E-08 | 5.400E-08 |
| 22149 | 1108 | S1108H | D | 7.6e-08 | 2.3e-08 | 2.9e-08 | 4.267E-08 | 2.902E-08 | 2.300E-08 | 7.600E-08 |
| 22150 | 1108 | S1108I | D | 5.8e-08 | 2.9e-08 | 3e-08 | 3.900E-08 | 1.646E-08 | 2.900E-08 | 5.800E-08 |
| 22151 | 1108 | S1108L | D | 2.8e-08 | 1.4e-07 | 4.8e-08 | 7.200E-08 | 5.973E-08 | 2.800E-08 | 1.400E-07 |
| 22152 | 1108 | S1108K | D | 6.9e-08 | 5.5e-08 | 2.1e-08 | 4.833E-08 | 2.468E-08 | 2.100E-08 | 6.900E-08 |
| 22153 | 1108 | S1108M | D | 2.3e-08 | 1e-07 | 2.5e-08 | 4.933E-08 | 4.389E-08 | 2.300E-08 | 1.000E-07 |
| 22154 | 1108 | S1108F | D | 1.1e-07 | 2.2e-08 | 2.4e-08 | 5.200E-08 | 5.024E-08 | 2.200E-08 | 1.100E-07 |
| 22155 | 1108 | S1108P | D | 5.2e-08 | 4.4e-08 | 3e-08 | 4.200E-08 | 1.114E-08 | 3.000E-08 | 5.200E-08 |
| 22156 | 1108 | S1108S | D | 4.9e-08 | 4.4e-08 | 3.6e-08 | 4.300E-08 | 6.557E-09 | 3.600E-08 | 4.900E-08 |
| 22157 | 1108 | S1108T | D | 1.3e-07 | 4.4e-08 | 5.7e-08 | 7.700E-08 | 4.636E-08 | 4.400E-08 | 1.300E-07 |
| 22158 | 1108 | S1108W | D | 2.4e-08 | 2.3e-08 | 7.5e-08 | 4.067E-08 | 2.974E-08 | 2.300E-08 | 7.500E-08 |
| 22159 | 1108 | S1108Y | D | 1.5e-08 | 3e-08 | 1e-07 | 4.833E-08 | 4.537E-08 | 1.500E-08 | 1.000E-07 |
| 22160 | 1108 | S1108V | D | 3.8e-08 | 1.5e-08 | 6.2e-08 | 3.833E-08 | 2.350E-08 | 1.500E-08 | 6.200E-08 |
| 22161 | 1109 | A1109A | D | 8e-08 | 4.2e-08 | 1.8e-08 | 4.667E-08 | 3.126E-08 | 1.800E-08 | 8.000E-08 |
| 22162 | 1109 | A1109R | D | 1.8e-08 | 3.6e-08 | 1.4e-08 | 2.267E-08 | 1.172E-08 | 1.400E-08 | 3.600E-08 |
| 22163 | 1109 | A1109N | D | 4.2e-08 | 6.2e-08 | 1.7e-08 | 4.033E-08 | 2.255E-08 | 1.700E-08 | 6.200E-08 |
| 22164 | 1109 | A1109D | D | 9.7e-08 | 2.9e-08 | 2.9e-08 | 5.167E-08 | 3.926E-08 | 2.900E-08 | 9.700E-08 |
| 22165 | 1109 | A1109C | D | 5.2e-08 | 3.5e-08 | 3.2e-08 | 3.967E-08 | 1.079E-08 | 3.200E-08 | 5.200E-08 |
| 22166 | 1109 | A1109E | D | 2.2e-08 | 1.4e-08 | 4.6e-08 | 2.733E-08 | 1.665E-08 | 1.400E-08 | 4.600E-08 |
| 22167 | 1109 | A1109Q | D | 2.5e-08 | 5.2e-08 | 5.9e-08 | 4.533E-08 | 1.795E-08 | 2.500E-08 | 5.900E-08 |
| 22168 | 1109 | A1109G | D | 8.7e-08 | 9e-08 | 3.6e-08 | 7.100E-08 | 3.035E-08 | 3.600E-08 | 9.000E-08 |
| 22169 | 1109 | A1109H | D | 6.9e-08 | 2e-08 | 8.2e-08 | 5.700E-08 | 3.270E-08 | 2.000E-08 | 8.200E-08 |
| 22170 | 1109 | A1109I | D | 6e-08 | 2.6e-08 | 4.2e-08 | 4.267E-08 | 1.701E-08 | 2.600E-08 | 6.000E-08 |
| 22171 | 1109 | A1109L | D | 2.8e-08 | 6.1e-08 | 5e-08 | 4.633E-08 | 1.680E-08 | 2.800E-08 | 6.100E-08 |
| 22172 | 1109 | A1109K | D | 1.8e-07 | 3.6e-08 | 3e-08 | 8.200E-08 | 8.492E-08 | 3.000E-08 | 1.800E-07 |
| 22173 | 1109 | A1109M | D | 6.1e-08 | 6.3e-08 | 4.8e-08 | 5.733E-08 | 8.145E-09 | 4.800E-08 | 6.300E-08 |
| 22174 | 1109 | A1109F | D | 2.9e-08 | 5.1e-08 | 2.8e-08 | 3.600E-08 | 1.300E-08 | 2.800E-08 | 5.100E-08 |
| 22175 | 1109 | A1109P | D | 6.6e-08 | 1.3e-07 | 6.7e-08 | 8.767E-08 | 3.667E-08 | 6.600E-08 | 1.300E-07 |
| 22176 | 1109 | A1109S | D | 1.5e-08 | 3.9e-08 | 1.6e-08 | 2.333E-08 | 1.358E-08 | 1.500E-08 | 3.900E-08 |
| 22177 | 1109 | A1109T | D | 8e-08 | 8.7e-08 | 6.1e-08 | 7.600E-08 | 1.345E-08 | 6.100E-08 | 8.700E-08 |
| 22178 | 1109 | A1109W | D | 4e-08 | 4.9e-08 | 1.5e-08 | 3.467E-08 | 1.762E-08 | 1.500E-08 | 4.900E-08 |
| 22179 | 1109 | A1109Y | D | 9.7e-09 | 1e-07 | 2.9e-08 | 4.623E-08 | 4.755E-08 | 9.700E-09 | 1.000E-07 |
| 22180 | 1109 | A1109V | D | 1.8e-08 | 7.1e-08 | 1.5e-08 | 3.467E-08 | 3.150E-08 | 1.500E-08 | 7.100E-08 |
| 22181 | 1110 | S1110A | D | 2.8e-08 | 4e-08 | 3.3e-08 | 3.367E-08 | 6.028E-09 | 2.800E-08 | 4.000E-08 |
| 22182 | 1110 | S1110R | D | 3.1e-08 | 7.8e-09 | 9.1e-08 | 4.327E-08 | 4.293E-08 | 7.800E-09 | 9.100E-08 |
| 22183 | 1110 | S1110N | D | 3.9e-08 | 6.2e-08 | 2.4e-08 | 4.167E-08 | 1.914E-08 | 2.400E-08 | 6.200E-08 |
| 22184 | 1110 | S1110D | D | 8.6e-08 | 5.5e-08 | 2.4e-07 | 1.270E-07 | 9.908E-08 | 5.500E-08 | 2.400E-07 |

| | | | | | | | | | | |
|---|---|---|---|---|---|---|---|---|---|---|
| 22185 | 1110 | S1110C | D | 3.1e-08 | 3.3e-08 | 5.9e-08 | 4.100E-08 | 1.562E-08 | 3.100E-08 | 5.900E-08 |
| 22186 | 1110 | S1110E | D | 2.9e-08 | 1.8e-08 | 1.5e-08 | 2.067E-08 | 7.371E-09 | 1.500E-08 | 2.900E-08 |
| 22187 | 1110 | S1110Q | D | 5.9e-08 | 6.1e-08 | 3.1e-08 | 5.033E-08 | 1.677E-08 | 3.100E-08 | 6.100E-08 |
| 22188 | 1110 | S1110G | D | 4.5e-08 | 5.8e-08 | 2.4e-08 | 4.233E-08 | 1.716E-08 | 2.400E-08 | 5.800E-08 |
| 22189 | 1110 | S1110H | D | 3.3e-08 | 6.3e-08 | 5e-08 | 4.867E-08 | 1.504E-08 | 3.300E-08 | 6.300E-08 |
| 22190 | 1110 | S1110I | D | 2.5e-08 | 1.2e-07 | 4.7e-08 | 6.400E-08 | 4.973E-08 | 2.500E-08 | 1.200E-07 |
| 22191 | 1110 | S1110L | D | 9.6e-08 | 3.7e-08 | 6e-08 | 6.433E-08 | 2.974E-08 | 3.700E-08 | 9.600E-08 |
| 22192 | 1110 | S1110K | D | 5.3e-08 | 7.7e-08 | 2.7e-08 | 5.233E-08 | 2.501E-08 | 2.700E-08 | 7.700E-08 |
| 22193 | 1110 | S1110M | D | 5.9e-08 | 7e-08 | 4.9e-08 | 5.933E-08 | 1.050E-08 | 4.900E-08 | 7.000E-08 |
| 22194 | 1110 | S1110F | D | 3.9e-08 | 5.2e-08 | 4.4e-08 | 4.500E-08 | 6.557E-09 | 3.900E-08 | 5.200E-08 |
| 22195 | 1110 | S1110P | D | 4.4e-08 | 6.2e-08 | 2.2e-08 | 4.267E-08 | 2.003E-08 | 2.200E-08 | 6.200E-08 |
| 22196 | 1110 | S1110S | D | 1.6e-08 | 5.7e-08 | 5.2e-08 | 4.167E-08 | 2.237E-08 | 1.600E-08 | 5.700E-08 |
| 22197 | 1110 | S1110T | D | 3.8e-08 | 5.1e-08 | 2.3e-08 | 3.733E-08 | 1.401E-08 | 2.300E-08 | 5.100E-08 |
| 22198 | 1110 | S1110W | D | 6.5e-08 | 1.2e-07 | 5.9e-08 | 8.133E-08 | 3.362E-08 | 5.900E-08 | 1.200E-07 |
| 22199 | 1110 | S1110Y | D | 3.5e-08 | 4.1e-08 | 2.3e-08 | 3.300E-08 | 9.165E-09 | 2.300E-08 | 4.100E-08 |
| 22200 | 1110 | S1110V | D | 1.2e-08 | 3.4e-08 | 4.9e-08 | 3.167E-08 | 1.861E-08 | 1.200E-08 | 4.900E-08 |
| 22201 | 1111 | F1111A | D | 2.3e-08 | 6.2e-08 | 2.2e-08 | 3.567E-08 | 2.281E-08 | 2.200E-08 | 6.200E-08 |
| 22202 | 1111 | F1111R | D | 3.5e-08 | 1e-07 | 9.6e-09 | 4.820E-08 | 4.662E-08 | 9.600E-09 | 1.000E-07 |
| 22203 | 1111 | F1111N | D | 1.3e-07 | 2.8e-08 | 2.8e-08 | 6.200E-08 | 5.889E-08 | 2.800E-08 | 1.300E-07 |
| 22204 | 1111 | F1111D | D | 1.9e-08 | 1.1e-08 | 6.8e-08 | 3.267E-08 | 3.086E-08 | 1.100E-08 | 6.800E-08 |
| 22205 | 1111 | F1111C | D | 5.2e-08 | 1.9e-08 | 8.4e-08 | 5.167E-08 | 3.250E-08 | 1.900E-08 | 8.400E-08 |
| 22206 | 1111 | F1111E | D | 3.3e-08 | 2.2e-08 | 1.3e-07 | 6.167E-08 | 5.943E-08 | 2.200E-08 | 1.300E-07 |
| 22207 | 1111 | F1111Q | D | 5.2e-08 | 1.9e-08 | 9.2e-08 | 5.433E-08 | 3.656E-08 | 1.900E-08 | 9.200E-08 |
| 22208 | 1111 | F1111G | D | 3e-08 | 6.4e-08 | 3.8e-08 | 4.400E-08 | 1.778E-08 | 3.000E-08 | 6.400E-08 |
| 22209 | 1111 | F1111H | D | 6.8e-08 | 3.1e-08 | 1.2e-07 | 7.300E-08 | 4.471E-08 | 3.100E-08 | 1.200E-07 |
| 22210 | 1111 | F1111I | D | 8.4e-08 | 3.5e-08 | 2.8e-08 | 4.900E-08 | 3.051E-08 | 2.800E-08 | 8.400E-08 |
| 22211 | 1111 | F1111L | D | 5.6e-08 | 5.6e-08 | 4e-08 | 5.067E-08 | 9.238E-09 | 4.000E-08 | 5.600E-08 |
| 22212 | 1111 | F1111K | D | 9.2e-08 | 2.3e-08 | 3.8e-08 | 5.100E-08 | 3.629E-08 | 2.300E-08 | 9.200E-08 |
| 22213 | 1111 | F1111M | D | 6.8e-08 | 5.8e-08 | 8.8e-08 | 7.133E-08 | 1.528E-08 | 5.800E-08 | 8.800E-08 |
| 22214 | 1111 | F1111F | D | 3.2e-08 | 8.3e-08 | 5.1e-08 | 5.533E-08 | 2.577E-08 | 3.200E-08 | 8.300E-08 |
| 22215 | 1111 | F1111P | D | 4.4e-08 | 3.2e-08 | 7.7e-08 | 5.100E-08 | 2.330E-08 | 3.200E-08 | 7.700E-08 |
| 22216 | 1111 | F1111S | D | 5.2e-08 | 4.5e-08 | 3.5e-08 | 4.400E-08 | 8.544E-09 | 3.500E-08 | 5.200E-08 |
| 22217 | 1111 | F1111T | D | 3.4e-08 | 5.6e-08 | 3.4e-08 | 4.133E-08 | 1.270E-08 | 3.400E-08 | 5.600E-08 |
| 22218 | 1111 | F1111W | D | 3.9e-08 | 2.1e-08 | 3.6e-08 | 3.200E-08 | 9.644E-09 | 2.100E-08 | 3.900E-08 |
| 22219 | 1111 | F1111Y | D | 2.2e-08 | 3.6e-08 | 2.9e-08 | 2.900E-08 | 7.000E-09 | 2.200E-08 | 3.600E-08 |
| 22220 | 1111 | F1111V | D | 7e-08 | 3.9e-08 | 1.7e-08 | 4.200E-08 | 2.663E-08 | 1.700E-08 | 7.000E-08 |
| 22221 | 1112 | L1112A | D | 2.4e-08 | 1.1e-07 | 8.5e-08 | 7.300E-08 | 4.424E-08 | 2.400E-08 | 1.100E-07 |
| 22222 | 1112 | L1112R | D | 3.1e-08 | 3.8e-08 | 3.3e-08 | 3.400E-08 | 3.606E-09 | 3.100E-08 | 3.800E-08 |
| 22223 | 1112 | L1112N | D | 4.2e-08 | 2.5e-08 | 4.7e-08 | 3.800E-08 | 1.153E-08 | 2.500E-08 | 4.700E-08 |
| 22224 | 1112 | L1112D | D | 5.2e-08 | 2e-08 | 3e-08 | 3.400E-08 | 1.637E-08 | 2.000E-08 | 5.200E-08 |
| 22225 | 1112 | L1112C | D | 1.7e-08 | 4.6e-08 | 6e-08 | 4.100E-08 | 2.193E-08 | 1.700E-08 | 6.000E-08 |
| 22226 | 1112 | L1112E | D | 1.8e-08 | 6.6e-08 | 5.6e-08 | 4.667E-08 | 2.532E-08 | 1.800E-08 | 6.600E-08 |
| 22227 | 1112 | L1112Q | D | 4.4e-08 | 9.8e-08 | 1.9e-08 | 5.367E-08 | 4.038E-08 | 1.900E-08 | 9.800E-08 |
| 22228 | 1112 | L1112G | D | 3.7e-08 | 1.8e-07 | 9.2e-08 | 1.030E-07 | 7.213E-08 | 3.700E-08 | 1.800E-07 |
| 22229 | 1112 | L1112H | D | 4.7e-08 | 1.6e-08 | 1.1e-08 | 2.467E-08 | 1.950E-08 | 1.100E-08 | 4.700E-08 |
| 22230 | 1112 | L1112I | D | 5.6e-08 | 7.5e-08 | 6.9e-08 | 6.667E-08 | 9.713E-09 | 5.600E-08 | 7.500E-08 |
| 22231 | 1112 | L1112L | D | 4.2e-08 | 2.8e-08 | 2.7e-08 | 3.233E-08 | 8.386E-09 | 2.700E-08 | 4.200E-08 |
| 22232 | 1112 | L1112K | D | 3.7e-08 | 1.6e-08 | 2.1e-08 | 2.467E-08 | 1.097E-08 | 1.600E-08 | 3.700E-08 |
| 22233 | 1112 | L1112M | D | 1.2e-08 | 8.6e-08 | 3.4e-08 | 4.400E-08 | 3.800E-08 | 1.200E-08 | 8.600E-08 |
| 22234 | 1112 | L1112F | D | 1.5e-07 | 3.2e-08 | 3.4e-08 | 7.200E-08 | 6.756E-08 | 3.200E-08 | 1.500E-07 |
| 22235 | 1112 | L1112P | D | 3.3e-08 | 4.6e-08 | 5.9e-08 | 4.600E-08 | 1.300E-08 | 3.300E-08 | 5.900E-08 |
| 22236 | 1112 | L1112S | D | 2.2e-08 | 4.3e-08 | 4.5e-08 | 3.667E-08 | 1.274E-08 | 2.200E-08 | 4.500E-08 |
| 22237 | 1112 | L1112T | D | 9.2e-08 | 8.2e-08 | 6.5e-08 | 7.967E-08 | 1.365E-08 | 6.500E-08 | 9.200E-08 |
| 22238 | 1112 | L1112W | D | 3.7e-08 | 6.1e-08 | 4.5e-08 | 4.767E-08 | 1.222E-08 | 3.700E-08 | 6.100E-08 |
| 22239 | 1112 | L1112Y | D | 2.4e-08 | 2.7e-08 | 1.4e-08 | 2.167E-08 | 6.807E-09 | 1.400E-08 | 2.700E-08 |
| 22240 | 1112 | L1112V | D | 7.9e-08 | 1.4e-08 | 2.1e-08 | 3.800E-08 | 3.568E-08 | 1.400E-08 | 7.900E-08 |
| 22241 | 1113 | Y1113A | D | 4.6e-08 | 2.4e-08 | 2.2e-08 | 3.067E-08 | 1.332E-08 | 2.200E-08 | 4.600E-08 |
| 22242 | 1113 | Y1113R | D | 5.3e-08 | 7.4e-08 | 3.1e-08 | 5.267E-08 | 2.150E-08 | 3.100E-08 | 7.400E-08 |
| 22243 | 1113 | Y1113N | D | 3.9e-08 | 2.3e-08 | 2.1e-08 | 2.767E-08 | 9.866E-09 | 2.100E-08 | 3.900E-08 |
| 22244 | 1113 | Y1113D | D | 6.8e-08 | 1.8e-08 | 2.8e-08 | 3.800E-08 | 2.646E-08 | 1.800E-08 | 6.800E-08 |
| 22245 | 1113 | Y1113C | D | 6.3e-08 | 6.8e-08 | 1.8e-08 | 4.967E-08 | 2.754E-08 | 1.800E-08 | 6.800E-08 |
| 22246 | 1113 | Y1113E | D | 1.1e-07 | 5.8e-08 | 7.8e-08 | 8.200E-08 | 2.623E-08 | 5.800E-08 | 1.100E-07 |
| 22247 | 1113 | Y1113Q | D | 1.6e-08 | 5e-08 | 2.6e-08 | 3.067E-08 | 1.747E-08 | 1.600E-08 | 5.000E-08 |
| 22248 | 1113 | Y1113G | D | 3.1e-08 | 4.7e-08 | 3.2e-08 | 3.667E-08 | 8.963E-09 | 3.100E-08 | 4.700E-08 |
| 22249 | 1113 | Y1113H | D | 3.5e-08 | 4.4e-08 | 2.9e-08 | 3.600E-08 | 7.550E-09 | 2.900E-08 | 4.400E-08 |
| 22250 | 1113 | Y1113I | D | 5e-08 | 4.8e-08 | 2.2e-08 | 4.000E-08 | 1.562E-08 | 2.200E-08 | 5.000E-08 |
| 22251 | 1113 | Y1113L | D | 7.3e-08 | 8.2e-08 | 4.9e-08 | 6.800E-08 | 1.706E-08 | 4.900E-08 | 8.200E-08 |
| 22252 | 1113 | Y1113K | D | 1e-07 | 1e-08 | 1.5e-08 | 4.167E-08 | 5.058E-08 | 1.000E-08 | 1.000E-07 |
| 22253 | 1113 | Y1113M | D | 6.9e-08 | 2.5e-08 | 2.7e-08 | 4.033E-08 | 2.485E-08 | 2.500E-08 | 6.900E-08 |
| 22254 | 1113 | Y1113F | D | 3.8e-08 | 3.5e-08 | 3.1e-08 | 3.467E-08 | 3.512E-09 | 3.100E-08 | 3.800E-08 |
| 22255 | 1113 | Y1113P | D | 6.6e-08 | 3.3e-08 | 5.7e-08 | 5.200E-08 | 1.706E-08 | 3.300E-08 | 6.600E-08 |
| 22256 | 1113 | Y1113S | D | 2.3e-08 | 2.3e-08 | 1.3e-07 | 5.867E-08 | 6.178E-08 | 2.300E-08 | 1.300E-07 |
| 22257 | 1113 | Y1113T | D | 2.4e-08 | 6.1e-08 | 3.6e-08 | 4.033E-08 | 1.888E-08 | 2.400E-08 | 6.100E-08 |
| 22258 | 1113 | Y1113W | D | 4.3e-08 | 3e-08 | 6.9e-09 | 2.663E-08 | 1.828E-08 | 6.900E-09 | 4.300E-08 |
| 22259 | 1113 | Y1113Y | D | 6.7e-08 | 4.8e-08 | 7.6e-08 | 6.367E-08 | 1.429E-08 | 4.800E-08 | 7.600E-08 |
| 22260 | 1113 | Y1113V | D | 3.1e-08 | 2.4e-08 | 3.1e-08 | 2.867E-08 | 4.041E-09 | 2.400E-08 | 3.100E-08 |
| 22261 | 1114 | S1114A | D | 7.8e-08 | 5.3e-08 | 8.7e-08 | 7.267E-08 | 1.762E-08 | 5.300E-08 | 8.700E-08 |
| 22262 | 1114 | S1114R | D | 3.4e-08 | 3.3e-08 | 7.7e-08 | 4.800E-08 | 2.512E-08 | 3.300E-08 | 7.700E-08 |
| 22263 | 1114 | S1114N | D | 4.2e-08 | 1.2e-07 | 3.8e-08 | 6.667E-08 | 4.623E-08 | 3.800E-08 | 1.200E-07 |
| 22264 | 1114 | S1114D | D | 3.8e-08 | 7.1e-09 | 5.5e-08 | 3.337E-08 | 2.428E-08 | 7.100E-09 | 5.500E-08 |
| 22265 | 1114 | S1114C | D | 4.7e-08 | 6.2e-08 | 3.8e-08 | 4.900E-08 | 1.212E-08 | 3.800E-08 | 6.200E-08 |
| 22266 | 1114 | S1114E | D | 7.2e-08 | 1.8e-08 | 3.6e-08 | 4.200E-08 | 2.750E-08 | 1.800E-08 | 7.200E-08 |
| 22267 | 1114 | S1114Q | D | 2.9e-08 | 2.9e-08 | 1e-07 | 5.267E-08 | 4.099E-08 | 2.900E-08 | 1.000E-07 |
| 22268 | 1114 | S1114G | D | 2.8e-08 | 2.7e-08 | 2.9e-08 | 2.800E-08 | 1.000E-09 | 2.700E-08 | 2.900E-08 |
| 22269 | 1114 | S1114H | D | 9e-08 | 6.4e-08 | 2.9e-08 | 6.100E-08 | 3.061E-08 | 2.900E-08 | 9.000E-08 |

| | | | | | | | | | | |
|---|---|---|---|---|---|---|---|---|---|---|
| 22270 | 1114 | S1114I | D | 5.1e-08 | 6.8e-08 | 3e-08 | 4.967E-08 | 1.904E-08 | 3.000E-08 | 6.800E-08 |
| 22271 | 1114 | S1114L | D | 1.3e-07 | 2.4e-08 | 1.9e-08 | 5.767E-08 | 6.269E-08 | 1.900E-08 | 1.300E-07 |
| 22272 | 1114 | S1114K | D | 7.8e-08 | 5e-08 | 7.3e-08 | 6.700E-08 | 1.493E-08 | 5.000E-08 | 7.800E-08 |
| 22273 | 1114 | S1114M | D | 4.7e-08 | 6e-08 | 1.1e-07 | 7.233E-08 | 3.326E-08 | 4.700E-08 | 1.100E-07 |
| 22274 | 1114 | S1114F | D | 4.7e-08 | 3.1e-08 | 6.4e-08 | 4.733E-08 | 1.650E-08 | 3.100E-08 | 6.400E-08 |
| 22275 | 1114 | S1114P | D | 9.3e-08 | 1.2e-08 | 4.2e-08 | 4.900E-08 | 4.095E-08 | 1.200E-08 | 9.300E-08 |
| 22276 | 1114 | S1114S | D | 3.6e-08 | 4.2e-08 | 9.7e-08 | 5.833E-08 | 3.362E-08 | 3.600E-08 | 9.700E-08 |
| 22277 | 1114 | S1114T | D | 9.6e-08 | 2.9e-08 | 4.2e-08 | 5.567E-08 | 3.553E-08 | 2.900E-08 | 9.600E-08 |
| 22278 | 1114 | S1114W | D | 2.5e-08 | 2.8e-08 | 8.6e-08 | 4.633E-08 | 3.439E-08 | 2.500E-08 | 8.600E-08 |
| 22279 | 1114 | S1114Y | D | 4.6e-08 | 7e-08 | 2.9e-08 | 4.833E-08 | 2.060E-08 | 2.900E-08 | 7.000E-08 |
| 22280 | 1114 | S1114V | D | 7.4e-08 | 2.3e-08 | 3.5e-08 | 4.400E-08 | 2.666E-08 | 2.300E-08 | 7.400E-08 |
| 22281 | 1115 | G1115A | D | 4.8e-08 | 7.3e-08 | 3.3e-08 | 5.133E-08 | 2.021E-08 | 3.300E-08 | 7.300E-08 |
| 22282 | 1115 | G1115R | D | 1.8e-08 | 4.3e-08 | 2.9e-08 | 3.000E-08 | 1.253E-08 | 1.800E-08 | 4.300E-08 |
| 22283 | 1115 | G1115N | D | 2.1e-08 | 3.3e-08 | 1.9e-08 | 2.433E-08 | 7.572E-09 | 1.900E-08 | 3.300E-08 |
| 22284 | 1115 | G1115D | D | 7.7e-08 | 6.3e-08 | 1.3e-07 | 9.000E-08 | 3.534E-08 | 6.300E-08 | 1.300E-07 |
| 22285 | 1115 | G1115C | D | 7.8e-08 | 3.4e-08 | 5.2e-08 | 5.467E-08 | 2.212E-08 | 3.400E-08 | 7.800E-08 |
| 22286 | 1115 | G1115E | D | 7e-08 | 3.6e-08 | 3.9e-08 | 4.833E-08 | 1.882E-08 | 3.600E-08 | 7.000E-08 |
| 22287 | 1115 | G1115Q | D | 5.5e-08 | 6e-08 | 4.2e-08 | 5.233E-08 | 9.292E-09 | 4.200E-08 | 6.000E-08 |
| 22288 | 1115 | G1115G | D | 4.3e-08 | 1.7e-08 | 4.9e-08 | 3.633E-08 | 1.701E-08 | 1.700E-08 | 4.900E-08 |
| 22289 | 1115 | G1115H | D | 7.3e-08 | 2.5e-08 | 3.7e-08 | 4.500E-08 | 2.498E-08 | 2.500E-08 | 7.300E-08 |
| 22290 | 1115 | G1115I | D | 2.4e-08 | 3.8e-08 | 7.6e-08 | 4.600E-08 | 2.691E-08 | 2.400E-08 | 7.600E-08 |
| 22291 | 1115 | G1115L | D | 1.9e-08 | 3.4e-08 | 5.1e-08 | 3.467E-08 | 1.601E-08 | 1.900E-08 | 5.100E-08 |
| 22292 | 1115 | G1115K | D | 7.3e-08 | 8.7e-08 | 3.6e-08 | 6.533E-08 | 2.635E-08 | 3.600E-08 | 8.700E-08 |
| 22293 | 1115 | G1115M | D | 7.2e-08 | 3.3e-08 | 4.4e-08 | 4.967E-08 | 2.011E-08 | 3.300E-08 | 7.200E-08 |
| 22294 | 1115 | G1115F | D | 1.5e-08 | 5.9e-08 | 6.3e-08 | 4.567E-08 | 2.663E-08 | 1.500E-08 | 6.300E-08 |
| 22295 | 1115 | G1115P | D | 5e-08 | 3e-08 | 4.7e-08 | 4.233E-08 | 1.079E-08 | 3.000E-08 | 5.000E-08 |
| 22296 | 1115 | G1115S | D | 6.5e-08 | 5e-08 | 2.6e-08 | 4.700E-08 | 1.967E-08 | 2.600E-08 | 6.500E-08 |
| 22297 | 1115 | G1115T | D | 5.5e-08 | 4.4e-08 | 5.3e-08 | 5.067E-08 | 5.859E-09 | 4.400E-08 | 5.500E-08 |
| 22298 | 1115 | G1115W | D | 5.2e-08 | 4.1e-08 | 2.8e-08 | 4.033E-08 | 1.201E-08 | 2.800E-08 | 5.200E-08 |
| 22299 | 1115 | G1115Y | D | 3.4e-08 | 7.8e-08 | 2e-08 | 4.400E-08 | 3.027E-08 | 2.000E-08 | 7.800E-08 |
| 22300 | 1115 | G1115V | D | 1.6e-07 | 7.8e-08 | 4.4e-08 | 9.400E-08 | 5.963E-08 | 4.400E-08 | 1.600E-07 |
| 22301 | 1116 | V1116A | D | 7.7e-08 | 4.9e-08 | 7.6e-08 | 6.733E-08 | 1.589E-08 | 4.900E-08 | 7.700E-08 |
| 22302 | 1116 | V1116R | D | 2.6e-08 | 2e-08 | 2.9e-08 | 2.500E-08 | 4.583E-09 | 2.000E-08 | 2.900E-08 |
| 22303 | 1116 | V1116N | D | 4e-08 | 3e-08 | 2e-08 | 3.000E-08 | 1.000E-08 | 2.000E-08 | 4.000E-08 |
| 22304 | 1116 | V1116D | D | 3.1e-08 | 4.3e-08 | 6.2e-08 | 4.533E-08 | 1.563E-08 | 3.100E-08 | 6.200E-08 |
| 22305 | 1116 | V1116C | D | 5.6e-08 | 3.1e-08 | 5.1e-08 | 4.600E-08 | 1.323E-08 | 3.100E-08 | 5.600E-08 |
| 22306 | 1116 | V1116E | D | 1.8e-08 | 4.5e-08 | 6.7e-08 | 4.333E-08 | 2.454E-08 | 1.800E-08 | 6.700E-08 |
| 22307 | 1116 | V1116Q | D | 4.5e-08 | 6.4e-08 | 5.5e-08 | 5.467E-08 | 9.504E-09 | 4.500E-08 | 6.400E-08 |
| 22308 | 1116 | V1116G | D | 6.5e-08 | 1.3e-08 | 1.4e-07 | 7.267E-08 | 6.385E-08 | 1.300E-08 | 1.400E-07 |
| 22309 | 1116 | V1116H | D | 3.3e-08 | 3.4e-08 | 7.8e-08 | 4.833E-08 | 2.570E-08 | 3.300E-08 | 7.800E-08 |
| 22310 | 1116 | V1116I | D | 3.4e-08 | 2.7e-08 | 3.3e-08 | 3.133E-08 | 3.786E-09 | 2.700E-08 | 3.400E-08 |
| 22311 | 1116 | V1116L | D | 6.9e-08 | 1.9e-08 | 4.5e-08 | 4.433E-08 | 2.501E-08 | 1.900E-08 | 6.900E-08 |
| 22312 | 1116 | V1116K | D | 2.5e-08 | 1.6e-08 | 7.4e-08 | 3.833E-08 | 3.121E-08 | 1.600E-08 | 7.400E-08 |
| 22313 | 1116 | V1116M | D | 1.7e-08 | 1.2e-08 | 1e-07 | 4.300E-08 | 4.943E-08 | 1.200E-08 | 1.000E-07 |
| 22314 | 1116 | V1116F | D | 3.5e-08 | 7.3e-08 | 5.2e-08 | 5.333E-08 | 1.904E-08 | 3.500E-08 | 7.300E-08 |
| 22315 | 1116 | V1116P | D | 2.7e-08 | 5.7e-08 | 2.3e-08 | 3.567E-08 | 1.858E-08 | 2.300E-08 | 5.700E-08 |
| 22316 | 1116 | V1116S | D | 2.1e-08 | 2.9e-08 | 3.6e-08 | 2.867E-08 | 7.506E-09 | 2.100E-08 | 3.600E-08 |
| 22317 | 1116 | V1116T | D | 1e-07 | 2.8e-08 | 1.1e-08 | 4.633E-08 | 4.725E-08 | 1.100E-08 | 1.000E-07 |
| 22318 | 1116 | V1116W | D | 4.3e-08 | 1.4e-08 | 6.6e-08 | 4.100E-08 | 2.606E-08 | 1.400E-08 | 6.600E-08 |
| 22319 | 1116 | V1116Y | D | 2.2e-08 | 1.9e-08 | 8.7e-08 | 4.267E-08 | 3.842E-08 | 1.900E-08 | 8.700E-08 |
| 22320 | 1116 | V1116V | D | 4.5e-08 | 3e-08 | 6.1e-08 | 4.533E-08 | 1.550E-08 | 3.000E-08 | 6.100E-08 |
| 22321 | 1117 | P1117A | D | 4.6e-08 | 1.9e-08 | 4.6e-08 | 3.700E-08 | 1.559E-08 | 1.900E-08 | 4.600E-08 |
| 22322 | 1117 | P1117R | D | 3.5e-08 | 2.1e-08 | 3.8e-08 | 3.133E-08 | 9.074E-09 | 2.100E-08 | 3.800E-08 |
| 22323 | 1117 | P1117N | D | 2.8e-08 | 6.2e-08 | 3.8e-08 | 4.267E-08 | 1.747E-08 | 2.800E-08 | 6.200E-08 |
| 22324 | 1117 | P1117D | D | 6.8e-08 | 2.7e-08 | 5.8e-08 | 5.100E-08 | 2.138E-08 | 2.700E-08 | 6.800E-08 |
| 22325 | 1117 | P1117C | D | 1.8e-08 | 2.1e-08 | 5.1e-08 | 3.000E-08 | 1.825E-08 | 1.800E-08 | 5.100E-08 |
| 22326 | 1117 | P1117E | D | 7.3e-08 | 1e-07 | 9.4e-08 | 8.900E-08 | 1.418E-08 | 7.300E-08 | 1.000E-07 |
| 22327 | 1117 | P1117Q | D | 4.2e-08 | 3.1e-08 | 1e-07 | 5.767E-08 | 3.707E-08 | 3.100E-08 | 1.000E-07 |
| 22328 | 1117 | P1117G | D | 1.5e-07 | 1.9e-08 | 3.2e-08 | 6.700E-08 | 7.217E-08 | 1.900E-08 | 1.500E-07 |
| 22329 | 1117 | P1117H | D | 7.4e-08 | 1.6e-08 | 6.8e-08 | 5.267E-08 | 3.190E-08 | 1.600E-08 | 7.400E-08 |
| 22330 | 1117 | P1117I | D | 2.9e-08 | 4.2e-08 | 3.1e-08 | 3.400E-08 | 7.000E-09 | 2.900E-08 | 4.200E-08 |
| 22331 | 1117 | P1117L | D | 4e-08 | 3.7e-08 | 3.6e-08 | 3.767E-08 | 2.082E-09 | 3.600E-08 | 4.000E-08 |
| 22332 | 1117 | P1117K | D | 3.1e-08 | 8.5e-08 | 1.5e-07 | 8.867E-08 | 5.958E-08 | 3.100E-08 | 1.500E-07 |
| 22333 | 1117 | P1117M | D | 3.8e-08 | 6e-08 | 8.8e-08 | 6.200E-08 | 2.506E-08 | 3.800E-08 | 8.800E-08 |
| 22334 | 1117 | P1117F | D | 1.1e-08 | 2.2e-08 | 2.7e-08 | 2.000E-08 | 8.185E-09 | 1.100E-08 | 2.700E-08 |
| 22335 | 1117 | P1117P | D | 2.5e-08 | 5.6e-08 | 4e-08 | 4.033E-08 | 1.550E-08 | 2.500E-08 | 5.600E-08 |
| 22336 | 1117 | P1117S | D | 2.4e-08 | 5.2e-08 | 2.4e-08 | 3.333E-08 | 1.617E-08 | 2.400E-08 | 5.200E-08 |
| 22337 | 1117 | P1117T | D | 2.7e-08 | 3.8e-08 | 3.1e-08 | 3.200E-08 | 5.568E-09 | 2.700E-08 | 3.800E-08 |
| 22338 | 1117 | P1117W | D | 1.2e-07 | 3e-08 | 1.1e-08 | 5.367E-08 | 5.823E-08 | 1.100E-08 | 1.200E-07 |
| 22339 | 1117 | P1117Y | D | 4.9e-08 | 3.9e-08 | 4.6e-08 | 4.467E-08 | 5.132E-09 | 3.900E-08 | 4.900E-08 |
| 22340 | 1117 | P1117V | D | 9.9e-08 | 4.7e-08 | 2.7e-08 | 5.767E-08 | 3.717E-08 | 2.700E-08 | 9.900E-08 |
| 22341 | 1118 | S1118A | D | 2.7e-08 | 3.8e-08 | 1.2e-08 | 2.567E-08 | 1.305E-08 | 1.200E-08 | 3.800E-08 |
| 22342 | 1118 | S1118R | D | 9.2e-08 | 2.2e-08 | 2.9e-08 | 4.767E-08 | 3.855E-08 | 2.200E-08 | 9.200E-08 |
| 22343 | 1118 | S1118N | D | 5.4e-08 | 3.5e-08 | 7.2e-09 | 3.207E-08 | 2.354E-08 | 7.200E-09 | 5.400E-08 |
| 22344 | 1118 | S1118D | D | 3.2e-08 | 1.1e-07 | 6.6e-08 | 6.933E-08 | 3.911E-08 | 3.200E-08 | 1.100E-07 |
| 22345 | 1118 | S1118C | D | 1.6e-08 | 4e-08 | 2.1e-08 | 2.567E-08 | 1.266E-08 | 1.600E-08 | 4.000E-08 |
| 22346 | 1118 | S1118E | D | 4.4e-08 | 2.7e-08 | 1.1e-07 | 6.033E-08 | 4.384E-08 | 2.700E-08 | 1.100E-07 |
| 22347 | 1118 | S1118Q | D | 1.2e-07 | 3.9e-08 | 4.8e-08 | 6.900E-08 | 4.440E-08 | 3.900E-08 | 1.200E-07 |
| 22348 | 1118 | S1118G | D | 6.6e-08 | 7.6e-08 | 5.5e-08 | 6.567E-08 | 1.050E-08 | 5.500E-08 | 7.600E-08 |
| 22349 | 1118 | S1118H | D | 1.3e-08 | 1.3e-07 | 6.9e-08 | 7.067E-08 | 5.852E-08 | 1.300E-08 | 1.300E-07 |
| 22350 | 1118 | S1118I | D | 2.8e-08 | 2e-08 | 3.4e-08 | 2.733E-08 | 7.024E-09 | 2.000E-08 | 3.400E-08 |
| 22351 | 1118 | S1118L | D | 3.1e-08 | 1.8e-08 | 6.1e-08 | 3.667E-08 | 2.205E-08 | 1.800E-08 | 6.100E-08 |
| 22352 | 1118 | S1118K | D | 4.2e-08 | 6.2e-08 | 3.4e-08 | 4.600E-08 | 1.442E-08 | 3.400E-08 | 6.200E-08 |
| 22353 | 1118 | S1118M | D | 6.3e-08 | 1.5e-08 | 4.1e-08 | 3.967E-08 | 2.403E-08 | 1.500E-08 | 6.300E-08 |
| 22354 | 1118 | S1118F | D | 1.8e-08 | 2.6e-08 | 6.4e-08 | 3.600E-08 | 2.458E-08 | 1.800E-08 | 6.400E-08 |

| | | | | | | | | | | |
|---|---|---|---|---|---|---|---|---|---|---|
| 22355 | 1118 | S1118P | D | 1.1e-07 | 9.6e-08 | 4.3e-08 | 8.300E-08 | 3.534E-08 | 4.300E-08 | 1.100E-07 |
| 22356 | 1118 | S1118S | D | 2.2e-08 | 3.9e-08 | 2.8e-08 | 2.967E-08 | 8.622E-09 | 2.200E-08 | 3.900E-08 |
| 22357 | 1118 | S1118T | D | 7e-08 | 2e-08 | 6.2e-08 | 5.067E-08 | 2.686E-08 | 2.000E-08 | 7.000E-08 |
| 22358 | 1118 | S1118W | D | 6.3e-08 | 2.8e-08 | 5.2e-08 | 4.767E-08 | 1.790E-08 | 2.800E-08 | 6.300E-08 |
| 22359 | 1118 | S1118Y | D | 3.4e-08 | 2e-08 | 5.1e-08 | 3.500E-08 | 1.552E-08 | 2.000E-08 | 5.100E-08 |
| 22360 | 1118 | S1118V | D | 3.5e-08 | 3.3e-08 | 3e-08 | 3.267E-08 | 2.517E-09 | 3.000E-08 | 3.500E-08 |
| 22361 | 1119 | R1119A | D | 3.7e-08 | 8.1e-08 | 3.1e-08 | 4.967E-08 | 2.730E-08 | 3.100E-08 | 8.100E-08 |
| 22362 | 1119 | R1119R | D | 3.7e-08 | 8.8e-08 | 6.1e-08 | 6.200E-08 | 2.551E-08 | 3.700E-08 | 8.800E-08 |
| 22363 | 1119 | R1119N | D | 4.2e-08 | 6.6e-08 | 5.2e-08 | 5.333E-08 | 1.206E-08 | 4.200E-08 | 6.600E-08 |
| 22364 | 1119 | R1119D | D | 4.8e-08 | 2.6e-08 | 3.8e-08 | 3.733E-08 | 1.102E-08 | 2.600E-08 | 4.800E-08 |
| 22365 | 1119 | R1119C | D | 2e-08 | 3.5e-08 | 4.6e-08 | 3.367E-08 | 1.305E-08 | 2.000E-08 | 4.600E-08 |
| 22366 | 1119 | R1119E | D | 1.5e-08 | 6e-08 | 3.5e-08 | 3.667E-08 | 2.255E-08 | 1.500E-08 | 6.000E-08 |
| 22367 | 1119 | R1119Q | D | 6.7e-08 | 5.1e-08 | 5.1e-08 | 5.633E-08 | 9.238E-09 | 5.100E-08 | 6.700E-08 |
| 22368 | 1119 | R1119G | D | 3.5e-08 | 3.6e-08 | 1.5e-08 | 2.867E-08 | 1.185E-08 | 1.500E-08 | 3.600E-08 |
| 22369 | 1119 | R1119H | D | 2.8e-08 | 4.2e-08 | 3.5e-08 | 3.500E-08 | 7.000E-09 | 2.800E-08 | 4.200E-08 |
| 22370 | 1119 | R1119I | D | 7.8e-08 | 3e-08 | 5.4e-08 | 5.400E-08 | 2.400E-08 | 3.000E-08 | 7.800E-08 |
| 22371 | 1119 | R1119L | D | 1.5e-07 | 7e-08 | 4.8e-08 | 8.933E-08 | 5.368E-08 | 4.800E-08 | 1.500E-07 |
| 22372 | 1119 | R1119K | D | 3.4e-08 | 3.7e-08 | 4.4e-08 | 3.833E-08 | 5.132E-09 | 3.400E-08 | 4.400E-08 |
| 22373 | 1119 | R1119M | D | 4.7e-08 | 3.6e-08 | 5e-08 | 4.433E-08 | 7.371E-09 | 3.600E-08 | 5.000E-08 |
| 22374 | 1119 | R1119F | D | 4.8e-08 | 5.1e-08 | 5e-08 | 4.967E-08 | 1.528E-09 | 4.800E-08 | 5.100E-08 |
| 22375 | 1119 | R1119P | D | 3.4e-08 | 5.9e-08 | 6.6e-08 | 5.300E-08 | 1.682E-08 | 3.400E-08 | 6.600E-08 |
| 22376 | 1119 | R1119S | D | 5.4e-08 | 6e-08 | 2.1e-08 | 4.500E-08 | 2.100E-08 | 2.100E-08 | 6.000E-08 |
| 22377 | 1119 | R1119T | D | 4.5e-08 | 2.6e-08 | 1.3e-07 | 6.700E-08 | 5.538E-08 | 2.600E-08 | 1.300E-07 |
| 22378 | 1119 | R1119W | D | 6e-08 | 4e-08 | 6e-08 | 5.333E-08 | 1.155E-08 | 4.000E-08 | 6.000E-08 |
| 22379 | 1119 | R1119Y | D | 2.7e-08 | 4.2e-08 | 4e-08 | 3.633E-08 | 8.145E-09 | 2.700E-08 | 4.200E-08 |
| 22380 | 1119 | R1119V | D | 3.5e-08 | 4.7e-08 | 3.6e-08 | 3.933E-08 | 6.658E-09 | 3.500E-08 | 4.700E-08 |
| 22381 | 1120 | F1120A | D | 4.3e-08 | 8.2e-08 | 5.9e-08 | 6.133E-08 | 1.960E-08 | 4.300E-08 | 8.200E-08 |
| 22382 | 1120 | F1120R | D | 5.5e-08 | 7.3e-08 | 2.2e-08 | 5.000E-08 | 2.587E-08 | 2.200E-08 | 7.300E-08 |
| 22383 | 1120 | F1120N | D | 4.3e-08 | 2.6e-08 | 7.2e-08 | 4.700E-08 | 2.326E-08 | 2.600E-08 | 7.200E-08 |
| 22384 | 1120 | F1120D | D | 4.2e-08 | 3.6e-08 | 4.4e-08 | 4.067E-08 | 4.163E-09 | 3.600E-08 | 4.400E-08 |
| 22385 | 1120 | F1120C | D | 3.9e-08 | 3.7e-08 | 4.8e-08 | 4.133E-08 | 5.859E-09 | 3.700E-08 | 4.800E-08 |
| 22386 | 1120 | F1120E | D | 5.3e-08 | 4.6e-08 | 3e-08 | 4.300E-08 | 1.179E-08 | 3.000E-08 | 5.300E-08 |
| 22387 | 1120 | F1120Q | D | 1.7e-08 | 6.4e-08 | 1.6e-08 | 3.233E-08 | 2.743E-08 | 1.600E-08 | 6.400E-08 |
| 22388 | 1120 | F1120G | D | 8.9e-08 | 9.4e-08 | 1.8e-08 | 6.700E-08 | 4.251E-08 | 1.800E-08 | 9.400E-08 |
| 22389 | 1120 | F1120H | D | 7.2e-08 | 2.2e-08 | 9.9e-09 | 3.463E-08 | 3.292E-08 | 9.900E-09 | 7.200E-08 |
| 22390 | 1120 | F1120I | D | 1.2e-07 | 4e-08 | 6.4e-08 | 7.467E-08 | 4.105E-08 | 4.000E-08 | 1.200E-07 |
| 22391 | 1120 | F1120L | D | 2.2e-08 | 2.3e-08 | 3.1e-08 | 2.533E-08 | 4.933E-09 | 2.200E-08 | 3.100E-08 |
| 22392 | 1120 | F1120K | D | 5.9e-08 | 4.4e-08 | 7.1e-08 | 5.800E-08 | 1.353E-08 | 4.400E-08 | 7.100E-08 |
| 22393 | 1120 | F1120M | D | 5.8e-08 | 4.7e-08 | 9.1e-08 | 6.533E-08 | 2.290E-08 | 4.700E-08 | 9.100E-08 |
| 22394 | 1120 | F1120F | D | 2.3e-08 | 2.4e-08 | 2.7e-08 | 2.467E-08 | 2.082E-09 | 2.300E-08 | 2.700E-08 |
| 22395 | 1120 | F1120P | D | 3.8e-08 | 3.7e-08 | 9.3e-08 | 5.600E-08 | 3.205E-08 | 3.700E-08 | 9.300E-08 |
| 22396 | 1120 | F1120S | D | 1.3e-08 | 5e-08 | 2.9e-08 | 3.067E-08 | 1.856E-08 | 1.300E-08 | 5.000E-08 |
| 22397 | 1120 | F1120T | D | 7e-08 | 4.8e-08 | 3.8e-08 | 5.200E-08 | 1.637E-08 | 3.800E-08 | 7.000E-08 |
| 22398 | 1120 | F1120W | D | 1.2e-07 | 5.7e-08 | 4.2e-08 | 7.300E-08 | 4.139E-08 | 4.200E-08 | 1.200E-07 |
| 22399 | 1120 | F1120Y | D | 7.6e-08 | 3.9e-08 | 3.7e-08 | 5.067E-08 | 2.196E-08 | 3.700E-08 | 7.600E-08 |
| 22400 | 1120 | F1120V | D | 8.5e-08 | 3.4e-08 | 6.4e-08 | 6.100E-08 | 2.563E-08 | 3.400E-08 | 8.500E-08 |
| 22401 | 1121 | S1121A | D | 3.9e-08 | 3.2e-08 | 3.5e-08 | 3.533E-08 | 3.512E-09 | 3.200E-08 | 3.900E-08 |
| 22402 | 1121 | S1121R | D | 4.1e-08 | 4.1e-08 | 6.7e-08 | 4.967E-08 | 1.501E-08 | 4.100E-08 | 6.700E-08 |
| 22403 | 1121 | S1121N | D | 3.6e-08 | 2.3e-08 | 1.6e-08 | 2.500E-08 | 1.015E-08 | 1.600E-08 | 3.600E-08 |
| 22404 | 1121 | S1121D | D | 5.5e-08 | 1.1e-07 | 1e-08 | 5.833E-08 | 5.008E-08 | 1.000E-08 | 1.100E-07 |
| 22405 | 1121 | S1121C | D | 9.6e-08 | 5.5e-08 | 2.9e-08 | 6.000E-08 | 3.378E-08 | 2.900E-08 | 9.600E-08 |
| 22406 | 1121 | S1121E | D | 2.4e-08 | 3.7e-08 | 3.7e-08 | 3.267E-08 | 7.506E-09 | 2.400E-08 | 3.700E-08 |
| 22407 | 1121 | S1121Q | D | 2.4e-08 | 4.9e-08 | 2.4e-07 | 1.043E-07 | 1.182E-07 | 2.400E-08 | 2.400E-07 |
| 22408 | 1121 | S1121G | D | 1.4e-07 | 2.1e-08 | 2.6e-08 | 6.233E-08 | 6.731E-08 | 2.100E-08 | 1.400E-07 |
| 22409 | 1121 | S1121H | D | 6.6e-08 | 3.2e-08 | 2.7e-08 | 4.167E-08 | 2.122E-08 | 2.700E-08 | 6.600E-08 |
| 22410 | 1121 | S1121I | D | 1.5e-08 | 1.5e-08 | 8.3e-08 | 3.767E-08 | 3.926E-08 | 1.500E-08 | 8.300E-08 |
| 22411 | 1121 | S1121L | D | 4.1e-08 | 1.1e-07 | 4.7e-08 | 6.600E-08 | 3.822E-08 | 4.100E-08 | 1.100E-07 |
| 22412 | 1121 | S1121K | D | 2.2e-08 | 5.3e-08 | 3.9e-08 | 3.800E-08 | 1.552E-08 | 2.200E-08 | 5.300E-08 |
| 22413 | 1121 | S1121M | D | 9.7e-08 | 6.8e-08 | 6.9e-08 | 7.800E-08 | 1.646E-08 | 6.800E-08 | 9.700E-08 |
| 22414 | 1121 | S1121F | D | 3.6e-08 | 8.6e-08 | 9.9e-08 | 7.367E-08 | 3.326E-08 | 3.600E-08 | 9.900E-08 |
| 22415 | 1121 | S1121P | D | 7e-08 | 2.6e-08 | 4e-08 | 4.533E-08 | 2.248E-08 | 2.600E-08 | 7.000E-08 |
| 22416 | 1121 | S1121S | D | 3.2e-08 | 6.5e-08 | 9.7e-08 | 6.467E-08 | 3.250E-08 | 3.200E-08 | 9.700E-08 |
| 22417 | 1121 | S1121T | D | 3.2e-08 | 2.9e-08 | 2.3e-08 | 2.800E-08 | 4.583E-09 | 2.300E-08 | 3.200E-08 |
| 22418 | 1121 | S1121W | D | 6.7e-08 | 4.4e-08 | 3e-08 | 4.700E-08 | 1.868E-08 | 3.000E-08 | 6.700E-08 |
| 22419 | 1121 | S1121Y | D | 9.1e-08 | 6.2e-08 | 4.1e-08 | 6.467E-08 | 2.511E-08 | 4.100E-08 | 9.100E-08 |
| 22420 | 1121 | S1121V | D | 4.3e-08 | 2.8e-08 | 5.4e-08 | 4.167E-08 | 1.305E-08 | 2.800E-08 | 5.400E-08 |
| 22421 | 1122 | G1122A | D | 3.8e-08 | 2.3e-08 | 2.5e-08 | 2.867E-08 | 8.145E-09 | 2.300E-08 | 3.800E-08 |
| 22422 | 1122 | G1122R | D | 4.2e-08 | 6.6e-08 | 3.5e-08 | 4.767E-08 | 1.626E-08 | 3.500E-08 | 6.600E-08 |
| 22423 | 1122 | G1122N | D | 2.1e-08 | 1.3e-08 | 4.8e-08 | 2.733E-08 | 1.834E-08 | 1.300E-08 | 4.800E-08 |
| 22424 | 1122 | G1122D | D | 3.7e-08 | 6.3e-08 | 5e-08 | 5.000E-08 | 1.300E-08 | 3.700E-08 | 6.300E-08 |
| 22425 | 1122 | G1122C | D | 3.9e-08 | 1.6e-08 | 2.7e-08 | 2.733E-08 | 1.150E-08 | 1.600E-08 | 3.900E-08 |
| 22426 | 1122 | G1122E | D | 3.6e-08 | 8.1e-08 | 3.4e-08 | 5.033E-08 | 2.658E-08 | 3.400E-08 | 8.100E-08 |
| 22427 | 1122 | G1122Q | D | 2.7e-08 | 4.1e-08 | 1.8e-08 | 2.867E-08 | 1.159E-08 | 1.800E-08 | 4.100E-08 |
| 22428 | 1122 | G1122G | D | 5.2e-08 | 9.8e-08 | 9.7e-08 | 8.233E-08 | 2.627E-08 | 5.200E-08 | 9.800E-08 |
| 22429 | 1122 | G1122H | D | 4.4e-08 | 4.6e-08 | 4.5e-08 | 4.500E-08 | 1.000E-09 | 4.400E-08 | 4.600E-08 |
| 22430 | 1122 | G1122I | D | 3.3e-08 | 3.5e-08 | 7.9e-08 | 4.900E-08 | 2.600E-08 | 3.300E-08 | 7.900E-08 |
| 22431 | 1122 | G1122L | D | 2.5e-08 | 1.2e-07 | 8.3e-08 | 7.600E-08 | 4.789E-08 | 2.500E-08 | 1.200E-07 |
| 22432 | 1122 | G1122K | D | 2.4e-08 | 4.6e-08 | 4.7e-08 | 3.900E-08 | 1.300E-08 | 2.400E-08 | 4.700E-08 |
| 22433 | 1122 | G1122M | D | 7.4e-08 | 2.4e-08 | 6.3e-08 | 5.367E-08 | 2.627E-08 | 2.400E-08 | 7.400E-08 |
| 22434 | 1122 | G1122F | D | 6.2e-08 | 6.2e-08 | 1.7e-08 | 4.700E-08 | 2.598E-08 | 1.700E-08 | 6.200E-08 |
| 22435 | 1122 | G1122P | D | 4.8e-08 | 2.1e-08 | 8e-08 | 4.967E-08 | 2.954E-08 | 2.100E-08 | 8.000E-08 |
| 22436 | 1122 | G1122S | D | 1.4e-08 | 2e-08 | 4.5e-08 | 2.633E-08 | 1.644E-08 | 1.400E-08 | 4.500E-08 |
| 22437 | 1122 | G1122T | D | 2e-08 | 3.5e-08 | 1.8e-07 | 7.833E-08 | 8.836E-08 | 2.000E-08 | 1.800E-07 |
| 22438 | 1122 | G1122W | D | 4.1e-08 | 3.2e-08 | 4.7e-08 | 4.000E-08 | 7.550E-09 | 3.200E-08 | 4.700E-08 |
| 22439 | 1122 | G1122Y | D | 4.4e-08 | 2.5e-08 | 3.1e-08 | 3.333E-08 | 9.713E-09 | 2.500E-08 | 4.400E-08 |

| | | | | | | | | | | |
|---|---|---|---|---|---|---|---|---|---|---|
| 22440 | 1122 | G1122V | D | 4.4e-08 | 2.4e-08 | 1.1e-07 | 5.933E-08 | 4.500E-08 | 2.400E-08 | 1.100E-07 |
| 22441 | 1123 | S1123A | D | 1.5e-08 | 2.4e-08 | 3e-08 | 2.300E-08 | 7.550E-09 | 1.500E-08 | 3.000E-08 |
| 22442 | 1123 | S1123R | D | 3.4e-08 | 4.8e-08 | 1.8e-08 | 3.333E-08 | 1.501E-08 | 1.800E-08 | 4.800E-08 |
| 22443 | 1123 | S1123N | D | 3.6e-08 | 6.6e-08 | 3.9e-08 | 4.700E-08 | 1.652E-08 | 3.600E-08 | 6.600E-08 |
| 22444 | 1123 | S1123D | D | 6.6e-08 | 8.6e-09 | 2.3e-08 | 3.253E-08 | 2.986E-08 | 8.600E-09 | 6.600E-08 |
| 22445 | 1123 | S1123C | D | 6.9e-08 | 6e-08 | 1.6e-08 | 4.833E-08 | 2.836E-08 | 1.600E-08 | 6.900E-08 |
| 22446 | 1123 | S1123E | D | 6.8e-08 | 9.3e-08 | 2.6e-08 | 6.233E-08 | 3.386E-08 | 2.600E-08 | 9.300E-08 |
| 22447 | 1123 | S1123Q | D | 1.2e-07 | 3.3e-08 | 3.3e-08 | 6.200E-08 | 5.023E-08 | 3.300E-08 | 1.200E-07 |
| 22448 | 1123 | S1123G | D | 6.8e-08 | 4.2e-08 | 5.9e-08 | 5.633E-08 | 1.320E-08 | 4.200E-08 | 6.800E-08 |
| 22449 | 1123 | S1123H | D | 4.9e-08 | 2.8e-08 | 2.5e-08 | 3.400E-08 | 1.308E-08 | 2.500E-08 | 4.900E-08 |
| 22450 | 1123 | S1123I | D | 1.1e-08 | 4.7e-08 | 3.3e-08 | 3.033E-08 | 1.815E-08 | 1.100E-08 | 4.700E-08 |
| 22451 | 1123 | S1123L | D | 2.7e-08 | 2.2e-08 | 3.9e-08 | 2.933E-08 | 8.737E-09 | 2.200E-08 | 3.900E-08 |
| 22452 | 1123 | S1123K | D | 6.4e-08 | 3.7e-08 | 3.2e-08 | 4.433E-08 | 1.721E-08 | 3.200E-08 | 6.400E-08 |
| 22453 | 1123 | S1123M | D | 4.5e-08 | 6.6e-08 | 9e-08 | 6.700E-08 | 2.252E-08 | 4.500E-08 | 9.000E-08 |
| 22454 | 1123 | S1123F | D | 4e-08 | 7.9e-08 | 2e-08 | 4.633E-08 | 3.001E-08 | 2.000E-08 | 7.900E-08 |
| 22455 | 1123 | S1123P | D | 6.9e-08 | 5.4e-08 | 4.1e-08 | 5.467E-08 | 1.401E-08 | 4.100E-08 | 6.900E-08 |
| 22456 | 1123 | S1123S | D | 2.5e-08 | 1.9e-08 | 4.5e-08 | 2.967E-08 | 1.361E-08 | 1.900E-08 | 4.500E-08 |
| 22457 | 1123 | S1123T | D | 6.4e-08 | 1.7e-08 | 3.9e-08 | 4.000E-08 | 2.352E-08 | 1.700E-08 | 6.400E-08 |
| 22458 | 1123 | S1123W | D | 1.8e-08 | 2.2e-09 | 6.6e-08 | 2.873E-08 | 3.323E-08 | 2.200E-09 | 6.600E-08 |
| 22459 | 1123 | S1123Y | D | 2.6e-08 | 1.3e-08 | 2.2e-08 | 2.033E-08 | 6.658E-09 | 1.300E-08 | 2.600E-08 |
| 22460 | 1123 | S1123V | D | 2.3e-08 | 1e-07 | 3.6e-08 | 5.300E-08 | 4.122E-08 | 2.300E-08 | 1.000E-07 |
| 22461 | 1124 | R1124A | D | 7.3e-08 | 1.1e-07 | 2e-08 | 6.767E-08 | 4.524E-08 | 2.000E-08 | 1.100E-07 |
| 22462 | 1124 | R1124R | D | 4.6e-08 | 3.8e-08 | 2.6e-08 | 3.667E-08 | 1.007E-08 | 2.600E-08 | 4.600E-08 |
| 22463 | 1124 | R1124N | D | 2.6e-08 | 5.6e-08 | 2.9e-08 | 3.700E-08 | 1.652E-08 | 2.600E-08 | 5.600E-08 |
| 22464 | 1124 | R1124D | D | 5.8e-08 | 7.9e-08 | 3.2e-08 | 5.633E-08 | 2.354E-08 | 3.200E-08 | 7.900E-08 |
| 22465 | 1124 | R1124C | D | 7.3e-08 | 1.4e-08 | 5.9e-08 | 4.867E-08 | 3.083E-08 | 1.400E-08 | 7.300E-08 |
| 22466 | 1124 | R1124E | D | 1e-07 | 4.4e-08 | 4.7e-08 | 6.367E-08 | 3.150E-08 | 4.400E-08 | 1.000E-07 |
| 22467 | 1124 | R1124Q | D | 9.2e-08 | 8.9e-08 | 6.3e-08 | 8.133E-08 | 1.595E-08 | 6.300E-08 | 9.200E-08 |
| 22468 | 1124 | R1124G | D | 1.7e-08 | 5.2e-08 | 5.1e-08 | 4.000E-08 | 1.992E-08 | 1.700E-08 | 5.200E-08 |
| 22469 | 1124 | R1124H | D | 2.7e-08 | 1.2e-07 | 2.3e-08 | 5.667E-08 | 5.488E-08 | 2.300E-08 | 1.200E-07 |
| 22470 | 1124 | R1124I | D | 6.7e-08 | 4e-08 | 8.1e-08 | 6.267E-08 | 2.084E-08 | 4.000E-08 | 8.100E-08 |
| 22471 | 1124 | R1124L | D | 6.1e-08 | 8.3e-08 | 5.7e-08 | 6.700E-08 | 1.400E-08 | 5.700E-08 | 8.300E-08 |
| 22472 | 1124 | R1124K | D | 5.5e-08 | 4.6e-08 | 2.7e-08 | 4.267E-08 | 1.429E-08 | 2.700E-08 | 5.500E-08 |
| 22473 | 1124 | R1124M | D | 3.3e-08 | 9.9e-08 | 2.3e-08 | 5.167E-08 | 4.130E-08 | 2.300E-08 | 9.900E-08 |
| 22474 | 1124 | R1124F | D | 2.9e-08 | 8.4e-08 | 2.2e-08 | 4.500E-08 | 3.396E-08 | 2.200E-08 | 8.400E-08 |
| 22475 | 1124 | R1124P | D | 5.5e-08 | 3.4e-08 | 7.7e-08 | 5.533E-08 | 2.150E-08 | 3.400E-08 | 7.700E-08 |
| 22476 | 1124 | R1124S | D | 2.8e-08 | 5.9e-08 | 3.3e-08 | 4.000E-08 | 1.664E-08 | 2.800E-08 | 5.900E-08 |
| 22477 | 1124 | R1124T | D | 4.7e-08 | 4.9e-08 | 3.2e-08 | 4.267E-08 | 9.292E-09 | 3.200E-08 | 4.900E-08 |
| 22478 | 1124 | R1124W | D | 4.5e-08 | 2.9e-08 | 5e-08 | 4.133E-08 | 1.097E-08 | 2.900E-08 | 5.000E-08 |
| 22479 | 1124 | R1124Y | D | 2.2e-08 | 1.1e-07 | 3.8e-08 | 5.667E-08 | 4.688E-08 | 2.200E-08 | 1.100E-07 |
| 22480 | 1124 | R1124V | D | 7.2e-08 | 2.7e-08 | 6.1e-08 | 5.333E-08 | 2.346E-08 | 2.700E-08 | 7.200E-08 |
| 22481 | 1125 | S1125A | D | 5.8e-08 | 3.4e-08 | 1.5e-08 | 3.567E-08 | 2.155E-08 | 1.500E-08 | 5.800E-08 |
| 22482 | 1125 | S1125R | D | 1.1e-07 | 9.4e-08 | 3e-08 | 7.800E-08 | 4.233E-08 | 3.000E-08 | 1.100E-07 |
| 22483 | 1125 | S1125N | D | 8.7e-08 | 9e-08 | 5.9e-08 | 7.867E-08 | 1.710E-08 | 5.900E-08 | 9.000E-08 |
| 22484 | 1125 | S1125D | D | 1.2e-08 | 1.4e-07 | 3e-08 | 6.067E-08 | 6.929E-08 | 1.200E-08 | 1.400E-07 |
| 22485 | 1125 | S1125C | D | 1.6e-07 | 3.4e-08 | 6.1e-08 | 8.500E-08 | 6.634E-08 | 3.400E-08 | 1.600E-07 |
| 22486 | 1125 | S1125E | D | 8.9e-08 | 3.5e-08 | 2e-08 | 4.800E-08 | 3.629E-08 | 2.000E-08 | 8.900E-08 |
| 22487 | 1125 | S1125Q | D | 4.2e-08 | 7.6e-08 | 7.9e-08 | 6.567E-08 | 2.055E-08 | 4.200E-08 | 7.900E-08 |
| 22488 | 1125 | S1125G | D | 1e-07 | 3.3e-08 | 7.9e-08 | 7.067E-08 | 3.427E-08 | 3.300E-08 | 1.000E-07 |
| 22489 | 1125 | S1125H | D | 3e-08 | 4.6e-08 | 1.4e-08 | 3.000E-08 | 1.600E-08 | 1.400E-08 | 4.600E-08 |
| 22490 | 1125 | S1125I | D | 4.7e-08 | 2.9e-08 | 5.6e-08 | 4.400E-08 | 1.375E-08 | 2.900E-08 | 5.600E-08 |
| 22491 | 1125 | S1125L | D | 6.9e-08 | 5.5e-08 | 4.4e-08 | 5.600E-08 | 1.253E-08 | 4.400E-08 | 6.900E-08 |
| 22492 | 1125 | S1125K | D | 4.1e-08 | 5.7e-08 | 4.7e-08 | 4.833E-08 | 8.083E-09 | 4.100E-08 | 5.700E-08 |
| 22493 | 1125 | S1125M | D | 4.1e-08 | 1.3e-08 | 2.2e-08 | 2.533E-08 | 1.429E-08 | 1.300E-08 | 4.100E-08 |
| 22494 | 1125 | S1125F | D | 3.6e-08 | 9.5e-08 | 4.1e-08 | 5.733E-08 | 3.272E-08 | 3.600E-08 | 9.500E-08 |
| 22495 | 1125 | S1125P | D | 2.6e-08 | 3.2e-08 | 3.4e-08 | 3.067E-08 | 4.163E-09 | 2.600E-08 | 3.400E-08 |
| 22496 | 1125 | S1125S | D | 2.5e-08 | 1.3e-08 | 5.2e-08 | 3.000E-08 | 1.997E-08 | 1.300E-08 | 5.200E-08 |
| 22497 | 1125 | S1125T | D | 6.2e-08 | 4.9e-08 | 1.2e-07 | 7.700E-08 | 3.780E-08 | 4.900E-08 | 1.200E-07 |
| 22498 | 1125 | S1125W | D | 1.7e-07 | 2.5e-08 | 2.5e-08 | 7.333E-08 | 8.372E-08 | 2.500E-08 | 1.700E-07 |
| 22499 | 1125 | S1125Y | D | 1.5e-08 | 2.9e-08 | 9e-08 | 4.467E-08 | 3.988E-08 | 1.500E-08 | 9.000E-08 |
| 22500 | 1125 | S1125V | D | 5.3e-08 | 4e-08 | 3.1e-08 | 4.133E-08 | 1.106E-08 | 3.100E-08 | 5.300E-08 |
| 22501 | 1126 | G1126A | D | 8.8e-08 | 7.8e-08 | 2.5e-08 | 6.367E-08 | 3.386E-08 | 2.500E-08 | 8.800E-08 |
| 22502 | 1126 | G1126R | D | 7.4e-08 | 1.1e-07 | 2.8e-08 | 7.067E-08 | 4.110E-08 | 2.800E-08 | 1.100E-07 |
| 22503 | 1126 | G1126N | D | 6.5e-08 | 5e-08 | 6.5e-08 | 6.000E-08 | 8.660E-09 | 5.000E-08 | 6.500E-08 |
| 22504 | 1126 | G1126D | D | 3.8e-08 | 2.9e-08 | 4.6e-08 | 3.767E-08 | 8.505E-09 | 2.900E-08 | 4.600E-08 |
| 22505 | 1126 | G1126C | D | 4.9e-08 | 4.9e-08 | 8.5e-08 | 6.100E-08 | 2.078E-08 | 4.900E-08 | 8.500E-08 |
| 22506 | 1126 | G1126E | D | 8.6e-08 | 8.8e-08 | 3e-08 | 6.800E-08 | 3.292E-08 | 3.000E-08 | 8.800E-08 |
| 22507 | 1126 | G1126Q | D | 1.6e-08 | 1.7e-08 | 8.1e-08 | 3.800E-08 | 3.724E-08 | 1.600E-08 | 8.100E-08 |
| 22508 | 1126 | G1126G | D | 9.1e-08 | 2.8e-08 | 1.8e-08 | 4.567E-08 | 3.958E-08 | 1.800E-08 | 9.100E-08 |
| 22509 | 1126 | G1126H | D | 2.1e-08 | 3.1e-08 | 4.9e-08 | 3.367E-08 | 1.419E-08 | 2.100E-08 | 4.900E-08 |
| 22510 | 1126 | G1126I | D | 5.1e-08 | 5.2e-08 | 2.9e-08 | 4.400E-08 | 1.300E-08 | 2.900E-08 | 5.200E-08 |
| 22511 | 1126 | G1126L | D | 6.8e-08 | 5.6e-08 | 2.2e-08 | 4.867E-08 | 2.386E-08 | 2.200E-08 | 6.800E-08 |
| 22512 | 1126 | G1126K | D | 5.9e-08 | 9.2e-08 | 4.5e-08 | 6.533E-08 | 2.413E-08 | 4.500E-08 | 9.200E-08 |
| 22513 | 1126 | G1126M | D | 3.4e-08 | 2.7e-08 | 2.6e-08 | 2.900E-08 | 4.359E-09 | 2.600E-08 | 3.400E-08 |
| 22514 | 1126 | G1126F | D | 1e-08 | 3.3e-08 | 6.8e-08 | 3.700E-08 | 2.921E-08 | 1.000E-08 | 6.800E-08 |
| 22515 | 1126 | G1126P | D | 4.1e-08 | 4.2e-08 | 3.2e-08 | 3.833E-08 | 5.508E-09 | 3.200E-08 | 4.200E-08 |
| 22516 | 1126 | G1126S | D | 2.4e-08 | 4.2e-08 | 4.6e-08 | 3.733E-08 | 1.172E-08 | 2.400E-08 | 4.600E-08 |
| 22517 | 1126 | G1126T | D | 3.7e-08 | 2.2e-08 | 2.5e-08 | 2.800E-08 | 7.937E-09 | 2.200E-08 | 3.700E-08 |
| 22518 | 1126 | G1126W | D | 1.6e-08 | 3.7e-08 | 2.4e-08 | 2.567E-08 | 1.060E-08 | 1.600E-08 | 3.700E-08 |
| 22519 | 1126 | G1126Y | D | 6.3e-08 | 1.4e-07 | 1.3e-08 | 7.200E-08 | 6.398E-08 | 1.300E-08 | 1.400E-07 |
| 22520 | 1126 | G1126V | D | 5.6e-08 | 3.9e-08 | 1.3e-08 | 3.600E-08 | 2.166E-08 | 1.300E-08 | 5.600E-08 |
| 22521 | 1127 | T1127A | D | 6e-08 | 1.4e-08 | 5.2e-08 | 4.200E-08 | 2.458E-08 | 1.400E-08 | 6.000E-08 |
| 22522 | 1127 | T1127R | D | 9.7e-08 | 2.7e-08 | 1.6e-08 | 4.667E-08 | 4.394E-08 | 1.600E-08 | 9.700E-08 |
| 22523 | 1127 | T1127N | D | 1.8e-08 | 4.5e-08 | 2.1e-08 | 2.800E-08 | 1.480E-08 | 1.800E-08 | 4.500E-08 |
| 22524 | 1127 | T1127D | D | 1e-07 | 3.4e-08 | 4.5e-08 | 5.967E-08 | 3.536E-08 | 3.400E-08 | 1.000E-07 |

| | | | | | | | | | | |
|---|---|---|---|---|---|---|---|---|---|---|
| 22525 | 1127 | T1127C | D | 3.7e-08 | 2.5e-08 | 1.5e-08 | 2.567E-08 | 1.102E-08 | 1.500E-08 | 3.700E-08 |
| 22526 | 1127 | T1127E | D | 4e-08 | 6.7e-08 | 1.6e-08 | 4.100E-08 | 2.551E-08 | 1.600E-08 | 6.700E-08 |
| 22527 | 1127 | T1127Q | D | 1.9e-08 | 4e-08 | 3.4e-08 | 3.100E-08 | 1.082E-08 | 1.900E-08 | 4.000E-08 |
| 22528 | 1127 | T1127G | D | 4.1e-08 | 7e-08 | 4.9e-08 | 5.333E-08 | 1.498E-08 | 4.100E-08 | 7.000E-08 |
| 22529 | 1127 | T1127H | D | 2.3e-08 | 2.4e-08 | 4.3e-08 | 3.000E-08 | 1.127E-08 | 2.300E-08 | 4.300E-08 |
| 22530 | 1127 | T1127I | D | 9.8e-09 | 7.3e-08 | 3.1e-08 | 3.793E-08 | 3.217E-08 | 9.800E-09 | 7.300E-08 |
| 22531 | 1127 | T1127L | D | 1.9e-08 | 1.5e-08 | 5e-08 | 2.800E-08 | 1.916E-08 | 1.500E-08 | 5.000E-08 |
| 22532 | 1127 | T1127K | D | 1e-07 | 7.5e-08 | 3.7e-08 | 7.067E-08 | 3.172E-08 | 3.700E-08 | 1.000E-07 |
| 22533 | 1127 | T1127M | D | 6.2e-08 | 8e-08 | 2.6e-08 | 5.600E-08 | 2.750E-08 | 2.600E-08 | 8.000E-08 |
| 22534 | 1127 | T1127F | D | 5.3e-08 | 1.2e-08 | 6.1e-08 | 4.200E-08 | 2.629E-08 | 1.200E-08 | 6.100E-08 |
| 22535 | 1127 | T1127P | D | 2.4e-08 | 3.5e-08 | 9.8e-08 | 5.233E-08 | 3.993E-08 | 2.400E-08 | 9.800E-08 |
| 22536 | 1127 | T1127S | D | 3.9e-08 | 4.2e-08 | 6.3e-08 | 4.800E-08 | 1.308E-08 | 3.900E-08 | 6.300E-08 |
| 22537 | 1127 | T1127T | D | 1.1e-07 | 3.9e-08 | 6.5e-08 | 7.133E-08 | 3.592E-08 | 3.900E-08 | 1.100E-07 |
| 22538 | 1127 | T1127W | D | 4.1e-08 | 5.2e-08 | 1.2e-08 | 3.500E-08 | 2.066E-08 | 1.200E-08 | 5.200E-08 |
| 22539 | 1127 | T1127Y | D | 3.2e-08 | 3.7e-08 | 1.3e-07 | 6.633E-08 | 5.519E-08 | 3.200E-08 | 1.300E-07 |
| 22540 | 1127 | T1127V | D | 2.9e-08 | 9.1e-08 | 2.6e-08 | 4.867E-08 | 3.669E-08 | 2.600E-08 | 9.100E-08 |
| 22541 | 1128 | D1128A | D | 1.1e-07 | 2.9e-08 | 2.9e-08 | 5.600E-08 | 4.677E-08 | 2.900E-08 | 1.100E-07 |
| 22542 | 1128 | D1128R | D | 7.7e-08 | 4.5e-08 | 8.1e-08 | 6.767E-08 | 1.973E-08 | 4.500E-08 | 8.100E-08 |
| 22543 | 1128 | D1128N | D | 1e-07 | 7.5e-08 | 7.2e-08 | 8.233E-08 | 1.537E-08 | 7.200E-08 | 1.000E-07 |
| 22544 | 1128 | D1128D | D | 2.5e-08 | 1e-08 | 1.9e-08 | 1.800E-08 | 7.550E-09 | 1.000E-08 | 2.500E-08 |
| 22545 | 1128 | D1128C | D | 9.8e-08 | 1.2e-07 | 3.7e-08 | 8.500E-08 | 4.300E-08 | 3.700E-08 | 1.200E-07 |
| 22546 | 1128 | D1128E | D | 4e-08 | 2.1e-08 | 5.8e-08 | 3.967E-08 | 1.850E-08 | 2.100E-08 | 5.800E-08 |
| 22547 | 1128 | D1128Q | D | 3.1e-08 | 7.5e-08 | 1.1e-07 | 7.200E-08 | 3.959E-08 | 3.100E-08 | 1.100E-07 |
| 22548 | 1128 | D1128G | D | 1.3e-08 | 5.9e-08 | 8.9e-08 | 5.367E-08 | 3.828E-08 | 1.300E-08 | 8.900E-08 |
| 22549 | 1128 | D1128H | D | 9.6e-09 | 1.7e-08 | 3.4e-08 | 2.020E-08 | 1.251E-08 | 9.600E-09 | 3.400E-08 |
| 22550 | 1128 | D1128I | D | 5.3e-08 | 5.5e-08 | 2.3e-08 | 4.367E-08 | 1.793E-08 | 2.300E-08 | 5.500E-08 |
| 22551 | 1128 | D1128L | D | 4.5e-08 | 1.9e-07 | 4.7e-08 | 9.400E-08 | 8.314E-08 | 4.500E-08 | 1.900E-07 |
| 22552 | 1128 | D1128K | D | 9e-08 | 5.6e-08 | 3.1e-08 | 5.900E-08 | 2.961E-08 | 3.100E-08 | 9.000E-08 |
| 22553 | 1128 | D1128M | D | 4.3e-08 | 2.1e-08 | 4.1e-08 | 3.500E-08 | 1.217E-08 | 2.100E-08 | 4.300E-08 |
| 22554 | 1128 | D1128F | D | 5e-08 | 3.9e-08 | 4e-08 | 4.300E-08 | 6.083E-09 | 3.900E-08 | 5.000E-08 |
| 22555 | 1128 | D1128P | D | 1.9e-08 | 2.8e-08 | 2.9e-08 | 2.533E-08 | 5.508E-09 | 1.900E-08 | 2.900E-08 |
| 22556 | 1128 | D1128S | D | 4.8e-08 | 6.5e-08 | 5.3e-08 | 5.533E-08 | 8.737E-09 | 4.800E-08 | 6.500E-08 |
| 22557 | 1128 | D1128T | D | 1.1e-07 | 3.5e-08 | 3.1e-08 | 5.867E-08 | 4.450E-08 | 3.100E-08 | 1.100E-07 |
| 22558 | 1128 | D1128W | D | 6.5e-08 | 1.3e-07 | 6.7e-08 | 8.733E-08 | 3.696E-08 | 6.500E-08 | 1.300E-07 |
| 22559 | 1128 | D1128Y | D | 4.6e-08 | 2e-08 | 4.5e-08 | 3.700E-08 | 1.473E-08 | 2.000E-08 | 4.600E-08 |
| 22560 | 1128 | D1128V | D | 2.8e-08 | 5.9e-08 | 1.1e-07 | 6.567E-08 | 4.140E-08 | 2.800E-08 | 1.100E-07 |
| 22561 | 1129 | F1129A | D | 4.3e-08 | 4e-08 | 7.6e-08 | 5.300E-08 | 1.997E-08 | 4.000E-08 | 7.600E-08 |
| 22562 | 1129 | F1129R | D | 4.5e-08 | 7e-08 | 3.2e-08 | 4.900E-08 | 1.931E-08 | 3.200E-08 | 7.000E-08 |
| 22563 | 1129 | F1129N | D | 6.1e-08 | 4.3e-08 | 4.8e-08 | 5.067E-08 | 9.292E-09 | 4.300E-08 | 6.100E-08 |
| 22564 | 1129 | F1129D | D | 5.9e-08 | 4.4e-08 | 2.8e-08 | 4.367E-08 | 1.550E-08 | 2.800E-08 | 5.900E-08 |
| 22565 | 1129 | F1129C | D | 6.9e-08 | 6.6e-08 | 1e-07 | 7.833E-08 | 1.882E-08 | 6.600E-08 | 1.000E-07 |
| 22566 | 1129 | F1129E | D | 1.7e-07 | 2e-08 | 4.5e-08 | 7.833E-08 | 8.036E-08 | 2.000E-08 | 1.700E-07 |
| 22567 | 1129 | F1129Q | D | 7e-08 | 9.5e-09 | 3.1e-08 | 3.683E-08 | 3.067E-08 | 9.500E-09 | 7.000E-08 |
| 22568 | 1129 | F1129G | D | 1.6e-08 | 1.9e-08 | 3.2e-07 | 1.183E-07 | 1.747E-07 | 1.600E-08 | 3.200E-07 |
| 22569 | 1129 | F1129H | D | 2.7e-08 | 5.1e-08 | 7.6e-08 | 5.133E-08 | 2.450E-08 | 2.700E-08 | 7.600E-08 |
| 22570 | 1129 | F1129I | D | 4.4e-08 | 2.8e-08 | 9.2e-08 | 5.467E-08 | 3.331E-08 | 2.800E-08 | 9.200E-08 |
| 22571 | 1129 | F1129L | D | 2.7e-08 | 2.9e-08 | 5.8e-08 | 3.800E-08 | 1.735E-08 | 2.700E-08 | 5.800E-08 |
| 22572 | 1129 | F1129K | D | 3.9e-08 | 1.7e-08 | 4.7e-08 | 3.433E-08 | 1.553E-08 | 1.700E-08 | 4.700E-08 |
| 22573 | 1129 | F1129M | D | 6.9e-08 | 3.6e-08 | 6.9e-08 | 5.800E-08 | 1.905E-08 | 3.600E-08 | 6.900E-08 |
| 22574 | 1129 | F1129F | D | 5.4e-08 | 1.4e-08 | 3e-08 | 3.267E-08 | 2.013E-08 | 1.400E-08 | 5.400E-08 |
| 22575 | 1129 | F1129P | D | 9.1e-08 | 5.5e-08 | 1.1e-08 | 5.233E-08 | 4.007E-08 | 1.100E-08 | 9.100E-08 |
| 22576 | 1129 | F1129S | D | 2.2e-08 | 2.4e-08 | 3.6e-08 | 2.733E-08 | 7.572E-09 | 2.200E-08 | 3.600E-08 |
| 22577 | 1129 | F1129T | D | 4.5e-08 | 2e-08 | 2.9e-08 | 3.133E-08 | 1.266E-08 | 2.000E-08 | 4.500E-08 |
| 22578 | 1129 | F1129W | D | 2e-07 | 1.3e-08 | 1e-07 | 1.043E-07 | 9.358E-08 | 1.300E-08 | 2.000E-07 |
| 22579 | 1129 | F1129Y | D | 5.6e-08 | 6.3e-08 | 3.5e-08 | 5.133E-08 | 1.457E-08 | 3.500E-08 | 6.300E-08 |
| 22580 | 1129 | F1129V | D | 6e-08 | 2.4e-08 | 3.9e-08 | 4.100E-08 | 1.808E-08 | 2.400E-08 | 6.000E-08 |
| 22581 | 1130 | T1130A | D | 2.8e-08 | 4.2e-08 | 5.4e-08 | 4.133E-08 | 1.301E-08 | 2.800E-08 | 5.400E-08 |
| 22582 | 1130 | T1130R | D | 5.7e-08 | 1.2e-07 | 5e-08 | 7.567E-08 | 3.855E-08 | 5.000E-08 | 1.200E-07 |
| 22583 | 1130 | T1130N | D | 3.9e-08 | 3.1e-08 | 1.9e-08 | 2.967E-08 | 1.007E-08 | 1.900E-08 | 3.900E-08 |
| 22584 | 1130 | T1130D | D | 6.5e-08 | 3e-08 | 2.9e-08 | 4.133E-08 | 2.050E-08 | 2.900E-08 | 6.500E-08 |
| 22585 | 1130 | T1130C | D | 3e-08 | 8.2e-08 | 7.7e-08 | 6.300E-08 | 2.869E-08 | 3.000E-08 | 8.200E-08 |
| 22586 | 1130 | T1130E | D | 1.1e-08 | 5.2e-08 | 6.9e-08 | 4.400E-08 | 2.982E-08 | 1.100E-08 | 6.900E-08 |
| 22587 | 1130 | T1130Q | D | 6.3e-08 | 2.7e-08 | 2.6e-08 | 3.867E-08 | 2.108E-08 | 2.600E-08 | 6.300E-08 |
| 22588 | 1130 | T1130G | D | 1.1e-07 | 5.6e-08 | 2.8e-08 | 6.467E-08 | 4.168E-08 | 2.800E-08 | 1.100E-07 |
| 22589 | 1130 | T1130H | D | 5.1e-08 | 7.7e-08 | 1.1e-07 | 7.933E-08 | 2.957E-08 | 5.100E-08 | 1.100E-07 |
| 22590 | 1130 | T1130I | D | 3.4e-08 | 7.5e-08 | 6.4e-08 | 5.767E-08 | 2.122E-08 | 3.400E-08 | 7.500E-08 |
| 22591 | 1130 | T1130L | D | 1.3e-07 | 3.5e-08 | 3.6e-08 | 6.700E-08 | 5.456E-08 | 3.500E-08 | 1.300E-07 |
| 22592 | 1130 | T1130K | D | 2.2e-07 | 3.9e-08 | 7.2e-08 | 1.103E-07 | 9.640E-08 | 3.900E-08 | 2.200E-07 |
| 22593 | 1130 | T1130M | D | 2.9e-08 | 3.2e-08 | 1.1e-07 | 5.700E-08 | 4.592E-08 | 2.900E-08 | 1.100E-07 |
| 22594 | 1130 | T1130F | D | 2.6e-08 | 4.2e-09 | 3.2e-08 | 2.073E-08 | 1.463E-08 | 4.200E-09 | 3.200E-08 |
| 22595 | 1130 | T1130P | D | 2e-08 | 3.9e-08 | 4.7e-08 | 3.533E-08 | 1.387E-08 | 2.000E-08 | 4.700E-08 |
| 22596 | 1130 | T1130S | D | 6e-08 | 1.3e-08 | 2.4e-08 | 3.233E-08 | 2.458E-08 | 1.300E-08 | 6.000E-08 |
| 22597 | 1130 | T1130T | D | 5.7e-08 | 7.2e-08 | 6.4e-08 | 6.433E-08 | 7.506E-09 | 5.700E-08 | 7.200E-08 |
| 22598 | 1130 | T1130W | D | 3.2e-08 | 4.6e-08 | 5.2e-08 | 4.333E-08 | 1.026E-08 | 3.200E-08 | 5.200E-08 |
| 22599 | 1130 | T1130Y | D | 6.1e-08 | 2.3e-08 | 2e-08 | 3.467E-08 | 2.285E-08 | 2.000E-08 | 6.100E-08 |
| 22600 | 1130 | T1130V | D | 1.1e-07 | 8.8e-08 | 4.2e-08 | 8.000E-08 | 3.470E-08 | 4.200E-08 | 1.100E-07 |
| 22601 | 1131 | L1131A | D | 6.5e-08 | 4.9e-08 | 3.5e-08 | 4.967E-08 | 1.501E-08 | 3.500E-08 | 6.500E-08 |
| 22602 | 1131 | L1131R | D | 5.7e-08 | 7.9e-08 | 5.2e-08 | 6.267E-08 | 1.436E-08 | 5.200E-08 | 7.900E-08 |
| 22603 | 1131 | L1131N | D | 5.9e-08 | 2.9e-08 | 3.7e-08 | 4.167E-08 | 1.553E-08 | 2.900E-08 | 5.900E-08 |
| 22604 | 1131 | L1131D | D | 5.6e-08 | 6.4e-08 | 2.5e-08 | 4.833E-08 | 2.060E-08 | 2.500E-08 | 6.400E-08 |
| 22605 | 1131 | L1131C | D | 2.5e-08 | 4.3e-08 | 2e-08 | 2.933E-08 | 1.210E-08 | 2.000E-08 | 4.300E-08 |
| 22606 | 1131 | L1131E | D | 1.7e-08 | 6.5e-08 | 5.3e-08 | 4.500E-08 | 2.498E-08 | 1.700E-08 | 6.500E-08 |
| 22607 | 1131 | L1131Q | D | 3.4e-08 | 4.8e-08 | 4.1e-08 | 4.100E-08 | 7.000E-09 | 3.400E-08 | 4.800E-08 |
| 22608 | 1131 | L1131G | D | 4.3e-08 | 1.1e-07 | 3.2e-08 | 6.167E-08 | 4.222E-08 | 3.200E-08 | 1.100E-07 |
| 22609 | 1131 | L1131H | D | 8e-08 | 2.7e-08 | 3.5e-08 | 4.733E-08 | 2.857E-08 | 2.700E-08 | 8.000E-08 |

| | | | | | | | | | | |
|---|---|---|---|---|---|---|---|---|---|---|
| 22610 | 1131 | L1131I | D | 2.3e-08 | 9.7e-08 | 3.2e-08 | 5.067E-08 | 4.038E-08 | 2.300E-08 | 9.700E-08 |
| 22611 | 1131 | L1131L | D | 1.7e-08 | 5.2e-08 | 3.5e-08 | 3.467E-08 | 1.750E-08 | 1.700E-08 | 5.200E-08 |
| 22612 | 1131 | L1131K | D | 2.3e-08 | 3.7e-08 | 4.8e-08 | 3.600E-08 | 1.253E-08 | 2.300E-08 | 4.800E-08 |
| 22613 | 1131 | L1131M | D | 3.2e-08 | 3.8e-08 | 8.3e-08 | 5.100E-08 | 2.787E-08 | 3.200E-08 | 8.300E-08 |
| 22614 | 1131 | L1131F | D | 6.6e-08 | 3.8e-08 | 1.9e-08 | 4.100E-08 | 2.364E-08 | 1.900E-08 | 6.600E-08 |
| 22615 | 1131 | L1131P | D | 4.4e-08 | 1.5e-08 | 3.5e-08 | 3.133E-08 | 1.484E-08 | 1.500E-08 | 4.400E-08 |
| 22616 | 1131 | L1131S | D | 3.7e-08 | 3.3e-08 | 4.4e-08 | 3.800E-08 | 5.568E-09 | 3.300E-08 | 4.400E-08 |
| 22617 | 1131 | L1131T | D | 3.6e-08 | 4.7e-08 | 1e-07 | 6.100E-08 | 3.422E-08 | 3.600E-08 | 1.000E-07 |
| 22618 | 1131 | L1131W | D | 7.1e-08 | 4.7e-08 | 5.5e-08 | 5.767E-08 | 1.222E-08 | 4.700E-08 | 7.100E-08 |
| 22619 | 1131 | L1131Y | D | 2.8e-08 | 4.5e-08 | 6.6e-08 | 4.633E-08 | 1.904E-08 | 2.800E-08 | 6.600E-08 |
| 22620 | 1131 | L1131V | D | 6.2e-08 | 2.8e-08 | 4.3e-08 | 4.433E-08 | 1.704E-08 | 2.800E-08 | 6.200E-08 |
| 22621 | 1132 | T1132A | D | 7e-08 | 7.8e-08 | 1.3e-08 | 5.367E-08 | 3.544E-08 | 1.300E-08 | 7.800E-08 |
| 22622 | 1132 | T1132R | D | 1.5e-08 | 4.2e-08 | 5.9e-08 | 3.867E-08 | 2.219E-08 | 1.500E-08 | 5.900E-08 |
| 22623 | 1132 | T1132N | D | 1.3e-07 | 4e-08 | 8.6e-08 | 8.533E-08 | 4.500E-08 | 4.000E-08 | 1.300E-07 |
| 22624 | 1132 | T1132D | D | 6.4e-08 | 8.9e-08 | 4.1e-08 | 6.467E-08 | 2.401E-08 | 4.100E-08 | 8.900E-08 |
| 22625 | 1132 | T1132C | D | 6.2e-08 | 3.6e-08 | 1.2e-08 | 3.667E-08 | 2.501E-08 | 1.200E-08 | 6.200E-08 |
| 22626 | 1132 | T1132E | D | 1.1e-07 | 5.4e-08 | 3.2e-08 | 6.533E-08 | 4.022E-08 | 3.200E-08 | 1.100E-07 |
| 22627 | 1132 | T1132Q | D | 7.2e-08 | 7.6e-08 | 4.4e-08 | 6.400E-08 | 1.744E-08 | 4.400E-08 | 7.600E-08 |
| 22628 | 1132 | T1132G | D | 6.2e-08 | 9.4e-08 | 7.6e-08 | 7.733E-08 | 1.604E-08 | 6.200E-08 | 9.400E-08 |
| 22629 | 1132 | T1132H | D | 8.2e-08 | 5.2e-08 | 1.1e-08 | 4.833E-08 | 3.564E-08 | 1.100E-08 | 8.200E-08 |
| 22630 | 1132 | T1132I | D | 2.8e-08 | 1.1e-07 | 4.7e-08 | 6.167E-08 | 4.292E-08 | 2.800E-08 | 1.100E-07 |
| 22631 | 1132 | T1132L | D | 6.2e-08 | 5.5e-08 | 4.3e-08 | 5.333E-08 | 9.609E-09 | 4.300E-08 | 6.200E-08 |
| 22632 | 1132 | T1132K | D | 2e-08 | 4.7e-08 | 5.7e-08 | 4.133E-08 | 1.914E-08 | 2.000E-08 | 5.700E-08 |
| 22633 | 1132 | T1132M | D | 4.5e-08 | 3.8e-08 | 8.7e-08 | 5.667E-08 | 2.650E-08 | 3.800E-08 | 8.700E-08 |
| 22634 | 1132 | T1132F | D | 2.8e-08 | 2.2e-08 | 4.9e-08 | 3.300E-08 | 1.418E-08 | 2.200E-08 | 4.900E-08 |
| 22635 | 1132 | T1132P | D | 2.7e-08 | 4e-08 | 1e-07 | 5.567E-08 | 3.894E-08 | 2.700E-08 | 1.000E-07 |
| 22636 | 1132 | T1132S | D | 1.4e-08 | 8.4e-09 | 9.8e-08 | 4.013E-08 | 5.019E-08 | 8.400E-09 | 9.800E-08 |
| 22637 | 1132 | T1132T | D | 6.3e-08 | 6.9e-08 | 3.3e-08 | 5.500E-08 | 1.929E-08 | 3.300E-08 | 6.900E-08 |
| 22638 | 1132 | T1132W | D | 5.8e-08 | 4.3e-08 | 4.4e-08 | 4.833E-08 | 8.386E-09 | 4.300E-08 | 5.800E-08 |
| 22639 | 1132 | T1132Y | D | 6.7e-08 | 4e-08 | 5e-08 | 5.233E-08 | 1.365E-08 | 4.000E-08 | 6.700E-08 |
| 22640 | 1132 | T1132V | D | 1.8e-08 | 1.1e-07 | 3.3e-08 | 5.367E-08 | 4.936E-08 | 1.800E-08 | 1.100E-07 |
| 22641 | 1133 | I1133A | D | 4.2e-08 | 5.5e-08 | 6.7e-08 | 5.467E-08 | 1.250E-08 | 4.200E-08 | 6.700E-08 |
| 22642 | 1133 | I1133R | D | 4.1e-08 | 2.8e-08 | 4.4e-08 | 3.767E-08 | 8.505E-09 | 2.800E-08 | 4.400E-08 |
| 22643 | 1133 | I1133N | D | 2.2e-08 | 3.4e-08 | 2.7e-08 | 2.767E-08 | 6.028E-09 | 2.200E-08 | 3.400E-08 |
| 22644 | 1133 | I1133D | D | 3.1e-08 | 2.7e-08 | 6.2e-08 | 4.000E-08 | 1.916E-08 | 2.700E-08 | 6.200E-08 |
| 22645 | 1133 | I1133C | D | 4.7e-08 | 3.9e-08 | 4.1e-08 | 4.233E-08 | 4.163E-09 | 3.900E-08 | 4.700E-08 |
| 22646 | 1133 | I1133E | D | 5.5e-08 | 5.1e-08 | 5.6e-08 | 5.400E-08 | 2.646E-09 | 5.100E-08 | 5.600E-08 |
| 22647 | 1133 | I1133Q | D | 2.2e-07 | 3.3e-08 | 4.8e-08 | 1.003E-07 | 1.039E-07 | 3.300E-08 | 2.200E-07 |
| 22648 | 1133 | I1133G | D | 4.9e-08 | 2.4e-08 | 4.3e-08 | 3.867E-08 | 1.305E-08 | 2.400E-08 | 4.900E-08 |
| 22649 | 1133 | I1133H | D | 4.2e-08 | 4.5e-08 | 7.5e-08 | 5.400E-08 | 1.825E-08 | 4.200E-08 | 7.500E-08 |
| 22650 | 1133 | I1133I | D | 5.8e-08 | 2.7e-08 | 5.7e-08 | 4.733E-08 | 1.762E-08 | 2.700E-08 | 5.800E-08 |
| 22651 | 1133 | I1133L | D | 5.4e-08 | 4.6e-08 | 1.9e-08 | 3.967E-08 | 1.834E-08 | 1.900E-08 | 5.400E-08 |
| 22652 | 1133 | I1133K | D | 3.3e-08 | 1.3e-07 | 4.3e-08 | 6.867E-08 | 5.335E-08 | 3.300E-08 | 1.300E-07 |
| 22653 | 1133 | I1133M | D | 3.9e-08 | 7.6e-08 | 5.3e-08 | 5.600E-08 | 1.868E-08 | 3.900E-08 | 7.600E-08 |
| 22654 | 1133 | I1133F | D | 5e-08 | 4.9e-08 | 4.4e-08 | 4.767E-08 | 3.215E-09 | 4.400E-08 | 5.000E-08 |
| 22655 | 1133 | I1133P | D | 6.4e-08 | 4e-08 | 6.1e-08 | 5.500E-08 | 1.308E-08 | 4.000E-08 | 6.400E-08 |
| 22656 | 1133 | I1133S | D | 5e-08 | 1.5e-08 | 1.4e-08 | 2.633E-08 | 2.050E-08 | 1.400E-08 | 5.000E-08 |
| 22657 | 1133 | I1133T | D | 1.8e-08 | 2.8e-08 | 3.5e-08 | 2.700E-08 | 8.544E-09 | 1.800E-08 | 3.500E-08 |
| 22658 | 1133 | I1133W | D | 3.8e-08 | 2.5e-08 | 2e-08 | 2.767E-08 | 9.292E-09 | 2.000E-08 | 3.800E-08 |
| 22659 | 1133 | I1133Y | D | 1.4e-08 | 2.2e-08 | 4.5e-08 | 2.700E-08 | 1.609E-08 | 1.400E-08 | 4.500E-08 |
| 22660 | 1133 | I1133V | D | 3.8e-08 | 4.8e-08 | 3e-08 | 3.867E-08 | 9.018E-09 | 3.000E-08 | 4.800E-08 |
| 22661 | 1134 | S1134A | D | 4.5e-08 | 1.5e-08 | 4.3e-08 | 3.433E-08 | 1.677E-08 | 1.500E-08 | 4.500E-08 |
| 22662 | 1134 | S1134R | D | 2.6e-08 | 2.7e-08 | 4e-08 | 3.100E-08 | 7.810E-09 | 2.600E-08 | 4.000E-08 |
| 22663 | 1134 | S1134N | D | 2.3e-08 | 5.7e-08 | 4.5e-08 | 4.167E-08 | 1.724E-08 | 2.300E-08 | 5.700E-08 |
| 22664 | 1134 | S1134D | D | 3.9e-08 | 6.4e-08 | 2e-08 | 4.100E-08 | 2.207E-08 | 2.000E-08 | 6.400E-08 |
| 22665 | 1134 | S1134C | D | 7.1e-09 | 1.5e-07 | 3.4e-08 | 6.370E-08 | 7.594E-08 | 7.100E-09 | 1.500E-07 |
| 22666 | 1134 | S1134E | D | 2.9e-08 | 7.9e-08 | 5.6e-08 | 5.467E-08 | 2.503E-08 | 2.900E-08 | 7.900E-08 |
| 22667 | 1134 | S1134Q | D | 5.7e-08 | 6.6e-08 | 4.4e-08 | 5.567E-08 | 1.106E-08 | 4.400E-08 | 6.600E-08 |
| 22668 | 1134 | S1134G | D | 3.9e-08 | 3.5e-08 | 3.2e-08 | 3.533E-08 | 3.512E-09 | 3.200E-08 | 3.900E-08 |
| 22669 | 1134 | S1134H | D | 3.3e-08 | 6.6e-08 | 2.9e-08 | 4.267E-08 | 2.031E-08 | 2.900E-08 | 6.600E-08 |
| 22670 | 1134 | S1134I | D | 1.5e-08 | 2.8e-08 | 1.1e-07 | 5.100E-08 | 5.151E-08 | 1.500E-08 | 1.100E-07 |
| 22671 | 1134 | S1134L | D | 1.4e-08 | 4.9e-08 | 9.3e-08 | 5.200E-08 | 3.959E-08 | 1.400E-08 | 9.300E-08 |
| 22672 | 1134 | S1134K | D | 2.2e-08 | 4.6e-08 | 2.5e-08 | 3.100E-08 | 1.308E-08 | 2.200E-08 | 4.600E-08 |
| 22673 | 1134 | S1134M | D | 2.3e-08 | 3e-08 | 3.1e-08 | 2.800E-08 | 4.359E-09 | 2.300E-08 | 3.100E-08 |
| 22674 | 1134 | S1134F | D | 5.3e-08 | 4.7e-08 | 3.5e-08 | 4.500E-08 | 9.165E-09 | 3.500E-08 | 5.300E-08 |
| 22675 | 1134 | S1134P | D | 5.3e-08 | 1.6e-08 | 2.4e-08 | 3.100E-08 | 1.947E-08 | 1.600E-08 | 5.300E-08 |
| 22676 | 1134 | S1134S | D | 6.1e-08 | 1.9e-08 | 1.4e-08 | 3.133E-08 | 2.581E-08 | 1.400E-08 | 6.100E-08 |
| 22677 | 1134 | S1134T | D | 3e-08 | 2.4e-08 | 2e-08 | 2.467E-08 | 5.033E-09 | 2.000E-08 | 3.000E-08 |
| 22678 | 1134 | S1134W | D | 1.2e-07 | 2.4e-08 | 7.1e-08 | 7.167E-08 | 4.800E-08 | 2.400E-08 | 1.200E-07 |
| 22679 | 1134 | S1134Y | D | 4.1e-08 | 3.9e-08 | 4.5e-08 | 4.167E-08 | 3.055E-09 | 3.900E-08 | 4.500E-08 |
| 22680 | 1134 | S1134V | D | 3.8e-08 | 9.1e-08 | 6.8e-08 | 6.567E-08 | 2.658E-08 | 3.800E-08 | 9.100E-08 |
| 22681 | 1135 | S1135A | D | 5e-08 | 3.4e-08 | 7.6e-08 | 5.333E-08 | 2.120E-08 | 3.400E-08 | 7.600E-08 |
| 22682 | 1135 | S1135R | D | 4.9e-08 | 4e-08 | 7.8e-08 | 5.567E-08 | 1.986E-08 | 4.000E-08 | 7.800E-08 |
| 22683 | 1135 | S1135N | D | 5.9e-08 | 6.4e-08 | 1.4e-07 | 8.767E-08 | 4.539E-08 | 5.900E-08 | 1.400E-07 |
| 22684 | 1135 | S1135D | D | 7.2e-08 | 5.2e-08 | 4.5e-08 | 5.633E-08 | 1.401E-08 | 4.500E-08 | 7.200E-08 |
| 22685 | 1135 | S1135C | D | 3.5e-08 | 3.7e-08 | 6.4e-08 | 4.533E-08 | 1.620E-08 | 3.500E-08 | 6.400E-08 |
| 22686 | 1135 | S1135E | D | 1.8e-08 | 3.4e-08 | 4.7e-08 | 3.300E-08 | 1.453E-08 | 1.800E-08 | 4.700E-08 |
| 22687 | 1135 | S1135Q | D | 2.9e-08 | 3.8e-08 | 3.9e-08 | 3.533E-08 | 5.508E-09 | 2.900E-08 | 3.900E-08 |
| 22688 | 1135 | S1135G | D | 2.8e-08 | 5.3e-08 | 5e-08 | 4.367E-08 | 1.365E-08 | 2.800E-08 | 5.300E-08 |
| 22689 | 1135 | S1135H | D | 1.4e-08 | 2.6e-08 | 4.2e-08 | 2.733E-08 | 1.405E-08 | 1.400E-08 | 4.200E-08 |
| 22690 | 1135 | S1135I | D | 5.2e-08 | 3.5e-08 | 6.5e-08 | 5.067E-08 | 1.504E-08 | 3.500E-08 | 6.500E-08 |
| 22691 | 1135 | S1135L | D | 6.9e-08 | 1.1e-07 | 4e-08 | 7.300E-08 | 3.517E-08 | 4.000E-08 | 1.100E-07 |
| 22692 | 1135 | S1135K | D | 5.9e-08 | 6.8e-08 | 2.5e-08 | 5.067E-08 | 2.268E-08 | 2.500E-08 | 6.800E-08 |
| 22693 | 1135 | S1135M | D | 4.8e-08 | 5.9e-08 | 4e-08 | 4.900E-08 | 9.539E-09 | 4.000E-08 | 5.900E-08 |
| 22694 | 1135 | S1135F | D | 4.1e-08 | 4.2e-08 | 5.2e-08 | 4.500E-08 | 6.083E-09 | 4.100E-08 | 5.200E-08 |

| | | | | | | | | | | |
|---|---|---|---|---|---|---|---|---|---|---|
| 22695 | 1135 | S1135P | D | 3.3e-08 | 6.2e-08 | 2.1e-08 | 3.867E-08 | 2.108E-08 | 2.100E-08 | 6.200E-08 |
| 22696 | 1135 | S1135S | D | 4.2e-08 | 3.7e-08 | 1.6e-08 | 3.167E-08 | 1.380E-08 | 1.600E-08 | 4.200E-08 |
| 22697 | 1135 | S1135T | D | 3.3e-08 | 3.1e-08 | 7.5e-08 | 4.633E-08 | 2.485E-08 | 3.100E-08 | 7.500E-08 |
| 22698 | 1135 | S1135W | D | 8.4e-08 | 3.8e-08 | 5.7e-08 | 5.967E-08 | 2.312E-08 | 3.800E-08 | 8.400E-08 |
| 22699 | 1135 | S1135Y | D | 5.4e-08 | 8.8e-09 | 2.9e-08 | 3.060E-08 | 2.264E-08 | 8.800E-09 | 5.400E-08 |
| 22700 | 1135 | S1135V | D | 2e-08 | 9e-08 | 7.2e-08 | 6.067E-08 | 3.635E-08 | 2.000E-08 | 9.000E-08 |
| 22701 | 1136 | L1136A | D | 9.4e-08 | 9.7e-08 | 2.2e-08 | 7.100E-08 | 4.246E-08 | 2.200E-08 | 9.700E-08 |
| 22702 | 1136 | L1136R | D | 1.2e-08 | 3.2e-08 | 1.1e-07 | 5.133E-08 | 5.178E-08 | 1.200E-08 | 1.100E-07 |
| 22703 | 1136 | L1136N | D | 5.2e-08 | 7.1e-08 | 5.5e-08 | 5.933E-08 | 1.021E-08 | 5.200E-08 | 7.100E-08 |
| 22704 | 1136 | L1136D | D | 4.1e-08 | 1.1e-08 | 7.5e-08 | 4.233E-08 | 3.202E-08 | 1.100E-08 | 7.500E-08 |
| 22705 | 1136 | L1136C | D | 2.2e-08 | 3.1e-08 | 1.3e-08 | 2.200E-08 | 9.000E-09 | 1.300E-08 | 3.100E-08 |
| 22706 | 1136 | L1136E | D | 2.7e-08 | 4.6e-08 | 5.5e-08 | 4.267E-08 | 1.429E-08 | 2.700E-08 | 5.500E-08 |
| 22707 | 1136 | L1136Q | D | 6.1e-08 | 6.4e-08 | 2.4e-08 | 4.967E-08 | 2.228E-08 | 2.400E-08 | 6.400E-08 |
| 22708 | 1136 | L1136G | D | 2.9e-08 | 4.2e-08 | 5.5e-08 | 4.200E-08 | 1.300E-08 | 2.900E-08 | 5.500E-08 |
| 22709 | 1136 | L1136H | D | 2e-08 | 2.6e-08 | 3.5e-08 | 2.700E-08 | 7.550E-09 | 2.000E-08 | 3.500E-08 |
| 22710 | 1136 | L1136I | D | 1.5e-08 | 1.6e-08 | 8.2e-08 | 3.767E-08 | 3.840E-08 | 1.500E-08 | 8.200E-08 |
| 22711 | 1136 | L1136L | D | 6e-08 | 8.7e-08 | 3.9e-08 | 6.200E-08 | 2.406E-08 | 3.900E-08 | 8.700E-08 |
| 22712 | 1136 | L1136K | D | 5.2e-08 | 3.8e-08 | 3.4e-08 | 4.133E-08 | 9.452E-09 | 3.400E-08 | 5.200E-08 |
| 22713 | 1136 | L1136M | D | 5.5e-08 | 4e-08 | 3.5e-08 | 4.333E-08 | 1.041E-08 | 3.500E-08 | 5.500E-08 |
| 22714 | 1136 | L1136F | D | 4.7e-08 | 9.3e-08 | 2.5e-08 | 5.500E-08 | 3.470E-08 | 2.500E-08 | 9.300E-08 |
| 22715 | 1136 | L1136P | D | 4.7e-08 | 3.2e-08 | 3.7e-08 | 3.867E-08 | 7.638E-09 | 3.200E-08 | 4.700E-08 |
| 22716 | 1136 | L1136S | D | 5.6e-08 | 7.5e-08 | 1.3e-08 | 4.800E-08 | 3.176E-08 | 1.300E-08 | 7.500E-08 |
| 22717 | 1136 | L1136T | D | 3.9e-08 | 3.2e-08 | 1.1e-07 | 6.033E-08 | 4.315E-08 | 3.200E-08 | 1.100E-07 |
| 22718 | 1136 | L1136W | D | 3.5e-08 | 2.2e-08 | 2.4e-08 | 2.700E-08 | 7.000E-09 | 2.200E-08 | 3.500E-08 |
| 22719 | 1136 | L1136Y | D | 1.5e-08 | 1.8e-08 | 4.5e-08 | 2.600E-08 | 1.652E-08 | 1.500E-08 | 4.500E-08 |
| 22720 | 1136 | L1136V | D | 3.6e-08 | 4.7e-08 | 2.8e-08 | 3.700E-08 | 9.539E-09 | 2.800E-08 | 4.700E-08 |
| 22721 | 1137 | Q1137A | D | 5.3e-08 | 2.4e-08 | 3.6e-08 | 3.767E-08 | 1.457E-08 | 2.400E-08 | 5.300E-08 |
| 22722 | 1137 | Q1137R | D | 4.6e-08 | 5.8e-08 | 6.6e-08 | 5.667E-08 | 1.007E-08 | 4.600E-08 | 6.600E-08 |
| 22723 | 1137 | Q1137N | D | 1.2e-07 | 2.1e-08 | 4.6e-08 | 6.233E-08 | 5.148E-08 | 2.100E-08 | 1.200E-07 |
| 22724 | 1137 | Q1137D | D | 5.3e-08 | 5e-08 | 5.2e-08 | 5.167E-08 | 1.528E-09 | 5.000E-08 | 5.300E-08 |
| 22725 | 1137 | Q1137C | D | 2.6e-08 | 5e-08 | 4.1e-08 | 3.900E-08 | 1.212E-08 | 2.600E-08 | 5.000E-08 |
| 22726 | 1137 | Q1137E | D | 3.5e-08 | 3.3e-08 | 5.7e-08 | 4.167E-08 | 1.332E-08 | 3.300E-08 | 5.700E-08 |
| 22727 | 1137 | Q1137Q | D | 4.4e-08 | 3.1e-08 | 2.4e-08 | 3.300E-08 | 1.015E-08 | 2.400E-08 | 4.400E-08 |
| 22728 | 1137 | Q1137G | D | 8e-08 | 2.8e-08 | 5.8e-08 | 5.533E-08 | 2.610E-08 | 2.800E-08 | 8.000E-08 |
| 22729 | 1137 | Q1137H | D | 3.5e-08 | 4.2e-08 | 5.1e-08 | 4.267E-08 | 8.021E-09 | 3.500E-08 | 5.100E-08 |
| 22730 | 1137 | Q1137I | D | 6.6e-08 | 3.3e-08 | 9.5e-08 | 6.467E-08 | 3.102E-08 | 3.300E-08 | 9.500E-08 |
| 22731 | 1137 | Q1137L | D | 2.7e-08 | 2.8e-08 | 5.4e-08 | 3.633E-08 | 1.531E-08 | 2.700E-08 | 5.400E-08 |
| 22732 | 1137 | Q1137K | D | 2.2e-08 | 4.8e-08 | 2e-08 | 3.000E-08 | 1.562E-08 | 2.000E-08 | 4.800E-08 |
| 22733 | 1137 | Q1137M | D | 4.1e-08 | 2e-08 | 3.1e-08 | 3.067E-08 | 1.050E-08 | 2.000E-08 | 4.100E-08 |
| 22734 | 1137 | Q1137F | D | 4.3e-08 | 6.2e-08 | 2.1e-08 | 4.200E-08 | 2.052E-08 | 2.100E-08 | 6.200E-08 |
| 22735 | 1137 | Q1137P | D | 4.7e-08 | 7.8e-08 | 4.2e-08 | 5.567E-08 | 1.950E-08 | 4.200E-08 | 7.800E-08 |
| 22736 | 1137 | Q1137S | D | 4e-08 | 3.4e-08 | 3.1e-08 | 3.500E-08 | 4.583E-09 | 3.100E-08 | 4.000E-08 |
| 22737 | 1137 | Q1137T | D | 6.2e-08 | 6.1e-08 | 5.2e-08 | 5.833E-08 | 5.508E-09 | 5.200E-08 | 6.200E-08 |
| 22738 | 1137 | Q1137W | D | 2.1e-08 | 5.8e-08 | 6.5e-08 | 4.800E-08 | 2.364E-08 | 2.100E-08 | 6.500E-08 |
| 22739 | 1137 | Q1137Y | D | 8.3e-08 | 2.1e-08 | 5.4e-08 | 5.267E-08 | 3.102E-08 | 2.100E-08 | 8.300E-08 |
| 22740 | 1137 | Q1137V | D | 7e-08 | 3.5e-08 | 3.2e-08 | 4.567E-08 | 2.113E-08 | 3.200E-08 | 7.000E-08 |
| 22741 | 1138 | P1138A | D | 2e-08 | 8e-08 | 7.4e-09 | 3.580E-08 | 3.879E-08 | 7.400E-09 | 8.000E-08 |
| 22742 | 1138 | P1138R | D | 2.2e-07 | 3.8e-08 | 2.6e-08 | 9.467E-08 | 1.087E-07 | 2.600E-08 | 2.200E-07 |
| 22743 | 1138 | P1138N | D | 3.8e-08 | 2e-08 | 3.6e-08 | 3.133E-08 | 9.866E-09 | 2.000E-08 | 3.800E-08 |
| 22744 | 1138 | P1138D | D | 3.2e-08 | 6.9e-08 | 7.3e-08 | 5.800E-08 | 2.261E-08 | 3.200E-08 | 7.300E-08 |
| 22745 | 1138 | P1138C | D | 3.2e-08 | 3e-08 | 4.2e-08 | 3.467E-08 | 6.429E-09 | 3.000E-08 | 4.200E-08 |
| 22746 | 1138 | P1138E | D | 3.8e-08 | 1.2e-08 | 1.4e-07 | 6.333E-08 | 6.766E-08 | 1.200E-08 | 1.400E-07 |
| 22747 | 1138 | P1138Q | D | 5.2e-08 | 7.8e-08 | 1.3e-08 | 4.767E-08 | 3.272E-08 | 1.300E-08 | 7.800E-08 |
| 22748 | 1138 | P1138G | D | 2.3e-08 | 3.6e-08 | 1.7e-08 | 2.533E-08 | 9.713E-09 | 1.700E-08 | 3.600E-08 |
| 22749 | 1138 | P1138H | D | 4.4e-08 | 3.1e-08 | 1.6e-08 | 3.033E-08 | 1.401E-08 | 1.600E-08 | 4.400E-08 |
| 22750 | 1138 | P1138I | D | 3.1e-08 | 2.7e-08 | 3.1e-08 | 2.967E-08 | 2.309E-09 | 2.700E-08 | 3.100E-08 |
| 22751 | 1138 | P1138L | D | 1.4e-08 | 2.7e-08 | 5.7e-08 | 3.267E-08 | 2.205E-08 | 1.400E-08 | 5.700E-08 |
| 22752 | 1138 | P1138K | D | 8e-08 | 7.8e-08 | 4.6e-08 | 6.800E-08 | 1.908E-08 | 4.600E-08 | 8.000E-08 |
| 22753 | 1138 | P1138M | D | 4.4e-08 | 4.4e-08 | 1.3e-08 | 3.367E-08 | 1.790E-08 | 1.300E-08 | 4.400E-08 |
| 22754 | 1138 | P1138F | D | 7.9e-08 | 2.1e-08 | 4.2e-08 | 4.733E-08 | 2.937E-08 | 2.100E-08 | 7.900E-08 |
| 22755 | 1138 | P1138P | D | 2.5e-08 | 3.9e-08 | 3.2e-08 | 3.200E-08 | 7.000E-09 | 2.500E-08 | 3.900E-08 |
| 22756 | 1138 | P1138S | D | 4.5e-08 | 3.9e-08 | 4.5e-08 | 4.300E-08 | 3.464E-09 | 3.900E-08 | 4.500E-08 |
| 22757 | 1138 | P1138T | D | 2.7e-08 | 1.5e-08 | 5.7e-08 | 3.300E-08 | 2.163E-08 | 1.500E-08 | 5.700E-08 |
| 22758 | 1138 | P1138W | D | 9e-08 | 2.7e-08 | 6.5e-08 | 6.067E-08 | 3.172E-08 | 2.700E-08 | 9.000E-08 |
| 22759 | 1138 | P1138Y | D | 2.8e-08 | 4.1e-08 | 3.5e-08 | 3.467E-08 | 6.506E-09 | 2.800E-08 | 4.100E-08 |
| 22760 | 1138 | P1138V | D | 6e-08 | 2e-08 | 1.1e-08 | 3.033E-08 | 2.608E-08 | 1.100E-08 | 6.000E-08 |
| 22761 | 1139 | E1139A | D | 5.2e-08 | 7.8e-08 | 6.9e-08 | 6.633E-08 | 1.320E-08 | 5.200E-08 | 7.800E-08 |
| 22762 | 1139 | E1139R | D | 2.7e-08 | 9.4e-08 | 7.7e-08 | 6.600E-08 | 3.483E-08 | 2.700E-08 | 9.400E-08 |
| 22763 | 1139 | E1139N | D | 1e-07 | 2.4e-08 | 7.7e-08 | 6.700E-08 | 3.897E-08 | 2.400E-08 | 1.000E-07 |
| 22764 | 1139 | E1139D | D | 5.3e-08 | 4.1e-08 | 6e-08 | 5.133E-08 | 9.609E-09 | 4.100E-08 | 6.000E-08 |
| 22765 | 1139 | E1139C | D | 3.6e-08 | 1.7e-08 | 1.8e-08 | 2.367E-08 | 1.069E-08 | 1.700E-08 | 3.600E-08 |
| 22766 | 1139 | E1139E | D | 4.2e-08 | 1.5e-07 | 6.4e-08 | 8.533E-08 | 5.707E-08 | 4.200E-08 | 1.500E-07 |
| 22767 | 1139 | E1139Q | D | 4.8e-08 | 8.8e-08 | 3.8e-08 | 5.800E-08 | 2.646E-08 | 3.800E-08 | 8.800E-08 |
| 22768 | 1139 | E1139G | D | 6.1e-08 | 1.2e-07 | 3.7e-08 | 7.267E-08 | 4.271E-08 | 3.700E-08 | 1.200E-07 |
| 22769 | 1139 | E1139H | D | 3.5e-08 | 3.9e-08 | 6.7e-08 | 4.700E-08 | 1.744E-08 | 3.500E-08 | 6.700E-08 |
| 22770 | 1139 | E1139I | D | 3.8e-08 | 3.4e-08 | 3.7e-08 | 3.633E-08 | 2.082E-09 | 3.400E-08 | 3.800E-08 |
| 22771 | 1139 | E1139L | D | 1.1e-08 | 1.4e-08 | 4.5e-08 | 2.333E-08 | 1.882E-08 | 1.100E-08 | 4.500E-08 |
| 22772 | 1139 | E1139K | D | 8.5e-08 | 2.7e-08 | 1.8e-07 | 9.733E-08 | 7.724E-08 | 2.700E-08 | 1.800E-07 |
| 22773 | 1139 | E1139M | D | 2e-08 | 1.1e-07 | 2.3e-08 | 5.100E-08 | 5.112E-08 | 2.000E-08 | 1.100E-07 |
| 22774 | 1139 | E1139F | D | 4e-08 | 5.9e-08 | 5.3e-08 | 5.067E-08 | 9.713E-09 | 4.000E-08 | 5.900E-08 |
| 22775 | 1139 | E1139P | D | 2.2e-08 | 5.7e-08 | 3.4e-08 | 3.767E-08 | 1.779E-08 | 2.200E-08 | 5.700E-08 |
| 22776 | 1139 | E1139S | D | 3.1e-08 | 9.7e-08 | 4.2e-08 | 5.667E-08 | 3.536E-08 | 3.100E-08 | 9.700E-08 |
| 22777 | 1139 | E1139T | D | 4.9e-08 | 2.9e-08 | 1.9e-08 | 3.233E-08 | 1.528E-08 | 1.900E-08 | 4.900E-08 |
| 22778 | 1139 | E1139W | D | 1.6e-08 | 1.4e-07 | 1.9e-08 | 5.833E-08 | 7.074E-08 | 1.600E-08 | 1.400E-07 |
| 22779 | 1139 | E1139Y | D | 2.2e-08 | 3.7e-08 | 3.3e-08 | 3.067E-08 | 7.767E-09 | 2.200E-08 | 3.700E-08 |

| | | | | | | | | | | |
|---|---|---|---|---|---|---|---|---|---|---|
| 22780 | 1139 | E1139V | D | 6.9e-08 | 2.5e-08 | 8.4e-09 | 3.413E-08 | 3.132E-08 | 8.400E-09 | 6.900E-08 |
| 22781 | 1140 | D1140A | D | 8.4e-08 | 5.3e-08 | 1.6e-07 | 9.900E-08 | 5.505E-08 | 5.300E-08 | 1.600E-07 |
| 22782 | 1140 | D1140R | D | 3e-08 | 3.7e-08 | 6e-08 | 4.233E-08 | 1.570E-08 | 3.000E-08 | 6.000E-08 |
| 22783 | 1140 | D1140N | D | 2.1e-08 | 7.8e-09 | 2.1e-08 | 1.660E-08 | 7.621E-09 | 7.800E-09 | 2.100E-08 |
| 22784 | 1140 | D1140D | D | 3.3e-08 | 5.7e-08 | 4.6e-08 | 4.533E-08 | 1.201E-08 | 3.300E-08 | 5.700E-08 |
| 22785 | 1140 | D1140C | D | 1.9e-08 | 4.6e-08 | 6.9e-08 | 4.467E-08 | 2.503E-08 | 1.900E-08 | 6.900E-08 |
| 22786 | 1140 | D1140E | D | 1.6e-08 | 7.6e-08 | 5.5e-08 | 4.900E-08 | 3.045E-08 | 1.600E-08 | 7.600E-08 |
| 22787 | 1140 | D1140Q | D | 2.7e-08 | 3.5e-08 | 6.1e-08 | 4.100E-08 | 1.778E-08 | 2.700E-08 | 6.100E-08 |
| 22788 | 1140 | D1140G | D | 4.4e-08 | 2.8e-08 | 5.5e-08 | 4.233E-08 | 1.358E-08 | 2.800E-08 | 5.500E-08 |
| 22789 | 1140 | D1140H | D | 2.8e-08 | 2.5e-08 | 3.2e-08 | 2.833E-08 | 3.512E-09 | 2.500E-08 | 3.200E-08 |
| 22790 | 1140 | D1140I | D | 2.8e-08 | 1.8e-08 | 6.1e-08 | 3.567E-08 | 2.250E-08 | 1.800E-08 | 6.100E-08 |
| 22791 | 1140 | D1140L | D | 3.9e-08 | 6.7e-08 | 4.1e-08 | 4.900E-08 | 1.562E-08 | 3.900E-08 | 6.700E-08 |
| 22792 | 1140 | D1140K | D | 7.6e-08 | 4.3e-08 | 8.7e-08 | 6.867E-08 | 2.290E-08 | 4.300E-08 | 8.700E-08 |
| 22793 | 1140 | D1140M | D | 3.3e-08 | 6.8e-08 | 1.9e-08 | 4.000E-08 | 2.524E-08 | 1.900E-08 | 6.800E-08 |
| 22794 | 1140 | D1140F | D | 2.4e-08 | 7.5e-08 | 5e-08 | 4.967E-08 | 2.550E-08 | 2.400E-08 | 7.500E-08 |
| 22795 | 1140 | D1140P | D | 2.1e-08 | 6.8e-08 | 4.4e-08 | 4.433E-08 | 2.350E-08 | 2.100E-08 | 6.800E-08 |
| 22796 | 1140 | D1140S | D | 5.3e-08 | 2.6e-08 | 5.6e-08 | 4.500E-08 | 1.652E-08 | 2.600E-08 | 5.600E-08 |
| 22797 | 1140 | D1140T | D | 6.8e-08 | 4.6e-08 | 5.4e-08 | 5.600E-08 | 1.114E-08 | 4.600E-08 | 6.800E-08 |
| 22798 | 1140 | D1140W | D | 8.5e-08 | 3.5e-08 | 7.8e-08 | 6.600E-08 | 2.707E-08 | 3.500E-08 | 8.500E-08 |
| 22799 | 1140 | D1140Y | D | 4.5e-08 | 1.6e-08 | 4e-08 | 3.367E-08 | 1.550E-08 | 1.600E-08 | 4.500E-08 |
| 22800 | 1140 | D1140V | D | 2.1e-08 | 3.9e-08 | 5.4e-08 | 3.800E-08 | 1.652E-08 | 2.100E-08 | 5.400E-08 |
| 22801 | 1141 | F1141A | D | 6.2e-08 | 9.2e-08 | 1.6e-08 | 5.667E-08 | 3.828E-08 | 1.600E-08 | 9.200E-08 |
| 22802 | 1141 | F1141R | D | 3.6e-08 | 1.7e-08 | 1.5e-08 | 2.267E-08 | 1.159E-08 | 1.500E-08 | 3.600E-08 |
| 22803 | 1141 | F1141N | D | 4.6e-08 | 2.2e-08 | 1.7e-08 | 2.833E-08 | 1.550E-08 | 1.700E-08 | 4.600E-08 |
| 22804 | 1141 | F1141D | D | 1.7e-07 | 3.4e-08 | 7.7e-08 | 9.367E-08 | 6.951E-08 | 3.400E-08 | 1.700E-07 |
| 22805 | 1141 | F1141C | D | 4.7e-08 | 1.4e-07 | 6.8e-08 | 8.500E-08 | 4.877E-08 | 4.700E-08 | 1.400E-07 |
| 22806 | 1141 | F1141E | D | 1.8e-08 | 1.4e-08 | 2.6e-08 | 1.933E-08 | 6.110E-09 | 1.400E-08 | 2.600E-08 |
| 22807 | 1141 | F1141Q | D | 4.1e-08 | 7.1e-08 | 3.1e-08 | 4.767E-08 | 2.082E-08 | 3.100E-08 | 7.100E-08 |
| 22808 | 1141 | F1141G | D | 6.8e-08 | 3.3e-08 | 3.8e-08 | 4.633E-08 | 1.893E-08 | 3.300E-08 | 6.800E-08 |
| 22809 | 1141 | F1141H | D | 5.1e-08 | 5.8e-08 | 5.4e-08 | 5.433E-08 | 3.512E-09 | 5.100E-08 | 5.800E-08 |
| 22810 | 1141 | F1141I | D | 3e-08 | 6.2e-08 | 8.7e-08 | 5.967E-08 | 2.857E-08 | 3.000E-08 | 8.700E-08 |
| 22811 | 1141 | F1141L | D | 5.9e-08 | 4.3e-08 | 2.3e-08 | 4.167E-08 | 1.804E-08 | 2.300E-08 | 5.900E-08 |
| 22812 | 1141 | F1141K | D | 4.6e-08 | 6.8e-08 | 2.8e-08 | 4.733E-08 | 2.003E-08 | 2.800E-08 | 6.800E-08 |
| 22813 | 1141 | F1141M | D | 4e-08 | 3.3e-08 | 2e-08 | 3.100E-08 | 1.015E-08 | 2.000E-08 | 4.000E-08 |
| 22814 | 1141 | F1141F | D | 3.3e-08 | 2e-08 | 3.3e-08 | 2.867E-08 | 7.506E-09 | 2.000E-08 | 3.300E-08 |
| 22815 | 1141 | F1141P | D | 1.7e-08 | 7.6e-08 | 7.5e-08 | 5.600E-08 | 3.378E-08 | 1.700E-08 | 7.600E-08 |
| 22816 | 1141 | F1141S | D | 6.7e-08 | 2.4e-08 | 2.4e-08 | 3.833E-08 | 2.483E-08 | 2.400E-08 | 6.700E-08 |
| 22817 | 1141 | F1141T | D | 3.7e-08 | 3.5e-08 | 1.9e-07 | 8.733E-08 | 8.892E-08 | 3.500E-08 | 1.900E-07 |
| 22818 | 1141 | F1141W | D | 1.1e-07 | 3.5e-08 | 5.5e-08 | 6.667E-08 | 3.884E-08 | 3.500E-08 | 1.100E-07 |
| 22819 | 1141 | F1141Y | D | 6e-08 | 2e-08 | 4.2e-08 | 4.067E-08 | 2.003E-08 | 2.000E-08 | 6.000E-08 |
| 22820 | 1141 | F1141V | D | 1.4e-07 | 4.4e-08 | 3e-08 | 7.133E-08 | 5.988E-08 | 3.000E-08 | 1.400E-07 |
| 22821 | 1142 | A1142A | D | 5.5e-08 | 8e-08 | 6.5e-08 | 6.667E-08 | 1.258E-08 | 5.500E-08 | 8.000E-08 |
| 22822 | 1142 | A1142R | D | 7.5e-08 | 2e-08 | 5e-08 | 4.833E-08 | 2.754E-08 | 2.000E-08 | 7.500E-08 |
| 22823 | 1142 | A1142N | D | 4.3e-08 | 4.4e-08 | 1.8e-08 | 3.500E-08 | 1.473E-08 | 1.800E-08 | 4.400E-08 |
| 22824 | 1142 | A1142D | D | 7e-08 | 5.6e-08 | 3.1e-08 | 5.233E-08 | 1.976E-08 | 3.100E-08 | 7.000E-08 |
| 22825 | 1142 | A1142C | D | 2e-08 | 9.1e-08 | 7.1e-08 | 6.067E-08 | 3.661E-08 | 2.000E-08 | 9.100E-08 |
| 22826 | 1142 | A1142E | D | 2.2e-08 | 4.3e-08 | 4.7e-08 | 3.733E-08 | 1.343E-08 | 2.200E-08 | 4.700E-08 |
| 22827 | 1142 | A1142Q | D | 2.5e-08 | 6e-08 | 4.3e-08 | 4.267E-08 | 1.750E-08 | 2.500E-08 | 6.000E-08 |
| 22828 | 1142 | A1142G | D | 6e-08 | 1.8e-08 | 7.3e-08 | 5.033E-08 | 2.875E-08 | 1.800E-08 | 7.300E-08 |
| 22829 | 1142 | A1142H | D | 3.3e-08 | 9.8e-09 | 4.7e-08 | 2.993E-08 | 1.879E-08 | 9.800E-09 | 4.700E-08 |
| 22830 | 1142 | A1142I | D | 1.1e-07 | 5e-08 | 5e-08 | 7.000E-08 | 3.464E-08 | 5.000E-08 | 1.100E-07 |
| 22831 | 1142 | A1142L | D | 3.7e-08 | 2.6e-08 | 3e-08 | 3.100E-08 | 5.568E-09 | 2.600E-08 | 3.700E-08 |
| 22832 | 1142 | A1142K | D | 5.1e-08 | 2.5e-08 | 3.8e-08 | 3.800E-08 | 1.300E-08 | 2.500E-08 | 5.100E-08 |
| 22833 | 1142 | A1142M | D | 6.4e-08 | 5e-08 | 3.3e-08 | 4.900E-08 | 1.552E-08 | 3.300E-08 | 6.400E-08 |
| 22834 | 1142 | A1142F | D | 8.2e-08 | 6.6e-08 | 1.9e-07 | 1.127E-07 | 6.745E-08 | 6.600E-08 | 1.900E-07 |
| 22835 | 1142 | A1142P | D | 2.9e-08 | 1.7e-08 | 1.8e-08 | 2.133E-08 | 6.658E-09 | 1.700E-08 | 2.900E-08 |
| 22836 | 1142 | A1142S | D | 3.3e-08 | 3.7e-08 | 4.2e-08 | 3.733E-08 | 4.509E-09 | 3.300E-08 | 4.200E-08 |
| 22837 | 1142 | A1142T | D | 1.4e-08 | 2.3e-08 | 5.6e-08 | 3.100E-08 | 2.211E-08 | 1.400E-08 | 5.600E-08 |
| 22838 | 1142 | A1142W | D | 2.8e-08 | 1.5e-08 | 1.7e-07 | 7.100E-08 | 8.598E-08 | 1.500E-08 | 1.700E-07 |
| 22839 | 1142 | A1142Y | D | 6.7e-08 | 2.8e-08 | 9e-08 | 6.167E-08 | 3.134E-08 | 2.800E-08 | 9.000E-08 |
| 22840 | 1142 | A1142V | D | 3.3e-08 | 2.4e-08 | 1.1e-07 | 5.567E-08 | 4.727E-08 | 2.400E-08 | 1.100E-07 |
| 22841 | 1143 | T1143A | D | 1.9e-07 | 1.7e-08 | 4e-08 | 8.233E-08 | 9.395E-08 | 1.700E-08 | 1.900E-07 |
| 22842 | 1143 | T1143R | D | 5.4e-08 | 2.9e-08 | 2.6e-08 | 3.633E-08 | 1.537E-08 | 2.600E-08 | 5.400E-08 |
| 22843 | 1143 | T1143N | D | 7e-08 | 1.7e-08 | 4.2e-08 | 4.300E-08 | 2.651E-08 | 1.700E-08 | 7.000E-08 |
| 22844 | 1143 | T1143D | D | 6.5e-08 | 4.4e-08 | 1e-07 | 6.967E-08 | 2.829E-08 | 4.400E-08 | 1.000E-07 |
| 22845 | 1143 | T1143C | D | 1.4e-08 | 3.6e-08 | 6e-08 | 3.667E-08 | 2.301E-08 | 1.400E-08 | 6.000E-08 |
| 22846 | 1143 | T1143E | D | 4.3e-08 | 8.3e-08 | 9.8e-09 | 4.527E-08 | 3.665E-08 | 9.800E-09 | 8.300E-08 |
| 22847 | 1143 | T1143Q | D | 3.1e-08 | 1.1e-07 | 4.2e-08 | 6.100E-08 | 4.279E-08 | 3.100E-08 | 1.100E-07 |
| 22848 | 1143 | T1143G | D | 7.4e-08 | 8e-08 | 5.9e-08 | 7.100E-08 | 1.082E-08 | 5.900E-08 | 8.000E-08 |
| 22849 | 1143 | T1143H | D | 3.7e-08 | 6.3e-08 | 3.4e-08 | 4.467E-08 | 1.595E-08 | 3.400E-08 | 6.300E-08 |
| 22850 | 1143 | T1143I | D | 8.9e-08 | 1.1e-07 | 2.1e-08 | 7.333E-08 | 4.652E-08 | 2.100E-08 | 1.100E-07 |
| 22851 | 1143 | T1143L | D | 6.3e-08 | 1.6e-08 | 5.3e-08 | 4.400E-08 | 2.476E-08 | 1.600E-08 | 6.300E-08 |
| 22852 | 1143 | T1143K | D | 1.5e-08 | 1.3e-08 | 2.4e-08 | 1.733E-08 | 5.859E-09 | 1.300E-08 | 2.400E-08 |
| 22853 | 1143 | T1143M | D | 1.1e-07 | 2.6e-08 | 1.3e-08 | 4.967E-08 | 5.265E-08 | 1.300E-08 | 1.100E-07 |
| 22854 | 1143 | T1143F | D | 2.1e-08 | 5.7e-08 | 2.7e-08 | 3.500E-08 | 1.929E-08 | 2.100E-08 | 5.700E-08 |
| 22855 | 1143 | T1143P | D | 5.7e-08 | 2.1e-08 | 2.8e-08 | 3.533E-08 | 1.909E-08 | 2.100E-08 | 5.700E-08 |
| 22856 | 1143 | T1143S | D | 1.4e-08 | 3.8e-08 | 1.3e-08 | 2.167E-08 | 1.415E-08 | 1.300E-08 | 3.800E-08 |
| 22857 | 1143 | T1143T | D | 4.8e-08 | 6.2e-08 | 8.8e-08 | 6.600E-08 | 2.030E-08 | 4.800E-08 | 8.800E-08 |
| 22858 | 1143 | T1143W | D | 9.8e-08 | 1.8e-08 | 5e-08 | 5.533E-08 | 4.027E-08 | 1.800E-08 | 9.800E-08 |
| 22859 | 1143 | T1143Y | D | 4.1e-08 | 6.4e-08 | 5.3e-08 | 5.267E-08 | 1.150E-08 | 4.100E-08 | 6.400E-08 |
| 22860 | 1143 | T1143V | D | 3.5e-08 | 1.5e-07 | 6.8e-08 | 8.433E-08 | 5.921E-08 | 3.500E-08 | 1.500E-07 |
| 22861 | 1144 | Y1144A | D | 3.9e-08 | 9.5e-08 | 3e-08 | 5.467E-08 | 3.522E-08 | 3.000E-08 | 9.500E-08 |
| 22862 | 1144 | Y1144R | D | 2.7e-08 | 1.5e-07 | 6.5e-08 | 8.067E-08 | 6.298E-08 | 2.700E-08 | 1.500E-07 |
| 22863 | 1144 | Y1144N | D | 5.1e-08 | 6.4e-08 | 3.1e-08 | 4.867E-08 | 1.662E-08 | 3.100E-08 | 6.400E-08 |
| 22864 | 1144 | Y1144D | D | 1.3e-08 | 4e-08 | 6.5e-08 | 3.933E-08 | 2.601E-08 | 1.300E-08 | 6.500E-08 |

| | | | | | | | | | | |
|---|---|---|---|---|---|---|---|---|---|---|
| 22865 | 1144 | Y1144C | D | 4.2e-08 | 5.1e-08 | 3.9e-08 | 4.400E-08 | 6.245E-09 | 3.900E-08 | 5.100E-08 |
| 22866 | 1144 | Y1144E | D | 1.1e-08 | 3.5e-08 | 1.1e-07 | 5.200E-08 | 5.164E-08 | 1.100E-08 | 1.100E-07 |
| 22867 | 1144 | Y1144Q | D | 5.3e-08 | 7.3e-08 | 8.5e-08 | 7.033E-08 | 1.617E-08 | 5.300E-08 | 8.500E-08 |
| 22868 | 1144 | Y1144G | D | 1.5e-07 | 7e-08 | 3.7e-08 | 8.567E-08 | 5.811E-08 | 3.700E-08 | 1.500E-07 |
| 22869 | 1144 | Y1144H | D | 5.1e-08 | 8.7e-08 | 4.3e-08 | 6.033E-08 | 2.344E-08 | 4.300E-08 | 8.700E-08 |
| 22870 | 1144 | Y1144I | D | 1.3e-08 | 4.9e-08 | 2.9e-08 | 3.033E-08 | 1.804E-08 | 1.300E-08 | 4.900E-08 |
| 22871 | 1144 | Y1144L | D | 2.9e-08 | 4e-08 | 2.9e-08 | 3.267E-08 | 6.351E-09 | 2.900E-08 | 4.000E-08 |
| 22872 | 1144 | Y1144K | D | 9.7e-09 | 2.3e-08 | 5.9e-08 | 3.057E-08 | 2.551E-08 | 9.700E-09 | 5.900E-08 |
| 22873 | 1144 | Y1144M | D | 1.1e-07 | 6.2e-08 | 8.9e-08 | 8.700E-08 | 2.406E-08 | 6.200E-08 | 1.100E-07 |
| 22874 | 1144 | Y1144F | D | 1.2e-07 | 2.4e-08 | 5.5e-08 | 6.633E-08 | 4.899E-08 | 2.400E-08 | 1.200E-07 |
| 22875 | 1144 | Y1144P | D | 5e-08 | 3.2e-08 | 4.7e-08 | 4.300E-08 | 9.644E-09 | 3.200E-08 | 5.000E-08 |
| 22876 | 1144 | Y1144S | D | 7.9e-08 | 9.3e-08 | 2.5e-08 | 6.567E-08 | 3.591E-08 | 2.500E-08 | 9.300E-08 |
| 22877 | 1144 | Y1144T | D | 7.9e-08 | 1.8e-08 | 1.1e-07 | 6.900E-08 | 4.681E-08 | 1.800E-08 | 1.100E-07 |
| 22878 | 1144 | Y1144W | D | 4.1e-08 | 7e-08 | 8.4e-08 | 6.500E-08 | 2.193E-08 | 4.100E-08 | 8.400E-08 |
| 22879 | 1144 | Y1144Y | D | 1.1e-08 | 2e-08 | 4.3e-08 | 2.467E-08 | 1.650E-08 | 1.100E-08 | 4.300E-08 |
| 22880 | 1144 | Y1144V | D | 3.4e-08 | 2.7e-08 | 2.5e-08 | 2.867E-08 | 4.726E-09 | 2.500E-08 | 3.400E-08 |
| 22881 | 1145 | Y1145A | D | 3.5e-08 | 3.4e-08 | 2.9e-08 | 3.267E-08 | 3.215E-09 | 2.900E-08 | 3.500E-08 |
| 22882 | 1145 | Y1145R | D | 3.8e-08 | 2.6e-08 | 5.2e-08 | 3.867E-08 | 1.301E-08 | 2.600E-08 | 5.200E-08 |
| 22883 | 1145 | Y1145N | D | 5.4e-08 | 1.4e-08 | 3.5e-08 | 3.433E-08 | 2.001E-08 | 1.400E-08 | 5.400E-08 |
| 22884 | 1145 | Y1145D | D | 8.4e-08 | 3.1e-08 | 1.2e-07 | 7.833E-08 | 4.477E-08 | 3.100E-08 | 1.200E-07 |
| 22885 | 1145 | Y1145C | D | 3.6e-08 | 4.8e-08 | 1.6e-08 | 3.333E-08 | 1.617E-08 | 1.600E-08 | 4.800E-08 |
| 22886 | 1145 | Y1145E | D | 5.8e-08 | 8.7e-08 | 3.9e-08 | 6.133E-08 | 2.417E-08 | 3.900E-08 | 8.700E-08 |
| 22887 | 1145 | Y1145Q | D | 3.8e-08 | 5.4e-08 | 6.5e-08 | 5.233E-08 | 1.358E-08 | 3.800E-08 | 6.500E-08 |
| 22888 | 1145 | Y1145G | D | 2.1e-08 | 3.3e-08 | 1.1e-07 | 5.467E-08 | 4.829E-08 | 2.100E-08 | 1.100E-07 |
| 22889 | 1145 | Y1145H | D | 3.8e-08 | 4.5e-08 | 5.1e-08 | 4.467E-08 | 6.506E-09 | 3.800E-08 | 5.100E-08 |
| 22890 | 1145 | Y1145I | D | 6.8e-08 | 3.5e-08 | 6.2e-08 | 5.500E-08 | 1.758E-08 | 3.500E-08 | 6.800E-08 |
| 22891 | 1145 | Y1145L | D | 2.6e-08 | 1.5e-08 | 6.4e-08 | 3.500E-08 | 2.571E-08 | 1.500E-08 | 6.400E-08 |
| 22892 | 1145 | Y1145K | D | 2.5e-08 | 3.6e-08 | 5.7e-08 | 3.933E-08 | 1.626E-08 | 2.500E-08 | 5.700E-08 |
| 22893 | 1145 | Y1145M | D | 4.6e-08 | 3.2e-08 | 6.7e-08 | 4.833E-08 | 1.762E-08 | 3.200E-08 | 6.700E-08 |
| 22894 | 1145 | Y1145F | D | 9.2e-08 | 2.8e-08 | 3.1e-08 | 5.033E-08 | 3.612E-08 | 2.800E-08 | 9.200E-08 |
| 22895 | 1145 | Y1145P | D | 2.1e-08 | 7.3e-08 | 7.5e-08 | 5.633E-08 | 3.062E-08 | 2.100E-08 | 7.500E-08 |
| 22896 | 1145 | Y1145S | D | 5.2e-08 | 3.3e-08 | 3.9e-08 | 4.133E-08 | 9.713E-09 | 3.300E-08 | 5.200E-08 |
| 22897 | 1145 | Y1145T | D | 2.3e-08 | 4.3e-08 | 7.8e-08 | 4.800E-08 | 2.784E-08 | 2.300E-08 | 7.800E-08 |
| 22898 | 1145 | Y1145W | D | 5.5e-08 | 3.4e-08 | 5.1e-08 | 4.667E-08 | 1.115E-08 | 3.400E-08 | 5.500E-08 |
| 22899 | 1145 | Y1145Y | D | 1.4e-08 | 1.6e-08 | 1.6e-08 | 1.533E-08 | 1.155E-09 | 1.400E-08 | 1.600E-08 |
| 22900 | 1145 | Y1145V | D | 3.2e-08 | 4e-08 | 9.9e-08 | 5.700E-08 | 3.659E-08 | 3.200E-08 | 9.900E-08 |
| 22901 | 1146 | C1146A | D | 3.2e-08 | 3.4e-08 | 1.8e-08 | 2.800E-08 | 8.718E-09 | 1.800E-08 | 3.400E-08 |
| 22902 | 1146 | C1146R | D | 3.3e-08 | 6.1e-08 | 5.1e-08 | 4.833E-08 | 1.419E-08 | 3.300E-08 | 6.100E-08 |
| 22903 | 1146 | C1146N | D | 3.2e-08 | 1.1e-07 | 1e-07 | 8.067E-08 | 4.244E-08 | 3.200E-08 | 1.100E-07 |
| 22904 | 1146 | C1146D | D | 4e-08 | 2.7e-08 | 4.1e-08 | 3.600E-08 | 7.810E-09 | 2.700E-08 | 4.100E-08 |
| 22905 | 1146 | C1146C | D | 1.1e-07 | 8.4e-08 | 2.5e-08 | 7.300E-08 | 4.355E-08 | 2.500E-08 | 1.100E-07 |
| 22906 | 1146 | C1146E | D | 2.7e-08 | 5.3e-08 | 2.8e-08 | 3.600E-08 | 1.473E-08 | 2.700E-08 | 5.300E-08 |
| 22907 | 1146 | C1146Q | D | 4.5e-08 | 4.6e-08 | 7.4e-08 | 5.500E-08 | 1.646E-08 | 4.500E-08 | 7.400E-08 |
| 22908 | 1146 | C1146G | D | 3e-08 | 4.4e-08 | 1.7e-07 | 8.133E-08 | 7.711E-08 | 3.000E-08 | 1.700E-07 |
| 22909 | 1146 | C1146H | D | 5.2e-08 | 2.1e-08 | 4.8e-08 | 4.033E-08 | 1.686E-08 | 2.100E-08 | 5.200E-08 |
| 22910 | 1146 | C1146I | D | 5.7e-08 | 4.8e-08 | 3.2e-08 | 4.567E-08 | 1.266E-08 | 3.200E-08 | 5.700E-08 |
| 22911 | 1146 | C1146L | D | 3.2e-08 | 2.5e-08 | 2.4e-08 | 2.700E-08 | 4.359E-09 | 2.400E-08 | 3.200E-08 |
| 22912 | 1146 | C1146K | D | 3e-08 | 5.1e-08 | 6.3e-08 | 4.800E-08 | 1.670E-08 | 3.000E-08 | 6.300E-08 |
| 22913 | 1146 | C1146M | D | 2.6e-08 | 4.2e-08 | 2.8e-08 | 3.200E-08 | 8.718E-09 | 2.600E-08 | 4.200E-08 |
| 22914 | 1146 | C1146F | D | 7e-08 | 1e-07 | 3.8e-08 | 6.933E-08 | 3.101E-08 | 3.800E-08 | 1.000E-07 |
| 22915 | 1146 | C1146P | D | 3.1e-08 | 2.3e-08 | 6.8e-08 | 4.067E-08 | 2.401E-08 | 2.300E-08 | 6.800E-08 |
| 22916 | 1146 | C1146S | D | 7e-08 | 7.2e-08 | 2.3e-08 | 5.500E-08 | 2.773E-08 | 2.300E-08 | 7.200E-08 |
| 22917 | 1146 | C1146T | D | 2e-08 | 5.7e-08 | 5e-08 | 4.233E-08 | 1.966E-08 | 2.000E-08 | 5.700E-08 |
| 22918 | 1146 | C1146W | D | 8.4e-08 | 6e-08 | 4e-08 | 6.133E-08 | 2.203E-08 | 4.000E-08 | 8.400E-08 |
| 22919 | 1146 | C1146Y | D | 6.6e-08 | 5.2e-08 | 2.4e-08 | 4.733E-08 | 2.139E-08 | 2.400E-08 | 6.600E-08 |
| 22920 | 1146 | C1146V | D | 3.4e-08 | 1.6e-07 | 2.6e-08 | 7.333E-08 | 7.516E-08 | 2.600E-08 | 1.600E-07 |
| 22921 | 1147 | Q1147A | D | 2.9e-08 | 3.9e-08 | 5.7e-08 | 4.167E-08 | 1.419E-08 | 2.900E-08 | 5.700E-08 |
| 22922 | 1147 | Q1147R | D | 4.3e-08 | 4.1e-08 | 8.3e-08 | 5.567E-08 | 2.369E-08 | 4.100E-08 | 8.300E-08 |
| 22923 | 1147 | Q1147N | D | 5.3e-08 | 2.2e-08 | 2.3e-08 | 3.267E-08 | 1.762E-08 | 2.200E-08 | 5.300E-08 |
| 22924 | 1147 | Q1147D | D | 1.8e-08 | 4.8e-08 | 2.5e-08 | 3.033E-08 | 1.570E-08 | 1.800E-08 | 4.800E-08 |
| 22925 | 1147 | Q1147C | D | 2.7e-08 | 5.7e-08 | 5.8e-08 | 4.733E-08 | 1.762E-08 | 2.700E-08 | 5.800E-08 |
| 22926 | 1147 | Q1147E | D | 6.5e-08 | 1.5e-08 | 4.2e-08 | 4.067E-08 | 2.503E-08 | 1.500E-08 | 6.500E-08 |
| 22927 | 1147 | Q1147Q | D | 4.6e-08 | 5.8e-08 | 6.4e-08 | 5.600E-08 | 9.165E-09 | 4.600E-08 | 6.400E-08 |
| 22928 | 1147 | Q1147G | D | 8.4e-08 | 1.1e-07 | 9.8e-08 | 9.733E-08 | 1.301E-08 | 8.400E-08 | 1.100E-07 |
| 22929 | 1147 | Q1147H | D | 4.9e-08 | 4.1e-08 | 1.7e-08 | 3.567E-08 | 1.665E-08 | 1.700E-08 | 4.900E-08 |
| 22930 | 1147 | Q1147I | D | 4.1e-08 | 4.2e-08 | 2e-08 | 3.433E-08 | 1.242E-08 | 2.000E-08 | 4.200E-08 |
| 22931 | 1147 | Q1147L | D | 1.5e-08 | 1.4e-07 | 3.7e-08 | 6.400E-08 | 6.673E-08 | 1.500E-08 | 1.400E-07 |
| 22932 | 1147 | Q1147K | D | 1e-07 | 1e-07 | 5.1e-08 | 8.367E-08 | 2.829E-08 | 5.100E-08 | 1.000E-07 |
| 22933 | 1147 | Q1147M | D | 9.1e-08 | 8.5e-08 | 6.2e-08 | 7.933E-08 | 1.531E-08 | 6.200E-08 | 9.100E-08 |
| 22934 | 1147 | Q1147F | D | 5.9e-08 | 5.2e-08 | 8.2e-08 | 6.433E-08 | 1.570E-08 | 5.200E-08 | 8.200E-08 |
| 22935 | 1147 | Q1147P | D | 8.4e-08 | 2.7e-08 | 2.1e-08 | 4.400E-08 | 3.477E-08 | 2.100E-08 | 8.400E-08 |
| 22936 | 1147 | Q1147S | D | 9.6e-09 | 4.1e-08 | 5.1e-08 | 3.387E-08 | 2.160E-08 | 9.600E-09 | 5.100E-08 |
| 22937 | 1147 | Q1147T | D | 4.3e-08 | 9.8e-08 | 4.8e-08 | 6.300E-08 | 3.041E-08 | 4.300E-08 | 9.800E-08 |
| 22938 | 1147 | Q1147W | D | 3.1e-08 | 4.3e-08 | 3.8e-08 | 3.733E-08 | 6.028E-09 | 3.100E-08 | 4.300E-08 |
| 22939 | 1147 | Q1147Y | D | 5.8e-08 | 5.2e-08 | 9.6e-08 | 6.867E-08 | 2.386E-08 | 5.200E-08 | 9.600E-08 |
| 22940 | 1147 | Q1147V | D | 6.8e-08 | 1.5e-08 | 4.3e-08 | 4.200E-08 | 2.651E-08 | 1.500E-08 | 6.800E-08 |
| 22941 | 1148 | Q1148A | D | 1.1e-07 | 4.5e-08 | 4.8e-08 | 6.767E-08 | 3.669E-08 | 4.500E-08 | 1.100E-07 |
| 22942 | 1148 | Q1148R | D | 6.4e-08 | 4.1e-08 | 3e-08 | 4.500E-08 | 1.735E-08 | 3.000E-08 | 6.400E-08 |
| 22943 | 1148 | Q1148N | D | 3.1e-08 | 3.4e-08 | 6.2e-08 | 4.233E-08 | 1.710E-08 | 3.100E-08 | 6.200E-08 |
| 22944 | 1148 | Q1148D | D | 5.2e-08 | 1.3e-08 | 2.3e-08 | 2.933E-08 | 2.026E-08 | 1.300E-08 | 5.200E-08 |
| 22945 | 1148 | Q1148C | D | 5e-08 | 4.8e-08 | 3.2e-08 | 4.333E-08 | 9.866E-09 | 3.200E-08 | 5.000E-08 |
| 22946 | 1148 | Q1148E | D | 3.7e-08 | 5.5e-08 | 1.3e-08 | 3.500E-08 | 2.107E-08 | 1.300E-08 | 5.500E-08 |
| 22947 | 1148 | Q1148Q | D | 6.1e-08 | 4.4e-08 | 8.8e-09 | 3.793E-08 | 2.662E-08 | 8.800E-09 | 6.100E-08 |
| 22948 | 1148 | Q1148G | D | 3e-08 | 6.3e-08 | 7.6e-08 | 5.633E-08 | 2.371E-08 | 3.000E-08 | 7.600E-08 |
| 22949 | 1148 | Q1148H | D | 5.7e-08 | 1.2e-08 | 5.2e-08 | 4.033E-08 | 2.466E-08 | 1.200E-08 | 5.700E-08 |

| | | | | | | | | | | |
|---|---|---|---|---|---|---|---|---|---|---|
| 22950 | 1148 | Q1148I | D | 7.5e-08 | 8.8e-08 | 7e-08 | 7.767E-08 | 9.292E-09 | 7.000E-08 | 8.800E-08 |
| 22951 | 1148 | Q1148L | D | 4.6e-08 | 5.9e-08 | 3.7e-08 | 4.733E-08 | 1.106E-08 | 3.700E-08 | 5.900E-08 |
| 22952 | 1148 | Q1148K | D | 3.2e-08 | 4.2e-08 | 2.6e-08 | 3.333E-08 | 8.083E-09 | 2.600E-08 | 4.200E-08 |
| 22953 | 1148 | Q1148M | D | 5e-08 | 6.5e-08 | 1.3e-08 | 4.267E-08 | 2.676E-08 | 1.300E-08 | 6.500E-08 |
| 22954 | 1148 | Q1148F | D | 4.3e-08 | 5.3e-08 | 4.1e-08 | 4.567E-08 | 6.429E-09 | 4.100E-08 | 5.300E-08 |
| 22955 | 1148 | Q1148P | D | 1.1e-08 | 3.4e-08 | 9.2e-08 | 4.567E-08 | 4.174E-08 | 1.100E-08 | 9.200E-08 |
| 22956 | 1148 | Q1148S | D | 5.8e-08 | 2.9e-08 | 4.1e-08 | 4.267E-08 | 1.457E-08 | 2.900E-08 | 5.800E-08 |
| 22957 | 1148 | Q1148T | D | 3.8e-08 | 7.6e-08 | 5.7e-08 | 5.700E-08 | 1.900E-08 | 3.800E-08 | 7.600E-08 |
| 22958 | 1148 | Q1148W | D | 1e-07 | 6.2e-08 | 6.7e-08 | 7.633E-08 | 2.065E-08 | 6.200E-08 | 1.000E-07 |
| 22959 | 1148 | Q1148Y | D | 3.5e-08 | 5.8e-08 | 5.9e-08 | 5.067E-08 | 1.358E-08 | 3.500E-08 | 5.900E-08 |
| 22960 | 1148 | Q1148V | D | 8.3e-08 | 1.9e-08 | 1.5e-07 | 8.400E-08 | 6.551E-08 | 1.900E-08 | 1.500E-07 |
| 22961 | 1149 | H1149A | D | 5.6e-08 | 1.4e-07 | 1.8e-08 | 7.133E-08 | 6.243E-08 | 1.800E-08 | 1.400E-07 |
| 22962 | 1149 | H1149R | D | 3.6e-08 | 4e-08 | 7.6e-08 | 5.067E-08 | 2.203E-08 | 3.600E-08 | 7.600E-08 |
| 22963 | 1149 | H1149N | D | 6.6e-08 | 1e-07 | 6.4e-08 | 7.667E-08 | 2.023E-08 | 6.400E-08 | 1.000E-07 |
| 22964 | 1149 | H1149D | D | 8.8e-08 | 3.4e-08 | 3.9e-08 | 5.367E-08 | 2.984E-08 | 3.400E-08 | 8.800E-08 |
| 22965 | 1149 | H1149C | D | 4.2e-08 | 6.6e-08 | 3.1e-08 | 4.633E-08 | 1.790E-08 | 3.100E-08 | 6.600E-08 |
| 22966 | 1149 | H1149E | D | 1.6e-08 | 4.1e-08 | 3.6e-08 | 3.100E-08 | 1.323E-08 | 1.600E-08 | 4.100E-08 |
| 22967 | 1149 | H1149Q | D | 2e-08 | 4.2e-08 | 3.7e-08 | 3.300E-08 | 1.153E-08 | 2.000E-08 | 4.200E-08 |
| 22968 | 1149 | H1149G | D | 2.7e-08 | 8.1e-08 | 2.3e-08 | 4.367E-08 | 3.239E-08 | 2.300E-08 | 8.100E-08 |
| 22969 | 1149 | H1149H | D | 2.3e-08 | 2.8e-08 | 6.3e-08 | 3.800E-08 | 2.179E-08 | 2.300E-08 | 6.300E-08 |
| 22970 | 1149 | H1149I | D | 7.5e-08 | 2.7e-08 | 1.2e-08 | 3.800E-08 | 3.291E-08 | 1.200E-08 | 7.500E-08 |
| 22971 | 1149 | H1149L | D | 2e-08 | 4.1e-08 | 5.6e-08 | 3.900E-08 | 1.808E-08 | 2.000E-08 | 5.600E-08 |
| 22972 | 1149 | H1149K | D | 4.1e-08 | 2.6e-08 | 5e-08 | 3.900E-08 | 1.212E-08 | 2.600E-08 | 5.000E-08 |
| 22973 | 1149 | H1149M | D | 5.2e-08 | 6.6e-08 | 4.6e-08 | 5.467E-08 | 1.026E-08 | 4.600E-08 | 6.600E-08 |
| 22974 | 1149 | H1149F | D | 3.7e-08 | 5.1e-08 | 2.1e-08 | 3.633E-08 | 1.501E-08 | 2.100E-08 | 5.100E-08 |
| 22975 | 1149 | H1149P | D | 4.1e-08 | 4.3e-08 | 5.3e-08 | 4.567E-08 | 6.429E-09 | 4.100E-08 | 5.300E-08 |
| 22976 | 1149 | H1149S | D | 3.8e-08 | 4.1e-08 | 2.1e-08 | 3.333E-08 | 1.079E-08 | 2.100E-08 | 4.100E-08 |
| 22977 | 1149 | H1149T | D | 5.1e-08 | 5.3e-08 | 2.3e-08 | 4.233E-08 | 1.677E-08 | 2.300E-08 | 5.300E-08 |
| 22978 | 1149 | H1149W | D | 5.8e-08 | 7e-08 | 1.4e-08 | 4.733E-08 | 2.948E-08 | 1.400E-08 | 7.000E-08 |
| 22979 | 1149 | H1149Y | D | 2.2e-08 | 2.2e-08 | 4.8e-08 | 3.067E-08 | 1.501E-08 | 2.200E-08 | 4.800E-08 |
| 22980 | 1149 | H1149V | D | 2e-08 | 6.6e-08 | 6.7e-08 | 5.100E-08 | 2.685E-08 | 2.000E-08 | 6.700E-08 |
| 22981 | 1150 | Y1150A | D | 1.1e-07 | 3.2e-08 | 5.5e-08 | 6.567E-08 | 4.008E-08 | 3.200E-08 | 1.100E-07 |
| 22982 | 1150 | Y1150R | D | 4.2e-08 | 6.5e-08 | 3.8e-08 | 4.833E-08 | 1.457E-08 | 3.800E-08 | 6.500E-08 |
| 22983 | 1150 | Y1150N | D | 2.5e-08 | 3.3e-08 | 7.6e-08 | 4.467E-08 | 2.743E-08 | 2.500E-08 | 7.600E-08 |
| 22984 | 1150 | Y1150D | D | 3.5e-08 | 4.9e-08 | 1e-07 | 6.133E-08 | 3.421E-08 | 3.500E-08 | 1.000E-07 |
| 22985 | 1150 | Y1150C | D | 3.3e-08 | 4.3e-08 | 1.7e-08 | 3.100E-08 | 1.311E-08 | 1.700E-08 | 4.300E-08 |
| 22986 | 1150 | Y1150E | D | 3.9e-08 | 3.8e-08 | 1.8e-08 | 3.167E-08 | 1.185E-08 | 1.800E-08 | 3.900E-08 |
| 22987 | 1150 | Y1150Q | D | 5.4e-08 | 9.7e-08 | 5.2e-08 | 6.767E-08 | 2.542E-08 | 5.200E-08 | 9.700E-08 |
| 22988 | 1150 | Y1150G | D | 8.4e-08 | 3.7e-08 | 7.2e-08 | 6.433E-08 | 2.442E-08 | 3.700E-08 | 8.400E-08 |
| 22989 | 1150 | Y1150H | D | 2.7e-08 | 4.7e-08 | 2.9e-08 | 3.433E-08 | 1.102E-08 | 2.700E-08 | 4.700E-08 |
| 22990 | 1150 | Y1150I | D | 5.2e-08 | 3.5e-08 | 2.9e-08 | 3.867E-08 | 1.193E-08 | 2.900E-08 | 5.200E-08 |
| 22991 | 1150 | Y1150L | D | 3.8e-08 | 1.3e-08 | 3.8e-08 | 2.967E-08 | 1.443E-08 | 1.300E-08 | 3.800E-08 |
| 22992 | 1150 | Y1150K | D | 1.1e-07 | 2.3e-08 | 3.1e-08 | 5.467E-08 | 4.809E-08 | 2.300E-08 | 1.100E-07 |
| 22993 | 1150 | Y1150M | D | 4.3e-08 | 2e-08 | 3.3e-08 | 3.200E-08 | 1.153E-08 | 2.000E-08 | 4.300E-08 |
| 22994 | 1150 | Y1150F | D | 6.2e-08 | 6.1e-08 | 1.2e-07 | 8.100E-08 | 3.378E-08 | 6.100E-08 | 1.200E-07 |
| 22995 | 1150 | Y1150P | D | 2.7e-08 | 2.8e-08 | 4e-08 | 3.167E-08 | 7.234E-09 | 2.700E-08 | 4.000E-08 |
| 22996 | 1150 | Y1150S | D | 5.1e-08 | 4e-08 | 6e-08 | 5.033E-08 | 1.002E-08 | 4.000E-08 | 6.000E-08 |
| 22997 | 1150 | Y1150T | D | 6e-08 | 8.9e-08 | 2.8e-08 | 5.900E-08 | 3.051E-08 | 2.800E-08 | 8.900E-08 |
| 22998 | 1150 | Y1150W | D | 4.4e-08 | 3.8e-08 | 3.6e-08 | 3.933E-08 | 4.163E-09 | 3.600E-08 | 4.400E-08 |
| 22999 | 1150 | Y1150Y | D | 7.3e-08 | 4.4e-08 | 2.7e-08 | 4.800E-08 | 2.326E-08 | 2.700E-08 | 7.300E-08 |
| 23000 | 1150 | Y1150V | D | 8.3e-08 | 3.8e-08 | 4.4e-08 | 5.500E-08 | 2.443E-08 | 3.800E-08 | 8.300E-08 |
| 23001 | 1151 | T1151A | D | 7e-08 | 3.9e-08 | 1.1e-07 | 7.300E-08 | 3.559E-08 | 3.900E-08 | 1.100E-07 |
| 23002 | 1151 | T1151R | D | 2.3e-08 | 1.6e-08 | 1.2e-07 | 5.300E-08 | 5.813E-08 | 1.600E-08 | 1.200E-07 |
| 23003 | 1151 | T1151N | D | 5.6e-08 | 6.1e-08 | 5.7e-08 | 5.800E-08 | 2.646E-09 | 5.600E-08 | 6.100E-08 |
| 23004 | 1151 | T1151D | D | 2e-08 | 2.4e-08 | 6.4e-08 | 3.600E-08 | 2.433E-08 | 2.000E-08 | 6.400E-08 |
| 23005 | 1151 | T1151C | D | 9.2e-08 | 3.2e-08 | 8.8e-09 | 4.427E-08 | 4.293E-08 | 8.800E-09 | 9.200E-08 |
| 23006 | 1151 | T1151E | D | 1e-07 | 3.5e-08 | 1.4e-07 | 9.167E-08 | 5.299E-08 | 3.500E-08 | 1.400E-07 |
| 23007 | 1151 | T1151Q | D | 6.7e-08 | 4.3e-08 | 3.1e-08 | 4.700E-08 | 1.833E-08 | 3.100E-08 | 6.700E-08 |
| 23008 | 1151 | T1151G | D | 1.5e-08 | 1.4e-08 | 1e-07 | 4.300E-08 | 4.937E-08 | 1.400E-08 | 1.000E-07 |
| 23009 | 1151 | T1151H | D | 5.7e-08 | 6.6e-08 | 8.4e-08 | 6.900E-08 | 1.375E-08 | 5.700E-08 | 8.400E-08 |
| 23010 | 1151 | T1151I | D | 3e-08 | 6.2e-08 | 2.8e-08 | 4.000E-08 | 1.908E-08 | 2.800E-08 | 6.200E-08 |
| 23011 | 1151 | T1151L | D | 6.4e-08 | 6.3e-08 | 5.2e-08 | 5.967E-08 | 6.658E-09 | 5.200E-08 | 6.400E-08 |
| 23012 | 1151 | T1151K | D | 5.7e-08 | 5.7e-08 | 7.3e-08 | 6.233E-08 | 9.238E-09 | 5.700E-08 | 7.300E-08 |
| 23013 | 1151 | T1151M | D | 4.8e-08 | 4.7e-08 | 3.2e-08 | 4.233E-08 | 8.963E-09 | 3.200E-08 | 4.800E-08 |
| 23014 | 1151 | T1151F | D | 5.9e-08 | 6.3e-08 | 3.3e-08 | 5.167E-08 | 1.629E-08 | 3.300E-08 | 6.300E-08 |
| 23015 | 1151 | T1151P | D | 3.1e-08 | 2.8e-08 | 2.6e-08 | 2.833E-08 | 2.517E-09 | 2.600E-08 | 3.100E-08 |
| 23016 | 1151 | T1151S | D | 7.2e-08 | 4.1e-08 | 4.5e-08 | 5.267E-08 | 1.686E-08 | 4.100E-08 | 7.200E-08 |
| 23017 | 1151 | T1151T | D | 6.3e-08 | 4.5e-08 | 7e-08 | 5.933E-08 | 1.290E-08 | 4.500E-08 | 7.000E-08 |
| 23018 | 1151 | T1151W | D | 3.6e-08 | 4.6e-08 | 3.6e-08 | 3.933E-08 | 5.774E-09 | 3.600E-08 | 4.600E-08 |
| 23019 | 1151 | T1151Y | D | 2.3e-08 | 1.2e-07 | 2.4e-08 | 5.567E-08 | 5.572E-08 | 2.300E-08 | 1.200E-07 |
| 23020 | 1151 | T1151V | D | 5.6e-08 | 3.4e-08 | 3.3e-08 | 4.100E-08 | 1.300E-08 | 3.300E-08 | 5.600E-08 |
| 23021 | 1152 | T1152A | D | 2.4e-08 | 5e-08 | 4.3e-08 | 3.900E-08 | 1.345E-08 | 2.400E-08 | 5.000E-08 |
| 23022 | 1152 | T1152R | D | 6.7e-08 | 2.7e-08 | 5.8e-08 | 5.067E-08 | 2.098E-08 | 2.700E-08 | 6.700E-08 |
| 23023 | 1152 | T1152N | D | 5.9e-08 | 7.8e-08 | 1.9e-08 | 5.200E-08 | 3.012E-08 | 1.900E-08 | 7.800E-08 |
| 23024 | 1152 | T1152D | D | 7.5e-08 | 6.1e-08 | 3e-08 | 5.533E-08 | 2.303E-08 | 3.000E-08 | 7.500E-08 |
| 23025 | 1152 | T1152C | D | 5.1e-08 | 9.9e-09 | 2.2e-08 | 2.763E-08 | 2.112E-08 | 9.900E-09 | 5.100E-08 |
| 23026 | 1152 | T1152E | D | 5.2e-08 | 5.9e-08 | 9.3e-08 | 6.800E-08 | 2.193E-08 | 5.200E-08 | 9.300E-08 |
| 23027 | 1152 | T1152Q | D | 3.1e-08 | 1.7e-08 | 3e-08 | 2.600E-08 | 7.810E-09 | 1.700E-08 | 3.100E-08 |
| 23028 | 1152 | T1152G | D | 5.8e-08 | 5.2e-08 | 2.2e-07 | 1.100E-07 | 9.531E-08 | 5.200E-08 | 2.200E-07 |
| 23029 | 1152 | T1152H | D | 5.2e-08 | 2.3e-08 | 2.4e-08 | 3.300E-08 | 1.646E-08 | 2.300E-08 | 5.200E-08 |
| 23030 | 1152 | T1152I | D | 4.5e-08 | 2.1e-08 | 2e-08 | 2.867E-08 | 1.415E-08 | 2.000E-08 | 4.500E-08 |
| 23031 | 1152 | T1152L | D | 9e-08 | 4.2e-08 | 6.9e-08 | 6.700E-08 | 2.406E-08 | 4.200E-08 | 9.000E-08 |
| 23032 | 1152 | T1152K | D | 4.6e-08 | 3.2e-08 | 2.9e-08 | 3.567E-08 | 9.074E-09 | 2.900E-08 | 4.600E-08 |
| 23033 | 1152 | T1152M | D | 2.6e-08 | 4.4e-08 | 3.8e-08 | 3.600E-08 | 9.165E-09 | 2.600E-08 | 4.400E-08 |
| 23034 | 1152 | T1152F | D | 5.1e-08 | 3.6e-08 | 5.9e-08 | 4.867E-08 | 1.168E-08 | 3.600E-08 | 5.900E-08 |

| | | | | | | | | | | |
|---|---|---|---|---|---|---|---|---|---|---|
| 23035 | 1152 | T1152P | D | 1e-08 | 6.4e-08 | 4.4e-08 | 3.933E-08 | 2.730E-08 | 1.000E-08 | 6.400E-08 |
| 23036 | 1152 | T1152S | D | 1.7e-08 | 4e-08 | 4.4e-08 | 3.367E-08 | 1.457E-08 | 1.700E-08 | 4.400E-08 |
| 23037 | 1152 | T1152T | D | 3.6e-08 | 2.8e-08 | 4.9e-08 | 3.767E-08 | 1.060E-08 | 2.800E-08 | 4.900E-08 |
| 23038 | 1152 | T1152W | D | 3.3e-08 | 6.2e-08 | 2.8e-08 | 4.100E-08 | 1.836E-08 | 2.800E-08 | 6.200E-08 |
| 23039 | 1152 | T1152Y | D | 3.2e-08 | 5.1e-08 | 5.6e-08 | 4.633E-08 | 1.266E-08 | 3.200E-08 | 5.600E-08 |
| 23040 | 1152 | T1152V | D | 3.2e-08 | 3.1e-08 | 2.9e-08 | 3.067E-08 | 1.528E-09 | 2.900E-08 | 3.200E-08 |
| 23041 | 1153 | P1153A | D | 7.5e-08 | 3.5e-08 | 8.7e-08 | 6.567E-08 | 2.723E-08 | 3.500E-08 | 8.700E-08 |
| 23042 | 1153 | P1153R | D | 5.1e-08 | 2.1e-08 | 2.7e-08 | 3.300E-08 | 1.587E-08 | 2.100E-08 | 5.100E-08 |
| 23043 | 1153 | P1153N | D | 2.9e-08 | 2.5e-08 | 5.2e-08 | 3.533E-08 | 1.457E-08 | 2.500E-08 | 5.200E-08 |
| 23044 | 1153 | P1153D | D | 5e-08 | 7.2e-08 | 2.8e-08 | 5.000E-08 | 2.200E-08 | 2.800E-08 | 7.200E-08 |
| 23045 | 1153 | P1153C | D | 1.5e-08 | 3.1e-08 | 5.3e-08 | 3.300E-08 | 1.908E-08 | 1.500E-08 | 5.300E-08 |
| 23046 | 1153 | P1153E | D | 2.1e-08 | 1.1e-07 | 4.9e-08 | 6.000E-08 | 4.551E-08 | 2.100E-08 | 1.100E-07 |
| 23047 | 1153 | P1153Q | D | 1.4e-08 | 1.8e-08 | 6.2e-08 | 3.133E-08 | 2.663E-08 | 1.400E-08 | 6.200E-08 |
| 23048 | 1153 | P1153G | D | 9e-08 | 2e-08 | 1.1e-07 | 7.333E-08 | 4.726E-08 | 2.000E-08 | 1.100E-07 |
| 23049 | 1153 | P1153H | D | 1.3e-07 | 6.3e-08 | 2.3e-08 | 7.200E-08 | 5.406E-08 | 2.300E-08 | 1.300E-07 |
| 23050 | 1153 | P1153I | D | 2.7e-08 | 5e-08 | 8.6e-08 | 5.433E-08 | 2.974E-08 | 2.700E-08 | 8.600E-08 |
| 23051 | 1153 | P1153L | D | 3e-08 | 6.4e-08 | 4.3e-08 | 4.567E-08 | 1.716E-08 | 3.000E-08 | 6.400E-08 |
| 23052 | 1153 | P1153K | D | 4.7e-08 | 1.8e-08 | 5.7e-08 | 4.067E-08 | 2.026E-08 | 1.800E-08 | 5.700E-08 |
| 23053 | 1153 | P1153M | D | 4e-08 | 9.4e-08 | 2.3e-08 | 5.233E-08 | 3.707E-08 | 2.300E-08 | 9.400E-08 |
| 23054 | 1153 | P1153F | D | 7.2e-08 | 2e-08 | 1.5e-07 | 8.067E-08 | 6.543E-08 | 2.000E-08 | 1.500E-07 |
| 23055 | 1153 | P1153P | D | 1.7e-08 | 3e-08 | 1.8e-07 | 7.567E-08 | 9.059E-08 | 1.700E-08 | 1.800E-07 |
| 23056 | 1153 | P1153S | D | 2.3e-08 | 4.6e-08 | 7.5e-08 | 4.800E-08 | 2.606E-08 | 2.300E-08 | 7.500E-08 |
| 23057 | 1153 | P1153T | D | 2.6e-08 | 4.2e-08 | 3e-08 | 3.267E-08 | 8.327E-09 | 2.600E-08 | 4.200E-08 |
| 23058 | 1153 | P1153W | D | 6.1e-08 | 1.2e-07 | 6.5e-08 | 8.200E-08 | 3.297E-08 | 6.100E-08 | 1.200E-07 |
| 23059 | 1153 | P1153Y | D | 5.1e-08 | 1.3e-08 | 4.5e-08 | 3.633E-08 | 2.043E-08 | 1.300E-08 | 5.100E-08 |
| 23060 | 1153 | P1153V | D | 4.5e-08 | 6.2e-08 | 7.1e-08 | 5.933E-08 | 1.320E-08 | 4.500E-08 | 7.100E-08 |
| 23061 | 1154 | P1154A | D | 1e-08 | 5.3e-08 | 2.9e-08 | 3.067E-08 | 2.155E-08 | 1.000E-08 | 5.300E-08 |
| 23062 | 1154 | P1154R | D | 3.4e-08 | 7.9e-08 | 5.2e-08 | 5.500E-08 | 2.265E-08 | 3.400E-08 | 7.900E-08 |
| 23063 | 1154 | P1154N | D | 4.3e-08 | 6.4e-08 | 4.2e-08 | 4.967E-08 | 1.242E-08 | 4.200E-08 | 6.400E-08 |
| 23064 | 1154 | P1154D | D | 1.7e-08 | 5.3e-08 | 3.6e-08 | 3.533E-08 | 1.801E-08 | 1.700E-08 | 5.300E-08 |
| 23065 | 1154 | P1154C | D | 2.8e-08 | 4e-08 | 7.6e-08 | 4.800E-08 | 2.498E-08 | 2.800E-08 | 7.600E-08 |
| 23066 | 1154 | P1154E | D | 2.6e-08 | 1.8e-08 | 6.2e-08 | 3.533E-08 | 2.344E-08 | 1.800E-08 | 6.200E-08 |
| 23067 | 1154 | P1154Q | D | 5.6e-08 | 7.1e-08 | 3.8e-08 | 5.500E-08 | 1.652E-08 | 3.800E-08 | 7.100E-08 |
| 23068 | 1154 | P1154G | D | 9.3e-08 | 5.6e-08 | 2.3e-08 | 5.733E-08 | 3.502E-08 | 2.300E-08 | 9.300E-08 |
| 23069 | 1154 | P1154H | D | 2.9e-08 | 4.2e-08 | 3.8e-08 | 3.633E-08 | 6.658E-09 | 2.900E-08 | 4.200E-08 |
| 23070 | 1154 | P1154I | D | 4e-08 | 1.9e-07 | 1.8e-08 | 8.267E-08 | 9.360E-08 | 1.800E-08 | 1.900E-07 |
| 23071 | 1154 | P1154L | D | 3.5e-08 | 5.4e-08 | 5.8e-08 | 4.900E-08 | 1.229E-08 | 3.500E-08 | 5.800E-08 |
| 23072 | 1154 | P1154K | D | 2.6e-08 | 4.6e-08 | 2e-08 | 3.067E-08 | 1.361E-08 | 2.000E-08 | 4.600E-08 |
| 23073 | 1154 | P1154M | D | 9.4e-08 | 7.7e-08 | 5.9e-08 | 7.667E-08 | 1.750E-08 | 5.900E-08 | 9.400E-08 |
| 23074 | 1154 | P1154F | D | 7.5e-08 | 8.4e-08 | 3.5e-08 | 6.467E-08 | 2.608E-08 | 3.500E-08 | 8.400E-08 |
| 23075 | 1154 | P1154P | D | 7.7e-08 | 3e-08 | 1.6e-07 | 8.900E-08 | 6.583E-08 | 3.000E-08 | 1.600E-07 |
| 23076 | 1154 | P1154S | D | 3.9e-08 | 8.5e-08 | 4.3e-08 | 5.567E-08 | 2.548E-08 | 3.900E-08 | 8.500E-08 |
| 23077 | 1154 | P1154T | D | 3.8e-08 | 1.4e-08 | 3.2e-08 | 2.800E-08 | 1.249E-08 | 1.400E-08 | 3.800E-08 |
| 23078 | 1154 | P1154W | D | 1.5e-07 | 5.4e-08 | 2e-08 | 7.467E-08 | 6.742E-08 | 2.000E-08 | 1.500E-07 |
| 23079 | 1154 | P1154Y | D | 6e-08 | 5.2e-08 | 3.4e-08 | 4.867E-08 | 1.332E-08 | 3.400E-08 | 6.000E-08 |
| 23080 | 1154 | P1154V | D | 4.8e-08 | 4.8e-08 | 4.1e-08 | 4.567E-08 | 4.041E-09 | 4.100E-08 | 4.800E-08 |
| 23081 | 1155 | T1155A | D | 1.7e-08 | 8e-08 | 8.2e-08 | 5.967E-08 | 3.696E-08 | 1.700E-08 | 8.200E-08 |
| 23082 | 1155 | T1155R | D | 5e-08 | 6.5e-08 | 7.4e-08 | 6.300E-08 | 1.212E-08 | 5.000E-08 | 7.400E-08 |
| 23083 | 1155 | T1155N | D | 1.4e-08 | 8.6e-08 | 9.9e-08 | 6.633E-08 | 4.579E-08 | 1.400E-08 | 9.900E-08 |
| 23084 | 1155 | T1155D | D | 4.3e-08 | 1.3e-07 | 5.4e-08 | 7.567E-08 | 4.737E-08 | 4.300E-08 | 1.300E-07 |
| 23085 | 1155 | T1155C | D | 4.6e-08 | 1.3e-07 | 1.2e-08 | 6.267E-08 | 6.074E-08 | 1.200E-08 | 1.300E-07 |
| 23086 | 1155 | T1155E | D | 5.6e-08 | 9e-09 | 3.3e-08 | 3.267E-08 | 2.350E-08 | 9.000E-09 | 5.600E-08 |
| 23087 | 1155 | T1155Q | D | 5.6e-08 | 2.5e-08 | 7.2e-08 | 5.100E-08 | 2.390E-08 | 2.500E-08 | 7.200E-08 |
| 23088 | 1155 | T1155G | D | 6.5e-08 | 8.6e-08 | 5e-08 | 6.700E-08 | 1.808E-08 | 5.000E-08 | 8.600E-08 |
| 23089 | 1155 | T1155H | D | 7.2e-08 | 6e-08 | 4.7e-08 | 5.967E-08 | 1.250E-08 | 4.700E-08 | 7.200E-08 |
| 23090 | 1155 | T1155I | D | 5.6e-08 | 1.1e-07 | 4.5e-08 | 7.033E-08 | 3.479E-08 | 4.500E-08 | 1.100E-07 |
| 23091 | 1155 | T1155L | D | 6.2e-08 | 2.6e-08 | 3.2e-08 | 4.000E-08 | 1.929E-08 | 2.600E-08 | 6.200E-08 |
| 23092 | 1155 | T1155K | D | 1e-07 | 2.4e-08 | 1.3e-08 | 4.567E-08 | 4.737E-08 | 1.300E-08 | 1.000E-07 |
| 23093 | 1155 | T1155M | D | 5.7e-08 | 3.3e-08 | 3.1e-08 | 4.033E-08 | 1.447E-08 | 3.100E-08 | 5.700E-08 |
| 23094 | 1155 | T1155F | D | 5.4e-08 | 5.1e-08 | 9e-09 | 3.800E-08 | 2.516E-08 | 9.000E-09 | 5.400E-08 |
| 23095 | 1155 | T1155P | D | 2.8e-08 | 2e-08 | 8.9e-08 | 4.567E-08 | 3.774E-08 | 2.000E-08 | 8.900E-08 |
| 23096 | 1155 | T1155S | D | 5.9e-08 | 2.6e-08 | 4.1e-08 | 4.200E-08 | 1.652E-08 | 2.600E-08 | 5.900E-08 |
| 23097 | 1155 | T1155T | D | 3.6e-08 | 4.9e-08 | 3e-08 | 3.833E-08 | 9.713E-09 | 3.000E-08 | 4.900E-08 |
| 23098 | 1155 | T1155W | D | 2.8e-08 | 4.4e-08 | 1e-07 | 5.733E-08 | 3.781E-08 | 2.800E-08 | 1.000E-07 |
| 23099 | 1155 | T1155Y | D | 1.7e-08 | 2.3e-08 | 7e-08 | 3.667E-08 | 2.902E-08 | 1.700E-08 | 7.000E-08 |
| 23100 | 1155 | T1155V | D | 1.8e-08 | 2.5e-08 | 7.8e-08 | 4.033E-08 | 3.281E-08 | 1.800E-08 | 7.800E-08 |
| 23101 | 1156 | F1156A | D | 3.3e-08 | 2.3e-08 | 2e-08 | 2.533E-08 | 6.807E-09 | 2.000E-08 | 3.300E-08 |
| 23102 | 1156 | F1156R | D | 3.1e-08 | 3.1e-08 | 4.9e-08 | 3.700E-08 | 1.039E-08 | 3.100E-08 | 4.900E-08 |
| 23103 | 1156 | F1156N | D | 3.8e-08 | 1.3e-08 | 2.6e-08 | 2.567E-08 | 1.250E-08 | 1.300E-08 | 3.800E-08 |
| 23104 | 1156 | F1156D | D | 2.4e-08 | 6.8e-08 | 4.8e-08 | 4.667E-08 | 2.203E-08 | 2.400E-08 | 6.800E-08 |
| 23105 | 1156 | F1156C | D | 3.2e-08 | 7e-08 | 7e-08 | 5.733E-08 | 2.194E-08 | 3.200E-08 | 7.000E-08 |
| 23106 | 1156 | F1156E | D | 1.1e-07 | 3.2e-08 | 2.4e-08 | 5.533E-08 | 4.751E-08 | 2.400E-08 | 1.100E-07 |
| 23107 | 1156 | F1156Q | D | 1e-07 | 4.5e-08 | 5.4e-08 | 6.633E-08 | 2.950E-08 | 4.500E-08 | 1.000E-07 |
| 23108 | 1156 | F1156G | D | 4.1e-08 | 4.2e-08 | 9.8e-09 | 3.093E-08 | 1.831E-08 | 9.800E-09 | 4.200E-08 |
| 23109 | 1156 | F1156H | D | 4.7e-08 | 3.3e-08 | 4.1e-08 | 4.033E-08 | 7.024E-09 | 3.300E-08 | 4.700E-08 |
| 23110 | 1156 | F1156I | D | 5.4e-08 | 5.5e-08 | 2.1e-08 | 4.333E-08 | 1.935E-08 | 2.100E-08 | 5.500E-08 |
| 23111 | 1156 | F1156L | D | 8.4e-08 | 2.5e-08 | 3.1e-08 | 4.667E-08 | 3.247E-08 | 2.500E-08 | 8.400E-08 |
| 23112 | 1156 | F1156K | D | 3.2e-08 | 3.9e-08 | 2.4e-08 | 3.167E-08 | 7.506E-09 | 2.400E-08 | 3.900E-08 |
| 23113 | 1156 | F1156M | D | 8.5e-08 | 1.7e-08 | 2.3e-08 | 4.167E-08 | 3.765E-08 | 1.700E-08 | 8.500E-08 |
| 23114 | 1156 | F1156F | D | 4.2e-08 | 5.4e-08 | 2.3e-08 | 3.967E-08 | 1.563E-08 | 2.300E-08 | 5.400E-08 |
| 23115 | 1156 | F1156P | D | 5.8e-08 | 6.2e-08 | 4.5e-08 | 5.500E-08 | 8.888E-09 | 4.500E-08 | 6.200E-08 |
| 23116 | 1156 | F1156S | D | 6.1e-08 | 1.2e-08 | 3.9e-08 | 3.733E-08 | 2.454E-08 | 1.200E-08 | 6.100E-08 |
| 23117 | 1156 | F1156T | D | 5.3e-08 | 2.1e-08 | 7.2e-09 | 2.707E-08 | 2.349E-08 | 7.200E-09 | 5.300E-08 |
| 23118 | 1156 | F1156W | D | 2.2e-08 | 6.8e-08 | 6.6e-08 | 5.200E-08 | 2.600E-08 | 2.200E-08 | 6.800E-08 |
| 23119 | 1156 | F1156Y | D | 2.1e-08 | 7.3e-08 | 4.3e-08 | 4.567E-08 | 2.610E-08 | 2.100E-08 | 7.300E-08 |

| | | | | | | | | | | |
|---|---|---|---|---|---|---|---|---|---|---|
| 23120 | 1156 | F1156V | D | 5e-08 | 4.2e-08 | 2.2e-08 | 3.800E-08 | 1.442E-08 | 2.200E-08 | 5.000E-08 |
| 23121 | 1157 | G1157A | D | 2.5e-08 | 1e-07 | 1.7e-08 | 4.733E-08 | 4.579E-08 | 1.700E-08 | 1.000E-07 |
| 23122 | 1157 | G1157R | D | 7.3e-08 | 6.9e-08 | 9.6e-08 | 7.933E-08 | 1.457E-08 | 6.900E-08 | 9.600E-08 |
| 23123 | 1157 | G1157N | D | 5.4e-08 | 4.5e-08 | 4.8e-08 | 4.900E-08 | 4.583E-09 | 4.500E-08 | 5.400E-08 |
| 23124 | 1157 | G1157D | D | 6.5e-08 | 6.3e-09 | 6e-08 | 4.377E-08 | 3.254E-08 | 6.300E-09 | 6.500E-08 |
| 23125 | 1157 | G1157C | D | 1.2e-07 | 1.7e-08 | 4.7e-08 | 6.133E-08 | 5.297E-08 | 1.700E-08 | 1.200E-07 |
| 23126 | 1157 | G1157E | D | 1.7e-07 | 5.5e-08 | 8.9e-08 | 1.047E-07 | 5.908E-08 | 5.500E-08 | 1.700E-07 |
| 23127 | 1157 | G1157Q | D | 4e-08 | 1.3e-08 | 4.2e-08 | 3.167E-08 | 1.620E-08 | 1.300E-08 | 4.200E-08 |
| 23128 | 1157 | G1157G | D | 3.8e-08 | 2.7e-08 | 3.8e-08 | 3.433E-08 | 6.351E-09 | 2.700E-08 | 3.800E-08 |
| 23129 | 1157 | G1157H | D | 3e-08 | 1.2e-08 | 4.9e-08 | 3.033E-08 | 1.850E-08 | 1.200E-08 | 4.900E-08 |
| 23130 | 1157 | G1157I | D | 4.9e-08 | 3.1e-08 | 5.9e-08 | 4.633E-08 | 1.419E-08 | 3.100E-08 | 5.900E-08 |
| 23131 | 1157 | G1157L | D | 5.4e-08 | 5.5e-08 | 3.5e-08 | 4.800E-08 | 1.127E-08 | 3.500E-08 | 5.500E-08 |
| 23132 | 1157 | G1157K | D | 2.5e-08 | 4.7e-08 | 1.9e-08 | 3.033E-08 | 1.474E-08 | 1.900E-08 | 4.700E-08 |
| 23133 | 1157 | G1157M | D | 5.4e-08 | 6.7e-08 | 5.3e-08 | 5.800E-08 | 7.810E-09 | 5.300E-08 | 6.700E-08 |
| 23134 | 1157 | G1157F | D | 7e-08 | 5.3e-08 | 7.8e-08 | 6.700E-08 | 1.277E-08 | 5.300E-08 | 7.800E-08 |
| 23135 | 1157 | G1157P | D | 8.4e-08 | 3.7e-08 | 3.1e-08 | 5.067E-08 | 2.902E-08 | 3.100E-08 | 8.400E-08 |
| 23136 | 1157 | G1157S | D | 3.3e-08 | 1.5e-08 | 6.6e-08 | 3.800E-08 | 2.587E-08 | 1.500E-08 | 6.600E-08 |
| 23137 | 1157 | G1157T | D | 4.8e-08 | 2.9e-08 | 5.6e-08 | 4.433E-08 | 1.387E-08 | 2.900E-08 | 5.600E-08 |
| 23138 | 1157 | G1157W | D | 3.7e-08 | 8.5e-08 | 8.1e-08 | 6.767E-08 | 2.663E-08 | 3.700E-08 | 8.500E-08 |
| 23139 | 1157 | G1157Y | D | 4.6e-08 | 5.2e-08 | 3.3e-08 | 4.367E-08 | 9.713E-09 | 3.300E-08 | 5.200E-08 |
| 23140 | 1157 | G1157V | D | 9.1e-08 | 1.8e-07 | 4.4e-08 | 1.050E-07 | 6.907E-08 | 4.400E-08 | 1.800E-07 |
| 23141 | 1158 | Q1158A | D | 1.5e-08 | 4.6e-08 | 3.1e-08 | 3.067E-08 | 1.550E-08 | 1.500E-08 | 4.600E-08 |
| 23142 | 1158 | Q1158R | D | 6.3e-08 | 1e-07 | 9.2e-08 | 8.500E-08 | 1.947E-08 | 6.300E-08 | 1.000E-07 |
| 23143 | 1158 | Q1158N | D | 4.7e-08 | 3.3e-08 | 7.9e-08 | 5.300E-08 | 2.358E-08 | 3.300E-08 | 7.900E-08 |
| 23144 | 1158 | Q1158D | D | 2e-08 | 5.7e-08 | 4.5e-08 | 4.067E-08 | 1.888E-08 | 2.000E-08 | 5.700E-08 |
| 23145 | 1158 | Q1158C | D | 3e-08 | 2.9e-08 | 1.1e-07 | 5.633E-08 | 4.648E-08 | 2.900E-08 | 1.100E-07 |
| 23146 | 1158 | Q1158E | D | 6.5e-08 | 5.8e-08 | 9e-08 | 7.100E-08 | 1.682E-08 | 5.800E-08 | 9.000E-08 |
| 23147 | 1158 | Q1158Q | D | 4.2e-08 | 7.2e-08 | 7.1e-08 | 6.167E-08 | 1.704E-08 | 4.200E-08 | 7.200E-08 |
| 23148 | 1158 | Q1158G | D | 2.5e-08 | 1.3e-08 | 7.7e-08 | 3.833E-08 | 3.402E-08 | 1.300E-08 | 7.700E-08 |
| 23149 | 1158 | Q1158H | D | 1.9e-08 | 4.9e-08 | 5.8e-08 | 4.200E-08 | 2.042E-08 | 1.900E-08 | 5.800E-08 |
| 23150 | 1158 | Q1158I | D | 5.2e-08 | 5.3e-08 | 3.1e-08 | 4.533E-08 | 1.242E-08 | 3.100E-08 | 5.300E-08 |
| 23151 | 1158 | Q1158L | D | 7e-08 | 2.3e-08 | 1.7e-08 | 3.667E-08 | 2.902E-08 | 1.700E-08 | 7.000E-08 |
| 23152 | 1158 | Q1158K | D | 2.2e-08 | 4.6e-08 | 5.5e-08 | 4.100E-08 | 1.706E-08 | 2.200E-08 | 5.500E-08 |
| 23153 | 1158 | Q1158M | D | 1.3e-08 | 3.8e-08 | 1.6e-08 | 2.233E-08 | 1.365E-08 | 1.300E-08 | 3.800E-08 |
| 23154 | 1158 | Q1158F | D | 9.7e-08 | 5.7e-08 | 1.5e-08 | 5.633E-08 | 4.100E-08 | 1.500E-08 | 9.700E-08 |
| 23155 | 1158 | Q1158P | D | 6.7e-08 | 1.7e-08 | 2.4e-08 | 3.600E-08 | 2.707E-08 | 1.700E-08 | 6.700E-08 |
| 23156 | 1158 | Q1158S | D | 5.9e-08 | 4.6e-08 | 1.3e-07 | 7.833E-08 | 4.521E-08 | 4.600E-08 | 1.300E-07 |
| 23157 | 1158 | Q1158T | D | 6.4e-08 | 1.8e-08 | 4.6e-08 | 4.267E-08 | 2.318E-08 | 1.800E-08 | 6.400E-08 |
| 23158 | 1158 | Q1158W | D | 3.5e-08 | 3.4e-08 | 7.2e-08 | 4.700E-08 | 2.166E-08 | 3.400E-08 | 7.200E-08 |
| 23159 | 1158 | Q1158Y | D | 3.1e-08 | 2e-08 | 2.8e-08 | 2.633E-08 | 5.686E-09 | 2.000E-08 | 3.100E-08 |
| 23160 | 1158 | Q1158V | D | 1.1e-07 | 1.4e-08 | 2.3e-08 | 4.900E-08 | 5.302E-08 | 1.400E-08 | 1.100E-07 |
| 23161 | 1159 | G1159A | D | 1.9e-08 | 7.9e-08 | 9.5e-08 | 6.433E-08 | 4.007E-08 | 1.900E-08 | 9.500E-08 |
| 23162 | 1159 | G1159R | D | 3.7e-08 | 6.3e-08 | 1.8e-08 | 3.933E-08 | 2.259E-08 | 1.800E-08 | 6.300E-08 |
| 23163 | 1159 | G1159N | D | 4.1e-08 | 5.4e-08 | 8.3e-08 | 5.933E-08 | 2.150E-08 | 4.100E-08 | 8.300E-08 |
| 23164 | 1159 | G1159D | D | 2.4e-08 | 2.8e-08 | 4.9e-08 | 3.367E-08 | 1.343E-08 | 2.400E-08 | 4.900E-08 |
| 23165 | 1159 | G1159C | D | 4.8e-08 | 9.1e-08 | 4.5e-08 | 6.133E-08 | 2.574E-08 | 4.500E-08 | 9.100E-08 |
| 23166 | 1159 | G1159E | D | 9.6e-08 | 1.2e-08 | 3.8e-08 | 4.867E-08 | 4.300E-08 | 1.200E-08 | 9.600E-08 |
| 23167 | 1159 | G1159Q | D | 9.4e-09 | 4.5e-08 | 3.8e-08 | 3.080E-08 | 1.886E-08 | 9.400E-09 | 4.500E-08 |
| 23168 | 1159 | G1159G | D | 6.3e-08 | 2.5e-08 | 4.4e-08 | 4.400E-08 | 1.900E-08 | 2.500E-08 | 6.300E-08 |
| 23169 | 1159 | G1159H | D | 4.3e-08 | 4.8e-08 | 3.1e-08 | 4.067E-08 | 8.737E-09 | 3.100E-08 | 4.800E-08 |
| 23170 | 1159 | G1159I | D | 3.9e-08 | 3.5e-08 | 1.8e-08 | 3.067E-08 | 1.115E-08 | 1.800E-08 | 3.900E-08 |
| 23171 | 1159 | G1159L | D | 5.6e-08 | 1.2e-08 | 5e-08 | 3.933E-08 | 2.386E-08 | 1.200E-08 | 5.600E-08 |
| 23172 | 1159 | G1159K | D | 4.5e-08 | 5.9e-08 | 3.7e-08 | 4.700E-08 | 1.114E-08 | 3.700E-08 | 5.900E-08 |
| 23173 | 1159 | G1159M | D | 1.3e-08 | 7e-08 | 2.2e-08 | 3.500E-08 | 3.064E-08 | 1.300E-08 | 7.000E-08 |
| 23174 | 1159 | G1159F | D | 1.2e-07 | 3.1e-08 | 1.2e-07 | 9.033E-08 | 5.138E-08 | 3.100E-08 | 1.200E-07 |
| 23175 | 1159 | G1159P | D | 7e-08 | 5.4e-08 | 1.4e-08 | 4.600E-08 | 2.884E-08 | 1.400E-08 | 7.000E-08 |
| 23176 | 1159 | G1159S | D | 1.9e-08 | 1.1e-07 | 4.8e-08 | 5.900E-08 | 4.649E-08 | 1.900E-08 | 1.100E-07 |
| 23177 | 1159 | G1159T | D | 4.5e-08 | 2.6e-08 | 7.4e-08 | 4.833E-08 | 2.417E-08 | 2.600E-08 | 7.400E-08 |
| 23178 | 1159 | G1159W | D | 3.5e-08 | 2.2e-08 | 4.8e-08 | 3.500E-08 | 1.300E-08 | 2.200E-08 | 4.800E-08 |
| 23179 | 1159 | G1159Y | D | 3.6e-08 | 4.4e-08 | 3.5e-08 | 3.833E-08 | 4.933E-09 | 3.500E-08 | 4.400E-08 |
| 23180 | 1159 | G1159V | D | 2.6e-08 | 3e-08 | 4.9e-08 | 3.500E-08 | 1.229E-08 | 2.600E-08 | 4.900E-08 |
| 23181 | 1160 | T1160A | D | 3.9e-08 | 6.1e-08 | 7.4e-08 | 5.800E-08 | 1.769E-08 | 3.900E-08 | 7.400E-08 |
| 23182 | 1160 | T1160R | D | 4.1e-08 | 1.4e-07 | 1.7e-08 | 6.600E-08 | 6.520E-08 | 1.700E-08 | 1.400E-07 |
| 23183 | 1160 | T1160N | D | 9.2e-08 | 2.8e-08 | 5.7e-08 | 5.900E-08 | 3.205E-08 | 2.800E-08 | 9.200E-08 |
| 23184 | 1160 | T1160D | D | 3.5e-08 | 3.2e-08 | 5.1e-08 | 3.933E-08 | 1.021E-08 | 3.200E-08 | 5.100E-08 |
| 23185 | 1160 | T1160C | D | 9.9e-08 | 3.6e-08 | 2.9e-08 | 5.467E-08 | 3.855E-08 | 2.900E-08 | 9.900E-08 |
| 23186 | 1160 | T1160E | D | 8.4e-08 | 1.2e-07 | 1.9e-07 | 1.313E-07 | 5.390E-08 | 8.400E-08 | 1.900E-07 |
| 23187 | 1160 | T1160Q | D | 2e-08 | 3e-08 | 1.3e-08 | 2.100E-08 | 8.544E-09 | 1.300E-08 | 3.000E-08 |
| 23188 | 1160 | T1160G | D | 4e-08 | 8.1e-08 | 6.2e-08 | 6.100E-08 | 2.052E-08 | 4.000E-08 | 8.100E-08 |
| 23189 | 1160 | T1160H | D | 4.5e-08 | 7.9e-08 | 1.4e-08 | 4.600E-08 | 3.251E-08 | 1.400E-08 | 7.900E-08 |
| 23190 | 1160 | T1160I | D | 1e-07 | 5.4e-08 | 1e-07 | 8.467E-08 | 2.656E-08 | 5.400E-08 | 1.000E-07 |
| 23191 | 1160 | T1160L | D | 1.4e-08 | 4.8e-08 | 3.1e-08 | 3.100E-08 | 1.700E-08 | 1.400E-08 | 4.800E-08 |
| 23192 | 1160 | T1160K | D | 5.4e-08 | 5.8e-08 | 4.2e-08 | 5.133E-08 | 8.327E-09 | 4.200E-08 | 5.800E-08 |
| 23193 | 1160 | T1160M | D | 5.5e-08 | 6.3e-08 | 1.8e-08 | 4.533E-08 | 2.401E-08 | 1.800E-08 | 6.300E-08 |
| 23194 | 1160 | T1160F | D | 3.6e-08 | 2.8e-08 | 2.3e-08 | 2.900E-08 | 6.557E-09 | 2.300E-08 | 3.600E-08 |
| 23195 | 1160 | T1160P | D | 3.4e-08 | 7.5e-08 | 7.2e-08 | 6.033E-08 | 2.285E-08 | 3.400E-08 | 7.500E-08 |
| 23196 | 1160 | T1160S | D | 6e-08 | 1.8e-08 | 1.3e-07 | 6.933E-08 | 5.658E-08 | 1.800E-08 | 1.300E-07 |
| 23197 | 1160 | T1160T | D | 7.7e-08 | 4.6e-08 | 9.7e-08 | 7.333E-08 | 2.570E-08 | 4.600E-08 | 9.700E-08 |
| 23198 | 1160 | T1160W | D | 3.3e-08 | 2.5e-08 | 1e-08 | 2.267E-08 | 1.168E-08 | 1.000E-08 | 3.300E-08 |
| 23199 | 1160 | T1160Y | D | 8.7e-08 | 1.2e-08 | 2.3e-08 | 4.067E-08 | 4.050E-08 | 1.200E-08 | 8.700E-08 |
| 23200 | 1160 | T1160V | D | 4.7e-08 | 3.9e-08 | 3.5e-08 | 4.033E-08 | 6.110E-09 | 3.500E-08 | 4.700E-08 |
| 23201 | 1161 | K1161A | D | 9.3e-08 | 2.9e-08 | 2.4e-08 | 4.867E-08 | 3.848E-08 | 2.400E-08 | 9.300E-08 |
| 23202 | 1161 | K1161R | D | 5.6e-08 | 5.3e-08 | 4.6e-08 | 5.167E-08 | 5.132E-09 | 4.600E-08 | 5.600E-08 |
| 23203 | 1161 | K1161N | D | 4.3e-08 | 6.8e-08 | 4.4e-08 | 5.167E-08 | 1.415E-08 | 4.300E-08 | 6.800E-08 |
| 23204 | 1161 | K1161D | D | 5.2e-08 | 5.3e-08 | 6.6e-08 | 5.700E-08 | 7.810E-09 | 5.200E-08 | 6.600E-08 |

| | | | | | | | | | | |
|---|---|---|---|---|---|---|---|---|---|---|
| 23205 | 1161 | K1161C | D | 5.2e-08 | 7.6e-08 | 1.8e-08 | 4.867E-08 | 2.914E-08 | 1.800E-08 | 7.600E-08 |
| 23206 | 1161 | K1161E | D | 2.6e-08 | 2.1e-08 | 2.7e-08 | 2.467E-08 | 3.215E-09 | 2.100E-08 | 2.700E-08 |
| 23207 | 1161 | K1161Q | D | 1.6e-07 | 7e-08 | 5.7e-08 | 9.567E-08 | 5.609E-08 | 5.700E-08 | 1.600E-07 |
| 23208 | 1161 | K1161G | D | 6.5e-08 | 4.8e-08 | 6.6e-08 | 5.967E-08 | 1.012E-08 | 4.800E-08 | 6.600E-08 |
| 23209 | 1161 | K1161H | D | 8.6e-08 | 7.4e-08 | 3.2e-08 | 6.400E-08 | 2.835E-08 | 3.200E-08 | 8.600E-08 |
| 23210 | 1161 | K1161I | D | 1.1e-08 | 3.6e-07 | 1e-07 | 1.570E-07 | 1.813E-07 | 1.100E-08 | 3.600E-07 |
| 23211 | 1161 | K1161L | D | 8.4e-08 | 4.2e-08 | 2.6e-08 | 5.067E-08 | 2.996E-08 | 2.600E-08 | 8.400E-08 |
| 23212 | 1161 | K1161K | D | 6.5e-08 | 7.9e-08 | 5.4e-08 | 6.600E-08 | 1.253E-08 | 5.400E-08 | 7.900E-08 |
| 23213 | 1161 | K1161M | D | 3.7e-08 | 6.1e-08 | 2.4e-08 | 4.067E-08 | 1.877E-08 | 2.400E-08 | 6.100E-08 |
| 23214 | 1161 | K1161F | D | 2.2e-08 | 2.3e-08 | 1.3e-07 | 5.833E-08 | 6.207E-08 | 2.200E-08 | 1.300E-07 |
| 23215 | 1161 | K1161P | D | 3.3e-08 | 1.3e-07 | 4.5e-08 | 6.933E-08 | 5.288E-08 | 3.300E-08 | 1.300E-07 |
| 23216 | 1161 | K1161S | D | 4.1e-08 | 3.4e-08 | 6.4e-08 | 4.633E-08 | 1.570E-08 | 3.400E-08 | 6.400E-08 |
| 23217 | 1161 | K1161T | D | 8e-08 | 4e-08 | 1.1e-07 | 7.667E-08 | 3.512E-08 | 4.000E-08 | 1.100E-07 |
| 23218 | 1161 | K1161W | D | 8.8e-08 | 7.8e-08 | 5.7e-08 | 7.433E-08 | 1.582E-08 | 5.700E-08 | 8.800E-08 |
| 23219 | 1161 | K1161Y | D | 1.7e-08 | 1.9e-08 | 3e-08 | 2.200E-08 | 7.000E-09 | 1.700E-08 | 3.000E-08 |
| 23220 | 1161 | K1161V | D | 1.8e-08 | 5.7e-08 | 1.8e-08 | 3.100E-08 | 2.252E-08 | 1.800E-08 | 5.700E-08 |
| 23221 | 1162 | V1162A | D | 9.2e-08 | 7.2e-08 | 1.7e-08 | 6.033E-08 | 3.884E-08 | 1.700E-08 | 9.200E-08 |
| 23222 | 1162 | V1162R | D | 4.5e-08 | 9.7e-08 | 2.2e-08 | 5.467E-08 | 3.842E-08 | 2.200E-08 | 9.700E-08 |
| 23223 | 1162 | V1162N | D | 8.4e-08 | 3e-08 | 2e-08 | 4.467E-08 | 3.443E-08 | 2.000E-08 | 8.400E-08 |
| 23224 | 1162 | V1162D | D | 8.3e-08 | 3.7e-08 | 3.5e-08 | 5.167E-08 | 2.715E-08 | 3.500E-08 | 8.300E-08 |
| 23225 | 1162 | V1162C | D | 4.5e-08 | 9.8e-08 | 2.6e-08 | 5.633E-08 | 3.731E-08 | 2.600E-08 | 9.800E-08 |
| 23226 | 1162 | V1162E | D | 6.7e-08 | 1.4e-07 | 3.5e-08 | 8.067E-08 | 5.382E-08 | 3.500E-08 | 1.400E-07 |
| 23227 | 1162 | V1162Q | D | 5.2e-08 | 3.6e-08 | 6.6e-08 | 5.133E-08 | 1.501E-08 | 3.600E-08 | 6.600E-08 |
| 23228 | 1162 | V1162G | D | 7.8e-08 | 7.9e-08 | 2.9e-08 | 6.200E-08 | 2.858E-08 | 2.900E-08 | 7.900E-08 |
| 23229 | 1162 | V1162H | D | 3.1e-08 | 1.1e-07 | 4.7e-08 | 6.267E-08 | 4.177E-08 | 3.100E-08 | 1.100E-07 |
| 23230 | 1162 | V1162I | D | 5.1e-08 | 7.5e-08 | 2e-08 | 4.867E-08 | 2.757E-08 | 2.000E-08 | 7.500E-08 |
| 23231 | 1162 | V1162L | D | 1e-07 | 1.7e-08 | 4.7e-08 | 5.467E-08 | 4.203E-08 | 1.700E-08 | 1.000E-07 |
| 23232 | 1162 | V1162K | D | 6.6e-08 | 3.5e-08 | 8.8e-08 | 6.300E-08 | 2.663E-08 | 3.500E-08 | 8.800E-08 |
| 23233 | 1162 | V1162M | D | 7.5e-08 | 1.5e-08 | 3e-08 | 4.000E-08 | 3.122E-08 | 1.500E-08 | 7.500E-08 |
| 23234 | 1162 | V1162F | D | 2.3e-08 | 1e-07 | 4.8e-08 | 5.700E-08 | 3.928E-08 | 2.300E-08 | 1.000E-07 |
| 23235 | 1162 | V1162P | D | 1.7e-08 | 4e-08 | 3.3e-08 | 3.000E-08 | 1.179E-08 | 1.700E-08 | 4.000E-08 |
| 23236 | 1162 | V1162S | D | 2.8e-08 | 7.6e-08 | 4.8e-08 | 5.067E-08 | 2.411E-08 | 2.800E-08 | 7.600E-08 |
| 23237 | 1162 | V1162T | D | 7.6e-08 | 4.8e-08 | 2.5e-08 | 4.967E-08 | 2.554E-08 | 2.500E-08 | 7.600E-08 |
| 23238 | 1162 | V1162W | D | 4.5e-08 | 2.8e-08 | 3.5e-08 | 3.600E-08 | 8.544E-09 | 2.800E-08 | 4.500E-08 |
| 23239 | 1162 | V1162Y | D | 6.1e-08 | 1.3e-08 | 3e-08 | 3.467E-08 | 2.434E-08 | 1.300E-08 | 6.100E-08 |
| 23240 | 1162 | V1162V | D | 5.8e-08 | 3.9e-08 | 4.4e-08 | 4.700E-08 | 9.849E-09 | 3.900E-08 | 5.800E-08 |
| 23241 | 1163 | E1163A | D | 1.5e-08 | 5.9e-08 | 4.2e-08 | 3.867E-08 | 2.219E-08 | 1.500E-08 | 5.900E-08 |
| 23242 | 1163 | E1163R | D | 5.8e-08 | 4.8e-08 | 4.2e-08 | 4.933E-08 | 8.083E-09 | 4.200E-08 | 5.800E-08 |
| 23243 | 1163 | E1163N | D | 3.6e-08 | 9.6e-08 | 1.3e-07 | 8.733E-08 | 4.760E-08 | 3.600E-08 | 1.300E-07 |
| 23244 | 1163 | E1163D | D | 3.4e-08 | 2.9e-08 | 8.6e-08 | 4.967E-08 | 3.156E-08 | 2.900E-08 | 8.600E-08 |
| 23245 | 1163 | E1163C | D | 2.4e-08 | 6.2e-08 | 2.6e-08 | 3.733E-08 | 2.139E-08 | 2.400E-08 | 6.200E-08 |
| 23246 | 1163 | E1163E | D | 4.1e-08 | 2.5e-08 | 6.1e-08 | 4.233E-08 | 1.804E-08 | 2.500E-08 | 6.100E-08 |
| 23247 | 1163 | E1163Q | D | 4.8e-08 | 7.4e-08 | 5.6e-08 | 5.933E-08 | 1.332E-08 | 4.800E-08 | 7.400E-08 |
| 23248 | 1163 | E1163G | D | 1.2e-07 | 4.1e-08 | 1.7e-08 | 5.933E-08 | 5.389E-08 | 1.700E-08 | 1.200E-07 |
| 23249 | 1163 | E1163H | D | 5.9e-08 | 3e-08 | 2.8e-08 | 3.900E-08 | 1.735E-08 | 2.800E-08 | 5.900E-08 |
| 23250 | 1163 | E1163I | D | 2.6e-08 | 1.4e-08 | 3.7e-08 | 2.567E-08 | 1.150E-08 | 1.400E-08 | 3.700E-08 |
| 23251 | 1163 | E1163L | D | 4.3e-08 | 1.9e-08 | 5.6e-08 | 3.933E-08 | 1.877E-08 | 1.900E-08 | 5.600E-08 |
| 23252 | 1163 | E1163K | D | 1.6e-07 | 1.1e-07 | 1.3e-07 | 1.333E-07 | 2.517E-08 | 1.100E-07 | 1.600E-07 |
| 23253 | 1163 | E1163M | D | 6.3e-08 | 4.6e-08 | 5.4e-08 | 5.433E-08 | 8.505E-09 | 4.600E-08 | 6.300E-08 |
| 23254 | 1163 | E1163F | D | 1.8e-08 | 2.5e-08 | 7.4e-08 | 3.900E-08 | 3.051E-08 | 1.800E-08 | 7.400E-08 |
| 23255 | 1163 | E1163P | D | 3.6e-08 | 4.7e-08 | 4.3e-08 | 4.200E-08 | 5.568E-09 | 3.600E-08 | 4.700E-08 |
| 23256 | 1163 | E1163S | D | 3e-08 | 4.3e-08 | 1.7e-08 | 3.000E-08 | 1.300E-08 | 1.700E-08 | 4.300E-08 |
| 23257 | 1163 | E1163T | D | 2.2e-08 | 2.8e-08 | 3.2e-08 | 2.733E-08 | 5.033E-09 | 2.200E-08 | 3.200E-08 |
| 23258 | 1163 | E1163W | D | 7e-08 | 5.6e-08 | 3.4e-08 | 5.333E-08 | 1.815E-08 | 3.400E-08 | 7.000E-08 |
| 23259 | 1163 | E1163Y | D | 3.2e-08 | 3e-08 | 3.3e-08 | 3.167E-08 | 1.528E-09 | 3.000E-08 | 3.300E-08 |
| 23260 | 1163 | E1163V | D | 1.1e-08 | 4.4e-08 | 5.9e-08 | 3.800E-08 | 2.456E-08 | 1.100E-08 | 5.900E-08 |
| 23261 | 1164 | I1164A | D | 4.4e-08 | 5.3e-08 | 4.9e-08 | 4.867E-08 | 4.509E-09 | 4.400E-08 | 5.300E-08 |
| 23262 | 1164 | I1164R | D | 2.6e-08 | 4.3e-08 | 2e-08 | 2.967E-08 | 1.193E-08 | 2.000E-08 | 4.300E-08 |
| 23263 | 1164 | I1164N | D | 6.5e-08 | 9.7e-08 | 4.5e-08 | 6.900E-08 | 2.623E-08 | 4.500E-08 | 9.700E-08 |
| 23264 | 1164 | I1164D | D | 3.4e-08 | 4.7e-08 | 7.2e-08 | 5.100E-08 | 1.931E-08 | 3.400E-08 | 7.200E-08 |
| 23265 | 1164 | I1164C | D | 4.3e-08 | 4.8e-08 | 3e-08 | 4.033E-08 | 9.292E-09 | 3.000E-08 | 4.800E-08 |
| 23266 | 1164 | I1164E | D | 3.2e-08 | 3.8e-08 | 7.9e-08 | 4.967E-08 | 2.558E-08 | 3.200E-08 | 7.900E-08 |
| 23267 | 1164 | I1164Q | D | 2e-08 | 2.2e-08 | 2.4e-08 | 2.200E-08 | 2.000E-09 | 2.000E-08 | 2.400E-08 |
| 23268 | 1164 | I1164G | D | 1.9e-08 | 1.3e-07 | 3.7e-08 | 6.200E-08 | 5.957E-08 | 1.900E-08 | 1.300E-07 |
| 23269 | 1164 | I1164H | D | 4.2e-08 | 1.5e-08 | 5.3e-08 | 3.667E-08 | 1.955E-08 | 1.500E-08 | 5.300E-08 |
| 23270 | 1164 | I1164I | D | 3.4e-08 | 5.2e-08 | 3.7e-08 | 4.100E-08 | 9.644E-09 | 3.400E-08 | 5.200E-08 |
| 23271 | 1164 | I1164L | D | 6.7e-08 | 2.7e-08 | 8.6e-08 | 6.000E-08 | 3.012E-08 | 2.700E-08 | 8.600E-08 |
| 23272 | 1164 | I1164K | D | 7.1e-08 | 7.2e-08 | 2.9e-08 | 5.733E-08 | 2.454E-08 | 2.900E-08 | 7.200E-08 |
| 23273 | 1164 | I1164M | D | 2.9e-08 | 4.7e-08 | 2.5e-08 | 3.367E-08 | 1.172E-08 | 2.500E-08 | 4.700E-08 |
| 23274 | 1164 | I1164F | D | 1.4e-08 | 4.7e-08 | 1.9e-08 | 2.667E-08 | 1.779E-08 | 1.400E-08 | 4.700E-08 |
| 23275 | 1164 | I1164P | D | 5.5e-08 | 1.2e-08 | 2.6e-08 | 3.100E-08 | 2.193E-08 | 1.200E-08 | 5.500E-08 |
| 23276 | 1164 | I1164S | D | 5.5e-08 | 2e-08 | 4.6e-08 | 4.033E-08 | 1.818E-08 | 2.000E-08 | 5.500E-08 |
| 23277 | 1164 | I1164T | D | 3.9e-08 | 7.5e-08 | 3.3e-08 | 4.900E-08 | 2.272E-08 | 3.300E-08 | 7.500E-08 |
| 23278 | 1164 | I1164W | D | 3.8e-08 | 2.8e-08 | 1.2e-08 | 2.600E-08 | 1.311E-08 | 1.200E-08 | 3.800E-08 |
| 23279 | 1164 | I1164Y | D | 3.1e-08 | 4.6e-08 | 3.7e-08 | 3.800E-08 | 7.550E-09 | 3.100E-08 | 4.600E-08 |
| 23280 | 1164 | I1164V | D | 1.2e-07 | 3.6e-08 | 1.8e-08 | 5.800E-08 | 5.444E-08 | 1.800E-08 | 1.200E-07 |
| 23281 | 1165 | K1165A | D | 2e-08 | 4.9e-08 | 5.8e-08 | 4.233E-08 | 1.986E-08 | 2.000E-08 | 5.800E-08 |
| 23282 | 1165 | K1165R | D | 9.4e-08 | 3.7e-08 | 1.3e-07 | 8.700E-08 | 4.689E-08 | 3.700E-08 | 1.300E-07 |
| 23283 | 1165 | K1165N | D | 2.4e-08 | 3.8e-08 | 9.6e-08 | 5.267E-08 | 3.818E-08 | 2.400E-08 | 9.600E-08 |
| 23284 | 1165 | K1165D | D | 1.4e-08 | 2e-07 | 4.3e-08 | 8.567E-08 | 1.001E-07 | 1.400E-08 | 2.000E-07 |
| 23285 | 1165 | K1165C | D | 5.8e-08 | 3.9e-08 | 3.4e-08 | 4.367E-08 | 1.266E-08 | 3.400E-08 | 5.800E-08 |
| 23286 | 1165 | K1165E | D | 1.8e-07 | 6.4e-08 | 2.1e-08 | 8.833E-08 | 8.225E-08 | 2.100E-08 | 1.800E-07 |
| 23287 | 1165 | K1165Q | D | 6.6e-08 | 5e-08 | 3.6e-08 | 5.067E-08 | 1.501E-08 | 3.600E-08 | 6.600E-08 |
| 23288 | 1165 | K1165G | D | 7.2e-08 | 9e-08 | 4.5e-08 | 6.900E-08 | 2.265E-08 | 4.500E-08 | 9.000E-08 |
| 23289 | 1165 | K1165H | D | 4.8e-08 | 5.4e-08 | 8.5e-08 | 6.233E-08 | 1.986E-08 | 4.800E-08 | 8.500E-08 |

| | | | | | | | | | | |
|---|---|---|---|---|---|---|---|---|---|---|
| 23290 | 1165 | K1165I | D | 1.5e-08 | 7.7e-09 | 7.5e-08 | 3.257E-08 | 3.693E-08 | 7.700E-09 | 7.500E-08 |
| 23291 | 1165 | K1165L | D | 5.1e-08 | 3.4e-08 | 8.9e-08 | 5.800E-08 | 2.816E-08 | 3.400E-08 | 8.900E-08 |
| 23292 | 1165 | K1165K | D | 3.1e-08 | 3.2e-08 | 3.5e-08 | 3.267E-08 | 2.082E-09 | 3.100E-08 | 3.500E-08 |
| 23293 | 1165 | K1165M | D | 3.9e-08 | 1.1e-07 | 3.3e-08 | 6.067E-08 | 4.283E-08 | 3.300E-08 | 1.100E-07 |
| 23294 | 1165 | K1165F | D | 4.1e-08 | 5.7e-08 | 9.9e-08 | 6.567E-08 | 2.996E-08 | 4.100E-08 | 9.900E-08 |
| 23295 | 1165 | K1165P | D | 1.7e-08 | 3.4e-08 | 6.7e-08 | 3.933E-08 | 2.542E-08 | 1.700E-08 | 6.700E-08 |
| 23296 | 1165 | K1165S | D | 3e-08 | 4.2e-08 | 7e-08 | 4.733E-08 | 2.053E-08 | 3.000E-08 | 7.000E-08 |
| 23297 | 1165 | K1165T | D | 2e-08 | 5.4e-08 | 1.1e-07 | 6.133E-08 | 4.545E-08 | 2.000E-08 | 1.100E-07 |
| 23298 | 1165 | K1165W | D | 2.6e-08 | 6.2e-08 | 2.3e-08 | 3.700E-08 | 2.170E-08 | 2.300E-08 | 6.200E-08 |
| 23299 | 1165 | K1165Y | D | 8.1e-08 | 4.5e-08 | 5e-08 | 5.867E-08 | 1.950E-08 | 4.500E-08 | 8.100E-08 |
| 23300 | 1165 | K1165V | D | 1.9e-08 | 3.5e-08 | 5.1e-08 | 3.500E-08 | 1.600E-08 | 1.900E-08 | 5.100E-08 |
| 23301 | 1166 | R1166A | D | 6.3e-08 | 3.8e-08 | 1.5e-08 | 3.867E-08 | 2.401E-08 | 1.500E-08 | 6.300E-08 |
| 23302 | 1166 | R1166R | D | 9.2e-08 | 1.4e-07 | 4.9e-08 | 9.367E-08 | 4.552E-08 | 4.900E-08 | 1.400E-07 |
| 23303 | 1166 | R1166N | D | 4.9e-08 | 6.1e-08 | 1.3e-07 | 8.000E-08 | 4.371E-08 | 4.900E-08 | 1.300E-07 |
| 23304 | 1166 | R1166D | D | 3.4e-08 | 6e-08 | 1.5e-08 | 3.633E-08 | 2.259E-08 | 1.500E-08 | 6.000E-08 |
| 23305 | 1166 | R1166C | D | 1e-07 | 3.4e-08 | 8.4e-08 | 7.267E-08 | 3.443E-08 | 3.400E-08 | 1.000E-07 |
| 23306 | 1166 | R1166E | D | 5.9e-08 | 4.4e-08 | 2.6e-08 | 4.300E-08 | 1.652E-08 | 2.600E-08 | 5.900E-08 |
| 23307 | 1166 | R1166Q | D | 5.5e-08 | 6.1e-08 | 3.7e-08 | 5.100E-08 | 1.249E-08 | 3.700E-08 | 6.100E-08 |
| 23308 | 1166 | R1166G | D | 7.6e-08 | 3.2e-08 | 2.8e-08 | 4.533E-08 | 2.663E-08 | 2.800E-08 | 7.600E-08 |
| 23309 | 1166 | R1166H | D | 8.1e-08 | 5e-08 | 6e-08 | 6.367E-08 | 1.582E-08 | 5.000E-08 | 8.100E-08 |
| 23310 | 1166 | R1166I | D | 2.7e-08 | 1.9e-08 | 5.6e-08 | 3.400E-08 | 1.947E-08 | 1.900E-08 | 5.600E-08 |
| 23311 | 1166 | R1166L | D | 9.8e-08 | 2e-08 | 1.5e-08 | 4.433E-08 | 4.654E-08 | 1.500E-08 | 9.800E-08 |
| 23312 | 1166 | R1166K | D | 3e-08 | 7.5e-08 | 5.5e-08 | 5.333E-08 | 2.255E-08 | 3.000E-08 | 7.500E-08 |
| 23313 | 1166 | R1166M | D | 3.9e-08 | 3.7e-08 | 4.7e-08 | 4.100E-08 | 5.292E-09 | 3.700E-08 | 4.700E-08 |
| 23314 | 1166 | R1166F | D | 1.8e-08 | 4.1e-08 | 8.4e-08 | 4.767E-08 | 3.350E-08 | 1.800E-08 | 8.400E-08 |
| 23315 | 1166 | R1166P | D | 4.7e-08 | 4.2e-08 | 7.5e-08 | 5.467E-08 | 1.779E-08 | 4.200E-08 | 7.500E-08 |
| 23316 | 1166 | R1166S | D | 3.7e-08 | 4.8e-08 | 3.5e-08 | 4.000E-08 | 7.000E-09 | 3.500E-08 | 4.800E-08 |
| 23317 | 1166 | R1166T | D | 3.5e-08 | 1e-07 | 2.5e-08 | 5.333E-08 | 4.072E-08 | 2.500E-08 | 1.000E-07 |
| 23318 | 1166 | R1166W | D | 5.8e-08 | 7.2e-08 | 4.3e-08 | 5.767E-08 | 1.450E-08 | 4.300E-08 | 7.200E-08 |
| 23319 | 1166 | R1166Y | D | 4.4e-08 | 2.1e-08 | 4.1e-08 | 3.533E-08 | 1.250E-08 | 2.100E-08 | 4.400E-08 |
| 23320 | 1166 | R1166V | D | 7.8e-08 | 9.4e-09 | 6.5e-08 | 5.080E-08 | 3.644E-08 | 9.400E-09 | 7.800E-08 |
| 23321 | 1167 | T1167A | D | 6e-08 | 2.3e-08 | 3.7e-08 | 4.000E-08 | 1.868E-08 | 2.300E-08 | 6.000E-08 |
| 23322 | 1167 | T1167R | D | 4.4e-08 | 2.8e-08 | 8.8e-08 | 5.333E-08 | 3.107E-08 | 2.800E-08 | 8.800E-08 |
| 23323 | 1167 | T1167N | D | 4.3e-08 | 5.3e-08 | 3.8e-08 | 4.467E-08 | 7.638E-09 | 3.800E-08 | 5.300E-08 |
| 23324 | 1167 | T1167D | D | 8.1e-08 | 4.5e-08 | 4.4e-08 | 5.667E-08 | 2.108E-08 | 4.400E-08 | 8.100E-08 |
| 23325 | 1167 | T1167C | D | 6.7e-08 | 5.1e-08 | 4.2e-08 | 5.333E-08 | 1.266E-08 | 4.200E-08 | 6.700E-08 |
| 23326 | 1167 | T1167E | D | 3.8e-08 | 5.3e-08 | 6.3e-09 | 3.243E-08 | 2.384E-08 | 6.300E-09 | 5.300E-08 |
| 23327 | 1167 | T1167Q | D | 1.3e-07 | 7.1e-08 | 3.9e-08 | 8.000E-08 | 4.616E-08 | 3.900E-08 | 1.300E-07 |
| 23328 | 1167 | T1167G | D | 4.6e-08 | 7.1e-08 | 2.7e-08 | 4.800E-08 | 2.207E-08 | 2.700E-08 | 7.100E-08 |
| 23329 | 1167 | T1167H | D | 4.7e-08 | 3.3e-08 | 4.3e-08 | 4.100E-08 | 7.211E-09 | 3.300E-08 | 4.700E-08 |
| 23330 | 1167 | T1167I | D | 4.5e-08 | 3.3e-08 | 1.2e-07 | 6.600E-08 | 4.715E-08 | 3.300E-08 | 1.200E-07 |
| 23331 | 1167 | T1167L | D | 1.3e-08 | 5.6e-08 | 2.9e-08 | 3.267E-08 | 2.173E-08 | 1.300E-08 | 5.600E-08 |
| 23332 | 1167 | T1167K | D | 1.4e-07 | 7.3e-08 | 8.9e-08 | 1.007E-07 | 3.499E-08 | 7.300E-08 | 1.400E-07 |
| 23333 | 1167 | T1167M | D | 3.4e-08 | 2.7e-08 | 1.2e-07 | 6.033E-08 | 5.179E-08 | 2.700E-08 | 1.200E-07 |
| 23334 | 1167 | T1167F | D | 5.1e-08 | 5.3e-08 | 4.2e-08 | 4.867E-08 | 5.859E-09 | 4.200E-08 | 5.300E-08 |
| 23335 | 1167 | T1167P | D | 3.6e-08 | 3.5e-08 | 9.8e-08 | 5.633E-08 | 3.609E-08 | 3.500E-08 | 9.800E-08 |
| 23336 | 1167 | T1167S | D | 8.8e-08 | 5.6e-08 | 7.1e-08 | 7.167E-08 | 1.601E-08 | 5.600E-08 | 8.800E-08 |
| 23337 | 1167 | T1167T | D | 6.2e-08 | 4.8e-08 | 4.5e-08 | 5.167E-08 | 9.074E-09 | 4.500E-08 | 6.200E-08 |
| 23338 | 1167 | T1167W | D | 2.1e-08 | 1.4e-08 | 4.7e-08 | 2.733E-08 | 1.739E-08 | 1.400E-08 | 4.700E-08 |
| 23339 | 1167 | T1167Y | D | 3.1e-08 | 6.8e-08 | 4.4e-08 | 4.767E-08 | 1.877E-08 | 3.100E-08 | 6.800E-08 |
| 23340 | 1167 | T1167V | D | 9.1e-08 | 2.4e-08 | 1.1e-07 | 7.500E-08 | 4.518E-08 | 2.400E-08 | 1.100E-07 |
| 23341 | 1168 | V1168A | D | 1.9e-08 | 3.9e-08 | 3.8e-08 | 3.200E-08 | 1.127E-08 | 1.900E-08 | 3.900E-08 |
| 23342 | 1168 | V1168R | D | 9.4e-08 | 7.7e-08 | 5.5e-08 | 7.533E-08 | 1.955E-08 | 5.500E-08 | 9.400E-08 |
| 23343 | 1168 | V1168N | D | 3.5e-08 | 5.9e-08 | 1.7e-07 | 8.800E-08 | 7.202E-08 | 3.500E-08 | 1.700E-07 |
| 23344 | 1168 | V1168D | D | 3.9e-08 | 4.8e-08 | 7e-08 | 5.233E-08 | 1.595E-08 | 3.900E-08 | 7.000E-08 |
| 23345 | 1168 | V1168C | D | 4.7e-08 | 2.5e-08 | 1.1e-07 | 6.067E-08 | 4.412E-08 | 2.500E-08 | 1.100E-07 |
| 23346 | 1168 | V1168E | D | 5.4e-08 | 3.6e-08 | 1.7e-08 | 3.567E-08 | 1.850E-08 | 1.700E-08 | 5.400E-08 |
| 23347 | 1168 | V1168Q | D | 1.2e-07 | 1.9e-08 | 4.7e-08 | 6.200E-08 | 5.214E-08 | 1.900E-08 | 1.200E-07 |
| 23348 | 1168 | V1168G | D | 3.2e-08 | 1.1e-07 | 1.8e-08 | 5.333E-08 | 4.957E-08 | 1.800E-08 | 1.100E-07 |
| 23349 | 1168 | V1168H | D | 5.8e-08 | 2.3e-08 | 1.1e-07 | 6.367E-08 | 4.378E-08 | 2.300E-08 | 1.100E-07 |
| 23350 | 1168 | V1168I | D | 6.1e-08 | 2.2e-08 | 7.2e-08 | 5.167E-08 | 2.627E-08 | 2.200E-08 | 7.200E-08 |
| 23351 | 1168 | V1168L | D | 5.6e-08 | 4e-08 | 2.9e-08 | 4.167E-08 | 1.358E-08 | 2.900E-08 | 5.600E-08 |
| 23352 | 1168 | V1168K | D | 1.7e-08 | 7.9e-08 | 9.2e-08 | 6.267E-08 | 4.008E-08 | 1.700E-08 | 9.200E-08 |
| 23353 | 1168 | V1168M | D | 4.1e-08 | 5.2e-08 | 1.1e-07 | 6.767E-08 | 3.707E-08 | 4.100E-08 | 1.100E-07 |
| 23354 | 1168 | V1168F | D | 9.1e-08 | 6.1e-08 | 2.5e-08 | 5.900E-08 | 3.305E-08 | 2.500E-08 | 9.100E-08 |
| 23355 | 1168 | V1168P | D | 5.2e-08 | 3.2e-08 | 8e-08 | 5.467E-08 | 2.411E-08 | 3.200E-08 | 8.000E-08 |
| 23356 | 1168 | V1168S | D | 2.4e-08 | 8.5e-08 | 5.6e-08 | 5.500E-08 | 3.051E-08 | 2.400E-08 | 8.500E-08 |
| 23357 | 1168 | V1168T | D | 1.6e-08 | 5.6e-08 | 4.9e-08 | 4.033E-08 | 2.136E-08 | 1.600E-08 | 5.600E-08 |
| 23358 | 1168 | V1168W | D | 5.4e-08 | 3.5e-08 | 4.4e-08 | 4.433E-08 | 9.504E-09 | 3.500E-08 | 5.400E-08 |
| 23359 | 1168 | V1168Y | D | 1e-07 | 5.7e-08 | 5e-08 | 6.900E-08 | 2.707E-08 | 5.000E-08 | 1.000E-07 |
| 23360 | 1168 | V1168V | D | 4.6e-08 | 3.5e-08 | 5.2e-08 | 4.433E-08 | 8.622E-09 | 3.500E-08 | 5.200E-08 |
| 23361 | 1169 | A1169A | D | 5.9e-08 | 3.5e-08 | 2.6e-08 | 4.000E-08 | 1.706E-08 | 2.600E-08 | 5.900E-08 |
| 23362 | 1169 | A1169R | D | 7e-08 | 3.7e-08 | 1.2e-07 | 7.567E-08 | 4.179E-08 | 3.700E-08 | 1.200E-07 |
| 23363 | 1169 | A1169N | D | 8.1e-09 | 6.1e-08 | 4.7e-08 | 3.870E-08 | 2.741E-08 | 8.100E-09 | 6.100E-08 |
| 23364 | 1169 | A1169D | D | 3.3e-08 | 5.5e-08 | 1.5e-08 | 3.433E-08 | 2.003E-08 | 1.500E-08 | 5.500E-08 |
| 23365 | 1169 | A1169C | D | 4.9e-08 | 2.9e-08 | 1.1e-07 | 6.267E-08 | 4.219E-08 | 2.900E-08 | 1.100E-07 |
| 23366 | 1169 | A1169E | D | 5.2e-08 | 1.6e-08 | 2.2e-08 | 3.000E-08 | 1.929E-08 | 1.600E-08 | 5.200E-08 |
| 23367 | 1169 | A1169Q | D | 6.8e-08 | 7.1e-08 | 6.1e-08 | 6.667E-08 | 5.132E-09 | 6.100E-08 | 7.100E-08 |
| 23368 | 1169 | A1169G | D | 2.9e-08 | 1.6e-08 | 8.2e-08 | 4.233E-08 | 3.496E-08 | 1.600E-08 | 8.200E-08 |
| 23369 | 1169 | A1169H | D | 6.6e-08 | 4.4e-08 | 4.9e-08 | 5.300E-08 | 1.153E-08 | 4.400E-08 | 6.600E-08 |
| 23370 | 1169 | A1169I | D | 3.3e-08 | 1.1e-08 | 2.7e-08 | 2.367E-08 | 1.137E-08 | 1.100E-08 | 3.300E-08 |
| 23371 | 1169 | A1169L | D | 1.5e-07 | 3e-08 | 3.1e-08 | 7.033E-08 | 6.900E-08 | 3.000E-08 | 1.500E-07 |
| 23372 | 1169 | A1169K | D | 2.1e-08 | 2.6e-08 | 4.7e-08 | 3.133E-08 | 1.380E-08 | 2.100E-08 | 4.700E-08 |
| 23373 | 1169 | A1169M | D | 2.6e-08 | 1.6e-08 | 5e-08 | 3.067E-08 | 1.747E-08 | 1.600E-08 | 5.000E-08 |
| 23374 | 1169 | A1169F | D | 2.3e-08 | 1.1e-08 | 2.3e-08 | 1.900E-08 | 6.928E-09 | 1.100E-08 | 2.300E-08 |

| | | | | | | | | | | |
|---|---|---|---|---|---|---|---|---|---|---|
| 23375 | 1169 | A1169P | D | 1.1e-07 | 4.8e-08 | 2.4e-08 | 6.067E-08 | 4.438E-08 | 2.400E-08 | 1.100E-07 |
| 23376 | 1169 | A1169S | D | 4.9e-08 | 8.9e-08 | 4.4e-08 | 6.067E-08 | 2.466E-08 | 4.400E-08 | 8.900E-08 |
| 23377 | 1169 | A1169T | D | 6.9e-08 | 1.4e-08 | 4.4e-08 | 4.233E-08 | 2.754E-08 | 1.400E-08 | 6.900E-08 |
| 23378 | 1169 | A1169W | D | 1.8e-08 | 9.6e-08 | 4.4e-08 | 5.267E-08 | 3.972E-08 | 1.800E-08 | 9.600E-08 |
| 23379 | 1169 | A1169Y | D | 2.1e-08 | 2.6e-08 | 3.9e-08 | 2.867E-08 | 9.292E-09 | 2.100E-08 | 3.900E-08 |
| 23380 | 1169 | A1169V | D | 3.2e-08 | 8.8e-08 | 3.3e-08 | 5.100E-08 | 3.205E-08 | 3.200E-08 | 8.800E-08 |
| 23381 | 1170 | A1170A | D | 4.8e-08 | 6.8e-08 | 5.8e-08 | 5.800E-08 | 1.000E-08 | 4.800E-08 | 6.800E-08 |
| 23382 | 1170 | A1170R | D | 6.9e-08 | 3.4e-08 | 7e-08 | 5.767E-08 | 2.050E-08 | 3.400E-08 | 7.000E-08 |
| 23383 | 1170 | A1170N | D | 1.3e-07 | 6.2e-08 | 1.1e-07 | 1.007E-07 | 3.495E-08 | 6.200E-08 | 1.300E-07 |
| 23384 | 1170 | A1170D | D | 3.6e-08 | 1.4e-07 | 7.7e-08 | 8.433E-08 | 5.239E-08 | 3.600E-08 | 1.400E-07 |
| 23385 | 1170 | A1170C | D | 6e-08 | 7.3e-08 | 3.1e-08 | 5.467E-08 | 2.150E-08 | 3.100E-08 | 7.300E-08 |
| 23386 | 1170 | A1170E | D | 1.1e-08 | 6.4e-08 | 2.6e-08 | 3.367E-08 | 2.732E-08 | 1.100E-08 | 6.400E-08 |
| 23387 | 1170 | A1170Q | D | 9.7e-08 | 3.9e-08 | 4.2e-08 | 5.933E-08 | 3.265E-08 | 3.900E-08 | 9.700E-08 |
| 23388 | 1170 | A1170G | D | 2.1e-08 | 6.6e-08 | 1.4e-07 | 7.567E-08 | 6.009E-08 | 2.100E-08 | 1.400E-07 |
| 23389 | 1170 | A1170H | D | 1.6e-08 | 4.3e-08 | 1.8e-08 | 2.567E-08 | 1.504E-08 | 1.600E-08 | 4.300E-08 |
| 23390 | 1170 | A1170I | D | 4.2e-08 | 5.8e-08 | 2.3e-08 | 4.100E-08 | 1.752E-08 | 2.300E-08 | 5.800E-08 |
| 23391 | 1170 | A1170L | D | 2.1e-08 | 3.9e-08 | 5.2e-08 | 3.733E-08 | 1.557E-08 | 2.100E-08 | 5.200E-08 |
| 23392 | 1170 | A1170K | D | 8.5e-08 | 7.8e-09 | 6.4e-08 | 5.227E-08 | 3.992E-08 | 7.800E-09 | 8.500E-08 |
| 23393 | 1170 | A1170M | D | 2.8e-08 | 8.7e-08 | 2.4e-08 | 4.633E-08 | 3.528E-08 | 2.400E-08 | 8.700E-08 |
| 23394 | 1170 | A1170F | D | 6.5e-08 | 3.6e-08 | 5.4e-08 | 5.167E-08 | 1.464E-08 | 3.600E-08 | 6.500E-08 |
| 23395 | 1170 | A1170P | D | 3.9e-08 | 4.4e-08 | 5.1e-08 | 4.467E-08 | 6.028E-09 | 3.900E-08 | 5.100E-08 |
| 23396 | 1170 | A1170S | D | 5.4e-08 | 1.4e-07 | 4.3e-08 | 7.900E-08 | 5.311E-08 | 4.300E-08 | 1.400E-07 |
| 23397 | 1170 | A1170T | D | 1.2e-08 | 7.7e-08 | 2.1e-07 | 9.967E-08 | 1.009E-07 | 1.200E-08 | 2.100E-07 |
| 23398 | 1170 | A1170W | D | 9.7e-08 | 8.9e-08 | 1.4e-07 | 1.087E-07 | 2.743E-08 | 8.900E-08 | 1.400E-07 |
| 23399 | 1170 | A1170Y | D | 2.7e-08 | 4.8e-08 | 4.6e-08 | 4.033E-08 | 1.159E-08 | 2.700E-08 | 4.800E-08 |
| 23400 | 1170 | A1170V | D | 6.3e-08 | 3.6e-08 | 4.7e-08 | 4.867E-08 | 1.358E-08 | 3.600E-08 | 6.300E-08 |
| 23401 | 1171 | P1171A | D | 2.4e-08 | 7.2e-08 | 1.2e-08 | 3.600E-08 | 3.175E-08 | 1.200E-08 | 7.200E-08 |
| 23402 | 1171 | P1171R | D | 8.8e-09 | 4.6e-08 | 5.7e-08 | 3.727E-08 | 2.526E-08 | 8.800E-09 | 5.700E-08 |
| 23403 | 1171 | P1171N | D | 4e-08 | 3.8e-08 | 6.2e-08 | 4.667E-08 | 1.332E-08 | 3.800E-08 | 6.200E-08 |
| 23404 | 1171 | P1171D | D | 5.7e-08 | 1.8e-08 | 3.2e-08 | 3.567E-08 | 1.976E-08 | 1.800E-08 | 5.700E-08 |
| 23405 | 1171 | P1171C | D | 2.7e-08 | 1e-07 | 2.5e-08 | 5.067E-08 | 4.274E-08 | 2.500E-08 | 1.000E-07 |
| 23406 | 1171 | P1171E | D | 2.2e-08 | 6.1e-08 | 3.3e-08 | 3.867E-08 | 2.011E-08 | 2.200E-08 | 6.100E-08 |
| 23407 | 1171 | P1171Q | D | 6.3e-08 | 3.1e-08 | 3.6e-08 | 4.333E-08 | 1.721E-08 | 3.100E-08 | 6.300E-08 |
| 23408 | 1171 | P1171G | D | 3.4e-08 | 5.7e-08 | 5.6e-08 | 4.900E-08 | 1.300E-08 | 3.400E-08 | 5.700E-08 |
| 23409 | 1171 | P1171H | D | 1.6e-08 | 1.9e-08 | 3.4e-08 | 2.300E-08 | 9.644E-09 | 1.600E-08 | 3.400E-08 |
| 23410 | 1171 | P1171I | D | 4.4e-08 | 3.1e-08 | 3.9e-08 | 3.800E-08 | 6.557E-09 | 3.100E-08 | 4.400E-08 |
| 23411 | 1171 | P1171L | D | 7.5e-08 | 1.4e-08 | 2.5e-08 | 3.800E-08 | 3.251E-08 | 1.400E-08 | 7.500E-08 |
| 23412 | 1171 | P1171K | D | 2.9e-08 | 7.2e-08 | 8.1e-08 | 6.067E-08 | 2.779E-08 | 2.900E-08 | 8.100E-08 |
| 23413 | 1171 | P1171M | D | 4.3e-08 | 1.4e-08 | 4.4e-08 | 3.367E-08 | 1.704E-08 | 1.400E-08 | 4.400E-08 |
| 23414 | 1171 | P1171F | D | 2.5e-08 | 2.5e-08 | 1.3e-07 | 6.000E-08 | 6.062E-08 | 2.500E-08 | 1.300E-07 |
| 23415 | 1171 | P1171P | D | 9.8e-08 | 6e-08 | 9.1e-08 | 8.300E-08 | 2.022E-08 | 6.000E-08 | 9.800E-08 |
| 23416 | 1171 | P1171S | D | 8.2e-09 | 2.2e-08 | 3.8e-08 | 2.273E-08 | 1.491E-08 | 8.200E-09 | 3.800E-08 |
| 23417 | 1171 | P1171T | D | 5.1e-08 | 2.3e-08 | 7.9e-08 | 5.100E-08 | 2.800E-08 | 2.300E-08 | 7.900E-08 |
| 23418 | 1171 | P1171W | D | 5e-08 | 5.4e-08 | 3.3e-08 | 4.567E-08 | 1.115E-08 | 3.300E-08 | 5.400E-08 |
| 23419 | 1171 | P1171Y | D | 5.3e-08 | 1.1e-07 | 2.6e-08 | 6.300E-08 | 4.288E-08 | 2.600E-08 | 1.100E-07 |
| 23420 | 1171 | P1171V | D | 4.2e-08 | 1.3e-08 | 1.6e-08 | 2.367E-08 | 1.595E-08 | 1.300E-08 | 4.200E-08 |
| 23421 | 1172 | S1172A | D | 1.6e-08 | 5.4e-08 | 5.4e-08 | 4.133E-08 | 2.194E-08 | 1.600E-08 | 5.400E-08 |
| 23422 | 1172 | S1172R | D | 1.8e-08 | 1.7e-08 | 4.5e-08 | 2.667E-08 | 1.589E-08 | 1.700E-08 | 4.500E-08 |
| 23423 | 1172 | S1172N | D | 1e-07 | 2.4e-08 | 4.2e-08 | 5.533E-08 | 3.972E-08 | 2.400E-08 | 1.000E-07 |
| 23424 | 1172 | S1172D | D | 3.8e-08 | 3.6e-08 | 3.2e-08 | 3.533E-08 | 3.055E-09 | 3.200E-08 | 3.800E-08 |
| 23425 | 1172 | S1172C | D | 2.8e-08 | 3e-08 | 2e-08 | 2.600E-08 | 5.292E-09 | 2.000E-08 | 3.000E-08 |
| 23426 | 1172 | S1172E | D | 5.8e-08 | 9.2e-08 | 1.2e-07 | 9.000E-08 | 3.105E-08 | 5.800E-08 | 1.200E-07 |
| 23427 | 1172 | S1172Q | D | 3.1e-08 | 6.3e-08 | 4.1e-08 | 4.500E-08 | 1.637E-08 | 3.100E-08 | 6.300E-08 |
| 23428 | 1172 | S1172G | D | 3e-08 | 3.2e-08 | 1e-07 | 5.400E-08 | 3.985E-08 | 3.000E-08 | 1.000E-07 |
| 23429 | 1172 | S1172H | D | 2.2e-08 | 2.7e-08 | 1.3e-08 | 2.067E-08 | 7.095E-09 | 1.300E-08 | 2.700E-08 |
| 23430 | 1172 | S1172I | D | 3.3e-08 | 3.9e-08 | 8.5e-08 | 5.233E-08 | 2.845E-08 | 3.300E-08 | 8.500E-08 |
| 23431 | 1172 | S1172L | D | 3e-08 | 4.3e-08 | 4.2e-08 | 3.833E-08 | 7.234E-09 | 3.000E-08 | 4.300E-08 |
| 23432 | 1172 | S1172K | D | 5e-08 | 8.7e-08 | 2.4e-08 | 5.367E-08 | 3.166E-08 | 2.400E-08 | 8.700E-08 |
| 23433 | 1172 | S1172M | D | 4.3e-08 | 9.6e-08 | 3.7e-08 | 5.867E-08 | 3.247E-08 | 3.700E-08 | 9.600E-08 |
| 23434 | 1172 | S1172F | D | 8.5e-08 | 7.5e-08 | 8.5e-08 | 8.167E-08 | 5.774E-09 | 7.500E-08 | 8.500E-08 |
| 23435 | 1172 | S1172P | D | 1.2e-08 | 3.4e-08 | 5e-08 | 3.200E-08 | 1.908E-08 | 1.200E-08 | 5.000E-08 |
| 23436 | 1172 | S1172S | D | 9.3e-08 | 1.9e-08 | 2.3e-08 | 4.500E-08 | 4.162E-08 | 1.900E-08 | 9.300E-08 |
| 23437 | 1172 | S1172T | D | 3.4e-08 | 3.2e-08 | 2.6e-08 | 3.067E-08 | 4.163E-09 | 2.600E-08 | 3.400E-08 |
| 23438 | 1172 | S1172W | D | 8e-09 | 3.3e-08 | 8.9e-08 | 4.333E-08 | 4.148E-08 | 8.000E-09 | 8.900E-08 |
| 23439 | 1172 | S1172Y | D | 2.4e-08 | 7.6e-08 | 4.7e-08 | 4.900E-08 | 2.606E-08 | 2.400E-08 | 7.600E-08 |
| 23440 | 1172 | S1172V | D | 5.9e-08 | 4.1e-08 | 3.6e-08 | 4.533E-08 | 1.210E-08 | 3.600E-08 | 5.900E-08 |
| 23441 | 1173 | V1173A | D | 4.6e-08 | 1e-08 | 5e-08 | 3.533E-08 | 2.203E-08 | 1.000E-08 | 5.000E-08 |
| 23442 | 1173 | V1173R | D | 6.6e-08 | 2.7e-08 | 2.1e-08 | 3.800E-08 | 2.443E-08 | 2.100E-08 | 6.600E-08 |
| 23443 | 1173 | V1173N | D | 7.7e-08 | 6.6e-08 | 5.3e-08 | 6.533E-08 | 1.201E-08 | 5.300E-08 | 7.700E-08 |
| 23444 | 1173 | V1173D | D | 4.9e-08 | 3.2e-08 | 4.1e-08 | 4.067E-08 | 8.505E-09 | 3.200E-08 | 4.900E-08 |
| 23445 | 1173 | V1173C | D | 5e-08 | 6.2e-08 | 7.3e-08 | 6.167E-08 | 1.150E-08 | 5.000E-08 | 7.300E-08 |
| 23446 | 1173 | V1173E | D | 2.5e-08 | 3.5e-08 | 4.4e-08 | 3.467E-08 | 9.504E-09 | 2.500E-08 | 4.400E-08 |
| 23447 | 1173 | V1173Q | D | 2.9e-08 | 2.7e-08 | 4.2e-08 | 3.267E-08 | 8.145E-09 | 2.700E-08 | 4.200E-08 |
| 23448 | 1173 | V1173G | D | 2.6e-08 | 3.6e-08 | 1.6e-08 | 2.600E-08 | 1.000E-08 | 1.600E-08 | 3.600E-08 |
| 23449 | 1173 | V1173H | D | 1e-07 | 3.4e-08 | 1e-07 | 7.800E-08 | 3.811E-08 | 3.400E-08 | 1.000E-07 |
| 23450 | 1173 | V1173I | D | 3.2e-08 | 7.8e-08 | 1.1e-07 | 7.333E-08 | 3.921E-08 | 3.200E-08 | 1.100E-07 |
| 23451 | 1173 | V1173L | D | 3.4e-08 | 5.6e-08 | 2.5e-08 | 3.833E-08 | 1.595E-08 | 2.500E-08 | 5.600E-08 |
| 23452 | 1173 | V1173K | D | 7.2e-08 | 3.5e-08 | 4.5e-08 | 5.067E-08 | 1.914E-08 | 3.500E-08 | 7.200E-08 |
| 23453 | 1173 | V1173M | D | 5.4e-08 | 3.3e-08 | 4.8e-08 | 4.500E-08 | 1.082E-08 | 3.300E-08 | 5.400E-08 |
| 23454 | 1173 | V1173F | D | 1.9e-08 | 2.3e-08 | 8.4e-08 | 4.200E-08 | 3.643E-08 | 1.900E-08 | 8.400E-08 |
| 23455 | 1173 | V1173P | D | 9.9e-08 | 6.2e-08 | 3.2e-08 | 6.433E-08 | 3.356E-08 | 3.200E-08 | 9.900E-08 |
| 23456 | 1173 | V1173S | D | 1.3e-07 | 1.1e-07 | 1e-07 | 1.133E-07 | 1.528E-08 | 1.000E-07 | 1.300E-07 |
| 23457 | 1173 | V1173T | D | 4.2e-08 | 1.9e-08 | 5.5e-08 | 3.867E-08 | 1.823E-08 | 1.900E-08 | 5.500E-08 |
| 23458 | 1173 | V1173W | D | 1.9e-08 | 2.4e-08 | 3.3e-08 | 2.533E-08 | 7.095E-09 | 1.900E-08 | 3.300E-08 |
| 23459 | 1173 | V1173Y | D | 1.3e-07 | 3.5e-08 | 4.8e-08 | 7.100E-08 | 5.151E-08 | 3.500E-08 | 1.300E-07 |

| | | | | | | | | | | |
|---|---|---|---|---|---|---|---|---|---|---|
| 23460 | 1173 | V1173V | D | 4.5e-08 | 6.7e-08 | 5.8e-08 | 5.667E-08 | 1.106E-08 | 4.500E-08 | 6.700E-08 |
| 23461 | 1174 | F1174A | D | 8.3e-08 | 1.1e-07 | 7.2e-08 | 8.833E-08 | 1.955E-08 | 7.200E-08 | 1.100E-07 |
| 23462 | 1174 | F1174R | D | 7.3e-08 | 7.4e-08 | 7.4e-08 | 7.367E-08 | 5.774E-10 | 7.300E-08 | 7.400E-08 |
| 23463 | 1174 | F1174N | D | 7.8e-08 | 2.1e-08 | 5.5e-08 | 5.133E-08 | 2.868E-08 | 2.100E-08 | 7.800E-08 |
| 23464 | 1174 | F1174D | D | 3.1e-08 | 5.4e-08 | 9.6e-08 | 6.033E-08 | 3.296E-08 | 3.100E-08 | 9.600E-08 |
| 23465 | 1174 | F1174C | D | 1.5e-07 | 2.9e-08 | 2.4e-08 | 6.767E-08 | 7.135E-08 | 2.400E-08 | 1.500E-07 |
| 23466 | 1174 | F1174E | D | 4.2e-08 | 2.5e-08 | 5e-08 | 3.900E-08 | 1.277E-08 | 2.500E-08 | 5.000E-08 |
| 23467 | 1174 | F1174Q | D | 4.8e-08 | 9.6e-09 | 3.6e-08 | 3.120E-08 | 1.964E-08 | 9.600E-09 | 4.800E-08 |
| 23468 | 1174 | F1174G | D | 5e-08 | 8.6e-08 | 3.8e-08 | 5.800E-08 | 2.498E-08 | 3.800E-08 | 8.600E-08 |
| 23469 | 1174 | F1174H | D | 2.6e-08 | 2.7e-08 | 2.4e-08 | 2.567E-08 | 1.528E-09 | 2.400E-08 | 2.700E-08 |
| 23470 | 1174 | F1174I | D | 3.3e-08 | 2.4e-08 | 5.5e-08 | 3.733E-08 | 1.595E-08 | 2.400E-08 | 5.500E-08 |
| 23471 | 1174 | F1174L | D | 6.3e-08 | 4.1e-08 | 2.4e-08 | 4.267E-08 | 1.955E-08 | 2.400E-08 | 6.300E-08 |
| 23472 | 1174 | F1174K | D | 2.4e-08 | 5.5e-08 | 2e-08 | 3.300E-08 | 1.916E-08 | 2.000E-08 | 5.500E-08 |
| 23473 | 1174 | F1174M | D | 4.7e-08 | 7e-08 | 4.7e-08 | 5.467E-08 | 1.328E-08 | 4.700E-08 | 7.000E-08 |
| 23474 | 1174 | F1174F | D | 6.7e-08 | 2.7e-08 | 3.7e-08 | 4.367E-08 | 2.082E-08 | 2.700E-08 | 6.700E-08 |
| 23475 | 1174 | F1174P | D | 7.9e-08 | 1.1e-08 | 3.7e-08 | 4.233E-08 | 3.431E-08 | 1.100E-08 | 7.900E-08 |
| 23476 | 1174 | F1174S | D | 1.1e-08 | 1.8e-07 | 9.5e-08 | 9.533E-08 | 8.450E-08 | 1.100E-08 | 1.800E-07 |
| 23477 | 1174 | F1174T | D | 4.9e-08 | 4e-08 | 2.3e-08 | 3.733E-08 | 1.320E-08 | 2.300E-08 | 4.900E-08 |
| 23478 | 1174 | F1174W | D | 3e-08 | 3.8e-08 | 6.3e-08 | 4.367E-08 | 1.721E-08 | 3.000E-08 | 6.300E-08 |
| 23479 | 1174 | F1174Y | D | 5.1e-08 | 1.7e-08 | 1.3e-07 | 6.600E-08 | 5.797E-08 | 1.700E-08 | 1.300E-07 |
| 23480 | 1174 | F1174V | D | 2.1e-08 | 4e-08 | 6.1e-08 | 4.067E-08 | 2.001E-08 | 2.100E-08 | 6.100E-08 |
| 23481 | 1175 | I1175A | D | 6.1e-08 | 3.4e-08 | 4e-08 | 4.500E-08 | 1.418E-08 | 3.400E-08 | 6.100E-08 |
| 23482 | 1175 | I1175R | D | 3.5e-08 | 1.5e-08 | 7e-08 | 4.000E-08 | 2.784E-08 | 1.500E-08 | 7.000E-08 |
| 23483 | 1175 | I1175N | D | 4.2e-08 | 2.4e-08 | 4.4e-08 | 3.667E-08 | 1.102E-08 | 2.400E-08 | 4.400E-08 |
| 23484 | 1175 | I1175D | D | 1.4e-08 | 1.3e-08 | 1.8e-08 | 1.500E-08 | 2.646E-09 | 1.300E-08 | 1.800E-08 |
| 23485 | 1175 | I1175C | D | 3.7e-08 | 2.7e-08 | 5e-08 | 3.800E-08 | 1.153E-08 | 2.700E-08 | 5.000E-08 |
| 23486 | 1175 | I1175E | D | 3e-08 | 8.8e-08 | 4.9e-08 | 5.567E-08 | 2.957E-08 | 3.000E-08 | 8.800E-08 |
| 23487 | 1175 | I1175Q | D | 5.8e-09 | 1.1e-07 | 2.8e-08 | 4.793E-08 | 5.489E-08 | 5.800E-09 | 1.100E-07 |
| 23488 | 1175 | I1175G | D | 3.2e-08 | 3.9e-08 | 4.3e-08 | 3.800E-08 | 5.568E-09 | 3.200E-08 | 4.300E-08 |
| 23489 | 1175 | I1175H | D | 2.5e-08 | 3.8e-08 | 4.4e-08 | 3.567E-08 | 9.713E-09 | 2.500E-08 | 4.400E-08 |
| 23490 | 1175 | I1175I | D | 8.3e-09 | 6.9e-08 | 7.5e-08 | 5.077E-08 | 3.690E-08 | 8.300E-09 | 7.500E-08 |
| 23491 | 1175 | I1175L | D | 1.4e-08 | 6.9e-08 | 6.9e-08 | 5.067E-08 | 3.175E-08 | 1.400E-08 | 6.900E-08 |
| 23492 | 1175 | I1175K | D | 4.2e-08 | 3e-08 | 1.8e-08 | 3.000E-08 | 1.200E-08 | 1.800E-08 | 4.200E-08 |
| 23493 | 1175 | I1175M | D | 9.8e-08 | 4.7e-08 | 4.8e-08 | 6.433E-08 | 2.916E-08 | 4.700E-08 | 9.800E-08 |
| 23494 | 1175 | I1175F | D | 4.8e-08 | 1.9e-08 | 3.4e-08 | 3.367E-08 | 1.450E-08 | 1.900E-08 | 4.800E-08 |
| 23495 | 1175 | I1175P | D | 7.4e-08 | 5.5e-08 | 2.9e-08 | 5.267E-08 | 2.259E-08 | 2.900E-08 | 7.400E-08 |
| 23496 | 1175 | I1175S | D | 1.8e-08 | 1.1e-07 | 7.4e-08 | 6.733E-08 | 4.636E-08 | 1.800E-08 | 1.100E-07 |
| 23497 | 1175 | I1175T | D | 7.7e-08 | 9.2e-08 | 4.7e-08 | 7.200E-08 | 2.291E-08 | 4.700E-08 | 9.200E-08 |
| 23498 | 1175 | I1175W | D | 3.3e-08 | 3.2e-08 | 1e-07 | 5.500E-08 | 3.897E-08 | 3.200E-08 | 1.000E-07 |
| 23499 | 1175 | I1175Y | D | 4.4e-08 | 5e-08 | 1.7e-08 | 3.700E-08 | 1.758E-08 | 1.700E-08 | 5.000E-08 |
| 23500 | 1175 | I1175V | D | 3.2e-08 | 1.2e-08 | 4.3e-08 | 2.900E-08 | 1.572E-08 | 1.200E-08 | 4.300E-08 |
| 23501 | 1176 | F1176A | D | 4.9e-08 | 1.1e-07 | 7.2e-08 | 7.700E-08 | 3.081E-08 | 4.900E-08 | 1.100E-07 |
| 23502 | 1176 | F1176R | D | 6.1e-08 | 3.3e-08 | 7.8e-08 | 5.733E-08 | 2.272E-08 | 3.300E-08 | 7.800E-08 |
| 23503 | 1176 | F1176N | D | 3.1e-08 | 3.5e-08 | 3.1e-08 | 3.233E-08 | 2.309E-09 | 3.100E-08 | 3.500E-08 |
| 23504 | 1176 | F1176D | D | 4.7e-08 | 5.9e-08 | 6.7e-08 | 5.767E-08 | 1.007E-08 | 4.700E-08 | 6.700E-08 |
| 23505 | 1176 | F1176C | D | 3.9e-08 | 2.1e-08 | 7.2e-08 | 4.400E-08 | 2.587E-08 | 2.100E-08 | 7.200E-08 |
| 23506 | 1176 | F1176E | D | 5e-08 | 3.8e-08 | 5.2e-08 | 4.667E-08 | 7.572E-09 | 3.800E-08 | 5.200E-08 |
| 23507 | 1176 | F1176Q | D | 2.6e-08 | 6.6e-08 | 6.5e-08 | 5.233E-08 | 2.281E-08 | 2.600E-08 | 6.600E-08 |
| 23508 | 1176 | F1176G | D | 7e-08 | 9.4e-08 | 4.9e-08 | 7.100E-08 | 2.252E-08 | 4.900E-08 | 9.400E-08 |
| 23509 | 1176 | F1176H | D | 3.8e-08 | 3.9e-08 | 3.6e-08 | 3.767E-08 | 1.528E-09 | 3.600E-08 | 3.900E-08 |
| 23510 | 1176 | F1176I | D | 4.9e-08 | 1.9e-08 | 3.2e-08 | 3.333E-08 | 1.504E-08 | 1.900E-08 | 4.900E-08 |
| 23511 | 1176 | F1176L | D | 4.2e-08 | 3.4e-08 | 1.9e-08 | 3.167E-08 | 1.168E-08 | 1.900E-08 | 4.200E-08 |
| 23512 | 1176 | F1176K | D | 3.8e-08 | 2.9e-08 | 2.9e-08 | 3.200E-08 | 5.196E-09 | 2.900E-08 | 3.800E-08 |
| 23513 | 1176 | F1176M | D | 1.2e-07 | 2.3e-08 | 4.7e-08 | 6.333E-08 | 5.052E-08 | 2.300E-08 | 1.200E-07 |
| 23514 | 1176 | F1176F | D | 2.1e-08 | 3.7e-08 | 7.9e-09 | 2.197E-08 | 1.457E-08 | 7.900E-09 | 3.700E-08 |
| 23515 | 1176 | F1176P | D | 2.2e-08 | 4.9e-08 | 5e-08 | 4.033E-08 | 1.589E-08 | 2.200E-08 | 5.000E-08 |
| 23516 | 1176 | F1176S | D | 4.5e-08 | 4.2e-08 | 5.6e-08 | 4.767E-08 | 7.371E-09 | 4.200E-08 | 5.600E-08 |
| 23517 | 1176 | F1176T | D | 1.7e-08 | 5.5e-08 | 6.3e-08 | 4.500E-08 | 2.458E-08 | 1.700E-08 | 6.300E-08 |
| 23518 | 1176 | F1176W | D | 4.8e-08 | 3.2e-08 | 3.5e-08 | 3.833E-08 | 8.505E-09 | 3.200E-08 | 4.800E-08 |
| 23519 | 1176 | F1176Y | D | 3.2e-08 | 4.1e-08 | 5.1e-08 | 4.133E-08 | 9.504E-09 | 3.200E-08 | 5.100E-08 |
| 23520 | 1176 | F1176V | D | 2.4e-08 | 4.5e-08 | 1.2e-07 | 6.300E-08 | 5.047E-08 | 2.400E-08 | 1.200E-07 |
| 23521 | 1177 | P1177A | D | 3.9e-08 | 1.2e-07 | 8.6e-08 | 8.167E-08 | 4.067E-08 | 3.900E-08 | 1.200E-07 |
| 23522 | 1177 | P1177R | D | 6.1e-08 | 3.3e-08 | 1.2e-08 | 3.533E-08 | 2.458E-08 | 1.200E-08 | 6.100E-08 |
| 23523 | 1177 | P1177N | D | 6.5e-08 | 5.2e-08 | 2.1e-08 | 4.600E-08 | 2.261E-08 | 2.100E-08 | 6.500E-08 |
| 23524 | 1177 | P1177D | D | 4.3e-08 | 6.9e-08 | 3.7e-08 | 4.967E-08 | 1.701E-08 | 3.700E-08 | 6.900E-08 |
| 23525 | 1177 | P1177C | D | 8.2e-08 | 2.3e-08 | 4.2e-08 | 4.900E-08 | 3.012E-08 | 2.300E-08 | 8.200E-08 |
| 23526 | 1177 | P1177E | D | 4.3e-08 | 3.1e-08 | 5.9e-08 | 4.433E-08 | 1.405E-08 | 3.100E-08 | 5.900E-08 |
| 23527 | 1177 | P1177Q | D | 1.7e-08 | 1.3e-08 | 3.8e-08 | 2.267E-08 | 1.343E-08 | 1.300E-08 | 3.800E-08 |
| 23528 | 1177 | P1177G | D | 2.3e-08 | 7.2e-08 | 1.3e-08 | 3.600E-08 | 3.158E-08 | 1.300E-08 | 7.200E-08 |
| 23529 | 1177 | P1177H | D | 2.6e-08 | 2.4e-08 | 2.4e-08 | 2.467E-08 | 1.155E-09 | 2.400E-08 | 2.600E-08 |
| 23530 | 1177 | P1177I | D | 5e-08 | 2.8e-08 | 3.3e-08 | 3.700E-08 | 1.153E-08 | 2.800E-08 | 5.000E-08 |
| 23531 | 1177 | P1177L | D | 6.1e-08 | 1.3e-07 | 4.6e-08 | 7.900E-08 | 4.480E-08 | 4.600E-08 | 1.300E-07 |
| 23532 | 1177 | P1177K | D | 2.5e-08 | 8.9e-08 | 4.7e-08 | 5.367E-08 | 3.252E-08 | 2.500E-08 | 8.900E-08 |
| 23533 | 1177 | P1177M | D | 7.6e-08 | 7.7e-08 | 4.5e-08 | 6.600E-08 | 1.819E-08 | 4.500E-08 | 7.700E-08 |
| 23534 | 1177 | P1177F | D | 4.8e-08 | 5.3e-08 | 3.9e-08 | 4.667E-08 | 7.095E-09 | 3.900E-08 | 5.300E-08 |
| 23535 | 1177 | P1177P | D | 1.8e-08 | 5.1e-08 | 4.7e-08 | 3.867E-08 | 1.801E-08 | 1.800E-08 | 5.100E-08 |
| 23536 | 1177 | P1177S | D | 9.2e-08 | 5.3e-08 | 1.8e-08 | 5.433E-08 | 3.702E-08 | 1.800E-08 | 9.200E-08 |
| 23537 | 1177 | P1177T | D | 2.4e-08 | 2.7e-08 | 7.1e-08 | 4.067E-08 | 2.631E-08 | 2.400E-08 | 7.100E-08 |
| 23538 | 1177 | P1177W | D | 7e-08 | 2.9e-08 | 3.7e-08 | 4.533E-08 | 2.173E-08 | 2.900E-08 | 7.000E-08 |
| 23539 | 1177 | P1177Y | D | 4e-08 | 2.4e-08 | 7e-08 | 4.467E-08 | 2.335E-08 | 2.400E-08 | 7.000E-08 |
| 23540 | 1177 | P1177V | D | 2.5e-08 | 1e-07 | 5.7e-08 | 6.067E-08 | 3.763E-08 | 2.500E-08 | 1.000E-07 |
| 23541 | 1178 | P1178A | D | 6.4e-08 | 6.2e-08 | 5.7e-08 | 6.100E-08 | 3.606E-09 | 5.700E-08 | 6.400E-08 |
| 23542 | 1178 | P1178R | D | 1.2e-07 | 2.7e-08 | 2.2e-08 | 5.633E-08 | 5.519E-08 | 2.200E-08 | 1.200E-07 |
| 23543 | 1178 | P1178N | D | 3.7e-09 | 3e-08 | 6.7e-08 | 3.357E-08 | 3.180E-08 | 3.700E-09 | 6.700E-08 |
| 23544 | 1178 | P1178D | D | 6e-08 | 8e-08 | 2.9e-08 | 5.633E-08 | 2.570E-08 | 2.900E-08 | 8.000E-08 |

| | | | | | | | | | | |
|---|---|---|---|---|---|---|---|---|---|---|
| 23545 | 1178 | P1178C | D | 3.9e-08 | 3.5e-08 | 1.8e-07 | 8.467E-08 | 8.259E-08 | 3.500E-08 | 1.800E-07 |
| 23546 | 1178 | P1178E | D | 2e-08 | 7e-08 | 3.2e-08 | 4.067E-08 | 2.610E-08 | 2.000E-08 | 7.000E-08 |
| 23547 | 1178 | P1178Q | D | 1.4e-07 | 3.6e-08 | 7e-08 | 8.200E-08 | 5.303E-08 | 3.600E-08 | 1.400E-07 |
| 23548 | 1178 | P1178G | D | 3.8e-08 | 7.7e-08 | 1.5e-07 | 8.833E-08 | 5.685E-08 | 3.800E-08 | 1.500E-07 |
| 23549 | 1178 | P1178H | D | 3.8e-08 | 2e-07 | 1.2e-07 | 1.193E-07 | 8.100E-08 | 3.800E-08 | 2.000E-07 |
| 23550 | 1178 | P1178I | D | 6.7e-08 | 2.4e-08 | 1.2e-07 | 7.033E-08 | 4.809E-08 | 2.400E-08 | 1.200E-07 |
| 23551 | 1178 | P1178L | D | 4.8e-08 | 3.4e-08 | 8.6e-08 | 5.600E-08 | 2.691E-08 | 3.400E-08 | 8.600E-08 |
| 23552 | 1178 | P1178K | D | 3e-08 | 9e-08 | 6.3e-08 | 6.100E-08 | 3.005E-08 | 3.000E-08 | 9.000E-08 |
| 23553 | 1178 | P1178M | D | 1.8e-07 | 7.9e-08 | 5.9e-08 | 1.060E-07 | 6.486E-08 | 5.900E-08 | 1.800E-07 |
| 23554 | 1178 | P1178F | D | 7.4e-08 | 2.5e-08 | 8.3e-08 | 6.067E-08 | 3.121E-08 | 2.500E-08 | 8.300E-08 |
| 23555 | 1178 | P1178P | D | 2.5e-08 | 7.2e-08 | 3.9e-08 | 4.533E-08 | 2.413E-08 | 2.500E-08 | 7.200E-08 |
| 23556 | 1178 | P1178S | D | 3.9e-08 | 6.6e-09 | 6.9e-08 | 3.820E-08 | 3.121E-08 | 6.600E-09 | 6.900E-08 |
| 23557 | 1178 | P1178T | D | 2.8e-08 | 4.1e-08 | 7.1e-08 | 4.667E-08 | 2.205E-08 | 2.800E-08 | 7.100E-08 |
| 23558 | 1178 | P1178W | D | 5.5e-08 | 1.2e-07 | 1.3e-08 | 6.267E-08 | 5.391E-08 | 1.300E-08 | 1.200E-07 |
| 23559 | 1178 | P1178Y | D | 3.5e-08 | 2e-08 | 3.1e-08 | 2.867E-08 | 7.767E-09 | 2.000E-08 | 3.500E-08 |
| 23560 | 1178 | P1178V | D | 6.7e-08 | 4.6e-08 | 1.7e-08 | 4.333E-08 | 2.511E-08 | 1.700E-08 | 6.700E-08 |
| 23561 | 1179 | S1179A | D | 4.8e-08 | 4e-08 | 8.9e-08 | 5.900E-08 | 2.629E-08 | 4.000E-08 | 8.900E-08 |
| 23562 | 1179 | S1179R | D | 7.3e-08 | 2.1e-08 | 1.8e-08 | 3.733E-08 | 3.092E-08 | 1.800E-08 | 7.300E-08 |
| 23563 | 1179 | S1179N | D | 1.3e-07 | 6.3e-08 | 4.8e-08 | 8.033E-08 | 4.366E-08 | 4.800E-08 | 1.300E-07 |
| 23564 | 1179 | S1179D | D | 5.1e-08 | 2.4e-08 | 1.4e-08 | 2.967E-08 | 1.914E-08 | 1.400E-08 | 5.100E-08 |
| 23565 | 1179 | S1179C | D | 4.8e-08 | 3e-08 | 1.9e-08 | 3.233E-08 | 1.464E-08 | 1.900E-08 | 4.800E-08 |
| 23566 | 1179 | S1179E | D | 1.3e-07 | 3e-08 | 7.7e-08 | 7.900E-08 | 5.003E-08 | 3.000E-08 | 1.300E-07 |
| 23567 | 1179 | S1179Q | D | 1e-07 | 7.2e-08 | 5.4e-08 | 7.533E-08 | 2.318E-08 | 5.400E-08 | 1.000E-07 |
| 23568 | 1179 | S1179G | D | 3.2e-08 | 2.6e-08 | 2.7e-08 | 2.833E-08 | 3.215E-09 | 2.600E-08 | 3.200E-08 |
| 23569 | 1179 | S1179H | D | 2.5e-08 | 5.1e-08 | 1.3e-07 | 6.867E-08 | 5.468E-08 | 2.500E-08 | 1.300E-07 |
| 23570 | 1179 | S1179I | D | 1.5e-07 | 2e-08 | 2.1e-08 | 6.367E-08 | 7.477E-08 | 2.000E-08 | 1.500E-07 |
| 23571 | 1179 | S1179L | D | 6.1e-08 | 3.1e-08 | 8.8e-08 | 6.000E-08 | 2.851E-08 | 3.100E-08 | 8.800E-08 |
| 23572 | 1179 | S1179K | D | 5.4e-08 | 5.1e-08 | 5.6e-08 | 5.367E-08 | 2.517E-09 | 5.100E-08 | 5.600E-08 |
| 23573 | 1179 | S1179M | D | 2.9e-08 | 7.5e-08 | 3.5e-08 | 4.633E-08 | 2.501E-08 | 2.900E-08 | 7.500E-08 |
| 23574 | 1179 | S1179F | D | 8.2e-08 | 7.6e-08 | 4.4e-08 | 6.733E-08 | 2.043E-08 | 4.400E-08 | 8.200E-08 |
| 23575 | 1179 | S1179P | D | 4.9e-08 | 1.3e-07 | 3e-08 | 6.967E-08 | 5.311E-08 | 3.000E-08 | 1.300E-07 |
| 23576 | 1179 | S1179S | D | 2.5e-08 | 3e-08 | 4.6e-08 | 3.367E-08 | 1.097E-08 | 2.500E-08 | 4.600E-08 |
| 23577 | 1179 | S1179T | D | 1e-07 | 7.8e-08 | 3.7e-08 | 7.167E-08 | 3.197E-08 | 3.700E-08 | 1.000E-07 |
| 23578 | 1179 | S1179W | D | 7.6e-09 | 1.6e-07 | 3.7e-08 | 6.820E-08 | 8.085E-08 | 7.600E-09 | 1.600E-07 |
| 23579 | 1179 | S1179Y | D | 2.2e-08 | 5.8e-08 | 5.7e-08 | 4.567E-08 | 2.050E-08 | 2.200E-08 | 5.800E-08 |
| 23580 | 1179 | S1179V | D | 2.6e-08 | 3.3e-08 | 3.5e-08 | 3.133E-08 | 4.726E-09 | 2.600E-08 | 3.500E-08 |
| 23581 | 1180 | D1180A | D | 1.6e-08 | 4.6e-08 | 2.5e-08 | 2.900E-08 | 1.539E-08 | 1.600E-08 | 4.600E-08 |
| 23582 | 1180 | D1180R | D | 3.6e-07 | 3e-08 | 6.5e-08 | 1.517E-07 | 1.813E-07 | 3.000E-08 | 3.600E-07 |
| 23583 | 1180 | D1180N | D | 5.2e-08 | 5.8e-08 | 4.9e-08 | 5.300E-08 | 4.583E-09 | 4.900E-08 | 5.800E-08 |
| 23584 | 1180 | D1180D | D | 2.9e-08 | 4.1e-08 | 1.7e-07 | 8.000E-08 | 7.817E-08 | 2.900E-08 | 1.700E-07 |
| 23585 | 1180 | D1180C | D | 1.6e-08 | 7.4e-08 | 1.2e-07 | 7.000E-08 | 5.212E-08 | 1.600E-08 | 1.200E-07 |
| 23586 | 1180 | D1180E | D | 3.1e-08 | 4.9e-08 | 4.1e-08 | 4.033E-08 | 9.018E-09 | 3.100E-08 | 4.900E-08 |
| 23587 | 1180 | D1180Q | D | 5.3e-08 | 1.3e-07 | 2.8e-08 | 7.033E-08 | 5.316E-08 | 2.800E-08 | 1.300E-07 |
| 23588 | 1180 | D1180G | D | 3.9e-08 | 5.4e-08 | 6.2e-08 | 5.167E-08 | 1.168E-08 | 3.900E-08 | 6.200E-08 |
| 23589 | 1180 | D1180H | D | 4e-08 | 8.3e-08 | 4.2e-08 | 5.500E-08 | 2.427E-08 | 4.000E-08 | 8.300E-08 |
| 23590 | 1180 | D1180I | D | 4e-08 | 4.3e-08 | 5.2e-08 | 4.500E-08 | 6.245E-09 | 4.000E-08 | 5.200E-08 |
| 23591 | 1180 | D1180L | D | 1.8e-08 | 8.9e-08 | 5.5e-08 | 5.400E-08 | 3.551E-08 | 1.800E-08 | 8.900E-08 |
| 23592 | 1180 | D1180K | D | 1.3e-07 | 2.1e-08 | 2.8e-08 | 5.967E-08 | 6.101E-08 | 2.100E-08 | 1.300E-07 |
| 23593 | 1180 | D1180M | D | 1e-07 | 4.2e-08 | 3.9e-08 | 6.033E-08 | 3.439E-08 | 3.900E-08 | 1.000E-07 |
| 23594 | 1180 | D1180F | D | 1.7e-08 | 6e-08 | 2.8e-08 | 3.500E-08 | 2.234E-08 | 1.700E-08 | 6.000E-08 |
| 23595 | 1180 | D1180P | D | 2.9e-08 | 1.9e-08 | 6.3e-08 | 3.700E-08 | 2.307E-08 | 1.900E-08 | 6.300E-08 |
| 23596 | 1180 | D1180S | D | 3.6e-08 | 1e-07 | 5.9e-08 | 6.500E-08 | 3.242E-08 | 3.600E-08 | 1.000E-07 |
| 23597 | 1180 | D1180T | D | 4.8e-08 | 4.9e-08 | 5e-08 | 4.900E-08 | 1.000E-09 | 4.800E-08 | 5.000E-08 |
| 23598 | 1180 | D1180W | D | 3.5e-08 | 1.2e-07 | 5.6e-08 | 7.033E-08 | 4.428E-08 | 3.500E-08 | 1.200E-07 |
| 23599 | 1180 | D1180Y | D | 1.6e-08 | 1.6e-08 | 7.6e-08 | 3.600E-08 | 3.464E-08 | 1.600E-08 | 7.600E-08 |
| 23600 | 1180 | D1180V | D | 5.8e-09 | 5e-08 | 2.3e-08 | 2.627E-08 | 2.228E-08 | 5.800E-09 | 5.000E-08 |
| 23601 | 1181 | E1181A | D | 3.2e-08 | 2.2e-08 | 4.8e-08 | 3.400E-08 | 1.311E-08 | 2.200E-08 | 4.800E-08 |
| 23602 | 1181 | E1181R | D | 2e-08 | 4.1e-08 | 2.2e-08 | 2.767E-08 | 1.159E-08 | 2.000E-08 | 4.100E-08 |
| 23603 | 1181 | E1181N | D | 1.7e-08 | 2.2e-08 | 5.1e-08 | 3.000E-08 | 1.836E-08 | 1.700E-08 | 5.100E-08 |
| 23604 | 1181 | E1181D | D | 3.8e-08 | 5e-08 | 3.6e-08 | 4.133E-08 | 7.572E-09 | 3.600E-08 | 5.000E-08 |
| 23605 | 1181 | E1181C | D | 6.8e-08 | 4.9e-08 | 6.1e-09 | 4.103E-08 | 3.171E-08 | 6.100E-09 | 6.800E-08 |
| 23606 | 1181 | E1181E | D | 2.8e-08 | 7.7e-08 | 7.1e-08 | 5.867E-08 | 2.673E-08 | 2.800E-08 | 7.700E-08 |
| 23607 | 1181 | E1181Q | D | 3.2e-08 | 4.4e-08 | 3.4e-08 | 3.667E-08 | 6.429E-09 | 3.200E-08 | 4.400E-08 |
| 23608 | 1181 | E1181G | D | 2.3e-08 | 5e-08 | 1.5e-07 | 7.433E-08 | 6.691E-08 | 2.300E-08 | 1.500E-07 |
| 23609 | 1181 | E1181H | D | 7.3e-08 | 5.2e-08 | 2e-08 | 4.833E-08 | 2.669E-08 | 2.000E-08 | 7.300E-08 |
| 23610 | 1181 | E1181I | D | 4e-08 | 6.3e-08 | 2.6e-08 | 4.300E-08 | 1.868E-08 | 2.600E-08 | 6.300E-08 |
| 23611 | 1181 | E1181L | D | 4.8e-08 | 8.1e-08 | 3e-08 | 5.300E-08 | 2.587E-08 | 3.000E-08 | 8.100E-08 |
| 23612 | 1181 | E1181K | D | 1.1e-07 | 2.9e-08 | 5.4e-08 | 6.433E-08 | 4.148E-08 | 2.900E-08 | 1.100E-07 |
| 23613 | 1181 | E1181M | D | 8.6e-08 | 2.6e-08 | 2.4e-08 | 4.533E-08 | 3.523E-08 | 2.400E-08 | 8.600E-08 |
| 23614 | 1181 | E1181F | D | 1.3e-07 | 6.8e-08 | 8.2e-08 | 9.333E-08 | 3.252E-08 | 6.800E-08 | 1.300E-07 |
| 23615 | 1181 | E1181P | D | 8.7e-08 | 2e-08 | 3.1e-08 | 4.600E-08 | 3.593E-08 | 2.000E-08 | 8.700E-08 |
| 23616 | 1181 | E1181S | D | 3.5e-08 | 5.5e-08 | 3.4e-08 | 4.133E-08 | 1.185E-08 | 3.400E-08 | 5.500E-08 |
| 23617 | 1181 | E1181T | D | 1.1e-07 | 4e-08 | 4.7e-08 | 6.567E-08 | 3.855E-08 | 4.000E-08 | 1.100E-07 |
| 23618 | 1181 | E1181W | D | 8.4e-09 | 7.5e-08 | 9.1e-08 | 5.813E-08 | 4.381E-08 | 8.400E-09 | 9.100E-08 |
| 23619 | 1181 | E1181Y | D | 6e-08 | 8.3e-08 | 1e-08 | 5.100E-08 | 3.732E-08 | 1.000E-08 | 8.300E-08 |
| 23620 | 1181 | E1181V | D | 3.9e-08 | 6.9e-08 | 3.7e-08 | 4.833E-08 | 1.793E-08 | 3.700E-08 | 6.900E-08 |
| 23621 | 1182 | Q1182A | D | 1.8e-07 | 2e-08 | 5.5e-08 | 8.500E-08 | 8.411E-08 | 2.000E-08 | 1.800E-07 |
| 23622 | 1182 | Q1182R | D | 1.4e-07 | 6.4e-08 | 3.8e-08 | 8.067E-08 | 5.300E-08 | 3.800E-08 | 1.400E-07 |
| 23623 | 1182 | Q1182N | D | 4.1e-08 | 1.7e-08 | 3.9e-08 | 3.233E-08 | 1.332E-08 | 1.700E-08 | 4.100E-08 |
| 23624 | 1182 | Q1182D | D | 4.2e-08 | 5.2e-08 | 1.6e-08 | 3.667E-08 | 1.858E-08 | 1.600E-08 | 5.200E-08 |
| 23625 | 1182 | Q1182C | D | 4.5e-08 | 3.7e-08 | 7.7e-08 | 5.300E-08 | 2.117E-08 | 3.700E-08 | 7.700E-08 |
| 23626 | 1182 | Q1182E | D | 6.9e-08 | 2.5e-08 | 9.7e-08 | 6.367E-08 | 3.630E-08 | 2.500E-08 | 9.700E-08 |
| 23627 | 1182 | Q1182Q | D | 3.7e-08 | 4.9e-08 | 1.4e-08 | 3.333E-08 | 1.779E-08 | 1.400E-08 | 4.900E-08 |
| 23628 | 1182 | Q1182G | D | 5.5e-08 | 3.3e-08 | 4.6e-08 | 4.467E-08 | 1.106E-08 | 3.300E-08 | 5.500E-08 |
| 23629 | 1182 | Q1182H | D | 7.5e-08 | 6.5e-08 | 3.2e-08 | 5.733E-08 | 2.250E-08 | 3.200E-08 | 7.500E-08 |

| | | | | | | | | | | |
|---|---|---|---|---|---|---|---|---|---|---|
| 23630 | 1182 | Q1182I | D | 7.6e-08 | 4.2e-08 | 6.5e-08 | 6.100E-08 | 1.735E-08 | 4.200E-08 | 7.600E-08 |
| 23631 | 1182 | Q1182L | D | 4.9e-08 | 4.4e-08 | 1.2e-07 | 7.100E-08 | 4.251E-08 | 4.400E-08 | 1.200E-07 |
| 23632 | 1182 | Q1182K | D | 1.4e-08 | 2.4e-08 | 6.4e-08 | 3.400E-08 | 2.646E-08 | 1.400E-08 | 6.400E-08 |
| 23633 | 1182 | Q1182M | D | 1.7e-08 | 4.9e-08 | 4.7e-08 | 3.767E-08 | 1.793E-08 | 1.700E-08 | 4.900E-08 |
| 23634 | 1182 | Q1182F | D | 1.2e-07 | 3.9e-08 | 4.1e-08 | 6.667E-08 | 4.620E-08 | 3.900E-08 | 1.200E-07 |
| 23635 | 1182 | Q1182P | D | 6.4e-08 | 3.3e-08 | 5.2e-08 | 4.967E-08 | 1.563E-08 | 3.300E-08 | 6.400E-08 |
| 23636 | 1182 | Q1182S | D | 3.2e-08 | 4.2e-08 | 3.6e-08 | 3.667E-08 | 5.033E-09 | 3.200E-08 | 4.200E-08 |
| 23637 | 1182 | Q1182T | D | 5.8e-08 | 3.4e-08 | 6.3e-08 | 5.167E-08 | 1.550E-08 | 3.400E-08 | 6.300E-08 |
| 23638 | 1182 | Q1182W | D | 3.7e-08 | 4.8e-08 | 5e-08 | 4.500E-08 | 7.000E-09 | 3.700E-08 | 5.000E-08 |
| 23639 | 1182 | Q1182Y | D | 3.8e-08 | 1.4e-08 | 4.2e-08 | 3.133E-08 | 1.514E-08 | 1.400E-08 | 4.200E-08 |
| 23640 | 1182 | Q1182V | D | 4.7e-08 | 2.2e-08 | 4.4e-08 | 3.767E-08 | 1.365E-08 | 2.200E-08 | 4.700E-08 |
| 23641 | 1183 | L1183A | D | 3.3e-08 | 7.2e-08 | 3.9e-08 | 4.800E-08 | 2.100E-08 | 3.300E-08 | 7.200E-08 |
| 23642 | 1183 | L1183R | D | 4.2e-08 | 4.3e-08 | 3e-08 | 3.833E-08 | 7.234E-09 | 3.000E-08 | 4.300E-08 |
| 23643 | 1183 | L1183N | D | 4.6e-08 | 3e-08 | 8.5e-08 | 5.367E-08 | 2.829E-08 | 3.000E-08 | 8.500E-08 |
| 23644 | 1183 | L1183D | D | 6.4e-08 | 7.9e-08 | 1.1e-07 | 8.433E-08 | 2.346E-08 | 6.400E-08 | 1.100E-07 |
| 23645 | 1183 | L1183C | D | 2.3e-08 | 6.8e-08 | 9.9e-08 | 6.333E-08 | 3.821E-08 | 2.300E-08 | 9.900E-08 |
| 23646 | 1183 | L1183E | D | 5.7e-08 | 2.8e-07 | 4.4e-08 | 1.270E-07 | 1.327E-07 | 4.400E-08 | 2.800E-07 |
| 23647 | 1183 | L1183Q | D | 9.4e-08 | 5.3e-08 | 6.5e-08 | 7.067E-08 | 2.108E-08 | 5.300E-08 | 9.400E-08 |
| 23648 | 1183 | L1183G | D | 5.4e-08 | 1.5e-08 | 5.2e-08 | 4.033E-08 | 2.196E-08 | 1.500E-08 | 5.400E-08 |
| 23649 | 1183 | L1183H | D | 2.6e-08 | 5.1e-08 | 1.4e-07 | 7.233E-08 | 5.992E-08 | 2.600E-08 | 1.400E-07 |
| 23650 | 1183 | L1183I | D | 2.8e-08 | 2.5e-08 | 6.1e-08 | 3.800E-08 | 1.997E-08 | 2.500E-08 | 6.100E-08 |
| 23651 | 1183 | L1183L | D | 9.8e-08 | 1.6e-08 | 1.9e-08 | 4.433E-08 | 4.650E-08 | 1.600E-08 | 9.800E-08 |
| 23652 | 1183 | L1183K | D | 2.8e-08 | 2.7e-08 | 6.3e-08 | 3.933E-08 | 2.050E-08 | 2.700E-08 | 6.300E-08 |
| 23653 | 1183 | L1183M | D | 2e-08 | 4e-08 | 3.2e-08 | 3.067E-08 | 1.007E-08 | 2.000E-08 | 4.000E-08 |
| 23654 | 1183 | L1183F | D | 7.9e-08 | 6.3e-08 | 4.1e-08 | 6.100E-08 | 1.908E-08 | 4.100E-08 | 7.900E-08 |
| 23655 | 1183 | L1183P | D | 4.1e-08 | 5.7e-08 | 1.4e-08 | 3.733E-08 | 2.173E-08 | 1.400E-08 | 5.700E-08 |
| 23656 | 1183 | L1183S | D | 1.2e-08 | 3.9e-08 | 5e-08 | 3.367E-08 | 1.955E-08 | 1.200E-08 | 5.000E-08 |
| 23657 | 1183 | L1183T | D | 2.3e-08 | 4.5e-08 | 6.1e-08 | 4.300E-08 | 1.908E-08 | 2.300E-08 | 6.100E-08 |
| 23658 | 1183 | L1183W | D | 4.8e-08 | 3.1e-08 | 8.1e-08 | 5.333E-08 | 2.542E-08 | 3.100E-08 | 8.100E-08 |
| 23659 | 1183 | L1183Y | D | 6.4e-08 | 2.6e-08 | 1.4e-08 | 3.467E-08 | 2.610E-08 | 1.400E-08 | 6.400E-08 |
| 23660 | 1183 | L1183V | D | 1e-07 | 3.9e-08 | 7.9e-08 | 7.267E-08 | 3.099E-08 | 3.900E-08 | 1.000E-07 |
| 23661 | 1184 | K1184A | D | 3.1e-08 | 7.7e-08 | 5.3e-08 | 5.367E-08 | 2.301E-08 | 3.100E-08 | 7.700E-08 |
| 23662 | 1184 | K1184R | D | 2.9e-08 | 7.4e-08 | 1e-07 | 6.767E-08 | 3.592E-08 | 2.900E-08 | 1.000E-07 |
| 23663 | 1184 | K1184N | D | 1.8e-08 | 3.3e-08 | 9.3e-08 | 4.800E-08 | 3.969E-08 | 1.800E-08 | 9.300E-08 |
| 23664 | 1184 | K1184D | D | 2.3e-08 | 5.9e-08 | 4.4e-08 | 4.200E-08 | 1.808E-08 | 2.300E-08 | 5.900E-08 |
| 23665 | 1184 | K1184C | D | 1.2e-07 | 5.7e-08 | 9.7e-08 | 9.133E-08 | 3.188E-08 | 5.700E-08 | 1.200E-07 |
| 23666 | 1184 | K1184E | D | 1.2e-08 | 3.9e-08 | 3.2e-08 | 2.767E-08 | 1.401E-08 | 1.200E-08 | 3.900E-08 |
| 23667 | 1184 | K1184Q | D | 1.6e-07 | 4.1e-08 | 3.6e-08 | 7.900E-08 | 7.019E-08 | 3.600E-08 | 1.600E-07 |
| 23668 | 1184 | K1184G | D | 4.9e-08 | 2.8e-08 | 5.7e-08 | 4.467E-08 | 1.498E-08 | 2.800E-08 | 5.700E-08 |
| 23669 | 1184 | K1184H | D | 1.9e-08 | 3.3e-08 | 2.2e-08 | 2.467E-08 | 7.371E-09 | 1.900E-08 | 3.300E-08 |
| 23670 | 1184 | K1184I | D | 3.1e-08 | 6.3e-08 | 8.4e-08 | 5.933E-08 | 2.669E-08 | 3.100E-08 | 8.400E-08 |
| 23671 | 1184 | K1184L | D | 1.1e-07 | 5.1e-08 | 3.6e-08 | 6.567E-08 | 3.912E-08 | 3.600E-08 | 1.100E-07 |
| 23672 | 1184 | K1184K | D | 3.6e-08 | 1.5e-08 | 5.5e-08 | 3.533E-08 | 2.001E-08 | 1.500E-08 | 5.500E-08 |
| 23673 | 1184 | K1184M | D | 4.4e-08 | 5.6e-08 | 6.7e-08 | 5.567E-08 | 1.150E-08 | 4.400E-08 | 6.700E-08 |
| 23674 | 1184 | K1184F | D | 6.2e-08 | 2.9e-08 | 5e-08 | 4.700E-08 | 1.670E-08 | 2.900E-08 | 6.200E-08 |
| 23675 | 1184 | K1184P | D | 8e-08 | 3.8e-08 | 3e-08 | 4.933E-08 | 2.686E-08 | 3.000E-08 | 8.000E-08 |
| 23676 | 1184 | K1184S | D | 1e-07 | 7.2e-08 | 4.6e-08 | 7.267E-08 | 2.701E-08 | 4.600E-08 | 1.000E-07 |
| 23677 | 1184 | K1184T | D | 3.6e-08 | 1.9e-08 | 1.1e-07 | 5.500E-08 | 4.838E-08 | 1.900E-08 | 1.100E-07 |
| 23678 | 1184 | K1184W | D | 3.7e-08 | 5.2e-08 | 8.4e-08 | 5.767E-08 | 2.401E-08 | 3.700E-08 | 8.400E-08 |
| 23679 | 1184 | K1184Y | D | 4.9e-08 | 2.1e-08 | 2.1e-08 | 3.033E-08 | 1.617E-08 | 2.100E-08 | 4.900E-08 |
| 23680 | 1184 | K1184V | D | 1.2e-07 | 1.4e-08 | 1.8e-08 | 5.067E-08 | 6.008E-08 | 1.400E-08 | 1.200E-07 |
| 23681 | 1185 | S1185A | D | 5.5e-08 | 5.1e-08 | 3.8e-08 | 4.800E-08 | 8.888E-09 | 3.800E-08 | 5.500E-08 |
| 23682 | 1185 | S1185R | D | 5.4e-08 | 5.2e-08 | 2.7e-08 | 4.433E-08 | 1.504E-08 | 2.700E-08 | 5.400E-08 |
| 23683 | 1185 | S1185N | D | 4.1e-08 | 6.6e-08 | 6.8e-08 | 5.833E-08 | 1.504E-08 | 4.100E-08 | 6.800E-08 |
| 23684 | 1185 | S1185D | D | 1.6e-08 | 5.7e-08 | 2.8e-08 | 3.367E-08 | 2.108E-08 | 1.600E-08 | 5.700E-08 |
| 23685 | 1185 | S1185C | D | 4.3e-08 | 5.1e-08 | 1.1e-08 | 3.500E-08 | 2.117E-08 | 1.100E-08 | 5.100E-08 |
| 23686 | 1185 | S1185E | D | 8.9e-08 | 7.3e-08 | 2.5e-08 | 6.233E-08 | 3.331E-08 | 2.500E-08 | 8.900E-08 |
| 23687 | 1185 | S1185Q | D | 3.6e-08 | 5.4e-08 | 5.8e-08 | 4.933E-08 | 1.172E-08 | 3.600E-08 | 5.800E-08 |
| 23688 | 1185 | S1185G | D | 9.1e-08 | 2.9e-08 | 2.3e-07 | 1.167E-07 | 1.029E-07 | 2.900E-08 | 2.300E-07 |
| 23689 | 1185 | S1185H | D | 4.7e-08 | 4.1e-08 | 2.1e-08 | 3.633E-08 | 1.361E-08 | 2.100E-08 | 4.700E-08 |
| 23690 | 1185 | S1185I | D | 8.8e-08 | 4e-08 | 6.1e-08 | 6.300E-08 | 2.406E-08 | 4.000E-08 | 8.800E-08 |
| 23691 | 1185 | S1185L | D | 6.4e-08 | 1e-07 | 8e-08 | 8.133E-08 | 1.804E-08 | 6.400E-08 | 1.000E-07 |
| 23692 | 1185 | S1185K | D | 4.3e-08 | 2.3e-08 | 5.6e-08 | 4.067E-08 | 1.662E-08 | 2.300E-08 | 5.600E-08 |
| 23693 | 1185 | S1185M | D | 3.9e-08 | 1.4e-08 | 3.1e-08 | 2.800E-08 | 1.277E-08 | 1.400E-08 | 3.900E-08 |
| 23694 | 1185 | S1185F | D | 2.5e-08 | 2.9e-08 | 1.7e-08 | 2.367E-08 | 6.110E-09 | 1.700E-08 | 2.900E-08 |
| 23695 | 1185 | S1185P | D | 3.2e-08 | 8.3e-08 | 1.3e-08 | 4.267E-08 | 3.620E-08 | 1.300E-08 | 8.300E-08 |
| 23696 | 1185 | S1185S | D | 5.1e-08 | 4.8e-08 | 5.8e-08 | 5.233E-08 | 5.132E-09 | 4.800E-08 | 5.800E-08 |
| 23697 | 1185 | S1185T | D | 3.5e-08 | 1.2e-07 | 4.6e-08 | 6.700E-08 | 4.623E-08 | 3.500E-08 | 1.200E-07 |
| 23698 | 1185 | S1185W | D | 1.4e-07 | 7.6e-08 | 2.8e-08 | 8.133E-08 | 5.619E-08 | 2.800E-08 | 1.400E-07 |
| 23699 | 1185 | S1185Y | D | 3.7e-08 | 1.9e-08 | 7.7e-08 | 4.433E-08 | 2.969E-08 | 1.900E-08 | 7.700E-08 |
| 23700 | 1185 | S1185V | D | 6.5e-08 | 1.5e-08 | 5.4e-08 | 4.467E-08 | 2.627E-08 | 1.500E-08 | 6.500E-08 |
| 23701 | 1186 | G1186A | D | 6.6e-08 | 5.1e-08 | 2.9e-08 | 4.867E-08 | 1.861E-08 | 2.900E-08 | 6.600E-08 |
| 23702 | 1186 | G1186R | D | 5.7e-08 | 4e-08 | 5.6e-08 | 5.100E-08 | 9.539E-09 | 4.000E-08 | 5.700E-08 |
| 23703 | 1186 | G1186N | D | 3.7e-08 | 4.3e-08 | 5.7e-08 | 4.567E-08 | 1.026E-08 | 3.700E-08 | 5.700E-08 |
| 23704 | 1186 | G1186D | D | 6.8e-08 | 1.4e-08 | 4.3e-08 | 4.167E-08 | 2.702E-08 | 1.400E-08 | 6.800E-08 |
| 23705 | 1186 | G1186C | D | 3.2e-08 | 4.1e-08 | 6.4e-08 | 4.567E-08 | 1.650E-08 | 3.200E-08 | 6.400E-08 |
| 23706 | 1186 | G1186E | D | 5.5e-08 | 4.4e-08 | 9e-08 | 6.300E-08 | 2.402E-08 | 4.400E-08 | 9.000E-08 |
| 23707 | 1186 | G1186Q | D | 2.9e-08 | 2.2e-08 | 5e-08 | 3.367E-08 | 1.457E-08 | 2.200E-08 | 5.000E-08 |
| 23708 | 1186 | G1186G | D | 3.6e-08 | 8.7e-08 | 4.8e-08 | 5.700E-08 | 2.666E-08 | 3.600E-08 | 8.700E-08 |
| 23709 | 1186 | G1186H | D | 9.3e-08 | 6.4e-08 | 7.2e-08 | 7.633E-08 | 1.498E-08 | 6.400E-08 | 9.300E-08 |
| 23710 | 1186 | G1186I | D | 3.5e-08 | 8.5e-08 | 4.2e-08 | 5.400E-08 | 2.707E-08 | 3.500E-08 | 8.500E-08 |
| 23711 | 1186 | G1186L | D | 2.4e-08 | 6.2e-08 | 7.1e-08 | 5.233E-08 | 2.495E-08 | 2.400E-08 | 7.100E-08 |
| 23712 | 1186 | G1186K | D | 4.9e-08 | 8e-08 | 4.6e-08 | 5.833E-08 | 1.882E-08 | 4.600E-08 | 8.000E-08 |
| 23713 | 1186 | G1186M | D | 3.2e-08 | 2.8e-08 | 2.4e-08 | 2.800E-08 | 4.000E-09 | 2.400E-08 | 3.200E-08 |
| 23714 | 1186 | G1186F | D | 4.2e-08 | 8.1e-08 | 4.4e-08 | 5.567E-08 | 2.196E-08 | 4.200E-08 | 8.100E-08 |

| | | | | | | | | | | |
|---|---|---|---|---|---|---|---|---|---|---|
| 23715 | 1186 | G1186P | D | 7.5e-08 | 3.8e-08 | 4.4e-08 | 5.233E-08 | 1.986E-08 | 3.800E-08 | 7.500E-08 |
| 23716 | 1186 | G1186S | D | 3.1e-08 | 7.6e-08 | 3.5e-08 | 4.733E-08 | 2.491E-08 | 3.100E-08 | 7.600E-08 |
| 23717 | 1186 | G1186T | D | 6.8e-08 | 1.7e-08 | 5.9e-08 | 4.800E-08 | 2.722E-08 | 1.700E-08 | 6.800E-08 |
| 23718 | 1186 | G1186W | D | 8.1e-09 | 1.7e-08 | 4.6e-08 | 2.370E-08 | 1.982E-08 | 8.100E-09 | 4.600E-08 |
| 23719 | 1186 | G1186Y | D | 5.1e-08 | 4.5e-08 | 1.5e-07 | 8.200E-08 | 5.897E-08 | 4.500E-08 | 1.500E-07 |
| 23720 | 1186 | G1186V | D | 5e-08 | 8.6e-08 | 3e-08 | 5.533E-08 | 2.838E-08 | 3.000E-08 | 8.600E-08 |
| 23721 | 1187 | T1187A | D | 7.6e-08 | 1e-08 | 2e-08 | 3.533E-08 | 3.557E-08 | 1.000E-08 | 7.600E-08 |
| 23722 | 1187 | T1187R | D | 3.4e-08 | 1.4e-08 | 5e-08 | 3.267E-08 | 1.804E-08 | 1.400E-08 | 5.000E-08 |
| 23723 | 1187 | T1187N | D | 5.4e-08 | 1.8e-08 | 3.6e-08 | 3.600E-08 | 1.800E-08 | 1.800E-08 | 5.400E-08 |
| 23724 | 1187 | T1187D | D | 1.8e-08 | 3.7e-08 | 6.1e-08 | 3.867E-08 | 2.155E-08 | 1.800E-08 | 6.100E-08 |
| 23725 | 1187 | T1187C | D | 2e-08 | 7.6e-08 | 3.6e-08 | 4.400E-08 | 2.884E-08 | 2.000E-08 | 7.600E-08 |
| 23726 | 1187 | T1187E | D | 7e-08 | 3.7e-08 | 6.5e-08 | 5.733E-08 | 1.779E-08 | 3.700E-08 | 7.000E-08 |
| 23727 | 1187 | T1187Q | D | 3.3e-08 | 1.8e-08 | 6.6e-08 | 3.900E-08 | 2.456E-08 | 1.800E-08 | 6.600E-08 |
| 23728 | 1187 | T1187G | D | 7.3e-09 | 1.1e-08 | 3.2e-08 | 1.677E-08 | 1.332E-08 | 7.300E-09 | 3.200E-08 |
| 23729 | 1187 | T1187H | D | 2.7e-08 | 5.9e-08 | 2.5e-08 | 3.700E-08 | 1.908E-08 | 2.500E-08 | 5.900E-08 |
| 23730 | 1187 | T1187I | D | 7.9e-08 | 3.6e-08 | 7.9e-08 | 6.467E-08 | 2.483E-08 | 3.600E-08 | 7.900E-08 |
| 23731 | 1187 | T1187L | D | 4.7e-08 | 5.6e-08 | 4.2e-08 | 4.833E-08 | 7.095E-09 | 4.200E-08 | 5.600E-08 |
| 23732 | 1187 | T1187K | D | 1.2e-07 | 4.6e-08 | 1.3e-07 | 9.867E-08 | 4.588E-08 | 4.600E-08 | 1.300E-07 |
| 23733 | 1187 | T1187M | D | 9.5e-09 | 2.3e-08 | 5.4e-08 | 2.883E-08 | 2.282E-08 | 9.500E-09 | 5.400E-08 |
| 23734 | 1187 | T1187F | D | 4.5e-08 | 1e-08 | 4e-08 | 3.167E-08 | 1.893E-08 | 1.000E-08 | 4.500E-08 |
| 23735 | 1187 | T1187P | D | 4.2e-08 | 2.3e-08 | 3.6e-08 | 3.367E-08 | 9.713E-09 | 2.300E-08 | 4.200E-08 |
| 23736 | 1187 | T1187S | D | 5.4e-08 | 5.8e-08 | 8.9e-08 | 6.700E-08 | 1.916E-08 | 5.400E-08 | 8.900E-08 |
| 23737 | 1187 | T1187T | D | 3.8e-08 | 1.6e-08 | 6.4e-08 | 3.933E-08 | 2.403E-08 | 1.600E-08 | 6.400E-08 |
| 23738 | 1187 | T1187W | D | 9.8e-08 | 4.5e-08 | 4.9e-08 | 6.400E-08 | 2.951E-08 | 4.500E-08 | 9.800E-08 |
| 23739 | 1187 | T1187Y | D | 4.5e-08 | 1.8e-08 | 5.9e-08 | 4.067E-08 | 2.084E-08 | 1.800E-08 | 5.900E-08 |
| 23740 | 1187 | T1187V | D | 2e-08 | 2.9e-08 | 1.8e-08 | 2.233E-08 | 5.859E-09 | 1.800E-08 | 2.900E-08 |
| 23741 | 1188 | A1188A | D | 2.7e-08 | 6.1e-08 | 4.6e-08 | 4.467E-08 | 1.704E-08 | 2.700E-08 | 6.100E-08 |
| 23742 | 1188 | A1188R | D | 2e-08 | 3.4e-08 | 1.2e-07 | 5.800E-08 | 5.415E-08 | 2.000E-08 | 1.200E-07 |
| 23743 | 1188 | A1188N | D | 5.1e-08 | 6.9e-08 | 6.7e-08 | 6.233E-08 | 9.866E-09 | 5.100E-08 | 6.900E-08 |
| 23744 | 1188 | A1188D | D | 4.8e-08 | 1.7e-07 | 2.5e-08 | 8.100E-08 | 7.793E-08 | 2.500E-08 | 1.700E-07 |
| 23745 | 1188 | A1188C | D | 2.2e-08 | 8.9e-08 | 8.5e-08 | 6.533E-08 | 3.758E-08 | 2.200E-08 | 8.900E-08 |
| 23746 | 1188 | A1188E | D | 3.8e-08 | 2.8e-08 | 2.1e-08 | 2.900E-08 | 8.544E-09 | 2.100E-08 | 3.800E-08 |
| 23747 | 1188 | A1188Q | D | 1.1e-07 | 1.5e-08 | 2.4e-08 | 4.967E-08 | 5.244E-08 | 1.500E-08 | 1.100E-07 |
| 23748 | 1188 | A1188G | D | 5.3e-08 | 3.3e-08 | 4.4e-08 | 4.333E-08 | 1.002E-08 | 3.300E-08 | 5.300E-08 |
| 23749 | 1188 | A1188H | D | 5.3e-08 | 1.6e-07 | 4.1e-08 | 8.467E-08 | 6.552E-08 | 4.100E-08 | 1.600E-07 |
| 23750 | 1188 | A1188I | D | 3.1e-08 | 2e-08 | 4.9e-08 | 3.333E-08 | 1.464E-08 | 2.000E-08 | 4.900E-08 |
| 23751 | 1188 | A1188L | D | 6e-08 | 1.8e-08 | 5.6e-08 | 4.467E-08 | 2.318E-08 | 1.800E-08 | 6.000E-08 |
| 23752 | 1188 | A1188K | D | 3.9e-08 | 4.5e-08 | 3.1e-08 | 3.833E-08 | 7.024E-09 | 3.100E-08 | 4.500E-08 |
| 23753 | 1188 | A1188M | D | 4.2e-08 | 5.5e-08 | 1.7e-07 | 8.900E-08 | 7.045E-08 | 4.200E-08 | 1.700E-07 |
| 23754 | 1188 | A1188F | D | 1e-07 | 7.9e-08 | 7e-08 | 8.300E-08 | 1.539E-08 | 7.000E-08 | 1.000E-07 |
| 23755 | 1188 | A1188P | D | 2.6e-08 | 2.5e-08 | 3.7e-08 | 2.933E-08 | 6.658E-09 | 2.500E-08 | 3.700E-08 |
| 23756 | 1188 | A1188S | D | 2.8e-08 | 5.3e-08 | 3.8e-08 | 3.967E-08 | 1.258E-08 | 2.800E-08 | 5.300E-08 |
| 23757 | 1188 | A1188T | D | 4.5e-08 | 4.3e-08 | 2.4e-08 | 3.733E-08 | 1.159E-08 | 2.400E-08 | 4.500E-08 |
| 23758 | 1188 | A1188W | D | 5.5e-08 | 4.1e-08 | 5e-08 | 4.867E-08 | 7.095E-09 | 4.100E-08 | 5.500E-08 |
| 23759 | 1188 | A1188Y | D | 4.1e-08 | 2.3e-08 | 2.4e-08 | 2.933E-08 | 1.012E-08 | 2.300E-08 | 4.100E-08 |
| 23760 | 1188 | A1188V | D | 1.8e-08 | 1.3e-07 | 2.9e-08 | 5.900E-08 | 6.173E-08 | 1.800E-08 | 1.300E-07 |
| 23761 | 1189 | S1189A | D | 5.2e-08 | 1.3e-08 | 8.1e-08 | 4.867E-08 | 3.412E-08 | 1.300E-08 | 8.100E-08 |
| 23762 | 1189 | S1189R | D | 5.1e-08 | 7.4e-08 | 7.6e-08 | 6.700E-08 | 1.389E-08 | 5.100E-08 | 7.600E-08 |
| 23763 | 1189 | S1189N | D | 4.9e-08 | 3.2e-08 | 8.2e-08 | 5.433E-08 | 2.542E-08 | 3.200E-08 | 8.200E-08 |
| 23764 | 1189 | S1189D | D | 3.3e-08 | 1.6e-08 | 5.5e-08 | 3.467E-08 | 1.955E-08 | 1.600E-08 | 5.500E-08 |
| 23765 | 1189 | S1189C | D | 1.5e-07 | 8.4e-08 | 5.3e-08 | 9.567E-08 | 4.954E-08 | 5.300E-08 | 1.500E-07 |
| 23766 | 1189 | S1189E | D | 6.2e-08 | 1.2e-07 | 6.7e-08 | 8.300E-08 | 3.214E-08 | 6.200E-08 | 1.200E-07 |
| 23767 | 1189 | S1189Q | D | 3.9e-08 | 7e-08 | 2e-08 | 4.300E-08 | 2.524E-08 | 2.000E-08 | 7.000E-08 |
| 23768 | 1189 | S1189G | D | 5.5e-08 | 3.7e-08 | 1.2e-07 | 7.067E-08 | 4.366E-08 | 3.700E-08 | 1.200E-07 |
| 23769 | 1189 | S1189H | D | 8e-08 | 6.2e-08 | 3.4e-08 | 5.867E-08 | 2.318E-08 | 3.400E-08 | 8.000E-08 |
| 23770 | 1189 | S1189I | D | 4.2e-08 | 2.8e-08 | 5.8e-08 | 4.267E-08 | 1.501E-08 | 2.800E-08 | 5.800E-08 |
| 23771 | 1189 | S1189L | D | 4.1e-08 | 1.8e-07 | 3.2e-08 | 8.433E-08 | 8.297E-08 | 3.200E-08 | 1.800E-07 |
| 23772 | 1189 | S1189K | D | 3.3e-08 | 9.8e-08 | 1.2e-07 | 8.367E-08 | 4.524E-08 | 3.300E-08 | 1.200E-07 |
| 23773 | 1189 | S1189M | D | 7.5e-09 | 3.5e-08 | 1e-07 | 4.750E-08 | 4.750E-08 | 7.500E-09 | 1.000E-07 |
| 23774 | 1189 | S1189F | D | 3.6e-08 | 2.5e-08 | 1.2e-07 | 6.033E-08 | 5.196E-08 | 2.500E-08 | 1.200E-07 |
| 23775 | 1189 | S1189P | D | 9.9e-08 | 7e-08 | 4e-08 | 6.967E-08 | 2.950E-08 | 4.000E-08 | 9.900E-08 |
| 23776 | 1189 | S1189S | D | 2e-08 | 4.9e-08 | 6e-08 | 4.300E-08 | 2.066E-08 | 2.000E-08 | 6.000E-08 |
| 23777 | 1189 | S1189T | D | 1e-07 | 1.2e-08 | 4e-08 | 5.067E-08 | 4.496E-08 | 1.200E-08 | 1.000E-07 |
| 23778 | 1189 | S1189W | D | 7.6e-08 | 2.9e-08 | 6.4e-08 | 5.633E-08 | 2.442E-08 | 2.900E-08 | 7.600E-08 |
| 23779 | 1189 | S1189Y | D | 2.4e-08 | 5.2e-08 | 2.3e-08 | 3.300E-08 | 1.646E-08 | 2.300E-08 | 5.200E-08 |
| 23780 | 1189 | S1189V | D | 1.6e-08 | 3.6e-08 | 4.9e-08 | 3.367E-08 | 1.662E-08 | 1.600E-08 | 4.900E-08 |
| 23781 | 1190 | V1190A | D | 3.8e-08 | 3.1e-08 | 7.8e-08 | 4.900E-08 | 2.536E-08 | 3.100E-08 | 7.800E-08 |
| 23782 | 1190 | V1190R | D | 6.2e-08 | 5.5e-08 | 3.7e-08 | 5.133E-08 | 1.290E-08 | 3.700E-08 | 6.200E-08 |
| 23783 | 1190 | V1190N | D | 3.4e-08 | 5.3e-08 | 5.4e-08 | 4.700E-08 | 1.127E-08 | 3.400E-08 | 5.400E-08 |
| 23784 | 1190 | V1190D | D | 9e-08 | 4.3e-08 | 6.7e-08 | 6.667E-08 | 2.350E-08 | 4.300E-08 | 9.000E-08 |
| 23785 | 1190 | V1190C | D | 4.7e-08 | 1.1e-07 | 1.9e-08 | 5.867E-08 | 4.661E-08 | 1.900E-08 | 1.100E-07 |
| 23786 | 1190 | V1190E | D | 4e-08 | 3.1e-08 | 4.9e-08 | 4.000E-08 | 9.000E-09 | 3.100E-08 | 4.900E-08 |
| 23787 | 1190 | V1190Q | D | 1.3e-08 | 4.4e-08 | 4.1e-08 | 3.267E-08 | 1.710E-08 | 1.300E-08 | 4.400E-08 |
| 23788 | 1190 | V1190G | D | 4.2e-08 | 1.6e-07 | 3.4e-08 | 7.867E-08 | 7.055E-08 | 3.400E-08 | 1.600E-07 |
| 23789 | 1190 | V1190H | D | 7.7e-08 | 3.5e-08 | 5.4e-08 | 5.533E-08 | 2.103E-08 | 3.500E-08 | 7.700E-08 |
| 23790 | 1190 | V1190I | D | 2.9e-08 | 8.6e-08 | 2.4e-08 | 4.633E-08 | 3.444E-08 | 2.400E-08 | 8.600E-08 |
| 23791 | 1190 | V1190L | D | 7.7e-08 | 6.8e-08 | 8.3e-08 | 7.600E-08 | 7.550E-09 | 6.800E-08 | 8.300E-08 |
| 23792 | 1190 | V1190K | D | 4.4e-08 | 4.7e-08 | 3.7e-08 | 4.267E-08 | 5.132E-09 | 3.700E-08 | 4.700E-08 |
| 23793 | 1190 | V1190M | D | 5.7e-08 | 4.9e-08 | 8.2e-08 | 6.267E-08 | 1.721E-08 | 4.900E-08 | 8.200E-08 |
| 23794 | 1190 | V1190F | D | 3.7e-08 | 3.1e-08 | 2.2e-08 | 3.000E-08 | 7.550E-09 | 2.200E-08 | 3.700E-08 |
| 23795 | 1190 | V1190P | D | 6.4e-08 | 2.7e-08 | 5.7e-08 | 4.933E-08 | 1.966E-08 | 2.700E-08 | 6.400E-08 |
| 23796 | 1190 | V1190S | D | 1.3e-08 | 9e-08 | 2.2e-08 | 4.167E-08 | 4.210E-08 | 1.300E-08 | 9.000E-08 |
| 23797 | 1190 | V1190T | D | 4.5e-08 | 3.8e-08 | 4.3e-08 | 4.200E-08 | 3.606E-09 | 3.800E-08 | 4.500E-08 |
| 23798 | 1190 | V1190W | D | 2.2e-08 | 8.1e-08 | 5.4e-08 | 5.233E-08 | 2.954E-08 | 2.200E-08 | 8.100E-08 |
| 23799 | 1190 | V1190Y | D | 4.6e-08 | 8.4e-08 | 5.5e-08 | 6.167E-08 | 1.986E-08 | 4.600E-08 | 8.400E-08 |

| | | | | | | | | | | |
|---|---|---|---|---|---|---|---|---|---|---|
| 23800 | 1190 | V1190V | D | 4.5e-08 | 1.8e-08 | 6.8e-08 | 4.367E-08 | 2.503E-08 | 1.800E-08 | 6.800E-08 |
| 23801 | 1191 | V1191A | D | 4.4e-08 | 1.3e-08 | 4.1e-08 | 3.267E-08 | 1.710E-08 | 1.300E-08 | 4.400E-08 |
| 23802 | 1191 | V1191R | D | 4.6e-08 | 1.5e-07 | 8.3e-08 | 9.300E-08 | 5.272E-08 | 4.600E-08 | 1.500E-07 |
| 23803 | 1191 | V1191N | D | 5.1e-08 | 6.1e-08 | 4.2e-08 | 5.133E-08 | 9.504E-09 | 4.200E-08 | 6.100E-08 |
| 23804 | 1191 | V1191D | D | 2.5e-08 | 7.3e-08 | 4.7e-08 | 4.833E-08 | 2.403E-08 | 2.500E-08 | 7.300E-08 |
| 23805 | 1191 | V1191C | D | 3.3e-08 | 4.9e-08 | 8.5e-08 | 5.567E-08 | 2.663E-08 | 3.300E-08 | 8.500E-08 |
| 23806 | 1191 | V1191E | D | 2.3e-08 | 4e-08 | 4.4e-08 | 3.567E-08 | 1.115E-08 | 2.300E-08 | 4.400E-08 |
| 23807 | 1191 | V1191Q | D | 8.7e-08 | 8.8e-08 | 4.6e-08 | 7.367E-08 | 2.397E-08 | 4.600E-08 | 8.800E-08 |
| 23808 | 1191 | V1191G | D | 1.6e-07 | 7.9e-08 | 3.5e-08 | 9.133E-08 | 6.341E-08 | 3.500E-08 | 1.600E-07 |
| 23809 | 1191 | V1191H | D | 4.3e-08 | 1.8e-08 | 3.6e-08 | 3.233E-08 | 1.290E-08 | 1.800E-08 | 4.300E-08 |
| 23810 | 1191 | V1191I | D | 7.6e-08 | 2.5e-08 | 2.7e-08 | 4.267E-08 | 2.888E-08 | 2.500E-08 | 7.600E-08 |
| 23811 | 1191 | V1191L | D | 2.9e-08 | 3.8e-08 | 2.8e-08 | 3.167E-08 | 5.508E-09 | 2.800E-08 | 3.800E-08 |
| 23812 | 1191 | V1191K | D | 1.2e-07 | 3.5e-08 | 1.9e-08 | 5.800E-08 | 5.429E-08 | 1.900E-08 | 1.200E-07 |
| 23813 | 1191 | V1191M | D | 4.2e-08 | 9.9e-09 | 3.1e-08 | 2.763E-08 | 1.631E-08 | 9.900E-09 | 4.200E-08 |
| 23814 | 1191 | V1191F | D | 3.9e-08 | 3e-08 | 6e-08 | 4.300E-08 | 1.539E-08 | 3.000E-08 | 6.000E-08 |
| 23815 | 1191 | V1191P | D | 5e-08 | 3.7e-08 | 1.3e-08 | 3.333E-08 | 1.877E-08 | 1.300E-08 | 5.000E-08 |
| 23816 | 1191 | V1191S | D | 1.3e-08 | 6.2e-08 | 5.5e-08 | 4.333E-08 | 2.650E-08 | 1.300E-08 | 6.200E-08 |
| 23817 | 1191 | V1191T | D | 2.7e-07 | 4.2e-08 | 2.7e-08 | 1.130E-07 | 1.362E-07 | 2.700E-08 | 2.700E-07 |
| 23818 | 1191 | V1191W | D | 4.3e-08 | 5e-08 | 3.8e-08 | 4.367E-08 | 6.028E-09 | 3.800E-08 | 5.000E-08 |
| 23819 | 1191 | V1191Y | D | 2.7e-08 | 7.5e-08 | 3.1e-08 | 4.433E-08 | 2.663E-08 | 2.700E-08 | 7.500E-08 |
| 23820 | 1191 | V1191V | D | 1.8e-08 | 2e-08 | 7.5e-08 | 3.767E-08 | 3.235E-08 | 1.800E-08 | 7.500E-08 |
| 23821 | 1192 | C1192A | D | 2.8e-08 | 2.7e-08 | 5.8e-08 | 3.767E-08 | 1.762E-08 | 2.700E-08 | 5.800E-08 |
| 23822 | 1192 | C1192R | D | 8.6e-08 | 2.7e-08 | 5.4e-08 | 5.567E-08 | 2.954E-08 | 2.700E-08 | 8.600E-08 |
| 23823 | 1192 | C1192N | D | 2.4e-08 | 2.2e-08 | 1e-07 | 4.867E-08 | 4.447E-08 | 2.200E-08 | 1.000E-07 |
| 23824 | 1192 | C1192D | D | 1.5e-08 | 2.7e-08 | 5.7e-08 | 3.300E-08 | 2.163E-08 | 1.500E-08 | 5.700E-08 |
| 23825 | 1192 | C1192C | D | 2.4e-08 | 3.8e-08 | 6.8e-08 | 4.333E-08 | 2.248E-08 | 2.400E-08 | 6.800E-08 |
| 23826 | 1192 | C1192E | D | 5.4e-08 | 1.3e-08 | 2.2e-08 | 2.967E-08 | 2.155E-08 | 1.300E-08 | 5.400E-08 |
| 23827 | 1192 | C1192Q | D | 3.5e-08 | 4.4e-08 | 2.5e-08 | 3.467E-08 | 9.504E-09 | 2.500E-08 | 4.400E-08 |
| 23828 | 1192 | C1192G | D | 5.9e-08 | 3.4e-08 | 6.6e-08 | 5.300E-08 | 1.682E-08 | 3.400E-08 | 6.600E-08 |
| 23829 | 1192 | C1192H | D | 1.1e-07 | 4.5e-08 | 1.9e-08 | 5.800E-08 | 4.687E-08 | 1.900E-08 | 1.100E-07 |
| 23830 | 1192 | C1192I | D | 3.8e-08 | 4.6e-08 | 7.5e-08 | 5.300E-08 | 1.947E-08 | 3.800E-08 | 7.500E-08 |
| 23831 | 1192 | C1192L | D | 3.7e-08 | 3.9e-08 | 1.8e-08 | 3.133E-08 | 1.159E-08 | 1.800E-08 | 3.900E-08 |
| 23832 | 1192 | C1192K | D | 2.4e-08 | 8.5e-08 | 2.7e-08 | 4.533E-08 | 3.439E-08 | 2.400E-08 | 8.500E-08 |
| 23833 | 1192 | C1192M | D | 2.7e-08 | 5.7e-08 | 1.1e-07 | 6.467E-08 | 4.203E-08 | 2.700E-08 | 1.100E-07 |
| 23834 | 1192 | C1192F | D | 5.2e-08 | 3.1e-08 | 3.6e-08 | 3.967E-08 | 1.097E-08 | 3.100E-08 | 5.200E-08 |
| 23835 | 1192 | C1192P | D | 4e-08 | 9.8e-08 | 4.5e-08 | 6.100E-08 | 3.214E-08 | 4.000E-08 | 9.800E-08 |
| 23836 | 1192 | C1192S | D | 8.4e-08 | 1.2e-07 | 1.6e-07 | 1.213E-07 | 3.802E-08 | 8.400E-08 | 1.600E-07 |
| 23837 | 1192 | C1192T | D | 4.2e-08 | 5.7e-08 | 4.6e-08 | 4.833E-08 | 7.767E-09 | 4.200E-08 | 5.700E-08 |
| 23838 | 1192 | C1192W | D | 6.7e-08 | 3.9e-08 | 2.7e-08 | 4.433E-08 | 2.053E-08 | 2.700E-08 | 6.700E-08 |
| 23839 | 1192 | C1192Y | D | 2.8e-08 | 7.8e-08 | 1e-07 | 6.867E-08 | 3.690E-08 | 2.800E-08 | 1.000E-07 |
| 23840 | 1192 | C1192V | D | 3.3e-08 | 2.5e-08 | 3.9e-08 | 3.233E-08 | 7.024E-09 | 2.500E-08 | 3.900E-08 |
| 23841 | 1193 | L1193A | D | 7.1e-08 | 3.6e-08 | 6.1e-08 | 5.600E-08 | 1.803E-08 | 3.600E-08 | 7.100E-08 |
| 23842 | 1193 | L1193R | D | 4.7e-08 | 3.2e-08 | 7e-08 | 4.967E-08 | 1.914E-08 | 3.200E-08 | 7.000E-08 |
| 23843 | 1193 | L1193N | D | 5.1e-08 | 7.9e-08 | 2.3e-08 | 5.100E-08 | 2.800E-08 | 2.300E-08 | 7.900E-08 |
| 23844 | 1193 | L1193D | D | 3.4e-08 | 4.9e-08 | 8.7e-08 | 5.667E-08 | 2.732E-08 | 3.400E-08 | 8.700E-08 |
| 23845 | 1193 | L1193C | D | 1.7e-08 | 2.3e-08 | 2.5e-08 | 2.167E-08 | 4.163E-09 | 1.700E-08 | 2.500E-08 |
| 23846 | 1193 | L1193E | D | 3.4e-08 | 1.4e-07 | 9.3e-09 | 6.110E-08 | 6.944E-08 | 9.300E-09 | 1.400E-07 |
| 23847 | 1193 | L1193Q | D | 1.7e-08 | 7e-08 | 4.7e-08 | 4.467E-08 | 2.658E-08 | 1.700E-08 | 7.000E-08 |
| 23848 | 1193 | L1193G | D | 9.8e-08 | 9.4e-08 | 5.6e-08 | 8.267E-08 | 2.318E-08 | 5.600E-08 | 9.800E-08 |
| 23849 | 1193 | L1193H | D | 2.2e-08 | 9e-08 | 3.1e-08 | 4.767E-08 | 3.694E-08 | 2.200E-08 | 9.000E-08 |
| 23850 | 1193 | L1193I | D | 4.8e-08 | 4.5e-08 | 3.9e-08 | 4.400E-08 | 4.583E-09 | 3.900E-08 | 4.800E-08 |
| 23851 | 1193 | L1193L | D | 3.4e-08 | 3.4e-08 | 2.2e-08 | 3.000E-08 | 6.928E-09 | 2.200E-08 | 3.400E-08 |
| 23852 | 1193 | L1193K | D | 2.4e-08 | 1.6e-08 | 7.8e-08 | 3.933E-08 | 3.372E-08 | 1.600E-08 | 7.800E-08 |
| 23853 | 1193 | L1193M | D | 7.4e-08 | 5.4e-08 | 3.3e-08 | 5.367E-08 | 2.050E-08 | 3.300E-08 | 7.400E-08 |
| 23854 | 1193 | L1193F | D | 5.6e-08 | 5.4e-08 | 5.1e-08 | 5.367E-08 | 2.517E-09 | 5.100E-08 | 5.600E-08 |
| 23855 | 1193 | L1193P | D | 6.5e-08 | 4.5e-08 | 1.5e-07 | 8.667E-08 | 5.575E-08 | 4.500E-08 | 1.500E-07 |
| 23856 | 1193 | L1193S | D | 2.8e-08 | 3.7e-08 | 5.3e-08 | 3.933E-08 | 1.266E-08 | 2.800E-08 | 5.300E-08 |
| 23857 | 1193 | L1193T | D | 4.8e-08 | 4.5e-08 | 5e-08 | 4.767E-08 | 2.517E-09 | 4.500E-08 | 5.000E-08 |
| 23858 | 1193 | L1193W | D | 3e-08 | 3.2e-08 | 5.8e-08 | 4.000E-08 | 1.562E-08 | 3.000E-08 | 5.800E-08 |
| 23859 | 1193 | L1193Y | D | 5e-08 | 2.8e-08 | 5.2e-08 | 4.333E-08 | 1.332E-08 | 2.800E-08 | 5.200E-08 |
| 23860 | 1193 | L1193V | D | 5.2e-08 | 3.2e-08 | 2.6e-08 | 3.667E-08 | 1.361E-08 | 2.600E-08 | 5.200E-08 |
| 23861 | 1194 | L1194A | D | 5.6e-08 | 4.4e-08 | 1.4e-08 | 3.800E-08 | 2.163E-08 | 1.400E-08 | 5.600E-08 |
| 23862 | 1194 | L1194R | D | 1.1e-07 | 1e-07 | 8.9e-08 | 9.967E-08 | 1.050E-08 | 8.900E-08 | 1.100E-07 |
| 23863 | 1194 | L1194N | D | 3.1e-08 | 2.5e-08 | 2.3e-08 | 2.633E-08 | 4.163E-09 | 2.300E-08 | 3.100E-08 |
| 23864 | 1194 | L1194D | D | 5.7e-08 | 3.8e-08 | 5.8e-08 | 5.100E-08 | 1.127E-08 | 3.800E-08 | 5.800E-08 |
| 23865 | 1194 | L1194C | D | 2.9e-08 | 2.5e-08 | 1.6e-08 | 2.333E-08 | 6.658E-09 | 1.600E-08 | 2.900E-08 |
| 23866 | 1194 | L1194E | D | 5e-08 | 3.6e-08 | 4e-08 | 4.200E-08 | 7.211E-09 | 3.600E-08 | 5.000E-08 |
| 23867 | 1194 | L1194Q | D | 2.8e-08 | 4e-08 | 5.1e-08 | 3.967E-08 | 1.150E-08 | 2.800E-08 | 5.100E-08 |
| 23868 | 1194 | L1194G | D | 3.9e-08 | 9.7e-08 | 8.2e-08 | 7.267E-08 | 3.011E-08 | 3.900E-08 | 9.700E-08 |
| 23869 | 1194 | L1194H | D | 1.1e-07 | 7.7e-08 | 6.4e-08 | 8.367E-08 | 2.371E-08 | 6.400E-08 | 1.100E-07 |
| 23870 | 1194 | L1194I | D | 2.7e-08 | 7e-08 | 5.2e-08 | 4.967E-08 | 2.159E-08 | 2.700E-08 | 7.000E-08 |
| 23871 | 1194 | L1194L | D | 4e-08 | 5e-08 | 2.1e-08 | 3.700E-08 | 1.473E-08 | 2.100E-08 | 5.000E-08 |
| 23872 | 1194 | L1194K | D | 4.5e-08 | 3.6e-08 | 2.6e-08 | 3.567E-08 | 9.504E-09 | 2.600E-08 | 4.500E-08 |
| 23873 | 1194 | L1194M | D | 6e-08 | 4.8e-08 | 1.7e-08 | 4.167E-08 | 2.219E-08 | 1.700E-08 | 6.000E-08 |
| 23874 | 1194 | L1194F | D | 7.7e-08 | 2.6e-08 | 5e-08 | 5.100E-08 | 2.551E-08 | 2.600E-08 | 7.700E-08 |
| 23875 | 1194 | L1194P | D | 6.5e-08 | 6e-08 | 4e-08 | 5.500E-08 | 1.323E-08 | 4.000E-08 | 6.500E-08 |
| 23876 | 1194 | L1194S | D | 2.5e-08 | 1.8e-08 | 1.6e-07 | 6.767E-08 | 8.004E-08 | 1.800E-08 | 1.600E-07 |
| 23877 | 1194 | L1194T | D | 2e-08 | 3.4e-08 | 1.4e-07 | 6.467E-08 | 6.562E-08 | 2.000E-08 | 1.400E-07 |
| 23878 | 1194 | L1194W | D | 6.2e-08 | 4.3e-08 | 3.7e-08 | 4.733E-08 | 1.305E-08 | 3.700E-08 | 6.200E-08 |
| 23879 | 1194 | L1194Y | D | 5.2e-08 | 5.7e-08 | 3.4e-08 | 4.767E-08 | 1.210E-08 | 3.400E-08 | 5.700E-08 |
| 23880 | 1194 | L1194V | D | 4.7e-08 | 1.2e-07 | 5.4e-08 | 7.367E-08 | 4.028E-08 | 4.700E-08 | 1.200E-07 |
| 23881 | 1195 | N1195A | D | 2.1e-08 | 5.1e-08 | 6.1e-08 | 4.433E-08 | 2.082E-08 | 2.100E-08 | 6.100E-08 |
| 23882 | 1195 | N1195R | D | 3e-08 | 2e-08 | 4.7e-08 | 3.233E-08 | 1.365E-08 | 2.000E-08 | 4.700E-08 |
| 23883 | 1195 | N1195N | D | 6e-08 | 1.2e-07 | 2.1e-08 | 6.700E-08 | 4.987E-08 | 2.100E-08 | 1.200E-07 |
| 23884 | 1195 | N1195D | D | 2.5e-08 | 6.6e-08 | 3.1e-08 | 4.067E-08 | 2.214E-08 | 2.500E-08 | 6.600E-08 |

| | | | | | | | | | | |
|---|---|---|---|---|---|---|---|---|---|---|
| 23885 | 1195 | N1195C | D | 6.9e-08 | 4.3e-08 | 3.7e-08 | 4.967E-08 | 1.701E-08 | 3.700E-08 | 6.900E-08 |
| 23886 | 1195 | N1195E | D | 4.8e-08 | 3.7e-08 | 5e-08 | 4.500E-08 | 7.000E-09 | 3.700E-08 | 5.000E-08 |
| 23887 | 1195 | N1195Q | D | 3.9e-08 | 4.8e-08 | 2.7e-08 | 3.800E-08 | 1.054E-08 | 2.700E-08 | 4.800E-08 |
| 23888 | 1195 | N1195G | D | 2e-08 | 2.7e-08 | 4.1e-08 | 2.933E-08 | 1.069E-08 | 2.000E-08 | 4.100E-08 |
| 23889 | 1195 | N1195H | D | 3.2e-08 | 1.3e-08 | 6.7e-08 | 3.733E-08 | 2.739E-08 | 1.300E-08 | 6.700E-08 |
| 23890 | 1195 | N1195I | D | 1.9e-08 | 7.3e-08 | 3.3e-08 | 4.167E-08 | 2.802E-08 | 1.900E-08 | 7.300E-08 |
| 23891 | 1195 | N1195L | D | 8.2e-08 | 6.3e-08 | 1.2e-07 | 8.833E-08 | 2.902E-08 | 6.300E-08 | 1.200E-07 |
| 23892 | 1195 | N1195K | D | 4.3e-08 | 1.4e-08 | 1.1e-07 | 5.567E-08 | 4.924E-08 | 1.400E-08 | 1.100E-07 |
| 23893 | 1195 | N1195M | D | 4.3e-08 | 2e-08 | 6.3e-08 | 4.200E-08 | 2.152E-08 | 2.000E-08 | 6.300E-08 |
| 23894 | 1195 | N1195F | D | 7.4e-08 | 1.1e-07 | 8.1e-08 | 8.833E-08 | 1.909E-08 | 7.400E-08 | 1.100E-07 |
| 23895 | 1195 | N1195P | D | 1.8e-08 | 1.5e-08 | 5.1e-08 | 2.800E-08 | 1.997E-08 | 1.500E-08 | 5.100E-08 |
| 23896 | 1195 | N1195S | D | 4.8e-08 | 5.3e-08 | 6.2e-08 | 5.433E-08 | 7.095E-09 | 4.800E-08 | 6.200E-08 |
| 23897 | 1195 | N1195T | D | 4.1e-08 | 4.7e-08 | 6.4e-08 | 5.067E-08 | 1.193E-08 | 4.100E-08 | 6.400E-08 |
| 23898 | 1195 | N1195W | D | 3.1e-08 | 5.6e-08 | 1.1e-07 | 6.567E-08 | 4.038E-08 | 3.100E-08 | 1.100E-07 |
| 23899 | 1195 | N1195Y | D | 2.4e-08 | 5.4e-08 | 7.4e-08 | 5.067E-08 | 2.517E-08 | 2.400E-08 | 7.400E-08 |
| 23900 | 1195 | N1195V | D | 3.1e-08 | 1.7e-08 | 3e-08 | 2.600E-08 | 7.810E-09 | 1.700E-08 | 3.100E-08 |
| 23901 | 1196 | N1196A | D | 5.7e-08 | 5.3e-08 | 1.4e-08 | 4.133E-08 | 2.376E-08 | 1.400E-08 | 5.700E-08 |
| 23902 | 1196 | N1196R | D | 9e-08 | 3.6e-08 | 2.9e-08 | 5.167E-08 | 3.338E-08 | 2.900E-08 | 9.000E-08 |
| 23903 | 1196 | N1196N | D | 8.2e-08 | 4.8e-08 | 3.7e-08 | 5.567E-08 | 2.346E-08 | 3.700E-08 | 8.200E-08 |
| 23904 | 1196 | N1196D | D | 4e-08 | 9e-09 | 5.2e-08 | 3.367E-08 | 2.219E-08 | 9.000E-09 | 5.200E-08 |
| 23905 | 1196 | N1196C | D | 3.5e-08 | 5.3e-08 | 2.2e-08 | 3.667E-08 | 1.557E-08 | 2.200E-08 | 5.300E-08 |
| 23906 | 1196 | N1196E | D | 2.4e-08 | 2.7e-08 | 4e-08 | 3.033E-08 | 8.505E-09 | 2.400E-08 | 4.000E-08 |
| 23907 | 1196 | N1196Q | D | 7.1e-08 | 2.9e-08 | 4.3e-08 | 4.767E-08 | 2.139E-08 | 2.900E-08 | 7.100E-08 |
| 23908 | 1196 | N1196G | D | 4.2e-08 | 5e-08 | 2.8e-08 | 4.000E-08 | 1.114E-08 | 2.800E-08 | 5.000E-08 |
| 23909 | 1196 | N1196H | D | 7.7e-08 | 3.7e-08 | 1.3e-08 | 4.233E-08 | 3.233E-08 | 1.300E-08 | 7.700E-08 |
| 23910 | 1196 | N1196I | D | 3.8e-08 | 4.6e-08 | 2.6e-08 | 3.667E-08 | 1.007E-08 | 2.600E-08 | 4.600E-08 |
| 23911 | 1196 | N1196L | D | 7.1e-08 | 6.4e-08 | 4.8e-08 | 6.100E-08 | 1.179E-08 | 4.800E-08 | 7.100E-08 |
| 23912 | 1196 | N1196K | D | 2.8e-08 | 5.4e-08 | 6.4e-08 | 4.867E-08 | 1.858E-08 | 2.800E-08 | 6.400E-08 |
| 23913 | 1196 | N1196M | D | 4e-08 | 3e-08 | 3.6e-08 | 3.533E-08 | 5.033E-09 | 3.000E-08 | 4.000E-08 |
| 23914 | 1196 | N1196F | D | 2.1e-08 | 1.1e-07 | 1.1e-07 | 8.033E-08 | 5.138E-08 | 2.100E-08 | 1.100E-07 |
| 23915 | 1196 | N1196P | D | 6.3e-08 | 3.7e-08 | 5.2e-08 | 5.067E-08 | 1.305E-08 | 3.700E-08 | 6.300E-08 |
| 23916 | 1196 | N1196S | D | 8e-08 | 2.9e-08 | 6.1e-08 | 5.667E-08 | 2.577E-08 | 2.900E-08 | 8.000E-08 |
| 23917 | 1196 | N1196T | D | 3.1e-08 | 1.9e-08 | 5.7e-08 | 3.567E-08 | 1.943E-08 | 1.900E-08 | 5.700E-08 |
| 23918 | 1196 | N1196W | D | 8.9e-08 | 6e-08 | 3.6e-08 | 6.167E-08 | 2.654E-08 | 3.600E-08 | 8.900E-08 |
| 23919 | 1196 | N1196Y | D | 1.9e-08 | 2.6e-08 | 2.2e-08 | 2.233E-08 | 3.512E-09 | 1.900E-08 | 2.600E-08 |
| 23920 | 1196 | N1196V | D | 5e-08 | 5.3e-08 | 1.9e-08 | 4.067E-08 | 1.882E-08 | 1.900E-08 | 5.300E-08 |
| 23921 | 1197 | F1197A | D | 3.9e-08 | 4.6e-08 | 2.3e-08 | 3.600E-08 | 1.179E-08 | 2.300E-08 | 4.600E-08 |
| 23922 | 1197 | F1197R | D | 9.5e-08 | 1.2e-07 | 7.8e-08 | 9.767E-08 | 2.113E-08 | 7.800E-08 | 1.200E-07 |
| 23923 | 1197 | F1197N | D | 3.4e-08 | 1.2e-07 | 4.5e-08 | 6.633E-08 | 4.680E-08 | 3.400E-08 | 1.200E-07 |
| 23924 | 1197 | F1197D | D | 3.3e-08 | 4.3e-08 | 3.6e-08 | 3.733E-08 | 5.132E-09 | 3.300E-08 | 4.300E-08 |
| 23925 | 1197 | F1197C | D | 9.6e-09 | 3.3e-08 | 4.3e-08 | 2.853E-08 | 1.714E-08 | 9.600E-09 | 4.300E-08 |
| 23926 | 1197 | F1197E | D | 3.3e-08 | 5.1e-08 | 2.8e-08 | 3.733E-08 | 1.210E-08 | 2.800E-08 | 5.100E-08 |
| 23927 | 1197 | F1197Q | D | 3.8e-08 | 5.2e-08 | 1.2e-08 | 3.400E-08 | 2.030E-08 | 1.200E-08 | 5.200E-08 |
| 23928 | 1197 | F1197G | D | 3.9e-08 | 1.7e-08 | 6e-08 | 3.867E-08 | 2.150E-08 | 1.700E-08 | 6.000E-08 |
| 23929 | 1197 | F1197H | D | 3.8e-08 | 4.6e-08 | 3.2e-08 | 3.867E-08 | 7.024E-09 | 3.200E-08 | 4.600E-08 |
| 23930 | 1197 | F1197I | D | 4e-08 | 1.2e-07 | 4e-08 | 6.667E-08 | 4.619E-08 | 4.000E-08 | 1.200E-07 |
| 23931 | 1197 | F1197L | D | 4.9e-08 | 3e-08 | 4e-08 | 3.967E-08 | 9.504E-09 | 3.000E-08 | 4.900E-08 |
| 23932 | 1197 | F1197K | D | 1e-07 | 2e-08 | 7.7e-08 | 6.567E-08 | 4.119E-08 | 2.000E-08 | 1.000E-07 |
| 23933 | 1197 | F1197M | D | 3.6e-08 | 4e-08 | 1.9e-08 | 3.167E-08 | 1.115E-08 | 1.900E-08 | 4.000E-08 |
| 23934 | 1197 | F1197F | D | 2.7e-08 | 2.5e-08 | 4.4e-08 | 3.200E-08 | 1.044E-08 | 2.500E-08 | 4.400E-08 |
| 23935 | 1197 | F1197P | D | 4.9e-08 | 5.4e-08 | 3.9e-08 | 4.733E-08 | 7.638E-09 | 3.900E-08 | 5.400E-08 |
| 23936 | 1197 | F1197S | D | 4.9e-08 | 6.8e-08 | 1.7e-08 | 4.467E-08 | 2.577E-08 | 1.700E-08 | 6.800E-08 |
| 23937 | 1197 | F1197T | D | 4.3e-08 | 2.1e-08 | 7.5e-08 | 4.633E-08 | 2.715E-08 | 2.100E-08 | 7.500E-08 |
| 23938 | 1197 | F1197W | D | 6e-08 | 4.2e-08 | 3.6e-08 | 4.600E-08 | 1.249E-08 | 3.600E-08 | 6.000E-08 |
| 23939 | 1197 | F1197Y | D | 5.7e-08 | 1.2e-07 | 4.9e-08 | 7.533E-08 | 3.889E-08 | 4.900E-08 | 1.200E-07 |
| 23940 | 1197 | F1197V | D | 3.2e-08 | 1.1e-07 | 6.5e-08 | 6.900E-08 | 3.915E-08 | 3.200E-08 | 1.100E-07 |
| 23941 | 1198 | Y1198A | D | 4.3e-08 | 3.3e-08 | 7.1e-08 | 4.900E-08 | 1.970E-08 | 3.300E-08 | 7.100E-08 |
| 23942 | 1198 | Y1198R | D | 4.7e-08 | 7.6e-08 | 1.9e-08 | 4.733E-08 | 2.850E-08 | 1.900E-08 | 7.600E-08 |
| 23943 | 1198 | Y1198N | D | 1.2e-07 | 1.6e-08 | 7.1e-08 | 6.900E-08 | 5.203E-08 | 1.600E-08 | 1.200E-07 |
| 23944 | 1198 | Y1198D | D | 2.5e-08 | 5e-08 | 1.7e-08 | 3.067E-08 | 1.721E-08 | 1.700E-08 | 5.000E-08 |
| 23945 | 1198 | Y1198C | D | 3.2e-08 | 1.3e-07 | 2.8e-08 | 6.333E-08 | 5.777E-08 | 2.800E-08 | 1.300E-07 |
| 23946 | 1198 | Y1198E | D | 2.5e-08 | 1.9e-08 | 2.4e-08 | 2.267E-08 | 3.215E-09 | 1.900E-08 | 2.500E-08 |
| 23947 | 1198 | Y1198Q | D | 9.6e-08 | 2.7e-08 | 8.2e-08 | 6.833E-08 | 3.647E-08 | 2.700E-08 | 9.600E-08 |
| 23948 | 1198 | Y1198G | D | 2e-08 | 5.8e-08 | 4.4e-08 | 4.067E-08 | 1.922E-08 | 2.000E-08 | 5.800E-08 |
| 23949 | 1198 | Y1198H | D | 6e-08 | 7.2e-08 | 4.3e-08 | 5.833E-08 | 1.457E-08 | 4.300E-08 | 7.200E-08 |
| 23950 | 1198 | Y1198I | D | 1.5e-08 | 6e-08 | 5.7e-08 | 4.400E-08 | 2.516E-08 | 1.500E-08 | 6.000E-08 |
| 23951 | 1198 | Y1198L | D | 4.2e-08 | 3.4e-08 | 8.3e-08 | 5.300E-08 | 2.629E-08 | 3.400E-08 | 8.300E-08 |
| 23952 | 1198 | Y1198K | D | 7.9e-08 | 1.7e-08 | 6.7e-08 | 5.433E-08 | 3.288E-08 | 1.700E-08 | 7.900E-08 |
| 23953 | 1198 | Y1198M | D | 4.6e-08 | 2.6e-08 | 4.5e-08 | 3.900E-08 | 1.127E-08 | 2.600E-08 | 4.600E-08 |
| 23954 | 1198 | Y1198F | D | 6e-08 | 1e-07 | 7.8e-08 | 7.933E-08 | 2.003E-08 | 6.000E-08 | 1.000E-07 |
| 23955 | 1198 | Y1198P | D | 3.6e-08 | 5.3e-08 | 2.2e-08 | 3.700E-08 | 1.552E-08 | 2.200E-08 | 5.300E-08 |
| 23956 | 1198 | Y1198S | D | 5.2e-08 | 2.1e-08 | 3.4e-08 | 3.567E-08 | 1.557E-08 | 2.100E-08 | 5.200E-08 |
| 23957 | 1198 | Y1198T | D | 1e-07 | 4.2e-08 | 4.2e-08 | 6.133E-08 | 3.349E-08 | 4.200E-08 | 1.000E-07 |
| 23958 | 1198 | Y1198W | D | 3.5e-08 | 3.4e-08 | 8.5e-08 | 5.133E-08 | 2.916E-08 | 3.400E-08 | 8.500E-08 |
| 23959 | 1198 | Y1198Y | D | 6.1e-08 | 5.7e-08 | 3.2e-08 | 5.000E-08 | 1.572E-08 | 3.200E-08 | 6.100E-08 |
| 23960 | 1198 | Y1198V | D | 7.9e-08 | 2.7e-08 | 6e-08 | 5.533E-08 | 2.631E-08 | 2.700E-08 | 7.900E-08 |
| 23961 | 1199 | P1199A | D | 3.9e-08 | 4.1e-08 | 4.3e-08 | 4.100E-08 | 2.000E-09 | 3.900E-08 | 4.300E-08 |
| 23962 | 1199 | P1199R | D | 3.1e-08 | 9.3e-08 | 3.1e-08 | 5.167E-08 | 3.580E-08 | 3.100E-08 | 9.300E-08 |
| 23963 | 1199 | P1199N | D | 2.1e-08 | 8e-08 | 7.6e-09 | 3.620E-08 | 3.852E-08 | 7.600E-09 | 8.000E-08 |
| 23964 | 1199 | P1199D | D | 7.9e-08 | 5.7e-08 | 2.1e-08 | 5.233E-08 | 2.928E-08 | 2.100E-08 | 7.900E-08 |
| 23965 | 1199 | P1199C | D | 7.8e-08 | 7.6e-08 | 4e-08 | 6.467E-08 | 2.139E-08 | 4.000E-08 | 7.800E-08 |
| 23966 | 1199 | P1199E | D | 1.2e-07 | 8.6e-08 | 2e-08 | 7.533E-08 | 5.085E-08 | 2.000E-08 | 1.200E-07 |
| 23967 | 1199 | P1199Q | D | 8.4e-08 | 9.1e-08 | 5.3e-08 | 7.600E-08 | 2.022E-08 | 5.300E-08 | 9.100E-08 |
| 23968 | 1199 | P1199G | D | 1e-07 | 4.6e-08 | 5.6e-08 | 6.733E-08 | 2.873E-08 | 4.600E-08 | 1.000E-07 |
| 23969 | 1199 | P1199H | D | 1.7e-08 | 4.2e-08 | 5e-08 | 3.633E-08 | 1.721E-08 | 1.700E-08 | 5.000E-08 |

| | | | | | | | | | | |
|---|---|---|---|---|---|---|---|---|---|---|
| 23970 | 1199 | P1199I | D | 2.6e-08 | 8.9e-09 | 2.2e-07 | 8.497E-08 | 1.173E-07 | 8.900E-09 | 2.200E-07 |
| 23971 | 1199 | P1199L | D | 2.1e-08 | 1.3e-07 | 1e-07 | 8.367E-08 | 5.631E-08 | 2.100E-08 | 1.300E-07 |
| 23972 | 1199 | P1199K | D | 3e-08 | 5.2e-08 | 3e-08 | 3.733E-08 | 1.270E-08 | 3.000E-08 | 5.200E-08 |
| 23973 | 1199 | P1199M | D | 2.3e-08 | 7.6e-08 | 4.5e-08 | 4.800E-08 | 2.663E-08 | 2.300E-08 | 7.600E-08 |
| 23974 | 1199 | P1199F | D | 5.5e-08 | 7.2e-08 | 6.6e-08 | 6.433E-08 | 8.622E-09 | 5.500E-08 | 7.200E-08 |
| 23975 | 1199 | P1199P | D | 9.8e-08 | 1.4e-08 | 3.4e-08 | 4.867E-08 | 4.388E-08 | 1.400E-08 | 9.800E-08 |
| 23976 | 1199 | P1199S | D | 5.8e-08 | 7.4e-08 | 2.1e-08 | 5.100E-08 | 2.718E-08 | 2.100E-08 | 7.400E-08 |
| 23977 | 1199 | P1199T | D | 7e-08 | 5.8e-08 | 7.3e-08 | 6.700E-08 | 7.937E-09 | 5.800E-08 | 7.300E-08 |
| 23978 | 1199 | P1199W | D | 3.9e-08 | 4.3e-08 | 4.7e-08 | 4.300E-08 | 4.000E-09 | 3.900E-08 | 4.700E-08 |
| 23979 | 1199 | P1199Y | D | 6.3e-08 | 5.2e-08 | 7.8e-08 | 6.433E-08 | 1.305E-08 | 5.200E-08 | 7.800E-08 |
| 23980 | 1199 | P1199V | D | 7.7e-08 | 5.1e-08 | 5.1e-08 | 5.967E-08 | 1.501E-08 | 5.100E-08 | 7.700E-08 |
| 23981 | 1200 | R1200A | D | 2.7e-08 | 2.3e-08 | 4.8e-08 | 3.267E-08 | 1.343E-08 | 2.300E-08 | 4.800E-08 |
| 23982 | 1200 | R1200R | D | 5.3e-08 | 1.6e-08 | 2.5e-08 | 3.133E-08 | 1.930E-08 | 1.600E-08 | 5.300E-08 |
| 23983 | 1200 | R1200N | D | 2.6e-08 | 4.2e-08 | 5.5e-08 | 4.100E-08 | 1.453E-08 | 2.600E-08 | 5.500E-08 |
| 23984 | 1200 | R1200D | D | 4.1e-08 | 3.2e-08 | 5.4e-08 | 4.233E-08 | 1.106E-08 | 3.200E-08 | 5.400E-08 |
| 23985 | 1200 | R1200C | D | 4.1e-08 | 4.9e-08 | 5.5e-08 | 4.833E-08 | 7.024E-09 | 4.100E-08 | 5.500E-08 |
| 23986 | 1200 | R1200E | D | 6.1e-08 | 1.6e-08 | 2.4e-08 | 3.367E-08 | 2.401E-08 | 1.600E-08 | 6.100E-08 |
| 23987 | 1200 | R1200Q | D | 2.1e-08 | 3.4e-08 | 3.2e-08 | 2.900E-08 | 7.000E-09 | 2.100E-08 | 3.400E-08 |
| 23988 | 1200 | R1200G | D | 9.1e-08 | 6.2e-08 | 9.6e-08 | 8.300E-08 | 1.836E-08 | 6.200E-08 | 9.600E-08 |
| 23989 | 1200 | R1200H | D | 3.3e-08 | 3.1e-08 | 1.8e-08 | 2.733E-08 | 8.145E-09 | 1.800E-08 | 3.300E-08 |
| 23990 | 1200 | R1200I | D | 3.2e-08 | 5.3e-08 | 3.1e-08 | 3.867E-08 | 1.242E-08 | 3.100E-08 | 5.300E-08 |
| 23991 | 1200 | R1200L | D | 1.8e-08 | 2.2e-08 | 1e-07 | 4.667E-08 | 4.623E-08 | 1.800E-08 | 1.000E-07 |
| 23992 | 1200 | R1200K | D | 1.8e-08 | 2.4e-08 | 9e-08 | 4.400E-08 | 3.995E-08 | 1.800E-08 | 9.000E-08 |
| 23993 | 1200 | R1200M | D | 2e-08 | 2.8e-08 | 2.6e-08 | 2.467E-08 | 4.163E-09 | 2.000E-08 | 2.800E-08 |
| 23994 | 1200 | R1200F | D | 2.4e-08 | 6e-08 | 5.1e-08 | 4.500E-08 | 1.873E-08 | 2.400E-08 | 6.000E-08 |
| 23995 | 1200 | R1200P | D | 3.2e-08 | 8.2e-08 | 5e-08 | 5.467E-08 | 2.532E-08 | 3.200E-08 | 8.200E-08 |
| 23996 | 1200 | R1200S | D | 4.2e-08 | 4e-08 | 5.8e-08 | 4.667E-08 | 9.866E-09 | 4.000E-08 | 5.800E-08 |
| 23997 | 1200 | R1200T | D | 3e-08 | 2.9e-08 | 2.8e-08 | 2.900E-08 | 1.000E-09 | 2.800E-08 | 3.000E-08 |
| 23998 | 1200 | R1200W | D | 6.4e-08 | 2.7e-08 | 2e-08 | 3.700E-08 | 2.364E-08 | 2.000E-08 | 6.400E-08 |
| 23999 | 1200 | R1200Y | D | 8.4e-08 | 2e-08 | 2.7e-08 | 4.367E-08 | 3.510E-08 | 2.000E-08 | 8.400E-08 |
| 24000 | 1200 | R1200V | D | 1e-07 | 3.4e-08 | 4.4e-08 | 5.933E-08 | 3.557E-08 | 3.400E-08 | 1.000E-07 |
| 24001 | 1201 | E1201A | D | 3.1e-08 | 2e-08 | 1.3e-07 | 6.033E-08 | 6.058E-08 | 2.000E-08 | 1.300E-07 |
| 24002 | 1201 | E1201R | D | 6.8e-08 | 2.6e-08 | 4.9e-08 | 4.767E-08 | 2.103E-08 | 2.600E-08 | 6.800E-08 |
| 24003 | 1201 | E1201N | D | 5e-08 | 1.1e-07 | 3.5e-08 | 6.500E-08 | 3.969E-08 | 3.500E-08 | 1.100E-07 |
| 24004 | 1201 | E1201D | D | 2.6e-08 | 3.8e-08 | 4.4e-08 | 3.600E-08 | 9.165E-09 | 2.600E-08 | 4.400E-08 |
| 24005 | 1201 | E1201C | D | 2.8e-08 | 2e-08 | 9.3e-08 | 4.700E-08 | 4.004E-08 | 2.000E-08 | 9.300E-08 |
| 24006 | 1201 | E1201E | D | 2.7e-08 | 9.8e-08 | 2.3e-08 | 4.933E-08 | 4.219E-08 | 2.300E-08 | 9.800E-08 |
| 24007 | 1201 | E1201Q | D | 4e-08 | 2.1e-08 | 5.6e-08 | 3.900E-08 | 1.752E-08 | 2.100E-08 | 5.600E-08 |
| 24008 | 1201 | E1201G | D | 5e-08 | 4.1e-08 | 5.6e-08 | 4.900E-08 | 7.550E-09 | 4.100E-08 | 5.600E-08 |
| 24009 | 1201 | E1201H | D | 5.4e-08 | 1.1e-07 | 3.9e-08 | 6.767E-08 | 3.742E-08 | 3.900E-08 | 1.100E-07 |
| 24010 | 1201 | E1201I | D | 5.8e-08 | 1.8e-08 | 6.2e-08 | 4.600E-08 | 2.433E-08 | 1.800E-08 | 6.200E-08 |
| 24011 | 1201 | E1201L | D | 9.3e-08 | 4.9e-08 | 6e-08 | 6.733E-08 | 2.290E-08 | 4.900E-08 | 9.300E-08 |
| 24012 | 1201 | E1201K | D | 4.5e-08 | 1.9e-08 | 6.9e-09 | 2.363E-08 | 1.947E-08 | 6.900E-09 | 4.500E-08 |
| 24013 | 1201 | E1201M | D | 2.3e-08 | 4.1e-08 | 3.4e-08 | 3.267E-08 | 9.074E-09 | 2.300E-08 | 4.100E-08 |
| 24014 | 1201 | E1201F | D | 1.8e-08 | 6.3e-08 | 6.5e-08 | 4.867E-08 | 2.658E-08 | 1.800E-08 | 6.500E-08 |
| 24015 | 1201 | E1201P | D | 8.2e-08 | 3.8e-08 | 4.9e-08 | 5.633E-08 | 2.290E-08 | 3.800E-08 | 8.200E-08 |
| 24016 | 1201 | E1201S | D | 2.6e-08 | 2.8e-08 | 4.2e-08 | 3.200E-08 | 8.718E-09 | 2.600E-08 | 4.200E-08 |
| 24017 | 1201 | E1201T | D | 4.2e-08 | 5.4e-08 | 2e-07 | 9.867E-08 | 8.796E-08 | 4.200E-08 | 2.000E-07 |
| 24018 | 1201 | E1201W | D | 2e-08 | 4e-08 | 1.8e-07 | 8.000E-08 | 8.718E-08 | 2.000E-08 | 1.800E-07 |
| 24019 | 1201 | E1201Y | D | 2e-07 | 2.3e-08 | 4.4e-08 | 8.900E-08 | 9.670E-08 | 2.300E-08 | 2.000E-07 |
| 24020 | 1201 | E1201V | D | 1.3e-07 | 4.2e-08 | 8.2e-08 | 8.467E-08 | 4.406E-08 | 4.200E-08 | 1.300E-07 |
| 24021 | 1202 | A1202A | D | 5.1e-08 | 7.9e-08 | 6.2e-08 | 6.400E-08 | 1.411E-08 | 5.100E-08 | 7.900E-08 |
| 24022 | 1202 | A1202R | D | 4.2e-08 | 1e-08 | 2.8e-08 | 2.667E-08 | 1.604E-08 | 1.000E-08 | 4.200E-08 |
| 24023 | 1202 | A1202N | D | 1.4e-08 | 2.2e-08 | 1.5e-08 | 1.700E-08 | 4.359E-09 | 1.400E-08 | 2.200E-08 |
| 24024 | 1202 | A1202D | D | 5e-08 | 1.5e-08 | 7.2e-08 | 4.567E-08 | 2.875E-08 | 1.500E-08 | 7.200E-08 |
| 24025 | 1202 | A1202C | D | 2.7e-08 | 3.1e-08 | 1.4e-08 | 2.400E-08 | 8.888E-09 | 1.400E-08 | 3.100E-08 |
| 24026 | 1202 | A1202E | D | 2.2e-08 | 6.6e-08 | 9.5e-08 | 6.100E-08 | 3.676E-08 | 2.200E-08 | 9.500E-08 |
| 24027 | 1202 | A1202Q | D | 5.6e-08 | 7e-08 | 6.9e-08 | 6.500E-08 | 7.810E-09 | 5.600E-08 | 7.000E-08 |
| 24028 | 1202 | A1202G | D | 4.2e-08 | 3.3e-08 | 2e-08 | 3.167E-08 | 1.106E-08 | 2.000E-08 | 4.200E-08 |
| 24029 | 1202 | A1202H | D | 1.3e-08 | 2.3e-08 | 5.6e-08 | 3.067E-08 | 2.250E-08 | 1.300E-08 | 5.600E-08 |
| 24030 | 1202 | A1202I | D | 2.6e-08 | 1.8e-08 | 2.2e-08 | 2.200E-08 | 4.000E-09 | 1.800E-08 | 2.600E-08 |
| 24031 | 1202 | A1202L | D | 4.9e-08 | 4.7e-08 | 3.5e-08 | 4.367E-08 | 7.572E-09 | 3.500E-08 | 4.900E-08 |
| 24032 | 1202 | A1202K | D | 1.7e-08 | 4.2e-08 | 4.1e-08 | 3.333E-08 | 1.415E-08 | 1.700E-08 | 4.200E-08 |
| 24033 | 1202 | A1202M | D | 3.4e-08 | 7.4e-08 | 1.9e-08 | 4.233E-08 | 2.843E-08 | 1.900E-08 | 7.400E-08 |
| 24034 | 1202 | A1202F | D | 2.6e-08 | 2.9e-08 | 6.8e-08 | 4.100E-08 | 2.343E-08 | 2.600E-08 | 6.800E-08 |
| 24035 | 1202 | A1202P | D | 3.2e-08 | 3.8e-08 | 3.1e-08 | 3.367E-08 | 3.786E-09 | 3.100E-08 | 3.800E-08 |
| 24036 | 1202 | A1202S | D | 2.7e-08 | 2.1e-08 | 7.9e-09 | 1.863E-08 | 9.767E-09 | 7.900E-09 | 2.700E-08 |
| 24037 | 1202 | A1202T | D | 6e-08 | 4.6e-08 | 5.9e-08 | 5.500E-08 | 7.810E-09 | 4.600E-08 | 6.000E-08 |
| 24038 | 1202 | A1202W | D | 1.9e-08 | 3.7e-08 | 1.2e-08 | 2.267E-08 | 1.290E-08 | 1.200E-08 | 3.700E-08 |
| 24039 | 1202 | A1202Y | D | 6.2e-08 | 4.5e-08 | 3e-08 | 4.567E-08 | 1.601E-08 | 3.000E-08 | 6.200E-08 |
| 24040 | 1202 | A1202V | D | 5.9e-08 | 4.2e-08 | 4.9e-08 | 5.000E-08 | 8.544E-09 | 4.200E-08 | 5.900E-08 |
| 24041 | 1203 | K1203A | D | 1.5e-07 | 3.4e-08 | 3.7e-08 | 7.367E-08 | 6.612E-08 | 3.400E-08 | 1.500E-07 |
| 24042 | 1203 | K1203R | D | 2.1e-08 | 3.2e-08 | 4e-08 | 3.100E-08 | 9.539E-09 | 2.100E-08 | 4.000E-08 |
| 24043 | 1203 | K1203N | D | 1.8e-08 | 5.4e-08 | 1.9e-08 | 3.033E-08 | 2.050E-08 | 1.800E-08 | 5.400E-08 |
| 24044 | 1203 | K1203D | D | 4e-08 | 1.4e-08 | 5.4e-08 | 3.600E-08 | 2.030E-08 | 1.400E-08 | 5.400E-08 |
| 24045 | 1203 | K1203C | D | 3.5e-08 | 3e-08 | 1.9e-08 | 2.800E-08 | 8.185E-09 | 1.900E-08 | 3.500E-08 |
| 24046 | 1203 | K1203E | D | 8.6e-08 | 1.8e-08 | 2.9e-08 | 4.433E-08 | 3.650E-08 | 1.800E-08 | 8.600E-08 |
| 24047 | 1203 | K1203Q | D | 3.7e-08 | 5.1e-08 | 6.8e-08 | 5.200E-08 | 1.552E-08 | 3.700E-08 | 6.800E-08 |
| 24048 | 1203 | K1203G | D | 5.9e-08 | 4.2e-08 | 1.6e-08 | 3.900E-08 | 2.166E-08 | 1.600E-08 | 5.900E-08 |
| 24049 | 1203 | K1203H | D | 3.1e-08 | 4.2e-08 | 1.7e-08 | 3.000E-08 | 1.253E-08 | 1.700E-08 | 4.200E-08 |
| 24050 | 1203 | K1203I | D | 4.9e-08 | 2.3e-08 | 2e-07 | 9.067E-08 | 9.557E-08 | 2.300E-08 | 2.000E-07 |
| 24051 | 1203 | K1203L | D | 1e-07 | 5.3e-08 | 5.1e-08 | 6.800E-08 | 2.773E-08 | 5.100E-08 | 1.000E-07 |
| 24052 | 1203 | K1203K | D | 9.8e-08 | 7.7e-08 | 4.2e-08 | 7.233E-08 | 2.829E-08 | 4.200E-08 | 9.800E-08 |
| 24053 | 1203 | K1203M | D | 1.9e-08 | 2.4e-08 | 2.5e-08 | 2.267E-08 | 3.215E-09 | 1.900E-08 | 2.500E-08 |
| 24054 | 1203 | K1203F | D | 7.7e-08 | 5.8e-08 | 8.3e-08 | 7.267E-08 | 1.305E-08 | 5.800E-08 | 8.300E-08 |

| | | | | | | | | | | |
|---|---|---|---|---|---|---|---|---|---|---|
| 24055 | 1203 | K1203P | D | 1.9e-08 | 3.3e-08 | 3.1e-08 | 2.767E-08 | 7.572E-09 | 1.900E-08 | 3.300E-08 |
| 24056 | 1203 | K1203S | D | 1.2e-07 | 3.1e-08 | 2.2e-08 | 5.767E-08 | 5.417E-08 | 2.200E-08 | 1.200E-07 |
| 24057 | 1203 | K1203T | D | 3.7e-08 | 6.4e-08 | 4.5e-08 | 4.867E-08 | 1.387E-08 | 3.700E-08 | 6.400E-08 |
| 24058 | 1203 | K1203W | D | 4.6e-08 | 5e-08 | 8.7e-08 | 6.100E-08 | 2.261E-08 | 4.600E-08 | 8.700E-08 |
| 24059 | 1203 | K1203Y | D | 3.9e-08 | 7.6e-08 | 4e-08 | 5.167E-08 | 2.108E-08 | 3.900E-08 | 7.600E-08 |
| 24060 | 1203 | K1203V | D | 5.3e-08 | 6.3e-08 | 4e-08 | 5.200E-08 | 1.153E-08 | 4.000E-08 | 6.300E-08 |
| 24061 | 1204 | V1204A | D | 1.9e-08 | 5.7e-08 | 1.1e-08 | 2.900E-08 | 2.458E-08 | 1.100E-08 | 5.700E-08 |
| 24062 | 1204 | V1204R | D | 3.4e-08 | 5.9e-08 | 2e-08 | 3.767E-08 | 1.976E-08 | 2.000E-08 | 5.900E-08 |
| 24063 | 1204 | V1204N | D | 7e-08 | 2e-08 | 3e-08 | 4.000E-08 | 2.646E-08 | 2.000E-08 | 7.000E-08 |
| 24064 | 1204 | V1204D | D | 9.1e-08 | 3.1e-08 | 7.9e-08 | 6.700E-08 | 3.175E-08 | 3.100E-08 | 9.100E-08 |
| 24065 | 1204 | V1204C | D | 3.9e-08 | 3.8e-08 | 4.3e-08 | 4.000E-08 | 2.646E-09 | 3.800E-08 | 4.300E-08 |
| 24066 | 1204 | V1204E | D | 5.1e-08 | 2.3e-08 | 5.1e-08 | 4.167E-08 | 1.617E-08 | 2.300E-08 | 5.100E-08 |
| 24067 | 1204 | V1204Q | D | 1e-07 | 4.3e-08 | 6.5e-08 | 6.933E-08 | 2.875E-08 | 4.300E-08 | 1.000E-07 |
| 24068 | 1204 | V1204G | D | 8e-08 | 3.7e-08 | 8.9e-08 | 6.867E-08 | 2.779E-08 | 3.700E-08 | 8.900E-08 |
| 24069 | 1204 | V1204H | D | 2.6e-08 | 3e-08 | 4.6e-08 | 3.400E-08 | 1.058E-08 | 2.600E-08 | 4.600E-08 |
| 24070 | 1204 | V1204I | D | 2e-08 | 3.8e-08 | 2.4e-08 | 2.733E-08 | 9.452E-09 | 2.000E-08 | 3.800E-08 |
| 24071 | 1204 | V1204L | D | 2.8e-08 | 4e-08 | 1.7e-08 | 2.833E-08 | 1.150E-08 | 1.700E-08 | 4.000E-08 |
| 24072 | 1204 | V1204K | D | 3.9e-08 | 1.3e-07 | 5.9e-08 | 7.600E-08 | 4.782E-08 | 3.900E-08 | 1.300E-07 |
| 24073 | 1204 | V1204M | D | 6.7e-08 | 6.4e-08 | 1.9e-08 | 5.000E-08 | 2.689E-08 | 1.900E-08 | 6.700E-08 |
| 24074 | 1204 | V1204F | D | 2.5e-08 | 2.4e-07 | 4e-08 | 1.017E-07 | 1.200E-07 | 2.500E-08 | 2.400E-07 |
| 24075 | 1204 | V1204P | D | 3.2e-08 | 3.7e-08 | 3.2e-08 | 3.367E-08 | 2.887E-09 | 3.200E-08 | 3.700E-08 |
| 24076 | 1204 | V1204S | D | 4.5e-08 | 5.9e-08 | 4.1e-08 | 4.833E-08 | 9.452E-09 | 4.100E-08 | 5.900E-08 |
| 24077 | 1204 | V1204T | D | 6.1e-08 | 5.4e-08 | 4.3e-08 | 5.267E-08 | 9.074E-09 | 4.300E-08 | 6.100E-08 |
| 24078 | 1204 | V1204W | D | 3.8e-08 | 4.2e-08 | 9.6e-08 | 5.867E-08 | 3.239E-08 | 3.800E-08 | 9.600E-08 |
| 24079 | 1204 | V1204Y | D | 4.9e-08 | 5.9e-08 | 3.8e-08 | 4.867E-08 | 1.050E-08 | 3.800E-08 | 5.900E-08 |
| 24080 | 1204 | V1204V | D | 2.2e-08 | 6.7e-08 | 2.2e-08 | 3.700E-08 | 2.598E-08 | 2.200E-08 | 6.700E-08 |
| 24081 | 1205 | Q1205A | D | 1.8e-08 | 1.2e-07 | 8.4e-08 | 7.400E-08 | 5.173E-08 | 1.800E-08 | 1.200E-07 |
| 24082 | 1205 | Q1205R | D | 1.1e-07 | 3.2e-08 | 4.3e-08 | 6.167E-08 | 4.222E-08 | 3.200E-08 | 1.100E-07 |
| 24083 | 1205 | Q1205N | D | 7e-08 | 3.4e-08 | 1.1e-07 | 7.133E-08 | 3.802E-08 | 3.400E-08 | 1.100E-07 |
| 24084 | 1205 | Q1205D | D | 2.5e-08 | 4e-08 | 3.2e-08 | 3.233E-08 | 7.506E-09 | 2.500E-08 | 4.000E-08 |
| 24085 | 1205 | Q1205C | D | 2.4e-08 | 5.9e-08 | 8.9e-08 | 5.733E-08 | 3.253E-08 | 2.400E-08 | 8.900E-08 |
| 24086 | 1205 | Q1205E | D | 2.5e-08 | 2.5e-08 | 1.2e-08 | 2.067E-08 | 7.506E-09 | 1.200E-08 | 2.500E-08 |
| 24087 | 1205 | Q1205Q | D | 7e-08 | 3.2e-08 | 8.2e-08 | 6.133E-08 | 2.610E-08 | 3.200E-08 | 8.200E-08 |
| 24088 | 1205 | Q1205G | D | 3.1e-08 | 3.7e-08 | 3.8e-08 | 3.533E-08 | 3.786E-09 | 3.100E-08 | 3.800E-08 |
| 24089 | 1205 | Q1205H | D | 4.4e-08 | 1e-07 | 3.9e-08 | 6.100E-08 | 3.387E-08 | 3.900E-08 | 1.000E-07 |
| 24090 | 1205 | Q1205I | D | 1.5e-07 | 4.7e-08 | 4.3e-08 | 8.000E-08 | 6.065E-08 | 4.300E-08 | 1.500E-07 |
| 24091 | 1205 | Q1205L | D | 4.3e-08 | 6.3e-08 | 3.8e-08 | 4.800E-08 | 1.323E-08 | 3.800E-08 | 6.300E-08 |
| 24092 | 1205 | Q1205K | D | 6.2e-08 | 3.5e-08 | 1.4e-08 | 3.700E-08 | 2.406E-08 | 1.400E-08 | 6.200E-08 |
| 24093 | 1205 | Q1205M | D | 7.6e-08 | 5e-08 | 6.3e-08 | 6.300E-08 | 1.300E-08 | 5.000E-08 | 7.600E-08 |
| 24094 | 1205 | Q1205F | D | 2.5e-08 | 1.4e-08 | 1.6e-08 | 1.833E-08 | 5.859E-09 | 1.400E-08 | 2.500E-08 |
| 24095 | 1205 | Q1205P | D | 2.9e-08 | 3.5e-08 | 7.6e-08 | 4.667E-08 | 2.558E-08 | 2.900E-08 | 7.600E-08 |
| 24096 | 1205 | Q1205S | D | 3.7e-08 | 8.8e-08 | 9.8e-09 | 4.493E-08 | 3.970E-08 | 9.800E-09 | 8.800E-08 |
| 24097 | 1205 | Q1205T | D | 2.8e-08 | 4.5e-08 | 6.7e-08 | 4.667E-08 | 1.955E-08 | 2.800E-08 | 6.700E-08 |
| 24098 | 1205 | Q1205W | D | 4.4e-08 | 4.9e-08 | 3e-08 | 4.100E-08 | 9.849E-09 | 3.000E-08 | 4.900E-08 |
| 24099 | 1205 | Q1205Y | D | 1e-07 | 6.1e-08 | 1.6e-08 | 5.900E-08 | 4.204E-08 | 1.600E-08 | 1.000E-07 |
| 24100 | 1205 | Q1205V | D | 5.2e-08 | 6.8e-08 | 8.9e-08 | 6.967E-08 | 1.856E-08 | 5.200E-08 | 8.900E-08 |
| 24101 | 1206 | W1206A | D | 2.9e-08 | 1.7e-08 | 5.4e-08 | 3.333E-08 | 1.888E-08 | 1.700E-08 | 5.400E-08 |
| 24102 | 1206 | W1206R | D | 1e-07 | 5e-08 | 7.5e-08 | 7.500E-08 | 2.500E-08 | 5.000E-08 | 1.000E-07 |
| 24103 | 1206 | W1206N | D | 3.8e-08 | 3.9e-08 | 1.3e-07 | 6.900E-08 | 5.283E-08 | 3.800E-08 | 1.300E-07 |
| 24104 | 1206 | W1206D | D | 6e-08 | 1.7e-08 | 6e-08 | 4.567E-08 | 2.483E-08 | 1.700E-08 | 6.000E-08 |
| 24105 | 1206 | W1206C | D | 5.1e-08 | 4.7e-08 | 6.9e-08 | 5.567E-08 | 1.172E-08 | 4.700E-08 | 6.900E-08 |
| 24106 | 1206 | W1206E | D | 7.5e-08 | 4.4e-08 | 2.8e-08 | 4.900E-08 | 2.390E-08 | 2.800E-08 | 7.500E-08 |
| 24107 | 1206 | W1206Q | D | 2.3e-08 | 1.9e-08 | 2.5e-08 | 2.233E-08 | 3.055E-09 | 1.900E-08 | 2.500E-08 |
| 24108 | 1206 | W1206G | D | 5.2e-08 | 5.8e-08 | 1.6e-08 | 4.200E-08 | 2.272E-08 | 1.600E-08 | 5.800E-08 |
| 24109 | 1206 | W1206H | D | 5e-08 | 3.5e-08 | 5.8e-08 | 4.767E-08 | 1.168E-08 | 3.500E-08 | 5.800E-08 |
| 24110 | 1206 | W1206I | D | 2.6e-08 | 4.6e-08 | 6.2e-08 | 4.467E-08 | 1.804E-08 | 2.600E-08 | 6.200E-08 |
| 24111 | 1206 | W1206L | D | 7.3e-08 | 2.8e-08 | 4e-08 | 4.700E-08 | 2.330E-08 | 2.800E-08 | 7.300E-08 |
| 24112 | 1206 | W1206K | D | 5.4e-08 | 5e-08 | 9.3e-09 | 3.777E-08 | 2.473E-08 | 9.300E-09 | 5.400E-08 |
| 24113 | 1206 | W1206M | D | 3.6e-08 | 6.1e-08 | 4.2e-08 | 4.633E-08 | 1.305E-08 | 3.600E-08 | 6.100E-08 |
| 24114 | 1206 | W1206F | D | 6e-08 | 2.4e-08 | 3.2e-08 | 3.867E-08 | 1.890E-08 | 2.400E-08 | 6.000E-08 |
| 24115 | 1206 | W1206P | D | 1.9e-08 | 1.6e-08 | 2.3e-08 | 1.933E-08 | 3.512E-09 | 1.600E-08 | 2.300E-08 |
| 24116 | 1206 | W1206S | D | 1.9e-08 | 3.7e-08 | 6e-08 | 3.867E-08 | 2.055E-08 | 1.900E-08 | 6.000E-08 |
| 24117 | 1206 | W1206T | D | 8.5e-08 | 6.3e-08 | 2e-08 | 5.600E-08 | 3.306E-08 | 2.000E-08 | 8.500E-08 |
| 24118 | 1206 | W1206W | D | 3.6e-08 | 2.5e-08 | 1.3e-07 | 6.367E-08 | 5.771E-08 | 2.500E-08 | 1.300E-07 |
| 24119 | 1206 | W1206Y | D | 6.9e-08 | 6.7e-08 | 2.8e-08 | 5.467E-08 | 2.312E-08 | 2.800E-08 | 6.900E-08 |
| 24120 | 1206 | W1206V | D | 4.8e-08 | 3.8e-08 | 1e-07 | 6.200E-08 | 3.329E-08 | 3.800E-08 | 1.000E-07 |
| 24121 | 1207 | K1207A | D | 4.8e-08 | 1.6e-08 | 5.8e-08 | 4.067E-08 | 2.194E-08 | 1.600E-08 | 5.800E-08 |
| 24122 | 1207 | K1207R | D | 8.6e-08 | 3.5e-08 | 1.7e-08 | 4.600E-08 | 3.579E-08 | 1.700E-08 | 8.600E-08 |
| 24123 | 1207 | K1207N | D | 2.8e-08 | 2.2e-08 | 6.2e-08 | 3.733E-08 | 2.157E-08 | 2.200E-08 | 6.200E-08 |
| 24124 | 1207 | K1207D | D | 4.4e-08 | 1.8e-08 | 1.9e-08 | 2.700E-08 | 1.473E-08 | 1.800E-08 | 4.400E-08 |
| 24125 | 1207 | K1207C | D | 2.5e-08 | 8.7e-08 | 4.2e-08 | 5.133E-08 | 3.204E-08 | 2.500E-08 | 8.700E-08 |
| 24126 | 1207 | K1207E | D | 7.2e-08 | 3.4e-08 | 3.6e-08 | 4.733E-08 | 2.139E-08 | 3.400E-08 | 7.200E-08 |
| 24127 | 1207 | K1207Q | D | 3.2e-08 | 3.2e-08 | 6.2e-08 | 4.200E-08 | 1.732E-08 | 3.200E-08 | 6.200E-08 |
| 24128 | 1207 | K1207G | D | 5.2e-08 | 4.2e-08 | 4.7e-08 | 4.700E-08 | 5.000E-09 | 4.200E-08 | 5.200E-08 |
| 24129 | 1207 | K1207H | D | 9.3e-08 | 3.7e-08 | 4.3e-08 | 5.767E-08 | 3.075E-08 | 3.700E-08 | 9.300E-08 |
| 24130 | 1207 | K1207I | D | 9.7e-08 | 5.4e-08 | 4e-08 | 6.367E-08 | 2.970E-08 | 4.000E-08 | 9.700E-08 |
| 24131 | 1207 | K1207L | D | 5.9e-08 | 3.2e-08 | 6.4e-08 | 5.167E-08 | 1.721E-08 | 3.200E-08 | 6.400E-08 |
| 24132 | 1207 | K1207K | D | 6.8e-08 | 3.8e-08 | 3.6e-08 | 4.733E-08 | 1.793E-08 | 3.600E-08 | 6.800E-08 |
| 24133 | 1207 | K1207M | D | 5.7e-08 | 2.3e-08 | 3.4e-08 | 3.800E-08 | 1.735E-08 | 2.300E-08 | 5.700E-08 |
| 24134 | 1207 | K1207F | D | 2.5e-08 | 2.4e-08 | 1.8e-08 | 2.233E-08 | 3.786E-09 | 1.800E-08 | 2.500E-08 |
| 24135 | 1207 | K1207P | D | 6.1e-08 | 2.6e-08 | 8e-08 | 5.567E-08 | 2.739E-08 | 2.600E-08 | 8.000E-08 |
| 24136 | 1207 | K1207S | D | 3.6e-08 | 3.6e-08 | 3.1e-08 | 3.433E-08 | 2.887E-09 | 3.100E-08 | 3.600E-08 |
| 24137 | 1207 | K1207T | D | 1.4e-08 | 3.6e-08 | 4.4e-08 | 3.133E-08 | 1.553E-08 | 1.400E-08 | 4.400E-08 |
| 24138 | 1207 | K1207W | D | 7.4e-08 | 5.5e-08 | 1.9e-07 | 1.063E-07 | 7.308E-08 | 5.500E-08 | 1.900E-07 |
| 24139 | 1207 | K1207Y | D | 3.9e-08 | 5e-08 | 6.9e-08 | 5.267E-08 | 1.518E-08 | 3.900E-08 | 6.900E-08 |

| | | | | | | | | | | |
|---|---|---|---|---|---|---|---|---|---|---|
| 24140 | 1207 | K1207V | D | 5.5e-08 | 1.1e-07 | 5.9e-08 | 7.467E-08 | 3.066E-08 | 5.500E-08 | 1.100E-07 |
| 24141 | 1208 | V1208A | D | 5.7e-08 | 1.8e-07 | 1.6e-08 | 8.433E-08 | 8.535E-08 | 1.600E-08 | 1.800E-07 |
| 24142 | 1208 | V1208R | D | 9.6e-08 | 2e-08 | 1.8e-08 | 4.467E-08 | 4.447E-08 | 1.800E-08 | 9.600E-08 |
| 24143 | 1208 | V1208N | D | 2e-08 | 2.9e-08 | 4.1e-08 | 3.000E-08 | 1.054E-08 | 2.000E-08 | 4.100E-08 |
| 24144 | 1208 | V1208D | D | 5.2e-08 | 2.5e-08 | 6.8e-08 | 4.833E-08 | 2.173E-08 | 2.500E-08 | 6.800E-08 |
| 24145 | 1208 | V1208C | D | 1.4e-07 | 4.5e-08 | 1.4e-08 | 6.633E-08 | 6.565E-08 | 1.400E-08 | 1.400E-07 |
| 24146 | 1208 | V1208E | D | 2.9e-08 | 4.5e-08 | 2.7e-08 | 3.367E-08 | 9.866E-09 | 2.700E-08 | 4.500E-08 |
| 24147 | 1208 | V1208Q | D | 3.4e-08 | 6.3e-08 | 9.5e-09 | 3.550E-08 | 2.678E-08 | 9.500E-09 | 6.300E-08 |
| 24148 | 1208 | V1208G | D | 7.5e-08 | 1.4e-08 | 6.1e-08 | 5.000E-08 | 3.195E-08 | 1.400E-08 | 7.500E-08 |
| 24149 | 1208 | V1208H | D | 1.6e-08 | 2.1e-08 | 5.8e-08 | 3.167E-08 | 2.294E-08 | 1.600E-08 | 5.800E-08 |
| 24150 | 1208 | V1208I | D | 6.6e-08 | 7.4e-08 | 2.8e-08 | 5.600E-08 | 2.458E-08 | 2.800E-08 | 7.400E-08 |
| 24151 | 1208 | V1208L | D | 3.9e-08 | 7.5e-08 | 6.6e-09 | 4.020E-08 | 3.422E-08 | 6.600E-09 | 7.500E-08 |
| 24152 | 1208 | V1208K | D | 2.2e-08 | 1.2e-07 | 4.5e-08 | 6.233E-08 | 5.125E-08 | 2.200E-08 | 1.200E-07 |
| 24153 | 1208 | V1208M | D | 4e-08 | 1.8e-08 | 2.1e-08 | 2.633E-08 | 1.193E-08 | 1.800E-08 | 4.000E-08 |
| 24154 | 1208 | V1208F | D | 2.7e-08 | 1.7e-08 | 1.7e-08 | 2.033E-08 | 5.774E-09 | 1.700E-08 | 2.700E-08 |
| 24155 | 1208 | V1208P | D | 3.8e-08 | 2.8e-08 | 5.2e-08 | 3.933E-08 | 1.206E-08 | 2.800E-08 | 5.200E-08 |
| 24156 | 1208 | V1208S | D | 4.2e-08 | 6e-08 | 3e-08 | 4.400E-08 | 1.510E-08 | 3.000E-08 | 6.000E-08 |
| 24157 | 1208 | V1208T | D | 4e-08 | 3.9e-08 | 6.5e-08 | 4.800E-08 | 1.473E-08 | 3.900E-08 | 6.500E-08 |
| 24158 | 1208 | V1208W | D | 1e-07 | 5.4e-08 | 3.5e-08 | 6.300E-08 | 3.342E-08 | 3.500E-08 | 1.000E-07 |
| 24159 | 1208 | V1208Y | D | 9e-08 | 3.8e-08 | 9.7e-08 | 7.500E-08 | 3.223E-08 | 3.800E-08 | 9.700E-08 |
| 24160 | 1208 | V1208V | D | 8.3e-08 | 1.4e-08 | 6.2e-08 | 5.300E-08 | 3.537E-08 | 1.400E-08 | 8.300E-08 |
| 24161 | 1209 | D1209A | D | 4.6e-08 | 1.1e-07 | 7.4e-08 | 7.667E-08 | 3.208E-08 | 4.600E-08 | 1.100E-07 |
| 24162 | 1209 | D1209R | D | 4.5e-08 | 5.1e-08 | 1.5e-07 | 8.200E-08 | 5.897E-08 | 4.500E-08 | 1.500E-07 |
| 24163 | 1209 | D1209N | D | 9.7e-08 | 3.2e-08 | 1.9e-07 | 1.063E-07 | 7.941E-08 | 3.200E-08 | 1.900E-07 |
| 24164 | 1209 | D1209D | D | 3.1e-08 | 4.2e-08 | 4.5e-08 | 3.933E-08 | 7.371E-09 | 3.100E-08 | 4.500E-08 |
| 24165 | 1209 | D1209C | D | 4.5e-08 | 1.6e-08 | 3.2e-08 | 3.100E-08 | 1.453E-08 | 1.600E-08 | 4.500E-08 |
| 24166 | 1209 | D1209E | D | 5.2e-08 | 4.3e-08 | 5.2e-08 | 4.900E-08 | 5.196E-09 | 4.300E-08 | 5.200E-08 |
| 24167 | 1209 | D1209Q | D | 3.8e-08 | 2.4e-08 | 1e-07 | 5.400E-08 | 4.045E-08 | 2.400E-08 | 1.000E-07 |
| 24168 | 1209 | D1209G | D | 2.3e-08 | 2.2e-08 | 3.1e-08 | 2.533E-08 | 4.933E-09 | 2.200E-08 | 3.100E-08 |
| 24169 | 1209 | D1209H | D | 4.5e-08 | 2.6e-08 | 4.1e-08 | 3.733E-08 | 1.002E-08 | 2.600E-08 | 4.500E-08 |
| 24170 | 1209 | D1209I | D | 4.1e-08 | 7.4e-08 | 3.3e-08 | 4.933E-08 | 2.173E-08 | 3.300E-08 | 7.400E-08 |
| 24171 | 1209 | D1209L | D | 1.5e-08 | 6.8e-08 | 8e-08 | 5.433E-08 | 3.459E-08 | 1.500E-08 | 8.000E-08 |
| 24172 | 1209 | D1209K | D | 3e-08 | 5.9e-08 | 1.3e-07 | 7.300E-08 | 5.145E-08 | 3.000E-08 | 1.300E-07 |
| 24173 | 1209 | D1209M | D | 3.1e-08 | 2.7e-08 | 4.5e-08 | 3.433E-08 | 9.452E-09 | 2.700E-08 | 4.500E-08 |
| 24174 | 1209 | D1209F | D | 5.7e-08 | 3.2e-08 | 6.1e-08 | 5.000E-08 | 1.572E-08 | 3.200E-08 | 6.100E-08 |
| 24175 | 1209 | D1209P | D | 4.1e-08 | 5.7e-08 | 9.9e-08 | 6.567E-08 | 2.996E-08 | 4.100E-08 | 9.900E-08 |
| 24176 | 1209 | D1209S | D | 3.4e-08 | 1.3e-08 | 3.5e-08 | 2.733E-08 | 1.242E-08 | 1.300E-08 | 3.500E-08 |
| 24177 | 1209 | D1209T | D | 1.3e-08 | 5.7e-08 | 2.5e-08 | 3.167E-08 | 2.274E-08 | 1.300E-08 | 5.700E-08 |
| 24178 | 1209 | D1209W | D | 6.5e-08 | 3.4e-08 | 9e-08 | 6.300E-08 | 2.805E-08 | 3.400E-08 | 9.000E-08 |
| 24179 | 1209 | D1209Y | D | 3.8e-08 | 5.7e-08 | 2.5e-08 | 4.000E-08 | 1.609E-08 | 2.500E-08 | 5.700E-08 |
| 24180 | 1209 | D1209V | D | 6e-08 | 1.1e-07 | 3.4e-08 | 6.800E-08 | 3.863E-08 | 3.400E-08 | 1.100E-07 |
| 24181 | 1210 | N1210A | D | 1.9e-08 | 6.7e-08 | 6.4e-08 | 5.000E-08 | 2.689E-08 | 1.900E-08 | 6.700E-08 |
| 24182 | 1210 | N1210R | D | 1e-08 | 3.1e-08 | 6.6e-08 | 3.567E-08 | 2.829E-08 | 1.000E-08 | 6.600E-08 |
| 24183 | 1210 | N1210N | D | 3.5e-08 | 2.7e-08 | 1.7e-08 | 2.633E-08 | 9.018E-09 | 1.700E-08 | 3.500E-08 |
| 24184 | 1210 | N1210D | D | 2.7e-08 | 2.6e-08 | 9.1e-08 | 4.800E-08 | 3.724E-08 | 2.600E-08 | 9.100E-08 |
| 24185 | 1210 | N1210C | D | 9.1e-08 | 5.3e-08 | 4.2e-08 | 6.200E-08 | 2.571E-08 | 4.200E-08 | 9.100E-08 |
| 24186 | 1210 | N1210E | D | 4.5e-08 | 5e-08 | 3.8e-08 | 4.433E-08 | 6.028E-09 | 3.800E-08 | 5.000E-08 |
| 24187 | 1210 | N1210Q | D | 9.6e-08 | 7.9e-08 | 2.7e-08 | 6.733E-08 | 3.595E-08 | 2.700E-08 | 9.600E-08 |
| 24188 | 1210 | N1210G | D | 5.2e-08 | 5.2e-09 | 1.9e-07 | 8.240E-08 | 9.608E-08 | 5.200E-09 | 1.900E-07 |
| 24189 | 1210 | N1210H | D | 1.2e-07 | 1.6e-08 | 6.4e-08 | 6.667E-08 | 5.205E-08 | 1.600E-08 | 1.200E-07 |
| 24190 | 1210 | N1210I | D | 1.8e-08 | 3e-08 | 8.6e-08 | 4.467E-08 | 3.630E-08 | 1.800E-08 | 8.600E-08 |
| 24191 | 1210 | N1210L | D | 5.4e-08 | 1.4e-08 | 5e-08 | 3.933E-08 | 2.203E-08 | 1.400E-08 | 5.400E-08 |
| 24192 | 1210 | N1210K | D | 8.3e-08 | 5.7e-08 | 3.3e-08 | 5.767E-08 | 2.501E-08 | 3.300E-08 | 8.300E-08 |
| 24193 | 1210 | N1210M | D | 6.9e-08 | 3.3e-08 | 7.3e-08 | 5.833E-08 | 2.203E-08 | 3.300E-08 | 7.300E-08 |
| 24194 | 1210 | N1210F | D | 2.1e-08 | 4e-08 | 2.9e-08 | 3.000E-08 | 9.539E-09 | 2.100E-08 | 4.000E-08 |
| 24195 | 1210 | N1210P | D | 3.9e-08 | 1.4e-08 | 3.7e-08 | 3.000E-08 | 1.389E-08 | 1.400E-08 | 3.900E-08 |
| 24196 | 1210 | N1210S | D | 1.1e-07 | 4.3e-08 | 1.9e-08 | 5.733E-08 | 4.716E-08 | 1.900E-08 | 1.100E-07 |
| 24197 | 1210 | N1210T | D | 4.3e-08 | 9.6e-08 | 3.4e-08 | 5.767E-08 | 3.350E-08 | 3.400E-08 | 9.600E-08 |
| 24198 | 1210 | N1210W | D | 2.1e-08 | 5.3e-08 | 3.1e-08 | 3.500E-08 | 1.637E-08 | 2.100E-08 | 5.300E-08 |
| 24199 | 1210 | N1210Y | D | 6.3e-08 | 4.1e-08 | 4e-08 | 4.800E-08 | 1.300E-08 | 4.000E-08 | 6.300E-08 |
| 24200 | 1210 | N1210V | D | 4.3e-08 | 2.5e-08 | 1.7e-08 | 2.833E-08 | 1.332E-08 | 1.700E-08 | 4.300E-08 |
| 24201 | 1211 | A1211A | D | 9.2e-08 | 6.1e-08 | 3.9e-08 | 6.400E-08 | 2.663E-08 | 3.900E-08 | 9.200E-08 |
| 24202 | 1211 | A1211R | D | 4.4e-08 | 1.1e-07 | 4.7e-08 | 6.700E-08 | 3.727E-08 | 4.400E-08 | 1.100E-07 |
| 24203 | 1211 | A1211N | D | 2.4e-08 | 4.3e-08 | 5.7e-08 | 4.133E-08 | 1.656E-08 | 2.400E-08 | 5.700E-08 |
| 24204 | 1211 | A1211D | D | 1.8e-08 | 4.7e-08 | 2.3e-08 | 2.933E-08 | 1.550E-08 | 1.800E-08 | 4.700E-08 |
| 24205 | 1211 | A1211C | D | 6.6e-08 | 8.5e-08 | 6.7e-08 | 7.267E-08 | 1.069E-08 | 6.600E-08 | 8.500E-08 |
| 24206 | 1211 | A1211E | D | 3.5e-08 | 7.8e-08 | 2.1e-08 | 4.467E-08 | 2.970E-08 | 2.100E-08 | 7.800E-08 |
| 24207 | 1211 | A1211Q | D | 6.9e-08 | 5.6e-08 | 3e-08 | 5.167E-08 | 1.986E-08 | 3.000E-08 | 6.900E-08 |
| 24208 | 1211 | A1211G | D | 4.2e-08 | 2.3e-08 | 2.5e-08 | 3.000E-08 | 1.044E-08 | 2.300E-08 | 4.200E-08 |
| 24209 | 1211 | A1211H | D | 1.6e-08 | 6.8e-08 | 3.9e-08 | 4.100E-08 | 2.606E-08 | 1.600E-08 | 6.800E-08 |
| 24210 | 1211 | A1211I | D | 2.8e-08 | 5.1e-08 | 5.5e-08 | 4.467E-08 | 1.457E-08 | 2.800E-08 | 5.500E-08 |
| 24211 | 1211 | A1211L | D | 1.3e-07 | 3.5e-08 | 4.1e-08 | 6.867E-08 | 5.320E-08 | 3.500E-08 | 1.300E-07 |
| 24212 | 1211 | A1211K | D | 1.9e-08 | 3.1e-08 | 3.4e-08 | 2.800E-08 | 7.937E-09 | 1.900E-08 | 3.400E-08 |
| 24213 | 1211 | A1211M | D | 3.7e-08 | 5.5e-08 | 5.4e-08 | 4.867E-08 | 1.012E-08 | 3.700E-08 | 5.500E-08 |
| 24214 | 1211 | A1211F | D | 6.2e-08 | 2.9e-08 | 6.7e-09 | 3.257E-08 | 2.782E-08 | 6.700E-09 | 6.200E-08 |
| 24215 | 1211 | A1211P | D | 2.6e-08 | 3.6e-08 | 1.8e-07 | 8.067E-08 | 8.617E-08 | 2.600E-08 | 1.800E-07 |
| 24216 | 1211 | A1211S | D | 4.3e-08 | 2.7e-08 | 2.5e-08 | 3.167E-08 | 9.866E-09 | 2.500E-08 | 4.300E-08 |
| 24217 | 1211 | A1211T | D | 8e-08 | 6.9e-08 | 5.1e-08 | 6.667E-08 | 1.464E-08 | 5.100E-08 | 8.000E-08 |
| 24218 | 1211 | A1211W | D | 6.2e-08 | 7.4e-08 | 1.9e-08 | 5.167E-08 | 2.892E-08 | 1.900E-08 | 7.400E-08 |
| 24219 | 1211 | A1211Y | D | 3.4e-08 | 2.9e-08 | 2.6e-08 | 2.967E-08 | 4.041E-09 | 2.600E-08 | 3.400E-08 |
| 24220 | 1211 | A1211V | D | 8.4e-08 | 2.1e-08 | 3.3e-08 | 4.600E-08 | 3.345E-08 | 2.100E-08 | 8.400E-08 |
| 24221 | 1212 | L1212A | D | 1.4e-08 | 1e-07 | 2.5e-08 | 4.633E-08 | 4.680E-08 | 1.400E-08 | 1.000E-07 |
| 24222 | 1212 | L1212R | D | 4.5e-08 | 5.7e-08 | 2.4e-08 | 4.200E-08 | 1.670E-08 | 2.400E-08 | 5.700E-08 |
| 24223 | 1212 | L1212N | D | 9.1e-08 | 3.7e-08 | 5.1e-08 | 5.967E-08 | 2.802E-08 | 3.700E-08 | 9.100E-08 |
| 24224 | 1212 | L1212D | D | 3.7e-08 | 3.5e-08 | 1.8e-08 | 3.000E-08 | 1.044E-08 | 1.800E-08 | 3.700E-08 |

| | | | | | | | | | | |
|---|---|---|---|---|---|---|---|---|---|---|
| 24225 | 1212 | L1212C | D | 1.7e-08 | 5.6e-08 | 1e-08 | 2.767E-08 | 2.479E-08 | 1.000E-08 | 5.600E-08 |
| 24226 | 1212 | L1212E | D | 3.5e-08 | 8.9e-08 | 8.5e-08 | 6.967E-08 | 3.009E-08 | 3.500E-08 | 8.900E-08 |
| 24227 | 1212 | L1212Q | D | 7.9e-08 | 6.1e-08 | 1.4e-07 | 9.333E-08 | 4.140E-08 | 6.100E-08 | 1.400E-07 |
| 24228 | 1212 | L1212G | D | 5.9e-08 | 3.3e-08 | 2.4e-08 | 3.867E-08 | 1.818E-08 | 2.400E-08 | 5.900E-08 |
| 24229 | 1212 | L1212H | D | 2.8e-08 | 6.3e-08 | 2e-08 | 3.700E-08 | 2.287E-08 | 2.000E-08 | 6.300E-08 |
| 24230 | 1212 | L1212I | D | 3.4e-08 | 1.6e-08 | 1.8e-08 | 2.267E-08 | 9.866E-09 | 1.600E-08 | 3.400E-08 |
| 24231 | 1212 | L1212L | D | 3.5e-08 | 7.6e-08 | 8.4e-08 | 6.500E-08 | 2.629E-08 | 3.500E-08 | 8.400E-08 |
| 24232 | 1212 | L1212K | D | 1.1e-07 | 8.7e-08 | 1.1e-07 | 1.023E-07 | 1.328E-08 | 8.700E-08 | 1.100E-07 |
| 24233 | 1212 | L1212M | D | 4.2e-08 | 3.8e-08 | 3.3e-08 | 3.767E-08 | 4.509E-09 | 3.300E-08 | 4.200E-08 |
| 24234 | 1212 | L1212F | D | 1.5e-08 | 1.3e-08 | 3e-08 | 1.933E-08 | 9.292E-09 | 1.300E-08 | 3.000E-08 |
| 24235 | 1212 | L1212P | D | 9.6e-08 | 8.3e-08 | 4.1e-08 | 7.333E-08 | 2.875E-08 | 4.100E-08 | 9.600E-08 |
| 24236 | 1212 | L1212S | D | 1.5e-08 | 1.8e-08 | 2.2e-08 | 1.833E-08 | 3.512E-09 | 1.500E-08 | 2.200E-08 |
| 24237 | 1212 | L1212T | D | 2.9e-08 | 1.7e-07 | 4.6e-08 | 8.167E-08 | 7.697E-08 | 2.900E-08 | 1.700E-07 |
| 24238 | 1212 | L1212W | D | 5.1e-08 | 1.5e-08 | 4.4e-08 | 3.667E-08 | 1.909E-08 | 1.500E-08 | 5.100E-08 |
| 24239 | 1212 | L1212Y | D | 2.8e-08 | 3.2e-08 | 1.3e-08 | 2.433E-08 | 1.002E-08 | 1.300E-08 | 3.200E-08 |
| 24240 | 1212 | L1212V | D | 1.6e-07 | 6.2e-08 | 2.1e-08 | 8.100E-08 | 7.142E-08 | 2.100E-08 | 1.600E-07 |
| 24241 | 1213 | Q1213A | D | 6.5e-08 | 2.4e-08 | 5.6e-08 | 4.833E-08 | 2.155E-08 | 2.400E-08 | 6.500E-08 |
| 24242 | 1213 | Q1213R | D | 1.3e-08 | 8.4e-08 | 9.9e-08 | 6.533E-08 | 4.594E-08 | 1.300E-08 | 9.900E-08 |
| 24243 | 1213 | Q1213N | D | 4.7e-08 | 1.4e-07 | 1.9e-08 | 6.867E-08 | 6.334E-08 | 1.900E-08 | 1.400E-07 |
| 24244 | 1213 | Q1213D | D | 3.8e-08 | 5e-08 | 3.6e-08 | 4.133E-08 | 7.572E-09 | 3.600E-08 | 5.000E-08 |
| 24245 | 1213 | Q1213C | D | 3.6e-08 | 6.7e-08 | 1e-07 | 6.767E-08 | 3.201E-08 | 3.600E-08 | 1.000E-07 |
| 24246 | 1213 | Q1213E | D | 5e-08 | 2.1e-08 | 5.5e-08 | 4.200E-08 | 1.836E-08 | 2.100E-08 | 5.500E-08 |
| 24247 | 1213 | Q1213Q | D | 8e-08 | 4.5e-08 | 1.7e-08 | 4.733E-08 | 3.156E-08 | 1.700E-08 | 8.000E-08 |
| 24248 | 1213 | Q1213G | D | 8.3e-08 | 6.2e-08 | 3.6e-08 | 6.033E-08 | 2.354E-08 | 3.600E-08 | 8.300E-08 |
| 24249 | 1213 | Q1213H | D | 4.6e-08 | 3.3e-08 | 2.7e-08 | 3.533E-08 | 9.713E-09 | 2.700E-08 | 4.600E-08 |
| 24250 | 1213 | Q1213I | D | 9.8e-08 | 2.2e-08 | 5.8e-08 | 5.933E-08 | 3.802E-08 | 2.200E-08 | 9.800E-08 |
| 24251 | 1213 | Q1213L | D | 8e-08 | 2.3e-08 | 1.1e-07 | 7.100E-08 | 4.419E-08 | 2.300E-08 | 1.100E-07 |
| 24252 | 1213 | Q1213K | D | 4e-08 | 2.9e-08 | 5.4e-08 | 4.100E-08 | 1.253E-08 | 2.900E-08 | 5.400E-08 |
| 24253 | 1213 | Q1213M | D | 6.2e-08 | 5.6e-08 | 4.3e-08 | 5.367E-08 | 9.713E-09 | 4.300E-08 | 6.200E-08 |
| 24254 | 1213 | Q1213F | D | 3.3e-08 | 7.1e-08 | 5.6e-08 | 5.333E-08 | 1.914E-08 | 3.300E-08 | 7.100E-08 |
| 24255 | 1213 | Q1213P | D | 2.5e-08 | 4.4e-08 | 4.9e-08 | 3.933E-08 | 1.266E-08 | 2.500E-08 | 4.900E-08 |
| 24256 | 1213 | Q1213S | D | 7.2e-08 | 5.1e-08 | 5.4e-08 | 5.900E-08 | 1.136E-08 | 5.100E-08 | 7.200E-08 |
| 24257 | 1213 | Q1213T | D | 1.1e-07 | 4e-08 | 5.9e-08 | 6.967E-08 | 3.620E-08 | 4.000E-08 | 1.100E-07 |
| 24258 | 1213 | Q1213W | D | 6.6e-08 | 5.8e-09 | 2.5e-08 | 3.227E-08 | 3.075E-08 | 5.800E-09 | 6.600E-08 |
| 24259 | 1213 | Q1213Y | D | 1.7e-08 | 5.4e-08 | 9.2e-08 | 5.433E-08 | 3.750E-08 | 1.700E-08 | 9.200E-08 |
| 24260 | 1213 | Q1213V | D | 5.3e-08 | 3.8e-08 | 6.8e-08 | 5.300E-08 | 1.500E-08 | 3.800E-08 | 6.800E-08 |
| 24261 | 1214 | S1214A | D | 5e-08 | 3.4e-08 | 4.5e-08 | 4.300E-08 | 8.185E-09 | 3.400E-08 | 5.000E-08 |
| 24262 | 1214 | S1214R | D | 7.3e-08 | 9.8e-08 | 2.6e-08 | 6.567E-08 | 3.656E-08 | 2.600E-08 | 9.800E-08 |
| 24263 | 1214 | S1214N | D | 1.9e-08 | 8.2e-08 | 7.4e-08 | 5.833E-08 | 3.430E-08 | 1.900E-08 | 8.200E-08 |
| 24264 | 1214 | S1214D | D | 6.7e-08 | 2.6e-08 | 2.1e-08 | 3.800E-08 | 2.524E-08 | 2.100E-08 | 6.700E-08 |
| 24265 | 1214 | S1214C | D | 4.9e-08 | 5.1e-08 | 4e-08 | 4.667E-08 | 5.859E-09 | 4.000E-08 | 5.100E-08 |
| 24266 | 1214 | S1214E | D | 3.4e-08 | 2.2e-08 | 2.6e-08 | 2.733E-08 | 6.110E-09 | 2.200E-08 | 3.400E-08 |
| 24267 | 1214 | S1214Q | D | 5.2e-08 | 4.4e-08 | 4.8e-08 | 4.800E-08 | 4.000E-09 | 4.400E-08 | 5.200E-08 |
| 24268 | 1214 | S1214G | D | 7.1e-08 | 5.2e-08 | 1.3e-08 | 4.533E-08 | 2.957E-08 | 1.300E-08 | 7.100E-08 |
| 24269 | 1214 | S1214H | D | 3.3e-08 | 1.9e-08 | 3.1e-08 | 2.767E-08 | 7.572E-09 | 1.900E-08 | 3.300E-08 |
| 24270 | 1214 | S1214I | D | 4e-08 | 6.1e-08 | 2.7e-08 | 4.267E-08 | 1.716E-08 | 2.700E-08 | 6.100E-08 |
| 24271 | 1214 | S1214L | D | 6.2e-08 | 4.7e-08 | 3.2e-08 | 4.700E-08 | 1.500E-08 | 3.200E-08 | 6.200E-08 |
| 24272 | 1214 | S1214K | D | 3.2e-08 | 1.3e-07 | 6.5e-08 | 7.567E-08 | 4.986E-08 | 3.200E-08 | 1.300E-07 |
| 24273 | 1214 | S1214M | D | 6.8e-08 | 3.7e-08 | 7.1e-08 | 5.867E-08 | 1.882E-08 | 3.700E-08 | 7.100E-08 |
| 24274 | 1214 | S1214F | D | 1.4e-08 | 5.3e-08 | 5e-08 | 3.900E-08 | 2.170E-08 | 1.400E-08 | 5.300E-08 |
| 24275 | 1214 | S1214P | D | 1.1e-07 | 4.1e-08 | 1.7e-08 | 5.600E-08 | 4.828E-08 | 1.700E-08 | 1.100E-07 |
| 24276 | 1214 | S1214S | D | 1.2e-08 | 1.9e-08 | 2.8e-08 | 1.967E-08 | 8.021E-09 | 1.200E-08 | 2.800E-08 |
| 24277 | 1214 | S1214T | D | 4.6e-08 | 5.1e-08 | 3.6e-08 | 4.433E-08 | 7.638E-09 | 3.600E-08 | 5.100E-08 |
| 24278 | 1214 | S1214W | D | 3.2e-08 | 2.5e-08 | 1.4e-08 | 2.367E-08 | 9.074E-09 | 1.400E-08 | 3.200E-08 |
| 24279 | 1214 | S1214Y | D | 3.6e-08 | 2.6e-08 | 4.7e-08 | 3.633E-08 | 1.050E-08 | 2.600E-08 | 4.700E-08 |
| 24280 | 1214 | S1214V | D | 5e-08 | 2e-08 | 4.5e-08 | 3.833E-08 | 1.607E-08 | 2.000E-08 | 5.000E-08 |
| 24281 | 1215 | G1215A | D | 8.5e-08 | 7.3e-08 | 5.8e-08 | 7.200E-08 | 1.353E-08 | 5.800E-08 | 8.500E-08 |
| 24282 | 1215 | G1215R | D | 2.4e-08 | 6e-08 | 2e-08 | 3.467E-08 | 2.203E-08 | 2.000E-08 | 6.000E-08 |
| 24283 | 1215 | G1215N | D | 3.9e-08 | 1.9e-08 | 1.6e-08 | 2.467E-08 | 1.250E-08 | 1.600E-08 | 3.900E-08 |
| 24284 | 1215 | G1215D | D | 4e-08 | 5.1e-08 | 5.3e-08 | 4.800E-08 | 7.000E-09 | 4.000E-08 | 5.300E-08 |
| 24285 | 1215 | G1215C | D | 4.4e-08 | 2.6e-08 | 5.1e-08 | 4.033E-08 | 1.290E-08 | 2.600E-08 | 5.100E-08 |
| 24286 | 1215 | G1215E | D | 2.9e-08 | 5e-08 | 4.7e-08 | 4.200E-08 | 1.136E-08 | 2.900E-08 | 5.000E-08 |
| 24287 | 1215 | G1215Q | D | 1.2e-08 | 6.7e-08 | 8.1e-08 | 5.333E-08 | 3.647E-08 | 1.200E-08 | 8.100E-08 |
| 24288 | 1215 | G1215G | D | 5.3e-08 | 6.7e-08 | 5.7e-08 | 5.900E-08 | 7.211E-09 | 5.300E-08 | 6.700E-08 |
| 24289 | 1215 | G1215H | D | 1.7e-08 | 3.5e-08 | 6.2e-08 | 3.800E-08 | 2.265E-08 | 1.700E-08 | 6.200E-08 |
| 24290 | 1215 | G1215I | D | 3.9e-08 | 6.4e-08 | 2.5e-08 | 4.267E-08 | 1.976E-08 | 2.500E-08 | 6.400E-08 |
| 24291 | 1215 | G1215L | D | 2.9e-08 | 9.1e-08 | 3.7e-08 | 5.233E-08 | 3.372E-08 | 2.900E-08 | 9.100E-08 |
| 24292 | 1215 | G1215K | D | 3.3e-08 | 5.9e-08 | 5.1e-08 | 4.767E-08 | 1.332E-08 | 3.300E-08 | 5.900E-08 |
| 24293 | 1215 | G1215M | D | 2e-08 | 3.9e-08 | 4e-08 | 3.300E-08 | 1.127E-08 | 2.000E-08 | 4.000E-08 |
| 24294 | 1215 | G1215F | D | 3.4e-08 | 4.4e-08 | 3.2e-08 | 3.667E-08 | 6.429E-09 | 3.200E-08 | 4.400E-08 |
| 24295 | 1215 | G1215P | D | 3.5e-08 | 7.4e-08 | 3.8e-08 | 4.900E-08 | 2.170E-08 | 3.500E-08 | 7.400E-08 |
| 24296 | 1215 | G1215S | D | 1.1e-07 | 9.5e-08 | 2.4e-08 | 7.633E-08 | 4.594E-08 | 2.400E-08 | 1.100E-07 |
| 24297 | 1215 | G1215T | D | 6.3e-08 | 6.6e-08 | 9.3e-08 | 7.400E-08 | 1.652E-08 | 6.300E-08 | 9.300E-08 |
| 24298 | 1215 | G1215W | D | 5.1e-08 | 2.6e-08 | 5.3e-08 | 4.333E-08 | 1.504E-08 | 2.600E-08 | 5.300E-08 |
| 24299 | 1215 | G1215Y | D | 5.7e-08 | 1e-07 | 1.5e-08 | 5.733E-08 | 4.250E-08 | 1.500E-08 | 1.000E-07 |
| 24300 | 1215 | G1215V | D | 3.3e-08 | 1.3e-08 | 1.5e-08 | 2.033E-08 | 1.102E-08 | 1.300E-08 | 3.300E-08 |
| 24301 | 1216 | N1216A | D | 2.8e-08 | 2.9e-08 | 3.7e-08 | 3.133E-08 | 4.933E-09 | 2.800E-08 | 3.700E-08 |
| 24302 | 1216 | N1216R | D | 9.9e-08 | 9.1e-08 | 4.8e-08 | 7.933E-08 | 2.743E-08 | 4.800E-08 | 9.900E-08 |
| 24303 | 1216 | N1216N | D | 3.3e-08 | 3.1e-08 | 4.6e-08 | 3.667E-08 | 8.145E-09 | 3.100E-08 | 4.600E-08 |
| 24304 | 1216 | N1216D | D | 3.5e-08 | 5.5e-08 | 3.7e-08 | 4.233E-08 | 1.102E-08 | 3.500E-08 | 5.500E-08 |
| 24305 | 1216 | N1216C | D | 4.4e-08 | 3.6e-08 | 9e-08 | 5.667E-08 | 2.914E-08 | 3.600E-08 | 9.000E-08 |
| 24306 | 1216 | N1216E | D | 3.7e-08 | 4.8e-08 | 1.9e-08 | 3.467E-08 | 1.464E-08 | 1.900E-08 | 4.800E-08 |
| 24307 | 1216 | N1216Q | D | 2.5e-08 | 2.4e-08 | 6.1e-08 | 3.667E-08 | 2.108E-08 | 2.400E-08 | 6.100E-08 |
| 24308 | 1216 | N1216G | D | 1.5e-08 | 1.4e-07 | 6.1e-08 | 7.200E-08 | 6.322E-08 | 1.500E-08 | 1.400E-07 |
| 24309 | 1216 | N1216H | D | 3.1e-08 | 3.8e-08 | 4.5e-08 | 3.800E-08 | 7.000E-09 | 3.100E-08 | 4.500E-08 |

| | | | | | | | | | | |
|---|---|---|---|---|---|---|---|---|---|---|
| 24310 | 1216 | N1216I | D | 2.5e-08 | 4e-08 | 1.9e-08 | 2.800E-08 | 1.082E-08 | 1.900E-08 | 4.000E-08 |
| 24311 | 1216 | N1216L | D | 1.6e-08 | 1.7e-08 | 2.3e-08 | 1.867E-08 | 3.786E-09 | 1.600E-08 | 2.300E-08 |
| 24312 | 1216 | N1216K | D | 8.1e-08 | 7.4e-08 | 3.4e-08 | 6.300E-08 | 2.536E-08 | 3.400E-08 | 8.100E-08 |
| 24313 | 1216 | N1216M | D | 5.4e-08 | 1.1e-07 | 2.3e-08 | 6.233E-08 | 4.409E-08 | 2.300E-08 | 1.100E-07 |
| 24314 | 1216 | N1216F | D | 2.5e-08 | 2.4e-08 | 3.7e-08 | 2.867E-08 | 7.234E-09 | 2.400E-08 | 3.700E-08 |
| 24315 | 1216 | N1216P | D | 1.7e-08 | 1.1e-07 | 6.1e-08 | 6.267E-08 | 4.652E-08 | 1.700E-08 | 1.100E-07 |
| 24316 | 1216 | N1216S | D | 5.7e-08 | 1.6e-07 | 4.1e-08 | 8.600E-08 | 6.458E-08 | 4.100E-08 | 1.600E-07 |
| 24317 | 1216 | N1216T | D | 2.3e-08 | 6e-08 | 3.1e-08 | 3.800E-08 | 1.947E-08 | 2.300E-08 | 6.000E-08 |
| 24318 | 1216 | N1216W | D | 4.1e-08 | 6.7e-08 | 7.3e-08 | 6.033E-08 | 1.701E-08 | 4.100E-08 | 7.300E-08 |
| 24319 | 1216 | N1216Y | D | 6.9e-08 | 2.3e-08 | 7e-09 | 3.300E-08 | 3.219E-08 | 7.000E-09 | 6.900E-08 |
| 24320 | 1216 | N1216V | D | 4.8e-08 | 1.5e-07 | 6.6e-08 | 8.800E-08 | 5.444E-08 | 4.800E-08 | 1.500E-07 |
| 24321 | 1217 | S1217A | D | 4.5e-08 | 3.4e-08 | 1.9e-08 | 3.267E-08 | 1.305E-08 | 1.900E-08 | 4.500E-08 |
| 24322 | 1217 | S1217R | D | 6.3e-08 | 7.3e-08 | 7.5e-08 | 7.033E-08 | 6.429E-09 | 6.300E-08 | 7.500E-08 |
| 24323 | 1217 | S1217N | D | 3.1e-08 | 4.5e-08 | 5.9e-08 | 4.500E-08 | 1.400E-08 | 3.100E-08 | 5.900E-08 |
| 24324 | 1217 | S1217D | D | 6.7e-08 | 1.5e-07 | 3.8e-08 | 8.500E-08 | 5.813E-08 | 3.800E-08 | 1.500E-07 |
| 24325 | 1217 | S1217C | D | 5.1e-08 | 3.5e-08 | 1.9e-08 | 3.500E-08 | 1.600E-08 | 1.900E-08 | 5.100E-08 |
| 24326 | 1217 | S1217E | D | 3.7e-08 | 3e-08 | 7.9e-08 | 4.867E-08 | 2.650E-08 | 3.000E-08 | 7.900E-08 |
| 24327 | 1217 | S1217Q | D | 2.1e-08 | 6.6e-08 | 4.6e-08 | 4.433E-08 | 2.255E-08 | 2.100E-08 | 6.600E-08 |
| 24328 | 1217 | S1217G | D | 3e-08 | 2.2e-08 | 1.7e-08 | 2.300E-08 | 6.557E-09 | 1.700E-08 | 3.000E-08 |
| 24329 | 1217 | S1217H | D | 2.2e-08 | 7.3e-08 | 2.4e-08 | 3.967E-08 | 2.888E-08 | 2.200E-08 | 7.300E-08 |
| 24330 | 1217 | S1217I | D | 5.5e-08 | 3.3e-08 | 7.3e-08 | 5.367E-08 | 2.003E-08 | 3.300E-08 | 7.300E-08 |
| 24331 | 1217 | S1217L | D | 5.4e-08 | 7.2e-08 | 3e-08 | 5.200E-08 | 2.107E-08 | 3.000E-08 | 7.200E-08 |
| 24332 | 1217 | S1217K | D | 1.9e-07 | 1.8e-08 | 1.3e-07 | 1.127E-07 | 8.730E-08 | 1.800E-08 | 1.900E-07 |
| 24333 | 1217 | S1217M | D | 1.4e-08 | 9.2e-08 | 5.5e-08 | 5.367E-08 | 3.902E-08 | 1.400E-08 | 9.200E-08 |
| 24334 | 1217 | S1217F | D | 4.5e-08 | 2.7e-08 | 1.3e-08 | 2.833E-08 | 1.604E-08 | 1.300E-08 | 4.500E-08 |
| 24335 | 1217 | S1217P | D | 4.9e-08 | 3.2e-08 | 1.9e-08 | 3.333E-08 | 1.504E-08 | 1.900E-08 | 4.900E-08 |
| 24336 | 1217 | S1217S | D | 4.2e-08 | 8e-08 | 5.4e-08 | 5.867E-08 | 1.943E-08 | 4.200E-08 | 8.000E-08 |
| 24337 | 1217 | S1217T | D | 3.7e-08 | 1.5e-08 | 4e-08 | 3.067E-08 | 1.365E-08 | 1.500E-08 | 4.000E-08 |
| 24338 | 1217 | S1217W | D | 6.8e-08 | 2.1e-08 | 2.1e-08 | 3.667E-08 | 2.714E-08 | 2.100E-08 | 6.800E-08 |
| 24339 | 1217 | S1217Y | D | 3.2e-08 | 3.8e-08 | 2.7e-08 | 3.233E-08 | 5.508E-09 | 2.700E-08 | 3.800E-08 |
| 24340 | 1217 | S1217V | D | 7.6e-08 | 3.9e-08 | 4.1e-08 | 5.200E-08 | 2.081E-08 | 3.900E-08 | 7.600E-08 |
| 24341 | 1218 | Q1218A | D | 1.2e-07 | 4.1e-08 | 6.6e-08 | 7.567E-08 | 4.038E-08 | 4.100E-08 | 1.200E-07 |
| 24342 | 1218 | Q1218R | D | 4.9e-08 | 3.5e-08 | 2.9e-08 | 3.767E-08 | 1.026E-08 | 2.900E-08 | 4.900E-08 |
| 24343 | 1218 | Q1218N | D | 4.5e-08 | 4.1e-08 | 2.4e-08 | 3.667E-08 | 1.115E-08 | 2.400E-08 | 4.500E-08 |
| 24344 | 1218 | Q1218D | D | 2.2e-08 | 4.4e-08 | 9.1e-08 | 5.233E-08 | 3.525E-08 | 2.200E-08 | 9.100E-08 |
| 24345 | 1218 | Q1218C | D | 3.6e-08 | 4e-08 | 8.7e-08 | 5.433E-08 | 2.836E-08 | 3.600E-08 | 8.700E-08 |
| 24346 | 1218 | Q1218E | D | 6.2e-08 | 2.4e-08 | 6.5e-08 | 5.033E-08 | 2.285E-08 | 2.400E-08 | 6.500E-08 |
| 24347 | 1218 | Q1218Q | D | 6.5e-08 | 4.3e-08 | 6.1e-08 | 5.633E-08 | 1.172E-08 | 4.300E-08 | 6.500E-08 |
| 24348 | 1218 | Q1218G | D | 8.4e-08 | 5e-08 | 3.3e-08 | 5.567E-08 | 2.597E-08 | 3.300E-08 | 8.400E-08 |
| 24349 | 1218 | Q1218H | D | 4.6e-08 | 2.7e-08 | 4.8e-08 | 4.033E-08 | 1.159E-08 | 2.700E-08 | 4.800E-08 |
| 24350 | 1218 | Q1218I | D | 2.9e-08 | 5.3e-08 | 6.6e-08 | 4.933E-08 | 1.877E-08 | 2.900E-08 | 6.600E-08 |
| 24351 | 1218 | Q1218L | D | 2.7e-08 | 4.3e-08 | 1.9e-08 | 2.967E-08 | 1.222E-08 | 1.900E-08 | 4.300E-08 |
| 24352 | 1218 | Q1218K | D | 4.2e-08 | 6.1e-08 | 2.2e-08 | 4.167E-08 | 1.950E-08 | 2.200E-08 | 6.100E-08 |
| 24353 | 1218 | Q1218M | D | 1.2e-08 | 3.4e-08 | 5.5e-08 | 3.367E-08 | 2.150E-08 | 1.200E-08 | 5.500E-08 |
| 24354 | 1218 | Q1218F | D | 3.6e-08 | 4.6e-08 | 7.1e-08 | 5.100E-08 | 1.803E-08 | 3.600E-08 | 7.100E-08 |
| 24355 | 1218 | Q1218P | D | 6.7e-08 | 5.1e-08 | 7.6e-08 | 6.467E-08 | 1.266E-08 | 5.100E-08 | 7.600E-08 |
| 24356 | 1218 | Q1218S | D | 5.2e-08 | 6.1e-08 | 5e-08 | 5.433E-08 | 5.859E-09 | 5.000E-08 | 6.100E-08 |
| 24357 | 1218 | Q1218T | D | 1.2e-08 | 5.7e-08 | 2.8e-08 | 3.233E-08 | 2.281E-08 | 1.200E-08 | 5.700E-08 |
| 24358 | 1218 | Q1218W | D | 5.6e-08 | 1e-07 | 3.3e-08 | 6.300E-08 | 3.404E-08 | 3.300E-08 | 1.000E-07 |
| 24359 | 1218 | Q1218Y | D | 2.7e-08 | 1.8e-08 | 7.2e-08 | 3.900E-08 | 2.893E-08 | 1.800E-08 | 7.200E-08 |
| 24360 | 1218 | Q1218V | D | 3.4e-08 | 5.4e-08 | 2.6e-08 | 3.800E-08 | 1.442E-08 | 2.600E-08 | 5.400E-08 |
| 24361 | 1219 | E1219A | D | 5.1e-08 | 2.1e-08 | 8.3e-08 | 5.167E-08 | 3.101E-08 | 2.100E-08 | 8.300E-08 |
| 24362 | 1219 | E1219R | D | 3.5e-08 | 3.8e-08 | 1.5e-07 | 7.433E-08 | 6.555E-08 | 3.500E-08 | 1.500E-07 |
| 24363 | 1219 | E1219N | D | 2.9e-08 | 4.7e-08 | 8e-08 | 5.200E-08 | 2.587E-08 | 2.900E-08 | 8.000E-08 |
| 24364 | 1219 | E1219D | D | 7.2e-08 | 3.5e-08 | 4.4e-08 | 5.033E-08 | 1.930E-08 | 3.500E-08 | 7.200E-08 |
| 24365 | 1219 | E1219C | D | 4.6e-08 | 7.5e-08 | 4.9e-08 | 5.667E-08 | 1.595E-08 | 4.600E-08 | 7.500E-08 |
| 24366 | 1219 | E1219E | D | 1.8e-08 | 7.1e-08 | 3.8e-08 | 4.233E-08 | 2.676E-08 | 1.800E-08 | 7.100E-08 |
| 24367 | 1219 | E1219Q | D | 2e-08 | 9.3e-08 | 9e-08 | 6.767E-08 | 4.131E-08 | 2.000E-08 | 9.300E-08 |
| 24368 | 1219 | E1219G | D | 7.5e-08 | 3.4e-08 | 2.2e-08 | 4.367E-08 | 2.779E-08 | 2.200E-08 | 7.500E-08 |
| 24369 | 1219 | E1219H | D | 4.4e-08 | 6.6e-08 | 7.6e-08 | 6.200E-08 | 1.637E-08 | 4.400E-08 | 7.600E-08 |
| 24370 | 1219 | E1219I | D | 5.2e-08 | 3.7e-08 | 5.5e-08 | 4.800E-08 | 9.644E-09 | 3.700E-08 | 5.500E-08 |
| 24371 | 1219 | E1219L | D | 5.9e-08 | 8.2e-08 | 4.3e-08 | 6.133E-08 | 1.960E-08 | 4.300E-08 | 8.200E-08 |
| 24372 | 1219 | E1219K | D | 4.6e-08 | 2.6e-08 | 3.5e-08 | 3.567E-08 | 1.002E-08 | 2.600E-08 | 4.600E-08 |
| 24373 | 1219 | E1219M | D | 4.1e-08 | 3.7e-08 | 3.8e-08 | 3.867E-08 | 2.082E-09 | 3.700E-08 | 4.100E-08 |
| 24374 | 1219 | E1219F | D | 5.3e-08 | 1e-07 | 1.1e-07 | 8.767E-08 | 3.044E-08 | 5.300E-08 | 1.100E-07 |
| 24375 | 1219 | E1219P | D | 7e-08 | 2.5e-08 | 3.5e-08 | 4.333E-08 | 2.363E-08 | 2.500E-08 | 7.000E-08 |
| 24376 | 1219 | E1219S | D | 3.6e-08 | 5.1e-08 | 2.9e-08 | 3.867E-08 | 1.124E-08 | 2.900E-08 | 5.100E-08 |
| 24377 | 1219 | E1219T | D | 1e-08 | 1.3e-07 | 4.3e-08 | 6.100E-08 | 6.199E-08 | 1.000E-08 | 1.300E-07 |
| 24378 | 1219 | E1219W | D | 4.3e-08 | 1.1e-07 | 1.3e-08 | 5.533E-08 | 4.966E-08 | 1.300E-08 | 1.100E-07 |
| 24379 | 1219 | E1219Y | D | 3.7e-08 | 6e-08 | 1e-08 | 3.567E-08 | 2.503E-08 | 1.000E-08 | 6.000E-08 |
| 24380 | 1219 | E1219V | D | 5.8e-08 | 1.1e-08 | 5.4e-08 | 4.100E-08 | 2.606E-08 | 1.100E-08 | 5.800E-08 |
| 24381 | 1220 | S1220A | D | 7.3e-08 | 3e-08 | 2.5e-08 | 4.267E-08 | 2.639E-08 | 2.500E-08 | 7.300E-08 |
| 24382 | 1220 | S1220R | D | 2.8e-08 | 5.6e-08 | 6.4e-08 | 4.933E-08 | 1.890E-08 | 2.800E-08 | 6.400E-08 |
| 24383 | 1220 | S1220N | D | 1.3e-08 | 4e-08 | 7.2e-08 | 4.167E-08 | 2.954E-08 | 1.300E-08 | 7.200E-08 |
| 24384 | 1220 | S1220D | D | 2.2e-08 | 1.3e-08 | 2.8e-08 | 2.100E-08 | 7.550E-09 | 1.300E-08 | 2.800E-08 |
| 24385 | 1220 | S1220C | D | 5.5e-08 | 6.8e-08 | 1.3e-08 | 4.533E-08 | 2.875E-08 | 1.300E-08 | 6.800E-08 |
| 24386 | 1220 | S1220E | D | 8e-08 | 2.7e-08 | 1.7e-08 | 4.133E-08 | 3.386E-08 | 1.700E-08 | 8.000E-08 |
| 24387 | 1220 | S1220Q | D | 5.1e-08 | 6.5e-08 | 2.3e-07 | 1.153E-07 | 9.955E-08 | 5.100E-08 | 2.300E-07 |
| 24388 | 1220 | S1220G | D | 2.3e-08 | 4.1e-08 | 3e-08 | 3.133E-08 | 9.074E-09 | 2.300E-08 | 4.100E-08 |
| 24389 | 1220 | S1220H | D | 1.5e-08 | 4.8e-08 | 8.1e-08 | 4.800E-08 | 3.300E-08 | 1.500E-08 | 8.100E-08 |
| 24390 | 1220 | S1220I | D | 5.3e-08 | 2.3e-08 | 4.7e-08 | 4.100E-08 | 1.587E-08 | 2.300E-08 | 5.300E-08 |
| 24391 | 1220 | S1220L | D | 8.2e-08 | 2.1e-08 | 4e-08 | 4.767E-08 | 3.121E-08 | 2.100E-08 | 8.200E-08 |
| 24392 | 1220 | S1220K | D | 3e-08 | 2.4e-08 | 5.6e-08 | 3.667E-08 | 1.701E-08 | 2.400E-08 | 5.600E-08 |
| 24393 | 1220 | S1220M | D | 9e-08 | 1.9e-08 | 3.5e-08 | 4.800E-08 | 3.724E-08 | 1.900E-08 | 9.000E-08 |
| 24394 | 1220 | S1220F | D | 3.5e-08 | 4.3e-08 | 3.3e-08 | 3.700E-08 | 5.292E-09 | 3.300E-08 | 4.300E-08 |

| | | | | | | | | | | |
|---|---|---|---|---|---|---|---|---|---|---|
| 24395 | 1220 | S1220P | D | 6.1e-08 | 5.6e-08 | 3.2e-08 | 4.967E-08 | 1.550E-08 | 3.200E-08 | 6.100E-08 |
| 24396 | 1220 | S1220S | D | 4.7e-08 | 5.7e-08 | 3.4e-08 | 4.600E-08 | 1.153E-08 | 3.400E-08 | 5.700E-08 |
| 24397 | 1220 | S1220T | D | 2e-08 | 4.7e-08 | 4.7e-08 | 3.800E-08 | 1.559E-08 | 2.000E-08 | 4.700E-08 |
| 24398 | 1220 | S1220W | D | 5.3e-08 | 6.2e-08 | 1.2e-08 | 4.233E-08 | 2.665E-08 | 1.200E-08 | 6.200E-08 |
| 24399 | 1220 | S1220Y | D | 4.7e-08 | 8.9e-08 | 9.2e-08 | 7.600E-08 | 2.516E-08 | 4.700E-08 | 9.200E-08 |
| 24400 | 1220 | S1220V | D | 6.5e-08 | 6.7e-08 | 5.8e-08 | 6.333E-08 | 4.726E-09 | 5.800E-08 | 6.700E-08 |
| 24401 | 1221 | V1221A | D | 2.7e-08 | 6.5e-08 | 7.1e-08 | 5.433E-08 | 2.386E-08 | 2.700E-08 | 7.100E-08 |
| 24402 | 1221 | V1221R | D | 4.4e-08 | 8.6e-08 | 4.7e-08 | 5.900E-08 | 2.343E-08 | 4.400E-08 | 8.600E-08 |
| 24403 | 1221 | V1221N | D | 6.4e-08 | 2.7e-08 | 5.5e-08 | 4.867E-08 | 1.930E-08 | 2.700E-08 | 6.400E-08 |
| 24404 | 1221 | V1221D | D | 7.3e-08 | 5.3e-08 | 2.4e-08 | 5.000E-08 | 2.464E-08 | 2.400E-08 | 7.300E-08 |
| 24405 | 1221 | V1221C | D | 1.6e-08 | 3.8e-08 | 2.5e-08 | 2.633E-08 | 1.106E-08 | 1.600E-08 | 3.800E-08 |
| 24406 | 1221 | V1221E | D | 7.8e-08 | 3.3e-08 | 5.8e-08 | 5.633E-08 | 2.255E-08 | 3.300E-08 | 7.800E-08 |
| 24407 | 1221 | V1221Q | D | 2.5e-08 | 4.6e-08 | 4.8e-08 | 3.967E-08 | 1.274E-08 | 2.500E-08 | 4.800E-08 |
| 24408 | 1221 | V1221G | D | 4.6e-08 | 3.3e-08 | 3e-08 | 3.633E-08 | 8.505E-09 | 3.000E-08 | 4.600E-08 |
| 24409 | 1221 | V1221H | D | 3.8e-08 | 1.4e-07 | 4.1e-08 | 7.300E-08 | 5.804E-08 | 3.800E-08 | 1.400E-07 |
| 24410 | 1221 | V1221I | D | 5.1e-08 | 4.8e-08 | 4.2e-08 | 4.700E-08 | 4.583E-09 | 4.200E-08 | 5.100E-08 |
| 24411 | 1221 | V1221L | D | 3.3e-08 | 6e-08 | 4.2e-08 | 4.500E-08 | 1.375E-08 | 3.300E-08 | 6.000E-08 |
| 24412 | 1221 | V1221K | D | 3.7e-08 | 1.4e-08 | 4.1e-08 | 3.067E-08 | 1.457E-08 | 1.400E-08 | 4.100E-08 |
| 24413 | 1221 | V1221M | D | 8.9e-08 | 8.2e-08 | 5.2e-08 | 7.433E-08 | 1.966E-08 | 5.200E-08 | 8.900E-08 |
| 24414 | 1221 | V1221F | D | 3.7e-08 | 2e-08 | 4.1e-08 | 3.267E-08 | 1.115E-08 | 2.000E-08 | 4.100E-08 |
| 24415 | 1221 | V1221P | D | 1.2e-07 | 1.6e-08 | 8.4e-08 | 7.333E-08 | 5.281E-08 | 1.600E-08 | 1.200E-07 |
| 24416 | 1221 | V1221S | D | 7.6e-08 | 6e-08 | 5.7e-08 | 6.433E-08 | 1.021E-08 | 5.700E-08 | 7.600E-08 |
| 24417 | 1221 | V1221T | D | 3.1e-08 | 2.2e-08 | 1.6e-08 | 2.300E-08 | 7.550E-09 | 1.600E-08 | 3.100E-08 |
| 24418 | 1221 | V1221W | D | 5.6e-08 | 4.5e-08 | 1.1e-08 | 3.733E-08 | 2.346E-08 | 1.100E-08 | 5.600E-08 |
| 24419 | 1221 | V1221Y | D | 4.4e-08 | 3e-08 | 5.2e-08 | 4.200E-08 | 1.114E-08 | 3.000E-08 | 5.200E-08 |
| 24420 | 1221 | V1221V | D | 2e-08 | 4.1e-08 | 1.2e-08 | 2.433E-08 | 1.498E-08 | 1.200E-08 | 4.100E-08 |
| 24421 | 1222 | T1222A | D | 2e-08 | 8e-08 | 8.8e-08 | 6.267E-08 | 3.717E-08 | 2.000E-08 | 8.800E-08 |
| 24422 | 1222 | T1222R | D | 6.1e-08 | 9.9e-08 | 4.5e-08 | 6.833E-08 | 2.774E-08 | 4.500E-08 | 9.900E-08 |
| 24423 | 1222 | T1222N | D | 3.9e-08 | 1.4e-07 | 2.6e-08 | 6.833E-08 | 6.240E-08 | 2.600E-08 | 1.400E-07 |
| 24424 | 1222 | T1222D | D | 4.2e-08 | 1.7e-08 | 4.9e-08 | 3.600E-08 | 1.682E-08 | 1.700E-08 | 4.900E-08 |
| 24425 | 1222 | T1222C | D | 2.4e-07 | 8e-08 | 2.9e-08 | 1.163E-07 | 1.101E-07 | 2.900E-08 | 2.400E-07 |
| 24426 | 1222 | T1222E | D | 2.1e-08 | 7.5e-08 | 2.2e-08 | 3.933E-08 | 3.089E-08 | 2.100E-08 | 7.500E-08 |
| 24427 | 1222 | T1222Q | D | 8.2e-08 | 3.8e-08 | 5e-08 | 5.667E-08 | 2.274E-08 | 3.800E-08 | 8.200E-08 |
| 24428 | 1222 | T1222G | D | 6.2e-08 | 3.6e-08 | 4.9e-08 | 4.900E-08 | 1.300E-08 | 3.600E-08 | 6.200E-08 |
| 24429 | 1222 | T1222H | D | 6.1e-08 | 2.9e-08 | 3.5e-08 | 4.167E-08 | 1.701E-08 | 2.900E-08 | 6.100E-08 |
| 24430 | 1222 | T1222I | D | 4.3e-08 | 7.5e-08 | 5.1e-08 | 5.633E-08 | 1.665E-08 | 4.300E-08 | 7.500E-08 |
| 24431 | 1222 | T1222L | D | 6.1e-08 | 5.1e-08 | 2.5e-08 | 4.567E-08 | 1.858E-08 | 2.500E-08 | 6.100E-08 |
| 24432 | 1222 | T1222K | D | 1.3e-08 | 2.4e-08 | 3.2e-08 | 2.300E-08 | 9.539E-09 | 1.300E-08 | 3.200E-08 |
| 24433 | 1222 | T1222M | D | 3.4e-08 | 2e-08 | 5.7e-08 | 3.700E-08 | 1.868E-08 | 2.000E-08 | 5.700E-08 |
| 24434 | 1222 | T1222F | D | 6.5e-08 | 1.3e-08 | 5.6e-08 | 4.467E-08 | 2.779E-08 | 1.300E-08 | 6.500E-08 |
| 24435 | 1222 | T1222P | D | 6.9e-08 | 6.6e-08 | 2.4e-08 | 5.300E-08 | 2.516E-08 | 2.400E-08 | 6.900E-08 |
| 24436 | 1222 | T1222S | D | 5.8e-08 | 7.5e-08 | 3.2e-08 | 5.500E-08 | 2.166E-08 | 3.200E-08 | 7.500E-08 |
| 24437 | 1222 | T1222T | D | 8.9e-08 | 9.7e-09 | 3.8e-08 | 4.557E-08 | 4.019E-08 | 9.700E-09 | 8.900E-08 |
| 24438 | 1222 | T1222W | D | 1.1e-08 | 2.7e-08 | 4.5e-08 | 2.767E-08 | 1.701E-08 | 1.100E-08 | 4.500E-08 |
| 24439 | 1222 | T1222Y | D | 4.2e-08 | 1e-07 | 2.9e-08 | 5.700E-08 | 3.780E-08 | 2.900E-08 | 1.000E-07 |
| 24440 | 1222 | T1222V | D | 2.7e-08 | 2.4e-08 | 3.5e-08 | 2.867E-08 | 5.686E-09 | 2.400E-08 | 3.500E-08 |
| 24441 | 1223 | E1223A | D | 1e-07 | 2.6e-08 | 4.6e-08 | 5.733E-08 | 3.828E-08 | 2.600E-08 | 1.000E-07 |
| 24442 | 1223 | E1223R | D | 3.1e-08 | 4.5e-08 | 3e-08 | 3.533E-08 | 8.386E-09 | 3.000E-08 | 4.500E-08 |
| 24443 | 1223 | E1223N | D | 3.2e-08 | 1.1e-07 | 9.5e-09 | 5.050E-08 | 5.274E-08 | 9.500E-09 | 1.100E-07 |
| 24444 | 1223 | E1223D | D | 3e-08 | 6e-08 | 3.3e-08 | 4.100E-08 | 1.652E-08 | 3.000E-08 | 6.000E-08 |
| 24445 | 1223 | E1223C | D | 4.2e-08 | 2.3e-08 | 7.7e-08 | 4.733E-08 | 2.739E-08 | 2.300E-08 | 7.700E-08 |
| 24446 | 1223 | E1223E | D | 2.6e-08 | 4.8e-08 | 7.3e-08 | 4.900E-08 | 2.352E-08 | 2.600E-08 | 7.300E-08 |
| 24447 | 1223 | E1223Q | D | 9e-08 | 5.2e-08 | 6.4e-08 | 6.867E-08 | 1.943E-08 | 5.200E-08 | 9.000E-08 |
| 24448 | 1223 | E1223G | D | 2.6e-08 | 1e-07 | 2.1e-08 | 4.900E-08 | 4.424E-08 | 2.100E-08 | 1.000E-07 |
| 24449 | 1223 | E1223H | D | 4.5e-08 | 1.2e-07 | 2.2e-08 | 6.233E-08 | 5.125E-08 | 2.200E-08 | 1.200E-07 |
| 24450 | 1223 | E1223I | D | 6.6e-08 | 3.5e-08 | 4.2e-08 | 4.767E-08 | 1.626E-08 | 3.500E-08 | 6.600E-08 |
| 24451 | 1223 | E1223L | D | 3.7e-08 | 1.2e-07 | 3.9e-09 | 5.363E-08 | 5.981E-08 | 3.900E-09 | 1.200E-07 |
| 24452 | 1223 | E1223K | D | 4.1e-08 | 8.2e-08 | 1.1e-07 | 7.767E-08 | 3.470E-08 | 4.100E-08 | 1.100E-07 |
| 24453 | 1223 | E1223M | D | 2.7e-08 | 2.1e-08 | 4e-08 | 2.933E-08 | 9.713E-09 | 2.100E-08 | 4.000E-08 |
| 24454 | 1223 | E1223F | D | 6.1e-08 | 3.8e-08 | 3e-08 | 4.300E-08 | 1.609E-08 | 3.000E-08 | 6.100E-08 |
| 24455 | 1223 | E1223P | D | 3.5e-08 | 1.6e-08 | 5.5e-08 | 3.533E-08 | 1.950E-08 | 1.600E-08 | 5.500E-08 |
| 24456 | 1223 | E1223S | D | 4e-08 | 3.3e-08 | 7e-08 | 4.767E-08 | 1.966E-08 | 3.300E-08 | 7.000E-08 |
| 24457 | 1223 | E1223T | D | 3.1e-08 | 9.8e-08 | 6.4e-08 | 6.433E-08 | 3.350E-08 | 3.100E-08 | 9.800E-08 |
| 24458 | 1223 | E1223W | D | 6.2e-08 | 4.6e-08 | 6.6e-08 | 5.800E-08 | 1.058E-08 | 4.600E-08 | 6.600E-08 |
| 24459 | 1223 | E1223Y | D | 4.2e-08 | 6.7e-08 | 5.9e-08 | 5.600E-08 | 1.277E-08 | 4.200E-08 | 6.700E-08 |
| 24460 | 1223 | E1223V | D | 5.5e-08 | 6.3e-08 | 3.4e-08 | 5.067E-08 | 1.498E-08 | 3.400E-08 | 6.300E-08 |
| 24461 | 1224 | Q1224A | D | 5e-08 | 4e-08 | 6.6e-08 | 5.200E-08 | 1.311E-08 | 4.000E-08 | 6.600E-08 |
| 24462 | 1224 | Q1224R | D | 1.1e-07 | 6.1e-08 | 6.5e-08 | 7.867E-08 | 2.721E-08 | 6.100E-08 | 1.100E-07 |
| 24463 | 1224 | Q1224N | D | 1.7e-08 | 4.4e-08 | 5.9e-08 | 4.000E-08 | 2.128E-08 | 1.700E-08 | 5.900E-08 |
| 24464 | 1224 | Q1224D | D | 3.3e-08 | 3.4e-08 | 4.6e-08 | 3.767E-08 | 7.234E-09 | 3.300E-08 | 4.600E-08 |
| 24465 | 1224 | Q1224C | D | 4.9e-08 | 5.9e-08 | 2e-08 | 4.267E-08 | 2.026E-08 | 2.000E-08 | 5.900E-08 |
| 24466 | 1224 | Q1224E | D | 5.1e-08 | 6.6e-08 | 3.4e-08 | 5.033E-08 | 1.601E-08 | 3.400E-08 | 6.600E-08 |
| 24467 | 1224 | Q1224Q | D | 2e-08 | 2e-08 | 4.7e-08 | 2.900E-08 | 1.559E-08 | 2.000E-08 | 4.700E-08 |
| 24468 | 1224 | Q1224G | D | 4.1e-08 | 5.5e-08 | 3.1e-08 | 4.233E-08 | 1.206E-08 | 3.100E-08 | 5.500E-08 |
| 24469 | 1224 | Q1224H | D | 3e-08 | 5.6e-08 | 4.1e-08 | 4.233E-08 | 1.305E-08 | 3.000E-08 | 5.600E-08 |
| 24470 | 1224 | Q1224I | D | 3.6e-08 | 2.2e-08 | 8.2e-08 | 4.667E-08 | 3.139E-08 | 2.200E-08 | 8.200E-08 |
| 24471 | 1224 | Q1224L | D | 5.1e-08 | 2.5e-08 | 5.5e-08 | 4.367E-08 | 1.629E-08 | 2.500E-08 | 5.500E-08 |
| 24472 | 1224 | Q1224K | D | 7.2e-08 | 3.4e-08 | 3.5e-08 | 4.700E-08 | 2.166E-08 | 3.400E-08 | 7.200E-08 |
| 24473 | 1224 | Q1224M | D | 2.8e-08 | 3.1e-08 | 3.7e-08 | 3.200E-08 | 4.583E-09 | 2.800E-08 | 3.700E-08 |
| 24474 | 1224 | Q1224F | D | 9.7e-08 | 9.9e-09 | 1.3e-07 | 7.897E-08 | 6.205E-08 | 9.900E-09 | 1.300E-07 |
| 24475 | 1224 | Q1224P | D | 1.8e-08 | 2.2e-08 | 3.1e-08 | 2.367E-08 | 6.658E-09 | 1.800E-08 | 3.100E-08 |
| 24476 | 1224 | Q1224S | D | 1.5e-08 | 2.4e-08 | 2.9e-08 | 2.267E-08 | 7.095E-09 | 1.500E-08 | 2.900E-08 |
| 24477 | 1224 | Q1224T | D | 4.9e-08 | 3.7e-08 | 3.8e-08 | 4.133E-08 | 6.658E-09 | 3.700E-08 | 4.900E-08 |
| 24478 | 1224 | Q1224W | D | 5.9e-08 | 2.2e-08 | 2.4e-08 | 3.500E-08 | 2.081E-08 | 2.200E-08 | 5.900E-08 |
| 24479 | 1224 | Q1224Y | D | 4.9e-08 | 2.2e-08 | 3.3e-08 | 3.467E-08 | 1.358E-08 | 2.200E-08 | 4.900E-08 |

| | | | | | | | | | | |
|---|---|---|---|---|---|---|---|---|---|---|
| 24480 | 1224 | Q1224V | D | 1.1e-07 | 1.9e-08 | 3.3e-08 | 5.400E-08 | 4.900E-08 | 1.900E-08 | 1.100E-07 |
| 24481 | 1225 | D1225A | D | 5.2e-08 | 6.5e-08 | 6.6e-08 | 6.100E-08 | 7.810E-09 | 5.200E-08 | 6.600E-08 |
| 24482 | 1225 | D1225R | D | 6e-08 | 3.9e-08 | 1.6e-08 | 3.833E-08 | 2.201E-08 | 1.600E-08 | 6.000E-08 |
| 24483 | 1225 | D1225N | D | 2.2e-07 | 2.2e-08 | 8.4e-08 | 1.087E-07 | 1.013E-07 | 2.200E-08 | 2.200E-07 |
| 24484 | 1225 | D1225D | D | 3.9e-08 | 1.9e-08 | 4e-08 | 3.267E-08 | 1.185E-08 | 1.900E-08 | 4.000E-08 |
| 24485 | 1225 | D1225C | D | 5.2e-08 | 1.9e-08 | 1e-07 | 5.700E-08 | 4.073E-08 | 1.900E-08 | 1.000E-07 |
| 24486 | 1225 | D1225E | D | 7.2e-08 | 5.8e-08 | 7.2e-08 | 6.733E-08 | 8.083E-09 | 5.800E-08 | 7.200E-08 |
| 24487 | 1225 | D1225Q | D | 6.9e-08 | 3.2e-08 | 4e-08 | 4.700E-08 | 1.947E-08 | 3.200E-08 | 6.900E-08 |
| 24488 | 1225 | D1225G | D | 5.6e-08 | 1.1e-07 | 4.8e-08 | 7.133E-08 | 3.372E-08 | 4.800E-08 | 1.100E-07 |
| 24489 | 1225 | D1225H | D | 3.3e-08 | 9.8e-08 | 8.6e-09 | 4.653E-08 | 4.621E-08 | 8.600E-09 | 9.800E-08 |
| 24490 | 1225 | D1225I | D | 4.8e-08 | 8.2e-08 | 6.7e-08 | 6.567E-08 | 1.704E-08 | 4.800E-08 | 8.200E-08 |
| 24491 | 1225 | D1225L | D | 1.3e-08 | 1.4e-07 | 2.5e-08 | 5.933E-08 | 7.012E-08 | 1.300E-08 | 1.400E-07 |
| 24492 | 1225 | D1225K | D | 3.3e-08 | 2.5e-08 | 1.6e-07 | 7.267E-08 | 7.574E-08 | 2.500E-08 | 1.600E-07 |
| 24493 | 1225 | D1225M | D | 4.2e-08 | 1.3e-07 | 1.9e-08 | 6.367E-08 | 5.859E-08 | 1.900E-08 | 1.300E-07 |
| 24494 | 1225 | D1225F | D | 3.7e-08 | 2.4e-08 | 2.1e-08 | 2.733E-08 | 8.505E-09 | 2.100E-08 | 3.700E-08 |
| 24495 | 1225 | D1225P | D | 7.8e-08 | 4.1e-08 | 5e-08 | 5.633E-08 | 1.930E-08 | 4.100E-08 | 7.800E-08 |
| 24496 | 1225 | D1225S | D | 8.5e-08 | 4.7e-08 | 5.6e-08 | 6.267E-08 | 1.986E-08 | 4.700E-08 | 8.500E-08 |
| 24497 | 1225 | D1225T | D | 4.5e-08 | 2.5e-08 | 1.5e-08 | 2.833E-08 | 1.528E-08 | 1.500E-08 | 4.500E-08 |
| 24498 | 1225 | D1225W | D | 3.8e-08 | 3.5e-08 | 8.1e-08 | 5.133E-08 | 2.574E-08 | 3.500E-08 | 8.100E-08 |
| 24499 | 1225 | D1225Y | D | 1.4e-08 | 3.1e-08 | 5.3e-08 | 3.267E-08 | 1.955E-08 | 1.400E-08 | 5.300E-08 |
| 24500 | 1225 | D1225V | D | 6.7e-08 | 4.3e-08 | 1.3e-07 | 8.000E-08 | 4.493E-08 | 4.300E-08 | 1.300E-07 |
| 24501 | 1226 | S1226A | D | 5.1e-08 | 2.7e-08 | 6.3e-08 | 4.700E-08 | 1.833E-08 | 2.700E-08 | 6.300E-08 |
| 24502 | 1226 | S1226R | D | 3.5e-08 | 3.5e-08 | 3.1e-08 | 3.367E-08 | 2.309E-09 | 3.100E-08 | 3.500E-08 |
| 24503 | 1226 | S1226N | D | 6.3e-08 | 4.4e-08 | 3.9e-08 | 4.867E-08 | 1.266E-08 | 3.900E-08 | 6.300E-08 |
| 24504 | 1226 | S1226D | D | 5.5e-08 | 1.3e-07 | 1.4e-07 | 1.083E-07 | 4.646E-08 | 5.500E-08 | 1.400E-07 |
| 24505 | 1226 | S1226C | D | 3.8e-08 | 2.2e-08 | 2e-08 | 2.667E-08 | 9.866E-09 | 2.000E-08 | 3.800E-08 |
| 24506 | 1226 | S1226E | D | 4.3e-08 | 1.7e-07 | 1.7e-08 | 7.667E-08 | 8.187E-08 | 1.700E-08 | 1.700E-07 |
| 24507 | 1226 | S1226Q | D | 1.9e-08 | 9e-08 | 7.2e-08 | 6.033E-08 | 3.691E-08 | 1.900E-08 | 9.000E-08 |
| 24508 | 1226 | S1226G | D | 3.2e-08 | 4e-08 | 3.4e-08 | 3.533E-08 | 4.163E-09 | 3.200E-08 | 4.000E-08 |
| 24509 | 1226 | S1226H | D | 1.1e-07 | 3.4e-08 | 5.7e-08 | 6.700E-08 | 3.897E-08 | 3.400E-08 | 1.100E-07 |
| 24510 | 1226 | S1226I | D | 5.6e-08 | 3.5e-08 | 3.4e-08 | 4.167E-08 | 1.242E-08 | 3.400E-08 | 5.600E-08 |
| 24511 | 1226 | S1226L | D | 5.8e-08 | 3.4e-08 | 2.8e-08 | 4.000E-08 | 1.587E-08 | 2.800E-08 | 5.800E-08 |
| 24512 | 1226 | S1226K | D | 1.1e-08 | 5.4e-08 | 1.8e-08 | 2.767E-08 | 2.307E-08 | 1.100E-08 | 5.400E-08 |
| 24513 | 1226 | S1226M | D | 3.3e-08 | 1.2e-08 | 1.6e-08 | 2.033E-08 | 1.115E-08 | 1.200E-08 | 3.300E-08 |
| 24514 | 1226 | S1226F | D | 5e-08 | 2.2e-08 | 3.5e-08 | 3.567E-08 | 1.401E-08 | 2.200E-08 | 5.000E-08 |
| 24515 | 1226 | S1226P | D | 3.3e-08 | 1.1e-07 | 3.2e-08 | 5.833E-08 | 4.475E-08 | 3.200E-08 | 1.100E-07 |
| 24516 | 1226 | S1226S | D | 2.1e-08 | 4.4e-08 | 4.7e-08 | 3.733E-08 | 1.422E-08 | 2.100E-08 | 4.700E-08 |
| 24517 | 1226 | S1226T | D | 2.5e-08 | 3.4e-08 | 5.3e-08 | 3.733E-08 | 1.429E-08 | 2.500E-08 | 5.300E-08 |
| 24518 | 1226 | S1226W | D | 3.7e-08 | 7.8e-08 | 4.3e-08 | 5.267E-08 | 2.214E-08 | 3.700E-08 | 7.800E-08 |
| 24519 | 1226 | S1226Y | D | 5.2e-08 | 3.2e-08 | 1.2e-07 | 6.800E-08 | 4.613E-08 | 3.200E-08 | 1.200E-07 |
| 24520 | 1226 | S1226V | D | 4.8e-08 | 9.6e-08 | 5.1e-08 | 6.500E-08 | 2.689E-08 | 4.800E-08 | 9.600E-08 |
| 24521 | 1227 | K1227A | D | 2.3e-08 | 2e-08 | 1.9e-08 | 2.067E-08 | 2.082E-09 | 1.900E-08 | 2.300E-08 |
| 24522 | 1227 | K1227R | D | 4.9e-08 | 3.7e-08 | 6.8e-08 | 5.133E-08 | 1.563E-08 | 3.700E-08 | 6.800E-08 |
| 24523 | 1227 | K1227N | D | 4.8e-08 | 5e-09 | 3.5e-08 | 2.933E-08 | 2.205E-08 | 5.000E-09 | 4.800E-08 |
| 24524 | 1227 | K1227D | D | 5.1e-08 | 4.9e-08 | 4.5e-08 | 4.833E-08 | 3.055E-09 | 4.500E-08 | 5.100E-08 |
| 24525 | 1227 | K1227C | D | 4.9e-08 | 4.8e-08 | 2.2e-08 | 3.967E-08 | 1.531E-08 | 2.200E-08 | 4.900E-08 |
| 24526 | 1227 | K1227E | D | 1.2e-08 | 3.6e-08 | 4.4e-08 | 3.067E-08 | 1.665E-08 | 1.200E-08 | 4.400E-08 |
| 24527 | 1227 | K1227Q | D | 8e-08 | 2.3e-08 | 2.7e-08 | 4.333E-08 | 3.182E-08 | 2.300E-08 | 8.000E-08 |
| 24528 | 1227 | K1227G | D | 5e-08 | 5.5e-08 | 3e-08 | 4.500E-08 | 1.323E-08 | 3.000E-08 | 5.500E-08 |
| 24529 | 1227 | K1227H | D | 1.8e-08 | 6.3e-08 | 5.6e-08 | 4.567E-08 | 2.421E-08 | 1.800E-08 | 6.300E-08 |
| 24530 | 1227 | K1227I | D | 8.2e-08 | 4.8e-08 | 4.9e-08 | 5.967E-08 | 1.935E-08 | 4.800E-08 | 8.200E-08 |
| 24531 | 1227 | K1227L | D | 3.7e-08 | 5.2e-08 | 1.6e-07 | 8.300E-08 | 6.710E-08 | 3.700E-08 | 1.600E-07 |
| 24532 | 1227 | K1227K | D | 4.1e-08 | 6.4e-08 | 3e-08 | 4.500E-08 | 1.735E-08 | 3.000E-08 | 6.400E-08 |
| 24533 | 1227 | K1227M | D | 7.4e-08 | 2.4e-08 | 4.2e-08 | 4.667E-08 | 2.532E-08 | 2.400E-08 | 7.400E-08 |
| 24534 | 1227 | K1227F | D | 4.5e-08 | 8e-08 | 3.4e-08 | 5.300E-08 | 2.402E-08 | 3.400E-08 | 8.000E-08 |
| 24535 | 1227 | K1227P | D | 3.9e-08 | 4.9e-08 | 5e-08 | 4.600E-08 | 6.083E-09 | 3.900E-08 | 5.000E-08 |
| 24536 | 1227 | K1227S | D | 1.9e-08 | 6.2e-08 | 7.3e-08 | 5.133E-08 | 2.854E-08 | 1.900E-08 | 7.300E-08 |
| 24537 | 1227 | K1227T | D | 2.1e-08 | 2e-08 | 2e-08 | 2.033E-08 | 5.774E-10 | 2.000E-08 | 2.100E-08 |
| 24538 | 1227 | K1227W | D | 2.6e-08 | 5.1e-08 | 8.2e-08 | 5.300E-08 | 2.805E-08 | 2.600E-08 | 8.200E-08 |
| 24539 | 1227 | K1227Y | D | 9.1e-08 | 3.1e-08 | 4e-08 | 5.400E-08 | 3.236E-08 | 3.100E-08 | 9.100E-08 |
| 24540 | 1227 | K1227V | D | 1.2e-07 | 2.1e-08 | 2.7e-08 | 5.600E-08 | 5.551E-08 | 2.100E-08 | 1.200E-07 |
| 24541 | 1228 | D1228A | D | 2.6e-08 | 4.2e-08 | 4.5e-08 | 3.767E-08 | 1.021E-08 | 2.600E-08 | 4.500E-08 |
| 24542 | 1228 | D1228R | D | 3e-08 | 1.9e-08 | 5.6e-08 | 3.500E-08 | 1.900E-08 | 1.900E-08 | 5.600E-08 |
| 24543 | 1228 | D1228N | D | 3e-08 | 4.9e-08 | 3.5e-08 | 3.800E-08 | 9.849E-09 | 3.000E-08 | 4.900E-08 |
| 24544 | 1228 | D1228D | D | 9.1e-08 | 3.7e-08 | 7.4e-08 | 6.733E-08 | 2.761E-08 | 3.700E-08 | 9.100E-08 |
| 24545 | 1228 | D1228C | D | 6.2e-08 | 5.9e-08 | 5.6e-08 | 5.900E-08 | 3.000E-09 | 5.600E-08 | 6.200E-08 |
| 24546 | 1228 | D1228E | D | 4.6e-08 | 5.3e-08 | 3.5e-08 | 4.467E-08 | 9.074E-09 | 3.500E-08 | 5.300E-08 |
| 24547 | 1228 | D1228Q | D | 9.5e-09 | 6.5e-08 | 8.6e-08 | 5.350E-08 | 3.953E-08 | 9.500E-09 | 8.600E-08 |
| 24548 | 1228 | D1228G | D | 6.3e-08 | 3.7e-08 | 1.2e-07 | 7.333E-08 | 4.245E-08 | 3.700E-08 | 1.200E-07 |
| 24549 | 1228 | D1228H | D | 2.2e-08 | 2.3e-08 | 8.2e-08 | 4.233E-08 | 3.436E-08 | 2.200E-08 | 8.200E-08 |
| 24550 | 1228 | D1228I | D | 6.7e-08 | 7.4e-08 | 7.6e-08 | 7.233E-08 | 4.726E-09 | 6.700E-08 | 7.600E-08 |
| 24551 | 1228 | D1228L | D | 3.2e-08 | 6.6e-08 | 2.2e-08 | 4.000E-08 | 2.307E-08 | 2.200E-08 | 6.600E-08 |
| 24552 | 1228 | D1228K | D | 2e-08 | 4.6e-08 | 6e-08 | 4.200E-08 | 2.030E-08 | 2.000E-08 | 6.000E-08 |
| 24553 | 1228 | D1228M | D | 6.4e-08 | 7.8e-08 | 6.4e-08 | 6.867E-08 | 8.083E-09 | 6.400E-08 | 7.800E-08 |
| 24554 | 1228 | D1228F | D | 4.3e-08 | 5e-08 | 4.6e-08 | 4.633E-08 | 3.512E-09 | 4.300E-08 | 5.000E-08 |
| 24555 | 1228 | D1228P | D | 6.1e-08 | 5e-08 | 6.5e-08 | 5.867E-08 | 7.767E-09 | 5.000E-08 | 6.500E-08 |
| 24556 | 1228 | D1228S | D | 2.8e-08 | 3e-08 | 2.2e-08 | 2.667E-08 | 4.163E-09 | 2.200E-08 | 3.000E-08 |
| 24557 | 1228 | D1228T | D | 5.6e-08 | 3.6e-08 | 5.2e-08 | 4.800E-08 | 1.058E-08 | 3.600E-08 | 5.600E-08 |
| 24558 | 1228 | D1228W | D | 5.2e-08 | 2e-08 | 2.4e-08 | 3.200E-08 | 1.744E-08 | 2.000E-08 | 5.200E-08 |
| 24559 | 1228 | D1228Y | D | 1.3e-07 | 8.9e-08 | 3e-08 | 8.300E-08 | 5.027E-08 | 3.000E-08 | 1.300E-07 |
| 24560 | 1228 | D1228V | D | 1.4e-07 | 5.8e-08 | 2.7e-08 | 7.500E-08 | 5.839E-08 | 2.700E-08 | 1.400E-07 |
| 24561 | 1229 | S1229A | D | 7.1e-08 | 4.6e-08 | 3.7e-07 | 1.623E-07 | 1.803E-07 | 4.600E-08 | 3.700E-07 |
| 24562 | 1229 | S1229R | D | 3.2e-08 | 4.4e-08 | 3.9e-08 | 3.833E-08 | 6.028E-09 | 3.200E-08 | 4.400E-08 |
| 24563 | 1229 | S1229N | D | 5.4e-08 | 3.2e-08 | 2.6e-08 | 3.733E-08 | 1.474E-08 | 2.600E-08 | 5.400E-08 |
| 24564 | 1229 | S1229D | D | 6.2e-08 | 5.5e-08 | 4e-08 | 5.233E-08 | 1.124E-08 | 4.000E-08 | 6.200E-08 |

| | | | | | | | | | | |
|---|---|---|---|---|---|---|---|---|---|---|
| 24565 | 1229 | S1229C | D | 1.6e-08 | 1.2e-07 | 8.8e-08 | 7.467E-08 | 5.327E-08 | 1.600E-08 | 1.200E-07 |
| 24566 | 1229 | S1229E | D | 2.7e-08 | 6.1e-08 | 3.8e-08 | 4.200E-08 | 1.735E-08 | 2.700E-08 | 6.100E-08 |
| 24567 | 1229 | S1229Q | D | 1.7e-08 | 6.7e-08 | 3.5e-08 | 3.967E-08 | 2.532E-08 | 1.700E-08 | 6.700E-08 |
| 24568 | 1229 | S1229G | D | 6.2e-08 | 6e-08 | 3.4e-08 | 5.200E-08 | 1.562E-08 | 3.400E-08 | 6.200E-08 |
| 24569 | 1229 | S1229H | D | 6.6e-08 | 7.1e-08 | 7.2e-08 | 6.967E-08 | 3.215E-09 | 6.600E-08 | 7.200E-08 |
| 24570 | 1229 | S1229I | D | 4.5e-08 | 3.7e-08 | 4.2e-08 | 4.133E-08 | 4.041E-09 | 3.700E-08 | 4.500E-08 |
| 24571 | 1229 | S1229L | D | 4.1e-08 | 6.9e-08 | 1.7e-07 | 9.333E-08 | 6.786E-08 | 4.100E-08 | 1.700E-07 |
| 24572 | 1229 | S1229K | D | 1.4e-07 | 7.5e-08 | 3.9e-08 | 8.467E-08 | 5.119E-08 | 3.900E-08 | 1.400E-07 |
| 24573 | 1229 | S1229M | D | 3e-08 | 8.7e-08 | 1.5e-08 | 4.400E-08 | 3.799E-08 | 1.500E-08 | 8.700E-08 |
| 24574 | 1229 | S1229F | D | 5.9e-08 | 2.3e-08 | 9.4e-08 | 5.867E-08 | 3.550E-08 | 2.300E-08 | 9.400E-08 |
| 24575 | 1229 | S1229P | D | 1.8e-08 | 3.8e-08 | 2.5e-08 | 2.700E-08 | 1.015E-08 | 1.800E-08 | 3.800E-08 |
| 24576 | 1229 | S1229S | D | 2.8e-08 | 4.2e-08 | 3.9e-08 | 3.633E-08 | 7.371E-09 | 2.800E-08 | 4.200E-08 |
| 24577 | 1229 | S1229T | D | 4.2e-08 | 3.6e-08 | 5.3e-08 | 4.367E-08 | 8.622E-09 | 3.600E-08 | 5.300E-08 |
| 24578 | 1229 | S1229W | D | 1.2e-08 | 2.3e-08 | 4.1e-08 | 2.533E-08 | 1.464E-08 | 1.200E-08 | 4.100E-08 |
| 24579 | 1229 | S1229Y | D | 7.9e-08 | 5.8e-08 | 9.3e-08 | 7.667E-08 | 1.762E-08 | 5.800E-08 | 9.300E-08 |
| 24580 | 1229 | S1229V | D | 2.4e-08 | 2.9e-08 | 3.5e-08 | 2.933E-08 | 5.508E-09 | 2.400E-08 | 3.500E-08 |
| 24581 | 1230 | T1230A | D | 7.2e-08 | 2e-08 | 8.6e-08 | 5.933E-08 | 3.478E-08 | 2.000E-08 | 8.600E-08 |
| 24582 | 1230 | T1230R | D | 4.6e-08 | 3.7e-08 | 5.9e-08 | 4.733E-08 | 1.106E-08 | 3.700E-08 | 5.900E-08 |
| 24583 | 1230 | T1230N | D | 2.5e-08 | 7.5e-08 | 5.2e-08 | 5.067E-08 | 2.503E-08 | 2.500E-08 | 7.500E-08 |
| 24584 | 1230 | T1230D | D | 3.4e-08 | 5.8e-08 | 5.3e-08 | 4.833E-08 | 1.266E-08 | 3.400E-08 | 5.800E-08 |
| 24585 | 1230 | T1230C | D | 5.4e-08 | 1.2e-07 | 4e-08 | 7.133E-08 | 4.272E-08 | 4.000E-08 | 1.200E-07 |
| 24586 | 1230 | T1230E | D | 3.8e-08 | 1.1e-07 | 2.7e-08 | 5.833E-08 | 4.508E-08 | 2.700E-08 | 1.100E-07 |
| 24587 | 1230 | T1230Q | D | 5.3e-08 | 3.9e-08 | 3.9e-08 | 4.367E-08 | 8.083E-09 | 3.900E-08 | 5.300E-08 |
| 24588 | 1230 | T1230G | D | 4.4e-08 | 5.6e-08 | 7.6e-08 | 5.867E-08 | 1.617E-08 | 4.400E-08 | 7.600E-08 |
| 24589 | 1230 | T1230H | D | 3.3e-08 | 5.9e-08 | 3e-08 | 4.067E-08 | 1.595E-08 | 3.000E-08 | 5.900E-08 |
| 24590 | 1230 | T1230I | D | 6.7e-08 | 5e-08 | 3.9e-08 | 5.200E-08 | 1.411E-08 | 3.900E-08 | 6.700E-08 |
| 24591 | 1230 | T1230L | D | 8.1e-08 | 3.8e-08 | 2.5e-08 | 4.800E-08 | 2.931E-08 | 2.500E-08 | 8.100E-08 |
| 24592 | 1230 | T1230K | D | 2.7e-08 | 4.2e-08 | 3.1e-08 | 3.333E-08 | 7.767E-09 | 2.700E-08 | 4.200E-08 |
| 24593 | 1230 | T1230M | D | 3.5e-08 | 4.2e-08 | 4.1e-08 | 3.933E-08 | 3.786E-09 | 3.500E-08 | 4.200E-08 |
| 24594 | 1230 | T1230F | D | 5.9e-08 | 3.9e-08 | 6e-08 | 5.267E-08 | 1.185E-08 | 3.900E-08 | 6.000E-08 |
| 24595 | 1230 | T1230P | D | 2.8e-08 | 8.6e-08 | 7.5e-08 | 6.300E-08 | 3.081E-08 | 2.800E-08 | 8.600E-08 |
| 24596 | 1230 | T1230S | D | 7e-08 | 2.3e-08 | 2.7e-08 | 4.000E-08 | 2.606E-08 | 2.300E-08 | 7.000E-08 |
| 24597 | 1230 | T1230T | D | 3.2e-08 | 6.4e-08 | 4.5e-08 | 4.700E-08 | 1.609E-08 | 3.200E-08 | 6.400E-08 |
| 24598 | 1230 | T1230W | D | 6e-08 | 4.2e-08 | 3.3e-08 | 4.500E-08 | 1.375E-08 | 3.300E-08 | 6.000E-08 |
| 24599 | 1230 | T1230Y | D | 5.5e-08 | 4e-08 | 9.4e-08 | 6.300E-08 | 2.787E-08 | 4.000E-08 | 9.400E-08 |
| 24600 | 1230 | T1230V | D | 2.8e-08 | 3.4e-08 | 1.2e-08 | 2.467E-08 | 1.137E-08 | 1.200E-08 | 3.400E-08 |
| 24601 | 1231 | Y1231A | D | 2.3e-08 | 3.1e-08 | 3.1e-08 | 2.833E-08 | 4.619E-09 | 2.300E-08 | 3.100E-08 |
| 24602 | 1231 | Y1231R | D | 8.8e-08 | 7.3e-08 | 3e-08 | 6.367E-08 | 3.011E-08 | 3.000E-08 | 8.800E-08 |
| 24603 | 1231 | Y1231N | D | 4.3e-08 | 1.5e-08 | 4.9e-08 | 3.567E-08 | 1.815E-08 | 1.500E-08 | 4.900E-08 |
| 24604 | 1231 | Y1231D | D | 5.2e-08 | 3.1e-08 | 6.6e-08 | 4.967E-08 | 1.762E-08 | 3.100E-08 | 6.600E-08 |
| 24605 | 1231 | Y1231C | D | 3.6e-08 | 7.2e-08 | 2.7e-08 | 4.500E-08 | 2.381E-08 | 2.700E-08 | 7.200E-08 |
| 24606 | 1231 | Y1231E | D | 6.4e-08 | 4.7e-08 | 3.5e-08 | 4.867E-08 | 1.457E-08 | 3.500E-08 | 6.400E-08 |
| 24607 | 1231 | Y1231Q | D | 3.1e-08 | 2.5e-08 | 1.1e-07 | 5.533E-08 | 4.744E-08 | 2.500E-08 | 1.100E-07 |
| 24608 | 1231 | Y1231G | D | 2.7e-08 | 3.2e-08 | 2.1e-08 | 2.667E-08 | 5.508E-09 | 2.100E-08 | 3.200E-08 |
| 24609 | 1231 | Y1231H | D | 2.5e-08 | 9.3e-09 | 6.4e-08 | 3.277E-08 | 2.816E-08 | 9.300E-09 | 6.400E-08 |
| 24610 | 1231 | Y1231I | D | 1.4e-07 | 1.6e-08 | 2.4e-08 | 6.000E-08 | 6.940E-08 | 1.600E-08 | 1.400E-07 |
| 24611 | 1231 | Y1231L | D | 6.6e-08 | 2e-08 | 1.6e-08 | 3.400E-08 | 2.778E-08 | 1.600E-08 | 6.600E-08 |
| 24612 | 1231 | Y1231K | D | 4.3e-08 | 7.3e-08 | 7.2e-08 | 6.267E-08 | 1.704E-08 | 4.300E-08 | 7.300E-08 |
| 24613 | 1231 | Y1231M | D | 8e-08 | 3.4e-08 | 3.1e-08 | 4.833E-08 | 2.747E-08 | 3.100E-08 | 8.000E-08 |
| 24614 | 1231 | Y1231F | D | 6.5e-08 | 2.1e-08 | 3e-08 | 3.867E-08 | 2.325E-08 | 2.100E-08 | 6.500E-08 |
| 24615 | 1231 | Y1231P | D | 3.4e-08 | 4.4e-08 | 5.3e-08 | 4.367E-08 | 9.504E-09 | 3.400E-08 | 5.300E-08 |
| 24616 | 1231 | Y1231S | D | 1.2e-07 | 3.2e-08 | 2.9e-08 | 6.033E-08 | 5.169E-08 | 2.900E-08 | 1.200E-07 |
| 24617 | 1231 | Y1231T | D | 2e-08 | 4.3e-08 | 3.5e-08 | 3.267E-08 | 1.168E-08 | 2.000E-08 | 4.300E-08 |
| 24618 | 1231 | Y1231W | D | 7.7e-08 | 3.3e-08 | 8.7e-08 | 6.567E-08 | 2.873E-08 | 3.300E-08 | 8.700E-08 |
| 24619 | 1231 | Y1231Y | D | 7.1e-08 | 3.6e-08 | 4.8e-08 | 5.167E-08 | 1.779E-08 | 3.600E-08 | 7.100E-08 |
| 24620 | 1231 | Y1231V | D | 4.6e-08 | 8.3e-08 | 2.7e-08 | 5.200E-08 | 2.848E-08 | 2.700E-08 | 8.300E-08 |
| 24621 | 1232 | S1232A | D | 2.5e-08 | 3.6e-08 | 3.1e-08 | 3.067E-08 | 5.508E-09 | 2.500E-08 | 3.600E-08 |
| 24622 | 1232 | S1232R | D | 7.3e-08 | 7.6e-08 | 2.5e-08 | 5.800E-08 | 2.862E-08 | 2.500E-08 | 7.600E-08 |
| 24623 | 1232 | S1232N | D | 3.3e-08 | 6.6e-08 | 4.9e-08 | 4.933E-08 | 1.650E-08 | 3.300E-08 | 6.600E-08 |
| 24624 | 1232 | S1232D | D | 7.1e-08 | 4.4e-08 | 1.1e-07 | 7.500E-08 | 3.318E-08 | 4.400E-08 | 1.100E-07 |
| 24625 | 1232 | S1232C | D | 1.2e-07 | 7.9e-08 | 3.6e-08 | 7.833E-08 | 4.200E-08 | 3.600E-08 | 1.200E-07 |
| 24626 | 1232 | S1232E | D | 1.7e-08 | 3.4e-08 | 3.8e-08 | 2.967E-08 | 1.115E-08 | 1.700E-08 | 3.800E-08 |
| 24627 | 1232 | S1232Q | D | 3e-08 | 1.9e-08 | 6.1e-08 | 3.667E-08 | 2.178E-08 | 1.900E-08 | 6.100E-08 |
| 24628 | 1232 | S1232G | D | 6e-08 | 2.5e-08 | 1.8e-08 | 3.433E-08 | 2.250E-08 | 1.800E-08 | 6.000E-08 |
| 24629 | 1232 | S1232H | D | 4.3e-08 | 6.2e-08 | 6.1e-08 | 5.533E-08 | 1.069E-08 | 4.300E-08 | 6.200E-08 |
| 24630 | 1232 | S1232I | D | 1.7e-08 | 3.6e-08 | 4.5e-08 | 3.267E-08 | 1.429E-08 | 1.700E-08 | 4.500E-08 |
| 24631 | 1232 | S1232L | D | 4.8e-08 | 4.6e-08 | 6.5e-08 | 5.300E-08 | 1.044E-08 | 4.600E-08 | 6.500E-08 |
| 24632 | 1232 | S1232K | D | 4.4e-08 | 7.4e-08 | 1.9e-08 | 4.567E-08 | 2.754E-08 | 1.900E-08 | 7.400E-08 |
| 24633 | 1232 | S1232M | D | 6.9e-08 | 9.3e-08 | 4e-08 | 6.733E-08 | 2.654E-08 | 4.000E-08 | 9.300E-08 |
| 24634 | 1232 | S1232F | D | 6e-08 | 3.6e-08 | 5.1e-08 | 4.900E-08 | 1.212E-08 | 3.600E-08 | 6.000E-08 |
| 24635 | 1232 | S1232P | D | 1.3e-07 | 1.1e-07 | 5e-08 | 9.667E-08 | 4.163E-08 | 5.000E-08 | 1.300E-07 |
| 24636 | 1232 | S1232S | D | 2.6e-08 | 5.3e-08 | 5e-08 | 4.300E-08 | 1.480E-08 | 2.600E-08 | 5.300E-08 |
| 24637 | 1232 | S1232T | D | 3e-08 | 6.4e-08 | 1.8e-08 | 3.733E-08 | 2.386E-08 | 1.800E-08 | 6.400E-08 |
| 24638 | 1232 | S1232W | D | 7.9e-08 | 4.9e-08 | 3.3e-08 | 5.367E-08 | 2.335E-08 | 3.300E-08 | 7.900E-08 |
| 24639 | 1232 | S1232Y | D | 5.8e-08 | 1.2e-07 | 5.6e-08 | 7.800E-08 | 3.639E-08 | 5.600E-08 | 1.200E-07 |
| 24640 | 1232 | S1232V | D | 5.6e-08 | 1.1e-08 | 6.6e-08 | 4.433E-08 | 2.930E-08 | 1.100E-08 | 6.600E-08 |
| 24641 | 1233 | L1233A | D | 5.7e-08 | 2e-08 | 1.7e-08 | 3.133E-08 | 2.228E-08 | 1.700E-08 | 5.700E-08 |
| 24642 | 1233 | L1233R | D | 1.6e-07 | 5.1e-08 | 6.6e-08 | 9.233E-08 | 5.908E-08 | 5.100E-08 | 1.600E-07 |
| 24643 | 1233 | L1233N | D | 1.3e-08 | 9.5e-09 | 2e-08 | 1.417E-08 | 5.346E-09 | 9.500E-09 | 2.000E-08 |
| 24644 | 1233 | L1233D | D | 6.1e-08 | 1.9e-08 | 8.4e-08 | 5.467E-08 | 3.296E-08 | 1.900E-08 | 8.400E-08 |
| 24645 | 1233 | L1233C | D | 3.7e-08 | 5.9e-08 | 3.7e-08 | 4.433E-08 | 1.270E-08 | 3.700E-08 | 5.900E-08 |
| 24646 | 1233 | L1233E | D | 6.6e-08 | 7.1e-08 | 5.6e-08 | 6.433E-08 | 7.638E-09 | 5.600E-08 | 7.100E-08 |
| 24647 | 1233 | L1233Q | D | 2.8e-08 | 7.8e-08 | 6.7e-08 | 5.767E-08 | 2.627E-08 | 2.800E-08 | 7.800E-08 |
| 24648 | 1233 | L1233G | D | 3e-08 | 6.2e-08 | 3.5e-08 | 4.233E-08 | 1.721E-08 | 3.000E-08 | 6.200E-08 |
| 24649 | 1233 | L1233H | D | 3.3e-08 | 5.1e-08 | 1.4e-07 | 7.467E-08 | 5.729E-08 | 3.300E-08 | 1.400E-07 |

| | | | | | | | | | | |
|---|---|---|---|---|---|---|---|---|---|---|
| 24650 | 1233 | L1233I | D | 2.2e-08 | 7.8e-08 | 4.6e-08 | 4.867E-08 | 2.810E-08 | 2.200E-08 | 7.800E-08 |
| 24651 | 1233 | L1233L | D | 5.6e-08 | 3.4e-08 | 4.4e-08 | 4.467E-08 | 1.102E-08 | 3.400E-08 | 5.600E-08 |
| 24652 | 1233 | L1233K | D | 8.2e-08 | 6.7e-08 | 3.9e-08 | 6.267E-08 | 2.183E-08 | 3.900E-08 | 8.200E-08 |
| 24653 | 1233 | L1233M | D | 5.7e-08 | 5.7e-08 | 4.7e-08 | 5.367E-08 | 5.774E-09 | 4.700E-08 | 5.700E-08 |
| 24654 | 1233 | L1233F | D | 9.4e-08 | 1.3e-08 | 7.6e-08 | 6.100E-08 | 4.253E-08 | 1.300E-08 | 9.400E-08 |
| 24655 | 1233 | L1233P | D | 4.9e-08 | 1.9e-08 | 9.5e-08 | 5.433E-08 | 3.828E-08 | 1.900E-08 | 9.500E-08 |
| 24656 | 1233 | L1233S | D | 9.3e-08 | 6.6e-08 | 3e-08 | 6.300E-08 | 3.161E-08 | 3.000E-08 | 9.300E-08 |
| 24657 | 1233 | L1233T | D | 1.4e-08 | 4.4e-08 | 4.8e-08 | 3.533E-08 | 1.858E-08 | 1.400E-08 | 4.800E-08 |
| 24658 | 1233 | L1233W | D | 1.1e-08 | 6.8e-08 | 6.7e-08 | 4.867E-08 | 3.262E-08 | 1.100E-08 | 6.800E-08 |
| 24659 | 1233 | L1233Y | D | 7.7e-08 | 6.2e-08 | 6.7e-08 | 6.867E-08 | 7.638E-09 | 6.200E-08 | 7.700E-08 |
| 24660 | 1233 | L1233V | D | 3.3e-08 | 2.1e-08 | 2.7e-08 | 2.700E-08 | 6.000E-09 | 2.100E-08 | 3.300E-08 |
| 24661 | 1234 | S1234A | D | 7.9e-08 | 3.2e-08 | 4.8e-08 | 5.300E-08 | 2.390E-08 | 3.200E-08 | 7.900E-08 |
| 24662 | 1234 | S1234R | D | 8.6e-08 | 2.7e-08 | 3.5e-08 | 4.933E-08 | 3.201E-08 | 2.700E-08 | 8.600E-08 |
| 24663 | 1234 | S1234N | D | 3.2e-08 | 5e-08 | 4.1e-08 | 4.100E-08 | 9.000E-09 | 3.200E-08 | 5.000E-08 |
| 24664 | 1234 | S1234D | D | 7.2e-08 | 9e-08 | 5.4e-08 | 7.200E-08 | 1.800E-08 | 5.400E-08 | 9.000E-08 |
| 24665 | 1234 | S1234C | D | 6.4e-08 | 5e-08 | 7.5e-08 | 6.300E-08 | 1.253E-08 | 5.000E-08 | 7.500E-08 |
| 24666 | 1234 | S1234E | D | 1.2e-08 | 7.6e-08 | 3.4e-08 | 4.067E-08 | 3.252E-08 | 1.200E-08 | 7.600E-08 |
| 24667 | 1234 | S1234Q | D | 6.2e-08 | 7.6e-08 | 2.2e-08 | 5.333E-08 | 2.802E-08 | 2.200E-08 | 7.600E-08 |
| 24668 | 1234 | S1234G | D | 2.1e-08 | 7.6e-08 | 3e-08 | 4.233E-08 | 2.950E-08 | 2.100E-08 | 7.600E-08 |
| 24669 | 1234 | S1234H | D | 9.6e-08 | 2.7e-08 | 3.9e-08 | 5.400E-08 | 3.686E-08 | 2.700E-08 | 9.600E-08 |
| 24670 | 1234 | S1234I | D | 2.7e-08 | 2.5e-08 | 1.4e-08 | 2.200E-08 | 7.000E-09 | 1.400E-08 | 2.700E-08 |
| 24671 | 1234 | S1234L | D | 5.4e-08 | 1.8e-08 | 6e-08 | 4.400E-08 | 2.272E-08 | 1.800E-08 | 6.000E-08 |
| 24672 | 1234 | S1234K | D | 3.3e-08 | 2.9e-08 | 2.1e-08 | 2.767E-08 | 6.110E-09 | 2.100E-08 | 3.300E-08 |
| 24673 | 1234 | S1234M | D | 5.2e-08 | 1.7e-08 | 2e-07 | 8.967E-08 | 9.714E-08 | 1.700E-08 | 2.000E-07 |
| 24674 | 1234 | S1234F | D | 2.5e-08 | 5e-08 | 7.3e-08 | 4.933E-08 | 2.401E-08 | 2.500E-08 | 7.300E-08 |
| 24675 | 1234 | S1234P | D | 8.2e-08 | 5.2e-08 | 6e-08 | 6.467E-08 | 1.553E-08 | 5.200E-08 | 8.200E-08 |
| 24676 | 1234 | S1234S | D | 3.7e-08 | 3.3e-08 | 2.7e-08 | 3.233E-08 | 5.033E-09 | 2.700E-08 | 3.700E-08 |
| 24677 | 1234 | S1234T | D | 4.4e-08 | 6.4e-08 | 4.8e-08 | 5.200E-08 | 1.058E-08 | 4.400E-08 | 6.400E-08 |
| 24678 | 1234 | S1234W | D | 3.2e-08 | 7.2e-08 | 2.4e-08 | 4.267E-08 | 2.572E-08 | 2.400E-08 | 7.200E-08 |
| 24679 | 1234 | S1234Y | D | 3.7e-08 | 7.8e-08 | 1e-07 | 7.167E-08 | 3.197E-08 | 3.700E-08 | 1.000E-07 |
| 24680 | 1234 | S1234V | D | 1.4e-07 | 5e-08 | 6.2e-08 | 8.400E-08 | 4.887E-08 | 5.000E-08 | 1.400E-07 |
| 24681 | 1235 | S1235A | D | 5.5e-08 | 5e-08 | 2.9e-08 | 4.467E-08 | 1.380E-08 | 2.900E-08 | 5.500E-08 |
| 24682 | 1235 | S1235R | D | 9.7e-08 | 6.4e-08 | 2.9e-08 | 6.333E-08 | 3.400E-08 | 2.900E-08 | 9.700E-08 |
| 24683 | 1235 | S1235N | D | 4e-08 | 2.6e-08 | 2.9e-08 | 3.167E-08 | 7.371E-09 | 2.600E-08 | 4.000E-08 |
| 24684 | 1235 | S1235D | D | 2.7e-08 | 2.3e-08 | 1.9e-08 | 2.300E-08 | 4.000E-09 | 1.900E-08 | 2.700E-08 |
| 24685 | 1235 | S1235C | D | 3.5e-08 | 2.2e-08 | 3.1e-08 | 2.933E-08 | 6.658E-09 | 2.200E-08 | 3.500E-08 |
| 24686 | 1235 | S1235E | D | 5.8e-08 | 3.2e-08 | 9.6e-09 | 3.320E-08 | 2.422E-08 | 9.600E-09 | 5.800E-08 |
| 24687 | 1235 | S1235Q | D | 4.9e-08 | 2.6e-08 | 2.7e-08 | 3.400E-08 | 1.300E-08 | 2.600E-08 | 4.900E-08 |
| 24688 | 1235 | S1235G | D | 6.1e-08 | 3e-08 | 7.8e-08 | 5.633E-08 | 2.434E-08 | 3.000E-08 | 7.800E-08 |
| 24689 | 1235 | S1235H | D | 4.1e-08 | 8.1e-08 | 1.6e-08 | 4.600E-08 | 3.279E-08 | 1.600E-08 | 8.100E-08 |
| 24690 | 1235 | S1235I | D | 8e-08 | 2.5e-08 | 6.9e-08 | 5.800E-08 | 2.910E-08 | 2.500E-08 | 8.000E-08 |
| 24691 | 1235 | S1235L | D | 2.1e-08 | 6e-08 | 1.5e-08 | 3.200E-08 | 2.443E-08 | 1.500E-08 | 6.000E-08 |
| 24692 | 1235 | S1235K | D | 1.7e-08 | 1.2e-07 | 3.9e-08 | 5.867E-08 | 5.424E-08 | 1.700E-08 | 1.200E-07 |
| 24693 | 1235 | S1235M | D | 3.6e-08 | 6.1e-08 | 2.2e-08 | 3.967E-08 | 1.976E-08 | 2.200E-08 | 6.100E-08 |
| 24694 | 1235 | S1235F | D | 1.1e-07 | 2.4e-08 | 1.8e-08 | 5.067E-08 | 5.147E-08 | 1.800E-08 | 1.100E-07 |
| 24695 | 1235 | S1235P | D | 6.2e-08 | 3.6e-08 | 3.4e-08 | 4.400E-08 | 1.562E-08 | 3.400E-08 | 6.200E-08 |
| 24696 | 1235 | S1235S | D | 9.4e-08 | 7.1e-08 | 1.9e-08 | 6.133E-08 | 3.842E-08 | 1.900E-08 | 9.400E-08 |
| 24697 | 1235 | S1235T | D | 2.8e-08 | 6.8e-08 | 2.2e-08 | 3.933E-08 | 2.501E-08 | 2.200E-08 | 6.800E-08 |
| 24698 | 1235 | S1235W | D | 7.3e-08 | 1.8e-08 | 3.8e-08 | 4.300E-08 | 2.784E-08 | 1.800E-08 | 7.300E-08 |
| 24699 | 1235 | S1235Y | D | 1.1e-07 | 4.9e-08 | 6.4e-08 | 7.433E-08 | 3.179E-08 | 4.900E-08 | 1.100E-07 |
| 24700 | 1235 | S1235V | D | 2.8e-08 | 3.5e-08 | 2.9e-08 | 3.067E-08 | 3.786E-09 | 2.800E-08 | 3.500E-08 |
| 24701 | 1236 | T1236A | D | 4.3e-08 | 3e-08 | 7e-08 | 4.767E-08 | 2.040E-08 | 3.000E-08 | 7.000E-08 |
| 24702 | 1236 | T1236R | D | 4.6e-08 | 3.2e-08 | 3.3e-08 | 3.700E-08 | 7.810E-09 | 3.200E-08 | 4.600E-08 |
| 24703 | 1236 | T1236N | D | 4.7e-08 | 2.2e-08 | 7e-08 | 4.633E-08 | 2.401E-08 | 2.200E-08 | 7.000E-08 |
| 24704 | 1236 | T1236D | D | 3.2e-08 | 8.9e-08 | 7.9e-08 | 6.667E-08 | 3.044E-08 | 3.200E-08 | 8.900E-08 |
| 24705 | 1236 | T1236C | D | 1.5e-08 | 2.3e-08 | 2.6e-08 | 2.133E-08 | 5.686E-09 | 1.500E-08 | 2.600E-08 |
| 24706 | 1236 | T1236E | D | 3.4e-08 | 4.1e-08 | 1e-07 | 5.833E-08 | 3.625E-08 | 3.400E-08 | 1.000E-07 |
| 24707 | 1236 | T1236Q | D | 4.3e-08 | 2.3e-08 | 3.8e-08 | 3.467E-08 | 1.041E-08 | 2.300E-08 | 4.300E-08 |
| 24708 | 1236 | T1236G | D | 2.6e-08 | 1.5e-08 | 7.6e-08 | 3.900E-08 | 3.251E-08 | 1.500E-08 | 7.600E-08 |
| 24709 | 1236 | T1236H | D | 5.2e-08 | 3.5e-08 | 2.7e-08 | 3.800E-08 | 1.277E-08 | 2.700E-08 | 5.200E-08 |
| 24710 | 1236 | T1236I | D | 5.5e-08 | 1.1e-08 | 3.9e-08 | 3.500E-08 | 2.227E-08 | 1.100E-08 | 5.500E-08 |
| 24711 | 1236 | T1236L | D | 3.9e-08 | 6.8e-08 | 1.6e-08 | 4.100E-08 | 2.606E-08 | 1.600E-08 | 6.800E-08 |
| 24712 | 1236 | T1236K | D | 3.4e-08 | 3.8e-08 | 2.1e-08 | 3.100E-08 | 8.888E-09 | 2.100E-08 | 3.800E-08 |
| 24713 | 1236 | T1236M | D | 2.9e-08 | 7.3e-08 | 3.5e-08 | 4.567E-08 | 2.386E-08 | 2.900E-08 | 7.300E-08 |
| 24714 | 1236 | T1236F | D | 3.4e-08 | 1.4e-08 | 4.7e-08 | 3.167E-08 | 1.662E-08 | 1.400E-08 | 4.700E-08 |
| 24715 | 1236 | T1236P | D | 2.9e-08 | 7.2e-08 | 6.3e-08 | 5.467E-08 | 2.268E-08 | 2.900E-08 | 7.200E-08 |
| 24716 | 1236 | T1236S | D | 6e-08 | 1.6e-07 | 5.5e-08 | 9.167E-08 | 5.923E-08 | 5.500E-08 | 1.600E-07 |
| 24717 | 1236 | T1236T | D | 1.2e-08 | 8.5e-08 | 7.9e-08 | 5.867E-08 | 4.053E-08 | 1.200E-08 | 8.500E-08 |
| 24718 | 1236 | T1236W | D | 6.2e-08 | 4e-08 | 7.3e-08 | 5.833E-08 | 1.680E-08 | 4.000E-08 | 7.300E-08 |
| 24719 | 1236 | T1236Y | D | 1.4e-07 | 2.5e-08 | 3.5e-08 | 6.667E-08 | 6.371E-08 | 2.500E-08 | 1.400E-07 |
| 24720 | 1236 | T1236V | D | 3.3e-08 | 1.9e-08 | 4.9e-08 | 3.367E-08 | 1.501E-08 | 1.900E-08 | 4.900E-08 |
| 24721 | 1237 | L1237A | D | 3.1e-08 | 9.3e-08 | 4.2e-08 | 5.533E-08 | 3.308E-08 | 3.100E-08 | 9.300E-08 |
| 24722 | 1237 | L1237R | D | 7.9e-08 | 5.5e-08 | 3.7e-08 | 5.700E-08 | 2.107E-08 | 3.700E-08 | 7.900E-08 |
| 24723 | 1237 | L1237N | D | 4.9e-08 | 1.3e-07 | 5e-08 | 7.633E-08 | 4.648E-08 | 4.900E-08 | 1.300E-07 |
| 24724 | 1237 | L1237D | D | 2.3e-08 | 3.8e-08 | 4.1e-08 | 3.400E-08 | 9.644E-09 | 2.300E-08 | 4.100E-08 |
| 24725 | 1237 | L1237C | D | 2.2e-08 | 1.1e-08 | 6e-08 | 3.100E-08 | 2.571E-08 | 1.100E-08 | 6.000E-08 |
| 24726 | 1237 | L1237E | D | 1.8e-08 | 4e-08 | 6.1e-08 | 3.967E-08 | 2.150E-08 | 1.800E-08 | 6.100E-08 |
| 24727 | 1237 | L1237Q | D | 1.7e-08 | 2.1e-08 | 4e-08 | 2.600E-08 | 1.229E-08 | 1.700E-08 | 4.000E-08 |
| 24728 | 1237 | L1237G | D | 3.4e-08 | 1.2e-07 | 3.9e-08 | 6.433E-08 | 4.827E-08 | 3.400E-08 | 1.200E-07 |
| 24729 | 1237 | L1237H | D | 6.8e-08 | 3.4e-08 | 9.6e-08 | 6.600E-08 | 3.105E-08 | 3.400E-08 | 9.600E-08 |
| 24730 | 1237 | L1237I | D | 4.4e-08 | 6.2e-08 | 2.4e-08 | 4.333E-08 | 1.901E-08 | 2.400E-08 | 6.200E-08 |
| 24731 | 1237 | L1237L | D | 8.2e-08 | 5.2e-08 | 2.6e-08 | 5.333E-08 | 2.802E-08 | 2.600E-08 | 8.200E-08 |
| 24732 | 1237 | L1237K | D | 3.2e-08 | 2.9e-08 | 4.8e-08 | 3.633E-08 | 1.021E-08 | 2.900E-08 | 4.800E-08 |
| 24733 | 1237 | L1237M | D | 3.3e-08 | 4.7e-08 | 1.5e-08 | 3.167E-08 | 1.604E-08 | 1.500E-08 | 4.700E-08 |
| 24734 | 1237 | L1237F | D | 9.7e-08 | 6.3e-08 | 1.1e-07 | 9.000E-08 | 2.427E-08 | 6.300E-08 | 1.100E-07 |

| | | | | | | | | | | |
|---|---|---|---|---|---|---|---|---|---|---|
| 24735 | 1237 | L1237P | D | 1.2e-07 | 4.6e-08 | 7.1e-08 | 7.900E-08 | 3.764E-08 | 4.600E-08 | 1.200E-07 |
| 24736 | 1237 | L1237S | D | 2.5e-08 | 3.8e-08 | 2.6e-08 | 2.967E-08 | 7.234E-09 | 2.500E-08 | 3.800E-08 |
| 24737 | 1237 | L1237T | D | 5.2e-08 | 3.2e-08 | 3.1e-08 | 3.833E-08 | 1.185E-08 | 3.100E-08 | 5.200E-08 |
| 24738 | 1237 | L1237W | D | 5.6e-08 | 5.3e-08 | 5.2e-08 | 5.367E-08 | 2.082E-09 | 5.200E-08 | 5.600E-08 |
| 24739 | 1237 | L1237Y | D | 1.7e-08 | 2.8e-08 | 2.8e-08 | 2.433E-08 | 6.351E-09 | 1.700E-08 | 2.800E-08 |
| 24740 | 1237 | L1237V | D | 2.7e-08 | 2.7e-08 | 4.6e-08 | 3.333E-08 | 1.097E-08 | 2.700E-08 | 4.600E-08 |
| 24741 | 1238 | T1238A | D | 2.8e-08 | 1.1e-07 | 3.6e-08 | 5.800E-08 | 4.521E-08 | 2.800E-08 | 1.100E-07 |
| 24742 | 1238 | T1238R | D | 1.5e-08 | 7.6e-08 | 7.7e-08 | 5.600E-08 | 3.551E-08 | 1.500E-08 | 7.700E-08 |
| 24743 | 1238 | T1238N | D | 5.8e-08 | 1e-07 | 4.8e-08 | 6.867E-08 | 2.759E-08 | 4.800E-08 | 1.000E-07 |
| 24744 | 1238 | T1238D | D | 6.4e-08 | 9.9e-08 | 2.2e-08 | 6.167E-08 | 3.855E-08 | 2.200E-08 | 9.900E-08 |
| 24745 | 1238 | T1238C | D | 4.8e-08 | 5.2e-08 | 3.5e-08 | 4.500E-08 | 8.888E-09 | 3.500E-08 | 5.200E-08 |
| 24746 | 1238 | T1238E | D | 4.3e-08 | 2.7e-08 | 6e-08 | 4.333E-08 | 1.650E-08 | 2.700E-08 | 6.000E-08 |
| 24747 | 1238 | T1238Q | D | 4.3e-08 | 5.4e-08 | 9.9e-08 | 6.533E-08 | 2.967E-08 | 4.300E-08 | 9.900E-08 |
| 24748 | 1238 | T1238G | D | 4.4e-08 | 2.6e-08 | 1.2e-08 | 2.733E-08 | 1.604E-08 | 1.200E-08 | 4.400E-08 |
| 24749 | 1238 | T1238H | D | 4.5e-08 | 2.7e-08 | 2.2e-08 | 3.133E-08 | 1.210E-08 | 2.200E-08 | 4.500E-08 |
| 24750 | 1238 | T1238I | D | 8.2e-08 | 5.3e-08 | 3e-08 | 5.500E-08 | 2.606E-08 | 3.000E-08 | 8.200E-08 |
| 24751 | 1238 | T1238L | D | 1.2e-08 | 4.8e-08 | 4.4e-08 | 3.467E-08 | 1.973E-08 | 1.200E-08 | 4.800E-08 |
| 24752 | 1238 | T1238K | D | 1.4e-08 | 3e-08 | 4.4e-08 | 2.933E-08 | 1.501E-08 | 1.400E-08 | 4.400E-08 |
| 24753 | 1238 | T1238M | D | 5.2e-08 | 3.9e-08 | 6e-08 | 5.033E-08 | 1.060E-08 | 3.900E-08 | 6.000E-08 |
| 24754 | 1238 | T1238F | D | 4.6e-08 | 3e-08 | 4.6e-08 | 4.067E-08 | 9.238E-09 | 3.000E-08 | 4.600E-08 |
| 24755 | 1238 | T1238P | D | 2.8e-08 | 3.9e-08 | 5.8e-08 | 4.167E-08 | 1.518E-08 | 2.800E-08 | 5.800E-08 |
| 24756 | 1238 | T1238S | D | 2e-08 | 3.4e-08 | 6.2e-08 | 3.867E-08 | 2.139E-08 | 2.000E-08 | 6.200E-08 |
| 24757 | 1238 | T1238T | D | 5.9e-08 | 1.9e-08 | 3e-08 | 3.600E-08 | 2.066E-08 | 1.900E-08 | 5.900E-08 |
| 24758 | 1238 | T1238W | D | 5.6e-08 | 1.9e-08 | 4.9e-08 | 4.133E-08 | 1.966E-08 | 1.900E-08 | 5.600E-08 |
| 24759 | 1238 | T1238Y | D | 8.2e-08 | 6e-08 | 3.2e-08 | 5.800E-08 | 2.506E-08 | 3.200E-08 | 8.200E-08 |
| 24760 | 1238 | T1238V | D | 2.4e-08 | 2.8e-08 | 1.6e-08 | 2.267E-08 | 6.110E-09 | 1.600E-08 | 2.800E-08 |
| 24761 | 1239 | L1239A | D | 4.3e-08 | 4.1e-08 | 8e-08 | 5.467E-08 | 2.196E-08 | 4.100E-08 | 8.000E-08 |
| 24762 | 1239 | L1239R | D | 3e-08 | 1.9e-08 | 8.3e-08 | 4.400E-08 | 3.422E-08 | 1.900E-08 | 8.300E-08 |
| 24763 | 1239 | L1239N | D | 6.1e-08 | 4.9e-08 | 5.2e-08 | 5.400E-08 | 6.245E-09 | 4.900E-08 | 6.100E-08 |
| 24764 | 1239 | L1239D | D | 4.4e-08 | 4.9e-08 | 4.4e-08 | 4.567E-08 | 2.887E-09 | 4.400E-08 | 4.900E-08 |
| 24765 | 1239 | L1239C | D | 6.9e-08 | 4.4e-08 | 2.1e-08 | 4.467E-08 | 2.401E-08 | 2.100E-08 | 6.900E-08 |
| 24766 | 1239 | L1239E | D | 6.2e-08 | 1.9e-08 | 1.3e-08 | 3.133E-08 | 2.673E-08 | 1.300E-08 | 6.200E-08 |
| 24767 | 1239 | L1239Q | D | 2e-08 | 4.3e-08 | 4e-08 | 3.433E-08 | 1.250E-08 | 2.000E-08 | 4.300E-08 |
| 24768 | 1239 | L1239G | D | 8e-08 | 3.2e-08 | 3.8e-08 | 5.000E-08 | 2.615E-08 | 3.200E-08 | 8.000E-08 |
| 24769 | 1239 | L1239H | D | 1.5e-07 | 3.6e-08 | 8.3e-08 | 8.967E-08 | 5.729E-08 | 3.600E-08 | 1.500E-07 |
| 24770 | 1239 | L1239I | D | 2.8e-08 | 2.9e-08 | 2.5e-08 | 2.733E-08 | 2.082E-09 | 2.500E-08 | 2.900E-08 |
| 24771 | 1239 | L1239L | D | 1.9e-08 | 2.8e-08 | 2.3e-08 | 2.333E-08 | 4.509E-09 | 1.900E-08 | 2.800E-08 |
| 24772 | 1239 | L1239K | D | 5e-08 | 4.6e-08 | 5.8e-08 | 5.133E-08 | 6.110E-09 | 4.600E-08 | 5.800E-08 |
| 24773 | 1239 | L1239M | D | 1.7e-08 | 1e-07 | 2.4e-08 | 4.700E-08 | 4.603E-08 | 1.700E-08 | 1.000E-07 |
| 24774 | 1239 | L1239F | D | 4e-08 | 5.5e-08 | 8.4e-08 | 5.967E-08 | 2.237E-08 | 4.000E-08 | 8.400E-08 |
| 24775 | 1239 | L1239P | D | 3.6e-08 | 4e-08 | 2.1e-08 | 3.233E-08 | 1.002E-08 | 2.100E-08 | 4.000E-08 |
| 24776 | 1239 | L1239S | D | 9.4e-08 | 3.9e-08 | 2.1e-08 | 5.133E-08 | 3.803E-08 | 2.100E-08 | 9.400E-08 |
| 24777 | 1239 | L1239T | D | 3.6e-08 | 6.3e-08 | 1.9e-08 | 3.933E-08 | 2.219E-08 | 1.900E-08 | 6.300E-08 |
| 24778 | 1239 | L1239W | D | 4.2e-08 | 9.2e-08 | 7.6e-08 | 7.000E-08 | 2.553E-08 | 4.200E-08 | 9.200E-08 |
| 24779 | 1239 | L1239Y | D | 2.2e-08 | 3.1e-08 | 6.5e-08 | 3.933E-08 | 2.268E-08 | 2.200E-08 | 6.500E-08 |
| 24780 | 1239 | L1239V | D | 1.3e-08 | 7e-08 | 2.7e-08 | 3.667E-08 | 2.970E-08 | 1.300E-08 | 7.000E-08 |
| 24781 | 1240 | S1240A | D | 5.1e-08 | 3.6e-08 | 6.8e-08 | 5.167E-08 | 1.601E-08 | 3.600E-08 | 6.800E-08 |
| 24782 | 1240 | S1240R | D | 5.4e-08 | 2.2e-08 | 7.2e-08 | 4.933E-08 | 2.532E-08 | 2.200E-08 | 7.200E-08 |
| 24783 | 1240 | S1240N | D | 4e-08 | 3e-08 | 2.7e-08 | 3.233E-08 | 6.807E-09 | 2.700E-08 | 4.000E-08 |
| 24784 | 1240 | S1240D | D | 3.4e-08 | 4.1e-08 | 1.3e-08 | 2.933E-08 | 1.457E-08 | 1.300E-08 | 4.100E-08 |
| 24785 | 1240 | S1240C | D | 2.8e-08 | 4.2e-08 | 8.7e-09 | 2.623E-08 | 1.672E-08 | 8.700E-09 | 4.200E-08 |
| 24786 | 1240 | S1240E | D | 2.6e-08 | 5.2e-08 | 7.6e-08 | 5.133E-08 | 2.501E-08 | 2.600E-08 | 7.600E-08 |
| 24787 | 1240 | S1240Q | D | 2.5e-08 | 4.5e-08 | 3.2e-08 | 3.400E-08 | 1.015E-08 | 2.500E-08 | 4.500E-08 |
| 24788 | 1240 | S1240G | D | 2.7e-08 | 1e-07 | 4e-08 | 5.567E-08 | 3.894E-08 | 2.700E-08 | 1.000E-07 |
| 24789 | 1240 | S1240H | D | 7.3e-08 | 7.7e-08 | 6.5e-08 | 7.167E-08 | 6.110E-09 | 6.500E-08 | 7.700E-08 |
| 24790 | 1240 | S1240I | D | 5.3e-08 | 1.5e-08 | 5.7e-08 | 4.167E-08 | 2.318E-08 | 1.500E-08 | 5.700E-08 |
| 24791 | 1240 | S1240L | D | 5.4e-08 | 4.1e-08 | 5.5e-08 | 5.000E-08 | 7.810E-09 | 4.100E-08 | 5.500E-08 |
| 24792 | 1240 | S1240K | D | 1.2e-07 | 1.8e-08 | 5.7e-08 | 6.500E-08 | 5.147E-08 | 1.800E-08 | 1.200E-07 |
| 24793 | 1240 | S1240M | D | 4.2e-08 | 3.7e-08 | 3.7e-08 | 3.867E-08 | 2.887E-09 | 3.700E-08 | 4.200E-08 |
| 24794 | 1240 | S1240F | D | 8.1e-08 | 4.8e-08 | 9.8e-09 | 4.627E-08 | 3.563E-08 | 9.800E-09 | 8.100E-08 |
| 24795 | 1240 | S1240P | D | 1.7e-08 | 3e-08 | 8.7e-08 | 4.467E-08 | 3.723E-08 | 1.700E-08 | 8.700E-08 |
| 24796 | 1240 | S1240S | D | 6.6e-08 | 2e-08 | 3.4e-08 | 4.000E-08 | 2.358E-08 | 2.000E-08 | 6.600E-08 |
| 24797 | 1240 | S1240T | D | 3.6e-08 | 6.6e-08 | 6.3e-08 | 5.500E-08 | 1.652E-08 | 3.600E-08 | 6.600E-08 |
| 24798 | 1240 | S1240W | D | 3.6e-08 | 3.6e-08 | 3.3e-08 | 3.500E-08 | 1.732E-09 | 3.300E-08 | 3.600E-08 |
| 24799 | 1240 | S1240Y | D | 3.2e-08 | 3.2e-08 | 2e-08 | 2.800E-08 | 6.928E-09 | 2.000E-08 | 3.200E-08 |
| 24800 | 1240 | S1240V | D | 1.8e-08 | 7.1e-08 | 3.1e-08 | 4.000E-08 | 2.762E-08 | 1.800E-08 | 7.100E-08 |
| 24801 | 1241 | K1241A | D | 1.3e-08 | 3.4e-08 | 9.3e-08 | 4.667E-08 | 4.148E-08 | 1.300E-08 | 9.300E-08 |
| 24802 | 1241 | K1241R | D | 1.9e-08 | 4.4e-08 | 3.8e-08 | 3.367E-08 | 1.305E-08 | 1.900E-08 | 4.400E-08 |
| 24803 | 1241 | K1241N | D | 3.5e-08 | 1.1e-08 | 2.9e-08 | 2.500E-08 | 1.249E-08 | 1.100E-08 | 3.500E-08 |
| 24804 | 1241 | K1241D | D | 2.2e-08 | 1.9e-08 | 7.5e-08 | 3.867E-08 | 3.150E-08 | 1.900E-08 | 7.500E-08 |
| 24805 | 1241 | K1241C | D | 2.2e-08 | 2.2e-08 | 8.2e-08 | 4.200E-08 | 3.464E-08 | 2.200E-08 | 8.200E-08 |
| 24806 | 1241 | K1241E | D | 2.5e-08 | 6.7e-08 | 3.5e-08 | 4.233E-08 | 2.194E-08 | 2.500E-08 | 6.700E-08 |
| 24807 | 1241 | K1241Q | D | 3.4e-08 | 4.7e-08 | 3.6e-08 | 3.900E-08 | 7.000E-09 | 3.400E-08 | 4.700E-08 |
| 24808 | 1241 | K1241G | D | 1.5e-08 | 6.7e-08 | 4.1e-08 | 4.100E-08 | 2.600E-08 | 1.500E-08 | 6.700E-08 |
| 24809 | 1241 | K1241H | D | 4.2e-08 | 6.1e-08 | 8.7e-08 | 6.333E-08 | 2.259E-08 | 4.200E-08 | 8.700E-08 |
| 24810 | 1241 | K1241I | D | 6.9e-08 | 1.2e-07 | 1.1e-08 | 6.667E-08 | 5.454E-08 | 1.100E-08 | 1.200E-07 |
| 24811 | 1241 | K1241L | D | 2.8e-08 | 1.1e-07 | 1.4e-07 | 9.267E-08 | 5.798E-08 | 2.800E-08 | 1.400E-07 |
| 24812 | 1241 | K1241K | D | 3.7e-08 | 2.4e-08 | 3.5e-08 | 3.200E-08 | 7.000E-09 | 2.400E-08 | 3.700E-08 |
| 24813 | 1241 | K1241M | D | 5.7e-08 | 3.3e-08 | 1e-07 | 6.333E-08 | 3.395E-08 | 3.300E-08 | 1.000E-07 |
| 24814 | 1241 | K1241F | D | 6.7e-08 | 2.2e-08 | 5.8e-08 | 4.900E-08 | 2.381E-08 | 2.200E-08 | 6.700E-08 |
| 24815 | 1241 | K1241P | D | 5.6e-08 | 1.2e-07 | 1.5e-08 | 6.367E-08 | 5.292E-08 | 1.500E-08 | 1.200E-07 |
| 24816 | 1241 | K1241S | D | 9.7e-08 | 6.8e-08 | 1.9e-08 | 6.133E-08 | 3.943E-08 | 1.900E-08 | 9.700E-08 |
| 24817 | 1241 | K1241T | D | 2.3e-08 | 1.4e-07 | 7e-08 | 7.767E-08 | 5.888E-08 | 2.300E-08 | 1.400E-07 |
| 24818 | 1241 | K1241W | D | 1.9e-08 | 7.7e-08 | 4.2e-08 | 4.600E-08 | 2.921E-08 | 1.900E-08 | 7.700E-08 |
| 24819 | 1241 | K1241Y | D | 5.4e-08 | 3.1e-08 | 5.3e-08 | 4.600E-08 | 1.300E-08 | 3.100E-08 | 5.400E-08 |

| | | | | | | | | | | |
|---|---|---|---|---|---|---|---|---|---|---|
| 24820 | 1241 | K1241V | D | 5e-08 | 2.2e-08 | 6.6e-08 | 4.600E-08 | 2.227E-08 | 2.200E-08 | 6.600E-08 |
| 24821 | 1242 | A1242A | D | 6.9e-08 | 9.5e-08 | 4.9e-08 | 7.100E-08 | 2.307E-08 | 4.900E-08 | 9.500E-08 |
| 24822 | 1242 | A1242R | D | 3e-08 | 6.5e-08 | 4.5e-08 | 4.667E-08 | 1.756E-08 | 3.000E-08 | 6.500E-08 |
| 24823 | 1242 | A1242N | D | 1.2e-08 | 1.6e-08 | 9.3e-09 | 1.243E-08 | 3.371E-09 | 9.300E-09 | 1.600E-08 |
| 24824 | 1242 | A1242D | D | 6.6e-08 | 7.6e-08 | 3.6e-08 | 5.933E-08 | 2.082E-08 | 3.600E-08 | 7.600E-08 |
| 24825 | 1242 | A1242C | D | 1.2e-07 | 8.9e-09 | 1.5e-08 | 4.797E-08 | 6.246E-08 | 8.900E-09 | 1.200E-07 |
| 24826 | 1242 | A1242E | D | 2.8e-08 | 6.6e-08 | 6.8e-08 | 5.400E-08 | 2.254E-08 | 2.800E-08 | 6.800E-08 |
| 24827 | 1242 | A1242Q | D | 5e-08 | 6.8e-08 | 8.2e-08 | 6.667E-08 | 1.604E-08 | 5.000E-08 | 8.200E-08 |
| 24828 | 1242 | A1242G | D | 7.9e-08 | 4.4e-08 | 2.4e-08 | 4.900E-08 | 2.784E-08 | 2.400E-08 | 7.900E-08 |
| 24829 | 1242 | A1242H | D | 2.7e-08 | 6.3e-08 | 7e-08 | 5.333E-08 | 2.307E-08 | 2.700E-08 | 7.000E-08 |
| 24830 | 1242 | A1242I | D | 3.4e-08 | 9.8e-08 | 1.1e-07 | 8.067E-08 | 4.086E-08 | 3.400E-08 | 1.100E-07 |
| 24831 | 1242 | A1242L | D | 9.1e-08 | 3.4e-08 | 3.5e-08 | 5.333E-08 | 3.262E-08 | 3.400E-08 | 9.100E-08 |
| 24832 | 1242 | A1242K | D | 3.5e-08 | 7.6e-08 | 7e-08 | 6.033E-08 | 2.214E-08 | 3.500E-08 | 7.600E-08 |
| 24833 | 1242 | A1242M | D | 3.4e-08 | 8.4e-08 | 2.9e-08 | 4.900E-08 | 3.041E-08 | 2.900E-08 | 8.400E-08 |
| 24834 | 1242 | A1242F | D | 5.4e-08 | 4.9e-08 | 9e-08 | 6.433E-08 | 2.237E-08 | 4.900E-08 | 9.000E-08 |
| 24835 | 1242 | A1242P | D | 1.9e-08 | 1.1e-07 | 2.6e-08 | 5.167E-08 | 5.064E-08 | 1.900E-08 | 1.100E-07 |
| 24836 | 1242 | A1242S | D | 6.5e-08 | 1.1e-07 | 2.1e-08 | 6.533E-08 | 4.450E-08 | 2.100E-08 | 1.100E-07 |
| 24837 | 1242 | A1242T | D | 1e-08 | 1e-08 | 2e-08 | 1.333E-08 | 5.774E-09 | 1.000E-08 | 2.000E-08 |
| 24838 | 1242 | A1242W | D | 1.1e-08 | 3.5e-08 | 4.2e-08 | 2.933E-08 | 1.626E-08 | 1.100E-08 | 4.200E-08 |
| 24839 | 1242 | A1242Y | D | 3.4e-08 | 2.6e-08 | 2.9e-08 | 2.967E-08 | 4.041E-09 | 2.600E-08 | 3.400E-08 |
| 24840 | 1242 | A1242V | D | 1.1e-07 | 8.1e-08 | 1.7e-08 | 6.933E-08 | 4.759E-08 | 1.700E-08 | 1.100E-07 |
| 24841 | 1243 | D1243A | D | 2.6e-08 | 7e-08 | 3.5e-08 | 4.367E-08 | 2.325E-08 | 2.600E-08 | 7.000E-08 |
| 24842 | 1243 | D1243R | D | 5.2e-08 | 4.5e-08 | 9.2e-08 | 6.300E-08 | 2.536E-08 | 4.500E-08 | 9.200E-08 |
| 24843 | 1243 | D1243N | D | 1.7e-08 | 8.4e-08 | 7.5e-08 | 5.867E-08 | 3.636E-08 | 1.700E-08 | 8.400E-08 |
| 24844 | 1243 | D1243D | D | 2.8e-08 | 8.8e-08 | 4.5e-08 | 5.367E-08 | 3.092E-08 | 2.800E-08 | 8.800E-08 |
| 24845 | 1243 | D1243C | D | 1e-07 | 3e-08 | 1e-07 | 7.667E-08 | 4.041E-08 | 3.000E-08 | 1.000E-07 |
| 24846 | 1243 | D1243E | D | 8.4e-08 | 2.1e-08 | 2e-08 | 4.167E-08 | 3.667E-08 | 2.000E-08 | 8.400E-08 |
| 24847 | 1243 | D1243Q | D | 4.3e-08 | 4.2e-08 | 7.9e-08 | 5.467E-08 | 2.108E-08 | 4.200E-08 | 7.900E-08 |
| 24848 | 1243 | D1243G | D | 2.2e-08 | 2.4e-08 | 7.1e-08 | 3.900E-08 | 2.773E-08 | 2.200E-08 | 7.100E-08 |
| 24849 | 1243 | D1243H | D | 5.2e-08 | 1.2e-07 | 6e-08 | 7.733E-08 | 3.717E-08 | 5.200E-08 | 1.200E-07 |
| 24850 | 1243 | D1243I | D | 6.8e-08 | 8.2e-09 | 5e-08 | 4.207E-08 | 3.068E-08 | 8.200E-09 | 6.800E-08 |
| 24851 | 1243 | D1243L | D | 2.6e-08 | 5.3e-08 | 3e-08 | 3.633E-08 | 1.457E-08 | 2.600E-08 | 5.300E-08 |
| 24852 | 1243 | D1243K | D | 2.1e-07 | 5e-08 | 1.2e-08 | 9.067E-08 | 1.051E-07 | 1.200E-08 | 2.100E-07 |
| 24853 | 1243 | D1243M | D | 1.2e-07 | 8e-08 | 5.6e-08 | 8.533E-08 | 3.233E-08 | 5.600E-08 | 1.200E-07 |
| 24854 | 1243 | D1243F | D | 9.4e-08 | 5.1e-08 | 8.2e-08 | 7.567E-08 | 2.219E-08 | 5.100E-08 | 9.400E-08 |
| 24855 | 1243 | D1243P | D | 5.2e-08 | 6.1e-08 | 8.4e-08 | 6.567E-08 | 1.650E-08 | 5.200E-08 | 8.400E-08 |
| 24856 | 1243 | D1243S | D | 9.8e-08 | 1.4e-08 | 3.4e-08 | 4.867E-08 | 4.388E-08 | 1.400E-08 | 9.800E-08 |
| 24857 | 1243 | D1243T | D | 3.2e-08 | 1.3e-08 | 1.8e-08 | 2.100E-08 | 9.849E-09 | 1.300E-08 | 3.200E-08 |
| 24858 | 1243 | D1243W | D | 4.9e-08 | 3.4e-08 | 3.4e-08 | 3.900E-08 | 8.660E-09 | 3.400E-08 | 4.900E-08 |
| 24859 | 1243 | D1243Y | D | 1.2e-07 | 7.9e-08 | 7.5e-08 | 9.133E-08 | 2.491E-08 | 7.500E-08 | 1.200E-07 |
| 24860 | 1243 | D1243V | D | 1.9e-08 | 7.5e-08 | 4.2e-08 | 4.533E-08 | 2.815E-08 | 1.900E-08 | 7.500E-08 |
| 24861 | 1244 | Y1244A | D | 4.2e-08 | 5.7e-08 | 1.1e-07 | 6.967E-08 | 3.573E-08 | 4.200E-08 | 1.100E-07 |
| 24862 | 1244 | Y1244R | D | 2.5e-08 | 2.4e-08 | 7.1e-08 | 4.000E-08 | 2.685E-08 | 2.400E-08 | 7.100E-08 |
| 24863 | 1244 | Y1244N | D | 3.1e-08 | 2.6e-08 | 2.9e-08 | 2.867E-08 | 2.517E-09 | 2.600E-08 | 3.100E-08 |
| 24864 | 1244 | Y1244D | D | 6.8e-08 | 1e-07 | 1.2e-07 | 9.600E-08 | 2.623E-08 | 6.800E-08 | 1.200E-07 |
| 24865 | 1244 | Y1244C | D | 3.7e-08 | 8.4e-08 | 4.6e-08 | 5.567E-08 | 2.495E-08 | 3.700E-08 | 8.400E-08 |
| 24866 | 1244 | Y1244E | D | 4.5e-08 | 1.2e-08 | 3.1e-08 | 2.933E-08 | 1.656E-08 | 1.200E-08 | 4.500E-08 |
| 24867 | 1244 | Y1244Q | D | 8.5e-09 | 3.2e-08 | 1.7e-08 | 1.917E-08 | 1.190E-08 | 8.500E-09 | 3.200E-08 |
| 24868 | 1244 | Y1244G | D | 2.2e-08 | 3.3e-08 | 8.5e-08 | 4.667E-08 | 3.365E-08 | 2.200E-08 | 8.500E-08 |
| 24869 | 1244 | Y1244H | D | 7.9e-08 | 1.8e-07 | 2.3e-08 | 9.400E-08 | 7.957E-08 | 2.300E-08 | 1.800E-07 |
| 24870 | 1244 | Y1244I | D | 2.4e-08 | 6.8e-08 | 5.4e-08 | 4.867E-08 | 2.248E-08 | 2.400E-08 | 6.800E-08 |
| 24871 | 1244 | Y1244L | D | 3.1e-08 | 1e-08 | 9.2e-08 | 4.433E-08 | 4.259E-08 | 1.000E-08 | 9.200E-08 |
| 24872 | 1244 | Y1244K | D | 9.5e-08 | 4.8e-08 | 3.5e-08 | 5.933E-08 | 3.156E-08 | 3.500E-08 | 9.500E-08 |
| 24873 | 1244 | Y1244M | D | 2.2e-07 | 3.2e-08 | 4.3e-08 | 9.833E-08 | 1.055E-07 | 3.200E-08 | 2.200E-07 |
| 24874 | 1244 | Y1244F | D | 2.7e-08 | 2.1e-08 | 2.6e-08 | 2.467E-08 | 3.215E-09 | 2.100E-08 | 2.700E-08 |
| 24875 | 1244 | Y1244P | D | 2.4e-08 | 3.3e-08 | 1.4e-08 | 2.367E-08 | 9.504E-09 | 1.400E-08 | 3.300E-08 |
| 24876 | 1244 | Y1244S | D | 3.4e-08 | 8.4e-08 | 1.1e-07 | 7.600E-08 | 3.863E-08 | 3.400E-08 | 1.100E-07 |
| 24877 | 1244 | Y1244T | D | 2.8e-08 | 3.6e-08 | 3.6e-08 | 3.333E-08 | 4.619E-09 | 2.800E-08 | 3.600E-08 |
| 24878 | 1244 | Y1244W | D | 6.4e-08 | 3.6e-08 | 5.4e-08 | 5.133E-08 | 1.419E-08 | 3.600E-08 | 6.400E-08 |
| 24879 | 1244 | Y1244Y | D | 4.5e-08 | 3.5e-08 | 9.2e-08 | 5.733E-08 | 3.044E-08 | 3.500E-08 | 9.200E-08 |
| 24880 | 1244 | Y1244V | D | 5.2e-08 | 5.3e-08 | 2.1e-08 | 4.200E-08 | 1.819E-08 | 2.100E-08 | 5.300E-08 |
| 24881 | 1245 | E1245A | D | 5.4e-08 | 7.5e-08 | 5.1e-08 | 6.000E-08 | 1.308E-08 | 5.100E-08 | 7.500E-08 |
| 24882 | 1245 | E1245R | D | 3e-08 | 5.2e-08 | 3.5e-08 | 3.900E-08 | 1.153E-08 | 3.000E-08 | 5.200E-08 |
| 24883 | 1245 | E1245N | D | 4.6e-08 | 1.7e-08 | 2.5e-08 | 2.933E-08 | 1.498E-08 | 1.700E-08 | 4.600E-08 |
| 24884 | 1245 | E1245D | D | 2.8e-08 | 1.9e-08 | 1.9e-08 | 2.200E-08 | 5.196E-09 | 1.900E-08 | 2.800E-08 |
| 24885 | 1245 | E1245C | D | 3.3e-08 | 4.2e-08 | 1.8e-08 | 3.100E-08 | 1.212E-08 | 1.800E-08 | 4.200E-08 |
| 24886 | 1245 | E1245E | D | 2.2e-08 | 1.6e-07 | 3.1e-08 | 7.100E-08 | 7.721E-08 | 2.200E-08 | 1.600E-07 |
| 24887 | 1245 | E1245Q | D | 3.2e-08 | 6.7e-08 | 5.7e-09 | 3.490E-08 | 3.075E-08 | 5.700E-09 | 6.700E-08 |
| 24888 | 1245 | E1245G | D | 9e-08 | 2.1e-08 | 3.3e-08 | 4.800E-08 | 3.686E-08 | 2.100E-08 | 9.000E-08 |
| 24889 | 1245 | E1245H | D | 7.9e-08 | 1.4e-07 | 2.6e-08 | 8.167E-08 | 5.705E-08 | 2.600E-08 | 1.400E-07 |
| 24890 | 1245 | E1245I | D | 1.9e-08 | 8.4e-09 | 4.9e-08 | 2.547E-08 | 2.106E-08 | 8.400E-09 | 4.900E-08 |
| 24891 | 1245 | E1245L | D | 4.6e-08 | 2.4e-08 | 2.6e-08 | 3.200E-08 | 1.217E-08 | 2.400E-08 | 4.600E-08 |
| 24892 | 1245 | E1245K | D | 5.2e-08 | 4e-08 | 5e-08 | 4.733E-08 | 6.429E-09 | 4.000E-08 | 5.200E-08 |
| 24893 | 1245 | E1245M | D | 4.5e-08 | 1.1e-07 | 2.6e-08 | 6.033E-08 | 4.405E-08 | 2.600E-08 | 1.100E-07 |
| 24894 | 1245 | E1245F | D | 3.7e-08 | 7.2e-08 | 5.2e-08 | 5.367E-08 | 1.756E-08 | 3.700E-08 | 7.200E-08 |
| 24895 | 1245 | E1245P | D | 2.7e-08 | 3.2e-08 | 3.6e-08 | 3.167E-08 | 4.509E-09 | 2.700E-08 | 3.600E-08 |
| 24896 | 1245 | E1245S | D | 4.4e-08 | 4e-08 | 3.4e-08 | 3.933E-08 | 5.033E-09 | 3.400E-08 | 4.400E-08 |
| 24897 | 1245 | E1245T | D | 3.2e-08 | 4.5e-08 | 1.4e-07 | 7.233E-08 | 5.896E-08 | 3.200E-08 | 1.400E-07 |
| 24898 | 1245 | E1245W | D | 2.5e-08 | 9.5e-08 | 8.9e-08 | 6.967E-08 | 3.880E-08 | 2.500E-08 | 9.500E-08 |
| 24899 | 1245 | E1245Y | D | 9.1e-08 | 4.4e-08 | 3.3e-08 | 5.600E-08 | 3.081E-08 | 3.300E-08 | 9.100E-08 |
| 24900 | 1245 | E1245V | D | 5.4e-08 | 3.9e-08 | 5.1e-08 | 4.800E-08 | 7.937E-09 | 3.900E-08 | 5.400E-08 |
| 24901 | 1246 | K1246A | D | 5.6e-08 | 2.7e-08 | 4.3e-08 | 4.200E-08 | 1.453E-08 | 2.700E-08 | 5.600E-08 |
| 24902 | 1246 | K1246R | D | 4.6e-08 | 4.4e-08 | 4.5e-08 | 4.500E-08 | 1.000E-09 | 4.400E-08 | 4.600E-08 |
| 24903 | 1246 | K1246N | D | 6.6e-08 | 5.5e-08 | 2.9e-08 | 5.000E-08 | 1.900E-08 | 2.900E-08 | 6.600E-08 |
| 24904 | 1246 | K1246D | D | 1.5e-07 | 4.8e-08 | 9.7e-08 | 9.833E-08 | 5.101E-08 | 4.800E-08 | 1.500E-07 |

| | | | | | | | | | | |
|---|---|---|---|---|---|---|---|---|---|---|
| 24905 | 1246 | K1246C | D | 4e-08 | 3.7e-08 | 7.5e-08 | 5.067E-08 | 2.113E-08 | 3.700E-08 | 7.500E-08 |
| 24906 | 1246 | K1246E | D | 6.8e-08 | 5.7e-08 | 3.9e-08 | 5.467E-08 | 1.464E-08 | 3.900E-08 | 6.800E-08 |
| 24907 | 1246 | K1246Q | D | 3.5e-08 | 1e-07 | 3.7e-08 | 5.733E-08 | 3.696E-08 | 3.500E-08 | 1.000E-07 |
| 24908 | 1246 | K1246G | D | 5.6e-08 | 3e-08 | 7.5e-08 | 5.367E-08 | 2.259E-08 | 3.000E-08 | 7.500E-08 |
| 24909 | 1246 | K1246H | D | 5.1e-08 | 5.8e-08 | 2.6e-08 | 4.500E-08 | 1.682E-08 | 2.600E-08 | 5.800E-08 |
| 24910 | 1246 | K1246I | D | 4.7e-08 | 5.4e-08 | 2.9e-08 | 4.333E-08 | 1.290E-08 | 2.900E-08 | 5.400E-08 |
| 24911 | 1246 | K1246L | D | 5.2e-08 | 4.9e-08 | 4.9e-08 | 5.000E-08 | 1.732E-09 | 4.900E-08 | 5.200E-08 |
| 24912 | 1246 | K1246K | D | 4e-08 | 5.3e-08 | 3.6e-08 | 4.300E-08 | 8.888E-09 | 3.600E-08 | 5.300E-08 |
| 24913 | 1246 | K1246M | D | 2.9e-08 | 4.9e-08 | 2.2e-08 | 3.333E-08 | 1.401E-08 | 2.200E-08 | 4.900E-08 |
| 24914 | 1246 | K1246F | D | 3.2e-08 | 6.8e-08 | 5e-08 | 5.000E-08 | 1.800E-08 | 3.200E-08 | 6.800E-08 |
| 24915 | 1246 | K1246P | D | 8.6e-08 | 1.7e-08 | 3e-08 | 4.433E-08 | 3.667E-08 | 1.700E-08 | 8.600E-08 |
| 24916 | 1246 | K1246S | D | 1.2e-08 | 4.2e-08 | 6.5e-08 | 3.967E-08 | 2.658E-08 | 1.200E-08 | 6.500E-08 |
| 24917 | 1246 | K1246T | D | 2.9e-08 | 4.1e-08 | 8.2e-08 | 5.067E-08 | 2.779E-08 | 2.900E-08 | 8.200E-08 |
| 24918 | 1246 | K1246W | D | 2.9e-08 | 7.5e-08 | 7.5e-08 | 5.967E-08 | 2.656E-08 | 2.900E-08 | 7.500E-08 |
| 24919 | 1246 | K1246Y | D | 4.2e-08 | 9.9e-09 | 1.4e-07 | 6.397E-08 | 6.777E-08 | 9.900E-09 | 1.400E-07 |
| 24920 | 1246 | K1246V | D | 1.5e-08 | 6.3e-08 | 2e-08 | 3.267E-08 | 2.639E-08 | 1.500E-08 | 6.300E-08 |
| 24921 | 1247 | H1247A | D | 2e-08 | 2.2e-08 | 3.5e-08 | 2.567E-08 | 8.145E-09 | 2.000E-08 | 3.500E-08 |
| 24922 | 1247 | H1247R | D | 3.2e-08 | 1.4e-07 | 6.7e-08 | 7.967E-08 | 5.510E-08 | 3.200E-08 | 1.400E-07 |
| 24923 | 1247 | H1247N | D | 2.3e-08 | 6.9e-08 | 3.8e-08 | 4.333E-08 | 2.346E-08 | 2.300E-08 | 6.900E-08 |
| 24924 | 1247 | H1247D | D | 4.3e-08 | 8.8e-08 | 1.2e-08 | 4.767E-08 | 3.821E-08 | 1.200E-08 | 8.800E-08 |
| 24925 | 1247 | H1247C | D | 8e-09 | 5.2e-08 | 3.4e-08 | 3.133E-08 | 2.212E-08 | 8.000E-09 | 5.200E-08 |
| 24926 | 1247 | H1247E | D | 5.2e-08 | 5.4e-08 | 3.8e-08 | 4.800E-08 | 8.718E-09 | 3.800E-08 | 5.400E-08 |
| 24927 | 1247 | H1247Q | D | 4.7e-08 | 4.7e-08 | 1.2e-07 | 7.133E-08 | 4.215E-08 | 4.700E-08 | 1.200E-07 |
| 24928 | 1247 | H1247G | D | 7.6e-08 | 2.3e-08 | 5.2e-08 | 5.033E-08 | 2.654E-08 | 2.300E-08 | 7.600E-08 |
| 24929 | 1247 | H1247H | D | 9.5e-08 | 6.1e-08 | 2.4e-08 | 6.000E-08 | 3.551E-08 | 2.400E-08 | 9.500E-08 |
| 24930 | 1247 | H1247I | D | 7.5e-08 | 4.5e-08 | 4.1e-08 | 5.367E-08 | 1.858E-08 | 4.100E-08 | 7.500E-08 |
| 24931 | 1247 | H1247L | D | 1.6e-07 | 2.6e-07 | 4.4e-08 | 1.547E-07 | 1.081E-07 | 4.400E-08 | 2.600E-07 |
| 24932 | 1247 | H1247K | D | 4e-08 | 1.1e-08 | 2.7e-08 | 2.600E-08 | 1.453E-08 | 1.100E-08 | 4.000E-08 |
| 24933 | 1247 | H1247M | D | 3.5e-08 | 5.6e-08 | 1.8e-08 | 3.633E-08 | 1.904E-08 | 1.800E-08 | 5.600E-08 |
| 24934 | 1247 | H1247F | D | 3.5e-08 | 1.5e-08 | 7.1e-08 | 4.033E-08 | 2.838E-08 | 1.500E-08 | 7.100E-08 |
| 24935 | 1247 | H1247P | D | 5.1e-08 | 6.1e-08 | 4.6e-08 | 5.267E-08 | 7.638E-09 | 4.600E-08 | 6.100E-08 |
| 24936 | 1247 | H1247S | D | 3.2e-08 | 8.1e-08 | 4.3e-08 | 5.200E-08 | 2.571E-08 | 3.200E-08 | 8.100E-08 |
| 24937 | 1247 | H1247T | D | 4.8e-08 | 7.7e-08 | 8.5e-08 | 7.000E-08 | 1.947E-08 | 4.800E-08 | 8.500E-08 |
| 24938 | 1247 | H1247W | D | 7.1e-08 | 1.7e-08 | 1.8e-08 | 3.533E-08 | 3.089E-08 | 1.700E-08 | 7.100E-08 |
| 24939 | 1247 | H1247Y | D | 4.6e-08 | 4.7e-08 | 4.8e-08 | 4.700E-08 | 1.000E-09 | 4.600E-08 | 4.800E-08 |
| 24940 | 1247 | H1247V | D | 1.9e-08 | 1.8e-08 | 1.1e-07 | 4.900E-08 | 5.283E-08 | 1.800E-08 | 1.100E-07 |
| 24941 | 1248 | K1248A | D | 1.3e-07 | 5.1e-08 | 3.4e-08 | 7.167E-08 | 5.123E-08 | 3.400E-08 | 1.300E-07 |
| 24942 | 1248 | K1248R | D | 3e-08 | 7.2e-08 | 3.4e-08 | 4.533E-08 | 2.318E-08 | 3.000E-08 | 7.200E-08 |
| 24943 | 1248 | K1248N | D | 4.3e-08 | 7.2e-08 | 2.3e-08 | 4.600E-08 | 2.464E-08 | 2.300E-08 | 7.200E-08 |
| 24944 | 1248 | K1248D | D | 5.6e-08 | 5.1e-08 | 5e-08 | 5.233E-08 | 3.215E-09 | 5.000E-08 | 5.600E-08 |
| 24945 | 1248 | K1248C | D | 7.3e-08 | 4.8e-08 | 3.3e-08 | 5.133E-08 | 2.021E-08 | 3.300E-08 | 7.300E-08 |
| 24946 | 1248 | K1248E | D | 6.5e-08 | 1.9e-08 | 3e-08 | 3.800E-08 | 2.402E-08 | 1.900E-08 | 6.500E-08 |
| 24947 | 1248 | K1248Q | D | 5.5e-08 | 1.4e-07 | 4.5e-08 | 8.000E-08 | 5.220E-08 | 4.500E-08 | 1.400E-07 |
| 24948 | 1248 | K1248G | D | 5.6e-08 | 4.3e-08 | 1.5e-08 | 3.800E-08 | 2.095E-08 | 1.500E-08 | 5.600E-08 |
| 24949 | 1248 | K1248H | D | 5e-08 | 4.4e-08 | 4.2e-08 | 4.533E-08 | 4.163E-09 | 4.200E-08 | 5.000E-08 |
| 24950 | 1248 | K1248I | D | 2.2e-08 | 7.2e-09 | 1.9e-08 | 1.607E-08 | 7.824E-09 | 7.200E-09 | 2.200E-08 |
| 24951 | 1248 | K1248L | D | 9.6e-08 | 2.8e-08 | 4.9e-08 | 5.767E-08 | 3.482E-08 | 2.800E-08 | 9.600E-08 |
| 24952 | 1248 | K1248K | D | 8.6e-08 | 5.8e-08 | 6.7e-08 | 7.033E-08 | 1.429E-08 | 5.800E-08 | 8.600E-08 |
| 24953 | 1248 | K1248M | D | 5.1e-08 | 5.1e-08 | 4.1e-08 | 4.767E-08 | 5.774E-09 | 4.100E-08 | 5.100E-08 |
| 24954 | 1248 | K1248F | D | 4e-08 | 3.6e-08 | 4.6e-08 | 4.067E-08 | 5.033E-09 | 3.600E-08 | 4.600E-08 |
| 24955 | 1248 | K1248P | D | 1.1e-07 | 3.7e-08 | 6.4e-08 | 7.033E-08 | 3.691E-08 | 3.700E-08 | 1.100E-07 |
| 24956 | 1248 | K1248S | D | 1.1e-07 | 2.9e-08 | 5.6e-08 | 6.500E-08 | 4.124E-08 | 2.900E-08 | 1.100E-07 |
| 24957 | 1248 | K1248T | D | 2.7e-08 | 3.1e-08 | 4.8e-08 | 3.533E-08 | 1.115E-08 | 2.700E-08 | 4.800E-08 |
| 24958 | 1248 | K1248W | D | 4.8e-08 | 8.5e-08 | 7.1e-08 | 6.800E-08 | 1.868E-08 | 4.800E-08 | 8.500E-08 |
| 24959 | 1248 | K1248Y | D | 3.1e-08 | 5.5e-08 | 1.2e-08 | 3.267E-08 | 2.155E-08 | 1.200E-08 | 5.500E-08 |
| 24960 | 1248 | K1248V | D | 8.2e-08 | 2.7e-08 | 5.4e-08 | 5.433E-08 | 2.750E-08 | 2.700E-08 | 8.200E-08 |
| 24961 | 1249 | V1249A | D | 2.5e-08 | 4.8e-08 | 1.9e-08 | 3.067E-08 | 1.531E-08 | 1.900E-08 | 4.800E-08 |
| 24962 | 1249 | V1249R | D | 4e-08 | 4e-08 | 4.3e-08 | 4.100E-08 | 1.732E-09 | 4.000E-08 | 4.300E-08 |
| 24963 | 1249 | V1249N | D | 1.5e-07 | 1.7e-08 | 5.6e-08 | 7.433E-08 | 6.837E-08 | 1.700E-08 | 1.500E-07 |
| 24964 | 1249 | V1249D | D | 3.7e-08 | 4.3e-08 | 2.8e-08 | 3.600E-08 | 7.550E-09 | 2.800E-08 | 4.300E-08 |
| 24965 | 1249 | V1249C | D | 4.7e-08 | 2.8e-08 | 2.4e-08 | 3.300E-08 | 1.229E-08 | 2.400E-08 | 4.700E-08 |
| 24966 | 1249 | V1249E | D | 3.9e-08 | 5.4e-08 | 6.9e-08 | 5.400E-08 | 1.500E-08 | 3.900E-08 | 6.900E-08 |
| 24967 | 1249 | V1249Q | D | 8.9e-08 | 3.9e-08 | 6.3e-08 | 6.367E-08 | 2.501E-08 | 3.900E-08 | 8.900E-08 |
| 24968 | 1249 | V1249G | D | 5.2e-08 | 4.3e-08 | 3e-08 | 4.167E-08 | 1.106E-08 | 3.000E-08 | 5.200E-08 |
| 24969 | 1249 | V1249H | D | 5.9e-08 | 4.8e-08 | 2.8e-08 | 4.500E-08 | 1.572E-08 | 2.800E-08 | 5.900E-08 |
| 24970 | 1249 | V1249I | D | 4.2e-08 | 6.6e-08 | 1.4e-08 | 4.067E-08 | 2.603E-08 | 1.400E-08 | 6.600E-08 |
| 24971 | 1249 | V1249L | D | 1.1e-07 | 2e-08 | 9.6e-08 | 7.533E-08 | 4.843E-08 | 2.000E-08 | 1.100E-07 |
| 24972 | 1249 | V1249K | D | 4.8e-08 | 1.9e-08 | 6e-08 | 4.233E-08 | 2.108E-08 | 1.900E-08 | 6.000E-08 |
| 24973 | 1249 | V1249M | D | 1.3e-07 | 1.9e-08 | 1.5e-07 | 9.967E-08 | 7.057E-08 | 1.900E-08 | 1.500E-07 |
| 24974 | 1249 | V1249F | D | 2e-08 | 2.4e-08 | 1.3e-07 | 5.800E-08 | 6.239E-08 | 2.000E-08 | 1.300E-07 |
| 24975 | 1249 | V1249P | D | 5.3e-08 | 1.9e-08 | 1.5e-08 | 2.900E-08 | 2.088E-08 | 1.500E-08 | 5.300E-08 |
| 24976 | 1249 | V1249S | D | 9.9e-08 | 6.4e-08 | 3.6e-08 | 6.633E-08 | 3.156E-08 | 3.600E-08 | 9.900E-08 |
| 24977 | 1249 | V1249T | D | 3.5e-08 | 6e-08 | 4.9e-08 | 4.800E-08 | 1.253E-08 | 3.500E-08 | 6.000E-08 |
| 24978 | 1249 | V1249W | D | 4e-08 | 5.6e-08 | 2.3e-08 | 3.967E-08 | 1.650E-08 | 2.300E-08 | 5.600E-08 |
| 24979 | 1249 | V1249Y | D | 8.1e-08 | 2.2e-08 | 3.1e-08 | 4.467E-08 | 3.179E-08 | 2.200E-08 | 8.100E-08 |
| 24980 | 1249 | V1249V | D | 3.4e-08 | 1.2e-07 | 4.5e-08 | 6.633E-08 | 4.680E-08 | 3.400E-08 | 1.200E-07 |
| 24981 | 1250 | Y1250A | D | 2.2e-08 | 6e-08 | 1.1e-07 | 6.400E-08 | 4.414E-08 | 2.200E-08 | 1.100E-07 |
| 24982 | 1250 | Y1250R | D | 5.4e-08 | 4.1e-08 | 6.1e-08 | 5.200E-08 | 1.015E-08 | 4.100E-08 | 6.100E-08 |
| 24983 | 1250 | Y1250N | D | 4.9e-08 | 2.1e-08 | 3.1e-08 | 3.367E-08 | 1.419E-08 | 2.100E-08 | 4.900E-08 |
| 24984 | 1250 | Y1250D | D | 3.8e-08 | 1e-07 | 1.1e-07 | 8.267E-08 | 3.900E-08 | 3.800E-08 | 1.100E-07 |
| 24985 | 1250 | Y1250C | D | 2.7e-08 | 1e-07 | 4.2e-08 | 5.633E-08 | 3.855E-08 | 2.700E-08 | 1.000E-07 |
| 24986 | 1250 | Y1250E | D | 8.1e-08 | 2e-08 | 1.6e-08 | 3.900E-08 | 3.643E-08 | 1.600E-08 | 8.100E-08 |
| 24987 | 1250 | Y1250Q | D | 2.7e-08 | 3.7e-08 | 4.5e-08 | 3.633E-08 | 9.018E-09 | 2.700E-08 | 4.500E-08 |
| 24988 | 1250 | Y1250G | D | 3.4e-08 | 7.7e-08 | 1.1e-08 | 4.067E-08 | 3.350E-08 | 1.100E-08 | 7.700E-08 |
| 24989 | 1250 | Y1250H | D | 6.7e-08 | 8.1e-08 | 2.6e-08 | 5.800E-08 | 2.858E-08 | 2.600E-08 | 8.100E-08 |

| | | | | | | | | | | |
|---|---|---|---|---|---|---|---|---|---|---|
| 24990 | 1250 | Y1250I | D | 2.9e-08 | 2.8e-08 | 3.8e-08 | 3.167E-08 | 5.508E-09 | 2.800E-08 | 3.800E-08 |
| 24991 | 1250 | Y1250L | D | 1.5e-08 | 4.6e-08 | 5e-08 | 3.700E-08 | 1.916E-08 | 1.500E-08 | 5.000E-08 |
| 24992 | 1250 | Y1250K | D | 7e-08 | 2.9e-08 | 2.9e-08 | 4.267E-08 | 2.367E-08 | 2.900E-08 | 7.000E-08 |
| 24993 | 1250 | Y1250M | D | 6e-08 | 4.1e-08 | 1e-07 | 6.700E-08 | 3.012E-08 | 4.100E-08 | 1.000E-07 |
| 24994 | 1250 | Y1250F | D | 2.7e-08 | 7.9e-08 | 7.8e-08 | 6.133E-08 | 2.974E-08 | 2.700E-08 | 7.900E-08 |
| 24995 | 1250 | Y1250P | D | 1.9e-08 | 6.5e-08 | 2.9e-08 | 3.767E-08 | 2.419E-08 | 1.900E-08 | 6.500E-08 |
| 24996 | 1250 | Y1250S | D | 4.9e-08 | 2.1e-08 | 7.5e-08 | 4.833E-08 | 2.701E-08 | 2.100E-08 | 7.500E-08 |
| 24997 | 1250 | Y1250T | D | 4.3e-08 | 4.7e-08 | 2.1e-08 | 3.700E-08 | 1.400E-08 | 2.100E-08 | 4.700E-08 |
| 24998 | 1250 | Y1250W | D | 4.1e-08 | 2.6e-08 | 5.3e-08 | 4.000E-08 | 1.353E-08 | 2.600E-08 | 5.300E-08 |
| 24999 | 1250 | Y1250Y | D | 6.1e-08 | 5.3e-08 | 1.7e-08 | 4.367E-08 | 2.344E-08 | 1.700E-08 | 6.100E-08 |
| 25000 | 1250 | Y1250V | D | 2.3e-08 | 5.8e-08 | 6.1e-08 | 4.733E-08 | 2.113E-08 | 2.300E-08 | 6.100E-08 |
| 25001 | 1251 | A1251A | D | 3.7e-08 | 4.7e-08 | 4e-08 | 4.133E-08 | 5.132E-09 | 3.700E-08 | 4.700E-08 |
| 25002 | 1251 | A1251R | D | 7.2e-08 | 9.7e-08 | 1.1e-07 | 9.300E-08 | 1.931E-08 | 7.200E-08 | 1.100E-07 |
| 25003 | 1251 | A1251N | D | 5.2e-08 | 3.9e-08 | 2.3e-08 | 3.800E-08 | 1.453E-08 | 2.300E-08 | 5.200E-08 |
| 25004 | 1251 | A1251D | D | 1.1e-08 | 3.8e-08 | 5.8e-08 | 3.567E-08 | 2.359E-08 | 1.100E-08 | 5.800E-08 |
| 25005 | 1251 | A1251C | D | 1.5e-08 | 1.6e-08 | 4.1e-08 | 2.400E-08 | 1.473E-08 | 1.500E-08 | 4.100E-08 |
| 25006 | 1251 | A1251E | D | 4.7e-08 | 2.7e-08 | 3e-08 | 3.467E-08 | 1.079E-08 | 2.700E-08 | 4.700E-08 |
| 25007 | 1251 | A1251Q | D | 6.3e-08 | 5.6e-08 | 7.3e-08 | 6.400E-08 | 8.544E-09 | 5.600E-08 | 7.300E-08 |
| 25008 | 1251 | A1251G | D | 2.7e-08 | 5.4e-08 | 2.3e-08 | 3.467E-08 | 1.686E-08 | 2.300E-08 | 5.400E-08 |
| 25009 | 1251 | A1251H | D | 6.3e-08 | 5.7e-08 | 3.2e-08 | 5.067E-08 | 1.644E-08 | 3.200E-08 | 6.300E-08 |
| 25010 | 1251 | A1251I | D | 1.4e-08 | 1.7e-08 | 7.7e-08 | 3.600E-08 | 3.554E-08 | 1.400E-08 | 7.700E-08 |
| 25011 | 1251 | A1251L | D | 2.5e-08 | 2.8e-08 | 7.6e-08 | 4.300E-08 | 2.862E-08 | 2.500E-08 | 7.600E-08 |
| 25012 | 1251 | A1251K | D | 4.7e-08 | 2.2e-08 | 4.2e-08 | 3.700E-08 | 1.323E-08 | 2.200E-08 | 4.700E-08 |
| 25013 | 1251 | A1251M | D | 5.1e-08 | 3.9e-08 | 2.3e-08 | 3.767E-08 | 1.405E-08 | 2.300E-08 | 5.100E-08 |
| 25014 | 1251 | A1251F | D | 5.5e-08 | 2e-08 | 2.3e-08 | 3.267E-08 | 1.940E-08 | 2.000E-08 | 5.500E-08 |
| 25015 | 1251 | A1251P | D | 1.7e-08 | 2.6e-08 | 7e-08 | 3.767E-08 | 2.836E-08 | 1.700E-08 | 7.000E-08 |
| 25016 | 1251 | A1251S | D | 4.2e-08 | 3.5e-08 | 4.1e-08 | 3.933E-08 | 3.786E-09 | 3.500E-08 | 4.200E-08 |
| 25017 | 1251 | A1251T | D | 1.6e-08 | 1.1e-07 | 1.5e-08 | 4.700E-08 | 5.456E-08 | 1.500E-08 | 1.100E-07 |
| 25018 | 1251 | A1251W | D | 4e-08 | 2.7e-08 | 4.9e-08 | 3.867E-08 | 1.106E-08 | 2.700E-08 | 4.900E-08 |
| 25019 | 1251 | A1251Y | D | 4.2e-08 | 2.4e-08 | 7.6e-08 | 4.733E-08 | 2.641E-08 | 2.400E-08 | 7.600E-08 |
| 25020 | 1251 | A1251V | D | 6.4e-08 | 2e-08 | 6.5e-08 | 4.967E-08 | 2.570E-08 | 2.000E-08 | 6.500E-08 |
| 25021 | 1252 | C1252A | D | 5e-08 | 6.7e-08 | 3.3e-08 | 5.000E-08 | 1.700E-08 | 3.300E-08 | 6.700E-08 |
| 25022 | 1252 | C1252R | D | 4.4e-08 | 6e-08 | 2.4e-08 | 4.267E-08 | 1.804E-08 | 2.400E-08 | 6.000E-08 |
| 25023 | 1252 | C1252N | D | 5.6e-08 | 5.5e-08 | 4e-08 | 5.033E-08 | 8.963E-09 | 4.000E-08 | 5.600E-08 |
| 25024 | 1252 | C1252D | D | 2.4e-08 | 1.8e-08 | 9.3e-08 | 4.500E-08 | 4.168E-08 | 1.800E-08 | 9.300E-08 |
| 25025 | 1252 | C1252C | D | 3.7e-08 | 6.3e-08 | 5e-08 | 5.000E-08 | 1.300E-08 | 3.700E-08 | 6.300E-08 |
| 25026 | 1252 | C1252E | D | 3e-08 | 1.1e-07 | 4e-08 | 6.000E-08 | 4.359E-08 | 3.000E-08 | 1.100E-07 |
| 25027 | 1252 | C1252Q | D | 6.2e-08 | 8.6e-08 | 2.3e-08 | 5.700E-08 | 3.180E-08 | 2.300E-08 | 8.600E-08 |
| 25028 | 1252 | C1252G | D | 1.8e-08 | 4.1e-08 | 4.6e-08 | 3.500E-08 | 1.493E-08 | 1.800E-08 | 4.600E-08 |
| 25029 | 1252 | C1252H | D | 1.3e-08 | 2.1e-08 | 1.9e-08 | 1.767E-08 | 4.163E-09 | 1.300E-08 | 2.100E-08 |
| 25030 | 1252 | C1252I | D | 3.3e-08 | 4.4e-08 | 1e-07 | 5.900E-08 | 3.593E-08 | 3.300E-08 | 1.000E-07 |
| 25031 | 1252 | C1252L | D | 3.1e-08 | 9.2e-08 | 3.8e-08 | 5.367E-08 | 3.338E-08 | 3.100E-08 | 9.200E-08 |
| 25032 | 1252 | C1252K | D | 3.8e-08 | 5.3e-08 | 3.4e-08 | 4.167E-08 | 1.002E-08 | 3.400E-08 | 5.300E-08 |
| 25033 | 1252 | C1252M | D | 3.7e-08 | 2.7e-08 | 2.5e-08 | 2.967E-08 | 6.429E-09 | 2.500E-08 | 3.700E-08 |
| 25034 | 1252 | C1252F | D | 1.6e-08 | 7.2e-08 | 4.8e-08 | 4.533E-08 | 2.810E-08 | 1.600E-08 | 7.200E-08 |
| 25035 | 1252 | C1252P | D | 2e-08 | 1.2e-08 | 2.9e-08 | 2.033E-08 | 8.505E-09 | 1.200E-08 | 2.900E-08 |
| 25036 | 1252 | C1252S | D | 3e-08 | 3.7e-08 | 4.4e-08 | 3.700E-08 | 7.000E-09 | 3.000E-08 | 4.400E-08 |
| 25037 | 1252 | C1252T | D | 3.4e-08 | 4.7e-08 | 8.9e-08 | 5.667E-08 | 2.875E-08 | 3.400E-08 | 8.900E-08 |
| 25038 | 1252 | C1252W | D | 1.5e-07 | 4.1e-08 | 1.8e-08 | 6.967E-08 | 7.051E-08 | 1.800E-08 | 1.500E-07 |
| 25039 | 1252 | C1252Y | D | 6.3e-08 | 9.7e-08 | 4.7e-08 | 6.900E-08 | 2.553E-08 | 4.700E-08 | 9.700E-08 |
| 25040 | 1252 | C1252V | D | 1.2e-07 | 4.3e-08 | 2.3e-08 | 6.200E-08 | 5.122E-08 | 2.300E-08 | 1.200E-07 |
| 25041 | 1253 | E1253A | D | 4.2e-08 | 5.9e-08 | 4.7e-08 | 4.933E-08 | 8.737E-09 | 4.200E-08 | 5.900E-08 |
| 25042 | 1253 | E1253R | D | 5.4e-08 | 3.8e-08 | 1.8e-08 | 3.667E-08 | 1.804E-08 | 1.800E-08 | 5.400E-08 |
| 25043 | 1253 | E1253N | D | 2.4e-08 | 6.4e-08 | 3.8e-08 | 4.200E-08 | 2.030E-08 | 2.400E-08 | 6.400E-08 |
| 25044 | 1253 | E1253D | D | 4.8e-08 | 3.4e-08 | 7.6e-08 | 5.267E-08 | 2.139E-08 | 3.400E-08 | 7.600E-08 |
| 25045 | 1253 | E1253C | D | 4.3e-08 | 8.5e-08 | 7.6e-09 | 4.520E-08 | 3.875E-08 | 7.600E-09 | 8.500E-08 |
| 25046 | 1253 | E1253E | D | 6.4e-08 | 4.9e-08 | 5.6e-08 | 5.633E-08 | 7.506E-09 | 4.900E-08 | 6.400E-08 |
| 25047 | 1253 | E1253Q | D | 6.9e-08 | 3.7e-08 | 7e-08 | 5.867E-08 | 1.877E-08 | 3.700E-08 | 7.000E-08 |
| 25048 | 1253 | E1253G | D | 3.7e-08 | 1.9e-08 | 4e-08 | 3.200E-08 | 1.136E-08 | 1.900E-08 | 4.000E-08 |
| 25049 | 1253 | E1253H | D | 6.4e-08 | 3.3e-08 | 3.4e-08 | 4.367E-08 | 1.762E-08 | 3.300E-08 | 6.400E-08 |
| 25050 | 1253 | E1253I | D | 3.8e-08 | 7.9e-08 | 3.1e-08 | 4.933E-08 | 2.593E-08 | 3.100E-08 | 7.900E-08 |
| 25051 | 1253 | E1253L | D | 6.9e-08 | 2.4e-08 | 8.6e-08 | 5.967E-08 | 3.204E-08 | 2.400E-08 | 8.600E-08 |
| 25052 | 1253 | E1253K | D | 9.3e-08 | 2.6e-08 | 4e-08 | 5.300E-08 | 3.534E-08 | 2.600E-08 | 9.300E-08 |
| 25053 | 1253 | E1253M | D | 5.1e-08 | 2.1e-08 | 1.5e-08 | 2.900E-08 | 1.929E-08 | 1.500E-08 | 5.100E-08 |
| 25054 | 1253 | E1253F | D | 5.1e-08 | 5.2e-08 | 5.3e-08 | 5.200E-08 | 1.000E-09 | 5.100E-08 | 5.300E-08 |
| 25055 | 1253 | E1253P | D | 2.1e-08 | 4.3e-08 | 3.1e-08 | 3.167E-08 | 1.102E-08 | 2.100E-08 | 4.300E-08 |
| 25056 | 1253 | E1253S | D | 6.2e-08 | 6.7e-08 | 7.4e-08 | 6.767E-08 | 6.028E-09 | 6.200E-08 | 7.400E-08 |
| 25057 | 1253 | E1253T | D | 6.5e-08 | 3.5e-08 | 7.5e-08 | 5.833E-08 | 2.082E-08 | 3.500E-08 | 7.500E-08 |
| 25058 | 1253 | E1253W | D | 1.2e-08 | 9.6e-08 | 6.3e-08 | 5.700E-08 | 4.232E-08 | 1.200E-08 | 9.600E-08 |
| 25059 | 1253 | E1253Y | D | 5.5e-08 | 2.3e-08 | 5e-08 | 4.267E-08 | 1.721E-08 | 2.300E-08 | 5.500E-08 |
| 25060 | 1253 | E1253V | D | 8.9e-08 | 5.1e-08 | 4.3e-08 | 6.100E-08 | 2.458E-08 | 4.300E-08 | 8.900E-08 |
| 25061 | 1254 | V1254A | D | 4.4e-08 | 1.2e-08 | 1.6e-07 | 7.200E-08 | 7.787E-08 | 1.200E-08 | 1.600E-07 |
| 25062 | 1254 | V1254R | D | 4e-08 | 4.3e-08 | 3.1e-08 | 3.800E-08 | 6.245E-09 | 3.100E-08 | 4.300E-08 |
| 25063 | 1254 | V1254N | D | 6.5e-08 | 3.2e-08 | 9.4e-08 | 6.367E-08 | 3.102E-08 | 3.200E-08 | 9.400E-08 |
| 25064 | 1254 | V1254D | D | 2.7e-08 | 1.8e-08 | 3.9e-08 | 2.800E-08 | 1.054E-08 | 1.800E-08 | 3.900E-08 |
| 25065 | 1254 | V1254C | D | 3.4e-08 | 7.5e-08 | 1.5e-08 | 4.133E-08 | 3.066E-08 | 1.500E-08 | 7.500E-08 |
| 25066 | 1254 | V1254E | D | 4.3e-08 | 2.4e-08 | 2.8e-08 | 3.167E-08 | 1.002E-08 | 2.400E-08 | 4.300E-08 |
| 25067 | 1254 | V1254Q | D | 2.6e-08 | 3.9e-08 | 2.1e-08 | 2.867E-08 | 9.292E-09 | 2.100E-08 | 3.900E-08 |
| 25068 | 1254 | V1254G | D | 7.3e-08 | 1.2e-07 | 2.3e-08 | 7.200E-08 | 4.851E-08 | 2.300E-08 | 1.200E-07 |
| 25069 | 1254 | V1254H | D | 3.1e-08 | 3.3e-08 | 1.9e-08 | 2.767E-08 | 7.572E-09 | 1.900E-08 | 3.300E-08 |
| 25070 | 1254 | V1254I | D | 1.9e-08 | 6.7e-08 | 1e-07 | 6.200E-08 | 4.073E-08 | 1.900E-08 | 1.000E-07 |
| 25071 | 1254 | V1254L | D | 1.4e-07 | 7.5e-08 | 3.2e-08 | 8.233E-08 | 5.437E-08 | 3.200E-08 | 1.400E-07 |
| 25072 | 1254 | V1254K | D | 9e-08 | 5.5e-08 | 6.7e-08 | 7.067E-08 | 1.779E-08 | 5.500E-08 | 9.000E-08 |
| 25073 | 1254 | V1254M | D | 5.2e-08 | 3.2e-08 | 1.6e-08 | 3.333E-08 | 1.804E-08 | 1.600E-08 | 5.200E-08 |
| 25074 | 1254 | V1254F | D | 4.1e-08 | 3.8e-08 | 5e-08 | 4.300E-08 | 6.245E-09 | 3.800E-08 | 5.000E-08 |

| | | | | | | | | | | |
|---|---|---|---|---|---|---|---|---|---|---|
| 25075 | 1254 | V1254P | D | 5.3e-08 | 3e-08 | 4.7e-08 | 4.333E-08 | 1.193E-08 | 3.000E-08 | 5.300E-08 |
| 25076 | 1254 | V1254S | D | 4e-08 | 3e-08 | 3.8e-08 | 3.600E-08 | 5.292E-09 | 3.000E-08 | 4.000E-08 |
| 25077 | 1254 | V1254T | D | 6.6e-08 | 5.1e-08 | 4.6e-08 | 5.433E-08 | 1.041E-08 | 4.600E-08 | 6.600E-08 |
| 25078 | 1254 | V1254W | D | 6.5e-08 | 1e-08 | 6.6e-08 | 4.700E-08 | 3.205E-08 | 1.000E-08 | 6.600E-08 |
| 25079 | 1254 | V1254Y | D | 7.1e-08 | 3.1e-08 | 1.4e-07 | 8.067E-08 | 5.514E-08 | 3.100E-08 | 1.400E-07 |
| 25080 | 1254 | V1254V | D | 3.2e-08 | 6e-08 | 4e-08 | 4.400E-08 | 1.442E-08 | 3.200E-08 | 6.000E-08 |
| 25081 | 1255 | T1255A | D | 4.6e-08 | 6.4e-08 | 2.9e-08 | 4.633E-08 | 1.750E-08 | 2.900E-08 | 6.400E-08 |
| 25082 | 1255 | T1255R | D | 2.1e-08 | 4e-08 | 4.5e-08 | 3.533E-08 | 1.266E-08 | 2.100E-08 | 4.500E-08 |
| 25083 | 1255 | T1255N | D | 3.8e-08 | 6.3e-08 | 1.5e-07 | 8.367E-08 | 5.879E-08 | 3.800E-08 | 1.500E-07 |
| 25084 | 1255 | T1255D | D | 5.2e-08 | 6.1e-08 | 2.5e-08 | 4.600E-08 | 1.873E-08 | 2.500E-08 | 6.100E-08 |
| 25085 | 1255 | T1255C | D | 5.2e-08 | 3.5e-08 | 1.1e-07 | 6.567E-08 | 3.932E-08 | 3.500E-08 | 1.100E-07 |
| 25086 | 1255 | T1255E | D | 8e-08 | 6.6e-08 | 7.1e-08 | 7.233E-08 | 7.095E-09 | 6.600E-08 | 8.000E-08 |
| 25087 | 1255 | T1255Q | D | 2.5e-08 | 1.7e-08 | 6.6e-08 | 3.600E-08 | 2.629E-08 | 1.700E-08 | 6.600E-08 |
| 25088 | 1255 | T1255G | D | 8.1e-09 | 5.6e-08 | 1.8e-07 | 8.137E-08 | 8.871E-08 | 8.100E-09 | 1.800E-07 |
| 25089 | 1255 | T1255H | D | 2.1e-08 | 1.4e-08 | 3.3e-08 | 2.267E-08 | 9.609E-09 | 1.400E-08 | 3.300E-08 |
| 25090 | 1255 | T1255I | D | 4.6e-08 | 5.3e-08 | 7.9e-08 | 5.933E-08 | 1.739E-08 | 4.600E-08 | 7.900E-08 |
| 25091 | 1255 | T1255L | D | 1.3e-07 | 3.5e-08 | 4.9e-08 | 7.133E-08 | 5.129E-08 | 3.500E-08 | 1.300E-07 |
| 25092 | 1255 | T1255K | D | 3.4e-08 | 7.3e-08 | 6.9e-08 | 5.867E-08 | 2.146E-08 | 3.400E-08 | 7.300E-08 |
| 25093 | 1255 | T1255M | D | 9.5e-09 | 1.1e-07 | 1.2e-07 | 7.983E-08 | 6.112E-08 | 9.500E-09 | 1.200E-07 |
| 25094 | 1255 | T1255F | D | 4.6e-08 | 5.5e-08 | 1.7e-08 | 3.933E-08 | 1.986E-08 | 1.700E-08 | 5.500E-08 |
| 25095 | 1255 | T1255P | D | 5.6e-08 | 8.4e-09 | 5.7e-08 | 4.047E-08 | 2.778E-08 | 8.400E-09 | 5.700E-08 |
| 25096 | 1255 | T1255S | D | 2.3e-08 | 2.4e-08 | 5.8e-08 | 3.500E-08 | 1.992E-08 | 2.300E-08 | 5.800E-08 |
| 25097 | 1255 | T1255T | D | 5e-08 | 2.7e-08 | 7.5e-08 | 5.067E-08 | 2.401E-08 | 2.700E-08 | 7.500E-08 |
| 25098 | 1255 | T1255W | D | 3.7e-08 | 8.6e-08 | 4.8e-08 | 5.700E-08 | 2.571E-08 | 3.700E-08 | 8.600E-08 |
| 25099 | 1255 | T1255Y | D | 2.6e-08 | 3.4e-08 | 1.2e-07 | 6.000E-08 | 5.212E-08 | 2.600E-08 | 1.200E-07 |
| 25100 | 1255 | T1255V | D | 6.9e-08 | 3.8e-08 | 6.7e-08 | 5.800E-08 | 1.735E-08 | 3.800E-08 | 6.900E-08 |
| 25101 | 1256 | H1256A | D | 2.9e-08 | 6.5e-08 | 6.3e-08 | 5.233E-08 | 2.023E-08 | 2.900E-08 | 6.500E-08 |
| 25102 | 1256 | H1256R | D | 2.6e-08 | 4.8e-08 | 5.9e-08 | 4.433E-08 | 1.680E-08 | 2.600E-08 | 5.900E-08 |
| 25103 | 1256 | H1256N | D | 5.2e-08 | 5.1e-08 | 8.9e-08 | 6.400E-08 | 2.166E-08 | 5.100E-08 | 8.900E-08 |
| 25104 | 1256 | H1256D | D | 4.6e-08 | 3e-08 | 5.6e-08 | 4.400E-08 | 1.311E-08 | 3.000E-08 | 5.600E-08 |
| 25105 | 1256 | H1256C | D | 3.3e-08 | 1.4e-08 | 7.9e-08 | 4.200E-08 | 3.342E-08 | 1.400E-08 | 7.900E-08 |
| 25106 | 1256 | H1256E | D | 3.1e-08 | 3.4e-08 | 1.8e-08 | 2.767E-08 | 8.505E-09 | 1.800E-08 | 3.400E-08 |
| 25107 | 1256 | H1256Q | D | 4.6e-08 | 5.9e-08 | 6.8e-08 | 5.767E-08 | 1.106E-08 | 4.600E-08 | 6.800E-08 |
| 25108 | 1256 | H1256G | D | 1.1e-07 | 9.8e-08 | 8.9e-08 | 9.900E-08 | 1.054E-08 | 8.900E-08 | 1.100E-07 |
| 25109 | 1256 | H1256H | D | 5.1e-08 | 3.4e-08 | 1.3e-08 | 3.267E-08 | 1.904E-08 | 1.300E-08 | 5.100E-08 |
| 25110 | 1256 | H1256I | D | 5.8e-08 | 2.4e-08 | 4.1e-08 | 4.100E-08 | 1.700E-08 | 2.400E-08 | 5.800E-08 |
| 25111 | 1256 | H1256L | D | 4.8e-08 | 4e-08 | 2.6e-08 | 3.800E-08 | 1.114E-08 | 2.600E-08 | 4.800E-08 |
| 25112 | 1256 | H1256K | D | 3.4e-08 | 6.4e-08 | 8e-08 | 5.933E-08 | 2.335E-08 | 3.400E-08 | 8.000E-08 |
| 25113 | 1256 | H1256M | D | 8.1e-08 | 5.2e-08 | 8.8e-08 | 7.367E-08 | 1.909E-08 | 5.200E-08 | 8.800E-08 |
| 25114 | 1256 | H1256F | D | 3.6e-08 | 9e-08 | 5.5e-08 | 6.033E-08 | 2.739E-08 | 3.600E-08 | 9.000E-08 |
| 25115 | 1256 | H1256P | D | 5.6e-08 | 2.3e-08 | 5.2e-08 | 4.367E-08 | 1.801E-08 | 2.300E-08 | 5.600E-08 |
| 25116 | 1256 | H1256S | D | 5e-08 | 3.8e-08 | 6.9e-08 | 5.233E-08 | 1.563E-08 | 3.800E-08 | 6.900E-08 |
| 25117 | 1256 | H1256T | D | 1.4e-07 | 5.4e-08 | 3.4e-08 | 7.600E-08 | 5.632E-08 | 3.400E-08 | 1.400E-07 |
| 25118 | 1256 | H1256W | D | 8.4e-08 | 4.9e-08 | 1.8e-08 | 5.033E-08 | 3.302E-08 | 1.800E-08 | 8.400E-08 |
| 25119 | 1256 | H1256Y | D | 3.2e-08 | 1.1e-07 | 8.3e-08 | 7.500E-08 | 3.961E-08 | 3.200E-08 | 1.100E-07 |
| 25120 | 1256 | H1256V | D | 4.2e-08 | 5.1e-08 | 4.3e-08 | 4.533E-08 | 4.933E-09 | 4.200E-08 | 5.100E-08 |
| 25121 | 1257 | Q1257A | D | 1.3e-07 | 4.1e-08 | 4.7e-08 | 7.267E-08 | 4.974E-08 | 4.100E-08 | 1.300E-07 |
| 25122 | 1257 | Q1257R | D | 1.2e-07 | 3e-08 | 2.4e-08 | 5.800E-08 | 5.378E-08 | 2.400E-08 | 1.200E-07 |
| 25123 | 1257 | Q1257N | D | 3.1e-08 | 1.1e-07 | 3.1e-08 | 5.733E-08 | 4.561E-08 | 3.100E-08 | 1.100E-07 |
| 25124 | 1257 | Q1257D | D | 1.1e-08 | 5.1e-08 | 4.2e-08 | 3.467E-08 | 2.098E-08 | 1.100E-08 | 5.100E-08 |
| 25125 | 1257 | Q1257C | D | 1.7e-08 | 1.3e-07 | 8.1e-08 | 7.600E-08 | 5.667E-08 | 1.700E-08 | 1.300E-07 |
| 25126 | 1257 | Q1257E | D | 4.8e-08 | 1.5e-08 | 4.2e-08 | 3.500E-08 | 1.758E-08 | 1.500E-08 | 4.800E-08 |
| 25127 | 1257 | Q1257Q | D | 6.4e-08 | 3.7e-08 | 9.6e-09 | 3.687E-08 | 2.720E-08 | 9.600E-09 | 6.400E-08 |
| 25128 | 1257 | Q1257G | D | 5.7e-08 | 9.3e-08 | 1.2e-07 | 9.000E-08 | 3.161E-08 | 5.700E-08 | 1.200E-07 |
| 25129 | 1257 | Q1257H | D | 1.3e-07 | 4.5e-08 | 3.7e-08 | 7.067E-08 | 5.154E-08 | 3.700E-08 | 1.300E-07 |
| 25130 | 1257 | Q1257I | D | 7.7e-08 | 1e-08 | 2.5e-08 | 3.733E-08 | 3.516E-08 | 1.000E-08 | 7.700E-08 |
| 25131 | 1257 | Q1257L | D | 4.2e-08 | 6.3e-08 | 9.1e-08 | 6.533E-08 | 2.458E-08 | 4.200E-08 | 9.100E-08 |
| 25132 | 1257 | Q1257K | D | 4.2e-08 | 4.2e-08 | 4.7e-08 | 4.367E-08 | 2.887E-09 | 4.200E-08 | 4.700E-08 |
| 25133 | 1257 | Q1257M | D | 3e-08 | 2e-08 | 2.2e-08 | 2.400E-08 | 5.292E-09 | 2.000E-08 | 3.000E-08 |
| 25134 | 1257 | Q1257F | D | 5.5e-08 | 5.1e-08 | 3.8e-08 | 4.800E-08 | 8.888E-09 | 3.800E-08 | 5.500E-08 |
| 25135 | 1257 | Q1257P | D | 4.2e-08 | 8.6e-08 | 1e-07 | 7.600E-08 | 3.027E-08 | 4.200E-08 | 1.000E-07 |
| 25136 | 1257 | Q1257S | D | 4e-08 | 2e-08 | 5.2e-08 | 3.733E-08 | 1.617E-08 | 2.000E-08 | 5.200E-08 |
| 25137 | 1257 | Q1257T | D | 4.5e-08 | 2.4e-08 | 2.1e-08 | 3.000E-08 | 1.308E-08 | 2.100E-08 | 4.500E-08 |
| 25138 | 1257 | Q1257W | D | 4.2e-08 | 2.9e-08 | 3.1e-08 | 3.400E-08 | 7.000E-09 | 2.900E-08 | 4.200E-08 |
| 25139 | 1257 | Q1257Y | D | 1.5e-07 | 5.3e-08 | 2.4e-08 | 7.567E-08 | 6.599E-08 | 2.400E-08 | 1.500E-07 |
| 25140 | 1257 | Q1257V | D | 6.6e-08 | 8.3e-08 | 8.4e-09 | 5.247E-08 | 3.910E-08 | 8.400E-09 | 8.300E-08 |
| 25141 | 1258 | G1258A | D | 1.8e-08 | 2.9e-08 | 4.9e-08 | 3.200E-08 | 1.572E-08 | 1.800E-08 | 4.900E-08 |
| 25142 | 1258 | G1258R | D | 4e-08 | 7.5e-08 | 2.4e-08 | 4.633E-08 | 2.608E-08 | 2.400E-08 | 7.500E-08 |
| 25143 | 1258 | G1258N | D | 4.4e-08 | 4e-08 | 4.3e-08 | 4.233E-08 | 2.082E-09 | 4.000E-08 | 4.400E-08 |
| 25144 | 1258 | G1258D | D | 3e-08 | 2.5e-08 | 5.6e-08 | 3.700E-08 | 1.664E-08 | 2.500E-08 | 5.600E-08 |
| 25145 | 1258 | G1258C | D | 2.2e-08 | 1.9e-08 | 3.8e-08 | 2.633E-08 | 1.021E-08 | 1.900E-08 | 3.800E-08 |
| 25146 | 1258 | G1258E | D | 4.8e-08 | 6.4e-08 | 2.1e-08 | 4.433E-08 | 2.173E-08 | 2.100E-08 | 6.400E-08 |
| 25147 | 1258 | G1258Q | D | 1.7e-08 | 1.9e-08 | 4.2e-08 | 2.600E-08 | 1.389E-08 | 1.700E-08 | 4.200E-08 |
| 25148 | 1258 | G1258G | D | 2.6e-08 | 4.8e-08 | 4.3e-08 | 3.900E-08 | 1.153E-08 | 2.600E-08 | 4.800E-08 |
| 25149 | 1258 | G1258H | D | 5.5e-08 | 2.5e-08 | 4.5e-08 | 4.167E-08 | 1.528E-08 | 2.500E-08 | 5.500E-08 |
| 25150 | 1258 | G1258I | D | 3.3e-08 | 6.9e-08 | 4.7e-08 | 4.967E-08 | 1.815E-08 | 3.300E-08 | 6.900E-08 |
| 25151 | 1258 | G1258L | D | 1.8e-07 | 1.3e-08 | 3.9e-08 | 7.733E-08 | 8.986E-08 | 1.300E-08 | 1.800E-07 |
| 25152 | 1258 | G1258K | D | 6.2e-08 | 1.4e-08 | 8.7e-08 | 5.433E-08 | 3.710E-08 | 1.400E-08 | 8.700E-08 |
| 25153 | 1258 | G1258M | D | 4.1e-08 | 9.1e-08 | 3.9e-08 | 5.700E-08 | 2.946E-08 | 3.900E-08 | 9.100E-08 |
| 25154 | 1258 | G1258F | D | 4.7e-08 | 4.8e-08 | 5.3e-08 | 4.933E-08 | 3.215E-09 | 4.700E-08 | 5.300E-08 |
| 25155 | 1258 | G1258P | D | 5.1e-08 | 5.9e-08 | 6.3e-08 | 5.767E-08 | 6.110E-09 | 5.100E-08 | 6.300E-08 |
| 25156 | 1258 | G1258S | D | 4.2e-08 | 5.6e-08 | 3.6e-08 | 4.467E-08 | 1.026E-08 | 3.600E-08 | 5.600E-08 |
| 25157 | 1258 | G1258T | D | 1.3e-08 | 2.5e-08 | 3.5e-08 | 2.433E-08 | 1.102E-08 | 1.300E-08 | 3.500E-08 |
| 25158 | 1258 | G1258W | D | 7.6e-08 | 2.8e-08 | 3.3e-08 | 4.567E-08 | 2.639E-08 | 2.800E-08 | 7.600E-08 |
| 25159 | 1258 | G1258Y | D | 4.9e-08 | 2.4e-08 | 4.2e-08 | 3.833E-08 | 1.290E-08 | 2.400E-08 | 4.900E-08 |

| | | | | | | | | | | |
|---|---|---|---|---|---|---|---|---|---|---|
| 25160 | 1258 | G1258V | D | 8.9e-08 | 8e-08 | 8.6e-08 | 8.500E-08 | 4.583E-09 | 8.000E-08 | 8.900E-08 |
| 25161 | 1259 | L1259A | D | 7.3e-08 | 2.3e-08 | 1.9e-07 | 9.533E-08 | 8.571E-08 | 2.300E-08 | 1.900E-07 |
| 25162 | 1259 | L1259R | D | 4.5e-08 | 4.2e-08 | 1.1e-07 | 6.567E-08 | 3.842E-08 | 4.200E-08 | 1.100E-07 |
| 25163 | 1259 | L1259N | D | 1.2e-08 | 1.1e-08 | 2.4e-08 | 1.567E-08 | 7.234E-09 | 1.100E-08 | 2.400E-08 |
| 25164 | 1259 | L1259D | D | 3.3e-08 | 1.2e-08 | 2.2e-08 | 2.233E-08 | 1.050E-08 | 1.200E-08 | 3.300E-08 |
| 25165 | 1259 | L1259C | D | 2.7e-08 | 9.3e-08 | 3.7e-08 | 5.233E-08 | 3.557E-08 | 2.700E-08 | 9.300E-08 |
| 25166 | 1259 | L1259E | D | 2.2e-08 | 4.3e-08 | 3.1e-08 | 3.200E-08 | 1.054E-08 | 2.200E-08 | 4.300E-08 |
| 25167 | 1259 | L1259Q | D | 4.6e-08 | 4.8e-08 | 5.5e-08 | 4.967E-08 | 4.726E-09 | 4.600E-08 | 5.500E-08 |
| 25168 | 1259 | L1259G | D | 4.9e-08 | 2.1e-08 | 3.3e-08 | 3.433E-08 | 1.405E-08 | 2.100E-08 | 4.900E-08 |
| 25169 | 1259 | L1259H | D | 2.4e-08 | 4.4e-08 | 3.3e-08 | 3.367E-08 | 1.002E-08 | 2.400E-08 | 4.400E-08 |
| 25170 | 1259 | L1259I | D | 2.2e-08 | 8.1e-08 | 8.7e-08 | 6.333E-08 | 3.592E-08 | 2.200E-08 | 8.700E-08 |
| 25171 | 1259 | L1259L | D | 5.7e-08 | 9.3e-09 | 6.3e-08 | 4.310E-08 | 2.942E-08 | 9.300E-09 | 6.300E-08 |
| 25172 | 1259 | L1259K | D | 6.1e-08 | 5.5e-08 | 1.2e-08 | 4.267E-08 | 2.673E-08 | 1.200E-08 | 6.100E-08 |
| 25173 | 1259 | L1259M | D | 6.1e-08 | 2.1e-08 | 4.3e-08 | 4.167E-08 | 2.003E-08 | 2.100E-08 | 6.100E-08 |
| 25174 | 1259 | L1259F | D | 3.2e-08 | 2.7e-08 | 3.8e-08 | 3.233E-08 | 5.508E-09 | 2.700E-08 | 3.800E-08 |
| 25175 | 1259 | L1259P | D | 2.9e-08 | 2.5e-08 | 6.9e-08 | 4.100E-08 | 2.433E-08 | 2.500E-08 | 6.900E-08 |
| 25176 | 1259 | L1259S | D | 3.5e-08 | 2.2e-08 | 1.7e-08 | 2.467E-08 | 9.292E-09 | 1.700E-08 | 3.500E-08 |
| 25177 | 1259 | L1259T | D | 2.5e-08 | 1.6e-08 | 2.7e-08 | 2.267E-08 | 5.859E-09 | 1.600E-08 | 2.700E-08 |
| 25178 | 1259 | L1259W | D | 4.4e-08 | 1e-07 | 3.6e-08 | 6.000E-08 | 3.487E-08 | 3.600E-08 | 1.000E-07 |
| 25179 | 1259 | L1259Y | D | 2e-08 | 3.9e-08 | 2.9e-08 | 2.933E-08 | 9.504E-09 | 2.000E-08 | 3.900E-08 |
| 25180 | 1259 | L1259V | D | 5.2e-08 | 4.1e-08 | 3.6e-08 | 4.300E-08 | 8.185E-09 | 3.600E-08 | 5.200E-08 |
| 25181 | 1260 | S1260A | D | 3.4e-08 | 4.3e-08 | 3.4e-08 | 3.700E-08 | 5.196E-09 | 3.400E-08 | 4.300E-08 |
| 25182 | 1260 | S1260R | D | 5.4e-08 | 1.1e-07 | 3.4e-08 | 6.600E-08 | 3.940E-08 | 3.400E-08 | 1.100E-07 |
| 25183 | 1260 | S1260N | D | 1.6e-08 | 5.5e-08 | 4.1e-08 | 3.733E-08 | 1.976E-08 | 1.600E-08 | 5.500E-08 |
| 25184 | 1260 | S1260D | D | 2.6e-08 | 1.7e-08 | 6.1e-08 | 3.467E-08 | 2.325E-08 | 1.700E-08 | 6.100E-08 |
| 25185 | 1260 | S1260C | D | 3.4e-08 | 1.3e-08 | 2.7e-08 | 2.467E-08 | 1.069E-08 | 1.300E-08 | 3.400E-08 |
| 25186 | 1260 | S1260E | D | 6e-08 | 5.4e-08 | 3.2e-08 | 4.867E-08 | 1.474E-08 | 3.200E-08 | 6.000E-08 |
| 25187 | 1260 | S1260Q | D | 5e-08 | 1.6e-08 | 2.3e-08 | 2.967E-08 | 1.795E-08 | 1.600E-08 | 5.000E-08 |
| 25188 | 1260 | S1260G | D | 7.4e-08 | 1.9e-08 | 3.8e-08 | 4.367E-08 | 2.793E-08 | 1.900E-08 | 7.400E-08 |
| 25189 | 1260 | S1260H | D | 2.4e-08 | 4.2e-08 | 8.6e-08 | 5.067E-08 | 3.190E-08 | 2.400E-08 | 8.600E-08 |
| 25190 | 1260 | S1260I | D | 5.3e-08 | 7.1e-08 | 3e-08 | 5.133E-08 | 2.055E-08 | 3.000E-08 | 7.100E-08 |
| 25191 | 1260 | S1260L | D | 7.5e-08 | 5.2e-08 | 2.7e-08 | 5.133E-08 | 2.401E-08 | 2.700E-08 | 7.500E-08 |
| 25192 | 1260 | S1260K | D | 3.7e-08 | 3.7e-08 | 6.5e-08 | 4.633E-08 | 1.617E-08 | 3.700E-08 | 6.500E-08 |
| 25193 | 1260 | S1260M | D | 6.5e-08 | 2.1e-08 | 2e-08 | 3.533E-08 | 2.570E-08 | 2.000E-08 | 6.500E-08 |
| 25194 | 1260 | S1260F | D | 7.6e-08 | 5.4e-08 | 2e-08 | 5.000E-08 | 2.821E-08 | 2.000E-08 | 7.600E-08 |
| 25195 | 1260 | S1260P | D | 6.1e-08 | 2.2e-08 | 5.9e-08 | 4.733E-08 | 2.196E-08 | 2.200E-08 | 6.100E-08 |
| 25196 | 1260 | S1260S | D | 1.7e-08 | 6.7e-08 | 7.1e-08 | 5.167E-08 | 3.009E-08 | 1.700E-08 | 7.100E-08 |
| 25197 | 1260 | S1260T | D | 6.6e-08 | 3.5e-08 | 9.3e-08 | 6.467E-08 | 2.902E-08 | 3.500E-08 | 9.300E-08 |
| 25198 | 1260 | S1260W | D | 1.8e-08 | 5.3e-08 | 2.2e-08 | 3.100E-08 | 1.916E-08 | 1.800E-08 | 5.300E-08 |
| 25199 | 1260 | S1260Y | D | 6.3e-08 | 5.3e-08 | 4.8e-08 | 5.467E-08 | 7.638E-09 | 4.800E-08 | 6.300E-08 |
| 25200 | 1260 | S1260V | D | 4.7e-08 | 9e-08 | 2.5e-08 | 5.400E-08 | 3.306E-08 | 2.500E-08 | 9.000E-08 |
| 25201 | 1261 | S1261A | D | 4.9e-08 | 7.2e-08 | 7.6e-08 | 6.567E-08 | 1.457E-08 | 4.900E-08 | 7.600E-08 |
| 25202 | 1261 | S1261R | D | 7.7e-08 | 4.5e-08 | 5.5e-08 | 5.900E-08 | 1.637E-08 | 4.500E-08 | 7.700E-08 |
| 25203 | 1261 | S1261N | D | 4.8e-08 | 2.7e-08 | 5.9e-08 | 4.467E-08 | 1.626E-08 | 2.700E-08 | 5.900E-08 |
| 25204 | 1261 | S1261D | D | 1.5e-08 | 5.6e-08 | 2.6e-08 | 3.233E-08 | 2.122E-08 | 1.500E-08 | 5.600E-08 |
| 25205 | 1261 | S1261C | D | 3.5e-08 | 2.8e-08 | 1.2e-07 | 6.100E-08 | 5.122E-08 | 2.800E-08 | 1.200E-07 |
| 25206 | 1261 | S1261E | D | 4.5e-08 | 5.6e-08 | 5.6e-08 | 5.233E-08 | 6.351E-09 | 4.500E-08 | 5.600E-08 |
| 25207 | 1261 | S1261Q | D | 6.1e-08 | 1e-08 | 3.5e-08 | 3.533E-08 | 2.550E-08 | 1.000E-08 | 6.100E-08 |
| 25208 | 1261 | S1261G | D | 8.7e-08 | 2.1e-08 | 2.4e-08 | 4.400E-08 | 3.727E-08 | 2.100E-08 | 8.700E-08 |
| 25209 | 1261 | S1261H | D | 9.1e-08 | 1.6e-07 | 4.8e-08 | 9.967E-08 | 5.650E-08 | 4.800E-08 | 1.600E-07 |
| 25210 | 1261 | S1261I | D | 3.7e-08 | 3.6e-08 | 5.5e-08 | 4.267E-08 | 1.069E-08 | 3.600E-08 | 5.500E-08 |
| 25211 | 1261 | S1261L | D | 6.1e-08 | 9.6e-08 | 2.2e-07 | 1.257E-07 | 8.355E-08 | 6.100E-08 | 2.200E-07 |
| 25212 | 1261 | S1261K | D | 6.3e-08 | 9.7e-08 | 7.6e-08 | 7.867E-08 | 1.716E-08 | 6.300E-08 | 9.700E-08 |
| 25213 | 1261 | S1261M | D | 2.9e-08 | 1.2e-07 | 5.1e-08 | 6.667E-08 | 4.748E-08 | 2.900E-08 | 1.200E-07 |
| 25214 | 1261 | S1261F | D | 3.1e-08 | 1.2e-07 | 4.4e-08 | 6.500E-08 | 4.807E-08 | 3.100E-08 | 1.200E-07 |
| 25215 | 1261 | S1261P | D | 2.5e-08 | 3.4e-08 | 2.3e-08 | 2.733E-08 | 5.859E-09 | 2.300E-08 | 3.400E-08 |
| 25216 | 1261 | S1261S | D | 3e-08 | 3.7e-08 | 3.2e-08 | 3.300E-08 | 3.606E-09 | 3.000E-08 | 3.700E-08 |
| 25217 | 1261 | S1261T | D | 2.8e-08 | 4.7e-08 | 3e-08 | 3.500E-08 | 1.044E-08 | 2.800E-08 | 4.700E-08 |
| 25218 | 1261 | S1261W | D | 4.2e-08 | 4.8e-08 | 2.4e-08 | 3.800E-08 | 1.249E-08 | 2.400E-08 | 4.800E-08 |
| 25219 | 1261 | S1261Y | D | 2e-08 | 4e-08 | 2.3e-08 | 2.767E-08 | 1.079E-08 | 2.000E-08 | 4.000E-08 |
| 25220 | 1261 | S1261V | D | 3.4e-08 | 1.7e-08 | 2.6e-08 | 2.567E-08 | 8.505E-09 | 1.700E-08 | 3.400E-08 |
| 25221 | 1262 | P1262A | D | 5.1e-08 | 3e-08 | 2.7e-08 | 3.600E-08 | 1.308E-08 | 2.700E-08 | 5.100E-08 |
| 25222 | 1262 | P1262R | D | 2.3e-08 | 2.4e-08 | 1.8e-08 | 2.167E-08 | 3.215E-09 | 1.800E-08 | 2.400E-08 |
| 25223 | 1262 | P1262N | D | 6.6e-08 | 4.2e-08 | 1.7e-07 | 9.267E-08 | 6.804E-08 | 4.200E-08 | 1.700E-07 |
| 25224 | 1262 | P1262D | D | 6.2e-08 | 8.8e-08 | 3.2e-08 | 6.067E-08 | 2.802E-08 | 3.200E-08 | 8.800E-08 |
| 25225 | 1262 | P1262C | D | 6.4e-08 | 3.5e-08 | 4.9e-08 | 4.933E-08 | 1.450E-08 | 3.500E-08 | 6.400E-08 |
| 25226 | 1262 | P1262E | D | 4.9e-08 | 5.6e-08 | 3.4e-08 | 4.633E-08 | 1.124E-08 | 3.400E-08 | 5.600E-08 |
| 25227 | 1262 | P1262Q | D | 3.7e-08 | 1.9e-08 | 5.6e-08 | 3.733E-08 | 1.850E-08 | 1.900E-08 | 5.600E-08 |
| 25228 | 1262 | P1262G | D | 4.7e-08 | 1.4e-08 | 1.4e-08 | 2.500E-08 | 1.905E-08 | 1.400E-08 | 4.700E-08 |
| 25229 | 1262 | P1262H | D | 3.8e-08 | 4.6e-08 | 3.5e-08 | 3.967E-08 | 5.686E-09 | 3.500E-08 | 4.600E-08 |
| 25230 | 1262 | P1262I | D | 7.1e-08 | 5e-08 | 4.4e-08 | 5.500E-08 | 1.418E-08 | 4.400E-08 | 7.100E-08 |
| 25231 | 1262 | P1262L | D | 4.2e-08 | 6.2e-08 | 1.1e-07 | 7.133E-08 | 3.495E-08 | 4.200E-08 | 1.100E-07 |
| 25232 | 1262 | P1262K | D | 2.3e-08 | 5.1e-08 | 1.4e-08 | 2.933E-08 | 1.930E-08 | 1.400E-08 | 5.100E-08 |
| 25233 | 1262 | P1262M | D | 1.3e-08 | 8.1e-08 | 9.2e-08 | 6.200E-08 | 4.279E-08 | 1.300E-08 | 9.200E-08 |
| 25234 | 1262 | P1262F | D | 4e-08 | 8.1e-08 | 4.1e-08 | 5.400E-08 | 2.339E-08 | 4.000E-08 | 8.100E-08 |
| 25235 | 1262 | P1262P | D | 3e-08 | 1.8e-08 | 2.5e-08 | 2.433E-08 | 6.028E-09 | 1.800E-08 | 3.000E-08 |
| 25236 | 1262 | P1262S | D | 1.7e-08 | 5.7e-08 | 4.8e-08 | 4.067E-08 | 2.098E-08 | 1.700E-08 | 5.700E-08 |
| 25237 | 1262 | P1262T | D | 3.1e-08 | 3.7e-08 | 4.9e-08 | 3.900E-08 | 9.165E-09 | 3.100E-08 | 4.900E-08 |
| 25238 | 1262 | P1262W | D | 4.3e-08 | 3.4e-08 | 3.4e-08 | 3.700E-08 | 5.196E-09 | 3.400E-08 | 4.300E-08 |
| 25239 | 1262 | P1262Y | D | 9.3e-08 | 5.5e-08 | 2.4e-08 | 5.733E-08 | 3.456E-08 | 2.400E-08 | 9.300E-08 |
| 25240 | 1262 | P1262V | D | 4e-08 | 5.3e-08 | 4.1e-08 | 4.467E-08 | 7.234E-09 | 4.000E-08 | 5.300E-08 |
| 25241 | 1263 | V1263A | D | 2e-08 | 3.5e-08 | 1.3e-07 | 6.167E-08 | 5.965E-08 | 2.000E-08 | 1.300E-07 |
| 25242 | 1263 | V1263R | D | 3.2e-08 | 4.1e-08 | 3.1e-08 | 3.467E-08 | 5.508E-09 | 3.100E-08 | 4.100E-08 |
| 25243 | 1263 | V1263N | D | 4.8e-08 | 5.7e-08 | 2e-08 | 4.167E-08 | 1.930E-08 | 2.000E-08 | 5.700E-08 |
| 25244 | 1263 | V1263D | D | 5.1e-08 | 2.5e-08 | 2e-08 | 3.200E-08 | 1.664E-08 | 2.000E-08 | 5.100E-08 |

| | | | | | | | | | | |
|---|---|---|---|---|---|---|---|---|---|---|
| 25245 | 1263 | V1263C | D | 1.6e-08 | 4.9e-08 | 3.7e-08 | 3.400E-08 | 1.670E-08 | 1.600E-08 | 4.900E-08 |
| 25246 | 1263 | V1263E | D | 8.6e-08 | 5e-08 | 3.5e-08 | 5.700E-08 | 2.621E-08 | 3.500E-08 | 8.600E-08 |
| 25247 | 1263 | V1263Q | D | 4.8e-08 | 9.4e-08 | 4.5e-08 | 6.233E-08 | 2.747E-08 | 4.500E-08 | 9.400E-08 |
| 25248 | 1263 | V1263G | D | 6.2e-08 | 2.2e-08 | 4.5e-08 | 4.300E-08 | 2.007E-08 | 2.200E-08 | 6.200E-08 |
| 25249 | 1263 | V1263H | D | 2.2e-08 | 8.6e-08 | 1e-07 | 6.933E-08 | 4.159E-08 | 2.200E-08 | 1.000E-07 |
| 25250 | 1263 | V1263I | D | 2.4e-08 | 4.9e-08 | 3.6e-08 | 3.633E-08 | 1.250E-08 | 2.400E-08 | 4.900E-08 |
| 25251 | 1263 | V1263L | D | 3.5e-08 | 2.9e-08 | 7.3e-08 | 4.567E-08 | 2.386E-08 | 2.900E-08 | 7.300E-08 |
| 25252 | 1263 | V1263K | D | 4.1e-08 | 2.9e-08 | 6.7e-08 | 4.567E-08 | 1.943E-08 | 2.900E-08 | 6.700E-08 |
| 25253 | 1263 | V1263M | D | 2.2e-08 | 4e-08 | 6.1e-08 | 4.100E-08 | 1.952E-08 | 2.200E-08 | 6.100E-08 |
| 25254 | 1263 | V1263F | D | 5.2e-08 | 9.8e-08 | 4.9e-08 | 6.633E-08 | 2.747E-08 | 4.900E-08 | 9.800E-08 |
| 25255 | 1263 | V1263P | D | 8e-08 | 2.9e-08 | 2.3e-08 | 4.400E-08 | 3.132E-08 | 2.300E-08 | 8.000E-08 |
| 25256 | 1263 | V1263S | D | 3.5e-08 | 4.7e-08 | 2.4e-08 | 3.533E-08 | 1.150E-08 | 2.400E-08 | 4.700E-08 |
| 25257 | 1263 | V1263T | D | 3.7e-08 | 6.1e-08 | 2.4e-08 | 4.067E-08 | 1.877E-08 | 2.400E-08 | 6.100E-08 |
| 25258 | 1263 | V1263W | D | 2.7e-08 | 7.3e-08 | 5.1e-08 | 5.033E-08 | 2.301E-08 | 2.700E-08 | 7.300E-08 |
| 25259 | 1263 | V1263Y | D | 6.7e-08 | 2.9e-08 | 9.2e-08 | 6.267E-08 | 3.172E-08 | 2.900E-08 | 9.200E-08 |
| 25260 | 1263 | V1263V | D | 4.6e-08 | 2.7e-08 | 1.5e-08 | 2.933E-08 | 1.563E-08 | 1.500E-08 | 4.600E-08 |
| 25261 | 1264 | T1264A | D | 5.1e-08 | 3.3e-08 | 2.6e-08 | 3.667E-08 | 1.290E-08 | 2.600E-08 | 5.100E-08 |
| 25262 | 1264 | T1264R | D | 1.7e-08 | 3.1e-08 | 1.6e-08 | 2.133E-08 | 8.386E-09 | 1.600E-08 | 3.100E-08 |
| 25263 | 1264 | T1264N | D | 1.5e-08 | 1.5e-08 | 4.7e-08 | 2.567E-08 | 1.848E-08 | 1.500E-08 | 4.700E-08 |
| 25264 | 1264 | T1264D | D | 1.9e-08 | 1.3e-07 | 5.8e-08 | 6.900E-08 | 5.631E-08 | 1.900E-08 | 1.300E-07 |
| 25265 | 1264 | T1264C | D | 1.4e-08 | 5.1e-08 | 7.2e-08 | 4.567E-08 | 2.937E-08 | 1.400E-08 | 7.200E-08 |
| 25266 | 1264 | T1264E | D | 5.9e-08 | 8e-08 | 3.9e-08 | 5.933E-08 | 2.050E-08 | 3.900E-08 | 8.000E-08 |
| 25267 | 1264 | T1264Q | D | 4.1e-08 | 4.9e-08 | 1.5e-08 | 3.500E-08 | 1.778E-08 | 1.500E-08 | 4.900E-08 |
| 25268 | 1264 | T1264G | D | 1.1e-07 | 2e-08 | 1.7e-07 | 1.000E-07 | 7.550E-08 | 2.000E-08 | 1.700E-07 |
| 25269 | 1264 | T1264H | D | 6.3e-08 | 4.1e-08 | 4.3e-08 | 4.900E-08 | 1.217E-08 | 4.100E-08 | 6.300E-08 |
| 25270 | 1264 | T1264I | D | 6.2e-08 | 1.1e-07 | 4.5e-08 | 7.233E-08 | 3.371E-08 | 4.500E-08 | 1.100E-07 |
| 25271 | 1264 | T1264L | D | 8.4e-08 | 3.4e-08 | 8.6e-08 | 6.800E-08 | 2.946E-08 | 3.400E-08 | 8.600E-08 |
| 25272 | 1264 | T1264K | D | 3.8e-08 | 4.8e-08 | 3.9e-08 | 4.167E-08 | 5.508E-09 | 3.800E-08 | 4.800E-08 |
| 25273 | 1264 | T1264M | D | 3.2e-08 | 1.1e-07 | 3.1e-08 | 5.767E-08 | 4.532E-08 | 3.100E-08 | 1.100E-07 |
| 25274 | 1264 | T1264F | D | 8.4e-08 | 4.4e-08 | 5.5e-08 | 6.100E-08 | 2.066E-08 | 4.400E-08 | 8.400E-08 |
| 25275 | 1264 | T1264P | D | 9.9e-08 | 4.8e-08 | 5.6e-08 | 6.767E-08 | 2.743E-08 | 4.800E-08 | 9.900E-08 |
| 25276 | 1264 | T1264S | D | 4.5e-08 | 1.9e-08 | 4.4e-08 | 3.600E-08 | 1.473E-08 | 1.900E-08 | 4.500E-08 |
| 25277 | 1264 | T1264T | D | 5.2e-08 | 7.4e-08 | 5.8e-08 | 6.133E-08 | 1.137E-08 | 5.200E-08 | 7.400E-08 |
| 25278 | 1264 | T1264W | D | 3.7e-08 | 3.4e-08 | 6.6e-08 | 4.567E-08 | 1.767E-08 | 3.400E-08 | 6.600E-08 |
| 25279 | 1264 | T1264Y | D | 9.9e-09 | 3.8e-08 | 1.3e-07 | 5.930E-08 | 6.282E-08 | 9.900E-09 | 1.300E-07 |
| 25280 | 1264 | T1264V | D | 3.4e-08 | 3.3e-08 | 4.9e-08 | 3.867E-08 | 8.963E-09 | 3.300E-08 | 4.900E-08 |
| 25281 | 1265 | K1265A | D | 5.7e-08 | 2.4e-08 | 8.6e-08 | 5.567E-08 | 3.102E-08 | 2.400E-08 | 8.600E-08 |
| 25282 | 1265 | K1265R | D | 2.7e-08 | 3.1e-08 | 1.1e-07 | 5.600E-08 | 4.681E-08 | 2.700E-08 | 1.100E-07 |
| 25283 | 1265 | K1265N | D | 4e-08 | 2.5e-08 | 7.9e-08 | 4.800E-08 | 2.787E-08 | 2.500E-08 | 7.900E-08 |
| 25284 | 1265 | K1265D | D | 4.6e-08 | 2.5e-08 | 4.3e-08 | 3.800E-08 | 1.136E-08 | 2.500E-08 | 4.600E-08 |
| 25285 | 1265 | K1265C | D | 4.5e-08 | 1.6e-08 | 2e-08 | 2.700E-08 | 1.572E-08 | 1.600E-08 | 4.500E-08 |
| 25286 | 1265 | K1265E | D | 3.9e-08 | 3.8e-08 | 5.7e-08 | 4.467E-08 | 1.069E-08 | 3.800E-08 | 5.700E-08 |
| 25287 | 1265 | K1265Q | D | 3e-08 | 2.2e-08 | 2.4e-08 | 2.533E-08 | 4.163E-09 | 2.200E-08 | 3.000E-08 |
| 25288 | 1265 | K1265G | D | 7.6e-08 | 1.7e-08 | 4.6e-08 | 4.633E-08 | 2.950E-08 | 1.700E-08 | 7.600E-08 |
| 25289 | 1265 | K1265H | D | 5.3e-08 | 3.4e-08 | 1e-07 | 6.233E-08 | 3.398E-08 | 3.400E-08 | 1.000E-07 |
| 25290 | 1265 | K1265I | D | 4.5e-08 | 9.5e-08 | 5e-08 | 6.333E-08 | 2.754E-08 | 4.500E-08 | 9.500E-08 |
| 25291 | 1265 | K1265L | D | 4.3e-08 | 3.5e-08 | 7.3e-08 | 5.033E-08 | 2.003E-08 | 3.500E-08 | 7.300E-08 |
| 25292 | 1265 | K1265K | D | 7.5e-08 | 4e-08 | 3.5e-08 | 5.000E-08 | 2.179E-08 | 3.500E-08 | 7.500E-08 |
| 25293 | 1265 | K1265M | D | 3.2e-08 | 7.1e-08 | 3.8e-08 | 4.700E-08 | 2.100E-08 | 3.200E-08 | 7.100E-08 |
| 25294 | 1265 | K1265F | D | 2.8e-08 | 4.4e-08 | 5.4e-08 | 4.200E-08 | 1.311E-08 | 2.800E-08 | 5.400E-08 |
| 25295 | 1265 | K1265P | D | 6.2e-08 | 7.8e-08 | 3.3e-08 | 5.767E-08 | 2.281E-08 | 3.300E-08 | 7.800E-08 |
| 25296 | 1265 | K1265S | D | 8.8e-08 | 3.4e-08 | 1.8e-08 | 4.667E-08 | 3.668E-08 | 1.800E-08 | 8.800E-08 |
| 25297 | 1265 | K1265T | D | 6e-08 | 1.5e-08 | 2.9e-08 | 3.467E-08 | 2.303E-08 | 1.500E-08 | 6.000E-08 |
| 25298 | 1265 | K1265W | D | 1.2e-08 | 5.8e-08 | 4.5e-08 | 3.833E-08 | 2.371E-08 | 1.200E-08 | 5.800E-08 |
| 25299 | 1265 | K1265Y | D | 5.6e-08 | 5.2e-08 | 6.1e-08 | 5.633E-08 | 4.509E-09 | 5.200E-08 | 6.100E-08 |
| 25300 | 1265 | K1265V | D | 1.1e-07 | 1.6e-08 | 5.8e-08 | 6.133E-08 | 4.709E-08 | 1.600E-08 | 1.100E-07 |
| 25301 | 1266 | S1266A | D | 5.9e-08 | 5.6e-08 | 4e-08 | 5.167E-08 | 1.021E-08 | 4.000E-08 | 5.900E-08 |
| 25302 | 1266 | S1266R | D | 1.1e-07 | 3e-08 | 2e-08 | 5.333E-08 | 4.933E-08 | 2.000E-08 | 1.100E-07 |
| 25303 | 1266 | S1266N | D | 2.5e-08 | 8.9e-08 | 5.9e-08 | 5.767E-08 | 3.202E-08 | 2.500E-08 | 8.900E-08 |
| 25304 | 1266 | S1266D | D | 7.9e-08 | 3.3e-08 | 2.2e-08 | 4.467E-08 | 3.024E-08 | 2.200E-08 | 7.900E-08 |
| 25305 | 1266 | S1266C | D | 3e-08 | 2.5e-08 | 3e-08 | 2.833E-08 | 2.887E-09 | 2.500E-08 | 3.000E-08 |
| 25306 | 1266 | S1266E | D | 4.2e-08 | 2.3e-08 | 3.5e-08 | 3.333E-08 | 9.609E-09 | 2.300E-08 | 4.200E-08 |
| 25307 | 1266 | S1266Q | D | 1.8e-08 | 5.2e-08 | 7.5e-08 | 4.833E-08 | 2.868E-08 | 1.800E-08 | 7.500E-08 |
| 25308 | 1266 | S1266G | D | 9.4e-08 | 1.7e-08 | 1.8e-08 | 4.300E-08 | 4.417E-08 | 1.700E-08 | 9.400E-08 |
| 25309 | 1266 | S1266H | D | 2.7e-08 | 2e-07 | 2.5e-08 | 8.400E-08 | 1.005E-07 | 2.500E-08 | 2.000E-07 |
| 25310 | 1266 | S1266I | D | 3.5e-08 | 1.2e-07 | 3.5e-08 | 6.333E-08 | 4.907E-08 | 3.500E-08 | 1.200E-07 |
| 25311 | 1266 | S1266L | D | 5.9e-08 | 2.1e-08 | 2e-08 | 3.333E-08 | 2.223E-08 | 2.000E-08 | 5.900E-08 |
| 25312 | 1266 | S1266K | D | 5.8e-08 | 1.4e-08 | 3.5e-08 | 3.567E-08 | 2.201E-08 | 1.400E-08 | 5.800E-08 |
| 25313 | 1266 | S1266M | D | 5.8e-08 | 1.8e-08 | 7e-08 | 4.867E-08 | 2.723E-08 | 1.800E-08 | 7.000E-08 |
| 25314 | 1266 | S1266F | D | 2.2e-08 | 4.4e-08 | 7.1e-08 | 4.567E-08 | 2.454E-08 | 2.200E-08 | 7.100E-08 |
| 25315 | 1266 | S1266P | D | 2.6e-08 | 4.4e-08 | 1.9e-08 | 2.967E-08 | 1.290E-08 | 1.900E-08 | 4.400E-08 |
| 25316 | 1266 | S1266S | D | 3.9e-08 | 4.7e-08 | 3.1e-08 | 3.900E-08 | 8.000E-09 | 3.100E-08 | 4.700E-08 |
| 25317 | 1266 | S1266T | D | 3e-08 | 7e-08 | 9.5e-09 | 3.650E-08 | 3.077E-08 | 9.500E-09 | 7.000E-08 |
| 25318 | 1266 | S1266W | D | 3.2e-08 | 1.8e-08 | 5.4e-08 | 3.467E-08 | 1.815E-08 | 1.800E-08 | 5.400E-08 |
| 25319 | 1266 | S1266Y | D | 5e-08 | 2e-08 | 2.4e-08 | 3.133E-08 | 1.629E-08 | 2.000E-08 | 5.000E-08 |
| 25320 | 1266 | S1266V | D | 7e-08 | 1.1e-07 | 1.8e-08 | 6.600E-08 | 4.613E-08 | 1.800E-08 | 1.100E-07 |
| 25321 | 1267 | F1267A | D | 1.6e-07 | 1.8e-08 | 3.8e-08 | 7.200E-08 | 7.686E-08 | 1.800E-08 | 1.600E-07 |
| 25322 | 1267 | F1267R | D | 4.5e-08 | 6.6e-08 | 2e-08 | 4.367E-08 | 2.303E-08 | 2.000E-08 | 6.600E-08 |
| 25323 | 1267 | F1267N | D | 3e-08 | 3.2e-08 | 4.4e-08 | 3.533E-08 | 7.572E-09 | 3.000E-08 | 4.400E-08 |
| 25324 | 1267 | F1267D | D | 5.5e-08 | 5.2e-08 | 5.9e-08 | 5.533E-08 | 3.512E-09 | 5.200E-08 | 5.900E-08 |
| 25325 | 1267 | F1267C | D | 1.4e-07 | 1e-07 | 3.6e-08 | 9.200E-08 | 5.246E-08 | 3.600E-08 | 1.400E-07 |
| 25326 | 1267 | F1267E | D | 4.2e-08 | 7.4e-08 | 5.9e-08 | 5.833E-08 | 1.601E-08 | 4.200E-08 | 7.400E-08 |
| 25327 | 1267 | F1267Q | D | 8.1e-08 | 3.4e-08 | 7.6e-08 | 6.367E-08 | 2.581E-08 | 3.400E-08 | 8.100E-08 |
| 25328 | 1267 | F1267G | D | 8.8e-08 | 2.4e-08 | 1.7e-08 | 4.300E-08 | 3.913E-08 | 1.700E-08 | 8.800E-08 |
| 25329 | 1267 | F1267H | D | 2.9e-08 | 1.4e-08 | 8.4e-08 | 4.233E-08 | 3.686E-08 | 1.400E-08 | 8.400E-08 |

| | | | | | | | | | | |
|---|---|---|---|---|---|---|---|---|---|---|
| 25330 | 1267 | F1267I | D | 7.4e-08 | 3.5e-08 | 3.2e-08 | 4.700E-08 | 2.343E-08 | 3.200E-08 | 7.400E-08 |
| 25331 | 1267 | F1267L | D | 8.7e-08 | 8.2e-08 | 4.2e-08 | 7.033E-08 | 2.466E-08 | 4.200E-08 | 8.700E-08 |
| 25332 | 1267 | F1267K | D | 5.6e-08 | 6.4e-08 | 3.7e-08 | 5.233E-08 | 1.387E-08 | 3.700E-08 | 6.400E-08 |
| 25333 | 1267 | F1267M | D | 4.1e-08 | 2.6e-08 | 2.2e-08 | 2.967E-08 | 1.002E-08 | 2.200E-08 | 4.100E-08 |
| 25334 | 1267 | F1267F | D | 5.7e-08 | 4e-08 | 7.5e-08 | 5.733E-08 | 1.750E-08 | 4.000E-08 | 7.500E-08 |
| 25335 | 1267 | F1267P | D | 3.5e-08 | 3.4e-08 | 3e-08 | 3.300E-08 | 2.646E-09 | 3.000E-08 | 3.500E-08 |
| 25336 | 1267 | F1267S | D | 3.9e-08 | 9.7e-08 | 2.4e-08 | 5.333E-08 | 3.855E-08 | 2.400E-08 | 9.700E-08 |
| 25337 | 1267 | F1267T | D | 4.5e-08 | 8.1e-08 | 2.2e-08 | 4.933E-08 | 2.974E-08 | 2.200E-08 | 8.100E-08 |
| 25338 | 1267 | F1267W | D | 1.6e-08 | 4.9e-08 | 6.4e-08 | 4.300E-08 | 2.456E-08 | 1.600E-08 | 6.400E-08 |
| 25339 | 1267 | F1267Y | D | 2.9e-08 | 5.1e-08 | 4.4e-08 | 4.133E-08 | 1.124E-08 | 2.900E-08 | 5.100E-08 |
| 25340 | 1267 | F1267V | D | 1.6e-08 | 4e-08 | 1.7e-08 | 2.433E-08 | 1.358E-08 | 1.600E-08 | 4.000E-08 |
| 25341 | 1268 | N1268A | D | 3.4e-08 | 7.9e-08 | 2.5e-08 | 4.600E-08 | 2.893E-08 | 2.500E-08 | 7.900E-08 |
| 25342 | 1268 | N1268R | D | 5.6e-08 | 7.1e-08 | 7e-08 | 6.567E-08 | 8.386E-09 | 5.600E-08 | 7.100E-08 |
| 25343 | 1268 | N1268N | D | 3.6e-08 | 3.2e-08 | 8.2e-08 | 5.000E-08 | 2.778E-08 | 3.200E-08 | 8.200E-08 |
| 25344 | 1268 | N1268D | D | 7.8e-09 | 3.5e-08 | 2.6e-08 | 2.293E-08 | 1.386E-08 | 7.800E-09 | 3.500E-08 |
| 25345 | 1268 | N1268C | D | 8.4e-08 | 1.5e-07 | 3.2e-08 | 8.867E-08 | 5.914E-08 | 3.200E-08 | 1.500E-07 |
| 25346 | 1268 | N1268E | D | 1e-07 | 1.5e-08 | 5.9e-08 | 5.800E-08 | 4.251E-08 | 1.500E-08 | 1.000E-07 |
| 25347 | 1268 | N1268Q | D | 5.9e-08 | 4.9e-08 | 1.5e-08 | 4.100E-08 | 2.307E-08 | 1.500E-08 | 5.900E-08 |
| 25348 | 1268 | N1268G | D | 2.2e-08 | 4.1e-08 | 5.9e-08 | 4.067E-08 | 1.850E-08 | 2.200E-08 | 5.900E-08 |
| 25349 | 1268 | N1268H | D | 3.9e-08 | 6.3e-08 | 7e-08 | 5.733E-08 | 1.626E-08 | 3.900E-08 | 7.000E-08 |
| 25350 | 1268 | N1268I | D | 3.3e-08 | 7.6e-08 | 2.9e-08 | 4.600E-08 | 2.606E-08 | 2.900E-08 | 7.600E-08 |
| 25351 | 1268 | N1268L | D | 2.7e-08 | 7.8e-08 | 5.4e-08 | 5.300E-08 | 2.551E-08 | 2.700E-08 | 7.800E-08 |
| 25352 | 1268 | N1268K | D | 2.5e-08 | 5.2e-08 | 1.2e-08 | 2.967E-08 | 2.040E-08 | 1.200E-08 | 5.200E-08 |
| 25353 | 1268 | N1268M | D | 7.8e-08 | 6.8e-08 | 3.4e-08 | 6.000E-08 | 2.307E-08 | 3.400E-08 | 7.800E-08 |
| 25354 | 1268 | N1268F | D | 2e-08 | 4.1e-08 | 1.8e-08 | 2.633E-08 | 1.274E-08 | 1.800E-08 | 4.100E-08 |
| 25355 | 1268 | N1268P | D | 1e-07 | 7.4e-08 | 9.4e-08 | 8.933E-08 | 1.361E-08 | 7.400E-08 | 1.000E-07 |
| 25356 | 1268 | N1268S | D | 2.3e-08 | 4.3e-08 | 2.9e-08 | 3.167E-08 | 1.026E-08 | 2.300E-08 | 4.300E-08 |
| 25357 | 1268 | N1268T | D | 9e-08 | 8.4e-08 | 1.1e-07 | 9.467E-08 | 1.361E-08 | 8.400E-08 | 1.100E-07 |
| 25358 | 1268 | N1268W | D | 4.1e-08 | 3.8e-08 | 3.9e-08 | 3.933E-08 | 1.528E-09 | 3.800E-08 | 4.100E-08 |
| 25359 | 1268 | N1268Y | D | 8.9e-08 | 2.9e-08 | 1.6e-07 | 9.267E-08 | 6.558E-08 | 2.900E-08 | 1.600E-07 |
| 25360 | 1268 | N1268V | D | 6.2e-08 | 4.2e-08 | 1.4e-07 | 8.133E-08 | 5.178E-08 | 4.200E-08 | 1.400E-07 |
| 25361 | 1269 | R1269A | D | 8.5e-08 | 4e-08 | 6.6e-08 | 6.367E-08 | 2.259E-08 | 4.000E-08 | 8.500E-08 |
| 25362 | 1269 | R1269R | D | 7.1e-08 | 2.7e-08 | 2.9e-08 | 4.233E-08 | 2.485E-08 | 2.700E-08 | 7.100E-08 |
| 25363 | 1269 | R1269N | D | 1.2e-07 | 3.5e-08 | 4e-08 | 6.500E-08 | 4.770E-08 | 3.500E-08 | 1.200E-07 |
| 25364 | 1269 | R1269D | D | 2e-08 | 3.8e-08 | 5.9e-08 | 3.900E-08 | 1.952E-08 | 2.000E-08 | 5.900E-08 |
| 25365 | 1269 | R1269C | D | 2.1e-08 | 6.9e-08 | 3.2e-08 | 4.067E-08 | 2.515E-08 | 2.100E-08 | 6.900E-08 |
| 25366 | 1269 | R1269E | D | 7.9e-08 | 2.3e-08 | 6.9e-08 | 5.700E-08 | 2.987E-08 | 2.300E-08 | 7.900E-08 |
| 25367 | 1269 | R1269Q | D | 2.8e-08 | 1.5e-08 | 1.6e-08 | 1.967E-08 | 7.234E-09 | 1.500E-08 | 2.800E-08 |
| 25368 | 1269 | R1269G | D | 2.3e-08 | 2.2e-08 | 6.1e-08 | 3.533E-08 | 2.223E-08 | 2.200E-08 | 6.100E-08 |
| 25369 | 1269 | R1269H | D | 3.3e-08 | 6.5e-08 | 5.8e-08 | 5.200E-08 | 1.682E-08 | 3.300E-08 | 6.500E-08 |
| 25370 | 1269 | R1269I | D | 4.1e-08 | 1.4e-08 | 4.4e-08 | 3.300E-08 | 1.652E-08 | 1.400E-08 | 4.400E-08 |
| 25371 | 1269 | R1269L | D | 6.2e-08 | 3e-08 | 7.2e-08 | 5.467E-08 | 2.194E-08 | 3.000E-08 | 7.200E-08 |
| 25372 | 1269 | R1269K | D | 3.8e-08 | 2.5e-08 | 4.7e-08 | 3.667E-08 | 1.106E-08 | 2.500E-08 | 4.700E-08 |
| 25373 | 1269 | R1269M | D | 3.9e-08 | 6.2e-08 | 6.3e-08 | 5.467E-08 | 1.358E-08 | 3.900E-08 | 6.300E-08 |
| 25374 | 1269 | R1269F | D | 2.8e-08 | 6.6e-08 | 5.7e-08 | 5.033E-08 | 1.986E-08 | 2.800E-08 | 6.600E-08 |
| 25375 | 1269 | R1269P | D | 3.2e-08 | 3.6e-08 | 5.5e-08 | 4.100E-08 | 1.229E-08 | 3.200E-08 | 5.500E-08 |
| 25376 | 1269 | R1269S | D | 9e-08 | 1.3e-08 | 3.5e-08 | 4.600E-08 | 3.966E-08 | 1.300E-08 | 9.000E-08 |
| 25377 | 1269 | R1269T | D | 8.7e-08 | 8.6e-08 | 3.1e-08 | 6.800E-08 | 3.205E-08 | 3.100E-08 | 8.700E-08 |
| 25378 | 1269 | R1269W | D | 1.8e-08 | 5.9e-08 | 7.4e-08 | 5.033E-08 | 2.899E-08 | 1.800E-08 | 7.400E-08 |
| 25379 | 1269 | R1269Y | D | 6.6e-08 | 3.7e-08 | 2.6e-08 | 4.300E-08 | 2.066E-08 | 2.600E-08 | 6.600E-08 |
| 25380 | 1269 | R1269V | D | 2.7e-08 | 5.3e-08 | 3.4e-08 | 3.800E-08 | 1.345E-08 | 2.700E-08 | 5.300E-08 |
| 25381 | 1270 | G1270A | D | 3.6e-08 | 4.1e-08 | 5.2e-08 | 4.300E-08 | 8.185E-09 | 3.600E-08 | 5.200E-08 |
| 25382 | 1270 | G1270R | D | 2.7e-08 | 6.7e-08 | 3.5e-08 | 4.300E-08 | 2.117E-08 | 2.700E-08 | 6.700E-08 |
| 25383 | 1270 | G1270N | D | 2.9e-07 | 3.5e-08 | 5.9e-08 | 1.280E-07 | 1.408E-07 | 3.500E-08 | 2.900E-07 |
| 25384 | 1270 | G1270D | D | 5.8e-08 | 4.6e-08 | 1.2e-07 | 7.467E-08 | 3.972E-08 | 4.600E-08 | 1.200E-07 |
| 25385 | 1270 | G1270C | D | 2e-08 | 2.5e-08 | 4.5e-08 | 3.000E-08 | 1.323E-08 | 2.000E-08 | 4.500E-08 |
| 25386 | 1270 | G1270E | D | 2.9e-08 | 1.6e-08 | 3e-08 | 2.500E-08 | 7.810E-09 | 1.600E-08 | 3.000E-08 |
| 25387 | 1270 | G1270Q | D | 3.8e-08 | 1.2e-08 | 1e-07 | 5.000E-08 | 4.521E-08 | 1.200E-08 | 1.000E-07 |
| 25388 | 1270 | G1270G | D | 4.6e-08 | 3.5e-08 | 3.5e-08 | 3.867E-08 | 6.351E-09 | 3.500E-08 | 4.600E-08 |
| 25389 | 1270 | G1270H | D | 3.8e-08 | 2.4e-08 | 2e-08 | 2.733E-08 | 9.452E-09 | 2.000E-08 | 3.800E-08 |
| 25390 | 1270 | G1270I | D | 1.6e-08 | 4.2e-08 | 2.6e-08 | 2.800E-08 | 1.311E-08 | 1.600E-08 | 4.200E-08 |
| 25391 | 1270 | G1270L | D | 1.8e-07 | 5.6e-08 | 2.8e-08 | 8.800E-08 | 8.089E-08 | 2.800E-08 | 1.800E-07 |
| 25392 | 1270 | G1270K | D | 3.3e-08 | 8.9e-08 | 3.2e-08 | 5.133E-08 | 3.262E-08 | 3.200E-08 | 8.900E-08 |
| 25393 | 1270 | G1270M | D | 3.7e-08 | 4.7e-08 | 3.5e-08 | 3.967E-08 | 6.429E-09 | 3.500E-08 | 4.700E-08 |
| 25394 | 1270 | G1270F | D | 3.9e-08 | 2.1e-08 | 1.1e-07 | 5.667E-08 | 4.706E-08 | 2.100E-08 | 1.100E-07 |
| 25395 | 1270 | G1270P | D | 3e-08 | 4.9e-08 | 5e-08 | 4.300E-08 | 1.127E-08 | 3.000E-08 | 5.000E-08 |
| 25396 | 1270 | G1270S | D | 1.1e-07 | 6.6e-08 | 6.8e-08 | 8.133E-08 | 2.485E-08 | 6.600E-08 | 1.100E-07 |
| 25397 | 1270 | G1270T | D | 1.1e-07 | 6.1e-08 | 8.2e-08 | 8.433E-08 | 2.458E-08 | 6.100E-08 | 1.100E-07 |
| 25398 | 1270 | G1270W | D | 2.8e-08 | 2.4e-08 | 3.1e-08 | 2.767E-08 | 3.512E-09 | 2.400E-08 | 3.100E-08 |
| 25399 | 1270 | G1270Y | D | 5e-08 | 1.1e-07 | 1.5e-07 | 1.033E-07 | 5.033E-08 | 5.000E-08 | 1.500E-07 |
| 25400 | 1270 | G1270V | D | 4.8e-08 | 2.3e-08 | 8.9e-08 | 5.333E-08 | 3.332E-08 | 2.300E-08 | 8.900E-08 |
| 25401 | 1271 | E1271A | D | 8.1e-08 | 4.6e-08 | 2.1e-08 | 4.933E-08 | 3.014E-08 | 2.100E-08 | 8.100E-08 |
| 25402 | 1271 | E1271R | D | 6.7e-08 | 5.1e-08 | 1.6e-08 | 4.467E-08 | 2.608E-08 | 1.600E-08 | 6.700E-08 |
| 25403 | 1271 | E1271N | D | 2.2e-08 | 3.6e-08 | 1.3e-08 | 2.367E-08 | 1.159E-08 | 1.300E-08 | 3.600E-08 |
| 25404 | 1271 | E1271D | D | 3.4e-08 | 4.1e-08 | 5.5e-08 | 4.333E-08 | 1.069E-08 | 3.400E-08 | 5.500E-08 |
| 25405 | 1271 | E1271C | D | 2.6e-08 | 3.1e-08 | 4e-08 | 3.233E-08 | 7.095E-09 | 2.600E-08 | 4.000E-08 |
| 25406 | 1271 | E1271E | D | 3.6e-08 | 3.9e-08 | 1.5e-07 | 7.500E-08 | 6.497E-08 | 3.600E-08 | 1.500E-07 |
| 25407 | 1271 | E1271Q | D | 9.3e-08 | 7.5e-08 | 5.6e-08 | 7.467E-08 | 1.850E-08 | 5.600E-08 | 9.300E-08 |
| 25408 | 1271 | E1271G | D | 1e-08 | 2.6e-08 | 2e-08 | 1.867E-08 | 8.083E-09 | 1.000E-08 | 2.600E-08 |
| 25409 | 1271 | E1271H | D | 8.4e-08 | 5.2e-08 | 9.2e-08 | 7.600E-08 | 2.117E-08 | 5.200E-08 | 9.200E-08 |
| 25410 | 1271 | E1271I | D | 1.1e-08 | 5.3e-08 | 1.8e-07 | 8.133E-08 | 8.799E-08 | 1.100E-08 | 1.800E-07 |
| 25411 | 1271 | E1271L | D | 3.6e-08 | 3.6e-08 | 5.4e-08 | 4.200E-08 | 1.039E-08 | 3.600E-08 | 5.400E-08 |
| 25412 | 1271 | E1271K | D | 1.7e-08 | 9.4e-08 | 7.3e-08 | 6.133E-08 | 3.980E-08 | 1.700E-08 | 9.400E-08 |
| 25413 | 1271 | E1271M | D | 5e-08 | 4.5e-08 | 4.1e-08 | 4.533E-08 | 4.509E-09 | 4.100E-08 | 5.000E-08 |
| 25414 | 1271 | E1271F | D | 3.6e-08 | 5.2e-08 | 5e-08 | 4.600E-08 | 8.718E-09 | 3.600E-08 | 5.200E-08 |

| | | | | | | | | | | |
|---|---|---|---|---|---|---|---|---|---|---|
| 25415 | 1271 | E1271P | D | 7.4e-08 | 1.3e-07 | 3.1e-08 | 7.833E-08 | 4.964E-08 | 3.100E-08 | 1.300E-07 |
| 25416 | 1271 | E1271S | D | 5.8e-08 | 4.9e-08 | 5.1e-08 | 5.267E-08 | 4.726E-09 | 4.900E-08 | 5.800E-08 |
| 25417 | 1271 | E1271T | D | 4e-08 | 5e-08 | 3.2e-08 | 4.067E-08 | 9.018E-09 | 3.200E-08 | 5.000E-08 |
| 25418 | 1271 | E1271W | D | 5.9e-08 | 6.4e-08 | 5.9e-08 | 6.067E-08 | 2.887E-09 | 5.900E-08 | 6.400E-08 |
| 25419 | 1271 | E1271Y | D | 3.2e-08 | 5e-08 | 3.9e-08 | 4.033E-08 | 9.074E-09 | 3.200E-08 | 5.000E-08 |
| 25420 | 1271 | E1271V | D | 1.5e-07 | 3.8e-08 | 3e-08 | 7.267E-08 | 6.709E-08 | 3.000E-08 | 1.500E-07 |
| 25421 | 1272 | C1272A | D | 2.4e-08 | 9e-08 | 2.9e-08 | 4.767E-08 | 3.675E-08 | 2.400E-08 | 9.000E-08 |
| 25422 | 1272 | C1272R | D | 3.5e-08 | 9.9e-09 | 6.8e-08 | 3.763E-08 | 2.914E-08 | 9.900E-09 | 6.800E-08 |
| 25423 | 1272 | C1272N | D | 7.4e-08 | 6.6e-08 | 3.9e-08 | 5.967E-08 | 1.834E-08 | 3.900E-08 | 7.400E-08 |
| 25424 | 1272 | C1272D | D | 2.2e-08 | 3.4e-08 | 2.7e-08 | 2.767E-08 | 6.028E-09 | 2.200E-08 | 3.400E-08 |
| 25425 | 1272 | C1272C | D | 4.8e-08 | 5.6e-08 | 5.8e-08 | 5.400E-08 | 5.292E-09 | 4.800E-08 | 5.800E-08 |
| 25426 | 1272 | C1272E | D | 2.6e-08 | 3.1e-08 | 2.1e-08 | 2.600E-08 | 5.000E-09 | 2.100E-08 | 3.100E-08 |
| 25427 | 1272 | C1272Q | D | 4.2e-08 | 5e-08 | 4.6e-08 | 4.600E-08 | 4.000E-09 | 4.200E-08 | 5.000E-08 |
| 25428 | 1272 | C1272G | D | 3.4e-08 | 5.9e-08 | 6.4e-08 | 5.233E-08 | 1.607E-08 | 3.400E-08 | 6.400E-08 |
| 25429 | 1272 | C1272H | D | 2.5e-08 | 1.3e-08 | 9e-08 | 4.267E-08 | 4.143E-08 | 1.300E-08 | 9.000E-08 |
| 25430 | 1272 | C1272I | D | 2.7e-08 | 4e-08 | 1.8e-08 | 2.833E-08 | 1.106E-08 | 1.800E-08 | 4.000E-08 |
| 25431 | 1272 | C1272L | D | 3.9e-08 | 3.4e-08 | 4e-08 | 3.767E-08 | 3.215E-09 | 3.400E-08 | 4.000E-08 |
| 25432 | 1272 | C1272K | D | 4.1e-08 | 3.6e-08 | 1.4e-07 | 7.233E-08 | 5.865E-08 | 3.600E-08 | 1.400E-07 |
| 25433 | 1272 | C1272M | D | 5.5e-08 | 3.5e-08 | 2.9e-08 | 3.967E-08 | 1.361E-08 | 2.900E-08 | 5.500E-08 |
| 25434 | 1272 | C1272F | D | 1.3e-08 | 8.2e-08 | 3.9e-08 | 4.467E-08 | 3.485E-08 | 1.300E-08 | 8.200E-08 |
| 25435 | 1272 | C1272P | D | 3.8e-08 | 2.4e-08 | 1.5e-08 | 2.567E-08 | 1.159E-08 | 1.500E-08 | 3.800E-08 |
| 25436 | 1272 | C1272S | D | 1.3e-07 | 5.2e-08 | 1.4e-08 | 6.533E-08 | 5.914E-08 | 1.400E-08 | 1.300E-07 |
| 25437 | 1272 | C1272T | D | 3.1e-08 | 1.8e-08 | 3e-08 | 2.633E-08 | 7.234E-09 | 1.800E-08 | 3.100E-08 |
| 25438 | 1272 | C1272W | D | 5.2e-08 | 3.2e-08 | 2.5e-08 | 3.633E-08 | 1.401E-08 | 2.500E-08 | 5.200E-08 |
| 25439 | 1272 | C1272Y | D | 4.4e-08 | 1.1e-07 | 2e-08 | 5.800E-08 | 4.660E-08 | 2.000E-08 | 1.100E-07 |
| 25440 | 1272 | C1272V | D | 5.9e-08 | 2.5e-08 | 6.3e-08 | 4.900E-08 | 2.088E-08 | 2.500E-08 | 6.300E-08 |
| 25441 | 1273 | E1273A | E | 2.9e-08 | 4.1e-08 | 2.9e-08 | 3.300E-08 | 6.928E-09 | 2.900E-08 | 4.100E-08 |
| 25442 | 1273 | E1273R | E | 3.8e-08 | 5.3e-08 | 2.4e-08 | 3.833E-08 | 1.450E-08 | 2.400E-08 | 5.300E-08 |
| 25443 | 1273 | E1273N | E | 9.2e-08 | 1e-07 | 5.6e-08 | 8.267E-08 | 2.344E-08 | 5.600E-08 | 1.000E-07 |
| 25444 | 1273 | E1273D | E | 1.8e-08 | 5.8e-08 | 3.6e-08 | 3.733E-08 | 2.003E-08 | 1.800E-08 | 5.800E-08 |
| 25445 | 1273 | E1273C | E | 4.5e-08 | 9.7e-08 | 4.6e-08 | 6.267E-08 | 2.974E-08 | 4.500E-08 | 9.700E-08 |
| 25446 | 1273 | E1273E | E | 3.5e-08 | 2.5e-08 | 5.5e-08 | 3.833E-08 | 1.528E-08 | 2.500E-08 | 5.500E-08 |
| 25447 | 1273 | E1273Q | E | 2.8e-08 | 3.4e-08 | 6.6e-08 | 4.267E-08 | 2.043E-08 | 2.800E-08 | 6.600E-08 |
| 25448 | 1273 | E1273G | E | 5.3e-08 | 2.3e-08 | 1.1e-08 | 2.900E-08 | 2.163E-08 | 1.100E-08 | 5.300E-08 |
| 25449 | 1273 | E1273H | E | 5.6e-08 | 3.5e-08 | 4e-08 | 4.367E-08 | 1.097E-08 | 3.500E-08 | 5.600E-08 |
| 25450 | 1273 | E1273I | E | 3.3e-08 | 2.6e-08 | 1.7e-08 | 2.533E-08 | 8.021E-09 | 1.700E-08 | 3.300E-08 |
| 25451 | 1273 | E1273L | E | 8e-08 | 3.8e-08 | 5.1e-08 | 5.633E-08 | 2.150E-08 | 3.800E-08 | 8.000E-08 |
| 25452 | 1273 | E1273K | E | 2.6e-08 | 4.1e-08 | 7.8e-08 | 4.833E-08 | 2.676E-08 | 2.600E-08 | 7.800E-08 |
| 25453 | 1273 | E1273M | E | 6.5e-08 | 3.3e-08 | 5.5e-08 | 5.100E-08 | 1.637E-08 | 3.300E-08 | 6.500E-08 |
| 25454 | 1273 | E1273F | E | 4.6e-08 | 7.2e-08 | 1.2e-07 | 7.933E-08 | 3.754E-08 | 4.600E-08 | 1.200E-07 |
| 25455 | 1273 | E1273P | E | 6.1e-08 | 2.7e-08 | 3.9e-08 | 4.233E-08 | 1.724E-08 | 2.700E-08 | 6.100E-08 |
| 25456 | 1273 | E1273S | E | 7.1e-08 | 2.8e-08 | 5.8e-08 | 5.233E-08 | 2.205E-08 | 2.800E-08 | 7.100E-08 |
| 25457 | 1273 | E1273T | E | 8.9e-08 | 2.6e-08 | 4.5e-08 | 5.333E-08 | 3.232E-08 | 2.600E-08 | 8.900E-08 |
| 25458 | 1273 | E1273W | E | 4.8e-08 | 6.4e-08 | 6.6e-08 | 5.933E-08 | 9.866E-09 | 4.800E-08 | 6.600E-08 |
| 25459 | 1273 | E1273Y | E | 1.2e-08 | 3e-08 | 8.2e-08 | 4.133E-08 | 3.635E-08 | 1.200E-08 | 8.200E-08 |
| 25460 | 1273 | E1273V | E | 9.5e-08 | 2.3e-08 | 1.2e-07 | 7.933E-08 | 5.036E-08 | 2.300E-08 | 1.200E-07 |
| 25461 | 1274 | V1274A | E | 2.2e-08 | 1.2e-07 | 3.6e-08 | 5.933E-08 | 5.300E-08 | 2.200E-08 | 1.200E-07 |
| 25462 | 1274 | V1274R | E | 6.3e-08 | 1.2e-08 | 6.4e-08 | 4.633E-08 | 2.974E-08 | 1.200E-08 | 6.400E-08 |
| 25463 | 1274 | V1274N | E | 1.6e-08 | 4.3e-08 | 2e-08 | 2.633E-08 | 1.457E-08 | 1.600E-08 | 4.300E-08 |
| 25464 | 1274 | V1274D | E | 4.9e-08 | 4.7e-08 | 6.8e-08 | 5.467E-08 | 1.159E-08 | 4.700E-08 | 6.800E-08 |
| 25465 | 1274 | V1274C | E | 6.3e-08 | 6.1e-08 | 3.8e-08 | 5.400E-08 | 1.389E-08 | 3.800E-08 | 6.300E-08 |
| 25466 | 1274 | V1274E | E | 7.1e-08 | 7.3e-09 | 2.6e-08 | 3.477E-08 | 3.274E-08 | 7.300E-09 | 7.100E-08 |
| 25467 | 1274 | V1274Q | E | 5e-08 | 2.9e-08 | 4.4e-08 | 4.100E-08 | 1.082E-08 | 2.900E-08 | 5.000E-08 |
| 25468 | 1274 | V1274G | E | 2.8e-08 | 2.2e-08 | 5.4e-08 | 3.467E-08 | 1.701E-08 | 2.200E-08 | 5.400E-08 |
| 25469 | 1274 | V1274H | E | 5e-08 | 2.7e-08 | 2.8e-08 | 3.500E-08 | 1.300E-08 | 2.700E-08 | 5.000E-08 |
| 25470 | 1274 | V1274I | E | 3.4e-08 | 2.9e-08 | 9e-08 | 5.100E-08 | 3.387E-08 | 2.900E-08 | 9.000E-08 |
| 25471 | 1274 | V1274L | E | 3.8e-08 | 4.6e-08 | 3.6e-08 | 4.000E-08 | 5.292E-09 | 3.600E-08 | 4.600E-08 |
| 25472 | 1274 | V1274K | E | 5.1e-08 | 8.7e-09 | 9.2e-08 | 5.057E-08 | 4.165E-08 | 8.700E-09 | 9.200E-08 |
| 25473 | 1274 | V1274M | E | 4.5e-08 | 5.6e-08 | 6.5e-08 | 5.533E-08 | 1.002E-08 | 4.500E-08 | 6.500E-08 |
| 25474 | 1274 | V1274F | E | 4e-08 | 3.6e-08 | 4.2e-08 | 3.933E-08 | 3.055E-09 | 3.600E-08 | 4.200E-08 |
| 25475 | 1274 | V1274P | E | 2.5e-08 | 3.5e-08 | 6.2e-09 | 2.207E-08 | 1.462E-08 | 6.200E-09 | 3.500E-08 |
| 25476 | 1274 | V1274S | E | 5.1e-08 | 8e-08 | 3.3e-08 | 5.467E-08 | 2.371E-08 | 3.300E-08 | 8.000E-08 |
| 25477 | 1274 | V1274T | E | 3.2e-08 | 8.3e-08 | 5.1e-08 | 5.533E-08 | 2.577E-08 | 3.200E-08 | 8.300E-08 |
| 25478 | 1274 | V1274W | E | 6.7e-08 | 1.6e-08 | 6.6e-08 | 4.967E-08 | 2.916E-08 | 1.600E-08 | 6.700E-08 |
| 25479 | 1274 | V1274Y | E | 5.2e-08 | 1.2e-08 | 2.3e-08 | 2.900E-08 | 2.066E-08 | 1.200E-08 | 5.200E-08 |
| 25480 | 1274 | V1274V | E | 3.9e-08 | 4.3e-08 | 3.8e-08 | 4.000E-08 | 2.646E-09 | 3.800E-08 | 4.300E-08 |
| 25481 | 1275 | Q1275A | E | 2.8e-08 | 4.2e-08 | 7.7e-08 | 4.900E-08 | 2.524E-08 | 2.800E-08 | 7.700E-08 |
| 25482 | 1275 | Q1275R | E | 3.2e-08 | 7.5e-08 | 7.1e-08 | 5.933E-08 | 2.376E-08 | 3.200E-08 | 7.500E-08 |
| 25483 | 1275 | Q1275N | E | 1.3e-07 | 4.4e-08 | 4.7e-08 | 7.367E-08 | 4.881E-08 | 4.400E-08 | 1.300E-07 |
| 25484 | 1275 | Q1275D | E | 5e-08 | 6.2e-08 | 9e-08 | 6.733E-08 | 2.053E-08 | 5.000E-08 | 9.000E-08 |
| 25485 | 1275 | Q1275C | E | 9.8e-08 | 9.8e-08 | 2.5e-08 | 7.367E-08 | 4.215E-08 | 2.500E-08 | 9.800E-08 |
| 25486 | 1275 | Q1275E | E | 5.6e-08 | 4.5e-08 | 7e-08 | 5.700E-08 | 1.253E-08 | 4.500E-08 | 7.000E-08 |
| 25487 | 1275 | Q1275Q | E | 5.8e-09 | 5.6e-08 | 5.1e-08 | 3.760E-08 | 2.765E-08 | 5.800E-09 | 5.600E-08 |
| 25488 | 1275 | Q1275G | E | 1.2e-07 | 6.5e-08 | 3.3e-08 | 7.267E-08 | 4.400E-08 | 3.300E-08 | 1.200E-07 |
| 25489 | 1275 | Q1275H | E | 2.5e-08 | 4.4e-08 | 2.6e-08 | 3.167E-08 | 1.069E-08 | 2.500E-08 | 4.400E-08 |
| 25490 | 1275 | Q1275I | E | 6.9e-08 | 5.2e-08 | 2.9e-08 | 5.000E-08 | 2.007E-08 | 2.900E-08 | 6.900E-08 |
| 25491 | 1275 | Q1275L | E | 2.3e-08 | 4.6e-08 | 1.9e-08 | 2.933E-08 | 1.457E-08 | 1.900E-08 | 4.600E-08 |
| 25492 | 1275 | Q1275K | E | 3.6e-08 | 2.4e-08 | 7.5e-08 | 4.500E-08 | 2.666E-08 | 2.400E-08 | 7.500E-08 |
| 25493 | 1275 | Q1275M | E | 6.7e-08 | 4e-08 | 9.4e-08 | 6.700E-08 | 2.700E-08 | 4.000E-08 | 9.400E-08 |
| 25494 | 1275 | Q1275F | E | 3e-08 | 7.9e-08 | 2.7e-08 | 4.533E-08 | 2.919E-08 | 2.700E-08 | 7.900E-08 |
| 25495 | 1275 | Q1275P | E | 3e-08 | 5.2e-08 | 6.2e-08 | 4.800E-08 | 1.637E-08 | 3.000E-08 | 6.200E-08 |
| 25496 | 1275 | Q1275S | E | 6.3e-08 | 2.9e-08 | 7.2e-08 | 5.467E-08 | 2.268E-08 | 2.900E-08 | 7.200E-08 |
| 25497 | 1275 | Q1275T | E | 4.1e-08 | 2.1e-08 | 9.3e-09 | 2.377E-08 | 1.603E-08 | 9.300E-09 | 4.100E-08 |
| 25498 | 1275 | Q1275W | E | 1.5e-07 | 8.7e-08 | 4.7e-08 | 9.467E-08 | 5.193E-08 | 4.700E-08 | 1.500E-07 |
| 25499 | 1275 | Q1275Y | E | 4.1e-08 | 5.6e-08 | 3e-08 | 4.233E-08 | 1.305E-08 | 3.000E-08 | 5.600E-08 |

| | | | | | | | | | | |
|---|---|---|---|---|---|---|---|---|---|---|
| 25500 | 1275 | Q1275V | E | 8.5e-08 | 4.9e-08 | 2.7e-07 | 1.347E-07 | 1.186E-07 | 4.900E-08 | 2.700E-07 |
| 25501 | 1276 | L1276A | E | 2.8e-08 | 3.3e-08 | 2.2e-08 | 2.767E-08 | 5.508E-09 | 2.200E-08 | 3.300E-08 |
| 25502 | 1276 | L1276R | E | 3.5e-08 | 1.4e-07 | 5.4e-08 | 7.633E-08 | 5.595E-08 | 3.500E-08 | 1.400E-07 |
| 25503 | 1276 | L1276N | E | 4.8e-08 | 1.8e-08 | 5.8e-08 | 4.133E-08 | 2.082E-08 | 1.800E-08 | 5.800E-08 |
| 25504 | 1276 | L1276D | E | 3.8e-08 | 1.6e-07 | 8.4e-08 | 9.400E-08 | 6.161E-08 | 3.800E-08 | 1.600E-07 |
| 25505 | 1276 | L1276C | E | 2.4e-08 | 6.4e-08 | 1.7e-08 | 3.500E-08 | 2.536E-08 | 1.700E-08 | 6.400E-08 |
| 25506 | 1276 | L1276E | E | 1.6e-08 | 1.1e-07 | 2.8e-08 | 5.133E-08 | 5.116E-08 | 1.600E-08 | 1.100E-07 |
| 25507 | 1276 | L1276Q | E | 1.4e-08 | 6.7e-08 | 8e-08 | 5.367E-08 | 3.496E-08 | 1.400E-08 | 8.000E-08 |
| 25508 | 1276 | L1276G | E | 1.8e-08 | 6.4e-08 | 9.6e-09 | 3.053E-08 | 2.929E-08 | 9.600E-09 | 6.400E-08 |
| 25509 | 1276 | L1276H | E | 4.7e-08 | 3.7e-08 | 5.8e-08 | 4.733E-08 | 1.050E-08 | 3.700E-08 | 5.800E-08 |
| 25510 | 1276 | L1276I | E | 4.2e-08 | 6.6e-08 | 6.6e-08 | 5.800E-08 | 1.386E-08 | 4.200E-08 | 6.600E-08 |
| 25511 | 1276 | L1276L | E | 5.5e-08 | 4.5e-08 | 2.8e-08 | 4.267E-08 | 1.365E-08 | 2.800E-08 | 5.500E-08 |
| 25512 | 1276 | L1276K | E | 2.9e-08 | 4.4e-08 | 2.8e-08 | 3.367E-08 | 8.963E-09 | 2.800E-08 | 4.400E-08 |
| 25513 | 1276 | L1276M | E | 7.2e-08 | 2e-08 | 4.4e-08 | 4.533E-08 | 2.603E-08 | 2.000E-08 | 7.200E-08 |
| 25514 | 1276 | L1276F | E | 2.2e-08 | 3.4e-08 | 7e-08 | 4.200E-08 | 2.498E-08 | 2.200E-08 | 7.000E-08 |
| 25515 | 1276 | L1276P | E | 1.1e-07 | 1.6e-07 | 5.3e-08 | 1.077E-07 | 5.354E-08 | 5.300E-08 | 1.600E-07 |
| 25516 | 1276 | L1276S | E | 4.6e-08 | 5e-08 | 3.2e-08 | 4.267E-08 | 9.452E-09 | 3.200E-08 | 5.000E-08 |
| 25517 | 1276 | L1276T | E | 2.3e-08 | 1.7e-08 | 9.3e-08 | 4.433E-08 | 4.225E-08 | 1.700E-08 | 9.300E-08 |
| 25518 | 1276 | L1276W | E | 1.2e-07 | 2.4e-08 | 4.5e-08 | 6.300E-08 | 5.047E-08 | 2.400E-08 | 1.200E-07 |
| 25519 | 1276 | L1276Y | E | 6.6e-08 | 6.3e-08 | 1.4e-08 | 4.767E-08 | 2.919E-08 | 1.400E-08 | 6.600E-08 |
| 25520 | 1276 | L1276V | E | 4.4e-08 | 2.8e-08 | 5.1e-08 | 4.100E-08 | 1.179E-08 | 2.800E-08 | 5.100E-08 |
| 25521 | 1277 | V1277A | E | 4e-08 | 3.8e-08 | 1.5e-08 | 3.100E-08 | 1.389E-08 | 1.500E-08 | 4.000E-08 |
| 25522 | 1277 | V1277R | E | 3.8e-08 | 9.7e-08 | 2e-08 | 5.167E-08 | 4.028E-08 | 2.000E-08 | 9.700E-08 |
| 25523 | 1277 | V1277N | E | 3.3e-08 | 1e-07 | 2.7e-08 | 5.333E-08 | 4.053E-08 | 2.700E-08 | 1.000E-07 |
| 25524 | 1277 | V1277D | E | 8.5e-08 | 3.5e-08 | 7.9e-08 | 6.633E-08 | 2.730E-08 | 3.500E-08 | 8.500E-08 |
| 25525 | 1277 | V1277C | E | 4.3e-08 | 3.4e-08 | 4e-08 | 3.900E-08 | 4.583E-09 | 3.400E-08 | 4.300E-08 |
| 25526 | 1277 | V1277E | E | 2e-08 | 1.3e-08 | 1.3e-07 | 5.433E-08 | 6.562E-08 | 1.300E-08 | 1.300E-07 |
| 25527 | 1277 | V1277Q | E | 6.4e-08 | 8.4e-09 | 1.8e-08 | 3.013E-08 | 2.972E-08 | 8.400E-09 | 6.400E-08 |
| 25528 | 1277 | V1277G | E | 5.1e-08 | 2e-08 | 3e-08 | 3.367E-08 | 1.582E-08 | 2.000E-08 | 5.100E-08 |
| 25529 | 1277 | V1277H | E | 7.3e-08 | 2.7e-08 | 7.9e-08 | 5.967E-08 | 2.845E-08 | 2.700E-08 | 7.900E-08 |
| 25530 | 1277 | V1277I | E | 5.9e-08 | 5.6e-08 | 7.4e-08 | 6.300E-08 | 9.644E-09 | 5.600E-08 | 7.400E-08 |
| 25531 | 1277 | V1277L | E | 6.7e-08 | 7.5e-08 | 2.1e-08 | 5.433E-08 | 2.914E-08 | 2.100E-08 | 7.500E-08 |
| 25532 | 1277 | V1277K | E | 9.1e-08 | 2.6e-08 | 4.3e-08 | 5.333E-08 | 3.371E-08 | 2.600E-08 | 9.100E-08 |
| 25533 | 1277 | V1277M | E | 9.6e-08 | 8.8e-08 | 3.4e-08 | 7.267E-08 | 3.372E-08 | 3.400E-08 | 9.600E-08 |
| 25534 | 1277 | V1277F | E | 7.2e-08 | 4.3e-08 | 2.1e-08 | 4.533E-08 | 2.558E-08 | 2.100E-08 | 7.200E-08 |
| 25535 | 1277 | V1277P | E | 2.6e-08 | 2.3e-08 | 5.7e-08 | 3.533E-08 | 1.882E-08 | 2.300E-08 | 5.700E-08 |
| 25536 | 1277 | V1277S | E | 5.5e-08 | 2.9e-08 | 1.9e-08 | 3.433E-08 | 1.858E-08 | 1.900E-08 | 5.500E-08 |
| 25537 | 1277 | V1277T | E | 7.1e-08 | 4.6e-08 | 2.2e-08 | 4.633E-08 | 2.450E-08 | 2.200E-08 | 7.100E-08 |
| 25538 | 1277 | V1277W | E | 7.5e-08 | 8.5e-09 | 3.2e-08 | 3.850E-08 | 3.372E-08 | 8.500E-09 | 7.500E-08 |
| 25539 | 1277 | V1277Y | E | 7.7e-08 | 4.2e-08 | 6e-08 | 5.967E-08 | 1.750E-08 | 4.200E-08 | 7.700E-08 |
| 25540 | 1277 | V1277V | E | 8e-08 | 4.2e-08 | 3.9e-08 | 5.367E-08 | 2.285E-08 | 3.900E-08 | 8.000E-08 |
| 25541 | 1278 | E1278A | E | 8.6e-08 | 6.1e-08 | 4e-08 | 6.233E-08 | 2.303E-08 | 4.000E-08 | 8.600E-08 |
| 25542 | 1278 | E1278R | E | 4.4e-08 | 3.6e-08 | 8.8e-08 | 5.600E-08 | 2.800E-08 | 3.600E-08 | 8.800E-08 |
| 25543 | 1278 | E1278N | E | 3.9e-08 | 5.9e-08 | 9.7e-08 | 6.500E-08 | 2.946E-08 | 3.900E-08 | 9.700E-08 |
| 25544 | 1278 | E1278D | E | 6.7e-08 | 2.8e-08 | 5.8e-08 | 5.100E-08 | 2.042E-08 | 2.800E-08 | 6.700E-08 |
| 25545 | 1278 | E1278C | E | 1.1e-07 | 5.2e-08 | 5.6e-08 | 7.267E-08 | 3.239E-08 | 5.200E-08 | 1.100E-07 |
| 25546 | 1278 | E1278E | E | 2.8e-08 | 4.7e-08 | 1e-07 | 5.833E-08 | 3.731E-08 | 2.800E-08 | 1.000E-07 |
| 25547 | 1278 | E1278Q | E | 3.7e-08 | 3.4e-08 | 6.8e-08 | 4.633E-08 | 1.882E-08 | 3.400E-08 | 6.800E-08 |
| 25548 | 1278 | E1278G | E | 9.4e-08 | 2.8e-08 | 1.5e-08 | 4.567E-08 | 4.236E-08 | 1.500E-08 | 9.400E-08 |
| 25549 | 1278 | E1278H | E | 6.7e-08 | 4.1e-08 | 8.3e-08 | 6.367E-08 | 2.120E-08 | 4.100E-08 | 8.300E-08 |
| 25550 | 1278 | E1278I | E | 4.2e-08 | 6.7e-08 | 2.8e-08 | 4.567E-08 | 1.976E-08 | 2.800E-08 | 6.700E-08 |
| 25551 | 1278 | E1278L | E | 7.5e-08 | 7.5e-08 | 6e-08 | 7.000E-08 | 8.660E-09 | 6.000E-08 | 7.500E-08 |
| 25552 | 1278 | E1278K | E | 8e-08 | 5.8e-08 | 1.8e-08 | 5.200E-08 | 3.143E-08 | 1.800E-08 | 8.000E-08 |
| 25553 | 1278 | E1278M | E | 1.6e-08 | 7.6e-08 | 7.2e-08 | 5.467E-08 | 3.355E-08 | 1.600E-08 | 7.600E-08 |
| 25554 | 1278 | E1278F | E | 1.9e-08 | 1e-08 | 1.2e-07 | 4.967E-08 | 6.108E-08 | 1.000E-08 | 1.200E-07 |
| 25555 | 1278 | E1278P | E | 2.9e-08 | 5.6e-08 | 5.7e-08 | 4.733E-08 | 1.589E-08 | 2.900E-08 | 5.700E-08 |
| 25556 | 1278 | E1278S | E | 2.5e-08 | 9.2e-08 | 2.5e-08 | 4.733E-08 | 3.868E-08 | 2.500E-08 | 9.200E-08 |
| 25557 | 1278 | E1278T | E | 3.6e-08 | 1.7e-07 | 4.7e-08 | 8.433E-08 | 7.439E-08 | 3.600E-08 | 1.700E-07 |
| 25558 | 1278 | E1278W | E | 3.2e-08 | 3.6e-08 | 1.1e-07 | 5.933E-08 | 4.392E-08 | 3.200E-08 | 1.100E-07 |
| 25559 | 1278 | E1278Y | E | 6.6e-08 | 6.9e-08 | 5.4e-08 | 6.300E-08 | 7.937E-09 | 5.400E-08 | 6.900E-08 |
| 25560 | 1278 | E1278V | E | 6e-08 | 1.1e-07 | 4.2e-08 | 7.067E-08 | 3.523E-08 | 4.200E-08 | 1.100E-07 |
| 25561 | 1279 | S1279A | E | 7.7e-08 | 5.3e-08 | 2.7e-08 | 5.233E-08 | 2.501E-08 | 2.700E-08 | 7.700E-08 |
| 25562 | 1279 | S1279R | E | 2.3e-08 | 6.8e-08 | 3.3e-08 | 4.133E-08 | 2.363E-08 | 2.300E-08 | 6.800E-08 |
| 25563 | 1279 | S1279N | E | 4.6e-08 | 1.3e-07 | 5.2e-08 | 7.600E-08 | 4.686E-08 | 4.600E-08 | 1.300E-07 |
| 25564 | 1279 | S1279D | E | 1e-08 | 6.4e-08 | 1.4e-08 | 2.933E-08 | 3.009E-08 | 1.000E-08 | 6.400E-08 |
| 25565 | 1279 | S1279C | E | 4e-08 | 8.3e-08 | 2.5e-08 | 4.933E-08 | 3.011E-08 | 2.500E-08 | 8.300E-08 |
| 25566 | 1279 | S1279E | E | 3.5e-08 | 6.6e-08 | 4.4e-08 | 4.833E-08 | 1.595E-08 | 3.500E-08 | 6.600E-08 |
| 25567 | 1279 | S1279Q | E | 5.5e-08 | 4.5e-08 | 5.6e-08 | 5.200E-08 | 6.083E-09 | 4.500E-08 | 5.600E-08 |
| 25568 | 1279 | S1279G | E | 3.2e-08 | 2.8e-08 | 4.1e-08 | 3.367E-08 | 6.658E-09 | 2.800E-08 | 4.100E-08 |
| 25569 | 1279 | S1279H | E | 6.2e-08 | 6.7e-08 | 1.4e-07 | 8.967E-08 | 4.366E-08 | 6.200E-08 | 1.400E-07 |
| 25570 | 1279 | S1279I | E | 1.6e-08 | 1.1e-08 | 8.8e-08 | 3.833E-08 | 4.309E-08 | 1.100E-08 | 8.800E-08 |
| 25571 | 1279 | S1279L | E | 2e-08 | 3.6e-08 | 2e-08 | 2.533E-08 | 9.238E-09 | 2.000E-08 | 3.600E-08 |
| 25572 | 1279 | S1279K | E | 4.2e-08 | 2.4e-08 | 2.3e-08 | 2.967E-08 | 1.069E-08 | 2.300E-08 | 4.200E-08 |
| 25573 | 1279 | S1279M | E | 2.6e-08 | 9.8e-09 | 4.9e-08 | 2.827E-08 | 1.970E-08 | 9.800E-09 | 4.900E-08 |
| 25574 | 1279 | S1279F | E | 5.3e-08 | 5.4e-08 | 4.1e-08 | 4.933E-08 | 7.234E-09 | 4.100E-08 | 5.400E-08 |
| 25575 | 1279 | S1279P | E | 3.4e-08 | 4.4e-08 | 4.9e-08 | 4.233E-08 | 7.638E-09 | 3.400E-08 | 4.900E-08 |
| 25576 | 1279 | S1279S | E | 6.1e-08 | 4.8e-08 | 5.5e-08 | 5.467E-08 | 6.506E-09 | 4.800E-08 | 6.100E-08 |
| 25577 | 1279 | S1279T | E | 2.5e-08 | 3.8e-08 | 1.2e-07 | 6.100E-08 | 5.151E-08 | 2.500E-08 | 1.200E-07 |
| 25578 | 1279 | S1279W | E | 4.6e-08 | 2.7e-08 | 7e-08 | 4.767E-08 | 2.155E-08 | 2.700E-08 | 7.000E-08 |
| 25579 | 1279 | S1279Y | E | 4e-08 | 5.4e-08 | 3.9e-08 | 4.433E-08 | 8.386E-09 | 3.900E-08 | 5.400E-08 |
| 25580 | 1279 | S1279V | E | 5.8e-08 | 5.3e-08 | 1.1e-07 | 7.367E-08 | 3.156E-08 | 5.300E-08 | 1.100E-07 |
| 25581 | 1280 | G1280A | E | 2.5e-08 | 5.5e-08 | 2.5e-08 | 3.500E-08 | 1.732E-08 | 2.500E-08 | 5.500E-08 |
| 25582 | 1280 | G1280R | E | 3.5e-08 | 6.6e-08 | 6.7e-08 | 5.600E-08 | 1.819E-08 | 3.500E-08 | 6.700E-08 |
| 25583 | 1280 | G1280N | E | 7e-08 | 3.2e-08 | 7.4e-08 | 5.867E-08 | 2.318E-08 | 3.200E-08 | 7.400E-08 |
| 25584 | 1280 | G1280D | E | 4.2e-08 | 1.4e-07 | 7.5e-08 | 8.567E-08 | 4.986E-08 | 4.200E-08 | 1.400E-07 |

| | | | | | | | | | | |
|---|---|---|---|---|---|---|---|---|---|---|
| 25585 | 1280 | G1280C | E | 3e-08 | 7.4e-08 | 2.7e-08 | 4.367E-08 | 2.631E-08 | 2.700E-08 | 7.400E-08 |
| 25586 | 1280 | G1280E | E | 4e-08 | 5.5e-08 | 1.1e-08 | 3.533E-08 | 2.237E-08 | 1.100E-08 | 5.500E-08 |
| 25587 | 1280 | G1280Q | E | 2.5e-08 | 5.1e-08 | 5.2e-08 | 4.267E-08 | 1.531E-08 | 2.500E-08 | 5.200E-08 |
| 25588 | 1280 | G1280G | E | 3.1e-08 | 4.4e-08 | 4.3e-08 | 3.933E-08 | 7.234E-09 | 3.100E-08 | 4.400E-08 |
| 25589 | 1280 | G1280H | E | 3.9e-08 | 2.9e-08 | 6e-08 | 4.267E-08 | 1.582E-08 | 2.900E-08 | 6.000E-08 |
| 25590 | 1280 | G1280I | E | 1.8e-07 | 4.7e-08 | 2.3e-08 | 8.333E-08 | 8.457E-08 | 2.300E-08 | 1.800E-07 |
| 25591 | 1280 | G1280L | E | 5.8e-08 | 2.4e-08 | 5.9e-08 | 4.700E-08 | 1.992E-08 | 2.400E-08 | 5.900E-08 |
| 25592 | 1280 | G1280K | E | 3.3e-08 | 2.7e-08 | 5.5e-08 | 3.833E-08 | 1.474E-08 | 2.700E-08 | 5.500E-08 |
| 25593 | 1280 | G1280M | E | 2.7e-08 | 5.5e-08 | 2.5e-08 | 3.567E-08 | 1.677E-08 | 2.500E-08 | 5.500E-08 |
| 25594 | 1280 | G1280F | E | 3.5e-08 | 2.4e-08 | 8.8e-08 | 4.900E-08 | 3.422E-08 | 2.400E-08 | 8.800E-08 |
| 25595 | 1280 | G1280P | E | 2.6e-08 | 1.8e-08 | 4e-08 | 2.800E-08 | 1.114E-08 | 1.800E-08 | 4.000E-08 |
| 25596 | 1280 | G1280S | E | 3.9e-08 | 2.7e-08 | 3e-08 | 3.200E-08 | 6.245E-09 | 2.700E-08 | 3.900E-08 |
| 25597 | 1280 | G1280T | E | 8.7e-08 | 6.4e-08 | 7.7e-08 | 7.600E-08 | 1.153E-08 | 6.400E-08 | 8.700E-08 |
| 25598 | 1280 | G1280W | E | 8.7e-08 | 5.4e-08 | 2.1e-08 | 5.400E-08 | 3.300E-08 | 2.100E-08 | 8.700E-08 |
| 25599 | 1280 | G1280Y | E | 3e-08 | 2.7e-08 | 7.4e-08 | 4.367E-08 | 2.631E-08 | 2.700E-08 | 7.400E-08 |
| 25600 | 1280 | G1280V | E | 2.6e-08 | 9.5e-08 | 3.7e-08 | 5.267E-08 | 3.707E-08 | 2.600E-08 | 9.500E-08 |
| 25601 | 1281 | G1281A | E | 2.4e-08 | 2.1e-08 | 2.7e-08 | 2.400E-08 | 3.000E-09 | 2.100E-08 | 2.700E-08 |
| 25602 | 1281 | G1281R | E | 8.9e-08 | 5.3e-08 | 4.7e-08 | 6.300E-08 | 2.272E-08 | 4.700E-08 | 8.900E-08 |
| 25603 | 1281 | G1281N | E | 4.6e-08 | 4.1e-08 | 5e-08 | 4.567E-08 | 4.509E-09 | 4.100E-08 | 5.000E-08 |
| 25604 | 1281 | G1281D | E | 3.9e-08 | 4.4e-08 | 2.9e-08 | 3.733E-08 | 7.638E-09 | 2.900E-08 | 4.400E-08 |
| 25605 | 1281 | G1281C | E | 3.6e-08 | 5.7e-08 | 3.4e-08 | 4.233E-08 | 1.274E-08 | 3.400E-08 | 5.700E-08 |
| 25606 | 1281 | G1281E | E | 5e-08 | 2.8e-08 | 5e-08 | 4.267E-08 | 1.270E-08 | 2.800E-08 | 5.000E-08 |
| 25607 | 1281 | G1281Q | E | 2.1e-08 | 9.4e-08 | 6.5e-08 | 6.000E-08 | 3.676E-08 | 2.100E-08 | 9.400E-08 |
| 25608 | 1281 | G1281G | E | 2.3e-08 | 6.2e-08 | 5.9e-08 | 4.800E-08 | 2.170E-08 | 2.300E-08 | 6.200E-08 |
| 25609 | 1281 | G1281H | E | 4.5e-08 | 2.9e-08 | 2.3e-08 | 3.233E-08 | 1.137E-08 | 2.300E-08 | 4.500E-08 |
| 25610 | 1281 | G1281I | E | 1.5e-08 | 7.4e-08 | 3.5e-08 | 4.133E-08 | 3.001E-08 | 1.500E-08 | 7.400E-08 |
| 25611 | 1281 | G1281L | E | 4.1e-08 | 1.1e-07 | 8.1e-08 | 7.733E-08 | 3.465E-08 | 4.100E-08 | 1.100E-07 |
| 25612 | 1281 | G1281K | E | 3.4e-08 | 9.2e-08 | 4.5e-08 | 5.700E-08 | 3.081E-08 | 3.400E-08 | 9.200E-08 |
| 25613 | 1281 | G1281M | E | 1.1e-07 | 9.9e-08 | 8.1e-08 | 9.667E-08 | 1.464E-08 | 8.100E-08 | 1.100E-07 |
| 25614 | 1281 | G1281F | E | 3.9e-08 | 5.1e-08 | 4.5e-08 | 4.500E-08 | 6.000E-09 | 3.900E-08 | 5.100E-08 |
| 25615 | 1281 | G1281P | E | 1.9e-08 | 2.8e-08 | 8.8e-08 | 4.500E-08 | 3.751E-08 | 1.900E-08 | 8.800E-08 |
| 25616 | 1281 | G1281S | E | 4e-08 | 2.3e-08 | 4.9e-08 | 3.733E-08 | 1.320E-08 | 2.300E-08 | 4.900E-08 |
| 25617 | 1281 | G1281T | E | 5.5e-08 | 6.1e-08 | 1.8e-08 | 4.467E-08 | 2.329E-08 | 1.800E-08 | 6.100E-08 |
| 25618 | 1281 | G1281W | E | 4.7e-08 | 8.1e-08 | 2.8e-08 | 5.200E-08 | 2.685E-08 | 2.800E-08 | 8.100E-08 |
| 25619 | 1281 | G1281Y | E | 3.1e-08 | 4.4e-08 | 4.3e-08 | 3.933E-08 | 7.234E-09 | 3.100E-08 | 4.400E-08 |
| 25620 | 1281 | G1281V | E | 2.9e-08 | 1.3e-08 | 4.4e-08 | 2.867E-08 | 1.550E-08 | 1.300E-08 | 4.400E-08 |
| 25621 | 1282 | G1282A | E | 9.9e-08 | 5.4e-08 | 4.5e-08 | 6.600E-08 | 2.893E-08 | 4.500E-08 | 9.900E-08 |
| 25622 | 1282 | G1282R | E | 4.1e-08 | 1.3e-08 | 4.9e-08 | 3.433E-08 | 1.890E-08 | 1.300E-08 | 4.900E-08 |
| 25623 | 1282 | G1282N | E | 9.4e-08 | 2.6e-08 | 1.3e-08 | 4.433E-08 | 4.350E-08 | 1.300E-08 | 9.400E-08 |
| 25624 | 1282 | G1282D | E | 3.7e-08 | 3.9e-08 | 5.7e-08 | 4.433E-08 | 1.102E-08 | 3.700E-08 | 5.700E-08 |
| 25625 | 1282 | G1282C | E | 1.2e-07 | 5e-08 | 9.3e-08 | 8.767E-08 | 3.530E-08 | 5.000E-08 | 1.200E-07 |
| 25626 | 1282 | G1282E | E | 1.5e-08 | 3e-08 | 1.7e-08 | 2.067E-08 | 8.145E-09 | 1.500E-08 | 3.000E-08 |
| 25627 | 1282 | G1282Q | E | 3.5e-08 | 3.2e-08 | 3e-08 | 3.233E-08 | 2.517E-09 | 3.000E-08 | 3.500E-08 |
| 25628 | 1282 | G1282G | E | 3.3e-08 | 3.7e-08 | 2.8e-08 | 3.267E-08 | 4.509E-09 | 2.800E-08 | 3.700E-08 |
| 25629 | 1282 | G1282H | E | 4.6e-08 | 3.4e-08 | 3e-08 | 3.667E-08 | 8.327E-09 | 3.000E-08 | 4.600E-08 |
| 25630 | 1282 | G1282I | E | 3.3e-08 | 2.8e-08 | 1.6e-08 | 2.567E-08 | 8.737E-09 | 1.600E-08 | 3.300E-08 |
| 25631 | 1282 | G1282L | E | 5e-08 | 1.2e-07 | 2.9e-08 | 6.633E-08 | 4.765E-08 | 2.900E-08 | 1.200E-07 |
| 25632 | 1282 | G1282K | E | 8.2e-08 | 1.8e-08 | 7.8e-08 | 5.933E-08 | 3.585E-08 | 1.800E-08 | 8.200E-08 |
| 25633 | 1282 | G1282M | E | 6.1e-08 | 4.9e-08 | 3.9e-08 | 4.967E-08 | 1.102E-08 | 3.900E-08 | 6.100E-08 |
| 25634 | 1282 | G1282F | E | 3.1e-08 | 3e-08 | 4.6e-08 | 3.567E-08 | 8.963E-09 | 3.000E-08 | 4.600E-08 |
| 25635 | 1282 | G1282P | E | 3e-08 | 1.1e-08 | 7.2e-08 | 3.767E-08 | 3.121E-08 | 1.100E-08 | 7.200E-08 |
| 25636 | 1282 | G1282S | E | 2.1e-08 | 4.2e-08 | 3.9e-08 | 3.400E-08 | 1.136E-08 | 2.100E-08 | 4.200E-08 |
| 25637 | 1282 | G1282T | E | 4.2e-08 | 3.3e-08 | 6.1e-08 | 4.533E-08 | 1.429E-08 | 3.300E-08 | 6.100E-08 |
| 25638 | 1282 | G1282W | E | 2e-08 | 2.4e-08 | 4.1e-08 | 2.833E-08 | 1.115E-08 | 2.000E-08 | 4.100E-08 |
| 25639 | 1282 | G1282Y | E | 8.2e-08 | 6.1e-08 | 4.1e-08 | 6.133E-08 | 2.050E-08 | 4.100E-08 | 8.200E-08 |
| 25640 | 1282 | G1282V | E | 1.8e-08 | 6e-08 | 8.5e-08 | 5.433E-08 | 3.386E-08 | 1.800E-08 | 8.500E-08 |
| 25641 | 1283 | L1283A | E | 2.9e-08 | 3.6e-08 | 2.4e-08 | 2.967E-08 | 6.028E-09 | 2.400E-08 | 3.600E-08 |
| 25642 | 1283 | L1283R | E | 6.4e-08 | 5.1e-08 | 1.4e-08 | 4.300E-08 | 2.594E-08 | 1.400E-08 | 6.400E-08 |
| 25643 | 1283 | L1283N | E | 2.4e-08 | 5.8e-08 | 6.1e-08 | 4.767E-08 | 2.055E-08 | 2.400E-08 | 6.100E-08 |
| 25644 | 1283 | L1283D | E | 2.9e-08 | 9.6e-08 | 1.1e-07 | 7.833E-08 | 4.329E-08 | 2.900E-08 | 1.100E-07 |
| 25645 | 1283 | L1283C | E | 3.2e-08 | 3e-08 | 3.3e-08 | 3.167E-08 | 1.528E-09 | 3.000E-08 | 3.300E-08 |
| 25646 | 1283 | L1283E | E | 2.3e-08 | 8.8e-08 | 2.7e-08 | 4.600E-08 | 3.643E-08 | 2.300E-08 | 8.800E-08 |
| 25647 | 1283 | L1283Q | E | 4.9e-08 | 4.9e-08 | 2.2e-08 | 4.000E-08 | 1.559E-08 | 2.200E-08 | 4.900E-08 |
| 25648 | 1283 | L1283G | E | 6.3e-08 | 7.2e-08 | 5.5e-08 | 6.333E-08 | 8.505E-09 | 5.500E-08 | 7.200E-08 |
| 25649 | 1283 | L1283H | E | 8.5e-08 | 6.7e-08 | 4.9e-08 | 6.700E-08 | 1.800E-08 | 4.900E-08 | 8.500E-08 |
| 25650 | 1283 | L1283I | E | 4.5e-08 | 9.1e-08 | 1.6e-07 | 9.867E-08 | 5.788E-08 | 4.500E-08 | 1.600E-07 |
| 25651 | 1283 | L1283L | E | 5.8e-08 | 1e-07 | 3.3e-08 | 6.367E-08 | 3.386E-08 | 3.300E-08 | 1.000E-07 |
| 25652 | 1283 | L1283K | E | 5e-08 | 5e-08 | 6.9e-08 | 5.633E-08 | 1.097E-08 | 5.000E-08 | 6.900E-08 |
| 25653 | 1283 | L1283M | E | 2.2e-08 | 1.6e-08 | 4.1e-08 | 2.633E-08 | 1.305E-08 | 1.600E-08 | 4.100E-08 |
| 25654 | 1283 | L1283F | E | 4.7e-08 | 2.5e-08 | 6.8e-08 | 4.667E-08 | 2.150E-08 | 2.500E-08 | 6.800E-08 |
| 25655 | 1283 | L1283P | E | 1.1e-08 | 3.1e-08 | 5e-08 | 3.067E-08 | 1.950E-08 | 1.100E-08 | 5.000E-08 |
| 25656 | 1283 | L1283S | E | 2.1e-08 | 5.4e-08 | 5e-08 | 4.167E-08 | 1.801E-08 | 2.100E-08 | 5.400E-08 |
| 25657 | 1283 | L1283T | E | 5.1e-08 | 2.4e-08 | 6.4e-08 | 4.633E-08 | 2.040E-08 | 2.400E-08 | 6.400E-08 |
| 25658 | 1283 | L1283W | E | 7.6e-08 | 6.7e-08 | 1.1e-07 | 8.433E-08 | 2.268E-08 | 6.700E-08 | 1.100E-07 |
| 25659 | 1283 | L1283Y | E | 3.7e-08 | 4e-08 | 5.4e-08 | 4.367E-08 | 9.074E-09 | 3.700E-08 | 5.400E-08 |
| 25660 | 1283 | L1283V | E | 5.6e-08 | 1.7e-08 | 3.1e-08 | 3.467E-08 | 1.976E-08 | 1.700E-08 | 5.600E-08 |
| 25661 | 1284 | V1284A | E | 2.9e-08 | 2.8e-08 | 6e-08 | 3.900E-08 | 1.819E-08 | 2.800E-08 | 6.000E-08 |
| 25662 | 1284 | V1284R | E | 6.9e-08 | 2.9e-08 | 8.5e-08 | 6.100E-08 | 2.884E-08 | 2.900E-08 | 8.500E-08 |
| 25663 | 1284 | V1284N | E | 5.6e-08 | 5.4e-08 | 7.7e-08 | 6.233E-08 | 1.274E-08 | 5.400E-08 | 7.700E-08 |
| 25664 | 1284 | V1284D | E | 6.8e-08 | 9.5e-08 | 3.3e-08 | 6.533E-08 | 3.109E-08 | 3.300E-08 | 9.500E-08 |
| 25665 | 1284 | V1284C | E | 4.6e-08 | 5.6e-08 | 4e-08 | 4.733E-08 | 8.083E-09 | 4.000E-08 | 5.600E-08 |
| 25666 | 1284 | V1284E | E | 9.6e-08 | 4.1e-08 | 2.1e-08 | 5.267E-08 | 3.884E-08 | 2.100E-08 | 9.600E-08 |
| 25667 | 1284 | V1284Q | E | 2.1e-08 | 5.9e-08 | 4.1e-08 | 4.033E-08 | 1.901E-08 | 2.100E-08 | 5.900E-08 |
| 25668 | 1284 | V1284G | E | 7.4e-08 | 4.9e-08 | 3.1e-08 | 5.133E-08 | 2.159E-08 | 3.100E-08 | 7.400E-08 |
| 25669 | 1284 | V1284H | E | 1e-07 | 2.4e-08 | 6e-08 | 6.133E-08 | 3.802E-08 | 2.400E-08 | 1.000E-07 |

| | | | | | | | | | | |
|---|---|---|---|---|---|---|---|---|---|---|
| 25670 | 1284 | V1284I | E | 3.7e-08 | 7.2e-08 | 6e-08 | 5.633E-08 | 1.779E-08 | 3.700E-08 | 7.200E-08 |
| 25671 | 1284 | V1284L | E | 2.3e-08 | 2.3e-08 | 5e-08 | 3.200E-08 | 1.559E-08 | 2.300E-08 | 5.000E-08 |
| 25672 | 1284 | V1284K | E | 2.3e-08 | 4.5e-08 | 2.7e-08 | 3.167E-08 | 1.172E-08 | 2.300E-08 | 4.500E-08 |
| 25673 | 1284 | V1284M | E | 2.9e-08 | 4.5e-08 | 6.2e-08 | 4.533E-08 | 1.650E-08 | 2.900E-08 | 6.200E-08 |
| 25674 | 1284 | V1284F | E | 1.3e-07 | 2.9e-08 | 1.1e-07 | 8.967E-08 | 5.348E-08 | 2.900E-08 | 1.300E-07 |
| 25675 | 1284 | V1284P | E | 5.8e-08 | 2.9e-08 | 9.8e-08 | 6.167E-08 | 3.465E-08 | 2.900E-08 | 9.800E-08 |
| 25676 | 1284 | V1284S | E | 2.4e-08 | 4.2e-08 | 6.9e-08 | 4.500E-08 | 2.265E-08 | 2.400E-08 | 6.900E-08 |
| 25677 | 1284 | V1284T | E | 3.9e-08 | 7.7e-08 | 1.8e-08 | 4.467E-08 | 2.991E-08 | 1.800E-08 | 7.700E-08 |
| 25678 | 1284 | V1284W | E | 8.8e-08 | 4.6e-08 | 4.3e-08 | 5.900E-08 | 2.516E-08 | 4.300E-08 | 8.800E-08 |
| 25679 | 1284 | V1284Y | E | 1.1e-07 | 1e-07 | 9.2e-08 | 1.007E-07 | 9.018E-09 | 9.200E-08 | 1.100E-07 |
| 25680 | 1284 | V1284V | E | 5.4e-08 | 5.8e-08 | 6.6e-08 | 5.933E-08 | 6.110E-09 | 5.400E-08 | 6.600E-08 |
| 25681 | 1285 | Q1285A | E | 9.9e-09 | 5.6e-08 | 5.9e-08 | 4.163E-08 | 2.752E-08 | 9.900E-09 | 5.900E-08 |
| 25682 | 1285 | Q1285R | E | 5.4e-08 | 2.9e-08 | 7.9e-09 | 3.030E-08 | 2.308E-08 | 7.900E-09 | 5.400E-08 |
| 25683 | 1285 | Q1285N | E | 4.1e-08 | 1.8e-08 | 2.8e-08 | 2.900E-08 | 1.153E-08 | 1.800E-08 | 4.100E-08 |
| 25684 | 1285 | Q1285D | E | 5.4e-08 | 5.5e-08 | 3.3e-08 | 4.733E-08 | 1.242E-08 | 3.300E-08 | 5.500E-08 |
| 25685 | 1285 | Q1285C | E | 4e-08 | 4.7e-08 | 3.4e-08 | 4.033E-08 | 6.506E-09 | 3.400E-08 | 4.700E-08 |
| 25686 | 1285 | Q1285E | E | 5.5e-08 | 4.4e-08 | 2.6e-08 | 4.167E-08 | 1.464E-08 | 2.600E-08 | 5.500E-08 |
| 25687 | 1285 | Q1285Q | E | 3.8e-08 | 1.2e-08 | 8.8e-08 | 4.600E-08 | 3.863E-08 | 1.200E-08 | 8.800E-08 |
| 25688 | 1285 | Q1285G | E | 1.2e-07 | 3.5e-08 | 9.1e-08 | 8.200E-08 | 4.321E-08 | 3.500E-08 | 1.200E-07 |
| 25689 | 1285 | Q1285H | E | 1.2e-07 | 6.6e-08 | 8.2e-09 | 6.473E-08 | 5.591E-08 | 8.200E-09 | 1.200E-07 |
| 25690 | 1285 | Q1285I | E | 3e-08 | 4.8e-08 | 6.8e-08 | 4.867E-08 | 1.901E-08 | 3.000E-08 | 6.800E-08 |
| 25691 | 1285 | Q1285L | E | 1.1e-07 | 1.8e-08 | 1.8e-08 | 4.867E-08 | 5.312E-08 | 1.800E-08 | 1.100E-07 |
| 25692 | 1285 | Q1285K | E | 3.6e-08 | 9.9e-09 | 2.4e-08 | 2.330E-08 | 1.306E-08 | 9.900E-09 | 3.600E-08 |
| 25693 | 1285 | Q1285M | E | 3.4e-08 | 9.2e-08 | 2.4e-08 | 5.000E-08 | 3.672E-08 | 2.400E-08 | 9.200E-08 |
| 25694 | 1285 | Q1285F | E | 3.6e-08 | 3.3e-08 | 6e-08 | 4.300E-08 | 1.480E-08 | 3.300E-08 | 6.000E-08 |
| 25695 | 1285 | Q1285P | E | 4.7e-08 | 1.4e-08 | 6.7e-08 | 4.267E-08 | 2.676E-08 | 1.400E-08 | 6.700E-08 |
| 25696 | 1285 | Q1285S | E | 5.3e-08 | 8.4e-08 | 2.4e-08 | 5.367E-08 | 3.001E-08 | 2.400E-08 | 8.400E-08 |
| 25697 | 1285 | Q1285T | E | 6.8e-08 | 1.7e-08 | 5.3e-08 | 4.600E-08 | 2.621E-08 | 1.700E-08 | 6.800E-08 |
| 25698 | 1285 | Q1285W | E | 4.4e-08 | 3.9e-08 | 3e-08 | 3.767E-08 | 7.095E-09 | 3.000E-08 | 4.400E-08 |
| 25699 | 1285 | Q1285Y | E | 2e-08 | 5.2e-08 | 4.9e-08 | 4.033E-08 | 1.767E-08 | 2.000E-08 | 5.200E-08 |
| 25700 | 1285 | Q1285V | E | 8.4e-08 | 5.1e-08 | 2.7e-08 | 5.400E-08 | 2.862E-08 | 2.700E-08 | 8.400E-08 |
| 25701 | 1286 | P1286A | E | 2.5e-08 | 4.2e-08 | 1.4e-08 | 2.700E-08 | 1.411E-08 | 1.400E-08 | 4.200E-08 |
| 25702 | 1286 | P1286R | E | 1.7e-08 | 2.5e-08 | 2.5e-08 | 2.233E-08 | 4.619E-09 | 1.700E-08 | 2.500E-08 |
| 25703 | 1286 | P1286N | E | 2e-08 | 5.6e-08 | 1.1e-08 | 2.900E-08 | 2.381E-08 | 1.100E-08 | 5.600E-08 |
| 25704 | 1286 | P1286D | E | 5.4e-08 | 1.8e-08 | 7.1e-08 | 4.767E-08 | 2.706E-08 | 1.800E-08 | 7.100E-08 |
| 25705 | 1286 | P1286C | E | 3.7e-08 | 6.1e-08 | 3.2e-08 | 4.333E-08 | 1.550E-08 | 3.200E-08 | 6.100E-08 |
| 25706 | 1286 | P1286E | E | 1.1e-07 | 5.5e-08 | 8.3e-08 | 8.267E-08 | 2.750E-08 | 5.500E-08 | 1.100E-07 |
| 25707 | 1286 | P1286Q | E | 4.1e-08 | 4.3e-08 | 7.4e-08 | 5.267E-08 | 1.850E-08 | 4.100E-08 | 7.400E-08 |
| 25708 | 1286 | P1286G | E | 2.9e-08 | 7.4e-08 | 1.5e-08 | 3.933E-08 | 3.083E-08 | 1.500E-08 | 7.400E-08 |
| 25709 | 1286 | P1286H | E | 1.1e-07 | 4.4e-08 | 2.5e-08 | 5.967E-08 | 4.461E-08 | 2.500E-08 | 1.100E-07 |
| 25710 | 1286 | P1286I | E | 5.7e-08 | 4.7e-08 | 2.9e-08 | 4.433E-08 | 1.419E-08 | 2.900E-08 | 5.700E-08 |
| 25711 | 1286 | P1286L | E | 6.4e-08 | 5.1e-08 | 1.4e-07 | 8.500E-08 | 4.807E-08 | 5.100E-08 | 1.400E-07 |
| 25712 | 1286 | P1286K | E | 2.7e-08 | 3.5e-08 | 3e-08 | 3.067E-08 | 4.041E-09 | 2.700E-08 | 3.500E-08 |
| 25713 | 1286 | P1286M | E | 2.5e-08 | 1.6e-08 | 1.7e-08 | 1.933E-08 | 4.933E-09 | 1.600E-08 | 2.500E-08 |
| 25714 | 1286 | P1286F | E | 1.6e-07 | 4.8e-08 | 5.2e-08 | 8.667E-08 | 6.354E-08 | 4.800E-08 | 1.600E-07 |
| 25715 | 1286 | P1286P | E | 1.3e-08 | 3.3e-08 | 4e-08 | 2.867E-08 | 1.401E-08 | 1.300E-08 | 4.000E-08 |
| 25716 | 1286 | P1286S | E | 2.5e-08 | 3.7e-08 | 3.5e-08 | 3.233E-08 | 6.429E-09 | 2.500E-08 | 3.700E-08 |
| 25717 | 1286 | P1286T | E | 3.7e-08 | 3.3e-08 | 2.6e-08 | 3.200E-08 | 5.568E-09 | 2.600E-08 | 3.700E-08 |
| 25718 | 1286 | P1286W | E | 8e-08 | 1.2e-08 | 2.4e-08 | 3.867E-08 | 3.630E-08 | 1.200E-08 | 8.000E-08 |
| 25719 | 1286 | P1286Y | E | 4.3e-08 | 2.7e-08 | 5e-08 | 4.000E-08 | 1.179E-08 | 2.700E-08 | 5.000E-08 |
| 25720 | 1286 | P1286V | E | 3e-08 | 4.2e-08 | 3.3e-08 | 3.500E-08 | 6.245E-09 | 3.000E-08 | 4.200E-08 |
| 25721 | 1287 | G1287A | E | 2.4e-08 | 1e-07 | 4.3e-08 | 5.567E-08 | 3.955E-08 | 2.400E-08 | 1.000E-07 |
| 25722 | 1287 | G1287R | E | 2.2e-08 | 4.7e-08 | 4.1e-08 | 3.667E-08 | 1.305E-08 | 2.200E-08 | 4.700E-08 |
| 25723 | 1287 | G1287N | E | 5.4e-08 | 5.6e-08 | 3.5e-08 | 4.833E-08 | 1.159E-08 | 3.500E-08 | 5.600E-08 |
| 25724 | 1287 | G1287D | E | 2.8e-08 | 4.5e-08 | 5.8e-08 | 4.367E-08 | 1.504E-08 | 2.800E-08 | 5.800E-08 |
| 25725 | 1287 | G1287C | E | 5.2e-08 | 6.3e-08 | 1.4e-08 | 4.300E-08 | 2.571E-08 | 1.400E-08 | 6.300E-08 |
| 25726 | 1287 | G1287E | E | 4.2e-08 | 3.1e-08 | 1.7e-08 | 3.000E-08 | 1.253E-08 | 1.700E-08 | 4.200E-08 |
| 25727 | 1287 | G1287Q | E | 1.4e-08 | 6.8e-09 | 6.6e-08 | 2.893E-08 | 3.230E-08 | 6.800E-09 | 6.600E-08 |
| 25728 | 1287 | G1287G | E | 5.8e-08 | 1.3e-07 | 2.1e-08 | 6.967E-08 | 5.543E-08 | 2.100E-08 | 1.300E-07 |
| 25729 | 1287 | G1287H | E | 2.4e-08 | 3.6e-08 | 1.3e-08 | 2.433E-08 | 1.150E-08 | 1.300E-08 | 3.600E-08 |
| 25730 | 1287 | G1287I | E | 1.1e-07 | 8e-08 | 4.5e-08 | 7.833E-08 | 3.253E-08 | 4.500E-08 | 1.100E-07 |
| 25731 | 1287 | G1287L | E | 3.1e-08 | 6.6e-08 | 5.2e-08 | 4.967E-08 | 1.762E-08 | 3.100E-08 | 6.600E-08 |
| 25732 | 1287 | G1287K | E | 1.4e-07 | 9e-08 | 8.6e-08 | 1.053E-07 | 3.009E-08 | 8.600E-08 | 1.400E-07 |
| 25733 | 1287 | G1287M | E | 2.1e-08 | 2e-08 | 5e-08 | 3.033E-08 | 1.704E-08 | 2.000E-08 | 5.000E-08 |
| 25734 | 1287 | G1287F | E | 2.5e-08 | 6e-08 | 3.6e-08 | 4.033E-08 | 1.790E-08 | 2.500E-08 | 6.000E-08 |
| 25735 | 1287 | G1287P | E | 5.1e-08 | 2e-08 | 1.1e-08 | 2.733E-08 | 2.098E-08 | 1.100E-08 | 5.100E-08 |
| 25736 | 1287 | G1287S | E | 5.4e-08 | 1.3e-08 | 4.8e-08 | 3.833E-08 | 2.214E-08 | 1.300E-08 | 5.400E-08 |
| 25737 | 1287 | G1287T | E | 1.6e-07 | 2.9e-08 | 4.1e-08 | 7.667E-08 | 7.242E-08 | 2.900E-08 | 1.600E-07 |
| 25738 | 1287 | G1287W | E | 6.7e-08 | 4.8e-08 | 1.8e-08 | 4.433E-08 | 2.470E-08 | 1.800E-08 | 6.700E-08 |
| 25739 | 1287 | G1287Y | E | 2.3e-08 | 6.4e-08 | 2.1e-08 | 3.600E-08 | 2.427E-08 | 2.100E-08 | 6.400E-08 |
| 25740 | 1287 | G1287V | E | 5.9e-08 | 6.4e-08 | 8e-08 | 6.767E-08 | 1.097E-08 | 5.900E-08 | 8.000E-08 |
| 25741 | 1288 | G1288A | E | 6.1e-08 | 6.9e-08 | 1.6e-08 | 4.867E-08 | 2.857E-08 | 1.600E-08 | 6.900E-08 |
| 25742 | 1288 | G1288R | E | 2.6e-08 | 3e-08 | 6.1e-08 | 3.900E-08 | 1.916E-08 | 2.600E-08 | 6.100E-08 |
| 25743 | 1288 | G1288N | E | 1e-08 | 3.7e-08 | 8.1e-08 | 4.267E-08 | 3.584E-08 | 1.000E-08 | 8.100E-08 |
| 25744 | 1288 | G1288D | E | 3.6e-08 | 8.4e-08 | 1.1e-07 | 7.667E-08 | 3.754E-08 | 3.600E-08 | 1.100E-07 |
| 25745 | 1288 | G1288C | E | 4.1e-08 | 3.5e-08 | 7.1e-08 | 4.900E-08 | 1.929E-08 | 3.500E-08 | 7.100E-08 |
| 25746 | 1288 | G1288E | E | 2.3e-08 | 3.4e-08 | 4e-08 | 3.233E-08 | 8.622E-09 | 2.300E-08 | 4.000E-08 |
| 25747 | 1288 | G1288Q | E | 5.7e-08 | 6e-08 | 8.9e-08 | 6.867E-08 | 1.767E-08 | 5.700E-08 | 8.900E-08 |
| 25748 | 1288 | G1288G | E | 2.7e-08 | 3.9e-08 | 4e-08 | 3.533E-08 | 7.234E-09 | 2.700E-08 | 4.000E-08 |
| 25749 | 1288 | G1288H | E | 1.6e-08 | 4.2e-08 | 6.2e-08 | 4.000E-08 | 2.307E-08 | 1.600E-08 | 6.200E-08 |
| 25750 | 1288 | G1288I | E | 3.8e-08 | 1.9e-08 | 5.2e-08 | 3.633E-08 | 1.656E-08 | 1.900E-08 | 5.200E-08 |
| 25751 | 1288 | G1288L | E | 4.2e-08 | 1.9e-08 | 7.7e-08 | 4.600E-08 | 2.921E-08 | 1.900E-08 | 7.700E-08 |
| 25752 | 1288 | G1288K | E | 1e-07 | 4.4e-08 | 7.6e-08 | 7.333E-08 | 2.810E-08 | 4.400E-08 | 1.000E-07 |
| 25753 | 1288 | G1288M | E | 3.7e-08 | 6.9e-08 | 1.5e-07 | 8.533E-08 | 5.824E-08 | 3.700E-08 | 1.500E-07 |
| 25754 | 1288 | G1288F | E | 2.8e-08 | 4.4e-08 | 6.3e-08 | 4.500E-08 | 1.752E-08 | 2.800E-08 | 6.300E-08 |

| | | | | | | | | | | |
|---|---|---|---|---|---|---|---|---|---|---|
| 25755 | 1288 | G1288P | E | 3.6e-08 | 3.9e-08 | 5.2e-08 | 4.233E-08 | 8.505E-09 | 3.600E-08 | 5.200E-08 |
| 25756 | 1288 | G1288S | E | 3.7e-08 | 6.5e-08 | 5.1e-08 | 5.100E-08 | 1.400E-08 | 3.700E-08 | 6.500E-08 |
| 25757 | 1288 | G1288T | E | 4.4e-08 | 2.8e-08 | 1.5e-07 | 7.400E-08 | 6.630E-08 | 2.800E-08 | 1.500E-07 |
| 25758 | 1288 | G1288W | E | 1.5e-07 | 8.1e-08 | 3.1e-08 | 8.733E-08 | 5.975E-08 | 3.100E-08 | 1.500E-07 |
| 25759 | 1288 | G1288Y | E | 5.8e-08 | 6.2e-08 | 5.6e-08 | 5.867E-08 | 3.055E-09 | 5.600E-08 | 6.200E-08 |
| 25760 | 1288 | G1288V | E | 2.9e-08 | 7.5e-08 | 4.6e-08 | 5.000E-08 | 2.326E-08 | 2.900E-08 | 7.500E-08 |
| 25761 | 1289 | S1289A | E | 2.5e-08 | 6e-08 | 9.4e-08 | 5.967E-08 | 3.450E-08 | 2.500E-08 | 9.400E-08 |
| 25762 | 1289 | S1289R | E | 1.1e-07 | 6.1e-08 | 1.5e-08 | 6.200E-08 | 4.751E-08 | 1.500E-08 | 1.100E-07 |
| 25763 | 1289 | S1289N | E | 4.2e-08 | 2e-08 | 4e-08 | 3.400E-08 | 1.217E-08 | 2.000E-08 | 4.200E-08 |
| 25764 | 1289 | S1289D | E | 5.3e-08 | 1.9e-08 | 5.7e-08 | 4.300E-08 | 2.088E-08 | 1.900E-08 | 5.700E-08 |
| 25765 | 1289 | S1289C | E | 1.9e-08 | 7.7e-08 | 4.2e-08 | 4.600E-08 | 2.921E-08 | 1.900E-08 | 7.700E-08 |
| 25766 | 1289 | S1289E | E | 1.9e-08 | 3.1e-08 | 2.8e-08 | 2.600E-08 | 6.245E-09 | 1.900E-08 | 3.100E-08 |
| 25767 | 1289 | S1289Q | E | 1e-07 | 9.1e-08 | 4.9e-08 | 8.000E-08 | 2.722E-08 | 4.900E-08 | 1.000E-07 |
| 25768 | 1289 | S1289G | E | 4.6e-08 | 5.1e-08 | 5.3e-08 | 5.000E-08 | 3.606E-09 | 4.600E-08 | 5.300E-08 |
| 25769 | 1289 | S1289H | E | 3e-08 | 2.4e-08 | 3.9e-08 | 3.100E-08 | 7.550E-09 | 2.400E-08 | 3.900E-08 |
| 25770 | 1289 | S1289I | E | 5.1e-08 | 3e-08 | 5.9e-08 | 4.667E-08 | 1.498E-08 | 3.000E-08 | 5.900E-08 |
| 25771 | 1289 | S1289L | E | 3e-08 | 4.1e-08 | 2.3e-08 | 3.133E-08 | 9.074E-09 | 2.300E-08 | 4.100E-08 |
| 25772 | 1289 | S1289K | E | 3.9e-08 | 2.3e-08 | 5.5e-08 | 3.900E-08 | 1.600E-08 | 2.300E-08 | 5.500E-08 |
| 25773 | 1289 | S1289M | E | 1.2e-07 | 3.6e-08 | 7.6e-08 | 7.733E-08 | 4.202E-08 | 3.600E-08 | 1.200E-07 |
| 25774 | 1289 | S1289F | E | 7.5e-08 | 3.2e-08 | 6.5e-08 | 5.733E-08 | 2.250E-08 | 3.200E-08 | 7.500E-08 |
| 25775 | 1289 | S1289P | E | 4.1e-08 | 4.5e-08 | 1e-07 | 6.200E-08 | 3.297E-08 | 4.100E-08 | 1.000E-07 |
| 25776 | 1289 | S1289S | E | 1.7e-08 | 1.3e-08 | 1e-07 | 4.333E-08 | 4.912E-08 | 1.300E-08 | 1.000E-07 |
| 25777 | 1289 | S1289T | E | 9.6e-08 | 1.1e-07 | 4e-08 | 8.200E-08 | 3.704E-08 | 4.000E-08 | 1.100E-07 |
| 25778 | 1289 | S1289W | E | 2.2e-08 | 5.3e-08 | 1.3e-08 | 2.933E-08 | 2.098E-08 | 1.300E-08 | 5.300E-08 |
| 25779 | 1289 | S1289Y | E | 9.1e-08 | 3.5e-08 | 8.7e-08 | 7.100E-08 | 3.124E-08 | 3.500E-08 | 9.100E-08 |
| 25780 | 1289 | S1289V | E | 2.5e-08 | 9.2e-08 | 2.2e-08 | 4.633E-08 | 3.958E-08 | 2.200E-08 | 9.200E-08 |
| 25781 | 1290 | L1290A | E | 2.7e-08 | 1.9e-08 | 2.5e-08 | 2.367E-08 | 4.163E-09 | 1.900E-08 | 2.700E-08 |
| 25782 | 1290 | L1290R | E | 5.6e-08 | 4.8e-08 | 3.3e-08 | 4.567E-08 | 1.168E-08 | 3.300E-08 | 5.600E-08 |
| 25783 | 1290 | L1290N | E | 6.2e-08 | 7e-08 | 7.9e-08 | 7.033E-08 | 8.505E-09 | 6.200E-08 | 7.900E-08 |
| 25784 | 1290 | L1290D | E | 3.9e-08 | 4.3e-08 | 3.8e-08 | 4.000E-08 | 2.646E-09 | 3.800E-08 | 4.300E-08 |
| 25785 | 1290 | L1290C | E | 8.2e-08 | 6.8e-08 | 4.4e-08 | 6.467E-08 | 1.922E-08 | 4.400E-08 | 8.200E-08 |
| 25786 | 1290 | L1290E | E | 9.3e-08 | 2.5e-08 | 5.9e-08 | 5.900E-08 | 3.400E-08 | 2.500E-08 | 9.300E-08 |
| 25787 | 1290 | L1290Q | E | 5.4e-08 | 5.6e-08 | 7e-08 | 6.000E-08 | 8.718E-09 | 5.400E-08 | 7.000E-08 |
| 25788 | 1290 | L1290G | E | 4.4e-08 | 8.9e-08 | 4.3e-08 | 5.867E-08 | 2.627E-08 | 4.300E-08 | 8.900E-08 |
| 25789 | 1290 | L1290H | E | 9.7e-08 | 5.1e-08 | 3.1e-08 | 5.967E-08 | 3.384E-08 | 3.100E-08 | 9.700E-08 |
| 25790 | 1290 | L1290I | E | 1.7e-08 | 4.4e-08 | 3.3e-08 | 3.133E-08 | 1.358E-08 | 1.700E-08 | 4.400E-08 |
| 25791 | 1290 | L1290L | E | 7e-08 | 4.9e-08 | 4.9e-08 | 5.600E-08 | 1.212E-08 | 4.900E-08 | 7.000E-08 |
| 25792 | 1290 | L1290K | E | 8.1e-08 | 2.9e-08 | 6.6e-08 | 5.867E-08 | 2.676E-08 | 2.900E-08 | 8.100E-08 |
| 25793 | 1290 | L1290M | E | 7.7e-08 | 2.1e-08 | 4.3e-08 | 4.700E-08 | 2.821E-08 | 2.100E-08 | 7.700E-08 |
| 25794 | 1290 | L1290F | E | 4.3e-08 | 9.9e-08 | 6e-08 | 6.733E-08 | 2.871E-08 | 4.300E-08 | 9.900E-08 |
| 25795 | 1290 | L1290P | E | 7e-08 | 1.5e-07 | 2.7e-08 | 8.233E-08 | 6.242E-08 | 2.700E-08 | 1.500E-07 |
| 25796 | 1290 | L1290S | E | 5.9e-08 | 3.1e-08 | 1.7e-08 | 3.567E-08 | 2.139E-08 | 1.700E-08 | 5.900E-08 |
| 25797 | 1290 | L1290T | E | 5.2e-08 | 1.1e-07 | 3.1e-08 | 6.433E-08 | 4.092E-08 | 3.100E-08 | 1.100E-07 |
| 25798 | 1290 | L1290W | E | 4.7e-08 | 3.3e-08 | 6.1e-08 | 4.700E-08 | 1.400E-08 | 3.300E-08 | 6.100E-08 |
| 25799 | 1290 | L1290Y | E | 2.4e-08 | 3.3e-08 | 3.8e-08 | 3.167E-08 | 7.095E-09 | 2.400E-08 | 3.800E-08 |
| 25800 | 1290 | L1290V | E | 5.4e-08 | 5.4e-08 | 1.4e-08 | 4.067E-08 | 2.309E-08 | 1.400E-08 | 5.400E-08 |
| 25801 | 1291 | R1291A | E | 5.4e-08 | 4.6e-08 | 1.2e-07 | 7.333E-08 | 4.061E-08 | 4.600E-08 | 1.200E-07 |
| 25802 | 1291 | R1291R | E | 3.8e-08 | 6.2e-08 | 8.8e-08 | 6.267E-08 | 2.501E-08 | 3.800E-08 | 8.800E-08 |
| 25803 | 1291 | R1291N | E | 2.5e-08 | 4.3e-08 | 5.2e-08 | 4.000E-08 | 1.375E-08 | 2.500E-08 | 5.200E-08 |
| 25804 | 1291 | R1291D | E | 4.1e-08 | 4.8e-08 | 2.9e-08 | 3.933E-08 | 9.609E-09 | 2.900E-08 | 4.800E-08 |
| 25805 | 1291 | R1291C | E | 3.2e-08 | 7.4e-08 | 4.4e-08 | 5.000E-08 | 2.163E-08 | 3.200E-08 | 7.400E-08 |
| 25806 | 1291 | R1291E | E | 9.4e-09 | 1.1e-08 | 4.2e-08 | 2.080E-08 | 1.838E-08 | 9.400E-09 | 4.200E-08 |
| 25807 | 1291 | R1291Q | E | 2.9e-08 | 5.5e-08 | 4.9e-08 | 4.433E-08 | 1.361E-08 | 2.900E-08 | 5.500E-08 |
| 25808 | 1291 | R1291G | E | 8e-08 | 2e-08 | 1.9e-08 | 3.967E-08 | 3.493E-08 | 1.900E-08 | 8.000E-08 |
| 25809 | 1291 | R1291H | E | 2.5e-08 | 3.3e-08 | 1.6e-08 | 2.467E-08 | 8.505E-09 | 1.600E-08 | 3.300E-08 |
| 25810 | 1291 | R1291I | E | 4.7e-08 | 5.4e-08 | 2.9e-08 | 4.333E-08 | 1.290E-08 | 2.900E-08 | 5.400E-08 |
| 25811 | 1291 | R1291L | E | 6.6e-08 | 5.1e-08 | 8.7e-08 | 6.800E-08 | 1.808E-08 | 5.100E-08 | 8.700E-08 |
| 25812 | 1291 | R1291K | E | 2.8e-08 | 3.5e-08 | 3.9e-08 | 3.400E-08 | 5.568E-09 | 2.800E-08 | 3.900E-08 |
| 25813 | 1291 | R1291M | E | 2e-07 | 2.4e-08 | 2.4e-08 | 8.267E-08 | 1.016E-07 | 2.400E-08 | 2.000E-07 |
| 25814 | 1291 | R1291F | E | 7.6e-08 | 2.1e-08 | 4.5e-08 | 4.733E-08 | 2.757E-08 | 2.100E-08 | 7.600E-08 |
| 25815 | 1291 | R1291P | E | 6.1e-08 | 1.2e-08 | 9.8e-09 | 2.760E-08 | 2.895E-08 | 9.800E-09 | 6.100E-08 |
| 25816 | 1291 | R1291S | E | 8.7e-08 | 7.4e-08 | 7.5e-08 | 7.867E-08 | 7.234E-09 | 7.400E-08 | 8.700E-08 |
| 25817 | 1291 | R1291T | E | 4.2e-08 | 3.3e-08 | 3.2e-08 | 3.567E-08 | 5.508E-09 | 3.200E-08 | 4.200E-08 |
| 25818 | 1291 | R1291W | E | 3.5e-08 | 3.2e-08 | 1.9e-08 | 2.867E-08 | 8.505E-09 | 1.900E-08 | 3.500E-08 |
| 25819 | 1291 | R1291Y | E | 6.9e-08 | 1.3e-08 | 1.4e-07 | 7.400E-08 | 6.365E-08 | 1.300E-08 | 1.400E-07 |
| 25820 | 1291 | R1291V | E | 5.8e-08 | 3.4e-08 | 7.9e-08 | 5.700E-08 | 2.252E-08 | 3.400E-08 | 7.900E-08 |
| 25821 | 1292 | L1292A | E | 3.7e-08 | 5.2e-08 | 2.8e-08 | 3.900E-08 | 1.212E-08 | 2.800E-08 | 5.200E-08 |
| 25822 | 1292 | L1292R | E | 3.1e-08 | 7.2e-08 | 3.2e-08 | 4.500E-08 | 2.339E-08 | 3.100E-08 | 7.200E-08 |
| 25823 | 1292 | L1292N | E | 2e-08 | 3.4e-08 | 1.1e-08 | 2.167E-08 | 1.159E-08 | 1.100E-08 | 3.400E-08 |
| 25824 | 1292 | L1292D | E | 3.6e-08 | 6.6e-08 | 8.6e-08 | 6.267E-08 | 2.517E-08 | 3.600E-08 | 8.600E-08 |
| 25825 | 1292 | L1292C | E | 4.5e-08 | 3.3e-08 | 3.2e-08 | 3.667E-08 | 7.234E-09 | 3.200E-08 | 4.500E-08 |
| 25826 | 1292 | L1292E | E | 1.2e-07 | 3.4e-08 | 1.8e-08 | 5.733E-08 | 5.486E-08 | 1.800E-08 | 1.200E-07 |
| 25827 | 1292 | L1292Q | E | 1.4e-07 | 1.9e-08 | 8.8e-08 | 8.233E-08 | 6.070E-08 | 1.900E-08 | 1.400E-07 |
| 25828 | 1292 | L1292G | E | 2.2e-08 | 4.4e-08 | 3.2e-08 | 3.267E-08 | 1.102E-08 | 2.200E-08 | 4.400E-08 |
| 25829 | 1292 | L1292H | E | 7.4e-08 | 6.3e-08 | 1.2e-07 | 8.567E-08 | 3.024E-08 | 6.300E-08 | 1.200E-07 |
| 25830 | 1292 | L1292I | E | 3.3e-08 | 1.5e-08 | 5e-08 | 3.267E-08 | 1.750E-08 | 1.500E-08 | 5.000E-08 |
| 25831 | 1292 | L1292L | E | 3.3e-08 | 6.8e-08 | 2e-08 | 4.033E-08 | 2.483E-08 | 2.000E-08 | 6.800E-08 |
| 25832 | 1292 | L1292K | E | 4.2e-08 | 2.1e-08 | 4.5e-08 | 3.600E-08 | 1.308E-08 | 2.100E-08 | 4.500E-08 |
| 25833 | 1292 | L1292M | E | 5.9e-08 | 8.6e-09 | 3.9e-08 | 3.553E-08 | 2.538E-08 | 8.600E-09 | 5.900E-08 |
| 25834 | 1292 | L1292F | E | 4.1e-08 | 2.1e-08 | 5.7e-08 | 3.967E-08 | 1.804E-08 | 2.100E-08 | 5.700E-08 |
| 25835 | 1292 | L1292P | E | 7.7e-08 | 6.3e-08 | 4.3e-08 | 6.100E-08 | 1.709E-08 | 4.300E-08 | 7.700E-08 |
| 25836 | 1292 | L1292S | E | 4.5e-08 | 5e-08 | 3.3e-08 | 4.267E-08 | 8.737E-09 | 3.300E-08 | 5.000E-08 |
| 25837 | 1292 | L1292T | E | 4.2e-08 | 5.1e-08 | 3.3e-08 | 4.200E-08 | 9.000E-09 | 3.300E-08 | 5.100E-08 |
| 25838 | 1292 | L1292W | E | 7.5e-08 | 1.1e-08 | 4.7e-08 | 4.433E-08 | 3.208E-08 | 1.100E-08 | 7.500E-08 |
| 25839 | 1292 | L1292Y | E | 4.3e-08 | 4e-08 | 6.8e-08 | 5.033E-08 | 1.537E-08 | 4.000E-08 | 6.800E-08 |

| | | | | | | | | | | |
|---|---|---|---|---|---|---|---|---|---|---|
| 25840 | 1292 | L1292V | E | 4.4e-08 | 7.8e-08 | 4.7e-08 | 5.633E-08 | 1.882E-08 | 4.400E-08 | 7.800E-08 |
| 25841 | 1293 | S1293A | E | 7.3e-08 | 4e-08 | 1.1e-07 | 7.433E-08 | 3.502E-08 | 4.000E-08 | 1.100E-07 |
| 25842 | 1293 | S1293R | E | 1.2e-08 | 6.2e-08 | 6.5e-08 | 4.633E-08 | 2.977E-08 | 1.200E-08 | 6.500E-08 |
| 25843 | 1293 | S1293N | E | 2.6e-08 | 5.7e-08 | 1.6e-08 | 3.300E-08 | 2.138E-08 | 1.600E-08 | 5.700E-08 |
| 25844 | 1293 | S1293D | E | 5e-08 | 3.3e-08 | 2e-08 | 3.433E-08 | 1.504E-08 | 2.000E-08 | 5.000E-08 |
| 25845 | 1293 | S1293C | E | 2.1e-08 | 4.2e-08 | 5.3e-08 | 3.867E-08 | 1.626E-08 | 2.100E-08 | 5.300E-08 |
| 25846 | 1293 | S1293E | E | 1.9e-08 | 6.9e-08 | 7.2e-08 | 5.333E-08 | 2.977E-08 | 1.900E-08 | 7.200E-08 |
| 25847 | 1293 | S1293Q | E | 7e-08 | 3e-08 | 3.3e-08 | 4.433E-08 | 2.228E-08 | 3.000E-08 | 7.000E-08 |
| 25848 | 1293 | S1293G | E | 4.8e-08 | 7e-08 | 3.7e-08 | 5.167E-08 | 1.680E-08 | 3.700E-08 | 7.000E-08 |
| 25849 | 1293 | S1293H | E | 3.2e-08 | 4.6e-08 | 5.3e-08 | 4.367E-08 | 1.069E-08 | 3.200E-08 | 5.300E-08 |
| 25850 | 1293 | S1293I | E | 2.2e-08 | 1.8e-08 | 1.9e-08 | 1.967E-08 | 2.082E-09 | 1.800E-08 | 2.200E-08 |
| 25851 | 1293 | S1293L | E | 1.2e-08 | 3.9e-08 | 5.2e-08 | 3.433E-08 | 2.040E-08 | 1.200E-08 | 5.200E-08 |
| 25852 | 1293 | S1293K | E | 4.4e-08 | 3.1e-08 | 7.7e-08 | 5.067E-08 | 2.371E-08 | 3.100E-08 | 7.700E-08 |
| 25853 | 1293 | S1293M | E | 4e-08 | 3.5e-08 | 4e-08 | 3.833E-08 | 2.887E-09 | 3.500E-08 | 4.000E-08 |
| 25854 | 1293 | S1293F | E | 4.4e-08 | 5e-08 | 4.5e-08 | 4.633E-08 | 3.215E-09 | 4.400E-08 | 5.000E-08 |
| 25855 | 1293 | S1293P | E | 5.9e-08 | 5.4e-08 | 2.1e-08 | 4.467E-08 | 2.065E-08 | 2.100E-08 | 5.900E-08 |
| 25856 | 1293 | S1293S | E | 5.6e-08 | 3e-08 | 5.7e-08 | 4.767E-08 | 1.531E-08 | 3.000E-08 | 5.700E-08 |
| 25857 | 1293 | S1293T | E | 3.5e-08 | 4e-08 | 7.6e-08 | 5.033E-08 | 2.237E-08 | 3.500E-08 | 7.600E-08 |
| 25858 | 1293 | S1293W | E | 1.6e-08 | 2.4e-08 | 3e-08 | 2.333E-08 | 7.024E-09 | 1.600E-08 | 3.000E-08 |
| 25859 | 1293 | S1293Y | E | 1.3e-08 | 7e-08 | 2.2e-08 | 3.500E-08 | 3.064E-08 | 1.300E-08 | 7.000E-08 |
| 25860 | 1293 | S1293V | E | 9.4e-08 | 4.2e-08 | 3.2e-08 | 5.600E-08 | 3.329E-08 | 3.200E-08 | 9.400E-08 |
| 25861 | 1294 | C1294A | E | 3.3e-08 | 6.5e-08 | 3.1e-08 | 4.300E-08 | 1.908E-08 | 3.100E-08 | 6.500E-08 |
| 25862 | 1294 | C1294R | E | 5.4e-08 | 3.9e-08 | 7.3e-08 | 5.533E-08 | 1.704E-08 | 3.900E-08 | 7.300E-08 |
| 25863 | 1294 | C1294N | E | 1.2e-08 | 5.4e-08 | 3.2e-08 | 3.267E-08 | 2.101E-08 | 1.200E-08 | 5.400E-08 |
| 25864 | 1294 | C1294D | E | 7.5e-08 | 5.3e-08 | 3.8e-08 | 5.533E-08 | 1.861E-08 | 3.800E-08 | 7.500E-08 |
| 25865 | 1294 | C1294C | E | 1.1e-08 | 2.4e-08 | 5e-08 | 2.833E-08 | 1.986E-08 | 1.100E-08 | 5.000E-08 |
| 25866 | 1294 | C1294E | E | 8.1e-08 | 6.1e-08 | 3.4e-08 | 5.867E-08 | 2.359E-08 | 3.400E-08 | 8.100E-08 |
| 25867 | 1294 | C1294Q | E | 3.8e-08 | 5e-08 | 4.4e-08 | 4.400E-08 | 6.000E-09 | 3.800E-08 | 5.000E-08 |
| 25868 | 1294 | C1294G | E | 4.8e-08 | 3e-08 | 5.4e-08 | 4.400E-08 | 1.249E-08 | 3.000E-08 | 5.400E-08 |
| 25869 | 1294 | C1294H | E | 5.5e-08 | 2.2e-08 | 4.2e-08 | 3.967E-08 | 1.662E-08 | 2.200E-08 | 5.500E-08 |
| 25870 | 1294 | C1294I | E | 5.8e-08 | 1.3e-07 | 3.8e-08 | 7.533E-08 | 4.839E-08 | 3.800E-08 | 1.300E-07 |
| 25871 | 1294 | C1294L | E | 8.6e-08 | 2.7e-08 | 8.7e-08 | 6.667E-08 | 3.436E-08 | 2.700E-08 | 8.700E-08 |
| 25872 | 1294 | C1294K | E | 6.8e-08 | 4e-08 | 3.3e-08 | 4.700E-08 | 1.852E-08 | 3.300E-08 | 6.800E-08 |
| 25873 | 1294 | C1294M | E | 4.2e-08 | 2.3e-08 | 4.8e-08 | 3.767E-08 | 1.305E-08 | 2.300E-08 | 4.800E-08 |
| 25874 | 1294 | C1294F | E | 3.7e-08 | 5.4e-08 | 2.3e-08 | 3.800E-08 | 1.552E-08 | 2.300E-08 | 5.400E-08 |
| 25875 | 1294 | C1294P | E | 1.3e-07 | 3.7e-08 | 8.9e-09 | 5.863E-08 | 6.338E-08 | 8.900E-09 | 1.300E-07 |
| 25876 | 1294 | C1294S | E | 3.6e-08 | 1.9e-08 | 3.2e-08 | 2.900E-08 | 8.888E-09 | 1.900E-08 | 3.600E-08 |
| 25877 | 1294 | C1294T | E | 5e-08 | 2.6e-08 | 5.8e-08 | 4.467E-08 | 1.665E-08 | 2.600E-08 | 5.800E-08 |
| 25878 | 1294 | C1294W | E | 4.9e-08 | 7.9e-08 | 7.3e-08 | 6.700E-08 | 1.587E-08 | 4.900E-08 | 7.900E-08 |
| 25879 | 1294 | C1294Y | E | 1.6e-08 | 5.4e-08 | 6.3e-08 | 4.433E-08 | 2.495E-08 | 1.600E-08 | 6.300E-08 |
| 25880 | 1294 | C1294V | E | 4.2e-08 | 5.7e-08 | 4.2e-08 | 4.700E-08 | 8.660E-09 | 4.200E-08 | 5.700E-08 |
| 25881 | 1295 | A1295A | E | 4.4e-08 | 5.1e-08 | 3.5e-08 | 4.333E-08 | 8.021E-09 | 3.500E-08 | 5.100E-08 |
| 25882 | 1295 | A1295R | E | 2.7e-08 | 1.2e-07 | 7.5e-08 | 7.400E-08 | 4.651E-08 | 2.700E-08 | 1.200E-07 |
| 25883 | 1295 | A1295N | E | 2.9e-08 | 1.8e-07 | 2.6e-08 | 7.833E-08 | 8.806E-08 | 2.600E-08 | 1.800E-07 |
| 25884 | 1295 | A1295D | E | 6.4e-08 | 5.5e-08 | 7.9e-08 | 6.600E-08 | 1.212E-08 | 5.500E-08 | 7.900E-08 |
| 25885 | 1295 | A1295C | E | 3e-07 | 5.7e-08 | 6.8e-09 | 1.213E-07 | 1.568E-07 | 6.800E-09 | 3.000E-07 |
| 25886 | 1295 | A1295E | E | 4.2e-08 | 1.5e-08 | 1.2e-07 | 5.900E-08 | 5.453E-08 | 1.500E-08 | 1.200E-07 |
| 25887 | 1295 | A1295Q | E | 6.2e-08 | 4.4e-08 | 2.9e-08 | 4.500E-08 | 1.652E-08 | 2.900E-08 | 6.200E-08 |
| 25888 | 1295 | A1295G | E | 5.1e-08 | 2.3e-08 | 1.9e-08 | 3.100E-08 | 1.744E-08 | 1.900E-08 | 5.100E-08 |
| 25889 | 1295 | A1295H | E | 4.4e-08 | 8.6e-09 | 6.3e-08 | 3.853E-08 | 2.761E-08 | 8.600E-09 | 6.300E-08 |
| 25890 | 1295 | A1295I | E | 4.4e-08 | 2.8e-08 | 2.5e-08 | 3.233E-08 | 1.021E-08 | 2.500E-08 | 4.400E-08 |
| 25891 | 1295 | A1295L | E | 4.2e-08 | 3.2e-08 | 4.8e-08 | 4.067E-08 | 8.083E-09 | 3.200E-08 | 4.800E-08 |
| 25892 | 1295 | A1295K | E | 2.4e-08 | 8.4e-08 | 2.4e-08 | 4.400E-08 | 3.464E-08 | 2.400E-08 | 8.400E-08 |
| 25893 | 1295 | A1295M | E | 2.8e-07 | 8.1e-08 | 5.2e-08 | 1.377E-07 | 1.241E-07 | 5.200E-08 | 2.800E-07 |
| 25894 | 1295 | A1295F | E | 3.9e-08 | 3.8e-08 | 3.3e-08 | 3.667E-08 | 3.215E-09 | 3.300E-08 | 3.900E-08 |
| 25895 | 1295 | A1295P | E | 9.8e-08 | 5.2e-08 | 5.4e-08 | 6.800E-08 | 2.600E-08 | 5.200E-08 | 9.800E-08 |
| 25896 | 1295 | A1295S | E | 3.2e-08 | 8.9e-08 | 6.4e-08 | 6.167E-08 | 2.857E-08 | 3.200E-08 | 8.900E-08 |
| 25897 | 1295 | A1295T | E | 2.7e-08 | 7e-08 | 2.9e-08 | 4.200E-08 | 2.427E-08 | 2.700E-08 | 7.000E-08 |
| 25898 | 1295 | A1295W | E | 8.3e-08 | 3.5e-08 | 2.8e-08 | 4.867E-08 | 2.994E-08 | 2.800E-08 | 8.300E-08 |
| 25899 | 1295 | A1295Y | E | 2.7e-08 | 4.1e-08 | 2e-08 | 2.933E-08 | 1.069E-08 | 2.000E-08 | 4.100E-08 |
| 25900 | 1295 | A1295V | E | 2.5e-08 | 4.8e-08 | 4.2e-08 | 3.833E-08 | 1.193E-08 | 2.500E-08 | 4.800E-08 |
| 25901 | 1296 | A1296A | E | 1.7e-08 | 5.6e-08 | 5.4e-08 | 4.233E-08 | 2.196E-08 | 1.700E-08 | 5.600E-08 |
| 25902 | 1296 | A1296R | E | 2.6e-08 | 5.7e-08 | 2.1e-08 | 3.467E-08 | 1.950E-08 | 2.100E-08 | 5.700E-08 |
| 25903 | 1296 | A1296N | E | 1.3e-07 | 6.1e-08 | 1.1e-07 | 1.003E-07 | 3.550E-08 | 6.100E-08 | 1.300E-07 |
| 25904 | 1296 | A1296D | E | 2e-08 | 3.4e-08 | 9.6e-09 | 2.120E-08 | 1.224E-08 | 9.600E-09 | 3.400E-08 |
| 25905 | 1296 | A1296C | E | 6.8e-08 | 2.8e-08 | 2e-08 | 3.867E-08 | 2.572E-08 | 2.000E-08 | 6.800E-08 |
| 25906 | 1296 | A1296E | E | 1.9e-08 | 7.8e-09 | 1.1e-07 | 4.560E-08 | 5.605E-08 | 7.800E-09 | 1.100E-07 |
| 25907 | 1296 | A1296Q | E | 3.9e-08 | 3.6e-08 | 9.2e-08 | 5.567E-08 | 3.150E-08 | 3.600E-08 | 9.200E-08 |
| 25908 | 1296 | A1296G | E | 4.7e-08 | 3.4e-08 | 6.9e-08 | 5.000E-08 | 1.769E-08 | 3.400E-08 | 6.900E-08 |
| 25909 | 1296 | A1296H | E | 5.1e-08 | 5e-08 | 4.1e-08 | 4.733E-08 | 5.508E-09 | 4.100E-08 | 5.100E-08 |
| 25910 | 1296 | A1296I | E | 4.3e-08 | 1.7e-07 | 3.2e-08 | 8.167E-08 | 7.670E-08 | 3.200E-08 | 1.700E-07 |
| 25911 | 1296 | A1296L | E | 7.5e-08 | 2.9e-08 | 1.8e-08 | 4.067E-08 | 3.024E-08 | 1.800E-08 | 7.500E-08 |
| 25912 | 1296 | A1296K | E | 2.9e-08 | 6.2e-08 | 4.1e-08 | 4.400E-08 | 1.670E-08 | 2.900E-08 | 6.200E-08 |
| 25913 | 1296 | A1296M | E | 2.9e-08 | 3.6e-08 | 4.5e-08 | 3.667E-08 | 8.021E-09 | 2.900E-08 | 4.500E-08 |
| 25914 | 1296 | A1296F | E | 6.4e-08 | 5.1e-08 | 4.9e-08 | 5.467E-08 | 8.145E-09 | 4.900E-08 | 6.400E-08 |
| 25915 | 1296 | A1296P | E | 3.9e-08 | 2.9e-08 | 1.3e-07 | 6.600E-08 | 5.565E-08 | 2.900E-08 | 1.300E-07 |
| 25916 | 1296 | A1296S | E | 3.5e-08 | 3.5e-08 | 3.9e-08 | 3.633E-08 | 2.309E-09 | 3.500E-08 | 3.900E-08 |
| 25917 | 1296 | A1296T | E | 1.8e-08 | 4e-08 | 4.8e-08 | 3.533E-08 | 1.553E-08 | 1.800E-08 | 4.800E-08 |
| 25918 | 1296 | A1296W | E | 3.8e-08 | 2.4e-08 | 4.5e-08 | 3.567E-08 | 1.069E-08 | 2.400E-08 | 4.500E-08 |
| 25919 | 1296 | A1296Y | E | 1.6e-08 | 4.4e-08 | 3.8e-08 | 3.267E-08 | 1.474E-08 | 1.600E-08 | 4.400E-08 |
| 25920 | 1296 | A1296V | E | 2.9e-08 | 3.5e-08 | 6.7e-08 | 4.367E-08 | 2.043E-08 | 2.900E-08 | 6.700E-08 |
| 25921 | 1297 | S1297A | E | 8.4e-08 | 4e-08 | 2.7e-08 | 5.033E-08 | 2.987E-08 | 2.700E-08 | 8.400E-08 |
| 25922 | 1297 | S1297R | E | 3.4e-08 | 1.4e-08 | 7.5e-08 | 4.100E-08 | 3.110E-08 | 1.400E-08 | 7.500E-08 |
| 25923 | 1297 | S1297N | E | 2.4e-08 | 3e-08 | 3.7e-08 | 3.033E-08 | 6.506E-09 | 2.400E-08 | 3.700E-08 |
| 25924 | 1297 | S1297D | E | 5.1e-08 | 7.4e-08 | 1.6e-08 | 4.700E-08 | 2.921E-08 | 1.600E-08 | 7.400E-08 |

| | | | | | | | | | | |
|---|---|---|---|---|---|---|---|---|---|---|
| 25925 | 1297 | S1297C | E | 1.1e-07 | 8e-08 | 4.4e-08 | 7.800E-08 | 3.305E-08 | 4.400E-08 | 1.100E-07 |
| 25926 | 1297 | S1297E | E | 1.5e-07 | 4.5e-08 | 9.6e-08 | 9.700E-08 | 5.251E-08 | 4.500E-08 | 1.500E-07 |
| 25927 | 1297 | S1297Q | E | 1.8e-08 | 5.7e-08 | 4.6e-08 | 4.033E-08 | 2.011E-08 | 1.800E-08 | 5.700E-08 |
| 25928 | 1297 | S1297G | E | 5.2e-08 | 2.3e-08 | 5.5e-08 | 4.333E-08 | 1.767E-08 | 2.300E-08 | 5.500E-08 |
| 25929 | 1297 | S1297H | E | 6.4e-08 | 2.8e-08 | 4.8e-08 | 4.667E-08 | 1.804E-08 | 2.800E-08 | 6.400E-08 |
| 25930 | 1297 | S1297I | E | 7.6e-08 | 2.2e-08 | 5.6e-08 | 5.133E-08 | 2.730E-08 | 2.200E-08 | 7.600E-08 |
| 25931 | 1297 | S1297L | E | 2e-08 | 2.4e-08 | 4.9e-08 | 3.100E-08 | 1.572E-08 | 2.000E-08 | 4.900E-08 |
| 25932 | 1297 | S1297K | E | 4.8e-08 | 1.2e-07 | 8e-08 | 8.267E-08 | 3.607E-08 | 4.800E-08 | 1.200E-07 |
| 25933 | 1297 | S1297M | E | 4.6e-08 | 1.4e-08 | 2.9e-08 | 2.967E-08 | 1.601E-08 | 1.400E-08 | 4.600E-08 |
| 25934 | 1297 | S1297F | E | 3.9e-08 | 3.2e-08 | 4.7e-08 | 3.933E-08 | 7.506E-09 | 3.200E-08 | 4.700E-08 |
| 25935 | 1297 | S1297P | E | 3.5e-08 | 3.5e-08 | 1.8e-08 | 2.933E-08 | 9.815E-09 | 1.800E-08 | 3.500E-08 |
| 25936 | 1297 | S1297S | E | 1.1e-07 | 8.4e-08 | 7.7e-08 | 9.033E-08 | 1.739E-08 | 7.700E-08 | 1.100E-07 |
| 25937 | 1297 | S1297T | E | 1.9e-08 | 3.9e-08 | 8.2e-08 | 4.667E-08 | 3.219E-08 | 1.900E-08 | 8.200E-08 |
| 25938 | 1297 | S1297W | E | 3.9e-08 | 3.5e-08 | 4e-08 | 3.800E-08 | 2.646E-09 | 3.500E-08 | 4.000E-08 |
| 25939 | 1297 | S1297Y | E | 3.8e-08 | 6.2e-08 | 5.3e-08 | 5.100E-08 | 1.212E-08 | 3.800E-08 | 6.200E-08 |
| 25940 | 1297 | S1297V | E | 1.7e-08 | 2.4e-08 | 5.5e-08 | 3.200E-08 | 2.022E-08 | 1.700E-08 | 5.500E-08 |
| 25941 | 1298 | G1298A | E | 3.6e-08 | 1.8e-07 | 3.6e-08 | 8.400E-08 | 8.314E-08 | 3.600E-08 | 1.800E-07 |
| 25942 | 1298 | G1298R | E | 5.7e-08 | 2.3e-08 | 3.3e-08 | 3.767E-08 | 1.747E-08 | 2.300E-08 | 5.700E-08 |
| 25943 | 1298 | G1298N | E | 4.5e-08 | 2.8e-08 | 1.4e-08 | 2.900E-08 | 1.552E-08 | 1.400E-08 | 4.500E-08 |
| 25944 | 1298 | G1298D | E | 6.7e-08 | 3.1e-08 | 1.4e-08 | 3.733E-08 | 2.706E-08 | 1.400E-08 | 6.700E-08 |
| 25945 | 1298 | G1298C | E | 1.7e-08 | 3.4e-08 | 5.5e-08 | 3.533E-08 | 1.904E-08 | 1.700E-08 | 5.500E-08 |
| 25946 | 1298 | G1298E | E | 7.9e-08 | 6.1e-08 | 9e-09 | 4.967E-08 | 3.635E-08 | 9.000E-09 | 7.900E-08 |
| 25947 | 1298 | G1298Q | E | 4.2e-08 | 1.6e-07 | 1.6e-08 | 7.267E-08 | 7.674E-08 | 1.600E-08 | 1.600E-07 |
| 25948 | 1298 | G1298G | E | 4.5e-08 | 8.1e-08 | 2.8e-08 | 5.133E-08 | 2.706E-08 | 2.800E-08 | 8.100E-08 |
| 25949 | 1298 | G1298H | E | 1.1e-07 | 6.1e-08 | 2.7e-08 | 6.600E-08 | 4.173E-08 | 2.700E-08 | 1.100E-07 |
| 25950 | 1298 | G1298I | E | 8.6e-08 | 3e-08 | 1.9e-08 | 4.500E-08 | 3.593E-08 | 1.900E-08 | 8.600E-08 |
| 25951 | 1298 | G1298L | E | 4.4e-08 | 7.4e-08 | 5.6e-08 | 5.800E-08 | 1.510E-08 | 4.400E-08 | 7.400E-08 |
| 25952 | 1298 | G1298K | E | 5.6e-08 | 3.6e-08 | 2.8e-08 | 4.000E-08 | 1.442E-08 | 2.800E-08 | 5.600E-08 |
| 25953 | 1298 | G1298M | E | 8.1e-08 | 2.8e-08 | 2.8e-08 | 4.567E-08 | 3.060E-08 | 2.800E-08 | 8.100E-08 |
| 25954 | 1298 | G1298F | E | 6.5e-08 | 7.1e-08 | 1.2e-07 | 8.533E-08 | 3.017E-08 | 6.500E-08 | 1.200E-07 |
| 25955 | 1298 | G1298P | E | 4.6e-08 | 4.3e-08 | 6.3e-08 | 5.067E-08 | 1.079E-08 | 4.300E-08 | 6.300E-08 |
| 25956 | 1298 | G1298S | E | 2.6e-08 | 3.4e-08 | 3.5e-08 | 3.167E-08 | 4.933E-09 | 2.600E-08 | 3.500E-08 |
| 25957 | 1298 | G1298T | E | 2.3e-08 | 5.3e-08 | 2.8e-08 | 3.467E-08 | 1.607E-08 | 2.300E-08 | 5.300E-08 |
| 25958 | 1298 | G1298W | E | 6.9e-08 | 6.1e-08 | 3.3e-08 | 5.433E-08 | 1.890E-08 | 3.300E-08 | 6.900E-08 |
| 25959 | 1298 | G1298Y | E | 2e-08 | 1.7e-08 | 1.1e-08 | 1.600E-08 | 4.583E-09 | 1.100E-08 | 2.000E-08 |
| 25960 | 1298 | G1298V | E | 6.7e-08 | 4.9e-08 | 1.4e-07 | 8.533E-08 | 4.819E-08 | 4.900E-08 | 1.400E-07 |
| 25961 | 1299 | F1299A | E | 4.6e-08 | 5.3e-08 | 2.6e-08 | 4.167E-08 | 1.401E-08 | 2.600E-08 | 5.300E-08 |
| 25962 | 1299 | F1299R | E | 5.3e-08 | 5e-08 | 7e-08 | 5.767E-08 | 1.079E-08 | 5.000E-08 | 7.000E-08 |
| 25963 | 1299 | F1299N | E | 5.2e-08 | 4e-08 | 6.8e-09 | 3.293E-08 | 2.341E-08 | 6.800E-09 | 5.200E-08 |
| 25964 | 1299 | F1299D | E | 3.2e-08 | 7.5e-08 | 7.1e-08 | 5.933E-08 | 2.376E-08 | 3.200E-08 | 7.500E-08 |
| 25965 | 1299 | F1299C | E | 5.1e-08 | 4.4e-08 | 3.4e-08 | 4.300E-08 | 8.544E-09 | 3.400E-08 | 5.100E-08 |
| 25966 | 1299 | F1299E | E | 1.3e-07 | 2.4e-08 | 8.6e-08 | 8.000E-08 | 5.325E-08 | 2.400E-08 | 1.300E-07 |
| 25967 | 1299 | F1299Q | E | 4.6e-08 | 3.5e-08 | 6.4e-08 | 4.833E-08 | 1.464E-08 | 3.500E-08 | 6.400E-08 |
| 25968 | 1299 | F1299G | E | 2.1e-08 | 4.8e-08 | 1.4e-07 | 6.967E-08 | 6.239E-08 | 2.100E-08 | 1.400E-07 |
| 25969 | 1299 | F1299H | E | 1.4e-08 | 2.8e-08 | 3e-08 | 2.400E-08 | 8.718E-09 | 1.400E-08 | 3.000E-08 |
| 25970 | 1299 | F1299I | E | 2.8e-08 | 1.3e-07 | 3e-08 | 6.267E-08 | 5.832E-08 | 2.800E-08 | 1.300E-07 |
| 25971 | 1299 | F1299L | E | 1.9e-08 | 5.2e-08 | 7.4e-08 | 4.833E-08 | 2.768E-08 | 1.900E-08 | 7.400E-08 |
| 25972 | 1299 | F1299K | E | 8.3e-08 | 4.4e-08 | 3.1e-08 | 5.267E-08 | 2.706E-08 | 3.100E-08 | 8.300E-08 |
| 25973 | 1299 | F1299M | E | 1.2e-07 | 1.1e-07 | 2.5e-08 | 8.500E-08 | 5.220E-08 | 2.500E-08 | 1.200E-07 |
| 25974 | 1299 | F1299F | E | 6.8e-08 | 4.4e-08 | 3.8e-08 | 5.000E-08 | 1.587E-08 | 3.800E-08 | 6.800E-08 |
| 25975 | 1299 | F1299P | E | 2.5e-08 | 7.3e-08 | 8.2e-08 | 6.000E-08 | 3.064E-08 | 2.500E-08 | 8.200E-08 |
| 25976 | 1299 | F1299S | E | 8.4e-08 | 1.7e-08 | 7.3e-08 | 5.800E-08 | 3.593E-08 | 1.700E-08 | 8.400E-08 |
| 25977 | 1299 | F1299T | E | 7.1e-08 | 3.6e-08 | 3.7e-08 | 4.800E-08 | 1.992E-08 | 3.600E-08 | 7.100E-08 |
| 25978 | 1299 | F1299W | E | 6.4e-08 | 1e-08 | 6.7e-08 | 4.700E-08 | 3.208E-08 | 1.000E-08 | 6.700E-08 |
| 25979 | 1299 | F1299Y | E | 2.3e-08 | 4e-08 | 4.5e-08 | 3.600E-08 | 1.153E-08 | 2.300E-08 | 4.500E-08 |
| 25980 | 1299 | F1299V | E | 4.3e-08 | 3.3e-08 | 3.4e-08 | 3.667E-08 | 5.508E-09 | 3.300E-08 | 4.300E-08 |
| 25981 | 1300 | N1300A | E | 6.1e-08 | 3.1e-08 | 4.6e-08 | 4.600E-08 | 1.500E-08 | 3.100E-08 | 6.100E-08 |
| 25982 | 1300 | N1300R | E | 3.1e-08 | 2.2e-08 | 2.5e-08 | 2.600E-08 | 4.583E-09 | 2.200E-08 | 3.100E-08 |
| 25983 | 1300 | N1300N | E | 1.7e-08 | 5.5e-08 | 2e-08 | 3.067E-08 | 2.113E-08 | 1.700E-08 | 5.500E-08 |
| 25984 | 1300 | N1300D | E | 1.8e-08 | 1.7e-08 | 2.1e-08 | 1.867E-08 | 2.082E-09 | 1.700E-08 | 2.100E-08 |
| 25985 | 1300 | N1300C | E | 5.7e-08 | 2e-08 | 7.3e-08 | 5.000E-08 | 2.718E-08 | 2.000E-08 | 7.300E-08 |
| 25986 | 1300 | N1300E | E | 5.6e-08 | 2e-08 | 2e-08 | 3.200E-08 | 2.078E-08 | 2.000E-08 | 5.600E-08 |
| 25987 | 1300 | N1300Q | E | 3.2e-08 | 3e-08 | 2.7e-08 | 2.967E-08 | 2.517E-09 | 2.700E-08 | 3.200E-08 |
| 25988 | 1300 | N1300G | E | 3.7e-08 | 6.2e-08 | 3.7e-08 | 4.533E-08 | 1.443E-08 | 3.700E-08 | 6.200E-08 |
| 25989 | 1300 | N1300H | E | 4.4e-08 | 5.6e-08 | 4.4e-08 | 4.800E-08 | 6.928E-09 | 4.400E-08 | 5.600E-08 |
| 25990 | 1300 | N1300I | E | 8.5e-08 | 2.8e-08 | 5.2e-08 | 5.500E-08 | 2.862E-08 | 2.800E-08 | 8.500E-08 |
| 25991 | 1300 | N1300L | E | 5.3e-08 | 4.8e-08 | 4.4e-08 | 4.833E-08 | 4.509E-09 | 4.400E-08 | 5.300E-08 |
| 25992 | 1300 | N1300K | E | 6.5e-08 | 4.7e-08 | 5e-08 | 5.400E-08 | 9.644E-09 | 4.700E-08 | 6.500E-08 |
| 25993 | 1300 | N1300M | E | 3.1e-08 | 1.4e-08 | 7.7e-08 | 4.067E-08 | 3.259E-08 | 1.400E-08 | 7.700E-08 |
| 25994 | 1300 | N1300F | E | 7.4e-08 | 2.6e-08 | 1.8e-08 | 3.933E-08 | 3.029E-08 | 1.800E-08 | 7.400E-08 |
| 25995 | 1300 | N1300P | E | 1.5e-07 | 2.1e-08 | 3.6e-08 | 6.900E-08 | 7.055E-08 | 2.100E-08 | 1.500E-07 |
| 25996 | 1300 | N1300S | E | 3.7e-08 | 3.7e-08 | 3e-08 | 3.467E-08 | 4.041E-09 | 3.000E-08 | 3.700E-08 |
| 25997 | 1300 | N1300T | E | 4.5e-08 | 3.4e-08 | 2.1e-08 | 3.333E-08 | 1.201E-08 | 2.100E-08 | 4.500E-08 |
| 25998 | 1300 | N1300W | E | 3.6e-08 | 2.9e-08 | 2.8e-08 | 3.100E-08 | 4.359E-09 | 2.800E-08 | 3.600E-08 |
| 25999 | 1300 | N1300Y | E | 3e-08 | 2.7e-08 | 4.9e-09 | 2.063E-08 | 1.371E-08 | 4.900E-09 | 3.000E-08 |
| 26000 | 1300 | N1300V | E | 2.1e-08 | 3.9e-08 | 1.9e-08 | 2.633E-08 | 1.102E-08 | 1.900E-08 | 3.900E-08 |
| 26001 | 1301 | I1301A | E | 1.8e-08 | 3.5e-08 | 2.3e-08 | 2.533E-08 | 8.737E-09 | 1.800E-08 | 3.500E-08 |
| 26002 | 1301 | I1301R | E | 3.3e-08 | 5e-08 | 2.3e-07 | 1.043E-07 | 1.092E-07 | 3.300E-08 | 2.300E-07 |
| 26003 | 1301 | I1301N | E | 9.9e-08 | 4.7e-08 | 2.4e-08 | 5.667E-08 | 3.842E-08 | 2.400E-08 | 9.900E-08 |
| 26004 | 1301 | I1301D | E | 2.7e-08 | 1.8e-08 | 5e-08 | 3.167E-08 | 1.650E-08 | 1.800E-08 | 5.000E-08 |
| 26005 | 1301 | I1301C | E | 1.8e-08 | 1.4e-08 | 1.4e-07 | 5.733E-08 | 7.162E-08 | 1.400E-08 | 1.400E-07 |
| 26006 | 1301 | I1301E | E | 1.1e-07 | 4.9e-08 | 5.4e-08 | 7.100E-08 | 3.387E-08 | 4.900E-08 | 1.100E-07 |
| 26007 | 1301 | I1301Q | E | 1.9e-08 | 2.4e-08 | 4.7e-08 | 3.000E-08 | 1.493E-08 | 1.900E-08 | 4.700E-08 |
| 26008 | 1301 | I1301G | E | 3.2e-08 | 6.7e-08 | 3.8e-08 | 4.567E-08 | 1.872E-08 | 3.200E-08 | 6.700E-08 |
| 26009 | 1301 | I1301H | E | 4.6e-08 | 6e-08 | 7.2e-08 | 5.933E-08 | 1.301E-08 | 4.600E-08 | 7.200E-08 |

| | | | | | | | | | | |
|---|---|---|---|---|---|---|---|---|---|---|
| 26010 | 1301 | I1301I | E | 3.3e-08 | 8.1e-09 | 3.6e-08 | 2.570E-08 | 1.532E-08 | 8.100E-09 | 3.600E-08 |
| 26011 | 1301 | I1301L | E | 9.8e-09 | 3.4e-08 | 1.1e-08 | 1.827E-08 | 1.364E-08 | 9.800E-09 | 3.400E-08 |
| 26012 | 1301 | I1301K | E | 5.6e-08 | 7.9e-08 | 4.2e-08 | 5.900E-08 | 1.868E-08 | 4.200E-08 | 7.900E-08 |
| 26013 | 1301 | I1301M | E | 2.4e-08 | 6.9e-08 | 5.9e-08 | 5.067E-08 | 2.363E-08 | 2.400E-08 | 6.900E-08 |
| 26014 | 1301 | I1301F | E | 5.3e-08 | 3.7e-08 | 3.1e-08 | 4.033E-08 | 1.137E-08 | 3.100E-08 | 5.300E-08 |
| 26015 | 1301 | I1301P | E | 9.5e-08 | 8.8e-08 | 2.2e-08 | 6.833E-08 | 4.028E-08 | 2.200E-08 | 9.500E-08 |
| 26016 | 1301 | I1301S | E | 3.9e-08 | 1.1e-07 | 4.4e-08 | 6.433E-08 | 3.963E-08 | 3.900E-08 | 1.100E-07 |
| 26017 | 1301 | I1301T | E | 2.3e-08 | 5.2e-08 | 3.2e-08 | 3.567E-08 | 1.484E-08 | 2.300E-08 | 5.200E-08 |
| 26018 | 1301 | I1301W | E | 5.3e-08 | 8.8e-08 | 1.5e-08 | 5.200E-08 | 3.651E-08 | 1.500E-08 | 8.800E-08 |
| 26019 | 1301 | I1301Y | E | 6.6e-08 | 2.4e-08 | 1e-07 | 6.333E-08 | 3.807E-08 | 2.400E-08 | 1.000E-07 |
| 26020 | 1301 | I1301V | E | 4.6e-09 | 7.7e-08 | 6.3e-08 | 4.820E-08 | 3.840E-08 | 4.600E-09 | 7.700E-08 |
| 26021 | 1302 | K1302A | E | 3.2e-08 | 3.4e-08 | 5.2e-08 | 3.933E-08 | 1.102E-08 | 3.200E-08 | 5.200E-08 |
| 26022 | 1302 | K1302R | E | 2e-08 | 6.2e-08 | 7.7e-08 | 5.300E-08 | 2.955E-08 | 2.000E-08 | 7.700E-08 |
| 26023 | 1302 | K1302N | E | 2.7e-08 | 2.2e-08 | 2.7e-08 | 2.533E-08 | 2.887E-09 | 2.200E-08 | 2.700E-08 |
| 26024 | 1302 | K1302D | E | 5.3e-08 | 1.7e-08 | 4.3e-08 | 3.767E-08 | 1.858E-08 | 1.700E-08 | 5.300E-08 |
| 26025 | 1302 | K1302C | E | 7.2e-08 | 1.5e-08 | 5.8e-08 | 4.833E-08 | 2.970E-08 | 1.500E-08 | 7.200E-08 |
| 26026 | 1302 | K1302E | E | 2.2e-08 | 8.2e-08 | 2e-08 | 4.133E-08 | 3.523E-08 | 2.000E-08 | 8.200E-08 |
| 26027 | 1302 | K1302Q | E | 5.5e-08 | 1.2e-08 | 3.4e-08 | 3.367E-08 | 2.150E-08 | 1.200E-08 | 5.500E-08 |
| 26028 | 1302 | K1302G | E | 4.5e-08 | 9.8e-08 | 2.7e-08 | 5.667E-08 | 3.691E-08 | 2.700E-08 | 9.800E-08 |
| 26029 | 1302 | K1302H | E | 1.1e-07 | 3.4e-08 | 2.6e-08 | 5.667E-08 | 4.636E-08 | 2.600E-08 | 1.100E-07 |
| 26030 | 1302 | K1302I | E | 5.6e-08 | 1.7e-08 | 3.1e-08 | 3.467E-08 | 1.976E-08 | 1.700E-08 | 5.600E-08 |
| 26031 | 1302 | K1302L | E | 4e-08 | 9.1e-08 | 6.2e-08 | 6.433E-08 | 2.558E-08 | 4.000E-08 | 9.100E-08 |
| 26032 | 1302 | K1302K | E | 2.4e-08 | 1.1e-07 | 7.1e-08 | 6.833E-08 | 4.306E-08 | 2.400E-08 | 1.100E-07 |
| 26033 | 1302 | K1302M | E | 2.5e-08 | 5.9e-08 | 7.6e-08 | 5.333E-08 | 2.597E-08 | 2.500E-08 | 7.600E-08 |
| 26034 | 1302 | K1302F | E | 4e-08 | 4.3e-08 | 8.1e-08 | 5.467E-08 | 2.285E-08 | 4.000E-08 | 8.100E-08 |
| 26035 | 1302 | K1302P | E | 4.8e-08 | 1.1e-07 | 2.7e-08 | 6.167E-08 | 4.315E-08 | 2.700E-08 | 1.100E-07 |
| 26036 | 1302 | K1302S | E | 3.7e-08 | 4.3e-08 | 2.8e-08 | 3.600E-08 | 7.550E-09 | 2.800E-08 | 4.300E-08 |
| 26037 | 1302 | K1302T | E | 1.1e-08 | 5.7e-08 | 3.9e-08 | 3.567E-08 | 2.318E-08 | 1.100E-08 | 5.700E-08 |
| 26038 | 1302 | K1302W | E | 3.2e-08 | 6e-08 | 1e-07 | 6.400E-08 | 3.418E-08 | 3.200E-08 | 1.000E-07 |
| 26039 | 1302 | K1302Y | E | 3.2e-08 | 6.6e-08 | 6e-08 | 5.267E-08 | 1.815E-08 | 3.200E-08 | 6.600E-08 |
| 26040 | 1302 | K1302V | E | 8.1e-08 | 2e-08 | 3.8e-08 | 4.633E-08 | 3.134E-08 | 2.000E-08 | 8.100E-08 |
| 26041 | 1303 | D1303A | E | 5.2e-08 | 2.5e-08 | 2.5e-08 | 3.400E-08 | 1.559E-08 | 2.500E-08 | 5.200E-08 |
| 26042 | 1303 | D1303R | E | 8e-08 | 5e-08 | 4.6e-08 | 5.867E-08 | 1.858E-08 | 4.600E-08 | 8.000E-08 |
| 26043 | 1303 | D1303N | E | 4.8e-08 | 4.3e-08 | 7.5e-08 | 5.533E-08 | 1.721E-08 | 4.300E-08 | 7.500E-08 |
| 26044 | 1303 | D1303D | E | 4.1e-08 | 4.7e-08 | 7.3e-08 | 5.367E-08 | 1.701E-08 | 4.100E-08 | 7.300E-08 |
| 26045 | 1303 | D1303C | E | 4.5e-08 | 4.6e-08 | 6.9e-08 | 5.333E-08 | 1.358E-08 | 4.500E-08 | 6.900E-08 |
| 26046 | 1303 | D1303E | E | 3.4e-08 | 6.7e-08 | 4.9e-08 | 5.000E-08 | 1.652E-08 | 3.400E-08 | 6.700E-08 |
| 26047 | 1303 | D1303Q | E | 4.8e-08 | 8.3e-08 | 8.5e-08 | 7.200E-08 | 2.081E-08 | 4.800E-08 | 8.500E-08 |
| 26048 | 1303 | D1303G | E | 5.2e-08 | 5e-08 | 8.1e-08 | 6.100E-08 | 1.735E-08 | 5.000E-08 | 8.100E-08 |
| 26049 | 1303 | D1303H | E | 5e-08 | 2.7e-08 | 3.3e-08 | 3.667E-08 | 1.193E-08 | 2.700E-08 | 5.000E-08 |
| 26050 | 1303 | D1303I | E | 2.1e-08 | 6.7e-08 | 6.7e-08 | 5.167E-08 | 2.656E-08 | 2.100E-08 | 6.700E-08 |
| 26051 | 1303 | D1303L | E | 2.2e-08 | 8.7e-08 | 5.7e-08 | 5.533E-08 | 3.253E-08 | 2.200E-08 | 8.700E-08 |
| 26052 | 1303 | D1303K | E | 1.9e-08 | 2.2e-08 | 2.8e-08 | 2.300E-08 | 4.583E-09 | 1.900E-08 | 2.800E-08 |
| 26053 | 1303 | D1303M | E | 5.4e-08 | 8.5e-08 | 3.1e-08 | 5.667E-08 | 2.710E-08 | 3.100E-08 | 8.500E-08 |
| 26054 | 1303 | D1303F | E | 1.2e-08 | 2.5e-08 | 4.2e-08 | 2.633E-08 | 1.504E-08 | 1.200E-08 | 4.200E-08 |
| 26055 | 1303 | D1303P | E | 7.4e-09 | 9.2e-08 | 9.5e-09 | 3.630E-08 | 4.825E-08 | 7.400E-09 | 9.200E-08 |
| 26056 | 1303 | D1303S | E | 1.9e-08 | 3.9e-08 | 3.7e-08 | 3.167E-08 | 1.102E-08 | 1.900E-08 | 3.900E-08 |
| 26057 | 1303 | D1303T | E | 4.6e-08 | 5.4e-08 | 3.2e-08 | 4.400E-08 | 1.114E-08 | 3.200E-08 | 5.400E-08 |
| 26058 | 1303 | D1303W | E | 8.6e-09 | 4.9e-08 | 3.1e-08 | 2.953E-08 | 2.024E-08 | 8.600E-09 | 4.900E-08 |
| 26059 | 1303 | D1303Y | E | 4.8e-08 | 3.8e-08 | 4.3e-08 | 4.300E-08 | 5.000E-09 | 3.800E-08 | 4.800E-08 |
| 26060 | 1303 | D1303V | E | 5.6e-08 | 2.8e-08 | 4.7e-08 | 4.367E-08 | 1.429E-08 | 2.800E-08 | 5.600E-08 |
| 26061 | 1304 | T1304A | E | 2.7e-08 | 2.9e-08 | 4.5e-08 | 3.367E-08 | 9.866E-09 | 2.700E-08 | 4.500E-08 |
| 26062 | 1304 | T1304R | E | 3.2e-08 | 3e-08 | 2.7e-08 | 2.967E-08 | 2.517E-09 | 2.700E-08 | 3.200E-08 |
| 26063 | 1304 | T1304N | E | 3.5e-08 | 5.7e-08 | 4.5e-08 | 4.567E-08 | 1.102E-08 | 3.500E-08 | 5.700E-08 |
| 26064 | 1304 | T1304D | E | 1.6e-08 | 4.5e-08 | 3.9e-08 | 3.333E-08 | 1.531E-08 | 1.600E-08 | 4.500E-08 |
| 26065 | 1304 | T1304C | E | 5.8e-08 | 6.5e-08 | 9.3e-08 | 7.200E-08 | 1.852E-08 | 5.800E-08 | 9.300E-08 |
| 26066 | 1304 | T1304E | E | 1.4e-08 | 5.6e-08 | 6.1e-08 | 4.367E-08 | 2.581E-08 | 1.400E-08 | 6.100E-08 |
| 26067 | 1304 | T1304Q | E | 6.3e-08 | 9.7e-08 | 3.3e-08 | 6.433E-08 | 3.202E-08 | 3.300E-08 | 9.700E-08 |
| 26068 | 1304 | T1304G | E | 1.3e-08 | 2.1e-07 | 3.1e-08 | 8.467E-08 | 1.089E-07 | 1.300E-08 | 2.100E-07 |
| 26069 | 1304 | T1304H | E | 7.7e-08 | 4.5e-08 | 7.3e-08 | 6.500E-08 | 1.744E-08 | 4.500E-08 | 7.700E-08 |
| 26070 | 1304 | T1304I | E | 1.7e-08 | 2.8e-08 | 5.5e-08 | 3.333E-08 | 1.955E-08 | 1.700E-08 | 5.500E-08 |
| 26071 | 1304 | T1304L | E | 5.9e-08 | 3.2e-08 | 1.5e-07 | 8.033E-08 | 6.183E-08 | 3.200E-08 | 1.500E-07 |
| 26072 | 1304 | T1304K | E | 4e-08 | 8.6e-08 | 5.2e-08 | 5.933E-08 | 2.386E-08 | 4.000E-08 | 8.600E-08 |
| 26073 | 1304 | T1304M | E | 5.3e-08 | 6.2e-08 | 3.3e-08 | 4.933E-08 | 1.484E-08 | 3.300E-08 | 6.200E-08 |
| 26074 | 1304 | T1304F | E | 3.6e-08 | 3.9e-08 | 9.1e-08 | 5.533E-08 | 3.092E-08 | 3.600E-08 | 9.100E-08 |
| 26075 | 1304 | T1304P | E | 4e-08 | 7.5e-08 | 3.7e-08 | 5.067E-08 | 2.113E-08 | 3.700E-08 | 7.500E-08 |
| 26076 | 1304 | T1304S | E | 8.5e-08 | 3.5e-08 | 3.4e-08 | 5.133E-08 | 2.916E-08 | 3.400E-08 | 8.500E-08 |
| 26077 | 1304 | T1304T | E | 5.1e-08 | 7.7e-08 | 3.6e-08 | 5.467E-08 | 2.074E-08 | 3.600E-08 | 7.700E-08 |
| 26078 | 1304 | T1304W | E | 7.3e-08 | 3.6e-08 | 5.1e-08 | 5.333E-08 | 1.861E-08 | 3.600E-08 | 7.300E-08 |
| 26079 | 1304 | T1304Y | E | 1.8e-08 | 1.2e-08 | 4.5e-08 | 2.500E-08 | 1.758E-08 | 1.200E-08 | 4.500E-08 |
| 26080 | 1304 | T1304V | E | 7.4e-08 | 3.3e-08 | 6.7e-08 | 5.800E-08 | 2.193E-08 | 3.300E-08 | 7.400E-08 |
| 26081 | 1305 | Y1305A | E | 1.2e-07 | 6.8e-08 | 6.1e-08 | 8.300E-08 | 3.223E-08 | 6.100E-08 | 1.200E-07 |
| 26082 | 1305 | Y1305R | E | 9.1e-08 | 1.8e-08 | 4.9e-08 | 5.267E-08 | 3.664E-08 | 1.800E-08 | 9.100E-08 |
| 26083 | 1305 | Y1305N | E | 4.2e-08 | 4.4e-08 | 9.8e-08 | 6.133E-08 | 3.177E-08 | 4.200E-08 | 9.800E-08 |
| 26084 | 1305 | Y1305D | E | 7.2e-08 | 3.5e-08 | 8.4e-08 | 6.367E-08 | 2.554E-08 | 3.500E-08 | 8.400E-08 |
| 26085 | 1305 | Y1305C | E | 7.5e-08 | 1.7e-08 | 8.1e-08 | 5.767E-08 | 3.535E-08 | 1.700E-08 | 8.100E-08 |
| 26086 | 1305 | Y1305E | E | 1.3e-07 | 2.2e-08 | 2.3e-08 | 5.833E-08 | 6.207E-08 | 2.200E-08 | 1.300E-07 |
| 26087 | 1305 | Y1305Q | E | 3.7e-08 | 6.9e-08 | 2.9e-08 | 4.500E-08 | 2.117E-08 | 2.900E-08 | 6.900E-08 |
| 26088 | 1305 | Y1305G | E | 5e-08 | 4.6e-08 | 5.9e-08 | 5.167E-08 | 6.658E-09 | 4.600E-08 | 5.900E-08 |
| 26089 | 1305 | Y1305H | E | 2e-08 | 8.7e-08 | 2.4e-08 | 4.367E-08 | 3.758E-08 | 2.000E-08 | 8.700E-08 |
| 26090 | 1305 | Y1305I | E | 1.8e-08 | 3.7e-08 | 3.1e-08 | 2.867E-08 | 9.713E-09 | 1.800E-08 | 3.700E-08 |
| 26091 | 1305 | Y1305L | E | 2.6e-08 | 4.5e-08 | 9e-08 | 5.367E-08 | 3.287E-08 | 2.600E-08 | 9.000E-08 |
| 26092 | 1305 | Y1305K | E | 4.1e-08 | 1.4e-08 | 4.1e-08 | 3.200E-08 | 1.559E-08 | 1.400E-08 | 4.100E-08 |
| 26093 | 1305 | Y1305M | E | 1.3e-08 | 6.2e-08 | 3.7e-08 | 3.733E-08 | 2.450E-08 | 1.300E-08 | 6.200E-08 |
| 26094 | 1305 | Y1305F | E | 7.2e-08 | 5.5e-08 | 1.1e-08 | 4.600E-08 | 3.148E-08 | 1.100E-08 | 7.200E-08 |

| | | | | | | | | | | |
|---|---|---|---|---|---|---|---|---|---|---|
| 26095 | 1305 | Y1305P | E | 6.1e-08 | 3.1e-08 | 4.1e-08 | 4.433E-08 | 1.528E-08 | 3.100E-08 | 6.100E-08 |
| 26096 | 1305 | Y1305S | E | 1.5e-08 | 5.6e-08 | 8.4e-08 | 5.167E-08 | 3.470E-08 | 1.500E-08 | 8.400E-08 |
| 26097 | 1305 | Y1305T | E | 3.7e-08 | 2.3e-08 | 9.6e-08 | 5.200E-08 | 3.874E-08 | 2.300E-08 | 9.600E-08 |
| 26098 | 1305 | Y1305W | E | 4.6e-08 | 1.8e-08 | 6.2e-08 | 4.200E-08 | 2.227E-08 | 1.800E-08 | 6.200E-08 |
| 26099 | 1305 | Y1305Y | E | 2.2e-08 | 3.2e-08 | 2e-08 | 2.467E-08 | 6.429E-09 | 2.000E-08 | 3.200E-08 |
| 26100 | 1305 | Y1305V | E | 3.5e-08 | 1.9e-07 | 5.1e-08 | 9.200E-08 | 8.525E-08 | 3.500E-08 | 1.900E-07 |
| 26101 | 1306 | I1306A | E | 3.1e-08 | 6.1e-08 | 5.1e-08 | 4.767E-08 | 1.528E-08 | 3.100E-08 | 6.100E-08 |
| 26102 | 1306 | I1306R | E | 4.7e-08 | 9.8e-08 | 8e-08 | 7.500E-08 | 2.587E-08 | 4.700E-08 | 9.800E-08 |
| 26103 | 1306 | I1306N | E | 1.5e-08 | 1.7e-07 | 5.7e-08 | 8.067E-08 | 8.016E-08 | 1.500E-08 | 1.700E-07 |
| 26104 | 1306 | I1306D | E | 3.6e-08 | 1.3e-08 | 1.1e-08 | 2.000E-08 | 1.389E-08 | 1.100E-08 | 3.600E-08 |
| 26105 | 1306 | I1306C | E | 2.4e-08 | 6e-08 | 8.3e-08 | 5.567E-08 | 2.974E-08 | 2.400E-08 | 8.300E-08 |
| 26106 | 1306 | I1306E | E | 5.9e-08 | 1.3e-07 | 3.3e-08 | 7.400E-08 | 5.021E-08 | 3.300E-08 | 1.300E-07 |
| 26107 | 1306 | I1306Q | E | 4.7e-08 | 1.4e-08 | 3.7e-08 | 3.267E-08 | 1.692E-08 | 1.400E-08 | 4.700E-08 |
| 26108 | 1306 | I1306G | E | 1.2e-07 | 2.1e-08 | 2.8e-08 | 5.633E-08 | 5.525E-08 | 2.100E-08 | 1.200E-07 |
| 26109 | 1306 | I1306H | E | 1.1e-07 | 2.9e-08 | 7.8e-08 | 7.233E-08 | 4.080E-08 | 2.900E-08 | 1.100E-07 |
| 26110 | 1306 | I1306I | E | 5e-08 | 1.1e-07 | 6.8e-08 | 7.600E-08 | 3.079E-08 | 5.000E-08 | 1.100E-07 |
| 26111 | 1306 | I1306L | E | 2.4e-08 | 1.4e-08 | 5.2e-08 | 3.000E-08 | 1.970E-08 | 1.400E-08 | 5.200E-08 |
| 26112 | 1306 | I1306K | E | 3.9e-08 | 6e-08 | 3e-08 | 4.300E-08 | 1.539E-08 | 3.000E-08 | 6.000E-08 |
| 26113 | 1306 | I1306M | E | 8.5e-08 | 4.5e-08 | 7.4e-09 | 4.580E-08 | 3.881E-08 | 7.400E-09 | 8.500E-08 |
| 26114 | 1306 | I1306F | E | 1.8e-07 | 1.9e-08 | 3.5e-08 | 7.800E-08 | 8.870E-08 | 1.900E-08 | 1.800E-07 |
| 26115 | 1306 | I1306P | E | 2.9e-08 | 4.7e-08 | 5.4e-08 | 4.333E-08 | 1.290E-08 | 2.900E-08 | 5.400E-08 |
| 26116 | 1306 | I1306S | E | 3.2e-08 | 5.5e-08 | 1.6e-08 | 3.433E-08 | 1.960E-08 | 1.600E-08 | 5.500E-08 |
| 26117 | 1306 | I1306T | E | 4.2e-08 | 1.5e-08 | 2e-08 | 2.567E-08 | 1.436E-08 | 1.500E-08 | 4.200E-08 |
| 26118 | 1306 | I1306W | E | 9.8e-08 | 3.6e-08 | 3e-08 | 5.467E-08 | 3.765E-08 | 3.000E-08 | 9.800E-08 |
| 26119 | 1306 | I1306Y | E | 1.4e-08 | 3.1e-08 | 2.4e-08 | 2.300E-08 | 8.544E-09 | 1.400E-08 | 3.100E-08 |
| 26120 | 1306 | I1306V | E | 4.4e-08 | 1.1e-08 | 5.6e-08 | 3.700E-08 | 2.330E-08 | 1.100E-08 | 5.600E-08 |
| 26121 | 1307 | H1307A | E | 4.5e-08 | 5.4e-08 | 4.9e-08 | 4.933E-08 | 4.509E-09 | 4.500E-08 | 5.400E-08 |
| 26122 | 1307 | H1307R | E | 6.1e-08 | 6e-08 | 1.1e-08 | 4.400E-08 | 2.858E-08 | 1.100E-08 | 6.100E-08 |
| 26123 | 1307 | H1307N | E | 5.9e-08 | 4.6e-08 | 5.6e-08 | 5.367E-08 | 6.807E-09 | 4.600E-08 | 5.900E-08 |
| 26124 | 1307 | H1307D | E | 3.7e-08 | 3.4e-08 | 7.8e-08 | 4.967E-08 | 2.458E-08 | 3.400E-08 | 7.800E-08 |
| 26125 | 1307 | H1307C | E | 2e-08 | 5e-08 | 7.4e-08 | 4.800E-08 | 2.706E-08 | 2.000E-08 | 7.400E-08 |
| 26126 | 1307 | H1307E | E | 3.4e-08 | 2.8e-08 | 2.6e-08 | 2.933E-08 | 4.163E-09 | 2.600E-08 | 3.400E-08 |
| 26127 | 1307 | H1307Q | E | 2.8e-08 | 6.1e-08 | 1.8e-08 | 3.567E-08 | 2.250E-08 | 1.800E-08 | 6.100E-08 |
| 26128 | 1307 | H1307G | E | 2.8e-08 | 7.4e-08 | 4e-08 | 4.733E-08 | 2.386E-08 | 2.800E-08 | 7.400E-08 |
| 26129 | 1307 | H1307H | E | 5.8e-08 | 2.5e-08 | 5e-08 | 4.433E-08 | 1.721E-08 | 2.500E-08 | 5.800E-08 |
| 26130 | 1307 | H1307I | E | 4.8e-08 | 4.4e-08 | 9.9e-08 | 6.367E-08 | 3.066E-08 | 4.400E-08 | 9.900E-08 |
| 26131 | 1307 | H1307L | E | 2.4e-08 | 5.3e-08 | 3.5e-08 | 3.733E-08 | 1.464E-08 | 2.400E-08 | 5.300E-08 |
| 26132 | 1307 | H1307K | E | 8.9e-08 | 6.8e-08 | 3.1e-08 | 6.267E-08 | 2.937E-08 | 3.100E-08 | 8.900E-08 |
| 26133 | 1307 | H1307M | E | 2.3e-08 | 5.6e-08 | 4.6e-08 | 4.167E-08 | 1.692E-08 | 2.300E-08 | 5.600E-08 |
| 26134 | 1307 | H1307F | E | 8.3e-08 | 3.7e-08 | 1.1e-07 | 7.667E-08 | 3.691E-08 | 3.700E-08 | 1.100E-07 |
| 26135 | 1307 | H1307P | E | 1.9e-08 | 9.3e-08 | 3.6e-08 | 4.933E-08 | 3.876E-08 | 1.900E-08 | 9.300E-08 |
| 26136 | 1307 | H1307S | E | 7.1e-08 | 4.4e-08 | 1.1e-07 | 7.500E-08 | 3.318E-08 | 4.400E-08 | 1.100E-07 |
| 26137 | 1307 | H1307T | E | 3.5e-08 | 1.3e-07 | 1.9e-07 | 1.183E-07 | 7.816E-08 | 3.500E-08 | 1.900E-07 |
| 26138 | 1307 | H1307W | E | 3.9e-08 | 3.3e-08 | 2.1e-08 | 3.100E-08 | 9.165E-09 | 2.100E-08 | 3.900E-08 |
| 26139 | 1307 | H1307Y | E | 1.5e-07 | 6.5e-08 | 5e-08 | 8.833E-08 | 5.393E-08 | 5.000E-08 | 1.500E-07 |
| 26140 | 1307 | H1307V | E | 2.8e-08 | 1.8e-08 | 1.6e-08 | 2.067E-08 | 6.429E-09 | 1.600E-08 | 2.800E-08 |
| 26141 | 1308 | W1308A | E | 1.2e-08 | 5.1e-08 | 4.8e-08 | 3.700E-08 | 2.170E-08 | 1.200E-08 | 5.100E-08 |
| 26142 | 1308 | W1308R | E | 3e-08 | 2.2e-08 | 9.3e-08 | 4.833E-08 | 3.889E-08 | 2.200E-08 | 9.300E-08 |
| 26143 | 1308 | W1308N | E | 5.3e-08 | 2.5e-08 | 2e-08 | 3.267E-08 | 1.779E-08 | 2.000E-08 | 5.300E-08 |
| 26144 | 1308 | W1308D | E | 1.4e-08 | 6.6e-08 | 1.8e-08 | 3.267E-08 | 2.894E-08 | 1.400E-08 | 6.600E-08 |
| 26145 | 1308 | W1308C | E | 6.5e-08 | 4.3e-08 | 7.4e-08 | 6.067E-08 | 1.595E-08 | 4.300E-08 | 7.400E-08 |
| 26146 | 1308 | W1308E | E | 2e-08 | 5.4e-08 | 1.8e-08 | 3.067E-08 | 2.023E-08 | 1.800E-08 | 5.400E-08 |
| 26147 | 1308 | W1308Q | E | 5.6e-08 | 1.3e-08 | 6.3e-08 | 4.400E-08 | 2.707E-08 | 1.300E-08 | 6.300E-08 |
| 26148 | 1308 | W1308G | E | 1.3e-08 | 4.3e-08 | 4e-08 | 3.200E-08 | 1.652E-08 | 1.300E-08 | 4.300E-08 |
| 26149 | 1308 | W1308H | E | 3.1e-08 | 2.8e-08 | 1.9e-08 | 2.600E-08 | 6.245E-09 | 1.900E-08 | 3.100E-08 |
| 26150 | 1308 | W1308I | E | 8.6e-08 | 6.2e-08 | 2.8e-08 | 5.867E-08 | 2.914E-08 | 2.800E-08 | 8.600E-08 |
| 26151 | 1308 | W1308L | E | 7e-08 | 9e-09 | 7.2e-08 | 5.033E-08 | 3.581E-08 | 9.000E-09 | 7.200E-08 |
| 26152 | 1308 | W1308K | E | 5.6e-08 | 4.9e-08 | 5.1e-08 | 5.200E-08 | 3.606E-09 | 4.900E-08 | 5.600E-08 |
| 26153 | 1308 | W1308M | E | 1.6e-08 | 3.5e-08 | 1.1e-08 | 2.067E-08 | 1.266E-08 | 1.100E-08 | 3.500E-08 |
| 26154 | 1308 | W1308F | E | 3.2e-08 | 1.1e-07 | 1.2e-08 | 5.133E-08 | 5.178E-08 | 1.200E-08 | 1.100E-07 |
| 26155 | 1308 | W1308P | E | 3.8e-08 | 1.2e-08 | 6.2e-08 | 3.733E-08 | 2.501E-08 | 1.200E-08 | 6.200E-08 |
| 26156 | 1308 | W1308S | E | 7.1e-08 | 1.4e-08 | 2.3e-08 | 3.600E-08 | 3.064E-08 | 1.400E-08 | 7.100E-08 |
| 26157 | 1308 | W1308T | E | 1.4e-08 | 5.8e-08 | 2.6e-08 | 3.267E-08 | 2.274E-08 | 1.400E-08 | 5.800E-08 |
| 26158 | 1308 | W1308W | E | 2.6e-08 | 8.1e-08 | 9.9e-08 | 6.867E-08 | 3.803E-08 | 2.600E-08 | 9.900E-08 |
| 26159 | 1308 | W1308Y | E | 2.7e-08 | 8.3e-08 | 1.3e-08 | 4.100E-08 | 3.704E-08 | 1.300E-08 | 8.300E-08 |
| 26160 | 1308 | W1308V | E | 3.4e-08 | 5.9e-08 | 4.3e-08 | 4.533E-08 | 1.266E-08 | 3.400E-08 | 5.900E-08 |
| 26161 | 1309 | V1309A | E | 6.6e-08 | 6.5e-08 | 2.9e-08 | 5.333E-08 | 2.108E-08 | 2.900E-08 | 6.600E-08 |
| 26162 | 1309 | V1309R | E | 5.6e-08 | 1.8e-08 | 7.7e-08 | 5.033E-08 | 2.991E-08 | 1.800E-08 | 7.700E-08 |
| 26163 | 1309 | V1309N | E | 4.9e-08 | 5.9e-08 | 8.1e-08 | 6.300E-08 | 1.637E-08 | 4.900E-08 | 8.100E-08 |
| 26164 | 1309 | V1309D | E | 2.7e-08 | 5.5e-08 | 1.1e-07 | 6.400E-08 | 4.223E-08 | 2.700E-08 | 1.100E-07 |
| 26165 | 1309 | V1309C | E | 4.1e-08 | 4.1e-08 | 5e-08 | 4.400E-08 | 5.196E-09 | 4.100E-08 | 5.000E-08 |
| 26166 | 1309 | V1309E | E | 2.4e-08 | 3.2e-08 | 3.2e-08 | 2.933E-08 | 4.619E-09 | 2.400E-08 | 3.200E-08 |
| 26167 | 1309 | V1309Q | E | 2.4e-08 | 5e-08 | 6.5e-08 | 4.633E-08 | 2.074E-08 | 2.400E-08 | 6.500E-08 |
| 26168 | 1309 | V1309G | E | 3.2e-08 | 8.3e-08 | 2.7e-08 | 4.733E-08 | 3.099E-08 | 2.700E-08 | 8.300E-08 |
| 26169 | 1309 | V1309H | E | 3.4e-08 | 3e-08 | 3.1e-08 | 3.167E-08 | 2.082E-09 | 3.000E-08 | 3.400E-08 |
| 26170 | 1309 | V1309I | E | 3.8e-08 | 1.4e-08 | 5e-08 | 3.400E-08 | 1.833E-08 | 1.400E-08 | 5.000E-08 |
| 26171 | 1309 | V1309L | E | 7.9e-08 | 2.7e-08 | 5.1e-08 | 5.233E-08 | 2.603E-08 | 2.700E-08 | 7.900E-08 |
| 26172 | 1309 | V1309K | E | 4.1e-08 | 6.3e-08 | 2.2e-08 | 4.200E-08 | 2.052E-08 | 2.200E-08 | 6.300E-08 |
| 26173 | 1309 | V1309M | E | 2e-08 | 3.4e-08 | 2e-08 | 2.467E-08 | 8.083E-09 | 2.000E-08 | 3.400E-08 |
| 26174 | 1309 | V1309F | E | 9e-08 | 3.4e-08 | 5.3e-08 | 5.900E-08 | 2.848E-08 | 3.400E-08 | 9.000E-08 |
| 26175 | 1309 | V1309P | E | 3.8e-08 | 2.9e-08 | 4.1e-08 | 3.600E-08 | 6.245E-09 | 2.900E-08 | 4.100E-08 |
| 26176 | 1309 | V1309S | E | 4.6e-08 | 1.8e-08 | 4.3e-08 | 3.567E-08 | 1.537E-08 | 1.800E-08 | 4.600E-08 |
| 26177 | 1309 | V1309T | E | 2e-08 | 4.6e-08 | 2.7e-08 | 3.100E-08 | 1.345E-08 | 2.000E-08 | 4.600E-08 |
| 26178 | 1309 | V1309W | E | 1.8e-08 | 3.5e-08 | 7.1e-08 | 4.133E-08 | 2.706E-08 | 1.800E-08 | 7.100E-08 |
| 26179 | 1309 | V1309Y | E | 3.5e-08 | 4.3e-08 | 5.3e-08 | 4.367E-08 | 9.018E-09 | 3.500E-08 | 5.300E-08 |

| | | | | | | | | | | |
|---|---|---|---|---|---|---|---|---|---|---|
| 26180 | 1309 | V1309V | E | 2.2e-08 | 3.5e-08 | 5.7e-08 | 3.800E-08 | 1.769E-08 | 2.200E-08 | 5.700E-08 |
| 26181 | 1310 | R1310A | E | 5.9e-08 | 7.2e-08 | 5.5e-08 | 6.200E-08 | 8.888E-09 | 5.500E-08 | 7.200E-08 |
| 26182 | 1310 | R1310R | E | 1.1e-07 | 3.3e-08 | 8.1e-08 | 7.467E-08 | 3.889E-08 | 3.300E-08 | 1.100E-07 |
| 26183 | 1310 | R1310N | E | 2.3e-08 | 4.1e-08 | 3.1e-08 | 3.167E-08 | 9.018E-09 | 2.300E-08 | 4.100E-08 |
| 26184 | 1310 | R1310D | E | 2.9e-08 | 6.5e-08 | 6.2e-08 | 5.200E-08 | 1.997E-08 | 2.900E-08 | 6.500E-08 |
| 26185 | 1310 | R1310C | E | 2.6e-08 | 9.1e-08 | 1.5e-08 | 4.400E-08 | 4.107E-08 | 1.500E-08 | 9.100E-08 |
| 26186 | 1310 | R1310E | E | 6e-08 | 4.7e-08 | 2e-08 | 4.233E-08 | 2.040E-08 | 2.000E-08 | 6.000E-08 |
| 26187 | 1310 | R1310Q | E | 5.4e-08 | 1.4e-07 | 6.3e-08 | 8.567E-08 | 4.727E-08 | 5.400E-08 | 1.400E-07 |
| 26188 | 1310 | R1310G | E | 3.5e-08 | 6.8e-08 | 9.1e-08 | 6.467E-08 | 2.815E-08 | 3.500E-08 | 9.100E-08 |
| 26189 | 1310 | R1310H | E | 3.1e-08 | 4.6e-08 | 3.5e-08 | 3.733E-08 | 7.767E-09 | 3.100E-08 | 4.600E-08 |
| 26190 | 1310 | R1310I | E | 3.4e-08 | 2.8e-08 | 1.3e-07 | 6.400E-08 | 5.724E-08 | 2.800E-08 | 1.300E-07 |
| 26191 | 1310 | R1310L | E | 4.2e-08 | 4.8e-08 | 5.1e-08 | 4.700E-08 | 4.583E-09 | 4.200E-08 | 5.100E-08 |
| 26192 | 1310 | R1310K | E | 2.4e-08 | 5e-08 | 2.1e-08 | 3.167E-08 | 1.595E-08 | 2.100E-08 | 5.000E-08 |
| 26193 | 1310 | R1310M | E | 8e-08 | 1.5e-08 | 2.6e-08 | 4.033E-08 | 3.479E-08 | 1.500E-08 | 8.000E-08 |
| 26194 | 1310 | R1310F | E | 4.6e-08 | 4.2e-08 | 6.5e-08 | 5.100E-08 | 1.229E-08 | 4.200E-08 | 6.500E-08 |
| 26195 | 1310 | R1310P | E | 3.8e-08 | 2.7e-08 | 3.4e-08 | 3.300E-08 | 5.568E-09 | 2.700E-08 | 3.800E-08 |
| 26196 | 1310 | R1310S | E | 3e-08 | 5.1e-08 | 3.3e-08 | 3.800E-08 | 1.136E-08 | 3.000E-08 | 5.100E-08 |
| 26197 | 1310 | R1310T | E | 3.2e-08 | 9.7e-08 | 5.9e-08 | 6.267E-08 | 3.265E-08 | 3.200E-08 | 9.700E-08 |
| 26198 | 1310 | R1310W | E | 9e-08 | 3.7e-08 | 3.4e-08 | 5.367E-08 | 3.150E-08 | 3.400E-08 | 9.000E-08 |
| 26199 | 1310 | R1310Y | E | 3.2e-08 | 1.1e-08 | 4.4e-08 | 2.900E-08 | 1.670E-08 | 1.100E-08 | 4.400E-08 |
| 26200 | 1310 | R1310V | E | 7.3e-08 | 6.8e-08 | 2.9e-08 | 5.667E-08 | 2.409E-08 | 2.900E-08 | 7.300E-08 |
| 26201 | 1311 | Q1311A | E | 2.3e-08 | 3.3e-08 | 4.3e-08 | 3.300E-08 | 1.000E-08 | 2.300E-08 | 4.300E-08 |
| 26202 | 1311 | Q1311R | E | 2.8e-08 | 3.2e-08 | 3.5e-08 | 3.167E-08 | 3.512E-09 | 2.800E-08 | 3.500E-08 |
| 26203 | 1311 | Q1311N | E | 6.9e-08 | 1.2e-07 | 3.1e-08 | 7.333E-08 | 4.466E-08 | 3.100E-08 | 1.200E-07 |
| 26204 | 1311 | Q1311D | E | 2.7e-08 | 2.3e-08 | 1.3e-07 | 6.000E-08 | 6.065E-08 | 2.300E-08 | 1.300E-07 |
| 26205 | 1311 | Q1311C | E | 9.1e-08 | 1.7e-08 | 9.2e-08 | 6.667E-08 | 4.302E-08 | 1.700E-08 | 9.200E-08 |
| 26206 | 1311 | Q1311E | E | 1.3e-07 | 2.7e-08 | 4.7e-08 | 6.800E-08 | 5.462E-08 | 2.700E-08 | 1.300E-07 |
| 26207 | 1311 | Q1311Q | E | 1.2e-07 | 6.3e-08 | 4.6e-08 | 7.633E-08 | 3.876E-08 | 4.600E-08 | 1.200E-07 |
| 26208 | 1311 | Q1311G | E | 2.1e-08 | 4.5e-08 | 3.3e-08 | 3.300E-08 | 1.200E-08 | 2.100E-08 | 4.500E-08 |
| 26209 | 1311 | Q1311H | E | 1.9e-08 | 4.5e-08 | 4.9e-08 | 3.767E-08 | 1.629E-08 | 1.900E-08 | 4.900E-08 |
| 26210 | 1311 | Q1311I | E | 2.2e-08 | 3.2e-08 | 2.4e-08 | 2.600E-08 | 5.292E-09 | 2.200E-08 | 3.200E-08 |
| 26211 | 1311 | Q1311L | E | 7.7e-08 | 5.7e-08 | 2.8e-08 | 5.400E-08 | 2.464E-08 | 2.800E-08 | 7.700E-08 |
| 26212 | 1311 | Q1311K | E | 5.3e-08 | 1.9e-08 | 2.2e-08 | 3.133E-08 | 1.882E-08 | 1.900E-08 | 5.300E-08 |
| 26213 | 1311 | Q1311M | E | 3.8e-08 | 4.5e-08 | 1.4e-08 | 3.233E-08 | 1.626E-08 | 1.400E-08 | 4.500E-08 |
| 26214 | 1311 | Q1311F | E | 7.4e-08 | 8.3e-08 | 1.7e-08 | 5.800E-08 | 3.579E-08 | 1.700E-08 | 8.300E-08 |
| 26215 | 1311 | Q1311P | E | 2.4e-08 | 5.3e-08 | 2.9e-08 | 3.533E-08 | 1.550E-08 | 2.400E-08 | 5.300E-08 |
| 26216 | 1311 | Q1311S | E | 5.8e-08 | 5.8e-08 | 5e-08 | 5.533E-08 | 4.619E-09 | 5.000E-08 | 5.800E-08 |
| 26217 | 1311 | Q1311T | E | 3.5e-08 | 3.7e-08 | 5.8e-08 | 4.333E-08 | 1.274E-08 | 3.500E-08 | 5.800E-08 |
| 26218 | 1311 | Q1311W | E | 7.3e-08 | 7e-08 | 1.8e-08 | 5.367E-08 | 3.092E-08 | 1.800E-08 | 7.300E-08 |
| 26219 | 1311 | Q1311Y | E | 1.5e-07 | 9.5e-08 | 3.4e-08 | 9.300E-08 | 5.803E-08 | 3.400E-08 | 1.500E-07 |
| 26220 | 1311 | Q1311V | E | 3.6e-08 | 2.3e-08 | 3.3e-08 | 3.067E-08 | 6.807E-09 | 2.300E-08 | 3.600E-08 |
| 26221 | 1312 | A1312A | E | 4.7e-08 | 3.5e-08 | 2.1e-08 | 3.433E-08 | 1.301E-08 | 2.100E-08 | 4.700E-08 |
| 26222 | 1312 | A1312R | E | 3.6e-08 | 3e-08 | 9.1e-08 | 5.233E-08 | 3.362E-08 | 3.000E-08 | 9.100E-08 |
| 26223 | 1312 | A1312N | E | 8.1e-08 | 2e-08 | 5.5e-08 | 5.200E-08 | 3.061E-08 | 2.000E-08 | 8.100E-08 |
| 26224 | 1312 | A1312D | E | 8.9e-08 | 3e-08 | 1.3e-07 | 8.300E-08 | 5.027E-08 | 3.000E-08 | 1.300E-07 |
| 26225 | 1312 | A1312C | E | 5.2e-08 | 1.8e-08 | 4.1e-08 | 3.700E-08 | 1.735E-08 | 1.800E-08 | 5.200E-08 |
| 26226 | 1312 | A1312E | E | 8e-08 | 4.2e-08 | 5.3e-08 | 5.833E-08 | 1.955E-08 | 4.200E-08 | 8.000E-08 |
| 26227 | 1312 | A1312Q | E | 5.3e-08 | 2.8e-08 | 1.3e-08 | 3.133E-08 | 2.021E-08 | 1.300E-08 | 5.300E-08 |
| 26228 | 1312 | A1312G | E | 3.9e-08 | 1.3e-07 | 7.4e-09 | 5.880E-08 | 6.365E-08 | 7.400E-09 | 1.300E-07 |
| 26229 | 1312 | A1312H | E | 5.6e-08 | 6.6e-08 | 3.6e-08 | 5.267E-08 | 1.528E-08 | 3.600E-08 | 6.600E-08 |
| 26230 | 1312 | A1312I | E | 5.2e-08 | 3.1e-08 | 4e-08 | 4.100E-08 | 1.054E-08 | 3.100E-08 | 5.200E-08 |
| 26231 | 1312 | A1312L | E | 5.4e-08 | 1e-07 | 4.9e-08 | 6.767E-08 | 2.811E-08 | 4.900E-08 | 1.000E-07 |
| 26232 | 1312 | A1312K | E | 4.4e-08 | 1.8e-08 | 7.1e-08 | 4.433E-08 | 2.650E-08 | 1.800E-08 | 7.100E-08 |
| 26233 | 1312 | A1312M | E | 3.5e-08 | 4.1e-08 | 1e-07 | 5.867E-08 | 3.592E-08 | 3.500E-08 | 1.000E-07 |
| 26234 | 1312 | A1312F | E | 5.1e-08 | 1.7e-08 | 3.6e-08 | 3.467E-08 | 1.704E-08 | 1.700E-08 | 5.100E-08 |
| 26235 | 1312 | A1312P | E | 3.8e-08 | 5.5e-08 | 7.5e-08 | 5.600E-08 | 1.852E-08 | 3.800E-08 | 7.500E-08 |
| 26236 | 1312 | A1312S | E | 1.4e-07 | 8.3e-08 | 4.6e-08 | 8.967E-08 | 4.735E-08 | 4.600E-08 | 1.400E-07 |
| 26237 | 1312 | A1312T | E | 4e-08 | 7.9e-08 | 1.1e-07 | 7.633E-08 | 3.508E-08 | 4.000E-08 | 1.100E-07 |
| 26238 | 1312 | A1312W | E | 4.4e-08 | 2.3e-08 | 5e-08 | 3.900E-08 | 1.418E-08 | 2.300E-08 | 5.000E-08 |
| 26239 | 1312 | A1312Y | E | 3.4e-08 | 1.6e-08 | 9.1e-08 | 4.700E-08 | 3.915E-08 | 1.600E-08 | 9.100E-08 |
| 26240 | 1312 | A1312V | E | 3.6e-08 | 2.5e-08 | 5.5e-08 | 3.867E-08 | 1.518E-08 | 2.500E-08 | 5.500E-08 |
| 26241 | 1313 | P1313A | E | 1.1e-08 | 4.5e-08 | 4.4e-08 | 3.333E-08 | 1.935E-08 | 1.100E-08 | 4.500E-08 |
| 26242 | 1313 | P1313R | E | 6e-08 | 5.3e-08 | 5.1e-08 | 5.467E-08 | 4.726E-09 | 5.100E-08 | 6.000E-08 |
| 26243 | 1313 | P1313N | E | 2.5e-08 | 1.3e-08 | 6.6e-08 | 3.467E-08 | 2.779E-08 | 1.300E-08 | 6.600E-08 |
| 26244 | 1313 | P1313D | E | 2.2e-08 | 4.5e-08 | 1.5e-07 | 7.233E-08 | 6.824E-08 | 2.200E-08 | 1.500E-07 |
| 26245 | 1313 | P1313C | E | 3.4e-08 | 8.6e-08 | 4.9e-08 | 5.633E-08 | 2.676E-08 | 3.400E-08 | 8.600E-08 |
| 26246 | 1313 | P1313E | E | 3.9e-08 | 3.3e-08 | 5.5e-08 | 4.233E-08 | 1.137E-08 | 3.300E-08 | 5.500E-08 |
| 26247 | 1313 | P1313Q | E | 6.3e-08 | 8.3e-09 | 2.1e-08 | 3.077E-08 | 2.863E-08 | 8.300E-09 | 6.300E-08 |
| 26248 | 1313 | P1313G | E | 3.1e-08 | 2.7e-08 | 6.6e-08 | 4.133E-08 | 2.146E-08 | 2.700E-08 | 6.600E-08 |
| 26249 | 1313 | P1313H | E | 4.5e-08 | 1e-07 | 4.6e-08 | 6.367E-08 | 3.147E-08 | 4.500E-08 | 1.000E-07 |
| 26250 | 1313 | P1313I | E | 3.4e-08 | 2.8e-08 | 1.8e-08 | 2.667E-08 | 8.083E-09 | 1.800E-08 | 3.400E-08 |
| 26251 | 1313 | P1313L | E | 3.1e-08 | 6.6e-08 | 4.4e-08 | 4.700E-08 | 1.769E-08 | 3.100E-08 | 6.600E-08 |
| 26252 | 1313 | P1313K | E | 4.2e-08 | 7.7e-08 | 7.2e-08 | 6.367E-08 | 1.893E-08 | 4.200E-08 | 7.700E-08 |
| 26253 | 1313 | P1313M | E | 5.1e-08 | 4.2e-08 | 2e-08 | 3.767E-08 | 1.595E-08 | 2.000E-08 | 5.100E-08 |
| 26254 | 1313 | P1313F | E | 1.7e-08 | 1.2e-07 | 3.8e-08 | 5.833E-08 | 5.443E-08 | 1.700E-08 | 1.200E-07 |
| 26255 | 1313 | P1313P | E | 5.1e-08 | 2.9e-08 | 4.3e-08 | 4.100E-08 | 1.114E-08 | 2.900E-08 | 5.100E-08 |
| 26256 | 1313 | P1313S | E | 7.3e-08 | 7.5e-08 | 7.3e-08 | 7.367E-08 | 1.155E-09 | 7.300E-08 | 7.500E-08 |
| 26257 | 1313 | P1313T | E | 1.6e-08 | 5.9e-08 | 6.9e-08 | 4.800E-08 | 2.816E-08 | 1.600E-08 | 6.900E-08 |
| 26258 | 1313 | P1313W | E | 2.2e-08 | 6.1e-08 | 7.1e-08 | 5.133E-08 | 2.589E-08 | 2.200E-08 | 7.100E-08 |
| 26259 | 1313 | P1313Y | E | 3.6e-08 | 2.3e-08 | 6.9e-08 | 4.267E-08 | 2.371E-08 | 2.300E-08 | 6.900E-08 |
| 26260 | 1313 | P1313V | E | 8.2e-08 | 8.1e-08 | 6.3e-08 | 7.533E-08 | 1.069E-08 | 6.300E-08 | 8.200E-08 |
| 26261 | 1314 | G1314A | E | 8.2e-08 | 1.7e-08 | 3.4e-08 | 4.433E-08 | 3.371E-08 | 1.700E-08 | 8.200E-08 |
| 26262 | 1314 | G1314R | E | 1.8e-08 | 3.2e-08 | 5.5e-08 | 3.500E-08 | 1.868E-08 | 1.800E-08 | 5.500E-08 |
| 26263 | 1314 | G1314N | E | 2.6e-08 | 1e-07 | 2.7e-08 | 5.100E-08 | 4.244E-08 | 2.600E-08 | 1.000E-07 |
| 26264 | 1314 | G1314D | E | 1.2e-07 | 5e-08 | 2.9e-08 | 6.633E-08 | 4.765E-08 | 2.900E-08 | 1.200E-07 |

| | | | | | | | | | | |
|---|---|---|---|---|---|---|---|---|---|---|
| 26265 | 1314 | G1314C | E | 5.1e-08 | 8.4e-08 | 2.9e-08 | 5.467E-08 | 2.768E-08 | 2.900E-08 | 8.400E-08 |
| 26266 | 1314 | G1314E | E | 2.4e-08 | 2.4e-08 | 3.5e-08 | 2.767E-08 | 6.351E-09 | 2.400E-08 | 3.500E-08 |
| 26267 | 1314 | G1314Q | E | 1.3e-08 | 2.4e-08 | 5.8e-08 | 3.167E-08 | 2.346E-08 | 1.300E-08 | 5.800E-08 |
| 26268 | 1314 | G1314G | E | 3.3e-08 | 4.1e-08 | 2.8e-08 | 3.400E-08 | 6.557E-09 | 2.800E-08 | 4.100E-08 |
| 26269 | 1314 | G1314H | E | 4.9e-08 | 7.1e-08 | 3.3e-08 | 5.100E-08 | 1.908E-08 | 3.300E-08 | 7.100E-08 |
| 26270 | 1314 | G1314I | E | 5.1e-08 | 2e-08 | 8.9e-09 | 2.663E-08 | 2.182E-08 | 8.900E-09 | 5.100E-08 |
| 26271 | 1314 | G1314L | E | 2.6e-08 | 3.9e-08 | 3.5e-08 | 3.333E-08 | 6.658E-09 | 2.600E-08 | 3.900E-08 |
| 26272 | 1314 | G1314K | E | 7.7e-08 | 2.3e-08 | 6.1e-08 | 5.367E-08 | 2.774E-08 | 2.300E-08 | 7.700E-08 |
| 26273 | 1314 | G1314M | E | 6e-08 | 5.4e-08 | 3.7e-08 | 5.033E-08 | 1.193E-08 | 3.700E-08 | 6.000E-08 |
| 26274 | 1314 | G1314F | E | 3.7e-08 | 2.3e-08 | 2.3e-08 | 2.767E-08 | 8.083E-09 | 2.300E-08 | 3.700E-08 |
| 26275 | 1314 | G1314P | E | 2.5e-08 | 5e-08 | 5.1e-08 | 4.200E-08 | 1.473E-08 | 2.500E-08 | 5.100E-08 |
| 26276 | 1314 | G1314S | E | 6.9e-08 | 1.7e-08 | 4.7e-08 | 4.433E-08 | 2.610E-08 | 1.700E-08 | 6.900E-08 |
| 26277 | 1314 | G1314T | E | 2.4e-08 | 2.7e-08 | 3e-08 | 2.700E-08 | 3.000E-09 | 2.400E-08 | 3.000E-08 |
| 26278 | 1314 | G1314W | E | 1.8e-08 | 5e-08 | 3.9e-08 | 3.567E-08 | 1.626E-08 | 1.800E-08 | 5.000E-08 |
| 26279 | 1314 | G1314Y | E | 3.1e-08 | 3.4e-08 | 3.9e-08 | 3.467E-08 | 4.041E-09 | 3.100E-08 | 3.900E-08 |
| 26280 | 1314 | G1314V | E | 2.2e-08 | 3.4e-08 | 1.9e-08 | 2.500E-08 | 7.937E-09 | 1.900E-08 | 3.400E-08 |
| 26281 | 1315 | K1315A | E | 4.5e-08 | 5.8e-08 | 6.5e-08 | 5.600E-08 | 1.015E-08 | 4.500E-08 | 6.500E-08 |
| 26282 | 1315 | K1315R | E | 1.5e-08 | 2e-08 | 8.1e-08 | 3.867E-08 | 3.675E-08 | 1.500E-08 | 8.100E-08 |
| 26283 | 1315 | K1315N | E | 4.3e-08 | 1.4e-07 | 4.6e-08 | 7.633E-08 | 5.516E-08 | 4.300E-08 | 1.400E-07 |
| 26284 | 1315 | K1315D | E | 7.8e-08 | 4.4e-08 | 1.8e-07 | 1.007E-07 | 7.078E-08 | 4.400E-08 | 1.800E-07 |
| 26285 | 1315 | K1315C | E | 5.4e-08 | 2.8e-08 | 2.3e-08 | 3.500E-08 | 1.664E-08 | 2.300E-08 | 5.400E-08 |
| 26286 | 1315 | K1315E | E | 6.2e-08 | 4.3e-08 | 2.6e-08 | 4.367E-08 | 1.801E-08 | 2.600E-08 | 6.200E-08 |
| 26287 | 1315 | K1315Q | E | 3.4e-08 | 7.1e-08 | 3e-08 | 4.500E-08 | 2.261E-08 | 3.000E-08 | 7.100E-08 |
| 26288 | 1315 | K1315G | E | 1.8e-08 | 3.9e-08 | 1.3e-08 | 2.333E-08 | 1.380E-08 | 1.300E-08 | 3.900E-08 |
| 26289 | 1315 | K1315H | E | 6.6e-08 | 6.1e-08 | 1.8e-08 | 4.833E-08 | 2.639E-08 | 1.800E-08 | 6.600E-08 |
| 26290 | 1315 | K1315I | E | 8.3e-08 | 2.1e-08 | 1.2e-08 | 3.867E-08 | 3.866E-08 | 1.200E-08 | 8.300E-08 |
| 26291 | 1315 | K1315L | E | 4.5e-08 | 4.3e-08 | 1.2e-07 | 6.933E-08 | 4.389E-08 | 4.300E-08 | 1.200E-07 |
| 26292 | 1315 | K1315K | E | 5.4e-08 | 6.2e-08 | 1.2e-08 | 4.267E-08 | 2.686E-08 | 1.200E-08 | 6.200E-08 |
| 26293 | 1315 | K1315M | E | 2.1e-08 | 2.2e-08 | 3.6e-08 | 2.633E-08 | 8.386E-09 | 2.100E-08 | 3.600E-08 |
| 26294 | 1315 | K1315F | E | 2.8e-08 | 4.4e-08 | 3.1e-08 | 3.433E-08 | 8.505E-09 | 2.800E-08 | 4.400E-08 |
| 26295 | 1315 | K1315P | E | 3.9e-08 | 7e-08 | 2.7e-08 | 4.533E-08 | 2.219E-08 | 2.700E-08 | 7.000E-08 |
| 26296 | 1315 | K1315S | E | 9.8e-08 | 9.1e-08 | 5.7e-08 | 8.200E-08 | 2.193E-08 | 5.700E-08 | 9.800E-08 |
| 26297 | 1315 | K1315T | E | 3.1e-08 | 1e-07 | 4.1e-08 | 5.733E-08 | 3.729E-08 | 3.100E-08 | 1.000E-07 |
| 26298 | 1315 | K1315W | E | 1e-08 | 7.5e-08 | 4.4e-08 | 4.300E-08 | 3.251E-08 | 1.000E-08 | 7.500E-08 |
| 26299 | 1315 | K1315Y | E | 4.3e-08 | 4.9e-08 | 1.3e-07 | 7.400E-08 | 4.859E-08 | 4.300E-08 | 1.300E-07 |
| 26300 | 1315 | K1315V | E | 3.8e-08 | 4.5e-08 | 3.3e-08 | 3.867E-08 | 6.028E-09 | 3.300E-08 | 4.500E-08 |
| 26301 | 1316 | G1316A | E | 7.2e-08 | 3.8e-08 | 4e-08 | 5.000E-08 | 1.908E-08 | 3.800E-08 | 7.200E-08 |
| 26302 | 1316 | G1316R | E | 4.4e-08 | 2.1e-08 | 2.5e-08 | 3.000E-08 | 1.229E-08 | 2.100E-08 | 4.400E-08 |
| 26303 | 1316 | G1316N | E | 5e-08 | 3.3e-08 | 1.2e-08 | 3.167E-08 | 1.904E-08 | 1.200E-08 | 5.000E-08 |
| 26304 | 1316 | G1316D | E | 1.8e-08 | 1e-07 | 4.1e-08 | 5.300E-08 | 4.230E-08 | 1.800E-08 | 1.000E-07 |
| 26305 | 1316 | G1316C | E | 1.6e-08 | 3.7e-08 | 1.3e-07 | 6.100E-08 | 6.067E-08 | 1.600E-08 | 1.300E-07 |
| 26306 | 1316 | G1316E | E | 9.3e-08 | 1.8e-08 | 1.7e-08 | 4.267E-08 | 4.359E-08 | 1.700E-08 | 9.300E-08 |
| 26307 | 1316 | G1316Q | E | 7e-08 | 4.5e-08 | 4.7e-08 | 5.400E-08 | 1.389E-08 | 4.500E-08 | 7.000E-08 |
| 26308 | 1316 | G1316G | E | 4.2e-08 | 2.9e-08 | 1.8e-08 | 2.967E-08 | 1.201E-08 | 1.800E-08 | 4.200E-08 |
| 26309 | 1316 | G1316H | E | 1.8e-08 | 5.5e-08 | 4.7e-08 | 4.000E-08 | 1.947E-08 | 1.800E-08 | 5.500E-08 |
| 26310 | 1316 | G1316I | E | 7.5e-08 | 1.1e-07 | 4e-08 | 7.500E-08 | 3.500E-08 | 4.000E-08 | 1.100E-07 |
| 26311 | 1316 | G1316L | E | 4.9e-08 | 5e-08 | 4.6e-08 | 4.833E-08 | 2.082E-09 | 4.600E-08 | 5.000E-08 |
| 26312 | 1316 | G1316K | E | 1.1e-07 | 4.7e-08 | 1.2e-08 | 5.633E-08 | 4.966E-08 | 1.200E-08 | 1.100E-07 |
| 26313 | 1316 | G1316M | E | 7.4e-08 | 7.9e-08 | 2.6e-08 | 5.967E-08 | 2.926E-08 | 2.600E-08 | 7.900E-08 |
| 26314 | 1316 | G1316F | E | 3.7e-08 | 2.9e-08 | 4.9e-09 | 2.363E-08 | 1.671E-08 | 4.900E-09 | 3.700E-08 |
| 26315 | 1316 | G1316P | E | 2.8e-08 | 3.9e-07 | 2.4e-08 | 1.473E-07 | 2.102E-07 | 2.400E-08 | 3.900E-07 |
| 26316 | 1316 | G1316S | E | 5.4e-08 | 5.1e-08 | 6.1e-08 | 5.533E-08 | 5.132E-09 | 5.100E-08 | 6.100E-08 |
| 26317 | 1316 | G1316T | E | 3.8e-08 | 8.2e-08 | 6.3e-08 | 6.100E-08 | 2.207E-08 | 3.800E-08 | 8.200E-08 |
| 26318 | 1316 | G1316W | E | 5.1e-08 | 3.4e-08 | 3.3e-08 | 3.933E-08 | 1.012E-08 | 3.300E-08 | 5.100E-08 |
| 26319 | 1316 | G1316Y | E | 7.1e-08 | 1.6e-07 | 2e-07 | 1.437E-07 | 6.603E-08 | 7.100E-08 | 2.000E-07 |
| 26320 | 1316 | G1316V | E | 1e-07 | 1.2e-07 | 5e-08 | 9.000E-08 | 3.606E-08 | 5.000E-08 | 1.200E-07 |
| 26321 | 1317 | L1317A | E | 6.4e-08 | 4.6e-08 | 1.3e-08 | 4.100E-08 | 2.587E-08 | 1.300E-08 | 6.400E-08 |
| 26322 | 1317 | L1317R | E | 7.4e-08 | 9.3e-08 | 4e-08 | 6.900E-08 | 2.685E-08 | 4.000E-08 | 9.300E-08 |
| 26323 | 1317 | L1317N | E | 5.2e-08 | 6.8e-08 | 6.3e-08 | 6.100E-08 | 8.185E-09 | 5.200E-08 | 6.800E-08 |
| 26324 | 1317 | L1317D | E | 3.8e-08 | 1.5e-08 | 5.3e-08 | 3.533E-08 | 1.914E-08 | 1.500E-08 | 5.300E-08 |
| 26325 | 1317 | L1317C | E | 3.4e-08 | 1.3e-07 | 6.1e-08 | 7.500E-08 | 4.951E-08 | 3.400E-08 | 1.300E-07 |
| 26326 | 1317 | L1317E | E | 7e-08 | 3.9e-08 | 4.2e-08 | 5.033E-08 | 1.710E-08 | 3.900E-08 | 7.000E-08 |
| 26327 | 1317 | L1317Q | E | 7.7e-08 | 3.9e-08 | 1.6e-07 | 9.200E-08 | 6.188E-08 | 3.900E-08 | 1.600E-07 |
| 26328 | 1317 | L1317G | E | 1.5e-08 | 2.8e-08 | 4.6e-08 | 2.967E-08 | 1.557E-08 | 1.500E-08 | 4.600E-08 |
| 26329 | 1317 | L1317H | E | 3.2e-08 | 4.2e-08 | 6.4e-08 | 4.600E-08 | 1.637E-08 | 3.200E-08 | 6.400E-08 |
| 26330 | 1317 | L1317I | E | 5.8e-08 | 6.1e-08 | 3.6e-08 | 5.167E-08 | 1.365E-08 | 3.600E-08 | 6.100E-08 |
| 26331 | 1317 | L1317L | E | 3.2e-08 | 1e-07 | 3e-08 | 5.400E-08 | 3.985E-08 | 3.000E-08 | 1.000E-07 |
| 26332 | 1317 | L1317K | E | 3.9e-08 | 4.4e-08 | 9.8e-08 | 6.033E-08 | 3.272E-08 | 3.900E-08 | 9.800E-08 |
| 26333 | 1317 | L1317M | E | 6.8e-08 | 1.2e-07 | 2.8e-08 | 7.200E-08 | 4.613E-08 | 2.800E-08 | 1.200E-07 |
| 26334 | 1317 | L1317F | E | 4.9e-08 | 1.5e-07 | 6.2e-08 | 8.700E-08 | 5.495E-08 | 4.900E-08 | 1.500E-07 |
| 26335 | 1317 | L1317P | E | 1.9e-07 | 8.1e-08 | 8.3e-08 | 1.180E-07 | 6.236E-08 | 8.100E-08 | 1.900E-07 |
| 26336 | 1317 | L1317S | E | 7.5e-08 | 6.3e-08 | 2.8e-08 | 5.533E-08 | 2.442E-08 | 2.800E-08 | 7.500E-08 |
| 26337 | 1317 | L1317T | E | 4.1e-08 | 2.1e-08 | 5.1e-08 | 3.767E-08 | 1.528E-08 | 2.100E-08 | 5.100E-08 |
| 26338 | 1317 | L1317W | E | 7.7e-08 | 4.1e-08 | 4.8e-08 | 5.533E-08 | 1.909E-08 | 4.100E-08 | 7.700E-08 |
| 26339 | 1317 | L1317Y | E | 4.8e-08 | 4.1e-08 | 9e-08 | 5.967E-08 | 2.650E-08 | 4.100E-08 | 9.000E-08 |
| 26340 | 1317 | L1317V | E | 2.8e-08 | 4.3e-08 | 8.3e-08 | 5.133E-08 | 2.843E-08 | 2.800E-08 | 8.300E-08 |
| 26341 | 1318 | E1318A | E | 3.1e-08 | 1.9e-08 | 9.3e-08 | 4.767E-08 | 3.972E-08 | 1.900E-08 | 9.300E-08 |
| 26342 | 1318 | E1318R | E | 2.7e-08 | 3.4e-08 | 1.6e-08 | 2.567E-08 | 9.074E-09 | 1.600E-08 | 3.400E-08 |
| 26343 | 1318 | E1318N | E | 4.5e-08 | 2.2e-08 | 3.4e-08 | 3.367E-08 | 1.150E-08 | 2.200E-08 | 4.500E-08 |
| 26344 | 1318 | E1318D | E | 4e-08 | 5.9e-08 | 2.1e-08 | 4.000E-08 | 1.900E-08 | 2.100E-08 | 5.900E-08 |
| 26345 | 1318 | E1318C | E | 2.4e-08 | 3e-08 | 2.4e-08 | 2.600E-08 | 3.464E-09 | 2.400E-08 | 3.000E-08 |
| 26346 | 1318 | E1318E | E | 6.1e-08 | 5.7e-08 | 6.3e-08 | 6.033E-08 | 3.055E-09 | 5.700E-08 | 6.300E-08 |
| 26347 | 1318 | E1318Q | E | 2.5e-08 | 2.3e-08 | 3.6e-08 | 2.800E-08 | 7.000E-09 | 2.300E-08 | 3.600E-08 |
| 26348 | 1318 | E1318G | E | 4.5e-08 | 2.4e-08 | 7.6e-08 | 4.833E-08 | 2.616E-08 | 2.400E-08 | 7.600E-08 |
| 26349 | 1318 | E1318H | E | 4.9e-08 | 2.8e-08 | 3.5e-08 | 3.733E-08 | 1.069E-08 | 2.800E-08 | 4.900E-08 |

| | | | | | | | | | | |
|---|---|---|---|---|---|---|---|---|---|---|
| 26350 | 1318 | E1318I | E | 2e-08 | 4.5e-08 | 4.7e-08 | 3.733E-08 | 1.504E-08 | 2.000E-08 | 4.700E-08 |
| 26351 | 1318 | E1318L | E | 3.5e-08 | 5.5e-08 | 1.4e-07 | 7.667E-08 | 5.575E-08 | 3.500E-08 | 1.400E-07 |
| 26352 | 1318 | E1318K | E | 7e-08 | 2.8e-08 | 3.6e-08 | 4.467E-08 | 2.230E-08 | 2.800E-08 | 7.000E-08 |
| 26353 | 1318 | E1318M | E | 5.2e-08 | 2.8e-08 | 9.7e-08 | 5.900E-08 | 3.503E-08 | 2.800E-08 | 9.700E-08 |
| 26354 | 1318 | E1318F | E | 6.8e-08 | 7.5e-08 | 7.1e-08 | 7.133E-08 | 3.512E-09 | 6.800E-08 | 7.500E-08 |
| 26355 | 1318 | E1318P | E | 4.2e-08 | 3.3e-08 | 5.8e-08 | 4.433E-08 | 1.266E-08 | 3.300E-08 | 5.800E-08 |
| 26356 | 1318 | E1318S | E | 3.5e-08 | 1e-07 | 5.1e-09 | 4.670E-08 | 4.852E-08 | 5.100E-09 | 1.000E-07 |
| 26357 | 1318 | E1318T | E | 5.4e-08 | 3.9e-08 | 3.7e-08 | 4.333E-08 | 9.292E-09 | 3.700E-08 | 5.400E-08 |
| 26358 | 1318 | E1318W | E | 6.3e-08 | 3.7e-08 | 2.5e-08 | 4.167E-08 | 1.943E-08 | 2.500E-08 | 6.300E-08 |
| 26359 | 1318 | E1318Y | E | 2.2e-08 | 2.2e-08 | 5.2e-08 | 3.200E-08 | 1.732E-08 | 2.200E-08 | 5.200E-08 |
| 26360 | 1318 | E1318V | E | 1.8e-08 | 1.5e-08 | 6e-08 | 3.100E-08 | 2.516E-08 | 1.500E-08 | 6.000E-08 |
| 26361 | 1319 | W1319A | E | 3.2e-08 | 3.2e-08 | 4.6e-08 | 3.667E-08 | 8.083E-09 | 3.200E-08 | 4.600E-08 |
| 26362 | 1319 | W1319R | E | 8.6e-08 | 5.3e-08 | 2.7e-08 | 5.533E-08 | 2.957E-08 | 2.700E-08 | 8.600E-08 |
| 26363 | 1319 | W1319N | E | 4.9e-08 | 4e-08 | 4.7e-08 | 4.533E-08 | 4.726E-09 | 4.000E-08 | 4.900E-08 |
| 26364 | 1319 | W1319D | E | 2.4e-08 | 2.1e-08 | 2.6e-07 | 1.017E-07 | 1.371E-07 | 2.100E-08 | 2.600E-07 |
| 26365 | 1319 | W1319C | E | 1.4e-07 | 3.8e-08 | 3.4e-08 | 7.067E-08 | 6.008E-08 | 3.400E-08 | 1.400E-07 |
| 26366 | 1319 | W1319E | E | 9.1e-08 | 3.8e-08 | 7.4e-08 | 6.767E-08 | 2.706E-08 | 3.800E-08 | 9.100E-08 |
| 26367 | 1319 | W1319Q | E | 3.9e-08 | 5.2e-08 | 2.8e-08 | 3.967E-08 | 1.201E-08 | 2.800E-08 | 5.200E-08 |
| 26368 | 1319 | W1319G | E | 3.7e-08 | 3.7e-08 | 2.9e-08 | 3.433E-08 | 4.619E-09 | 2.900E-08 | 3.700E-08 |
| 26369 | 1319 | W1319H | E | 1.6e-08 | 3.5e-08 | 4.8e-08 | 3.300E-08 | 1.609E-08 | 1.600E-08 | 4.800E-08 |
| 26370 | 1319 | W1319I | E | 4.7e-08 | 1.9e-08 | 4.7e-08 | 3.767E-08 | 1.617E-08 | 1.900E-08 | 4.700E-08 |
| 26371 | 1319 | W1319L | E | 2.9e-08 | 3.3e-08 | 1.3e-07 | 6.400E-08 | 5.719E-08 | 2.900E-08 | 1.300E-07 |
| 26372 | 1319 | W1319K | E | 3.2e-08 | 3.2e-08 | 1.8e-08 | 2.733E-08 | 8.083E-09 | 1.800E-08 | 3.200E-08 |
| 26373 | 1319 | W1319M | E | 3.4e-08 | 1.4e-08 | 4e-08 | 2.933E-08 | 1.361E-08 | 1.400E-08 | 4.000E-08 |
| 26374 | 1319 | W1319F | E | 2e-08 | 4.9e-08 | 2.6e-08 | 3.167E-08 | 1.531E-08 | 2.000E-08 | 4.900E-08 |
| 26375 | 1319 | W1319P | E | 4.5e-08 | 3.7e-08 | 2.9e-08 | 3.700E-08 | 8.000E-09 | 2.900E-08 | 4.500E-08 |
| 26376 | 1319 | W1319S | E | 4.2e-08 | 7.3e-08 | 4.4e-08 | 5.300E-08 | 1.735E-08 | 4.200E-08 | 7.300E-08 |
| 26377 | 1319 | W1319T | E | 3.9e-08 | 1.3e-08 | 3.8e-08 | 3.000E-08 | 1.473E-08 | 1.300E-08 | 3.900E-08 |
| 26378 | 1319 | W1319W | E | 7.9e-08 | 4.9e-08 | 6.7e-09 | 4.490E-08 | 3.632E-08 | 6.700E-09 | 7.900E-08 |
| 26379 | 1319 | W1319Y | E | 6.5e-08 | 5.7e-08 | 8e-08 | 6.733E-08 | 1.168E-08 | 5.700E-08 | 8.000E-08 |
| 26380 | 1319 | W1319V | E | 4.1e-08 | 3.8e-08 | 1.6e-08 | 3.167E-08 | 1.365E-08 | 1.600E-08 | 4.100E-08 |
| 26381 | 1320 | V1320A | E | 3e-08 | 1.8e-08 | 4.5e-08 | 3.100E-08 | 1.353E-08 | 1.800E-08 | 4.500E-08 |
| 26382 | 1320 | V1320R | E | 7.9e-08 | 5.4e-08 | 2e-08 | 5.100E-08 | 2.961E-08 | 2.000E-08 | 7.900E-08 |
| 26383 | 1320 | V1320N | E | 1.9e-08 | 1.5e-07 | 4.7e-08 | 7.200E-08 | 6.899E-08 | 1.900E-08 | 1.500E-07 |
| 26384 | 1320 | V1320D | E | 4.4e-08 | 4.6e-08 | 1.5e-08 | 3.500E-08 | 1.735E-08 | 1.500E-08 | 4.600E-08 |
| 26385 | 1320 | V1320C | E | 6.4e-08 | 4e-08 | 1.6e-08 | 4.000E-08 | 2.400E-08 | 1.600E-08 | 6.400E-08 |
| 26386 | 1320 | V1320E | E | 3e-08 | 2.8e-08 | 9.5e-08 | 5.100E-08 | 3.812E-08 | 2.800E-08 | 9.500E-08 |
| 26387 | 1320 | V1320Q | E | 5.3e-08 | 6.3e-08 | 2.6e-08 | 4.733E-08 | 1.914E-08 | 2.600E-08 | 6.300E-08 |
| 26388 | 1320 | V1320G | E | 2.3e-08 | 2.5e-08 | 2.5e-08 | 2.433E-08 | 1.155E-09 | 2.300E-08 | 2.500E-08 |
| 26389 | 1320 | V1320H | E | 5.4e-08 | 2.5e-08 | 4.8e-08 | 4.233E-08 | 1.531E-08 | 2.500E-08 | 5.400E-08 |
| 26390 | 1320 | V1320I | E | 2.9e-08 | 4.2e-08 | 7e-08 | 4.700E-08 | 2.095E-08 | 2.900E-08 | 7.000E-08 |
| 26391 | 1320 | V1320L | E | 3.7e-08 | 2.8e-08 | 4e-08 | 3.500E-08 | 6.245E-09 | 2.800E-08 | 4.000E-08 |
| 26392 | 1320 | V1320K | E | 3.1e-08 | 2.1e-08 | 3.3e-08 | 2.833E-08 | 6.429E-09 | 2.100E-08 | 3.300E-08 |
| 26393 | 1320 | V1320M | E | 3.1e-08 | 1.3e-08 | 1.4e-08 | 1.933E-08 | 1.012E-08 | 1.300E-08 | 3.100E-08 |
| 26394 | 1320 | V1320F | E | 6.5e-08 | 6.3e-08 | 4.3e-08 | 5.700E-08 | 1.217E-08 | 4.300E-08 | 6.500E-08 |
| 26395 | 1320 | V1320P | E | 4.2e-08 | 3.2e-08 | 3.6e-08 | 3.667E-08 | 5.033E-09 | 3.200E-08 | 4.200E-08 |
| 26396 | 1320 | V1320S | E | 1.7e-08 | 2e-08 | 7.2e-08 | 3.633E-08 | 3.092E-08 | 1.700E-08 | 7.200E-08 |
| 26397 | 1320 | V1320T | E | 6.4e-08 | 2.2e-08 | 2.6e-08 | 3.733E-08 | 2.318E-08 | 2.200E-08 | 6.400E-08 |
| 26398 | 1320 | V1320W | E | 5.8e-08 | 3.3e-08 | 1.8e-08 | 3.633E-08 | 2.021E-08 | 1.800E-08 | 5.800E-08 |
| 26399 | 1320 | V1320Y | E | 3.1e-08 | 4.1e-08 | 6.3e-08 | 4.500E-08 | 1.637E-08 | 3.100E-08 | 6.300E-08 |
| 26400 | 1320 | V1320V | E | 2.7e-08 | 6.7e-08 | 2.8e-08 | 4.067E-08 | 2.281E-08 | 2.700E-08 | 6.700E-08 |
| 26401 | 1321 | A1321A | E | 5e-08 | 4.7e-08 | 1.9e-08 | 3.867E-08 | 1.710E-08 | 1.900E-08 | 5.000E-08 |
| 26402 | 1321 | A1321R | E | 5.4e-08 | 2.4e-08 | 1.9e-08 | 3.233E-08 | 1.893E-08 | 1.900E-08 | 5.400E-08 |
| 26403 | 1321 | A1321N | E | 5.7e-08 | 5.2e-08 | 5.5e-08 | 5.467E-08 | 2.517E-09 | 5.200E-08 | 5.700E-08 |
| 26404 | 1321 | A1321D | E | 4.5e-08 | 4.6e-08 | 1.5e-08 | 3.533E-08 | 1.762E-08 | 1.500E-08 | 4.600E-08 |
| 26405 | 1321 | A1321C | E | 4.3e-08 | 2.4e-08 | 1.6e-08 | 2.767E-08 | 1.387E-08 | 1.600E-08 | 4.300E-08 |
| 26406 | 1321 | A1321E | E | 3.8e-08 | 2.1e-08 | 6.4e-08 | 4.100E-08 | 2.166E-08 | 2.100E-08 | 6.400E-08 |
| 26407 | 1321 | A1321Q | E | 5e-08 | 3.7e-08 | 3e-08 | 3.900E-08 | 1.015E-08 | 3.000E-08 | 5.000E-08 |
| 26408 | 1321 | A1321G | E | 1.6e-08 | 1.6e-08 | 3.5e-08 | 2.233E-08 | 1.097E-08 | 1.600E-08 | 3.500E-08 |
| 26409 | 1321 | A1321H | E | 1.3e-07 | 4.2e-08 | 1.8e-08 | 6.333E-08 | 5.897E-08 | 1.800E-08 | 1.300E-07 |
| 26410 | 1321 | A1321I | E | 5.4e-08 | 4.2e-08 | 2.7e-08 | 4.100E-08 | 1.353E-08 | 2.700E-08 | 5.400E-08 |
| 26411 | 1321 | A1321L | E | 2.7e-08 | 4e-08 | 4.1e-08 | 3.600E-08 | 7.810E-09 | 2.700E-08 | 4.100E-08 |
| 26412 | 1321 | A1321K | E | 4e-08 | 1.7e-08 | 5.6e-08 | 3.767E-08 | 1.960E-08 | 1.700E-08 | 5.600E-08 |
| 26413 | 1321 | A1321M | E | 5.7e-08 | 4.3e-08 | 4.8e-08 | 4.933E-08 | 7.095E-09 | 4.300E-08 | 5.700E-08 |
| 26414 | 1321 | A1321F | E | 7.5e-08 | 3.8e-08 | 3e-08 | 4.767E-08 | 2.401E-08 | 3.000E-08 | 7.500E-08 |
| 26415 | 1321 | A1321P | E | 3.5e-08 | 8.3e-08 | 9.7e-09 | 4.257E-08 | 3.723E-08 | 9.700E-09 | 8.300E-08 |
| 26416 | 1321 | A1321S | E | 9.8e-08 | 4e-08 | 5.1e-08 | 6.300E-08 | 3.081E-08 | 4.000E-08 | 9.800E-08 |
| 26417 | 1321 | A1321T | E | 1.1e-07 | 2.4e-08 | 6.7e-08 | 6.700E-08 | 4.300E-08 | 2.400E-08 | 1.100E-07 |
| 26418 | 1321 | A1321W | E | 9e-08 | 2.2e-08 | 6.8e-08 | 6.000E-08 | 3.470E-08 | 2.200E-08 | 9.000E-08 |
| 26419 | 1321 | A1321Y | E | 2.3e-08 | 5.8e-08 | 4e-08 | 4.033E-08 | 1.750E-08 | 2.300E-08 | 5.800E-08 |
| 26420 | 1321 | A1321V | E | 2.5e-08 | 7.4e-08 | 3.2e-08 | 4.367E-08 | 2.650E-08 | 2.500E-08 | 7.400E-08 |
| 26421 | 1322 | R1322A | E | 2.8e-08 | 4.9e-08 | 7.8e-08 | 5.167E-08 | 2.511E-08 | 2.800E-08 | 7.800E-08 |
| 26422 | 1322 | R1322R | E | 3.6e-08 | 1.5e-07 | 5.6e-08 | 8.067E-08 | 6.087E-08 | 3.600E-08 | 1.500E-07 |
| 26423 | 1322 | R1322N | E | 3e-08 | 4.5e-08 | 3.3e-08 | 3.600E-08 | 7.937E-09 | 3.000E-08 | 4.500E-08 |
| 26424 | 1322 | R1322D | E | 1.1e-08 | 8.2e-08 | 3.3e-08 | 4.200E-08 | 3.635E-08 | 1.100E-08 | 8.200E-08 |
| 26425 | 1322 | R1322C | E | 3.5e-08 | 1e-07 | 8.4e-08 | 7.300E-08 | 3.387E-08 | 3.500E-08 | 1.000E-07 |
| 26426 | 1322 | R1322E | E | 3.3e-08 | 6.7e-08 | 4.5e-08 | 4.833E-08 | 1.724E-08 | 3.300E-08 | 6.700E-08 |
| 26427 | 1322 | R1322Q | E | 3.6e-08 | 4.7e-08 | 4e-08 | 4.100E-08 | 5.568E-09 | 3.600E-08 | 4.700E-08 |
| 26428 | 1322 | R1322G | E | 8.3e-08 | 2.8e-08 | 3.5e-08 | 4.867E-08 | 2.994E-08 | 2.800E-08 | 8.300E-08 |
| 26429 | 1322 | R1322H | E | 2.9e-08 | 1.6e-08 | 4.3e-08 | 2.933E-08 | 1.350E-08 | 1.600E-08 | 4.300E-08 |
| 26430 | 1322 | R1322I | E | 3.6e-08 | 7.7e-08 | 1.7e-08 | 4.333E-08 | 3.066E-08 | 1.700E-08 | 7.700E-08 |
| 26431 | 1322 | R1322L | E | 2.8e-08 | 4.9e-08 | 3.4e-08 | 3.700E-08 | 1.082E-08 | 2.800E-08 | 4.900E-08 |
| 26432 | 1322 | R1322K | E | 4.1e-08 | 3.2e-08 | 2.5e-08 | 3.267E-08 | 8.021E-09 | 2.500E-08 | 4.100E-08 |
| 26433 | 1322 | R1322M | E | 2.1e-08 | 6.9e-08 | 3.8e-08 | 4.267E-08 | 2.434E-08 | 2.100E-08 | 6.900E-08 |
| 26434 | 1322 | R1322F | E | 3.6e-08 | 2.1e-08 | 7.3e-08 | 4.333E-08 | 2.676E-08 | 2.100E-08 | 7.300E-08 |

| | | | | | | | | | | |
|---|---|---|---|---|---|---|---|---|---|---|
| 26435 | 1322 | R1322P | E | 1.8e-08 | 2.9e-08 | 7e-08 | 3.900E-08 | 2.740E-08 | 1.800E-08 | 7.000E-08 |
| 26436 | 1322 | R1322S | E | 2.1e-08 | 6.4e-08 | 6.5e-08 | 5.000E-08 | 2.512E-08 | 2.100E-08 | 6.500E-08 |
| 26437 | 1322 | R1322T | E | 6.9e-08 | 1.7e-08 | 3.1e-08 | 3.900E-08 | 2.691E-08 | 1.700E-08 | 6.900E-08 |
| 26438 | 1322 | R1322W | E | 3.4e-08 | 5.2e-08 | 3.2e-08 | 3.933E-08 | 1.102E-08 | 3.200E-08 | 5.200E-08 |
| 26439 | 1322 | R1322Y | E | 8.6e-08 | 4.3e-08 | 8.1e-08 | 7.000E-08 | 2.352E-08 | 4.300E-08 | 8.600E-08 |
| 26440 | 1322 | R1322V | E | 3.4e-08 | 6.4e-08 | 1.7e-08 | 3.833E-08 | 2.380E-08 | 1.700E-08 | 6.400E-08 |
| 26441 | 1323 | I1323A | E | 3.5e-08 | 1.5e-07 | 8.2e-08 | 8.900E-08 | 5.782E-08 | 3.500E-08 | 1.500E-07 |
| 26442 | 1323 | I1323R | E | 9.1e-08 | 2.7e-08 | 6.2e-08 | 6.000E-08 | 3.205E-08 | 2.700E-08 | 9.100E-08 |
| 26443 | 1323 | I1323N | E | 5.7e-08 | 1e-08 | 2.4e-08 | 3.033E-08 | 2.413E-08 | 1.000E-08 | 5.700E-08 |
| 26444 | 1323 | I1323D | E | 1.5e-07 | 3.2e-08 | 7e-08 | 8.400E-08 | 6.023E-08 | 3.200E-08 | 1.500E-07 |
| 26445 | 1323 | I1323C | E | 2.8e-08 | 6.1e-08 | 1.2e-07 | 6.967E-08 | 4.661E-08 | 2.800E-08 | 1.200E-07 |
| 26446 | 1323 | I1323E | E | 1e-07 | 3.7e-08 | 1.5e-08 | 5.067E-08 | 4.412E-08 | 1.500E-08 | 1.000E-07 |
| 26447 | 1323 | I1323Q | E | 2.3e-08 | 4.5e-08 | 6.7e-08 | 4.500E-08 | 2.200E-08 | 2.300E-08 | 6.700E-08 |
| 26448 | 1323 | I1323G | E | 3.1e-08 | 1.9e-08 | 4.9e-08 | 3.300E-08 | 1.510E-08 | 1.900E-08 | 4.900E-08 |
| 26449 | 1323 | I1323H | E | 6.2e-08 | 3.4e-08 | 4.7e-08 | 4.767E-08 | 1.401E-08 | 3.400E-08 | 6.200E-08 |
| 26450 | 1323 | I1323I | E | 2.6e-08 | 4.1e-08 | 3.6e-08 | 3.433E-08 | 7.638E-09 | 2.600E-08 | 4.100E-08 |
| 26451 | 1323 | I1323L | E | 1e-08 | 8.1e-08 | 3.2e-08 | 4.100E-08 | 3.635E-08 | 1.000E-08 | 8.100E-08 |
| 26452 | 1323 | I1323K | E | 3.8e-08 | 8.2e-08 | 6.7e-08 | 6.233E-08 | 2.237E-08 | 3.800E-08 | 8.200E-08 |
| 26453 | 1323 | I1323M | E | 5.4e-08 | 3.6e-08 | 8e-08 | 5.667E-08 | 2.212E-08 | 3.600E-08 | 8.000E-08 |
| 26454 | 1323 | I1323F | E | 7.4e-08 | 4.3e-08 | 3.9e-08 | 5.200E-08 | 1.916E-08 | 3.900E-08 | 7.400E-08 |
| 26455 | 1323 | I1323P | E | 1.1e-07 | 5.6e-08 | 5.6e-08 | 7.400E-08 | 3.118E-08 | 5.600E-08 | 1.100E-07 |
| 26456 | 1323 | I1323S | E | 6.5e-08 | 2.1e-08 | 2.5e-08 | 3.700E-08 | 2.433E-08 | 2.100E-08 | 6.500E-08 |
| 26457 | 1323 | I1323T | E | 6.6e-08 | 4e-08 | 3.4e-08 | 4.667E-08 | 1.701E-08 | 3.400E-08 | 6.600E-08 |
| 26458 | 1323 | I1323W | E | 1.9e-08 | 3.4e-08 | 3.7e-08 | 3.000E-08 | 9.644E-09 | 1.900E-08 | 3.700E-08 |
| 26459 | 1323 | I1323Y | E | 2.2e-08 | 4.6e-08 | 2e-08 | 2.933E-08 | 1.447E-08 | 2.000E-08 | 4.600E-08 |
| 26460 | 1323 | I1323V | E | 8.2e-08 | 7.1e-08 | 5.2e-08 | 6.833E-08 | 1.518E-08 | 5.200E-08 | 8.200E-08 |
| 26461 | 1324 | Y1324A | E | 9.4e-08 | 6.3e-08 | 3.7e-08 | 6.467E-08 | 2.854E-08 | 3.700E-08 | 9.400E-08 |
| 26462 | 1324 | Y1324R | E | 4.2e-08 | 6.5e-08 | 6e-08 | 5.567E-08 | 1.210E-08 | 4.200E-08 | 6.500E-08 |
| 26463 | 1324 | Y1324N | E | 2.5e-08 | 3e-08 | 4e-08 | 3.167E-08 | 7.638E-09 | 2.500E-08 | 4.000E-08 |
| 26464 | 1324 | Y1324D | E | 2.4e-08 | 5.1e-08 | 4e-08 | 3.833E-08 | 1.358E-08 | 2.400E-08 | 5.100E-08 |
| 26465 | 1324 | Y1324C | E | 1e-07 | 6.3e-08 | 3.6e-08 | 6.633E-08 | 3.213E-08 | 3.600E-08 | 1.000E-07 |
| 26466 | 1324 | Y1324E | E | 2.8e-08 | 5.9e-08 | 2.6e-08 | 3.767E-08 | 1.850E-08 | 2.600E-08 | 5.900E-08 |
| 26467 | 1324 | Y1324Q | E | 5.6e-08 | 2.3e-08 | 4.1e-08 | 4.000E-08 | 1.652E-08 | 2.300E-08 | 5.600E-08 |
| 26468 | 1324 | Y1324G | E | 4.1e-08 | 1.4e-07 | 5.2e-08 | 7.767E-08 | 5.426E-08 | 4.100E-08 | 1.400E-07 |
| 26469 | 1324 | Y1324H | E | 9e-08 | 6.1e-08 | 1.9e-08 | 5.667E-08 | 3.570E-08 | 1.900E-08 | 9.000E-08 |
| 26470 | 1324 | Y1324I | E | 2.8e-08 | 4.8e-08 | 3.5e-08 | 3.700E-08 | 1.015E-08 | 2.800E-08 | 4.800E-08 |
| 26471 | 1324 | Y1324L | E | 1.1e-07 | 4.9e-08 | 4.8e-08 | 6.900E-08 | 3.551E-08 | 4.800E-08 | 1.100E-07 |
| 26472 | 1324 | Y1324K | E | 3.2e-08 | 6.1e-08 | 3.6e-08 | 4.300E-08 | 1.572E-08 | 3.200E-08 | 6.100E-08 |
| 26473 | 1324 | Y1324M | E | 1.9e-07 | 5.5e-08 | 2.8e-08 | 9.100E-08 | 8.679E-08 | 2.800E-08 | 1.900E-07 |
| 26474 | 1324 | Y1324F | E | 4.4e-08 | 2.6e-08 | 1.7e-08 | 2.900E-08 | 1.375E-08 | 1.700E-08 | 4.400E-08 |
| 26475 | 1324 | Y1324P | E | 4.8e-08 | 3.7e-07 | 7.8e-08 | 1.653E-07 | 1.779E-07 | 4.800E-08 | 3.700E-07 |
| 26476 | 1324 | Y1324S | E | 1.4e-07 | 1.8e-08 | 2.5e-08 | 6.100E-08 | 6.851E-08 | 1.800E-08 | 1.400E-07 |
| 26477 | 1324 | Y1324T | E | 1.6e-07 | 3.1e-08 | 4.3e-08 | 7.800E-08 | 7.127E-08 | 3.100E-08 | 1.600E-07 |
| 26478 | 1324 | Y1324W | E | 7.1e-08 | 4.1e-08 | 1.1e-07 | 7.400E-08 | 3.460E-08 | 4.100E-08 | 1.100E-07 |
| 26479 | 1324 | Y1324Y | E | 7.1e-08 | 4.1e-08 | 4.3e-08 | 5.167E-08 | 1.677E-08 | 4.100E-08 | 7.100E-08 |
| 26480 | 1324 | Y1324V | E | 3.9e-08 | 3.1e-08 | 2.4e-08 | 3.133E-08 | 7.506E-09 | 2.400E-08 | 3.900E-08 |
| 26481 | 1325 | P1325A | E | 3.1e-08 | 1.1e-07 | 9e-08 | 7.700E-08 | 4.107E-08 | 3.100E-08 | 1.100E-07 |
| 26482 | 1325 | P1325R | E | 2.7e-08 | 4.7e-08 | 2.6e-08 | 3.333E-08 | 1.185E-08 | 2.600E-08 | 4.700E-08 |
| 26483 | 1325 | P1325N | E | 1.8e-08 | 7e-09 | 5.3e-08 | 2.600E-08 | 2.402E-08 | 7.000E-09 | 5.300E-08 |
| 26484 | 1325 | P1325D | E | 5.8e-08 | 5.5e-08 | 4.2e-08 | 5.167E-08 | 8.505E-09 | 4.200E-08 | 5.800E-08 |
| 26485 | 1325 | P1325C | E | 2e-08 | 1.5e-08 | 4.2e-08 | 2.567E-08 | 1.436E-08 | 1.500E-08 | 4.200E-08 |
| 26486 | 1325 | P1325E | E | 3.4e-08 | 5.2e-08 | 2.1e-08 | 3.567E-08 | 1.557E-08 | 2.100E-08 | 5.200E-08 |
| 26487 | 1325 | P1325Q | E | 7.7e-08 | 3e-08 | 4.2e-08 | 4.967E-08 | 2.442E-08 | 3.000E-08 | 7.700E-08 |
| 26488 | 1325 | P1325G | E | 5.3e-08 | 3.4e-08 | 3.3e-08 | 4.000E-08 | 1.127E-08 | 3.300E-08 | 5.300E-08 |
| 26489 | 1325 | P1325H | E | 5.3e-08 | 1.1e-08 | 2.7e-08 | 3.033E-08 | 2.120E-08 | 1.100E-08 | 5.300E-08 |
| 26490 | 1325 | P1325I | E | 6.4e-08 | 6e-08 | 3.1e-08 | 5.167E-08 | 1.801E-08 | 3.100E-08 | 6.400E-08 |
| 26491 | 1325 | P1325L | E | 1.3e-08 | 1.6e-08 | 3.6e-08 | 2.167E-08 | 1.250E-08 | 1.300E-08 | 3.600E-08 |
| 26492 | 1325 | P1325K | E | 3.6e-08 | 3.6e-08 | 3e-08 | 3.400E-08 | 3.464E-09 | 3.000E-08 | 3.600E-08 |
| 26493 | 1325 | P1325M | E | 1.1e-07 | 4.8e-08 | 6e-08 | 7.267E-08 | 3.288E-08 | 4.800E-08 | 1.100E-07 |
| 26494 | 1325 | P1325F | E | 2.2e-08 | 4.5e-08 | 5e-08 | 3.900E-08 | 1.493E-08 | 2.200E-08 | 5.000E-08 |
| 26495 | 1325 | P1325P | E | 5.1e-08 | 1.8e-08 | 2.9e-08 | 3.267E-08 | 1.680E-08 | 1.800E-08 | 5.100E-08 |
| 26496 | 1325 | P1325S | E | 3.1e-08 | 3.6e-08 | 6.3e-08 | 4.333E-08 | 1.721E-08 | 3.100E-08 | 6.300E-08 |
| 26497 | 1325 | P1325T | E | 2.2e-08 | 5.4e-08 | 2e-08 | 3.200E-08 | 1.908E-08 | 2.000E-08 | 5.400E-08 |
| 26498 | 1325 | P1325W | E | 4.5e-08 | 2e-08 | 2.7e-08 | 3.067E-08 | 1.290E-08 | 2.000E-08 | 4.500E-08 |
| 26499 | 1325 | P1325Y | E | 6.9e-08 | 1.1e-07 | 2.2e-08 | 6.700E-08 | 4.403E-08 | 2.200E-08 | 1.100E-07 |
| 26500 | 1325 | P1325V | E | 4.3e-08 | 5.8e-08 | 3.9e-08 | 4.667E-08 | 1.002E-08 | 3.900E-08 | 5.800E-08 |
| 26501 | 1326 | T1326A | E | 3.1e-08 | 1e-07 | 2.4e-08 | 5.167E-08 | 4.200E-08 | 2.400E-08 | 1.000E-07 |
| 26502 | 1326 | T1326R | E | 2.1e-08 | 7.8e-08 | 1.2e-08 | 3.700E-08 | 3.579E-08 | 1.200E-08 | 7.800E-08 |
| 26503 | 1326 | T1326N | E | 3.5e-08 | 8.3e-08 | 3.8e-08 | 5.200E-08 | 2.689E-08 | 3.500E-08 | 8.300E-08 |
| 26504 | 1326 | T1326D | E | 6.3e-08 | 3.8e-08 | 5.6e-08 | 5.233E-08 | 1.290E-08 | 3.800E-08 | 6.300E-08 |
| 26505 | 1326 | T1326C | E | 2.4e-08 | 2.6e-07 | 2.2e-08 | 1.020E-07 | 1.368E-07 | 2.200E-08 | 2.600E-07 |
| 26506 | 1326 | T1326E | E | 5.5e-08 | 3.4e-08 | 1.6e-07 | 8.300E-08 | 6.751E-08 | 3.400E-08 | 1.600E-07 |
| 26507 | 1326 | T1326Q | E | 2.5e-08 | 2.5e-08 | 6.6e-08 | 3.867E-08 | 2.367E-08 | 2.500E-08 | 6.600E-08 |
| 26508 | 1326 | T1326G | E | 5.2e-08 | 4.6e-08 | 5.3e-08 | 5.033E-08 | 3.786E-09 | 4.600E-08 | 5.300E-08 |
| 26509 | 1326 | T1326H | E | 3.3e-08 | 5.4e-08 | 7.4e-08 | 5.367E-08 | 2.050E-08 | 3.300E-08 | 7.400E-08 |
| 26510 | 1326 | T1326I | E | 4.6e-08 | 4.1e-08 | 9.2e-08 | 5.967E-08 | 2.811E-08 | 4.100E-08 | 9.200E-08 |
| 26511 | 1326 | T1326L | E | 5e-08 | 3e-08 | 1.5e-08 | 3.167E-08 | 1.756E-08 | 1.500E-08 | 5.000E-08 |
| 26512 | 1326 | T1326K | E | 2.2e-08 | 3e-08 | 2.3e-08 | 2.500E-08 | 4.359E-09 | 2.200E-08 | 3.000E-08 |
| 26513 | 1326 | T1326M | E | 2.4e-08 | 5.3e-08 | 6.5e-08 | 4.733E-08 | 2.108E-08 | 2.400E-08 | 6.500E-08 |
| 26514 | 1326 | T1326F | E | 2.6e-08 | 3.6e-08 | 1.1e-07 | 5.733E-08 | 4.588E-08 | 2.600E-08 | 1.100E-07 |
| 26515 | 1326 | T1326P | E | 5.1e-08 | 3e-08 | 6.6e-08 | 4.900E-08 | 1.808E-08 | 3.000E-08 | 6.600E-08 |
| 26516 | 1326 | T1326S | E | 3.3e-08 | 2.7e-08 | 1.4e-07 | 6.667E-08 | 6.358E-08 | 2.700E-08 | 1.400E-07 |
| 26517 | 1326 | T1326T | E | 6.1e-08 | 1.4e-07 | 2.3e-08 | 7.467E-08 | 5.969E-08 | 2.300E-08 | 1.400E-07 |
| 26518 | 1326 | T1326W | E | 7.5e-08 | 5e-08 | 2.9e-08 | 5.133E-08 | 2.303E-08 | 2.900E-08 | 7.500E-08 |
| 26519 | 1326 | T1326Y | E | 4.7e-08 | 1.9e-08 | 8.3e-08 | 4.967E-08 | 3.208E-08 | 1.900E-08 | 8.300E-08 |

| | | | | | | | | | | |
|---|---|---|---|---|---|---|---|---|---|---|
| 26520 | 1326 | T1326V | E | 2.4e-08 | 1.5e-08 | 2.3e-08 | 2.067E-08 | 4.933E-09 | 1.500E-08 | 2.400E-08 |
| 26521 | 1327 | N1327A | E | 3.9e-08 | 5.3e-08 | 2.2e-08 | 3.800E-08 | 1.552E-08 | 2.200E-08 | 5.300E-08 |
| 26522 | 1327 | N1327R | E | 9.4e-08 | 1.3e-07 | 6.3e-08 | 9.567E-08 | 3.353E-08 | 6.300E-08 | 1.300E-07 |
| 26523 | 1327 | N1327N | E | 2e-08 | 3.3e-08 | 7.6e-08 | 4.300E-08 | 2.931E-08 | 2.000E-08 | 7.600E-08 |
| 26524 | 1327 | N1327D | E | 2.7e-08 | 2.9e-08 | 3.9e-08 | 3.167E-08 | 6.429E-09 | 2.700E-08 | 3.900E-08 |
| 26525 | 1327 | N1327C | E | 5.7e-08 | 7.3e-08 | 7e-08 | 6.667E-08 | 8.505E-09 | 5.700E-08 | 7.300E-08 |
| 26526 | 1327 | N1327E | E | 2.6e-08 | 4.5e-08 | 3.1e-08 | 3.400E-08 | 9.849E-09 | 2.600E-08 | 4.500E-08 |
| 26527 | 1327 | N1327Q | E | 2.6e-08 | 1.7e-08 | 7.8e-08 | 4.033E-08 | 3.293E-08 | 1.700E-08 | 7.800E-08 |
| 26528 | 1327 | N1327G | E | 9.7e-08 | 3.6e-08 | 3.9e-08 | 5.733E-08 | 3.439E-08 | 3.600E-08 | 9.700E-08 |
| 26529 | 1327 | N1327H | E | 3.4e-08 | 2.4e-08 | 3.4e-08 | 3.067E-08 | 5.774E-09 | 2.400E-08 | 3.400E-08 |
| 26530 | 1327 | N1327I | E | 2.9e-08 | 3e-08 | 3.4e-08 | 3.100E-08 | 2.646E-09 | 2.900E-08 | 3.400E-08 |
| 26531 | 1327 | N1327L | E | 4e-08 | 2.5e-08 | 2.5e-08 | 3.000E-08 | 8.660E-09 | 2.500E-08 | 4.000E-08 |
| 26532 | 1327 | N1327K | E | 5e-08 | 4.4e-08 | 2.7e-08 | 4.033E-08 | 1.193E-08 | 2.700E-08 | 5.000E-08 |
| 26533 | 1327 | N1327M | E | 2.6e-08 | 8.4e-08 | 5e-08 | 5.333E-08 | 2.914E-08 | 2.600E-08 | 8.400E-08 |
| 26534 | 1327 | N1327F | E | 5.5e-08 | 5.6e-08 | 1.4e-08 | 4.167E-08 | 2.397E-08 | 1.400E-08 | 5.600E-08 |
| 26535 | 1327 | N1327P | E | 8.9e-08 | 2.6e-08 | 2.4e-08 | 4.633E-08 | 3.696E-08 | 2.400E-08 | 8.900E-08 |
| 26536 | 1327 | N1327S | E | 6.2e-08 | 3.7e-08 | 4.6e-08 | 4.833E-08 | 1.266E-08 | 3.700E-08 | 6.200E-08 |
| 26537 | 1327 | N1327T | E | 3.1e-08 | 2.8e-08 | 3.2e-08 | 3.033E-08 | 2.082E-09 | 2.800E-08 | 3.200E-08 |
| 26538 | 1327 | N1327W | E | 3.2e-08 | 3.5e-08 | 2.8e-08 | 3.167E-08 | 3.512E-09 | 2.800E-08 | 3.500E-08 |
| 26539 | 1327 | N1327Y | E | 1.2e-08 | 3.1e-08 | 8.1e-08 | 4.133E-08 | 3.564E-08 | 1.200E-08 | 8.100E-08 |
| 26540 | 1327 | N1327V | E | 2.8e-08 | 4.4e-08 | 2.5e-08 | 3.233E-08 | 1.021E-08 | 2.500E-08 | 4.400E-08 |
| 26541 | 1328 | G1328A | E | 1.1e-07 | 2.2e-08 | 6.2e-08 | 6.467E-08 | 4.406E-08 | 2.200E-08 | 1.100E-07 |
| 26542 | 1328 | G1328R | E | 1.1e-07 | 3.4e-08 | 9.1e-08 | 7.833E-08 | 3.955E-08 | 3.400E-08 | 1.100E-07 |
| 26543 | 1328 | G1328N | E | 4.2e-08 | 3e-08 | 3.6e-08 | 3.600E-08 | 6.000E-09 | 3.000E-08 | 4.200E-08 |
| 26544 | 1328 | G1328D | E | 2.9e-08 | 4.1e-08 | 8.7e-08 | 5.233E-08 | 3.062E-08 | 2.900E-08 | 8.700E-08 |
| 26545 | 1328 | G1328C | E | 2.1e-08 | 2.2e-08 | 5.2e-08 | 3.167E-08 | 1.762E-08 | 2.100E-08 | 5.200E-08 |
| 26546 | 1328 | G1328E | E | 4e-08 | 5.6e-08 | 4.3e-08 | 4.633E-08 | 8.505E-09 | 4.000E-08 | 5.600E-08 |
| 26547 | 1328 | G1328Q | E | 6.8e-08 | 6.3e-09 | 4e-08 | 3.810E-08 | 3.089E-08 | 6.300E-09 | 6.800E-08 |
| 26548 | 1328 | G1328G | E | 4.1e-08 | 5.8e-08 | 4.7e-08 | 4.867E-08 | 8.622E-09 | 4.100E-08 | 5.800E-08 |
| 26549 | 1328 | G1328H | E | 2.6e-08 | 4.5e-08 | 4.2e-08 | 3.767E-08 | 1.021E-08 | 2.600E-08 | 4.500E-08 |
| 26550 | 1328 | G1328I | E | 6.8e-08 | 7.6e-08 | 1e-07 | 8.133E-08 | 1.665E-08 | 6.800E-08 | 1.000E-07 |
| 26551 | 1328 | G1328L | E | 6.1e-08 | 1.2e-08 | 7.6e-08 | 4.967E-08 | 3.347E-08 | 1.200E-08 | 7.600E-08 |
| 26552 | 1328 | G1328K | E | 1.4e-07 | 3.8e-08 | 3.3e-08 | 7.033E-08 | 6.038E-08 | 3.300E-08 | 1.400E-07 |
| 26553 | 1328 | G1328M | E | 4.8e-08 | 3.8e-08 | 8.2e-08 | 5.600E-08 | 2.307E-08 | 3.800E-08 | 8.200E-08 |
| 26554 | 1328 | G1328F | E | 6.9e-08 | 7.3e-08 | 2.8e-08 | 5.667E-08 | 2.491E-08 | 2.800E-08 | 7.300E-08 |
| 26555 | 1328 | G1328P | E | 3.6e-08 | 4.9e-08 | 3.1e-08 | 3.867E-08 | 9.292E-09 | 3.100E-08 | 4.900E-08 |
| 26556 | 1328 | G1328S | E | 7.6e-08 | 2.8e-08 | 4.2e-08 | 4.867E-08 | 2.468E-08 | 2.800E-08 | 7.600E-08 |
| 26557 | 1328 | G1328T | E | 3.1e-08 | 4.6e-08 | 2.2e-08 | 3.300E-08 | 1.212E-08 | 2.200E-08 | 4.600E-08 |
| 26558 | 1328 | G1328W | E | 4.2e-08 | 9.5e-09 | 6.5e-08 | 3.883E-08 | 2.789E-08 | 9.500E-09 | 6.500E-08 |
| 26559 | 1328 | G1328Y | E | 8.3e-09 | 2.5e-08 | 5.3e-08 | 2.877E-08 | 2.259E-08 | 8.300E-09 | 5.300E-08 |
| 26560 | 1328 | G1328V | E | 4.2e-08 | 3.3e-08 | 8.8e-08 | 5.433E-08 | 2.950E-08 | 3.300E-08 | 8.800E-08 |
| 26561 | 1329 | Y1329A | E | 3.2e-08 | 4.2e-08 | 2.8e-08 | 3.400E-08 | 7.211E-09 | 2.800E-08 | 4.200E-08 |
| 26562 | 1329 | Y1329R | E | 2.1e-08 | 2.8e-08 | 7.6e-09 | 1.887E-08 | 1.037E-08 | 7.600E-09 | 2.800E-08 |
| 26563 | 1329 | Y1329N | E | 2.6e-08 | 7e-08 | 3.9e-08 | 4.500E-08 | 2.261E-08 | 2.600E-08 | 7.000E-08 |
| 26564 | 1329 | Y1329D | E | 1.9e-08 | 3.9e-08 | 1.5e-07 | 6.933E-08 | 7.057E-08 | 1.900E-08 | 1.500E-07 |
| 26565 | 1329 | Y1329C | E | 2.7e-08 | 2.8e-08 | 3e-08 | 2.833E-08 | 1.528E-09 | 2.700E-08 | 3.000E-08 |
| 26566 | 1329 | Y1329E | E | 9.9e-08 | 1.4e-07 | 1.1e-07 | 1.163E-07 | 2.122E-08 | 9.900E-08 | 1.400E-07 |
| 26567 | 1329 | Y1329Q | E | 4.9e-08 | 7.3e-08 | 1.2e-08 | 4.467E-08 | 3.073E-08 | 1.200E-08 | 7.300E-08 |
| 26568 | 1329 | Y1329G | E | 5.2e-08 | 5.5e-08 | 7.6e-08 | 6.100E-08 | 1.308E-08 | 5.200E-08 | 7.600E-08 |
| 26569 | 1329 | Y1329H | E | 5.4e-08 | 6e-08 | 1.3e-08 | 4.233E-08 | 2.558E-08 | 1.300E-08 | 6.000E-08 |
| 26570 | 1329 | Y1329I | E | 1.1e-07 | 2.8e-08 | 4.3e-08 | 6.033E-08 | 4.366E-08 | 2.800E-08 | 1.100E-07 |
| 26571 | 1329 | Y1329L | E | 2.8e-08 | 2.3e-08 | 6.5e-08 | 3.867E-08 | 2.294E-08 | 2.300E-08 | 6.500E-08 |
| 26572 | 1329 | Y1329K | E | 8e-08 | 3e-08 | 2.1e-07 | 1.067E-07 | 9.292E-08 | 3.000E-08 | 2.100E-07 |
| 26573 | 1329 | Y1329M | E | 3.9e-08 | 5.8e-08 | 1.3e-07 | 7.567E-08 | 4.800E-08 | 3.900E-08 | 1.300E-07 |
| 26574 | 1329 | Y1329F | E | 3.4e-08 | 1.3e-07 | 3.6e-08 | 6.667E-08 | 5.486E-08 | 3.400E-08 | 1.300E-07 |
| 26575 | 1329 | Y1329P | E | 4.2e-08 | 7.3e-08 | 2.4e-08 | 4.633E-08 | 2.479E-08 | 2.400E-08 | 7.300E-08 |
| 26576 | 1329 | Y1329S | E | 1.5e-08 | 3e-08 | 5.6e-08 | 3.367E-08 | 2.074E-08 | 1.500E-08 | 5.600E-08 |
| 26577 | 1329 | Y1329T | E | 1.7e-07 | 3.9e-08 | 1.5e-08 | 7.467E-08 | 8.343E-08 | 1.500E-08 | 1.700E-07 |
| 26578 | 1329 | Y1329W | E | 2.2e-08 | 3.4e-08 | 2.1e-08 | 2.567E-08 | 7.234E-09 | 2.100E-08 | 3.400E-08 |
| 26579 | 1329 | Y1329Y | E | 5e-08 | 2.1e-08 | 4e-08 | 3.700E-08 | 1.473E-08 | 2.100E-08 | 5.000E-08 |
| 26580 | 1329 | Y1329V | E | 3.4e-08 | 4.9e-08 | 6.5e-08 | 4.933E-08 | 1.550E-08 | 3.400E-08 | 6.500E-08 |
| 26581 | 1330 | T1330A | E | 3.2e-08 | 5.3e-08 | 6.8e-08 | 5.100E-08 | 1.808E-08 | 3.200E-08 | 6.800E-08 |
| 26582 | 1330 | T1330R | E | 4e-08 | 8e-08 | 6.5e-08 | 6.167E-08 | 2.021E-08 | 4.000E-08 | 8.000E-08 |
| 26583 | 1330 | T1330N | E | 6.2e-08 | 6.6e-08 | 3.4e-08 | 5.400E-08 | 1.744E-08 | 3.400E-08 | 6.600E-08 |
| 26584 | 1330 | T1330D | E | 4.8e-08 | 4.7e-08 | 3.2e-08 | 4.233E-08 | 8.963E-09 | 3.200E-08 | 4.800E-08 |
| 26585 | 1330 | T1330C | E | 3.9e-08 | 1.3e-08 | 4.3e-08 | 3.167E-08 | 1.629E-08 | 1.300E-08 | 4.300E-08 |
| 26586 | 1330 | T1330E | E | 8e-08 | 4.1e-08 | 3.5e-08 | 5.200E-08 | 2.443E-08 | 3.500E-08 | 8.000E-08 |
| 26587 | 1330 | T1330Q | E | 5.2e-08 | 7e-08 | 7.9e-08 | 6.700E-08 | 1.375E-08 | 5.200E-08 | 7.900E-08 |
| 26588 | 1330 | T1330G | E | 6.5e-08 | 4.5e-08 | 5.2e-08 | 5.400E-08 | 1.015E-08 | 4.500E-08 | 6.500E-08 |
| 26589 | 1330 | T1330H | E | 6.5e-08 | 2.6e-08 | 5.8e-08 | 4.967E-08 | 2.079E-08 | 2.600E-08 | 6.500E-08 |
| 26590 | 1330 | T1330I | E | 7e-08 | 3.7e-08 | 1.2e-08 | 3.967E-08 | 2.909E-08 | 1.200E-08 | 7.000E-08 |
| 26591 | 1330 | T1330L | E | 7.7e-08 | 4.4e-08 | 7.6e-08 | 6.567E-08 | 1.877E-08 | 4.400E-08 | 7.700E-08 |
| 26592 | 1330 | T1330K | E | 1.1e-07 | 2.3e-08 | 1.5e-08 | 4.933E-08 | 5.269E-08 | 1.500E-08 | 1.100E-07 |
| 26593 | 1330 | T1330M | E | 6.3e-08 | 7.4e-08 | 4.5e-08 | 6.067E-08 | 1.464E-08 | 4.500E-08 | 7.400E-08 |
| 26594 | 1330 | T1330F | E | 3.2e-08 | 2.9e-08 | 2.1e-08 | 2.733E-08 | 5.686E-09 | 2.100E-08 | 3.200E-08 |
| 26595 | 1330 | T1330P | E | 1e-07 | 6.6e-08 | 1.8e-08 | 6.133E-08 | 4.120E-08 | 1.800E-08 | 1.000E-07 |
| 26596 | 1330 | T1330S | E | 7.9e-08 | 3.9e-08 | 6.8e-08 | 6.200E-08 | 2.066E-08 | 3.900E-08 | 7.900E-08 |
| 26597 | 1330 | T1330T | E | 3.5e-08 | 5.9e-08 | 8.2e-08 | 5.867E-08 | 2.350E-08 | 3.500E-08 | 8.200E-08 |
| 26598 | 1330 | T1330W | E | 2.1e-08 | 3.3e-08 | 3.3e-08 | 2.900E-08 | 6.928E-09 | 2.100E-08 | 3.300E-08 |
| 26599 | 1330 | T1330Y | E | 4.6e-08 | 4.2e-08 | 1.2e-07 | 6.933E-08 | 4.392E-08 | 4.200E-08 | 1.200E-07 |
| 26600 | 1330 | T1330V | E | 6.8e-08 | 2.9e-08 | 3e-08 | 4.233E-08 | 2.223E-08 | 2.900E-08 | 6.800E-08 |
| 26601 | 1331 | R1331A | E | 5.9e-08 | 3.6e-08 | 1e-07 | 6.500E-08 | 3.242E-08 | 3.600E-08 | 1.000E-07 |
| 26602 | 1331 | R1331R | E | 3.2e-08 | 3.4e-08 | 2e-08 | 2.867E-08 | 7.572E-09 | 2.000E-08 | 3.400E-08 |
| 26603 | 1331 | R1331N | E | 5.3e-08 | 6.7e-08 | 1.8e-08 | 4.600E-08 | 2.524E-08 | 1.800E-08 | 6.700E-08 |
| 26604 | 1331 | R1331D | E | 2.6e-08 | 7.7e-08 | 4.5e-08 | 4.933E-08 | 2.577E-08 | 2.600E-08 | 7.700E-08 |

| | | | | | | | | | | |
|---|---|---|---|---|---|---|---|---|---|---|
| 26605 | 1331 | R1331C | E | 5.3e-08 | 7.5e-08 | 4.9e-08 | 5.900E-08 | 1.400E-08 | 4.900E-08 | 7.500E-08 |
| 26606 | 1331 | R1331E | E | 6.8e-08 | 2.3e-08 | 7.2e-08 | 5.433E-08 | 2.721E-08 | 2.300E-08 | 7.200E-08 |
| 26607 | 1331 | R1331Q | E | 1.1e-07 | 4.6e-08 | 5.8e-08 | 7.133E-08 | 3.402E-08 | 4.600E-08 | 1.100E-07 |
| 26608 | 1331 | R1331G | E | 4.2e-08 | 3.1e-08 | 2.4e-08 | 3.233E-08 | 9.074E-09 | 2.400E-08 | 4.200E-08 |
| 26609 | 1331 | R1331H | E | 3.5e-08 | 6.1e-08 | 1.3e-08 | 3.633E-08 | 2.403E-08 | 1.300E-08 | 6.100E-08 |
| 26610 | 1331 | R1331I | E | 5.1e-08 | 1.2e-07 | 7.9e-08 | 8.333E-08 | 3.470E-08 | 5.100E-08 | 1.200E-07 |
| 26611 | 1331 | R1331L | E | 6.7e-09 | 3.1e-08 | 4.1e-08 | 2.623E-08 | 1.764E-08 | 6.700E-09 | 4.100E-08 |
| 26612 | 1331 | R1331K | E | 5.9e-08 | 5.2e-08 | 5.7e-08 | 5.600E-08 | 3.606E-09 | 5.200E-08 | 5.900E-08 |
| 26613 | 1331 | R1331M | E | 2e-07 | 6.5e-08 | 3.9e-08 | 1.013E-07 | 8.643E-08 | 3.900E-08 | 2.000E-07 |
| 26614 | 1331 | R1331F | E | 5.1e-08 | 4e-08 | 2.9e-08 | 4.000E-08 | 1.100E-08 | 2.900E-08 | 5.100E-08 |
| 26615 | 1331 | R1331P | E | 1.1e-07 | 1.6e-07 | 7.2e-08 | 1.140E-07 | 4.414E-08 | 7.200E-08 | 1.600E-07 |
| 26616 | 1331 | R1331S | E | 2.5e-08 | 1.4e-07 | 8.3e-08 | 8.267E-08 | 5.750E-08 | 2.500E-08 | 1.400E-07 |
| 26617 | 1331 | R1331T | E | 3.3e-08 | 6e-08 | 1e-07 | 6.433E-08 | 3.371E-08 | 3.300E-08 | 1.000E-07 |
| 26618 | 1331 | R1331W | E | 4.4e-08 | 1.3e-08 | 9.4e-08 | 5.033E-08 | 4.087E-08 | 1.300E-08 | 9.400E-08 |
| 26619 | 1331 | R1331Y | E | 5.1e-08 | 2.9e-08 | 4.2e-08 | 4.067E-08 | 1.106E-08 | 2.900E-08 | 5.100E-08 |
| 26620 | 1331 | R1331V | E | 6.5e-08 | 6.7e-08 | 8.3e-08 | 7.167E-08 | 9.866E-09 | 6.500E-08 | 8.300E-08 |
| 26621 | 1332 | Y1332A | E | 3.5e-08 | 6.8e-08 | 1.9e-08 | 4.067E-08 | 2.499E-08 | 1.900E-08 | 6.800E-08 |
| 26622 | 1332 | Y1332R | E | 5.4e-08 | 3.4e-08 | 4e-08 | 4.267E-08 | 1.026E-08 | 3.400E-08 | 5.400E-08 |
| 26623 | 1332 | Y1332N | E | 4.1e-08 | 5.9e-08 | 3.1e-08 | 4.367E-08 | 1.419E-08 | 3.100E-08 | 5.900E-08 |
| 26624 | 1332 | Y1332D | E | 2.2e-08 | 3.4e-08 | 5.3e-08 | 3.633E-08 | 1.563E-08 | 2.200E-08 | 5.300E-08 |
| 26625 | 1332 | Y1332C | E | 7.1e-08 | 6.8e-08 | 8.2e-09 | 4.907E-08 | 3.542E-08 | 8.200E-09 | 7.100E-08 |
| 26626 | 1332 | Y1332E | E | 2.4e-08 | 6.1e-08 | 4.7e-08 | 4.400E-08 | 1.868E-08 | 2.400E-08 | 6.100E-08 |
| 26627 | 1332 | Y1332Q | E | 1.4e-08 | 2.9e-08 | 8.2e-08 | 4.167E-08 | 3.573E-08 | 1.400E-08 | 8.200E-08 |
| 26628 | 1332 | Y1332G | E | 4.7e-08 | 1.9e-08 | 8.3e-08 | 4.967E-08 | 3.208E-08 | 1.900E-08 | 8.300E-08 |
| 26629 | 1332 | Y1332H | E | 1e-07 | 2.5e-08 | 4.9e-08 | 5.800E-08 | 3.830E-08 | 2.500E-08 | 1.000E-07 |
| 26630 | 1332 | Y1332I | E | 3e-08 | 7.3e-08 | 2.5e-08 | 4.267E-08 | 2.639E-08 | 2.500E-08 | 7.300E-08 |
| 26631 | 1332 | Y1332L | E | 5e-08 | 8.5e-08 | 4.4e-08 | 5.967E-08 | 2.214E-08 | 4.400E-08 | 8.500E-08 |
| 26632 | 1332 | Y1332K | E | 2.8e-08 | 4.8e-08 | 4.4e-08 | 4.000E-08 | 1.058E-08 | 2.800E-08 | 4.800E-08 |
| 26633 | 1332 | Y1332M | E | 9.6e-08 | 8e-08 | 3.1e-08 | 6.900E-08 | 3.387E-08 | 3.100E-08 | 9.600E-08 |
| 26634 | 1332 | Y1332F | E | 4.1e-08 | 3.8e-08 | 2.7e-08 | 3.533E-08 | 7.371E-09 | 2.700E-08 | 4.100E-08 |
| 26635 | 1332 | Y1332P | E | 3.1e-08 | 2.8e-08 | 2.7e-08 | 2.867E-08 | 2.082E-09 | 2.700E-08 | 3.100E-08 |
| 26636 | 1332 | Y1332S | E | 3.6e-08 | 6.3e-08 | 8.4e-08 | 6.100E-08 | 2.406E-08 | 3.600E-08 | 8.400E-08 |
| 26637 | 1332 | Y1332T | E | 7e-08 | 8.6e-08 | 9.6e-09 | 5.520E-08 | 4.029E-08 | 9.600E-09 | 8.600E-08 |
| 26638 | 1332 | Y1332W | E | 3.6e-08 | 6.1e-08 | 8.8e-08 | 6.167E-08 | 2.601E-08 | 3.600E-08 | 8.800E-08 |
| 26639 | 1332 | Y1332Y | E | 5.4e-08 | 3.4e-08 | 2.4e-08 | 3.733E-08 | 1.528E-08 | 2.400E-08 | 5.400E-08 |
| 26640 | 1332 | Y1332V | E | 4.8e-08 | 6.7e-08 | 2.8e-08 | 4.767E-08 | 1.950E-08 | 2.800E-08 | 6.700E-08 |
| 26641 | 1333 | A1333A | E | 5.2e-08 | 3.5e-08 | 1.8e-08 | 3.500E-08 | 1.700E-08 | 1.800E-08 | 5.200E-08 |
| 26642 | 1333 | A1333R | E | 3.6e-08 | 2.1e-08 | 5.8e-08 | 3.833E-08 | 1.861E-08 | 2.100E-08 | 5.800E-08 |
| 26643 | 1333 | A1333N | E | 1.2e-08 | 1.1e-07 | 4.6e-08 | 5.600E-08 | 4.976E-08 | 1.200E-08 | 1.100E-07 |
| 26644 | 1333 | A1333D | E | 4.2e-08 | 9.3e-08 | 3.9e-08 | 5.800E-08 | 3.035E-08 | 3.900E-08 | 9.300E-08 |
| 26645 | 1333 | A1333C | E | 1.5e-07 | 7.4e-08 | 2.1e-08 | 8.167E-08 | 6.484E-08 | 2.100E-08 | 1.500E-07 |
| 26646 | 1333 | A1333E | E | 2.8e-08 | 6.4e-08 | 2.1e-08 | 3.767E-08 | 2.307E-08 | 2.100E-08 | 6.400E-08 |
| 26647 | 1333 | A1333Q | E | 4.7e-08 | 1.9e-08 | 1.1e-07 | 5.867E-08 | 4.661E-08 | 1.900E-08 | 1.100E-07 |
| 26648 | 1333 | A1333G | E | 3.3e-08 | 5.5e-08 | 6.8e-08 | 5.200E-08 | 1.769E-08 | 3.300E-08 | 6.800E-08 |
| 26649 | 1333 | A1333H | E | 7e-08 | 4.8e-08 | 6.6e-08 | 6.133E-08 | 1.172E-08 | 4.800E-08 | 7.000E-08 |
| 26650 | 1333 | A1333I | E | 1.8e-08 | 3.1e-08 | 3.6e-08 | 2.833E-08 | 9.292E-09 | 1.800E-08 | 3.600E-08 |
| 26651 | 1333 | A1333L | E | 3.7e-08 | 4.9e-08 | 7.6e-08 | 5.400E-08 | 1.997E-08 | 3.700E-08 | 7.600E-08 |
| 26652 | 1333 | A1333K | E | 3.6e-08 | 2.3e-08 | 3.5e-08 | 3.133E-08 | 7.234E-09 | 2.300E-08 | 3.600E-08 |
| 26653 | 1333 | A1333M | E | 6.8e-08 | 1.4e-08 | 3.8e-08 | 4.000E-08 | 2.706E-08 | 1.400E-08 | 6.800E-08 |
| 26654 | 1333 | A1333F | E | 3.9e-08 | 7.2e-08 | 2.8e-08 | 4.633E-08 | 2.290E-08 | 2.800E-08 | 7.200E-08 |
| 26655 | 1333 | A1333P | E | 2e-08 | 3e-08 | 3e-08 | 2.667E-08 | 5.774E-09 | 2.000E-08 | 3.000E-08 |
| 26656 | 1333 | A1333S | E | 2.3e-08 | 2e-08 | 7.7e-08 | 4.000E-08 | 3.208E-08 | 2.000E-08 | 7.700E-08 |
| 26657 | 1333 | A1333T | E | 4.7e-08 | 5.7e-08 | 2.6e-08 | 4.333E-08 | 1.582E-08 | 2.600E-08 | 5.700E-08 |
| 26658 | 1333 | A1333W | E | 8.5e-08 | 6.2e-08 | 4.2e-08 | 6.300E-08 | 2.152E-08 | 4.200E-08 | 8.500E-08 |
| 26659 | 1333 | A1333Y | E | 6.6e-08 | 6.3e-08 | 3.5e-08 | 5.467E-08 | 1.710E-08 | 3.500E-08 | 6.600E-08 |
| 26660 | 1333 | A1333V | E | 2.6e-08 | 5.4e-08 | 4.7e-08 | 4.233E-08 | 1.457E-08 | 2.600E-08 | 5.400E-08 |
| 26661 | 1334 | D1334A | E | 8.4e-08 | 4.9e-08 | 9.4e-08 | 7.567E-08 | 2.363E-08 | 4.900E-08 | 9.400E-08 |
| 26662 | 1334 | D1334R | E | 8.6e-08 | 3.1e-08 | 4.4e-08 | 5.367E-08 | 2.875E-08 | 3.100E-08 | 8.600E-08 |
| 26663 | 1334 | D1334N | E | 1.5e-07 | 9.5e-08 | 3.1e-08 | 9.200E-08 | 5.956E-08 | 3.100E-08 | 1.500E-07 |
| 26664 | 1334 | D1334D | E | 2.8e-08 | 2e-08 | 3.3e-08 | 2.700E-08 | 6.557E-09 | 2.000E-08 | 3.300E-08 |
| 26665 | 1334 | D1334C | E | 3.3e-08 | 1.7e-08 | 7.6e-08 | 4.200E-08 | 3.051E-08 | 1.700E-08 | 7.600E-08 |
| 26666 | 1334 | D1334E | E | 4.9e-08 | 1e-07 | 1.3e-07 | 9.300E-08 | 4.095E-08 | 4.900E-08 | 1.300E-07 |
| 26667 | 1334 | D1334Q | E | 4.5e-08 | 5.7e-08 | 3.8e-08 | 4.667E-08 | 9.609E-09 | 3.800E-08 | 5.700E-08 |
| 26668 | 1334 | D1334G | E | 4.3e-08 | 5.9e-08 | 4.4e-08 | 4.867E-08 | 8.963E-09 | 4.300E-08 | 5.900E-08 |
| 26669 | 1334 | D1334H | E | 6.4e-08 | 3e-08 | 2.9e-08 | 4.100E-08 | 1.992E-08 | 2.900E-08 | 6.400E-08 |
| 26670 | 1334 | D1334I | E | 1.1e-08 | 9.3e-08 | 4.6e-08 | 5.000E-08 | 4.115E-08 | 1.100E-08 | 9.300E-08 |
| 26671 | 1334 | D1334L | E | 7.2e-08 | 6.8e-08 | 2.9e-08 | 5.633E-08 | 2.376E-08 | 2.900E-08 | 7.200E-08 |
| 26672 | 1334 | D1334K | E | 7.9e-08 | 2.2e-08 | 9.6e-08 | 6.567E-08 | 3.876E-08 | 2.200E-08 | 9.600E-08 |
| 26673 | 1334 | D1334M | E | 3.4e-08 | 1.2e-07 | 2.6e-08 | 6.000E-08 | 5.212E-08 | 2.600E-08 | 1.200E-07 |
| 26674 | 1334 | D1334F | E | 8.5e-08 | 7.8e-08 | 4.2e-08 | 6.833E-08 | 2.307E-08 | 4.200E-08 | 8.500E-08 |
| 26675 | 1334 | D1334P | E | 2.7e-08 | 2.9e-08 | 5.4e-08 | 3.667E-08 | 1.504E-08 | 2.700E-08 | 5.400E-08 |
| 26676 | 1334 | D1334S | E | 5.6e-08 | 5.3e-08 | 1.7e-08 | 4.200E-08 | 2.170E-08 | 1.700E-08 | 5.600E-08 |
| 26677 | 1334 | D1334T | E | 3.5e-08 | 4e-08 | 8.2e-08 | 5.233E-08 | 2.581E-08 | 3.500E-08 | 8.200E-08 |
| 26678 | 1334 | D1334W | E | 2.3e-08 | 4.4e-08 | 3.9e-08 | 3.533E-08 | 1.097E-08 | 2.300E-08 | 4.400E-08 |
| 26679 | 1334 | D1334Y | E | 5.9e-08 | 8.1e-08 | 3.5e-08 | 5.833E-08 | 2.301E-08 | 3.500E-08 | 8.100E-08 |
| 26680 | 1334 | D1334V | E | 6.3e-08 | 3.7e-08 | 2.1e-07 | 1.033E-07 | 9.329E-08 | 3.700E-08 | 2.100E-07 |
| 26681 | 1335 | S1335A | E | 2.9e-08 | 4.5e-08 | 2.1e-08 | 3.167E-08 | 1.222E-08 | 2.100E-08 | 4.500E-08 |
| 26682 | 1335 | S1335R | E | 5.2e-08 | 5.3e-08 | 2.5e-08 | 4.333E-08 | 1.589E-08 | 2.500E-08 | 5.300E-08 |
| 26683 | 1335 | S1335N | E | 3.8e-08 | 3.3e-08 | 7e-08 | 4.700E-08 | 2.007E-08 | 3.300E-08 | 7.000E-08 |
| 26684 | 1335 | S1335D | E | 6.3e-08 | 5.7e-08 | 2.2e-08 | 4.733E-08 | 2.214E-08 | 2.200E-08 | 6.300E-08 |
| 26685 | 1335 | S1335C | E | 2e-07 | 2.1e-08 | 1.8e-08 | 7.967E-08 | 1.042E-07 | 1.800E-08 | 2.000E-07 |
| 26686 | 1335 | S1335E | E | 1.9e-08 | 3.3e-08 | 1.3e-07 | 6.067E-08 | 6.045E-08 | 1.900E-08 | 1.300E-07 |
| 26687 | 1335 | S1335Q | E | 6.1e-08 | 7.8e-08 | 4.8e-08 | 6.233E-08 | 1.504E-08 | 4.800E-08 | 7.800E-08 |
| 26688 | 1335 | S1335G | E | 1e-07 | 3.4e-08 | 3.3e-08 | 5.567E-08 | 3.840E-08 | 3.300E-08 | 1.000E-07 |
| 26689 | 1335 | S1335H | E | 9.1e-08 | 4.5e-08 | 3e-08 | 5.533E-08 | 3.179E-08 | 3.000E-08 | 9.100E-08 |

| | | | | | | | | | | |
|---|---|---|---|---|---|---|---|---|---|---|
| 26690 | 1335 | S1335I | E | 1.8e-08 | 5e-08 | 2.4e-08 | 3.067E-08 | 1.701E-08 | 1.800E-08 | 5.000E-08 |
| 26691 | 1335 | S1335L | E | 2.8e-08 | 1.3e-07 | 3.3e-08 | 6.367E-08 | 5.750E-08 | 2.800E-08 | 1.300E-07 |
| 26692 | 1335 | S1335K | E | 7.1e-08 | 6e-08 | 1.4e-08 | 4.833E-08 | 3.024E-08 | 1.400E-08 | 7.100E-08 |
| 26693 | 1335 | S1335M | E | 7.2e-08 | 3.7e-08 | 4.8e-08 | 5.233E-08 | 1.790E-08 | 3.700E-08 | 7.200E-08 |
| 26694 | 1335 | S1335F | E | 8.7e-08 | 2.7e-08 | 2.9e-08 | 4.767E-08 | 3.408E-08 | 2.700E-08 | 8.700E-08 |
| 26695 | 1335 | S1335P | E | 8.7e-08 | 2.4e-07 | 6.6e-08 | 1.310E-07 | 9.498E-08 | 6.600E-08 | 2.400E-07 |
| 26696 | 1335 | S1335S | E | 7.6e-08 | 4.1e-08 | 8.1e-08 | 6.600E-08 | 2.179E-08 | 4.100E-08 | 8.100E-08 |
| 26697 | 1335 | S1335T | E | 1.7e-08 | 6.4e-08 | 1.4e-08 | 3.167E-08 | 2.804E-08 | 1.400E-08 | 6.400E-08 |
| 26698 | 1335 | S1335W | E | 3.4e-08 | 2.5e-08 | 4.8e-08 | 3.567E-08 | 1.159E-08 | 2.500E-08 | 4.800E-08 |
| 26699 | 1335 | S1335Y | E | 2.5e-08 | 7.7e-08 | 2.5e-08 | 4.233E-08 | 3.002E-08 | 2.500E-08 | 7.700E-08 |
| 26700 | 1335 | S1335V | E | 5.6e-08 | 5.2e-08 | 2e-08 | 4.267E-08 | 1.973E-08 | 2.000E-08 | 5.600E-08 |
| 26701 | 1336 | V1336A | E | 4.9e-08 | 3.9e-08 | 4.3e-08 | 4.367E-08 | 5.033E-09 | 3.900E-08 | 4.900E-08 |
| 26702 | 1336 | V1336R | E | 3.5e-08 | 1.8e-08 | 1.7e-08 | 2.333E-08 | 1.012E-08 | 1.700E-08 | 3.500E-08 |
| 26703 | 1336 | V1336N | E | 4.9e-08 | 7.2e-08 | 4.8e-08 | 5.633E-08 | 1.358E-08 | 4.800E-08 | 7.200E-08 |
| 26704 | 1336 | V1336D | E | 4.8e-08 | 2.4e-08 | 4.2e-08 | 3.800E-08 | 1.249E-08 | 2.400E-08 | 4.800E-08 |
| 26705 | 1336 | V1336C | E | 1.9e-08 | 2e-08 | 1.8e-08 | 1.900E-08 | 1.000E-09 | 1.800E-08 | 2.000E-08 |
| 26706 | 1336 | V1336E | E | 2.7e-08 | 3e-08 | 5e-08 | 3.567E-08 | 1.250E-08 | 2.700E-08 | 5.000E-08 |
| 26707 | 1336 | V1336Q | E | 6.9e-08 | 3.2e-08 | 3.8e-08 | 4.633E-08 | 1.986E-08 | 3.200E-08 | 6.900E-08 |
| 26708 | 1336 | V1336G | E | 7e-08 | 8.6e-08 | 8.3e-08 | 7.967E-08 | 8.505E-09 | 7.000E-08 | 8.600E-08 |
| 26709 | 1336 | V1336H | E | 5e-08 | 2.9e-08 | 7.9e-09 | 2.897E-08 | 2.105E-08 | 7.900E-09 | 5.000E-08 |
| 26710 | 1336 | V1336I | E | 8.4e-08 | 9.6e-08 | 2.6e-08 | 6.867E-08 | 3.743E-08 | 2.600E-08 | 9.600E-08 |
| 26711 | 1336 | V1336L | E | 4.8e-08 | 5.5e-08 | 4.2e-08 | 4.833E-08 | 6.506E-09 | 4.200E-08 | 5.500E-08 |
| 26712 | 1336 | V1336K | E | 5.6e-08 | 9e-08 | 3.9e-08 | 6.167E-08 | 2.597E-08 | 3.900E-08 | 9.000E-08 |
| 26713 | 1336 | V1336M | E | 2e-08 | 3.9e-08 | 3.2e-08 | 3.033E-08 | 9.609E-09 | 2.000E-08 | 3.900E-08 |
| 26714 | 1336 | V1336F | E | 4.4e-08 | 7e-08 | 3.6e-08 | 5.000E-08 | 1.778E-08 | 3.600E-08 | 7.000E-08 |
| 26715 | 1336 | V1336P | E | 7e-08 | 1.1e-07 | 4.5e-08 | 7.500E-08 | 3.279E-08 | 4.500E-08 | 1.100E-07 |
| 26716 | 1336 | V1336S | E | 2.2e-08 | 4.6e-08 | 4.1e-08 | 3.633E-08 | 1.266E-08 | 2.200E-08 | 4.600E-08 |
| 26717 | 1336 | V1336T | E | 3.4e-08 | 6.4e-08 | 1.8e-08 | 3.867E-08 | 2.335E-08 | 1.800E-08 | 6.400E-08 |
| 26718 | 1336 | V1336W | E | 5.9e-08 | 9e-09 | 5.4e-08 | 4.067E-08 | 2.754E-08 | 9.000E-09 | 5.900E-08 |
| 26719 | 1336 | V1336Y | E | 8.4e-08 | 3.5e-08 | 4.6e-08 | 5.500E-08 | 2.571E-08 | 3.500E-08 | 8.400E-08 |
| 26720 | 1336 | V1336V | E | 1.1e-07 | 1.1e-07 | 4.7e-08 | 8.900E-08 | 3.637E-08 | 4.700E-08 | 1.100E-07 |
| 26721 | 1337 | K1337A | E | 2.9e-08 | 5.8e-08 | 1.1e-07 | 6.567E-08 | 4.104E-08 | 2.900E-08 | 1.100E-07 |
| 26722 | 1337 | K1337R | E | 3.5e-08 | 4.2e-08 | 4.6e-08 | 4.100E-08 | 5.568E-09 | 3.500E-08 | 4.600E-08 |
| 26723 | 1337 | K1337N | E | 6.7e-08 | 5e-08 | 2.4e-08 | 4.700E-08 | 2.166E-08 | 2.400E-08 | 6.700E-08 |
| 26724 | 1337 | K1337D | E | 5.7e-08 | 4.2e-08 | 2.3e-08 | 4.067E-08 | 1.704E-08 | 2.300E-08 | 5.700E-08 |
| 26725 | 1337 | K1337C | E | 8.8e-08 | 3.3e-08 | 2.8e-08 | 4.967E-08 | 3.329E-08 | 2.800E-08 | 8.800E-08 |
| 26726 | 1337 | K1337E | E | 4.1e-08 | 9e-08 | 2.8e-08 | 5.300E-08 | 3.270E-08 | 2.800E-08 | 9.000E-08 |
| 26727 | 1337 | K1337Q | E | 5.2e-08 | 2.7e-08 | 1.5e-07 | 7.633E-08 | 6.501E-08 | 2.700E-08 | 1.500E-07 |
| 26728 | 1337 | K1337G | E | 4.5e-08 | 3.5e-08 | 1.2e-07 | 6.667E-08 | 4.646E-08 | 3.500E-08 | 1.200E-07 |
| 26729 | 1337 | K1337H | E | 2.4e-08 | 9.1e-08 | 4.1e-08 | 5.200E-08 | 3.483E-08 | 2.400E-08 | 9.100E-08 |
| 26730 | 1337 | K1337I | E | 2.9e-08 | 2.2e-08 | 5.3e-08 | 3.467E-08 | 1.626E-08 | 2.200E-08 | 5.300E-08 |
| 26731 | 1337 | K1337L | E | 2.6e-08 | 8.8e-08 | 4e-08 | 5.133E-08 | 3.252E-08 | 2.600E-08 | 8.800E-08 |
| 26732 | 1337 | K1337K | E | 4.6e-08 | 4.4e-08 | 4.9e-08 | 4.633E-08 | 2.517E-09 | 4.400E-08 | 4.900E-08 |
| 26733 | 1337 | K1337M | E | 2.5e-08 | 3e-08 | 1.6e-08 | 2.367E-08 | 7.095E-09 | 1.600E-08 | 3.000E-08 |
| 26734 | 1337 | K1337F | E | 8.3e-08 | 7.4e-08 | 1.3e-07 | 9.567E-08 | 3.007E-08 | 7.400E-08 | 1.300E-07 |
| 26735 | 1337 | K1337P | E | 7e-08 | 2.3e-08 | 5.2e-08 | 4.833E-08 | 2.371E-08 | 2.300E-08 | 7.000E-08 |
| 26736 | 1337 | K1337S | E | 1.2e-08 | 1.4e-08 | 5.9e-08 | 2.833E-08 | 2.658E-08 | 1.200E-08 | 5.900E-08 |
| 26737 | 1337 | K1337T | E | 2.7e-08 | 8.5e-08 | 3.5e-08 | 4.900E-08 | 3.143E-08 | 2.700E-08 | 8.500E-08 |
| 26738 | 1337 | K1337W | E | 6.4e-08 | 3.4e-08 | 2.3e-08 | 4.033E-08 | 2.122E-08 | 2.300E-08 | 6.400E-08 |
| 26739 | 1337 | K1337Y | E | 1.7e-08 | 1.4e-08 | 4.5e-08 | 2.533E-08 | 1.710E-08 | 1.400E-08 | 4.500E-08 |
| 26740 | 1337 | K1337V | E | 5e-08 | 2.6e-08 | 3.4e-08 | 3.667E-08 | 1.222E-08 | 2.600E-08 | 5.000E-08 |
| 26741 | 1338 | G1338A | E | 2.4e-08 | 3.4e-08 | 2.5e-08 | 2.767E-08 | 5.508E-09 | 2.400E-08 | 3.400E-08 |
| 26742 | 1338 | G1338R | E | 2.9e-08 | 1.2e-07 | 4e-08 | 6.300E-08 | 4.967E-08 | 2.900E-08 | 1.200E-07 |
| 26743 | 1338 | G1338N | E | 5.3e-08 | 2.6e-08 | 3.9e-08 | 3.933E-08 | 1.350E-08 | 2.600E-08 | 5.300E-08 |
| 26744 | 1338 | G1338D | E | 5.4e-08 | 5.3e-08 | 6.7e-08 | 5.800E-08 | 7.810E-09 | 5.300E-08 | 6.700E-08 |
| 26745 | 1338 | G1338C | E | 1.6e-08 | 6.2e-08 | 5.2e-08 | 4.333E-08 | 2.419E-08 | 1.600E-08 | 6.200E-08 |
| 26746 | 1338 | G1338E | E | 3.2e-08 | 4.8e-08 | 7.3e-08 | 5.100E-08 | 2.066E-08 | 3.200E-08 | 7.300E-08 |
| 26747 | 1338 | G1338Q | E | 3.6e-08 | 3e-08 | 4.3e-08 | 3.633E-08 | 6.506E-09 | 3.000E-08 | 4.300E-08 |
| 26748 | 1338 | G1338G | E | 6.8e-08 | 4.7e-08 | 2.3e-08 | 4.600E-08 | 2.252E-08 | 2.300E-08 | 6.800E-08 |
| 26749 | 1338 | G1338H | E | 1.3e-08 | 1.3e-08 | 1.4e-08 | 1.333E-08 | 5.774E-10 | 1.300E-08 | 1.400E-08 |
| 26750 | 1338 | G1338I | E | 2.9e-08 | 5e-08 | 1.3e-07 | 6.967E-08 | 5.329E-08 | 2.900E-08 | 1.300E-07 |
| 26751 | 1338 | G1338L | E | 4.8e-08 | 4.8e-08 | 5.4e-08 | 5.000E-08 | 3.464E-09 | 4.800E-08 | 5.400E-08 |
| 26752 | 1338 | G1338K | E | 5.4e-08 | 1.4e-08 | 1e-07 | 5.600E-08 | 4.303E-08 | 1.400E-08 | 1.000E-07 |
| 26753 | 1338 | G1338M | E | 1.5e-08 | 1.4e-08 | 1e-07 | 4.300E-08 | 4.937E-08 | 1.400E-08 | 1.000E-07 |
| 26754 | 1338 | G1338F | E | 2.1e-08 | 3e-08 | 1.6e-08 | 2.233E-08 | 7.095E-09 | 1.600E-08 | 3.000E-08 |
| 26755 | 1338 | G1338P | E | 1.5e-07 | 9.9e-08 | 4.3e-08 | 9.733E-08 | 5.352E-08 | 4.300E-08 | 1.500E-07 |
| 26756 | 1338 | G1338S | E | 3.6e-08 | 2e-08 | 2e-08 | 2.533E-08 | 9.238E-09 | 2.000E-08 | 3.600E-08 |
| 26757 | 1338 | G1338T | E | 3.9e-08 | 6.1e-08 | 4.3e-08 | 4.767E-08 | 1.172E-08 | 3.900E-08 | 6.100E-08 |
| 26758 | 1338 | G1338W | E | 4e-08 | 1.8e-08 | 3.5e-08 | 3.100E-08 | 1.153E-08 | 1.800E-08 | 4.000E-08 |
| 26759 | 1338 | G1338Y | E | 5.8e-08 | 4e-08 | 7.6e-08 | 5.800E-08 | 1.800E-08 | 4.000E-08 | 7.600E-08 |
| 26760 | 1338 | G1338V | E | 6.3e-08 | 1.1e-07 | 2e-08 | 6.433E-08 | 4.501E-08 | 2.000E-08 | 1.100E-07 |
| 26761 | 1339 | R1339A | E | 2.3e-08 | 3.6e-08 | 2.4e-08 | 2.767E-08 | 7.234E-09 | 2.300E-08 | 3.600E-08 |
| 26762 | 1339 | R1339R | E | 3.9e-08 | 9.7e-08 | 5.5e-08 | 6.367E-08 | 2.996E-08 | 3.900E-08 | 9.700E-08 |
| 26763 | 1339 | R1339N | E | 4.7e-08 | 3.4e-08 | 5.8e-08 | 4.633E-08 | 1.201E-08 | 3.400E-08 | 5.800E-08 |
| 26764 | 1339 | R1339D | E | 4.7e-08 | 1.7e-07 | 1.3e-07 | 1.157E-07 | 6.274E-08 | 4.700E-08 | 1.700E-07 |
| 26765 | 1339 | R1339C | E | 5.7e-08 | 6.5e-08 | 3.5e-08 | 5.233E-08 | 1.553E-08 | 3.500E-08 | 6.500E-08 |
| 26766 | 1339 | R1339E | E | 7.6e-08 | 4.5e-08 | 7.2e-08 | 6.433E-08 | 1.686E-08 | 4.500E-08 | 7.600E-08 |
| 26767 | 1339 | R1339Q | E | 2.5e-08 | 3.8e-08 | 8.9e-08 | 5.067E-08 | 3.383E-08 | 2.500E-08 | 8.900E-08 |
| 26768 | 1339 | R1339G | E | 6.8e-08 | 1.6e-08 | 3.7e-08 | 4.033E-08 | 2.616E-08 | 1.600E-08 | 6.800E-08 |
| 26769 | 1339 | R1339H | E | 1.1e-07 | 5.4e-08 | 4.3e-08 | 6.900E-08 | 3.593E-08 | 4.300E-08 | 1.100E-07 |
| 26770 | 1339 | R1339I | E | 1.5e-07 | 4.4e-08 | 3.1e-08 | 7.500E-08 | 6.528E-08 | 3.100E-08 | 1.500E-07 |
| 26771 | 1339 | R1339L | E | 3.9e-08 | 7.9e-08 | 1.9e-08 | 4.567E-08 | 3.055E-08 | 1.900E-08 | 7.900E-08 |
| 26772 | 1339 | R1339K | E | 3.6e-08 | 3.4e-08 | 1.1e-07 | 6.000E-08 | 4.331E-08 | 3.400E-08 | 1.100E-07 |
| 26773 | 1339 | R1339M | E | 2.9e-08 | 6.2e-08 | 5.5e-08 | 4.867E-08 | 1.739E-08 | 2.900E-08 | 6.200E-08 |
| 26774 | 1339 | R1339F | E | 3.5e-08 | 4.6e-08 | 5.2e-08 | 4.433E-08 | 8.622E-09 | 3.500E-08 | 5.200E-08 |

| | | | | | | | | | | |
|---|---|---|---|---|---|---|---|---|---|---|
| 26775 | 1339 | R1339P | E | 2.7e-08 | 2.8e-08 | 6.8e-08 | 4.100E-08 | 2.339E-08 | 2.700E-08 | 6.800E-08 |
| 26776 | 1339 | R1339S | E | 4.3e-08 | 4.2e-08 | 3.7e-08 | 4.067E-08 | 3.215E-09 | 3.700E-08 | 4.300E-08 |
| 26777 | 1339 | R1339T | E | 1.7e-08 | 3.3e-08 | 3.5e-08 | 2.833E-08 | 9.866E-09 | 1.700E-08 | 3.500E-08 |
| 26778 | 1339 | R1339W | E | 1.8e-08 | 5.7e-08 | 7.8e-08 | 5.100E-08 | 3.045E-08 | 1.800E-08 | 7.800E-08 |
| 26779 | 1339 | R1339Y | E | 5.2e-08 | 4.9e-08 | 4e-08 | 4.700E-08 | 6.245E-09 | 4.000E-08 | 5.200E-08 |
| 26780 | 1339 | R1339V | E | 4.8e-08 | 3e-08 | 2.9e-08 | 3.567E-08 | 1.069E-08 | 2.900E-08 | 4.800E-08 |
| 26781 | 1340 | F1340A | E | 4.3e-08 | 4.8e-08 | 7.8e-08 | 5.633E-08 | 1.893E-08 | 4.300E-08 | 7.800E-08 |
| 26782 | 1340 | F1340R | E | 5.7e-08 | 5.2e-08 | 5.7e-08 | 5.533E-08 | 2.887E-09 | 5.200E-08 | 5.700E-08 |
| 26783 | 1340 | F1340N | E | 3.3e-08 | 5.9e-08 | 3.6e-08 | 4.267E-08 | 1.422E-08 | 3.300E-08 | 5.900E-08 |
| 26784 | 1340 | F1340D | E | 2.9e-08 | 3.8e-08 | 5e-08 | 3.900E-08 | 1.054E-08 | 2.900E-08 | 5.000E-08 |
| 26785 | 1340 | F1340C | E | 3.3e-08 | 3.7e-08 | 9.2e-08 | 5.400E-08 | 3.297E-08 | 3.300E-08 | 9.200E-08 |
| 26786 | 1340 | F1340E | E | 7.1e-08 | 4.6e-08 | 2e-08 | 4.567E-08 | 2.550E-08 | 2.000E-08 | 7.100E-08 |
| 26787 | 1340 | F1340Q | E | 5.5e-08 | 6.3e-08 | 6.2e-08 | 6.000E-08 | 4.359E-09 | 5.500E-08 | 6.300E-08 |
| 26788 | 1340 | F1340G | E | 4.5e-08 | 1.1e-07 | 4.1e-08 | 6.533E-08 | 3.873E-08 | 4.100E-08 | 1.100E-07 |
| 26789 | 1340 | F1340H | E | 7.2e-08 | 4.1e-08 | 8.4e-08 | 6.567E-08 | 2.219E-08 | 4.100E-08 | 8.400E-08 |
| 26790 | 1340 | F1340I | E | 4.4e-08 | 2.2e-08 | 5.3e-08 | 3.967E-08 | 1.595E-08 | 2.200E-08 | 5.300E-08 |
| 26791 | 1340 | F1340L | E | 7.7e-08 | 4.1e-08 | 2.1e-08 | 4.633E-08 | 2.838E-08 | 2.100E-08 | 7.700E-08 |
| 26792 | 1340 | F1340K | E | 4.6e-08 | 1.3e-07 | 5.4e-08 | 7.667E-08 | 4.636E-08 | 4.600E-08 | 1.300E-07 |
| 26793 | 1340 | F1340M | E | 2.8e-08 | 9.1e-08 | 6.3e-08 | 6.067E-08 | 3.156E-08 | 2.800E-08 | 9.100E-08 |
| 26794 | 1340 | F1340F | E | 5.2e-08 | 5.2e-08 | 5.1e-08 | 5.167E-08 | 5.774E-10 | 5.100E-08 | 5.200E-08 |
| 26795 | 1340 | F1340P | E | 3.3e-08 | 2.8e-08 | 3e-08 | 3.033E-08 | 2.517E-09 | 2.800E-08 | 3.300E-08 |
| 26796 | 1340 | F1340S | E | 7.5e-08 | 1.6e-07 | 1.2e-07 | 1.183E-07 | 4.252E-08 | 7.500E-08 | 1.600E-07 |
| 26797 | 1340 | F1340T | E | 9.5e-08 | 6.6e-08 | 3.7e-08 | 6.600E-08 | 2.900E-08 | 3.700E-08 | 9.500E-08 |
| 26798 | 1340 | F1340W | E | 1.5e-08 | 1.2e-07 | 4.2e-08 | 5.900E-08 | 5.453E-08 | 1.500E-08 | 1.200E-07 |
| 26799 | 1340 | F1340Y | E | 4.6e-08 | 5.4e-08 | 2.7e-08 | 4.233E-08 | 1.387E-08 | 2.700E-08 | 5.400E-08 |
| 26800 | 1340 | F1340V | E | 9.6e-08 | 2.7e-08 | 4.6e-08 | 5.633E-08 | 3.564E-08 | 2.700E-08 | 9.600E-08 |
| 26801 | 1341 | T1341A | E | 3.7e-08 | 4.2e-08 | 2.6e-08 | 3.500E-08 | 8.185E-09 | 2.600E-08 | 4.200E-08 |
| 26802 | 1341 | T1341R | E | 2e-08 | 2.9e-08 | 2.4e-08 | 2.433E-08 | 4.509E-09 | 2.000E-08 | 2.900E-08 |
| 26803 | 1341 | T1341N | E | 6.2e-08 | 2.8e-08 | 3.8e-08 | 4.267E-08 | 1.747E-08 | 2.800E-08 | 6.200E-08 |
| 26804 | 1341 | T1341D | E | 6.9e-08 | 1.3e-07 | 1.3e-08 | 7.067E-08 | 5.852E-08 | 1.300E-08 | 1.300E-07 |
| 26805 | 1341 | T1341C | E | 4.7e-08 | 5.7e-08 | 5.7e-08 | 5.367E-08 | 5.774E-09 | 4.700E-08 | 5.700E-08 |
| 26806 | 1341 | T1341E | E | 4.6e-08 | 1.9e-08 | 3.7e-08 | 3.400E-08 | 1.375E-08 | 1.900E-08 | 4.600E-08 |
| 26807 | 1341 | T1341Q | E | 7.2e-08 | 5.4e-08 | 7e-08 | 6.533E-08 | 9.866E-09 | 5.400E-08 | 7.200E-08 |
| 26808 | 1341 | T1341G | E | 4.1e-08 | 5.5e-08 | 1.1e-08 | 3.567E-08 | 2.248E-08 | 1.100E-08 | 5.500E-08 |
| 26809 | 1341 | T1341H | E | 4.7e-08 | 2.2e-08 | 8.9e-09 | 2.597E-08 | 1.936E-08 | 8.900E-09 | 4.700E-08 |
| 26810 | 1341 | T1341I | E | 9.5e-08 | 1.4e-07 | 1.6e-08 | 8.367E-08 | 6.277E-08 | 1.600E-08 | 1.400E-07 |
| 26811 | 1341 | T1341L | E | 5.2e-08 | 3.5e-08 | 1.3e-07 | 7.233E-08 | 5.066E-08 | 3.500E-08 | 1.300E-07 |
| 26812 | 1341 | T1341K | E | 1.1e-07 | 2.2e-08 | 2.6e-08 | 5.267E-08 | 4.969E-08 | 2.200E-08 | 1.100E-07 |
| 26813 | 1341 | T1341M | E | 2.3e-08 | 6.5e-08 | 2.5e-08 | 3.767E-08 | 2.369E-08 | 2.300E-08 | 6.500E-08 |
| 26814 | 1341 | T1341F | E | 9.8e-08 | 1.6e-08 | 4.9e-08 | 5.433E-08 | 4.126E-08 | 1.600E-08 | 9.800E-08 |
| 26815 | 1341 | T1341P | E | 4.1e-08 | 4.9e-08 | 1.6e-08 | 3.533E-08 | 1.721E-08 | 1.600E-08 | 4.900E-08 |
| 26816 | 1341 | T1341S | E | 6e-08 | 7.3e-08 | 3.3e-08 | 5.533E-08 | 2.040E-08 | 3.300E-08 | 7.300E-08 |
| 26817 | 1341 | T1341T | E | 4.8e-08 | 2.2e-08 | 6.5e-08 | 4.500E-08 | 2.166E-08 | 2.200E-08 | 6.500E-08 |
| 26818 | 1341 | T1341W | E | 4.9e-08 | 1.7e-08 | 4.8e-08 | 3.800E-08 | 1.819E-08 | 1.700E-08 | 4.900E-08 |
| 26819 | 1341 | T1341Y | E | 5.3e-08 | 2.8e-08 | 2e-07 | 9.367E-08 | 9.293E-08 | 2.800E-08 | 2.000E-07 |
| 26820 | 1341 | T1341V | E | 1e-07 | 9.6e-09 | 6e-08 | 5.653E-08 | 4.530E-08 | 9.600E-09 | 1.000E-07 |
| 26821 | 1342 | I1342A | E | 1e-07 | 3.1e-08 | 6.2e-08 | 6.433E-08 | 3.456E-08 | 3.100E-08 | 1.000E-07 |
| 26822 | 1342 | I1342R | E | 5.1e-08 | 5.8e-08 | 1.3e-08 | 4.067E-08 | 2.421E-08 | 1.300E-08 | 5.800E-08 |
| 26823 | 1342 | I1342N | E | 1.4e-08 | 9.6e-08 | 7.6e-08 | 6.200E-08 | 4.276E-08 | 1.400E-08 | 9.600E-08 |
| 26824 | 1342 | I1342D | E | 5.7e-08 | 4.1e-08 | 9e-08 | 6.267E-08 | 2.499E-08 | 4.100E-08 | 9.000E-08 |
| 26825 | 1342 | I1342C | E | 5.6e-08 | 2.3e-08 | 5e-08 | 4.300E-08 | 1.758E-08 | 2.300E-08 | 5.600E-08 |
| 26826 | 1342 | I1342E | E | 3.6e-08 | 4.2e-08 | 1e-08 | 2.933E-08 | 1.701E-08 | 1.000E-08 | 4.200E-08 |
| 26827 | 1342 | I1342Q | E | 2.1e-08 | 1e-07 | 2.2e-08 | 4.767E-08 | 4.532E-08 | 2.100E-08 | 1.000E-07 |
| 26828 | 1342 | I1342G | E | 3.9e-08 | 2.1e-08 | 1.1e-07 | 5.667E-08 | 4.706E-08 | 2.100E-08 | 1.100E-07 |
| 26829 | 1342 | I1342H | E | 1.2e-08 | 1.4e-07 | 4.6e-08 | 6.600E-08 | 6.630E-08 | 1.200E-08 | 1.400E-07 |
| 26830 | 1342 | I1342I | E | 9.7e-08 | 7e-08 | 4.5e-08 | 7.067E-08 | 2.601E-08 | 4.500E-08 | 9.700E-08 |
| 26831 | 1342 | I1342L | E | 1.5e-08 | 2.9e-08 | 1.5e-08 | 1.967E-08 | 8.083E-09 | 1.500E-08 | 2.900E-08 |
| 26832 | 1342 | I1342K | E | 5.8e-08 | 2e-07 | 2.1e-08 | 9.300E-08 | 9.449E-08 | 2.100E-08 | 2.000E-07 |
| 26833 | 1342 | I1342M | E | 4e-08 | 4.1e-08 | 1.5e-08 | 3.200E-08 | 1.473E-08 | 1.500E-08 | 4.100E-08 |
| 26834 | 1342 | I1342F | E | 3.1e-08 | 1e-07 | 6.3e-08 | 6.467E-08 | 3.453E-08 | 3.100E-08 | 1.000E-07 |
| 26835 | 1342 | I1342P | E | 5.2e-08 | 5.1e-08 | 9.8e-08 | 6.700E-08 | 2.685E-08 | 5.100E-08 | 9.800E-08 |
| 26836 | 1342 | I1342S | E | 3.8e-08 | 5.9e-08 | 3.3e-08 | 4.333E-08 | 1.380E-08 | 3.300E-08 | 5.900E-08 |
| 26837 | 1342 | I1342T | E | 3.7e-08 | 2.6e-08 | 7.7e-08 | 4.667E-08 | 2.684E-08 | 2.600E-08 | 7.700E-08 |
| 26838 | 1342 | I1342W | E | 2.6e-08 | 5.8e-08 | 6.4e-08 | 4.933E-08 | 2.043E-08 | 2.600E-08 | 6.400E-08 |
| 26839 | 1342 | I1342Y | E | 3.1e-08 | 3.2e-08 | 7.9e-08 | 4.733E-08 | 2.743E-08 | 3.100E-08 | 7.900E-08 |
| 26840 | 1342 | I1342V | E | 4.4e-08 | 2.8e-08 | 3.5e-08 | 3.567E-08 | 8.021E-09 | 2.800E-08 | 4.400E-08 |
| 26841 | 1343 | S1343A | E | 4.5e-08 | 1.5e-07 | 8.6e-08 | 9.367E-08 | 5.292E-08 | 4.500E-08 | 1.500E-07 |
| 26842 | 1343 | S1343R | E | 1.9e-08 | 1.9e-08 | 4.3e-08 | 2.700E-08 | 1.386E-08 | 1.900E-08 | 4.300E-08 |
| 26843 | 1343 | S1343N | E | 6.6e-08 | 8.1e-08 | 2.8e-08 | 5.833E-08 | 2.732E-08 | 2.800E-08 | 8.100E-08 |
| 26844 | 1343 | S1343D | E | 6.7e-08 | 2.6e-08 | 2.5e-08 | 3.933E-08 | 2.397E-08 | 2.500E-08 | 6.700E-08 |
| 26845 | 1343 | S1343C | E | 2.9e-08 | 5.4e-08 | 1e-07 | 6.100E-08 | 3.601E-08 | 2.900E-08 | 1.000E-07 |
| 26846 | 1343 | S1343E | E | 3.6e-08 | 1.6e-07 | 2.6e-08 | 7.400E-08 | 7.465E-08 | 2.600E-08 | 1.600E-07 |
| 26847 | 1343 | S1343Q | E | 3.4e-08 | 1.1e-07 | 4.9e-08 | 6.433E-08 | 4.025E-08 | 3.400E-08 | 1.100E-07 |
| 26848 | 1343 | S1343G | E | 6.1e-08 | 3.2e-08 | 3e-08 | 4.100E-08 | 1.735E-08 | 3.000E-08 | 6.100E-08 |
| 26849 | 1343 | S1343H | E | 2.7e-08 | 2.7e-08 | 3.1e-08 | 2.833E-08 | 2.309E-09 | 2.700E-08 | 3.100E-08 |
| 26850 | 1343 | S1343I | E | 7.1e-08 | 6.7e-08 | 1e-07 | 7.933E-08 | 1.801E-08 | 6.700E-08 | 1.000E-07 |
| 26851 | 1343 | S1343L | E | 3.4e-08 | 1.5e-08 | 2.8e-08 | 2.567E-08 | 9.713E-09 | 1.500E-08 | 3.400E-08 |
| 26852 | 1343 | S1343K | E | 3.9e-08 | 1.3e-07 | 3.4e-08 | 6.767E-08 | 5.404E-08 | 3.400E-08 | 1.300E-07 |
| 26853 | 1343 | S1343M | E | 2.8e-08 | 3.1e-08 | 4.6e-08 | 3.500E-08 | 9.644E-09 | 2.800E-08 | 4.600E-08 |
| 26854 | 1343 | S1343F | E | 2.1e-08 | 3.6e-08 | 2.5e-07 | 1.023E-07 | 1.281E-07 | 2.100E-08 | 2.500E-07 |
| 26855 | 1343 | S1343P | E | 1.7e-07 | 3.1e-08 | 1.1e-07 | 1.037E-07 | 6.972E-08 | 3.100E-08 | 1.700E-07 |
| 26856 | 1343 | S1343S | E | 4.6e-08 | 5.1e-08 | 4.7e-09 | 3.390E-08 | 2.541E-08 | 4.700E-09 | 5.100E-08 |
| 26857 | 1343 | S1343T | E | 9.3e-08 | 6.6e-08 | 3.7e-08 | 6.533E-08 | 2.801E-08 | 3.700E-08 | 9.300E-08 |
| 26858 | 1343 | S1343W | E | 8.9e-09 | 1.5e-08 | 2.2e-08 | 1.530E-08 | 6.555E-09 | 8.900E-09 | 2.200E-08 |
| 26859 | 1343 | S1343Y | E | 5.3e-08 | 4.3e-08 | 7.3e-08 | 5.633E-08 | 1.528E-08 | 4.300E-08 | 7.300E-08 |

| | | | | | | | | | | |
|---|---|---|---|---|---|---|---|---|---|---|
| 26860 | 1343 | S1343V | E | 3.3e-08 | 1.8e-08 | 1.7e-07 | 7.367E-08 | 8.376E-08 | 1.800E-08 | 1.700E-07 |
| 26861 | 1344 | A1344A | E | 4.2e-08 | 6.7e-08 | 3.3e-08 | 4.733E-08 | 1.762E-08 | 3.300E-08 | 6.700E-08 |
| 26862 | 1344 | A1344R | E | 6.1e-08 | 1.2e-08 | 2.4e-08 | 3.233E-08 | 2.554E-08 | 1.200E-08 | 6.100E-08 |
| 26863 | 1344 | A1344N | E | 6.3e-08 | 4.5e-08 | 2e-08 | 4.267E-08 | 2.159E-08 | 2.000E-08 | 6.300E-08 |
| 26864 | 1344 | A1344D | E | 6.7e-08 | 3.1e-08 | 6.2e-09 | 3.473E-08 | 3.057E-08 | 6.200E-09 | 6.700E-08 |
| 26865 | 1344 | A1344C | E | 3.5e-08 | 1.5e-07 | 1.9e-08 | 6.800E-08 | 7.146E-08 | 1.900E-08 | 1.500E-07 |
| 26866 | 1344 | A1344E | E | 7e-08 | 1.3e-07 | 7.6e-08 | 9.200E-08 | 3.305E-08 | 7.000E-08 | 1.300E-07 |
| 26867 | 1344 | A1344Q | E | 6.6e-08 | 4e-08 | 3.9e-08 | 4.833E-08 | 1.531E-08 | 3.900E-08 | 6.600E-08 |
| 26868 | 1344 | A1344G | E | 7.4e-08 | 1.2e-07 | 4.4e-08 | 7.933E-08 | 3.828E-08 | 4.400E-08 | 1.200E-07 |
| 26869 | 1344 | A1344H | E | 3.6e-08 | 5.9e-08 | 3.3e-08 | 4.267E-08 | 1.422E-08 | 3.300E-08 | 5.900E-08 |
| 26870 | 1344 | A1344I | E | 4.5e-08 | 1.5e-08 | 3.4e-08 | 3.133E-08 | 1.518E-08 | 1.500E-08 | 4.500E-08 |
| 26871 | 1344 | A1344L | E | 6.5e-08 | 7.2e-08 | 5.1e-08 | 6.267E-08 | 1.069E-08 | 5.100E-08 | 7.200E-08 |
| 26872 | 1344 | A1344K | E | 4.1e-08 | 2.8e-08 | 7.5e-08 | 4.800E-08 | 2.427E-08 | 2.800E-08 | 7.500E-08 |
| 26873 | 1344 | A1344M | E | 4.6e-08 | 2.5e-08 | 4.6e-08 | 3.900E-08 | 1.212E-08 | 2.500E-08 | 4.600E-08 |
| 26874 | 1344 | A1344F | E | 3.1e-08 | 3e-08 | 7.5e-08 | 4.533E-08 | 2.570E-08 | 3.000E-08 | 7.500E-08 |
| 26875 | 1344 | A1344P | E | 2.1e-08 | 6.2e-08 | 1e-07 | 6.100E-08 | 3.951E-08 | 2.100E-08 | 1.000E-07 |
| 26876 | 1344 | A1344S | E | 2.9e-08 | 6.5e-08 | 4.8e-08 | 4.733E-08 | 1.801E-08 | 2.900E-08 | 6.500E-08 |
| 26877 | 1344 | A1344T | E | 1.6e-08 | 4.6e-08 | 2.3e-08 | 2.833E-08 | 1.570E-08 | 1.600E-08 | 4.600E-08 |
| 26878 | 1344 | A1344W | E | 1.6e-07 | 5.9e-08 | 1e-07 | 1.063E-07 | 5.080E-08 | 5.900E-08 | 1.600E-07 |
| 26879 | 1344 | A1344Y | E | 5e-08 | 7e-08 | 4.4e-08 | 5.467E-08 | 1.361E-08 | 4.400E-08 | 7.000E-08 |
| 26880 | 1344 | A1344V | E | 7.1e-08 | 2.3e-08 | 6.9e-08 | 5.433E-08 | 2.715E-08 | 2.300E-08 | 7.100E-08 |
| 26881 | 1345 | D1345A | E | 5.2e-08 | 5.1e-08 | 2.7e-08 | 4.333E-08 | 1.415E-08 | 2.700E-08 | 5.200E-08 |
| 26882 | 1345 | D1345R | E | 3.3e-08 | 8.5e-08 | 5.1e-08 | 5.633E-08 | 2.641E-08 | 3.300E-08 | 8.500E-08 |
| 26883 | 1345 | D1345N | E | 7.7e-08 | 9.7e-08 | 4.2e-08 | 7.200E-08 | 2.784E-08 | 4.200E-08 | 9.700E-08 |
| 26884 | 1345 | D1345D | E | 1e-08 | 6.2e-08 | 2.3e-07 | 1.007E-07 | 1.150E-07 | 1.000E-08 | 2.300E-07 |
| 26885 | 1345 | D1345C | E | 3.4e-08 | 5.8e-08 | 1.7e-08 | 3.633E-08 | 2.060E-08 | 1.700E-08 | 5.800E-08 |
| 26886 | 1345 | D1345E | E | 8.3e-08 | 1.1e-08 | 3.1e-08 | 4.167E-08 | 3.717E-08 | 1.100E-08 | 8.300E-08 |
| 26887 | 1345 | D1345Q | E | 4e-08 | 3.5e-08 | 4.2e-08 | 3.900E-08 | 3.606E-09 | 3.500E-08 | 4.200E-08 |
| 26888 | 1345 | D1345G | E | 9.5e-08 | 5.5e-08 | 3.1e-08 | 6.033E-08 | 3.233E-08 | 3.100E-08 | 9.500E-08 |
| 26889 | 1345 | D1345H | E | 1.8e-08 | 1.5e-08 | 2.5e-08 | 1.933E-08 | 5.132E-09 | 1.500E-08 | 2.500E-08 |
| 26890 | 1345 | D1345I | E | 2.8e-08 | 6.9e-08 | 3.7e-08 | 4.467E-08 | 2.155E-08 | 2.800E-08 | 6.900E-08 |
| 26891 | 1345 | D1345L | E | 1.8e-08 | 2.4e-08 | 7.4e-08 | 3.867E-08 | 3.075E-08 | 1.800E-08 | 7.400E-08 |
| 26892 | 1345 | D1345K | E | 7.5e-08 | 2.1e-08 | 7e-08 | 5.533E-08 | 2.984E-08 | 2.100E-08 | 7.500E-08 |
| 26893 | 1345 | D1345M | E | 5.1e-08 | 2.9e-08 | 1.6e-08 | 3.200E-08 | 1.769E-08 | 1.600E-08 | 5.100E-08 |
| 26894 | 1345 | D1345F | E | 4.2e-08 | 3.7e-08 | 2.4e-08 | 3.433E-08 | 9.292E-09 | 2.400E-08 | 4.200E-08 |
| 26895 | 1345 | D1345P | E | 1.8e-08 | 3e-08 | 9.6e-08 | 4.800E-08 | 4.200E-08 | 1.800E-08 | 9.600E-08 |
| 26896 | 1345 | D1345S | E | 2.8e-08 | 3.3e-08 | 3.1e-08 | 3.067E-08 | 2.517E-09 | 2.800E-08 | 3.300E-08 |
| 26897 | 1345 | D1345T | E | 9.5e-08 | 4.9e-09 | 3.5e-08 | 4.497E-08 | 4.587E-08 | 4.900E-09 | 9.500E-08 |
| 26898 | 1345 | D1345W | E | 4.4e-08 | 2.8e-08 | 1.4e-08 | 2.867E-08 | 1.501E-08 | 1.400E-08 | 4.400E-08 |
| 26899 | 1345 | D1345Y | E | 6.5e-08 | 5.7e-08 | 6.8e-08 | 6.333E-08 | 5.686E-09 | 5.700E-08 | 6.800E-08 |
| 26900 | 1345 | D1345V | E | 6.4e-08 | 1.4e-07 | 2e-08 | 7.467E-08 | 6.071E-08 | 2.000E-08 | 1.400E-07 |
| 26901 | 1346 | T1346A | E | 6.2e-08 | 1.1e-07 | 1.2e-08 | 6.133E-08 | 4.900E-08 | 1.200E-08 | 1.100E-07 |
| 26902 | 1346 | T1346R | E | 5.7e-08 | 2.1e-08 | 7.6e-08 | 5.133E-08 | 2.793E-08 | 2.100E-08 | 7.600E-08 |
| 26903 | 1346 | T1346N | E | 7.6e-08 | 6e-08 | 4.3e-08 | 5.967E-08 | 1.650E-08 | 4.300E-08 | 7.600E-08 |
| 26904 | 1346 | T1346D | E | 3.6e-08 | 3.3e-08 | 1.2e-07 | 6.300E-08 | 4.939E-08 | 3.300E-08 | 1.200E-07 |
| 26905 | 1346 | T1346C | E | 4.4e-08 | 3e-08 | 3.5e-08 | 3.633E-08 | 7.095E-09 | 3.000E-08 | 4.400E-08 |
| 26906 | 1346 | T1346E | E | 3.7e-08 | 1.7e-08 | 6.6e-08 | 4.000E-08 | 2.464E-08 | 1.700E-08 | 6.600E-08 |
| 26907 | 1346 | T1346Q | E | 5.2e-08 | 4.8e-08 | 3.2e-08 | 4.400E-08 | 1.058E-08 | 3.200E-08 | 5.200E-08 |
| 26908 | 1346 | T1346G | E | 3.3e-08 | 5.2e-08 | 3.1e-08 | 3.867E-08 | 1.159E-08 | 3.100E-08 | 5.200E-08 |
| 26909 | 1346 | T1346H | E | 3.8e-08 | 5.7e-08 | 7.5e-08 | 5.667E-08 | 1.850E-08 | 3.800E-08 | 7.500E-08 |
| 26910 | 1346 | T1346I | E | 5e-08 | 5.1e-08 | 1e-07 | 6.700E-08 | 2.858E-08 | 5.000E-08 | 1.000E-07 |
| 26911 | 1346 | T1346L | E | 6.1e-08 | 2e-08 | 6.5e-08 | 4.867E-08 | 2.491E-08 | 2.000E-08 | 6.500E-08 |
| 26912 | 1346 | T1346K | E | 3.9e-08 | 1.7e-08 | 7.3e-08 | 4.300E-08 | 2.821E-08 | 1.700E-08 | 7.300E-08 |
| 26913 | 1346 | T1346M | E | 8.7e-08 | 6e-08 | 4.4e-08 | 6.367E-08 | 2.173E-08 | 4.400E-08 | 8.700E-08 |
| 26914 | 1346 | T1346F | E | 2e-08 | 2.8e-08 | 1.1e-07 | 5.267E-08 | 4.981E-08 | 2.000E-08 | 1.100E-07 |
| 26915 | 1346 | T1346P | E | 6.1e-08 | 6.3e-08 | 2.7e-08 | 5.033E-08 | 2.023E-08 | 2.700E-08 | 6.300E-08 |
| 26916 | 1346 | T1346S | E | 3e-08 | 3.1e-08 | 1.6e-08 | 2.567E-08 | 8.386E-09 | 1.600E-08 | 3.100E-08 |
| 26917 | 1346 | T1346T | E | 2.6e-08 | 7.3e-08 | 5.1e-08 | 5.000E-08 | 2.352E-08 | 2.600E-08 | 7.300E-08 |
| 26918 | 1346 | T1346W | E | 1.3e-07 | 1.7e-07 | 7.5e-08 | 1.250E-07 | 4.770E-08 | 7.500E-08 | 1.700E-07 |
| 26919 | 1346 | T1346Y | E | 8.3e-08 | 7.6e-08 | 5.3e-08 | 7.067E-08 | 1.570E-08 | 5.300E-08 | 8.300E-08 |
| 26920 | 1346 | T1346V | E | 9.6e-08 | 4.5e-08 | 4.5e-08 | 6.200E-08 | 2.944E-08 | 4.500E-08 | 9.600E-08 |
| 26921 | 1347 | S1347A | E | 4.9e-08 | 7.3e-08 | 1.8e-07 | 1.007E-07 | 6.974E-08 | 4.900E-08 | 1.800E-07 |
| 26922 | 1347 | S1347R | E | 2.4e-08 | 3.2e-08 | 2.4e-08 | 2.667E-08 | 4.619E-09 | 2.400E-08 | 3.200E-08 |
| 26923 | 1347 | S1347N | E | 1.1e-07 | 2.4e-08 | 1.7e-07 | 1.013E-07 | 7.338E-08 | 2.400E-08 | 1.700E-07 |
| 26924 | 1347 | S1347D | E | 4.4e-08 | 5.1e-08 | 2.6e-08 | 4.033E-08 | 1.290E-08 | 2.600E-08 | 5.100E-08 |
| 26925 | 1347 | S1347C | E | 5.3e-08 | 1.3e-07 | 3.1e-08 | 7.133E-08 | 5.198E-08 | 3.100E-08 | 1.300E-07 |
| 26926 | 1347 | S1347E | E | 1.9e-08 | 4.9e-08 | 5e-08 | 3.933E-08 | 1.762E-08 | 1.900E-08 | 5.000E-08 |
| 26927 | 1347 | S1347Q | E | 2e-08 | 2e-08 | 6.8e-08 | 3.600E-08 | 2.771E-08 | 2.000E-08 | 6.800E-08 |
| 26928 | 1347 | S1347G | E | 4.6e-08 | 2.1e-08 | 3.6e-08 | 3.433E-08 | 1.258E-08 | 2.100E-08 | 4.600E-08 |
| 26929 | 1347 | S1347H | E | 3.7e-08 | 3.6e-08 | 8.6e-08 | 5.300E-08 | 2.858E-08 | 3.600E-08 | 8.600E-08 |
| 26930 | 1347 | S1347I | E | 5.5e-08 | 1.9e-08 | 4.4e-08 | 3.933E-08 | 1.845E-08 | 1.900E-08 | 5.500E-08 |
| 26931 | 1347 | S1347L | E | 4.1e-08 | 3.6e-08 | 4.9e-08 | 4.200E-08 | 6.557E-09 | 3.600E-08 | 4.900E-08 |
| 26932 | 1347 | S1347K | E | 4.9e-08 | 2.6e-08 | 2.7e-08 | 3.400E-08 | 1.300E-08 | 2.600E-08 | 4.900E-08 |
| 26933 | 1347 | S1347M | E | 5.3e-08 | 4.5e-08 | 3.2e-08 | 4.333E-08 | 1.060E-08 | 3.200E-08 | 5.300E-08 |
| 26934 | 1347 | S1347F | E | 9.1e-08 | 6e-08 | 2.7e-08 | 5.933E-08 | 3.201E-08 | 2.700E-08 | 9.100E-08 |
| 26935 | 1347 | S1347P | E | 1.6e-08 | 5e-08 | 8e-08 | 4.867E-08 | 3.202E-08 | 1.600E-08 | 8.000E-08 |
| 26936 | 1347 | S1347S | E | 7.9e-08 | 1.2e-07 | 2.8e-08 | 7.567E-08 | 4.609E-08 | 2.800E-08 | 1.200E-07 |
| 26937 | 1347 | S1347T | E | 1.6e-08 | 2.2e-08 | 1.1e-07 | 4.933E-08 | 5.262E-08 | 1.600E-08 | 1.100E-07 |
| 26938 | 1347 | S1347W | E | 4.4e-08 | 5.7e-08 | 3.6e-08 | 4.567E-08 | 1.060E-08 | 3.600E-08 | 5.700E-08 |
| 26939 | 1347 | S1347Y | E | 8.7e-08 | 4.8e-08 | 4.6e-08 | 6.033E-08 | 2.312E-08 | 4.600E-08 | 8.700E-08 |
| 26940 | 1347 | S1347V | E | 6e-08 | 6.8e-08 | 2.5e-08 | 5.100E-08 | 2.287E-08 | 2.500E-08 | 6.800E-08 |
| 26941 | 1348 | K1348A | E | 6.4e-09 | 4.8e-08 | 3e-08 | 2.813E-08 | 2.086E-08 | 6.400E-09 | 4.800E-08 |
| 26942 | 1348 | K1348R | E | 1.4e-08 | 9e-08 | 3.1e-08 | 4.500E-08 | 3.989E-08 | 1.400E-08 | 9.000E-08 |
| 26943 | 1348 | K1348N | E | 2.5e-08 | 4.7e-08 | 1.4e-08 | 2.867E-08 | 1.680E-08 | 1.400E-08 | 4.700E-08 |
| 26944 | 1348 | K1348D | E | 1.1e-08 | 2.1e-08 | 9.5e-08 | 4.233E-08 | 4.588E-08 | 1.100E-08 | 9.500E-08 |

| | | | | | | | | | | |
|---|---|---|---|---|---|---|---|---|---|---|
| 26945 | 1348 | K1348C | E | 1.8e-07 | 6.9e-08 | 1.6e-07 | 1.363E-07 | 5.916E-08 | 6.900E-08 | 1.800E-07 |
| 26946 | 1348 | K1348E | E | 2.2e-08 | 6.5e-08 | 4.5e-08 | 4.400E-08 | 2.152E-08 | 2.200E-08 | 6.500E-08 |
| 26947 | 1348 | K1348Q | E | 2.9e-08 | 3.1e-08 | 2.5e-08 | 2.833E-08 | 3.055E-09 | 2.500E-08 | 3.100E-08 |
| 26948 | 1348 | K1348G | E | 3.7e-08 | 5.2e-08 | 3.7e-08 | 4.200E-08 | 8.660E-09 | 3.700E-08 | 5.200E-08 |
| 26949 | 1348 | K1348H | E | 2.4e-08 | 2e-08 | 6e-08 | 3.467E-08 | 2.203E-08 | 2.000E-08 | 6.000E-08 |
| 26950 | 1348 | K1348I | E | 9.4e-08 | 3.7e-08 | 3e-08 | 5.367E-08 | 3.510E-08 | 3.000E-08 | 9.400E-08 |
| 26951 | 1348 | K1348L | E | 3.5e-08 | 2.9e-08 | 4.2e-08 | 3.533E-08 | 6.506E-09 | 2.900E-08 | 4.200E-08 |
| 26952 | 1348 | K1348K | E | 4.1e-08 | 9.4e-08 | 7.1e-08 | 6.867E-08 | 2.658E-08 | 4.100E-08 | 9.400E-08 |
| 26953 | 1348 | K1348M | E | 1e-07 | 3.8e-08 | 4.7e-08 | 6.167E-08 | 3.350E-08 | 3.800E-08 | 1.000E-07 |
| 26954 | 1348 | K1348F | E | 1.7e-08 | 3.9e-08 | 3.6e-08 | 3.067E-08 | 1.193E-08 | 1.700E-08 | 3.900E-08 |
| 26955 | 1348 | K1348P | E | 8.7e-08 | 4e-08 | 2.1e-08 | 4.933E-08 | 3.398E-08 | 2.100E-08 | 8.700E-08 |
| 26956 | 1348 | K1348S | E | 2.9e-08 | 3.5e-08 | 3.7e-08 | 3.367E-08 | 4.163E-09 | 2.900E-08 | 3.700E-08 |
| 26957 | 1348 | K1348T | E | 1.6e-08 | 3.1e-08 | 1.1e-07 | 5.233E-08 | 5.050E-08 | 1.600E-08 | 1.100E-07 |
| 26958 | 1348 | K1348W | E | 3.7e-08 | 2.9e-08 | 5.7e-08 | 4.100E-08 | 1.442E-08 | 2.900E-08 | 5.700E-08 |
| 26959 | 1348 | K1348Y | E | 2e-08 | 2.6e-08 | 6.6e-08 | 3.733E-08 | 2.501E-08 | 2.000E-08 | 6.600E-08 |
| 26960 | 1348 | K1348V | E | 5.2e-08 | 3.2e-08 | 5.5e-08 | 4.633E-08 | 1.250E-08 | 3.200E-08 | 5.500E-08 |
| 26961 | 1349 | N1349A | E | 5.7e-08 | 2.2e-08 | 4.3e-08 | 4.067E-08 | 1.762E-08 | 2.200E-08 | 5.700E-08 |
| 26962 | 1349 | N1349R | E | 9.3e-08 | 5.6e-08 | 5.8e-08 | 6.900E-08 | 2.081E-08 | 5.600E-08 | 9.300E-08 |
| 26963 | 1349 | N1349N | E | 7.4e-08 | 4e-08 | 5.8e-08 | 5.733E-08 | 1.701E-08 | 4.000E-08 | 7.400E-08 |
| 26964 | 1349 | N1349D | E | 2.3e-08 | 3.6e-08 | 7.4e-08 | 4.433E-08 | 2.650E-08 | 2.300E-08 | 7.400E-08 |
| 26965 | 1349 | N1349C | E | 7e-08 | 9e-08 | 3.2e-08 | 6.400E-08 | 2.946E-08 | 3.200E-08 | 9.000E-08 |
| 26966 | 1349 | N1349E | E | 1e-07 | 2.5e-08 | 1.3e-07 | 8.500E-08 | 5.408E-08 | 2.500E-08 | 1.300E-07 |
| 26967 | 1349 | N1349Q | E | 4.1e-08 | 6.4e-08 | 5.1e-08 | 5.200E-08 | 1.153E-08 | 4.100E-08 | 6.400E-08 |
| 26968 | 1349 | N1349G | E | 6e-08 | 2.9e-08 | 5.1e-08 | 4.667E-08 | 1.595E-08 | 2.900E-08 | 6.000E-08 |
| 26969 | 1349 | N1349H | E | 6.9e-08 | 2.3e-08 | 3.5e-08 | 4.233E-08 | 2.386E-08 | 2.300E-08 | 6.900E-08 |
| 26970 | 1349 | N1349I | E | 1.3e-08 | 1.7e-08 | 3e-08 | 2.000E-08 | 8.888E-09 | 1.300E-08 | 3.000E-08 |
| 26971 | 1349 | N1349L | E | 9.7e-08 | 3.9e-08 | 9.3e-08 | 7.633E-08 | 3.239E-08 | 3.900E-08 | 9.700E-08 |
| 26972 | 1349 | N1349K | E | 3.8e-08 | 4.9e-08 | 4.4e-08 | 4.367E-08 | 5.508E-09 | 3.800E-08 | 4.900E-08 |
| 26973 | 1349 | N1349M | E | 3.8e-08 | 1e-07 | 1.9e-07 | 1.093E-07 | 7.643E-08 | 3.800E-08 | 1.900E-07 |
| 26974 | 1349 | N1349F | E | 5.4e-08 | 3.4e-08 | 3.2e-08 | 4.000E-08 | 1.217E-08 | 3.200E-08 | 5.400E-08 |
| 26975 | 1349 | N1349P | E | 4.5e-08 | 3.3e-08 | 2.2e-07 | 9.933E-08 | 1.047E-07 | 3.300E-08 | 2.200E-07 |
| 26976 | 1349 | N1349S | E | 2.3e-08 | 5.7e-08 | 5.4e-08 | 4.467E-08 | 1.882E-08 | 2.300E-08 | 5.700E-08 |
| 26977 | 1349 | N1349T | E | 6.5e-08 | 4e-08 | 6.6e-08 | 5.700E-08 | 1.473E-08 | 4.000E-08 | 6.600E-08 |
| 26978 | 1349 | N1349W | E | 3.3e-08 | 2.6e-08 | 1.1e-07 | 5.633E-08 | 4.661E-08 | 2.600E-08 | 1.100E-07 |
| 26979 | 1349 | N1349Y | E | 2.1e-08 | 9.3e-08 | 4e-08 | 5.133E-08 | 3.731E-08 | 2.100E-08 | 9.300E-08 |
| 26980 | 1349 | N1349V | E | 1.6e-07 | 4.4e-08 | 3.9e-08 | 8.100E-08 | 6.846E-08 | 3.900E-08 | 1.600E-07 |
| 26981 | 1350 | T1350A | E | 3.3e-08 | 3.4e-08 | 6.3e-08 | 4.333E-08 | 1.704E-08 | 3.300E-08 | 6.300E-08 |
| 26982 | 1350 | T1350R | E | 8.8e-08 | 3.6e-08 | 3.7e-08 | 5.367E-08 | 2.974E-08 | 3.600E-08 | 8.800E-08 |
| 26983 | 1350 | T1350N | E | 1.2e-07 | 3.7e-08 | 3.2e-08 | 6.300E-08 | 4.943E-08 | 3.200E-08 | 1.200E-07 |
| 26984 | 1350 | T1350D | E | 3.9e-08 | 4.6e-08 | 2.7e-08 | 3.733E-08 | 9.609E-09 | 2.700E-08 | 4.600E-08 |
| 26985 | 1350 | T1350C | E | 3.3e-08 | 4.5e-08 | 4.4e-08 | 4.067E-08 | 6.658E-09 | 3.300E-08 | 4.500E-08 |
| 26986 | 1350 | T1350E | E | 7.9e-08 | 4.1e-08 | 8.6e-08 | 6.867E-08 | 2.421E-08 | 4.100E-08 | 8.600E-08 |
| 26987 | 1350 | T1350Q | E | 2.7e-08 | 1.5e-07 | 3.1e-08 | 6.933E-08 | 6.989E-08 | 2.700E-08 | 1.500E-07 |
| 26988 | 1350 | T1350G | E | 6.1e-08 | 5.8e-08 | 4.7e-08 | 5.533E-08 | 7.371E-09 | 4.700E-08 | 6.100E-08 |
| 26989 | 1350 | T1350H | E | 7.3e-08 | 5.6e-08 | 5.2e-08 | 6.033E-08 | 1.115E-08 | 5.200E-08 | 7.300E-08 |
| 26990 | 1350 | T1350I | E | 2.9e-08 | 5.6e-08 | 9.7e-08 | 6.067E-08 | 3.424E-08 | 2.900E-08 | 9.700E-08 |
| 26991 | 1350 | T1350L | E | 6.2e-08 | 7.2e-08 | 6.3e-08 | 6.567E-08 | 5.508E-09 | 6.200E-08 | 7.200E-08 |
| 26992 | 1350 | T1350K | E | 9.4e-08 | 6.8e-08 | 1.9e-08 | 6.033E-08 | 3.808E-08 | 1.900E-08 | 9.400E-08 |
| 26993 | 1350 | T1350M | E | 8.4e-08 | 5.2e-08 | 4.2e-08 | 5.933E-08 | 2.194E-08 | 4.200E-08 | 8.400E-08 |
| 26994 | 1350 | T1350F | E | 4.4e-08 | 3.9e-08 | 7.8e-08 | 5.367E-08 | 2.122E-08 | 3.900E-08 | 7.800E-08 |
| 26995 | 1350 | T1350P | E | 8.4e-08 | 6.1e-08 | 4.4e-08 | 6.300E-08 | 2.007E-08 | 4.400E-08 | 8.400E-08 |
| 26996 | 1350 | T1350S | E | 7e-08 | 3.8e-08 | 4.8e-08 | 5.200E-08 | 1.637E-08 | 3.800E-08 | 7.000E-08 |
| 26997 | 1350 | T1350T | E | 2.2e-08 | 3.4e-08 | 1.1e-07 | 5.533E-08 | 4.772E-08 | 2.200E-08 | 1.100E-07 |
| 26998 | 1350 | T1350W | E | 4.8e-08 | 1.2e-07 | 4.7e-08 | 7.167E-08 | 4.186E-08 | 4.700E-08 | 1.200E-07 |
| 26999 | 1350 | T1350Y | E | 8.4e-08 | 8.6e-08 | 1.7e-08 | 6.233E-08 | 3.927E-08 | 1.700E-08 | 8.600E-08 |
| 27000 | 1350 | T1350V | E | 1.6e-07 | 6e-08 | 2.8e-08 | 8.267E-08 | 6.886E-08 | 2.800E-08 | 1.600E-07 |
| 27001 | 1351 | A1351A | E | 2.5e-08 | 2.3e-08 | 4.4e-08 | 3.067E-08 | 1.159E-08 | 2.300E-08 | 4.400E-08 |
| 27002 | 1351 | A1351R | E | 5.6e-08 | 3.3e-08 | 6.2e-08 | 5.033E-08 | 1.531E-08 | 3.300E-08 | 6.200E-08 |
| 27003 | 1351 | A1351N | E | 3.2e-08 | 4.4e-08 | 3e-08 | 3.533E-08 | 7.572E-09 | 3.000E-08 | 4.400E-08 |
| 27004 | 1351 | A1351D | E | 4e-08 | 5.4e-08 | 1.7e-08 | 3.700E-08 | 1.868E-08 | 1.700E-08 | 5.400E-08 |
| 27005 | 1351 | A1351C | E | 4.5e-08 | 3.7e-08 | 5.7e-08 | 4.633E-08 | 1.007E-08 | 3.700E-08 | 5.700E-08 |
| 27006 | 1351 | A1351E | E | 6.2e-08 | 2.3e-08 | 5.5e-08 | 4.667E-08 | 2.079E-08 | 2.300E-08 | 6.200E-08 |
| 27007 | 1351 | A1351Q | E | 5.6e-08 | 4.1e-08 | 6.9e-08 | 5.533E-08 | 1.401E-08 | 4.100E-08 | 6.900E-08 |
| 27008 | 1351 | A1351G | E | 6.7e-08 | 2.8e-08 | 4.9e-08 | 4.800E-08 | 1.952E-08 | 2.800E-08 | 6.700E-08 |
| 27009 | 1351 | A1351H | E | 8.3e-08 | 4.7e-08 | 7.8e-08 | 6.933E-08 | 1.950E-08 | 4.700E-08 | 8.300E-08 |
| 27010 | 1351 | A1351I | E | 1.1e-08 | 1.4e-07 | 5.8e-08 | 6.967E-08 | 6.529E-08 | 1.100E-08 | 1.400E-07 |
| 27011 | 1351 | A1351L | E | 5.5e-08 | 7.6e-08 | 3.8e-08 | 5.633E-08 | 1.904E-08 | 3.800E-08 | 7.600E-08 |
| 27012 | 1351 | A1351K | E | 3.9e-08 | 4.8e-08 | 1.1e-07 | 6.567E-08 | 3.866E-08 | 3.900E-08 | 1.100E-07 |
| 27013 | 1351 | A1351M | E | 1.1e-07 | 3.8e-08 | 3.9e-08 | 6.233E-08 | 4.128E-08 | 3.800E-08 | 1.100E-07 |
| 27014 | 1351 | A1351F | E | 6.7e-08 | 4.5e-08 | 3.3e-08 | 4.833E-08 | 1.724E-08 | 3.300E-08 | 6.700E-08 |
| 27015 | 1351 | A1351P | E | 5.3e-08 | 3.3e-08 | 4.5e-08 | 4.367E-08 | 1.007E-08 | 3.300E-08 | 5.300E-08 |
| 27016 | 1351 | A1351S | E | 3.4e-08 | 3.3e-08 | 4.6e-08 | 3.767E-08 | 7.234E-09 | 3.300E-08 | 4.600E-08 |
| 27017 | 1351 | A1351T | E | 1.1e-07 | 1.1e-07 | 3.7e-08 | 8.567E-08 | 4.215E-08 | 3.700E-08 | 1.100E-07 |
| 27018 | 1351 | A1351W | E | 5.3e-08 | 6.9e-08 | 5e-08 | 5.733E-08 | 1.021E-08 | 5.000E-08 | 6.900E-08 |
| 27019 | 1351 | A1351Y | E | 4.8e-08 | 2.5e-08 | 2.2e-08 | 3.167E-08 | 1.422E-08 | 2.200E-08 | 4.800E-08 |
| 27020 | 1351 | A1351V | E | 2.3e-08 | 2.2e-08 | 6.7e-08 | 3.733E-08 | 2.570E-08 | 2.200E-08 | 6.700E-08 |
| 27021 | 1352 | Y1352A | E | 4.5e-08 | 6.7e-08 | 3.7e-08 | 4.967E-08 | 1.553E-08 | 3.700E-08 | 6.700E-08 |
| 27022 | 1352 | Y1352R | E | 2.4e-08 | 4.5e-08 | 1.3e-07 | 6.633E-08 | 5.613E-08 | 2.400E-08 | 1.300E-07 |
| 27023 | 1352 | Y1352N | E | 3.6e-08 | 6.3e-08 | 9.4e-08 | 6.433E-08 | 2.902E-08 | 3.600E-08 | 9.400E-08 |
| 27024 | 1352 | Y1352D | E | 9.6e-08 | 1.5e-08 | 4.6e-08 | 5.233E-08 | 4.087E-08 | 1.500E-08 | 9.600E-08 |
| 27025 | 1352 | Y1352C | E | 9.6e-08 | 6.6e-08 | 3e-08 | 6.400E-08 | 3.305E-08 | 3.000E-08 | 9.600E-08 |
| 27026 | 1352 | Y1352E | E | 2.2e-08 | 5.3e-08 | 2.3e-08 | 3.267E-08 | 1.762E-08 | 2.200E-08 | 5.300E-08 |
| 27027 | 1352 | Y1352Q | E | 3e-08 | 6.6e-08 | 6.3e-08 | 5.300E-08 | 1.997E-08 | 3.000E-08 | 6.600E-08 |
| 27028 | 1352 | Y1352G | E | 7.3e-08 | 5.5e-08 | 5.4e-08 | 6.067E-08 | 1.069E-08 | 5.400E-08 | 7.300E-08 |
| 27029 | 1352 | Y1352H | E | 4.1e-08 | 1.4e-08 | 3e-08 | 2.833E-08 | 1.358E-08 | 1.400E-08 | 4.100E-08 |

| | | | | | | | | | | |
|---|---|---|---|---|---|---|---|---|---|---|
| 27030 | 1352 | Y1352I | E | 3.9e-08 | 1.5e-07 | 4.6e-08 | 7.833E-08 | 6.216E-08 | 3.900E-08 | 1.500E-07 |
| 27031 | 1352 | Y1352L | E | 3.1e-08 | 7.6e-08 | 6.7e-08 | 5.800E-08 | 2.381E-08 | 3.100E-08 | 7.600E-08 |
| 27032 | 1352 | Y1352K | E | 9.5e-08 | 3.8e-08 | 8.3e-08 | 7.200E-08 | 3.005E-08 | 3.800E-08 | 9.500E-08 |
| 27033 | 1352 | Y1352M | E | 1.3e-07 | 5.9e-08 | 8.2e-08 | 9.033E-08 | 3.623E-08 | 5.900E-08 | 1.300E-07 |
| 27034 | 1352 | Y1352F | E | 6.3e-08 | 5.5e-08 | 3.2e-08 | 5.000E-08 | 1.609E-08 | 3.200E-08 | 6.300E-08 |
| 27035 | 1352 | Y1352P | E | 2e-08 | 3.3e-08 | 4.5e-08 | 3.267E-08 | 1.250E-08 | 2.000E-08 | 4.500E-08 |
| 27036 | 1352 | Y1352S | E | 3.2e-08 | 8.3e-08 | 1.6e-07 | 9.167E-08 | 6.444E-08 | 3.200E-08 | 1.600E-07 |
| 27037 | 1352 | Y1352T | E | 7.2e-08 | 1.9e-08 | 2.1e-08 | 3.733E-08 | 3.004E-08 | 1.900E-08 | 7.200E-08 |
| 27038 | 1352 | Y1352W | E | 4.4e-08 | 3.2e-08 | 4e-08 | 3.867E-08 | 6.110E-09 | 3.200E-08 | 4.400E-08 |
| 27039 | 1352 | Y1352Y | E | 6.7e-08 | 5.7e-08 | 3.5e-08 | 5.300E-08 | 1.637E-08 | 3.500E-08 | 6.700E-08 |
| 27040 | 1352 | Y1352V | E | 2.9e-08 | 3.6e-08 | 2.1e-08 | 2.867E-08 | 7.506E-09 | 2.100E-08 | 3.600E-08 |
| 27041 | 1353 | L1353A | E | 4.4e-08 | 3.6e-08 | 6.1e-08 | 4.700E-08 | 1.277E-08 | 3.600E-08 | 6.100E-08 |
| 27042 | 1353 | L1353R | E | 3.9e-08 | 2.5e-08 | 3.3e-08 | 3.233E-08 | 7.024E-09 | 2.500E-08 | 3.900E-08 |
| 27043 | 1353 | L1353N | E | 1.3e-07 | 5.7e-08 | 2.8e-08 | 7.167E-08 | 5.256E-08 | 2.800E-08 | 1.300E-07 |
| 27044 | 1353 | L1353D | E | 2.2e-08 | 3.6e-08 | 4.4e-08 | 3.400E-08 | 1.114E-08 | 2.200E-08 | 4.400E-08 |
| 27045 | 1353 | L1353C | E | 1.5e-08 | 5.8e-08 | 5.7e-08 | 4.333E-08 | 2.454E-08 | 1.500E-08 | 5.800E-08 |
| 27046 | 1353 | L1353E | E | 5.2e-08 | 7.5e-08 | 6.7e-08 | 6.467E-08 | 1.168E-08 | 5.200E-08 | 7.500E-08 |
| 27047 | 1353 | L1353Q | E | 2.7e-08 | 2.8e-08 | 3.5e-08 | 3.000E-08 | 4.359E-09 | 2.700E-08 | 3.500E-08 |
| 27048 | 1353 | L1353G | E | 4.1e-08 | 6.1e-08 | 3.5e-08 | 4.567E-08 | 1.361E-08 | 3.500E-08 | 6.100E-08 |
| 27049 | 1353 | L1353H | E | 4.8e-08 | 1.9e-08 | 1.4e-08 | 2.700E-08 | 1.836E-08 | 1.400E-08 | 4.800E-08 |
| 27050 | 1353 | L1353I | E | 1.6e-08 | 6.3e-08 | 1.5e-08 | 3.133E-08 | 2.743E-08 | 1.500E-08 | 6.300E-08 |
| 27051 | 1353 | L1353L | E | 4.8e-08 | 6.2e-08 | 2.4e-08 | 4.467E-08 | 1.922E-08 | 2.400E-08 | 6.200E-08 |
| 27052 | 1353 | L1353K | E | 7.7e-08 | 6.2e-08 | 2.7e-08 | 5.533E-08 | 2.566E-08 | 2.700E-08 | 7.700E-08 |
| 27053 | 1353 | L1353M | E | 2.6e-08 | 2.2e-08 | 2.6e-08 | 2.467E-08 | 2.309E-09 | 2.200E-08 | 2.600E-08 |
| 27054 | 1353 | L1353F | E | 5.3e-08 | 5.9e-08 | 2.4e-08 | 4.533E-08 | 1.872E-08 | 2.400E-08 | 5.900E-08 |
| 27055 | 1353 | L1353P | E | 4.5e-08 | 5.8e-08 | 5.5e-08 | 5.267E-08 | 6.807E-09 | 4.500E-08 | 5.800E-08 |
| 27056 | 1353 | L1353S | E | 4.3e-08 | 9.1e-09 | 5.2e-08 | 3.470E-08 | 2.262E-08 | 9.100E-09 | 5.200E-08 |
| 27057 | 1353 | L1353T | E | 1.1e-07 | 1e-08 | 1.3e-08 | 4.433E-08 | 5.689E-08 | 1.000E-08 | 1.100E-07 |
| 27058 | 1353 | L1353W | E | 3.8e-08 | 7.8e-08 | 6.3e-08 | 5.967E-08 | 2.021E-08 | 3.800E-08 | 7.800E-08 |
| 27059 | 1353 | L1353Y | E | 1.4e-08 | 4.7e-08 | 5.1e-08 | 3.733E-08 | 2.031E-08 | 1.400E-08 | 5.100E-08 |
| 27060 | 1353 | L1353V | E | 7.9e-08 | 6.3e-08 | 1.3e-07 | 9.067E-08 | 3.499E-08 | 6.300E-08 | 1.300E-07 |
| 27061 | 1354 | Q1354A | E | 8e-08 | 4.3e-08 | 5.8e-08 | 6.033E-08 | 1.861E-08 | 4.300E-08 | 8.000E-08 |
| 27062 | 1354 | Q1354R | E | 2.4e-08 | 2.8e-08 | 2.7e-08 | 2.633E-08 | 2.082E-09 | 2.400E-08 | 2.800E-08 |
| 27063 | 1354 | Q1354N | E | 4.3e-08 | 2.8e-08 | 3e-08 | 3.367E-08 | 8.145E-09 | 2.800E-08 | 4.300E-08 |
| 27064 | 1354 | Q1354D | E | 3.7e-08 | 2.1e-08 | 4.5e-08 | 3.433E-08 | 1.222E-08 | 2.100E-08 | 4.500E-08 |
| 27065 | 1354 | Q1354C | E | 6.6e-08 | 6.6e-08 | 1.1e-07 | 8.067E-08 | 2.540E-08 | 6.600E-08 | 1.100E-07 |
| 27066 | 1354 | Q1354E | E | 4.7e-08 | 4.9e-08 | 6.9e-08 | 5.500E-08 | 1.217E-08 | 4.700E-08 | 6.900E-08 |
| 27067 | 1354 | Q1354Q | E | 3.6e-08 | 6.4e-08 | 8.8e-08 | 6.267E-08 | 2.603E-08 | 3.600E-08 | 8.800E-08 |
| 27068 | 1354 | Q1354G | E | 9.8e-08 | 2.9e-08 | 5.3e-08 | 6.000E-08 | 3.503E-08 | 2.900E-08 | 9.800E-08 |
| 27069 | 1354 | Q1354H | E | 4.3e-08 | 7.4e-08 | 2.5e-08 | 4.733E-08 | 2.479E-08 | 2.500E-08 | 7.400E-08 |
| 27070 | 1354 | Q1354I | E | 8.2e-08 | 4e-08 | 5.1e-08 | 5.767E-08 | 2.178E-08 | 4.000E-08 | 8.200E-08 |
| 27071 | 1354 | Q1354L | E | 7e-08 | 4.5e-08 | 1e-07 | 7.167E-08 | 2.754E-08 | 4.500E-08 | 1.000E-07 |
| 27072 | 1354 | Q1354K | E | 5.6e-08 | 2.9e-08 | 3.4e-08 | 3.967E-08 | 1.436E-08 | 2.900E-08 | 5.600E-08 |
| 27073 | 1354 | Q1354M | E | 2.8e-08 | 8.7e-08 | 5.3e-08 | 5.600E-08 | 2.961E-08 | 2.800E-08 | 8.700E-08 |
| 27074 | 1354 | Q1354F | E | 2.8e-08 | 5e-08 | 4.5e-08 | 4.100E-08 | 1.153E-08 | 2.800E-08 | 5.000E-08 |
| 27075 | 1354 | Q1354P | E | 1.9e-08 | 5.8e-08 | 6.7e-08 | 4.800E-08 | 2.551E-08 | 1.900E-08 | 6.700E-08 |
| 27076 | 1354 | Q1354S | E | 4.7e-08 | 5.7e-08 | 1.8e-08 | 4.067E-08 | 2.026E-08 | 1.800E-08 | 5.700E-08 |
| 27077 | 1354 | Q1354T | E | 5.2e-08 | 8.3e-08 | 2.1e-08 | 5.200E-08 | 3.100E-08 | 2.100E-08 | 8.300E-08 |
| 27078 | 1354 | Q1354W | E | 3.5e-08 | 6.4e-08 | 2.9e-08 | 4.267E-08 | 1.872E-08 | 2.900E-08 | 6.400E-08 |
| 27079 | 1354 | Q1354Y | E | 1e-08 | 3e-08 | 1.6e-08 | 1.867E-08 | 1.026E-08 | 1.000E-08 | 3.000E-08 |
| 27080 | 1354 | Q1354V | E | 1.3e-08 | 6e-08 | 3.9e-08 | 3.733E-08 | 2.354E-08 | 1.300E-08 | 6.000E-08 |
| 27081 | 1355 | M1355A | E | 4.4e-08 | 4.4e-08 | 4.5e-08 | 4.433E-08 | 5.774E-10 | 4.400E-08 | 4.500E-08 |
| 27082 | 1355 | M1355R | E | 3.8e-08 | 2e-08 | 3.9e-08 | 3.233E-08 | 1.069E-08 | 2.000E-08 | 3.900E-08 |
| 27083 | 1355 | M1355N | E | 4.5e-08 | 4.2e-08 | 8.1e-08 | 5.600E-08 | 2.170E-08 | 4.200E-08 | 8.100E-08 |
| 27084 | 1355 | M1355D | E | 2.9e-08 | 2.4e-08 | 4.4e-08 | 3.233E-08 | 1.041E-08 | 2.400E-08 | 4.400E-08 |
| 27085 | 1355 | M1355C | E | 2.3e-08 | 3.1e-08 | 4.2e-08 | 3.200E-08 | 9.539E-09 | 2.300E-08 | 4.200E-08 |
| 27086 | 1355 | M1355E | E | 5.6e-08 | 4.1e-08 | 7.1e-08 | 5.600E-08 | 1.500E-08 | 4.100E-08 | 7.100E-08 |
| 27087 | 1355 | M1355Q | E | 3.1e-08 | 6.8e-08 | 9.6e-08 | 6.500E-08 | 3.260E-08 | 3.100E-08 | 9.600E-08 |
| 27088 | 1355 | M1355G | E | 6.9e-08 | 2.5e-08 | 8.1e-08 | 5.833E-08 | 2.948E-08 | 2.500E-08 | 8.100E-08 |
| 27089 | 1355 | M1355H | E | 3.3e-08 | 4.5e-08 | 5.3e-08 | 4.367E-08 | 1.007E-08 | 3.300E-08 | 5.300E-08 |
| 27090 | 1355 | M1355I | E | 1.1e-07 | 8e-08 | 4.3e-08 | 7.767E-08 | 3.356E-08 | 4.300E-08 | 1.100E-07 |
| 27091 | 1355 | M1355L | E | 2.3e-08 | 1.3e-08 | 1.8e-08 | 1.800E-08 | 5.000E-09 | 1.300E-08 | 2.300E-08 |
| 27092 | 1355 | M1355K | E | 8.1e-08 | 3.2e-08 | 1.1e-07 | 7.433E-08 | 3.943E-08 | 3.200E-08 | 1.100E-07 |
| 27093 | 1355 | M1355M | E | 5.5e-08 | 8.3e-08 | 4e-08 | 5.933E-08 | 2.183E-08 | 4.000E-08 | 8.300E-08 |
| 27094 | 1355 | M1355F | E | 2.8e-08 | 4.8e-08 | 1.3e-07 | 6.867E-08 | 5.405E-08 | 2.800E-08 | 1.300E-07 |
| 27095 | 1355 | M1355P | E | 6.3e-08 | 6.8e-08 | 2.3e-08 | 5.133E-08 | 2.466E-08 | 2.300E-08 | 6.800E-08 |
| 27096 | 1355 | M1355S | E | 1.5e-08 | 5e-08 | 6.1e-08 | 4.200E-08 | 2.402E-08 | 1.500E-08 | 6.100E-08 |
| 27097 | 1355 | M1355T | E | 5.2e-08 | 4.4e-08 | 3.9e-08 | 4.500E-08 | 6.557E-09 | 3.900E-08 | 5.200E-08 |
| 27098 | 1355 | M1355W | E | 4.3e-08 | 5.2e-08 | 2.1e-08 | 3.867E-08 | 1.595E-08 | 2.100E-08 | 5.200E-08 |
| 27099 | 1355 | M1355Y | E | 2.8e-08 | 1.8e-08 | 4e-08 | 2.867E-08 | 1.102E-08 | 1.800E-08 | 4.000E-08 |
| 27100 | 1355 | M1355V | E | 6e-08 | 3.1e-08 | 3.4e-08 | 4.167E-08 | 1.595E-08 | 3.100E-08 | 6.000E-08 |
| 27101 | 1356 | N1356A | E | 3.5e-08 | 4.2e-08 | 4.7e-08 | 4.133E-08 | 6.028E-09 | 3.500E-08 | 4.700E-08 |
| 27102 | 1356 | N1356R | E | 2.5e-08 | 2.1e-08 | 1.1e-07 | 5.200E-08 | 5.027E-08 | 2.100E-08 | 1.100E-07 |
| 27103 | 1356 | N1356N | E | 6.8e-08 | 1.5e-08 | 4.1e-08 | 4.133E-08 | 2.650E-08 | 1.500E-08 | 6.800E-08 |
| 27104 | 1356 | N1356D | E | 1.9e-08 | 7.9e-08 | 8.2e-08 | 6.000E-08 | 3.554E-08 | 1.900E-08 | 8.200E-08 |
| 27105 | 1356 | N1356C | E | 5.5e-08 | 3.9e-08 | 4.5e-08 | 4.633E-08 | 8.083E-09 | 3.900E-08 | 5.500E-08 |
| 27106 | 1356 | N1356E | E | 1.4e-07 | 6.7e-08 | 2.9e-08 | 7.867E-08 | 5.641E-08 | 2.900E-08 | 1.400E-07 |
| 27107 | 1356 | N1356Q | E | 1.2e-07 | 2.7e-08 | 8.2e-08 | 7.633E-08 | 4.676E-08 | 2.700E-08 | 1.200E-07 |
| 27108 | 1356 | N1356G | E | 3.9e-08 | 5.3e-08 | 4.3e-08 | 4.500E-08 | 7.211E-09 | 3.900E-08 | 5.300E-08 |
| 27109 | 1356 | N1356H | E | 3.5e-08 | 2.8e-08 | 5.7e-08 | 4.000E-08 | 1.513E-08 | 2.800E-08 | 5.700E-08 |
| 27110 | 1356 | N1356I | E | 3.6e-08 | 6.3e-08 | 4e-08 | 4.633E-08 | 1.457E-08 | 3.600E-08 | 6.300E-08 |
| 27111 | 1356 | N1356L | E | 3.4e-08 | 3.1e-08 | 6.5e-08 | 4.333E-08 | 1.882E-08 | 3.100E-08 | 6.500E-08 |
| 27112 | 1356 | N1356K | E | 6.5e-08 | 6.4e-08 | 1.9e-07 | 1.063E-07 | 7.246E-08 | 6.400E-08 | 1.900E-07 |
| 27113 | 1356 | N1356M | E | 3.5e-08 | 4.7e-08 | 4.6e-08 | 4.267E-08 | 6.658E-09 | 3.500E-08 | 4.700E-08 |
| 27114 | 1356 | N1356F | E | 2.1e-08 | 2e-08 | 5.1e-08 | 3.067E-08 | 1.762E-08 | 2.000E-08 | 5.100E-08 |

| | | | | | | | | | | |
|---|---|---|---|---|---|---|---|---|---|---|
| 27115 | 1356 | N1356P | E | 4.4e-08 | 3.3e-08 | 7.9e-08 | 5.200E-08 | 2.402E-08 | 3.300E-08 | 7.900E-08 |
| 27116 | 1356 | N1356S | E | 6.7e-08 | 2.6e-08 | 3.6e-08 | 4.300E-08 | 2.138E-08 | 2.600E-08 | 6.700E-08 |
| 27117 | 1356 | N1356T | E | 9.2e-09 | 5.7e-08 | 8.2e-08 | 4.940E-08 | 3.699E-08 | 9.200E-09 | 8.200E-08 |
| 27118 | 1356 | N1356W | E | 2e-08 | 5.9e-08 | 1.5e-08 | 3.133E-08 | 2.409E-08 | 1.500E-08 | 5.900E-08 |
| 27119 | 1356 | N1356Y | E | 2.2e-08 | 2e-08 | 5.5e-08 | 3.233E-08 | 1.966E-08 | 2.000E-08 | 5.500E-08 |
| 27120 | 1356 | N1356V | E | 3.9e-08 | 5.1e-08 | 1.9e-08 | 3.633E-08 | 1.617E-08 | 1.900E-08 | 5.100E-08 |
| 27121 | 1357 | S1357A | E | 4.5e-08 | 2e-08 | 3.9e-08 | 3.467E-08 | 1.305E-08 | 2.000E-08 | 4.500E-08 |
| 27122 | 1357 | S1357R | E | 2.8e-08 | 1.3e-07 | 5.5e-08 | 7.100E-08 | 5.285E-08 | 2.800E-08 | 1.300E-07 |
| 27123 | 1357 | S1357N | E | 2.7e-08 | 5.9e-08 | 3.8e-08 | 4.133E-08 | 1.626E-08 | 2.700E-08 | 5.900E-08 |
| 27124 | 1357 | S1357D | E | 1e-07 | 2.5e-07 | 3.4e-08 | 1.280E-07 | 1.107E-07 | 3.400E-08 | 2.500E-07 |
| 27125 | 1357 | S1357C | E | 4.8e-08 | 4.8e-08 | 8.1e-08 | 5.900E-08 | 1.905E-08 | 4.800E-08 | 8.100E-08 |
| 27126 | 1357 | S1357E | E | 2.9e-08 | 8e-08 | 2.1e-08 | 4.333E-08 | 3.201E-08 | 2.100E-08 | 8.000E-08 |
| 27127 | 1357 | S1357Q | E | 2.4e-08 | 4.2e-08 | 6.9e-08 | 4.500E-08 | 2.265E-08 | 2.400E-08 | 6.900E-08 |
| 27128 | 1357 | S1357G | E | 9.4e-08 | 4e-08 | 5.9e-08 | 6.433E-08 | 2.739E-08 | 4.000E-08 | 9.400E-08 |
| 27129 | 1357 | S1357H | E | 8.2e-08 | 2.2e-08 | 4.1e-08 | 4.833E-08 | 3.066E-08 | 2.200E-08 | 8.200E-08 |
| 27130 | 1357 | S1357I | E | 1.4e-08 | 2.6e-08 | 2.8e-08 | 2.267E-08 | 7.572E-09 | 1.400E-08 | 2.800E-08 |
| 27131 | 1357 | S1357L | E | 6.7e-08 | 7.4e-08 | 4e-08 | 6.033E-08 | 1.795E-08 | 4.000E-08 | 7.400E-08 |
| 27132 | 1357 | S1357K | E | 2.6e-08 | 7.1e-08 | 4.4e-08 | 4.700E-08 | 2.265E-08 | 2.600E-08 | 7.100E-08 |
| 27133 | 1357 | S1357M | E | 3.3e-08 | 6.8e-08 | 2.5e-08 | 4.200E-08 | 2.287E-08 | 2.500E-08 | 6.800E-08 |
| 27134 | 1357 | S1357F | E | 1.1e-07 | 8.5e-08 | 4.5e-08 | 8.000E-08 | 3.279E-08 | 4.500E-08 | 1.100E-07 |
| 27135 | 1357 | S1357P | E | 1.5e-08 | 5.9e-08 | 1.2e-07 | 6.467E-08 | 5.273E-08 | 1.500E-08 | 1.200E-07 |
| 27136 | 1357 | S1357S | E | 3.2e-08 | 8.2e-08 | 7.3e-08 | 6.233E-08 | 2.665E-08 | 3.200E-08 | 8.200E-08 |
| 27137 | 1357 | S1357T | E | 5e-08 | 1.2e-08 | 2.6e-08 | 2.933E-08 | 1.922E-08 | 1.200E-08 | 5.000E-08 |
| 27138 | 1357 | S1357W | E | 2.2e-08 | 4.9e-08 | 4.1e-08 | 3.733E-08 | 1.387E-08 | 2.200E-08 | 4.900E-08 |
| 27139 | 1357 | S1357Y | E | 4e-08 | 4.3e-08 | 5.7e-08 | 4.667E-08 | 9.074E-09 | 4.000E-08 | 5.700E-08 |
| 27140 | 1357 | S1357V | E | 4.5e-08 | 5.1e-08 | 6.4e-08 | 5.333E-08 | 9.713E-09 | 4.500E-08 | 6.400E-08 |
| 27141 | 1358 | L1358A | E | 4.4e-08 | 5.3e-08 | 3.2e-08 | 4.300E-08 | 1.054E-08 | 3.200E-08 | 5.300E-08 |
| 27142 | 1358 | L1358R | E | 7.5e-08 | 9.3e-08 | 5.5e-08 | 7.433E-08 | 1.901E-08 | 5.500E-08 | 9.300E-08 |
| 27143 | 1358 | L1358N | E | 6.1e-08 | 3.4e-08 | 3.2e-08 | 4.233E-08 | 1.620E-08 | 3.200E-08 | 6.100E-08 |
| 27144 | 1358 | L1358D | E | 6.4e-08 | 4e-08 | 5.3e-08 | 5.233E-08 | 1.201E-08 | 4.000E-08 | 6.400E-08 |
| 27145 | 1358 | L1358C | E | 2.3e-08 | 6.3e-08 | 5.3e-08 | 4.633E-08 | 2.082E-08 | 2.300E-08 | 6.300E-08 |
| 27146 | 1358 | L1358E | E | 3.7e-08 | 6.3e-08 | 5.1e-08 | 5.033E-08 | 1.301E-08 | 3.700E-08 | 6.300E-08 |
| 27147 | 1358 | L1358Q | E | 9.8e-09 | 1.1e-07 | 2.8e-08 | 4.927E-08 | 5.338E-08 | 9.800E-09 | 1.100E-07 |
| 27148 | 1358 | L1358G | E | 3e-08 | 3.4e-08 | 6.1e-08 | 4.167E-08 | 1.686E-08 | 3.000E-08 | 6.100E-08 |
| 27149 | 1358 | L1358H | E | 2.5e-08 | 2.3e-08 | 8e-08 | 4.267E-08 | 3.235E-08 | 2.300E-08 | 8.000E-08 |
| 27150 | 1358 | L1358I | E | 7.4e-08 | 3.2e-08 | 8.9e-08 | 6.500E-08 | 2.955E-08 | 3.200E-08 | 8.900E-08 |
| 27151 | 1358 | L1358L | E | 4.3e-08 | 8e-08 | 9e-08 | 7.100E-08 | 2.476E-08 | 4.300E-08 | 9.000E-08 |
| 27152 | 1358 | L1358K | E | 6.1e-08 | 2.1e-08 | 9e-08 | 5.733E-08 | 3.465E-08 | 2.100E-08 | 9.000E-08 |
| 27153 | 1358 | L1358M | E | 2.2e-08 | 2.3e-07 | 5.2e-08 | 1.013E-07 | 1.124E-07 | 2.200E-08 | 2.300E-07 |
| 27154 | 1358 | L1358F | E | 4.7e-08 | 3.1e-08 | 5.7e-08 | 4.500E-08 | 1.311E-08 | 3.100E-08 | 5.700E-08 |
| 27155 | 1358 | L1358P | E | 1.7e-08 | 3e-08 | 6.3e-08 | 3.667E-08 | 2.371E-08 | 1.700E-08 | 6.300E-08 |
| 27156 | 1358 | L1358S | E | 4.3e-08 | 1.3e-07 | 2.5e-08 | 6.600E-08 | 5.615E-08 | 2.500E-08 | 1.300E-07 |
| 27157 | 1358 | L1358T | E | 5.9e-08 | 4.6e-08 | 1.2e-07 | 7.500E-08 | 3.951E-08 | 4.600E-08 | 1.200E-07 |
| 27158 | 1358 | L1358W | E | 6.7e-08 | 2.7e-08 | 4.3e-08 | 4.567E-08 | 2.013E-08 | 2.700E-08 | 6.700E-08 |
| 27159 | 1358 | L1358Y | E | 5e-08 | 5.2e-08 | 2.7e-08 | 4.300E-08 | 1.389E-08 | 2.700E-08 | 5.200E-08 |
| 27160 | 1358 | L1358V | E | 4.2e-08 | 2.8e-08 | 3.7e-08 | 3.567E-08 | 7.095E-09 | 2.800E-08 | 4.200E-08 |
| 27161 | 1359 | R1359A | E | 6.9e-08 | 4.2e-08 | 6e-08 | 5.700E-08 | 1.375E-08 | 4.200E-08 | 6.900E-08 |
| 27162 | 1359 | R1359R | E | 5e-08 | 3.4e-08 | 6.2e-08 | 4.867E-08 | 1.405E-08 | 3.400E-08 | 6.200E-08 |
| 27163 | 1359 | R1359N | E | 1.1e-07 | 3.8e-08 | 3.5e-08 | 6.100E-08 | 4.246E-08 | 3.500E-08 | 1.100E-07 |
| 27164 | 1359 | R1359D | E | 5.7e-08 | 8.1e-08 | 2.5e-08 | 5.433E-08 | 2.810E-08 | 2.500E-08 | 8.100E-08 |
| 27165 | 1359 | R1359C | E | 6.6e-08 | 1.6e-08 | 3e-08 | 3.733E-08 | 2.579E-08 | 1.600E-08 | 6.600E-08 |
| 27166 | 1359 | R1359E | E | 3.3e-08 | 1.8e-08 | 1.1e-08 | 2.067E-08 | 1.124E-08 | 1.100E-08 | 3.300E-08 |
| 27167 | 1359 | R1359Q | E | 2.2e-08 | 1.1e-07 | 4.3e-08 | 5.833E-08 | 4.596E-08 | 2.200E-08 | 1.100E-07 |
| 27168 | 1359 | R1359G | E | 4.7e-08 | 1.6e-08 | 2.1e-08 | 2.800E-08 | 1.664E-08 | 1.600E-08 | 4.700E-08 |
| 27169 | 1359 | R1359H | E | 4.4e-08 | 6.2e-08 | 2.9e-08 | 4.500E-08 | 1.652E-08 | 2.900E-08 | 6.200E-08 |
| 27170 | 1359 | R1359I | E | 2.2e-08 | 8.8e-08 | 4.9e-08 | 5.300E-08 | 3.318E-08 | 2.200E-08 | 8.800E-08 |
| 27171 | 1359 | R1359L | E | 5.4e-08 | 6e-08 | 3.2e-08 | 4.867E-08 | 1.474E-08 | 3.200E-08 | 6.000E-08 |
| 27172 | 1359 | R1359K | E | 8.1e-08 | 2.4e-08 | 6e-08 | 5.500E-08 | 2.883E-08 | 2.400E-08 | 8.100E-08 |
| 27173 | 1359 | R1359M | E | 2.1e-08 | 3.5e-08 | 8.4e-08 | 4.667E-08 | 3.308E-08 | 2.100E-08 | 8.400E-08 |
| 27174 | 1359 | R1359F | E | 2.6e-08 | 2.6e-08 | 1.3e-08 | 2.167E-08 | 7.506E-09 | 1.300E-08 | 2.600E-08 |
| 27175 | 1359 | R1359P | E | 8.3e-08 | 2.5e-08 | 7.2e-08 | 6.000E-08 | 3.081E-08 | 2.500E-08 | 8.300E-08 |
| 27176 | 1359 | R1359S | E | 1e-07 | 2.8e-08 | 6.4e-08 | 6.400E-08 | 3.600E-08 | 2.800E-08 | 1.000E-07 |
| 27177 | 1359 | R1359T | E | 3.1e-08 | 6.5e-08 | 6.9e-08 | 5.500E-08 | 2.088E-08 | 3.100E-08 | 6.900E-08 |
| 27178 | 1359 | R1359W | E | 4.4e-08 | 4.5e-08 | 3.6e-08 | 4.167E-08 | 4.933E-09 | 3.600E-08 | 4.500E-08 |
| 27179 | 1359 | R1359Y | E | 2.1e-07 | 2.8e-08 | 1.2e-07 | 1.193E-07 | 9.100E-08 | 2.800E-08 | 2.100E-07 |
| 27180 | 1359 | R1359V | E | 3.7e-08 | 7.9e-08 | 1.1e-07 | 7.533E-08 | 3.664E-08 | 3.700E-08 | 1.100E-07 |
| 27181 | 1360 | A1360A | E | 5.5e-08 | 7.8e-08 | 2.8e-08 | 5.367E-08 | 2.503E-08 | 2.800E-08 | 7.800E-08 |
| 27182 | 1360 | A1360R | E | 1.4e-08 | 4.3e-08 | 6.2e-08 | 3.967E-08 | 2.417E-08 | 1.400E-08 | 6.200E-08 |
| 27183 | 1360 | A1360N | E | 1.7e-08 | 5.3e-08 | 4.5e-08 | 3.833E-08 | 1.890E-08 | 1.700E-08 | 5.300E-08 |
| 27184 | 1360 | A1360D | E | 5.8e-08 | 2.4e-08 | 5.7e-08 | 4.633E-08 | 1.935E-08 | 2.400E-08 | 5.800E-08 |
| 27185 | 1360 | A1360C | E | 1.6e-08 | 6.2e-08 | 6.1e-08 | 4.633E-08 | 2.627E-08 | 1.600E-08 | 6.200E-08 |
| 27186 | 1360 | A1360E | E | 6.5e-08 | 1.7e-08 | 4.9e-08 | 4.367E-08 | 2.444E-08 | 1.700E-08 | 6.500E-08 |
| 27187 | 1360 | A1360Q | E | 4.3e-08 | 7.2e-08 | 3.2e-08 | 4.900E-08 | 2.066E-08 | 3.200E-08 | 7.200E-08 |
| 27188 | 1360 | A1360G | E | 5.4e-08 | 3.8e-08 | 7.3e-08 | 5.500E-08 | 1.752E-08 | 3.800E-08 | 7.300E-08 |
| 27189 | 1360 | A1360H | E | 2.4e-08 | 1.7e-08 | 7.7e-08 | 3.933E-08 | 3.281E-08 | 1.700E-08 | 7.700E-08 |
| 27190 | 1360 | A1360I | E | 2.9e-08 | 1.5e-07 | 3.4e-08 | 7.100E-08 | 6.846E-08 | 2.900E-08 | 1.500E-07 |
| 27191 | 1360 | A1360L | E | 3.3e-08 | 5.8e-08 | 2.9e-08 | 4.000E-08 | 1.572E-08 | 2.900E-08 | 5.800E-08 |
| 27192 | 1360 | A1360K | E | 8.8e-08 | 4.4e-08 | 2.1e-08 | 5.100E-08 | 3.404E-08 | 2.100E-08 | 8.800E-08 |
| 27193 | 1360 | A1360M | E | 5.4e-08 | 7.2e-08 | 1e-07 | 7.533E-08 | 2.318E-08 | 5.400E-08 | 1.000E-07 |
| 27194 | 1360 | A1360F | E | 5e-08 | 6.6e-08 | 3.7e-08 | 5.100E-08 | 1.453E-08 | 3.700E-08 | 6.600E-08 |
| 27195 | 1360 | A1360P | E | 8.1e-08 | 6.3e-08 | 3.4e-08 | 5.933E-08 | 2.371E-08 | 3.400E-08 | 8.100E-08 |
| 27196 | 1360 | A1360S | E | 9.6e-08 | 5.6e-08 | 3.4e-08 | 6.200E-08 | 3.143E-08 | 3.400E-08 | 9.600E-08 |
| 27197 | 1360 | A1360T | E | 3.5e-08 | 1.4e-07 | 1e-07 | 9.167E-08 | 5.299E-08 | 3.500E-08 | 1.400E-07 |
| 27198 | 1360 | A1360W | E | 6.8e-08 | 5.3e-08 | 4.7e-08 | 5.600E-08 | 1.082E-08 | 4.700E-08 | 6.800E-08 |
| 27199 | 1360 | A1360Y | E | 2.6e-08 | 5.3e-08 | 2.8e-08 | 3.567E-08 | 1.504E-08 | 2.600E-08 | 5.300E-08 |

| | | | | | | | | | | |
|---|---|---|---|---|---|---|---|---|---|---|
| 27200 | 1360 | A1360V | E | 3.3e-08 | 7.6e-08 | 1.2e-07 | 7.633E-08 | 4.350E-08 | 3.300E-08 | 1.200E-07 |
| 27201 | 1361 | E1361A | E | 1.9e-08 | 3.4e-08 | 4.1e-08 | 3.133E-08 | 1.124E-08 | 1.900E-08 | 4.100E-08 |
| 27202 | 1361 | E1361R | E | 5.5e-08 | 2.2e-08 | 7.6e-08 | 5.100E-08 | 2.722E-08 | 2.200E-08 | 7.600E-08 |
| 27203 | 1361 | E1361N | E | 3.8e-08 | 5.6e-08 | 4.5e-08 | 4.633E-08 | 9.074E-09 | 3.800E-08 | 5.600E-08 |
| 27204 | 1361 | E1361D | E | 1.3e-07 | 5.9e-08 | 4.9e-08 | 7.933E-08 | 4.416E-08 | 4.900E-08 | 1.300E-07 |
| 27205 | 1361 | E1361C | E | 5.4e-08 | 2.2e-08 | 7e-09 | 2.767E-08 | 2.401E-08 | 7.000E-09 | 5.400E-08 |
| 27206 | 1361 | E1361E | E | 7.3e-08 | 4.2e-08 | 5.8e-08 | 5.767E-08 | 1.550E-08 | 4.200E-08 | 7.300E-08 |
| 27207 | 1361 | E1361Q | E | 1.1e-07 | 3.4e-08 | 5e-08 | 6.467E-08 | 4.007E-08 | 3.400E-08 | 1.100E-07 |
| 27208 | 1361 | E1361G | E | 1.1e-08 | 4.8e-08 | 2.6e-08 | 2.833E-08 | 1.861E-08 | 1.100E-08 | 4.800E-08 |
| 27209 | 1361 | E1361H | E | 5.2e-08 | 3e-08 | 3.6e-08 | 3.933E-08 | 1.137E-08 | 3.000E-08 | 5.200E-08 |
| 27210 | 1361 | E1361I | E | 1e-07 | 1.2e-07 | 5.8e-08 | 9.267E-08 | 3.164E-08 | 5.800E-08 | 1.200E-07 |
| 27211 | 1361 | E1361L | E | 6.8e-08 | 7.3e-08 | 4.6e-08 | 6.233E-08 | 1.436E-08 | 4.600E-08 | 7.300E-08 |
| 27212 | 1361 | E1361K | E | 2.7e-08 | 7e-08 | 1e-07 | 6.567E-08 | 3.669E-08 | 2.700E-08 | 1.000E-07 |
| 27213 | 1361 | E1361M | E | 5.9e-08 | 4.7e-08 | 5.5e-08 | 5.367E-08 | 6.110E-09 | 4.700E-08 | 5.900E-08 |
| 27214 | 1361 | E1361F | E | 4.2e-08 | 4e-08 | 4.6e-08 | 4.267E-08 | 3.055E-09 | 4.000E-08 | 4.600E-08 |
| 27215 | 1361 | E1361P | E | 3.1e-08 | 1.7e-08 | 1.7e-08 | 2.167E-08 | 8.083E-09 | 1.700E-08 | 3.100E-08 |
| 27216 | 1361 | E1361S | E | 4.2e-08 | 1.2e-08 | 4.8e-08 | 3.400E-08 | 1.929E-08 | 1.200E-08 | 4.800E-08 |
| 27217 | 1361 | E1361T | E | 5e-08 | 8.4e-08 | 2.7e-08 | 5.367E-08 | 2.868E-08 | 2.700E-08 | 8.400E-08 |
| 27218 | 1361 | E1361W | E | 4e-08 | 2.6e-08 | 7.3e-08 | 4.633E-08 | 2.413E-08 | 2.600E-08 | 7.300E-08 |
| 27219 | 1361 | E1361Y | E | 1.1e-07 | 7.3e-08 | 3.7e-08 | 7.333E-08 | 3.650E-08 | 3.700E-08 | 1.100E-07 |
| 27220 | 1361 | E1361V | E | 3.4e-08 | 2.9e-08 | 3e-08 | 3.100E-08 | 2.646E-09 | 2.900E-08 | 3.400E-08 |
| 27221 | 1362 | D1362A | E | 4.3e-08 | 8.3e-08 | 6.9e-08 | 6.500E-08 | 2.030E-08 | 4.300E-08 | 8.300E-08 |
| 27222 | 1362 | D1362R | E | 6e-08 | 8.2e-08 | 4.1e-08 | 6.100E-08 | 2.052E-08 | 4.100E-08 | 8.200E-08 |
| 27223 | 1362 | D1362N | E | 8.2e-08 | 1.7e-08 | 1.1e-07 | 6.967E-08 | 4.771E-08 | 1.700E-08 | 1.100E-07 |
| 27224 | 1362 | D1362D | E | 8.4e-08 | 2.9e-08 | 1.1e-07 | 7.433E-08 | 4.136E-08 | 2.900E-08 | 1.100E-07 |
| 27225 | 1362 | D1362C | E | 2.1e-08 | 5.7e-08 | 5.7e-08 | 4.500E-08 | 2.078E-08 | 2.100E-08 | 5.700E-08 |
| 27226 | 1362 | D1362E | E | 4.3e-08 | 5.6e-08 | 3.6e-08 | 4.500E-08 | 1.015E-08 | 3.600E-08 | 5.600E-08 |
| 27227 | 1362 | D1362Q | E | 7.6e-08 | 5.4e-08 | 2.7e-08 | 5.233E-08 | 2.454E-08 | 2.700E-08 | 7.600E-08 |
| 27228 | 1362 | D1362G | E | 8.3e-08 | 9e-08 | 6e-08 | 7.767E-08 | 1.570E-08 | 6.000E-08 | 9.000E-08 |
| 27229 | 1362 | D1362H | E | 3.1e-08 | 3.3e-08 | 2.3e-08 | 2.900E-08 | 5.292E-09 | 2.300E-08 | 3.300E-08 |
| 27230 | 1362 | D1362I | E | 5.1e-08 | 2.8e-08 | 7.8e-08 | 5.233E-08 | 2.503E-08 | 2.800E-08 | 7.800E-08 |
| 27231 | 1362 | D1362L | E | 1.5e-07 | 8.2e-08 | 1.4e-07 | 1.240E-07 | 3.672E-08 | 8.200E-08 | 1.500E-07 |
| 27232 | 1362 | D1362K | E | 5.3e-08 | 1.9e-08 | 3.9e-08 | 3.700E-08 | 1.709E-08 | 1.900E-08 | 5.300E-08 |
| 27233 | 1362 | D1362M | E | 5.4e-08 | 1.9e-08 | 1.8e-08 | 3.033E-08 | 2.050E-08 | 1.800E-08 | 5.400E-08 |
| 27234 | 1362 | D1362F | E | 2.4e-08 | 5.6e-08 | 4.6e-08 | 4.200E-08 | 1.637E-08 | 2.400E-08 | 5.600E-08 |
| 27235 | 1362 | D1362P | E | 3.5e-08 | 1.1e-07 | 2.7e-08 | 5.733E-08 | 4.579E-08 | 2.700E-08 | 1.100E-07 |
| 27236 | 1362 | D1362S | E | 1.4e-08 | 4.4e-08 | 7.2e-08 | 4.333E-08 | 2.901E-08 | 1.400E-08 | 7.200E-08 |
| 27237 | 1362 | D1362T | E | 5.7e-08 | 2.6e-08 | 3.5e-08 | 3.933E-08 | 1.595E-08 | 2.600E-08 | 5.700E-08 |
| 27238 | 1362 | D1362W | E | 5.7e-08 | 6.6e-08 | 5.6e-08 | 5.967E-08 | 5.508E-09 | 5.600E-08 | 6.600E-08 |
| 27239 | 1362 | D1362Y | E | 8.1e-08 | 3.2e-08 | 1.9e-07 | 1.010E-07 | 8.088E-08 | 3.200E-08 | 1.900E-07 |
| 27240 | 1362 | D1362V | E | 1.4e-08 | 2.9e-08 | 4.1e-08 | 2.800E-08 | 1.353E-08 | 1.400E-08 | 4.100E-08 |
| 27241 | 1363 | T1363A | E | 1.2e-07 | 3.4e-08 | 6.1e-08 | 7.167E-08 | 4.398E-08 | 3.400E-08 | 1.200E-07 |
| 27242 | 1363 | T1363R | E | 2.5e-08 | 1.9e-08 | 6.5e-08 | 3.633E-08 | 2.501E-08 | 1.900E-08 | 6.500E-08 |
| 27243 | 1363 | T1363N | E | 6e-08 | 1e-08 | 3.3e-08 | 3.433E-08 | 2.503E-08 | 1.000E-08 | 6.000E-08 |
| 27244 | 1363 | T1363D | E | 3.7e-08 | 3.6e-08 | 4.1e-08 | 3.800E-08 | 2.646E-09 | 3.600E-08 | 4.100E-08 |
| 27245 | 1363 | T1363C | E | 7.7e-08 | 2.9e-08 | 2.6e-08 | 4.400E-08 | 2.862E-08 | 2.600E-08 | 7.700E-08 |
| 27246 | 1363 | T1363E | E | 7.6e-08 | 3.6e-08 | 9.9e-08 | 7.033E-08 | 3.188E-08 | 3.600E-08 | 9.900E-08 |
| 27247 | 1363 | T1363Q | E | 2.6e-08 | 1.1e-07 | 2.4e-08 | 5.333E-08 | 4.908E-08 | 2.400E-08 | 1.100E-07 |
| 27248 | 1363 | T1363G | E | 6.1e-08 | 4.1e-08 | 4.7e-08 | 4.967E-08 | 1.026E-08 | 4.100E-08 | 6.100E-08 |
| 27249 | 1363 | T1363H | E | 1.1e-07 | 4.2e-08 | 4.9e-08 | 6.700E-08 | 3.740E-08 | 4.200E-08 | 1.100E-07 |
| 27250 | 1363 | T1363I | E | 4.2e-08 | 4e-08 | 5.5e-08 | 4.567E-08 | 8.145E-09 | 4.000E-08 | 5.500E-08 |
| 27251 | 1363 | T1363L | E | 9.5e-08 | 1.3e-08 | 2e-08 | 4.267E-08 | 4.546E-08 | 1.300E-08 | 9.500E-08 |
| 27252 | 1363 | T1363K | E | 2.5e-08 | 2.7e-08 | 6.4e-08 | 3.867E-08 | 2.196E-08 | 2.500E-08 | 6.400E-08 |
| 27253 | 1363 | T1363M | E | 3e-08 | 2.3e-08 | 3.1e-08 | 2.800E-08 | 4.359E-09 | 2.300E-08 | 3.100E-08 |
| 27254 | 1363 | T1363F | E | 1.4e-07 | 1e-07 | 3.5e-08 | 9.167E-08 | 5.299E-08 | 3.500E-08 | 1.400E-07 |
| 27255 | 1363 | T1363P | E | 5.3e-08 | 4.7e-08 | 8.8e-08 | 6.267E-08 | 2.214E-08 | 4.700E-08 | 8.800E-08 |
| 27256 | 1363 | T1363S | E | 6e-08 | 2.2e-08 | 3.7e-08 | 3.967E-08 | 1.914E-08 | 2.200E-08 | 6.000E-08 |
| 27257 | 1363 | T1363T | E | 4.7e-08 | 4.1e-08 | 6.7e-08 | 5.167E-08 | 1.361E-08 | 4.100E-08 | 6.700E-08 |
| 27258 | 1363 | T1363W | E | 7.7e-08 | 1.1e-07 | 5.1e-08 | 7.933E-08 | 2.957E-08 | 5.100E-08 | 1.100E-07 |
| 27259 | 1363 | T1363Y | E | 8.2e-08 | 2.9e-08 | 3.3e-08 | 4.800E-08 | 2.951E-08 | 2.900E-08 | 8.200E-08 |
| 27260 | 1363 | T1363V | E | 5.4e-08 | 4e-08 | 4.9e-08 | 4.767E-08 | 7.095E-09 | 4.000E-08 | 5.400E-08 |
| 27261 | 1364 | A1364A | E | 5.5e-08 | 6.4e-08 | 4.6e-08 | 5.500E-08 | 9.000E-09 | 4.600E-08 | 6.400E-08 |
| 27262 | 1364 | A1364R | E | 8.5e-09 | 3.8e-08 | 1.6e-07 | 6.883E-08 | 8.032E-08 | 8.500E-09 | 1.600E-07 |
| 27263 | 1364 | A1364N | E | 1.1e-08 | 4e-08 | 5.8e-08 | 3.633E-08 | 2.371E-08 | 1.100E-08 | 5.800E-08 |
| 27264 | 1364 | A1364D | E | 3.8e-08 | 2.2e-08 | 3.7e-08 | 3.233E-08 | 8.963E-09 | 2.200E-08 | 3.800E-08 |
| 27265 | 1364 | A1364C | E | 3.3e-08 | 5.3e-08 | 4.3e-08 | 4.300E-08 | 1.000E-08 | 3.300E-08 | 5.300E-08 |
| 27266 | 1364 | A1364E | E | 3.5e-08 | 2.1e-08 | 5e-08 | 3.533E-08 | 1.450E-08 | 2.100E-08 | 5.000E-08 |
| 27267 | 1364 | A1364Q | E | 3.4e-08 | 8.3e-09 | 2.9e-08 | 2.377E-08 | 1.363E-08 | 8.300E-09 | 3.400E-08 |
| 27268 | 1364 | A1364G | E | 1.9e-08 | 1.2e-08 | 1.3e-07 | 5.367E-08 | 6.620E-08 | 1.200E-08 | 1.300E-07 |
| 27269 | 1364 | A1364H | E | 4.7e-08 | 3e-08 | 5.3e-08 | 4.333E-08 | 1.193E-08 | 3.000E-08 | 5.300E-08 |
| 27270 | 1364 | A1364I | E | 3.7e-08 | 3.9e-08 | 4.3e-08 | 3.967E-08 | 3.055E-09 | 3.700E-08 | 4.300E-08 |
| 27271 | 1364 | A1364L | E | 6.1e-08 | 6e-08 | 5.5e-08 | 5.867E-08 | 3.215E-09 | 5.500E-08 | 6.100E-08 |
| 27272 | 1364 | A1364K | E | 3.5e-08 | 1.6e-07 | 3.4e-08 | 7.633E-08 | 7.246E-08 | 3.400E-08 | 1.600E-07 |
| 27273 | 1364 | A1364M | E | 1.6e-08 | 5.2e-08 | 7.5e-08 | 4.767E-08 | 2.974E-08 | 1.600E-08 | 7.500E-08 |
| 27274 | 1364 | A1364F | E | 4.3e-08 | 2e-08 | 5.2e-08 | 3.833E-08 | 1.650E-08 | 2.000E-08 | 5.200E-08 |
| 27275 | 1364 | A1364P | E | 4.5e-08 | 4.5e-08 | 2.6e-08 | 3.867E-08 | 1.097E-08 | 2.600E-08 | 4.500E-08 |
| 27276 | 1364 | A1364S | E | 1e-07 | 1e-07 | 1.5e-08 | 7.167E-08 | 4.907E-08 | 1.500E-08 | 1.000E-07 |
| 27277 | 1364 | A1364T | E | 3.3e-08 | 5.5e-08 | 3.4e-08 | 4.067E-08 | 1.242E-08 | 3.300E-08 | 5.500E-08 |
| 27278 | 1364 | A1364W | E | 4.3e-08 | 8.4e-08 | 8.8e-08 | 7.167E-08 | 2.491E-08 | 4.300E-08 | 8.800E-08 |
| 27279 | 1364 | A1364Y | E | 3.2e-08 | 2.3e-08 | 2.6e-08 | 2.700E-08 | 4.583E-09 | 2.300E-08 | 3.200E-08 |
| 27280 | 1364 | A1364V | E | 3.4e-08 | 2.2e-08 | 4.6e-08 | 3.400E-08 | 1.200E-08 | 2.200E-08 | 4.600E-08 |
| 27281 | 1365 | V1365A | E | 5.3e-08 | 3.2e-08 | 7.9e-08 | 5.467E-08 | 2.354E-08 | 3.200E-08 | 7.900E-08 |
| 27282 | 1365 | V1365R | E | 2.7e-08 | 9.9e-08 | 3.6e-08 | 5.400E-08 | 3.923E-08 | 2.700E-08 | 9.900E-08 |
| 27283 | 1365 | V1365N | E | 3.2e-08 | 5e-08 | 2.6e-08 | 3.600E-08 | 1.249E-08 | 2.600E-08 | 5.000E-08 |
| 27284 | 1365 | V1365D | E | 1.4e-08 | 2.5e-08 | 4.1e-08 | 2.667E-08 | 1.358E-08 | 1.400E-08 | 4.100E-08 |

| | | | | | | | | | | |
|---|---|---|---|---|---|---|---|---|---|---|
| 27285 | 1365 | V1365C | E | 4.4e-08 | 9e-08 | 3e-08 | 5.467E-08 | 3.139E-08 | 3.000E-08 | 9.000E-08 |
| 27286 | 1365 | V1365E | E | 2.8e-08 | 9.3e-09 | 1.7e-08 | 1.810E-08 | 9.398E-09 | 9.300E-09 | 2.800E-08 |
| 27287 | 1365 | V1365Q | E | 5.9e-08 | 1.4e-08 | 6.8e-08 | 4.700E-08 | 2.893E-08 | 1.400E-08 | 6.800E-08 |
| 27288 | 1365 | V1365G | E | 1.5e-08 | 5.6e-08 | 2.7e-08 | 3.267E-08 | 2.108E-08 | 1.500E-08 | 5.600E-08 |
| 27289 | 1365 | V1365H | E | 4.1e-08 | 7.2e-08 | 4.2e-08 | 5.167E-08 | 1.762E-08 | 4.100E-08 | 7.200E-08 |
| 27290 | 1365 | V1365I | E | 2.8e-08 | 4.2e-08 | 8.8e-08 | 5.267E-08 | 3.139E-08 | 2.800E-08 | 8.800E-08 |
| 27291 | 1365 | V1365L | E | 7.8e-08 | 3.4e-08 | 1.8e-08 | 4.333E-08 | 3.107E-08 | 1.800E-08 | 7.800E-08 |
| 27292 | 1365 | V1365K | E | 2.4e-08 | 2.2e-08 | 4.6e-08 | 3.067E-08 | 1.332E-08 | 2.200E-08 | 4.600E-08 |
| 27293 | 1365 | V1365M | E | 7.4e-09 | 2.9e-08 | 2.8e-08 | 2.147E-08 | 1.219E-08 | 7.400E-09 | 2.900E-08 |
| 27294 | 1365 | V1365F | E | 2.5e-08 | 2.8e-08 | 3.3e-08 | 2.867E-08 | 4.041E-09 | 2.500E-08 | 3.300E-08 |
| 27295 | 1365 | V1365P | E | 5.1e-08 | 4.1e-08 | 9.2e-09 | 3.373E-08 | 2.183E-08 | 9.200E-09 | 5.100E-08 |
| 27296 | 1365 | V1365S | E | 2.5e-08 | 5.6e-08 | 1.4e-08 | 3.167E-08 | 2.178E-08 | 1.400E-08 | 5.600E-08 |
| 27297 | 1365 | V1365T | E | 1.4e-07 | 2.7e-08 | 2.5e-08 | 6.400E-08 | 6.583E-08 | 2.500E-08 | 1.400E-07 |
| 27298 | 1365 | V1365W | E | 1.8e-08 | 3e-08 | 2.7e-08 | 2.500E-08 | 6.245E-09 | 1.800E-08 | 3.000E-08 |
| 27299 | 1365 | V1365Y | E | 2e-08 | 6.6e-08 | 3.7e-08 | 4.100E-08 | 2.326E-08 | 2.000E-08 | 6.600E-08 |
| 27300 | 1365 | V1365V | E | 6.6e-08 | 6e-08 | 3.5e-08 | 5.367E-08 | 1.644E-08 | 3.500E-08 | 6.600E-08 |
| 27301 | 1366 | Y1366A | E | 4.2e-08 | 6.2e-08 | 8.3e-08 | 6.233E-08 | 2.050E-08 | 4.200E-08 | 8.300E-08 |
| 27302 | 1366 | Y1366R | E | 5.7e-08 | 5.4e-08 | 9e-08 | 6.700E-08 | 1.997E-08 | 5.400E-08 | 9.000E-08 |
| 27303 | 1366 | Y1366N | E | 1.5e-08 | 3.2e-08 | 8e-08 | 4.233E-08 | 3.371E-08 | 1.500E-08 | 8.000E-08 |
| 27304 | 1366 | Y1366D | E | 2.8e-08 | 5.5e-08 | 4e-08 | 4.100E-08 | 1.353E-08 | 2.800E-08 | 5.500E-08 |
| 27305 | 1366 | Y1366C | E | 6.7e-08 | 5.7e-08 | 2.8e-08 | 5.067E-08 | 2.026E-08 | 2.800E-08 | 6.700E-08 |
| 27306 | 1366 | Y1366E | E | 7.4e-08 | 8.9e-09 | 2.6e-08 | 3.630E-08 | 3.375E-08 | 8.900E-09 | 7.400E-08 |
| 27307 | 1366 | Y1366Q | E | 6.5e-08 | 5.6e-08 | 2.3e-08 | 4.800E-08 | 2.211E-08 | 2.300E-08 | 6.500E-08 |
| 27308 | 1366 | Y1366G | E | 1.5e-08 | 5.9e-08 | 6.7e-08 | 4.700E-08 | 2.800E-08 | 1.500E-08 | 6.700E-08 |
| 27309 | 1366 | Y1366H | E | 3.9e-08 | 6.5e-08 | 1.7e-08 | 4.033E-08 | 2.403E-08 | 1.700E-08 | 6.500E-08 |
| 27310 | 1366 | Y1366I | E | 2.5e-08 | 5.4e-08 | 2.1e-08 | 3.333E-08 | 1.801E-08 | 2.100E-08 | 5.400E-08 |
| 27311 | 1366 | Y1366L | E | 3.7e-08 | 3e-07 | 5.5e-08 | 1.307E-07 | 1.469E-07 | 3.700E-08 | 3.000E-07 |
| 27312 | 1366 | Y1366K | E | 2.3e-08 | 3.8e-08 | 9.1e-08 | 5.067E-08 | 3.573E-08 | 2.300E-08 | 9.100E-08 |
| 27313 | 1366 | Y1366M | E | 2.7e-08 | 3.6e-08 | 5.8e-08 | 4.033E-08 | 1.595E-08 | 2.700E-08 | 5.800E-08 |
| 27314 | 1366 | Y1366F | E | 5.5e-08 | 2.6e-08 | 3.3e-08 | 3.800E-08 | 1.513E-08 | 2.600E-08 | 5.500E-08 |
| 27315 | 1366 | Y1366P | E | 3.1e-08 | 6.3e-08 | 5.8e-08 | 5.067E-08 | 1.721E-08 | 3.100E-08 | 6.300E-08 |
| 27316 | 1366 | Y1366S | E | 7.8e-08 | 9.8e-08 | 8.5e-08 | 8.700E-08 | 1.015E-08 | 7.800E-08 | 9.800E-08 |
| 27317 | 1366 | Y1366T | E | 3e-08 | 5.1e-08 | 2.7e-08 | 3.600E-08 | 1.308E-08 | 2.700E-08 | 5.100E-08 |
| 27318 | 1366 | Y1366W | E | 5.6e-08 | 7.6e-08 | 2.2e-07 | 1.173E-07 | 8.947E-08 | 5.600E-08 | 2.200E-07 |
| 27319 | 1366 | Y1366Y | E | 3.5e-08 | 5.2e-08 | 3.2e-08 | 3.967E-08 | 1.079E-08 | 3.200E-08 | 5.200E-08 |
| 27320 | 1366 | Y1366V | E | 2.1e-08 | 1.6e-07 | 1.2e-07 | 1.003E-07 | 7.156E-08 | 2.100E-08 | 1.600E-07 |
| 27321 | 1367 | Y1367A | E | 2.5e-08 | 6e-08 | 4.9e-08 | 4.467E-08 | 1.790E-08 | 2.500E-08 | 6.000E-08 |
| 27322 | 1367 | Y1367R | E | 2.4e-08 | 5.7e-08 | 4.6e-08 | 4.233E-08 | 1.680E-08 | 2.400E-08 | 5.700E-08 |
| 27323 | 1367 | Y1367N | E | 5.6e-08 | 1.5e-08 | 3.6e-08 | 3.567E-08 | 2.050E-08 | 1.500E-08 | 5.600E-08 |
| 27324 | 1367 | Y1367D | E | 3.9e-08 | 4.6e-08 | 5.8e-08 | 4.767E-08 | 9.609E-09 | 3.900E-08 | 5.800E-08 |
| 27325 | 1367 | Y1367C | E | 3e-08 | 2.8e-08 | 3.3e-08 | 3.033E-08 | 2.517E-09 | 2.800E-08 | 3.300E-08 |
| 27326 | 1367 | Y1367E | E | 6.3e-08 | 5.8e-08 | 4.2e-08 | 5.433E-08 | 1.097E-08 | 4.200E-08 | 6.300E-08 |
| 27327 | 1367 | Y1367Q | E | 3.7e-08 | 6.3e-08 | 1.8e-08 | 3.933E-08 | 2.259E-08 | 1.800E-08 | 6.300E-08 |
| 27328 | 1367 | Y1367G | E | 8.4e-08 | 4.6e-08 | 6e-08 | 6.333E-08 | 1.922E-08 | 4.600E-08 | 8.400E-08 |
| 27329 | 1367 | Y1367H | E | 8.1e-08 | 5.5e-09 | 1.2e-07 | 6.883E-08 | 5.821E-08 | 5.500E-09 | 1.200E-07 |
| 27330 | 1367 | Y1367I | E | 6.1e-08 | 6.3e-08 | 5.9e-08 | 6.100E-08 | 2.000E-09 | 5.900E-08 | 6.300E-08 |
| 27331 | 1367 | Y1367L | E | 6.7e-08 | 2.5e-08 | 9.3e-08 | 6.167E-08 | 3.431E-08 | 2.500E-08 | 9.300E-08 |
| 27332 | 1367 | Y1367K | E | 1.5e-08 | 3.9e-08 | 3.5e-08 | 2.967E-08 | 1.286E-08 | 1.500E-08 | 3.900E-08 |
| 27333 | 1367 | Y1367M | E | 2.1e-08 | 1.7e-08 | 3.1e-08 | 2.300E-08 | 7.211E-09 | 1.700E-08 | 3.100E-08 |
| 27334 | 1367 | Y1367F | E | 2.6e-08 | 6.3e-08 | 1e-07 | 6.300E-08 | 3.700E-08 | 2.600E-08 | 1.000E-07 |
| 27335 | 1367 | Y1367P | E | 2.6e-08 | 6.9e-08 | 2.5e-08 | 4.000E-08 | 2.512E-08 | 2.500E-08 | 6.900E-08 |
| 27336 | 1367 | Y1367S | E | 1.4e-07 | 3.5e-08 | 5.2e-08 | 7.567E-08 | 5.636E-08 | 3.500E-08 | 1.400E-07 |
| 27337 | 1367 | Y1367T | E | 1.9e-08 | 1.1e-07 | 1.5e-08 | 4.800E-08 | 5.373E-08 | 1.500E-08 | 1.100E-07 |
| 27338 | 1367 | Y1367W | E | 4e-08 | 2.8e-08 | 7.4e-08 | 4.733E-08 | 2.386E-08 | 2.800E-08 | 7.400E-08 |
| 27339 | 1367 | Y1367Y | E | 1.1e-07 | 3.3e-08 | 2e-08 | 5.433E-08 | 4.864E-08 | 2.000E-08 | 1.100E-07 |
| 27340 | 1367 | Y1367V | E | 8.1e-08 | 6.8e-08 | 4.8e-08 | 6.567E-08 | 1.662E-08 | 4.800E-08 | 8.100E-08 |
| 27341 | 1368 | C1368A | E | 1e-07 | 4.1e-08 | 3e-08 | 5.700E-08 | 3.764E-08 | 3.000E-08 | 1.000E-07 |
| 27342 | 1368 | C1368R | E | 4.2e-08 | 2.7e-08 | 7.1e-08 | 4.667E-08 | 2.237E-08 | 2.700E-08 | 7.100E-08 |
| 27343 | 1368 | C1368N | E | 7.5e-08 | 3.5e-08 | 6.5e-08 | 5.833E-08 | 2.082E-08 | 3.500E-08 | 7.500E-08 |
| 27344 | 1368 | C1368D | E | 3.7e-08 | 1.4e-07 | 3.3e-08 | 7.000E-08 | 6.065E-08 | 3.300E-08 | 1.400E-07 |
| 27345 | 1368 | C1368C | E | 9.5e-09 | 9.5e-08 | 2.6e-08 | 4.350E-08 | 4.536E-08 | 9.500E-09 | 9.500E-08 |
| 27346 | 1368 | C1368E | E | 1.8e-08 | 2.7e-08 | 3.8e-08 | 2.767E-08 | 1.002E-08 | 1.800E-08 | 3.800E-08 |
| 27347 | 1368 | C1368Q | E | 5.7e-08 | 4e-08 | 5.8e-08 | 5.167E-08 | 1.012E-08 | 4.000E-08 | 5.800E-08 |
| 27348 | 1368 | C1368G | E | 6e-08 | 7.1e-08 | 6.9e-08 | 6.667E-08 | 5.859E-09 | 6.000E-08 | 7.100E-08 |
| 27349 | 1368 | C1368H | E | 5.6e-08 | 2.8e-08 | 4.2e-08 | 4.200E-08 | 1.400E-08 | 2.800E-08 | 5.600E-08 |
| 27350 | 1368 | C1368I | E | 4.4e-08 | 2.2e-08 | 1e-07 | 5.533E-08 | 4.022E-08 | 2.200E-08 | 1.000E-07 |
| 27351 | 1368 | C1368L | E | 6.3e-08 | 3.7e-08 | 3.7e-08 | 4.567E-08 | 1.501E-08 | 3.700E-08 | 6.300E-08 |
| 27352 | 1368 | C1368K | E | 7.1e-08 | 4.2e-08 | 4.6e-08 | 5.300E-08 | 1.572E-08 | 4.200E-08 | 7.100E-08 |
| 27353 | 1368 | C1368M | E | 1.6e-08 | 8.7e-08 | 5.2e-08 | 5.167E-08 | 3.550E-08 | 1.600E-08 | 8.700E-08 |
| 27354 | 1368 | C1368F | E | 2.2e-08 | 2.7e-08 | 8.9e-08 | 4.600E-08 | 3.732E-08 | 2.200E-08 | 8.900E-08 |
| 27355 | 1368 | C1368P | E | 3.6e-08 | 3.7e-08 | 1.5e-08 | 2.933E-08 | 1.242E-08 | 1.500E-08 | 3.700E-08 |
| 27356 | 1368 | C1368S | E | 1.5e-07 | 6.5e-08 | 3.8e-08 | 8.433E-08 | 5.845E-08 | 3.800E-08 | 1.500E-07 |
| 27357 | 1368 | C1368T | E | 1.8e-08 | 1.2e-07 | 1.5e-07 | 9.600E-08 | 6.920E-08 | 1.800E-08 | 1.500E-07 |
| 27358 | 1368 | C1368W | E | 3.7e-08 | 3.3e-08 | 1.1e-07 | 6.000E-08 | 4.335E-08 | 3.300E-08 | 1.100E-07 |
| 27359 | 1368 | C1368Y | E | 5.1e-08 | 4.2e-08 | 7e-08 | 5.433E-08 | 1.429E-08 | 4.200E-08 | 7.000E-08 |
| 27360 | 1368 | C1368V | E | 1.4e-08 | 3.5e-08 | 1.7e-08 | 2.200E-08 | 1.136E-08 | 1.400E-08 | 3.500E-08 |
| 27361 | 1369 | S1369A | E | 2.8e-08 | 1.6e-08 | 4.4e-08 | 2.933E-08 | 1.405E-08 | 1.600E-08 | 4.400E-08 |
| 27362 | 1369 | S1369R | E | 4.7e-08 | 8.1e-08 | 3.5e-08 | 5.433E-08 | 2.386E-08 | 3.500E-08 | 8.100E-08 |
| 27363 | 1369 | S1369N | E | 3.3e-08 | 1.4e-07 | 3e-08 | 6.767E-08 | 6.266E-08 | 3.000E-08 | 1.400E-07 |
| 27364 | 1369 | S1369D | E | 5.7e-08 | 7.8e-08 | 5.5e-08 | 6.333E-08 | 1.274E-08 | 5.500E-08 | 7.800E-08 |
| 27365 | 1369 | S1369C | E | 3.5e-08 | 2e-08 | 6.3e-08 | 3.933E-08 | 2.183E-08 | 2.000E-08 | 6.300E-08 |
| 27366 | 1369 | S1369E | E | 3.3e-08 | 2e-08 | 5.9e-08 | 3.733E-08 | 1.986E-08 | 2.000E-08 | 5.900E-08 |
| 27367 | 1369 | S1369Q | E | 5.6e-08 | 5.6e-08 | 6.6e-08 | 5.933E-08 | 5.774E-09 | 5.600E-08 | 6.600E-08 |
| 27368 | 1369 | S1369G | E | 5.5e-08 | 4.8e-08 | 4.9e-08 | 5.067E-08 | 3.786E-09 | 4.800E-08 | 5.500E-08 |
| 27369 | 1369 | S1369H | E | 3.9e-08 | 1e-07 | 9.7e-08 | 7.867E-08 | 3.439E-08 | 3.900E-08 | 1.000E-07 |

| | | | | | | | | | | |
|---|---|---|---|---|---|---|---|---|---|---|
| 27370 | 1369 | S1369I | E | 2.1e-08 | 3e-08 | 5.2e-08 | 3.433E-08 | 1.595E-08 | 2.100E-08 | 5.200E-08 |
| 27371 | 1369 | S1369L | E | 6.1e-08 | 4.5e-08 | 4.1e-08 | 4.900E-08 | 1.058E-08 | 4.100E-08 | 6.100E-08 |
| 27372 | 1369 | S1369K | E | 1.6e-08 | 5.4e-08 | 5.2e-08 | 4.067E-08 | 2.139E-08 | 1.600E-08 | 5.400E-08 |
| 27373 | 1369 | S1369M | E | 1.6e-07 | 3e-08 | 3.9e-08 | 7.633E-08 | 7.260E-08 | 3.000E-08 | 1.600E-07 |
| 27374 | 1369 | S1369F | E | 1e-07 | 1.4e-08 | 2.7e-08 | 4.700E-08 | 4.636E-08 | 1.400E-08 | 1.000E-07 |
| 27375 | 1369 | S1369P | E | 9.6e-08 | 1.2e-07 | 9.7e-08 | 1.043E-07 | 1.358E-08 | 9.600E-08 | 1.200E-07 |
| 27376 | 1369 | S1369S | E | 4.8e-08 | 1.4e-07 | 3.8e-08 | 7.533E-08 | 5.623E-08 | 3.800E-08 | 1.400E-07 |
| 27377 | 1369 | S1369T | E | 3.8e-08 | 6.2e-08 | 3.5e-08 | 4.500E-08 | 1.480E-08 | 3.500E-08 | 6.200E-08 |
| 27378 | 1369 | S1369W | E | 5.2e-08 | 2.6e-08 | 6.3e-08 | 4.700E-08 | 1.900E-08 | 2.600E-08 | 6.300E-08 |
| 27379 | 1369 | S1369Y | E | 4.8e-08 | 6e-08 | 7e-08 | 5.933E-08 | 1.102E-08 | 4.800E-08 | 7.000E-08 |
| 27380 | 1369 | S1369V | E | 2.8e-08 | 2.8e-08 | 6.5e-08 | 4.033E-08 | 2.136E-08 | 2.800E-08 | 6.500E-08 |
| 27381 | 1370 | R1370A | E | 2.2e-08 | 4.7e-08 | 1.5e-08 | 2.800E-08 | 1.682E-08 | 1.500E-08 | 4.700E-08 |
| 27382 | 1370 | R1370R | E | 4.4e-08 | 4e-08 | 8.3e-08 | 5.567E-08 | 2.376E-08 | 4.000E-08 | 8.300E-08 |
| 27383 | 1370 | R1370N | E | 2.5e-08 | 5.1e-08 | 3.1e-08 | 3.567E-08 | 1.361E-08 | 2.500E-08 | 5.100E-08 |
| 27384 | 1370 | R1370D | E | 7.3e-08 | 3.4e-08 | 2.5e-08 | 4.400E-08 | 2.551E-08 | 2.500E-08 | 7.300E-08 |
| 27385 | 1370 | R1370C | E | 3.4e-08 | 3.8e-08 | 7.2e-08 | 4.800E-08 | 2.088E-08 | 3.400E-08 | 7.200E-08 |
| 27386 | 1370 | R1370E | E | 5.3e-08 | 7.6e-08 | 3e-08 | 5.300E-08 | 2.300E-08 | 3.000E-08 | 7.600E-08 |
| 27387 | 1370 | R1370Q | E | 7.1e-08 | 5.7e-08 | 7.2e-08 | 6.667E-08 | 8.386E-09 | 5.700E-08 | 7.200E-08 |
| 27388 | 1370 | R1370G | E | 1.8e-08 | 2.5e-08 | 2.8e-08 | 2.367E-08 | 5.132E-09 | 1.800E-08 | 2.800E-08 |
| 27389 | 1370 | R1370H | E | 3.5e-08 | 2.9e-08 | 4.7e-08 | 3.700E-08 | 9.165E-09 | 2.900E-08 | 4.700E-08 |
| 27390 | 1370 | R1370I | E | 6.5e-08 | 4.4e-08 | 2.4e-08 | 4.433E-08 | 2.050E-08 | 2.400E-08 | 6.500E-08 |
| 27391 | 1370 | R1370L | E | 5.2e-08 | 5e-08 | 2.7e-08 | 4.300E-08 | 1.389E-08 | 2.700E-08 | 5.200E-08 |
| 27392 | 1370 | R1370K | E | 7.4e-08 | 4.4e-08 | 4.8e-08 | 5.533E-08 | 1.629E-08 | 4.400E-08 | 7.400E-08 |
| 27393 | 1370 | R1370M | E | 5.7e-08 | 5.1e-08 | 6.7e-08 | 5.833E-08 | 8.083E-09 | 5.100E-08 | 6.700E-08 |
| 27394 | 1370 | R1370F | E | 2.3e-08 | 1.9e-08 | 2.8e-08 | 2.333E-08 | 4.509E-09 | 1.900E-08 | 2.800E-08 |
| 27395 | 1370 | R1370P | E | 5.9e-08 | 4.9e-08 | 2.6e-08 | 4.467E-08 | 1.692E-08 | 2.600E-08 | 5.900E-08 |
| 27396 | 1370 | R1370S | E | 6e-08 | 7.2e-08 | 7.2e-09 | 4.640E-08 | 3.447E-08 | 7.200E-09 | 7.200E-08 |
| 27397 | 1370 | R1370T | E | 3.2e-08 | 5.1e-08 | 6e-08 | 4.767E-08 | 1.429E-08 | 3.200E-08 | 6.000E-08 |
| 27398 | 1370 | R1370W | E | 6.2e-08 | 4e-08 | 2.8e-08 | 4.333E-08 | 1.724E-08 | 2.800E-08 | 6.200E-08 |
| 27399 | 1370 | R1370Y | E | 1.1e-07 | 5.2e-08 | 4.7e-08 | 6.967E-08 | 3.502E-08 | 4.700E-08 | 1.100E-07 |
| 27400 | 1370 | R1370V | E | 2.9e-08 | 4.1e-08 | 3.6e-08 | 3.533E-08 | 6.028E-09 | 2.900E-08 | 4.100E-08 |
| 27401 | 1371 | W1371A | E | 4e-08 | 3.9e-08 | 1e-08 | 2.967E-08 | 1.704E-08 | 1.000E-08 | 4.000E-08 |
| 27402 | 1371 | W1371R | E | 6.2e-08 | 7.6e-08 | 8.8e-08 | 7.533E-08 | 1.301E-08 | 6.200E-08 | 8.800E-08 |
| 27403 | 1371 | W1371N | E | 1.2e-08 | 2.8e-08 | 3.1e-08 | 2.367E-08 | 1.021E-08 | 1.200E-08 | 3.100E-08 |
| 27404 | 1371 | W1371D | E | 6e-08 | 2.9e-08 | 2.1e-08 | 3.667E-08 | 2.060E-08 | 2.100E-08 | 6.000E-08 |
| 27405 | 1371 | W1371C | E | 4.9e-08 | 8.3e-08 | 8.7e-08 | 7.300E-08 | 2.088E-08 | 4.900E-08 | 8.700E-08 |
| 27406 | 1371 | W1371E | E | 5.4e-08 | 2.2e-08 | 1.4e-07 | 7.200E-08 | 6.102E-08 | 2.200E-08 | 1.400E-07 |
| 27407 | 1371 | W1371Q | E | 5e-08 | 6.1e-08 | 3.1e-08 | 4.733E-08 | 1.518E-08 | 3.100E-08 | 6.100E-08 |
| 27408 | 1371 | W1371G | E | 1e-07 | 7.6e-08 | 1.9e-08 | 6.500E-08 | 4.161E-08 | 1.900E-08 | 1.000E-07 |
| 27409 | 1371 | W1371H | E | 2.8e-08 | 3.9e-08 | 4.5e-08 | 3.733E-08 | 8.622E-09 | 2.800E-08 | 4.500E-08 |
| 27410 | 1371 | W1371I | E | 4e-08 | 4.6e-08 | 3.9e-08 | 4.167E-08 | 3.786E-09 | 3.900E-08 | 4.600E-08 |
| 27411 | 1371 | W1371L | E | 1.7e-08 | 3.3e-08 | 6.2e-08 | 3.733E-08 | 2.281E-08 | 1.700E-08 | 6.200E-08 |
| 27412 | 1371 | W1371K | E | 6.2e-08 | 4.8e-08 | 2.1e-08 | 4.367E-08 | 2.084E-08 | 2.100E-08 | 6.200E-08 |
| 27413 | 1371 | W1371M | E | 6e-08 | 3.2e-08 | 3.2e-08 | 4.133E-08 | 1.617E-08 | 3.200E-08 | 6.000E-08 |
| 27414 | 1371 | W1371F | E | 3.4e-08 | 7.3e-08 | 3.6e-08 | 4.767E-08 | 2.196E-08 | 3.400E-08 | 7.300E-08 |
| 27415 | 1371 | W1371P | E | 4.4e-08 | 3.3e-08 | 1.4e-08 | 3.033E-08 | 1.518E-08 | 1.400E-08 | 4.400E-08 |
| 27416 | 1371 | W1371S | E | 4.6e-08 | 4.5e-08 | 5.4e-08 | 4.833E-08 | 4.933E-09 | 4.500E-08 | 5.400E-08 |
| 27417 | 1371 | W1371T | E | 4.3e-08 | 4e-08 | 3e-08 | 3.767E-08 | 6.807E-09 | 3.000E-08 | 4.300E-08 |
| 27418 | 1371 | W1371W | E | 5.6e-08 | 3.8e-08 | 7e-08 | 5.467E-08 | 1.604E-08 | 3.800E-08 | 7.000E-08 |
| 27419 | 1371 | W1371Y | E | 2.4e-08 | 6.6e-08 | 5e-08 | 4.667E-08 | 2.120E-08 | 2.400E-08 | 6.600E-08 |
| 27420 | 1371 | W1371V | E | 1.3e-08 | 5e-08 | 3.2e-08 | 3.167E-08 | 1.850E-08 | 1.300E-08 | 5.000E-08 |
| 27421 | 1372 | G1372A | E | 5.1e-08 | 9.6e-08 | 4.5e-08 | 6.400E-08 | 2.787E-08 | 4.500E-08 | 9.600E-08 |
| 27422 | 1372 | G1372R | E | 4.5e-08 | 3.8e-08 | 3.6e-08 | 3.967E-08 | 4.726E-09 | 3.600E-08 | 4.500E-08 |
| 27423 | 1372 | G1372N | E | 9.3e-08 | 3.9e-08 | 5.5e-08 | 6.233E-08 | 2.774E-08 | 3.900E-08 | 9.300E-08 |
| 27424 | 1372 | G1372D | E | 4e-08 | 1.6e-08 | 2.6e-08 | 2.733E-08 | 1.206E-08 | 1.600E-08 | 4.000E-08 |
| 27425 | 1372 | G1372C | E | 3.3e-07 | 4.8e-08 | 3e-08 | 1.360E-07 | 1.682E-07 | 3.000E-08 | 3.300E-07 |
| 27426 | 1372 | G1372E | E | 3.1e-08 | 7e-08 | 3.9e-08 | 4.667E-08 | 2.060E-08 | 3.100E-08 | 7.000E-08 |
| 27427 | 1372 | G1372Q | E | 2.8e-08 | 2e-08 | 5e-08 | 3.267E-08 | 1.553E-08 | 2.000E-08 | 5.000E-08 |
| 27428 | 1372 | G1372G | E | 5.5e-08 | 4.1e-08 | 8.5e-08 | 6.033E-08 | 2.248E-08 | 4.100E-08 | 8.500E-08 |
| 27429 | 1372 | G1372H | E | 1e-07 | 2e-08 | 4.6e-08 | 5.533E-08 | 4.081E-08 | 2.000E-08 | 1.000E-07 |
| 27430 | 1372 | G1372I | E | 3e-08 | 2e-08 | 4.7e-08 | 3.233E-08 | 1.365E-08 | 2.000E-08 | 4.700E-08 |
| 27431 | 1372 | G1372L | E | 5.1e-08 | 2.8e-08 | 7.1e-08 | 5.000E-08 | 2.152E-08 | 2.800E-08 | 7.100E-08 |
| 27432 | 1372 | G1372K | E | 2.9e-08 | 3.6e-08 | 6.6e-08 | 4.367E-08 | 1.966E-08 | 2.900E-08 | 6.600E-08 |
| 27433 | 1372 | G1372M | E | 2.3e-08 | 1.9e-08 | 6.2e-08 | 3.467E-08 | 2.376E-08 | 1.900E-08 | 6.200E-08 |
| 27434 | 1372 | G1372F | E | 4.7e-08 | 5.3e-08 | 2.2e-08 | 4.067E-08 | 1.644E-08 | 2.200E-08 | 5.300E-08 |
| 27435 | 1372 | G1372P | E | 9.2e-09 | 3.3e-08 | 4.8e-08 | 3.007E-08 | 1.957E-08 | 9.200E-09 | 4.800E-08 |
| 27436 | 1372 | G1372S | E | 3.5e-08 | 1.8e-08 | 5e-08 | 3.433E-08 | 1.601E-08 | 1.800E-08 | 5.000E-08 |
| 27437 | 1372 | G1372T | E | 1.4e-07 | 6.9e-08 | 1.2e-07 | 1.097E-07 | 3.661E-08 | 6.900E-08 | 1.400E-07 |
| 27438 | 1372 | G1372W | E | 4.1e-08 | 8.2e-08 | 3.9e-08 | 5.400E-08 | 2.427E-08 | 3.900E-08 | 8.200E-08 |
| 27439 | 1372 | G1372Y | E | 2.8e-08 | 2.9e-08 | 1.3e-07 | 6.233E-08 | 5.860E-08 | 2.800E-08 | 1.300E-07 |
| 27440 | 1372 | G1372V | E | 5.6e-08 | 6.8e-08 | 1.5e-08 | 4.633E-08 | 2.779E-08 | 1.500E-08 | 6.800E-08 |
| 27441 | 1373 | G1373A | E | 2.4e-08 | 4.4e-08 | 2.5e-08 | 3.100E-08 | 1.127E-08 | 2.400E-08 | 4.400E-08 |
| 27442 | 1373 | G1373R | E | 3.2e-08 | 9.6e-08 | 3.2e-08 | 5.333E-08 | 3.695E-08 | 3.200E-08 | 9.600E-08 |
| 27443 | 1373 | G1373N | E | 3.5e-08 | 4.1e-08 | 3.7e-08 | 3.767E-08 | 3.055E-09 | 3.500E-08 | 4.100E-08 |
| 27444 | 1373 | G1373D | E | 4.7e-08 | 8.1e-08 | 4e-08 | 5.600E-08 | 2.193E-08 | 4.000E-08 | 8.100E-08 |
| 27445 | 1373 | G1373C | E | 4.6e-08 | 4e-08 | 1.8e-08 | 3.467E-08 | 1.474E-08 | 1.800E-08 | 4.600E-08 |
| 27446 | 1373 | G1373E | E | 6.3e-08 | 6.5e-08 | 3.3e-08 | 5.367E-08 | 1.793E-08 | 3.300E-08 | 6.500E-08 |
| 27447 | 1373 | G1373Q | E | 4.4e-08 | 4.6e-08 | 2.2e-08 | 3.733E-08 | 1.332E-08 | 2.200E-08 | 4.600E-08 |
| 27448 | 1373 | G1373G | E | 2.6e-08 | 2.7e-08 | 6.7e-09 | 1.990E-08 | 1.144E-08 | 6.700E-09 | 2.700E-08 |
| 27449 | 1373 | G1373H | E | 4.8e-08 | 4.3e-08 | 8.9e-08 | 6.000E-08 | 2.524E-08 | 4.300E-08 | 8.900E-08 |
| 27450 | 1373 | G1373I | E | 4.9e-08 | 5.5e-08 | 2.2e-08 | 4.200E-08 | 1.758E-08 | 2.200E-08 | 5.500E-08 |
| 27451 | 1373 | G1373L | E | 8.4e-08 | 4.6e-08 | 3.3e-08 | 5.433E-08 | 2.650E-08 | 3.300E-08 | 8.400E-08 |
| 27452 | 1373 | G1373K | E | 3.5e-08 | 5.8e-08 | 5.6e-08 | 4.967E-08 | 1.274E-08 | 3.500E-08 | 5.800E-08 |
| 27453 | 1373 | G1373M | E | 3.2e-08 | 1.2e-07 | 2.4e-08 | 5.867E-08 | 5.327E-08 | 2.400E-08 | 1.200E-07 |
| 27454 | 1373 | G1373F | E | 1.5e-08 | 8.3e-08 | 2.5e-08 | 4.100E-08 | 3.672E-08 | 1.500E-08 | 8.300E-08 |

| | | | | | | | | | | |
|---|---|---|---|---|---|---|---|---|---|---|
| 27455 | 1373 | G1373P | E | 5.6e-08 | 2e-08 | 7.1e-08 | 4.900E-08 | 2.621E-08 | 2.000E-08 | 7.100E-08 |
| 27456 | 1373 | G1373S | E | 1.6e-08 | 5.1e-08 | 4.8e-08 | 3.833E-08 | 1.940E-08 | 1.600E-08 | 5.100E-08 |
| 27457 | 1373 | G1373T | E | 7.1e-08 | 3.8e-08 | 3.6e-08 | 4.833E-08 | 1.966E-08 | 3.600E-08 | 7.100E-08 |
| 27458 | 1373 | G1373W | E | 5.5e-08 | 3.8e-08 | 7.7e-08 | 5.667E-08 | 1.955E-08 | 3.800E-08 | 7.700E-08 |
| 27459 | 1373 | G1373Y | E | 5e-08 | 1.9e-08 | 4.4e-08 | 3.767E-08 | 1.644E-08 | 1.900E-08 | 5.000E-08 |
| 27460 | 1373 | G1373V | E | 1.1e-07 | 2.4e-08 | 5.6e-08 | 6.333E-08 | 4.347E-08 | 2.400E-08 | 1.100E-07 |
| 27461 | 1374 | D1374A | E | 4.7e-08 | 4e-08 | 2.5e-08 | 3.733E-08 | 1.124E-08 | 2.500E-08 | 4.700E-08 |
| 27462 | 1374 | D1374R | E | 1.5e-07 | 6.2e-08 | 2.5e-08 | 7.900E-08 | 6.421E-08 | 2.500E-08 | 1.500E-07 |
| 27463 | 1374 | D1374N | E | 1.5e-08 | 4.7e-08 | 2.8e-08 | 3.000E-08 | 1.609E-08 | 1.500E-08 | 4.700E-08 |
| 27464 | 1374 | D1374D | E | 7.3e-08 | 3.9e-08 | 4.3e-08 | 5.167E-08 | 1.858E-08 | 3.900E-08 | 7.300E-08 |
| 27465 | 1374 | D1374C | E | 1.4e-07 | 1.2e-08 | 4e-08 | 6.400E-08 | 6.729E-08 | 1.200E-08 | 1.400E-07 |
| 27466 | 1374 | D1374E | E | 2.9e-08 | 7.3e-08 | 3.4e-08 | 4.533E-08 | 2.409E-08 | 2.900E-08 | 7.300E-08 |
| 27467 | 1374 | D1374Q | E | 1.3e-08 | 5.7e-08 | 5.4e-08 | 4.133E-08 | 2.458E-08 | 1.300E-08 | 5.700E-08 |
| 27468 | 1374 | D1374G | E | 5.1e-08 | 7.2e-08 | 1.9e-08 | 4.733E-08 | 2.669E-08 | 1.900E-08 | 7.200E-08 |
| 27469 | 1374 | D1374H | E | 3.5e-08 | 5e-08 | 8.6e-08 | 5.700E-08 | 2.621E-08 | 3.500E-08 | 8.600E-08 |
| 27470 | 1374 | D1374I | E | 3.1e-08 | 7.9e-08 | 6.2e-09 | 3.873E-08 | 3.701E-08 | 6.200E-09 | 7.900E-08 |
| 27471 | 1374 | D1374L | E | 2.2e-07 | 9.2e-08 | 3.2e-08 | 1.147E-07 | 9.603E-08 | 3.200E-08 | 2.200E-07 |
| 27472 | 1374 | D1374K | E | 4e-08 | 4.9e-08 | 3.6e-08 | 4.167E-08 | 6.658E-09 | 3.600E-08 | 4.900E-08 |
| 27473 | 1374 | D1374M | E | 5.5e-08 | 5.2e-08 | 2.8e-08 | 4.500E-08 | 1.480E-08 | 2.800E-08 | 5.500E-08 |
| 27474 | 1374 | D1374F | E | 4.9e-08 | 7.8e-08 | 3e-08 | 5.233E-08 | 2.417E-08 | 3.000E-08 | 7.800E-08 |
| 27475 | 1374 | D1374P | E | 4.4e-08 | 2.4e-08 | 4.2e-08 | 3.667E-08 | 1.102E-08 | 2.400E-08 | 4.400E-08 |
| 27476 | 1374 | D1374S | E | 3.6e-08 | 8.9e-08 | 4.5e-08 | 5.667E-08 | 2.836E-08 | 3.600E-08 | 8.900E-08 |
| 27477 | 1374 | D1374T | E | 6.8e-08 | 9.5e-08 | 5.7e-08 | 7.333E-08 | 1.955E-08 | 5.700E-08 | 9.500E-08 |
| 27478 | 1374 | D1374W | E | 3.3e-08 | 3.6e-08 | 3.6e-08 | 3.500E-08 | 1.732E-09 | 3.300E-08 | 3.600E-08 |
| 27479 | 1374 | D1374Y | E | 1e-07 | 6.3e-08 | 4.9e-08 | 7.067E-08 | 2.635E-08 | 4.900E-08 | 1.000E-07 |
| 27480 | 1374 | D1374V | E | 5.2e-08 | 9.6e-08 | 4e-08 | 6.267E-08 | 2.948E-08 | 4.000E-08 | 9.600E-08 |
| 27481 | 1375 | G1375A | E | 2.8e-08 | 4.6e-08 | 6.7e-08 | 4.700E-08 | 1.952E-08 | 2.800E-08 | 6.700E-08 |
| 27482 | 1375 | G1375R | E | 4.4e-08 | 8e-08 | 4.1e-08 | 5.500E-08 | 2.170E-08 | 4.100E-08 | 8.000E-08 |
| 27483 | 1375 | G1375N | E | 9.8e-09 | 9.9e-08 | 2.3e-08 | 4.393E-08 | 4.814E-08 | 9.800E-09 | 9.900E-08 |
| 27484 | 1375 | G1375D | E | 5.1e-08 | 1.5e-08 | 2e-08 | 2.867E-08 | 1.950E-08 | 1.500E-08 | 5.100E-08 |
| 27485 | 1375 | G1375C | E | 3e-08 | 4e-08 | 3.4e-08 | 3.467E-08 | 5.033E-09 | 3.000E-08 | 4.000E-08 |
| 27486 | 1375 | G1375E | E | 7.7e-08 | 2e-08 | 2.1e-08 | 3.933E-08 | 3.262E-08 | 2.000E-08 | 7.700E-08 |
| 27487 | 1375 | G1375Q | E | 2.9e-08 | 3.8e-08 | 8.4e-08 | 5.033E-08 | 2.950E-08 | 2.900E-08 | 8.400E-08 |
| 27488 | 1375 | G1375G | E | 6.9e-08 | 5e-08 | 9.4e-08 | 7.100E-08 | 2.207E-08 | 5.000E-08 | 9.400E-08 |
| 27489 | 1375 | G1375H | E | 5e-08 | 5.8e-08 | 5.4e-08 | 5.400E-08 | 4.000E-09 | 5.000E-08 | 5.800E-08 |
| 27490 | 1375 | G1375I | E | 1e-07 | 5.1e-08 | 1.1e-07 | 8.700E-08 | 3.158E-08 | 5.100E-08 | 1.100E-07 |
| 27491 | 1375 | G1375L | E | 3.3e-08 | 2.6e-08 | 2.2e-08 | 2.700E-08 | 5.568E-09 | 2.200E-08 | 3.300E-08 |
| 27492 | 1375 | G1375K | E | 6.5e-08 | 4.3e-08 | 7.2e-08 | 6.000E-08 | 1.513E-08 | 4.300E-08 | 7.200E-08 |
| 27493 | 1375 | G1375M | E | 2.7e-08 | 7e-08 | 7.4e-08 | 5.700E-08 | 2.606E-08 | 2.700E-08 | 7.400E-08 |
| 27494 | 1375 | G1375F | E | 2.3e-08 | 4.2e-08 | 3.7e-08 | 3.400E-08 | 9.849E-09 | 2.300E-08 | 4.200E-08 |
| 27495 | 1375 | G1375P | E | 3.5e-08 | 1e-07 | 7.3e-08 | 6.933E-08 | 3.265E-08 | 3.500E-08 | 1.000E-07 |
| 27496 | 1375 | G1375S | E | 1.7e-08 | 3.3e-08 | 4.8e-08 | 3.267E-08 | 1.550E-08 | 1.700E-08 | 4.800E-08 |
| 27497 | 1375 | G1375T | E | 1.8e-08 | 1.3e-08 | 1.9e-08 | 1.667E-08 | 3.215E-09 | 1.300E-08 | 1.900E-08 |
| 27498 | 1375 | G1375W | E | 2.2e-08 | 2.8e-08 | 4.9e-08 | 3.300E-08 | 1.418E-08 | 2.200E-08 | 4.900E-08 |
| 27499 | 1375 | G1375Y | E | 3.5e-08 | 1.2e-07 | 1.8e-08 | 5.767E-08 | 5.465E-08 | 1.800E-08 | 1.200E-07 |
| 27500 | 1375 | G1375V | E | 2.6e-08 | 4.1e-08 | 1.5e-07 | 7.233E-08 | 6.768E-08 | 2.600E-08 | 1.500E-07 |
| 27501 | 1376 | F1376A | E | 2.9e-08 | 7.7e-09 | 4.6e-08 | 2.757E-08 | 1.919E-08 | 7.700E-09 | 4.600E-08 |
| 27502 | 1376 | F1376R | E | 4.3e-08 | 6e-08 | 5.9e-08 | 5.400E-08 | 9.539E-09 | 4.300E-08 | 6.000E-08 |
| 27503 | 1376 | F1376N | E | 2.3e-08 | 3.3e-08 | 9.5e-08 | 5.033E-08 | 3.900E-08 | 2.300E-08 | 9.500E-08 |
| 27504 | 1376 | F1376D | E | 1.3e-07 | 9.4e-08 | 3e-08 | 8.467E-08 | 5.065E-08 | 3.000E-08 | 1.300E-07 |
| 27505 | 1376 | F1376C | E | 4.4e-08 | 3.1e-08 | 6.3e-08 | 4.600E-08 | 1.609E-08 | 3.100E-08 | 6.300E-08 |
| 27506 | 1376 | F1376E | E | 4.5e-08 | 7.9e-08 | 2.5e-08 | 4.967E-08 | 2.730E-08 | 2.500E-08 | 7.900E-08 |
| 27507 | 1376 | F1376Q | E | 2.4e-08 | 6.3e-08 | 4.8e-08 | 4.500E-08 | 1.967E-08 | 2.400E-08 | 6.300E-08 |
| 27508 | 1376 | F1376G | E | 4.4e-08 | 6.9e-08 | 4.7e-08 | 5.333E-08 | 1.365E-08 | 4.400E-08 | 6.900E-08 |
| 27509 | 1376 | F1376H | E | 4.5e-08 | 5.7e-08 | 5.5e-08 | 5.233E-08 | 6.429E-09 | 4.500E-08 | 5.700E-08 |
| 27510 | 1376 | F1376I | E | 5.7e-08 | 5.1e-08 | 8.4e-08 | 6.400E-08 | 1.758E-08 | 5.100E-08 | 8.400E-08 |
| 27511 | 1376 | F1376L | E | 3.1e-08 | 4.9e-08 | 2.3e-07 | 1.033E-07 | 1.101E-07 | 3.100E-08 | 2.300E-07 |
| 27512 | 1376 | F1376K | E | 4.3e-08 | 6.3e-08 | 3.7e-08 | 4.767E-08 | 1.361E-08 | 3.700E-08 | 6.300E-08 |
| 27513 | 1376 | F1376M | E | 1.5e-07 | 4e-08 | 2.6e-08 | 7.200E-08 | 6.791E-08 | 2.600E-08 | 1.500E-07 |
| 27514 | 1376 | F1376F | E | 2.3e-08 | 5.3e-08 | 2.8e-08 | 3.467E-08 | 1.607E-08 | 2.300E-08 | 5.300E-08 |
| 27515 | 1376 | F1376P | E | 8.6e-08 | 5.1e-08 | 2e-08 | 5.233E-08 | 3.302E-08 | 2.000E-08 | 8.600E-08 |
| 27516 | 1376 | F1376S | E | 1.6e-07 | 4e-07 | 6.2e-08 | 2.073E-07 | 1.739E-07 | 6.200E-08 | 4.000E-07 |
| 27517 | 1376 | F1376T | E | 3.4e-08 | 7e-08 | 6.8e-08 | 5.733E-08 | 2.023E-08 | 3.400E-08 | 7.000E-08 |
| 27518 | 1376 | F1376W | E | 1.4e-08 | 8.5e-08 | 4.7e-08 | 4.867E-08 | 3.553E-08 | 1.400E-08 | 8.500E-08 |
| 27519 | 1376 | F1376Y | E | 1.7e-08 | 5.7e-08 | 6.3e-08 | 4.567E-08 | 2.501E-08 | 1.700E-08 | 6.300E-08 |
| 27520 | 1376 | F1376V | E | 1.1e-07 | 2.4e-08 | 5.9e-08 | 6.433E-08 | 4.325E-08 | 2.400E-08 | 1.100E-07 |
| 27521 | 1377 | Y1377A | E | 4.2e-08 | 3.3e-08 | 1.8e-08 | 3.100E-08 | 1.212E-08 | 1.800E-08 | 4.200E-08 |
| 27522 | 1377 | Y1377R | E | 2.6e-08 | 2.8e-08 | 1.2e-07 | 5.800E-08 | 5.370E-08 | 2.600E-08 | 1.200E-07 |
| 27523 | 1377 | Y1377N | E | 4.8e-08 | 3e-08 | 2.6e-08 | 3.467E-08 | 1.172E-08 | 2.600E-08 | 4.800E-08 |
| 27524 | 1377 | Y1377D | E | 2.7e-08 | 1.7e-08 | 5.5e-08 | 3.300E-08 | 1.970E-08 | 1.700E-08 | 5.500E-08 |
| 27525 | 1377 | Y1377C | E | 1.8e-08 | 2.7e-08 | 4.8e-08 | 3.100E-08 | 1.539E-08 | 1.800E-08 | 4.800E-08 |
| 27526 | 1377 | Y1377E | E | 8.8e-08 | 3.4e-08 | 7.9e-08 | 6.700E-08 | 2.893E-08 | 3.400E-08 | 8.800E-08 |
| 27527 | 1377 | Y1377Q | E | 1.5e-07 | 4.6e-08 | 4.7e-08 | 8.100E-08 | 5.976E-08 | 4.600E-08 | 1.500E-07 |
| 27528 | 1377 | Y1377G | E | 4.8e-08 | 2.8e-08 | 4.1e-08 | 3.900E-08 | 1.015E-08 | 2.800E-08 | 4.800E-08 |
| 27529 | 1377 | Y1377H | E | 8.7e-08 | 6.5e-08 | 5.7e-08 | 6.967E-08 | 1.553E-08 | 5.700E-08 | 8.700E-08 |
| 27530 | 1377 | Y1377I | E | 4.3e-08 | 2.1e-08 | 4.1e-08 | 3.500E-08 | 1.217E-08 | 2.100E-08 | 4.300E-08 |
| 27531 | 1377 | Y1377L | E | 2.6e-08 | 3.6e-08 | 2.3e-08 | 2.833E-08 | 6.807E-09 | 2.300E-08 | 3.600E-08 |
| 27532 | 1377 | Y1377K | E | 5.6e-08 | 1.6e-08 | 5.9e-08 | 4.367E-08 | 2.401E-08 | 1.600E-08 | 5.900E-08 |
| 27533 | 1377 | Y1377M | E | 4.4e-08 | 4.9e-08 | 7.2e-08 | 5.500E-08 | 1.493E-08 | 4.400E-08 | 7.200E-08 |
| 27534 | 1377 | Y1377F | E | 7.4e-08 | 2.5e-08 | 3.5e-08 | 4.467E-08 | 2.589E-08 | 2.500E-08 | 7.400E-08 |
| 27535 | 1377 | Y1377P | E | 6.3e-08 | 2.1e-08 | 5.9e-08 | 4.767E-08 | 2.318E-08 | 2.100E-08 | 6.300E-08 |
| 27536 | 1377 | Y1377S | E | 6e-08 | 6.4e-08 | 1.7e-08 | 4.700E-08 | 2.606E-08 | 1.700E-08 | 6.400E-08 |
| 27537 | 1377 | Y1377T | E | 4.6e-08 | 1.2e-07 | 5.1e-08 | 7.233E-08 | 4.136E-08 | 4.600E-08 | 1.200E-07 |
| 27538 | 1377 | Y1377W | E | 5.3e-08 | 3.7e-08 | 4.5e-08 | 4.500E-08 | 8.000E-09 | 3.700E-08 | 5.300E-08 |
| 27539 | 1377 | Y1377Y | E | 3e-08 | 5.2e-08 | 2e-08 | 3.400E-08 | 1.637E-08 | 2.000E-08 | 5.200E-08 |

| | | | | | | | | | | |
|---|---|---|---|---|---|---|---|---|---|---|
| 27540 | 1377 | Y1377V | E | 3.3e-08 | 4.6e-08 | 4.4e-08 | 4.100E-08 | 7.000E-09 | 3.300E-08 | 4.600E-08 |
| 27541 | 1378 | A1378A | E | 2.4e-08 | 3.4e-08 | 2.5e-08 | 2.767E-08 | 5.508E-09 | 2.400E-08 | 3.400E-08 |
| 27542 | 1378 | A1378R | E | 2.6e-08 | 2.6e-08 | 2.2e-08 | 2.467E-08 | 2.309E-09 | 2.200E-08 | 2.600E-08 |
| 27543 | 1378 | A1378N | E | 3.7e-08 | 4e-08 | 6.6e-08 | 4.767E-08 | 1.595E-08 | 3.700E-08 | 6.600E-08 |
| 27544 | 1378 | A1378D | E | 1.2e-07 | 1e-07 | 2.7e-08 | 8.233E-08 | 4.895E-08 | 2.700E-08 | 1.200E-07 |
| 27545 | 1378 | A1378C | E | 2.6e-08 | 1.7e-08 | 2.3e-08 | 2.200E-08 | 4.583E-09 | 1.700E-08 | 2.600E-08 |
| 27546 | 1378 | A1378E | E | 3e-08 | 2.3e-08 | 4.9e-08 | 3.400E-08 | 1.345E-08 | 2.300E-08 | 4.900E-08 |
| 27547 | 1378 | A1378Q | E | 1.3e-07 | 3.3e-08 | 4.8e-08 | 7.033E-08 | 5.221E-08 | 3.300E-08 | 1.300E-07 |
| 27548 | 1378 | A1378G | E | 3.6e-08 | 1e-07 | 4.7e-08 | 6.100E-08 | 3.422E-08 | 3.600E-08 | 1.000E-07 |
| 27549 | 1378 | A1378H | E | 8.7e-08 | 5.1e-08 | 8.6e-08 | 7.467E-08 | 2.050E-08 | 5.100E-08 | 8.700E-08 |
| 27550 | 1378 | A1378I | E | 7e-08 | 3.2e-08 | 9.9e-09 | 3.730E-08 | 3.040E-08 | 9.900E-09 | 7.000E-08 |
| 27551 | 1378 | A1378L | E | 2.8e-08 | 1.3e-07 | 6.5e-09 | 5.483E-08 | 6.598E-08 | 6.500E-09 | 1.300E-07 |
| 27552 | 1378 | A1378K | E | 6e-08 | 7.2e-08 | 4e-08 | 5.733E-08 | 1.617E-08 | 4.000E-08 | 7.200E-08 |
| 27553 | 1378 | A1378M | E | 7.2e-08 | 1.8e-08 | 1.8e-08 | 3.600E-08 | 3.118E-08 | 1.800E-08 | 7.200E-08 |
| 27554 | 1378 | A1378F | E | 4.4e-08 | 5.6e-08 | 9.4e-08 | 6.467E-08 | 2.610E-08 | 4.400E-08 | 9.400E-08 |
| 27555 | 1378 | A1378P | E | 1.8e-08 | 2.5e-08 | 2.8e-08 | 2.367E-08 | 5.132E-09 | 1.800E-08 | 2.800E-08 |
| 27556 | 1378 | A1378S | E | 4.2e-08 | 9.9e-08 | 2.5e-08 | 5.533E-08 | 3.876E-08 | 2.500E-08 | 9.900E-08 |
| 27557 | 1378 | A1378T | E | 5.2e-08 | 5.1e-08 | 3.1e-08 | 4.467E-08 | 1.185E-08 | 3.100E-08 | 5.200E-08 |
| 27558 | 1378 | A1378W | E | 3.8e-08 | 2.9e-08 | 5e-08 | 3.900E-08 | 1.054E-08 | 2.900E-08 | 5.000E-08 |
| 27559 | 1378 | A1378Y | E | 4.4e-08 | 1.6e-07 | 7.7e-08 | 9.367E-08 | 5.977E-08 | 4.400E-08 | 1.600E-07 |
| 27560 | 1378 | A1378V | E | 1.9e-08 | 2.9e-08 | 5.2e-08 | 3.333E-08 | 1.692E-08 | 1.900E-08 | 5.200E-08 |
| 27561 | 1379 | M1379A | E | 6.4e-08 | 2.9e-08 | 3.5e-08 | 4.267E-08 | 1.872E-08 | 2.900E-08 | 6.400E-08 |
| 27562 | 1379 | M1379R | E | 4.3e-08 | 6.3e-08 | 4.7e-08 | 5.100E-08 | 1.058E-08 | 4.300E-08 | 6.300E-08 |
| 27563 | 1379 | M1379N | E | 5.4e-08 | 8.2e-08 | 9.4e-08 | 7.667E-08 | 2.053E-08 | 5.400E-08 | 9.400E-08 |
| 27564 | 1379 | M1379D | E | 4e-08 | 1.7e-08 | 5.7e-08 | 3.800E-08 | 2.007E-08 | 1.700E-08 | 5.700E-08 |
| 27565 | 1379 | M1379C | E | 7.4e-08 | 7.9e-08 | 3.3e-08 | 6.200E-08 | 2.524E-08 | 3.300E-08 | 7.900E-08 |
| 27566 | 1379 | M1379E | E | 3.2e-08 | 6.6e-08 | 2.5e-08 | 4.100E-08 | 2.193E-08 | 2.500E-08 | 6.600E-08 |
| 27567 | 1379 | M1379Q | E | 2.8e-08 | 4.4e-08 | 4.5e-08 | 3.900E-08 | 9.539E-09 | 2.800E-08 | 4.500E-08 |
| 27568 | 1379 | M1379G | E | 4e-08 | 3.7e-08 | 3.6e-08 | 3.767E-08 | 2.082E-09 | 3.600E-08 | 4.000E-08 |
| 27569 | 1379 | M1379H | E | 4.5e-08 | 1.2e-08 | 2.8e-08 | 2.833E-08 | 1.650E-08 | 1.200E-08 | 4.500E-08 |
| 27570 | 1379 | M1379I | E | 2.5e-08 | 5.5e-08 | 2.9e-08 | 3.633E-08 | 1.629E-08 | 2.500E-08 | 5.500E-08 |
| 27571 | 1379 | M1379L | E | 5.2e-08 | 8.5e-08 | 3.7e-08 | 5.800E-08 | 2.456E-08 | 3.700E-08 | 8.500E-08 |
| 27572 | 1379 | M1379K | E | 4.1e-08 | 2.8e-08 | 4.8e-08 | 3.900E-08 | 1.015E-08 | 2.800E-08 | 4.800E-08 |
| 27573 | 1379 | M1379M | E | 8.4e-08 | 3.5e-08 | 2.6e-08 | 4.833E-08 | 3.121E-08 | 2.600E-08 | 8.400E-08 |
| 27574 | 1379 | M1379F | E | 6.2e-08 | 2.4e-08 | 2e-08 | 3.533E-08 | 2.318E-08 | 2.000E-08 | 6.200E-08 |
| 27575 | 1379 | M1379P | E | 3.6e-08 | 5.2e-08 | 2.7e-08 | 3.833E-08 | 1.266E-08 | 2.700E-08 | 5.200E-08 |
| 27576 | 1379 | M1379S | E | 7.4e-08 | 5.7e-08 | 5.2e-08 | 6.100E-08 | 1.153E-08 | 5.200E-08 | 7.400E-08 |
| 27577 | 1379 | M1379T | E | 1.7e-08 | 6.4e-08 | 3.2e-08 | 3.767E-08 | 2.401E-08 | 1.700E-08 | 6.400E-08 |
| 27578 | 1379 | M1379W | E | 6.5e-08 | 5.5e-08 | 9e-08 | 7.000E-08 | 1.803E-08 | 5.500E-08 | 9.000E-08 |
| 27579 | 1379 | M1379Y | E | 2.9e-08 | 4.7e-08 | 6.2e-08 | 4.600E-08 | 1.652E-08 | 2.900E-08 | 6.200E-08 |
| 27580 | 1379 | M1379V | E | 2.6e-08 | 4.2e-08 | 2.8e-08 | 3.200E-08 | 8.718E-09 | 2.600E-08 | 4.200E-08 |
| 27581 | 1380 | D1380A | E | 6e-08 | 4.2e-08 | 4.2e-08 | 4.800E-08 | 1.039E-08 | 4.200E-08 | 6.000E-08 |
| 27582 | 1380 | D1380R | E | 7.4e-08 | 3.5e-08 | 1.8e-08 | 4.233E-08 | 2.871E-08 | 1.800E-08 | 7.400E-08 |
| 27583 | 1380 | D1380N | E | 9.9e-08 | 7.9e-08 | 7.9e-08 | 8.567E-08 | 1.155E-08 | 7.900E-08 | 9.900E-08 |
| 27584 | 1380 | D1380D | E | 6.3e-08 | 4.3e-08 | 2.7e-08 | 4.433E-08 | 1.804E-08 | 2.700E-08 | 6.300E-08 |
| 27585 | 1380 | D1380C | E | 7.6e-08 | 1e-08 | 1.3e-08 | 3.300E-08 | 3.727E-08 | 1.000E-08 | 7.600E-08 |
| 27586 | 1380 | D1380E | E | 6.5e-08 | 1.2e-07 | 3.2e-08 | 7.233E-08 | 4.446E-08 | 3.200E-08 | 1.200E-07 |
| 27587 | 1380 | D1380Q | E | 3.7e-08 | 2e-08 | 6.6e-08 | 4.100E-08 | 2.326E-08 | 2.000E-08 | 6.600E-08 |
| 27588 | 1380 | D1380G | E | 1.1e-07 | 4.7e-08 | 3.7e-08 | 6.467E-08 | 3.958E-08 | 3.700E-08 | 1.100E-07 |
| 27589 | 1380 | D1380H | E | 2e-08 | 3.1e-08 | 2.1e-08 | 2.400E-08 | 6.083E-09 | 2.000E-08 | 3.100E-08 |
| 27590 | 1380 | D1380I | E | 1.1e-07 | 4.2e-08 | 1.6e-07 | 1.040E-07 | 5.923E-08 | 4.200E-08 | 1.600E-07 |
| 27591 | 1380 | D1380L | E | 1.1e-07 | 6.2e-08 | 3.6e-08 | 6.933E-08 | 3.754E-08 | 3.600E-08 | 1.100E-07 |
| 27592 | 1380 | D1380K | E | 3.1e-08 | 3e-08 | 2.2e-08 | 2.767E-08 | 4.933E-09 | 2.200E-08 | 3.100E-08 |
| 27593 | 1380 | D1380M | E | 7.2e-08 | 1.1e-07 | 5.7e-08 | 7.967E-08 | 2.732E-08 | 5.700E-08 | 1.100E-07 |
| 27594 | 1380 | D1380F | E | 3.9e-08 | 5.5e-08 | 1.9e-08 | 3.767E-08 | 1.804E-08 | 1.900E-08 | 5.500E-08 |
| 27595 | 1380 | D1380P | E | 5e-08 | 5.9e-08 | 5.1e-08 | 5.333E-08 | 4.933E-09 | 5.000E-08 | 5.900E-08 |
| 27596 | 1380 | D1380S | E | 2.9e-08 | 7.1e-08 | 5e-08 | 5.000E-08 | 2.100E-08 | 2.900E-08 | 7.100E-08 |
| 27597 | 1380 | D1380T | E | 3.1e-08 | 1.9e-08 | 2.5e-08 | 2.500E-08 | 6.000E-09 | 1.900E-08 | 3.100E-08 |
| 27598 | 1380 | D1380W | E | 7.3e-08 | 6.2e-08 | 1.2e-08 | 4.900E-08 | 3.251E-08 | 1.200E-08 | 7.300E-08 |
| 27599 | 1380 | D1380Y | E | 5.7e-09 | 5.9e-08 | 3.1e-08 | 3.190E-08 | 2.666E-08 | 5.700E-09 | 5.900E-08 |
| 27600 | 1380 | D1380V | E | 2.4e-08 | 1e-08 | 3.5e-08 | 2.300E-08 | 1.253E-08 | 1.000E-08 | 3.500E-08 |
| 27601 | 1381 | Y1381A | E | 3.5e-08 | 4.1e-08 | 3.6e-08 | 3.733E-08 | 3.215E-09 | 3.500E-08 | 4.100E-08 |
| 27602 | 1381 | Y1381R | E | 5.1e-08 | 4.5e-08 | 4.4e-08 | 4.667E-08 | 3.786E-09 | 4.400E-08 | 5.100E-08 |
| 27603 | 1381 | Y1381N | E | 1.7e-08 | 2.7e-08 | 2.8e-08 | 2.400E-08 | 6.083E-09 | 1.700E-08 | 2.800E-08 |
| 27604 | 1381 | Y1381D | E | 5.3e-08 | 5.8e-08 | 3.7e-08 | 4.933E-08 | 1.097E-08 | 3.700E-08 | 5.800E-08 |
| 27605 | 1381 | Y1381C | E | 5.4e-08 | 6.2e-08 | 2.8e-08 | 4.800E-08 | 1.778E-08 | 2.800E-08 | 6.200E-08 |
| 27606 | 1381 | Y1381E | E | 2.3e-08 | 3.3e-08 | 5.1e-08 | 3.567E-08 | 1.419E-08 | 2.300E-08 | 5.100E-08 |
| 27607 | 1381 | Y1381Q | E | 5.9e-08 | 8.4e-08 | 5e-08 | 6.433E-08 | 1.762E-08 | 5.000E-08 | 8.400E-08 |
| 27608 | 1381 | Y1381G | E | 6.3e-08 | 4.4e-08 | 1.6e-08 | 4.100E-08 | 2.364E-08 | 1.600E-08 | 6.300E-08 |
| 27609 | 1381 | Y1381H | E | 5.4e-08 | 6.6e-08 | 3e-08 | 5.000E-08 | 1.833E-08 | 3.000E-08 | 6.600E-08 |
| 27610 | 1381 | Y1381I | E | 4.3e-08 | 4.8e-08 | 4.6e-08 | 4.567E-08 | 2.517E-09 | 4.300E-08 | 4.800E-08 |
| 27611 | 1381 | Y1381L | E | 2.8e-08 | 2.1e-08 | 5.2e-08 | 3.367E-08 | 1.626E-08 | 2.100E-08 | 5.200E-08 |
| 27612 | 1381 | Y1381K | E | 1.6e-07 | 3.9e-08 | 2.6e-08 | 7.500E-08 | 7.390E-08 | 2.600E-08 | 1.600E-07 |
| 27613 | 1381 | Y1381M | E | 2.2e-08 | 4.9e-08 | 2.1e-08 | 3.067E-08 | 1.589E-08 | 2.100E-08 | 4.900E-08 |
| 27614 | 1381 | Y1381F | E | 1.2e-08 | 3.9e-08 | 4.1e-08 | 3.067E-08 | 1.620E-08 | 1.200E-08 | 4.100E-08 |
| 27615 | 1381 | Y1381P | E | 3e-08 | 2.9e-08 | 4e-08 | 3.300E-08 | 6.083E-09 | 2.900E-08 | 4.000E-08 |
| 27616 | 1381 | Y1381S | E | 1.5e-08 | 6.4e-08 | 3.2e-08 | 3.700E-08 | 2.488E-08 | 1.500E-08 | 6.400E-08 |
| 27617 | 1381 | Y1381T | E | 1.1e-07 | 3.5e-08 | 5.3e-08 | 6.600E-08 | 3.915E-08 | 3.500E-08 | 1.100E-07 |
| 27618 | 1381 | Y1381W | E | 1.1e-07 | 2.4e-08 | 4.7e-08 | 6.033E-08 | 4.452E-08 | 2.400E-08 | 1.100E-07 |
| 27619 | 1381 | Y1381Y | E | 1.6e-08 | 1e-07 | 9.1e-08 | 6.900E-08 | 4.612E-08 | 1.600E-08 | 1.000E-07 |
| 27620 | 1381 | Y1381V | E | 2.4e-08 | 1.9e-08 | 5.4e-08 | 3.233E-08 | 1.893E-08 | 1.900E-08 | 5.400E-08 |
| 27621 | 1382 | W1382A | E | 4.8e-08 | 3e-08 | 2.4e-08 | 3.400E-08 | 1.249E-08 | 2.400E-08 | 4.800E-08 |
| 27622 | 1382 | W1382R | E | 2.2e-08 | 1.2e-07 | 5.4e-08 | 6.533E-08 | 4.997E-08 | 2.200E-08 | 1.200E-07 |
| 27623 | 1382 | W1382N | E | 5.9e-08 | 7.4e-08 | 1.6e-07 | 9.767E-08 | 5.450E-08 | 5.900E-08 | 1.600E-07 |
| 27624 | 1382 | W1382D | E | 4.5e-08 | 9.3e-08 | 4.3e-08 | 6.033E-08 | 2.831E-08 | 4.300E-08 | 9.300E-08 |

| | | | | | | | | | | |
|---|---|---|---|---|---|---|---|---|---|---|
| 27625 | 1382 | W1382C | E | 5.5e-08 | 6.5e-08 | 1.3e-07 | 8.333E-08 | 4.072E-08 | 5.500E-08 | 1.300E-07 |
| 27626 | 1382 | W1382E | E | 7.6e-08 | 8.4e-08 | 4.8e-08 | 6.933E-08 | 1.890E-08 | 4.800E-08 | 8.400E-08 |
| 27627 | 1382 | W1382Q | E | 6.6e-08 | 2.7e-08 | 2.1e-08 | 3.800E-08 | 2.443E-08 | 2.100E-08 | 6.600E-08 |
| 27628 | 1382 | W1382G | E | 6.3e-09 | 8.8e-08 | 8.6e-08 | 6.010E-08 | 4.660E-08 | 6.300E-09 | 8.800E-08 |
| 27629 | 1382 | W1382H | E | 1.7e-08 | 5.6e-08 | 2.8e-08 | 3.367E-08 | 2.011E-08 | 1.700E-08 | 5.600E-08 |
| 27630 | 1382 | W1382I | E | 4.5e-08 | 5.5e-08 | 4.5e-08 | 4.833E-08 | 5.774E-09 | 4.500E-08 | 5.500E-08 |
| 27631 | 1382 | W1382L | E | 3e-08 | 3.5e-08 | 3.1e-08 | 3.200E-08 | 2.646E-09 | 3.000E-08 | 3.500E-08 |
| 27632 | 1382 | W1382K | E | 3.7e-08 | 8.7e-08 | 3.9e-08 | 5.433E-08 | 2.831E-08 | 3.700E-08 | 8.700E-08 |
| 27633 | 1382 | W1382M | E | 4.7e-08 | 2.3e-08 | 1.3e-08 | 2.767E-08 | 1.747E-08 | 1.300E-08 | 4.700E-08 |
| 27634 | 1382 | W1382F | E | 5.9e-08 | 6.7e-08 | 3.8e-08 | 5.467E-08 | 1.498E-08 | 3.800E-08 | 6.700E-08 |
| 27635 | 1382 | W1382P | E | 2.4e-08 | 6.7e-08 | 3.9e-08 | 4.333E-08 | 2.183E-08 | 2.400E-08 | 6.700E-08 |
| 27636 | 1382 | W1382S | E | 3.5e-08 | 3.2e-08 | 3.5e-08 | 3.400E-08 | 1.732E-09 | 3.200E-08 | 3.500E-08 |
| 27637 | 1382 | W1382T | E | 2.9e-08 | 6.7e-08 | 4.7e-08 | 4.767E-08 | 1.901E-08 | 2.900E-08 | 6.700E-08 |
| 27638 | 1382 | W1382W | E | 2.9e-08 | 2.5e-08 | 2.8e-08 | 2.733E-08 | 2.082E-09 | 2.500E-08 | 2.900E-08 |
| 27639 | 1382 | W1382Y | E | 1.9e-08 | 5.4e-08 | 4.2e-08 | 3.833E-08 | 1.779E-08 | 1.900E-08 | 5.400E-08 |
| 27640 | 1382 | W1382V | E | 2.9e-08 | 3e-08 | 1.3e-07 | 6.300E-08 | 5.803E-08 | 2.900E-08 | 1.300E-07 |
| 27641 | 1383 | G1383A | E | 2.6e-08 | 5.3e-08 | 3e-08 | 3.633E-08 | 1.457E-08 | 2.600E-08 | 5.300E-08 |
| 27642 | 1383 | G1383R | E | 2.2e-08 | 6e-08 | 4.7e-08 | 4.300E-08 | 1.931E-08 | 2.200E-08 | 6.000E-08 |
| 27643 | 1383 | G1383N | E | 3.9e-08 | 4.3e-08 | 7.3e-08 | 5.167E-08 | 1.858E-08 | 3.900E-08 | 7.300E-08 |
| 27644 | 1383 | G1383D | E | 2.1e-08 | 4.4e-08 | 3e-08 | 3.167E-08 | 1.159E-08 | 2.100E-08 | 4.400E-08 |
| 27645 | 1383 | G1383C | E | 4e-08 | 1.1e-07 | 2.3e-08 | 5.767E-08 | 4.611E-08 | 2.300E-08 | 1.100E-07 |
| 27646 | 1383 | G1383E | E | 2.7e-08 | 4.7e-08 | 4.7e-08 | 4.033E-08 | 1.155E-08 | 2.700E-08 | 4.700E-08 |
| 27647 | 1383 | G1383Q | E | 4.5e-08 | 2.8e-08 | 7e-08 | 4.767E-08 | 2.113E-08 | 2.800E-08 | 7.000E-08 |
| 27648 | 1383 | G1383G | E | 7.3e-08 | 9.2e-08 | 3.9e-08 | 6.800E-08 | 2.685E-08 | 3.900E-08 | 9.200E-08 |
| 27649 | 1383 | G1383H | E | 6.5e-08 | 2.9e-08 | 1.2e-08 | 3.533E-08 | 2.706E-08 | 1.200E-08 | 6.500E-08 |
| 27650 | 1383 | G1383I | E | 4.7e-08 | 4.9e-08 | 1.4e-08 | 3.667E-08 | 1.966E-08 | 1.400E-08 | 4.900E-08 |
| 27651 | 1383 | G1383L | E | 1.6e-07 | 5.4e-08 | 7.1e-08 | 9.500E-08 | 5.693E-08 | 5.400E-08 | 1.600E-07 |
| 27652 | 1383 | G1383K | E | 8.4e-08 | 1.6e-08 | 3e-08 | 4.333E-08 | 3.591E-08 | 1.600E-08 | 8.400E-08 |
| 27653 | 1383 | G1383M | E | 4.7e-08 | 1.6e-08 | 7.1e-08 | 4.467E-08 | 2.757E-08 | 1.600E-08 | 7.100E-08 |
| 27654 | 1383 | G1383F | E | 6e-08 | 4.3e-08 | 2.9e-08 | 4.400E-08 | 1.552E-08 | 2.900E-08 | 6.000E-08 |
| 27655 | 1383 | G1383P | E | 1.5e-08 | 5.1e-08 | 1e-07 | 5.533E-08 | 4.267E-08 | 1.500E-08 | 1.000E-07 |
| 27656 | 1383 | G1383S | E | 5.4e-08 | 2.8e-08 | 4.7e-08 | 4.300E-08 | 1.345E-08 | 2.800E-08 | 5.400E-08 |
| 27657 | 1383 | G1383T | E | 7.5e-08 | 5.7e-08 | 3.6e-08 | 5.600E-08 | 1.952E-08 | 3.600E-08 | 7.500E-08 |
| 27658 | 1383 | G1383W | E | 7.2e-08 | 2.7e-08 | 2.5e-08 | 4.133E-08 | 2.658E-08 | 2.500E-08 | 7.200E-08 |
| 27659 | 1383 | G1383Y | E | 4.7e-08 | 4.1e-08 | 3.8e-08 | 4.200E-08 | 4.583E-09 | 3.800E-08 | 4.700E-08 |
| 27660 | 1383 | G1383V | E | 6.4e-08 | 5.9e-08 | 1.6e-07 | 9.433E-08 | 5.692E-08 | 5.900E-08 | 1.600E-07 |
| 27661 | 1384 | Q1384A | E | 3.5e-08 | 3.3e-08 | 2.8e-08 | 3.200E-08 | 3.606E-09 | 2.800E-08 | 3.500E-08 |
| 27662 | 1384 | Q1384R | E | 3.5e-08 | 1e-07 | 3.6e-08 | 5.700E-08 | 3.724E-08 | 3.500E-08 | 1.000E-07 |
| 27663 | 1384 | Q1384N | E | 7e-08 | 5.3e-08 | 9.3e-08 | 7.200E-08 | 2.007E-08 | 5.300E-08 | 9.300E-08 |
| 27664 | 1384 | Q1384D | E | 2.1e-08 | 5.2e-08 | 5.2e-08 | 4.167E-08 | 1.790E-08 | 2.100E-08 | 5.200E-08 |
| 27665 | 1384 | Q1384C | E | 5.2e-08 | 2e-08 | 5.8e-08 | 4.333E-08 | 2.043E-08 | 2.000E-08 | 5.800E-08 |
| 27666 | 1384 | Q1384E | E | 2.7e-08 | 2.1e-08 | 9e-08 | 4.600E-08 | 3.822E-08 | 2.100E-08 | 9.000E-08 |
| 27667 | 1384 | Q1384Q | E | 2.6e-08 | 3e-08 | 6e-08 | 3.867E-08 | 1.858E-08 | 2.600E-08 | 6.000E-08 |
| 27668 | 1384 | Q1384G | E | 1.2e-08 | 5.1e-08 | 4.5e-08 | 3.600E-08 | 2.100E-08 | 1.200E-08 | 5.100E-08 |
| 27669 | 1384 | Q1384H | E | 2.9e-08 | 5.4e-08 | 2.9e-08 | 3.733E-08 | 1.443E-08 | 2.900E-08 | 5.400E-08 |
| 27670 | 1384 | Q1384I | E | 4.1e-08 | 9.9e-08 | 2.8e-08 | 5.600E-08 | 3.780E-08 | 2.800E-08 | 9.900E-08 |
| 27671 | 1384 | Q1384L | E | 2.9e-08 | 4e-08 | 7.5e-08 | 4.800E-08 | 2.402E-08 | 2.900E-08 | 7.500E-08 |
| 27672 | 1384 | Q1384K | E | 3e-08 | 5.9e-08 | 5.5e-08 | 4.800E-08 | 1.572E-08 | 3.000E-08 | 5.900E-08 |
| 27673 | 1384 | Q1384M | E | 4.4e-08 | 4.9e-08 | 4.1e-08 | 4.467E-08 | 4.041E-09 | 4.100E-08 | 4.900E-08 |
| 27674 | 1384 | Q1384F | E | 3.4e-08 | 5.2e-08 | 3.9e-08 | 4.167E-08 | 9.292E-09 | 3.400E-08 | 5.200E-08 |
| 27675 | 1384 | Q1384P | E | 6.9e-08 | 1.4e-08 | 4.5e-08 | 4.267E-08 | 2.757E-08 | 1.400E-08 | 6.900E-08 |
| 27676 | 1384 | Q1384S | E | 5.6e-08 | 2.9e-08 | 1.9e-08 | 3.467E-08 | 1.914E-08 | 1.900E-08 | 5.600E-08 |
| 27677 | 1384 | Q1384T | E | 5.2e-08 | 2.9e-08 | 4.6e-08 | 4.233E-08 | 1.193E-08 | 2.900E-08 | 5.200E-08 |
| 27678 | 1384 | Q1384W | E | 4.5e-08 | 3.4e-08 | 2.7e-08 | 3.533E-08 | 9.074E-09 | 2.700E-08 | 4.500E-08 |
| 27679 | 1384 | Q1384Y | E | 6.7e-08 | 6e-08 | 8.2e-08 | 6.967E-08 | 1.124E-08 | 6.000E-08 | 8.200E-08 |
| 27680 | 1384 | Q1384V | E | 5.8e-08 | 3.9e-08 | 3e-08 | 4.233E-08 | 1.429E-08 | 3.000E-08 | 5.800E-08 |
| 27681 | 1385 | G1385A | E | 8.9e-08 | 1e-07 | 6.2e-08 | 8.367E-08 | 1.955E-08 | 6.200E-08 | 1.000E-07 |
| 27682 | 1385 | G1385R | E | 6.5e-08 | 2.6e-08 | 7.4e-08 | 5.500E-08 | 2.551E-08 | 2.600E-08 | 7.400E-08 |
| 27683 | 1385 | G1385N | E | 5.1e-08 | 4.8e-08 | 7.6e-08 | 5.833E-08 | 1.537E-08 | 4.800E-08 | 7.600E-08 |
| 27684 | 1385 | G1385D | E | 4.2e-08 | 4.6e-08 | 3.2e-08 | 4.000E-08 | 7.211E-09 | 3.200E-08 | 4.600E-08 |
| 27685 | 1385 | G1385C | E | 4.9e-08 | 5.1e-08 | 2.2e-08 | 4.067E-08 | 1.620E-08 | 2.200E-08 | 5.100E-08 |
| 27686 | 1385 | G1385E | E | 6.3e-08 | 4.5e-08 | 4.3e-08 | 5.033E-08 | 1.102E-08 | 4.300E-08 | 6.300E-08 |
| 27687 | 1385 | G1385Q | E | 4.3e-08 | 4.7e-08 | 5.3e-08 | 4.767E-08 | 5.033E-09 | 4.300E-08 | 5.300E-08 |
| 27688 | 1385 | G1385G | E | 4.2e-08 | 3.2e-08 | 6.9e-08 | 4.767E-08 | 1.914E-08 | 3.200E-08 | 6.900E-08 |
| 27689 | 1385 | G1385H | E | 4e-08 | 6.9e-08 | 5.3e-08 | 5.400E-08 | 1.453E-08 | 4.000E-08 | 6.900E-08 |
| 27690 | 1385 | G1385I | E | 8.6e-08 | 1.7e-08 | 8.7e-08 | 6.333E-08 | 4.013E-08 | 1.700E-08 | 8.700E-08 |
| 27691 | 1385 | G1385L | E | 5e-08 | 4.4e-08 | 5e-08 | 4.800E-08 | 3.464E-09 | 4.400E-08 | 5.000E-08 |
| 27692 | 1385 | G1385K | E | 1.2e-07 | 4.5e-08 | 4.3e-08 | 6.933E-08 | 4.389E-08 | 4.300E-08 | 1.200E-07 |
| 27693 | 1385 | G1385M | E | 2.5e-08 | 3.9e-08 | 6.1e-08 | 4.167E-08 | 1.815E-08 | 2.500E-08 | 6.100E-08 |
| 27694 | 1385 | G1385F | E | 6.4e-08 | 2.8e-08 | 7.6e-08 | 5.600E-08 | 2.498E-08 | 2.800E-08 | 7.600E-08 |
| 27695 | 1385 | G1385P | E | 2.8e-08 | 3.6e-08 | 1.1e-07 | 5.800E-08 | 4.521E-08 | 2.800E-08 | 1.100E-07 |
| 27696 | 1385 | G1385S | E | 5.5e-08 | 9.5e-08 | 9e-08 | 8.000E-08 | 2.179E-08 | 5.500E-08 | 9.500E-08 |
| 27697 | 1385 | G1385T | E | 3.2e-08 | 2e-08 | 7e-08 | 4.067E-08 | 2.610E-08 | 2.000E-08 | 7.000E-08 |
| 27698 | 1385 | G1385W | E | 5.1e-08 | 1.3e-07 | 1.9e-08 | 6.667E-08 | 5.713E-08 | 1.900E-08 | 1.300E-07 |
| 27699 | 1385 | G1385Y | E | 5e-08 | 1.1e-08 | 2.6e-08 | 2.900E-08 | 1.967E-08 | 1.100E-08 | 5.000E-08 |
| 27700 | 1385 | G1385V | E | 7.1e-08 | 1.2e-08 | 1.8e-08 | 3.367E-08 | 3.247E-08 | 1.200E-08 | 7.100E-08 |
| 27701 | 1386 | T1386A | E | 1e-07 | 5.5e-08 | 1.2e-07 | 9.167E-08 | 3.329E-08 | 5.500E-08 | 1.200E-07 |
| 27702 | 1386 | T1386R | E | 5.3e-08 | 4.9e-08 | 8.5e-08 | 6.233E-08 | 1.973E-08 | 4.900E-08 | 8.500E-08 |
| 27703 | 1386 | T1386N | E | 3.5e-08 | 1.6e-07 | 6.4e-08 | 8.633E-08 | 6.542E-08 | 3.500E-08 | 1.600E-07 |
| 27704 | 1386 | T1386D | E | 3.7e-08 | 2.6e-08 | 1e-07 | 5.433E-08 | 3.993E-08 | 2.600E-08 | 1.000E-07 |
| 27705 | 1386 | T1386C | E | 5.3e-08 | 9.8e-08 | 2.8e-08 | 5.967E-08 | 3.547E-08 | 2.800E-08 | 9.800E-08 |
| 27706 | 1386 | T1386E | E | 3.8e-08 | 4.6e-08 | 7.6e-08 | 5.333E-08 | 2.003E-08 | 3.800E-08 | 7.600E-08 |
| 27707 | 1386 | T1386Q | E | 4.9e-08 | 2.2e-07 | 3.4e-08 | 1.010E-07 | 1.033E-07 | 3.400E-08 | 2.200E-07 |
| 27708 | 1386 | T1386G | E | 5.4e-08 | 2.1e-08 | 7.2e-08 | 4.900E-08 | 2.587E-08 | 2.100E-08 | 7.200E-08 |
| 27709 | 1386 | T1386H | E | 7.3e-08 | 1.3e-08 | 3.9e-08 | 4.167E-08 | 3.009E-08 | 1.300E-08 | 7.300E-08 |

| | | | | | | | | | | |
|---|---|---|---|---|---|---|---|---|---|---|
| 27710 | 1386 | T1386I | E | 6.8e-08 | 5.2e-08 | 3.3e-08 | 5.100E-08 | 1.752E-08 | 3.300E-08 | 6.800E-08 |
| 27711 | 1386 | T1386L | E | 2.9e-08 | 1.4e-08 | 3.2e-08 | 2.500E-08 | 9.644E-09 | 1.400E-08 | 3.200E-08 |
| 27712 | 1386 | T1386K | E | 2.6e-08 | 4.1e-08 | 1.9e-08 | 2.867E-08 | 1.124E-08 | 1.900E-08 | 4.100E-08 |
| 27713 | 1386 | T1386M | E | 6.3e-08 | 2.4e-08 | 5.5e-08 | 4.733E-08 | 2.060E-08 | 2.400E-08 | 6.300E-08 |
| 27714 | 1386 | T1386F | E | 3.9e-08 | 5.3e-08 | 3.1e-08 | 4.100E-08 | 1.114E-08 | 3.100E-08 | 5.300E-08 |
| 27715 | 1386 | T1386P | E | 7.4e-08 | 3.6e-08 | 8.5e-08 | 6.500E-08 | 2.571E-08 | 3.600E-08 | 8.500E-08 |
| 27716 | 1386 | T1386S | E | 8.1e-08 | 7.8e-08 | 7.1e-08 | 7.667E-08 | 5.132E-09 | 7.100E-08 | 8.100E-08 |
| 27717 | 1386 | T1386T | E | 3.6e-08 | 6.3e-08 | 6.6e-08 | 5.500E-08 | 1.652E-08 | 3.600E-08 | 6.600E-08 |
| 27718 | 1386 | T1386W | E | 1.9e-08 | 1.3e-08 | 2.6e-08 | 1.933E-08 | 6.506E-09 | 1.300E-08 | 2.600E-08 |
| 27719 | 1386 | T1386Y | E | 1.2e-07 | 4.4e-08 | 3.8e-08 | 6.733E-08 | 4.571E-08 | 3.800E-08 | 1.200E-07 |
| 27720 | 1386 | T1386V | E | 4.5e-08 | 5.6e-08 | 3.2e-08 | 4.433E-08 | 1.201E-08 | 3.200E-08 | 5.600E-08 |
| 27721 | 1387 | L1387A | E | 1.4e-08 | 4.2e-08 | 2.8e-08 | 2.800E-08 | 1.400E-08 | 1.400E-08 | 4.200E-08 |
| 27722 | 1387 | L1387R | E | 2.7e-08 | 7.3e-08 | 3.3e-08 | 4.433E-08 | 2.501E-08 | 2.700E-08 | 7.300E-08 |
| 27723 | 1387 | L1387N | E | 5.5e-08 | 3e-08 | 1.7e-07 | 8.500E-08 | 7.467E-08 | 3.000E-08 | 1.700E-07 |
| 27724 | 1387 | L1387D | E | 3.5e-08 | 6.7e-08 | 2.5e-08 | 4.233E-08 | 2.194E-08 | 2.500E-08 | 6.700E-08 |
| 27725 | 1387 | L1387C | E | 1.9e-08 | 4.6e-08 | 3.4e-08 | 3.300E-08 | 1.353E-08 | 1.900E-08 | 4.600E-08 |
| 27726 | 1387 | L1387E | E | 2.5e-08 | 7e-08 | 5.7e-08 | 5.067E-08 | 2.316E-08 | 2.500E-08 | 7.000E-08 |
| 27727 | 1387 | L1387Q | E | 1.9e-08 | 2e-08 | 2.9e-08 | 2.267E-08 | 5.508E-09 | 1.900E-08 | 2.900E-08 |
| 27728 | 1387 | L1387G | E | 1.4e-07 | 5.8e-08 | 2.9e-08 | 7.567E-08 | 5.757E-08 | 2.900E-08 | 1.400E-07 |
| 27729 | 1387 | L1387H | E | 1.6e-08 | 4.6e-08 | 2.5e-08 | 2.900E-08 | 1.539E-08 | 1.600E-08 | 4.600E-08 |
| 27730 | 1387 | L1387I | E | 1.6e-07 | 4.4e-08 | 2.1e-08 | 7.500E-08 | 7.451E-08 | 2.100E-08 | 1.600E-07 |
| 27731 | 1387 | L1387L | E | 8.5e-08 | 3.2e-08 | 3.8e-08 | 5.167E-08 | 2.902E-08 | 3.200E-08 | 8.500E-08 |
| 27732 | 1387 | L1387K | E | 3.5e-08 | 5.1e-08 | 3e-08 | 3.867E-08 | 1.097E-08 | 3.000E-08 | 5.100E-08 |
| 27733 | 1387 | L1387M | E | 4.5e-08 | 5.9e-08 | 3.7e-08 | 4.700E-08 | 1.114E-08 | 3.700E-08 | 5.900E-08 |
| 27734 | 1387 | L1387F | E | 2.4e-08 | 6.8e-08 | 4.7e-08 | 4.633E-08 | 2.201E-08 | 2.400E-08 | 6.800E-08 |
| 27735 | 1387 | L1387P | E | 4.1e-08 | 1.4e-08 | 7e-08 | 4.167E-08 | 2.801E-08 | 1.400E-08 | 7.000E-08 |
| 27736 | 1387 | L1387S | E | 4.4e-08 | 9e-08 | 9.4e-08 | 7.600E-08 | 2.778E-08 | 4.400E-08 | 9.400E-08 |
| 27737 | 1387 | L1387T | E | 8.3e-08 | 7.8e-08 | 5.5e-08 | 7.200E-08 | 1.493E-08 | 5.500E-08 | 8.300E-08 |
| 27738 | 1387 | L1387W | E | 3.8e-08 | 2.9e-08 | 1.4e-08 | 2.700E-08 | 1.212E-08 | 1.400E-08 | 3.800E-08 |
| 27739 | 1387 | L1387Y | E | 4.1e-08 | 3.1e-08 | 7.9e-09 | 2.663E-08 | 1.698E-08 | 7.900E-09 | 4.100E-08 |
| 27740 | 1387 | L1387V | E | 1.4e-08 | 3.2e-08 | 5.5e-08 | 3.367E-08 | 2.055E-08 | 1.400E-08 | 5.500E-08 |
| 27741 | 1388 | V1388A | E | 2.2e-07 | 2.5e-08 | 2.9e-08 | 9.133E-08 | 1.114E-07 | 2.500E-08 | 2.200E-07 |
| 27742 | 1388 | V1388R | E | 5.6e-08 | 8.9e-09 | 4.5e-08 | 3.663E-08 | 2.464E-08 | 8.900E-09 | 5.600E-08 |
| 27743 | 1388 | V1388N | E | 5.8e-08 | 2.4e-08 | 2.8e-08 | 3.667E-08 | 1.858E-08 | 2.400E-08 | 5.800E-08 |
| 27744 | 1388 | V1388D | E | 8.5e-08 | 5.6e-08 | 6.5e-08 | 6.867E-08 | 1.484E-08 | 5.600E-08 | 8.500E-08 |
| 27745 | 1388 | V1388C | E | 6.8e-08 | 7.4e-08 | 5e-08 | 6.400E-08 | 1.249E-08 | 5.000E-08 | 7.400E-08 |
| 27746 | 1388 | V1388E | E | 5.9e-08 | 1e-07 | 2.2e-08 | 6.033E-08 | 3.902E-08 | 2.200E-08 | 1.000E-07 |
| 27747 | 1388 | V1388Q | E | 5.2e-08 | 1.3e-08 | 1.2e-07 | 6.167E-08 | 5.415E-08 | 1.300E-08 | 1.200E-07 |
| 27748 | 1388 | V1388G | E | 1.5e-07 | 4.2e-08 | 2.2e-08 | 7.133E-08 | 6.886E-08 | 2.200E-08 | 1.500E-07 |
| 27749 | 1388 | V1388H | E | 5.8e-08 | 3.7e-08 | 3.2e-08 | 4.233E-08 | 1.380E-08 | 3.200E-08 | 5.800E-08 |
| 27750 | 1388 | V1388I | E | 3.8e-08 | 4.7e-08 | 2.6e-08 | 3.700E-08 | 1.054E-08 | 2.600E-08 | 4.700E-08 |
| 27751 | 1388 | V1388L | E | 2.1e-08 | 3.2e-08 | 8.5e-08 | 4.600E-08 | 3.422E-08 | 2.100E-08 | 8.500E-08 |
| 27752 | 1388 | V1388K | E | 5.5e-08 | 8e-08 | 4.1e-08 | 5.867E-08 | 1.976E-08 | 4.100E-08 | 8.000E-08 |
| 27753 | 1388 | V1388M | E | 3.7e-08 | 1.5e-08 | 3.9e-08 | 3.033E-08 | 1.332E-08 | 1.500E-08 | 3.900E-08 |
| 27754 | 1388 | V1388F | E | 9.4e-08 | 5.2e-08 | 5.4e-08 | 6.667E-08 | 2.369E-08 | 5.200E-08 | 9.400E-08 |
| 27755 | 1388 | V1388P | E | 4.3e-08 | 6.7e-08 | 3.7e-08 | 4.900E-08 | 1.587E-08 | 3.700E-08 | 6.700E-08 |
| 27756 | 1388 | V1388S | E | 5.3e-08 | 4.4e-08 | 8.2e-08 | 5.967E-08 | 1.986E-08 | 4.400E-08 | 8.200E-08 |
| 27757 | 1388 | V1388T | E | 2.5e-08 | 1.3e-07 | 5.6e-08 | 7.033E-08 | 5.395E-08 | 2.500E-08 | 1.300E-07 |
| 27758 | 1388 | V1388W | E | 4.1e-08 | 2e-08 | 1.3e-08 | 2.467E-08 | 1.457E-08 | 1.300E-08 | 4.100E-08 |
| 27759 | 1388 | V1388Y | E | 2.3e-08 | 9e-08 | 5.6e-08 | 5.633E-08 | 3.350E-08 | 2.300E-08 | 9.000E-08 |
| 27760 | 1388 | V1388V | E | 2.9e-08 | 4.1e-08 | 1.6e-08 | 2.867E-08 | 1.250E-08 | 1.600E-08 | 4.100E-08 |
| 27761 | 1389 | T1389A | E | 7.7e-08 | 8.1e-08 | 7.1e-08 | 7.633E-08 | 5.033E-09 | 7.100E-08 | 8.100E-08 |
| 27762 | 1389 | T1389R | E | 1.5e-08 | 2.7e-08 | 6.7e-08 | 3.633E-08 | 2.723E-08 | 1.500E-08 | 6.700E-08 |
| 27763 | 1389 | T1389N | E | 2.1e-08 | 1.6e-08 | 3.5e-08 | 2.400E-08 | 9.849E-09 | 1.600E-08 | 3.500E-08 |
| 27764 | 1389 | T1389D | E | 4.6e-08 | 4.3e-08 | 3.5e-08 | 4.133E-08 | 5.686E-09 | 3.500E-08 | 4.600E-08 |
| 27765 | 1389 | T1389C | E | 5.2e-08 | 2.4e-08 | 1.9e-08 | 3.167E-08 | 1.779E-08 | 1.900E-08 | 5.200E-08 |
| 27766 | 1389 | T1389E | E | 1.9e-08 | 5e-08 | 3.1e-08 | 3.333E-08 | 1.563E-08 | 1.900E-08 | 5.000E-08 |
| 27767 | 1389 | T1389Q | E | 9.1e-08 | 6e-08 | 3.4e-08 | 6.167E-08 | 2.854E-08 | 3.400E-08 | 9.100E-08 |
| 27768 | 1389 | T1389G | E | 4.7e-08 | 5.8e-08 | 3.2e-08 | 4.567E-08 | 1.305E-08 | 3.200E-08 | 5.800E-08 |
| 27769 | 1389 | T1389H | E | 4e-08 | 5.9e-08 | 2.8e-08 | 4.233E-08 | 1.563E-08 | 2.800E-08 | 5.900E-08 |
| 27770 | 1389 | T1389I | E | 1.7e-07 | 3.9e-08 | 2.6e-08 | 7.833E-08 | 7.965E-08 | 2.600E-08 | 1.700E-07 |
| 27771 | 1389 | T1389L | E | 1.9e-08 | 1.6e-08 | 6.1e-08 | 3.200E-08 | 2.516E-08 | 1.600E-08 | 6.100E-08 |
| 27772 | 1389 | T1389K | E | 4.2e-08 | 1.6e-08 | 4.3e-08 | 3.367E-08 | 1.531E-08 | 1.600E-08 | 4.300E-08 |
| 27773 | 1389 | T1389M | E | 7.2e-08 | 1e-07 | 6.9e-08 | 8.033E-08 | 1.710E-08 | 6.900E-08 | 1.000E-07 |
| 27774 | 1389 | T1389F | E | 1.1e-07 | 1.5e-08 | 2.1e-08 | 4.867E-08 | 5.320E-08 | 1.500E-08 | 1.100E-07 |
| 27775 | 1389 | T1389P | E | 3.7e-08 | 1.3e-08 | 6.6e-08 | 3.867E-08 | 2.654E-08 | 1.300E-08 | 6.600E-08 |
| 27776 | 1389 | T1389S | E | 2.1e-08 | 5.2e-08 | 1.3e-08 | 2.867E-08 | 2.060E-08 | 1.300E-08 | 5.200E-08 |
| 27777 | 1389 | T1389T | E | 3.1e-08 | 6.6e-08 | 4.4e-08 | 4.700E-08 | 1.769E-08 | 3.100E-08 | 6.600E-08 |
| 27778 | 1389 | T1389W | E | 4e-08 | 9.6e-08 | 6e-08 | 6.533E-08 | 2.838E-08 | 4.000E-08 | 9.600E-08 |
| 27779 | 1389 | T1389Y | E | 4.6e-08 | 5.8e-08 | 4.1e-08 | 4.833E-08 | 8.737E-09 | 4.100E-08 | 5.800E-08 |
| 27780 | 1389 | T1389V | E | 9.5e-08 | 5e-08 | 7.7e-08 | 7.400E-08 | 2.265E-08 | 5.000E-08 | 9.500E-08 |
| 27781 | 1390 | V1390A | E | 3.9e-08 | 2.4e-08 | 7.1e-08 | 4.467E-08 | 2.401E-08 | 2.400E-08 | 7.100E-08 |
| 27782 | 1390 | V1390R | E | 9.6e-08 | 6.2e-08 | 3.5e-08 | 6.433E-08 | 3.057E-08 | 3.500E-08 | 9.600E-08 |
| 27783 | 1390 | V1390N | E | 6.5e-08 | 2.9e-08 | 2.3e-08 | 3.900E-08 | 2.272E-08 | 2.300E-08 | 6.500E-08 |
| 27784 | 1390 | V1390D | E | 6.9e-08 | 2.4e-08 | 2.9e-08 | 4.067E-08 | 2.466E-08 | 2.400E-08 | 6.900E-08 |
| 27785 | 1390 | V1390C | E | 2.2e-08 | 3.6e-08 | 2.6e-08 | 2.800E-08 | 7.211E-09 | 2.200E-08 | 3.600E-08 |
| 27786 | 1390 | V1390E | E | 5.6e-08 | 5.8e-08 | 2.5e-08 | 4.633E-08 | 1.850E-08 | 2.500E-08 | 5.800E-08 |
| 27787 | 1390 | V1390Q | E | 1.9e-08 | 4.4e-08 | 4.4e-08 | 3.567E-08 | 1.443E-08 | 1.900E-08 | 4.400E-08 |
| 27788 | 1390 | V1390G | E | 8.8e-09 | 3.3e-08 | 6.8e-08 | 3.660E-08 | 2.976E-08 | 8.800E-09 | 6.800E-08 |
| 27789 | 1390 | V1390H | E | 4.9e-08 | 3.2e-08 | 3.9e-08 | 4.000E-08 | 8.544E-09 | 3.200E-08 | 4.900E-08 |
| 27790 | 1390 | V1390I | E | 9.5e-08 | 7.7e-09 | 3.8e-08 | 4.690E-08 | 4.433E-08 | 7.700E-09 | 9.500E-08 |
| 27791 | 1390 | V1390L | E | 1.7e-08 | 6.8e-08 | 1.6e-07 | 8.167E-08 | 7.247E-08 | 1.700E-08 | 1.600E-07 |
| 27792 | 1390 | V1390K | E | 9.8e-08 | 2.8e-08 | 3.2e-08 | 5.267E-08 | 3.931E-08 | 2.800E-08 | 9.800E-08 |
| 27793 | 1390 | V1390M | E | 2.4e-08 | 5.5e-08 | 4.7e-08 | 4.200E-08 | 1.609E-08 | 2.400E-08 | 5.500E-08 |
| 27794 | 1390 | V1390F | E | 4e-08 | 7.3e-08 | 3.4e-08 | 4.900E-08 | 2.100E-08 | 3.400E-08 | 7.300E-08 |

| | | | | | | | | | | |
|---|---|---|---|---|---|---|---|---|---|---|
| 27795 | 1390 | V1390P | E | 4.7e-08 | 4e-08 | 2.7e-08 | 3.800E-08 | 1.015E-08 | 2.700E-08 | 4.700E-08 |
| 27796 | 1390 | V1390S | E | 4e-08 | 6.2e-08 | 6e-08 | 5.400E-08 | 1.217E-08 | 4.000E-08 | 6.200E-08 |
| 27797 | 1390 | V1390T | E | 2.2e-08 | 9.4e-08 | 8e-08 | 6.533E-08 | 3.818E-08 | 2.200E-08 | 9.400E-08 |
| 27798 | 1390 | V1390W | E | 1.4e-08 | 7.8e-08 | 3.6e-08 | 4.267E-08 | 3.252E-08 | 1.400E-08 | 7.800E-08 |
| 27799 | 1390 | V1390Y | E | 8.7e-08 | 1.1e-07 | 3.8e-08 | 7.833E-08 | 3.677E-08 | 3.800E-08 | 1.100E-07 |
| 27800 | 1390 | V1390V | E | 1.3e-07 | 1.1e-07 | 3.7e-08 | 9.233E-08 | 4.895E-08 | 3.700E-08 | 1.300E-07 |
| 27801 | 1391 | S1391A | E | 4e-08 | 3.1e-08 | 8.4e-09 | 2.647E-08 | 1.628E-08 | 8.400E-09 | 4.000E-08 |
| 27802 | 1391 | S1391R | E | 2.1e-08 | 3.3e-08 | 1.3e-07 | 6.133E-08 | 5.977E-08 | 2.100E-08 | 1.300E-07 |
| 27803 | 1391 | S1391N | E | 4.3e-08 | 8.2e-08 | 7.7e-08 | 6.733E-08 | 2.122E-08 | 4.300E-08 | 8.200E-08 |
| 27804 | 1391 | S1391D | E | 6.3e-08 | 4.9e-08 | 4.4e-08 | 5.200E-08 | 9.849E-09 | 4.400E-08 | 6.300E-08 |
| 27805 | 1391 | S1391C | E | 7.2e-08 | 3.2e-08 | 4.1e-08 | 4.833E-08 | 2.098E-08 | 3.200E-08 | 7.200E-08 |
| 27806 | 1391 | S1391E | E | 2.1e-08 | 6.6e-08 | 7.1e-08 | 5.267E-08 | 2.754E-08 | 2.100E-08 | 7.100E-08 |
| 27807 | 1391 | S1391Q | E | 2.9e-08 | 3.5e-08 | 4e-08 | 3.467E-08 | 5.508E-09 | 2.900E-08 | 4.000E-08 |
| 27808 | 1391 | S1391G | E | 6e-08 | 1.3e-07 | 1.8e-07 | 1.233E-07 | 6.028E-08 | 6.000E-08 | 1.800E-07 |
| 27809 | 1391 | S1391H | E | 7.3e-08 | 4.5e-08 | 2.4e-08 | 4.733E-08 | 2.458E-08 | 2.400E-08 | 7.300E-08 |
| 27810 | 1391 | S1391I | E | 4.1e-08 | 5.3e-08 | 1e-08 | 3.467E-08 | 2.219E-08 | 1.000E-08 | 5.300E-08 |
| 27811 | 1391 | S1391L | E | 2.7e-08 | 7.3e-08 | 2.8e-08 | 4.267E-08 | 2.627E-08 | 2.700E-08 | 7.300E-08 |
| 27812 | 1391 | S1391K | E | 1.4e-07 | 6.3e-08 | 1.4e-07 | 1.143E-07 | 4.446E-08 | 6.300E-08 | 1.400E-07 |
| 27813 | 1391 | S1391M | E | 5e-08 | 3.5e-08 | 6.4e-08 | 4.967E-08 | 1.450E-08 | 3.500E-08 | 6.400E-08 |
| 27814 | 1391 | S1391F | E | 1.9e-07 | 6.2e-08 | 5.6e-08 | 1.027E-07 | 7.569E-08 | 5.600E-08 | 1.900E-07 |
| 27815 | 1391 | S1391P | E | 1.2e-08 | 3.2e-08 | 1.5e-08 | 1.967E-08 | 1.079E-08 | 1.200E-08 | 3.200E-08 |
| 27816 | 1391 | S1391S | E | 4.2e-08 | 1.1e-08 | 7.8e-08 | 4.367E-08 | 3.353E-08 | 1.100E-08 | 7.800E-08 |
| 27817 | 1391 | S1391T | E | 1.9e-08 | 5.3e-08 | 4.2e-08 | 3.800E-08 | 1.735E-08 | 1.900E-08 | 5.300E-08 |
| 27818 | 1391 | S1391W | E | 1.4e-07 | 3e-08 | 1.5e-08 | 6.167E-08 | 6.825E-08 | 1.500E-08 | 1.400E-07 |
| 27819 | 1391 | S1391Y | E | 5.1e-08 | 8.2e-08 | 9.5e-08 | 7.600E-08 | 2.261E-08 | 5.100E-08 | 9.500E-08 |
| 27820 | 1391 | S1391V | E | 2.5e-08 | 1.9e-08 | 1.1e-07 | 5.133E-08 | 5.090E-08 | 1.900E-08 | 1.100E-07 |
| 27821 | 1392 | S1392A | E | 3.8e-08 | 5.2e-08 | 6.2e-08 | 5.067E-08 | 1.206E-08 | 3.800E-08 | 6.200E-08 |
| 27822 | 1392 | S1392R | E | 5e-08 | 2.4e-08 | 3.2e-08 | 3.533E-08 | 1.332E-08 | 2.400E-08 | 5.000E-08 |
| 27823 | 1392 | S1392N | E | 6.2e-08 | 3.6e-08 | 4.6e-08 | 4.800E-08 | 1.311E-08 | 3.600E-08 | 6.200E-08 |
| 27824 | 1392 | S1392D | E | 9.6e-08 | 2.8e-08 | 1.7e-08 | 4.700E-08 | 4.279E-08 | 1.700E-08 | 9.600E-08 |
| 27825 | 1392 | S1392C | E | 4.1e-08 | 7e-08 | 5.5e-08 | 5.533E-08 | 1.450E-08 | 4.100E-08 | 7.000E-08 |
| 27826 | 1392 | S1392E | E | 7.3e-08 | 1.3e-08 | 1.3e-07 | 7.200E-08 | 5.851E-08 | 1.300E-08 | 1.300E-07 |
| 27827 | 1392 | S1392Q | E | 2.7e-08 | 5.7e-08 | 3.6e-08 | 4.000E-08 | 1.539E-08 | 2.700E-08 | 5.700E-08 |
| 27828 | 1392 | S1392G | E | 1.1e-07 | 1.1e-07 | 4.7e-08 | 8.900E-08 | 3.637E-08 | 4.700E-08 | 1.100E-07 |
| 27829 | 1392 | S1392H | E | 6.8e-08 | 6.3e-08 | 3.5e-08 | 5.533E-08 | 1.779E-08 | 3.500E-08 | 6.800E-08 |
| 27830 | 1392 | S1392I | E | 5.8e-08 | 2.9e-08 | 4.6e-08 | 4.433E-08 | 1.457E-08 | 2.900E-08 | 5.800E-08 |
| 27831 | 1392 | S1392L | E | 5.1e-08 | 1e-08 | 1.3e-07 | 6.367E-08 | 6.099E-08 | 1.000E-08 | 1.300E-07 |
| 27832 | 1392 | S1392K | E | 2.6e-08 | 7.7e-08 | 1.5e-08 | 3.933E-08 | 3.308E-08 | 1.500E-08 | 7.700E-08 |
| 27833 | 1392 | S1392M | E | 8.7e-08 | 4.3e-08 | 1.4e-07 | 9.000E-08 | 4.857E-08 | 4.300E-08 | 1.400E-07 |
| 27834 | 1392 | S1392F | E | 1.5e-08 | 1.3e-08 | 7e-08 | 3.267E-08 | 3.235E-08 | 1.300E-08 | 7.000E-08 |
| 27835 | 1392 | S1392P | E | 3.6e-08 | 3.7e-08 | 2.7e-08 | 3.333E-08 | 5.508E-09 | 2.700E-08 | 3.700E-08 |
| 27836 | 1392 | S1392S | E | 5.2e-08 | 2.2e-08 | 4.4e-08 | 3.933E-08 | 1.553E-08 | 2.200E-08 | 5.200E-08 |
| 27837 | 1392 | S1392T | E | 5.6e-08 | 2.7e-08 | 9.7e-08 | 6.000E-08 | 3.517E-08 | 2.700E-08 | 9.700E-08 |
| 27838 | 1392 | S1392W | E | 6.9e-08 | 7.2e-08 | 5.6e-08 | 6.567E-08 | 8.505E-09 | 5.600E-08 | 7.200E-08 |
| 27839 | 1392 | S1392Y | E | 1.4e-08 | 3.2e-08 | 9.7e-08 | 4.767E-08 | 4.366E-08 | 1.400E-08 | 9.700E-08 |
| 27840 | 1392 | S1392V | E | 2.5e-08 | 2.5e-08 | 5e-08 | 3.333E-08 | 1.443E-08 | 2.500E-08 | 5.000E-08 |
| 27841 | 1393 | A1393A | E | 5.7e-08 | 3e-08 | 4.8e-08 | 4.500E-08 | 1.375E-08 | 3.000E-08 | 5.700E-08 |
| 27842 | 1393 | A1393R | E | 1.7e-08 | 5e-09 | 3.5e-08 | 1.900E-08 | 1.510E-08 | 5.000E-09 | 3.500E-08 |
| 27843 | 1393 | A1393N | E | 1.7e-08 | 2e-08 | 7.3e-08 | 3.667E-08 | 3.150E-08 | 1.700E-08 | 7.300E-08 |
| 27844 | 1393 | A1393D | E | 4e-08 | 3e-08 | 1.1e-07 | 6.000E-08 | 4.359E-08 | 3.000E-08 | 1.100E-07 |
| 27845 | 1393 | A1393C | E | 4.8e-08 | 2.5e-08 | 6.1e-08 | 4.467E-08 | 1.823E-08 | 2.500E-08 | 6.100E-08 |
| 27846 | 1393 | A1393E | E | 9.1e-08 | 2.3e-08 | 5.6e-08 | 5.667E-08 | 3.400E-08 | 2.300E-08 | 9.100E-08 |
| 27847 | 1393 | A1393Q | E | 7.3e-08 | 2.6e-08 | 4.5e-08 | 4.800E-08 | 2.364E-08 | 2.600E-08 | 7.300E-08 |
| 27848 | 1393 | A1393G | E | 4.3e-08 | 3.1e-08 | 6.4e-08 | 4.600E-08 | 1.670E-08 | 3.100E-08 | 6.400E-08 |
| 27849 | 1393 | A1393H | E | 1.7e-08 | 4.5e-08 | 2.3e-08 | 2.833E-08 | 1.474E-08 | 1.700E-08 | 4.500E-08 |
| 27850 | 1393 | A1393I | E | 1.1e-07 | 9.2e-08 | 4.9e-08 | 8.367E-08 | 3.134E-08 | 4.900E-08 | 1.100E-07 |
| 27851 | 1393 | A1393L | E | 6e-08 | 9.1e-08 | 3.6e-08 | 6.233E-08 | 2.757E-08 | 3.600E-08 | 9.100E-08 |
| 27852 | 1393 | A1393K | E | 4e-08 | 4.6e-08 | 3.9e-08 | 4.167E-08 | 3.786E-09 | 3.900E-08 | 4.600E-08 |
| 27853 | 1393 | A1393M | E | 3.6e-08 | 2.4e-08 | 3.1e-08 | 3.033E-08 | 6.028E-09 | 2.400E-08 | 3.600E-08 |
| 27854 | 1393 | A1393F | E | 9.1e-09 | 4.8e-08 | 9.3e-08 | 5.003E-08 | 4.199E-08 | 9.100E-09 | 9.300E-08 |
| 27855 | 1393 | A1393P | E | 2.7e-08 | 2.6e-08 | 3e-08 | 2.767E-08 | 2.082E-09 | 2.600E-08 | 3.000E-08 |
| 27856 | 1393 | A1393S | E | 6.4e-08 | 3.1e-08 | 4.3e-08 | 4.600E-08 | 1.670E-08 | 3.100E-08 | 6.400E-08 |
| 27857 | 1393 | A1393T | E | 7.6e-08 | 3.8e-08 | 7.8e-08 | 6.400E-08 | 2.254E-08 | 3.800E-08 | 7.800E-08 |
| 27858 | 1393 | A1393W | E | 2.6e-08 | 2.6e-08 | 2.9e-08 | 2.700E-08 | 1.732E-09 | 2.600E-08 | 2.900E-08 |
| 27859 | 1393 | A1393Y | E | 7.2e-08 | 1.6e-08 | 7.2e-08 | 5.333E-08 | 3.233E-08 | 1.600E-08 | 7.200E-08 |
| 27860 | 1393 | A1393V | E | 2.1e-08 | 8e-08 | 1.6e-08 | 3.900E-08 | 3.559E-08 | 1.600E-08 | 8.000E-08 |
| 27861 | 1394 | S1394A | E | 1.8e-08 | 4e-08 | 7.6e-08 | 4.467E-08 | 2.928E-08 | 1.800E-08 | 7.600E-08 |
| 27862 | 1394 | S1394R | E | 3.7e-08 | 5.7e-08 | 4.4e-08 | 4.600E-08 | 1.015E-08 | 3.700E-08 | 5.700E-08 |
| 27863 | 1394 | S1394N | E | 6.5e-08 | 2.5e-08 | 4.2e-08 | 4.400E-08 | 2.007E-08 | 2.500E-08 | 6.500E-08 |
| 27864 | 1394 | S1394D | E | 8.6e-08 | 7.1e-09 | 4.6e-08 | 4.637E-08 | 3.945E-08 | 7.100E-09 | 8.600E-08 |
| 27865 | 1394 | S1394C | E | 3.7e-08 | 2e-08 | 6.7e-08 | 4.133E-08 | 2.380E-08 | 2.000E-08 | 6.700E-08 |
| 27866 | 1394 | S1394E | E | 7.3e-08 | 2.4e-08 | 5.6e-08 | 5.100E-08 | 2.488E-08 | 2.400E-08 | 7.300E-08 |
| 27867 | 1394 | S1394Q | E | 6e-08 | 3.5e-08 | 5.1e-08 | 4.867E-08 | 1.266E-08 | 3.500E-08 | 6.000E-08 |
| 27868 | 1394 | S1394G | E | 9.4e-08 | 2e-08 | 4.9e-08 | 5.433E-08 | 3.729E-08 | 2.000E-08 | 9.400E-08 |
| 27869 | 1394 | S1394H | E | 1.8e-08 | 1.6e-08 | 2.1e-08 | 1.833E-08 | 2.517E-09 | 1.600E-08 | 2.100E-08 |
| 27870 | 1394 | S1394I | E | 5e-08 | 6.4e-08 | 4.2e-08 | 5.200E-08 | 1.114E-08 | 4.200E-08 | 6.400E-08 |
| 27871 | 1394 | S1394L | E | 2.2e-08 | 4.3e-08 | 4.3e-08 | 3.600E-08 | 1.212E-08 | 2.200E-08 | 4.300E-08 |
| 27872 | 1394 | S1394K | E | 8.2e-08 | 7e-08 | 5.4e-08 | 6.867E-08 | 1.405E-08 | 5.400E-08 | 8.200E-08 |
| 27873 | 1394 | S1394M | E | 3e-08 | 6.2e-08 | 4.5e-08 | 4.567E-08 | 1.601E-08 | 3.000E-08 | 6.200E-08 |
| 27874 | 1394 | S1394F | E | 5e-08 | 5.3e-08 | 6.9e-08 | 5.733E-08 | 1.021E-08 | 5.000E-08 | 6.900E-08 |
| 27875 | 1394 | S1394P | E | 4.7e-08 | 1.8e-08 | 1.6e-08 | 2.700E-08 | 1.735E-08 | 1.600E-08 | 4.700E-08 |
| 27876 | 1394 | S1394S | E | 4.4e-08 | 2.2e-08 | 5.6e-08 | 4.067E-08 | 1.724E-08 | 2.200E-08 | 5.600E-08 |
| 27877 | 1394 | S1394T | E | 4.4e-08 | 2.2e-08 | 2.1e-08 | 2.900E-08 | 1.300E-08 | 2.100E-08 | 4.400E-08 |
| 27878 | 1394 | S1394W | E | 3.8e-08 | 5.4e-08 | 7.9e-08 | 5.700E-08 | 2.066E-08 | 3.800E-08 | 7.900E-08 |
| 27879 | 1394 | S1394Y | E | 4.2e-08 | 1.1e-07 | 1.1e-07 | 8.733E-08 | 3.926E-08 | 4.200E-08 | 1.100E-07 |

| | | | | | | | | | | |
|---|---|---|---|---|---|---|---|---|---|---|
| 27880 | 1394 | S1394V | E | 1e-07 | 3.9e-08 | 4.4e-08 | 6.100E-08 | 3.387E-08 | 3.900E-08 | 1.000E-07 |
| 27881 | 1395 | T1395A | E | 4.7e-08 | 3.2e-08 | 3.7e-08 | 3.867E-08 | 7.638E-09 | 3.200E-08 | 4.700E-08 |
| 27882 | 1395 | T1395R | E | 3.9e-08 | 1.1e-07 | 1.9e-08 | 5.600E-08 | 4.782E-08 | 1.900E-08 | 1.100E-07 |
| 27883 | 1395 | T1395N | E | 3.9e-08 | 1e-07 | 9.3e-08 | 7.733E-08 | 3.338E-08 | 3.900E-08 | 1.000E-07 |
| 27884 | 1395 | T1395D | E | 4.4e-08 | 4e-08 | 5.7e-08 | 4.700E-08 | 8.888E-09 | 4.000E-08 | 5.700E-08 |
| 27885 | 1395 | T1395C | E | 2.2e-08 | 2.9e-08 | 4.9e-08 | 3.333E-08 | 1.401E-08 | 2.200E-08 | 4.900E-08 |
| 27886 | 1395 | T1395E | E | 4.1e-08 | 4.5e-08 | 5.8e-08 | 4.800E-08 | 8.888E-09 | 4.100E-08 | 5.800E-08 |
| 27887 | 1395 | T1395Q | E | 1e-07 | 5.8e-08 | 8.3e-08 | 8.033E-08 | 2.113E-08 | 5.800E-08 | 1.000E-07 |
| 27888 | 1395 | T1395G | E | 5.2e-08 | 3.2e-08 | 4.2e-08 | 4.200E-08 | 1.000E-08 | 3.200E-08 | 5.200E-08 |
| 27889 | 1395 | T1395H | E | 4.6e-08 | 1.4e-07 | 2.7e-08 | 7.100E-08 | 6.051E-08 | 2.700E-08 | 1.400E-07 |
| 27890 | 1395 | T1395I | E | 2.8e-08 | 5.6e-08 | 3.6e-08 | 4.000E-08 | 1.442E-08 | 2.800E-08 | 5.600E-08 |
| 27891 | 1395 | T1395L | E | 1.4e-08 | 6.8e-08 | 6.2e-08 | 4.800E-08 | 2.960E-08 | 1.400E-08 | 6.800E-08 |
| 27892 | 1395 | T1395K | E | 3.4e-08 | 9.8e-08 | 3.5e-08 | 5.567E-08 | 3.667E-08 | 3.400E-08 | 9.800E-08 |
| 27893 | 1395 | T1395M | E | 7e-08 | 2.4e-08 | 6.4e-08 | 5.267E-08 | 2.501E-08 | 2.400E-08 | 7.000E-08 |
| 27894 | 1395 | T1395F | E | 6.1e-08 | 2.5e-08 | 4e-08 | 4.200E-08 | 1.808E-08 | 2.500E-08 | 6.100E-08 |
| 27895 | 1395 | T1395P | E | 1.1e-07 | 4.4e-08 | 3.4e-08 | 6.267E-08 | 4.130E-08 | 3.400E-08 | 1.100E-07 |
| 27896 | 1395 | T1395S | E | 1e-08 | 3.4e-08 | 5e-08 | 3.133E-08 | 2.013E-08 | 1.000E-08 | 5.000E-08 |
| 27897 | 1395 | T1395T | E | 4.4e-08 | 4.5e-08 | 1e-07 | 6.300E-08 | 3.205E-08 | 4.400E-08 | 1.000E-07 |
| 27898 | 1395 | T1395W | E | 3.3e-08 | 8.1e-08 | 5e-08 | 5.467E-08 | 2.434E-08 | 3.300E-08 | 8.100E-08 |
| 27899 | 1395 | T1395Y | E | 2.5e-08 | 2.7e-08 | 8.2e-08 | 4.467E-08 | 3.235E-08 | 2.500E-08 | 8.200E-08 |
| 27900 | 1395 | T1395V | E | 4.4e-08 | 8.6e-08 | 4.9e-08 | 5.967E-08 | 2.294E-08 | 4.400E-08 | 8.600E-08 |
| 27901 | 1396 | K1396A | E | 3e-08 | 3.4e-08 | 4.8e-08 | 3.733E-08 | 9.452E-09 | 3.000E-08 | 4.800E-08 |
| 27902 | 1396 | K1396R | E | 7.7e-08 | 7.1e-08 | 6.9e-08 | 7.233E-08 | 4.163E-09 | 6.900E-08 | 7.700E-08 |
| 27903 | 1396 | K1396N | E | 3.8e-08 | 8.6e-08 | 2.7e-08 | 5.033E-08 | 3.137E-08 | 2.700E-08 | 8.600E-08 |
| 27904 | 1396 | K1396D | E | 4e-08 | 9.8e-08 | 1.2e-07 | 8.600E-08 | 4.133E-08 | 4.000E-08 | 1.200E-07 |
| 27905 | 1396 | K1396C | E | 7.4e-08 | 8e-08 | 2.2e-08 | 5.867E-08 | 3.190E-08 | 2.200E-08 | 8.000E-08 |
| 27906 | 1396 | K1396E | E | 2.5e-08 | 2.6e-08 | 8.3e-08 | 4.467E-08 | 3.320E-08 | 2.500E-08 | 8.300E-08 |
| 27907 | 1396 | K1396Q | E | 6.5e-08 | 2.1e-08 | 8e-08 | 5.533E-08 | 3.066E-08 | 2.100E-08 | 8.000E-08 |
| 27908 | 1396 | K1396G | E | 2.8e-08 | 2.1e-08 | 4.2e-08 | 3.033E-08 | 1.069E-08 | 2.100E-08 | 4.200E-08 |
| 27909 | 1396 | K1396H | E | 5.6e-08 | 1.9e-08 | 3.7e-08 | 3.733E-08 | 1.850E-08 | 1.900E-08 | 5.600E-08 |
| 27910 | 1396 | K1396I | E | 4.6e-08 | 5.4e-08 | 1.3e-07 | 7.667E-08 | 4.636E-08 | 4.600E-08 | 1.300E-07 |
| 27911 | 1396 | K1396L | E | 4.8e-08 | 5.6e-08 | 1.2e-07 | 7.467E-08 | 3.946E-08 | 4.800E-08 | 1.200E-07 |
| 27912 | 1396 | K1396K | E | 3.8e-08 | 4.2e-08 | 4.5e-08 | 4.167E-08 | 3.512E-09 | 3.800E-08 | 4.500E-08 |
| 27913 | 1396 | K1396M | E | 5.3e-08 | 1.3e-08 | 2.8e-08 | 3.133E-08 | 2.021E-08 | 1.300E-08 | 5.300E-08 |
| 27914 | 1396 | K1396F | E | 6.8e-08 | 3e-08 | 2.8e-08 | 4.200E-08 | 2.254E-08 | 2.800E-08 | 6.800E-08 |
| 27915 | 1396 | K1396P | E | 6.9e-08 | 5.1e-08 | 9.2e-08 | 7.067E-08 | 2.055E-08 | 5.100E-08 | 9.200E-08 |
| 27916 | 1396 | K1396S | E | 4.8e-08 | 4.5e-08 | 1e-07 | 6.433E-08 | 3.092E-08 | 4.500E-08 | 1.000E-07 |
| 27917 | 1396 | K1396T | E | 2.8e-08 | 1.1e-07 | 1.8e-08 | 5.200E-08 | 5.048E-08 | 1.800E-08 | 1.100E-07 |
| 27918 | 1396 | K1396W | E | 4.6e-08 | 2.5e-08 | 4.4e-08 | 3.833E-08 | 1.159E-08 | 2.500E-08 | 4.600E-08 |
| 27919 | 1396 | K1396Y | E | 1.2e-08 | 1.8e-08 | 2.2e-08 | 1.733E-08 | 5.033E-09 | 1.200E-08 | 2.200E-08 |
| 27920 | 1396 | K1396V | E | 2.2e-08 | 1.6e-08 | 4.9e-08 | 2.900E-08 | 1.758E-08 | 1.600E-08 | 4.900E-08 |
| 27921 | 1397 | G1397A | E | 3e-08 | 2.1e-08 | 1.4e-08 | 2.167E-08 | 8.021E-09 | 1.400E-08 | 3.000E-08 |
| 27922 | 1397 | G1397R | E | 2.2e-08 | 5.8e-08 | 3.6e-08 | 3.867E-08 | 1.815E-08 | 2.200E-08 | 5.800E-08 |
| 27923 | 1397 | G1397N | E | 3.2e-08 | 3.1e-08 | 4.2e-08 | 3.500E-08 | 6.083E-09 | 3.100E-08 | 4.200E-08 |
| 27924 | 1397 | G1397D | E | 4.2e-08 | 8.9e-09 | 9.5e-08 | 4.863E-08 | 4.343E-08 | 8.900E-09 | 9.500E-08 |
| 27925 | 1397 | G1397C | E | 7.2e-08 | 7.5e-08 | 4.7e-08 | 6.467E-08 | 1.537E-08 | 4.700E-08 | 7.500E-08 |
| 27926 | 1397 | G1397E | E | 5.1e-08 | 4.4e-08 | 2.8e-08 | 4.100E-08 | 1.179E-08 | 2.800E-08 | 5.100E-08 |
| 27927 | 1397 | G1397Q | E | 1.8e-08 | 4.5e-08 | 3.4e-08 | 3.233E-08 | 1.358E-08 | 1.800E-08 | 4.500E-08 |
| 27928 | 1397 | G1397G | E | 2.3e-08 | 1.4e-08 | 6.3e-08 | 3.333E-08 | 2.608E-08 | 1.400E-08 | 6.300E-08 |
| 27929 | 1397 | G1397H | E | 1.1e-07 | 4e-08 | 1.1e-07 | 8.667E-08 | 4.041E-08 | 4.000E-08 | 1.100E-07 |
| 27930 | 1397 | G1397I | E | 5.3e-08 | 5.1e-08 | 4.8e-08 | 5.067E-08 | 2.517E-09 | 4.800E-08 | 5.300E-08 |
| 27931 | 1397 | G1397L | E | 4.5e-08 | 2.4e-08 | 1.5e-07 | 7.300E-08 | 6.751E-08 | 2.400E-08 | 1.500E-07 |
| 27932 | 1397 | G1397K | E | 6.2e-08 | 2.5e-08 | 3.6e-08 | 4.100E-08 | 1.900E-08 | 2.500E-08 | 6.200E-08 |
| 27933 | 1397 | G1397M | E | 7.4e-08 | 4.4e-08 | 1.4e-08 | 4.400E-08 | 3.000E-08 | 1.400E-08 | 7.400E-08 |
| 27934 | 1397 | G1397F | E | 3e-08 | 5.4e-08 | 5.8e-08 | 4.733E-08 | 1.514E-08 | 3.000E-08 | 5.800E-08 |
| 27935 | 1397 | G1397P | E | 3.3e-08 | 5.7e-08 | 2.7e-08 | 3.900E-08 | 1.587E-08 | 2.700E-08 | 5.700E-08 |
| 27936 | 1397 | G1397S | E | 5.4e-08 | 6.2e-08 | 1.9e-08 | 4.500E-08 | 2.287E-08 | 1.900E-08 | 6.200E-08 |
| 27937 | 1397 | G1397T | E | 1.8e-08 | 5e-08 | 5.8e-08 | 4.200E-08 | 2.117E-08 | 1.800E-08 | 5.800E-08 |
| 27938 | 1397 | G1397W | E | 6.7e-08 | 3.8e-08 | 5.4e-08 | 5.300E-08 | 1.453E-08 | 3.800E-08 | 6.700E-08 |
| 27939 | 1397 | G1397Y | E | 5.8e-08 | 2.6e-08 | 3.9e-08 | 4.100E-08 | 1.609E-08 | 2.600E-08 | 5.800E-08 |
| 27940 | 1397 | G1397V | E | 4.4e-08 | 1.6e-08 | 8.9e-08 | 4.967E-08 | 3.683E-08 | 1.600E-08 | 8.900E-08 |
| 27941 | 1398 | P1398A | E | 5.9e-08 | 3e-08 | 3.1e-08 | 4.000E-08 | 1.646E-08 | 3.000E-08 | 5.900E-08 |
| 27942 | 1398 | P1398R | E | 4.8e-08 | 3.9e-08 | 8.1e-08 | 5.600E-08 | 2.211E-08 | 3.900E-08 | 8.100E-08 |
| 27943 | 1398 | P1398N | E | 3.6e-08 | 2.3e-08 | 4.4e-08 | 3.433E-08 | 1.060E-08 | 2.300E-08 | 4.400E-08 |
| 27944 | 1398 | P1398D | E | 1.7e-07 | 1.6e-08 | 2.5e-08 | 7.033E-08 | 8.643E-08 | 1.600E-08 | 1.700E-07 |
| 27945 | 1398 | P1398C | E | 2.6e-08 | 3.3e-08 | 1.5e-07 | 6.967E-08 | 6.966E-08 | 2.600E-08 | 1.500E-07 |
| 27946 | 1398 | P1398E | E | 3.8e-08 | 2.9e-08 | 7.7e-08 | 4.800E-08 | 2.551E-08 | 2.900E-08 | 7.700E-08 |
| 27947 | 1398 | P1398Q | E | 2.2e-08 | 1.1e-07 | 5.8e-08 | 6.333E-08 | 4.424E-08 | 2.200E-08 | 1.100E-07 |
| 27948 | 1398 | P1398G | E | 1.6e-08 | 1.2e-07 | 1.3e-08 | 4.967E-08 | 6.093E-08 | 1.300E-08 | 1.200E-07 |
| 27949 | 1398 | P1398H | E | 4.8e-08 | 1.2e-08 | 3.7e-08 | 3.233E-08 | 1.845E-08 | 1.200E-08 | 4.800E-08 |
| 27950 | 1398 | P1398I | E | 1.9e-08 | 4.3e-08 | 1.8e-08 | 2.667E-08 | 1.415E-08 | 1.800E-08 | 4.300E-08 |
| 27951 | 1398 | P1398L | E | 1.1e-07 | 2.1e-08 | 3.9e-08 | 5.667E-08 | 4.706E-08 | 2.100E-08 | 1.100E-07 |
| 27952 | 1398 | P1398K | E | 3.2e-08 | 8.7e-08 | 5e-08 | 5.633E-08 | 2.804E-08 | 3.200E-08 | 8.700E-08 |
| 27953 | 1398 | P1398M | E | 2.1e-08 | 1.9e-08 | 2.4e-08 | 2.133E-08 | 2.517E-09 | 1.900E-08 | 2.400E-08 |
| 27954 | 1398 | P1398F | E | 1.4e-07 | 1.5e-08 | 2.5e-08 | 6.000E-08 | 6.946E-08 | 1.500E-08 | 1.400E-07 |
| 27955 | 1398 | P1398P | E | 7.3e-08 | 7.2e-08 | 4.4e-08 | 6.300E-08 | 1.646E-08 | 4.400E-08 | 7.300E-08 |
| 27956 | 1398 | P1398S | E | 9.2e-08 | 2.1e-08 | 5.1e-08 | 5.467E-08 | 3.564E-08 | 2.100E-08 | 9.200E-08 |
| 27957 | 1398 | P1398T | E | 1.1e-07 | 4.1e-08 | 3.7e-08 | 6.267E-08 | 4.104E-08 | 3.700E-08 | 1.100E-07 |
| 27958 | 1398 | P1398W | E | 4.9e-08 | 4.5e-08 | 3.3e-08 | 4.233E-08 | 8.327E-09 | 3.300E-08 | 4.900E-08 |
| 27959 | 1398 | P1398Y | E | 1.8e-08 | 6.2e-08 | 3.5e-08 | 3.833E-08 | 2.219E-08 | 1.800E-08 | 6.200E-08 |
| 27960 | 1398 | P1398V | E | 4.2e-08 | 5.6e-08 | 5.6e-08 | 5.133E-08 | 8.083E-09 | 4.200E-08 | 5.600E-08 |
| 27961 | 1399 | S1399A | E | 8.3e-08 | 1.4e-08 | 2.1e-07 | 1.023E-07 | 9.942E-08 | 1.400E-08 | 2.100E-07 |
| 27962 | 1399 | S1399R | E | 2.6e-08 | 1e-07 | 1.5e-08 | 4.700E-08 | 4.623E-08 | 1.500E-08 | 1.000E-07 |
| 27963 | 1399 | S1399N | E | 1.2e-08 | 3.6e-08 | 4.2e-08 | 3.000E-08 | 1.587E-08 | 1.200E-08 | 4.200E-08 |
| 27964 | 1399 | S1399D | E | 8.1e-08 | 2.1e-08 | 2.6e-08 | 4.267E-08 | 3.329E-08 | 2.100E-08 | 8.100E-08 |

| | | | | | | | | | | |
|---|---|---|---|---|---|---|---|---|---|---|
| 27965 | 1399 | S1399C | E | 2.7e-08 | 3.9e-08 | 4.9e-08 | 3.833E-08 | 1.102E-08 | 2.700E-08 | 4.900E-08 |
| 27966 | 1399 | S1399E | E | 3.8e-08 | 2.6e-08 | 1.2e-08 | 2.533E-08 | 1.301E-08 | 1.200E-08 | 3.800E-08 |
| 27967 | 1399 | S1399Q | E | 3.5e-08 | 3.4e-08 | 6.9e-08 | 4.600E-08 | 1.992E-08 | 3.400E-08 | 6.900E-08 |
| 27968 | 1399 | S1399G | E | 8.4e-08 | 1e-07 | 2.4e-08 | 6.933E-08 | 4.007E-08 | 2.400E-08 | 1.000E-07 |
| 27969 | 1399 | S1399H | E | 2.5e-08 | 1.3e-07 | 4.9e-08 | 6.800E-08 | 5.502E-08 | 2.500E-08 | 1.300E-07 |
| 27970 | 1399 | S1399I | E | 7.9e-08 | 1.4e-08 | 3e-08 | 4.100E-08 | 3.387E-08 | 1.400E-08 | 7.900E-08 |
| 27971 | 1399 | S1399L | E | 2.5e-08 | 1.3e-07 | 7.8e-08 | 7.767E-08 | 5.250E-08 | 2.500E-08 | 1.300E-07 |
| 27972 | 1399 | S1399K | E | 6.2e-08 | 4.8e-08 | 5e-08 | 5.333E-08 | 7.572E-09 | 4.800E-08 | 6.200E-08 |
| 27973 | 1399 | S1399M | E | 4.4e-08 | 3e-08 | 3.8e-08 | 3.733E-08 | 7.024E-09 | 3.000E-08 | 4.400E-08 |
| 27974 | 1399 | S1399F | E | 5.3e-08 | 1.5e-08 | 5.5e-08 | 4.100E-08 | 2.254E-08 | 1.500E-08 | 5.500E-08 |
| 27975 | 1399 | S1399P | E | 4.3e-08 | 1.3e-08 | 2.3e-08 | 2.633E-08 | 1.528E-08 | 1.300E-08 | 4.300E-08 |
| 27976 | 1399 | S1399S | E | 5.4e-08 | 2.7e-08 | 3e-08 | 3.700E-08 | 1.480E-08 | 2.700E-08 | 5.400E-08 |
| 27977 | 1399 | S1399T | E | 1.7e-08 | 2.6e-08 | 3.8e-08 | 2.700E-08 | 1.054E-08 | 1.700E-08 | 3.800E-08 |
| 27978 | 1399 | S1399W | E | 4.5e-08 | 3e-08 | 2.6e-08 | 3.367E-08 | 1.002E-08 | 2.600E-08 | 4.500E-08 |
| 27979 | 1399 | S1399Y | E | 3.2e-08 | 4.3e-08 | 2.8e-08 | 3.433E-08 | 7.767E-09 | 2.800E-08 | 4.300E-08 |
| 27980 | 1399 | S1399V | E | 4.5e-08 | 5.5e-08 | 1.2e-07 | 7.333E-08 | 4.072E-08 | 4.500E-08 | 1.200E-07 |
| 27981 | 1400 | V1400A | E | 5.6e-08 | 3.5e-08 | 2.5e-08 | 3.867E-08 | 1.582E-08 | 2.500E-08 | 5.600E-08 |
| 27982 | 1400 | V1400R | E | 5.9e-08 | 6.2e-08 | 8.8e-08 | 6.967E-08 | 1.595E-08 | 5.900E-08 | 8.800E-08 |
| 27983 | 1400 | V1400N | E | 6.6e-08 | 3.7e-08 | 5.9e-08 | 5.400E-08 | 1.513E-08 | 3.700E-08 | 6.600E-08 |
| 27984 | 1400 | V1400D | E | 2e-08 | 4.5e-08 | 1.2e-07 | 6.167E-08 | 5.204E-08 | 2.000E-08 | 1.200E-07 |
| 27985 | 1400 | V1400C | E | 3.8e-08 | 8.4e-08 | 4.2e-08 | 5.467E-08 | 2.548E-08 | 3.800E-08 | 8.400E-08 |
| 27986 | 1400 | V1400E | E | 2.3e-08 | 2e-08 | 1.8e-08 | 2.033E-08 | 2.517E-09 | 1.800E-08 | 2.300E-08 |
| 27987 | 1400 | V1400Q | E | 2.4e-08 | 3.7e-08 | 1.2e-07 | 6.033E-08 | 5.208E-08 | 2.400E-08 | 1.200E-07 |
| 27988 | 1400 | V1400G | E | 4.6e-08 | 3.2e-08 | 5.7e-08 | 4.500E-08 | 1.253E-08 | 3.200E-08 | 5.700E-08 |
| 27989 | 1400 | V1400H | E | 4.2e-08 | 3.3e-08 | 6.1e-08 | 4.533E-08 | 1.429E-08 | 3.300E-08 | 6.100E-08 |
| 27990 | 1400 | V1400I | E | 4.6e-08 | 2.5e-08 | 3.4e-08 | 3.500E-08 | 1.054E-08 | 2.500E-08 | 4.600E-08 |
| 27991 | 1400 | V1400L | E | 2.7e-08 | 3.1e-08 | 2.3e-08 | 2.700E-08 | 4.000E-09 | 2.300E-08 | 3.100E-08 |
| 27992 | 1400 | V1400K | E | 5e-08 | 4.2e-08 | 1.3e-07 | 7.400E-08 | 4.866E-08 | 4.200E-08 | 1.300E-07 |
| 27993 | 1400 | V1400M | E | 5.8e-08 | 4.2e-08 | 6.9e-08 | 5.633E-08 | 1.358E-08 | 4.200E-08 | 6.900E-08 |
| 27994 | 1400 | V1400F | E | 5.5e-08 | 5.4e-08 | 5.7e-08 | 5.533E-08 | 1.528E-09 | 5.400E-08 | 5.700E-08 |
| 27995 | 1400 | V1400P | E | 3.1e-08 | 6.8e-08 | 5.6e-08 | 5.167E-08 | 1.888E-08 | 3.100E-08 | 6.800E-08 |
| 27996 | 1400 | V1400S | E | 2e-08 | 4.8e-08 | 5.9e-08 | 4.233E-08 | 2.011E-08 | 2.000E-08 | 5.900E-08 |
| 27997 | 1400 | V1400T | E | 2e-08 | 2.9e-08 | 7.9e-08 | 4.267E-08 | 3.179E-08 | 2.000E-08 | 7.900E-08 |
| 27998 | 1400 | V1400W | E | 7.4e-08 | 7.8e-08 | 1e-07 | 8.400E-08 | 1.400E-08 | 7.400E-08 | 1.000E-07 |
| 27999 | 1400 | V1400Y | E | 3.4e-08 | 4.2e-08 | 2.3e-08 | 3.300E-08 | 9.539E-09 | 2.300E-08 | 4.200E-08 |
| 28000 | 1400 | V1400V | E | 5.4e-08 | 8.6e-08 | 4.6e-08 | 6.200E-08 | 2.117E-08 | 4.600E-08 | 8.600E-08 |
| 28001 | 1401 | F1401A | E | 5.7e-08 | 4.3e-08 | 6e-08 | 5.333E-08 | 9.074E-09 | 4.300E-08 | 6.000E-08 |
| 28002 | 1401 | F1401R | E | 5.3e-08 | 3.1e-08 | 3.4e-08 | 3.933E-08 | 1.193E-08 | 3.100E-08 | 5.300E-08 |
| 28003 | 1401 | F1401N | E | 2.3e-08 | 5.4e-08 | 6.6e-08 | 4.767E-08 | 2.219E-08 | 2.300E-08 | 6.600E-08 |
| 28004 | 1401 | F1401D | E | 5.1e-08 | 3.5e-08 | 2.8e-08 | 3.800E-08 | 1.179E-08 | 2.800E-08 | 5.100E-08 |
| 28005 | 1401 | F1401C | E | 3.4e-08 | 1.8e-08 | 5.5e-08 | 3.567E-08 | 1.856E-08 | 1.800E-08 | 5.500E-08 |
| 28006 | 1401 | F1401E | E | 2.2e-08 | 4.4e-08 | 2e-08 | 2.867E-08 | 1.332E-08 | 2.000E-08 | 4.400E-08 |
| 28007 | 1401 | F1401Q | E | 2.4e-08 | 1.1e-07 | 2.4e-08 | 5.267E-08 | 4.965E-08 | 2.400E-08 | 1.100E-07 |
| 28008 | 1401 | F1401G | E | 8.5e-08 | 7.6e-08 | 4.5e-08 | 6.867E-08 | 2.098E-08 | 4.500E-08 | 8.500E-08 |
| 28009 | 1401 | F1401H | E | 5.2e-08 | 1.4e-08 | 7.1e-08 | 4.567E-08 | 2.902E-08 | 1.400E-08 | 7.100E-08 |
| 28010 | 1401 | F1401I | E | 5.3e-09 | 6e-08 | 3.9e-08 | 3.477E-08 | 2.759E-08 | 5.300E-09 | 6.000E-08 |
| 28011 | 1401 | F1401L | E | 9.7e-09 | 2.7e-08 | 1.7e-08 | 1.790E-08 | 8.685E-09 | 9.700E-09 | 2.700E-08 |
| 28012 | 1401 | F1401K | E | 6.9e-08 | 1.6e-08 | 5.2e-08 | 4.567E-08 | 2.706E-08 | 1.600E-08 | 6.900E-08 |
| 28013 | 1401 | F1401M | E | 3.1e-08 | 4.8e-08 | 3.3e-08 | 3.733E-08 | 9.292E-09 | 3.100E-08 | 4.800E-08 |
| 28014 | 1401 | F1401F | E | 6.7e-08 | 1.8e-08 | 3.6e-08 | 4.033E-08 | 2.479E-08 | 1.800E-08 | 6.700E-08 |
| 28015 | 1401 | F1401P | E | 2.5e-08 | 4.3e-08 | 6.3e-08 | 4.367E-08 | 1.901E-08 | 2.500E-08 | 6.300E-08 |
| 28016 | 1401 | F1401S | E | 5.1e-08 | 4.6e-08 | 1e-07 | 6.567E-08 | 2.984E-08 | 4.600E-08 | 1.000E-07 |
| 28017 | 1401 | F1401T | E | 3.8e-08 | 4.1e-08 | 2.8e-08 | 3.567E-08 | 6.807E-09 | 2.800E-08 | 4.100E-08 |
| 28018 | 1401 | F1401W | E | 4.8e-08 | 3.8e-08 | 1.3e-08 | 3.300E-08 | 1.803E-08 | 1.300E-08 | 4.800E-08 |
| 28019 | 1401 | F1401Y | E | 4.4e-08 | 1.3e-07 | 9.8e-08 | 9.067E-08 | 4.347E-08 | 4.400E-08 | 1.300E-07 |
| 28020 | 1401 | F1401V | E | 2.7e-08 | 3e-08 | 1.5e-08 | 2.400E-08 | 7.937E-09 | 1.500E-08 | 3.000E-08 |
| 28021 | 1402 | P1402A | E | 3e-08 | 8.1e-08 | 3.9e-08 | 5.000E-08 | 2.722E-08 | 3.000E-08 | 8.100E-08 |
| 28022 | 1402 | P1402R | E | 7e-08 | 1.2e-07 | 7.1e-08 | 8.700E-08 | 2.858E-08 | 7.000E-08 | 1.200E-07 |
| 28023 | 1402 | P1402N | E | 3.2e-08 | 3.8e-08 | 8.3e-08 | 5.100E-08 | 2.787E-08 | 3.200E-08 | 8.300E-08 |
| 28024 | 1402 | P1402D | E | 5.9e-08 | 4.4e-08 | 2.4e-08 | 4.233E-08 | 1.756E-08 | 2.400E-08 | 5.900E-08 |
| 28025 | 1402 | P1402C | E | 3.7e-08 | 4.9e-08 | 6e-08 | 4.867E-08 | 1.150E-08 | 3.700E-08 | 6.000E-08 |
| 28026 | 1402 | P1402E | E | 2.8e-08 | 3.5e-08 | 2e-08 | 2.767E-08 | 7.506E-09 | 2.000E-08 | 3.500E-08 |
| 28027 | 1402 | P1402Q | E | 1.5e-08 | 1.6e-08 | 3.7e-08 | 2.267E-08 | 1.242E-08 | 1.500E-08 | 3.700E-08 |
| 28028 | 1402 | P1402G | E | 4.9e-08 | 2.9e-08 | 6.2e-08 | 4.667E-08 | 1.662E-08 | 2.900E-08 | 6.200E-08 |
| 28029 | 1402 | P1402H | E | 4.7e-08 | 4.9e-08 | 1.1e-07 | 6.867E-08 | 3.581E-08 | 4.700E-08 | 1.100E-07 |
| 28030 | 1402 | P1402I | E | 1.7e-08 | 2.9e-08 | 1.6e-08 | 2.067E-08 | 7.234E-09 | 1.600E-08 | 2.900E-08 |
| 28031 | 1402 | P1402L | E | 5.9e-08 | 3.5e-08 | 4.1e-08 | 4.500E-08 | 1.249E-08 | 3.500E-08 | 5.900E-08 |
| 28032 | 1402 | P1402K | E | 2.3e-08 | 6.1e-08 | 4.6e-08 | 4.333E-08 | 1.914E-08 | 2.300E-08 | 6.100E-08 |
| 28033 | 1402 | P1402M | E | 7e-08 | 2.2e-08 | 6.1e-08 | 5.100E-08 | 2.551E-08 | 2.200E-08 | 7.000E-08 |
| 28034 | 1402 | P1402F | E | 9.3e-08 | 8.1e-08 | 3.1e-08 | 6.833E-08 | 3.288E-08 | 3.100E-08 | 9.300E-08 |
| 28035 | 1402 | P1402P | E | 4e-08 | 3.2e-08 | 1.8e-08 | 3.000E-08 | 1.114E-08 | 1.800E-08 | 4.000E-08 |
| 28036 | 1402 | P1402S | E | 1.4e-08 | 5e-08 | 4.7e-08 | 3.700E-08 | 1.997E-08 | 1.400E-08 | 5.000E-08 |
| 28037 | 1402 | P1402T | E | 4.3e-08 | 3.1e-08 | 1.4e-08 | 2.933E-08 | 1.457E-08 | 1.400E-08 | 4.300E-08 |
| 28038 | 1402 | P1402W | E | 3.3e-08 | 4.6e-08 | 2.8e-08 | 3.567E-08 | 9.292E-09 | 2.800E-08 | 4.600E-08 |
| 28039 | 1402 | P1402Y | E | 3.3e-08 | 3.7e-08 | 3.5e-08 | 3.500E-08 | 2.000E-09 | 3.300E-08 | 3.700E-08 |
| 28040 | 1402 | P1402V | E | 5.2e-08 | 1.1e-07 | 4.2e-08 | 6.800E-08 | 3.672E-08 | 4.200E-08 | 1.100E-07 |
| 28041 | 1403 | L1403A | E | 4.9e-08 | 3.6e-08 | 6.1e-08 | 4.867E-08 | 1.250E-08 | 3.600E-08 | 6.100E-08 |
| 28042 | 1403 | L1403R | E | 4.3e-08 | 6.6e-08 | 2.4e-08 | 4.433E-08 | 2.103E-08 | 2.400E-08 | 6.600E-08 |
| 28043 | 1403 | L1403N | E | 9.4e-08 | 1.1e-07 | 1.2e-08 | 7.200E-08 | 5.257E-08 | 1.200E-08 | 1.100E-07 |
| 28044 | 1403 | L1403D | E | 3.8e-08 | 7.1e-08 | 1.1e-07 | 7.300E-08 | 3.604E-08 | 3.800E-08 | 1.100E-07 |
| 28045 | 1403 | L1403C | E | 5.6e-08 | 7.5e-08 | 4.4e-08 | 5.833E-08 | 1.563E-08 | 4.400E-08 | 7.500E-08 |
| 28046 | 1403 | L1403E | E | 9.1e-08 | 1.1e-07 | 3e-08 | 7.700E-08 | 4.180E-08 | 3.000E-08 | 1.100E-07 |
| 28047 | 1403 | L1403Q | E | 4.2e-08 | 9e-08 | 1.5e-08 | 4.900E-08 | 3.799E-08 | 1.500E-08 | 9.000E-08 |
| 28048 | 1403 | L1403G | E | 3.2e-08 | 5.8e-08 | 3.7e-08 | 4.233E-08 | 1.380E-08 | 3.200E-08 | 5.800E-08 |
| 28049 | 1403 | L1403H | E | 3.8e-08 | 7e-08 | 3.3e-08 | 4.700E-08 | 2.007E-08 | 3.300E-08 | 7.000E-08 |

| | | | | | | | | | | |
|---|---|---|---|---|---|---|---|---|---|---|
| 28050 | 1403 | L1403I | E | 5e-08 | 7.7e-08 | 5.2e-08 | 5.967E-08 | 1.504E-08 | 5.000E-08 | 7.700E-08 |
| 28051 | 1403 | L1403L | E | 6.6e-08 | 4.8e-08 | 2.9e-08 | 4.767E-08 | 1.850E-08 | 2.900E-08 | 6.600E-08 |
| 28052 | 1403 | L1403K | E | 2.7e-08 | 3.9e-08 | 2.3e-08 | 2.967E-08 | 8.327E-09 | 2.300E-08 | 3.900E-08 |
| 28053 | 1403 | L1403M | E | 6.9e-08 | 6.7e-08 | 3.4e-08 | 5.667E-08 | 1.966E-08 | 3.400E-08 | 6.900E-08 |
| 28054 | 1403 | L1403F | E | 1.5e-08 | 3.9e-08 | 4e-08 | 3.133E-08 | 1.415E-08 | 1.500E-08 | 4.000E-08 |
| 28055 | 1403 | L1403P | E | 1.4e-07 | 6.2e-08 | 4.6e-08 | 8.267E-08 | 5.029E-08 | 4.600E-08 | 1.400E-07 |
| 28056 | 1403 | L1403S | E | 1.3e-07 | 5.1e-08 | 9.7e-08 | 9.267E-08 | 3.968E-08 | 5.100E-08 | 1.300E-07 |
| 28057 | 1403 | L1403T | E | 2.2e-08 | 4.2e-08 | 3.6e-08 | 3.333E-08 | 1.026E-08 | 2.200E-08 | 4.200E-08 |
| 28058 | 1403 | L1403W | E | 4.3e-08 | 7.8e-08 | 5.8e-08 | 5.967E-08 | 1.756E-08 | 4.300E-08 | 7.800E-08 |
| 28059 | 1403 | L1403Y | E | 8.2e-08 | 7.3e-08 | 4.8e-08 | 6.767E-08 | 1.762E-08 | 4.800E-08 | 8.200E-08 |
| 28060 | 1403 | L1403V | E | 6.3e-08 | 1e-07 | 1.2e-07 | 9.433E-08 | 2.892E-08 | 6.300E-08 | 1.200E-07 |
| 28061 | 1404 | A1404A | E | 4.5e-08 | 1.2e-08 | 3.1e-08 | 2.933E-08 | 1.656E-08 | 1.200E-08 | 4.500E-08 |
| 28062 | 1404 | A1404R | E | 7.1e-08 | 3.1e-08 | 3.1e-08 | 4.433E-08 | 2.309E-08 | 3.100E-08 | 7.100E-08 |
| 28063 | 1404 | A1404N | E | 4.7e-08 | 1.4e-07 | 4.1e-08 | 7.600E-08 | 5.551E-08 | 4.100E-08 | 1.400E-07 |
| 28064 | 1404 | A1404D | E | 7.1e-08 | 5.1e-08 | 4e-08 | 5.400E-08 | 1.572E-08 | 4.000E-08 | 7.100E-08 |
| 28065 | 1404 | A1404C | E | 6.2e-08 | 6e-08 | 7.2e-08 | 6.467E-08 | 6.429E-09 | 6.000E-08 | 7.200E-08 |
| 28066 | 1404 | A1404E | E | 6e-09 | 3.3e-08 | 2.6e-08 | 2.167E-08 | 1.401E-08 | 6.000E-09 | 3.300E-08 |
| 28067 | 1404 | A1404Q | E | 5.7e-08 | 5.9e-08 | 3.4e-08 | 5.000E-08 | 1.389E-08 | 3.400E-08 | 5.900E-08 |
| 28068 | 1404 | A1404G | E | 3.6e-08 | 8.8e-08 | 6.9e-08 | 6.433E-08 | 2.631E-08 | 3.600E-08 | 8.800E-08 |
| 28069 | 1404 | A1404H | E | 2.1e-08 | 1.2e-07 | 7.4e-08 | 7.167E-08 | 4.954E-08 | 2.100E-08 | 1.200E-07 |
| 28070 | 1404 | A1404I | E | 3.2e-08 | 2.5e-08 | 2.8e-08 | 2.833E-08 | 3.512E-09 | 2.500E-08 | 3.200E-08 |
| 28071 | 1404 | A1404L | E | 3e-08 | 1.4e-07 | 3.9e-08 | 6.967E-08 | 6.108E-08 | 3.000E-08 | 1.400E-07 |
| 28072 | 1404 | A1404K | E | 5.4e-08 | 5.3e-08 | 1e-07 | 6.900E-08 | 2.685E-08 | 5.300E-08 | 1.000E-07 |
| 28073 | 1404 | A1404M | E | 1.2e-07 | 8.4e-08 | 3.6e-08 | 8.000E-08 | 4.214E-08 | 3.600E-08 | 1.200E-07 |
| 28074 | 1404 | A1404F | E | 4.3e-08 | 4.1e-08 | 1.3e-07 | 7.133E-08 | 5.082E-08 | 4.100E-08 | 1.300E-07 |
| 28075 | 1404 | A1404P | E | 4.3e-08 | 2.7e-08 | 3.3e-08 | 3.433E-08 | 8.083E-09 | 2.700E-08 | 4.300E-08 |
| 28076 | 1404 | A1404S | E | 8.5e-09 | 5.3e-08 | 1e-07 | 5.383E-08 | 4.576E-08 | 8.500E-09 | 1.000E-07 |
| 28077 | 1404 | A1404T | E | 7.1e-09 | 4.9e-08 | 1.1e-07 | 5.537E-08 | 5.174E-08 | 7.100E-09 | 1.100E-07 |
| 28078 | 1404 | A1404W | E | 2.8e-08 | 2e-08 | 3.3e-08 | 2.700E-08 | 6.557E-09 | 2.000E-08 | 3.300E-08 |
| 28079 | 1404 | A1404Y | E | 4e-08 | 3.1e-08 | 5.1e-08 | 4.067E-08 | 1.002E-08 | 3.100E-08 | 5.100E-08 |
| 28080 | 1404 | A1404V | E | 2.6e-08 | 3.2e-08 | 1.1e-07 | 5.600E-08 | 4.686E-08 | 2.600E-08 | 1.100E-07 |
| 28081 | 1405 | P1405A | E | 1.2e-07 | 3.8e-08 | 2.2e-08 | 6.000E-08 | 5.257E-08 | 2.200E-08 | 1.200E-07 |
| 28082 | 1405 | P1405R | E | 5.2e-08 | 3.8e-08 | 3.1e-08 | 4.033E-08 | 1.069E-08 | 3.100E-08 | 5.200E-08 |
| 28083 | 1405 | P1405N | E | 5.1e-08 | 1.1e-07 | 4.4e-08 | 6.833E-08 | 3.625E-08 | 4.400E-08 | 1.100E-07 |
| 28084 | 1405 | P1405D | E | 1.4e-07 | 1.8e-08 | 8.5e-08 | 8.100E-08 | 6.110E-08 | 1.800E-08 | 1.400E-07 |
| 28085 | 1405 | P1405C | E | 5.9e-08 | 4.3e-08 | 5.4e-08 | 5.200E-08 | 8.185E-09 | 4.300E-08 | 5.900E-08 |
| 28086 | 1405 | P1405E | E | 1.8e-08 | 2.7e-08 | 5.9e-08 | 3.467E-08 | 2.155E-08 | 1.800E-08 | 5.900E-08 |
| 28087 | 1405 | P1405Q | E | 5e-08 | 9.2e-08 | 2.9e-08 | 5.700E-08 | 3.208E-08 | 2.900E-08 | 9.200E-08 |
| 28088 | 1405 | P1405G | E | 2.7e-08 | 1.6e-07 | 3.2e-08 | 7.300E-08 | 7.539E-08 | 2.700E-08 | 1.600E-07 |
| 28089 | 1405 | P1405H | E | 2.3e-08 | 3.6e-08 | 4.5e-08 | 3.467E-08 | 1.106E-08 | 2.300E-08 | 4.500E-08 |
| 28090 | 1405 | P1405I | E | 4.9e-08 | 6.3e-08 | 5.2e-08 | 5.467E-08 | 7.371E-09 | 4.900E-08 | 6.300E-08 |
| 28091 | 1405 | P1405L | E | 2.2e-08 | 3e-08 | 3.4e-08 | 2.867E-08 | 6.110E-09 | 2.200E-08 | 3.400E-08 |
| 28092 | 1405 | P1405K | E | 5e-08 | 1.3e-07 | 5.6e-08 | 7.867E-08 | 4.456E-08 | 5.000E-08 | 1.300E-07 |
| 28093 | 1405 | P1405M | E | 3.2e-08 | 3.9e-08 | 2.3e-08 | 3.133E-08 | 8.021E-09 | 2.300E-08 | 3.900E-08 |
| 28094 | 1405 | P1405F | E | 4.6e-08 | 5.8e-08 | 4.4e-08 | 4.933E-08 | 7.572E-09 | 4.400E-08 | 5.800E-08 |
| 28095 | 1405 | P1405P | E | 3.7e-08 | 4e-08 | 1.2e-07 | 6.567E-08 | 4.708E-08 | 3.700E-08 | 1.200E-07 |
| 28096 | 1405 | P1405S | E | 2.5e-08 | 3.4e-08 | 4.3e-08 | 3.400E-08 | 9.000E-09 | 2.500E-08 | 4.300E-08 |
| 28097 | 1405 | P1405T | E | 2.9e-08 | 2.9e-08 | 5.9e-08 | 3.900E-08 | 1.732E-08 | 2.900E-08 | 5.900E-08 |
| 28098 | 1405 | P1405W | E | 1.1e-07 | 4.3e-08 | 6.5e-08 | 7.267E-08 | 3.415E-08 | 4.300E-08 | 1.100E-07 |
| 28099 | 1405 | P1405Y | E | 7.2e-08 | 5e-08 | 7.7e-08 | 6.633E-08 | 1.436E-08 | 5.000E-08 | 7.700E-08 |
| 28100 | 1405 | P1405V | E | 2.3e-08 | 1.1e-07 | 5.4e-08 | 6.233E-08 | 4.409E-08 | 2.300E-08 | 1.100E-07 |
| 28101 | 1406 | S1406A | E | 7.9e-08 | 5.6e-08 | 1.4e-07 | 9.167E-08 | 4.341E-08 | 5.600E-08 | 1.400E-07 |
| 28102 | 1406 | S1406R | E | 5.5e-08 | 1.5e-07 | 3.2e-08 | 7.900E-08 | 6.255E-08 | 3.200E-08 | 1.500E-07 |
| 28103 | 1406 | S1406N | E | 6.4e-08 | 1.5e-08 | 6.2e-08 | 4.700E-08 | 2.773E-08 | 1.500E-08 | 6.400E-08 |
| 28104 | 1406 | S1406D | E | 4.3e-08 | 4.6e-08 | 4e-08 | 4.300E-08 | 3.000E-09 | 4.000E-08 | 4.600E-08 |
| 28105 | 1406 | S1406C | E | 3.1e-08 | 6.6e-08 | 8.1e-08 | 5.933E-08 | 2.566E-08 | 3.100E-08 | 8.100E-08 |
| 28106 | 1406 | S1406E | E | 5.8e-08 | 6.6e-08 | 8.8e-09 | 4.427E-08 | 3.097E-08 | 8.800E-09 | 6.600E-08 |
| 28107 | 1406 | S1406Q | E | 8.6e-08 | 4.3e-08 | 5.6e-08 | 6.167E-08 | 2.205E-08 | 4.300E-08 | 8.600E-08 |
| 28108 | 1406 | S1406G | E | 7.1e-08 | 1.3e-08 | 2.1e-08 | 3.500E-08 | 3.143E-08 | 1.300E-08 | 7.100E-08 |
| 28109 | 1406 | S1406H | E | 3.7e-08 | 6.1e-08 | 5.4e-08 | 5.067E-08 | 1.234E-08 | 3.700E-08 | 6.100E-08 |
| 28110 | 1406 | S1406I | E | 1e-07 | 1.4e-07 | 4.1e-08 | 9.367E-08 | 4.980E-08 | 4.100E-08 | 1.400E-07 |
| 28111 | 1406 | S1406L | E | 6.5e-08 | 4.7e-08 | 1.2e-07 | 7.733E-08 | 3.803E-08 | 4.700E-08 | 1.200E-07 |
| 28112 | 1406 | S1406K | E | 5.2e-08 | 5.3e-08 | 8.8e-08 | 6.433E-08 | 2.050E-08 | 5.200E-08 | 8.800E-08 |
| 28113 | 1406 | S1406M | E | 2.6e-08 | 4.2e-08 | 3.9e-08 | 3.567E-08 | 8.505E-09 | 2.600E-08 | 4.200E-08 |
| 28114 | 1406 | S1406F | E | 3.8e-08 | 5.3e-08 | 2.9e-08 | 4.000E-08 | 1.212E-08 | 2.900E-08 | 5.300E-08 |
| 28115 | 1406 | S1406P | E | 9.8e-08 | 2.6e-08 | 6e-08 | 6.133E-08 | 3.602E-08 | 2.600E-08 | 9.800E-08 |
| 28116 | 1406 | S1406S | E | 2.2e-08 | 3.1e-08 | 1.5e-08 | 2.267E-08 | 8.021E-09 | 1.500E-08 | 3.100E-08 |
| 28117 | 1406 | S1406T | E | 3.7e-08 | 9.9e-08 | 2.9e-08 | 5.500E-08 | 3.831E-08 | 2.900E-08 | 9.900E-08 |
| 28118 | 1406 | S1406W | E | 8.7e-08 | 2.7e-08 | 1.2e-07 | 7.800E-08 | 4.715E-08 | 2.700E-08 | 1.200E-07 |
| 28119 | 1406 | S1406Y | E | 1.1e-07 | 4.1e-08 | 2.8e-08 | 5.967E-08 | 4.407E-08 | 2.800E-08 | 1.100E-07 |
| 28120 | 1406 | S1406V | E | 4.5e-08 | 7.3e-08 | 3.5e-08 | 5.100E-08 | 1.970E-08 | 3.500E-08 | 7.300E-08 |
| 28121 | 1407 | S1407A | E | 4.4e-08 | 1.8e-08 | 4e-08 | 3.400E-08 | 1.400E-08 | 1.800E-08 | 4.400E-08 |
| 28122 | 1407 | S1407R | E | 5.6e-08 | 3.6e-08 | 6.4e-08 | 5.200E-08 | 1.442E-08 | 3.600E-08 | 6.400E-08 |
| 28123 | 1407 | S1407N | E | 1.8e-08 | 1.2e-08 | 6.4e-08 | 3.133E-08 | 2.845E-08 | 1.200E-08 | 6.400E-08 |
| 28124 | 1407 | S1407D | E | 4.4e-08 | 3.3e-08 | 6.7e-08 | 4.800E-08 | 1.735E-08 | 3.300E-08 | 6.700E-08 |
| 28125 | 1407 | S1407C | E | 7.5e-08 | 2e-08 | 5.2e-08 | 4.900E-08 | 2.762E-08 | 2.000E-08 | 7.500E-08 |
| 28126 | 1407 | S1407E | E | 3e-08 | 5.3e-08 | 2.3e-08 | 3.533E-08 | 1.570E-08 | 2.300E-08 | 5.300E-08 |
| 28127 | 1407 | S1407Q | E | 6e-08 | 2.3e-08 | 5.2e-08 | 4.500E-08 | 1.947E-08 | 2.300E-08 | 6.000E-08 |
| 28128 | 1407 | S1407G | E | 5.9e-08 | 1.1e-07 | 2.1e-08 | 6.333E-08 | 4.466E-08 | 2.100E-08 | 1.100E-07 |
| 28129 | 1407 | S1407H | E | 5.9e-08 | 8.4e-08 | 6.1e-08 | 6.800E-08 | 1.389E-08 | 5.900E-08 | 8.400E-08 |
| 28130 | 1407 | S1407I | E | 5.7e-08 | 5.3e-08 | 8.1e-08 | 6.367E-08 | 1.514E-08 | 5.300E-08 | 8.100E-08 |
| 28131 | 1407 | S1407L | E | 7.9e-08 | 5.9e-08 | 3.9e-08 | 5.900E-08 | 2.000E-08 | 3.900E-08 | 7.900E-08 |
| 28132 | 1407 | S1407K | E | 5.7e-08 | 5.4e-08 | 7.4e-08 | 6.167E-08 | 1.079E-08 | 5.400E-08 | 7.400E-08 |
| 28133 | 1407 | S1407M | E | 5.8e-08 | 2.9e-08 | 3.8e-08 | 4.167E-08 | 1.484E-08 | 2.900E-08 | 5.800E-08 |
| 28134 | 1407 | S1407F | E | 5.1e-08 | 3.3e-08 | 4.3e-08 | 4.233E-08 | 9.018E-09 | 3.300E-08 | 5.100E-08 |

| | | | | | | | | | | |
|---|---|---|---|---|---|---|---|---|---|---|
| 28135 | 1407 | S1407P | E | 1.9e-08 | 1.5e-08 | 2.1e-08 | 1.833E-08 | 3.055E-09 | 1.500E-08 | 2.100E-08 |
| 28136 | 1407 | S1407S | E | 4.7e-08 | 1.5e-07 | 2.2e-08 | 7.300E-08 | 6.785E-08 | 2.200E-08 | 1.500E-07 |
| 28137 | 1407 | S1407T | E | 1.9e-08 | 5e-08 | 3.8e-08 | 3.567E-08 | 1.563E-08 | 1.900E-08 | 5.000E-08 |
| 28138 | 1407 | S1407W | E | 1.5e-08 | 4.5e-08 | 6.7e-08 | 4.233E-08 | 2.610E-08 | 1.500E-08 | 6.700E-08 |
| 28139 | 1407 | S1407Y | E | 4.1e-08 | 6.5e-08 | 1.9e-08 | 4.167E-08 | 2.301E-08 | 1.900E-08 | 6.500E-08 |
| 28140 | 1407 | S1407V | E | 2.7e-08 | 6e-08 | 1e-07 | 6.233E-08 | 3.656E-08 | 2.700E-08 | 1.000E-07 |
| 28141 | 1408 | K1408A | E | 3.5e-08 | 7.5e-08 | 5.3e-08 | 5.433E-08 | 2.003E-08 | 3.500E-08 | 7.500E-08 |
| 28142 | 1408 | K1408R | E | 5.7e-08 | 4.6e-08 | 5.2e-08 | 5.167E-08 | 5.508E-09 | 4.600E-08 | 5.700E-08 |
| 28143 | 1408 | K1408N | E | 5.3e-08 | 4.6e-08 | 1.9e-08 | 3.933E-08 | 1.795E-08 | 1.900E-08 | 5.300E-08 |
| 28144 | 1408 | K1408D | E | 3.2e-08 | 9.4e-08 | 5.3e-08 | 5.967E-08 | 3.153E-08 | 3.200E-08 | 9.400E-08 |
| 28145 | 1408 | K1408C | E | 4.3e-08 | 9.6e-08 | 3.6e-08 | 5.833E-08 | 3.281E-08 | 3.600E-08 | 9.600E-08 |
| 28146 | 1408 | K1408E | E | 3.2e-08 | 3.2e-08 | 3.7e-08 | 3.367E-08 | 2.887E-09 | 3.200E-08 | 3.700E-08 |
| 28147 | 1408 | K1408Q | E | 1.8e-08 | 4.5e-08 | 7.3e-08 | 4.533E-08 | 2.750E-08 | 1.800E-08 | 7.300E-08 |
| 28148 | 1408 | K1408G | E | 6.4e-08 | 6.2e-08 | 5.4e-08 | 6.000E-08 | 5.292E-09 | 5.400E-08 | 6.400E-08 |
| 28149 | 1408 | K1408H | E | 2.9e-08 | 7.2e-08 | 2.3e-08 | 4.133E-08 | 2.673E-08 | 2.300E-08 | 7.200E-08 |
| 28150 | 1408 | K1408I | E | 6.9e-08 | 6.3e-08 | 3.1e-08 | 5.433E-08 | 2.043E-08 | 3.100E-08 | 6.900E-08 |
| 28151 | 1408 | K1408L | E | 9.5e-08 | 4e-08 | 2e-08 | 5.167E-08 | 3.884E-08 | 2.000E-08 | 9.500E-08 |
| 28152 | 1408 | K1408K | E | 4.9e-08 | 2.1e-08 | 4.4e-08 | 3.800E-08 | 1.493E-08 | 2.100E-08 | 4.900E-08 |
| 28153 | 1408 | K1408M | E | 9.7e-08 | 2.2e-08 | 3.6e-08 | 5.167E-08 | 3.988E-08 | 2.200E-08 | 9.700E-08 |
| 28154 | 1408 | K1408F | E | 3.4e-08 | 2.6e-08 | 5.8e-08 | 3.933E-08 | 1.665E-08 | 2.600E-08 | 5.800E-08 |
| 28155 | 1408 | K1408P | E | 4.1e-08 | 2.1e-08 | 6.4e-08 | 4.200E-08 | 2.152E-08 | 2.100E-08 | 6.400E-08 |
| 28156 | 1408 | K1408S | E | 1.1e-07 | 3.4e-08 | 5.9e-08 | 6.767E-08 | 3.873E-08 | 3.400E-08 | 1.100E-07 |
| 28157 | 1408 | K1408T | E | 1.1e-08 | 3.7e-08 | 2.4e-08 | 2.400E-08 | 1.300E-08 | 1.100E-08 | 3.700E-08 |
| 28158 | 1408 | K1408W | E | 2.9e-08 | 2.3e-08 | 1.9e-08 | 2.367E-08 | 5.033E-09 | 1.900E-08 | 2.900E-08 |
| 28159 | 1408 | K1408Y | E | 6.7e-08 | 1.1e-07 | 5.2e-08 | 7.633E-08 | 3.011E-08 | 5.200E-08 | 1.100E-07 |
| 28160 | 1408 | K1408V | E | 7.1e-08 | 7.1e-08 | 3e-08 | 5.733E-08 | 2.367E-08 | 3.000E-08 | 7.100E-08 |
| 28161 | 1409 | S1409A | E | 4.3e-08 | 8.3e-08 | 4.2e-08 | 5.600E-08 | 2.339E-08 | 4.200E-08 | 8.300E-08 |
| 28162 | 1409 | S1409R | E | 6.8e-08 | 3.1e-08 | 5.9e-08 | 5.267E-08 | 1.930E-08 | 3.100E-08 | 6.800E-08 |
| 28163 | 1409 | S1409N | E | 2.7e-08 | 2.4e-08 | 1.1e-07 | 5.367E-08 | 4.881E-08 | 2.400E-08 | 1.100E-07 |
| 28164 | 1409 | S1409D | E | 6.9e-08 | 2.3e-08 | 1.8e-08 | 3.667E-08 | 2.811E-08 | 1.800E-08 | 6.900E-08 |
| 28165 | 1409 | S1409C | E | 2.2e-08 | 2.1e-08 | 4.5e-08 | 2.933E-08 | 1.358E-08 | 2.100E-08 | 4.500E-08 |
| 28166 | 1409 | S1409E | E | 2.4e-08 | 1.2e-07 | 4.4e-08 | 6.267E-08 | 5.065E-08 | 2.400E-08 | 1.200E-07 |
| 28167 | 1409 | S1409Q | E | 2.5e-08 | 7e-08 | 8.8e-09 | 3.460E-08 | 3.171E-08 | 8.800E-09 | 7.000E-08 |
| 28168 | 1409 | S1409G | E | 6.5e-08 | 8.6e-08 | 2.3e-08 | 5.800E-08 | 3.208E-08 | 2.300E-08 | 8.600E-08 |
| 28169 | 1409 | S1409H | E | 4e-08 | 4.2e-08 | 2.8e-08 | 3.667E-08 | 7.572E-09 | 2.800E-08 | 4.200E-08 |
| 28170 | 1409 | S1409I | E | 3e-08 | 5.2e-08 | 4.2e-08 | 4.133E-08 | 1.102E-08 | 3.000E-08 | 5.200E-08 |
| 28171 | 1409 | S1409L | E | 1.8e-07 | 4.4e-08 | 3.9e-08 | 8.767E-08 | 8.000E-08 | 3.900E-08 | 1.800E-07 |
| 28172 | 1409 | S1409K | E | 4.5e-08 | 6.4e-08 | 6.5e-08 | 5.800E-08 | 1.127E-08 | 4.500E-08 | 6.500E-08 |
| 28173 | 1409 | S1409M | E | 2.9e-08 | 4.4e-08 | 6.3e-08 | 4.533E-08 | 1.704E-08 | 2.900E-08 | 6.300E-08 |
| 28174 | 1409 | S1409F | E | 1.4e-08 | 1.3e-08 | 2.8e-08 | 1.833E-08 | 8.386E-09 | 1.300E-08 | 2.800E-08 |
| 28175 | 1409 | S1409P | E | 1.9e-08 | 5.3e-08 | 7.4e-08 | 4.867E-08 | 2.775E-08 | 1.900E-08 | 7.400E-08 |
| 28176 | 1409 | S1409S | E | 3.7e-08 | 2.5e-08 | 9.5e-08 | 5.233E-08 | 3.743E-08 | 2.500E-08 | 9.500E-08 |
| 28177 | 1409 | S1409T | E | 1.3e-08 | 7.7e-08 | 2e-08 | 3.667E-08 | 3.510E-08 | 1.300E-08 | 7.700E-08 |
| 28178 | 1409 | S1409W | E | 7.7e-08 | 4.7e-08 | 5e-08 | 5.800E-08 | 1.652E-08 | 4.700E-08 | 7.700E-08 |
| 28179 | 1409 | S1409Y | E | 6.9e-08 | 2.2e-08 | 6.5e-08 | 5.200E-08 | 2.606E-08 | 2.200E-08 | 6.900E-08 |
| 28180 | 1409 | S1409V | E | 3.8e-08 | 4.8e-08 | 7.5e-08 | 5.367E-08 | 1.914E-08 | 3.800E-08 | 7.500E-08 |
| 28181 | 1410 | T1410A | E | 2.9e-08 | 2.4e-08 | 6.4e-08 | 3.900E-08 | 2.179E-08 | 2.400E-08 | 6.400E-08 |
| 28182 | 1410 | T1410R | E | 4.4e-08 | 2.9e-08 | 1.8e-08 | 3.033E-08 | 1.305E-08 | 1.800E-08 | 4.400E-08 |
| 28183 | 1410 | T1410N | E | 9e-08 | 7.4e-08 | 1.7e-08 | 6.033E-08 | 3.837E-08 | 1.700E-08 | 9.000E-08 |
| 28184 | 1410 | T1410D | E | 5.5e-08 | 3e-08 | 4.2e-08 | 4.233E-08 | 1.250E-08 | 3.000E-08 | 5.500E-08 |
| 28185 | 1410 | T1410C | E | 1.7e-08 | 6.2e-08 | 6.5e-08 | 4.800E-08 | 2.689E-08 | 1.700E-08 | 6.500E-08 |
| 28186 | 1410 | T1410E | E | 8.4e-08 | 1.7e-08 | 4.3e-08 | 4.800E-08 | 3.378E-08 | 1.700E-08 | 8.400E-08 |
| 28187 | 1410 | T1410Q | E | 3.7e-08 | 1.6e-08 | 4.6e-08 | 3.300E-08 | 1.539E-08 | 1.600E-08 | 4.600E-08 |
| 28188 | 1410 | T1410G | E | 1.1e-07 | 3.4e-08 | 4.8e-08 | 6.400E-08 | 4.045E-08 | 3.400E-08 | 1.100E-07 |
| 28189 | 1410 | T1410H | E | 1.9e-08 | 2.9e-08 | 4.4e-08 | 3.067E-08 | 1.258E-08 | 1.900E-08 | 4.400E-08 |
| 28190 | 1410 | T1410I | E | 2.1e-07 | 4.3e-08 | 2.1e-08 | 9.133E-08 | 1.034E-07 | 2.100E-08 | 2.100E-07 |
| 28191 | 1410 | T1410L | E | 9.1e-08 | 2.4e-08 | 7.4e-08 | 6.300E-08 | 3.483E-08 | 2.400E-08 | 9.100E-08 |
| 28192 | 1410 | T1410K | E | 1.5e-08 | 4e-08 | 6.2e-08 | 3.900E-08 | 2.352E-08 | 1.500E-08 | 6.200E-08 |
| 28193 | 1410 | T1410M | E | 6.3e-08 | 3.7e-08 | 6.1e-08 | 5.367E-08 | 1.447E-08 | 3.700E-08 | 6.300E-08 |
| 28194 | 1410 | T1410F | E | 3.4e-08 | 2.6e-08 | 5.3e-08 | 3.767E-08 | 1.387E-08 | 2.600E-08 | 5.300E-08 |
| 28195 | 1410 | T1410P | E | 2.5e-08 | 6.9e-08 | 2.3e-08 | 3.900E-08 | 2.600E-08 | 2.300E-08 | 6.900E-08 |
| 28196 | 1410 | T1410S | E | 2.2e-08 | 3.9e-08 | 1.6e-08 | 2.567E-08 | 1.193E-08 | 1.600E-08 | 3.900E-08 |
| 28197 | 1410 | T1410T | E | 4.2e-08 | 4.1e-08 | 4.8e-08 | 4.367E-08 | 3.786E-09 | 4.100E-08 | 4.800E-08 |
| 28198 | 1410 | T1410W | E | 1.4e-08 | 5.9e-08 | 5.1e-08 | 4.133E-08 | 2.401E-08 | 1.400E-08 | 5.900E-08 |
| 28199 | 1410 | T1410Y | E | 5.4e-08 | 2.2e-08 | 5.7e-08 | 4.433E-08 | 1.940E-08 | 2.200E-08 | 5.700E-08 |
| 28200 | 1410 | T1410V | E | 3.8e-08 | 5.8e-08 | 6.3e-08 | 5.300E-08 | 1.323E-08 | 3.800E-08 | 6.300E-08 |
| 28201 | 1411 | S1411A | E | 8.1e-09 | 3.4e-08 | 2.6e-08 | 2.270E-08 | 1.326E-08 | 8.100E-09 | 3.400E-08 |
| 28202 | 1411 | S1411R | E | 9.5e-08 | 7.7e-08 | 5.2e-08 | 7.467E-08 | 2.159E-08 | 5.200E-08 | 9.500E-08 |
| 28203 | 1411 | S1411N | E | 5.5e-08 | 2.9e-08 | 3.3e-08 | 3.900E-08 | 1.400E-08 | 2.900E-08 | 5.500E-08 |
| 28204 | 1411 | S1411D | E | 1.2e-08 | 3.1e-08 | 4.4e-08 | 2.900E-08 | 1.609E-08 | 1.200E-08 | 4.400E-08 |
| 28205 | 1411 | S1411C | E | 5.8e-08 | 5.8e-08 | 4e-08 | 5.200E-08 | 1.039E-08 | 4.000E-08 | 5.800E-08 |
| 28206 | 1411 | S1411E | E | 3.1e-08 | 8.7e-08 | 2e-07 | 1.060E-07 | 8.609E-08 | 3.100E-08 | 2.000E-07 |
| 28207 | 1411 | S1411Q | E | 1.8e-08 | 4.9e-08 | 3.1e-08 | 3.267E-08 | 1.557E-08 | 1.800E-08 | 4.900E-08 |
| 28208 | 1411 | S1411G | E | 3e-08 | 7.1e-08 | 2.4e-08 | 4.167E-08 | 2.558E-08 | 2.400E-08 | 7.100E-08 |
| 28209 | 1411 | S1411H | E | 3.5e-08 | 1e-08 | 5.3e-08 | 3.267E-08 | 2.159E-08 | 1.000E-08 | 5.300E-08 |
| 28210 | 1411 | S1411I | E | 8.1e-08 | 6.2e-08 | 4.5e-08 | 6.267E-08 | 1.801E-08 | 4.500E-08 | 8.100E-08 |
| 28211 | 1411 | S1411L | E | 2.5e-08 | 3.9e-08 | 8.9e-08 | 5.100E-08 | 3.365E-08 | 2.500E-08 | 8.900E-08 |
| 28212 | 1411 | S1411K | E | 1.8e-08 | 3.6e-08 | 2e-08 | 2.467E-08 | 9.866E-09 | 1.800E-08 | 3.600E-08 |
| 28213 | 1411 | S1411M | E | 2.2e-08 | 1e-07 | 4.3e-08 | 5.500E-08 | 4.036E-08 | 2.200E-08 | 1.000E-07 |
| 28214 | 1411 | S1411F | E | 1.3e-07 | 8.1e-08 | 3.3e-08 | 8.133E-08 | 4.850E-08 | 3.300E-08 | 1.300E-07 |
| 28215 | 1411 | S1411P | E | 4.7e-08 | 8.5e-08 | 2.9e-08 | 5.367E-08 | 2.859E-08 | 2.900E-08 | 8.500E-08 |
| 28216 | 1411 | S1411S | E | 6.3e-08 | 4.4e-08 | 7.8e-08 | 6.167E-08 | 1.704E-08 | 4.400E-08 | 7.800E-08 |
| 28217 | 1411 | S1411T | E | 6.3e-08 | 2.5e-08 | 5.3e-08 | 4.700E-08 | 1.970E-08 | 2.500E-08 | 6.300E-08 |
| 28218 | 1411 | S1411W | E | 1e-07 | 2.2e-08 | 4e-08 | 5.400E-08 | 4.084E-08 | 2.200E-08 | 1.000E-07 |
| 28219 | 1411 | S1411Y | E | 5.1e-08 | 9e-08 | 6.7e-08 | 6.933E-08 | 1.960E-08 | 5.100E-08 | 9.000E-08 |

| | | | | | | | | | | |
|---|---|---|---|---|---|---|---|---|---|---|
| 28220 | 1411 | S1411V | E | 1.1e-07 | 4.1e-08 | 6.8e-08 | 7.300E-08 | 3.477E-08 | 4.100E-08 | 1.100E-07 |
| 28221 | 1412 | G1412A | E | 2.1e-08 | 6.1e-08 | 1.8e-08 | 3.333E-08 | 2.401E-08 | 1.800E-08 | 6.100E-08 |
| 28222 | 1412 | G1412R | E | 2.1e-08 | 3.5e-08 | 7e-08 | 4.200E-08 | 2.524E-08 | 2.100E-08 | 7.000E-08 |
| 28223 | 1412 | G1412N | E | 3.4e-08 | 2.5e-08 | 4.5e-08 | 3.467E-08 | 1.002E-08 | 2.500E-08 | 4.500E-08 |
| 28224 | 1412 | G1412D | E | 3.4e-08 | 4.5e-08 | 3.5e-08 | 3.800E-08 | 6.083E-09 | 3.400E-08 | 4.500E-08 |
| 28225 | 1412 | G1412C | E | 2.4e-08 | 3.1e-08 | 3.7e-08 | 3.067E-08 | 6.506E-09 | 2.400E-08 | 3.700E-08 |
| 28226 | 1412 | G1412E | E | 6e-08 | 6.4e-08 | 3.1e-08 | 5.167E-08 | 1.801E-08 | 3.100E-08 | 6.400E-08 |
| 28227 | 1412 | G1412Q | E | 5e-08 | 5.3e-08 | 5.3e-08 | 5.200E-08 | 1.732E-09 | 5.000E-08 | 5.300E-08 |
| 28228 | 1412 | G1412G | E | 5.7e-08 | 6.8e-08 | 1.5e-08 | 4.667E-08 | 2.797E-08 | 1.500E-08 | 6.800E-08 |
| 28229 | 1412 | G1412H | E | 4.9e-08 | 9.9e-08 | 1.7e-08 | 5.500E-08 | 4.133E-08 | 1.700E-08 | 9.900E-08 |
| 28230 | 1412 | G1412I | E | 3.5e-08 | 5.7e-08 | 4.8e-08 | 4.667E-08 | 1.106E-08 | 3.500E-08 | 5.700E-08 |
| 28231 | 1412 | G1412L | E | 5.4e-08 | 6.3e-08 | 2.4e-08 | 4.700E-08 | 2.042E-08 | 2.400E-08 | 6.300E-08 |
| 28232 | 1412 | G1412K | E | 5.6e-08 | 5e-08 | 5.2e-08 | 5.267E-08 | 3.055E-09 | 5.000E-08 | 5.600E-08 |
| 28233 | 1412 | G1412M | E | 3.6e-08 | 1.7e-08 | 4.7e-08 | 3.333E-08 | 1.518E-08 | 1.700E-08 | 4.700E-08 |
| 28234 | 1412 | G1412F | E | 1e-07 | 2.8e-08 | 3.8e-08 | 5.533E-08 | 3.900E-08 | 2.800E-08 | 1.000E-07 |
| 28235 | 1412 | G1412P | E | 6e-08 | 3.4e-08 | 4.2e-08 | 4.533E-08 | 1.332E-08 | 3.400E-08 | 6.000E-08 |
| 28236 | 1412 | G1412S | E | 4.5e-08 | 1.3e-08 | 5.5e-08 | 3.767E-08 | 2.194E-08 | 1.300E-08 | 5.500E-08 |
| 28237 | 1412 | G1412T | E | 2.2e-08 | 7.5e-08 | 1e-07 | 6.567E-08 | 3.983E-08 | 2.200E-08 | 1.000E-07 |
| 28238 | 1412 | G1412W | E | 2.4e-08 | 4.4e-08 | 5.9e-08 | 4.233E-08 | 1.756E-08 | 2.400E-08 | 5.900E-08 |
| 28239 | 1412 | G1412Y | E | 4.4e-08 | 2.8e-08 | 8.6e-08 | 5.267E-08 | 2.996E-08 | 2.800E-08 | 8.600E-08 |
| 28240 | 1412 | G1412V | E | 3.4e-08 | 1.1e-07 | 2.1e-07 | 1.180E-07 | 8.827E-08 | 3.400E-08 | 2.100E-07 |
| 28241 | 1413 | G1413A | E | 1.7e-08 | 7.1e-08 | 2.3e-08 | 3.700E-08 | 2.960E-08 | 1.700E-08 | 7.100E-08 |
| 28242 | 1413 | G1413R | E | 3.5e-08 | 4.6e-08 | 4.4e-08 | 4.167E-08 | 5.859E-09 | 3.500E-08 | 4.600E-08 |
| 28243 | 1413 | G1413N | E | 9.7e-08 | 3.4e-08 | 4.9e-08 | 6.000E-08 | 3.291E-08 | 3.400E-08 | 9.700E-08 |
| 28244 | 1413 | G1413D | E | 6.4e-08 | 7.9e-08 | 6.4e-08 | 6.900E-08 | 8.660E-09 | 6.400E-08 | 7.900E-08 |
| 28245 | 1413 | G1413C | E | 2.9e-08 | 2.7e-08 | 4e-08 | 3.200E-08 | 7.000E-09 | 2.700E-08 | 4.000E-08 |
| 28246 | 1413 | G1413E | E | 5e-08 | 2.9e-08 | 1.1e-07 | 6.300E-08 | 4.204E-08 | 2.900E-08 | 1.100E-07 |
| 28247 | 1413 | G1413Q | E | 6.9e-08 | 3.9e-08 | 5.8e-08 | 5.533E-08 | 1.518E-08 | 3.900E-08 | 6.900E-08 |
| 28248 | 1413 | G1413G | E | 3.6e-08 | 1.5e-07 | 6.5e-08 | 8.367E-08 | 5.925E-08 | 3.600E-08 | 1.500E-07 |
| 28249 | 1413 | G1413H | E | 2.1e-08 | 8e-08 | 7.6e-08 | 5.900E-08 | 3.297E-08 | 2.100E-08 | 8.000E-08 |
| 28250 | 1413 | G1413I | E | 4.7e-08 | 7.9e-08 | 3.3e-08 | 5.300E-08 | 2.358E-08 | 3.300E-08 | 7.900E-08 |
| 28251 | 1413 | G1413L | E | 2.4e-08 | 6.3e-08 | 8.8e-08 | 5.833E-08 | 3.225E-08 | 2.400E-08 | 8.800E-08 |
| 28252 | 1413 | G1413K | E | 1.8e-08 | 3e-08 | 2e-08 | 2.267E-08 | 6.429E-09 | 1.800E-08 | 3.000E-08 |
| 28253 | 1413 | G1413M | E | 2.2e-08 | 3.3e-08 | 7.3e-08 | 4.267E-08 | 2.684E-08 | 2.200E-08 | 7.300E-08 |
| 28254 | 1413 | G1413F | E | 5.3e-08 | 1.7e-08 | 1.7e-08 | 2.900E-08 | 2.078E-08 | 1.700E-08 | 5.300E-08 |
| 28255 | 1413 | G1413P | E | 2.7e-08 | 4.2e-08 | 3.5e-08 | 3.467E-08 | 7.506E-09 | 2.700E-08 | 4.200E-08 |
| 28256 | 1413 | G1413S | E | 4.3e-08 | 2.5e-08 | 8.1e-08 | 4.967E-08 | 2.859E-08 | 2.500E-08 | 8.100E-08 |
| 28257 | 1413 | G1413T | E | 3.6e-08 | 2.6e-08 | 4.1e-08 | 3.433E-08 | 7.638E-09 | 2.600E-08 | 4.100E-08 |
| 28258 | 1413 | G1413W | E | 4.7e-08 | 9.4e-08 | 3.8e-08 | 5.967E-08 | 3.007E-08 | 3.800E-08 | 9.400E-08 |
| 28259 | 1413 | G1413Y | E | 1e-08 | 6.9e-08 | 8e-08 | 5.300E-08 | 3.764E-08 | 1.000E-08 | 8.000E-08 |
| 28260 | 1413 | G1413V | E | 3.4e-08 | 3.1e-08 | 3.1e-08 | 3.200E-08 | 1.732E-09 | 3.100E-08 | 3.400E-08 |
| 28261 | 1414 | T1414A | E | 7e-08 | 2.7e-08 | 1.2e-08 | 3.633E-08 | 3.011E-08 | 1.200E-08 | 7.000E-08 |
| 28262 | 1414 | T1414R | E | 4.5e-08 | 2.5e-08 | 1.7e-08 | 2.900E-08 | 1.442E-08 | 1.700E-08 | 4.500E-08 |
| 28263 | 1414 | T1414N | E | 4.9e-08 | 1e-07 | 1.9e-08 | 5.600E-08 | 4.095E-08 | 1.900E-08 | 1.000E-07 |
| 28264 | 1414 | T1414D | E | 3.5e-08 | 1.4e-07 | 3.2e-08 | 6.900E-08 | 6.151E-08 | 3.200E-08 | 1.400E-07 |
| 28265 | 1414 | T1414C | E | 4.4e-08 | 1.5e-08 | 1.3e-08 | 2.400E-08 | 1.735E-08 | 1.300E-08 | 4.400E-08 |
| 28266 | 1414 | T1414E | E | 2.9e-08 | 2e-08 | 2.5e-08 | 2.467E-08 | 4.509E-09 | 2.000E-08 | 2.900E-08 |
| 28267 | 1414 | T1414Q | E | 2.7e-08 | 1.9e-08 | 9.2e-08 | 4.600E-08 | 4.004E-08 | 1.900E-08 | 9.200E-08 |
| 28268 | 1414 | T1414G | E | 1.7e-08 | 1e-07 | 3.2e-08 | 4.967E-08 | 4.423E-08 | 1.700E-08 | 1.000E-07 |
| 28269 | 1414 | T1414H | E | 2.4e-08 | 2.4e-08 | 6.2e-08 | 3.667E-08 | 2.194E-08 | 2.400E-08 | 6.200E-08 |
| 28270 | 1414 | T1414I | E | 5.2e-08 | 2.1e-08 | 3.1e-08 | 3.467E-08 | 1.582E-08 | 2.100E-08 | 5.200E-08 |
| 28271 | 1414 | T1414L | E | 7.1e-08 | 1.9e-08 | 3.6e-08 | 4.200E-08 | 2.651E-08 | 1.900E-08 | 7.100E-08 |
| 28272 | 1414 | T1414K | E | 1.1e-08 | 4.9e-08 | 3.9e-08 | 3.300E-08 | 1.970E-08 | 1.100E-08 | 4.900E-08 |
| 28273 | 1414 | T1414M | E | 1.8e-08 | 2.5e-08 | 2.7e-08 | 2.333E-08 | 4.726E-09 | 1.800E-08 | 2.700E-08 |
| 28274 | 1414 | T1414F | E | 2.7e-08 | 4.2e-08 | 1.8e-08 | 2.900E-08 | 1.212E-08 | 1.800E-08 | 4.200E-08 |
| 28275 | 1414 | T1414P | E | 7.9e-08 | 1.8e-08 | 4.6e-08 | 4.767E-08 | 3.053E-08 | 1.800E-08 | 7.900E-08 |
| 28276 | 1414 | T1414S | E | 2.5e-08 | 7.7e-08 | 1.6e-08 | 3.933E-08 | 3.293E-08 | 1.600E-08 | 7.700E-08 |
| 28277 | 1414 | T1414T | E | 2.8e-08 | 4.7e-08 | 5.1e-08 | 4.200E-08 | 1.229E-08 | 2.800E-08 | 5.100E-08 |
| 28278 | 1414 | T1414W | E | 9.6e-08 | 7e-08 | 4.9e-08 | 7.167E-08 | 2.354E-08 | 4.900E-08 | 9.600E-08 |
| 28279 | 1414 | T1414Y | E | 2.2e-08 | 2.3e-08 | 3.6e-08 | 2.700E-08 | 7.810E-09 | 2.200E-08 | 3.600E-08 |
| 28280 | 1414 | T1414V | E | 3.4e-08 | 5.5e-08 | 6e-08 | 4.967E-08 | 1.380E-08 | 3.400E-08 | 6.000E-08 |
| 28281 | 1415 | A1415A | E | 2.8e-08 | 6.1e-08 | 5.5e-08 | 4.800E-08 | 1.758E-08 | 2.800E-08 | 6.100E-08 |
| 28282 | 1415 | A1415R | E | 4.2e-08 | 7.6e-08 | 2.3e-08 | 4.700E-08 | 2.685E-08 | 2.300E-08 | 7.600E-08 |
| 28283 | 1415 | A1415N | E | 8.6e-08 | 3.3e-08 | 9e-08 | 6.967E-08 | 3.182E-08 | 3.300E-08 | 9.000E-08 |
| 28284 | 1415 | A1415D | E | 6.2e-09 | 4e-08 | 5.3e-08 | 3.307E-08 | 2.416E-08 | 6.200E-09 | 5.300E-08 |
| 28285 | 1415 | A1415C | E | 9.9e-09 | 9.3e-08 | 2.9e-08 | 4.397E-08 | 4.352E-08 | 9.900E-09 | 9.300E-08 |
| 28286 | 1415 | A1415E | E | 5.9e-08 | 3.5e-08 | 4e-08 | 4.467E-08 | 1.266E-08 | 3.500E-08 | 5.900E-08 |
| 28287 | 1415 | A1415Q | E | 3.8e-08 | 4.9e-08 | 5.6e-08 | 4.767E-08 | 9.074E-09 | 3.800E-08 | 5.600E-08 |
| 28288 | 1415 | A1415G | E | 7.3e-08 | 2.4e-07 | 4e-08 | 1.177E-07 | 1.072E-07 | 4.000E-08 | 2.400E-07 |
| 28289 | 1415 | A1415H | E | 9.4e-08 | 2.3e-08 | 4.4e-08 | 5.367E-08 | 3.647E-08 | 2.300E-08 | 9.400E-08 |
| 28290 | 1415 | A1415I | E | 2.2e-08 | 3.7e-08 | 2.1e-08 | 2.667E-08 | 8.963E-09 | 2.100E-08 | 3.700E-08 |
| 28291 | 1415 | A1415L | E | 4.5e-08 | 5.8e-08 | 3.7e-08 | 4.667E-08 | 1.060E-08 | 3.700E-08 | 5.800E-08 |
| 28292 | 1415 | A1415K | E | 3.7e-08 | 6.8e-08 | 2e-08 | 4.167E-08 | 2.434E-08 | 2.000E-08 | 6.800E-08 |
| 28293 | 1415 | A1415M | E | 4.2e-08 | 2.6e-08 | 4.3e-08 | 3.700E-08 | 9.539E-09 | 2.600E-08 | 4.300E-08 |
| 28294 | 1415 | A1415F | E | 1.9e-08 | 3.6e-08 | 4.9e-08 | 3.467E-08 | 1.504E-08 | 1.900E-08 | 4.900E-08 |
| 28295 | 1415 | A1415P | E | 1.3e-08 | 3.6e-08 | 3.1e-08 | 2.667E-08 | 1.210E-08 | 1.300E-08 | 3.600E-08 |
| 28296 | 1415 | A1415S | E | 1.9e-08 | 5.1e-08 | 3e-08 | 3.333E-08 | 1.626E-08 | 1.900E-08 | 5.100E-08 |
| 28297 | 1415 | A1415T | E | 3.5e-08 | 5.7e-08 | 2.6e-08 | 3.933E-08 | 1.595E-08 | 2.600E-08 | 5.700E-08 |
| 28298 | 1415 | A1415W | E | 3.4e-08 | 3.1e-08 | 5.8e-08 | 4.100E-08 | 1.480E-08 | 3.100E-08 | 5.800E-08 |
| 28299 | 1415 | A1415Y | E | 5.4e-08 | 9.1e-08 | 2.3e-08 | 5.600E-08 | 3.404E-08 | 2.300E-08 | 9.100E-08 |
| 28300 | 1415 | A1415V | E | 5.7e-08 | 7.4e-08 | 6.2e-08 | 6.433E-08 | 8.737E-09 | 5.700E-08 | 7.400E-08 |
| 28301 | 1416 | A1416A | E | 3.6e-08 | 1.1e-07 | 1.7e-08 | 5.433E-08 | 4.914E-08 | 1.700E-08 | 1.100E-07 |
| 28302 | 1416 | A1416R | E | 6.3e-08 | 5.7e-08 | 5.3e-08 | 5.767E-08 | 5.033E-09 | 5.300E-08 | 6.300E-08 |
| 28303 | 1416 | A1416N | E | 5.4e-08 | 2.3e-08 | 4.5e-08 | 4.067E-08 | 1.595E-08 | 2.300E-08 | 5.400E-08 |
| 28304 | 1416 | A1416D | E | 5.1e-08 | 4.4e-08 | 3.4e-08 | 4.300E-08 | 8.544E-09 | 3.400E-08 | 5.100E-08 |

| | | | | | | | | | | |
|---|---|---|---|---|---|---|---|---|---|---|
| 28305 | 1416 | A1416C | E | 4e-08 | 5e-08 | 4.4e-08 | 4.467E-08 | 5.033E-09 | 4.000E-08 | 5.000E-08 |
| 28306 | 1416 | A1416E | E | 6e-08 | 3.7e-08 | 6.3e-08 | 5.333E-08 | 1.422E-08 | 3.700E-08 | 6.300E-08 |
| 28307 | 1416 | A1416Q | E | 5.3e-08 | 6.5e-08 | 4.5e-08 | 5.433E-08 | 1.007E-08 | 4.500E-08 | 6.500E-08 |
| 28308 | 1416 | A1416G | E | 4.8e-08 | 4.9e-08 | 2.1e-08 | 3.933E-08 | 1.589E-08 | 2.100E-08 | 4.900E-08 |
| 28309 | 1416 | A1416H | E | 4e-08 | 7.7e-08 | 6.4e-08 | 6.033E-08 | 1.877E-08 | 4.000E-08 | 7.700E-08 |
| 28310 | 1416 | A1416I | E | 5.2e-08 | 2.1e-08 | 4.4e-08 | 3.900E-08 | 1.609E-08 | 2.100E-08 | 5.200E-08 |
| 28311 | 1416 | A1416L | E | 3e-08 | 2.5e-08 | 1.1e-07 | 5.500E-08 | 4.770E-08 | 2.500E-08 | 1.100E-07 |
| 28312 | 1416 | A1416K | E | 2.9e-08 | 4.3e-08 | 7.1e-08 | 4.767E-08 | 2.139E-08 | 2.900E-08 | 7.100E-08 |
| 28313 | 1416 | A1416M | E | 5.6e-08 | 4.3e-08 | 1.9e-08 | 3.933E-08 | 1.877E-08 | 1.900E-08 | 5.600E-08 |
| 28314 | 1416 | A1416F | E | 8e-08 | 4.1e-08 | 5.4e-08 | 5.833E-08 | 1.986E-08 | 4.100E-08 | 8.000E-08 |
| 28315 | 1416 | A1416P | E | 5.1e-08 | 2.4e-08 | 8e-08 | 5.167E-08 | 2.801E-08 | 2.400E-08 | 8.000E-08 |
| 28316 | 1416 | A1416S | E | 2.5e-08 | 3.9e-08 | 2.7e-08 | 3.033E-08 | 7.572E-09 | 2.500E-08 | 3.900E-08 |
| 28317 | 1416 | A1416T | E | 1.2e-07 | 2.8e-08 | 3.2e-08 | 6.000E-08 | 5.200E-08 | 2.800E-08 | 1.200E-07 |
| 28318 | 1416 | A1416W | E | 1.9e-08 | 2e-08 | 3.3e-08 | 2.400E-08 | 7.810E-09 | 1.900E-08 | 3.300E-08 |
| 28319 | 1416 | A1416Y | E | 3.9e-08 | 5.7e-08 | 4.4e-08 | 4.667E-08 | 9.292E-09 | 3.900E-08 | 5.700E-08 |
| 28320 | 1416 | A1416V | E | 1e-07 | 1.5e-07 | 5.5e-08 | 1.017E-07 | 4.752E-08 | 5.500E-08 | 1.500E-07 |
| 28321 | 1417 | L1417A | E | 8.7e-09 | 7e-08 | 9e-08 | 5.623E-08 | 4.236E-08 | 8.700E-09 | 9.000E-08 |
| 28322 | 1417 | L1417R | E | 4.7e-08 | 2.9e-08 | 6.1e-08 | 4.567E-08 | 1.604E-08 | 2.900E-08 | 6.100E-08 |
| 28323 | 1417 | L1417N | E | 1.5e-07 | 7.2e-08 | 4.5e-08 | 8.900E-08 | 5.453E-08 | 4.500E-08 | 1.500E-07 |
| 28324 | 1417 | L1417D | E | 1.3e-07 | 2.6e-08 | 3.9e-08 | 6.500E-08 | 5.667E-08 | 2.600E-08 | 1.300E-07 |
| 28325 | 1417 | L1417C | E | 4.7e-08 | 3e-08 | 4.9e-08 | 4.200E-08 | 1.044E-08 | 3.000E-08 | 4.900E-08 |
| 28326 | 1417 | L1417E | E | 6.3e-08 | 2.1e-08 | 2.4e-08 | 3.600E-08 | 2.343E-08 | 2.100E-08 | 6.300E-08 |
| 28327 | 1417 | L1417Q | E | 8.5e-08 | 1.9e-08 | 4.4e-08 | 4.933E-08 | 3.332E-08 | 1.900E-08 | 8.500E-08 |
| 28328 | 1417 | L1417G | E | 6.6e-08 | 3.7e-08 | 4.7e-08 | 5.000E-08 | 1.473E-08 | 3.700E-08 | 6.600E-08 |
| 28329 | 1417 | L1417H | E | 5.3e-08 | 3.3e-08 | 4.8e-08 | 4.467E-08 | 1.041E-08 | 3.300E-08 | 5.300E-08 |
| 28330 | 1417 | L1417I | E | 6.3e-08 | 1.3e-08 | 6.2e-08 | 4.600E-08 | 2.858E-08 | 1.300E-08 | 6.300E-08 |
| 28331 | 1417 | L1417L | E | 2.3e-08 | 8.4e-09 | 5.5e-08 | 2.880E-08 | 2.384E-08 | 8.400E-09 | 5.500E-08 |
| 28332 | 1417 | L1417K | E | 5.3e-08 | 1.1e-07 | 5.1e-08 | 7.133E-08 | 3.350E-08 | 5.100E-08 | 1.100E-07 |
| 28333 | 1417 | L1417M | E | 4.6e-08 | 4.5e-08 | 2.7e-08 | 3.933E-08 | 1.069E-08 | 2.700E-08 | 4.600E-08 |
| 28334 | 1417 | L1417F | E | 2.2e-08 | 3.6e-08 | 3.6e-08 | 3.133E-08 | 8.083E-09 | 2.200E-08 | 3.600E-08 |
| 28335 | 1417 | L1417P | E | 1.6e-08 | 8.5e-08 | 4.7e-08 | 4.933E-08 | 3.456E-08 | 1.600E-08 | 8.500E-08 |
| 28336 | 1417 | L1417S | E | 1.8e-07 | 3.4e-08 | 1e-07 | 1.047E-07 | 7.311E-08 | 3.400E-08 | 1.800E-07 |
| 28337 | 1417 | L1417T | E | 5.2e-08 | 5e-08 | 1e-08 | 3.733E-08 | 2.369E-08 | 1.000E-08 | 5.200E-08 |
| 28338 | 1417 | L1417W | E | 3.4e-08 | 9.4e-08 | 3.6e-08 | 5.467E-08 | 3.408E-08 | 3.400E-08 | 9.400E-08 |
| 28339 | 1417 | L1417Y | E | 1.8e-08 | 3.9e-08 | 3.2e-08 | 2.967E-08 | 1.069E-08 | 1.800E-08 | 3.900E-08 |
| 28340 | 1417 | L1417V | E | 4e-08 | 1.1e-07 | 7.6e-08 | 7.533E-08 | 3.500E-08 | 4.000E-08 | 1.100E-07 |
| 28341 | 1418 | G1418A | E | 3.4e-08 | 4.9e-08 | 7.6e-08 | 5.300E-08 | 2.128E-08 | 3.400E-08 | 7.600E-08 |
| 28342 | 1418 | G1418R | E | 1.1e-07 | 5.8e-08 | 5.2e-08 | 7.333E-08 | 3.190E-08 | 5.200E-08 | 1.100E-07 |
| 28343 | 1418 | G1418N | E | 3e-08 | 2.3e-08 | 5.9e-08 | 3.733E-08 | 1.909E-08 | 2.300E-08 | 5.900E-08 |
| 28344 | 1418 | G1418D | E | 1.4e-08 | 3.1e-08 | 6e-08 | 3.500E-08 | 2.326E-08 | 1.400E-08 | 6.000E-08 |
| 28345 | 1418 | G1418C | E | 3.5e-08 | 2.9e-08 | 7.9e-08 | 4.767E-08 | 2.730E-08 | 2.900E-08 | 7.900E-08 |
| 28346 | 1418 | G1418E | E | 5e-08 | 1.7e-08 | 4.1e-08 | 3.600E-08 | 1.706E-08 | 1.700E-08 | 5.000E-08 |
| 28347 | 1418 | G1418Q | E | 6.2e-08 | 3.6e-08 | 3.1e-08 | 4.300E-08 | 1.664E-08 | 3.100E-08 | 6.200E-08 |
| 28348 | 1418 | G1418G | E | 3.2e-08 | 8.3e-09 | 3.3e-08 | 2.443E-08 | 1.398E-08 | 8.300E-09 | 3.300E-08 |
| 28349 | 1418 | G1418H | E | 1.8e-08 | 2.7e-08 | 5.1e-08 | 3.200E-08 | 1.706E-08 | 1.800E-08 | 5.100E-08 |
| 28350 | 1418 | G1418I | E | 5.3e-08 | 3.2e-08 | 1.1e-07 | 6.500E-08 | 4.036E-08 | 3.200E-08 | 1.100E-07 |
| 28351 | 1418 | G1418L | E | 3e-08 | 8.1e-08 | 3.3e-08 | 4.800E-08 | 2.862E-08 | 3.000E-08 | 8.100E-08 |
| 28352 | 1418 | G1418K | E | 7.7e-08 | 4.4e-08 | 4.6e-08 | 5.567E-08 | 1.850E-08 | 4.400E-08 | 7.700E-08 |
| 28353 | 1418 | G1418M | E | 6e-08 | 1.7e-08 | 2.7e-08 | 3.467E-08 | 2.250E-08 | 1.700E-08 | 6.000E-08 |
| 28354 | 1418 | G1418F | E | 5.2e-08 | 5.3e-08 | 2e-08 | 4.167E-08 | 1.877E-08 | 2.000E-08 | 5.300E-08 |
| 28355 | 1418 | G1418P | E | 9.9e-08 | 2e-08 | 4.3e-08 | 5.400E-08 | 4.063E-08 | 2.000E-08 | 9.900E-08 |
| 28356 | 1418 | G1418S | E | 3.1e-08 | 1.2e-08 | 7.2e-08 | 3.833E-08 | 3.066E-08 | 1.200E-08 | 7.200E-08 |
| 28357 | 1418 | G1418T | E | 2.2e-08 | 4.4e-08 | 5.8e-08 | 4.133E-08 | 1.815E-08 | 2.200E-08 | 5.800E-08 |
| 28358 | 1418 | G1418W | E | 3.8e-08 | 6e-08 | 2.9e-08 | 4.233E-08 | 1.595E-08 | 2.900E-08 | 6.000E-08 |
| 28359 | 1418 | G1418Y | E | 3.4e-08 | 5.5e-08 | 3.8e-08 | 4.233E-08 | 1.115E-08 | 3.400E-08 | 5.500E-08 |
| 28360 | 1418 | G1418V | E | 6.7e-08 | 5.4e-08 | 4.7e-08 | 5.600E-08 | 1.015E-08 | 4.700E-08 | 6.700E-08 |
| 28361 | 1419 | C1419A | E | 4.8e-08 | 9.2e-08 | 4.3e-08 | 6.100E-08 | 2.696E-08 | 4.300E-08 | 9.200E-08 |
| 28362 | 1419 | C1419R | E | 2e-08 | 3.7e-08 | 5e-08 | 3.567E-08 | 1.504E-08 | 2.000E-08 | 5.000E-08 |
| 28363 | 1419 | C1419N | E | 3e-08 | 1.7e-08 | 3.3e-08 | 2.667E-08 | 8.505E-09 | 1.700E-08 | 3.300E-08 |
| 28364 | 1419 | C1419D | E | 3.4e-08 | 1.4e-08 | 2.1e-08 | 2.300E-08 | 1.015E-08 | 1.400E-08 | 3.400E-08 |
| 28365 | 1419 | C1419C | E | 2.1e-08 | 7.8e-08 | 2.1e-08 | 4.000E-08 | 3.291E-08 | 2.100E-08 | 7.800E-08 |
| 28366 | 1419 | C1419E | E | 1.6e-07 | 2.3e-08 | 1.3e-08 | 6.533E-08 | 8.214E-08 | 1.300E-08 | 1.600E-07 |
| 28367 | 1419 | C1419Q | E | 2.9e-08 | 7.7e-08 | 6.1e-08 | 5.567E-08 | 2.444E-08 | 2.900E-08 | 7.700E-08 |
| 28368 | 1419 | C1419G | E | 6.9e-08 | 6.6e-08 | 2e-08 | 5.167E-08 | 2.747E-08 | 2.000E-08 | 6.900E-08 |
| 28369 | 1419 | C1419H | E | 3.3e-08 | 6.1e-08 | 3.4e-08 | 4.267E-08 | 1.589E-08 | 3.300E-08 | 6.100E-08 |
| 28370 | 1419 | C1419I | E | 9.4e-09 | 6.7e-08 | 1.7e-08 | 3.113E-08 | 3.129E-08 | 9.400E-09 | 6.700E-08 |
| 28371 | 1419 | C1419L | E | 1.3e-07 | 2.7e-08 | 1.6e-08 | 5.767E-08 | 6.288E-08 | 1.600E-08 | 1.300E-07 |
| 28372 | 1419 | C1419K | E | 1.8e-07 | 4.8e-08 | 3.3e-08 | 8.700E-08 | 8.089E-08 | 3.300E-08 | 1.800E-07 |
| 28373 | 1419 | C1419M | E | 1.6e-08 | 5e-08 | 1.8e-08 | 2.800E-08 | 1.908E-08 | 1.600E-08 | 5.000E-08 |
| 28374 | 1419 | C1419F | E | 4.7e-08 | 3.8e-08 | 7.5e-08 | 5.333E-08 | 1.930E-08 | 3.800E-08 | 7.500E-08 |
| 28375 | 1419 | C1419P | E | 2.7e-08 | 3.2e-07 | 6.5e-08 | 1.373E-07 | 1.593E-07 | 2.700E-08 | 3.200E-07 |
| 28376 | 1419 | C1419S | E | 4.6e-08 | 2.4e-08 | 1.1e-07 | 6.000E-08 | 4.468E-08 | 2.400E-08 | 1.100E-07 |
| 28377 | 1419 | C1419T | E | 1.1e-07 | 2.1e-08 | 3.4e-08 | 5.500E-08 | 4.807E-08 | 2.100E-08 | 1.100E-07 |
| 28378 | 1419 | C1419W | E | 4e-08 | 3.1e-08 | 4.5e-08 | 3.867E-08 | 7.095E-09 | 3.100E-08 | 4.500E-08 |
| 28379 | 1419 | C1419Y | E | 2.4e-08 | 4.5e-08 | 3e-08 | 3.300E-08 | 1.082E-08 | 2.400E-08 | 4.500E-08 |
| 28380 | 1419 | C1419V | E | 5.9e-08 | 1.7e-08 | 6e-08 | 4.533E-08 | 2.454E-08 | 1.700E-08 | 6.000E-08 |
| 28381 | 1420 | L1420A | E | 4.1e-08 | 5.1e-08 | 6.5e-08 | 5.233E-08 | 1.206E-08 | 4.100E-08 | 6.500E-08 |
| 28382 | 1420 | L1420R | E | 9.1e-08 | 2.8e-08 | 5.7e-08 | 5.867E-08 | 3.153E-08 | 2.800E-08 | 9.100E-08 |
| 28383 | 1420 | L1420N | E | 3.8e-08 | 3.6e-08 | 5.5e-08 | 4.300E-08 | 1.044E-08 | 3.600E-08 | 5.500E-08 |
| 28384 | 1420 | L1420D | E | 4.1e-08 | 6.7e-08 | 1e-07 | 6.933E-08 | 2.957E-08 | 4.100E-08 | 1.000E-07 |
| 28385 | 1420 | L1420C | E | 7.6e-08 | 5.5e-08 | 8.5e-08 | 7.200E-08 | 1.539E-08 | 5.500E-08 | 8.500E-08 |
| 28386 | 1420 | L1420E | E | 4.3e-08 | 8.4e-08 | 4e-08 | 5.567E-08 | 2.458E-08 | 4.000E-08 | 8.400E-08 |
| 28387 | 1420 | L1420Q | E | 4.8e-08 | 2.6e-08 | 5.2e-08 | 4.200E-08 | 1.400E-08 | 2.600E-08 | 5.200E-08 |
| 28388 | 1420 | L1420G | E | 5.2e-08 | 4.7e-08 | 5.7e-08 | 5.200E-08 | 5.000E-09 | 4.700E-08 | 5.700E-08 |
| 28389 | 1420 | L1420H | E | 3.3e-08 | 6.3e-08 | 4.4e-08 | 4.667E-08 | 1.518E-08 | 3.300E-08 | 6.300E-08 |

| | | | | | | | | | | |
|---|---|---|---|---|---|---|---|---|---|---|
| 28390 | 1420 | L1420I | E | 3.9e-08 | 2.7e-08 | 1.3e-07 | 6.533E-08 | 5.632E-08 | 2.700E-08 | 1.300E-07 |
| 28391 | 1420 | L1420L | E | 5.9e-08 | 2.1e-08 | 2.7e-08 | 3.567E-08 | 2.043E-08 | 2.100E-08 | 5.900E-08 |
| 28392 | 1420 | L1420K | E | 1.5e-08 | 7.8e-08 | 5.7e-08 | 5.000E-08 | 3.208E-08 | 1.500E-08 | 7.800E-08 |
| 28393 | 1420 | L1420M | E | 6.6e-08 | 3.6e-08 | 7.7e-08 | 5.967E-08 | 2.122E-08 | 3.600E-08 | 7.700E-08 |
| 28394 | 1420 | L1420F | E | 3e-08 | 2.9e-08 | 4.2e-08 | 3.367E-08 | 7.234E-09 | 2.900E-08 | 4.200E-08 |
| 28395 | 1420 | L1420P | E | 2.7e-08 | 5.9e-08 | 5.5e-08 | 4.700E-08 | 1.744E-08 | 2.700E-08 | 5.900E-08 |
| 28396 | 1420 | L1420S | E | 2.6e-08 | 1.7e-07 | 2.1e-08 | 7.233E-08 | 8.462E-08 | 2.100E-08 | 1.700E-07 |
| 28397 | 1420 | L1420T | E | 5.3e-08 | 6e-08 | 4.1e-08 | 5.133E-08 | 9.609E-09 | 4.100E-08 | 6.000E-08 |
| 28398 | 1420 | L1420W | E | 3e-08 | 3e-08 | 2.9e-08 | 2.967E-08 | 5.774E-10 | 2.900E-08 | 3.000E-08 |
| 28399 | 1420 | L1420Y | E | 6.3e-08 | 1.2e-07 | 3.1e-08 | 7.133E-08 | 4.508E-08 | 3.100E-08 | 1.200E-07 |
| 28400 | 1420 | L1420V | E | 1.8e-08 | 6.5e-08 | 5e-08 | 4.433E-08 | 2.401E-08 | 1.800E-08 | 6.500E-08 |
| 28401 | 1421 | V1421A | E | 1.2e-07 | 8.9e-08 | 6.9e-08 | 9.267E-08 | 2.570E-08 | 6.900E-08 | 1.200E-07 |
| 28402 | 1421 | V1421R | E | 5.9e-08 | 3.6e-08 | 4.1e-08 | 4.533E-08 | 1.210E-08 | 3.600E-08 | 5.900E-08 |
| 28403 | 1421 | V1421N | E | 2.6e-08 | 1.7e-08 | 5.4e-08 | 3.233E-08 | 1.930E-08 | 1.700E-08 | 5.400E-08 |
| 28404 | 1421 | V1421D | E | 7.9e-08 | 2.8e-08 | 2.7e-08 | 4.467E-08 | 2.974E-08 | 2.700E-08 | 7.900E-08 |
| 28405 | 1421 | V1421C | E | 5.7e-08 | 2.8e-08 | 1.5e-07 | 7.833E-08 | 6.374E-08 | 2.800E-08 | 1.500E-07 |
| 28406 | 1421 | V1421E | E | 2.4e-08 | 6.4e-08 | 3.9e-08 | 4.233E-08 | 2.021E-08 | 2.400E-08 | 6.400E-08 |
| 28407 | 1421 | V1421Q | E | 7.3e-08 | 5e-08 | 8e-08 | 6.767E-08 | 1.570E-08 | 5.000E-08 | 8.000E-08 |
| 28408 | 1421 | V1421G | E | 9e-08 | 4.1e-08 | 4.2e-08 | 5.767E-08 | 2.801E-08 | 4.100E-08 | 9.000E-08 |
| 28409 | 1421 | V1421H | E | 5.1e-08 | 4.6e-08 | 1.2e-07 | 7.233E-08 | 4.136E-08 | 4.600E-08 | 1.200E-07 |
| 28410 | 1421 | V1421I | E | 8.3e-08 | 5.4e-08 | 7.3e-08 | 7.000E-08 | 1.473E-08 | 5.400E-08 | 8.300E-08 |
| 28411 | 1421 | V1421L | E | 4.9e-08 | 5.1e-08 | 1.8e-08 | 3.933E-08 | 1.850E-08 | 1.800E-08 | 5.100E-08 |
| 28412 | 1421 | V1421K | E | 2.8e-08 | 4.4e-08 | 2e-08 | 3.067E-08 | 1.222E-08 | 2.000E-08 | 4.400E-08 |
| 28413 | 1421 | V1421M | E | 1.5e-08 | 4e-08 | 2e-08 | 2.500E-08 | 1.323E-08 | 1.500E-08 | 4.000E-08 |
| 28414 | 1421 | V1421F | E | 3.6e-08 | 5.8e-08 | 4e-08 | 4.467E-08 | 1.172E-08 | 3.600E-08 | 5.800E-08 |
| 28415 | 1421 | V1421P | E | 5.3e-08 | 7.4e-08 | 3.9e-08 | 5.533E-08 | 1.762E-08 | 3.900E-08 | 7.400E-08 |
| 28416 | 1421 | V1421S | E | 1.4e-07 | 6e-08 | 5.5e-08 | 8.500E-08 | 4.770E-08 | 5.500E-08 | 1.400E-07 |
| 28417 | 1421 | V1421T | E | 6.9e-08 | 4.1e-08 | 4.6e-08 | 5.200E-08 | 1.493E-08 | 4.100E-08 | 6.900E-08 |
| 28418 | 1421 | V1421W | E | 1.2e-08 | 2.5e-08 | 1.4e-08 | 1.700E-08 | 7.000E-09 | 1.200E-08 | 2.500E-08 |
| 28419 | 1421 | V1421Y | E | 9.5e-08 | 3.1e-08 | 6.1e-08 | 6.233E-08 | 3.202E-08 | 3.100E-08 | 9.500E-08 |
| 28420 | 1421 | V1421V | E | 1.6e-08 | 7e-08 | 6.4e-08 | 5.000E-08 | 2.960E-08 | 1.600E-08 | 7.000E-08 |
| 28421 | 1422 | K1422A | E | 1.2e-07 | 3.3e-08 | 2.6e-08 | 5.967E-08 | 5.237E-08 | 2.600E-08 | 1.200E-07 |
| 28422 | 1422 | K1422R | E | 2.2e-08 | 8.4e-08 | 6.1e-08 | 5.567E-08 | 3.134E-08 | 2.200E-08 | 8.400E-08 |
| 28423 | 1422 | K1422N | E | 3.8e-08 | 7.9e-09 | 6.1e-08 | 3.563E-08 | 2.663E-08 | 7.900E-09 | 6.100E-08 |
| 28424 | 1422 | K1422D | E | 7.1e-08 | 5e-08 | 1e-07 | 7.367E-08 | 2.511E-08 | 5.000E-08 | 1.000E-07 |
| 28425 | 1422 | K1422C | E | 4.4e-08 | 3.1e-08 | 2e-08 | 3.167E-08 | 1.201E-08 | 2.000E-08 | 4.400E-08 |
| 28426 | 1422 | K1422E | E | 6.4e-08 | 3.8e-08 | 2.5e-08 | 4.233E-08 | 1.986E-08 | 2.500E-08 | 6.400E-08 |
| 28427 | 1422 | K1422Q | E | 4.6e-08 | 3.8e-08 | 4.5e-08 | 4.300E-08 | 4.359E-09 | 3.800E-08 | 4.600E-08 |
| 28428 | 1422 | K1422G | E | 2.5e-08 | 2e-08 | 8.3e-08 | 4.267E-08 | 3.502E-08 | 2.000E-08 | 8.300E-08 |
| 28429 | 1422 | K1422H | E | 8.6e-08 | 1e-07 | 6.9e-08 | 8.500E-08 | 1.552E-08 | 6.900E-08 | 1.000E-07 |
| 28430 | 1422 | K1422I | E | 2.6e-08 | 4.3e-08 | 6.5e-08 | 4.467E-08 | 1.955E-08 | 2.600E-08 | 6.500E-08 |
| 28431 | 1422 | K1422L | E | 3.8e-08 | 6.5e-08 | 4.1e-08 | 4.800E-08 | 1.480E-08 | 3.800E-08 | 6.500E-08 |
| 28432 | 1422 | K1422K | E | 4.7e-08 | 5.5e-08 | 5.2e-08 | 5.133E-08 | 4.041E-09 | 4.700E-08 | 5.500E-08 |
| 28433 | 1422 | K1422M | E | 9.4e-08 | 2.3e-08 | 5.4e-08 | 5.700E-08 | 3.559E-08 | 2.300E-08 | 9.400E-08 |
| 28434 | 1422 | K1422F | E | 3.8e-08 | 1.9e-08 | 5.8e-08 | 3.833E-08 | 1.950E-08 | 1.900E-08 | 5.800E-08 |
| 28435 | 1422 | K1422P | E | 5.8e-08 | 1.7e-08 | 5.3e-08 | 4.267E-08 | 2.237E-08 | 1.700E-08 | 5.800E-08 |
| 28436 | 1422 | K1422S | E | 4.8e-08 | 2.2e-08 | 3.3e-08 | 3.433E-08 | 1.305E-08 | 2.200E-08 | 4.800E-08 |
| 28437 | 1422 | K1422T | E | 1.5e-08 | 3.9e-08 | 5e-08 | 3.467E-08 | 1.790E-08 | 1.500E-08 | 5.000E-08 |
| 28438 | 1422 | K1422W | E | 3.8e-08 | 6.6e-08 | 2.9e-08 | 4.433E-08 | 1.930E-08 | 2.900E-08 | 6.600E-08 |
| 28439 | 1422 | K1422Y | E | 4.5e-08 | 8.7e-08 | 4e-08 | 5.733E-08 | 2.581E-08 | 4.000E-08 | 8.700E-08 |
| 28440 | 1422 | K1422V | E | 6e-08 | 9.9e-09 | 4.7e-08 | 3.897E-08 | 2.600E-08 | 9.900E-09 | 6.000E-08 |
| 28441 | 1423 | D1423A | E | 2e-08 | 6.1e-08 | 1.4e-08 | 3.167E-08 | 2.558E-08 | 1.400E-08 | 6.100E-08 |
| 28442 | 1423 | D1423R | E | 7.6e-08 | 2.2e-08 | 4.9e-08 | 4.900E-08 | 2.700E-08 | 2.200E-08 | 7.600E-08 |
| 28443 | 1423 | D1423N | E | 3.8e-08 | 2.2e-08 | 1.1e-07 | 5.667E-08 | 4.688E-08 | 2.200E-08 | 1.100E-07 |
| 28444 | 1423 | D1423D | E | 3.6e-08 | 3.7e-08 | 4.6e-08 | 3.967E-08 | 5.508E-09 | 3.600E-08 | 4.600E-08 |
| 28445 | 1423 | D1423C | E | 4.6e-08 | 4.2e-08 | 5.4e-08 | 4.733E-08 | 6.110E-09 | 4.200E-08 | 5.400E-08 |
| 28446 | 1423 | D1423E | E | 1.5e-08 | 5e-08 | 2.8e-08 | 3.100E-08 | 1.769E-08 | 1.500E-08 | 5.000E-08 |
| 28447 | 1423 | D1423Q | E | 4e-08 | 3.3e-08 | 3.4e-08 | 3.567E-08 | 3.786E-09 | 3.300E-08 | 4.000E-08 |
| 28448 | 1423 | D1423G | E | 3.7e-08 | 2.9e-08 | 2.7e-08 | 3.100E-08 | 5.292E-09 | 2.700E-08 | 3.700E-08 |
| 28449 | 1423 | D1423H | E | 6.7e-08 | 5.1e-08 | 2.8e-08 | 4.867E-08 | 1.960E-08 | 2.800E-08 | 6.700E-08 |
| 28450 | 1423 | D1423I | E | 3.1e-08 | 3.9e-08 | 6.2e-08 | 4.400E-08 | 1.609E-08 | 3.100E-08 | 6.200E-08 |
| 28451 | 1423 | D1423L | E | 3.3e-08 | 1.3e-07 | 3.6e-08 | 6.633E-08 | 5.516E-08 | 3.300E-08 | 1.300E-07 |
| 28452 | 1423 | D1423K | E | 3.3e-08 | 1e-07 | 5.6e-08 | 6.300E-08 | 3.404E-08 | 3.300E-08 | 1.000E-07 |
| 28453 | 1423 | D1423M | E | 9.6e-08 | 6.1e-08 | 4.6e-08 | 6.767E-08 | 2.566E-08 | 4.600E-08 | 9.600E-08 |
| 28454 | 1423 | D1423F | E | 2.8e-08 | 8.5e-08 | 1e-07 | 7.100E-08 | 3.799E-08 | 2.800E-08 | 1.000E-07 |
| 28455 | 1423 | D1423P | E | 7e-08 | 1.3e-07 | 1.6e-08 | 7.200E-08 | 5.703E-08 | 1.600E-08 | 1.300E-07 |
| 28456 | 1423 | D1423S | E | 6e-08 | 5.5e-08 | 2.6e-08 | 4.700E-08 | 1.836E-08 | 2.600E-08 | 6.000E-08 |
| 28457 | 1423 | D1423T | E | 1.2e-08 | 2.6e-08 | 2.5e-08 | 2.100E-08 | 7.810E-09 | 1.200E-08 | 2.600E-08 |
| 28458 | 1423 | D1423W | E | 1.1e-07 | 2.1e-08 | 2.7e-08 | 5.267E-08 | 4.974E-08 | 2.100E-08 | 1.100E-07 |
| 28459 | 1423 | D1423Y | E | 3.2e-08 | 1.9e-08 | 5.1e-08 | 3.400E-08 | 1.609E-08 | 1.900E-08 | 5.100E-08 |
| 28460 | 1423 | D1423V | E | 1.1e-08 | 7.1e-08 | 5.2e-08 | 4.467E-08 | 3.066E-08 | 1.100E-08 | 7.100E-08 |
| 28461 | 1424 | Y1424A | E | 8.3e-08 | 2.7e-08 | 5.2e-08 | 5.400E-08 | 2.805E-08 | 2.700E-08 | 8.300E-08 |
| 28462 | 1424 | Y1424R | E | 3.5e-08 | 8e-08 | 6.4e-08 | 5.967E-08 | 2.281E-08 | 3.500E-08 | 8.000E-08 |
| 28463 | 1424 | Y1424N | E | 2.6e-08 | 3e-08 | 5.6e-08 | 3.733E-08 | 1.629E-08 | 2.600E-08 | 5.600E-08 |
| 28464 | 1424 | Y1424D | E | 6e-08 | 2.4e-08 | 3e-08 | 3.800E-08 | 1.929E-08 | 2.400E-08 | 6.000E-08 |
| 28465 | 1424 | Y1424C | E | 2.8e-08 | 9.7e-08 | 1.8e-08 | 4.767E-08 | 4.302E-08 | 1.800E-08 | 9.700E-08 |
| 28466 | 1424 | Y1424E | E | 3.1e-08 | 3.2e-08 | 6.2e-08 | 4.167E-08 | 1.762E-08 | 3.100E-08 | 6.200E-08 |
| 28467 | 1424 | Y1424Q | E | 5.8e-08 | 1.9e-08 | 5.9e-08 | 4.533E-08 | 2.281E-08 | 1.900E-08 | 5.900E-08 |
| 28468 | 1424 | Y1424G | E | 6.1e-08 | 3.5e-08 | 7.1e-08 | 5.567E-08 | 1.858E-08 | 3.500E-08 | 7.100E-08 |
| 28469 | 1424 | Y1424H | E | 4.9e-08 | 2.1e-08 | 3.6e-08 | 3.533E-08 | 1.401E-08 | 2.100E-08 | 4.900E-08 |
| 28470 | 1424 | Y1424I | E | 1.8e-08 | 7.5e-08 | 3.4e-08 | 4.233E-08 | 2.940E-08 | 1.800E-08 | 7.500E-08 |
| 28471 | 1424 | Y1424L | E | 2.8e-08 | 3.2e-08 | 8.4e-08 | 4.800E-08 | 3.124E-08 | 2.800E-08 | 8.400E-08 |
| 28472 | 1424 | Y1424K | E | 3.5e-08 | 2.4e-08 | 4.2e-08 | 3.367E-08 | 9.074E-09 | 2.400E-08 | 4.200E-08 |
| 28473 | 1424 | Y1424M | E | 4e-08 | 9.3e-08 | 7.5e-08 | 6.933E-08 | 2.695E-08 | 4.000E-08 | 9.300E-08 |
| 28474 | 1424 | Y1424F | E | 2.4e-08 | 4.4e-08 | 7.3e-08 | 4.700E-08 | 2.464E-08 | 2.400E-08 | 7.300E-08 |

| | | | | | | | | | | |
|---|---|---|---|---|---|---|---|---|---|---|
| 28475 | 1424 | Y1424P | E | 1.7e-08 | 9.4e-09 | 4.8e-08 | 2.480E-08 | 2.045E-08 | 9.400E-09 | 4.800E-08 |
| 28476 | 1424 | Y1424S | E | 1.5e-08 | 4e-08 | 1.5e-08 | 2.333E-08 | 1.443E-08 | 1.500E-08 | 4.000E-08 |
| 28477 | 1424 | Y1424T | E | 8.3e-08 | 6.4e-08 | 2e-08 | 5.567E-08 | 3.232E-08 | 2.000E-08 | 8.300E-08 |
| 28478 | 1424 | Y1424W | E | 2.1e-08 | 5.1e-08 | 1e-07 | 5.733E-08 | 3.988E-08 | 2.100E-08 | 1.000E-07 |
| 28479 | 1424 | Y1424Y | E | 2.5e-08 | 5e-08 | 2.6e-08 | 3.367E-08 | 1.415E-08 | 2.500E-08 | 5.000E-08 |
| 28480 | 1424 | Y1424V | E | 4.5e-08 | 4.2e-08 | 3.7e-08 | 4.133E-08 | 4.041E-09 | 3.700E-08 | 4.500E-08 |
| 28481 | 1425 | F1425A | E | 1.2e-07 | 1.6e-08 | 2.8e-08 | 5.467E-08 | 5.690E-08 | 1.600E-08 | 1.200E-07 |
| 28482 | 1425 | F1425R | E | 2.5e-08 | 7.2e-08 | 3.8e-08 | 4.500E-08 | 2.427E-08 | 2.500E-08 | 7.200E-08 |
| 28483 | 1425 | F1425N | E | 4.3e-08 | 2.9e-08 | 2.9e-08 | 3.367E-08 | 8.083E-09 | 2.900E-08 | 4.300E-08 |
| 28484 | 1425 | F1425D | E | 1.2e-08 | 5.6e-08 | 2.8e-08 | 3.200E-08 | 2.227E-08 | 1.200E-08 | 5.600E-08 |
| 28485 | 1425 | F1425C | E | 6.7e-08 | 7.4e-08 | 6.6e-08 | 6.900E-08 | 4.359E-09 | 6.600E-08 | 7.400E-08 |
| 28486 | 1425 | F1425E | E | 4e-08 | 7.2e-08 | 3.8e-08 | 5.000E-08 | 1.908E-08 | 3.800E-08 | 7.200E-08 |
| 28487 | 1425 | F1425Q | E | 2.4e-08 | 4.2e-08 | 2e-08 | 2.867E-08 | 1.172E-08 | 2.000E-08 | 4.200E-08 |
| 28488 | 1425 | F1425G | E | 5.5e-08 | 1e-07 | 3.3e-08 | 6.267E-08 | 3.415E-08 | 3.300E-08 | 1.000E-07 |
| 28489 | 1425 | F1425H | E | 4.8e-08 | 5.1e-08 | 9e-09 | 3.600E-08 | 2.343E-08 | 9.000E-09 | 5.100E-08 |
| 28490 | 1425 | F1425I | E | 4e-08 | 3e-08 | 5.3e-08 | 4.100E-08 | 1.153E-08 | 3.000E-08 | 5.300E-08 |
| 28491 | 1425 | F1425L | E | 8.9e-08 | 7.9e-08 | 3.1e-08 | 6.633E-08 | 3.101E-08 | 3.100E-08 | 8.900E-08 |
| 28492 | 1425 | F1425K | E | 3.3e-08 | 5e-08 | 5.7e-08 | 4.667E-08 | 1.234E-08 | 3.300E-08 | 5.700E-08 |
| 28493 | 1425 | F1425M | E | 4.1e-08 | 3.8e-08 | 1.4e-07 | 7.300E-08 | 5.804E-08 | 3.800E-08 | 1.400E-07 |
| 28494 | 1425 | F1425F | E | 1.2e-07 | 6.3e-08 | 8.3e-08 | 8.867E-08 | 2.892E-08 | 6.300E-08 | 1.200E-07 |
| 28495 | 1425 | F1425P | E | 8.9e-08 | 3.6e-08 | 1.3e-08 | 4.600E-08 | 3.897E-08 | 1.300E-08 | 8.900E-08 |
| 28496 | 1425 | F1425S | E | 2.4e-08 | 1.9e-08 | 2.8e-08 | 2.367E-08 | 4.509E-09 | 1.900E-08 | 2.800E-08 |
| 28497 | 1425 | F1425T | E | 1.2e-08 | 7.1e-08 | 2.4e-08 | 3.567E-08 | 3.118E-08 | 1.200E-08 | 7.100E-08 |
| 28498 | 1425 | F1425W | E | 3.5e-08 | 2.7e-08 | 5.7e-08 | 3.967E-08 | 1.553E-08 | 2.700E-08 | 5.700E-08 |
| 28499 | 1425 | F1425Y | E | 3.3e-08 | 3.1e-08 | 3.4e-08 | 3.267E-08 | 1.528E-09 | 3.100E-08 | 3.400E-08 |
| 28500 | 1425 | F1425V | E | 6.6e-08 | 2.8e-08 | 5.3e-08 | 4.900E-08 | 1.931E-08 | 2.800E-08 | 6.600E-08 |
| 28501 | 1426 | P1426A | E | 2.1e-08 | 2.8e-08 | 1.3e-08 | 2.067E-08 | 7.506E-09 | 1.300E-08 | 2.800E-08 |
| 28502 | 1426 | P1426R | E | 7.4e-08 | 1.8e-08 | 6.7e-08 | 5.300E-08 | 3.051E-08 | 1.800E-08 | 7.400E-08 |
| 28503 | 1426 | P1426N | E | 2.7e-08 | 5.1e-08 | 3.8e-08 | 3.867E-08 | 1.201E-08 | 2.700E-08 | 5.100E-08 |
| 28504 | 1426 | P1426D | E | 3.3e-08 | 5.1e-08 | 3.4e-08 | 3.933E-08 | 1.012E-08 | 3.300E-08 | 5.100E-08 |
| 28505 | 1426 | P1426C | E | 1.8e-08 | 5.4e-08 | 3.4e-08 | 3.533E-08 | 1.804E-08 | 1.800E-08 | 5.400E-08 |
| 28506 | 1426 | P1426E | E | 5.5e-08 | 2.2e-08 | 5.8e-08 | 4.500E-08 | 1.997E-08 | 2.200E-08 | 5.800E-08 |
| 28507 | 1426 | P1426Q | E | 3.9e-08 | 9.1e-08 | 5.1e-08 | 6.033E-08 | 2.723E-08 | 3.900E-08 | 9.100E-08 |
| 28508 | 1426 | P1426G | E | 4.4e-08 | 3e-08 | 2.2e-08 | 3.200E-08 | 1.114E-08 | 2.200E-08 | 4.400E-08 |
| 28509 | 1426 | P1426H | E | 4.8e-08 | 2.9e-08 | 6.6e-08 | 4.767E-08 | 1.850E-08 | 2.900E-08 | 6.600E-08 |
| 28510 | 1426 | P1426I | E | 3.5e-08 | 1.8e-08 | 8.5e-09 | 2.050E-08 | 1.343E-08 | 8.500E-09 | 3.500E-08 |
| 28511 | 1426 | P1426L | E | 4.2e-08 | 5.2e-08 | 3.1e-08 | 4.167E-08 | 1.050E-08 | 3.100E-08 | 5.200E-08 |
| 28512 | 1426 | P1426K | E | 2.1e-08 | 3.4e-08 | 4.6e-08 | 3.367E-08 | 1.250E-08 | 2.100E-08 | 4.600E-08 |
| 28513 | 1426 | P1426M | E | 2.9e-08 | 2.2e-08 | 7.5e-08 | 4.200E-08 | 2.879E-08 | 2.200E-08 | 7.500E-08 |
| 28514 | 1426 | P1426F | E | 6.8e-08 | 4.6e-08 | 2.1e-08 | 4.500E-08 | 2.352E-08 | 2.100E-08 | 6.800E-08 |
| 28515 | 1426 | P1426P | E | 2.3e-08 | 4.9e-08 | 3.5e-08 | 3.567E-08 | 1.301E-08 | 2.300E-08 | 4.900E-08 |
| 28516 | 1426 | P1426S | E | 4.9e-08 | 5.8e-08 | 7.4e-08 | 6.033E-08 | 1.266E-08 | 4.900E-08 | 7.400E-08 |
| 28517 | 1426 | P1426T | E | 6.9e-08 | 3.4e-08 | 4.4e-08 | 4.900E-08 | 1.803E-08 | 3.400E-08 | 6.900E-08 |
| 28518 | 1426 | P1426W | E | 1.4e-08 | 2.4e-08 | 6.4e-08 | 3.400E-08 | 2.646E-08 | 1.400E-08 | 6.400E-08 |
| 28519 | 1426 | P1426Y | E | 8.6e-08 | 4.1e-08 | 7.7e-08 | 6.800E-08 | 2.381E-08 | 4.100E-08 | 8.600E-08 |
| 28520 | 1426 | P1426V | E | 3.3e-08 | 3.6e-08 | 6e-08 | 4.300E-08 | 1.480E-08 | 3.300E-08 | 6.000E-08 |
| 28521 | 1427 | E1427A | E | 4.3e-08 | 4.1e-08 | 6.2e-08 | 4.867E-08 | 1.159E-08 | 4.100E-08 | 6.200E-08 |
| 28522 | 1427 | E1427R | E | 2.5e-08 | 2.2e-08 | 2.4e-08 | 2.367E-08 | 1.528E-09 | 2.200E-08 | 2.500E-08 |
| 28523 | 1427 | E1427N | E | 4.3e-08 | 2.1e-08 | 1.7e-08 | 2.700E-08 | 1.400E-08 | 1.700E-08 | 4.300E-08 |
| 28524 | 1427 | E1427D | E | 5.8e-08 | 7.3e-08 | 1.3e-07 | 8.700E-08 | 3.799E-08 | 5.800E-08 | 1.300E-07 |
| 28525 | 1427 | E1427C | E | 4.3e-08 | 2.6e-08 | 3.4e-08 | 3.433E-08 | 8.505E-09 | 2.600E-08 | 4.300E-08 |
| 28526 | 1427 | E1427E | E | 2.2e-08 | 5.6e-08 | 8.2e-08 | 5.333E-08 | 3.009E-08 | 2.200E-08 | 8.200E-08 |
| 28527 | 1427 | E1427Q | E | 1.6e-08 | 4.5e-08 | 6.3e-08 | 4.133E-08 | 2.371E-08 | 1.600E-08 | 6.300E-08 |
| 28528 | 1427 | E1427G | E | 7.1e-08 | 3.3e-08 | 7.7e-08 | 6.033E-08 | 2.386E-08 | 3.300E-08 | 7.700E-08 |
| 28529 | 1427 | E1427H | E | 6e-08 | 3.3e-08 | 2.8e-08 | 4.033E-08 | 1.721E-08 | 2.800E-08 | 6.000E-08 |
| 28530 | 1427 | E1427I | E | 4.5e-08 | 2.8e-08 | 2.3e-08 | 3.200E-08 | 1.153E-08 | 2.300E-08 | 4.500E-08 |
| 28531 | 1427 | E1427L | E | 5.6e-08 | 8.7e-08 | 2.7e-08 | 5.667E-08 | 3.001E-08 | 2.700E-08 | 8.700E-08 |
| 28532 | 1427 | E1427K | E | 4.6e-08 | 2.1e-08 | 2.2e-08 | 2.967E-08 | 1.415E-08 | 2.100E-08 | 4.600E-08 |
| 28533 | 1427 | E1427M | E | 4.8e-08 | 6.3e-08 | 3.1e-08 | 4.733E-08 | 1.601E-08 | 3.100E-08 | 6.300E-08 |
| 28534 | 1427 | E1427F | E | 6e-08 | 8.1e-08 | 2.9e-08 | 5.667E-08 | 2.616E-08 | 2.900E-08 | 8.100E-08 |
| 28535 | 1427 | E1427P | E | 4e-08 | 4.1e-08 | 2.8e-08 | 3.633E-08 | 7.234E-09 | 2.800E-08 | 4.100E-08 |
| 28536 | 1427 | E1427S | E | 1.9e-08 | 2.7e-08 | 5.9e-08 | 3.500E-08 | 2.117E-08 | 1.900E-08 | 5.900E-08 |
| 28537 | 1427 | E1427T | E | 2.6e-08 | 8e-08 | 4.2e-08 | 4.933E-08 | 2.774E-08 | 2.600E-08 | 8.000E-08 |
| 28538 | 1427 | E1427W | E | 2.3e-08 | 9.4e-08 | 6.1e-08 | 5.933E-08 | 3.553E-08 | 2.300E-08 | 9.400E-08 |
| 28539 | 1427 | E1427Y | E | 2.1e-08 | 3.3e-08 | 5.8e-08 | 3.733E-08 | 1.888E-08 | 2.100E-08 | 5.800E-08 |
| 28540 | 1427 | E1427V | E | 7.6e-08 | 1e-07 | 6.9e-08 | 8.167E-08 | 1.626E-08 | 6.900E-08 | 1.000E-07 |
| 28541 | 1428 | P1428A | E | 6.9e-08 | 2.7e-08 | 1.4e-07 | 7.867E-08 | 5.712E-08 | 2.700E-08 | 1.400E-07 |
| 28542 | 1428 | P1428R | E | 4e-08 | 4e-08 | 4.1e-08 | 4.033E-08 | 5.774E-10 | 4.000E-08 | 4.100E-08 |
| 28543 | 1428 | P1428N | E | 5.6e-09 | 7.5e-08 | 2.6e-08 | 3.553E-08 | 3.567E-08 | 5.600E-09 | 7.500E-08 |
| 28544 | 1428 | P1428D | E | 2.5e-08 | 2.1e-08 | 2e-07 | 8.200E-08 | 1.022E-07 | 2.100E-08 | 2.000E-07 |
| 28545 | 1428 | P1428C | E | 6.3e-08 | 8.9e-08 | 4.7e-08 | 6.633E-08 | 2.120E-08 | 4.700E-08 | 8.900E-08 |
| 28546 | 1428 | P1428E | E | 4e-08 | 6.4e-08 | 5.4e-08 | 5.267E-08 | 1.206E-08 | 4.000E-08 | 6.400E-08 |
| 28547 | 1428 | P1428Q | E | 4.3e-08 | 2.7e-08 | 8.6e-08 | 5.200E-08 | 3.051E-08 | 2.700E-08 | 8.600E-08 |
| 28548 | 1428 | P1428G | E | 2.9e-08 | 5.3e-08 | 2.2e-08 | 3.467E-08 | 1.626E-08 | 2.200E-08 | 5.300E-08 |
| 28549 | 1428 | P1428H | E | 5.4e-08 | 3.4e-08 | 2.8e-08 | 3.867E-08 | 1.361E-08 | 2.800E-08 | 5.400E-08 |
| 28550 | 1428 | P1428I | E | 4.2e-08 | 9.5e-08 | 5.9e-08 | 6.533E-08 | 2.706E-08 | 4.200E-08 | 9.500E-08 |
| 28551 | 1428 | P1428L | E | 6.3e-08 | 3.5e-08 | 6.1e-08 | 5.300E-08 | 1.562E-08 | 3.500E-08 | 6.300E-08 |
| 28552 | 1428 | P1428K | E | 3.4e-08 | 6.2e-08 | 3.2e-08 | 4.267E-08 | 1.677E-08 | 3.200E-08 | 6.200E-08 |
| 28553 | 1428 | P1428M | E | 1.8e-08 | 7.8e-08 | 2.3e-08 | 3.967E-08 | 3.329E-08 | 1.800E-08 | 7.800E-08 |
| 28554 | 1428 | P1428F | E | 7.3e-08 | 3.5e-08 | 7.3e-08 | 6.033E-08 | 2.194E-08 | 3.500E-08 | 7.300E-08 |
| 28555 | 1428 | P1428P | E | 2.3e-08 | 4.7e-08 | 5.4e-08 | 4.133E-08 | 1.626E-08 | 2.300E-08 | 5.400E-08 |
| 28556 | 1428 | P1428S | E | 4.1e-08 | 6.8e-08 | 2.5e-08 | 4.467E-08 | 2.173E-08 | 2.500E-08 | 6.800E-08 |
| 28557 | 1428 | P1428T | E | 2.8e-08 | 7.1e-08 | 2.7e-08 | 4.200E-08 | 2.512E-08 | 2.700E-08 | 7.100E-08 |
| 28558 | 1428 | P1428W | E | 1.4e-07 | 3.4e-08 | 2.7e-08 | 6.700E-08 | 6.332E-08 | 2.700E-08 | 1.400E-07 |
| 28559 | 1428 | P1428Y | E | 2.8e-08 | 4e-08 | 5.2e-08 | 4.000E-08 | 1.200E-08 | 2.800E-08 | 5.200E-08 |

| | | | | | | | | | | |
|---|---|---|---|---|---|---|---|---|---|---|
| 28560 | 1428 | P1428V | E | 4.9e-08 | 1.1e-08 | 2.4e-08 | 2.800E-08 | 1.931E-08 | 1.100E-08 | 4.900E-08 |
| 28561 | 1429 | V1429A | E | 3.7e-08 | 5.5e-08 | 4.2e-08 | 4.467E-08 | 9.292E-09 | 3.700E-08 | 5.500E-08 |
| 28562 | 1429 | V1429R | E | 7.2e-08 | 2.4e-08 | 1.5e-08 | 3.700E-08 | 3.064E-08 | 1.500E-08 | 7.200E-08 |
| 28563 | 1429 | V1429N | E | 2.4e-08 | 6.3e-08 | 6.2e-08 | 4.967E-08 | 2.223E-08 | 2.400E-08 | 6.300E-08 |
| 28564 | 1429 | V1429D | E | 4.8e-08 | 8.9e-08 | 5e-08 | 6.233E-08 | 2.312E-08 | 4.800E-08 | 8.900E-08 |
| 28565 | 1429 | V1429C | E | 1.3e-07 | 4.6e-08 | 7.3e-08 | 8.300E-08 | 4.288E-08 | 4.600E-08 | 1.300E-07 |
| 28566 | 1429 | V1429E | E | 5.3e-08 | 5.2e-08 | 2.5e-08 | 4.333E-08 | 1.589E-08 | 2.500E-08 | 5.300E-08 |
| 28567 | 1429 | V1429Q | E | 1.1e-08 | 5.5e-08 | 1.9e-08 | 2.833E-08 | 2.344E-08 | 1.100E-08 | 5.500E-08 |
| 28568 | 1429 | V1429G | E | 2.4e-08 | 3.6e-08 | 3.2e-08 | 3.067E-08 | 6.110E-09 | 2.400E-08 | 3.600E-08 |
| 28569 | 1429 | V1429H | E | 2.3e-08 | 2.2e-08 | 4.6e-08 | 3.033E-08 | 1.358E-08 | 2.200E-08 | 4.600E-08 |
| 28570 | 1429 | V1429I | E | 1.1e-07 | 4.1e-08 | 7.4e-08 | 7.500E-08 | 3.451E-08 | 4.100E-08 | 1.100E-07 |
| 28571 | 1429 | V1429L | E | 4.4e-08 | 8.1e-08 | 4.4e-08 | 5.633E-08 | 2.136E-08 | 4.400E-08 | 8.100E-08 |
| 28572 | 1429 | V1429K | E | 8.8e-08 | 6.1e-08 | 1.9e-08 | 5.600E-08 | 3.477E-08 | 1.900E-08 | 8.800E-08 |
| 28573 | 1429 | V1429M | E | 3.1e-08 | 6.6e-08 | 3.9e-08 | 4.533E-08 | 1.834E-08 | 3.100E-08 | 6.600E-08 |
| 28574 | 1429 | V1429F | E | 5.5e-08 | 3e-08 | 4.5e-08 | 4.333E-08 | 1.258E-08 | 3.000E-08 | 5.500E-08 |
| 28575 | 1429 | V1429P | E | 2.3e-08 | 3.7e-08 | 2e-08 | 2.667E-08 | 9.074E-09 | 2.000E-08 | 3.700E-08 |
| 28576 | 1429 | V1429S | E | 6.6e-08 | 1.2e-08 | 6.2e-08 | 4.667E-08 | 3.009E-08 | 1.200E-08 | 6.600E-08 |
| 28577 | 1429 | V1429T | E | 2.6e-08 | 4.5e-08 | 1.4e-08 | 2.833E-08 | 1.563E-08 | 1.400E-08 | 4.500E-08 |
| 28578 | 1429 | V1429W | E | 5.1e-08 | 8.1e-08 | 3.4e-08 | 5.533E-08 | 2.380E-08 | 3.400E-08 | 8.100E-08 |
| 28579 | 1429 | V1429Y | E | 2.9e-08 | 1.8e-08 | 5.3e-08 | 3.333E-08 | 1.790E-08 | 1.800E-08 | 5.300E-08 |
| 28580 | 1429 | V1429V | E | 5.6e-08 | 6.7e-08 | 3.6e-08 | 5.300E-08 | 1.572E-08 | 3.600E-08 | 6.700E-08 |
| 28581 | 1430 | T1430A | E | 1.3e-08 | 2.5e-07 | 8.9e-08 | 1.173E-07 | 1.210E-07 | 1.300E-08 | 2.500E-07 |
| 28582 | 1430 | T1430R | E | 1.6e-08 | 3.8e-08 | 2.3e-08 | 2.567E-08 | 1.124E-08 | 1.600E-08 | 3.800E-08 |
| 28583 | 1430 | T1430N | E | 7.2e-08 | 1.5e-08 | 3.2e-08 | 3.967E-08 | 2.926E-08 | 1.500E-08 | 7.200E-08 |
| 28584 | 1430 | T1430D | E | 3.1e-08 | 2.5e-08 | 1.3e-07 | 6.200E-08 | 5.897E-08 | 2.500E-08 | 1.300E-07 |
| 28585 | 1430 | T1430C | E | 4.5e-08 | 3.9e-08 | 3.2e-08 | 3.867E-08 | 6.506E-09 | 3.200E-08 | 4.500E-08 |
| 28586 | 1430 | T1430E | E | 2.2e-08 | 8.6e-08 | 3.2e-08 | 4.667E-08 | 3.443E-08 | 2.200E-08 | 8.600E-08 |
| 28587 | 1430 | T1430Q | E | 5e-08 | 1.8e-08 | 3e-08 | 3.267E-08 | 1.617E-08 | 1.800E-08 | 5.000E-08 |
| 28588 | 1430 | T1430G | E | 3.1e-08 | 1.5e-08 | 4.5e-08 | 3.033E-08 | 1.501E-08 | 1.500E-08 | 4.500E-08 |
| 28589 | 1430 | T1430H | E | 3.2e-08 | 6.3e-08 | 4e-08 | 4.500E-08 | 1.609E-08 | 3.200E-08 | 6.300E-08 |
| 28590 | 1430 | T1430I | E | 4.4e-08 | 1.9e-08 | 5e-08 | 3.767E-08 | 1.644E-08 | 1.900E-08 | 5.000E-08 |
| 28591 | 1430 | T1430L | E | 6.2e-08 | 7e-08 | 3.9e-08 | 5.700E-08 | 1.609E-08 | 3.900E-08 | 7.000E-08 |
| 28592 | 1430 | T1430K | E | 9.5e-08 | 5.9e-08 | 2.5e-08 | 5.967E-08 | 3.500E-08 | 2.500E-08 | 9.500E-08 |
| 28593 | 1430 | T1430M | E | 2.6e-08 | 3.8e-08 | 9.1e-08 | 5.167E-08 | 3.459E-08 | 2.600E-08 | 9.100E-08 |
| 28594 | 1430 | T1430F | E | 8.6e-08 | 9.5e-08 | 4.5e-08 | 7.533E-08 | 2.665E-08 | 4.500E-08 | 9.500E-08 |
| 28595 | 1430 | T1430P | E | 4.4e-08 | 1.5e-08 | 7.4e-08 | 4.433E-08 | 2.950E-08 | 1.500E-08 | 7.400E-08 |
| 28596 | 1430 | T1430S | E | 5e-08 | 6.7e-08 | 5.4e-08 | 5.700E-08 | 8.888E-09 | 5.000E-08 | 6.700E-08 |
| 28597 | 1430 | T1430T | E | 5.1e-08 | 3.7e-08 | 2.5e-08 | 3.767E-08 | 1.301E-08 | 2.500E-08 | 5.100E-08 |
| 28598 | 1430 | T1430W | E | 3e-08 | 4.7e-08 | 2.1e-08 | 3.267E-08 | 1.320E-08 | 2.100E-08 | 4.700E-08 |
| 28599 | 1430 | T1430Y | E | 3.8e-08 | 3.3e-08 | 5.6e-08 | 4.233E-08 | 1.210E-08 | 3.300E-08 | 5.600E-08 |
| 28600 | 1430 | T1430V | E | 1.3e-07 | 8.7e-08 | 6.3e-08 | 9.333E-08 | 3.395E-08 | 6.300E-08 | 1.300E-07 |
| 28601 | 1431 | V1431A | E | 5.1e-08 | 1.8e-08 | 1.7e-08 | 2.867E-08 | 1.935E-08 | 1.700E-08 | 5.100E-08 |
| 28602 | 1431 | V1431R | E | 2.2e-08 | 2.7e-08 | 4e-08 | 2.967E-08 | 9.292E-09 | 2.200E-08 | 4.000E-08 |
| 28603 | 1431 | V1431N | E | 9.2e-08 | 4.6e-08 | 4.2e-08 | 6.000E-08 | 2.778E-08 | 4.200E-08 | 9.200E-08 |
| 28604 | 1431 | V1431D | E | 3e-08 | 4.1e-08 | 2.5e-08 | 3.200E-08 | 8.185E-09 | 2.500E-08 | 4.100E-08 |
| 28605 | 1431 | V1431C | E | 4.6e-08 | 1.2e-07 | 8.5e-09 | 5.817E-08 | 5.674E-08 | 8.500E-09 | 1.200E-07 |
| 28606 | 1431 | V1431E | E | 3.4e-08 | 1.8e-08 | 5.7e-08 | 3.633E-08 | 1.960E-08 | 1.800E-08 | 5.700E-08 |
| 28607 | 1431 | V1431Q | E | 6.5e-08 | 7.8e-08 | 3e-08 | 5.767E-08 | 2.483E-08 | 3.000E-08 | 7.800E-08 |
| 28608 | 1431 | V1431G | E | 2e-08 | 2.1e-08 | 8e-08 | 4.033E-08 | 3.436E-08 | 2.000E-08 | 8.000E-08 |
| 28609 | 1431 | V1431H | E | 7.1e-08 | 3.3e-08 | 7.6e-08 | 6.000E-08 | 2.352E-08 | 3.300E-08 | 7.600E-08 |
| 28610 | 1431 | V1431I | E | 8.5e-09 | 1.1e-07 | 8.3e-08 | 6.717E-08 | 5.257E-08 | 8.500E-09 | 1.100E-07 |
| 28611 | 1431 | V1431L | E | 6.5e-08 | 5.1e-08 | 5.2e-08 | 5.600E-08 | 7.810E-09 | 5.100E-08 | 6.500E-08 |
| 28612 | 1431 | V1431K | E | 1.7e-08 | 1e-07 | 4.2e-08 | 5.300E-08 | 4.258E-08 | 1.700E-08 | 1.000E-07 |
| 28613 | 1431 | V1431M | E | 5e-08 | 5.5e-08 | 5e-08 | 5.167E-08 | 2.887E-09 | 5.000E-08 | 5.500E-08 |
| 28614 | 1431 | V1431F | E | 9.9e-08 | 6.1e-08 | 1.6e-08 | 5.867E-08 | 4.155E-08 | 1.600E-08 | 9.900E-08 |
| 28615 | 1431 | V1431P | E | 5.9e-08 | 6.1e-08 | 5.2e-08 | 5.733E-08 | 4.726E-09 | 5.200E-08 | 6.100E-08 |
| 28616 | 1431 | V1431S | E | 2.1e-08 | 4.1e-08 | 3.9e-08 | 3.367E-08 | 1.102E-08 | 2.100E-08 | 4.100E-08 |
| 28617 | 1431 | V1431T | E | 7.4e-08 | 4.9e-08 | 4e-08 | 5.433E-08 | 1.762E-08 | 4.000E-08 | 7.400E-08 |
| 28618 | 1431 | V1431W | E | 2.2e-08 | 4.2e-08 | 5.8e-08 | 4.067E-08 | 1.804E-08 | 2.200E-08 | 5.800E-08 |
| 28619 | 1431 | V1431Y | E | 6.8e-08 | 1.6e-08 | 2.7e-08 | 3.700E-08 | 2.740E-08 | 1.600E-08 | 6.800E-08 |
| 28620 | 1431 | V1431V | E | 2.3e-08 | 6e-08 | 2e-08 | 3.433E-08 | 2.228E-08 | 2.000E-08 | 6.000E-08 |
| 28621 | 1432 | S1432A | E | 4e-08 | 1.8e-07 | 5.8e-08 | 9.267E-08 | 7.617E-08 | 4.000E-08 | 1.800E-07 |
| 28622 | 1432 | S1432R | E | 2.3e-08 | 6e-08 | 4.8e-08 | 4.367E-08 | 1.888E-08 | 2.300E-08 | 6.000E-08 |
| 28623 | 1432 | S1432N | E | 1.7e-08 | 3e-08 | 7.8e-08 | 4.167E-08 | 3.213E-08 | 1.700E-08 | 7.800E-08 |
| 28624 | 1432 | S1432D | E | 3.1e-08 | 5.1e-08 | 4.2e-08 | 4.133E-08 | 1.002E-08 | 3.100E-08 | 5.100E-08 |
| 28625 | 1432 | S1432C | E | 3.2e-08 | 2.2e-08 | 5.8e-08 | 3.733E-08 | 1.858E-08 | 2.200E-08 | 5.800E-08 |
| 28626 | 1432 | S1432E | E | 9.9e-08 | 3.5e-08 | 9.5e-08 | 7.633E-08 | 3.585E-08 | 3.500E-08 | 9.900E-08 |
| 28627 | 1432 | S1432Q | E | 6.8e-08 | 4.6e-08 | 4.2e-08 | 5.200E-08 | 1.400E-08 | 4.200E-08 | 6.800E-08 |
| 28628 | 1432 | S1432G | E | 1.9e-08 | 3.4e-08 | 5.4e-08 | 3.567E-08 | 1.756E-08 | 1.900E-08 | 5.400E-08 |
| 28629 | 1432 | S1432H | E | 5.7e-08 | 2e-08 | 2.6e-08 | 3.433E-08 | 1.986E-08 | 2.000E-08 | 5.700E-08 |
| 28630 | 1432 | S1432I | E | 5.9e-08 | 4.8e-08 | 1e-07 | 6.900E-08 | 2.740E-08 | 4.800E-08 | 1.000E-07 |
| 28631 | 1432 | S1432L | E | 1.7e-08 | 3.3e-08 | 5.8e-08 | 3.600E-08 | 2.066E-08 | 1.700E-08 | 5.800E-08 |
| 28632 | 1432 | S1432K | E | 7e-08 | 4.6e-08 | 8.8e-08 | 6.800E-08 | 2.107E-08 | 4.600E-08 | 8.800E-08 |
| 28633 | 1432 | S1432M | E | 2.4e-08 | 4.2e-08 | 3.8e-08 | 3.467E-08 | 9.452E-09 | 2.400E-08 | 4.200E-08 |
| 28634 | 1432 | S1432F | E | 8.5e-08 | 6.1e-08 | 1.3e-07 | 9.200E-08 | 3.503E-08 | 6.100E-08 | 1.300E-07 |
| 28635 | 1432 | S1432P | E | 2.6e-08 | 7.9e-08 | 2.5e-08 | 4.333E-08 | 3.089E-08 | 2.500E-08 | 7.900E-08 |
| 28636 | 1432 | S1432S | E | 4.9e-08 | 1.9e-08 | 4.4e-08 | 3.733E-08 | 1.607E-08 | 1.900E-08 | 4.900E-08 |
| 28637 | 1432 | S1432T | E | 1e-07 | 5.7e-08 | 3e-08 | 6.233E-08 | 3.530E-08 | 3.000E-08 | 1.000E-07 |
| 28638 | 1432 | S1432W | E | 4.2e-08 | 2.9e-08 | 3.6e-08 | 3.567E-08 | 6.506E-09 | 2.900E-08 | 4.200E-08 |
| 28639 | 1432 | S1432Y | E | 4e-08 | 1.8e-08 | 5.2e-08 | 3.667E-08 | 1.724E-08 | 1.800E-08 | 5.200E-08 |
| 28640 | 1432 | S1432V | E | 2.7e-08 | 4.5e-08 | 6.2e-08 | 4.467E-08 | 1.750E-08 | 2.700E-08 | 6.200E-08 |
| 28641 | 1433 | W1433A | E | 1.9e-08 | 1.2e-07 | 1.7e-08 | 5.200E-08 | 5.890E-08 | 1.700E-08 | 1.200E-07 |
| 28642 | 1433 | W1433R | E | 3.2e-08 | 6.3e-08 | 7.9e-08 | 5.800E-08 | 2.390E-08 | 3.200E-08 | 7.900E-08 |
| 28643 | 1433 | W1433N | E | 3.1e-08 | 6.5e-08 | 6.2e-08 | 5.267E-08 | 1.882E-08 | 3.100E-08 | 6.500E-08 |
| 28644 | 1433 | W1433D | E | 1.1e-07 | 6.5e-08 | 7.9e-08 | 8.467E-08 | 2.303E-08 | 6.500E-08 | 1.100E-07 |

| | | | | | | | | | | |
|---|---|---|---|---|---|---|---|---|---|---|
| 28645 | 1433 | W1433C | E | 4.4e-08 | 4.1e-08 | 4e-08 | 4.167E-08 | 2.082E-09 | 4.000E-08 | 4.400E-08 |
| 28646 | 1433 | W1433E | E | 5.4e-08 | 8e-08 | 6.3e-08 | 6.567E-08 | 1.320E-08 | 5.400E-08 | 8.000E-08 |
| 28647 | 1433 | W1433Q | E | 5.9e-08 | 6.1e-08 | 2.9e-08 | 4.967E-08 | 1.793E-08 | 2.900E-08 | 6.100E-08 |
| 28648 | 1433 | W1433G | E | 1.7e-08 | 4.4e-08 | 6.2e-08 | 4.100E-08 | 2.265E-08 | 1.700E-08 | 6.200E-08 |
| 28649 | 1433 | W1433H | E | 3.2e-08 | 5.2e-08 | 7.7e-09 | 3.057E-08 | 2.218E-08 | 7.700E-09 | 5.200E-08 |
| 28650 | 1433 | W1433I | E | 2.2e-08 | 3.5e-08 | 2.7e-08 | 2.800E-08 | 6.557E-09 | 2.200E-08 | 3.500E-08 |
| 28651 | 1433 | W1433L | E | 8.4e-08 | 1.5e-07 | 9e-08 | 1.080E-07 | 3.650E-08 | 8.400E-08 | 1.500E-07 |
| 28652 | 1433 | W1433K | E | 5.1e-08 | 7.5e-08 | 1.9e-08 | 4.833E-08 | 2.810E-08 | 1.900E-08 | 7.500E-08 |
| 28653 | 1433 | W1433M | E | 5.3e-08 | 3e-08 | 4.5e-08 | 4.267E-08 | 1.168E-08 | 3.000E-08 | 5.300E-08 |
| 28654 | 1433 | W1433F | E | 1.9e-08 | 7.8e-08 | 6.2e-08 | 5.300E-08 | 3.051E-08 | 1.900E-08 | 7.800E-08 |
| 28655 | 1433 | W1433P | E | 8e-08 | 2.7e-08 | 2.1e-08 | 4.267E-08 | 3.247E-08 | 2.100E-08 | 8.000E-08 |
| 28656 | 1433 | W1433S | E | 2e-08 | 1.5e-08 | 4.7e-08 | 2.733E-08 | 1.721E-08 | 1.500E-08 | 4.700E-08 |
| 28657 | 1433 | W1433T | E | 4.3e-08 | 5.9e-08 | 7.4e-08 | 5.867E-08 | 1.550E-08 | 4.300E-08 | 7.400E-08 |
| 28658 | 1433 | W1433W | E | 3.5e-08 | 3.4e-08 | 6.3e-08 | 4.400E-08 | 1.646E-08 | 3.400E-08 | 6.300E-08 |
| 28659 | 1433 | W1433Y | E | 6.7e-08 | 6.3e-08 | 6.3e-08 | 6.433E-08 | 2.309E-09 | 6.300E-08 | 6.700E-08 |
| 28660 | 1433 | W1433V | E | 4.7e-08 | 5.3e-08 | 6.9e-08 | 5.633E-08 | 1.137E-08 | 4.700E-08 | 6.900E-08 |
| 28661 | 1434 | N1434A | E | 4e-08 | 1.6e-08 | 9e-08 | 4.867E-08 | 3.775E-08 | 1.600E-08 | 9.000E-08 |
| 28662 | 1434 | N1434R | E | 1.6e-08 | 5.6e-08 | 5.2e-08 | 4.133E-08 | 2.203E-08 | 1.600E-08 | 5.600E-08 |
| 28663 | 1434 | N1434N | E | 5.1e-08 | 2.4e-08 | 3.2e-08 | 3.567E-08 | 1.387E-08 | 2.400E-08 | 5.100E-08 |
| 28664 | 1434 | N1434D | E | 3.3e-08 | 1.3e-08 | 1.8e-08 | 2.133E-08 | 1.041E-08 | 1.300E-08 | 3.300E-08 |
| 28665 | 1434 | N1434C | E | 1.8e-08 | 1.6e-08 | 3.1e-08 | 2.167E-08 | 8.145E-09 | 1.600E-08 | 3.100E-08 |
| 28666 | 1434 | N1434E | E | 5.1e-08 | 6.2e-08 | 1.2e-08 | 4.167E-08 | 2.627E-08 | 1.200E-08 | 6.200E-08 |
| 28667 | 1434 | N1434Q | E | 7.1e-08 | 3.4e-08 | 6.1e-08 | 5.533E-08 | 1.914E-08 | 3.400E-08 | 7.100E-08 |
| 28668 | 1434 | N1434G | E | 1.1e-07 | 3.5e-08 | 6.8e-08 | 7.100E-08 | 3.759E-08 | 3.500E-08 | 1.100E-07 |
| 28669 | 1434 | N1434H | E | 3.9e-08 | 2.4e-08 | 3.1e-08 | 3.133E-08 | 7.506E-09 | 2.400E-08 | 3.900E-08 |
| 28670 | 1434 | N1434I | E | 5.2e-08 | 4.8e-08 | 6.3e-08 | 5.433E-08 | 7.767E-09 | 4.800E-08 | 6.300E-08 |
| 28671 | 1434 | N1434L | E | 4.9e-08 | 1.3e-07 | 4.2e-08 | 7.367E-08 | 4.891E-08 | 4.200E-08 | 1.300E-07 |
| 28672 | 1434 | N1434K | E | 7.7e-08 | 9.1e-08 | 4.8e-08 | 7.200E-08 | 2.193E-08 | 4.800E-08 | 9.100E-08 |
| 28673 | 1434 | N1434M | E | 3.5e-08 | 6.8e-08 | 4.4e-08 | 4.900E-08 | 1.706E-08 | 3.500E-08 | 6.800E-08 |
| 28674 | 1434 | N1434F | E | 3.6e-08 | 3.3e-08 | 4.8e-08 | 3.900E-08 | 7.937E-09 | 3.300E-08 | 4.800E-08 |
| 28675 | 1434 | N1434P | E | 3.6e-08 | 1.7e-08 | 5.9e-08 | 3.733E-08 | 2.103E-08 | 1.700E-08 | 5.900E-08 |
| 28676 | 1434 | N1434S | E | 7.1e-08 | 4e-08 | 7.7e-08 | 6.267E-08 | 1.986E-08 | 4.000E-08 | 7.700E-08 |
| 28677 | 1434 | N1434T | E | 5.7e-08 | 5.8e-08 | 2.3e-08 | 4.600E-08 | 1.992E-08 | 2.300E-08 | 5.800E-08 |
| 28678 | 1434 | N1434W | E | 5.7e-08 | 5.8e-08 | 2.2e-08 | 4.567E-08 | 2.050E-08 | 2.200E-08 | 5.800E-08 |
| 28679 | 1434 | N1434Y | E | 3.5e-08 | 5.3e-08 | 4.4e-08 | 4.400E-08 | 9.000E-09 | 3.500E-08 | 5.300E-08 |
| 28680 | 1434 | N1434V | E | 3.5e-08 | 3.8e-08 | 4.3e-08 | 3.867E-08 | 4.041E-09 | 3.500E-08 | 4.300E-08 |
| 28681 | 1435 | S1435A | E | 2.6e-08 | 7.3e-08 | 1.3e-07 | 7.633E-08 | 5.208E-08 | 2.600E-08 | 1.300E-07 |
| 28682 | 1435 | S1435R | E | 4.7e-08 | 7.8e-08 | 2.2e-08 | 4.900E-08 | 2.805E-08 | 2.200E-08 | 7.800E-08 |
| 28683 | 1435 | S1435N | E | 4.5e-08 | 2.3e-08 | 4e-08 | 3.600E-08 | 1.153E-08 | 2.300E-08 | 4.500E-08 |
| 28684 | 1435 | S1435D | E | 4e-08 | 6.3e-08 | 2.9e-08 | 4.400E-08 | 1.735E-08 | 2.900E-08 | 6.300E-08 |
| 28685 | 1435 | S1435C | E | 4e-08 | 2e-08 | 4.7e-08 | 3.567E-08 | 1.401E-08 | 2.000E-08 | 4.700E-08 |
| 28686 | 1435 | S1435E | E | 2e-08 | 6.2e-09 | 9.7e-08 | 4.107E-08 | 4.893E-08 | 6.200E-09 | 9.700E-08 |
| 28687 | 1435 | S1435Q | E | 2.8e-08 | 4.4e-08 | 4e-08 | 3.733E-08 | 8.327E-09 | 2.800E-08 | 4.400E-08 |
| 28688 | 1435 | S1435G | E | 5.8e-08 | 7.1e-08 | 4.9e-08 | 5.933E-08 | 1.106E-08 | 4.900E-08 | 7.100E-08 |
| 28689 | 1435 | S1435H | E | 3e-08 | 4.3e-08 | 3.5e-08 | 3.600E-08 | 6.557E-09 | 3.000E-08 | 4.300E-08 |
| 28690 | 1435 | S1435I | E | 2.8e-08 | 7.4e-08 | 2.3e-08 | 4.167E-08 | 2.811E-08 | 2.300E-08 | 7.400E-08 |
| 28691 | 1435 | S1435L | E | 2e-08 | 3.7e-08 | 5.5e-08 | 3.733E-08 | 1.750E-08 | 2.000E-08 | 5.500E-08 |
| 28692 | 1435 | S1435K | E | 3.8e-08 | 4.1e-08 | 3.3e-08 | 3.733E-08 | 4.041E-09 | 3.300E-08 | 4.100E-08 |
| 28693 | 1435 | S1435M | E | 6.6e-08 | 3.6e-08 | 8.1e-08 | 6.100E-08 | 2.291E-08 | 3.600E-08 | 8.100E-08 |
| 28694 | 1435 | S1435F | E | 3.6e-08 | 2.4e-08 | 4e-08 | 3.333E-08 | 8.327E-09 | 2.400E-08 | 4.000E-08 |
| 28695 | 1435 | S1435P | E | 7.5e-08 | 4.9e-08 | 4.1e-08 | 5.500E-08 | 1.778E-08 | 4.100E-08 | 7.500E-08 |
| 28696 | 1435 | S1435S | E | 2.1e-08 | 3.8e-08 | 3.5e-08 | 3.133E-08 | 9.074E-09 | 2.100E-08 | 3.800E-08 |
| 28697 | 1435 | S1435T | E | 1.2e-07 | 5.7e-08 | 3e-08 | 6.900E-08 | 4.618E-08 | 3.000E-08 | 1.200E-07 |
| 28698 | 1435 | S1435W | E | 5.9e-08 | 1e-07 | 2.4e-08 | 6.100E-08 | 3.804E-08 | 2.400E-08 | 1.000E-07 |
| 28699 | 1435 | S1435Y | E | 3.9e-08 | 2e-08 | 1.8e-08 | 2.567E-08 | 1.159E-08 | 1.800E-08 | 3.900E-08 |
| 28700 | 1435 | S1435V | E | 4.9e-08 | 3.8e-08 | 3.1e-08 | 3.933E-08 | 9.074E-09 | 3.100E-08 | 4.900E-08 |
| 28701 | 1436 | G1436A | E | 3.4e-08 | 6.3e-08 | 6.9e-08 | 5.533E-08 | 1.872E-08 | 3.400E-08 | 6.900E-08 |
| 28702 | 1436 | G1436R | E | 3.6e-08 | 3.9e-08 | 1.2e-08 | 2.900E-08 | 1.480E-08 | 1.200E-08 | 3.900E-08 |
| 28703 | 1436 | G1436N | E | 4.4e-08 | 2.5e-08 | 6.4e-08 | 4.433E-08 | 1.950E-08 | 2.500E-08 | 6.400E-08 |
| 28704 | 1436 | G1436D | E | 6.3e-08 | 2.9e-08 | 2.9e-08 | 4.033E-08 | 1.963E-08 | 2.900E-08 | 6.300E-08 |
| 28705 | 1436 | G1436C | E | 6.9e-08 | 5.3e-08 | 6.3e-08 | 6.167E-08 | 8.083E-09 | 5.300E-08 | 6.900E-08 |
| 28706 | 1436 | G1436E | E | 7.6e-08 | 2.9e-08 | 9.2e-08 | 6.567E-08 | 3.275E-08 | 2.900E-08 | 9.200E-08 |
| 28707 | 1436 | G1436Q | E | 4.3e-08 | 1.3e-08 | 5.2e-08 | 3.600E-08 | 2.042E-08 | 1.300E-08 | 5.200E-08 |
| 28708 | 1436 | G1436G | E | 2e-08 | 5e-08 | 2.9e-08 | 3.300E-08 | 1.539E-08 | 2.000E-08 | 5.000E-08 |
| 28709 | 1436 | G1436H | E | 3.9e-08 | 7.2e-08 | 6.9e-08 | 6.000E-08 | 1.825E-08 | 3.900E-08 | 7.200E-08 |
| 28710 | 1436 | G1436I | E | 1.8e-08 | 3.7e-08 | 2.6e-08 | 2.700E-08 | 9.539E-09 | 1.800E-08 | 3.700E-08 |
| 28711 | 1436 | G1436L | E | 1.9e-08 | 1.5e-08 | 3.3e-08 | 2.233E-08 | 9.452E-09 | 1.500E-08 | 3.300E-08 |
| 28712 | 1436 | G1436K | E | 8.6e-08 | 4.8e-08 | 2.5e-08 | 5.300E-08 | 3.081E-08 | 2.500E-08 | 8.600E-08 |
| 28713 | 1436 | G1436M | E | 7.1e-08 | 5e-08 | 2.7e-08 | 4.933E-08 | 2.201E-08 | 2.700E-08 | 7.100E-08 |
| 28714 | 1436 | G1436F | E | 3.4e-08 | 4.3e-08 | 5.6e-08 | 4.433E-08 | 1.106E-08 | 3.400E-08 | 5.600E-08 |
| 28715 | 1436 | G1436P | E | 4.7e-08 | 9.2e-08 | 3.2e-08 | 5.700E-08 | 3.122E-08 | 3.200E-08 | 9.200E-08 |
| 28716 | 1436 | G1436S | E | 1.6e-08 | 1.1e-07 | 8.8e-08 | 7.133E-08 | 4.917E-08 | 1.600E-08 | 1.100E-07 |
| 28717 | 1436 | G1436T | E | 4e-08 | 1e-08 | 6.6e-08 | 3.867E-08 | 2.802E-08 | 1.000E-08 | 6.600E-08 |
| 28718 | 1436 | G1436W | E | 5.3e-08 | 3e-08 | 6.4e-08 | 4.900E-08 | 1.735E-08 | 3.000E-08 | 6.400E-08 |
| 28719 | 1436 | G1436Y | E | 5.1e-08 | 1.9e-08 | 3.3e-08 | 3.433E-08 | 1.604E-08 | 1.900E-08 | 5.100E-08 |
| 28720 | 1436 | G1436V | E | 5.3e-08 | 3.7e-08 | 6.8e-08 | 5.267E-08 | 1.550E-08 | 3.700E-08 | 6.800E-08 |
| 28721 | 1437 | A1437A | E | 5.8e-08 | 2.2e-08 | 3.9e-08 | 3.967E-08 | 1.801E-08 | 2.200E-08 | 5.800E-08 |
| 28722 | 1437 | A1437R | E | 3.4e-08 | 3.5e-08 | 3e-08 | 3.300E-08 | 2.646E-09 | 3.000E-08 | 3.500E-08 |
| 28723 | 1437 | A1437N | E | 1e-07 | 4.9e-08 | 6e-08 | 6.967E-08 | 2.684E-08 | 4.900E-08 | 1.000E-07 |
| 28724 | 1437 | A1437D | E | 6.7e-08 | 7e-08 | 2.6e-08 | 5.433E-08 | 2.458E-08 | 2.600E-08 | 7.000E-08 |
| 28725 | 1437 | A1437C | E | 1.5e-08 | 3.5e-08 | 2.3e-08 | 2.433E-08 | 1.007E-08 | 1.500E-08 | 3.500E-08 |
| 28726 | 1437 | A1437E | E | 2.1e-08 | 6e-08 | 4.1e-08 | 4.067E-08 | 1.950E-08 | 2.100E-08 | 6.000E-08 |
| 28727 | 1437 | A1437Q | E | 2.7e-08 | 4.1e-08 | 1.2e-07 | 6.267E-08 | 5.014E-08 | 2.700E-08 | 1.200E-07 |
| 28728 | 1437 | A1437G | E | 3e-08 | 3.2e-08 | 2.7e-08 | 2.967E-08 | 2.517E-09 | 2.700E-08 | 3.200E-08 |
| 28729 | 1437 | A1437H | E | 5e-08 | 3.9e-08 | 9.9e-08 | 6.267E-08 | 3.194E-08 | 3.900E-08 | 9.900E-08 |

| | | | | | | | | | | |
|---|---|---|---|---|---|---|---|---|---|---|
| 28730 | 1437 | A1437I | E | 1.5e-08 | 4.4e-08 | 8e-08 | 4.633E-08 | 3.256E-08 | 1.500E-08 | 8.000E-08 |
| 28731 | 1437 | A1437L | E | 8.1e-08 | 1.5e-08 | 1e-07 | 6.533E-08 | 4.461E-08 | 1.500E-08 | 1.000E-07 |
| 28732 | 1437 | A1437K | E | 2.9e-08 | 5.7e-08 | 5.3e-08 | 4.633E-08 | 1.514E-08 | 2.900E-08 | 5.700E-08 |
| 28733 | 1437 | A1437M | E | 2.4e-08 | 2.6e-08 | 6.4e-08 | 3.800E-08 | 2.254E-08 | 2.400E-08 | 6.400E-08 |
| 28734 | 1437 | A1437F | E | 7.2e-08 | 7.5e-08 | 8.1e-08 | 7.600E-08 | 4.583E-09 | 7.200E-08 | 8.100E-08 |
| 28735 | 1437 | A1437P | E | 1.6e-08 | 4.7e-08 | 6.1e-08 | 4.133E-08 | 2.303E-08 | 1.600E-08 | 6.100E-08 |
| 28736 | 1437 | A1437S | E | 3.8e-08 | 7.8e-08 | 7.6e-08 | 6.400E-08 | 2.254E-08 | 3.800E-08 | 7.800E-08 |
| 28737 | 1437 | A1437T | E | 6.2e-08 | 6.2e-08 | 3.6e-08 | 5.333E-08 | 1.501E-08 | 3.600E-08 | 6.200E-08 |
| 28738 | 1437 | A1437W | E | 3.2e-08 | 2.8e-08 | 6.4e-08 | 4.133E-08 | 1.973E-08 | 2.800E-08 | 6.400E-08 |
| 28739 | 1437 | A1437Y | E | 1.1e-07 | 1e-07 | 3.1e-08 | 8.033E-08 | 4.302E-08 | 3.100E-08 | 1.100E-07 |
| 28740 | 1437 | A1437V | E | 5.2e-08 | 1.5e-08 | 8.2e-08 | 4.967E-08 | 3.356E-08 | 1.500E-08 | 8.200E-08 |
| 28741 | 1438 | L1438A | E | 3.7e-08 | 1.7e-08 | 7.4e-08 | 4.267E-08 | 2.892E-08 | 1.700E-08 | 7.400E-08 |
| 28742 | 1438 | L1438R | E | 2.5e-08 | 9e-09 | 2.9e-08 | 2.100E-08 | 1.058E-08 | 9.000E-09 | 2.900E-08 |
| 28743 | 1438 | L1438N | E | 4.3e-08 | 3.3e-08 | 5.5e-08 | 4.367E-08 | 1.102E-08 | 3.300E-08 | 5.500E-08 |
| 28744 | 1438 | L1438D | E | 2.7e-08 | 1.3e-07 | 2.8e-08 | 6.167E-08 | 5.918E-08 | 2.700E-08 | 1.300E-07 |
| 28745 | 1438 | L1438C | E | 4.7e-08 | 6.3e-08 | 3.3e-08 | 4.767E-08 | 1.501E-08 | 3.300E-08 | 6.300E-08 |
| 28746 | 1438 | L1438E | E | 2.4e-08 | 7.6e-08 | 4.4e-08 | 4.800E-08 | 2.623E-08 | 2.400E-08 | 7.600E-08 |
| 28747 | 1438 | L1438Q | E | 7.2e-08 | 2.8e-08 | 3.9e-08 | 4.633E-08 | 2.290E-08 | 2.800E-08 | 7.200E-08 |
| 28748 | 1438 | L1438G | E | 8e-08 | 7.1e-08 | 5.1e-08 | 6.733E-08 | 1.484E-08 | 5.100E-08 | 8.000E-08 |
| 28749 | 1438 | L1438H | E | 1.1e-07 | 8.4e-08 | 3.2e-08 | 7.533E-08 | 3.972E-08 | 3.200E-08 | 1.100E-07 |
| 28750 | 1438 | L1438I | E | 5.5e-08 | 1.8e-08 | 3.4e-08 | 3.567E-08 | 1.856E-08 | 1.800E-08 | 5.500E-08 |
| 28751 | 1438 | L1438L | E | 6.4e-08 | 7.4e-08 | 4.3e-08 | 6.033E-08 | 1.582E-08 | 4.300E-08 | 7.400E-08 |
| 28752 | 1438 | L1438K | E | 1e-07 | 8.5e-08 | 7e-08 | 8.500E-08 | 1.500E-08 | 7.000E-08 | 1.000E-07 |
| 28753 | 1438 | L1438M | E | 1.8e-07 | 6.2e-08 | 1.9e-08 | 8.700E-08 | 8.336E-08 | 1.900E-08 | 1.800E-07 |
| 28754 | 1438 | L1438F | E | 1.8e-08 | 1.2e-07 | 8.9e-08 | 7.567E-08 | 5.229E-08 | 1.800E-08 | 1.200E-07 |
| 28755 | 1438 | L1438P | E | 6.3e-08 | 3.4e-08 | 3.3e-08 | 4.333E-08 | 1.704E-08 | 3.300E-08 | 6.300E-08 |
| 28756 | 1438 | L1438S | E | 4.9e-08 | 3.8e-08 | 2.4e-08 | 3.700E-08 | 1.253E-08 | 2.400E-08 | 4.900E-08 |
| 28757 | 1438 | L1438T | E | 2.6e-08 | 7.2e-08 | 3.8e-08 | 4.533E-08 | 2.386E-08 | 2.600E-08 | 7.200E-08 |
| 28758 | 1438 | L1438W | E | 4.3e-08 | 4.9e-08 | 4.6e-08 | 4.600E-08 | 3.000E-09 | 4.300E-08 | 4.900E-08 |
| 28759 | 1438 | L1438Y | E | 7.1e-08 | 5.1e-08 | 4.3e-08 | 5.500E-08 | 1.442E-08 | 4.300E-08 | 7.100E-08 |
| 28760 | 1438 | L1438V | E | 1.5e-07 | 4.4e-08 | 2.1e-08 | 7.167E-08 | 6.881E-08 | 2.100E-08 | 1.500E-07 |
| 28761 | 1439 | T1439A | E | 2.5e-08 | 3.2e-08 | 4.1e-08 | 3.267E-08 | 8.021E-09 | 2.500E-08 | 4.100E-08 |
| 28762 | 1439 | T1439R | E | 2.2e-08 | 3.6e-08 | 6.5e-08 | 4.100E-08 | 2.193E-08 | 2.200E-08 | 6.500E-08 |
| 28763 | 1439 | T1439N | E | 3.1e-08 | 1.8e-07 | 5.3e-08 | 8.800E-08 | 8.043E-08 | 3.100E-08 | 1.800E-07 |
| 28764 | 1439 | T1439D | E | 1.6e-08 | 5.9e-08 | 3.2e-08 | 3.567E-08 | 2.173E-08 | 1.600E-08 | 5.900E-08 |
| 28765 | 1439 | T1439C | E | 7.1e-08 | 2.7e-08 | 3.3e-08 | 4.367E-08 | 2.386E-08 | 2.700E-08 | 7.100E-08 |
| 28766 | 1439 | T1439E | E | 4.9e-08 | 1.1e-07 | 4.6e-08 | 6.833E-08 | 3.612E-08 | 4.600E-08 | 1.100E-07 |
| 28767 | 1439 | T1439Q | E | 4e-08 | 8.7e-08 | 4.6e-08 | 5.767E-08 | 2.558E-08 | 4.000E-08 | 8.700E-08 |
| 28768 | 1439 | T1439G | E | 8e-08 | 6.9e-08 | 3.1e-08 | 6.000E-08 | 2.571E-08 | 3.100E-08 | 8.000E-08 |
| 28769 | 1439 | T1439H | E | 4.4e-08 | 2.1e-08 | 6.5e-08 | 4.333E-08 | 2.201E-08 | 2.100E-08 | 6.500E-08 |
| 28770 | 1439 | T1439I | E | 7.1e-08 | 2.6e-08 | 3.7e-08 | 4.467E-08 | 2.346E-08 | 2.600E-08 | 7.100E-08 |
| 28771 | 1439 | T1439L | E | 1.1e-07 | 3.6e-08 | 2.2e-07 | 1.220E-07 | 9.259E-08 | 3.600E-08 | 2.200E-07 |
| 28772 | 1439 | T1439K | E | 2.1e-08 | 1.6e-08 | 1.2e-07 | 5.233E-08 | 5.865E-08 | 1.600E-08 | 1.200E-07 |
| 28773 | 1439 | T1439M | E | 4.6e-08 | 5.8e-08 | 3.4e-08 | 4.600E-08 | 1.200E-08 | 3.400E-08 | 5.800E-08 |
| 28774 | 1439 | T1439F | E | 2.8e-08 | 6e-08 | 3.7e-08 | 4.167E-08 | 1.650E-08 | 2.800E-08 | 6.000E-08 |
| 28775 | 1439 | T1439P | E | 3.2e-08 | 3e-08 | 4e-08 | 3.400E-08 | 5.292E-09 | 3.000E-08 | 4.000E-08 |
| 28776 | 1439 | T1439S | E | 6.3e-08 | 3.8e-08 | 6.5e-08 | 5.533E-08 | 1.504E-08 | 3.800E-08 | 6.500E-08 |
| 28777 | 1439 | T1439T | E | 3.5e-08 | 1.4e-08 | 4.1e-08 | 3.000E-08 | 1.418E-08 | 1.400E-08 | 4.100E-08 |
| 28778 | 1439 | T1439W | E | 2.1e-08 | 4.3e-08 | 8.9e-08 | 5.100E-08 | 3.470E-08 | 2.100E-08 | 8.900E-08 |
| 28779 | 1439 | T1439Y | E | 4.9e-08 | 6.4e-08 | 5.5e-08 | 5.600E-08 | 7.550E-09 | 4.900E-08 | 6.400E-08 |
| 28780 | 1439 | T1439V | E | 2.7e-08 | 1.3e-08 | 9.8e-08 | 4.600E-08 | 4.557E-08 | 1.300E-08 | 9.800E-08 |
| 28781 | 1440 | S1440A | E | 6.2e-08 | 7.7e-08 | 2.3e-08 | 5.400E-08 | 2.787E-08 | 2.300E-08 | 7.700E-08 |
| 28782 | 1440 | S1440R | E | 5e-08 | 2.6e-08 | 3.8e-08 | 3.800E-08 | 1.200E-08 | 2.600E-08 | 5.000E-08 |
| 28783 | 1440 | S1440N | E | 8.3e-08 | 2.9e-08 | 5.2e-08 | 5.467E-08 | 2.710E-08 | 2.900E-08 | 8.300E-08 |
| 28784 | 1440 | S1440D | E | 2e-08 | 2.8e-08 | 2.7e-08 | 2.500E-08 | 4.359E-09 | 2.000E-08 | 2.800E-08 |
| 28785 | 1440 | S1440C | E | 6.2e-08 | 2.1e-08 | 8.3e-08 | 5.533E-08 | 3.153E-08 | 2.100E-08 | 8.300E-08 |
| 28786 | 1440 | S1440E | E | 3.8e-08 | 1.7e-08 | 5.1e-08 | 3.533E-08 | 1.716E-08 | 1.700E-08 | 5.100E-08 |
| 28787 | 1440 | S1440Q | E | 4.1e-08 | 1.8e-08 | 2.8e-08 | 2.900E-08 | 1.153E-08 | 1.800E-08 | 4.100E-08 |
| 28788 | 1440 | S1440G | E | 3e-08 | 5.6e-08 | 6.9e-08 | 5.167E-08 | 1.986E-08 | 3.000E-08 | 6.900E-08 |
| 28789 | 1440 | S1440H | E | 2.2e-08 | 1.3e-07 | 6.4e-08 | 7.200E-08 | 5.444E-08 | 2.200E-08 | 1.300E-07 |
| 28790 | 1440 | S1440I | E | 1.8e-08 | 2.3e-08 | 4.4e-08 | 2.833E-08 | 1.380E-08 | 1.800E-08 | 4.400E-08 |
| 28791 | 1440 | S1440L | E | 4.5e-08 | 2.5e-08 | 7.7e-08 | 4.900E-08 | 2.623E-08 | 2.500E-08 | 7.700E-08 |
| 28792 | 1440 | S1440K | E | 6.1e-08 | 6.8e-08 | 6.4e-08 | 6.433E-08 | 3.512E-09 | 6.100E-08 | 6.800E-08 |
| 28793 | 1440 | S1440M | E | 2.3e-08 | 8.1e-08 | 2.8e-08 | 4.400E-08 | 3.214E-08 | 2.300E-08 | 8.100E-08 |
| 28794 | 1440 | S1440F | E | 3.7e-08 | 3.1e-08 | 2.1e-08 | 2.967E-08 | 8.083E-09 | 2.100E-08 | 3.700E-08 |
| 28795 | 1440 | S1440P | E | 6.1e-08 | 2.1e-08 | 3e-08 | 3.733E-08 | 2.098E-08 | 2.100E-08 | 6.100E-08 |
| 28796 | 1440 | S1440S | E | 2.1e-08 | 1e-07 | 9.3e-08 | 7.133E-08 | 4.373E-08 | 2.100E-08 | 1.000E-07 |
| 28797 | 1440 | S1440T | E | 1.4e-07 | 1.4e-08 | 3.5e-08 | 6.300E-08 | 6.751E-08 | 1.400E-08 | 1.400E-07 |
| 28798 | 1440 | S1440W | E | 8.9e-08 | 3.9e-08 | 1.2e-08 | 4.667E-08 | 3.907E-08 | 1.200E-08 | 8.900E-08 |
| 28799 | 1440 | S1440Y | E | 6.3e-08 | 8.5e-08 | 8.6e-08 | 7.800E-08 | 1.300E-08 | 6.300E-08 | 8.600E-08 |
| 28800 | 1440 | S1440V | E | 1.9e-08 | 1.1e-07 | 7.9e-08 | 6.933E-08 | 4.626E-08 | 1.900E-08 | 1.100E-07 |
| 28801 | 1441 | G1441A | E | 3.2e-08 | 5.9e-08 | 7e-08 | 5.367E-08 | 1.955E-08 | 3.200E-08 | 7.000E-08 |
| 28802 | 1441 | G1441R | E | 1.2e-07 | 1.1e-07 | 1.6e-07 | 1.300E-07 | 2.646E-08 | 1.100E-07 | 1.600E-07 |
| 28803 | 1441 | G1441N | E | 2.9e-08 | 3.4e-08 | 3.4e-08 | 3.233E-08 | 2.887E-09 | 2.900E-08 | 3.400E-08 |
| 28804 | 1441 | G1441D | E | 5.2e-08 | 2.9e-08 | 1.2e-07 | 6.700E-08 | 4.732E-08 | 2.900E-08 | 1.200E-07 |
| 28805 | 1441 | G1441C | E | 3.3e-08 | 6e-08 | 2.2e-08 | 3.833E-08 | 1.955E-08 | 2.200E-08 | 6.000E-08 |
| 28806 | 1441 | G1441E | E | 5.7e-08 | 3.2e-08 | 5.9e-08 | 4.933E-08 | 1.504E-08 | 3.200E-08 | 5.900E-08 |
| 28807 | 1441 | G1441Q | E | 3.3e-08 | 1.6e-07 | 3.1e-08 | 7.467E-08 | 7.391E-08 | 3.100E-08 | 1.600E-07 |
| 28808 | 1441 | G1441G | E | 2.5e-08 | 4.1e-08 | 6.1e-08 | 4.233E-08 | 1.804E-08 | 2.500E-08 | 6.100E-08 |
| 28809 | 1441 | G1441H | E | 5.8e-08 | 6.2e-08 | 8e-08 | 6.667E-08 | 1.172E-08 | 5.800E-08 | 8.000E-08 |
| 28810 | 1441 | G1441I | E | 1.3e-07 | 2e-08 | 3.2e-08 | 6.067E-08 | 6.034E-08 | 2.000E-08 | 1.300E-07 |
| 28811 | 1441 | G1441L | E | 1.3e-07 | 7e-08 | 6.2e-08 | 8.733E-08 | 3.717E-08 | 6.200E-08 | 1.300E-07 |
| 28812 | 1441 | G1441K | E | 4.6e-08 | 1.2e-07 | 4.1e-08 | 6.900E-08 | 4.424E-08 | 4.100E-08 | 1.200E-07 |
| 28813 | 1441 | G1441M | E | 5.3e-08 | 5.4e-08 | 4.8e-08 | 5.167E-08 | 3.215E-09 | 4.800E-08 | 5.400E-08 |
| 28814 | 1441 | G1441F | E | 5.7e-08 | 4.4e-08 | 1.9e-08 | 4.000E-08 | 1.931E-08 | 1.900E-08 | 5.700E-08 |

| | | | | | | | | | | |
|---|---|---|---|---|---|---|---|---|---|---|
| 28815 | 1441 | G1441P | E | 5.5e-08 | 2.9e-08 | 5e-08 | 4.467E-08 | 1.380E-08 | 2.900E-08 | 5.500E-08 |
| 28816 | 1441 | G1441S | E | 3.5e-08 | 5.4e-08 | 3.5e-08 | 4.133E-08 | 1.097E-08 | 3.500E-08 | 5.400E-08 |
| 28817 | 1441 | G1441T | E | 1.1e-08 | 4.4e-08 | 5.5e-08 | 3.667E-08 | 2.290E-08 | 1.100E-08 | 5.500E-08 |
| 28818 | 1441 | G1441W | E | 6.2e-08 | 3.1e-08 | 2.7e-08 | 4.000E-08 | 1.916E-08 | 2.700E-08 | 6.200E-08 |
| 28819 | 1441 | G1441Y | E | 6.5e-08 | 4.4e-08 | 8.2e-08 | 6.367E-08 | 1.904E-08 | 4.400E-08 | 8.200E-08 |
| 28820 | 1441 | G1441V | E | 1.1e-08 | 1.9e-08 | 7.9e-08 | 3.633E-08 | 3.717E-08 | 1.100E-08 | 7.900E-08 |
| 28821 | 1442 | V1442A | E | 1.5e-08 | 7.5e-09 | 5.4e-08 | 2.550E-08 | 2.496E-08 | 7.500E-09 | 5.400E-08 |
| 28822 | 1442 | V1442R | E | 2.3e-08 | 1.8e-08 | 5.9e-08 | 3.333E-08 | 2.237E-08 | 1.800E-08 | 5.900E-08 |
| 28823 | 1442 | V1442N | E | 5.9e-08 | 1e-07 | 8.1e-08 | 8.000E-08 | 2.052E-08 | 5.900E-08 | 1.000E-07 |
| 28824 | 1442 | V1442D | E | 5.9e-08 | 3.9e-08 | 2.9e-08 | 4.233E-08 | 1.528E-08 | 2.900E-08 | 5.900E-08 |
| 28825 | 1442 | V1442C | E | 3e-08 | 6.2e-08 | 5.6e-08 | 4.933E-08 | 1.701E-08 | 3.000E-08 | 6.200E-08 |
| 28826 | 1442 | V1442E | E | 2.6e-08 | 1.1e-08 | 2.3e-08 | 2.000E-08 | 7.937E-09 | 1.100E-08 | 2.600E-08 |
| 28827 | 1442 | V1442Q | E | 3e-08 | 4.3e-08 | 3.2e-08 | 3.500E-08 | 7.000E-09 | 3.000E-08 | 4.300E-08 |
| 28828 | 1442 | V1442G | E | 1.1e-08 | 1.5e-08 | 5e-08 | 2.533E-08 | 2.146E-08 | 1.100E-08 | 5.000E-08 |
| 28829 | 1442 | V1442H | E | 3.9e-08 | 3.9e-08 | 5.8e-08 | 4.533E-08 | 1.097E-08 | 3.900E-08 | 5.800E-08 |
| 28830 | 1442 | V1442I | E | 3.3e-08 | 5.3e-08 | 6.4e-08 | 5.000E-08 | 1.572E-08 | 3.300E-08 | 6.400E-08 |
| 28831 | 1442 | V1442L | E | 6.8e-08 | 1.9e-08 | 5e-08 | 4.567E-08 | 2.479E-08 | 1.900E-08 | 6.800E-08 |
| 28832 | 1442 | V1442K | E | 2.5e-08 | 1.1e-07 | 8.1e-09 | 4.770E-08 | 5.461E-08 | 8.100E-09 | 1.100E-07 |
| 28833 | 1442 | V1442M | E | 4.3e-08 | 2.2e-08 | 4.2e-08 | 3.567E-08 | 1.185E-08 | 2.200E-08 | 4.300E-08 |
| 28834 | 1442 | V1442F | E | 4e-08 | 2.3e-08 | 2.8e-08 | 3.033E-08 | 8.737E-09 | 2.300E-08 | 4.000E-08 |
| 28835 | 1442 | V1442P | E | 4.6e-08 | 5.7e-08 | 1.1e-08 | 3.800E-08 | 2.402E-08 | 1.100E-08 | 5.700E-08 |
| 28836 | 1442 | V1442S | E | 2.1e-08 | 2.9e-08 | 3.3e-08 | 2.767E-08 | 6.110E-09 | 2.100E-08 | 3.300E-08 |
| 28837 | 1442 | V1442T | E | 3.3e-08 | 4.5e-08 | 3.3e-08 | 3.700E-08 | 6.928E-09 | 3.300E-08 | 4.500E-08 |
| 28838 | 1442 | V1442W | E | 7.6e-09 | 3.7e-08 | 4.2e-08 | 2.887E-08 | 1.859E-08 | 7.600E-09 | 4.200E-08 |
| 28839 | 1442 | V1442Y | E | 6.4e-08 | 9.1e-08 | 4.8e-08 | 6.767E-08 | 2.173E-08 | 4.800E-08 | 9.100E-08 |
| 28840 | 1442 | V1442V | E | 4.7e-08 | 3e-08 | 3.3e-08 | 3.667E-08 | 9.074E-09 | 3.000E-08 | 4.700E-08 |
| 28841 | 1443 | H1443A | E | 2.4e-08 | 1.3e-07 | 4.2e-08 | 6.533E-08 | 5.672E-08 | 2.400E-08 | 1.300E-07 |
| 28842 | 1443 | H1443R | E | 4.4e-08 | 6.1e-08 | 1e-08 | 3.833E-08 | 2.597E-08 | 1.000E-08 | 6.100E-08 |
| 28843 | 1443 | H1443N | E | 6.4e-08 | 7.4e-08 | 5.2e-08 | 6.333E-08 | 1.102E-08 | 5.200E-08 | 7.400E-08 |
| 28844 | 1443 | H1443D | E | 3.6e-08 | 1.3e-08 | 5.4e-08 | 3.433E-08 | 2.055E-08 | 1.300E-08 | 5.400E-08 |
| 28845 | 1443 | H1443C | E | 5.9e-08 | 3.4e-08 | 9e-08 | 6.100E-08 | 2.805E-08 | 3.400E-08 | 9.000E-08 |
| 28846 | 1443 | H1443E | E | 9.3e-08 | 5.9e-08 | 6.1e-08 | 7.100E-08 | 1.908E-08 | 5.900E-08 | 9.300E-08 |
| 28847 | 1443 | H1443Q | E | 6.2e-08 | 6e-08 | 5.9e-08 | 6.033E-08 | 1.528E-09 | 5.900E-08 | 6.200E-08 |
| 28848 | 1443 | H1443G | E | 3.2e-08 | 3.8e-08 | 1.3e-07 | 6.667E-08 | 5.493E-08 | 3.200E-08 | 1.300E-07 |
| 28849 | 1443 | H1443H | E | 3.8e-08 | 3.3e-08 | 2.5e-08 | 3.200E-08 | 6.557E-09 | 2.500E-08 | 3.800E-08 |
| 28850 | 1443 | H1443I | E | 4.2e-08 | 6.6e-08 | 1.6e-08 | 4.133E-08 | 2.501E-08 | 1.600E-08 | 6.600E-08 |
| 28851 | 1443 | H1443L | E | 9.7e-08 | 5.2e-08 | 4.1e-08 | 6.333E-08 | 2.967E-08 | 4.100E-08 | 9.700E-08 |
| 28852 | 1443 | H1443K | E | 4.8e-08 | 3.8e-08 | 3.3e-08 | 3.967E-08 | 7.638E-09 | 3.300E-08 | 4.800E-08 |
| 28853 | 1443 | H1443M | E | 5.6e-08 | 2.4e-08 | 2.8e-08 | 3.600E-08 | 1.744E-08 | 2.400E-08 | 5.600E-08 |
| 28854 | 1443 | H1443F | E | 2.6e-08 | 9.8e-08 | 1.4e-07 | 8.800E-08 | 5.765E-08 | 2.600E-08 | 1.400E-07 |
| 28855 | 1443 | H1443P | E | 2.8e-08 | 3.9e-08 | 8.3e-08 | 5.000E-08 | 2.910E-08 | 2.800E-08 | 8.300E-08 |
| 28856 | 1443 | H1443S | E | 7.6e-08 | 3.9e-08 | 7.7e-08 | 6.400E-08 | 2.166E-08 | 3.900E-08 | 7.700E-08 |
| 28857 | 1443 | H1443T | E | 9.7e-08 | 2.8e-08 | 7.4e-08 | 6.633E-08 | 3.513E-08 | 2.800E-08 | 9.700E-08 |
| 28858 | 1443 | H1443W | E | 3.8e-08 | 3.5e-08 | 3.4e-08 | 3.567E-08 | 2.082E-09 | 3.400E-08 | 3.800E-08 |
| 28859 | 1443 | H1443Y | E | 2e-08 | 6.4e-08 | 6.5e-08 | 4.967E-08 | 2.570E-08 | 2.000E-08 | 6.500E-08 |
| 28860 | 1443 | H1443V | E | 6.1e-08 | 4.3e-08 | 3.8e-08 | 4.733E-08 | 1.210E-08 | 3.800E-08 | 6.100E-08 |
| 28861 | 1444 | T1444A | E | 2.8e-08 | 3.1e-08 | 8.3e-08 | 4.733E-08 | 3.092E-08 | 2.800E-08 | 8.300E-08 |
| 28862 | 1444 | T1444R | E | 4.7e-08 | 6.4e-08 | 8.1e-08 | 6.400E-08 | 1.700E-08 | 4.700E-08 | 8.100E-08 |
| 28863 | 1444 | T1444N | E | 2.5e-08 | 6.3e-08 | 1.9e-08 | 3.567E-08 | 2.386E-08 | 1.900E-08 | 6.300E-08 |
| 28864 | 1444 | T1444D | E | 2.3e-08 | 9.1e-08 | 2.9e-08 | 4.767E-08 | 3.765E-08 | 2.300E-08 | 9.100E-08 |
| 28865 | 1444 | T1444C | E | 2e-08 | 3.5e-08 | 4.7e-08 | 3.400E-08 | 1.353E-08 | 2.000E-08 | 4.700E-08 |
| 28866 | 1444 | T1444E | E | 5.9e-08 | 8.1e-08 | 2.4e-08 | 5.467E-08 | 2.875E-08 | 2.400E-08 | 8.100E-08 |
| 28867 | 1444 | T1444Q | E | 2.2e-08 | 4.3e-08 | 3.5e-08 | 3.333E-08 | 1.060E-08 | 2.200E-08 | 4.300E-08 |
| 28868 | 1444 | T1444G | E | 5.2e-08 | 3.2e-08 | 7.5e-08 | 5.300E-08 | 2.152E-08 | 3.200E-08 | 7.500E-08 |
| 28869 | 1444 | T1444H | E | 5.8e-08 | 3.4e-08 | 3e-08 | 4.067E-08 | 1.514E-08 | 3.000E-08 | 5.800E-08 |
| 28870 | 1444 | T1444I | E | 4.4e-08 | 1.6e-07 | 5.5e-08 | 8.633E-08 | 6.403E-08 | 4.400E-08 | 1.600E-07 |
| 28871 | 1444 | T1444L | E | 6.7e-08 | 1.7e-07 | 5.4e-08 | 9.700E-08 | 6.355E-08 | 5.400E-08 | 1.700E-07 |
| 28872 | 1444 | T1444K | E | 3.7e-08 | 8.7e-08 | 3.9e-08 | 5.433E-08 | 2.831E-08 | 3.700E-08 | 8.700E-08 |
| 28873 | 1444 | T1444M | E | 3.6e-08 | 2.7e-08 | 4.2e-08 | 3.500E-08 | 7.550E-09 | 2.700E-08 | 4.200E-08 |
| 28874 | 1444 | T1444F | E | 7.7e-08 | 3.7e-08 | 4.4e-08 | 5.267E-08 | 2.136E-08 | 3.700E-08 | 7.700E-08 |
| 28875 | 1444 | T1444P | E | 4.9e-08 | 3.3e-08 | 4.2e-08 | 4.133E-08 | 8.021E-09 | 3.300E-08 | 4.900E-08 |
| 28876 | 1444 | T1444S | E | 4.6e-08 | 8.5e-08 | 6.4e-08 | 6.500E-08 | 1.952E-08 | 4.600E-08 | 8.500E-08 |
| 28877 | 1444 | T1444T | E | 6.1e-08 | 4.3e-08 | 2.1e-08 | 4.167E-08 | 2.003E-08 | 2.100E-08 | 6.100E-08 |
| 28878 | 1444 | T1444W | E | 7e-08 | 4.8e-08 | 4.7e-08 | 5.500E-08 | 1.300E-08 | 4.700E-08 | 7.000E-08 |
| 28879 | 1444 | T1444Y | E | 4.6e-08 | 4.8e-08 | 1.6e-07 | 8.467E-08 | 6.525E-08 | 4.600E-08 | 1.600E-07 |
| 28880 | 1444 | T1444V | E | 7.3e-09 | 3.5e-08 | 4.7e-08 | 2.977E-08 | 2.036E-08 | 7.300E-09 | 4.700E-08 |
| 28881 | 1445 | F1445A | E | 3.6e-08 | 6.5e-08 | 7.2e-08 | 5.767E-08 | 1.909E-08 | 3.600E-08 | 7.200E-08 |
| 28882 | 1445 | F1445R | E | 1.3e-08 | 2.3e-08 | 2.6e-08 | 2.067E-08 | 6.807E-09 | 1.300E-08 | 2.600E-08 |
| 28883 | 1445 | F1445N | E | 3.4e-08 | 2.5e-08 | 3.5e-08 | 3.133E-08 | 5.508E-09 | 2.500E-08 | 3.500E-08 |
| 28884 | 1445 | F1445D | E | 1.4e-07 | 2.4e-08 | 5.5e-08 | 7.300E-08 | 6.006E-08 | 2.400E-08 | 1.400E-07 |
| 28885 | 1445 | F1445C | E | 2.4e-08 | 3.3e-08 | 6e-08 | 3.900E-08 | 1.873E-08 | 2.400E-08 | 6.000E-08 |
| 28886 | 1445 | F1445E | E | 3.5e-08 | 7.2e-08 | 8.1e-08 | 6.267E-08 | 2.438E-08 | 3.500E-08 | 8.100E-08 |
| 28887 | 1445 | F1445Q | E | 1.4e-07 | 2e-08 | 2.5e-08 | 6.167E-08 | 6.788E-08 | 2.000E-08 | 1.400E-07 |
| 28888 | 1445 | F1445G | E | 1.9e-07 | 3.2e-08 | 3e-08 | 8.400E-08 | 9.180E-08 | 3.000E-08 | 1.900E-07 |
| 28889 | 1445 | F1445H | E | 3e-08 | 2.1e-08 | 3.2e-08 | 2.767E-08 | 5.859E-09 | 2.100E-08 | 3.200E-08 |
| 28890 | 1445 | F1445I | E | 3.8e-08 | 1.7e-08 | 8.4e-08 | 4.633E-08 | 3.427E-08 | 1.700E-08 | 8.400E-08 |
| 28891 | 1445 | F1445L | E | 3.4e-08 | 1.6e-07 | 3.5e-08 | 7.633E-08 | 7.246E-08 | 3.400E-08 | 1.600E-07 |
| 28892 | 1445 | F1445K | E | 1.4e-08 | 3.9e-08 | 4e-08 | 3.100E-08 | 1.473E-08 | 1.400E-08 | 4.000E-08 |
| 28893 | 1445 | F1445M | E | 2.5e-08 | 6.8e-08 | 5e-08 | 4.767E-08 | 2.159E-08 | 2.500E-08 | 6.800E-08 |
| 28894 | 1445 | F1445F | E | 2.1e-08 | 1.4e-08 | 1.5e-08 | 1.667E-08 | 3.786E-09 | 1.400E-08 | 2.100E-08 |
| 28895 | 1445 | F1445P | E | 4.3e-08 | 7e-08 | 1.3e-08 | 4.200E-08 | 2.851E-08 | 1.300E-08 | 7.000E-08 |
| 28896 | 1445 | F1445S | E | 4.1e-08 | 3.8e-08 | 3.5e-08 | 3.800E-08 | 3.000E-09 | 3.500E-08 | 4.100E-08 |
| 28897 | 1445 | F1445T | E | 3.1e-08 | 7.8e-08 | 3.1e-08 | 4.667E-08 | 2.714E-08 | 3.100E-08 | 7.800E-08 |
| 28898 | 1445 | F1445W | E | 6e-08 | 3.5e-08 | 3.7e-08 | 4.400E-08 | 1.389E-08 | 3.500E-08 | 6.000E-08 |
| 28899 | 1445 | F1445Y | E | 2.1e-08 | 5.9e-08 | 3e-08 | 3.667E-08 | 1.986E-08 | 2.100E-08 | 5.900E-08 |

| | | | | | | | | | | |
|---|---|---|---|---|---|---|---|---|---|---|
| 28900 | 1445 | F1445V | E | 3.6e-08 | 2.8e-08 | 4.8e-08 | 3.733E-08 | 1.007E-08 | 2.800E-08 | 4.800E-08 |
| 28901 | 1446 | P1446A | E | 9.2e-08 | 3.6e-08 | 1.1e-08 | 4.633E-08 | 4.148E-08 | 1.100E-08 | 9.200E-08 |
| 28902 | 1446 | P1446R | E | 9e-08 | 2.1e-08 | 4.9e-08 | 5.333E-08 | 3.470E-08 | 2.100E-08 | 9.000E-08 |
| 28903 | 1446 | P1446N | E | 4.8e-08 | 4.5e-08 | 4.9e-08 | 4.733E-08 | 2.082E-09 | 4.500E-08 | 4.900E-08 |
| 28904 | 1446 | P1446D | E | 3.1e-08 | 2.5e-08 | 2.4e-07 | 9.867E-08 | 1.224E-07 | 2.500E-08 | 2.400E-07 |
| 28905 | 1446 | P1446C | E | 2.2e-08 | 6.9e-08 | 4.1e-08 | 4.400E-08 | 2.364E-08 | 2.200E-08 | 6.900E-08 |
| 28906 | 1446 | P1446E | E | 3.3e-08 | 5.5e-08 | 3.4e-08 | 4.067E-08 | 1.242E-08 | 3.300E-08 | 5.500E-08 |
| 28907 | 1446 | P1446Q | E | 6.8e-08 | 7.9e-08 | 2.9e-08 | 5.867E-08 | 2.627E-08 | 2.900E-08 | 7.900E-08 |
| 28908 | 1446 | P1446G | E | 5.5e-08 | 7e-08 | 2e-08 | 4.833E-08 | 2.566E-08 | 2.000E-08 | 7.000E-08 |
| 28909 | 1446 | P1446H | E | 3.9e-08 | 2.8e-08 | 1.9e-08 | 2.867E-08 | 1.002E-08 | 1.900E-08 | 3.900E-08 |
| 28910 | 1446 | P1446I | E | 1.4e-08 | 1.8e-07 | 4.9e-08 | 8.100E-08 | 8.750E-08 | 1.400E-08 | 1.800E-07 |
| 28911 | 1446 | P1446L | E | 5.7e-08 | 6.9e-08 | 7.6e-08 | 6.733E-08 | 9.609E-09 | 5.700E-08 | 7.600E-08 |
| 28912 | 1446 | P1446K | E | 6.3e-08 | 2.8e-08 | 3.3e-08 | 4.133E-08 | 1.893E-08 | 2.800E-08 | 6.300E-08 |
| 28913 | 1446 | P1446M | E | 5e-08 | 1.2e-08 | 7.7e-09 | 2.323E-08 | 2.328E-08 | 7.700E-09 | 5.000E-08 |
| 28914 | 1446 | P1446F | E | 7.7e-08 | 4e-08 | 2.4e-08 | 4.700E-08 | 2.718E-08 | 2.400E-08 | 7.700E-08 |
| 28915 | 1446 | P1446P | E | 3.5e-08 | 1.8e-08 | 6e-08 | 3.767E-08 | 2.113E-08 | 1.800E-08 | 6.000E-08 |
| 28916 | 1446 | P1446S | E | 5e-08 | 3.9e-08 | 5.9e-08 | 4.933E-08 | 1.002E-08 | 3.900E-08 | 5.900E-08 |
| 28917 | 1446 | P1446T | E | 7.6e-08 | 3.5e-08 | 1.5e-08 | 4.200E-08 | 3.110E-08 | 1.500E-08 | 7.600E-08 |
| 28918 | 1446 | P1446W | E | 5.7e-08 | 9e-08 | 6e-08 | 6.900E-08 | 1.825E-08 | 5.700E-08 | 9.000E-08 |
| 28919 | 1446 | P1446Y | E | 3.2e-08 | 1.2e-07 | 2.5e-08 | 5.900E-08 | 5.294E-08 | 2.500E-08 | 1.200E-07 |
| 28920 | 1446 | P1446V | E | 7.2e-08 | 2.7e-08 | 1.5e-08 | 3.800E-08 | 3.005E-08 | 1.500E-08 | 7.200E-08 |
| 28921 | 1447 | A1447A | E | 6e-08 | 3.9e-08 | 4.1e-08 | 4.667E-08 | 1.159E-08 | 3.900E-08 | 6.000E-08 |
| 28922 | 1447 | A1447R | E | 2.1e-08 | 3e-08 | 3.5e-08 | 2.867E-08 | 7.095E-09 | 2.100E-08 | 3.500E-08 |
| 28923 | 1447 | A1447N | E | 4.4e-08 | 6.9e-08 | 5.6e-08 | 5.633E-08 | 1.250E-08 | 4.400E-08 | 6.900E-08 |
| 28924 | 1447 | A1447D | E | 4.6e-08 | 7.3e-08 | 3.8e-08 | 5.233E-08 | 1.834E-08 | 3.800E-08 | 7.300E-08 |
| 28925 | 1447 | A1447C | E | 5.3e-08 | 6.8e-09 | 1.1e-07 | 5.660E-08 | 5.169E-08 | 6.800E-09 | 1.100E-07 |
| 28926 | 1447 | A1447E | E | 1.7e-08 | 1.1e-07 | 5.6e-08 | 6.100E-08 | 4.670E-08 | 1.700E-08 | 1.100E-07 |
| 28927 | 1447 | A1447Q | E | 3.6e-08 | 9.2e-09 | 5.2e-08 | 3.240E-08 | 2.163E-08 | 9.200E-09 | 5.200E-08 |
| 28928 | 1447 | A1447G | E | 2.3e-08 | 2.3e-08 | 4.7e-08 | 3.100E-08 | 1.386E-08 | 2.300E-08 | 4.700E-08 |
| 28929 | 1447 | A1447H | E | 5.1e-08 | 2.9e-08 | 3.1e-08 | 3.700E-08 | 1.217E-08 | 2.900E-08 | 5.100E-08 |
| 28930 | 1447 | A1447I | E | 6.1e-08 | 3.2e-08 | 3.7e-08 | 4.333E-08 | 1.550E-08 | 3.200E-08 | 6.100E-08 |
| 28931 | 1447 | A1447L | E | 4.7e-08 | 3.4e-08 | 1.6e-08 | 3.233E-08 | 1.557E-08 | 1.600E-08 | 4.700E-08 |
| 28932 | 1447 | A1447K | E | 7.3e-08 | 1.3e-07 | 5.5e-08 | 8.600E-08 | 3.915E-08 | 5.500E-08 | 1.300E-07 |
| 28933 | 1447 | A1447M | E | 4.3e-08 | 4.9e-08 | 3.5e-08 | 4.233E-08 | 7.024E-09 | 3.500E-08 | 4.900E-08 |
| 28934 | 1447 | A1447F | E | 2.7e-08 | 6.5e-08 | 3.7e-08 | 4.300E-08 | 1.970E-08 | 2.700E-08 | 6.500E-08 |
| 28935 | 1447 | A1447P | E | 7.5e-08 | 3.6e-08 | 2.4e-08 | 4.500E-08 | 2.666E-08 | 2.400E-08 | 7.500E-08 |
| 28936 | 1447 | A1447S | E | 3.8e-08 | 7.3e-08 | 8.8e-08 | 6.633E-08 | 2.566E-08 | 3.800E-08 | 8.800E-08 |
| 28937 | 1447 | A1447T | E | 1.4e-08 | 3.7e-08 | 8.2e-08 | 4.433E-08 | 3.459E-08 | 1.400E-08 | 8.200E-08 |
| 28938 | 1447 | A1447W | E | 3.5e-08 | 9.4e-08 | 1.9e-08 | 4.933E-08 | 3.950E-08 | 1.900E-08 | 9.400E-08 |
| 28939 | 1447 | A1447Y | E | 4.7e-08 | 3.5e-08 | 3.2e-08 | 3.800E-08 | 7.937E-09 | 3.200E-08 | 4.700E-08 |
| 28940 | 1447 | A1447V | E | 1.5e-08 | 6.4e-08 | 3.9e-08 | 3.933E-08 | 2.450E-08 | 1.500E-08 | 6.400E-08 |
| 28941 | 1448 | V1448A | E | 7.7e-08 | 3.6e-08 | 3.3e-08 | 4.867E-08 | 2.458E-08 | 3.300E-08 | 7.700E-08 |
| 28942 | 1448 | V1448R | E | 2e-08 | 4.1e-08 | 3.8e-08 | 3.300E-08 | 1.136E-08 | 2.000E-08 | 4.100E-08 |
| 28943 | 1448 | V1448N | E | 2.3e-08 | 3.9e-08 | 1e-07 | 5.400E-08 | 4.063E-08 | 2.300E-08 | 1.000E-07 |
| 28944 | 1448 | V1448D | E | 3e-08 | 6.6e-08 | 7.3e-08 | 5.633E-08 | 2.307E-08 | 3.000E-08 | 7.300E-08 |
| 28945 | 1448 | V1448C | E | 2.9e-08 | 1.9e-08 | 3.8e-08 | 2.867E-08 | 9.504E-09 | 1.900E-08 | 3.800E-08 |
| 28946 | 1448 | V1448E | E | 1.7e-08 | 4.5e-08 | 5.2e-08 | 3.800E-08 | 1.852E-08 | 1.700E-08 | 5.200E-08 |
| 28947 | 1448 | V1448Q | E | 2.6e-08 | 1.1e-08 | 8.6e-08 | 4.100E-08 | 3.969E-08 | 1.100E-08 | 8.600E-08 |
| 28948 | 1448 | V1448G | E | 7.8e-08 | 4.6e-08 | 2.2e-08 | 4.867E-08 | 2.810E-08 | 2.200E-08 | 7.800E-08 |
| 28949 | 1448 | V1448H | E | 4.2e-08 | 1.3e-07 | 4.5e-08 | 7.233E-08 | 4.996E-08 | 4.200E-08 | 1.300E-07 |
| 28950 | 1448 | V1448I | E | 4.2e-08 | 5.4e-08 | 2.4e-08 | 4.000E-08 | 1.510E-08 | 2.400E-08 | 5.400E-08 |
| 28951 | 1448 | V1448L | E | 3.2e-08 | 6.4e-08 | 1.3e-07 | 7.533E-08 | 4.997E-08 | 3.200E-08 | 1.300E-07 |
| 28952 | 1448 | V1448K | E | 2.9e-08 | 1.1e-08 | 5.3e-08 | 3.100E-08 | 2.107E-08 | 1.100E-08 | 5.300E-08 |
| 28953 | 1448 | V1448M | E | 1.6e-07 | 4.7e-08 | 5.3e-08 | 8.667E-08 | 6.358E-08 | 4.700E-08 | 1.600E-07 |
| 28954 | 1448 | V1448F | E | 5e-08 | 3.5e-08 | 6e-08 | 4.833E-08 | 1.258E-08 | 3.500E-08 | 6.000E-08 |
| 28955 | 1448 | V1448P | E | 1e-08 | 2.8e-08 | 3.8e-08 | 2.533E-08 | 1.419E-08 | 1.000E-08 | 3.800E-08 |
| 28956 | 1448 | V1448S | E | 1.6e-08 | 2.4e-08 | 2.4e-08 | 2.133E-08 | 4.619E-09 | 1.600E-08 | 2.400E-08 |
| 28957 | 1448 | V1448T | E | 2.4e-08 | 3.2e-08 | 1.7e-08 | 2.433E-08 | 7.506E-09 | 1.700E-08 | 3.200E-08 |
| 28958 | 1448 | V1448W | E | 2.4e-08 | 3.1e-08 | 1.7e-08 | 2.400E-08 | 7.000E-09 | 1.700E-08 | 3.100E-08 |
| 28959 | 1448 | V1448Y | E | 3.4e-08 | 2.7e-08 | 5.6e-08 | 3.900E-08 | 1.513E-08 | 2.700E-08 | 5.600E-08 |
| 28960 | 1448 | V1448V | E | 5.9e-08 | 4.6e-08 | 6.5e-08 | 5.667E-08 | 9.713E-09 | 4.600E-08 | 6.500E-08 |
| 28961 | 1449 | L1449A | E | 4e-08 | 2.8e-08 | 5.8e-08 | 4.200E-08 | 1.510E-08 | 2.800E-08 | 5.800E-08 |
| 28962 | 1449 | L1449R | E | 4.7e-08 | 1.8e-08 | 3.7e-08 | 3.400E-08 | 1.473E-08 | 1.800E-08 | 4.700E-08 |
| 28963 | 1449 | L1449N | E | 1.2e-07 | 1.5e-07 | 1.1e-07 | 1.267E-07 | 2.082E-08 | 1.100E-07 | 1.500E-07 |
| 28964 | 1449 | L1449D | E | 9.7e-08 | 2.5e-08 | 2.2e-08 | 4.800E-08 | 4.246E-08 | 2.200E-08 | 9.700E-08 |
| 28965 | 1449 | L1449C | E | 5e-08 | 3.2e-08 | 4.3e-08 | 4.167E-08 | 9.074E-09 | 3.200E-08 | 5.000E-08 |
| 28966 | 1449 | L1449E | E | 2.7e-08 | 2.9e-08 | 4e-08 | 3.200E-08 | 7.000E-09 | 2.700E-08 | 4.000E-08 |
| 28967 | 1449 | L1449Q | E | 4.1e-08 | 6.7e-08 | 7.6e-08 | 6.133E-08 | 1.818E-08 | 4.100E-08 | 7.600E-08 |
| 28968 | 1449 | L1449G | E | 4.6e-08 | 2.9e-08 | 2.5e-08 | 3.333E-08 | 1.115E-08 | 2.500E-08 | 4.600E-08 |
| 28969 | 1449 | L1449H | E | 1.3e-07 | 1.4e-08 | 8.3e-08 | 7.567E-08 | 5.835E-08 | 1.400E-08 | 1.300E-07 |
| 28970 | 1449 | L1449I | E | 2.8e-08 | 8e-08 | 5e-08 | 5.267E-08 | 2.610E-08 | 2.800E-08 | 8.000E-08 |
| 28971 | 1449 | L1449L | E | 2.7e-08 | 1.2e-07 | 6.2e-08 | 6.967E-08 | 4.697E-08 | 2.700E-08 | 1.200E-07 |
| 28972 | 1449 | L1449K | E | 4.9e-08 | 3.4e-08 | 5.5e-08 | 4.600E-08 | 1.082E-08 | 3.400E-08 | 5.500E-08 |
| 28973 | 1449 | L1449M | E | 1.8e-08 | 5e-08 | 6.3e-08 | 4.367E-08 | 2.316E-08 | 1.800E-08 | 6.300E-08 |
| 28974 | 1449 | L1449F | E | 1.6e-08 | 6.8e-08 | 5.9e-08 | 4.767E-08 | 2.779E-08 | 1.600E-08 | 6.800E-08 |
| 28975 | 1449 | L1449P | E | 3e-08 | 5.1e-08 | 4.9e-08 | 4.333E-08 | 1.159E-08 | 3.000E-08 | 5.100E-08 |
| 28976 | 1449 | L1449S | E | 8.3e-08 | 4.3e-08 | 4.9e-08 | 5.833E-08 | 2.157E-08 | 4.300E-08 | 8.300E-08 |
| 28977 | 1449 | L1449T | E | 6.5e-08 | 4.7e-08 | 1.1e-07 | 7.400E-08 | 3.245E-08 | 4.700E-08 | 1.100E-07 |
| 28978 | 1449 | L1449W | E | 2.4e-08 | 4.4e-08 | 4.3e-08 | 3.700E-08 | 1.127E-08 | 2.400E-08 | 4.400E-08 |
| 28979 | 1449 | L1449Y | E | 1.1e-08 | 7.5e-08 | 2.9e-08 | 3.833E-08 | 3.301E-08 | 1.100E-08 | 7.500E-08 |
| 28980 | 1449 | L1449V | E | 3.5e-08 | 1.2e-07 | 7.9e-08 | 7.800E-08 | 4.251E-08 | 3.500E-08 | 1.200E-07 |
| 28981 | 1450 | Q1450A | E | 4.8e-08 | 7.1e-08 | 3.3e-08 | 5.067E-08 | 1.914E-08 | 3.300E-08 | 7.100E-08 |
| 28982 | 1450 | Q1450R | E | 1.9e-08 | 6.8e-08 | 5.8e-08 | 4.833E-08 | 2.589E-08 | 1.900E-08 | 6.800E-08 |
| 28983 | 1450 | Q1450N | E | 6e-08 | 3.3e-08 | 5.3e-08 | 4.867E-08 | 1.401E-08 | 3.300E-08 | 6.000E-08 |
| 28984 | 1450 | Q1450D | E | 6.6e-08 | 1.1e-07 | 2e-08 | 6.533E-08 | 4.500E-08 | 2.000E-08 | 1.100E-07 |

| | | | | | | | | | | |
|---|---|---|---|---|---|---|---|---|---|---|
| 28985 | 1450 | Q1450C | E | 6.8e-08 | 2.1e-08 | 2.5e-08 | 3.800E-08 | 2.606E-08 | 2.100E-08 | 6.800E-08 |
| 28986 | 1450 | Q1450E | E | 7.5e-09 | 5.1e-08 | 7.8e-08 | 4.550E-08 | 3.557E-08 | 7.500E-09 | 7.800E-08 |
| 28987 | 1450 | Q1450Q | E | 2.9e-08 | 5.4e-08 | 1.4e-08 | 3.233E-08 | 2.021E-08 | 1.400E-08 | 5.400E-08 |
| 28988 | 1450 | Q1450G | E | 7.8e-08 | 3.6e-08 | 4.5e-08 | 5.300E-08 | 2.211E-08 | 3.600E-08 | 7.800E-08 |
| 28989 | 1450 | Q1450H | E | 2e-08 | 2.9e-08 | 2.1e-08 | 2.333E-08 | 4.933E-09 | 2.000E-08 | 2.900E-08 |
| 28990 | 1450 | Q1450I | E | 5.3e-08 | 6.9e-08 | 6e-08 | 6.067E-08 | 8.021E-09 | 5.300E-08 | 6.900E-08 |
| 28991 | 1450 | Q1450L | E | 2.1e-08 | 8.5e-08 | 2.5e-08 | 4.367E-08 | 3.585E-08 | 2.100E-08 | 8.500E-08 |
| 28992 | 1450 | Q1450K | E | 4.9e-08 | 2.6e-08 | 3.5e-08 | 3.667E-08 | 1.159E-08 | 2.600E-08 | 4.900E-08 |
| 28993 | 1450 | Q1450M | E | 4.5e-08 | 4.7e-08 | 3.5e-08 | 4.233E-08 | 6.429E-09 | 3.500E-08 | 4.700E-08 |
| 28994 | 1450 | Q1450F | E | 3.9e-08 | 6.2e-08 | 4.2e-08 | 4.767E-08 | 1.250E-08 | 3.900E-08 | 6.200E-08 |
| 28995 | 1450 | Q1450P | E | 5e-08 | 2.7e-08 | 1.5e-08 | 3.067E-08 | 1.779E-08 | 1.500E-08 | 5.000E-08 |
| 28996 | 1450 | Q1450S | E | 4.9e-08 | 2e-08 | 3.2e-08 | 3.367E-08 | 1.457E-08 | 2.000E-08 | 4.900E-08 |
| 28997 | 1450 | Q1450T | E | 1.1e-07 | 3.1e-08 | 4.5e-08 | 6.200E-08 | 4.215E-08 | 3.100E-08 | 1.100E-07 |
| 28998 | 1450 | Q1450W | E | 2.3e-08 | 4.2e-08 | 2.7e-08 | 3.067E-08 | 1.002E-08 | 2.300E-08 | 4.200E-08 |
| 28999 | 1450 | Q1450Y | E | 2.9e-08 | 4.4e-08 | 7.6e-08 | 4.967E-08 | 2.401E-08 | 2.900E-08 | 7.600E-08 |
| 29000 | 1450 | Q1450V | E | 1.1e-07 | 2.5e-08 | 2.2e-08 | 5.233E-08 | 4.996E-08 | 2.200E-08 | 1.100E-07 |
| 29001 | 1451 | S1451A | E | 2.5e-08 | 3.1e-08 | 6.8e-08 | 4.133E-08 | 2.329E-08 | 2.500E-08 | 6.800E-08 |
| 29002 | 1451 | S1451R | E | 2.5e-08 | 1.6e-08 | 4.4e-08 | 2.833E-08 | 1.429E-08 | 1.600E-08 | 4.400E-08 |
| 29003 | 1451 | S1451N | E | 6.8e-08 | 2e-08 | 2.1e-08 | 3.633E-08 | 2.743E-08 | 2.000E-08 | 6.800E-08 |
| 29004 | 1451 | S1451D | E | 1.5e-07 | 3e-08 | 5.9e-08 | 7.967E-08 | 6.261E-08 | 3.000E-08 | 1.500E-07 |
| 29005 | 1451 | S1451C | E | 7.9e-08 | 3.5e-08 | 1e-07 | 7.133E-08 | 3.317E-08 | 3.500E-08 | 1.000E-07 |
| 29006 | 1451 | S1451E | E | 6.6e-08 | 4.1e-08 | 5.1e-08 | 5.267E-08 | 1.258E-08 | 4.100E-08 | 6.600E-08 |
| 29007 | 1451 | S1451Q | E | 2.6e-08 | 7.9e-08 | 4.5e-08 | 5.000E-08 | 2.685E-08 | 2.600E-08 | 7.900E-08 |
| 29008 | 1451 | S1451G | E | 3.1e-08 | 7.1e-08 | 7e-08 | 5.733E-08 | 2.281E-08 | 3.100E-08 | 7.100E-08 |
| 29009 | 1451 | S1451H | E | 3.8e-08 | 2.9e-08 | 2.5e-08 | 3.067E-08 | 6.658E-09 | 2.500E-08 | 3.800E-08 |
| 29010 | 1451 | S1451I | E | 1e-07 | 6.2e-08 | 4.2e-08 | 6.800E-08 | 2.946E-08 | 4.200E-08 | 1.000E-07 |
| 29011 | 1451 | S1451L | E | 5.5e-08 | 2.8e-08 | 1.4e-07 | 7.433E-08 | 5.845E-08 | 2.800E-08 | 1.400E-07 |
| 29012 | 1451 | S1451K | E | 4.3e-08 | 2.9e-08 | 4e-08 | 3.733E-08 | 7.371E-09 | 2.900E-08 | 4.300E-08 |
| 29013 | 1451 | S1451M | E | 2.9e-08 | 5.8e-08 | 5.2e-08 | 4.633E-08 | 1.531E-08 | 2.900E-08 | 5.800E-08 |
| 29014 | 1451 | S1451F | E | 4.3e-07 | 3.2e-08 | 4.6e-08 | 1.693E-07 | 2.259E-07 | 3.200E-08 | 4.300E-07 |
| 29015 | 1451 | S1451P | E | 3.4e-08 | 2.3e-08 | 4.3e-08 | 3.333E-08 | 1.002E-08 | 2.300E-08 | 4.300E-08 |
| 29016 | 1451 | S1451S | E | 5.5e-08 | 1.9e-08 | 4.3e-08 | 3.900E-08 | 1.833E-08 | 1.900E-08 | 5.500E-08 |
| 29017 | 1451 | S1451T | E | 6.1e-08 | 2.8e-08 | 3.9e-08 | 4.267E-08 | 1.680E-08 | 2.800E-08 | 6.100E-08 |
| 29018 | 1451 | S1451W | E | 5.6e-08 | 4.6e-08 | 4.6e-08 | 4.933E-08 | 5.774E-09 | 4.600E-08 | 5.600E-08 |
| 29019 | 1451 | S1451Y | E | 3.4e-08 | 4.9e-08 | 5.4e-08 | 4.567E-08 | 1.041E-08 | 3.400E-08 | 5.400E-08 |
| 29020 | 1451 | S1451V | E | 4.1e-08 | 1e-07 | 1.9e-08 | 5.333E-08 | 4.188E-08 | 1.900E-08 | 1.000E-07 |
| 29021 | 1452 | S1452A | E | 2.9e-08 | 2.4e-08 | 2.9e-08 | 2.733E-08 | 2.887E-09 | 2.400E-08 | 2.900E-08 |
| 29022 | 1452 | S1452R | E | 4.4e-08 | 4.8e-08 | 6.6e-08 | 5.267E-08 | 1.172E-08 | 4.400E-08 | 6.600E-08 |
| 29023 | 1452 | S1452N | E | 1.1e-08 | 2.5e-08 | 3.2e-08 | 2.267E-08 | 1.069E-08 | 1.100E-08 | 3.200E-08 |
| 29024 | 1452 | S1452D | E | 3.2e-08 | 4.7e-08 | 1.7e-08 | 3.200E-08 | 1.500E-08 | 1.700E-08 | 4.700E-08 |
| 29025 | 1452 | S1452C | E | 6.6e-08 | 6.3e-08 | 3.7e-08 | 5.533E-08 | 1.595E-08 | 3.700E-08 | 6.600E-08 |
| 29026 | 1452 | S1452E | E | 2.2e-08 | 2.7e-08 | 2.7e-08 | 2.533E-08 | 2.887E-09 | 2.200E-08 | 2.700E-08 |
| 29027 | 1452 | S1452Q | E | 2.1e-08 | 3.2e-08 | 2.9e-08 | 2.733E-08 | 5.686E-09 | 2.100E-08 | 3.200E-08 |
| 29028 | 1452 | S1452G | E | 5.9e-08 | 2.9e-08 | 3.7e-08 | 4.167E-08 | 1.553E-08 | 2.900E-08 | 5.900E-08 |
| 29029 | 1452 | S1452H | E | 3.9e-08 | 5.6e-08 | 4.4e-08 | 4.633E-08 | 8.737E-09 | 3.900E-08 | 5.600E-08 |
| 29030 | 1452 | S1452I | E | 2.1e-07 | 4.3e-08 | 6.5e-08 | 1.060E-07 | 9.074E-08 | 4.300E-08 | 2.100E-07 |
| 29031 | 1452 | S1452L | E | 1.1e-08 | 1.4e-08 | 6.4e-08 | 2.967E-08 | 2.977E-08 | 1.100E-08 | 6.400E-08 |
| 29032 | 1452 | S1452K | E | 4.5e-08 | 5.4e-08 | 8e-08 | 5.967E-08 | 1.818E-08 | 4.500E-08 | 8.000E-08 |
| 29033 | 1452 | S1452M | E | 8.5e-08 | 1.2e-08 | 4.2e-08 | 4.633E-08 | 3.669E-08 | 1.200E-08 | 8.500E-08 |
| 29034 | 1452 | S1452F | E | 1.9e-08 | 2e-08 | 2.8e-08 | 2.233E-08 | 4.933E-09 | 1.900E-08 | 2.800E-08 |
| 29035 | 1452 | S1452P | E | 3.1e-08 | 9.5e-08 | 4.4e-08 | 5.667E-08 | 3.383E-08 | 3.100E-08 | 9.500E-08 |
| 29036 | 1452 | S1452S | E | 2.5e-08 | 1.2e-08 | 1.2e-07 | 5.233E-08 | 5.896E-08 | 1.200E-08 | 1.200E-07 |
| 29037 | 1452 | S1452T | E | 2.7e-08 | 4e-08 | 6.2e-08 | 4.300E-08 | 1.769E-08 | 2.700E-08 | 6.200E-08 |
| 29038 | 1452 | S1452W | E | 1.5e-08 | 5.4e-08 | 8.7e-08 | 5.200E-08 | 3.604E-08 | 1.500E-08 | 8.700E-08 |
| 29039 | 1452 | S1452Y | E | 4.2e-08 | 7.2e-08 | 6.1e-08 | 5.833E-08 | 1.518E-08 | 4.200E-08 | 7.200E-08 |
| 29040 | 1452 | S1452V | E | 5.8e-08 | 2.1e-08 | 2.6e-08 | 3.500E-08 | 2.007E-08 | 2.100E-08 | 5.800E-08 |
| 29041 | 1453 | G1453A | E | 1.1e-07 | 5e-08 | 2.9e-08 | 6.300E-08 | 4.204E-08 | 2.900E-08 | 1.100E-07 |
| 29042 | 1453 | G1453R | E | 5.2e-08 | 3.5e-08 | 3.6e-08 | 4.100E-08 | 9.539E-09 | 3.500E-08 | 5.200E-08 |
| 29043 | 1453 | G1453N | E | 6.7e-08 | 4.1e-08 | 4.5e-08 | 5.100E-08 | 1.400E-08 | 4.100E-08 | 6.700E-08 |
| 29044 | 1453 | G1453D | E | 3.3e-08 | 7.3e-08 | 5.8e-08 | 5.467E-08 | 2.021E-08 | 3.300E-08 | 7.300E-08 |
| 29045 | 1453 | G1453C | E | 6.5e-08 | 5.1e-08 | 4.2e-08 | 5.267E-08 | 1.159E-08 | 4.200E-08 | 6.500E-08 |
| 29046 | 1453 | G1453E | E | 2.4e-08 | 5.9e-08 | 2.9e-08 | 3.733E-08 | 1.893E-08 | 2.400E-08 | 5.900E-08 |
| 29047 | 1453 | G1453Q | E | 3e-08 | 4e-08 | 5.6e-08 | 4.200E-08 | 1.311E-08 | 3.000E-08 | 5.600E-08 |
| 29048 | 1453 | G1453G | E | 3.9e-08 | 5e-08 | 8.4e-08 | 5.767E-08 | 2.346E-08 | 3.900E-08 | 8.400E-08 |
| 29049 | 1453 | G1453H | E | 5.9e-08 | 1e-08 | 2.7e-08 | 3.200E-08 | 2.488E-08 | 1.000E-08 | 5.900E-08 |
| 29050 | 1453 | G1453I | E | 4.4e-08 | 2.2e-08 | 3.4e-08 | 3.333E-08 | 1.102E-08 | 2.200E-08 | 4.400E-08 |
| 29051 | 1453 | G1453L | E | 7.4e-08 | 4e-08 | 3.3e-08 | 4.900E-08 | 2.193E-08 | 3.300E-08 | 7.400E-08 |
| 29052 | 1453 | G1453K | E | 2.9e-08 | 5e-08 | 7.4e-08 | 5.100E-08 | 2.252E-08 | 2.900E-08 | 7.400E-08 |
| 29053 | 1453 | G1453M | E | 1.5e-08 | 1.3e-07 | 2e-08 | 5.500E-08 | 6.500E-08 | 1.500E-08 | 1.300E-07 |
| 29054 | 1453 | G1453F | E | 7.1e-08 | 4e-08 | 3.4e-08 | 4.833E-08 | 1.986E-08 | 3.400E-08 | 7.100E-08 |
| 29055 | 1453 | G1453P | E | 4e-08 | 5.5e-08 | 1.1e-07 | 6.833E-08 | 3.686E-08 | 4.000E-08 | 1.100E-07 |
| 29056 | 1453 | G1453S | E | 1.8e-08 | 2.9e-08 | 6.9e-08 | 3.867E-08 | 2.684E-08 | 1.800E-08 | 6.900E-08 |
| 29057 | 1453 | G1453T | E | 2.6e-08 | 1.2e-08 | 7.3e-08 | 3.700E-08 | 3.195E-08 | 1.200E-08 | 7.300E-08 |
| 29058 | 1453 | G1453W | E | 3.6e-08 | 4.6e-08 | 9.9e-08 | 6.033E-08 | 3.386E-08 | 3.600E-08 | 9.900E-08 |
| 29059 | 1453 | G1453Y | E | 9.2e-08 | 2.5e-08 | 2.1e-08 | 4.600E-08 | 3.989E-08 | 2.100E-08 | 9.200E-08 |
| 29060 | 1453 | G1453V | E | 3.3e-08 | 6e-08 | 8.2e-08 | 5.833E-08 | 2.454E-08 | 3.300E-08 | 8.200E-08 |
| 29061 | 1454 | L1454A | E | 2.9e-08 | 6.8e-08 | 3.9e-08 | 4.533E-08 | 2.026E-08 | 2.900E-08 | 6.800E-08 |
| 29062 | 1454 | L1454R | E | 2.7e-08 | 6e-08 | 1.9e-08 | 3.533E-08 | 2.173E-08 | 1.900E-08 | 6.000E-08 |
| 29063 | 1454 | L1454N | E | 5.1e-08 | 3.3e-08 | 2.5e-08 | 3.633E-08 | 1.332E-08 | 2.500E-08 | 5.100E-08 |
| 29064 | 1454 | L1454D | E | 4.1e-08 | 3.1e-08 | 5e-08 | 4.067E-08 | 9.504E-09 | 3.100E-08 | 5.000E-08 |
| 29065 | 1454 | L1454C | E | 3.6e-08 | 8.2e-08 | 3.7e-08 | 5.167E-08 | 2.627E-08 | 3.600E-08 | 8.200E-08 |
| 29066 | 1454 | L1454E | E | 3.5e-08 | 1.7e-08 | 1e-07 | 5.067E-08 | 4.366E-08 | 1.700E-08 | 1.000E-07 |
| 29067 | 1454 | L1454Q | E | 3.5e-08 | 3.5e-08 | 8e-08 | 5.000E-08 | 2.598E-08 | 3.500E-08 | 8.000E-08 |
| 29068 | 1454 | L1454G | E | 6.4e-08 | 5.9e-08 | 4.7e-08 | 5.667E-08 | 8.737E-09 | 4.700E-08 | 6.400E-08 |
| 29069 | 1454 | L1454H | E | 3.7e-08 | 8e-08 | 4.8e-08 | 5.500E-08 | 2.234E-08 | 3.700E-08 | 8.000E-08 |

| | | | | | | | | | | |
|---|---|---|---|---|---|---|---|---|---|---|
| 29070 | 1454 | L1454I | E | 4.6e-08 | 1.3e-07 | 3e-08 | 6.867E-08 | 5.372E-08 | 3.000E-08 | 1.300E-07 |
| 29071 | 1454 | L1454L | E | 1.6e-08 | 2.5e-08 | 9.5e-08 | 4.533E-08 | 4.325E-08 | 1.600E-08 | 9.500E-08 |
| 29072 | 1454 | L1454K | E | 6.2e-08 | 3.8e-08 | 4.4e-08 | 4.800E-08 | 1.249E-08 | 3.800E-08 | 6.200E-08 |
| 29073 | 1454 | L1454M | E | 6.3e-08 | 2.3e-08 | 4.2e-08 | 4.267E-08 | 2.001E-08 | 2.300E-08 | 6.300E-08 |
| 29074 | 1454 | L1454F | E | 3.2e-08 | 7.9e-08 | 8e-08 | 6.367E-08 | 2.743E-08 | 3.200E-08 | 8.000E-08 |
| 29075 | 1454 | L1454P | E | 3.1e-08 | 2.7e-08 | 1.1e-07 | 5.600E-08 | 4.681E-08 | 2.700E-08 | 1.100E-07 |
| 29076 | 1454 | L1454S | E | 6.4e-08 | 3.4e-08 | 2.2e-08 | 4.000E-08 | 2.163E-08 | 2.200E-08 | 6.400E-08 |
| 29077 | 1454 | L1454T | E | 5e-08 | 1.1e-07 | 6e-08 | 7.333E-08 | 3.215E-08 | 5.000E-08 | 1.100E-07 |
| 29078 | 1454 | L1454W | E | 6.3e-08 | 1.2e-08 | 2.5e-08 | 3.333E-08 | 2.650E-08 | 1.200E-08 | 6.300E-08 |
| 29079 | 1454 | L1454Y | E | 1.9e-08 | 2.9e-08 | 2e-08 | 2.267E-08 | 5.508E-09 | 1.900E-08 | 2.900E-08 |
| 29080 | 1454 | L1454V | E | 4.6e-08 | 2e-08 | 6.3e-08 | 4.300E-08 | 2.166E-08 | 2.000E-08 | 6.300E-08 |
| 29081 | 1455 | Y1455A | E | 4e-08 | 4.1e-08 | 6.4e-08 | 4.833E-08 | 1.358E-08 | 4.000E-08 | 6.400E-08 |
| 29082 | 1455 | Y1455R | E | 3.9e-08 | 2.5e-08 | 1.8e-07 | 8.133E-08 | 8.573E-08 | 2.500E-08 | 1.800E-07 |
| 29083 | 1455 | Y1455N | E | 5.2e-08 | 5.9e-08 | 5.3e-08 | 5.467E-08 | 3.786E-09 | 5.200E-08 | 5.900E-08 |
| 29084 | 1455 | Y1455D | E | 3.2e-08 | 2.8e-08 | 2.1e-08 | 2.700E-08 | 5.568E-09 | 2.100E-08 | 3.200E-08 |
| 29085 | 1455 | Y1455C | E | 2.9e-08 | 9.9e-08 | 1.4e-08 | 4.733E-08 | 4.537E-08 | 1.400E-08 | 9.900E-08 |
| 29086 | 1455 | Y1455E | E | 6.1e-08 | 3.9e-08 | 3.5e-08 | 4.500E-08 | 1.400E-08 | 3.500E-08 | 6.100E-08 |
| 29087 | 1455 | Y1455Q | E | 4.2e-08 | 3.8e-08 | 1e-08 | 3.000E-08 | 1.744E-08 | 1.000E-08 | 4.200E-08 |
| 29088 | 1455 | Y1455G | E | 6.3e-08 | 2.9e-08 | 3.7e-08 | 4.300E-08 | 1.778E-08 | 2.900E-08 | 6.300E-08 |
| 29089 | 1455 | Y1455H | E | 5.7e-08 | 2.8e-08 | 4e-08 | 4.167E-08 | 1.457E-08 | 2.800E-08 | 5.700E-08 |
| 29090 | 1455 | Y1455I | E | 5e-08 | 4.9e-08 | 1.3e-07 | 7.633E-08 | 4.648E-08 | 4.900E-08 | 1.300E-07 |
| 29091 | 1455 | Y1455L | E | 1.6e-08 | 6.3e-08 | 2.4e-08 | 3.433E-08 | 2.515E-08 | 1.600E-08 | 6.300E-08 |
| 29092 | 1455 | Y1455K | E | 9.7e-08 | 2.6e-08 | 9.3e-08 | 7.200E-08 | 3.989E-08 | 2.600E-08 | 9.700E-08 |
| 29093 | 1455 | Y1455M | E | 1.1e-07 | 2.2e-08 | 1.3e-07 | 8.733E-08 | 5.746E-08 | 2.200E-08 | 1.300E-07 |
| 29094 | 1455 | Y1455F | E | 3.2e-08 | 3e-08 | 5e-08 | 3.733E-08 | 1.102E-08 | 3.000E-08 | 5.000E-08 |
| 29095 | 1455 | Y1455P | E | 7.1e-08 | 3.3e-08 | 5.8e-08 | 5.400E-08 | 1.931E-08 | 3.300E-08 | 7.100E-08 |
| 29096 | 1455 | Y1455S | E | 3.7e-08 | 7.4e-08 | 5.9e-08 | 5.667E-08 | 1.861E-08 | 3.700E-08 | 7.400E-08 |
| 29097 | 1455 | Y1455T | E | 8e-08 | 6e-08 | 3.2e-08 | 5.733E-08 | 2.411E-08 | 3.200E-08 | 8.000E-08 |
| 29098 | 1455 | Y1455W | E | 2.1e-08 | 4.5e-08 | 4.8e-08 | 3.800E-08 | 1.480E-08 | 2.100E-08 | 4.800E-08 |
| 29099 | 1455 | Y1455Y | E | 6.6e-08 | 1.8e-08 | 3e-08 | 3.800E-08 | 2.498E-08 | 1.800E-08 | 6.600E-08 |
| 29100 | 1455 | Y1455V | E | 5.9e-08 | 3.2e-08 | 2e-08 | 3.700E-08 | 1.997E-08 | 2.000E-08 | 5.900E-08 |
| 29101 | 1456 | S1456A | E | 2.8e-08 | 5.2e-08 | 2.9e-08 | 3.633E-08 | 1.358E-08 | 2.800E-08 | 5.200E-08 |
| 29102 | 1456 | S1456R | E | 7.4e-08 | 2.5e-08 | 5.4e-08 | 5.100E-08 | 2.464E-08 | 2.500E-08 | 7.400E-08 |
| 29103 | 1456 | S1456N | E | 1.1e-07 | 1.2e-07 | 1.5e-08 | 8.167E-08 | 5.795E-08 | 1.500E-08 | 1.200E-07 |
| 29104 | 1456 | S1456D | E | 5.4e-08 | 2.9e-08 | 3.5e-08 | 3.933E-08 | 1.305E-08 | 2.900E-08 | 5.400E-08 |
| 29105 | 1456 | S1456C | E | 5e-08 | 2.3e-08 | 1.5e-08 | 2.933E-08 | 1.834E-08 | 1.500E-08 | 5.000E-08 |
| 29106 | 1456 | S1456E | E | 3.1e-08 | 1.5e-08 | 4e-08 | 2.867E-08 | 1.266E-08 | 1.500E-08 | 4.000E-08 |
| 29107 | 1456 | S1456Q | E | 6.6e-08 | 2.2e-08 | 6.2e-08 | 5.000E-08 | 2.433E-08 | 2.200E-08 | 6.600E-08 |
| 29108 | 1456 | S1456G | E | 1e-07 | 4.4e-08 | 5.4e-08 | 6.600E-08 | 2.987E-08 | 4.400E-08 | 1.000E-07 |
| 29109 | 1456 | S1456H | E | 1.3e-08 | 2.3e-08 | 5.5e-08 | 3.033E-08 | 2.194E-08 | 1.300E-08 | 5.500E-08 |
| 29110 | 1456 | S1456I | E | 4e-08 | 9.8e-08 | 4.9e-08 | 6.233E-08 | 3.121E-08 | 4.000E-08 | 9.800E-08 |
| 29111 | 1456 | S1456L | E | 4.2e-08 | 4.6e-08 | 3.8e-08 | 4.200E-08 | 4.000E-09 | 3.800E-08 | 4.600E-08 |
| 29112 | 1456 | S1456K | E | 6.5e-08 | 7.9e-08 | 3.1e-08 | 5.833E-08 | 2.468E-08 | 3.100E-08 | 7.900E-08 |
| 29113 | 1456 | S1456M | E | 7.7e-08 | 1.2e-07 | 8.7e-08 | 9.467E-08 | 2.250E-08 | 7.700E-08 | 1.200E-07 |
| 29114 | 1456 | S1456F | E | 7.9e-08 | 6.1e-08 | 6.8e-08 | 6.933E-08 | 9.074E-09 | 6.100E-08 | 7.900E-08 |
| 29115 | 1456 | S1456P | E | 2.4e-08 | 2.8e-08 | 3.2e-08 | 2.800E-08 | 4.000E-09 | 2.400E-08 | 3.200E-08 |
| 29116 | 1456 | S1456S | E | 4.3e-08 | 4.5e-08 | 2.8e-08 | 3.867E-08 | 9.292E-09 | 2.800E-08 | 4.500E-08 |
| 29117 | 1456 | S1456T | E | 2e-08 | 4.2e-08 | 3.1e-08 | 3.100E-08 | 1.100E-08 | 2.000E-08 | 4.200E-08 |
| 29118 | 1456 | S1456W | E | 6.4e-08 | 5.6e-08 | 6.5e-08 | 6.167E-08 | 4.933E-09 | 5.600E-08 | 6.500E-08 |
| 29119 | 1456 | S1456Y | E | 8.9e-08 | 4.4e-08 | 1.6e-08 | 4.967E-08 | 3.683E-08 | 1.600E-08 | 8.900E-08 |
| 29120 | 1456 | S1456V | E | 3e-08 | 5.8e-08 | 3.4e-08 | 4.067E-08 | 1.514E-08 | 3.000E-08 | 5.800E-08 |
| 29121 | 1457 | L1457A | E | 2.6e-08 | 3e-08 | 8.1e-08 | 4.567E-08 | 3.066E-08 | 2.600E-08 | 8.100E-08 |
| 29122 | 1457 | L1457R | E | 2.6e-08 | 3.3e-08 | 6.8e-08 | 4.233E-08 | 2.250E-08 | 2.600E-08 | 6.800E-08 |
| 29123 | 1457 | L1457N | E | 1.8e-08 | 1.1e-07 | 1.7e-08 | 4.833E-08 | 5.341E-08 | 1.700E-08 | 1.100E-07 |
| 29124 | 1457 | L1457D | E | 2.8e-08 | 5.7e-08 | 4.5e-08 | 4.333E-08 | 1.457E-08 | 2.800E-08 | 5.700E-08 |
| 29125 | 1457 | L1457C | E | 6.1e-08 | 6.1e-09 | 2.9e-08 | 3.203E-08 | 2.758E-08 | 6.100E-09 | 6.100E-08 |
| 29126 | 1457 | L1457E | E | 5.8e-08 | 2.4e-08 | 2.7e-08 | 3.633E-08 | 1.882E-08 | 2.400E-08 | 5.800E-08 |
| 29127 | 1457 | L1457Q | E | 6.8e-08 | 2.5e-08 | 3.5e-08 | 4.267E-08 | 2.250E-08 | 2.500E-08 | 6.800E-08 |
| 29128 | 1457 | L1457G | E | 1.3e-07 | 2.9e-08 | 3.1e-08 | 6.333E-08 | 5.774E-08 | 2.900E-08 | 1.300E-07 |
| 29129 | 1457 | L1457H | E | 1.5e-08 | 2.8e-08 | 9.8e-08 | 4.700E-08 | 4.464E-08 | 1.500E-08 | 9.800E-08 |
| 29130 | 1457 | L1457I | E | 6.3e-08 | 3.2e-08 | 5.9e-08 | 5.133E-08 | 1.686E-08 | 3.200E-08 | 6.300E-08 |
| 29131 | 1457 | L1457L | E | 1.8e-08 | 2.6e-08 | 1.5e-08 | 1.967E-08 | 5.686E-09 | 1.500E-08 | 2.600E-08 |
| 29132 | 1457 | L1457K | E | 1.1e-07 | 3.7e-08 | 6.1e-08 | 6.933E-08 | 3.721E-08 | 3.700E-08 | 1.100E-07 |
| 29133 | 1457 | L1457M | E | 2.1e-08 | 8.8e-08 | 4.1e-08 | 5.000E-08 | 3.439E-08 | 2.100E-08 | 8.800E-08 |
| 29134 | 1457 | L1457F | E | 8e-08 | 3.5e-08 | 4.3e-08 | 5.267E-08 | 2.401E-08 | 3.500E-08 | 8.000E-08 |
| 29135 | 1457 | L1457P | E | 3.5e-08 | 2.5e-08 | 4.3e-08 | 3.433E-08 | 9.018E-09 | 2.500E-08 | 4.300E-08 |
| 29136 | 1457 | L1457S | E | 3.5e-08 | 9.2e-08 | 1.8e-08 | 4.833E-08 | 3.876E-08 | 1.800E-08 | 9.200E-08 |
| 29137 | 1457 | L1457T | E | 8.3e-08 | 5e-08 | 2.4e-08 | 5.233E-08 | 2.957E-08 | 2.400E-08 | 8.300E-08 |
| 29138 | 1457 | L1457W | E | 2.2e-08 | 3.3e-08 | 5.7e-08 | 3.733E-08 | 1.790E-08 | 2.200E-08 | 5.700E-08 |
| 29139 | 1457 | L1457Y | E | 3e-08 | 8.8e-08 | 1.3e-07 | 8.267E-08 | 5.021E-08 | 3.000E-08 | 1.300E-07 |
| 29140 | 1457 | L1457V | E | 5.1e-08 | 8.3e-08 | 5e-08 | 6.133E-08 | 1.877E-08 | 5.000E-08 | 8.300E-08 |
| 29141 | 1458 | S1458A | E | 1.3e-07 | 1.9e-08 | 7.4e-08 | 7.433E-08 | 5.550E-08 | 1.900E-08 | 1.300E-07 |
| 29142 | 1458 | S1458R | E | 3.3e-08 | 9.3e-08 | 5.1e-08 | 5.900E-08 | 3.079E-08 | 3.300E-08 | 9.300E-08 |
| 29143 | 1458 | S1458N | E | 3.7e-08 | 6.7e-08 | 6.4e-08 | 5.600E-08 | 1.652E-08 | 3.700E-08 | 6.700E-08 |
| 29144 | 1458 | S1458D | E | 3.8e-08 | 5.2e-08 | 9.8e-08 | 6.267E-08 | 3.139E-08 | 3.800E-08 | 9.800E-08 |
| 29145 | 1458 | S1458C | E | 1.8e-08 | 2.6e-08 | 4.3e-08 | 2.900E-08 | 1.277E-08 | 1.800E-08 | 4.300E-08 |
| 29146 | 1458 | S1458E | E | 2e-08 | 2.8e-08 | 2.4e-08 | 2.400E-08 | 4.000E-09 | 2.000E-08 | 2.800E-08 |
| 29147 | 1458 | S1458Q | E | 3.4e-08 | 6.1e-08 | 3e-08 | 4.167E-08 | 1.686E-08 | 3.000E-08 | 6.100E-08 |
| 29148 | 1458 | S1458G | E | 1.2e-07 | 4.3e-08 | 3.1e-08 | 6.467E-08 | 4.829E-08 | 3.100E-08 | 1.200E-07 |
| 29149 | 1458 | S1458H | E | 2.5e-08 | 8.6e-08 | 2e-08 | 4.367E-08 | 3.675E-08 | 2.000E-08 | 8.600E-08 |
| 29150 | 1458 | S1458I | E | 4.2e-08 | 2.5e-08 | 3e-08 | 3.233E-08 | 8.737E-09 | 2.500E-08 | 4.200E-08 |
| 29151 | 1458 | S1458L | E | 6.8e-08 | 2.5e-08 | 9.1e-08 | 6.133E-08 | 3.350E-08 | 2.500E-08 | 9.100E-08 |
| 29152 | 1458 | S1458K | E | 8.3e-08 | 3.3e-08 | 3.3e-08 | 4.967E-08 | 2.887E-08 | 3.300E-08 | 8.300E-08 |
| 29153 | 1458 | S1458M | E | 4.5e-08 | 3.8e-08 | 3.6e-08 | 3.967E-08 | 4.726E-09 | 3.600E-08 | 4.500E-08 |
| 29154 | 1458 | S1458F | E | 4.7e-08 | 1.4e-07 | 6.3e-08 | 8.333E-08 | 4.972E-08 | 4.700E-08 | 1.400E-07 |

| | | | | | | | | | | |
|---|---|---|---|---|---|---|---|---|---|---|
| 29155 | 1458 | S1458P | E | 3.1e-08 | 1.1e-07 | 7.5e-08 | 7.200E-08 | 3.959E-08 | 3.100E-08 | 1.100E-07 |
| 29156 | 1458 | S1458S | E | 6.5e-08 | 1.8e-08 | 2.4e-08 | 3.567E-08 | 2.558E-08 | 1.800E-08 | 6.500E-08 |
| 29157 | 1458 | S1458T | E | 3.1e-08 | 1.7e-07 | 3e-08 | 7.700E-08 | 8.054E-08 | 3.000E-08 | 1.700E-07 |
| 29158 | 1458 | S1458W | E | 3e-08 | 6.6e-08 | 4e-08 | 4.533E-08 | 1.858E-08 | 3.000E-08 | 6.600E-08 |
| 29159 | 1458 | S1458Y | E | 3.1e-08 | 6.3e-08 | 1.5e-08 | 3.633E-08 | 2.444E-08 | 1.500E-08 | 6.300E-08 |
| 29160 | 1458 | S1458V | E | 1.2e-07 | 5e-08 | 5.4e-08 | 7.467E-08 | 3.931E-08 | 5.000E-08 | 1.200E-07 |
| 29161 | 1459 | S1459A | E | 2.8e-08 | 3.1e-08 | 1.3e-08 | 2.400E-08 | 9.644E-09 | 1.300E-08 | 3.100E-08 |
| 29162 | 1459 | S1459R | E | 8.5e-08 | 5.2e-08 | 1.9e-07 | 1.090E-07 | 7.206E-08 | 5.200E-08 | 1.900E-07 |
| 29163 | 1459 | S1459N | E | 2.6e-08 | 8.4e-08 | 1.5e-08 | 4.167E-08 | 3.707E-08 | 1.500E-08 | 8.400E-08 |
| 29164 | 1459 | S1459D | E | 9.2e-08 | 2.3e-08 | 3.5e-08 | 5.000E-08 | 3.686E-08 | 2.300E-08 | 9.200E-08 |
| 29165 | 1459 | S1459C | E | 3.3e-08 | 9.2e-09 | 9.3e-08 | 4.507E-08 | 4.318E-08 | 9.200E-09 | 9.300E-08 |
| 29166 | 1459 | S1459E | E | 5.8e-08 | 4.3e-08 | 5.4e-08 | 5.167E-08 | 7.767E-09 | 4.300E-08 | 5.800E-08 |
| 29167 | 1459 | S1459Q | E | 3.9e-08 | 2.8e-08 | 2.2e-08 | 2.967E-08 | 8.622E-09 | 2.200E-08 | 3.900E-08 |
| 29168 | 1459 | S1459G | E | 3.4e-08 | 7.9e-08 | 2.1e-08 | 4.467E-08 | 3.044E-08 | 2.100E-08 | 7.900E-08 |
| 29169 | 1459 | S1459H | E | 2.2e-08 | 3e-08 | 5.4e-08 | 3.533E-08 | 1.665E-08 | 2.200E-08 | 5.400E-08 |
| 29170 | 1459 | S1459I | E | 4.1e-08 | 3.5e-08 | 4.7e-08 | 4.100E-08 | 6.000E-09 | 3.500E-08 | 4.700E-08 |
| 29171 | 1459 | S1459L | E | 2.4e-08 | 1.4e-08 | 4.7e-08 | 2.833E-08 | 1.692E-08 | 1.400E-08 | 4.700E-08 |
| 29172 | 1459 | S1459K | E | 3.2e-08 | 6.4e-08 | 7.3e-08 | 5.633E-08 | 2.155E-08 | 3.200E-08 | 7.300E-08 |
| 29173 | 1459 | S1459M | E | 9e-09 | 5.7e-08 | 3.3e-08 | 3.300E-08 | 2.400E-08 | 9.000E-09 | 5.700E-08 |
| 29174 | 1459 | S1459F | E | 2.4e-08 | 1.1e-08 | 6.3e-08 | 3.267E-08 | 2.706E-08 | 1.100E-08 | 6.300E-08 |
| 29175 | 1459 | S1459P | E | 3.5e-08 | 5.3e-08 | 5.8e-08 | 4.867E-08 | 1.210E-08 | 3.500E-08 | 5.800E-08 |
| 29176 | 1459 | S1459S | E | 4.2e-08 | 4.4e-08 | 5.5e-08 | 4.700E-08 | 7.000E-09 | 4.200E-08 | 5.500E-08 |
| 29177 | 1459 | S1459T | E | 1.4e-07 | 5.1e-08 | 1.7e-08 | 6.933E-08 | 6.352E-08 | 1.700E-08 | 1.400E-07 |
| 29178 | 1459 | S1459W | E | 1e-07 | 7e-08 | 3.9e-08 | 6.967E-08 | 3.050E-08 | 3.900E-08 | 1.000E-07 |
| 29179 | 1459 | S1459Y | E | 8.2e-08 | 3.5e-08 | 1.6e-08 | 4.433E-08 | 3.398E-08 | 1.600E-08 | 8.200E-08 |
| 29180 | 1459 | S1459V | E | 2.6e-08 | 4.2e-08 | 3.2e-08 | 3.333E-08 | 8.083E-09 | 2.600E-08 | 4.200E-08 |
| 29181 | 1460 | V1460A | E | 3.8e-08 | 2.2e-08 | 6.2e-08 | 4.067E-08 | 2.013E-08 | 2.200E-08 | 6.200E-08 |
| 29182 | 1460 | V1460R | E | 1.3e-07 | 9.7e-08 | 9.4e-08 | 1.070E-07 | 1.997E-08 | 9.400E-08 | 1.300E-07 |
| 29183 | 1460 | V1460N | E | 5e-08 | 4.9e-08 | 4.5e-08 | 4.800E-08 | 2.646E-09 | 4.500E-08 | 5.000E-08 |
| 29184 | 1460 | V1460D | E | 3.1e-08 | 1.3e-07 | 2.1e-08 | 6.067E-08 | 6.025E-08 | 2.100E-08 | 1.300E-07 |
| 29185 | 1460 | V1460C | E | 1.3e-08 | 6.5e-08 | 3.1e-08 | 3.633E-08 | 2.641E-08 | 1.300E-08 | 6.500E-08 |
| 29186 | 1460 | V1460E | E | 4.8e-08 | 1e-07 | 1.7e-07 | 1.060E-07 | 6.122E-08 | 4.800E-08 | 1.700E-07 |
| 29187 | 1460 | V1460Q | E | 5.1e-09 | 4.9e-08 | 3.3e-08 | 2.903E-08 | 2.222E-08 | 5.100E-09 | 4.900E-08 |
| 29188 | 1460 | V1460G | E | 4.7e-08 | 3.4e-08 | 3.1e-08 | 3.733E-08 | 8.505E-09 | 3.100E-08 | 4.700E-08 |
| 29189 | 1460 | V1460H | E | 2.1e-08 | 8.2e-08 | 3.3e-08 | 4.533E-08 | 3.232E-08 | 2.100E-08 | 8.200E-08 |
| 29190 | 1460 | V1460I | E | 8.2e-08 | 2.4e-08 | 1.8e-08 | 4.133E-08 | 3.535E-08 | 1.800E-08 | 8.200E-08 |
| 29191 | 1460 | V1460L | E | 8.6e-08 | 6.1e-08 | 4.1e-08 | 6.267E-08 | 2.255E-08 | 4.100E-08 | 8.600E-08 |
| 29192 | 1460 | V1460K | E | 3.3e-08 | 3.3e-08 | 4.1e-08 | 3.567E-08 | 4.619E-09 | 3.300E-08 | 4.100E-08 |
| 29193 | 1460 | V1460M | E | 2.9e-08 | 8.6e-08 | 3.1e-08 | 4.867E-08 | 3.235E-08 | 2.900E-08 | 8.600E-08 |
| 29194 | 1460 | V1460F | E | 2.2e-08 | 1.7e-08 | 1e-07 | 4.633E-08 | 4.654E-08 | 1.700E-08 | 1.000E-07 |
| 29195 | 1460 | V1460P | E | 6.3e-08 | 3.3e-08 | 5.6e-08 | 5.067E-08 | 1.570E-08 | 3.300E-08 | 6.300E-08 |
| 29196 | 1460 | V1460S | E | 3.4e-08 | 2e-08 | 1.8e-08 | 2.400E-08 | 8.718E-09 | 1.800E-08 | 3.400E-08 |
| 29197 | 1460 | V1460T | E | 2.4e-08 | 7.1e-08 | 1.2e-08 | 3.567E-08 | 3.118E-08 | 1.200E-08 | 7.100E-08 |
| 29198 | 1460 | V1460W | E | 6.3e-08 | 7.9e-08 | 2.5e-08 | 5.567E-08 | 2.774E-08 | 2.500E-08 | 7.900E-08 |
| 29199 | 1460 | V1460Y | E | 1.7e-08 | 9e-08 | 6.8e-08 | 5.833E-08 | 3.745E-08 | 1.700E-08 | 9.000E-08 |
| 29200 | 1460 | V1460V | E | 4e-08 | 2.3e-08 | 4.1e-08 | 3.467E-08 | 1.012E-08 | 2.300E-08 | 4.100E-08 |
| 29201 | 1461 | V1461A | E | 2.6e-08 | 2.1e-08 | 2.7e-08 | 2.467E-08 | 3.215E-09 | 2.100E-08 | 2.700E-08 |
| 29202 | 1461 | V1461R | E | 5.2e-08 | 2.4e-08 | 1.6e-08 | 3.067E-08 | 1.890E-08 | 1.600E-08 | 5.200E-08 |
| 29203 | 1461 | V1461N | E | 9e-08 | 7.7e-08 | 3.9e-08 | 6.867E-08 | 2.650E-08 | 3.900E-08 | 9.000E-08 |
| 29204 | 1461 | V1461D | E | 2e-08 | 5.3e-08 | 6.2e-08 | 4.500E-08 | 2.211E-08 | 2.000E-08 | 6.200E-08 |
| 29205 | 1461 | V1461C | E | 5.5e-08 | 3e-08 | 6e-08 | 4.833E-08 | 1.607E-08 | 3.000E-08 | 6.000E-08 |
| 29206 | 1461 | V1461E | E | 3.3e-08 | 7.5e-08 | 8.3e-08 | 6.367E-08 | 2.686E-08 | 3.300E-08 | 8.300E-08 |
| 29207 | 1461 | V1461Q | E | 1.6e-07 | 3.6e-08 | 5e-08 | 8.200E-08 | 6.791E-08 | 3.600E-08 | 1.600E-07 |
| 29208 | 1461 | V1461G | E | 3.5e-08 | 2.8e-08 | 3.9e-08 | 3.400E-08 | 5.568E-09 | 2.800E-08 | 3.900E-08 |
| 29209 | 1461 | V1461H | E | 5.7e-08 | 6.4e-08 | 2.5e-08 | 4.867E-08 | 2.079E-08 | 2.500E-08 | 6.400E-08 |
| 29210 | 1461 | V1461I | E | 3.6e-08 | 9.2e-08 | 7.3e-08 | 6.700E-08 | 2.848E-08 | 3.600E-08 | 9.200E-08 |
| 29211 | 1461 | V1461L | E | 3.1e-08 | 5.1e-08 | 8.6e-08 | 5.600E-08 | 2.784E-08 | 3.100E-08 | 8.600E-08 |
| 29212 | 1461 | V1461K | E | 2.1e-08 | 3.5e-08 | 4.3e-08 | 3.300E-08 | 1.114E-08 | 2.100E-08 | 4.300E-08 |
| 29213 | 1461 | V1461M | E | 8.8e-08 | 4.5e-08 | 3.1e-08 | 5.467E-08 | 2.970E-08 | 3.100E-08 | 8.800E-08 |
| 29214 | 1461 | V1461F | E | 4.2e-08 | 3.1e-08 | 3.9e-08 | 3.733E-08 | 5.686E-09 | 3.100E-08 | 4.200E-08 |
| 29215 | 1461 | V1461P | E | 5.4e-08 | 1.5e-08 | 1e-08 | 2.633E-08 | 2.409E-08 | 1.000E-08 | 5.400E-08 |
| 29216 | 1461 | V1461S | E | 1.8e-08 | 3.3e-08 | 4.7e-08 | 3.267E-08 | 1.450E-08 | 1.800E-08 | 4.700E-08 |
| 29217 | 1461 | V1461T | E | 1.7e-08 | 2e-08 | 4.7e-08 | 2.800E-08 | 1.652E-08 | 1.700E-08 | 4.700E-08 |
| 29218 | 1461 | V1461W | E | 4e-08 | 3.6e-08 | 1.5e-08 | 3.033E-08 | 1.343E-08 | 1.500E-08 | 4.000E-08 |
| 29219 | 1461 | V1461Y | E | 1.1e-07 | 2.9e-08 | 4.9e-08 | 6.267E-08 | 4.219E-08 | 2.900E-08 | 1.100E-07 |
| 29220 | 1461 | V1461V | E | 8.7e-08 | 6.2e-08 | 2.6e-08 | 5.833E-08 | 3.066E-08 | 2.600E-08 | 8.700E-08 |
| 29221 | 1462 | T1462A | E | 1.9e-08 | 5.9e-08 | 1.4e-08 | 3.067E-08 | 2.466E-08 | 1.400E-08 | 5.900E-08 |
| 29222 | 1462 | T1462R | E | 7.3e-08 | 2e-08 | 3.2e-08 | 4.167E-08 | 2.779E-08 | 2.000E-08 | 7.300E-08 |
| 29223 | 1462 | T1462N | E | 7e-08 | 1.7e-08 | 2.2e-08 | 3.633E-08 | 2.926E-08 | 1.700E-08 | 7.000E-08 |
| 29224 | 1462 | T1462D | E | 7.4e-08 | 4.2e-08 | 4.7e-08 | 5.433E-08 | 1.721E-08 | 4.200E-08 | 7.400E-08 |
| 29225 | 1462 | T1462C | E | 2e-08 | 4.2e-08 | 4.9e-08 | 3.700E-08 | 1.513E-08 | 2.000E-08 | 4.900E-08 |
| 29226 | 1462 | T1462E | E | 1.1e-07 | 5.4e-08 | 2.8e-08 | 6.400E-08 | 4.190E-08 | 2.800E-08 | 1.100E-07 |
| 29227 | 1462 | T1462Q | E | 3.5e-08 | 5.7e-08 | 1e-08 | 3.400E-08 | 2.352E-08 | 1.000E-08 | 5.700E-08 |
| 29228 | 1462 | T1462G | E | 7.3e-08 | 3.3e-08 | 1.2e-07 | 7.533E-08 | 4.355E-08 | 3.300E-08 | 1.200E-07 |
| 29229 | 1462 | T1462H | E | 2.6e-08 | 4.7e-08 | 4.1e-08 | 3.800E-08 | 1.082E-08 | 2.600E-08 | 4.700E-08 |
| 29230 | 1462 | T1462I | E | 3.5e-08 | 4e-08 | 6.2e-08 | 4.567E-08 | 1.436E-08 | 3.500E-08 | 6.200E-08 |
| 29231 | 1462 | T1462L | E | 4e-08 | 3.7e-08 | 2.7e-08 | 3.467E-08 | 6.807E-09 | 2.700E-08 | 4.000E-08 |
| 29232 | 1462 | T1462K | E | 3.4e-08 | 4.7e-08 | 5.7e-08 | 4.600E-08 | 1.153E-08 | 3.400E-08 | 5.700E-08 |
| 29233 | 1462 | T1462M | E | 9.8e-08 | 4.2e-08 | 3.4e-08 | 5.800E-08 | 3.487E-08 | 3.400E-08 | 9.800E-08 |
| 29234 | 1462 | T1462F | E | 2.9e-08 | 9.5e-08 | 9.8e-08 | 7.400E-08 | 3.900E-08 | 2.900E-08 | 9.800E-08 |
| 29235 | 1462 | T1462P | E | 1.1e-08 | 3.9e-08 | 3.5e-08 | 2.833E-08 | 1.514E-08 | 1.100E-08 | 3.900E-08 |
| 29236 | 1462 | T1462S | E | 2.8e-08 | 1.7e-08 | 4.3e-08 | 2.933E-08 | 1.305E-08 | 1.700E-08 | 4.300E-08 |
| 29237 | 1462 | T1462T | E | 1.9e-08 | 4.8e-08 | 1.4e-07 | 6.900E-08 | 6.317E-08 | 1.900E-08 | 1.400E-07 |
| 29238 | 1462 | T1462W | E | 1.6e-08 | 2e-08 | 5.9e-08 | 3.167E-08 | 2.376E-08 | 1.600E-08 | 5.900E-08 |
| 29239 | 1462 | T1462Y | E | 5.1e-08 | 7.3e-08 | 2.7e-08 | 5.033E-08 | 2.301E-08 | 2.700E-08 | 7.300E-08 |

| | | | | | | | | | | |
|---|---|---|---|---|---|---|---|---|---|---|
| 29240 | 1462 | T1462V | E | 5.8e-08 | 1e-08 | 7.2e-08 | 4.667E-08 | 3.252E-08 | 1.000E-08 | 7.200E-08 |
| 29241 | 1463 | V1463A | E | 8.1e-08 | 9.3e-08 | 4.7e-08 | 7.367E-08 | 2.386E-08 | 4.700E-08 | 9.300E-08 |
| 29242 | 1463 | V1463R | E | 1.4e-08 | 2.6e-08 | 2.8e-08 | 2.267E-08 | 7.572E-09 | 1.400E-08 | 2.800E-08 |
| 29243 | 1463 | V1463N | E | 1.4e-08 | 3.3e-08 | 8.9e-08 | 4.533E-08 | 3.899E-08 | 1.400E-08 | 8.900E-08 |
| 29244 | 1463 | V1463D | E | 5.5e-08 | 2.6e-08 | 4.3e-08 | 4.133E-08 | 1.457E-08 | 2.600E-08 | 5.500E-08 |
| 29245 | 1463 | V1463C | E | 3.6e-08 | 4.6e-08 | 1.4e-08 | 3.200E-08 | 1.637E-08 | 1.400E-08 | 4.600E-08 |
| 29246 | 1463 | V1463E | E | 5.4e-08 | 4.3e-08 | 4.3e-08 | 4.667E-08 | 6.351E-09 | 4.300E-08 | 5.400E-08 |
| 29247 | 1463 | V1463Q | E | 5e-08 | 2.7e-08 | 5.6e-08 | 4.433E-08 | 1.531E-08 | 2.700E-08 | 5.600E-08 |
| 29248 | 1463 | V1463G | E | 3.9e-08 | 2.7e-08 | 2e-08 | 2.867E-08 | 9.609E-09 | 2.000E-08 | 3.900E-08 |
| 29249 | 1463 | V1463H | E | 7.5e-08 | 5.1e-08 | 2.4e-08 | 5.000E-08 | 2.551E-08 | 2.400E-08 | 7.500E-08 |
| 29250 | 1463 | V1463I | E | 3.9e-08 | 5.9e-08 | 8e-08 | 5.933E-08 | 2.050E-08 | 3.900E-08 | 8.000E-08 |
| 29251 | 1463 | V1463L | E | 2.1e-08 | 2.5e-08 | 5.4e-08 | 3.333E-08 | 1.801E-08 | 2.100E-08 | 5.400E-08 |
| 29252 | 1463 | V1463K | E | 4.5e-08 | 3.2e-08 | 6.7e-08 | 4.800E-08 | 1.769E-08 | 3.200E-08 | 6.700E-08 |
| 29253 | 1463 | V1463M | E | 4.1e-08 | 3.2e-08 | 1.5e-07 | 7.433E-08 | 6.568E-08 | 3.200E-08 | 1.500E-07 |
| 29254 | 1463 | V1463F | E | 1.7e-08 | 6.9e-08 | 9.9e-08 | 6.167E-08 | 4.149E-08 | 1.700E-08 | 9.900E-08 |
| 29255 | 1463 | V1463P | E | 7.2e-08 | 1.1e-07 | 6.8e-08 | 8.333E-08 | 2.318E-08 | 6.800E-08 | 1.100E-07 |
| 29256 | 1463 | V1463S | E | 4.3e-08 | 3.5e-08 | 3.8e-08 | 3.867E-08 | 4.041E-09 | 3.500E-08 | 4.300E-08 |
| 29257 | 1463 | V1463T | E | 7.4e-08 | 4.6e-08 | 4.2e-08 | 5.400E-08 | 1.744E-08 | 4.200E-08 | 7.400E-08 |
| 29258 | 1463 | V1463W | E | 3.4e-08 | 6.9e-08 | 6.6e-08 | 5.633E-08 | 1.940E-08 | 3.400E-08 | 6.900E-08 |
| 29259 | 1463 | V1463Y | E | 5.3e-08 | 3.3e-08 | 1.3e-07 | 7.200E-08 | 5.122E-08 | 3.300E-08 | 1.300E-07 |
| 29260 | 1463 | V1463V | E | 2e-07 | 2.7e-08 | 1.3e-08 | 8.000E-08 | 1.042E-07 | 1.300E-08 | 2.000E-07 |
| 29261 | 1464 | P1464A | E | 3.9e-08 | 7.1e-08 | 6.7e-08 | 5.900E-08 | 1.744E-08 | 3.900E-08 | 7.100E-08 |
| 29262 | 1464 | P1464R | E | 1.7e-07 | 4.6e-08 | 5.6e-08 | 9.067E-08 | 6.889E-08 | 4.600E-08 | 1.700E-07 |
| 29263 | 1464 | P1464N | E | 3.5e-08 | 3.6e-08 | 6.6e-08 | 4.567E-08 | 1.762E-08 | 3.500E-08 | 6.600E-08 |
| 29264 | 1464 | P1464D | E | 1.5e-08 | 3.5e-08 | 3.7e-08 | 2.900E-08 | 1.217E-08 | 1.500E-08 | 3.700E-08 |
| 29265 | 1464 | P1464C | E | 2.1e-08 | 4.4e-08 | 3.8e-08 | 3.433E-08 | 1.193E-08 | 2.100E-08 | 4.400E-08 |
| 29266 | 1464 | P1464E | E | 9.8e-08 | 5e-08 | 2.3e-08 | 5.700E-08 | 3.799E-08 | 2.300E-08 | 9.800E-08 |
| 29267 | 1464 | P1464Q | E | 2.9e-08 | 5.9e-08 | 5.4e-08 | 4.733E-08 | 1.607E-08 | 2.900E-08 | 5.900E-08 |
| 29268 | 1464 | P1464G | E | 1.3e-08 | 7.5e-08 | 9.1e-08 | 5.967E-08 | 4.120E-08 | 1.300E-08 | 9.100E-08 |
| 29269 | 1464 | P1464H | E | 4.5e-08 | 2.9e-08 | 5.3e-08 | 4.233E-08 | 1.222E-08 | 2.900E-08 | 5.300E-08 |
| 29270 | 1464 | P1464I | E | 1e-07 | 9.5e-08 | 4.2e-08 | 7.900E-08 | 3.214E-08 | 4.200E-08 | 1.000E-07 |
| 29271 | 1464 | P1464L | E | 1.7e-08 | 3.1e-08 | 2.3e-08 | 2.367E-08 | 7.024E-09 | 1.700E-08 | 3.100E-08 |
| 29272 | 1464 | P1464K | E | 1.4e-07 | 2.9e-08 | 4.9e-08 | 7.267E-08 | 5.916E-08 | 2.900E-08 | 1.400E-07 |
| 29273 | 1464 | P1464M | E | 1.8e-08 | 2.8e-08 | 3.7e-08 | 2.767E-08 | 9.504E-09 | 1.800E-08 | 3.700E-08 |
| 29274 | 1464 | P1464F | E | 4.8e-08 | 5.4e-08 | 4.7e-08 | 4.967E-08 | 3.786E-09 | 4.700E-08 | 5.400E-08 |
| 29275 | 1464 | P1464P | E | 1.2e-07 | 1e-07 | 6.9e-08 | 9.633E-08 | 2.570E-08 | 6.900E-08 | 1.200E-07 |
| 29276 | 1464 | P1464S | E | 1.7e-08 | 8.6e-08 | 1.6e-07 | 8.767E-08 | 7.151E-08 | 1.700E-08 | 1.600E-07 |
| 29277 | 1464 | P1464T | E | 3e-08 | 2.3e-08 | 4.4e-08 | 3.233E-08 | 1.069E-08 | 2.300E-08 | 4.400E-08 |
| 29278 | 1464 | P1464W | E | 3.6e-08 | 1.7e-08 | 5.5e-08 | 3.600E-08 | 1.900E-08 | 1.700E-08 | 5.500E-08 |
| 29279 | 1464 | P1464Y | E | 2.5e-08 | 4.7e-08 | 3.3e-08 | 3.500E-08 | 1.114E-08 | 2.500E-08 | 4.700E-08 |
| 29280 | 1464 | P1464V | E | 4.1e-08 | 3.8e-08 | 5.6e-08 | 4.500E-08 | 9.644E-09 | 3.800E-08 | 5.600E-08 |
| 29281 | 1465 | S1465A | E | 6.4e-08 | 6.1e-08 | 3.7e-08 | 5.400E-08 | 1.480E-08 | 3.700E-08 | 6.400E-08 |
| 29282 | 1465 | S1465R | E | 2.6e-08 | 8e-08 | 1.1e-07 | 7.200E-08 | 4.257E-08 | 2.600E-08 | 1.100E-07 |
| 29283 | 1465 | S1465N | E | 3.2e-08 | 3.9e-08 | 1e-07 | 5.700E-08 | 3.740E-08 | 3.200E-08 | 1.000E-07 |
| 29284 | 1465 | S1465D | E | 6.4e-08 | 7.4e-08 | 2.9e-08 | 5.567E-08 | 2.363E-08 | 2.900E-08 | 7.400E-08 |
| 29285 | 1465 | S1465C | E | 5.8e-08 | 6.2e-08 | 2.9e-08 | 4.967E-08 | 1.801E-08 | 2.900E-08 | 6.200E-08 |
| 29286 | 1465 | S1465E | E | 3.4e-08 | 8.1e-08 | 5.5e-08 | 5.667E-08 | 2.354E-08 | 3.400E-08 | 8.100E-08 |
| 29287 | 1465 | S1465Q | E | 6.1e-08 | 4.2e-08 | 6.3e-08 | 5.533E-08 | 1.159E-08 | 4.200E-08 | 6.300E-08 |
| 29288 | 1465 | S1465G | E | 6.6e-08 | 6.7e-08 | 4.1e-08 | 5.800E-08 | 1.473E-08 | 4.100E-08 | 6.700E-08 |
| 29289 | 1465 | S1465H | E | 7e-08 | 2.4e-08 | 7.4e-08 | 5.600E-08 | 2.778E-08 | 2.400E-08 | 7.400E-08 |
| 29290 | 1465 | S1465I | E | 3.3e-08 | 2.1e-08 | 8.8e-08 | 4.733E-08 | 3.573E-08 | 2.100E-08 | 8.800E-08 |
| 29291 | 1465 | S1465L | E | 4.3e-08 | 5.3e-08 | 6e-08 | 5.200E-08 | 8.544E-09 | 4.300E-08 | 6.000E-08 |
| 29292 | 1465 | S1465K | E | 3.8e-08 | 3.9e-08 | 4.9e-08 | 4.200E-08 | 6.083E-09 | 3.800E-08 | 4.900E-08 |
| 29293 | 1465 | S1465M | E | 5.3e-08 | 7.5e-08 | 5.4e-08 | 6.067E-08 | 1.242E-08 | 5.300E-08 | 7.500E-08 |
| 29294 | 1465 | S1465F | E | 3.5e-08 | 2.5e-08 | 3.8e-08 | 3.267E-08 | 6.807E-09 | 2.500E-08 | 3.800E-08 |
| 29295 | 1465 | S1465P | E | 2.9e-08 | 7.4e-08 | 2.1e-08 | 4.133E-08 | 2.857E-08 | 2.100E-08 | 7.400E-08 |
| 29296 | 1465 | S1465S | E | 4.1e-08 | 1.3e-07 | 3.8e-08 | 6.967E-08 | 5.227E-08 | 3.800E-08 | 1.300E-07 |
| 29297 | 1465 | S1465T | E | 5.3e-08 | 4e-08 | 4.2e-08 | 4.500E-08 | 7.000E-09 | 4.000E-08 | 5.300E-08 |
| 29298 | 1465 | S1465W | E | 5.4e-08 | 9.8e-08 | 4.6e-08 | 6.600E-08 | 2.800E-08 | 4.600E-08 | 9.800E-08 |
| 29299 | 1465 | S1465Y | E | 2.3e-08 | 3.1e-08 | 4.2e-08 | 3.200E-08 | 9.539E-09 | 2.300E-08 | 4.200E-08 |
| 29300 | 1465 | S1465V | E | 2.9e-08 | 3.2e-08 | 1.1e-07 | 5.700E-08 | 4.592E-08 | 2.900E-08 | 1.100E-07 |
| 29301 | 1466 | S1466A | E | 5e-08 | 4e-08 | 3.6e-08 | 4.200E-08 | 7.211E-09 | 3.600E-08 | 5.000E-08 |
| 29302 | 1466 | S1466R | E | 4.2e-08 | 9.8e-08 | 4.8e-08 | 6.267E-08 | 3.075E-08 | 4.200E-08 | 9.800E-08 |
| 29303 | 1466 | S1466N | E | 3.1e-08 | 9.2e-08 | 3.3e-08 | 5.200E-08 | 3.466E-08 | 3.100E-08 | 9.200E-08 |
| 29304 | 1466 | S1466D | E | 2.5e-08 | 3.9e-08 | 4.6e-08 | 3.667E-08 | 1.069E-08 | 2.500E-08 | 4.600E-08 |
| 29305 | 1466 | S1466C | E | 2.8e-08 | 5.3e-08 | 6.2e-08 | 4.767E-08 | 1.762E-08 | 2.800E-08 | 6.200E-08 |
| 29306 | 1466 | S1466E | E | 3.4e-08 | 5.4e-08 | 5.8e-08 | 4.867E-08 | 1.286E-08 | 3.400E-08 | 5.800E-08 |
| 29307 | 1466 | S1466Q | E | 3.3e-08 | 2.6e-08 | 4.4e-08 | 3.433E-08 | 9.074E-09 | 2.600E-08 | 4.400E-08 |
| 29308 | 1466 | S1466G | E | 3.3e-08 | 5.7e-08 | 2.3e-07 | 1.067E-07 | 1.075E-07 | 3.300E-08 | 2.300E-07 |
| 29309 | 1466 | S1466H | E | 2.1e-07 | 2.7e-08 | 8.4e-08 | 1.070E-07 | 9.364E-08 | 2.700E-08 | 2.100E-07 |
| 29310 | 1466 | S1466I | E | 7.9e-08 | 3.5e-08 | 3.9e-08 | 5.100E-08 | 2.433E-08 | 3.500E-08 | 7.900E-08 |
| 29311 | 1466 | S1466L | E | 2.7e-08 | 2.3e-08 | 4.5e-08 | 3.167E-08 | 1.172E-08 | 2.300E-08 | 4.500E-08 |
| 29312 | 1466 | S1466K | E | 1.3e-08 | 2.8e-08 | 4.4e-08 | 2.833E-08 | 1.550E-08 | 1.300E-08 | 4.400E-08 |
| 29313 | 1466 | S1466M | E | 2.2e-08 | 3.5e-08 | 5.2e-08 | 3.633E-08 | 1.504E-08 | 2.200E-08 | 5.200E-08 |
| 29314 | 1466 | S1466F | E | 6.1e-08 | 1.8e-08 | 1.3e-08 | 3.067E-08 | 2.639E-08 | 1.300E-08 | 6.100E-08 |
| 29315 | 1466 | S1466P | E | 5.9e-08 | 2.2e-08 | 4.2e-08 | 4.100E-08 | 1.852E-08 | 2.200E-08 | 5.900E-08 |
| 29316 | 1466 | S1466S | E | 3e-08 | 5.3e-08 | 5.1e-08 | 4.467E-08 | 1.274E-08 | 3.000E-08 | 5.300E-08 |
| 29317 | 1466 | S1466T | E | 7.3e-08 | 7.6e-08 | 3e-08 | 5.967E-08 | 2.574E-08 | 3.000E-08 | 7.600E-08 |
| 29318 | 1466 | S1466W | E | 2.4e-08 | 4.9e-08 | 6.3e-08 | 4.533E-08 | 1.976E-08 | 2.400E-08 | 6.300E-08 |
| 29319 | 1466 | S1466Y | E | 5.1e-08 | 4.3e-08 | 8.2e-08 | 5.867E-08 | 2.060E-08 | 4.300E-08 | 8.200E-08 |
| 29320 | 1466 | S1466V | E | 1.4e-07 | 3.1e-08 | 2.8e-08 | 6.633E-08 | 6.381E-08 | 2.800E-08 | 1.400E-07 |
| 29321 | 1467 | S1467A | E | 7.4e-08 | 6.2e-08 | 1.5e-08 | 5.033E-08 | 3.118E-08 | 1.500E-08 | 7.400E-08 |
| 29322 | 1467 | S1467R | E | 3.4e-08 | 3.6e-08 | 5.2e-08 | 4.067E-08 | 9.866E-09 | 3.400E-08 | 5.200E-08 |
| 29323 | 1467 | S1467N | E | 6.1e-08 | 8.8e-08 | 2.1e-07 | 1.197E-07 | 7.939E-08 | 6.100E-08 | 2.100E-07 |
| 29324 | 1467 | S1467D | E | 4.7e-08 | 4.3e-08 | 6.5e-08 | 5.167E-08 | 1.172E-08 | 4.300E-08 | 6.500E-08 |

| | | | | | | | | | | |
|---|---|---|---|---|---|---|---|---|---|---|
| 29325 | 1467 | S1467C | E | 4.3e-08 | 1e-07 | 6e-08 | 6.767E-08 | 2.926E-08 | 4.300E-08 | 1.000E-07 |
| 29326 | 1467 | S1467E | E | 5.2e-08 | 2.3e-08 | 3.2e-08 | 3.567E-08 | 1.484E-08 | 2.300E-08 | 5.200E-08 |
| 29327 | 1467 | S1467Q | E | 5.9e-08 | 4.4e-08 | 3.5e-08 | 4.600E-08 | 1.212E-08 | 3.500E-08 | 5.900E-08 |
| 29328 | 1467 | S1467G | E | 3e-08 | 4.5e-08 | 1.9e-08 | 3.133E-08 | 1.305E-08 | 1.900E-08 | 4.500E-08 |
| 29329 | 1467 | S1467H | E | 5.3e-08 | 2.7e-08 | 9.2e-09 | 2.973E-08 | 2.203E-08 | 9.200E-09 | 5.300E-08 |
| 29330 | 1467 | S1467I | E | 3e-08 | 6e-08 | 6.7e-08 | 5.233E-08 | 1.966E-08 | 3.000E-08 | 6.700E-08 |
| 29331 | 1467 | S1467L | E | 3.6e-08 | 1.4e-08 | 2.5e-08 | 2.500E-08 | 1.100E-08 | 1.400E-08 | 3.600E-08 |
| 29332 | 1467 | S1467K | E | 3.8e-08 | 5.5e-08 | 4.9e-08 | 4.733E-08 | 8.622E-09 | 3.800E-08 | 5.500E-08 |
| 29333 | 1467 | S1467M | E | 2e-08 | 1.7e-08 | 7.1e-08 | 3.600E-08 | 3.035E-08 | 1.700E-08 | 7.100E-08 |
| 29334 | 1467 | S1467F | E | 1.1e-07 | 2.4e-08 | 4.4e-08 | 5.933E-08 | 4.500E-08 | 2.400E-08 | 1.100E-07 |
| 29335 | 1467 | S1467P | E | 1.8e-08 | 5.4e-08 | 4.5e-08 | 3.900E-08 | 1.873E-08 | 1.800E-08 | 5.400E-08 |
| 29336 | 1467 | S1467S | E | 7.7e-08 | 4e-08 | 2.1e-08 | 4.600E-08 | 2.848E-08 | 2.100E-08 | 7.700E-08 |
| 29337 | 1467 | S1467T | E | 1.7e-08 | 7.3e-08 | 4.5e-08 | 4.500E-08 | 2.800E-08 | 1.700E-08 | 7.300E-08 |
| 29338 | 1467 | S1467W | E | 3e-08 | 5.5e-08 | 6.7e-08 | 5.067E-08 | 1.888E-08 | 3.000E-08 | 6.700E-08 |
| 29339 | 1467 | S1467Y | E | 7.8e-08 | 1.3e-07 | 7.6e-08 | 9.467E-08 | 3.062E-08 | 7.600E-08 | 1.300E-07 |
| 29340 | 1467 | S1467V | E | 4.7e-08 | 4.3e-08 | 6.7e-08 | 5.233E-08 | 1.286E-08 | 4.300E-08 | 6.700E-08 |
| 29341 | 1468 | L1468A | E | 1e-08 | 3.5e-08 | 8.5e-09 | 1.783E-08 | 1.489E-08 | 8.500E-09 | 3.500E-08 |
| 29342 | 1468 | L1468R | E | 3.2e-08 | 5.1e-08 | 8.3e-08 | 5.533E-08 | 2.577E-08 | 3.200E-08 | 8.300E-08 |
| 29343 | 1468 | L1468N | E | 6.1e-08 | 8e-08 | 8.2e-08 | 7.433E-08 | 1.159E-08 | 6.100E-08 | 8.200E-08 |
| 29344 | 1468 | L1468D | E | 5.4e-08 | 6.3e-08 | 4e-08 | 5.233E-08 | 1.159E-08 | 4.000E-08 | 6.300E-08 |
| 29345 | 1468 | L1468C | E | 2.3e-08 | 7.4e-08 | 2.6e-08 | 4.100E-08 | 2.862E-08 | 2.300E-08 | 7.400E-08 |
| 29346 | 1468 | L1468E | E | 2.7e-08 | 6.7e-08 | 6.4e-08 | 5.267E-08 | 2.228E-08 | 2.700E-08 | 6.700E-08 |
| 29347 | 1468 | L1468Q | E | 3.7e-08 | 2e-08 | 1e-07 | 5.233E-08 | 4.215E-08 | 2.000E-08 | 1.000E-07 |
| 29348 | 1468 | L1468G | E | 5.3e-08 | 1.9e-08 | 6.2e-08 | 4.467E-08 | 2.268E-08 | 1.900E-08 | 6.200E-08 |
| 29349 | 1468 | L1468H | E | 1.3e-08 | 2.4e-08 | 2.4e-08 | 2.033E-08 | 6.351E-09 | 1.300E-08 | 2.400E-08 |
| 29350 | 1468 | L1468I | E | 2.6e-08 | 1.6e-08 | 4.5e-08 | 2.900E-08 | 1.473E-08 | 1.600E-08 | 4.500E-08 |
| 29351 | 1468 | L1468L | E | 3.6e-08 | 7e-09 | 1.6e-07 | 6.767E-08 | 8.127E-08 | 7.000E-09 | 1.600E-07 |
| 29352 | 1468 | L1468K | E | 9.7e-09 | 1.4e-08 | 7e-08 | 3.123E-08 | 3.364E-08 | 9.700E-09 | 7.000E-08 |
| 29353 | 1468 | L1468M | E | 2.9e-08 | 6.3e-08 | 7.9e-08 | 5.700E-08 | 2.553E-08 | 2.900E-08 | 7.900E-08 |
| 29354 | 1468 | L1468F | E | 3.2e-08 | 3.3e-08 | 3.8e-08 | 3.433E-08 | 3.215E-09 | 3.200E-08 | 3.800E-08 |
| 29355 | 1468 | L1468P | E | 5.6e-08 | 7.2e-08 | 4.7e-08 | 5.833E-08 | 1.266E-08 | 4.700E-08 | 7.200E-08 |
| 29356 | 1468 | L1468S | E | 4.4e-08 | 4.2e-08 | 3e-08 | 3.867E-08 | 7.572E-09 | 3.000E-08 | 4.400E-08 |
| 29357 | 1468 | L1468T | E | 2.6e-08 | 4.7e-08 | 5.6e-08 | 4.300E-08 | 1.539E-08 | 2.600E-08 | 5.600E-08 |
| 29358 | 1468 | L1468W | E | 3.9e-08 | 7.4e-08 | 4e-08 | 5.100E-08 | 1.992E-08 | 3.900E-08 | 7.400E-08 |
| 29359 | 1468 | L1468Y | E | 2.2e-08 | 3.1e-08 | 1.5e-08 | 2.267E-08 | 8.021E-09 | 1.500E-08 | 3.100E-08 |
| 29360 | 1468 | L1468V | E | 5.7e-08 | 3.2e-08 | 7.4e-08 | 5.433E-08 | 2.113E-08 | 3.200E-08 | 7.400E-08 |
| 29361 | 1469 | G1469A | E | 2.6e-08 | 5.3e-08 | 1.2e-08 | 3.033E-08 | 2.084E-08 | 1.200E-08 | 5.300E-08 |
| 29362 | 1469 | G1469R | E | 3.2e-08 | 7.3e-08 | 4.1e-08 | 4.867E-08 | 2.155E-08 | 3.200E-08 | 7.300E-08 |
| 29363 | 1469 | G1469N | E | 1.1e-07 | 8.9e-08 | 3.1e-08 | 7.667E-08 | 4.092E-08 | 3.100E-08 | 1.100E-07 |
| 29364 | 1469 | G1469D | E | 6.2e-08 | 2.4e-08 | 1.5e-07 | 7.867E-08 | 6.463E-08 | 2.400E-08 | 1.500E-07 |
| 29365 | 1469 | G1469C | E | 6.8e-08 | 2.5e-08 | 5.7e-08 | 5.000E-08 | 2.234E-08 | 2.500E-08 | 6.800E-08 |
| 29366 | 1469 | G1469E | E | 6.9e-08 | 3.5e-08 | 3.9e-08 | 4.767E-08 | 1.858E-08 | 3.500E-08 | 6.900E-08 |
| 29367 | 1469 | G1469Q | E | 4e-08 | 9.9e-08 | 9.9e-09 | 4.963E-08 | 4.532E-08 | 9.900E-09 | 9.900E-08 |
| 29368 | 1469 | G1469G | E | 5.4e-08 | 2.4e-08 | 4.5e-08 | 4.100E-08 | 1.539E-08 | 2.400E-08 | 5.400E-08 |
| 29369 | 1469 | G1469H | E | 2.1e-08 | 4.6e-08 | 3.8e-08 | 3.500E-08 | 1.277E-08 | 2.100E-08 | 4.600E-08 |
| 29370 | 1469 | G1469I | E | 8.4e-08 | 2.9e-08 | 1.2e-07 | 7.767E-08 | 4.583E-08 | 2.900E-08 | 1.200E-07 |
| 29371 | 1469 | G1469L | E | 8.4e-08 | 2.7e-08 | 3.1e-08 | 4.733E-08 | 3.182E-08 | 2.700E-08 | 8.400E-08 |
| 29372 | 1469 | G1469K | E | 1.8e-08 | 6e-08 | 2.1e-08 | 3.300E-08 | 2.343E-08 | 1.800E-08 | 6.000E-08 |
| 29373 | 1469 | G1469M | E | 6.2e-08 | 1.7e-08 | 7.9e-08 | 5.267E-08 | 3.204E-08 | 1.700E-08 | 7.900E-08 |
| 29374 | 1469 | G1469F | E | 1.8e-08 | 1.7e-08 | 8.1e-08 | 3.867E-08 | 3.667E-08 | 1.700E-08 | 8.100E-08 |
| 29375 | 1469 | G1469P | E | 5.2e-08 | 5.5e-08 | 3.1e-08 | 4.600E-08 | 1.308E-08 | 3.100E-08 | 5.500E-08 |
| 29376 | 1469 | G1469S | E | 2.2e-08 | 4.7e-08 | 4.3e-08 | 3.733E-08 | 1.343E-08 | 2.200E-08 | 4.700E-08 |
| 29377 | 1469 | G1469T | E | 3.5e-08 | 5.3e-08 | 3.3e-08 | 4.033E-08 | 1.102E-08 | 3.300E-08 | 5.300E-08 |
| 29378 | 1469 | G1469W | E | 9.2e-09 | 1.5e-08 | 4e-08 | 2.140E-08 | 1.637E-08 | 9.200E-09 | 4.000E-08 |
| 29379 | 1469 | G1469Y | E | 1.4e-08 | 5.1e-08 | 3.4e-08 | 3.300E-08 | 1.852E-08 | 1.400E-08 | 5.100E-08 |
| 29380 | 1469 | G1469V | E | 4.1e-08 | 2.9e-08 | 4.3e-08 | 3.767E-08 | 7.572E-09 | 2.900E-08 | 4.300E-08 |
| 29381 | 1470 | T1470A | E | 3.7e-08 | 3.9e-08 | 4.4e-08 | 4.000E-08 | 3.606E-09 | 3.700E-08 | 4.400E-08 |
| 29382 | 1470 | T1470R | E | 2.6e-08 | 8.1e-08 | 2e-08 | 4.233E-08 | 3.362E-08 | 2.000E-08 | 8.100E-08 |
| 29383 | 1470 | T1470N | E | 1.7e-08 | 1e-07 | 2.9e-08 | 4.867E-08 | 4.486E-08 | 1.700E-08 | 1.000E-07 |
| 29384 | 1470 | T1470D | E | 4.3e-08 | 1.2e-07 | 4.1e-08 | 6.800E-08 | 4.504E-08 | 4.100E-08 | 1.200E-07 |
| 29385 | 1470 | T1470C | E | 5e-08 | 2.5e-08 | 6.1e-08 | 4.533E-08 | 1.845E-08 | 2.500E-08 | 6.100E-08 |
| 29386 | 1470 | T1470E | E | 5e-08 | 4.2e-08 | 5.2e-08 | 4.800E-08 | 5.292E-09 | 4.200E-08 | 5.200E-08 |
| 29387 | 1470 | T1470Q | E | 6.9e-08 | 4.3e-08 | 6.6e-08 | 5.933E-08 | 1.422E-08 | 4.300E-08 | 6.900E-08 |
| 29388 | 1470 | T1470G | E | 8.6e-08 | 8.7e-08 | 4e-08 | 7.100E-08 | 2.685E-08 | 4.000E-08 | 8.700E-08 |
| 29389 | 1470 | T1470H | E | 4.4e-08 | 3e-08 | 2.5e-08 | 3.300E-08 | 9.849E-09 | 2.500E-08 | 4.400E-08 |
| 29390 | 1470 | T1470I | E | 2.7e-08 | 3.9e-08 | 2.4e-08 | 3.000E-08 | 7.937E-09 | 2.400E-08 | 3.900E-08 |
| 29391 | 1470 | T1470L | E | 4.7e-08 | 2.2e-08 | 4.8e-08 | 3.900E-08 | 1.473E-08 | 2.200E-08 | 4.800E-08 |
| 29392 | 1470 | T1470K | E | 5e-08 | 4.2e-08 | 3.5e-08 | 4.233E-08 | 7.506E-09 | 3.500E-08 | 5.000E-08 |
| 29393 | 1470 | T1470M | E | 3.2e-08 | 5.1e-08 | 9e-08 | 5.767E-08 | 2.957E-08 | 3.200E-08 | 9.000E-08 |
| 29394 | 1470 | T1470F | E | 4e-08 | 3.9e-08 | 4.6e-08 | 4.167E-08 | 3.786E-09 | 3.900E-08 | 4.600E-08 |
| 29395 | 1470 | T1470P | E | 4.4e-08 | 2.5e-08 | 4.9e-08 | 3.933E-08 | 1.266E-08 | 2.500E-08 | 4.900E-08 |
| 29396 | 1470 | T1470S | E | 9.3e-08 | 4.9e-08 | 5.1e-08 | 6.433E-08 | 2.485E-08 | 4.900E-08 | 9.300E-08 |
| 29397 | 1470 | T1470T | E | 1.4e-08 | 6.1e-08 | 5.9e-08 | 4.467E-08 | 2.658E-08 | 1.400E-08 | 6.100E-08 |
| 29398 | 1470 | T1470W | E | 3.8e-08 | 1.9e-08 | 4.4e-08 | 3.367E-08 | 1.305E-08 | 1.900E-08 | 4.400E-08 |
| 29399 | 1470 | T1470Y | E | 1.6e-08 | 3.1e-08 | 8e-08 | 4.233E-08 | 3.347E-08 | 1.600E-08 | 8.000E-08 |
| 29400 | 1470 | T1470V | E | 5.7e-08 | 4.9e-08 | 2.6e-08 | 4.400E-08 | 1.609E-08 | 2.600E-08 | 5.700E-08 |
| 29401 | 1471 | Q1471A | E | 3e-08 | 2.3e-08 | 7.8e-08 | 4.367E-08 | 2.994E-08 | 2.300E-08 | 7.800E-08 |
| 29402 | 1471 | Q1471R | E | 5.6e-08 | 3e-08 | 1.8e-07 | 8.867E-08 | 8.016E-08 | 3.000E-08 | 1.800E-07 |
| 29403 | 1471 | Q1471N | E | 8.4e-08 | 3.3e-08 | 4.4e-08 | 5.367E-08 | 2.684E-08 | 3.300E-08 | 8.400E-08 |
| 29404 | 1471 | Q1471D | E | 4.2e-08 | 5.4e-08 | 7.5e-08 | 5.700E-08 | 1.670E-08 | 4.200E-08 | 7.500E-08 |
| 29405 | 1471 | Q1471C | E | 5.9e-08 | 4.3e-08 | 3.8e-08 | 4.667E-08 | 1.097E-08 | 3.800E-08 | 5.900E-08 |
| 29406 | 1471 | Q1471E | E | 4e-08 | 9e-08 | 5.7e-08 | 6.233E-08 | 2.542E-08 | 4.000E-08 | 9.000E-08 |
| 29407 | 1471 | Q1471Q | E | 5.5e-08 | 3.8e-08 | 2.9e-08 | 4.067E-08 | 1.320E-08 | 2.900E-08 | 5.500E-08 |
| 29408 | 1471 | Q1471G | E | 2.5e-08 | 9e-08 | 4e-08 | 5.167E-08 | 3.403E-08 | 2.500E-08 | 9.000E-08 |
| 29409 | 1471 | Q1471H | E | 6.3e-08 | 6.8e-08 | 5.4e-08 | 6.167E-08 | 7.095E-09 | 5.400E-08 | 6.800E-08 |

| | | | | | | | | | | |
|---|---|---|---|---|---|---|---|---|---|---|
| 29410 | 1471 | Q1471I | E | 4.7e-08 | 2.2e-08 | 7.2e-08 | 4.700E-08 | 2.500E-08 | 2.200E-08 | 7.200E-08 |
| 29411 | 1471 | Q1471L | E | 1.8e-08 | 4.1e-08 | 6.5e-08 | 4.133E-08 | 2.350E-08 | 1.800E-08 | 6.500E-08 |
| 29412 | 1471 | Q1471K | E | 3.1e-08 | 2e-08 | 1.4e-08 | 2.167E-08 | 8.622E-09 | 1.400E-08 | 3.100E-08 |
| 29413 | 1471 | Q1471M | E | 6.9e-08 | 2.7e-08 | 1.9e-08 | 3.833E-08 | 2.686E-08 | 1.900E-08 | 6.900E-08 |
| 29414 | 1471 | Q1471F | E | 4e-08 | 1.2e-07 | 1.8e-08 | 5.933E-08 | 5.368E-08 | 1.800E-08 | 1.200E-07 |
| 29415 | 1471 | Q1471P | E | 1.2e-07 | 4e-08 | 5.2e-08 | 7.067E-08 | 4.314E-08 | 4.000E-08 | 1.200E-07 |
| 29416 | 1471 | Q1471S | E | 2.2e-08 | 7.9e-08 | 7.2e-08 | 5.767E-08 | 3.109E-08 | 2.200E-08 | 7.900E-08 |
| 29417 | 1471 | Q1471T | E | 2.5e-08 | 7.9e-08 | 5e-08 | 5.133E-08 | 2.702E-08 | 2.500E-08 | 7.900E-08 |
| 29418 | 1471 | Q1471W | E | 9.7e-08 | 6.6e-08 | 7e-08 | 7.767E-08 | 1.686E-08 | 6.600E-08 | 9.700E-08 |
| 29419 | 1471 | Q1471Y | E | 4.4e-08 | 1.6e-08 | 7.1e-08 | 4.367E-08 | 2.750E-08 | 1.600E-08 | 7.100E-08 |
| 29420 | 1471 | Q1471V | E | 2.5e-08 | 6.1e-08 | 1.5e-07 | 7.867E-08 | 6.435E-08 | 2.500E-08 | 1.500E-07 |
| 29421 | 1472 | T1472A | E | 1.3e-07 | 2.9e-08 | 1.8e-07 | 1.130E-07 | 7.692E-08 | 2.900E-08 | 1.800E-07 |
| 29422 | 1472 | T1472R | E | 1.9e-08 | 6e-08 | 2.5e-08 | 3.467E-08 | 2.214E-08 | 1.900E-08 | 6.000E-08 |
| 29423 | 1472 | T1472N | E | 1.8e-08 | 5.8e-08 | 5.4e-08 | 4.333E-08 | 2.203E-08 | 1.800E-08 | 5.800E-08 |
| 29424 | 1472 | T1472D | E | 4.8e-08 | 7e-08 | 2.2e-08 | 4.667E-08 | 2.403E-08 | 2.200E-08 | 7.000E-08 |
| 29425 | 1472 | T1472C | E | 5.5e-08 | 1.2e-07 | 2.4e-08 | 6.633E-08 | 4.899E-08 | 2.400E-08 | 1.200E-07 |
| 29426 | 1472 | T1472E | E | 2.6e-08 | 5.7e-08 | 4e-08 | 4.100E-08 | 1.552E-08 | 2.600E-08 | 5.700E-08 |
| 29427 | 1472 | T1472Q | E | 1.5e-08 | 1.4e-07 | 5.2e-08 | 6.900E-08 | 6.421E-08 | 1.500E-08 | 1.400E-07 |
| 29428 | 1472 | T1472G | E | 3.8e-08 | 5e-08 | 2.7e-08 | 3.833E-08 | 1.150E-08 | 2.700E-08 | 5.000E-08 |
| 29429 | 1472 | T1472H | E | 2.3e-08 | 3.4e-08 | 4.3e-08 | 3.333E-08 | 1.002E-08 | 2.300E-08 | 4.300E-08 |
| 29430 | 1472 | T1472I | E | 9e-09 | 1.1e-08 | 1e-07 | 4.000E-08 | 5.197E-08 | 9.000E-09 | 1.000E-07 |
| 29431 | 1472 | T1472L | E | 4.7e-08 | 2.5e-08 | 2.5e-08 | 3.233E-08 | 1.270E-08 | 2.500E-08 | 4.700E-08 |
| 29432 | 1472 | T1472K | E | 3.6e-08 | 9.5e-08 | 7.3e-08 | 6.800E-08 | 2.982E-08 | 3.600E-08 | 9.500E-08 |
| 29433 | 1472 | T1472M | E | 8.8e-08 | 4.8e-08 | 5e-08 | 6.200E-08 | 2.254E-08 | 4.800E-08 | 8.800E-08 |
| 29434 | 1472 | T1472F | E | 1.2e-08 | 2.7e-08 | 3.1e-08 | 2.333E-08 | 1.002E-08 | 1.200E-08 | 3.100E-08 |
| 29435 | 1472 | T1472P | E | 1.1e-07 | 1e-07 | 7.2e-08 | 9.400E-08 | 1.970E-08 | 7.200E-08 | 1.100E-07 |
| 29436 | 1472 | T1472S | E | 1.6e-08 | 4e-08 | 6e-08 | 3.867E-08 | 2.203E-08 | 1.600E-08 | 6.000E-08 |
| 29437 | 1472 | T1472T | E | 6.4e-08 | 3.8e-08 | 8.7e-08 | 6.300E-08 | 2.452E-08 | 3.800E-08 | 8.700E-08 |
| 29438 | 1472 | T1472W | E | 2e-07 | 1.5e-08 | 4.7e-08 | 8.733E-08 | 9.888E-08 | 1.500E-08 | 2.000E-07 |
| 29439 | 1472 | T1472Y | E | 7.9e-08 | 3.6e-08 | 1.3e-07 | 8.167E-08 | 4.706E-08 | 3.600E-08 | 1.300E-07 |
| 29440 | 1472 | T1472V | E | 1.5e-07 | 2.9e-08 | 1.3e-08 | 6.400E-08 | 7.491E-08 | 1.300E-08 | 1.500E-07 |
| 29441 | 1473 | Y1473A | E | 3.9e-08 | 1e-07 | 3.3e-08 | 5.733E-08 | 3.707E-08 | 3.300E-08 | 1.000E-07 |
| 29442 | 1473 | Y1473R | E | 6.4e-08 | 4e-08 | 2.7e-08 | 4.367E-08 | 1.877E-08 | 2.700E-08 | 6.400E-08 |
| 29443 | 1473 | Y1473N | E | 5.4e-08 | 5.5e-08 | 8.9e-08 | 6.600E-08 | 1.992E-08 | 5.400E-08 | 8.900E-08 |
| 29444 | 1473 | Y1473D | E | 4.7e-08 | 3.1e-08 | 4.9e-08 | 4.233E-08 | 9.866E-09 | 3.100E-08 | 4.900E-08 |
| 29445 | 1473 | Y1473C | E | 3.4e-08 | 4.2e-08 | 7.7e-08 | 5.100E-08 | 2.287E-08 | 3.400E-08 | 7.700E-08 |
| 29446 | 1473 | Y1473E | E | 7e-08 | 4.7e-08 | 1.3e-07 | 8.233E-08 | 4.285E-08 | 4.700E-08 | 1.300E-07 |
| 29447 | 1473 | Y1473Q | E | 8.3e-08 | 1.3e-08 | 5.2e-08 | 4.933E-08 | 3.508E-08 | 1.300E-08 | 8.300E-08 |
| 29448 | 1473 | Y1473G | E | 4e-08 | 3.2e-08 | 5.5e-08 | 4.233E-08 | 1.168E-08 | 3.200E-08 | 5.500E-08 |
| 29449 | 1473 | Y1473H | E | 3.9e-08 | 9.7e-08 | 3.3e-08 | 5.633E-08 | 3.535E-08 | 3.300E-08 | 9.700E-08 |
| 29450 | 1473 | Y1473I | E | 8e-08 | 3.4e-08 | 5.6e-08 | 5.667E-08 | 2.301E-08 | 3.400E-08 | 8.000E-08 |
| 29451 | 1473 | Y1473L | E | 6.7e-08 | 2.7e-08 | 7.6e-08 | 5.667E-08 | 2.608E-08 | 2.700E-08 | 7.600E-08 |
| 29452 | 1473 | Y1473K | E | 5.2e-08 | 9.6e-08 | 5.4e-08 | 6.733E-08 | 2.485E-08 | 5.200E-08 | 9.600E-08 |
| 29453 | 1473 | Y1473M | E | 3.7e-08 | 1.3e-07 | 7.8e-08 | 8.167E-08 | 4.661E-08 | 3.700E-08 | 1.300E-07 |
| 29454 | 1473 | Y1473F | E | 1e-07 | 4.2e-08 | 2e-08 | 5.400E-08 | 4.133E-08 | 2.000E-08 | 1.000E-07 |
| 29455 | 1473 | Y1473P | E | 2.3e-08 | 2.6e-08 | 1.2e-07 | 5.633E-08 | 5.516E-08 | 2.300E-08 | 1.200E-07 |
| 29456 | 1473 | Y1473S | E | 3.5e-08 | 1e-07 | 4.4e-08 | 5.967E-08 | 3.522E-08 | 3.500E-08 | 1.000E-07 |
| 29457 | 1473 | Y1473T | E | 2.6e-08 | 4.2e-08 | 5.7e-08 | 4.167E-08 | 1.550E-08 | 2.600E-08 | 5.700E-08 |
| 29458 | 1473 | Y1473W | E | 3.6e-08 | 3.5e-08 | 8.2e-08 | 5.100E-08 | 2.685E-08 | 3.500E-08 | 8.200E-08 |
| 29459 | 1473 | Y1473Y | E | 5.3e-08 | 1.8e-08 | 9.3e-08 | 5.467E-08 | 3.753E-08 | 1.800E-08 | 9.300E-08 |
| 29460 | 1473 | Y1473V | E | 1.8e-08 | 4.7e-08 | 4.7e-08 | 3.733E-08 | 1.674E-08 | 1.800E-08 | 4.700E-08 |
| 29461 | 1474 | I1474A | E | 3.6e-08 | 1.9e-08 | 3.3e-08 | 2.933E-08 | 9.074E-09 | 1.900E-08 | 3.600E-08 |
| 29462 | 1474 | I1474R | E | 4e-08 | 4.2e-08 | 9.1e-08 | 5.767E-08 | 2.888E-08 | 4.000E-08 | 9.100E-08 |
| 29463 | 1474 | I1474N | E | 3e-08 | 8.1e-08 | 3e-08 | 4.700E-08 | 2.944E-08 | 3.000E-08 | 8.100E-08 |
| 29464 | 1474 | I1474D | E | 6.9e-08 | 5.2e-08 | 3.1e-08 | 5.067E-08 | 1.904E-08 | 3.100E-08 | 6.900E-08 |
| 29465 | 1474 | I1474C | E | 5.5e-08 | 5.8e-08 | 5.8e-08 | 5.700E-08 | 1.732E-09 | 5.500E-08 | 5.800E-08 |
| 29466 | 1474 | I1474E | E | 1.9e-08 | 2.1e-08 | 2.7e-08 | 2.233E-08 | 4.163E-09 | 1.900E-08 | 2.700E-08 |
| 29467 | 1474 | I1474Q | E | 1.3e-08 | 2.2e-07 | 2.5e-08 | 8.600E-08 | 1.162E-07 | 1.300E-08 | 2.200E-07 |
| 29468 | 1474 | I1474G | E | 7.1e-08 | 4.7e-08 | 2.1e-08 | 4.633E-08 | 2.501E-08 | 2.100E-08 | 7.100E-08 |
| 29469 | 1474 | I1474H | E | 1.3e-08 | 9.8e-08 | 3.6e-08 | 4.900E-08 | 4.397E-08 | 1.300E-08 | 9.800E-08 |
| 29470 | 1474 | I1474I | E | 2.8e-08 | 2.7e-08 | 8.6e-08 | 4.700E-08 | 3.378E-08 | 2.700E-08 | 8.600E-08 |
| 29471 | 1474 | I1474L | E | 8.7e-08 | 3.3e-08 | 6.4e-08 | 6.133E-08 | 2.710E-08 | 3.300E-08 | 8.700E-08 |
| 29472 | 1474 | I1474K | E | 2.2e-08 | 1.4e-07 | 5.3e-08 | 7.167E-08 | 6.117E-08 | 2.200E-08 | 1.400E-07 |
| 29473 | 1474 | I1474M | E | 2.4e-08 | 3.5e-08 | 1.5e-08 | 2.467E-08 | 1.002E-08 | 1.500E-08 | 3.500E-08 |
| 29474 | 1474 | I1474F | E | 4.1e-08 | 5.1e-08 | 3.2e-08 | 4.133E-08 | 9.504E-09 | 3.200E-08 | 5.100E-08 |
| 29475 | 1474 | I1474P | E | 1.2e-07 | 2.2e-08 | 2.4e-08 | 5.533E-08 | 5.601E-08 | 2.200E-08 | 1.200E-07 |
| 29476 | 1474 | I1474S | E | 1.9e-08 | 1.1e-07 | 4e-08 | 5.633E-08 | 4.765E-08 | 1.900E-08 | 1.100E-07 |
| 29477 | 1474 | I1474T | E | 5.6e-08 | 2.7e-08 | 2.9e-08 | 3.733E-08 | 1.620E-08 | 2.700E-08 | 5.600E-08 |
| 29478 | 1474 | I1474W | E | 9.1e-08 | 3.5e-08 | 7.9e-08 | 6.833E-08 | 2.948E-08 | 3.500E-08 | 9.100E-08 |
| 29479 | 1474 | I1474Y | E | 4.9e-08 | 3.5e-08 | 4.8e-08 | 4.400E-08 | 7.810E-09 | 3.500E-08 | 4.900E-08 |
| 29480 | 1474 | I1474V | E | 3.9e-08 | 1.6e-08 | 5.7e-08 | 3.733E-08 | 2.055E-08 | 1.600E-08 | 5.700E-08 |
| 29481 | 1475 | C1475A | E | 4e-08 | 7.2e-08 | 3.4e-08 | 4.867E-08 | 2.043E-08 | 3.400E-08 | 7.200E-08 |
| 29482 | 1475 | C1475R | E | 1.9e-08 | 4.6e-08 | 1.3e-07 | 6.500E-08 | 5.789E-08 | 1.900E-08 | 1.300E-07 |
| 29483 | 1475 | C1475N | E | 4.1e-08 | 4.7e-08 | 3.4e-08 | 4.067E-08 | 6.506E-09 | 3.400E-08 | 4.700E-08 |
| 29484 | 1475 | C1475D | E | 3.2e-08 | 8.1e-08 | 1.4e-08 | 4.233E-08 | 3.467E-08 | 1.400E-08 | 8.100E-08 |
| 29485 | 1475 | C1475C | E | 8.9e-08 | 2.5e-08 | 6.4e-08 | 5.933E-08 | 3.225E-08 | 2.500E-08 | 8.900E-08 |
| 29486 | 1475 | C1475E | E | 9.3e-08 | 2.5e-08 | 4.7e-08 | 5.500E-08 | 3.470E-08 | 2.500E-08 | 9.300E-08 |
| 29487 | 1475 | C1475Q | E | 6.5e-08 | 6e-08 | 1.8e-08 | 4.767E-08 | 2.581E-08 | 1.800E-08 | 6.500E-08 |
| 29488 | 1475 | C1475G | E | 3.9e-08 | 4.9e-08 | 3.2e-08 | 4.000E-08 | 8.544E-09 | 3.200E-08 | 4.900E-08 |
| 29489 | 1475 | C1475H | E | 7.9e-09 | 3.6e-08 | 7.6e-08 | 3.997E-08 | 3.422E-08 | 7.900E-09 | 7.600E-08 |
| 29490 | 1475 | C1475I | E | 3.5e-08 | 2e-08 | 3.7e-08 | 3.067E-08 | 9.292E-09 | 2.000E-08 | 3.700E-08 |
| 29491 | 1475 | C1475L | E | 1.6e-08 | 6.2e-08 | 3e-08 | 3.600E-08 | 2.358E-08 | 1.600E-08 | 6.200E-08 |
| 29492 | 1475 | C1475K | E | 3.2e-08 | 3.1e-08 | 1.5e-08 | 2.600E-08 | 9.539E-09 | 1.500E-08 | 3.200E-08 |
| 29493 | 1475 | C1475M | E | 4.6e-08 | 3e-08 | 2.6e-08 | 3.400E-08 | 1.058E-08 | 2.600E-08 | 4.600E-08 |
| 29494 | 1475 | C1475F | E | 4.1e-08 | 1.5e-08 | 3.7e-08 | 3.100E-08 | 1.400E-08 | 1.500E-08 | 4.100E-08 |

| | | | | | | | | | | |
|---|---|---|---|---|---|---|---|---|---|---|
| 29495 | 1475 | C1475P | E | 7e-08 | 1.4e-08 | 3.4e-08 | 3.933E-08 | 2.838E-08 | 1.400E-08 | 7.000E-08 |
| 29496 | 1475 | C1475S | E | 3.6e-08 | 4.4e-08 | 6.3e-08 | 4.767E-08 | 1.387E-08 | 3.600E-08 | 6.300E-08 |
| 29497 | 1475 | C1475T | E | 8.2e-08 | 5.2e-08 | 6.5e-08 | 6.633E-08 | 1.504E-08 | 5.200E-08 | 8.200E-08 |
| 29498 | 1475 | C1475W | E | 2.1e-08 | 3.7e-08 | 8.4e-08 | 4.733E-08 | 3.275E-08 | 2.100E-08 | 8.400E-08 |
| 29499 | 1475 | C1475Y | E | 3.9e-08 | 9.1e-08 | 7.3e-08 | 6.767E-08 | 2.641E-08 | 3.900E-08 | 9.100E-08 |
| 29500 | 1475 | C1475V | E | 6.3e-09 | 9e-08 | 4.5e-08 | 4.710E-08 | 4.189E-08 | 6.300E-09 | 9.000E-08 |
| 29501 | 1476 | N1476A | E | 4e-08 | 4.8e-08 | 6.9e-08 | 5.233E-08 | 1.498E-08 | 4.000E-08 | 6.900E-08 |
| 29502 | 1476 | N1476R | E | 6.9e-08 | 7.6e-08 | 1.4e-08 | 5.300E-08 | 3.396E-08 | 1.400E-08 | 7.600E-08 |
| 29503 | 1476 | N1476N | E | 3.4e-08 | 2.7e-08 | 2.2e-08 | 2.767E-08 | 6.028E-09 | 2.200E-08 | 3.400E-08 |
| 29504 | 1476 | N1476D | E | 3.4e-08 | 7.8e-08 | 2.3e-08 | 4.500E-08 | 2.910E-08 | 2.300E-08 | 7.800E-08 |
| 29505 | 1476 | N1476C | E | 3.1e-08 | 2.7e-08 | 5.5e-08 | 3.767E-08 | 1.514E-08 | 2.700E-08 | 5.500E-08 |
| 29506 | 1476 | N1476E | E | 6.7e-08 | 1.4e-07 | 7.1e-08 | 9.267E-08 | 4.104E-08 | 6.700E-08 | 1.400E-07 |
| 29507 | 1476 | N1476Q | E | 7.4e-08 | 1.5e-08 | 4.6e-08 | 4.500E-08 | 2.951E-08 | 1.500E-08 | 7.400E-08 |
| 29508 | 1476 | N1476G | E | 4.9e-08 | 4.3e-08 | 4.4e-08 | 4.533E-08 | 3.215E-09 | 4.300E-08 | 4.900E-08 |
| 29509 | 1476 | N1476H | E | 5.1e-08 | 3.7e-08 | 5.5e-08 | 4.767E-08 | 9.452E-09 | 3.700E-08 | 5.500E-08 |
| 29510 | 1476 | N1476I | E | 6.1e-08 | 5.4e-08 | 2e-08 | 4.500E-08 | 2.193E-08 | 2.000E-08 | 6.100E-08 |
| 29511 | 1476 | N1476L | E | 7.5e-08 | 1.5e-08 | 5.3e-08 | 4.767E-08 | 3.035E-08 | 1.500E-08 | 7.500E-08 |
| 29512 | 1476 | N1476K | E | 6.6e-08 | 2.5e-08 | 7.1e-08 | 5.400E-08 | 2.524E-08 | 2.500E-08 | 7.100E-08 |
| 29513 | 1476 | N1476M | E | 2.1e-08 | 9.1e-08 | 6.8e-08 | 6.000E-08 | 3.568E-08 | 2.100E-08 | 9.100E-08 |
| 29514 | 1476 | N1476F | E | 4.4e-08 | 4.5e-08 | 5.6e-08 | 4.833E-08 | 6.658E-09 | 4.400E-08 | 5.600E-08 |
| 29515 | 1476 | N1476P | E | 3.2e-08 | 2.7e-08 | 4.8e-08 | 3.567E-08 | 1.097E-08 | 2.700E-08 | 4.800E-08 |
| 29516 | 1476 | N1476S | E | 1.3e-08 | 8.7e-08 | 3.2e-08 | 4.400E-08 | 3.843E-08 | 1.300E-08 | 8.700E-08 |
| 29517 | 1476 | N1476T | E | 2.9e-08 | 2.6e-08 | 5.2e-08 | 3.567E-08 | 1.422E-08 | 2.600E-08 | 5.200E-08 |
| 29518 | 1476 | N1476W | E | 4.2e-08 | 7.2e-08 | 4.1e-08 | 5.167E-08 | 1.762E-08 | 4.100E-08 | 7.200E-08 |
| 29519 | 1476 | N1476Y | E | 5e-08 | 3.6e-08 | 3.6e-08 | 4.067E-08 | 8.083E-09 | 3.600E-08 | 5.000E-08 |
| 29520 | 1476 | N1476V | E | 4.8e-08 | 2.1e-08 | 1.5e-08 | 2.800E-08 | 1.758E-08 | 1.500E-08 | 4.800E-08 |
| 29521 | 1477 | V1477A | E | 5.5e-08 | 1.8e-08 | 2.3e-08 | 3.200E-08 | 2.007E-08 | 1.800E-08 | 5.500E-08 |
| 29522 | 1477 | V1477R | E | 8.5e-08 | 6.8e-08 | 7.5e-08 | 7.600E-08 | 8.544E-09 | 6.800E-08 | 8.500E-08 |
| 29523 | 1477 | V1477N | E | 7.5e-08 | 5.2e-08 | 3.3e-08 | 5.333E-08 | 2.103E-08 | 3.300E-08 | 7.500E-08 |
| 29524 | 1477 | V1477D | E | 4.5e-08 | 1.7e-08 | 3.8e-08 | 3.333E-08 | 1.457E-08 | 1.700E-08 | 4.500E-08 |
| 29525 | 1477 | V1477C | E | 1.4e-08 | 2.2e-08 | 1.7e-08 | 1.767E-08 | 4.041E-09 | 1.400E-08 | 2.200E-08 |
| 29526 | 1477 | V1477E | E | 1.8e-08 | 3.3e-08 | 2.6e-08 | 2.567E-08 | 7.506E-09 | 1.800E-08 | 3.300E-08 |
| 29527 | 1477 | V1477Q | E | 3.5e-08 | 2e-08 | 2.4e-08 | 2.633E-08 | 7.767E-09 | 2.000E-08 | 3.500E-08 |
| 29528 | 1477 | V1477G | E | 3.5e-08 | 5.7e-08 | 2.3e-08 | 3.833E-08 | 1.724E-08 | 2.300E-08 | 5.700E-08 |
| 29529 | 1477 | V1477H | E | 1.9e-08 | 9.7e-08 | 2.8e-08 | 4.800E-08 | 4.267E-08 | 1.900E-08 | 9.700E-08 |
| 29530 | 1477 | V1477I | E | 5.3e-08 | 2.3e-08 | 8.6e-08 | 5.400E-08 | 3.151E-08 | 2.300E-08 | 8.600E-08 |
| 29531 | 1477 | V1477L | E | 5e-08 | 4.4e-08 | 5.1e-08 | 4.833E-08 | 3.786E-09 | 4.400E-08 | 5.100E-08 |
| 29532 | 1477 | V1477K | E | 4.2e-08 | 4.3e-08 | 9.1e-08 | 5.867E-08 | 2.801E-08 | 4.200E-08 | 9.100E-08 |
| 29533 | 1477 | V1477M | E | 3.9e-08 | 3.1e-08 | 1e-07 | 5.667E-08 | 3.774E-08 | 3.100E-08 | 1.000E-07 |
| 29534 | 1477 | V1477F | E | 8.1e-09 | 8e-08 | 4.5e-08 | 4.437E-08 | 3.595E-08 | 8.100E-09 | 8.000E-08 |
| 29535 | 1477 | V1477P | E | 2e-08 | 8.6e-08 | 3.3e-08 | 4.633E-08 | 3.496E-08 | 2.000E-08 | 8.600E-08 |
| 29536 | 1477 | V1477S | E | 9.8e-08 | 5.5e-08 | 1.2e-08 | 5.500E-08 | 4.300E-08 | 1.200E-08 | 9.800E-08 |
| 29537 | 1477 | V1477T | E | 4.5e-08 | 4.9e-08 | 5.3e-08 | 4.900E-08 | 4.000E-09 | 4.500E-08 | 5.300E-08 |
| 29538 | 1477 | V1477W | E | 3.8e-08 | 1.7e-08 | 7.7e-08 | 4.400E-08 | 3.045E-08 | 1.700E-08 | 7.700E-08 |
| 29539 | 1477 | V1477Y | E | 2.8e-08 | 4.4e-08 | 3.6e-08 | 3.600E-08 | 8.000E-09 | 2.800E-08 | 4.400E-08 |
| 29540 | 1477 | V1477V | E | 3.9e-08 | 5e-08 | 5.7e-08 | 4.867E-08 | 9.074E-09 | 3.900E-08 | 5.700E-08 |
| 29541 | 1478 | N1478A | E | 9.4e-08 | 6.5e-08 | 1.3e-08 | 5.733E-08 | 4.104E-08 | 1.300E-08 | 9.400E-08 |
| 29542 | 1478 | N1478R | E | 5.9e-08 | 3.7e-08 | 1.9e-08 | 3.833E-08 | 2.003E-08 | 1.900E-08 | 5.900E-08 |
| 29543 | 1478 | N1478N | E | 2.8e-08 | 5.3e-08 | 4.2e-08 | 4.100E-08 | 1.253E-08 | 2.800E-08 | 5.300E-08 |
| 29544 | 1478 | N1478D | E | 3.7e-08 | 1.8e-08 | 2.4e-08 | 2.633E-08 | 9.713E-09 | 1.800E-08 | 3.700E-08 |
| 29545 | 1478 | N1478C | E | 5.6e-08 | 4.7e-08 | 4.5e-08 | 4.933E-08 | 5.859E-09 | 4.500E-08 | 5.600E-08 |
| 29546 | 1478 | N1478E | E | 1.5e-08 | 3.6e-08 | 1.8e-08 | 2.300E-08 | 1.136E-08 | 1.500E-08 | 3.600E-08 |
| 29547 | 1478 | N1478Q | E | 3.8e-08 | 2.6e-08 | 6.6e-08 | 4.333E-08 | 2.053E-08 | 2.600E-08 | 6.600E-08 |
| 29548 | 1478 | N1478G | E | 4.8e-08 | 2.9e-08 | 5.8e-08 | 4.500E-08 | 1.473E-08 | 2.900E-08 | 5.800E-08 |
| 29549 | 1478 | N1478H | E | 5e-08 | 3.1e-08 | 2.1e-08 | 3.400E-08 | 1.473E-08 | 2.100E-08 | 5.000E-08 |
| 29550 | 1478 | N1478I | E | 2.8e-08 | 6.2e-08 | 8.2e-08 | 5.733E-08 | 2.730E-08 | 2.800E-08 | 8.200E-08 |
| 29551 | 1478 | N1478L | E | 6.3e-08 | 5.6e-08 | 1.5e-07 | 8.967E-08 | 5.237E-08 | 5.600E-08 | 1.500E-07 |
| 29552 | 1478 | N1478K | E | 9.4e-09 | 3e-08 | 1.2e-07 | 5.313E-08 | 5.882E-08 | 9.400E-09 | 1.200E-07 |
| 29553 | 1478 | N1478M | E | 6.1e-08 | 6.1e-08 | 1.9e-08 | 4.700E-08 | 2.425E-08 | 1.900E-08 | 6.100E-08 |
| 29554 | 1478 | N1478F | E | 6e-08 | 2.3e-08 | 5.9e-08 | 4.733E-08 | 2.108E-08 | 2.300E-08 | 6.000E-08 |
| 29555 | 1478 | N1478P | E | 2.9e-08 | 4.9e-08 | 1.1e-08 | 2.967E-08 | 1.901E-08 | 1.100E-08 | 4.900E-08 |
| 29556 | 1478 | N1478S | E | 4e-08 | 7.1e-08 | 6.2e-08 | 5.767E-08 | 1.595E-08 | 4.000E-08 | 7.100E-08 |
| 29557 | 1478 | N1478T | E | 3.3e-08 | 7.2e-08 | 2.9e-08 | 4.467E-08 | 2.376E-08 | 2.900E-08 | 7.200E-08 |
| 29558 | 1478 | N1478W | E | 4e-08 | 6.3e-08 | 5.2e-08 | 5.167E-08 | 1.150E-08 | 4.000E-08 | 6.300E-08 |
| 29559 | 1478 | N1478Y | E | 1.4e-08 | 1.2e-07 | 6.1e-08 | 6.500E-08 | 5.311E-08 | 1.400E-08 | 1.200E-07 |
| 29560 | 1478 | N1478V | E | 1.1e-07 | 2.1e-08 | 5.5e-08 | 6.200E-08 | 4.491E-08 | 2.100E-08 | 1.100E-07 |
| 29561 | 1479 | H1479A | E | 3.7e-08 | 1e-07 | 1.6e-08 | 5.100E-08 | 4.371E-08 | 1.600E-08 | 1.000E-07 |
| 29562 | 1479 | H1479R | E | 5.2e-08 | 2.9e-08 | 3e-08 | 3.700E-08 | 1.300E-08 | 2.900E-08 | 5.200E-08 |
| 29563 | 1479 | H1479N | E | 4.2e-08 | 1.5e-08 | 5.8e-08 | 3.833E-08 | 2.173E-08 | 1.500E-08 | 5.800E-08 |
| 29564 | 1479 | H1479D | E | 2.6e-08 | 2.5e-08 | 9.7e-08 | 4.933E-08 | 4.128E-08 | 2.500E-08 | 9.700E-08 |
| 29565 | 1479 | H1479C | E | 3e-08 | 4.8e-08 | 8.8e-08 | 5.533E-08 | 2.969E-08 | 3.000E-08 | 8.800E-08 |
| 29566 | 1479 | H1479E | E | 5.1e-08 | 5.3e-08 | 1.9e-08 | 4.100E-08 | 1.908E-08 | 1.900E-08 | 5.300E-08 |
| 29567 | 1479 | H1479Q | E | 5.9e-08 | 4.7e-08 | 7e-08 | 5.867E-08 | 1.150E-08 | 4.700E-08 | 7.000E-08 |
| 29568 | 1479 | H1479G | E | 3.5e-08 | 9.7e-08 | 2.8e-08 | 5.333E-08 | 3.798E-08 | 2.800E-08 | 9.700E-08 |
| 29569 | 1479 | H1479H | E | 4.3e-08 | 3.2e-08 | 2.6e-08 | 3.367E-08 | 8.622E-09 | 2.600E-08 | 4.300E-08 |
| 29570 | 1479 | H1479I | E | 3e-08 | 6.3e-08 | 4.7e-08 | 4.667E-08 | 1.650E-08 | 3.000E-08 | 6.300E-08 |
| 29571 | 1479 | H1479L | E | 2.4e-08 | 4.8e-08 | 4.1e-08 | 3.767E-08 | 1.234E-08 | 2.400E-08 | 4.800E-08 |
| 29572 | 1479 | H1479K | E | 4.3e-08 | 3.3e-08 | 5.5e-08 | 4.367E-08 | 1.102E-08 | 3.300E-08 | 5.500E-08 |
| 29573 | 1479 | H1479M | E | 4.4e-08 | 2.5e-08 | 2.7e-08 | 3.200E-08 | 1.044E-08 | 2.500E-08 | 4.400E-08 |
| 29574 | 1479 | H1479F | E | 7.3e-08 | 3.1e-08 | 5.9e-08 | 5.433E-08 | 2.139E-08 | 3.100E-08 | 7.300E-08 |
| 29575 | 1479 | H1479P | E | 2.8e-08 | 4e-08 | 6.7e-08 | 4.500E-08 | 1.997E-08 | 2.800E-08 | 6.700E-08 |
| 29576 | 1479 | H1479S | E | 3.3e-08 | 4.6e-08 | 5.4e-08 | 4.433E-08 | 1.060E-08 | 3.300E-08 | 5.400E-08 |
| 29577 | 1479 | H1479T | E | 8.4e-08 | 3.6e-08 | 3.3e-08 | 5.100E-08 | 2.862E-08 | 3.300E-08 | 8.400E-08 |
| 29578 | 1479 | H1479W | E | 5.5e-08 | 1.4e-07 | 5.3e-08 | 8.267E-08 | 4.966E-08 | 5.300E-08 | 1.400E-07 |
| 29579 | 1479 | H1479Y | E | 1.2e-08 | 6.3e-08 | 8.5e-08 | 5.333E-08 | 3.745E-08 | 1.200E-08 | 8.500E-08 |

| | | | | | | | | | | |
|---|---|---|---|---|---|---|---|---|---|---|
| 29580 | 1479 | H1479V | E | 6.2e-08 | 2.1e-08 | 4.6e-08 | 4.300E-08 | 2.066E-08 | 2.100E-08 | 6.200E-08 |
| 29581 | 1480 | K1480A | E | 5e-08 | 6.3e-08 | 1.1e-07 | 7.433E-08 | 3.156E-08 | 5.000E-08 | 1.100E-07 |
| 29582 | 1480 | K1480R | E | 3.4e-08 | 6.7e-08 | 1.1e-07 | 7.033E-08 | 3.811E-08 | 3.400E-08 | 1.100E-07 |
| 29583 | 1480 | K1480N | E | 5.3e-08 | 3.3e-08 | 7.2e-08 | 5.267E-08 | 1.950E-08 | 3.300E-08 | 7.200E-08 |
| 29584 | 1480 | K1480D | E | 4e-08 | 2.4e-08 | 5.8e-08 | 4.067E-08 | 1.701E-08 | 2.400E-08 | 5.800E-08 |
| 29585 | 1480 | K1480C | E | 3.8e-08 | 2.9e-08 | 7.8e-08 | 4.833E-08 | 2.608E-08 | 2.900E-08 | 7.800E-08 |
| 29586 | 1480 | K1480E | E | 6.4e-08 | 2e-08 | 4.8e-08 | 4.400E-08 | 2.227E-08 | 2.000E-08 | 6.400E-08 |
| 29587 | 1480 | K1480Q | E | 4.2e-08 | 2.6e-08 | 2.3e-08 | 3.033E-08 | 1.021E-08 | 2.300E-08 | 4.200E-08 |
| 29588 | 1480 | K1480G | E | 5.7e-08 | 1.1e-07 | 7.5e-08 | 8.067E-08 | 2.695E-08 | 5.700E-08 | 1.100E-07 |
| 29589 | 1480 | K1480H | E | 1.4e-07 | 6.9e-08 | 7.9e-08 | 9.600E-08 | 3.843E-08 | 6.900E-08 | 1.400E-07 |
| 29590 | 1480 | K1480I | E | 3.4e-08 | 2.7e-08 | 4.3e-08 | 3.467E-08 | 8.021E-09 | 2.700E-08 | 4.300E-08 |
| 29591 | 1480 | K1480L | E | 7.1e-08 | 5.1e-08 | 1.8e-08 | 4.667E-08 | 2.676E-08 | 1.800E-08 | 7.100E-08 |
| 29592 | 1480 | K1480K | E | 2.7e-08 | 3.2e-08 | 6.9e-08 | 4.267E-08 | 2.294E-08 | 2.700E-08 | 6.900E-08 |
| 29593 | 1480 | K1480M | E | 5.2e-08 | 6e-08 | 6.6e-08 | 5.933E-08 | 7.024E-09 | 5.200E-08 | 6.600E-08 |
| 29594 | 1480 | K1480F | E | 2.2e-08 | 6.2e-08 | 6.1e-08 | 4.833E-08 | 2.281E-08 | 2.200E-08 | 6.200E-08 |
| 29595 | 1480 | K1480P | E | 1.4e-08 | 4.5e-08 | 3.1e-08 | 3.000E-08 | 1.552E-08 | 1.400E-08 | 4.500E-08 |
| 29596 | 1480 | K1480S | E | 2.2e-08 | 2e-08 | 3.4e-08 | 2.533E-08 | 7.572E-09 | 2.000E-08 | 3.400E-08 |
| 29597 | 1480 | K1480T | E | 1e-07 | 2.1e-08 | 2.3e-08 | 4.800E-08 | 4.504E-08 | 2.100E-08 | 1.000E-07 |
| 29598 | 1480 | K1480W | E | 3.2e-08 | 7.7e-08 | 3.4e-08 | 4.767E-08 | 2.542E-08 | 3.200E-08 | 7.700E-08 |
| 29599 | 1480 | K1480Y | E | 5.4e-08 | 1.7e-08 | 1.1e-07 | 6.033E-08 | 4.682E-08 | 1.700E-08 | 1.100E-07 |
| 29600 | 1480 | K1480V | E | 4.5e-08 | 2.9e-08 | 6.4e-08 | 4.600E-08 | 1.752E-08 | 2.900E-08 | 6.400E-08 |
| 29601 | 1481 | P1481A | E | 4e-08 | 1.2e-08 | 2e-08 | 2.400E-08 | 1.442E-08 | 1.200E-08 | 4.000E-08 |
| 29602 | 1481 | P1481R | E | 9.8e-09 | 6.2e-08 | 4.6e-08 | 3.927E-08 | 2.674E-08 | 9.800E-09 | 6.200E-08 |
| 29603 | 1481 | P1481N | E | 5e-08 | 1.5e-08 | 7.6e-08 | 4.700E-08 | 3.061E-08 | 1.500E-08 | 7.600E-08 |
| 29604 | 1481 | P1481D | E | 2e-08 | 5.3e-08 | 5.4e-08 | 4.233E-08 | 1.935E-08 | 2.000E-08 | 5.400E-08 |
| 29605 | 1481 | P1481C | E | 5e-08 | 5.4e-08 | 2.7e-08 | 4.367E-08 | 1.457E-08 | 2.700E-08 | 5.400E-08 |
| 29606 | 1481 | P1481E | E | 1.2e-08 | 6.3e-08 | 8.2e-08 | 5.233E-08 | 3.620E-08 | 1.200E-08 | 8.200E-08 |
| 29607 | 1481 | P1481Q | E | 3.6e-08 | 2.9e-08 | 1.3e-08 | 2.600E-08 | 1.179E-08 | 1.300E-08 | 3.600E-08 |
| 29608 | 1481 | P1481G | E | 4.7e-08 | 8.2e-08 | 3.6e-08 | 5.500E-08 | 2.402E-08 | 3.600E-08 | 8.200E-08 |
| 29609 | 1481 | P1481H | E | 6.3e-08 | 1.3e-08 | 3.8e-08 | 3.800E-08 | 2.500E-08 | 1.300E-08 | 6.300E-08 |
| 29610 | 1481 | P1481I | E | 4.8e-08 | 7.3e-08 | 9.9e-08 | 7.333E-08 | 2.550E-08 | 4.800E-08 | 9.900E-08 |
| 29611 | 1481 | P1481L | E | 2e-08 | 3.2e-08 | 3.8e-08 | 3.000E-08 | 9.165E-09 | 2.000E-08 | 3.800E-08 |
| 29612 | 1481 | P1481K | E | 5.3e-08 | 3.6e-08 | 8.6e-08 | 5.833E-08 | 2.542E-08 | 3.600E-08 | 8.600E-08 |
| 29613 | 1481 | P1481M | E | 2.9e-08 | 9.7e-08 | 9.5e-09 | 4.517E-08 | 4.594E-08 | 9.500E-09 | 9.700E-08 |
| 29614 | 1481 | P1481F | E | 3.5e-08 | 1.4e-08 | 3.7e-08 | 2.867E-08 | 1.274E-08 | 1.400E-08 | 3.700E-08 |
| 29615 | 1481 | P1481P | E | 1.4e-07 | 1.7e-07 | 2.1e-08 | 1.103E-07 | 7.881E-08 | 2.100E-08 | 1.700E-07 |
| 29616 | 1481 | P1481S | E | 1.9e-08 | 3e-08 | 4.4e-08 | 3.100E-08 | 1.253E-08 | 1.900E-08 | 4.400E-08 |
| 29617 | 1481 | P1481T | E | 9e-08 | 7.4e-08 | 3.1e-08 | 6.500E-08 | 3.051E-08 | 3.100E-08 | 9.000E-08 |
| 29618 | 1481 | P1481W | E | 4e-08 | 3.4e-08 | 9.1e-08 | 5.500E-08 | 3.132E-08 | 3.400E-08 | 9.100E-08 |
| 29619 | 1481 | P1481Y | E | 1.2e-08 | 5.7e-08 | 8.9e-08 | 5.267E-08 | 3.868E-08 | 1.200E-08 | 8.900E-08 |
| 29620 | 1481 | P1481V | E | 4.3e-08 | 5.8e-08 | 3.1e-08 | 4.400E-08 | 1.353E-08 | 3.100E-08 | 5.800E-08 |
| 29621 | 1482 | S1482A | E | 3.3e-08 | 8.8e-08 | 3.1e-08 | 5.067E-08 | 3.235E-08 | 3.100E-08 | 8.800E-08 |
| 29622 | 1482 | S1482R | E | 4.9e-08 | 7.6e-08 | 3.1e-08 | 5.200E-08 | 2.265E-08 | 3.100E-08 | 7.600E-08 |
| 29623 | 1482 | S1482N | E | 1.3e-08 | 2e-08 | 4.3e-08 | 2.533E-08 | 1.570E-08 | 1.300E-08 | 4.300E-08 |
| 29624 | 1482 | S1482D | E | 1.2e-08 | 6e-08 | 5.4e-08 | 4.200E-08 | 2.615E-08 | 1.200E-08 | 6.000E-08 |
| 29625 | 1482 | S1482C | E | 2.5e-08 | 5.9e-08 | 2.3e-08 | 3.567E-08 | 2.023E-08 | 2.300E-08 | 5.900E-08 |
| 29626 | 1482 | S1482E | E | 2.7e-08 | 1.8e-08 | 5.5e-08 | 3.333E-08 | 1.930E-08 | 1.800E-08 | 5.500E-08 |
| 29627 | 1482 | S1482Q | E | 3e-08 | 2.1e-08 | 1e-07 | 5.033E-08 | 4.325E-08 | 2.100E-08 | 1.000E-07 |
| 29628 | 1482 | S1482G | E | 2.8e-08 | 7.1e-08 | 1.6e-08 | 3.833E-08 | 2.892E-08 | 1.600E-08 | 7.100E-08 |
| 29629 | 1482 | S1482H | E | 6.3e-08 | 4.5e-08 | 1.7e-08 | 4.167E-08 | 2.318E-08 | 1.700E-08 | 6.300E-08 |
| 29630 | 1482 | S1482I | E | 2.8e-08 | 3.4e-08 | 1.6e-08 | 2.600E-08 | 9.165E-09 | 1.600E-08 | 3.400E-08 |
| 29631 | 1482 | S1482L | E | 7e-08 | 2.7e-08 | 3.4e-08 | 4.367E-08 | 2.307E-08 | 2.700E-08 | 7.000E-08 |
| 29632 | 1482 | S1482K | E | 7.7e-08 | 3.2e-08 | 2.5e-08 | 4.467E-08 | 2.822E-08 | 2.500E-08 | 7.700E-08 |
| 29633 | 1482 | S1482M | E | 1.3e-08 | 7.3e-08 | 2.2e-08 | 3.600E-08 | 3.236E-08 | 1.300E-08 | 7.300E-08 |
| 29634 | 1482 | S1482F | E | 5.5e-08 | 1.1e-07 | 3.9e-08 | 6.800E-08 | 3.724E-08 | 3.900E-08 | 1.100E-07 |
| 29635 | 1482 | S1482P | E | 3.1e-08 | 1.1e-07 | 1.3e-07 | 9.033E-08 | 5.235E-08 | 3.100E-08 | 1.300E-07 |
| 29636 | 1482 | S1482S | E | 3.8e-08 | 3.6e-08 | 1e-07 | 5.800E-08 | 3.639E-08 | 3.600E-08 | 1.000E-07 |
| 29637 | 1482 | S1482T | E | 2.3e-08 | 5.7e-08 | 2.6e-08 | 3.533E-08 | 1.882E-08 | 2.300E-08 | 5.700E-08 |
| 29638 | 1482 | S1482W | E | 4.5e-08 | 2.6e-08 | 2.2e-08 | 3.100E-08 | 1.229E-08 | 2.200E-08 | 4.500E-08 |
| 29639 | 1482 | S1482Y | E | 3.9e-08 | 1.9e-08 | 1.5e-08 | 2.433E-08 | 1.286E-08 | 1.500E-08 | 3.900E-08 |
| 29640 | 1482 | S1482V | E | 2.1e-08 | 5.9e-08 | 3.6e-08 | 3.867E-08 | 1.914E-08 | 2.100E-08 | 5.900E-08 |
| 29641 | 1483 | N1483A | E | 7.7e-08 | 5.3e-08 | 2.6e-08 | 5.200E-08 | 2.551E-08 | 2.600E-08 | 7.700E-08 |
| 29642 | 1483 | N1483R | E | 5.2e-08 | 4.1e-08 | 8e-08 | 5.767E-08 | 2.011E-08 | 4.100E-08 | 8.000E-08 |
| 29643 | 1483 | N1483N | E | 5.4e-08 | 1.8e-08 | 8.8e-08 | 5.333E-08 | 3.500E-08 | 1.800E-08 | 8.800E-08 |
| 29644 | 1483 | N1483D | E | 4.8e-08 | 7.5e-08 | 4.9e-08 | 5.733E-08 | 1.531E-08 | 4.800E-08 | 7.500E-08 |
| 29645 | 1483 | N1483C | E | 2.4e-08 | 4.8e-08 | 5.4e-08 | 4.200E-08 | 1.587E-08 | 2.400E-08 | 5.400E-08 |
| 29646 | 1483 | N1483E | E | 1.9e-08 | 1.6e-08 | 5.5e-08 | 3.000E-08 | 2.170E-08 | 1.600E-08 | 5.500E-08 |
| 29647 | 1483 | N1483Q | E | 6e-08 | 3.9e-08 | 5.1e-08 | 5.000E-08 | 1.054E-08 | 3.900E-08 | 6.000E-08 |
| 29648 | 1483 | N1483G | E | 3.8e-08 | 2.1e-08 | 5.7e-08 | 3.867E-08 | 1.801E-08 | 2.100E-08 | 5.700E-08 |
| 29649 | 1483 | N1483H | E | 4.2e-08 | 4e-08 | 6.9e-08 | 5.033E-08 | 1.620E-08 | 4.000E-08 | 6.900E-08 |
| 29650 | 1483 | N1483I | E | 2.4e-08 | 2.4e-08 | 8.2e-08 | 4.333E-08 | 3.349E-08 | 2.400E-08 | 8.200E-08 |
| 29651 | 1483 | N1483L | E | 3e-08 | 3.1e-08 | 3.9e-08 | 3.333E-08 | 4.933E-09 | 3.000E-08 | 3.900E-08 |
| 29652 | 1483 | N1483K | E | 5.8e-08 | 6.1e-08 | 2.9e-08 | 4.933E-08 | 1.767E-08 | 2.900E-08 | 6.100E-08 |
| 29653 | 1483 | N1483M | E | 3.6e-08 | 1.9e-08 | 6.3e-08 | 3.933E-08 | 2.219E-08 | 1.900E-08 | 6.300E-08 |
| 29654 | 1483 | N1483F | E | 3.8e-08 | 1.3e-07 | 2.6e-08 | 6.467E-08 | 5.690E-08 | 2.600E-08 | 1.300E-07 |
| 29655 | 1483 | N1483P | E | 7e-08 | 4.3e-08 | 8.4e-08 | 6.567E-08 | 2.084E-08 | 4.300E-08 | 8.400E-08 |
| 29656 | 1483 | N1483S | E | 8.1e-08 | 2.3e-08 | 1.4e-08 | 3.933E-08 | 3.636E-08 | 1.400E-08 | 8.100E-08 |
| 29657 | 1483 | N1483T | E | 7.8e-08 | 3.3e-08 | 1.9e-08 | 4.333E-08 | 3.083E-08 | 1.900E-08 | 7.800E-08 |
| 29658 | 1483 | N1483W | E | 2.9e-08 | 4.4e-08 | 4.8e-08 | 4.033E-08 | 1.002E-08 | 2.900E-08 | 4.800E-08 |
| 29659 | 1483 | N1483Y | E | 8.7e-08 | 6.9e-08 | 2.3e-08 | 5.967E-08 | 3.301E-08 | 2.300E-08 | 8.700E-08 |
| 29660 | 1483 | N1483V | E | 4.9e-08 | 1.9e-07 | 5.4e-08 | 9.767E-08 | 8.000E-08 | 4.900E-08 | 1.900E-07 |
| 29661 | 1484 | T1484A | E | 4.7e-08 | 4.4e-08 | 3e-08 | 4.033E-08 | 9.074E-09 | 3.000E-08 | 4.700E-08 |
| 29662 | 1484 | T1484R | E | 4.1e-08 | 5.8e-08 | 5.6e-08 | 5.167E-08 | 9.292E-09 | 4.100E-08 | 5.800E-08 |
| 29663 | 1484 | T1484N | E | 5.9e-08 | 4.1e-08 | 4.3e-08 | 4.767E-08 | 9.866E-09 | 4.100E-08 | 5.900E-08 |
| 29664 | 1484 | T1484D | E | 2.8e-08 | 2.2e-08 | 4e-08 | 3.000E-08 | 9.165E-09 | 2.200E-08 | 4.000E-08 |

| | | | | | | | | | | |
|---|---|---|---|---|---|---|---|---|---|---|
| 29665 | 1484 | T1484C | E | 4.7e-08 | 4.4e-08 | 8.5e-08 | 5.867E-08 | 2.285E-08 | 4.400E-08 | 8.500E-08 |
| 29666 | 1484 | T1484E | E | 4.5e-08 | 2.8e-08 | 2.4e-08 | 3.233E-08 | 1.115E-08 | 2.400E-08 | 4.500E-08 |
| 29667 | 1484 | T1484Q | E | 3.7e-08 | 6e-08 | 6.5e-08 | 5.400E-08 | 1.493E-08 | 3.700E-08 | 6.500E-08 |
| 29668 | 1484 | T1484G | E | 5.5e-08 | 6.3e-08 | 3.9e-08 | 5.233E-08 | 1.222E-08 | 3.900E-08 | 6.300E-08 |
| 29669 | 1484 | T1484H | E | 5.2e-08 | 2.3e-08 | 4.1e-08 | 3.867E-08 | 1.464E-08 | 2.300E-08 | 5.200E-08 |
| 29670 | 1484 | T1484I | E | 4.3e-08 | 6.6e-08 | 1.7e-08 | 4.200E-08 | 2.452E-08 | 1.700E-08 | 6.600E-08 |
| 29671 | 1484 | T1484L | E | 3.6e-08 | 5e-08 | 8.7e-08 | 5.767E-08 | 2.635E-08 | 3.600E-08 | 8.700E-08 |
| 29672 | 1484 | T1484K | E | 3.4e-08 | 7e-08 | 5.7e-08 | 5.367E-08 | 1.823E-08 | 3.400E-08 | 7.000E-08 |
| 29673 | 1484 | T1484M | E | 5.9e-08 | 6.7e-08 | 4.9e-08 | 5.833E-08 | 9.018E-09 | 4.900E-08 | 6.700E-08 |
| 29674 | 1484 | T1484F | E | 2.6e-08 | 4.3e-08 | 1.2e-08 | 2.700E-08 | 1.552E-08 | 1.200E-08 | 4.300E-08 |
| 29675 | 1484 | T1484P | E | 4.3e-08 | 3.1e-08 | 1.2e-07 | 6.467E-08 | 4.829E-08 | 3.100E-08 | 1.200E-07 |
| 29676 | 1484 | T1484S | E | 4e-08 | 1.6e-08 | 4.6e-08 | 3.400E-08 | 1.587E-08 | 1.600E-08 | 4.600E-08 |
| 29677 | 1484 | T1484T | E | 5e-09 | 1.2e-07 | 3.4e-08 | 5.300E-08 | 5.981E-08 | 5.000E-09 | 1.200E-07 |
| 29678 | 1484 | T1484W | E | 5.1e-08 | 3.9e-08 | 3.8e-08 | 4.267E-08 | 7.234E-09 | 3.800E-08 | 5.100E-08 |
| 29679 | 1484 | T1484Y | E | 2.2e-08 | 8.4e-08 | 2.7e-08 | 4.433E-08 | 3.444E-08 | 2.200E-08 | 8.400E-08 |
| 29680 | 1484 | T1484V | E | 2.1e-08 | 4e-08 | 4.2e-08 | 3.433E-08 | 1.159E-08 | 2.100E-08 | 4.200E-08 |
| 29681 | 1485 | K1485A | E | 7.2e-08 | 1.6e-08 | 5.2e-08 | 4.667E-08 | 2.838E-08 | 1.600E-08 | 7.200E-08 |
| 29682 | 1485 | K1485R | E | 3.3e-08 | 6.1e-08 | 3.3e-08 | 4.233E-08 | 1.617E-08 | 3.300E-08 | 6.100E-08 |
| 29683 | 1485 | K1485N | E | 5.9e-08 | 5.7e-08 | 3.6e-08 | 5.067E-08 | 1.274E-08 | 3.600E-08 | 5.900E-08 |
| 29684 | 1485 | K1485D | E | 4.3e-08 | 3e-08 | 7.7e-08 | 5.000E-08 | 2.427E-08 | 3.000E-08 | 7.700E-08 |
| 29685 | 1485 | K1485C | E | 5.5e-08 | 1.2e-07 | 4.9e-08 | 7.467E-08 | 3.937E-08 | 4.900E-08 | 1.200E-07 |
| 29686 | 1485 | K1485E | E | 4.9e-08 | 6.2e-08 | 2.1e-08 | 4.400E-08 | 2.095E-08 | 2.100E-08 | 6.200E-08 |
| 29687 | 1485 | K1485Q | E | 3.5e-08 | 2.8e-08 | 1.7e-08 | 2.667E-08 | 9.074E-09 | 1.700E-08 | 3.500E-08 |
| 29688 | 1485 | K1485G | E | 2.4e-08 | 2.6e-08 | 1.9e-08 | 2.300E-08 | 3.606E-09 | 1.900E-08 | 2.600E-08 |
| 29689 | 1485 | K1485H | E | 2.4e-08 | 1.8e-07 | 5e-08 | 8.467E-08 | 8.358E-08 | 2.400E-08 | 1.800E-07 |
| 29690 | 1485 | K1485I | E | 3.3e-08 | 4.4e-08 | 3.7e-08 | 3.800E-08 | 5.568E-09 | 3.300E-08 | 4.400E-08 |
| 29691 | 1485 | K1485L | E | 3.1e-08 | 3.7e-08 | 7.7e-08 | 4.833E-08 | 2.501E-08 | 3.100E-08 | 7.700E-08 |
| 29692 | 1485 | K1485K | E | 5.4e-08 | 5.1e-08 | 4.4e-08 | 4.967E-08 | 5.132E-09 | 4.400E-08 | 5.400E-08 |
| 29693 | 1485 | K1485M | E | 7.9e-08 | 2.4e-08 | 4.9e-08 | 5.067E-08 | 2.754E-08 | 2.400E-08 | 7.900E-08 |
| 29694 | 1485 | K1485F | E | 4.4e-08 | 7.5e-08 | 5.5e-08 | 5.800E-08 | 1.572E-08 | 4.400E-08 | 7.500E-08 |
| 29695 | 1485 | K1485P | E | 6.6e-08 | 1.5e-08 | 3.2e-08 | 3.767E-08 | 2.597E-08 | 1.500E-08 | 6.600E-08 |
| 29696 | 1485 | K1485S | E | 4.6e-08 | 8.3e-08 | 3.1e-08 | 5.333E-08 | 2.676E-08 | 3.100E-08 | 8.300E-08 |
| 29697 | 1485 | K1485T | E | 6.3e-08 | 3.5e-08 | 1.2e-07 | 7.267E-08 | 4.332E-08 | 3.500E-08 | 1.200E-07 |
| 29698 | 1485 | K1485W | E | 3.1e-08 | 2.5e-08 | 5e-08 | 3.533E-08 | 1.305E-08 | 2.500E-08 | 5.000E-08 |
| 29699 | 1485 | K1485Y | E | 3.4e-08 | 6.1e-08 | 9e-08 | 6.167E-08 | 2.801E-08 | 3.400E-08 | 9.000E-08 |
| 29700 | 1485 | K1485V | E | 3.3e-08 | 6.5e-08 | 2.2e-08 | 4.000E-08 | 2.234E-08 | 2.200E-08 | 6.500E-08 |
| 29701 | 1486 | V1486A | E | 4e-08 | 5.6e-08 | 6.8e-08 | 5.467E-08 | 1.405E-08 | 4.000E-08 | 6.800E-08 |
| 29702 | 1486 | V1486R | E | 4.1e-08 | 3.1e-08 | 1e-07 | 5.733E-08 | 3.729E-08 | 3.100E-08 | 1.000E-07 |
| 29703 | 1486 | V1486N | E | 1.5e-08 | 5.9e-08 | 5e-08 | 4.133E-08 | 2.325E-08 | 1.500E-08 | 5.900E-08 |
| 29704 | 1486 | V1486D | E | 4e-08 | 6.4e-08 | 2.8e-08 | 4.400E-08 | 1.833E-08 | 2.800E-08 | 6.400E-08 |
| 29705 | 1486 | V1486C | E | 9.3e-08 | 3.1e-08 | 1.7e-08 | 4.700E-08 | 4.045E-08 | 1.700E-08 | 9.300E-08 |
| 29706 | 1486 | V1486E | E | 7.3e-08 | 9e-08 | 5.1e-08 | 7.133E-08 | 1.955E-08 | 5.100E-08 | 9.000E-08 |
| 29707 | 1486 | V1486Q | E | 2.8e-08 | 4.4e-08 | 1.7e-08 | 2.967E-08 | 1.358E-08 | 1.700E-08 | 4.400E-08 |
| 29708 | 1486 | V1486G | E | 6.9e-08 | 3.4e-08 | 5e-08 | 5.100E-08 | 1.752E-08 | 3.400E-08 | 6.900E-08 |
| 29709 | 1486 | V1486H | E | 2.4e-08 | 2e-08 | 3.6e-08 | 2.667E-08 | 8.327E-09 | 2.000E-08 | 3.600E-08 |
| 29710 | 1486 | V1486I | E | 3.3e-08 | 9.2e-09 | 2.9e-08 | 2.373E-08 | 1.274E-08 | 9.200E-09 | 3.300E-08 |
| 29711 | 1486 | V1486L | E | 8.3e-08 | 1.9e-08 | 6.8e-08 | 5.667E-08 | 3.347E-08 | 1.900E-08 | 8.300E-08 |
| 29712 | 1486 | V1486K | E | 3.6e-08 | 4.8e-08 | 3e-08 | 3.800E-08 | 9.165E-09 | 3.000E-08 | 4.800E-08 |
| 29713 | 1486 | V1486M | E | 2e-08 | 4.1e-08 | 9.2e-08 | 5.100E-08 | 3.703E-08 | 2.000E-08 | 9.200E-08 |
| 29714 | 1486 | V1486F | E | 4.7e-08 | 2.4e-08 | 6.2e-08 | 4.433E-08 | 1.914E-08 | 2.400E-08 | 6.200E-08 |
| 29715 | 1486 | V1486P | E | 5.3e-08 | 3.8e-08 | 4.8e-08 | 4.633E-08 | 7.638E-09 | 3.800E-08 | 5.300E-08 |
| 29716 | 1486 | V1486S | E | 1.7e-08 | 9.2e-09 | 2.3e-08 | 1.640E-08 | 6.920E-09 | 9.200E-09 | 2.300E-08 |
| 29717 | 1486 | V1486T | E | 1.2e-07 | 6.1e-08 | 7e-08 | 8.367E-08 | 3.179E-08 | 6.100E-08 | 1.200E-07 |
| 29718 | 1486 | V1486W | E | 4.2e-08 | 3.9e-08 | 1.3e-07 | 7.033E-08 | 5.169E-08 | 3.900E-08 | 1.300E-07 |
| 29719 | 1486 | V1486Y | E | 1e-07 | 5.3e-08 | 2.9e-08 | 6.067E-08 | 3.612E-08 | 2.900E-08 | 1.000E-07 |
| 29720 | 1486 | V1486V | E | 2.8e-08 | 4.6e-08 | 4e-08 | 3.800E-08 | 9.165E-09 | 2.800E-08 | 4.600E-08 |
| 29721 | 1487 | D1487A | E | 7.6e-08 | 2.2e-08 | 4.2e-08 | 4.667E-08 | 2.730E-08 | 2.200E-08 | 7.600E-08 |
| 29722 | 1487 | D1487R | E | 5.4e-08 | 5.8e-08 | 4.7e-08 | 5.300E-08 | 5.568E-09 | 4.700E-08 | 5.800E-08 |
| 29723 | 1487 | D1487N | E | 5.3e-08 | 4.3e-08 | 1e-07 | 6.533E-08 | 3.044E-08 | 4.300E-08 | 1.000E-07 |
| 29724 | 1487 | D1487D | E | 6.2e-08 | 5.8e-08 | 4.4e-08 | 5.467E-08 | 9.452E-09 | 4.400E-08 | 6.200E-08 |
| 29725 | 1487 | D1487C | E | 3e-08 | 1.1e-07 | 7e-08 | 7.000E-08 | 4.000E-08 | 3.000E-08 | 1.100E-07 |
| 29726 | 1487 | D1487E | E | 9.4e-08 | 8.1e-08 | 3.1e-08 | 6.867E-08 | 3.326E-08 | 3.100E-08 | 9.400E-08 |
| 29727 | 1487 | D1487Q | E | 7.3e-08 | 1.6e-08 | 5e-08 | 4.633E-08 | 2.868E-08 | 1.600E-08 | 7.300E-08 |
| 29728 | 1487 | D1487G | E | 1.4e-07 | 5e-08 | 3.8e-08 | 7.600E-08 | 5.575E-08 | 3.800E-08 | 1.400E-07 |
| 29729 | 1487 | D1487H | E | 2.4e-08 | 2e-08 | 4.8e-08 | 3.067E-08 | 1.514E-08 | 2.000E-08 | 4.800E-08 |
| 29730 | 1487 | D1487I | E | 1.9e-08 | 5.7e-08 | 1.1e-07 | 6.200E-08 | 4.571E-08 | 1.900E-08 | 1.100E-07 |
| 29731 | 1487 | D1487L | E | 1.1e-07 | 3.1e-08 | 3.7e-08 | 5.933E-08 | 4.398E-08 | 3.100E-08 | 1.100E-07 |
| 29732 | 1487 | D1487K | E | 3.5e-08 | 2.2e-08 | 2.7e-08 | 2.800E-08 | 6.557E-09 | 2.200E-08 | 3.500E-08 |
| 29733 | 1487 | D1487M | E | 4.3e-08 | 1.1e-08 | 6e-08 | 3.800E-08 | 2.488E-08 | 1.100E-08 | 6.000E-08 |
| 29734 | 1487 | D1487F | E | 7.9e-08 | 3.3e-08 | 4.7e-08 | 5.300E-08 | 2.358E-08 | 3.300E-08 | 7.900E-08 |
| 29735 | 1487 | D1487P | E | 2.4e-08 | 3.7e-08 | 1.1e-08 | 2.400E-08 | 1.300E-08 | 1.100E-08 | 3.700E-08 |
| 29736 | 1487 | D1487S | E | 2.4e-08 | 7e-08 | 6.3e-08 | 5.233E-08 | 2.479E-08 | 2.400E-08 | 7.000E-08 |
| 29737 | 1487 | D1487T | E | 2.3e-08 | 3.7e-08 | 3.1e-08 | 3.033E-08 | 7.024E-09 | 2.300E-08 | 3.700E-08 |
| 29738 | 1487 | D1487W | E | 3.9e-08 | 9.5e-08 | 4.3e-08 | 5.900E-08 | 3.124E-08 | 3.900E-08 | 9.500E-08 |
| 29739 | 1487 | D1487Y | E | 3.6e-08 | 2.9e-07 | 9.4e-08 | 1.400E-07 | 1.331E-07 | 3.600E-08 | 2.900E-07 |
| 29740 | 1487 | D1487V | E | 7e-08 | 5.8e-08 | 1.7e-08 | 4.833E-08 | 2.779E-08 | 1.700E-08 | 7.000E-08 |
| 29741 | 1488 | K1488A | E | 2.9e-08 | 1.9e-08 | 2.8e-08 | 2.533E-08 | 5.508E-09 | 1.900E-08 | 2.900E-08 |
| 29742 | 1488 | K1488R | E | 2.9e-08 | 5.8e-08 | 6.1e-08 | 4.933E-08 | 1.767E-08 | 2.900E-08 | 6.100E-08 |
| 29743 | 1488 | K1488N | E | 5e-08 | 3.8e-08 | 8.8e-09 | 3.227E-08 | 2.119E-08 | 8.800E-09 | 5.000E-08 |
| 29744 | 1488 | K1488D | E | 4.1e-08 | 2e-08 | 5.4e-08 | 3.833E-08 | 1.716E-08 | 2.000E-08 | 5.400E-08 |
| 29745 | 1488 | K1488C | E | 3.3e-08 | 2.6e-08 | 2.8e-08 | 2.900E-08 | 3.606E-09 | 2.600E-08 | 3.300E-08 |
| 29746 | 1488 | K1488E | E | 5.4e-08 | 4.8e-08 | 6.1e-08 | 5.433E-08 | 6.506E-09 | 4.800E-08 | 6.100E-08 |
| 29747 | 1488 | K1488Q | E | 1e-07 | 2.5e-08 | 3e-08 | 5.167E-08 | 4.193E-08 | 2.500E-08 | 1.000E-07 |
| 29748 | 1488 | K1488G | E | 2.8e-08 | 3.7e-08 | 7.9e-08 | 4.800E-08 | 2.722E-08 | 2.800E-08 | 7.900E-08 |
| 29749 | 1488 | K1488H | E | 2.1e-08 | 2.1e-08 | 2.2e-08 | 2.133E-08 | 5.774E-10 | 2.100E-08 | 2.200E-08 |

| | | | | | | | | | | |
|---|---|---|---|---|---|---|---|---|---|---|
| 29750 | 1488 | K1488I | E | 1.6e-08 | 3.3e-08 | 3.1e-08 | 2.667E-08 | 9.292E-09 | 1.600E-08 | 3.300E-08 |
| 29751 | 1488 | K1488L | E | 4.9e-08 | 2.8e-08 | 6.5e-08 | 4.733E-08 | 1.856E-08 | 2.800E-08 | 6.500E-08 |
| 29752 | 1488 | K1488K | E | 1.6e-08 | 2.4e-08 | 2.8e-08 | 2.267E-08 | 6.110E-09 | 1.600E-08 | 2.800E-08 |
| 29753 | 1488 | K1488M | E | 8.6e-08 | 4.5e-08 | 7.8e-08 | 6.967E-08 | 2.173E-08 | 4.500E-08 | 8.600E-08 |
| 29754 | 1488 | K1488F | E | 7.9e-08 | 6.1e-08 | 5.6e-08 | 6.533E-08 | 1.210E-08 | 5.600E-08 | 7.900E-08 |
| 29755 | 1488 | K1488P | E | 5e-08 | 1.5e-07 | 9.1e-08 | 9.700E-08 | 5.027E-08 | 5.000E-08 | 1.500E-07 |
| 29756 | 1488 | K1488S | E | 2.9e-08 | 2.1e-08 | 5.2e-08 | 3.400E-08 | 1.609E-08 | 2.100E-08 | 5.200E-08 |
| 29757 | 1488 | K1488T | E | 3.8e-08 | 8.3e-09 | 5.8e-08 | 3.477E-08 | 2.501E-08 | 8.300E-09 | 5.800E-08 |
| 29758 | 1488 | K1488W | E | 4e-08 | 6.1e-08 | 3.7e-08 | 4.600E-08 | 1.308E-08 | 3.700E-08 | 6.100E-08 |
| 29759 | 1488 | K1488Y | E | 2.7e-08 | 1.5e-08 | 3e-08 | 2.400E-08 | 7.937E-09 | 1.500E-08 | 3.000E-08 |
| 29760 | 1488 | K1488V | E | 1.7e-08 | 1.3e-07 | 3.4e-08 | 6.033E-08 | 6.093E-08 | 1.700E-08 | 1.300E-07 |
| 29761 | 1489 | K1489A | E | 4.1e-08 | 5.2e-08 | 4.4e-08 | 4.567E-08 | 5.686E-09 | 4.100E-08 | 5.200E-08 |
| 29762 | 1489 | K1489R | E | 3.9e-08 | 7.9e-08 | 1.4e-08 | 4.400E-08 | 3.279E-08 | 1.400E-08 | 7.900E-08 |
| 29763 | 1489 | K1489N | E | 4.2e-08 | 4.6e-08 | 8.2e-09 | 3.207E-08 | 2.077E-08 | 8.200E-09 | 4.600E-08 |
| 29764 | 1489 | K1489D | E | 7e-08 | 1.6e-07 | 1e-08 | 8.000E-08 | 7.550E-08 | 1.000E-08 | 1.600E-07 |
| 29765 | 1489 | K1489C | E | 1.9e-08 | 9e-08 | 7.5e-08 | 6.133E-08 | 3.742E-08 | 1.900E-08 | 9.000E-08 |
| 29766 | 1489 | K1489E | E | 4.5e-08 | 2.3e-08 | 1.7e-08 | 2.833E-08 | 1.474E-08 | 1.700E-08 | 4.500E-08 |
| 29767 | 1489 | K1489Q | E | 2.6e-08 | 2.7e-08 | 2.4e-08 | 2.567E-08 | 1.528E-09 | 2.400E-08 | 2.700E-08 |
| 29768 | 1489 | K1489G | E | 3.4e-08 | 5.5e-08 | 5.4e-08 | 4.767E-08 | 1.185E-08 | 3.400E-08 | 5.500E-08 |
| 29769 | 1489 | K1489H | E | 2.6e-08 | 1.8e-08 | 4e-08 | 2.800E-08 | 1.114E-08 | 1.800E-08 | 4.000E-08 |
| 29770 | 1489 | K1489I | E | 7.6e-08 | 2.2e-07 | 1e-08 | 1.020E-07 | 1.074E-07 | 1.000E-08 | 2.200E-07 |
| 29771 | 1489 | K1489L | E | 1.6e-08 | 3.1e-08 | 6.6e-08 | 3.767E-08 | 2.566E-08 | 1.600E-08 | 6.600E-08 |
| 29772 | 1489 | K1489K | E | 4.7e-08 | 2.5e-08 | 4.1e-08 | 3.767E-08 | 1.137E-08 | 2.500E-08 | 4.700E-08 |
| 29773 | 1489 | K1489M | E | 8.5e-08 | 3.2e-08 | 7.8e-08 | 6.500E-08 | 2.879E-08 | 3.200E-08 | 8.500E-08 |
| 29774 | 1489 | K1489F | E | 3.1e-08 | 5.2e-08 | 7.8e-08 | 5.367E-08 | 2.354E-08 | 3.100E-08 | 7.800E-08 |
| 29775 | 1489 | K1489P | E | 1.3e-07 | 6.7e-08 | 4.6e-08 | 8.100E-08 | 4.371E-08 | 4.600E-08 | 1.300E-07 |
| 29776 | 1489 | K1489S | E | 6e-08 | 6.7e-08 | 1.7e-08 | 4.800E-08 | 2.707E-08 | 1.700E-08 | 6.700E-08 |
| 29777 | 1489 | K1489T | E | 4.5e-08 | 9.4e-08 | 3.2e-08 | 5.700E-08 | 3.270E-08 | 3.200E-08 | 9.400E-08 |
| 29778 | 1489 | K1489W | E | 2.9e-08 | 6.2e-08 | 1.1e-08 | 3.400E-08 | 2.587E-08 | 1.100E-08 | 6.200E-08 |
| 29779 | 1489 | K1489Y | E | 4.6e-08 | 2.7e-08 | 6.3e-08 | 4.533E-08 | 1.801E-08 | 2.700E-08 | 6.300E-08 |
| 29780 | 1489 | K1489V | E | 4.7e-08 | 3.4e-08 | 3.3e-08 | 3.800E-08 | 7.810E-09 | 3.300E-08 | 4.700E-08 |
| 29781 | 1490 | V1490A | E | 1.1e-08 | 3.5e-08 | 5.9e-08 | 3.500E-08 | 2.400E-08 | 1.100E-08 | 5.900E-08 |
| 29782 | 1490 | V1490R | E | 4.3e-08 | 7.1e-09 | 2.9e-08 | 2.637E-08 | 1.809E-08 | 7.100E-09 | 4.300E-08 |
| 29783 | 1490 | V1490N | E | 1.1e-07 | 4.3e-08 | 9.5e-08 | 8.267E-08 | 3.516E-08 | 4.300E-08 | 1.100E-07 |
| 29784 | 1490 | V1490D | E | 2.9e-08 | 6e-08 | 5.1e-08 | 4.667E-08 | 1.595E-08 | 2.900E-08 | 6.000E-08 |
| 29785 | 1490 | V1490C | E | 1.6e-08 | 3e-08 | 3.8e-08 | 2.800E-08 | 1.114E-08 | 1.600E-08 | 3.800E-08 |
| 29786 | 1490 | V1490E | E | 2.1e-08 | 1.1e-07 | 2.6e-08 | 5.233E-08 | 5.000E-08 | 2.100E-08 | 1.100E-07 |
| 29787 | 1490 | V1490Q | E | 7e-08 | 2.4e-08 | 2e-08 | 3.800E-08 | 2.778E-08 | 2.000E-08 | 7.000E-08 |
| 29788 | 1490 | V1490G | E | 4.9e-08 | 9.4e-08 | 1.3e-08 | 5.200E-08 | 4.058E-08 | 1.300E-08 | 9.400E-08 |
| 29789 | 1490 | V1490H | E | 1.2e-08 | 8.5e-08 | 4.2e-08 | 4.633E-08 | 3.669E-08 | 1.200E-08 | 8.500E-08 |
| 29790 | 1490 | V1490I | E | 5.1e-08 | 5.5e-08 | 7.5e-08 | 6.033E-08 | 1.286E-08 | 5.100E-08 | 7.500E-08 |
| 29791 | 1490 | V1490L | E | 3.9e-08 | 2.8e-08 | 7.4e-08 | 4.700E-08 | 2.402E-08 | 2.800E-08 | 7.400E-08 |
| 29792 | 1490 | V1490K | E | 1.2e-07 | 4.8e-08 | 6e-08 | 7.600E-08 | 3.857E-08 | 4.800E-08 | 1.200E-07 |
| 29793 | 1490 | V1490M | E | 6e-08 | 1.4e-08 | 1.8e-08 | 3.067E-08 | 2.548E-08 | 1.400E-08 | 6.000E-08 |
| 29794 | 1490 | V1490F | E | 1.4e-08 | 5.2e-08 | 7e-08 | 4.533E-08 | 2.859E-08 | 1.400E-08 | 7.000E-08 |
| 29795 | 1490 | V1490P | E | 1.6e-08 | 4.1e-08 | 3e-08 | 2.900E-08 | 1.253E-08 | 1.600E-08 | 4.100E-08 |
| 29796 | 1490 | V1490S | E | 6.1e-08 | 3.8e-08 | 5.6e-08 | 5.167E-08 | 1.210E-08 | 3.800E-08 | 6.100E-08 |
| 29797 | 1490 | V1490T | E | 1.6e-08 | 2.6e-08 | 5.2e-08 | 3.133E-08 | 1.858E-08 | 1.600E-08 | 5.200E-08 |
| 29798 | 1490 | V1490W | E | 1.7e-08 | 7.6e-08 | 2.1e-07 | 1.010E-07 | 9.890E-08 | 1.700E-08 | 2.100E-07 |
| 29799 | 1490 | V1490Y | E | 5.8e-08 | 3.5e-08 | 5.9e-08 | 5.067E-08 | 1.358E-08 | 3.500E-08 | 5.900E-08 |
| 29800 | 1490 | V1490V | E | 7.3e-08 | 1.8e-08 | 4.7e-08 | 4.600E-08 | 2.751E-08 | 1.800E-08 | 7.300E-08 |
| 29801 | 1491 | E1491A | E | 5.5e-08 | 5.5e-08 | 6.4e-08 | 5.800E-08 | 5.196E-09 | 5.500E-08 | 6.400E-08 |
| 29802 | 1491 | E1491R | E | 4.3e-08 | 3e-08 | 1.2e-08 | 2.833E-08 | 1.557E-08 | 1.200E-08 | 4.300E-08 |
| 29803 | 1491 | E1491N | E | 5.3e-08 | 5.5e-08 | 5e-08 | 5.267E-08 | 2.517E-09 | 5.000E-08 | 5.500E-08 |
| 29804 | 1491 | E1491D | E | 1.8e-08 | 2.5e-08 | 1.9e-08 | 2.067E-08 | 3.786E-09 | 1.800E-08 | 2.500E-08 |
| 29805 | 1491 | E1491C | E | 3.3e-08 | 7.3e-08 | 8.4e-08 | 6.333E-08 | 2.684E-08 | 3.300E-08 | 8.400E-08 |
| 29806 | 1491 | E1491E | E | 7.9e-08 | 3.4e-08 | 1.7e-08 | 4.333E-08 | 3.204E-08 | 1.700E-08 | 7.900E-08 |
| 29807 | 1491 | E1491Q | E | 3.6e-08 | 6.7e-08 | 2e-08 | 4.100E-08 | 2.390E-08 | 2.000E-08 | 6.700E-08 |
| 29808 | 1491 | E1491G | E | 5.2e-08 | 3.9e-08 | 3.2e-08 | 4.100E-08 | 1.015E-08 | 3.200E-08 | 5.200E-08 |
| 29809 | 1491 | E1491H | E | 4.2e-08 | 4e-08 | 1.2e-08 | 3.133E-08 | 1.677E-08 | 1.200E-08 | 4.200E-08 |
| 29810 | 1491 | E1491I | E | 3.3e-08 | 2.4e-08 | 6.8e-08 | 4.167E-08 | 2.325E-08 | 2.400E-08 | 6.800E-08 |
| 29811 | 1491 | E1491L | E | 3.7e-08 | 3.4e-08 | 4.5e-08 | 3.867E-08 | 5.686E-09 | 3.400E-08 | 4.500E-08 |
| 29812 | 1491 | E1491K | E | 2.6e-08 | 6e-08 | 1.3e-08 | 3.300E-08 | 2.427E-08 | 1.300E-08 | 6.000E-08 |
| 29813 | 1491 | E1491M | E | 1.8e-07 | 6e-08 | 6.4e-08 | 1.013E-07 | 6.816E-08 | 6.000E-08 | 1.800E-07 |
| 29814 | 1491 | E1491F | E | 4.9e-08 | 2.2e-08 | 8.7e-08 | 5.267E-08 | 3.265E-08 | 2.200E-08 | 8.700E-08 |
| 29815 | 1491 | E1491P | E | 3.8e-08 | 6.8e-08 | 5.3e-08 | 5.300E-08 | 1.500E-08 | 3.800E-08 | 6.800E-08 |
| 29816 | 1491 | E1491S | E | 5.7e-08 | 2.4e-08 | 1.1e-07 | 6.367E-08 | 4.339E-08 | 2.400E-08 | 1.100E-07 |
| 29817 | 1491 | E1491T | E | 1.5e-07 | 5.3e-08 | 8.3e-08 | 9.533E-08 | 4.966E-08 | 5.300E-08 | 1.500E-07 |
| 29818 | 1491 | E1491W | E | 3.2e-08 | 5.9e-08 | 3.2e-08 | 4.100E-08 | 1.559E-08 | 3.200E-08 | 5.900E-08 |
| 29819 | 1491 | E1491Y | E | 2.1e-08 | 3.3e-08 | 1e-07 | 5.133E-08 | 4.257E-08 | 2.100E-08 | 1.000E-07 |
| 29820 | 1491 | E1491V | E | 6.6e-08 | 6.1e-08 | 4.3e-08 | 5.667E-08 | 1.210E-08 | 4.300E-08 | 6.600E-08 |
| 29821 | 1492 | P1492A | E | 2.1e-08 | 8.3e-08 | 2.8e-08 | 4.400E-08 | 3.396E-08 | 2.100E-08 | 8.300E-08 |
| 29822 | 1492 | P1492R | E | 1.9e-08 | 2.3e-08 | 2.2e-08 | 2.133E-08 | 2.082E-09 | 1.900E-08 | 2.300E-08 |
| 29823 | 1492 | P1492N | E | 5.8e-08 | 4.3e-08 | 6e-08 | 5.367E-08 | 9.292E-09 | 4.300E-08 | 6.000E-08 |
| 29824 | 1492 | P1492D | E | 1.1e-07 | 1e-08 | 2.3e-08 | 4.767E-08 | 5.437E-08 | 1.000E-08 | 1.100E-07 |
| 29825 | 1492 | P1492C | E | 2.8e-08 | 1.7e-08 | 9.9e-08 | 4.800E-08 | 4.451E-08 | 1.700E-08 | 9.900E-08 |
| 29826 | 1492 | P1492E | E | 4.4e-08 | 2.1e-08 | 4.7e-08 | 3.733E-08 | 1.422E-08 | 2.100E-08 | 4.700E-08 |
| 29827 | 1492 | P1492Q | E | 4.3e-08 | 5.6e-08 | 5.9e-08 | 5.267E-08 | 8.505E-09 | 4.300E-08 | 5.900E-08 |
| 29828 | 1492 | P1492G | E | 7.3e-08 | 4.9e-08 | 5.1e-08 | 5.767E-08 | 1.332E-08 | 4.900E-08 | 7.300E-08 |
| 29829 | 1492 | P1492H | E | 3.1e-08 | 5.3e-08 | 3.9e-08 | 4.100E-08 | 1.114E-08 | 3.100E-08 | 5.300E-08 |
| 29830 | 1492 | P1492I | E | 9.5e-08 | 1.2e-07 | 4.5e-08 | 8.667E-08 | 3.819E-08 | 4.500E-08 | 1.200E-07 |
| 29831 | 1492 | P1492L | E | 2.1e-08 | 6.2e-08 | 6.2e-08 | 4.833E-08 | 2.367E-08 | 2.100E-08 | 6.200E-08 |
| 29832 | 1492 | P1492K | E | 5.8e-08 | 5.1e-08 | 4.4e-08 | 5.100E-08 | 7.000E-09 | 4.400E-08 | 5.800E-08 |
| 29833 | 1492 | P1492M | E | 6.5e-08 | 1.6e-08 | 3.8e-08 | 3.967E-08 | 2.454E-08 | 1.600E-08 | 6.500E-08 |
| 29834 | 1492 | P1492F | E | 4.6e-08 | 1.9e-07 | 2.5e-08 | 8.700E-08 | 8.982E-08 | 2.500E-08 | 1.900E-07 |

| | | | | | | | | | | |
|---|---|---|---|---|---|---|---|---|---|---|
| 29835 | 1492 | P1492P | E | 2.1e-08 | 9.3e-08 | 7.5e-08 | 6.300E-08 | 3.747E-08 | 2.100E-08 | 9.300E-08 |
| 29836 | 1492 | P1492S | E | 8e-08 | 4.6e-08 | 7.9e-09 | 4.463E-08 | 3.607E-08 | 7.900E-09 | 8.000E-08 |
| 29837 | 1492 | P1492T | E | 1.4e-08 | 4.1e-08 | 4.6e-08 | 3.367E-08 | 1.721E-08 | 1.400E-08 | 4.600E-08 |
| 29838 | 1492 | P1492W | E | 4.3e-08 | 4.6e-08 | 4.7e-08 | 4.533E-08 | 2.082E-09 | 4.300E-08 | 4.700E-08 |
| 29839 | 1492 | P1492Y | E | 5.7e-08 | 6.2e-08 | 1.6e-08 | 4.500E-08 | 2.524E-08 | 1.600E-08 | 6.200E-08 |
| 29840 | 1492 | P1492V | E | 1.2e-07 | 1.8e-08 | 4.7e-08 | 6.167E-08 | 5.256E-08 | 1.800E-08 | 1.200E-07 |

Table 4: A scalable Her2-Trastuzumab-Pertuzumab binding affinity landscape. In this table, **ResID** represents the amino acid residue number of PDB entry 6OGE, while **Muta** represents the site-specific missense mutation introduced into the 1492 amino acid residues in PDB entry 6OGE, **Min**, **Max**, **Mean** and **Std** represent the minimum, the maximum, the average and the standard deviation of the $K_d$ values calculated using Prodigy, respectively.

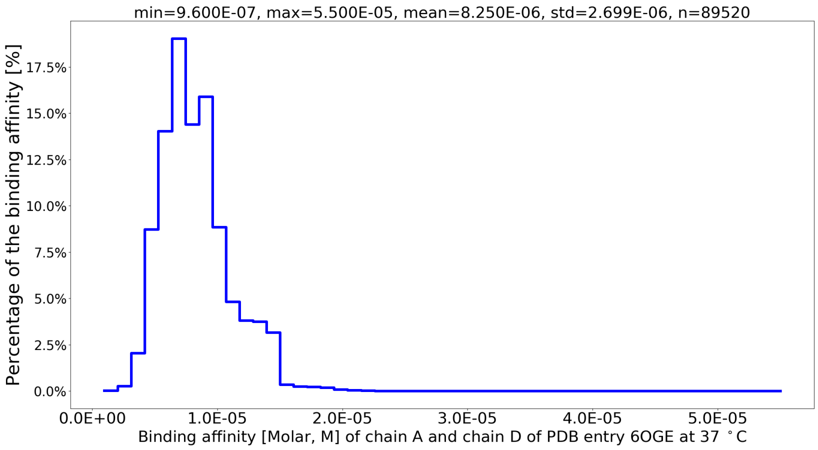


Figure 3: A histogram of the distribution pattern of the Her2-Trastuzumab-Pertuzumab binding affinities between chain A and chain D (listed in Table 1) of PDB entry 6OGE with one site-specific missense mutation.

| No. | ResID | Muta | ChainID | $K_{d1}$ (M) | $K_{d2}$ (M) | $K_{d3}$ (M) | Mean (M) | Std (M) | Min (M) | Max (M) |
|---|---|---|---|---|---|---|---|---|---|---|
| 1 | 1 | T1A | A | 7.8e-06 | 7.8e-06 | 7.8e-06 | 7.800E-06 | 0.000E+00 | 7.800E-06 | 7.800E-06 |
| 2 | 1 | T1R | A | 1.2e-05 | 1.2e-05 | 1.2e-05 | 1.200E-05 | 0.000E+00 | 1.200E-05 | 1.200E-05 |
| 3 | 1 | T1N | A | 5.6e-06 | 5.6e-06 | 5.6e-06 | 5.600E-06 | 0.000E+00 | 5.600E-06 | 5.600E-06 |
| 4 | 1 | T1D | A | 1.3e-05 | 1.3e-05 | 1.3e-05 | 1.300E-05 | 0.000E+00 | 1.300E-05 | 1.300E-05 |
| 5 | 1 | T1C | A | 4e-06 | 4e-06 | 4e-06 | 4.000E-06 | 0.000E+00 | 4.000E-06 | 4.000E-06 |
| 6 | 1 | T1E | A | 6.7e-06 | 6.7e-06 | 6.7e-06 | 6.700E-06 | 0.000E+00 | 6.700E-06 | 6.700E-06 |
| 7 | 1 | T1Q | A | 7.1e-06 | 7.1e-06 | 7.1e-06 | 7.100E-06 | 0.000E+00 | 7.100E-06 | 7.100E-06 |
| 8 | 1 | T1G | A | 1.1e-05 | 1.1e-05 | 1.1e-05 | 1.100E-05 | 2.075E-21 | 1.100E-05 | 1.100E-05 |
| 9 | 1 | T1H | A | 6.4e-06 | 6.4e-06 | 6.4e-06 | 6.400E-06 | 0.000E+00 | 6.400E-06 | 6.400E-06 |
| 10 | 1 | T1I | A | 7.3e-06 | 7.3e-06 | 7.3e-06 | 7.300E-06 | 0.000E+00 | 7.300E-06 | 7.300E-06 |
| 11 | 1 | T1L | A | 1.3e-05 | 1.3e-05 | 1.3e-05 | 1.300E-05 | 0.000E+00 | 1.300E-05 | 1.300E-05 |
| 12 | 1 | T1K | A | 8.2e-06 | 8.2e-06 | 8.2e-06 | 8.200E-06 | 0.000E+00 | 8.200E-06 | 8.200E-06 |
| 13 | 1 | T1M | A | 9.8e-06 | 9.8e-06 | 9.8e-06 | 9.800E-06 | 0.000E+00 | 9.800E-06 | 9.800E-06 |
| 14 | 1 | T1F | A | 1.2e-05 | 1.2e-05 | 1.2e-05 | 1.200E-05 | 0.000E+00 | 1.200E-05 | 1.200E-05 |
| 15 | 1 | T1P | A | 6.3e-06 | 6.3e-06 | 6.3e-06 | 6.300E-06 | 0.000E+00 | 6.300E-06 | 6.300E-06 |
| 16 | 1 | T1S | A | 5e-06 | 5e-06 | 5e-06 | 5.000E-06 | 0.000E+00 | 5.000E-06 | 5.000E-06 |
| 17 | 1 | T1T | A | 4.9e-06 | 4.9e-06 | 4.9e-06 | 4.900E-06 | 0.000E+00 | 4.900E-06 | 4.900E-06 |
| 18 | 1 | T1W | A | 8.7e-06 | 8.7e-06 | 8.7e-06 | 8.700E-06 | 0.000E+00 | 8.700E-06 | 8.700E-06 |
| 19 | 1 | T1Y | A | 1.4e-05 | 1.4e-05 | 1.4e-05 | 1.400E-05 | 0.000E+00 | 1.400E-05 | 1.400E-05 |
| 20 | 1 | T1V | A | 4.9e-06 | 4.9e-06 | 4.9e-06 | 4.900E-06 | 0.000E+00 | 4.900E-06 | 4.900E-06 |
| 21 | 2 | Q2A | A | 1.1e-05 | 1.1e-05 | 1.1e-05 | 1.100E-05 | 2.075E-21 | 1.100E-05 | 1.100E-05 |
| 22 | 2 | Q2R | A | 1.3e-05 | 1.3e-05 | 1.3e-05 | 1.300E-05 | 0.000E+00 | 1.300E-05 | 1.300E-05 |
| 23 | 2 | Q2N | A | 5.6e-06 | 5.6e-06 | 5.6e-06 | 5.600E-06 | 0.000E+00 | 5.600E-06 | 5.600E-06 |
| 24 | 2 | Q2D | A | 9.1e-06 | 9.1e-06 | 9.1e-06 | 9.100E-06 | 0.000E+00 | 9.100E-06 | 9.100E-06 |
| 25 | 2 | Q2C | A | 8.6e-06 | 1.1e-05 | 1.1e-05 | 1.020E-05 | 1.386E-06 | 8.600E-06 | 1.100E-05 |
| 26 | 2 | Q2E | A | 6.3e-06 | 1.3e-05 | 1.1e-05 | 1.010E-05 | 3.439E-06 | 6.300E-06 | 1.300E-05 |
| 27 | 2 | Q2Q | A | 1.3e-05 | 4.5e-06 | 9.3e-06 | 8.933E-06 | 4.262E-06 | 4.500E-06 | 1.300E-05 |
| 28 | 2 | Q2G | A | 9.8e-06 | 1.4e-05 | 5.8e-06 | 9.867E-06 | 4.100E-06 | 5.800E-06 | 1.400E-05 |
| 29 | 2 | Q2H | A | 6.9e-06 | 1.2e-05 | 1.2e-05 | 1.030E-05 | 2.944E-06 | 6.900E-06 | 1.200E-05 |
| 30 | 2 | Q2I | A | 6e-06 | 7.2e-06 | 6e-06 | 6.400E-06 | 6.928E-07 | 6.000E-06 | 7.200E-06 |
| 31 | 2 | Q2L | A | 7.9e-06 | 5.6e-06 | 7.9e-06 | 7.133E-06 | 1.328E-06 | 5.600E-06 | 7.900E-06 |
| 32 | 2 | Q2K | A | 1e-05 | 1e-05 | 8.9e-06 | 9.633E-06 | 6.351E-07 | 8.900E-06 | 1.000E-05 |
| 33 | 2 | Q2M | A | 1.1e-05 | 1.1e-05 | 8.2e-06 | 1.007E-05 | 1.617E-06 | 8.200E-06 | 1.100E-05 |
| 34 | 2 | Q2F | A | 5.2e-06 | 1.4e-05 | 5.2e-06 | 8.133E-06 | 5.081E-06 | 5.200E-06 | 1.400E-05 |
| 35 | 2 | Q2P | A | 7.4e-06 | 7.5e-06 | 7.4e-06 | 7.433E-06 | 5.774E-08 | 7.400E-06 | 7.500E-06 |
| 36 | 2 | Q2S | A | 1.5e-05 | 1.5e-05 | 1.5e-05 | 1.500E-05 | 0.000E+00 | 1.500E-05 | 1.500E-05 |
| 37 | 2 | Q2T | A | 7.2e-06 | 5.9e-06 | 7e-06 | 6.700E-06 | 7.000E-07 | 5.900E-06 | 7.200E-06 |
| 38 | 2 | Q2W | A | 1.4e-05 | 1e-05 | 7.7e-06 | 1.057E-05 | 3.188E-06 | 7.700E-06 | 1.400E-05 |
| 39 | 2 | Q2Y | A | 4e-06 | 8.4e-06 | 8.7e-06 | 7.033E-06 | 2.631E-06 | 4.000E-06 | 8.700E-06 |
| 40 | 2 | Q2V | A | 1.1e-05 | 6.9e-06 | 6.9e-06 | 8.267E-06 | 2.367E-06 | 6.900E-06 | 1.100E-05 |
| 41 | 3 | V3A | A | 6.3e-06 | 6.3e-06 | 1.1e-05 | 7.867E-06 | 2.714E-06 | 6.300E-06 | 1.100E-05 |
| 42 | 3 | V3R | A | 8.1e-06 | 8.3e-06 | 8.3e-06 | 8.233E-06 | 1.155E-07 | 8.100E-06 | 8.300E-06 |
| 43 | 3 | V3N | A | 5.9e-06 | 5.9e-06 | 5.9e-06 | 5.900E-06 | 0.000E+00 | 5.900E-06 | 5.900E-06 |
| 44 | 3 | V3D | A | 6.5e-06 | 6.5e-06 | 6.5e-06 | 6.500E-06 | 0.000E+00 | 6.500E-06 | 6.500E-06 |
| 45 | 3 | V3C | A | 1.5e-05 | 1.5e-05 | 5.8e-06 | 1.193E-05 | 5.312E-06 | 5.800E-06 | 1.500E-05 |
| 46 | 3 | V3E | A | 5e-06 | 5e-06 | 7.2e-06 | 5.733E-06 | 1.270E-06 | 5.000E-06 | 7.200E-06 |
| 47 | 3 | V3Q | A | 1e-05 | 1e-05 | 8.5e-06 | 9.500E-06 | 8.660E-07 | 8.500E-06 | 1.000E-05 |
| 48 | 3 | V3G | A | 7.1e-06 | 7.1e-06 | 7.1e-06 | 7.100E-06 | 0.000E+00 | 7.100E-06 | 7.100E-06 |
| 49 | 3 | V3H | A | 5.6e-06 | 9.6e-06 | 7.7e-06 | 7.633E-06 | 2.001E-06 | 5.600E-06 | 9.600E-06 |
| 50 | 3 | V3I | A | 7.7e-06 | 7.8e-06 | 7.7e-06 | 7.733E-06 | 5.774E-08 | 7.700E-06 | 7.800E-06 |
| 51 | 3 | V3L | A | 1.3e-05 | 6.4e-06 | 7.2e-06 | 8.867E-06 | 3.602E-06 | 6.400E-06 | 1.300E-05 |
| 52 | 3 | V3K | A | 1.6e-05 | 6.5e-06 | 8e-06 | 1.017E-05 | 5.107E-06 | 6.500E-06 | 1.600E-05 |
| 53 | 3 | V3M | A | 7.8e-06 | 4.5e-06 | 5.9e-06 | 6.067E-06 | 1.656E-06 | 4.500E-06 | 7.800E-06 |
| 54 | 3 | V3F | A | 9.3e-06 | 7e-06 | 8.7e-06 | 8.333E-06 | 1.193E-06 | 7.000E-06 | 9.300E-06 |
| 55 | 3 | V3P | A | 7.5e-06 | 5.7e-06 | 5.7e-06 | 6.300E-06 | 1.039E-06 | 5.700E-06 | 7.500E-06 |
| 56 | 3 | V3S | A | 7.3e-06 | 7.3e-06 | 4.7e-06 | 6.433E-06 | 1.501E-06 | 4.700E-06 | 7.300E-06 |
| 57 | 3 | V3T | A | 3.7e-06 | 7e-06 | 7.2e-06 | 5.967E-06 | 1.966E-06 | 3.700E-06 | 7.200E-06 |
| 58 | 3 | V3W | A | 8.8e-06 | 8.8e-06 | 4.7e-06 | 7.433E-06 | 2.367E-06 | 4.700E-06 | 8.800E-06 |
| 59 | 3 | V3Y | A | 9.5e-06 | 8.9e-06 | 6.9e-06 | 8.433E-06 | 1.361E-06 | 6.900E-06 | 9.500E-06 |
| 60 | 3 | V3V | A | 1e-05 | 7.8e-06 | 1.3e-05 | 1.027E-05 | 2.610E-06 | 7.800E-06 | 1.300E-05 |
| 61 | 4 | C4A | A | 1.1e-05 | 1.5e-05 | 8.4e-06 | 1.147E-05 | 3.325E-06 | 8.400E-06 | 1.500E-05 |
| 62 | 4 | C4R | A | 4.5e-06 | 4.2e-06 | 6.4e-06 | 5.033E-06 | 1.193E-06 | 4.200E-06 | 6.400E-06 |
| 63 | 4 | C4N | A | 8.3e-06 | 7.7e-06 | 8.9e-06 | 8.300E-06 | 6.000E-07 | 7.700E-06 | 8.900E-06 |
| 64 | 4 | C4D | A | 6e-06 | 9.7e-06 | 4.5e-06 | 6.733E-06 | 2.676E-06 | 4.500E-06 | 9.700E-06 |
| 65 | 4 | C4C | A | 1.5e-05 | 1.5e-05 | 1.5e-05 | 1.500E-05 | 0.000E+00 | 1.500E-05 | 1.500E-05 |
| 66 | 4 | C4E | A | 8.7e-06 | 7e-06 | 5.5e-06 | 7.067E-06 | 1.601E-06 | 5.500E-06 | 8.700E-06 |
| 67 | 4 | C4Q | A | 6.2e-06 | 9.1e-06 | 7.2e-06 | 7.500E-06 | 1.473E-06 | 6.200E-06 | 9.100E-06 |
| 68 | 4 | C4G | A | 7.8e-06 | 1.1e-05 | 7e-06 | 8.600E-06 | 2.117E-06 | 7.000E-06 | 1.100E-05 |
| 69 | 4 | C4H | A | 1.4e-05 | 7.2e-06 | 8.5e-06 | 9.900E-06 | 3.610E-06 | 7.200E-06 | 1.400E-05 |
| 70 | 4 | C4I | A | 5.9e-06 | 5.9e-06 | 9.1e-06 | 6.967E-06 | 1.848E-06 | 5.900E-06 | 9.100E-06 |
| 71 | 4 | C4L | A | 4.5e-06 | 6.6e-06 | 8.1e-06 | 6.400E-06 | 1.808E-06 | 4.500E-06 | 8.100E-06 |
| 72 | 4 | C4K | A | 1e-05 | 1e-05 | 8.2e-06 | 9.400E-06 | 1.039E-06 | 8.200E-06 | 1.000E-05 |
| 73 | 4 | C4M | A | 1.3e-05 | 8.1e-06 | 8e-06 | 9.700E-06 | 2.858E-06 | 8.000E-06 | 1.300E-05 |
| 74 | 4 | C4F | A | 8.7e-06 | 6.3e-06 | 6.4e-06 | 7.133E-06 | 1.358E-06 | 6.300E-06 | 8.700E-06 |
| 75 | 4 | C4P | A | 5.7e-06 | 7e-06 | 8e-06 | 6.900E-06 | 1.153E-06 | 5.700E-06 | 8.000E-06 |
| 76 | 4 | C4S | A | 6.9e-06 | 3.9e-06 | 9.4e-06 | 6.733E-06 | 2.754E-06 | 3.900E-06 | 9.400E-06 |
| 77 | 4 | C4T | A | 8.8e-06 | 6.1e-06 | 9.7e-06 | 8.200E-06 | 1.873E-06 | 6.100E-06 | 9.700E-06 |
| 78 | 4 | C4W | A | 9.3e-06 | 4.2e-06 | 8e-06 | 7.167E-06 | 2.650E-06 | 4.200E-06 | 9.300E-06 |
| 79 | 4 | C4Y | A | 6.2e-06 | 1.2e-05 | 5.4e-06 | 7.867E-06 | 3.602E-06 | 5.400E-06 | 1.200E-05 |
| 80 | 4 | C4V | A | 7.9e-06 | 9e-06 | 6.5e-06 | 7.800E-06 | 1.253E-06 | 6.500E-06 | 9.000E-06 |
| 81 | 5 | T5A | A | 9e-06 | 8.9e-06 | 1e-05 | 9.300E-06 | 6.083E-07 | 8.900E-06 | 1.000E-05 |
| 82 | 5 | T5R | A | 5.4e-06 | 7.7e-06 | 1.6e-05 | 9.700E-06 | 5.576E-06 | 5.400E-06 | 1.600E-05 |
| 83 | 5 | T5N | A | 4.7e-06 | 7.4e-06 | 6e-06 | 6.033E-06 | 1.350E-06 | 4.700E-06 | 7.400E-06 |
| 84 | 5 | T5D | A | 1e-05 | 4.3e-06 | 4.3e-06 | 6.200E-06 | 3.291E-06 | 4.300E-06 | 1.000E-05 |

| | | | | | | | | | | |
|---|---|---|---|---|---|---|---|---|---|---|
| 85 | 5 | T5C | A | 6.3e-06 | 8.1e-06 | 5.3e-06 | 6.567E-06 | 1.419E-06 | 5.300E-06 | 8.100E-06 |
| 86 | 5 | T5E | A | 9.7e-06 | 6.8e-06 | 1.3e-05 | 9.833E-06 | 3.102E-06 | 6.800E-06 | 1.300E-05 |
| 87 | 5 | T5Q | A | 6.6e-06 | 4.9e-06 | 7.6e-06 | 6.367E-06 | 1.365E-06 | 4.900E-06 | 7.600E-06 |
| 88 | 5 | T5G | A | 6.2e-06 | 6.6e-06 | 7.2e-06 | 6.667E-06 | 5.033E-07 | 6.200E-06 | 7.200E-06 |
| 89 | 5 | T5H | A | 4.6e-06 | 5.5e-06 | 6.8e-06 | 5.633E-06 | 1.106E-06 | 4.600E-06 | 6.800E-06 |
| 90 | 5 | T5I | A | 1e-05 | 5.1e-06 | 6.6e-06 | 7.233E-06 | 2.511E-06 | 5.100E-06 | 1.000E-05 |
| 91 | 5 | T5L | A | 9e-06 | 9e-06 | 7.3e-06 | 8.433E-06 | 9.815E-07 | 7.300E-06 | 9.000E-06 |
| 92 | 5 | T5K | A | 6.3e-06 | 9.7e-06 | 1.2e-05 | 9.333E-06 | 2.868E-06 | 6.300E-06 | 1.200E-05 |
| 93 | 5 | T5M | A | 1e-05 | 6.8e-06 | 9.9e-06 | 8.900E-06 | 1.819E-06 | 6.800E-06 | 1.000E-05 |
| 94 | 5 | T5F | A | 1.9e-05 | 1.3e-05 | 1.3e-05 | 1.500E-05 | 3.464E-06 | 1.300E-05 | 1.900E-05 |
| 95 | 5 | T5P | A | 7.2e-06 | 7e-06 | 1.3e-05 | 9.067E-06 | 3.408E-06 | 7.000E-06 | 1.300E-05 |
| 96 | 5 | T5S | A | 1.2e-05 | 4.6e-06 | 8.4e-06 | 8.333E-06 | 3.700E-06 | 4.600E-06 | 1.200E-05 |
| 97 | 5 | T5T | A | 8.3e-06 | 1.2e-05 | 1.2e-05 | 1.077E-05 | 2.136E-06 | 8.300E-06 | 1.200E-05 |
| 98 | 5 | T5W | A | 7.7e-06 | 6.2e-06 | 1.3e-05 | 8.967E-06 | 3.573E-06 | 6.200E-06 | 1.300E-05 |
| 99 | 5 | T5Y | A | 7.3e-06 | 7.3e-06 | 6.8e-06 | 7.133E-06 | 2.887E-07 | 6.800E-06 | 7.300E-06 |
| 100 | 5 | T5V | A | 5.4e-06 | 9.6e-06 | 1.4e-05 | 9.667E-06 | 4.300E-06 | 5.400E-06 | 1.400E-05 |
| 101 | 6 | G6A | A | 9.9e-06 | 9.8e-06 | 4.4e-06 | 8.033E-06 | 3.147E-06 | 4.400E-06 | 9.900E-06 |
| 102 | 6 | G6R | A | 8.5e-06 | 5.4e-06 | 9.1e-06 | 7.667E-06 | 1.986E-06 | 5.400E-06 | 9.100E-06 |
| 103 | 6 | G6N | A | 9.3e-06 | 7.4e-06 | 6.4e-06 | 7.700E-06 | 1.473E-06 | 6.400E-06 | 9.300E-06 |
| 104 | 6 | G6D | A | 9.8e-06 | 8.9e-06 | 1.3e-05 | 1.057E-05 | 2.155E-06 | 8.900E-06 | 1.300E-05 |
| 105 | 6 | G6C | A | 9.2e-06 | 4.2e-06 | 6e-06 | 6.467E-06 | 2.532E-06 | 4.200E-06 | 9.200E-06 |
| 106 | 6 | G6E | A | 5e-06 | 8.4e-06 | 1.1e-05 | 8.133E-06 | 3.009E-06 | 5.000E-06 | 1.100E-05 |
| 107 | 6 | G6Q | A | 5.9e-06 | 6.9e-06 | 5.2e-06 | 6.000E-06 | 8.544E-07 | 5.200E-06 | 6.900E-06 |
| 108 | 6 | G6G | A | 1.2e-05 | 5e-06 | 3.6e-06 | 6.867E-06 | 4.500E-06 | 3.600E-06 | 1.200E-05 |
| 109 | 6 | G6H | A | 8.7e-06 | 1.7e-05 | 4.3e-06 | 1.000E-05 | 6.449E-06 | 4.300E-06 | 1.700E-05 |
| 110 | 6 | G6I | A | 9.9e-06 | 7.3e-06 | 1.3e-05 | 1.007E-05 | 2.854E-06 | 7.300E-06 | 1.300E-05 |
| 111 | 6 | G6L | A | 9.8e-06 | 7.1e-06 | 1.1e-05 | 9.300E-06 | 1.997E-06 | 7.100E-06 | 1.100E-05 |
| 112 | 6 | G6K | A | 4.3e-06 | 1.3e-05 | 3.7e-06 | 7.000E-06 | 5.205E-06 | 3.700E-06 | 1.300E-05 |
| 113 | 6 | G6M | A | 9.6e-06 | 4.9e-06 | 1.2e-05 | 8.833E-06 | 3.612E-06 | 4.900E-06 | 1.200E-05 |
| 114 | 6 | G6F | A | 6.9e-06 | 1.3e-05 | 6.6e-06 | 8.833E-06 | 3.612E-06 | 6.600E-06 | 1.300E-05 |
| 115 | 6 | G6P | A | 8.3e-06 | 8.7e-06 | 5.2e-06 | 7.400E-06 | 1.916E-06 | 5.200E-06 | 8.700E-06 |
| 116 | 6 | G6S | A | 8.7e-06 | 1e-05 | 7.4e-06 | 8.700E-06 | 1.300E-06 | 7.400E-06 | 1.000E-05 |
| 117 | 6 | G6T | A | 9.3e-06 | 8.8e-06 | 8.7e-06 | 8.933E-06 | 3.215E-07 | 8.700E-06 | 9.300E-06 |
| 118 | 6 | G6W | A | 5.3e-06 | 1.5e-05 | 4.7e-06 | 8.333E-06 | 5.781E-06 | 4.700E-06 | 1.500E-05 |
| 119 | 6 | G6Y | A | 8.6e-06 | 6.8e-06 | 5.6e-06 | 7.000E-06 | 1.510E-06 | 5.600E-06 | 8.600E-06 |
| 120 | 6 | G6V | A | 4.2e-06 | 4.7e-06 | 8.4e-06 | 5.767E-06 | 2.294E-06 | 4.200E-06 | 8.400E-06 |
| 121 | 7 | T7A | A | 5.9e-06 | 6.3e-06 | 9.4e-06 | 7.200E-06 | 1.916E-06 | 5.900E-06 | 9.400E-06 |
| 122 | 7 | T7R | A | 7.6e-06 | 1.1e-05 | 6.4e-06 | 8.333E-06 | 2.386E-06 | 6.400E-06 | 1.100E-05 |
| 123 | 7 | T7N | A | 5.8e-06 | 9.5e-06 | 6e-06 | 7.100E-06 | 2.081E-06 | 5.800E-06 | 9.500E-06 |
| 124 | 7 | T7D | A | 4.5e-06 | 9e-06 | 4.6e-06 | 6.033E-06 | 2.570E-06 | 4.500E-06 | 9.000E-06 |
| 125 | 7 | T7C | A | 8.6e-06 | 8.2e-06 | 6.3e-06 | 7.700E-06 | 1.229E-06 | 6.300E-06 | 8.600E-06 |
| 126 | 7 | T7E | A | 6.6e-06 | 7.4e-06 | 6.3e-06 | 6.767E-06 | 5.686E-07 | 6.300E-06 | 7.400E-06 |
| 127 | 7 | T7Q | A | 8.3e-06 | 7.2e-06 | 1.1e-05 | 8.833E-06 | 1.955E-06 | 7.200E-06 | 1.100E-05 |
| 128 | 7 | T7G | A | 8.7e-06 | 7.4e-06 | 7.5e-06 | 7.867E-06 | 7.234E-07 | 7.400E-06 | 8.700E-06 |
| 129 | 7 | T7H | A | 1.8e-05 | 5.1e-06 | 5.1e-06 | 9.400E-06 | 7.448E-06 | 5.100E-06 | 1.800E-05 |
| 130 | 7 | T7I | A | 9.9e-06 | 1e-05 | 6.6e-06 | 8.833E-06 | 1.935E-06 | 6.600E-06 | 1.000E-05 |
| 131 | 7 | T7L | A | 5.3e-06 | 5.4e-06 | 7.1e-06 | 5.933E-06 | 1.012E-06 | 5.300E-06 | 7.100E-06 |
| 132 | 7 | T7K | A | 5.3e-06 | 6.8e-06 | 4.4e-06 | 5.500E-06 | 1.212E-06 | 4.400E-06 | 6.800E-06 |
| 133 | 7 | T7M | A | 6.2e-06 | 6.6e-06 | 6.8e-06 | 6.533E-06 | 3.055E-07 | 6.200E-06 | 6.800E-06 |
| 134 | 7 | T7F | A | 6.9e-06 | 5.9e-06 | 7.1e-06 | 6.633E-06 | 6.429E-07 | 5.900E-06 | 7.100E-06 |
| 135 | 7 | T7P | A | 1.1e-05 | 1.1e-05 | 5.7e-06 | 9.233E-06 | 3.060E-06 | 5.700E-06 | 1.100E-05 |
| 136 | 7 | T7S | A | 5.5e-06 | 8e-06 | 6.7e-06 | 6.733E-06 | 1.250E-06 | 5.500E-06 | 8.000E-06 |
| 137 | 7 | T7T | A | 7.9e-06 | 1e-05 | 6.9e-06 | 8.267E-06 | 1.582E-06 | 6.900E-06 | 1.000E-05 |
| 138 | 7 | T7W | A | 9.1e-06 | 1.2e-05 | 8.3e-06 | 9.800E-06 | 1.947E-06 | 8.300E-06 | 1.200E-05 |
| 139 | 7 | T7Y | A | 4.5e-06 | 5e-06 | 7.5e-06 | 5.667E-06 | 1.607E-06 | 4.500E-06 | 7.500E-06 |
| 140 | 7 | T7V | A | 7.8e-06 | 8.3e-06 | 5.3e-06 | 7.133E-06 | 1.607E-06 | 5.300E-06 | 8.300E-06 |
| 141 | 8 | D8A | A | 4.3e-06 | 9.2e-06 | 6.4e-06 | 6.633E-06 | 2.458E-06 | 4.300E-06 | 9.200E-06 |
| 142 | 8 | D8R | A | 7.8e-06 | 9.3e-06 | 9.3e-06 | 8.800E-06 | 8.660E-07 | 7.800E-06 | 9.300E-06 |
| 143 | 8 | D8N | A | 1.1e-05 | 3.6e-06 | 8.9e-06 | 7.833E-06 | 3.814E-06 | 3.600E-06 | 1.100E-05 |
| 144 | 8 | D8D | A | 9e-06 | 6.9e-06 | 1.1e-05 | 8.967E-06 | 2.050E-06 | 6.900E-06 | 1.100E-05 |
| 145 | 8 | D8C | A | 1.2e-05 | 1e-05 | 9e-06 | 1.033E-05 | 1.528E-06 | 9.000E-06 | 1.200E-05 |
| 146 | 8 | D8E | A | 6.8e-06 | 7.5e-06 | 9.2e-06 | 7.833E-06 | 1.234E-06 | 6.800E-06 | 9.200E-06 |
| 147 | 8 | D8Q | A | 1e-05 | 8.7e-06 | 6.8e-06 | 8.500E-06 | 1.609E-06 | 6.800E-06 | 1.000E-05 |
| 148 | 8 | D8G | A | 1.5e-05 | 1.3e-05 | 8.3e-06 | 1.210E-05 | 3.439E-06 | 8.300E-06 | 1.500E-05 |
| 149 | 8 | D8H | A | 3e-06 | 1.4e-05 | 1.5e-05 | 1.067E-05 | 6.658E-06 | 3.000E-06 | 1.500E-05 |
| 150 | 8 | D8I | A | 8.3e-06 | 1e-05 | 5.1e-06 | 7.800E-06 | 2.488E-06 | 5.100E-06 | 1.000E-05 |
| 151 | 8 | D8L | A | 4.5e-06 | 1.3e-05 | 9.5e-06 | 9.000E-06 | 4.272E-06 | 4.500E-06 | 1.300E-05 |
| 152 | 8 | D8K | A | 8.5e-06 | 5.5e-06 | 9.8e-06 | 7.933E-06 | 2.205E-06 | 5.500E-06 | 9.800E-06 |
| 153 | 8 | D8M | A | 4.8e-06 | 6.5e-06 | 9.1e-06 | 6.800E-06 | 2.166E-06 | 4.800E-06 | 9.100E-06 |
| 154 | 8 | D8F | A | 6.6e-06 | 4.5e-06 | 4.7e-06 | 5.267E-06 | 1.159E-06 | 4.500E-06 | 6.600E-06 |
| 155 | 8 | D8P | A | 1e-05 | 6.7e-06 | 8.7e-06 | 8.467E-06 | 1.662E-06 | 6.700E-06 | 1.000E-05 |
| 156 | 8 | D8S | A | 7.2e-06 | 1.2e-05 | 9e-06 | 9.400E-06 | 2.425E-06 | 7.200E-06 | 1.200E-05 |
| 157 | 8 | D8T | A | 8.2e-06 | 5.4e-06 | 9.3e-06 | 7.633E-06 | 2.011E-06 | 5.400E-06 | 9.300E-06 |
| 158 | 8 | D8W | A | 6.6e-06 | 7.1e-06 | 1.3e-05 | 8.900E-06 | 3.559E-06 | 6.600E-06 | 1.300E-05 |
| 159 | 8 | D8Y | A | 9.5e-06 | 1e-05 | 9.2e-06 | 9.567E-06 | 4.041E-07 | 9.200E-06 | 1.000E-05 |
| 160 | 8 | D8V | A | 6.4e-06 | 8.6e-06 | 6.8e-06 | 7.267E-06 | 1.172E-06 | 6.400E-06 | 8.600E-06 |
| 161 | 9 | M9A | A | 1.1e-05 | 4.8e-06 | 9.5e-06 | 8.433E-06 | 3.235E-06 | 4.800E-06 | 1.100E-05 |
| 162 | 9 | M9R | A | 9.4e-06 | 1.2e-05 | 6.7e-06 | 9.367E-06 | 2.650E-06 | 6.700E-06 | 1.200E-05 |
| 163 | 9 | M9N | A | 1.2e-05 | 8.9e-06 | 4.6e-06 | 8.500E-06 | 3.716E-06 | 4.600E-06 | 1.200E-05 |
| 164 | 9 | M9D | A | 5.9e-06 | 8.4e-06 | 1.1e-05 | 8.433E-06 | 2.550E-06 | 5.900E-06 | 1.100E-05 |
| 165 | 9 | M9C | A | 1.1e-05 | 7.2e-06 | 1.2e-05 | 1.007E-05 | 2.532E-06 | 7.200E-06 | 1.200E-05 |
| 166 | 9 | M9E | A | 1e-05 | 4.6e-06 | 8.1e-06 | 7.567E-06 | 2.739E-06 | 4.600E-06 | 1.000E-05 |
| 167 | 9 | M9Q | A | 3.7e-06 | 8.7e-06 | 1.1e-05 | 7.800E-06 | 3.732E-06 | 3.700E-06 | 1.100E-05 |
| 168 | 9 | M9G | A | 1.3e-05 | 6e-06 | 1.3e-05 | 1.067E-05 | 4.041E-06 | 6.000E-06 | 1.300E-05 |
| 169 | 9 | M9H | A | 6.3e-06 | 6.9e-06 | 4.7e-06 | 5.967E-06 | 1.137E-06 | 4.700E-06 | 6.900E-06 |

| | | | | | | | | | | |
|---|---|---|---|---|---|---|---|---|---|---|
| 170 | 9 | M9I | A | 8e-06 | 5.4e-06 | 8.8e-06 | 7.400E-06 | 1.778E-06 | 5.400E-06 | 8.800E-06 |
| 171 | 9 | M9L | A | 7.9e-06 | 1e-05 | 4.6e-06 | 7.500E-06 | 2.722E-06 | 4.600E-06 | 1.000E-05 |
| 172 | 9 | M9K | A | 5.8e-06 | 6.6e-06 | 6.8e-06 | 6.400E-06 | 5.292E-07 | 5.800E-06 | 6.800E-06 |
| 173 | 9 | M9M | A | 6e-06 | 6e-06 | 9.1e-06 | 7.033E-06 | 1.790E-06 | 6.000E-06 | 9.100E-06 |
| 174 | 9 | M9F | A | 4.9e-06 | 1.1e-05 | 6.1e-06 | 7.333E-06 | 3.232E-06 | 4.900E-06 | 1.100E-05 |
| 175 | 9 | M9P | A | 5.6e-06 | 9.9e-06 | 6.4e-06 | 7.300E-06 | 2.287E-06 | 5.600E-06 | 9.900E-06 |
| 176 | 9 | M9S | A | 6.8e-06 | 5.8e-06 | 5.1e-06 | 5.900E-06 | 8.544E-07 | 5.100E-06 | 6.800E-06 |
| 177 | 9 | M9T | A | 8.6e-06 | 7e-06 | 6.1e-06 | 7.233E-06 | 1.266E-06 | 6.100E-06 | 8.600E-06 |
| 178 | 9 | M9W | A | 4.6e-06 | 5.7e-06 | 6.4e-06 | 5.567E-06 | 9.074E-07 | 4.600E-06 | 6.400E-06 |
| 179 | 9 | M9Y | A | 7.3e-06 | 8e-06 | 1e-05 | 8.433E-06 | 1.401E-06 | 7.300E-06 | 1.000E-05 |
| 180 | 9 | M9V | A | 9.6e-06 | 9.8e-06 | 6.9e-06 | 8.767E-06 | 1.620E-06 | 6.900E-06 | 9.800E-06 |
| 181 | 10 | K10A | A | 8.5e-06 | 5.9e-06 | 5e-06 | 6.467E-06 | 1.818E-06 | 5.000E-06 | 8.500E-06 |
| 182 | 10 | K10R | A | 1.1e-05 | 8.8e-06 | 1.3e-05 | 1.093E-05 | 2.101E-06 | 8.800E-06 | 1.300E-05 |
| 183 | 10 | K10N | A | 5.6e-06 | 8.3e-06 | 6.8e-06 | 6.900E-06 | 1.353E-06 | 5.600E-06 | 8.300E-06 |
| 184 | 10 | K10D | A | 6.4e-06 | 8.6e-06 | 8.7e-06 | 7.900E-06 | 1.300E-06 | 6.400E-06 | 8.700E-06 |
| 185 | 10 | K10C | A | 4.3e-06 | 6.7e-06 | 4.3e-06 | 5.100E-06 | 1.386E-06 | 4.300E-06 | 6.700E-06 |
| 186 | 10 | K10E | A | 1.4e-05 | 5.2e-06 | 6.9e-06 | 8.700E-06 | 4.668E-06 | 5.200E-06 | 1.400E-05 |
| 187 | 10 | K10Q | A | 7.8e-06 | 1.1e-05 | 1.3e-05 | 1.060E-05 | 2.623E-06 | 7.800E-06 | 1.300E-05 |
| 188 | 10 | K10G | A | 6.1e-06 | 9.8e-06 | 7.4e-06 | 7.767E-06 | 1.877E-06 | 6.100E-06 | 9.800E-06 |
| 189 | 10 | K10H | A | 7e-06 | 8e-06 | 9.3e-06 | 8.100E-06 | 1.153E-06 | 7.000E-06 | 9.300E-06 |
| 190 | 10 | K10I | A | 8.6e-06 | 8.3e-06 | 4.8e-06 | 7.233E-06 | 2.113E-06 | 4.800E-06 | 8.600E-06 |
| 191 | 10 | K10L | A | 1e-05 | 7.9e-06 | 9.6e-06 | 9.167E-06 | 1.115E-06 | 7.900E-06 | 1.000E-05 |
| 192 | 10 | K10K | A | 8.7e-06 | 9e-06 | 5.9e-06 | 7.867E-06 | 1.710E-06 | 5.900E-06 | 9.000E-06 |
| 193 | 10 | K10M | A | 5.4e-06 | 7.1e-06 | 1e-05 | 7.500E-06 | 2.326E-06 | 5.400E-06 | 1.000E-05 |
| 194 | 10 | K10F | A | 1e-05 | 5.2e-06 | 1.1e-05 | 8.733E-06 | 3.101E-06 | 5.200E-06 | 1.100E-05 |
| 195 | 10 | K10P | A | 9.2e-06 | 1.1e-05 | 6e-06 | 8.733E-06 | 2.532E-06 | 6.000E-06 | 1.100E-05 |
| 196 | 10 | K10S | A | 6.2e-06 | 7.3e-06 | 8.6e-06 | 7.367E-06 | 1.201E-06 | 6.200E-06 | 8.600E-06 |
| 197 | 10 | K10T | A | 9.4e-06 | 1.2e-05 | 1.3e-05 | 1.147E-05 | 1.858E-06 | 9.400E-06 | 1.300E-05 |
| 198 | 10 | K10W | A | 5.9e-06 | 5e-06 | 9.1e-06 | 6.667E-06 | 2.155E-06 | 5.000E-06 | 9.100E-06 |
| 199 | 10 | K10Y | A | 1.4e-05 | 9.4e-06 | 4.9e-06 | 9.433E-06 | 4.550E-06 | 4.900E-06 | 1.400E-05 |
| 200 | 10 | K10V | A | 3.5e-06 | 7.8e-06 | 4.8e-06 | 5.367E-06 | 2.205E-06 | 3.500E-06 | 7.800E-06 |
| 201 | 11 | L11A | A | 1.3e-05 | 1.1e-05 | 1.4e-05 | 1.267E-05 | 1.528E-06 | 1.100E-05 | 1.400E-05 |
| 202 | 11 | L11R | A | 6.7e-06 | 5.8e-06 | 9.3e-06 | 7.267E-06 | 1.818E-06 | 5.800E-06 | 9.300E-06 |
| 203 | 11 | L11N | A | 6.3e-06 | 8.7e-06 | 7.9e-06 | 7.633E-06 | 1.222E-06 | 6.300E-06 | 8.700E-06 |
| 204 | 11 | L11D | A | 6.5e-06 | 9.6e-06 | 1.2e-05 | 9.367E-06 | 2.757E-06 | 6.500E-06 | 1.200E-05 |
| 205 | 11 | L11C | A | 9.4e-06 | 4.6e-06 | 5.9e-06 | 6.633E-06 | 2.483E-06 | 4.600E-06 | 9.400E-06 |
| 206 | 11 | L11E | A | 1.2e-05 | 8.9e-06 | 5.2e-06 | 8.700E-06 | 3.404E-06 | 5.200E-06 | 1.200E-05 |
| 207 | 11 | L11Q | A | 8.8e-06 | 3.7e-06 | 9.4e-06 | 7.300E-06 | 3.132E-06 | 3.700E-06 | 9.400E-06 |
| 208 | 11 | L11G | A | 8.7e-06 | 5.4e-06 | 6.8e-06 | 6.967E-06 | 1.656E-06 | 5.400E-06 | 8.700E-06 |
| 209 | 11 | L11H | A | 6.5e-06 | 5.4e-06 | 6.2e-06 | 6.033E-06 | 5.686E-07 | 5.400E-06 | 6.500E-06 |
| 210 | 11 | L11I | A | 6.4e-06 | 5.1e-06 | 8.5e-06 | 6.667E-06 | 1.716E-06 | 5.100E-06 | 8.500E-06 |
| 211 | 11 | L11L | A | 8.3e-06 | 1.3e-05 | 1.2e-05 | 1.110E-05 | 2.476E-06 | 8.300E-06 | 1.300E-05 |
| 212 | 11 | L11K | A | 1.5e-05 | 1e-05 | 4.4e-06 | 9.800E-06 | 5.303E-06 | 4.400E-06 | 1.500E-05 |
| 213 | 11 | L11M | A | 7e-06 | 1.2e-05 | 9.4e-06 | 9.467E-06 | 2.501E-06 | 7.000E-06 | 1.200E-05 |
| 214 | 11 | L11F | A | 7e-06 | 6.1e-06 | 1.1e-05 | 8.033E-06 | 2.608E-06 | 6.100E-06 | 1.100E-05 |
| 215 | 11 | L11P | A | 8.1e-06 | 6.5e-06 | 9.3e-06 | 7.967E-06 | 1.405E-06 | 6.500E-06 | 9.300E-06 |
| 216 | 11 | L11S | A | 7.1e-06 | 6e-06 | 5.9e-06 | 6.333E-06 | 6.658E-07 | 5.900E-06 | 7.100E-06 |
| 217 | 11 | L11T | A | 6.9e-06 | 9.9e-06 | 5.8e-06 | 7.533E-06 | 2.122E-06 | 5.800E-06 | 9.900E-06 |
| 218 | 11 | L11W | A | 9.1e-06 | 5.9e-06 | 1.3e-05 | 9.333E-06 | 3.556E-06 | 5.900E-06 | 1.300E-05 |
| 219 | 11 | L11Y | A | 9.6e-06 | 7.8e-06 | 1.1e-05 | 9.467E-06 | 1.604E-06 | 7.800E-06 | 1.100E-05 |
| 220 | 11 | L11V | A | 6.1e-06 | 6.9e-06 | 1e-05 | 7.667E-06 | 2.060E-06 | 6.100E-06 | 1.000E-05 |
| 221 | 12 | R12A | A | 6.3e-06 | 1.1e-05 | 3.5e-06 | 6.933E-06 | 3.790E-06 | 3.500E-06 | 1.100E-05 |
| 222 | 12 | R12R | A | 6.6e-06 | 9.4e-06 | 9.1e-06 | 8.367E-06 | 1.537E-06 | 6.600E-06 | 9.400E-06 |
| 223 | 12 | R12N | A | 9.2e-06 | 5.2e-06 | 6.4e-06 | 6.933E-06 | 2.053E-06 | 5.200E-06 | 9.200E-06 |
| 224 | 12 | R12D | A | 6.8e-06 | 8.8e-06 | 5.8e-06 | 7.133E-06 | 1.528E-06 | 5.800E-06 | 8.800E-06 |
| 225 | 12 | R12C | A | 6.9e-06 | 7.3e-06 | 1e-05 | 8.067E-06 | 1.686E-06 | 6.900E-06 | 1.000E-05 |
| 226 | 12 | R12E | A | 4e-06 | 6.6e-06 | 7.2e-06 | 5.933E-06 | 1.701E-06 | 4.000E-06 | 7.200E-06 |
| 227 | 12 | R12Q | A | 6.5e-06 | 7.9e-06 | 6.9e-06 | 7.100E-06 | 7.211E-07 | 6.500E-06 | 7.900E-06 |
| 228 | 12 | R12G | A | 8.9e-06 | 8.6e-06 | 1.2e-05 | 9.833E-06 | 1.882E-06 | 8.600E-06 | 1.200E-05 |
| 229 | 12 | R12H | A | 8.6e-06 | 8.3e-06 | 7.3e-06 | 8.067E-06 | 6.807E-07 | 7.300E-06 | 8.600E-06 |
| 230 | 12 | R12I | A | 1e-05 | 8.1e-06 | 7.4e-06 | 8.500E-06 | 1.345E-06 | 7.400E-06 | 1.000E-05 |
| 231 | 12 | R12L | A | 9.6e-06 | 9.6e-06 | 8.9e-06 | 9.367E-06 | 4.041E-07 | 8.900E-06 | 9.600E-06 |
| 232 | 12 | R12K | A | 8.7e-06 | 6.3e-06 | 6e-06 | 7.000E-06 | 1.480E-06 | 6.000E-06 | 8.700E-06 |
| 233 | 12 | R12M | A | 6.7e-06 | 9.3e-06 | 8.1e-06 | 8.033E-06 | 1.301E-06 | 6.700E-06 | 9.300E-06 |
| 234 | 12 | R12F | A | 6.8e-06 | 5.1e-06 | 1.4e-05 | 8.633E-06 | 4.725E-06 | 5.100E-06 | 1.400E-05 |
| 235 | 12 | R12P | A | 1.4e-05 | 5.7e-06 | 8.2e-06 | 9.300E-06 | 4.258E-06 | 5.700E-06 | 1.400E-05 |
| 236 | 12 | R12S | A | 7.5e-06 | 4.7e-06 | 7.6e-06 | 6.600E-06 | 1.646E-06 | 4.700E-06 | 7.600E-06 |
| 237 | 12 | R12T | A | 5.5e-06 | 7.4e-06 | 9.7e-06 | 7.533E-06 | 2.103E-06 | 5.500E-06 | 9.700E-06 |
| 238 | 12 | R12W | A | 6.5e-06 | 8.2e-06 | 1.1e-05 | 8.567E-06 | 2.272E-06 | 6.500E-06 | 1.100E-05 |
| 239 | 12 | R12Y | A | 1.9e-05 | 7.9e-06 | 9e-06 | 1.197E-05 | 6.116E-06 | 7.900E-06 | 1.900E-05 |
| 240 | 12 | R12V | A | 7.7e-06 | 1e-05 | 6.9e-06 | 8.200E-06 | 1.609E-06 | 6.900E-06 | 1.000E-05 |
| 241 | 13 | L13A | A | 1e-05 | 9.7e-06 | 8.1e-06 | 9.267E-06 | 1.021E-06 | 8.100E-06 | 1.000E-05 |
| 242 | 13 | L13R | A | 7.3e-06 | 8.8e-06 | 5.9e-06 | 7.333E-06 | 1.450E-06 | 5.900E-06 | 8.800E-06 |
| 243 | 13 | L13N | A | 6.1e-06 | 5.8e-06 | 8.8e-06 | 6.900E-06 | 1.652E-06 | 5.800E-06 | 8.800E-06 |
| 244 | 13 | L13D | A | 1.3e-05 | 6e-06 | 8e-06 | 9.000E-06 | 3.606E-06 | 6.000E-06 | 1.300E-05 |
| 245 | 13 | L13C | A | 3.5e-06 | 6.9e-06 | 9.4e-06 | 6.600E-06 | 2.961E-06 | 3.500E-06 | 9.400E-06 |
| 246 | 13 | L13E | A | 8.4e-06 | 1.1e-05 | 8.8e-06 | 9.400E-06 | 1.400E-06 | 8.400E-06 | 1.100E-05 |
| 247 | 13 | L13Q | A | 5.6e-06 | 8.8e-06 | 5.2e-06 | 6.533E-06 | 1.973E-06 | 5.200E-06 | 8.800E-06 |
| 248 | 13 | L13G | A | 1.1e-05 | 3.9e-06 | 8.4e-06 | 7.767E-06 | 3.592E-06 | 3.900E-06 | 1.100E-05 |
| 249 | 13 | L13H | A | 7.3e-06 | 7e-06 | 1.2e-05 | 8.767E-06 | 2.804E-06 | 7.000E-06 | 1.200E-05 |
| 250 | 13 | L13I | A | 6.9e-06 | 1.5e-05 | 4.4e-06 | 8.767E-06 | 5.541E-06 | 4.400E-06 | 1.500E-05 |
| 251 | 13 | L13L | A | 1.2e-05 | 1e-05 | 1.1e-05 | 1.100E-05 | 1.000E-06 | 1.000E-05 | 1.200E-05 |
| 252 | 13 | L13K | A | 9.2e-06 | 1.3e-05 | 4.7e-06 | 8.967E-06 | 4.155E-06 | 4.700E-06 | 1.300E-05 |
| 253 | 13 | L13M | A | 3.4e-06 | 9.1e-06 | 1.2e-05 | 8.167E-06 | 4.375E-06 | 3.400E-06 | 1.200E-05 |
| 254 | 13 | L13F | A | 9.3e-06 | 9.6e-06 | 1.3e-05 | 1.063E-05 | 2.055E-06 | 9.300E-06 | 1.300E-05 |

| | | | | | | | | | | |
|---|---|---|---|---|---|---|---|---|---|---|
| 255 | 13 | L13P | A | 6.2e-06 | 1.4e-05 | 8.4e-06 | 9.533E-06 | 4.022E-06 | 6.200E-06 | 1.400E-05 |
| 256 | 13 | L13S | A | 6.1e-06 | 1.1e-05 | 8.7e-06 | 8.600E-06 | 2.452E-06 | 6.100E-06 | 1.100E-05 |
| 257 | 13 | L13T | A | 6e-06 | 6.1e-06 | 3.2e-06 | 5.100E-06 | 1.646E-06 | 3.200E-06 | 6.100E-06 |
| 258 | 13 | L13W | A | 7.5e-06 | 6.5e-06 | 7.5e-06 | 7.167E-06 | 5.774E-07 | 6.500E-06 | 7.500E-06 |
| 259 | 13 | L13Y | A | 5.7e-06 | 6.8e-06 | 6.8e-06 | 6.433E-06 | 6.351E-07 | 5.700E-06 | 6.800E-06 |
| 260 | 13 | L13V | A | 4.5e-06 | 6.2e-06 | 1.1e-05 | 7.233E-06 | 3.371E-06 | 4.500E-06 | 1.100E-05 |
| 261 | 14 | P14A | A | 8.5e-06 | 6.8e-06 | 8.9e-06 | 8.067E-06 | 1.115E-06 | 6.800E-06 | 8.900E-06 |
| 262 | 14 | P14R | A | 6.8e-06 | 5.6e-06 | 4.6e-06 | 5.667E-06 | 1.102E-06 | 4.600E-06 | 6.800E-06 |
| 263 | 14 | P14N | A | 9.4e-06 | 1.3e-05 | 7.8e-06 | 1.007E-05 | 2.663E-06 | 7.800E-06 | 1.300E-05 |
| 264 | 14 | P14D | A | 9.9e-06 | 8.7e-06 | 8.3e-06 | 8.967E-06 | 8.327E-07 | 8.300E-06 | 9.900E-06 |
| 265 | 14 | P14C | A | 4.7e-06 | 1.2e-05 | 8.9e-06 | 8.533E-06 | 3.664E-06 | 4.700E-06 | 1.200E-05 |
| 266 | 14 | P14E | A | 4.8e-06 | 1e-05 | 8.1e-06 | 7.633E-06 | 2.631E-06 | 4.800E-06 | 1.000E-05 |
| 267 | 14 | P14Q | A | 7.8e-06 | 8.6e-06 | 8.2e-06 | 8.200E-06 | 4.000E-07 | 7.800E-06 | 8.600E-06 |
| 268 | 14 | P14G | A | 6.2e-06 | 6.2e-06 | 6.3e-06 | 6.233E-06 | 5.774E-08 | 6.200E-06 | 6.300E-06 |
| 269 | 14 | P14H | A | 9.5e-06 | 6.3e-06 | 9.6e-06 | 8.467E-06 | 1.877E-06 | 6.300E-06 | 9.600E-06 |
| 270 | 14 | P14I | A | 7.7e-06 | 7e-06 | 9.5e-06 | 8.067E-06 | 1.290E-06 | 7.000E-06 | 9.500E-06 |
| 271 | 14 | P14L | A | 8.3e-06 | 9.1e-06 | 5.4e-06 | 7.600E-06 | 1.947E-06 | 5.400E-06 | 9.100E-06 |
| 272 | 14 | P14K | A | 6.5e-06 | 8.4e-06 | 8.5e-06 | 7.800E-06 | 1.127E-06 | 6.500E-06 | 8.500E-06 |
| 273 | 14 | P14M | A | 7.7e-06 | 7.4e-06 | 5.1e-06 | 6.733E-06 | 1.422E-06 | 5.100E-06 | 7.700E-06 |
| 274 | 14 | P14F | A | 6e-06 | 8e-06 | 6.5e-06 | 6.833E-06 | 1.041E-06 | 6.000E-06 | 8.000E-06 |
| 275 | 14 | P14P | A | 1.1e-05 | 6.7e-06 | 3.2e-06 | 6.967E-06 | 3.907E-06 | 3.200E-06 | 1.100E-05 |
| 276 | 14 | P14S | A | 1.6e-05 | 6.6e-06 | 5.6e-06 | 9.400E-06 | 5.738E-06 | 5.600E-06 | 1.600E-05 |
| 277 | 14 | P14T | A | 6.6e-06 | 8.4e-06 | 1.2e-05 | 9.000E-06 | 2.750E-06 | 6.600E-06 | 1.200E-05 |
| 278 | 14 | P14W | A | 9.3e-06 | 6.4e-06 | 6.3e-06 | 7.333E-06 | 1.704E-06 | 6.300E-06 | 9.300E-06 |
| 279 | 14 | P14Y | A | 6.6e-06 | 8.8e-06 | 6.7e-06 | 7.367E-06 | 1.242E-06 | 6.600E-06 | 8.800E-06 |
| 280 | 14 | P14V | A | 7.2e-06 | 9.1e-06 | 9.2e-06 | 8.500E-06 | 1.127E-06 | 7.200E-06 | 9.200E-06 |
| 281 | 15 | A15A | A | 9.2e-06 | 8.2e-06 | 6.1e-06 | 7.833E-06 | 1.582E-06 | 6.100E-06 | 9.200E-06 |
| 282 | 15 | A15R | A | 6.7e-06 | 9.2e-06 | 5.6e-06 | 7.167E-06 | 1.845E-06 | 5.600E-06 | 9.200E-06 |
| 283 | 15 | A15N | A | 7e-06 | 9.4e-06 | 8.9e-06 | 8.433E-06 | 1.266E-06 | 7.000E-06 | 9.400E-06 |
| 284 | 15 | A15D | A | 8e-06 | 8.2e-06 | 4.3e-06 | 6.833E-06 | 2.196E-06 | 4.300E-06 | 8.200E-06 |
| 285 | 15 | A15C | A | 8.9e-06 | 7.3e-06 | 1.3e-05 | 9.733E-06 | 2.940E-06 | 7.300E-06 | 1.300E-05 |
| 286 | 15 | A15E | A | 6.3e-06 | 6e-06 | 1e-05 | 7.433E-06 | 2.228E-06 | 6.000E-06 | 1.000E-05 |
| 287 | 15 | A15Q | A | 4.7e-06 | 8.2e-06 | 5.6e-06 | 6.167E-06 | 1.818E-06 | 4.700E-06 | 8.200E-06 |
| 288 | 15 | A15G | A | 1.6e-05 | 7.8e-06 | 1.5e-05 | 1.293E-05 | 4.474E-06 | 7.800E-06 | 1.600E-05 |
| 289 | 15 | A15H | A | 9.8e-06 | 8.7e-06 | 7.5e-06 | 8.667E-06 | 1.150E-06 | 7.500E-06 | 9.800E-06 |
| 290 | 15 | A15I | A | 6.9e-06 | 7.5e-06 | 8.5e-06 | 7.633E-06 | 8.083E-07 | 6.900E-06 | 8.500E-06 |
| 291 | 15 | A15L | A | 6.9e-06 | 9e-06 | 1.3e-05 | 9.633E-06 | 3.099E-06 | 6.900E-06 | 1.300E-05 |
| 292 | 15 | A15K | A | 7.5e-06 | 1.1e-05 | 7.4e-06 | 8.633E-06 | 2.050E-06 | 7.400E-06 | 1.100E-05 |
| 293 | 15 | A15M | A | 8.8e-06 | 8.8e-06 | 8.4e-06 | 8.667E-06 | 2.309E-07 | 8.400E-06 | 8.800E-06 |
| 294 | 15 | A15F | A | 1.6e-05 | 9.2e-06 | 5e-06 | 1.007E-05 | 5.551E-06 | 5.000E-06 | 1.600E-05 |
| 295 | 15 | A15P | A | 1.3e-05 | 9.7e-06 | 7.6e-06 | 1.010E-05 | 2.722E-06 | 7.600E-06 | 1.300E-05 |
| 296 | 15 | A15S | A | 1.1e-05 | 8.9e-06 | 9e-06 | 9.633E-06 | 1.185E-06 | 8.900E-06 | 1.100E-05 |
| 297 | 15 | A15T | A | 1.3e-05 | 9.5e-06 | 5.9e-06 | 9.467E-06 | 3.550E-06 | 5.900E-06 | 1.300E-05 |
| 298 | 15 | A15W | A | 7.6e-06 | 9.2e-06 | 9.8e-06 | 8.867E-06 | 1.137E-06 | 7.600E-06 | 9.800E-06 |
| 299 | 15 | A15Y | A | 8e-06 | 5.8e-06 | 1.1e-05 | 8.267E-06 | 2.610E-06 | 5.800E-06 | 1.100E-05 |
| 300 | 15 | A15V | A | 7e-06 | 1.1e-05 | 8.7e-06 | 8.900E-06 | 2.007E-06 | 7.000E-06 | 1.100E-05 |
| 301 | 16 | S16A | A | 7.4e-06 | 6.2e-06 | 8.4e-06 | 7.333E-06 | 1.102E-06 | 6.200E-06 | 8.400E-06 |
| 302 | 16 | S16R | A | 9.1e-06 | 9.1e-06 | 1e-05 | 9.400E-06 | 5.196E-07 | 9.100E-06 | 1.000E-05 |
| 303 | 16 | S16N | A | 7.5e-06 | 1e-05 | 4.1e-06 | 7.200E-06 | 2.961E-06 | 4.100E-06 | 1.000E-05 |
| 304 | 16 | S16D | A | 6e-06 | 6.4e-06 | 7.3e-06 | 6.567E-06 | 6.658E-07 | 6.000E-06 | 7.300E-06 |
| 305 | 16 | S16C | A | 9.6e-06 | 5.8e-06 | 1.3e-05 | 9.467E-06 | 3.602E-06 | 5.800E-06 | 1.300E-05 |
| 306 | 16 | S16E | A | 5.6e-06 | 1.4e-05 | 8.9e-06 | 9.500E-06 | 4.232E-06 | 5.600E-06 | 1.400E-05 |
| 307 | 16 | S16Q | A | 4.7e-06 | 7.4e-06 | 1.4e-05 | 8.700E-06 | 4.784E-06 | 4.700E-06 | 1.400E-05 |
| 308 | 16 | S16G | A | 9.1e-06 | 5.4e-06 | 1.4e-05 | 9.500E-06 | 4.314E-06 | 5.400E-06 | 1.400E-05 |
| 309 | 16 | S16H | A | 1.2e-05 | 1.4e-05 | 7e-06 | 1.100E-05 | 3.606E-06 | 7.000E-06 | 1.400E-05 |
| 310 | 16 | S16I | A | 9.7e-06 | 9.1e-06 | 6.1e-06 | 8.300E-06 | 1.929E-06 | 6.100E-06 | 9.700E-06 |
| 311 | 16 | S16L | A | 7e-06 | 9.8e-06 | 8.4e-06 | 8.400E-06 | 1.400E-06 | 7.000E-06 | 9.800E-06 |
| 312 | 16 | S16K | A | 5.7e-06 | 4.9e-06 | 8.7e-06 | 6.433E-06 | 2.003E-06 | 4.900E-06 | 8.700E-06 |
| 313 | 16 | S16M | A | 1e-05 | 9.4e-06 | 8.4e-06 | 9.267E-06 | 8.083E-07 | 8.400E-06 | 1.000E-05 |
| 314 | 16 | S16F | A | 1.2e-05 | 8.3e-06 | 7.8e-06 | 9.367E-06 | 2.294E-06 | 7.800E-06 | 1.200E-05 |
| 315 | 16 | S16P | A | 4.8e-06 | 6.5e-06 | 8.5e-06 | 6.600E-06 | 1.852E-06 | 4.800E-06 | 8.500E-06 |
| 316 | 16 | S16S | A | 8.2e-06 | 6.2e-06 | 8.4e-06 | 7.600E-06 | 1.217E-06 | 6.200E-06 | 8.400E-06 |
| 317 | 16 | S16T | A | 6.7e-06 | 8.2e-06 | 9e-06 | 7.967E-06 | 1.168E-06 | 6.700E-06 | 9.000E-06 |
| 318 | 16 | S16W | A | 4.8e-06 | 1.3e-05 | 8.9e-06 | 8.900E-06 | 4.100E-06 | 4.800E-06 | 1.300E-05 |
| 319 | 16 | S16Y | A | 4.9e-06 | 5.1e-06 | 5.5e-06 | 5.167E-06 | 3.055E-07 | 4.900E-06 | 5.500E-06 |
| 320 | 16 | S16V | A | 1e-05 | 6.4e-06 | 9.3e-06 | 8.567E-06 | 1.909E-06 | 6.400E-06 | 1.000E-05 |
| 321 | 17 | P17A | A | 7.9e-06 | 9.9e-06 | 5.5e-06 | 7.767E-06 | 2.203E-06 | 5.500E-06 | 9.900E-06 |
| 322 | 17 | P17R | A | 9.3e-06 | 1.2e-05 | 8.1e-06 | 9.800E-06 | 1.997E-06 | 8.100E-06 | 1.200E-05 |
| 323 | 17 | P17N | A | 1.6e-05 | 9.2e-06 | 6.3e-06 | 1.050E-05 | 4.979E-06 | 6.300E-06 | 1.600E-05 |
| 324 | 17 | P17D | A | 8.4e-06 | 9.3e-06 | 9.4e-06 | 9.033E-06 | 5.508E-07 | 8.400E-06 | 9.400E-06 |
| 325 | 17 | P17C | A | 7.8e-06 | 9e-06 | 3.8e-06 | 6.867E-06 | 2.723E-06 | 3.800E-06 | 9.000E-06 |
| 326 | 17 | P17E | A | 8.6e-06 | 6.9e-06 | 6e-06 | 7.167E-06 | 1.320E-06 | 6.000E-06 | 8.600E-06 |
| 327 | 17 | P17Q | A | 1.2e-05 | 6.8e-06 | 8.2e-06 | 9.000E-06 | 2.691E-06 | 6.800E-06 | 1.200E-05 |
| 328 | 17 | P17G | A | 1.3e-05 | 9.1e-06 | 9.5e-06 | 1.053E-05 | 2.146E-06 | 9.100E-06 | 1.300E-05 |
| 329 | 17 | P17H | A | 6.5e-06 | 9.9e-06 | 1.2e-05 | 9.467E-06 | 2.775E-06 | 6.500E-06 | 1.200E-05 |
| 330 | 17 | P17I | A | 6.7e-06 | 9.3e-06 | 1.2e-05 | 9.333E-06 | 2.650E-06 | 6.700E-06 | 1.200E-05 |
| 331 | 17 | P17L | A | 7.5e-06 | 9.6e-06 | 1e-05 | 9.033E-06 | 1.343E-06 | 7.500E-06 | 1.000E-05 |
| 332 | 17 | P17K | A | 9.8e-06 | 1e-05 | 8.6e-06 | 9.467E-06 | 7.572E-07 | 8.600E-06 | 1.000E-05 |
| 333 | 17 | P17M | A | 6.3e-06 | 7.5e-06 | 9.7e-06 | 7.833E-06 | 1.724E-06 | 6.300E-06 | 9.700E-06 |
| 334 | 17 | P17F | A | 1.4e-05 | 1.1e-05 | 8.3e-06 | 1.110E-05 | 2.851E-06 | 8.300E-06 | 1.400E-05 |
| 335 | 17 | P17P | A | 1.5e-05 | 6e-06 | 9.1e-06 | 1.003E-05 | 4.572E-06 | 6.000E-06 | 1.500E-05 |
| 336 | 17 | P17S | A | 1.3e-05 | 6.5e-06 | 8.2e-06 | 9.233E-06 | 3.371E-06 | 6.500E-06 | 1.300E-05 |
| 337 | 17 | P17T | A | 9.7e-06 | 9.3e-06 | 4.5e-06 | 7.833E-06 | 2.894E-06 | 4.500E-06 | 9.700E-06 |
| 338 | 17 | P17W | A | 9.8e-06 | 7.8e-06 | 6.4e-06 | 8.000E-06 | 1.709E-06 | 6.400E-06 | 9.800E-06 |
| 339 | 17 | P17Y | A | 6.2e-06 | 6.6e-06 | 6.3e-06 | 6.367E-06 | 2.082E-07 | 6.200E-06 | 6.600E-06 |

| | | | | | | | | | | |
|---|---|---|---|---|---|---|---|---|---|---|
| 340 | 17 | P17V | A | 9.5e-06 | 7.8e-06 | 9.5e-06 | 8.933E-06 | 9.815E-07 | 7.800E-06 | 9.500E-06 |
| 341 | 18 | E18A | A | 5e-06 | 8.2e-06 | 7.7e-06 | 6.967E-06 | 1.721E-06 | 5.000E-06 | 8.200E-06 |
| 342 | 18 | E18R | A | 8.7e-06 | 4.9e-06 | 1e-05 | 7.867E-06 | 2.650E-06 | 4.900E-06 | 1.000E-05 |
| 343 | 18 | E18N | A | 9.4e-06 | 8.6e-06 | 8.3e-06 | 8.767E-06 | 5.686E-07 | 8.300E-06 | 9.400E-06 |
| 344 | 18 | E18D | A | 9.1e-06 | 7.5e-06 | 8.1e-06 | 8.233E-06 | 8.083E-07 | 7.500E-06 | 9.100E-06 |
| 345 | 18 | E18C | A | 1.2e-05 | 1e-05 | 4.2e-06 | 8.733E-06 | 4.051E-06 | 4.200E-06 | 1.200E-05 |
| 346 | 18 | E18E | A | 9.4e-06 | 5.9e-06 | 5.3e-06 | 6.867E-06 | 2.214E-06 | 5.300E-06 | 9.400E-06 |
| 347 | 18 | E18Q | A | 8.5e-06 | 4.5e-06 | 6.1e-06 | 6.367E-06 | 2.013E-06 | 4.500E-06 | 8.500E-06 |
| 348 | 18 | E18G | A | 6.5e-06 | 7.2e-06 | 3.4e-06 | 5.700E-06 | 2.022E-06 | 3.400E-06 | 7.200E-06 |
| 349 | 18 | E18H | A | 5.3e-06 | 6.7e-06 | 6e-06 | 6.000E-06 | 7.000E-07 | 5.300E-06 | 6.700E-06 |
| 350 | 18 | E18I | A | 1.9e-05 | 4.3e-06 | 6.7e-06 | 1.000E-05 | 7.886E-06 | 4.300E-06 | 1.900E-05 |
| 351 | 18 | E18L | A | 6.9e-06 | 9.1e-06 | 6.5e-06 | 7.500E-06 | 1.400E-06 | 6.500E-06 | 9.100E-06 |
| 352 | 18 | E18K | A | 4.7e-06 | 8.7e-06 | 7.7e-06 | 7.033E-06 | 2.082E-06 | 4.700E-06 | 8.700E-06 |
| 353 | 18 | E18M | A | 7.9e-06 | 7.1e-06 | 5.1e-06 | 6.700E-06 | 1.442E-06 | 5.100E-06 | 7.900E-06 |
| 354 | 18 | E18F | A | 8.9e-06 | 6e-06 | 8e-06 | 7.633E-06 | 1.484E-06 | 6.000E-06 | 8.900E-06 |
| 355 | 18 | E18P | A | 6.3e-06 | 9.2e-06 | 1e-05 | 8.500E-06 | 1.947E-06 | 6.300E-06 | 1.000E-05 |
| 356 | 18 | E18S | A | 1.4e-05 | 7e-06 | 7.4e-06 | 9.467E-06 | 3.931E-06 | 7.000E-06 | 1.400E-05 |
| 357 | 18 | E18T | A | 8.8e-06 | 8.7e-06 | 9.2e-06 | 8.900E-06 | 2.646E-07 | 8.700E-06 | 9.200E-06 |
| 358 | 18 | E18W | A | 6.7e-06 | 8.8e-06 | 1.2e-05 | 9.167E-06 | 2.669E-06 | 6.700E-06 | 1.200E-05 |
| 359 | 18 | E18Y | A | 5.9e-06 | 1.1e-05 | 7.4e-06 | 8.100E-06 | 2.621E-06 | 5.900E-06 | 1.100E-05 |
| 360 | 18 | E18V | A | 8.7e-06 | 1.1e-05 | 6.5e-06 | 8.733E-06 | 2.250E-06 | 6.500E-06 | 1.100E-05 |
| 361 | 19 | T19A | A | 5.8e-06 | 1e-05 | 1.3e-05 | 9.600E-06 | 3.617E-06 | 5.800E-06 | 1.300E-05 |
| 362 | 19 | T19R | A | 7.2e-06 | 6.8e-06 | 1.1e-05 | 8.333E-06 | 2.318E-06 | 6.800E-06 | 1.100E-05 |
| 363 | 19 | T19N | A | 6.4e-06 | 4.5e-06 | 6.5e-06 | 5.800E-06 | 1.127E-06 | 4.500E-06 | 6.500E-06 |
| 364 | 19 | T19D | A | 9.3e-06 | 5.4e-06 | 8.9e-06 | 7.867E-06 | 2.146E-06 | 5.400E-06 | 9.300E-06 |
| 365 | 19 | T19C | A | 7.5e-06 | 5.3e-06 | 5.5e-06 | 6.100E-06 | 1.217E-06 | 5.300E-06 | 7.500E-06 |
| 366 | 19 | T19E | A | 8.5e-06 | 9.3e-06 | 1e-05 | 9.267E-06 | 7.506E-07 | 8.500E-06 | 1.000E-05 |
| 367 | 19 | T19Q | A | 1.3e-05 | 9.6e-06 | 9.5e-06 | 1.070E-05 | 1.992E-06 | 9.500E-06 | 1.300E-05 |
| 368 | 19 | T19G | A | 4.8e-06 | 5.7e-06 | 8.4e-06 | 6.300E-06 | 1.873E-06 | 4.800E-06 | 8.400E-06 |
| 369 | 19 | T19H | A | 8.9e-06 | 5.9e-06 | 9.5e-06 | 8.100E-06 | 1.929E-06 | 5.900E-06 | 9.500E-06 |
| 370 | 19 | T19I | A | 9.7e-06 | 8.3e-06 | 7.2e-06 | 8.400E-06 | 1.253E-06 | 7.200E-06 | 9.700E-06 |
| 371 | 19 | T19L | A | 8.8e-06 | 9.3e-06 | 6e-06 | 8.033E-06 | 1.779E-06 | 6.000E-06 | 9.300E-06 |
| 372 | 19 | T19K | A | 8.4e-06 | 4.8e-06 | 8.9e-06 | 7.367E-06 | 2.237E-06 | 4.800E-06 | 8.900E-06 |
| 373 | 19 | T19M | A | 5.2e-06 | 4.8e-06 | 5.3e-06 | 5.100E-06 | 2.646E-07 | 4.800E-06 | 5.300E-06 |
| 374 | 19 | T19F | A | 1e-05 | 1.3e-05 | 4.3e-06 | 9.100E-06 | 4.419E-06 | 4.300E-06 | 1.300E-05 |
| 375 | 19 | T19P | A | 6.6e-06 | 1e-05 | 6.6e-06 | 7.733E-06 | 1.963E-06 | 6.600E-06 | 1.000E-05 |
| 376 | 19 | T19S | A | 1.3e-05 | 4.2e-06 | 1e-05 | 9.067E-06 | 4.474E-06 | 4.200E-06 | 1.300E-05 |
| 377 | 19 | T19T | A | 9.4e-06 | 6.2e-06 | 1.3e-05 | 9.533E-06 | 3.402E-06 | 6.200E-06 | 1.300E-05 |
| 378 | 19 | T19W | A | 3.2e-06 | 8e-06 | 6.4e-06 | 5.867E-06 | 2.444E-06 | 3.200E-06 | 8.000E-06 |
| 379 | 19 | T19Y | A | 8.2e-06 | 1.2e-05 | 6.3e-06 | 8.833E-06 | 2.902E-06 | 6.300E-06 | 1.200E-05 |
| 380 | 19 | T19V | A | 9.3e-06 | 9.7e-06 | 1.1e-05 | 1.000E-05 | 8.888E-07 | 9.300E-06 | 1.100E-05 |
| 381 | 20 | H20A | A | 9.1e-06 | 7.5e-06 | 7.5e-06 | 8.033E-06 | 9.238E-07 | 7.500E-06 | 9.100E-06 |
| 382 | 20 | H20R | A | 8.3e-06 | 6.3e-06 | 1.4e-05 | 9.533E-06 | 3.995E-06 | 6.300E-06 | 1.400E-05 |
| 383 | 20 | H20N | A | 5.4e-06 | 6.6e-06 | 6.7e-06 | 6.233E-06 | 7.234E-07 | 5.400E-06 | 6.700E-06 |
| 384 | 20 | H20D | A | 7.5e-06 | 6.4e-06 | 8.2e-06 | 7.367E-06 | 9.074E-07 | 6.400E-06 | 8.200E-06 |
| 385 | 20 | H20C | A | 1.1e-05 | 4.6e-06 | 8.6e-06 | 8.067E-06 | 3.233E-06 | 4.600E-06 | 1.100E-05 |
| 386 | 20 | H20E | A | 7.4e-06 | 1.3e-05 | 1.9e-05 | 1.313E-05 | 5.801E-06 | 7.400E-06 | 1.900E-05 |
| 387 | 20 | H20Q | A | 3.3e-06 | 1.4e-05 | 1.3e-05 | 1.010E-05 | 5.910E-06 | 3.300E-06 | 1.400E-05 |
| 388 | 20 | H20G | A | 7.9e-06 | 1.4e-05 | 5.4e-06 | 9.100E-06 | 4.424E-06 | 5.400E-06 | 1.400E-05 |
| 389 | 20 | H20H | A | 1.3e-05 | 9.3e-06 | 1e-05 | 1.077E-05 | 1.966E-06 | 9.300E-06 | 1.300E-05 |
| 390 | 20 | H20I | A | 6.4e-06 | 8.9e-06 | 7.6e-06 | 7.633E-06 | 1.250E-06 | 6.400E-06 | 8.900E-06 |
| 391 | 20 | H20L | A | 7.4e-06 | 8.8e-06 | 7.5e-06 | 7.900E-06 | 7.810E-07 | 7.400E-06 | 8.800E-06 |
| 392 | 20 | H20K | A | 1e-05 | 6.4e-06 | 7e-06 | 7.800E-06 | 1.929E-06 | 6.400E-06 | 1.000E-05 |
| 393 | 20 | H20M | A | 1.1e-05 | 9e-06 | 7.3e-06 | 9.100E-06 | 1.852E-06 | 7.300E-06 | 1.100E-05 |
| 394 | 20 | H20F | A | 6.5e-06 | 6.2e-06 | 1.2e-05 | 8.233E-06 | 3.265E-06 | 6.200E-06 | 1.200E-05 |
| 395 | 20 | H20P | A | 8.8e-06 | 6.3e-06 | 6.5e-06 | 7.200E-06 | 1.389E-06 | 6.300E-06 | 8.800E-06 |
| 396 | 20 | H20S | A | 1.1e-05 | 1e-05 | 1.3e-05 | 1.133E-05 | 1.528E-06 | 1.000E-05 | 1.300E-05 |
| 397 | 20 | H20T | A | 6.8e-06 | 5.2e-06 | 7.1e-06 | 6.367E-06 | 1.021E-06 | 5.200E-06 | 7.100E-06 |
| 398 | 20 | H20W | A | 5.8e-06 | 5.2e-06 | 8.5e-06 | 6.500E-06 | 1.758E-06 | 5.200E-06 | 8.500E-06 |
| 399 | 20 | H20Y | A | 1.1e-05 | 1.3e-05 | 6.1e-06 | 1.003E-05 | 3.550E-06 | 6.100E-06 | 1.300E-05 |
| 400 | 20 | H20V | A | 5.2e-06 | 1.6e-05 | 9.9e-06 | 1.037E-05 | 5.415E-06 | 5.200E-06 | 1.600E-05 |
| 401 | 21 | L21A | A | 7.6e-06 | 1.4e-05 | 1e-05 | 1.053E-05 | 3.233E-06 | 7.600E-06 | 1.400E-05 |
| 402 | 21 | L21R | A | 1.1e-05 | 5.6e-06 | 9.6e-06 | 8.733E-06 | 2.802E-06 | 5.600E-06 | 1.100E-05 |
| 403 | 21 | L21N | A | 6.2e-06 | 8.2e-06 | 4.5e-06 | 6.300E-06 | 1.852E-06 | 4.500E-06 | 8.200E-06 |
| 404 | 21 | L21D | A | 1e-05 | 9.2e-06 | 1.1e-05 | 1.007E-05 | 9.018E-07 | 9.200E-06 | 1.100E-05 |
| 405 | 21 | L21C | A | 1.1e-05 | 9.6e-06 | 8.1e-06 | 9.567E-06 | 1.450E-06 | 8.100E-06 | 1.100E-05 |
| 406 | 21 | L21E | A | 8.1e-06 | 6.7e-06 | 1e-05 | 8.267E-06 | 1.656E-06 | 6.700E-06 | 1.000E-05 |
| 407 | 21 | L21Q | A | 8.4e-06 | 6.3e-06 | 5.4e-06 | 6.700E-06 | 1.539E-06 | 5.400E-06 | 8.400E-06 |
| 408 | 21 | L21G | A | 6.3e-06 | 6.6e-06 | 6.5e-06 | 6.467E-06 | 1.528E-07 | 6.300E-06 | 6.600E-06 |
| 409 | 21 | L21H | A | 5.4e-06 | 9.2e-06 | 4e-06 | 6.200E-06 | 2.691E-06 | 4.000E-06 | 9.200E-06 |
| 410 | 21 | L21I | A | 5e-06 | 7.8e-06 | 7.9e-06 | 6.900E-06 | 1.646E-06 | 5.000E-06 | 7.900E-06 |
| 411 | 21 | L21L | A | 7.2e-06 | 6.2e-06 | 9.3e-06 | 7.567E-06 | 1.582E-06 | 6.200E-06 | 9.300E-06 |
| 412 | 21 | L21K | A | 1.3e-05 | 8.2e-06 | 1.3e-05 | 1.140E-05 | 2.771E-06 | 8.200E-06 | 1.300E-05 |
| 413 | 21 | L21M | A | 6e-06 | 1.3e-05 | 8.8e-06 | 9.267E-06 | 3.523E-06 | 6.000E-06 | 1.300E-05 |
| 414 | 21 | L21F | A | 7.4e-06 | 1.1e-05 | 9.5e-06 | 9.300E-06 | 1.808E-06 | 7.400E-06 | 1.100E-05 |
| 415 | 21 | L21P | A | 9.5e-06 | 8.2e-06 | 1.1e-05 | 9.567E-06 | 1.401E-06 | 8.200E-06 | 1.100E-05 |
| 416 | 21 | L21S | A | 5.2e-06 | 1.8e-05 | 3.3e-06 | 8.833E-06 | 7.995E-06 | 3.300E-06 | 1.800E-05 |
| 417 | 21 | L21T | A | 6.6e-06 | 6.2e-06 | 1.2e-05 | 8.267E-06 | 3.239E-06 | 6.200E-06 | 1.200E-05 |
| 418 | 21 | L21W | A | 5.1e-06 | 6.5e-06 | 3.1e-06 | 4.900E-06 | 1.709E-06 | 3.100E-06 | 6.500E-06 |
| 419 | 21 | L21Y | A | 7.5e-06 | 5.6e-06 | 3.5e-06 | 5.533E-06 | 2.001E-06 | 3.500E-06 | 7.500E-06 |
| 420 | 21 | L21V | A | 6.7e-06 | 1e-05 | 7.2e-06 | 7.967E-06 | 1.779E-06 | 6.700E-06 | 1.000E-05 |
| 421 | 22 | D22A | A | 8.1e-06 | 1.2e-05 | 5.7e-06 | 8.600E-06 | 3.180E-06 | 5.700E-06 | 1.200E-05 |
| 422 | 22 | D22R | A | 4.2e-06 | 8.6e-06 | 6.5e-06 | 6.433E-06 | 2.201E-06 | 4.200E-06 | 8.600E-06 |
| 423 | 22 | D22N | A | 5.4e-06 | 1e-05 | 9.5e-06 | 8.300E-06 | 2.524E-06 | 5.400E-06 | 1.000E-05 |
| 424 | 22 | D22D | A | 8.7e-06 | 6.7e-06 | 4.9e-06 | 6.767E-06 | 1.901E-06 | 4.900E-06 | 8.700E-06 |

| | | | | | | | | | | |
|---|---|---|---|---|---|---|---|---|---|---|
| 425 | 22 | D22C | A | 9.3e-06 | 6.3e-06 | 8.8e-06 | 8.133E-06 | 1.607E-06 | 6.300E-06 | 9.300E-06 |
| 426 | 22 | D22E | A | 7.3e-06 | 7.6e-06 | 4.8e-06 | 6.567E-06 | 1.537E-06 | 4.800E-06 | 7.600E-06 |
| 427 | 22 | D22Q | A | 8.7e-06 | 8.3e-06 | 7.9e-06 | 8.300E-06 | 4.000E-07 | 7.900E-06 | 8.700E-06 |
| 428 | 22 | D22G | A | 8e-06 | 7.3e-06 | 7.4e-06 | 7.567E-06 | 3.786E-07 | 7.300E-06 | 8.000E-06 |
| 429 | 22 | D22H | A | 8.9e-06 | 1.3e-05 | 8.8e-06 | 1.023E-05 | 2.397E-06 | 8.800E-06 | 1.300E-05 |
| 430 | 22 | D22I | A | 7.3e-06 | 6.5e-06 | 1.2e-05 | 8.600E-06 | 2.972E-06 | 6.500E-06 | 1.200E-05 |
| 431 | 22 | D22L | A | 4.8e-06 | 4.2e-06 | 4.5e-06 | 4.500E-06 | 3.000E-07 | 4.200E-06 | 4.800E-06 |
| 432 | 22 | D22K | A | 8.6e-06 | 1.2e-05 | 7.3e-06 | 9.300E-06 | 2.427E-06 | 7.300E-06 | 1.200E-05 |
| 433 | 22 | D22M | A | 7.3e-06 | 8.6e-06 | 8.5e-06 | 8.133E-06 | 7.234E-07 | 7.300E-06 | 8.600E-06 |
| 434 | 22 | D22F | A | 8.9e-06 | 8.9e-06 | 1.2e-05 | 9.933E-06 | 1.790E-06 | 8.900E-06 | 1.200E-05 |
| 435 | 22 | D22P | A | 8.6e-06 | 6.8e-06 | 9.8e-06 | 8.400E-06 | 1.510E-06 | 6.800E-06 | 9.800E-06 |
| 436 | 22 | D22S | A | 6.1e-06 | 9.2e-06 | 1.5e-05 | 1.010E-05 | 4.518E-06 | 6.100E-06 | 1.500E-05 |
| 437 | 22 | D22T | A | 1.1e-05 | 7.5e-06 | 1e-05 | 9.500E-06 | 1.803E-06 | 7.500E-06 | 1.100E-05 |
| 438 | 22 | D22W | A | 5.1e-06 | 3.7e-06 | 5.9e-06 | 4.900E-06 | 1.114E-06 | 3.700E-06 | 5.900E-06 |
| 439 | 22 | D22Y | A | 8.6e-06 | 4.6e-06 | 1.3e-05 | 8.733E-06 | 4.202E-06 | 4.600E-06 | 1.300E-05 |
| 440 | 22 | D22V | A | 3.7e-06 | 9.8e-06 | 1.4e-05 | 9.167E-06 | 5.179E-06 | 3.700E-06 | 1.400E-05 |
| 441 | 23 | M23A | A | 7.9e-06 | 9.7e-06 | 9e-06 | 8.867E-06 | 9.074E-07 | 7.900E-06 | 9.700E-06 |
| 442 | 23 | M23R | A | 5.1e-06 | 8.2e-06 | 1.1e-05 | 8.100E-06 | 2.951E-06 | 5.100E-06 | 1.100E-05 |
| 443 | 23 | M23N | A | 4.8e-06 | 5.9e-06 | 6.6e-06 | 5.767E-06 | 9.074E-07 | 4.800E-06 | 6.600E-06 |
| 444 | 23 | M23D | A | 5.2e-06 | 7e-06 | 9.4e-06 | 7.200E-06 | 2.107E-06 | 5.200E-06 | 9.400E-06 |
| 445 | 23 | M23C | A | 9.4e-06 | 1e-05 | 5.3e-06 | 8.233E-06 | 2.558E-06 | 5.300E-06 | 1.000E-05 |
| 446 | 23 | M23E | A | 5.2e-06 | 4.8e-06 | 7.4e-06 | 5.800E-06 | 1.400E-06 | 4.800E-06 | 7.400E-06 |
| 447 | 23 | M23Q | A | 5.5e-06 | 6.4e-06 | 1.4e-05 | 8.633E-06 | 4.669E-06 | 5.500E-06 | 1.400E-05 |
| 448 | 23 | M23G | A | 8e-06 | 8.6e-06 | 6.1e-06 | 7.567E-06 | 1.305E-06 | 6.100E-06 | 8.600E-06 |
| 449 | 23 | M23H | A | 1.5e-05 | 7.8e-06 | 9.5e-06 | 1.077E-05 | 3.763E-06 | 7.800E-06 | 1.500E-05 |
| 450 | 23 | M23I | A | 1.1e-05 | 8.8e-06 | 1.3e-05 | 1.093E-05 | 2.101E-06 | 8.800E-06 | 1.300E-05 |
| 451 | 23 | M23L | A | 5.3e-06 | 6.5e-06 | 6e-06 | 5.933E-06 | 6.028E-07 | 5.300E-06 | 6.500E-06 |
| 452 | 23 | M23K | A | 8.8e-06 | 8e-06 | 6.6e-06 | 7.800E-06 | 1.114E-06 | 6.600E-06 | 8.800E-06 |
| 453 | 23 | M23M | A | 1.2e-05 | 5.1e-06 | 5e-06 | 7.367E-06 | 4.013E-06 | 5.000E-06 | 1.200E-05 |
| 454 | 23 | M23F | A | 8.6e-06 | 5.7e-06 | 8e-06 | 7.433E-06 | 1.531E-06 | 5.700E-06 | 8.600E-06 |
| 455 | 23 | M23P | A | 7.2e-06 | 5.2e-06 | 7.5e-06 | 6.633E-06 | 1.250E-06 | 5.200E-06 | 7.500E-06 |
| 456 | 23 | M23S | A | 1.1e-05 | 5.5e-06 | 7.2e-06 | 7.900E-06 | 2.816E-06 | 5.500E-06 | 1.100E-05 |
| 457 | 23 | M23T | A | 5.6e-06 | 7e-06 | 7.7e-06 | 6.767E-06 | 1.069E-06 | 5.600E-06 | 7.700E-06 |
| 458 | 23 | M23W | A | 1e-05 | 8.1e-06 | 6.4e-06 | 8.167E-06 | 1.801E-06 | 6.400E-06 | 1.000E-05 |
| 459 | 23 | M23Y | A | 7.7e-06 | 7.9e-06 | 8.4e-06 | 8.000E-06 | 3.606E-07 | 7.700E-06 | 8.400E-06 |
| 460 | 23 | M23V | A | 1.1e-05 | 1.1e-05 | 5.9e-06 | 9.300E-06 | 2.944E-06 | 5.900E-06 | 1.100E-05 |
| 461 | 24 | L24A | A | 7e-06 | 6.3e-06 | 8.3e-06 | 7.200E-06 | 1.015E-06 | 6.300E-06 | 8.300E-06 |
| 462 | 24 | L24R | A | 6.8e-06 | 6.2e-06 | 9.6e-06 | 7.533E-06 | 1.815E-06 | 6.200E-06 | 9.600E-06 |
| 463 | 24 | L24N | A | 9.7e-06 | 1e-05 | 1.1e-05 | 1.023E-05 | 6.807E-07 | 9.700E-06 | 1.100E-05 |
| 464 | 24 | L24D | A | 3.2e-06 | 9.7e-06 | 1e-05 | 7.633E-06 | 3.842E-06 | 3.200E-06 | 1.000E-05 |
| 465 | 24 | L24C | A | 6.5e-06 | 1.3e-05 | 1.3e-05 | 1.083E-05 | 3.753E-06 | 6.500E-06 | 1.300E-05 |
| 466 | 24 | L24E | A | 9.4e-06 | 1.3e-05 | 5.8e-06 | 9.400E-06 | 3.600E-06 | 5.800E-06 | 1.300E-05 |
| 467 | 24 | L24Q | A | 8e-06 | 6.9e-06 | 6e-06 | 6.967E-06 | 1.002E-06 | 6.000E-06 | 8.000E-06 |
| 468 | 24 | L24G | A | 6.1e-06 | 7.5e-06 | 9.9e-06 | 7.833E-06 | 1.922E-06 | 6.100E-06 | 9.900E-06 |
| 469 | 24 | L24H | A | 1.4e-05 | 9.9e-06 | 1e-05 | 1.130E-05 | 2.339E-06 | 9.900E-06 | 1.400E-05 |
| 470 | 24 | L24I | A | 9.2e-06 | 6.3e-06 | 1.4e-05 | 9.833E-06 | 3.889E-06 | 6.300E-06 | 1.400E-05 |
| 471 | 24 | L24L | A | 6.3e-06 | 7.1e-06 | 6.6e-06 | 6.667E-06 | 4.041E-07 | 6.300E-06 | 7.100E-06 |
| 472 | 24 | L24K | A | 4.9e-06 | 6.3e-06 | 9.3e-06 | 6.833E-06 | 2.248E-06 | 4.900E-06 | 9.300E-06 |
| 473 | 24 | L24M | A | 6.6e-06 | 8.3e-06 | 1.5e-05 | 9.967E-06 | 4.441E-06 | 6.600E-06 | 1.500E-05 |
| 474 | 24 | L24F | A | 5.9e-06 | 9.9e-06 | 8.7e-06 | 8.167E-06 | 2.053E-06 | 5.900E-06 | 9.900E-06 |
| 475 | 24 | L24P | A | 6.2e-06 | 9.8e-06 | 7.4e-06 | 7.800E-06 | 1.833E-06 | 6.200E-06 | 9.800E-06 |
| 476 | 24 | L24S | A | 1.3e-05 | 9.2e-06 | 1.1e-05 | 1.107E-05 | 1.901E-06 | 9.200E-06 | 1.300E-05 |
| 477 | 24 | L24T | A | 9.3e-06 | 8.6e-06 | 6.5e-06 | 8.133E-06 | 1.457E-06 | 6.500E-06 | 9.300E-06 |
| 478 | 24 | L24W | A | 4.9e-06 | 7.6e-06 | 8.6e-06 | 7.033E-06 | 1.914E-06 | 4.900E-06 | 8.600E-06 |
| 479 | 24 | L24Y | A | 7.8e-06 | 9.8e-06 | 9.8e-06 | 9.133E-06 | 1.155E-06 | 7.800E-06 | 9.800E-06 |
| 480 | 24 | L24V | A | 9.6e-06 | 6.7e-06 | 1.9e-05 | 1.177E-05 | 6.430E-06 | 6.700E-06 | 1.900E-05 |
| 481 | 25 | R25A | A | 6.7e-06 | 7.6e-06 | 9.5e-06 | 7.933E-06 | 1.429E-06 | 6.700E-06 | 9.500E-06 |
| 482 | 25 | R25R | A | 6.7e-06 | 6e-06 | 5.7e-06 | 6.133E-06 | 5.132E-07 | 5.700E-06 | 6.700E-06 |
| 483 | 25 | R25N | A | 1.1e-05 | 9e-06 | 6.2e-06 | 8.733E-06 | 2.411E-06 | 6.200E-06 | 1.100E-05 |
| 484 | 25 | R25D | A | 5.1e-06 | 7.9e-06 | 7.4e-06 | 6.800E-06 | 1.493E-06 | 5.100E-06 | 7.900E-06 |
| 485 | 25 | R25C | A | 6.2e-06 | 6.2e-06 | 5.5e-06 | 5.967E-06 | 4.041E-07 | 5.500E-06 | 6.200E-06 |
| 486 | 25 | R25E | A | 1e-05 | 6.6e-06 | 7.4e-06 | 8.000E-06 | 1.778E-06 | 6.600E-06 | 1.000E-05 |
| 487 | 25 | R25Q | A | 8.9e-06 | 6.5e-06 | 1.2e-05 | 9.133E-06 | 2.757E-06 | 6.500E-06 | 1.200E-05 |
| 488 | 25 | R25G | A | 6.8e-06 | 7.1e-06 | 5.2e-06 | 6.367E-06 | 1.021E-06 | 5.200E-06 | 7.100E-06 |
| 489 | 25 | R25H | A | 4.6e-06 | 4.4e-06 | 9.4e-06 | 6.133E-06 | 2.831E-06 | 4.400E-06 | 9.400E-06 |
| 490 | 25 | R25I | A | 8e-06 | 1.2e-05 | 5.6e-06 | 8.533E-06 | 3.233E-06 | 5.600E-06 | 1.200E-05 |
| 491 | 25 | R25L | A | 4.4e-06 | 6.2e-06 | 1.3e-05 | 7.867E-06 | 4.536E-06 | 4.400E-06 | 1.300E-05 |
| 492 | 25 | R25K | A | 6.5e-06 | 1.2e-05 | 7.4e-06 | 8.633E-06 | 2.950E-06 | 6.500E-06 | 1.200E-05 |
| 493 | 25 | R25M | A | 5.7e-06 | 5.4e-06 | 7.1e-06 | 6.067E-06 | 9.074E-07 | 5.400E-06 | 7.100E-06 |
| 494 | 25 | R25F | A | 7.7e-06 | 7.3e-06 | 9.8e-06 | 8.267E-06 | 1.343E-06 | 7.300E-06 | 9.800E-06 |
| 495 | 25 | R25P | A | 5.8e-06 | 1e-05 | 9.4e-06 | 8.400E-06 | 2.272E-06 | 5.800E-06 | 1.000E-05 |
| 496 | 25 | R25S | A | 1.1e-05 | 2.9e-06 | 9.8e-06 | 7.900E-06 | 4.371E-06 | 2.900E-06 | 1.100E-05 |
| 497 | 25 | R25T | A | 6.9e-06 | 1.3e-05 | 5.6e-06 | 8.500E-06 | 3.951E-06 | 5.600E-06 | 1.300E-05 |
| 498 | 25 | R25W | A | 1.1e-05 | 8.3e-06 | 6.4e-06 | 8.567E-06 | 2.312E-06 | 6.400E-06 | 1.100E-05 |
| 499 | 25 | R25Y | A | 5.2e-06 | 7.9e-06 | 9.1e-06 | 7.400E-06 | 1.997E-06 | 5.200E-06 | 9.100E-06 |
| 500 | 25 | R25V | A | 4.5e-06 | 6.1e-06 | 6.3e-06 | 5.633E-06 | 9.866E-07 | 4.500E-06 | 6.300E-06 |
| 501 | 26 | H26A | A | 7.1e-06 | 3.9e-06 | 8.7e-06 | 6.567E-06 | 2.444E-06 | 3.900E-06 | 8.700E-06 |
| 502 | 26 | H26R | A | 5.5e-06 | 9.9e-06 | 8.6e-06 | 8.000E-06 | 2.261E-06 | 5.500E-06 | 9.900E-06 |
| 503 | 26 | H26N | A | 8.5e-06 | 9.6e-06 | 5.7e-06 | 7.933E-06 | 2.011E-06 | 5.700E-06 | 9.600E-06 |
| 504 | 26 | H26D | A | 1.1e-05 | 9.6e-06 | 1e-05 | 1.020E-05 | 7.211E-07 | 9.600E-06 | 1.100E-05 |
| 505 | 26 | H26C | A | 1.2e-05 | 1.4e-05 | 7.3e-06 | 1.110E-05 | 3.439E-06 | 7.300E-06 | 1.400E-05 |
| 506 | 26 | H26E | A | 8.4e-06 | 1e-05 | 8.9e-06 | 9.100E-06 | 8.185E-07 | 8.400E-06 | 1.000E-05 |
| 507 | 26 | H26Q | A | 6.6e-06 | 7e-06 | 6.6e-06 | 6.733E-06 | 2.309E-07 | 6.600E-06 | 7.000E-06 |
| 508 | 26 | H26G | A | 8.1e-06 | 6.8e-06 | 8.2e-06 | 7.700E-06 | 7.810E-07 | 6.800E-06 | 8.200E-06 |
| 509 | 26 | H26H | A | 8.7e-06 | 1.1e-05 | 7.5e-06 | 9.067E-06 | 1.779E-06 | 7.500E-06 | 1.100E-05 |

| | | | | | | | | | | |
|---|---|---|---|---|---|---|---|---|---|---|
| 510 | 26 | H26I | A | 1e-05 | 1e-05 | 9.7e-06 | 9.900E-06 | 1.732E-07 | 9.700E-06 | 1.000E-05 |
| 511 | 26 | H26L | A | 1.2e-05 | 1.1e-05 | 6.9e-06 | 9.967E-06 | 2.702E-06 | 6.900E-06 | 1.200E-05 |
| 512 | 26 | H26K | A | 8e-06 | 1.4e-05 | 1.1e-05 | 1.100E-05 | 3.000E-06 | 8.000E-06 | 1.400E-05 |
| 513 | 26 | H26M | A | 6.4e-06 | 1.1e-05 | 4.4e-06 | 7.267E-06 | 3.384E-06 | 4.400E-06 | 1.100E-05 |
| 514 | 26 | H26F | A | 6e-06 | 9.1e-06 | 5.7e-06 | 6.933E-06 | 1.882E-06 | 5.700E-06 | 9.100E-06 |
| 515 | 26 | H26P | A | 9.6e-06 | 1.3e-05 | 7.5e-06 | 1.003E-05 | 2.775E-06 | 7.500E-06 | 1.300E-05 |
| 516 | 26 | H26S | A | 8.4e-06 | 1.6e-05 | 8.9e-06 | 1.110E-05 | 4.251E-06 | 8.400E-06 | 1.600E-05 |
| 517 | 26 | H26T | A | 6.2e-06 | 4.5e-06 | 8.3e-06 | 6.333E-06 | 1.904E-06 | 4.500E-06 | 8.300E-06 |
| 518 | 26 | H26W | A | 8e-06 | 1e-05 | 1.6e-05 | 1.133E-05 | 4.163E-06 | 8.000E-06 | 1.600E-05 |
| 519 | 26 | H26Y | A | 6.2e-06 | 7.4e-06 | 7e-06 | 6.867E-06 | 6.110E-07 | 6.200E-06 | 7.400E-06 |
| 520 | 26 | H26V | A | 9.7e-06 | 1.3e-05 | 1e-05 | 1.090E-05 | 1.825E-06 | 9.700E-06 | 1.300E-05 |
| 521 | 27 | L27A | A | 9.5e-06 | 6.5e-06 | 7.3e-06 | 7.767E-06 | 1.553E-06 | 6.500E-06 | 9.500E-06 |
| 522 | 27 | L27R | A | 4.8e-06 | 1.4e-05 | 1.3e-05 | 1.060E-05 | 5.048E-06 | 4.800E-06 | 1.400E-05 |
| 523 | 27 | L27N | A | 4.8e-06 | 7.7e-06 | 9.6e-06 | 7.367E-06 | 2.417E-06 | 4.800E-06 | 9.600E-06 |
| 524 | 27 | L27D | A | 1.2e-05 | 1.2e-05 | 8.3e-06 | 1.077E-05 | 2.136E-06 | 8.300E-06 | 1.200E-05 |
| 525 | 27 | L27C | A | 1.2e-05 | 1.4e-05 | 1.1e-05 | 1.233E-05 | 1.528E-06 | 1.100E-05 | 1.400E-05 |
| 526 | 27 | L27E | A | 4.6e-06 | 9.8e-06 | 8.8e-06 | 7.733E-06 | 2.759E-06 | 4.600E-06 | 9.800E-06 |
| 527 | 27 | L27Q | A | 7.2e-06 | 8.3e-06 | 9.4e-06 | 8.300E-06 | 1.100E-06 | 7.200E-06 | 9.400E-06 |
| 528 | 27 | L27G | A | 1.2e-05 | 8.9e-06 | 8e-06 | 9.633E-06 | 2.098E-06 | 8.000E-06 | 1.200E-05 |
| 529 | 27 | L27H | A | 5.2e-06 | 6.3e-06 | 7.2e-06 | 6.233E-06 | 1.002E-06 | 5.200E-06 | 7.200E-06 |
| 530 | 27 | L27I | A | 6.5e-06 | 6.9e-06 | 7e-06 | 6.800E-06 | 2.646E-07 | 6.500E-06 | 7.000E-06 |
| 531 | 27 | L27L | A | 7.4e-06 | 9.7e-06 | 6.4e-06 | 7.833E-06 | 1.692E-06 | 6.400E-06 | 9.700E-06 |
| 532 | 27 | L27K | A | 6.4e-06 | 6e-06 | 7.2e-06 | 6.533E-06 | 6.110E-07 | 6.000E-06 | 7.200E-06 |
| 533 | 27 | L27M | A | 9.5e-06 | 7.2e-06 | 1.3e-05 | 9.900E-06 | 2.921E-06 | 7.200E-06 | 1.300E-05 |
| 534 | 27 | L27F | A | 1.3e-05 | 1.3e-05 | 6.8e-06 | 1.093E-05 | 3.580E-06 | 6.800E-06 | 1.300E-05 |
| 535 | 27 | L27P | A | 9.3e-06 | 7.5e-06 | 5e-06 | 7.267E-06 | 2.159E-06 | 5.000E-06 | 9.300E-06 |
| 536 | 27 | L27S | A | 9.5e-06 | 5.5e-06 | 5e-06 | 6.667E-06 | 2.466E-06 | 5.000E-06 | 9.500E-06 |
| 537 | 27 | L27T | A | 6.7e-06 | 4.5e-06 | 8.8e-06 | 6.667E-06 | 2.150E-06 | 4.500E-06 | 8.800E-06 |
| 538 | 27 | L27W | A | 5.7e-06 | 9e-06 | 6.1e-06 | 6.933E-06 | 1.801E-06 | 5.700E-06 | 9.000E-06 |
| 539 | 27 | L27Y | A | 4.2e-06 | 6e-06 | 8.7e-06 | 6.300E-06 | 2.265E-06 | 4.200E-06 | 8.700E-06 |
| 540 | 27 | L27V | A | 8.6e-06 | 1e-05 | 8.8e-06 | 9.133E-06 | 7.572E-07 | 8.600E-06 | 1.000E-05 |
| 541 | 28 | Y28A | A | 1.4e-05 | 1.1e-05 | 9.3e-06 | 1.143E-05 | 2.380E-06 | 9.300E-06 | 1.400E-05 |
| 542 | 28 | Y28R | A | 6.8e-06 | 9.3e-06 | 1e-05 | 8.700E-06 | 1.682E-06 | 6.800E-06 | 1.000E-05 |
| 543 | 28 | Y28N | A | 6.6e-06 | 6.2e-06 | 1.4e-05 | 8.933E-06 | 4.392E-06 | 6.200E-06 | 1.400E-05 |
| 544 | 28 | Y28D | A | 6.2e-06 | 1.4e-05 | 9.7e-06 | 9.967E-06 | 3.907E-06 | 6.200E-06 | 1.400E-05 |
| 545 | 28 | Y28C | A | 8.7e-06 | 7.2e-06 | 7.2e-06 | 7.700E-06 | 8.660E-07 | 7.200E-06 | 8.700E-06 |
| 546 | 28 | Y28E | A | 1.2e-05 | 5.7e-06 | 6e-06 | 7.900E-06 | 3.554E-06 | 5.700E-06 | 1.200E-05 |
| 547 | 28 | Y28Q | A | 7.2e-06 | 8.5e-06 | 8.7e-06 | 8.133E-06 | 8.145E-07 | 7.200E-06 | 8.700E-06 |
| 548 | 28 | Y28G | A | 5.4e-06 | 7.3e-06 | 7.9e-06 | 6.867E-06 | 1.305E-06 | 5.400E-06 | 7.900E-06 |
| 549 | 28 | Y28H | A | 9.6e-06 | 7e-06 | 6.9e-06 | 7.833E-06 | 1.531E-06 | 6.900E-06 | 9.600E-06 |
| 550 | 28 | Y28I | A | 9e-06 | 9.2e-06 | 9.4e-06 | 9.200E-06 | 2.000E-07 | 9.000E-06 | 9.400E-06 |
| 551 | 28 | Y28L | A | 1e-05 | 9.2e-06 | 1.3e-05 | 1.073E-05 | 2.003E-06 | 9.200E-06 | 1.300E-05 |
| 552 | 28 | Y28K | A | 8.2e-06 | 8.5e-06 | 7.2e-06 | 7.967E-06 | 6.807E-07 | 7.200E-06 | 8.500E-06 |
| 553 | 28 | Y28M | A | 6.2e-06 | 9.2e-06 | 1.1e-05 | 8.800E-06 | 2.425E-06 | 6.200E-06 | 1.100E-05 |
| 554 | 28 | Y28F | A | 6.9e-06 | 1e-05 | 1.1e-05 | 9.300E-06 | 2.138E-06 | 6.900E-06 | 1.100E-05 |
| 555 | 28 | Y28P | A | 1.2e-05 | 3.2e-06 | 8.1e-06 | 7.767E-06 | 4.409E-06 | 3.200E-06 | 1.200E-05 |
| 556 | 28 | Y28S | A | 5.1e-06 | 4.7e-06 | 4e-06 | 4.600E-06 | 5.568E-07 | 4.000E-06 | 5.100E-06 |
| 557 | 28 | Y28T | A | 1.5e-05 | 1.4e-05 | 1.3e-05 | 1.400E-05 | 1.000E-06 | 1.300E-05 | 1.500E-05 |
| 558 | 28 | Y28W | A | 6.3e-06 | 8.9e-06 | 1.1e-05 | 8.733E-06 | 2.354E-06 | 6.300E-06 | 1.100E-05 |
| 559 | 28 | Y28Y | A | 1.1e-05 | 7.6e-06 | 6.5e-06 | 8.367E-06 | 2.346E-06 | 6.500E-06 | 1.100E-05 |
| 560 | 28 | Y28V | A | 1.1e-05 | 9e-06 | 8.4e-06 | 9.467E-06 | 1.361E-06 | 8.400E-06 | 1.100E-05 |
| 561 | 29 | Q29A | A | 9e-06 | 1.2e-05 | 8.6e-06 | 9.867E-06 | 1.858E-06 | 8.600E-06 | 1.200E-05 |
| 562 | 29 | Q29R | A | 1.1e-05 | 7.9e-06 | 4.7e-06 | 7.867E-06 | 3.150E-06 | 4.700E-06 | 1.100E-05 |
| 563 | 29 | Q29N | A | 8.5e-06 | 5.7e-06 | 8.3e-06 | 7.500E-06 | 1.562E-06 | 5.700E-06 | 8.500E-06 |
| 564 | 29 | Q29D | A | 7.1e-06 | 9e-06 | 5.5e-06 | 7.200E-06 | 1.752E-06 | 5.500E-06 | 9.000E-06 |
| 565 | 29 | Q29C | A | 6.9e-06 | 9.1e-06 | 8.7e-06 | 8.233E-06 | 1.172E-06 | 6.900E-06 | 9.100E-06 |
| 566 | 29 | Q29E | A | 9.8e-06 | 7e-06 | 8.6e-06 | 8.467E-06 | 1.405E-06 | 7.000E-06 | 9.800E-06 |
| 567 | 29 | Q29Q | A | 5.3e-06 | 5.8e-06 | 1e-05 | 7.033E-06 | 2.581E-06 | 5.300E-06 | 1.000E-05 |
| 568 | 29 | Q29G | A | 5.2e-06 | 8.8e-06 | 1.3e-05 | 9.000E-06 | 3.904E-06 | 5.200E-06 | 1.300E-05 |
| 569 | 29 | Q29H | A | 8e-06 | 9.8e-06 | 9e-06 | 8.933E-06 | 9.018E-07 | 8.000E-06 | 9.800E-06 |
| 570 | 29 | Q29I | A | 6.7e-06 | 5.8e-06 | 1.1e-05 | 7.833E-06 | 2.779E-06 | 5.800E-06 | 1.100E-05 |
| 571 | 29 | Q29L | A | 1.1e-05 | 7.1e-06 | 1.8e-05 | 1.203E-05 | 5.523E-06 | 7.100E-06 | 1.800E-05 |
| 572 | 29 | Q29K | A | 5.1e-06 | 3.7e-06 | 1.1e-05 | 6.600E-06 | 3.874E-06 | 3.700E-06 | 1.100E-05 |
| 573 | 29 | Q29M | A | 2.8e-05 | 6e-06 | 9.9e-06 | 1.463E-05 | 1.174E-05 | 6.000E-06 | 2.800E-05 |
| 574 | 29 | Q29F | A | 1e-05 | 5.5e-06 | 5.1e-06 | 6.867E-06 | 2.721E-06 | 5.100E-06 | 1.000E-05 |
| 575 | 29 | Q29P | A | 7.4e-06 | 8.6e-06 | 9.2e-06 | 8.400E-06 | 9.165E-07 | 7.400E-06 | 9.200E-06 |
| 576 | 29 | Q29S | A | 1.2e-05 | 1.2e-05 | 7.8e-06 | 1.060E-05 | 2.425E-06 | 7.800E-06 | 1.200E-05 |
| 577 | 29 | Q29T | A | 4e-06 | 8.9e-06 | 5.6e-06 | 6.167E-06 | 2.499E-06 | 4.000E-06 | 8.900E-06 |
| 578 | 29 | Q29W | A | 1e-05 | 7.9e-06 | 4.3e-06 | 7.400E-06 | 2.883E-06 | 4.300E-06 | 1.000E-05 |
| 579 | 29 | Q29Y | A | 8e-06 | 9.5e-06 | 1e-05 | 9.167E-06 | 1.041E-06 | 8.000E-06 | 1.000E-05 |
| 580 | 29 | Q29V | A | 6.4e-06 | 5.4e-06 | 1.1e-05 | 7.600E-06 | 2.987E-06 | 5.400E-06 | 1.100E-05 |
| 581 | 30 | G30A | A | 9.5e-06 | 4.6e-06 | 6.4e-06 | 6.833E-06 | 2.479E-06 | 4.600E-06 | 9.500E-06 |
| 582 | 30 | G30R | A | 7.2e-06 | 9.1e-06 | 8.1e-06 | 8.133E-06 | 9.504E-07 | 7.200E-06 | 9.100E-06 |
| 583 | 30 | G30N | A | 7.6e-06 | 7.1e-06 | 8.2e-06 | 7.633E-06 | 5.508E-07 | 7.100E-06 | 8.200E-06 |
| 584 | 30 | G30D | A | 9.1e-06 | 6.7e-06 | 5.2e-06 | 7.000E-06 | 1.967E-06 | 5.200E-06 | 9.100E-06 |
| 585 | 30 | G30C | A | 7.7e-06 | 6.3e-06 | 4.9e-06 | 6.300E-06 | 1.400E-06 | 4.900E-06 | 7.700E-06 |
| 586 | 30 | G30E | A | 5.9e-06 | 9.9e-06 | 7.6e-06 | 7.800E-06 | 2.007E-06 | 5.900E-06 | 9.900E-06 |
| 587 | 30 | G30Q | A | 5.9e-06 | 4.6e-06 | 9.8e-06 | 6.767E-06 | 2.706E-06 | 4.600E-06 | 9.800E-06 |
| 588 | 30 | G30G | A | 7.8e-06 | 1.1e-05 | 9.6e-06 | 9.467E-06 | 1.604E-06 | 7.800E-06 | 1.100E-05 |
| 589 | 30 | G30H | A | 1.4e-05 | 7.7e-06 | 9.1e-06 | 1.027E-05 | 3.308E-06 | 7.700E-06 | 1.400E-05 |
| 590 | 30 | G30I | A | 9e-06 | 8.9e-06 | 6.8e-06 | 8.233E-06 | 1.242E-06 | 6.800E-06 | 9.000E-06 |
| 591 | 30 | G30L | A | 9.7e-06 | 9.2e-06 | 6.7e-06 | 8.533E-06 | 1.607E-06 | 6.700E-06 | 9.700E-06 |
| 592 | 30 | G30K | A | 8e-06 | 9.1e-06 | 8.4e-06 | 8.500E-06 | 5.568E-07 | 8.000E-06 | 9.100E-06 |
| 593 | 30 | G30M | A | 6.1e-06 | 1e-05 | 9.5e-06 | 8.533E-06 | 2.122E-06 | 6.100E-06 | 1.000E-05 |
| 594 | 30 | G30F | A | 9.9e-06 | 5.2e-06 | 9.9e-06 | 8.333E-06 | 2.714E-06 | 5.200E-06 | 9.900E-06 |

| | | | | | | | | | | |
|---|---|---|---|---|---|---|---|---|---|---|
| 595 | 30 | G30P | A | 7e-06 | 1.1e-05 | 4.8e-06 | 7.600E-06 | 3.143E-06 | 4.800E-06 | 1.100E-05 |
| 596 | 30 | G30S | A | 9e-06 | 5.6e-06 | 9.9e-06 | 8.167E-06 | 2.268E-06 | 5.600E-06 | 9.900E-06 |
| 597 | 30 | G30T | A | 1.1e-05 | 8.8e-06 | 8.8e-06 | 9.533E-06 | 1.270E-06 | 8.800E-06 | 1.100E-05 |
| 598 | 30 | G30W | A | 7.9e-06 | 6.9e-06 | 7.8e-06 | 7.533E-06 | 5.508E-07 | 6.900E-06 | 7.900E-06 |
| 599 | 30 | G30Y | A | 9.2e-06 | 1.2e-05 | 4.8e-06 | 8.667E-06 | 3.630E-06 | 4.800E-06 | 1.200E-05 |
| 600 | 30 | G30V | A | 1.2e-05 | 7.5e-06 | 7.8e-06 | 9.100E-06 | 2.516E-06 | 7.500E-06 | 1.200E-05 |
| 601 | 31 | C31A | A | 5.8e-06 | 1.3e-05 | 9.4e-06 | 9.400E-06 | 3.600E-06 | 5.800E-06 | 1.300E-05 |
| 602 | 31 | C31R | A | 9.4e-06 | 1.1e-05 | 9.5e-06 | 9.967E-06 | 8.963E-07 | 9.400E-06 | 1.100E-05 |
| 603 | 31 | C31N | A | 9.1e-06 | 8e-06 | 1.1e-05 | 9.367E-06 | 1.518E-06 | 8.000E-06 | 1.100E-05 |
| 604 | 31 | C31D | A | 6e-06 | 6.4e-06 | 6.4e-06 | 6.267E-06 | 2.309E-07 | 6.000E-06 | 6.400E-06 |
| 605 | 31 | C31C | A | 7.5e-06 | 5.7e-06 | 1.2e-05 | 8.400E-06 | 3.245E-06 | 5.700E-06 | 1.200E-05 |
| 606 | 31 | C31E | A | 9.4e-06 | 9.4e-06 | 1.1e-05 | 9.933E-06 | 9.238E-07 | 9.400E-06 | 1.100E-05 |
| 607 | 31 | C31Q | A | 1.2e-05 | 8e-06 | 9.7e-06 | 9.900E-06 | 2.007E-06 | 8.000E-06 | 1.200E-05 |
| 608 | 31 | C31G | A | 6.1e-06 | 1e-05 | 7.3e-06 | 7.800E-06 | 1.997E-06 | 6.100E-06 | 1.000E-05 |
| 609 | 31 | C31H | A | 4.6e-06 | 8.8e-06 | 5.9e-06 | 6.433E-06 | 2.150E-06 | 4.600E-06 | 8.800E-06 |
| 610 | 31 | C31I | A | 1.4e-05 | 7.7e-06 | 5.9e-06 | 9.200E-06 | 4.253E-06 | 5.900E-06 | 1.400E-05 |
| 611 | 31 | C31L | A | 8.7e-06 | 1.1e-05 | 4.7e-06 | 8.133E-06 | 3.188E-06 | 4.700E-06 | 1.100E-05 |
| 612 | 31 | C31K | A | 9.1e-06 | 6.7e-06 | 4.5e-06 | 6.767E-06 | 2.301E-06 | 4.500E-06 | 9.100E-06 |
| 613 | 31 | C31M | A | 8.9e-06 | 8.1e-06 | 5.4e-06 | 7.467E-06 | 1.834E-06 | 5.400E-06 | 8.900E-06 |
| 614 | 31 | C31F | A | 6.4e-06 | 4.8e-06 | 1.3e-05 | 8.067E-06 | 4.347E-06 | 4.800E-06 | 1.300E-05 |
| 615 | 31 | C31P | A | 9.1e-06 | 4.8e-06 | 5.4e-06 | 6.433E-06 | 2.329E-06 | 4.800E-06 | 9.100E-06 |
| 616 | 31 | C31S | A | 5.5e-06 | 1e-05 | 6.5e-06 | 7.333E-06 | 2.363E-06 | 5.500E-06 | 1.000E-05 |
| 617 | 31 | C31T | A | 7.1e-06 | 6.9e-06 | 7.3e-06 | 7.100E-06 | 2.000E-07 | 6.900E-06 | 7.300E-06 |
| 618 | 31 | C31W | A | 8.7e-06 | 7.4e-06 | 8.2e-06 | 8.100E-06 | 6.557E-07 | 7.400E-06 | 8.700E-06 |
| 619 | 31 | C31Y | A | 5.6e-06 | 5.5e-06 | 4.3e-06 | 5.133E-06 | 7.234E-07 | 4.300E-06 | 5.600E-06 |
| 620 | 31 | C31V | A | 8.3e-06 | 8.5e-06 | 9e-06 | 8.600E-06 | 3.606E-07 | 8.300E-06 | 9.000E-06 |
| 621 | 32 | Q32A | A | 7e-06 | 8.6e-06 | 7.2e-06 | 7.600E-06 | 8.718E-07 | 7.000E-06 | 8.600E-06 |
| 622 | 32 | Q32R | A | 5e-06 | 8.1e-06 | 5.4e-06 | 6.167E-06 | 1.686E-06 | 5.000E-06 | 8.100E-06 |
| 623 | 32 | Q32N | A | 5.4e-06 | 7.3e-06 | 9.7e-06 | 7.467E-06 | 2.155E-06 | 5.400E-06 | 9.700E-06 |
| 624 | 32 | Q32D | A | 1e-05 | 8.2e-06 | 1.1e-05 | 9.733E-06 | 1.419E-06 | 8.200E-06 | 1.100E-05 |
| 625 | 32 | Q32C | A | 6e-06 | 4.5e-06 | 8.9e-06 | 6.467E-06 | 2.237E-06 | 4.500E-06 | 8.900E-06 |
| 626 | 32 | Q32E | A | 4.8e-06 | 8.2e-06 | 6.6e-06 | 6.533E-06 | 1.701E-06 | 4.800E-06 | 8.200E-06 |
| 627 | 32 | Q32Q | A | 6.6e-06 | 8e-06 | 7.8e-06 | 7.467E-06 | 7.572E-07 | 6.600E-06 | 8.000E-06 |
| 628 | 32 | Q32G | A | 8.7e-06 | 7.3e-06 | 1.4e-05 | 1.000E-05 | 3.534E-06 | 7.300E-06 | 1.400E-05 |
| 629 | 32 | Q32H | A | 4.9e-06 | 8.5e-06 | 5.5e-06 | 6.300E-06 | 1.929E-06 | 4.900E-06 | 8.500E-06 |
| 630 | 32 | Q32I | A | 4.4e-06 | 8.8e-06 | 1e-05 | 7.733E-06 | 2.948E-06 | 4.400E-06 | 1.000E-05 |
| 631 | 32 | Q32L | A | 7.5e-06 | 7.1e-06 | 5e-06 | 6.533E-06 | 1.343E-06 | 5.000E-06 | 7.500E-06 |
| 632 | 32 | Q32K | A | 5.6e-06 | 8.7e-06 | 6.5e-06 | 6.933E-06 | 1.595E-06 | 5.600E-06 | 8.700E-06 |
| 633 | 32 | Q32M | A | 7.3e-06 | 7.3e-06 | 8.8e-06 | 7.800E-06 | 8.660E-07 | 7.300E-06 | 8.800E-06 |
| 634 | 32 | Q32F | A | 6.3e-06 | 8e-06 | 9e-06 | 7.767E-06 | 1.365E-06 | 6.300E-06 | 9.000E-06 |
| 635 | 32 | Q32P | A | 7.1e-06 | 1e-05 | 1.3e-05 | 1.003E-05 | 2.950E-06 | 7.100E-06 | 1.300E-05 |
| 636 | 32 | Q32S | A | 4.9e-06 | 6.4e-06 | 4.9e-06 | 5.400E-06 | 8.660E-07 | 4.900E-06 | 6.400E-06 |
| 637 | 32 | Q32T | A | 5.6e-06 | 9.2e-06 | 9.5e-06 | 8.100E-06 | 2.170E-06 | 5.600E-06 | 9.500E-06 |
| 638 | 32 | Q32W | A | 6.4e-06 | 1e-05 | 9.8e-06 | 8.733E-06 | 2.023E-06 | 6.400E-06 | 1.000E-05 |
| 639 | 32 | Q32Y | A | 7.3e-06 | 4.4e-06 | 5.5e-06 | 5.733E-06 | 1.464E-06 | 4.400E-06 | 7.300E-06 |
| 640 | 32 | Q32V | A | 7.1e-06 | 4.9e-06 | 9.4e-06 | 7.133E-06 | 2.250E-06 | 4.900E-06 | 9.400E-06 |
| 641 | 33 | V33A | A | 6.2e-06 | 1.1e-05 | 9.4e-06 | 8.867E-06 | 2.444E-06 | 6.200E-06 | 1.100E-05 |
| 642 | 33 | V33R | A | 4.3e-06 | 1.5e-05 | 9.4e-06 | 9.567E-06 | 5.352E-06 | 4.300E-06 | 1.500E-05 |
| 643 | 33 | V33N | A | 5.3e-06 | 1.3e-05 | 1.4e-05 | 1.077E-05 | 4.761E-06 | 5.300E-06 | 1.400E-05 |
| 644 | 33 | V33D | A | 1.1e-05 | 9e-06 | 4.1e-06 | 8.033E-06 | 3.550E-06 | 4.100E-06 | 1.100E-05 |
| 645 | 33 | V33C | A | 9.1e-06 | 7.5e-06 | 6.8e-06 | 7.800E-06 | 1.179E-06 | 6.800E-06 | 9.100E-06 |
| 646 | 33 | V33E | A | 6.4e-06 | 9.5e-06 | 5.5e-06 | 7.133E-06 | 2.098E-06 | 5.500E-06 | 9.500E-06 |
| 647 | 33 | V33Q | A | 8.2e-06 | 6.3e-06 | 1e-05 | 8.167E-06 | 1.850E-06 | 6.300E-06 | 1.000E-05 |
| 648 | 33 | V33G | A | 5.4e-06 | 8.1e-06 | 7.8e-06 | 7.100E-06 | 1.480E-06 | 5.400E-06 | 8.100E-06 |
| 649 | 33 | V33H | A | 9.1e-06 | 5.5e-06 | 7.4e-06 | 7.333E-06 | 1.801E-06 | 5.500E-06 | 9.100E-06 |
| 650 | 33 | V33I | A | 7.2e-06 | 7e-06 | 6.7e-06 | 6.967E-06 | 2.517E-07 | 6.700E-06 | 7.200E-06 |
| 651 | 33 | V33L | A | 7.6e-06 | 9.2e-06 | 1.3e-05 | 9.933E-06 | 2.774E-06 | 7.600E-06 | 1.300E-05 |
| 652 | 33 | V33K | A | 4.5e-06 | 1.3e-05 | 5.1e-06 | 7.533E-06 | 4.744E-06 | 4.500E-06 | 1.300E-05 |
| 653 | 33 | V33M | A | 9.9e-06 | 8.1e-06 | 9.8e-06 | 9.267E-06 | 1.012E-06 | 8.100E-06 | 9.900E-06 |
| 654 | 33 | V33F | A | 7e-06 | 1.1e-05 | 4.9e-06 | 7.633E-06 | 3.099E-06 | 4.900E-06 | 1.100E-05 |
| 655 | 33 | V33P | A | 6.5e-06 | 7.2e-06 | 7.5e-06 | 7.067E-06 | 5.132E-07 | 6.500E-06 | 7.500E-06 |
| 656 | 33 | V33S | A | 4.5e-06 | 1e-05 | 7.7e-06 | 7.400E-06 | 2.762E-06 | 4.500E-06 | 1.000E-05 |
| 657 | 33 | V33T | A | 7.4e-06 | 1.2e-05 | 6.1e-06 | 8.500E-06 | 3.100E-06 | 6.100E-06 | 1.200E-05 |
| 658 | 33 | V33W | A | 7.4e-06 | 6.2e-06 | 9.9e-06 | 7.833E-06 | 1.888E-06 | 6.200E-06 | 9.900E-06 |
| 659 | 33 | V33Y | A | 1e-05 | 5.4e-06 | 9e-06 | 8.133E-06 | 2.419E-06 | 5.400E-06 | 1.000E-05 |
| 660 | 33 | V33V | A | 6.9e-06 | 9.1e-06 | 6.8e-06 | 7.600E-06 | 1.300E-06 | 6.800E-06 | 9.100E-06 |
| 661 | 34 | V34A | A | 9.2e-06 | 7e-06 | 6.6e-06 | 7.600E-06 | 1.400E-06 | 6.600E-06 | 9.200E-06 |
| 662 | 34 | V34R | A | 1.5e-05 | 1e-05 | 6.8e-06 | 1.060E-05 | 4.133E-06 | 6.800E-06 | 1.500E-05 |
| 663 | 34 | V34N | A | 1.3e-05 | 1e-05 | 3.7e-06 | 8.900E-06 | 4.747E-06 | 3.700E-06 | 1.300E-05 |
| 664 | 34 | V34D | A | 6.7e-06 | 5.4e-06 | 6.3e-06 | 6.133E-06 | 6.658E-07 | 5.400E-06 | 6.700E-06 |
| 665 | 34 | V34C | A | 9.6e-06 | 5.1e-06 | 6.3e-06 | 7.000E-06 | 2.330E-06 | 5.100E-06 | 9.600E-06 |
| 666 | 34 | V34E | A | 5.8e-06 | 6.7e-06 | 8e-06 | 6.833E-06 | 1.106E-06 | 5.800E-06 | 8.000E-06 |
| 667 | 34 | V34Q | A | 8.1e-06 | 6.8e-06 | 7.5e-06 | 7.467E-06 | 6.506E-07 | 6.800E-06 | 8.100E-06 |
| 668 | 34 | V34G | A | 7.2e-06 | 4.8e-06 | 1.3e-05 | 8.333E-06 | 4.216E-06 | 4.800E-06 | 1.300E-05 |
| 669 | 34 | V34H | A | 6.1e-06 | 1e-05 | 1.1e-05 | 9.033E-06 | 2.589E-06 | 6.100E-06 | 1.100E-05 |
| 670 | 34 | V34I | A | 6.3e-06 | 5.4e-06 | 8.5e-06 | 6.733E-06 | 1.595E-06 | 5.400E-06 | 8.500E-06 |
| 671 | 34 | V34L | A | 7.9e-06 | 6.2e-06 | 1.4e-05 | 9.367E-06 | 4.102E-06 | 6.200E-06 | 1.400E-05 |
| 672 | 34 | V34K | A | 5.5e-06 | 8.4e-06 | 1e-05 | 7.967E-06 | 2.281E-06 | 5.500E-06 | 1.000E-05 |
| 673 | 34 | V34M | A | 9.4e-06 | 1e-05 | 1.1e-05 | 1.013E-05 | 8.083E-07 | 9.400E-06 | 1.100E-05 |
| 674 | 34 | V34F | A | 1.4e-05 | 7.4e-06 | 8.2e-06 | 9.867E-06 | 3.602E-06 | 7.400E-06 | 1.400E-05 |
| 675 | 34 | V34P | A | 3.7e-06 | 7.9e-06 | 1.3e-05 | 8.200E-06 | 4.657E-06 | 3.700E-06 | 1.300E-05 |
| 676 | 34 | V34S | A | 1.4e-05 | 7.1e-06 | 1e-05 | 1.037E-05 | 3.465E-06 | 7.100E-06 | 1.400E-05 |
| 677 | 34 | V34T | A | 8.2e-06 | 1.2e-05 | 1.1e-05 | 1.040E-05 | 1.970E-06 | 8.200E-06 | 1.200E-05 |
| 678 | 34 | V34W | A | 5.3e-06 | 1.3e-05 | 6.9e-06 | 8.400E-06 | 4.063E-06 | 5.300E-06 | 1.300E-05 |
| 679 | 34 | V34Y | A | 4.3e-06 | 1.1e-05 | 5.7e-06 | 7.000E-06 | 3.534E-06 | 4.300E-06 | 1.100E-05 |

| | | | | | | | | | | |
|---|---|---|---|---|---|---|---|---|---|---|
| 680 | 34 | V34V | A | 6.2e-06 | 6.7e-06 | 1.3e-05 | 8.633E-06 | 3.790E-06 | 6.200E-06 | 1.300E-05 |
| 681 | 35 | Q35A | A | 1.2e-05 | 1.2e-05 | 1.1e-05 | 1.167E-05 | 5.774E-07 | 1.100E-05 | 1.200E-05 |
| 682 | 35 | Q35R | A | 6.6e-06 | 7.8e-06 | 7.1e-06 | 7.167E-06 | 6.028E-07 | 6.600E-06 | 7.800E-06 |
| 683 | 35 | Q35N | A | 8.2e-06 | 9.8e-06 | 7.1e-06 | 8.367E-06 | 1.358E-06 | 7.100E-06 | 9.800E-06 |
| 684 | 35 | Q35D | A | 5.7e-06 | 9.2e-06 | 1.3e-05 | 9.300E-06 | 3.651E-06 | 5.700E-06 | 1.300E-05 |
| 685 | 35 | Q35C | A | 6.6e-06 | 1.3e-05 | 1.2e-05 | 1.053E-05 | 3.443E-06 | 6.600E-06 | 1.300E-05 |
| 686 | 35 | Q35E | A | 4.4e-06 | 6.4e-06 | 1.3e-05 | 7.933E-06 | 4.500E-06 | 4.400E-06 | 1.300E-05 |
| 687 | 35 | Q35Q | A | 6.3e-06 | 9e-06 | 9.3e-06 | 8.200E-06 | 1.652E-06 | 6.300E-06 | 9.300E-06 |
| 688 | 35 | Q35G | A | 1.3e-05 | 5.9e-06 | 9.4e-06 | 9.433E-06 | 3.550E-06 | 5.900E-06 | 1.300E-05 |
| 689 | 35 | Q35H | A | 6.7e-06 | 1.2e-05 | 4.3e-06 | 7.667E-06 | 3.940E-06 | 4.300E-06 | 1.200E-05 |
| 690 | 35 | Q35I | A | 1e-05 | 9.8e-06 | 5.5e-06 | 8.433E-06 | 2.542E-06 | 5.500E-06 | 1.000E-05 |
| 691 | 35 | Q35L | A | 9.7e-06 | 6.6e-06 | 4.2e-06 | 6.833E-06 | 2.757E-06 | 4.200E-06 | 9.700E-06 |
| 692 | 35 | Q35K | A | 4.5e-06 | 4.9e-06 | 8.1e-06 | 5.833E-06 | 1.973E-06 | 4.500E-06 | 8.100E-06 |
| 693 | 35 | Q35M | A | 1.2e-05 | 6.6e-06 | 7.6e-06 | 8.733E-06 | 2.873E-06 | 6.600E-06 | 1.200E-05 |
| 694 | 35 | Q35F | A | 5.2e-06 | 1.1e-05 | 1e-05 | 8.733E-06 | 3.101E-06 | 5.200E-06 | 1.100E-05 |
| 695 | 35 | Q35P | A | 1.1e-05 | 6.8e-06 | 7.2e-06 | 8.333E-06 | 2.318E-06 | 6.800E-06 | 1.100E-05 |
| 696 | 35 | Q35S | A | 6e-06 | 5.7e-06 | 6.8e-06 | 6.167E-06 | 5.686E-07 | 5.700E-06 | 6.800E-06 |
| 697 | 35 | Q35T | A | 1.3e-05 | 7.8e-06 | 4.7e-06 | 8.500E-06 | 4.194E-06 | 4.700E-06 | 1.300E-05 |
| 698 | 35 | Q35W | A | 9.9e-06 | 5e-06 | 9.4e-06 | 8.100E-06 | 2.696E-06 | 5.000E-06 | 9.900E-06 |
| 699 | 35 | Q35Y | A | 1.3e-05 | 5.3e-06 | 7.1e-06 | 8.467E-06 | 4.028E-06 | 5.300E-06 | 1.300E-05 |
| 700 | 35 | Q35V | A | 1.2e-05 | 9.5e-06 | 8.9e-06 | 1.013E-05 | 1.644E-06 | 8.900E-06 | 1.200E-05 |
| 701 | 36 | G36A | A | 6e-06 | 6.5e-06 | 1.5e-05 | 9.167E-06 | 5.058E-06 | 6.000E-06 | 1.500E-05 |
| 702 | 36 | G36R | A | 4.6e-06 | 9.9e-06 | 1.3e-05 | 9.167E-06 | 4.248E-06 | 4.600E-06 | 1.300E-05 |
| 703 | 36 | G36N | A | 1.3e-05 | 5.9e-06 | 6.7e-06 | 8.533E-06 | 3.889E-06 | 5.900E-06 | 1.300E-05 |
| 704 | 36 | G36D | A | 5e-06 | 5.5e-06 | 5.4e-06 | 5.300E-06 | 2.646E-07 | 5.000E-06 | 5.500E-06 |
| 705 | 36 | G36C | A | 1e-05 | 7.6e-06 | 9.8e-06 | 9.133E-06 | 1.332E-06 | 7.600E-06 | 1.000E-05 |
| 706 | 36 | G36E | A | 1.3e-05 | 7.7e-06 | 6.2e-06 | 8.967E-06 | 3.573E-06 | 6.200E-06 | 1.300E-05 |
| 707 | 36 | G36Q | A | 6e-06 | 7.9e-06 | 7.9e-06 | 7.267E-06 | 1.097E-06 | 6.000E-06 | 7.900E-06 |
| 708 | 36 | G36G | A | 1e-05 | 5.6e-06 | 6.9e-06 | 7.500E-06 | 2.261E-06 | 5.600E-06 | 1.000E-05 |
| 709 | 36 | G36H | A | 7.2e-06 | 8.9e-06 | 4.1e-06 | 6.733E-06 | 2.434E-06 | 4.100E-06 | 8.900E-06 |
| 710 | 36 | G36I | A | 8e-06 | 8.8e-06 | 8.3e-06 | 8.367E-06 | 4.041E-07 | 8.000E-06 | 8.800E-06 |
| 711 | 36 | G36L | A | 1.3e-05 | 7e-06 | 6e-06 | 8.667E-06 | 3.786E-06 | 6.000E-06 | 1.300E-05 |
| 712 | 36 | G36K | A | 1.3e-05 | 1.2e-05 | 6.7e-06 | 1.057E-05 | 3.386E-06 | 6.700E-06 | 1.300E-05 |
| 713 | 36 | G36M | A | 1.1e-05 | 4.5e-06 | 8.1e-06 | 7.867E-06 | 3.256E-06 | 4.500E-06 | 1.100E-05 |
| 714 | 36 | G36F | A | 1.3e-05 | 5.7e-06 | 5.6e-06 | 8.100E-06 | 4.244E-06 | 5.600E-06 | 1.300E-05 |
| 715 | 36 | G36P | A | 1.4e-05 | 1.3e-05 | 5.1e-06 | 1.070E-05 | 4.875E-06 | 5.100E-06 | 1.400E-05 |
| 716 | 36 | G36S | A | 5.3e-06 | 7.9e-06 | 6.4e-06 | 6.533E-06 | 1.305E-06 | 5.300E-06 | 7.900E-06 |
| 717 | 36 | G36T | A | 5.2e-06 | 5.5e-06 | 6.9e-06 | 5.867E-06 | 9.074E-07 | 5.200E-06 | 6.900E-06 |
| 718 | 36 | G36W | A | 8.3e-06 | 6.5e-06 | 9.1e-06 | 7.967E-06 | 1.332E-06 | 6.500E-06 | 9.100E-06 |
| 719 | 36 | G36Y | A | 1.4e-05 | 6.9e-06 | 5.3e-06 | 8.733E-06 | 4.631E-06 | 5.300E-06 | 1.400E-05 |
| 720 | 36 | G36V | A | 1.1e-05 | 8.6e-06 | 8.2e-06 | 9.267E-06 | 1.514E-06 | 8.200E-06 | 1.100E-05 |
| 721 | 37 | N37A | A | 8.2e-06 | 1.7e-05 | 9.6e-06 | 1.160E-05 | 4.729E-06 | 8.200E-06 | 1.700E-05 |
| 722 | 37 | N37R | A | 5.7e-06 | 6.9e-06 | 7.1e-06 | 6.567E-06 | 7.572E-07 | 5.700E-06 | 7.100E-06 |
| 723 | 37 | N37N | A | 5.5e-06 | 8.9e-06 | 6.3e-06 | 6.900E-06 | 1.778E-06 | 5.500E-06 | 8.900E-06 |
| 724 | 37 | N37D | A | 6.2e-06 | 9.2e-06 | 4.9e-06 | 6.767E-06 | 2.205E-06 | 4.900E-06 | 9.200E-06 |
| 725 | 37 | N37C | A | 5.6e-06 | 7.2e-06 | 5.8e-06 | 6.200E-06 | 8.718E-07 | 5.600E-06 | 7.200E-06 |
| 726 | 37 | N37E | A | 9.2e-06 | 1e-05 | 4.1e-06 | 7.767E-06 | 3.201E-06 | 4.100E-06 | 1.000E-05 |
| 727 | 37 | N37Q | A | 5.9e-06 | 1.2e-05 | 3.2e-06 | 7.033E-06 | 4.508E-06 | 3.200E-06 | 1.200E-05 |
| 728 | 37 | N37G | A | 8.6e-06 | 4.6e-06 | 6.9e-06 | 6.700E-06 | 2.007E-06 | 4.600E-06 | 8.600E-06 |
| 729 | 37 | N37H | A | 6.4e-06 | 8e-06 | 8.6e-06 | 7.667E-06 | 1.137E-06 | 6.400E-06 | 8.600E-06 |
| 730 | 37 | N37I | A | 8.6e-06 | 6.6e-06 | 4.9e-06 | 6.700E-06 | 1.852E-06 | 4.900E-06 | 8.600E-06 |
| 731 | 37 | N37L | A | 7.9e-06 | 1.2e-05 | 6.8e-06 | 8.900E-06 | 2.740E-06 | 6.800E-06 | 1.200E-05 |
| 732 | 37 | N37K | A | 2.4e-05 | 7.3e-06 | 8e-06 | 1.310E-05 | 9.446E-06 | 7.300E-06 | 2.400E-05 |
| 733 | 37 | N37M | A | 1e-05 | 7.6e-06 | 9.4e-06 | 9.000E-06 | 1.249E-06 | 7.600E-06 | 1.000E-05 |
| 734 | 37 | N37F | A | 1.1e-05 | 9.4e-06 | 9.5e-06 | 9.967E-06 | 8.963E-07 | 9.400E-06 | 1.100E-05 |
| 735 | 37 | N37P | A | 5.5e-06 | 1.8e-05 | 4.6e-06 | 9.367E-06 | 7.490E-06 | 4.600E-06 | 1.800E-05 |
| 736 | 37 | N37S | A | 1.2e-05 | 5.3e-06 | 1e-05 | 9.100E-06 | 3.439E-06 | 5.300E-06 | 1.200E-05 |
| 737 | 37 | N37T | A | 9.3e-06 | 6e-06 | 7.1e-06 | 7.467E-06 | 1.680E-06 | 6.000E-06 | 9.300E-06 |
| 738 | 37 | N37W | A | 9e-06 | 5.4e-06 | 1.2e-05 | 8.800E-06 | 3.305E-06 | 5.400E-06 | 1.200E-05 |
| 739 | 37 | N37Y | A | 9.4e-06 | 4.3e-06 | 9.2e-06 | 7.633E-06 | 2.888E-06 | 4.300E-06 | 9.400E-06 |
| 740 | 37 | N37V | A | 4.3e-06 | 5.6e-06 | 6.8e-06 | 5.567E-06 | 1.250E-06 | 4.300E-06 | 6.800E-06 |
| 741 | 38 | L38A | A | 6e-06 | 4.9e-06 | 7.1e-06 | 6.000E-06 | 1.100E-06 | 4.900E-06 | 7.100E-06 |
| 742 | 38 | L38R | A | 9.1e-06 | 8.8e-06 | 9e-06 | 8.967E-06 | 1.528E-07 | 8.800E-06 | 9.100E-06 |
| 743 | 38 | L38N | A | 1.1e-05 | 8.2e-06 | 1e-05 | 9.733E-06 | 1.419E-06 | 8.200E-06 | 1.100E-05 |
| 744 | 38 | L38D | A | 6.9e-06 | 9.2e-06 | 5.6e-06 | 7.233E-06 | 1.823E-06 | 5.600E-06 | 9.200E-06 |
| 745 | 38 | L38C | A | 6.6e-06 | 9.1e-06 | 9.4e-06 | 8.367E-06 | 1.537E-06 | 6.600E-06 | 9.400E-06 |
| 746 | 38 | L38E | A | 4e-06 | 5.3e-06 | 9.8e-06 | 6.367E-06 | 3.044E-06 | 4.000E-06 | 9.800E-06 |
| 747 | 38 | L38Q | A | 1.1e-05 | 4.9e-06 | 4.6e-06 | 6.833E-06 | 3.612E-06 | 4.600E-06 | 1.100E-05 |
| 748 | 38 | L38G | A | 6e-06 | 6.9e-06 | 1.3e-05 | 8.633E-06 | 3.808E-06 | 6.000E-06 | 1.300E-05 |
| 749 | 38 | L38H | A | 6e-06 | 1.2e-05 | 6e-06 | 8.000E-06 | 3.464E-06 | 6.000E-06 | 1.200E-05 |
| 750 | 38 | L38I | A | 7.2e-06 | 6.1e-06 | 1e-05 | 7.767E-06 | 2.011E-06 | 6.100E-06 | 1.000E-05 |
| 751 | 38 | L38L | A | 1.3e-05 | 8.4e-06 | 6.7e-06 | 9.367E-06 | 3.259E-06 | 6.700E-06 | 1.300E-05 |
| 752 | 38 | L38K | A | 6.9e-06 | 8.1e-06 | 6.7e-06 | 7.233E-06 | 7.572E-07 | 6.700E-06 | 8.100E-06 |
| 753 | 38 | L38M | A | 7.4e-06 | 4.9e-06 | 9.1e-06 | 7.133E-06 | 2.113E-06 | 4.900E-06 | 9.100E-06 |
| 754 | 38 | L38F | A | 1e-05 | 7.6e-06 | 7.6e-06 | 8.400E-06 | 1.386E-06 | 7.600E-06 | 1.000E-05 |
| 755 | 38 | L38P | A | 8.6e-06 | 5.4e-06 | 6.6e-06 | 6.867E-06 | 1.617E-06 | 5.400E-06 | 8.600E-06 |
| 756 | 38 | L38S | A | 6.5e-06 | 1.2e-05 | 1e-05 | 9.500E-06 | 2.784E-06 | 6.500E-06 | 1.200E-05 |
| 757 | 38 | L38T | A | 9.3e-06 | 8.7e-06 | 5.8e-06 | 7.933E-06 | 1.872E-06 | 5.800E-06 | 9.300E-06 |
| 758 | 38 | L38W | A | 8.3e-06 | 6e-06 | 1.1e-05 | 8.433E-06 | 2.503E-06 | 6.000E-06 | 1.100E-05 |
| 759 | 38 | L38Y | A | 6.8e-06 | 7.4e-06 | 8.3e-06 | 7.500E-06 | 7.550E-07 | 6.800E-06 | 8.300E-06 |
| 760 | 38 | L38V | A | 5.1e-06 | 5.5e-06 | 9.4e-06 | 6.667E-06 | 2.376E-06 | 5.100E-06 | 9.400E-06 |
| 761 | 39 | E39A | A | 7.5e-06 | 5.3e-06 | 7.4e-06 | 6.733E-06 | 1.242E-06 | 5.300E-06 | 7.500E-06 |
| 762 | 39 | E39R | A | 9.1e-06 | 8.8e-06 | 7e-06 | 8.300E-06 | 1.136E-06 | 7.000E-06 | 9.100E-06 |
| 763 | 39 | E39N | A | 8.8e-06 | 9.5e-06 | 8.5e-06 | 8.933E-06 | 5.132E-07 | 8.500E-06 | 9.500E-06 |
| 764 | 39 | E39D | A | 1.5e-05 | 1.2e-05 | 1.4e-05 | 1.367E-05 | 1.528E-06 | 1.200E-05 | 1.500E-05 |

| | | | | | | | | | | |
|---|---|---|---|---|---|---|---|---|---|---|
| 765 | 39 | E39C | A | 9.5e-06 | 7.6e-06 | 6.2e-06 | 7.767E-06 | 1.656E-06 | 6.200E-06 | 9.500E-06 |
| 766 | 39 | E39E | A | 5.2e-06 | 7.1e-06 | 6.2e-06 | 6.167E-06 | 9.504E-07 | 5.200E-06 | 7.100E-06 |
| 767 | 39 | E39Q | A | 6.7e-06 | 9.1e-06 | 5.7e-06 | 7.167E-06 | 1.747E-06 | 5.700E-06 | 9.100E-06 |
| 768 | 39 | E39G | A | 7.2e-06 | 1e-05 | 9.1e-06 | 8.767E-06 | 1.429E-06 | 7.200E-06 | 1.000E-05 |
| 769 | 39 | E39H | A | 8.9e-06 | 7.5e-06 | 1.1e-05 | 9.133E-06 | 1.762E-06 | 7.500E-06 | 1.100E-05 |
| 770 | 39 | E39I | A | 8.4e-06 | 6.5e-06 | 8.5e-06 | 7.800E-06 | 1.127E-06 | 6.500E-06 | 8.500E-06 |
| 771 | 39 | E39L | A | 1e-05 | 6.9e-06 | 6.9e-06 | 7.933E-06 | 1.790E-06 | 6.900E-06 | 1.000E-05 |
| 772 | 39 | E39K | A | 1.6e-05 | 4.3e-06 | 8.3e-06 | 9.533E-06 | 5.947E-06 | 4.300E-06 | 1.600E-05 |
| 773 | 39 | E39M | A | 6.5e-06 | 8.7e-06 | 8.7e-06 | 7.967E-06 | 1.270E-06 | 6.500E-06 | 8.700E-06 |
| 774 | 39 | E39F | A | 8.6e-06 | 6.4e-06 | 1.4e-05 | 9.667E-06 | 3.911E-06 | 6.400E-06 | 1.400E-05 |
| 775 | 39 | E39P | A | 1.6e-05 | 7.3e-06 | 6.9e-06 | 1.007E-05 | 5.142E-06 | 6.900E-06 | 1.600E-05 |
| 776 | 39 | E39S | A | 8.1e-06 | 4.6e-06 | 8.9e-06 | 7.200E-06 | 2.287E-06 | 4.600E-06 | 8.900E-06 |
| 777 | 39 | E39T | A | 9.3e-06 | 6.1e-06 | 7.7e-06 | 7.700E-06 | 1.600E-06 | 6.100E-06 | 9.300E-06 |
| 778 | 39 | E39W | A | 6.9e-06 | 8.6e-06 | 9.4e-06 | 8.300E-06 | 1.277E-06 | 6.900E-06 | 9.400E-06 |
| 779 | 39 | E39Y | A | 7.3e-06 | 5e-06 | 4.3e-06 | 5.533E-06 | 1.570E-06 | 4.300E-06 | 7.300E-06 |
| 780 | 39 | E39V | A | 5.3e-06 | 6.6e-06 | 6.8e-06 | 6.233E-06 | 8.145E-07 | 5.300E-06 | 6.800E-06 |
| 781 | 40 | L40A | A | 8.8e-06 | 8.8e-06 | 4.2e-06 | 7.267E-06 | 2.656E-06 | 4.200E-06 | 8.800E-06 |
| 782 | 40 | L40R | A | 6.3e-06 | 1.2e-05 | 4.1e-06 | 7.467E-06 | 4.077E-06 | 4.100E-06 | 1.200E-05 |
| 783 | 40 | L40N | A | 4.5e-06 | 7.9e-06 | 9.2e-06 | 7.200E-06 | 2.427E-06 | 4.500E-06 | 9.200E-06 |
| 784 | 40 | L40D | A | 7.4e-06 | 4.7e-06 | 6.8e-06 | 6.300E-06 | 1.418E-06 | 4.700E-06 | 7.400E-06 |
| 785 | 40 | L40C | A | 8.1e-06 | 6.5e-06 | 7.6e-06 | 7.400E-06 | 8.185E-07 | 6.500E-06 | 8.100E-06 |
| 786 | 40 | L40E | A | 6.5e-06 | 5.7e-06 | 9.2e-06 | 7.133E-06 | 1.834E-06 | 5.700E-06 | 9.200E-06 |
| 787 | 40 | L40Q | A | 9.7e-06 | 1.5e-05 | 6.7e-06 | 1.047E-05 | 4.203E-06 | 6.700E-06 | 1.500E-05 |
| 788 | 40 | L40G | A | 7.1e-06 | 8.7e-06 | 8.2e-06 | 8.000E-06 | 8.185E-07 | 7.100E-06 | 8.700E-06 |
| 789 | 40 | L40H | A | 1e-05 | 7.4e-06 | 1.1e-05 | 9.467E-06 | 1.858E-06 | 7.400E-06 | 1.100E-05 |
| 790 | 40 | L40I | A | 9e-06 | 6.9e-06 | 1e-05 | 8.633E-06 | 1.582E-06 | 6.900E-06 | 1.000E-05 |
| 791 | 40 | L40L | A | 4.5e-06 | 9.4e-06 | 6.4e-06 | 6.767E-06 | 2.470E-06 | 4.500E-06 | 9.400E-06 |
| 792 | 40 | L40K | A | 7.2e-06 | 1.1e-05 | 7.8e-06 | 8.667E-06 | 2.043E-06 | 7.200E-06 | 1.100E-05 |
| 793 | 40 | L40M | A | 5.9e-06 | 1.1e-05 | 8e-06 | 8.300E-06 | 2.563E-06 | 5.900E-06 | 1.100E-05 |
| 794 | 40 | L40F | A | 6e-06 | 6.7e-06 | 8.4e-06 | 7.033E-06 | 1.234E-06 | 6.000E-06 | 8.400E-06 |
| 795 | 40 | L40P | A | 9.7e-06 | 7.8e-06 | 8.8e-06 | 8.767E-06 | 9.504E-07 | 7.800E-06 | 9.700E-06 |
| 796 | 40 | L40S | A | 5.5e-06 | 8.3e-06 | 5.7e-06 | 6.500E-06 | 1.562E-06 | 5.500E-06 | 8.300E-06 |
| 797 | 40 | L40T | A | 8.4e-06 | 9.5e-06 | 1e-05 | 9.300E-06 | 8.185E-07 | 8.400E-06 | 1.000E-05 |
| 798 | 40 | L40W | A | 9.2e-06 | 1.4e-05 | 8.3e-06 | 1.050E-05 | 3.064E-06 | 8.300E-06 | 1.400E-05 |
| 799 | 40 | L40Y | A | 7.5e-06 | 6.8e-06 | 6.7e-06 | 7.000E-06 | 4.359E-07 | 6.700E-06 | 7.500E-06 |
| 800 | 40 | L40V | A | 1.3e-05 | 6.2e-06 | 6.7e-06 | 8.633E-06 | 3.790E-06 | 6.200E-06 | 1.300E-05 |
| 801 | 41 | T41A | A | 7.3e-06 | 1.3e-05 | 1e-05 | 1.010E-05 | 2.851E-06 | 7.300E-06 | 1.300E-05 |
| 802 | 41 | T41R | A | 8.7e-06 | 8.3e-06 | 7.1e-06 | 8.033E-06 | 8.327E-07 | 7.100E-06 | 8.700E-06 |
| 803 | 41 | T41N | A | 7e-06 | 8.5e-06 | 5e-06 | 6.833E-06 | 1.756E-06 | 5.000E-06 | 8.500E-06 |
| 804 | 41 | T41D | A | 4.1e-06 | 4.9e-06 | 4.9e-06 | 4.633E-06 | 4.619E-07 | 4.100E-06 | 4.900E-06 |
| 805 | 41 | T41C | A | 5.2e-06 | 9.1e-06 | 1.4e-05 | 9.433E-06 | 4.409E-06 | 5.200E-06 | 1.400E-05 |
| 806 | 41 | T41E | A | 7.6e-06 | 9.5e-06 | 8.9e-06 | 8.667E-06 | 9.713E-07 | 7.600E-06 | 9.500E-06 |
| 807 | 41 | T41Q | A | 1e-05 | 1e-05 | 8.7e-06 | 9.567E-06 | 7.506E-07 | 8.700E-06 | 1.000E-05 |
| 808 | 41 | T41G | A | 1e-05 | 8.5e-06 | 1.3e-05 | 1.050E-05 | 2.291E-06 | 8.500E-06 | 1.300E-05 |
| 809 | 41 | T41H | A | 6.1e-06 | 4.9e-06 | 6.3e-06 | 5.767E-06 | 7.572E-07 | 4.900E-06 | 6.300E-06 |
| 810 | 41 | T41I | A | 7.5e-06 | 9.5e-06 | 1.3e-05 | 1.000E-05 | 2.784E-06 | 7.500E-06 | 1.300E-05 |
| 811 | 41 | T41L | A | 5.5e-06 | 8.6e-06 | 5.2e-06 | 6.433E-06 | 1.882E-06 | 5.200E-06 | 8.600E-06 |
| 812 | 41 | T41K | A | 7e-06 | 6.7e-06 | 1.1e-05 | 8.233E-06 | 2.401E-06 | 6.700E-06 | 1.100E-05 |
| 813 | 41 | T41M | A | 9e-06 | 7.1e-06 | 9.5e-06 | 8.533E-06 | 1.266E-06 | 7.100E-06 | 9.500E-06 |
| 814 | 41 | T41F | A | 6.5e-06 | 6.8e-06 | 1.3e-05 | 8.767E-06 | 3.669E-06 | 6.500E-06 | 1.300E-05 |
| 815 | 41 | T41P | A | 1.1e-05 | 6.6e-06 | 8.4e-06 | 8.667E-06 | 2.212E-06 | 6.600E-06 | 1.100E-05 |
| 816 | 41 | T41S | A | 5.2e-06 | 1.2e-05 | 7.9e-06 | 8.367E-06 | 3.424E-06 | 5.200E-06 | 1.200E-05 |
| 817 | 41 | T41T | A | 6.5e-06 | 5.3e-06 | 7.5e-06 | 6.433E-06 | 1.102E-06 | 5.300E-06 | 7.500E-06 |
| 818 | 41 | T41W | A | 8.7e-06 | 7.3e-06 | 8.7e-06 | 8.233E-06 | 8.083E-07 | 7.300E-06 | 8.700E-06 |
| 819 | 41 | T41Y | A | 1e-05 | 4.4e-06 | 7.2e-06 | 7.200E-06 | 2.800E-06 | 4.400E-06 | 1.000E-05 |
| 820 | 41 | T41V | A | 1.4e-05 | 1.2e-05 | 1.3e-05 | 1.300E-05 | 1.000E-06 | 1.200E-05 | 1.400E-05 |
| 821 | 42 | Y42A | A | 9.1e-06 | 1e-05 | 1.1e-05 | 1.003E-05 | 9.504E-07 | 9.100E-06 | 1.100E-05 |
| 822 | 42 | Y42R | A | 8e-06 | 7.3e-06 | 1.1e-05 | 8.767E-06 | 1.966E-06 | 7.300E-06 | 1.100E-05 |
| 823 | 42 | Y42N | A | 7.5e-06 | 1e-05 | 7.6e-06 | 8.367E-06 | 1.415E-06 | 7.500E-06 | 1.000E-05 |
| 824 | 42 | Y42D | A | 6.9e-06 | 8.8e-06 | 5e-06 | 6.900E-06 | 1.900E-06 | 5.000E-06 | 8.800E-06 |
| 825 | 42 | Y42C | A | 9.8e-06 | 6e-06 | 1e-05 | 8.600E-06 | 2.254E-06 | 6.000E-06 | 1.000E-05 |
| 826 | 42 | Y42E | A | 1.1e-05 | 8.3e-06 | 9.4e-06 | 9.567E-06 | 1.358E-06 | 8.300E-06 | 1.100E-05 |
| 827 | 42 | Y42Q | A | 9.2e-06 | 8.6e-06 | 4.9e-06 | 7.567E-06 | 2.329E-06 | 4.900E-06 | 9.200E-06 |
| 828 | 42 | Y42G | A | 8.8e-06 | 6.5e-06 | 9.7e-06 | 8.333E-06 | 1.650E-06 | 6.500E-06 | 9.700E-06 |
| 829 | 42 | Y42H | A | 8.7e-06 | 5.3e-06 | 1e-05 | 8.000E-06 | 2.427E-06 | 5.300E-06 | 1.000E-05 |
| 830 | 42 | Y42I | A | 1.1e-05 | 1.6e-05 | 6.4e-06 | 1.113E-05 | 4.801E-06 | 6.400E-06 | 1.600E-05 |
| 831 | 42 | Y42L | A | 9.1e-06 | 4.6e-06 | 7.8e-06 | 7.167E-06 | 2.316E-06 | 4.600E-06 | 9.100E-06 |
| 832 | 42 | Y42K | A | 5.9e-06 | 9.4e-06 | 8.1e-06 | 7.800E-06 | 1.769E-06 | 5.900E-06 | 9.400E-06 |
| 833 | 42 | Y42M | A | 6.6e-06 | 9.3e-06 | 7.1e-06 | 7.667E-06 | 1.436E-06 | 6.600E-06 | 9.300E-06 |
| 834 | 42 | Y42F | A | 1.5e-05 | 6.2e-06 | 7.5e-06 | 9.567E-06 | 4.750E-06 | 6.200E-06 | 1.500E-05 |
| 835 | 42 | Y42P | A | 4.8e-06 | 3.6e-06 | 1e-05 | 6.133E-06 | 3.402E-06 | 3.600E-06 | 1.000E-05 |
| 836 | 42 | Y42S | A | 7.5e-06 | 9.3e-06 | 6.8e-06 | 7.867E-06 | 1.290E-06 | 6.800E-06 | 9.300E-06 |
| 837 | 42 | Y42T | A | 7.2e-06 | 4.8e-06 | 5.3e-06 | 5.767E-06 | 1.266E-06 | 4.800E-06 | 7.200E-06 |
| 838 | 42 | Y42W | A | 1e-05 | 1.1e-05 | 6.6e-06 | 9.200E-06 | 2.307E-06 | 6.600E-06 | 1.100E-05 |
| 839 | 42 | Y42Y | A | 1.2e-05 | 5.6e-06 | 1.7e-05 | 1.153E-05 | 5.714E-06 | 5.600E-06 | 1.700E-05 |
| 840 | 42 | Y42V | A | 8.6e-06 | 7.3e-06 | 8.5e-06 | 8.133E-06 | 7.234E-07 | 7.300E-06 | 8.600E-06 |
| 841 | 43 | L43A | A | 8.6e-06 | 8.3e-06 | 7.2e-06 | 8.033E-06 | 7.371E-07 | 7.200E-06 | 8.600E-06 |
| 842 | 43 | L43R | A | 7e-06 | 5.5e-06 | 1.1e-05 | 7.833E-06 | 2.843E-06 | 5.500E-06 | 1.100E-05 |
| 843 | 43 | L43N | A | 1e-05 | 1.3e-05 | 8.5e-06 | 1.050E-05 | 2.291E-06 | 8.500E-06 | 1.300E-05 |
| 844 | 43 | L43D | A | 8.1e-06 | 9.1e-06 | 6.6e-06 | 7.933E-06 | 1.258E-06 | 6.600E-06 | 9.100E-06 |
| 845 | 43 | L43C | A | 7.1e-06 | 8.6e-06 | 1.1e-05 | 8.900E-06 | 1.967E-06 | 7.100E-06 | 1.100E-05 |
| 846 | 43 | L43E | A | 8.3e-06 | 1.3e-05 | 1.3e-05 | 1.143E-05 | 2.714E-06 | 8.300E-06 | 1.300E-05 |
| 847 | 43 | L43Q | A | 7.1e-06 | 4.6e-06 | 9.3e-06 | 7.000E-06 | 2.352E-06 | 4.600E-06 | 9.300E-06 |
| 848 | 43 | L43G | A | 1.7e-05 | 5.6e-06 | 1e-05 | 1.087E-05 | 5.749E-06 | 5.600E-06 | 1.700E-05 |
| 849 | 43 | L43H | A | 6.6e-06 | 7.1e-06 | 5.8e-06 | 6.500E-06 | 6.557E-07 | 5.800E-06 | 7.100E-06 |

| | | | | | | | | | | |
|---|---|---|---|---|---|---|---|---|---|---|
| 850 | 43 | L43I | A | 9.9e-06 | 8.6e-06 | 5.6e-06 | 8.033E-06 | 2.205E-06 | 5.600E-06 | 9.900E-06 |
| 851 | 43 | L43L | A | 1.1e-05 | 5.3e-06 | 7.2e-06 | 7.833E-06 | 2.902E-06 | 5.300E-06 | 1.100E-05 |
| 852 | 43 | L43K | A | 6.4e-06 | 7.9e-06 | 7e-06 | 7.100E-06 | 7.550E-07 | 6.400E-06 | 7.900E-06 |
| 853 | 43 | L43M | A | 9.6e-06 | 1.1e-05 | 8.9e-06 | 9.833E-06 | 1.069E-06 | 8.900E-06 | 1.100E-05 |
| 854 | 43 | L43F | A | 9.2e-06 | 9.8e-06 | 1.5e-05 | 1.133E-05 | 3.190E-06 | 9.200E-06 | 1.500E-05 |
| 855 | 43 | L43P | A | 1.1e-05 | 6.6e-06 | 6.3e-06 | 7.967E-06 | 2.631E-06 | 6.300E-06 | 1.100E-05 |
| 856 | 43 | L43S | A | 8.5e-06 | 9.1e-06 | 7.6e-06 | 8.400E-06 | 7.550E-07 | 7.600E-06 | 9.100E-06 |
| 857 | 43 | L43T | A | 4.2e-06 | 8.7e-06 | 8.8e-06 | 7.233E-06 | 2.627E-06 | 4.200E-06 | 8.800E-06 |
| 858 | 43 | L43W | A | 5.4e-06 | 6.7e-06 | 1e-05 | 7.367E-06 | 2.371E-06 | 5.400E-06 | 1.000E-05 |
| 859 | 43 | L43Y | A | 1.4e-05 | 1.1e-05 | 9.6e-06 | 1.153E-05 | 2.248E-06 | 9.600E-06 | 1.400E-05 |
| 860 | 43 | L43V | A | 5.8e-06 | 6e-06 | 1.2e-05 | 7.933E-06 | 3.523E-06 | 5.800E-06 | 1.200E-05 |
| 861 | 44 | P44A | A | 1.2e-05 | 5.1e-06 | 4.4e-06 | 7.167E-06 | 4.200E-06 | 4.400E-06 | 1.200E-05 |
| 862 | 44 | P44R | A | 7.4e-06 | 6.3e-06 | 6.2e-06 | 6.633E-06 | 6.658E-07 | 6.200E-06 | 7.400E-06 |
| 863 | 44 | P44N | A | 6.9e-06 | 8.1e-06 | 6.4e-06 | 7.133E-06 | 8.737E-07 | 6.400E-06 | 8.100E-06 |
| 864 | 44 | P44D | A | 7.8e-06 | 7.1e-06 | 6.7e-06 | 7.200E-06 | 5.568E-07 | 6.700E-06 | 7.800E-06 |
| 865 | 44 | P44C | A | 9.4e-06 | 1.5e-05 | 5.5e-06 | 9.967E-06 | 4.775E-06 | 5.500E-06 | 1.500E-05 |
| 866 | 44 | P44E | A | 8.7e-06 | 5.8e-06 | 4.5e-06 | 6.333E-06 | 2.150E-06 | 4.500E-06 | 8.700E-06 |
| 867 | 44 | P44Q | A | 4.3e-06 | 6.5e-06 | 1.2e-05 | 7.600E-06 | 3.966E-06 | 4.300E-06 | 1.200E-05 |
| 868 | 44 | P44G | A | 6.6e-06 | 8.7e-06 | 1.1e-05 | 8.767E-06 | 2.201E-06 | 6.600E-06 | 1.100E-05 |
| 869 | 44 | P44H | A | 5.4e-06 | 6.2e-06 | 6.7e-06 | 6.100E-06 | 6.557E-07 | 5.400E-06 | 6.700E-06 |
| 870 | 44 | P44I | A | 8.2e-06 | 6.3e-06 | 5.6e-06 | 6.700E-06 | 1.345E-06 | 5.600E-06 | 8.200E-06 |
| 871 | 44 | P44L | A | 1.2e-05 | 8.2e-06 | 4.4e-06 | 8.200E-06 | 3.800E-06 | 4.400E-06 | 1.200E-05 |
| 872 | 44 | P44K | A | 9.7e-06 | 1.1e-05 | 4.9e-06 | 8.533E-06 | 3.213E-06 | 4.900E-06 | 1.100E-05 |
| 873 | 44 | P44M | A | 8.9e-06 | 1e-05 | 1e-05 | 9.633E-06 | 6.351E-07 | 8.900E-06 | 1.000E-05 |
| 874 | 44 | P44F | A | 4.5e-06 | 8.8e-06 | 1e-05 | 7.767E-06 | 2.892E-06 | 4.500E-06 | 1.000E-05 |
| 875 | 44 | P44P | A | 3.9e-06 | 7.6e-06 | 7.5e-06 | 6.333E-06 | 2.108E-06 | 3.900E-06 | 7.600E-06 |
| 876 | 44 | P44S | A | 6.3e-06 | 3.1e-06 | 8.7e-06 | 6.033E-06 | 2.810E-06 | 3.100E-06 | 8.700E-06 |
| 877 | 44 | P44T | A | 5e-06 | 9.8e-06 | 8.4e-06 | 7.733E-06 | 2.468E-06 | 5.000E-06 | 9.800E-06 |
| 878 | 44 | P44W | A | 5.1e-06 | 4.8e-06 | 3.3e-06 | 4.400E-06 | 9.644E-07 | 3.300E-06 | 5.100E-06 |
| 879 | 44 | P44Y | A | 1.2e-05 | 6e-06 | 7.2e-06 | 8.400E-06 | 3.175E-06 | 6.000E-06 | 1.200E-05 |
| 880 | 44 | P44V | A | 4.9e-06 | 6.8e-06 | 6.3e-06 | 6.000E-06 | 9.849E-07 | 4.900E-06 | 6.800E-06 |
| 881 | 45 | T45A | A | 5e-06 | 1e-05 | 1e-05 | 8.333E-06 | 2.887E-06 | 5.000E-06 | 1.000E-05 |
| 882 | 45 | T45R | A | 9.8e-06 | 1.4e-05 | 7.3e-06 | 1.037E-05 | 3.386E-06 | 7.300E-06 | 1.400E-05 |
| 883 | 45 | T45N | A | 6e-06 | 9.5e-06 | 6.3e-06 | 7.267E-06 | 1.940E-06 | 6.000E-06 | 9.500E-06 |
| 884 | 45 | T45D | A | 1.1e-05 | 6.5e-06 | 9.6e-06 | 9.033E-06 | 2.303E-06 | 6.500E-06 | 1.100E-05 |
| 885 | 45 | T45C | A | 4.9e-06 | 4.9e-06 | 6.6e-06 | 5.467E-06 | 9.815E-07 | 4.900E-06 | 6.600E-06 |
| 886 | 45 | T45E | A | 9.9e-06 | 6.5e-06 | 1.1e-05 | 9.133E-06 | 2.346E-06 | 6.500E-06 | 1.100E-05 |
| 887 | 45 | T45Q | A | 1.3e-05 | 6.7e-06 | 8.3e-06 | 9.333E-06 | 3.275E-06 | 6.700E-06 | 1.300E-05 |
| 888 | 45 | T45G | A | 8.8e-06 | 1e-05 | 1.2e-05 | 1.027E-05 | 1.617E-06 | 8.800E-06 | 1.200E-05 |
| 889 | 45 | T45H | A | 5.6e-06 | 6.7e-06 | 7.8e-06 | 6.700E-06 | 1.100E-06 | 5.600E-06 | 7.800E-06 |
| 890 | 45 | T45I | A | 7.6e-06 | 1.3e-05 | 1.4e-05 | 1.153E-05 | 3.443E-06 | 7.600E-06 | 1.400E-05 |
| 891 | 45 | T45L | A | 7.3e-06 | 9.3e-06 | 9.6e-06 | 8.733E-06 | 1.250E-06 | 7.300E-06 | 9.600E-06 |
| 892 | 45 | T45K | A | 7.1e-06 | 7.7e-06 | 1.4e-05 | 9.600E-06 | 3.822E-06 | 7.100E-06 | 1.400E-05 |
| 893 | 45 | T45M | A | 1.1e-05 | 1.2e-05 | 8.8e-06 | 1.060E-05 | 1.637E-06 | 8.800E-06 | 1.200E-05 |
| 894 | 45 | T45F | A | 9.2e-06 | 1e-05 | 5.3e-06 | 8.167E-06 | 2.515E-06 | 5.300E-06 | 1.000E-05 |
| 895 | 45 | T45P | A | 9.6e-06 | 6.6e-06 | 1.2e-05 | 9.400E-06 | 2.706E-06 | 6.600E-06 | 1.200E-05 |
| 896 | 45 | T45S | A | 5.8e-06 | 5e-06 | 9e-06 | 6.600E-06 | 2.117E-06 | 5.000E-06 | 9.000E-06 |
| 897 | 45 | T45T | A | 1.2e-05 | 5.8e-06 | 5.7e-06 | 7.833E-06 | 3.609E-06 | 5.700E-06 | 1.200E-05 |
| 898 | 45 | T45W | A | 5.1e-06 | 6.3e-06 | 8.8e-06 | 6.733E-06 | 1.888E-06 | 5.100E-06 | 8.800E-06 |
| 899 | 45 | T45Y | A | 9.3e-06 | 7e-06 | 5.9e-06 | 7.400E-06 | 1.735E-06 | 5.900E-06 | 9.300E-06 |
| 900 | 45 | T45V | A | 4.2e-06 | 1.1e-05 | 8.6e-06 | 7.933E-06 | 3.449E-06 | 4.200E-06 | 1.100E-05 |
| 901 | 46 | N46A | A | 9.2e-06 | 1.5e-05 | 1.1e-05 | 1.173E-05 | 2.969E-06 | 9.200E-06 | 1.500E-05 |
| 902 | 46 | N46R | A | 1e-05 | 9.3e-06 | 4.9e-06 | 8.067E-06 | 2.765E-06 | 4.900E-06 | 1.000E-05 |
| 903 | 46 | N46N | A | 1.3e-05 | 1e-05 | 7.6e-06 | 1.020E-05 | 2.706E-06 | 7.600E-06 | 1.300E-05 |
| 904 | 46 | N46D | A | 5.5e-06 | 7.9e-06 | 5.8e-06 | 6.400E-06 | 1.308E-06 | 5.500E-06 | 7.900E-06 |
| 905 | 46 | N46C | A | 6.9e-06 | 9.8e-06 | 4.9e-06 | 7.200E-06 | 2.464E-06 | 4.900E-06 | 9.800E-06 |
| 906 | 46 | N46E | A | 7.2e-06 | 8e-06 | 8.8e-06 | 8.000E-06 | 8.000E-07 | 7.200E-06 | 8.800E-06 |
| 907 | 46 | N46Q | A | 7.6e-06 | 9.5e-06 | 9.6e-06 | 8.900E-06 | 1.127E-06 | 7.600E-06 | 9.600E-06 |
| 908 | 46 | N46G | A | 1.2e-05 | 9.5e-06 | 9.7e-06 | 1.040E-05 | 1.389E-06 | 9.500E-06 | 1.200E-05 |
| 909 | 46 | N46H | A | 5.4e-06 | 8.9e-06 | 7e-06 | 7.100E-06 | 1.752E-06 | 5.400E-06 | 8.900E-06 |
| 910 | 46 | N46I | A | 7.5e-06 | 7.2e-06 | 6.8e-06 | 7.167E-06 | 3.512E-07 | 6.800E-06 | 7.500E-06 |
| 911 | 46 | N46L | A | 1.1e-05 | 7.6e-06 | 6.1e-06 | 8.233E-06 | 2.511E-06 | 6.100E-06 | 1.100E-05 |
| 912 | 46 | N46K | A | 8e-06 | 1e-05 | 6.1e-06 | 8.033E-06 | 1.950E-06 | 6.100E-06 | 1.000E-05 |
| 913 | 46 | N46M | A | 1.3e-05 | 4.7e-06 | 6.8e-06 | 8.167E-06 | 4.315E-06 | 4.700E-06 | 1.300E-05 |
| 914 | 46 | N46F | A | 5.3e-06 | 9.5e-06 | 1.3e-05 | 9.267E-06 | 3.855E-06 | 5.300E-06 | 1.300E-05 |
| 915 | 46 | N46P | A | 8.5e-06 | 7.4e-06 | 4.8e-06 | 6.900E-06 | 1.900E-06 | 4.800E-06 | 8.500E-06 |
| 916 | 46 | N46S | A | 6.9e-06 | 6.4e-06 | 9.9e-06 | 7.733E-06 | 1.893E-06 | 6.400E-06 | 9.900E-06 |
| 917 | 46 | N46T | A | 1.4e-05 | 5.6e-06 | 1.2e-05 | 1.053E-05 | 4.388E-06 | 5.600E-06 | 1.400E-05 |
| 918 | 46 | N46W | A | 4.9e-06 | 8e-06 | 3.5e-06 | 5.467E-06 | 2.303E-06 | 3.500E-06 | 8.000E-06 |
| 919 | 46 | N46Y | A | 9.3e-06 | 6.6e-06 | 4.6e-06 | 6.833E-06 | 2.359E-06 | 4.600E-06 | 9.300E-06 |
| 920 | 46 | N46V | A | 6.6e-06 | 1.1e-05 | 9.2e-06 | 8.933E-06 | 2.212E-06 | 6.600E-06 | 1.100E-05 |
| 921 | 47 | A47A | A | 9.4e-06 | 1.3e-05 | 7.7e-06 | 1.003E-05 | 2.706E-06 | 7.700E-06 | 1.300E-05 |
| 922 | 47 | A47R | A | 1.1e-05 | 6.4e-06 | 6.8e-06 | 8.067E-06 | 2.548E-06 | 6.400E-06 | 1.100E-05 |
| 923 | 47 | A47N | A | 8.2e-06 | 5.3e-06 | 6.3e-06 | 6.600E-06 | 1.473E-06 | 5.300E-06 | 8.200E-06 |
| 924 | 47 | A47D | A | 9.2e-06 | 9.7e-06 | 8.8e-06 | 9.233E-06 | 4.509E-07 | 8.800E-06 | 9.700E-06 |
| 925 | 47 | A47C | A | 7.6e-06 | 4.9e-06 | 7.5e-06 | 6.667E-06 | 1.531E-06 | 4.900E-06 | 7.600E-06 |
| 926 | 47 | A47E | A | 6e-06 | 4.6e-06 | 5.2e-06 | 5.267E-06 | 7.024E-07 | 4.600E-06 | 6.000E-06 |
| 927 | 47 | A47Q | A | 9.9e-06 | 7.2e-06 | 7e-06 | 8.033E-06 | 1.620E-06 | 7.000E-06 | 9.900E-06 |
| 928 | 47 | A47G | A | 6.8e-06 | 1.4e-05 | 7.6e-06 | 9.467E-06 | 3.946E-06 | 6.800E-06 | 1.400E-05 |
| 929 | 47 | A47H | A | 7e-06 | 9.4e-06 | 4.3e-06 | 6.900E-06 | 2.551E-06 | 4.300E-06 | 9.400E-06 |
| 930 | 47 | A47I | A | 6.8e-06 | 1.3e-05 | 4.4e-06 | 8.067E-06 | 4.438E-06 | 4.400E-06 | 1.300E-05 |
| 931 | 47 | A47L | A | 1.2e-05 | 7.9e-06 | 8.5e-06 | 9.467E-06 | 2.214E-06 | 7.900E-06 | 1.200E-05 |
| 932 | 47 | A47K | A | 5.8e-06 | 5.4e-06 | 6.6e-06 | 5.933E-06 | 6.110E-07 | 5.400E-06 | 6.600E-06 |
| 933 | 47 | A47M | A | 1.6e-05 | 9.2e-06 | 1.3e-05 | 1.273E-05 | 3.408E-06 | 9.200E-06 | 1.600E-05 |
| 934 | 47 | A47F | A | 5.6e-06 | 1.1e-05 | 4.5e-06 | 7.033E-06 | 3.479E-06 | 4.500E-06 | 1.100E-05 |

| | | | | | | | | | | |
|---|---|---|---|---|---|---|---|---|---|---|
| 935 | 47 | A47P | A | 6.7e-06 | 5.6e-06 | 1.1e-05 | 7.767E-06 | 2.854E-06 | 5.600E-06 | 1.100E-05 |
| 936 | 47 | A47S | A | 6.3e-06 | 6.5e-06 | 6.8e-06 | 6.533E-06 | 2.517E-07 | 6.300E-06 | 6.800E-06 |
| 937 | 47 | A47T | A | 6.7e-06 | 6.5e-06 | 7.2e-06 | 6.800E-06 | 3.606E-07 | 6.500E-06 | 7.200E-06 |
| 938 | 47 | A47W | A | 1.4e-05 | 6.7e-06 | 6.7e-06 | 9.133E-06 | 4.215E-06 | 6.700E-06 | 1.400E-05 |
| 939 | 47 | A47Y | A | 5.6e-06 | 6.3e-06 | 8e-06 | 6.633E-06 | 1.234E-06 | 5.600E-06 | 8.000E-06 |
| 940 | 47 | A47V | A | 8.2e-06 | 7.6e-06 | 1.1e-05 | 8.933E-06 | 1.815E-06 | 7.600E-06 | 1.100E-05 |
| 941 | 48 | S48A | A | 4.6e-06 | 1.1e-05 | 5.5e-06 | 7.033E-06 | 3.465E-06 | 4.600E-06 | 1.100E-05 |
| 942 | 48 | S48R | A | 8.5e-06 | 6.4e-06 | 1.3e-05 | 9.300E-06 | 3.372E-06 | 6.400E-06 | 1.300E-05 |
| 943 | 48 | S48N | A | 4.6e-06 | 7.2e-06 | 6.8e-06 | 6.200E-06 | 1.400E-06 | 4.600E-06 | 7.200E-06 |
| 944 | 48 | S48D | A | 1.1e-05 | 8.1e-06 | 4.1e-06 | 7.733E-06 | 3.465E-06 | 4.100E-06 | 1.100E-05 |
| 945 | 48 | S48C | A | 6.7e-06 | 9.3e-06 | 1e-05 | 8.667E-06 | 1.739E-06 | 6.700E-06 | 1.000E-05 |
| 946 | 48 | S48E | A | 5.9e-06 | 1.4e-05 | 6.9e-06 | 8.933E-06 | 4.416E-06 | 5.900E-06 | 1.400E-05 |
| 947 | 48 | S48Q | A | 8.8e-06 | 1.1e-05 | 7.9e-06 | 9.233E-06 | 1.595E-06 | 7.900E-06 | 1.100E-05 |
| 948 | 48 | S48G | A | 7.2e-06 | 7.7e-06 | 9.5e-06 | 8.133E-06 | 1.210E-06 | 7.200E-06 | 9.500E-06 |
| 949 | 48 | S48H | A | 9.3e-06 | 9.8e-06 | 5.7e-06 | 8.267E-06 | 2.237E-06 | 5.700E-06 | 9.800E-06 |
| 950 | 48 | S48I | A | 7.2e-06 | 8.3e-06 | 1.4e-05 | 9.833E-06 | 3.650E-06 | 7.200E-06 | 1.400E-05 |
| 951 | 48 | S48L | A | 1.4e-05 | 9.5e-06 | 1.6e-05 | 1.317E-05 | 3.329E-06 | 9.500E-06 | 1.600E-05 |
| 952 | 48 | S48K | A | 7.3e-06 | 9e-06 | 1.2e-05 | 9.433E-06 | 2.380E-06 | 7.300E-06 | 1.200E-05 |
| 953 | 48 | S48M | A | 7e-06 | 8e-06 | 9.9e-06 | 8.300E-06 | 1.473E-06 | 7.000E-06 | 9.900E-06 |
| 954 | 48 | S48F | A | 4.8e-06 | 7.1e-06 | 6.4e-06 | 6.100E-06 | 1.179E-06 | 4.800E-06 | 7.100E-06 |
| 955 | 48 | S48P | A | 1.1e-05 | 6.5e-06 | 1.2e-05 | 9.833E-06 | 2.930E-06 | 6.500E-06 | 1.200E-05 |
| 956 | 48 | S48S | A | 5.7e-06 | 6.3e-06 | 5.8e-06 | 5.933E-06 | 3.215E-07 | 5.700E-06 | 6.300E-06 |
| 957 | 48 | S48T | A | 4.1e-06 | 6.4e-06 | 9.8e-06 | 6.767E-06 | 2.868E-06 | 4.100E-06 | 9.800E-06 |
| 958 | 48 | S48W | A | 5.6e-06 | 9.1e-06 | 9e-06 | 7.900E-06 | 1.992E-06 | 5.600E-06 | 9.100E-06 |
| 959 | 48 | S48Y | A | 6.8e-06 | 1.1e-05 | 8.8e-06 | 8.867E-06 | 2.101E-06 | 6.800E-06 | 1.100E-05 |
| 960 | 48 | S48V | A | 6.7e-06 | 7.9e-06 | 7.7e-06 | 7.433E-06 | 6.429E-07 | 6.700E-06 | 7.900E-06 |
| 961 | 49 | L49A | A | 6.4e-06 | 5.7e-06 | 7e-06 | 6.367E-06 | 6.506E-07 | 5.700E-06 | 7.000E-06 |
| 962 | 49 | L49R | A | 5.2e-06 | 5.3e-06 | 7.4e-06 | 5.967E-06 | 1.242E-06 | 5.200E-06 | 7.400E-06 |
| 963 | 49 | L49N | A | 5.2e-06 | 1.2e-05 | 1.9e-05 | 1.207E-05 | 6.900E-06 | 5.200E-06 | 1.900E-05 |
| 964 | 49 | L49D | A | 5.4e-06 | 5.5e-06 | 6.6e-06 | 5.833E-06 | 6.658E-07 | 5.400E-06 | 6.600E-06 |
| 965 | 49 | L49C | A | 6.8e-06 | 1e-05 | 8.5e-06 | 8.433E-06 | 1.601E-06 | 6.800E-06 | 1.000E-05 |
| 966 | 49 | L49E | A | 6.1e-06 | 7.7e-06 | 9.4e-06 | 7.733E-06 | 1.650E-06 | 6.100E-06 | 9.400E-06 |
| 967 | 49 | L49Q | A | 6.1e-06 | 9.3e-06 | 6.1e-06 | 7.167E-06 | 1.848E-06 | 6.100E-06 | 9.300E-06 |
| 968 | 49 | L49G | A | 7.4e-06 | 3.4e-06 | 5.9e-06 | 5.567E-06 | 2.021E-06 | 3.400E-06 | 7.400E-06 |
| 969 | 49 | L49H | A | 4.1e-06 | 7.9e-06 | 6.8e-06 | 6.267E-06 | 1.955E-06 | 4.100E-06 | 7.900E-06 |
| 970 | 49 | L49I | A | 6e-06 | 6.1e-06 | 5.4e-06 | 5.833E-06 | 3.786E-07 | 5.400E-06 | 6.100E-06 |
| 971 | 49 | L49L | A | 9.7e-06 | 8.5e-06 | 9.2e-06 | 9.133E-06 | 6.028E-07 | 8.500E-06 | 9.700E-06 |
| 972 | 49 | L49K | A | 1e-05 | 9.7e-06 | 8.8e-06 | 9.500E-06 | 6.245E-07 | 8.800E-06 | 1.000E-05 |
| 973 | 49 | L49M | A | 9.4e-06 | 5.1e-06 | 6.5e-06 | 7.000E-06 | 2.193E-06 | 5.100E-06 | 9.400E-06 |
| 974 | 49 | L49F | A | 1.5e-05 | 6.6e-06 | 8.8e-06 | 1.013E-05 | 4.356E-06 | 6.600E-06 | 1.500E-05 |
| 975 | 49 | L49P | A | 5.4e-06 | 9.8e-06 | 9e-06 | 8.067E-06 | 2.344E-06 | 5.400E-06 | 9.800E-06 |
| 976 | 49 | L49S | A | 5.9e-06 | 6.7e-06 | 7.9e-06 | 6.833E-06 | 1.007E-06 | 5.900E-06 | 7.900E-06 |
| 977 | 49 | L49T | A | 1.2e-05 | 1e-05 | 7e-06 | 9.667E-06 | 2.517E-06 | 7.000E-06 | 1.200E-05 |
| 978 | 49 | L49W | A | 6.1e-06 | 6.9e-06 | 1.9e-05 | 1.067E-05 | 7.228E-06 | 6.100E-06 | 1.900E-05 |
| 979 | 49 | L49Y | A | 8.2e-06 | 5.7e-06 | 1e-05 | 7.967E-06 | 2.159E-06 | 5.700E-06 | 1.000E-05 |
| 980 | 49 | L49V | A | 9.7e-06 | 9.5e-06 | 5.3e-06 | 8.167E-06 | 2.485E-06 | 5.300E-06 | 9.700E-06 |
| 981 | 50 | S50A | A | 9.2e-06 | 1.1e-05 | 8.1e-06 | 9.433E-06 | 1.464E-06 | 8.100E-06 | 1.100E-05 |
| 982 | 50 | S50R | A | 5.5e-06 | 7e-06 | 9.5e-06 | 7.333E-06 | 2.021E-06 | 5.500E-06 | 9.500E-06 |
| 983 | 50 | S50N | A | 9.6e-06 | 7.1e-06 | 1.4e-05 | 1.023E-05 | 3.493E-06 | 7.100E-06 | 1.400E-05 |
| 984 | 50 | S50D | A | 5.8e-06 | 1.4e-05 | 7.3e-06 | 9.033E-06 | 4.366E-06 | 5.800E-06 | 1.400E-05 |
| 985 | 50 | S50C | A | 1.3e-05 | 1.2e-05 | 1.1e-05 | 1.200E-05 | 1.000E-06 | 1.100E-05 | 1.300E-05 |
| 986 | 50 | S50E | A | 8.7e-06 | 6.3e-06 | 7.7e-06 | 7.567E-06 | 1.206E-06 | 6.300E-06 | 8.700E-06 |
| 987 | 50 | S50Q | A | 1.2e-05 | 1.2e-05 | 5.9e-06 | 9.967E-06 | 3.522E-06 | 5.900E-06 | 1.200E-05 |
| 988 | 50 | S50G | A | 9.4e-06 | 6.8e-06 | 1e-05 | 8.733E-06 | 1.701E-06 | 6.800E-06 | 1.000E-05 |
| 989 | 50 | S50H | A | 8.1e-06 | 5.5e-06 | 1.4e-05 | 9.200E-06 | 4.355E-06 | 5.500E-06 | 1.400E-05 |
| 990 | 50 | S50I | A | 8e-06 | 7.6e-06 | 6e-06 | 7.200E-06 | 1.058E-06 | 6.000E-06 | 8.000E-06 |
| 991 | 50 | S50L | A | 6.7e-06 | 6.9e-06 | 8.5e-06 | 7.367E-06 | 9.866E-07 | 6.700E-06 | 8.500E-06 |
| 992 | 50 | S50K | A | 1.1e-05 | 1.3e-05 | 9.1e-06 | 1.103E-05 | 1.950E-06 | 9.100E-06 | 1.300E-05 |
| 993 | 50 | S50M | A | 8.1e-06 | 8.1e-06 | 1.1e-05 | 9.067E-06 | 1.674E-06 | 8.100E-06 | 1.100E-05 |
| 994 | 50 | S50F | A | 8.5e-06 | 8e-06 | 9.8e-06 | 8.767E-06 | 9.292E-07 | 8.000E-06 | 9.800E-06 |
| 995 | 50 | S50P | A | 4.8e-06 | 1.2e-05 | 1.2e-05 | 9.600E-06 | 4.157E-06 | 4.800E-06 | 1.200E-05 |
| 996 | 50 | S50S | A | 6.1e-06 | 8.4e-06 | 8.9e-06 | 7.800E-06 | 1.493E-06 | 6.100E-06 | 8.900E-06 |
| 997 | 50 | S50T | A | 8.7e-06 | 7.8e-06 | 5.3e-06 | 7.267E-06 | 1.762E-06 | 5.300E-06 | 8.700E-06 |
| 998 | 50 | S50W | A | 1.1e-05 | 5.6e-06 | 7.6e-06 | 8.067E-06 | 2.730E-06 | 5.600E-06 | 1.100E-05 |
| 999 | 50 | S50Y | A | 9.8e-06 | 1.3e-05 | 6.2e-06 | 9.667E-06 | 3.402E-06 | 6.200E-06 | 1.300E-05 |
| 1000 | 50 | S50V | A | 9.3e-06 | 8.1e-06 | 7.8e-06 | 8.400E-06 | 7.937E-07 | 7.800E-06 | 9.300E-06 |
| 1001 | 51 | F51A | A | 9.3e-06 | 9e-06 | 4.1e-06 | 7.467E-06 | 2.919E-06 | 4.100E-06 | 9.300E-06 |
| 1002 | 51 | F51R | A | 6.9e-06 | 7.3e-06 | 6.6e-06 | 6.933E-06 | 3.512E-07 | 6.600E-06 | 7.300E-06 |
| 1003 | 51 | F51N | A | 1e-05 | 4.7e-06 | 8.4e-06 | 7.700E-06 | 2.718E-06 | 4.700E-06 | 1.000E-05 |
| 1004 | 51 | F51D | A | 1.3e-05 | 1.1e-05 | 1e-05 | 1.133E-05 | 1.528E-06 | 1.000E-05 | 1.300E-05 |
| 1005 | 51 | F51C | A | 1.2e-05 | 9.8e-06 | 1e-05 | 1.060E-05 | 1.217E-06 | 9.800E-06 | 1.200E-05 |
| 1006 | 51 | F51E | A | 7.7e-06 | 4.5e-06 | 4.5e-06 | 5.567E-06 | 1.848E-06 | 4.500E-06 | 7.700E-06 |
| 1007 | 51 | F51Q | A | 7.1e-06 | 4.8e-06 | 1e-05 | 7.300E-06 | 2.606E-06 | 4.800E-06 | 1.000E-05 |
| 1008 | 51 | F51G | A | 9.4e-06 | 1.3e-05 | 4.6e-06 | 9.000E-06 | 4.214E-06 | 4.600E-06 | 1.300E-05 |
| 1009 | 51 | F51H | A | 1.1e-05 | 8.7e-06 | 4.7e-06 | 8.133E-06 | 3.188E-06 | 4.700E-06 | 1.100E-05 |
| 1010 | 51 | F51I | A | 6.9e-06 | 9.9e-06 | 1.4e-05 | 1.027E-05 | 3.564E-06 | 6.900E-06 | 1.400E-05 |
| 1011 | 51 | F51L | A | 1.2e-05 | 9.1e-06 | 1.3e-05 | 1.137E-05 | 2.026E-06 | 9.100E-06 | 1.300E-05 |
| 1012 | 51 | F51K | A | 8e-06 | 1e-05 | 1.7e-05 | 1.167E-05 | 4.726E-06 | 8.000E-06 | 1.700E-05 |
| 1013 | 51 | F51M | A | 6.7e-06 | 5.7e-06 | 4.4e-06 | 5.600E-06 | 1.153E-06 | 4.400E-06 | 6.700E-06 |
| 1014 | 51 | F51F | A | 8e-06 | 9e-06 | 6e-06 | 7.667E-06 | 1.528E-06 | 6.000E-06 | 9.000E-06 |
| 1015 | 51 | F51P | A | 9e-06 | 1.1e-05 | 9.4e-06 | 9.800E-06 | 1.058E-06 | 9.000E-06 | 1.100E-05 |
| 1016 | 51 | F51S | A | 9.6e-06 | 8.5e-06 | 6.9e-06 | 8.333E-06 | 1.358E-06 | 6.900E-06 | 9.600E-06 |
| 1017 | 51 | F51T | A | 7.9e-06 | 7.3e-06 | 1.2e-05 | 9.067E-06 | 2.558E-06 | 7.300E-06 | 1.200E-05 |
| 1018 | 51 | F51W | A | 6.2e-06 | 8.6e-06 | 6.7e-06 | 7.167E-06 | 1.266E-06 | 6.200E-06 | 8.600E-06 |
| 1019 | 51 | F51Y | A | 1.3e-05 | 6.6e-06 | 7e-06 | 8.867E-06 | 3.585E-06 | 6.600E-06 | 1.300E-05 |

| | | | | | | | | | | |
|---|---|---|---|---|---|---|---|---|---|---|
| 1020 | 51 | F51V | A | 9.7e-06 | 4.7e-06 | 7.4e-06 | 7.267E-06 | 2.503E-06 | 4.700E-06 | 9.700E-06 |
| 1021 | 52 | L52A | A | 6.4e-06 | 8.1e-06 | 5.8e-06 | 6.767E-06 | 1.193E-06 | 5.800E-06 | 8.100E-06 |
| 1022 | 52 | L52R | A | 6.8e-06 | 1.1e-05 | 7.9e-06 | 8.567E-06 | 2.178E-06 | 6.800E-06 | 1.100E-05 |
| 1023 | 52 | L52N | A | 9.8e-06 | 1.2e-05 | 1.3e-05 | 1.160E-05 | 1.637E-06 | 9.800E-06 | 1.300E-05 |
| 1024 | 52 | L52D | A | 5.5e-06 | 6.6e-06 | 8.3e-06 | 6.800E-06 | 1.411E-06 | 5.500E-06 | 8.300E-06 |
| 1025 | 52 | L52C | A | 5e-06 | 6.6e-06 | 1.2e-05 | 7.867E-06 | 3.668E-06 | 5.000E-06 | 1.200E-05 |
| 1026 | 52 | L52E | A | 1.2e-05 | 7.4e-06 | 1e-05 | 9.800E-06 | 2.307E-06 | 7.400E-06 | 1.200E-05 |
| 1027 | 52 | L52Q | A | 5.7e-06 | 4.8e-06 | 8.9e-06 | 6.467E-06 | 2.155E-06 | 4.800E-06 | 8.900E-06 |
| 1028 | 52 | L52G | A | 8e-06 | 6.6e-06 | 1.9e-05 | 1.120E-05 | 6.791E-06 | 6.600E-06 | 1.900E-05 |
| 1029 | 52 | L52H | A | 1e-05 | 1.1e-05 | 1.3e-05 | 1.133E-05 | 1.528E-06 | 1.000E-05 | 1.300E-05 |
| 1030 | 52 | L52I | A | 7.6e-06 | 9.5e-06 | 1.2e-05 | 9.700E-06 | 2.207E-06 | 7.600E-06 | 1.200E-05 |
| 1031 | 52 | L52L | A | 9.4e-06 | 8.3e-06 | 1e-05 | 9.233E-06 | 8.622E-07 | 8.300E-06 | 1.000E-05 |
| 1032 | 52 | L52K | A | 9.5e-06 | 1.1e-05 | 9.8e-06 | 1.010E-05 | 7.937E-07 | 9.500E-06 | 1.100E-05 |
| 1033 | 52 | L52M | A | 5.2e-06 | 1.5e-05 | 7.2e-06 | 9.133E-06 | 5.178E-06 | 5.200E-06 | 1.500E-05 |
| 1034 | 52 | L52F | A | 1.2e-05 | 1e-05 | 8.5e-06 | 1.017E-05 | 1.756E-06 | 8.500E-06 | 1.200E-05 |
| 1035 | 52 | L52P | A | 4.9e-06 | 7e-06 | 8.8e-06 | 6.900E-06 | 1.952E-06 | 4.900E-06 | 8.800E-06 |
| 1036 | 52 | L52S | A | 6.6e-06 | 3.5e-06 | 6.6e-06 | 5.567E-06 | 1.790E-06 | 3.500E-06 | 6.600E-06 |
| 1037 | 52 | L52T | A | 6.7e-06 | 1.2e-05 | 5.1e-06 | 7.933E-06 | 3.612E-06 | 5.100E-06 | 1.200E-05 |
| 1038 | 52 | L52W | A | 9.2e-06 | 7.2e-06 | 4.1e-06 | 6.833E-06 | 2.570E-06 | 4.100E-06 | 9.200E-06 |
| 1039 | 52 | L52Y | A | 6.5e-06 | 6.3e-06 | 8.9e-06 | 7.233E-06 | 1.447E-06 | 6.300E-06 | 8.900E-06 |
| 1040 | 52 | L52V | A | 7.6e-06 | 1e-05 | 4.8e-06 | 7.467E-06 | 2.603E-06 | 4.800E-06 | 1.000E-05 |
| 1041 | 53 | Q53A | A | 9.1e-06 | 9.7e-06 | 9.1e-06 | 9.300E-06 | 3.464E-07 | 9.100E-06 | 9.700E-06 |
| 1042 | 53 | Q53R | A | 7.5e-06 | 6.4e-06 | 1.1e-05 | 8.300E-06 | 2.402E-06 | 6.400E-06 | 1.100E-05 |
| 1043 | 53 | Q53N | A | 6.7e-06 | 6.1e-06 | 4.1e-06 | 5.633E-06 | 1.361E-06 | 4.100E-06 | 6.700E-06 |
| 1044 | 53 | Q53D | A | 1.4e-05 | 5.8e-06 | 5.4e-06 | 8.400E-06 | 4.854E-06 | 5.400E-06 | 1.400E-05 |
| 1045 | 53 | Q53C | A | 7.5e-06 | 1e-05 | 9.9e-06 | 9.133E-06 | 1.415E-06 | 7.500E-06 | 1.000E-05 |
| 1046 | 53 | Q53E | A | 7.1e-06 | 9.7e-06 | 8.8e-06 | 8.533E-06 | 1.320E-06 | 7.100E-06 | 9.700E-06 |
| 1047 | 53 | Q53Q | A | 7.7e-06 | 6.1e-06 | 7e-06 | 6.933E-06 | 8.021E-07 | 6.100E-06 | 7.700E-06 |
| 1048 | 53 | Q53G | A | 6.9e-06 | 6.8e-06 | 5.1e-06 | 6.267E-06 | 1.012E-06 | 5.100E-06 | 6.900E-06 |
| 1049 | 53 | Q53H | A | 4.9e-06 | 6.7e-06 | 9.7e-06 | 7.100E-06 | 2.425E-06 | 4.900E-06 | 9.700E-06 |
| 1050 | 53 | Q53I | A | 1e-05 | 8.3e-06 | 1.2e-05 | 1.010E-05 | 1.852E-06 | 8.300E-06 | 1.200E-05 |
| 1051 | 53 | Q53L | A | 1.1e-05 | 3e-06 | 7.5e-06 | 7.167E-06 | 4.010E-06 | 3.000E-06 | 1.100E-05 |
| 1052 | 53 | Q53K | A | 8.3e-06 | 7.1e-06 | 1e-05 | 8.467E-06 | 1.457E-06 | 7.100E-06 | 1.000E-05 |
| 1053 | 53 | Q53M | A | 6.6e-06 | 9.5e-06 | 8.5e-06 | 8.200E-06 | 1.473E-06 | 6.600E-06 | 9.500E-06 |
| 1054 | 53 | Q53F | A | 6.8e-06 | 7.2e-06 | 5.7e-06 | 6.567E-06 | 7.767E-07 | 5.700E-06 | 7.200E-06 |
| 1055 | 53 | Q53P | A | 6.8e-06 | 5.8e-06 | 1.3e-05 | 8.533E-06 | 3.900E-06 | 5.800E-06 | 1.300E-05 |
| 1056 | 53 | Q53S | A | 1e-05 | 1e-05 | 8.4e-06 | 9.467E-06 | 9.238E-07 | 8.400E-06 | 1.000E-05 |
| 1057 | 53 | Q53T | A | 8.6e-06 | 1.1e-05 | 7.4e-06 | 9.000E-06 | 1.833E-06 | 7.400E-06 | 1.100E-05 |
| 1058 | 53 | Q53W | A | 8.9e-06 | 6.2e-06 | 9.7e-06 | 8.267E-06 | 1.834E-06 | 6.200E-06 | 9.700E-06 |
| 1059 | 53 | Q53Y | A | 6.5e-06 | 1.2e-05 | 5e-06 | 7.833E-06 | 3.686E-06 | 5.000E-06 | 1.200E-05 |
| 1060 | 53 | Q53V | A | 7.7e-06 | 7.1e-06 | 1e-05 | 8.267E-06 | 1.531E-06 | 7.100E-06 | 1.000E-05 |
| 1061 | 54 | D54A | A | 9e-06 | 6.4e-06 | 9.3e-06 | 8.233E-06 | 1.595E-06 | 6.400E-06 | 9.300E-06 |
| 1062 | 54 | D54R | A | 1.1e-05 | 6.6e-06 | 9.4e-06 | 9.000E-06 | 2.227E-06 | 6.600E-06 | 1.100E-05 |
| 1063 | 54 | D54N | A | 1.3e-05 | 8.1e-06 | 6.3e-06 | 9.133E-06 | 3.467E-06 | 6.300E-06 | 1.300E-05 |
| 1064 | 54 | D54D | A | 1.3e-05 | 1.6e-05 | 5.6e-06 | 1.153E-05 | 5.353E-06 | 5.600E-06 | 1.600E-05 |
| 1065 | 54 | D54C | A | 8.8e-06 | 6.4e-06 | 8.8e-06 | 8.000E-06 | 1.386E-06 | 6.400E-06 | 8.800E-06 |
| 1066 | 54 | D54E | A | 9.2e-06 | 6.8e-06 | 1.5e-05 | 1.033E-05 | 4.216E-06 | 6.800E-06 | 1.500E-05 |
| 1067 | 54 | D54Q | A | 9.3e-06 | 1.1e-05 | 4.5e-06 | 8.267E-06 | 3.371E-06 | 4.500E-06 | 1.100E-05 |
| 1068 | 54 | D54G | A | 8.1e-06 | 4.6e-06 | 6.6e-06 | 6.433E-06 | 1.756E-06 | 4.600E-06 | 8.100E-06 |
| 1069 | 54 | D54H | A | 9.6e-06 | 9.8e-06 | 6.2e-06 | 8.533E-06 | 2.023E-06 | 6.200E-06 | 9.800E-06 |
| 1070 | 54 | D54I | A | 6.2e-06 | 4.8e-06 | 9.6e-06 | 6.867E-06 | 2.468E-06 | 4.800E-06 | 9.600E-06 |
| 1071 | 54 | D54L | A | 8.6e-06 | 5.1e-06 | 1.3e-05 | 8.900E-06 | 3.959E-06 | 5.100E-06 | 1.300E-05 |
| 1072 | 54 | D54K | A | 8.5e-06 | 9.1e-06 | 4.8e-06 | 7.467E-06 | 2.329E-06 | 4.800E-06 | 9.100E-06 |
| 1073 | 54 | D54M | A | 1e-05 | 7.8e-06 | 4.2e-06 | 7.333E-06 | 2.928E-06 | 4.200E-06 | 1.000E-05 |
| 1074 | 54 | D54F | A | 9.9e-06 | 7.9e-06 | 1.2e-05 | 9.933E-06 | 2.050E-06 | 7.900E-06 | 1.200E-05 |
| 1075 | 54 | D54P | A | 9.6e-06 | 6.1e-06 | 9.8e-06 | 8.500E-06 | 2.081E-06 | 6.100E-06 | 9.800E-06 |
| 1076 | 54 | D54S | A | 6.7e-06 | 6.2e-06 | 6.6e-06 | 6.500E-06 | 2.646E-07 | 6.200E-06 | 6.700E-06 |
| 1077 | 54 | D54T | A | 6.4e-06 | 1.2e-05 | 8.1e-06 | 8.833E-06 | 2.871E-06 | 6.400E-06 | 1.200E-05 |
| 1078 | 54 | D54W | A | 1.3e-05 | 7.3e-06 | 7.9e-06 | 9.400E-06 | 3.132E-06 | 7.300E-06 | 1.300E-05 |
| 1079 | 54 | D54Y | A | 5.2e-06 | 5.5e-06 | 1.3e-05 | 7.900E-06 | 4.419E-06 | 5.200E-06 | 1.300E-05 |
| 1080 | 54 | D54V | A | 9.4e-06 | 6.2e-06 | 6e-06 | 7.200E-06 | 1.908E-06 | 6.000E-06 | 9.400E-06 |
| 1081 | 55 | I55A | A | 5.1e-06 | 9.6e-06 | 6.9e-06 | 7.200E-06 | 2.265E-06 | 5.100E-06 | 9.600E-06 |
| 1082 | 55 | I55R | A | 6.6e-06 | 6.9e-06 | 8.7e-06 | 7.400E-06 | 1.136E-06 | 6.600E-06 | 8.700E-06 |
| 1083 | 55 | I55N | A | 1.1e-05 | 6.5e-06 | 4.8e-06 | 7.433E-06 | 3.204E-06 | 4.800E-06 | 1.100E-05 |
| 1084 | 55 | I55D | A | 6.7e-06 | 4.9e-06 | 6.6e-06 | 6.067E-06 | 1.012E-06 | 4.900E-06 | 6.700E-06 |
| 1085 | 55 | I55C | A | 9.3e-06 | 8.8e-06 | 1e-05 | 9.367E-06 | 6.028E-07 | 8.800E-06 | 1.000E-05 |
| 1086 | 55 | I55E | A | 1.3e-05 | 9.4e-06 | 1e-05 | 1.080E-05 | 1.929E-06 | 9.400E-06 | 1.300E-05 |
| 1087 | 55 | I55Q | A | 6.5e-06 | 9.3e-06 | 7.7e-06 | 7.833E-06 | 1.405E-06 | 6.500E-06 | 9.300E-06 |
| 1088 | 55 | I55G | A | 8.9e-06 | 8.7e-06 | 1e-05 | 9.200E-06 | 7.000E-07 | 8.700E-06 | 1.000E-05 |
| 1089 | 55 | I55H | A | 8.7e-06 | 8.9e-06 | 7.8e-06 | 8.467E-06 | 5.859E-07 | 7.800E-06 | 8.900E-06 |
| 1090 | 55 | I55I | A | 8.1e-06 | 5.9e-06 | 7.5e-06 | 7.167E-06 | 1.137E-06 | 5.900E-06 | 8.100E-06 |
| 1091 | 55 | I55L | A | 6.8e-06 | 9.9e-06 | 6.4e-06 | 7.700E-06 | 1.916E-06 | 6.400E-06 | 9.900E-06 |
| 1092 | 55 | I55K | A | 6.4e-06 | 7.4e-06 | 9.3e-06 | 7.700E-06 | 1.473E-06 | 6.400E-06 | 9.300E-06 |
| 1093 | 55 | I55M | A | 9.8e-06 | 5.4e-06 | 4e-06 | 6.400E-06 | 3.027E-06 | 4.000E-06 | 9.800E-06 |
| 1094 | 55 | I55F | A | 6.5e-06 | 7.7e-06 | 8.1e-06 | 7.433E-06 | 8.327E-07 | 6.500E-06 | 8.100E-06 |
| 1095 | 55 | I55P | A | 5.4e-06 | 4.9e-06 | 1.2e-05 | 7.433E-06 | 3.963E-06 | 4.900E-06 | 1.200E-05 |
| 1096 | 55 | I55S | A | 4.4e-06 | 8.1e-06 | 9e-06 | 7.167E-06 | 2.438E-06 | 4.400E-06 | 9.000E-06 |
| 1097 | 55 | I55T | A | 6.5e-06 | 6.3e-06 | 5.5e-06 | 6.100E-06 | 5.292E-07 | 5.500E-06 | 6.500E-06 |
| 1098 | 55 | I55W | A | 8.9e-06 | 1.1e-05 | 3.4e-06 | 7.767E-06 | 3.925E-06 | 3.400E-06 | 1.100E-05 |
| 1099 | 55 | I55Y | A | 7.4e-06 | 1.2e-05 | 7.5e-06 | 8.967E-06 | 2.627E-06 | 7.400E-06 | 1.200E-05 |
| 1100 | 55 | I55V | A | 6e-06 | 8.4e-06 | 8.8e-06 | 7.733E-06 | 1.514E-06 | 6.000E-06 | 8.800E-06 |
| 1101 | 56 | Q56A | A | 1e-05 | 6.6e-06 | 1.1e-05 | 9.200E-06 | 2.307E-06 | 6.600E-06 | 1.100E-05 |
| 1102 | 56 | Q56R | A | 9.4e-06 | 6.8e-06 | 5.9e-06 | 7.367E-06 | 1.818E-06 | 5.900E-06 | 9.400E-06 |
| 1103 | 56 | Q56N | A | 1e-05 | 9e-06 | 7.1e-06 | 8.700E-06 | 1.473E-06 | 7.100E-06 | 1.000E-05 |
| 1104 | 56 | Q56D | A | 9.7e-06 | 5.9e-06 | 1.4e-05 | 9.867E-06 | 4.053E-06 | 5.900E-06 | 1.400E-05 |

| | | | | | | | | | | |
|---|---|---|---|---|---|---|---|---|---|---|
| 1105 | 56 | Q56C | A | 1e-05 | 4.9e-06 | 1e-05 | 8.300E-06 | 2.944E-06 | 4.900E-06 | 1.000E-05 |
| 1106 | 56 | Q56E | A | 6.6e-06 | 5.4e-06 | 6e-06 | 6.000E-06 | 6.000E-07 | 5.400E-06 | 6.600E-06 |
| 1107 | 56 | Q56Q | A | 8.8e-06 | 7.7e-06 | 7.5e-06 | 8.000E-06 | 7.000E-07 | 7.500E-06 | 8.800E-06 |
| 1108 | 56 | Q56G | A | 4.9e-06 | 4.9e-06 | 1e-05 | 6.600E-06 | 2.944E-06 | 4.900E-06 | 1.000E-05 |
| 1109 | 56 | Q56H | A | 1e-05 | 5.1e-06 | 8.2e-06 | 7.767E-06 | 2.479E-06 | 5.100E-06 | 1.000E-05 |
| 1110 | 56 | Q56I | A | 1e-05 | 1.2e-05 | 8.2e-06 | 1.007E-05 | 1.901E-06 | 8.200E-06 | 1.200E-05 |
| 1111 | 56 | Q56L | A | 6.8e-06 | 5.5e-06 | 6.8e-06 | 6.367E-06 | 7.506E-07 | 5.500E-06 | 6.800E-06 |
| 1112 | 56 | Q56K | A | 8.4e-06 | 5.5e-06 | 3.2e-06 | 5.700E-06 | 2.606E-06 | 3.200E-06 | 8.400E-06 |
| 1113 | 56 | Q56M | A | 1.1e-05 | 9.6e-06 | 8.3e-06 | 9.633E-06 | 1.350E-06 | 8.300E-06 | 1.100E-05 |
| 1114 | 56 | Q56F | A | 1e-05 | 8.5e-06 | 9.4e-06 | 9.300E-06 | 7.550E-07 | 8.500E-06 | 1.000E-05 |
| 1115 | 56 | Q56P | A | 1.3e-05 | 8.2e-06 | 9.4e-06 | 1.020E-05 | 2.498E-06 | 8.200E-06 | 1.300E-05 |
| 1116 | 56 | Q56S | A | 6.4e-06 | 4e-06 | 9.9e-06 | 6.767E-06 | 2.967E-06 | 4.000E-06 | 9.900E-06 |
| 1117 | 56 | Q56T | A | 4e-06 | 7.1e-06 | 7.5e-06 | 6.200E-06 | 1.916E-06 | 4.000E-06 | 7.500E-06 |
| 1118 | 56 | Q56W | A | 7.6e-06 | 4.9e-06 | 7.1e-06 | 6.533E-06 | 1.436E-06 | 4.900E-06 | 7.600E-06 |
| 1119 | 56 | Q56Y | A | 6.8e-06 | 6.4e-06 | 4.9e-06 | 6.033E-06 | 1.002E-06 | 4.900E-06 | 6.800E-06 |
| 1120 | 56 | Q56V | A | 6.7e-06 | 4.6e-06 | 5.8e-06 | 5.700E-06 | 1.054E-06 | 4.600E-06 | 6.700E-06 |
| 1121 | 57 | E57A | A | 7e-06 | 4.1e-06 | 8.1e-06 | 6.400E-06 | 2.066E-06 | 4.100E-06 | 8.100E-06 |
| 1122 | 57 | E57R | A | 4.8e-06 | 6e-06 | 7e-06 | 5.933E-06 | 1.102E-06 | 4.800E-06 | 7.000E-06 |
| 1123 | 57 | E57N | A | 4.5e-06 | 8e-06 | 8.4e-06 | 6.967E-06 | 2.146E-06 | 4.500E-06 | 8.400E-06 |
| 1124 | 57 | E57D | A | 6e-06 | 9.8e-06 | 9.1e-06 | 8.300E-06 | 2.022E-06 | 6.000E-06 | 9.800E-06 |
| 1125 | 57 | E57C | A | 6.2e-06 | 4.2e-06 | 6.2e-06 | 5.533E-06 | 1.155E-06 | 4.200E-06 | 6.200E-06 |
| 1126 | 57 | E57E | A | 1.2e-05 | 4.9e-06 | 4.5e-06 | 7.133E-06 | 4.219E-06 | 4.500E-06 | 1.200E-05 |
| 1127 | 57 | E57Q | A | 6.7e-06 | 4.9e-06 | 5.7e-06 | 5.767E-06 | 9.018E-07 | 4.900E-06 | 6.700E-06 |
| 1128 | 57 | E57G | A | 8.2e-06 | 1.5e-05 | 7.1e-06 | 1.010E-05 | 4.279E-06 | 7.100E-06 | 1.500E-05 |
| 1129 | 57 | E57H | A | 4.5e-06 | 6.3e-06 | 6.4e-06 | 5.733E-06 | 1.069E-06 | 4.500E-06 | 6.400E-06 |
| 1130 | 57 | E57I | A | 6.5e-06 | 9.3e-06 | 5.3e-06 | 7.033E-06 | 2.053E-06 | 5.300E-06 | 9.300E-06 |
| 1131 | 57 | E57L | A | 9.4e-06 | 3.5e-06 | 8.3e-06 | 7.067E-06 | 3.137E-06 | 3.500E-06 | 9.400E-06 |
| 1132 | 57 | E57K | A | 7.6e-06 | 8e-06 | 1.1e-05 | 8.867E-06 | 1.858E-06 | 7.600E-06 | 1.100E-05 |
| 1133 | 57 | E57M | A | 1.8e-05 | 1.4e-05 | 5.2e-06 | 1.240E-05 | 6.548E-06 | 5.200E-06 | 1.800E-05 |
| 1134 | 57 | E57F | A | 1.4e-05 | 8e-06 | 6.5e-06 | 9.500E-06 | 3.969E-06 | 6.500E-06 | 1.400E-05 |
| 1135 | 57 | E57P | A | 1e-05 | 6.8e-06 | 9.4e-06 | 8.733E-06 | 1.701E-06 | 6.800E-06 | 1.000E-05 |
| 1136 | 57 | E57S | A | 8.4e-06 | 6.2e-06 | 6e-06 | 6.867E-06 | 1.332E-06 | 6.000E-06 | 8.400E-06 |
| 1137 | 57 | E57T | A | 7.5e-06 | 6.3e-06 | 7.1e-06 | 6.967E-06 | 6.110E-07 | 6.300E-06 | 7.500E-06 |
| 1138 | 57 | E57W | A | 6.3e-06 | 7.2e-06 | 8.9e-06 | 7.467E-06 | 1.320E-06 | 6.300E-06 | 8.900E-06 |
| 1139 | 57 | E57Y | A | 6.9e-06 | 9.9e-06 | 7.2e-06 | 8.000E-06 | 1.652E-06 | 6.900E-06 | 9.900E-06 |
| 1140 | 57 | E57V | A | 9.2e-06 | 1.2e-05 | 7.1e-06 | 9.433E-06 | 2.458E-06 | 7.100E-06 | 1.200E-05 |
| 1141 | 58 | V58A | A | 6.7e-06 | 5.2e-06 | 1.3e-05 | 8.300E-06 | 4.139E-06 | 5.200E-06 | 1.300E-05 |
| 1142 | 58 | V58R | A | 7.5e-06 | 6e-06 | 6.7e-06 | 6.733E-06 | 7.506E-07 | 6.000E-06 | 7.500E-06 |
| 1143 | 58 | V58N | A | 5.2e-06 | 1e-05 | 5.8e-06 | 7.000E-06 | 2.615E-06 | 5.200E-06 | 1.000E-05 |
| 1144 | 58 | V58D | A | 5.8e-06 | 1.5e-05 | 5.3e-06 | 8.700E-06 | 5.462E-06 | 5.300E-06 | 1.500E-05 |
| 1145 | 58 | V58C | A | 1e-05 | 5.2e-06 | 8.6e-06 | 7.933E-06 | 2.468E-06 | 5.200E-06 | 1.000E-05 |
| 1146 | 58 | V58E | A | 8.7e-06 | 6.8e-06 | 9.9e-06 | 8.467E-06 | 1.563E-06 | 6.800E-06 | 9.900E-06 |
| 1147 | 58 | V58Q | A | 9.9e-06 | 8.7e-06 | 4.5e-06 | 7.700E-06 | 2.835E-06 | 4.500E-06 | 9.900E-06 |
| 1148 | 58 | V58G | A | 1e-05 | 6.8e-06 | 7.1e-06 | 7.967E-06 | 1.767E-06 | 6.800E-06 | 1.000E-05 |
| 1149 | 58 | V58H | A | 8.9e-06 | 8.7e-06 | 7.9e-06 | 8.500E-06 | 5.292E-07 | 7.900E-06 | 8.900E-06 |
| 1150 | 58 | V58I | A | 1.2e-05 | 3.9e-06 | 6.3e-06 | 7.400E-06 | 4.161E-06 | 3.900E-06 | 1.200E-05 |
| 1151 | 58 | V58L | A | 9.2e-06 | 5.6e-06 | 5.7e-06 | 6.833E-06 | 2.050E-06 | 5.600E-06 | 9.200E-06 |
| 1152 | 58 | V58K | A | 8.8e-06 | 7.4e-06 | 8.9e-06 | 8.367E-06 | 8.386E-07 | 7.400E-06 | 8.900E-06 |
| 1153 | 58 | V58M | A | 4e-06 | 6.5e-06 | 1.2e-05 | 7.500E-06 | 4.093E-06 | 4.000E-06 | 1.200E-05 |
| 1154 | 58 | V58F | A | 1e-05 | 7.1e-06 | 5.3e-06 | 7.467E-06 | 2.371E-06 | 5.300E-06 | 1.000E-05 |
| 1155 | 58 | V58P | A | 7.3e-06 | 1e-05 | 7.2e-06 | 8.167E-06 | 1.589E-06 | 7.200E-06 | 1.000E-05 |
| 1156 | 58 | V58S | A | 8.7e-06 | 9.1e-06 | 7.2e-06 | 8.333E-06 | 1.002E-06 | 7.200E-06 | 9.100E-06 |
| 1157 | 58 | V58T | A | 8.1e-06 | 9.3e-06 | 7.4e-06 | 8.267E-06 | 9.609E-07 | 7.400E-06 | 9.300E-06 |
| 1158 | 58 | V58W | A | 7.1e-06 | 7.6e-06 | 9e-06 | 7.900E-06 | 9.849E-07 | 7.100E-06 | 9.000E-06 |
| 1159 | 58 | V58Y | A | 7.6e-06 | 8.9e-06 | 8.5e-06 | 8.333E-06 | 6.658E-07 | 7.600E-06 | 8.900E-06 |
| 1160 | 58 | V58V | A | 1.1e-05 | 6.5e-06 | 1.3e-05 | 1.017E-05 | 3.329E-06 | 6.500E-06 | 1.300E-05 |
| 1161 | 59 | Q59A | A | 9.1e-06 | 5.2e-06 | 9.5e-06 | 7.933E-06 | 2.376E-06 | 5.200E-06 | 9.500E-06 |
| 1162 | 59 | Q59R | A | 4.9e-06 | 7.5e-06 | 7e-06 | 6.467E-06 | 1.380E-06 | 4.900E-06 | 7.500E-06 |
| 1163 | 59 | Q59N | A | 1e-05 | 4.1e-06 | 5.3e-06 | 6.467E-06 | 3.118E-06 | 4.100E-06 | 1.000E-05 |
| 1164 | 59 | Q59D | A | 7.1e-06 | 6.9e-06 | 9.3e-06 | 7.767E-06 | 1.332E-06 | 6.900E-06 | 9.300E-06 |
| 1165 | 59 | Q59C | A | 1.1e-05 | 9.1e-06 | 6.5e-06 | 8.867E-06 | 2.259E-06 | 6.500E-06 | 1.100E-05 |
| 1166 | 59 | Q59E | A | 7.8e-06 | 9e-06 | 4.1e-06 | 6.967E-06 | 2.554E-06 | 4.100E-06 | 9.000E-06 |
| 1167 | 59 | Q59Q | A | 6e-06 | 7e-06 | 9.9e-06 | 7.633E-06 | 2.026E-06 | 6.000E-06 | 9.900E-06 |
| 1168 | 59 | Q59G | A | 6.7e-06 | 9.1e-06 | 5.5e-06 | 7.100E-06 | 1.833E-06 | 5.500E-06 | 9.100E-06 |
| 1169 | 59 | Q59H | A | 6.9e-06 | 1.2e-05 | 6.5e-06 | 8.467E-06 | 3.066E-06 | 6.500E-06 | 1.200E-05 |
| 1170 | 59 | Q59I | A | 1e-05 | 9.3e-06 | 1.2e-05 | 1.043E-05 | 1.401E-06 | 9.300E-06 | 1.200E-05 |
| 1171 | 59 | Q59L | A | 5.3e-06 | 4.9e-06 | 6.4e-06 | 5.533E-06 | 7.767E-07 | 4.900E-06 | 6.400E-06 |
| 1172 | 59 | Q59K | A | 5.6e-06 | 4.7e-06 | 7.2e-06 | 5.833E-06 | 1.266E-06 | 4.700E-06 | 7.200E-06 |
| 1173 | 59 | Q59M | A | 5.7e-06 | 1.4e-05 | 1.2e-05 | 1.057E-05 | 4.332E-06 | 5.700E-06 | 1.400E-05 |
| 1174 | 59 | Q59F | A | 1.3e-05 | 7e-06 | 7.4e-06 | 9.133E-06 | 3.355E-06 | 7.000E-06 | 1.300E-05 |
| 1175 | 59 | Q59P | A | 7.1e-06 | 1e-05 | 5.6e-06 | 7.567E-06 | 2.237E-06 | 5.600E-06 | 1.000E-05 |
| 1176 | 59 | Q59S | A | 5.4e-06 | 8.9e-06 | 8.6e-06 | 7.633E-06 | 1.940E-06 | 5.400E-06 | 8.900E-06 |
| 1177 | 59 | Q59T | A | 8.7e-06 | 1.1e-05 | 8.3e-06 | 9.333E-06 | 1.457E-06 | 8.300E-06 | 1.100E-05 |
| 1178 | 59 | Q59W | A | 1.1e-05 | 6.1e-06 | 1.1e-05 | 9.367E-06 | 2.829E-06 | 6.100E-06 | 1.100E-05 |
| 1179 | 59 | Q59Y | A | 1.3e-05 | 8.6e-06 | 1.3e-05 | 1.153E-05 | 2.540E-06 | 8.600E-06 | 1.300E-05 |
| 1180 | 59 | Q59V | A | 5.5e-06 | 1.3e-05 | 9.8e-06 | 9.433E-06 | 3.763E-06 | 5.500E-06 | 1.300E-05 |
| 1181 | 60 | G60A | A | 5.6e-06 | 6.1e-06 | 8.8e-06 | 6.833E-06 | 1.721E-06 | 5.600E-06 | 8.800E-06 |
| 1182 | 60 | G60R | A | 1.2e-05 | 4.3e-06 | 8.8e-06 | 8.367E-06 | 3.868E-06 | 4.300E-06 | 1.200E-05 |
| 1183 | 60 | G60N | A | 5.6e-06 | 9.4e-06 | 3.8e-06 | 6.267E-06 | 2.859E-06 | 3.800E-06 | 9.400E-06 |
| 1184 | 60 | G60D | A | 8.1e-06 | 6.9e-06 | 1.3e-05 | 9.333E-06 | 3.232E-06 | 6.900E-06 | 1.300E-05 |
| 1185 | 60 | G60C | A | 9.9e-06 | 1.5e-05 | 5.7e-06 | 1.020E-05 | 4.657E-06 | 5.700E-06 | 1.500E-05 |
| 1186 | 60 | G60E | A | 4.3e-06 | 5e-06 | 1.4e-05 | 7.767E-06 | 5.410E-06 | 4.300E-06 | 1.400E-05 |
| 1187 | 60 | G60Q | A | 5.6e-06 | 8.4e-06 | 7.3e-06 | 7.100E-06 | 1.411E-06 | 5.600E-06 | 8.400E-06 |
| 1188 | 60 | G60G | A | 1.5e-05 | 5.4e-06 | 7.5e-06 | 9.300E-06 | 5.047E-06 | 5.400E-06 | 1.500E-05 |
| 1189 | 60 | G60H | A | 1.2e-05 | 6.8e-06 | 6.6e-06 | 8.467E-06 | 3.062E-06 | 6.600E-06 | 1.200E-05 |

| | | | | | | | | | | |
|---|---|---|---|---|---|---|---|---|---|---|
| 1190 | 60 | G60I | A | 7.5e-06 | 1.3e-05 | 1e-05 | 1.017E-05 | 2.754E-06 | 7.500E-06 | 1.300E-05 |
| 1191 | 60 | G60L | A | 8.4e-06 | 4.4e-06 | 7.5e-06 | 6.767E-06 | 2.098E-06 | 4.400E-06 | 8.400E-06 |
| 1192 | 60 | G60K | A | 7.9e-06 | 6.6e-06 | 6.7e-06 | 7.067E-06 | 7.234E-07 | 6.600E-06 | 7.900E-06 |
| 1193 | 60 | G60M | A | 8.7e-06 | 9.3e-06 | 8.4e-06 | 8.800E-06 | 4.583E-07 | 8.400E-06 | 9.300E-06 |
| 1194 | 60 | G60F | A | 5.9e-06 | 8.5e-06 | 6.1e-06 | 6.833E-06 | 1.447E-06 | 5.900E-06 | 8.500E-06 |
| 1195 | 60 | G60P | A | 8.4e-06 | 6e-06 | 7.3e-06 | 7.233E-06 | 1.201E-06 | 6.000E-06 | 8.400E-06 |
| 1196 | 60 | G60S | A | 4.7e-06 | 1.1e-05 | 6.7e-06 | 7.467E-06 | 3.219E-06 | 4.700E-06 | 1.100E-05 |
| 1197 | 60 | G60T | A | 9.6e-06 | 1.4e-05 | 8.3e-06 | 1.063E-05 | 2.987E-06 | 8.300E-06 | 1.400E-05 |
| 1198 | 60 | G60W | A | 7.4e-06 | 8.6e-06 | 1.1e-05 | 9.000E-06 | 1.833E-06 | 7.400E-06 | 1.100E-05 |
| 1199 | 60 | G60Y | A | 5.8e-06 | 5.7e-06 | 9.7e-06 | 7.067E-06 | 2.281E-06 | 5.700E-06 | 9.700E-06 |
| 1200 | 60 | G60V | A | 1e-05 | 8.2e-06 | 8.2e-06 | 8.800E-06 | 1.039E-06 | 8.200E-06 | 1.000E-05 |
| 1201 | 61 | Y61A | A | 1.3e-05 | 6.5e-06 | 1.2e-05 | 1.050E-05 | 3.500E-06 | 6.500E-06 | 1.300E-05 |
| 1202 | 61 | Y61R | A | 1.1e-05 | 9.7e-06 | 1e-05 | 1.023E-05 | 6.807E-07 | 9.700E-06 | 1.100E-05 |
| 1203 | 61 | Y61N | A | 6.9e-06 | 9.3e-06 | 7.4e-06 | 7.867E-06 | 1.266E-06 | 6.900E-06 | 9.300E-06 |
| 1204 | 61 | Y61D | A | 8.3e-06 | 9e-06 | 7.8e-06 | 8.367E-06 | 6.028E-07 | 7.800E-06 | 9.000E-06 |
| 1205 | 61 | Y61C | A | 9.2e-06 | 6.8e-06 | 1e-05 | 8.667E-06 | 1.665E-06 | 6.800E-06 | 1.000E-05 |
| 1206 | 61 | Y61E | A | 6.7e-06 | 5.8e-06 | 8.8e-06 | 7.100E-06 | 1.539E-06 | 5.800E-06 | 8.800E-06 |
| 1207 | 61 | Y61Q | A | 7.9e-06 | 7.6e-06 | 5.6e-06 | 7.033E-06 | 1.250E-06 | 5.600E-06 | 7.900E-06 |
| 1208 | 61 | Y61G | A | 1.5e-05 | 1.2e-05 | 8.4e-06 | 1.180E-05 | 3.305E-06 | 8.400E-06 | 1.500E-05 |
| 1209 | 61 | Y61H | A | 4.6e-06 | 1.4e-05 | 8e-06 | 8.867E-06 | 4.760E-06 | 4.600E-06 | 1.400E-05 |
| 1210 | 61 | Y61I | A | 7.1e-06 | 9.4e-06 | 8.6e-06 | 8.367E-06 | 1.168E-06 | 7.100E-06 | 9.400E-06 |
| 1211 | 61 | Y61L | A | 1.3e-05 | 6.9e-06 | 1.3e-05 | 1.097E-05 | 3.522E-06 | 6.900E-06 | 1.300E-05 |
| 1212 | 61 | Y61K | A | 7.9e-06 | 8.4e-06 | 6.7e-06 | 7.667E-06 | 8.737E-07 | 6.700E-06 | 8.400E-06 |
| 1213 | 61 | Y61M | A | 9.3e-06 | 6.1e-06 | 9.5e-06 | 8.300E-06 | 1.908E-06 | 6.100E-06 | 9.500E-06 |
| 1214 | 61 | Y61F | A | 6.3e-06 | 6.2e-06 | 5.3e-06 | 5.933E-06 | 5.508E-07 | 5.300E-06 | 6.300E-06 |
| 1215 | 61 | Y61P | A | 8.4e-06 | 8.6e-06 | 9.8e-06 | 8.933E-06 | 7.572E-07 | 8.400E-06 | 9.800E-06 |
| 1216 | 61 | Y61S | A | 4.2e-06 | 6.4e-06 | 6.6e-06 | 5.733E-06 | 1.332E-06 | 4.200E-06 | 6.600E-06 |
| 1217 | 61 | Y61T | A | 8.3e-06 | 8.7e-06 | 7.6e-06 | 8.200E-06 | 5.568E-07 | 7.600E-06 | 8.700E-06 |
| 1218 | 61 | Y61W | A | 1.1e-05 | 8.8e-06 | 9.4e-06 | 9.733E-06 | 1.137E-06 | 8.800E-06 | 1.100E-05 |
| 1219 | 61 | Y61Y | A | 7.7e-06 | 9.5e-06 | 1.2e-05 | 9.733E-06 | 2.159E-06 | 7.700E-06 | 1.200E-05 |
| 1220 | 61 | Y61V | A | 4.9e-06 | 4.7e-06 | 6.4e-06 | 5.333E-06 | 9.292E-07 | 4.700E-06 | 6.400E-06 |
| 1221 | 62 | V62A | A | 1.1e-05 | 9.1e-06 | 6.5e-06 | 8.867E-06 | 2.259E-06 | 6.500E-06 | 1.100E-05 |
| 1222 | 62 | V62R | A | 1.1e-05 | 9.6e-06 | 6.1e-06 | 8.900E-06 | 2.524E-06 | 6.100E-06 | 1.100E-05 |
| 1223 | 62 | V62N | A | 7.9e-06 | 4.7e-06 | 4.9e-06 | 5.833E-06 | 1.793E-06 | 4.700E-06 | 7.900E-06 |
| 1224 | 62 | V62D | A | 1.3e-05 | 1.2e-05 | 9.6e-06 | 1.153E-05 | 1.747E-06 | 9.600E-06 | 1.300E-05 |
| 1225 | 62 | V62C | A | 5e-06 | 7.6e-06 | 7.7e-06 | 6.767E-06 | 1.531E-06 | 5.000E-06 | 7.700E-06 |
| 1226 | 62 | V62E | A | 1e-05 | 4.9e-06 | 6.5e-06 | 7.133E-06 | 2.608E-06 | 4.900E-06 | 1.000E-05 |
| 1227 | 62 | V62Q | A | 8.2e-06 | 1.1e-05 | 9.8e-06 | 9.667E-06 | 1.405E-06 | 8.200E-06 | 1.100E-05 |
| 1228 | 62 | V62G | A | 1.1e-05 | 8.3e-06 | 7.1e-06 | 8.800E-06 | 1.997E-06 | 7.100E-06 | 1.100E-05 |
| 1229 | 62 | V62H | A | 6e-06 | 7.3e-06 | 9.7e-06 | 7.667E-06 | 1.877E-06 | 6.000E-06 | 9.700E-06 |
| 1230 | 62 | V62I | A | 9.1e-06 | 1e-05 | 6.1e-06 | 8.400E-06 | 2.042E-06 | 6.100E-06 | 1.000E-05 |
| 1231 | 62 | V62L | A | 5.8e-06 | 4.4e-06 | 1e-05 | 6.733E-06 | 2.914E-06 | 4.400E-06 | 1.000E-05 |
| 1232 | 62 | V62K | A | 7.5e-06 | 1e-05 | 6.1e-06 | 7.867E-06 | 1.976E-06 | 6.100E-06 | 1.000E-05 |
| 1233 | 62 | V62M | A | 8.7e-06 | 4.2e-06 | 5.7e-06 | 6.200E-06 | 2.291E-06 | 4.200E-06 | 8.700E-06 |
| 1234 | 62 | V62F | A | 8.7e-06 | 7.8e-06 | 8.6e-06 | 8.367E-06 | 4.933E-07 | 7.800E-06 | 8.700E-06 |
| 1235 | 62 | V62P | A | 6.2e-06 | 6.5e-06 | 9.7e-06 | 7.467E-06 | 1.940E-06 | 6.200E-06 | 9.700E-06 |
| 1236 | 62 | V62S | A | 1.1e-05 | 8e-06 | 6.4e-06 | 8.467E-06 | 2.335E-06 | 6.400E-06 | 1.100E-05 |
| 1237 | 62 | V62T | A | 8.6e-06 | 6e-06 | 9.8e-06 | 8.133E-06 | 1.943E-06 | 6.000E-06 | 9.800E-06 |
| 1238 | 62 | V62W | A | 1e-05 | 8.1e-06 | 7.1e-06 | 8.400E-06 | 1.473E-06 | 7.100E-06 | 1.000E-05 |
| 1239 | 62 | V62Y | A | 7.3e-06 | 4.5e-06 | 1e-05 | 7.267E-06 | 2.750E-06 | 4.500E-06 | 1.000E-05 |
| 1240 | 62 | V62V | A | 4.5e-06 | 1e-05 | 8.1e-06 | 7.533E-06 | 2.793E-06 | 4.500E-06 | 1.000E-05 |
| 1241 | 63 | L63A | A | 1.2e-05 | 1.3e-05 | 9.6e-06 | 1.153E-05 | 1.747E-06 | 9.600E-06 | 1.300E-05 |
| 1242 | 63 | L63R | A | 8.9e-06 | 4.1e-06 | 5.2e-06 | 6.067E-06 | 2.515E-06 | 4.100E-06 | 8.900E-06 |
| 1243 | 63 | L63N | A | 6e-06 | 1.2e-05 | 1.4e-05 | 1.067E-05 | 4.163E-06 | 6.000E-06 | 1.400E-05 |
| 1244 | 63 | L63D | A | 4.5e-06 | 1.5e-05 | 1.3e-05 | 1.083E-05 | 5.575E-06 | 4.500E-06 | 1.500E-05 |
| 1245 | 63 | L63C | A | 1e-05 | 7.5e-06 | 6.6e-06 | 8.033E-06 | 1.762E-06 | 6.600E-06 | 1.000E-05 |
| 1246 | 63 | L63E | A | 9.2e-06 | 7.5e-06 | 1e-05 | 8.900E-06 | 1.277E-06 | 7.500E-06 | 1.000E-05 |
| 1247 | 63 | L63Q | A | 5.9e-06 | 8.6e-06 | 8.7e-06 | 7.733E-06 | 1.589E-06 | 5.900E-06 | 8.700E-06 |
| 1248 | 63 | L63G | A | 1e-05 | 7.9e-06 | 8.1e-06 | 8.667E-06 | 1.159E-06 | 7.900E-06 | 1.000E-05 |
| 1249 | 63 | L63H | A | 4.4e-06 | 6.4e-06 | 9.3e-06 | 6.700E-06 | 2.464E-06 | 4.400E-06 | 9.300E-06 |
| 1250 | 63 | L63I | A | 5.6e-06 | 6e-06 | 8.9e-06 | 6.833E-06 | 1.801E-06 | 5.600E-06 | 8.900E-06 |
| 1251 | 63 | L63L | A | 7.8e-06 | 6.7e-06 | 1.4e-05 | 9.500E-06 | 3.936E-06 | 6.700E-06 | 1.400E-05 |
| 1252 | 63 | L63K | A | 4.5e-06 | 9.3e-06 | 7.9e-06 | 7.233E-06 | 2.468E-06 | 4.500E-06 | 9.300E-06 |
| 1253 | 63 | L63M | A | 7.5e-06 | 9.9e-06 | 9.6e-06 | 9.000E-06 | 1.308E-06 | 7.500E-06 | 9.900E-06 |
| 1254 | 63 | L63F | A | 7.3e-06 | 4.9e-06 | 6.1e-06 | 6.100E-06 | 1.200E-06 | 4.900E-06 | 7.300E-06 |
| 1255 | 63 | L63P | A | 5.5e-06 | 7.3e-06 | 4.6e-06 | 5.800E-06 | 1.375E-06 | 4.600E-06 | 7.300E-06 |
| 1256 | 63 | L63S | A | 5.6e-06 | 5.7e-06 | 9.3e-06 | 6.867E-06 | 2.108E-06 | 5.600E-06 | 9.300E-06 |
| 1257 | 63 | L63T | A | 9.2e-06 | 8.8e-06 | 6.6e-06 | 8.200E-06 | 1.400E-06 | 6.600E-06 | 9.200E-06 |
| 1258 | 63 | L63W | A | 8.3e-06 | 9.7e-06 | 1e-05 | 9.333E-06 | 9.074E-07 | 8.300E-06 | 1.000E-05 |
| 1259 | 63 | L63Y | A | 7.4e-06 | 7.5e-06 | 9.9e-06 | 8.267E-06 | 1.415E-06 | 7.400E-06 | 9.900E-06 |
| 1260 | 63 | L63V | A | 1.5e-05 | 4.1e-06 | 9.2e-06 | 9.433E-06 | 5.454E-06 | 4.100E-06 | 1.500E-05 |
| 1261 | 64 | I64A | A | 9.1e-06 | 6.7e-06 | 7.5e-06 | 7.767E-06 | 1.222E-06 | 6.700E-06 | 9.100E-06 |
| 1262 | 64 | I64R | A | 1.2e-05 | 1.3e-05 | 1e-05 | 1.167E-05 | 1.528E-06 | 1.000E-05 | 1.300E-05 |
| 1263 | 64 | I64N | A | 8.4e-06 | 6.4e-06 | 8.8e-06 | 7.867E-06 | 1.286E-06 | 6.400E-06 | 8.800E-06 |
| 1264 | 64 | I64D | A | 1.4e-05 | 9.9e-06 | 8.1e-06 | 1.067E-05 | 3.024E-06 | 8.100E-06 | 1.400E-05 |
| 1265 | 64 | I64C | A | 6.9e-06 | 1e-05 | 8e-06 | 8.300E-06 | 1.572E-06 | 6.900E-06 | 1.000E-05 |
| 1266 | 64 | I64E | A | 8.8e-06 | 1.1e-05 | 5.8e-06 | 8.533E-06 | 2.610E-06 | 5.800E-06 | 1.100E-05 |
| 1267 | 64 | I64Q | A | 1.1e-05 | 9.3e-06 | 4.2e-06 | 8.167E-06 | 3.539E-06 | 4.200E-06 | 1.100E-05 |
| 1268 | 64 | I64G | A | 6.9e-06 | 4.8e-06 | 5.3e-06 | 5.667E-06 | 1.097E-06 | 4.800E-06 | 6.900E-06 |
| 1269 | 64 | I64H | A | 9.7e-06 | 1.4e-05 | 5.7e-06 | 9.800E-06 | 4.151E-06 | 5.700E-06 | 1.400E-05 |
| 1270 | 64 | I64I | A | 6.9e-06 | 6.5e-06 | 4.5e-06 | 5.967E-06 | 1.286E-06 | 4.500E-06 | 6.900E-06 |
| 1271 | 64 | I64L | A | 9.1e-06 | 1.2e-05 | 8.5e-06 | 9.867E-06 | 1.872E-06 | 8.500E-06 | 1.200E-05 |
| 1272 | 64 | I64K | A | 9e-06 | 6.2e-06 | 3e-06 | 6.067E-06 | 3.002E-06 | 3.000E-06 | 9.000E-06 |
| 1273 | 64 | I64M | A | 9e-06 | 8.3e-06 | 8.6e-06 | 8.633E-06 | 3.512E-07 | 8.300E-06 | 9.000E-06 |
| 1274 | 64 | I64F | A | 4.5e-06 | 8.7e-06 | 5.2e-06 | 6.133E-06 | 2.250E-06 | 4.500E-06 | 8.700E-06 |

| | | | | | | | | | | |
|---|---|---|---|---|---|---|---|---|---|---|
| 1275 | 64 | I64P | A | 9.7e-06 | 9.9e-06 | 8.2e-06 | 9.267E-06 | 9.292E-07 | 8.200E-06 | 9.900E-06 |
| 1276 | 64 | I64S | A | 7.2e-06 | 7.6e-06 | 8.2e-06 | 7.667E-06 | 5.033E-07 | 7.200E-06 | 8.200E-06 |
| 1277 | 64 | I64T | A | 1.1e-05 | 2e-05 | 5.6e-06 | 1.220E-05 | 7.275E-06 | 5.600E-06 | 2.000E-05 |
| 1278 | 64 | I64W | A | 7.3e-06 | 6.9e-06 | 6e-06 | 6.733E-06 | 6.658E-07 | 6.000E-06 | 7.300E-06 |
| 1279 | 64 | I64Y | A | 8.4e-06 | 7.8e-06 | 8.1e-06 | 8.100E-06 | 3.000E-07 | 7.800E-06 | 8.400E-06 |
| 1280 | 64 | I64V | A | 1.4e-05 | 6.9e-06 | 8.7e-06 | 9.867E-06 | 3.691E-06 | 6.900E-06 | 1.400E-05 |
| 1281 | 65 | A65A | A | 3.8e-06 | 5.3e-06 | 1.4e-05 | 7.700E-06 | 5.507E-06 | 3.800E-06 | 1.400E-05 |
| 1282 | 65 | A65R | A | 8.1e-06 | 8.8e-06 | 1.5e-05 | 1.063E-05 | 3.798E-06 | 8.100E-06 | 1.500E-05 |
| 1283 | 65 | A65N | A | 8.5e-06 | 9.1e-06 | 5.1e-06 | 7.567E-06 | 2.157E-06 | 5.100E-06 | 9.100E-06 |
| 1284 | 65 | A65D | A | 8.4e-06 | 8.6e-06 | 6.4e-06 | 7.800E-06 | 1.217E-06 | 6.400E-06 | 8.600E-06 |
| 1285 | 65 | A65C | A | 1.3e-05 | 1.3e-05 | 1.1e-05 | 1.233E-05 | 1.155E-06 | 1.100E-05 | 1.300E-05 |
| 1286 | 65 | A65E | A | 4.9e-06 | 5.5e-06 | 7.7e-06 | 6.033E-06 | 1.474E-06 | 4.900E-06 | 7.700E-06 |
| 1287 | 65 | A65Q | A | 6.1e-06 | 1.2e-05 | 7.4e-06 | 8.500E-06 | 3.100E-06 | 6.100E-06 | 1.200E-05 |
| 1288 | 65 | A65G | A | 4.5e-06 | 7.6e-06 | 6.3e-06 | 6.133E-06 | 1.557E-06 | 4.500E-06 | 7.600E-06 |
| 1289 | 65 | A65H | A | 7.9e-06 | 9e-06 | 5.1e-06 | 7.333E-06 | 2.011E-06 | 5.100E-06 | 9.000E-06 |
| 1290 | 65 | A65I | A | 9.7e-06 | 6.8e-06 | 7.8e-06 | 8.100E-06 | 1.473E-06 | 6.800E-06 | 9.700E-06 |
| 1291 | 65 | A65L | A | 9.8e-06 | 9.1e-06 | 1.1e-05 | 9.967E-06 | 9.609E-07 | 9.100E-06 | 1.100E-05 |
| 1292 | 65 | A65K | A | 8.4e-06 | 8.1e-06 | 9.4e-06 | 8.633E-06 | 6.807E-07 | 8.100E-06 | 9.400E-06 |
| 1293 | 65 | A65M | A | 7.1e-06 | 1.7e-05 | 8.7e-06 | 1.093E-05 | 5.314E-06 | 7.100E-06 | 1.700E-05 |
| 1294 | 65 | A65F | A | 8e-06 | 6.5e-06 | 5.6e-06 | 6.700E-06 | 1.212E-06 | 5.600E-06 | 8.000E-06 |
| 1295 | 65 | A65P | A | 5.5e-06 | 6.2e-06 | 9.3e-06 | 7.000E-06 | 2.022E-06 | 5.500E-06 | 9.300E-06 |
| 1296 | 65 | A65S | A | 7.4e-06 | 1.2e-05 | 1.8e-05 | 1.247E-05 | 5.315E-06 | 7.400E-06 | 1.800E-05 |
| 1297 | 65 | A65T | A | 1.2e-05 | 8.8e-06 | 8.2e-06 | 9.667E-06 | 2.043E-06 | 8.200E-06 | 1.200E-05 |
| 1298 | 65 | A65W | A | 8.3e-06 | 1.4e-05 | 1.3e-05 | 1.177E-05 | 3.044E-06 | 8.300E-06 | 1.400E-05 |
| 1299 | 65 | A65Y | A | 6e-06 | 9.1e-06 | 7.7e-06 | 7.600E-06 | 1.552E-06 | 6.000E-06 | 9.100E-06 |
| 1300 | 65 | A65V | A | 6.2e-06 | 6.9e-06 | 1.1e-05 | 8.033E-06 | 2.593E-06 | 6.200E-06 | 1.100E-05 |
| 1301 | 66 | H66A | A | 4.9e-06 | 7.3e-06 | 9.4e-06 | 7.200E-06 | 2.252E-06 | 4.900E-06 | 9.400E-06 |
| 1302 | 66 | H66R | A | 1.4e-05 | 7.5e-06 | 4.5e-06 | 8.667E-06 | 4.856E-06 | 4.500E-06 | 1.400E-05 |
| 1303 | 66 | H66N | A | 4.8e-06 | 9.5e-06 | 9.8e-06 | 8.033E-06 | 2.804E-06 | 4.800E-06 | 9.800E-06 |
| 1304 | 66 | H66D | A | 6e-06 | 9.6e-06 | 6.8e-06 | 7.467E-06 | 1.890E-06 | 6.000E-06 | 9.600E-06 |
| 1305 | 66 | H66C | A | 4.6e-06 | 7e-06 | 5.1e-06 | 5.567E-06 | 1.266E-06 | 4.600E-06 | 7.000E-06 |
| 1306 | 66 | H66E | A | 5e-06 | 7.8e-06 | 9.2e-06 | 7.333E-06 | 2.139E-06 | 5.000E-06 | 9.200E-06 |
| 1307 | 66 | H66Q | A | 8.2e-06 | 8.8e-06 | 4.8e-06 | 7.267E-06 | 2.157E-06 | 4.800E-06 | 8.800E-06 |
| 1308 | 66 | H66G | A | 4.9e-06 | 5.5e-06 | 5e-06 | 5.133E-06 | 3.215E-07 | 4.900E-06 | 5.500E-06 |
| 1309 | 66 | H66H | A | 1e-05 | 6.9e-06 | 1.4e-05 | 1.030E-05 | 3.559E-06 | 6.900E-06 | 1.400E-05 |
| 1310 | 66 | H66I | A | 1e-05 | 6.6e-06 | 1.3e-05 | 9.867E-06 | 3.202E-06 | 6.600E-06 | 1.300E-05 |
| 1311 | 66 | H66L | A | 1.6e-05 | 6.1e-06 | 6.2e-06 | 9.433E-06 | 5.687E-06 | 6.100E-06 | 1.600E-05 |
| 1312 | 66 | H66K | A | 7.6e-06 | 5.6e-06 | 6.9e-06 | 6.700E-06 | 1.015E-06 | 5.600E-06 | 7.600E-06 |
| 1313 | 66 | H66M | A | 6.8e-06 | 7.8e-06 | 8.5e-06 | 7.700E-06 | 8.544E-07 | 6.800E-06 | 8.500E-06 |
| 1314 | 66 | H66F | A | 5.8e-06 | 1.1e-05 | 1.2e-05 | 9.600E-06 | 3.329E-06 | 5.800E-06 | 1.200E-05 |
| 1315 | 66 | H66P | A | 9.5e-06 | 7.6e-06 | 8.2e-06 | 8.433E-06 | 9.713E-07 | 7.600E-06 | 9.500E-06 |
| 1316 | 66 | H66S | A | 6.8e-06 | 1.3e-05 | 5e-06 | 8.267E-06 | 4.197E-06 | 5.000E-06 | 1.300E-05 |
| 1317 | 66 | H66T | A | 7.3e-06 | 8.8e-06 | 1e-05 | 8.700E-06 | 1.353E-06 | 7.300E-06 | 1.000E-05 |
| 1318 | 66 | H66W | A | 9e-06 | 5.9e-06 | 1.3e-05 | 9.300E-06 | 3.559E-06 | 5.900E-06 | 1.300E-05 |
| 1319 | 66 | H66Y | A | 1.2e-05 | 9.6e-06 | 5.4e-06 | 9.000E-06 | 3.341E-06 | 5.400E-06 | 1.200E-05 |
| 1320 | 66 | H66V | A | 7.8e-06 | 1.1e-05 | 1e-05 | 9.600E-06 | 1.637E-06 | 7.800E-06 | 1.100E-05 |
| 1321 | 67 | N67A | A | 9.6e-06 | 7e-06 | 1.5e-05 | 1.053E-05 | 4.081E-06 | 7.000E-06 | 1.500E-05 |
| 1322 | 67 | N67R | A | 1.3e-05 | 6.6e-06 | 4.6e-06 | 8.067E-06 | 4.388E-06 | 4.600E-06 | 1.300E-05 |
| 1323 | 67 | N67N | A | 5.6e-06 | 5.6e-06 | 8e-06 | 6.400E-06 | 1.386E-06 | 5.600E-06 | 8.000E-06 |
| 1324 | 67 | N67D | A | 4.9e-06 | 1e-05 | 4.3e-06 | 6.400E-06 | 3.132E-06 | 4.300E-06 | 1.000E-05 |
| 1325 | 67 | N67C | A | 1.1e-05 | 6.7e-06 | 8.5e-06 | 8.733E-06 | 2.159E-06 | 6.700E-06 | 1.100E-05 |
| 1326 | 67 | N67E | A | 9e-06 | 7.4e-06 | 6e-06 | 7.467E-06 | 1.501E-06 | 6.000E-06 | 9.000E-06 |
| 1327 | 67 | N67Q | A | 7e-06 | 8.9e-06 | 9.3e-06 | 8.400E-06 | 1.229E-06 | 7.000E-06 | 9.300E-06 |
| 1328 | 67 | N67G | A | 8.3e-06 | 1.3e-05 | 1.2e-05 | 1.110E-05 | 2.476E-06 | 8.300E-06 | 1.300E-05 |
| 1329 | 67 | N67H | A | 1.1e-05 | 5.5e-06 | 8.2e-06 | 8.233E-06 | 2.750E-06 | 5.500E-06 | 1.100E-05 |
| 1330 | 67 | N67I | A | 9.6e-06 | 8e-06 | 5.3e-06 | 7.633E-06 | 2.173E-06 | 5.300E-06 | 9.600E-06 |
| 1331 | 67 | N67L | A | 6.8e-06 | 4.8e-06 | 6.6e-06 | 6.067E-06 | 1.102E-06 | 4.800E-06 | 6.800E-06 |
| 1332 | 67 | N67K | A | 1.4e-05 | 5.7e-06 | 6.8e-06 | 8.833E-06 | 4.508E-06 | 5.700E-06 | 1.400E-05 |
| 1333 | 67 | N67M | A | 5.4e-06 | 5.7e-06 | 4.5e-06 | 5.200E-06 | 6.245E-07 | 4.500E-06 | 5.700E-06 |
| 1334 | 67 | N67F | A | 1.6e-05 | 7.5e-06 | 6.6e-06 | 1.003E-05 | 5.187E-06 | 6.600E-06 | 1.600E-05 |
| 1335 | 67 | N67P | A | 6.5e-06 | 5.2e-06 | 1e-05 | 7.233E-06 | 2.483E-06 | 5.200E-06 | 1.000E-05 |
| 1336 | 67 | N67S | A | 9.6e-06 | 1.1e-05 | 1e-05 | 1.020E-05 | 7.211E-07 | 9.600E-06 | 1.100E-05 |
| 1337 | 67 | N67T | A | 8.7e-06 | 7e-06 | 3.7e-06 | 6.467E-06 | 2.542E-06 | 3.700E-06 | 8.700E-06 |
| 1338 | 67 | N67W | A | 9.3e-06 | 8.7e-06 | 9.5e-06 | 9.167E-06 | 4.163E-07 | 8.700E-06 | 9.500E-06 |
| 1339 | 67 | N67Y | A | 7.4e-06 | 8.6e-06 | 7.8e-06 | 7.933E-06 | 6.110E-07 | 7.400E-06 | 8.600E-06 |
| 1340 | 67 | N67V | A | 6.9e-06 | 9.1e-06 | 9.5e-06 | 8.500E-06 | 1.400E-06 | 6.900E-06 | 9.500E-06 |
| 1341 | 68 | Q68A | A | 1.4e-05 | 8.3e-06 | 8.7e-06 | 1.033E-05 | 3.182E-06 | 8.300E-06 | 1.400E-05 |
| 1342 | 68 | Q68R | A | 9.9e-06 | 7.1e-06 | 1.3e-05 | 1.000E-05 | 2.951E-06 | 7.100E-06 | 1.300E-05 |
| 1343 | 68 | Q68N | A | 1.2e-05 | 1.3e-05 | 4.8e-06 | 9.933E-06 | 4.474E-06 | 4.800E-06 | 1.300E-05 |
| 1344 | 68 | Q68D | A | 5.5e-06 | 1.6e-05 | 1.2e-05 | 1.117E-05 | 5.299E-06 | 5.500E-06 | 1.600E-05 |
| 1345 | 68 | Q68C | A | 8.6e-06 | 9.6e-06 | 8.6e-06 | 8.933E-06 | 5.774E-07 | 8.600E-06 | 9.600E-06 |
| 1346 | 68 | Q68E | A | 1.3e-05 | 7.1e-06 | 6e-06 | 8.700E-06 | 3.764E-06 | 6.000E-06 | 1.300E-05 |
| 1347 | 68 | Q68Q | A | 7.6e-06 | 3.3e-06 | 7.1e-06 | 6.000E-06 | 2.352E-06 | 3.300E-06 | 7.600E-06 |
| 1348 | 68 | Q68G | A | 5.1e-06 | 4.8e-06 | 6.6e-06 | 5.500E-06 | 9.644E-07 | 4.800E-06 | 6.600E-06 |
| 1349 | 68 | Q68H | A | 8.6e-06 | 7.9e-06 | 4.1e-06 | 6.867E-06 | 2.421E-06 | 4.100E-06 | 8.600E-06 |
| 1350 | 68 | Q68I | A | 1.1e-05 | 6.4e-06 | 6e-06 | 7.800E-06 | 2.778E-06 | 6.000E-06 | 1.100E-05 |
| 1351 | 68 | Q68L | A | 6.3e-06 | 1.1e-05 | 5e-06 | 7.433E-06 | 3.156E-06 | 5.000E-06 | 1.100E-05 |
| 1352 | 68 | Q68K | A | 8.3e-06 | 9.8e-06 | 1e-05 | 9.367E-06 | 9.292E-07 | 8.300E-06 | 1.000E-05 |
| 1353 | 68 | Q68M | A | 3.5e-06 | 1.1e-05 | 8.7e-06 | 7.733E-06 | 3.842E-06 | 3.500E-06 | 1.100E-05 |
| 1354 | 68 | Q68F | A | 7.4e-06 | 3.8e-06 | 1e-05 | 7.067E-06 | 3.113E-06 | 3.800E-06 | 1.000E-05 |
| 1355 | 68 | Q68P | A | 7.4e-06 | 1.4e-05 | 9.6e-06 | 1.033E-05 | 3.361E-06 | 7.400E-06 | 1.400E-05 |
| 1356 | 68 | Q68S | A | 6.2e-06 | 7.9e-06 | 9e-06 | 7.700E-06 | 1.411E-06 | 6.200E-06 | 9.000E-06 |
| 1357 | 68 | Q68T | A | 8.4e-06 | 1e-05 | 6.2e-06 | 8.200E-06 | 1.908E-06 | 6.200E-06 | 1.000E-05 |
| 1358 | 68 | Q68W | A | 9.1e-06 | 5.8e-06 | 8.4e-06 | 7.767E-06 | 1.739E-06 | 5.800E-06 | 9.100E-06 |
| 1359 | 68 | Q68Y | A | 7.7e-06 | 6.4e-06 | 6.9e-06 | 7.000E-06 | 6.557E-07 | 6.400E-06 | 7.700E-06 |

| | | | | | | | | | | |
|---|---|---|---|---|---|---|---|---|---|---|
| 1360 | 68 | Q68V | A | 1.4e-05 | 7.4e-06 | 5.2e-06 | 8.867E-06 | 4.580E-06 | 5.200E-06 | 1.400E-05 |
| 1361 | 69 | V69A | A | 5.3e-06 | 8.2e-06 | 6.9e-06 | 6.800E-06 | 1.453E-06 | 5.300E-06 | 8.200E-06 |
| 1362 | 69 | V69R | A | 5e-06 | 8.2e-06 | 6.5e-06 | 6.567E-06 | 1.601E-06 | 5.000E-06 | 8.200E-06 |
| 1363 | 69 | V69N | A | 6.4e-06 | 1e-05 | 7.7e-06 | 8.033E-06 | 1.823E-06 | 6.400E-06 | 1.000E-05 |
| 1364 | 69 | V69D | A | 1.1e-05 | 7.3e-06 | 9.2e-06 | 9.167E-06 | 1.850E-06 | 7.300E-06 | 1.100E-05 |
| 1365 | 69 | V69C | A | 6.6e-06 | 1.2e-05 | 1.1e-05 | 9.867E-06 | 2.873E-06 | 6.600E-06 | 1.200E-05 |
| 1366 | 69 | V69E | A | 6.3e-06 | 8.6e-06 | 8e-06 | 7.633E-06 | 1.193E-06 | 6.300E-06 | 8.600E-06 |
| 1367 | 69 | V69Q | A | 3.8e-06 | 4.4e-06 | 4.3e-06 | 4.167E-06 | 3.215E-07 | 3.800E-06 | 4.400E-06 |
| 1368 | 69 | V69G | A | 8.3e-06 | 1.4e-05 | 6.9e-06 | 9.733E-06 | 3.761E-06 | 6.900E-06 | 1.400E-05 |
| 1369 | 69 | V69H | A | 4.4e-06 | 9.3e-06 | 7.4e-06 | 7.033E-06 | 2.470E-06 | 4.400E-06 | 9.300E-06 |
| 1370 | 69 | V69I | A | 5.9e-06 | 6.4e-06 | 1.6e-05 | 9.433E-06 | 5.692E-06 | 5.900E-06 | 1.600E-05 |
| 1371 | 69 | V69L | A | 1e-05 | 5.3e-06 | 5.7e-06 | 7.000E-06 | 2.606E-06 | 5.300E-06 | 1.000E-05 |
| 1372 | 69 | V69K | A | 7.3e-06 | 6.2e-06 | 1.4e-05 | 9.167E-06 | 4.222E-06 | 6.200E-06 | 1.400E-05 |
| 1373 | 69 | V69M | A | 9e-06 | 7.3e-06 | 9.9e-06 | 8.733E-06 | 1.320E-06 | 7.300E-06 | 9.900E-06 |
| 1374 | 69 | V69F | A | 5.5e-06 | 9.1e-06 | 1.1e-05 | 8.533E-06 | 2.793E-06 | 5.500E-06 | 1.100E-05 |
| 1375 | 69 | V69P | A | 9e-06 | 5.3e-06 | 1.1e-05 | 8.433E-06 | 2.892E-06 | 5.300E-06 | 1.100E-05 |
| 1376 | 69 | V69S | A | 3.9e-06 | 8.7e-06 | 8.5e-06 | 7.033E-06 | 2.715E-06 | 3.900E-06 | 8.700E-06 |
| 1377 | 69 | V69T | A | 5.2e-06 | 6.6e-06 | 7.3e-06 | 6.367E-06 | 1.069E-06 | 5.200E-06 | 7.300E-06 |
| 1378 | 69 | V69W | A | 9.7e-06 | 5.6e-06 | 8.6e-06 | 7.967E-06 | 2.122E-06 | 5.600E-06 | 9.700E-06 |
| 1379 | 69 | V69Y | A | 1e-05 | 5.2e-06 | 6.2e-06 | 7.133E-06 | 2.532E-06 | 5.200E-06 | 1.000E-05 |
| 1380 | 69 | V69V | A | 1.3e-05 | 9.3e-06 | 4.7e-06 | 9.000E-06 | 4.158E-06 | 4.700E-06 | 1.300E-05 |
| 1381 | 70 | R70A | A | 8.6e-06 | 7.4e-06 | 6.3e-06 | 7.433E-06 | 1.150E-06 | 6.300E-06 | 8.600E-06 |
| 1382 | 70 | R70R | A | 8.4e-06 | 9.6e-06 | 8.5e-06 | 8.833E-06 | 6.658E-07 | 8.400E-06 | 9.600E-06 |
| 1383 | 70 | R70N | A | 1e-05 | 6.2e-06 | 7.3e-06 | 7.833E-06 | 1.955E-06 | 6.200E-06 | 1.000E-05 |
| 1384 | 70 | R70D | A | 1.1e-05 | 6.9e-06 | 8e-06 | 8.633E-06 | 2.122E-06 | 6.900E-06 | 1.100E-05 |
| 1385 | 70 | R70C | A | 6.8e-06 | 9.2e-06 | 9.4e-06 | 8.467E-06 | 1.447E-06 | 6.800E-06 | 9.400E-06 |
| 1386 | 70 | R70E | A | 7.1e-06 | 8.4e-06 | 6.1e-06 | 7.200E-06 | 1.153E-06 | 6.100E-06 | 8.400E-06 |
| 1387 | 70 | R70Q | A | 1.3e-05 | 8.9e-06 | 6.4e-06 | 9.433E-06 | 3.332E-06 | 6.400E-06 | 1.300E-05 |
| 1388 | 70 | R70G | A | 6.6e-06 | 7e-06 | 8.7e-06 | 7.433E-06 | 1.115E-06 | 6.600E-06 | 8.700E-06 |
| 1389 | 70 | R70H | A | 6.3e-06 | 1.3e-05 | 7.4e-06 | 8.900E-06 | 3.593E-06 | 6.300E-06 | 1.300E-05 |
| 1390 | 70 | R70I | A | 1.5e-05 | 9e-06 | 8e-06 | 1.067E-05 | 3.786E-06 | 8.000E-06 | 1.500E-05 |
| 1391 | 70 | R70L | A | 6.6e-06 | 1e-05 | 4.4e-06 | 7.000E-06 | 2.821E-06 | 4.400E-06 | 1.000E-05 |
| 1392 | 70 | R70K | A | 1e-05 | 6.7e-06 | 9.5e-06 | 8.733E-06 | 1.779E-06 | 6.700E-06 | 1.000E-05 |
| 1393 | 70 | R70M | A | 4.1e-06 | 7.2e-06 | 9.7e-06 | 7.000E-06 | 2.805E-06 | 4.100E-06 | 9.700E-06 |
| 1394 | 70 | R70F | A | 9.3e-06 | 5.5e-06 | 7.7e-06 | 7.500E-06 | 1.908E-06 | 5.500E-06 | 9.300E-06 |
| 1395 | 70 | R70P | A | 1.1e-05 | 7.8e-06 | 7.7e-06 | 8.833E-06 | 1.877E-06 | 7.700E-06 | 1.100E-05 |
| 1396 | 70 | R70S | A | 5e-06 | 1.1e-05 | 4.8e-06 | 6.933E-06 | 3.523E-06 | 4.800E-06 | 1.100E-05 |
| 1397 | 70 | R70T | A | 6e-06 | 5.6e-06 | 6.2e-06 | 5.933E-06 | 3.055E-07 | 5.600E-06 | 6.200E-06 |
| 1398 | 70 | R70W | A | 9.1e-06 | 5.2e-06 | 6e-06 | 6.767E-06 | 2.060E-06 | 5.200E-06 | 9.100E-06 |
| 1399 | 70 | R70Y | A | 8e-06 | 5.8e-06 | 6.8e-06 | 6.867E-06 | 1.102E-06 | 5.800E-06 | 8.000E-06 |
| 1400 | 70 | R70V | A | 9.6e-06 | 6.8e-06 | 7.1e-06 | 7.833E-06 | 1.537E-06 | 6.800E-06 | 9.600E-06 |
| 1401 | 71 | Q71A | A | 7.1e-06 | 4e-06 | 7e-06 | 6.033E-06 | 1.762E-06 | 4.000E-06 | 7.100E-06 |
| 1402 | 71 | Q71R | A | 8.3e-06 | 1.3e-05 | 8e-06 | 9.767E-06 | 2.804E-06 | 8.000E-06 | 1.300E-05 |
| 1403 | 71 | Q71N | A | 1.1e-05 | 1e-05 | 6.9e-06 | 9.300E-06 | 2.138E-06 | 6.900E-06 | 1.100E-05 |
| 1404 | 71 | Q71D | A | 8.5e-06 | 1.2e-05 | 9e-06 | 9.833E-06 | 1.893E-06 | 8.500E-06 | 1.200E-05 |
| 1405 | 71 | Q71C | A | 4.6e-06 | 8.1e-06 | 7.8e-06 | 6.833E-06 | 1.940E-06 | 4.600E-06 | 8.100E-06 |
| 1406 | 71 | Q71E | A | 1.1e-05 | 9.1e-06 | 5.6e-06 | 8.567E-06 | 2.739E-06 | 5.600E-06 | 1.100E-05 |
| 1407 | 71 | Q71Q | A | 6.7e-06 | 4.6e-06 | 7e-06 | 6.100E-06 | 1.308E-06 | 4.600E-06 | 7.000E-06 |
| 1408 | 71 | Q71G | A | 8.2e-06 | 9.3e-06 | 7.7e-06 | 8.400E-06 | 8.185E-07 | 7.700E-06 | 9.300E-06 |
| 1409 | 71 | Q71H | A | 7.2e-06 | 1.3e-05 | 1.5e-05 | 1.173E-05 | 4.051E-06 | 7.200E-06 | 1.500E-05 |
| 1410 | 71 | Q71I | A | 1.1e-05 | 5.8e-06 | 5.3e-06 | 7.367E-06 | 3.156E-06 | 5.300E-06 | 1.100E-05 |
| 1411 | 71 | Q71L | A | 4.6e-06 | 1.3e-05 | 1.2e-05 | 9.867E-06 | 4.588E-06 | 4.600E-06 | 1.300E-05 |
| 1412 | 71 | Q71K | A | 7.6e-06 | 9.2e-06 | 9e-06 | 8.600E-06 | 8.718E-07 | 7.600E-06 | 9.200E-06 |
| 1413 | 71 | Q71M | A | 5.4e-06 | 7.2e-06 | 7.5e-06 | 6.700E-06 | 1.136E-06 | 5.400E-06 | 7.500E-06 |
| 1414 | 71 | Q71F | A | 4.5e-06 | 9.1e-06 | 6.3e-06 | 6.633E-06 | 2.318E-06 | 4.500E-06 | 9.100E-06 |
| 1415 | 71 | Q71P | A | 7.2e-06 | 7e-06 | 8.6e-06 | 7.600E-06 | 8.718E-07 | 7.000E-06 | 8.600E-06 |
| 1416 | 71 | Q71S | A | 1.3e-05 | 9.7e-06 | 1.3e-05 | 1.190E-05 | 1.905E-06 | 9.700E-06 | 1.300E-05 |
| 1417 | 71 | Q71T | A | 5.8e-06 | 1e-05 | 7.9e-06 | 7.900E-06 | 2.100E-06 | 5.800E-06 | 1.000E-05 |
| 1418 | 71 | Q71W | A | 9.7e-06 | 7.1e-06 | 4.5e-06 | 7.100E-06 | 2.600E-06 | 4.500E-06 | 9.700E-06 |
| 1419 | 71 | Q71Y | A | 6.5e-06 | 9.8e-06 | 6.6e-06 | 7.633E-06 | 1.877E-06 | 6.500E-06 | 9.800E-06 |
| 1420 | 71 | Q71V | A | 6.4e-06 | 6.7e-06 | 8.6e-06 | 7.233E-06 | 1.193E-06 | 6.400E-06 | 8.600E-06 |
| 1421 | 72 | V72A | A | 6.9e-06 | 7.4e-06 | 4.5e-06 | 6.267E-06 | 1.550E-06 | 4.500E-06 | 7.400E-06 |
| 1422 | 72 | V72R | A | 7.7e-06 | 1.1e-05 | 1.2e-05 | 1.023E-05 | 2.250E-06 | 7.700E-06 | 1.200E-05 |
| 1423 | 72 | V72N | A | 9.4e-06 | 1e-05 | 9.4e-06 | 9.600E-06 | 3.464E-07 | 9.400E-06 | 1.000E-05 |
| 1424 | 72 | V72D | A | 7.9e-06 | 9.2e-06 | 8.9e-06 | 8.667E-06 | 6.807E-07 | 7.900E-06 | 9.200E-06 |
| 1425 | 72 | V72C | A | 9.4e-06 | 7.4e-06 | 1.3e-05 | 9.933E-06 | 2.838E-06 | 7.400E-06 | 1.300E-05 |
| 1426 | 72 | V72E | A | 3.8e-06 | 9.8e-06 | 8.1e-06 | 7.233E-06 | 3.092E-06 | 3.800E-06 | 9.800E-06 |
| 1427 | 72 | V72Q | A | 7.9e-06 | 8.4e-06 | 6.8e-06 | 7.700E-06 | 8.185E-07 | 6.800E-06 | 8.400E-06 |
| 1428 | 72 | V72G | A | 1.5e-05 | 1.2e-05 | 1.1e-05 | 1.267E-05 | 2.082E-06 | 1.100E-05 | 1.500E-05 |
| 1429 | 72 | V72H | A | 8.5e-06 | 6.8e-06 | 4.7e-06 | 6.667E-06 | 1.904E-06 | 4.700E-06 | 8.500E-06 |
| 1430 | 72 | V72I | A | 6.3e-06 | 8.6e-06 | 7.7e-06 | 7.533E-06 | 1.159E-06 | 6.300E-06 | 8.600E-06 |
| 1431 | 72 | V72L | A | 9.4e-06 | 6.5e-06 | 5.5e-06 | 7.133E-06 | 2.026E-06 | 5.500E-06 | 9.400E-06 |
| 1432 | 72 | V72K | A | 9.1e-06 | 9.4e-06 | 7.7e-06 | 8.733E-06 | 9.074E-07 | 7.700E-06 | 9.400E-06 |
| 1433 | 72 | V72M | A | 9.2e-06 | 6.5e-06 | 8.7e-06 | 8.133E-06 | 1.436E-06 | 6.500E-06 | 9.200E-06 |
| 1434 | 72 | V72F | A | 8.7e-06 | 7.3e-06 | 6e-06 | 7.333E-06 | 1.350E-06 | 6.000E-06 | 8.700E-06 |
| 1435 | 72 | V72P | A | 1e-05 | 1e-05 | 5.8e-06 | 8.600E-06 | 2.425E-06 | 5.800E-06 | 1.000E-05 |
| 1436 | 72 | V72S | A | 1.4e-05 | 1e-05 | 5.2e-06 | 9.733E-06 | 4.406E-06 | 5.200E-06 | 1.400E-05 |
| 1437 | 72 | V72T | A | 4.4e-06 | 8.2e-06 | 1.3e-05 | 8.533E-06 | 4.310E-06 | 4.400E-06 | 1.300E-05 |
| 1438 | 72 | V72W | A | 1.1e-05 | 1e-05 | 5.1e-06 | 8.700E-06 | 3.158E-06 | 5.100E-06 | 1.100E-05 |
| 1439 | 72 | V72Y | A | 1e-05 | 9.8e-06 | 6.4e-06 | 8.733E-06 | 2.023E-06 | 6.400E-06 | 1.000E-05 |
| 1440 | 72 | V72V | A | 1.3e-05 | 5.2e-06 | 6.9e-06 | 8.367E-06 | 4.102E-06 | 5.200E-06 | 1.300E-05 |
| 1441 | 73 | P73A | A | 1e-05 | 9.8e-06 | 7.3e-06 | 9.033E-06 | 1.504E-06 | 7.300E-06 | 1.000E-05 |
| 1442 | 73 | P73R | A | 8.3e-06 | 8.5e-06 | 7.2e-06 | 8.000E-06 | 7.000E-07 | 7.200E-06 | 8.500E-06 |
| 1443 | 73 | P73N | A | 9e-06 | 1.1e-05 | 4.9e-06 | 8.300E-06 | 3.110E-06 | 4.900E-06 | 1.100E-05 |
| 1444 | 73 | P73D | A | 6.2e-06 | 1.3e-05 | 4.2e-06 | 7.800E-06 | 4.613E-06 | 4.200E-06 | 1.300E-05 |

| | | | | | | | | | | |
|---|---|---|---|---|---|---|---|---|---|---|
| 1445 | 73 | P73C | A | 6.6e-06 | 8.5e-06 | 7.2e-06 | 7.433E-06 | 9.713E-07 | 6.600E-06 | 8.500E-06 |
| 1446 | 73 | P73E | A | 1.3e-05 | 7.8e-06 | 8.7e-06 | 9.833E-06 | 2.779E-06 | 7.800E-06 | 1.300E-05 |
| 1447 | 73 | P73Q | A | 7e-06 | 4e-06 | 7.8e-06 | 6.267E-06 | 2.003E-06 | 4.000E-06 | 7.800E-06 |
| 1448 | 73 | P73G | A | 5.7e-06 | 7.6e-06 | 5.9e-06 | 6.400E-06 | 1.044E-06 | 5.700E-06 | 7.600E-06 |
| 1449 | 73 | P73H | A | 1.2e-05 | 9.7e-06 | 9.5e-06 | 1.040E-05 | 1.389E-06 | 9.500E-06 | 1.200E-05 |
| 1450 | 73 | P73I | A | 1.1e-05 | 1.1e-05 | 6.2e-06 | 9.400E-06 | 2.771E-06 | 6.200E-06 | 1.100E-05 |
| 1451 | 73 | P73L | A | 9.1e-06 | 7.8e-06 | 8e-06 | 8.300E-06 | 7.000E-07 | 7.800E-06 | 9.100E-06 |
| 1452 | 73 | P73K | A | 9.5e-06 | 7.8e-06 | 6.9e-06 | 8.067E-06 | 1.320E-06 | 6.900E-06 | 9.500E-06 |
| 1453 | 73 | P73M | A | 2.9e-06 | 1.1e-05 | 6.8e-06 | 6.900E-06 | 4.051E-06 | 2.900E-06 | 1.100E-05 |
| 1454 | 73 | P73F | A | 6.3e-06 | 8.9e-06 | 9.8e-06 | 8.333E-06 | 1.818E-06 | 6.300E-06 | 9.800E-06 |
| 1455 | 73 | P73P | A | 6.6e-06 | 9.3e-06 | 8.5e-06 | 8.133E-06 | 1.387E-06 | 6.600E-06 | 9.300E-06 |
| 1456 | 73 | P73S | A | 5.9e-06 | 8.1e-06 | 8.5e-06 | 7.500E-06 | 1.400E-06 | 5.900E-06 | 8.500E-06 |
| 1457 | 73 | P73T | A | 7.7e-06 | 6.4e-06 | 6e-06 | 6.700E-06 | 8.888E-07 | 6.000E-06 | 7.700E-06 |
| 1458 | 73 | P73W | A | 9.9e-06 | 9.5e-06 | 9.5e-06 | 9.633E-06 | 2.309E-07 | 9.500E-06 | 9.900E-06 |
| 1459 | 73 | P73Y | A | 5e-06 | 9.4e-06 | 6.4e-06 | 6.933E-06 | 2.248E-06 | 5.000E-06 | 9.400E-06 |
| 1460 | 73 | P73V | A | 5e-06 | 4.3e-06 | 5.8e-06 | 5.033E-06 | 7.506E-07 | 4.300E-06 | 5.800E-06 |
| 1461 | 74 | L74A | A | 6.6e-06 | 8.8e-06 | 7.3e-06 | 7.567E-06 | 1.124E-06 | 6.600E-06 | 8.800E-06 |
| 1462 | 74 | L74R | A | 1.2e-05 | 6.3e-06 | 3.4e-06 | 7.233E-06 | 4.375E-06 | 3.400E-06 | 1.200E-05 |
| 1463 | 74 | L74N | A | 8.9e-06 | 5.9e-06 | 1.3e-05 | 9.267E-06 | 3.564E-06 | 5.900E-06 | 1.300E-05 |
| 1464 | 74 | L74D | A | 7.1e-06 | 1.1e-05 | 6.1e-06 | 8.067E-06 | 2.589E-06 | 6.100E-06 | 1.100E-05 |
| 1465 | 74 | L74C | A | 9.2e-06 | 5.4e-06 | 9.9e-06 | 8.167E-06 | 2.421E-06 | 5.400E-06 | 9.900E-06 |
| 1466 | 74 | L74E | A | 9.7e-06 | 4.9e-06 | 3.9e-06 | 6.167E-06 | 3.101E-06 | 3.900E-06 | 9.700E-06 |
| 1467 | 74 | L74Q | A | 9.9e-06 | 7.4e-06 | 9.7e-06 | 9.000E-06 | 1.389E-06 | 7.400E-06 | 9.900E-06 |
| 1468 | 74 | L74G | A | 9.5e-06 | 4.9e-06 | 7.3e-06 | 7.233E-06 | 2.301E-06 | 4.900E-06 | 9.500E-06 |
| 1469 | 74 | L74H | A | 1.2e-05 | 1.1e-05 | 1.2e-05 | 1.167E-05 | 5.774E-07 | 1.100E-05 | 1.200E-05 |
| 1470 | 74 | L74I | A | 1.3e-05 | 5e-06 | 6.5e-06 | 8.167E-06 | 4.252E-06 | 5.000E-06 | 1.300E-05 |
| 1471 | 74 | L74L | A | 7.7e-06 | 7.3e-06 | 1e-05 | 8.333E-06 | 1.457E-06 | 7.300E-06 | 1.000E-05 |
| 1472 | 74 | L74K | A | 4.6e-06 | 1.1e-05 | 3.3e-06 | 6.300E-06 | 4.122E-06 | 3.300E-06 | 1.100E-05 |
| 1473 | 74 | L74M | A | 1.1e-05 | 7.3e-06 | 9.4e-06 | 9.233E-06 | 1.856E-06 | 7.300E-06 | 1.100E-05 |
| 1474 | 74 | L74F | A | 1.2e-05 | 9.9e-06 | 6.6e-06 | 9.500E-06 | 2.722E-06 | 6.600E-06 | 1.200E-05 |
| 1475 | 74 | L74P | A | 1e-05 | 7.2e-06 | 7.9e-06 | 8.367E-06 | 1.457E-06 | 7.200E-06 | 1.000E-05 |
| 1476 | 74 | L74S | A | 5.9e-06 | 4.8e-06 | 9.2e-06 | 6.633E-06 | 2.290E-06 | 4.800E-06 | 9.200E-06 |
| 1477 | 74 | L74T | A | 8.7e-06 | 1e-05 | 9.7e-06 | 9.467E-06 | 6.807E-07 | 8.700E-06 | 1.000E-05 |
| 1478 | 74 | L74W | A | 6.3e-06 | 1e-05 | 7.1e-06 | 7.800E-06 | 1.947E-06 | 6.300E-06 | 1.000E-05 |
| 1479 | 74 | L74Y | A | 8.6e-06 | 8.8e-06 | 9.4e-06 | 8.933E-06 | 4.163E-07 | 8.600E-06 | 9.400E-06 |
| 1480 | 74 | L74V | A | 7.4e-06 | 8.7e-06 | 7.3e-06 | 7.800E-06 | 7.810E-07 | 7.300E-06 | 8.700E-06 |
| 1481 | 75 | Q75A | A | 7.2e-06 | 8e-06 | 9.4e-06 | 8.200E-06 | 1.114E-06 | 7.200E-06 | 9.400E-06 |
| 1482 | 75 | Q75R | A | 6.5e-06 | 7e-06 | 9.9e-06 | 7.800E-06 | 1.836E-06 | 6.500E-06 | 9.900E-06 |
| 1483 | 75 | Q75N | A | 6.1e-06 | 9.9e-06 | 8e-06 | 8.000E-06 | 1.900E-06 | 6.100E-06 | 9.900E-06 |
| 1484 | 75 | Q75D | A | 1e-05 | 8.1e-06 | 1.1e-05 | 9.700E-06 | 1.473E-06 | 8.100E-06 | 1.100E-05 |
| 1485 | 75 | Q75C | A | 9.3e-06 | 7.5e-06 | 1.2e-05 | 9.600E-06 | 2.265E-06 | 7.500E-06 | 1.200E-05 |
| 1486 | 75 | Q75E | A | 9.7e-06 | 8.7e-06 | 9.5e-06 | 9.300E-06 | 5.292E-07 | 8.700E-06 | 9.700E-06 |
| 1487 | 75 | Q75Q | A | 1.2e-05 | 1.2e-05 | 1.4e-05 | 1.267E-05 | 1.155E-06 | 1.200E-05 | 1.400E-05 |
| 1488 | 75 | Q75G | A | 6.7e-06 | 9.9e-06 | 8e-06 | 8.200E-06 | 1.609E-06 | 6.700E-06 | 9.900E-06 |
| 1489 | 75 | Q75H | A | 5.8e-06 | 1.1e-05 | 6.2e-06 | 7.667E-06 | 2.894E-06 | 5.800E-06 | 1.100E-05 |
| 1490 | 75 | Q75I | A | 5.1e-06 | 8.2e-06 | 3.4e-06 | 5.567E-06 | 2.434E-06 | 3.400E-06 | 8.200E-06 |
| 1491 | 75 | Q75L | A | 5.5e-06 | 8.8e-06 | 1e-05 | 8.100E-06 | 2.330E-06 | 5.500E-06 | 1.000E-05 |
| 1492 | 75 | Q75K | A | 8.3e-06 | 3.1e-06 | 9.1e-06 | 6.833E-06 | 3.258E-06 | 3.100E-06 | 9.100E-06 |
| 1493 | 75 | Q75M | A | 1.9e-05 | 5e-06 | 5.1e-06 | 9.700E-06 | 8.054E-06 | 5.000E-06 | 1.900E-05 |
| 1494 | 75 | Q75F | A | 8.5e-06 | 7.1e-06 | 9.4e-06 | 8.333E-06 | 1.159E-06 | 7.100E-06 | 9.400E-06 |
| 1495 | 75 | Q75P | A | 7e-06 | 1.2e-05 | 6.1e-06 | 8.367E-06 | 3.179E-06 | 6.100E-06 | 1.200E-05 |
| 1496 | 75 | Q75S | A | 5.4e-06 | 6.5e-06 | 4.9e-06 | 5.600E-06 | 8.185E-07 | 4.900E-06 | 6.500E-06 |
| 1497 | 75 | Q75T | A | 1e-05 | 1e-05 | 7.1e-06 | 9.033E-06 | 1.674E-06 | 7.100E-06 | 1.000E-05 |
| 1498 | 75 | Q75W | A | 7.8e-06 | 6.1e-06 | 1.1e-05 | 8.300E-06 | 2.488E-06 | 6.100E-06 | 1.100E-05 |
| 1499 | 75 | Q75Y | A | 5.3e-06 | 7.6e-06 | 4.5e-06 | 5.800E-06 | 1.609E-06 | 4.500E-06 | 7.600E-06 |
| 1500 | 75 | Q75V | A | 6.9e-06 | 9.7e-06 | 5.2e-06 | 7.267E-06 | 2.272E-06 | 5.200E-06 | 9.700E-06 |
| 1501 | 76 | R76A | A | 1.7e-05 | 7e-06 | 6.6e-06 | 1.020E-05 | 5.892E-06 | 6.600E-06 | 1.700E-05 |
| 1502 | 76 | R76R | A | 7.7e-06 | 6.6e-06 | 9.4e-06 | 7.900E-06 | 1.411E-06 | 6.600E-06 | 9.400E-06 |
| 1503 | 76 | R76N | A | 4.1e-06 | 7.7e-06 | 8.9e-06 | 6.900E-06 | 2.498E-06 | 4.100E-06 | 8.900E-06 |
| 1504 | 76 | R76D | A | 6.3e-06 | 1.2e-05 | 6.1e-06 | 8.133E-06 | 3.350E-06 | 6.100E-06 | 1.200E-05 |
| 1505 | 76 | R76C | A | 8.1e-06 | 4.8e-06 | 1.3e-05 | 8.633E-06 | 4.126E-06 | 4.800E-06 | 1.300E-05 |
| 1506 | 76 | R76E | A | 1.4e-05 | 1.1e-05 | 7.8e-06 | 1.093E-05 | 3.101E-06 | 7.800E-06 | 1.400E-05 |
| 1507 | 76 | R76Q | A | 7.5e-06 | 6.4e-06 | 5.8e-06 | 6.567E-06 | 8.622E-07 | 5.800E-06 | 7.500E-06 |
| 1508 | 76 | R76G | A | 1.1e-05 | 1.1e-05 | 4.9e-06 | 8.967E-06 | 3.522E-06 | 4.900E-06 | 1.100E-05 |
| 1509 | 76 | R76H | A | 8.3e-06 | 4.7e-06 | 9.8e-06 | 7.600E-06 | 2.621E-06 | 4.700E-06 | 9.800E-06 |
| 1510 | 76 | R76I | A | 7e-06 | 1.4e-05 | 9.6e-06 | 1.020E-05 | 3.538E-06 | 7.000E-06 | 1.400E-05 |
| 1511 | 76 | R76L | A | 7.2e-06 | 9.2e-06 | 9.1e-06 | 8.500E-06 | 1.127E-06 | 7.200E-06 | 9.200E-06 |
| 1512 | 76 | R76K | A | 8.1e-06 | 1.1e-05 | 5.1e-06 | 8.067E-06 | 2.950E-06 | 5.100E-06 | 1.100E-05 |
| 1513 | 76 | R76M | A | 1.5e-05 | 6.5e-06 | 5.6e-06 | 9.033E-06 | 5.187E-06 | 5.600E-06 | 1.500E-05 |
| 1514 | 76 | R76F | A | 8.1e-06 | 7e-06 | 8.1e-06 | 7.733E-06 | 6.351E-07 | 7.000E-06 | 8.100E-06 |
| 1515 | 76 | R76P | A | 9.6e-06 | 1.2e-05 | 5.5e-06 | 9.033E-06 | 3.287E-06 | 5.500E-06 | 1.200E-05 |
| 1516 | 76 | R76S | A | 4.3e-06 | 9.7e-06 | 4.6e-06 | 6.200E-06 | 3.035E-06 | 4.300E-06 | 9.700E-06 |
| 1517 | 76 | R76T | A | 6.3e-06 | 7.2e-06 | 6e-06 | 6.500E-06 | 6.245E-07 | 6.000E-06 | 7.200E-06 |
| 1518 | 76 | R76W | A | 5e-06 | 1.3e-05 | 7e-06 | 8.333E-06 | 4.163E-06 | 5.000E-06 | 1.300E-05 |
| 1519 | 76 | R76Y | A | 6.7e-06 | 9.3e-06 | 6.6e-06 | 7.533E-06 | 1.531E-06 | 6.600E-06 | 9.300E-06 |
| 1520 | 76 | R76V | A | 5.2e-06 | 8.2e-06 | 4.5e-06 | 5.967E-06 | 1.966E-06 | 4.500E-06 | 8.200E-06 |
| 1521 | 77 | L77A | A | 8.7e-06 | 9.9e-06 | 6.5e-06 | 8.367E-06 | 1.724E-06 | 6.500E-06 | 9.900E-06 |
| 1522 | 77 | L77R | A | 7.1e-06 | 5.4e-06 | 8e-06 | 6.833E-06 | 1.320E-06 | 5.400E-06 | 8.000E-06 |
| 1523 | 77 | L77N | A | 9.2e-06 | 7.9e-06 | 9.3e-06 | 8.800E-06 | 7.810E-07 | 7.900E-06 | 9.300E-06 |
| 1524 | 77 | L77D | A | 7.8e-06 | 5.7e-06 | 1.3e-05 | 8.833E-06 | 3.758E-06 | 5.700E-06 | 1.300E-05 |
| 1525 | 77 | L77C | A | 6.1e-06 | 7.9e-06 | 8.3e-06 | 7.433E-06 | 1.172E-06 | 6.100E-06 | 8.300E-06 |
| 1526 | 77 | L77E | A | 7.5e-06 | 1.1e-05 | 8.9e-06 | 9.133E-06 | 1.762E-06 | 7.500E-06 | 1.100E-05 |
| 1527 | 77 | L77Q | A | 6.7e-06 | 9.1e-06 | 8.3e-06 | 8.033E-06 | 1.222E-06 | 6.700E-06 | 9.100E-06 |
| 1528 | 77 | L77G | A | 6.9e-06 | 1e-05 | 5.6e-06 | 7.500E-06 | 2.261E-06 | 5.600E-06 | 1.000E-05 |
| 1529 | 77 | L77H | A | 9e-06 | 9.6e-06 | 5.6e-06 | 8.067E-06 | 2.157E-06 | 5.600E-06 | 9.600E-06 |

| | | | | | | | | | | |
|---|---|---|---|---|---|---|---|---|---|---|
| 1530 | 77 | L77I | A | 1.3e-05 | 8.4e-06 | 5e-06 | 8.800E-06 | 4.015E-06 | 5.000E-06 | 1.300E-05 |
| 1531 | 77 | L77L | A | 6.7e-06 | 7.5e-06 | 5.8e-06 | 6.667E-06 | 8.505E-07 | 5.800E-06 | 7.500E-06 |
| 1532 | 77 | L77K | A | 1.2e-05 | 5.1e-06 | 5.5e-06 | 7.533E-06 | 3.873E-06 | 5.100E-06 | 1.200E-05 |
| 1533 | 77 | L77M | A | 6.3e-06 | 6e-06 | 4.3e-06 | 5.533E-06 | 1.079E-06 | 4.300E-06 | 6.300E-06 |
| 1534 | 77 | L77F | A | 8.4e-06 | 5.9e-06 | 7.4e-06 | 7.233E-06 | 1.258E-06 | 5.900E-06 | 8.400E-06 |
| 1535 | 77 | L77P | A | 8.6e-06 | 1.3e-05 | 8.9e-06 | 1.017E-05 | 2.458E-06 | 8.600E-06 | 1.300E-05 |
| 1536 | 77 | L77S | A | 4.8e-06 | 6.9e-06 | 9.4e-06 | 7.033E-06 | 2.303E-06 | 4.800E-06 | 9.400E-06 |
| 1537 | 77 | L77T | A | 1.2e-05 | 1e-05 | 1.8e-05 | 1.333E-05 | 4.163E-06 | 1.000E-05 | 1.800E-05 |
| 1538 | 77 | L77W | A | 4.4e-06 | 9e-06 | 6e-06 | 6.467E-06 | 2.335E-06 | 4.400E-06 | 9.000E-06 |
| 1539 | 77 | L77Y | A | 9e-06 | 1.1e-05 | 6e-06 | 8.667E-06 | 2.517E-06 | 6.000E-06 | 1.100E-05 |
| 1540 | 77 | L77V | A | 6e-06 | 9e-06 | 9.7e-06 | 8.233E-06 | 1.966E-06 | 6.000E-06 | 9.700E-06 |
| 1541 | 78 | R78A | A | 8.2e-06 | 7.6e-06 | 1.3e-05 | 9.600E-06 | 2.960E-06 | 7.600E-06 | 1.300E-05 |
| 1542 | 78 | R78R | A | 6.2e-06 | 7.8e-06 | 3.6e-06 | 5.867E-06 | 2.120E-06 | 3.600E-06 | 7.800E-06 |
| 1543 | 78 | R78N | A | 7.8e-06 | 5e-06 | 6e-06 | 6.267E-06 | 1.419E-06 | 5.000E-06 | 7.800E-06 |
| 1544 | 78 | R78D | A | 4.3e-06 | 7.9e-06 | 1.3e-05 | 8.400E-06 | 4.371E-06 | 4.300E-06 | 1.300E-05 |
| 1545 | 78 | R78C | A | 7.8e-06 | 6.3e-06 | 8.7e-06 | 7.600E-06 | 1.212E-06 | 6.300E-06 | 8.700E-06 |
| 1546 | 78 | R78E | A | 5.1e-06 | 1e-05 | 9.8e-06 | 8.300E-06 | 2.773E-06 | 5.100E-06 | 1.000E-05 |
| 1547 | 78 | R78Q | A | 5.1e-06 | 7e-06 | 6.6e-06 | 6.233E-06 | 1.002E-06 | 5.100E-06 | 7.000E-06 |
| 1548 | 78 | R78G | A | 7.4e-06 | 7.3e-06 | 7e-06 | 7.233E-06 | 2.082E-07 | 7.000E-06 | 7.400E-06 |
| 1549 | 78 | R78H | A | 1e-05 | 1e-05 | 1.5e-05 | 1.167E-05 | 2.887E-06 | 1.000E-05 | 1.500E-05 |
| 1550 | 78 | R78I | A | 5e-06 | 1.9e-05 | 4.8e-06 | 9.600E-06 | 8.141E-06 | 4.800E-06 | 1.900E-05 |
| 1551 | 78 | R78L | A | 4.6e-06 | 4.8e-06 | 4.5e-06 | 4.633E-06 | 1.528E-07 | 4.500E-06 | 4.800E-06 |
| 1552 | 78 | R78K | A | 7.7e-06 | 4.4e-06 | 5.2e-06 | 5.767E-06 | 1.721E-06 | 4.400E-06 | 7.700E-06 |
| 1553 | 78 | R78M | A | 6.7e-06 | 6.7e-06 | 6.2e-06 | 6.533E-06 | 2.887E-07 | 6.200E-06 | 6.700E-06 |
| 1554 | 78 | R78F | A | 9.7e-06 | 6.8e-06 | 8.1e-06 | 8.200E-06 | 1.453E-06 | 6.800E-06 | 9.700E-06 |
| 1555 | 78 | R78P | A | 7.2e-06 | 9.5e-06 | 7.5e-06 | 8.067E-06 | 1.250E-06 | 7.200E-06 | 9.500E-06 |
| 1556 | 78 | R78S | A | 6.2e-06 | 6.9e-06 | 6.8e-06 | 6.633E-06 | 3.786E-07 | 6.200E-06 | 6.900E-06 |
| 1557 | 78 | R78T | A | 1.2e-05 | 1.1e-05 | 1.1e-05 | 1.133E-05 | 5.774E-07 | 1.100E-05 | 1.200E-05 |
| 1558 | 78 | R78W | A | 3.3e-06 | 9e-06 | 6.7e-06 | 6.333E-06 | 2.868E-06 | 3.300E-06 | 9.000E-06 |
| 1559 | 78 | R78Y | A | 1.4e-05 | 1.1e-05 | 1.2e-05 | 1.233E-05 | 1.528E-06 | 1.100E-05 | 1.400E-05 |
| 1560 | 78 | R78V | A | 1e-05 | 1.1e-05 | 1.4e-05 | 1.167E-05 | 2.082E-06 | 1.000E-05 | 1.400E-05 |
| 1561 | 79 | I79A | A | 1.1e-05 | 6.7e-06 | 4.5e-06 | 7.400E-06 | 3.306E-06 | 4.500E-06 | 1.100E-05 |
| 1562 | 79 | I79R | A | 6.3e-06 | 8.9e-06 | 5.2e-06 | 6.800E-06 | 1.900E-06 | 5.200E-06 | 8.900E-06 |
| 1563 | 79 | I79N | A | 1.4e-05 | 1.3e-05 | 1.3e-05 | 1.333E-05 | 5.774E-07 | 1.300E-05 | 1.400E-05 |
| 1564 | 79 | I79D | A | 5.1e-06 | 8.6e-06 | 7.4e-06 | 7.033E-06 | 1.779E-06 | 5.100E-06 | 8.600E-06 |
| 1565 | 79 | I79C | A | 1e-05 | 8.8e-06 | 9.8e-06 | 9.533E-06 | 6.429E-07 | 8.800E-06 | 1.000E-05 |
| 1566 | 79 | I79E | A | 9.2e-06 | 1e-05 | 6.5e-06 | 8.567E-06 | 1.834E-06 | 6.500E-06 | 1.000E-05 |
| 1567 | 79 | I79Q | A | 7.8e-06 | 6.3e-06 | 9.8e-06 | 7.967E-06 | 1.756E-06 | 6.300E-06 | 9.800E-06 |
| 1568 | 79 | I79G | A | 1e-05 | 7.5e-06 | 4.1e-06 | 7.200E-06 | 2.961E-06 | 4.100E-06 | 1.000E-05 |
| 1569 | 79 | I79H | A | 9.2e-06 | 9.1e-06 | 8.3e-06 | 8.867E-06 | 4.933E-07 | 8.300E-06 | 9.200E-06 |
| 1570 | 79 | I79I | A | 9e-06 | 8.3e-06 | 1.1e-05 | 9.433E-06 | 1.401E-06 | 8.300E-06 | 1.100E-05 |
| 1571 | 79 | I79L | A | 4.5e-06 | 3.9e-06 | 4.5e-06 | 4.300E-06 | 3.464E-07 | 3.900E-06 | 4.500E-06 |
| 1572 | 79 | I79K | A | 7.8e-06 | 8.6e-06 | 7.2e-06 | 7.867E-06 | 7.024E-07 | 7.200E-06 | 8.600E-06 |
| 1573 | 79 | I79M | A | 1.3e-05 | 1.1e-05 | 5.6e-06 | 9.867E-06 | 3.828E-06 | 5.600E-06 | 1.300E-05 |
| 1574 | 79 | I79F | A | 9.3e-06 | 6.3e-06 | 4.9e-06 | 6.833E-06 | 2.248E-06 | 4.900E-06 | 9.300E-06 |
| 1575 | 79 | I79P | A | 4.5e-06 | 7.8e-06 | 1.2e-05 | 8.100E-06 | 3.759E-06 | 4.500E-06 | 1.200E-05 |
| 1576 | 79 | I79S | A | 6.8e-06 | 8.6e-06 | 7.3e-06 | 7.567E-06 | 9.292E-07 | 6.800E-06 | 8.600E-06 |
| 1577 | 79 | I79T | A | 7.1e-06 | 5.6e-06 | 8.4e-06 | 7.033E-06 | 1.401E-06 | 5.600E-06 | 8.400E-06 |
| 1578 | 79 | I79W | A | 9.6e-06 | 1.1e-05 | 1.8e-05 | 1.287E-05 | 4.500E-06 | 9.600E-06 | 1.800E-05 |
| 1579 | 79 | I79Y | A | 1e-05 | 9.3e-06 | 9.4e-06 | 9.567E-06 | 3.786E-07 | 9.300E-06 | 1.000E-05 |
| 1580 | 79 | I79V | A | 9.4e-06 | 9.7e-06 | 1.3e-05 | 1.070E-05 | 1.997E-06 | 9.400E-06 | 1.300E-05 |
| 1581 | 80 | V80A | A | 6.8e-06 | 5.1e-06 | 5.8e-06 | 5.900E-06 | 8.544E-07 | 5.100E-06 | 6.800E-06 |
| 1582 | 80 | V80R | A | 8.4e-06 | 6.6e-06 | 4.7e-06 | 6.567E-06 | 1.850E-06 | 4.700E-06 | 8.400E-06 |
| 1583 | 80 | V80N | A | 1.2e-05 | 7.7e-06 | 7e-06 | 8.900E-06 | 2.707E-06 | 7.000E-06 | 1.200E-05 |
| 1584 | 80 | V80D | A | 4.4e-06 | 8.9e-06 | 1.1e-05 | 8.100E-06 | 3.372E-06 | 4.400E-06 | 1.100E-05 |
| 1585 | 80 | V80C | A | 8.5e-06 | 1.1e-05 | 4.8e-06 | 8.100E-06 | 3.119E-06 | 4.800E-06 | 1.100E-05 |
| 1586 | 80 | V80E | A | 6.9e-06 | 7.9e-06 | 8.3e-06 | 7.700E-06 | 7.211E-07 | 6.900E-06 | 8.300E-06 |
| 1587 | 80 | V80Q | A | 5.7e-06 | 8.1e-06 | 4.9e-06 | 6.233E-06 | 1.665E-06 | 4.900E-06 | 8.100E-06 |
| 1588 | 80 | V80G | A | 1.1e-05 | 1.3e-05 | 6.7e-06 | 1.023E-05 | 3.219E-06 | 6.700E-06 | 1.300E-05 |
| 1589 | 80 | V80H | A | 6.2e-06 | 7.2e-06 | 9.5e-06 | 7.633E-06 | 1.692E-06 | 6.200E-06 | 9.500E-06 |
| 1590 | 80 | V80I | A | 9.7e-06 | 1.4e-05 | 9.6e-06 | 1.110E-05 | 2.512E-06 | 9.600E-06 | 1.400E-05 |
| 1591 | 80 | V80L | A | 3.8e-06 | 5e-06 | 5.8e-06 | 4.867E-06 | 1.007E-06 | 3.800E-06 | 5.800E-06 |
| 1592 | 80 | V80K | A | 4.9e-06 | 9.7e-06 | 9.5e-06 | 8.033E-06 | 2.715E-06 | 4.900E-06 | 9.700E-06 |
| 1593 | 80 | V80M | A | 9.4e-06 | 6.5e-06 | 9.2e-06 | 8.367E-06 | 1.620E-06 | 6.500E-06 | 9.400E-06 |
| 1594 | 80 | V80F | A | 9.3e-06 | 6.9e-06 | 6.6e-06 | 7.600E-06 | 1.480E-06 | 6.600E-06 | 9.300E-06 |
| 1595 | 80 | V80P | A | 1.4e-05 | 8.4e-06 | 9.7e-06 | 1.070E-05 | 2.931E-06 | 8.400E-06 | 1.400E-05 |
| 1596 | 80 | V80S | A | 9.4e-06 | 6.1e-06 | 9.7e-06 | 8.400E-06 | 1.997E-06 | 6.100E-06 | 9.700E-06 |
| 1597 | 80 | V80T | A | 3.6e-06 | 5e-06 | 3.7e-06 | 4.100E-06 | 7.810E-07 | 3.600E-06 | 5.000E-06 |
| 1598 | 80 | V80W | A | 6.7e-06 | 7.5e-06 | 4.7e-06 | 6.300E-06 | 1.442E-06 | 4.700E-06 | 7.500E-06 |
| 1599 | 80 | V80Y | A | 1.3e-05 | 8.2e-06 | 6.1e-06 | 9.100E-06 | 3.537E-06 | 6.100E-06 | 1.300E-05 |
| 1600 | 80 | V80V | A | 7.6e-06 | 5.9e-06 | 9.5e-06 | 7.667E-06 | 1.801E-06 | 5.900E-06 | 9.500E-06 |
| 1601 | 81 | R81A | A | 7.9e-06 | 5.6e-06 | 8.4e-06 | 7.300E-06 | 1.493E-06 | 5.600E-06 | 8.400E-06 |
| 1602 | 81 | R81R | A | 8.7e-06 | 1.1e-05 | 6.3e-06 | 8.667E-06 | 2.350E-06 | 6.300E-06 | 1.100E-05 |
| 1603 | 81 | R81N | A | 1.3e-05 | 8e-06 | 8.9e-06 | 9.967E-06 | 2.665E-06 | 8.000E-06 | 1.300E-05 |
| 1604 | 81 | R81D | A | 8.9e-06 | 8.6e-06 | 5.4e-06 | 7.633E-06 | 1.940E-06 | 5.400E-06 | 8.900E-06 |
| 1605 | 81 | R81C | A | 7.9e-06 | 5.2e-06 | 6.9e-06 | 6.667E-06 | 1.365E-06 | 5.200E-06 | 7.900E-06 |
| 1606 | 81 | R81E | A | 9e-06 | 1e-05 | 8e-06 | 9.000E-06 | 1.000E-06 | 8.000E-06 | 1.000E-05 |
| 1607 | 81 | R81Q | A | 1e-05 | 8.3e-06 | 3.8e-06 | 7.367E-06 | 3.204E-06 | 3.800E-06 | 1.000E-05 |
| 1608 | 81 | R81G | A | 1.2e-05 | 3e-06 | 9.3e-06 | 8.100E-06 | 4.618E-06 | 3.000E-06 | 1.200E-05 |
| 1609 | 81 | R81H | A | 7.3e-06 | 9.1e-06 | 5.1e-06 | 7.167E-06 | 2.003E-06 | 5.100E-06 | 9.100E-06 |
| 1610 | 81 | R81I | A | 6.8e-06 | 6.6e-06 | 9e-06 | 7.467E-06 | 1.332E-06 | 6.600E-06 | 9.000E-06 |
| 1611 | 81 | R81L | A | 5.8e-06 | 6.8e-06 | 9.5e-06 | 7.367E-06 | 1.914E-06 | 5.800E-06 | 9.500E-06 |
| 1612 | 81 | R81K | A | 6.4e-06 | 5.1e-06 | 7.1e-06 | 6.200E-06 | 1.015E-06 | 5.100E-06 | 7.100E-06 |
| 1613 | 81 | R81M | A | 7e-06 | 6.8e-06 | 8.6e-06 | 7.467E-06 | 9.866E-07 | 6.800E-06 | 8.600E-06 |
| 1614 | 81 | R81F | A | 9.8e-06 | 1.4e-05 | 4.4e-06 | 9.400E-06 | 4.812E-06 | 4.400E-06 | 1.400E-05 |

| | | | | | | | | | | |
|---|---|---|---|---|---|---|---|---|---|---|
| 1615 | 81 | R81P | A | 1.1e-05 | 6.8e-06 | 3.6e-06 | 7.133E-06 | 3.711E-06 | 3.600E-06 | 1.100E-05 |
| 1616 | 81 | R81S | A | 7.9e-06 | 6.1e-06 | 9.4e-06 | 7.800E-06 | 1.652E-06 | 6.100E-06 | 9.400E-06 |
| 1617 | 81 | R81T | A | 9.3e-06 | 6.8e-06 | 4.1e-06 | 6.733E-06 | 2.601E-06 | 4.100E-06 | 9.300E-06 |
| 1618 | 81 | R81W | A | 1.1e-05 | 8.5e-06 | 6.7e-06 | 8.733E-06 | 2.159E-06 | 6.700E-06 | 1.100E-05 |
| 1619 | 81 | R81Y | A | 7.5e-06 | 7e-06 | 1.1e-05 | 8.500E-06 | 2.179E-06 | 7.000E-06 | 1.100E-05 |
| 1620 | 81 | R81V | A | 7.1e-06 | 5.4e-06 | 6.6e-06 | 6.367E-06 | 8.737E-07 | 5.400E-06 | 7.100E-06 |
| 1621 | 82 | G82A | A | 7.6e-06 | 1e-05 | 6.7e-06 | 8.100E-06 | 1.706E-06 | 6.700E-06 | 1.000E-05 |
| 1622 | 82 | G82R | A | 7.8e-06 | 5.3e-06 | 5e-06 | 6.033E-06 | 1.537E-06 | 5.000E-06 | 7.800E-06 |
| 1623 | 82 | G82N | A | 8e-06 | 5.6e-06 | 1.1e-05 | 8.200E-06 | 2.706E-06 | 5.600E-06 | 1.100E-05 |
| 1624 | 82 | G82D | A | 7.5e-06 | 5.8e-06 | 5.6e-06 | 6.300E-06 | 1.044E-06 | 5.600E-06 | 7.500E-06 |
| 1625 | 82 | G82C | A | 9.7e-06 | 7.6e-06 | 5.5e-06 | 7.600E-06 | 2.100E-06 | 5.500E-06 | 9.700E-06 |
| 1626 | 82 | G82E | A | 1.9e-05 | 9.2e-06 | 9.1e-06 | 1.243E-05 | 5.687E-06 | 9.100E-06 | 1.900E-05 |
| 1627 | 82 | G82Q | A | 9.8e-06 | 7e-06 | 5.2e-06 | 7.333E-06 | 2.318E-06 | 5.200E-06 | 9.800E-06 |
| 1628 | 82 | G82G | A | 9.4e-06 | 6.3e-06 | 1e-05 | 8.567E-06 | 1.986E-06 | 6.300E-06 | 1.000E-05 |
| 1629 | 82 | G82H | A | 6.8e-06 | 6.2e-06 | 1.1e-05 | 8.000E-06 | 2.615E-06 | 6.200E-06 | 1.100E-05 |
| 1630 | 82 | G82I | A | 9.1e-06 | 8.5e-06 | 8.9e-06 | 8.833E-06 | 3.055E-07 | 8.500E-06 | 9.100E-06 |
| 1631 | 82 | G82L | A | 6.4e-06 | 4.8e-06 | 1.3e-05 | 8.067E-06 | 4.347E-06 | 4.800E-06 | 1.300E-05 |
| 1632 | 82 | G82K | A | 5e-06 | 6.3e-06 | 7.4e-06 | 6.233E-06 | 1.201E-06 | 5.000E-06 | 7.400E-06 |
| 1633 | 82 | G82M | A | 1.2e-05 | 6.4e-06 | 5.7e-06 | 8.033E-06 | 3.453E-06 | 5.700E-06 | 1.200E-05 |
| 1634 | 82 | G82F | A | 6e-06 | 1e-05 | 1.1e-05 | 9.000E-06 | 2.646E-06 | 6.000E-06 | 1.100E-05 |
| 1635 | 82 | G82P | A | 2.7e-06 | 7.7e-06 | 1e-05 | 6.800E-06 | 3.732E-06 | 2.700E-06 | 1.000E-05 |
| 1636 | 82 | G82S | A | 9.9e-06 | 8.7e-06 | 1.1e-05 | 9.867E-06 | 1.150E-06 | 8.700E-06 | 1.100E-05 |
| 1637 | 82 | G82T | A | 6.3e-06 | 1e-05 | 9.7e-06 | 8.667E-06 | 2.055E-06 | 6.300E-06 | 1.000E-05 |
| 1638 | 82 | G82W | A | 8.2e-06 | 6.2e-06 | 1.2e-05 | 8.800E-06 | 2.946E-06 | 6.200E-06 | 1.200E-05 |
| 1639 | 82 | G82Y | A | 1.2e-05 | 1.5e-05 | 6.3e-06 | 1.110E-05 | 4.419E-06 | 6.300E-06 | 1.500E-05 |
| 1640 | 82 | G82V | A | 4.9e-06 | 5e-06 | 3e-06 | 4.300E-06 | 1.127E-06 | 3.000E-06 | 5.000E-06 |
| 1641 | 83 | T83A | A | 1e-05 | 1.4e-05 | 1.2e-05 | 1.200E-05 | 2.000E-06 | 1.000E-05 | 1.400E-05 |
| 1642 | 83 | T83R | A | 9.3e-06 | 6e-06 | 6.4e-06 | 7.233E-06 | 1.801E-06 | 6.000E-06 | 9.300E-06 |
| 1643 | 83 | T83N | A | 4.3e-06 | 7.3e-06 | 5e-06 | 5.533E-06 | 1.570E-06 | 4.300E-06 | 7.300E-06 |
| 1644 | 83 | T83D | A | 1.3e-05 | 7e-06 | 5.8e-06 | 8.600E-06 | 3.857E-06 | 5.800E-06 | 1.300E-05 |
| 1645 | 83 | T83C | A | 7.7e-06 | 9.4e-06 | 1.1e-05 | 9.367E-06 | 1.650E-06 | 7.700E-06 | 1.100E-05 |
| 1646 | 83 | T83E | A | 1.3e-05 | 8.8e-06 | 5.8e-06 | 9.200E-06 | 3.617E-06 | 5.800E-06 | 1.300E-05 |
| 1647 | 83 | T83Q | A | 8.9e-06 | 6.1e-06 | 1.2e-05 | 9.000E-06 | 2.951E-06 | 6.100E-06 | 1.200E-05 |
| 1648 | 83 | T83G | A | 8e-06 | 1.3e-05 | 8.6e-06 | 9.867E-06 | 2.730E-06 | 8.000E-06 | 1.300E-05 |
| 1649 | 83 | T83H | A | 4.7e-06 | 1.2e-05 | 5.3e-06 | 7.333E-06 | 4.053E-06 | 4.700E-06 | 1.200E-05 |
| 1650 | 83 | T83I | A | 1.1e-05 | 4.3e-06 | 7e-06 | 7.433E-06 | 3.371E-06 | 4.300E-06 | 1.100E-05 |
| 1651 | 83 | T83L | A | 4.9e-06 | 1.4e-05 | 1e-05 | 9.633E-06 | 4.561E-06 | 4.900E-06 | 1.400E-05 |
| 1652 | 83 | T83K | A | 9.4e-06 | 7.2e-06 | 6.5e-06 | 7.700E-06 | 1.513E-06 | 6.500E-06 | 9.400E-06 |
| 1653 | 83 | T83M | A | 1.4e-05 | 7.8e-06 | 1e-05 | 1.060E-05 | 3.143E-06 | 7.800E-06 | 1.400E-05 |
| 1654 | 83 | T83F | A | 5.1e-06 | 6.1e-06 | 1.2e-05 | 7.733E-06 | 3.729E-06 | 5.100E-06 | 1.200E-05 |
| 1655 | 83 | T83P | A | 1.2e-05 | 8.7e-06 | 5.5e-06 | 8.733E-06 | 3.250E-06 | 5.500E-06 | 1.200E-05 |
| 1656 | 83 | T83S | A | 1.3e-05 | 1.1e-05 | 1.2e-05 | 1.200E-05 | 1.000E-06 | 1.100E-05 | 1.300E-05 |
| 1657 | 83 | T83T | A | 1.5e-05 | 1.1e-05 | 8.7e-06 | 1.157E-05 | 3.188E-06 | 8.700E-06 | 1.500E-05 |
| 1658 | 83 | T83W | A | 6.8e-06 | 1.7e-05 | 4.4e-06 | 9.400E-06 | 6.690E-06 | 4.400E-06 | 1.700E-05 |
| 1659 | 83 | T83Y | A | 9.4e-06 | 9.2e-06 | 1.2e-05 | 1.020E-05 | 1.562E-06 | 9.200E-06 | 1.200E-05 |
| 1660 | 83 | T83V | A | 1.4e-05 | 5.3e-06 | 1.3e-05 | 1.077E-05 | 4.761E-06 | 5.300E-06 | 1.400E-05 |
| 1661 | 84 | Q84A | A | 5.2e-06 | 8.4e-06 | 6.3e-06 | 6.633E-06 | 1.626E-06 | 5.200E-06 | 8.400E-06 |
| 1662 | 84 | Q84R | A | 6.8e-06 | 7.6e-06 | 7.2e-06 | 7.200E-06 | 4.000E-07 | 6.800E-06 | 7.600E-06 |
| 1663 | 84 | Q84N | A | 6.4e-06 | 6.9e-06 | 3e-06 | 5.433E-06 | 2.122E-06 | 3.000E-06 | 6.900E-06 |
| 1664 | 84 | Q84D | A | 9.6e-06 | 1.1e-05 | 9.6e-06 | 1.007E-05 | 8.083E-07 | 9.600E-06 | 1.100E-05 |
| 1665 | 84 | Q84C | A | 8.3e-06 | 6.3e-06 | 4.8e-06 | 6.467E-06 | 1.756E-06 | 4.800E-06 | 8.300E-06 |
| 1666 | 84 | Q84E | A | 3.9e-06 | 1.2e-05 | 9.2e-06 | 8.367E-06 | 4.114E-06 | 3.900E-06 | 1.200E-05 |
| 1667 | 84 | Q84Q | A | 8.9e-06 | 7e-06 | 7.7e-06 | 7.867E-06 | 9.609E-07 | 7.000E-06 | 8.900E-06 |
| 1668 | 84 | Q84G | A | 9.7e-06 | 9.7e-06 | 1e-05 | 9.800E-06 | 1.732E-07 | 9.700E-06 | 1.000E-05 |
| 1669 | 84 | Q84H | A | 1.2e-05 | 1.2e-05 | 8.8e-06 | 1.093E-05 | 1.848E-06 | 8.800E-06 | 1.200E-05 |
| 1670 | 84 | Q84I | A | 6.2e-06 | 3.7e-06 | 8.1e-06 | 6.000E-06 | 2.207E-06 | 3.700E-06 | 8.100E-06 |
| 1671 | 84 | Q84L | A | 4.9e-06 | 5.9e-06 | 8.8e-06 | 6.533E-06 | 2.026E-06 | 4.900E-06 | 8.800E-06 |
| 1672 | 84 | Q84K | A | 7.4e-06 | 6.6e-06 | 6.3e-06 | 6.767E-06 | 5.686E-07 | 6.300E-06 | 7.400E-06 |
| 1673 | 84 | Q84M | A | 5.8e-06 | 5.5e-06 | 6.2e-06 | 5.833E-06 | 3.512E-07 | 5.500E-06 | 6.200E-06 |
| 1674 | 84 | Q84F | A | 7e-06 | 1e-05 | 8.6e-06 | 8.533E-06 | 1.501E-06 | 7.000E-06 | 1.000E-05 |
| 1675 | 84 | Q84P | A | 9.9e-06 | 1.3e-05 | 9.1e-06 | 1.067E-05 | 2.060E-06 | 9.100E-06 | 1.300E-05 |
| 1676 | 84 | Q84S | A | 9.5e-06 | 1e-05 | 9.1e-06 | 9.533E-06 | 4.509E-07 | 9.100E-06 | 1.000E-05 |
| 1677 | 84 | Q84T | A | 1.1e-05 | 6.6e-06 | 1e-05 | 9.200E-06 | 2.307E-06 | 6.600E-06 | 1.100E-05 |
| 1678 | 84 | Q84W | A | 9e-06 | 1.2e-05 | 6.2e-06 | 9.067E-06 | 2.901E-06 | 6.200E-06 | 1.200E-05 |
| 1679 | 84 | Q84Y | A | 6.4e-06 | 6.6e-06 | 6e-06 | 6.333E-06 | 3.055E-07 | 6.000E-06 | 6.600E-06 |
| 1680 | 84 | Q84V | A | 4.8e-06 | 6e-06 | 7.9e-06 | 6.233E-06 | 1.563E-06 | 4.800E-06 | 7.900E-06 |
| 1681 | 85 | L85A | A | 8.5e-06 | 5.9e-06 | 9.2e-06 | 7.867E-06 | 1.739E-06 | 5.900E-06 | 9.200E-06 |
| 1682 | 85 | L85R | A | 4.1e-06 | 1.2e-05 | 1.4e-05 | 1.003E-05 | 5.235E-06 | 4.100E-06 | 1.400E-05 |
| 1683 | 85 | L85N | A | 5.5e-06 | 1.3e-05 | 8.4e-06 | 8.967E-06 | 3.782E-06 | 5.500E-06 | 1.300E-05 |
| 1684 | 85 | L85D | A | 4.7e-06 | 9.5e-06 | 8.9e-06 | 7.700E-06 | 2.615E-06 | 4.700E-06 | 9.500E-06 |
| 1685 | 85 | L85C | A | 4.5e-06 | 8.8e-06 | 1.1e-05 | 8.100E-06 | 3.306E-06 | 4.500E-06 | 1.100E-05 |
| 1686 | 85 | L85E | A | 9.1e-06 | 6.9e-06 | 8.5e-06 | 8.167E-06 | 1.137E-06 | 6.900E-06 | 9.100E-06 |
| 1687 | 85 | L85Q | A | 9.4e-06 | 1e-05 | 9.5e-06 | 9.633E-06 | 3.215E-07 | 9.400E-06 | 1.000E-05 |
| 1688 | 85 | L85G | A | 9.6e-06 | 9.9e-06 | 8.5e-06 | 9.333E-06 | 7.371E-07 | 8.500E-06 | 9.900E-06 |
| 1689 | 85 | L85H | A | 5.8e-06 | 9.1e-06 | 1e-05 | 8.300E-06 | 2.211E-06 | 5.800E-06 | 1.000E-05 |
| 1690 | 85 | L85I | A | 1.4e-05 | 1e-05 | 6.4e-06 | 1.013E-05 | 3.802E-06 | 6.400E-06 | 1.400E-05 |
| 1691 | 85 | L85L | A | 6.7e-06 | 8e-06 | 7.9e-06 | 7.533E-06 | 7.234E-07 | 6.700E-06 | 8.000E-06 |
| 1692 | 85 | L85K | A | 8.2e-06 | 5.3e-06 | 6.7e-06 | 6.733E-06 | 1.450E-06 | 5.300E-06 | 8.200E-06 |
| 1693 | 85 | L85M | A | 6.4e-06 | 6.8e-06 | 1e-05 | 7.733E-06 | 1.973E-06 | 6.400E-06 | 1.000E-05 |
| 1694 | 85 | L85F | A | 7.1e-06 | 7.6e-06 | 1.3e-05 | 9.233E-06 | 3.272E-06 | 7.100E-06 | 1.300E-05 |
| 1695 | 85 | L85P | A | 8.6e-06 | 7.1e-06 | 7e-06 | 7.567E-06 | 8.963E-07 | 7.000E-06 | 8.600E-06 |
| 1696 | 85 | L85S | A | 6.1e-06 | 7.6e-06 | 1e-05 | 7.900E-06 | 1.967E-06 | 6.100E-06 | 1.000E-05 |
| 1697 | 85 | L85T | A | 2e-05 | 7.6e-06 | 9.7e-06 | 1.243E-05 | 6.637E-06 | 7.600E-06 | 2.000E-05 |
| 1698 | 85 | L85W | A | 9.3e-06 | 7.4e-06 | 9.4e-06 | 8.700E-06 | 1.127E-06 | 7.400E-06 | 9.400E-06 |
| 1699 | 85 | L85Y | A | 5.1e-06 | 6e-06 | 8.2e-06 | 6.433E-06 | 1.595E-06 | 5.100E-06 | 8.200E-06 |

| | | | | | | | | | | |
|---|---|---|---|---|---|---|---|---|---|---|
| 1700 | 85 | L85V | A | 1.4e-05 | 9e-06 | 1e-05 | 1.100E-05 | 2.646E-06 | 9.000E-06 | 1.400E-05 |
| 1701 | 86 | F86A | A | 8.9e-06 | 4.7e-06 | 1.3e-05 | 8.867E-06 | 4.150E-06 | 4.700E-06 | 1.300E-05 |
| 1702 | 86 | F86R | A | 9.3e-06 | 9.7e-06 | 9.3e-06 | 9.433E-06 | 2.309E-07 | 9.300E-06 | 9.700E-06 |
| 1703 | 86 | F86N | A | 5.9e-06 | 8.5e-06 | 6.1e-06 | 6.833E-06 | 1.447E-06 | 5.900E-06 | 8.500E-06 |
| 1704 | 86 | F86D | A | 1.2e-05 | 8.8e-06 | 6.5e-06 | 9.100E-06 | 2.762E-06 | 6.500E-06 | 1.200E-05 |
| 1705 | 86 | F86C | A | 9.9e-06 | 7.6e-06 | 1e-05 | 9.167E-06 | 1.358E-06 | 7.600E-06 | 1.000E-05 |
| 1706 | 86 | F86E | A | 6.7e-06 | 8.4e-06 | 4.9e-06 | 6.667E-06 | 1.750E-06 | 4.900E-06 | 8.400E-06 |
| 1707 | 86 | F86Q | A | 7e-06 | 1e-05 | 8.9e-06 | 8.633E-06 | 1.518E-06 | 7.000E-06 | 1.000E-05 |
| 1708 | 86 | F86G | A | 1.1e-05 | 7e-06 | 8.8e-06 | 8.933E-06 | 2.003E-06 | 7.000E-06 | 1.100E-05 |
| 1709 | 86 | F86H | A | 8.6e-06 | 9.3e-06 | 9.6e-06 | 9.167E-06 | 5.132E-07 | 8.600E-06 | 9.600E-06 |
| 1710 | 86 | F86I | A | 9.1e-06 | 1.4e-05 | 8.9e-06 | 1.067E-05 | 2.888E-06 | 8.900E-06 | 1.400E-05 |
| 1711 | 86 | F86L | A | 9.4e-06 | 4.7e-06 | 9e-06 | 7.700E-06 | 2.606E-06 | 4.700E-06 | 9.400E-06 |
| 1712 | 86 | F86K | A | 4.6e-06 | 1.4e-05 | 1.4e-05 | 1.087E-05 | 5.427E-06 | 4.600E-06 | 1.400E-05 |
| 1713 | 86 | F86M | A | 9.4e-06 | 1.3e-05 | 5.9e-06 | 9.433E-06 | 3.550E-06 | 5.900E-06 | 1.300E-05 |
| 1714 | 86 | F86F | A | 4.9e-06 | 7.5e-06 | 7.1e-06 | 6.500E-06 | 1.400E-06 | 4.900E-06 | 7.500E-06 |
| 1715 | 86 | F86P | A | 6.4e-06 | 1.1e-05 | 5.9e-06 | 7.767E-06 | 2.811E-06 | 5.900E-06 | 1.100E-05 |
| 1716 | 86 | F86S | A | 7.4e-06 | 5e-06 | 9.8e-06 | 7.400E-06 | 2.400E-06 | 5.000E-06 | 9.800E-06 |
| 1717 | 86 | F86T | A | 9.1e-06 | 4.9e-06 | 5.2e-06 | 6.400E-06 | 2.343E-06 | 4.900E-06 | 9.100E-06 |
| 1718 | 86 | F86W | A | 8.6e-06 | 7.5e-06 | 8.8e-06 | 8.300E-06 | 7.000E-07 | 7.500E-06 | 8.800E-06 |
| 1719 | 86 | F86Y | A | 8.1e-06 | 1.6e-05 | 1.4e-05 | 1.270E-05 | 4.107E-06 | 8.100E-06 | 1.600E-05 |
| 1720 | 86 | F86V | A | 9.7e-06 | 8.3e-06 | 1.2e-05 | 1.000E-05 | 1.868E-06 | 8.300E-06 | 1.200E-05 |
| 1721 | 87 | E87A | A | 7.4e-06 | 1.3e-05 | 7e-06 | 9.133E-06 | 3.355E-06 | 7.000E-06 | 1.300E-05 |
| 1722 | 87 | E87R | A | 9.8e-06 | 8e-06 | 6.2e-06 | 8.000E-06 | 1.800E-06 | 6.200E-06 | 9.800E-06 |
| 1723 | 87 | E87N | A | 7.8e-06 | 6.4e-06 | 1e-05 | 8.067E-06 | 1.815E-06 | 6.400E-06 | 1.000E-05 |
| 1724 | 87 | E87D | A | 9.8e-06 | 1e-05 | 9.7e-06 | 9.833E-06 | 1.528E-07 | 9.700E-06 | 1.000E-05 |
| 1725 | 87 | E87C | A | 6e-06 | 5.7e-06 | 9.9e-06 | 7.200E-06 | 2.343E-06 | 5.700E-06 | 9.900E-06 |
| 1726 | 87 | E87E | A | 6.4e-06 | 8.2e-06 | 7.8e-06 | 7.467E-06 | 9.452E-07 | 6.400E-06 | 8.200E-06 |
| 1727 | 87 | E87Q | A | 8.9e-06 | 8.2e-06 | 6.6e-06 | 7.900E-06 | 1.179E-06 | 6.600E-06 | 8.900E-06 |
| 1728 | 87 | E87G | A | 7.3e-06 | 7.7e-06 | 4.9e-06 | 6.633E-06 | 1.514E-06 | 4.900E-06 | 7.700E-06 |
| 1729 | 87 | E87H | A | 8.6e-06 | 6.5e-06 | 8.6e-06 | 7.900E-06 | 1.212E-06 | 6.500E-06 | 8.600E-06 |
| 1730 | 87 | E87I | A | 1.7e-05 | 1.2e-05 | 9.8e-06 | 1.293E-05 | 3.690E-06 | 9.800E-06 | 1.700E-05 |
| 1731 | 87 | E87L | A | 6.2e-06 | 6.6e-06 | 5.6e-06 | 6.133E-06 | 5.033E-07 | 5.600E-06 | 6.600E-06 |
| 1732 | 87 | E87K | A | 1e-05 | 6.7e-06 | 7.1e-06 | 7.933E-06 | 1.801E-06 | 6.700E-06 | 1.000E-05 |
| 1733 | 87 | E87M | A | 7.9e-06 | 1.1e-05 | 6.1e-06 | 8.333E-06 | 2.479E-06 | 6.100E-06 | 1.100E-05 |
| 1734 | 87 | E87F | A | 9.5e-06 | 1e-05 | 7.5e-06 | 9.000E-06 | 1.323E-06 | 7.500E-06 | 1.000E-05 |
| 1735 | 87 | E87P | A | 8.8e-06 | 1.1e-05 | 5.3e-06 | 8.367E-06 | 2.875E-06 | 5.300E-06 | 1.100E-05 |
| 1736 | 87 | E87S | A | 1.1e-05 | 9.7e-06 | 4.3e-06 | 8.333E-06 | 3.553E-06 | 4.300E-06 | 1.100E-05 |
| 1737 | 87 | E87T | A | 6.1e-06 | 8.3e-06 | 7.3e-06 | 7.233E-06 | 1.102E-06 | 6.100E-06 | 8.300E-06 |
| 1738 | 87 | E87W | A | 1.1e-05 | 6.1e-06 | 6.3e-06 | 7.800E-06 | 2.773E-06 | 6.100E-06 | 1.100E-05 |
| 1739 | 87 | E87Y | A | 6.4e-06 | 1.3e-05 | 8.2e-06 | 9.200E-06 | 3.412E-06 | 6.400E-06 | 1.300E-05 |
| 1740 | 87 | E87V | A | 8.2e-06 | 8.2e-06 | 1.3e-05 | 9.800E-06 | 2.771E-06 | 8.200E-06 | 1.300E-05 |
| 1741 | 88 | D88A | A | 6.9e-06 | 6.9e-06 | 9.2e-06 | 7.667E-06 | 1.328E-06 | 6.900E-06 | 9.200E-06 |
| 1742 | 88 | D88R | A | 7.6e-06 | 3.1e-06 | 9.9e-06 | 6.867E-06 | 3.459E-06 | 3.100E-06 | 9.900E-06 |
| 1743 | 88 | D88N | A | 8.4e-06 | 7.7e-06 | 9.4e-06 | 8.500E-06 | 8.544E-07 | 7.700E-06 | 9.400E-06 |
| 1744 | 88 | D88D | A | 5.8e-06 | 7.2e-06 | 6.5e-06 | 6.500E-06 | 7.000E-07 | 5.800E-06 | 7.200E-06 |
| 1745 | 88 | D88C | A | 9.1e-06 | 6.4e-06 | 4.4e-06 | 6.633E-06 | 2.359E-06 | 4.400E-06 | 9.100E-06 |
| 1746 | 88 | D88E | A | 8.1e-06 | 8.6e-06 | 7.1e-06 | 7.933E-06 | 7.638E-07 | 7.100E-06 | 8.600E-06 |
| 1747 | 88 | D88Q | A | 6.3e-06 | 1.2e-05 | 1.3e-05 | 1.043E-05 | 3.614E-06 | 6.300E-06 | 1.300E-05 |
| 1748 | 88 | D88G | A | 6.1e-06 | 8e-06 | 6.9e-06 | 7.000E-06 | 9.539E-07 | 6.100E-06 | 8.000E-06 |
| 1749 | 88 | D88H | A | 8e-06 | 8.6e-06 | 1.2e-05 | 9.533E-06 | 2.157E-06 | 8.000E-06 | 1.200E-05 |
| 1750 | 88 | D88I | A | 9.5e-06 | 5.8e-06 | 7.3e-06 | 7.533E-06 | 1.861E-06 | 5.800E-06 | 9.500E-06 |
| 1751 | 88 | D88L | A | 1.1e-05 | 1.8e-05 | 9.6e-06 | 1.287E-05 | 4.500E-06 | 9.600E-06 | 1.800E-05 |
| 1752 | 88 | D88K | A | 1.6e-05 | 1.8e-05 | 6.8e-06 | 1.360E-05 | 5.973E-06 | 6.800E-06 | 1.800E-05 |
| 1753 | 88 | D88M | A | 9.8e-06 | 6.3e-06 | 8.5e-06 | 8.200E-06 | 1.769E-06 | 6.300E-06 | 9.800E-06 |
| 1754 | 88 | D88F | A | 8.9e-06 | 5.2e-06 | 6.1e-06 | 6.733E-06 | 1.930E-06 | 5.200E-06 | 8.900E-06 |
| 1755 | 88 | D88P | A | 1e-05 | 9.3e-06 | 1.1e-05 | 1.010E-05 | 8.544E-07 | 9.300E-06 | 1.100E-05 |
| 1756 | 88 | D88S | A | 6.7e-06 | 5.3e-06 | 9.3e-06 | 7.100E-06 | 2.030E-06 | 5.300E-06 | 9.300E-06 |
| 1757 | 88 | D88T | A | 4.7e-06 | 6.4e-06 | 7.3e-06 | 6.133E-06 | 1.320E-06 | 4.700E-06 | 7.300E-06 |
| 1758 | 88 | D88W | A | 6.9e-06 | 6.6e-06 | 1.3e-05 | 8.833E-06 | 3.612E-06 | 6.600E-06 | 1.300E-05 |
| 1759 | 88 | D88Y | A | 9e-06 | 4.8e-06 | 8.6e-06 | 7.467E-06 | 2.318E-06 | 4.800E-06 | 9.000E-06 |
| 1760 | 88 | D88V | A | 9.5e-06 | 4.2e-06 | 8.9e-06 | 7.533E-06 | 2.902E-06 | 4.200E-06 | 9.500E-06 |
| 1761 | 89 | N89A | A | 9.7e-06 | 9.7e-06 | 6.4e-06 | 8.600E-06 | 1.905E-06 | 6.400E-06 | 9.700E-06 |
| 1762 | 89 | N89R | A | 1.3e-05 | 5.5e-06 | 1.3e-05 | 1.050E-05 | 4.330E-06 | 5.500E-06 | 1.300E-05 |
| 1763 | 89 | N89N | A | 8e-06 | 7.5e-06 | 8e-06 | 7.833E-06 | 2.887E-07 | 7.500E-06 | 8.000E-06 |
| 1764 | 89 | N89D | A | 7.3e-06 | 6.4e-06 | 6.6e-06 | 6.767E-06 | 4.726E-07 | 6.400E-06 | 7.300E-06 |
| 1765 | 89 | N89C | A | 9.2e-06 | 8.9e-06 | 6.9e-06 | 8.333E-06 | 1.250E-06 | 6.900E-06 | 9.200E-06 |
| 1766 | 89 | N89E | A | 7.1e-06 | 6.6e-06 | 6.1e-06 | 6.600E-06 | 5.000E-07 | 6.100E-06 | 7.100E-06 |
| 1767 | 89 | N89Q | A | 1.1e-05 | 5.6e-06 | 5.6e-06 | 7.400E-06 | 3.118E-06 | 5.600E-06 | 1.100E-05 |
| 1768 | 89 | N89G | A | 9.9e-06 | 9.3e-06 | 7.3e-06 | 8.833E-06 | 1.361E-06 | 7.300E-06 | 9.900E-06 |
| 1769 | 89 | N89H | A | 8.5e-06 | 1.1e-05 | 8.7e-06 | 9.400E-06 | 1.389E-06 | 8.500E-06 | 1.100E-05 |
| 1770 | 89 | N89I | A | 1.5e-05 | 6.7e-06 | 5.7e-06 | 9.133E-06 | 5.105E-06 | 5.700E-06 | 1.500E-05 |
| 1771 | 89 | N89L | A | 7.3e-06 | 6.9e-06 | 8.2e-06 | 7.467E-06 | 6.658E-07 | 6.900E-06 | 8.200E-06 |
| 1772 | 89 | N89K | A | 9.8e-06 | 5.2e-06 | 6.2e-06 | 7.067E-06 | 2.419E-06 | 5.200E-06 | 9.800E-06 |
| 1773 | 89 | N89M | A | 6.2e-06 | 3.3e-06 | 3.6e-06 | 4.367E-06 | 1.595E-06 | 3.300E-06 | 6.200E-06 |
| 1774 | 89 | N89F | A | 1.1e-05 | 7e-06 | 1.2e-05 | 1.000E-05 | 2.646E-06 | 7.000E-06 | 1.200E-05 |
| 1775 | 89 | N89P | A | 6.6e-06 | 1.1e-05 | 2.7e-06 | 6.767E-06 | 4.153E-06 | 2.700E-06 | 1.100E-05 |
| 1776 | 89 | N89S | A | 5.6e-06 | 8.9e-06 | 5.4e-06 | 6.633E-06 | 1.966E-06 | 5.400E-06 | 8.900E-06 |
| 1777 | 89 | N89T | A | 2.3e-06 | 9.9e-06 | 6.2e-06 | 6.133E-06 | 3.800E-06 | 2.300E-06 | 9.900E-06 |
| 1778 | 89 | N89W | A | 4.5e-06 | 7.1e-06 | 4e-06 | 5.200E-06 | 1.664E-06 | 4.000E-06 | 7.100E-06 |
| 1779 | 89 | N89Y | A | 9.5e-06 | 5.2e-06 | 9.1e-06 | 7.933E-06 | 2.376E-06 | 5.200E-06 | 9.500E-06 |
| 1780 | 89 | N89V | A | 1.1e-05 | 6.4e-06 | 8.3e-06 | 8.567E-06 | 2.312E-06 | 6.400E-06 | 1.100E-05 |
| 1781 | 90 | Y90A | A | 6.9e-06 | 5.1e-06 | 8.7e-06 | 6.900E-06 | 1.800E-06 | 5.100E-06 | 8.700E-06 |
| 1782 | 90 | Y90R | A | 5.1e-06 | 4.6e-06 | 1.2e-05 | 7.233E-06 | 4.136E-06 | 4.600E-06 | 1.200E-05 |
| 1783 | 90 | Y90N | A | 5.9e-06 | 5.9e-06 | 9.8e-06 | 7.200E-06 | 2.252E-06 | 5.900E-06 | 9.800E-06 |
| 1784 | 90 | Y90D | A | 8.2e-06 | 5.1e-06 | 1.2e-05 | 8.433E-06 | 3.456E-06 | 5.100E-06 | 1.200E-05 |

| | | | | | | | | | | |
|---|---|---|---|---|---|---|---|---|---|---|
| 1785 | 90 | Y90C | A | 1.3e-05 | 1.1e-05 | 4.5e-06 | 9.500E-06 | 4.444E-06 | 4.500E-06 | 1.300E-05 |
| 1786 | 90 | Y90E | A | 5.8e-06 | 1.2e-05 | 1e-05 | 9.267E-06 | 3.164E-06 | 5.800E-06 | 1.200E-05 |
| 1787 | 90 | Y90Q | A | 8e-06 | 5.4e-06 | 9.8e-06 | 7.733E-06 | 2.212E-06 | 5.400E-06 | 9.800E-06 |
| 1788 | 90 | Y90G | A | 8.8e-06 | 6.9e-06 | 9.4e-06 | 8.367E-06 | 1.305E-06 | 6.900E-06 | 9.400E-06 |
| 1789 | 90 | Y90H | A | 9.6e-06 | 2.3e-06 | 6.6e-06 | 6.167E-06 | 3.669E-06 | 2.300E-06 | 9.600E-06 |
| 1790 | 90 | Y90I | A | 9.9e-06 | 1.3e-05 | 1.3e-05 | 1.197E-05 | 1.790E-06 | 9.900E-06 | 1.300E-05 |
| 1791 | 90 | Y90L | A | 9.6e-06 | 8.5e-06 | 1e-05 | 9.367E-06 | 7.767E-07 | 8.500E-06 | 1.000E-05 |
| 1792 | 90 | Y90K | A | 5.6e-06 | 1.4e-05 | 5.4e-06 | 8.333E-06 | 4.908E-06 | 5.400E-06 | 1.400E-05 |
| 1793 | 90 | Y90M | A | 7.2e-06 | 5.3e-06 | 7.7e-06 | 6.733E-06 | 1.266E-06 | 5.300E-06 | 7.700E-06 |
| 1794 | 90 | Y90F | A | 9e-06 | 9.4e-06 | 1e-05 | 9.467E-06 | 5.033E-07 | 9.000E-06 | 1.000E-05 |
| 1795 | 90 | Y90P | A | 7.9e-06 | 9.6e-06 | 9.1e-06 | 8.867E-06 | 8.737E-07 | 7.900E-06 | 9.600E-06 |
| 1796 | 90 | Y90S | A | 1.1e-05 | 6.2e-06 | 4.8e-06 | 7.333E-06 | 3.252E-06 | 4.800E-06 | 1.100E-05 |
| 1797 | 90 | Y90T | A | 4.4e-06 | 8.4e-06 | 9.6e-06 | 7.467E-06 | 2.723E-06 | 4.400E-06 | 9.600E-06 |
| 1798 | 90 | Y90W | A | 6.5e-06 | 1e-05 | 7.6e-06 | 8.033E-06 | 1.790E-06 | 6.500E-06 | 1.000E-05 |
| 1799 | 90 | Y90Y | A | 6.5e-06 | 5.8e-06 | 6.2e-06 | 6.167E-06 | 3.512E-07 | 5.800E-06 | 6.500E-06 |
| 1800 | 90 | Y90V | A | 4.9e-06 | 1.2e-05 | 1.3e-05 | 9.967E-06 | 4.416E-06 | 4.900E-06 | 1.300E-05 |
| 1801 | 91 | A91A | A | 6.8e-06 | 9.4e-06 | 1.2e-05 | 9.400E-06 | 2.600E-06 | 6.800E-06 | 1.200E-05 |
| 1802 | 91 | A91R | A | 1.1e-05 | 1.5e-05 | 1e-05 | 1.200E-05 | 2.646E-06 | 1.000E-05 | 1.500E-05 |
| 1803 | 91 | A91N | A | 6.4e-06 | 1e-05 | 4.3e-06 | 6.900E-06 | 2.883E-06 | 4.300E-06 | 1.000E-05 |
| 1804 | 91 | A91D | A | 5e-06 | 8.5e-06 | 8.5e-06 | 7.333E-06 | 2.021E-06 | 5.000E-06 | 8.500E-06 |
| 1805 | 91 | A91C | A | 1.2e-05 | 8e-06 | 5.5e-06 | 8.500E-06 | 3.279E-06 | 5.500E-06 | 1.200E-05 |
| 1806 | 91 | A91E | A | 5e-06 | 8.4e-06 | 6.6e-06 | 6.667E-06 | 1.701E-06 | 5.000E-06 | 8.400E-06 |
| 1807 | 91 | A91Q | A | 1.1e-05 | 5.3e-06 | 6.8e-06 | 7.700E-06 | 2.955E-06 | 5.300E-06 | 1.100E-05 |
| 1808 | 91 | A91G | A | 7.7e-06 | 7.3e-06 | 9.5e-06 | 8.167E-06 | 1.172E-06 | 7.300E-06 | 9.500E-06 |
| 1809 | 91 | A91H | A | 1e-05 | 7.7e-06 | 6.7e-06 | 8.133E-06 | 1.692E-06 | 6.700E-06 | 1.000E-05 |
| 1810 | 91 | A91I | A | 4.4e-06 | 6.2e-06 | 4.6e-06 | 5.067E-06 | 9.866E-07 | 4.400E-06 | 6.200E-06 |
| 1811 | 91 | A91L | A | 7.5e-06 | 6.3e-06 | 6.9e-06 | 6.900E-06 | 6.000E-07 | 6.300E-06 | 7.500E-06 |
| 1812 | 91 | A91K | A | 1.1e-05 | 5.7e-06 | 8.6e-06 | 8.433E-06 | 2.654E-06 | 5.700E-06 | 1.100E-05 |
| 1813 | 91 | A91M | A | 6.7e-06 | 4.6e-06 | 5.2e-06 | 5.500E-06 | 1.082E-06 | 4.600E-06 | 6.700E-06 |
| 1814 | 91 | A91F | A | 4.7e-06 | 7.7e-06 | 5.4e-06 | 5.933E-06 | 1.570E-06 | 4.700E-06 | 7.700E-06 |
| 1815 | 91 | A91P | A | 6.1e-06 | 1.6e-05 | 7e-06 | 9.700E-06 | 5.474E-06 | 6.100E-06 | 1.600E-05 |
| 1816 | 91 | A91S | A | 6.3e-06 | 9e-06 | 9.3e-06 | 8.200E-06 | 1.652E-06 | 6.300E-06 | 9.300E-06 |
| 1817 | 91 | A91T | A | 9.8e-06 | 1.2e-05 | 6.2e-06 | 9.333E-06 | 2.928E-06 | 6.200E-06 | 1.200E-05 |
| 1818 | 91 | A91W | A | 1e-05 | 6.4e-06 | 6.6e-06 | 7.667E-06 | 2.023E-06 | 6.400E-06 | 1.000E-05 |
| 1819 | 91 | A91Y | A | 1.5e-05 | 6.1e-06 | 4.6e-06 | 8.567E-06 | 5.622E-06 | 4.600E-06 | 1.500E-05 |
| 1820 | 91 | A91V | A | 7.2e-06 | 8.5e-06 | 1e-05 | 8.567E-06 | 1.401E-06 | 7.200E-06 | 1.000E-05 |
| 1821 | 92 | L92A | A | 7.4e-06 | 5e-06 | 6e-06 | 6.133E-06 | 1.206E-06 | 5.000E-06 | 7.400E-06 |
| 1822 | 92 | L92R | A | 6.7e-06 | 5.7e-06 | 5.2e-06 | 5.867E-06 | 7.638E-07 | 5.200E-06 | 6.700E-06 |
| 1823 | 92 | L92N | A | 5.1e-06 | 6.1e-06 | 1.3e-05 | 8.067E-06 | 4.302E-06 | 5.100E-06 | 1.300E-05 |
| 1824 | 92 | L92D | A | 1.9e-05 | 7.8e-06 | 6.9e-06 | 1.123E-05 | 6.741E-06 | 6.900E-06 | 1.900E-05 |
| 1825 | 92 | L92C | A | 6.7e-06 | 7.1e-06 | 9.8e-06 | 7.867E-06 | 1.686E-06 | 6.700E-06 | 9.800E-06 |
| 1826 | 92 | L92E | A | 7.8e-06 | 8.2e-06 | 5.1e-06 | 7.033E-06 | 1.686E-06 | 5.100E-06 | 8.200E-06 |
| 1827 | 92 | L92Q | A | 6.4e-06 | 6.1e-06 | 9.1e-06 | 7.200E-06 | 1.652E-06 | 6.100E-06 | 9.100E-06 |
| 1828 | 92 | L92G | A | 9.8e-06 | 5.7e-06 | 6.7e-06 | 7.400E-06 | 2.138E-06 | 5.700E-06 | 9.800E-06 |
| 1829 | 92 | L92H | A | 1.2e-05 | 4.9e-06 | 3.7e-06 | 6.867E-06 | 4.486E-06 | 3.700E-06 | 1.200E-05 |
| 1830 | 92 | L92I | A | 8.8e-06 | 8.4e-06 | 1e-05 | 9.067E-06 | 8.327E-07 | 8.400E-06 | 1.000E-05 |
| 1831 | 92 | L92L | A | 7.7e-06 | 7e-06 | 5.8e-06 | 6.833E-06 | 9.609E-07 | 5.800E-06 | 7.700E-06 |
| 1832 | 92 | L92K | A | 7.8e-06 | 1e-05 | 9.3e-06 | 9.033E-06 | 1.124E-06 | 7.800E-06 | 1.000E-05 |
| 1833 | 92 | L92M | A | 5.3e-06 | 5.8e-06 | 7.4e-06 | 6.167E-06 | 1.097E-06 | 5.300E-06 | 7.400E-06 |
| 1834 | 92 | L92F | A | 5.6e-06 | 5.5e-06 | 5e-06 | 5.367E-06 | 3.215E-07 | 5.000E-06 | 5.600E-06 |
| 1835 | 92 | L92P | A | 1.3e-05 | 1e-05 | 7e-06 | 1.000E-05 | 3.000E-06 | 7.000E-06 | 1.300E-05 |
| 1836 | 92 | L92S | A | 8.8e-06 | 7e-06 | 1.2e-05 | 9.267E-06 | 2.532E-06 | 7.000E-06 | 1.200E-05 |
| 1837 | 92 | L92T | A | 6.9e-06 | 7.5e-06 | 7e-06 | 7.133E-06 | 3.215E-07 | 6.900E-06 | 7.500E-06 |
| 1838 | 92 | L92W | A | 6.1e-06 | 8.4e-06 | 8.8e-06 | 7.767E-06 | 1.457E-06 | 6.100E-06 | 8.800E-06 |
| 1839 | 92 | L92Y | A | 5.3e-06 | 8.1e-06 | 6.2e-06 | 6.533E-06 | 1.429E-06 | 5.300E-06 | 8.100E-06 |
| 1840 | 92 | L92V | A | 8.2e-06 | 7.2e-06 | 2e-05 | 1.180E-05 | 7.119E-06 | 7.200E-06 | 2.000E-05 |
| 1841 | 93 | A93A | A | 5.9e-06 | 4.6e-06 | 1.3e-05 | 7.833E-06 | 4.521E-06 | 4.600E-06 | 1.300E-05 |
| 1842 | 93 | A93R | A | 1.4e-05 | 5.2e-06 | 6.7e-06 | 8.633E-06 | 4.708E-06 | 5.200E-06 | 1.400E-05 |
| 1843 | 93 | A93N | A | 5.3e-06 | 1e-05 | 6.9e-06 | 7.400E-06 | 2.390E-06 | 5.300E-06 | 1.000E-05 |
| 1844 | 93 | A93D | A | 6.2e-06 | 1e-05 | 5.2e-06 | 7.133E-06 | 2.532E-06 | 5.200E-06 | 1.000E-05 |
| 1845 | 93 | A93C | A | 8.4e-06 | 9e-06 | 9.4e-06 | 8.933E-06 | 5.033E-07 | 8.400E-06 | 9.400E-06 |
| 1846 | 93 | A93E | A | 4.5e-06 | 6.9e-06 | 7.1e-06 | 6.167E-06 | 1.447E-06 | 4.500E-06 | 7.100E-06 |
| 1847 | 93 | A93Q | A | 7.2e-06 | 3.8e-06 | 9.8e-06 | 6.933E-06 | 3.009E-06 | 3.800E-06 | 9.800E-06 |
| 1848 | 93 | A93G | A | 9.5e-06 | 1e-05 | 6.5e-06 | 8.667E-06 | 1.893E-06 | 6.500E-06 | 1.000E-05 |
| 1849 | 93 | A93H | A | 7.4e-06 | 1e-05 | 9.3e-06 | 8.900E-06 | 1.345E-06 | 7.400E-06 | 1.000E-05 |
| 1850 | 93 | A93I | A | 6.8e-06 | 5.6e-06 | 1.3e-05 | 8.467E-06 | 3.972E-06 | 5.600E-06 | 1.300E-05 |
| 1851 | 93 | A93L | A | 6.4e-06 | 1.3e-05 | 7.3e-06 | 8.900E-06 | 3.579E-06 | 6.400E-06 | 1.300E-05 |
| 1852 | 93 | A93K | A | 7.1e-06 | 4.8e-06 | 5.6e-06 | 5.833E-06 | 1.168E-06 | 4.800E-06 | 7.100E-06 |
| 1853 | 93 | A93M | A | 7.3e-06 | 1.3e-05 | 1.3e-05 | 1.110E-05 | 3.291E-06 | 7.300E-06 | 1.300E-05 |
| 1854 | 93 | A93F | A | 8.9e-06 | 6.6e-06 | 1.4e-05 | 9.833E-06 | 3.787E-06 | 6.600E-06 | 1.400E-05 |
| 1855 | 93 | A93P | A | 1.1e-05 | 5.1e-06 | 4.8e-06 | 6.967E-06 | 3.496E-06 | 4.800E-06 | 1.100E-05 |
| 1856 | 93 | A93S | A | 7.2e-06 | 7.6e-06 | 5.3e-06 | 6.700E-06 | 1.229E-06 | 5.300E-06 | 7.600E-06 |
| 1857 | 93 | A93T | A | 8.1e-06 | 8.2e-06 | 5.1e-06 | 7.133E-06 | 1.762E-06 | 5.100E-06 | 8.200E-06 |
| 1858 | 93 | A93W | A | 1e-05 | 6.3e-06 | 9.6e-06 | 8.633E-06 | 2.031E-06 | 6.300E-06 | 1.000E-05 |
| 1859 | 93 | A93Y | A | 1.3e-05 | 9e-06 | 6.2e-06 | 9.400E-06 | 3.418E-06 | 6.200E-06 | 1.300E-05 |
| 1860 | 93 | A93V | A | 7.6e-06 | 9.8e-06 | 9.1e-06 | 8.833E-06 | 1.124E-06 | 7.600E-06 | 9.800E-06 |
| 1861 | 94 | V94A | A | 9.2e-06 | 8.1e-06 | 7.5e-06 | 8.267E-06 | 8.622E-07 | 7.500E-06 | 9.200E-06 |
| 1862 | 94 | V94R | A | 9e-06 | 4.2e-06 | 5.9e-06 | 6.367E-06 | 2.434E-06 | 4.200E-06 | 9.000E-06 |
| 1863 | 94 | V94N | A | 5.2e-06 | 6.2e-06 | 9.1e-06 | 6.833E-06 | 2.026E-06 | 5.200E-06 | 9.100E-06 |
| 1864 | 94 | V94D | A | 5.9e-06 | 8.5e-06 | 1.1e-05 | 8.467E-06 | 2.550E-06 | 5.900E-06 | 1.100E-05 |
| 1865 | 94 | V94C | A | 9.2e-06 | 9.4e-06 | 5.7e-06 | 8.100E-06 | 2.081E-06 | 5.700E-06 | 9.400E-06 |
| 1866 | 94 | V94E | A | 8.9e-06 | 5.6e-06 | 9e-06 | 7.833E-06 | 1.935E-06 | 5.600E-06 | 9.000E-06 |
| 1867 | 94 | V94Q | A | 8.5e-06 | 1.3e-05 | 1e-05 | 1.050E-05 | 2.291E-06 | 8.500E-06 | 1.300E-05 |
| 1868 | 94 | V94G | A | 1.1e-05 | 9.2e-06 | 9.9e-06 | 1.003E-05 | 9.074E-07 | 9.200E-06 | 1.100E-05 |
| 1869 | 94 | V94H | A | 5.5e-06 | 6.8e-06 | 7.1e-06 | 6.467E-06 | 8.505E-07 | 5.500E-06 | 7.100E-06 |

| | | | | | | | | | | |
|---|---|---|---|---|---|---|---|---|---|---|
| 1870 | 94 | V94I | A | 1.3e-05 | 1e-05 | 7.9e-06 | 1.030E-05 | 2.563E-06 | 7.900E-06 | 1.300E-05 |
| 1871 | 94 | V94L | A | 1e-05 | 9.3e-06 | 1.4e-05 | 1.110E-05 | 2.536E-06 | 9.300E-06 | 1.400E-05 |
| 1872 | 94 | V94K | A | 6.4e-06 | 6.5e-06 | 1.9e-05 | 1.063E-05 | 7.246E-06 | 6.400E-06 | 1.900E-05 |
| 1873 | 94 | V94M | A | 6.8e-06 | 1.4e-05 | 1.6e-05 | 1.227E-05 | 4.839E-06 | 6.800E-06 | 1.600E-05 |
| 1874 | 94 | V94F | A | 1.4e-05 | 7e-06 | 7.2e-06 | 9.400E-06 | 3.985E-06 | 7.000E-06 | 1.400E-05 |
| 1875 | 94 | V94P | A | 4.9e-06 | 5e-06 | 7e-06 | 5.633E-06 | 1.185E-06 | 4.900E-06 | 7.000E-06 |
| 1876 | 94 | V94S | A | 6e-06 | 9.1e-06 | 6.2e-06 | 7.100E-06 | 1.735E-06 | 6.000E-06 | 9.100E-06 |
| 1877 | 94 | V94T | A | 6.7e-06 | 1.1e-05 | 6.6e-06 | 8.100E-06 | 2.512E-06 | 6.600E-06 | 1.100E-05 |
| 1878 | 94 | V94W | A | 1.1e-05 | 1.1e-05 | 8.8e-06 | 1.027E-05 | 1.270E-06 | 8.800E-06 | 1.100E-05 |
| 1879 | 94 | V94Y | A | 6.2e-06 | 8.2e-06 | 8.6e-06 | 7.667E-06 | 1.286E-06 | 6.200E-06 | 8.600E-06 |
| 1880 | 94 | V94V | A | 4.9e-06 | 3.6e-06 | 6.4e-06 | 4.967E-06 | 1.401E-06 | 3.600E-06 | 6.400E-06 |
| 1881 | 95 | L95A | A | 9.7e-06 | 6.7e-06 | 1.3e-05 | 9.800E-06 | 3.151E-06 | 6.700E-06 | 1.300E-05 |
| 1882 | 95 | L95R | A | 4.8e-06 | 7e-06 | 5.8e-06 | 5.867E-06 | 1.102E-06 | 4.800E-06 | 7.000E-06 |
| 1883 | 95 | L95N | A | 6.2e-06 | 8.6e-06 | 6.8e-06 | 7.200E-06 | 1.249E-06 | 6.200E-06 | 8.600E-06 |
| 1884 | 95 | L95D | A | 7.8e-06 | 6.5e-06 | 6.6e-06 | 6.967E-06 | 7.234E-07 | 6.500E-06 | 7.800E-06 |
| 1885 | 95 | L95C | A | 1e-05 | 6.3e-06 | 3.8e-06 | 6.700E-06 | 3.119E-06 | 3.800E-06 | 1.000E-05 |
| 1886 | 95 | L95E | A | 8.5e-06 | 8.1e-06 | 8.2e-06 | 8.267E-06 | 2.082E-07 | 8.100E-06 | 8.500E-06 |
| 1887 | 95 | L95Q | A | 6.5e-06 | 4.4e-06 | 8.4e-06 | 6.433E-06 | 2.001E-06 | 4.400E-06 | 8.400E-06 |
| 1888 | 95 | L95G | A | 8.6e-06 | 6.5e-06 | 1.1e-05 | 8.700E-06 | 2.252E-06 | 6.500E-06 | 1.100E-05 |
| 1889 | 95 | L95H | A | 4.7e-06 | 9.9e-06 | 5.8e-06 | 6.800E-06 | 2.740E-06 | 4.700E-06 | 9.900E-06 |
| 1890 | 95 | L95I | A | 9.9e-06 | 9.5e-06 | 7.4e-06 | 8.933E-06 | 1.343E-06 | 7.400E-06 | 9.900E-06 |
| 1891 | 95 | L95L | A | 5.2e-06 | 7e-06 | 8.7e-06 | 6.967E-06 | 1.750E-06 | 5.200E-06 | 8.700E-06 |
| 1892 | 95 | L95K | A | 8.5e-06 | 1e-05 | 9.7e-06 | 9.400E-06 | 7.937E-07 | 8.500E-06 | 1.000E-05 |
| 1893 | 95 | L95M | A | 9.9e-06 | 7.4e-06 | 9.7e-06 | 9.000E-06 | 1.389E-06 | 7.400E-06 | 9.900E-06 |
| 1894 | 95 | L95F | A | 8e-06 | 8.3e-06 | 1e-05 | 8.767E-06 | 1.079E-06 | 8.000E-06 | 1.000E-05 |
| 1895 | 95 | L95P | A | 6.9e-06 | 8.4e-06 | 5.6e-06 | 6.967E-06 | 1.401E-06 | 5.600E-06 | 8.400E-06 |
| 1896 | 95 | L95S | A | 1.1e-05 | 4.8e-06 | 8.6e-06 | 8.133E-06 | 3.126E-06 | 4.800E-06 | 1.100E-05 |
| 1897 | 95 | L95T | A | 8.9e-06 | 9.6e-06 | 6.3e-06 | 8.267E-06 | 1.739E-06 | 6.300E-06 | 9.600E-06 |
| 1898 | 95 | L95W | A | 4.4e-06 | 1e-05 | 1.1e-05 | 8.467E-06 | 3.557E-06 | 4.400E-06 | 1.100E-05 |
| 1899 | 95 | L95Y | A | 1.2e-05 | 6.5e-06 | 6.4e-06 | 8.300E-06 | 3.205E-06 | 6.400E-06 | 1.200E-05 |
| 1900 | 95 | L95V | A | 6.4e-06 | 7.1e-06 | 6.4e-06 | 6.633E-06 | 4.041E-07 | 6.400E-06 | 7.100E-06 |
| 1901 | 96 | D96A | A | 1e-05 | 6.8e-06 | 1.3e-05 | 9.933E-06 | 3.101E-06 | 6.800E-06 | 1.300E-05 |
| 1902 | 96 | D96R | A | 5.8e-06 | 6.9e-06 | 8e-06 | 6.900E-06 | 1.100E-06 | 5.800E-06 | 8.000E-06 |
| 1903 | 96 | D96N | A | 7.3e-06 | 1.3e-05 | 9e-06 | 9.767E-06 | 2.926E-06 | 7.300E-06 | 1.300E-05 |
| 1904 | 96 | D96D | A | 7.9e-06 | 8.3e-06 | 1.3e-05 | 9.733E-06 | 2.836E-06 | 7.900E-06 | 1.300E-05 |
| 1905 | 96 | D96C | A | 8.6e-06 | 1.3e-05 | 7.1e-06 | 9.567E-06 | 3.066E-06 | 7.100E-06 | 1.300E-05 |
| 1906 | 96 | D96E | A | 6.3e-06 | 1.3e-05 | 7.2e-06 | 8.833E-06 | 3.636E-06 | 6.300E-06 | 1.300E-05 |
| 1907 | 96 | D96Q | A | 7.7e-06 | 1.3e-05 | 6.2e-06 | 8.967E-06 | 3.573E-06 | 6.200E-06 | 1.300E-05 |
| 1908 | 96 | D96G | A | 5.5e-06 | 1.5e-05 | 5.8e-06 | 8.767E-06 | 5.400E-06 | 5.500E-06 | 1.500E-05 |
| 1909 | 96 | D96H | A | 6.6e-06 | 9.1e-06 | 6.4e-06 | 7.367E-06 | 1.504E-06 | 6.400E-06 | 9.100E-06 |
| 1910 | 96 | D96I | A | 7e-06 | 9.9e-06 | 8.4e-06 | 8.433E-06 | 1.450E-06 | 7.000E-06 | 9.900E-06 |
| 1911 | 96 | D96L | A | 6.7e-06 | 9.7e-06 | 9.3e-06 | 8.567E-06 | 1.629E-06 | 6.700E-06 | 9.700E-06 |
| 1912 | 96 | D96K | A | 1e-05 | 1.4e-05 | 6.3e-06 | 1.010E-05 | 3.851E-06 | 6.300E-06 | 1.400E-05 |
| 1913 | 96 | D96M | A | 8e-06 | 7.6e-06 | 5.9e-06 | 7.167E-06 | 1.115E-06 | 5.900E-06 | 8.000E-06 |
| 1914 | 96 | D96F | A | 8e-06 | 6.5e-06 | 1e-05 | 8.167E-06 | 1.756E-06 | 6.500E-06 | 1.000E-05 |
| 1915 | 96 | D96P | A | 9.3e-06 | 1e-05 | 6.5e-06 | 8.600E-06 | 1.852E-06 | 6.500E-06 | 1.000E-05 |
| 1916 | 96 | D96S | A | 8.1e-06 | 1.1e-05 | 9.8e-06 | 9.633E-06 | 1.457E-06 | 8.100E-06 | 1.100E-05 |
| 1917 | 96 | D96T | A | 5.2e-06 | 6.2e-06 | 7.4e-06 | 6.267E-06 | 1.102E-06 | 5.200E-06 | 7.400E-06 |
| 1918 | 96 | D96W | A | 8.4e-06 | 5.6e-06 | 7.5e-06 | 7.167E-06 | 1.429E-06 | 5.600E-06 | 8.400E-06 |
| 1919 | 96 | D96Y | A | 5.1e-06 | 8.6e-06 | 8.7e-06 | 7.467E-06 | 2.050E-06 | 5.100E-06 | 8.700E-06 |
| 1920 | 96 | D96V | A | 6.1e-06 | 8.1e-06 | 1.4e-05 | 9.400E-06 | 4.107E-06 | 6.100E-06 | 1.400E-05 |
| 1921 | 97 | N97A | A | 9.9e-06 | 5.9e-06 | 1e-05 | 8.600E-06 | 2.339E-06 | 5.900E-06 | 1.000E-05 |
| 1922 | 97 | N97R | A | 6.9e-06 | 9e-06 | 3.1e-06 | 6.333E-06 | 2.991E-06 | 3.100E-06 | 9.000E-06 |
| 1923 | 97 | N97N | A | 6.2e-06 | 9.1e-06 | 9.1e-06 | 8.133E-06 | 1.674E-06 | 6.200E-06 | 9.100E-06 |
| 1924 | 97 | N97D | A | 6.3e-06 | 7.9e-06 | 4.7e-06 | 6.300E-06 | 1.600E-06 | 4.700E-06 | 7.900E-06 |
| 1925 | 97 | N97C | A | 8.5e-06 | 6.6e-06 | 6.7e-06 | 7.267E-06 | 1.069E-06 | 6.600E-06 | 8.500E-06 |
| 1926 | 97 | N97E | A | 1.3e-05 | 9.5e-06 | 7.2e-06 | 9.900E-06 | 2.921E-06 | 7.200E-06 | 1.300E-05 |
| 1927 | 97 | N97Q | A | 4.5e-06 | 6.8e-06 | 9.9e-06 | 7.067E-06 | 2.710E-06 | 4.500E-06 | 9.900E-06 |
| 1928 | 97 | N97G | A | 3.9e-06 | 6.2e-06 | 8.1e-06 | 6.067E-06 | 2.103E-06 | 3.900E-06 | 8.100E-06 |
| 1929 | 97 | N97H | A | 5e-06 | 9.6e-06 | 5.4e-06 | 6.667E-06 | 2.548E-06 | 5.000E-06 | 9.600E-06 |
| 1930 | 97 | N97I | A | 7.3e-06 | 1e-05 | 8.9e-06 | 8.733E-06 | 1.358E-06 | 7.300E-06 | 1.000E-05 |
| 1931 | 97 | N97L | A | 7.5e-06 | 7.9e-06 | 8e-06 | 7.800E-06 | 2.646E-07 | 7.500E-06 | 8.000E-06 |
| 1932 | 97 | N97K | A | 9.5e-06 | 9.4e-06 | 8.4e-06 | 9.100E-06 | 6.083E-07 | 8.400E-06 | 9.500E-06 |
| 1933 | 97 | N97M | A | 1.5e-05 | 6.2e-06 | 8.5e-06 | 9.900E-06 | 4.564E-06 | 6.200E-06 | 1.500E-05 |
| 1934 | 97 | N97F | A | 7.6e-06 | 7.3e-06 | 1e-05 | 8.300E-06 | 1.480E-06 | 7.300E-06 | 1.000E-05 |
| 1935 | 97 | N97P | A | 4.6e-06 | 8.4e-06 | 7.3e-06 | 6.767E-06 | 1.955E-06 | 4.600E-06 | 8.400E-06 |
| 1936 | 97 | N97S | A | 8.2e-06 | 9e-06 | 1.4e-05 | 1.040E-05 | 3.143E-06 | 8.200E-06 | 1.400E-05 |
| 1937 | 97 | N97T | A | 5.3e-06 | 1.5e-05 | 6.8e-06 | 9.033E-06 | 5.221E-06 | 5.300E-06 | 1.500E-05 |
| 1938 | 97 | N97W | A | 5.5e-06 | 5.8e-06 | 7.3e-06 | 6.200E-06 | 9.644E-07 | 5.500E-06 | 7.300E-06 |
| 1939 | 97 | N97Y | A | 1.2e-05 | 1e-05 | 1.1e-05 | 1.100E-05 | 1.000E-06 | 1.000E-05 | 1.200E-05 |
| 1940 | 97 | N97V | A | 8.2e-06 | 3.3e-06 | 8.1e-06 | 6.533E-06 | 2.801E-06 | 3.300E-06 | 8.200E-06 |
| 1941 | 98 | G98A | A | 4.5e-06 | 8e-06 | 7.6e-06 | 6.700E-06 | 1.916E-06 | 4.500E-06 | 8.000E-06 |
| 1942 | 98 | G98R | A | 1.3e-05 | 9.9e-06 | 6.8e-06 | 9.900E-06 | 3.100E-06 | 6.800E-06 | 1.300E-05 |
| 1943 | 98 | G98N | A | 8.5e-06 | 9.3e-06 | 7.3e-06 | 8.367E-06 | 1.007E-06 | 7.300E-06 | 9.300E-06 |
| 1944 | 98 | G98D | A | 1e-05 | 1.2e-05 | 6.6e-06 | 9.533E-06 | 2.730E-06 | 6.600E-06 | 1.200E-05 |
| 1945 | 98 | G98C | A | 5.7e-06 | 4.5e-06 | 8.6e-06 | 6.267E-06 | 2.108E-06 | 4.500E-06 | 8.600E-06 |
| 1946 | 98 | G98E | A | 7.8e-06 | 1e-05 | 7.8e-06 | 8.533E-06 | 1.270E-06 | 7.800E-06 | 1.000E-05 |
| 1947 | 98 | G98Q | A | 9.5e-06 | 8.1e-06 | 1e-05 | 9.200E-06 | 9.849E-07 | 8.100E-06 | 1.000E-05 |
| 1948 | 98 | G98G | A | 1.2e-05 | 1.3e-05 | 9.3e-06 | 1.143E-05 | 1.914E-06 | 9.300E-06 | 1.300E-05 |
| 1949 | 98 | G98H | A | 6e-06 | 8.1e-06 | 3.9e-06 | 6.000E-06 | 2.100E-06 | 3.900E-06 | 8.100E-06 |
| 1950 | 98 | G98I | A | 7.5e-06 | 7.5e-06 | 1.2e-05 | 9.000E-06 | 2.598E-06 | 7.500E-06 | 1.200E-05 |
| 1951 | 98 | G98L | A | 1.1e-05 | 7e-06 | 9.4e-06 | 9.133E-06 | 2.013E-06 | 7.000E-06 | 1.100E-05 |
| 1952 | 98 | G98K | A | 9.1e-06 | 6.5e-06 | 5.7e-06 | 7.100E-06 | 1.778E-06 | 5.700E-06 | 9.100E-06 |
| 1953 | 98 | G98M | A | 5e-06 | 6.2e-06 | 5.6e-06 | 5.600E-06 | 6.000E-07 | 5.000E-06 | 6.200E-06 |
| 1954 | 98 | G98F | A | 3.5e-06 | 7.2e-06 | 9.5e-06 | 6.733E-06 | 3.027E-06 | 3.500E-06 | 9.500E-06 |

| | | | | | | | | | | |
|---|---|---|---|---|---|---|---|---|---|---|
| 1955 | 98 | G98P | A | 5.4e-06 | 6.7e-06 | 6.6e-06 | 6.233E-06 | 7.234E-07 | 5.400E-06 | 6.700E-06 |
| 1956 | 98 | G98S | A | 9.4e-06 | 4.5e-06 | 6e-06 | 6.633E-06 | 2.511E-06 | 4.500E-06 | 9.400E-06 |
| 1957 | 98 | G98T | A | 4.2e-06 | 1.4e-05 | 8.2e-06 | 8.800E-06 | 4.927E-06 | 4.200E-06 | 1.400E-05 |
| 1958 | 98 | G98W | A | 5.8e-06 | 1.4e-05 | 5.1e-06 | 8.300E-06 | 4.949E-06 | 5.100E-06 | 1.400E-05 |
| 1959 | 98 | G98Y | A | 8.9e-06 | 3.9e-06 | 6e-06 | 6.267E-06 | 2.511E-06 | 3.900E-06 | 8.900E-06 |
| 1960 | 98 | G98V | A | 6.9e-06 | 8.8e-06 | 8.4e-06 | 8.033E-06 | 1.002E-06 | 6.900E-06 | 8.800E-06 |
| 1961 | 99 | D99A | A | 1.1e-05 | 1.5e-05 | 6.1e-06 | 1.070E-05 | 4.458E-06 | 6.100E-06 | 1.500E-05 |
| 1962 | 99 | D99R | A | 1.9e-05 | 9.8e-06 | 6.7e-06 | 1.183E-05 | 6.397E-06 | 6.700E-06 | 1.900E-05 |
| 1963 | 99 | D99N | A | 6.5e-06 | 7.3e-06 | 6.7e-06 | 6.833E-06 | 4.163E-07 | 6.500E-06 | 7.300E-06 |
| 1964 | 99 | D99D | A | 6.9e-06 | 5.7e-06 | 7e-06 | 6.533E-06 | 7.234E-07 | 5.700E-06 | 7.000E-06 |
| 1965 | 99 | D99C | A | 7e-06 | 8.7e-06 | 4.6e-06 | 6.767E-06 | 2.060E-06 | 4.600E-06 | 8.700E-06 |
| 1966 | 99 | D99E | A | 8.4e-06 | 8.6e-06 | 8.5e-06 | 8.500E-06 | 1.000E-07 | 8.400E-06 | 8.600E-06 |
| 1967 | 99 | D99Q | A | 6.7e-06 | 7.5e-06 | 7.8e-06 | 7.333E-06 | 5.686E-07 | 6.700E-06 | 7.800E-06 |
| 1968 | 99 | D99G | A | 8.6e-06 | 1e-05 | 9.2e-06 | 9.267E-06 | 7.024E-07 | 8.600E-06 | 1.000E-05 |
| 1969 | 99 | D99H | A | 9.8e-06 | 8.1e-06 | 1e-05 | 9.300E-06 | 1.044E-06 | 8.100E-06 | 1.000E-05 |
| 1970 | 99 | D99I | A | 9.7e-06 | 6.7e-06 | 8.9e-06 | 8.433E-06 | 1.553E-06 | 6.700E-06 | 9.700E-06 |
| 1971 | 99 | D99L | A | 7.3e-06 | 9.5e-06 | 4.8e-06 | 7.200E-06 | 2.352E-06 | 4.800E-06 | 9.500E-06 |
| 1972 | 99 | D99K | A | 1.2e-05 | 1e-05 | 6.6e-06 | 9.533E-06 | 2.730E-06 | 6.600E-06 | 1.200E-05 |
| 1973 | 99 | D99M | A | 5.4e-06 | 5.1e-06 | 4.9e-06 | 5.133E-06 | 2.517E-07 | 4.900E-06 | 5.400E-06 |
| 1974 | 99 | D99F | A | 4.8e-06 | 7.9e-06 | 8.2e-06 | 6.967E-06 | 1.882E-06 | 4.800E-06 | 8.200E-06 |
| 1975 | 99 | D99P | A | 8e-06 | 9.9e-06 | 9.9e-06 | 9.267E-06 | 1.097E-06 | 8.000E-06 | 9.900E-06 |
| 1976 | 99 | D99S | A | 1.1e-05 | 8e-06 | 6.5e-06 | 8.500E-06 | 2.291E-06 | 6.500E-06 | 1.100E-05 |
| 1977 | 99 | D99T | A | 6.9e-06 | 7.7e-06 | 1.3e-05 | 9.200E-06 | 3.315E-06 | 6.900E-06 | 1.300E-05 |
| 1978 | 99 | D99W | A | 9.2e-06 | 8.8e-06 | 5.6e-06 | 7.867E-06 | 1.973E-06 | 5.600E-06 | 9.200E-06 |
| 1979 | 99 | D99Y | A | 5.8e-06 | 6.4e-06 | 5.5e-06 | 5.900E-06 | 4.583E-07 | 5.500E-06 | 6.400E-06 |
| 1980 | 99 | D99V | A | 6.8e-06 | 7.2e-06 | 8.9e-06 | 7.633E-06 | 1.115E-06 | 6.800E-06 | 8.900E-06 |
| 1981 | 100 | P100A | A | 8.3e-06 | 6.6e-06 | 9.3e-06 | 8.067E-06 | 1.365E-06 | 6.600E-06 | 9.300E-06 |
| 1982 | 100 | P100R | A | 1.4e-05 | 8.6e-06 | 3.8e-06 | 8.800E-06 | 5.103E-06 | 3.800E-06 | 1.400E-05 |
| 1983 | 100 | P100N | A | 1.4e-05 | 1.1e-05 | 5.3e-06 | 1.010E-05 | 4.419E-06 | 5.300E-06 | 1.400E-05 |
| 1984 | 100 | P100D | A | 8.4e-06 | 7.6e-06 | 1.1e-05 | 9.000E-06 | 1.778E-06 | 7.600E-06 | 1.100E-05 |
| 1985 | 100 | P100C | A | 7.1e-06 | 1.2e-05 | 1.2e-05 | 1.037E-05 | 2.829E-06 | 7.100E-06 | 1.200E-05 |
| 1986 | 100 | P100E | A | 1.6e-05 | 1.1e-05 | 1.2e-05 | 1.300E-05 | 2.646E-06 | 1.100E-05 | 1.600E-05 |
| 1987 | 100 | P100Q | A | 9.7e-06 | 6.6e-06 | 1e-05 | 8.767E-06 | 1.882E-06 | 6.600E-06 | 1.000E-05 |
| 1988 | 100 | P100G | A | 6.3e-06 | 7.6e-06 | 8.1e-06 | 7.333E-06 | 9.292E-07 | 6.300E-06 | 8.100E-06 |
| 1989 | 100 | P100H | A | 6.4e-06 | 8.2e-06 | 7.9e-06 | 7.500E-06 | 9.644E-07 | 6.400E-06 | 8.200E-06 |
| 1990 | 100 | P100I | A | 1e-05 | 8e-06 | 7.2e-06 | 8.400E-06 | 1.442E-06 | 7.200E-06 | 1.000E-05 |
| 1991 | 100 | P100L | A | 6.8e-06 | 8.1e-06 | 6.1e-06 | 7.000E-06 | 1.015E-06 | 6.100E-06 | 8.100E-06 |
| 1992 | 100 | P100K | A | 6.7e-06 | 5.5e-06 | 6.2e-06 | 6.133E-06 | 6.028E-07 | 5.500E-06 | 6.700E-06 |
| 1993 | 100 | P100M | A | 9.5e-06 | 1.3e-05 | 6.7e-06 | 9.733E-06 | 3.156E-06 | 6.700E-06 | 1.300E-05 |
| 1994 | 100 | P100F | A | 6.8e-06 | 6.9e-06 | 7.2e-06 | 6.967E-06 | 2.082E-07 | 6.800E-06 | 7.200E-06 |
| 1995 | 100 | P100P | A | 1.1e-05 | 6.2e-06 | 1.3e-05 | 1.007E-05 | 3.495E-06 | 6.200E-06 | 1.300E-05 |
| 1996 | 100 | P100S | A | 9.6e-06 | 3.5e-06 | 8.6e-06 | 7.233E-06 | 3.272E-06 | 3.500E-06 | 9.600E-06 |
| 1997 | 100 | P100T | A | 7.2e-06 | 9.1e-06 | 1.1e-05 | 9.100E-06 | 1.900E-06 | 7.200E-06 | 1.100E-05 |
| 1998 | 100 | P100W | A | 9.6e-06 | 5.9e-06 | 9.5e-06 | 8.333E-06 | 2.108E-06 | 5.900E-06 | 9.600E-06 |
| 1999 | 100 | P100Y | A | 1.3e-05 | 6.8e-06 | 6.5e-06 | 8.767E-06 | 3.669E-06 | 6.500E-06 | 1.300E-05 |
| 2000 | 100 | P100V | A | 8.9e-06 | 8.8e-06 | 8.9e-06 | 8.867E-06 | 5.774E-08 | 8.800E-06 | 8.900E-06 |
| 2001 | 101 | L101A | A | 1e-05 | 1e-05 | 9.2e-06 | 9.733E-06 | 4.619E-07 | 9.200E-06 | 1.000E-05 |
| 2002 | 101 | L101R | A | 7.3e-06 | 6.2e-06 | 4.4e-06 | 5.967E-06 | 1.464E-06 | 4.400E-06 | 7.300E-06 |
| 2003 | 101 | L101N | A | 9.8e-06 | 9.4e-06 | 1e-05 | 9.733E-06 | 3.055E-07 | 9.400E-06 | 1.000E-05 |
| 2004 | 101 | L101D | A | 8.3e-06 | 5.3e-06 | 6.9e-06 | 6.833E-06 | 1.501E-06 | 5.300E-06 | 8.300E-06 |
| 2005 | 101 | L101C | A | 6.5e-06 | 9.1e-06 | 6.1e-06 | 7.233E-06 | 1.629E-06 | 6.100E-06 | 9.100E-06 |
| 2006 | 101 | L101E | A | 8.4e-06 | 1.1e-05 | 9.1e-06 | 9.500E-06 | 1.345E-06 | 8.400E-06 | 1.100E-05 |
| 2007 | 101 | L101Q | A | 6.2e-06 | 1e-05 | 1.3e-05 | 9.733E-06 | 3.408E-06 | 6.200E-06 | 1.300E-05 |
| 2008 | 101 | L101G | A | 7.7e-06 | 8e-06 | 5.6e-06 | 7.100E-06 | 1.308E-06 | 5.600E-06 | 8.000E-06 |
| 2009 | 101 | L101H | A | 6.3e-06 | 3.6e-06 | 1.3e-05 | 7.633E-06 | 4.840E-06 | 3.600E-06 | 1.300E-05 |
| 2010 | 101 | L101I | A | 1.2e-05 | 8.8e-06 | 1.3e-05 | 1.127E-05 | 2.194E-06 | 8.800E-06 | 1.300E-05 |
| 2011 | 101 | L101L | A | 1.1e-05 | 6.2e-06 | 9.6e-06 | 8.933E-06 | 2.468E-06 | 6.200E-06 | 1.100E-05 |
| 2012 | 101 | L101K | A | 8.3e-06 | 1.2e-05 | 6.5e-06 | 8.933E-06 | 2.804E-06 | 6.500E-06 | 1.200E-05 |
| 2013 | 101 | L101M | A | 5.8e-06 | 8e-06 | 1e-05 | 7.933E-06 | 2.101E-06 | 5.800E-06 | 1.000E-05 |
| 2014 | 101 | L101F | A | 5.5e-06 | 1.4e-05 | 8.9e-06 | 9.467E-06 | 4.278E-06 | 5.500E-06 | 1.400E-05 |
| 2015 | 101 | L101P | A | 1e-05 | 8.1e-06 | 1.3e-05 | 1.037E-05 | 2.470E-06 | 8.100E-06 | 1.300E-05 |
| 2016 | 101 | L101S | A | 7.3e-06 | 9.3e-06 | 9.4e-06 | 8.667E-06 | 1.185E-06 | 7.300E-06 | 9.400E-06 |
| 2017 | 101 | L101T | A | 1.2e-05 | 7.6e-06 | 1.5e-05 | 1.153E-05 | 3.722E-06 | 7.600E-06 | 1.500E-05 |
| 2018 | 101 | L101W | A | 6.6e-06 | 4.8e-06 | 8.6e-06 | 6.667E-06 | 1.901E-06 | 4.800E-06 | 8.600E-06 |
| 2019 | 101 | L101Y | A | 8.5e-06 | 5.1e-06 | 4.2e-06 | 5.933E-06 | 2.268E-06 | 4.200E-06 | 8.500E-06 |
| 2020 | 101 | L101V | A | 6.4e-06 | 1.3e-05 | 9.1e-06 | 9.500E-06 | 3.318E-06 | 6.400E-06 | 1.300E-05 |
| 2021 | 102 | N102A | A | 7.4e-06 | 8.4e-06 | 1.4e-05 | 9.933E-06 | 3.557E-06 | 7.400E-06 | 1.400E-05 |
| 2022 | 102 | N102R | A | 7.8e-06 | 6.8e-06 | 7.3e-06 | 7.300E-06 | 5.000E-07 | 6.800E-06 | 7.800E-06 |
| 2023 | 102 | N102N | A | 6.8e-06 | 4.7e-06 | 5e-06 | 5.500E-06 | 1.136E-06 | 4.700E-06 | 6.800E-06 |
| 2024 | 102 | N102D | A | 6.9e-06 | 1.2e-05 | 7.8e-06 | 8.900E-06 | 2.722E-06 | 6.900E-06 | 1.200E-05 |
| 2025 | 102 | N102C | A | 6e-06 | 5.1e-06 | 6.9e-06 | 6.000E-06 | 9.000E-07 | 5.100E-06 | 6.900E-06 |
| 2026 | 102 | N102E | A | 6.7e-06 | 9.6e-06 | 9.9e-06 | 8.733E-06 | 1.767E-06 | 6.700E-06 | 9.900E-06 |
| 2027 | 102 | N102Q | A | 9.5e-06 | 9.4e-06 | 9.7e-06 | 9.533E-06 | 1.528E-07 | 9.400E-06 | 9.700E-06 |
| 2028 | 102 | N102G | A | 5.9e-06 | 5.8e-06 | 9.4e-06 | 7.033E-06 | 2.050E-06 | 5.800E-06 | 9.400E-06 |
| 2029 | 102 | N102H | A | 4.9e-06 | 1.3e-05 | 9.3e-06 | 9.067E-06 | 4.055E-06 | 4.900E-06 | 1.300E-05 |
| 2030 | 102 | N102I | A | 6.9e-06 | 9.1e-06 | 7.5e-06 | 7.833E-06 | 1.137E-06 | 6.900E-06 | 9.100E-06 |
| 2031 | 102 | N102L | A | 6.8e-06 | 1.1e-05 | 9e-06 | 8.933E-06 | 2.101E-06 | 6.800E-06 | 1.100E-05 |
| 2032 | 102 | N102K | A | 6.8e-06 | 9e-06 | 6.1e-06 | 7.300E-06 | 1.513E-06 | 6.100E-06 | 9.000E-06 |
| 2033 | 102 | N102M | A | 6.4e-06 | 6.5e-06 | 9.3e-06 | 7.400E-06 | 1.646E-06 | 6.400E-06 | 9.300E-06 |
| 2034 | 102 | N102F | A | 6.1e-06 | 1.1e-05 | 9.3e-06 | 8.800E-06 | 2.488E-06 | 6.100E-06 | 1.100E-05 |
| 2035 | 102 | N102P | A | 8.8e-06 | 6.9e-06 | 1.1e-05 | 8.900E-06 | 2.052E-06 | 6.900E-06 | 1.100E-05 |
| 2036 | 102 | N102S | A | 7e-06 | 1e-05 | 8e-06 | 8.333E-06 | 1.528E-06 | 7.000E-06 | 1.000E-05 |
| 2037 | 102 | N102T | A | 5.4e-06 | 8e-06 | 1.3e-05 | 8.800E-06 | 3.863E-06 | 5.400E-06 | 1.300E-05 |
| 2038 | 102 | N102W | A | 1.3e-05 | 9.9e-06 | 6.4e-06 | 9.767E-06 | 3.302E-06 | 6.400E-06 | 1.300E-05 |
| 2039 | 102 | N102Y | A | 1.3e-05 | 9.4e-06 | 1e-05 | 1.080E-05 | 1.929E-06 | 9.400E-06 | 1.300E-05 |

| | | | | | | | | | | |
|---|---|---|---|---|---|---|---|---|---|---|
| 2040 | 102 | N102V | A | 7.3e-06 | 1.3e-05 | 6.8e-06 | 9.033E-06 | 3.444E-06 | 6.800E-06 | 1.300E-05 |
| 2041 | 103 | N103A | A | 5.6e-06 | 6.2e-06 | 8.7e-06 | 6.833E-06 | 1.644E-06 | 5.600E-06 | 8.700E-06 |
| 2042 | 103 | N103R | A | 6.4e-06 | 1.4e-05 | 7.6e-06 | 9.333E-06 | 4.086E-06 | 6.400E-06 | 1.400E-05 |
| 2043 | 103 | N103N | A | 6.4e-06 | 7.4e-06 | 5.8e-06 | 6.533E-06 | 8.083E-07 | 5.800E-06 | 7.400E-06 |
| 2044 | 103 | N103D | A | 6.9e-06 | 9.6e-06 | 5.4e-06 | 7.300E-06 | 2.128E-06 | 5.400E-06 | 9.600E-06 |
| 2045 | 103 | N103C | A | 9.1e-06 | 3.7e-06 | 5.7e-06 | 6.167E-06 | 2.730E-06 | 3.700E-06 | 9.100E-06 |
| 2046 | 103 | N103E | A | 8.6e-06 | 4.7e-06 | 5.7e-06 | 6.333E-06 | 2.026E-06 | 4.700E-06 | 8.600E-06 |
| 2047 | 103 | N103Q | A | 8.8e-06 | 1.1e-05 | 1e-05 | 9.933E-06 | 1.102E-06 | 8.800E-06 | 1.100E-05 |
| 2048 | 103 | N103G | A | 7.4e-06 | 5.8e-06 | 6.2e-06 | 6.467E-06 | 8.327E-07 | 5.800E-06 | 7.400E-06 |
| 2049 | 103 | N103H | A | 5.8e-06 | 4.1e-06 | 9.5e-06 | 6.467E-06 | 2.761E-06 | 4.100E-06 | 9.500E-06 |
| 2050 | 103 | N103I | A | 9e-06 | 7.2e-06 | 7.1e-06 | 7.767E-06 | 1.069E-06 | 7.100E-06 | 9.000E-06 |
| 2051 | 103 | N103L | A | 1.2e-05 | 4.8e-06 | 4.3e-06 | 7.033E-06 | 4.309E-06 | 4.300E-06 | 1.200E-05 |
| 2052 | 103 | N103K | A | 9.1e-06 | 9.8e-06 | 6.8e-06 | 8.567E-06 | 1.570E-06 | 6.800E-06 | 9.800E-06 |
| 2053 | 103 | N103M | A | 4.6e-06 | 9.1e-06 | 8.6e-06 | 7.433E-06 | 2.466E-06 | 4.600E-06 | 9.100E-06 |
| 2054 | 103 | N103F | A | 5e-06 | 3.7e-06 | 7.5e-06 | 5.400E-06 | 1.931E-06 | 3.700E-06 | 7.500E-06 |
| 2055 | 103 | N103P | A | 3.9e-06 | 1.4e-05 | 6.4e-06 | 8.100E-06 | 5.260E-06 | 3.900E-06 | 1.400E-05 |
| 2056 | 103 | N103S | A | 7.4e-06 | 1.4e-05 | 5.2e-06 | 8.867E-06 | 4.580E-06 | 5.200E-06 | 1.400E-05 |
| 2057 | 103 | N103T | A | 4.7e-06 | 1.3e-05 | 6.4e-06 | 8.033E-06 | 4.384E-06 | 4.700E-06 | 1.300E-05 |
| 2058 | 103 | N103W | A | 6.9e-06 | 7.9e-06 | 4.9e-06 | 6.567E-06 | 1.528E-06 | 4.900E-06 | 7.900E-06 |
| 2059 | 103 | N103Y | A | 6.4e-06 | 1.1e-05 | 1.4e-05 | 1.047E-05 | 3.828E-06 | 6.400E-06 | 1.400E-05 |
| 2060 | 103 | N103V | A | 1.8e-05 | 5.8e-06 | 7.9e-06 | 1.057E-05 | 6.523E-06 | 5.800E-06 | 1.800E-05 |
| 2061 | 104 | T104A | A | 1.3e-05 | 9.1e-06 | 9.2e-06 | 1.043E-05 | 2.223E-06 | 9.100E-06 | 1.300E-05 |
| 2062 | 104 | T104R | A | 8.2e-06 | 7.7e-06 | 7.2e-06 | 7.700E-06 | 5.000E-07 | 7.200E-06 | 8.200E-06 |
| 2063 | 104 | T104N | A | 7.8e-06 | 6.7e-06 | 6.3e-06 | 6.933E-06 | 7.767E-07 | 6.300E-06 | 7.800E-06 |
| 2064 | 104 | T104D | A | 5.4e-06 | 6.5e-06 | 6.8e-06 | 6.233E-06 | 7.371E-07 | 5.400E-06 | 6.800E-06 |
| 2065 | 104 | T104C | A | 9.2e-06 | 1e-05 | 6.8e-06 | 8.667E-06 | 1.665E-06 | 6.800E-06 | 1.000E-05 |
| 2066 | 104 | T104E | A | 1.1e-05 | 4.4e-06 | 1.3e-05 | 9.467E-06 | 4.500E-06 | 4.400E-06 | 1.300E-05 |
| 2067 | 104 | T104Q | A | 1.3e-05 | 9.8e-06 | 5.5e-06 | 9.433E-06 | 3.763E-06 | 5.500E-06 | 1.300E-05 |
| 2068 | 104 | T104G | A | 6.6e-06 | 8.5e-06 | 9.7e-06 | 8.267E-06 | 1.563E-06 | 6.600E-06 | 9.700E-06 |
| 2069 | 104 | T104H | A | 6.1e-06 | 7.2e-06 | 6.5e-06 | 6.600E-06 | 5.568E-07 | 6.100E-06 | 7.200E-06 |
| 2070 | 104 | T104I | A | 6.4e-06 | 6.5e-06 | 9.6e-06 | 7.500E-06 | 1.819E-06 | 6.400E-06 | 9.600E-06 |
| 2071 | 104 | T104L | A | 8.1e-06 | 1e-05 | 6e-06 | 8.033E-06 | 2.001E-06 | 6.000E-06 | 1.000E-05 |
| 2072 | 104 | T104K | A | 1.4e-05 | 1.3e-05 | 7.5e-06 | 1.150E-05 | 3.500E-06 | 7.500E-06 | 1.400E-05 |
| 2073 | 104 | T104M | A | 1e-05 | 6.2e-06 | 9.4e-06 | 8.533E-06 | 2.043E-06 | 6.200E-06 | 1.000E-05 |
| 2074 | 104 | T104F | A | 5e-06 | 8.6e-06 | 5.3e-06 | 6.300E-06 | 1.997E-06 | 5.000E-06 | 8.600E-06 |
| 2075 | 104 | T104P | A | 9.2e-06 | 9e-06 | 7.4e-06 | 8.533E-06 | 9.866E-07 | 7.400E-06 | 9.200E-06 |
| 2076 | 104 | T104S | A | 1e-05 | 1.1e-05 | 8.3e-06 | 9.767E-06 | 1.365E-06 | 8.300E-06 | 1.100E-05 |
| 2077 | 104 | T104T | A | 8.5e-06 | 1e-05 | 1.1e-05 | 9.833E-06 | 1.258E-06 | 8.500E-06 | 1.100E-05 |
| 2078 | 104 | T104W | A | 1.1e-05 | 8e-06 | 1.3e-05 | 1.067E-05 | 2.517E-06 | 8.000E-06 | 1.300E-05 |
| 2079 | 104 | T104Y | A | 5.4e-06 | 9.6e-06 | 6.6e-06 | 7.200E-06 | 2.163E-06 | 5.400E-06 | 9.600E-06 |
| 2080 | 104 | T104V | A | 9.3e-06 | 1.3e-05 | 8.7e-06 | 1.033E-05 | 2.329E-06 | 8.700E-06 | 1.300E-05 |
| 2081 | 105 | T105A | A | 6.5e-06 | 1.2e-05 | 9.7e-06 | 9.400E-06 | 2.762E-06 | 6.500E-06 | 1.200E-05 |
| 2082 | 105 | T105R | A | 6.5e-06 | 8.6e-06 | 7.9e-06 | 7.667E-06 | 1.069E-06 | 6.500E-06 | 8.600E-06 |
| 2083 | 105 | T105N | A | 7.1e-06 | 9.1e-06 | 7.6e-06 | 7.933E-06 | 1.041E-06 | 7.100E-06 | 9.100E-06 |
| 2084 | 105 | T105D | A | 5.3e-06 | 9.1e-06 | 7.7e-06 | 7.367E-06 | 1.922E-06 | 5.300E-06 | 9.100E-06 |
| 2085 | 105 | T105C | A | 6.6e-06 | 1e-05 | 9.7e-06 | 8.767E-06 | 1.882E-06 | 6.600E-06 | 1.000E-05 |
| 2086 | 105 | T105E | A | 8.3e-06 | 1.1e-05 | 1.1e-05 | 1.010E-05 | 1.559E-06 | 8.300E-06 | 1.100E-05 |
| 2087 | 105 | T105Q | A | 9.6e-06 | 8.5e-06 | 8.6e-06 | 8.900E-06 | 6.083E-07 | 8.500E-06 | 9.600E-06 |
| 2088 | 105 | T105G | A | 8.8e-06 | 6.5e-06 | 6.9e-06 | 7.400E-06 | 1.229E-06 | 6.500E-06 | 8.800E-06 |
| 2089 | 105 | T105H | A | 1.1e-05 | 8.3e-06 | 3.3e-06 | 7.533E-06 | 3.907E-06 | 3.300E-06 | 1.100E-05 |
| 2090 | 105 | T105I | A | 6.2e-06 | 8.1e-06 | 1.1e-05 | 8.433E-06 | 2.417E-06 | 6.200E-06 | 1.100E-05 |
| 2091 | 105 | T105L | A | 7e-06 | 8.3e-06 | 1.4e-05 | 9.767E-06 | 3.723E-06 | 7.000E-06 | 1.400E-05 |
| 2092 | 105 | T105K | A | 8.1e-06 | 5.9e-06 | 1.2e-05 | 8.667E-06 | 3.089E-06 | 5.900E-06 | 1.200E-05 |
| 2093 | 105 | T105M | A | 4.5e-06 | 6.1e-06 | 7e-06 | 5.867E-06 | 1.266E-06 | 4.500E-06 | 7.000E-06 |
| 2094 | 105 | T105F | A | 9.3e-06 | 1.1e-05 | 9.2e-06 | 9.833E-06 | 1.012E-06 | 9.200E-06 | 1.100E-05 |
| 2095 | 105 | T105P | A | 8.3e-06 | 7.6e-06 | 7.3e-06 | 7.733E-06 | 5.132E-07 | 7.300E-06 | 8.300E-06 |
| 2096 | 105 | T105S | A | 8.2e-06 | 4.3e-06 | 1.2e-05 | 8.167E-06 | 3.850E-06 | 4.300E-06 | 1.200E-05 |
| 2097 | 105 | T105T | A | 4.6e-06 | 6.2e-06 | 9e-06 | 6.600E-06 | 2.227E-06 | 4.600E-06 | 9.000E-06 |
| 2098 | 105 | T105W | A | 1e-05 | 9.8e-06 | 7.3e-06 | 9.033E-06 | 1.504E-06 | 7.300E-06 | 1.000E-05 |
| 2099 | 105 | T105Y | A | 7.7e-06 | 7.9e-06 | 1e-05 | 8.533E-06 | 1.274E-06 | 7.700E-06 | 1.000E-05 |
| 2100 | 105 | T105V | A | 9.4e-06 | 6.9e-06 | 4.6e-06 | 6.967E-06 | 2.401E-06 | 4.600E-06 | 9.400E-06 |
| 2101 | 106 | P106A | A | 5.2e-06 | 8.2e-06 | 1.3e-05 | 8.800E-06 | 3.934E-06 | 5.200E-06 | 1.300E-05 |
| 2102 | 106 | P106R | A | 7.6e-06 | 3.3e-06 | 1.1e-05 | 7.300E-06 | 3.859E-06 | 3.300E-06 | 1.100E-05 |
| 2103 | 106 | P106N | A | 6.2e-06 | 8.2e-06 | 5.9e-06 | 6.767E-06 | 1.250E-06 | 5.900E-06 | 8.200E-06 |
| 2104 | 106 | P106D | A | 7.4e-06 | 8.6e-06 | 6.8e-06 | 7.600E-06 | 9.165E-07 | 6.800E-06 | 8.600E-06 |
| 2105 | 106 | P106C | A | 4.8e-06 | 6.6e-06 | 5.3e-06 | 5.567E-06 | 9.292E-07 | 4.800E-06 | 6.600E-06 |
| 2106 | 106 | P106E | A | 6.1e-06 | 1.2e-05 | 9.2e-06 | 9.100E-06 | 2.951E-06 | 6.100E-06 | 1.200E-05 |
| 2107 | 106 | P106Q | A | 8.6e-06 | 7.2e-06 | 5.6e-06 | 7.133E-06 | 1.501E-06 | 5.600E-06 | 8.600E-06 |
| 2108 | 106 | P106G | A | 7.6e-06 | 4.4e-06 | 7.9e-06 | 6.633E-06 | 1.940E-06 | 4.400E-06 | 7.900E-06 |
| 2109 | 106 | P106H | A | 1.2e-05 | 1.2e-05 | 4.4e-06 | 9.467E-06 | 4.388E-06 | 4.400E-06 | 1.200E-05 |
| 2110 | 106 | P106I | A | 6.4e-06 | 6.2e-06 | 7.7e-06 | 6.767E-06 | 8.145E-07 | 6.200E-06 | 7.700E-06 |
| 2111 | 106 | P106L | A | 9.3e-06 | 1.1e-05 | 6.4e-06 | 8.900E-06 | 2.326E-06 | 6.400E-06 | 1.100E-05 |
| 2112 | 106 | P106K | A | 5.3e-06 | 1.4e-05 | 5e-06 | 8.100E-06 | 5.112E-06 | 5.000E-06 | 1.400E-05 |
| 2113 | 106 | P106M | A | 7.1e-06 | 8.5e-06 | 9e-06 | 8.200E-06 | 9.849E-07 | 7.100E-06 | 9.000E-06 |
| 2114 | 106 | P106F | A | 1.5e-05 | 5.8e-06 | 9.3e-06 | 1.003E-05 | 4.644E-06 | 5.800E-06 | 1.500E-05 |
| 2115 | 106 | P106P | A | 9.1e-06 | 3.5e-06 | 6.5e-06 | 6.367E-06 | 2.802E-06 | 3.500E-06 | 9.100E-06 |
| 2116 | 106 | P106S | A | 5.7e-06 | 7.5e-06 | 5.1e-06 | 6.100E-06 | 1.249E-06 | 5.100E-06 | 7.500E-06 |
| 2117 | 106 | P106T | A | 8.6e-06 | 4.5e-06 | 1e-05 | 7.700E-06 | 2.858E-06 | 4.500E-06 | 1.000E-05 |
| 2118 | 106 | P106W | A | 1.2e-05 | 6.2e-06 | 6.1e-06 | 8.100E-06 | 3.378E-06 | 6.100E-06 | 1.200E-05 |
| 2119 | 106 | P106Y | A | 9.8e-06 | 6.1e-06 | 4.9e-06 | 6.933E-06 | 2.554E-06 | 4.900E-06 | 9.800E-06 |
| 2120 | 106 | P106V | A | 9.9e-06 | 6.5e-06 | 9.8e-06 | 8.733E-06 | 1.935E-06 | 6.500E-06 | 9.900E-06 |
| 2121 | 107 | V107A | A | 5.8e-06 | 6e-06 | 1.9e-05 | 1.027E-05 | 7.564E-06 | 5.800E-06 | 1.900E-05 |
| 2122 | 107 | V107R | A | 8.8e-06 | 5e-06 | 8.3e-06 | 7.367E-06 | 2.065E-06 | 5.000E-06 | 8.800E-06 |
| 2123 | 107 | V107N | A | 9.9e-06 | 9.5e-06 | 1.1e-05 | 1.013E-05 | 7.767E-07 | 9.500E-06 | 1.100E-05 |
| 2124 | 107 | V107D | A | 9.4e-06 | 5.8e-06 | 6.9e-06 | 7.367E-06 | 1.845E-06 | 5.800E-06 | 9.400E-06 |

| | | | | | | | | | | |
|---|---|---|---|---|---|---|---|---|---|---|
| 2125 | 107 | V107C | A | 5.1e-06 | 1.2e-05 | 1.1e-05 | 9.367E-06 | 3.729E-06 | 5.100E-06 | 1.200E-05 |
| 2126 | 107 | V107E | A | 7.4e-06 | 1.1e-05 | 6.1e-06 | 8.167E-06 | 2.538E-06 | 6.100E-06 | 1.100E-05 |
| 2127 | 107 | V107Q | A | 9.4e-06 | 9.4e-06 | 1.3e-05 | 1.060E-05 | 2.078E-06 | 9.400E-06 | 1.300E-05 |
| 2128 | 107 | V107G | A | 8.3e-06 | 1e-05 | 1e-05 | 9.433E-06 | 9.815E-07 | 8.300E-06 | 1.000E-05 |
| 2129 | 107 | V107H | A | 4.9e-06 | 5.3e-06 | 6.6e-06 | 5.600E-06 | 8.888E-07 | 4.900E-06 | 6.600E-06 |
| 2130 | 107 | V107I | A | 7e-06 | 8.9e-06 | 1e-05 | 8.633E-06 | 1.518E-06 | 7.000E-06 | 1.000E-05 |
| 2131 | 107 | V107L | A | 7.1e-06 | 4.7e-06 | 6.4e-06 | 6.067E-06 | 1.234E-06 | 4.700E-06 | 7.100E-06 |
| 2132 | 107 | V107K | A | 9.5e-06 | 6.1e-06 | 4.6e-06 | 6.733E-06 | 2.511E-06 | 4.600E-06 | 9.500E-06 |
| 2133 | 107 | V107M | A | 8.7e-06 | 5.5e-06 | 1.3e-05 | 9.067E-06 | 3.763E-06 | 5.500E-06 | 1.300E-05 |
| 2134 | 107 | V107F | A | 6.2e-06 | 1.1e-05 | 1.1e-05 | 9.400E-06 | 2.771E-06 | 6.200E-06 | 1.100E-05 |
| 2135 | 107 | V107P | A | 1.3e-05 | 9.3e-06 | 1e-05 | 1.077E-05 | 1.966E-06 | 9.300E-06 | 1.300E-05 |
| 2136 | 107 | V107S | A | 7.4e-06 | 7.9e-06 | 1e-05 | 8.433E-06 | 1.380E-06 | 7.400E-06 | 1.000E-05 |
| 2137 | 107 | V107T | A | 9.1e-06 | 5.4e-06 | 4.2e-06 | 6.233E-06 | 2.554E-06 | 4.200E-06 | 9.100E-06 |
| 2138 | 107 | V107W | A | 4.4e-06 | 1.3e-05 | 8.5e-06 | 8.633E-06 | 4.302E-06 | 4.400E-06 | 1.300E-05 |
| 2139 | 107 | V107Y | A | 6.5e-06 | 9.6e-06 | 7.8e-06 | 7.967E-06 | 1.557E-06 | 6.500E-06 | 9.600E-06 |
| 2140 | 107 | V107V | A | 1.3e-05 | 9.4e-06 | 1.8e-05 | 1.347E-05 | 4.319E-06 | 9.400E-06 | 1.800E-05 |
| 2141 | 108 | T108A | A | 7.6e-06 | 5.3e-06 | 1.1e-05 | 7.967E-06 | 2.868E-06 | 5.300E-06 | 1.100E-05 |
| 2142 | 108 | T108R | A | 7.4e-06 | 9.7e-06 | 1.3e-05 | 1.003E-05 | 2.815E-06 | 7.400E-06 | 1.300E-05 |
| 2143 | 108 | T108N | A | 8.3e-06 | 7.1e-06 | 1.3e-05 | 9.467E-06 | 3.118E-06 | 7.100E-06 | 1.300E-05 |
| 2144 | 108 | T108D | A | 1.1e-05 | 6.5e-06 | 1e-05 | 9.167E-06 | 2.363E-06 | 6.500E-06 | 1.100E-05 |
| 2145 | 108 | T108C | A | 1.3e-05 | 4.5e-06 | 8.6e-06 | 8.700E-06 | 4.251E-06 | 4.500E-06 | 1.300E-05 |
| 2146 | 108 | T108E | A | 1.8e-05 | 7.4e-06 | 9.7e-06 | 1.170E-05 | 5.576E-06 | 7.400E-06 | 1.800E-05 |
| 2147 | 108 | T108Q | A | 6.8e-06 | 6.7e-06 | 7.7e-06 | 7.067E-06 | 5.508E-07 | 6.700E-06 | 7.700E-06 |
| 2148 | 108 | T108G | A | 6.9e-06 | 9.4e-06 | 1e-05 | 8.767E-06 | 1.644E-06 | 6.900E-06 | 1.000E-05 |
| 2149 | 108 | T108H | A | 9.9e-06 | 8.5e-06 | 5.7e-06 | 8.033E-06 | 2.139E-06 | 5.700E-06 | 9.900E-06 |
| 2150 | 108 | T108I | A | 6.8e-06 | 9.1e-06 | 1.3e-05 | 9.633E-06 | 3.134E-06 | 6.800E-06 | 1.300E-05 |
| 2151 | 108 | T108L | A | 9.9e-06 | 1.5e-05 | 7.1e-06 | 1.067E-05 | 4.005E-06 | 7.100E-06 | 1.500E-05 |
| 2152 | 108 | T108K | A | 1e-05 | 9.8e-06 | 1.2e-05 | 1.060E-05 | 1.217E-06 | 9.800E-06 | 1.200E-05 |
| 2153 | 108 | T108M | A | 6.4e-06 | 6.8e-06 | 7e-06 | 6.733E-06 | 3.055E-07 | 6.400E-06 | 7.000E-06 |
| 2154 | 108 | T108F | A | 7.5e-06 | 5.4e-06 | 9.5e-06 | 7.467E-06 | 2.050E-06 | 5.400E-06 | 9.500E-06 |
| 2155 | 108 | T108P | A | 6.7e-06 | 5.3e-06 | 1.3e-05 | 8.333E-06 | 4.102E-06 | 5.300E-06 | 1.300E-05 |
| 2156 | 108 | T108S | A | 6e-06 | 7.8e-06 | 8.8e-06 | 7.533E-06 | 1.419E-06 | 6.000E-06 | 8.800E-06 |
| 2157 | 108 | T108T | A | 7.4e-06 | 9e-06 | 1.1e-05 | 9.133E-06 | 1.804E-06 | 7.400E-06 | 1.100E-05 |
| 2158 | 108 | T108W | A | 9.7e-06 | 9.7e-06 | 5e-06 | 8.133E-06 | 2.714E-06 | 5.000E-06 | 9.700E-06 |
| 2159 | 108 | T108Y | A | 4.4e-06 | 4.5e-06 | 4.4e-06 | 4.433E-06 | 5.774E-08 | 4.400E-06 | 4.500E-06 |
| 2160 | 108 | T108V | A | 4.9e-06 | 6.9e-06 | 8.7e-06 | 6.833E-06 | 1.901E-06 | 4.900E-06 | 8.700E-06 |
| 2161 | 109 | G109A | A | 9.5e-06 | 6.5e-06 | 8e-06 | 8.000E-06 | 1.500E-06 | 6.500E-06 | 9.500E-06 |
| 2162 | 109 | G109R | A | 1.2e-05 | 6.5e-06 | 5e-06 | 7.833E-06 | 3.686E-06 | 5.000E-06 | 1.200E-05 |
| 2163 | 109 | G109N | A | 7e-06 | 3.6e-06 | 6.6e-06 | 5.733E-06 | 1.858E-06 | 3.600E-06 | 7.000E-06 |
| 2164 | 109 | G109D | A | 1.2e-05 | 9.9e-06 | 1.3e-05 | 1.163E-05 | 1.582E-06 | 9.900E-06 | 1.300E-05 |
| 2165 | 109 | G109C | A | 8.2e-06 | 1.4e-05 | 8.6e-06 | 1.027E-05 | 3.239E-06 | 8.200E-06 | 1.400E-05 |
| 2166 | 109 | G109E | A | 6e-06 | 5.6e-06 | 6.5e-06 | 6.033E-06 | 4.509E-07 | 5.600E-06 | 6.500E-06 |
| 2167 | 109 | G109Q | A | 8.6e-06 | 7.6e-06 | 7.2e-06 | 7.800E-06 | 7.211E-07 | 7.200E-06 | 8.600E-06 |
| 2168 | 109 | G109G | A | 9.2e-06 | 1.2e-05 | 1.2e-05 | 1.107E-05 | 1.617E-06 | 9.200E-06 | 1.200E-05 |
| 2169 | 109 | G109H | A | 5.9e-06 | 8.2e-06 | 1.1e-05 | 8.367E-06 | 2.554E-06 | 5.900E-06 | 1.100E-05 |
| 2170 | 109 | G109I | A | 1.1e-05 | 8.2e-06 | 4.5e-06 | 7.900E-06 | 3.260E-06 | 4.500E-06 | 1.100E-05 |
| 2171 | 109 | G109L | A | 5.1e-06 | 1e-05 | 6.8e-06 | 7.300E-06 | 2.488E-06 | 5.100E-06 | 1.000E-05 |
| 2172 | 109 | G109K | A | 7.8e-06 | 6.9e-06 | 5.1e-06 | 6.600E-06 | 1.375E-06 | 5.100E-06 | 7.800E-06 |
| 2173 | 109 | G109M | A | 7e-06 | 4.7e-06 | 1.3e-05 | 8.233E-06 | 4.285E-06 | 4.700E-06 | 1.300E-05 |
| 2174 | 109 | G109F | A | 9.7e-06 | 3.6e-06 | 9.4e-06 | 7.567E-06 | 3.439E-06 | 3.600E-06 | 9.700E-06 |
| 2175 | 109 | G109P | A | 9.1e-06 | 7.3e-06 | 8.6e-06 | 8.333E-06 | 9.292E-07 | 7.300E-06 | 9.100E-06 |
| 2176 | 109 | G109S | A | 1.3e-05 | 5.6e-06 | 6.4e-06 | 8.333E-06 | 4.061E-06 | 5.600E-06 | 1.300E-05 |
| 2177 | 109 | G109T | A | 9.9e-06 | 5.4e-06 | 5.9e-06 | 7.067E-06 | 2.466E-06 | 5.400E-06 | 9.900E-06 |
| 2178 | 109 | G109W | A | 1e-05 | 4.7e-06 | 1.2e-05 | 8.900E-06 | 3.772E-06 | 4.700E-06 | 1.200E-05 |
| 2179 | 109 | G109Y | A | 6.6e-06 | 5.9e-06 | 9.3e-06 | 7.267E-06 | 1.795E-06 | 5.900E-06 | 9.300E-06 |
| 2180 | 109 | G109V | A | 8.5e-06 | 3.3e-06 | 1e-05 | 7.267E-06 | 3.516E-06 | 3.300E-06 | 1.000E-05 |
| 2181 | 110 | A110A | A | 7.5e-06 | 8.6e-06 | 6.5e-06 | 7.533E-06 | 1.050E-06 | 6.500E-06 | 8.600E-06 |
| 2182 | 110 | A110R | A | 7.9e-06 | 5.9e-06 | 9.8e-06 | 7.867E-06 | 1.950E-06 | 5.900E-06 | 9.800E-06 |
| 2183 | 110 | A110N | A | 6.5e-06 | 8.1e-06 | 1.1e-05 | 8.533E-06 | 2.281E-06 | 6.500E-06 | 1.100E-05 |
| 2184 | 110 | A110D | A | 5.4e-06 | 7.1e-06 | 4.2e-06 | 5.567E-06 | 1.457E-06 | 4.200E-06 | 7.100E-06 |
| 2185 | 110 | A110C | A | 8.2e-06 | 4.8e-06 | 8.1e-06 | 7.033E-06 | 1.935E-06 | 4.800E-06 | 8.200E-06 |
| 2186 | 110 | A110E | A | 1.5e-05 | 8.7e-06 | 8.6e-06 | 1.077E-05 | 3.667E-06 | 8.600E-06 | 1.500E-05 |
| 2187 | 110 | A110Q | A | 5.8e-06 | 4.5e-06 | 1.2e-05 | 7.433E-06 | 4.008E-06 | 4.500E-06 | 1.200E-05 |
| 2188 | 110 | A110G | A | 1.2e-05 | 7.8e-06 | 4.7e-06 | 8.167E-06 | 3.664E-06 | 4.700E-06 | 1.200E-05 |
| 2189 | 110 | A110H | A | 8.4e-06 | 1.2e-05 | 8.2e-06 | 9.533E-06 | 2.139E-06 | 8.200E-06 | 1.200E-05 |
| 2190 | 110 | A110I | A | 5.8e-06 | 8.2e-06 | 6.1e-06 | 6.700E-06 | 1.308E-06 | 5.800E-06 | 8.200E-06 |
| 2191 | 110 | A110L | A | 6.1e-06 | 8.3e-06 | 6.8e-06 | 7.067E-06 | 1.124E-06 | 6.100E-06 | 8.300E-06 |
| 2192 | 110 | A110K | A | 8.2e-06 | 8.5e-06 | 6.4e-06 | 7.700E-06 | 1.136E-06 | 6.400E-06 | 8.500E-06 |
| 2193 | 110 | A110M | A | 9.6e-06 | 8.6e-06 | 7.7e-06 | 8.633E-06 | 9.504E-07 | 7.700E-06 | 9.600E-06 |
| 2194 | 110 | A110F | A | 6.9e-06 | 7.4e-06 | 7e-06 | 7.100E-06 | 2.646E-07 | 6.900E-06 | 7.400E-06 |
| 2195 | 110 | A110P | A | 1.3e-05 | 9.9e-06 | 7.4e-06 | 1.010E-05 | 2.805E-06 | 7.400E-06 | 1.300E-05 |
| 2196 | 110 | A110S | A | 9.8e-06 | 5.8e-06 | 1.3e-05 | 9.533E-06 | 3.607E-06 | 5.800E-06 | 1.300E-05 |
| 2197 | 110 | A110T | A | 7.2e-06 | 7.5e-06 | 6.6e-06 | 7.100E-06 | 4.583E-07 | 6.600E-06 | 7.500E-06 |
| 2198 | 110 | A110W | A | 8.5e-06 | 6.8e-06 | 6.3e-06 | 7.200E-06 | 1.153E-06 | 6.300E-06 | 8.500E-06 |
| 2199 | 110 | A110Y | A | 8e-06 | 4.8e-06 | 9.8e-06 | 7.533E-06 | 2.532E-06 | 4.800E-06 | 9.800E-06 |
| 2200 | 110 | A110V | A | 7.6e-06 | 5.9e-06 | 1.3e-05 | 8.833E-06 | 3.707E-06 | 5.900E-06 | 1.300E-05 |
| 2201 | 111 | S111A | A | 5.4e-06 | 7.1e-06 | 5.7e-06 | 6.067E-06 | 9.074E-07 | 5.400E-06 | 7.100E-06 |
| 2202 | 111 | S111R | A | 8e-06 | 5.7e-06 | 9.3e-06 | 7.667E-06 | 1.823E-06 | 5.700E-06 | 9.300E-06 |
| 2203 | 111 | S111N | A | 9.6e-06 | 8.2e-06 | 9.5e-06 | 9.100E-06 | 7.810E-07 | 8.200E-06 | 9.600E-06 |
| 2204 | 111 | S111D | A | 4.6e-06 | 5.9e-06 | 7.1e-06 | 5.867E-06 | 1.250E-06 | 4.600E-06 | 7.100E-06 |
| 2205 | 111 | S111C | A | 8e-06 | 6.8e-06 | 7.3e-06 | 7.367E-06 | 6.028E-07 | 6.800E-06 | 8.000E-06 |
| 2206 | 111 | S111E | A | 4.6e-06 | 1.3e-05 | 9.7e-06 | 9.100E-06 | 4.232E-06 | 4.600E-06 | 1.300E-05 |
| 2207 | 111 | S111Q | A | 6.3e-06 | 7.1e-06 | 8.8e-06 | 7.400E-06 | 1.277E-06 | 6.300E-06 | 8.800E-06 |
| 2208 | 111 | S111G | A | 1.5e-05 | 6.3e-06 | 1.4e-05 | 1.177E-05 | 4.761E-06 | 6.300E-06 | 1.500E-05 |
| 2209 | 111 | S111H | A | 9.8e-06 | 9.9e-06 | 1.3e-05 | 1.090E-05 | 1.819E-06 | 9.800E-06 | 1.300E-05 |

| | | | | | | | | | | |
|---|---|---|---|---|---|---|---|---|---|---|
| 2210 | 111 | S111I | A | 6.4e-06 | 9e-06 | 1e-05 | 8.467E-06 | 1.858E-06 | 6.400E-06 | 1.000E-05 |
| 2211 | 111 | S111L | A | 9.6e-06 | 1.3e-05 | 6.3e-06 | 9.633E-06 | 3.350E-06 | 6.300E-06 | 1.300E-05 |
| 2212 | 111 | S111K | A | 5.9e-06 | 9.9e-06 | 6e-06 | 7.267E-06 | 2.281E-06 | 5.900E-06 | 9.900E-06 |
| 2213 | 111 | S111M | A | 6.8e-06 | 9.4e-06 | 9.9e-06 | 8.700E-06 | 1.664E-06 | 6.800E-06 | 9.900E-06 |
| 2214 | 111 | S111F | A | 7.5e-06 | 4.8e-06 | 6.3e-06 | 6.200E-06 | 1.353E-06 | 4.800E-06 | 7.500E-06 |
| 2215 | 111 | S111P | A | 4.8e-06 | 7.7e-06 | 5e-06 | 5.833E-06 | 1.620E-06 | 4.800E-06 | 7.700E-06 |
| 2216 | 111 | S111S | A | 1.4e-05 | 1.1e-05 | 1e-05 | 1.167E-05 | 2.082E-06 | 1.000E-05 | 1.400E-05 |
| 2217 | 111 | S111T | A | 3.6e-06 | 8.3e-06 | 1.5e-05 | 8.967E-06 | 5.729E-06 | 3.600E-06 | 1.500E-05 |
| 2218 | 111 | S111W | A | 8.4e-06 | 1e-05 | 9.2e-06 | 9.200E-06 | 8.000E-07 | 8.400E-06 | 1.000E-05 |
| 2219 | 111 | S111Y | A | 5.4e-06 | 1.1e-05 | 1.3e-05 | 9.800E-06 | 3.940E-06 | 5.400E-06 | 1.300E-05 |
| 2220 | 111 | S111V | A | 9.9e-06 | 9.4e-06 | 1.1e-05 | 1.010E-05 | 8.185E-07 | 9.400E-06 | 1.100E-05 |
| 2221 | 112 | P112A | A | 7.6e-06 | 9.5e-06 | 8.8e-06 | 8.633E-06 | 9.609E-07 | 7.600E-06 | 9.500E-06 |
| 2222 | 112 | P112R | A | 7.3e-06 | 5.9e-06 | 6.9e-06 | 6.700E-06 | 7.211E-07 | 5.900E-06 | 7.300E-06 |
| 2223 | 112 | P112N | A | 8.6e-06 | 5.6e-06 | 9.7e-06 | 7.967E-06 | 2.122E-06 | 5.600E-06 | 9.700E-06 |
| 2224 | 112 | P112D | A | 4.6e-06 | 7.9e-06 | 7.1e-06 | 6.533E-06 | 1.721E-06 | 4.600E-06 | 7.900E-06 |
| 2225 | 112 | P112C | A | 6.1e-06 | 4.3e-06 | 1.2e-05 | 7.467E-06 | 4.028E-06 | 4.300E-06 | 1.200E-05 |
| 2226 | 112 | P112E | A | 1e-05 | 9.4e-06 | 9.5e-06 | 9.633E-06 | 3.215E-07 | 9.400E-06 | 1.000E-05 |
| 2227 | 112 | P112Q | A | 6.4e-06 | 4.4e-06 | 5.3e-06 | 5.367E-06 | 1.002E-06 | 4.400E-06 | 6.400E-06 |
| 2228 | 112 | P112G | A | 8.9e-06 | 1.1e-05 | 9.1e-06 | 9.667E-06 | 1.159E-06 | 8.900E-06 | 1.100E-05 |
| 2229 | 112 | P112H | A | 5.2e-06 | 7.9e-06 | 9.4e-06 | 7.500E-06 | 2.128E-06 | 5.200E-06 | 9.400E-06 |
| 2230 | 112 | P112I | A | 6.4e-06 | 6.4e-06 | 1.1e-05 | 7.933E-06 | 2.656E-06 | 6.400E-06 | 1.100E-05 |
| 2231 | 112 | P112L | A | 9.4e-06 | 6.7e-06 | 8e-06 | 8.033E-06 | 1.350E-06 | 6.700E-06 | 9.400E-06 |
| 2232 | 112 | P112K | A | 1.1e-05 | 1.1e-05 | 9.4e-06 | 1.047E-05 | 9.238E-07 | 9.400E-06 | 1.100E-05 |
| 2233 | 112 | P112M | A | 1e-05 | 7.4e-06 | 6.4e-06 | 7.933E-06 | 1.858E-06 | 6.400E-06 | 1.000E-05 |
| 2234 | 112 | P112F | A | 7e-06 | 7.7e-06 | 9.8e-06 | 8.167E-06 | 1.457E-06 | 7.000E-06 | 9.800E-06 |
| 2235 | 112 | P112P | A | 8.4e-06 | 7.6e-06 | 4.9e-06 | 6.967E-06 | 1.834E-06 | 4.900E-06 | 8.400E-06 |
| 2236 | 112 | P112S | A | 6.8e-06 | 5.3e-06 | 6.7e-06 | 6.267E-06 | 8.386E-07 | 5.300E-06 | 6.800E-06 |
| 2237 | 112 | P112T | A | 1e-05 | 1.3e-05 | 5.1e-06 | 9.367E-06 | 3.988E-06 | 5.100E-06 | 1.300E-05 |
| 2238 | 112 | P112W | A | 7.2e-06 | 4.7e-06 | 6.5e-06 | 6.133E-06 | 1.290E-06 | 4.700E-06 | 7.200E-06 |
| 2239 | 112 | P112Y | A | 8.7e-06 | 9.6e-06 | 8.6e-06 | 8.967E-06 | 5.508E-07 | 8.600E-06 | 9.600E-06 |
| 2240 | 112 | P112V | A | 8.7e-06 | 1e-05 | 6.5e-06 | 8.400E-06 | 1.769E-06 | 6.500E-06 | 1.000E-05 |
| 2241 | 113 | G113A | A | 9.6e-06 | 5.1e-06 | 7.5e-06 | 7.400E-06 | 2.252E-06 | 5.100E-06 | 9.600E-06 |
| 2242 | 113 | G113R | A | 7.3e-06 | 6e-06 | 8.7e-06 | 7.333E-06 | 1.350E-06 | 6.000E-06 | 8.700E-06 |
| 2243 | 113 | G113N | A | 9.7e-06 | 1.3e-05 | 5.5e-06 | 9.400E-06 | 3.759E-06 | 5.500E-06 | 1.300E-05 |
| 2244 | 113 | G113D | A | 6.8e-06 | 1.4e-05 | 7.7e-06 | 9.500E-06 | 3.923E-06 | 6.800E-06 | 1.400E-05 |
| 2245 | 113 | G113C | A | 1e-05 | 5.2e-06 | 8.1e-06 | 7.767E-06 | 2.417E-06 | 5.200E-06 | 1.000E-05 |
| 2246 | 113 | G113E | A | 8.2e-06 | 5.8e-06 | 8.1e-06 | 7.367E-06 | 1.358E-06 | 5.800E-06 | 8.200E-06 |
| 2247 | 113 | G113Q | A | 6.9e-06 | 6.8e-06 | 7.5e-06 | 7.067E-06 | 3.786E-07 | 6.800E-06 | 7.500E-06 |
| 2248 | 113 | G113G | A | 7.6e-06 | 6.6e-06 | 6.9e-06 | 7.033E-06 | 5.132E-07 | 6.600E-06 | 7.600E-06 |
| 2249 | 113 | G113H | A | 8.1e-06 | 6.7e-06 | 5.2e-06 | 6.667E-06 | 1.450E-06 | 5.200E-06 | 8.100E-06 |
| 2250 | 113 | G113I | A | 6.6e-06 | 7.6e-06 | 1.1e-05 | 8.400E-06 | 2.307E-06 | 6.600E-06 | 1.100E-05 |
| 2251 | 113 | G113L | A | 4.3e-06 | 7.1e-06 | 8.8e-06 | 6.733E-06 | 2.272E-06 | 4.300E-06 | 8.800E-06 |
| 2252 | 113 | G113K | A | 9.2e-06 | 1.8e-05 | 4.8e-06 | 1.067E-05 | 6.721E-06 | 4.800E-06 | 1.800E-05 |
| 2253 | 113 | G113M | A | 1e-05 | 6.3e-06 | 6.4e-06 | 7.567E-06 | 2.108E-06 | 6.300E-06 | 1.000E-05 |
| 2254 | 113 | G113F | A | 9.9e-06 | 9.2e-06 | 9.8e-06 | 9.633E-06 | 3.786E-07 | 9.200E-06 | 9.900E-06 |
| 2255 | 113 | G113P | A | 9.7e-06 | 5.2e-06 | 5.9e-06 | 6.933E-06 | 2.421E-06 | 5.200E-06 | 9.700E-06 |
| 2256 | 113 | G113S | A | 7.6e-06 | 1.9e-05 | 7.4e-06 | 1.133E-05 | 6.640E-06 | 7.400E-06 | 1.900E-05 |
| 2257 | 113 | G113T | A | 8.7e-06 | 6.9e-06 | 1e-05 | 8.533E-06 | 1.557E-06 | 6.900E-06 | 1.000E-05 |
| 2258 | 113 | G113W | A | 6.9e-06 | 9.8e-06 | 7.3e-06 | 8.000E-06 | 1.572E-06 | 6.900E-06 | 9.800E-06 |
| 2259 | 113 | G113Y | A | 6.4e-06 | 5.6e-06 | 1.1e-05 | 7.667E-06 | 2.914E-06 | 5.600E-06 | 1.100E-05 |
| 2260 | 113 | G113V | A | 6.5e-06 | 1.2e-05 | 6.6e-06 | 8.367E-06 | 3.147E-06 | 6.500E-06 | 1.200E-05 |
| 2261 | 114 | G114A | A | 4.3e-06 | 7.9e-06 | 5.8e-06 | 6.000E-06 | 1.808E-06 | 4.300E-06 | 7.900E-06 |
| 2262 | 114 | G114R | A | 8.1e-06 | 4.3e-06 | 5.9e-06 | 6.100E-06 | 1.908E-06 | 4.300E-06 | 8.100E-06 |
| 2263 | 114 | G114N | A | 6.9e-06 | 8.8e-06 | 1.2e-05 | 9.233E-06 | 2.577E-06 | 6.900E-06 | 1.200E-05 |
| 2264 | 114 | G114D | A | 6.1e-06 | 7e-06 | 5.1e-06 | 6.067E-06 | 9.504E-07 | 5.100E-06 | 7.000E-06 |
| 2265 | 114 | G114C | A | 1.2e-05 | 6.7e-06 | 5.5e-06 | 8.067E-06 | 3.459E-06 | 5.500E-06 | 1.200E-05 |
| 2266 | 114 | G114E | A | 8.2e-06 | 9.5e-06 | 5e-06 | 7.567E-06 | 2.316E-06 | 5.000E-06 | 9.500E-06 |
| 2267 | 114 | G114Q | A | 9.1e-06 | 9.8e-06 | 9.3e-06 | 9.400E-06 | 3.606E-07 | 9.100E-06 | 9.800E-06 |
| 2268 | 114 | G114G | A | 6.9e-06 | 1e-05 | 9.2e-06 | 8.700E-06 | 1.609E-06 | 6.900E-06 | 1.000E-05 |
| 2269 | 114 | G114H | A | 1.2e-05 | 5.1e-06 | 1e-05 | 9.033E-06 | 3.550E-06 | 5.100E-06 | 1.200E-05 |
| 2270 | 114 | G114I | A | 6.3e-06 | 8.3e-06 | 6e-06 | 6.867E-06 | 1.250E-06 | 6.000E-06 | 8.300E-06 |
| 2271 | 114 | G114L | A | 8.9e-06 | 6.3e-06 | 6.8e-06 | 7.333E-06 | 1.380E-06 | 6.300E-06 | 8.900E-06 |
| 2272 | 114 | G114K | A | 1.3e-05 | 5.1e-06 | 1.6e-05 | 1.137E-05 | 5.631E-06 | 5.100E-06 | 1.600E-05 |
| 2273 | 114 | G114M | A | 9.7e-06 | 8.9e-06 | 4.2e-06 | 7.600E-06 | 2.972E-06 | 4.200E-06 | 9.700E-06 |
| 2274 | 114 | G114F | A | 5.4e-06 | 4.1e-06 | 9.4e-06 | 6.300E-06 | 2.762E-06 | 4.100E-06 | 9.400E-06 |
| 2275 | 114 | G114P | A | 6.5e-06 | 5.9e-06 | 6.5e-06 | 6.300E-06 | 3.464E-07 | 5.900E-06 | 6.500E-06 |
| 2276 | 114 | G114S | A | 7.3e-06 | 6.3e-06 | 7.9e-06 | 7.167E-06 | 8.083E-07 | 6.300E-06 | 7.900E-06 |
| 2277 | 114 | G114T | A | 9.9e-06 | 6.2e-06 | 9.8e-06 | 8.633E-06 | 2.108E-06 | 6.200E-06 | 9.900E-06 |
| 2278 | 114 | G114W | A | 8.3e-06 | 1.3e-05 | 9.3e-06 | 1.020E-05 | 2.476E-06 | 8.300E-06 | 1.300E-05 |
| 2279 | 114 | G114Y | A | 4.9e-06 | 9.3e-06 | 9.9e-06 | 8.033E-06 | 2.730E-06 | 4.900E-06 | 9.900E-06 |
| 2280 | 114 | G114V | A | 1.2e-05 | 1.2e-05 | 8.6e-06 | 1.087E-05 | 1.963E-06 | 8.600E-06 | 1.200E-05 |
| 2281 | 115 | L115A | A | 9.1e-06 | 8.5e-06 | 7.3e-06 | 8.300E-06 | 9.165E-07 | 7.300E-06 | 9.100E-06 |
| 2282 | 115 | L115R | A | 9e-06 | 5.8e-06 | 4.5e-06 | 6.433E-06 | 2.316E-06 | 4.500E-06 | 9.000E-06 |
| 2283 | 115 | L115N | A | 9e-06 | 6.4e-06 | 4.7e-06 | 6.700E-06 | 2.166E-06 | 4.700E-06 | 9.000E-06 |
| 2284 | 115 | L115D | A | 1e-05 | 1.1e-05 | 1.2e-05 | 1.100E-05 | 1.000E-06 | 1.000E-05 | 1.200E-05 |
| 2285 | 115 | L115C | A | 7.6e-06 | 8.8e-06 | 8.4e-06 | 8.267E-06 | 6.110E-07 | 7.600E-06 | 8.800E-06 |
| 2286 | 115 | L115E | A | 1.8e-05 | 5.5e-06 | 1.7e-05 | 1.350E-05 | 6.946E-06 | 5.500E-06 | 1.800E-05 |
| 2287 | 115 | L115Q | A | 5.1e-06 | 8.5e-06 | 7.4e-06 | 7.000E-06 | 1.735E-06 | 5.100E-06 | 8.500E-06 |
| 2288 | 115 | L115G | A | 1.1e-05 | 4.9e-06 | 5.8e-06 | 7.233E-06 | 3.293E-06 | 4.900E-06 | 1.100E-05 |
| 2289 | 115 | L115H | A | 3.9e-06 | 1.1e-05 | 6.7e-06 | 7.200E-06 | 3.576E-06 | 3.900E-06 | 1.100E-05 |
| 2290 | 115 | L115I | A | 1.2e-05 | 9.8e-06 | 8.6e-06 | 1.013E-05 | 1.724E-06 | 8.600E-06 | 1.200E-05 |
| 2291 | 115 | L115L | A | 1e-05 | 9.7e-06 | 4.2e-06 | 7.967E-06 | 3.265E-06 | 4.200E-06 | 1.000E-05 |
| 2292 | 115 | L115K | A | 7.7e-06 | 8.7e-06 | 9.7e-06 | 8.700E-06 | 1.000E-06 | 7.700E-06 | 9.700E-06 |
| 2293 | 115 | L115M | A | 6.6e-06 | 1.3e-05 | 4.7e-06 | 8.100E-06 | 4.349E-06 | 4.700E-06 | 1.300E-05 |
| 2294 | 115 | L115F | A | 9.7e-06 | 5.6e-06 | 1e-05 | 8.433E-06 | 2.458E-06 | 5.600E-06 | 1.000E-05 |

| | | | | | | | | | | |
|---|---|---|---|---|---|---|---|---|---|---|
| 2295 | 115 | L115P | A | 1.3e-05 | 1.3e-05 | 1.4e-05 | 1.333E-05 | 5.774E-07 | 1.300E-05 | 1.400E-05 |
| 2296 | 115 | L115S | A | 9.5e-06 | 1.4e-05 | 6.3e-06 | 9.933E-06 | 3.868E-06 | 6.300E-06 | 1.400E-05 |
| 2297 | 115 | L115T | A | 1.2e-05 | 5.6e-06 | 1.1e-05 | 9.533E-06 | 3.443E-06 | 5.600E-06 | 1.200E-05 |
| 2298 | 115 | L115W | A | 8.3e-06 | 8.8e-06 | 4.8e-06 | 7.300E-06 | 2.179E-06 | 4.800E-06 | 8.800E-06 |
| 2299 | 115 | L115Y | A | 5.9e-06 | 1.1e-05 | 6.5e-06 | 7.800E-06 | 2.787E-06 | 5.900E-06 | 1.100E-05 |
| 2300 | 115 | L115V | A | 6.3e-06 | 8.6e-06 | 1e-05 | 8.300E-06 | 1.868E-06 | 6.300E-06 | 1.000E-05 |
| 2301 | 116 | R116A | A | 6.5e-06 | 1.2e-05 | 1.7e-05 | 1.183E-05 | 5.252E-06 | 6.500E-06 | 1.700E-05 |
| 2302 | 116 | R116R | A | 5.3e-06 | 9.2e-06 | 7.7e-06 | 7.400E-06 | 1.967E-06 | 5.300E-06 | 9.200E-06 |
| 2303 | 116 | R116N | A | 9.2e-06 | 8.1e-06 | 5.3e-06 | 7.533E-06 | 2.011E-06 | 5.300E-06 | 9.200E-06 |
| 2304 | 116 | R116D | A | 1.2e-05 | 8.8e-06 | 9.5e-06 | 1.010E-05 | 1.682E-06 | 8.800E-06 | 1.200E-05 |
| 2305 | 116 | R116C | A | 6.9e-06 | 9e-06 | 7.6e-06 | 7.833E-06 | 1.069E-06 | 6.900E-06 | 9.000E-06 |
| 2306 | 116 | R116E | A | 7.4e-06 | 8.6e-06 | 4.6e-06 | 6.867E-06 | 2.053E-06 | 4.600E-06 | 8.600E-06 |
| 2307 | 116 | R116Q | A | 8.7e-06 | 6.8e-06 | 1.1e-05 | 8.833E-06 | 2.103E-06 | 6.800E-06 | 1.100E-05 |
| 2308 | 116 | R116G | A | 5.4e-06 | 1.3e-05 | 9.2e-06 | 9.200E-06 | 3.800E-06 | 5.400E-06 | 1.300E-05 |
| 2309 | 116 | R116H | A | 9.3e-06 | 6.6e-06 | 1.4e-05 | 9.967E-06 | 3.745E-06 | 6.600E-06 | 1.400E-05 |
| 2310 | 116 | R116I | A | 6.7e-06 | 4.3e-06 | 1e-05 | 7.000E-06 | 2.862E-06 | 4.300E-06 | 1.000E-05 |
| 2311 | 116 | R116L | A | 6.3e-06 | 9.3e-06 | 5.9e-06 | 7.167E-06 | 1.858E-06 | 5.900E-06 | 9.300E-06 |
| 2312 | 116 | R116K | A | 9.4e-06 | 9.5e-06 | 9.9e-06 | 9.600E-06 | 2.646E-07 | 9.400E-06 | 9.900E-06 |
| 2313 | 116 | R116M | A | 1.4e-05 | 1.4e-05 | 6.3e-06 | 1.143E-05 | 4.446E-06 | 6.300E-06 | 1.400E-05 |
| 2314 | 116 | R116F | A | 1.1e-05 | 7.5e-06 | 6.6e-06 | 8.367E-06 | 2.325E-06 | 6.600E-06 | 1.100E-05 |
| 2315 | 116 | R116P | A | 6.5e-06 | 4.9e-06 | 5.3e-06 | 5.567E-06 | 8.327E-07 | 4.900E-06 | 6.500E-06 |
| 2316 | 116 | R116S | A | 4.3e-06 | 6.7e-06 | 6.8e-06 | 5.933E-06 | 1.415E-06 | 4.300E-06 | 6.800E-06 |
| 2317 | 116 | R116T | A | 7e-06 | 7e-06 | 6.1e-06 | 6.700E-06 | 5.196E-07 | 6.100E-06 | 7.000E-06 |
| 2318 | 116 | R116W | A | 6.7e-06 | 7e-06 | 7.2e-06 | 6.967E-06 | 2.517E-07 | 6.700E-06 | 7.200E-06 |
| 2319 | 116 | R116Y | A | 7.1e-06 | 5.3e-06 | 9.6e-06 | 7.333E-06 | 2.159E-06 | 5.300E-06 | 9.600E-06 |
| 2320 | 116 | R116V | A | 7.5e-06 | 6.8e-06 | 5.3e-06 | 6.533E-06 | 1.124E-06 | 5.300E-06 | 7.500E-06 |
| 2321 | 117 | E117A | A | 9.4e-06 | 6.9e-06 | 5.1e-06 | 7.133E-06 | 2.159E-06 | 5.100E-06 | 9.400E-06 |
| 2322 | 117 | E117R | A | 5.7e-06 | 1e-05 | 8e-06 | 7.900E-06 | 2.152E-06 | 5.700E-06 | 1.000E-05 |
| 2323 | 117 | E117N | A | 1.2e-05 | 8.8e-06 | 8.5e-06 | 9.767E-06 | 1.940E-06 | 8.500E-06 | 1.200E-05 |
| 2324 | 117 | E117D | A | 7.6e-06 | 1.2e-05 | 5.4e-06 | 8.333E-06 | 3.361E-06 | 5.400E-06 | 1.200E-05 |
| 2325 | 117 | E117C | A | 6.9e-06 | 1.4e-05 | 6.1e-06 | 9.000E-06 | 4.349E-06 | 6.100E-06 | 1.400E-05 |
| 2326 | 117 | E117E | A | 6.4e-06 | 8.8e-06 | 1.4e-05 | 9.733E-06 | 3.885E-06 | 6.400E-06 | 1.400E-05 |
| 2327 | 117 | E117Q | A | 7.7e-06 | 1.1e-05 | 6.5e-06 | 8.400E-06 | 2.330E-06 | 6.500E-06 | 1.100E-05 |
| 2328 | 117 | E117G | A | 7.6e-06 | 1.6e-05 | 1.4e-05 | 1.253E-05 | 4.388E-06 | 7.600E-06 | 1.600E-05 |
| 2329 | 117 | E117H | A | 4.7e-06 | 6.2e-06 | 1.3e-05 | 7.967E-06 | 4.423E-06 | 4.700E-06 | 1.300E-05 |
| 2330 | 117 | E117I | A | 7.7e-06 | 9.7e-06 | 5.3e-06 | 7.567E-06 | 2.203E-06 | 5.300E-06 | 9.700E-06 |
| 2331 | 117 | E117L | A | 8.1e-06 | 1.3e-05 | 1.1e-05 | 1.070E-05 | 2.464E-06 | 8.100E-06 | 1.300E-05 |
| 2332 | 117 | E117K | A | 7.1e-06 | 6.2e-06 | 8.6e-06 | 7.300E-06 | 1.212E-06 | 6.200E-06 | 8.600E-06 |
| 2333 | 117 | E117M | A | 9.6e-06 | 4.6e-06 | 5.1e-06 | 6.433E-06 | 2.754E-06 | 4.600E-06 | 9.600E-06 |
| 2334 | 117 | E117F | A | 9.3e-06 | 6.3e-06 | 9.4e-06 | 8.333E-06 | 1.762E-06 | 6.300E-06 | 9.400E-06 |
| 2335 | 117 | E117P | A | 9.1e-06 | 4.4e-06 | 8.1e-06 | 7.200E-06 | 2.476E-06 | 4.400E-06 | 9.100E-06 |
| 2336 | 117 | E117S | A | 7.8e-06 | 9e-06 | 1.4e-05 | 1.027E-05 | 3.288E-06 | 7.800E-06 | 1.400E-05 |
| 2337 | 117 | E117T | A | 5.9e-06 | 1.2e-05 | 1.2e-05 | 9.967E-06 | 3.522E-06 | 5.900E-06 | 1.200E-05 |
| 2338 | 117 | E117W | A | 7.9e-06 | 8.9e-06 | 8.6e-06 | 8.467E-06 | 5.132E-07 | 7.900E-06 | 8.900E-06 |
| 2339 | 117 | E117Y | A | 9.7e-06 | 7.2e-06 | 1e-05 | 8.967E-06 | 1.537E-06 | 7.200E-06 | 1.000E-05 |
| 2340 | 117 | E117V | A | 1.5e-05 | 8e-06 | 8.3e-06 | 1.043E-05 | 3.958E-06 | 8.000E-06 | 1.500E-05 |
| 2341 | 118 | L118A | A | 6.1e-06 | 8.8e-06 | 7.5e-06 | 7.467E-06 | 1.350E-06 | 6.100E-06 | 8.800E-06 |
| 2342 | 118 | L118R | A | 4.6e-06 | 8.8e-06 | 6.1e-06 | 6.500E-06 | 2.128E-06 | 4.600E-06 | 8.800E-06 |
| 2343 | 118 | L118N | A | 6.3e-06 | 8.9e-06 | 9.8e-06 | 8.333E-06 | 1.818E-06 | 6.300E-06 | 9.800E-06 |
| 2344 | 118 | L118D | A | 6.4e-06 | 5.8e-06 | 6.4e-06 | 6.200E-06 | 3.464E-07 | 5.800E-06 | 6.400E-06 |
| 2345 | 118 | L118C | A | 6.3e-06 | 9e-06 | 7.6e-06 | 7.633E-06 | 1.350E-06 | 6.300E-06 | 9.000E-06 |
| 2346 | 118 | L118E | A | 6.5e-06 | 1e-05 | 8.2e-06 | 8.233E-06 | 1.750E-06 | 6.500E-06 | 1.000E-05 |
| 2347 | 118 | L118Q | A | 8.8e-06 | 1.3e-05 | 4.4e-06 | 8.733E-06 | 4.300E-06 | 4.400E-06 | 1.300E-05 |
| 2348 | 118 | L118G | A | 6.1e-06 | 6.7e-06 | 5.7e-06 | 6.167E-06 | 5.033E-07 | 5.700E-06 | 6.700E-06 |
| 2349 | 118 | L118H | A | 5.1e-06 | 9.5e-06 | 1.2e-05 | 8.867E-06 | 3.493E-06 | 5.100E-06 | 1.200E-05 |
| 2350 | 118 | L118I | A | 1.1e-05 | 8.1e-06 | 8.2e-06 | 9.100E-06 | 1.646E-06 | 8.100E-06 | 1.100E-05 |
| 2351 | 118 | L118L | A | 1.1e-05 | 5.1e-06 | 1.2e-05 | 9.367E-06 | 3.729E-06 | 5.100E-06 | 1.200E-05 |
| 2352 | 118 | L118K | A | 7.6e-06 | 7.9e-06 | 1e-05 | 8.500E-06 | 1.308E-06 | 7.600E-06 | 1.000E-05 |
| 2353 | 118 | L118M | A | 9.5e-06 | 9.3e-06 | 9.6e-06 | 9.467E-06 | 1.528E-07 | 9.300E-06 | 9.600E-06 |
| 2354 | 118 | L118F | A | 1.1e-05 | 6.2e-06 | 4.5e-06 | 7.233E-06 | 3.371E-06 | 4.500E-06 | 1.100E-05 |
| 2355 | 118 | L118P | A | 7.6e-06 | 1.1e-05 | 8.6e-06 | 9.067E-06 | 1.747E-06 | 7.600E-06 | 1.100E-05 |
| 2356 | 118 | L118S | A | 9.7e-06 | 6.5e-06 | 7.8e-06 | 8.000E-06 | 1.609E-06 | 6.500E-06 | 9.700E-06 |
| 2357 | 118 | L118T | A | 9.5e-06 | 5.9e-06 | 9.9e-06 | 8.433E-06 | 2.203E-06 | 5.900E-06 | 9.900E-06 |
| 2358 | 118 | L118W | A | 6.7e-06 | 4.3e-06 | 1e-05 | 7.000E-06 | 2.862E-06 | 4.300E-06 | 1.000E-05 |
| 2359 | 118 | L118Y | A | 1.2e-05 | 7.1e-06 | 1.2e-05 | 1.037E-05 | 2.829E-06 | 7.100E-06 | 1.200E-05 |
| 2360 | 118 | L118V | A | 3e-06 | 6.2e-06 | 5.9e-06 | 5.033E-06 | 1.767E-06 | 3.000E-06 | 6.200E-06 |
| 2361 | 119 | Q119A | A | 6.3e-06 | 7.2e-06 | 1.1e-05 | 8.167E-06 | 2.495E-06 | 6.300E-06 | 1.100E-05 |
| 2362 | 119 | Q119R | A | 5.3e-06 | 1.2e-05 | 6.1e-06 | 7.800E-06 | 3.659E-06 | 5.300E-06 | 1.200E-05 |
| 2363 | 119 | Q119N | A | 8.2e-06 | 6.8e-06 | 8.1e-06 | 7.700E-06 | 7.810E-07 | 6.800E-06 | 8.200E-06 |
| 2364 | 119 | Q119D | A | 9.6e-06 | 6.3e-06 | 6.8e-06 | 7.567E-06 | 1.779E-06 | 6.300E-06 | 9.600E-06 |
| 2365 | 119 | Q119C | A | 9e-06 | 7.6e-06 | 7.5e-06 | 8.033E-06 | 8.386E-07 | 7.500E-06 | 9.000E-06 |
| 2366 | 119 | Q119E | A | 6.3e-06 | 6.2e-06 | 9.7e-06 | 7.400E-06 | 1.992E-06 | 6.200E-06 | 9.700E-06 |
| 2367 | 119 | Q119Q | A | 6.4e-06 | 1.1e-05 | 4.5e-06 | 7.300E-06 | 3.342E-06 | 4.500E-06 | 1.100E-05 |
| 2368 | 119 | Q119G | A | 1.4e-05 | 7.3e-06 | 1.2e-05 | 1.110E-05 | 3.439E-06 | 7.300E-06 | 1.400E-05 |
| 2369 | 119 | Q119H | A | 8e-06 | 7.4e-06 | 6.5e-06 | 7.300E-06 | 7.550E-07 | 6.500E-06 | 8.000E-06 |
| 2370 | 119 | Q119I | A | 8.8e-06 | 6.9e-06 | 1.2e-05 | 9.233E-06 | 2.577E-06 | 6.900E-06 | 1.200E-05 |
| 2371 | 119 | Q119L | A | 9.1e-06 | 1.2e-05 | 9.7e-06 | 1.027E-05 | 1.531E-06 | 9.100E-06 | 1.200E-05 |
| 2372 | 119 | Q119K | A | 9.6e-06 | 9.7e-06 | 1.1e-05 | 1.010E-05 | 7.810E-07 | 9.600E-06 | 1.100E-05 |
| 2373 | 119 | Q119M | A | 4.7e-06 | 3.6e-06 | 4.5e-06 | 4.267E-06 | 5.859E-07 | 3.600E-06 | 4.700E-06 |
| 2374 | 119 | Q119F | A | 4.2e-06 | 6.2e-06 | 7.2e-06 | 5.867E-06 | 1.528E-06 | 4.200E-06 | 7.200E-06 |
| 2375 | 119 | Q119P | A | 1.3e-05 | 7.2e-06 | 1.4e-05 | 1.140E-05 | 3.672E-06 | 7.200E-06 | 1.400E-05 |
| 2376 | 119 | Q119S | A | 1.1e-05 | 6.6e-06 | 6.5e-06 | 8.033E-06 | 2.570E-06 | 6.500E-06 | 1.100E-05 |
| 2377 | 119 | Q119T | A | 4.4e-06 | 1.4e-05 | 1.4e-05 | 1.080E-05 | 5.543E-06 | 4.400E-06 | 1.400E-05 |
| 2378 | 119 | Q119W | A | 1e-05 | 1.3e-05 | 7.4e-06 | 1.013E-05 | 2.802E-06 | 7.400E-06 | 1.300E-05 |
| 2379 | 119 | Q119Y | A | 5.7e-06 | 1e-05 | 9.4e-06 | 8.367E-06 | 2.329E-06 | 5.700E-06 | 1.000E-05 |

| | | | | | | | | | | |
|---|---|---|---|---|---|---|---|---|---|---|
| 2380 | 119 | Q119V | A | 7.1e-06 | 1.4e-05 | 8.8e-06 | 9.967E-06 | 3.595E-06 | 7.100E-06 | 1.400E-05 |
| 2381 | 120 | L120A | A | 7.2e-06 | 6.7e-06 | 6.4e-06 | 6.767E-06 | 4.041E-07 | 6.400E-06 | 7.200E-06 |
| 2382 | 120 | L120R | A | 5.8e-06 | 6.3e-06 | 4.4e-06 | 5.500E-06 | 9.849E-07 | 4.400E-06 | 6.300E-06 |
| 2383 | 120 | L120N | A | 7.6e-06 | 7.6e-06 | 1.3e-05 | 9.400E-06 | 3.118E-06 | 7.600E-06 | 1.300E-05 |
| 2384 | 120 | L120D | A | 4.6e-06 | 9.5e-06 | 9.2e-06 | 7.767E-06 | 2.747E-06 | 4.600E-06 | 9.500E-06 |
| 2385 | 120 | L120C | A | 9.2e-06 | 9.1e-06 | 1.5e-05 | 1.110E-05 | 3.378E-06 | 9.100E-06 | 1.500E-05 |
| 2386 | 120 | L120E | A | 1.1e-05 | 1.1e-05 | 1.1e-05 | 1.100E-05 | 2.075E-21 | 1.100E-05 | 1.100E-05 |
| 2387 | 120 | L120Q | A | 1.1e-05 | 6.7e-06 | 1.6e-05 | 1.123E-05 | 4.654E-06 | 6.700E-06 | 1.600E-05 |
| 2388 | 120 | L120G | A | 1e-05 | 6.8e-06 | 7.5e-06 | 8.100E-06 | 1.682E-06 | 6.800E-06 | 1.000E-05 |
| 2389 | 120 | L120H | A | 8.5e-06 | 5.6e-06 | 8.9e-06 | 7.667E-06 | 1.801E-06 | 5.600E-06 | 8.900E-06 |
| 2390 | 120 | L120I | A | 8.8e-06 | 1.7e-05 | 7.4e-06 | 1.107E-05 | 5.186E-06 | 7.400E-06 | 1.700E-05 |
| 2391 | 120 | L120L | A | 1e-05 | 9e-06 | 1.9e-05 | 1.267E-05 | 5.508E-06 | 9.000E-06 | 1.900E-05 |
| 2392 | 120 | L120K | A | 1e-05 | 7.3e-06 | 1.4e-05 | 1.043E-05 | 3.371E-06 | 7.300E-06 | 1.400E-05 |
| 2393 | 120 | L120M | A | 6.2e-06 | 5.2e-06 | 8.1e-06 | 6.500E-06 | 1.473E-06 | 5.200E-06 | 8.100E-06 |
| 2394 | 120 | L120F | A | 1e-05 | 6.7e-06 | 6.2e-06 | 7.633E-06 | 2.065E-06 | 6.200E-06 | 1.000E-05 |
| 2395 | 120 | L120P | A | 3.9e-06 | 8.4e-06 | 8e-06 | 6.767E-06 | 2.491E-06 | 3.900E-06 | 8.400E-06 |
| 2396 | 120 | L120S | A | 1e-05 | 7.6e-06 | 1.3e-05 | 1.020E-05 | 2.706E-06 | 7.600E-06 | 1.300E-05 |
| 2397 | 120 | L120T | A | 6.6e-06 | 3.3e-06 | 5.8e-06 | 5.233E-06 | 1.721E-06 | 3.300E-06 | 6.600E-06 |
| 2398 | 120 | L120W | A | 4.5e-06 | 8.6e-06 | 8.5e-06 | 7.200E-06 | 2.339E-06 | 4.500E-06 | 8.600E-06 |
| 2399 | 120 | L120Y | A | 7.4e-06 | 9.1e-06 | 7.3e-06 | 7.933E-06 | 1.012E-06 | 7.300E-06 | 9.100E-06 |
| 2400 | 120 | L120V | A | 7e-06 | 7e-06 | 6.1e-06 | 6.700E-06 | 5.196E-07 | 6.100E-06 | 7.000E-06 |
| 2401 | 121 | R121A | A | 1.1e-05 | 1.2e-05 | 4.9e-06 | 9.300E-06 | 3.843E-06 | 4.900E-06 | 1.200E-05 |
| 2402 | 121 | R121R | A | 9.4e-06 | 1e-05 | 8.8e-06 | 9.400E-06 | 6.000E-07 | 8.800E-06 | 1.000E-05 |
| 2403 | 121 | R121N | A | 4.4e-06 | 6.6e-06 | 8e-06 | 6.333E-06 | 1.815E-06 | 4.400E-06 | 8.000E-06 |
| 2404 | 121 | R121D | A | 6.4e-06 | 7.8e-06 | 9.1e-06 | 7.767E-06 | 1.350E-06 | 6.400E-06 | 9.100E-06 |
| 2405 | 121 | R121C | A | 5.6e-06 | 8.7e-06 | 1e-05 | 8.100E-06 | 2.261E-06 | 5.600E-06 | 1.000E-05 |
| 2406 | 121 | R121E | A | 7e-06 | 4.6e-06 | 7.3e-06 | 6.300E-06 | 1.480E-06 | 4.600E-06 | 7.300E-06 |
| 2407 | 121 | R121Q | A | 6.3e-06 | 9.1e-06 | 6.4e-06 | 7.267E-06 | 1.589E-06 | 6.300E-06 | 9.100E-06 |
| 2408 | 121 | R121G | A | 9.3e-06 | 6.2e-06 | 1e-05 | 8.500E-06 | 2.022E-06 | 6.200E-06 | 1.000E-05 |
| 2409 | 121 | R121H | A | 6.3e-06 | 4.9e-06 | 1e-05 | 7.067E-06 | 2.635E-06 | 4.900E-06 | 1.000E-05 |
| 2410 | 121 | R121I | A | 7.5e-06 | 7e-06 | 1.3e-05 | 9.167E-06 | 3.329E-06 | 7.000E-06 | 1.300E-05 |
| 2411 | 121 | R121L | A | 7.7e-06 | 5.4e-06 | 8.7e-06 | 7.267E-06 | 1.692E-06 | 5.400E-06 | 8.700E-06 |
| 2412 | 121 | R121K | A | 4.6e-06 | 3.4e-06 | 1.1e-05 | 6.333E-06 | 4.086E-06 | 3.400E-06 | 1.100E-05 |
| 2413 | 121 | R121M | A | 9.4e-06 | 9.5e-06 | 5.8e-06 | 8.233E-06 | 2.108E-06 | 5.800E-06 | 9.500E-06 |
| 2414 | 121 | R121F | A | 8.5e-06 | 5.7e-06 | 8.4e-06 | 7.533E-06 | 1.589E-06 | 5.700E-06 | 8.500E-06 |
| 2415 | 121 | R121P | A | 6.4e-06 | 9.4e-06 | 9.2e-06 | 8.333E-06 | 1.677E-06 | 6.400E-06 | 9.400E-06 |
| 2416 | 121 | R121S | A | 6.7e-06 | 5.5e-06 | 8.6e-06 | 6.933E-06 | 1.563E-06 | 5.500E-06 | 8.600E-06 |
| 2417 | 121 | R121T | A | 6.5e-06 | 7.4e-06 | 8.8e-06 | 7.567E-06 | 1.159E-06 | 6.500E-06 | 8.800E-06 |
| 2418 | 121 | R121W | A | 4.3e-06 | 1.3e-05 | 5e-06 | 7.433E-06 | 4.834E-06 | 4.300E-06 | 1.300E-05 |
| 2419 | 121 | R121Y | A | 6.3e-06 | 1.3e-05 | 8.4e-06 | 9.233E-06 | 3.427E-06 | 6.300E-06 | 1.300E-05 |
| 2420 | 121 | R121V | A | 1.1e-05 | 9.6e-06 | 1e-05 | 1.020E-05 | 7.211E-07 | 9.600E-06 | 1.100E-05 |
| 2421 | 122 | S122A | A | 1e-05 | 1e-05 | 3.7e-06 | 7.900E-06 | 3.637E-06 | 3.700E-06 | 1.000E-05 |
| 2422 | 122 | S122R | A | 7e-06 | 4.4e-06 | 7.2e-06 | 6.200E-06 | 1.562E-06 | 4.400E-06 | 7.200E-06 |
| 2423 | 122 | S122N | A | 1.7e-05 | 8e-06 | 8.6e-06 | 1.120E-05 | 5.032E-06 | 8.000E-06 | 1.700E-05 |
| 2424 | 122 | S122D | A | 9.9e-06 | 3.8e-06 | 4.9e-06 | 6.200E-06 | 3.251E-06 | 3.800E-06 | 9.900E-06 |
| 2425 | 122 | S122C | A | 4.4e-06 | 7.9e-06 | 9.6e-06 | 7.300E-06 | 2.651E-06 | 4.400E-06 | 9.600E-06 |
| 2426 | 122 | S122E | A | 8.9e-06 | 6.7e-06 | 1e-05 | 8.533E-06 | 1.680E-06 | 6.700E-06 | 1.000E-05 |
| 2427 | 122 | S122Q | A | 5.9e-06 | 9e-06 | 7.7e-06 | 7.533E-06 | 1.557E-06 | 5.900E-06 | 9.000E-06 |
| 2428 | 122 | S122G | A | 1.4e-05 | 8e-06 | 8.8e-06 | 1.027E-05 | 3.258E-06 | 8.000E-06 | 1.400E-05 |
| 2429 | 122 | S122H | A | 8.5e-06 | 6.5e-06 | 8.8e-06 | 7.933E-06 | 1.250E-06 | 6.500E-06 | 8.800E-06 |
| 2430 | 122 | S122I | A | 5.9e-06 | 9.5e-06 | 6.4e-06 | 7.267E-06 | 1.950E-06 | 5.900E-06 | 9.500E-06 |
| 2431 | 122 | S122L | A | 8.6e-06 | 9.6e-06 | 4.8e-06 | 7.667E-06 | 2.532E-06 | 4.800E-06 | 9.600E-06 |
| 2432 | 122 | S122K | A | 1.1e-05 | 7.8e-06 | 4.7e-06 | 7.833E-06 | 3.150E-06 | 4.700E-06 | 1.100E-05 |
| 2433 | 122 | S122M | A | 9.5e-06 | 5.7e-06 | 5.8e-06 | 7.000E-06 | 2.166E-06 | 5.700E-06 | 9.500E-06 |
| 2434 | 122 | S122F | A | 1e-05 | 6e-06 | 7.4e-06 | 7.800E-06 | 2.030E-06 | 6.000E-06 | 1.000E-05 |
| 2435 | 122 | S122P | A | 9.8e-06 | 1.1e-05 | 6.8e-06 | 9.200E-06 | 2.163E-06 | 6.800E-06 | 1.100E-05 |
| 2436 | 122 | S122S | A | 1.3e-05 | 1.2e-05 | 4.2e-06 | 9.733E-06 | 4.818E-06 | 4.200E-06 | 1.300E-05 |
| 2437 | 122 | S122T | A | 7.1e-06 | 9.5e-06 | 5.2e-06 | 7.267E-06 | 2.155E-06 | 5.200E-06 | 9.500E-06 |
| 2438 | 122 | S122W | A | 1.6e-05 | 7.2e-06 | 1.4e-05 | 1.240E-05 | 4.613E-06 | 7.200E-06 | 1.600E-05 |
| 2439 | 122 | S122Y | A | 6.3e-06 | 6.4e-06 | 8.2e-06 | 6.967E-06 | 1.069E-06 | 6.300E-06 | 8.200E-06 |
| 2440 | 122 | S122V | A | 9.9e-06 | 4.6e-06 | 7.4e-06 | 7.300E-06 | 2.651E-06 | 4.600E-06 | 9.900E-06 |
| 2441 | 123 | L123A | A | 1e-05 | 8.5e-06 | 6.8e-06 | 8.433E-06 | 1.601E-06 | 6.800E-06 | 1.000E-05 |
| 2442 | 123 | L123R | A | 3.4e-06 | 8.5e-06 | 7.1e-06 | 6.333E-06 | 2.635E-06 | 3.400E-06 | 8.500E-06 |
| 2443 | 123 | L123N | A | 6.1e-06 | 9.6e-06 | 8.3e-06 | 8.000E-06 | 1.769E-06 | 6.100E-06 | 9.600E-06 |
| 2444 | 123 | L123D | A | 6.1e-06 | 7.5e-06 | 6.2e-06 | 6.600E-06 | 7.810E-07 | 6.100E-06 | 7.500E-06 |
| 2445 | 123 | L123C | A | 6.1e-06 | 9.3e-06 | 1.2e-05 | 9.133E-06 | 2.954E-06 | 6.100E-06 | 1.200E-05 |
| 2446 | 123 | L123E | A | 1.5e-05 | 1.1e-05 | 9.4e-06 | 1.180E-05 | 2.884E-06 | 9.400E-06 | 1.500E-05 |
| 2447 | 123 | L123Q | A | 9.7e-06 | 9.3e-06 | 5e-06 | 8.000E-06 | 2.606E-06 | 5.000E-06 | 9.700E-06 |
| 2448 | 123 | L123G | A | 9.8e-06 | 1.1e-05 | 1.4e-05 | 1.160E-05 | 2.163E-06 | 9.800E-06 | 1.400E-05 |
| 2449 | 123 | L123H | A | 6.9e-06 | 1.4e-05 | 7.3e-06 | 9.400E-06 | 3.989E-06 | 6.900E-06 | 1.400E-05 |
| 2450 | 123 | L123I | A | 7.3e-06 | 1e-05 | 9.1e-06 | 8.800E-06 | 1.375E-06 | 7.300E-06 | 1.000E-05 |
| 2451 | 123 | L123L | A | 1.3e-05 | 9.9e-06 | 9.2e-06 | 1.070E-05 | 2.022E-06 | 9.200E-06 | 1.300E-05 |
| 2452 | 123 | L123K | A | 8.5e-06 | 4.6e-06 | 6.5e-06 | 6.533E-06 | 1.950E-06 | 4.600E-06 | 8.500E-06 |
| 2453 | 123 | L123M | A | 5.2e-06 | 6e-06 | 4.6e-06 | 5.267E-06 | 7.024E-07 | 4.600E-06 | 6.000E-06 |
| 2454 | 123 | L123F | A | 4.9e-06 | 4.4e-06 | 1e-05 | 6.433E-06 | 3.099E-06 | 4.400E-06 | 1.000E-05 |
| 2455 | 123 | L123P | A | 9.6e-06 | 1.2e-05 | 1.2e-05 | 1.120E-05 | 1.386E-06 | 9.600E-06 | 1.200E-05 |
| 2456 | 123 | L123S | A | 1.3e-05 | 9.1e-06 | 6.7e-06 | 9.600E-06 | 3.180E-06 | 6.700E-06 | 1.300E-05 |
| 2457 | 123 | L123T | A | 6.1e-06 | 8.2e-06 | 1.1e-05 | 8.433E-06 | 2.458E-06 | 6.100E-06 | 1.100E-05 |
| 2458 | 123 | L123W | A | 6.6e-06 | 6.6e-06 | 5.8e-06 | 6.333E-06 | 4.619E-07 | 5.800E-06 | 6.600E-06 |
| 2459 | 123 | L123Y | A | 9.5e-06 | 6.6e-06 | 7.7e-06 | 7.933E-06 | 1.464E-06 | 6.600E-06 | 9.500E-06 |
| 2460 | 123 | L123V | A | 7.2e-06 | 7.7e-06 | 9.3e-06 | 8.067E-06 | 1.097E-06 | 7.200E-06 | 9.300E-06 |
| 2461 | 124 | T124A | A | 9.1e-06 | 9.6e-06 | 8.1e-06 | 8.933E-06 | 7.638E-07 | 8.100E-06 | 9.600E-06 |
| 2462 | 124 | T124R | A | 9.7e-06 | 1.1e-05 | 8e-06 | 9.567E-06 | 1.504E-06 | 8.000E-06 | 1.100E-05 |
| 2463 | 124 | T124N | A | 9.2e-06 | 9.5e-06 | 7.1e-06 | 8.600E-06 | 1.308E-06 | 7.100E-06 | 9.500E-06 |
| 2464 | 124 | T124D | A | 7.1e-06 | 8.9e-06 | 5.4e-06 | 7.133E-06 | 1.750E-06 | 5.400E-06 | 8.900E-06 |

| | | | | | | | | | | |
|---|---|---|---|---|---|---|---|---|---|---|
| 2465 | 124 | T124C | A | 6.5e-06 | 9e-06 | 7.2e-06 | 7.567E-06 | 1.290E-06 | 6.500E-06 | 9.000E-06 |
| 2466 | 124 | T124E | A | 1e-05 | 1e-05 | 9.7e-06 | 9.900E-06 | 1.732E-07 | 9.700E-06 | 1.000E-05 |
| 2467 | 124 | T124Q | A | 1.1e-05 | 9.8e-06 | 8.3e-06 | 9.700E-06 | 1.353E-06 | 8.300E-06 | 1.100E-05 |
| 2468 | 124 | T124G | A | 8.3e-06 | 8.4e-06 | 7.7e-06 | 8.133E-06 | 3.786E-07 | 7.700E-06 | 8.400E-06 |
| 2469 | 124 | T124H | A | 1.4e-05 | 8.4e-06 | 9.7e-06 | 1.070E-05 | 2.931E-06 | 8.400E-06 | 1.400E-05 |
| 2470 | 124 | T124I | A | 9.4e-06 | 6e-06 | 7.3e-06 | 7.567E-06 | 1.716E-06 | 6.000E-06 | 9.400E-06 |
| 2471 | 124 | T124L | A | 6.1e-06 | 6.3e-06 | 1.3e-05 | 8.467E-06 | 3.927E-06 | 6.100E-06 | 1.300E-05 |
| 2472 | 124 | T124K | A | 7.3e-06 | 7.5e-06 | 7.3e-06 | 7.367E-06 | 1.155E-07 | 7.300E-06 | 7.500E-06 |
| 2473 | 124 | T124M | A | 6.2e-06 | 3.9e-06 | 1e-05 | 6.700E-06 | 3.081E-06 | 3.900E-06 | 1.000E-05 |
| 2474 | 124 | T124F | A | 9.3e-06 | 6.2e-06 | 9.4e-06 | 8.300E-06 | 1.819E-06 | 6.200E-06 | 9.400E-06 |
| 2475 | 124 | T124P | A | 5.4e-06 | 9.8e-06 | 7.5e-06 | 7.567E-06 | 2.201E-06 | 5.400E-06 | 9.800E-06 |
| 2476 | 124 | T124S | A | 6.7e-06 | 5.6e-06 | 8.4e-06 | 6.900E-06 | 1.411E-06 | 5.600E-06 | 8.400E-06 |
| 2477 | 124 | T124T | A | 4.7e-06 | 8.9e-06 | 8.9e-06 | 7.500E-06 | 2.425E-06 | 4.700E-06 | 8.900E-06 |
| 2478 | 124 | T124W | A | 7.8e-06 | 9.1e-06 | 1.1e-05 | 9.300E-06 | 1.609E-06 | 7.800E-06 | 1.100E-05 |
| 2479 | 124 | T124Y | A | 1.1e-05 | 6.5e-06 | 7.2e-06 | 8.233E-06 | 2.421E-06 | 6.500E-06 | 1.100E-05 |
| 2480 | 124 | T124V | A | 6.5e-06 | 7e-06 | 7.8e-06 | 7.100E-06 | 6.557E-07 | 6.500E-06 | 7.800E-06 |
| 2481 | 125 | E125A | A | 8e-06 | 5.5e-06 | 1.3e-05 | 8.833E-06 | 3.819E-06 | 5.500E-06 | 1.300E-05 |
| 2482 | 125 | E125R | A | 8.5e-06 | 1.2e-05 | 8.9e-06 | 9.800E-06 | 1.916E-06 | 8.500E-06 | 1.200E-05 |
| 2483 | 125 | E125N | A | 7.7e-06 | 6.8e-06 | 6.7e-06 | 7.067E-06 | 5.508E-07 | 6.700E-06 | 7.700E-06 |
| 2484 | 125 | E125D | A | 5.2e-06 | 7.3e-06 | 9.2e-06 | 7.233E-06 | 2.001E-06 | 5.200E-06 | 9.200E-06 |
| 2485 | 125 | E125C | A | 1.1e-05 | 6.1e-06 | 7.1e-06 | 8.067E-06 | 2.589E-06 | 6.100E-06 | 1.100E-05 |
| 2486 | 125 | E125E | A | 4.6e-06 | 4e-06 | 1.4e-05 | 7.533E-06 | 5.608E-06 | 4.000E-06 | 1.400E-05 |
| 2487 | 125 | E125Q | A | 6.4e-06 | 7.9e-06 | 6.7e-06 | 7.000E-06 | 7.937E-07 | 6.400E-06 | 7.900E-06 |
| 2488 | 125 | E125G | A | 9.9e-06 | 8.2e-06 | 6.3e-06 | 8.133E-06 | 1.801E-06 | 6.300E-06 | 9.900E-06 |
| 2489 | 125 | E125H | A | 9e-06 | 7.6e-06 | 7.9e-06 | 8.167E-06 | 7.371E-07 | 7.600E-06 | 9.000E-06 |
| 2490 | 125 | E125I | A | 5.5e-06 | 8.4e-06 | 1e-05 | 7.967E-06 | 2.281E-06 | 5.500E-06 | 1.000E-05 |
| 2491 | 125 | E125L | A | 6e-06 | 4.6e-06 | 7e-06 | 5.867E-06 | 1.206E-06 | 4.600E-06 | 7.000E-06 |
| 2492 | 125 | E125K | A | 6.7e-06 | 9.8e-06 | 8.9e-06 | 8.467E-06 | 1.595E-06 | 6.700E-06 | 9.800E-06 |
| 2493 | 125 | E125M | A | 6.4e-06 | 8.3e-06 | 8e-06 | 7.567E-06 | 1.021E-06 | 6.400E-06 | 8.300E-06 |
| 2494 | 125 | E125F | A | 1.2e-05 | 6.3e-06 | 5.8e-06 | 8.033E-06 | 3.444E-06 | 5.800E-06 | 1.200E-05 |
| 2495 | 125 | E125P | A | 5.7e-06 | 1.4e-05 | 9.8e-06 | 9.833E-06 | 4.150E-06 | 5.700E-06 | 1.400E-05 |
| 2496 | 125 | E125S | A | 5e-06 | 5.8e-06 | 8.8e-06 | 6.533E-06 | 2.003E-06 | 5.000E-06 | 8.800E-06 |
| 2497 | 125 | E125T | A | 5.9e-06 | 7.5e-06 | 6.7e-06 | 6.700E-06 | 8.000E-07 | 5.900E-06 | 7.500E-06 |
| 2498 | 125 | E125W | A | 4.2e-06 | 5.3e-06 | 9e-06 | 6.167E-06 | 2.515E-06 | 4.200E-06 | 9.000E-06 |
| 2499 | 125 | E125Y | A | 7.2e-06 | 1.2e-05 | 6.8e-06 | 8.667E-06 | 2.894E-06 | 6.800E-06 | 1.200E-05 |
| 2500 | 125 | E125V | A | 7.8e-06 | 8.9e-06 | 7.3e-06 | 8.000E-06 | 8.185E-07 | 7.300E-06 | 8.900E-06 |
| 2501 | 126 | I126A | A | 7e-06 | 6e-06 | 6.1e-06 | 6.367E-06 | 5.508E-07 | 6.000E-06 | 7.000E-06 |
| 2502 | 126 | I126R | A | 5.6e-06 | 6.7e-06 | 5e-06 | 5.767E-06 | 8.622E-07 | 5.000E-06 | 6.700E-06 |
| 2503 | 126 | I126N | A | 9.8e-06 | 1e-05 | 4.8e-06 | 8.200E-06 | 2.946E-06 | 4.800E-06 | 1.000E-05 |
| 2504 | 126 | I126D | A | 1e-05 | 1.1e-05 | 7.7e-06 | 9.567E-06 | 1.692E-06 | 7.700E-06 | 1.100E-05 |
| 2505 | 126 | I126C | A | 5.8e-06 | 8.8e-06 | 6.8e-06 | 7.133E-06 | 1.528E-06 | 5.800E-06 | 8.800E-06 |
| 2506 | 126 | I126E | A | 1e-05 | 9.8e-06 | 6e-06 | 8.600E-06 | 2.254E-06 | 6.000E-06 | 1.000E-05 |
| 2507 | 126 | I126Q | A | 8.9e-06 | 4.3e-06 | 9.8e-06 | 7.667E-06 | 2.950E-06 | 4.300E-06 | 9.800E-06 |
| 2508 | 126 | I126G | A | 7e-06 | 7.4e-06 | 6e-06 | 6.800E-06 | 7.211E-07 | 6.000E-06 | 7.400E-06 |
| 2509 | 126 | I126H | A | 1.1e-05 | 6.4e-06 | 8.7e-06 | 8.700E-06 | 2.300E-06 | 6.400E-06 | 1.100E-05 |
| 2510 | 126 | I126I | A | 1.3e-05 | 4.7e-06 | 6.5e-06 | 8.067E-06 | 4.366E-06 | 4.700E-06 | 1.300E-05 |
| 2511 | 126 | I126L | A | 6.9e-06 | 1.6e-05 | 1.5e-05 | 1.263E-05 | 4.990E-06 | 6.900E-06 | 1.600E-05 |
| 2512 | 126 | I126K | A | 7.6e-06 | 7.5e-06 | 4.7e-06 | 6.600E-06 | 1.646E-06 | 4.700E-06 | 7.600E-06 |
| 2513 | 126 | I126M | A | 8e-06 | 8.7e-06 | 8.1e-06 | 8.267E-06 | 3.786E-07 | 8.000E-06 | 8.700E-06 |
| 2514 | 126 | I126F | A | 7e-06 | 8e-06 | 1e-05 | 8.333E-06 | 1.528E-06 | 7.000E-06 | 1.000E-05 |
| 2515 | 126 | I126P | A | 8.5e-06 | 4.5e-06 | 9.3e-06 | 7.433E-06 | 2.572E-06 | 4.500E-06 | 9.300E-06 |
| 2516 | 126 | I126S | A | 1.3e-05 | 9.5e-06 | 7e-06 | 9.833E-06 | 3.014E-06 | 7.000E-06 | 1.300E-05 |
| 2517 | 126 | I126T | A | 9.1e-06 | 6.8e-06 | 7e-06 | 7.633E-06 | 1.274E-06 | 6.800E-06 | 9.100E-06 |
| 2518 | 126 | I126W | A | 5.9e-06 | 6.2e-06 | 1.1e-05 | 7.700E-06 | 2.862E-06 | 5.900E-06 | 1.100E-05 |
| 2519 | 126 | I126Y | A | 7.5e-06 | 6.2e-06 | 8.3e-06 | 7.333E-06 | 1.060E-06 | 6.200E-06 | 8.300E-06 |
| 2520 | 126 | I126V | A | 8.6e-06 | 6e-06 | 5.9e-06 | 6.833E-06 | 1.531E-06 | 5.900E-06 | 8.600E-06 |
| 2521 | 127 | L127A | A | 1.2e-05 | 8.3e-06 | 4e-06 | 8.100E-06 | 4.004E-06 | 4.000E-06 | 1.200E-05 |
| 2522 | 127 | L127R | A | 6.7e-06 | 8.2e-06 | 6.8e-06 | 7.233E-06 | 8.386E-07 | 6.700E-06 | 8.200E-06 |
| 2523 | 127 | L127N | A | 9.7e-06 | 5e-06 | 8.6e-06 | 7.767E-06 | 2.458E-06 | 5.000E-06 | 9.700E-06 |
| 2524 | 127 | L127D | A | 6.7e-06 | 9.2e-06 | 1.1e-05 | 8.967E-06 | 2.159E-06 | 6.700E-06 | 1.100E-05 |
| 2525 | 127 | L127C | A | 5.1e-06 | 1.3e-05 | 1.1e-05 | 9.700E-06 | 4.107E-06 | 5.100E-06 | 1.300E-05 |
| 2526 | 127 | L127E | A | 6.5e-06 | 1.3e-05 | 5e-06 | 8.167E-06 | 4.252E-06 | 5.000E-06 | 1.300E-05 |
| 2527 | 127 | L127Q | A | 9e-06 | 5.8e-06 | 1.5e-05 | 9.933E-06 | 4.670E-06 | 5.800E-06 | 1.500E-05 |
| 2528 | 127 | L127G | A | 7.9e-06 | 8.4e-06 | 6.2e-06 | 7.500E-06 | 1.153E-06 | 6.200E-06 | 8.400E-06 |
| 2529 | 127 | L127H | A | 5.3e-06 | 9.5e-06 | 9e-06 | 7.933E-06 | 2.294E-06 | 5.300E-06 | 9.500E-06 |
| 2530 | 127 | L127I | A | 8.5e-06 | 9.9e-06 | 5.1e-06 | 7.833E-06 | 2.468E-06 | 5.100E-06 | 9.900E-06 |
| 2531 | 127 | L127L | A | 4.8e-06 | 5.9e-06 | 8.4e-06 | 6.367E-06 | 1.845E-06 | 4.800E-06 | 8.400E-06 |
| 2532 | 127 | L127K | A | 8.7e-06 | 8.4e-06 | 9.8e-06 | 8.967E-06 | 7.371E-07 | 8.400E-06 | 9.800E-06 |
| 2533 | 127 | L127M | A | 6.8e-06 | 8.3e-06 | 1e-05 | 8.367E-06 | 1.601E-06 | 6.800E-06 | 1.000E-05 |
| 2534 | 127 | L127F | A | 1.4e-05 | 6.7e-06 | 9e-06 | 9.900E-06 | 3.732E-06 | 6.700E-06 | 1.400E-05 |
| 2535 | 127 | L127P | A | 6.7e-06 | 5e-06 | 1.3e-05 | 8.233E-06 | 4.215E-06 | 5.000E-06 | 1.300E-05 |
| 2536 | 127 | L127S | A | 6e-06 | 9.9e-06 | 6.2e-06 | 7.367E-06 | 2.196E-06 | 6.000E-06 | 9.900E-06 |
| 2537 | 127 | L127T | A | 5.9e-06 | 1.2e-05 | 9e-06 | 8.967E-06 | 3.050E-06 | 5.900E-06 | 1.200E-05 |
| 2538 | 127 | L127W | A | 7.8e-06 | 7.2e-06 | 5.7e-06 | 6.900E-06 | 1.082E-06 | 5.700E-06 | 7.800E-06 |
| 2539 | 127 | L127Y | A | 6.5e-06 | 7.1e-06 | 5.8e-06 | 6.467E-06 | 6.506E-07 | 5.800E-06 | 7.100E-06 |
| 2540 | 127 | L127V | A | 1.1e-05 | 1.2e-05 | 3.3e-06 | 8.767E-06 | 4.761E-06 | 3.300E-06 | 1.200E-05 |
| 2541 | 128 | K128A | A | 1e-05 | 7.3e-06 | 1.3e-05 | 1.010E-05 | 2.851E-06 | 7.300E-06 | 1.300E-05 |
| 2542 | 128 | K128R | A | 8.3e-06 | 4.5e-06 | 1.1e-05 | 7.933E-06 | 3.265E-06 | 4.500E-06 | 1.100E-05 |
| 2543 | 128 | K128N | A | 8.8e-06 | 7.5e-06 | 9.4e-06 | 8.567E-06 | 9.713E-07 | 7.500E-06 | 9.400E-06 |
| 2544 | 128 | K128D | A | 1.4e-05 | 8.4e-06 | 8.7e-06 | 1.037E-05 | 3.150E-06 | 8.400E-06 | 1.400E-05 |
| 2545 | 128 | K128C | A | 5.6e-06 | 9.3e-06 | 8.5e-06 | 7.800E-06 | 1.947E-06 | 5.600E-06 | 9.300E-06 |
| 2546 | 128 | K128E | A | 9.5e-06 | 1e-05 | 7.8e-06 | 9.100E-06 | 1.153E-06 | 7.800E-06 | 1.000E-05 |
| 2547 | 128 | K128Q | A | 8.4e-06 | 6e-06 | 8.1e-06 | 7.500E-06 | 1.308E-06 | 6.000E-06 | 8.400E-06 |
| 2548 | 128 | K128G | A | 8.9e-06 | 5.7e-06 | 1.1e-05 | 8.533E-06 | 2.669E-06 | 5.700E-06 | 1.100E-05 |
| 2549 | 128 | K128H | A | 1.1e-05 | 6.4e-06 | 6.3e-06 | 7.900E-06 | 2.685E-06 | 6.300E-06 | 1.100E-05 |

| | | | | | | | | | | |
|---|---|---|---|---|---|---|---|---|---|---|
| 2550 | 128 | K128I | A | 8.6e-06 | 2e-05 | 7.4e-06 | 1.200E-05 | 6.954E-06 | 7.400E-06 | 2.000E-05 |
| 2551 | 128 | K128L | A | 2.1e-05 | 5.9e-06 | 1e-05 | 1.230E-05 | 7.808E-06 | 5.900E-06 | 2.100E-05 |
| 2552 | 128 | K128K | A | 1.3e-05 | 7.2e-06 | 5.9e-06 | 8.700E-06 | 3.780E-06 | 5.900E-06 | 1.300E-05 |
| 2553 | 128 | K128M | A | 8.5e-06 | 1e-05 | 9.1e-06 | 9.200E-06 | 7.550E-07 | 8.500E-06 | 1.000E-05 |
| 2554 | 128 | K128F | A | 5.1e-06 | 5.5e-06 | 1.1e-05 | 7.200E-06 | 3.297E-06 | 5.100E-06 | 1.100E-05 |
| 2555 | 128 | K128P | A | 9e-06 | 6.4e-06 | 5.8e-06 | 7.067E-06 | 1.701E-06 | 5.800E-06 | 9.000E-06 |
| 2556 | 128 | K128S | A | 7.9e-06 | 5.4e-06 | 4.8e-06 | 6.033E-06 | 1.644E-06 | 4.800E-06 | 7.900E-06 |
| 2557 | 128 | K128T | A | 1.3e-05 | 9.2e-06 | 6.2e-06 | 9.467E-06 | 3.408E-06 | 6.200E-06 | 1.300E-05 |
| 2558 | 128 | K128W | A | 8.8e-06 | 9.5e-06 | 5.3e-06 | 7.867E-06 | 2.250E-06 | 5.300E-06 | 9.500E-06 |
| 2559 | 128 | K128Y | A | 4.6e-06 | 4.2e-06 | 7.6e-06 | 5.467E-06 | 1.858E-06 | 4.200E-06 | 7.600E-06 |
| 2560 | 128 | K128V | A | 1.3e-05 | 7.2e-06 | 8.6e-06 | 9.600E-06 | 3.027E-06 | 7.200E-06 | 1.300E-05 |
| 2561 | 129 | G129A | A | 4.9e-06 | 3.6e-06 | 6e-06 | 4.833E-06 | 1.201E-06 | 3.600E-06 | 6.000E-06 |
| 2562 | 129 | G129R | A | 1.1e-05 | 6.7e-06 | 6.7e-06 | 8.133E-06 | 2.483E-06 | 6.700E-06 | 1.100E-05 |
| 2563 | 129 | G129N | A | 9.2e-06 | 1.1e-05 | 1.1e-05 | 1.040E-05 | 1.039E-06 | 9.200E-06 | 1.100E-05 |
| 2564 | 129 | G129D | A | 7.5e-06 | 9.2e-06 | 9.9e-06 | 8.867E-06 | 1.234E-06 | 7.500E-06 | 9.900E-06 |
| 2565 | 129 | G129C | A | 7e-06 | 7.6e-06 | 8.3e-06 | 7.633E-06 | 6.506E-07 | 7.000E-06 | 8.300E-06 |
| 2566 | 129 | G129E | A | 7.5e-06 | 1.3e-05 | 8e-06 | 9.500E-06 | 3.041E-06 | 7.500E-06 | 1.300E-05 |
| 2567 | 129 | G129Q | A | 1.2e-05 | 1.4e-05 | 6.7e-06 | 1.090E-05 | 3.772E-06 | 6.700E-06 | 1.400E-05 |
| 2568 | 129 | G129G | A | 8.3e-06 | 6.4e-06 | 1.1e-05 | 8.567E-06 | 2.312E-06 | 6.400E-06 | 1.100E-05 |
| 2569 | 129 | G129H | A | 5.9e-06 | 6.6e-06 | 4e-06 | 5.500E-06 | 1.345E-06 | 4.000E-06 | 6.600E-06 |
| 2570 | 129 | G129I | A | 9.6e-06 | 1.1e-05 | 8.5e-06 | 9.700E-06 | 1.253E-06 | 8.500E-06 | 1.100E-05 |
| 2571 | 129 | G129L | A | 7.1e-06 | 8.2e-06 | 6.7e-06 | 7.333E-06 | 7.767E-07 | 6.700E-06 | 8.200E-06 |
| 2572 | 129 | G129K | A | 9.1e-06 | 6.1e-06 | 1.4e-05 | 9.733E-06 | 3.988E-06 | 6.100E-06 | 1.400E-05 |
| 2573 | 129 | G129M | A | 4.5e-06 | 6.2e-06 | 9.2e-06 | 6.633E-06 | 2.380E-06 | 4.500E-06 | 9.200E-06 |
| 2574 | 129 | G129F | A | 8.6e-06 | 1.6e-05 | 9.3e-06 | 1.130E-05 | 4.085E-06 | 8.600E-06 | 1.600E-05 |
| 2575 | 129 | G129P | A | 7.2e-06 | 6.8e-06 | 6.1e-06 | 6.700E-06 | 5.568E-07 | 6.100E-06 | 7.200E-06 |
| 2576 | 129 | G129S | A | 8.6e-06 | 9.5e-06 | 6.5e-06 | 8.200E-06 | 1.539E-06 | 6.500E-06 | 9.500E-06 |
| 2577 | 129 | G129T | A | 7.2e-06 | 9.1e-06 | 1.1e-05 | 9.100E-06 | 1.900E-06 | 7.200E-06 | 1.100E-05 |
| 2578 | 129 | G129W | A | 8.4e-06 | 1.5e-05 | 7.3e-06 | 1.023E-05 | 4.165E-06 | 7.300E-06 | 1.500E-05 |
| 2579 | 129 | G129Y | A | 7.2e-06 | 6.7e-06 | 7.7e-06 | 7.200E-06 | 5.000E-07 | 6.700E-06 | 7.700E-06 |
| 2580 | 129 | G129V | A | 9.3e-06 | 6.2e-06 | 8.7e-06 | 8.067E-06 | 1.644E-06 | 6.200E-06 | 9.300E-06 |
| 2581 | 130 | G130A | A | 7.4e-06 | 1.4e-05 | 4.8e-06 | 8.733E-06 | 4.743E-06 | 4.800E-06 | 1.400E-05 |
| 2582 | 130 | G130R | A | 5e-06 | 4.5e-06 | 1.4e-05 | 7.833E-06 | 5.346E-06 | 4.500E-06 | 1.400E-05 |
| 2583 | 130 | G130N | A | 1e-05 | 9.4e-06 | 1.1e-05 | 1.013E-05 | 8.083E-07 | 9.400E-06 | 1.100E-05 |
| 2584 | 130 | G130D | A | 8.8e-06 | 1.3e-05 | 1.3e-05 | 1.160E-05 | 2.425E-06 | 8.800E-06 | 1.300E-05 |
| 2585 | 130 | G130C | A | 1.2e-05 | 4.5e-06 | 7e-06 | 7.833E-06 | 3.819E-06 | 4.500E-06 | 1.200E-05 |
| 2586 | 130 | G130E | A | 9e-06 | 6.1e-06 | 9.4e-06 | 8.167E-06 | 1.801E-06 | 6.100E-06 | 9.400E-06 |
| 2587 | 130 | G130Q | A | 1.4e-05 | 6.2e-06 | 7.8e-06 | 9.333E-06 | 4.120E-06 | 6.200E-06 | 1.400E-05 |
| 2588 | 130 | G130G | A | 7.6e-06 | 6.8e-06 | 6e-06 | 6.800E-06 | 8.000E-07 | 6.000E-06 | 7.600E-06 |
| 2589 | 130 | G130H | A | 1.2e-05 | 6.2e-06 | 5.9e-06 | 8.033E-06 | 3.439E-06 | 5.900E-06 | 1.200E-05 |
| 2590 | 130 | G130I | A | 8.8e-06 | 1e-05 | 5.9e-06 | 8.233E-06 | 2.108E-06 | 5.900E-06 | 1.000E-05 |
| 2591 | 130 | G130L | A | 1e-05 | 7.3e-06 | 1.3e-05 | 1.010E-05 | 2.851E-06 | 7.300E-06 | 1.300E-05 |
| 2592 | 130 | G130K | A | 1.1e-05 | 4e-06 | 1.1e-05 | 8.667E-06 | 4.041E-06 | 4.000E-06 | 1.100E-05 |
| 2593 | 130 | G130M | A | 1.1e-05 | 5.4e-06 | 1.3e-05 | 9.800E-06 | 3.940E-06 | 5.400E-06 | 1.300E-05 |
| 2594 | 130 | G130F | A | 7.9e-06 | 1e-05 | 1.1e-05 | 9.633E-06 | 1.582E-06 | 7.900E-06 | 1.100E-05 |
| 2595 | 130 | G130P | A | 5.1e-06 | 7.1e-06 | 8.6e-06 | 6.933E-06 | 1.756E-06 | 5.100E-06 | 8.600E-06 |
| 2596 | 130 | G130S | A | 6.6e-06 | 1e-05 | 7.7e-06 | 8.100E-06 | 1.735E-06 | 6.600E-06 | 1.000E-05 |
| 2597 | 130 | G130T | A | 9.5e-06 | 8.2e-06 | 7.7e-06 | 8.467E-06 | 9.292E-07 | 7.700E-06 | 9.500E-06 |
| 2598 | 130 | G130W | A | 6.6e-06 | 1.1e-05 | 7e-06 | 8.200E-06 | 2.433E-06 | 6.600E-06 | 1.100E-05 |
| 2599 | 130 | G130Y | A | 6.5e-06 | 1.1e-05 | 6.6e-06 | 8.033E-06 | 2.570E-06 | 6.500E-06 | 1.100E-05 |
| 2600 | 130 | G130V | A | 9.5e-06 | 9.2e-06 | 6e-06 | 8.233E-06 | 1.940E-06 | 6.000E-06 | 9.500E-06 |
| 2601 | 131 | V131A | A | 6.2e-06 | 6.1e-06 | 9.2e-06 | 7.167E-06 | 1.762E-06 | 6.100E-06 | 9.200E-06 |
| 2602 | 131 | V131R | A | 1.4e-05 | 8.2e-06 | 6.3e-06 | 9.500E-06 | 4.011E-06 | 6.300E-06 | 1.400E-05 |
| 2603 | 131 | V131N | A | 6.7e-06 | 7.5e-06 | 1.6e-05 | 1.007E-05 | 5.154E-06 | 6.700E-06 | 1.600E-05 |
| 2604 | 131 | V131D | A | 6.8e-06 | 5.8e-06 | 9.8e-06 | 7.467E-06 | 2.082E-06 | 5.800E-06 | 9.800E-06 |
| 2605 | 131 | V131C | A | 8.3e-06 | 1.2e-05 | 7e-06 | 9.100E-06 | 2.594E-06 | 7.000E-06 | 1.200E-05 |
| 2606 | 131 | V131E | A | 8.9e-06 | 6.3e-06 | 9.1e-06 | 8.100E-06 | 1.562E-06 | 6.300E-06 | 9.100E-06 |
| 2607 | 131 | V131Q | A | 5.6e-06 | 1.1e-05 | 6.2e-06 | 7.600E-06 | 2.960E-06 | 5.600E-06 | 1.100E-05 |
| 2608 | 131 | V131G | A | 4.5e-06 | 7.5e-06 | 8.4e-06 | 6.800E-06 | 2.042E-06 | 4.500E-06 | 8.400E-06 |
| 2609 | 131 | V131H | A | 6.7e-06 | 9.4e-06 | 2.1e-05 | 1.237E-05 | 7.598E-06 | 6.700E-06 | 2.100E-05 |
| 2610 | 131 | V131I | A | 1e-05 | 8.9e-06 | 1e-05 | 9.633E-06 | 6.351E-07 | 8.900E-06 | 1.000E-05 |
| 2611 | 131 | V131L | A | 7.6e-06 | 1e-05 | 9.4e-06 | 9.000E-06 | 1.249E-06 | 7.600E-06 | 1.000E-05 |
| 2612 | 131 | V131K | A | 1e-05 | 1.1e-05 | 1.1e-05 | 1.067E-05 | 5.774E-07 | 1.000E-05 | 1.100E-05 |
| 2613 | 131 | V131M | A | 7.5e-06 | 9.4e-06 | 5.1e-06 | 7.333E-06 | 2.155E-06 | 5.100E-06 | 9.400E-06 |
| 2614 | 131 | V131F | A | 6.5e-06 | 4.6e-06 | 5.7e-06 | 5.600E-06 | 9.539E-07 | 4.600E-06 | 6.500E-06 |
| 2615 | 131 | V131P | A | 6.9e-06 | 6e-06 | 9.7e-06 | 7.533E-06 | 1.930E-06 | 6.000E-06 | 9.700E-06 |
| 2616 | 131 | V131S | A | 1.1e-05 | 5.4e-06 | 4.1e-06 | 6.833E-06 | 3.667E-06 | 4.100E-06 | 1.100E-05 |
| 2617 | 131 | V131T | A | 1.2e-05 | 9.5e-06 | 6.2e-06 | 9.233E-06 | 2.909E-06 | 6.200E-06 | 1.200E-05 |
| 2618 | 131 | V131W | A | 1e-05 | 8.8e-06 | 5.9e-06 | 8.233E-06 | 2.108E-06 | 5.900E-06 | 1.000E-05 |
| 2619 | 131 | V131Y | A | 9.3e-06 | 4.5e-06 | 5.4e-06 | 6.400E-06 | 2.551E-06 | 4.500E-06 | 9.300E-06 |
| 2620 | 131 | V131V | A | 7.4e-06 | 8.9e-06 | 7e-06 | 7.767E-06 | 1.002E-06 | 7.000E-06 | 8.900E-06 |
| 2621 | 132 | L132A | A | 1.3e-05 | 3.8e-06 | 1e-05 | 8.933E-06 | 4.692E-06 | 3.800E-06 | 1.300E-05 |
| 2622 | 132 | L132R | A | 1.4e-05 | 5.7e-06 | 7.7e-06 | 9.133E-06 | 4.332E-06 | 5.700E-06 | 1.400E-05 |
| 2623 | 132 | L132N | A | 4e-06 | 6.4e-06 | 5.5e-06 | 5.300E-06 | 1.212E-06 | 4.000E-06 | 6.400E-06 |
| 2624 | 132 | L132D | A | 4.7e-06 | 4.2e-06 | 5.7e-06 | 4.867E-06 | 7.638E-07 | 4.200E-06 | 5.700E-06 |
| 2625 | 132 | L132C | A | 8.6e-06 | 7.2e-06 | 9.5e-06 | 8.433E-06 | 1.159E-06 | 7.200E-06 | 9.500E-06 |
| 2626 | 132 | L132E | A | 1e-05 | 4.6e-06 | 7.5e-06 | 7.367E-06 | 2.702E-06 | 4.600E-06 | 1.000E-05 |
| 2627 | 132 | L132Q | A | 7.6e-06 | 8.5e-06 | 6.2e-06 | 7.433E-06 | 1.159E-06 | 6.200E-06 | 8.500E-06 |
| 2628 | 132 | L132G | A | 9.6e-06 | 8.9e-06 | 1e-05 | 9.500E-06 | 5.568E-07 | 8.900E-06 | 1.000E-05 |
| 2629 | 132 | L132H | A | 6.7e-06 | 9.2e-06 | 1.4e-05 | 9.967E-06 | 3.710E-06 | 6.700E-06 | 1.400E-05 |
| 2630 | 132 | L132I | A | 7.9e-06 | 1.2e-05 | 1.3e-05 | 1.097E-05 | 2.702E-06 | 7.900E-06 | 1.300E-05 |
| 2631 | 132 | L132L | A | 5e-06 | 7.3e-06 | 6.7e-06 | 6.333E-06 | 1.193E-06 | 5.000E-06 | 7.300E-06 |
| 2632 | 132 | L132K | A | 9.7e-06 | 1.1e-05 | 5.5e-06 | 8.733E-06 | 2.875E-06 | 5.500E-06 | 1.100E-05 |
| 2633 | 132 | L132M | A | 8.9e-06 | 4.3e-06 | 1.1e-05 | 8.067E-06 | 3.427E-06 | 4.300E-06 | 1.100E-05 |
| 2634 | 132 | L132F | A | 4.4e-06 | 5.1e-06 | 9.5e-06 | 6.333E-06 | 2.765E-06 | 4.400E-06 | 9.500E-06 |

| | | | | | | | | | | |
|---|---|---|---|---|---|---|---|---|---|---|
| 2635 | 132 | L132P | A | 6.5e-06 | 6e-06 | 7.5e-06 | 6.667E-06 | 7.638E-07 | 6.000E-06 | 7.500E-06 |
| 2636 | 132 | L132S | A | 7.1e-06 | 5.2e-06 | 7.3e-06 | 6.533E-06 | 1.159E-06 | 5.200E-06 | 7.300E-06 |
| 2637 | 132 | L132T | A | 1.4e-05 | 6.2e-06 | 8.8e-06 | 9.667E-06 | 3.972E-06 | 6.200E-06 | 1.400E-05 |
| 2638 | 132 | L132W | A | 9e-06 | 9.1e-06 | 1.4e-05 | 1.070E-05 | 2.858E-06 | 9.000E-06 | 1.400E-05 |
| 2639 | 132 | L132Y | A | 1.3e-05 | 6.5e-06 | 1.2e-05 | 1.050E-05 | 3.500E-06 | 6.500E-06 | 1.300E-05 |
| 2640 | 132 | L132V | A | 1e-05 | 1.1e-05 | 7.9e-06 | 9.633E-06 | 1.582E-06 | 7.900E-06 | 1.100E-05 |
| 2641 | 133 | I133A | A | 9.2e-06 | 1e-05 | 9.8e-06 | 9.667E-06 | 4.163E-07 | 9.200E-06 | 1.000E-05 |
| 2642 | 133 | I133R | A | 6.2e-06 | 7.5e-06 | 6.9e-06 | 6.867E-06 | 6.506E-07 | 6.200E-06 | 7.500E-06 |
| 2643 | 133 | I133N | A | 1.2e-05 | 4.8e-06 | 1.2e-05 | 9.600E-06 | 4.157E-06 | 4.800E-06 | 1.200E-05 |
| 2644 | 133 | I133D | A | 6.9e-06 | 8.4e-06 | 6.9e-06 | 7.400E-06 | 8.660E-07 | 6.900E-06 | 8.400E-06 |
| 2645 | 133 | I133C | A | 9.8e-06 | 1.6e-05 | 6.1e-06 | 1.063E-05 | 5.002E-06 | 6.100E-06 | 1.600E-05 |
| 2646 | 133 | I133E | A | 8.3e-06 | 1.1e-05 | 6.1e-06 | 8.467E-06 | 2.454E-06 | 6.100E-06 | 1.100E-05 |
| 2647 | 133 | I133Q | A | 9.3e-06 | 6.4e-06 | 1.2e-05 | 9.233E-06 | 2.801E-06 | 6.400E-06 | 1.200E-05 |
| 2648 | 133 | I133G | A | 1e-05 | 1.3e-05 | 5.5e-06 | 9.500E-06 | 3.775E-06 | 5.500E-06 | 1.300E-05 |
| 2649 | 133 | I133H | A | 9.7e-06 | 4.9e-06 | 6.1e-06 | 6.900E-06 | 2.498E-06 | 4.900E-06 | 9.700E-06 |
| 2650 | 133 | I133I | A | 6.5e-06 | 6.9e-06 | 6.9e-06 | 6.767E-06 | 2.309E-07 | 6.500E-06 | 6.900E-06 |
| 2651 | 133 | I133L | A | 7.3e-06 | 7.2e-06 | 9.7e-06 | 8.067E-06 | 1.415E-06 | 7.200E-06 | 9.700E-06 |
| 2652 | 133 | I133K | A | 1.5e-05 | 9.6e-06 | 9.1e-06 | 1.123E-05 | 3.272E-06 | 9.100E-06 | 1.500E-05 |
| 2653 | 133 | I133M | A | 6e-06 | 8.8e-06 | 6.3e-06 | 7.033E-06 | 1.537E-06 | 6.000E-06 | 8.800E-06 |
| 2654 | 133 | I133F | A | 7.8e-06 | 8.2e-06 | 9.7e-06 | 8.567E-06 | 1.002E-06 | 7.800E-06 | 9.700E-06 |
| 2655 | 133 | I133P | A | 7.9e-06 | 7.5e-06 | 7.7e-06 | 7.700E-06 | 2.000E-07 | 7.500E-06 | 7.900E-06 |
| 2656 | 133 | I133S | A | 1.2e-05 | 8.4e-06 | 9.7e-06 | 1.003E-05 | 1.823E-06 | 8.400E-06 | 1.200E-05 |
| 2657 | 133 | I133T | A | 6.2e-06 | 6.4e-06 | 4.7e-06 | 5.767E-06 | 9.292E-07 | 4.700E-06 | 6.400E-06 |
| 2658 | 133 | I133W | A | 9.6e-06 | 5.9e-06 | 8.1e-06 | 7.867E-06 | 1.861E-06 | 5.900E-06 | 9.600E-06 |
| 2659 | 133 | I133Y | A | 9.2e-06 | 4.6e-06 | 6.5e-06 | 6.767E-06 | 2.312E-06 | 4.600E-06 | 9.200E-06 |
| 2660 | 133 | I133V | A | 1e-05 | 8.5e-06 | 5.7e-06 | 8.067E-06 | 2.183E-06 | 5.700E-06 | 1.000E-05 |
| 2661 | 134 | Q134A | A | 1.2e-05 | 6.2e-06 | 8.3e-06 | 8.833E-06 | 2.937E-06 | 6.200E-06 | 1.200E-05 |
| 2662 | 134 | Q134R | A | 4.9e-06 | 1.4e-05 | 1e-05 | 9.633E-06 | 4.561E-06 | 4.900E-06 | 1.400E-05 |
| 2663 | 134 | Q134N | A | 1.2e-05 | 1.3e-05 | 1e-05 | 1.167E-05 | 1.528E-06 | 1.000E-05 | 1.300E-05 |
| 2664 | 134 | Q134D | A | 9e-06 | 1.5e-05 | 8e-06 | 1.067E-05 | 3.786E-06 | 8.000E-06 | 1.500E-05 |
| 2665 | 134 | Q134C | A | 9.4e-06 | 6.4e-06 | 9.9e-06 | 8.567E-06 | 1.893E-06 | 6.400E-06 | 9.900E-06 |
| 2666 | 134 | Q134E | A | 4e-06 | 7e-06 | 5.8e-06 | 5.600E-06 | 1.510E-06 | 4.000E-06 | 7.000E-06 |
| 2667 | 134 | Q134Q | A | 7.5e-06 | 6.3e-06 | 8e-06 | 7.267E-06 | 8.737E-07 | 6.300E-06 | 8.000E-06 |
| 2668 | 134 | Q134G | A | 9.3e-06 | 6.8e-06 | 1.1e-05 | 9.033E-06 | 2.113E-06 | 6.800E-06 | 1.100E-05 |
| 2669 | 134 | Q134H | A | 7e-06 | 7e-06 | 6.4e-06 | 6.800E-06 | 3.464E-07 | 6.400E-06 | 7.000E-06 |
| 2670 | 134 | Q134I | A | 9.7e-06 | 5.1e-06 | 9.3e-06 | 8.033E-06 | 2.548E-06 | 5.100E-06 | 9.700E-06 |
| 2671 | 134 | Q134L | A | 8.1e-06 | 9.7e-06 | 4.7e-06 | 7.500E-06 | 2.553E-06 | 4.700E-06 | 9.700E-06 |
| 2672 | 134 | Q134K | A | 9.5e-06 | 1.1e-05 | 5.9e-06 | 8.800E-06 | 2.621E-06 | 5.900E-06 | 1.100E-05 |
| 2673 | 134 | Q134M | A | 9.2e-06 | 9.6e-06 | 7.8e-06 | 8.867E-06 | 9.452E-07 | 7.800E-06 | 9.600E-06 |
| 2674 | 134 | Q134F | A | 8.3e-06 | 6.8e-06 | 9.4e-06 | 8.167E-06 | 1.305E-06 | 6.800E-06 | 9.400E-06 |
| 2675 | 134 | Q134P | A | 1.4e-05 | 9.8e-06 | 7e-06 | 1.027E-05 | 3.523E-06 | 7.000E-06 | 1.400E-05 |
| 2676 | 134 | Q134S | A | 5.8e-06 | 7.3e-06 | 5.8e-06 | 6.300E-06 | 8.660E-07 | 5.800E-06 | 7.300E-06 |
| 2677 | 134 | Q134T | A | 1e-05 | 8.8e-06 | 6.7e-06 | 8.500E-06 | 1.670E-06 | 6.700E-06 | 1.000E-05 |
| 2678 | 134 | Q134W | A | 4.1e-06 | 7.7e-06 | 1.6e-05 | 9.267E-06 | 6.103E-06 | 4.100E-06 | 1.600E-05 |
| 2679 | 134 | Q134Y | A | 6.5e-06 | 3.5e-06 | 7.7e-06 | 5.900E-06 | 2.163E-06 | 3.500E-06 | 7.700E-06 |
| 2680 | 134 | Q134V | A | 6.5e-06 | 1e-05 | 1.3e-05 | 9.833E-06 | 3.253E-06 | 6.500E-06 | 1.300E-05 |
| 2681 | 135 | R135A | A | 7.6e-06 | 1.6e-05 | 6.1e-06 | 9.900E-06 | 5.336E-06 | 6.100E-06 | 1.600E-05 |
| 2682 | 135 | R135R | A | 9.9e-06 | 1.4e-05 | 9.7e-06 | 1.120E-05 | 2.427E-06 | 9.700E-06 | 1.400E-05 |
| 2683 | 135 | R135N | A | 7.2e-06 | 6.1e-06 | 6.2e-06 | 6.500E-06 | 6.083E-07 | 6.100E-06 | 7.200E-06 |
| 2684 | 135 | R135D | A | 7.3e-06 | 9e-06 | 7.2e-06 | 7.833E-06 | 1.012E-06 | 7.200E-06 | 9.000E-06 |
| 2685 | 135 | R135C | A | 9.3e-06 | 3.1e-06 | 7.3e-06 | 6.567E-06 | 3.164E-06 | 3.100E-06 | 9.300E-06 |
| 2686 | 135 | R135E | A | 8e-06 | 9.8e-06 | 4.2e-06 | 7.333E-06 | 2.859E-06 | 4.200E-06 | 9.800E-06 |
| 2687 | 135 | R135Q | A | 7.1e-06 | 6.3e-06 | 4.3e-06 | 5.900E-06 | 1.442E-06 | 4.300E-06 | 7.100E-06 |
| 2688 | 135 | R135G | A | 9.4e-06 | 1e-05 | 9.5e-06 | 9.633E-06 | 3.215E-07 | 9.400E-06 | 1.000E-05 |
| 2689 | 135 | R135H | A | 8.3e-06 | 6.7e-06 | 8.4e-06 | 7.800E-06 | 9.539E-07 | 6.700E-06 | 8.400E-06 |
| 2690 | 135 | R135I | A | 7.9e-06 | 5.4e-06 | 6.3e-06 | 6.533E-06 | 1.266E-06 | 5.400E-06 | 7.900E-06 |
| 2691 | 135 | R135L | A | 2e-05 | 9.8e-06 | 1.1e-05 | 1.360E-05 | 5.575E-06 | 9.800E-06 | 2.000E-05 |
| 2692 | 135 | R135K | A | 7.3e-06 | 5.3e-06 | 9.5e-06 | 7.367E-06 | 2.101E-06 | 5.300E-06 | 9.500E-06 |
| 2693 | 135 | R135M | A | 5.1e-06 | 9.1e-06 | 5.8e-06 | 6.667E-06 | 2.136E-06 | 5.100E-06 | 9.100E-06 |
| 2694 | 135 | R135F | A | 6.9e-06 | 8.2e-06 | 6.3e-06 | 7.133E-06 | 9.713E-07 | 6.300E-06 | 8.200E-06 |
| 2695 | 135 | R135P | A | 4.6e-06 | 6.8e-06 | 9.3e-06 | 6.900E-06 | 2.352E-06 | 4.600E-06 | 9.300E-06 |
| 2696 | 135 | R135S | A | 7.1e-06 | 9.3e-06 | 6.6e-06 | 7.667E-06 | 1.436E-06 | 6.600E-06 | 9.300E-06 |
| 2697 | 135 | R135T | A | 8.8e-06 | 4.5e-06 | 7e-06 | 6.767E-06 | 2.159E-06 | 4.500E-06 | 8.800E-06 |
| 2698 | 135 | R135W | A | 5.9e-06 | 8.6e-06 | 5.1e-06 | 6.533E-06 | 1.834E-06 | 5.100E-06 | 8.600E-06 |
| 2699 | 135 | R135Y | A | 9.7e-06 | 1.3e-05 | 8e-06 | 1.023E-05 | 2.542E-06 | 8.000E-06 | 1.300E-05 |
| 2700 | 135 | R135V | A | 6.5e-06 | 7.4e-06 | 6.3e-06 | 6.733E-06 | 5.859E-07 | 6.300E-06 | 7.400E-06 |
| 2701 | 136 | N136A | A | 7e-06 | 1.8e-05 | 7.2e-06 | 1.073E-05 | 6.294E-06 | 7.000E-06 | 1.800E-05 |
| 2702 | 136 | N136R | A | 8.9e-06 | 7.7e-06 | 6.9e-06 | 7.833E-06 | 1.007E-06 | 6.900E-06 | 8.900E-06 |
| 2703 | 136 | N136N | A | 8.6e-06 | 5.7e-06 | 8.9e-06 | 7.733E-06 | 1.767E-06 | 5.700E-06 | 8.900E-06 |
| 2704 | 136 | N136D | A | 7.2e-06 | 1.4e-05 | 4.4e-06 | 8.533E-06 | 4.937E-06 | 4.400E-06 | 1.400E-05 |
| 2705 | 136 | N136C | A | 7.1e-06 | 7.9e-06 | 6.5e-06 | 7.167E-06 | 7.024E-07 | 6.500E-06 | 7.900E-06 |
| 2706 | 136 | N136E | A | 5.8e-06 | 5.3e-06 | 7.4e-06 | 6.167E-06 | 1.097E-06 | 5.300E-06 | 7.400E-06 |
| 2707 | 136 | N136Q | A | 6.4e-06 | 9.9e-06 | 9.9e-06 | 8.733E-06 | 2.021E-06 | 6.400E-06 | 9.900E-06 |
| 2708 | 136 | N136G | A | 1.2e-05 | 6.9e-06 | 1.3e-05 | 1.063E-05 | 3.272E-06 | 6.900E-06 | 1.300E-05 |
| 2709 | 136 | N136H | A | 4.8e-06 | 6e-06 | 1.3e-05 | 7.933E-06 | 4.429E-06 | 4.800E-06 | 1.300E-05 |
| 2710 | 136 | N136I | A | 1.3e-05 | 6.5e-06 | 7.3e-06 | 8.933E-06 | 3.544E-06 | 6.500E-06 | 1.300E-05 |
| 2711 | 136 | N136L | A | 1.2e-05 | 9.4e-06 | 1.1e-05 | 1.080E-05 | 1.311E-06 | 9.400E-06 | 1.200E-05 |
| 2712 | 136 | N136K | A | 8e-06 | 7.3e-06 | 9.9e-06 | 8.400E-06 | 1.345E-06 | 7.300E-06 | 9.900E-06 |
| 2713 | 136 | N136M | A | 3.2e-06 | 6.7e-06 | 6.1e-06 | 5.333E-06 | 1.872E-06 | 3.200E-06 | 6.700E-06 |
| 2714 | 136 | N136F | A | 7.1e-06 | 5.7e-06 | 6.4e-06 | 6.400E-06 | 7.000E-07 | 5.700E-06 | 7.100E-06 |
| 2715 | 136 | N136P | A | 1.3e-05 | 5.2e-06 | 1.3e-05 | 1.040E-05 | 4.503E-06 | 5.200E-06 | 1.300E-05 |
| 2716 | 136 | N136S | A | 7.4e-06 | 8.3e-06 | 1e-05 | 8.567E-06 | 1.320E-06 | 7.400E-06 | 1.000E-05 |
| 2717 | 136 | N136T | A | 6.8e-06 | 5.3e-06 | 1.1e-05 | 7.700E-06 | 2.955E-06 | 5.300E-06 | 1.100E-05 |
| 2718 | 136 | N136W | A | 6.9e-06 | 5.5e-06 | 5.5e-06 | 5.967E-06 | 8.083E-07 | 5.500E-06 | 6.900E-06 |
| 2719 | 136 | N136Y | A | 8e-06 | 9.1e-06 | 7.2e-06 | 8.100E-06 | 9.539E-07 | 7.200E-06 | 9.100E-06 |

| | | | | | | | | | | |
|---|---|---|---|---|---|---|---|---|---|---|
| 2720 | 136 | N136V | A | 5.2e-06 | 1.8e-05 | 8.8e-06 | 1.067E-05 | 6.601E-06 | 5.200E-06 | 1.800E-05 |
| 2721 | 137 | P137A | A | 7.5e-06 | 1.2e-05 | 5.2e-06 | 8.233E-06 | 3.459E-06 | 5.200E-06 | 1.200E-05 |
| 2722 | 137 | P137R | A | 7e-06 | 1.3e-05 | 1e-05 | 1.000E-05 | 3.000E-06 | 7.000E-06 | 1.300E-05 |
| 2723 | 137 | P137N | A | 7.2e-06 | 1.1e-05 | 6.6e-06 | 8.267E-06 | 2.386E-06 | 6.600E-06 | 1.100E-05 |
| 2724 | 137 | P137D | A | 7.3e-06 | 9.5e-06 | 6.6e-06 | 7.800E-06 | 1.513E-06 | 6.600E-06 | 9.500E-06 |
| 2725 | 137 | P137C | A | 5e-06 | 9.1e-06 | 7.9e-06 | 7.333E-06 | 2.108E-06 | 5.000E-06 | 9.100E-06 |
| 2726 | 137 | P137E | A | 8.6e-06 | 7.7e-06 | 8.9e-06 | 8.400E-06 | 6.245E-07 | 7.700E-06 | 8.900E-06 |
| 2727 | 137 | P137Q | A | 1.4e-05 | 5.9e-06 | 8.7e-06 | 9.533E-06 | 4.114E-06 | 5.900E-06 | 1.400E-05 |
| 2728 | 137 | P137G | A | 1.3e-05 | 4.8e-06 | 6.6e-06 | 8.133E-06 | 4.310E-06 | 4.800E-06 | 1.300E-05 |
| 2729 | 137 | P137H | A | 9e-06 | 1e-05 | 6.6e-06 | 8.533E-06 | 1.747E-06 | 6.600E-06 | 1.000E-05 |
| 2730 | 137 | P137I | A | 6e-06 | 5.8e-06 | 5.1e-06 | 5.633E-06 | 4.726E-07 | 5.100E-06 | 6.000E-06 |
| 2731 | 137 | P137L | A | 8.8e-06 | 9e-06 | 8e-06 | 8.600E-06 | 5.292E-07 | 8.000E-06 | 9.000E-06 |
| 2732 | 137 | P137K | A | 5.9e-06 | 1.1e-05 | 4.5e-06 | 7.133E-06 | 3.421E-06 | 4.500E-06 | 1.100E-05 |
| 2733 | 137 | P137M | A | 6e-06 | 1.2e-05 | 5.7e-06 | 7.900E-06 | 3.554E-06 | 5.700E-06 | 1.200E-05 |
| 2734 | 137 | P137F | A | 1.3e-05 | 8.5e-06 | 6.1e-06 | 9.200E-06 | 3.503E-06 | 6.100E-06 | 1.300E-05 |
| 2735 | 137 | P137P | A | 6.8e-06 | 6.7e-06 | 6.8e-06 | 6.767E-06 | 5.774E-08 | 6.700E-06 | 6.800E-06 |
| 2736 | 137 | P137S | A | 7.3e-06 | 9.1e-06 | 7e-06 | 7.800E-06 | 1.136E-06 | 7.000E-06 | 9.100E-06 |
| 2737 | 137 | P137T | A | 1.7e-05 | 6.1e-06 | 7.2e-06 | 1.010E-05 | 6.001E-06 | 6.100E-06 | 1.700E-05 |
| 2738 | 137 | P137W | A | 9e-06 | 8.7e-06 | 6.9e-06 | 8.200E-06 | 1.136E-06 | 6.900E-06 | 9.000E-06 |
| 2739 | 137 | P137Y | A | 6.8e-06 | 1.1e-05 | 7.9e-06 | 8.567E-06 | 2.178E-06 | 6.800E-06 | 1.100E-05 |
| 2740 | 137 | P137V | A | 7.3e-06 | 7.4e-06 | 8.9e-06 | 7.867E-06 | 8.963E-07 | 7.300E-06 | 8.900E-06 |
| 2741 | 138 | Q138A | A | 1.3e-05 | 5.4e-06 | 6.7e-06 | 8.367E-06 | 4.065E-06 | 5.400E-06 | 1.300E-05 |
| 2742 | 138 | Q138R | A | 7e-06 | 5.7e-06 | 9.2e-06 | 7.300E-06 | 1.769E-06 | 5.700E-06 | 9.200E-06 |
| 2743 | 138 | Q138N | A | 9.5e-06 | 7.6e-06 | 4.5e-06 | 7.200E-06 | 2.524E-06 | 4.500E-06 | 9.500E-06 |
| 2744 | 138 | Q138D | A | 4.5e-06 | 6.4e-06 | 6.3e-06 | 5.733E-06 | 1.069E-06 | 4.500E-06 | 6.400E-06 |
| 2745 | 138 | Q138C | A | 8.9e-06 | 9.4e-06 | 5.3e-06 | 7.867E-06 | 2.237E-06 | 5.300E-06 | 9.400E-06 |
| 2746 | 138 | Q138E | A | 4.7e-06 | 1e-05 | 1.3e-05 | 9.233E-06 | 4.203E-06 | 4.700E-06 | 1.300E-05 |
| 2747 | 138 | Q138Q | A | 9.2e-06 | 1e-05 | 9.5e-06 | 9.567E-06 | 4.041E-07 | 9.200E-06 | 1.000E-05 |
| 2748 | 138 | Q138G | A | 6.2e-06 | 9.2e-06 | 6e-06 | 7.133E-06 | 1.793E-06 | 6.000E-06 | 9.200E-06 |
| 2749 | 138 | Q138H | A | 5e-06 | 1.2e-05 | 5.9e-06 | 7.633E-06 | 3.808E-06 | 5.000E-06 | 1.200E-05 |
| 2750 | 138 | Q138I | A | 1e-05 | 8.7e-06 | 1.4e-05 | 1.090E-05 | 2.762E-06 | 8.700E-06 | 1.400E-05 |
| 2751 | 138 | Q138L | A | 8.3e-06 | 6.2e-06 | 1.2e-05 | 8.833E-06 | 2.937E-06 | 6.200E-06 | 1.200E-05 |
| 2752 | 138 | Q138K | A | 5.5e-06 | 7.3e-06 | 5.3e-06 | 6.033E-06 | 1.102E-06 | 5.300E-06 | 7.300E-06 |
| 2753 | 138 | Q138M | A | 7.1e-06 | 9.2e-06 | 1.4e-05 | 1.010E-05 | 3.537E-06 | 7.100E-06 | 1.400E-05 |
| 2754 | 138 | Q138F | A | 5.2e-06 | 5.7e-06 | 1e-05 | 6.967E-06 | 2.639E-06 | 5.200E-06 | 1.000E-05 |
| 2755 | 138 | Q138P | A | 7e-06 | 9.6e-06 | 5.1e-06 | 7.233E-06 | 2.259E-06 | 5.100E-06 | 9.600E-06 |
| 2756 | 138 | Q138S | A | 7e-06 | 5.1e-06 | 7.1e-06 | 6.400E-06 | 1.127E-06 | 5.100E-06 | 7.100E-06 |
| 2757 | 138 | Q138T | A | 1.1e-05 | 7.5e-06 | 6.9e-06 | 8.467E-06 | 2.214E-06 | 6.900E-06 | 1.100E-05 |
| 2758 | 138 | Q138W | A | 1.1e-05 | 5.6e-06 | 8.2e-06 | 8.267E-06 | 2.701E-06 | 5.600E-06 | 1.100E-05 |
| 2759 | 138 | Q138Y | A | 4.6e-06 | 1e-05 | 6e-06 | 6.867E-06 | 2.802E-06 | 4.600E-06 | 1.000E-05 |
| 2760 | 138 | Q138V | A | 1.4e-05 | 6.9e-06 | 9e-06 | 9.967E-06 | 3.647E-06 | 6.900E-06 | 1.400E-05 |
| 2761 | 139 | L139A | A | 9.2e-06 | 1.2e-05 | 4.1e-06 | 8.433E-06 | 4.005E-06 | 4.100E-06 | 1.200E-05 |
| 2762 | 139 | L139R | A | 7.5e-06 | 5.5e-06 | 7.2e-06 | 6.733E-06 | 1.079E-06 | 5.500E-06 | 7.500E-06 |
| 2763 | 139 | L139N | A | 9e-06 | 5.7e-06 | 1.3e-05 | 9.233E-06 | 3.656E-06 | 5.700E-06 | 1.300E-05 |
| 2764 | 139 | L139D | A | 1e-05 | 5.5e-06 | 9.2e-06 | 8.233E-06 | 2.401E-06 | 5.500E-06 | 1.000E-05 |
| 2765 | 139 | L139C | A | 6.6e-06 | 6.2e-06 | 8.8e-06 | 7.200E-06 | 1.400E-06 | 6.200E-06 | 8.800E-06 |
| 2766 | 139 | L139E | A | 4.1e-06 | 8.7e-06 | 8.1e-06 | 6.967E-06 | 2.501E-06 | 4.100E-06 | 8.700E-06 |
| 2767 | 139 | L139Q | A | 8.6e-06 | 6.3e-06 | 5.3e-06 | 6.733E-06 | 1.692E-06 | 5.300E-06 | 8.600E-06 |
| 2768 | 139 | L139G | A | 6.3e-06 | 7.8e-06 | 7e-06 | 7.033E-06 | 7.506E-07 | 6.300E-06 | 7.800E-06 |
| 2769 | 139 | L139H | A | 7.1e-06 | 6.8e-06 | 9.8e-06 | 7.900E-06 | 1.652E-06 | 6.800E-06 | 9.800E-06 |
| 2770 | 139 | L139I | A | 9.2e-06 | 1.4e-05 | 1.1e-05 | 1.140E-05 | 2.425E-06 | 9.200E-06 | 1.400E-05 |
| 2771 | 139 | L139L | A | 9.2e-06 | 7.3e-06 | 4.1e-06 | 6.867E-06 | 2.577E-06 | 4.100E-06 | 9.200E-06 |
| 2772 | 139 | L139K | A | 5.9e-06 | 3.8e-06 | 4.2e-06 | 4.633E-06 | 1.115E-06 | 3.800E-06 | 5.900E-06 |
| 2773 | 139 | L139M | A | 7.1e-06 | 9.9e-06 | 4e-06 | 7.000E-06 | 2.951E-06 | 4.000E-06 | 9.900E-06 |
| 2774 | 139 | L139F | A | 7.9e-06 | 5.8e-06 | 5.9e-06 | 6.533E-06 | 1.185E-06 | 5.800E-06 | 7.900E-06 |
| 2775 | 139 | L139P | A | 9.7e-06 | 1.2e-05 | 1.1e-05 | 1.090E-05 | 1.153E-06 | 9.700E-06 | 1.200E-05 |
| 2776 | 139 | L139S | A | 9.6e-06 | 5.6e-06 | 7.6e-06 | 7.600E-06 | 2.000E-06 | 5.600E-06 | 9.600E-06 |
| 2777 | 139 | L139T | A | 1.3e-05 | 6.4e-06 | 3.6e-06 | 7.667E-06 | 4.826E-06 | 3.600E-06 | 1.300E-05 |
| 2778 | 139 | L139W | A | 4.3e-06 | 5.1e-06 | 6.6e-06 | 5.333E-06 | 1.168E-06 | 4.300E-06 | 6.600E-06 |
| 2779 | 139 | L139Y | A | 4.6e-06 | 6.8e-06 | 6.4e-06 | 5.933E-06 | 1.172E-06 | 4.600E-06 | 6.800E-06 |
| 2780 | 139 | L139V | A | 1.3e-05 | 7.8e-06 | 6.1e-06 | 8.967E-06 | 3.595E-06 | 6.100E-06 | 1.300E-05 |
| 2781 | 140 | C140A | A | 5e-06 | 6.5e-06 | 1.1e-05 | 7.500E-06 | 3.122E-06 | 5.000E-06 | 1.100E-05 |
| 2782 | 140 | C140R | A | 1.2e-05 | 7.4e-06 | 1e-05 | 9.800E-06 | 2.307E-06 | 7.400E-06 | 1.200E-05 |
| 2783 | 140 | C140N | A | 6.8e-06 | 1.1e-05 | 1.4e-05 | 1.060E-05 | 3.617E-06 | 6.800E-06 | 1.400E-05 |
| 2784 | 140 | C140D | A | 5.4e-06 | 6.6e-06 | 1.1e-05 | 7.667E-06 | 2.948E-06 | 5.400E-06 | 1.100E-05 |
| 2785 | 140 | C140C | A | 8.7e-06 | 8.4e-06 | 8.6e-06 | 8.567E-06 | 1.528E-07 | 8.400E-06 | 8.700E-06 |
| 2786 | 140 | C140E | A | 6.2e-06 | 1.2e-05 | 1.4e-05 | 1.073E-05 | 4.051E-06 | 6.200E-06 | 1.400E-05 |
| 2787 | 140 | C140Q | A | 4.7e-06 | 1e-05 | 3.9e-06 | 6.200E-06 | 3.315E-06 | 3.900E-06 | 1.000E-05 |
| 2788 | 140 | C140G | A | 1.3e-05 | 6.7e-06 | 1.1e-05 | 1.023E-05 | 3.219E-06 | 6.700E-06 | 1.300E-05 |
| 2789 | 140 | C140H | A | 1.1e-05 | 6.2e-06 | 1.3e-05 | 1.007E-05 | 3.495E-06 | 6.200E-06 | 1.300E-05 |
| 2790 | 140 | C140I | A | 5.6e-06 | 7.6e-06 | 8.3e-06 | 7.167E-06 | 1.401E-06 | 5.600E-06 | 8.300E-06 |
| 2791 | 140 | C140L | A | 9.1e-06 | 5.1e-06 | 8.9e-06 | 7.700E-06 | 2.254E-06 | 5.100E-06 | 9.100E-06 |
| 2792 | 140 | C140K | A | 5.3e-06 | 1.2e-05 | 1.4e-05 | 1.043E-05 | 4.557E-06 | 5.300E-06 | 1.400E-05 |
| 2793 | 140 | C140M | A | 6.4e-06 | 1.3e-05 | 5.4e-06 | 8.267E-06 | 4.130E-06 | 5.400E-06 | 1.300E-05 |
| 2794 | 140 | C140F | A | 8.5e-06 | 6.7e-06 | 5e-06 | 6.733E-06 | 1.750E-06 | 5.000E-06 | 8.500E-06 |
| 2795 | 140 | C140P | A | 6.1e-06 | 9.4e-06 | 4.1e-06 | 6.533E-06 | 2.676E-06 | 4.100E-06 | 9.400E-06 |
| 2796 | 140 | C140S | A | 9.7e-06 | 1e-05 | 6.9e-06 | 8.867E-06 | 1.710E-06 | 6.900E-06 | 1.000E-05 |
| 2797 | 140 | C140T | A | 6.3e-06 | 4e-06 | 1.5e-05 | 8.433E-06 | 5.802E-06 | 4.000E-06 | 1.500E-05 |
| 2798 | 140 | C140W | A | 6.7e-06 | 6.3e-06 | 1.1e-05 | 8.000E-06 | 2.606E-06 | 6.300E-06 | 1.100E-05 |
| 2799 | 140 | C140Y | A | 6.5e-06 | 8.7e-06 | 5.8e-06 | 7.000E-06 | 1.513E-06 | 5.800E-06 | 8.700E-06 |
| 2800 | 140 | C140V | A | 6e-06 | 8.8e-06 | 1e-05 | 8.267E-06 | 2.053E-06 | 6.000E-06 | 1.000E-05 |
| 2801 | 141 | Y141A | A | 5.9e-06 | 1.3e-05 | 7e-06 | 8.633E-06 | 3.821E-06 | 5.900E-06 | 1.300E-05 |
| 2802 | 141 | Y141R | A | 8.6e-06 | 1.2e-05 | 9.2e-06 | 9.933E-06 | 1.815E-06 | 8.600E-06 | 1.200E-05 |
| 2803 | 141 | Y141N | A | 7.1e-06 | 1.1e-05 | 5.8e-06 | 7.967E-06 | 2.706E-06 | 5.800E-06 | 1.100E-05 |
| 2804 | 141 | Y141D | A | 6.5e-06 | 7.3e-06 | 4.8e-06 | 6.200E-06 | 1.277E-06 | 4.800E-06 | 7.300E-06 |

| | | | | | | | | | | |
|---|---|---|---|---|---|---|---|---|---|---|
| 2805 | 141 | Y141C | A | 7.8e-06 | 8.2e-06 | 3.9e-06 | 6.633E-06 | 2.376E-06 | 3.900E-06 | 8.200E-06 |
| 2806 | 141 | Y141E | A | 6.7e-06 | 4.9e-06 | 7e-06 | 6.200E-06 | 1.136E-06 | 4.900E-06 | 7.000E-06 |
| 2807 | 141 | Y141Q | A | 8.3e-06 | 8.7e-06 | 6.6e-06 | 7.867E-06 | 1.115E-06 | 6.600E-06 | 8.700E-06 |
| 2808 | 141 | Y141G | A | 9.7e-06 | 9.3e-06 | 6e-06 | 8.333E-06 | 2.031E-06 | 6.000E-06 | 9.700E-06 |
| 2809 | 141 | Y141H | A | 1e-05 | 8.5e-06 | 1.3e-05 | 1.050E-05 | 2.291E-06 | 8.500E-06 | 1.300E-05 |
| 2810 | 141 | Y141I | A | 6.9e-06 | 8.7e-06 | 9.6e-06 | 8.400E-06 | 1.375E-06 | 6.900E-06 | 9.600E-06 |
| 2811 | 141 | Y141L | A | 9.6e-06 | 8.8e-06 | 6.5e-06 | 8.300E-06 | 1.609E-06 | 6.500E-06 | 9.600E-06 |
| 2812 | 141 | Y141K | A | 9.7e-06 | 4.4e-06 | 8.7e-06 | 7.600E-06 | 2.816E-06 | 4.400E-06 | 9.700E-06 |
| 2813 | 141 | Y141M | A | 1.4e-05 | 8e-06 | 5.3e-06 | 9.100E-06 | 4.453E-06 | 5.300E-06 | 1.400E-05 |
| 2814 | 141 | Y141F | A | 1e-05 | 7.4e-06 | 4.4e-06 | 7.267E-06 | 2.802E-06 | 4.400E-06 | 1.000E-05 |
| 2815 | 141 | Y141P | A | 4.9e-06 | 1.1e-05 | 1e-05 | 8.633E-06 | 3.272E-06 | 4.900E-06 | 1.100E-05 |
| 2816 | 141 | Y141S | A | 1.2e-05 | 7.2e-06 | 6.3e-06 | 8.500E-06 | 3.064E-06 | 6.300E-06 | 1.200E-05 |
| 2817 | 141 | Y141T | A | 1.1e-05 | 7.1e-06 | 8.7e-06 | 8.933E-06 | 1.960E-06 | 7.100E-06 | 1.100E-05 |
| 2818 | 141 | Y141W | A | 1.1e-05 | 3.5e-06 | 9.1e-06 | 7.867E-06 | 3.899E-06 | 3.500E-06 | 1.100E-05 |
| 2819 | 141 | Y141Y | A | 5.1e-06 | 6.9e-06 | 7.6e-06 | 6.533E-06 | 1.290E-06 | 5.100E-06 | 7.600E-06 |
| 2820 | 141 | Y141V | A | 1.7e-05 | 9e-06 | 1.3e-05 | 1.300E-05 | 4.000E-06 | 9.000E-06 | 1.700E-05 |
| 2821 | 142 | Q142A | A | 7.8e-06 | 9.1e-06 | 6.9e-06 | 7.933E-06 | 1.106E-06 | 6.900E-06 | 9.100E-06 |
| 2822 | 142 | Q142R | A | 1.3e-05 | 4.8e-06 | 6.4e-06 | 8.067E-06 | 4.347E-06 | 4.800E-06 | 1.300E-05 |
| 2823 | 142 | Q142N | A | 8.5e-06 | 7.3e-06 | 1.1e-05 | 8.933E-06 | 1.888E-06 | 7.300E-06 | 1.100E-05 |
| 2824 | 142 | Q142D | A | 7.2e-06 | 5.9e-06 | 9.1e-06 | 7.400E-06 | 1.609E-06 | 5.900E-06 | 9.100E-06 |
| 2825 | 142 | Q142C | A | 7.2e-06 | 4.9e-06 | 7.1e-06 | 6.400E-06 | 1.300E-06 | 4.900E-06 | 7.200E-06 |
| 2826 | 142 | Q142E | A | 6.5e-06 | 6.8e-06 | 1.1e-05 | 8.100E-06 | 2.516E-06 | 6.500E-06 | 1.100E-05 |
| 2827 | 142 | Q142Q | A | 4e-06 | 8.7e-06 | 8.3e-06 | 7.000E-06 | 2.606E-06 | 4.000E-06 | 8.700E-06 |
| 2828 | 142 | Q142G | A | 1.2e-05 | 9.7e-06 | 5.1e-06 | 8.933E-06 | 3.513E-06 | 5.100E-06 | 1.200E-05 |
| 2829 | 142 | Q142H | A | 7.9e-06 | 1.1e-05 | 7e-06 | 8.633E-06 | 2.098E-06 | 7.000E-06 | 1.100E-05 |
| 2830 | 142 | Q142I | A | 7.3e-06 | 1e-05 | 8e-06 | 8.433E-06 | 1.401E-06 | 7.300E-06 | 1.000E-05 |
| 2831 | 142 | Q142L | A | 4.2e-06 | 8.4e-06 | 9.7e-06 | 7.433E-06 | 2.875E-06 | 4.200E-06 | 9.700E-06 |
| 2832 | 142 | Q142K | A | 1e-05 | 6.5e-06 | 6.5e-06 | 7.667E-06 | 2.021E-06 | 6.500E-06 | 1.000E-05 |
| 2833 | 142 | Q142M | A | 8.6e-06 | 7.1e-06 | 6.5e-06 | 7.400E-06 | 1.082E-06 | 6.500E-06 | 8.600E-06 |
| 2834 | 142 | Q142F | A | 9.1e-06 | 5.9e-06 | 1.4e-05 | 9.667E-06 | 4.080E-06 | 5.900E-06 | 1.400E-05 |
| 2835 | 142 | Q142P | A | 4.8e-06 | 1.1e-05 | 5.8e-06 | 7.200E-06 | 3.329E-06 | 4.800E-06 | 1.100E-05 |
| 2836 | 142 | Q142S | A | 6.7e-06 | 1.1e-05 | 8.9e-06 | 8.867E-06 | 2.150E-06 | 6.700E-06 | 1.100E-05 |
| 2837 | 142 | Q142T | A | 7.9e-06 | 1.2e-05 | 8.9e-06 | 9.600E-06 | 2.138E-06 | 7.900E-06 | 1.200E-05 |
| 2838 | 142 | Q142W | A | 6.9e-06 | 7.8e-06 | 1.2e-05 | 8.900E-06 | 2.722E-06 | 6.900E-06 | 1.200E-05 |
| 2839 | 142 | Q142Y | A | 6.1e-06 | 6.9e-06 | 4.6e-06 | 5.867E-06 | 1.168E-06 | 4.600E-06 | 6.900E-06 |
| 2840 | 142 | Q142V | A | 1.1e-05 | 6.2e-06 | 8.6e-06 | 8.600E-06 | 2.400E-06 | 6.200E-06 | 1.100E-05 |
| 2841 | 143 | D143A | A | 5.5e-06 | 6.2e-06 | 8.7e-06 | 6.800E-06 | 1.682E-06 | 5.500E-06 | 8.700E-06 |
| 2842 | 143 | D143R | A | 1.3e-05 | 5.8e-06 | 8.1e-06 | 8.967E-06 | 3.677E-06 | 5.800E-06 | 1.300E-05 |
| 2843 | 143 | D143N | A | 6.8e-06 | 5e-06 | 7.4e-06 | 6.400E-06 | 1.249E-06 | 5.000E-06 | 7.400E-06 |
| 2844 | 143 | D143D | A | 8.2e-06 | 6.2e-06 | 6e-06 | 6.800E-06 | 1.217E-06 | 6.000E-06 | 8.200E-06 |
| 2845 | 143 | D143C | A | 7.5e-06 | 8.1e-06 | 5.9e-06 | 7.167E-06 | 1.137E-06 | 5.900E-06 | 8.100E-06 |
| 2846 | 143 | D143E | A | 1.1e-05 | 7.5e-06 | 1e-05 | 9.500E-06 | 1.803E-06 | 7.500E-06 | 1.100E-05 |
| 2847 | 143 | D143Q | A | 8.3e-06 | 5.7e-06 | 8.6e-06 | 7.533E-06 | 1.595E-06 | 5.700E-06 | 8.600E-06 |
| 2848 | 143 | D143G | A | 1e-05 | 5.5e-06 | 1.3e-05 | 9.500E-06 | 3.775E-06 | 5.500E-06 | 1.300E-05 |
| 2849 | 143 | D143H | A | 7e-06 | 1.1e-05 | 6.1e-06 | 8.033E-06 | 2.608E-06 | 6.100E-06 | 1.100E-05 |
| 2850 | 143 | D143I | A | 8.9e-06 | 7.1e-06 | 9.9e-06 | 8.633E-06 | 1.419E-06 | 7.100E-06 | 9.900E-06 |
| 2851 | 143 | D143L | A | 6.8e-06 | 7.5e-06 | 5.3e-06 | 6.533E-06 | 1.124E-06 | 5.300E-06 | 7.500E-06 |
| 2852 | 143 | D143K | A | 9.5e-06 | 9.4e-06 | 6.2e-06 | 8.367E-06 | 1.877E-06 | 6.200E-06 | 9.500E-06 |
| 2853 | 143 | D143M | A | 5.3e-06 | 6.3e-06 | 6.5e-06 | 6.033E-06 | 6.429E-07 | 5.300E-06 | 6.500E-06 |
| 2854 | 143 | D143F | A | 4.6e-06 | 1.2e-05 | 5.9e-06 | 7.500E-06 | 3.951E-06 | 4.600E-06 | 1.200E-05 |
| 2855 | 143 | D143P | A | 6.7e-06 | 1.6e-05 | 9.8e-06 | 1.083E-05 | 4.735E-06 | 6.700E-06 | 1.600E-05 |
| 2856 | 143 | D143S | A | 1.3e-05 | 6.4e-06 | 9.5e-06 | 9.633E-06 | 3.302E-06 | 6.400E-06 | 1.300E-05 |
| 2857 | 143 | D143T | A | 7.9e-06 | 8.3e-06 | 6.9e-06 | 7.700E-06 | 7.211E-07 | 6.900E-06 | 8.300E-06 |
| 2858 | 143 | D143W | A | 8.9e-06 | 4.3e-06 | 8.1e-06 | 7.100E-06 | 2.458E-06 | 4.300E-06 | 8.900E-06 |
| 2859 | 143 | D143Y | A | 8.2e-06 | 1e-05 | 9.2e-06 | 9.133E-06 | 9.018E-07 | 8.200E-06 | 1.000E-05 |
| 2860 | 143 | D143V | A | 5.1e-06 | 9.9e-06 | 1.3e-05 | 9.333E-06 | 3.980E-06 | 5.100E-06 | 1.300E-05 |
| 2861 | 144 | T144A | A | 1e-05 | 5.4e-06 | 9.1e-06 | 8.167E-06 | 2.438E-06 | 5.400E-06 | 1.000E-05 |
| 2862 | 144 | T144R | A | 1e-05 | 1.1e-05 | 7.8e-06 | 9.600E-06 | 1.637E-06 | 7.800E-06 | 1.100E-05 |
| 2863 | 144 | T144N | A | 9.4e-06 | 1.2e-05 | 1.4e-05 | 1.180E-05 | 2.307E-06 | 9.400E-06 | 1.400E-05 |
| 2864 | 144 | T144D | A | 9.2e-06 | 1.1e-05 | 5.8e-06 | 8.667E-06 | 2.641E-06 | 5.800E-06 | 1.100E-05 |
| 2865 | 144 | T144C | A | 1e-05 | 7e-06 | 8.9e-06 | 8.633E-06 | 1.518E-06 | 7.000E-06 | 1.000E-05 |
| 2866 | 144 | T144E | A | 6.6e-06 | 1.4e-05 | 6.5e-06 | 9.033E-06 | 4.302E-06 | 6.500E-06 | 1.400E-05 |
| 2867 | 144 | T144Q | A | 7.1e-06 | 4.7e-06 | 5.4e-06 | 5.733E-06 | 1.234E-06 | 4.700E-06 | 7.100E-06 |
| 2868 | 144 | T144G | A | 9.4e-06 | 1.3e-05 | 1.1e-05 | 1.113E-05 | 1.804E-06 | 9.400E-06 | 1.300E-05 |
| 2869 | 144 | T144H | A | 1.3e-05 | 7e-06 | 8.2e-06 | 9.400E-06 | 3.175E-06 | 7.000E-06 | 1.300E-05 |
| 2870 | 144 | T144I | A | 9.8e-06 | 3.2e-06 | 6.4e-06 | 6.467E-06 | 3.301E-06 | 3.200E-06 | 9.800E-06 |
| 2871 | 144 | T144L | A | 9.5e-06 | 1e-05 | 9.2e-06 | 9.567E-06 | 4.041E-07 | 9.200E-06 | 1.000E-05 |
| 2872 | 144 | T144K | A | 7.5e-06 | 8e-06 | 9.2e-06 | 8.233E-06 | 8.737E-07 | 7.500E-06 | 9.200E-06 |
| 2873 | 144 | T144M | A | 5.7e-06 | 1.2e-05 | 4.8e-06 | 7.500E-06 | 3.923E-06 | 4.800E-06 | 1.200E-05 |
| 2874 | 144 | T144F | A | 6.6e-06 | 1.3e-05 | 9.6e-06 | 9.733E-06 | 3.202E-06 | 6.600E-06 | 1.300E-05 |
| 2875 | 144 | T144P | A | 9.6e-06 | 7e-06 | 6.6e-06 | 7.733E-06 | 1.629E-06 | 6.600E-06 | 9.600E-06 |
| 2876 | 144 | T144S | A | 6.2e-06 | 5.9e-06 | 1.4e-05 | 8.700E-06 | 4.592E-06 | 5.900E-06 | 1.400E-05 |
| 2877 | 144 | T144T | A | 6.8e-06 | 1.3e-05 | 6.6e-06 | 8.800E-06 | 3.639E-06 | 6.600E-06 | 1.300E-05 |
| 2878 | 144 | T144W | A | 9.1e-06 | 5.3e-06 | 1.5e-05 | 9.800E-06 | 4.888E-06 | 5.300E-06 | 1.500E-05 |
| 2879 | 144 | T144Y | A | 1.1e-05 | 1.3e-05 | 7.3e-06 | 1.043E-05 | 2.892E-06 | 7.300E-06 | 1.300E-05 |
| 2880 | 144 | T144V | A | 6.1e-06 | 8.7e-06 | 1.7e-05 | 1.060E-05 | 5.693E-06 | 6.100E-06 | 1.700E-05 |
| 2881 | 145 | I145A | A | 5.2e-06 | 8.6e-06 | 6.6e-06 | 6.800E-06 | 1.709E-06 | 5.200E-06 | 8.600E-06 |
| 2882 | 145 | I145R | A | 1.3e-05 | 8.5e-06 | 7.9e-06 | 9.800E-06 | 2.787E-06 | 7.900E-06 | 1.300E-05 |
| 2883 | 145 | I145N | A | 7.2e-06 | 1.2e-05 | 8.4e-06 | 9.200E-06 | 2.498E-06 | 7.200E-06 | 1.200E-05 |
| 2884 | 145 | I145D | A | 5.5e-06 | 4.3e-06 | 8.4e-06 | 6.067E-06 | 2.108E-06 | 4.300E-06 | 8.400E-06 |
| 2885 | 145 | I145C | A | 9.1e-06 | 9.6e-06 | 1.1e-05 | 9.900E-06 | 9.849E-07 | 9.100E-06 | 1.100E-05 |
| 2886 | 145 | I145E | A | 6.8e-06 | 8.7e-06 | 9.4e-06 | 8.300E-06 | 1.345E-06 | 6.800E-06 | 9.400E-06 |
| 2887 | 145 | I145Q | A | 6.2e-06 | 5.9e-06 | 1.2e-05 | 8.033E-06 | 3.439E-06 | 5.900E-06 | 1.200E-05 |
| 2888 | 145 | I145G | A | 6.5e-06 | 7.5e-06 | 1.4e-05 | 9.333E-06 | 4.072E-06 | 6.500E-06 | 1.400E-05 |
| 2889 | 145 | I145H | A | 7.3e-06 | 1.3e-05 | 9.7e-06 | 1.000E-05 | 2.862E-06 | 7.300E-06 | 1.300E-05 |

| | | | | | | | | | | |
|---|---|---|---|---|---|---|---|---|---|---|
| 2890 | 145 | I145I | A | 6.3e-06 | 1e-05 | 9.1e-06 | 8.467E-06 | 1.930E-06 | 6.300E-06 | 1.000E-05 |
| 2891 | 145 | I145L | A | 6.3e-06 | 7.9e-06 | 1.1e-05 | 8.400E-06 | 2.390E-06 | 6.300E-06 | 1.100E-05 |
| 2892 | 145 | I145K | A | 5e-06 | 9e-06 | 4.2e-06 | 6.067E-06 | 2.572E-06 | 4.200E-06 | 9.000E-06 |
| 2893 | 145 | I145M | A | 1e-05 | 5.3e-06 | 7e-06 | 7.433E-06 | 2.380E-06 | 5.300E-06 | 1.000E-05 |
| 2894 | 145 | I145F | A | 6e-06 | 8.8e-06 | 9e-06 | 7.933E-06 | 1.677E-06 | 6.000E-06 | 9.000E-06 |
| 2895 | 145 | I145P | A | 9.5e-06 | 7.1e-06 | 8.2e-06 | 8.267E-06 | 1.201E-06 | 7.100E-06 | 9.500E-06 |
| 2896 | 145 | I145S | A | 8.1e-06 | 7.6e-06 | 6.6e-06 | 7.433E-06 | 7.638E-07 | 6.600E-06 | 8.100E-06 |
| 2897 | 145 | I145T | A | 6.2e-06 | 4.1e-06 | 4.6e-06 | 4.967E-06 | 1.097E-06 | 4.100E-06 | 6.200E-06 |
| 2898 | 145 | I145W | A | 1.2e-05 | 8.2e-06 | 7.3e-06 | 9.167E-06 | 2.495E-06 | 7.300E-06 | 1.200E-05 |
| 2899 | 145 | I145Y | A | 4.1e-06 | 8.7e-06 | 1.1e-05 | 7.933E-06 | 3.513E-06 | 4.100E-06 | 1.100E-05 |
| 2900 | 145 | I145V | A | 6.8e-06 | 4.6e-06 | 7.6e-06 | 6.333E-06 | 1.553E-06 | 4.600E-06 | 7.600E-06 |
| 2901 | 146 | L146A | A | 7.6e-06 | 7.9e-06 | 8.6e-06 | 8.033E-06 | 5.132E-07 | 7.600E-06 | 8.600E-06 |
| 2902 | 146 | L146R | A | 9.5e-06 | 5.4e-06 | 4.9e-06 | 6.600E-06 | 2.524E-06 | 4.900E-06 | 9.500E-06 |
| 2903 | 146 | L146N | A | 9.3e-06 | 5.1e-06 | 6.1e-06 | 6.833E-06 | 2.194E-06 | 5.100E-06 | 9.300E-06 |
| 2904 | 146 | L146D | A | 8.6e-06 | 4.8e-06 | 4.8e-06 | 6.067E-06 | 2.194E-06 | 4.800E-06 | 8.600E-06 |
| 2905 | 146 | L146C | A | 7.1e-06 | 7.6e-06 | 6e-06 | 6.900E-06 | 8.185E-07 | 6.000E-06 | 7.600E-06 |
| 2906 | 146 | L146E | A | 1.4e-05 | 6.8e-06 | 6.1e-06 | 8.967E-06 | 4.373E-06 | 6.100E-06 | 1.400E-05 |
| 2907 | 146 | L146Q | A | 1.1e-05 | 3.3e-06 | 5.9e-06 | 6.733E-06 | 3.917E-06 | 3.300E-06 | 1.100E-05 |
| 2908 | 146 | L146G | A | 1e-05 | 7.8e-06 | 1.1e-05 | 9.600E-06 | 1.637E-06 | 7.800E-06 | 1.100E-05 |
| 2909 | 146 | L146H | A | 6.4e-06 | 1.1e-05 | 4.2e-06 | 7.200E-06 | 3.470E-06 | 4.200E-06 | 1.100E-05 |
| 2910 | 146 | L146I | A | 6.8e-06 | 1.1e-05 | 6.8e-06 | 8.200E-06 | 2.425E-06 | 6.800E-06 | 1.100E-05 |
| 2911 | 146 | L146L | A | 6.5e-06 | 7.8e-06 | 5.1e-06 | 6.467E-06 | 1.350E-06 | 5.100E-06 | 7.800E-06 |
| 2912 | 146 | L146K | A | 6.6e-06 | 5.4e-06 | 4.9e-06 | 5.633E-06 | 8.737E-07 | 4.900E-06 | 6.600E-06 |
| 2913 | 146 | L146M | A | 1.1e-05 | 7e-06 | 1.1e-05 | 9.667E-06 | 2.309E-06 | 7.000E-06 | 1.100E-05 |
| 2914 | 146 | L146F | A | 6.9e-06 | 8.7e-06 | 1.4e-05 | 9.867E-06 | 3.691E-06 | 6.900E-06 | 1.400E-05 |
| 2915 | 146 | L146P | A | 8.2e-06 | 6.8e-06 | 1.2e-05 | 9.000E-06 | 2.691E-06 | 6.800E-06 | 1.200E-05 |
| 2916 | 146 | L146S | A | 8e-06 | 5.4e-06 | 7.2e-06 | 6.867E-06 | 1.332E-06 | 5.400E-06 | 8.000E-06 |
| 2917 | 146 | L146T | A | 6.4e-06 | 5.4e-06 | 6.4e-06 | 6.067E-06 | 5.774E-07 | 5.400E-06 | 6.400E-06 |
| 2918 | 146 | L146W | A | 7.8e-06 | 1.1e-05 | 5.5e-06 | 8.100E-06 | 2.762E-06 | 5.500E-06 | 1.100E-05 |
| 2919 | 146 | L146Y | A | 9.2e-06 | 4.2e-06 | 8.6e-06 | 7.333E-06 | 2.730E-06 | 4.200E-06 | 9.200E-06 |
| 2920 | 146 | L146V | A | 1.4e-05 | 3.9e-06 | 7.3e-06 | 8.400E-06 | 5.139E-06 | 3.900E-06 | 1.400E-05 |
| 2921 | 147 | W147A | A | 7.5e-06 | 9.7e-06 | 9.9e-06 | 9.033E-06 | 1.332E-06 | 7.500E-06 | 9.900E-06 |
| 2922 | 147 | W147R | A | 9.8e-06 | 7.9e-06 | 1.1e-05 | 9.567E-06 | 1.563E-06 | 7.900E-06 | 1.100E-05 |
| 2923 | 147 | W147N | A | 6.7e-06 | 6.1e-06 | 5.6e-06 | 6.133E-06 | 5.508E-07 | 5.600E-06 | 6.700E-06 |
| 2924 | 147 | W147D | A | 8.6e-06 | 7.1e-06 | 7.4e-06 | 7.700E-06 | 7.937E-07 | 7.100E-06 | 8.600E-06 |
| 2925 | 147 | W147C | A | 6.1e-06 | 9.5e-06 | 7.3e-06 | 7.633E-06 | 1.724E-06 | 6.100E-06 | 9.500E-06 |
| 2926 | 147 | W147E | A | 6.5e-06 | 5.9e-06 | 6.4e-06 | 6.267E-06 | 3.215E-07 | 5.900E-06 | 6.500E-06 |
| 2927 | 147 | W147Q | A | 1.5e-05 | 6.7e-06 | 1e-05 | 1.057E-05 | 4.179E-06 | 6.700E-06 | 1.500E-05 |
| 2928 | 147 | W147G | A | 1.3e-05 | 6.6e-06 | 1.2e-05 | 1.053E-05 | 3.443E-06 | 6.600E-06 | 1.300E-05 |
| 2929 | 147 | W147H | A | 6.8e-06 | 5.7e-06 | 7.3e-06 | 6.600E-06 | 8.185E-07 | 5.700E-06 | 7.300E-06 |
| 2930 | 147 | W147I | A | 1e-05 | 9.9e-06 | 1e-05 | 9.967E-06 | 5.774E-08 | 9.900E-06 | 1.000E-05 |
| 2931 | 147 | W147L | A | 7.1e-06 | 1.1e-05 | 1.5e-05 | 1.103E-05 | 3.950E-06 | 7.100E-06 | 1.500E-05 |
| 2932 | 147 | W147K | A | 7.7e-06 | 9.4e-06 | 5.2e-06 | 7.433E-06 | 2.113E-06 | 5.200E-06 | 9.400E-06 |
| 2933 | 147 | W147M | A | 1.1e-05 | 8.8e-06 | 5.9e-06 | 8.567E-06 | 2.558E-06 | 5.900E-06 | 1.100E-05 |
| 2934 | 147 | W147F | A | 1.1e-05 | 1.1e-05 | 1e-05 | 1.067E-05 | 5.774E-07 | 1.000E-05 | 1.100E-05 |
| 2935 | 147 | W147P | A | 1.4e-05 | 7.8e-06 | 1.1e-05 | 1.093E-05 | 3.101E-06 | 7.800E-06 | 1.400E-05 |
| 2936 | 147 | W147S | A | 1.3e-05 | 2e-05 | 9.6e-06 | 1.420E-05 | 5.303E-06 | 9.600E-06 | 2.000E-05 |
| 2937 | 147 | W147T | A | 8.8e-06 | 7.4e-06 | 7.1e-06 | 7.767E-06 | 9.074E-07 | 7.100E-06 | 8.800E-06 |
| 2938 | 147 | W147W | A | 6.2e-06 | 1.3e-05 | 8.5e-06 | 9.233E-06 | 3.459E-06 | 6.200E-06 | 1.300E-05 |
| 2939 | 147 | W147Y | A | 7e-06 | 8.2e-06 | 1e-05 | 8.400E-06 | 1.510E-06 | 7.000E-06 | 1.000E-05 |
| 2940 | 147 | W147V | A | 5.3e-06 | 6.7e-06 | 6e-06 | 6.000E-06 | 7.000E-07 | 5.300E-06 | 6.700E-06 |
| 2941 | 148 | K148A | A | 1.2e-05 | 6.8e-06 | 8.1e-06 | 8.967E-06 | 2.706E-06 | 6.800E-06 | 1.200E-05 |
| 2942 | 148 | K148R | A | 5e-06 | 9.4e-06 | 8.6e-06 | 7.667E-06 | 2.344E-06 | 5.000E-06 | 9.400E-06 |
| 2943 | 148 | K148N | A | 1.2e-05 | 8.4e-06 | 7.9e-06 | 9.433E-06 | 2.237E-06 | 7.900E-06 | 1.200E-05 |
| 2944 | 148 | K148D | A | 7.8e-06 | 1.2e-05 | 8e-06 | 9.267E-06 | 2.369E-06 | 7.800E-06 | 1.200E-05 |
| 2945 | 148 | K148C | A | 8.8e-06 | 1.3e-05 | 9.4e-06 | 1.040E-05 | 2.272E-06 | 8.800E-06 | 1.300E-05 |
| 2946 | 148 | K148E | A | 9.3e-06 | 5.4e-06 | 1e-05 | 8.233E-06 | 2.479E-06 | 5.400E-06 | 1.000E-05 |
| 2947 | 148 | K148Q | A | 1.1e-05 | 8.2e-06 | 9.3e-06 | 9.500E-06 | 1.411E-06 | 8.200E-06 | 1.100E-05 |
| 2948 | 148 | K148G | A | 5.4e-06 | 5.6e-06 | 6.8e-06 | 5.933E-06 | 7.572E-07 | 5.400E-06 | 6.800E-06 |
| 2949 | 148 | K148H | A | 8.5e-06 | 9.3e-06 | 4.3e-06 | 7.367E-06 | 2.686E-06 | 4.300E-06 | 9.300E-06 |
| 2950 | 148 | K148I | A | 5.5e-06 | 1.3e-05 | 9.3e-06 | 9.267E-06 | 3.750E-06 | 5.500E-06 | 1.300E-05 |
| 2951 | 148 | K148L | A | 1e-05 | 9.4e-06 | 6.4e-06 | 8.600E-06 | 1.929E-06 | 6.400E-06 | 1.000E-05 |
| 2952 | 148 | K148K | A | 4.7e-06 | 1.5e-05 | 1.4e-05 | 1.123E-05 | 5.680E-06 | 4.700E-06 | 1.500E-05 |
| 2953 | 148 | K148M | A | 8.4e-06 | 6.8e-06 | 9e-06 | 8.067E-06 | 1.137E-06 | 6.800E-06 | 9.000E-06 |
| 2954 | 148 | K148F | A | 9.7e-06 | 7e-06 | 6.8e-06 | 7.833E-06 | 1.620E-06 | 6.800E-06 | 9.700E-06 |
| 2955 | 148 | K148P | A | 7.6e-06 | 1.3e-05 | 1.4e-05 | 1.153E-05 | 3.443E-06 | 7.600E-06 | 1.400E-05 |
| 2956 | 148 | K148S | A | 9.8e-06 | 1.2e-05 | 9.6e-06 | 1.047E-05 | 1.332E-06 | 9.600E-06 | 1.200E-05 |
| 2957 | 148 | K148T | A | 4.5e-06 | 1.1e-05 | 6.8e-06 | 7.433E-06 | 3.296E-06 | 4.500E-06 | 1.100E-05 |
| 2958 | 148 | K148W | A | 8.6e-06 | 5e-06 | 9.1e-06 | 7.567E-06 | 2.237E-06 | 5.000E-06 | 9.100E-06 |
| 2959 | 148 | K148Y | A | 1.1e-05 | 1.1e-05 | 6.5e-06 | 9.500E-06 | 2.598E-06 | 6.500E-06 | 1.100E-05 |
| 2960 | 148 | K148V | A | 6.7e-06 | 8.7e-06 | 6.9e-06 | 7.433E-06 | 1.102E-06 | 6.700E-06 | 8.700E-06 |
| 2961 | 149 | D149A | A | 8.8e-06 | 4.5e-06 | 1.4e-05 | 9.100E-06 | 4.757E-06 | 4.500E-06 | 1.400E-05 |
| 2962 | 149 | D149R | A | 7.5e-06 | 7.5e-06 | 1.4e-05 | 9.667E-06 | 3.753E-06 | 7.500E-06 | 1.400E-05 |
| 2963 | 149 | D149N | A | 1.2e-05 | 6.6e-06 | 1.4e-05 | 1.087E-05 | 3.828E-06 | 6.600E-06 | 1.400E-05 |
| 2964 | 149 | D149D | A | 6.3e-06 | 5.6e-06 | 9.5e-06 | 7.133E-06 | 2.079E-06 | 5.600E-06 | 9.500E-06 |
| 2965 | 149 | D149C | A | 5.6e-06 | 1e-05 | 9.5e-06 | 8.367E-06 | 2.409E-06 | 5.600E-06 | 1.000E-05 |
| 2966 | 149 | D149E | A | 5.4e-06 | 9.6e-06 | 6.3e-06 | 7.100E-06 | 2.211E-06 | 5.400E-06 | 9.600E-06 |
| 2967 | 149 | D149Q | A | 9.9e-06 | 5.2e-06 | 8.2e-06 | 7.767E-06 | 2.380E-06 | 5.200E-06 | 9.900E-06 |
| 2968 | 149 | D149G | A | 7.2e-06 | 7.3e-06 | 1.1e-05 | 8.500E-06 | 2.166E-06 | 7.200E-06 | 1.100E-05 |
| 2969 | 149 | D149H | A | 6.8e-06 | 3.4e-06 | 1e-05 | 6.733E-06 | 3.301E-06 | 3.400E-06 | 1.000E-05 |
| 2970 | 149 | D149I | A | 4.7e-06 | 6e-06 | 8.6e-06 | 6.433E-06 | 1.986E-06 | 4.700E-06 | 8.600E-06 |
| 2971 | 149 | D149L | A | 5.5e-06 | 8.2e-06 | 9.4e-06 | 7.700E-06 | 1.997E-06 | 5.500E-06 | 9.400E-06 |
| 2972 | 149 | D149K | A | 7.6e-06 | 3.9e-06 | 1.1e-05 | 7.500E-06 | 3.551E-06 | 3.900E-06 | 1.100E-05 |
| 2973 | 149 | D149M | A | 1.3e-05 | 4.5e-06 | 9.8e-06 | 9.100E-06 | 4.293E-06 | 4.500E-06 | 1.300E-05 |
| 2974 | 149 | D149F | A | 7.4e-06 | 5.2e-06 | 6.5e-06 | 6.367E-06 | 1.106E-06 | 5.200E-06 | 7.400E-06 |

| | | | | | | | | | | |
|---|---|---|---|---|---|---|---|---|---|---|
| 2975 | 149 | D149P | A | 1.2e-05 | 5.8e-06 | 6.5e-06 | 8.100E-06 | 3.396E-06 | 5.800E-06 | 1.200E-05 |
| 2976 | 149 | D149S | A | 9.1e-06 | 4.5e-06 | 9.8e-06 | 7.800E-06 | 2.879E-06 | 4.500E-06 | 9.800E-06 |
| 2977 | 149 | D149T | A | 6.8e-06 | 6.1e-06 | 6.7e-06 | 6.533E-06 | 3.786E-07 | 6.100E-06 | 6.800E-06 |
| 2978 | 149 | D149W | A | 7.9e-06 | 6.9e-06 | 9.7e-06 | 8.167E-06 | 1.419E-06 | 6.900E-06 | 9.700E-06 |
| 2979 | 149 | D149Y | A | 5.8e-06 | 1.2e-05 | 5.9e-06 | 7.900E-06 | 3.551E-06 | 5.800E-06 | 1.200E-05 |
| 2980 | 149 | D149V | A | 1.1e-05 | 1.2e-05 | 7.9e-06 | 1.030E-05 | 2.138E-06 | 7.900E-06 | 1.200E-05 |
| 2981 | 150 | I150A | A | 5.9e-06 | 7.1e-06 | 1.2e-05 | 8.333E-06 | 3.232E-06 | 5.900E-06 | 1.200E-05 |
| 2982 | 150 | I150R | A | 6.5e-06 | 9.8e-06 | 7.4e-06 | 7.900E-06 | 1.706E-06 | 6.500E-06 | 9.800E-06 |
| 2983 | 150 | I150N | A | 8.2e-06 | 9.5e-06 | 6.3e-06 | 8.000E-06 | 1.609E-06 | 6.300E-06 | 9.500E-06 |
| 2984 | 150 | I150D | A | 1e-05 | 1.1e-05 | 1.1e-05 | 1.067E-05 | 5.774E-07 | 1.000E-05 | 1.100E-05 |
| 2985 | 150 | I150C | A | 8.7e-06 | 1e-05 | 4.5e-06 | 7.733E-06 | 2.875E-06 | 4.500E-06 | 1.000E-05 |
| 2986 | 150 | I150E | A | 4.4e-06 | 6.4e-06 | 6.3e-06 | 5.700E-06 | 1.127E-06 | 4.400E-06 | 6.400E-06 |
| 2987 | 150 | I150Q | A | 9.3e-06 | 7e-06 | 7.6e-06 | 7.967E-06 | 1.193E-06 | 7.000E-06 | 9.300E-06 |
| 2988 | 150 | I150G | A | 4.9e-06 | 5.8e-06 | 1.2e-05 | 7.567E-06 | 3.866E-06 | 4.900E-06 | 1.200E-05 |
| 2989 | 150 | I150H | A | 1.3e-05 | 5.2e-06 | 8.5e-06 | 8.900E-06 | 3.915E-06 | 5.200E-06 | 1.300E-05 |
| 2990 | 150 | I150I | A | 6.4e-06 | 7.9e-06 | 9.6e-06 | 7.967E-06 | 1.601E-06 | 6.400E-06 | 9.600E-06 |
| 2991 | 150 | I150L | A | 1.1e-05 | 6.2e-06 | 7.4e-06 | 8.200E-06 | 2.498E-06 | 6.200E-06 | 1.100E-05 |
| 2992 | 150 | I150K | A | 5.7e-06 | 1.2e-05 | 8e-06 | 8.567E-06 | 3.188E-06 | 5.700E-06 | 1.200E-05 |
| 2993 | 150 | I150M | A | 1.1e-05 | 7.1e-06 | 9.9e-06 | 9.333E-06 | 2.011E-06 | 7.100E-06 | 1.100E-05 |
| 2994 | 150 | I150F | A | 6.7e-06 | 9.1e-06 | 1e-05 | 8.600E-06 | 1.706E-06 | 6.700E-06 | 1.000E-05 |
| 2995 | 150 | I150P | A | 1.1e-05 | 8.4e-06 | 6.3e-06 | 8.567E-06 | 2.354E-06 | 6.300E-06 | 1.100E-05 |
| 2996 | 150 | I150S | A | 6.6e-06 | 6.4e-06 | 7.6e-06 | 6.867E-06 | 6.429E-07 | 6.400E-06 | 7.600E-06 |
| 2997 | 150 | I150T | A | 8.4e-06 | 7.9e-06 | 5.8e-06 | 7.367E-06 | 1.380E-06 | 5.800E-06 | 8.400E-06 |
| 2998 | 150 | I150W | A | 9.9e-06 | 6.6e-06 | 9.8e-06 | 8.767E-06 | 1.877E-06 | 6.600E-06 | 9.900E-06 |
| 2999 | 150 | I150Y | A | 6.5e-06 | 9.4e-06 | 7.2e-06 | 7.700E-06 | 1.513E-06 | 6.500E-06 | 9.400E-06 |
| 3000 | 150 | I150V | A | 1.3e-05 | 6.5e-06 | 9e-06 | 9.500E-06 | 3.279E-06 | 6.500E-06 | 1.300E-05 |
| 3001 | 151 | F151A | A | 4.5e-06 | 6.5e-06 | 9.8e-06 | 6.933E-06 | 2.676E-06 | 4.500E-06 | 9.800E-06 |
| 3002 | 151 | F151R | A | 9.6e-06 | 6.7e-06 | 7.2e-06 | 7.833E-06 | 1.550E-06 | 6.700E-06 | 9.600E-06 |
| 3003 | 151 | F151N | A | 8.5e-06 | 7.7e-06 | 1.4e-05 | 1.007E-05 | 3.430E-06 | 7.700E-06 | 1.400E-05 |
| 3004 | 151 | F151D | A | 1.3e-05 | 8.6e-06 | 5.5e-06 | 9.033E-06 | 3.769E-06 | 5.500E-06 | 1.300E-05 |
| 3005 | 151 | F151C | A | 8.6e-06 | 5.2e-06 | 6.8e-06 | 6.867E-06 | 1.701E-06 | 5.200E-06 | 8.600E-06 |
| 3006 | 151 | F151E | A | 1.1e-05 | 6.9e-06 | 8.4e-06 | 8.767E-06 | 2.074E-06 | 6.900E-06 | 1.100E-05 |
| 3007 | 151 | F151Q | A | 6.4e-06 | 8.3e-06 | 1.1e-05 | 8.567E-06 | 2.312E-06 | 6.400E-06 | 1.100E-05 |
| 3008 | 151 | F151G | A | 9.2e-06 | 8.4e-06 | 6.6e-06 | 8.067E-06 | 1.332E-06 | 6.600E-06 | 9.200E-06 |
| 3009 | 151 | F151H | A | 6.1e-06 | 8e-06 | 6e-06 | 6.700E-06 | 1.127E-06 | 6.000E-06 | 8.000E-06 |
| 3010 | 151 | F151I | A | 9.1e-06 | 1.4e-05 | 9.3e-06 | 1.080E-05 | 2.773E-06 | 9.100E-06 | 1.400E-05 |
| 3011 | 151 | F151L | A | 1e-05 | 1.6e-05 | 9.2e-06 | 1.173E-05 | 3.717E-06 | 9.200E-06 | 1.600E-05 |
| 3012 | 151 | F151K | A | 5.3e-06 | 5.1e-06 | 7.9e-06 | 6.100E-06 | 1.562E-06 | 5.100E-06 | 7.900E-06 |
| 3013 | 151 | F151M | A | 1.1e-05 | 7e-06 | 4.8e-06 | 7.600E-06 | 3.143E-06 | 4.800E-06 | 1.100E-05 |
| 3014 | 151 | F151F | A | 9.1e-06 | 6.5e-06 | 9.5e-06 | 8.367E-06 | 1.629E-06 | 6.500E-06 | 9.500E-06 |
| 3015 | 151 | F151P | A | 7.5e-06 | 5.4e-06 | 9.7e-06 | 7.533E-06 | 2.150E-06 | 5.400E-06 | 9.700E-06 |
| 3016 | 151 | F151S | A | 9e-06 | 3.6e-06 | 1.1e-05 | 7.867E-06 | 3.828E-06 | 3.600E-06 | 1.100E-05 |
| 3017 | 151 | F151T | A | 9.9e-06 | 9.1e-06 | 8e-06 | 9.000E-06 | 9.539E-07 | 8.000E-06 | 9.900E-06 |
| 3018 | 151 | F151W | A | 5.8e-06 | 9.5e-06 | 4.8e-06 | 6.700E-06 | 2.476E-06 | 4.800E-06 | 9.500E-06 |
| 3019 | 151 | F151Y | A | 9.9e-06 | 5e-06 | 6.3e-06 | 7.067E-06 | 2.538E-06 | 5.000E-06 | 9.900E-06 |
| 3020 | 151 | F151V | A | 6.5e-06 | 1.2e-05 | 6.9e-06 | 8.467E-06 | 3.066E-06 | 6.500E-06 | 1.200E-05 |
| 3021 | 152 | H152A | A | 1e-05 | 9.1e-06 | 5.9e-06 | 8.333E-06 | 2.155E-06 | 5.900E-06 | 1.000E-05 |
| 3022 | 152 | H152R | A | 1.1e-05 | 7.5e-06 | 9.4e-06 | 9.300E-06 | 1.752E-06 | 7.500E-06 | 1.100E-05 |
| 3023 | 152 | H152N | A | 8.3e-06 | 7.8e-06 | 1.1e-05 | 9.033E-06 | 1.721E-06 | 7.800E-06 | 1.100E-05 |
| 3024 | 152 | H152D | A | 9.2e-06 | 6.4e-06 | 5.1e-06 | 6.900E-06 | 2.095E-06 | 5.100E-06 | 9.200E-06 |
| 3025 | 152 | H152C | A | 7.8e-06 | 7.1e-06 | 6.2e-06 | 7.033E-06 | 8.021E-07 | 6.200E-06 | 7.800E-06 |
| 3026 | 152 | H152E | A | 4.2e-06 | 9.9e-06 | 7.3e-06 | 7.133E-06 | 2.854E-06 | 4.200E-06 | 9.900E-06 |
| 3027 | 152 | H152Q | A | 5.1e-06 | 6.4e-06 | 1.2e-05 | 7.833E-06 | 3.667E-06 | 5.100E-06 | 1.200E-05 |
| 3028 | 152 | H152G | A | 8.1e-06 | 7e-06 | 7.9e-06 | 7.667E-06 | 5.859E-07 | 7.000E-06 | 8.100E-06 |
| 3029 | 152 | H152H | A | 5.1e-06 | 1.1e-05 | 7.5e-06 | 7.867E-06 | 2.967E-06 | 5.100E-06 | 1.100E-05 |
| 3030 | 152 | H152I | A | 5.5e-06 | 8.3e-06 | 7.2e-06 | 7.000E-06 | 1.411E-06 | 5.500E-06 | 8.300E-06 |
| 3031 | 152 | H152L | A | 1.5e-05 | 5.1e-06 | 9.1e-06 | 9.733E-06 | 4.980E-06 | 5.100E-06 | 1.500E-05 |
| 3032 | 152 | H152K | A | 6.8e-06 | 7.3e-06 | 7.1e-06 | 7.067E-06 | 2.517E-07 | 6.800E-06 | 7.300E-06 |
| 3033 | 152 | H152M | A | 5.9e-06 | 7.2e-06 | 5.9e-06 | 6.333E-06 | 7.506E-07 | 5.900E-06 | 7.200E-06 |
| 3034 | 152 | H152F | A | 9.8e-06 | 9e-06 | 7.2e-06 | 8.667E-06 | 1.332E-06 | 7.200E-06 | 9.800E-06 |
| 3035 | 152 | H152P | A | 9.6e-06 | 6.3e-06 | 8.2e-06 | 8.033E-06 | 1.656E-06 | 6.300E-06 | 9.600E-06 |
| 3036 | 152 | H152S | A | 5.2e-06 | 8.9e-06 | 1e-05 | 8.033E-06 | 2.515E-06 | 5.200E-06 | 1.000E-05 |
| 3037 | 152 | H152T | A | 4.4e-06 | 6.8e-06 | 6e-06 | 5.733E-06 | 1.222E-06 | 4.400E-06 | 6.800E-06 |
| 3038 | 152 | H152W | A | 9.6e-06 | 9.4e-06 | 3.1e-06 | 7.367E-06 | 3.696E-06 | 3.100E-06 | 9.600E-06 |
| 3039 | 152 | H152Y | A | 9.8e-06 | 6.6e-06 | 9.7e-06 | 8.700E-06 | 1.819E-06 | 6.600E-06 | 9.800E-06 |
| 3040 | 152 | H152V | A | 8.7e-06 | 7.9e-06 | 7e-06 | 7.867E-06 | 8.505E-07 | 7.000E-06 | 8.700E-06 |
| 3041 | 153 | K153A | A | 7e-06 | 1.3e-05 | 8.2e-06 | 9.400E-06 | 3.175E-06 | 7.000E-06 | 1.300E-05 |
| 3042 | 153 | K153R | A | 5.2e-06 | 4.5e-06 | 5.6e-06 | 5.100E-06 | 5.568E-07 | 4.500E-06 | 5.600E-06 |
| 3043 | 153 | K153N | A | 5.4e-06 | 1.2e-05 | 8.3e-06 | 8.567E-06 | 3.308E-06 | 5.400E-06 | 1.200E-05 |
| 3044 | 153 | K153D | A | 7e-06 | 8.9e-06 | 8.9e-06 | 8.267E-06 | 1.097E-06 | 7.000E-06 | 8.900E-06 |
| 3045 | 153 | K153C | A | 6.1e-06 | 4.2e-06 | 8.6e-06 | 6.300E-06 | 2.207E-06 | 4.200E-06 | 8.600E-06 |
| 3046 | 153 | K153E | A | 7.9e-06 | 6.4e-06 | 6.7e-06 | 7.000E-06 | 7.937E-07 | 6.400E-06 | 7.900E-06 |
| 3047 | 153 | K153Q | A | 6.9e-06 | 5.7e-06 | 5.4e-06 | 6.000E-06 | 7.937E-07 | 5.400E-06 | 6.900E-06 |
| 3048 | 153 | K153G | A | 1.4e-05 | 3.1e-06 | 1.1e-05 | 9.367E-06 | 5.631E-06 | 3.100E-06 | 1.400E-05 |
| 3049 | 153 | K153H | A | 9.4e-06 | 1.5e-05 | 5.5e-06 | 9.967E-06 | 4.775E-06 | 5.500E-06 | 1.500E-05 |
| 3050 | 153 | K153I | A | 5.2e-06 | 9.3e-06 | 7.1e-06 | 7.200E-06 | 2.052E-06 | 5.200E-06 | 9.300E-06 |
| 3051 | 153 | K153L | A | 1e-05 | 1.1e-05 | 9.5e-06 | 1.017E-05 | 7.638E-07 | 9.500E-06 | 1.100E-05 |
| 3052 | 153 | K153K | A | 7.6e-06 | 5.1e-06 | 7.2e-06 | 6.633E-06 | 1.343E-06 | 5.100E-06 | 7.600E-06 |
| 3053 | 153 | K153M | A | 5.9e-06 | 1e-05 | 5.8e-06 | 7.233E-06 | 2.397E-06 | 5.800E-06 | 1.000E-05 |
| 3054 | 153 | K153F | A | 5.7e-06 | 8.9e-06 | 9e-06 | 7.867E-06 | 1.877E-06 | 5.700E-06 | 9.000E-06 |
| 3055 | 153 | K153P | A | 5.3e-06 | 4e-06 | 8.9e-06 | 6.067E-06 | 2.538E-06 | 4.000E-06 | 8.900E-06 |
| 3056 | 153 | K153S | A | 6.6e-06 | 5.7e-06 | 9.8e-06 | 7.367E-06 | 2.155E-06 | 5.700E-06 | 9.800E-06 |
| 3057 | 153 | K153T | A | 1.2e-05 | 6e-06 | 8.5e-06 | 8.833E-06 | 3.014E-06 | 6.000E-06 | 1.200E-05 |
| 3058 | 153 | K153W | A | 1.4e-05 | 8.9e-06 | 4.4e-06 | 9.100E-06 | 4.803E-06 | 4.400E-06 | 1.400E-05 |
| 3059 | 153 | K153Y | A | 6.8e-06 | 1e-05 | 4.9e-06 | 7.233E-06 | 2.577E-06 | 4.900E-06 | 1.000E-05 |

| | | | | | | | | | | |
|---|---|---|---|---|---|---|---|---|---|---|
| 3060 | 153 | K153V | A | 6.8e-06 | 5.6e-06 | 8.4e-06 | 6.933E-06 | 1.405E-06 | 5.600E-06 | 8.400E-06 |
| 3061 | 154 | N154A | A | 1.4e-05 | 7.1e-06 | 1.8e-05 | 1.303E-05 | 5.514E-06 | 7.100E-06 | 1.800E-05 |
| 3062 | 154 | N154R | A | 1e-05 | 9.3e-06 | 9.1e-06 | 9.467E-06 | 4.726E-07 | 9.100E-06 | 1.000E-05 |
| 3063 | 154 | N154N | A | 7.1e-06 | 6.3e-06 | 1.3e-05 | 8.800E-06 | 3.659E-06 | 6.300E-06 | 1.300E-05 |
| 3064 | 154 | N154D | A | 7e-06 | 8e-06 | 5.4e-06 | 6.800E-06 | 1.311E-06 | 5.400E-06 | 8.000E-06 |
| 3065 | 154 | N154C | A | 6.9e-06 | 7.5e-06 | 5.8e-06 | 6.733E-06 | 8.622E-07 | 5.800E-06 | 7.500E-06 |
| 3066 | 154 | N154E | A | 7.9e-06 | 4.8e-06 | 1.3e-05 | 8.567E-06 | 4.140E-06 | 4.800E-06 | 1.300E-05 |
| 3067 | 154 | N154Q | A | 9.4e-06 | 9.6e-06 | 6.3e-06 | 8.433E-06 | 1.850E-06 | 6.300E-06 | 9.600E-06 |
| 3068 | 154 | N154G | A | 1.5e-05 | 5.7e-06 | 1.1e-05 | 1.057E-05 | 4.665E-06 | 5.700E-06 | 1.500E-05 |
| 3069 | 154 | N154H | A | 9.5e-06 | 1e-05 | 1.4e-05 | 1.117E-05 | 2.466E-06 | 9.500E-06 | 1.400E-05 |
| 3070 | 154 | N154I | A | 7.7e-06 | 1e-05 | 1.1e-05 | 9.567E-06 | 1.692E-06 | 7.700E-06 | 1.100E-05 |
| 3071 | 154 | N154L | A | 1.1e-05 | 6.7e-06 | 4.6e-06 | 7.433E-06 | 3.262E-06 | 4.600E-06 | 1.100E-05 |
| 3072 | 154 | N154K | A | 5.3e-06 | 1.4e-05 | 4e-06 | 7.767E-06 | 5.437E-06 | 4.000E-06 | 1.400E-05 |
| 3073 | 154 | N154M | A | 1e-05 | 4.8e-06 | 1.1e-05 | 8.600E-06 | 3.329E-06 | 4.800E-06 | 1.100E-05 |
| 3074 | 154 | N154F | A | 5.3e-06 | 8.1e-06 | 3.5e-06 | 5.633E-06 | 2.318E-06 | 3.500E-06 | 8.100E-06 |
| 3075 | 154 | N154P | A | 7.1e-06 | 1.4e-05 | 6.5e-06 | 9.200E-06 | 4.168E-06 | 6.500E-06 | 1.400E-05 |
| 3076 | 154 | N154S | A | 1.6e-05 | 1e-05 | 6.7e-06 | 1.090E-05 | 4.715E-06 | 6.700E-06 | 1.600E-05 |
| 3077 | 154 | N154T | A | 8.6e-06 | 5.6e-06 | 8.4e-06 | 7.533E-06 | 1.677E-06 | 5.600E-06 | 8.600E-06 |
| 3078 | 154 | N154W | A | 7.5e-06 | 1.1e-05 | 5.4e-06 | 7.967E-06 | 2.829E-06 | 5.400E-06 | 1.100E-05 |
| 3079 | 154 | N154Y | A | 6.4e-06 | 6e-06 | 6.5e-06 | 6.300E-06 | 2.646E-07 | 6.000E-06 | 6.500E-06 |
| 3080 | 154 | N154V | A | 8.4e-06 | 5.1e-06 | 7.8e-06 | 7.100E-06 | 1.758E-06 | 5.100E-06 | 8.400E-06 |
| 3081 | 155 | N155A | A | 6.4e-06 | 9.5e-06 | 1.4e-05 | 9.967E-06 | 3.821E-06 | 6.400E-06 | 1.400E-05 |
| 3082 | 155 | N155R | A | 1.3e-05 | 6.7e-06 | 9.2e-06 | 9.633E-06 | 3.172E-06 | 6.700E-06 | 1.300E-05 |
| 3083 | 155 | N155N | A | 1.3e-05 | 1e-05 | 5e-06 | 9.333E-06 | 4.041E-06 | 5.000E-06 | 1.300E-05 |
| 3084 | 155 | N155D | A | 7.1e-06 | 9.7e-06 | 6.4e-06 | 7.733E-06 | 1.739E-06 | 6.400E-06 | 9.700E-06 |
| 3085 | 155 | N155C | A | 1.1e-05 | 1.2e-05 | 4.5e-06 | 9.167E-06 | 4.072E-06 | 4.500E-06 | 1.200E-05 |
| 3086 | 155 | N155E | A | 8.6e-06 | 9.5e-06 | 9.3e-06 | 9.133E-06 | 4.726E-07 | 8.600E-06 | 9.500E-06 |
| 3087 | 155 | N155Q | A | 6.5e-06 | 9.6e-06 | 8.8e-06 | 8.300E-06 | 1.609E-06 | 6.500E-06 | 9.600E-06 |
| 3088 | 155 | N155G | A | 6.5e-06 | 8.9e-06 | 6.4e-06 | 7.267E-06 | 1.415E-06 | 6.400E-06 | 8.900E-06 |
| 3089 | 155 | N155H | A | 1.3e-05 | 6.8e-06 | 8.5e-06 | 9.433E-06 | 3.204E-06 | 6.800E-06 | 1.300E-05 |
| 3090 | 155 | N155I | A | 1.4e-05 | 1e-05 | 4.3e-06 | 9.433E-06 | 4.875E-06 | 4.300E-06 | 1.400E-05 |
| 3091 | 155 | N155L | A | 5.3e-06 | 1.3e-05 | 6.4e-06 | 8.233E-06 | 4.165E-06 | 5.300E-06 | 1.300E-05 |
| 3092 | 155 | N155K | A | 5.7e-06 | 8.1e-06 | 5.5e-06 | 6.433E-06 | 1.447E-06 | 5.500E-06 | 8.100E-06 |
| 3093 | 155 | N155M | A | 6.8e-06 | 8.3e-06 | 7e-06 | 7.367E-06 | 8.145E-07 | 6.800E-06 | 8.300E-06 |
| 3094 | 155 | N155F | A | 6.2e-06 | 6.2e-06 | 5.9e-06 | 6.100E-06 | 1.732E-07 | 5.900E-06 | 6.200E-06 |
| 3095 | 155 | N155P | A | 7.5e-06 | 6.1e-06 | 9e-06 | 7.533E-06 | 1.450E-06 | 6.100E-06 | 9.000E-06 |
| 3096 | 155 | N155S | A | 8.1e-06 | 1.2e-05 | 9.2e-06 | 9.767E-06 | 2.011E-06 | 8.100E-06 | 1.200E-05 |
| 3097 | 155 | N155T | A | 9.5e-06 | 6.9e-06 | 5.6e-06 | 7.333E-06 | 1.986E-06 | 5.600E-06 | 9.500E-06 |
| 3098 | 155 | N155W | A | 1.3e-05 | 7.9e-06 | 1.1e-05 | 1.063E-05 | 2.570E-06 | 7.900E-06 | 1.300E-05 |
| 3099 | 155 | N155Y | A | 1e-05 | 4.9e-06 | 5.6e-06 | 6.833E-06 | 2.765E-06 | 4.900E-06 | 1.000E-05 |
| 3100 | 155 | N155V | A | 6.7e-06 | 1e-05 | 7.4e-06 | 8.033E-06 | 1.739E-06 | 6.700E-06 | 1.000E-05 |
| 3101 | 156 | Q156A | A | 6e-06 | 9.9e-06 | 1e-05 | 8.633E-06 | 2.281E-06 | 6.000E-06 | 1.000E-05 |
| 3102 | 156 | Q156R | A | 5e-06 | 9.2e-06 | 4.6e-06 | 6.267E-06 | 2.548E-06 | 4.600E-06 | 9.200E-06 |
| 3103 | 156 | Q156N | A | 9.7e-06 | 9.4e-06 | 7.8e-06 | 8.967E-06 | 1.021E-06 | 7.800E-06 | 9.700E-06 |
| 3104 | 156 | Q156D | A | 4.9e-06 | 7.4e-06 | 9.7e-06 | 7.333E-06 | 2.401E-06 | 4.900E-06 | 9.700E-06 |
| 3105 | 156 | Q156C | A | 1e-05 | 7.2e-06 | 1.4e-05 | 1.040E-05 | 3.418E-06 | 7.200E-06 | 1.400E-05 |
| 3106 | 156 | Q156E | A | 6.7e-06 | 5.4e-06 | 6.8e-06 | 6.300E-06 | 7.810E-07 | 5.400E-06 | 6.800E-06 |
| 3107 | 156 | Q156Q | A | 6.4e-06 | 1.4e-05 | 7.9e-06 | 9.433E-06 | 4.025E-06 | 6.400E-06 | 1.400E-05 |
| 3108 | 156 | Q156G | A | 8.1e-06 | 1e-05 | 7.4e-06 | 8.500E-06 | 1.345E-06 | 7.400E-06 | 1.000E-05 |
| 3109 | 156 | Q156H | A | 5.7e-06 | 9.4e-06 | 6.9e-06 | 7.333E-06 | 1.888E-06 | 5.700E-06 | 9.400E-06 |
| 3110 | 156 | Q156I | A | 6e-06 | 1.2e-05 | 8.9e-06 | 8.967E-06 | 3.001E-06 | 6.000E-06 | 1.200E-05 |
| 3111 | 156 | Q156L | A | 8.2e-06 | 1.3e-05 | 9.9e-06 | 1.037E-05 | 2.434E-06 | 8.200E-06 | 1.300E-05 |
| 3112 | 156 | Q156K | A | 8.3e-06 | 5.4e-06 | 9e-06 | 7.567E-06 | 1.909E-06 | 5.400E-06 | 9.000E-06 |
| 3113 | 156 | Q156M | A | 8.1e-06 | 1.4e-05 | 1e-05 | 1.070E-05 | 3.012E-06 | 8.100E-06 | 1.400E-05 |
| 3114 | 156 | Q156F | A | 7.5e-06 | 1.3e-05 | 2.9e-06 | 7.800E-06 | 5.057E-06 | 2.900E-06 | 1.300E-05 |
| 3115 | 156 | Q156P | A | 6.9e-06 | 8.7e-06 | 1.3e-05 | 9.533E-06 | 3.134E-06 | 6.900E-06 | 1.300E-05 |
| 3116 | 156 | Q156S | A | 6.1e-06 | 1.2e-05 | 9.1e-06 | 9.067E-06 | 2.950E-06 | 6.100E-06 | 1.200E-05 |
| 3117 | 156 | Q156T | A | 3.9e-06 | 9e-06 | 7.3e-06 | 6.733E-06 | 2.597E-06 | 3.900E-06 | 9.000E-06 |
| 3118 | 156 | Q156W | A | 9.6e-06 | 2.2e-05 | 7.3e-06 | 1.297E-05 | 7.907E-06 | 7.300E-06 | 2.200E-05 |
| 3119 | 156 | Q156Y | A | 6.4e-06 | 1.5e-05 | 1.3e-05 | 1.147E-05 | 4.500E-06 | 6.400E-06 | 1.500E-05 |
| 3120 | 156 | Q156V | A | 6.5e-06 | 4.5e-06 | 1.3e-05 | 8.000E-06 | 4.444E-06 | 4.500E-06 | 1.300E-05 |
| 3121 | 157 | L157A | A | 5.6e-06 | 8.9e-06 | 6.5e-06 | 7.000E-06 | 1.706E-06 | 5.600E-06 | 8.900E-06 |
| 3122 | 157 | L157R | A | 4.7e-06 | 7e-06 | 6e-06 | 5.900E-06 | 1.153E-06 | 4.700E-06 | 7.000E-06 |
| 3123 | 157 | L157N | A | 4.5e-06 | 6.8e-06 | 8e-06 | 6.433E-06 | 1.779E-06 | 4.500E-06 | 8.000E-06 |
| 3124 | 157 | L157D | A | 1e-05 | 8.9e-06 | 9.3e-06 | 9.400E-06 | 5.568E-07 | 8.900E-06 | 1.000E-05 |
| 3125 | 157 | L157C | A | 9.6e-06 | 6.6e-06 | 1.4e-05 | 1.007E-05 | 3.722E-06 | 6.600E-06 | 1.400E-05 |
| 3126 | 157 | L157E | A | 1.2e-05 | 9.7e-06 | 1.1e-05 | 1.090E-05 | 1.153E-06 | 9.700E-06 | 1.200E-05 |
| 3127 | 157 | L157Q | A | 4.4e-06 | 8.4e-06 | 6.5e-06 | 6.433E-06 | 2.001E-06 | 4.400E-06 | 8.400E-06 |
| 3128 | 157 | L157G | A | 4.8e-06 | 9.3e-06 | 9.9e-06 | 8.000E-06 | 2.787E-06 | 4.800E-06 | 9.900E-06 |
| 3129 | 157 | L157H | A | 6.5e-06 | 6.5e-06 | 4.7e-06 | 5.900E-06 | 1.039E-06 | 4.700E-06 | 6.500E-06 |
| 3130 | 157 | L157I | A | 1e-05 | 8e-06 | 7.7e-06 | 8.567E-06 | 1.250E-06 | 7.700E-06 | 1.000E-05 |
| 3131 | 157 | L157L | A | 6.6e-06 | 7e-06 | 8.4e-06 | 7.333E-06 | 9.452E-07 | 6.600E-06 | 8.400E-06 |
| 3132 | 157 | L157K | A | 5.3e-06 | 8.2e-06 | 4.6e-06 | 6.033E-06 | 1.909E-06 | 4.600E-06 | 8.200E-06 |
| 3133 | 157 | L157M | A | 9.5e-06 | 9.4e-06 | 9.4e-06 | 9.433E-06 | 5.774E-08 | 9.400E-06 | 9.500E-06 |
| 3134 | 157 | L157F | A | 8.7e-06 | 8.3e-06 | 8.3e-06 | 8.433E-06 | 2.309E-07 | 8.300E-06 | 8.700E-06 |
| 3135 | 157 | L157P | A | 4.7e-06 | 4.5e-06 | 9.3e-06 | 6.167E-06 | 2.715E-06 | 4.500E-06 | 9.300E-06 |
| 3136 | 157 | L157S | A | 6.9e-06 | 1.2e-05 | 7.7e-06 | 8.867E-06 | 2.743E-06 | 6.900E-06 | 1.200E-05 |
| 3137 | 157 | L157T | A | 3.5e-06 | 8.3e-06 | 6.3e-06 | 6.033E-06 | 2.411E-06 | 3.500E-06 | 8.300E-06 |
| 3138 | 157 | L157W | A | 6.5e-06 | 6.5e-06 | 5e-06 | 6.000E-06 | 8.660E-07 | 5.000E-06 | 6.500E-06 |
| 3139 | 157 | L157Y | A | 6e-06 | 6.5e-06 | 6.4e-06 | 6.300E-06 | 2.646E-07 | 6.000E-06 | 6.500E-06 |
| 3140 | 157 | L157V | A | 6.6e-06 | 6.9e-06 | 8.9e-06 | 7.467E-06 | 1.250E-06 | 6.600E-06 | 8.900E-06 |
| 3141 | 158 | A158A | A | 4.6e-06 | 8.1e-06 | 4.9e-06 | 5.867E-06 | 1.940E-06 | 4.600E-06 | 8.100E-06 |
| 3142 | 158 | A158R | A | 2.8e-06 | 4.3e-06 | 5.5e-06 | 4.200E-06 | 1.353E-06 | 2.800E-06 | 5.500E-06 |
| 3143 | 158 | A158N | A | 1.1e-05 | 8.2e-06 | 5.2e-06 | 8.133E-06 | 2.901E-06 | 5.200E-06 | 1.100E-05 |
| 3144 | 158 | A158D | A | 1.1e-05 | 8.5e-06 | 1.2e-05 | 1.050E-05 | 1.803E-06 | 8.500E-06 | 1.200E-05 |

| | | | | | | | | | | |
|---|---|---|---|---|---|---|---|---|---|---|
| 3145 | 158 | A158C | A | 5.5e-06 | 9.6e-06 | 8.7e-06 | 7.933E-06 | 2.155E-06 | 5.500E-06 | 9.600E-06 |
| 3146 | 158 | A158E | A | 6.6e-06 | 7.8e-06 | 1e-05 | 8.133E-06 | 1.724E-06 | 6.600E-06 | 1.000E-05 |
| 3147 | 158 | A158Q | A | 1e-05 | 9.8e-06 | 9.3e-06 | 9.700E-06 | 3.606E-07 | 9.300E-06 | 1.000E-05 |
| 3148 | 158 | A158G | A | 7.8e-06 | 7.1e-06 | 1.4e-05 | 9.633E-06 | 3.798E-06 | 7.100E-06 | 1.400E-05 |
| 3149 | 158 | A158H | A | 6.9e-06 | 6.2e-06 | 5.6e-06 | 6.233E-06 | 6.506E-07 | 5.600E-06 | 6.900E-06 |
| 3150 | 158 | A158I | A | 9.6e-06 | 8.3e-06 | 6.1e-06 | 8.000E-06 | 1.769E-06 | 6.100E-06 | 9.600E-06 |
| 3151 | 158 | A158L | A | 8.3e-06 | 1.3e-05 | 9e-06 | 1.010E-05 | 2.536E-06 | 8.300E-06 | 1.300E-05 |
| 3152 | 158 | A158K | A | 8.4e-06 | 5.6e-06 | 8e-06 | 7.333E-06 | 1.514E-06 | 5.600E-06 | 8.400E-06 |
| 3153 | 158 | A158M | A | 6e-06 | 1e-05 | 6.8e-06 | 7.600E-06 | 2.117E-06 | 6.000E-06 | 1.000E-05 |
| 3154 | 158 | A158F | A | 6.7e-06 | 8.6e-06 | 8.6e-06 | 7.967E-06 | 1.097E-06 | 6.700E-06 | 8.600E-06 |
| 3155 | 158 | A158P | A | 6.2e-06 | 8e-06 | 6.9e-06 | 7.033E-06 | 9.074E-07 | 6.200E-06 | 8.000E-06 |
| 3156 | 158 | A158S | A | 1.4e-05 | 6.4e-06 | 5.6e-06 | 8.667E-06 | 4.636E-06 | 5.600E-06 | 1.400E-05 |
| 3157 | 158 | A158T | A | 6.5e-06 | 8.9e-06 | 1.4e-05 | 9.800E-06 | 3.830E-06 | 6.500E-06 | 1.400E-05 |
| 3158 | 158 | A158W | A | 1.1e-05 | 5.9e-06 | 1e-05 | 8.967E-06 | 2.702E-06 | 5.900E-06 | 1.100E-05 |
| 3159 | 158 | A158Y | A | 6.5e-06 | 5.9e-06 | 9.5e-06 | 7.300E-06 | 1.929E-06 | 5.900E-06 | 9.500E-06 |
| 3160 | 158 | A158V | A | 3.9e-06 | 4.1e-06 | 6.6e-06 | 4.867E-06 | 1.504E-06 | 3.900E-06 | 6.600E-06 |
| 3161 | 159 | L159A | A | 9.2e-06 | 4.7e-06 | 9.4e-06 | 7.767E-06 | 2.658E-06 | 4.700E-06 | 9.400E-06 |
| 3162 | 159 | L159R | A | 7.3e-06 | 7.1e-06 | 7.8e-06 | 7.400E-06 | 3.606E-07 | 7.100E-06 | 7.800E-06 |
| 3163 | 159 | L159N | A | 8.8e-06 | 5.3e-06 | 5.9e-06 | 6.667E-06 | 1.872E-06 | 5.300E-06 | 8.800E-06 |
| 3164 | 159 | L159D | A | 4.6e-06 | 5.3e-06 | 5.8e-06 | 5.233E-06 | 6.028E-07 | 4.600E-06 | 5.800E-06 |
| 3165 | 159 | L159C | A | 9.6e-06 | 6.1e-06 | 9.7e-06 | 8.467E-06 | 2.050E-06 | 6.100E-06 | 9.700E-06 |
| 3166 | 159 | L159E | A | 9.9e-06 | 1e-05 | 5.5e-06 | 8.467E-06 | 2.570E-06 | 5.500E-06 | 1.000E-05 |
| 3167 | 159 | L159Q | A | 8e-06 | 5.6e-06 | 1.3e-05 | 8.867E-06 | 3.775E-06 | 5.600E-06 | 1.300E-05 |
| 3168 | 159 | L159G | A | 1.5e-05 | 9.5e-06 | 7.5e-06 | 1.067E-05 | 3.884E-06 | 7.500E-06 | 1.500E-05 |
| 3169 | 159 | L159H | A | 1.1e-05 | 1.3e-05 | 1.2e-05 | 1.200E-05 | 1.000E-06 | 1.100E-05 | 1.300E-05 |
| 3170 | 159 | L159I | A | 1.2e-05 | 1.1e-05 | 8.4e-06 | 1.047E-05 | 1.858E-06 | 8.400E-06 | 1.200E-05 |
| 3171 | 159 | L159L | A | 1.2e-05 | 7e-06 | 6.6e-06 | 8.533E-06 | 3.009E-06 | 6.600E-06 | 1.200E-05 |
| 3172 | 159 | L159K | A | 9.9e-06 | 1.2e-05 | 4.6e-06 | 8.833E-06 | 3.814E-06 | 4.600E-06 | 1.200E-05 |
| 3173 | 159 | L159M | A | 8.8e-06 | 6.4e-06 | 6e-06 | 7.067E-06 | 1.514E-06 | 6.000E-06 | 8.800E-06 |
| 3174 | 159 | L159F | A | 1e-05 | 6.7e-06 | 1.4e-05 | 1.023E-05 | 3.656E-06 | 6.700E-06 | 1.400E-05 |
| 3175 | 159 | L159P | A | 8.2e-06 | 1.2e-05 | 8.2e-06 | 9.467E-06 | 2.194E-06 | 8.200E-06 | 1.200E-05 |
| 3176 | 159 | L159S | A | 8e-06 | 6.1e-06 | 5.7e-06 | 6.600E-06 | 1.229E-06 | 5.700E-06 | 8.000E-06 |
| 3177 | 159 | L159T | A | 1e-05 | 1.3e-05 | 8.1e-06 | 1.037E-05 | 2.470E-06 | 8.100E-06 | 1.300E-05 |
| 3178 | 159 | L159W | A | 5.8e-06 | 6.7e-06 | 6.6e-06 | 6.367E-06 | 4.933E-07 | 5.800E-06 | 6.700E-06 |
| 3179 | 159 | L159Y | A | 8.1e-06 | 5e-06 | 5.7e-06 | 6.267E-06 | 1.626E-06 | 5.000E-06 | 8.100E-06 |
| 3180 | 159 | L159V | A | 1.4e-05 | 8.1e-06 | 7.2e-06 | 9.767E-06 | 3.694E-06 | 7.200E-06 | 1.400E-05 |
| 3181 | 160 | T160A | A | 1e-05 | 1.1e-05 | 5.7e-06 | 8.900E-06 | 2.816E-06 | 5.700E-06 | 1.100E-05 |
| 3182 | 160 | T160R | A | 7.1e-06 | 9e-06 | 1.4e-05 | 1.003E-05 | 3.564E-06 | 7.100E-06 | 1.400E-05 |
| 3183 | 160 | T160N | A | 6.2e-06 | 7.4e-06 | 1e-05 | 7.867E-06 | 1.943E-06 | 6.200E-06 | 1.000E-05 |
| 3184 | 160 | T160D | A | 1.1e-05 | 1.3e-05 | 5.2e-06 | 9.733E-06 | 4.051E-06 | 5.200E-06 | 1.300E-05 |
| 3185 | 160 | T160C | A | 6.7e-06 | 8.9e-06 | 1e-05 | 8.533E-06 | 1.680E-06 | 6.700E-06 | 1.000E-05 |
| 3186 | 160 | T160E | A | 7.3e-06 | 9.3e-06 | 9.6e-06 | 8.733E-06 | 1.250E-06 | 7.300E-06 | 9.600E-06 |
| 3187 | 160 | T160Q | A | 6.3e-06 | 9.1e-06 | 1.3e-05 | 9.467E-06 | 3.365E-06 | 6.300E-06 | 1.300E-05 |
| 3188 | 160 | T160G | A | 6.2e-06 | 4.5e-06 | 1.3e-05 | 7.900E-06 | 4.498E-06 | 4.500E-06 | 1.300E-05 |
| 3189 | 160 | T160H | A | 8.7e-06 | 8.5e-06 | 9.6e-06 | 8.933E-06 | 5.859E-07 | 8.500E-06 | 9.600E-06 |
| 3190 | 160 | T160I | A | 9.1e-06 | 8.2e-06 | 8.2e-06 | 8.500E-06 | 5.196E-07 | 8.200E-06 | 9.100E-06 |
| 3191 | 160 | T160L | A | 8.7e-06 | 4.1e-06 | 9.7e-06 | 7.500E-06 | 2.987E-06 | 4.100E-06 | 9.700E-06 |
| 3192 | 160 | T160K | A | 6.8e-06 | 7.6e-06 | 4.7e-06 | 6.367E-06 | 1.498E-06 | 4.700E-06 | 7.600E-06 |
| 3193 | 160 | T160M | A | 5.7e-06 | 8.1e-06 | 8.9e-06 | 7.567E-06 | 1.665E-06 | 5.700E-06 | 8.900E-06 |
| 3194 | 160 | T160F | A | 8.3e-06 | 6.3e-06 | 1.3e-05 | 9.200E-06 | 3.439E-06 | 6.300E-06 | 1.300E-05 |
| 3195 | 160 | T160P | A | 7.3e-06 | 5.8e-06 | 1.4e-05 | 9.033E-06 | 4.366E-06 | 5.800E-06 | 1.400E-05 |
| 3196 | 160 | T160S | A | 5.3e-06 | 7.6e-06 | 6.9e-06 | 6.600E-06 | 1.179E-06 | 5.300E-06 | 7.600E-06 |
| 3197 | 160 | T160T | A | 1e-05 | 6.9e-06 | 6.8e-06 | 7.900E-06 | 1.819E-06 | 6.800E-06 | 1.000E-05 |
| 3198 | 160 | T160W | A | 5.7e-06 | 6e-06 | 9.9e-06 | 7.200E-06 | 2.343E-06 | 5.700E-06 | 9.900E-06 |
| 3199 | 160 | T160Y | A | 9.8e-06 | 6.8e-06 | 8.5e-06 | 8.367E-06 | 1.504E-06 | 6.800E-06 | 9.800E-06 |
| 3200 | 160 | T160V | A | 1.1e-05 | 6.9e-06 | 1e-05 | 9.300E-06 | 2.138E-06 | 6.900E-06 | 1.100E-05 |
| 3201 | 161 | L161A | A | 1.4e-05 | 7.9e-06 | 8e-06 | 9.967E-06 | 3.493E-06 | 7.900E-06 | 1.400E-05 |
| 3202 | 161 | L161R | A | 9.8e-06 | 9.2e-06 | 7e-06 | 8.667E-06 | 1.474E-06 | 7.000E-06 | 9.800E-06 |
| 3203 | 161 | L161N | A | 1.3e-05 | 8.6e-06 | 9.1e-06 | 1.023E-05 | 2.409E-06 | 8.600E-06 | 1.300E-05 |
| 3204 | 161 | L161D | A | 4.8e-06 | 7e-06 | 8.2e-06 | 6.667E-06 | 1.724E-06 | 4.800E-06 | 8.200E-06 |
| 3205 | 161 | L161C | A | 5.5e-06 | 5.5e-06 | 1.1e-05 | 7.333E-06 | 3.175E-06 | 5.500E-06 | 1.100E-05 |
| 3206 | 161 | L161E | A | 9.6e-06 | 9e-06 | 8.7e-06 | 9.100E-06 | 4.583E-07 | 8.700E-06 | 9.600E-06 |
| 3207 | 161 | L161Q | A | 1.1e-05 | 1e-05 | 8.3e-06 | 9.767E-06 | 1.365E-06 | 8.300E-06 | 1.100E-05 |
| 3208 | 161 | L161G | A | 9.8e-06 | 9.9e-06 | 1e-05 | 9.900E-06 | 1.000E-07 | 9.800E-06 | 1.000E-05 |
| 3209 | 161 | L161H | A | 7.1e-06 | 1.3e-05 | 9.3e-06 | 9.800E-06 | 2.982E-06 | 7.100E-06 | 1.300E-05 |
| 3210 | 161 | L161I | A | 1.3e-05 | 5.6e-06 | 9.3e-06 | 9.300E-06 | 3.700E-06 | 5.600E-06 | 1.300E-05 |
| 3211 | 161 | L161L | A | 9.9e-06 | 7.4e-06 | 1.3e-05 | 1.010E-05 | 2.805E-06 | 7.400E-06 | 1.300E-05 |
| 3212 | 161 | L161K | A | 8.9e-06 | 7e-06 | 7.1e-06 | 7.667E-06 | 1.069E-06 | 7.000E-06 | 8.900E-06 |
| 3213 | 161 | L161M | A | 5.7e-06 | 6.7e-06 | 7.5e-06 | 6.633E-06 | 9.018E-07 | 5.700E-06 | 7.500E-06 |
| 3214 | 161 | L161F | A | 8.3e-06 | 7.6e-06 | 1.4e-05 | 9.967E-06 | 3.510E-06 | 7.600E-06 | 1.400E-05 |
| 3215 | 161 | L161P | A | 6.9e-06 | 7.1e-06 | 6.3e-06 | 6.767E-06 | 4.163E-07 | 6.300E-06 | 7.100E-06 |
| 3216 | 161 | L161S | A | 9.4e-06 | 1.3e-05 | 7.5e-06 | 9.967E-06 | 2.793E-06 | 7.500E-06 | 1.300E-05 |
| 3217 | 161 | L161T | A | 8.3e-06 | 5.2e-06 | 9e-06 | 7.500E-06 | 2.022E-06 | 5.200E-06 | 9.000E-06 |
| 3218 | 161 | L161W | A | 4.5e-06 | 6.9e-06 | 7.3e-06 | 6.233E-06 | 1.514E-06 | 4.500E-06 | 7.300E-06 |
| 3219 | 161 | L161Y | A | 6.4e-06 | 6.1e-06 | 6.5e-06 | 6.333E-06 | 2.082E-07 | 6.100E-06 | 6.500E-06 |
| 3220 | 161 | L161V | A | 7.7e-06 | 5.5e-06 | 6.8e-06 | 6.667E-06 | 1.106E-06 | 5.500E-06 | 7.700E-06 |
| 3221 | 162 | I162A | A | 4.7e-06 | 9.5e-06 | 7.9e-06 | 7.367E-06 | 2.444E-06 | 4.700E-06 | 9.500E-06 |
| 3222 | 162 | I162R | A | 1.4e-05 | 7.3e-06 | 1e-05 | 1.043E-05 | 3.371E-06 | 7.300E-06 | 1.400E-05 |
| 3223 | 162 | I162N | A | 1.2e-05 | 6.5e-06 | 4.9e-06 | 7.800E-06 | 3.724E-06 | 4.900E-06 | 1.200E-05 |
| 3224 | 162 | I162D | A | 1.3e-05 | 8.7e-06 | 9.9e-06 | 1.053E-05 | 2.219E-06 | 8.700E-06 | 1.300E-05 |
| 3225 | 162 | I162C | A | 1.2e-05 | 6.6e-06 | 8.9e-06 | 9.167E-06 | 2.710E-06 | 6.600E-06 | 1.200E-05 |
| 3226 | 162 | I162E | A | 1.3e-05 | 5.5e-06 | 7e-06 | 8.500E-06 | 3.969E-06 | 5.500E-06 | 1.300E-05 |
| 3227 | 162 | I162Q | A | 9.9e-06 | 6.5e-06 | 8.3e-06 | 8.233E-06 | 1.701E-06 | 6.500E-06 | 9.900E-06 |
| 3228 | 162 | I162G | A | 9.2e-06 | 9.2e-06 | 1.1e-05 | 9.800E-06 | 1.039E-06 | 9.200E-06 | 1.100E-05 |
| 3229 | 162 | I162H | A | 8.8e-06 | 6.5e-06 | 6.4e-06 | 7.233E-06 | 1.358E-06 | 6.400E-06 | 8.800E-06 |

| | | | | | | | | | | |
|---|---|---|---|---|---|---|---|---|---|---|
| 3230 | 162 | I162I | A | 8.6e-06 | 6.9e-06 | 7.8e-06 | 7.767E-06 | 8.505E-07 | 6.900E-06 | 8.600E-06 |
| 3231 | 162 | I162L | A | 6.2e-06 | 9.7e-06 | 8.7e-06 | 8.200E-06 | 1.803E-06 | 6.200E-06 | 9.700E-06 |
| 3232 | 162 | I162K | A | 8.1e-06 | 8.1e-06 | 8.8e-06 | 8.333E-06 | 4.041E-07 | 8.100E-06 | 8.800E-06 |
| 3233 | 162 | I162M | A | 5.3e-06 | 6.4e-06 | 9.8e-06 | 7.167E-06 | 2.346E-06 | 5.300E-06 | 9.800E-06 |
| 3234 | 162 | I162F | A | 7.2e-06 | 5.3e-06 | 8.1e-06 | 6.867E-06 | 1.429E-06 | 5.300E-06 | 8.100E-06 |
| 3235 | 162 | I162P | A | 1e-05 | 6.1e-06 | 8e-06 | 8.033E-06 | 1.950E-06 | 6.100E-06 | 1.000E-05 |
| 3236 | 162 | I162S | A | 8.9e-06 | 6.5e-06 | 8.5e-06 | 7.967E-06 | 1.286E-06 | 6.500E-06 | 8.900E-06 |
| 3237 | 162 | I162T | A | 1.4e-05 | 1.2e-05 | 1e-05 | 1.200E-05 | 2.000E-06 | 1.000E-05 | 1.400E-05 |
| 3238 | 162 | I162W | A | 4.8e-06 | 5.6e-06 | 7.5e-06 | 5.967E-06 | 1.387E-06 | 4.800E-06 | 7.500E-06 |
| 3239 | 162 | I162Y | A | 9.4e-06 | 9.6e-06 | 7.6e-06 | 8.867E-06 | 1.102E-06 | 7.600E-06 | 9.600E-06 |
| 3240 | 162 | I162V | A | 5.4e-06 | 7.5e-06 | 5.8e-06 | 6.233E-06 | 1.115E-06 | 5.400E-06 | 7.500E-06 |
| 3241 | 163 | D163A | A | 1.3e-05 | 3.3e-06 | 1.4e-05 | 1.010E-05 | 5.910E-06 | 3.300E-06 | 1.400E-05 |
| 3242 | 163 | D163R | A | 7.5e-06 | 1e-05 | 7.2e-06 | 8.233E-06 | 1.537E-06 | 7.200E-06 | 1.000E-05 |
| 3243 | 163 | D163N | A | 1e-05 | 1.3e-05 | 6.5e-06 | 9.833E-06 | 3.253E-06 | 6.500E-06 | 1.300E-05 |
| 3244 | 163 | D163D | A | 6.7e-06 | 8.3e-06 | 7.7e-06 | 7.567E-06 | 8.083E-07 | 6.700E-06 | 8.300E-06 |
| 3245 | 163 | D163C | A | 8.3e-06 | 8e-06 | 1.3e-05 | 9.767E-06 | 2.804E-06 | 8.000E-06 | 1.300E-05 |
| 3246 | 163 | D163E | A | 6.3e-06 | 1.2e-05 | 5.9e-06 | 8.067E-06 | 3.412E-06 | 5.900E-06 | 1.200E-05 |
| 3247 | 163 | D163Q | A | 5.2e-06 | 7.3e-06 | 1e-05 | 7.500E-06 | 2.406E-06 | 5.200E-06 | 1.000E-05 |
| 3248 | 163 | D163G | A | 4.2e-06 | 5.5e-06 | 9.3e-06 | 6.333E-06 | 2.650E-06 | 4.200E-06 | 9.300E-06 |
| 3249 | 163 | D163H | A | 6.9e-06 | 6.9e-06 | 7.7e-06 | 7.167E-06 | 4.619E-07 | 6.900E-06 | 7.700E-06 |
| 3250 | 163 | D163I | A | 9.6e-06 | 1.3e-05 | 8.2e-06 | 1.027E-05 | 2.468E-06 | 8.200E-06 | 1.300E-05 |
| 3251 | 163 | D163L | A | 5.2e-06 | 9.3e-06 | 6.2e-06 | 6.900E-06 | 2.138E-06 | 5.200E-06 | 9.300E-06 |
| 3252 | 163 | D163K | A | 6e-06 | 9.8e-06 | 1.2e-05 | 9.267E-06 | 3.035E-06 | 6.000E-06 | 1.200E-05 |
| 3253 | 163 | D163M | A | 9.6e-06 | 1e-05 | 9.2e-06 | 9.600E-06 | 4.000E-07 | 9.200E-06 | 1.000E-05 |
| 3254 | 163 | D163F | A | 8.4e-06 | 4.7e-06 | 9.1e-06 | 7.400E-06 | 2.364E-06 | 4.700E-06 | 9.100E-06 |
| 3255 | 163 | D163P | A | 9.9e-06 | 9.4e-06 | 7.3e-06 | 8.867E-06 | 1.380E-06 | 7.300E-06 | 9.900E-06 |
| 3256 | 163 | D163S | A | 4.8e-06 | 4.8e-06 | 6.4e-06 | 5.333E-06 | 9.238E-07 | 4.800E-06 | 6.400E-06 |
| 3257 | 163 | D163T | A | 8.9e-06 | 6.3e-06 | 5.9e-06 | 7.033E-06 | 1.629E-06 | 5.900E-06 | 8.900E-06 |
| 3258 | 163 | D163W | A | 9.2e-06 | 5.3e-06 | 9.3e-06 | 7.933E-06 | 2.281E-06 | 5.300E-06 | 9.300E-06 |
| 3259 | 163 | D163Y | A | 6.5e-06 | 4.7e-06 | 6.5e-06 | 5.900E-06 | 1.039E-06 | 4.700E-06 | 6.500E-06 |
| 3260 | 163 | D163V | A | 9.6e-06 | 7.9e-06 | 9.8e-06 | 9.100E-06 | 1.044E-06 | 7.900E-06 | 9.800E-06 |
| 3261 | 164 | T164A | A | 6.5e-06 | 9.6e-06 | 1.5e-05 | 1.037E-05 | 4.302E-06 | 6.500E-06 | 1.500E-05 |
| 3262 | 164 | T164R | A | 8.5e-06 | 1.2e-05 | 6.7e-06 | 9.067E-06 | 2.695E-06 | 6.700E-06 | 1.200E-05 |
| 3263 | 164 | T164N | A | 4.1e-06 | 1e-05 | 8.9e-06 | 7.667E-06 | 3.137E-06 | 4.100E-06 | 1.000E-05 |
| 3264 | 164 | T164D | A | 7.4e-06 | 6.2e-06 | 1.3e-05 | 8.867E-06 | 3.630E-06 | 6.200E-06 | 1.300E-05 |
| 3265 | 164 | T164C | A | 1.1e-05 | 9.3e-06 | 5.2e-06 | 8.500E-06 | 2.982E-06 | 5.200E-06 | 1.100E-05 |
| 3266 | 164 | T164E | A | 9.9e-06 | 5.4e-06 | 1.4e-05 | 9.767E-06 | 4.302E-06 | 5.400E-06 | 1.400E-05 |
| 3267 | 164 | T164Q | A | 6.3e-06 | 1e-05 | 5.7e-06 | 7.333E-06 | 2.329E-06 | 5.700E-06 | 1.000E-05 |
| 3268 | 164 | T164G | A | 8.5e-06 | 8.6e-06 | 1.4e-05 | 1.037E-05 | 3.147E-06 | 8.500E-06 | 1.400E-05 |
| 3269 | 164 | T164H | A | 8.6e-06 | 6.4e-06 | 6.3e-06 | 7.100E-06 | 1.300E-06 | 6.300E-06 | 8.600E-06 |
| 3270 | 164 | T164I | A | 1.1e-05 | 7.1e-06 | 1e-05 | 9.367E-06 | 2.026E-06 | 7.100E-06 | 1.100E-05 |
| 3271 | 164 | T164L | A | 7e-06 | 8.9e-06 | 9e-06 | 8.300E-06 | 1.127E-06 | 7.000E-06 | 9.000E-06 |
| 3272 | 164 | T164K | A | 8.1e-06 | 8.4e-06 | 8.2e-06 | 8.233E-06 | 1.528E-07 | 8.100E-06 | 8.400E-06 |
| 3273 | 164 | T164M | A | 3.3e-06 | 6.6e-06 | 9.3e-06 | 6.400E-06 | 3.005E-06 | 3.300E-06 | 9.300E-06 |
| 3274 | 164 | T164F | A | 1.2e-05 | 8.5e-06 | 9.6e-06 | 1.003E-05 | 1.790E-06 | 8.500E-06 | 1.200E-05 |
| 3275 | 164 | T164P | A | 7.6e-06 | 6.7e-06 | 5e-06 | 6.433E-06 | 1.320E-06 | 5.000E-06 | 7.600E-06 |
| 3276 | 164 | T164S | A | 9.6e-06 | 6.3e-06 | 8.8e-06 | 8.233E-06 | 1.721E-06 | 6.300E-06 | 9.600E-06 |
| 3277 | 164 | T164T | A | 3.5e-06 | 7.7e-06 | 1.1e-05 | 7.400E-06 | 3.759E-06 | 3.500E-06 | 1.100E-05 |
| 3278 | 164 | T164W | A | 1.5e-05 | 1.2e-05 | 1.1e-05 | 1.267E-05 | 2.082E-06 | 1.100E-05 | 1.500E-05 |
| 3279 | 164 | T164Y | A | 1.3e-05 | 7.8e-06 | 4.6e-06 | 8.467E-06 | 4.239E-06 | 4.600E-06 | 1.300E-05 |
| 3280 | 164 | T164V | A | 7.8e-06 | 1.5e-05 | 7e-06 | 9.933E-06 | 4.406E-06 | 7.000E-06 | 1.500E-05 |
| 3281 | 165 | N165A | A | 7.4e-06 | 1e-05 | 1.6e-05 | 1.113E-05 | 4.411E-06 | 7.400E-06 | 1.600E-05 |
| 3282 | 165 | N165R | A | 8.3e-06 | 6.7e-06 | 9.2e-06 | 8.067E-06 | 1.266E-06 | 6.700E-06 | 9.200E-06 |
| 3283 | 165 | N165N | A | 7.8e-06 | 8.5e-06 | 6.7e-06 | 7.667E-06 | 9.074E-07 | 6.700E-06 | 8.500E-06 |
| 3284 | 165 | N165D | A | 1.5e-05 | 8.9e-06 | 1.2e-05 | 1.197E-05 | 3.050E-06 | 8.900E-06 | 1.500E-05 |
| 3285 | 165 | N165C | A | 1.1e-05 | 8.9e-06 | 7.2e-06 | 9.033E-06 | 1.904E-06 | 7.200E-06 | 1.100E-05 |
| 3286 | 165 | N165E | A | 1.2e-05 | 1.4e-05 | 9.6e-06 | 1.187E-05 | 2.203E-06 | 9.600E-06 | 1.400E-05 |
| 3287 | 165 | N165Q | A | 1e-05 | 9.5e-06 | 8.5e-06 | 9.333E-06 | 7.638E-07 | 8.500E-06 | 1.000E-05 |
| 3288 | 165 | N165G | A | 4.5e-06 | 6.8e-06 | 8.9e-06 | 6.733E-06 | 2.201E-06 | 4.500E-06 | 8.900E-06 |
| 3289 | 165 | N165H | A | 1.3e-05 | 1e-05 | 6.8e-06 | 9.933E-06 | 3.101E-06 | 6.800E-06 | 1.300E-05 |
| 3290 | 165 | N165I | A | 6.4e-06 | 1e-05 | 5.3e-06 | 7.233E-06 | 2.458E-06 | 5.300E-06 | 1.000E-05 |
| 3291 | 165 | N165L | A | 5.4e-06 | 9e-06 | 1.2e-05 | 8.800E-06 | 3.305E-06 | 5.400E-06 | 1.200E-05 |
| 3292 | 165 | N165K | A | 6.4e-06 | 9.2e-06 | 1e-05 | 8.533E-06 | 1.890E-06 | 6.400E-06 | 1.000E-05 |
| 3293 | 165 | N165M | A | 1.4e-05 | 5.2e-06 | 9.2e-06 | 9.467E-06 | 4.406E-06 | 5.200E-06 | 1.400E-05 |
| 3294 | 165 | N165F | A | 8.5e-06 | 6.3e-06 | 8.1e-06 | 7.633E-06 | 1.172E-06 | 6.300E-06 | 8.500E-06 |
| 3295 | 165 | N165P | A | 1e-05 | 5.7e-06 | 6.6e-06 | 7.433E-06 | 2.268E-06 | 5.700E-06 | 1.000E-05 |
| 3296 | 165 | N165S | A | 8.9e-06 | 1.4e-05 | 8.2e-06 | 1.037E-05 | 3.166E-06 | 8.200E-06 | 1.400E-05 |
| 3297 | 165 | N165T | A | 7.5e-06 | 5.5e-06 | 9.8e-06 | 7.600E-06 | 2.152E-06 | 5.500E-06 | 9.800E-06 |
| 3298 | 165 | N165W | A | 6.9e-06 | 9.3e-06 | 7e-06 | 7.733E-06 | 1.358E-06 | 6.900E-06 | 9.300E-06 |
| 3299 | 165 | N165Y | A | 9.2e-06 | 5.1e-06 | 1.3e-05 | 9.100E-06 | 3.951E-06 | 5.100E-06 | 1.300E-05 |
| 3300 | 165 | N165V | A | 6.5e-06 | 5.2e-06 | 5.8e-06 | 5.833E-06 | 6.506E-07 | 5.200E-06 | 6.500E-06 |
| 3301 | 166 | R166A | A | 8.4e-06 | 8.6e-06 | 5.3e-06 | 7.433E-06 | 1.850E-06 | 5.300E-06 | 8.600E-06 |
| 3302 | 166 | R166R | A | 6.7e-06 | 5.4e-06 | 1.2e-05 | 8.033E-06 | 3.496E-06 | 5.400E-06 | 1.200E-05 |
| 3303 | 166 | R166N | A | 6.9e-06 | 1.2e-05 | 5.4e-06 | 8.100E-06 | 3.460E-06 | 5.400E-06 | 1.200E-05 |
| 3304 | 166 | R166D | A | 9.8e-06 | 7.8e-06 | 1.1e-05 | 9.533E-06 | 1.617E-06 | 7.800E-06 | 1.100E-05 |
| 3305 | 166 | R166C | A | 5.8e-06 | 4.2e-06 | 6.2e-06 | 5.400E-06 | 1.058E-06 | 4.200E-06 | 6.200E-06 |
| 3306 | 166 | R166E | A | 8e-06 | 6.1e-06 | 6.4e-06 | 6.833E-06 | 1.021E-06 | 6.100E-06 | 8.000E-06 |
| 3307 | 166 | R166Q | A | 9.1e-06 | 7.2e-06 | 7.6e-06 | 7.967E-06 | 1.002E-06 | 7.200E-06 | 9.100E-06 |
| 3308 | 166 | R166G | A | 1.2e-05 | 1.5e-05 | 9.6e-06 | 1.220E-05 | 2.706E-06 | 9.600E-06 | 1.500E-05 |
| 3309 | 166 | R166H | A | 3.7e-06 | 6.7e-06 | 5.5e-06 | 5.300E-06 | 1.510E-06 | 3.700E-06 | 6.700E-06 |
| 3310 | 166 | R166I | A | 9.9e-06 | 9.8e-06 | 9.3e-06 | 9.667E-06 | 3.215E-07 | 9.300E-06 | 9.900E-06 |
| 3311 | 166 | R166L | A | 6.8e-06 | 7.1e-06 | 6.5e-06 | 6.800E-06 | 3.000E-07 | 6.500E-06 | 7.100E-06 |
| 3312 | 166 | R166K | A | 8.5e-06 | 7.4e-06 | 9e-06 | 8.300E-06 | 8.185E-07 | 7.400E-06 | 9.000E-06 |
| 3313 | 166 | R166M | A | 4.8e-06 | 8.5e-06 | 8.5e-06 | 7.267E-06 | 2.136E-06 | 4.800E-06 | 8.500E-06 |
| 3314 | 166 | R166F | A | 4.2e-06 | 9.5e-06 | 8e-06 | 7.233E-06 | 2.732E-06 | 4.200E-06 | 9.500E-06 |

| | | | | | | | | | | |
|---|---|---|---|---|---|---|---|---|---|---|
| 3315 | 166 | R166P | A | 4.6e-06 | 8.3e-06 | 6.7e-06 | 6.533E-06 | 1.856E-06 | 4.600E-06 | 8.300E-06 |
| 3316 | 166 | R166S | A | 7.1e-06 | 6.2e-06 | 3.9e-06 | 5.733E-06 | 1.650E-06 | 3.900E-06 | 7.100E-06 |
| 3317 | 166 | R166T | A | 7.4e-06 | 8.4e-06 | 8e-06 | 7.933E-06 | 5.033E-07 | 7.400E-06 | 8.400E-06 |
| 3318 | 166 | R166W | A | 8.2e-06 | 6.5e-06 | 5.1e-06 | 6.600E-06 | 1.552E-06 | 5.100E-06 | 8.200E-06 |
| 3319 | 166 | R166Y | A | 1.1e-05 | 1.2e-05 | 5.3e-06 | 9.433E-06 | 3.614E-06 | 5.300E-06 | 1.200E-05 |
| 3320 | 166 | R166V | A | 5.4e-06 | 1e-05 | 6.3e-06 | 7.233E-06 | 2.438E-06 | 5.400E-06 | 1.000E-05 |
| 3321 | 167 | S167A | A | 7.3e-06 | 1e-05 | 4.6e-06 | 7.300E-06 | 2.700E-06 | 4.600E-06 | 1.000E-05 |
| 3322 | 167 | S167R | A | 6.9e-06 | 7.2e-06 | 1.4e-05 | 9.367E-06 | 4.015E-06 | 6.900E-06 | 1.400E-05 |
| 3323 | 167 | S167N | A | 9e-06 | 5e-06 | 8.7e-06 | 7.567E-06 | 2.228E-06 | 5.000E-06 | 9.000E-06 |
| 3324 | 167 | S167D | A | 9.7e-06 | 1e-05 | 1.4e-05 | 1.123E-05 | 2.401E-06 | 9.700E-06 | 1.400E-05 |
| 3325 | 167 | S167C | A | 1.1e-05 | 9.8e-06 | 1e-05 | 1.027E-05 | 6.429E-07 | 9.800E-06 | 1.100E-05 |
| 3326 | 167 | S167E | A | 7e-06 | 1e-05 | 8.9e-06 | 8.633E-06 | 1.518E-06 | 7.000E-06 | 1.000E-05 |
| 3327 | 167 | S167Q | A | 6e-06 | 5.5e-06 | 9e-06 | 6.833E-06 | 1.893E-06 | 5.500E-06 | 9.000E-06 |
| 3328 | 167 | S167G | A | 7.7e-06 | 6.7e-06 | 1.2e-05 | 8.800E-06 | 2.816E-06 | 6.700E-06 | 1.200E-05 |
| 3329 | 167 | S167H | A | 7e-06 | 9e-06 | 7.5e-06 | 7.833E-06 | 1.041E-06 | 7.000E-06 | 9.000E-06 |
| 3330 | 167 | S167I | A | 8.3e-06 | 1.3e-05 | 9.2e-06 | 1.017E-05 | 2.495E-06 | 8.300E-06 | 1.300E-05 |
| 3331 | 167 | S167L | A | 5e-06 | 5.4e-06 | 6.5e-06 | 5.633E-06 | 7.767E-07 | 5.000E-06 | 6.500E-06 |
| 3332 | 167 | S167K | A | 6.8e-06 | 4.8e-06 | 6.5e-06 | 6.033E-06 | 1.079E-06 | 4.800E-06 | 6.800E-06 |
| 3333 | 167 | S167M | A | 7.5e-06 | 7.1e-06 | 1.4e-05 | 9.533E-06 | 3.873E-06 | 7.100E-06 | 1.400E-05 |
| 3334 | 167 | S167F | A | 7.4e-06 | 7.3e-06 | 7.7e-06 | 7.467E-06 | 2.082E-07 | 7.300E-06 | 7.700E-06 |
| 3335 | 167 | S167P | A | 8.8e-06 | 6.3e-06 | 6.4e-06 | 7.167E-06 | 1.415E-06 | 6.300E-06 | 8.800E-06 |
| 3336 | 167 | S167S | A | 5.2e-06 | 8e-06 | 6.9e-06 | 6.700E-06 | 1.411E-06 | 5.200E-06 | 8.000E-06 |
| 3337 | 167 | S167T | A | 6.8e-06 | 1.1e-05 | 6.8e-06 | 8.200E-06 | 2.425E-06 | 6.800E-06 | 1.100E-05 |
| 3338 | 167 | S167W | A | 8.7e-06 | 4.5e-06 | 5.4e-06 | 6.200E-06 | 2.211E-06 | 4.500E-06 | 8.700E-06 |
| 3339 | 167 | S167Y | A | 7.9e-06 | 5.7e-06 | 1.1e-05 | 8.200E-06 | 2.663E-06 | 5.700E-06 | 1.100E-05 |
| 3340 | 167 | S167V | A | 1.4e-05 | 8e-06 | 1.2e-05 | 1.133E-05 | 3.055E-06 | 8.000E-06 | 1.400E-05 |
| 3341 | 168 | R168A | A | 6.1e-06 | 4e-06 | 6e-06 | 5.367E-06 | 1.185E-06 | 4.000E-06 | 6.100E-06 |
| 3342 | 168 | R168R | A | 5.9e-06 | 8.3e-06 | 7.1e-06 | 7.100E-06 | 1.200E-06 | 5.900E-06 | 8.300E-06 |
| 3343 | 168 | R168N | A | 1.1e-05 | 7.7e-06 | 7.8e-06 | 8.833E-06 | 1.877E-06 | 7.700E-06 | 1.100E-05 |
| 3344 | 168 | R168D | A | 6.9e-06 | 6.5e-06 | 8e-06 | 7.133E-06 | 7.767E-07 | 6.500E-06 | 8.000E-06 |
| 3345 | 168 | R168C | A | 7.2e-06 | 7.6e-06 | 6.6e-06 | 7.133E-06 | 5.033E-07 | 6.600E-06 | 7.600E-06 |
| 3346 | 168 | R168E | A | 9.2e-06 | 9.2e-06 | 5.9e-06 | 8.100E-06 | 1.905E-06 | 5.900E-06 | 9.200E-06 |
| 3347 | 168 | R168Q | A | 8.2e-06 | 7.4e-06 | 6.6e-06 | 7.400E-06 | 8.000E-07 | 6.600E-06 | 8.200E-06 |
| 3348 | 168 | R168G | A | 1.3e-05 | 6.1e-06 | 6.7e-06 | 8.600E-06 | 3.822E-06 | 6.100E-06 | 1.300E-05 |
| 3349 | 168 | R168H | A | 4.6e-06 | 8.9e-06 | 1.3e-05 | 8.833E-06 | 4.200E-06 | 4.600E-06 | 1.300E-05 |
| 3350 | 168 | R168I | A | 1.4e-05 | 5.5e-06 | 4e-06 | 7.833E-06 | 5.393E-06 | 4.000E-06 | 1.400E-05 |
| 3351 | 168 | R168L | A | 1.4e-05 | 6.5e-06 | 1.2e-05 | 1.083E-05 | 3.884E-06 | 6.500E-06 | 1.400E-05 |
| 3352 | 168 | R168K | A | 6.9e-06 | 6.3e-06 | 3.9e-06 | 5.700E-06 | 1.587E-06 | 3.900E-06 | 6.900E-06 |
| 3353 | 168 | R168M | A | 9.6e-06 | 6.1e-06 | 8e-06 | 7.900E-06 | 1.752E-06 | 6.100E-06 | 9.600E-06 |
| 3354 | 168 | R168F | A | 1e-05 | 8.6e-06 | 1.3e-05 | 1.053E-05 | 2.248E-06 | 8.600E-06 | 1.300E-05 |
| 3355 | 168 | R168P | A | 1e-05 | 6.7e-06 | 5.2e-06 | 7.300E-06 | 2.456E-06 | 5.200E-06 | 1.000E-05 |
| 3356 | 168 | R168S | A | 6.8e-06 | 5.2e-06 | 5.4e-06 | 5.800E-06 | 8.718E-07 | 5.200E-06 | 6.800E-06 |
| 3357 | 168 | R168T | A | 6.6e-06 | 7.8e-06 | 5e-06 | 6.467E-06 | 1.405E-06 | 5.000E-06 | 7.800E-06 |
| 3358 | 168 | R168W | A | 1e-05 | 1.2e-05 | 1.1e-05 | 1.100E-05 | 1.000E-06 | 1.000E-05 | 1.200E-05 |
| 3359 | 168 | R168Y | A | 1.4e-05 | 4.7e-06 | 8.8e-06 | 9.167E-06 | 4.661E-06 | 4.700E-06 | 1.400E-05 |
| 3360 | 168 | R168V | A | 7e-06 | 6.4e-06 | 7.3e-06 | 6.900E-06 | 4.583E-07 | 6.400E-06 | 7.300E-06 |
| 3361 | 169 | A169A | A | 9.2e-06 | 6.7e-06 | 5.7e-06 | 7.200E-06 | 1.803E-06 | 5.700E-06 | 9.200E-06 |
| 3362 | 169 | A169R | A | 4.6e-06 | 6.4e-06 | 1.2e-05 | 7.667E-06 | 3.859E-06 | 4.600E-06 | 1.200E-05 |
| 3363 | 169 | A169N | A | 6.5e-06 | 7.5e-06 | 9.3e-06 | 7.767E-06 | 1.419E-06 | 6.500E-06 | 9.300E-06 |
| 3364 | 169 | A169D | A | 6.5e-06 | 5.1e-06 | 9.1e-06 | 6.900E-06 | 2.030E-06 | 5.100E-06 | 9.100E-06 |
| 3365 | 169 | A169C | A | 8.9e-06 | 8.5e-06 | 1e-05 | 9.133E-06 | 7.767E-07 | 8.500E-06 | 1.000E-05 |
| 3366 | 169 | A169E | A | 1e-05 | 1.2e-05 | 7.6e-06 | 9.867E-06 | 2.203E-06 | 7.600E-06 | 1.200E-05 |
| 3367 | 169 | A169Q | A | 9e-06 | 6.8e-06 | 7e-06 | 7.600E-06 | 1.217E-06 | 6.800E-06 | 9.000E-06 |
| 3368 | 169 | A169G | A | 6.3e-06 | 6.4e-06 | 6.4e-06 | 6.367E-06 | 5.774E-08 | 6.300E-06 | 6.400E-06 |
| 3369 | 169 | A169H | A | 9.5e-06 | 7.4e-06 | 6.2e-06 | 7.700E-06 | 1.670E-06 | 6.200E-06 | 9.500E-06 |
| 3370 | 169 | A169I | A | 8.7e-06 | 6.9e-06 | 5e-06 | 6.867E-06 | 1.850E-06 | 5.000E-06 | 8.700E-06 |
| 3371 | 169 | A169L | A | 6.5e-06 | 1.5e-05 | 5.6e-06 | 9.033E-06 | 5.187E-06 | 5.600E-06 | 1.500E-05 |
| 3372 | 169 | A169K | A | 1.4e-05 | 9e-06 | 5.5e-06 | 9.500E-06 | 4.272E-06 | 5.500E-06 | 1.400E-05 |
| 3373 | 169 | A169M | A | 1.1e-05 | 1.2e-05 | 6.4e-06 | 9.800E-06 | 2.987E-06 | 6.400E-06 | 1.200E-05 |
| 3374 | 169 | A169F | A | 8.6e-06 | 7e-06 | 8.2e-06 | 7.933E-06 | 8.327E-07 | 7.000E-06 | 8.600E-06 |
| 3375 | 169 | A169P | A | 6e-06 | 6.2e-06 | 1.4e-05 | 8.733E-06 | 4.562E-06 | 6.000E-06 | 1.400E-05 |
| 3376 | 169 | A169S | A | 9.5e-06 | 1.2e-05 | 7.5e-06 | 9.667E-06 | 2.255E-06 | 7.500E-06 | 1.200E-05 |
| 3377 | 169 | A169T | A | 9.6e-06 | 8.8e-06 | 5.5e-06 | 7.967E-06 | 2.173E-06 | 5.500E-06 | 9.600E-06 |
| 3378 | 169 | A169W | A | 4.3e-06 | 8.5e-06 | 7.1e-06 | 6.633E-06 | 2.139E-06 | 4.300E-06 | 8.500E-06 |
| 3379 | 169 | A169Y | A | 6.3e-06 | 5.5e-06 | 8.3e-06 | 6.700E-06 | 1.442E-06 | 5.500E-06 | 8.300E-06 |
| 3380 | 169 | A169V | A | 1.2e-05 | 5.4e-06 | 6.6e-06 | 8.000E-06 | 3.516E-06 | 5.400E-06 | 1.200E-05 |
| 3381 | 170 | C170A | A | 5.9e-06 | 9.2e-06 | 6.2e-06 | 7.100E-06 | 1.825E-06 | 5.900E-06 | 9.200E-06 |
| 3382 | 170 | C170R | A | 8.4e-06 | 8.1e-06 | 1.1e-05 | 9.167E-06 | 1.595E-06 | 8.100E-06 | 1.100E-05 |
| 3383 | 170 | C170N | A | 9.7e-06 | 9.1e-06 | 7.4e-06 | 8.733E-06 | 1.193E-06 | 7.400E-06 | 9.700E-06 |
| 3384 | 170 | C170D | A | 1.3e-05 | 9.5e-06 | 9.9e-06 | 1.080E-05 | 1.916E-06 | 9.500E-06 | 1.300E-05 |
| 3385 | 170 | C170C | A | 6.1e-06 | 7.9e-06 | 8.5e-06 | 7.500E-06 | 1.249E-06 | 6.100E-06 | 8.500E-06 |
| 3386 | 170 | C170E | A | 1.5e-05 | 1.1e-05 | 1.3e-05 | 1.300E-05 | 2.000E-06 | 1.100E-05 | 1.500E-05 |
| 3387 | 170 | C170Q | A | 1.2e-05 | 6.6e-06 | 6.6e-06 | 8.400E-06 | 3.118E-06 | 6.600E-06 | 1.200E-05 |
| 3388 | 170 | C170G | A | 1e-05 | 4.7e-06 | 9e-06 | 7.900E-06 | 2.816E-06 | 4.700E-06 | 1.000E-05 |
| 3389 | 170 | C170H | A | 6.2e-06 | 1.8e-05 | 9.2e-06 | 1.113E-05 | 6.133E-06 | 6.200E-06 | 1.800E-05 |
| 3390 | 170 | C170I | A | 8.9e-06 | 8e-06 | 5e-06 | 7.300E-06 | 2.042E-06 | 5.000E-06 | 8.900E-06 |
| 3391 | 170 | C170L | A | 7.5e-06 | 1.1e-05 | 7.9e-06 | 8.800E-06 | 1.916E-06 | 7.500E-06 | 1.100E-05 |
| 3392 | 170 | C170K | A | 8.5e-06 | 1e-05 | 1.1e-05 | 9.833E-06 | 1.258E-06 | 8.500E-06 | 1.100E-05 |
| 3393 | 170 | C170M | A | 1.1e-05 | 6.6e-06 | 7.1e-06 | 8.233E-06 | 2.409E-06 | 6.600E-06 | 1.100E-05 |
| 3394 | 170 | C170F | A | 5.2e-06 | 1.1e-05 | 1e-05 | 8.733E-06 | 3.101E-06 | 5.200E-06 | 1.100E-05 |
| 3395 | 170 | C170P | A | 1.1e-05 | 1e-05 | 1.1e-05 | 1.067E-05 | 5.774E-07 | 1.000E-05 | 1.100E-05 |
| 3396 | 170 | C170S | A | 6e-06 | 6.5e-06 | 1.3e-05 | 8.500E-06 | 3.905E-06 | 6.000E-06 | 1.300E-05 |
| 3397 | 170 | C170T | A | 1.2e-05 | 6.3e-06 | 7.9e-06 | 8.733E-06 | 2.940E-06 | 6.300E-06 | 1.200E-05 |
| 3398 | 170 | C170W | A | 1.2e-05 | 8.1e-06 | 1.2e-05 | 1.070E-05 | 2.252E-06 | 8.100E-06 | 1.200E-05 |
| 3399 | 170 | C170Y | A | 9.5e-06 | 3.2e-06 | 8.6e-06 | 7.100E-06 | 3.407E-06 | 3.200E-06 | 9.500E-06 |

| | | | | | | | | | | |
|---|---|---|---|---|---|---|---|---|---|---|
| 3400 | 170 | C170V | A | 8.2e-06 | 7.9e-06 | 6.6e-06 | 7.567E-06 | 8.505E-07 | 6.600E-06 | 8.200E-06 |
| 3401 | 171 | H171A | A | 6.3e-06 | 1e-05 | 8.7e-06 | 8.333E-06 | 1.877E-06 | 6.300E-06 | 1.000E-05 |
| 3402 | 171 | H171R | A | 1e-05 | 7.1e-06 | 9.8e-06 | 8.967E-06 | 1.620E-06 | 7.100E-06 | 1.000E-05 |
| 3403 | 171 | H171N | A | 7.2e-06 | 4.8e-06 | 9.8e-06 | 7.267E-06 | 2.501E-06 | 4.800E-06 | 9.800E-06 |
| 3404 | 171 | H171D | A | 3.9e-06 | 6.9e-06 | 6.5e-06 | 5.767E-06 | 1.629E-06 | 3.900E-06 | 6.900E-06 |
| 3405 | 171 | H171C | A | 6.6e-06 | 5.8e-06 | 9.2e-06 | 7.200E-06 | 1.778E-06 | 5.800E-06 | 9.200E-06 |
| 3406 | 171 | H171E | A | 8.7e-06 | 3.4e-06 | 1.2e-05 | 8.033E-06 | 4.339E-06 | 3.400E-06 | 1.200E-05 |
| 3407 | 171 | H171Q | A | 7e-06 | 8.3e-06 | 9.7e-06 | 8.333E-06 | 1.350E-06 | 7.000E-06 | 9.700E-06 |
| 3408 | 171 | H171G | A | 5.5e-06 | 9.5e-06 | 9.7e-06 | 8.233E-06 | 2.369E-06 | 5.500E-06 | 9.700E-06 |
| 3409 | 171 | H171H | A | 1.3e-05 | 6.6e-06 | 9.2e-06 | 9.600E-06 | 3.219E-06 | 6.600E-06 | 1.300E-05 |
| 3410 | 171 | H171I | A | 6.8e-06 | 9.7e-06 | 6.7e-06 | 7.733E-06 | 1.704E-06 | 6.700E-06 | 9.700E-06 |
| 3411 | 171 | H171L | A | 8.4e-06 | 7.6e-06 | 9.4e-06 | 8.467E-06 | 9.018E-07 | 7.600E-06 | 9.400E-06 |
| 3412 | 171 | H171K | A | 8.4e-06 | 7e-06 | 7.3e-06 | 7.567E-06 | 7.371E-07 | 7.000E-06 | 8.400E-06 |
| 3413 | 171 | H171M | A | 9e-06 | 5.8e-06 | 6.7e-06 | 7.167E-06 | 1.650E-06 | 5.800E-06 | 9.000E-06 |
| 3414 | 171 | H171F | A | 8e-06 | 6.5e-06 | 1.3e-05 | 9.167E-06 | 3.403E-06 | 6.500E-06 | 1.300E-05 |
| 3415 | 171 | H171P | A | 8e-06 | 9.1e-06 | 6.7e-06 | 7.933E-06 | 1.201E-06 | 6.700E-06 | 9.100E-06 |
| 3416 | 171 | H171S | A | 8e-06 | 1e-05 | 6.6e-06 | 8.200E-06 | 1.709E-06 | 6.600E-06 | 1.000E-05 |
| 3417 | 171 | H171T | A | 8.2e-06 | 8.4e-06 | 1.3e-05 | 9.867E-06 | 2.715E-06 | 8.200E-06 | 1.300E-05 |
| 3418 | 171 | H171W | A | 8e-06 | 7.2e-06 | 4.6e-06 | 6.600E-06 | 1.778E-06 | 4.600E-06 | 8.000E-06 |
| 3419 | 171 | H171Y | A | 7.6e-06 | 1.2e-05 | 7.5e-06 | 9.033E-06 | 2.570E-06 | 7.500E-06 | 1.200E-05 |
| 3420 | 171 | H171V | A | 6.5e-06 | 1.3e-05 | 5.8e-06 | 8.433E-06 | 3.970E-06 | 5.800E-06 | 1.300E-05 |
| 3421 | 172 | P172A | A | 1.3e-05 | 1.5e-05 | 1.2e-05 | 1.333E-05 | 1.528E-06 | 1.200E-05 | 1.500E-05 |
| 3422 | 172 | P172R | A | 7.6e-06 | 6.5e-06 | 1.5e-05 | 9.700E-06 | 4.623E-06 | 6.500E-06 | 1.500E-05 |
| 3423 | 172 | P172N | A | 9.7e-06 | 6.7e-06 | 8.9e-06 | 8.433E-06 | 1.553E-06 | 6.700E-06 | 9.700E-06 |
| 3424 | 172 | P172D | A | 5.2e-06 | 4.3e-06 | 1e-05 | 6.500E-06 | 3.064E-06 | 4.300E-06 | 1.000E-05 |
| 3425 | 172 | P172C | A | 8e-06 | 3.5e-06 | 4.5e-06 | 5.333E-06 | 2.363E-06 | 3.500E-06 | 8.000E-06 |
| 3426 | 172 | P172E | A | 1.1e-05 | 8.6e-06 | 7.5e-06 | 9.033E-06 | 1.790E-06 | 7.500E-06 | 1.100E-05 |
| 3427 | 172 | P172Q | A | 8e-06 | 1.4e-05 | 5.3e-06 | 9.100E-06 | 4.453E-06 | 5.300E-06 | 1.400E-05 |
| 3428 | 172 | P172G | A | 1.1e-05 | 9.6e-06 | 6.8e-06 | 9.133E-06 | 2.139E-06 | 6.800E-06 | 1.100E-05 |
| 3429 | 172 | P172H | A | 6.2e-06 | 6.8e-06 | 1e-05 | 7.667E-06 | 2.043E-06 | 6.200E-06 | 1.000E-05 |
| 3430 | 172 | P172I | A | 4.8e-06 | 1.1e-05 | 9.7e-06 | 8.500E-06 | 3.270E-06 | 4.800E-06 | 1.100E-05 |
| 3431 | 172 | P172L | A | 3e-06 | 8.6e-06 | 7.7e-06 | 6.433E-06 | 3.007E-06 | 3.000E-06 | 8.600E-06 |
| 3432 | 172 | P172K | A | 5.8e-06 | 4.7e-06 | 7.5e-06 | 6.000E-06 | 1.411E-06 | 4.700E-06 | 7.500E-06 |
| 3433 | 172 | P172M | A | 9.7e-06 | 5.5e-06 | 8.9e-06 | 8.033E-06 | 2.230E-06 | 5.500E-06 | 9.700E-06 |
| 3434 | 172 | P172F | A | 9.9e-06 | 4.5e-06 | 9.4e-06 | 7.933E-06 | 2.984E-06 | 4.500E-06 | 9.900E-06 |
| 3435 | 172 | P172P | A | 9.1e-06 | 6.8e-06 | 6.8e-06 | 7.567E-06 | 1.328E-06 | 6.800E-06 | 9.100E-06 |
| 3436 | 172 | P172S | A | 7e-06 | 1.2e-05 | 1.1e-05 | 1.000E-05 | 2.646E-06 | 7.000E-06 | 1.200E-05 |
| 3437 | 172 | P172T | A | 7.7e-06 | 1.3e-05 | 6.4e-06 | 9.033E-06 | 3.496E-06 | 6.400E-06 | 1.300E-05 |
| 3438 | 172 | P172W | A | 7.7e-06 | 7e-06 | 1e-05 | 8.233E-06 | 1.570E-06 | 7.000E-06 | 1.000E-05 |
| 3439 | 172 | P172Y | A | 9.3e-06 | 7.2e-06 | 1.3e-05 | 9.833E-06 | 2.937E-06 | 7.200E-06 | 1.300E-05 |
| 3440 | 172 | P172V | A | 8.2e-06 | 5.7e-06 | 1.5e-05 | 9.633E-06 | 4.813E-06 | 5.700E-06 | 1.500E-05 |
| 3441 | 173 | C173A | A | 6.3e-06 | 8e-06 | 5.2e-06 | 6.500E-06 | 1.411E-06 | 5.200E-06 | 8.000E-06 |
| 3442 | 173 | C173R | A | 1.2e-05 | 7.1e-06 | 4.2e-06 | 7.767E-06 | 3.943E-06 | 4.200E-06 | 1.200E-05 |
| 3443 | 173 | C173N | A | 9.2e-06 | 1e-05 | 8.7e-06 | 9.300E-06 | 6.557E-07 | 8.700E-06 | 1.000E-05 |
| 3444 | 173 | C173D | A | 7e-06 | 9.5e-06 | 4.6e-06 | 7.033E-06 | 2.450E-06 | 4.600E-06 | 9.500E-06 |
| 3445 | 173 | C173C | A | 4.5e-06 | 9.2e-06 | 7.8e-06 | 7.167E-06 | 2.413E-06 | 4.500E-06 | 9.200E-06 |
| 3446 | 173 | C173E | A | 7.3e-06 | 9.3e-06 | 9.8e-06 | 8.800E-06 | 1.323E-06 | 7.300E-06 | 9.800E-06 |
| 3447 | 173 | C173Q | A | 9.5e-06 | 9.9e-06 | 4.7e-06 | 8.033E-06 | 2.894E-06 | 4.700E-06 | 9.900E-06 |
| 3448 | 173 | C173G | A | 9.8e-06 | 1.7e-05 | 8.7e-06 | 1.183E-05 | 4.508E-06 | 8.700E-06 | 1.700E-05 |
| 3449 | 173 | C173H | A | 8.3e-06 | 7.5e-06 | 1e-05 | 8.600E-06 | 1.277E-06 | 7.500E-06 | 1.000E-05 |
| 3450 | 173 | C173I | A | 1.2e-05 | 8.5e-06 | 7.1e-06 | 9.200E-06 | 2.524E-06 | 7.100E-06 | 1.200E-05 |
| 3451 | 173 | C173L | A | 8.1e-06 | 5.7e-06 | 1e-05 | 7.933E-06 | 2.155E-06 | 5.700E-06 | 1.000E-05 |
| 3452 | 173 | C173K | A | 9.6e-06 | 1.2e-05 | 9.6e-06 | 1.040E-05 | 1.386E-06 | 9.600E-06 | 1.200E-05 |
| 3453 | 173 | C173M | A | 9.9e-06 | 8.6e-06 | 9.8e-06 | 9.433E-06 | 7.234E-07 | 8.600E-06 | 9.900E-06 |
| 3454 | 173 | C173F | A | 7.5e-06 | 1.2e-05 | 9.6e-06 | 9.700E-06 | 2.252E-06 | 7.500E-06 | 1.200E-05 |
| 3455 | 173 | C173P | A | 5.2e-06 | 1e-05 | 1.4e-05 | 9.733E-06 | 4.406E-06 | 5.200E-06 | 1.400E-05 |
| 3456 | 173 | C173S | A | 8.8e-06 | 1.3e-05 | 5.1e-06 | 8.967E-06 | 3.953E-06 | 5.100E-06 | 1.300E-05 |
| 3457 | 173 | C173T | A | 1.3e-05 | 5.9e-06 | 5.7e-06 | 8.200E-06 | 4.158E-06 | 5.700E-06 | 1.300E-05 |
| 3458 | 173 | C173W | A | 7.2e-06 | 7.2e-06 | 6.8e-06 | 7.067E-06 | 2.309E-07 | 6.800E-06 | 7.200E-06 |
| 3459 | 173 | C173Y | A | 1.1e-05 | 6.9e-06 | 1.2e-05 | 9.967E-06 | 2.702E-06 | 6.900E-06 | 1.200E-05 |
| 3460 | 173 | C173V | A | 1e-05 | 9.7e-06 | 9.1e-06 | 9.600E-06 | 4.583E-07 | 9.100E-06 | 1.000E-05 |
| 3461 | 174 | S174A | A | 6.5e-06 | 6.8e-06 | 9.5e-06 | 7.600E-06 | 1.652E-06 | 6.500E-06 | 9.500E-06 |
| 3462 | 174 | S174R | A | 7.2e-06 | 1e-05 | 5.2e-06 | 7.467E-06 | 2.411E-06 | 5.200E-06 | 1.000E-05 |
| 3463 | 174 | S174N | A | 5.1e-06 | 9.7e-06 | 8.2e-06 | 7.667E-06 | 2.346E-06 | 5.100E-06 | 9.700E-06 |
| 3464 | 174 | S174D | A | 1.2e-05 | 6e-06 | 7e-06 | 8.333E-06 | 3.215E-06 | 6.000E-06 | 1.200E-05 |
| 3465 | 174 | S174C | A | 7e-06 | 6.4e-06 | 5.8e-06 | 6.400E-06 | 6.000E-07 | 5.800E-06 | 7.000E-06 |
| 3466 | 174 | S174E | A | 1.4e-05 | 6.6e-06 | 5.4e-06 | 8.667E-06 | 4.658E-06 | 5.400E-06 | 1.400E-05 |
| 3467 | 174 | S174Q | A | 1.2e-05 | 9.3e-06 | 5.1e-06 | 8.800E-06 | 3.477E-06 | 5.100E-06 | 1.200E-05 |
| 3468 | 174 | S174G | A | 3.7e-06 | 9.5e-06 | 1.1e-05 | 8.067E-06 | 3.855E-06 | 3.700E-06 | 1.100E-05 |
| 3469 | 174 | S174H | A | 5.8e-06 | 1.2e-05 | 9.3e-06 | 9.033E-06 | 3.109E-06 | 5.800E-06 | 1.200E-05 |
| 3470 | 174 | S174I | A | 8.7e-06 | 1.5e-05 | 7e-06 | 1.023E-05 | 4.215E-06 | 7.000E-06 | 1.500E-05 |
| 3471 | 174 | S174L | A | 5.5e-06 | 6.6e-06 | 6.6e-06 | 6.233E-06 | 6.351E-07 | 5.500E-06 | 6.600E-06 |
| 3472 | 174 | S174K | A | 6.8e-06 | 8.3e-06 | 8.6e-06 | 7.900E-06 | 9.644E-07 | 6.800E-06 | 8.600E-06 |
| 3473 | 174 | S174M | A | 6.8e-06 | 6.9e-06 | 7.7e-06 | 7.133E-06 | 4.933E-07 | 6.800E-06 | 7.700E-06 |
| 3474 | 174 | S174F | A | 7.8e-06 | 1e-05 | 1.4e-05 | 1.060E-05 | 3.143E-06 | 7.800E-06 | 1.400E-05 |
| 3475 | 174 | S174P | A | 1.4e-05 | 7.8e-06 | 5.9e-06 | 9.233E-06 | 4.236E-06 | 5.900E-06 | 1.400E-05 |
| 3476 | 174 | S174S | A | 6.9e-06 | 5.7e-06 | 9.3e-06 | 7.300E-06 | 1.833E-06 | 5.700E-06 | 9.300E-06 |
| 3477 | 174 | S174T | A | 5.4e-06 | 7.5e-06 | 1e-05 | 7.633E-06 | 2.303E-06 | 5.400E-06 | 1.000E-05 |
| 3478 | 174 | S174W | A | 9.1e-06 | 6.4e-06 | 4.5e-06 | 6.667E-06 | 2.312E-06 | 4.500E-06 | 9.100E-06 |
| 3479 | 174 | S174Y | A | 1.2e-05 | 8.3e-06 | 8.9e-06 | 9.733E-06 | 1.986E-06 | 8.300E-06 | 1.200E-05 |
| 3480 | 174 | S174V | A | 8.2e-06 | 9e-06 | 9.3e-06 | 8.833E-06 | 5.686E-07 | 8.200E-06 | 9.300E-06 |
| 3481 | 175 | P175A | A | 1.1e-05 | 2e-05 | 9.8e-06 | 1.360E-05 | 5.575E-06 | 9.800E-06 | 2.000E-05 |
| 3482 | 175 | P175R | A | 7.5e-06 | 9.1e-06 | 9.9e-06 | 8.833E-06 | 1.222E-06 | 7.500E-06 | 9.900E-06 |
| 3483 | 175 | P175N | A | 3.1e-06 | 7.9e-06 | 8.5e-06 | 6.500E-06 | 2.960E-06 | 3.100E-06 | 8.500E-06 |
| 3484 | 175 | P175D | A | 9.7e-06 | 8.4e-06 | 7.2e-06 | 8.433E-06 | 1.250E-06 | 7.200E-06 | 9.700E-06 |

| | | | | | | | | | | |
|---|---|---|---|---|---|---|---|---|---|---|
| 3485 | 175 | P175C | A | 1.2e-05 | 7.4e-06 | 9.5e-06 | 9.633E-06 | 2.303E-06 | 7.400E-06 | 1.200E-05 |
| 3486 | 175 | P175E | A | 7.1e-06 | 7.6e-06 | 8.1e-06 | 7.600E-06 | 5.000E-07 | 7.100E-06 | 8.100E-06 |
| 3487 | 175 | P175Q | A | 8.8e-06 | 1.1e-05 | 7.8e-06 | 9.200E-06 | 1.637E-06 | 7.800E-06 | 1.100E-05 |
| 3488 | 175 | P175G | A | 7.3e-06 | 1.2e-05 | 4.9e-06 | 8.067E-06 | 3.612E-06 | 4.900E-06 | 1.200E-05 |
| 3489 | 175 | P175H | A | 7.9e-06 | 6e-06 | 8.7e-06 | 7.533E-06 | 1.387E-06 | 6.000E-06 | 8.700E-06 |
| 3490 | 175 | P175I | A | 9.2e-06 | 9.7e-06 | 1e-05 | 9.633E-06 | 4.041E-07 | 9.200E-06 | 1.000E-05 |
| 3491 | 175 | P175L | A | 7.4e-06 | 4.4e-06 | 6.1e-06 | 5.967E-06 | 1.504E-06 | 4.400E-06 | 7.400E-06 |
| 3492 | 175 | P175K | A | 5.9e-06 | 9.9e-06 | 7.3e-06 | 7.700E-06 | 2.030E-06 | 5.900E-06 | 9.900E-06 |
| 3493 | 175 | P175M | A | 9.1e-06 | 4.7e-06 | 9.5e-06 | 7.767E-06 | 2.663E-06 | 4.700E-06 | 9.500E-06 |
| 3494 | 175 | P175F | A | 5.2e-06 | 6.6e-06 | 6.2e-06 | 6.000E-06 | 7.211E-07 | 5.200E-06 | 6.600E-06 |
| 3495 | 175 | P175P | A | 7.9e-06 | 3.9e-06 | 9.6e-06 | 7.133E-06 | 2.926E-06 | 3.900E-06 | 9.600E-06 |
| 3496 | 175 | P175S | A | 5.8e-06 | 7.8e-06 | 7.8e-06 | 7.133E-06 | 1.155E-06 | 5.800E-06 | 7.800E-06 |
| 3497 | 175 | P175T | A | 9.8e-06 | 6.4e-06 | 1.1e-05 | 9.067E-06 | 2.386E-06 | 6.400E-06 | 1.100E-05 |
| 3498 | 175 | P175W | A | 5.1e-06 | 1.3e-05 | 8.7e-06 | 8.933E-06 | 3.955E-06 | 5.100E-06 | 1.300E-05 |
| 3499 | 175 | P175Y | A | 6.1e-06 | 5.8e-06 | 9.5e-06 | 7.133E-06 | 2.055E-06 | 5.800E-06 | 9.500E-06 |
| 3500 | 175 | P175V | A | 5.2e-06 | 7.1e-06 | 8.5e-06 | 6.933E-06 | 1.656E-06 | 5.200E-06 | 8.500E-06 |
| 3501 | 176 | M176A | A | 1.2e-05 | 5.6e-06 | 4.6e-06 | 7.400E-06 | 4.015E-06 | 4.600E-06 | 1.200E-05 |
| 3502 | 176 | M176R | A | 6.4e-06 | 6.2e-06 | 9.4e-06 | 7.333E-06 | 1.793E-06 | 6.200E-06 | 9.400E-06 |
| 3503 | 176 | M176N | A | 7.5e-06 | 1.1e-05 | 7.3e-06 | 8.600E-06 | 2.081E-06 | 7.300E-06 | 1.100E-05 |
| 3504 | 176 | M176D | A | 1.4e-05 | 1.3e-05 | 8.4e-06 | 1.180E-05 | 2.987E-06 | 8.400E-06 | 1.400E-05 |
| 3505 | 176 | M176C | A | 1e-05 | 9.8e-06 | 1.1e-05 | 1.027E-05 | 6.429E-07 | 9.800E-06 | 1.100E-05 |
| 3506 | 176 | M176E | A | 7.7e-06 | 8.4e-06 | 9.9e-06 | 8.667E-06 | 1.124E-06 | 7.700E-06 | 9.900E-06 |
| 3507 | 176 | M176Q | A | 6.9e-06 | 5.9e-06 | 1.4e-05 | 8.933E-06 | 4.416E-06 | 5.900E-06 | 1.400E-05 |
| 3508 | 176 | M176G | A | 8.7e-06 | 7.9e-06 | 1.3e-05 | 9.867E-06 | 2.743E-06 | 7.900E-06 | 1.300E-05 |
| 3509 | 176 | M176H | A | 6.7e-06 | 9.1e-06 | 4e-06 | 6.600E-06 | 2.551E-06 | 4.000E-06 | 9.100E-06 |
| 3510 | 176 | M176I | A | 7.5e-06 | 5.8e-06 | 8.4e-06 | 7.233E-06 | 1.320E-06 | 5.800E-06 | 8.400E-06 |
| 3511 | 176 | M176L | A | 1.2e-05 | 9.7e-06 | 8.1e-06 | 9.933E-06 | 1.960E-06 | 8.100E-06 | 1.200E-05 |
| 3512 | 176 | M176K | A | 7.6e-06 | 6.1e-06 | 8.7e-06 | 7.467E-06 | 1.305E-06 | 6.100E-06 | 8.700E-06 |
| 3513 | 176 | M176M | A | 1.8e-05 | 7.9e-06 | 8.9e-06 | 1.160E-05 | 5.565E-06 | 7.900E-06 | 1.800E-05 |
| 3514 | 176 | M176F | A | 8.4e-06 | 5.8e-06 | 5e-06 | 6.400E-06 | 1.778E-06 | 5.000E-06 | 8.400E-06 |
| 3515 | 176 | M176P | A | 9.8e-06 | 1.2e-05 | 1.3e-05 | 1.160E-05 | 1.637E-06 | 9.800E-06 | 1.300E-05 |
| 3516 | 176 | M176S | A | 4.2e-06 | 9e-06 | 6.3e-06 | 6.500E-06 | 2.406E-06 | 4.200E-06 | 9.000E-06 |
| 3517 | 176 | M176T | A | 1.2e-05 | 8.7e-06 | 6.3e-06 | 9.000E-06 | 2.862E-06 | 6.300E-06 | 1.200E-05 |
| 3518 | 176 | M176W | A | 1.1e-05 | 6.2e-06 | 7.9e-06 | 8.367E-06 | 2.434E-06 | 6.200E-06 | 1.100E-05 |
| 3519 | 176 | M176Y | A | 9.5e-06 | 5e-06 | 4.3e-06 | 6.267E-06 | 2.822E-06 | 4.300E-06 | 9.500E-06 |
| 3520 | 176 | M176V | A | 8.9e-06 | 5.3e-06 | 4.9e-06 | 6.367E-06 | 2.203E-06 | 4.900E-06 | 8.900E-06 |
| 3521 | 177 | C177A | A | 7.7e-06 | 6.8e-06 | 1.3e-05 | 9.167E-06 | 3.350E-06 | 6.800E-06 | 1.300E-05 |
| 3522 | 177 | C177R | A | 7.8e-06 | 6.8e-06 | 1.3e-05 | 9.200E-06 | 3.329E-06 | 6.800E-06 | 1.300E-05 |
| 3523 | 177 | C177N | A | 6.2e-06 | 1e-05 | 4.9e-06 | 7.033E-06 | 2.650E-06 | 4.900E-06 | 1.000E-05 |
| 3524 | 177 | C177D | A | 7.8e-06 | 1.5e-05 | 1.1e-05 | 1.127E-05 | 3.607E-06 | 7.800E-06 | 1.500E-05 |
| 3525 | 177 | C177C | A | 5.6e-06 | 4.4e-06 | 7.2e-06 | 5.733E-06 | 1.405E-06 | 4.400E-06 | 7.200E-06 |
| 3526 | 177 | C177E | A | 1e-05 | 9.1e-06 | 6.5e-06 | 8.533E-06 | 1.818E-06 | 6.500E-06 | 1.000E-05 |
| 3527 | 177 | C177Q | A | 6.8e-06 | 7.3e-06 | 9.9e-06 | 8.000E-06 | 1.664E-06 | 6.800E-06 | 9.900E-06 |
| 3528 | 177 | C177G | A | 7.4e-06 | 1e-05 | 9e-06 | 8.800E-06 | 1.311E-06 | 7.400E-06 | 1.000E-05 |
| 3529 | 177 | C177H | A | 1.4e-05 | 9.2e-06 | 7.1e-06 | 1.010E-05 | 3.537E-06 | 7.100E-06 | 1.400E-05 |
| 3530 | 177 | C177I | A | 5.7e-06 | 6.7e-06 | 7.3e-06 | 6.567E-06 | 8.083E-07 | 5.700E-06 | 7.300E-06 |
| 3531 | 177 | C177L | A | 5e-06 | 1.2e-05 | 1.3e-05 | 1.000E-05 | 4.359E-06 | 5.000E-06 | 1.300E-05 |
| 3532 | 177 | C177K | A | 9.6e-06 | 8.1e-06 | 5.9e-06 | 7.867E-06 | 1.861E-06 | 5.900E-06 | 9.600E-06 |
| 3533 | 177 | C177M | A | 1.2e-05 | 8.5e-06 | 5.1e-06 | 8.533E-06 | 3.450E-06 | 5.100E-06 | 1.200E-05 |
| 3534 | 177 | C177F | A | 1.4e-05 | 6e-06 | 1e-05 | 1.000E-05 | 4.000E-06 | 6.000E-06 | 1.400E-05 |
| 3535 | 177 | C177P | A | 8e-06 | 7.4e-06 | 9.6e-06 | 8.333E-06 | 1.137E-06 | 7.400E-06 | 9.600E-06 |
| 3536 | 177 | C177S | A | 1e-05 | 7.1e-06 | 1.3e-05 | 1.003E-05 | 2.950E-06 | 7.100E-06 | 1.300E-05 |
| 3537 | 177 | C177T | A | 6.9e-06 | 8.1e-06 | 4.3e-06 | 6.433E-06 | 1.943E-06 | 4.300E-06 | 8.100E-06 |
| 3538 | 177 | C177W | A | 1e-05 | 7.2e-06 | 9.3e-06 | 8.833E-06 | 1.457E-06 | 7.200E-06 | 1.000E-05 |
| 3539 | 177 | C177Y | A | 1.1e-05 | 1e-05 | 6.3e-06 | 9.100E-06 | 2.476E-06 | 6.300E-06 | 1.100E-05 |
| 3540 | 177 | C177V | A | 5.9e-06 | 5.6e-06 | 1.1e-05 | 7.500E-06 | 3.035E-06 | 5.600E-06 | 1.100E-05 |
| 3541 | 178 | K178A | A | 1.1e-05 | 4.9e-06 | 4.6e-06 | 6.833E-06 | 3.612E-06 | 4.600E-06 | 1.100E-05 |
| 3542 | 178 | K178R | A | 8.3e-06 | 5.8e-06 | 6.6e-06 | 6.900E-06 | 1.277E-06 | 5.800E-06 | 8.300E-06 |
| 3543 | 178 | K178N | A | 5.7e-06 | 1.3e-05 | 5.1e-06 | 7.933E-06 | 4.398E-06 | 5.100E-06 | 1.300E-05 |
| 3544 | 178 | K178D | A | 6.8e-06 | 6.4e-06 | 1.4e-05 | 9.067E-06 | 4.277E-06 | 6.400E-06 | 1.400E-05 |
| 3545 | 178 | K178C | A | 5.3e-06 | 6.7e-06 | 7.9e-06 | 6.633E-06 | 1.301E-06 | 5.300E-06 | 7.900E-06 |
| 3546 | 178 | K178E | A | 9e-06 | 6.2e-06 | 5.9e-06 | 7.033E-06 | 1.710E-06 | 5.900E-06 | 9.000E-06 |
| 3547 | 178 | K178Q | A | 8.2e-06 | 3.9e-06 | 6.6e-06 | 6.233E-06 | 2.173E-06 | 3.900E-06 | 8.200E-06 |
| 3548 | 178 | K178G | A | 1.1e-05 | 6.7e-06 | 9.4e-06 | 9.033E-06 | 2.173E-06 | 6.700E-06 | 1.100E-05 |
| 3549 | 178 | K178H | A | 8.3e-06 | 6.8e-06 | 1.3e-05 | 9.367E-06 | 3.235E-06 | 6.800E-06 | 1.300E-05 |
| 3550 | 178 | K178I | A | 1.3e-05 | 8e-06 | 1.1e-05 | 1.067E-05 | 2.517E-06 | 8.000E-06 | 1.300E-05 |
| 3551 | 178 | K178L | A | 1.7e-05 | 9e-06 | 5.8e-06 | 1.060E-05 | 5.769E-06 | 5.800E-06 | 1.700E-05 |
| 3552 | 178 | K178K | A | 4.7e-06 | 8.8e-06 | 6.5e-06 | 6.667E-06 | 2.055E-06 | 4.700E-06 | 8.800E-06 |
| 3553 | 178 | K178M | A | 4.3e-06 | 6.4e-06 | 1.2e-05 | 7.567E-06 | 3.980E-06 | 4.300E-06 | 1.200E-05 |
| 3554 | 178 | K178F | A | 7.8e-06 | 1.2e-05 | 8.6e-06 | 9.467E-06 | 2.230E-06 | 7.800E-06 | 1.200E-05 |
| 3555 | 178 | K178P | A | 3.9e-06 | 5.7e-06 | 7.2e-06 | 5.600E-06 | 1.652E-06 | 3.900E-06 | 7.200E-06 |
| 3556 | 178 | K178S | A | 6.7e-06 | 9.6e-06 | 1e-05 | 8.767E-06 | 1.801E-06 | 6.700E-06 | 1.000E-05 |
| 3557 | 178 | K178T | A | 7.7e-06 | 8.5e-06 | 4.1e-06 | 6.767E-06 | 2.344E-06 | 4.100E-06 | 8.500E-06 |
| 3558 | 178 | K178W | A | 8.9e-06 | 6.8e-06 | 6.4e-06 | 7.367E-06 | 1.343E-06 | 6.400E-06 | 8.900E-06 |
| 3559 | 178 | K178Y | A | 9.9e-06 | 6.8e-06 | 8.5e-06 | 8.400E-06 | 1.552E-06 | 6.800E-06 | 9.900E-06 |
| 3560 | 178 | K178V | A | 1.3e-05 | 7e-06 | 9.1e-06 | 9.700E-06 | 3.045E-06 | 7.000E-06 | 1.300E-05 |
| 3561 | 179 | G179A | A | 4.8e-06 | 1.3e-05 | 9.4e-06 | 9.067E-06 | 4.110E-06 | 4.800E-06 | 1.300E-05 |
| 3562 | 179 | G179R | A | 8.8e-06 | 6.1e-06 | 7.1e-06 | 7.333E-06 | 1.365E-06 | 6.100E-06 | 8.800E-06 |
| 3563 | 179 | G179N | A | 4.6e-06 | 8.5e-06 | 8.2e-06 | 7.100E-06 | 2.170E-06 | 4.600E-06 | 8.500E-06 |
| 3564 | 179 | G179D | A | 7.3e-06 | 6.6e-06 | 9.9e-06 | 7.933E-06 | 1.739E-06 | 6.600E-06 | 9.900E-06 |
| 3565 | 179 | G179C | A | 4e-06 | 4.3e-06 | 6.9e-06 | 5.067E-06 | 1.595E-06 | 4.000E-06 | 6.900E-06 |
| 3566 | 179 | G179E | A | 6.5e-06 | 9.9e-06 | 9.7e-06 | 8.700E-06 | 1.908E-06 | 6.500E-06 | 9.900E-06 |
| 3567 | 179 | G179Q | A | 6.7e-06 | 5.1e-06 | 8.9e-06 | 6.900E-06 | 1.908E-06 | 5.100E-06 | 8.900E-06 |
| 3568 | 179 | G179G | A | 6.6e-06 | 4.5e-06 | 7.7e-06 | 6.267E-06 | 1.626E-06 | 4.500E-06 | 7.700E-06 |
| 3569 | 179 | G179H | A | 6.4e-06 | 6.3e-06 | 1.4e-05 | 8.900E-06 | 4.417E-06 | 6.300E-06 | 1.400E-05 |

| | | | | | | | | | | |
|---|---|---|---|---|---|---|---|---|---|---|
| 3570 | 179 | G179I | A | 6.5e-06 | 7e-06 | 8e-06 | 7.167E-06 | 7.638E-07 | 6.500E-06 | 8.000E-06 |
| 3571 | 179 | G179L | A | 6.4e-06 | 6.4e-06 | 9.5e-06 | 7.433E-06 | 1.790E-06 | 6.400E-06 | 9.500E-06 |
| 3572 | 179 | G179K | A | 7.5e-06 | 8.2e-06 | 5.3e-06 | 7.000E-06 | 1.513E-06 | 5.300E-06 | 8.200E-06 |
| 3573 | 179 | G179M | A | 6.5e-06 | 8.7e-06 | 8.5e-06 | 7.900E-06 | 1.217E-06 | 6.500E-06 | 8.700E-06 |
| 3574 | 179 | G179F | A | 1.7e-05 | 6.4e-06 | 1.3e-05 | 1.213E-05 | 5.353E-06 | 6.400E-06 | 1.700E-05 |
| 3575 | 179 | G179P | A | 8.3e-06 | 9.2e-06 | 1.1e-05 | 9.500E-06 | 1.375E-06 | 8.300E-06 | 1.100E-05 |
| 3576 | 179 | G179S | A | 5.8e-06 | 8.3e-06 | 1.2e-05 | 8.700E-06 | 3.119E-06 | 5.800E-06 | 1.200E-05 |
| 3577 | 179 | G179T | A | 7.1e-06 | 9e-06 | 7.4e-06 | 7.833E-06 | 1.021E-06 | 7.100E-06 | 9.000E-06 |
| 3578 | 179 | G179W | A | 1.3e-05 | 1.1e-05 | 6.4e-06 | 1.013E-05 | 3.384E-06 | 6.400E-06 | 1.300E-05 |
| 3579 | 179 | G179Y | A | 7.3e-06 | 7.2e-06 | 6.7e-06 | 7.067E-06 | 3.215E-07 | 6.700E-06 | 7.300E-06 |
| 3580 | 179 | G179V | A | 6.4e-06 | 1.2e-05 | 6.4e-06 | 8.267E-06 | 3.233E-06 | 6.400E-06 | 1.200E-05 |
| 3581 | 180 | S180A | A | 5.2e-06 | 6.8e-06 | 5.7e-06 | 5.900E-06 | 8.185E-07 | 5.200E-06 | 6.800E-06 |
| 3582 | 180 | S180R | A | 8.9e-06 | 9.1e-06 | 9.7e-06 | 9.233E-06 | 4.163E-07 | 8.900E-06 | 9.700E-06 |
| 3583 | 180 | S180N | A | 9.3e-06 | 9.5e-06 | 5.1e-06 | 7.967E-06 | 2.485E-06 | 5.100E-06 | 9.500E-06 |
| 3584 | 180 | S180D | A | 4.7e-06 | 6.9e-06 | 1e-05 | 7.200E-06 | 2.663E-06 | 4.700E-06 | 1.000E-05 |
| 3585 | 180 | S180C | A | 6.8e-06 | 4.4e-06 | 6.2e-06 | 5.800E-06 | 1.249E-06 | 4.400E-06 | 6.800E-06 |
| 3586 | 180 | S180E | A | 6.3e-06 | 6.6e-06 | 8.1e-06 | 7.000E-06 | 9.644E-07 | 6.300E-06 | 8.100E-06 |
| 3587 | 180 | S180Q | A | 8.4e-06 | 7.9e-06 | 4.7e-06 | 7.000E-06 | 2.007E-06 | 4.700E-06 | 8.400E-06 |
| 3588 | 180 | S180G | A | 6.8e-06 | 9e-06 | 1e-05 | 8.600E-06 | 1.637E-06 | 6.800E-06 | 1.000E-05 |
| 3589 | 180 | S180H | A | 1e-05 | 1.1e-05 | 1.2e-05 | 1.100E-05 | 1.000E-06 | 1.000E-05 | 1.200E-05 |
| 3590 | 180 | S180I | A | 7.1e-06 | 6.9e-06 | 6.6e-06 | 6.867E-06 | 2.517E-07 | 6.600E-06 | 7.100E-06 |
| 3591 | 180 | S180L | A | 1.5e-05 | 7.2e-06 | 8.7e-06 | 1.030E-05 | 4.139E-06 | 7.200E-06 | 1.500E-05 |
| 3592 | 180 | S180K | A | 1e-05 | 9.4e-06 | 7.4e-06 | 8.933E-06 | 1.361E-06 | 7.400E-06 | 1.000E-05 |
| 3593 | 180 | S180M | A | 1.1e-05 | 1e-05 | 1.2e-05 | 1.100E-05 | 1.000E-06 | 1.000E-05 | 1.200E-05 |
| 3594 | 180 | S180F | A | 1.2e-05 | 6.5e-06 | 9.7e-06 | 9.400E-06 | 2.762E-06 | 6.500E-06 | 1.200E-05 |
| 3595 | 180 | S180P | A | 9.1e-06 | 9.5e-06 | 5.2e-06 | 7.933E-06 | 2.376E-06 | 5.200E-06 | 9.500E-06 |
| 3596 | 180 | S180S | A | 6.7e-06 | 9.6e-06 | 1.4e-05 | 1.010E-05 | 3.676E-06 | 6.700E-06 | 1.400E-05 |
| 3597 | 180 | S180T | A | 1.4e-05 | 6e-06 | 9.1e-06 | 9.700E-06 | 4.034E-06 | 6.000E-06 | 1.400E-05 |
| 3598 | 180 | S180W | A | 9.3e-06 | 9.3e-06 | 1e-05 | 9.533E-06 | 4.041E-07 | 9.300E-06 | 1.000E-05 |
| 3599 | 180 | S180Y | A | 1.7e-05 | 9.9e-06 | 8.1e-06 | 1.167E-05 | 4.706E-06 | 8.100E-06 | 1.700E-05 |
| 3600 | 180 | S180V | A | 6.1e-06 | 1e-05 | 1e-05 | 8.700E-06 | 2.252E-06 | 6.100E-06 | 1.000E-05 |
| 3601 | 181 | R181A | A | 1e-05 | 7e-06 | 8.1e-06 | 8.367E-06 | 1.518E-06 | 7.000E-06 | 1.000E-05 |
| 3602 | 181 | R181R | A | 1.4e-05 | 9.1e-06 | 1e-05 | 1.103E-05 | 2.608E-06 | 9.100E-06 | 1.400E-05 |
| 3603 | 181 | R181N | A | 7.7e-06 | 8e-06 | 1.3e-05 | 9.567E-06 | 2.977E-06 | 7.700E-06 | 1.300E-05 |
| 3604 | 181 | R181D | A | 5.8e-06 | 1e-05 | 7.2e-06 | 7.667E-06 | 2.139E-06 | 5.800E-06 | 1.000E-05 |
| 3605 | 181 | R181C | A | 3.8e-06 | 6.7e-06 | 4.8e-06 | 5.100E-06 | 1.473E-06 | 3.800E-06 | 6.700E-06 |
| 3606 | 181 | R181E | A | 5.1e-06 | 3.3e-06 | 6.3e-06 | 4.900E-06 | 1.510E-06 | 3.300E-06 | 6.300E-06 |
| 3607 | 181 | R181Q | A | 1.2e-05 | 1.1e-05 | 5.5e-06 | 9.500E-06 | 3.500E-06 | 5.500E-06 | 1.200E-05 |
| 3608 | 181 | R181G | A | 1.2e-05 | 1.9e-05 | 9.1e-06 | 1.337E-05 | 5.090E-06 | 9.100E-06 | 1.900E-05 |
| 3609 | 181 | R181H | A | 8.7e-06 | 4.5e-06 | 3e-06 | 5.400E-06 | 2.955E-06 | 3.000E-06 | 8.700E-06 |
| 3610 | 181 | R181I | A | 6.8e-06 | 6.9e-06 | 9.5e-06 | 7.733E-06 | 1.531E-06 | 6.800E-06 | 9.500E-06 |
| 3611 | 181 | R181L | A | 9.8e-06 | 5.9e-06 | 8.7e-06 | 8.133E-06 | 2.011E-06 | 5.900E-06 | 9.800E-06 |
| 3612 | 181 | R181K | A | 5.3e-06 | 7.6e-06 | 9.7e-06 | 7.533E-06 | 2.201E-06 | 5.300E-06 | 9.700E-06 |
| 3613 | 181 | R181M | A | 7.8e-06 | 8.2e-06 | 7.3e-06 | 7.767E-06 | 4.509E-07 | 7.300E-06 | 8.200E-06 |
| 3614 | 181 | R181F | A | 1.1e-05 | 9e-06 | 9.4e-06 | 9.800E-06 | 1.058E-06 | 9.000E-06 | 1.100E-05 |
| 3615 | 181 | R181P | A | 6.2e-06 | 4.8e-06 | 7.7e-06 | 6.233E-06 | 1.450E-06 | 4.800E-06 | 7.700E-06 |
| 3616 | 181 | R181S | A | 1.2e-05 | 1.3e-05 | 4.5e-06 | 9.833E-06 | 4.646E-06 | 4.500E-06 | 1.300E-05 |
| 3617 | 181 | R181T | A | 8.4e-06 | 6.3e-06 | 4.6e-06 | 6.433E-06 | 1.904E-06 | 4.600E-06 | 8.400E-06 |
| 3618 | 181 | R181W | A | 6.5e-06 | 9.2e-06 | 7.8e-06 | 7.833E-06 | 1.350E-06 | 6.500E-06 | 9.200E-06 |
| 3619 | 181 | R181Y | A | 1.3e-05 | 5.5e-06 | 8.3e-06 | 8.933E-06 | 3.790E-06 | 5.500E-06 | 1.300E-05 |
| 3620 | 181 | R181V | A | 8.3e-06 | 1.4e-05 | 1.4e-05 | 1.210E-05 | 3.291E-06 | 8.300E-06 | 1.400E-05 |
| 3621 | 182 | C182A | A | 1e-05 | 1e-05 | 4.8e-06 | 8.267E-06 | 3.002E-06 | 4.800E-06 | 1.000E-05 |
| 3622 | 182 | C182R | A | 8.4e-06 | 8.7e-06 | 1e-05 | 9.033E-06 | 8.505E-07 | 8.400E-06 | 1.000E-05 |
| 3623 | 182 | C182N | A | 7e-06 | 4.6e-06 | 7.8e-06 | 6.467E-06 | 1.665E-06 | 4.600E-06 | 7.800E-06 |
| 3624 | 182 | C182D | A | 4.9e-06 | 9e-06 | 4.6e-06 | 6.167E-06 | 2.458E-06 | 4.600E-06 | 9.000E-06 |
| 3625 | 182 | C182C | A | 7.6e-06 | 8.6e-06 | 1.2e-05 | 9.400E-06 | 2.307E-06 | 7.600E-06 | 1.200E-05 |
| 3626 | 182 | C182E | A | 1.3e-05 | 7.4e-06 | 8.3e-06 | 9.567E-06 | 3.007E-06 | 7.400E-06 | 1.300E-05 |
| 3627 | 182 | C182Q | A | 7.7e-06 | 5.8e-06 | 1.2e-05 | 8.500E-06 | 3.176E-06 | 5.800E-06 | 1.200E-05 |
| 3628 | 182 | C182G | A | 1.3e-05 | 4.7e-06 | 4.8e-06 | 7.500E-06 | 4.763E-06 | 4.700E-06 | 1.300E-05 |
| 3629 | 182 | C182H | A | 9.3e-06 | 1.3e-05 | 8.3e-06 | 1.020E-05 | 2.476E-06 | 8.300E-06 | 1.300E-05 |
| 3630 | 182 | C182I | A | 7.4e-06 | 5.5e-06 | 9.6e-06 | 7.500E-06 | 2.052E-06 | 5.500E-06 | 9.600E-06 |
| 3631 | 182 | C182L | A | 9.3e-06 | 9.4e-06 | 9.6e-06 | 9.433E-06 | 1.528E-07 | 9.300E-06 | 9.600E-06 |
| 3632 | 182 | C182K | A | 1.3e-05 | 7.5e-06 | 1.4e-05 | 1.150E-05 | 3.500E-06 | 7.500E-06 | 1.400E-05 |
| 3633 | 182 | C182M | A | 1.3e-05 | 6.7e-06 | 1.3e-05 | 1.090E-05 | 3.637E-06 | 6.700E-06 | 1.300E-05 |
| 3634 | 182 | C182F | A | 9.5e-06 | 6.8e-06 | 5.5e-06 | 7.267E-06 | 2.040E-06 | 5.500E-06 | 9.500E-06 |
| 3635 | 182 | C182P | A | 2e-05 | 5.4e-06 | 9.1e-06 | 1.150E-05 | 7.590E-06 | 5.400E-06 | 2.000E-05 |
| 3636 | 182 | C182S | A | 1e-05 | 8.4e-06 | 1.3e-05 | 1.047E-05 | 2.335E-06 | 8.400E-06 | 1.300E-05 |
| 3637 | 182 | C182T | A | 7.6e-06 | 4.8e-06 | 5.7e-06 | 6.033E-06 | 1.429E-06 | 4.800E-06 | 7.600E-06 |
| 3638 | 182 | C182W | A | 5.4e-06 | 4.6e-06 | 8.7e-06 | 6.233E-06 | 2.173E-06 | 4.600E-06 | 8.700E-06 |
| 3639 | 182 | C182Y | A | 9.9e-06 | 8.8e-06 | 9.7e-06 | 9.467E-06 | 5.859E-07 | 8.800E-06 | 9.900E-06 |
| 3640 | 182 | C182V | A | 4.5e-06 | 7e-06 | 4e-06 | 5.167E-06 | 1.607E-06 | 4.000E-06 | 7.000E-06 |
| 3641 | 183 | W183A | A | 9.5e-06 | 9.7e-06 | 7.5e-06 | 8.900E-06 | 1.217E-06 | 7.500E-06 | 9.700E-06 |
| 3642 | 183 | W183R | A | 9.3e-06 | 9.4e-06 | 7.1e-06 | 8.600E-06 | 1.300E-06 | 7.100E-06 | 9.400E-06 |
| 3643 | 183 | W183N | A | 9.4e-06 | 6.7e-06 | 9.2e-06 | 8.433E-06 | 1.504E-06 | 6.700E-06 | 9.400E-06 |
| 3644 | 183 | W183D | A | 7.9e-06 | 6.6e-06 | 9.9e-06 | 8.133E-06 | 1.662E-06 | 6.600E-06 | 9.900E-06 |
| 3645 | 183 | W183C | A | 1.1e-05 | 5.1e-06 | 8.5e-06 | 8.200E-06 | 2.961E-06 | 5.100E-06 | 1.100E-05 |
| 3646 | 183 | W183E | A | 1.2e-05 | 1e-05 | 9.9e-06 | 1.063E-05 | 1.185E-06 | 9.900E-06 | 1.200E-05 |
| 3647 | 183 | W183Q | A | 8.6e-06 | 7e-06 | 7.7e-06 | 7.767E-06 | 8.021E-07 | 7.000E-06 | 8.600E-06 |
| 3648 | 183 | W183G | A | 7.2e-06 | 9e-06 | 7.9e-06 | 8.033E-06 | 9.074E-07 | 7.200E-06 | 9.000E-06 |
| 3649 | 183 | W183H | A | 8.6e-06 | 1.1e-05 | 6.6e-06 | 8.733E-06 | 2.203E-06 | 6.600E-06 | 1.100E-05 |
| 3650 | 183 | W183I | A | 7e-06 | 8.3e-06 | 4.9e-06 | 6.733E-06 | 1.716E-06 | 4.900E-06 | 8.300E-06 |
| 3651 | 183 | W183L | A | 8.7e-06 | 9.6e-06 | 4.6e-06 | 7.633E-06 | 2.665E-06 | 4.600E-06 | 9.600E-06 |
| 3652 | 183 | W183K | A | 8.9e-06 | 6.2e-06 | 8.3e-06 | 7.800E-06 | 1.418E-06 | 6.200E-06 | 8.900E-06 |
| 3653 | 183 | W183M | A | 5.1e-06 | 5.6e-06 | 6.7e-06 | 5.800E-06 | 8.185E-07 | 5.100E-06 | 6.700E-06 |
| 3654 | 183 | W183F | A | 5.9e-06 | 1.2e-05 | 1.6e-05 | 1.130E-05 | 5.086E-06 | 5.900E-06 | 1.600E-05 |

| | | | | | | | | | | |
|---|---|---|---|---|---|---|---|---|---|---|
| 3655 | 183 | W183P | A | 6.8e-06 | 9.5e-06 | 3.9e-06 | 6.733E-06 | 2.801E-06 | 3.900E-06 | 9.500E-06 |
| 3656 | 183 | W183S | A | 8.9e-06 | 6.8e-06 | 7.2e-06 | 7.633E-06 | 1.115E-06 | 6.800E-06 | 8.900E-06 |
| 3657 | 183 | W183T | A | 7.1e-06 | 4.6e-06 | 1.1e-05 | 7.567E-06 | 3.225E-06 | 4.600E-06 | 1.100E-05 |
| 3658 | 183 | W183W | A | 5e-06 | 5.2e-06 | 5e-06 | 5.067E-06 | 1.155E-07 | 5.000E-06 | 5.200E-06 |
| 3659 | 183 | W183Y | A | 5.5e-06 | 1.3e-05 | 9e-06 | 9.167E-06 | 3.753E-06 | 5.500E-06 | 1.300E-05 |
| 3660 | 183 | W183V | A | 8.2e-06 | 7.2e-06 | 1.4e-05 | 9.800E-06 | 3.672E-06 | 7.200E-06 | 1.400E-05 |
| 3661 | 184 | G184A | A | 5.2e-06 | 1.1e-05 | 6.2e-06 | 7.467E-06 | 3.101E-06 | 5.200E-06 | 1.100E-05 |
| 3662 | 184 | G184R | A | 1.3e-05 | 5.7e-06 | 9.1e-06 | 9.267E-06 | 3.653E-06 | 5.700E-06 | 1.300E-05 |
| 3663 | 184 | G184N | A | 9.9e-06 | 1.5e-05 | 8.8e-06 | 1.123E-05 | 3.308E-06 | 8.800E-06 | 1.500E-05 |
| 3664 | 184 | G184D | A | 1.2e-05 | 8.6e-06 | 1.3e-05 | 1.120E-05 | 2.307E-06 | 8.600E-06 | 1.300E-05 |
| 3665 | 184 | G184C | A | 1.5e-05 | 9.2e-06 | 9e-06 | 1.107E-05 | 3.408E-06 | 9.000E-06 | 1.500E-05 |
| 3666 | 184 | G184E | A | 9.2e-06 | 6.4e-06 | 1.1e-05 | 8.867E-06 | 2.318E-06 | 6.400E-06 | 1.100E-05 |
| 3667 | 184 | G184Q | A | 7.8e-06 | 8.8e-06 | 6.2e-06 | 7.600E-06 | 1.311E-06 | 6.200E-06 | 8.800E-06 |
| 3668 | 184 | G184G | A | 1.3e-05 | 1.1e-05 | 1.2e-05 | 1.200E-05 | 1.000E-06 | 1.100E-05 | 1.300E-05 |
| 3669 | 184 | G184H | A | 1.1e-05 | 8.7e-06 | 1.7e-05 | 1.223E-05 | 4.285E-06 | 8.700E-06 | 1.700E-05 |
| 3670 | 184 | G184I | A | 1.4e-05 | 8.1e-06 | 9.1e-06 | 1.040E-05 | 3.158E-06 | 8.100E-06 | 1.400E-05 |
| 3671 | 184 | G184L | A | 7.5e-06 | 6.3e-06 | 9.9e-06 | 7.900E-06 | 1.833E-06 | 6.300E-06 | 9.900E-06 |
| 3672 | 184 | G184K | A | 4.8e-06 | 1.2e-05 | 1e-05 | 8.933E-06 | 3.717E-06 | 4.800E-06 | 1.200E-05 |
| 3673 | 184 | G184M | A | 8.3e-06 | 6.5e-06 | 1.3e-05 | 9.267E-06 | 3.356E-06 | 6.500E-06 | 1.300E-05 |
| 3674 | 184 | G184F | A | 8.4e-06 | 7.8e-06 | 1.4e-05 | 1.007E-05 | 3.420E-06 | 7.800E-06 | 1.400E-05 |
| 3675 | 184 | G184P | A | 3.2e-06 | 6.8e-06 | 1.3e-05 | 7.667E-06 | 4.957E-06 | 3.200E-06 | 1.300E-05 |
| 3676 | 184 | G184S | A | 8.9e-06 | 6.4e-06 | 1.5e-05 | 1.010E-05 | 4.424E-06 | 6.400E-06 | 1.500E-05 |
| 3677 | 184 | G184T | A | 6e-06 | 1.3e-05 | 7e-06 | 8.667E-06 | 3.786E-06 | 6.000E-06 | 1.300E-05 |
| 3678 | 184 | G184W | A | 6.1e-06 | 5.7e-06 | 1.4e-05 | 8.600E-06 | 4.681E-06 | 5.700E-06 | 1.400E-05 |
| 3679 | 184 | G184Y | A | 5.6e-06 | 5.6e-06 | 8.8e-06 | 6.667E-06 | 1.848E-06 | 5.600E-06 | 8.800E-06 |
| 3680 | 184 | G184V | A | 5.6e-06 | 8.1e-06 | 5.6e-06 | 6.433E-06 | 1.443E-06 | 5.600E-06 | 8.100E-06 |
| 3681 | 185 | E185A | A | 1.9e-05 | 8.5e-06 | 9.1e-06 | 1.220E-05 | 5.897E-06 | 8.500E-06 | 1.900E-05 |
| 3682 | 185 | E185R | A | 9e-06 | 5.6e-06 | 8.7e-06 | 7.767E-06 | 1.882E-06 | 5.600E-06 | 9.000E-06 |
| 3683 | 185 | E185N | A | 5.5e-06 | 5.6e-06 | 9.2e-06 | 6.767E-06 | 2.108E-06 | 5.500E-06 | 9.200E-06 |
| 3684 | 185 | E185D | A | 9.5e-06 | 4.1e-06 | 7.9e-06 | 7.167E-06 | 2.774E-06 | 4.100E-06 | 9.500E-06 |
| 3685 | 185 | E185C | A | 1.1e-05 | 6.1e-06 | 5.3e-06 | 7.467E-06 | 3.086E-06 | 5.300E-06 | 1.100E-05 |
| 3686 | 185 | E185E | A | 9.4e-06 | 9.5e-06 | 8.2e-06 | 9.033E-06 | 7.234E-07 | 8.200E-06 | 9.500E-06 |
| 3687 | 185 | E185Q | A | 9e-06 | 8.9e-06 | 1.1e-05 | 9.633E-06 | 1.185E-06 | 8.900E-06 | 1.100E-05 |
| 3688 | 185 | E185G | A | 6.7e-06 | 1.1e-05 | 2.1e-05 | 1.290E-05 | 7.337E-06 | 6.700E-06 | 2.100E-05 |
| 3689 | 185 | E185H | A | 5.7e-06 | 6.2e-06 | 5.7e-06 | 5.867E-06 | 2.887E-07 | 5.700E-06 | 6.200E-06 |
| 3690 | 185 | E185I | A | 1.2e-05 | 5.2e-06 | 1.4e-05 | 1.040E-05 | 4.613E-06 | 5.200E-06 | 1.400E-05 |
| 3691 | 185 | E185L | A | 6.2e-06 | 9.2e-06 | 7.7e-06 | 7.700E-06 | 1.500E-06 | 6.200E-06 | 9.200E-06 |
| 3692 | 185 | E185K | A | 1e-05 | 7.8e-06 | 9e-06 | 8.933E-06 | 1.102E-06 | 7.800E-06 | 1.000E-05 |
| 3693 | 185 | E185M | A | 9.4e-06 | 4.6e-06 | 8.2e-06 | 7.400E-06 | 2.498E-06 | 4.600E-06 | 9.400E-06 |
| 3694 | 185 | E185F | A | 1e-05 | 1.3e-05 | 1.4e-05 | 1.233E-05 | 2.082E-06 | 1.000E-05 | 1.400E-05 |
| 3695 | 185 | E185P | A | 8.4e-06 | 1.2e-05 | 8e-06 | 9.467E-06 | 2.203E-06 | 8.000E-06 | 1.200E-05 |
| 3696 | 185 | E185S | A | 1.1e-05 | 6.1e-06 | 1.2e-05 | 9.700E-06 | 3.158E-06 | 6.100E-06 | 1.200E-05 |
| 3697 | 185 | E185T | A | 7.9e-06 | 9.6e-06 | 8.8e-06 | 8.767E-06 | 8.505E-07 | 7.900E-06 | 9.600E-06 |
| 3698 | 185 | E185W | A | 7.1e-06 | 7.9e-06 | 1.4e-05 | 9.667E-06 | 3.774E-06 | 7.100E-06 | 1.400E-05 |
| 3699 | 185 | E185Y | A | 1.4e-05 | 8.8e-06 | 5.6e-06 | 9.467E-06 | 4.239E-06 | 5.600E-06 | 1.400E-05 |
| 3700 | 185 | E185V | A | 6.9e-06 | 1.2e-05 | 6.9e-06 | 8.600E-06 | 2.944E-06 | 6.900E-06 | 1.200E-05 |
| 3701 | 186 | S186A | A | 7.1e-06 | 5.6e-06 | 1.1e-05 | 7.900E-06 | 2.787E-06 | 5.600E-06 | 1.100E-05 |
| 3702 | 186 | S186R | A | 1.5e-05 | 5.7e-06 | 7.6e-06 | 9.433E-06 | 4.914E-06 | 5.700E-06 | 1.500E-05 |
| 3703 | 186 | S186N | A | 8.9e-06 | 1e-05 | 5.9e-06 | 8.267E-06 | 2.122E-06 | 5.900E-06 | 1.000E-05 |
| 3704 | 186 | S186D | A | 5.8e-06 | 8.4e-06 | 1.4e-05 | 9.400E-06 | 4.190E-06 | 5.800E-06 | 1.400E-05 |
| 3705 | 186 | S186C | A | 9.6e-06 | 8.8e-06 | 5.6e-06 | 8.000E-06 | 2.117E-06 | 5.600E-06 | 9.600E-06 |
| 3706 | 186 | S186E | A | 1.2e-05 | 8.2e-06 | 5.8e-06 | 8.667E-06 | 3.126E-06 | 5.800E-06 | 1.200E-05 |
| 3707 | 186 | S186Q | A | 1.3e-05 | 1.1e-05 | 6.5e-06 | 1.017E-05 | 3.329E-06 | 6.500E-06 | 1.300E-05 |
| 3708 | 186 | S186G | A | 1.1e-05 | 4e-06 | 5.6e-06 | 6.867E-06 | 3.668E-06 | 4.000E-06 | 1.100E-05 |
| 3709 | 186 | S186H | A | 8.8e-06 | 5.7e-06 | 9.1e-06 | 7.867E-06 | 1.882E-06 | 5.700E-06 | 9.100E-06 |
| 3710 | 186 | S186I | A | 5.5e-06 | 1.3e-05 | 1.1e-05 | 9.833E-06 | 3.884E-06 | 5.500E-06 | 1.300E-05 |
| 3711 | 186 | S186L | A | 6.4e-06 | 6.2e-06 | 1e-05 | 7.533E-06 | 2.139E-06 | 6.200E-06 | 1.000E-05 |
| 3712 | 186 | S186K | A | 8.4e-06 | 9.2e-06 | 7e-06 | 8.200E-06 | 1.114E-06 | 7.000E-06 | 9.200E-06 |
| 3713 | 186 | S186M | A | 1.1e-05 | 4.7e-06 | 9.4e-06 | 8.367E-06 | 3.275E-06 | 4.700E-06 | 1.100E-05 |
| 3714 | 186 | S186F | A | 7.9e-06 | 9.8e-06 | 7.3e-06 | 8.333E-06 | 1.305E-06 | 7.300E-06 | 9.800E-06 |
| 3715 | 186 | S186P | A | 9e-06 | 6.3e-06 | 8.8e-06 | 8.033E-06 | 1.504E-06 | 6.300E-06 | 9.000E-06 |
| 3716 | 186 | S186S | A | 2.3e-05 | 4.5e-06 | 6.8e-06 | 1.143E-05 | 1.008E-05 | 4.500E-06 | 2.300E-05 |
| 3717 | 186 | S186T | A | 8.9e-06 | 1.2e-05 | 7.5e-06 | 9.467E-06 | 2.303E-06 | 7.500E-06 | 1.200E-05 |
| 3718 | 186 | S186W | A | 3.7e-06 | 7.6e-06 | 1.2e-05 | 7.767E-06 | 4.153E-06 | 3.700E-06 | 1.200E-05 |
| 3719 | 186 | S186Y | A | 6.5e-06 | 5.9e-06 | 1e-05 | 7.467E-06 | 2.214E-06 | 5.900E-06 | 1.000E-05 |
| 3720 | 186 | S186V | A | 1e-05 | 7e-06 | 9.1e-06 | 8.700E-06 | 1.539E-06 | 7.000E-06 | 1.000E-05 |
| 3721 | 187 | S187A | A | 4.3e-06 | 4.5e-06 | 1.4e-05 | 7.600E-06 | 5.543E-06 | 4.300E-06 | 1.400E-05 |
| 3722 | 187 | S187R | A | 5.9e-06 | 9.9e-06 | 8.5e-06 | 8.100E-06 | 2.030E-06 | 5.900E-06 | 9.900E-06 |
| 3723 | 187 | S187N | A | 1.4e-05 | 7.5e-06 | 6.4e-06 | 9.300E-06 | 4.107E-06 | 6.400E-06 | 1.400E-05 |
| 3724 | 187 | S187D | A | 4.8e-06 | 6.8e-06 | 4.5e-06 | 5.367E-06 | 1.250E-06 | 4.500E-06 | 6.800E-06 |
| 3725 | 187 | S187C | A | 1.3e-05 | 7.2e-06 | 7.8e-06 | 9.333E-06 | 3.190E-06 | 7.200E-06 | 1.300E-05 |
| 3726 | 187 | S187E | A | 1.1e-05 | 7.8e-06 | 5.3e-06 | 8.033E-06 | 2.857E-06 | 5.300E-06 | 1.100E-05 |
| 3727 | 187 | S187Q | A | 8.8e-06 | 5.6e-06 | 9.3e-06 | 7.900E-06 | 2.007E-06 | 5.600E-06 | 9.300E-06 |
| 3728 | 187 | S187G | A | 6.4e-06 | 5.2e-06 | 6.9e-06 | 6.167E-06 | 8.737E-07 | 5.200E-06 | 6.900E-06 |
| 3729 | 187 | S187H | A | 8.5e-06 | 7.3e-06 | 9.3e-06 | 8.367E-06 | 1.007E-06 | 7.300E-06 | 9.300E-06 |
| 3730 | 187 | S187I | A | 6.5e-06 | 8.4e-06 | 5.2e-06 | 6.700E-06 | 1.609E-06 | 5.200E-06 | 8.400E-06 |
| 3731 | 187 | S187L | A | 5.7e-06 | 8.1e-06 | 1.4e-05 | 9.267E-06 | 4.271E-06 | 5.700E-06 | 1.400E-05 |
| 3732 | 187 | S187K | A | 6.3e-06 | 5.6e-06 | 1e-05 | 7.300E-06 | 2.364E-06 | 5.600E-06 | 1.000E-05 |
| 3733 | 187 | S187M | A | 9.6e-06 | 7.6e-06 | 5.1e-06 | 7.433E-06 | 2.255E-06 | 5.100E-06 | 9.600E-06 |
| 3734 | 187 | S187F | A | 4e-06 | 1e-05 | 4.8e-06 | 6.267E-06 | 3.258E-06 | 4.000E-06 | 1.000E-05 |
| 3735 | 187 | S187P | A | 7.1e-06 | 8e-06 | 1.4e-05 | 9.700E-06 | 3.751E-06 | 7.100E-06 | 1.400E-05 |
| 3736 | 187 | S187S | A | 1.2e-05 | 6.7e-06 | 1.3e-05 | 1.057E-05 | 3.386E-06 | 6.700E-06 | 1.300E-05 |
| 3737 | 187 | S187T | A | 8e-06 | 7.4e-06 | 8.5e-06 | 7.967E-06 | 5.508E-07 | 7.400E-06 | 8.500E-06 |
| 3738 | 187 | S187W | A | 8.2e-06 | 5.8e-06 | 3.4e-06 | 5.800E-06 | 2.400E-06 | 3.400E-06 | 8.200E-06 |
| 3739 | 187 | S187Y | A | 6.9e-06 | 1.3e-05 | 4.8e-06 | 8.233E-06 | 4.259E-06 | 4.800E-06 | 1.300E-05 |

| | | | | | | | | | | |
|---|---|---|---|---|---|---|---|---|---|---|
| 3740 | 187 | S187V | A | 9.3e-06 | 9.7e-06 | 7.3e-06 | 8.767E-06 | 1.286E-06 | 7.300E-06 | 9.700E-06 |
| 3741 | 188 | E188A | A | 6.2e-06 | 6.6e-06 | 1.5e-05 | 9.267E-06 | 4.969E-06 | 6.200E-06 | 1.500E-05 |
| 3742 | 188 | E188R | A | 1.1e-05 | 1.2e-05 | 4.9e-06 | 9.300E-06 | 3.843E-06 | 4.900E-06 | 1.200E-05 |
| 3743 | 188 | E188N | A | 9.5e-06 | 1.3e-05 | 8.2e-06 | 1.023E-05 | 2.483E-06 | 8.200E-06 | 1.300E-05 |
| 3744 | 188 | E188D | A | 6.7e-06 | 7.2e-06 | 7.3e-06 | 7.067E-06 | 3.215E-07 | 6.700E-06 | 7.300E-06 |
| 3745 | 188 | E188C | A | 6e-06 | 5.9e-06 | 9.3e-06 | 7.067E-06 | 1.935E-06 | 5.900E-06 | 9.300E-06 |
| 3746 | 188 | E188E | A | 9.3e-06 | 1e-05 | 1.2e-05 | 1.043E-05 | 1.401E-06 | 9.300E-06 | 1.200E-05 |
| 3747 | 188 | E188Q | A | 8.5e-06 | 9.7e-06 | 9.8e-06 | 9.333E-06 | 7.234E-07 | 8.500E-06 | 9.800E-06 |
| 3748 | 188 | E188G | A | 1e-05 | 9e-06 | 1.3e-05 | 1.067E-05 | 2.082E-06 | 9.000E-06 | 1.300E-05 |
| 3749 | 188 | E188H | A | 1.4e-05 | 8.1e-06 | 6.9e-06 | 9.667E-06 | 3.800E-06 | 6.900E-06 | 1.400E-05 |
| 3750 | 188 | E188I | A | 8.6e-06 | 7.6e-06 | 5.1e-06 | 7.100E-06 | 1.803E-06 | 5.100E-06 | 8.600E-06 |
| 3751 | 188 | E188L | A | 8.7e-06 | 5.8e-06 | 1.3e-05 | 9.167E-06 | 3.623E-06 | 5.800E-06 | 1.300E-05 |
| 3752 | 188 | E188K | A | 4.9e-06 | 6.1e-06 | 6.8e-06 | 5.933E-06 | 9.609E-07 | 4.900E-06 | 6.800E-06 |
| 3753 | 188 | E188M | A | 1.1e-05 | 5.2e-06 | 7.8e-06 | 8.000E-06 | 2.905E-06 | 5.200E-06 | 1.100E-05 |
| 3754 | 188 | E188F | A | 7.2e-06 | 9.6e-06 | 4.9e-06 | 7.233E-06 | 2.350E-06 | 4.900E-06 | 9.600E-06 |
| 3755 | 188 | E188P | A | 5.1e-06 | 7e-06 | 1.1e-05 | 7.700E-06 | 3.012E-06 | 5.100E-06 | 1.100E-05 |
| 3756 | 188 | E188S | A | 8.5e-06 | 7.2e-06 | 1.3e-05 | 9.567E-06 | 3.044E-06 | 7.200E-06 | 1.300E-05 |
| 3757 | 188 | E188T | A | 8.3e-06 | 9.2e-06 | 6.2e-06 | 7.900E-06 | 1.539E-06 | 6.200E-06 | 9.200E-06 |
| 3758 | 188 | E188W | A | 6e-06 | 8.3e-06 | 6.1e-06 | 6.800E-06 | 1.300E-06 | 6.000E-06 | 8.300E-06 |
| 3759 | 188 | E188Y | A | 4.6e-06 | 6.8e-06 | 1.1e-05 | 7.467E-06 | 3.252E-06 | 4.600E-06 | 1.100E-05 |
| 3760 | 188 | E188V | A | 9.4e-06 | 6.3e-06 | 9.7e-06 | 8.467E-06 | 1.882E-06 | 6.300E-06 | 9.700E-06 |
| 3761 | 189 | D189A | A | 6.4e-06 | 7.1e-06 | 7.4e-06 | 6.967E-06 | 5.132E-07 | 6.400E-06 | 7.400E-06 |
| 3762 | 189 | D189R | A | 4.8e-06 | 4.4e-06 | 7.9e-06 | 5.700E-06 | 1.916E-06 | 4.400E-06 | 7.900E-06 |
| 3763 | 189 | D189N | A | 8.3e-06 | 8.7e-06 | 9.4e-06 | 8.800E-06 | 5.568E-07 | 8.300E-06 | 9.400E-06 |
| 3764 | 189 | D189D | A | 6.3e-06 | 1.5e-05 | 6.9e-06 | 9.400E-06 | 4.859E-06 | 6.300E-06 | 1.500E-05 |
| 3765 | 189 | D189C | A | 5.8e-06 | 1.4e-05 | 4.7e-06 | 8.167E-06 | 5.082E-06 | 4.700E-06 | 1.400E-05 |
| 3766 | 189 | D189E | A | 8.9e-06 | 1.2e-05 | 1.2e-05 | 1.097E-05 | 1.790E-06 | 8.900E-06 | 1.200E-05 |
| 3767 | 189 | D189Q | A | 5.4e-06 | 1.3e-05 | 6.8e-06 | 8.400E-06 | 4.045E-06 | 5.400E-06 | 1.300E-05 |
| 3768 | 189 | D189G | A | 1.1e-05 | 1.1e-05 | 6.8e-06 | 9.600E-06 | 2.425E-06 | 6.800E-06 | 1.100E-05 |
| 3769 | 189 | D189H | A | 6.1e-06 | 9.8e-06 | 7e-06 | 7.633E-06 | 1.930E-06 | 6.100E-06 | 9.800E-06 |
| 3770 | 189 | D189I | A | 6.3e-06 | 8.5e-06 | 5e-06 | 6.600E-06 | 1.769E-06 | 5.000E-06 | 8.500E-06 |
| 3771 | 189 | D189L | A | 1.4e-05 | 1.1e-05 | 1.2e-05 | 1.233E-05 | 1.528E-06 | 1.100E-05 | 1.400E-05 |
| 3772 | 189 | D189K | A | 6.7e-06 | 6.8e-06 | 9.8e-06 | 7.767E-06 | 1.762E-06 | 6.700E-06 | 9.800E-06 |
| 3773 | 189 | D189M | A | 5e-06 | 6.1e-06 | 4.9e-06 | 5.333E-06 | 6.658E-07 | 4.900E-06 | 6.100E-06 |
| 3774 | 189 | D189F | A | 1.4e-05 | 1e-05 | 6.8e-06 | 1.027E-05 | 3.607E-06 | 6.800E-06 | 1.400E-05 |
| 3775 | 189 | D189P | A | 5.1e-06 | 9.1e-06 | 4.5e-06 | 6.233E-06 | 2.501E-06 | 4.500E-06 | 9.100E-06 |
| 3776 | 189 | D189S | A | 1.3e-05 | 1.3e-05 | 5.5e-06 | 1.050E-05 | 4.330E-06 | 5.500E-06 | 1.300E-05 |
| 3777 | 189 | D189T | A | 9.5e-06 | 7.7e-06 | 6e-06 | 7.733E-06 | 1.750E-06 | 6.000E-06 | 9.500E-06 |
| 3778 | 189 | D189W | A | 5.5e-06 | 9.9e-06 | 1.4e-05 | 9.800E-06 | 4.251E-06 | 5.500E-06 | 1.400E-05 |
| 3779 | 189 | D189Y | A | 1.5e-05 | 9.2e-06 | 8.1e-06 | 1.077E-05 | 3.707E-06 | 8.100E-06 | 1.500E-05 |
| 3780 | 189 | D189V | A | 9.9e-06 | 5.1e-06 | 9e-06 | 8.000E-06 | 2.551E-06 | 5.100E-06 | 9.900E-06 |
| 3781 | 190 | C190A | A | 7.7e-06 | 1e-05 | 9.4e-06 | 9.033E-06 | 1.193E-06 | 7.700E-06 | 1.000E-05 |
| 3782 | 190 | C190R | A | 1.2e-05 | 9.8e-06 | 1.3e-05 | 1.160E-05 | 1.637E-06 | 9.800E-06 | 1.300E-05 |
| 3783 | 190 | C190N | A | 7.1e-06 | 1.6e-05 | 8.1e-06 | 1.040E-05 | 4.875E-06 | 7.100E-06 | 1.600E-05 |
| 3784 | 190 | C190D | A | 7.4e-06 | 8.6e-06 | 1e-05 | 8.667E-06 | 1.301E-06 | 7.400E-06 | 1.000E-05 |
| 3785 | 190 | C190C | A | 4.6e-06 | 8.8e-06 | 7.9e-06 | 7.100E-06 | 2.211E-06 | 4.600E-06 | 8.800E-06 |
| 3786 | 190 | C190E | A | 8.4e-06 | 7.6e-06 | 7.2e-06 | 7.733E-06 | 6.110E-07 | 7.200E-06 | 8.400E-06 |
| 3787 | 190 | C190Q | A | 5.6e-06 | 4.6e-06 | 1e-05 | 6.733E-06 | 2.873E-06 | 4.600E-06 | 1.000E-05 |
| 3788 | 190 | C190G | A | 1e-05 | 6e-06 | 4.2e-06 | 6.733E-06 | 2.969E-06 | 4.200E-06 | 1.000E-05 |
| 3789 | 190 | C190H | A | 8.7e-06 | 6.7e-06 | 9.2e-06 | 8.200E-06 | 1.323E-06 | 6.700E-06 | 9.200E-06 |
| 3790 | 190 | C190I | A | 9.9e-06 | 9e-06 | 7.2e-06 | 8.700E-06 | 1.375E-06 | 7.200E-06 | 9.900E-06 |
| 3791 | 190 | C190L | A | 8.2e-06 | 1e-05 | 9.1e-06 | 9.100E-06 | 9.000E-07 | 8.200E-06 | 1.000E-05 |
| 3792 | 190 | C190K | A | 1.4e-05 | 1.3e-05 | 7.9e-06 | 1.163E-05 | 3.272E-06 | 7.900E-06 | 1.400E-05 |
| 3793 | 190 | C190M | A | 9.9e-06 | 6.3e-06 | 8.6e-06 | 8.267E-06 | 1.823E-06 | 6.300E-06 | 9.900E-06 |
| 3794 | 190 | C190F | A | 5.2e-06 | 1.3e-05 | 8.7e-06 | 8.967E-06 | 3.907E-06 | 5.200E-06 | 1.300E-05 |
| 3795 | 190 | C190P | A | 8.9e-06 | 7e-06 | 8e-06 | 7.967E-06 | 9.504E-07 | 7.000E-06 | 8.900E-06 |
| 3796 | 190 | C190S | A | 6.6e-06 | 8.6e-06 | 8.1e-06 | 7.767E-06 | 1.041E-06 | 6.600E-06 | 8.600E-06 |
| 3797 | 190 | C190T | A | 5.6e-06 | 7.7e-06 | 8.5e-06 | 7.267E-06 | 1.498E-06 | 5.600E-06 | 8.500E-06 |
| 3798 | 190 | C190W | A | 9.3e-06 | 6.7e-06 | 9.7e-06 | 8.567E-06 | 1.629E-06 | 6.700E-06 | 9.700E-06 |
| 3799 | 190 | C190Y | A | 8.6e-06 | 8.9e-06 | 8.1e-06 | 8.533E-06 | 4.041E-07 | 8.100E-06 | 8.900E-06 |
| 3800 | 190 | C190V | A | 5.1e-06 | 6.6e-06 | 6.9e-06 | 6.200E-06 | 9.644E-07 | 5.100E-06 | 6.900E-06 |
| 3801 | 191 | Q191A | A | 8.7e-06 | 8.4e-06 | 4.8e-06 | 7.300E-06 | 2.170E-06 | 4.800E-06 | 8.700E-06 |
| 3802 | 191 | Q191R | A | 6.4e-06 | 4.7e-06 | 1.3e-05 | 8.033E-06 | 4.384E-06 | 4.700E-06 | 1.300E-05 |
| 3803 | 191 | Q191N | A | 1e-05 | 8.9e-06 | 8.2e-06 | 9.033E-06 | 9.074E-07 | 8.200E-06 | 1.000E-05 |
| 3804 | 191 | Q191D | A | 1.3e-05 | 9e-06 | 1.3e-05 | 1.167E-05 | 2.309E-06 | 9.000E-06 | 1.300E-05 |
| 3805 | 191 | Q191C | A | 6.4e-06 | 6.3e-06 | 9e-06 | 7.233E-06 | 1.531E-06 | 6.300E-06 | 9.000E-06 |
| 3806 | 191 | Q191E | A | 8.9e-06 | 1.1e-05 | 5.2e-06 | 8.367E-06 | 2.937E-06 | 5.200E-06 | 1.100E-05 |
| 3807 | 191 | Q191Q | A | 8.7e-06 | 6.6e-06 | 8.1e-06 | 7.800E-06 | 1.082E-06 | 6.600E-06 | 8.700E-06 |
| 3808 | 191 | Q191G | A | 6.7e-06 | 7e-06 | 6.8e-06 | 6.833E-06 | 1.528E-07 | 6.700E-06 | 7.000E-06 |
| 3809 | 191 | Q191H | A | 6.8e-06 | 8.4e-06 | 8.5e-06 | 7.900E-06 | 9.539E-07 | 6.800E-06 | 8.500E-06 |
| 3810 | 191 | Q191I | A | 1.1e-05 | 9.5e-06 | 9.1e-06 | 9.867E-06 | 1.002E-06 | 9.100E-06 | 1.100E-05 |
| 3811 | 191 | Q191L | A | 1.5e-05 | 5.4e-06 | 9.3e-06 | 9.900E-06 | 4.828E-06 | 5.400E-06 | 1.500E-05 |
| 3812 | 191 | Q191K | A | 9e-06 | 1.4e-05 | 5.7e-06 | 9.567E-06 | 4.179E-06 | 5.700E-06 | 1.400E-05 |
| 3813 | 191 | Q191M | A | 8.6e-06 | 8.7e-06 | 5e-06 | 7.433E-06 | 2.108E-06 | 5.000E-06 | 8.700E-06 |
| 3814 | 191 | Q191F | A | 7.6e-06 | 5.7e-06 | 5.3e-06 | 6.200E-06 | 1.229E-06 | 5.300E-06 | 7.600E-06 |
| 3815 | 191 | Q191P | A | 7e-06 | 6.7e-06 | 1.2e-05 | 8.567E-06 | 2.977E-06 | 6.700E-06 | 1.200E-05 |
| 3816 | 191 | Q191S | A | 4e-06 | 1.1e-05 | 8.3e-06 | 7.767E-06 | 3.530E-06 | 4.000E-06 | 1.100E-05 |
| 3817 | 191 | Q191T | A | 8.7e-06 | 4.6e-06 | 9.8e-06 | 7.700E-06 | 2.740E-06 | 4.600E-06 | 9.800E-06 |
| 3818 | 191 | Q191W | A | 5.8e-06 | 6.6e-06 | 5.3e-06 | 5.900E-06 | 6.557E-07 | 5.300E-06 | 6.600E-06 |
| 3819 | 191 | Q191Y | A | 5.7e-06 | 5.3e-06 | 7.1e-06 | 6.033E-06 | 9.452E-07 | 5.300E-06 | 7.100E-06 |
| 3820 | 191 | Q191V | A | 9.7e-06 | 7.6e-06 | 1.4e-05 | 1.043E-05 | 3.262E-06 | 7.600E-06 | 1.400E-05 |
| 3821 | 192 | S192A | A | 1.1e-05 | 6.7e-06 | 8.6e-06 | 8.767E-06 | 2.155E-06 | 6.700E-06 | 1.100E-05 |
| 3822 | 192 | S192R | A | 7.8e-06 | 3.3e-06 | 1e-05 | 7.033E-06 | 3.415E-06 | 3.300E-06 | 1.000E-05 |
| 3823 | 192 | S192N | A | 6.8e-06 | 7.8e-06 | 5e-06 | 6.533E-06 | 1.419E-06 | 5.000E-06 | 7.800E-06 |
| 3824 | 192 | S192D | A | 6.4e-06 | 3.7e-06 | 1e-05 | 6.700E-06 | 3.161E-06 | 3.700E-06 | 1.000E-05 |

| | | | | | | | | | | |
|---|---|---|---|---|---|---|---|---|---|---|
| 3825 | 192 | S192C | A | 1.4e-05 | 4.2e-06 | 8.2e-06 | 8.800E-06 | 4.927E-06 | 4.200E-06 | 1.400E-05 |
| 3826 | 192 | S192E | A | 1.4e-05 | 8.5e-06 | 8.9e-06 | 1.047E-05 | 3.066E-06 | 8.500E-06 | 1.400E-05 |
| 3827 | 192 | S192Q | A | 8e-06 | 5.2e-06 | 6.1e-06 | 6.433E-06 | 1.429E-06 | 5.200E-06 | 8.000E-06 |
| 3828 | 192 | S192G | A | 6.3e-06 | 5.7e-06 | 7.9e-06 | 6.633E-06 | 1.137E-06 | 5.700E-06 | 7.900E-06 |
| 3829 | 192 | S192H | A | 1.2e-05 | 6.9e-06 | 9e-06 | 9.300E-06 | 2.563E-06 | 6.900E-06 | 1.200E-05 |
| 3830 | 192 | S192I | A | 1.3e-05 | 8.4e-06 | 1.1e-05 | 1.080E-05 | 2.307E-06 | 8.400E-06 | 1.300E-05 |
| 3831 | 192 | S192L | A | 1.3e-05 | 7.8e-06 | 1.1e-05 | 1.060E-05 | 2.623E-06 | 7.800E-06 | 1.300E-05 |
| 3832 | 192 | S192K | A | 6.8e-06 | 1.3e-05 | 6.4e-06 | 8.733E-06 | 3.700E-06 | 6.400E-06 | 1.300E-05 |
| 3833 | 192 | S192M | A | 1.3e-05 | 1.1e-05 | 6.8e-06 | 1.027E-05 | 3.164E-06 | 6.800E-06 | 1.300E-05 |
| 3834 | 192 | S192F | A | 8.5e-06 | 7.3e-06 | 8.9e-06 | 8.233E-06 | 8.327E-07 | 7.300E-06 | 8.900E-06 |
| 3835 | 192 | S192P | A | 1.1e-05 | 6.5e-06 | 8.9e-06 | 8.800E-06 | 2.252E-06 | 6.500E-06 | 1.100E-05 |
| 3836 | 192 | S192S | A | 1.3e-05 | 8.8e-06 | 1e-05 | 1.060E-05 | 2.163E-06 | 8.800E-06 | 1.300E-05 |
| 3837 | 192 | S192T | A | 4.3e-06 | 1.1e-05 | 9.3e-06 | 8.200E-06 | 3.483E-06 | 4.300E-06 | 1.100E-05 |
| 3838 | 192 | S192W | A | 8.4e-06 | 4.6e-06 | 8.7e-06 | 7.233E-06 | 2.285E-06 | 4.600E-06 | 8.700E-06 |
| 3839 | 192 | S192Y | A | 9e-06 | 9.5e-06 | 7e-06 | 8.500E-06 | 1.323E-06 | 7.000E-06 | 9.500E-06 |
| 3840 | 192 | S192V | A | 5.5e-06 | 9.3e-06 | 1.1e-05 | 8.600E-06 | 2.816E-06 | 5.500E-06 | 1.100E-05 |
| 3841 | 193 | L193A | A | 9.2e-06 | 1.3e-05 | 5e-06 | 9.067E-06 | 4.002E-06 | 5.000E-06 | 1.300E-05 |
| 3842 | 193 | L193R | A | 1.1e-05 | 8.3e-06 | 5.5e-06 | 8.267E-06 | 2.750E-06 | 5.500E-06 | 1.100E-05 |
| 3843 | 193 | L193N | A | 6.9e-06 | 1.2e-05 | 4.6e-06 | 7.833E-06 | 3.787E-06 | 4.600E-06 | 1.200E-05 |
| 3844 | 193 | L193D | A | 7.8e-06 | 4.9e-06 | 1.1e-05 | 7.900E-06 | 3.051E-06 | 4.900E-06 | 1.100E-05 |
| 3845 | 193 | L193C | A | 6.6e-06 | 7.8e-06 | 7.8e-06 | 7.400E-06 | 6.928E-07 | 6.600E-06 | 7.800E-06 |
| 3846 | 193 | L193E | A | 6.6e-06 | 9e-06 | 7.9e-06 | 7.833E-06 | 1.201E-06 | 6.600E-06 | 9.000E-06 |
| 3847 | 193 | L193Q | A | 9.3e-06 | 1.3e-05 | 8.1e-06 | 1.013E-05 | 2.554E-06 | 8.100E-06 | 1.300E-05 |
| 3848 | 193 | L193G | A | 8.7e-06 | 8.9e-06 | 7.1e-06 | 8.233E-06 | 9.866E-07 | 7.100E-06 | 8.900E-06 |
| 3849 | 193 | L193H | A | 9.6e-06 | 1.2e-05 | 6.9e-06 | 9.500E-06 | 2.551E-06 | 6.900E-06 | 1.200E-05 |
| 3850 | 193 | L193I | A | 7e-06 | 1.1e-05 | 1.1e-05 | 9.667E-06 | 2.309E-06 | 7.000E-06 | 1.100E-05 |
| 3851 | 193 | L193L | A | 6.8e-06 | 1.1e-05 | 1.2e-05 | 9.933E-06 | 2.759E-06 | 6.800E-06 | 1.200E-05 |
| 3852 | 193 | L193K | A | 9.1e-06 | 5.5e-06 | 6.1e-06 | 6.900E-06 | 1.929E-06 | 5.500E-06 | 9.100E-06 |
| 3853 | 193 | L193M | A | 9.8e-06 | 1.3e-05 | 7.1e-06 | 9.967E-06 | 2.954E-06 | 7.100E-06 | 1.300E-05 |
| 3854 | 193 | L193F | A | 5.9e-06 | 1.2e-05 | 5.4e-06 | 7.767E-06 | 3.675E-06 | 5.400E-06 | 1.200E-05 |
| 3855 | 193 | L193P | A | 1.1e-05 | 1.3e-05 | 7.3e-06 | 1.043E-05 | 2.892E-06 | 7.300E-06 | 1.300E-05 |
| 3856 | 193 | L193S | A | 6.3e-06 | 5.5e-06 | 8.2e-06 | 6.667E-06 | 1.387E-06 | 5.500E-06 | 8.200E-06 |
| 3857 | 193 | L193T | A | 8e-06 | 4.8e-06 | 9.1e-06 | 7.300E-06 | 2.234E-06 | 4.800E-06 | 9.100E-06 |
| 3858 | 193 | L193W | A | 8.2e-06 | 1.1e-05 | 8.1e-06 | 9.100E-06 | 1.646E-06 | 8.100E-06 | 1.100E-05 |
| 3859 | 193 | L193Y | A | 1e-05 | 8.9e-06 | 7.6e-06 | 8.833E-06 | 1.201E-06 | 7.600E-06 | 1.000E-05 |
| 3860 | 193 | L193V | A | 8.6e-06 | 6.4e-06 | 5.1e-06 | 6.700E-06 | 1.769E-06 | 5.100E-06 | 8.600E-06 |
| 3861 | 194 | T194A | A | 6.2e-06 | 1.3e-05 | 1.3e-05 | 1.073E-05 | 3.926E-06 | 6.200E-06 | 1.300E-05 |
| 3862 | 194 | T194R | A | 8.5e-06 | 9e-06 | 8.7e-06 | 8.733E-06 | 2.517E-07 | 8.500E-06 | 9.000E-06 |
| 3863 | 194 | T194N | A | 4.4e-06 | 1.7e-05 | 9.1e-06 | 1.017E-05 | 6.367E-06 | 4.400E-06 | 1.700E-05 |
| 3864 | 194 | T194D | A | 9.8e-06 | 1.2e-05 | 5.8e-06 | 9.200E-06 | 3.143E-06 | 5.800E-06 | 1.200E-05 |
| 3865 | 194 | T194C | A | 8.9e-06 | 9.8e-06 | 7.3e-06 | 8.667E-06 | 1.266E-06 | 7.300E-06 | 9.800E-06 |
| 3866 | 194 | T194E | A | 9e-06 | 1e-05 | 9.3e-06 | 9.433E-06 | 5.132E-07 | 9.000E-06 | 1.000E-05 |
| 3867 | 194 | T194Q | A | 5.3e-06 | 1.3e-05 | 4.7e-06 | 7.667E-06 | 4.629E-06 | 4.700E-06 | 1.300E-05 |
| 3868 | 194 | T194G | A | 5.6e-06 | 1.5e-05 | 9.8e-06 | 1.013E-05 | 4.709E-06 | 5.600E-06 | 1.500E-05 |
| 3869 | 194 | T194H | A | 1.4e-05 | 5.4e-06 | 8.6e-06 | 9.333E-06 | 4.347E-06 | 5.400E-06 | 1.400E-05 |
| 3870 | 194 | T194I | A | 5.6e-06 | 6.9e-06 | 9.4e-06 | 7.300E-06 | 1.931E-06 | 5.600E-06 | 9.400E-06 |
| 3871 | 194 | T194L | A | 5.5e-06 | 9.2e-06 | 8.2e-06 | 7.633E-06 | 1.914E-06 | 5.500E-06 | 9.200E-06 |
| 3872 | 194 | T194K | A | 9.4e-06 | 8.7e-06 | 8.3e-06 | 8.800E-06 | 5.568E-07 | 8.300E-06 | 9.400E-06 |
| 3873 | 194 | T194M | A | 9.3e-06 | 7.6e-06 | 7.7e-06 | 8.200E-06 | 9.539E-07 | 7.600E-06 | 9.300E-06 |
| 3874 | 194 | T194F | A | 9.1e-06 | 9.5e-06 | 5.3e-06 | 7.967E-06 | 2.318E-06 | 5.300E-06 | 9.500E-06 |
| 3875 | 194 | T194P | A | 1.4e-05 | 9.7e-06 | 1.1e-05 | 1.157E-05 | 2.205E-06 | 9.700E-06 | 1.400E-05 |
| 3876 | 194 | T194S | A | 8.6e-06 | 9.5e-06 | 9.1e-06 | 9.067E-06 | 4.509E-07 | 8.600E-06 | 9.500E-06 |
| 3877 | 194 | T194T | A | 6.5e-06 | 1.2e-05 | 7.5e-06 | 8.667E-06 | 2.930E-06 | 6.500E-06 | 1.200E-05 |
| 3878 | 194 | T194W | A | 7.6e-06 | 6.4e-06 | 7e-06 | 7.000E-06 | 6.000E-07 | 6.400E-06 | 7.600E-06 |
| 3879 | 194 | T194Y | A | 8e-06 | 7.7e-06 | 9e-06 | 8.233E-06 | 6.807E-07 | 7.700E-06 | 9.000E-06 |
| 3880 | 194 | T194V | A | 5e-06 | 5.8e-06 | 1e-05 | 6.933E-06 | 2.686E-06 | 5.000E-06 | 1.000E-05 |
| 3881 | 195 | R195A | A | 9.2e-06 | 6.7e-06 | 7.3e-06 | 7.733E-06 | 1.305E-06 | 6.700E-06 | 9.200E-06 |
| 3882 | 195 | R195R | A | 8e-06 | 8.1e-06 | 8.4e-06 | 8.167E-06 | 2.082E-07 | 8.000E-06 | 8.400E-06 |
| 3883 | 195 | R195N | A | 1.3e-05 | 6.4e-06 | 7.2e-06 | 8.867E-06 | 3.602E-06 | 6.400E-06 | 1.300E-05 |
| 3884 | 195 | R195D | A | 9.6e-06 | 6.2e-06 | 1.2e-05 | 9.267E-06 | 2.914E-06 | 6.200E-06 | 1.200E-05 |
| 3885 | 195 | R195C | A | 7.1e-06 | 6.6e-06 | 6e-06 | 6.567E-06 | 5.508E-07 | 6.000E-06 | 7.100E-06 |
| 3886 | 195 | R195E | A | 6.6e-06 | 8.5e-06 | 6.5e-06 | 7.200E-06 | 1.127E-06 | 6.500E-06 | 8.500E-06 |
| 3887 | 195 | R195Q | A | 9.5e-06 | 7.3e-06 | 4.3e-06 | 7.033E-06 | 2.610E-06 | 4.300E-06 | 9.500E-06 |
| 3888 | 195 | R195G | A | 1.4e-05 | 7.9e-06 | 5.3e-06 | 9.067E-06 | 4.466E-06 | 5.300E-06 | 1.400E-05 |
| 3889 | 195 | R195H | A | 1.1e-05 | 7.2e-06 | 6.2e-06 | 8.133E-06 | 2.532E-06 | 6.200E-06 | 1.100E-05 |
| 3890 | 195 | R195I | A | 1e-05 | 6.1e-06 | 7.7e-06 | 7.933E-06 | 1.960E-06 | 6.100E-06 | 1.000E-05 |
| 3891 | 195 | R195L | A | 5.5e-06 | 5.9e-06 | 7.1e-06 | 6.167E-06 | 8.327E-07 | 5.500E-06 | 7.100E-06 |
| 3892 | 195 | R195K | A | 4.6e-06 | 7e-06 | 6.7e-06 | 6.100E-06 | 1.308E-06 | 4.600E-06 | 7.000E-06 |
| 3893 | 195 | R195M | A | 8.1e-06 | 9.9e-06 | 7.7e-06 | 8.567E-06 | 1.172E-06 | 7.700E-06 | 9.900E-06 |
| 3894 | 195 | R195F | A | 9.1e-06 | 8.7e-06 | 6.9e-06 | 8.233E-06 | 1.172E-06 | 6.900E-06 | 9.100E-06 |
| 3895 | 195 | R195P | A | 6.9e-06 | 9.3e-06 | 1.3e-05 | 9.733E-06 | 3.073E-06 | 6.900E-06 | 1.300E-05 |
| 3896 | 195 | R195S | A | 1.2e-05 | 8.8e-06 | 8.2e-06 | 9.667E-06 | 2.043E-06 | 8.200E-06 | 1.200E-05 |
| 3897 | 195 | R195T | A | 1.3e-05 | 1.1e-05 | 7e-06 | 1.033E-05 | 3.055E-06 | 7.000E-06 | 1.300E-05 |
| 3898 | 195 | R195W | A | 9.8e-06 | 6.1e-06 | 6.9e-06 | 7.600E-06 | 1.947E-06 | 6.100E-06 | 9.800E-06 |
| 3899 | 195 | R195Y | A | 8.9e-06 | 5.5e-06 | 1.2e-05 | 8.800E-06 | 3.251E-06 | 5.500E-06 | 1.200E-05 |
| 3900 | 195 | R195V | A | 9.3e-06 | 9.5e-06 | 6.4e-06 | 8.400E-06 | 1.735E-06 | 6.400E-06 | 9.500E-06 |
| 3901 | 196 | T196A | A | 1.1e-05 | 7.7e-06 | 6.1e-06 | 8.267E-06 | 2.499E-06 | 6.100E-06 | 1.100E-05 |
| 3902 | 196 | T196R | A | 1.2e-05 | 1e-05 | 8.1e-06 | 1.003E-05 | 1.950E-06 | 8.100E-06 | 1.200E-05 |
| 3903 | 196 | T196N | A | 7e-06 | 5.8e-06 | 7.6e-06 | 6.800E-06 | 9.165E-07 | 5.800E-06 | 7.600E-06 |
| 3904 | 196 | T196D | A | 1.4e-05 | 6.1e-06 | 9.2e-06 | 9.767E-06 | 3.980E-06 | 6.100E-06 | 1.400E-05 |
| 3905 | 196 | T196C | A | 1.2e-05 | 6.5e-06 | 1.1e-05 | 9.833E-06 | 2.930E-06 | 6.500E-06 | 1.200E-05 |
| 3906 | 196 | T196E | A | 7.4e-06 | 9.7e-06 | 1.4e-05 | 1.037E-05 | 3.350E-06 | 7.400E-06 | 1.400E-05 |
| 3907 | 196 | T196Q | A | 9.6e-06 | 1.1e-05 | 9.4e-06 | 1.000E-05 | 8.718E-07 | 9.400E-06 | 1.100E-05 |
| 3908 | 196 | T196G | A | 1e-05 | 7.4e-06 | 6.1e-06 | 7.833E-06 | 1.986E-06 | 6.100E-06 | 1.000E-05 |
| 3909 | 196 | T196H | A | 5.3e-06 | 1.4e-05 | 1e-05 | 9.767E-06 | 4.355E-06 | 5.300E-06 | 1.400E-05 |

| | | | | | | | | | | |
|---|---|---|---|---|---|---|---|---|---|---|
| 3910 | 196 | T196I | A | 9.3e-06 | 7.6e-06 | 9.9e-06 | 8.933E-06 | 1.193E-06 | 7.600E-06 | 9.900E-06 |
| 3911 | 196 | T196L | A | 5.6e-06 | 1.4e-05 | 6.2e-06 | 8.600E-06 | 4.686E-06 | 5.600E-06 | 1.400E-05 |
| 3912 | 196 | T196K | A | 9e-06 | 5.4e-06 | 7.1e-06 | 7.167E-06 | 1.801E-06 | 5.400E-06 | 9.000E-06 |
| 3913 | 196 | T196M | A | 9.1e-06 | 7.4e-06 | 6.8e-06 | 7.767E-06 | 1.193E-06 | 6.800E-06 | 9.100E-06 |
| 3914 | 196 | T196F | A | 9.8e-06 | 7.1e-06 | 8.5e-06 | 8.467E-06 | 1.350E-06 | 7.100E-06 | 9.800E-06 |
| 3915 | 196 | T196P | A | 7.6e-06 | 1.1e-05 | 9e-06 | 9.200E-06 | 1.709E-06 | 7.600E-06 | 1.100E-05 |
| 3916 | 196 | T196S | A | 7.8e-06 | 9.2e-06 | 9.4e-06 | 8.800E-06 | 8.718E-07 | 7.800E-06 | 9.400E-06 |
| 3917 | 196 | T196T | A | 6.2e-06 | 5.6e-06 | 4.2e-06 | 5.333E-06 | 1.026E-06 | 4.200E-06 | 6.200E-06 |
| 3918 | 196 | T196W | A | 1e-05 | 1.2e-05 | 1.1e-05 | 1.100E-05 | 1.000E-06 | 1.000E-05 | 1.200E-05 |
| 3919 | 196 | T196Y | A | 5.2e-06 | 9.8e-06 | 8.1e-06 | 7.700E-06 | 2.326E-06 | 5.200E-06 | 9.800E-06 |
| 3920 | 196 | T196V | A | 9.1e-06 | 8.8e-06 | 6.3e-06 | 8.067E-06 | 1.537E-06 | 6.300E-06 | 9.100E-06 |
| 3921 | 197 | V197A | A | 6.1e-06 | 9.2e-06 | 7.2e-06 | 7.500E-06 | 1.572E-06 | 6.100E-06 | 9.200E-06 |
| 3922 | 197 | V197R | A | 3.3e-06 | 1.2e-05 | 8.5e-06 | 7.933E-06 | 4.378E-06 | 3.300E-06 | 1.200E-05 |
| 3923 | 197 | V197N | A | 9.1e-06 | 1.4e-05 | 1e-05 | 1.103E-05 | 2.608E-06 | 9.100E-06 | 1.400E-05 |
| 3924 | 197 | V197D | A | 9.6e-06 | 1.4e-05 | 5.9e-06 | 9.833E-06 | 4.055E-06 | 5.900E-06 | 1.400E-05 |
| 3925 | 197 | V197C | A | 5.4e-06 | 6.4e-06 | 8.6e-06 | 6.800E-06 | 1.637E-06 | 5.400E-06 | 8.600E-06 |
| 3926 | 197 | V197E | A | 9.9e-06 | 7.9e-06 | 5.9e-06 | 7.900E-06 | 2.000E-06 | 5.900E-06 | 9.900E-06 |
| 3927 | 197 | V197Q | A | 1e-05 | 7.2e-06 | 7.7e-06 | 8.300E-06 | 1.493E-06 | 7.200E-06 | 1.000E-05 |
| 3928 | 197 | V197G | A | 4.6e-06 | 6.9e-06 | 8.2e-06 | 6.567E-06 | 1.823E-06 | 4.600E-06 | 8.200E-06 |
| 3929 | 197 | V197H | A | 1.4e-05 | 1.1e-05 | 6.8e-06 | 1.060E-05 | 3.617E-06 | 6.800E-06 | 1.400E-05 |
| 3930 | 197 | V197I | A | 9.4e-06 | 1.3e-05 | 6.2e-06 | 9.533E-06 | 3.402E-06 | 6.200E-06 | 1.300E-05 |
| 3931 | 197 | V197L | A | 7.1e-06 | 1.2e-05 | 6.5e-06 | 8.533E-06 | 3.017E-06 | 6.500E-06 | 1.200E-05 |
| 3932 | 197 | V197K | A | 6.3e-06 | 1e-05 | 6.9e-06 | 7.733E-06 | 1.986E-06 | 6.300E-06 | 1.000E-05 |
| 3933 | 197 | V197M | A | 8.3e-06 | 7.5e-06 | 6.8e-06 | 7.533E-06 | 7.506E-07 | 6.800E-06 | 8.300E-06 |
| 3934 | 197 | V197F | A | 9.6e-06 | 6.3e-06 | 6.4e-06 | 7.433E-06 | 1.877E-06 | 6.300E-06 | 9.600E-06 |
| 3935 | 197 | V197P | A | 4.1e-06 | 9e-06 | 7.6e-06 | 6.900E-06 | 2.524E-06 | 4.100E-06 | 9.000E-06 |
| 3936 | 197 | V197S | A | 1.4e-05 | 8.3e-06 | 9.1e-06 | 1.047E-05 | 3.086E-06 | 8.300E-06 | 1.400E-05 |
| 3937 | 197 | V197T | A | 7e-06 | 6.6e-06 | 9.7e-06 | 7.767E-06 | 1.686E-06 | 6.600E-06 | 9.700E-06 |
| 3938 | 197 | V197W | A | 1.2e-05 | 6e-06 | 6.8e-06 | 8.267E-06 | 3.258E-06 | 6.000E-06 | 1.200E-05 |
| 3939 | 197 | V197Y | A | 7.6e-06 | 1.5e-05 | 8.5e-06 | 1.037E-05 | 4.038E-06 | 7.600E-06 | 1.500E-05 |
| 3940 | 197 | V197V | A | 8.4e-06 | 7.3e-06 | 4.9e-06 | 6.867E-06 | 1.790E-06 | 4.900E-06 | 8.400E-06 |
| 3941 | 198 | C198A | A | 1e-05 | 1.1e-05 | 9.1e-06 | 1.003E-05 | 9.504E-07 | 9.100E-06 | 1.100E-05 |
| 3942 | 198 | C198R | A | 7.1e-06 | 7.5e-06 | 5.3e-06 | 6.633E-06 | 1.172E-06 | 5.300E-06 | 7.500E-06 |
| 3943 | 198 | C198N | A | 5.5e-06 | 7.8e-06 | 1.3e-05 | 8.767E-06 | 3.842E-06 | 5.500E-06 | 1.300E-05 |
| 3944 | 198 | C198D | A | 6.4e-06 | 8.5e-06 | 8.3e-06 | 7.733E-06 | 1.159E-06 | 6.400E-06 | 8.500E-06 |
| 3945 | 198 | C198C | A | 5.4e-06 | 8.8e-06 | 8.3e-06 | 7.500E-06 | 1.836E-06 | 5.400E-06 | 8.800E-06 |
| 3946 | 198 | C198E | A | 9.8e-06 | 7.4e-06 | 7.2e-06 | 8.133E-06 | 1.447E-06 | 7.200E-06 | 9.800E-06 |
| 3947 | 198 | C198Q | A | 6.3e-06 | 9.6e-06 | 1.2e-05 | 9.300E-06 | 2.862E-06 | 6.300E-06 | 1.200E-05 |
| 3948 | 198 | C198G | A | 1.2e-05 | 9.8e-06 | 4.8e-06 | 8.867E-06 | 3.690E-06 | 4.800E-06 | 1.200E-05 |
| 3949 | 198 | C198H | A | 7.8e-06 | 7.3e-06 | 8.8e-06 | 7.967E-06 | 7.638E-07 | 7.300E-06 | 8.800E-06 |
| 3950 | 198 | C198I | A | 4.7e-06 | 3.7e-06 | 1.1e-05 | 6.467E-06 | 3.958E-06 | 3.700E-06 | 1.100E-05 |
| 3951 | 198 | C198L | A | 9.7e-06 | 7.8e-06 | 6.9e-06 | 8.133E-06 | 1.429E-06 | 6.900E-06 | 9.700E-06 |
| 3952 | 198 | C198K | A | 8e-06 | 6.7e-06 | 5.5e-06 | 6.733E-06 | 1.250E-06 | 5.500E-06 | 8.000E-06 |
| 3953 | 198 | C198M | A | 1.8e-05 | 4.7e-06 | 6e-06 | 9.567E-06 | 7.332E-06 | 4.700E-06 | 1.800E-05 |
| 3954 | 198 | C198F | A | 1.2e-05 | 1.8e-05 | 9.2e-06 | 1.307E-05 | 4.496E-06 | 9.200E-06 | 1.800E-05 |
| 3955 | 198 | C198P | A | 9.2e-06 | 6.9e-06 | 9.2e-06 | 8.433E-06 | 1.328E-06 | 6.900E-06 | 9.200E-06 |
| 3956 | 198 | C198S | A | 1.1e-05 | 9.2e-06 | 8.1e-06 | 9.433E-06 | 1.464E-06 | 8.100E-06 | 1.100E-05 |
| 3957 | 198 | C198T | A | 1.1e-05 | 4.2e-06 | 4.8e-06 | 6.667E-06 | 3.765E-06 | 4.200E-06 | 1.100E-05 |
| 3958 | 198 | C198W | A | 1.2e-05 | 7e-06 | 1.1e-05 | 1.000E-05 | 2.646E-06 | 7.000E-06 | 1.200E-05 |
| 3959 | 198 | C198Y | A | 4.9e-06 | 6.1e-06 | 9.6e-06 | 6.867E-06 | 2.442E-06 | 4.900E-06 | 9.600E-06 |
| 3960 | 198 | C198V | A | 5.8e-06 | 9.7e-06 | 4.7e-06 | 6.733E-06 | 2.627E-06 | 4.700E-06 | 9.700E-06 |
| 3961 | 199 | A199A | A | 7.2e-06 | 6e-06 | 4.6e-06 | 5.933E-06 | 1.301E-06 | 4.600E-06 | 7.200E-06 |
| 3962 | 199 | A199R | A | 5.2e-06 | 6.8e-06 | 5.4e-06 | 5.800E-06 | 8.718E-07 | 5.200E-06 | 6.800E-06 |
| 3963 | 199 | A199N | A | 1.1e-05 | 6.8e-06 | 7e-06 | 8.267E-06 | 2.369E-06 | 6.800E-06 | 1.100E-05 |
| 3964 | 199 | A199D | A | 9.1e-06 | 9.2e-06 | 1e-05 | 9.433E-06 | 4.933E-07 | 9.100E-06 | 1.000E-05 |
| 3965 | 199 | A199C | A | 7.1e-06 | 9.1e-06 | 4.1e-06 | 6.767E-06 | 2.517E-06 | 4.100E-06 | 9.100E-06 |
| 3966 | 199 | A199E | A | 1.3e-05 | 8.5e-06 | 7e-06 | 9.500E-06 | 3.122E-06 | 7.000E-06 | 1.300E-05 |
| 3967 | 199 | A199Q | A | 4.9e-06 | 9.1e-06 | 1e-05 | 8.000E-06 | 2.722E-06 | 4.900E-06 | 1.000E-05 |
| 3968 | 199 | A199G | A | 5.7e-06 | 9.7e-06 | 6.2e-06 | 7.200E-06 | 2.179E-06 | 5.700E-06 | 9.700E-06 |
| 3969 | 199 | A199H | A | 8.6e-06 | 8.5e-06 | 5.5e-06 | 7.533E-06 | 1.762E-06 | 5.500E-06 | 8.600E-06 |
| 3970 | 199 | A199I | A | 9.5e-06 | 8e-06 | 1e-05 | 9.167E-06 | 1.041E-06 | 8.000E-06 | 1.000E-05 |
| 3971 | 199 | A199L | A | 9.6e-06 | 6.1e-06 | 7.8e-06 | 7.833E-06 | 1.750E-06 | 6.100E-06 | 9.600E-06 |
| 3972 | 199 | A199K | A | 8.2e-06 | 6.8e-06 | 1e-05 | 8.333E-06 | 1.604E-06 | 6.800E-06 | 1.000E-05 |
| 3973 | 199 | A199M | A | 1.5e-05 | 3.5e-06 | 1e-05 | 9.500E-06 | 5.766E-06 | 3.500E-06 | 1.500E-05 |
| 3974 | 199 | A199F | A | 6.5e-06 | 4.6e-06 | 1.1e-05 | 7.367E-06 | 3.287E-06 | 4.600E-06 | 1.100E-05 |
| 3975 | 199 | A199P | A | 1.1e-05 | 1.3e-05 | 6.9e-06 | 1.030E-05 | 3.110E-06 | 6.900E-06 | 1.300E-05 |
| 3976 | 199 | A199S | A | 4.8e-06 | 8.3e-06 | 6.3e-06 | 6.467E-06 | 1.756E-06 | 4.800E-06 | 8.300E-06 |
| 3977 | 199 | A199T | A | 8.5e-06 | 1e-05 | 6.3e-06 | 8.267E-06 | 1.861E-06 | 6.300E-06 | 1.000E-05 |
| 3978 | 199 | A199W | A | 1e-05 | 7.7e-06 | 6.3e-06 | 8.000E-06 | 1.868E-06 | 6.300E-06 | 1.000E-05 |
| 3979 | 199 | A199Y | A | 4.7e-06 | 9.4e-06 | 6.4e-06 | 6.833E-06 | 2.380E-06 | 4.700E-06 | 9.400E-06 |
| 3980 | 199 | A199V | A | 1e-05 | 1e-05 | 1.1e-05 | 1.033E-05 | 5.774E-07 | 1.000E-05 | 1.100E-05 |
| 3981 | 200 | G200A | A | 7.5e-06 | 4.6e-06 | 9e-06 | 7.033E-06 | 2.237E-06 | 4.600E-06 | 9.000E-06 |
| 3982 | 200 | G200R | A | 1.2e-05 | 8.7e-06 | 1.1e-05 | 1.057E-05 | 1.692E-06 | 8.700E-06 | 1.200E-05 |
| 3983 | 200 | G200N | A | 4.7e-06 | 5.8e-06 | 7.8e-06 | 6.100E-06 | 1.572E-06 | 4.700E-06 | 7.800E-06 |
| 3984 | 200 | G200D | A | 8e-06 | 8.9e-06 | 5.1e-06 | 7.333E-06 | 1.986E-06 | 5.100E-06 | 8.900E-06 |
| 3985 | 200 | G200C | A | 5.6e-06 | 5.2e-06 | 5.5e-06 | 5.433E-06 | 2.082E-07 | 5.200E-06 | 5.600E-06 |
| 3986 | 200 | G200E | A | 1e-05 | 8.5e-06 | 6.7e-06 | 8.400E-06 | 1.652E-06 | 6.700E-06 | 1.000E-05 |
| 3987 | 200 | G200Q | A | 9.2e-06 | 4.6e-06 | 1.2e-05 | 8.600E-06 | 3.736E-06 | 4.600E-06 | 1.200E-05 |
| 3988 | 200 | G200G | A | 4.6e-06 | 9.3e-06 | 6.4e-06 | 6.767E-06 | 2.371E-06 | 4.600E-06 | 9.300E-06 |
| 3989 | 200 | G200H | A | 6.1e-06 | 1e-05 | 6.5e-06 | 7.533E-06 | 2.146E-06 | 6.100E-06 | 1.000E-05 |
| 3990 | 200 | G200I | A | 8.9e-06 | 6.5e-06 | 6.8e-06 | 7.400E-06 | 1.308E-06 | 6.500E-06 | 8.900E-06 |
| 3991 | 200 | G200L | A | 6.2e-06 | 1.3e-05 | 5e-06 | 8.067E-06 | 4.314E-06 | 5.000E-06 | 1.300E-05 |
| 3992 | 200 | G200K | A | 9.9e-06 | 7.7e-06 | 4.4e-06 | 7.333E-06 | 2.768E-06 | 4.400E-06 | 9.900E-06 |
| 3993 | 200 | G200M | A | 5.9e-06 | 9.1e-06 | 1.2e-05 | 9.000E-06 | 3.051E-06 | 5.900E-06 | 1.200E-05 |
| 3994 | 200 | G200F | A | 3.5e-06 | 1e-05 | 9.8e-06 | 7.767E-06 | 3.696E-06 | 3.500E-06 | 1.000E-05 |

| | | | | | | | | | | |
|---|---|---|---|---|---|---|---|---|---|---|
| 3995 | 200 | G200P | A | 7e-06 | 8.2e-06 | 7e-06 | 7.400E-06 | 6.928E-07 | 7.000E-06 | 8.200E-06 |
| 3996 | 200 | G200S | A | 6.2e-06 | 3.7e-06 | 9.9e-06 | 6.600E-06 | 3.119E-06 | 3.700E-06 | 9.900E-06 |
| 3997 | 200 | G200T | A | 9.2e-06 | 1.1e-05 | 8.6e-06 | 9.600E-06 | 1.249E-06 | 8.600E-06 | 1.100E-05 |
| 3998 | 200 | G200W | A | 4.7e-06 | 6.6e-06 | 1.2e-05 | 7.767E-06 | 3.787E-06 | 4.700E-06 | 1.200E-05 |
| 3999 | 200 | G200Y | A | 1.3e-05 | 5.1e-06 | 9.7e-06 | 9.267E-06 | 3.968E-06 | 5.100E-06 | 1.300E-05 |
| 4000 | 200 | G200V | A | 1.3e-05 | 4.6e-06 | 8.6e-06 | 8.733E-06 | 4.202E-06 | 4.600E-06 | 1.300E-05 |
| 4001 | 201 | G201A | A | 5.4e-06 | 7.9e-06 | 7.1e-06 | 6.800E-06 | 1.277E-06 | 5.400E-06 | 7.900E-06 |
| 4002 | 201 | G201R | A | 8.8e-06 | 7.8e-06 | 4.4e-06 | 7.000E-06 | 2.307E-06 | 4.400E-06 | 8.800E-06 |
| 4003 | 201 | G201N | A | 6.6e-06 | 8.3e-06 | 9e-06 | 7.967E-06 | 1.234E-06 | 6.600E-06 | 9.000E-06 |
| 4004 | 201 | G201D | A | 8.2e-06 | 6.9e-06 | 8.5e-06 | 7.867E-06 | 8.505E-07 | 6.900E-06 | 8.500E-06 |
| 4005 | 201 | G201C | A | 5.3e-06 | 7.9e-06 | 9.5e-06 | 7.567E-06 | 2.120E-06 | 5.300E-06 | 9.500E-06 |
| 4006 | 201 | G201E | A | 1.6e-05 | 6.3e-06 | 6.8e-06 | 9.700E-06 | 5.462E-06 | 6.300E-06 | 1.600E-05 |
| 4007 | 201 | G201Q | A | 9.4e-06 | 4.3e-06 | 1e-05 | 7.900E-06 | 3.132E-06 | 4.300E-06 | 1.000E-05 |
| 4008 | 201 | G201G | A | 1.1e-05 | 8.8e-06 | 1.3e-05 | 1.093E-05 | 2.101E-06 | 8.800E-06 | 1.300E-05 |
| 4009 | 201 | G201H | A | 1e-05 | 1e-05 | 1.1e-05 | 1.033E-05 | 5.774E-07 | 1.000E-05 | 1.100E-05 |
| 4010 | 201 | G201I | A | 4.3e-06 | 1.1e-05 | 9.2e-06 | 8.167E-06 | 3.467E-06 | 4.300E-06 | 1.100E-05 |
| 4011 | 201 | G201L | A | 6.9e-06 | 9.3e-06 | 8e-06 | 8.067E-06 | 1.201E-06 | 6.900E-06 | 9.300E-06 |
| 4012 | 201 | G201K | A | 1.1e-05 | 9.3e-06 | 7.9e-06 | 9.400E-06 | 1.552E-06 | 7.900E-06 | 1.100E-05 |
| 4013 | 201 | G201M | A | 4.4e-06 | 7.8e-06 | 8.2e-06 | 6.800E-06 | 2.088E-06 | 4.400E-06 | 8.200E-06 |
| 4014 | 201 | G201F | A | 1.1e-05 | 8.1e-06 | 1e-05 | 9.700E-06 | 1.473E-06 | 8.100E-06 | 1.100E-05 |
| 4015 | 201 | G201P | A | 1.4e-05 | 1.2e-05 | 9.6e-06 | 1.187E-05 | 2.203E-06 | 9.600E-06 | 1.400E-05 |
| 4016 | 201 | G201S | A | 9.6e-06 | 8.7e-06 | 8e-06 | 8.767E-06 | 8.021E-07 | 8.000E-06 | 9.600E-06 |
| 4017 | 201 | G201T | A | 1.2e-05 | 7.1e-06 | 5.7e-06 | 8.267E-06 | 3.308E-06 | 5.700E-06 | 1.200E-05 |
| 4018 | 201 | G201W | A | 8.3e-06 | 6.6e-06 | 8.2e-06 | 7.700E-06 | 9.539E-07 | 6.600E-06 | 8.300E-06 |
| 4019 | 201 | G201Y | A | 9.7e-06 | 6.7e-06 | 8.8e-06 | 8.400E-06 | 1.539E-06 | 6.700E-06 | 9.700E-06 |
| 4020 | 201 | G201V | A | 9.4e-06 | 9.4e-06 | 5.2e-06 | 8.000E-06 | 2.425E-06 | 5.200E-06 | 9.400E-06 |
| 4021 | 202 | C202A | A | 1e-05 | 1.2e-05 | 9.8e-06 | 1.060E-05 | 1.217E-06 | 9.800E-06 | 1.200E-05 |
| 4022 | 202 | C202R | A | 7.7e-06 | 5.3e-06 | 6.6e-06 | 6.533E-06 | 1.201E-06 | 5.300E-06 | 7.700E-06 |
| 4023 | 202 | C202N | A | 6.8e-06 | 8.5e-06 | 9.1e-06 | 8.133E-06 | 1.193E-06 | 6.800E-06 | 9.100E-06 |
| 4024 | 202 | C202D | A | 6.6e-06 | 8.4e-06 | 9.3e-06 | 8.100E-06 | 1.375E-06 | 6.600E-06 | 9.300E-06 |
| 4025 | 202 | C202C | A | 1e-05 | 9.6e-06 | 1.4e-05 | 1.120E-05 | 2.433E-06 | 9.600E-06 | 1.400E-05 |
| 4026 | 202 | C202E | A | 6.3e-06 | 7.6e-06 | 7.9e-06 | 7.267E-06 | 8.505E-07 | 6.300E-06 | 7.900E-06 |
| 4027 | 202 | C202Q | A | 1.4e-05 | 6e-06 | 9.7e-06 | 9.900E-06 | 4.004E-06 | 6.000E-06 | 1.400E-05 |
| 4028 | 202 | C202G | A | 9.4e-06 | 4.8e-06 | 1e-05 | 8.067E-06 | 2.845E-06 | 4.800E-06 | 1.000E-05 |
| 4029 | 202 | C202H | A | 7.8e-06 | 8.3e-06 | 4.3e-06 | 6.800E-06 | 2.179E-06 | 4.300E-06 | 8.300E-06 |
| 4030 | 202 | C202I | A | 8.6e-06 | 8.3e-06 | 1.8e-05 | 1.163E-05 | 5.516E-06 | 8.300E-06 | 1.800E-05 |
| 4031 | 202 | C202L | A | 5.4e-06 | 8.6e-06 | 9.9e-06 | 7.967E-06 | 2.316E-06 | 5.400E-06 | 9.900E-06 |
| 4032 | 202 | C202K | A | 7.3e-06 | 5.5e-06 | 1.3e-05 | 8.600E-06 | 3.915E-06 | 5.500E-06 | 1.300E-05 |
| 4033 | 202 | C202M | A | 1.2e-05 | 4.6e-06 | 8.5e-06 | 8.367E-06 | 3.702E-06 | 4.600E-06 | 1.200E-05 |
| 4034 | 202 | C202F | A | 6.9e-06 | 7.4e-06 | 5.2e-06 | 6.500E-06 | 1.153E-06 | 5.200E-06 | 7.400E-06 |
| 4035 | 202 | C202P | A | 4.9e-06 | 6.3e-06 | 4.8e-06 | 5.333E-06 | 8.386E-07 | 4.800E-06 | 6.300E-06 |
| 4036 | 202 | C202S | A | 8.8e-06 | 6.6e-06 | 1.3e-05 | 9.467E-06 | 3.252E-06 | 6.600E-06 | 1.300E-05 |
| 4037 | 202 | C202T | A | 1e-05 | 1.4e-05 | 8.8e-06 | 1.093E-05 | 2.723E-06 | 8.800E-06 | 1.400E-05 |
| 4038 | 202 | C202W | A | 1.3e-05 | 8.9e-06 | 5.6e-06 | 9.167E-06 | 3.707E-06 | 5.600E-06 | 1.300E-05 |
| 4039 | 202 | C202Y | A | 5.3e-06 | 6.4e-06 | 9.7e-06 | 7.133E-06 | 2.290E-06 | 5.300E-06 | 9.700E-06 |
| 4040 | 202 | C202V | A | 6.7e-06 | 1.1e-05 | 1.3e-05 | 1.023E-05 | 3.219E-06 | 6.700E-06 | 1.300E-05 |
| 4041 | 203 | A203A | A | 8.4e-06 | 9.4e-06 | 8.9e-06 | 8.900E-06 | 5.000E-07 | 8.400E-06 | 9.400E-06 |
| 4042 | 203 | A203R | A | 9.3e-06 | 8.2e-06 | 5.4e-06 | 7.633E-06 | 2.011E-06 | 5.400E-06 | 9.300E-06 |
| 4043 | 203 | A203N | A | 7.9e-06 | 6.6e-06 | 1.1e-05 | 8.500E-06 | 2.261E-06 | 6.600E-06 | 1.100E-05 |
| 4044 | 203 | A203D | A | 9.7e-06 | 7e-06 | 8.3e-06 | 8.333E-06 | 1.350E-06 | 7.000E-06 | 9.700E-06 |
| 4045 | 203 | A203C | A | 7.5e-06 | 5e-06 | 4.3e-06 | 5.600E-06 | 1.682E-06 | 4.300E-06 | 7.500E-06 |
| 4046 | 203 | A203E | A | 1.5e-05 | 5e-06 | 7.7e-06 | 9.233E-06 | 5.173E-06 | 5.000E-06 | 1.500E-05 |
| 4047 | 203 | A203Q | A | 5.9e-06 | 8.8e-06 | 4.9e-06 | 6.533E-06 | 2.026E-06 | 4.900E-06 | 8.800E-06 |
| 4048 | 203 | A203G | A | 1.2e-05 | 4.9e-06 | 6.4e-06 | 7.767E-06 | 3.742E-06 | 4.900E-06 | 1.200E-05 |
| 4049 | 203 | A203H | A | 4.6e-06 | 6.7e-06 | 1.1e-05 | 7.433E-06 | 3.262E-06 | 4.600E-06 | 1.100E-05 |
| 4050 | 203 | A203I | A | 8.7e-06 | 7.7e-06 | 8.7e-06 | 8.367E-06 | 5.774E-07 | 7.700E-06 | 8.700E-06 |
| 4051 | 203 | A203L | A | 1.3e-05 | 8.4e-06 | 8.6e-06 | 1.000E-05 | 2.600E-06 | 8.400E-06 | 1.300E-05 |
| 4052 | 203 | A203K | A | 1.1e-05 | 6e-06 | 1.4e-05 | 1.033E-05 | 4.041E-06 | 6.000E-06 | 1.400E-05 |
| 4053 | 203 | A203M | A | 1.3e-05 | 8.2e-06 | 6.7e-06 | 9.300E-06 | 3.291E-06 | 6.700E-06 | 1.300E-05 |
| 4054 | 203 | A203F | A | 9.9e-06 | 9.9e-06 | 1.4e-05 | 1.127E-05 | 2.367E-06 | 9.900E-06 | 1.400E-05 |
| 4055 | 203 | A203P | A | 9.4e-06 | 6.1e-06 | 6.2e-06 | 7.233E-06 | 1.877E-06 | 6.100E-06 | 9.400E-06 |
| 4056 | 203 | A203S | A | 5.4e-06 | 1.2e-05 | 7e-06 | 8.133E-06 | 3.443E-06 | 5.400E-06 | 1.200E-05 |
| 4057 | 203 | A203T | A | 9.9e-06 | 7.6e-06 | 4.2e-06 | 7.233E-06 | 2.868E-06 | 4.200E-06 | 9.900E-06 |
| 4058 | 203 | A203W | A | 5.1e-06 | 8.2e-06 | 1.1e-05 | 8.100E-06 | 2.951E-06 | 5.100E-06 | 1.100E-05 |
| 4059 | 203 | A203Y | A | 5.6e-06 | 1.1e-05 | 6.4e-06 | 7.667E-06 | 2.914E-06 | 5.600E-06 | 1.100E-05 |
| 4060 | 203 | A203V | A | 7.3e-06 | 8.1e-06 | 6.3e-06 | 7.233E-06 | 9.018E-07 | 6.300E-06 | 8.100E-06 |
| 4061 | 204 | R204A | A | 4.8e-06 | 6.6e-06 | 4.5e-06 | 5.300E-06 | 1.136E-06 | 4.500E-06 | 6.600E-06 |
| 4062 | 204 | R204R | A | 1e-05 | 6.5e-06 | 4.3e-06 | 6.933E-06 | 2.875E-06 | 4.300E-06 | 1.000E-05 |
| 4063 | 204 | R204N | A | 1.4e-05 | 6.8e-06 | 1.1e-05 | 1.060E-05 | 3.617E-06 | 6.800E-06 | 1.400E-05 |
| 4064 | 204 | R204D | A | 1e-05 | 7.8e-06 | 7.7e-06 | 8.500E-06 | 1.300E-06 | 7.700E-06 | 1.000E-05 |
| 4065 | 204 | R204C | A | 9.2e-06 | 8.2e-06 | 7.5e-06 | 8.300E-06 | 8.544E-07 | 7.500E-06 | 9.200E-06 |
| 4066 | 204 | R204E | A | 6.3e-06 | 1e-05 | 5.1e-06 | 7.133E-06 | 2.554E-06 | 5.100E-06 | 1.000E-05 |
| 4067 | 204 | R204Q | A | 7.4e-06 | 8.8e-06 | 1.5e-05 | 1.040E-05 | 4.045E-06 | 7.400E-06 | 1.500E-05 |
| 4068 | 204 | R204G | A | 6.6e-06 | 1.3e-05 | 5.5e-06 | 8.367E-06 | 4.050E-06 | 5.500E-06 | 1.300E-05 |
| 4069 | 204 | R204H | A | 9.4e-06 | 1.2e-05 | 9e-06 | 1.013E-05 | 1.629E-06 | 9.000E-06 | 1.200E-05 |
| 4070 | 204 | R204I | A | 5.7e-06 | 7.2e-06 | 7.5e-06 | 6.800E-06 | 9.644E-07 | 5.700E-06 | 7.500E-06 |
| 4071 | 204 | R204L | A | 4.7e-06 | 4.6e-06 | 6.2e-06 | 5.167E-06 | 8.963E-07 | 4.600E-06 | 6.200E-06 |
| 4072 | 204 | R204K | A | 1.3e-05 | 5.8e-06 | 8.4e-06 | 9.067E-06 | 3.646E-06 | 5.800E-06 | 1.300E-05 |
| 4073 | 204 | R204M | A | 8.7e-06 | 7.9e-06 | 6.9e-06 | 7.833E-06 | 9.018E-07 | 6.900E-06 | 8.700E-06 |
| 4074 | 204 | R204F | A | 7.3e-06 | 8.5e-06 | 9.4e-06 | 8.400E-06 | 1.054E-06 | 7.300E-06 | 9.400E-06 |
| 4075 | 204 | R204P | A | 9.5e-06 | 9e-06 | 7.2e-06 | 8.567E-06 | 1.210E-06 | 7.200E-06 | 9.500E-06 |
| 4076 | 204 | R204S | A | 9.4e-06 | 9.7e-06 | 6.4e-06 | 8.500E-06 | 1.825E-06 | 6.400E-06 | 9.700E-06 |
| 4077 | 204 | R204T | A | 7.9e-06 | 8.2e-06 | 6.2e-06 | 7.433E-06 | 1.079E-06 | 6.200E-06 | 8.200E-06 |
| 4078 | 204 | R204W | A | 6.7e-06 | 8.9e-06 | 5.7e-06 | 7.100E-06 | 1.637E-06 | 5.700E-06 | 8.900E-06 |
| 4079 | 204 | R204Y | A | 7.1e-06 | 6.6e-06 | 9.3e-06 | 7.667E-06 | 1.436E-06 | 6.600E-06 | 9.300E-06 |

| | | | | | | | | | | |
|---|---|---|---|---|---|---|---|---|---|---|
| 4080 | 204 | R204V | A | 6.5e-06 | 9.4e-06 | 9.4e-06 | 8.433E-06 | 1.674E-06 | 6.500E-06 | 9.400E-06 |
| 4081 | 205 | C205A | A | 5.9e-06 | 8.8e-06 | 1.3e-05 | 9.233E-06 | 3.570E-06 | 5.900E-06 | 1.300E-05 |
| 4082 | 205 | C205R | A | 7.4e-06 | 7.3e-06 | 1.4e-05 | 9.567E-06 | 3.840E-06 | 7.300E-06 | 1.400E-05 |
| 4083 | 205 | C205N | A | 1.1e-05 | 9.2e-06 | 6.3e-06 | 8.833E-06 | 2.371E-06 | 6.300E-06 | 1.100E-05 |
| 4084 | 205 | C205D | A | 8.8e-06 | 8.7e-06 | 1.9e-05 | 1.217E-05 | 5.918E-06 | 8.700E-06 | 1.900E-05 |
| 4085 | 205 | C205C | A | 1.4e-05 | 5.9e-06 | 4.2e-06 | 8.033E-06 | 5.237E-06 | 4.200E-06 | 1.400E-05 |
| 4086 | 205 | C205E | A | 8.3e-06 | 1e-05 | 1.4e-05 | 1.077E-05 | 2.926E-06 | 8.300E-06 | 1.400E-05 |
| 4087 | 205 | C205Q | A | 9.4e-06 | 7.1e-06 | 6.7e-06 | 7.733E-06 | 1.457E-06 | 6.700E-06 | 9.400E-06 |
| 4088 | 205 | C205G | A | 1.3e-05 | 9.7e-06 | 1.1e-05 | 1.123E-05 | 1.662E-06 | 9.700E-06 | 1.300E-05 |
| 4089 | 205 | C205H | A | 1.4e-05 | 4e-06 | 9.1e-06 | 9.033E-06 | 5.000E-06 | 4.000E-06 | 1.400E-05 |
| 4090 | 205 | C205I | A | 7.4e-06 | 1e-05 | 6.8e-06 | 8.067E-06 | 1.701E-06 | 6.800E-06 | 1.000E-05 |
| 4091 | 205 | C205L | A | 4.1e-06 | 7.6e-06 | 9.5e-06 | 7.067E-06 | 2.739E-06 | 4.100E-06 | 9.500E-06 |
| 4092 | 205 | C205K | A | 7.1e-06 | 5.5e-06 | 7e-06 | 6.533E-06 | 8.963E-07 | 5.500E-06 | 7.100E-06 |
| 4093 | 205 | C205M | A | 1e-05 | 7.2e-06 | 1.1e-05 | 9.400E-06 | 1.970E-06 | 7.200E-06 | 1.100E-05 |
| 4094 | 205 | C205F | A | 6.9e-06 | 6.1e-06 | 5.4e-06 | 6.133E-06 | 7.506E-07 | 5.400E-06 | 6.900E-06 |
| 4095 | 205 | C205P | A | 9.9e-06 | 4.5e-06 | 9.4e-06 | 7.933E-06 | 2.984E-06 | 4.500E-06 | 9.900E-06 |
| 4096 | 205 | C205S | A | 5.8e-06 | 8.4e-06 | 8.4e-06 | 7.533E-06 | 1.501E-06 | 5.800E-06 | 8.400E-06 |
| 4097 | 205 | C205T | A | 9.3e-06 | 5.4e-06 | 6.8e-06 | 7.167E-06 | 1.976E-06 | 5.400E-06 | 9.300E-06 |
| 4098 | 205 | C205W | A | 1e-05 | 9.2e-06 | 1e-05 | 9.733E-06 | 4.619E-07 | 9.200E-06 | 1.000E-05 |
| 4099 | 205 | C205Y | A | 1.2e-05 | 1.1e-05 | 9.8e-06 | 1.093E-05 | 1.102E-06 | 9.800E-06 | 1.200E-05 |
| 4100 | 205 | C205V | A | 1.4e-05 | 6.9e-06 | 5.3e-06 | 8.733E-06 | 4.631E-06 | 5.300E-06 | 1.400E-05 |
| 4101 | 206 | K206A | A | 8.3e-06 | 6.6e-06 | 5.7e-06 | 6.867E-06 | 1.320E-06 | 5.700E-06 | 8.300E-06 |
| 4102 | 206 | K206R | A | 6.4e-06 | 8.6e-06 | 9.5e-06 | 8.167E-06 | 1.595E-06 | 6.400E-06 | 9.500E-06 |
| 4103 | 206 | K206N | A | 1.3e-05 | 5.7e-06 | 7e-06 | 8.567E-06 | 3.894E-06 | 5.700E-06 | 1.300E-05 |
| 4104 | 206 | K206D | A | 1.1e-05 | 9.4e-06 | 1.1e-05 | 1.047E-05 | 9.238E-07 | 9.400E-06 | 1.100E-05 |
| 4105 | 206 | K206C | A | 5.4e-06 | 5.7e-06 | 4.3e-06 | 5.133E-06 | 7.371E-07 | 4.300E-06 | 5.700E-06 |
| 4106 | 206 | K206E | A | 9.3e-06 | 6.5e-06 | 6.7e-06 | 7.500E-06 | 1.562E-06 | 6.500E-06 | 9.300E-06 |
| 4107 | 206 | K206Q | A | 5.8e-06 | 4.3e-06 | 1.2e-05 | 7.367E-06 | 4.082E-06 | 4.300E-06 | 1.200E-05 |
| 4108 | 206 | K206G | A | 1.1e-05 | 9.5e-06 | 9.6e-06 | 1.003E-05 | 8.386E-07 | 9.500E-06 | 1.100E-05 |
| 4109 | 206 | K206H | A | 5.1e-06 | 5.2e-06 | 5.4e-06 | 5.233E-06 | 1.528E-07 | 5.100E-06 | 5.400E-06 |
| 4110 | 206 | K206I | A | 1.1e-05 | 6.3e-06 | 9.5e-06 | 8.933E-06 | 2.401E-06 | 6.300E-06 | 1.100E-05 |
| 4111 | 206 | K206L | A | 4.4e-06 | 7e-06 | 6.3e-06 | 5.900E-06 | 1.345E-06 | 4.400E-06 | 7.000E-06 |
| 4112 | 206 | K206K | A | 6e-06 | 4.6e-06 | 8.1e-06 | 6.233E-06 | 1.762E-06 | 4.600E-06 | 8.100E-06 |
| 4113 | 206 | K206M | A | 7.8e-06 | 9e-06 | 6.7e-06 | 7.833E-06 | 1.150E-06 | 6.700E-06 | 9.000E-06 |
| 4114 | 206 | K206F | A | 4.7e-06 | 1.5e-05 | 9e-06 | 9.567E-06 | 5.173E-06 | 4.700E-06 | 1.500E-05 |
| 4115 | 206 | K206P | A | 7.3e-06 | 1e-05 | 1.1e-05 | 9.433E-06 | 1.914E-06 | 7.300E-06 | 1.100E-05 |
| 4116 | 206 | K206S | A | 7.8e-06 | 9.7e-06 | 9.1e-06 | 8.867E-06 | 9.713E-07 | 7.800E-06 | 9.700E-06 |
| 4117 | 206 | K206T | A | 7.4e-06 | 5e-06 | 7.1e-06 | 6.500E-06 | 1.308E-06 | 5.000E-06 | 7.400E-06 |
| 4118 | 206 | K206W | A | 8.2e-06 | 1.3e-05 | 6.4e-06 | 9.200E-06 | 3.412E-06 | 6.400E-06 | 1.300E-05 |
| 4119 | 206 | K206Y | A | 5.7e-06 | 9.3e-06 | 1.1e-05 | 8.667E-06 | 2.706E-06 | 5.700E-06 | 1.100E-05 |
| 4120 | 206 | K206V | A | 7.6e-06 | 1e-05 | 4.6e-06 | 7.400E-06 | 2.706E-06 | 4.600E-06 | 1.000E-05 |
| 4121 | 207 | G207A | A | 9.1e-06 | 4.9e-06 | 7.5e-06 | 7.167E-06 | 2.120E-06 | 4.900E-06 | 9.100E-06 |
| 4122 | 207 | G207R | A | 1.4e-05 | 6.8e-06 | 1.3e-05 | 1.127E-05 | 3.900E-06 | 6.800E-06 | 1.400E-05 |
| 4123 | 207 | G207N | A | 9.4e-06 | 8.8e-06 | 5.7e-06 | 7.967E-06 | 1.986E-06 | 5.700E-06 | 9.400E-06 |
| 4124 | 207 | G207D | A | 1.3e-05 | 1e-05 | 6.4e-06 | 9.800E-06 | 3.305E-06 | 6.400E-06 | 1.300E-05 |
| 4125 | 207 | G207C | A | 9.4e-06 | 6e-06 | 4.6e-06 | 6.667E-06 | 2.468E-06 | 4.600E-06 | 9.400E-06 |
| 4126 | 207 | G207E | A | 7e-06 | 8.7e-06 | 4.9e-06 | 6.867E-06 | 1.904E-06 | 4.900E-06 | 8.700E-06 |
| 4127 | 207 | G207Q | A | 7.8e-06 | 1e-05 | 4.1e-06 | 7.300E-06 | 2.982E-06 | 4.100E-06 | 1.000E-05 |
| 4128 | 207 | G207G | A | 7.8e-06 | 6.4e-06 | 9.5e-06 | 7.900E-06 | 1.552E-06 | 6.400E-06 | 9.500E-06 |
| 4129 | 207 | G207H | A | 6.1e-06 | 8.8e-06 | 7.3e-06 | 7.400E-06 | 1.353E-06 | 6.100E-06 | 8.800E-06 |
| 4130 | 207 | G207I | A | 8.4e-06 | 9.3e-06 | 1.5e-05 | 1.090E-05 | 3.579E-06 | 8.400E-06 | 1.500E-05 |
| 4131 | 207 | G207L | A | 7.1e-06 | 7e-06 | 5.8e-06 | 6.633E-06 | 7.234E-07 | 5.800E-06 | 7.100E-06 |
| 4132 | 207 | G207K | A | 8.3e-06 | 1e-05 | 6.6e-06 | 8.300E-06 | 1.700E-06 | 6.600E-06 | 1.000E-05 |
| 4133 | 207 | G207M | A | 1e-05 | 8.7e-06 | 1.2e-05 | 1.023E-05 | 1.662E-06 | 8.700E-06 | 1.200E-05 |
| 4134 | 207 | G207F | A | 9.4e-06 | 5.1e-06 | 7.9e-06 | 7.467E-06 | 2.183E-06 | 5.100E-06 | 9.400E-06 |
| 4135 | 207 | G207P | A | 5.2e-06 | 6.4e-06 | 8.2e-06 | 6.600E-06 | 1.510E-06 | 5.200E-06 | 8.200E-06 |
| 4136 | 207 | G207S | A | 6.6e-06 | 3.6e-06 | 1.6e-05 | 8.733E-06 | 6.469E-06 | 3.600E-06 | 1.600E-05 |
| 4137 | 207 | G207T | A | 1e-05 | 7e-06 | 6.6e-06 | 7.867E-06 | 1.858E-06 | 6.600E-06 | 1.000E-05 |
| 4138 | 207 | G207W | A | 1.4e-05 | 4.9e-06 | 5e-06 | 7.967E-06 | 5.225E-06 | 4.900E-06 | 1.400E-05 |
| 4139 | 207 | G207Y | A | 6.6e-06 | 4.4e-06 | 5.4e-06 | 5.467E-06 | 1.102E-06 | 4.400E-06 | 6.600E-06 |
| 4140 | 207 | G207V | A | 5.2e-06 | 4.5e-06 | 6.4e-06 | 5.367E-06 | 9.609E-07 | 4.500E-06 | 6.400E-06 |
| 4141 | 208 | P208A | A | 8.9e-06 | 4.9e-06 | 9.3e-06 | 7.700E-06 | 2.433E-06 | 4.900E-06 | 9.300E-06 |
| 4142 | 208 | P208R | A | 4e-06 | 5.3e-06 | 8.8e-06 | 6.033E-06 | 2.483E-06 | 4.000E-06 | 8.800E-06 |
| 4143 | 208 | P208N | A | 4.7e-06 | 8.9e-06 | 9.3e-06 | 7.633E-06 | 2.548E-06 | 4.700E-06 | 9.300E-06 |
| 4144 | 208 | P208D | A | 1.1e-05 | 1.1e-05 | 6.2e-06 | 9.400E-06 | 2.771E-06 | 6.200E-06 | 1.100E-05 |
| 4145 | 208 | P208C | A | 1.4e-05 | 1.4e-05 | 4.9e-06 | 1.097E-05 | 5.254E-06 | 4.900E-06 | 1.400E-05 |
| 4146 | 208 | P208E | A | 7.1e-06 | 9.5e-06 | 9.1e-06 | 8.567E-06 | 1.286E-06 | 7.100E-06 | 9.500E-06 |
| 4147 | 208 | P208Q | A | 6e-06 | 9.3e-06 | 6.7e-06 | 7.333E-06 | 1.739E-06 | 6.000E-06 | 9.300E-06 |
| 4148 | 208 | P208G | A | 7.2e-06 | 4.9e-06 | 1.2e-05 | 8.033E-06 | 3.623E-06 | 4.900E-06 | 1.200E-05 |
| 4149 | 208 | P208H | A | 6.3e-06 | 6.5e-06 | 6.6e-06 | 6.467E-06 | 1.528E-07 | 6.300E-06 | 6.600E-06 |
| 4150 | 208 | P208I | A | 6.7e-06 | 9.7e-06 | 1e-05 | 8.800E-06 | 1.825E-06 | 6.700E-06 | 1.000E-05 |
| 4151 | 208 | P208L | A | 6.6e-06 | 3.2e-06 | 9e-06 | 6.267E-06 | 2.914E-06 | 3.200E-06 | 9.000E-06 |
| 4152 | 208 | P208K | A | 6.1e-06 | 6.5e-06 | 6.9e-06 | 6.500E-06 | 4.000E-07 | 6.100E-06 | 6.900E-06 |
| 4153 | 208 | P208M | A | 6.9e-06 | 6.3e-06 | 5.9e-06 | 6.367E-06 | 5.033E-07 | 5.900E-06 | 6.900E-06 |
| 4154 | 208 | P208F | A | 6.9e-06 | 6.9e-06 | 5.6e-06 | 6.467E-06 | 7.506E-07 | 5.600E-06 | 6.900E-06 |
| 4155 | 208 | P208P | A | 1.5e-05 | 7.8e-06 | 7.1e-06 | 9.967E-06 | 4.373E-06 | 7.100E-06 | 1.500E-05 |
| 4156 | 208 | P208S | A | 8.5e-06 | 6.5e-06 | 6e-06 | 7.000E-06 | 1.323E-06 | 6.000E-06 | 8.500E-06 |
| 4157 | 208 | P208T | A | 6.5e-06 | 7.1e-06 | 4.5e-06 | 6.033E-06 | 1.361E-06 | 4.500E-06 | 7.100E-06 |
| 4158 | 208 | P208W | A | 8.6e-06 | 6.2e-06 | 8.9e-06 | 7.900E-06 | 1.480E-06 | 6.200E-06 | 8.900E-06 |
| 4159 | 208 | P208Y | A | 9.4e-06 | 9.1e-06 | 9.9e-06 | 9.467E-06 | 4.041E-07 | 9.100E-06 | 9.900E-06 |
| 4160 | 208 | P208V | A | 1.1e-05 | 1.4e-05 | 5.9e-06 | 1.030E-05 | 4.095E-06 | 5.900E-06 | 1.400E-05 |
| 4161 | 209 | L209A | A | 5.3e-06 | 9.5e-06 | 8.7e-06 | 7.833E-06 | 2.230E-06 | 5.300E-06 | 9.500E-06 |
| 4162 | 209 | L209R | A | 7.3e-06 | 4.4e-06 | 8.3e-06 | 6.667E-06 | 2.026E-06 | 4.400E-06 | 8.300E-06 |
| 4163 | 209 | L209N | A | 5.9e-06 | 7.5e-06 | 5.9e-06 | 6.433E-06 | 9.238E-07 | 5.900E-06 | 7.500E-06 |
| 4164 | 209 | L209D | A | 6.7e-06 | 6.9e-06 | 9.8e-06 | 7.800E-06 | 1.735E-06 | 6.700E-06 | 9.800E-06 |

| | | | | | | | | | | |
|---|---|---|---|---|---|---|---|---|---|---|
| 4165 | 209 | L209C | A | 1.3e-05 | 1.2e-05 | 5e-06 | 1.000E-05 | 4.359E-06 | 5.000E-06 | 1.300E-05 |
| 4166 | 209 | L209E | A | 6.6e-06 | 9.4e-06 | 5.2e-06 | 7.067E-06 | 2.139E-06 | 5.200E-06 | 9.400E-06 |
| 4167 | 209 | L209Q | A | 6.9e-06 | 8.7e-06 | 1e-05 | 8.533E-06 | 1.557E-06 | 6.900E-06 | 1.000E-05 |
| 4168 | 209 | L209G | A | 1.1e-05 | 5.7e-06 | 6.8e-06 | 7.833E-06 | 2.797E-06 | 5.700E-06 | 1.100E-05 |
| 4169 | 209 | L209H | A | 7.8e-06 | 6.6e-06 | 5.3e-06 | 6.567E-06 | 1.250E-06 | 5.300E-06 | 7.800E-06 |
| 4170 | 209 | L209I | A | 1.4e-05 | 3.2e-06 | 1.3e-05 | 1.007E-05 | 5.968E-06 | 3.200E-06 | 1.400E-05 |
| 4171 | 209 | L209L | A | 1.2e-05 | 8.6e-06 | 6.7e-06 | 9.100E-06 | 2.685E-06 | 6.700E-06 | 1.200E-05 |
| 4172 | 209 | L209K | A | 6.9e-06 | 8.9e-06 | 3.6e-06 | 6.467E-06 | 2.676E-06 | 3.600E-06 | 8.900E-06 |
| 4173 | 209 | L209M | A | 1e-05 | 8.8e-06 | 9.1e-06 | 9.300E-06 | 6.245E-07 | 8.800E-06 | 1.000E-05 |
| 4174 | 209 | L209F | A | 9.2e-06 | 5.8e-06 | 7.1e-06 | 7.367E-06 | 1.716E-06 | 5.800E-06 | 9.200E-06 |
| 4175 | 209 | L209P | A | 6.5e-06 | 1.1e-05 | 6.3e-06 | 7.933E-06 | 2.658E-06 | 6.300E-06 | 1.100E-05 |
| 4176 | 209 | L209S | A | 8.5e-06 | 8.4e-06 | 5.9e-06 | 7.600E-06 | 1.473E-06 | 5.900E-06 | 8.500E-06 |
| 4177 | 209 | L209T | A | 5.1e-06 | 7.8e-06 | 1e-05 | 7.633E-06 | 2.454E-06 | 5.100E-06 | 1.000E-05 |
| 4178 | 209 | L209W | A | 1.5e-05 | 1.1e-05 | 1.3e-05 | 1.300E-05 | 2.000E-06 | 1.100E-05 | 1.500E-05 |
| 4179 | 209 | L209Y | A | 7e-06 | 9.3e-06 | 8.2e-06 | 8.167E-06 | 1.150E-06 | 7.000E-06 | 9.300E-06 |
| 4180 | 209 | L209V | A | 4.6e-06 | 1.2e-05 | 1.2e-05 | 9.533E-06 | 4.272E-06 | 4.600E-06 | 1.200E-05 |
| 4181 | 210 | P210A | A | 9.5e-06 | 1.2e-05 | 1e-05 | 1.050E-05 | 1.323E-06 | 9.500E-06 | 1.200E-05 |
| 4182 | 210 | P210R | A | 6e-06 | 9e-06 | 1e-05 | 8.333E-06 | 2.082E-06 | 6.000E-06 | 1.000E-05 |
| 4183 | 210 | P210N | A | 5.6e-06 | 8e-06 | 6.4e-06 | 6.667E-06 | 1.222E-06 | 5.600E-06 | 8.000E-06 |
| 4184 | 210 | P210D | A | 1e-05 | 1.2e-05 | 1e-05 | 1.067E-05 | 1.155E-06 | 1.000E-05 | 1.200E-05 |
| 4185 | 210 | P210C | A | 1.2e-05 | 1.2e-05 | 6.1e-06 | 1.003E-05 | 3.406E-06 | 6.100E-06 | 1.200E-05 |
| 4186 | 210 | P210E | A | 6.9e-06 | 8.1e-06 | 1.3e-05 | 9.333E-06 | 3.232E-06 | 6.900E-06 | 1.300E-05 |
| 4187 | 210 | P210Q | A | 9.8e-06 | 4.4e-06 | 5.9e-06 | 6.700E-06 | 2.787E-06 | 4.400E-06 | 9.800E-06 |
| 4188 | 210 | P210G | A | 7.8e-06 | 4.9e-06 | 7.9e-06 | 6.867E-06 | 1.704E-06 | 4.900E-06 | 7.900E-06 |
| 4189 | 210 | P210H | A | 1.1e-05 | 9.6e-06 | 8.5e-06 | 9.700E-06 | 1.253E-06 | 8.500E-06 | 1.100E-05 |
| 4190 | 210 | P210I | A | 8.5e-06 | 9e-06 | 6.2e-06 | 7.900E-06 | 1.493E-06 | 6.200E-06 | 9.000E-06 |
| 4191 | 210 | P210L | A | 7.7e-06 | 6.2e-06 | 9.3e-06 | 7.733E-06 | 1.550E-06 | 6.200E-06 | 9.300E-06 |
| 4192 | 210 | P210K | A | 8.7e-06 | 6.8e-06 | 1.3e-05 | 9.500E-06 | 3.176E-06 | 6.800E-06 | 1.300E-05 |
| 4193 | 210 | P210M | A | 6.3e-06 | 8.3e-06 | 9.5e-06 | 8.033E-06 | 1.617E-06 | 6.300E-06 | 9.500E-06 |
| 4194 | 210 | P210F | A | 9e-06 | 6.8e-06 | 1.3e-05 | 9.600E-06 | 3.143E-06 | 6.800E-06 | 1.300E-05 |
| 4195 | 210 | P210P | A | 6.9e-06 | 7.4e-06 | 1.2e-05 | 8.767E-06 | 2.811E-06 | 6.900E-06 | 1.200E-05 |
| 4196 | 210 | P210S | A | 7.8e-06 | 6.6e-06 | 7.6e-06 | 7.333E-06 | 6.429E-07 | 6.600E-06 | 7.800E-06 |
| 4197 | 210 | P210T | A | 8.7e-06 | 9.6e-06 | 5.4e-06 | 7.900E-06 | 2.211E-06 | 5.400E-06 | 9.600E-06 |
| 4198 | 210 | P210W | A | 1.1e-05 | 6e-06 | 7e-06 | 8.000E-06 | 2.646E-06 | 6.000E-06 | 1.100E-05 |
| 4199 | 210 | P210Y | A | 7e-06 | 1.2e-05 | 8.9e-06 | 9.300E-06 | 2.524E-06 | 7.000E-06 | 1.200E-05 |
| 4200 | 210 | P210V | A | 4.9e-06 | 1.4e-05 | 8.5e-06 | 9.133E-06 | 4.583E-06 | 4.900E-06 | 1.400E-05 |
| 4201 | 211 | T211A | A | 1.2e-05 | 8.2e-06 | 5.2e-06 | 8.467E-06 | 3.408E-06 | 5.200E-06 | 1.200E-05 |
| 4202 | 211 | T211R | A | 6e-06 | 9.2e-06 | 6.2e-06 | 7.133E-06 | 1.793E-06 | 6.000E-06 | 9.200E-06 |
| 4203 | 211 | T211N | A | 6.3e-06 | 1.1e-05 | 9.9e-06 | 9.067E-06 | 2.458E-06 | 6.300E-06 | 1.100E-05 |
| 4204 | 211 | T211D | A | 6.5e-06 | 1.1e-05 | 8.7e-06 | 8.733E-06 | 2.250E-06 | 6.500E-06 | 1.100E-05 |
| 4205 | 211 | T211C | A | 6.4e-06 | 1.1e-05 | 6.9e-06 | 8.100E-06 | 2.524E-06 | 6.400E-06 | 1.100E-05 |
| 4206 | 211 | T211E | A | 6.5e-06 | 1.1e-05 | 6.7e-06 | 8.067E-06 | 2.542E-06 | 6.500E-06 | 1.100E-05 |
| 4207 | 211 | T211Q | A | 6.9e-06 | 9.6e-06 | 5.2e-06 | 7.233E-06 | 2.219E-06 | 5.200E-06 | 9.600E-06 |
| 4208 | 211 | T211G | A | 9.8e-06 | 1.5e-05 | 4.8e-06 | 9.867E-06 | 5.100E-06 | 4.800E-06 | 1.500E-05 |
| 4209 | 211 | T211H | A | 6e-06 | 5.8e-06 | 4.9e-06 | 5.567E-06 | 5.859E-07 | 4.900E-06 | 6.000E-06 |
| 4210 | 211 | T211I | A | 6.4e-06 | 7.8e-06 | 9e-06 | 7.733E-06 | 1.301E-06 | 6.400E-06 | 9.000E-06 |
| 4211 | 211 | T211L | A | 7.7e-06 | 6.3e-06 | 7.9e-06 | 7.300E-06 | 8.718E-07 | 6.300E-06 | 7.900E-06 |
| 4212 | 211 | T211K | A | 8.2e-06 | 9.6e-06 | 5.5e-06 | 7.767E-06 | 2.084E-06 | 5.500E-06 | 9.600E-06 |
| 4213 | 211 | T211M | A | 9e-06 | 6.5e-06 | 4.6e-06 | 6.700E-06 | 2.207E-06 | 4.600E-06 | 9.000E-06 |
| 4214 | 211 | T211F | A | 7.9e-06 | 7.3e-06 | 4.6e-06 | 6.600E-06 | 1.758E-06 | 4.600E-06 | 7.900E-06 |
| 4215 | 211 | T211P | A | 6.7e-06 | 5.5e-06 | 9.7e-06 | 7.300E-06 | 2.163E-06 | 5.500E-06 | 9.700E-06 |
| 4216 | 211 | T211S | A | 1.3e-05 | 5.4e-06 | 7.2e-06 | 8.533E-06 | 3.972E-06 | 5.400E-06 | 1.300E-05 |
| 4217 | 211 | T211T | A | 1.4e-05 | 1.5e-05 | 1.3e-05 | 1.400E-05 | 1.000E-06 | 1.300E-05 | 1.500E-05 |
| 4218 | 211 | T211W | A | 9.1e-06 | 9.4e-06 | 8.9e-06 | 9.133E-06 | 2.517E-07 | 8.900E-06 | 9.400E-06 |
| 4219 | 211 | T211Y | A | 8e-06 | 1.5e-05 | 1e-05 | 1.100E-05 | 3.606E-06 | 8.000E-06 | 1.500E-05 |
| 4220 | 211 | T211V | A | 5.6e-06 | 7.6e-06 | 1e-05 | 7.733E-06 | 2.203E-06 | 5.600E-06 | 1.000E-05 |
| 4221 | 212 | D212A | A | 1e-05 | 9.3e-06 | 6.2e-06 | 8.500E-06 | 2.022E-06 | 6.200E-06 | 1.000E-05 |
| 4222 | 212 | D212R | A | 8.8e-06 | 7.2e-06 | 1.1e-05 | 9.000E-06 | 1.908E-06 | 7.200E-06 | 1.100E-05 |
| 4223 | 212 | D212N | A | 9e-06 | 9.6e-06 | 7.9e-06 | 8.833E-06 | 8.622E-07 | 7.900E-06 | 9.600E-06 |
| 4224 | 212 | D212D | A | 3.3e-06 | 7.6e-06 | 9e-06 | 6.633E-06 | 2.970E-06 | 3.300E-06 | 9.000E-06 |
| 4225 | 212 | D212C | A | 4.4e-06 | 1e-05 | 5.3e-06 | 6.567E-06 | 3.007E-06 | 4.400E-06 | 1.000E-05 |
| 4226 | 212 | D212E | A | 9.6e-06 | 9.3e-06 | 1.2e-05 | 1.030E-05 | 1.480E-06 | 9.300E-06 | 1.200E-05 |
| 4227 | 212 | D212Q | A | 7.9e-06 | 1.3e-05 | 7.8e-06 | 9.567E-06 | 2.974E-06 | 7.800E-06 | 1.300E-05 |
| 4228 | 212 | D212G | A | 7.3e-06 | 1.5e-05 | 9.2e-06 | 1.050E-05 | 4.011E-06 | 7.300E-06 | 1.500E-05 |
| 4229 | 212 | D212H | A | 6.6e-06 | 9.1e-06 | 8.6e-06 | 8.100E-06 | 1.323E-06 | 6.600E-06 | 9.100E-06 |
| 4230 | 212 | D212I | A | 3.6e-06 | 7.6e-06 | 6.7e-06 | 5.967E-06 | 2.098E-06 | 3.600E-06 | 7.600E-06 |
| 4231 | 212 | D212L | A | 6.1e-06 | 7.3e-06 | 1e-05 | 7.800E-06 | 1.997E-06 | 6.100E-06 | 1.000E-05 |
| 4232 | 212 | D212K | A | 1.1e-05 | 5.7e-06 | 6.5e-06 | 7.733E-06 | 2.857E-06 | 5.700E-06 | 1.100E-05 |
| 4233 | 212 | D212M | A | 1.4e-05 | 7.2e-06 | 6.8e-06 | 9.333E-06 | 4.046E-06 | 6.800E-06 | 1.400E-05 |
| 4234 | 212 | D212F | A | 5.5e-06 | 7.3e-06 | 1e-05 | 7.600E-06 | 2.265E-06 | 5.500E-06 | 1.000E-05 |
| 4235 | 212 | D212P | A | 6.6e-06 | 8.1e-06 | 7.6e-06 | 7.433E-06 | 7.638E-07 | 6.600E-06 | 8.100E-06 |
| 4236 | 212 | D212S | A | 5.9e-06 | 7.8e-06 | 5.5e-06 | 6.400E-06 | 1.229E-06 | 5.500E-06 | 7.800E-06 |
| 4237 | 212 | D212T | A | 6.2e-06 | 8.5e-06 | 5.7e-06 | 6.800E-06 | 1.493E-06 | 5.700E-06 | 8.500E-06 |
| 4238 | 212 | D212W | A | 8.2e-06 | 9.2e-06 | 2.1e-05 | 1.280E-05 | 7.119E-06 | 8.200E-06 | 2.100E-05 |
| 4239 | 212 | D212Y | A | 4.8e-06 | 8e-06 | 6.7e-06 | 6.500E-06 | 1.609E-06 | 4.800E-06 | 8.000E-06 |
| 4240 | 212 | D212V | A | 9e-06 | 7.9e-06 | 5.9e-06 | 7.600E-06 | 1.572E-06 | 5.900E-06 | 9.000E-06 |
| 4241 | 213 | C213A | A | 7.9e-06 | 8.8e-06 | 8.4e-06 | 8.367E-06 | 4.509E-07 | 7.900E-06 | 8.800E-06 |
| 4242 | 213 | C213R | A | 6.9e-06 | 5.8e-06 | 9.3e-06 | 7.333E-06 | 1.790E-06 | 5.800E-06 | 9.300E-06 |
| 4243 | 213 | C213N | A | 9.3e-06 | 7.4e-06 | 5.4e-06 | 7.367E-06 | 1.950E-06 | 5.400E-06 | 9.300E-06 |
| 4244 | 213 | C213D | A | 7.7e-06 | 7e-06 | 1.3e-05 | 9.233E-06 | 3.281E-06 | 7.000E-06 | 1.300E-05 |
| 4245 | 213 | C213C | A | 1.2e-05 | 9.5e-06 | 4.6e-06 | 8.700E-06 | 3.764E-06 | 4.600E-06 | 1.200E-05 |
| 4246 | 213 | C213E | A | 1.5e-05 | 1e-05 | 9.2e-06 | 1.140E-05 | 3.143E-06 | 9.200E-06 | 1.500E-05 |
| 4247 | 213 | C213Q | A | 6.6e-06 | 5.7e-06 | 6.6e-06 | 6.300E-06 | 5.196E-07 | 5.700E-06 | 6.600E-06 |
| 4248 | 213 | C213G | A | 6.9e-06 | 8.8e-06 | 9.2e-06 | 8.300E-06 | 1.229E-06 | 6.900E-06 | 9.200E-06 |
| 4249 | 213 | C213H | A | 7.8e-06 | 6.4e-06 | 6.7e-06 | 6.967E-06 | 7.371E-07 | 6.400E-06 | 7.800E-06 |

| | | | | | | | | | | |
|---|---|---|---|---|---|---|---|---|---|---|
| 4250 | 213 | C213I | A | 7.9e-06 | 8.4e-06 | 8.4e-06 | 8.233E-06 | 2.887E-07 | 7.900E-06 | 8.400E-06 |
| 4251 | 213 | C213L | A | 1e-05 | 8.9e-06 | 4.8e-06 | 7.900E-06 | 2.740E-06 | 4.800E-06 | 1.000E-05 |
| 4252 | 213 | C213K | A | 3.3e-06 | 3.9e-06 | 6e-06 | 4.400E-06 | 1.418E-06 | 3.300E-06 | 6.000E-06 |
| 4253 | 213 | C213M | A | 1.4e-05 | 5.4e-06 | 1e-05 | 9.800E-06 | 4.303E-06 | 5.400E-06 | 1.400E-05 |
| 4254 | 213 | C213F | A | 4.4e-06 | 1e-05 | 5.7e-06 | 6.700E-06 | 2.931E-06 | 4.400E-06 | 1.000E-05 |
| 4255 | 213 | C213P | A | 1.2e-05 | 9.6e-06 | 1e-05 | 1.053E-05 | 1.286E-06 | 9.600E-06 | 1.200E-05 |
| 4256 | 213 | C213S | A | 4.7e-06 | 4.8e-06 | 8.4e-06 | 5.967E-06 | 2.108E-06 | 4.700E-06 | 8.400E-06 |
| 4257 | 213 | C213T | A | 7.7e-06 | 7.1e-06 | 5.9e-06 | 6.900E-06 | 9.165E-07 | 5.900E-06 | 7.700E-06 |
| 4258 | 213 | C213W | A | 5.7e-06 | 6.6e-06 | 4.6e-06 | 5.633E-06 | 1.002E-06 | 4.600E-06 | 6.600E-06 |
| 4259 | 213 | C213Y | A | 6.4e-06 | 1.4e-05 | 6.2e-06 | 8.867E-06 | 4.447E-06 | 6.200E-06 | 1.400E-05 |
| 4260 | 213 | C213V | A | 7.4e-06 | 9.2e-06 | 5.8e-06 | 7.467E-06 | 1.701E-06 | 5.800E-06 | 9.200E-06 |
| 4261 | 214 | C214A | A | 7.9e-06 | 5.9e-06 | 1.2e-05 | 8.600E-06 | 3.110E-06 | 5.900E-06 | 1.200E-05 |
| 4262 | 214 | C214R | A | 1e-05 | 1e-05 | 5.5e-06 | 8.500E-06 | 2.598E-06 | 5.500E-06 | 1.000E-05 |
| 4263 | 214 | C214N | A | 1.5e-05 | 1.2e-05 | 1.2e-05 | 1.300E-05 | 1.732E-06 | 1.200E-05 | 1.500E-05 |
| 4264 | 214 | C214D | A | 9.9e-06 | 1.2e-05 | 9.9e-06 | 1.060E-05 | 1.212E-06 | 9.900E-06 | 1.200E-05 |
| 4265 | 214 | C214C | A | 7.9e-06 | 7.4e-06 | 1.3e-05 | 9.433E-06 | 3.099E-06 | 7.400E-06 | 1.300E-05 |
| 4266 | 214 | C214E | A | 9e-06 | 5.8e-06 | 1e-05 | 8.267E-06 | 2.194E-06 | 5.800E-06 | 1.000E-05 |
| 4267 | 214 | C214Q | A | 9.3e-06 | 6.8e-06 | 1.2e-05 | 9.367E-06 | 2.601E-06 | 6.800E-06 | 1.200E-05 |
| 4268 | 214 | C214G | A | 5.9e-06 | 9e-06 | 1.7e-05 | 1.063E-05 | 5.727E-06 | 5.900E-06 | 1.700E-05 |
| 4269 | 214 | C214H | A | 6.5e-06 | 6.1e-06 | 8e-06 | 6.867E-06 | 1.002E-06 | 6.100E-06 | 8.000E-06 |
| 4270 | 214 | C214I | A | 9.8e-06 | 9.7e-06 | 1.6e-05 | 1.183E-05 | 3.609E-06 | 9.700E-06 | 1.600E-05 |
| 4271 | 214 | C214L | A | 5.7e-06 | 6.7e-06 | 6.2e-06 | 6.200E-06 | 5.000E-07 | 5.700E-06 | 6.700E-06 |
| 4272 | 214 | C214K | A | 9.7e-06 | 1.4e-05 | 6.6e-06 | 1.010E-05 | 3.716E-06 | 6.600E-06 | 1.400E-05 |
| 4273 | 214 | C214M | A | 1.3e-05 | 4e-06 | 1e-05 | 9.000E-06 | 4.583E-06 | 4.000E-06 | 1.300E-05 |
| 4274 | 214 | C214F | A | 5.9e-06 | 5.3e-06 | 8.7e-06 | 6.633E-06 | 1.815E-06 | 5.300E-06 | 8.700E-06 |
| 4275 | 214 | C214P | A | 4.8e-06 | 1.4e-05 | 1.1e-05 | 9.933E-06 | 4.692E-06 | 4.800E-06 | 1.400E-05 |
| 4276 | 214 | C214S | A | 9.6e-06 | 6.4e-06 | 7.7e-06 | 7.900E-06 | 1.609E-06 | 6.400E-06 | 9.600E-06 |
| 4277 | 214 | C214T | A | 7.9e-06 | 6.2e-06 | 7.5e-06 | 7.200E-06 | 8.888E-07 | 6.200E-06 | 7.900E-06 |
| 4278 | 214 | C214W | A | 8.7e-06 | 6.7e-06 | 1.2e-05 | 9.133E-06 | 2.676E-06 | 6.700E-06 | 1.200E-05 |
| 4279 | 214 | C214Y | A | 9.9e-06 | 5.7e-06 | 6.7e-06 | 7.433E-06 | 2.194E-06 | 5.700E-06 | 9.900E-06 |
| 4280 | 214 | C214V | A | 1e-05 | 1.1e-05 | 9.5e-06 | 1.017E-05 | 7.638E-07 | 9.500E-06 | 1.100E-05 |
| 4281 | 215 | H215A | A | 6.2e-06 | 7.1e-06 | 9.7e-06 | 7.667E-06 | 1.818E-06 | 6.200E-06 | 9.700E-06 |
| 4282 | 215 | H215R | A | 8.7e-06 | 1.1e-05 | 8.4e-06 | 9.367E-06 | 1.422E-06 | 8.400E-06 | 1.100E-05 |
| 4283 | 215 | H215N | A | 8.7e-06 | 8.9e-06 | 9.9e-06 | 9.167E-06 | 6.429E-07 | 8.700E-06 | 9.900E-06 |
| 4284 | 215 | H215D | A | 5.3e-06 | 9.1e-06 | 1.4e-05 | 9.467E-06 | 4.362E-06 | 5.300E-06 | 1.400E-05 |
| 4285 | 215 | H215C | A | 8.7e-06 | 6.7e-06 | 5e-06 | 6.800E-06 | 1.852E-06 | 5.000E-06 | 8.700E-06 |
| 4286 | 215 | H215E | A | 6.1e-06 | 1.1e-05 | 8.7e-06 | 8.600E-06 | 2.452E-06 | 6.100E-06 | 1.100E-05 |
| 4287 | 215 | H215Q | A | 3.7e-06 | 5.8e-06 | 8.7e-06 | 6.067E-06 | 2.511E-06 | 3.700E-06 | 8.700E-06 |
| 4288 | 215 | H215G | A | 1.3e-05 | 9.2e-06 | 1.7e-05 | 1.307E-05 | 3.900E-06 | 9.200E-06 | 1.700E-05 |
| 4289 | 215 | H215H | A | 1e-05 | 6e-06 | 7.9e-06 | 7.967E-06 | 2.001E-06 | 6.000E-06 | 1.000E-05 |
| 4290 | 215 | H215I | A | 9.3e-06 | 1e-05 | 9.9e-06 | 9.733E-06 | 3.786E-07 | 9.300E-06 | 1.000E-05 |
| 4291 | 215 | H215L | A | 8.5e-06 | 5.4e-06 | 1.1e-05 | 8.300E-06 | 2.805E-06 | 5.400E-06 | 1.100E-05 |
| 4292 | 215 | H215K | A | 5.5e-06 | 5.7e-06 | 1.4e-05 | 8.400E-06 | 4.851E-06 | 5.500E-06 | 1.400E-05 |
| 4293 | 215 | H215M | A | 7.9e-06 | 6.5e-06 | 1.1e-05 | 8.467E-06 | 2.303E-06 | 6.500E-06 | 1.100E-05 |
| 4294 | 215 | H215F | A | 1.4e-05 | 1e-05 | 1e-05 | 1.133E-05 | 2.309E-06 | 1.000E-05 | 1.400E-05 |
| 4295 | 215 | H215P | A | 8.3e-06 | 9.2e-06 | 4.6e-06 | 7.367E-06 | 2.438E-06 | 4.600E-06 | 9.200E-06 |
| 4296 | 215 | H215S | A | 1.2e-05 | 1.4e-05 | 6.7e-06 | 1.090E-05 | 3.772E-06 | 6.700E-06 | 1.400E-05 |
| 4297 | 215 | H215T | A | 7e-06 | 8.4e-06 | 5.6e-06 | 7.000E-06 | 1.400E-06 | 5.600E-06 | 8.400E-06 |
| 4298 | 215 | H215W | A | 1.2e-05 | 9.5e-06 | 9.3e-06 | 1.027E-05 | 1.504E-06 | 9.300E-06 | 1.200E-05 |
| 4299 | 215 | H215Y | A | 9.9e-06 | 5.7e-06 | 7.9e-06 | 7.833E-06 | 2.101E-06 | 5.700E-06 | 9.900E-06 |
| 4300 | 215 | H215V | A | 9.4e-06 | 9.7e-06 | 1.3e-05 | 1.070E-05 | 1.997E-06 | 9.400E-06 | 1.300E-05 |
| 4301 | 216 | E216A | A | 1.4e-05 | 9.9e-06 | 8.4e-06 | 1.077E-05 | 2.899E-06 | 8.400E-06 | 1.400E-05 |
| 4302 | 216 | E216R | A | 7.2e-06 | 5.4e-06 | 8.1e-06 | 6.900E-06 | 1.375E-06 | 5.400E-06 | 8.100E-06 |
| 4303 | 216 | E216N | A | 1.1e-05 | 8.7e-06 | 7.9e-06 | 9.200E-06 | 1.609E-06 | 7.900E-06 | 1.100E-05 |
| 4304 | 216 | E216D | A | 8.2e-06 | 1.3e-05 | 6e-06 | 9.067E-06 | 3.580E-06 | 6.000E-06 | 1.300E-05 |
| 4305 | 216 | E216C | A | 6.4e-06 | 8.2e-06 | 9.5e-06 | 8.033E-06 | 1.557E-06 | 6.400E-06 | 9.500E-06 |
| 4306 | 216 | E216E | A | 9e-06 | 4.7e-06 | 1e-05 | 7.900E-06 | 2.816E-06 | 4.700E-06 | 1.000E-05 |
| 4307 | 216 | E216Q | A | 7.3e-06 | 5e-06 | 7.2e-06 | 6.500E-06 | 1.300E-06 | 5.000E-06 | 7.300E-06 |
| 4308 | 216 | E216G | A | 1.3e-05 | 6.9e-06 | 6.3e-06 | 8.733E-06 | 3.707E-06 | 6.300E-06 | 1.300E-05 |
| 4309 | 216 | E216H | A | 8.9e-06 | 1e-05 | 4.7e-06 | 7.867E-06 | 2.797E-06 | 4.700E-06 | 1.000E-05 |
| 4310 | 216 | E216I | A | 1e-05 | 7.2e-06 | 7.3e-06 | 8.167E-06 | 1.589E-06 | 7.200E-06 | 1.000E-05 |
| 4311 | 216 | E216L | A | 1e-05 | 1.2e-05 | 9e-06 | 1.033E-05 | 1.528E-06 | 9.000E-06 | 1.200E-05 |
| 4312 | 216 | E216K | A | 8e-06 | 9.1e-06 | 5.1e-06 | 7.400E-06 | 2.066E-06 | 5.100E-06 | 9.100E-06 |
| 4313 | 216 | E216M | A | 8.2e-06 | 1.2e-05 | 1.2e-05 | 1.073E-05 | 2.194E-06 | 8.200E-06 | 1.200E-05 |
| 4314 | 216 | E216F | A | 1.4e-05 | 9.4e-06 | 8.3e-06 | 1.057E-05 | 3.024E-06 | 8.300E-06 | 1.400E-05 |
| 4315 | 216 | E216P | A | 7.7e-06 | 6.5e-06 | 1.3e-05 | 9.067E-06 | 3.459E-06 | 6.500E-06 | 1.300E-05 |
| 4316 | 216 | E216S | A | 7.2e-06 | 9.7e-06 | 5e-06 | 7.300E-06 | 2.352E-06 | 5.000E-06 | 9.700E-06 |
| 4317 | 216 | E216T | A | 5.5e-06 | 1.2e-05 | 8e-06 | 8.500E-06 | 3.279E-06 | 5.500E-06 | 1.200E-05 |
| 4318 | 216 | E216W | A | 9.1e-06 | 9.1e-06 | 9e-06 | 9.067E-06 | 5.774E-08 | 9.000E-06 | 9.100E-06 |
| 4319 | 216 | E216Y | A | 6e-06 | 5.3e-06 | 9.9e-06 | 7.067E-06 | 2.479E-06 | 5.300E-06 | 9.900E-06 |
| 4320 | 216 | E216V | A | 9e-06 | 1.4e-05 | 9.5e-06 | 1.083E-05 | 2.754E-06 | 9.000E-06 | 1.400E-05 |
| 4321 | 217 | Q217A | A | 8.6e-06 | 4e-06 | 7.3e-06 | 6.633E-06 | 2.371E-06 | 4.000E-06 | 8.600E-06 |
| 4322 | 217 | Q217R | A | 4.3e-06 | 8.9e-06 | 1.3e-05 | 8.733E-06 | 4.352E-06 | 4.300E-06 | 1.300E-05 |
| 4323 | 217 | Q217N | A | 9.4e-06 | 9.9e-06 | 5.5e-06 | 8.267E-06 | 2.409E-06 | 5.500E-06 | 9.900E-06 |
| 4324 | 217 | Q217D | A | 8.4e-06 | 8e-06 | 7.4e-06 | 7.933E-06 | 5.033E-07 | 7.400E-06 | 8.400E-06 |
| 4325 | 217 | Q217C | A | 7e-06 | 6.1e-06 | 7.3e-06 | 6.800E-06 | 6.245E-07 | 6.100E-06 | 7.300E-06 |
| 4326 | 217 | Q217E | A | 5.4e-06 | 5.9e-06 | 5.3e-06 | 5.533E-06 | 3.215E-07 | 5.300E-06 | 5.900E-06 |
| 4327 | 217 | Q217Q | A | 6.4e-06 | 1.1e-05 | 6.4e-06 | 7.933E-06 | 2.656E-06 | 6.400E-06 | 1.100E-05 |
| 4328 | 217 | Q217G | A | 9.6e-06 | 8.7e-06 | 7.4e-06 | 8.567E-06 | 1.106E-06 | 7.400E-06 | 9.600E-06 |
| 4329 | 217 | Q217H | A | 1.1e-05 | 8e-06 | 7.4e-06 | 8.800E-06 | 1.929E-06 | 7.400E-06 | 1.100E-05 |
| 4330 | 217 | Q217I | A | 1e-05 | 6.4e-06 | 8.1e-06 | 8.167E-06 | 1.801E-06 | 6.400E-06 | 1.000E-05 |
| 4331 | 217 | Q217L | A | 6.5e-06 | 1.3e-05 | 1e-05 | 9.833E-06 | 3.253E-06 | 6.500E-06 | 1.300E-05 |
| 4332 | 217 | Q217K | A | 7.1e-06 | 6.4e-06 | 8.9e-06 | 7.467E-06 | 1.290E-06 | 6.400E-06 | 8.900E-06 |
| 4333 | 217 | Q217M | A | 1.1e-05 | 1.4e-05 | 9.1e-06 | 1.137E-05 | 2.470E-06 | 9.100E-06 | 1.400E-05 |
| 4334 | 217 | Q217F | A | 9.3e-06 | 9.8e-06 | 1.2e-05 | 1.037E-05 | 1.436E-06 | 9.300E-06 | 1.200E-05 |

| | | | | | | | | | | |
|---|---|---|---|---|---|---|---|---|---|---|
| 4335 | 217 | Q217P | A | 4.9e-06 | 1e-05 | 8.9e-06 | 7.933E-06 | 2.684E-06 | 4.900E-06 | 1.000E-05 |
| 4336 | 217 | Q217S | A | 9e-06 | 6.3e-06 | 6.7e-06 | 7.333E-06 | 1.457E-06 | 6.300E-06 | 9.000E-06 |
| 4337 | 217 | Q217T | A | 6.2e-06 | 7.8e-06 | 6.8e-06 | 6.933E-06 | 8.083E-07 | 6.200E-06 | 7.800E-06 |
| 4338 | 217 | Q217W | A | 6.7e-06 | 1.4e-05 | 6.4e-06 | 9.033E-06 | 4.304E-06 | 6.400E-06 | 1.400E-05 |
| 4339 | 217 | Q217Y | A | 6e-06 | 8.3e-06 | 6.8e-06 | 7.033E-06 | 1.168E-06 | 6.000E-06 | 8.300E-06 |
| 4340 | 217 | Q217V | A | 1.1e-05 | 7.4e-06 | 7.4e-06 | 8.600E-06 | 2.078E-06 | 7.400E-06 | 1.100E-05 |
| 4341 | 218 | C218A | A | 8.1e-06 | 8.6e-06 | 1.4e-05 | 1.023E-05 | 3.272E-06 | 8.100E-06 | 1.400E-05 |
| 4342 | 218 | C218R | A | 8.1e-06 | 8.2e-06 | 1.3e-05 | 9.767E-06 | 2.801E-06 | 8.100E-06 | 1.300E-05 |
| 4343 | 218 | C218N | A | 9e-06 | 7.5e-06 | 9.1e-06 | 8.533E-06 | 8.963E-07 | 7.500E-06 | 9.100E-06 |
| 4344 | 218 | C218D | A | 6e-06 | 5.5e-06 | 6.2e-06 | 5.900E-06 | 3.606E-07 | 5.500E-06 | 6.200E-06 |
| 4345 | 218 | C218C | A | 6.7e-06 | 9e-06 | 9.8e-06 | 8.500E-06 | 1.609E-06 | 6.700E-06 | 9.800E-06 |
| 4346 | 218 | C218E | A | 4.8e-06 | 8.1e-06 | 1e-05 | 7.633E-06 | 2.631E-06 | 4.800E-06 | 1.000E-05 |
| 4347 | 218 | C218Q | A | 9.6e-06 | 9.4e-06 | 1.1e-05 | 1.000E-05 | 8.718E-07 | 9.400E-06 | 1.100E-05 |
| 4348 | 218 | C218G | A | 8.7e-06 | 1.4e-05 | 6.2e-06 | 9.633E-06 | 3.983E-06 | 6.200E-06 | 1.400E-05 |
| 4349 | 218 | C218H | A | 8.4e-06 | 1.1e-05 | 4.6e-06 | 8.000E-06 | 3.219E-06 | 4.600E-06 | 1.100E-05 |
| 4350 | 218 | C218I | A | 8.7e-06 | 1.8e-05 | 1.3e-05 | 1.323E-05 | 4.654E-06 | 8.700E-06 | 1.800E-05 |
| 4351 | 218 | C218L | A | 7.8e-06 | 5.7e-06 | 1.4e-05 | 9.167E-06 | 4.315E-06 | 5.700E-06 | 1.400E-05 |
| 4352 | 218 | C218K | A | 7.9e-06 | 1.2e-05 | 1.4e-05 | 1.130E-05 | 3.110E-06 | 7.900E-06 | 1.400E-05 |
| 4353 | 218 | C218M | A | 7.4e-06 | 8.2e-06 | 6.2e-06 | 7.267E-06 | 1.007E-06 | 6.200E-06 | 8.200E-06 |
| 4354 | 218 | C218F | A | 8.1e-06 | 8.7e-06 | 6.5e-06 | 7.767E-06 | 1.137E-06 | 6.500E-06 | 8.700E-06 |
| 4355 | 218 | C218P | A | 6.3e-06 | 8.8e-06 | 9.3e-06 | 8.133E-06 | 1.607E-06 | 6.300E-06 | 9.300E-06 |
| 4356 | 218 | C218S | A | 1.1e-05 | 9.8e-06 | 1.1e-05 | 1.060E-05 | 6.928E-07 | 9.800E-06 | 1.100E-05 |
| 4357 | 218 | C218T | A | 6.6e-06 | 6.8e-06 | 9.1e-06 | 7.500E-06 | 1.389E-06 | 6.600E-06 | 9.100E-06 |
| 4358 | 218 | C218W | A | 5.4e-06 | 8.1e-06 | 1.2e-05 | 8.500E-06 | 3.318E-06 | 5.400E-06 | 1.200E-05 |
| 4359 | 218 | C218Y | A | 1.1e-05 | 7.8e-06 | 4.9e-06 | 7.900E-06 | 3.051E-06 | 4.900E-06 | 1.100E-05 |
| 4360 | 218 | C218V | A | 3.4e-06 | 1.2e-05 | 6.4e-06 | 7.267E-06 | 4.365E-06 | 3.400E-06 | 1.200E-05 |
| 4361 | 219 | A219A | A | 6.7e-06 | 1.2e-05 | 6.2e-06 | 8.300E-06 | 3.214E-06 | 6.200E-06 | 1.200E-05 |
| 4362 | 219 | A219R | A | 1.4e-05 | 1.9e-05 | 1.1e-05 | 1.467E-05 | 4.041E-06 | 1.100E-05 | 1.900E-05 |
| 4363 | 219 | A219N | A | 8.9e-06 | 3.8e-06 | 8e-06 | 6.900E-06 | 2.722E-06 | 3.800E-06 | 8.900E-06 |
| 4364 | 219 | A219D | A | 6.8e-06 | 5.8e-06 | 1.4e-05 | 8.867E-06 | 4.474E-06 | 5.800E-06 | 1.400E-05 |
| 4365 | 219 | A219C | A | 1.5e-05 | 1.3e-05 | 6.9e-06 | 1.163E-05 | 4.219E-06 | 6.900E-06 | 1.500E-05 |
| 4366 | 219 | A219E | A | 9.4e-06 | 9.7e-06 | 4.7e-06 | 7.933E-06 | 2.804E-06 | 4.700E-06 | 9.700E-06 |
| 4367 | 219 | A219Q | A | 7.1e-06 | 9.8e-06 | 5.3e-06 | 7.400E-06 | 2.265E-06 | 5.300E-06 | 9.800E-06 |
| 4368 | 219 | A219G | A | 6.5e-06 | 6.8e-06 | 5.1e-06 | 6.133E-06 | 9.074E-07 | 5.100E-06 | 6.800E-06 |
| 4369 | 219 | A219H | A | 6.2e-06 | 8.4e-06 | 1e-05 | 8.200E-06 | 1.908E-06 | 6.200E-06 | 1.000E-05 |
| 4370 | 219 | A219I | A | 1.4e-05 | 5.3e-06 | 1.2e-05 | 1.043E-05 | 4.557E-06 | 5.300E-06 | 1.400E-05 |
| 4371 | 219 | A219L | A | 9.1e-06 | 8.3e-06 | 1e-05 | 9.133E-06 | 8.505E-07 | 8.300E-06 | 1.000E-05 |
| 4372 | 219 | A219K | A | 8.2e-06 | 1.2e-05 | 6e-06 | 8.733E-06 | 3.035E-06 | 6.000E-06 | 1.200E-05 |
| 4373 | 219 | A219M | A | 1.1e-05 | 8.6e-06 | 7.1e-06 | 8.900E-06 | 1.967E-06 | 7.100E-06 | 1.100E-05 |
| 4374 | 219 | A219F | A | 8.1e-06 | 3.9e-06 | 7.4e-06 | 6.467E-06 | 2.250E-06 | 3.900E-06 | 8.100E-06 |
| 4375 | 219 | A219P | A | 6.4e-06 | 1.1e-05 | 1.7e-05 | 1.147E-05 | 5.315E-06 | 6.400E-06 | 1.700E-05 |
| 4376 | 219 | A219S | A | 1.3e-05 | 6.1e-06 | 8.4e-06 | 9.167E-06 | 3.513E-06 | 6.100E-06 | 1.300E-05 |
| 4377 | 219 | A219T | A | 9.1e-06 | 3.9e-06 | 7.1e-06 | 6.700E-06 | 2.623E-06 | 3.900E-06 | 9.100E-06 |
| 4378 | 219 | A219W | A | 8e-06 | 7.2e-06 | 6e-06 | 7.067E-06 | 1.007E-06 | 6.000E-06 | 8.000E-06 |
| 4379 | 219 | A219Y | A | 5.6e-06 | 6.6e-06 | 6.2e-06 | 6.133E-06 | 5.033E-07 | 5.600E-06 | 6.600E-06 |
| 4380 | 219 | A219V | A | 6.1e-06 | 6.4e-06 | 6.5e-06 | 6.333E-06 | 2.082E-07 | 6.100E-06 | 6.500E-06 |
| 4381 | 220 | A220A | A | 9.6e-06 | 5.2e-06 | 7e-06 | 7.267E-06 | 2.212E-06 | 5.200E-06 | 9.600E-06 |
| 4382 | 220 | A220R | A | 7.1e-06 | 5.4e-06 | 6.7e-06 | 6.400E-06 | 8.888E-07 | 5.400E-06 | 7.100E-06 |
| 4383 | 220 | A220N | A | 7.2e-06 | 1e-05 | 5.6e-06 | 7.600E-06 | 2.227E-06 | 5.600E-06 | 1.000E-05 |
| 4384 | 220 | A220D | A | 5.2e-06 | 1.3e-05 | 6.6e-06 | 8.267E-06 | 4.159E-06 | 5.200E-06 | 1.300E-05 |
| 4385 | 220 | A220C | A | 5.8e-06 | 8.8e-06 | 5.9e-06 | 6.833E-06 | 1.704E-06 | 5.800E-06 | 8.800E-06 |
| 4386 | 220 | A220E | A | 7.2e-06 | 8.2e-06 | 1.3e-05 | 9.467E-06 | 3.101E-06 | 7.200E-06 | 1.300E-05 |
| 4387 | 220 | A220Q | A | 5.4e-06 | 8.8e-06 | 9.5e-06 | 7.900E-06 | 2.193E-06 | 5.400E-06 | 9.500E-06 |
| 4388 | 220 | A220G | A | 8.5e-06 | 7.3e-06 | 6e-06 | 7.267E-06 | 1.250E-06 | 6.000E-06 | 8.500E-06 |
| 4389 | 220 | A220H | A | 9.8e-06 | 7e-06 | 7.8e-06 | 8.200E-06 | 1.442E-06 | 7.000E-06 | 9.800E-06 |
| 4390 | 220 | A220I | A | 6.6e-06 | 1.4e-05 | 1e-05 | 1.020E-05 | 3.704E-06 | 6.600E-06 | 1.400E-05 |
| 4391 | 220 | A220L | A | 9.7e-06 | 1e-05 | 5.5e-06 | 8.400E-06 | 2.516E-06 | 5.500E-06 | 1.000E-05 |
| 4392 | 220 | A220K | A | 1.1e-05 | 7.9e-06 | 6.9e-06 | 8.600E-06 | 2.138E-06 | 6.900E-06 | 1.100E-05 |
| 4393 | 220 | A220M | A | 1.3e-05 | 9.8e-06 | 9.8e-06 | 1.087E-05 | 1.848E-06 | 9.800E-06 | 1.300E-05 |
| 4394 | 220 | A220F | A | 1.2e-05 | 2e-05 | 5e-06 | 1.233E-05 | 7.506E-06 | 5.000E-06 | 2.000E-05 |
| 4395 | 220 | A220P | A | 9.5e-06 | 7.6e-06 | 9.7e-06 | 8.933E-06 | 1.159E-06 | 7.600E-06 | 9.700E-06 |
| 4396 | 220 | A220S | A | 6.5e-06 | 9.8e-06 | 1.3e-05 | 9.767E-06 | 3.250E-06 | 6.500E-06 | 1.300E-05 |
| 4397 | 220 | A220T | A | 1.3e-05 | 8.2e-06 | 6.7e-06 | 9.300E-06 | 3.291E-06 | 6.700E-06 | 1.300E-05 |
| 4398 | 220 | A220W | A | 1.2e-05 | 1.4e-05 | 4.4e-06 | 1.013E-05 | 5.065E-06 | 4.400E-06 | 1.400E-05 |
| 4399 | 220 | A220Y | A | 8.6e-06 | 1e-05 | 1.3e-05 | 1.053E-05 | 2.248E-06 | 8.600E-06 | 1.300E-05 |
| 4400 | 220 | A220V | A | 1e-05 | 5e-06 | 1.1e-05 | 8.667E-06 | 3.215E-06 | 5.000E-06 | 1.100E-05 |
| 4401 | 221 | G221A | A | 1.5e-05 | 1.2e-05 | 5e-06 | 1.067E-05 | 5.132E-06 | 5.000E-06 | 1.500E-05 |
| 4402 | 221 | G221R | A | 4.9e-06 | 7.4e-06 | 5.9e-06 | 6.067E-06 | 1.258E-06 | 4.900E-06 | 7.400E-06 |
| 4403 | 221 | G221N | A | 8.6e-06 | 5.3e-06 | 9.1e-06 | 7.667E-06 | 2.065E-06 | 5.300E-06 | 9.100E-06 |
| 4404 | 221 | G221D | A | 6.3e-06 | 1.4e-05 | 6.4e-06 | 8.900E-06 | 4.417E-06 | 6.300E-06 | 1.400E-05 |
| 4405 | 221 | G221C | A | 1.3e-05 | 7.8e-06 | 5.8e-06 | 8.867E-06 | 3.717E-06 | 5.800E-06 | 1.300E-05 |
| 4406 | 221 | G221E | A | 1.2e-05 | 5e-06 | 1.4e-05 | 1.033E-05 | 4.726E-06 | 5.000E-06 | 1.400E-05 |
| 4407 | 221 | G221Q | A | 5.2e-06 | 1.3e-05 | 9.6e-06 | 9.267E-06 | 3.911E-06 | 5.200E-06 | 1.300E-05 |
| 4408 | 221 | G221G | A | 9e-06 | 9.1e-06 | 4.5e-06 | 7.533E-06 | 2.627E-06 | 4.500E-06 | 9.100E-06 |
| 4409 | 221 | G221H | A | 7e-06 | 4.7e-06 | 7.3e-06 | 6.333E-06 | 1.422E-06 | 4.700E-06 | 7.300E-06 |
| 4410 | 221 | G221I | A | 1e-05 | 1e-05 | 7.3e-06 | 9.100E-06 | 1.559E-06 | 7.300E-06 | 1.000E-05 |
| 4411 | 221 | G221L | A | 5e-06 | 9e-06 | 4.8e-06 | 6.267E-06 | 2.369E-06 | 4.800E-06 | 9.000E-06 |
| 4412 | 221 | G221K | A | 7.5e-06 | 1.3e-05 | 6.4e-06 | 8.967E-06 | 3.536E-06 | 6.400E-06 | 1.300E-05 |
| 4413 | 221 | G221M | A | 7.6e-06 | 1.1e-05 | 5.7e-06 | 8.100E-06 | 2.685E-06 | 5.700E-06 | 1.100E-05 |
| 4414 | 221 | G221F | A | 8.2e-06 | 1.6e-05 | 1e-05 | 1.140E-05 | 4.084E-06 | 8.200E-06 | 1.600E-05 |
| 4415 | 221 | G221P | A | 9.3e-06 | 8.9e-06 | 1.1e-05 | 9.733E-06 | 1.115E-06 | 8.900E-06 | 1.100E-05 |
| 4416 | 221 | G221S | A | 1.3e-05 | 1.3e-05 | 9.2e-06 | 1.173E-05 | 2.194E-06 | 9.200E-06 | 1.300E-05 |
| 4417 | 221 | G221T | A | 7e-06 | 6.7e-06 | 5.8e-06 | 6.500E-06 | 6.245E-07 | 5.800E-06 | 7.000E-06 |
| 4418 | 221 | G221W | A | 2.6e-06 | 7.7e-06 | 6.6e-06 | 5.633E-06 | 2.684E-06 | 2.600E-06 | 7.700E-06 |
| 4419 | 221 | G221Y | A | 7.5e-06 | 1.3e-05 | 1e-05 | 1.017E-05 | 2.754E-06 | 7.500E-06 | 1.300E-05 |

| | | | | | | | | | | |
|---|---|---|---|---|---|---|---|---|---|---|
| 4420 | 221 | G221V | A | 7.4e-06 | 9.1e-06 | 6.1e-06 | 7.533E-06 | 1.504E-06 | 6.100E-06 | 9.100E-06 |
| 4421 | 222 | C222A | A | 6.6e-06 | 1e-05 | 8.8e-06 | 8.467E-06 | 1.724E-06 | 6.600E-06 | 1.000E-05 |
| 4422 | 222 | C222R | A | 6.2e-06 | 8.6e-06 | 1.3e-05 | 9.267E-06 | 3.449E-06 | 6.200E-06 | 1.300E-05 |
| 4423 | 222 | C222N | A | 7.8e-06 | 5.8e-06 | 1.3e-05 | 8.867E-06 | 3.717E-06 | 5.800E-06 | 1.300E-05 |
| 4424 | 222 | C222D | A | 6.2e-06 | 8e-06 | 4.7e-06 | 6.300E-06 | 1.652E-06 | 4.700E-06 | 8.000E-06 |
| 4425 | 222 | C222C | A | 8.4e-06 | 4.6e-06 | 6.8e-06 | 6.600E-06 | 1.908E-06 | 4.600E-06 | 8.400E-06 |
| 4426 | 222 | C222E | A | 8.8e-06 | 1.2e-05 | 5.1e-06 | 8.633E-06 | 3.453E-06 | 5.100E-06 | 1.200E-05 |
| 4427 | 222 | C222Q | A | 4.7e-06 | 8.4e-06 | 9.4e-06 | 7.500E-06 | 2.476E-06 | 4.700E-06 | 9.400E-06 |
| 4428 | 222 | C222G | A | 4.5e-06 | 1.4e-05 | 5.8e-06 | 8.100E-06 | 5.151E-06 | 4.500E-06 | 1.400E-05 |
| 4429 | 222 | C222H | A | 1.3e-05 | 5.8e-06 | 6.6e-06 | 8.467E-06 | 3.946E-06 | 5.800E-06 | 1.300E-05 |
| 4430 | 222 | C222I | A | 7.8e-06 | 1.3e-05 | 8.6e-06 | 9.800E-06 | 2.800E-06 | 7.800E-06 | 1.300E-05 |
| 4431 | 222 | C222L | A | 5.9e-06 | 4.5e-06 | 1.4e-05 | 8.133E-06 | 5.129E-06 | 4.500E-06 | 1.400E-05 |
| 4432 | 222 | C222K | A | 4.9e-06 | 1e-05 | 9.4e-06 | 8.100E-06 | 2.787E-06 | 4.900E-06 | 1.000E-05 |
| 4433 | 222 | C222M | A | 5.8e-06 | 6.6e-06 | 3.8e-06 | 5.400E-06 | 1.442E-06 | 3.800E-06 | 6.600E-06 |
| 4434 | 222 | C222F | A | 4.7e-06 | 9.1e-06 | 1.3e-05 | 8.933E-06 | 4.153E-06 | 4.700E-06 | 1.300E-05 |
| 4435 | 222 | C222P | A | 7.4e-06 | 9.6e-06 | 8.7e-06 | 8.567E-06 | 1.106E-06 | 7.400E-06 | 9.600E-06 |
| 4436 | 222 | C222S | A | 5.9e-06 | 7.5e-06 | 9.2e-06 | 7.533E-06 | 1.650E-06 | 5.900E-06 | 9.200E-06 |
| 4437 | 222 | C222T | A | 1.3e-05 | 1e-05 | 1.4e-05 | 1.233E-05 | 2.082E-06 | 1.000E-05 | 1.400E-05 |
| 4438 | 222 | C222W | A | 9.6e-06 | 9.1e-06 | 9.2e-06 | 9.300E-06 | 2.646E-07 | 9.100E-06 | 9.600E-06 |
| 4439 | 222 | C222Y | A | 7.1e-06 | 9.7e-06 | 9.2e-06 | 8.667E-06 | 1.380E-06 | 7.100E-06 | 9.700E-06 |
| 4440 | 222 | C222V | A | 4.2e-06 | 7.9e-06 | 8.4e-06 | 6.833E-06 | 2.294E-06 | 4.200E-06 | 8.400E-06 |
| 4441 | 223 | T223A | A | 9.4e-06 | 9.6e-06 | 8.6e-06 | 9.200E-06 | 5.292E-07 | 8.600E-06 | 9.600E-06 |
| 4442 | 223 | T223R | A | 5.1e-06 | 1.2e-05 | 9.3e-06 | 8.800E-06 | 3.477E-06 | 5.100E-06 | 1.200E-05 |
| 4443 | 223 | T223N | A | 7e-06 | 7.7e-06 | 8.6e-06 | 7.767E-06 | 8.021E-07 | 7.000E-06 | 8.600E-06 |
| 4444 | 223 | T223D | A | 8e-06 | 1e-05 | 1e-05 | 9.333E-06 | 1.155E-06 | 8.000E-06 | 1.000E-05 |
| 4445 | 223 | T223C | A | 1.1e-05 | 7.7e-06 | 5e-06 | 7.900E-06 | 3.005E-06 | 5.000E-06 | 1.100E-05 |
| 4446 | 223 | T223E | A | 9.3e-06 | 1.1e-05 | 1.1e-05 | 1.043E-05 | 9.815E-07 | 9.300E-06 | 1.100E-05 |
| 4447 | 223 | T223Q | A | 6.2e-06 | 5e-06 | 6.9e-06 | 6.033E-06 | 9.609E-07 | 5.000E-06 | 6.900E-06 |
| 4448 | 223 | T223G | A | 6.4e-06 | 1.1e-05 | 6.3e-06 | 7.900E-06 | 2.685E-06 | 6.300E-06 | 1.100E-05 |
| 4449 | 223 | T223H | A | 6.4e-06 | 1e-05 | 5.5e-06 | 7.300E-06 | 2.381E-06 | 5.500E-06 | 1.000E-05 |
| 4450 | 223 | T223I | A | 1.3e-05 | 1.5e-05 | 9.8e-06 | 1.260E-05 | 2.623E-06 | 9.800E-06 | 1.500E-05 |
| 4451 | 223 | T223L | A | 1.3e-05 | 6.9e-06 | 7e-06 | 8.967E-06 | 3.493E-06 | 6.900E-06 | 1.300E-05 |
| 4452 | 223 | T223K | A | 1.1e-05 | 9.7e-06 | 7.4e-06 | 9.367E-06 | 1.823E-06 | 7.400E-06 | 1.100E-05 |
| 4453 | 223 | T223M | A | 9.3e-06 | 1.3e-05 | 9.2e-06 | 1.050E-05 | 2.166E-06 | 9.200E-06 | 1.300E-05 |
| 4454 | 223 | T223F | A | 1.4e-05 | 6.1e-06 | 6.4e-06 | 8.833E-06 | 4.477E-06 | 6.100E-06 | 1.400E-05 |
| 4455 | 223 | T223P | A | 5.4e-06 | 1e-05 | 9.3e-06 | 8.233E-06 | 2.479E-06 | 5.400E-06 | 1.000E-05 |
| 4456 | 223 | T223S | A | 1.2e-05 | 6.5e-06 | 9.7e-06 | 9.400E-06 | 2.762E-06 | 6.500E-06 | 1.200E-05 |
| 4457 | 223 | T223T | A | 5e-06 | 6.9e-06 | 1.3e-05 | 8.300E-06 | 4.180E-06 | 5.000E-06 | 1.300E-05 |
| 4458 | 223 | T223W | A | 5.3e-06 | 9.3e-06 | 6.1e-06 | 6.900E-06 | 2.117E-06 | 5.300E-06 | 9.300E-06 |
| 4459 | 223 | T223Y | A | 8.3e-06 | 9.2e-06 | 5.4e-06 | 7.633E-06 | 1.986E-06 | 5.400E-06 | 9.200E-06 |
| 4460 | 223 | T223V | A | 9.3e-06 | 8e-06 | 8.3e-06 | 8.533E-06 | 6.807E-07 | 8.000E-06 | 9.300E-06 |
| 4461 | 224 | G224A | A | 6.9e-06 | 8.6e-06 | 8.6e-06 | 8.033E-06 | 9.815E-07 | 6.900E-06 | 8.600E-06 |
| 4462 | 224 | G224R | A | 6.4e-06 | 6.5e-06 | 6.6e-06 | 6.500E-06 | 1.000E-07 | 6.400E-06 | 6.600E-06 |
| 4463 | 224 | G224N | A | 7e-06 | 6.1e-06 | 7e-06 | 6.700E-06 | 5.196E-07 | 6.100E-06 | 7.000E-06 |
| 4464 | 224 | G224D | A | 9.8e-06 | 6.8e-06 | 9.3e-06 | 8.633E-06 | 1.607E-06 | 6.800E-06 | 9.800E-06 |
| 4465 | 224 | G224C | A | 7.4e-06 | 1.3e-05 | 5.1e-06 | 8.500E-06 | 4.063E-06 | 5.100E-06 | 1.300E-05 |
| 4466 | 224 | G224E | A | 8.8e-06 | 1e-05 | 1e-05 | 9.600E-06 | 6.928E-07 | 8.800E-06 | 1.000E-05 |
| 4467 | 224 | G224Q | A | 1.7e-05 | 9.7e-06 | 6.7e-06 | 1.113E-05 | 5.297E-06 | 6.700E-06 | 1.700E-05 |
| 4468 | 224 | G224G | A | 9.1e-06 | 5.7e-06 | 4.8e-06 | 6.533E-06 | 2.268E-06 | 4.800E-06 | 9.100E-06 |
| 4469 | 224 | G224H | A | 4.2e-06 | 7.4e-06 | 1.5e-05 | 8.867E-06 | 5.547E-06 | 4.200E-06 | 1.500E-05 |
| 4470 | 224 | G224I | A | 4.7e-06 | 1e-05 | 5.4e-06 | 6.700E-06 | 2.879E-06 | 4.700E-06 | 1.000E-05 |
| 4471 | 224 | G224L | A | 8.6e-06 | 9.8e-06 | 9.2e-06 | 9.200E-06 | 6.000E-07 | 8.600E-06 | 9.800E-06 |
| 4472 | 224 | G224K | A | 1e-05 | 7.9e-06 | 2.4e-05 | 1.397E-05 | 8.752E-06 | 7.900E-06 | 2.400E-05 |
| 4473 | 224 | G224M | A | 8.7e-06 | 6.9e-06 | 1.2e-05 | 9.200E-06 | 2.587E-06 | 6.900E-06 | 1.200E-05 |
| 4474 | 224 | G224F | A | 4.2e-06 | 1.1e-05 | 4.4e-06 | 6.533E-06 | 3.870E-06 | 4.200E-06 | 1.100E-05 |
| 4475 | 224 | G224P | A | 6.1e-06 | 6.7e-06 | 9.8e-06 | 7.533E-06 | 1.986E-06 | 6.100E-06 | 9.800E-06 |
| 4476 | 224 | G224S | A | 1.2e-05 | 5.6e-06 | 1.2e-05 | 9.867E-06 | 3.695E-06 | 5.600E-06 | 1.200E-05 |
| 4477 | 224 | G224T | A | 1e-05 | 1.3e-05 | 1.4e-05 | 1.233E-05 | 2.082E-06 | 1.000E-05 | 1.400E-05 |
| 4478 | 224 | G224W | A | 1e-05 | 1.2e-05 | 1.2e-05 | 1.133E-05 | 1.155E-06 | 1.000E-05 | 1.200E-05 |
| 4479 | 224 | G224Y | A | 7.5e-06 | 5.2e-06 | 8.8e-06 | 7.167E-06 | 1.823E-06 | 5.200E-06 | 8.800E-06 |
| 4480 | 224 | G224V | A | 5.3e-06 | 6.4e-06 | 9.9e-06 | 7.200E-06 | 2.402E-06 | 5.300E-06 | 9.900E-06 |
| 4481 | 225 | P225A | A | 7e-06 | 9.4e-06 | 1.1e-05 | 9.133E-06 | 2.013E-06 | 7.000E-06 | 1.100E-05 |
| 4482 | 225 | P225R | A | 7.3e-06 | 5.8e-06 | 6.2e-06 | 6.433E-06 | 7.767E-07 | 5.800E-06 | 7.300E-06 |
| 4483 | 225 | P225N | A | 1.4e-05 | 6.2e-06 | 8.6e-06 | 9.600E-06 | 3.995E-06 | 6.200E-06 | 1.400E-05 |
| 4484 | 225 | P225D | A | 4.6e-06 | 6.4e-06 | 8.1e-06 | 6.367E-06 | 1.750E-06 | 4.600E-06 | 8.100E-06 |
| 4485 | 225 | P225C | A | 8e-06 | 8.8e-06 | 7.8e-06 | 8.200E-06 | 5.292E-07 | 7.800E-06 | 8.800E-06 |
| 4486 | 225 | P225E | A | 8.6e-06 | 9.4e-06 | 5.9e-06 | 7.967E-06 | 1.834E-06 | 5.900E-06 | 9.400E-06 |
| 4487 | 225 | P225Q | A | 7.6e-06 | 7.9e-06 | 8.9e-06 | 8.133E-06 | 6.807E-07 | 7.600E-06 | 8.900E-06 |
| 4488 | 225 | P225G | A | 6.5e-06 | 5.5e-06 | 5.4e-06 | 5.800E-06 | 6.083E-07 | 5.400E-06 | 6.500E-06 |
| 4489 | 225 | P225H | A | 8e-06 | 6.5e-06 | 7.3e-06 | 7.267E-06 | 7.506E-07 | 6.500E-06 | 8.000E-06 |
| 4490 | 225 | P225I | A | 8.6e-06 | 1.1e-05 | 4.6e-06 | 8.067E-06 | 3.233E-06 | 4.600E-06 | 1.100E-05 |
| 4491 | 225 | P225L | A | 8e-06 | 6.2e-06 | 8.9e-06 | 7.700E-06 | 1.375E-06 | 6.200E-06 | 8.900E-06 |
| 4492 | 225 | P225K | A | 1.3e-05 | 7e-06 | 6.8e-06 | 8.933E-06 | 3.523E-06 | 6.800E-06 | 1.300E-05 |
| 4493 | 225 | P225M | A | 6.8e-06 | 6.3e-06 | 9.6e-06 | 7.567E-06 | 1.779E-06 | 6.300E-06 | 9.600E-06 |
| 4494 | 225 | P225F | A | 1.4e-05 | 6.3e-06 | 1.1e-05 | 1.043E-05 | 3.881E-06 | 6.300E-06 | 1.400E-05 |
| 4495 | 225 | P225P | A | 5.8e-06 | 4.7e-06 | 1.2e-05 | 7.500E-06 | 3.936E-06 | 4.700E-06 | 1.200E-05 |
| 4496 | 225 | P225S | A | 8.4e-06 | 9.7e-06 | 6.7e-06 | 8.267E-06 | 1.504E-06 | 6.700E-06 | 9.700E-06 |
| 4497 | 225 | P225T | A | 1.2e-05 | 7.1e-06 | 6.1e-06 | 8.400E-06 | 3.158E-06 | 6.100E-06 | 1.200E-05 |
| 4498 | 225 | P225W | A | 7.7e-06 | 7.8e-06 | 1e-05 | 8.500E-06 | 1.300E-06 | 7.700E-06 | 1.000E-05 |
| 4499 | 225 | P225Y | A | 6.6e-06 | 5.9e-06 | 4.3e-06 | 5.600E-06 | 1.179E-06 | 4.300E-06 | 6.600E-06 |
| 4500 | 225 | P225V | A | 1.1e-05 | 7e-06 | 1.4e-05 | 1.067E-05 | 3.512E-06 | 7.000E-06 | 1.400E-05 |
| 4501 | 226 | K226A | A | 1e-05 | 7.9e-06 | 1.3e-05 | 1.030E-05 | 2.563E-06 | 7.900E-06 | 1.300E-05 |
| 4502 | 226 | K226R | A | 6.1e-06 | 9e-06 | 7.1e-06 | 7.400E-06 | 1.473E-06 | 6.100E-06 | 9.000E-06 |
| 4503 | 226 | K226N | A | 1.5e-05 | 5e-06 | 6.6e-06 | 8.867E-06 | 5.372E-06 | 5.000E-06 | 1.500E-05 |
| 4504 | 226 | K226D | A | 8.2e-06 | 6.7e-06 | 8.7e-06 | 7.867E-06 | 1.041E-06 | 6.700E-06 | 8.700E-06 |

| | | | | | | | | | | |
|---|---|---|---|---|---|---|---|---|---|---|
| 4505 | 226 | K226C | A | 6.6e-06 | 8.4e-06 | 6.6e-06 | 7.200E-06 | 1.039E-06 | 6.600E-06 | 8.400E-06 |
| 4506 | 226 | K226E | A | 1.4e-05 | 6.8e-06 | 5e-06 | 8.600E-06 | 4.762E-06 | 5.000E-06 | 1.400E-05 |
| 4507 | 226 | K226Q | A | 3.8e-06 | 6.7e-06 | 6.8e-06 | 5.767E-06 | 1.704E-06 | 3.800E-06 | 6.800E-06 |
| 4508 | 226 | K226G | A | 1.3e-05 | 8.1e-06 | 5.9e-06 | 9.000E-06 | 3.635E-06 | 5.900E-06 | 1.300E-05 |
| 4509 | 226 | K226H | A | 8.8e-06 | 9.3e-06 | 1.2e-05 | 1.003E-05 | 1.721E-06 | 8.800E-06 | 1.200E-05 |
| 4510 | 226 | K226I | A | 1e-05 | 9e-06 | 5.4e-06 | 8.133E-06 | 2.419E-06 | 5.400E-06 | 1.000E-05 |
| 4511 | 226 | K226L | A | 9.8e-06 | 6.6e-06 | 6.2e-06 | 7.533E-06 | 1.973E-06 | 6.200E-06 | 9.800E-06 |
| 4512 | 226 | K226K | A | 5.8e-06 | 4.3e-06 | 1.4e-05 | 8.033E-06 | 5.221E-06 | 4.300E-06 | 1.400E-05 |
| 4513 | 226 | K226M | A | 7.7e-06 | 6.4e-06 | 6.8e-06 | 6.967E-06 | 6.658E-07 | 6.400E-06 | 7.700E-06 |
| 4514 | 226 | K226F | A | 5.3e-06 | 1e-05 | 7.7e-06 | 7.667E-06 | 2.350E-06 | 5.300E-06 | 1.000E-05 |
| 4515 | 226 | K226P | A | 6.9e-06 | 6.8e-06 | 6.1e-06 | 6.600E-06 | 4.359E-07 | 6.100E-06 | 6.900E-06 |
| 4516 | 226 | K226S | A | 8.5e-06 | 8.5e-06 | 6.5e-06 | 7.833E-06 | 1.155E-06 | 6.500E-06 | 8.500E-06 |
| 4517 | 226 | K226T | A | 6.3e-06 | 9.2e-06 | 8.3e-06 | 7.933E-06 | 1.484E-06 | 6.300E-06 | 9.200E-06 |
| 4518 | 226 | K226W | A | 7.9e-06 | 7.4e-06 | 6.5e-06 | 7.267E-06 | 7.095E-07 | 6.500E-06 | 7.900E-06 |
| 4519 | 226 | K226Y | A | 6.9e-06 | 8.3e-06 | 5.8e-06 | 7.000E-06 | 1.253E-06 | 5.800E-06 | 8.300E-06 |
| 4520 | 226 | K226V | A | 1e-05 | 5.4e-06 | 9e-06 | 8.133E-06 | 2.419E-06 | 5.400E-06 | 1.000E-05 |
| 4521 | 227 | H227A | A | 7.9e-06 | 6.6e-06 | 5e-06 | 6.500E-06 | 1.453E-06 | 5.000E-06 | 7.900E-06 |
| 4522 | 227 | H227R | A | 6.6e-06 | 1.2e-05 | 1.5e-05 | 1.120E-05 | 4.257E-06 | 6.600E-06 | 1.500E-05 |
| 4523 | 227 | H227N | A | 9.2e-06 | 1e-05 | 6.5e-06 | 8.567E-06 | 1.834E-06 | 6.500E-06 | 1.000E-05 |
| 4524 | 227 | H227D | A | 6.5e-06 | 6.8e-06 | 6.5e-06 | 6.600E-06 | 1.732E-07 | 6.500E-06 | 6.800E-06 |
| 4525 | 227 | H227C | A | 1e-05 | 1.2e-05 | 6.9e-06 | 9.633E-06 | 2.570E-06 | 6.900E-06 | 1.200E-05 |
| 4526 | 227 | H227E | A | 6.7e-06 | 7.4e-06 | 8.9e-06 | 7.667E-06 | 1.124E-06 | 6.700E-06 | 8.900E-06 |
| 4527 | 227 | H227Q | A | 5.7e-06 | 1.6e-05 | 7.8e-06 | 9.833E-06 | 5.443E-06 | 5.700E-06 | 1.600E-05 |
| 4528 | 227 | H227G | A | 4.8e-06 | 8.6e-06 | 1e-05 | 7.800E-06 | 2.691E-06 | 4.800E-06 | 1.000E-05 |
| 4529 | 227 | H227H | A | 6.4e-06 | 6.2e-06 | 5.8e-06 | 6.133E-06 | 3.055E-07 | 5.800E-06 | 6.400E-06 |
| 4530 | 227 | H227I | A | 1.4e-05 | 9.3e-06 | 1.4e-05 | 1.243E-05 | 2.714E-06 | 9.300E-06 | 1.400E-05 |
| 4531 | 227 | H227L | A | 5.9e-06 | 3.3e-06 | 9.7e-06 | 6.300E-06 | 3.219E-06 | 3.300E-06 | 9.700E-06 |
| 4532 | 227 | H227K | A | 1e-05 | 8.3e-06 | 3.9e-06 | 7.400E-06 | 3.148E-06 | 3.900E-06 | 1.000E-05 |
| 4533 | 227 | H227M | A | 5.5e-06 | 1.4e-05 | 8.5e-06 | 9.333E-06 | 4.311E-06 | 5.500E-06 | 1.400E-05 |
| 4534 | 227 | H227F | A | 5.4e-06 | 9.8e-06 | 9.6e-06 | 8.267E-06 | 2.485E-06 | 5.400E-06 | 9.800E-06 |
| 4535 | 227 | H227P | A | 6.9e-06 | 5.3e-06 | 6.8e-06 | 6.333E-06 | 8.963E-07 | 5.300E-06 | 6.900E-06 |
| 4536 | 227 | H227S | A | 5.6e-06 | 1.5e-05 | 9.3e-06 | 9.967E-06 | 4.735E-06 | 5.600E-06 | 1.500E-05 |
| 4537 | 227 | H227T | A | 1.2e-05 | 6.5e-06 | 5.4e-06 | 7.967E-06 | 3.536E-06 | 5.400E-06 | 1.200E-05 |
| 4538 | 227 | H227W | A | 7.5e-06 | 1.5e-05 | 1.5e-05 | 1.250E-05 | 4.330E-06 | 7.500E-06 | 1.500E-05 |
| 4539 | 227 | H227Y | A | 9e-06 | 6.2e-06 | 1.1e-05 | 8.733E-06 | 2.411E-06 | 6.200E-06 | 1.100E-05 |
| 4540 | 227 | H227V | A | 8.1e-06 | 1.2e-05 | 9.6e-06 | 9.900E-06 | 1.967E-06 | 8.100E-06 | 1.200E-05 |
| 4541 | 228 | S228A | A | 8.1e-06 | 3.3e-06 | 6.3e-06 | 5.900E-06 | 2.425E-06 | 3.300E-06 | 8.100E-06 |
| 4542 | 228 | S228R | A | 9.4e-06 | 6.7e-06 | 8.2e-06 | 8.100E-06 | 1.353E-06 | 6.700E-06 | 9.400E-06 |
| 4543 | 228 | S228N | A | 1.3e-05 | 1.2e-05 | 1.2e-05 | 1.233E-05 | 5.774E-07 | 1.200E-05 | 1.300E-05 |
| 4544 | 228 | S228D | A | 9.7e-06 | 1.1e-05 | 5.3e-06 | 8.667E-06 | 2.987E-06 | 5.300E-06 | 1.100E-05 |
| 4545 | 228 | S228C | A | 1.6e-05 | 1.5e-05 | 1e-05 | 1.367E-05 | 3.215E-06 | 1.000E-05 | 1.600E-05 |
| 4546 | 228 | S228E | A | 4.5e-06 | 7e-06 | 1.3e-05 | 8.167E-06 | 4.368E-06 | 4.500E-06 | 1.300E-05 |
| 4547 | 228 | S228Q | A | 9.4e-06 | 4.9e-06 | 1.2e-05 | 8.767E-06 | 3.592E-06 | 4.900E-06 | 1.200E-05 |
| 4548 | 228 | S228G | A | 7.9e-06 | 1.2e-05 | 5.6e-06 | 8.500E-06 | 3.242E-06 | 5.600E-06 | 1.200E-05 |
| 4549 | 228 | S228H | A | 9.3e-06 | 7.1e-06 | 7.6e-06 | 8.000E-06 | 1.153E-06 | 7.100E-06 | 9.300E-06 |
| 4550 | 228 | S228I | A | 4.7e-06 | 1.5e-05 | 1e-05 | 9.900E-06 | 5.151E-06 | 4.700E-06 | 1.500E-05 |
| 4551 | 228 | S228L | A | 6.6e-06 | 9.2e-06 | 5.2e-06 | 7.000E-06 | 2.030E-06 | 5.200E-06 | 9.200E-06 |
| 4552 | 228 | S228K | A | 8.3e-06 | 6.9e-06 | 8e-06 | 7.733E-06 | 7.371E-07 | 6.900E-06 | 8.300E-06 |
| 4553 | 228 | S228M | A | 8.8e-06 | 3.9e-06 | 6.4e-06 | 6.367E-06 | 2.450E-06 | 3.900E-06 | 8.800E-06 |
| 4554 | 228 | S228F | A | 1.1e-05 | 9.7e-06 | 8.4e-06 | 9.700E-06 | 1.300E-06 | 8.400E-06 | 1.100E-05 |
| 4555 | 228 | S228P | A | 4.3e-06 | 6.9e-06 | 8.1e-06 | 6.433E-06 | 1.943E-06 | 4.300E-06 | 8.100E-06 |
| 4556 | 228 | S228S | A | 7.9e-06 | 9.5e-06 | 1.1e-05 | 9.467E-06 | 1.550E-06 | 7.900E-06 | 1.100E-05 |
| 4557 | 228 | S228T | A | 5.6e-06 | 7e-06 | 6.2e-06 | 6.267E-06 | 7.024E-07 | 5.600E-06 | 7.000E-06 |
| 4558 | 228 | S228W | A | 8.5e-06 | 7.6e-06 | 6.7e-06 | 7.600E-06 | 9.000E-07 | 6.700E-06 | 8.500E-06 |
| 4559 | 228 | S228Y | A | 1.6e-05 | 7.9e-06 | 5e-06 | 9.633E-06 | 5.701E-06 | 5.000E-06 | 1.600E-05 |
| 4560 | 228 | S228V | A | 1.2e-05 | 8.8e-06 | 1e-05 | 1.027E-05 | 1.617E-06 | 8.800E-06 | 1.200E-05 |
| 4561 | 229 | D229A | A | 1e-05 | 1.7e-05 | 1.1e-05 | 1.267E-05 | 3.786E-06 | 1.000E-05 | 1.700E-05 |
| 4562 | 229 | D229R | A | 6.7e-06 | 7.4e-06 | 6.8e-06 | 6.967E-06 | 3.786E-07 | 6.700E-06 | 7.400E-06 |
| 4563 | 229 | D229N | A | 1.6e-05 | 5.4e-06 | 1.4e-05 | 1.180E-05 | 5.632E-06 | 5.400E-06 | 1.600E-05 |
| 4564 | 229 | D229D | A | 7.9e-06 | 1.1e-05 | 6.6e-06 | 8.500E-06 | 2.261E-06 | 6.600E-06 | 1.100E-05 |
| 4565 | 229 | D229C | A | 7e-06 | 6.5e-06 | 4.7e-06 | 6.067E-06 | 1.210E-06 | 4.700E-06 | 7.000E-06 |
| 4566 | 229 | D229E | A | 6.9e-06 | 1.1e-05 | 1.4e-05 | 1.063E-05 | 3.564E-06 | 6.900E-06 | 1.400E-05 |
| 4567 | 229 | D229Q | A | 8.4e-06 | 1.2e-05 | 8.1e-06 | 9.500E-06 | 2.170E-06 | 8.100E-06 | 1.200E-05 |
| 4568 | 229 | D229G | A | 8.8e-06 | 8.5e-06 | 9.4e-06 | 8.900E-06 | 4.583E-07 | 8.500E-06 | 9.400E-06 |
| 4569 | 229 | D229H | A | 8.3e-06 | 5.1e-06 | 7e-06 | 6.800E-06 | 1.609E-06 | 5.100E-06 | 8.300E-06 |
| 4570 | 229 | D229I | A | 8.8e-06 | 6.9e-06 | 7.2e-06 | 7.633E-06 | 1.021E-06 | 6.900E-06 | 8.800E-06 |
| 4571 | 229 | D229L | A | 6.9e-06 | 1e-05 | 1.1e-05 | 9.300E-06 | 2.138E-06 | 6.900E-06 | 1.100E-05 |
| 4572 | 229 | D229K | A | 8.1e-06 | 1.1e-05 | 5.4e-06 | 8.167E-06 | 2.801E-06 | 5.400E-06 | 1.100E-05 |
| 4573 | 229 | D229M | A | 9.6e-06 | 8e-06 | 6.8e-06 | 8.133E-06 | 1.405E-06 | 6.800E-06 | 9.600E-06 |
| 4574 | 229 | D229F | A | 6.6e-06 | 8.9e-06 | 5.1e-06 | 6.867E-06 | 1.914E-06 | 5.100E-06 | 8.900E-06 |
| 4575 | 229 | D229P | A | 1.3e-05 | 9.5e-06 | 7.4e-06 | 9.967E-06 | 2.829E-06 | 7.400E-06 | 1.300E-05 |
| 4576 | 229 | D229S | A | 1e-05 | 6.7e-06 | 1e-05 | 8.900E-06 | 1.905E-06 | 6.700E-06 | 1.000E-05 |
| 4577 | 229 | D229T | A | 6.9e-06 | 4.3e-06 | 9.3e-06 | 6.833E-06 | 2.501E-06 | 4.300E-06 | 9.300E-06 |
| 4578 | 229 | D229W | A | 7.5e-06 | 8.2e-06 | 5.1e-06 | 6.933E-06 | 1.626E-06 | 5.100E-06 | 8.200E-06 |
| 4579 | 229 | D229Y | A | 5.8e-06 | 1.1e-05 | 7.2e-06 | 8.000E-06 | 2.691E-06 | 5.800E-06 | 1.100E-05 |
| 4580 | 229 | D229V | A | 6.8e-06 | 7.4e-06 | 8.7e-06 | 7.633E-06 | 9.713E-07 | 6.800E-06 | 8.700E-06 |
| 4581 | 230 | C230A | A | 9.3e-06 | 9.1e-06 | 1e-05 | 9.467E-06 | 4.726E-07 | 9.100E-06 | 1.000E-05 |
| 4582 | 230 | C230R | A | 9.9e-06 | 4.8e-06 | 9e-06 | 7.900E-06 | 2.722E-06 | 4.800E-06 | 9.900E-06 |
| 4583 | 230 | C230N | A | 5e-06 | 5.5e-06 | 1e-05 | 6.833E-06 | 2.754E-06 | 5.000E-06 | 1.000E-05 |
| 4584 | 230 | C230D | A | 7.2e-06 | 9.3e-06 | 6.7e-06 | 7.733E-06 | 1.380E-06 | 6.700E-06 | 9.300E-06 |
| 4585 | 230 | C230C | A | 1.2e-05 | 4.8e-06 | 6.1e-06 | 7.633E-06 | 3.837E-06 | 4.800E-06 | 1.200E-05 |
| 4586 | 230 | C230E | A | 8.8e-06 | 5.1e-06 | 8.8e-06 | 7.567E-06 | 2.136E-06 | 5.100E-06 | 8.800E-06 |
| 4587 | 230 | C230Q | A | 6.5e-06 | 9.5e-06 | 1.4e-05 | 1.000E-05 | 3.775E-06 | 6.500E-06 | 1.400E-05 |
| 4588 | 230 | C230G | A | 4.6e-06 | 1.2e-05 | 4.8e-06 | 7.133E-06 | 4.216E-06 | 4.600E-06 | 1.200E-05 |
| 4589 | 230 | C230H | A | 6e-06 | 4.9e-06 | 6.1e-06 | 5.667E-06 | 6.658E-07 | 4.900E-06 | 6.100E-06 |

| | | | | | | | | | | |
|---|---|---|---|---|---|---|---|---|---|---|
| 4590 | 230 | C230I | A | 7.5e-06 | 1.4e-05 | 1.3e-05 | 1.150E-05 | 3.500E-06 | 7.500E-06 | 1.400E-05 |
| 4591 | 230 | C230L | A | 1.1e-05 | 1.2e-05 | 7.4e-06 | 1.013E-05 | 2.419E-06 | 7.400E-06 | 1.200E-05 |
| 4592 | 230 | C230K | A | 6.9e-06 | 1.2e-05 | 4.8e-06 | 7.900E-06 | 3.703E-06 | 4.800E-06 | 1.200E-05 |
| 4593 | 230 | C230M | A | 7.1e-06 | 1.1e-05 | 6.8e-06 | 8.300E-06 | 2.343E-06 | 6.800E-06 | 1.100E-05 |
| 4594 | 230 | C230F | A | 1.4e-05 | 9.3e-06 | 7.5e-06 | 1.027E-05 | 3.356E-06 | 7.500E-06 | 1.400E-05 |
| 4595 | 230 | C230P | A | 8.3e-06 | 9.3e-06 | 7.2e-06 | 8.267E-06 | 1.050E-06 | 7.200E-06 | 9.300E-06 |
| 4596 | 230 | C230S | A | 4.5e-06 | 5.3e-06 | 6.6e-06 | 5.467E-06 | 1.060E-06 | 4.500E-06 | 6.600E-06 |
| 4597 | 230 | C230T | A | 1.2e-05 | 1.3e-05 | 6.2e-06 | 1.040E-05 | 3.672E-06 | 6.200E-06 | 1.300E-05 |
| 4598 | 230 | C230W | A | 1.3e-05 | 1.3e-05 | 7.8e-06 | 1.127E-05 | 3.002E-06 | 7.800E-06 | 1.300E-05 |
| 4599 | 230 | C230Y | A | 8.9e-06 | 4.7e-06 | 6.7e-06 | 6.767E-06 | 2.101E-06 | 4.700E-06 | 8.900E-06 |
| 4600 | 230 | C230V | A | 7.8e-06 | 8.3e-06 | 6.8e-06 | 7.633E-06 | 7.638E-07 | 6.800E-06 | 8.300E-06 |
| 4601 | 231 | L231A | A | 4.5e-06 | 5.6e-06 | 4.5e-06 | 4.867E-06 | 6.351E-07 | 4.500E-06 | 5.600E-06 |
| 4602 | 231 | L231R | A | 8.3e-06 | 8.4e-06 | 8.7e-06 | 8.467E-06 | 2.082E-07 | 8.300E-06 | 8.700E-06 |
| 4603 | 231 | L231N | A | 8.2e-06 | 6.4e-06 | 4.3e-06 | 6.300E-06 | 1.952E-06 | 4.300E-06 | 8.200E-06 |
| 4604 | 231 | L231D | A | 1.2e-05 | 4.1e-06 | 5.6e-06 | 7.233E-06 | 4.196E-06 | 4.100E-06 | 1.200E-05 |
| 4605 | 231 | L231C | A | 6.9e-06 | 8.2e-06 | 6.8e-06 | 7.300E-06 | 7.810E-07 | 6.800E-06 | 8.200E-06 |
| 4606 | 231 | L231E | A | 8.1e-06 | 9.7e-06 | 1.3e-05 | 1.027E-05 | 2.499E-06 | 8.100E-06 | 1.300E-05 |
| 4607 | 231 | L231Q | A | 7.9e-06 | 8.2e-06 | 7e-06 | 7.700E-06 | 6.245E-07 | 7.000E-06 | 8.200E-06 |
| 4608 | 231 | L231G | A | 6.6e-06 | 9.3e-06 | 8.9e-06 | 8.267E-06 | 1.457E-06 | 6.600E-06 | 9.300E-06 |
| 4609 | 231 | L231H | A | 9e-06 | 1e-05 | 1.1e-05 | 1.000E-05 | 1.000E-06 | 9.000E-06 | 1.100E-05 |
| 4610 | 231 | L231I | A | 6.3e-06 | 8.2e-06 | 8.5e-06 | 7.667E-06 | 1.193E-06 | 6.300E-06 | 8.500E-06 |
| 4611 | 231 | L231L | A | 6.6e-06 | 1e-05 | 1e-05 | 8.867E-06 | 1.963E-06 | 6.600E-06 | 1.000E-05 |
| 4612 | 231 | L231K | A | 6.1e-06 | 8.3e-06 | 5.6e-06 | 6.667E-06 | 1.436E-06 | 5.600E-06 | 8.300E-06 |
| 4613 | 231 | L231M | A | 6.3e-06 | 9.7e-06 | 9.5e-06 | 8.500E-06 | 1.908E-06 | 6.300E-06 | 9.700E-06 |
| 4614 | 231 | L231F | A | 1.4e-05 | 7.3e-06 | 9.1e-06 | 1.013E-05 | 3.467E-06 | 7.300E-06 | 1.400E-05 |
| 4615 | 231 | L231P | A | 7.7e-06 | 3.8e-06 | 1.3e-05 | 8.167E-06 | 4.618E-06 | 3.800E-06 | 1.300E-05 |
| 4616 | 231 | L231S | A | 8.4e-06 | 1e-05 | 9.6e-06 | 9.333E-06 | 8.327E-07 | 8.400E-06 | 1.000E-05 |
| 4617 | 231 | L231T | A | 5.7e-06 | 4.8e-06 | 7.1e-06 | 5.867E-06 | 1.159E-06 | 4.800E-06 | 7.100E-06 |
| 4618 | 231 | L231W | A | 5.9e-06 | 4.8e-06 | 1.5e-05 | 8.567E-06 | 5.599E-06 | 4.800E-06 | 1.500E-05 |
| 4619 | 231 | L231Y | A | 6.2e-06 | 9.5e-06 | 8.1e-06 | 7.933E-06 | 1.656E-06 | 6.200E-06 | 9.500E-06 |
| 4620 | 231 | L231V | A | 5.6e-06 | 1.2e-05 | 7.4e-06 | 8.333E-06 | 3.301E-06 | 5.600E-06 | 1.200E-05 |
| 4621 | 232 | A232A | A | 5.2e-06 | 9.3e-06 | 5.5e-06 | 6.667E-06 | 2.285E-06 | 5.200E-06 | 9.300E-06 |
| 4622 | 232 | A232R | A | 5.7e-06 | 9.5e-06 | 1e-05 | 8.400E-06 | 2.352E-06 | 5.700E-06 | 1.000E-05 |
| 4623 | 232 | A232N | A | 8.4e-06 | 1.1e-05 | 9.7e-06 | 9.700E-06 | 1.300E-06 | 8.400E-06 | 1.100E-05 |
| 4624 | 232 | A232D | A | 8.8e-06 | 4.3e-06 | 7.4e-06 | 6.833E-06 | 2.303E-06 | 4.300E-06 | 8.800E-06 |
| 4625 | 232 | A232C | A | 1.4e-05 | 9.4e-06 | 1.3e-05 | 1.213E-05 | 2.419E-06 | 9.400E-06 | 1.400E-05 |
| 4626 | 232 | A232E | A | 6.5e-06 | 5e-06 | 9.8e-06 | 7.100E-06 | 2.456E-06 | 5.000E-06 | 9.800E-06 |
| 4627 | 232 | A232Q | A | 5.2e-06 | 1.2e-05 | 5.8e-06 | 7.667E-06 | 3.765E-06 | 5.200E-06 | 1.200E-05 |
| 4628 | 232 | A232G | A | 6e-06 | 6.8e-06 | 5.9e-06 | 6.233E-06 | 4.933E-07 | 5.900E-06 | 6.800E-06 |
| 4629 | 232 | A232H | A | 1.2e-05 | 4.8e-06 | 6.8e-06 | 7.867E-06 | 3.717E-06 | 4.800E-06 | 1.200E-05 |
| 4630 | 232 | A232I | A | 1.4e-05 | 9.4e-06 | 8.6e-06 | 1.067E-05 | 2.914E-06 | 8.600E-06 | 1.400E-05 |
| 4631 | 232 | A232L | A | 7.7e-06 | 7.1e-06 | 6e-06 | 6.933E-06 | 8.622E-07 | 6.000E-06 | 7.700E-06 |
| 4632 | 232 | A232K | A | 7.8e-06 | 6.6e-06 | 1.1e-05 | 8.467E-06 | 2.274E-06 | 6.600E-06 | 1.100E-05 |
| 4633 | 232 | A232M | A | 8.3e-06 | 9e-06 | 1.1e-05 | 9.433E-06 | 1.401E-06 | 8.300E-06 | 1.100E-05 |
| 4634 | 232 | A232F | A | 6.1e-06 | 6.9e-06 | 8.8e-06 | 7.267E-06 | 1.387E-06 | 6.100E-06 | 8.800E-06 |
| 4635 | 232 | A232P | A | 1.4e-05 | 9.9e-06 | 6.8e-06 | 1.023E-05 | 3.612E-06 | 6.800E-06 | 1.400E-05 |
| 4636 | 232 | A232S | A | 1.2e-05 | 5.1e-06 | 6.6e-06 | 7.900E-06 | 3.629E-06 | 5.100E-06 | 1.200E-05 |
| 4637 | 232 | A232T | A | 5.7e-06 | 9e-06 | 1.3e-05 | 9.233E-06 | 3.656E-06 | 5.700E-06 | 1.300E-05 |
| 4638 | 232 | A232W | A | 1.3e-05 | 1.1e-05 | 7e-06 | 1.033E-05 | 3.055E-06 | 7.000E-06 | 1.300E-05 |
| 4639 | 232 | A232Y | A | 9.2e-06 | 1.3e-05 | 6.4e-06 | 9.533E-06 | 3.313E-06 | 6.400E-06 | 1.300E-05 |
| 4640 | 232 | A232V | A | 8e-06 | 2.9e-06 | 7.2e-06 | 6.033E-06 | 2.743E-06 | 2.900E-06 | 8.000E-06 |
| 4641 | 233 | C233A | A | 9.1e-06 | 1.1e-05 | 7.4e-06 | 9.167E-06 | 1.801E-06 | 7.400E-06 | 1.100E-05 |
| 4642 | 233 | C233R | A | 5.3e-06 | 4.9e-06 | 7.6e-06 | 5.933E-06 | 1.457E-06 | 4.900E-06 | 7.600E-06 |
| 4643 | 233 | C233N | A | 5.5e-06 | 6.1e-06 | 5.5e-06 | 5.700E-06 | 3.464E-07 | 5.500E-06 | 6.100E-06 |
| 4644 | 233 | C233D | A | 1.2e-05 | 5.4e-06 | 9.3e-06 | 8.900E-06 | 3.318E-06 | 5.400E-06 | 1.200E-05 |
| 4645 | 233 | C233C | A | 8.9e-06 | 4.7e-06 | 1e-05 | 7.867E-06 | 2.797E-06 | 4.700E-06 | 1.000E-05 |
| 4646 | 233 | C233E | A | 8.4e-06 | 9.9e-06 | 1.1e-05 | 9.767E-06 | 1.305E-06 | 8.400E-06 | 1.100E-05 |
| 4647 | 233 | C233Q | A | 5.5e-06 | 9.5e-06 | 9.4e-06 | 8.133E-06 | 2.281E-06 | 5.500E-06 | 9.500E-06 |
| 4648 | 233 | C233G | A | 1.3e-05 | 5.6e-06 | 7e-06 | 8.533E-06 | 3.931E-06 | 5.600E-06 | 1.300E-05 |
| 4649 | 233 | C233H | A | 9.1e-06 | 5.8e-06 | 6.5e-06 | 7.133E-06 | 1.739E-06 | 5.800E-06 | 9.100E-06 |
| 4650 | 233 | C233I | A | 7.4e-06 | 6.6e-06 | 1.2e-05 | 8.667E-06 | 2.914E-06 | 6.600E-06 | 1.200E-05 |
| 4651 | 233 | C233L | A | 5.7e-06 | 1.1e-05 | 6.3e-06 | 7.667E-06 | 2.902E-06 | 5.700E-06 | 1.100E-05 |
| 4652 | 233 | C233K | A | 1.9e-05 | 6.4e-06 | 5.3e-06 | 1.023E-05 | 7.612E-06 | 5.300E-06 | 1.900E-05 |
| 4653 | 233 | C233M | A | 1.1e-05 | 8.8e-06 | 6.3e-06 | 8.700E-06 | 2.352E-06 | 6.300E-06 | 1.100E-05 |
| 4654 | 233 | C233F | A | 4.6e-06 | 8.8e-06 | 9.6e-06 | 7.667E-06 | 2.686E-06 | 4.600E-06 | 9.600E-06 |
| 4655 | 233 | C233P | A | 8.1e-06 | 1.8e-05 | 7e-06 | 1.103E-05 | 6.058E-06 | 7.000E-06 | 1.800E-05 |
| 4656 | 233 | C233S | A | 9.2e-06 | 7.2e-06 | 8.3e-06 | 8.233E-06 | 1.002E-06 | 7.200E-06 | 9.200E-06 |
| 4657 | 233 | C233T | A | 1e-05 | 7.9e-06 | 1.6e-05 | 1.130E-05 | 4.204E-06 | 7.900E-06 | 1.600E-05 |
| 4658 | 233 | C233W | A | 1.2e-05 | 8.5e-06 | 1e-05 | 1.017E-05 | 1.756E-06 | 8.500E-06 | 1.200E-05 |
| 4659 | 233 | C233Y | A | 1.2e-05 | 6.5e-06 | 4.7e-06 | 7.733E-06 | 3.803E-06 | 4.700E-06 | 1.200E-05 |
| 4660 | 233 | C233V | A | 8.6e-06 | 1.1e-05 | 9.1e-06 | 9.567E-06 | 1.266E-06 | 8.600E-06 | 1.100E-05 |
| 4661 | 234 | L234A | A | 9.6e-06 | 5.3e-06 | 7.9e-06 | 7.600E-06 | 2.166E-06 | 5.300E-06 | 9.600E-06 |
| 4662 | 234 | L234R | A | 6.7e-06 | 6.6e-06 | 7.9e-06 | 7.067E-06 | 7.234E-07 | 6.600E-06 | 7.900E-06 |
| 4663 | 234 | L234N | A | 8.5e-06 | 2e-05 | 6.5e-06 | 1.167E-05 | 7.286E-06 | 6.500E-06 | 2.000E-05 |
| 4664 | 234 | L234D | A | 1.2e-05 | 1.3e-05 | 1.1e-05 | 1.200E-05 | 1.000E-06 | 1.100E-05 | 1.300E-05 |
| 4665 | 234 | L234C | A | 1.2e-05 | 1.2e-05 | 3.8e-06 | 9.267E-06 | 4.734E-06 | 3.800E-06 | 1.200E-05 |
| 4666 | 234 | L234E | A | 6.3e-06 | 1e-05 | 6.7e-06 | 7.667E-06 | 2.031E-06 | 6.300E-06 | 1.000E-05 |
| 4667 | 234 | L234Q | A | 1.2e-05 | 1e-05 | 8.3e-06 | 1.010E-05 | 1.852E-06 | 8.300E-06 | 1.200E-05 |
| 4668 | 234 | L234G | A | 6.7e-06 | 1e-05 | 4.9e-06 | 7.200E-06 | 2.587E-06 | 4.900E-06 | 1.000E-05 |
| 4669 | 234 | L234H | A | 1.6e-05 | 1e-05 | 1e-05 | 1.200E-05 | 3.464E-06 | 1.000E-05 | 1.600E-05 |
| 4670 | 234 | L234I | A | 6.2e-06 | 4.6e-06 | 5.1e-06 | 5.300E-06 | 8.185E-07 | 4.600E-06 | 6.200E-06 |
| 4671 | 234 | L234L | A | 7.3e-06 | 7.6e-06 | 1.4e-05 | 9.633E-06 | 3.785E-06 | 7.300E-06 | 1.400E-05 |
| 4672 | 234 | L234K | A | 4.7e-06 | 5.7e-06 | 6.6e-06 | 5.667E-06 | 9.504E-07 | 4.700E-06 | 6.600E-06 |
| 4673 | 234 | L234M | A | 4.6e-06 | 1.7e-05 | 6.9e-06 | 9.500E-06 | 6.596E-06 | 4.600E-06 | 1.700E-05 |
| 4674 | 234 | L234F | A | 5.9e-06 | 5.4e-06 | 7.5e-06 | 6.267E-06 | 1.097E-06 | 5.400E-06 | 7.500E-06 |

| | | | | | | | | | | |
|---|---|---|---|---|---|---|---|---|---|---|
| 4675 | 234 | L234P | A | 1.2e-05 | 1.4e-05 | 1.2e-05 | 1.267E-05 | 1.155E-06 | 1.200E-05 | 1.400E-05 |
| 4676 | 234 | L234S | A | 6.1e-06 | 6.4e-06 | 8.5e-06 | 7.000E-06 | 1.308E-06 | 6.100E-06 | 8.500E-06 |
| 4677 | 234 | L234T | A | 3.6e-06 | 7.9e-06 | 4.3e-06 | 5.267E-06 | 2.307E-06 | 3.600E-06 | 7.900E-06 |
| 4678 | 234 | L234W | A | 3.7e-06 | 1e-05 | 7.2e-06 | 6.967E-06 | 3.156E-06 | 3.700E-06 | 1.000E-05 |
| 4679 | 234 | L234Y | A | 6.9e-06 | 1.3e-05 | 1e-05 | 9.967E-06 | 3.050E-06 | 6.900E-06 | 1.300E-05 |
| 4680 | 234 | L234V | A | 9.7e-06 | 1.1e-05 | 8.9e-06 | 9.867E-06 | 1.060E-06 | 8.900E-06 | 1.100E-05 |
| 4681 | 235 | H235A | A | 5.6e-06 | 8e-06 | 6.2e-06 | 6.600E-06 | 1.249E-06 | 5.600E-06 | 8.000E-06 |
| 4682 | 235 | H235R | A | 8.7e-06 | 1.4e-05 | 5.6e-06 | 9.433E-06 | 4.248E-06 | 5.600E-06 | 1.400E-05 |
| 4683 | 235 | H235N | A | 7.7e-06 | 9.7e-06 | 5.2e-06 | 7.533E-06 | 2.255E-06 | 5.200E-06 | 9.700E-06 |
| 4684 | 235 | H235D | A | 9.4e-06 | 4.2e-06 | 8.2e-06 | 7.267E-06 | 2.723E-06 | 4.200E-06 | 9.400E-06 |
| 4685 | 235 | H235C | A | 1.2e-05 | 1.1e-05 | 6e-06 | 9.667E-06 | 3.215E-06 | 6.000E-06 | 1.200E-05 |
| 4686 | 235 | H235E | A | 7.1e-06 | 8.1e-06 | 9.2e-06 | 8.133E-06 | 1.050E-06 | 7.100E-06 | 9.200E-06 |
| 4687 | 235 | H235Q | A | 1e-05 | 5.6e-06 | 9.9e-06 | 8.500E-06 | 2.512E-06 | 5.600E-06 | 1.000E-05 |
| 4688 | 235 | H235G | A | 1.5e-05 | 9.7e-06 | 1.4e-05 | 1.290E-05 | 2.816E-06 | 9.700E-06 | 1.500E-05 |
| 4689 | 235 | H235H | A | 1e-05 | 8.9e-06 | 1.2e-05 | 1.030E-05 | 1.572E-06 | 8.900E-06 | 1.200E-05 |
| 4690 | 235 | H235I | A | 9.8e-06 | 1.1e-05 | 7.9e-06 | 9.567E-06 | 1.563E-06 | 7.900E-06 | 1.100E-05 |
| 4691 | 235 | H235L | A | 6.6e-06 | 9.6e-06 | 7.9e-06 | 8.033E-06 | 1.504E-06 | 6.600E-06 | 9.600E-06 |
| 4692 | 235 | H235K | A | 5.9e-06 | 5.6e-06 | 8.8e-06 | 6.767E-06 | 1.767E-06 | 5.600E-06 | 8.800E-06 |
| 4693 | 235 | H235M | A | 6.5e-06 | 7.6e-06 | 1.5e-05 | 9.700E-06 | 4.623E-06 | 6.500E-06 | 1.500E-05 |
| 4694 | 235 | H235F | A | 7.1e-06 | 5.5e-06 | 9e-06 | 7.200E-06 | 1.752E-06 | 5.500E-06 | 9.000E-06 |
| 4695 | 235 | H235P | A | 5e-06 | 8.8e-06 | 4.3e-06 | 6.033E-06 | 2.421E-06 | 4.300E-06 | 8.800E-06 |
| 4696 | 235 | H235S | A | 6.4e-06 | 7e-06 | 8.8e-06 | 7.400E-06 | 1.249E-06 | 6.400E-06 | 8.800E-06 |
| 4697 | 235 | H235T | A | 1.1e-05 | 1.2e-05 | 1e-05 | 1.100E-05 | 1.000E-06 | 1.000E-05 | 1.200E-05 |
| 4698 | 235 | H235W | A | 7.8e-06 | 1e-05 | 7.7e-06 | 8.500E-06 | 1.300E-06 | 7.700E-06 | 1.000E-05 |
| 4699 | 235 | H235Y | A | 1.3e-05 | 9.4e-06 | 7.2e-06 | 9.867E-06 | 2.928E-06 | 7.200E-06 | 1.300E-05 |
| 4700 | 235 | H235V | A | 6.5e-06 | 9.3e-06 | 8.4e-06 | 8.067E-06 | 1.429E-06 | 6.500E-06 | 9.300E-06 |
| 4701 | 236 | F236A | A | 6.3e-06 | 5.8e-06 | 6.9e-06 | 6.333E-06 | 5.508E-07 | 5.800E-06 | 6.900E-06 |
| 4702 | 236 | F236R | A | 6.5e-06 | 5.1e-06 | 5.5e-06 | 5.700E-06 | 7.211E-07 | 5.100E-06 | 6.500E-06 |
| 4703 | 236 | F236N | A | 6.4e-06 | 1.1e-05 | 7.3e-06 | 8.233E-06 | 2.438E-06 | 6.400E-06 | 1.100E-05 |
| 4704 | 236 | F236D | A | 1.2e-05 | 1.3e-05 | 9.4e-06 | 1.147E-05 | 1.858E-06 | 9.400E-06 | 1.300E-05 |
| 4705 | 236 | F236C | A | 6.6e-06 | 7.2e-06 | 4.9e-06 | 6.233E-06 | 1.193E-06 | 4.900E-06 | 7.200E-06 |
| 4706 | 236 | F236E | A | 9.1e-06 | 7.8e-06 | 4.5e-06 | 7.133E-06 | 2.371E-06 | 4.500E-06 | 9.100E-06 |
| 4707 | 236 | F236Q | A | 8e-06 | 8e-06 | 5.7e-06 | 7.233E-06 | 1.328E-06 | 5.700E-06 | 8.000E-06 |
| 4708 | 236 | F236G | A | 1.2e-05 | 9e-06 | 5.1e-06 | 8.700E-06 | 3.460E-06 | 5.100E-06 | 1.200E-05 |
| 4709 | 236 | F236H | A | 4.6e-06 | 1.1e-05 | 8.7e-06 | 8.100E-06 | 3.242E-06 | 4.600E-06 | 1.100E-05 |
| 4710 | 236 | F236I | A | 7.8e-06 | 5e-06 | 4.9e-06 | 5.900E-06 | 1.646E-06 | 4.900E-06 | 7.800E-06 |
| 4711 | 236 | F236L | A | 5.2e-06 | 4.6e-06 | 6.4e-06 | 5.400E-06 | 9.165E-07 | 4.600E-06 | 6.400E-06 |
| 4712 | 236 | F236K | A | 8.7e-06 | 8.5e-06 | 1.1e-05 | 9.400E-06 | 1.389E-06 | 8.500E-06 | 1.100E-05 |
| 4713 | 236 | F236M | A | 1.5e-05 | 1e-05 | 5.5e-06 | 1.017E-05 | 4.752E-06 | 5.500E-06 | 1.500E-05 |
| 4714 | 236 | F236F | A | 9.7e-06 | 6.8e-06 | 9.5e-06 | 8.667E-06 | 1.620E-06 | 6.800E-06 | 9.700E-06 |
| 4715 | 236 | F236P | A | 1.2e-05 | 6.1e-06 | 6e-06 | 8.033E-06 | 3.436E-06 | 6.000E-06 | 1.200E-05 |
| 4716 | 236 | F236S | A | 6.7e-06 | 4.3e-06 | 9e-06 | 6.667E-06 | 2.350E-06 | 4.300E-06 | 9.000E-06 |
| 4717 | 236 | F236T | A | 4.8e-06 | 8.4e-06 | 9.7e-06 | 7.633E-06 | 2.538E-06 | 4.800E-06 | 9.700E-06 |
| 4718 | 236 | F236W | A | 7.7e-06 | 6.5e-06 | 8.4e-06 | 7.533E-06 | 9.609E-07 | 6.500E-06 | 8.400E-06 |
| 4719 | 236 | F236Y | A | 4.6e-06 | 9.2e-06 | 9e-06 | 7.600E-06 | 2.600E-06 | 4.600E-06 | 9.200E-06 |
| 4720 | 236 | F236V | A | 8.4e-06 | 5.8e-06 | 4.4e-06 | 6.200E-06 | 2.030E-06 | 4.400E-06 | 8.400E-06 |
| 4721 | 237 | N237A | A | 7.2e-06 | 1.2e-05 | 9.4e-06 | 9.533E-06 | 2.403E-06 | 7.200E-06 | 1.200E-05 |
| 4722 | 237 | N237R | A | 6.1e-06 | 6.7e-06 | 7.1e-06 | 6.633E-06 | 5.033E-07 | 6.100E-06 | 7.100E-06 |
| 4723 | 237 | N237N | A | 5.5e-06 | 6.3e-06 | 5.6e-06 | 5.800E-06 | 4.359E-07 | 5.500E-06 | 6.300E-06 |
| 4724 | 237 | N237D | A | 8.1e-06 | 1.2e-05 | 3.1e-06 | 7.733E-06 | 4.461E-06 | 3.100E-06 | 1.200E-05 |
| 4725 | 237 | N237C | A | 5.2e-06 | 1e-05 | 4.8e-06 | 6.667E-06 | 2.894E-06 | 4.800E-06 | 1.000E-05 |
| 4726 | 237 | N237E | A | 9.3e-06 | 7.5e-06 | 5.9e-06 | 7.567E-06 | 1.701E-06 | 5.900E-06 | 9.300E-06 |
| 4727 | 237 | N237Q | A | 1e-05 | 5.1e-06 | 1.1e-05 | 8.700E-06 | 3.158E-06 | 5.100E-06 | 1.100E-05 |
| 4728 | 237 | N237G | A | 5.2e-06 | 8.8e-06 | 7e-06 | 7.000E-06 | 1.800E-06 | 5.200E-06 | 8.800E-06 |
| 4729 | 237 | N237H | A | 6.1e-06 | 1e-05 | 8.5e-06 | 8.200E-06 | 1.967E-06 | 6.100E-06 | 1.000E-05 |
| 4730 | 237 | N237I | A | 5.9e-06 | 5.6e-06 | 8.2e-06 | 6.567E-06 | 1.422E-06 | 5.600E-06 | 8.200E-06 |
| 4731 | 237 | N237L | A | 1.2e-05 | 1.6e-05 | 1.1e-05 | 1.300E-05 | 2.646E-06 | 1.100E-05 | 1.600E-05 |
| 4732 | 237 | N237K | A | 9.9e-06 | 1.2e-05 | 8.2e-06 | 1.003E-05 | 1.904E-06 | 8.200E-06 | 1.200E-05 |
| 4733 | 237 | N237M | A | 4.3e-06 | 7.1e-06 | 5.8e-06 | 5.733E-06 | 1.401E-06 | 4.300E-06 | 7.100E-06 |
| 4734 | 237 | N237F | A | 4.4e-06 | 6.7e-06 | 7.2e-06 | 6.100E-06 | 1.493E-06 | 4.400E-06 | 7.200E-06 |
| 4735 | 237 | N237P | A | 8.1e-06 | 7e-06 | 6.9e-06 | 7.333E-06 | 6.658E-07 | 6.900E-06 | 8.100E-06 |
| 4736 | 237 | N237S | A | 7e-06 | 1.2e-05 | 1e-05 | 9.667E-06 | 2.517E-06 | 7.000E-06 | 1.200E-05 |
| 4737 | 237 | N237T | A | 1.1e-05 | 5.6e-06 | 1e-05 | 8.867E-06 | 2.873E-06 | 5.600E-06 | 1.100E-05 |
| 4738 | 237 | N237W | A | 9.1e-06 | 7.6e-06 | 9.6e-06 | 8.767E-06 | 1.041E-06 | 7.600E-06 | 9.600E-06 |
| 4739 | 237 | N237Y | A | 1.1e-05 | 1.3e-05 | 5.6e-06 | 9.867E-06 | 3.828E-06 | 5.600E-06 | 1.300E-05 |
| 4740 | 237 | N237V | A | 5.4e-06 | 1.1e-05 | 8.3e-06 | 8.233E-06 | 2.801E-06 | 5.400E-06 | 1.100E-05 |
| 4741 | 238 | H238A | A | 1.3e-05 | 7.8e-06 | 1.8e-05 | 1.293E-05 | 5.100E-06 | 7.800E-06 | 1.800E-05 |
| 4742 | 238 | H238R | A | 1.3e-05 | 4.8e-06 | 7e-06 | 8.267E-06 | 4.244E-06 | 4.800E-06 | 1.300E-05 |
| 4743 | 238 | H238N | A | 8.9e-06 | 5.5e-06 | 9.1e-06 | 7.833E-06 | 2.023E-06 | 5.500E-06 | 9.100E-06 |
| 4744 | 238 | H238D | A | 6.8e-06 | 7.1e-06 | 7.9e-06 | 7.267E-06 | 5.686E-07 | 6.800E-06 | 7.900E-06 |
| 4745 | 238 | H238C | A | 4.9e-06 | 7.6e-06 | 1.1e-05 | 7.833E-06 | 3.057E-06 | 4.900E-06 | 1.100E-05 |
| 4746 | 238 | H238E | A | 6.7e-06 | 1e-05 | 7.5e-06 | 8.067E-06 | 1.721E-06 | 6.700E-06 | 1.000E-05 |
| 4747 | 238 | H238Q | A | 1.4e-05 | 6.6e-06 | 7e-06 | 9.200E-06 | 4.162E-06 | 6.600E-06 | 1.400E-05 |
| 4748 | 238 | H238G | A | 8.8e-06 | 7.3e-06 | 7.8e-06 | 7.967E-06 | 7.638E-07 | 7.300E-06 | 8.800E-06 |
| 4749 | 238 | H238H | A | 9.1e-06 | 7.3e-06 | 5.4e-06 | 7.267E-06 | 1.850E-06 | 5.400E-06 | 9.100E-06 |
| 4750 | 238 | H238I | A | 5.8e-06 | 1e-05 | 9.2e-06 | 8.333E-06 | 2.230E-06 | 5.800E-06 | 1.000E-05 |
| 4751 | 238 | H238L | A | 1e-05 | 1.5e-05 | 5.8e-06 | 1.027E-05 | 4.606E-06 | 5.800E-06 | 1.500E-05 |
| 4752 | 238 | H238K | A | 6.8e-06 | 9.7e-06 | 9.9e-06 | 8.800E-06 | 1.735E-06 | 6.800E-06 | 9.900E-06 |
| 4753 | 238 | H238M | A | 6.3e-06 | 1.3e-05 | 1.5e-05 | 1.143E-05 | 4.557E-06 | 6.300E-06 | 1.500E-05 |
| 4754 | 238 | H238F | A | 4.7e-06 | 9.3e-06 | 1.1e-05 | 8.333E-06 | 3.259E-06 | 4.700E-06 | 1.100E-05 |
| 4755 | 238 | H238P | A | 6e-06 | 5.7e-06 | 5e-06 | 5.567E-06 | 5.132E-07 | 5.000E-06 | 6.000E-06 |
| 4756 | 238 | H238S | A | 5.7e-06 | 5.5e-06 | 8.4e-06 | 6.533E-06 | 1.620E-06 | 5.500E-06 | 8.400E-06 |
| 4757 | 238 | H238T | A | 6.9e-06 | 8.5e-06 | 9.7e-06 | 8.367E-06 | 1.405E-06 | 6.900E-06 | 9.700E-06 |
| 4758 | 238 | H238W | A | 1.3e-05 | 3.3e-06 | 1e-05 | 8.767E-06 | 4.966E-06 | 3.300E-06 | 1.300E-05 |
| 4759 | 238 | H238Y | A | 1.1e-05 | 8.9e-06 | 8.7e-06 | 9.533E-06 | 1.274E-06 | 8.700E-06 | 1.100E-05 |

| | | | | | | | | | | |
|---|---|---|---|---|---|---|---|---|---|---|
| 4760 | 238 | H238V | A | 7.1e-06 | 8.8e-06 | 6.2e-06 | 7.367E-06 | 1.320E-06 | 6.200E-06 | 8.800E-06 |
| 4761 | 239 | S239A | A | 1e-05 | 7.3e-06 | 8e-06 | 8.433E-06 | 1.401E-06 | 7.300E-06 | 1.000E-05 |
| 4762 | 239 | S239R | A | 1.1e-05 | 9.6e-06 | 6.7e-06 | 9.100E-06 | 2.193E-06 | 6.700E-06 | 1.100E-05 |
| 4763 | 239 | S239N | A | 9.2e-06 | 9.9e-06 | 9e-06 | 9.367E-06 | 4.726E-07 | 9.000E-06 | 9.900E-06 |
| 4764 | 239 | S239D | A | 6.9e-06 | 8.6e-06 | 9.1e-06 | 8.200E-06 | 1.153E-06 | 6.900E-06 | 9.100E-06 |
| 4765 | 239 | S239C | A | 1.2e-05 | 1.3e-05 | 1.4e-05 | 1.300E-05 | 1.000E-06 | 1.200E-05 | 1.400E-05 |
| 4766 | 239 | S239E | A | 9.2e-06 | 6.5e-06 | 9.3e-06 | 8.333E-06 | 1.589E-06 | 6.500E-06 | 9.300E-06 |
| 4767 | 239 | S239Q | A | 1e-05 | 4.6e-06 | 5.6e-06 | 6.733E-06 | 2.873E-06 | 4.600E-06 | 1.000E-05 |
| 4768 | 239 | S239G | A | 7.6e-06 | 5.2e-06 | 1.3e-05 | 8.600E-06 | 3.995E-06 | 5.200E-06 | 1.300E-05 |
| 4769 | 239 | S239H | A | 9.7e-06 | 7.5e-06 | 1e-05 | 9.067E-06 | 1.365E-06 | 7.500E-06 | 1.000E-05 |
| 4770 | 239 | S239I | A | 5.1e-06 | 6.7e-06 | 6.6e-06 | 6.133E-06 | 8.963E-07 | 5.100E-06 | 6.700E-06 |
| 4771 | 239 | S239L | A | 5.3e-06 | 1.2e-05 | 8.2e-06 | 8.500E-06 | 3.360E-06 | 5.300E-06 | 1.200E-05 |
| 4772 | 239 | S239K | A | 9.3e-06 | 4.9e-06 | 5.4e-06 | 6.533E-06 | 2.409E-06 | 4.900E-06 | 9.300E-06 |
| 4773 | 239 | S239M | A | 7.7e-06 | 5.1e-06 | 7.6e-06 | 6.800E-06 | 1.473E-06 | 5.100E-06 | 7.700E-06 |
| 4774 | 239 | S239F | A | 8.5e-06 | 5.4e-06 | 5e-06 | 6.300E-06 | 1.916E-06 | 5.000E-06 | 8.500E-06 |
| 4775 | 239 | S239P | A | 9.7e-06 | 9.9e-06 | 1.4e-05 | 1.120E-05 | 2.427E-06 | 9.700E-06 | 1.400E-05 |
| 4776 | 239 | S239S | A | 1.1e-05 | 7.5e-06 | 9.7e-06 | 9.400E-06 | 1.769E-06 | 7.500E-06 | 1.100E-05 |
| 4777 | 239 | S239T | A | 6.3e-06 | 8.1e-06 | 1.5e-05 | 9.800E-06 | 4.592E-06 | 6.300E-06 | 1.500E-05 |
| 4778 | 239 | S239W | A | 1.2e-05 | 6.8e-06 | 6.9e-06 | 8.567E-06 | 2.974E-06 | 6.800E-06 | 1.200E-05 |
| 4779 | 239 | S239Y | A | 7.5e-06 | 9.4e-06 | 6.6e-06 | 7.833E-06 | 1.429E-06 | 6.600E-06 | 9.400E-06 |
| 4780 | 239 | S239V | A | 1e-05 | 7.4e-06 | 5.8e-06 | 7.733E-06 | 2.120E-06 | 5.800E-06 | 1.000E-05 |
| 4781 | 240 | G240A | A | 5.9e-06 | 7.2e-06 | 5.5e-06 | 6.200E-06 | 8.888E-07 | 5.500E-06 | 7.200E-06 |
| 4782 | 240 | G240R | A | 9.6e-06 | 8.2e-06 | 7e-06 | 8.267E-06 | 1.301E-06 | 7.000E-06 | 9.600E-06 |
| 4783 | 240 | G240N | A | 1.1e-05 | 5.9e-06 | 4.1e-06 | 7.000E-06 | 3.579E-06 | 4.100E-06 | 1.100E-05 |
| 4784 | 240 | G240D | A | 8.2e-06 | 7.6e-06 | 1e-05 | 8.600E-06 | 1.249E-06 | 7.600E-06 | 1.000E-05 |
| 4785 | 240 | G240C | A | 6.1e-06 | 4.7e-06 | 7.3e-06 | 6.033E-06 | 1.301E-06 | 4.700E-06 | 7.300E-06 |
| 4786 | 240 | G240E | A | 6.5e-06 | 1.4e-05 | 7.9e-06 | 9.467E-06 | 3.988E-06 | 6.500E-06 | 1.400E-05 |
| 4787 | 240 | G240Q | A | 7.3e-06 | 1.5e-05 | 1e-05 | 1.077E-05 | 3.907E-06 | 7.300E-06 | 1.500E-05 |
| 4788 | 240 | G240G | A | 7.4e-06 | 6.6e-06 | 1.2e-05 | 8.667E-06 | 2.914E-06 | 6.600E-06 | 1.200E-05 |
| 4789 | 240 | G240H | A | 7.9e-06 | 6.6e-06 | 6.2e-06 | 6.900E-06 | 8.888E-07 | 6.200E-06 | 7.900E-06 |
| 4790 | 240 | G240I | A | 7e-06 | 4.4e-06 | 1.1e-05 | 7.467E-06 | 3.325E-06 | 4.400E-06 | 1.100E-05 |
| 4791 | 240 | G240L | A | 6.1e-06 | 1.1e-05 | 4.6e-06 | 7.233E-06 | 3.347E-06 | 4.600E-06 | 1.100E-05 |
| 4792 | 240 | G240K | A | 4.8e-06 | 5.3e-06 | 5.5e-06 | 5.200E-06 | 3.606E-07 | 4.800E-06 | 5.500E-06 |
| 4793 | 240 | G240M | A | 9.3e-06 | 1.3e-05 | 9.6e-06 | 1.063E-05 | 2.055E-06 | 9.300E-06 | 1.300E-05 |
| 4794 | 240 | G240F | A | 9.1e-06 | 7.1e-06 | 1.4e-05 | 1.007E-05 | 3.550E-06 | 7.100E-06 | 1.400E-05 |
| 4795 | 240 | G240P | A | 1.4e-05 | 7.6e-06 | 1.2e-05 | 1.120E-05 | 3.274E-06 | 7.600E-06 | 1.400E-05 |
| 4796 | 240 | G240S | A | 7.7e-06 | 6.4e-06 | 4e-06 | 6.033E-06 | 1.877E-06 | 4.000E-06 | 7.700E-06 |
| 4797 | 240 | G240T | A | 5.2e-06 | 9.5e-06 | 9.8e-06 | 8.167E-06 | 2.574E-06 | 5.200E-06 | 9.800E-06 |
| 4798 | 240 | G240W | A | 1.2e-05 | 9.3e-06 | 8.5e-06 | 9.933E-06 | 1.834E-06 | 8.500E-06 | 1.200E-05 |
| 4799 | 240 | G240Y | A | 6.5e-06 | 1.3e-05 | 7.2e-06 | 8.900E-06 | 3.568E-06 | 6.500E-06 | 1.300E-05 |
| 4800 | 240 | G240V | A | 6.8e-06 | 1e-05 | 5.1e-06 | 7.300E-06 | 2.488E-06 | 5.100E-06 | 1.000E-05 |
| 4801 | 241 | I241A | A | 9.2e-06 | 5.8e-06 | 8.8e-06 | 7.933E-06 | 1.858E-06 | 5.800E-06 | 9.200E-06 |
| 4802 | 241 | I241R | A | 1e-05 | 1e-05 | 9.3e-06 | 9.767E-06 | 4.041E-07 | 9.300E-06 | 1.000E-05 |
| 4803 | 241 | I241N | A | 9e-06 | 6.7e-06 | 9e-06 | 8.233E-06 | 1.328E-06 | 6.700E-06 | 9.000E-06 |
| 4804 | 241 | I241D | A | 6.6e-06 | 1.1e-05 | 9.7e-06 | 9.100E-06 | 2.261E-06 | 6.600E-06 | 1.100E-05 |
| 4805 | 241 | I241C | A | 1.4e-05 | 6.4e-06 | 6.5e-06 | 8.967E-06 | 4.359E-06 | 6.400E-06 | 1.400E-05 |
| 4806 | 241 | I241E | A | 5e-06 | 1.1e-05 | 1.1e-05 | 9.000E-06 | 3.464E-06 | 5.000E-06 | 1.100E-05 |
| 4807 | 241 | I241Q | A | 1.3e-05 | 1.4e-05 | 4.5e-06 | 1.050E-05 | 5.220E-06 | 4.500E-06 | 1.400E-05 |
| 4808 | 241 | I241G | A | 5.6e-06 | 1.1e-05 | 6.8e-06 | 7.800E-06 | 2.835E-06 | 5.600E-06 | 1.100E-05 |
| 4809 | 241 | I241H | A | 6.5e-06 | 6.4e-06 | 5.6e-06 | 6.167E-06 | 4.933E-07 | 5.600E-06 | 6.500E-06 |
| 4810 | 241 | I241I | A | 1.1e-05 | 1.1e-05 | 7.1e-06 | 9.700E-06 | 2.252E-06 | 7.100E-06 | 1.100E-05 |
| 4811 | 241 | I241L | A | 1.2e-05 | 5.7e-06 | 6.5e-06 | 8.067E-06 | 3.430E-06 | 5.700E-06 | 1.200E-05 |
| 4812 | 241 | I241K | A | 8.6e-06 | 9.3e-06 | 6.6e-06 | 8.167E-06 | 1.401E-06 | 6.600E-06 | 9.300E-06 |
| 4813 | 241 | I241M | A | 8.1e-06 | 1.4e-05 | 1.6e-05 | 1.270E-05 | 4.107E-06 | 8.100E-06 | 1.600E-05 |
| 4814 | 241 | I241F | A | 7.5e-06 | 7.5e-06 | 8.2e-06 | 7.733E-06 | 4.041E-07 | 7.500E-06 | 8.200E-06 |
| 4815 | 241 | I241P | A | 1e-05 | 6.9e-06 | 6.9e-06 | 7.933E-06 | 1.790E-06 | 6.900E-06 | 1.000E-05 |
| 4816 | 241 | I241S | A | 7.1e-06 | 8.1e-06 | 5.9e-06 | 7.033E-06 | 1.102E-06 | 5.900E-06 | 8.100E-06 |
| 4817 | 241 | I241T | A | 1.5e-05 | 6.7e-06 | 8.7e-06 | 1.013E-05 | 4.332E-06 | 6.700E-06 | 1.500E-05 |
| 4818 | 241 | I241W | A | 8e-06 | 7.5e-06 | 5.2e-06 | 6.900E-06 | 1.493E-06 | 5.200E-06 | 8.000E-06 |
| 4819 | 241 | I241Y | A | 1e-05 | 8.3e-06 | 6.4e-06 | 8.233E-06 | 1.801E-06 | 6.400E-06 | 1.000E-05 |
| 4820 | 241 | I241V | A | 9.8e-06 | 5.1e-06 | 1.3e-05 | 9.300E-06 | 3.974E-06 | 5.100E-06 | 1.300E-05 |
| 4821 | 242 | C242A | A | 1.4e-05 | 4.7e-06 | 8.2e-06 | 8.967E-06 | 4.697E-06 | 4.700E-06 | 1.400E-05 |
| 4822 | 242 | C242R | A | 5.5e-06 | 6.6e-06 | 8e-06 | 6.700E-06 | 1.253E-06 | 5.500E-06 | 8.000E-06 |
| 4823 | 242 | C242N | A | 6.7e-06 | 8.5e-06 | 6.2e-06 | 7.133E-06 | 1.210E-06 | 6.200E-06 | 8.500E-06 |
| 4824 | 242 | C242D | A | 8.3e-06 | 1.4e-05 | 5.7e-06 | 9.333E-06 | 4.245E-06 | 5.700E-06 | 1.400E-05 |
| 4825 | 242 | C242C | A | 7e-06 | 6.3e-06 | 8.6e-06 | 7.300E-06 | 1.179E-06 | 6.300E-06 | 8.600E-06 |
| 4826 | 242 | C242E | A | 8.7e-06 | 1.1e-05 | 6e-06 | 8.567E-06 | 2.503E-06 | 6.000E-06 | 1.100E-05 |
| 4827 | 242 | C242Q | A | 9.9e-06 | 6.8e-06 | 7.1e-06 | 7.933E-06 | 1.710E-06 | 6.800E-06 | 9.900E-06 |
| 4828 | 242 | C242G | A | 9.4e-06 | 6.1e-06 | 6.3e-06 | 7.267E-06 | 1.850E-06 | 6.100E-06 | 9.400E-06 |
| 4829 | 242 | C242H | A | 9.4e-06 | 7.7e-06 | 1.3e-05 | 1.003E-05 | 2.706E-06 | 7.700E-06 | 1.300E-05 |
| 4830 | 242 | C242I | A | 1.2e-05 | 5.3e-06 | 6.1e-06 | 7.800E-06 | 3.659E-06 | 5.300E-06 | 1.200E-05 |
| 4831 | 242 | C242L | A | 7.7e-06 | 7.5e-06 | 4.4e-06 | 6.533E-06 | 1.850E-06 | 4.400E-06 | 7.700E-06 |
| 4832 | 242 | C242K | A | 1.5e-05 | 1.1e-05 | 5.4e-06 | 1.047E-05 | 4.822E-06 | 5.400E-06 | 1.500E-05 |
| 4833 | 242 | C242M | A | 6.6e-06 | 6.2e-06 | 9e-06 | 7.267E-06 | 1.514E-06 | 6.200E-06 | 9.000E-06 |
| 4834 | 242 | C242F | A | 5e-06 | 6.9e-06 | 1.3e-05 | 8.300E-06 | 4.180E-06 | 5.000E-06 | 1.300E-05 |
| 4835 | 242 | C242P | A | 9e-06 | 4.4e-06 | 1e-05 | 7.800E-06 | 2.987E-06 | 4.400E-06 | 1.000E-05 |
| 4836 | 242 | C242S | A | 5.5e-06 | 1.3e-05 | 5.7e-06 | 8.067E-06 | 4.274E-06 | 5.500E-06 | 1.300E-05 |
| 4837 | 242 | C242T | A | 6.8e-06 | 7.3e-06 | 6.6e-06 | 6.900E-06 | 3.606E-07 | 6.600E-06 | 7.300E-06 |
| 4838 | 242 | C242W | A | 9.4e-06 | 5.3e-06 | 4.4e-06 | 6.367E-06 | 2.665E-06 | 4.400E-06 | 9.400E-06 |
| 4839 | 242 | C242Y | A | 1e-05 | 6.7e-06 | 5.9e-06 | 7.533E-06 | 2.173E-06 | 5.900E-06 | 1.000E-05 |
| 4840 | 242 | C242V | A | 7.3e-06 | 7.7e-06 | 1e-05 | 8.333E-06 | 1.457E-06 | 7.300E-06 | 1.000E-05 |
| 4841 | 243 | E243A | A | 4.8e-06 | 1e-05 | 1.1e-05 | 8.600E-06 | 3.329E-06 | 4.800E-06 | 1.100E-05 |
| 4842 | 243 | E243R | A | 7.8e-06 | 4.8e-06 | 5.3e-06 | 5.967E-06 | 1.607E-06 | 4.800E-06 | 7.800E-06 |
| 4843 | 243 | E243N | A | 8.2e-06 | 8.6e-06 | 8.7e-06 | 8.500E-06 | 2.646E-07 | 8.200E-06 | 8.700E-06 |
| 4844 | 243 | E243D | A | 7.2e-06 | 5.4e-06 | 9.6e-06 | 7.400E-06 | 2.107E-06 | 5.400E-06 | 9.600E-06 |

| | | | | | | | | | | |
|---|---|---|---|---|---|---|---|---|---|---|
| 4845 | 243 | E243C | A | 8e-06 | 5.8e-06 | 1.3e-05 | 8.933E-06 | 3.690E-06 | 5.800E-06 | 1.300E-05 |
| 4846 | 243 | E243E | A | 8.9e-06 | 7.3e-06 | 7.3e-06 | 7.833E-06 | 9.238E-07 | 7.300E-06 | 8.900E-06 |
| 4847 | 243 | E243Q | A | 9.6e-06 | 5.4e-06 | 7e-06 | 7.333E-06 | 2.120E-06 | 5.400E-06 | 9.600E-06 |
| 4848 | 243 | E243G | A | 9.6e-06 | 1e-05 | 1e-05 | 9.867E-06 | 2.309E-07 | 9.600E-06 | 1.000E-05 |
| 4849 | 243 | E243H | A | 1.2e-05 | 7e-06 | 8.3e-06 | 9.100E-06 | 2.594E-06 | 7.000E-06 | 1.200E-05 |
| 4850 | 243 | E243I | A | 8.6e-06 | 1.2e-05 | 6.5e-06 | 9.033E-06 | 2.775E-06 | 6.500E-06 | 1.200E-05 |
| 4851 | 243 | E243L | A | 1.7e-05 | 7.5e-06 | 4.3e-06 | 9.600E-06 | 6.605E-06 | 4.300E-06 | 1.700E-05 |
| 4852 | 243 | E243K | A | 1e-05 | 6.4e-06 | 1.4e-05 | 1.013E-05 | 3.802E-06 | 6.400E-06 | 1.400E-05 |
| 4853 | 243 | E243M | A | 1e-05 | 6.4e-06 | 9.7e-06 | 8.700E-06 | 1.997E-06 | 6.400E-06 | 1.000E-05 |
| 4854 | 243 | E243F | A | 1.5e-05 | 8.8e-06 | 8.1e-06 | 1.063E-05 | 3.798E-06 | 8.100E-06 | 1.500E-05 |
| 4855 | 243 | E243P | A | 7.1e-06 | 1.2e-05 | 8.7e-06 | 9.267E-06 | 2.499E-06 | 7.100E-06 | 1.200E-05 |
| 4856 | 243 | E243S | A | 9.7e-06 | 7.8e-06 | 7.3e-06 | 8.267E-06 | 1.266E-06 | 7.300E-06 | 9.700E-06 |
| 4857 | 243 | E243T | A | 9.6e-06 | 1e-05 | 1.3e-05 | 1.087E-05 | 1.858E-06 | 9.600E-06 | 1.300E-05 |
| 4858 | 243 | E243W | A | 1e-05 | 8.3e-06 | 8.5e-06 | 8.933E-06 | 9.292E-07 | 8.300E-06 | 1.000E-05 |
| 4859 | 243 | E243Y | A | 9e-06 | 5.9e-06 | 9e-06 | 7.967E-06 | 1.790E-06 | 5.900E-06 | 9.000E-06 |
| 4860 | 243 | E243V | A | 8.4e-06 | 1.6e-05 | 6.3e-06 | 1.023E-05 | 5.103E-06 | 6.300E-06 | 1.600E-05 |
| 4861 | 244 | L244A | A | 7.3e-06 | 7.2e-06 | 6.7e-06 | 7.067E-06 | 3.215E-07 | 6.700E-06 | 7.300E-06 |
| 4862 | 244 | L244R | A | 4.7e-06 | 8.6e-06 | 4.4e-06 | 5.900E-06 | 2.343E-06 | 4.400E-06 | 8.600E-06 |
| 4863 | 244 | L244N | A | 4.1e-06 | 6.5e-06 | 5.5e-06 | 5.367E-06 | 1.206E-06 | 4.100E-06 | 6.500E-06 |
| 4864 | 244 | L244D | A | 5.5e-06 | 7.2e-06 | 6.9e-06 | 6.533E-06 | 9.074E-07 | 5.500E-06 | 7.200E-06 |
| 4865 | 244 | L244C | A | 8e-06 | 1.3e-05 | 6.3e-06 | 9.100E-06 | 3.483E-06 | 6.300E-06 | 1.300E-05 |
| 4866 | 244 | L244E | A | 7.2e-06 | 9e-06 | 6.1e-06 | 7.433E-06 | 1.464E-06 | 6.100E-06 | 9.000E-06 |
| 4867 | 244 | L244Q | A | 1e-05 | 6.5e-06 | 6.2e-06 | 7.567E-06 | 2.113E-06 | 6.200E-06 | 1.000E-05 |
| 4868 | 244 | L244G | A | 8.3e-06 | 7.1e-06 | 1.3e-05 | 9.467E-06 | 3.118E-06 | 7.100E-06 | 1.300E-05 |
| 4869 | 244 | L244H | A | 1.8e-05 | 5.2e-06 | 8.9e-06 | 1.070E-05 | 6.587E-06 | 5.200E-06 | 1.800E-05 |
| 4870 | 244 | L244I | A | 1.6e-05 | 9e-06 | 9.5e-06 | 1.150E-05 | 3.905E-06 | 9.000E-06 | 1.600E-05 |
| 4871 | 244 | L244L | A | 1.2e-05 | 7.6e-06 | 7e-06 | 8.867E-06 | 2.730E-06 | 7.000E-06 | 1.200E-05 |
| 4872 | 244 | L244K | A | 8.4e-06 | 1.3e-05 | 6.6e-06 | 9.333E-06 | 3.301E-06 | 6.600E-06 | 1.300E-05 |
| 4873 | 244 | L244M | A | 1e-05 | 9.3e-06 | 5.5e-06 | 8.267E-06 | 2.421E-06 | 5.500E-06 | 1.000E-05 |
| 4874 | 244 | L244F | A | 4.8e-06 | 4.6e-06 | 6.4e-06 | 5.267E-06 | 9.866E-07 | 4.600E-06 | 6.400E-06 |
| 4875 | 244 | L244P | A | 5.1e-06 | 8.9e-06 | 7.1e-06 | 7.033E-06 | 1.901E-06 | 5.100E-06 | 8.900E-06 |
| 4876 | 244 | L244S | A | 4.7e-06 | 5.4e-06 | 9.3e-06 | 6.467E-06 | 2.479E-06 | 4.700E-06 | 9.300E-06 |
| 4877 | 244 | L244T | A | 7.8e-06 | 4.7e-06 | 9.7e-06 | 7.400E-06 | 2.524E-06 | 4.700E-06 | 9.700E-06 |
| 4878 | 244 | L244W | A | 8.3e-06 | 9.7e-06 | 9.1e-06 | 9.033E-06 | 7.024E-07 | 8.300E-06 | 9.700E-06 |
| 4879 | 244 | L244Y | A | 1.2e-05 | 7.1e-06 | 1.2e-05 | 1.037E-05 | 2.829E-06 | 7.100E-06 | 1.200E-05 |
| 4880 | 244 | L244V | A | 1e-05 | 8.3e-06 | 6.7e-06 | 8.333E-06 | 1.650E-06 | 6.700E-06 | 1.000E-05 |
| 4881 | 245 | H245A | A | 1.2e-05 | 8.1e-06 | 5.8e-06 | 8.633E-06 | 3.134E-06 | 5.800E-06 | 1.200E-05 |
| 4882 | 245 | H245R | A | 9.3e-06 | 5.6e-06 | 6.5e-06 | 7.133E-06 | 1.930E-06 | 5.600E-06 | 9.300E-06 |
| 4883 | 245 | H245N | A | 1e-05 | 1.1e-05 | 1e-05 | 1.033E-05 | 5.774E-07 | 1.000E-05 | 1.100E-05 |
| 4884 | 245 | H245D | A | 1e-05 | 7.3e-06 | 4.9e-06 | 7.400E-06 | 2.551E-06 | 4.900E-06 | 1.000E-05 |
| 4885 | 245 | H245C | A | 5.6e-06 | 1e-05 | 1.3e-05 | 9.533E-06 | 3.722E-06 | 5.600E-06 | 1.300E-05 |
| 4886 | 245 | H245E | A | 6.3e-06 | 1.1e-05 | 1.1e-05 | 9.433E-06 | 2.714E-06 | 6.300E-06 | 1.100E-05 |
| 4887 | 245 | H245Q | A | 6.3e-06 | 1.2e-05 | 6.2e-06 | 8.167E-06 | 3.320E-06 | 6.200E-06 | 1.200E-05 |
| 4888 | 245 | H245G | A | 6.1e-06 | 5.8e-06 | 8e-06 | 6.633E-06 | 1.193E-06 | 5.800E-06 | 8.000E-06 |
| 4889 | 245 | H245H | A | 1.4e-05 | 8.4e-06 | 7.8e-06 | 1.007E-05 | 3.420E-06 | 7.800E-06 | 1.400E-05 |
| 4890 | 245 | H245I | A | 6.4e-06 | 6.2e-06 | 1e-05 | 7.533E-06 | 2.139E-06 | 6.200E-06 | 1.000E-05 |
| 4891 | 245 | H245L | A | 6.2e-06 | 6e-06 | 8.9e-06 | 7.033E-06 | 1.620E-06 | 6.000E-06 | 8.900E-06 |
| 4892 | 245 | H245K | A | 9.5e-06 | 8.4e-06 | 8.7e-06 | 8.867E-06 | 5.686E-07 | 8.400E-06 | 9.500E-06 |
| 4893 | 245 | H245M | A | 5.6e-06 | 1.4e-05 | 6.9e-06 | 8.833E-06 | 4.521E-06 | 5.600E-06 | 1.400E-05 |
| 4894 | 245 | H245F | A | 4.8e-06 | 7.5e-06 | 6.9e-06 | 6.400E-06 | 1.418E-06 | 4.800E-06 | 7.500E-06 |
| 4895 | 245 | H245P | A | 7.7e-06 | 4.6e-06 | 5.1e-06 | 5.800E-06 | 1.664E-06 | 4.600E-06 | 7.700E-06 |
| 4896 | 245 | H245S | A | 6.3e-06 | 6.8e-06 | 4.8e-06 | 5.967E-06 | 1.041E-06 | 4.800E-06 | 6.800E-06 |
| 4897 | 245 | H245T | A | 1.8e-05 | 5.9e-06 | 8.1e-06 | 1.067E-05 | 6.445E-06 | 5.900E-06 | 1.800E-05 |
| 4898 | 245 | H245W | A | 9.6e-06 | 1.2e-05 | 7.4e-06 | 9.667E-06 | 2.301E-06 | 7.400E-06 | 1.200E-05 |
| 4899 | 245 | H245Y | A | 1.2e-05 | 9.9e-06 | 7e-06 | 9.633E-06 | 2.511E-06 | 7.000E-06 | 1.200E-05 |
| 4900 | 245 | H245V | A | 1.1e-05 | 5.5e-06 | 1.4e-05 | 1.017E-05 | 4.311E-06 | 5.500E-06 | 1.400E-05 |
| 4901 | 246 | C246A | A | 6.3e-06 | 5.9e-06 | 4.9e-06 | 5.700E-06 | 7.211E-07 | 4.900E-06 | 6.300E-06 |
| 4902 | 246 | C246R | A | 9.5e-06 | 6.2e-06 | 8.9e-06 | 8.200E-06 | 1.758E-06 | 6.200E-06 | 9.500E-06 |
| 4903 | 246 | C246N | A | 7.9e-06 | 6.4e-06 | 9.3e-06 | 7.867E-06 | 1.450E-06 | 6.400E-06 | 9.300E-06 |
| 4904 | 246 | C246D | A | 9.8e-06 | 1.1e-05 | 9.8e-06 | 1.020E-05 | 6.928E-07 | 9.800E-06 | 1.100E-05 |
| 4905 | 246 | C246C | A | 7.8e-06 | 8.2e-06 | 1.4e-05 | 1.000E-05 | 3.470E-06 | 7.800E-06 | 1.400E-05 |
| 4906 | 246 | C246E | A | 4.8e-06 | 1.3e-05 | 6.3e-06 | 8.033E-06 | 4.366E-06 | 4.800E-06 | 1.300E-05 |
| 4907 | 246 | C246Q | A | 4.7e-06 | 2.1e-05 | 7.1e-06 | 1.093E-05 | 8.800E-06 | 4.700E-06 | 2.100E-05 |
| 4908 | 246 | C246G | A | 7.1e-06 | 6.1e-06 | 5.5e-06 | 6.233E-06 | 8.083E-07 | 5.500E-06 | 7.100E-06 |
| 4909 | 246 | C246H | A | 8.4e-06 | 8.7e-06 | 8.8e-06 | 8.633E-06 | 2.082E-07 | 8.400E-06 | 8.800E-06 |
| 4910 | 246 | C246I | A | 4.9e-06 | 8e-06 | 9.2e-06 | 7.367E-06 | 2.219E-06 | 4.900E-06 | 9.200E-06 |
| 4911 | 246 | C246L | A | 9.4e-06 | 8.7e-06 | 9.5e-06 | 9.200E-06 | 4.359E-07 | 8.700E-06 | 9.500E-06 |
| 4912 | 246 | C246K | A | 5.2e-06 | 1.2e-05 | 6.4e-06 | 7.867E-06 | 3.630E-06 | 5.200E-06 | 1.200E-05 |
| 4913 | 246 | C246M | A | 6.9e-06 | 1.2e-05 | 9.4e-06 | 9.433E-06 | 2.550E-06 | 6.900E-06 | 1.200E-05 |
| 4914 | 246 | C246F | A | 6.6e-06 | 6.7e-06 | 7.7e-06 | 7.000E-06 | 6.083E-07 | 6.600E-06 | 7.700E-06 |
| 4915 | 246 | C246P | A | 1.3e-05 | 9.6e-06 | 4.8e-06 | 9.133E-06 | 4.120E-06 | 4.800E-06 | 1.300E-05 |
| 4916 | 246 | C246S | A | 8.5e-06 | 8.4e-06 | 9.4e-06 | 8.767E-06 | 5.508E-07 | 8.400E-06 | 9.400E-06 |
| 4917 | 246 | C246T | A | 7.2e-06 | 7.9e-06 | 5.9e-06 | 7.000E-06 | 1.015E-06 | 5.900E-06 | 7.900E-06 |
| 4918 | 246 | C246W | A | 9.9e-06 | 9.9e-06 | 9.7e-06 | 9.833E-06 | 1.155E-07 | 9.700E-06 | 9.900E-06 |
| 4919 | 246 | C246Y | A | 6.6e-06 | 1.3e-05 | 1e-05 | 9.867E-06 | 3.202E-06 | 6.600E-06 | 1.300E-05 |
| 4920 | 246 | C246V | A | 5.2e-06 | 9.7e-06 | 6e-06 | 6.967E-06 | 2.401E-06 | 5.200E-06 | 9.700E-06 |
| 4921 | 247 | P247A | A | 1.3e-05 | 8.5e-06 | 1.3e-05 | 1.150E-05 | 2.598E-06 | 8.500E-06 | 1.300E-05 |
| 4922 | 247 | P247R | A | 6.3e-06 | 6.1e-06 | 8.8e-06 | 7.067E-06 | 1.504E-06 | 6.100E-06 | 8.800E-06 |
| 4923 | 247 | P247N | A | 4.2e-06 | 7e-06 | 6.5e-06 | 5.900E-06 | 1.493E-06 | 4.200E-06 | 7.000E-06 |
| 4924 | 247 | P247D | A | 1.1e-05 | 5.1e-06 | 5.1e-06 | 7.067E-06 | 3.406E-06 | 5.100E-06 | 1.100E-05 |
| 4925 | 247 | P247C | A | 9.5e-06 | 5.1e-06 | 4.6e-06 | 6.400E-06 | 2.696E-06 | 4.600E-06 | 9.500E-06 |
| 4926 | 247 | P247E | A | 1e-05 | 5.8e-06 | 7.5e-06 | 7.767E-06 | 2.113E-06 | 5.800E-06 | 1.000E-05 |
| 4927 | 247 | P247Q | A | 9.3e-06 | 7.7e-06 | 8.8e-06 | 8.600E-06 | 8.185E-07 | 7.700E-06 | 9.300E-06 |
| 4928 | 247 | P247G | A | 7.8e-06 | 7.6e-06 | 4.9e-06 | 6.767E-06 | 1.620E-06 | 4.900E-06 | 7.800E-06 |
| 4929 | 247 | P247H | A | 1.1e-05 | 7.8e-06 | 9.8e-06 | 9.533E-06 | 1.617E-06 | 7.800E-06 | 1.100E-05 |

| | | | | | | | | | | |
|---|---|---|---|---|---|---|---|---|---|---|
| 4930 | 247 | P247I | A | 6e-06 | 1e-05 | 6.7e-06 | 7.567E-06 | 2.136E-06 | 6.000E-06 | 1.000E-05 |
| 4931 | 247 | P247L | A | 5e-06 | 6.3e-06 | 9e-06 | 6.767E-06 | 2.040E-06 | 5.000E-06 | 9.000E-06 |
| 4932 | 247 | P247K | A | 4.7e-06 | 8.8e-06 | 1e-05 | 7.833E-06 | 2.779E-06 | 4.700E-06 | 1.000E-05 |
| 4933 | 247 | P247M | A | 1.1e-05 | 7.7e-06 | 6.6e-06 | 8.433E-06 | 2.290E-06 | 6.600E-06 | 1.100E-05 |
| 4934 | 247 | P247F | A | 8.6e-06 | 5.8e-06 | 5.6e-06 | 6.667E-06 | 1.677E-06 | 5.600E-06 | 8.600E-06 |
| 4935 | 247 | P247P | A | 5.8e-06 | 9.4e-06 | 9.8e-06 | 8.333E-06 | 2.203E-06 | 5.800E-06 | 9.800E-06 |
| 4936 | 247 | P247S | A | 5e-06 | 1.3e-05 | 1.1e-05 | 9.667E-06 | 4.163E-06 | 5.000E-06 | 1.300E-05 |
| 4937 | 247 | P247T | A | 5.4e-06 | 9.6e-06 | 5.5e-06 | 6.833E-06 | 2.397E-06 | 5.400E-06 | 9.600E-06 |
| 4938 | 247 | P247W | A | 4.6e-06 | 1.8e-05 | 3.4e-06 | 8.667E-06 | 8.105E-06 | 3.400E-06 | 1.800E-05 |
| 4939 | 247 | P247Y | A | 8e-06 | 1.2e-05 | 7.2e-06 | 9.067E-06 | 2.572E-06 | 7.200E-06 | 1.200E-05 |
| 4940 | 247 | P247V | A | 8.7e-06 | 8.5e-06 | 8.6e-06 | 8.600E-06 | 1.000E-07 | 8.500E-06 | 8.700E-06 |
| 4941 | 248 | A248A | A | 8.9e-06 | 7.6e-06 | 5.6e-06 | 7.367E-06 | 1.662E-06 | 5.600E-06 | 8.900E-06 |
| 4942 | 248 | A248R | A | 2.1e-05 | 5.7e-06 | 5.3e-06 | 1.067E-05 | 8.951E-06 | 5.300E-06 | 2.100E-05 |
| 4943 | 248 | A248N | A | 7.2e-06 | 6.6e-06 | 6.5e-06 | 6.767E-06 | 3.786E-07 | 6.500E-06 | 7.200E-06 |
| 4944 | 248 | A248D | A | 8.9e-06 | 5e-06 | 4.7e-06 | 6.200E-06 | 2.343E-06 | 4.700E-06 | 8.900E-06 |
| 4945 | 248 | A248C | A | 6.4e-06 | 7.4e-06 | 1.1e-05 | 8.267E-06 | 2.419E-06 | 6.400E-06 | 1.100E-05 |
| 4946 | 248 | A248E | A | 6.4e-06 | 8.7e-06 | 1.1e-05 | 8.700E-06 | 2.300E-06 | 6.400E-06 | 1.100E-05 |
| 4947 | 248 | A248Q | A | 8.6e-06 | 6.4e-06 | 8.3e-06 | 7.767E-06 | 1.193E-06 | 6.400E-06 | 8.600E-06 |
| 4948 | 248 | A248G | A | 9.3e-06 | 1.2e-05 | 8.5e-06 | 9.933E-06 | 1.834E-06 | 8.500E-06 | 1.200E-05 |
| 4949 | 248 | A248H | A | 1e-05 | 1.2e-05 | 6.9e-06 | 9.633E-06 | 2.570E-06 | 6.900E-06 | 1.200E-05 |
| 4950 | 248 | A248I | A | 1.2e-05 | 1.1e-05 | 4.8e-06 | 9.267E-06 | 3.900E-06 | 4.800E-06 | 1.200E-05 |
| 4951 | 248 | A248L | A | 4.6e-06 | 8.5e-06 | 7.1e-06 | 6.733E-06 | 1.976E-06 | 4.600E-06 | 8.500E-06 |
| 4952 | 248 | A248K | A | 6.1e-06 | 7.9e-06 | 6.3e-06 | 6.767E-06 | 9.866E-07 | 6.100E-06 | 7.900E-06 |
| 4953 | 248 | A248M | A | 8.5e-06 | 7.1e-06 | 1.4e-05 | 9.867E-06 | 3.647E-06 | 7.100E-06 | 1.400E-05 |
| 4954 | 248 | A248F | A | 1e-05 | 1.1e-05 | 1.2e-05 | 1.100E-05 | 1.000E-06 | 1.000E-05 | 1.200E-05 |
| 4955 | 248 | A248P | A | 9.7e-06 | 8.7e-06 | 6.2e-06 | 8.200E-06 | 1.803E-06 | 6.200E-06 | 9.700E-06 |
| 4956 | 248 | A248S | A | 8.5e-06 | 1.3e-05 | 3.1e-06 | 8.200E-06 | 4.957E-06 | 3.100E-06 | 1.300E-05 |
| 4957 | 248 | A248T | A | 9.4e-06 | 5e-06 | 9.1e-06 | 7.833E-06 | 2.458E-06 | 5.000E-06 | 9.400E-06 |
| 4958 | 248 | A248W | A | 9.2e-06 | 6.4e-06 | 6.5e-06 | 7.367E-06 | 1.589E-06 | 6.400E-06 | 9.200E-06 |
| 4959 | 248 | A248Y | A | 4.8e-06 | 1.1e-05 | 9.8e-06 | 8.533E-06 | 3.288E-06 | 4.800E-06 | 1.100E-05 |
| 4960 | 248 | A248V | A | 7.9e-06 | 6.4e-06 | 1.3e-05 | 9.100E-06 | 3.460E-06 | 6.400E-06 | 1.300E-05 |
| 4961 | 249 | L249A | A | 1.4e-05 | 7e-06 | 7.8e-06 | 9.600E-06 | 3.831E-06 | 7.000E-06 | 1.400E-05 |
| 4962 | 249 | L249R | A | 1.8e-05 | 9.6e-06 | 8.5e-06 | 1.203E-05 | 5.196E-06 | 8.500E-06 | 1.800E-05 |
| 4963 | 249 | L249N | A | 6.9e-06 | 8.8e-06 | 9.5e-06 | 8.400E-06 | 1.345E-06 | 6.900E-06 | 9.500E-06 |
| 4964 | 249 | L249D | A | 6.6e-06 | 8.7e-06 | 7.7e-06 | 7.667E-06 | 1.050E-06 | 6.600E-06 | 8.700E-06 |
| 4965 | 249 | L249C | A | 9.4e-06 | 4.7e-06 | 7.7e-06 | 7.267E-06 | 2.380E-06 | 4.700E-06 | 9.400E-06 |
| 4966 | 249 | L249E | A | 6.8e-06 | 1.1e-05 | 1.3e-05 | 1.027E-05 | 3.164E-06 | 6.800E-06 | 1.300E-05 |
| 4967 | 249 | L249Q | A | 4.2e-06 | 6.6e-06 | 7.5e-06 | 6.100E-06 | 1.706E-06 | 4.200E-06 | 7.500E-06 |
| 4968 | 249 | L249G | A | 8.8e-06 | 7.4e-06 | 6.9e-06 | 7.700E-06 | 9.849E-07 | 6.900E-06 | 8.800E-06 |
| 4969 | 249 | L249H | A | 9.7e-06 | 8.3e-06 | 8.4e-06 | 8.800E-06 | 7.810E-07 | 8.300E-06 | 9.700E-06 |
| 4970 | 249 | L249I | A | 7.5e-06 | 7.4e-06 | 6.6e-06 | 7.167E-06 | 4.933E-07 | 6.600E-06 | 7.500E-06 |
| 4971 | 249 | L249L | A | 1.9e-05 | 6.6e-06 | 7.3e-06 | 1.097E-05 | 6.966E-06 | 6.600E-06 | 1.900E-05 |
| 4972 | 249 | L249K | A | 1e-05 | 6.9e-06 | 6.2e-06 | 7.700E-06 | 2.022E-06 | 6.200E-06 | 1.000E-05 |
| 4973 | 249 | L249M | A | 9.6e-06 | 6.6e-06 | 7.2e-06 | 7.800E-06 | 1.587E-06 | 6.600E-06 | 9.600E-06 |
| 4974 | 249 | L249F | A | 1.3e-05 | 1.2e-05 | 7e-06 | 1.067E-05 | 3.215E-06 | 7.000E-06 | 1.300E-05 |
| 4975 | 249 | L249P | A | 8.8e-06 | 1.2e-05 | 7e-06 | 9.267E-06 | 2.532E-06 | 7.000E-06 | 1.200E-05 |
| 4976 | 249 | L249S | A | 4.9e-06 | 6.8e-06 | 7.1e-06 | 6.267E-06 | 1.193E-06 | 4.900E-06 | 7.100E-06 |
| 4977 | 249 | L249T | A | 6.3e-06 | 7e-06 | 9e-06 | 7.433E-06 | 1.401E-06 | 6.300E-06 | 9.000E-06 |
| 4978 | 249 | L249W | A | 5.8e-06 | 1.3e-05 | 8.8e-06 | 9.200E-06 | 3.617E-06 | 5.800E-06 | 1.300E-05 |
| 4979 | 249 | L249Y | A | 4.9e-06 | 9.4e-06 | 9e-06 | 7.767E-06 | 2.491E-06 | 4.900E-06 | 9.400E-06 |
| 4980 | 249 | L249V | A | 1.3e-05 | 7.3e-06 | 6.6e-06 | 8.967E-06 | 3.510E-06 | 6.600E-06 | 1.300E-05 |
| 4981 | 250 | V250A | A | 1.5e-05 | 1.1e-05 | 4.4e-06 | 1.013E-05 | 5.353E-06 | 4.400E-06 | 1.500E-05 |
| 4982 | 250 | V250R | A | 3.6e-06 | 1.2e-05 | 6.2e-06 | 7.267E-06 | 4.300E-06 | 3.600E-06 | 1.200E-05 |
| 4983 | 250 | V250N | A | 1.1e-05 | 6.8e-06 | 8.1e-06 | 8.633E-06 | 2.150E-06 | 6.800E-06 | 1.100E-05 |
| 4984 | 250 | V250D | A | 5.3e-06 | 8.8e-06 | 6.6e-06 | 6.900E-06 | 1.769E-06 | 5.300E-06 | 8.800E-06 |
| 4985 | 250 | V250C | A | 1.2e-05 | 5.8e-06 | 5.1e-06 | 7.633E-06 | 3.798E-06 | 5.100E-06 | 1.200E-05 |
| 4986 | 250 | V250E | A | 8.8e-06 | 1e-05 | 9e-06 | 9.267E-06 | 6.429E-07 | 8.800E-06 | 1.000E-05 |
| 4987 | 250 | V250Q | A | 7.1e-06 | 6.4e-06 | 5.6e-06 | 6.367E-06 | 7.506E-07 | 5.600E-06 | 7.100E-06 |
| 4988 | 250 | V250G | A | 4.7e-06 | 7.5e-06 | 4.9e-06 | 5.700E-06 | 1.562E-06 | 4.700E-06 | 7.500E-06 |
| 4989 | 250 | V250H | A | 8.8e-06 | 4.9e-06 | 6.4e-06 | 6.700E-06 | 1.967E-06 | 4.900E-06 | 8.800E-06 |
| 4990 | 250 | V250I | A | 1e-05 | 6.6e-06 | 1.5e-05 | 1.053E-05 | 4.225E-06 | 6.600E-06 | 1.500E-05 |
| 4991 | 250 | V250L | A | 9.5e-06 | 8e-06 | 9.8e-06 | 9.100E-06 | 9.644E-07 | 8.000E-06 | 9.800E-06 |
| 4992 | 250 | V250K | A | 5.6e-06 | 9.8e-06 | 1.4e-05 | 9.800E-06 | 4.200E-06 | 5.600E-06 | 1.400E-05 |
| 4993 | 250 | V250M | A | 8.4e-06 | 5.6e-06 | 7.6e-06 | 7.200E-06 | 1.442E-06 | 5.600E-06 | 8.400E-06 |
| 4994 | 250 | V250F | A | 8.6e-06 | 5.1e-06 | 8.8e-06 | 7.500E-06 | 2.081E-06 | 5.100E-06 | 8.800E-06 |
| 4995 | 250 | V250P | A | 9.5e-06 | 7.6e-06 | 7e-06 | 8.033E-06 | 1.305E-06 | 7.000E-06 | 9.500E-06 |
| 4996 | 250 | V250S | A | 7.1e-06 | 1e-05 | 9.8e-06 | 8.967E-06 | 1.620E-06 | 7.100E-06 | 1.000E-05 |
| 4997 | 250 | V250T | A | 4.4e-06 | 8.5e-06 | 3.2e-06 | 5.367E-06 | 2.779E-06 | 3.200E-06 | 8.500E-06 |
| 4998 | 250 | V250W | A | 4.8e-06 | 9.6e-06 | 8.2e-06 | 7.533E-06 | 2.468E-06 | 4.800E-06 | 9.600E-06 |
| 4999 | 250 | V250Y | A | 1.3e-05 | 8e-06 | 1e-05 | 1.033E-05 | 2.517E-06 | 8.000E-06 | 1.300E-05 |
| 5000 | 250 | V250V | A | 7.2e-06 | 9.2e-06 | 7e-06 | 7.800E-06 | 1.217E-06 | 7.000E-06 | 9.200E-06 |
| 5001 | 251 | T251A | A | 8.8e-06 | 8.9e-06 | 8.9e-06 | 8.867E-06 | 5.774E-08 | 8.800E-06 | 8.900E-06 |
| 5002 | 251 | T251R | A | 9.8e-06 | 6.8e-06 | 7.4e-06 | 8.000E-06 | 1.587E-06 | 6.800E-06 | 9.800E-06 |
| 5003 | 251 | T251N | A | 1.4e-05 | 8.9e-06 | 8.5e-06 | 1.047E-05 | 3.066E-06 | 8.500E-06 | 1.400E-05 |
| 5004 | 251 | T251D | A | 5.7e-06 | 7.9e-06 | 7.3e-06 | 6.967E-06 | 1.137E-06 | 5.700E-06 | 7.900E-06 |
| 5005 | 251 | T251C | A | 9.6e-06 | 9e-06 | 8.2e-06 | 8.933E-06 | 7.024E-07 | 8.200E-06 | 9.600E-06 |
| 5006 | 251 | T251E | A | 8.6e-06 | 6.2e-06 | 7.8e-06 | 7.533E-06 | 1.222E-06 | 6.200E-06 | 8.600E-06 |
| 5007 | 251 | T251Q | A | 9.9e-06 | 6.7e-06 | 1.3e-05 | 9.867E-06 | 3.150E-06 | 6.700E-06 | 1.300E-05 |
| 5008 | 251 | T251G | A | 1.1e-05 | 8.4e-06 | 6.9e-06 | 8.767E-06 | 2.074E-06 | 6.900E-06 | 1.100E-05 |
| 5009 | 251 | T251H | A | 6.8e-06 | 6.8e-06 | 6.1e-06 | 6.567E-06 | 4.041E-07 | 6.100E-06 | 6.800E-06 |
| 5010 | 251 | T251I | A | 6.5e-06 | 5.3e-06 | 7.3e-06 | 6.367E-06 | 1.007E-06 | 5.300E-06 | 7.300E-06 |
| 5011 | 251 | T251L | A | 9.3e-06 | 1.6e-05 | 7e-06 | 1.077E-05 | 4.676E-06 | 7.000E-06 | 1.600E-05 |
| 5012 | 251 | T251K | A | 1e-05 | 8.8e-06 | 7.4e-06 | 8.733E-06 | 1.301E-06 | 7.400E-06 | 1.000E-05 |
| 5013 | 251 | T251M | A | 9.5e-06 | 6.6e-06 | 4.9e-06 | 7.000E-06 | 2.326E-06 | 4.900E-06 | 9.500E-06 |
| 5014 | 251 | T251F | A | 7.2e-06 | 8e-06 | 5.5e-06 | 6.900E-06 | 1.277E-06 | 5.500E-06 | 8.000E-06 |

| | | | | | | | | | | |
|---|---|---|---|---|---|---|---|---|---|---|
| 5015 | 251 | T251P | A | 6.8e-06 | 8.2e-06 | 6.1e-06 | 7.033E-06 | 1.069E-06 | 6.100E-06 | 8.200E-06 |
| 5016 | 251 | T251S | A | 3.1e-06 | 5.7e-06 | 6.8e-06 | 5.200E-06 | 1.900E-06 | 3.100E-06 | 6.800E-06 |
| 5017 | 251 | T251T | A | 6.3e-06 | 4.8e-06 | 5.8e-06 | 5.633E-06 | 7.638E-07 | 4.800E-06 | 6.300E-06 |
| 5018 | 251 | T251W | A | 9e-06 | 1.1e-05 | 6.6e-06 | 8.867E-06 | 2.203E-06 | 6.600E-06 | 1.100E-05 |
| 5019 | 251 | T251Y | A | 6.9e-06 | 5.9e-06 | 4.6e-06 | 5.800E-06 | 1.153E-06 | 4.600E-06 | 6.900E-06 |
| 5020 | 251 | T251V | A | 4.8e-06 | 6.9e-06 | 9.9e-06 | 7.200E-06 | 2.563E-06 | 4.800E-06 | 9.900E-06 |
| 5021 | 252 | Y252A | A | 7.2e-06 | 5e-06 | 5.4e-06 | 5.867E-06 | 1.172E-06 | 5.000E-06 | 7.200E-06 |
| 5022 | 252 | Y252R | A | 1.2e-05 | 1.4e-05 | 6.4e-06 | 1.080E-05 | 3.940E-06 | 6.400E-06 | 1.400E-05 |
| 5023 | 252 | Y252N | A | 1.1e-05 | 4.7e-06 | 7.8e-06 | 7.833E-06 | 3.150E-06 | 4.700E-06 | 1.100E-05 |
| 5024 | 252 | Y252D | A | 8.1e-06 | 1.4e-05 | 8.4e-06 | 1.017E-05 | 3.323E-06 | 8.100E-06 | 1.400E-05 |
| 5025 | 252 | Y252C | A | 4.8e-06 | 7e-06 | 5.5e-06 | 5.767E-06 | 1.124E-06 | 4.800E-06 | 7.000E-06 |
| 5026 | 252 | Y252E | A | 6.9e-06 | 6.3e-06 | 8e-06 | 7.067E-06 | 8.622E-07 | 6.300E-06 | 8.000E-06 |
| 5027 | 252 | Y252Q | A | 4.7e-06 | 4.4e-06 | 3.2e-06 | 4.100E-06 | 7.937E-07 | 3.200E-06 | 4.700E-06 |
| 5028 | 252 | Y252G | A | 3.2e-06 | 9.3e-06 | 8.4e-06 | 6.967E-06 | 3.293E-06 | 3.200E-06 | 9.300E-06 |
| 5029 | 252 | Y252H | A | 5.8e-06 | 1e-05 | 1.3e-05 | 9.600E-06 | 3.617E-06 | 5.800E-06 | 1.300E-05 |
| 5030 | 252 | Y252I | A | 1.5e-05 | 4.9e-06 | 1e-05 | 9.967E-06 | 5.050E-06 | 4.900E-06 | 1.500E-05 |
| 5031 | 252 | Y252L | A | 8.3e-06 | 1e-05 | 1.4e-05 | 1.077E-05 | 2.926E-06 | 8.300E-06 | 1.400E-05 |
| 5032 | 252 | Y252K | A | 8.5e-06 | 8.6e-06 | 3.2e-06 | 6.767E-06 | 3.089E-06 | 3.200E-06 | 8.600E-06 |
| 5033 | 252 | Y252M | A | 9.7e-06 | 5.3e-06 | 9.3e-06 | 8.100E-06 | 2.433E-06 | 5.300E-06 | 9.700E-06 |
| 5034 | 252 | Y252F | A | 5.6e-06 | 6.6e-06 | 7e-06 | 6.400E-06 | 7.211E-07 | 5.600E-06 | 7.000E-06 |
| 5035 | 252 | Y252P | A | 2e-05 | 9.7e-06 | 1.6e-05 | 1.523E-05 | 5.193E-06 | 9.700E-06 | 2.000E-05 |
| 5036 | 252 | Y252S | A | 9.8e-06 | 1.3e-05 | 8.2e-06 | 1.033E-05 | 2.444E-06 | 8.200E-06 | 1.300E-05 |
| 5037 | 252 | Y252T | A | 4.6e-06 | 1.2e-05 | 8e-06 | 8.200E-06 | 3.704E-06 | 4.600E-06 | 1.200E-05 |
| 5038 | 252 | Y252W | A | 7.7e-06 | 8e-06 | 1e-05 | 8.567E-06 | 1.250E-06 | 7.700E-06 | 1.000E-05 |
| 5039 | 252 | Y252Y | A | 8.7e-06 | 6.8e-06 | 9.2e-06 | 8.233E-06 | 1.266E-06 | 6.800E-06 | 9.200E-06 |
| 5040 | 252 | Y252V | A | 5.5e-06 | 1.3e-05 | 8.8e-06 | 9.100E-06 | 3.759E-06 | 5.500E-06 | 1.300E-05 |
| 5041 | 253 | N253A | A | 5.3e-06 | 9.4e-06 | 6.3e-06 | 7.000E-06 | 2.138E-06 | 5.300E-06 | 9.400E-06 |
| 5042 | 253 | N253R | A | 1.3e-05 | 1.1e-05 | 8.7e-06 | 1.090E-05 | 2.152E-06 | 8.700E-06 | 1.300E-05 |
| 5043 | 253 | N253N | A | 8.5e-06 | 8.6e-06 | 1.1e-05 | 9.367E-06 | 1.415E-06 | 8.500E-06 | 1.100E-05 |
| 5044 | 253 | N253D | A | 1e-05 | 6.7e-06 | 5.5e-06 | 7.400E-06 | 2.330E-06 | 5.500E-06 | 1.000E-05 |
| 5045 | 253 | N253C | A | 4.9e-06 | 5.2e-06 | 1.1e-05 | 7.033E-06 | 3.439E-06 | 4.900E-06 | 1.100E-05 |
| 5046 | 253 | N253E | A | 6.8e-06 | 9e-06 | 1e-05 | 8.600E-06 | 1.637E-06 | 6.800E-06 | 1.000E-05 |
| 5047 | 253 | N253Q | A | 1.1e-05 | 6.5e-06 | 9.9e-06 | 9.133E-06 | 2.346E-06 | 6.500E-06 | 1.100E-05 |
| 5048 | 253 | N253G | A | 4.9e-06 | 9.3e-06 | 7.8e-06 | 7.333E-06 | 2.237E-06 | 4.900E-06 | 9.300E-06 |
| 5049 | 253 | N253H | A | 7e-06 | 6e-06 | 6.2e-06 | 6.400E-06 | 5.292E-07 | 6.000E-06 | 7.000E-06 |
| 5050 | 253 | N253I | A | 9.7e-06 | 7.3e-06 | 6.5e-06 | 7.833E-06 | 1.665E-06 | 6.500E-06 | 9.700E-06 |
| 5051 | 253 | N253L | A | 8.3e-06 | 1e-05 | 8e-06 | 8.767E-06 | 1.079E-06 | 8.000E-06 | 1.000E-05 |
| 5052 | 253 | N253K | A | 1.4e-05 | 1.5e-05 | 1.4e-05 | 1.433E-05 | 5.774E-07 | 1.400E-05 | 1.500E-05 |
| 5053 | 253 | N253M | A | 1.1e-05 | 1.6e-05 | 8.5e-06 | 1.183E-05 | 3.819E-06 | 8.500E-06 | 1.600E-05 |
| 5054 | 253 | N253F | A | 1.4e-05 | 7e-06 | 4.8e-06 | 8.600E-06 | 4.804E-06 | 4.800E-06 | 1.400E-05 |
| 5055 | 253 | N253P | A | 8.5e-06 | 9.3e-06 | 1e-05 | 9.267E-06 | 7.506E-07 | 8.500E-06 | 1.000E-05 |
| 5056 | 253 | N253S | A | 9.1e-06 | 1.3e-05 | 1.6e-05 | 1.270E-05 | 3.460E-06 | 9.100E-06 | 1.600E-05 |
| 5057 | 253 | N253T | A | 9.5e-06 | 9.1e-06 | 6e-06 | 8.200E-06 | 1.916E-06 | 6.000E-06 | 9.500E-06 |
| 5058 | 253 | N253W | A | 6e-06 | 1.4e-05 | 5.4e-06 | 8.467E-06 | 4.801E-06 | 5.400E-06 | 1.400E-05 |
| 5059 | 253 | N253Y | A | 5.7e-06 | 4.3e-06 | 5.9e-06 | 5.300E-06 | 8.718E-07 | 4.300E-06 | 5.900E-06 |
| 5060 | 253 | N253V | A | 1.2e-05 | 6.4e-06 | 8.5e-06 | 8.967E-06 | 2.829E-06 | 6.400E-06 | 1.200E-05 |
| 5061 | 254 | T254A | A | 7.1e-06 | 9.3e-06 | 1.5e-05 | 1.047E-05 | 4.077E-06 | 7.100E-06 | 1.500E-05 |
| 5062 | 254 | T254R | A | 8.3e-06 | 1.2e-05 | 8.3e-06 | 9.533E-06 | 2.136E-06 | 8.300E-06 | 1.200E-05 |
| 5063 | 254 | T254N | A | 6e-06 | 1.9e-05 | 9.4e-06 | 1.147E-05 | 6.742E-06 | 6.000E-06 | 1.900E-05 |
| 5064 | 254 | T254D | A | 6.2e-06 | 1.4e-05 | 5.8e-06 | 8.667E-06 | 4.623E-06 | 5.800E-06 | 1.400E-05 |
| 5065 | 254 | T254C | A | 8e-06 | 4.9e-06 | 8.7e-06 | 7.200E-06 | 2.022E-06 | 4.900E-06 | 8.700E-06 |
| 5066 | 254 | T254E | A | 9.3e-06 | 1.1e-05 | 7.8e-06 | 9.367E-06 | 1.601E-06 | 7.800E-06 | 1.100E-05 |
| 5067 | 254 | T254Q | A | 7.2e-06 | 6.1e-06 | 8.1e-06 | 7.133E-06 | 1.002E-06 | 6.100E-06 | 8.100E-06 |
| 5068 | 254 | T254G | A | 2.1e-05 | 6.5e-06 | 1.2e-05 | 1.317E-05 | 7.320E-06 | 6.500E-06 | 2.100E-05 |
| 5069 | 254 | T254H | A | 7e-06 | 1.4e-05 | 8e-06 | 9.667E-06 | 3.786E-06 | 7.000E-06 | 1.400E-05 |
| 5070 | 254 | T254I | A | 6.9e-06 | 1.2e-05 | 1.2e-05 | 1.030E-05 | 2.944E-06 | 6.900E-06 | 1.200E-05 |
| 5071 | 254 | T254L | A | 1.3e-05 | 3e-06 | 7.5e-06 | 7.833E-06 | 5.008E-06 | 3.000E-06 | 1.300E-05 |
| 5072 | 254 | T254K | A | 9.2e-06 | 1.1e-05 | 1.1e-05 | 1.040E-05 | 1.039E-06 | 9.200E-06 | 1.100E-05 |
| 5073 | 254 | T254M | A | 9.4e-06 | 1.5e-05 | 9.1e-06 | 1.117E-05 | 3.323E-06 | 9.100E-06 | 1.500E-05 |
| 5074 | 254 | T254F | A | 8.4e-06 | 5.6e-06 | 1e-05 | 8.000E-06 | 2.227E-06 | 5.600E-06 | 1.000E-05 |
| 5075 | 254 | T254P | A | 5.8e-06 | 7.2e-06 | 6.6e-06 | 6.533E-06 | 7.024E-07 | 5.800E-06 | 7.200E-06 |
| 5076 | 254 | T254S | A | 7.2e-06 | 9.6e-06 | 9e-06 | 8.600E-06 | 1.249E-06 | 7.200E-06 | 9.600E-06 |
| 5077 | 254 | T254T | A | 6.4e-06 | 8e-06 | 8.3e-06 | 7.567E-06 | 1.021E-06 | 6.400E-06 | 8.300E-06 |
| 5078 | 254 | T254W | A | 8.3e-06 | 7.6e-06 | 6.5e-06 | 7.467E-06 | 9.074E-07 | 6.500E-06 | 8.300E-06 |
| 5079 | 254 | T254Y | A | 7.6e-06 | 6.9e-06 | 1.3e-05 | 9.167E-06 | 3.338E-06 | 6.900E-06 | 1.300E-05 |
| 5080 | 254 | T254V | A | 8.8e-06 | 9.1e-06 | 9.5e-06 | 9.133E-06 | 3.512E-07 | 8.800E-06 | 9.500E-06 |
| 5081 | 255 | D255A | A | 4.7e-06 | 1.1e-05 | 1e-05 | 8.567E-06 | 3.386E-06 | 4.700E-06 | 1.100E-05 |
| 5082 | 255 | D255R | A | 4.5e-06 | 5.5e-06 | 6.9e-06 | 5.633E-06 | 1.206E-06 | 4.500E-06 | 6.900E-06 |
| 5083 | 255 | D255N | A | 7.4e-06 | 1.2e-05 | 5.7e-06 | 8.367E-06 | 3.259E-06 | 5.700E-06 | 1.200E-05 |
| 5084 | 255 | D255D | A | 6.7e-06 | 1e-05 | 4.8e-06 | 7.167E-06 | 2.631E-06 | 4.800E-06 | 1.000E-05 |
| 5085 | 255 | D255C | A | 7.9e-06 | 6.7e-06 | 6.7e-06 | 7.100E-06 | 6.928E-07 | 6.700E-06 | 7.900E-06 |
| 5086 | 255 | D255E | A | 7.6e-06 | 1.2e-05 | 3.9e-06 | 7.833E-06 | 4.055E-06 | 3.900E-06 | 1.200E-05 |
| 5087 | 255 | D255Q | A | 9e-06 | 4.7e-06 | 9.8e-06 | 7.833E-06 | 2.743E-06 | 4.700E-06 | 9.800E-06 |
| 5088 | 255 | D255G | A | 9.2e-06 | 8.9e-06 | 7.9e-06 | 8.667E-06 | 6.807E-07 | 7.900E-06 | 9.200E-06 |
| 5089 | 255 | D255H | A | 7.9e-06 | 7.9e-06 | 4.6e-06 | 6.800E-06 | 1.905E-06 | 4.600E-06 | 7.900E-06 |
| 5090 | 255 | D255I | A | 1.3e-05 | 5e-06 | 8.3e-06 | 8.767E-06 | 4.020E-06 | 5.000E-06 | 1.300E-05 |
| 5091 | 255 | D255L | A | 6.5e-06 | 5.9e-06 | 6.2e-06 | 6.200E-06 | 3.000E-07 | 5.900E-06 | 6.500E-06 |
| 5092 | 255 | D255K | A | 3.8e-06 | 7.4e-06 | 5.3e-06 | 5.500E-06 | 1.808E-06 | 3.800E-06 | 7.400E-06 |
| 5093 | 255 | D255M | A | 7e-06 | 3.3e-06 | 1e-05 | 6.767E-06 | 3.356E-06 | 3.300E-06 | 1.000E-05 |
| 5094 | 255 | D255F | A | 9.6e-06 | 8.8e-06 | 6.4e-06 | 8.267E-06 | 1.665E-06 | 6.400E-06 | 9.600E-06 |
| 5095 | 255 | D255P | A | 5.8e-06 | 8.9e-06 | 4.4e-06 | 6.367E-06 | 2.303E-06 | 4.400E-06 | 8.900E-06 |
| 5096 | 255 | D255S | A | 8.5e-06 | 7.4e-06 | 9.2e-06 | 8.367E-06 | 9.074E-07 | 7.400E-06 | 9.200E-06 |
| 5097 | 255 | D255T | A | 4.8e-06 | 1.2e-05 | 6.5e-06 | 7.767E-06 | 3.763E-06 | 4.800E-06 | 1.200E-05 |
| 5098 | 255 | D255W | A | 5.2e-06 | 7.8e-06 | 1.2e-05 | 8.333E-06 | 3.431E-06 | 5.200E-06 | 1.200E-05 |
| 5099 | 255 | D255Y | A | 1.4e-05 | 5.9e-06 | 8.7e-06 | 9.533E-06 | 4.114E-06 | 5.900E-06 | 1.400E-05 |

| | | | | | | | | | | |
|---|---|---|---|---|---|---|---|---|---|---|
| 5100 | 255 | D255V | A | 8.4e-06 | 6.9e-06 | 7.9e-06 | 7.733E-06 | 7.638E-07 | 6.900E-06 | 8.400E-06 |
| 5101 | 256 | T256A | A | 6.9e-06 | 1e-05 | 1.4e-05 | 1.030E-05 | 3.559E-06 | 6.900E-06 | 1.400E-05 |
| 5102 | 256 | T256R | A | 6.7e-06 | 9.4e-06 | 6.4e-06 | 7.500E-06 | 1.652E-06 | 6.400E-06 | 9.400E-06 |
| 5103 | 256 | T256N | A | 4.6e-06 | 9.4e-06 | 6.8e-06 | 6.933E-06 | 2.403E-06 | 4.600E-06 | 9.400E-06 |
| 5104 | 256 | T256D | A | 7.5e-06 | 1.4e-05 | 9.8e-06 | 1.043E-05 | 3.296E-06 | 7.500E-06 | 1.400E-05 |
| 5105 | 256 | T256C | A | 9.2e-06 | 6.9e-06 | 8.4e-06 | 8.167E-06 | 1.168E-06 | 6.900E-06 | 9.200E-06 |
| 5106 | 256 | T256E | A | 6.7e-06 | 9.6e-06 | 1.7e-05 | 1.110E-05 | 5.311E-06 | 6.700E-06 | 1.700E-05 |
| 5107 | 256 | T256Q | A | 7.1e-06 | 1e-05 | 2e-05 | 1.237E-05 | 6.768E-06 | 7.100E-06 | 2.000E-05 |
| 5108 | 256 | T256G | A | 7.8e-06 | 2.7e-06 | 1.3e-05 | 7.833E-06 | 5.150E-06 | 2.700E-06 | 1.300E-05 |
| 5109 | 256 | T256H | A | 6.3e-06 | 4.9e-06 | 8.8e-06 | 6.667E-06 | 1.976E-06 | 4.900E-06 | 8.800E-06 |
| 5110 | 256 | T256I | A | 1e-05 | 1.1e-05 | 5.7e-06 | 8.900E-06 | 2.816E-06 | 5.700E-06 | 1.100E-05 |
| 5111 | 256 | T256L | A | 1.1e-05 | 1.5e-05 | 4.8e-06 | 1.027E-05 | 5.139E-06 | 4.800E-06 | 1.500E-05 |
| 5112 | 256 | T256K | A | 9.1e-06 | 8.4e-06 | 9.1e-06 | 8.867E-06 | 4.041E-07 | 8.400E-06 | 9.100E-06 |
| 5113 | 256 | T256M | A | 7.6e-06 | 9.3e-06 | 8.4e-06 | 8.433E-06 | 8.505E-07 | 7.600E-06 | 9.300E-06 |
| 5114 | 256 | T256F | A | 5.8e-06 | 6e-06 | 1.2e-05 | 7.933E-06 | 3.523E-06 | 5.800E-06 | 1.200E-05 |
| 5115 | 256 | T256P | A | 9.2e-06 | 7.3e-06 | 1e-05 | 8.833E-06 | 1.387E-06 | 7.300E-06 | 1.000E-05 |
| 5116 | 256 | T256S | A | 8.7e-06 | 9.7e-06 | 6.2e-06 | 8.200E-06 | 1.803E-06 | 6.200E-06 | 9.700E-06 |
| 5117 | 256 | T256T | A | 1e-05 | 9e-06 | 9.8e-06 | 9.600E-06 | 5.292E-07 | 9.000E-06 | 1.000E-05 |
| 5118 | 256 | T256W | A | 5.3e-06 | 1e-05 | 1.3e-05 | 9.433E-06 | 3.881E-06 | 5.300E-06 | 1.300E-05 |
| 5119 | 256 | T256Y | A | 1.1e-05 | 1.4e-05 | 9.9e-06 | 1.163E-05 | 2.122E-06 | 9.900E-06 | 1.400E-05 |
| 5120 | 256 | T256V | A | 9.6e-06 | 7.2e-06 | 5.9e-06 | 7.567E-06 | 1.877E-06 | 5.900E-06 | 9.600E-06 |
| 5121 | 257 | F257A | A | 1.2e-05 | 8.2e-06 | 9.2e-06 | 9.800E-06 | 1.970E-06 | 8.200E-06 | 1.200E-05 |
| 5122 | 257 | F257R | A | 6.6e-06 | 5.6e-06 | 5.7e-06 | 5.967E-06 | 5.508E-07 | 5.600E-06 | 6.600E-06 |
| 5123 | 257 | F257N | A | 7.9e-06 | 5.7e-06 | 6.9e-06 | 6.833E-06 | 1.102E-06 | 5.700E-06 | 7.900E-06 |
| 5124 | 257 | F257D | A | 5.6e-06 | 1.2e-05 | 7.2e-06 | 8.267E-06 | 3.331E-06 | 5.600E-06 | 1.200E-05 |
| 5125 | 257 | F257C | A | 1.1e-05 | 6.2e-06 | 6e-06 | 7.733E-06 | 2.831E-06 | 6.000E-06 | 1.100E-05 |
| 5126 | 257 | F257E | A | 3e-06 | 1.2e-05 | 7.5e-06 | 7.500E-06 | 4.500E-06 | 3.000E-06 | 1.200E-05 |
| 5127 | 257 | F257Q | A | 5.8e-06 | 9e-06 | 9.1e-06 | 7.967E-06 | 1.877E-06 | 5.800E-06 | 9.100E-06 |
| 5128 | 257 | F257G | A | 7.2e-06 | 7.2e-06 | 9.5e-06 | 7.967E-06 | 1.328E-06 | 7.200E-06 | 9.500E-06 |
| 5129 | 257 | F257H | A | 5.2e-06 | 7e-06 | 9.9e-06 | 7.367E-06 | 2.371E-06 | 5.200E-06 | 9.900E-06 |
| 5130 | 257 | F257I | A | 7.7e-06 | 6.2e-06 | 9.9e-06 | 7.933E-06 | 1.861E-06 | 6.200E-06 | 9.900E-06 |
| 5131 | 257 | F257L | A | 5.1e-06 | 6.5e-06 | 2e-05 | 1.053E-05 | 8.228E-06 | 5.100E-06 | 2.000E-05 |
| 5132 | 257 | F257K | A | 7e-06 | 8.2e-06 | 8.3e-06 | 7.833E-06 | 7.234E-07 | 7.000E-06 | 8.300E-06 |
| 5133 | 257 | F257M | A | 4.1e-06 | 7.5e-06 | 1.3e-05 | 8.200E-06 | 4.491E-06 | 4.100E-06 | 1.300E-05 |
| 5134 | 257 | F257F | A | 7.8e-06 | 7.9e-06 | 1e-05 | 8.567E-06 | 1.242E-06 | 7.800E-06 | 1.000E-05 |
| 5135 | 257 | F257P | A | 6.6e-06 | 8.6e-06 | 9.5e-06 | 8.233E-06 | 1.484E-06 | 6.600E-06 | 9.500E-06 |
| 5136 | 257 | F257S | A | 6.7e-06 | 9.9e-06 | 4e-06 | 6.867E-06 | 2.954E-06 | 4.000E-06 | 9.900E-06 |
| 5137 | 257 | F257T | A | 9.3e-06 | 6.4e-06 | 7e-06 | 7.567E-06 | 1.531E-06 | 6.400E-06 | 9.300E-06 |
| 5138 | 257 | F257W | A | 9.5e-06 | 5.7e-06 | 7.5e-06 | 7.567E-06 | 1.901E-06 | 5.700E-06 | 9.500E-06 |
| 5139 | 257 | F257Y | A | 5.9e-06 | 7.8e-06 | 4.2e-06 | 5.967E-06 | 1.801E-06 | 4.200E-06 | 7.800E-06 |
| 5140 | 257 | F257V | A | 9.3e-06 | 1e-05 | 9.4e-06 | 9.567E-06 | 3.786E-07 | 9.300E-06 | 1.000E-05 |
| 5141 | 258 | E258A | A | 7.4e-06 | 1.4e-05 | 5e-06 | 8.800E-06 | 4.660E-06 | 5.000E-06 | 1.400E-05 |
| 5142 | 258 | E258R | A | 8.1e-06 | 9.1e-06 | 1.6e-05 | 1.107E-05 | 4.302E-06 | 8.100E-06 | 1.600E-05 |
| 5143 | 258 | E258N | A | 6e-06 | 8.1e-06 | 8.4e-06 | 7.500E-06 | 1.308E-06 | 6.000E-06 | 8.400E-06 |
| 5144 | 258 | E258D | A | 6.8e-06 | 1.1e-05 | 9.5e-06 | 9.100E-06 | 2.128E-06 | 6.800E-06 | 1.100E-05 |
| 5145 | 258 | E258C | A | 6.3e-06 | 8.6e-06 | 1.3e-05 | 9.300E-06 | 3.404E-06 | 6.300E-06 | 1.300E-05 |
| 5146 | 258 | E258E | A | 8.1e-06 | 6.6e-06 | 8e-06 | 7.567E-06 | 8.386E-07 | 6.600E-06 | 8.100E-06 |
| 5147 | 258 | E258Q | A | 6.1e-06 | 1.2e-05 | 9.5e-06 | 9.200E-06 | 2.961E-06 | 6.100E-06 | 1.200E-05 |
| 5148 | 258 | E258G | A | 1.1e-05 | 6.8e-06 | 7.7e-06 | 8.500E-06 | 2.211E-06 | 6.800E-06 | 1.100E-05 |
| 5149 | 258 | E258H | A | 9.6e-06 | 7.4e-06 | 7.4e-06 | 8.133E-06 | 1.270E-06 | 7.400E-06 | 9.600E-06 |
| 5150 | 258 | E258I | A | 9.7e-06 | 8.9e-06 | 4.9e-06 | 7.833E-06 | 2.572E-06 | 4.900E-06 | 9.700E-06 |
| 5151 | 258 | E258L | A | 7.6e-06 | 1.2e-05 | 6.6e-06 | 8.733E-06 | 2.873E-06 | 6.600E-06 | 1.200E-05 |
| 5152 | 258 | E258K | A | 6.4e-06 | 8.4e-06 | 6.2e-06 | 7.000E-06 | 1.217E-06 | 6.200E-06 | 8.400E-06 |
| 5153 | 258 | E258M | A | 5e-06 | 1e-05 | 1.3e-05 | 9.333E-06 | 4.041E-06 | 5.000E-06 | 1.300E-05 |
| 5154 | 258 | E258F | A | 1.1e-05 | 8.6e-06 | 8.8e-06 | 9.467E-06 | 1.332E-06 | 8.600E-06 | 1.100E-05 |
| 5155 | 258 | E258P | A | 1e-05 | 9.6e-06 | 1.2e-05 | 1.053E-05 | 1.286E-06 | 9.600E-06 | 1.200E-05 |
| 5156 | 258 | E258S | A | 8.5e-06 | 7.5e-06 | 7.4e-06 | 7.800E-06 | 6.083E-07 | 7.400E-06 | 8.500E-06 |
| 5157 | 258 | E258T | A | 5.4e-06 | 6.6e-06 | 1.4e-05 | 8.667E-06 | 4.658E-06 | 5.400E-06 | 1.400E-05 |
| 5158 | 258 | E258W | A | 9.9e-06 | 6.9e-06 | 7e-06 | 7.933E-06 | 1.704E-06 | 6.900E-06 | 9.900E-06 |
| 5159 | 258 | E258Y | A | 6.4e-06 | 1.8e-05 | 4.2e-06 | 9.533E-06 | 7.414E-06 | 4.200E-06 | 1.800E-05 |
| 5160 | 258 | E258V | A | 6.5e-06 | 6.8e-06 | 1.1e-05 | 8.100E-06 | 2.516E-06 | 6.500E-06 | 1.100E-05 |
| 5161 | 259 | S259A | A | 1.1e-05 | 1.1e-05 | 5.6e-06 | 9.200E-06 | 3.118E-06 | 5.600E-06 | 1.100E-05 |
| 5162 | 259 | S259R | A | 1.1e-05 | 8.7e-06 | 3.7e-06 | 7.800E-06 | 3.732E-06 | 3.700E-06 | 1.100E-05 |
| 5163 | 259 | S259N | A | 9.2e-06 | 6.8e-06 | 1e-05 | 8.667E-06 | 1.665E-06 | 6.800E-06 | 1.000E-05 |
| 5164 | 259 | S259D | A | 6.4e-06 | 1.2e-05 | 9e-06 | 9.133E-06 | 2.802E-06 | 6.400E-06 | 1.200E-05 |
| 5165 | 259 | S259C | A | 4.4e-06 | 4.4e-06 | 7.5e-06 | 5.433E-06 | 1.790E-06 | 4.400E-06 | 7.500E-06 |
| 5166 | 259 | S259E | A | 7e-06 | 7.1e-06 | 6.1e-06 | 6.733E-06 | 5.508E-07 | 6.100E-06 | 7.100E-06 |
| 5167 | 259 | S259Q | A | 6.5e-06 | 9.4e-06 | 6.2e-06 | 7.367E-06 | 1.767E-06 | 6.200E-06 | 9.400E-06 |
| 5168 | 259 | S259G | A | 1.4e-05 | 6.9e-06 | 6.6e-06 | 9.167E-06 | 4.188E-06 | 6.600E-06 | 1.400E-05 |
| 5169 | 259 | S259H | A | 7e-06 | 7.7e-06 | 5.8e-06 | 6.833E-06 | 9.609E-07 | 5.800E-06 | 7.700E-06 |
| 5170 | 259 | S259I | A | 9.8e-06 | 9e-06 | 1.3e-05 | 1.060E-05 | 2.117E-06 | 9.000E-06 | 1.300E-05 |
| 5171 | 259 | S259L | A | 9.3e-06 | 8e-06 | 5.4e-06 | 7.567E-06 | 1.986E-06 | 5.400E-06 | 9.300E-06 |
| 5172 | 259 | S259K | A | 9.2e-06 | 8.4e-06 | 8.7e-06 | 8.767E-06 | 4.041E-07 | 8.400E-06 | 9.200E-06 |
| 5173 | 259 | S259M | A | 1.1e-05 | 1.1e-05 | 8.1e-06 | 1.003E-05 | 1.674E-06 | 8.100E-06 | 1.100E-05 |
| 5174 | 259 | S259F | A | 5.4e-06 | 6.7e-06 | 9e-06 | 7.033E-06 | 1.823E-06 | 5.400E-06 | 9.000E-06 |
| 5175 | 259 | S259P | A | 9.9e-06 | 1.2e-05 | 8.2e-06 | 1.003E-05 | 1.904E-06 | 8.200E-06 | 1.200E-05 |
| 5176 | 259 | S259S | A | 6.5e-06 | 8e-06 | 1.2e-05 | 8.833E-06 | 2.843E-06 | 6.500E-06 | 1.200E-05 |
| 5177 | 259 | S259T | A | 5.4e-06 | 6.7e-06 | 1.5e-05 | 9.033E-06 | 5.208E-06 | 5.400E-06 | 1.500E-05 |
| 5178 | 259 | S259W | A | 1.1e-05 | 1.4e-05 | 9e-06 | 1.133E-05 | 2.517E-06 | 9.000E-06 | 1.400E-05 |
| 5179 | 259 | S259Y | A | 7.2e-06 | 9.7e-06 | 1.1e-05 | 9.300E-06 | 1.931E-06 | 7.200E-06 | 1.100E-05 |
| 5180 | 259 | S259V | A | 8.6e-06 | 9.6e-06 | 3.6e-06 | 7.267E-06 | 3.215E-06 | 3.600E-06 | 9.600E-06 |
| 5181 | 260 | M260A | A | 5.8e-06 | 7e-06 | 1.7e-05 | 9.933E-06 | 6.149E-06 | 5.800E-06 | 1.700E-05 |
| 5182 | 260 | M260R | A | 1.3e-05 | 7.4e-06 | 9.1e-06 | 9.833E-06 | 2.871E-06 | 7.400E-06 | 1.300E-05 |
| 5183 | 260 | M260N | A | 9e-06 | 7.3e-06 | 6e-06 | 7.433E-06 | 1.504E-06 | 6.000E-06 | 9.000E-06 |
| 5184 | 260 | M260D | A | 7.9e-06 | 6.7e-06 | 5.4e-06 | 6.667E-06 | 1.250E-06 | 5.400E-06 | 7.900E-06 |

| | | | | | | | | | | |
|---|---|---|---|---|---|---|---|---|---|---|
| 5185 | 260 | M260C | A | 6.6e-06 | 1.6e-05 | 4.3e-06 | 8.967E-06 | 6.199E-06 | 4.300E-06 | 1.600E-05 |
| 5186 | 260 | M260E | A | 1.2e-05 | 7.2e-06 | 7.7e-06 | 8.967E-06 | 2.639E-06 | 7.200E-06 | 1.200E-05 |
| 5187 | 260 | M260Q | A | 9.2e-06 | 5.1e-06 | 6.7e-06 | 7.000E-06 | 2.066E-06 | 5.100E-06 | 9.200E-06 |
| 5188 | 260 | M260G | A | 9.1e-06 | 4.1e-06 | 9e-06 | 7.400E-06 | 2.858E-06 | 4.100E-06 | 9.100E-06 |
| 5189 | 260 | M260H | A | 8.7e-06 | 4.6e-06 | 1e-05 | 7.767E-06 | 2.818E-06 | 4.600E-06 | 1.000E-05 |
| 5190 | 260 | M260I | A | 9.8e-06 | 1.3e-05 | 6.2e-06 | 9.667E-06 | 3.402E-06 | 6.200E-06 | 1.300E-05 |
| 5191 | 260 | M260L | A | 6.2e-06 | 4.1e-06 | 7.3e-06 | 5.867E-06 | 1.626E-06 | 4.100E-06 | 7.300E-06 |
| 5192 | 260 | M260K | A | 8.1e-06 | 6.8e-06 | 6.6e-06 | 7.167E-06 | 8.145E-07 | 6.600E-06 | 8.100E-06 |
| 5193 | 260 | M260M | A | 8.5e-06 | 9.5e-06 | 1.3e-05 | 1.033E-05 | 2.363E-06 | 8.500E-06 | 1.300E-05 |
| 5194 | 260 | M260F | A | 8.5e-06 | 9e-06 | 6.6e-06 | 8.033E-06 | 1.266E-06 | 6.600E-06 | 9.000E-06 |
| 5195 | 260 | M260P | A | 7.9e-06 | 1.6e-05 | 7.5e-06 | 1.047E-05 | 4.796E-06 | 7.500E-06 | 1.600E-05 |
| 5196 | 260 | M260S | A | 4.3e-06 | 6.1e-06 | 1.3e-05 | 7.800E-06 | 4.592E-06 | 4.300E-06 | 1.300E-05 |
| 5197 | 260 | M260T | A | 1.2e-05 | 5.2e-06 | 1.1e-05 | 9.400E-06 | 3.672E-06 | 5.200E-06 | 1.200E-05 |
| 5198 | 260 | M260W | A | 8.3e-06 | 6.7e-06 | 8.5e-06 | 7.833E-06 | 9.866E-07 | 6.700E-06 | 8.500E-06 |
| 5199 | 260 | M260Y | A | 5.2e-06 | 7.1e-06 | 4.6e-06 | 5.633E-06 | 1.305E-06 | 4.600E-06 | 7.100E-06 |
| 5200 | 260 | M260V | A | 5.6e-06 | 1.1e-05 | 1.4e-05 | 1.020E-05 | 4.257E-06 | 5.600E-06 | 1.400E-05 |
| 5201 | 261 | P261A | A | 7.2e-06 | 8.7e-06 | 5.3e-06 | 7.067E-06 | 1.704E-06 | 5.300E-06 | 8.700E-06 |
| 5202 | 261 | P261R | A | 5.1e-06 | 8.6e-06 | 1e-05 | 7.900E-06 | 2.524E-06 | 5.100E-06 | 1.000E-05 |
| 5203 | 261 | P261N | A | 7.1e-06 | 5.7e-06 | 8.9e-06 | 7.233E-06 | 1.604E-06 | 5.700E-06 | 8.900E-06 |
| 5204 | 261 | P261D | A | 9.9e-06 | 8.3e-06 | 6e-06 | 8.067E-06 | 1.960E-06 | 6.000E-06 | 9.900E-06 |
| 5205 | 261 | P261C | A | 1.2e-05 | 6e-06 | 9.7e-06 | 9.233E-06 | 3.027E-06 | 6.000E-06 | 1.200E-05 |
| 5206 | 261 | P261E | A | 9.9e-06 | 8.8e-06 | 1.3e-05 | 1.057E-05 | 2.178E-06 | 8.800E-06 | 1.300E-05 |
| 5207 | 261 | P261Q | A | 7.5e-06 | 4.8e-06 | 1e-05 | 7.433E-06 | 2.601E-06 | 4.800E-06 | 1.000E-05 |
| 5208 | 261 | P261G | A | 8e-06 | 9.7e-06 | 7.1e-06 | 8.267E-06 | 1.320E-06 | 7.100E-06 | 9.700E-06 |
| 5209 | 261 | P261H | A | 1.2e-05 | 7.8e-06 | 6.4e-06 | 8.733E-06 | 2.914E-06 | 6.400E-06 | 1.200E-05 |
| 5210 | 261 | P261I | A | 9.7e-06 | 9.9e-06 | 5.9e-06 | 8.500E-06 | 2.254E-06 | 5.900E-06 | 9.900E-06 |
| 5211 | 261 | P261L | A | 1e-05 | 9.2e-06 | 1.3e-05 | 1.073E-05 | 2.003E-06 | 9.200E-06 | 1.300E-05 |
| 5212 | 261 | P261K | A | 4.8e-06 | 8.6e-06 | 1.1e-05 | 8.133E-06 | 3.126E-06 | 4.800E-06 | 1.100E-05 |
| 5213 | 261 | P261M | A | 5.3e-06 | 9.8e-06 | 4.6e-06 | 6.567E-06 | 2.822E-06 | 4.600E-06 | 9.800E-06 |
| 5214 | 261 | P261F | A | 1.2e-05 | 5.3e-06 | 1.2e-05 | 9.767E-06 | 3.868E-06 | 5.300E-06 | 1.200E-05 |
| 5215 | 261 | P261P | A | 9.6e-06 | 6.9e-06 | 6.8e-06 | 7.767E-06 | 1.589E-06 | 6.800E-06 | 9.600E-06 |
| 5216 | 261 | P261S | A | 7e-06 | 9.1e-06 | 5e-06 | 7.033E-06 | 2.050E-06 | 5.000E-06 | 9.100E-06 |
| 5217 | 261 | P261T | A | 6.6e-06 | 4.9e-06 | 7.3e-06 | 6.267E-06 | 1.234E-06 | 4.900E-06 | 7.300E-06 |
| 5218 | 261 | P261W | A | 4.7e-06 | 4.9e-06 | 8.3e-06 | 5.967E-06 | 2.023E-06 | 4.700E-06 | 8.300E-06 |
| 5219 | 261 | P261Y | A | 5.3e-06 | 6.9e-06 | 9.7e-06 | 7.300E-06 | 2.227E-06 | 5.300E-06 | 9.700E-06 |
| 5220 | 261 | P261V | A | 1.2e-05 | 8e-06 | 3.1e-06 | 7.700E-06 | 4.458E-06 | 3.100E-06 | 1.200E-05 |
| 5221 | 262 | N262A | A | 1.3e-05 | 1e-05 | 7.7e-06 | 1.023E-05 | 2.658E-06 | 7.700E-06 | 1.300E-05 |
| 5222 | 262 | N262R | A | 9.1e-06 | 1e-05 | 1e-05 | 9.700E-06 | 5.196E-07 | 9.100E-06 | 1.000E-05 |
| 5223 | 262 | N262N | A | 7.4e-06 | 9.4e-06 | 6.8e-06 | 7.867E-06 | 1.361E-06 | 6.800E-06 | 9.400E-06 |
| 5224 | 262 | N262D | A | 8.4e-06 | 7.5e-06 | 6.1e-06 | 7.333E-06 | 1.159E-06 | 6.100E-06 | 8.400E-06 |
| 5225 | 262 | N262C | A | 1.2e-05 | 1e-05 | 8.4e-06 | 1.013E-05 | 1.804E-06 | 8.400E-06 | 1.200E-05 |
| 5226 | 262 | N262E | A | 7.9e-06 | 9.7e-06 | 9.8e-06 | 9.133E-06 | 1.069E-06 | 7.900E-06 | 9.800E-06 |
| 5227 | 262 | N262Q | A | 6.8e-06 | 5.7e-06 | 9.9e-06 | 7.467E-06 | 2.178E-06 | 5.700E-06 | 9.900E-06 |
| 5228 | 262 | N262G | A | 1.2e-05 | 4.3e-06 | 9.3e-06 | 8.533E-06 | 3.907E-06 | 4.300E-06 | 1.200E-05 |
| 5229 | 262 | N262H | A | 1.1e-05 | 8.4e-06 | 7.3e-06 | 8.900E-06 | 1.900E-06 | 7.300E-06 | 1.100E-05 |
| 5230 | 262 | N262I | A | 7e-06 | 1.1e-05 | 9.9e-06 | 9.300E-06 | 2.066E-06 | 7.000E-06 | 1.100E-05 |
| 5231 | 262 | N262L | A | 6.2e-06 | 1.1e-05 | 7.9e-06 | 8.367E-06 | 2.434E-06 | 6.200E-06 | 1.100E-05 |
| 5232 | 262 | N262K | A | 7e-06 | 5.6e-06 | 1.1e-05 | 7.867E-06 | 2.802E-06 | 5.600E-06 | 1.100E-05 |
| 5233 | 262 | N262M | A | 4.8e-06 | 8.5e-06 | 4e-06 | 5.767E-06 | 2.401E-06 | 4.000E-06 | 8.500E-06 |
| 5234 | 262 | N262F | A | 5.4e-06 | 1.4e-05 | 4.9e-06 | 8.100E-06 | 5.116E-06 | 4.900E-06 | 1.400E-05 |
| 5235 | 262 | N262P | A | 9.3e-06 | 4.5e-06 | 1e-05 | 7.933E-06 | 2.994E-06 | 4.500E-06 | 1.000E-05 |
| 5236 | 262 | N262S | A | 7.1e-06 | 7.7e-06 | 1.2e-05 | 8.933E-06 | 2.673E-06 | 7.100E-06 | 1.200E-05 |
| 5237 | 262 | N262T | A | 5.1e-06 | 1.5e-05 | 8.2e-06 | 9.433E-06 | 5.064E-06 | 5.100E-06 | 1.500E-05 |
| 5238 | 262 | N262W | A | 6.6e-06 | 4e-06 | 1.4e-05 | 8.200E-06 | 5.188E-06 | 4.000E-06 | 1.400E-05 |
| 5239 | 262 | N262Y | A | 6.9e-06 | 9e-06 | 6.4e-06 | 7.433E-06 | 1.380E-06 | 6.400E-06 | 9.000E-06 |
| 5240 | 262 | N262V | A | 8.2e-06 | 9.9e-06 | 6.3e-06 | 8.133E-06 | 1.801E-06 | 6.300E-06 | 9.900E-06 |
| 5241 | 263 | P263A | A | 1.2e-05 | 8.3e-06 | 5.9e-06 | 8.733E-06 | 3.073E-06 | 5.900E-06 | 1.200E-05 |
| 5242 | 263 | P263R | A | 5.7e-06 | 6.4e-06 | 9.2e-06 | 7.100E-06 | 1.852E-06 | 5.700E-06 | 9.200E-06 |
| 5243 | 263 | P263N | A | 7.6e-06 | 1.2e-05 | 1.3e-05 | 1.087E-05 | 2.873E-06 | 7.600E-06 | 1.300E-05 |
| 5244 | 263 | P263D | A | 8.3e-06 | 9.8e-06 | 1.1e-05 | 9.700E-06 | 1.353E-06 | 8.300E-06 | 1.100E-05 |
| 5245 | 263 | P263C | A | 1.2e-05 | 9.9e-06 | 8e-06 | 9.967E-06 | 2.001E-06 | 8.000E-06 | 1.200E-05 |
| 5246 | 263 | P263E | A | 7.5e-06 | 9.1e-06 | 8.1e-06 | 8.233E-06 | 8.083E-07 | 7.500E-06 | 9.100E-06 |
| 5247 | 263 | P263Q | A | 5.5e-06 | 6.6e-06 | 7e-06 | 6.367E-06 | 7.767E-07 | 5.500E-06 | 7.000E-06 |
| 5248 | 263 | P263G | A | 3.8e-06 | 5.5e-06 | 3.1e-06 | 4.133E-06 | 1.234E-06 | 3.100E-06 | 5.500E-06 |
| 5249 | 263 | P263H | A | 5.7e-06 | 7e-06 | 5.1e-06 | 5.933E-06 | 9.713E-07 | 5.100E-06 | 7.000E-06 |
| 5250 | 263 | P263I | A | 1e-05 | 8e-06 | 1.3e-05 | 1.033E-05 | 2.517E-06 | 8.000E-06 | 1.300E-05 |
| 5251 | 263 | P263L | A | 8.8e-06 | 9.9e-06 | 1.4e-05 | 1.090E-05 | 2.740E-06 | 8.800E-06 | 1.400E-05 |
| 5252 | 263 | P263K | A | 5.2e-06 | 8.7e-06 | 9.3e-06 | 7.733E-06 | 2.214E-06 | 5.200E-06 | 9.300E-06 |
| 5253 | 263 | P263M | A | 7.4e-06 | 7.1e-06 | 9.2e-06 | 7.900E-06 | 1.136E-06 | 7.100E-06 | 9.200E-06 |
| 5254 | 263 | P263F | A | 6.9e-06 | 7e-06 | 6.5e-06 | 6.800E-06 | 2.646E-07 | 6.500E-06 | 7.000E-06 |
| 5255 | 263 | P263P | A | 7.1e-06 | 1.1e-05 | 5e-06 | 7.700E-06 | 3.045E-06 | 5.000E-06 | 1.100E-05 |
| 5256 | 263 | P263S | A | 8.8e-06 | 8.6e-06 | 1.1e-05 | 9.467E-06 | 1.332E-06 | 8.600E-06 | 1.100E-05 |
| 5257 | 263 | P263T | A | 5.8e-06 | 1e-05 | 6.1e-06 | 7.300E-06 | 2.343E-06 | 5.800E-06 | 1.000E-05 |
| 5258 | 263 | P263W | A | 7.4e-06 | 6.8e-06 | 1.1e-05 | 8.400E-06 | 2.272E-06 | 6.800E-06 | 1.100E-05 |
| 5259 | 263 | P263Y | A | 7.6e-06 | 9.8e-06 | 1.1e-05 | 9.467E-06 | 1.724E-06 | 7.600E-06 | 1.100E-05 |
| 5260 | 263 | P263V | A | 9.6e-06 | 1.1e-05 | 7.7e-06 | 9.433E-06 | 1.656E-06 | 7.700E-06 | 1.100E-05 |
| 5261 | 264 | E264A | A | 1.2e-05 | 1.2e-05 | 6.5e-06 | 1.017E-05 | 3.175E-06 | 6.500E-06 | 1.200E-05 |
| 5262 | 264 | E264R | A | 1.2e-05 | 8.2e-06 | 8.1e-06 | 9.433E-06 | 2.223E-06 | 8.100E-06 | 1.200E-05 |
| 5263 | 264 | E264N | A | 1.2e-05 | 9e-06 | 7.6e-06 | 9.533E-06 | 2.248E-06 | 7.600E-06 | 1.200E-05 |
| 5264 | 264 | E264D | A | 5.9e-06 | 1.3e-05 | 7.8e-06 | 8.900E-06 | 3.676E-06 | 5.900E-06 | 1.300E-05 |
| 5265 | 264 | E264C | A | 8.6e-06 | 9.9e-06 | 6.9e-06 | 8.467E-06 | 1.504E-06 | 6.900E-06 | 9.900E-06 |
| 5266 | 264 | E264E | A | 1.1e-05 | 9.1e-06 | 4.9e-06 | 8.333E-06 | 3.121E-06 | 4.900E-06 | 1.100E-05 |
| 5267 | 264 | E264Q | A | 8.4e-06 | 6.9e-06 | 5.7e-06 | 7.000E-06 | 1.353E-06 | 5.700E-06 | 8.400E-06 |
| 5268 | 264 | E264G | A | 6.2e-06 | 7.4e-06 | 1e-05 | 7.867E-06 | 1.943E-06 | 6.200E-06 | 1.000E-05 |
| 5269 | 264 | E264H | A | 5.2e-06 | 5.5e-06 | 7.7e-06 | 6.133E-06 | 1.365E-06 | 5.200E-06 | 7.700E-06 |

| | | | | | | | | | | |
|---|---|---|---|---|---|---|---|---|---|---|
| 5270 | 264 | E264I | A | 1.3e-05 | 8.9e-06 | 6.2e-06 | 9.367E-06 | 3.424E-06 | 6.200E-06 | 1.300E-05 |
| 5271 | 264 | E264L | A | 7.7e-06 | 4.9e-06 | 5.2e-06 | 5.933E-06 | 1.537E-06 | 4.900E-06 | 7.700E-06 |
| 5272 | 264 | E264K | A | 6.5e-06 | 8.8e-06 | 6.9e-06 | 7.400E-06 | 1.229E-06 | 6.500E-06 | 8.800E-06 |
| 5273 | 264 | E264M | A | 9.7e-06 | 1e-05 | 9.2e-06 | 9.633E-06 | 4.041E-07 | 9.200E-06 | 1.000E-05 |
| 5274 | 264 | E264F | A | 5.4e-06 | 8.1e-06 | 4.5e-06 | 6.000E-06 | 1.873E-06 | 4.500E-06 | 8.100E-06 |
| 5275 | 264 | E264P | A | 1.4e-05 | 8.5e-06 | 1.3e-05 | 1.183E-05 | 2.930E-06 | 8.500E-06 | 1.400E-05 |
| 5276 | 264 | E264S | A | 1e-05 | 6.6e-06 | 6.3e-06 | 7.633E-06 | 2.055E-06 | 6.300E-06 | 1.000E-05 |
| 5277 | 264 | E264T | A | 6.9e-06 | 1e-05 | 6.6e-06 | 7.833E-06 | 1.882E-06 | 6.600E-06 | 1.000E-05 |
| 5278 | 264 | E264W | A | 7.3e-06 | 8.2e-06 | 9.1e-06 | 8.200E-06 | 9.000E-07 | 7.300E-06 | 9.100E-06 |
| 5279 | 264 | E264Y | A | 8.2e-06 | 6.5e-06 | 3.7e-06 | 6.133E-06 | 2.272E-06 | 3.700E-06 | 8.200E-06 |
| 5280 | 264 | E264V | A | 1.1e-05 | 7.1e-06 | 6.9e-06 | 8.333E-06 | 2.312E-06 | 6.900E-06 | 1.100E-05 |
| 5281 | 265 | G265A | A | 5.7e-06 | 6.6e-06 | 1e-05 | 7.433E-06 | 2.268E-06 | 5.700E-06 | 1.000E-05 |
| 5282 | 265 | G265R | A | 6.5e-06 | 8.2e-06 | 4.9e-06 | 6.533E-06 | 1.650E-06 | 4.900E-06 | 8.200E-06 |
| 5283 | 265 | G265N | A | 1.1e-05 | 8.9e-06 | 9e-06 | 9.633E-06 | 1.185E-06 | 8.900E-06 | 1.100E-05 |
| 5284 | 265 | G265D | A | 7.1e-06 | 5.2e-06 | 9.8e-06 | 7.367E-06 | 2.312E-06 | 5.200E-06 | 9.800E-06 |
| 5285 | 265 | G265C | A | 5.2e-06 | 1.3e-05 | 4.7e-06 | 7.633E-06 | 4.654E-06 | 4.700E-06 | 1.300E-05 |
| 5286 | 265 | G265E | A | 1e-05 | 1.2e-05 | 8.6e-06 | 1.020E-05 | 1.709E-06 | 8.600E-06 | 1.200E-05 |
| 5287 | 265 | G265Q | A | 6.3e-06 | 1.3e-05 | 1.1e-05 | 1.010E-05 | 3.439E-06 | 6.300E-06 | 1.300E-05 |
| 5288 | 265 | G265G | A | 8.3e-06 | 1.2e-05 | 1e-05 | 1.010E-05 | 1.852E-06 | 8.300E-06 | 1.200E-05 |
| 5289 | 265 | G265H | A | 1e-05 | 5.8e-06 | 4.7e-06 | 6.833E-06 | 2.797E-06 | 4.700E-06 | 1.000E-05 |
| 5290 | 265 | G265I | A | 1.1e-05 | 6.4e-06 | 7.2e-06 | 8.200E-06 | 2.458E-06 | 6.400E-06 | 1.100E-05 |
| 5291 | 265 | G265L | A | 9.7e-06 | 7.2e-06 | 7.7e-06 | 8.200E-06 | 1.323E-06 | 7.200E-06 | 9.700E-06 |
| 5292 | 265 | G265K | A | 6e-06 | 1.7e-05 | 5.2e-06 | 9.400E-06 | 6.594E-06 | 5.200E-06 | 1.700E-05 |
| 5293 | 265 | G265M | A | 7.4e-06 | 4.5e-06 | 1.2e-05 | 7.967E-06 | 3.782E-06 | 4.500E-06 | 1.200E-05 |
| 5294 | 265 | G265F | A | 7.3e-06 | 6.2e-06 | 8e-06 | 7.167E-06 | 9.074E-07 | 6.200E-06 | 8.000E-06 |
| 5295 | 265 | G265P | A | 7.7e-06 | 9.6e-06 | 5.9e-06 | 7.733E-06 | 1.850E-06 | 5.900E-06 | 9.600E-06 |
| 5296 | 265 | G265S | A | 1e-05 | 5.8e-06 | 8.7e-06 | 8.167E-06 | 2.150E-06 | 5.800E-06 | 1.000E-05 |
| 5297 | 265 | G265T | A | 1.3e-05 | 6.4e-06 | 6.8e-06 | 8.733E-06 | 3.700E-06 | 6.400E-06 | 1.300E-05 |
| 5298 | 265 | G265W | A | 1.1e-05 | 1.1e-05 | 5.2e-06 | 9.067E-06 | 3.349E-06 | 5.200E-06 | 1.100E-05 |
| 5299 | 265 | G265Y | A | 4.6e-06 | 7e-06 | 8.9e-06 | 6.833E-06 | 2.155E-06 | 4.600E-06 | 8.900E-06 |
| 5300 | 265 | G265V | A | 8.1e-06 | 6.3e-06 | 1e-05 | 8.133E-06 | 1.850E-06 | 6.300E-06 | 1.000E-05 |
| 5301 | 266 | R266A | A | 5.9e-06 | 5.5e-06 | 6.2e-06 | 5.867E-06 | 3.512E-07 | 5.500E-06 | 6.200E-06 |
| 5302 | 266 | R266R | A | 1.1e-05 | 9.3e-06 | 9.4e-06 | 9.900E-06 | 9.539E-07 | 9.300E-06 | 1.100E-05 |
| 5303 | 266 | R266N | A | 9.5e-06 | 9.6e-06 | 1.1e-05 | 1.003E-05 | 8.386E-07 | 9.500E-06 | 1.100E-05 |
| 5304 | 266 | R266D | A | 1e-05 | 1e-05 | 6.2e-06 | 8.733E-06 | 2.194E-06 | 6.200E-06 | 1.000E-05 |
| 5305 | 266 | R266C | A | 5.8e-06 | 8.1e-06 | 8.8e-06 | 7.567E-06 | 1.570E-06 | 5.800E-06 | 8.800E-06 |
| 5306 | 266 | R266E | A | 5.7e-06 | 6e-06 | 1.1e-05 | 7.567E-06 | 2.977E-06 | 5.700E-06 | 1.100E-05 |
| 5307 | 266 | R266Q | A | 5.3e-06 | 8.6e-06 | 9.3e-06 | 7.733E-06 | 2.136E-06 | 5.300E-06 | 9.300E-06 |
| 5308 | 266 | R266G | A | 6.8e-06 | 8.3e-06 | 6.8e-06 | 7.300E-06 | 8.660E-07 | 6.800E-06 | 8.300E-06 |
| 5309 | 266 | R266H | A | 5.2e-06 | 9.5e-06 | 9.3e-06 | 8.000E-06 | 2.427E-06 | 5.200E-06 | 9.500E-06 |
| 5310 | 266 | R266I | A | 1.4e-05 | 7.5e-06 | 1.4e-05 | 1.183E-05 | 3.753E-06 | 7.500E-06 | 1.400E-05 |
| 5311 | 266 | R266L | A | 7.9e-06 | 6.8e-06 | 9.6e-06 | 8.100E-06 | 1.411E-06 | 6.800E-06 | 9.600E-06 |
| 5312 | 266 | R266K | A | 4.7e-06 | 9.3e-06 | 4.9e-06 | 6.300E-06 | 2.600E-06 | 4.700E-06 | 9.300E-06 |
| 5313 | 266 | R266M | A | 6.4e-06 | 8.7e-06 | 8.8e-06 | 7.967E-06 | 1.358E-06 | 6.400E-06 | 8.800E-06 |
| 5314 | 266 | R266F | A | 6.8e-06 | 2e-05 | 7e-06 | 1.127E-05 | 7.564E-06 | 6.800E-06 | 2.000E-05 |
| 5315 | 266 | R266P | A | 1.1e-05 | 7.2e-06 | 6.7e-06 | 8.300E-06 | 2.352E-06 | 6.700E-06 | 1.100E-05 |
| 5316 | 266 | R266S | A | 8e-06 | 8.3e-06 | 7.5e-06 | 7.933E-06 | 4.041E-07 | 7.500E-06 | 8.300E-06 |
| 5317 | 266 | R266T | A | 1.4e-05 | 5.4e-06 | 7.8e-06 | 9.067E-06 | 4.438E-06 | 5.400E-06 | 1.400E-05 |
| 5318 | 266 | R266W | A | 8.3e-06 | 4.5e-06 | 4.8e-06 | 5.867E-06 | 2.113E-06 | 4.500E-06 | 8.300E-06 |
| 5319 | 266 | R266Y | A | 9.2e-06 | 5.5e-06 | 1.1e-05 | 8.567E-06 | 2.804E-06 | 5.500E-06 | 1.100E-05 |
| 5320 | 266 | R266V | A | 1.3e-05 | 5.7e-06 | 7.5e-06 | 8.733E-06 | 3.803E-06 | 5.700E-06 | 1.300E-05 |
| 5321 | 267 | Y267A | A | 1e-05 | 8e-06 | 7.2e-06 | 8.400E-06 | 1.442E-06 | 7.200E-06 | 1.000E-05 |
| 5322 | 267 | Y267R | A | 7.5e-06 | 8.7e-06 | 1.4e-05 | 1.007E-05 | 3.459E-06 | 7.500E-06 | 1.400E-05 |
| 5323 | 267 | Y267N | A | 1.1e-05 | 8.7e-06 | 6.3e-06 | 8.667E-06 | 2.350E-06 | 6.300E-06 | 1.100E-05 |
| 5324 | 267 | Y267D | A | 9.1e-06 | 6.7e-06 | 1.5e-05 | 1.027E-05 | 4.271E-06 | 6.700E-06 | 1.500E-05 |
| 5325 | 267 | Y267C | A | 9.1e-06 | 1.2e-05 | 9.9e-06 | 1.033E-05 | 1.498E-06 | 9.100E-06 | 1.200E-05 |
| 5326 | 267 | Y267E | A | 8e-06 | 9.4e-06 | 7.1e-06 | 8.167E-06 | 1.159E-06 | 7.100E-06 | 9.400E-06 |
| 5327 | 267 | Y267Q | A | 5.4e-06 | 7.8e-06 | 1.2e-05 | 8.400E-06 | 3.341E-06 | 5.400E-06 | 1.200E-05 |
| 5328 | 267 | Y267G | A | 7e-06 | 1.2e-05 | 9.5e-06 | 9.500E-06 | 2.500E-06 | 7.000E-06 | 1.200E-05 |
| 5329 | 267 | Y267H | A | 6.3e-06 | 6.9e-06 | 1.1e-05 | 8.067E-06 | 2.558E-06 | 6.300E-06 | 1.100E-05 |
| 5330 | 267 | Y267I | A | 1.4e-05 | 6.3e-06 | 8.8e-06 | 9.700E-06 | 3.928E-06 | 6.300E-06 | 1.400E-05 |
| 5331 | 267 | Y267L | A | 9.3e-06 | 6.1e-06 | 6.5e-06 | 7.300E-06 | 1.744E-06 | 6.100E-06 | 9.300E-06 |
| 5332 | 267 | Y267K | A | 4.9e-06 | 5.1e-06 | 5.4e-06 | 5.133E-06 | 2.517E-07 | 4.900E-06 | 5.400E-06 |
| 5333 | 267 | Y267M | A | 4.9e-06 | 7.3e-06 | 1e-05 | 7.400E-06 | 2.551E-06 | 4.900E-06 | 1.000E-05 |
| 5334 | 267 | Y267F | A | 4.3e-06 | 9.4e-06 | 6e-06 | 6.567E-06 | 2.597E-06 | 4.300E-06 | 9.400E-06 |
| 5335 | 267 | Y267P | A | 6.7e-06 | 9.2e-06 | 1.3e-05 | 9.633E-06 | 3.172E-06 | 6.700E-06 | 1.300E-05 |
| 5336 | 267 | Y267S | A | 5.1e-06 | 6.6e-06 | 9.7e-06 | 7.133E-06 | 2.346E-06 | 5.100E-06 | 9.700E-06 |
| 5337 | 267 | Y267T | A | 9.1e-06 | 1e-05 | 7.3e-06 | 8.800E-06 | 1.375E-06 | 7.300E-06 | 1.000E-05 |
| 5338 | 267 | Y267W | A | 7.4e-06 | 8.5e-06 | 5.4e-06 | 7.100E-06 | 1.572E-06 | 5.400E-06 | 8.500E-06 |
| 5339 | 267 | Y267Y | A | 9.7e-06 | 6.2e-06 | 2.8e-06 | 6.233E-06 | 3.450E-06 | 2.800E-06 | 9.700E-06 |
| 5340 | 267 | Y267V | A | 5.1e-06 | 9.7e-06 | 9.7e-06 | 8.167E-06 | 2.656E-06 | 5.100E-06 | 9.700E-06 |
| 5341 | 268 | T268A | A | 6.3e-06 | 6.6e-06 | 6.2e-06 | 6.367E-06 | 2.082E-07 | 6.200E-06 | 6.600E-06 |
| 5342 | 268 | T268R | A | 1e-05 | 6.9e-06 | 8.4e-06 | 8.433E-06 | 1.550E-06 | 6.900E-06 | 1.000E-05 |
| 5343 | 268 | T268N | A | 5.7e-06 | 1.4e-05 | 6.5e-06 | 8.733E-06 | 4.579E-06 | 5.700E-06 | 1.400E-05 |
| 5344 | 268 | T268D | A | 7.2e-06 | 8.3e-06 | 8e-06 | 7.833E-06 | 5.686E-07 | 7.200E-06 | 8.300E-06 |
| 5345 | 268 | T268C | A | 1e-05 | 6.2e-06 | 1.1e-05 | 9.067E-06 | 2.532E-06 | 6.200E-06 | 1.100E-05 |
| 5346 | 268 | T268E | A | 1e-05 | 9.3e-06 | 5.3e-06 | 8.200E-06 | 2.536E-06 | 5.300E-06 | 1.000E-05 |
| 5347 | 268 | T268Q | A | 6.3e-06 | 8e-06 | 7.3e-06 | 7.200E-06 | 8.544E-07 | 6.300E-06 | 8.000E-06 |
| 5348 | 268 | T268G | A | 5.2e-06 | 9e-06 | 9.8e-06 | 8.000E-06 | 2.458E-06 | 5.200E-06 | 9.800E-06 |
| 5349 | 268 | T268H | A | 9.1e-06 | 8.1e-06 | 6.8e-06 | 8.000E-06 | 1.153E-06 | 6.800E-06 | 9.100E-06 |
| 5350 | 268 | T268I | A | 5e-06 | 7.1e-06 | 9.5e-06 | 7.200E-06 | 2.252E-06 | 5.000E-06 | 9.500E-06 |
| 5351 | 268 | T268L | A | 4.6e-06 | 4.8e-06 | 6.5e-06 | 5.300E-06 | 1.044E-06 | 4.600E-06 | 6.500E-06 |
| 5352 | 268 | T268K | A | 8.9e-06 | 8.3e-06 | 8.3e-06 | 8.500E-06 | 3.464E-07 | 8.300E-06 | 8.900E-06 |
| 5353 | 268 | T268M | A | 8.2e-06 | 9.1e-06 | 5.7e-06 | 7.667E-06 | 1.762E-06 | 5.700E-06 | 9.100E-06 |
| 5354 | 268 | T268F | A | 1.3e-05 | 5.8e-06 | 9.3e-06 | 9.367E-06 | 3.600E-06 | 5.800E-06 | 1.300E-05 |

| | | | | | | | | | | |
|---|---|---|---|---|---|---|---|---|---|---|
| 5355 | 268 | T268P | A | 4.4e-06 | 1.4e-05 | 1.3e-05 | 1.047E-05 | 5.278E-06 | 4.400E-06 | 1.400E-05 |
| 5356 | 268 | T268S | A | 6.4e-06 | 6.6e-06 | 7.7e-06 | 6.900E-06 | 7.000E-07 | 6.400E-06 | 7.700E-06 |
| 5357 | 268 | T268T | A | 9.3e-06 | 5.7e-06 | 1.1e-05 | 8.667E-06 | 2.706E-06 | 5.700E-06 | 1.100E-05 |
| 5358 | 268 | T268W | A | 9.2e-06 | 6.4e-06 | 5.4e-06 | 7.000E-06 | 1.970E-06 | 5.400E-06 | 9.200E-06 |
| 5359 | 268 | T268Y | A | 8.2e-06 | 7e-06 | 9.6e-06 | 8.267E-06 | 1.301E-06 | 7.000E-06 | 9.600E-06 |
| 5360 | 268 | T268V | A | 8.4e-06 | 9.7e-06 | 1.2e-05 | 1.003E-05 | 1.823E-06 | 8.400E-06 | 1.200E-05 |
| 5361 | 269 | F269A | A | 1e-05 | 7.6e-06 | 9e-06 | 8.867E-06 | 1.206E-06 | 7.600E-06 | 1.000E-05 |
| 5362 | 269 | F269R | A | 7.5e-06 | 8.3e-06 | 1.1e-05 | 8.933E-06 | 1.834E-06 | 7.500E-06 | 1.100E-05 |
| 5363 | 269 | F269N | A | 5.5e-06 | 7.6e-06 | 6e-06 | 6.367E-06 | 1.097E-06 | 5.500E-06 | 7.600E-06 |
| 5364 | 269 | F269D | A | 9.3e-06 | 6.9e-06 | 5.3e-06 | 7.167E-06 | 2.013E-06 | 5.300E-06 | 9.300E-06 |
| 5365 | 269 | F269C | A | 8.7e-06 | 9e-06 | 8.1e-06 | 8.600E-06 | 4.583E-07 | 8.100E-06 | 9.000E-06 |
| 5366 | 269 | F269E | A | 9.7e-06 | 3.5e-06 | 1e-05 | 7.733E-06 | 3.669E-06 | 3.500E-06 | 1.000E-05 |
| 5367 | 269 | F269Q | A | 1e-05 | 6.9e-06 | 9.2e-06 | 8.700E-06 | 1.609E-06 | 6.900E-06 | 1.000E-05 |
| 5368 | 269 | F269G | A | 8e-06 | 6.6e-06 | 6.6e-06 | 7.067E-06 | 8.083E-07 | 6.600E-06 | 8.000E-06 |
| 5369 | 269 | F269H | A | 6.7e-06 | 9.2e-06 | 9.3e-06 | 8.400E-06 | 1.473E-06 | 6.700E-06 | 9.300E-06 |
| 5370 | 269 | F269I | A | 9.7e-06 | 1.1e-05 | 5.7e-06 | 8.800E-06 | 2.762E-06 | 5.700E-06 | 1.100E-05 |
| 5371 | 269 | F269L | A | 8.4e-06 | 7.5e-06 | 8.8e-06 | 8.233E-06 | 6.658E-07 | 7.500E-06 | 8.800E-06 |
| 5372 | 269 | F269K | A | 1.7e-05 | 1e-05 | 8.1e-06 | 1.170E-05 | 4.687E-06 | 8.100E-06 | 1.700E-05 |
| 5373 | 269 | F269M | A | 6.9e-06 | 6.4e-06 | 9.3e-06 | 7.533E-06 | 1.550E-06 | 6.400E-06 | 9.300E-06 |
| 5374 | 269 | F269F | A | 6.7e-06 | 5.5e-06 | 1.1e-05 | 7.733E-06 | 2.892E-06 | 5.500E-06 | 1.100E-05 |
| 5375 | 269 | F269P | A | 5.9e-06 | 9.8e-06 | 1.2e-05 | 9.233E-06 | 3.089E-06 | 5.900E-06 | 1.200E-05 |
| 5376 | 269 | F269S | A | 6.1e-06 | 8.2e-06 | 4.4e-06 | 6.233E-06 | 1.904E-06 | 4.400E-06 | 8.200E-06 |
| 5377 | 269 | F269T | A | 7.6e-06 | 7.9e-06 | 6.9e-06 | 7.467E-06 | 5.132E-07 | 6.900E-06 | 7.900E-06 |
| 5378 | 269 | F269W | A | 7.3e-06 | 9.3e-06 | 1.1e-05 | 9.200E-06 | 1.852E-06 | 7.300E-06 | 1.100E-05 |
| 5379 | 269 | F269Y | A | 8.8e-06 | 6.9e-06 | 5.6e-06 | 7.100E-06 | 1.609E-06 | 5.600E-06 | 8.800E-06 |
| 5380 | 269 | F269V | A | 6.8e-06 | 4.5e-06 | 5.7e-06 | 5.667E-06 | 1.150E-06 | 4.500E-06 | 6.800E-06 |
| 5381 | 270 | G270A | A | 9.3e-06 | 1.2e-05 | 6.6e-06 | 9.300E-06 | 2.700E-06 | 6.600E-06 | 1.200E-05 |
| 5382 | 270 | G270R | A | 8.8e-06 | 6.6e-06 | 4.7e-06 | 6.700E-06 | 2.052E-06 | 4.700E-06 | 8.800E-06 |
| 5383 | 270 | G270N | A | 6.6e-06 | 1e-05 | 9.8e-06 | 8.800E-06 | 1.908E-06 | 6.600E-06 | 1.000E-05 |
| 5384 | 270 | G270D | A | 1.1e-05 | 1.1e-05 | 6.6e-06 | 9.533E-06 | 2.540E-06 | 6.600E-06 | 1.100E-05 |
| 5385 | 270 | G270C | A | 8.8e-06 | 1.3e-05 | 9.1e-06 | 1.030E-05 | 2.343E-06 | 8.800E-06 | 1.300E-05 |
| 5386 | 270 | G270E | A | 9.1e-06 | 6.9e-06 | 1.4e-05 | 1.000E-05 | 3.635E-06 | 6.900E-06 | 1.400E-05 |
| 5387 | 270 | G270Q | A | 8.6e-06 | 9.2e-06 | 6.9e-06 | 8.233E-06 | 1.193E-06 | 6.900E-06 | 9.200E-06 |
| 5388 | 270 | G270G | A | 8.9e-06 | 9.5e-06 | 1e-05 | 9.467E-06 | 5.508E-07 | 8.900E-06 | 1.000E-05 |
| 5389 | 270 | G270H | A | 4.3e-06 | 5e-06 | 5.5e-06 | 4.933E-06 | 6.028E-07 | 4.300E-06 | 5.500E-06 |
| 5390 | 270 | G270I | A | 1e-05 | 6.1e-06 | 1.5e-05 | 1.037E-05 | 4.461E-06 | 6.100E-06 | 1.500E-05 |
| 5391 | 270 | G270L | A | 8e-06 | 7.4e-06 | 6.1e-06 | 7.167E-06 | 9.713E-07 | 6.100E-06 | 8.000E-06 |
| 5392 | 270 | G270K | A | 7.5e-06 | 6e-06 | 5.6e-06 | 6.367E-06 | 1.002E-06 | 5.600E-06 | 7.500E-06 |
| 5393 | 270 | G270M | A | 1e-05 | 9.5e-06 | 8.4e-06 | 9.300E-06 | 8.185E-07 | 8.400E-06 | 1.000E-05 |
| 5394 | 270 | G270F | A | 5.3e-06 | 7.1e-06 | 9.1e-06 | 7.167E-06 | 1.901E-06 | 5.300E-06 | 9.100E-06 |
| 5395 | 270 | G270P | A | 5.6e-06 | 1.2e-05 | 6.8e-06 | 8.133E-06 | 3.402E-06 | 5.600E-06 | 1.200E-05 |
| 5396 | 270 | G270S | A | 1.1e-05 | 9.1e-06 | 8.5e-06 | 9.533E-06 | 1.305E-06 | 8.500E-06 | 1.100E-05 |
| 5397 | 270 | G270T | A | 9.7e-06 | 4.6e-06 | 7.5e-06 | 7.267E-06 | 2.558E-06 | 4.600E-06 | 9.700E-06 |
| 5398 | 270 | G270W | A | 8.3e-06 | 8.5e-06 | 6.2e-06 | 7.667E-06 | 1.274E-06 | 6.200E-06 | 8.500E-06 |
| 5399 | 270 | G270Y | A | 7.4e-06 | 6.7e-06 | 5.6e-06 | 6.567E-06 | 9.074E-07 | 5.600E-06 | 7.400E-06 |
| 5400 | 270 | G270V | A | 8.3e-06 | 1.4e-05 | 1.4e-05 | 1.210E-05 | 3.291E-06 | 8.300E-06 | 1.400E-05 |
| 5401 | 271 | A271A | A | 8.6e-06 | 1.5e-05 | 9e-06 | 1.087E-05 | 3.585E-06 | 8.600E-06 | 1.500E-05 |
| 5402 | 271 | A271R | A | 7.4e-06 | 1.1e-05 | 7.4e-06 | 8.600E-06 | 2.078E-06 | 7.400E-06 | 1.100E-05 |
| 5403 | 271 | A271N | A | 1.1e-05 | 5.8e-06 | 6.8e-06 | 7.867E-06 | 2.759E-06 | 5.800E-06 | 1.100E-05 |
| 5404 | 271 | A271D | A | 8.2e-06 | 1e-05 | 9.9e-06 | 9.367E-06 | 1.012E-06 | 8.200E-06 | 1.000E-05 |
| 5405 | 271 | A271C | A | 9.5e-06 | 8.8e-06 | 6.4e-06 | 8.233E-06 | 1.626E-06 | 6.400E-06 | 9.500E-06 |
| 5406 | 271 | A271E | A | 1.3e-05 | 5.7e-06 | 1.1e-05 | 9.900E-06 | 3.772E-06 | 5.700E-06 | 1.300E-05 |
| 5407 | 271 | A271Q | A | 6.2e-06 | 9.4e-06 | 6.1e-06 | 7.233E-06 | 1.877E-06 | 6.100E-06 | 9.400E-06 |
| 5408 | 271 | A271G | A | 8.5e-06 | 6.6e-06 | 9.5e-06 | 8.200E-06 | 1.473E-06 | 6.600E-06 | 9.500E-06 |
| 5409 | 271 | A271H | A | 8.6e-06 | 6.5e-06 | 1.1e-05 | 8.700E-06 | 2.252E-06 | 6.500E-06 | 1.100E-05 |
| 5410 | 271 | A271I | A | 9.4e-06 | 6.8e-06 | 7.5e-06 | 7.900E-06 | 1.345E-06 | 6.800E-06 | 9.400E-06 |
| 5411 | 271 | A271L | A | 7.1e-06 | 9.4e-06 | 4.9e-06 | 7.133E-06 | 2.250E-06 | 4.900E-06 | 9.400E-06 |
| 5412 | 271 | A271K | A | 5.7e-06 | 9.3e-06 | 8.2e-06 | 7.733E-06 | 1.845E-06 | 5.700E-06 | 9.300E-06 |
| 5413 | 271 | A271M | A | 7.5e-06 | 1.2e-05 | 8.7e-06 | 9.400E-06 | 2.330E-06 | 7.500E-06 | 1.200E-05 |
| 5414 | 271 | A271F | A | 4.7e-06 | 5.7e-06 | 7.6e-06 | 6.000E-06 | 1.473E-06 | 4.700E-06 | 7.600E-06 |
| 5415 | 271 | A271P | A | 8.8e-06 | 9e-06 | 7.3e-06 | 8.367E-06 | 9.292E-07 | 7.300E-06 | 9.000E-06 |
| 5416 | 271 | A271S | A | 2.8e-06 | 9.5e-06 | 6.3e-06 | 6.200E-06 | 3.351E-06 | 2.800E-06 | 9.500E-06 |
| 5417 | 271 | A271T | A | 4.9e-06 | 9.6e-06 | 3.2e-06 | 5.900E-06 | 3.315E-06 | 3.200E-06 | 9.600E-06 |
| 5418 | 271 | A271W | A | 8.3e-06 | 6e-06 | 7.8e-06 | 7.367E-06 | 1.210E-06 | 6.000E-06 | 8.300E-06 |
| 5419 | 271 | A271Y | A | 8.8e-06 | 4.6e-06 | 5.9e-06 | 6.433E-06 | 2.150E-06 | 4.600E-06 | 8.800E-06 |
| 5420 | 271 | A271V | A | 1.4e-05 | 5.5e-06 | 6.5e-06 | 8.667E-06 | 4.646E-06 | 5.500E-06 | 1.400E-05 |
| 5421 | 272 | S272A | A | 1.2e-05 | 8.5e-06 | 4.6e-06 | 8.367E-06 | 3.702E-06 | 4.600E-06 | 1.200E-05 |
| 5422 | 272 | S272R | A | 4.4e-06 | 7.2e-06 | 6.9e-06 | 6.167E-06 | 1.537E-06 | 4.400E-06 | 7.200E-06 |
| 5423 | 272 | S272N | A | 3.3e-06 | 5e-06 | 9.5e-06 | 5.933E-06 | 3.204E-06 | 3.300E-06 | 9.500E-06 |
| 5424 | 272 | S272D | A | 8.5e-06 | 8e-06 | 1e-05 | 8.833E-06 | 1.041E-06 | 8.000E-06 | 1.000E-05 |
| 5425 | 272 | S272C | A | 7.3e-06 | 4.9e-06 | 1.6e-05 | 9.400E-06 | 5.840E-06 | 4.900E-06 | 1.600E-05 |
| 5426 | 272 | S272E | A | 1e-05 | 8.9e-06 | 1.2e-05 | 1.030E-05 | 1.572E-06 | 8.900E-06 | 1.200E-05 |
| 5427 | 272 | S272Q | A | 4.9e-06 | 6.2e-06 | 1.7e-05 | 9.367E-06 | 6.643E-06 | 4.900E-06 | 1.700E-05 |
| 5428 | 272 | S272G | A | 9.7e-06 | 8.8e-06 | 9.5e-06 | 9.333E-06 | 4.726E-07 | 8.800E-06 | 9.700E-06 |
| 5429 | 272 | S272H | A | 6e-06 | 7.1e-06 | 6e-06 | 6.367E-06 | 6.351E-07 | 6.000E-06 | 7.100E-06 |
| 5430 | 272 | S272I | A | 1.1e-05 | 7e-06 | 8.4e-06 | 8.800E-06 | 2.030E-06 | 7.000E-06 | 1.100E-05 |
| 5431 | 272 | S272L | A | 5.1e-06 | 1e-05 | 9.9e-06 | 8.333E-06 | 2.801E-06 | 5.100E-06 | 1.000E-05 |
| 5432 | 272 | S272K | A | 2e-05 | 8.2e-06 | 1.1e-05 | 1.307E-05 | 6.165E-06 | 8.200E-06 | 2.000E-05 |
| 5433 | 272 | S272M | A | 6.7e-06 | 1.3e-05 | 5.3e-06 | 8.333E-06 | 4.102E-06 | 5.300E-06 | 1.300E-05 |
| 5434 | 272 | S272F | A | 9.5e-06 | 1.4e-05 | 1e-05 | 1.117E-05 | 2.466E-06 | 9.500E-06 | 1.400E-05 |
| 5435 | 272 | S272P | A | 2e-05 | 1e-05 | 9.3e-06 | 1.310E-05 | 5.986E-06 | 9.300E-06 | 2.000E-05 |
| 5436 | 272 | S272S | A | 6.7e-06 | 9.7e-06 | 9.1e-06 | 8.500E-06 | 1.587E-06 | 6.700E-06 | 9.700E-06 |
| 5437 | 272 | S272T | A | 5.3e-06 | 8.6e-06 | 9.5e-06 | 7.800E-06 | 2.211E-06 | 5.300E-06 | 9.500E-06 |
| 5438 | 272 | S272W | A | 7.1e-06 | 6.6e-06 | 1.6e-05 | 9.900E-06 | 5.289E-06 | 6.600E-06 | 1.600E-05 |
| 5439 | 272 | S272Y | A | 4.5e-06 | 1.1e-05 | 1e-05 | 8.500E-06 | 3.500E-06 | 4.500E-06 | 1.100E-05 |

| | | | | | | | | | | |
|---|---|---|---|---|---|---|---|---|---|---|
| 5440 | 272 | S272V | A | 9.5e-06 | 1.1e-05 | 6.3e-06 | 8.933E-06 | 2.401E-06 | 6.300E-06 | 1.100E-05 |
| 5441 | 273 | C273A | A | 1.5e-05 | 1.4e-05 | 8.5e-06 | 1.250E-05 | 3.500E-06 | 8.500E-06 | 1.500E-05 |
| 5442 | 273 | C273R | A | 4.9e-06 | 4.4e-06 | 1e-05 | 6.433E-06 | 3.099E-06 | 4.400E-06 | 1.000E-05 |
| 5443 | 273 | C273N | A | 4.4e-06 | 9.1e-06 | 6.8e-06 | 6.767E-06 | 2.350E-06 | 4.400E-06 | 9.100E-06 |
| 5444 | 273 | C273D | A | 7.7e-06 | 9.8e-06 | 9.7e-06 | 9.067E-06 | 1.185E-06 | 7.700E-06 | 9.800E-06 |
| 5445 | 273 | C273C | A | 8e-06 | 9.6e-06 | 5.1e-06 | 7.567E-06 | 2.281E-06 | 5.100E-06 | 9.600E-06 |
| 5446 | 273 | C273E | A | 7.7e-06 | 1.7e-05 | 9.2e-06 | 1.130E-05 | 4.993E-06 | 7.700E-06 | 1.700E-05 |
| 5447 | 273 | C273Q | A | 1.3e-05 | 8e-06 | 6.2e-06 | 9.067E-06 | 3.523E-06 | 6.200E-06 | 1.300E-05 |
| 5448 | 273 | C273G | A | 9.4e-06 | 4.5e-06 | 8e-06 | 7.300E-06 | 2.524E-06 | 4.500E-06 | 9.400E-06 |
| 5449 | 273 | C273H | A | 1.1e-05 | 6.7e-06 | 1.1e-05 | 9.567E-06 | 2.483E-06 | 6.700E-06 | 1.100E-05 |
| 5450 | 273 | C273I | A | 9.2e-06 | 6.1e-06 | 7.5e-06 | 7.600E-06 | 1.552E-06 | 6.100E-06 | 9.200E-06 |
| 5451 | 273 | C273L | A | 1.3e-05 | 1.5e-05 | 1.5e-05 | 1.433E-05 | 1.155E-06 | 1.300E-05 | 1.500E-05 |
| 5452 | 273 | C273K | A | 1.4e-05 | 6.5e-06 | 1e-05 | 1.017E-05 | 3.753E-06 | 6.500E-06 | 1.400E-05 |
| 5453 | 273 | C273M | A | 5.4e-06 | 6.6e-06 | 1.1e-05 | 7.667E-06 | 2.948E-06 | 5.400E-06 | 1.100E-05 |
| 5454 | 273 | C273F | A | 5.9e-06 | 6.7e-06 | 1.1e-05 | 7.867E-06 | 2.743E-06 | 5.900E-06 | 1.100E-05 |
| 5455 | 273 | C273P | A | 9.5e-06 | 1.2e-05 | 1.2e-05 | 1.117E-05 | 1.443E-06 | 9.500E-06 | 1.200E-05 |
| 5456 | 273 | C273S | A | 5.2e-06 | 5.7e-06 | 1.1e-05 | 7.300E-06 | 3.214E-06 | 5.200E-06 | 1.100E-05 |
| 5457 | 273 | C273T | A | 8.3e-06 | 5.7e-06 | 8.6e-06 | 7.533E-06 | 1.595E-06 | 5.700E-06 | 8.600E-06 |
| 5458 | 273 | C273W | A | 1.1e-05 | 9.7e-06 | 8.5e-06 | 9.733E-06 | 1.250E-06 | 8.500E-06 | 1.100E-05 |
| 5459 | 273 | C273Y | A | 1.2e-05 | 6.9e-06 | 1.7e-05 | 1.197E-05 | 5.050E-06 | 6.900E-06 | 1.700E-05 |
| 5460 | 273 | C273V | A | 9.4e-06 | 8.9e-06 | 5.4e-06 | 7.900E-06 | 2.179E-06 | 5.400E-06 | 9.400E-06 |
| 5461 | 274 | V274A | A | 5.8e-06 | 6.7e-06 | 6.7e-06 | 6.400E-06 | 5.196E-07 | 5.800E-06 | 6.700E-06 |
| 5462 | 274 | V274R | A | 9.6e-06 | 8.5e-06 | 1.2e-05 | 1.003E-05 | 1.790E-06 | 8.500E-06 | 1.200E-05 |
| 5463 | 274 | V274N | A | 6.9e-06 | 5e-06 | 5.8e-06 | 5.900E-06 | 9.539E-07 | 5.000E-06 | 6.900E-06 |
| 5464 | 274 | V274D | A | 7.5e-06 | 7.4e-06 | 9.8e-06 | 8.233E-06 | 1.358E-06 | 7.400E-06 | 9.800E-06 |
| 5465 | 274 | V274C | A | 4.3e-06 | 8.4e-06 | 4.5e-06 | 5.733E-06 | 2.312E-06 | 4.300E-06 | 8.400E-06 |
| 5466 | 274 | V274E | A | 1.7e-05 | 1.3e-05 | 1e-05 | 1.333E-05 | 3.512E-06 | 1.000E-05 | 1.700E-05 |
| 5467 | 274 | V274Q | A | 6.1e-06 | 4e-06 | 1.3e-05 | 7.700E-06 | 4.709E-06 | 4.000E-06 | 1.300E-05 |
| 5468 | 274 | V274G | A | 6e-06 | 6.8e-06 | 3.3e-06 | 5.367E-06 | 1.834E-06 | 3.300E-06 | 6.800E-06 |
| 5469 | 274 | V274H | A | 7.4e-06 | 4.9e-06 | 5.5e-06 | 5.933E-06 | 1.305E-06 | 4.900E-06 | 7.400E-06 |
| 5470 | 274 | V274I | A | 1.4e-05 | 1.4e-05 | 1.3e-05 | 1.367E-05 | 5.774E-07 | 1.300E-05 | 1.400E-05 |
| 5471 | 274 | V274L | A | 1.4e-05 | 6.1e-06 | 5.5e-06 | 8.533E-06 | 4.744E-06 | 5.500E-06 | 1.400E-05 |
| 5472 | 274 | V274K | A | 6e-06 | 1.3e-05 | 7.1e-06 | 8.700E-06 | 3.764E-06 | 6.000E-06 | 1.300E-05 |
| 5473 | 274 | V274M | A | 5.7e-06 | 7.6e-06 | 1.2e-05 | 8.433E-06 | 3.232E-06 | 5.700E-06 | 1.200E-05 |
| 5474 | 274 | V274F | A | 5.2e-06 | 9.7e-06 | 6.9e-06 | 7.267E-06 | 2.272E-06 | 5.200E-06 | 9.700E-06 |
| 5475 | 274 | V274P | A | 5.7e-06 | 1.4e-05 | 1.1e-05 | 1.023E-05 | 4.203E-06 | 5.700E-06 | 1.400E-05 |
| 5476 | 274 | V274S | A | 5.8e-06 | 9e-06 | 1.2e-05 | 8.933E-06 | 3.101E-06 | 5.800E-06 | 1.200E-05 |
| 5477 | 274 | V274T | A | 9.3e-06 | 6.8e-06 | 1e-05 | 8.700E-06 | 1.682E-06 | 6.800E-06 | 1.000E-05 |
| 5478 | 274 | V274W | A | 6.7e-06 | 1.2e-05 | 8.7e-06 | 9.133E-06 | 2.676E-06 | 6.700E-06 | 1.200E-05 |
| 5479 | 274 | V274Y | A | 1.1e-05 | 6.4e-06 | 5.3e-06 | 7.567E-06 | 3.024E-06 | 5.300E-06 | 1.100E-05 |
| 5480 | 274 | V274V | A | 8.4e-06 | 8.8e-06 | 6.3e-06 | 7.833E-06 | 1.343E-06 | 6.300E-06 | 8.800E-06 |
| 5481 | 275 | T275A | A | 1.1e-05 | 1.4e-05 | 6.8e-06 | 1.060E-05 | 3.617E-06 | 6.800E-06 | 1.400E-05 |
| 5482 | 275 | T275R | A | 7.5e-06 | 9.8e-06 | 8.6e-06 | 8.633E-06 | 1.150E-06 | 7.500E-06 | 9.800E-06 |
| 5483 | 275 | T275N | A | 1.1e-05 | 6.7e-06 | 6e-06 | 7.900E-06 | 2.707E-06 | 6.000E-06 | 1.100E-05 |
| 5484 | 275 | T275D | A | 5.5e-06 | 8.1e-06 | 9.1e-06 | 7.567E-06 | 1.858E-06 | 5.500E-06 | 9.100E-06 |
| 5485 | 275 | T275C | A | 6.9e-06 | 9.4e-06 | 1.1e-05 | 9.100E-06 | 2.066E-06 | 6.900E-06 | 1.100E-05 |
| 5486 | 275 | T275E | A | 1.3e-05 | 1.1e-05 | 7.9e-06 | 1.063E-05 | 2.570E-06 | 7.900E-06 | 1.300E-05 |
| 5487 | 275 | T275Q | A | 8.8e-06 | 4.7e-06 | 9.4e-06 | 7.633E-06 | 2.558E-06 | 4.700E-06 | 9.400E-06 |
| 5488 | 275 | T275G | A | 6.6e-06 | 1.1e-05 | 9.9e-06 | 9.167E-06 | 2.290E-06 | 6.600E-06 | 1.100E-05 |
| 5489 | 275 | T275H | A | 8.6e-06 | 8.6e-06 | 1.1e-05 | 9.400E-06 | 1.386E-06 | 8.600E-06 | 1.100E-05 |
| 5490 | 275 | T275I | A | 9.5e-06 | 7.2e-06 | 6.4e-06 | 7.700E-06 | 1.609E-06 | 6.400E-06 | 9.500E-06 |
| 5491 | 275 | T275L | A | 5.1e-06 | 6.7e-06 | 8.4e-06 | 6.733E-06 | 1.650E-06 | 5.100E-06 | 8.400E-06 |
| 5492 | 275 | T275K | A | 8.8e-06 | 7.8e-06 | 5.9e-06 | 7.500E-06 | 1.473E-06 | 5.900E-06 | 8.800E-06 |
| 5493 | 275 | T275M | A | 9e-06 | 8.4e-06 | 9.6e-06 | 9.000E-06 | 6.000E-07 | 8.400E-06 | 9.600E-06 |
| 5494 | 275 | T275F | A | 5.3e-06 | 9e-06 | 5.9e-06 | 6.733E-06 | 1.986E-06 | 5.300E-06 | 9.000E-06 |
| 5495 | 275 | T275P | A | 4.8e-06 | 8.1e-06 | 1e-05 | 7.633E-06 | 2.631E-06 | 4.800E-06 | 1.000E-05 |
| 5496 | 275 | T275S | A | 1.4e-05 | 4.2e-06 | 1.2e-05 | 1.007E-05 | 5.178E-06 | 4.200E-06 | 1.400E-05 |
| 5497 | 275 | T275T | A | 4.7e-06 | 1.1e-05 | 5.5e-06 | 7.067E-06 | 3.430E-06 | 4.700E-06 | 1.100E-05 |
| 5498 | 275 | T275W | A | 7.6e-06 | 5.1e-06 | 8.5e-06 | 7.067E-06 | 1.762E-06 | 5.100E-06 | 8.500E-06 |
| 5499 | 275 | T275Y | A | 6.2e-06 | 6.8e-06 | 1.1e-05 | 8.000E-06 | 2.615E-06 | 6.200E-06 | 1.100E-05 |
| 5500 | 275 | T275V | A | 9.7e-06 | 6.6e-06 | 9.9e-06 | 8.733E-06 | 1.850E-06 | 6.600E-06 | 9.900E-06 |
| 5501 | 276 | A276A | A | 6.8e-06 | 4.6e-06 | 1.1e-05 | 7.467E-06 | 3.252E-06 | 4.600E-06 | 1.100E-05 |
| 5502 | 276 | A276R | A | 6.9e-06 | 8.3e-06 | 6.3e-06 | 7.167E-06 | 1.026E-06 | 6.300E-06 | 8.300E-06 |
| 5503 | 276 | A276N | A | 6.7e-06 | 1.4e-05 | 8.3e-06 | 9.667E-06 | 3.837E-06 | 6.700E-06 | 1.400E-05 |
| 5504 | 276 | A276D | A | 8.6e-06 | 1e-05 | 1e-05 | 9.533E-06 | 8.083E-07 | 8.600E-06 | 1.000E-05 |
| 5505 | 276 | A276C | A | 1.1e-05 | 6.2e-06 | 4.1e-06 | 7.100E-06 | 3.537E-06 | 4.100E-06 | 1.100E-05 |
| 5506 | 276 | A276E | A | 9.6e-06 | 7.5e-06 | 1.2e-05 | 9.700E-06 | 2.252E-06 | 7.500E-06 | 1.200E-05 |
| 5507 | 276 | A276Q | A | 9e-06 | 7e-06 | 8.6e-06 | 8.200E-06 | 1.058E-06 | 7.000E-06 | 9.000E-06 |
| 5508 | 276 | A276G | A | 1e-05 | 5.4e-06 | 8.5e-06 | 7.967E-06 | 2.346E-06 | 5.400E-06 | 1.000E-05 |
| 5509 | 276 | A276H | A | 7.2e-06 | 6.4e-06 | 6.5e-06 | 6.700E-06 | 4.359E-07 | 6.400E-06 | 7.200E-06 |
| 5510 | 276 | A276I | A | 7e-06 | 9.7e-06 | 1e-05 | 8.900E-06 | 1.652E-06 | 7.000E-06 | 1.000E-05 |
| 5511 | 276 | A276L | A | 8.3e-06 | 5e-06 | 6.7e-06 | 6.667E-06 | 1.650E-06 | 5.000E-06 | 8.300E-06 |
| 5512 | 276 | A276K | A | 8.5e-06 | 1.1e-05 | 9e-06 | 9.500E-06 | 1.323E-06 | 8.500E-06 | 1.100E-05 |
| 5513 | 276 | A276M | A | 7.4e-06 | 5.5e-06 | 6.8e-06 | 6.567E-06 | 9.713E-07 | 5.500E-06 | 7.400E-06 |
| 5514 | 276 | A276F | A | 9.4e-06 | 1e-05 | 6.9e-06 | 8.767E-06 | 1.644E-06 | 6.900E-06 | 1.000E-05 |
| 5515 | 276 | A276P | A | 8.8e-06 | 1.2e-05 | 6.8e-06 | 9.200E-06 | 2.623E-06 | 6.800E-06 | 1.200E-05 |
| 5516 | 276 | A276S | A | 4.8e-06 | 1e-05 | 8.7e-06 | 7.833E-06 | 2.706E-06 | 4.800E-06 | 1.000E-05 |
| 5517 | 276 | A276T | A | 4.9e-06 | 8.7e-06 | 5.9e-06 | 6.500E-06 | 1.970E-06 | 4.900E-06 | 8.700E-06 |
| 5518 | 276 | A276W | A | 7e-06 | 6.5e-06 | 1.3e-05 | 8.833E-06 | 3.617E-06 | 6.500E-06 | 1.300E-05 |
| 5519 | 276 | A276Y | A | 7.7e-06 | 7.9e-06 | 9.2e-06 | 8.267E-06 | 8.145E-07 | 7.700E-06 | 9.200E-06 |
| 5520 | 276 | A276V | A | 5.5e-06 | 6.1e-06 | 1.3e-05 | 8.200E-06 | 4.168E-06 | 5.500E-06 | 1.300E-05 |
| 5521 | 277 | C277A | A | 5.1e-06 | 1.2e-05 | 6.5e-06 | 7.867E-06 | 3.647E-06 | 5.100E-06 | 1.200E-05 |
| 5522 | 277 | C277R | A | 4.4e-06 | 5.9e-06 | 1.1e-05 | 7.100E-06 | 3.460E-06 | 4.400E-06 | 1.100E-05 |
| 5523 | 277 | C277N | A | 1e-05 | 9.2e-06 | 7.4e-06 | 8.867E-06 | 1.332E-06 | 7.400E-06 | 1.000E-05 |
| 5524 | 277 | C277D | A | 7.7e-06 | 8.7e-06 | 4.9e-06 | 7.100E-06 | 1.970E-06 | 4.900E-06 | 8.700E-06 |

| | | | | | | | | | | |
|---|---|---|---|---|---|---|---|---|---|---|
| 5525 | 277 | C277C | A | 9.5e-06 | 9e-06 | 8.2e-06 | 8.900E-06 | 6.557E-07 | 8.200E-06 | 9.500E-06 |
| 5526 | 277 | C277E | A | 7.5e-06 | 8.2e-06 | 6.2e-06 | 7.300E-06 | 1.015E-06 | 6.200E-06 | 8.200E-06 |
| 5527 | 277 | C277Q | A | 3.2e-06 | 9.6e-06 | 4.9e-06 | 5.900E-06 | 3.315E-06 | 3.200E-06 | 9.600E-06 |
| 5528 | 277 | C277G | A | 9.6e-06 | 7.7e-06 | 9.6e-06 | 8.967E-06 | 1.097E-06 | 7.700E-06 | 9.600E-06 |
| 5529 | 277 | C277H | A | 8.6e-06 | 9.8e-06 | 8.1e-06 | 8.833E-06 | 8.737E-07 | 8.100E-06 | 9.800E-06 |
| 5530 | 277 | C277I | A | 6.6e-06 | 7.2e-06 | 5.2e-06 | 6.333E-06 | 1.026E-06 | 5.200E-06 | 7.200E-06 |
| 5531 | 277 | C277L | A | 7.3e-06 | 7.6e-06 | 5.5e-06 | 6.800E-06 | 1.136E-06 | 5.500E-06 | 7.600E-06 |
| 5532 | 277 | C277K | A | 1.2e-05 | 9.1e-06 | 7.4e-06 | 9.500E-06 | 2.326E-06 | 7.400E-06 | 1.200E-05 |
| 5533 | 277 | C277M | A | 1.2e-05 | 4.8e-06 | 4.9e-06 | 7.233E-06 | 4.128E-06 | 4.800E-06 | 1.200E-05 |
| 5534 | 277 | C277F | A | 7.3e-06 | 1e-05 | 9.2e-06 | 8.833E-06 | 1.387E-06 | 7.300E-06 | 1.000E-05 |
| 5535 | 277 | C277P | A | 7e-06 | 1.1e-05 | 6.1e-06 | 8.033E-06 | 2.608E-06 | 6.100E-06 | 1.100E-05 |
| 5536 | 277 | C277S | A | 5.4e-06 | 5.3e-06 | 6.9e-06 | 5.867E-06 | 8.963E-07 | 5.300E-06 | 6.900E-06 |
| 5537 | 277 | C277T | A | 9.4e-06 | 6.7e-06 | 7.8e-06 | 7.967E-06 | 1.358E-06 | 6.700E-06 | 9.400E-06 |
| 5538 | 277 | C277W | A | 7.9e-06 | 4.2e-06 | 6.5e-06 | 6.200E-06 | 1.868E-06 | 4.200E-06 | 7.900E-06 |
| 5539 | 277 | C277Y | A | 6.9e-06 | 1.4e-05 | 8.3e-06 | 9.733E-06 | 3.761E-06 | 6.900E-06 | 1.400E-05 |
| 5540 | 277 | C277V | A | 6.7e-06 | 1e-05 | 8.8e-06 | 8.500E-06 | 1.670E-06 | 6.700E-06 | 1.000E-05 |
| 5541 | 278 | P278A | A | 7.3e-06 | 5e-06 | 8.5e-06 | 6.933E-06 | 1.779E-06 | 5.000E-06 | 8.500E-06 |
| 5542 | 278 | P278R | A | 4.3e-06 | 7.4e-06 | 1.2e-05 | 7.900E-06 | 3.874E-06 | 4.300E-06 | 1.200E-05 |
| 5543 | 278 | P278N | A | 1e-05 | 4.3e-06 | 9.1e-06 | 7.800E-06 | 3.064E-06 | 4.300E-06 | 1.000E-05 |
| 5544 | 278 | P278D | A | 4.3e-06 | 5.9e-06 | 8.4e-06 | 6.200E-06 | 2.066E-06 | 4.300E-06 | 8.400E-06 |
| 5545 | 278 | P278C | A | 7.6e-06 | 9.5e-06 | 7.6e-06 | 8.233E-06 | 1.097E-06 | 7.600E-06 | 9.500E-06 |
| 5546 | 278 | P278E | A | 1e-05 | 1.4e-05 | 1.4e-05 | 1.267E-05 | 2.309E-06 | 1.000E-05 | 1.400E-05 |
| 5547 | 278 | P278Q | A | 8.8e-06 | 5.2e-06 | 7.2e-06 | 7.067E-06 | 1.804E-06 | 5.200E-06 | 8.800E-06 |
| 5548 | 278 | P278G | A | 1.4e-05 | 1.8e-05 | 6.7e-06 | 1.290E-05 | 5.730E-06 | 6.700E-06 | 1.800E-05 |
| 5549 | 278 | P278H | A | 7e-06 | 9.3e-06 | 7.7e-06 | 8.000E-06 | 1.179E-06 | 7.000E-06 | 9.300E-06 |
| 5550 | 278 | P278I | A | 8.2e-06 | 6.5e-06 | 1e-05 | 8.233E-06 | 1.750E-06 | 6.500E-06 | 1.000E-05 |
| 5551 | 278 | P278L | A | 1.4e-05 | 1.1e-05 | 6.1e-06 | 1.037E-05 | 3.988E-06 | 6.100E-06 | 1.400E-05 |
| 5552 | 278 | P278K | A | 6.7e-06 | 1.2e-05 | 6.4e-06 | 8.367E-06 | 3.150E-06 | 6.400E-06 | 1.200E-05 |
| 5553 | 278 | P278M | A | 6.7e-06 | 9.5e-06 | 9.2e-06 | 8.467E-06 | 1.537E-06 | 6.700E-06 | 9.500E-06 |
| 5554 | 278 | P278F | A | 6.4e-06 | 8.5e-06 | 9.9e-06 | 8.267E-06 | 1.762E-06 | 6.400E-06 | 9.900E-06 |
| 5555 | 278 | P278P | A | 9.4e-06 | 8.5e-06 | 4.7e-06 | 7.533E-06 | 2.495E-06 | 4.700E-06 | 9.400E-06 |
| 5556 | 278 | P278S | A | 8.3e-06 | 6.5e-06 | 5.7e-06 | 6.833E-06 | 1.332E-06 | 5.700E-06 | 8.300E-06 |
| 5557 | 278 | P278T | A | 9e-06 | 4.1e-06 | 9.5e-06 | 7.533E-06 | 2.984E-06 | 4.100E-06 | 9.500E-06 |
| 5558 | 278 | P278W | A | 8.3e-06 | 5.3e-06 | 9e-06 | 7.533E-06 | 1.966E-06 | 5.300E-06 | 9.000E-06 |
| 5559 | 278 | P278Y | A | 9.3e-06 | 9.7e-06 | 1.1e-05 | 1.000E-05 | 8.888E-07 | 9.300E-06 | 1.100E-05 |
| 5560 | 278 | P278V | A | 5.1e-06 | 6.6e-06 | 8.7e-06 | 6.800E-06 | 1.808E-06 | 5.100E-06 | 8.700E-06 |
| 5561 | 279 | Y279A | A | 7.8e-06 | 9.9e-06 | 9.5e-06 | 9.067E-06 | 1.115E-06 | 7.800E-06 | 9.900E-06 |
| 5562 | 279 | Y279R | A | 7.6e-06 | 6.4e-06 | 4.7e-06 | 6.233E-06 | 1.457E-06 | 4.700E-06 | 7.600E-06 |
| 5563 | 279 | Y279N | A | 4.5e-06 | 9.3e-06 | 1.1e-05 | 8.267E-06 | 3.371E-06 | 4.500E-06 | 1.100E-05 |
| 5564 | 279 | Y279D | A | 7.3e-06 | 9.8e-06 | 1.1e-05 | 9.367E-06 | 1.888E-06 | 7.300E-06 | 1.100E-05 |
| 5565 | 279 | Y279C | A | 8.4e-06 | 1e-05 | 7.9e-06 | 8.767E-06 | 1.097E-06 | 7.900E-06 | 1.000E-05 |
| 5566 | 279 | Y279E | A | 8.9e-06 | 8.8e-06 | 1.1e-05 | 9.567E-06 | 1.242E-06 | 8.800E-06 | 1.100E-05 |
| 5567 | 279 | Y279Q | A | 5.5e-06 | 8.6e-06 | 1e-05 | 8.033E-06 | 2.303E-06 | 5.500E-06 | 1.000E-05 |
| 5568 | 279 | Y279G | A | 6.2e-06 | 9e-06 | 4.6e-06 | 6.600E-06 | 2.227E-06 | 4.600E-06 | 9.000E-06 |
| 5569 | 279 | Y279H | A | 9.7e-06 | 5.3e-06 | 8.7e-06 | 7.900E-06 | 2.307E-06 | 5.300E-06 | 9.700E-06 |
| 5570 | 279 | Y279I | A | 1e-05 | 7e-06 | 6.5e-06 | 7.833E-06 | 1.893E-06 | 6.500E-06 | 1.000E-05 |
| 5571 | 279 | Y279L | A | 7.8e-06 | 8.2e-06 | 6e-06 | 7.333E-06 | 1.172E-06 | 6.000E-06 | 8.200E-06 |
| 5572 | 279 | Y279K | A | 7.2e-06 | 9e-06 | 5.9e-06 | 7.367E-06 | 1.557E-06 | 5.900E-06 | 9.000E-06 |
| 5573 | 279 | Y279M | A | 1.1e-05 | 6.8e-06 | 6.3e-06 | 8.033E-06 | 2.581E-06 | 6.300E-06 | 1.100E-05 |
| 5574 | 279 | Y279F | A | 9.1e-06 | 6.3e-06 | 9.8e-06 | 8.400E-06 | 1.852E-06 | 6.300E-06 | 9.800E-06 |
| 5575 | 279 | Y279P | A | 6.5e-06 | 8.3e-06 | 1.2e-05 | 8.933E-06 | 2.804E-06 | 6.500E-06 | 1.200E-05 |
| 5576 | 279 | Y279S | A | 1.4e-05 | 6.9e-06 | 6.6e-06 | 9.167E-06 | 4.188E-06 | 6.600E-06 | 1.400E-05 |
| 5577 | 279 | Y279T | A | 1.4e-05 | 6.7e-06 | 6.5e-06 | 9.067E-06 | 4.274E-06 | 6.500E-06 | 1.400E-05 |
| 5578 | 279 | Y279W | A | 9.5e-06 | 1.7e-05 | 8.3e-06 | 1.160E-05 | 4.715E-06 | 8.300E-06 | 1.700E-05 |
| 5579 | 279 | Y279Y | A | 9.7e-06 | 7.2e-06 | 9.8e-06 | 8.900E-06 | 1.473E-06 | 7.200E-06 | 9.800E-06 |
| 5580 | 279 | Y279V | A | 9.1e-06 | 6.3e-06 | 5.5e-06 | 6.967E-06 | 1.890E-06 | 5.500E-06 | 9.100E-06 |
| 5581 | 280 | N280A | A | 1.3e-05 | 8e-06 | 9.7e-06 | 1.023E-05 | 2.542E-06 | 8.000E-06 | 1.300E-05 |
| 5582 | 280 | N280R | A | 1.4e-05 | 6e-06 | 6.2e-06 | 8.733E-06 | 4.562E-06 | 6.000E-06 | 1.400E-05 |
| 5583 | 280 | N280N | A | 1e-05 | 6.3e-06 | 1.3e-05 | 9.767E-06 | 3.356E-06 | 6.300E-06 | 1.300E-05 |
| 5584 | 280 | N280D | A | 8.3e-06 | 9e-06 | 4.9e-06 | 7.400E-06 | 2.193E-06 | 4.900E-06 | 9.000E-06 |
| 5585 | 280 | N280C | A | 7.1e-06 | 1.3e-05 | 1e-05 | 1.003E-05 | 2.950E-06 | 7.100E-06 | 1.300E-05 |
| 5586 | 280 | N280E | A | 9.9e-06 | 6.9e-06 | 7.5e-06 | 8.100E-06 | 1.587E-06 | 6.900E-06 | 9.900E-06 |
| 5587 | 280 | N280Q | A | 6.5e-06 | 8.8e-06 | 7.5e-06 | 7.600E-06 | 1.153E-06 | 6.500E-06 | 8.800E-06 |
| 5588 | 280 | N280G | A | 6e-06 | 6.7e-06 | 8e-06 | 6.900E-06 | 1.015E-06 | 6.000E-06 | 8.000E-06 |
| 5589 | 280 | N280H | A | 7.7e-06 | 7.8e-06 | 6.4e-06 | 7.300E-06 | 7.810E-07 | 6.400E-06 | 7.800E-06 |
| 5590 | 280 | N280I | A | 8.2e-06 | 6.2e-06 | 4.6e-06 | 6.333E-06 | 1.804E-06 | 4.600E-06 | 8.200E-06 |
| 5591 | 280 | N280L | A | 4.9e-06 | 7.6e-06 | 9e-06 | 7.167E-06 | 2.084E-06 | 4.900E-06 | 9.000E-06 |
| 5592 | 280 | N280K | A | 6.7e-06 | 4e-06 | 8.6e-06 | 6.433E-06 | 2.312E-06 | 4.000E-06 | 8.600E-06 |
| 5593 | 280 | N280M | A | 1.3e-05 | 4.8e-06 | 8.8e-06 | 8.867E-06 | 4.100E-06 | 4.800E-06 | 1.300E-05 |
| 5594 | 280 | N280F | A | 7.9e-06 | 1.3e-05 | 6.1e-06 | 9.000E-06 | 3.579E-06 | 6.100E-06 | 1.300E-05 |
| 5595 | 280 | N280P | A | 7.5e-06 | 1.3e-05 | 1e-05 | 1.017E-05 | 2.754E-06 | 7.500E-06 | 1.300E-05 |
| 5596 | 280 | N280S | A | 7e-06 | 8.6e-06 | 6.2e-06 | 7.267E-06 | 1.222E-06 | 6.200E-06 | 8.600E-06 |
| 5597 | 280 | N280T | A | 1.2e-05 | 9.3e-06 | 8e-06 | 9.767E-06 | 2.040E-06 | 8.000E-06 | 1.200E-05 |
| 5598 | 280 | N280W | A | 5.7e-06 | 7.9e-06 | 9.2e-06 | 7.600E-06 | 1.769E-06 | 5.700E-06 | 9.200E-06 |
| 5599 | 280 | N280Y | A | 1.1e-05 | 8.8e-06 | 1.9e-05 | 1.293E-05 | 5.368E-06 | 8.800E-06 | 1.900E-05 |
| 5600 | 280 | N280V | A | 7.6e-06 | 7.2e-06 | 1.4e-05 | 9.600E-06 | 3.816E-06 | 7.200E-06 | 1.400E-05 |
| 5601 | 281 | Y281A | A | 7e-06 | 4.6e-06 | 4.6e-06 | 5.400E-06 | 1.386E-06 | 4.600E-06 | 7.000E-06 |
| 5602 | 281 | Y281R | A | 1.1e-05 | 9.3e-06 | 6e-06 | 8.767E-06 | 2.542E-06 | 6.000E-06 | 1.100E-05 |
| 5603 | 281 | Y281N | A | 7.2e-06 | 8.9e-06 | 6.8e-06 | 7.633E-06 | 1.115E-06 | 6.800E-06 | 8.900E-06 |
| 5604 | 281 | Y281D | A | 1.1e-05 | 1.2e-05 | 1.4e-05 | 1.233E-05 | 1.528E-06 | 1.100E-05 | 1.400E-05 |
| 5605 | 281 | Y281C | A | 6e-06 | 4.7e-06 | 8.5e-06 | 6.400E-06 | 1.931E-06 | 4.700E-06 | 8.500E-06 |
| 5606 | 281 | Y281E | A | 9.7e-06 | 5.7e-06 | 9.4e-06 | 8.267E-06 | 2.228E-06 | 5.700E-06 | 9.700E-06 |
| 5607 | 281 | Y281Q | A | 8.8e-06 | 7.2e-06 | 1.9e-05 | 1.167E-05 | 6.401E-06 | 7.200E-06 | 1.900E-05 |
| 5608 | 281 | Y281G | A | 5.7e-06 | 1e-05 | 5e-06 | 6.900E-06 | 2.707E-06 | 5.000E-06 | 1.000E-05 |
| 5609 | 281 | Y281H | A | 5.4e-06 | 1.2e-05 | 5.6e-06 | 7.667E-06 | 3.754E-06 | 5.400E-06 | 1.200E-05 |

| | | | | | | | | | | |
|---|---|---|---|---|---|---|---|---|---|---|
| 5610 | 281 | Y281I | A | 6.6e-06 | 9.7e-06 | 9.3e-06 | 8.533E-06 | 1.686E-06 | 6.600E-06 | 9.700E-06 |
| 5611 | 281 | Y281L | A | 5.1e-06 | 5.9e-06 | 7.9e-06 | 6.300E-06 | 1.442E-06 | 5.100E-06 | 7.900E-06 |
| 5612 | 281 | Y281K | A | 9e-06 | 9e-06 | 9.6e-06 | 9.200E-06 | 3.464E-07 | 9.000E-06 | 9.600E-06 |
| 5613 | 281 | Y281M | A | 8.4e-06 | 1.1e-05 | 1.3e-05 | 1.080E-05 | 2.307E-06 | 8.400E-06 | 1.300E-05 |
| 5614 | 281 | Y281F | A | 8.7e-06 | 8.9e-06 | 1.2e-05 | 9.867E-06 | 1.850E-06 | 8.700E-06 | 1.200E-05 |
| 5615 | 281 | Y281P | A | 7.5e-06 | 6.3e-06 | 7e-06 | 6.933E-06 | 6.028E-07 | 6.300E-06 | 7.500E-06 |
| 5616 | 281 | Y281S | A | 7.1e-06 | 1.1e-05 | 6.9e-06 | 8.333E-06 | 2.312E-06 | 6.900E-06 | 1.100E-05 |
| 5617 | 281 | Y281T | A | 5.6e-06 | 1.4e-05 | 7.6e-06 | 9.067E-06 | 4.388E-06 | 5.600E-06 | 1.400E-05 |
| 5618 | 281 | Y281W | A | 1.1e-05 | 7.5e-06 | 1e-05 | 9.500E-06 | 1.803E-06 | 7.500E-06 | 1.100E-05 |
| 5619 | 281 | Y281Y | A | 7.4e-06 | 5.5e-06 | 1.1e-05 | 7.967E-06 | 2.793E-06 | 5.500E-06 | 1.100E-05 |
| 5620 | 281 | Y281V | A | 6.2e-06 | 7.6e-06 | 6e-06 | 6.600E-06 | 8.718E-07 | 6.000E-06 | 7.600E-06 |
| 5621 | 282 | L282A | A | 9.5e-06 | 8.7e-06 | 7.1e-06 | 8.433E-06 | 1.222E-06 | 7.100E-06 | 9.500E-06 |
| 5622 | 282 | L282R | A | 5.3e-06 | 7.7e-06 | 9.1e-06 | 7.367E-06 | 1.922E-06 | 5.300E-06 | 9.100E-06 |
| 5623 | 282 | L282N | A | 1.2e-05 | 6.1e-06 | 5.4e-06 | 7.833E-06 | 3.625E-06 | 5.400E-06 | 1.200E-05 |
| 5624 | 282 | L282D | A | 5.1e-06 | 9.6e-06 | 6.7e-06 | 7.133E-06 | 2.281E-06 | 5.100E-06 | 9.600E-06 |
| 5625 | 282 | L282C | A | 7.1e-06 | 5.9e-06 | 6.8e-06 | 6.600E-06 | 6.245E-07 | 5.900E-06 | 7.100E-06 |
| 5626 | 282 | L282E | A | 6.4e-06 | 1e-05 | 9.6e-06 | 8.667E-06 | 1.973E-06 | 6.400E-06 | 1.000E-05 |
| 5627 | 282 | L282Q | A | 6.7e-06 | 6.1e-06 | 6e-06 | 6.267E-06 | 3.786E-07 | 6.000E-06 | 6.700E-06 |
| 5628 | 282 | L282G | A | 8.9e-06 | 8e-06 | 8.7e-06 | 8.533E-06 | 4.726E-07 | 8.000E-06 | 8.900E-06 |
| 5629 | 282 | L282H | A | 6.4e-06 | 9e-06 | 5.5e-06 | 6.967E-06 | 1.818E-06 | 5.500E-06 | 9.000E-06 |
| 5630 | 282 | L282I | A | 4.7e-06 | 9.3e-06 | 1e-05 | 8.000E-06 | 2.879E-06 | 4.700E-06 | 1.000E-05 |
| 5631 | 282 | L282L | A | 1.6e-05 | 1.3e-05 | 8.5e-06 | 1.250E-05 | 3.775E-06 | 8.500E-06 | 1.600E-05 |
| 5632 | 282 | L282K | A | 1.2e-05 | 7.4e-06 | 6.8e-06 | 8.733E-06 | 2.845E-06 | 6.800E-06 | 1.200E-05 |
| 5633 | 282 | L282M | A | 3e-06 | 6.3e-06 | 9.8e-06 | 6.367E-06 | 3.400E-06 | 3.000E-06 | 9.800E-06 |
| 5634 | 282 | L282F | A | 1.4e-05 | 1.4e-05 | 9.9e-06 | 1.263E-05 | 2.367E-06 | 9.900E-06 | 1.400E-05 |
| 5635 | 282 | L282P | A | 6.5e-06 | 7.2e-06 | 8.3e-06 | 7.333E-06 | 9.074E-07 | 6.500E-06 | 8.300E-06 |
| 5636 | 282 | L282S | A | 5.5e-06 | 7.1e-06 | 1e-05 | 7.533E-06 | 2.281E-06 | 5.500E-06 | 1.000E-05 |
| 5637 | 282 | L282T | A | 5.9e-06 | 9.7e-06 | 1e-05 | 8.533E-06 | 2.285E-06 | 5.900E-06 | 1.000E-05 |
| 5638 | 282 | L282W | A | 6.2e-06 | 6.6e-06 | 8.1e-06 | 6.967E-06 | 1.002E-06 | 6.200E-06 | 8.100E-06 |
| 5639 | 282 | L282Y | A | 8.4e-06 | 8.2e-06 | 7.7e-06 | 8.100E-06 | 3.606E-07 | 7.700E-06 | 8.400E-06 |
| 5640 | 282 | L282V | A | 8.5e-06 | 1.4e-05 | 9.8e-06 | 1.077E-05 | 2.875E-06 | 8.500E-06 | 1.400E-05 |
| 5641 | 283 | S283A | A | 7.2e-06 | 1.1e-05 | 6.5e-06 | 8.233E-06 | 2.421E-06 | 6.500E-06 | 1.100E-05 |
| 5642 | 283 | S283R | A | 1.2e-05 | 6.2e-06 | 9.6e-06 | 9.267E-06 | 2.914E-06 | 6.200E-06 | 1.200E-05 |
| 5643 | 283 | S283N | A | 1e-05 | 6.6e-06 | 9.9e-06 | 8.833E-06 | 1.935E-06 | 6.600E-06 | 1.000E-05 |
| 5644 | 283 | S283D | A | 5.7e-06 | 6.3e-06 | 6.7e-06 | 6.233E-06 | 5.033E-07 | 5.700E-06 | 6.700E-06 |
| 5645 | 283 | S283C | A | 1.4e-05 | 5.2e-06 | 1.5e-05 | 1.140E-05 | 5.393E-06 | 5.200E-06 | 1.500E-05 |
| 5646 | 283 | S283E | A | 1.1e-05 | 9.1e-06 | 9.6e-06 | 9.900E-06 | 9.849E-07 | 9.100E-06 | 1.100E-05 |
| 5647 | 283 | S283Q | A | 8.8e-06 | 1.2e-05 | 6.6e-06 | 9.133E-06 | 2.715E-06 | 6.600E-06 | 1.200E-05 |
| 5648 | 283 | S283G | A | 8.7e-06 | 5.1e-06 | 4.6e-06 | 6.133E-06 | 2.237E-06 | 4.600E-06 | 8.700E-06 |
| 5649 | 283 | S283H | A | 1.3e-05 | 7.5e-06 | 5.8e-06 | 8.767E-06 | 3.763E-06 | 5.800E-06 | 1.300E-05 |
| 5650 | 283 | S283I | A | 8.4e-06 | 5.7e-06 | 7.5e-06 | 7.200E-06 | 1.375E-06 | 5.700E-06 | 8.400E-06 |
| 5651 | 283 | S283L | A | 8e-06 | 7.8e-06 | 6.8e-06 | 7.533E-06 | 6.429E-07 | 6.800E-06 | 8.000E-06 |
| 5652 | 283 | S283K | A | 9.5e-06 | 6.6e-06 | 1.1e-05 | 9.033E-06 | 2.237E-06 | 6.600E-06 | 1.100E-05 |
| 5653 | 283 | S283M | A | 8.5e-06 | 5.9e-06 | 7.6e-06 | 7.333E-06 | 1.320E-06 | 5.900E-06 | 8.500E-06 |
| 5654 | 283 | S283F | A | 1e-05 | 1e-05 | 7.9e-06 | 9.300E-06 | 1.212E-06 | 7.900E-06 | 1.000E-05 |
| 5655 | 283 | S283P | A | 6.5e-06 | 9.5e-06 | 7.8e-06 | 7.933E-06 | 1.504E-06 | 6.500E-06 | 9.500E-06 |
| 5656 | 283 | S283S | A | 5.6e-06 | 6.7e-06 | 5e-06 | 5.767E-06 | 8.622E-07 | 5.000E-06 | 6.700E-06 |
| 5657 | 283 | S283T | A | 5.9e-06 | 5.7e-06 | 7.5e-06 | 6.367E-06 | 9.866E-07 | 5.700E-06 | 7.500E-06 |
| 5658 | 283 | S283W | A | 8.1e-06 | 1e-05 | 6.8e-06 | 8.300E-06 | 1.609E-06 | 6.800E-06 | 1.000E-05 |
| 5659 | 283 | S283Y | A | 9e-06 | 9.7e-06 | 7.3e-06 | 8.667E-06 | 1.234E-06 | 7.300E-06 | 9.700E-06 |
| 5660 | 283 | S283V | A | 7.1e-06 | 7.2e-06 | 5.1e-06 | 6.467E-06 | 1.185E-06 | 5.100E-06 | 7.200E-06 |
| 5661 | 284 | T284A | A | 1.3e-05 | 9.3e-06 | 1e-05 | 1.077E-05 | 1.966E-06 | 9.300E-06 | 1.300E-05 |
| 5662 | 284 | T284R | A | 8.6e-06 | 1e-05 | 4.9e-06 | 7.833E-06 | 2.635E-06 | 4.900E-06 | 1.000E-05 |
| 5663 | 284 | T284N | A | 6.2e-06 | 1.3e-05 | 7.5e-06 | 8.900E-06 | 3.610E-06 | 6.200E-06 | 1.300E-05 |
| 5664 | 284 | T284D | A | 4.6e-06 | 8.1e-06 | 8.4e-06 | 7.033E-06 | 2.113E-06 | 4.600E-06 | 8.400E-06 |
| 5665 | 284 | T284C | A | 1.3e-05 | 6.4e-06 | 6.6e-06 | 8.667E-06 | 3.754E-06 | 6.400E-06 | 1.300E-05 |
| 5666 | 284 | T284E | A | 1.5e-05 | 8.2e-06 | 9.2e-06 | 1.080E-05 | 3.672E-06 | 8.200E-06 | 1.500E-05 |
| 5667 | 284 | T284Q | A | 5.1e-06 | 4.3e-06 | 5.3e-06 | 4.900E-06 | 5.292E-07 | 4.300E-06 | 5.300E-06 |
| 5668 | 284 | T284G | A | 6.6e-06 | 1.2e-05 | 4.4e-06 | 7.667E-06 | 3.911E-06 | 4.400E-06 | 1.200E-05 |
| 5669 | 284 | T284H | A | 6.7e-06 | 6.6e-06 | 9.9e-06 | 7.733E-06 | 1.877E-06 | 6.600E-06 | 9.900E-06 |
| 5670 | 284 | T284I | A | 9.9e-06 | 1.3e-05 | 6.9e-06 | 9.933E-06 | 3.050E-06 | 6.900E-06 | 1.300E-05 |
| 5671 | 284 | T284L | A | 1.1e-05 | 1e-05 | 5.6e-06 | 8.867E-06 | 2.873E-06 | 5.600E-06 | 1.100E-05 |
| 5672 | 284 | T284K | A | 1.3e-05 | 7.4e-06 | 7.6e-06 | 9.333E-06 | 3.177E-06 | 7.400E-06 | 1.300E-05 |
| 5673 | 284 | T284M | A | 9.7e-06 | 7.4e-06 | 8.9e-06 | 8.667E-06 | 1.168E-06 | 7.400E-06 | 9.700E-06 |
| 5674 | 284 | T284F | A | 8e-06 | 8.4e-06 | 6.5e-06 | 7.633E-06 | 1.002E-06 | 6.500E-06 | 8.400E-06 |
| 5675 | 284 | T284P | A | 6.9e-06 | 9.2e-06 | 9.2e-06 | 8.433E-06 | 1.328E-06 | 6.900E-06 | 9.200E-06 |
| 5676 | 284 | T284S | A | 4.3e-06 | 8e-06 | 5.4e-06 | 5.900E-06 | 1.900E-06 | 4.300E-06 | 8.000E-06 |
| 5677 | 284 | T284T | A | 1.2e-05 | 7.3e-06 | 7.3e-06 | 8.867E-06 | 2.714E-06 | 7.300E-06 | 1.200E-05 |
| 5678 | 284 | T284W | A | 8.7e-06 | 4.9e-06 | 7.3e-06 | 6.967E-06 | 1.922E-06 | 4.900E-06 | 8.700E-06 |
| 5679 | 284 | T284Y | A | 7.4e-06 | 1.2e-05 | 8.6e-06 | 9.333E-06 | 2.386E-06 | 7.400E-06 | 1.200E-05 |
| 5680 | 284 | T284V | A | 9.3e-06 | 8e-06 | 8.3e-06 | 8.533E-06 | 6.807E-07 | 8.000E-06 | 9.300E-06 |
| 5681 | 285 | D285A | A | 1.3e-05 | 1e-05 | 5.5e-06 | 9.500E-06 | 3.775E-06 | 5.500E-06 | 1.300E-05 |
| 5682 | 285 | D285R | A | 1e-05 | 1.3e-05 | 7.1e-06 | 1.003E-05 | 2.950E-06 | 7.100E-06 | 1.300E-05 |
| 5683 | 285 | D285N | A | 5.6e-06 | 6.6e-06 | 8.9e-06 | 7.033E-06 | 1.692E-06 | 5.600E-06 | 8.900E-06 |
| 5684 | 285 | D285D | A | 7.3e-06 | 5.3e-06 | 8.6e-06 | 7.067E-06 | 1.662E-06 | 5.300E-06 | 8.600E-06 |
| 5685 | 285 | D285C | A | 1.3e-05 | 1.2e-05 | 1.3e-05 | 1.267E-05 | 5.774E-07 | 1.200E-05 | 1.300E-05 |
| 5686 | 285 | D285E | A | 6.2e-06 | 8.9e-06 | 8.1e-06 | 7.733E-06 | 1.387E-06 | 6.200E-06 | 8.900E-06 |
| 5687 | 285 | D285Q | A | 8.8e-06 | 5.1e-06 | 7.4e-06 | 7.100E-06 | 1.868E-06 | 5.100E-06 | 8.800E-06 |
| 5688 | 285 | D285G | A | 7.7e-06 | 4.5e-06 | 5.1e-06 | 5.767E-06 | 1.701E-06 | 4.500E-06 | 7.700E-06 |
| 5689 | 285 | D285H | A | 9.1e-06 | 5.9e-06 | 8.2e-06 | 7.733E-06 | 1.650E-06 | 5.900E-06 | 9.100E-06 |
| 5690 | 285 | D285I | A | 4e-06 | 4.6e-06 | 1.7e-05 | 8.533E-06 | 7.338E-06 | 4.000E-06 | 1.700E-05 |
| 5691 | 285 | D285L | A | 8.8e-06 | 6.2e-06 | 7.9e-06 | 7.633E-06 | 1.320E-06 | 6.200E-06 | 8.800E-06 |
| 5692 | 285 | D285K | A | 9.3e-06 | 7.8e-06 | 9e-06 | 8.700E-06 | 7.937E-07 | 7.800E-06 | 9.300E-06 |
| 5693 | 285 | D285M | A | 8.5e-06 | 9.8e-06 | 9.4e-06 | 9.233E-06 | 6.658E-07 | 8.500E-06 | 9.800E-06 |
| 5694 | 285 | D285F | A | 8.8e-06 | 6.2e-06 | 8.8e-06 | 7.933E-06 | 1.501E-06 | 6.200E-06 | 8.800E-06 |

| | | | | | | | | | | |
|---|---|---|---|---|---|---|---|---|---|---|
| 5695 | 285 | D285P | A | 1e-05 | 9e-06 | 6.4e-06 | 8.467E-06 | 1.858E-06 | 6.400E-06 | 1.000E-05 |
| 5696 | 285 | D285S | A | 7.1e-06 | 1e-05 | 1e-05 | 9.033E-06 | 1.674E-06 | 7.100E-06 | 1.000E-05 |
| 5697 | 285 | D285T | A | 1.3e-05 | 7.2e-06 | 8.3e-06 | 9.500E-06 | 3.081E-06 | 7.200E-06 | 1.300E-05 |
| 5698 | 285 | D285W | A | 6.9e-06 | 8.9e-06 | 4.4e-06 | 6.733E-06 | 2.255E-06 | 4.400E-06 | 8.900E-06 |
| 5699 | 285 | D285Y | A | 6.8e-06 | 7.3e-06 | 7.6e-06 | 7.233E-06 | 4.041E-07 | 6.800E-06 | 7.600E-06 |
| 5700 | 285 | D285V | A | 3.6e-06 | 8e-06 | 8.7e-06 | 6.767E-06 | 2.765E-06 | 3.600E-06 | 8.700E-06 |
| 5701 | 286 | V286A | A | 6.6e-06 | 1e-05 | 1e-05 | 8.867E-06 | 1.963E-06 | 6.600E-06 | 1.000E-05 |
| 5702 | 286 | V286R | A | 8.1e-06 | 5.8e-06 | 8e-06 | 7.300E-06 | 1.300E-06 | 5.800E-06 | 8.100E-06 |
| 5703 | 286 | V286N | A | 7.1e-06 | 6.7e-06 | 9.4e-06 | 7.733E-06 | 1.457E-06 | 6.700E-06 | 9.400E-06 |
| 5704 | 286 | V286D | A | 7.7e-06 | 1.1e-05 | 6.9e-06 | 8.533E-06 | 2.173E-06 | 6.900E-06 | 1.100E-05 |
| 5705 | 286 | V286C | A | 1e-05 | 4.8e-06 | 8.9e-06 | 7.900E-06 | 2.740E-06 | 4.800E-06 | 1.000E-05 |
| 5706 | 286 | V286E | A | 6.3e-06 | 9.7e-06 | 9.4e-06 | 8.467E-06 | 1.882E-06 | 6.300E-06 | 9.700E-06 |
| 5707 | 286 | V286Q | A | 9.8e-06 | 8.5e-06 | 6.9e-06 | 8.400E-06 | 1.453E-06 | 6.900E-06 | 9.800E-06 |
| 5708 | 286 | V286G | A | 7e-06 | 4.3e-06 | 8.1e-06 | 6.467E-06 | 1.955E-06 | 4.300E-06 | 8.100E-06 |
| 5709 | 286 | V286H | A | 7e-06 | 9.9e-06 | 6.8e-06 | 7.900E-06 | 1.735E-06 | 6.800E-06 | 9.900E-06 |
| 5710 | 286 | V286I | A | 6.2e-06 | 6.1e-06 | 4.6e-06 | 5.633E-06 | 8.963E-07 | 4.600E-06 | 6.200E-06 |
| 5711 | 286 | V286L | A | 8.2e-06 | 8.6e-06 | 5.6e-06 | 7.467E-06 | 1.629E-06 | 5.600E-06 | 8.600E-06 |
| 5712 | 286 | V286K | A | 8.2e-06 | 6.3e-06 | 6.7e-06 | 7.067E-06 | 1.002E-06 | 6.300E-06 | 8.200E-06 |
| 5713 | 286 | V286M | A | 6.2e-06 | 4.4e-06 | 6.9e-06 | 5.833E-06 | 1.290E-06 | 4.400E-06 | 6.900E-06 |
| 5714 | 286 | V286F | A | 8.4e-06 | 8.7e-06 | 9.4e-06 | 8.833E-06 | 5.132E-07 | 8.400E-06 | 9.400E-06 |
| 5715 | 286 | V286P | A | 1.4e-05 | 6.3e-06 | 5.4e-06 | 8.567E-06 | 4.727E-06 | 5.400E-06 | 1.400E-05 |
| 5716 | 286 | V286S | A | 6.6e-06 | 7.1e-06 | 7.9e-06 | 7.200E-06 | 6.557E-07 | 6.600E-06 | 7.900E-06 |
| 5717 | 286 | V286T | A | 4.8e-06 | 7.9e-06 | 1.1e-05 | 7.900E-06 | 3.100E-06 | 4.800E-06 | 1.100E-05 |
| 5718 | 286 | V286W | A | 6.8e-06 | 7.7e-06 | 9.8e-06 | 8.100E-06 | 1.539E-06 | 6.800E-06 | 9.800E-06 |
| 5719 | 286 | V286Y | A | 9e-06 | 7.6e-06 | 9.1e-06 | 8.567E-06 | 8.386E-07 | 7.600E-06 | 9.100E-06 |
| 5720 | 286 | V286V | A | 7.3e-06 | 6.2e-06 | 1.4e-05 | 9.167E-06 | 4.222E-06 | 6.200E-06 | 1.400E-05 |
| 5721 | 287 | G287A | A | 5.1e-06 | 1.3e-05 | 1.3e-05 | 1.037E-05 | 4.561E-06 | 5.100E-06 | 1.300E-05 |
| 5722 | 287 | G287R | A | 9.5e-06 | 7.5e-06 | 4.8e-06 | 7.267E-06 | 2.359E-06 | 4.800E-06 | 9.500E-06 |
| 5723 | 287 | G287N | A | 8.3e-06 | 7.8e-06 | 8.8e-06 | 8.300E-06 | 5.000E-07 | 7.800E-06 | 8.800E-06 |
| 5724 | 287 | G287D | A | 4.9e-06 | 6.7e-06 | 8.4e-06 | 6.667E-06 | 1.750E-06 | 4.900E-06 | 8.400E-06 |
| 5725 | 287 | G287C | A | 1.3e-05 | 1.2e-05 | 9.1e-06 | 1.137E-05 | 2.026E-06 | 9.100E-06 | 1.300E-05 |
| 5726 | 287 | G287E | A | 1.1e-05 | 9.8e-06 | 4.6e-06 | 8.467E-06 | 3.402E-06 | 4.600E-06 | 1.100E-05 |
| 5727 | 287 | G287Q | A | 4.9e-06 | 5.5e-06 | 1.5e-05 | 8.467E-06 | 5.666E-06 | 4.900E-06 | 1.500E-05 |
| 5728 | 287 | G287G | A | 6.9e-06 | 9.6e-06 | 8.5e-06 | 8.333E-06 | 1.358E-06 | 6.900E-06 | 9.600E-06 |
| 5729 | 287 | G287H | A | 8.9e-06 | 3e-06 | 7.9e-06 | 6.600E-06 | 3.158E-06 | 3.000E-06 | 8.900E-06 |
| 5730 | 287 | G287I | A | 4.3e-06 | 1.1e-05 | 1.2e-05 | 9.100E-06 | 4.187E-06 | 4.300E-06 | 1.200E-05 |
| 5731 | 287 | G287L | A | 1.1e-05 | 8.8e-06 | 6.8e-06 | 8.867E-06 | 2.101E-06 | 6.800E-06 | 1.100E-05 |
| 5732 | 287 | G287K | A | 7e-06 | 5.3e-06 | 7.4e-06 | 6.567E-06 | 1.115E-06 | 5.300E-06 | 7.400E-06 |
| 5733 | 287 | G287M | A | 6.2e-06 | 7.2e-06 | 5.3e-06 | 6.233E-06 | 9.504E-07 | 5.300E-06 | 7.200E-06 |
| 5734 | 287 | G287F | A | 1.2e-05 | 7.7e-06 | 6e-06 | 8.567E-06 | 3.092E-06 | 6.000E-06 | 1.200E-05 |
| 5735 | 287 | G287P | A | 3.7e-06 | 8.9e-06 | 9.1e-06 | 7.233E-06 | 3.062E-06 | 3.700E-06 | 9.100E-06 |
| 5736 | 287 | G287S | A | 7.1e-06 | 8.2e-06 | 4.9e-06 | 6.733E-06 | 1.680E-06 | 4.900E-06 | 8.200E-06 |
| 5737 | 287 | G287T | A | 1.1e-05 | 4.4e-06 | 8.6e-06 | 8.000E-06 | 3.341E-06 | 4.400E-06 | 1.100E-05 |
| 5738 | 287 | G287W | A | 9.5e-06 | 7e-06 | 8.2e-06 | 8.233E-06 | 1.250E-06 | 7.000E-06 | 9.500E-06 |
| 5739 | 287 | G287Y | A | 7e-06 | 9.9e-06 | 8.9e-06 | 8.600E-06 | 1.473E-06 | 7.000E-06 | 9.900E-06 |
| 5740 | 287 | G287V | A | 5.5e-06 | 8.4e-06 | 1.3e-05 | 8.967E-06 | 3.782E-06 | 5.500E-06 | 1.300E-05 |
| 5741 | 288 | S288A | A | 5.2e-06 | 1.5e-05 | 9.4e-06 | 9.867E-06 | 4.917E-06 | 5.200E-06 | 1.500E-05 |
| 5742 | 288 | S288R | A | 3.8e-06 | 7.7e-06 | 6.4e-06 | 5.967E-06 | 1.986E-06 | 3.800E-06 | 7.700E-06 |
| 5743 | 288 | S288N | A | 1e-05 | 6.5e-06 | 3.1e-06 | 6.533E-06 | 3.450E-06 | 3.100E-06 | 1.000E-05 |
| 5744 | 288 | S288D | A | 3.8e-06 | 5.5e-06 | 7e-06 | 5.433E-06 | 1.601E-06 | 3.800E-06 | 7.000E-06 |
| 5745 | 288 | S288C | A | 6.2e-06 | 1.3e-05 | 1.5e-05 | 1.140E-05 | 4.613E-06 | 6.200E-06 | 1.500E-05 |
| 5746 | 288 | S288E | A | 6.7e-06 | 5.3e-06 | 6.9e-06 | 6.300E-06 | 8.718E-07 | 5.300E-06 | 6.900E-06 |
| 5747 | 288 | S288Q | A | 9.6e-06 | 6.8e-06 | 3.2e-06 | 6.533E-06 | 3.208E-06 | 3.200E-06 | 9.600E-06 |
| 5748 | 288 | S288G | A | 6.6e-06 | 1.5e-05 | 7e-06 | 9.533E-06 | 4.738E-06 | 6.600E-06 | 1.500E-05 |
| 5749 | 288 | S288H | A | 7.5e-06 | 6.2e-06 | 9.3e-06 | 7.667E-06 | 1.557E-06 | 6.200E-06 | 9.300E-06 |
| 5750 | 288 | S288I | A | 8.3e-06 | 1e-05 | 5.9e-06 | 8.067E-06 | 2.060E-06 | 5.900E-06 | 1.000E-05 |
| 5751 | 288 | S288L | A | 1.4e-05 | 1.2e-05 | 7.6e-06 | 1.120E-05 | 3.274E-06 | 7.600E-06 | 1.400E-05 |
| 5752 | 288 | S288K | A | 9.6e-06 | 1.7e-05 | 9.5e-06 | 1.203E-05 | 4.302E-06 | 9.500E-06 | 1.700E-05 |
| 5753 | 288 | S288M | A | 1.1e-05 | 5.6e-06 | 7.7e-06 | 8.100E-06 | 2.722E-06 | 5.600E-06 | 1.100E-05 |
| 5754 | 288 | S288F | A | 7.6e-06 | 9e-06 | 1e-05 | 8.867E-06 | 1.206E-06 | 7.600E-06 | 1.000E-05 |
| 5755 | 288 | S288P | A | 1.5e-05 | 1.3e-05 | 1.1e-05 | 1.300E-05 | 2.000E-06 | 1.100E-05 | 1.500E-05 |
| 5756 | 288 | S288S | A | 9.8e-06 | 1.3e-05 | 1.5e-05 | 1.260E-05 | 2.623E-06 | 9.800E-06 | 1.500E-05 |
| 5757 | 288 | S288T | A | 1.1e-05 | 4.5e-06 | 6.5e-06 | 7.333E-06 | 3.329E-06 | 4.500E-06 | 1.100E-05 |
| 5758 | 288 | S288W | A | 8.7e-06 | 9.3e-06 | 1.2e-05 | 1.000E-05 | 1.758E-06 | 8.700E-06 | 1.200E-05 |
| 5759 | 288 | S288Y | A | 9.1e-06 | 6.6e-06 | 5.8e-06 | 7.167E-06 | 1.721E-06 | 5.800E-06 | 9.100E-06 |
| 5760 | 288 | S288V | A | 7e-06 | 1e-05 | 6.7e-06 | 7.900E-06 | 1.825E-06 | 6.700E-06 | 1.000E-05 |
| 5761 | 289 | C289A | A | 7.1e-06 | 7.3e-06 | 1.1e-05 | 8.467E-06 | 2.196E-06 | 7.100E-06 | 1.100E-05 |
| 5762 | 289 | C289R | A | 1.8e-05 | 6.5e-06 | 7.9e-06 | 1.080E-05 | 6.275E-06 | 6.500E-06 | 1.800E-05 |
| 5763 | 289 | C289N | A | 5.5e-06 | 1.2e-05 | 6.2e-06 | 7.900E-06 | 3.568E-06 | 5.500E-06 | 1.200E-05 |
| 5764 | 289 | C289D | A | 8.8e-06 | 7.8e-06 | 1.1e-05 | 9.200E-06 | 1.637E-06 | 7.800E-06 | 1.100E-05 |
| 5765 | 289 | C289C | A | 5.4e-06 | 6.8e-06 | 9.1e-06 | 7.100E-06 | 1.868E-06 | 5.400E-06 | 9.100E-06 |
| 5766 | 289 | C289E | A | 5.7e-06 | 6.5e-06 | 6.8e-06 | 6.333E-06 | 5.686E-07 | 5.700E-06 | 6.800E-06 |
| 5767 | 289 | C289Q | A | 1.2e-05 | 6.1e-06 | 1.4e-05 | 1.070E-05 | 4.107E-06 | 6.100E-06 | 1.400E-05 |
| 5768 | 289 | C289G | A | 1.2e-05 | 6.6e-06 | 1.2e-05 | 1.020E-05 | 3.118E-06 | 6.600E-06 | 1.200E-05 |
| 5769 | 289 | C289H | A | 1.5e-05 | 7.4e-06 | 7.9e-06 | 1.010E-05 | 4.251E-06 | 7.400E-06 | 1.500E-05 |
| 5770 | 289 | C289I | A | 6.3e-06 | 5.3e-06 | 1.2e-05 | 7.867E-06 | 3.614E-06 | 5.300E-06 | 1.200E-05 |
| 5771 | 289 | C289L | A | 1.1e-05 | 1.1e-05 | 6.7e-06 | 9.567E-06 | 2.483E-06 | 6.700E-06 | 1.100E-05 |
| 5772 | 289 | C289K | A | 9.8e-06 | 3.8e-06 | 7.2e-06 | 6.933E-06 | 3.009E-06 | 3.800E-06 | 9.800E-06 |
| 5773 | 289 | C289M | A | 1e-05 | 1.1e-05 | 6.9e-06 | 9.300E-06 | 2.138E-06 | 6.900E-06 | 1.100E-05 |
| 5774 | 289 | C289F | A | 6.5e-06 | 1.5e-05 | 8.4e-06 | 9.967E-06 | 4.461E-06 | 6.500E-06 | 1.500E-05 |
| 5775 | 289 | C289P | A | 7.5e-06 | 1.1e-05 | 1.1e-05 | 9.833E-06 | 2.021E-06 | 7.500E-06 | 1.100E-05 |
| 5776 | 289 | C289S | A | 9.4e-06 | 1.1e-05 | 1.3e-05 | 1.113E-05 | 1.804E-06 | 9.400E-06 | 1.300E-05 |
| 5777 | 289 | C289T | A | 1e-05 | 5e-06 | 8.9e-06 | 7.967E-06 | 2.627E-06 | 5.000E-06 | 1.000E-05 |
| 5778 | 289 | C289W | A | 5.2e-06 | 8.6e-06 | 9.2e-06 | 7.667E-06 | 2.157E-06 | 5.200E-06 | 9.200E-06 |
| 5779 | 289 | C289Y | A | 5.9e-06 | 1.1e-05 | 7.2e-06 | 8.033E-06 | 2.650E-06 | 5.900E-06 | 1.100E-05 |

| | | | | | | | | | | |
|---|---|---|---|---|---|---|---|---|---|---|
| 5780 | 289 | C289V | A | 6.2e-06 | 5.5e-06 | 4e-06 | 5.233E-06 | 1.124E-06 | 4.000E-06 | 6.200E-06 |
| 5781 | 290 | T290A | A | 8.7e-06 | 9.8e-06 | 9.6e-06 | 9.367E-06 | 5.859E-07 | 8.700E-06 | 9.800E-06 |
| 5782 | 290 | T290R | A | 1.1e-05 | 7.3e-06 | 1.2e-05 | 1.010E-05 | 2.476E-06 | 7.300E-06 | 1.200E-05 |
| 5783 | 290 | T290N | A | 8.5e-06 | 4.7e-06 | 1.3e-05 | 8.733E-06 | 4.155E-06 | 4.700E-06 | 1.300E-05 |
| 5784 | 290 | T290D | A | 1.1e-05 | 7.4e-06 | 1.6e-05 | 1.147E-05 | 4.319E-06 | 7.400E-06 | 1.600E-05 |
| 5785 | 290 | T290C | A | 8.4e-06 | 9.1e-06 | 1.3e-05 | 1.017E-05 | 2.479E-06 | 8.400E-06 | 1.300E-05 |
| 5786 | 290 | T290E | A | 1e-05 | 4.6e-06 | 1e-05 | 8.200E-06 | 3.118E-06 | 4.600E-06 | 1.000E-05 |
| 5787 | 290 | T290Q | A | 8.1e-06 | 1.3e-05 | 6.6e-06 | 9.233E-06 | 3.347E-06 | 6.600E-06 | 1.300E-05 |
| 5788 | 290 | T290G | A | 9.2e-06 | 9.2e-06 | 8.8e-06 | 9.067E-06 | 2.309E-07 | 8.800E-06 | 9.200E-06 |
| 5789 | 290 | T290H | A | 4.4e-06 | 5.9e-06 | 6.9e-06 | 5.733E-06 | 1.258E-06 | 4.400E-06 | 6.900E-06 |
| 5790 | 290 | T290I | A | 6e-06 | 6.8e-06 | 8.7e-06 | 7.167E-06 | 1.387E-06 | 6.000E-06 | 8.700E-06 |
| 5791 | 290 | T290L | A | 8.1e-06 | 7e-06 | 7.4e-06 | 7.500E-06 | 5.568E-07 | 7.000E-06 | 8.100E-06 |
| 5792 | 290 | T290K | A | 5.7e-06 | 1.1e-05 | 8.4e-06 | 8.367E-06 | 2.650E-06 | 5.700E-06 | 1.100E-05 |
| 5793 | 290 | T290M | A | 3.5e-06 | 9.8e-06 | 4.5e-06 | 5.933E-06 | 3.386E-06 | 3.500E-06 | 9.800E-06 |
| 5794 | 290 | T290F | A | 9e-06 | 5.3e-06 | 8e-06 | 7.433E-06 | 1.914E-06 | 5.300E-06 | 9.000E-06 |
| 5795 | 290 | T290P | A | 9e-06 | 8.9e-06 | 1.1e-05 | 9.633E-06 | 1.185E-06 | 8.900E-06 | 1.100E-05 |
| 5796 | 290 | T290S | A | 4.6e-06 | 4.2e-06 | 9.3e-06 | 6.033E-06 | 2.836E-06 | 4.200E-06 | 9.300E-06 |
| 5797 | 290 | T290T | A | 9.3e-06 | 8.5e-06 | 1e-05 | 9.267E-06 | 7.506E-07 | 8.500E-06 | 1.000E-05 |
| 5798 | 290 | T290W | A | 6.5e-06 | 1.2e-05 | 6.4e-06 | 8.300E-06 | 3.205E-06 | 6.400E-06 | 1.200E-05 |
| 5799 | 290 | T290Y | A | 1e-05 | 8e-06 | 8.6e-06 | 8.867E-06 | 1.026E-06 | 8.000E-06 | 1.000E-05 |
| 5800 | 290 | T290V | A | 6.2e-06 | 1e-05 | 4.7e-06 | 6.967E-06 | 2.732E-06 | 4.700E-06 | 1.000E-05 |
| 5801 | 291 | L291A | A | 8.7e-06 | 1.4e-05 | 9.4e-06 | 1.070E-05 | 2.879E-06 | 8.700E-06 | 1.400E-05 |
| 5802 | 291 | L291R | A | 6.6e-06 | 6.5e-06 | 6.9e-06 | 6.667E-06 | 2.082E-07 | 6.500E-06 | 6.900E-06 |
| 5803 | 291 | L291N | A | 4.7e-06 | 7.1e-06 | 5.5e-06 | 5.767E-06 | 1.222E-06 | 4.700E-06 | 7.100E-06 |
| 5804 | 291 | L291D | A | 9.7e-06 | 5.7e-06 | 7.5e-06 | 7.633E-06 | 2.003E-06 | 5.700E-06 | 9.700E-06 |
| 5805 | 291 | L291C | A | 8.1e-06 | 1.1e-05 | 7.3e-06 | 8.800E-06 | 1.947E-06 | 7.300E-06 | 1.100E-05 |
| 5806 | 291 | L291E | A | 8e-06 | 5.8e-06 | 9.1e-06 | 7.633E-06 | 1.680E-06 | 5.800E-06 | 9.100E-06 |
| 5807 | 291 | L291Q | A | 6.6e-06 | 1.2e-05 | 5.7e-06 | 8.100E-06 | 3.407E-06 | 5.700E-06 | 1.200E-05 |
| 5808 | 291 | L291G | A | 7.3e-06 | 6.2e-06 | 6.4e-06 | 6.633E-06 | 5.859E-07 | 6.200E-06 | 7.300E-06 |
| 5809 | 291 | L291H | A | 1.4e-05 | 9.5e-06 | 1.2e-05 | 1.183E-05 | 2.255E-06 | 9.500E-06 | 1.400E-05 |
| 5810 | 291 | L291I | A | 1e-05 | 8.3e-06 | 8.9e-06 | 9.067E-06 | 8.622E-07 | 8.300E-06 | 1.000E-05 |
| 5811 | 291 | L291L | A | 6.4e-06 | 6.4e-06 | 1e-05 | 7.600E-06 | 2.078E-06 | 6.400E-06 | 1.000E-05 |
| 5812 | 291 | L291K | A | 1.1e-05 | 6.3e-06 | 6.9e-06 | 8.067E-06 | 2.558E-06 | 6.300E-06 | 1.100E-05 |
| 5813 | 291 | L291M | A | 5.7e-06 | 6.7e-06 | 7.7e-06 | 6.700E-06 | 1.000E-06 | 5.700E-06 | 7.700E-06 |
| 5814 | 291 | L291F | A | 7.4e-06 | 4.8e-06 | 1.3e-05 | 8.400E-06 | 4.190E-06 | 4.800E-06 | 1.300E-05 |
| 5815 | 291 | L291P | A | 9.2e-06 | 7.9e-06 | 5e-06 | 7.367E-06 | 2.150E-06 | 5.000E-06 | 9.200E-06 |
| 5816 | 291 | L291S | A | 5.5e-06 | 1.1e-05 | 9.9e-06 | 8.800E-06 | 2.910E-06 | 5.500E-06 | 1.100E-05 |
| 5817 | 291 | L291T | A | 6.5e-06 | 6.7e-06 | 5.8e-06 | 6.333E-06 | 4.726E-07 | 5.800E-06 | 6.700E-06 |
| 5818 | 291 | L291W | A | 5.6e-06 | 1.2e-05 | 1.1e-05 | 9.533E-06 | 3.443E-06 | 5.600E-06 | 1.200E-05 |
| 5819 | 291 | L291Y | A | 8.4e-06 | 7.7e-06 | 8.2e-06 | 8.100E-06 | 3.606E-07 | 7.700E-06 | 8.400E-06 |
| 5820 | 291 | L291V | A | 4.2e-06 | 1e-05 | 9.5e-06 | 7.900E-06 | 3.214E-06 | 4.200E-06 | 1.000E-05 |
| 5821 | 292 | V292A | A | 7.2e-06 | 6.6e-06 | 6.6e-06 | 6.800E-06 | 3.464E-07 | 6.600E-06 | 7.200E-06 |
| 5822 | 292 | V292R | A | 6.1e-06 | 9.6e-06 | 8.4e-06 | 8.033E-06 | 1.779E-06 | 6.100E-06 | 9.600E-06 |
| 5823 | 292 | V292N | A | 4.9e-06 | 7.2e-06 | 7.6e-06 | 6.567E-06 | 1.457E-06 | 4.900E-06 | 7.600E-06 |
| 5824 | 292 | V292D | A | 1.1e-05 | 6.7e-06 | 4.6e-06 | 7.433E-06 | 3.262E-06 | 4.600E-06 | 1.100E-05 |
| 5825 | 292 | V292C | A | 9e-06 | 6e-06 | 7.8e-06 | 7.600E-06 | 1.510E-06 | 6.000E-06 | 9.000E-06 |
| 5826 | 292 | V292E | A | 1.4e-05 | 6.5e-06 | 4.2e-06 | 8.233E-06 | 5.125E-06 | 4.200E-06 | 1.400E-05 |
| 5827 | 292 | V292Q | A | 1.2e-05 | 4.7e-06 | 5.7e-06 | 7.467E-06 | 3.958E-06 | 4.700E-06 | 1.200E-05 |
| 5828 | 292 | V292G | A | 4.8e-06 | 6.2e-06 | 5.7e-06 | 5.567E-06 | 7.095E-07 | 4.800E-06 | 6.200E-06 |
| 5829 | 292 | V292H | A | 6.3e-06 | 3.7e-06 | 7.1e-06 | 5.700E-06 | 1.778E-06 | 3.700E-06 | 7.100E-06 |
| 5830 | 292 | V292I | A | 1e-05 | 1e-05 | 9.4e-06 | 9.800E-06 | 3.464E-07 | 9.400E-06 | 1.000E-05 |
| 5831 | 292 | V292L | A | 5e-06 | 6.2e-06 | 1.3e-05 | 8.067E-06 | 4.314E-06 | 5.000E-06 | 1.300E-05 |
| 5832 | 292 | V292K | A | 8.1e-06 | 9.1e-06 | 7.1e-06 | 8.100E-06 | 1.000E-06 | 7.100E-06 | 9.100E-06 |
| 5833 | 292 | V292M | A | 7.1e-06 | 1.4e-05 | 1.1e-05 | 1.070E-05 | 3.460E-06 | 7.100E-06 | 1.400E-05 |
| 5834 | 292 | V292F | A | 6.7e-06 | 7.2e-06 | 4.8e-06 | 6.233E-06 | 1.266E-06 | 4.800E-06 | 7.200E-06 |
| 5835 | 292 | V292P | A | 7.6e-06 | 5.3e-06 | 7.9e-06 | 6.933E-06 | 1.422E-06 | 5.300E-06 | 7.900E-06 |
| 5836 | 292 | V292S | A | 4.3e-06 | 5e-06 | 4.8e-06 | 4.700E-06 | 3.606E-07 | 4.300E-06 | 5.000E-06 |
| 5837 | 292 | V292T | A | 7e-06 | 5.9e-06 | 9.8e-06 | 7.567E-06 | 2.011E-06 | 5.900E-06 | 9.800E-06 |
| 5838 | 292 | V292W | A | 9.8e-06 | 6.1e-06 | 7.3e-06 | 7.733E-06 | 1.888E-06 | 6.100E-06 | 9.800E-06 |
| 5839 | 292 | V292Y | A | 1.4e-05 | 6.8e-06 | 7.7e-06 | 9.500E-06 | 3.923E-06 | 6.800E-06 | 1.400E-05 |
| 5840 | 292 | V292V | A | 6.2e-06 | 5.8e-06 | 5.8e-06 | 5.933E-06 | 2.309E-07 | 5.800E-06 | 6.200E-06 |
| 5841 | 293 | C293A | A | 8.5e-06 | 6.6e-06 | 8.6e-06 | 7.900E-06 | 1.127E-06 | 6.600E-06 | 8.600E-06 |
| 5842 | 293 | C293R | A | 1.1e-05 | 8.9e-06 | 6.8e-06 | 8.900E-06 | 2.100E-06 | 6.800E-06 | 1.100E-05 |
| 5843 | 293 | C293N | A | 6.2e-06 | 6.1e-06 | 8.8e-06 | 7.033E-06 | 1.531E-06 | 6.100E-06 | 8.800E-06 |
| 5844 | 293 | C293D | A | 1.3e-05 | 6.5e-06 | 7.3e-06 | 8.933E-06 | 3.544E-06 | 6.500E-06 | 1.300E-05 |
| 5845 | 293 | C293C | A | 5.3e-06 | 8.8e-06 | 7.3e-06 | 7.133E-06 | 1.756E-06 | 5.300E-06 | 8.800E-06 |
| 5846 | 293 | C293E | A | 5.6e-06 | 1e-05 | 1.4e-05 | 9.867E-06 | 4.202E-06 | 5.600E-06 | 1.400E-05 |
| 5847 | 293 | C293Q | A | 8.5e-06 | 1.2e-05 | 8.3e-06 | 9.600E-06 | 2.081E-06 | 8.300E-06 | 1.200E-05 |
| 5848 | 293 | C293G | A | 8.2e-06 | 9.8e-06 | 7.7e-06 | 8.567E-06 | 1.097E-06 | 7.700E-06 | 9.800E-06 |
| 5849 | 293 | C293H | A | 1.3e-05 | 1.1e-05 | 5.4e-06 | 9.800E-06 | 3.940E-06 | 5.400E-06 | 1.300E-05 |
| 5850 | 293 | C293I | A | 6.9e-06 | 1.1e-05 | 6.5e-06 | 8.133E-06 | 2.491E-06 | 6.500E-06 | 1.100E-05 |
| 5851 | 293 | C293L | A | 1.3e-05 | 3.8e-06 | 1.4e-05 | 1.027E-05 | 5.623E-06 | 3.800E-06 | 1.400E-05 |
| 5852 | 293 | C293K | A | 1.2e-05 | 9e-06 | 5.6e-06 | 8.867E-06 | 3.202E-06 | 5.600E-06 | 1.200E-05 |
| 5853 | 293 | C293M | A | 4.3e-06 | 5.9e-06 | 7.7e-06 | 5.967E-06 | 1.701E-06 | 4.300E-06 | 7.700E-06 |
| 5854 | 293 | C293F | A | 9.9e-06 | 8.7e-06 | 6.3e-06 | 8.300E-06 | 1.833E-06 | 6.300E-06 | 9.900E-06 |
| 5855 | 293 | C293P | A | 6e-06 | 1.3e-05 | 1e-05 | 9.667E-06 | 3.512E-06 | 6.000E-06 | 1.300E-05 |
| 5856 | 293 | C293S | A | 3.4e-06 | 9.6e-06 | 3.3e-06 | 5.433E-06 | 3.609E-06 | 3.300E-06 | 9.600E-06 |
| 5857 | 293 | C293T | A | 1e-05 | 7.5e-06 | 5.9e-06 | 7.800E-06 | 2.066E-06 | 5.900E-06 | 1.000E-05 |
| 5858 | 293 | C293W | A | 6e-06 | 7e-06 | 6.9e-06 | 6.633E-06 | 5.508E-07 | 6.000E-06 | 7.000E-06 |
| 5859 | 293 | C293Y | A | 7.8e-06 | 6.1e-06 | 1.1e-05 | 8.300E-06 | 2.488E-06 | 6.100E-06 | 1.100E-05 |
| 5860 | 293 | C293V | A | 9.8e-06 | 6.7e-06 | 5.9e-06 | 7.467E-06 | 2.060E-06 | 5.900E-06 | 9.800E-06 |
| 5861 | 294 | P294A | A | 4.9e-06 | 8.9e-06 | 4.2e-06 | 6.000E-06 | 2.536E-06 | 4.200E-06 | 8.900E-06 |
| 5862 | 294 | P294R | A | 6.7e-06 | 1e-05 | 8.6e-06 | 8.433E-06 | 1.656E-06 | 6.700E-06 | 1.000E-05 |
| 5863 | 294 | P294N | A | 9e-06 | 6.4e-06 | 5e-06 | 6.800E-06 | 2.030E-06 | 5.000E-06 | 9.000E-06 |
| 5864 | 294 | P294D | A | 4.7e-06 | 9.1e-06 | 7.7e-06 | 7.167E-06 | 2.248E-06 | 4.700E-06 | 9.100E-06 |

| | | | | | | | | | | |
|---|---|---|---|---|---|---|---|---|---|---|
| 5865 | 294 | P294C | A | 7.4e-06 | 1.2e-05 | 6.9e-06 | 8.767E-06 | 2.811E-06 | 6.900E-06 | 1.200E-05 |
| 5866 | 294 | P294E | A | 9.4e-06 | 1.1e-05 | 8.7e-06 | 9.700E-06 | 1.179E-06 | 8.700E-06 | 1.100E-05 |
| 5867 | 294 | P294Q | A | 7.2e-06 | 6.5e-06 | 8.5e-06 | 7.400E-06 | 1.015E-06 | 6.500E-06 | 8.500E-06 |
| 5868 | 294 | P294G | A | 6.1e-06 | 6.8e-06 | 6.6e-06 | 6.500E-06 | 3.606E-07 | 6.100E-06 | 6.800E-06 |
| 5869 | 294 | P294H | A | 1.3e-05 | 7.7e-06 | 8.9e-06 | 9.867E-06 | 2.779E-06 | 7.700E-06 | 1.300E-05 |
| 5870 | 294 | P294I | A | 7.2e-06 | 7.6e-06 | 1.1e-05 | 8.600E-06 | 2.088E-06 | 7.200E-06 | 1.100E-05 |
| 5871 | 294 | P294L | A | 1e-05 | 6.9e-06 | 8.8e-06 | 8.567E-06 | 1.563E-06 | 6.900E-06 | 1.000E-05 |
| 5872 | 294 | P294K | A | 8.5e-06 | 1.5e-05 | 9.9e-06 | 1.113E-05 | 3.421E-06 | 8.500E-06 | 1.500E-05 |
| 5873 | 294 | P294M | A | 7.4e-06 | 7.5e-06 | 7.5e-06 | 7.467E-06 | 5.774E-08 | 7.400E-06 | 7.500E-06 |
| 5874 | 294 | P294F | A | 8.8e-06 | 5.6e-06 | 6.6e-06 | 7.000E-06 | 1.637E-06 | 5.600E-06 | 8.800E-06 |
| 5875 | 294 | P294P | A | 6.9e-06 | 5e-06 | 1.4e-05 | 8.633E-06 | 4.744E-06 | 5.000E-06 | 1.400E-05 |
| 5876 | 294 | P294S | A | 5.2e-06 | 5.4e-06 | 7.4e-06 | 6.000E-06 | 1.217E-06 | 5.200E-06 | 7.400E-06 |
| 5877 | 294 | P294T | A | 8.5e-06 | 1.4e-05 | 9.5e-06 | 1.067E-05 | 2.930E-06 | 8.500E-06 | 1.400E-05 |
| 5878 | 294 | P294W | A | 8.6e-06 | 6.4e-06 | 6.1e-06 | 7.033E-06 | 1.365E-06 | 6.100E-06 | 8.600E-06 |
| 5879 | 294 | P294Y | A | 7.6e-06 | 6.3e-06 | 6.4e-06 | 6.767E-06 | 7.234E-07 | 6.300E-06 | 7.600E-06 |
| 5880 | 294 | P294V | A | 1.2e-05 | 9.5e-06 | 6.2e-06 | 9.233E-06 | 2.909E-06 | 6.200E-06 | 1.200E-05 |
| 5881 | 295 | L295A | A | 8.8e-06 | 1.2e-05 | 1e-05 | 1.027E-05 | 1.617E-06 | 8.800E-06 | 1.200E-05 |
| 5882 | 295 | L295R | A | 9.4e-06 | 6e-06 | 4.9e-06 | 6.767E-06 | 2.346E-06 | 4.900E-06 | 9.400E-06 |
| 5883 | 295 | L295N | A | 4.1e-06 | 7e-06 | 5.9e-06 | 5.667E-06 | 1.464E-06 | 4.100E-06 | 7.000E-06 |
| 5884 | 295 | L295D | A | 5.7e-06 | 6.9e-06 | 8.5e-06 | 7.033E-06 | 1.405E-06 | 5.700E-06 | 8.500E-06 |
| 5885 | 295 | L295C | A | 6.1e-06 | 7.3e-06 | 6e-06 | 6.467E-06 | 7.234E-07 | 6.000E-06 | 7.300E-06 |
| 5886 | 295 | L295E | A | 4.7e-06 | 1.3e-05 | 8.6e-06 | 8.767E-06 | 4.153E-06 | 4.700E-06 | 1.300E-05 |
| 5887 | 295 | L295Q | A | 7.5e-06 | 1e-05 | 8.5e-06 | 8.667E-06 | 1.258E-06 | 7.500E-06 | 1.000E-05 |
| 5888 | 295 | L295G | A | 6.1e-06 | 5.4e-06 | 9.3e-06 | 6.933E-06 | 2.079E-06 | 5.400E-06 | 9.300E-06 |
| 5889 | 295 | L295H | A | 9.1e-06 | 4.8e-06 | 1.1e-05 | 8.300E-06 | 3.176E-06 | 4.800E-06 | 1.100E-05 |
| 5890 | 295 | L295I | A | 5.6e-06 | 4.6e-06 | 8.9e-06 | 6.367E-06 | 2.250E-06 | 4.600E-06 | 8.900E-06 |
| 5891 | 295 | L295L | A | 6.9e-06 | 1.2e-05 | 6.5e-06 | 8.467E-06 | 3.066E-06 | 6.500E-06 | 1.200E-05 |
| 5892 | 295 | L295K | A | 9.5e-06 | 6.4e-06 | 6.2e-06 | 7.367E-06 | 1.850E-06 | 6.200E-06 | 9.500E-06 |
| 5893 | 295 | L295M | A | 7.1e-06 | 8.5e-06 | 9.2e-06 | 8.267E-06 | 1.069E-06 | 7.100E-06 | 9.200E-06 |
| 5894 | 295 | L295F | A | 9.5e-06 | 6.8e-06 | 8.9e-06 | 8.400E-06 | 1.418E-06 | 6.800E-06 | 9.500E-06 |
| 5895 | 295 | L295P | A | 6.6e-06 | 8.5e-06 | 1e-05 | 8.367E-06 | 1.704E-06 | 6.600E-06 | 1.000E-05 |
| 5896 | 295 | L295S | A | 9.9e-06 | 9.6e-06 | 7.7e-06 | 9.067E-06 | 1.193E-06 | 7.700E-06 | 9.900E-06 |
| 5897 | 295 | L295T | A | 8.9e-06 | 1e-05 | 5.4e-06 | 8.100E-06 | 2.402E-06 | 5.400E-06 | 1.000E-05 |
| 5898 | 295 | L295W | A | 7e-06 | 1e-05 | 6.9e-06 | 7.967E-06 | 1.762E-06 | 6.900E-06 | 1.000E-05 |
| 5899 | 295 | L295Y | A | 5.9e-06 | 9.2e-06 | 5e-06 | 6.700E-06 | 2.211E-06 | 5.000E-06 | 9.200E-06 |
| 5900 | 295 | L295V | A | 7.1e-06 | 5.2e-06 | 9.7e-06 | 7.333E-06 | 2.259E-06 | 5.200E-06 | 9.700E-06 |
| 5901 | 296 | H296A | A | 8.6e-06 | 7.4e-06 | 6.7e-06 | 7.567E-06 | 9.609E-07 | 6.700E-06 | 8.600E-06 |
| 5902 | 296 | H296R | A | 9.1e-06 | 1e-05 | 1e-05 | 9.700E-06 | 5.196E-07 | 9.100E-06 | 1.000E-05 |
| 5903 | 296 | H296N | A | 8e-06 | 9.6e-06 | 5.7e-06 | 7.767E-06 | 1.960E-06 | 5.700E-06 | 9.600E-06 |
| 5904 | 296 | H296D | A | 5.3e-06 | 8.4e-06 | 5.4e-06 | 6.367E-06 | 1.762E-06 | 5.300E-06 | 8.400E-06 |
| 5905 | 296 | H296C | A | 9.6e-06 | 6.1e-06 | 7.1e-06 | 7.600E-06 | 1.803E-06 | 6.100E-06 | 9.600E-06 |
| 5906 | 296 | H296E | A | 9.1e-06 | 9.2e-06 | 9.4e-06 | 9.233E-06 | 1.528E-07 | 9.100E-06 | 9.400E-06 |
| 5907 | 296 | H296Q | A | 1.2e-05 | 1.2e-05 | 8.7e-06 | 1.090E-05 | 1.905E-06 | 8.700E-06 | 1.200E-05 |
| 5908 | 296 | H296G | A | 6e-06 | 1.2e-05 | 9.4e-06 | 9.133E-06 | 3.009E-06 | 6.000E-06 | 1.200E-05 |
| 5909 | 296 | H296H | A | 1.1e-05 | 4.8e-06 | 6.6e-06 | 7.467E-06 | 3.190E-06 | 4.800E-06 | 1.100E-05 |
| 5910 | 296 | H296I | A | 8.3e-06 | 1e-05 | 7e-06 | 8.433E-06 | 1.504E-06 | 7.000E-06 | 1.000E-05 |
| 5911 | 296 | H296L | A | 6.6e-06 | 8.9e-06 | 5.4e-06 | 6.967E-06 | 1.779E-06 | 5.400E-06 | 8.900E-06 |
| 5912 | 296 | H296K | A | 6.8e-06 | 8.4e-06 | 5.7e-06 | 6.967E-06 | 1.358E-06 | 5.700E-06 | 8.400E-06 |
| 5913 | 296 | H296M | A | 1.5e-05 | 1.1e-05 | 6.8e-06 | 1.093E-05 | 4.100E-06 | 6.800E-06 | 1.500E-05 |
| 5914 | 296 | H296F | A | 1e-05 | 1e-05 | 5.4e-06 | 8.467E-06 | 2.656E-06 | 5.400E-06 | 1.000E-05 |
| 5915 | 296 | H296P | A | 8.4e-06 | 8.8e-06 | 9.1e-06 | 8.767E-06 | 3.512E-07 | 8.400E-06 | 9.100E-06 |
| 5916 | 296 | H296S | A | 6.7e-06 | 1.1e-05 | 1.1e-05 | 9.567E-06 | 2.483E-06 | 6.700E-06 | 1.100E-05 |
| 5917 | 296 | H296T | A | 6.5e-06 | 5.7e-06 | 5.7e-06 | 5.967E-06 | 4.619E-07 | 5.700E-06 | 6.500E-06 |
| 5918 | 296 | H296W | A | 7.3e-06 | 5.8e-06 | 1e-05 | 7.700E-06 | 2.128E-06 | 5.800E-06 | 1.000E-05 |
| 5919 | 296 | H296Y | A | 5.7e-06 | 1e-05 | 1.1e-05 | 8.900E-06 | 2.816E-06 | 5.700E-06 | 1.100E-05 |
| 5920 | 296 | H296V | A | 1.2e-05 | 1.2e-05 | 9e-06 | 1.100E-05 | 1.732E-06 | 9.000E-06 | 1.200E-05 |
| 5921 | 297 | N297A | A | 1.9e-05 | 8.2e-06 | 6e-06 | 1.107E-05 | 6.958E-06 | 6.000E-06 | 1.900E-05 |
| 5922 | 297 | N297R | A | 9.2e-06 | 6.7e-06 | 8.9e-06 | 8.267E-06 | 1.365E-06 | 6.700E-06 | 9.200E-06 |
| 5923 | 297 | N297N | A | 8.3e-06 | 1.2e-05 | 8.1e-06 | 9.467E-06 | 2.196E-06 | 8.100E-06 | 1.200E-05 |
| 5924 | 297 | N297D | A | 7.4e-06 | 7.3e-06 | 1.3e-05 | 9.233E-06 | 3.262E-06 | 7.300E-06 | 1.300E-05 |
| 5925 | 297 | N297C | A | 7.5e-06 | 1.1e-05 | 3.9e-06 | 7.467E-06 | 3.550E-06 | 3.900E-06 | 1.100E-05 |
| 5926 | 297 | N297E | A | 6.7e-06 | 8.5e-06 | 4.7e-06 | 6.633E-06 | 1.901E-06 | 4.700E-06 | 8.500E-06 |
| 5927 | 297 | N297Q | A | 8e-06 | 1e-05 | 9.1e-06 | 9.033E-06 | 1.002E-06 | 8.000E-06 | 1.000E-05 |
| 5928 | 297 | N297G | A | 1.4e-05 | 8.1e-06 | 6.1e-06 | 9.400E-06 | 4.107E-06 | 6.100E-06 | 1.400E-05 |
| 5929 | 297 | N297H | A | 5.8e-06 | 7.8e-06 | 1.1e-05 | 8.200E-06 | 2.623E-06 | 5.800E-06 | 1.100E-05 |
| 5930 | 297 | N297I | A | 1.1e-05 | 1.4e-05 | 1.2e-05 | 1.233E-05 | 1.528E-06 | 1.100E-05 | 1.400E-05 |
| 5931 | 297 | N297L | A | 6.7e-06 | 1.1e-05 | 6.3e-06 | 8.000E-06 | 2.606E-06 | 6.300E-06 | 1.100E-05 |
| 5932 | 297 | N297K | A | 9.5e-06 | 7.6e-06 | 5.4e-06 | 7.500E-06 | 2.052E-06 | 5.400E-06 | 9.500E-06 |
| 5933 | 297 | N297M | A | 6.5e-06 | 1.1e-05 | 4.4e-06 | 7.300E-06 | 3.372E-06 | 4.400E-06 | 1.100E-05 |
| 5934 | 297 | N297F | A | 7.7e-06 | 1.2e-05 | 1.2e-05 | 1.057E-05 | 2.483E-06 | 7.700E-06 | 1.200E-05 |
| 5935 | 297 | N297P | A | 4.3e-06 | 1.5e-05 | 6.2e-06 | 8.500E-06 | 5.709E-06 | 4.300E-06 | 1.500E-05 |
| 5936 | 297 | N297S | A | 8.3e-06 | 6.8e-06 | 5.6e-06 | 6.900E-06 | 1.353E-06 | 5.600E-06 | 8.300E-06 |
| 5937 | 297 | N297T | A | 8.7e-06 | 9.9e-06 | 8.8e-06 | 9.133E-06 | 6.658E-07 | 8.700E-06 | 9.900E-06 |
| 5938 | 297 | N297W | A | 1.1e-05 | 9.6e-06 | 8e-06 | 9.533E-06 | 1.501E-06 | 8.000E-06 | 1.100E-05 |
| 5939 | 297 | N297Y | A | 7.3e-06 | 9.4e-06 | 1.1e-05 | 9.233E-06 | 1.856E-06 | 7.300E-06 | 1.100E-05 |
| 5940 | 297 | N297V | A | 1e-05 | 8.6e-06 | 7e-06 | 8.533E-06 | 1.501E-06 | 7.000E-06 | 1.000E-05 |
| 5941 | 298 | Q298A | A | 1e-05 | 6.7e-06 | 5.1e-06 | 7.267E-06 | 2.499E-06 | 5.100E-06 | 1.000E-05 |
| 5942 | 298 | Q298R | A | 7.9e-06 | 7.5e-06 | 5.9e-06 | 7.100E-06 | 1.058E-06 | 5.900E-06 | 7.900E-06 |
| 5943 | 298 | Q298N | A | 1e-05 | 7.6e-06 | 5.9e-06 | 7.833E-06 | 2.060E-06 | 5.900E-06 | 1.000E-05 |
| 5944 | 298 | Q298D | A | 1.4e-05 | 9.8e-06 | 1.4e-05 | 1.260E-05 | 2.425E-06 | 9.800E-06 | 1.400E-05 |
| 5945 | 298 | Q298C | A | 6.8e-06 | 1.1e-05 | 1.1e-05 | 9.600E-06 | 2.425E-06 | 6.800E-06 | 1.100E-05 |
| 5946 | 298 | Q298E | A | 1.5e-05 | 7.2e-06 | 6.9e-06 | 9.700E-06 | 4.592E-06 | 6.900E-06 | 1.500E-05 |
| 5947 | 298 | Q298Q | A | 8.7e-06 | 1.1e-05 | 2.1e-05 | 1.357E-05 | 6.539E-06 | 8.700E-06 | 2.100E-05 |
| 5948 | 298 | Q298G | A | 1.6e-05 | 1.3e-05 | 9.7e-06 | 1.290E-05 | 3.151E-06 | 9.700E-06 | 1.600E-05 |
| 5949 | 298 | Q298H | A | 5.2e-06 | 5.8e-06 | 8.8e-06 | 6.600E-06 | 1.929E-06 | 5.200E-06 | 8.800E-06 |

| | | | | | | | | | | |
|---|---|---|---|---|---|---|---|---|---|---|
| 5950 | 298 | Q298I | A | 7.3e-06 | 1e-05 | 1.3e-05 | 1.010E-05 | 2.851E-06 | 7.300E-06 | 1.300E-05 |
| 5951 | 298 | Q298L | A | 9.9e-06 | 7.7e-06 | 8.4e-06 | 8.667E-06 | 1.124E-06 | 7.700E-06 | 9.900E-06 |
| 5952 | 298 | Q298K | A | 6.6e-06 | 7.8e-06 | 6.5e-06 | 6.967E-06 | 7.234E-07 | 6.500E-06 | 7.800E-06 |
| 5953 | 298 | Q298M | A | 1.2e-05 | 5.5e-06 | 1e-05 | 9.167E-06 | 3.329E-06 | 5.500E-06 | 1.200E-05 |
| 5954 | 298 | Q298F | A | 1.1e-05 | 5.3e-06 | 8.8e-06 | 8.367E-06 | 2.875E-06 | 5.300E-06 | 1.100E-05 |
| 5955 | 298 | Q298P | A | 9.7e-06 | 1.4e-05 | 1.5e-05 | 1.290E-05 | 2.816E-06 | 9.700E-06 | 1.500E-05 |
| 5956 | 298 | Q298S | A | 1.3e-05 | 9.5e-06 | 6.4e-06 | 9.633E-06 | 3.302E-06 | 6.400E-06 | 1.300E-05 |
| 5957 | 298 | Q298T | A | 1.3e-05 | 9.5e-06 | 8.9e-06 | 1.047E-05 | 2.214E-06 | 8.900E-06 | 1.300E-05 |
| 5958 | 298 | Q298W | A | 7.8e-06 | 1.1e-05 | 1.3e-05 | 1.060E-05 | 2.623E-06 | 7.800E-06 | 1.300E-05 |
| 5959 | 298 | Q298Y | A | 9.4e-06 | 6.5e-06 | 9.1e-06 | 8.333E-06 | 1.595E-06 | 6.500E-06 | 9.400E-06 |
| 5960 | 298 | Q298V | A | 6.8e-06 | 6.4e-06 | 1e-05 | 7.733E-06 | 1.973E-06 | 6.400E-06 | 1.000E-05 |
| 5961 | 299 | E299A | A | 9.5e-06 | 5.2e-06 | 9.6e-06 | 8.100E-06 | 2.512E-06 | 5.200E-06 | 9.600E-06 |
| 5962 | 299 | E299R | A | 6.9e-06 | 1.4e-05 | 8.6e-06 | 9.833E-06 | 3.707E-06 | 6.900E-06 | 1.400E-05 |
| 5963 | 299 | E299N | A | 8.6e-06 | 1.2e-05 | 3.9e-06 | 8.167E-06 | 4.067E-06 | 3.900E-06 | 1.200E-05 |
| 5964 | 299 | E299D | A | 7.7e-06 | 7.3e-06 | 4.5e-06 | 6.500E-06 | 1.744E-06 | 4.500E-06 | 7.700E-06 |
| 5965 | 299 | E299C | A | 7.3e-06 | 1.3e-05 | 7.3e-06 | 9.200E-06 | 3.291E-06 | 7.300E-06 | 1.300E-05 |
| 5966 | 299 | E299E | A | 8.1e-06 | 5e-06 | 1.3e-05 | 8.700E-06 | 4.034E-06 | 5.000E-06 | 1.300E-05 |
| 5967 | 299 | E299Q | A | 6.9e-06 | 9.1e-06 | 5.4e-06 | 7.133E-06 | 1.861E-06 | 5.400E-06 | 9.100E-06 |
| 5968 | 299 | E299G | A | 1e-05 | 9.6e-06 | 7.8e-06 | 9.133E-06 | 1.172E-06 | 7.800E-06 | 1.000E-05 |
| 5969 | 299 | E299H | A | 1e-05 | 8.7e-06 | 1e-05 | 9.567E-06 | 7.506E-07 | 8.700E-06 | 1.000E-05 |
| 5970 | 299 | E299I | A | 8.3e-06 | 9.5e-06 | 6e-06 | 7.933E-06 | 1.779E-06 | 6.000E-06 | 9.500E-06 |
| 5971 | 299 | E299L | A | 6.9e-06 | 7.7e-06 | 8.4e-06 | 7.667E-06 | 7.506E-07 | 6.900E-06 | 8.400E-06 |
| 5972 | 299 | E299K | A | 7.8e-06 | 1e-05 | 7.1e-06 | 8.300E-06 | 1.513E-06 | 7.100E-06 | 1.000E-05 |
| 5973 | 299 | E299M | A | 6.5e-06 | 1e-05 | 8.4e-06 | 8.300E-06 | 1.752E-06 | 6.500E-06 | 1.000E-05 |
| 5974 | 299 | E299F | A | 8.7e-06 | 6.9e-06 | 6.1e-06 | 7.233E-06 | 1.332E-06 | 6.100E-06 | 8.700E-06 |
| 5975 | 299 | E299P | A | 5.8e-06 | 6.4e-06 | 4.2e-06 | 5.467E-06 | 1.137E-06 | 4.200E-06 | 6.400E-06 |
| 5976 | 299 | E299S | A | 6.8e-06 | 5.2e-06 | 5e-06 | 5.667E-06 | 9.866E-07 | 5.000E-06 | 6.800E-06 |
| 5977 | 299 | E299T | A | 6.9e-06 | 1e-05 | 9.3e-06 | 8.733E-06 | 1.626E-06 | 6.900E-06 | 1.000E-05 |
| 5978 | 299 | E299W | A | 3.2e-06 | 9.7e-06 | 6.4e-06 | 6.433E-06 | 3.250E-06 | 3.200E-06 | 9.700E-06 |
| 5979 | 299 | E299Y | A | 6.1e-06 | 1.3e-05 | 1.1e-05 | 1.003E-05 | 3.550E-06 | 6.100E-06 | 1.300E-05 |
| 5980 | 299 | E299V | A | 9.5e-06 | 6.5e-06 | 4.9e-06 | 6.967E-06 | 2.335E-06 | 4.900E-06 | 9.500E-06 |
| 5981 | 300 | V300A | A | 2.1e-05 | 9e-06 | 8.5e-06 | 1.283E-05 | 7.077E-06 | 8.500E-06 | 2.100E-05 |
| 5982 | 300 | V300R | A | 4.3e-06 | 8.4e-06 | 9e-06 | 7.233E-06 | 2.558E-06 | 4.300E-06 | 9.000E-06 |
| 5983 | 300 | V300N | A | 1e-05 | 9.5e-06 | 1.1e-05 | 1.017E-05 | 7.638E-07 | 9.500E-06 | 1.100E-05 |
| 5984 | 300 | V300D | A | 9.1e-06 | 5.4e-06 | 9.4e-06 | 7.967E-06 | 2.228E-06 | 5.400E-06 | 9.400E-06 |
| 5985 | 300 | V300C | A | 6.5e-06 | 6.2e-06 | 8.2e-06 | 6.967E-06 | 1.079E-06 | 6.200E-06 | 8.200E-06 |
| 5986 | 300 | V300E | A | 5.7e-06 | 1e-05 | 9.3e-06 | 8.333E-06 | 2.307E-06 | 5.700E-06 | 1.000E-05 |
| 5987 | 300 | V300Q | A | 1.4e-05 | 1.3e-05 | 1e-05 | 1.233E-05 | 2.082E-06 | 1.000E-05 | 1.400E-05 |
| 5988 | 300 | V300G | A | 9e-06 | 5.3e-06 | 1.2e-05 | 8.767E-06 | 3.356E-06 | 5.300E-06 | 1.200E-05 |
| 5989 | 300 | V300H | A | 8.4e-06 | 5e-06 | 7.5e-06 | 6.967E-06 | 1.762E-06 | 5.000E-06 | 8.400E-06 |
| 5990 | 300 | V300I | A | 9e-06 | 9.1e-06 | 1e-05 | 9.367E-06 | 5.508E-07 | 9.000E-06 | 1.000E-05 |
| 5991 | 300 | V300L | A | 9.6e-06 | 8.7e-06 | 6.7e-06 | 8.333E-06 | 1.484E-06 | 6.700E-06 | 9.600E-06 |
| 5992 | 300 | V300K | A | 4.4e-06 | 1.1e-05 | 8.5e-06 | 7.967E-06 | 3.332E-06 | 4.400E-06 | 1.100E-05 |
| 5993 | 300 | V300M | A | 7.8e-06 | 9e-06 | 8.6e-06 | 8.467E-06 | 6.110E-07 | 7.800E-06 | 9.000E-06 |
| 5994 | 300 | V300F | A | 1.1e-05 | 5.7e-06 | 5.7e-06 | 7.467E-06 | 3.060E-06 | 5.700E-06 | 1.100E-05 |
| 5995 | 300 | V300P | A | 8.2e-06 | 5.7e-06 | 9.9e-06 | 7.933E-06 | 2.113E-06 | 5.700E-06 | 9.900E-06 |
| 5996 | 300 | V300S | A | 5.9e-06 | 5e-06 | 8e-06 | 6.300E-06 | 1.539E-06 | 5.000E-06 | 8.000E-06 |
| 5997 | 300 | V300T | A | 1.1e-05 | 1.2e-05 | 7.8e-06 | 1.027E-05 | 2.194E-06 | 7.800E-06 | 1.200E-05 |
| 5998 | 300 | V300W | A | 1.2e-05 | 6.9e-06 | 4.8e-06 | 7.900E-06 | 3.703E-06 | 4.800E-06 | 1.200E-05 |
| 5999 | 300 | V300Y | A | 9.9e-06 | 1e-05 | 7.2e-06 | 9.033E-06 | 1.589E-06 | 7.200E-06 | 1.000E-05 |
| 6000 | 300 | V300V | A | 9.8e-06 | 7e-06 | 7.2e-06 | 8.000E-06 | 1.562E-06 | 7.000E-06 | 9.800E-06 |
| 6001 | 301 | T301A | A | 9.3e-06 | 9.7e-06 | 6.6e-06 | 8.533E-06 | 1.686E-06 | 6.600E-06 | 9.700E-06 |
| 6002 | 301 | T301R | A | 1.4e-05 | 7.5e-06 | 9.7e-06 | 1.040E-05 | 3.306E-06 | 7.500E-06 | 1.400E-05 |
| 6003 | 301 | T301N | A | 4.9e-06 | 9.9e-06 | 4.1e-06 | 6.300E-06 | 3.143E-06 | 4.100E-06 | 9.900E-06 |
| 6004 | 301 | T301D | A | 1.1e-05 | 8.8e-06 | 9.6e-06 | 9.800E-06 | 1.114E-06 | 8.800E-06 | 1.100E-05 |
| 6005 | 301 | T301C | A | 4.3e-06 | 5.7e-06 | 6e-06 | 5.333E-06 | 9.074E-07 | 4.300E-06 | 6.000E-06 |
| 6006 | 301 | T301E | A | 7.5e-06 | 5.7e-06 | 7.9e-06 | 7.033E-06 | 1.172E-06 | 5.700E-06 | 7.900E-06 |
| 6007 | 301 | T301Q | A | 1.6e-05 | 1e-05 | 6.6e-06 | 1.087E-05 | 4.760E-06 | 6.600E-06 | 1.600E-05 |
| 6008 | 301 | T301G | A | 7.2e-06 | 1e-05 | 1.1e-05 | 9.400E-06 | 1.970E-06 | 7.200E-06 | 1.100E-05 |
| 6009 | 301 | T301H | A | 5e-06 | 7e-06 | 5.1e-06 | 5.700E-06 | 1.127E-06 | 5.000E-06 | 7.000E-06 |
| 6010 | 301 | T301I | A | 5.4e-06 | 6.9e-06 | 5.3e-06 | 5.867E-06 | 8.963E-07 | 5.300E-06 | 6.900E-06 |
| 6011 | 301 | T301L | A | 5.1e-06 | 7.1e-06 | 1e-05 | 7.400E-06 | 2.464E-06 | 5.100E-06 | 1.000E-05 |
| 6012 | 301 | T301K | A | 1.1e-05 | 7e-06 | 6.9e-06 | 8.300E-06 | 2.339E-06 | 6.900E-06 | 1.100E-05 |
| 6013 | 301 | T301M | A | 5.9e-06 | 1.1e-05 | 8.1e-06 | 8.333E-06 | 2.558E-06 | 5.900E-06 | 1.100E-05 |
| 6014 | 301 | T301F | A | 6.9e-06 | 1.1e-05 | 8.6e-06 | 8.833E-06 | 2.060E-06 | 6.900E-06 | 1.100E-05 |
| 6015 | 301 | T301P | A | 3.9e-06 | 7.4e-06 | 1.3e-05 | 8.100E-06 | 4.590E-06 | 3.900E-06 | 1.300E-05 |
| 6016 | 301 | T301S | A | 6.5e-06 | 1.6e-05 | 1.5e-05 | 1.250E-05 | 5.220E-06 | 6.500E-06 | 1.600E-05 |
| 6017 | 301 | T301T | A | 5.4e-06 | 8.8e-06 | 9.6e-06 | 7.933E-06 | 2.230E-06 | 5.400E-06 | 9.600E-06 |
| 6018 | 301 | T301W | A | 6.8e-06 | 5.6e-06 | 1.3e-05 | 8.467E-06 | 3.972E-06 | 5.600E-06 | 1.300E-05 |
| 6019 | 301 | T301Y | A | 3.5e-06 | 9.2e-06 | 6.7e-06 | 6.467E-06 | 2.857E-06 | 3.500E-06 | 9.200E-06 |
| 6020 | 301 | T301V | A | 6.1e-06 | 8.2e-06 | 7.4e-06 | 7.233E-06 | 1.060E-06 | 6.100E-06 | 8.200E-06 |
| 6021 | 302 | A302A | A | 9.9e-06 | 7e-06 | 6.6e-06 | 7.833E-06 | 1.801E-06 | 6.600E-06 | 9.900E-06 |
| 6022 | 302 | A302R | A | 1e-05 | 8.9e-06 | 5.4e-06 | 8.100E-06 | 2.402E-06 | 5.400E-06 | 1.000E-05 |
| 6023 | 302 | A302N | A | 6.2e-06 | 8.7e-06 | 1e-05 | 8.300E-06 | 1.931E-06 | 6.200E-06 | 1.000E-05 |
| 6024 | 302 | A302D | A | 9.4e-06 | 6.9e-06 | 8.1e-06 | 8.133E-06 | 1.250E-06 | 6.900E-06 | 9.400E-06 |
| 6025 | 302 | A302C | A | 8.3e-06 | 6.9e-06 | 6.6e-06 | 7.267E-06 | 9.074E-07 | 6.600E-06 | 8.300E-06 |
| 6026 | 302 | A302E | A | 8.6e-06 | 1.1e-05 | 6.7e-06 | 8.767E-06 | 2.155E-06 | 6.700E-06 | 1.100E-05 |
| 6027 | 302 | A302Q | A | 1e-05 | 1.1e-05 | 7.7e-06 | 9.567E-06 | 1.692E-06 | 7.700E-06 | 1.100E-05 |
| 6028 | 302 | A302G | A | 5.1e-06 | 1.1e-05 | 1.4e-05 | 1.003E-05 | 4.528E-06 | 5.100E-06 | 1.400E-05 |
| 6029 | 302 | A302H | A | 1e-05 | 1e-05 | 6.5e-06 | 8.833E-06 | 2.021E-06 | 6.500E-06 | 1.000E-05 |
| 6030 | 302 | A302I | A | 6.2e-06 | 5.8e-06 | 8.2e-06 | 6.733E-06 | 1.286E-06 | 5.800E-06 | 8.200E-06 |
| 6031 | 302 | A302L | A | 9.5e-06 | 9.6e-06 | 9.4e-06 | 9.500E-06 | 1.000E-07 | 9.400E-06 | 9.600E-06 |
| 6032 | 302 | A302K | A | 6.4e-06 | 9.4e-06 | 8.3e-06 | 8.033E-06 | 1.518E-06 | 6.400E-06 | 9.400E-06 |
| 6033 | 302 | A302M | A | 8.4e-06 | 1.1e-05 | 7.8e-06 | 9.067E-06 | 1.701E-06 | 7.800E-06 | 1.100E-05 |
| 6034 | 302 | A302F | A | 1.3e-05 | 1e-05 | 1e-05 | 1.100E-05 | 1.732E-06 | 1.000E-05 | 1.300E-05 |

| | | | | | | | | | | |
|---|---|---|---|---|---|---|---|---|---|---|
| 6035 | 302 | A302P | A | 2.1e-05 | 1e-05 | 1.3e-05 | 1.467E-05 | 5.686E-06 | 1.000E-05 | 2.100E-05 |
| 6036 | 302 | A302S | A | 7.2e-06 | 9.1e-06 | 4.4e-06 | 6.900E-06 | 2.364E-06 | 4.400E-06 | 9.100E-06 |
| 6037 | 302 | A302T | A | 8e-06 | 7.7e-06 | 1.1e-05 | 8.900E-06 | 1.825E-06 | 7.700E-06 | 1.100E-05 |
| 6038 | 302 | A302W | A | 7.4e-06 | 4.9e-06 | 4.3e-06 | 5.533E-06 | 1.644E-06 | 4.300E-06 | 7.400E-06 |
| 6039 | 302 | A302Y | A | 4.3e-06 | 7.3e-06 | 6.9e-06 | 6.167E-06 | 1.629E-06 | 4.300E-06 | 7.300E-06 |
| 6040 | 302 | A302V | A | 6.8e-06 | 8.1e-06 | 6.2e-06 | 7.033E-06 | 9.713E-07 | 6.200E-06 | 8.100E-06 |
| 6041 | 303 | E303A | A | 7.9e-06 | 4.5e-06 | 2.3e-06 | 4.900E-06 | 2.821E-06 | 2.300E-06 | 7.900E-06 |
| 6042 | 303 | E303R | A | 1.2e-05 | 6e-06 | 8.4e-06 | 8.800E-06 | 3.020E-06 | 6.000E-06 | 1.200E-05 |
| 6043 | 303 | E303N | A | 6.5e-06 | 7e-06 | 8.5e-06 | 7.333E-06 | 1.041E-06 | 6.500E-06 | 8.500E-06 |
| 6044 | 303 | E303D | A | 8.6e-06 | 5.3e-06 | 1.3e-05 | 8.967E-06 | 3.863E-06 | 5.300E-06 | 1.300E-05 |
| 6045 | 303 | E303C | A | 6.5e-06 | 5.9e-06 | 1.3e-05 | 8.467E-06 | 3.937E-06 | 5.900E-06 | 1.300E-05 |
| 6046 | 303 | E303E | A | 9.4e-06 | 6.3e-06 | 6.7e-06 | 7.467E-06 | 1.686E-06 | 6.300E-06 | 9.400E-06 |
| 6047 | 303 | E303Q | A | 6.2e-06 | 6.4e-06 | 1.4e-05 | 8.867E-06 | 4.447E-06 | 6.200E-06 | 1.400E-05 |
| 6048 | 303 | E303G | A | 9.9e-06 | 6.3e-06 | 8.9e-06 | 8.367E-06 | 1.858E-06 | 6.300E-06 | 9.900E-06 |
| 6049 | 303 | E303H | A | 1.3e-05 | 9.6e-06 | 4.7e-06 | 9.100E-06 | 4.173E-06 | 4.700E-06 | 1.300E-05 |
| 6050 | 303 | E303I | A | 5.9e-06 | 1.2e-05 | 6.9e-06 | 8.267E-06 | 3.272E-06 | 5.900E-06 | 1.200E-05 |
| 6051 | 303 | E303L | A | 7e-06 | 8.1e-06 | 7.5e-06 | 7.533E-06 | 5.508E-07 | 7.000E-06 | 8.100E-06 |
| 6052 | 303 | E303K | A | 9.9e-06 | 9.6e-06 | 5.3e-06 | 8.267E-06 | 2.574E-06 | 5.300E-06 | 9.900E-06 |
| 6053 | 303 | E303M | A | 6.8e-06 | 8.3e-06 | 8.7e-06 | 7.933E-06 | 1.002E-06 | 6.800E-06 | 8.700E-06 |
| 6054 | 303 | E303F | A | 6.1e-06 | 4.9e-06 | 6.1e-06 | 5.700E-06 | 6.928E-07 | 4.900E-06 | 6.100E-06 |
| 6055 | 303 | E303P | A | 3.9e-06 | 4.7e-06 | 8.8e-06 | 5.800E-06 | 2.629E-06 | 3.900E-06 | 8.800E-06 |
| 6056 | 303 | E303S | A | 7e-06 | 5.3e-06 | 9.7e-06 | 7.333E-06 | 2.219E-06 | 5.300E-06 | 9.700E-06 |
| 6057 | 303 | E303T | A | 7.3e-06 | 9.3e-06 | 6.7e-06 | 7.767E-06 | 1.361E-06 | 6.700E-06 | 9.300E-06 |
| 6058 | 303 | E303W | A | 1.1e-05 | 8.9e-06 | 9.2e-06 | 9.700E-06 | 1.136E-06 | 8.900E-06 | 1.100E-05 |
| 6059 | 303 | E303Y | A | 6.3e-06 | 4.7e-06 | 5.5e-06 | 5.500E-06 | 8.000E-07 | 4.700E-06 | 6.300E-06 |
| 6060 | 303 | E303V | A | 6.2e-06 | 8.1e-06 | 9.8e-06 | 8.033E-06 | 1.801E-06 | 6.200E-06 | 9.800E-06 |
| 6061 | 304 | D304A | A | 1.2e-05 | 5.6e-06 | 8.8e-06 | 8.800E-06 | 3.200E-06 | 5.600E-06 | 1.200E-05 |
| 6062 | 304 | D304R | A | 1.4e-05 | 6e-06 | 4.4e-06 | 8.133E-06 | 5.143E-06 | 4.400E-06 | 1.400E-05 |
| 6063 | 304 | D304N | A | 9e-06 | 9.3e-06 | 1.3e-05 | 1.043E-05 | 2.228E-06 | 9.000E-06 | 1.300E-05 |
| 6064 | 304 | D304D | A | 6.3e-06 | 4.7e-06 | 5.2e-06 | 5.400E-06 | 8.185E-07 | 4.700E-06 | 6.300E-06 |
| 6065 | 304 | D304C | A | 5.4e-06 | 8.2e-06 | 5.8e-06 | 6.467E-06 | 1.514E-06 | 5.400E-06 | 8.200E-06 |
| 6066 | 304 | D304E | A | 4.8e-06 | 4.8e-06 | 6.2e-06 | 5.267E-06 | 8.083E-07 | 4.800E-06 | 6.200E-06 |
| 6067 | 304 | D304Q | A | 8.3e-06 | 7.8e-06 | 1.3e-05 | 9.700E-06 | 2.869E-06 | 7.800E-06 | 1.300E-05 |
| 6068 | 304 | D304G | A | 1.3e-05 | 1.1e-05 | 5.5e-06 | 9.833E-06 | 3.884E-06 | 5.500E-06 | 1.300E-05 |
| 6069 | 304 | D304H | A | 8.3e-06 | 7e-06 | 1.2e-05 | 9.100E-06 | 2.594E-06 | 7.000E-06 | 1.200E-05 |
| 6070 | 304 | D304I | A | 6.6e-06 | 1.1e-05 | 6.4e-06 | 8.000E-06 | 2.600E-06 | 6.400E-06 | 1.100E-05 |
| 6071 | 304 | D304L | A | 3.2e-06 | 9.4e-06 | 7.5e-06 | 6.700E-06 | 3.176E-06 | 3.200E-06 | 9.400E-06 |
| 6072 | 304 | D304K | A | 6.9e-06 | 1e-05 | 8.6e-06 | 8.500E-06 | 1.552E-06 | 6.900E-06 | 1.000E-05 |
| 6073 | 304 | D304M | A | 8.2e-06 | 1.1e-05 | 7.7e-06 | 8.967E-06 | 1.779E-06 | 7.700E-06 | 1.100E-05 |
| 6074 | 304 | D304F | A | 8e-06 | 4.5e-06 | 8.2e-06 | 6.900E-06 | 2.081E-06 | 4.500E-06 | 8.200E-06 |
| 6075 | 304 | D304P | A | 9.1e-06 | 7.4e-06 | 9.7e-06 | 8.733E-06 | 1.193E-06 | 7.400E-06 | 9.700E-06 |
| 6076 | 304 | D304S | A | 1.2e-05 | 9.4e-06 | 7.8e-06 | 9.733E-06 | 2.120E-06 | 7.800E-06 | 1.200E-05 |
| 6077 | 304 | D304T | A | 8.3e-06 | 1.2e-05 | 4.6e-06 | 8.300E-06 | 3.700E-06 | 4.600E-06 | 1.200E-05 |
| 6078 | 304 | D304W | A | 5.9e-06 | 9.2e-06 | 7.6e-06 | 7.567E-06 | 1.650E-06 | 5.900E-06 | 9.200E-06 |
| 6079 | 304 | D304Y | A | 5.5e-06 | 9.2e-06 | 9.1e-06 | 7.933E-06 | 2.108E-06 | 5.500E-06 | 9.200E-06 |
| 6080 | 304 | D304V | A | 6.6e-06 | 5.8e-06 | 9.4e-06 | 7.267E-06 | 1.890E-06 | 5.800E-06 | 9.400E-06 |
| 6081 | 305 | G305A | A | 7.9e-06 | 6.2e-06 | 6.8e-06 | 6.967E-06 | 8.622E-07 | 6.200E-06 | 7.900E-06 |
| 6082 | 305 | G305R | A | 5.1e-06 | 6.7e-06 | 6.2e-06 | 6.000E-06 | 8.185E-07 | 5.100E-06 | 6.700E-06 |
| 6083 | 305 | G305N | A | 5.8e-06 | 1.1e-05 | 9.1e-06 | 8.633E-06 | 2.631E-06 | 5.800E-06 | 1.100E-05 |
| 6084 | 305 | G305D | A | 8.6e-06 | 7.4e-06 | 1.2e-05 | 9.333E-06 | 2.386E-06 | 7.400E-06 | 1.200E-05 |
| 6085 | 305 | G305C | A | 5.2e-06 | 1e-05 | 9.3e-06 | 8.167E-06 | 2.593E-06 | 5.200E-06 | 1.000E-05 |
| 6086 | 305 | G305E | A | 5.7e-06 | 4.7e-06 | 1.3e-05 | 7.800E-06 | 4.531E-06 | 4.700E-06 | 1.300E-05 |
| 6087 | 305 | G305Q | A | 6.5e-06 | 9.3e-06 | 8.4e-06 | 8.067E-06 | 1.429E-06 | 6.500E-06 | 9.300E-06 |
| 6088 | 305 | G305G | A | 1.3e-05 | 1.4e-05 | 5.2e-06 | 1.073E-05 | 4.818E-06 | 5.200E-06 | 1.400E-05 |
| 6089 | 305 | G305H | A | 1.4e-05 | 5.8e-06 | 1e-05 | 9.933E-06 | 4.100E-06 | 5.800E-06 | 1.400E-05 |
| 6090 | 305 | G305I | A | 6.4e-06 | 6.9e-06 | 3.4e-06 | 5.567E-06 | 1.893E-06 | 3.400E-06 | 6.900E-06 |
| 6091 | 305 | G305L | A | 1e-05 | 6.2e-06 | 4.1e-06 | 6.767E-06 | 2.991E-06 | 4.100E-06 | 1.000E-05 |
| 6092 | 305 | G305K | A | 4.6e-06 | 1e-05 | 8.9e-06 | 7.833E-06 | 2.854E-06 | 4.600E-06 | 1.000E-05 |
| 6093 | 305 | G305M | A | 7.1e-06 | 9.9e-06 | 1.2e-05 | 9.667E-06 | 2.458E-06 | 7.100E-06 | 1.200E-05 |
| 6094 | 305 | G305F | A | 5.1e-06 | 1e-05 | 1.3e-05 | 9.367E-06 | 3.988E-06 | 5.100E-06 | 1.300E-05 |
| 6095 | 305 | G305P | A | 5.2e-06 | 7e-06 | 7.2e-06 | 6.467E-06 | 1.102E-06 | 5.200E-06 | 7.200E-06 |
| 6096 | 305 | G305S | A | 9e-06 | 6.5e-06 | 9.2e-06 | 8.233E-06 | 1.504E-06 | 6.500E-06 | 9.200E-06 |
| 6097 | 305 | G305T | A | 7.6e-06 | 1.3e-05 | 9.2e-06 | 9.933E-06 | 2.774E-06 | 7.600E-06 | 1.300E-05 |
| 6098 | 305 | G305W | A | 8.5e-06 | 7.8e-06 | 7.7e-06 | 8.000E-06 | 4.359E-07 | 7.700E-06 | 8.500E-06 |
| 6099 | 305 | G305Y | A | 8e-06 | 9.1e-06 | 6.7e-06 | 7.933E-06 | 1.201E-06 | 6.700E-06 | 9.100E-06 |
| 6100 | 305 | G305V | A | 8.9e-06 | 1.4e-05 | 5.4e-06 | 9.433E-06 | 4.325E-06 | 5.400E-06 | 1.400E-05 |
| 6101 | 306 | T306A | A | 7e-06 | 9.8e-06 | 1e-05 | 8.933E-06 | 1.677E-06 | 7.000E-06 | 1.000E-05 |
| 6102 | 306 | T306R | A | 9.2e-06 | 5.7e-06 | 9.7e-06 | 8.200E-06 | 2.179E-06 | 5.700E-06 | 9.700E-06 |
| 6103 | 306 | T306N | A | 7.5e-06 | 8.5e-06 | 5.5e-06 | 7.167E-06 | 1.528E-06 | 5.500E-06 | 8.500E-06 |
| 6104 | 306 | T306D | A | 6.1e-06 | 1e-05 | 5.3e-06 | 7.133E-06 | 2.515E-06 | 5.300E-06 | 1.000E-05 |
| 6105 | 306 | T306C | A | 5.3e-06 | 5.3e-06 | 6.4e-06 | 5.667E-06 | 6.351E-07 | 5.300E-06 | 6.400E-06 |
| 6106 | 306 | T306E | A | 1.4e-05 | 6.9e-06 | 1e-05 | 1.030E-05 | 3.559E-06 | 6.900E-06 | 1.400E-05 |
| 6107 | 306 | T306Q | A | 5.3e-06 | 7e-06 | 9.1e-06 | 7.133E-06 | 1.904E-06 | 5.300E-06 | 9.100E-06 |
| 6108 | 306 | T306G | A | 7.6e-06 | 6.5e-06 | 8.6e-06 | 7.567E-06 | 1.050E-06 | 6.500E-06 | 8.600E-06 |
| 6109 | 306 | T306H | A | 9.3e-06 | 6.1e-06 | 8.2e-06 | 7.867E-06 | 1.626E-06 | 6.100E-06 | 9.300E-06 |
| 6110 | 306 | T306I | A | 7.1e-06 | 9.2e-06 | 5.2e-06 | 7.167E-06 | 2.001E-06 | 5.200E-06 | 9.200E-06 |
| 6111 | 306 | T306L | A | 9.8e-06 | 8.7e-06 | 6.4e-06 | 8.300E-06 | 1.735E-06 | 6.400E-06 | 9.800E-06 |
| 6112 | 306 | T306K | A | 1e-05 | 6.8e-06 | 6.8e-06 | 7.867E-06 | 1.848E-06 | 6.800E-06 | 1.000E-05 |
| 6113 | 306 | T306M | A | 6.2e-06 | 7.5e-06 | 1.3e-05 | 8.900E-06 | 3.610E-06 | 6.200E-06 | 1.300E-05 |
| 6114 | 306 | T306F | A | 7e-06 | 7.8e-06 | 1e-05 | 8.267E-06 | 1.553E-06 | 7.000E-06 | 1.000E-05 |
| 6115 | 306 | T306P | A | 6.6e-06 | 1.2e-05 | 9.9e-06 | 9.500E-06 | 2.722E-06 | 6.600E-06 | 1.200E-05 |
| 6116 | 306 | T306S | A | 1e-05 | 6.6e-06 | 8.9e-06 | 8.500E-06 | 1.735E-06 | 6.600E-06 | 1.000E-05 |
| 6117 | 306 | T306T | A | 8.3e-06 | 1.1e-05 | 2.4e-06 | 7.233E-06 | 4.398E-06 | 2.400E-06 | 1.100E-05 |
| 6118 | 306 | T306W | A | 6.8e-06 | 8.4e-06 | 7.7e-06 | 7.633E-06 | 8.021E-07 | 6.800E-06 | 8.400E-06 |
| 6119 | 306 | T306Y | A | 6.6e-06 | 6.3e-06 | 1.3e-05 | 8.633E-06 | 3.785E-06 | 6.300E-06 | 1.300E-05 |

| | | | | | | | | | | |
|---|---|---|---|---|---|---|---|---|---|---|
| 6120 | 306 | T306V | A | 7.9e-06 | 9.8e-06 | 1e-05 | 9.233E-06 | 1.159E-06 | 7.900E-06 | 1.000E-05 |
| 6121 | 307 | Q307A | A | 7.7e-06 | 5.8e-06 | 7.7e-06 | 7.067E-06 | 1.097E-06 | 5.800E-06 | 7.700E-06 |
| 6122 | 307 | Q307R | A | 6.9e-06 | 9e-06 | 9.5e-06 | 8.467E-06 | 1.380E-06 | 6.900E-06 | 9.500E-06 |
| 6123 | 307 | Q307N | A | 1.3e-05 | 1.2e-05 | 7.1e-06 | 1.070E-05 | 3.158E-06 | 7.100E-06 | 1.300E-05 |
| 6124 | 307 | Q307D | A | 3.6e-06 | 9.8e-06 | 9.8e-06 | 7.733E-06 | 3.580E-06 | 3.600E-06 | 9.800E-06 |
| 6125 | 307 | Q307C | A | 7.6e-06 | 3.1e-06 | 7e-06 | 5.900E-06 | 2.443E-06 | 3.100E-06 | 7.600E-06 |
| 6126 | 307 | Q307E | A | 1.2e-05 | 6.4e-06 | 6.7e-06 | 8.367E-06 | 3.150E-06 | 6.400E-06 | 1.200E-05 |
| 6127 | 307 | Q307Q | A | 8.1e-06 | 1.1e-05 | 5.7e-06 | 8.267E-06 | 2.654E-06 | 5.700E-06 | 1.100E-05 |
| 6128 | 307 | Q307G | A | 3.5e-06 | 9.4e-06 | 1.2e-05 | 8.300E-06 | 4.355E-06 | 3.500E-06 | 1.200E-05 |
| 6129 | 307 | Q307H | A | 6.9e-06 | 5.7e-06 | 3.9e-06 | 5.500E-06 | 1.510E-06 | 3.900E-06 | 6.900E-06 |
| 6130 | 307 | Q307I | A | 5.2e-06 | 1.2e-05 | 6.7e-06 | 7.967E-06 | 3.573E-06 | 5.200E-06 | 1.200E-05 |
| 6131 | 307 | Q307L | A | 7.9e-06 | 4.6e-06 | 7.4e-06 | 6.633E-06 | 1.779E-06 | 4.600E-06 | 7.900E-06 |
| 6132 | 307 | Q307K | A | 5.2e-06 | 5.7e-06 | 1.3e-05 | 7.967E-06 | 4.366E-06 | 5.200E-06 | 1.300E-05 |
| 6133 | 307 | Q307M | A | 4.8e-06 | 9.1e-06 | 5.8e-06 | 6.567E-06 | 2.250E-06 | 4.800E-06 | 9.100E-06 |
| 6134 | 307 | Q307F | A | 1.6e-05 | 9.3e-06 | 8.9e-06 | 1.140E-05 | 3.989E-06 | 8.900E-06 | 1.600E-05 |
| 6135 | 307 | Q307P | A | 1e-05 | 9.4e-06 | 7.7e-06 | 9.033E-06 | 1.193E-06 | 7.700E-06 | 1.000E-05 |
| 6136 | 307 | Q307S | A | 8.3e-06 | 6.2e-06 | 9.9e-06 | 8.133E-06 | 1.856E-06 | 6.200E-06 | 9.900E-06 |
| 6137 | 307 | Q307T | A | 7.5e-06 | 8e-06 | 8.7e-06 | 8.067E-06 | 6.028E-07 | 7.500E-06 | 8.700E-06 |
| 6138 | 307 | Q307W | A | 1.1e-05 | 9.1e-06 | 7.6e-06 | 9.233E-06 | 1.704E-06 | 7.600E-06 | 1.100E-05 |
| 6139 | 307 | Q307Y | A | 5.2e-06 | 1e-05 | 6e-06 | 7.067E-06 | 2.572E-06 | 5.200E-06 | 1.000E-05 |
| 6140 | 307 | Q307V | A | 3.5e-06 | 6e-06 | 7e-06 | 5.500E-06 | 1.803E-06 | 3.500E-06 | 7.000E-06 |
| 6141 | 308 | R308A | A | 8.3e-06 | 7.2e-06 | 1.4e-05 | 9.833E-06 | 3.650E-06 | 7.200E-06 | 1.400E-05 |
| 6142 | 308 | R308R | A | 4.8e-06 | 7.1e-06 | 4.8e-06 | 5.567E-06 | 1.328E-06 | 4.800E-06 | 7.100E-06 |
| 6143 | 308 | R308N | A | 4.1e-06 | 9.5e-06 | 7e-06 | 6.867E-06 | 2.702E-06 | 4.100E-06 | 9.500E-06 |
| 6144 | 308 | R308D | A | 9.6e-06 | 5e-06 | 5.2e-06 | 6.600E-06 | 2.600E-06 | 5.000E-06 | 9.600E-06 |
| 6145 | 308 | R308C | A | 4.3e-06 | 9.4e-06 | 9.5e-06 | 7.733E-06 | 2.974E-06 | 4.300E-06 | 9.500E-06 |
| 6146 | 308 | R308E | A | 6.4e-06 | 1.1e-05 | 6.7e-06 | 8.033E-06 | 2.574E-06 | 6.400E-06 | 1.100E-05 |
| 6147 | 308 | R308Q | A | 6.5e-06 | 1e-05 | 1.1e-05 | 9.167E-06 | 2.363E-06 | 6.500E-06 | 1.100E-05 |
| 6148 | 308 | R308G | A | 5.4e-06 | 9.7e-06 | 1.3e-05 | 9.367E-06 | 3.811E-06 | 5.400E-06 | 1.300E-05 |
| 6149 | 308 | R308H | A | 1.1e-05 | 7.9e-06 | 5.4e-06 | 8.100E-06 | 2.805E-06 | 5.400E-06 | 1.100E-05 |
| 6150 | 308 | R308I | A | 5.5e-06 | 8.9e-06 | 9.2e-06 | 7.867E-06 | 2.055E-06 | 5.500E-06 | 9.200E-06 |
| 6151 | 308 | R308L | A | 1e-05 | 7e-06 | 6.2e-06 | 7.733E-06 | 2.003E-06 | 6.200E-06 | 1.000E-05 |
| 6152 | 308 | R308K | A | 7.6e-06 | 1.5e-05 | 7e-06 | 9.867E-06 | 4.456E-06 | 7.000E-06 | 1.500E-05 |
| 6153 | 308 | R308M | A | 1e-05 | 9.4e-06 | 7.3e-06 | 8.900E-06 | 1.418E-06 | 7.300E-06 | 1.000E-05 |
| 6154 | 308 | R308F | A | 6.8e-06 | 5.4e-06 | 7.3e-06 | 6.500E-06 | 9.849E-07 | 5.400E-06 | 7.300E-06 |
| 6155 | 308 | R308P | A | 1.3e-05 | 9.6e-06 | 7e-06 | 9.867E-06 | 3.009E-06 | 7.000E-06 | 1.300E-05 |
| 6156 | 308 | R308S | A | 1.6e-05 | 7.4e-06 | 7.3e-06 | 1.023E-05 | 4.994E-06 | 7.300E-06 | 1.600E-05 |
| 6157 | 308 | R308T | A | 8.9e-06 | 8e-06 | 5.3e-06 | 7.400E-06 | 1.873E-06 | 5.300E-06 | 8.900E-06 |
| 6158 | 308 | R308W | A | 9.4e-06 | 7.3e-06 | 8.4e-06 | 8.367E-06 | 1.050E-06 | 7.300E-06 | 9.400E-06 |
| 6159 | 308 | R308Y | A | 5.6e-06 | 1.6e-05 | 1e-05 | 1.053E-05 | 5.220E-06 | 5.600E-06 | 1.600E-05 |
| 6160 | 308 | R308V | A | 7.3e-06 | 9.7e-06 | 6.8e-06 | 7.933E-06 | 1.550E-06 | 6.800E-06 | 9.700E-06 |
| 6161 | 309 | C309A | A | 8.3e-06 | 6.2e-06 | 8.8e-06 | 7.767E-06 | 1.380E-06 | 6.200E-06 | 8.800E-06 |
| 6162 | 309 | C309R | A | 5.8e-06 | 6.9e-06 | 7.4e-06 | 6.700E-06 | 8.185E-07 | 5.800E-06 | 7.400E-06 |
| 6163 | 309 | C309N | A | 6.6e-06 | 1.7e-05 | 1.3e-05 | 1.220E-05 | 5.246E-06 | 6.600E-06 | 1.700E-05 |
| 6164 | 309 | C309D | A | 8.8e-06 | 1e-05 | 7.6e-06 | 8.800E-06 | 1.200E-06 | 7.600E-06 | 1.000E-05 |
| 6165 | 309 | C309C | A | 6.2e-06 | 8.1e-06 | 7.8e-06 | 7.367E-06 | 1.021E-06 | 6.200E-06 | 8.100E-06 |
| 6166 | 309 | C309E | A | 8.5e-06 | 6.2e-06 | 7.9e-06 | 7.533E-06 | 1.193E-06 | 6.200E-06 | 8.500E-06 |
| 6167 | 309 | C309Q | A | 7.8e-06 | 9.4e-06 | 1e-05 | 9.067E-06 | 1.137E-06 | 7.800E-06 | 1.000E-05 |
| 6168 | 309 | C309G | A | 7.8e-06 | 1e-05 | 1.6e-05 | 1.127E-05 | 4.244E-06 | 7.800E-06 | 1.600E-05 |
| 6169 | 309 | C309H | A | 1e-05 | 4.9e-06 | 9.3e-06 | 8.067E-06 | 2.765E-06 | 4.900E-06 | 1.000E-05 |
| 6170 | 309 | C309I | A | 6.4e-06 | 1.5e-05 | 7.6e-06 | 9.667E-06 | 4.658E-06 | 6.400E-06 | 1.500E-05 |
| 6171 | 309 | C309L | A | 8.9e-06 | 7.6e-06 | 8.3e-06 | 8.267E-06 | 6.506E-07 | 7.600E-06 | 8.900E-06 |
| 6172 | 309 | C309K | A | 4.8e-06 | 4.5e-06 | 4.8e-06 | 4.700E-06 | 1.732E-07 | 4.500E-06 | 4.800E-06 |
| 6173 | 309 | C309M | A | 9.8e-06 | 1.1e-05 | 1.4e-05 | 1.160E-05 | 2.163E-06 | 9.800E-06 | 1.400E-05 |
| 6174 | 309 | C309F | A | 8.9e-06 | 9e-06 | 1.1e-05 | 9.633E-06 | 1.185E-06 | 8.900E-06 | 1.100E-05 |
| 6175 | 309 | C309P | A | 6.5e-06 | 8e-06 | 9.4e-06 | 7.967E-06 | 1.450E-06 | 6.500E-06 | 9.400E-06 |
| 6176 | 309 | C309S | A | 5e-06 | 4.7e-06 | 7.8e-06 | 5.833E-06 | 1.710E-06 | 4.700E-06 | 7.800E-06 |
| 6177 | 309 | C309T | A | 1.2e-05 | 5.1e-06 | 1.3e-05 | 1.003E-05 | 4.302E-06 | 5.100E-06 | 1.300E-05 |
| 6178 | 309 | C309W | A | 1.1e-05 | 6.1e-06 | 1.2e-05 | 9.700E-06 | 3.158E-06 | 6.100E-06 | 1.200E-05 |
| 6179 | 309 | C309Y | A | 1.1e-05 | 4.8e-06 | 5.7e-06 | 7.167E-06 | 3.350E-06 | 4.800E-06 | 1.100E-05 |
| 6180 | 309 | C309V | A | 1e-05 | 8.3e-06 | 5e-06 | 7.767E-06 | 2.542E-06 | 5.000E-06 | 1.000E-05 |
| 6181 | 310 | E310A | A | 9.2e-06 | 8.2e-06 | 9.9e-06 | 9.100E-06 | 8.544E-07 | 8.200E-06 | 9.900E-06 |
| 6182 | 310 | E310R | A | 9.7e-06 | 1e-05 | 4.2e-06 | 7.967E-06 | 3.265E-06 | 4.200E-06 | 1.000E-05 |
| 6183 | 310 | E310N | A | 6e-06 | 6e-06 | 5.8e-06 | 5.933E-06 | 1.155E-07 | 5.800E-06 | 6.000E-06 |
| 6184 | 310 | E310D | A | 5e-06 | 7.2e-06 | 6.6e-06 | 6.267E-06 | 1.137E-06 | 5.000E-06 | 7.200E-06 |
| 6185 | 310 | E310C | A | 1.4e-05 | 7.3e-06 | 1.2e-05 | 1.110E-05 | 3.439E-06 | 7.300E-06 | 1.400E-05 |
| 6186 | 310 | E310E | A | 1.1e-05 | 6.3e-06 | 9.2e-06 | 8.833E-06 | 2.371E-06 | 6.300E-06 | 1.100E-05 |
| 6187 | 310 | E310Q | A | 6.2e-06 | 6.3e-06 | 1e-05 | 7.500E-06 | 2.166E-06 | 6.200E-06 | 1.000E-05 |
| 6188 | 310 | E310G | A | 6.9e-06 | 1.5e-05 | 5.6e-06 | 9.167E-06 | 5.093E-06 | 5.600E-06 | 1.500E-05 |
| 6189 | 310 | E310H | A | 7.6e-06 | 9.1e-06 | 4.8e-06 | 7.167E-06 | 2.183E-06 | 4.800E-06 | 9.100E-06 |
| 6190 | 310 | E310I | A | 7.9e-06 | 9.3e-06 | 6.4e-06 | 7.867E-06 | 1.450E-06 | 6.400E-06 | 9.300E-06 |
| 6191 | 310 | E310L | A | 7.5e-06 | 6.9e-06 | 6.2e-06 | 6.867E-06 | 6.506E-07 | 6.200E-06 | 7.500E-06 |
| 6192 | 310 | E310K | A | 5.9e-06 | 8.7e-06 | 6.9e-06 | 7.167E-06 | 1.419E-06 | 5.900E-06 | 8.700E-06 |
| 6193 | 310 | E310M | A | 1e-05 | 6.4e-06 | 8e-06 | 8.133E-06 | 1.804E-06 | 6.400E-06 | 1.000E-05 |
| 6194 | 310 | E310F | A | 8.6e-06 | 7e-06 | 5.8e-06 | 7.133E-06 | 1.405E-06 | 5.800E-06 | 8.600E-06 |
| 6195 | 310 | E310P | A | 9.4e-06 | 7.9e-06 | 5.5e-06 | 7.600E-06 | 1.967E-06 | 5.500E-06 | 9.400E-06 |
| 6196 | 310 | E310S | A | 6.3e-06 | 6.7e-06 | 9.2e-06 | 7.400E-06 | 1.572E-06 | 6.300E-06 | 9.200E-06 |
| 6197 | 310 | E310T | A | 9.6e-06 | 6.6e-06 | 1.3e-05 | 9.733E-06 | 3.202E-06 | 6.600E-06 | 1.300E-05 |
| 6198 | 310 | E310W | A | 6.5e-06 | 4.5e-06 | 5e-06 | 5.333E-06 | 1.041E-06 | 4.500E-06 | 6.500E-06 |
| 6199 | 310 | E310Y | A | 8.8e-06 | 7.6e-06 | 5.9e-06 | 7.433E-06 | 1.457E-06 | 5.900E-06 | 8.800E-06 |
| 6200 | 310 | E310V | A | 7.6e-06 | 9.3e-06 | 9.5e-06 | 8.800E-06 | 1.044E-06 | 7.600E-06 | 9.500E-06 |
| 6201 | 311 | K311A | A | 7.8e-06 | 9.8e-06 | 5.4e-06 | 7.667E-06 | 2.203E-06 | 5.400E-06 | 9.800E-06 |
| 6202 | 311 | K311R | A | 1e-05 | 1.3e-05 | 7.2e-06 | 1.007E-05 | 2.901E-06 | 7.200E-06 | 1.300E-05 |
| 6203 | 311 | K311N | A | 5.9e-06 | 8.9e-06 | 5.7e-06 | 6.833E-06 | 1.793E-06 | 5.700E-06 | 8.900E-06 |
| 6204 | 311 | K311D | A | 7.7e-06 | 7.9e-06 | 7.1e-06 | 7.567E-06 | 4.163E-07 | 7.100E-06 | 7.900E-06 |

| | | | | | | | | | | |
|---|---|---|---|---|---|---|---|---|---|---|
| 6205 | 311 | K311C | A | 5e-06 | 5.4e-06 | 9e-06 | 6.467E-06 | 2.203E-06 | 5.000E-06 | 9.000E-06 |
| 6206 | 311 | K311E | A | 6.4e-06 | 7.8e-06 | 1e-05 | 8.067E-06 | 1.815E-06 | 6.400E-06 | 1.000E-05 |
| 6207 | 311 | K311Q | A | 4.9e-06 | 1e-05 | 1.3e-05 | 9.300E-06 | 4.095E-06 | 4.900E-06 | 1.300E-05 |
| 6208 | 311 | K311G | A | 9.8e-06 | 8.3e-06 | 1e-05 | 9.367E-06 | 9.292E-07 | 8.300E-06 | 1.000E-05 |
| 6209 | 311 | K311H | A | 7.5e-06 | 5.9e-06 | 5.1e-06 | 6.167E-06 | 1.222E-06 | 5.100E-06 | 7.500E-06 |
| 6210 | 311 | K311I | A | 7.3e-06 | 7.3e-06 | 8.9e-06 | 7.833E-06 | 9.238E-07 | 7.300E-06 | 8.900E-06 |
| 6211 | 311 | K311L | A | 1.2e-05 | 8.1e-06 | 4.5e-06 | 8.200E-06 | 3.751E-06 | 4.500E-06 | 1.200E-05 |
| 6212 | 311 | K311K | A | 7.3e-06 | 4.5e-06 | 9e-06 | 6.933E-06 | 2.272E-06 | 4.500E-06 | 9.000E-06 |
| 6213 | 311 | K311M | A | 6.5e-06 | 7.1e-06 | 9.8e-06 | 7.800E-06 | 1.758E-06 | 6.500E-06 | 9.800E-06 |
| 6214 | 311 | K311F | A | 6.8e-06 | 1e-05 | 4.8e-06 | 7.200E-06 | 2.623E-06 | 4.800E-06 | 1.000E-05 |
| 6215 | 311 | K311P | A | 6.3e-06 | 8.7e-06 | 8.4e-06 | 7.800E-06 | 1.308E-06 | 6.300E-06 | 8.700E-06 |
| 6216 | 311 | K311S | A | 4.7e-06 | 6.9e-06 | 1.4e-05 | 8.533E-06 | 4.860E-06 | 4.700E-06 | 1.400E-05 |
| 6217 | 311 | K311T | A | 7.3e-06 | 5.7e-06 | 1.2e-05 | 8.333E-06 | 3.275E-06 | 5.700E-06 | 1.200E-05 |
| 6218 | 311 | K311W | A | 5.7e-06 | 6.1e-06 | 1.7e-05 | 9.600E-06 | 6.412E-06 | 5.700E-06 | 1.700E-05 |
| 6219 | 311 | K311Y | A | 7.8e-06 | 9.5e-06 | 7.4e-06 | 8.233E-06 | 1.115E-06 | 7.400E-06 | 9.500E-06 |
| 6220 | 311 | K311V | A | 9e-06 | 7e-06 | 7.1e-06 | 7.700E-06 | 1.127E-06 | 7.000E-06 | 9.000E-06 |
| 6221 | 312 | C312A | A | 9.5e-06 | 6.2e-06 | 5.8e-06 | 7.167E-06 | 2.031E-06 | 5.800E-06 | 9.500E-06 |
| 6222 | 312 | C312R | A | 1e-05 | 4.7e-06 | 1.2e-05 | 8.900E-06 | 3.772E-06 | 4.700E-06 | 1.200E-05 |
| 6223 | 312 | C312N | A | 7.8e-06 | 1.4e-05 | 7.8e-06 | 9.867E-06 | 3.580E-06 | 7.800E-06 | 1.400E-05 |
| 6224 | 312 | C312D | A | 1.4e-05 | 5e-06 | 8.5e-06 | 9.167E-06 | 4.537E-06 | 5.000E-06 | 1.400E-05 |
| 6225 | 312 | C312C | A | 8.8e-06 | 8.5e-06 | 9.5e-06 | 8.933E-06 | 5.132E-07 | 8.500E-06 | 9.500E-06 |
| 6226 | 312 | C312E | A | 6.3e-06 | 1e-05 | 1.1e-05 | 9.100E-06 | 2.476E-06 | 6.300E-06 | 1.100E-05 |
| 6227 | 312 | C312Q | A | 7e-06 | 8.3e-06 | 7.3e-06 | 7.533E-06 | 6.807E-07 | 7.000E-06 | 8.300E-06 |
| 6228 | 312 | C312G | A | 4.5e-06 | 1.2e-05 | 7.5e-06 | 8.000E-06 | 3.775E-06 | 4.500E-06 | 1.200E-05 |
| 6229 | 312 | C312H | A | 1.4e-05 | 1.2e-05 | 5.3e-06 | 1.043E-05 | 4.557E-06 | 5.300E-06 | 1.400E-05 |
| 6230 | 312 | C312I | A | 5.8e-06 | 9.1e-06 | 5.9e-06 | 6.933E-06 | 1.877E-06 | 5.800E-06 | 9.100E-06 |
| 6231 | 312 | C312L | A | 7.7e-06 | 6.8e-06 | 5.7e-06 | 6.733E-06 | 1.002E-06 | 5.700E-06 | 7.700E-06 |
| 6232 | 312 | C312K | A | 8.1e-06 | 4.6e-06 | 4.2e-06 | 5.633E-06 | 2.146E-06 | 4.200E-06 | 8.100E-06 |
| 6233 | 312 | C312M | A | 6.4e-06 | 4.8e-06 | 7e-06 | 6.067E-06 | 1.137E-06 | 4.800E-06 | 7.000E-06 |
| 6234 | 312 | C312F | A | 9e-06 | 6.9e-06 | 1e-05 | 8.633E-06 | 1.582E-06 | 6.900E-06 | 1.000E-05 |
| 6235 | 312 | C312P | A | 1e-05 | 6.7e-06 | 7.9e-06 | 8.200E-06 | 1.670E-06 | 6.700E-06 | 1.000E-05 |
| 6236 | 312 | C312S | A | 1.4e-05 | 7.7e-06 | 4.9e-06 | 8.867E-06 | 4.661E-06 | 4.900E-06 | 1.400E-05 |
| 6237 | 312 | C312T | A | 1.1e-05 | 7.3e-06 | 1.3e-05 | 1.043E-05 | 2.892E-06 | 7.300E-06 | 1.300E-05 |
| 6238 | 312 | C312W | A | 8.2e-06 | 6.6e-06 | 5.9e-06 | 6.900E-06 | 1.179E-06 | 5.900E-06 | 8.200E-06 |
| 6239 | 312 | C312Y | A | 4.8e-06 | 6.3e-06 | 6.8e-06 | 5.967E-06 | 1.041E-06 | 4.800E-06 | 6.800E-06 |
| 6240 | 312 | C312V | A | 7e-06 | 9.4e-06 | 7.8e-06 | 8.067E-06 | 1.222E-06 | 7.000E-06 | 9.400E-06 |
| 6241 | 313 | S313A | A | 5.2e-06 | 1.1e-05 | 1e-05 | 8.733E-06 | 3.101E-06 | 5.200E-06 | 1.100E-05 |
| 6242 | 313 | S313R | A | 3.3e-06 | 1.5e-05 | 1e-05 | 9.433E-06 | 5.871E-06 | 3.300E-06 | 1.500E-05 |
| 6243 | 313 | S313N | A | 1e-05 | 6.6e-06 | 6.1e-06 | 7.567E-06 | 2.122E-06 | 6.100E-06 | 1.000E-05 |
| 6244 | 313 | S313D | A | 9e-06 | 8.9e-06 | 9.8e-06 | 9.233E-06 | 4.933E-07 | 8.900E-06 | 9.800E-06 |
| 6245 | 313 | S313C | A | 8.5e-06 | 7.2e-06 | 1.1e-05 | 8.900E-06 | 1.931E-06 | 7.200E-06 | 1.100E-05 |
| 6246 | 313 | S313E | A | 6.2e-06 | 6.1e-06 | 9.1e-06 | 7.133E-06 | 1.704E-06 | 6.100E-06 | 9.100E-06 |
| 6247 | 313 | S313Q | A | 9.2e-06 | 7.1e-06 | 1.1e-05 | 9.100E-06 | 1.952E-06 | 7.100E-06 | 1.100E-05 |
| 6248 | 313 | S313G | A | 1.2e-05 | 7.1e-06 | 9.1e-06 | 9.400E-06 | 2.464E-06 | 7.100E-06 | 1.200E-05 |
| 6249 | 313 | S313H | A | 9e-06 | 7.2e-06 | 7.8e-06 | 8.000E-06 | 9.165E-07 | 7.200E-06 | 9.000E-06 |
| 6250 | 313 | S313I | A | 4.7e-06 | 7.1e-06 | 8.4e-06 | 6.733E-06 | 1.877E-06 | 4.700E-06 | 8.400E-06 |
| 6251 | 313 | S313L | A | 1.3e-05 | 9e-06 | 1e-05 | 1.067E-05 | 2.082E-06 | 9.000E-06 | 1.300E-05 |
| 6252 | 313 | S313K | A | 4.9e-06 | 8.1e-06 | 1.4e-05 | 9.000E-06 | 4.616E-06 | 4.900E-06 | 1.400E-05 |
| 6253 | 313 | S313M | A | 6.8e-06 | 7.4e-06 | 1.4e-05 | 9.400E-06 | 3.995E-06 | 6.800E-06 | 1.400E-05 |
| 6254 | 313 | S313F | A | 6.1e-06 | 1.1e-05 | 9.2e-06 | 8.767E-06 | 2.479E-06 | 6.100E-06 | 1.100E-05 |
| 6255 | 313 | S313P | A | 3.4e-06 | 5.3e-06 | 4.8e-06 | 4.500E-06 | 9.849E-07 | 3.400E-06 | 5.300E-06 |
| 6256 | 313 | S313S | A | 8.5e-06 | 5.3e-06 | 7.1e-06 | 6.967E-06 | 1.604E-06 | 5.300E-06 | 8.500E-06 |
| 6257 | 313 | S313T | A | 5.8e-06 | 6.4e-06 | 1e-05 | 7.400E-06 | 2.272E-06 | 5.800E-06 | 1.000E-05 |
| 6258 | 313 | S313W | A | 6.7e-06 | 1.4e-05 | 9.6e-06 | 1.010E-05 | 3.676E-06 | 6.700E-06 | 1.400E-05 |
| 6259 | 313 | S313Y | A | 1.5e-05 | 7.8e-06 | 5.5e-06 | 9.433E-06 | 4.956E-06 | 5.500E-06 | 1.500E-05 |
| 6260 | 313 | S313V | A | 1.1e-05 | 1.2e-05 | 1.6e-05 | 1.300E-05 | 2.646E-06 | 1.100E-05 | 1.600E-05 |
| 6261 | 314 | K314A | A | 4.8e-06 | 7.1e-06 | 9.9e-06 | 7.267E-06 | 2.554E-06 | 4.800E-06 | 9.900E-06 |
| 6262 | 314 | K314R | A | 5e-06 | 7.8e-06 | 3.5e-06 | 5.433E-06 | 2.183E-06 | 3.500E-06 | 7.800E-06 |
| 6263 | 314 | K314N | A | 1.1e-05 | 6.3e-06 | 9.9e-06 | 9.067E-06 | 2.458E-06 | 6.300E-06 | 1.100E-05 |
| 6264 | 314 | K314D | A | 8.1e-06 | 4.7e-06 | 6e-06 | 6.267E-06 | 1.716E-06 | 4.700E-06 | 8.100E-06 |
| 6265 | 314 | K314C | A | 1e-05 | 1.1e-05 | 9.6e-06 | 1.020E-05 | 7.211E-07 | 9.600E-06 | 1.100E-05 |
| 6266 | 314 | K314E | A | 7.6e-06 | 6.8e-06 | 1.4e-05 | 9.467E-06 | 3.946E-06 | 6.800E-06 | 1.400E-05 |
| 6267 | 314 | K314Q | A | 8.4e-06 | 5.9e-06 | 1.1e-05 | 8.433E-06 | 2.550E-06 | 5.900E-06 | 1.100E-05 |
| 6268 | 314 | K314G | A | 9e-06 | 4.6e-06 | 7.7e-06 | 7.100E-06 | 2.261E-06 | 4.600E-06 | 9.000E-06 |
| 6269 | 314 | K314H | A | 1.4e-05 | 1e-05 | 7.7e-06 | 1.057E-05 | 3.188E-06 | 7.700E-06 | 1.400E-05 |
| 6270 | 314 | K314I | A | 6.4e-06 | 8.6e-06 | 6.5e-06 | 7.167E-06 | 1.242E-06 | 6.400E-06 | 8.600E-06 |
| 6271 | 314 | K314L | A | 1.4e-05 | 6.6e-06 | 7.3e-06 | 9.300E-06 | 4.085E-06 | 6.600E-06 | 1.400E-05 |
| 6272 | 314 | K314K | A | 6.7e-06 | 4.8e-06 | 5.8e-06 | 5.767E-06 | 9.504E-07 | 4.800E-06 | 6.700E-06 |
| 6273 | 314 | K314M | A | 1.1e-05 | 9.4e-06 | 1.2e-05 | 1.080E-05 | 1.311E-06 | 9.400E-06 | 1.200E-05 |
| 6274 | 314 | K314F | A | 4.5e-06 | 7.1e-06 | 4.7e-06 | 5.433E-06 | 1.447E-06 | 4.500E-06 | 7.100E-06 |
| 6275 | 314 | K314P | A | 6.2e-06 | 3.3e-06 | 9.4e-06 | 6.300E-06 | 3.051E-06 | 3.300E-06 | 9.400E-06 |
| 6276 | 314 | K314S | A | 9.1e-06 | 1.1e-05 | 1.1e-05 | 1.037E-05 | 1.097E-06 | 9.100E-06 | 1.100E-05 |
| 6277 | 314 | K314T | A | 1e-05 | 7.1e-06 | 8.8e-06 | 8.633E-06 | 1.457E-06 | 7.100E-06 | 1.000E-05 |
| 6278 | 314 | K314W | A | 1.2e-05 | 1.4e-05 | 8.8e-06 | 1.160E-05 | 2.623E-06 | 8.800E-06 | 1.400E-05 |
| 6279 | 314 | K314Y | A | 4.3e-06 | 7.7e-06 | 1.1e-05 | 7.667E-06 | 3.350E-06 | 4.300E-06 | 1.100E-05 |
| 6280 | 314 | K314V | A | 4.6e-06 | 1.2e-05 | 8.5e-06 | 8.367E-06 | 3.702E-06 | 4.600E-06 | 1.200E-05 |
| 6281 | 315 | P315A | A | 9.4e-06 | 1.2e-05 | 1e-05 | 1.047E-05 | 1.361E-06 | 9.400E-06 | 1.200E-05 |
| 6282 | 315 | P315R | A | 7.3e-06 | 8.3e-06 | 4.2e-06 | 6.600E-06 | 2.138E-06 | 4.200E-06 | 8.300E-06 |
| 6283 | 315 | P315N | A | 5.6e-06 | 5.8e-06 | 8.6e-06 | 6.667E-06 | 1.677E-06 | 5.600E-06 | 8.600E-06 |
| 6284 | 315 | P315D | A | 6.6e-06 | 6.2e-06 | 7.2e-06 | 6.667E-06 | 5.033E-07 | 6.200E-06 | 7.200E-06 |
| 6285 | 315 | P315C | A | 1.4e-05 | 5.1e-06 | 1.2e-05 | 1.037E-05 | 4.669E-06 | 5.100E-06 | 1.400E-05 |
| 6286 | 315 | P315E | A | 8.6e-06 | 9.6e-06 | 8.5e-06 | 8.900E-06 | 6.083E-07 | 8.500E-06 | 9.600E-06 |
| 6287 | 315 | P315Q | A | 3.3e-06 | 6.5e-06 | 5.3e-06 | 5.033E-06 | 1.617E-06 | 3.300E-06 | 6.500E-06 |
| 6288 | 315 | P315G | A | 7.4e-06 | 4.5e-06 | 9e-06 | 6.967E-06 | 2.281E-06 | 4.500E-06 | 9.000E-06 |
| 6289 | 315 | P315H | A | 9.4e-06 | 1.1e-05 | 3.3e-06 | 7.900E-06 | 4.063E-06 | 3.300E-06 | 1.100E-05 |

| | | | | | | | | | | |
|---|---|---|---|---|---|---|---|---|---|---|
| 6290 | 315 | P315I | A | 5.2e-06 | 6.5e-06 | 1.2e-05 | 7.900E-06 | 3.610E-06 | 5.200E-06 | 1.200E-05 |
| 6291 | 315 | P315L | A | 8.9e-06 | 8.8e-06 | 1.3e-05 | 1.023E-05 | 2.397E-06 | 8.800E-06 | 1.300E-05 |
| 6292 | 315 | P315K | A | 6.3e-06 | 1.1e-05 | 6.7e-06 | 8.000E-06 | 2.606E-06 | 6.300E-06 | 1.100E-05 |
| 6293 | 315 | P315M | A | 7.2e-06 | 6e-06 | 5.6e-06 | 6.267E-06 | 8.327E-07 | 5.600E-06 | 7.200E-06 |
| 6294 | 315 | P315F | A | 1.5e-05 | 8.5e-06 | 8.9e-06 | 1.080E-05 | 3.643E-06 | 8.500E-06 | 1.500E-05 |
| 6295 | 315 | P315P | A | 5.7e-06 | 6.8e-06 | 9.4e-06 | 7.300E-06 | 1.900E-06 | 5.700E-06 | 9.400E-06 |
| 6296 | 315 | P315S | A | 8.2e-06 | 6.9e-06 | 8.5e-06 | 7.867E-06 | 8.505E-07 | 6.900E-06 | 8.500E-06 |
| 6297 | 315 | P315T | A | 1.2e-05 | 6.9e-06 | 9.8e-06 | 9.567E-06 | 2.558E-06 | 6.900E-06 | 1.200E-05 |
| 6298 | 315 | P315W | A | 9.5e-06 | 4.9e-06 | 7.1e-06 | 7.167E-06 | 2.301E-06 | 4.900E-06 | 9.500E-06 |
| 6299 | 315 | P315Y | A | 6.5e-06 | 8.6e-06 | 6.8e-06 | 7.300E-06 | 1.136E-06 | 6.500E-06 | 8.600E-06 |
| 6300 | 315 | P315V | A | 1.2e-05 | 6.5e-06 | 1.9e-05 | 1.250E-05 | 6.265E-06 | 6.500E-06 | 1.900E-05 |
| 6301 | 316 | C316A | A | 1.1e-05 | 1.1e-05 | 6.6e-06 | 9.533E-06 | 2.540E-06 | 6.600E-06 | 1.100E-05 |
| 6302 | 316 | C316R | A | 1.3e-05 | 1.2e-05 | 7.1e-06 | 1.070E-05 | 3.158E-06 | 7.100E-06 | 1.300E-05 |
| 6303 | 316 | C316N | A | 8.2e-06 | 6.7e-06 | 7.3e-06 | 7.400E-06 | 7.550E-07 | 6.700E-06 | 8.200E-06 |
| 6304 | 316 | C316D | A | 7.8e-06 | 4.6e-06 | 6.8e-06 | 6.400E-06 | 1.637E-06 | 4.600E-06 | 7.800E-06 |
| 6305 | 316 | C316C | A | 7.5e-06 | 6.2e-06 | 1e-05 | 7.900E-06 | 1.931E-06 | 6.200E-06 | 1.000E-05 |
| 6306 | 316 | C316E | A | 6.2e-06 | 5.8e-06 | 1.1e-05 | 7.667E-06 | 2.894E-06 | 5.800E-06 | 1.100E-05 |
| 6307 | 316 | C316Q | A | 8.5e-06 | 7.5e-06 | 1.1e-05 | 9.000E-06 | 1.803E-06 | 7.500E-06 | 1.100E-05 |
| 6308 | 316 | C316G | A | 1.2e-05 | 9.1e-06 | 1e-05 | 1.037E-05 | 1.484E-06 | 9.100E-06 | 1.200E-05 |
| 6309 | 316 | C316H | A | 7.4e-06 | 9.2e-06 | 7.6e-06 | 8.067E-06 | 9.866E-07 | 7.400E-06 | 9.200E-06 |
| 6310 | 316 | C316I | A | 1e-05 | 5.2e-06 | 8.5e-06 | 7.900E-06 | 2.456E-06 | 5.200E-06 | 1.000E-05 |
| 6311 | 316 | C316L | A | 9.8e-06 | 9.8e-06 | 8.2e-06 | 9.267E-06 | 9.238E-07 | 8.200E-06 | 9.800E-06 |
| 6312 | 316 | C316K | A | 1.1e-05 | 8e-06 | 7.4e-06 | 8.800E-06 | 1.929E-06 | 7.400E-06 | 1.100E-05 |
| 6313 | 316 | C316M | A | 8.4e-06 | 1.1e-05 | 6.6e-06 | 8.667E-06 | 2.212E-06 | 6.600E-06 | 1.100E-05 |
| 6314 | 316 | C316F | A | 9.4e-06 | 1.5e-05 | 1.1e-05 | 1.180E-05 | 2.884E-06 | 9.400E-06 | 1.500E-05 |
| 6315 | 316 | C316P | A | 2.9e-06 | 1.3e-05 | 8.8e-06 | 8.233E-06 | 5.074E-06 | 2.900E-06 | 1.300E-05 |
| 6316 | 316 | C316S | A | 4.9e-06 | 4.3e-06 | 6.3e-06 | 5.167E-06 | 1.026E-06 | 4.300E-06 | 6.300E-06 |
| 6317 | 316 | C316T | A | 7.6e-06 | 9.6e-06 | 4.9e-06 | 7.367E-06 | 2.359E-06 | 4.900E-06 | 9.600E-06 |
| 6318 | 316 | C316W | A | 5.6e-06 | 7.3e-06 | 1.2e-05 | 8.300E-06 | 3.315E-06 | 5.600E-06 | 1.200E-05 |
| 6319 | 316 | C316Y | A | 6.1e-06 | 9.5e-06 | 6.3e-06 | 7.300E-06 | 1.908E-06 | 6.100E-06 | 9.500E-06 |
| 6320 | 316 | C316V | A | 1.3e-05 | 8.2e-06 | 7.7e-06 | 9.633E-06 | 2.926E-06 | 7.700E-06 | 1.300E-05 |
| 6321 | 317 | A317A | A | 9.4e-06 | 5.1e-06 | 9.2e-06 | 7.900E-06 | 2.427E-06 | 5.100E-06 | 9.400E-06 |
| 6322 | 317 | A317R | A | 7.1e-06 | 1.3e-05 | 8.5e-06 | 9.533E-06 | 3.083E-06 | 7.100E-06 | 1.300E-05 |
| 6323 | 317 | A317N | A | 9.4e-06 | 1.7e-05 | 1.2e-05 | 1.280E-05 | 3.863E-06 | 9.400E-06 | 1.700E-05 |
| 6324 | 317 | A317D | A | 1.3e-05 | 8.5e-06 | 6.8e-06 | 9.433E-06 | 3.204E-06 | 6.800E-06 | 1.300E-05 |
| 6325 | 317 | A317C | A | 9.6e-06 | 1.1e-05 | 1.1e-05 | 1.053E-05 | 8.083E-07 | 9.600E-06 | 1.100E-05 |
| 6326 | 317 | A317E | A | 1.8e-05 | 1.2e-05 | 7.2e-06 | 1.240E-05 | 5.411E-06 | 7.200E-06 | 1.800E-05 |
| 6327 | 317 | A317Q | A | 1.2e-05 | 6.9e-06 | 6.2e-06 | 8.367E-06 | 3.166E-06 | 6.200E-06 | 1.200E-05 |
| 6328 | 317 | A317G | A | 6.5e-06 | 3.6e-06 | 1e-05 | 6.700E-06 | 3.205E-06 | 3.600E-06 | 1.000E-05 |
| 6329 | 317 | A317H | A | 1.3e-05 | 7.3e-06 | 9.3e-06 | 9.867E-06 | 2.892E-06 | 7.300E-06 | 1.300E-05 |
| 6330 | 317 | A317I | A | 2.8e-05 | 9.1e-06 | 4.8e-06 | 1.397E-05 | 1.234E-05 | 4.800E-06 | 2.800E-05 |
| 6331 | 317 | A317L | A | 1e-05 | 6.3e-06 | 9.2e-06 | 8.500E-06 | 1.947E-06 | 6.300E-06 | 1.000E-05 |
| 6332 | 317 | A317K | A | 8.4e-06 | 7.5e-06 | 4.7e-06 | 6.867E-06 | 1.930E-06 | 4.700E-06 | 8.400E-06 |
| 6333 | 317 | A317M | A | 1.4e-05 | 8.7e-06 | 6.4e-06 | 9.700E-06 | 3.897E-06 | 6.400E-06 | 1.400E-05 |
| 6334 | 317 | A317F | A | 6e-06 | 5.2e-06 | 8.4e-06 | 6.533E-06 | 1.665E-06 | 5.200E-06 | 8.400E-06 |
| 6335 | 317 | A317P | A | 1.2e-05 | 6.6e-06 | 8.6e-06 | 9.067E-06 | 2.730E-06 | 6.600E-06 | 1.200E-05 |
| 6336 | 317 | A317S | A | 1.2e-05 | 8.3e-06 | 5e-06 | 8.433E-06 | 3.502E-06 | 5.000E-06 | 1.200E-05 |
| 6337 | 317 | A317T | A | 8.5e-06 | 9.7e-06 | 1.3e-05 | 1.040E-05 | 2.330E-06 | 8.500E-06 | 1.300E-05 |
| 6338 | 317 | A317W | A | 8.7e-06 | 5.1e-06 | 4.3e-06 | 6.033E-06 | 2.344E-06 | 4.300E-06 | 8.700E-06 |
| 6339 | 317 | A317Y | A | 8.4e-06 | 4.4e-06 | 5.7e-06 | 6.167E-06 | 2.040E-06 | 4.400E-06 | 8.400E-06 |
| 6340 | 317 | A317V | A | 1.1e-05 | 8.9e-06 | 8.7e-06 | 9.533E-06 | 1.274E-06 | 8.700E-06 | 1.100E-05 |
| 6341 | 318 | R318A | A | 7.2e-06 | 8.9e-06 | 1.4e-05 | 1.003E-05 | 3.539E-06 | 7.200E-06 | 1.400E-05 |
| 6342 | 318 | R318R | A | 6.4e-06 | 1.1e-05 | 8.1e-06 | 8.500E-06 | 2.326E-06 | 6.400E-06 | 1.100E-05 |
| 6343 | 318 | R318N | A | 4.4e-06 | 7.6e-06 | 9.7e-06 | 7.233E-06 | 2.669E-06 | 4.400E-06 | 9.700E-06 |
| 6344 | 318 | R318D | A | 9.4e-06 | 7e-06 | 5.5e-06 | 7.300E-06 | 1.967E-06 | 5.500E-06 | 9.400E-06 |
| 6345 | 318 | R318C | A | 5.3e-06 | 1.5e-05 | 8.4e-06 | 9.567E-06 | 4.954E-06 | 5.300E-06 | 1.500E-05 |
| 6346 | 318 | R318E | A | 8.8e-06 | 1e-05 | 6.1e-06 | 8.300E-06 | 1.997E-06 | 6.100E-06 | 1.000E-05 |
| 6347 | 318 | R318Q | A | 6.1e-06 | 1e-05 | 9.9e-06 | 8.667E-06 | 2.223E-06 | 6.100E-06 | 1.000E-05 |
| 6348 | 318 | R318G | A | 8.8e-06 | 8.6e-06 | 1.4e-05 | 1.047E-05 | 3.062E-06 | 8.600E-06 | 1.400E-05 |
| 6349 | 318 | R318H | A | 1.1e-05 | 8.8e-06 | 1.1e-05 | 1.027E-05 | 1.270E-06 | 8.800E-06 | 1.100E-05 |
| 6350 | 318 | R318I | A | 6.3e-06 | 6.5e-06 | 7e-06 | 6.600E-06 | 3.606E-07 | 6.300E-06 | 7.000E-06 |
| 6351 | 318 | R318L | A | 8.6e-06 | 5.2e-06 | 8.7e-06 | 7.500E-06 | 1.992E-06 | 5.200E-06 | 8.700E-06 |
| 6352 | 318 | R318K | A | 6.5e-06 | 9.6e-06 | 7.3e-06 | 7.800E-06 | 1.609E-06 | 6.500E-06 | 9.600E-06 |
| 6353 | 318 | R318M | A | 9.8e-06 | 9.6e-06 | 3.2e-06 | 7.533E-06 | 3.754E-06 | 3.200E-06 | 9.800E-06 |
| 6354 | 318 | R318F | A | 6.4e-06 | 8.2e-06 | 9.4e-06 | 8.000E-06 | 1.510E-06 | 6.400E-06 | 9.400E-06 |
| 6355 | 318 | R318P | A | 8.9e-06 | 1.2e-05 | 7e-06 | 9.300E-06 | 2.524E-06 | 7.000E-06 | 1.200E-05 |
| 6356 | 318 | R318S | A | 8.8e-06 | 8.7e-06 | 7.2e-06 | 8.233E-06 | 8.963E-07 | 7.200E-06 | 8.800E-06 |
| 6357 | 318 | R318T | A | 6.8e-06 | 1.3e-05 | 6.3e-06 | 8.700E-06 | 3.732E-06 | 6.300E-06 | 1.300E-05 |
| 6358 | 318 | R318W | A | 5.9e-06 | 1.3e-05 | 9.1e-06 | 9.333E-06 | 3.556E-06 | 5.900E-06 | 1.300E-05 |
| 6359 | 318 | R318Y | A | 4.3e-06 | 4.6e-06 | 1.1e-05 | 6.633E-06 | 3.785E-06 | 4.300E-06 | 1.100E-05 |
| 6360 | 318 | R318V | A | 7.5e-06 | 6.9e-06 | 6.6e-06 | 7.000E-06 | 4.583E-07 | 6.600E-06 | 7.500E-06 |
| 6361 | 319 | V319A | A | 4.7e-06 | 4.5e-06 | 6.7e-06 | 5.300E-06 | 1.217E-06 | 4.500E-06 | 6.700E-06 |
| 6362 | 319 | V319R | A | 2.2e-05 | 9.5e-06 | 5.2e-06 | 1.223E-05 | 8.727E-06 | 5.200E-06 | 2.200E-05 |
| 6363 | 319 | V319N | A | 6.6e-06 | 7.3e-06 | 6.6e-06 | 6.833E-06 | 4.041E-07 | 6.600E-06 | 7.300E-06 |
| 6364 | 319 | V319D | A | 6.9e-06 | 1.3e-05 | 9e-06 | 9.633E-06 | 3.099E-06 | 6.900E-06 | 1.300E-05 |
| 6365 | 319 | V319C | A | 9e-06 | 1.1e-05 | 9.9e-06 | 9.967E-06 | 1.002E-06 | 9.000E-06 | 1.100E-05 |
| 6366 | 319 | V319E | A | 1e-05 | 9.7e-06 | 7.6e-06 | 9.100E-06 | 1.308E-06 | 7.600E-06 | 1.000E-05 |
| 6367 | 319 | V319Q | A | 1e-05 | 6.1e-06 | 4.4e-06 | 6.833E-06 | 2.871E-06 | 4.400E-06 | 1.000E-05 |
| 6368 | 319 | V319G | A | 8.5e-06 | 3.9e-06 | 1.3e-05 | 8.467E-06 | 4.550E-06 | 3.900E-06 | 1.300E-05 |
| 6369 | 319 | V319H | A | 5e-06 | 1.2e-05 | 8.3e-06 | 8.433E-06 | 3.502E-06 | 5.000E-06 | 1.200E-05 |
| 6370 | 319 | V319I | A | 4.3e-06 | 1e-05 | 1.3e-05 | 9.100E-06 | 4.419E-06 | 4.300E-06 | 1.300E-05 |
| 6371 | 319 | V319L | A | 7e-06 | 1e-05 | 9.7e-06 | 8.900E-06 | 1.652E-06 | 7.000E-06 | 1.000E-05 |
| 6372 | 319 | V319K | A | 9.6e-06 | 1.1e-05 | 6.6e-06 | 9.067E-06 | 2.248E-06 | 6.600E-06 | 1.100E-05 |
| 6373 | 319 | V319M | A | 4.5e-06 | 8.2e-06 | 1.4e-05 | 8.900E-06 | 4.789E-06 | 4.500E-06 | 1.400E-05 |
| 6374 | 319 | V319F | A | 6.3e-06 | 9.7e-06 | 8.8e-06 | 8.267E-06 | 1.762E-06 | 6.300E-06 | 9.700E-06 |

| | | | | | | | | | | |
|---|---|---|---|---|---|---|---|---|---|---|
| 6375 | 319 | V319P | A | 9.6e-06 | 6.9e-06 | 1.2e-05 | 9.500E-06 | 2.551E-06 | 6.900E-06 | 1.200E-05 |
| 6376 | 319 | V319S | A | 6.8e-06 | 4.9e-06 | 9.3e-06 | 7.000E-06 | 2.207E-06 | 4.900E-06 | 9.300E-06 |
| 6377 | 319 | V319T | A | 7.4e-06 | 5.2e-06 | 5.1e-06 | 5.900E-06 | 1.300E-06 | 5.100E-06 | 7.400E-06 |
| 6378 | 319 | V319W | A | 5.1e-06 | 5e-06 | 6.7e-06 | 5.600E-06 | 9.539E-07 | 5.000E-06 | 6.700E-06 |
| 6379 | 319 | V319Y | A | 9.2e-06 | 7.5e-06 | 3.3e-06 | 6.667E-06 | 3.037E-06 | 3.300E-06 | 9.200E-06 |
| 6380 | 319 | V319V | A | 7.6e-06 | 9.2e-06 | 5e-06 | 7.267E-06 | 2.120E-06 | 5.000E-06 | 9.200E-06 |
| 6381 | 320 | C320A | A | 6.3e-06 | 5.6e-06 | 9.3e-06 | 7.067E-06 | 1.966E-06 | 5.600E-06 | 9.300E-06 |
| 6382 | 320 | C320R | A | 7.5e-06 | 7.6e-06 | 6.8e-06 | 7.300E-06 | 4.359E-07 | 6.800E-06 | 7.600E-06 |
| 6383 | 320 | C320N | A | 1e-05 | 4.5e-06 | 7.3e-06 | 7.267E-06 | 2.750E-06 | 4.500E-06 | 1.000E-05 |
| 6384 | 320 | C320D | A | 6.5e-06 | 8.4e-06 | 7.9e-06 | 7.600E-06 | 9.849E-07 | 6.500E-06 | 8.400E-06 |
| 6385 | 320 | C320C | A | 6.9e-06 | 1.1e-05 | 8.9e-06 | 8.933E-06 | 2.050E-06 | 6.900E-06 | 1.100E-05 |
| 6386 | 320 | C320E | A | 5.1e-06 | 7.8e-06 | 8.3e-06 | 7.067E-06 | 1.721E-06 | 5.100E-06 | 8.300E-06 |
| 6387 | 320 | C320Q | A | 9.3e-06 | 8e-06 | 9.2e-06 | 8.833E-06 | 7.234E-07 | 8.000E-06 | 9.300E-06 |
| 6388 | 320 | C320G | A | 5.1e-06 | 1.3e-05 | 4.7e-06 | 7.600E-06 | 4.681E-06 | 4.700E-06 | 1.300E-05 |
| 6389 | 320 | C320H | A | 6.7e-06 | 4.9e-06 | 8.4e-06 | 6.667E-06 | 1.750E-06 | 4.900E-06 | 8.400E-06 |
| 6390 | 320 | C320I | A | 7.5e-06 | 6.4e-06 | 9.9e-06 | 7.933E-06 | 1.790E-06 | 6.400E-06 | 9.900E-06 |
| 6391 | 320 | C320L | A | 5.9e-06 | 9.8e-06 | 5.7e-06 | 7.133E-06 | 2.312E-06 | 5.700E-06 | 9.800E-06 |
| 6392 | 320 | C320K | A | 3.9e-06 | 7.8e-06 | 5.9e-06 | 5.867E-06 | 1.950E-06 | 3.900E-06 | 7.800E-06 |
| 6393 | 320 | C320M | A | 1.2e-05 | 6.9e-06 | 9.6e-06 | 9.500E-06 | 2.551E-06 | 6.900E-06 | 1.200E-05 |
| 6394 | 320 | C320F | A | 6.5e-06 | 7.7e-06 | 1.1e-05 | 8.400E-06 | 2.330E-06 | 6.500E-06 | 1.100E-05 |
| 6395 | 320 | C320P | A | 1e-05 | 9.5e-06 | 3.7e-06 | 7.733E-06 | 3.502E-06 | 3.700E-06 | 1.000E-05 |
| 6396 | 320 | C320S | A | 6.5e-06 | 5.3e-06 | 6.7e-06 | 6.167E-06 | 7.572E-07 | 5.300E-06 | 6.700E-06 |
| 6397 | 320 | C320T | A | 1.1e-05 | 7.6e-06 | 4.5e-06 | 7.700E-06 | 3.251E-06 | 4.500E-06 | 1.100E-05 |
| 6398 | 320 | C320W | A | 7.9e-06 | 8.7e-06 | 8.1e-06 | 8.233E-06 | 4.163E-07 | 7.900E-06 | 8.700E-06 |
| 6399 | 320 | C320Y | A | 8.3e-06 | 6.5e-06 | 1.1e-05 | 8.600E-06 | 2.265E-06 | 6.500E-06 | 1.100E-05 |
| 6400 | 320 | C320V | A | 5.3e-06 | 3.2e-06 | 6.4e-06 | 4.967E-06 | 1.626E-06 | 3.200E-06 | 6.400E-06 |
| 6401 | 321 | Y321A | A | 6.5e-06 | 1.3e-05 | 6.2e-06 | 8.567E-06 | 3.842E-06 | 6.200E-06 | 1.300E-05 |
| 6402 | 321 | Y321R | A | 1.5e-05 | 7.1e-06 | 1e-05 | 1.070E-05 | 3.996E-06 | 7.100E-06 | 1.500E-05 |
| 6403 | 321 | Y321N | A | 9e-06 | 8.6e-06 | 1.4e-05 | 1.053E-05 | 3.009E-06 | 8.600E-06 | 1.400E-05 |
| 6404 | 321 | Y321D | A | 1.1e-05 | 1.2e-05 | 1e-05 | 1.100E-05 | 1.000E-06 | 1.000E-05 | 1.200E-05 |
| 6405 | 321 | Y321C | A | 1e-05 | 4.7e-06 | 9.7e-06 | 8.133E-06 | 2.977E-06 | 4.700E-06 | 1.000E-05 |
| 6406 | 321 | Y321E | A | 6.5e-06 | 7.2e-06 | 4.6e-06 | 6.100E-06 | 1.345E-06 | 4.600E-06 | 7.200E-06 |
| 6407 | 321 | Y321Q | A | 8.9e-06 | 9.1e-06 | 1.3e-05 | 1.033E-05 | 2.312E-06 | 8.900E-06 | 1.300E-05 |
| 6408 | 321 | Y321G | A | 1.1e-05 | 9.2e-06 | 5.4e-06 | 8.533E-06 | 2.859E-06 | 5.400E-06 | 1.100E-05 |
| 6409 | 321 | Y321H | A | 5.8e-06 | 1e-05 | 6.2e-06 | 7.333E-06 | 2.318E-06 | 5.800E-06 | 1.000E-05 |
| 6410 | 321 | Y321I | A | 1.2e-05 | 9.2e-06 | 6.2e-06 | 9.133E-06 | 2.901E-06 | 6.200E-06 | 1.200E-05 |
| 6411 | 321 | Y321L | A | 5.9e-06 | 5.7e-06 | 8.6e-06 | 6.733E-06 | 1.620E-06 | 5.700E-06 | 8.600E-06 |
| 6412 | 321 | Y321K | A | 1.1e-05 | 6.1e-06 | 1e-05 | 9.033E-06 | 2.589E-06 | 6.100E-06 | 1.100E-05 |
| 6413 | 321 | Y321M | A | 6.1e-06 | 6.8e-06 | 5.5e-06 | 6.133E-06 | 6.506E-07 | 5.500E-06 | 6.800E-06 |
| 6414 | 321 | Y321F | A | 9.3e-06 | 8.5e-06 | 4.7e-06 | 7.500E-06 | 2.458E-06 | 4.700E-06 | 9.300E-06 |
| 6415 | 321 | Y321P | A | 9.9e-06 | 1.4e-05 | 6.7e-06 | 1.020E-05 | 3.659E-06 | 6.700E-06 | 1.400E-05 |
| 6416 | 321 | Y321S | A | 1.3e-05 | 6.1e-06 | 8.7e-06 | 9.267E-06 | 3.485E-06 | 6.100E-06 | 1.300E-05 |
| 6417 | 321 | Y321T | A | 4.6e-06 | 1e-05 | 5.9e-06 | 6.833E-06 | 2.818E-06 | 4.600E-06 | 1.000E-05 |
| 6418 | 321 | Y321W | A | 7.7e-06 | 9.6e-06 | 8.9e-06 | 8.733E-06 | 9.609E-07 | 7.700E-06 | 9.600E-06 |
| 6419 | 321 | Y321Y | A | 7.9e-06 | 9.2e-06 | 1.4e-05 | 1.037E-05 | 3.213E-06 | 7.900E-06 | 1.400E-05 |
| 6420 | 321 | Y321V | A | 1.5e-05 | 6.2e-06 | 7e-06 | 9.400E-06 | 4.866E-06 | 6.200E-06 | 1.500E-05 |
| 6421 | 322 | G322A | A | 6.7e-06 | 6.5e-06 | 8e-06 | 7.067E-06 | 8.145E-07 | 6.500E-06 | 8.000E-06 |
| 6422 | 322 | G322R | A | 9.4e-06 | 9.4e-06 | 5.7e-06 | 8.167E-06 | 2.136E-06 | 5.700E-06 | 9.400E-06 |
| 6423 | 322 | G322N | A | 7.4e-06 | 1.2e-05 | 9.2e-06 | 9.533E-06 | 2.318E-06 | 7.400E-06 | 1.200E-05 |
| 6424 | 322 | G322D | A | 6.5e-06 | 1e-05 | 6.6e-06 | 7.700E-06 | 1.992E-06 | 6.500E-06 | 1.000E-05 |
| 6425 | 322 | G322C | A | 7.9e-06 | 5.6e-06 | 5.1e-06 | 6.200E-06 | 1.493E-06 | 5.100E-06 | 7.900E-06 |
| 6426 | 322 | G322E | A | 8.9e-06 | 1e-05 | 9.7e-06 | 9.533E-06 | 5.686E-07 | 8.900E-06 | 1.000E-05 |
| 6427 | 322 | G322Q | A | 7.6e-06 | 7.3e-06 | 9.1e-06 | 8.000E-06 | 9.644E-07 | 7.300E-06 | 9.100E-06 |
| 6428 | 322 | G322G | A | 6.5e-06 | 4.9e-06 | 6.2e-06 | 5.867E-06 | 8.505E-07 | 4.900E-06 | 6.500E-06 |
| 6429 | 322 | G322H | A | 1.6e-05 | 4.1e-06 | 6.8e-06 | 8.967E-06 | 6.239E-06 | 4.100E-06 | 1.600E-05 |
| 6430 | 322 | G322I | A | 9.4e-06 | 1.1e-05 | 7.8e-06 | 9.400E-06 | 1.600E-06 | 7.800E-06 | 1.100E-05 |
| 6431 | 322 | G322L | A | 9.5e-06 | 5.6e-06 | 8.3e-06 | 7.800E-06 | 1.997E-06 | 5.600E-06 | 9.500E-06 |
| 6432 | 322 | G322K | A | 6.8e-06 | 6.3e-06 | 4.8e-06 | 5.967E-06 | 1.041E-06 | 4.800E-06 | 6.800E-06 |
| 6433 | 322 | G322M | A | 6.3e-06 | 6.3e-06 | 1.2e-05 | 8.200E-06 | 3.291E-06 | 6.300E-06 | 1.200E-05 |
| 6434 | 322 | G322F | A | 4.1e-06 | 7.9e-06 | 1.5e-05 | 9.000E-06 | 5.533E-06 | 4.100E-06 | 1.500E-05 |
| 6435 | 322 | G322P | A | 5e-06 | 7.7e-06 | 6e-06 | 6.233E-06 | 1.365E-06 | 5.000E-06 | 7.700E-06 |
| 6436 | 322 | G322S | A | 6.9e-06 | 7.4e-06 | 8.8e-06 | 7.700E-06 | 9.849E-07 | 6.900E-06 | 8.800E-06 |
| 6437 | 322 | G322T | A | 6e-06 | 7.7e-06 | 6.5e-06 | 6.733E-06 | 8.737E-07 | 6.000E-06 | 7.700E-06 |
| 6438 | 322 | G322W | A | 1.4e-05 | 9.5e-06 | 7.1e-06 | 1.020E-05 | 3.503E-06 | 7.100E-06 | 1.400E-05 |
| 6439 | 322 | G322Y | A | 7.6e-06 | 9.4e-06 | 4.6e-06 | 7.200E-06 | 2.425E-06 | 4.600E-06 | 9.400E-06 |
| 6440 | 322 | G322V | A | 1e-05 | 5.2e-06 | 7.9e-06 | 7.700E-06 | 2.406E-06 | 5.200E-06 | 1.000E-05 |
| 6441 | 323 | L323A | A | 7.7e-06 | 1.4e-05 | 4.4e-06 | 8.700E-06 | 4.877E-06 | 4.400E-06 | 1.400E-05 |
| 6442 | 323 | L323R | A | 7e-06 | 5e-06 | 7.5e-06 | 6.500E-06 | 1.323E-06 | 5.000E-06 | 7.500E-06 |
| 6443 | 323 | L323N | A | 8.8e-06 | 7e-06 | 5.1e-06 | 6.967E-06 | 1.850E-06 | 5.100E-06 | 8.800E-06 |
| 6444 | 323 | L323D | A | 1.3e-05 | 5.9e-06 | 9.1e-06 | 9.333E-06 | 3.556E-06 | 5.900E-06 | 1.300E-05 |
| 6445 | 323 | L323C | A | 4.6e-06 | 8.6e-06 | 7.6e-06 | 6.933E-06 | 2.082E-06 | 4.600E-06 | 8.600E-06 |
| 6446 | 323 | L323E | A | 6.1e-06 | 6.7e-06 | 4.7e-06 | 5.833E-06 | 1.026E-06 | 4.700E-06 | 6.700E-06 |
| 6447 | 323 | L323Q | A | 1.2e-05 | 5.3e-06 | 8.2e-06 | 8.500E-06 | 3.360E-06 | 5.300E-06 | 1.200E-05 |
| 6448 | 323 | L323G | A | 5.5e-06 | 9.1e-06 | 1.2e-05 | 8.867E-06 | 3.256E-06 | 5.500E-06 | 1.200E-05 |
| 6449 | 323 | L323H | A | 6.3e-06 | 6.3e-06 | 4.9e-06 | 5.833E-06 | 8.083E-07 | 4.900E-06 | 6.300E-06 |
| 6450 | 323 | L323I | A | 5.7e-06 | 1.1e-05 | 7.6e-06 | 8.100E-06 | 2.685E-06 | 5.700E-06 | 1.100E-05 |
| 6451 | 323 | L323L | A | 8.2e-06 | 7.4e-06 | 5.8e-06 | 7.133E-06 | 1.222E-06 | 5.800E-06 | 8.200E-06 |
| 6452 | 323 | L323K | A | 6.3e-06 | 8e-06 | 6.5e-06 | 6.933E-06 | 9.292E-07 | 6.300E-06 | 8.000E-06 |
| 6453 | 323 | L323M | A | 6.6e-06 | 1.2e-05 | 8.2e-06 | 8.933E-06 | 2.774E-06 | 6.600E-06 | 1.200E-05 |
| 6454 | 323 | L323F | A | 7.4e-06 | 6.5e-06 | 8.3e-06 | 7.400E-06 | 9.000E-07 | 6.500E-06 | 8.300E-06 |
| 6455 | 323 | L323P | A | 6.3e-06 | 8.1e-06 | 6.3e-06 | 6.900E-06 | 1.039E-06 | 6.300E-06 | 8.100E-06 |
| 6456 | 323 | L323S | A | 7.8e-06 | 6.1e-06 | 8.4e-06 | 7.433E-06 | 1.193E-06 | 6.100E-06 | 8.400E-06 |
| 6457 | 323 | L323T | A | 6.5e-06 | 5.9e-06 | 9.3e-06 | 7.233E-06 | 1.815E-06 | 5.900E-06 | 9.300E-06 |
| 6458 | 323 | L323W | A | 7.3e-06 | 9e-06 | 7.8e-06 | 8.033E-06 | 8.737E-07 | 7.300E-06 | 9.000E-06 |
| 6459 | 323 | L323Y | A | 9.5e-06 | 1.2e-05 | 1.3e-05 | 1.150E-05 | 1.803E-06 | 9.500E-06 | 1.300E-05 |

| | | | | | | | | | | |
|---|---|---|---|---|---|---|---|---|---|---|
| 6460 | 323 | L323V | A | 6.6e-06 | 6.6e-06 | 6.7e-06 | 6.633E-06 | 5.774E-08 | 6.600E-06 | 6.700E-06 |
| 6461 | 324 | G324A | A | 4.7e-06 | 4.2e-06 | 9.6e-06 | 6.167E-06 | 2.984E-06 | 4.200E-06 | 9.600E-06 |
| 6462 | 324 | G324R | A | 1.1e-05 | 1.4e-05 | 6.8e-06 | 1.060E-05 | 3.617E-06 | 6.800E-06 | 1.400E-05 |
| 6463 | 324 | G324N | A | 4.3e-06 | 6.7e-06 | 8.1e-06 | 6.367E-06 | 1.922E-06 | 4.300E-06 | 8.100E-06 |
| 6464 | 324 | G324D | A | 7.1e-06 | 1.2e-05 | 4.8e-06 | 7.967E-06 | 3.677E-06 | 4.800E-06 | 1.200E-05 |
| 6465 | 324 | G324C | A | 4.6e-06 | 5.9e-06 | 8.5e-06 | 6.333E-06 | 1.986E-06 | 4.600E-06 | 8.500E-06 |
| 6466 | 324 | G324E | A | 3.9e-06 | 4.3e-06 | 8.3e-06 | 5.500E-06 | 2.433E-06 | 3.900E-06 | 8.300E-06 |
| 6467 | 324 | G324Q | A | 1.1e-05 | 7.1e-06 | 6e-06 | 8.033E-06 | 2.627E-06 | 6.000E-06 | 1.100E-05 |
| 6468 | 324 | G324G | A | 8.2e-06 | 1e-05 | 8.1e-06 | 8.767E-06 | 1.069E-06 | 8.100E-06 | 1.000E-05 |
| 6469 | 324 | G324H | A | 5.3e-06 | 7.5e-06 | 1.1e-05 | 7.933E-06 | 2.875E-06 | 5.300E-06 | 1.100E-05 |
| 6470 | 324 | G324I | A | 5.7e-06 | 9.8e-06 | 6.8e-06 | 7.433E-06 | 2.122E-06 | 5.700E-06 | 9.800E-06 |
| 6471 | 324 | G324L | A | 8.2e-06 | 7.8e-06 | 9e-06 | 8.333E-06 | 6.110E-07 | 7.800E-06 | 9.000E-06 |
| 6472 | 324 | G324K | A | 4.9e-06 | 9.5e-06 | 8.4e-06 | 7.600E-06 | 2.402E-06 | 4.900E-06 | 9.500E-06 |
| 6473 | 324 | G324M | A | 9.1e-06 | 1.2e-05 | 8.7e-06 | 9.933E-06 | 1.801E-06 | 8.700E-06 | 1.200E-05 |
| 6474 | 324 | G324F | A | 9.2e-06 | 7.8e-06 | 7.3e-06 | 8.100E-06 | 9.849E-07 | 7.300E-06 | 9.200E-06 |
| 6475 | 324 | G324P | A | 6.7e-06 | 7.1e-06 | 5.5e-06 | 6.433E-06 | 8.327E-07 | 5.500E-06 | 7.100E-06 |
| 6476 | 324 | G324S | A | 6.1e-06 | 1.1e-05 | 5.9e-06 | 7.667E-06 | 2.888E-06 | 5.900E-06 | 1.100E-05 |
| 6477 | 324 | G324T | A | 9.8e-06 | 9.2e-06 | 4e-06 | 7.667E-06 | 3.190E-06 | 4.000E-06 | 9.800E-06 |
| 6478 | 324 | G324W | A | 5.2e-06 | 8.4e-06 | 3.7e-06 | 5.767E-06 | 2.401E-06 | 3.700E-06 | 8.400E-06 |
| 6479 | 324 | G324Y | A | 8.3e-06 | 1.2e-05 | 6e-06 | 8.767E-06 | 3.027E-06 | 6.000E-06 | 1.200E-05 |
| 6480 | 324 | G324V | A | 8.3e-06 | 1.2e-05 | 4.8e-06 | 8.367E-06 | 3.600E-06 | 4.800E-06 | 1.200E-05 |
| 6481 | 325 | M325A | A | 8.3e-06 | 1e-05 | 6.6e-06 | 8.300E-06 | 1.700E-06 | 6.600E-06 | 1.000E-05 |
| 6482 | 325 | M325R | A | 7e-06 | 9.3e-06 | 3.4e-06 | 6.567E-06 | 2.974E-06 | 3.400E-06 | 9.300E-06 |
| 6483 | 325 | M325N | A | 1.1e-05 | 9.9e-06 | 4.7e-06 | 8.533E-06 | 3.365E-06 | 4.700E-06 | 1.100E-05 |
| 6484 | 325 | M325D | A | 7.2e-06 | 7.3e-06 | 4.7e-06 | 6.400E-06 | 1.473E-06 | 4.700E-06 | 7.300E-06 |
| 6485 | 325 | M325C | A | 7e-06 | 4.5e-06 | 9.5e-06 | 7.000E-06 | 2.500E-06 | 4.500E-06 | 9.500E-06 |
| 6486 | 325 | M325E | A | 5.9e-06 | 7.6e-06 | 5.9e-06 | 6.467E-06 | 9.815E-07 | 5.900E-06 | 7.600E-06 |
| 6487 | 325 | M325Q | A | 5.3e-06 | 6.2e-06 | 5.6e-06 | 5.700E-06 | 4.583E-07 | 5.300E-06 | 6.200E-06 |
| 6488 | 325 | M325G | A | 6.9e-06 | 1.6e-05 | 3e-06 | 8.633E-06 | 6.671E-06 | 3.000E-06 | 1.600E-05 |
| 6489 | 325 | M325H | A | 8.5e-06 | 1.7e-05 | 8.9e-06 | 1.147E-05 | 4.796E-06 | 8.500E-06 | 1.700E-05 |
| 6490 | 325 | M325I | A | 1.6e-05 | 5.5e-06 | 7e-06 | 9.500E-06 | 5.679E-06 | 5.500E-06 | 1.600E-05 |
| 6491 | 325 | M325L | A | 1.3e-05 | 9.1e-06 | 5.2e-06 | 9.100E-06 | 3.900E-06 | 5.200E-06 | 1.300E-05 |
| 6492 | 325 | M325K | A | 7.1e-06 | 1.1e-05 | 6.2e-06 | 8.100E-06 | 2.551E-06 | 6.200E-06 | 1.100E-05 |
| 6493 | 325 | M325M | A | 9.8e-06 | 1.2e-05 | 8.2e-06 | 1.000E-05 | 1.908E-06 | 8.200E-06 | 1.200E-05 |
| 6494 | 325 | M325F | A | 7.9e-06 | 1.5e-05 | 9.4e-06 | 1.077E-05 | 3.742E-06 | 7.900E-06 | 1.500E-05 |
| 6495 | 325 | M325P | A | 7.8e-06 | 1.2e-05 | 4.6e-06 | 8.133E-06 | 3.711E-06 | 4.600E-06 | 1.200E-05 |
| 6496 | 325 | M325S | A | 5.4e-06 | 6.2e-06 | 7.1e-06 | 6.233E-06 | 8.505E-07 | 5.400E-06 | 7.100E-06 |
| 6497 | 325 | M325T | A | 6.5e-06 | 5.8e-06 | 8.8e-06 | 7.033E-06 | 1.570E-06 | 5.800E-06 | 8.800E-06 |
| 6498 | 325 | M325W | A | 8e-06 | 1.1e-05 | 9.6e-06 | 9.533E-06 | 1.501E-06 | 8.000E-06 | 1.100E-05 |
| 6499 | 325 | M325Y | A | 7.1e-06 | 1.4e-05 | 4.5e-06 | 8.533E-06 | 4.910E-06 | 4.500E-06 | 1.400E-05 |
| 6500 | 325 | M325V | A | 5e-06 | 9.4e-06 | 5e-06 | 6.467E-06 | 2.540E-06 | 5.000E-06 | 9.400E-06 |
| 6501 | 326 | E326A | A | 8.6e-06 | 6.4e-06 | 4.5e-06 | 6.500E-06 | 2.052E-06 | 4.500E-06 | 8.600E-06 |
| 6502 | 326 | E326R | A | 9.2e-06 | 8.9e-06 | 1.1e-05 | 9.700E-06 | 1.136E-06 | 8.900E-06 | 1.100E-05 |
| 6503 | 326 | E326N | A | 9.6e-06 | 5.5e-06 | 6.9e-06 | 7.333E-06 | 2.084E-06 | 5.500E-06 | 9.600E-06 |
| 6504 | 326 | E326D | A | 6.4e-06 | 8.7e-06 | 1.2e-05 | 9.033E-06 | 2.815E-06 | 6.400E-06 | 1.200E-05 |
| 6505 | 326 | E326C | A | 7e-06 | 1.2e-05 | 9.8e-06 | 9.600E-06 | 2.506E-06 | 7.000E-06 | 1.200E-05 |
| 6506 | 326 | E326E | A | 1.2e-05 | 4.9e-06 | 5.7e-06 | 7.533E-06 | 3.889E-06 | 4.900E-06 | 1.200E-05 |
| 6507 | 326 | E326Q | A | 5.6e-06 | 8.4e-06 | 9.1e-06 | 7.700E-06 | 1.852E-06 | 5.600E-06 | 9.100E-06 |
| 6508 | 326 | E326G | A | 9.6e-06 | 1.1e-05 | 1e-05 | 1.020E-05 | 7.211E-07 | 9.600E-06 | 1.100E-05 |
| 6509 | 326 | E326H | A | 7e-06 | 5.5e-06 | 6.8e-06 | 6.433E-06 | 8.145E-07 | 5.500E-06 | 7.000E-06 |
| 6510 | 326 | E326I | A | 5.5e-06 | 9.8e-06 | 1.5e-05 | 1.010E-05 | 4.757E-06 | 5.500E-06 | 1.500E-05 |
| 6511 | 326 | E326L | A | 8.5e-06 | 6.1e-06 | 6.2e-06 | 6.933E-06 | 1.358E-06 | 6.100E-06 | 8.500E-06 |
| 6512 | 326 | E326K | A | 5.9e-06 | 6.1e-06 | 1.1e-05 | 7.667E-06 | 2.888E-06 | 5.900E-06 | 1.100E-05 |
| 6513 | 326 | E326M | A | 9.2e-06 | 9.8e-06 | 6.7e-06 | 8.567E-06 | 1.644E-06 | 6.700E-06 | 9.800E-06 |
| 6514 | 326 | E326F | A | 7.8e-06 | 5.8e-06 | 6.5e-06 | 6.700E-06 | 1.015E-06 | 5.800E-06 | 7.800E-06 |
| 6515 | 326 | E326P | A | 9.2e-06 | 8e-06 | 8.7e-06 | 8.633E-06 | 6.028E-07 | 8.000E-06 | 9.200E-06 |
| 6516 | 326 | E326S | A | 3.6e-06 | 4e-06 | 4.9e-06 | 4.167E-06 | 6.658E-07 | 3.600E-06 | 4.900E-06 |
| 6517 | 326 | E326T | A | 9.3e-06 | 9.6e-06 | 8.8e-06 | 9.233E-06 | 4.041E-07 | 8.800E-06 | 9.600E-06 |
| 6518 | 326 | E326W | A | 6.6e-06 | 1.3e-05 | 8.1e-06 | 9.233E-06 | 3.347E-06 | 6.600E-06 | 1.300E-05 |
| 6519 | 326 | E326Y | A | 7.7e-06 | 8.5e-06 | 5.2e-06 | 7.133E-06 | 1.721E-06 | 5.200E-06 | 8.500E-06 |
| 6520 | 326 | E326V | A | 1.1e-05 | 5.8e-06 | 9.3e-06 | 8.700E-06 | 2.651E-06 | 5.800E-06 | 1.100E-05 |
| 6521 | 327 | H327A | A | 9e-06 | 7.2e-06 | 1.1e-05 | 9.067E-06 | 1.901E-06 | 7.200E-06 | 1.100E-05 |
| 6522 | 327 | H327R | A | 8.7e-06 | 1.4e-05 | 1.3e-05 | 1.190E-05 | 2.816E-06 | 8.700E-06 | 1.400E-05 |
| 6523 | 327 | H327N | A | 6.9e-06 | 6.9e-06 | 6.1e-06 | 6.633E-06 | 4.619E-07 | 6.100E-06 | 6.900E-06 |
| 6524 | 327 | H327D | A | 7.6e-06 | 6.6e-06 | 1.1e-05 | 8.400E-06 | 2.307E-06 | 6.600E-06 | 1.100E-05 |
| 6525 | 327 | H327C | A | 6.5e-06 | 8.1e-06 | 8.7e-06 | 7.767E-06 | 1.137E-06 | 6.500E-06 | 8.700E-06 |
| 6526 | 327 | H327E | A | 5.6e-06 | 9.7e-06 | 1e-05 | 8.433E-06 | 2.458E-06 | 5.600E-06 | 1.000E-05 |
| 6527 | 327 | H327Q | A | 8e-06 | 6.4e-06 | 8.6e-06 | 7.667E-06 | 1.137E-06 | 6.400E-06 | 8.600E-06 |
| 6528 | 327 | H327G | A | 1.4e-05 | 1.3e-05 | 9.5e-06 | 1.217E-05 | 2.363E-06 | 9.500E-06 | 1.400E-05 |
| 6529 | 327 | H327H | A | 1.4e-05 | 6.2e-06 | 5.7e-06 | 8.633E-06 | 4.654E-06 | 5.700E-06 | 1.400E-05 |
| 6530 | 327 | H327I | A | 8.6e-06 | 6.3e-06 | 1e-05 | 8.300E-06 | 1.868E-06 | 6.300E-06 | 1.000E-05 |
| 6531 | 327 | H327L | A | 1.3e-05 | 9.7e-06 | 1e-05 | 1.090E-05 | 1.825E-06 | 9.700E-06 | 1.300E-05 |
| 6532 | 327 | H327K | A | 5.2e-06 | 6.4e-06 | 5.2e-06 | 5.600E-06 | 6.928E-07 | 5.200E-06 | 6.400E-06 |
| 6533 | 327 | H327M | A | 1.3e-05 | 7e-06 | 1.1e-05 | 1.033E-05 | 3.055E-06 | 7.000E-06 | 1.300E-05 |
| 6534 | 327 | H327F | A | 6.6e-06 | 8.9e-06 | 7.9e-06 | 7.800E-06 | 1.153E-06 | 6.600E-06 | 8.900E-06 |
| 6535 | 327 | H327P | A | 5.3e-06 | 4.9e-06 | 4.5e-06 | 4.900E-06 | 4.000E-07 | 4.500E-06 | 5.300E-06 |
| 6536 | 327 | H327S | A | 8.5e-06 | 8.2e-06 | 1.3e-05 | 9.900E-06 | 2.689E-06 | 8.200E-06 | 1.300E-05 |
| 6537 | 327 | H327T | A | 8.5e-06 | 7.8e-06 | 1e-05 | 8.767E-06 | 1.124E-06 | 7.800E-06 | 1.000E-05 |
| 6538 | 327 | H327W | A | 6e-06 | 7.8e-06 | 8e-06 | 7.267E-06 | 1.102E-06 | 6.000E-06 | 8.000E-06 |
| 6539 | 327 | H327Y | A | 5.9e-06 | 9.9e-06 | 7.1e-06 | 7.633E-06 | 2.053E-06 | 5.900E-06 | 9.900E-06 |
| 6540 | 327 | H327V | A | 6.1e-06 | 7.4e-06 | 1.5e-05 | 9.500E-06 | 4.807E-06 | 6.100E-06 | 1.500E-05 |
| 6541 | 328 | L328A | A | 6.2e-06 | 5.3e-06 | 9.4e-06 | 6.967E-06 | 2.155E-06 | 5.300E-06 | 9.400E-06 |
| 6542 | 328 | L328R | A | 9.2e-06 | 6.5e-06 | 8.9e-06 | 8.200E-06 | 1.480E-06 | 6.500E-06 | 9.200E-06 |
| 6543 | 328 | L328N | A | 7.2e-06 | 6.5e-06 | 1.2e-05 | 8.567E-06 | 2.994E-06 | 6.500E-06 | 1.200E-05 |
| 6544 | 328 | L328D | A | 8.4e-06 | 5.7e-06 | 5e-06 | 6.367E-06 | 1.795E-06 | 5.000E-06 | 8.400E-06 |

| | | | | | | | | | | |
|---|---|---|---|---|---|---|---|---|---|---|
| 6545 | 328 | L328C | A | 7.5e-06 | 6.3e-06 | 9.9e-06 | 7.900E-06 | 1.833E-06 | 6.300E-06 | 9.900E-06 |
| 6546 | 328 | L328E | A | 1.4e-05 | 7.6e-06 | 6.6e-06 | 9.400E-06 | 4.015E-06 | 6.600E-06 | 1.400E-05 |
| 6547 | 328 | L328Q | A | 8.6e-06 | 6.1e-06 | 6.2e-06 | 6.967E-06 | 1.415E-06 | 6.100E-06 | 8.600E-06 |
| 6548 | 328 | L328G | A | 6.9e-06 | 8.3e-06 | 1.5e-05 | 1.007E-05 | 4.329E-06 | 6.900E-06 | 1.500E-05 |
| 6549 | 328 | L328H | A | 1e-05 | 1.4e-05 | 6.3e-06 | 1.010E-05 | 3.851E-06 | 6.300E-06 | 1.400E-05 |
| 6550 | 328 | L328I | A | 9.5e-06 | 6.7e-06 | 5.9e-06 | 7.367E-06 | 1.890E-06 | 5.900E-06 | 9.500E-06 |
| 6551 | 328 | L328L | A | 8.4e-06 | 6.9e-06 | 1.3e-05 | 9.433E-06 | 3.179E-06 | 6.900E-06 | 1.300E-05 |
| 6552 | 328 | L328K | A | 1.4e-05 | 5.9e-06 | 7.7e-06 | 9.200E-06 | 4.253E-06 | 5.900E-06 | 1.400E-05 |
| 6553 | 328 | L328M | A | 9.3e-06 | 1e-05 | 9.3e-06 | 9.533E-06 | 4.041E-07 | 9.300E-06 | 1.000E-05 |
| 6554 | 328 | L328F | A | 7e-06 | 9.6e-06 | 9.3e-06 | 8.633E-06 | 1.422E-06 | 7.000E-06 | 9.600E-06 |
| 6555 | 328 | L328P | A | 4.2e-06 | 9.9e-06 | 6.7e-06 | 6.933E-06 | 2.857E-06 | 4.200E-06 | 9.900E-06 |
| 6556 | 328 | L328S | A | 9e-06 | 6.6e-06 | 1e-05 | 8.533E-06 | 1.747E-06 | 6.600E-06 | 1.000E-05 |
| 6557 | 328 | L328T | A | 8.9e-06 | 9e-06 | 5.8e-06 | 7.900E-06 | 1.819E-06 | 5.800E-06 | 9.000E-06 |
| 6558 | 328 | L328W | A | 9.6e-06 | 6.2e-06 | 5.9e-06 | 7.233E-06 | 2.055E-06 | 5.900E-06 | 9.600E-06 |
| 6559 | 328 | L328Y | A | 7.1e-06 | 6.6e-06 | 9e-06 | 7.567E-06 | 1.266E-06 | 6.600E-06 | 9.000E-06 |
| 6560 | 328 | L328V | A | 8e-06 | 1.4e-05 | 7.6e-06 | 9.867E-06 | 3.585E-06 | 7.600E-06 | 1.400E-05 |
| 6561 | 329 | R329A | A | 7.2e-06 | 7.7e-06 | 9e-06 | 7.967E-06 | 9.292E-07 | 7.200E-06 | 9.000E-06 |
| 6562 | 329 | R329R | A | 1.3e-05 | 6e-06 | 6.3e-06 | 8.433E-06 | 3.958E-06 | 6.000E-06 | 1.300E-05 |
| 6563 | 329 | R329N | A | 8.8e-06 | 1.1e-05 | 1.6e-05 | 1.193E-05 | 3.690E-06 | 8.800E-06 | 1.600E-05 |
| 6564 | 329 | R329D | A | 5.9e-06 | 4.9e-06 | 8.1e-06 | 6.300E-06 | 1.637E-06 | 4.900E-06 | 8.100E-06 |
| 6565 | 329 | R329C | A | 6.6e-06 | 8.1e-06 | 5.2e-06 | 6.633E-06 | 1.450E-06 | 5.200E-06 | 8.100E-06 |
| 6566 | 329 | R329E | A | 1.3e-05 | 6.5e-06 | 9.9e-06 | 9.800E-06 | 3.251E-06 | 6.500E-06 | 1.300E-05 |
| 6567 | 329 | R329Q | A | 8.2e-06 | 5.2e-06 | 1.1e-05 | 8.133E-06 | 2.901E-06 | 5.200E-06 | 1.100E-05 |
| 6568 | 329 | R329G | A | 8e-06 | 7.8e-06 | 6.9e-06 | 7.567E-06 | 5.859E-07 | 6.900E-06 | 8.000E-06 |
| 6569 | 329 | R329H | A | 7.8e-06 | 9.5e-06 | 7.4e-06 | 8.233E-06 | 1.115E-06 | 7.400E-06 | 9.500E-06 |
| 6570 | 329 | R329I | A | 6.9e-06 | 1.3e-05 | 1.2e-05 | 1.063E-05 | 3.272E-06 | 6.900E-06 | 1.300E-05 |
| 6571 | 329 | R329L | A | 6.2e-06 | 6.2e-06 | 1.1e-05 | 7.800E-06 | 2.771E-06 | 6.200E-06 | 1.100E-05 |
| 6572 | 329 | R329K | A | 1.2e-05 | 5.8e-06 | 8.5e-06 | 8.767E-06 | 3.109E-06 | 5.800E-06 | 1.200E-05 |
| 6573 | 329 | R329M | A | 8.5e-06 | 5.7e-06 | 5.8e-06 | 6.667E-06 | 1.589E-06 | 5.700E-06 | 8.500E-06 |
| 6574 | 329 | R329F | A | 9.2e-06 | 6e-06 | 9.2e-06 | 8.133E-06 | 1.848E-06 | 6.000E-06 | 9.200E-06 |
| 6575 | 329 | R329P | A | 4.9e-06 | 8.1e-06 | 1.2e-05 | 8.333E-06 | 3.556E-06 | 4.900E-06 | 1.200E-05 |
| 6576 | 329 | R329S | A | 5.1e-06 | 5.9e-06 | 1.4e-05 | 8.333E-06 | 4.924E-06 | 5.100E-06 | 1.400E-05 |
| 6577 | 329 | R329T | A | 9.1e-06 | 9.6e-06 | 7.9e-06 | 8.867E-06 | 8.737E-07 | 7.900E-06 | 9.600E-06 |
| 6578 | 329 | R329W | A | 7.5e-06 | 1.1e-05 | 6.2e-06 | 8.233E-06 | 2.483E-06 | 6.200E-06 | 1.100E-05 |
| 6579 | 329 | R329Y | A | 7.2e-06 | 6.2e-06 | 5.5e-06 | 6.300E-06 | 8.544E-07 | 5.500E-06 | 7.200E-06 |
| 6580 | 329 | R329V | A | 9.5e-06 | 8.9e-06 | 7.3e-06 | 8.567E-06 | 1.137E-06 | 7.300E-06 | 9.500E-06 |
| 6581 | 330 | E330A | A | 9.4e-06 | 1.1e-05 | 9.3e-06 | 9.900E-06 | 9.539E-07 | 9.300E-06 | 1.100E-05 |
| 6582 | 330 | E330R | A | 5.4e-06 | 1.3e-05 | 4.3e-06 | 7.567E-06 | 4.737E-06 | 4.300E-06 | 1.300E-05 |
| 6583 | 330 | E330N | A | 7.9e-06 | 9.8e-06 | 5.1e-06 | 7.600E-06 | 2.364E-06 | 5.100E-06 | 9.800E-06 |
| 6584 | 330 | E330D | A | 1.2e-05 | 9.1e-06 | 6.6e-06 | 9.233E-06 | 2.702E-06 | 6.600E-06 | 1.200E-05 |
| 6585 | 330 | E330C | A | 7.2e-06 | 1.8e-05 | 7.6e-06 | 1.093E-05 | 6.123E-06 | 7.200E-06 | 1.800E-05 |
| 6586 | 330 | E330E | A | 5.9e-06 | 7.3e-06 | 8.1e-06 | 7.100E-06 | 1.114E-06 | 5.900E-06 | 8.100E-06 |
| 6587 | 330 | E330Q | A | 7.4e-06 | 4.7e-06 | 9.5e-06 | 7.200E-06 | 2.406E-06 | 4.700E-06 | 9.500E-06 |
| 6588 | 330 | E330G | A | 4.9e-06 | 9e-06 | 7.8e-06 | 7.233E-06 | 2.108E-06 | 4.900E-06 | 9.000E-06 |
| 6589 | 330 | E330H | A | 8.8e-06 | 9.6e-06 | 5.1e-06 | 7.833E-06 | 2.401E-06 | 5.100E-06 | 9.600E-06 |
| 6590 | 330 | E330I | A | 5.6e-06 | 9.1e-06 | 7e-06 | 7.233E-06 | 1.762E-06 | 5.600E-06 | 9.100E-06 |
| 6591 | 330 | E330L | A | 8.5e-06 | 7.7e-06 | 6.6e-06 | 7.600E-06 | 9.539E-07 | 6.600E-06 | 8.500E-06 |
| 6592 | 330 | E330K | A | 5e-06 | 6.6e-06 | 8.6e-06 | 6.733E-06 | 1.804E-06 | 5.000E-06 | 8.600E-06 |
| 6593 | 330 | E330M | A | 7.1e-06 | 8.1e-06 | 5.5e-06 | 6.900E-06 | 1.311E-06 | 5.500E-06 | 8.100E-06 |
| 6594 | 330 | E330F | A | 7.2e-06 | 8.4e-06 | 1.1e-05 | 8.867E-06 | 1.943E-06 | 7.200E-06 | 1.100E-05 |
| 6595 | 330 | E330P | A | 8e-06 | 7.1e-06 | 5e-06 | 6.700E-06 | 1.539E-06 | 5.000E-06 | 8.000E-06 |
| 6596 | 330 | E330S | A | 9.1e-06 | 5.4e-06 | 3.9e-06 | 6.133E-06 | 2.676E-06 | 3.900E-06 | 9.100E-06 |
| 6597 | 330 | E330T | A | 8.3e-06 | 5.6e-06 | 5.4e-06 | 6.433E-06 | 1.620E-06 | 5.400E-06 | 8.300E-06 |
| 6598 | 330 | E330W | A | 8.8e-06 | 9.1e-06 | 4.8e-06 | 7.567E-06 | 2.401E-06 | 4.800E-06 | 9.100E-06 |
| 6599 | 330 | E330Y | A | 5.7e-06 | 9.8e-06 | 6.6e-06 | 7.367E-06 | 2.155E-06 | 5.700E-06 | 9.800E-06 |
| 6600 | 330 | E330V | A | 6e-06 | 9.6e-06 | 1.2e-05 | 9.200E-06 | 3.020E-06 | 6.000E-06 | 1.200E-05 |
| 6601 | 331 | V331A | A | 7e-06 | 9.6e-06 | 1.4e-05 | 1.020E-05 | 3.538E-06 | 7.000E-06 | 1.400E-05 |
| 6602 | 331 | V331R | A | 7.3e-06 | 1e-05 | 7.6e-06 | 8.300E-06 | 1.480E-06 | 7.300E-06 | 1.000E-05 |
| 6603 | 331 | V331N | A | 7.8e-06 | 5.6e-06 | 1.3e-05 | 8.800E-06 | 3.800E-06 | 5.600E-06 | 1.300E-05 |
| 6604 | 331 | V331D | A | 9.8e-06 | 5.9e-06 | 5.2e-06 | 6.967E-06 | 2.479E-06 | 5.200E-06 | 9.800E-06 |
| 6605 | 331 | V331C | A | 6e-06 | 1.2e-05 | 1e-05 | 9.333E-06 | 3.055E-06 | 6.000E-06 | 1.200E-05 |
| 6606 | 331 | V331E | A | 1.3e-05 | 6.3e-06 | 1.1e-05 | 1.010E-05 | 3.439E-06 | 6.300E-06 | 1.300E-05 |
| 6607 | 331 | V331Q | A | 9.7e-06 | 7.4e-06 | 8.2e-06 | 8.433E-06 | 1.168E-06 | 7.400E-06 | 9.700E-06 |
| 6608 | 331 | V331G | A | 8.6e-06 | 6.3e-06 | 9e-06 | 7.967E-06 | 1.457E-06 | 6.300E-06 | 9.000E-06 |
| 6609 | 331 | V331H | A | 6.5e-06 | 7.7e-06 | 1.3e-05 | 9.067E-06 | 3.459E-06 | 6.500E-06 | 1.300E-05 |
| 6610 | 331 | V331I | A | 8.3e-06 | 7.2e-06 | 7.1e-06 | 7.533E-06 | 6.658E-07 | 7.100E-06 | 8.300E-06 |
| 6611 | 331 | V331L | A | 1.2e-05 | 9.8e-06 | 8.5e-06 | 1.010E-05 | 1.769E-06 | 8.500E-06 | 1.200E-05 |
| 6612 | 331 | V331K | A | 8.4e-06 | 6.8e-06 | 6.6e-06 | 7.267E-06 | 9.866E-07 | 6.600E-06 | 8.400E-06 |
| 6613 | 331 | V331M | A | 1.3e-05 | 9e-06 | 7.7e-06 | 9.900E-06 | 2.762E-06 | 7.700E-06 | 1.300E-05 |
| 6614 | 331 | V331F | A | 3.7e-06 | 9.9e-06 | 5.6e-06 | 6.400E-06 | 3.176E-06 | 3.700E-06 | 9.900E-06 |
| 6615 | 331 | V331P | A | 8.6e-06 | 7.5e-06 | 8.8e-06 | 8.300E-06 | 7.000E-07 | 7.500E-06 | 8.800E-06 |
| 6616 | 331 | V331S | A | 5.8e-06 | 1e-05 | 1.1e-05 | 8.933E-06 | 2.759E-06 | 5.800E-06 | 1.100E-05 |
| 6617 | 331 | V331T | A | 8.6e-06 | 7.4e-06 | 1.3e-05 | 9.667E-06 | 2.948E-06 | 7.400E-06 | 1.300E-05 |
| 6618 | 331 | V331W | A | 4.5e-06 | 6.9e-06 | 1.6e-05 | 9.133E-06 | 6.067E-06 | 4.500E-06 | 1.600E-05 |
| 6619 | 331 | V331Y | A | 4.7e-06 | 5.5e-06 | 8.6e-06 | 6.267E-06 | 2.060E-06 | 4.700E-06 | 8.600E-06 |
| 6620 | 331 | V331V | A | 4.6e-06 | 9.5e-06 | 1.1e-05 | 8.367E-06 | 3.347E-06 | 4.600E-06 | 1.100E-05 |
| 6621 | 332 | R332A | A | 1.5e-05 | 1.1e-05 | 5.8e-06 | 1.060E-05 | 4.613E-06 | 5.800E-06 | 1.500E-05 |
| 6622 | 332 | R332R | A | 1.1e-05 | 7.9e-06 | 4.5e-06 | 7.800E-06 | 3.251E-06 | 4.500E-06 | 1.100E-05 |
| 6623 | 332 | R332N | A | 5.8e-06 | 9.1e-06 | 1.3e-05 | 9.300E-06 | 3.604E-06 | 5.800E-06 | 1.300E-05 |
| 6624 | 332 | R332D | A | 1.6e-05 | 1.1e-05 | 4e-06 | 1.033E-05 | 6.028E-06 | 4.000E-06 | 1.600E-05 |
| 6625 | 332 | R332C | A | 5.5e-06 | 1.3e-05 | 9e-06 | 9.167E-06 | 3.753E-06 | 5.500E-06 | 1.300E-05 |
| 6626 | 332 | R332E | A | 1.1e-05 | 1.1e-05 | 5.9e-06 | 9.300E-06 | 2.944E-06 | 5.900E-06 | 1.100E-05 |
| 6627 | 332 | R332Q | A | 6.4e-06 | 1e-05 | 8.7e-06 | 8.367E-06 | 1.823E-06 | 6.400E-06 | 1.000E-05 |
| 6628 | 332 | R332G | A | 1.1e-05 | 7.5e-06 | 9.3e-06 | 9.267E-06 | 1.750E-06 | 7.500E-06 | 1.100E-05 |
| 6629 | 332 | R332H | A | 4.5e-06 | 5.2e-06 | 1.8e-05 | 9.233E-06 | 7.600E-06 | 4.500E-06 | 1.800E-05 |

| | | | | | | | | | | |
|---|---|---|---|---|---|---|---|---|---|---|
| 6630 | 332 | R332I | A | 1.2e-05 | 8.1e-06 | 7.1e-06 | 9.067E-06 | 2.589E-06 | 7.100E-06 | 1.200E-05 |
| 6631 | 332 | R332L | A | 3.8e-06 | 1.2e-05 | 3e-06 | 6.267E-06 | 4.981E-06 | 3.000E-06 | 1.200E-05 |
| 6632 | 332 | R332K | A | 8.7e-06 | 9.5e-06 | 6.8e-06 | 8.333E-06 | 1.387E-06 | 6.800E-06 | 9.500E-06 |
| 6633 | 332 | R332M | A | 5.7e-06 | 5.7e-06 | 7.6e-06 | 6.333E-06 | 1.097E-06 | 5.700E-06 | 7.600E-06 |
| 6634 | 332 | R332F | A | 1.1e-05 | 6.6e-06 | 7.3e-06 | 8.300E-06 | 2.364E-06 | 6.600E-06 | 1.100E-05 |
| 6635 | 332 | R332P | A | 5.9e-06 | 7.9e-06 | 6.2e-06 | 6.667E-06 | 1.079E-06 | 5.900E-06 | 7.900E-06 |
| 6636 | 332 | R332S | A | 8.9e-06 | 7.8e-06 | 1.2e-05 | 9.567E-06 | 2.178E-06 | 7.800E-06 | 1.200E-05 |
| 6637 | 332 | R332T | A | 1.7e-05 | 8.7e-06 | 4.8e-06 | 1.017E-05 | 6.231E-06 | 4.800E-06 | 1.700E-05 |
| 6638 | 332 | R332W | A | 9.8e-06 | 8.7e-06 | 8.5e-06 | 9.000E-06 | 7.000E-07 | 8.500E-06 | 9.800E-06 |
| 6639 | 332 | R332Y | A | 4.6e-06 | 1.5e-05 | 9.3e-06 | 9.633E-06 | 5.208E-06 | 4.600E-06 | 1.500E-05 |
| 6640 | 332 | R332V | A | 1.1e-05 | 1e-05 | 6.5e-06 | 9.167E-06 | 2.363E-06 | 6.500E-06 | 1.100E-05 |
| 6641 | 333 | A333A | A | 7.2e-06 | 1.2e-05 | 5.1e-06 | 8.100E-06 | 3.537E-06 | 5.100E-06 | 1.200E-05 |
| 6642 | 333 | A333R | A | 6.6e-06 | 5.3e-06 | 6.6e-06 | 6.167E-06 | 7.506E-07 | 5.300E-06 | 6.600E-06 |
| 6643 | 333 | A333N | A | 6.6e-06 | 1.4e-05 | 9e-06 | 9.867E-06 | 3.775E-06 | 6.600E-06 | 1.400E-05 |
| 6644 | 333 | A333D | A | 5.4e-06 | 1.4e-05 | 9.9e-06 | 9.767E-06 | 4.302E-06 | 5.400E-06 | 1.400E-05 |
| 6645 | 333 | A333C | A | 8.3e-06 | 6e-06 | 1.4e-05 | 9.433E-06 | 4.119E-06 | 6.000E-06 | 1.400E-05 |
| 6646 | 333 | A333E | A | 6.2e-06 | 7.4e-06 | 7.3e-06 | 6.967E-06 | 6.658E-07 | 6.200E-06 | 7.400E-06 |
| 6647 | 333 | A333Q | A | 5.5e-06 | 5.1e-06 | 1.2e-05 | 7.533E-06 | 3.873E-06 | 5.100E-06 | 1.200E-05 |
| 6648 | 333 | A333G | A | 1.4e-05 | 1.2e-05 | 9e-06 | 1.167E-05 | 2.517E-06 | 9.000E-06 | 1.400E-05 |
| 6649 | 333 | A333H | A | 3e-06 | 1.2e-05 | 1.1e-05 | 8.667E-06 | 4.933E-06 | 3.000E-06 | 1.200E-05 |
| 6650 | 333 | A333I | A | 5.2e-06 | 6.5e-06 | 5.7e-06 | 5.800E-06 | 6.557E-07 | 5.200E-06 | 6.500E-06 |
| 6651 | 333 | A333L | A | 5.8e-06 | 8.5e-06 | 7.2e-06 | 7.167E-06 | 1.350E-06 | 5.800E-06 | 8.500E-06 |
| 6652 | 333 | A333K | A | 7.1e-06 | 8.9e-06 | 1.2e-05 | 9.333E-06 | 2.479E-06 | 7.100E-06 | 1.200E-05 |
| 6653 | 333 | A333M | A | 7.3e-06 | 6.9e-06 | 9.4e-06 | 7.867E-06 | 1.343E-06 | 6.900E-06 | 9.400E-06 |
| 6654 | 333 | A333F | A | 7.4e-06 | 6.2e-06 | 7.2e-06 | 6.933E-06 | 6.429E-07 | 6.200E-06 | 7.400E-06 |
| 6655 | 333 | A333P | A | 8.7e-06 | 5.3e-06 | 9.8e-06 | 7.933E-06 | 2.346E-06 | 5.300E-06 | 9.800E-06 |
| 6656 | 333 | A333S | A | 9.7e-06 | 8.3e-06 | 6.6e-06 | 8.200E-06 | 1.552E-06 | 6.600E-06 | 9.700E-06 |
| 6657 | 333 | A333T | A | 1.3e-05 | 9.7e-06 | 4.5e-06 | 9.067E-06 | 4.285E-06 | 4.500E-06 | 1.300E-05 |
| 6658 | 333 | A333W | A | 6.6e-06 | 4.7e-06 | 8.7e-06 | 6.667E-06 | 2.001E-06 | 4.700E-06 | 8.700E-06 |
| 6659 | 333 | A333Y | A | 6.8e-06 | 8.5e-06 | 9.6e-06 | 8.300E-06 | 1.411E-06 | 6.800E-06 | 9.600E-06 |
| 6660 | 333 | A333V | A | 8e-06 | 9.4e-06 | 9.3e-06 | 8.900E-06 | 7.810E-07 | 8.000E-06 | 9.400E-06 |
| 6661 | 334 | V334A | A | 7.1e-06 | 6.7e-06 | 9.4e-06 | 7.733E-06 | 1.457E-06 | 6.700E-06 | 9.400E-06 |
| 6662 | 334 | V334R | A | 1.4e-05 | 4.5e-06 | 6.2e-06 | 8.233E-06 | 5.066E-06 | 4.500E-06 | 1.400E-05 |
| 6663 | 334 | V334N | A | 5.1e-06 | 6.2e-06 | 1.1e-05 | 7.433E-06 | 3.137E-06 | 5.100E-06 | 1.100E-05 |
| 6664 | 334 | V334D | A | 1.3e-05 | 9.4e-06 | 5.7e-06 | 9.367E-06 | 3.650E-06 | 5.700E-06 | 1.300E-05 |
| 6665 | 334 | V334C | A | 7.3e-06 | 1.1e-05 | 8e-06 | 8.767E-06 | 1.966E-06 | 7.300E-06 | 1.100E-05 |
| 6666 | 334 | V334E | A | 7.6e-06 | 1.3e-05 | 5e-06 | 8.533E-06 | 4.081E-06 | 5.000E-06 | 1.300E-05 |
| 6667 | 334 | V334Q | A | 4.1e-06 | 4.4e-06 | 7e-06 | 5.167E-06 | 1.595E-06 | 4.100E-06 | 7.000E-06 |
| 6668 | 334 | V334G | A | 6.6e-06 | 1.1e-05 | 1.2e-05 | 9.867E-06 | 2.873E-06 | 6.600E-06 | 1.200E-05 |
| 6669 | 334 | V334H | A | 6.6e-06 | 8.6e-06 | 6.8e-06 | 7.333E-06 | 1.102E-06 | 6.600E-06 | 8.600E-06 |
| 6670 | 334 | V334I | A | 6.8e-06 | 7e-06 | 6.9e-06 | 6.900E-06 | 1.000E-07 | 6.800E-06 | 7.000E-06 |
| 6671 | 334 | V334L | A | 9.7e-06 | 1e-05 | 9.2e-06 | 9.633E-06 | 4.041E-07 | 9.200E-06 | 1.000E-05 |
| 6672 | 334 | V334K | A | 1e-05 | 9.6e-06 | 1e-05 | 9.867E-06 | 2.309E-07 | 9.600E-06 | 1.000E-05 |
| 6673 | 334 | V334M | A | 6.8e-06 | 5.7e-06 | 1.2e-05 | 8.167E-06 | 3.365E-06 | 5.700E-06 | 1.200E-05 |
| 6674 | 334 | V334F | A | 6.4e-06 | 5.6e-06 | 1.7e-05 | 9.667E-06 | 6.363E-06 | 5.600E-06 | 1.700E-05 |
| 6675 | 334 | V334P | A | 5.2e-06 | 9.2e-06 | 1e-05 | 8.133E-06 | 2.572E-06 | 5.200E-06 | 1.000E-05 |
| 6676 | 334 | V334S | A | 9e-06 | 6.2e-06 | 4.7e-06 | 6.633E-06 | 2.183E-06 | 4.700E-06 | 9.000E-06 |
| 6677 | 334 | V334T | A | 5e-06 | 7.5e-06 | 5.6e-06 | 6.033E-06 | 1.305E-06 | 5.000E-06 | 7.500E-06 |
| 6678 | 334 | V334W | A | 7.5e-06 | 1.3e-05 | 8.6e-06 | 9.700E-06 | 2.910E-06 | 7.500E-06 | 1.300E-05 |
| 6679 | 334 | V334Y | A | 6.6e-06 | 1.2e-05 | 4.1e-06 | 7.567E-06 | 4.038E-06 | 4.100E-06 | 1.200E-05 |
| 6680 | 334 | V334V | A | 9.3e-06 | 8.6e-06 | 5.5e-06 | 7.800E-06 | 2.022E-06 | 5.500E-06 | 9.300E-06 |
| 6681 | 335 | T335A | A | 4.1e-06 | 7.6e-06 | 8.5e-06 | 6.733E-06 | 2.325E-06 | 4.100E-06 | 8.500E-06 |
| 6682 | 335 | T335R | A | 1.7e-05 | 5e-06 | 4.8e-06 | 8.933E-06 | 6.987E-06 | 4.800E-06 | 1.700E-05 |
| 6683 | 335 | T335N | A | 9.4e-06 | 8.2e-06 | 1e-05 | 9.200E-06 | 9.165E-07 | 8.200E-06 | 1.000E-05 |
| 6684 | 335 | T335D | A | 5.9e-06 | 9.6e-06 | 6.3e-06 | 7.267E-06 | 2.031E-06 | 5.900E-06 | 9.600E-06 |
| 6685 | 335 | T335C | A | 6.5e-06 | 6.4e-06 | 5.3e-06 | 6.067E-06 | 6.658E-07 | 5.300E-06 | 6.500E-06 |
| 6686 | 335 | T335E | A | 1.4e-05 | 8.2e-06 | 1.1e-05 | 1.107E-05 | 2.901E-06 | 8.200E-06 | 1.400E-05 |
| 6687 | 335 | T335Q | A | 4.4e-06 | 4.7e-06 | 7.4e-06 | 5.500E-06 | 1.652E-06 | 4.400E-06 | 7.400E-06 |
| 6688 | 335 | T335G | A | 1.2e-05 | 1.3e-05 | 6.3e-06 | 1.043E-05 | 3.614E-06 | 6.300E-06 | 1.300E-05 |
| 6689 | 335 | T335H | A | 9.5e-06 | 9.3e-06 | 8.6e-06 | 9.133E-06 | 4.726E-07 | 8.600E-06 | 9.500E-06 |
| 6690 | 335 | T335I | A | 6.8e-06 | 6.6e-06 | 6.8e-06 | 6.733E-06 | 1.155E-07 | 6.600E-06 | 6.800E-06 |
| 6691 | 335 | T335L | A | 8.4e-06 | 7.6e-06 | 5.6e-06 | 7.200E-06 | 1.442E-06 | 5.600E-06 | 8.400E-06 |
| 6692 | 335 | T335K | A | 1.7e-05 | 6.8e-06 | 1.1e-05 | 1.160E-05 | 5.126E-06 | 6.800E-06 | 1.700E-05 |
| 6693 | 335 | T335M | A | 9.9e-06 | 9.3e-06 | 1.2e-05 | 1.040E-05 | 1.418E-06 | 9.300E-06 | 1.200E-05 |
| 6694 | 335 | T335F | A | 6.8e-06 | 4.9e-06 | 8.8e-06 | 6.833E-06 | 1.950E-06 | 4.900E-06 | 8.800E-06 |
| 6695 | 335 | T335P | A | 6.8e-06 | 9e-06 | 6.5e-06 | 7.433E-06 | 1.365E-06 | 6.500E-06 | 9.000E-06 |
| 6696 | 335 | T335S | A | 7.7e-06 | 6.8e-06 | 4.8e-06 | 6.433E-06 | 1.484E-06 | 4.800E-06 | 7.700E-06 |
| 6697 | 335 | T335T | A | 5.2e-06 | 6.4e-06 | 1.3e-05 | 8.200E-06 | 4.200E-06 | 5.200E-06 | 1.300E-05 |
| 6698 | 335 | T335W | A | 4.2e-06 | 7.8e-06 | 8.2e-06 | 6.733E-06 | 2.203E-06 | 4.200E-06 | 8.200E-06 |
| 6699 | 335 | T335Y | A | 9.3e-06 | 4.6e-06 | 9.2e-06 | 7.700E-06 | 2.685E-06 | 4.600E-06 | 9.300E-06 |
| 6700 | 335 | T335V | A | 8.8e-06 | 6.3e-06 | 9.6e-06 | 8.233E-06 | 1.721E-06 | 6.300E-06 | 9.600E-06 |
| 6701 | 336 | S336A | A | 6.2e-06 | 4.9e-06 | 3.6e-06 | 4.900E-06 | 1.300E-06 | 3.600E-06 | 6.200E-06 |
| 6702 | 336 | S336R | A | 5.3e-06 | 5.6e-06 | 9e-06 | 6.633E-06 | 2.055E-06 | 5.300E-06 | 9.000E-06 |
| 6703 | 336 | S336N | A | 9.5e-06 | 1.3e-05 | 6.7e-06 | 9.733E-06 | 3.156E-06 | 6.700E-06 | 1.300E-05 |
| 6704 | 336 | S336D | A | 7.6e-06 | 8.2e-06 | 8.9e-06 | 8.233E-06 | 6.506E-07 | 7.600E-06 | 8.900E-06 |
| 6705 | 336 | S336C | A | 9.5e-06 | 1.2e-05 | 1e-05 | 1.050E-05 | 1.323E-06 | 9.500E-06 | 1.200E-05 |
| 6706 | 336 | S336E | A | 9.6e-06 | 8.8e-06 | 7.9e-06 | 8.767E-06 | 8.505E-07 | 7.900E-06 | 9.600E-06 |
| 6707 | 336 | S336Q | A | 7.1e-06 | 8.2e-06 | 9.8e-06 | 8.367E-06 | 1.358E-06 | 7.100E-06 | 9.800E-06 |
| 6708 | 336 | S336G | A | 1.2e-05 | 1.2e-05 | 1e-05 | 1.133E-05 | 1.155E-06 | 1.000E-05 | 1.200E-05 |
| 6709 | 336 | S336H | A | 6.5e-06 | 7.7e-06 | 4.2e-06 | 6.133E-06 | 1.779E-06 | 4.200E-06 | 7.700E-06 |
| 6710 | 336 | S336I | A | 7.9e-06 | 1.3e-05 | 4.6e-06 | 8.500E-06 | 4.232E-06 | 4.600E-06 | 1.300E-05 |
| 6711 | 336 | S336L | A | 6.2e-06 | 8e-06 | 1e-05 | 8.067E-06 | 1.901E-06 | 6.200E-06 | 1.000E-05 |
| 6712 | 336 | S336K | A | 7.1e-06 | 7.1e-06 | 1e-05 | 8.067E-06 | 1.674E-06 | 7.100E-06 | 1.000E-05 |
| 6713 | 336 | S336M | A | 6.7e-06 | 9.7e-06 | 7.9e-06 | 8.100E-06 | 1.510E-06 | 6.700E-06 | 9.700E-06 |
| 6714 | 336 | S336F | A | 9.7e-06 | 1.3e-05 | 5.2e-06 | 9.300E-06 | 3.915E-06 | 5.200E-06 | 1.300E-05 |

| | | | | | | | | | | |
|---|---|---|---|---|---|---|---|---|---|---|
| 6715 | 336 | S336P | A | 3.7e-06 | 9.9e-06 | 1.3e-05 | 8.867E-06 | 4.735E-06 | 3.700E-06 | 1.300E-05 |
| 6716 | 336 | S336S | A | 8.7e-06 | 9.8e-06 | 4.4e-06 | 7.633E-06 | 2.854E-06 | 4.400E-06 | 9.800E-06 |
| 6717 | 336 | S336T | A | 1.2e-05 | 8.5e-06 | 6.4e-06 | 8.967E-06 | 2.829E-06 | 6.400E-06 | 1.200E-05 |
| 6718 | 336 | S336W | A | 7.6e-06 | 8.4e-06 | 5.7e-06 | 7.233E-06 | 1.387E-06 | 5.700E-06 | 8.400E-06 |
| 6719 | 336 | S336Y | A | 5.8e-06 | 8e-06 | 8.1e-06 | 7.300E-06 | 1.300E-06 | 5.800E-06 | 8.100E-06 |
| 6720 | 336 | S336V | A | 1.4e-05 | 1.6e-05 | 1.1e-05 | 1.367E-05 | 2.517E-06 | 1.100E-05 | 1.600E-05 |
| 6721 | 337 | A337A | A | 6.6e-06 | 9.2e-06 | 7.7e-06 | 7.833E-06 | 1.305E-06 | 6.600E-06 | 9.200E-06 |
| 6722 | 337 | A337R | A | 1.2e-05 | 7e-06 | 9.7e-06 | 9.567E-06 | 2.503E-06 | 7.000E-06 | 1.200E-05 |
| 6723 | 337 | A337N | A | 6.9e-06 | 7.5e-06 | 9.9e-06 | 8.100E-06 | 1.587E-06 | 6.900E-06 | 9.900E-06 |
| 6724 | 337 | A337D | A | 9.2e-06 | 6.7e-06 | 9.6e-06 | 8.500E-06 | 1.572E-06 | 6.700E-06 | 9.600E-06 |
| 6725 | 337 | A337C | A | 5.7e-06 | 5e-06 | 5.8e-06 | 5.500E-06 | 4.359E-07 | 5.000E-06 | 5.800E-06 |
| 6726 | 337 | A337E | A | 1.3e-05 | 6.6e-06 | 9.2e-06 | 9.600E-06 | 3.219E-06 | 6.600E-06 | 1.300E-05 |
| 6727 | 337 | A337Q | A | 8.3e-06 | 6.7e-06 | 7.7e-06 | 7.567E-06 | 8.083E-07 | 6.700E-06 | 8.300E-06 |
| 6728 | 337 | A337G | A | 9.5e-06 | 5.7e-06 | 7e-06 | 7.400E-06 | 1.931E-06 | 5.700E-06 | 9.500E-06 |
| 6729 | 337 | A337H | A | 6.8e-06 | 6.9e-06 | 9.4e-06 | 7.700E-06 | 1.473E-06 | 6.800E-06 | 9.400E-06 |
| 6730 | 337 | A337I | A | 6.4e-06 | 1.1e-05 | 6.1e-06 | 7.833E-06 | 2.747E-06 | 6.100E-06 | 1.100E-05 |
| 6731 | 337 | A337L | A | 1.2e-05 | 4.5e-06 | 8.4e-06 | 8.300E-06 | 3.751E-06 | 4.500E-06 | 1.200E-05 |
| 6732 | 337 | A337K | A | 9.9e-06 | 5.9e-06 | 6.4e-06 | 7.400E-06 | 2.179E-06 | 5.900E-06 | 9.900E-06 |
| 6733 | 337 | A337M | A | 9.7e-06 | 7e-06 | 6.8e-06 | 7.833E-06 | 1.620E-06 | 6.800E-06 | 9.700E-06 |
| 6734 | 337 | A337F | A | 9.9e-06 | 5.1e-06 | 1.5e-05 | 1.000E-05 | 4.951E-06 | 5.100E-06 | 1.500E-05 |
| 6735 | 337 | A337P | A | 1.1e-05 | 7.9e-06 | 4.9e-06 | 7.933E-06 | 3.050E-06 | 4.900E-06 | 1.100E-05 |
| 6736 | 337 | A337S | A | 7e-06 | 1.2e-05 | 1.1e-05 | 1.000E-05 | 2.646E-06 | 7.000E-06 | 1.200E-05 |
| 6737 | 337 | A337T | A | 6.9e-06 | 9.5e-06 | 6e-06 | 7.467E-06 | 1.818E-06 | 6.000E-06 | 9.500E-06 |
| 6738 | 337 | A337W | A | 9.8e-06 | 9.3e-06 | 5.8e-06 | 8.300E-06 | 2.179E-06 | 5.800E-06 | 9.800E-06 |
| 6739 | 337 | A337Y | A | 9.4e-06 | 7.4e-06 | 1.4e-05 | 1.027E-05 | 3.384E-06 | 7.400E-06 | 1.400E-05 |
| 6740 | 337 | A337V | A | 1.1e-05 | 1e-05 | 7.6e-06 | 9.533E-06 | 1.747E-06 | 7.600E-06 | 1.100E-05 |
| 6741 | 338 | N338A | A | 6.7e-06 | 9.1e-06 | 8.3e-06 | 8.033E-06 | 1.222E-06 | 6.700E-06 | 9.100E-06 |
| 6742 | 338 | N338R | A | 7.4e-06 | 7.2e-06 | 5.6e-06 | 6.733E-06 | 9.866E-07 | 5.600E-06 | 7.400E-06 |
| 6743 | 338 | N338N | A | 7e-06 | 9.7e-06 | 9.3e-06 | 8.667E-06 | 1.457E-06 | 7.000E-06 | 9.700E-06 |
| 6744 | 338 | N338D | A | 6.6e-06 | 9.4e-06 | 7.3e-06 | 7.767E-06 | 1.457E-06 | 6.600E-06 | 9.400E-06 |
| 6745 | 338 | N338C | A | 1.3e-05 | 7.4e-06 | 7.3e-06 | 9.233E-06 | 3.262E-06 | 7.300E-06 | 1.300E-05 |
| 6746 | 338 | N338E | A | 4.9e-06 | 6.4e-06 | 1.7e-05 | 9.433E-06 | 6.596E-06 | 4.900E-06 | 1.700E-05 |
| 6747 | 338 | N338Q | A | 1.1e-05 | 7.2e-06 | 9.4e-06 | 9.200E-06 | 1.908E-06 | 7.200E-06 | 1.100E-05 |
| 6748 | 338 | N338G | A | 8.5e-06 | 1.2e-05 | 9.1e-06 | 9.867E-06 | 1.872E-06 | 8.500E-06 | 1.200E-05 |
| 6749 | 338 | N338H | A | 6.9e-06 | 8.2e-06 | 5.1e-06 | 6.733E-06 | 1.557E-06 | 5.100E-06 | 8.200E-06 |
| 6750 | 338 | N338I | A | 6e-06 | 1.3e-05 | 6.8e-06 | 8.600E-06 | 3.831E-06 | 6.000E-06 | 1.300E-05 |
| 6751 | 338 | N338L | A | 8.5e-06 | 5.6e-06 | 6.4e-06 | 6.833E-06 | 1.498E-06 | 5.600E-06 | 8.500E-06 |
| 6752 | 338 | N338K | A | 9.5e-06 | 4.7e-06 | 1.3e-05 | 9.067E-06 | 4.167E-06 | 4.700E-06 | 1.300E-05 |
| 6753 | 338 | N338M | A | 1.2e-05 | 5.5e-06 | 5.9e-06 | 7.800E-06 | 3.643E-06 | 5.500E-06 | 1.200E-05 |
| 6754 | 338 | N338F | A | 1.5e-05 | 6.9e-06 | 4.1e-06 | 8.667E-06 | 5.661E-06 | 4.100E-06 | 1.500E-05 |
| 6755 | 338 | N338P | A | 5.8e-06 | 9.7e-06 | 9.4e-06 | 8.300E-06 | 2.170E-06 | 5.800E-06 | 9.700E-06 |
| 6756 | 338 | N338S | A | 1.2e-05 | 9.2e-06 | 6.9e-06 | 9.367E-06 | 2.554E-06 | 6.900E-06 | 1.200E-05 |
| 6757 | 338 | N338T | A | 9.7e-06 | 1e-05 | 9.9e-06 | 9.867E-06 | 1.528E-07 | 9.700E-06 | 1.000E-05 |
| 6758 | 338 | N338W | A | 6.2e-06 | 1e-05 | 6.1e-06 | 7.433E-06 | 2.223E-06 | 6.100E-06 | 1.000E-05 |
| 6759 | 338 | N338Y | A | 8.7e-06 | 9e-06 | 4.4e-06 | 7.367E-06 | 2.574E-06 | 4.400E-06 | 9.000E-06 |
| 6760 | 338 | N338V | A | 9.2e-06 | 1.3e-05 | 6.3e-06 | 9.500E-06 | 3.360E-06 | 6.300E-06 | 1.300E-05 |
| 6761 | 339 | I339A | A | 1.4e-05 | 9.5e-06 | 1.2e-05 | 1.183E-05 | 2.255E-06 | 9.500E-06 | 1.400E-05 |
| 6762 | 339 | I339R | A | 6.9e-06 | 7.1e-06 | 7.2e-06 | 7.067E-06 | 1.528E-07 | 6.900E-06 | 7.200E-06 |
| 6763 | 339 | I339N | A | 1.6e-05 | 7e-06 | 7.2e-06 | 1.007E-05 | 5.139E-06 | 7.000E-06 | 1.600E-05 |
| 6764 | 339 | I339D | A | 9e-06 | 6.5e-06 | 6.1e-06 | 7.200E-06 | 1.572E-06 | 6.100E-06 | 9.000E-06 |
| 6765 | 339 | I339C | A | 6.7e-06 | 7.4e-06 | 1e-05 | 8.033E-06 | 1.739E-06 | 6.700E-06 | 1.000E-05 |
| 6766 | 339 | I339E | A | 1.2e-05 | 5.6e-06 | 8.8e-06 | 8.800E-06 | 3.200E-06 | 5.600E-06 | 1.200E-05 |
| 6767 | 339 | I339Q | A | 8.3e-06 | 1.1e-05 | 1.2e-05 | 1.043E-05 | 1.914E-06 | 8.300E-06 | 1.200E-05 |
| 6768 | 339 | I339G | A | 8.2e-06 | 1.2e-05 | 1.1e-05 | 1.040E-05 | 1.970E-06 | 8.200E-06 | 1.200E-05 |
| 6769 | 339 | I339H | A | 9.8e-06 | 9.5e-06 | 7.6e-06 | 8.967E-06 | 1.193E-06 | 7.600E-06 | 9.800E-06 |
| 6770 | 339 | I339I | A | 4.7e-06 | 4.9e-06 | 9.5e-06 | 6.367E-06 | 2.715E-06 | 4.700E-06 | 9.500E-06 |
| 6771 | 339 | I339L | A | 7.5e-06 | 7.2e-06 | 4.6e-06 | 6.433E-06 | 1.595E-06 | 4.600E-06 | 7.500E-06 |
| 6772 | 339 | I339K | A | 7.5e-06 | 9.5e-06 | 9.5e-06 | 8.833E-06 | 1.155E-06 | 7.500E-06 | 9.500E-06 |
| 6773 | 339 | I339M | A | 4.6e-06 | 9e-06 | 9e-06 | 7.533E-06 | 2.540E-06 | 4.600E-06 | 9.000E-06 |
| 6774 | 339 | I339F | A | 9.8e-06 | 8.7e-06 | 9.3e-06 | 9.267E-06 | 5.508E-07 | 8.700E-06 | 9.800E-06 |
| 6775 | 339 | I339P | A | 4.9e-06 | 6.3e-06 | 6.4e-06 | 5.867E-06 | 8.386E-07 | 4.900E-06 | 6.400E-06 |
| 6776 | 339 | I339S | A | 6.6e-06 | 5.9e-06 | 8e-06 | 6.833E-06 | 1.069E-06 | 5.900E-06 | 8.000E-06 |
| 6777 | 339 | I339T | A | 1.1e-05 | 5.1e-06 | 1.3e-05 | 9.700E-06 | 4.107E-06 | 5.100E-06 | 1.300E-05 |
| 6778 | 339 | I339W | A | 8.4e-06 | 5.9e-06 | 6.5e-06 | 6.933E-06 | 1.305E-06 | 5.900E-06 | 8.400E-06 |
| 6779 | 339 | I339Y | A | 8.9e-06 | 7.2e-06 | 4.3e-06 | 6.800E-06 | 2.326E-06 | 4.300E-06 | 8.900E-06 |
| 6780 | 339 | I339V | A | 3.5e-06 | 1e-05 | 9.9e-06 | 7.800E-06 | 3.724E-06 | 3.500E-06 | 1.000E-05 |
| 6781 | 340 | Q340A | A | 9.5e-06 | 5.4e-06 | 9.7e-06 | 8.200E-06 | 2.427E-06 | 5.400E-06 | 9.700E-06 |
| 6782 | 340 | Q340R | A | 8e-06 | 1.3e-05 | 1.7e-05 | 1.267E-05 | 4.509E-06 | 8.000E-06 | 1.700E-05 |
| 6783 | 340 | Q340N | A | 7.6e-06 | 1e-05 | 9.8e-06 | 9.133E-06 | 1.332E-06 | 7.600E-06 | 1.000E-05 |
| 6784 | 340 | Q340D | A | 1.1e-05 | 6.5e-06 | 7.7e-06 | 8.400E-06 | 2.330E-06 | 6.500E-06 | 1.100E-05 |
| 6785 | 340 | Q340C | A | 5.4e-06 | 1.4e-05 | 6.6e-06 | 8.667E-06 | 4.658E-06 | 5.400E-06 | 1.400E-05 |
| 6786 | 340 | Q340E | A | 1.1e-05 | 8.7e-06 | 9.1e-06 | 9.600E-06 | 1.229E-06 | 8.700E-06 | 1.100E-05 |
| 6787 | 340 | Q340Q | A | 9.2e-06 | 7.9e-06 | 7.4e-06 | 8.167E-06 | 9.292E-07 | 7.400E-06 | 9.200E-06 |
| 6788 | 340 | Q340G | A | 5.9e-06 | 1.3e-05 | 7.7e-06 | 8.867E-06 | 3.691E-06 | 5.900E-06 | 1.300E-05 |
| 6789 | 340 | Q340H | A | 9.3e-06 | 8.7e-06 | 5.4e-06 | 7.800E-06 | 2.100E-06 | 5.400E-06 | 9.300E-06 |
| 6790 | 340 | Q340I | A | 1.3e-05 | 5.7e-06 | 1e-05 | 9.567E-06 | 3.669E-06 | 5.700E-06 | 1.300E-05 |
| 6791 | 340 | Q340L | A | 1.4e-05 | 6.4e-06 | 5.2e-06 | 8.533E-06 | 4.772E-06 | 5.200E-06 | 1.400E-05 |
| 6792 | 340 | Q340K | A | 1.9e-05 | 8.3e-06 | 6.2e-06 | 1.117E-05 | 6.865E-06 | 6.200E-06 | 1.900E-05 |
| 6793 | 340 | Q340M | A | 5e-06 | 5.2e-06 | 9.4e-06 | 6.533E-06 | 2.485E-06 | 5.000E-06 | 9.400E-06 |
| 6794 | 340 | Q340F | A | 5.9e-06 | 5.8e-06 | 4.7e-06 | 5.467E-06 | 6.658E-07 | 4.700E-06 | 5.900E-06 |
| 6795 | 340 | Q340P | A | 6.2e-06 | 1.3e-05 | 1.1e-05 | 1.007E-05 | 3.495E-06 | 6.200E-06 | 1.300E-05 |
| 6796 | 340 | Q340S | A | 9.4e-06 | 6.3e-06 | 7.8e-06 | 7.833E-06 | 1.550E-06 | 6.300E-06 | 9.400E-06 |
| 6797 | 340 | Q340T | A | 7.3e-06 | 7.5e-06 | 9e-06 | 7.933E-06 | 9.292E-07 | 7.300E-06 | 9.000E-06 |
| 6798 | 340 | Q340W | A | 7.5e-06 | 7.2e-06 | 7e-06 | 7.233E-06 | 2.517E-07 | 7.000E-06 | 7.500E-06 |
| 6799 | 340 | Q340Y | A | 5.6e-06 | 8.9e-06 | 9.3e-06 | 7.933E-06 | 2.031E-06 | 5.600E-06 | 9.300E-06 |

| | | | | | | | | | | |
|---|---|---|---|---|---|---|---|---|---|---|
| 6800 | 340 | Q340V | A | 8e-06 | 5.7e-06 | 6e-06 | 6.567E-06 | 1.250E-06 | 5.700E-06 | 8.000E-06 |
| 6801 | 341 | E341A | A | 5.3e-06 | 1.2e-05 | 1.4e-05 | 1.043E-05 | 4.557E-06 | 5.300E-06 | 1.400E-05 |
| 6802 | 341 | E341R | A | 5.8e-06 | 6.8e-06 | 9.5e-06 | 7.367E-06 | 1.914E-06 | 5.800E-06 | 9.500E-06 |
| 6803 | 341 | E341N | A | 8.3e-06 | 6.7e-06 | 1e-05 | 8.333E-06 | 1.650E-06 | 6.700E-06 | 1.000E-05 |
| 6804 | 341 | E341D | A | 7.6e-06 | 8e-06 | 4.8e-06 | 6.800E-06 | 1.744E-06 | 4.800E-06 | 8.000E-06 |
| 6805 | 341 | E341C | A | 8.1e-06 | 9.6e-06 | 7.3e-06 | 8.333E-06 | 1.168E-06 | 7.300E-06 | 9.600E-06 |
| 6806 | 341 | E341E | A | 8e-06 | 9.5e-06 | 9.7e-06 | 9.067E-06 | 9.292E-07 | 8.000E-06 | 9.700E-06 |
| 6807 | 341 | E341Q | A | 4.6e-06 | 6.1e-06 | 5.4e-06 | 5.367E-06 | 7.506E-07 | 4.600E-06 | 6.100E-06 |
| 6808 | 341 | E341G | A | 6.5e-06 | 1.1e-05 | 9e-06 | 8.833E-06 | 2.255E-06 | 6.500E-06 | 1.100E-05 |
| 6809 | 341 | E341H | A | 1.3e-05 | 8.9e-06 | 1.5e-05 | 1.230E-05 | 3.110E-06 | 8.900E-06 | 1.500E-05 |
| 6810 | 341 | E341I | A | 4.9e-06 | 8.8e-06 | 1.3e-05 | 8.900E-06 | 4.051E-06 | 4.900E-06 | 1.300E-05 |
| 6811 | 341 | E341L | A | 6.6e-06 | 1.1e-05 | 7.2e-06 | 8.267E-06 | 2.386E-06 | 6.600E-06 | 1.100E-05 |
| 6812 | 341 | E341K | A | 8.8e-06 | 5e-06 | 1e-05 | 7.933E-06 | 2.610E-06 | 5.000E-06 | 1.000E-05 |
| 6813 | 341 | E341M | A | 8.1e-06 | 9.8e-06 | 6.3e-06 | 8.067E-06 | 1.750E-06 | 6.300E-06 | 9.800E-06 |
| 6814 | 341 | E341F | A | 7.6e-06 | 7.3e-06 | 8.3e-06 | 7.733E-06 | 5.132E-07 | 7.300E-06 | 8.300E-06 |
| 6815 | 341 | E341P | A | 8.9e-06 | 8.7e-06 | 5.3e-06 | 7.633E-06 | 2.023E-06 | 5.300E-06 | 8.900E-06 |
| 6816 | 341 | E341S | A | 8.6e-06 | 7.8e-06 | 6.2e-06 | 7.533E-06 | 1.222E-06 | 6.200E-06 | 8.600E-06 |
| 6817 | 341 | E341T | A | 9.2e-06 | 8.4e-06 | 1.3e-05 | 1.020E-05 | 2.458E-06 | 8.400E-06 | 1.300E-05 |
| 6818 | 341 | E341W | A | 6.1e-06 | 5.7e-06 | 7.2e-06 | 6.333E-06 | 7.767E-07 | 5.700E-06 | 7.200E-06 |
| 6819 | 341 | E341Y | A | 1e-05 | 9.6e-06 | 7e-06 | 8.867E-06 | 1.629E-06 | 7.000E-06 | 1.000E-05 |
| 6820 | 341 | E341V | A | 9.8e-06 | 6.2e-06 | 1.2e-05 | 9.333E-06 | 2.928E-06 | 6.200E-06 | 1.200E-05 |
| 6821 | 342 | F342A | A | 6.9e-06 | 9.3e-06 | 6.8e-06 | 7.667E-06 | 1.415E-06 | 6.800E-06 | 9.300E-06 |
| 6822 | 342 | F342R | A | 8.3e-06 | 4.4e-06 | 1.1e-05 | 7.900E-06 | 3.318E-06 | 4.400E-06 | 1.100E-05 |
| 6823 | 342 | F342N | A | 1.3e-05 | 1.4e-05 | 1.5e-05 | 1.400E-05 | 1.000E-06 | 1.300E-05 | 1.500E-05 |
| 6824 | 342 | F342D | A | 9.8e-06 | 1.8e-05 | 6.2e-06 | 1.133E-05 | 6.048E-06 | 6.200E-06 | 1.800E-05 |
| 6825 | 342 | F342C | A | 7.3e-06 | 3.7e-06 | 1.3e-05 | 8.000E-06 | 4.689E-06 | 3.700E-06 | 1.300E-05 |
| 6826 | 342 | F342E | A | 5.5e-06 | 2.8e-06 | 7.7e-06 | 5.333E-06 | 2.454E-06 | 2.800E-06 | 7.700E-06 |
| 6827 | 342 | F342Q | A | 6e-06 | 6.3e-06 | 6.2e-06 | 6.167E-06 | 1.528E-07 | 6.000E-06 | 6.300E-06 |
| 6828 | 342 | F342G | A | 8.2e-06 | 8.5e-06 | 5.7e-06 | 7.467E-06 | 1.537E-06 | 5.700E-06 | 8.500E-06 |
| 6829 | 342 | F342H | A | 9.4e-06 | 9.9e-06 | 1.2e-05 | 1.043E-05 | 1.380E-06 | 9.400E-06 | 1.200E-05 |
| 6830 | 342 | F342I | A | 6.4e-06 | 8.2e-06 | 7.1e-06 | 7.233E-06 | 9.074E-07 | 6.400E-06 | 8.200E-06 |
| 6831 | 342 | F342L | A | 1.4e-05 | 7.7e-06 | 1e-05 | 1.057E-05 | 3.188E-06 | 7.700E-06 | 1.400E-05 |
| 6832 | 342 | F342K | A | 6.4e-06 | 6.2e-06 | 7.6e-06 | 6.733E-06 | 7.572E-07 | 6.200E-06 | 7.600E-06 |
| 6833 | 342 | F342M | A | 5.5e-06 | 7.5e-06 | 9.9e-06 | 7.633E-06 | 2.203E-06 | 5.500E-06 | 9.900E-06 |
| 6834 | 342 | F342F | A | 7.2e-06 | 5e-06 | 4.6e-06 | 5.600E-06 | 1.400E-06 | 4.600E-06 | 7.200E-06 |
| 6835 | 342 | F342P | A | 1.4e-05 | 5.7e-06 | 8.2e-06 | 9.300E-06 | 4.258E-06 | 5.700E-06 | 1.400E-05 |
| 6836 | 342 | F342S | A | 8.2e-06 | 5.9e-06 | 9.3e-06 | 7.800E-06 | 1.735E-06 | 5.900E-06 | 9.300E-06 |
| 6837 | 342 | F342T | A | 8.1e-06 | 9.3e-06 | 7.2e-06 | 8.200E-06 | 1.054E-06 | 7.200E-06 | 9.300E-06 |
| 6838 | 342 | F342W | A | 6.7e-06 | 6.5e-06 | 8.3e-06 | 7.167E-06 | 9.866E-07 | 6.500E-06 | 8.300E-06 |
| 6839 | 342 | F342Y | A | 6.7e-06 | 1.3e-05 | 9.9e-06 | 9.867E-06 | 3.150E-06 | 6.700E-06 | 1.300E-05 |
| 6840 | 342 | F342V | A | 6.5e-06 | 8.7e-06 | 8.8e-06 | 8.000E-06 | 1.300E-06 | 6.500E-06 | 8.800E-06 |
| 6841 | 343 | A343A | A | 9.4e-06 | 7.8e-06 | 6e-06 | 7.733E-06 | 1.701E-06 | 6.000E-06 | 9.400E-06 |
| 6842 | 343 | A343R | A | 6.7e-06 | 5.9e-06 | 5.3e-06 | 5.967E-06 | 7.024E-07 | 5.300E-06 | 6.700E-06 |
| 6843 | 343 | A343N | A | 7.7e-06 | 7.8e-06 | 4.6e-06 | 6.700E-06 | 1.819E-06 | 4.600E-06 | 7.800E-06 |
| 6844 | 343 | A343D | A | 6e-06 | 8.6e-06 | 6.5e-06 | 7.033E-06 | 1.380E-06 | 6.000E-06 | 8.600E-06 |
| 6845 | 343 | A343C | A | 1.4e-05 | 9.8e-06 | 7e-06 | 1.027E-05 | 3.523E-06 | 7.000E-06 | 1.400E-05 |
| 6846 | 343 | A343E | A | 1.4e-05 | 5.7e-06 | 8.4e-06 | 9.367E-06 | 4.234E-06 | 5.700E-06 | 1.400E-05 |
| 6847 | 343 | A343Q | A | 7.8e-06 | 1.2e-05 | 4.8e-06 | 8.200E-06 | 3.617E-06 | 4.800E-06 | 1.200E-05 |
| 6848 | 343 | A343G | A | 1.1e-05 | 7.6e-06 | 7.8e-06 | 8.800E-06 | 1.908E-06 | 7.600E-06 | 1.100E-05 |
| 6849 | 343 | A343H | A | 7.4e-06 | 8.6e-06 | 4.4e-06 | 6.800E-06 | 2.163E-06 | 4.400E-06 | 8.600E-06 |
| 6850 | 343 | A343I | A | 7.8e-06 | 7.4e-06 | 3.8e-06 | 6.333E-06 | 2.203E-06 | 3.800E-06 | 7.800E-06 |
| 6851 | 343 | A343L | A | 1.2e-05 | 9.3e-06 | 4.3e-06 | 8.533E-06 | 3.907E-06 | 4.300E-06 | 1.200E-05 |
| 6852 | 343 | A343K | A | 8.4e-06 | 8.8e-06 | 7e-06 | 8.067E-06 | 9.452E-07 | 7.000E-06 | 8.800E-06 |
| 6853 | 343 | A343M | A | 7.1e-06 | 1.2e-05 | 5.6e-06 | 8.233E-06 | 3.347E-06 | 5.600E-06 | 1.200E-05 |
| 6854 | 343 | A343F | A | 1.4e-05 | 9.4e-06 | 4.4e-06 | 9.267E-06 | 4.801E-06 | 4.400E-06 | 1.400E-05 |
| 6855 | 343 | A343P | A | 4.7e-06 | 5.9e-06 | 7.6e-06 | 6.067E-06 | 1.457E-06 | 4.700E-06 | 7.600E-06 |
| 6856 | 343 | A343S | A | 1.2e-05 | 5.3e-06 | 4.6e-06 | 7.300E-06 | 4.085E-06 | 4.600E-06 | 1.200E-05 |
| 6857 | 343 | A343T | A | 1e-05 | 7.3e-06 | 8.1e-06 | 8.467E-06 | 1.387E-06 | 7.300E-06 | 1.000E-05 |
| 6858 | 343 | A343W | A | 5.5e-06 | 9.6e-06 | 1.4e-05 | 9.700E-06 | 4.251E-06 | 5.500E-06 | 1.400E-05 |
| 6859 | 343 | A343Y | A | 7.4e-06 | 1.1e-05 | 9.4e-06 | 9.267E-06 | 1.804E-06 | 7.400E-06 | 1.100E-05 |
| 6860 | 343 | A343V | A | 5.1e-06 | 1.3e-05 | 9.9e-06 | 9.333E-06 | 3.980E-06 | 5.100E-06 | 1.300E-05 |
| 6861 | 344 | G344A | A | 1.4e-05 | 1.4e-05 | 7.8e-06 | 1.193E-05 | 3.580E-06 | 7.800E-06 | 1.400E-05 |
| 6862 | 344 | G344R | A | 8.1e-06 | 6.6e-06 | 8.4e-06 | 7.700E-06 | 9.644E-07 | 6.600E-06 | 8.400E-06 |
| 6863 | 344 | G344N | A | 5.3e-06 | 9.7e-06 | 7.5e-06 | 7.500E-06 | 2.200E-06 | 5.300E-06 | 9.700E-06 |
| 6864 | 344 | G344D | A | 7.8e-06 | 1e-05 | 8e-06 | 8.600E-06 | 1.217E-06 | 7.800E-06 | 1.000E-05 |
| 6865 | 344 | G344C | A | 6.7e-06 | 9.6e-06 | 5.7e-06 | 7.333E-06 | 2.026E-06 | 5.700E-06 | 9.600E-06 |
| 6866 | 344 | G344E | A | 6.3e-06 | 1e-05 | 6.1e-06 | 7.467E-06 | 2.196E-06 | 6.100E-06 | 1.000E-05 |
| 6867 | 344 | G344Q | A | 7.1e-06 | 4.7e-06 | 5.5e-06 | 5.767E-06 | 1.222E-06 | 4.700E-06 | 7.100E-06 |
| 6868 | 344 | G344G | A | 9.8e-06 | 1e-05 | 8.5e-06 | 9.433E-06 | 8.145E-07 | 8.500E-06 | 1.000E-05 |
| 6869 | 344 | G344H | A | 7.5e-06 | 6.5e-06 | 1.4e-05 | 9.333E-06 | 4.072E-06 | 6.500E-06 | 1.400E-05 |
| 6870 | 344 | G344I | A | 1.4e-05 | 5.6e-06 | 9e-06 | 9.533E-06 | 4.225E-06 | 5.600E-06 | 1.400E-05 |
| 6871 | 344 | G344L | A | 1.1e-05 | 8.6e-06 | 1.2e-05 | 1.053E-05 | 1.747E-06 | 8.600E-06 | 1.200E-05 |
| 6872 | 344 | G344K | A | 7.9e-06 | 7.9e-06 | 9e-06 | 8.267E-06 | 6.351E-07 | 7.900E-06 | 9.000E-06 |
| 6873 | 344 | G344M | A | 6.8e-06 | 9.4e-06 | 8.9e-06 | 8.367E-06 | 1.380E-06 | 6.800E-06 | 9.400E-06 |
| 6874 | 344 | G344F | A | 1e-05 | 1.1e-05 | 6.6e-06 | 9.200E-06 | 2.307E-06 | 6.600E-06 | 1.100E-05 |
| 6875 | 344 | G344P | A | 8.2e-06 | 8.9e-06 | 1e-05 | 9.033E-06 | 9.074E-07 | 8.200E-06 | 1.000E-05 |
| 6876 | 344 | G344S | A | 5.8e-06 | 4.8e-06 | 8.9e-06 | 6.500E-06 | 2.138E-06 | 4.800E-06 | 8.900E-06 |
| 6877 | 344 | G344T | A | 5.1e-06 | 6e-06 | 5.1e-06 | 5.400E-06 | 5.196E-07 | 5.100E-06 | 6.000E-06 |
| 6878 | 344 | G344W | A | 1e-05 | 5.4e-06 | 9.2e-06 | 8.200E-06 | 2.458E-06 | 5.400E-06 | 1.000E-05 |
| 6879 | 344 | G344Y | A | 6.6e-06 | 6.2e-06 | 8.1e-06 | 6.967E-06 | 1.002E-06 | 6.200E-06 | 8.100E-06 |
| 6880 | 344 | G344V | A | 8.3e-06 | 7e-06 | 7.7e-06 | 7.667E-06 | 6.506E-07 | 7.000E-06 | 8.300E-06 |
| 6881 | 345 | C345A | A | 9.7e-06 | 1.3e-05 | 1.3e-05 | 1.190E-05 | 1.905E-06 | 9.700E-06 | 1.300E-05 |
| 6882 | 345 | C345R | A | 4.5e-06 | 9.7e-06 | 6.8e-06 | 7.000E-06 | 2.606E-06 | 4.500E-06 | 9.700E-06 |
| 6883 | 345 | C345N | A | 8.8e-06 | 5e-06 | 9.6e-06 | 7.800E-06 | 2.458E-06 | 5.000E-06 | 9.600E-06 |
| 6884 | 345 | C345D | A | 7.4e-06 | 6.6e-06 | 4.5e-06 | 6.167E-06 | 1.498E-06 | 4.500E-06 | 7.400E-06 |

| | | | | | | | | | | |
|---|---|---|---|---|---|---|---|---|---|---|
| 6885 | 345 | C345C | A | 9.8e-06 | 1e-05 | 7.4e-06 | 9.067E-06 | 1.447E-06 | 7.400E-06 | 1.000E-05 |
| 6886 | 345 | C345E | A | 4.3e-06 | 6.4e-06 | 6.6e-06 | 5.767E-06 | 1.274E-06 | 4.300E-06 | 6.600E-06 |
| 6887 | 345 | C345Q | A | 8.9e-06 | 6e-06 | 7.4e-06 | 7.433E-06 | 1.450E-06 | 6.000E-06 | 8.900E-06 |
| 6888 | 345 | C345G | A | 9.8e-06 | 1e-05 | 5.7e-06 | 8.500E-06 | 2.427E-06 | 5.700E-06 | 1.000E-05 |
| 6889 | 345 | C345H | A | 7e-06 | 1.2e-05 | 1.3e-05 | 1.067E-05 | 3.215E-06 | 7.000E-06 | 1.300E-05 |
| 6890 | 345 | C345I | A | 7.3e-06 | 8e-06 | 9.9e-06 | 8.400E-06 | 1.345E-06 | 7.300E-06 | 9.900E-06 |
| 6891 | 345 | C345L | A | 1.2e-05 | 1.4e-05 | 8e-06 | 1.133E-05 | 3.055E-06 | 8.000E-06 | 1.400E-05 |
| 6892 | 345 | C345K | A | 7e-06 | 6.3e-06 | 7.5e-06 | 6.933E-06 | 6.028E-07 | 6.300E-06 | 7.500E-06 |
| 6893 | 345 | C345M | A | 7.8e-06 | 9.8e-06 | 7.5e-06 | 8.367E-06 | 1.250E-06 | 7.500E-06 | 9.800E-06 |
| 6894 | 345 | C345F | A | 1.3e-05 | 1.5e-05 | 1.1e-05 | 1.300E-05 | 2.000E-06 | 1.100E-05 | 1.500E-05 |
| 6895 | 345 | C345P | A | 3.9e-06 | 6.5e-06 | 1.2e-05 | 7.467E-06 | 4.136E-06 | 3.900E-06 | 1.200E-05 |
| 6896 | 345 | C345S | A | 5.5e-06 | 1e-05 | 7e-06 | 7.500E-06 | 2.291E-06 | 5.500E-06 | 1.000E-05 |
| 6897 | 345 | C345T | A | 8e-06 | 5.8e-06 | 1.4e-05 | 9.267E-06 | 4.244E-06 | 5.800E-06 | 1.400E-05 |
| 6898 | 345 | C345W | A | 1.5e-05 | 5.5e-06 | 1.4e-05 | 1.150E-05 | 5.220E-06 | 5.500E-06 | 1.500E-05 |
| 6899 | 345 | C345Y | A | 4.4e-06 | 8.9e-06 | 1.1e-05 | 8.100E-06 | 3.372E-06 | 4.400E-06 | 1.100E-05 |
| 6900 | 345 | C345V | A | 1.2e-05 | 1.3e-05 | 4.9e-06 | 9.967E-06 | 4.416E-06 | 4.900E-06 | 1.300E-05 |
| 6901 | 346 | K346A | A | 7e-06 | 9e-06 | 9.7e-06 | 8.567E-06 | 1.401E-06 | 7.000E-06 | 9.700E-06 |
| 6902 | 346 | K346R | A | 6.6e-06 | 7.1e-06 | 6.9e-06 | 6.867E-06 | 2.517E-07 | 6.600E-06 | 7.100E-06 |
| 6903 | 346 | K346N | A | 5.2e-06 | 7.1e-06 | 4.7e-06 | 5.667E-06 | 1.266E-06 | 4.700E-06 | 7.100E-06 |
| 6904 | 346 | K346D | A | 1.3e-05 | 1.2e-05 | 8.6e-06 | 1.120E-05 | 2.307E-06 | 8.600E-06 | 1.300E-05 |
| 6905 | 346 | K346C | A | 8.5e-06 | 6.8e-06 | 4.7e-06 | 6.667E-06 | 1.904E-06 | 4.700E-06 | 8.500E-06 |
| 6906 | 346 | K346E | A | 1.7e-05 | 8.4e-06 | 6.5e-06 | 1.063E-05 | 5.595E-06 | 6.500E-06 | 1.700E-05 |
| 6907 | 346 | K346Q | A | 1.2e-05 | 7.3e-06 | 1.3e-05 | 1.077E-05 | 3.044E-06 | 7.300E-06 | 1.300E-05 |
| 6908 | 346 | K346G | A | 8.8e-06 | 7.4e-06 | 7.3e-06 | 7.833E-06 | 8.386E-07 | 7.300E-06 | 8.800E-06 |
| 6909 | 346 | K346H | A | 1.7e-05 | 6.6e-06 | 5.3e-06 | 9.633E-06 | 6.413E-06 | 5.300E-06 | 1.700E-05 |
| 6910 | 346 | K346I | A | 6.9e-06 | 7.8e-06 | 1.2e-05 | 8.900E-06 | 2.722E-06 | 6.900E-06 | 1.200E-05 |
| 6911 | 346 | K346L | A | 5.5e-06 | 7.1e-06 | 4.8e-06 | 5.800E-06 | 1.179E-06 | 4.800E-06 | 7.100E-06 |
| 6912 | 346 | K346K | A | 7.5e-06 | 7.9e-06 | 1.2e-05 | 9.133E-06 | 2.491E-06 | 7.500E-06 | 1.200E-05 |
| 6913 | 346 | K346M | A | 6.2e-06 | 6.9e-06 | 5.1e-06 | 6.067E-06 | 9.074E-07 | 5.100E-06 | 6.900E-06 |
| 6914 | 346 | K346F | A | 8.3e-06 | 5.3e-06 | 1.2e-05 | 8.533E-06 | 3.356E-06 | 5.300E-06 | 1.200E-05 |
| 6915 | 346 | K346P | A | 1.1e-05 | 1.3e-05 | 8.1e-06 | 1.070E-05 | 2.464E-06 | 8.100E-06 | 1.300E-05 |
| 6916 | 346 | K346S | A | 8.7e-06 | 6.7e-06 | 5.5e-06 | 6.967E-06 | 1.617E-06 | 5.500E-06 | 8.700E-06 |
| 6917 | 346 | K346T | A | 1.2e-05 | 8.3e-06 | 9.2e-06 | 9.833E-06 | 1.930E-06 | 8.300E-06 | 1.200E-05 |
| 6918 | 346 | K346W | A | 3.5e-06 | 8.9e-06 | 9.2e-06 | 7.200E-06 | 3.208E-06 | 3.500E-06 | 9.200E-06 |
| 6919 | 346 | K346Y | A | 7.3e-06 | 6.6e-06 | 7.1e-06 | 7.000E-06 | 3.606E-07 | 6.600E-06 | 7.300E-06 |
| 6920 | 346 | K346V | A | 1.1e-05 | 1e-05 | 1e-05 | 1.033E-05 | 5.774E-07 | 1.000E-05 | 1.100E-05 |
| 6921 | 347 | K347A | A | 9.5e-06 | 9.7e-06 | 4.7e-06 | 7.967E-06 | 2.831E-06 | 4.700E-06 | 9.700E-06 |
| 6922 | 347 | K347R | A | 1.4e-05 | 8.4e-06 | 1e-05 | 1.080E-05 | 2.884E-06 | 8.400E-06 | 1.400E-05 |
| 6923 | 347 | K347N | A | 8.2e-06 | 1.1e-05 | 8.5e-06 | 9.233E-06 | 1.537E-06 | 8.200E-06 | 1.100E-05 |
| 6924 | 347 | K347D | A | 4.5e-06 | 5.1e-06 | 4.5e-06 | 4.700E-06 | 3.464E-07 | 4.500E-06 | 5.100E-06 |
| 6925 | 347 | K347C | A | 8.1e-06 | 7.1e-06 | 1.3e-05 | 9.400E-06 | 3.158E-06 | 7.100E-06 | 1.300E-05 |
| 6926 | 347 | K347E | A | 8.5e-06 | 4.7e-06 | 1e-05 | 7.733E-06 | 2.732E-06 | 4.700E-06 | 1.000E-05 |
| 6927 | 347 | K347Q | A | 7.2e-06 | 8.9e-06 | 6.4e-06 | 7.500E-06 | 1.277E-06 | 6.400E-06 | 8.900E-06 |
| 6928 | 347 | K347G | A | 9.5e-06 | 9e-06 | 6.6e-06 | 8.367E-06 | 1.550E-06 | 6.600E-06 | 9.500E-06 |
| 6929 | 347 | K347H | A | 9.6e-06 | 9.2e-06 | 9.7e-06 | 9.500E-06 | 2.646E-07 | 9.200E-06 | 9.700E-06 |
| 6930 | 347 | K347I | A | 1.3e-05 | 6.4e-06 | 6.7e-06 | 8.700E-06 | 3.727E-06 | 6.400E-06 | 1.300E-05 |
| 6931 | 347 | K347L | A | 8.1e-06 | 7.1e-06 | 9.8e-06 | 8.333E-06 | 1.365E-06 | 7.100E-06 | 9.800E-06 |
| 6932 | 347 | K347K | A | 9.3e-06 | 4.6e-06 | 5.9e-06 | 6.600E-06 | 2.427E-06 | 4.600E-06 | 9.300E-06 |
| 6933 | 347 | K347M | A | 6e-06 | 5.7e-06 | 5.9e-06 | 5.867E-06 | 1.528E-07 | 5.700E-06 | 6.000E-06 |
| 6934 | 347 | K347F | A | 7e-06 | 6.3e-06 | 1.2e-05 | 8.433E-06 | 3.109E-06 | 6.300E-06 | 1.200E-05 |
| 6935 | 347 | K347P | A | 7.6e-06 | 5.8e-06 | 1.4e-05 | 9.133E-06 | 4.310E-06 | 5.800E-06 | 1.400E-05 |
| 6936 | 347 | K347S | A | 9.1e-06 | 9e-06 | 9.4e-06 | 9.167E-06 | 2.082E-07 | 9.000E-06 | 9.400E-06 |
| 6937 | 347 | K347T | A | 1e-05 | 8.9e-06 | 4.3e-06 | 7.733E-06 | 3.024E-06 | 4.300E-06 | 1.000E-05 |
| 6938 | 347 | K347W | A | 6.9e-06 | 6.9e-06 | 1e-05 | 7.933E-06 | 1.790E-06 | 6.900E-06 | 1.000E-05 |
| 6939 | 347 | K347Y | A | 5e-06 | 6.2e-06 | 1.2e-05 | 7.733E-06 | 3.743E-06 | 5.000E-06 | 1.200E-05 |
| 6940 | 347 | K347V | A | 8.7e-06 | 9.1e-06 | 6.1e-06 | 7.967E-06 | 1.629E-06 | 6.100E-06 | 9.100E-06 |
| 6941 | 348 | I348A | A | 5.4e-06 | 5.7e-06 | 4.4e-06 | 5.167E-06 | 6.807E-07 | 4.400E-06 | 5.700E-06 |
| 6942 | 348 | I348R | A | 7.1e-06 | 1e-05 | 4.7e-06 | 7.267E-06 | 2.654E-06 | 4.700E-06 | 1.000E-05 |
| 6943 | 348 | I348N | A | 6.3e-06 | 9.6e-06 | 1.3e-05 | 9.633E-06 | 3.350E-06 | 6.300E-06 | 1.300E-05 |
| 6944 | 348 | I348D | A | 9.3e-06 | 6.5e-06 | 9.4e-06 | 8.400E-06 | 1.646E-06 | 6.500E-06 | 9.400E-06 |
| 6945 | 348 | I348C | A | 6.3e-06 | 1.1e-05 | 1.4e-05 | 1.043E-05 | 3.881E-06 | 6.300E-06 | 1.400E-05 |
| 6946 | 348 | I348E | A | 6.5e-06 | 1.3e-05 | 8.9e-06 | 9.467E-06 | 3.287E-06 | 6.500E-06 | 1.300E-05 |
| 6947 | 348 | I348Q | A | 9.5e-06 | 6.9e-06 | 1e-05 | 8.800E-06 | 1.664E-06 | 6.900E-06 | 1.000E-05 |
| 6948 | 348 | I348G | A | 5.1e-06 | 1.2e-05 | 1.1e-05 | 9.367E-06 | 3.729E-06 | 5.100E-06 | 1.200E-05 |
| 6949 | 348 | I348H | A | 9.3e-06 | 1.6e-05 | 1.4e-05 | 1.310E-05 | 3.439E-06 | 9.300E-06 | 1.600E-05 |
| 6950 | 348 | I348I | A | 1.1e-05 | 9.4e-06 | 9.5e-06 | 9.967E-06 | 8.963E-07 | 9.400E-06 | 1.100E-05 |
| 6951 | 348 | I348L | A | 8.4e-06 | 8.9e-06 | 6.3e-06 | 7.867E-06 | 1.380E-06 | 6.300E-06 | 8.900E-06 |
| 6952 | 348 | I348K | A | 6.9e-06 | 7.1e-06 | 5.9e-06 | 6.633E-06 | 6.429E-07 | 5.900E-06 | 7.100E-06 |
| 6953 | 348 | I348M | A | 1.7e-05 | 6.2e-06 | 6e-06 | 9.733E-06 | 6.294E-06 | 6.000E-06 | 1.700E-05 |
| 6954 | 348 | I348F | A | 7.2e-06 | 8.1e-06 | 5.6e-06 | 6.967E-06 | 1.266E-06 | 5.600E-06 | 8.100E-06 |
| 6955 | 348 | I348P | A | 1.3e-05 | 6e-06 | 6.5e-06 | 8.500E-06 | 3.905E-06 | 6.000E-06 | 1.300E-05 |
| 6956 | 348 | I348S | A | 1.6e-05 | 9.1e-06 | 9.9e-06 | 1.167E-05 | 3.774E-06 | 9.100E-06 | 1.600E-05 |
| 6957 | 348 | I348T | A | 8.4e-06 | 5.6e-06 | 6.7e-06 | 6.900E-06 | 1.411E-06 | 5.600E-06 | 8.400E-06 |
| 6958 | 348 | I348W | A | 1.1e-05 | 6.9e-06 | 5.1e-06 | 7.667E-06 | 3.024E-06 | 5.100E-06 | 1.100E-05 |
| 6959 | 348 | I348Y | A | 5.4e-06 | 6e-06 | 9.1e-06 | 6.833E-06 | 1.986E-06 | 5.400E-06 | 9.100E-06 |
| 6960 | 348 | I348V | A | 6.6e-06 | 5.3e-06 | 1.6e-05 | 9.300E-06 | 5.839E-06 | 5.300E-06 | 1.600E-05 |
| 6961 | 349 | F349A | A | 4.6e-06 | 1e-05 | 1.1e-05 | 8.533E-06 | 3.443E-06 | 4.600E-06 | 1.100E-05 |
| 6962 | 349 | F349R | A | 1.5e-05 | 1.1e-05 | 5e-06 | 1.033E-05 | 5.033E-06 | 5.000E-06 | 1.500E-05 |
| 6963 | 349 | F349N | A | 7.7e-06 | 4.4e-06 | 8.4e-06 | 6.833E-06 | 2.136E-06 | 4.400E-06 | 8.400E-06 |
| 6964 | 349 | F349D | A | 7.6e-06 | 6.6e-06 | 8.2e-06 | 7.467E-06 | 8.083E-07 | 6.600E-06 | 8.200E-06 |
| 6965 | 349 | F349C | A | 9.9e-06 | 9.7e-06 | 1.4e-05 | 1.120E-05 | 2.427E-06 | 9.700E-06 | 1.400E-05 |
| 6966 | 349 | F349E | A | 1.3e-05 | 9.7e-06 | 8.8e-06 | 1.050E-05 | 2.211E-06 | 8.800E-06 | 1.300E-05 |
| 6967 | 349 | F349Q | A | 8.7e-06 | 5.4e-06 | 9e-06 | 7.700E-06 | 1.997E-06 | 5.400E-06 | 9.000E-06 |
| 6968 | 349 | F349G | A | 1.3e-05 | 7.6e-06 | 8.5e-06 | 9.700E-06 | 2.893E-06 | 7.600E-06 | 1.300E-05 |
| 6969 | 349 | F349H | A | 1.1e-05 | 4.9e-06 | 6.1e-06 | 7.333E-06 | 3.232E-06 | 4.900E-06 | 1.100E-05 |

| | | | | | | | | | | |
|---|---|---|---|---|---|---|---|---|---|---|
| 6970 | 349 | F349I | A | 1.2e-05 | 9.4e-06 | 8.5e-06 | 9.967E-06 | 1.818E-06 | 8.500E-06 | 1.200E-05 |
| 6971 | 349 | F349L | A | 1.2e-05 | 6.6e-06 | 8.2e-06 | 8.933E-06 | 2.774E-06 | 6.600E-06 | 1.200E-05 |
| 6972 | 349 | F349K | A | 6.4e-06 | 5.9e-06 | 8.1e-06 | 6.800E-06 | 1.153E-06 | 5.900E-06 | 8.100E-06 |
| 6973 | 349 | F349M | A | 6.4e-06 | 1.1e-05 | 9.3e-06 | 8.900E-06 | 2.326E-06 | 6.400E-06 | 1.100E-05 |
| 6974 | 349 | F349F | A | 7.5e-06 | 1.2e-05 | 7.3e-06 | 8.933E-06 | 2.658E-06 | 7.300E-06 | 1.200E-05 |
| 6975 | 349 | F349P | A | 1.2e-05 | 1.2e-05 | 9.8e-06 | 1.127E-05 | 1.270E-06 | 9.800E-06 | 1.200E-05 |
| 6976 | 349 | F349S | A | 7.9e-06 | 9.8e-06 | 7.4e-06 | 8.367E-06 | 1.266E-06 | 7.400E-06 | 9.800E-06 |
| 6977 | 349 | F349T | A | 9.6e-06 | 6.4e-06 | 6.6e-06 | 7.533E-06 | 1.793E-06 | 6.400E-06 | 9.600E-06 |
| 6978 | 349 | F349W | A | 5.6e-06 | 8.7e-06 | 8.5e-06 | 7.600E-06 | 1.735E-06 | 5.600E-06 | 8.700E-06 |
| 6979 | 349 | F349Y | A | 8.8e-06 | 7.2e-06 | 6.9e-06 | 7.633E-06 | 1.021E-06 | 6.900E-06 | 8.800E-06 |
| 6980 | 349 | F349V | A | 4.8e-06 | 7.8e-06 | 1.5e-05 | 9.200E-06 | 5.242E-06 | 4.800E-06 | 1.500E-05 |
| 6981 | 350 | G350A | A | 8.4e-06 | 5.5e-06 | 6.7e-06 | 6.867E-06 | 1.457E-06 | 5.500E-06 | 8.400E-06 |
| 6982 | 350 | G350R | A | 5.9e-06 | 8.8e-06 | 9.4e-06 | 8.033E-06 | 1.872E-06 | 5.900E-06 | 9.400E-06 |
| 6983 | 350 | G350N | A | 6.8e-06 | 1.1e-05 | 7e-06 | 8.267E-06 | 2.369E-06 | 6.800E-06 | 1.100E-05 |
| 6984 | 350 | G350D | A | 6.2e-06 | 8.6e-06 | 1e-05 | 8.267E-06 | 1.922E-06 | 6.200E-06 | 1.000E-05 |
| 6985 | 350 | G350C | A | 8.5e-06 | 9e-06 | 1e-05 | 9.167E-06 | 7.638E-07 | 8.500E-06 | 1.000E-05 |
| 6986 | 350 | G350E | A | 8.5e-06 | 1.2e-05 | 6e-06 | 8.833E-06 | 3.014E-06 | 6.000E-06 | 1.200E-05 |
| 6987 | 350 | G350Q | A | 6.2e-06 | 5.1e-06 | 8.2e-06 | 6.500E-06 | 1.572E-06 | 5.100E-06 | 8.200E-06 |
| 6988 | 350 | G350G | A | 1.2e-05 | 9.4e-06 | 7.3e-06 | 9.567E-06 | 2.354E-06 | 7.300E-06 | 1.200E-05 |
| 6989 | 350 | G350H | A | 1.6e-05 | 7.2e-06 | 6e-06 | 9.733E-06 | 5.460E-06 | 6.000E-06 | 1.600E-05 |
| 6990 | 350 | G350I | A | 5.5e-06 | 6.2e-06 | 6.2e-06 | 5.967E-06 | 4.041E-07 | 5.500E-06 | 6.200E-06 |
| 6991 | 350 | G350L | A | 1e-05 | 4.4e-06 | 1e-05 | 8.133E-06 | 3.233E-06 | 4.400E-06 | 1.000E-05 |
| 6992 | 350 | G350K | A | 9.3e-06 | 4.4e-06 | 7e-06 | 6.900E-06 | 2.452E-06 | 4.400E-06 | 9.300E-06 |
| 6993 | 350 | G350M | A | 5.6e-06 | 6.4e-06 | 8.3e-06 | 6.767E-06 | 1.387E-06 | 5.600E-06 | 8.300E-06 |
| 6994 | 350 | G350F | A | 5.3e-06 | 8.9e-06 | 9.8e-06 | 8.000E-06 | 2.381E-06 | 5.300E-06 | 9.800E-06 |
| 6995 | 350 | G350P | A | 8.2e-06 | 4.4e-06 | 7.2e-06 | 6.600E-06 | 1.970E-06 | 4.400E-06 | 8.200E-06 |
| 6996 | 350 | G350S | A | 5.7e-06 | 6.6e-06 | 1.1e-05 | 7.767E-06 | 2.836E-06 | 5.700E-06 | 1.100E-05 |
| 6997 | 350 | G350T | A | 8.1e-06 | 1e-05 | 6e-06 | 8.033E-06 | 2.001E-06 | 6.000E-06 | 1.000E-05 |
| 6998 | 350 | G350W | A | 4.4e-06 | 1.1e-05 | 8.1e-06 | 7.833E-06 | 3.308E-06 | 4.400E-06 | 1.100E-05 |
| 6999 | 350 | G350Y | A | 9.8e-06 | 8.5e-06 | 7.8e-06 | 8.700E-06 | 1.015E-06 | 7.800E-06 | 9.800E-06 |
| 7000 | 350 | G350V | A | 6.6e-06 | 1.1e-05 | 8.8e-06 | 8.800E-06 | 2.200E-06 | 6.600E-06 | 1.100E-05 |
| 7001 | 351 | S351A | A | 9.9e-06 | 1.4e-05 | 8.7e-06 | 1.087E-05 | 2.779E-06 | 8.700E-06 | 1.400E-05 |
| 7002 | 351 | S351R | A | 1.4e-05 | 8.3e-06 | 8.3e-06 | 1.020E-05 | 3.291E-06 | 8.300E-06 | 1.400E-05 |
| 7003 | 351 | S351N | A | 8.9e-06 | 4.3e-06 | 5e-06 | 6.067E-06 | 2.479E-06 | 4.300E-06 | 8.900E-06 |
| 7004 | 351 | S351D | A | 8.2e-06 | 7.6e-06 | 6.9e-06 | 7.567E-06 | 6.506E-07 | 6.900E-06 | 8.200E-06 |
| 7005 | 351 | S351C | A | 8.6e-06 | 6.6e-06 | 5.9e-06 | 7.033E-06 | 1.401E-06 | 5.900E-06 | 8.600E-06 |
| 7006 | 351 | S351E | A | 9.8e-06 | 9.5e-06 | 1.4e-05 | 1.110E-05 | 2.516E-06 | 9.500E-06 | 1.400E-05 |
| 7007 | 351 | S351Q | A | 1e-05 | 4.6e-06 | 9.5e-06 | 8.033E-06 | 2.984E-06 | 4.600E-06 | 1.000E-05 |
| 7008 | 351 | S351G | A | 9.3e-06 | 9.5e-06 | 6.2e-06 | 8.333E-06 | 1.850E-06 | 6.200E-06 | 9.500E-06 |
| 7009 | 351 | S351H | A | 5.3e-06 | 6.1e-06 | 1.4e-05 | 8.467E-06 | 4.809E-06 | 5.300E-06 | 1.400E-05 |
| 7010 | 351 | S351I | A | 4.7e-06 | 6.9e-06 | 6.6e-06 | 6.067E-06 | 1.193E-06 | 4.700E-06 | 6.900E-06 |
| 7011 | 351 | S351L | A | 6e-06 | 6.5e-06 | 1.1e-05 | 7.833E-06 | 2.754E-06 | 6.000E-06 | 1.100E-05 |
| 7012 | 351 | S351K | A | 5.4e-06 | 5.5e-06 | 5.8e-06 | 5.567E-06 | 2.082E-07 | 5.400E-06 | 5.800E-06 |
| 7013 | 351 | S351M | A | 9.2e-06 | 7.5e-06 | 6.8e-06 | 7.833E-06 | 1.234E-06 | 6.800E-06 | 9.200E-06 |
| 7014 | 351 | S351F | A | 1.2e-05 | 9.3e-06 | 4.9e-06 | 8.733E-06 | 3.584E-06 | 4.900E-06 | 1.200E-05 |
| 7015 | 351 | S351P | A | 9.6e-06 | 8.6e-06 | 1.5e-05 | 1.107E-05 | 3.443E-06 | 8.600E-06 | 1.500E-05 |
| 7016 | 351 | S351S | A | 7.5e-06 | 5.9e-06 | 6.3e-06 | 6.567E-06 | 8.327E-07 | 5.900E-06 | 7.500E-06 |
| 7017 | 351 | S351T | A | 7.4e-06 | 1.2e-05 | 3.2e-06 | 7.533E-06 | 4.402E-06 | 3.200E-06 | 1.200E-05 |
| 7018 | 351 | S351W | A | 1.4e-05 | 1.1e-05 | 9.4e-06 | 1.147E-05 | 2.335E-06 | 9.400E-06 | 1.400E-05 |
| 7019 | 351 | S351Y | A | 9.3e-06 | 7.8e-06 | 6.9e-06 | 8.000E-06 | 1.212E-06 | 6.900E-06 | 9.300E-06 |
| 7020 | 351 | S351V | A | 1e-05 | 4.5e-06 | 6.3e-06 | 6.933E-06 | 2.804E-06 | 4.500E-06 | 1.000E-05 |
| 7021 | 352 | L352A | A | 9.5e-06 | 9.1e-06 | 1e-05 | 9.533E-06 | 4.509E-07 | 9.100E-06 | 1.000E-05 |
| 7022 | 352 | L352R | A | 5.7e-06 | 7e-06 | 4.1e-06 | 5.600E-06 | 1.453E-06 | 4.100E-06 | 7.000E-06 |
| 7023 | 352 | L352N | A | 1.4e-05 | 7.5e-06 | 8e-06 | 9.833E-06 | 3.617E-06 | 7.500E-06 | 1.400E-05 |
| 7024 | 352 | L352D | A | 6.5e-06 | 1.5e-05 | 6.6e-06 | 9.367E-06 | 4.879E-06 | 6.500E-06 | 1.500E-05 |
| 7025 | 352 | L352C | A | 8.9e-06 | 5.6e-06 | 1.3e-05 | 9.167E-06 | 3.707E-06 | 5.600E-06 | 1.300E-05 |
| 7026 | 352 | L352E | A | 8.2e-06 | 9.6e-06 | 6.1e-06 | 7.967E-06 | 1.762E-06 | 6.100E-06 | 9.600E-06 |
| 7027 | 352 | L352Q | A | 4.8e-06 | 1e-05 | 6.1e-06 | 6.967E-06 | 2.706E-06 | 4.800E-06 | 1.000E-05 |
| 7028 | 352 | L352G | A | 3.7e-06 | 9.3e-06 | 5.4e-06 | 6.133E-06 | 2.871E-06 | 3.700E-06 | 9.300E-06 |
| 7029 | 352 | L352H | A | 1.2e-05 | 1.1e-05 | 8.9e-06 | 1.063E-05 | 1.582E-06 | 8.900E-06 | 1.200E-05 |
| 7030 | 352 | L352I | A | 5.1e-06 | 7.9e-06 | 7.4e-06 | 6.800E-06 | 1.493E-06 | 5.100E-06 | 7.900E-06 |
| 7031 | 352 | L352L | A | 6.3e-06 | 1.2e-05 | 9.1e-06 | 9.133E-06 | 2.850E-06 | 6.300E-06 | 1.200E-05 |
| 7032 | 352 | L352K | A | 7.5e-06 | 7.3e-06 | 4.8e-06 | 6.533E-06 | 1.504E-06 | 4.800E-06 | 7.500E-06 |
| 7033 | 352 | L352M | A | 4.8e-06 | 1.3e-05 | 1.4e-05 | 1.060E-05 | 5.048E-06 | 4.800E-06 | 1.400E-05 |
| 7034 | 352 | L352F | A | 6.5e-06 | 1.4e-05 | 1.2e-05 | 1.083E-05 | 3.884E-06 | 6.500E-06 | 1.400E-05 |
| 7035 | 352 | L352P | A | 6.6e-06 | 6.1e-06 | 6.6e-06 | 6.433E-06 | 2.887E-07 | 6.100E-06 | 6.600E-06 |
| 7036 | 352 | L352S | A | 1.2e-05 | 1.2e-05 | 9.9e-06 | 1.130E-05 | 1.212E-06 | 9.900E-06 | 1.200E-05 |
| 7037 | 352 | L352T | A | 1.2e-05 | 6.4e-06 | 7.7e-06 | 8.700E-06 | 2.931E-06 | 6.400E-06 | 1.200E-05 |
| 7038 | 352 | L352W | A | 7.1e-06 | 9.1e-06 | 1.2e-05 | 9.400E-06 | 2.464E-06 | 7.100E-06 | 1.200E-05 |
| 7039 | 352 | L352Y | A | 3.8e-06 | 1.2e-05 | 9.2e-06 | 8.333E-06 | 4.168E-06 | 3.800E-06 | 1.200E-05 |
| 7040 | 352 | L352V | A | 7.7e-06 | 8.6e-06 | 8.1e-06 | 8.133E-06 | 4.509E-07 | 7.700E-06 | 8.600E-06 |
| 7041 | 353 | A353A | A | 5.9e-06 | 1.1e-05 | 9.4e-06 | 8.767E-06 | 2.608E-06 | 5.900E-06 | 1.100E-05 |
| 7042 | 353 | A353R | A | 4.9e-06 | 7e-06 | 9.6e-06 | 7.167E-06 | 2.354E-06 | 4.900E-06 | 9.600E-06 |
| 7043 | 353 | A353N | A | 8.1e-06 | 4.1e-06 | 9.4e-06 | 7.200E-06 | 2.762E-06 | 4.100E-06 | 9.400E-06 |
| 7044 | 353 | A353D | A | 1.1e-05 | 6.6e-06 | 1.1e-05 | 9.533E-06 | 2.540E-06 | 6.600E-06 | 1.100E-05 |
| 7045 | 353 | A353C | A | 7.3e-06 | 1.1e-05 | 3.5e-06 | 7.267E-06 | 3.750E-06 | 3.500E-06 | 1.100E-05 |
| 7046 | 353 | A353E | A | 7.5e-06 | 1e-05 | 5.1e-06 | 7.533E-06 | 2.450E-06 | 5.100E-06 | 1.000E-05 |
| 7047 | 353 | A353Q | A | 5.9e-06 | 9.2e-06 | 7e-06 | 7.367E-06 | 1.680E-06 | 5.900E-06 | 9.200E-06 |
| 7048 | 353 | A353G | A | 9.5e-06 | 1.1e-05 | 6.3e-06 | 8.933E-06 | 2.401E-06 | 6.300E-06 | 1.100E-05 |
| 7049 | 353 | A353H | A | 1.2e-05 | 9e-06 | 8.7e-06 | 9.900E-06 | 1.825E-06 | 8.700E-06 | 1.200E-05 |
| 7050 | 353 | A353I | A | 1.4e-05 | 6.4e-06 | 7.4e-06 | 9.267E-06 | 4.130E-06 | 6.400E-06 | 1.400E-05 |
| 7051 | 353 | A353L | A | 9.8e-06 | 7.7e-06 | 9.3e-06 | 8.933E-06 | 1.097E-06 | 7.700E-06 | 9.800E-06 |
| 7052 | 353 | A353K | A | 1.4e-05 | 9.1e-06 | 8.4e-06 | 1.050E-05 | 3.051E-06 | 8.400E-06 | 1.400E-05 |
| 7053 | 353 | A353M | A | 1e-05 | 7.4e-06 | 7.6e-06 | 8.333E-06 | 1.447E-06 | 7.400E-06 | 1.000E-05 |
| 7054 | 353 | A353F | A | 7.8e-06 | 8.2e-06 | 6.8e-06 | 7.600E-06 | 7.211E-07 | 6.800E-06 | 8.200E-06 |

| | | | | | | | | | | |
|---|---|---|---|---|---|---|---|---|---|---|
| 7055 | 353 | A353P | A | 8.4e-06 | 1.9e-05 | 1e-05 | 1.247E-05 | 5.714E-06 | 8.400E-06 | 1.900E-05 |
| 7056 | 353 | A353S | A | 5.3e-06 | 1.3e-05 | 4.5e-06 | 7.600E-06 | 4.694E-06 | 4.500E-06 | 1.300E-05 |
| 7057 | 353 | A353T | A | 4.5e-06 | 9.4e-06 | 1.2e-05 | 8.633E-06 | 3.808E-06 | 4.500E-06 | 1.200E-05 |
| 7058 | 353 | A353W | A | 6.7e-06 | 5.8e-06 | 6.3e-06 | 6.267E-06 | 4.509E-07 | 5.800E-06 | 6.700E-06 |
| 7059 | 353 | A353Y | A | 5.7e-06 | 8.1e-06 | 4.4e-06 | 6.067E-06 | 1.877E-06 | 4.400E-06 | 8.100E-06 |
| 7060 | 353 | A353V | A | 9.2e-06 | 9.1e-06 | 8.3e-06 | 8.867E-06 | 4.933E-07 | 8.300E-06 | 9.200E-06 |
| 7061 | 354 | F354A | A | 8.5e-06 | 9.4e-06 | 1.4e-05 | 1.063E-05 | 2.950E-06 | 8.500E-06 | 1.400E-05 |
| 7062 | 354 | F354R | A | 8e-06 | 1e-05 | 6.7e-06 | 8.233E-06 | 1.662E-06 | 6.700E-06 | 1.000E-05 |
| 7063 | 354 | F354N | A | 6.5e-06 | 8.4e-06 | 1.2e-05 | 8.967E-06 | 2.793E-06 | 6.500E-06 | 1.200E-05 |
| 7064 | 354 | F354D | A | 8.1e-06 | 9.3e-06 | 7e-06 | 8.133E-06 | 1.150E-06 | 7.000E-06 | 9.300E-06 |
| 7065 | 354 | F354C | A | 5.8e-06 | 1.4e-05 | 6.8e-06 | 8.867E-06 | 4.474E-06 | 5.800E-06 | 1.400E-05 |
| 7066 | 354 | F354E | A | 1.1e-05 | 6.5e-06 | 5e-06 | 7.500E-06 | 3.122E-06 | 5.000E-06 | 1.100E-05 |
| 7067 | 354 | F354Q | A | 9.2e-06 | 7.3e-06 | 8.1e-06 | 8.200E-06 | 9.539E-07 | 7.300E-06 | 9.200E-06 |
| 7068 | 354 | F354G | A | 7.2e-06 | 6.6e-06 | 7.6e-06 | 7.133E-06 | 5.033E-07 | 6.600E-06 | 7.600E-06 |
| 7069 | 354 | F354H | A | 1.4e-05 | 7.3e-06 | 9.6e-06 | 1.030E-05 | 3.404E-06 | 7.300E-06 | 1.400E-05 |
| 7070 | 354 | F354I | A | 5.1e-06 | 1.1e-05 | 6.3e-06 | 7.467E-06 | 3.118E-06 | 5.100E-06 | 1.100E-05 |
| 7071 | 354 | F354L | A | 8.5e-06 | 7.4e-06 | 6.7e-06 | 7.533E-06 | 9.074E-07 | 6.700E-06 | 8.500E-06 |
| 7072 | 354 | F354K | A | 6.6e-06 | 4.7e-06 | 5.8e-06 | 5.700E-06 | 9.539E-07 | 4.700E-06 | 6.600E-06 |
| 7073 | 354 | F354M | A | 6.1e-06 | 1e-05 | 7.5e-06 | 7.867E-06 | 1.976E-06 | 6.100E-06 | 1.000E-05 |
| 7074 | 354 | F354F | A | 6.8e-06 | 4.7e-06 | 9.9e-06 | 7.133E-06 | 2.616E-06 | 4.700E-06 | 9.900E-06 |
| 7075 | 354 | F354P | A | 1.1e-05 | 1e-05 | 5.6e-06 | 8.867E-06 | 2.873E-06 | 5.600E-06 | 1.100E-05 |
| 7076 | 354 | F354S | A | 8.3e-06 | 6.7e-06 | 7.3e-06 | 7.433E-06 | 8.083E-07 | 6.700E-06 | 8.300E-06 |
| 7077 | 354 | F354T | A | 6.5e-06 | 6.3e-06 | 9.5e-06 | 7.433E-06 | 1.793E-06 | 6.300E-06 | 9.500E-06 |
| 7078 | 354 | F354W | A | 1.3e-05 | 1.1e-05 | 8.6e-06 | 1.087E-05 | 2.203E-06 | 8.600E-06 | 1.300E-05 |
| 7079 | 354 | F354Y | A | 9.2e-06 | 6.7e-06 | 7.4e-06 | 7.767E-06 | 1.290E-06 | 6.700E-06 | 9.200E-06 |
| 7080 | 354 | F354V | A | 6.6e-06 | 1.4e-05 | 6.6e-06 | 9.067E-06 | 4.272E-06 | 6.600E-06 | 1.400E-05 |
| 7081 | 355 | L355A | A | 4.3e-06 | 8.1e-06 | 5.3e-06 | 5.900E-06 | 1.970E-06 | 4.300E-06 | 8.100E-06 |
| 7082 | 355 | L355R | A | 7.5e-06 | 6.4e-06 | 7.6e-06 | 7.167E-06 | 6.658E-07 | 6.400E-06 | 7.600E-06 |
| 7083 | 355 | L355N | A | 8.4e-06 | 5e-06 | 1.2e-05 | 8.467E-06 | 3.500E-06 | 5.000E-06 | 1.200E-05 |
| 7084 | 355 | L355D | A | 9.8e-06 | 1.2e-05 | 1.1e-05 | 1.093E-05 | 1.102E-06 | 9.800E-06 | 1.200E-05 |
| 7085 | 355 | L355C | A | 6.2e-06 | 5.6e-06 | 8.8e-06 | 6.867E-06 | 1.701E-06 | 5.600E-06 | 8.800E-06 |
| 7086 | 355 | L355E | A | 1e-05 | 8.8e-06 | 3.8e-06 | 7.533E-06 | 3.288E-06 | 3.800E-06 | 1.000E-05 |
| 7087 | 355 | L355Q | A | 9.9e-06 | 1.4e-05 | 6e-06 | 9.967E-06 | 4.000E-06 | 6.000E-06 | 1.400E-05 |
| 7088 | 355 | L355G | A | 1.6e-05 | 5.6e-06 | 5.9e-06 | 9.167E-06 | 5.920E-06 | 5.600E-06 | 1.600E-05 |
| 7089 | 355 | L355H | A | 5.9e-06 | 4.5e-06 | 6e-06 | 5.467E-06 | 8.386E-07 | 4.500E-06 | 6.000E-06 |
| 7090 | 355 | L355I | A | 5.3e-06 | 3.8e-06 | 6.2e-06 | 5.100E-06 | 1.212E-06 | 3.800E-06 | 6.200E-06 |
| 7091 | 355 | L355L | A | 5.2e-06 | 7.6e-06 | 9.6e-06 | 7.467E-06 | 2.203E-06 | 5.200E-06 | 9.600E-06 |
| 7092 | 355 | L355K | A | 7.8e-06 | 5.4e-06 | 6.2e-06 | 6.467E-06 | 1.222E-06 | 5.400E-06 | 7.800E-06 |
| 7093 | 355 | L355M | A | 7e-06 | 5.9e-06 | 6.9e-06 | 6.600E-06 | 6.083E-07 | 5.900E-06 | 7.000E-06 |
| 7094 | 355 | L355F | A | 6.3e-06 | 7.3e-06 | 7.6e-06 | 7.067E-06 | 6.807E-07 | 6.300E-06 | 7.600E-06 |
| 7095 | 355 | L355P | A | 9.7e-06 | 6.9e-06 | 5.5e-06 | 7.367E-06 | 2.139E-06 | 5.500E-06 | 9.700E-06 |
| 7096 | 355 | L355S | A | 5.9e-06 | 7e-06 | 6.8e-06 | 6.567E-06 | 5.859E-07 | 5.900E-06 | 7.000E-06 |
| 7097 | 355 | L355T | A | 9.9e-06 | 6.6e-06 | 7.4e-06 | 7.967E-06 | 1.721E-06 | 6.600E-06 | 9.900E-06 |
| 7098 | 355 | L355W | A | 1.1e-05 | 6.2e-06 | 7e-06 | 8.067E-06 | 2.572E-06 | 6.200E-06 | 1.100E-05 |
| 7099 | 355 | L355Y | A | 1e-05 | 6e-06 | 5.2e-06 | 7.067E-06 | 2.572E-06 | 5.200E-06 | 1.000E-05 |
| 7100 | 355 | L355V | A | 5.3e-06 | 6.5e-06 | 9.4e-06 | 7.067E-06 | 2.108E-06 | 5.300E-06 | 9.400E-06 |
| 7101 | 356 | P356A | A | 8.4e-06 | 1.1e-05 | 9.4e-06 | 9.600E-06 | 1.311E-06 | 8.400E-06 | 1.100E-05 |
| 7102 | 356 | P356R | A | 1.1e-05 | 1.2e-05 | 9.9e-06 | 1.097E-05 | 1.050E-06 | 9.900E-06 | 1.200E-05 |
| 7103 | 356 | P356N | A | 8.3e-06 | 1.1e-05 | 5.7e-06 | 8.333E-06 | 2.650E-06 | 5.700E-06 | 1.100E-05 |
| 7104 | 356 | P356D | A | 9e-06 | 1e-05 | 1e-05 | 9.667E-06 | 5.774E-07 | 9.000E-06 | 1.000E-05 |
| 7105 | 356 | P356C | A | 9.1e-06 | 5.4e-06 | 6.2e-06 | 6.900E-06 | 1.947E-06 | 5.400E-06 | 9.100E-06 |
| 7106 | 356 | P356E | A | 6.5e-06 | 1.3e-05 | 8e-06 | 9.167E-06 | 3.403E-06 | 6.500E-06 | 1.300E-05 |
| 7107 | 356 | P356Q | A | 8.7e-06 | 1.2e-05 | 1e-05 | 1.023E-05 | 1.662E-06 | 8.700E-06 | 1.200E-05 |
| 7108 | 356 | P356G | A | 5.3e-06 | 7e-06 | 5.1e-06 | 5.800E-06 | 1.044E-06 | 5.100E-06 | 7.000E-06 |
| 7109 | 356 | P356H | A | 8.2e-06 | 1.1e-05 | 4.8e-06 | 8.000E-06 | 3.105E-06 | 4.800E-06 | 1.100E-05 |
| 7110 | 356 | P356I | A | 9.4e-06 | 1.1e-05 | 6.8e-06 | 9.067E-06 | 2.120E-06 | 6.800E-06 | 1.100E-05 |
| 7111 | 356 | P356L | A | 8.5e-06 | 9.5e-06 | 8.6e-06 | 8.867E-06 | 5.508E-07 | 8.500E-06 | 9.500E-06 |
| 7112 | 356 | P356K | A | 9.3e-06 | 8.1e-06 | 9.4e-06 | 8.933E-06 | 7.234E-07 | 8.100E-06 | 9.400E-06 |
| 7113 | 356 | P356M | A | 5.1e-06 | 7.3e-06 | 8.6e-06 | 7.000E-06 | 1.769E-06 | 5.100E-06 | 8.600E-06 |
| 7114 | 356 | P356F | A | 5.5e-06 | 1.4e-05 | 7.2e-06 | 8.900E-06 | 4.498E-06 | 5.500E-06 | 1.400E-05 |
| 7115 | 356 | P356P | A | 7.6e-06 | 5e-06 | 8.2e-06 | 6.933E-06 | 1.701E-06 | 5.000E-06 | 8.200E-06 |
| 7116 | 356 | P356S | A | 4.5e-06 | 5.6e-06 | 6.2e-06 | 5.433E-06 | 8.622E-07 | 4.500E-06 | 6.200E-06 |
| 7117 | 356 | P356T | A | 9.2e-06 | 5.5e-06 | 6e-06 | 6.900E-06 | 2.007E-06 | 5.500E-06 | 9.200E-06 |
| 7118 | 356 | P356W | A | 9e-06 | 7.2e-06 | 5.4e-06 | 7.200E-06 | 1.800E-06 | 5.400E-06 | 9.000E-06 |
| 7119 | 356 | P356Y | A | 9.8e-06 | 6.4e-06 | 9.9e-06 | 8.700E-06 | 1.992E-06 | 6.400E-06 | 9.900E-06 |
| 7120 | 356 | P356V | A | 7.1e-06 | 5.7e-06 | 9.2e-06 | 7.333E-06 | 1.762E-06 | 5.700E-06 | 9.200E-06 |
| 7121 | 357 | E357A | A | 7.2e-06 | 6.3e-06 | 1e-05 | 7.833E-06 | 1.930E-06 | 6.300E-06 | 1.000E-05 |
| 7122 | 357 | E357R | A | 8.6e-06 | 5.2e-06 | 9.1e-06 | 7.633E-06 | 2.122E-06 | 5.200E-06 | 9.100E-06 |
| 7123 | 357 | E357N | A | 6.9e-06 | 9.9e-06 | 8.8e-06 | 8.533E-06 | 1.518E-06 | 6.900E-06 | 9.900E-06 |
| 7124 | 357 | E357D | A | 7e-06 | 8.1e-06 | 8.2e-06 | 7.767E-06 | 6.658E-07 | 7.000E-06 | 8.200E-06 |
| 7125 | 357 | E357C | A | 5.3e-06 | 9e-06 | 6.2e-06 | 6.833E-06 | 1.930E-06 | 5.300E-06 | 9.000E-06 |
| 7126 | 357 | E357E | A | 1e-05 | 1.2e-05 | 8.5e-06 | 1.017E-05 | 1.756E-06 | 8.500E-06 | 1.200E-05 |
| 7127 | 357 | E357Q | A | 6.6e-06 | 6.2e-06 | 6.1e-06 | 6.300E-06 | 2.646E-07 | 6.100E-06 | 6.600E-06 |
| 7128 | 357 | E357G | A | 1e-05 | 1.3e-05 | 1.1e-05 | 1.133E-05 | 1.528E-06 | 1.000E-05 | 1.300E-05 |
| 7129 | 357 | E357H | A | 6.9e-06 | 9.5e-06 | 6.1e-06 | 7.500E-06 | 1.778E-06 | 6.100E-06 | 9.500E-06 |
| 7130 | 357 | E357I | A | 1e-05 | 6.4e-06 | 9.3e-06 | 8.567E-06 | 1.909E-06 | 6.400E-06 | 1.000E-05 |
| 7131 | 357 | E357L | A | 5.6e-06 | 1e-05 | 1e-05 | 8.533E-06 | 2.540E-06 | 5.600E-06 | 1.000E-05 |
| 7132 | 357 | E357K | A | 1e-05 | 9.4e-06 | 6.5e-06 | 8.633E-06 | 1.872E-06 | 6.500E-06 | 1.000E-05 |
| 7133 | 357 | E357M | A | 5.4e-06 | 7.9e-06 | 9.4e-06 | 7.567E-06 | 2.021E-06 | 5.400E-06 | 9.400E-06 |
| 7134 | 357 | E357F | A | 2.1e-05 | 1.4e-05 | 8.4e-06 | 1.447E-05 | 6.313E-06 | 8.400E-06 | 2.100E-05 |
| 7135 | 357 | E357P | A | 1.4e-05 | 4.4e-06 | 7.1e-06 | 8.500E-06 | 4.951E-06 | 4.400E-06 | 1.400E-05 |
| 7136 | 357 | E357S | A | 6.2e-06 | 8e-06 | 5.5e-06 | 6.567E-06 | 1.290E-06 | 5.500E-06 | 8.000E-06 |
| 7137 | 357 | E357T | A | 1e-05 | 1.4e-05 | 1.4e-05 | 1.267E-05 | 2.309E-06 | 1.000E-05 | 1.400E-05 |
| 7138 | 357 | E357W | A | 6.1e-06 | 6.1e-06 | 4.1e-06 | 5.433E-06 | 1.155E-06 | 4.100E-06 | 6.100E-06 |
| 7139 | 357 | E357Y | A | 1.2e-05 | 1e-05 | 7.5e-06 | 9.833E-06 | 2.255E-06 | 7.500E-06 | 1.200E-05 |

| | | | | | | | | | | |
|---|---|---|---|---|---|---|---|---|---|---|
| 7140 | 357 | E357V | A | 8.3e-06 | 8.9e-06 | 9.9e-06 | 9.033E-06 | 8.083E-07 | 8.300E-06 | 9.900E-06 |
| 7141 | 358 | S358A | A | 9.6e-06 | 1.2e-05 | 6.9e-06 | 9.500E-06 | 2.551E-06 | 6.900E-06 | 1.200E-05 |
| 7142 | 358 | S358R | A | 1.1e-05 | 9e-06 | 7.2e-06 | 9.067E-06 | 1.901E-06 | 7.200E-06 | 1.100E-05 |
| 7143 | 358 | S358N | A | 7.2e-06 | 7.1e-06 | 7.9e-06 | 7.400E-06 | 4.359E-07 | 7.100E-06 | 7.900E-06 |
| 7144 | 358 | S358D | A | 6.9e-06 | 7.1e-06 | 5.6e-06 | 6.533E-06 | 8.145E-07 | 5.600E-06 | 7.100E-06 |
| 7145 | 358 | S358C | A | 6.2e-06 | 1.4e-05 | 8.5e-06 | 9.567E-06 | 4.008E-06 | 6.200E-06 | 1.400E-05 |
| 7146 | 358 | S358E | A | 8e-06 | 9.1e-06 | 6.1e-06 | 7.733E-06 | 1.518E-06 | 6.100E-06 | 9.100E-06 |
| 7147 | 358 | S358Q | A | 1.2e-05 | 6e-06 | 5.8e-06 | 7.933E-06 | 3.523E-06 | 5.800E-06 | 1.200E-05 |
| 7148 | 358 | S358G | A | 1e-05 | 1.1e-05 | 9.6e-06 | 1.020E-05 | 7.211E-07 | 9.600E-06 | 1.100E-05 |
| 7149 | 358 | S358H | A | 1.7e-05 | 7.7e-06 | 7e-06 | 1.057E-05 | 5.582E-06 | 7.000E-06 | 1.700E-05 |
| 7150 | 358 | S358I | A | 6e-06 | 7.2e-06 | 8.3e-06 | 7.167E-06 | 1.150E-06 | 6.000E-06 | 8.300E-06 |
| 7151 | 358 | S358L | A | 5.5e-06 | 6.9e-06 | 8.1e-06 | 6.833E-06 | 1.301E-06 | 5.500E-06 | 8.100E-06 |
| 7152 | 358 | S358K | A | 8.3e-06 | 6.1e-06 | 5.5e-06 | 6.633E-06 | 1.474E-06 | 5.500E-06 | 8.300E-06 |
| 7153 | 358 | S358M | A | 6.6e-06 | 9.1e-06 | 1e-05 | 8.567E-06 | 1.762E-06 | 6.600E-06 | 1.000E-05 |
| 7154 | 358 | S358F | A | 9.1e-06 | 6.1e-06 | 9.6e-06 | 8.267E-06 | 1.893E-06 | 6.100E-06 | 9.600E-06 |
| 7155 | 358 | S358P | A | 1.4e-05 | 5.7e-06 | 1.5e-05 | 1.157E-05 | 5.105E-06 | 5.700E-06 | 1.500E-05 |
| 7156 | 358 | S358S | A | 7.7e-06 | 5.9e-06 | 1.1e-05 | 8.200E-06 | 2.587E-06 | 5.900E-06 | 1.100E-05 |
| 7157 | 358 | S358T | A | 8.1e-06 | 6.8e-06 | 6e-06 | 6.967E-06 | 1.060E-06 | 6.000E-06 | 8.100E-06 |
| 7158 | 358 | S358W | A | 1.1e-05 | 9.5e-06 | 1.1e-05 | 1.050E-05 | 8.660E-07 | 9.500E-06 | 1.100E-05 |
| 7159 | 358 | S358Y | A | 4.8e-06 | 6.6e-06 | 8.2e-06 | 6.533E-06 | 1.701E-06 | 4.800E-06 | 8.200E-06 |
| 7160 | 358 | S358V | A | 1.3e-05 | 7e-06 | 9.7e-06 | 9.900E-06 | 3.005E-06 | 7.000E-06 | 1.300E-05 |
| 7161 | 359 | F359A | A | 1.5e-05 | 9.2e-06 | 7e-06 | 1.040E-05 | 4.133E-06 | 7.000E-06 | 1.500E-05 |
| 7162 | 359 | F359R | A | 7.1e-06 | 1.3e-05 | 7.5e-06 | 9.200E-06 | 3.297E-06 | 7.100E-06 | 1.300E-05 |
| 7163 | 359 | F359N | A | 7.6e-06 | 8.9e-06 | 1.2e-05 | 9.500E-06 | 2.261E-06 | 7.600E-06 | 1.200E-05 |
| 7164 | 359 | F359D | A | 5.9e-06 | 7.4e-06 | 7.7e-06 | 7.000E-06 | 9.644E-07 | 5.900E-06 | 7.700E-06 |
| 7165 | 359 | F359C | A | 9.2e-06 | 7.2e-06 | 5.4e-06 | 7.267E-06 | 1.901E-06 | 5.400E-06 | 9.200E-06 |
| 7166 | 359 | F359E | A | 5.3e-06 | 9.2e-06 | 7.7e-06 | 7.400E-06 | 1.967E-06 | 5.300E-06 | 9.200E-06 |
| 7167 | 359 | F359Q | A | 6.5e-06 | 8.4e-06 | 3.7e-06 | 6.200E-06 | 2.364E-06 | 3.700E-06 | 8.400E-06 |
| 7168 | 359 | F359G | A | 6.4e-06 | 9e-06 | 1.1e-05 | 8.800E-06 | 2.307E-06 | 6.400E-06 | 1.100E-05 |
| 7169 | 359 | F359H | A | 1.2e-05 | 1.1e-05 | 6.1e-06 | 9.700E-06 | 3.158E-06 | 6.100E-06 | 1.200E-05 |
| 7170 | 359 | F359I | A | 1e-05 | 8.2e-06 | 7.4e-06 | 8.533E-06 | 1.332E-06 | 7.400E-06 | 1.000E-05 |
| 7171 | 359 | F359L | A | 7.5e-06 | 1.1e-05 | 7.1e-06 | 8.533E-06 | 2.146E-06 | 7.100E-06 | 1.100E-05 |
| 7172 | 359 | F359K | A | 6.3e-06 | 1.1e-05 | 9.2e-06 | 8.833E-06 | 2.371E-06 | 6.300E-06 | 1.100E-05 |
| 7173 | 359 | F359M | A | 5.7e-06 | 6.9e-06 | 5.5e-06 | 6.033E-06 | 7.572E-07 | 5.500E-06 | 6.900E-06 |
| 7174 | 359 | F359F | A | 9.8e-06 | 7.2e-06 | 6.2e-06 | 7.733E-06 | 1.858E-06 | 6.200E-06 | 9.800E-06 |
| 7175 | 359 | F359P | A | 7.8e-06 | 6.5e-06 | 1.2e-05 | 8.767E-06 | 2.875E-06 | 6.500E-06 | 1.200E-05 |
| 7176 | 359 | F359S | A | 9.4e-06 | 7e-06 | 9.9e-06 | 8.767E-06 | 1.550E-06 | 7.000E-06 | 9.900E-06 |
| 7177 | 359 | F359T | A | 9.8e-06 | 3.4e-06 | 5.3e-06 | 6.167E-06 | 3.287E-06 | 3.400E-06 | 9.800E-06 |
| 7178 | 359 | F359W | A | 6.6e-06 | 6.6e-06 | 1.2e-05 | 8.400E-06 | 3.118E-06 | 6.600E-06 | 1.200E-05 |
| 7179 | 359 | F359Y | A | 8.2e-06 | 8e-06 | 1e-05 | 8.733E-06 | 1.102E-06 | 8.000E-06 | 1.000E-05 |
| 7180 | 359 | F359V | A | 6.8e-06 | 5e-06 | 7.5e-06 | 6.433E-06 | 1.290E-06 | 5.000E-06 | 7.500E-06 |
| 7181 | 360 | D360A | A | 7e-06 | 7e-06 | 9.1e-06 | 7.700E-06 | 1.212E-06 | 7.000E-06 | 9.100E-06 |
| 7182 | 360 | D360R | A | 6.7e-06 | 7.9e-06 | 1.5e-05 | 9.867E-06 | 4.486E-06 | 6.700E-06 | 1.500E-05 |
| 7183 | 360 | D360N | A | 1.1e-05 | 9.9e-06 | 9.6e-06 | 1.017E-05 | 7.371E-07 | 9.600E-06 | 1.100E-05 |
| 7184 | 360 | D360D | A | 9.7e-06 | 8.5e-06 | 6.1e-06 | 8.100E-06 | 1.833E-06 | 6.100E-06 | 9.700E-06 |
| 7185 | 360 | D360C | A | 8.5e-06 | 1.1e-05 | 8.1e-06 | 9.200E-06 | 1.572E-06 | 8.100E-06 | 1.100E-05 |
| 7186 | 360 | D360E | A | 4.8e-06 | 7.9e-06 | 8.7e-06 | 7.133E-06 | 2.060E-06 | 4.800E-06 | 8.700E-06 |
| 7187 | 360 | D360Q | A | 5.9e-06 | 1.3e-05 | 7.1e-06 | 8.667E-06 | 3.800E-06 | 5.900E-06 | 1.300E-05 |
| 7188 | 360 | D360G | A | 7.1e-06 | 9.9e-06 | 1.1e-05 | 9.333E-06 | 2.011E-06 | 7.100E-06 | 1.100E-05 |
| 7189 | 360 | D360H | A | 9e-06 | 8.7e-06 | 8.8e-06 | 8.833E-06 | 1.528E-07 | 8.700E-06 | 9.000E-06 |
| 7190 | 360 | D360I | A | 6.5e-06 | 8.5e-06 | 9.3e-06 | 8.100E-06 | 1.442E-06 | 6.500E-06 | 9.300E-06 |
| 7191 | 360 | D360L | A | 5.1e-06 | 6.9e-06 | 8.8e-06 | 6.933E-06 | 1.850E-06 | 5.100E-06 | 8.800E-06 |
| 7192 | 360 | D360K | A | 4.8e-06 | 7.9e-06 | 6.5e-06 | 6.400E-06 | 1.552E-06 | 4.800E-06 | 7.900E-06 |
| 7193 | 360 | D360M | A | 5.2e-06 | 1e-05 | 7.7e-06 | 7.633E-06 | 2.401E-06 | 5.200E-06 | 1.000E-05 |
| 7194 | 360 | D360F | A | 5.6e-06 | 5.4e-06 | 8.5e-06 | 6.500E-06 | 1.735E-06 | 5.400E-06 | 8.500E-06 |
| 7195 | 360 | D360P | A | 1e-05 | 5.3e-06 | 1.1e-05 | 8.767E-06 | 3.044E-06 | 5.300E-06 | 1.100E-05 |
| 7196 | 360 | D360S | A | 6.9e-06 | 7.1e-06 | 7e-06 | 7.000E-06 | 1.000E-07 | 6.900E-06 | 7.100E-06 |
| 7197 | 360 | D360T | A | 6.6e-06 | 6.7e-06 | 9.8e-06 | 7.700E-06 | 1.819E-06 | 6.600E-06 | 9.800E-06 |
| 7198 | 360 | D360W | A | 4e-06 | 4.9e-06 | 9.5e-06 | 6.133E-06 | 2.950E-06 | 4.000E-06 | 9.500E-06 |
| 7199 | 360 | D360Y | A | 9e-06 | 5.8e-06 | 1.2e-05 | 8.933E-06 | 3.101E-06 | 5.800E-06 | 1.200E-05 |
| 7200 | 360 | D360V | A | 5.5e-06 | 8.7e-06 | 1.2e-05 | 8.733E-06 | 3.250E-06 | 5.500E-06 | 1.200E-05 |
| 7201 | 361 | G361A | A | 1.1e-05 | 1.4e-05 | 8.2e-06 | 1.107E-05 | 2.901E-06 | 8.200E-06 | 1.400E-05 |
| 7202 | 361 | G361R | A | 8e-06 | 6.8e-06 | 7.6e-06 | 7.467E-06 | 6.110E-07 | 6.800E-06 | 8.000E-06 |
| 7203 | 361 | G361N | A | 7.6e-06 | 1.1e-05 | 6e-06 | 8.200E-06 | 2.553E-06 | 6.000E-06 | 1.100E-05 |
| 7204 | 361 | G361D | A | 7.5e-06 | 5.4e-06 | 4.7e-06 | 5.867E-06 | 1.457E-06 | 4.700E-06 | 7.500E-06 |
| 7205 | 361 | G361C | A | 4.6e-06 | 1.9e-05 | 1.2e-05 | 1.187E-05 | 7.201E-06 | 4.600E-06 | 1.900E-05 |
| 7206 | 361 | G361E | A | 9.8e-06 | 9.8e-06 | 7.5e-06 | 9.033E-06 | 1.328E-06 | 7.500E-06 | 9.800E-06 |
| 7207 | 361 | G361Q | A | 8.7e-06 | 1.1e-05 | 9e-06 | 9.567E-06 | 1.250E-06 | 8.700E-06 | 1.100E-05 |
| 7208 | 361 | G361G | A | 1.2e-05 | 1.2e-05 | 9e-06 | 1.100E-05 | 1.732E-06 | 9.000E-06 | 1.200E-05 |
| 7209 | 361 | G361H | A | 8.8e-06 | 1.1e-05 | 6.6e-06 | 8.800E-06 | 2.200E-06 | 6.600E-06 | 1.100E-05 |
| 7210 | 361 | G361I | A | 5.9e-06 | 7.7e-06 | 5.3e-06 | 6.300E-06 | 1.249E-06 | 5.300E-06 | 7.700E-06 |
| 7211 | 361 | G361L | A | 9.5e-06 | 8.3e-06 | 7.9e-06 | 8.567E-06 | 8.327E-07 | 7.900E-06 | 9.500E-06 |
| 7212 | 361 | G361K | A | 7.9e-06 | 1.2e-05 | 1.3e-05 | 1.097E-05 | 2.702E-06 | 7.900E-06 | 1.300E-05 |
| 7213 | 361 | G361M | A | 8.4e-06 | 6.4e-06 | 5.5e-06 | 6.767E-06 | 1.484E-06 | 5.500E-06 | 8.400E-06 |
| 7214 | 361 | G361F | A | 1.1e-05 | 9.7e-06 | 1.3e-05 | 1.123E-05 | 1.662E-06 | 9.700E-06 | 1.300E-05 |
| 7215 | 361 | G361P | A | 4.7e-06 | 9.2e-06 | 8.7e-06 | 7.533E-06 | 2.466E-06 | 4.700E-06 | 9.200E-06 |
| 7216 | 361 | G361S | A | 9.5e-06 | 4.7e-06 | 5.6e-06 | 6.600E-06 | 2.551E-06 | 4.700E-06 | 9.500E-06 |
| 7217 | 361 | G361T | A | 1.1e-05 | 6.4e-06 | 5.2e-06 | 7.533E-06 | 3.062E-06 | 5.200E-06 | 1.100E-05 |
| 7218 | 361 | G361W | A | 4.8e-06 | 8e-06 | 8.3e-06 | 7.033E-06 | 1.940E-06 | 4.800E-06 | 8.300E-06 |
| 7219 | 361 | G361Y | A | 4.7e-06 | 7.4e-06 | 6.8e-06 | 6.300E-06 | 1.418E-06 | 4.700E-06 | 7.400E-06 |
| 7220 | 361 | G361V | A | 5.3e-06 | 8.3e-06 | 5.2e-06 | 6.267E-06 | 1.762E-06 | 5.200E-06 | 8.300E-06 |
| 7221 | 362 | D362A | A | 9.4e-06 | 4.6e-06 | 6.8e-06 | 6.933E-06 | 2.403E-06 | 4.600E-06 | 9.400E-06 |
| 7222 | 362 | D362R | A | 1.4e-05 | 8.2e-06 | 6.5e-06 | 9.567E-06 | 3.932E-06 | 6.500E-06 | 1.400E-05 |
| 7223 | 362 | D362N | A | 6.7e-06 | 4.7e-06 | 7.9e-06 | 6.433E-06 | 1.617E-06 | 4.700E-06 | 7.900E-06 |
| 7224 | 362 | D362D | A | 8.7e-06 | 1.2e-05 | 1e-05 | 1.023E-05 | 1.662E-06 | 8.700E-06 | 1.200E-05 |

| | | | | | | | | | | |
|---|---|---|---|---|---|---|---|---|---|---|
| 7225 | 362 | D362C | A | 1.2e-05 | 6.7e-06 | 1.1e-05 | 9.900E-06 | 2.816E-06 | 6.700E-06 | 1.200E-05 |
| 7226 | 362 | D362E | A | 1.2e-05 | 9.9e-06 | 7.5e-06 | 9.800E-06 | 2.252E-06 | 7.500E-06 | 1.200E-05 |
| 7227 | 362 | D362Q | A | 4.4e-06 | 3.6e-06 | 6.1e-06 | 4.700E-06 | 1.277E-06 | 3.600E-06 | 6.100E-06 |
| 7228 | 362 | D362G | A | 3.1e-06 | 1.2e-05 | 1.1e-05 | 8.700E-06 | 4.875E-06 | 3.100E-06 | 1.200E-05 |
| 7229 | 362 | D362H | A | 6.4e-06 | 6e-06 | 4.5e-06 | 5.633E-06 | 1.002E-06 | 4.500E-06 | 6.400E-06 |
| 7230 | 362 | D362I | A | 8.3e-06 | 7.8e-06 | 7e-06 | 7.700E-06 | 6.557E-07 | 7.000E-06 | 8.300E-06 |
| 7231 | 362 | D362L | A | 9.5e-06 | 9e-06 | 8.6e-06 | 9.033E-06 | 4.509E-07 | 8.600E-06 | 9.500E-06 |
| 7232 | 362 | D362K | A | 1.1e-05 | 1.4e-05 | 1.3e-05 | 1.267E-05 | 1.528E-06 | 1.100E-05 | 1.400E-05 |
| 7233 | 362 | D362M | A | 5.1e-06 | 3.5e-06 | 1.3e-05 | 7.200E-06 | 5.086E-06 | 3.500E-06 | 1.300E-05 |
| 7234 | 362 | D362F | A | 5.4e-06 | 4.9e-06 | 8.4e-06 | 6.233E-06 | 1.893E-06 | 4.900E-06 | 8.400E-06 |
| 7235 | 362 | D362P | A | 9.2e-06 | 6.6e-06 | 6.1e-06 | 7.300E-06 | 1.664E-06 | 6.100E-06 | 9.200E-06 |
| 7236 | 362 | D362S | A | 6.4e-06 | 9.1e-06 | 6.8e-06 | 7.433E-06 | 1.457E-06 | 6.400E-06 | 9.100E-06 |
| 7237 | 362 | D362T | A | 5.8e-06 | 9.6e-06 | 1e-05 | 8.467E-06 | 2.318E-06 | 5.800E-06 | 1.000E-05 |
| 7238 | 362 | D362W | A | 7e-06 | 8.5e-06 | 8.5e-06 | 8.000E-06 | 8.660E-07 | 7.000E-06 | 8.500E-06 |
| 7239 | 362 | D362Y | A | 9.2e-06 | 5.6e-06 | 6.4e-06 | 7.067E-06 | 1.890E-06 | 5.600E-06 | 9.200E-06 |
| 7240 | 362 | D362V | A | 7.8e-06 | 9.7e-06 | 6.6e-06 | 8.033E-06 | 1.563E-06 | 6.600E-06 | 9.700E-06 |
| 7241 | 363 | P363A | A | 7.6e-06 | 7.3e-06 | 9.6e-06 | 8.167E-06 | 1.250E-06 | 7.300E-06 | 9.600E-06 |
| 7242 | 363 | P363R | A | 9.7e-06 | 6.7e-06 | 9.7e-06 | 8.700E-06 | 1.732E-06 | 6.700E-06 | 9.700E-06 |
| 7243 | 363 | P363N | A | 7.3e-06 | 4.9e-06 | 5.6e-06 | 5.933E-06 | 1.234E-06 | 4.900E-06 | 7.300E-06 |
| 7244 | 363 | P363D | A | 5.4e-06 | 1.4e-05 | 9.4e-06 | 9.600E-06 | 4.303E-06 | 5.400E-06 | 1.400E-05 |
| 7245 | 363 | P363C | A | 9.3e-06 | 2.8e-06 | 5.1e-06 | 5.733E-06 | 3.296E-06 | 2.800E-06 | 9.300E-06 |
| 7246 | 363 | P363E | A | 9.8e-06 | 7e-06 | 7.6e-06 | 8.133E-06 | 1.474E-06 | 7.000E-06 | 9.800E-06 |
| 7247 | 363 | P363Q | A | 8.7e-06 | 6e-06 | 1.1e-05 | 8.567E-06 | 2.503E-06 | 6.000E-06 | 1.100E-05 |
| 7248 | 363 | P363G | A | 7.3e-06 | 8.3e-06 | 8.3e-06 | 7.967E-06 | 5.774E-07 | 7.300E-06 | 8.300E-06 |
| 7249 | 363 | P363H | A | 8e-06 | 9.2e-06 | 5.1e-06 | 7.433E-06 | 2.108E-06 | 5.100E-06 | 9.200E-06 |
| 7250 | 363 | P363I | A | 5e-06 | 1.3e-05 | 9.2e-06 | 9.067E-06 | 4.002E-06 | 5.000E-06 | 1.300E-05 |
| 7251 | 363 | P363L | A | 5.9e-06 | 7.8e-06 | 7.5e-06 | 7.067E-06 | 1.021E-06 | 5.900E-06 | 7.800E-06 |
| 7252 | 363 | P363K | A | 8.5e-06 | 6.6e-06 | 1.1e-05 | 8.700E-06 | 2.207E-06 | 6.600E-06 | 1.100E-05 |
| 7253 | 363 | P363M | A | 8.5e-06 | 6.8e-06 | 5.3e-06 | 6.867E-06 | 1.601E-06 | 5.300E-06 | 8.500E-06 |
| 7254 | 363 | P363F | A | 4.7e-06 | 6e-06 | 8.3e-06 | 6.333E-06 | 1.823E-06 | 4.700E-06 | 8.300E-06 |
| 7255 | 363 | P363P | A | 6.2e-06 | 5.9e-06 | 8.4e-06 | 6.833E-06 | 1.365E-06 | 5.900E-06 | 8.400E-06 |
| 7256 | 363 | P363S | A | 9.7e-06 | 6.3e-06 | 9e-06 | 8.333E-06 | 1.795E-06 | 6.300E-06 | 9.700E-06 |
| 7257 | 363 | P363T | A | 8.5e-06 | 8.4e-06 | 1.1e-05 | 9.300E-06 | 1.473E-06 | 8.400E-06 | 1.100E-05 |
| 7258 | 363 | P363W | A | 6.6e-06 | 5.4e-06 | 8.9e-06 | 6.967E-06 | 1.779E-06 | 5.400E-06 | 8.900E-06 |
| 7259 | 363 | P363Y | A | 6.7e-06 | 1.4e-05 | 1.1e-05 | 1.057E-05 | 3.669E-06 | 6.700E-06 | 1.400E-05 |
| 7260 | 363 | P363V | A | 6.1e-06 | 9.8e-06 | 5.8e-06 | 7.233E-06 | 2.228E-06 | 5.800E-06 | 9.800E-06 |
| 7261 | 364 | A364A | A | 5.8e-06 | 5.7e-06 | 4.4e-06 | 5.300E-06 | 7.810E-07 | 4.400E-06 | 5.800E-06 |
| 7262 | 364 | A364R | A | 1.2e-05 | 8.6e-06 | 9.7e-06 | 1.010E-05 | 1.735E-06 | 8.600E-06 | 1.200E-05 |
| 7263 | 364 | A364N | A | 5.8e-06 | 1.2e-05 | 6.3e-06 | 8.033E-06 | 3.444E-06 | 5.800E-06 | 1.200E-05 |
| 7264 | 364 | A364D | A | 1e-05 | 9e-06 | 6.2e-06 | 8.400E-06 | 1.970E-06 | 6.200E-06 | 1.000E-05 |
| 7265 | 364 | A364C | A | 6.9e-06 | 8.8e-06 | 8.5e-06 | 8.067E-06 | 1.021E-06 | 6.900E-06 | 8.800E-06 |
| 7266 | 364 | A364E | A | 7.4e-06 | 1.4e-05 | 8.1e-06 | 9.833E-06 | 3.625E-06 | 7.400E-06 | 1.400E-05 |
| 7267 | 364 | A364Q | A | 6.7e-06 | 1.2e-05 | 7.4e-06 | 8.700E-06 | 2.879E-06 | 6.700E-06 | 1.200E-05 |
| 7268 | 364 | A364G | A | 6.7e-06 | 8.1e-06 | 7.1e-06 | 7.300E-06 | 7.211E-07 | 6.700E-06 | 8.100E-06 |
| 7269 | 364 | A364H | A | 9.2e-06 | 8.1e-06 | 9.8e-06 | 9.033E-06 | 8.622E-07 | 8.100E-06 | 9.800E-06 |
| 7270 | 364 | A364I | A | 5.3e-06 | 8.3e-06 | 1.3e-05 | 8.867E-06 | 3.881E-06 | 5.300E-06 | 1.300E-05 |
| 7271 | 364 | A364L | A | 1.3e-05 | 5.2e-06 | 4.8e-06 | 7.667E-06 | 4.623E-06 | 4.800E-06 | 1.300E-05 |
| 7272 | 364 | A364K | A | 5.1e-06 | 6.7e-06 | 7.6e-06 | 6.467E-06 | 1.266E-06 | 5.100E-06 | 7.600E-06 |
| 7273 | 364 | A364M | A | 8.1e-06 | 9.2e-06 | 9.3e-06 | 8.867E-06 | 6.658E-07 | 8.100E-06 | 9.300E-06 |
| 7274 | 364 | A364F | A | 9.6e-06 | 9.7e-06 | 6.8e-06 | 8.700E-06 | 1.646E-06 | 6.800E-06 | 9.700E-06 |
| 7275 | 364 | A364P | A | 5.3e-06 | 6.7e-06 | 9.1e-06 | 7.033E-06 | 1.922E-06 | 5.300E-06 | 9.100E-06 |
| 7276 | 364 | A364S | A | 8.6e-06 | 1.2e-05 | 6.5e-06 | 9.033E-06 | 2.775E-06 | 6.500E-06 | 1.200E-05 |
| 7277 | 364 | A364T | A | 5.4e-06 | 8.5e-06 | 9.4e-06 | 7.767E-06 | 2.098E-06 | 5.400E-06 | 9.400E-06 |
| 7278 | 364 | A364W | A | 8.4e-06 | 7.7e-06 | 5e-06 | 7.033E-06 | 1.795E-06 | 5.000E-06 | 8.400E-06 |
| 7279 | 364 | A364Y | A | 9.3e-06 | 1.4e-05 | 6.7e-06 | 1.000E-05 | 3.700E-06 | 6.700E-06 | 1.400E-05 |
| 7280 | 364 | A364V | A | 8.4e-06 | 7e-06 | 8.6e-06 | 8.000E-06 | 8.718E-07 | 7.000E-06 | 8.600E-06 |
| 7281 | 365 | S365A | A | 6.8e-06 | 6.8e-06 | 6e-06 | 6.533E-06 | 4.619E-07 | 6.000E-06 | 6.800E-06 |
| 7282 | 365 | S365R | A | 8.7e-06 | 9.2e-06 | 9.7e-06 | 9.200E-06 | 5.000E-07 | 8.700E-06 | 9.700E-06 |
| 7283 | 365 | S365N | A | 9.2e-06 | 7.1e-06 | 8.6e-06 | 8.300E-06 | 1.082E-06 | 7.100E-06 | 9.200E-06 |
| 7284 | 365 | S365D | A | 1.4e-05 | 4.7e-06 | 4.6e-06 | 7.767E-06 | 5.398E-06 | 4.600E-06 | 1.400E-05 |
| 7285 | 365 | S365C | A | 1.1e-05 | 7.1e-06 | 9.1e-06 | 9.067E-06 | 1.950E-06 | 7.100E-06 | 1.100E-05 |
| 7286 | 365 | S365E | A | 1.1e-05 | 7.6e-06 | 8.1e-06 | 8.900E-06 | 1.836E-06 | 7.600E-06 | 1.100E-05 |
| 7287 | 365 | S365Q | A | 6.8e-06 | 9.6e-06 | 6.1e-06 | 7.500E-06 | 1.852E-06 | 6.100E-06 | 9.600E-06 |
| 7288 | 365 | S365G | A | 9.6e-06 | 8.7e-06 | 1.3e-05 | 1.043E-05 | 2.268E-06 | 8.700E-06 | 1.300E-05 |
| 7289 | 365 | S365H | A | 9.3e-06 | 1.2e-05 | 6.9e-06 | 9.400E-06 | 2.551E-06 | 6.900E-06 | 1.200E-05 |
| 7290 | 365 | S365I | A | 1.1e-05 | 1.5e-05 | 7.6e-06 | 1.120E-05 | 3.704E-06 | 7.600E-06 | 1.500E-05 |
| 7291 | 365 | S365L | A | 1.4e-05 | 7.4e-06 | 5.6e-06 | 9.000E-06 | 4.423E-06 | 5.600E-06 | 1.400E-05 |
| 7292 | 365 | S365K | A | 4.6e-06 | 6.3e-06 | 4.6e-06 | 5.167E-06 | 9.815E-07 | 4.600E-06 | 6.300E-06 |
| 7293 | 365 | S365M | A | 9.5e-06 | 1e-05 | 1.3e-05 | 1.083E-05 | 1.893E-06 | 9.500E-06 | 1.300E-05 |
| 7294 | 365 | S365F | A | 5.5e-06 | 1e-05 | 7.1e-06 | 7.533E-06 | 2.281E-06 | 5.500E-06 | 1.000E-05 |
| 7295 | 365 | S365P | A | 4.2e-06 | 8e-06 | 6.5e-06 | 6.233E-06 | 1.914E-06 | 4.200E-06 | 8.000E-06 |
| 7296 | 365 | S365S | A | 2.1e-05 | 1.2e-05 | 6.7e-06 | 1.323E-05 | 7.229E-06 | 6.700E-06 | 2.100E-05 |
| 7297 | 365 | S365T | A | 9.9e-06 | 7.4e-06 | 6.7e-06 | 8.000E-06 | 1.682E-06 | 6.700E-06 | 9.900E-06 |
| 7298 | 365 | S365W | A | 1.1e-05 | 6.2e-06 | 1.2e-05 | 9.733E-06 | 3.101E-06 | 6.200E-06 | 1.200E-05 |
| 7299 | 365 | S365Y | A | 9.8e-06 | 7.7e-06 | 6.7e-06 | 8.067E-06 | 1.582E-06 | 6.700E-06 | 9.800E-06 |
| 7300 | 365 | S365V | A | 7.5e-06 | 5.2e-06 | 7.5e-06 | 6.733E-06 | 1.328E-06 | 5.200E-06 | 7.500E-06 |
| 7301 | 366 | N366A | A | 9e-06 | 9.1e-06 | 5.4e-06 | 7.833E-06 | 2.108E-06 | 5.400E-06 | 9.100E-06 |
| 7302 | 366 | N366R | A | 9.7e-06 | 7.3e-06 | 1.1e-05 | 9.333E-06 | 1.877E-06 | 7.300E-06 | 1.100E-05 |
| 7303 | 366 | N366N | A | 6.6e-06 | 1.2e-05 | 1e-05 | 9.533E-06 | 2.730E-06 | 6.600E-06 | 1.200E-05 |
| 7304 | 366 | N366D | A | 6.9e-06 | 6.6e-06 | 4.6e-06 | 6.033E-06 | 1.250E-06 | 4.600E-06 | 6.900E-06 |
| 7305 | 366 | N366C | A | 8e-06 | 9.2e-06 | 1.1e-05 | 9.400E-06 | 1.510E-06 | 8.000E-06 | 1.100E-05 |
| 7306 | 366 | N366E | A | 6.3e-06 | 7.4e-06 | 1.2e-05 | 8.567E-06 | 3.024E-06 | 6.300E-06 | 1.200E-05 |
| 7307 | 366 | N366Q | A | 7.2e-06 | 7e-06 | 8.4e-06 | 7.533E-06 | 7.572E-07 | 7.000E-06 | 8.400E-06 |
| 7308 | 366 | N366G | A | 9.2e-06 | 1.5e-05 | 7.1e-06 | 1.043E-05 | 4.092E-06 | 7.100E-06 | 1.500E-05 |
| 7309 | 366 | N366H | A | 5.6e-06 | 4.4e-06 | 8.8e-06 | 6.267E-06 | 2.274E-06 | 4.400E-06 | 8.800E-06 |

| | | | | | | | | | | |
|---|---|---|---|---|---|---|---|---|---|---|
| 7310 | 366 | N366I | A | 4.5e-06 | 9.3e-06 | 7.6e-06 | 7.133E-06 | 2.434E-06 | 4.500E-06 | 9.300E-06 |
| 7311 | 366 | N366L | A | 1.1e-05 | 6.4e-06 | 7.4e-06 | 8.267E-06 | 2.419E-06 | 6.400E-06 | 1.100E-05 |
| 7312 | 366 | N366K | A | 1.3e-05 | 4.5e-06 | 6.9e-06 | 8.133E-06 | 4.382E-06 | 4.500E-06 | 1.300E-05 |
| 7313 | 366 | N366M | A | 8.2e-06 | 6.7e-06 | 1e-05 | 8.300E-06 | 1.652E-06 | 6.700E-06 | 1.000E-05 |
| 7314 | 366 | N366F | A | 7.8e-06 | 6.2e-06 | 6.2e-06 | 6.733E-06 | 9.238E-07 | 6.200E-06 | 7.800E-06 |
| 7315 | 366 | N366P | A | 1.1e-05 | 7.2e-06 | 6.6e-06 | 8.267E-06 | 2.386E-06 | 6.600E-06 | 1.100E-05 |
| 7316 | 366 | N366S | A | 6.4e-06 | 1.2e-05 | 4.9e-06 | 7.767E-06 | 3.742E-06 | 4.900E-06 | 1.200E-05 |
| 7317 | 366 | N366T | A | 1.5e-05 | 6.5e-06 | 9.9e-06 | 1.047E-05 | 4.278E-06 | 6.500E-06 | 1.500E-05 |
| 7318 | 366 | N366W | A | 6.3e-06 | 8.7e-06 | 4.8e-06 | 6.600E-06 | 1.967E-06 | 4.800E-06 | 8.700E-06 |
| 7319 | 366 | N366Y | A | 6.6e-06 | 6.8e-06 | 8.3e-06 | 7.233E-06 | 9.292E-07 | 6.600E-06 | 8.300E-06 |
| 7320 | 366 | N366V | A | 1.2e-05 | 8.2e-06 | 7.1e-06 | 9.100E-06 | 2.571E-06 | 7.100E-06 | 1.200E-05 |
| 7321 | 367 | T367A | A | 6.8e-06 | 5.9e-06 | 9.2e-06 | 7.300E-06 | 1.706E-06 | 5.900E-06 | 9.200E-06 |
| 7322 | 367 | T367R | A | 5.2e-06 | 9.8e-06 | 6.8e-06 | 7.267E-06 | 2.335E-06 | 5.200E-06 | 9.800E-06 |
| 7323 | 367 | T367N | A | 6.3e-06 | 4.8e-06 | 9.9e-06 | 7.000E-06 | 2.621E-06 | 4.800E-06 | 9.900E-06 |
| 7324 | 367 | T367D | A | 9e-06 | 5.3e-06 | 9.6e-06 | 7.967E-06 | 2.329E-06 | 5.300E-06 | 9.600E-06 |
| 7325 | 367 | T367C | A | 6e-06 | 5.6e-06 | 6.6e-06 | 6.067E-06 | 5.033E-07 | 5.600E-06 | 6.600E-06 |
| 7326 | 367 | T367E | A | 1.2e-05 | 1.1e-05 | 5.8e-06 | 9.600E-06 | 3.329E-06 | 5.800E-06 | 1.200E-05 |
| 7327 | 367 | T367Q | A | 7.7e-06 | 4.7e-06 | 6.6e-06 | 6.333E-06 | 1.518E-06 | 4.700E-06 | 7.700E-06 |
| 7328 | 367 | T367G | A | 8.2e-06 | 9.9e-06 | 9.5e-06 | 9.200E-06 | 8.888E-07 | 8.200E-06 | 9.900E-06 |
| 7329 | 367 | T367H | A | 1.3e-05 | 1e-05 | 4.9e-06 | 9.300E-06 | 4.095E-06 | 4.900E-06 | 1.300E-05 |
| 7330 | 367 | T367I | A | 8.3e-06 | 4.9e-06 | 4.8e-06 | 6.000E-06 | 1.992E-06 | 4.800E-06 | 8.300E-06 |
| 7331 | 367 | T367L | A | 7e-06 | 4.4e-06 | 8.1e-06 | 6.500E-06 | 1.900E-06 | 4.400E-06 | 8.100E-06 |
| 7332 | 367 | T367K | A | 1.3e-05 | 9.5e-06 | 6.5e-06 | 9.667E-06 | 3.253E-06 | 6.500E-06 | 1.300E-05 |
| 7333 | 367 | T367M | A | 5.6e-06 | 5.5e-06 | 8.2e-06 | 6.433E-06 | 1.531E-06 | 5.500E-06 | 8.200E-06 |
| 7334 | 367 | T367F | A | 7.1e-06 | 8.4e-06 | 7.4e-06 | 7.633E-06 | 6.807E-07 | 7.100E-06 | 8.400E-06 |
| 7335 | 367 | T367P | A | 1.3e-05 | 7.6e-06 | 9.3e-06 | 9.967E-06 | 2.761E-06 | 7.600E-06 | 1.300E-05 |
| 7336 | 367 | T367S | A | 8.9e-06 | 9.6e-06 | 1.1e-05 | 9.833E-06 | 1.069E-06 | 8.900E-06 | 1.100E-05 |
| 7337 | 367 | T367T | A | 1.3e-05 | 9e-06 | 9.4e-06 | 1.047E-05 | 2.203E-06 | 9.000E-06 | 1.300E-05 |
| 7338 | 367 | T367W | A | 8.1e-06 | 9.2e-06 | 5.3e-06 | 7.533E-06 | 2.011E-06 | 5.300E-06 | 9.200E-06 |
| 7339 | 367 | T367Y | A | 9.9e-06 | 1.3e-05 | 5.5e-06 | 9.467E-06 | 3.769E-06 | 5.500E-06 | 1.300E-05 |
| 7340 | 367 | T367V | A | 1.9e-05 | 1e-05 | 4.6e-06 | 1.120E-05 | 7.275E-06 | 4.600E-06 | 1.900E-05 |
| 7341 | 368 | A368A | A | 8.5e-06 | 1.4e-05 | 7.1e-06 | 9.867E-06 | 3.647E-06 | 7.100E-06 | 1.400E-05 |
| 7342 | 368 | A368R | A | 1.4e-05 | 8.7e-06 | 1.4e-05 | 1.223E-05 | 3.060E-06 | 8.700E-06 | 1.400E-05 |
| 7343 | 368 | A368N | A | 3.1e-06 | 1.3e-05 | 6e-06 | 7.367E-06 | 5.090E-06 | 3.100E-06 | 1.300E-05 |
| 7344 | 368 | A368D | A | 9.8e-06 | 1.1e-05 | 6.3e-06 | 9.033E-06 | 2.442E-06 | 6.300E-06 | 1.100E-05 |
| 7345 | 368 | A368C | A | 9.4e-06 | 5.1e-06 | 9.5e-06 | 8.000E-06 | 2.512E-06 | 5.100E-06 | 9.500E-06 |
| 7346 | 368 | A368E | A | 9.9e-06 | 1.3e-05 | 4.9e-06 | 9.267E-06 | 4.087E-06 | 4.900E-06 | 1.300E-05 |
| 7347 | 368 | A368Q | A | 9.6e-06 | 5.9e-06 | 7.9e-06 | 7.800E-06 | 1.852E-06 | 5.900E-06 | 9.600E-06 |
| 7348 | 368 | A368G | A | 1.2e-05 | 9.4e-06 | 8.8e-06 | 1.007E-05 | 1.701E-06 | 8.800E-06 | 1.200E-05 |
| 7349 | 368 | A368H | A | 9.8e-06 | 5.9e-06 | 4e-06 | 6.567E-06 | 2.957E-06 | 4.000E-06 | 9.800E-06 |
| 7350 | 368 | A368I | A | 9.8e-06 | 7.7e-06 | 6.6e-06 | 8.033E-06 | 1.626E-06 | 6.600E-06 | 9.800E-06 |
| 7351 | 368 | A368L | A | 7.6e-06 | 1.2e-05 | 1.3e-05 | 1.087E-05 | 2.873E-06 | 7.600E-06 | 1.300E-05 |
| 7352 | 368 | A368K | A | 3.2e-06 | 1.3e-05 | 7.8e-06 | 8.000E-06 | 4.903E-06 | 3.200E-06 | 1.300E-05 |
| 7353 | 368 | A368M | A | 7.4e-06 | 4.1e-06 | 7.2e-06 | 6.233E-06 | 1.850E-06 | 4.100E-06 | 7.400E-06 |
| 7354 | 368 | A368F | A | 5.4e-06 | 5.1e-06 | 7.7e-06 | 6.067E-06 | 1.422E-06 | 5.100E-06 | 7.700E-06 |
| 7355 | 368 | A368P | A | 9.3e-06 | 5.7e-06 | 1.1e-05 | 8.667E-06 | 2.706E-06 | 5.700E-06 | 1.100E-05 |
| 7356 | 368 | A368S | A | 7.7e-06 | 8.1e-06 | 9.2e-06 | 8.333E-06 | 7.767E-07 | 7.700E-06 | 9.200E-06 |
| 7357 | 368 | A368T | A | 7.6e-06 | 7.9e-06 | 4.6e-06 | 6.700E-06 | 1.825E-06 | 4.600E-06 | 7.900E-06 |
| 7358 | 368 | A368W | A | 9.5e-06 | 9e-06 | 9.1e-06 | 9.200E-06 | 2.646E-07 | 9.000E-06 | 9.500E-06 |
| 7359 | 368 | A368Y | A | 4.8e-06 | 8.8e-06 | 5.5e-06 | 6.367E-06 | 2.136E-06 | 4.800E-06 | 8.800E-06 |
| 7360 | 368 | A368V | A | 6.9e-06 | 6.9e-06 | 5.4e-06 | 6.400E-06 | 8.660E-07 | 5.400E-06 | 6.900E-06 |
| 7361 | 369 | P369A | A | 5.3e-06 | 9.3e-06 | 4e-06 | 6.200E-06 | 2.762E-06 | 4.000E-06 | 9.300E-06 |
| 7362 | 369 | P369R | A | 6.1e-06 | 7.3e-06 | 8.6e-06 | 7.333E-06 | 1.250E-06 | 6.100E-06 | 8.600E-06 |
| 7363 | 369 | P369N | A | 8.8e-06 | 7.1e-06 | 5.8e-06 | 7.233E-06 | 1.504E-06 | 5.800E-06 | 8.800E-06 |
| 7364 | 369 | P369D | A | 3.2e-06 | 5.2e-06 | 9.2e-06 | 5.867E-06 | 3.055E-06 | 3.200E-06 | 9.200E-06 |
| 7365 | 369 | P369C | A | 5.9e-06 | 6e-06 | 6.5e-06 | 6.133E-06 | 3.215E-07 | 5.900E-06 | 6.500E-06 |
| 7366 | 369 | P369E | A | 6.7e-06 | 9.4e-06 | 7.2e-06 | 7.767E-06 | 1.436E-06 | 6.700E-06 | 9.400E-06 |
| 7367 | 369 | P369Q | A | 9.6e-06 | 4.7e-06 | 6.1e-06 | 6.800E-06 | 2.524E-06 | 4.700E-06 | 9.600E-06 |
| 7368 | 369 | P369G | A | 8.8e-06 | 7.6e-06 | 6.3e-06 | 7.567E-06 | 1.250E-06 | 6.300E-06 | 8.800E-06 |
| 7369 | 369 | P369H | A | 8.9e-06 | 8.4e-06 | 1.3e-05 | 1.010E-05 | 2.524E-06 | 8.400E-06 | 1.300E-05 |
| 7370 | 369 | P369I | A | 9.1e-06 | 9.3e-06 | 8.9e-06 | 9.100E-06 | 2.000E-07 | 8.900E-06 | 9.300E-06 |
| 7371 | 369 | P369L | A | 6.6e-06 | 6.9e-06 | 8e-06 | 7.167E-06 | 7.371E-07 | 6.600E-06 | 8.000E-06 |
| 7372 | 369 | P369K | A | 4.8e-06 | 9.1e-06 | 1e-05 | 7.967E-06 | 2.779E-06 | 4.800E-06 | 1.000E-05 |
| 7373 | 369 | P369M | A | 1.3e-05 | 1.1e-05 | 4.3e-06 | 9.433E-06 | 4.557E-06 | 4.300E-06 | 1.300E-05 |
| 7374 | 369 | P369F | A | 6e-06 | 8.7e-06 | 4.9e-06 | 6.533E-06 | 1.955E-06 | 4.900E-06 | 8.700E-06 |
| 7375 | 369 | P369P | A | 9.9e-06 | 7.5e-06 | 6.9e-06 | 8.100E-06 | 1.587E-06 | 6.900E-06 | 9.900E-06 |
| 7376 | 369 | P369S | A | 7.4e-06 | 1.1e-05 | 1.3e-05 | 1.047E-05 | 2.838E-06 | 7.400E-06 | 1.300E-05 |
| 7377 | 369 | P369T | A | 1.1e-05 | 8.9e-06 | 5.5e-06 | 8.467E-06 | 2.775E-06 | 5.500E-06 | 1.100E-05 |
| 7378 | 369 | P369W | A | 6.1e-06 | 7.4e-06 | 8.2e-06 | 7.233E-06 | 1.060E-06 | 6.100E-06 | 8.200E-06 |
| 7379 | 369 | P369Y | A | 1e-05 | 1.3e-05 | 1e-05 | 1.100E-05 | 1.732E-06 | 1.000E-05 | 1.300E-05 |
| 7380 | 369 | P369V | A | 6.2e-06 | 7.1e-06 | 8.6e-06 | 7.300E-06 | 1.212E-06 | 6.200E-06 | 8.600E-06 |
| 7381 | 370 | L370A | A | 5e-06 | 9.3e-06 | 8.7e-06 | 7.667E-06 | 2.329E-06 | 5.000E-06 | 9.300E-06 |
| 7382 | 370 | L370R | A | 5.8e-06 | 1.3e-05 | 9.5e-06 | 9.433E-06 | 3.600E-06 | 5.800E-06 | 1.300E-05 |
| 7383 | 370 | L370N | A | 1e-05 | 7e-06 | 5e-06 | 7.333E-06 | 2.517E-06 | 5.000E-06 | 1.000E-05 |
| 7384 | 370 | L370D | A | 1.4e-05 | 4.8e-06 | 9.5e-06 | 9.433E-06 | 4.600E-06 | 4.800E-06 | 1.400E-05 |
| 7385 | 370 | L370C | A | 5.6e-06 | 6.7e-06 | 4.8e-06 | 5.700E-06 | 9.539E-07 | 4.800E-06 | 6.700E-06 |
| 7386 | 370 | L370E | A | 7.6e-06 | 9.8e-06 | 5e-06 | 7.467E-06 | 2.403E-06 | 5.000E-06 | 9.800E-06 |
| 7387 | 370 | L370Q | A | 7.2e-06 | 5.5e-06 | 3.9e-06 | 5.533E-06 | 1.650E-06 | 3.900E-06 | 7.200E-06 |
| 7388 | 370 | L370G | A | 1.1e-05 | 9.3e-06 | 7.7e-06 | 9.333E-06 | 1.650E-06 | 7.700E-06 | 1.100E-05 |
| 7389 | 370 | L370H | A | 9e-06 | 7.1e-06 | 8e-06 | 8.033E-06 | 9.504E-07 | 7.100E-06 | 9.000E-06 |
| 7390 | 370 | L370I | A | 5.4e-06 | 5.9e-06 | 6.6e-06 | 5.967E-06 | 6.028E-07 | 5.400E-06 | 6.600E-06 |
| 7391 | 370 | L370L | A | 1.1e-05 | 6.3e-06 | 6.8e-06 | 8.033E-06 | 2.581E-06 | 6.300E-06 | 1.100E-05 |
| 7392 | 370 | L370K | A | 1.6e-05 | 5.9e-06 | 5.2e-06 | 9.033E-06 | 6.043E-06 | 5.200E-06 | 1.600E-05 |
| 7393 | 370 | L370M | A | 1.1e-05 | 6.5e-06 | 6.8e-06 | 8.100E-06 | 2.516E-06 | 6.500E-06 | 1.100E-05 |
| 7394 | 370 | L370F | A | 7.1e-06 | 7.1e-06 | 4.1e-06 | 6.100E-06 | 1.732E-06 | 4.100E-06 | 7.100E-06 |

| | | | | | | | | | | |
|---|---|---|---|---|---|---|---|---|---|---|
| 7395 | 370 | L370P | A | 1.2e-05 | 6.9e-06 | 1.4e-05 | 1.097E-05 | 3.661E-06 | 6.900E-06 | 1.400E-05 |
| 7396 | 370 | L370S | A | 1e-05 | 9e-06 | 1.3e-05 | 1.067E-05 | 2.082E-06 | 9.000E-06 | 1.300E-05 |
| 7397 | 370 | L370T | A | 7.2e-06 | 8.2e-06 | 1.2e-05 | 9.133E-06 | 2.532E-06 | 7.200E-06 | 1.200E-05 |
| 7398 | 370 | L370W | A | 5.6e-06 | 1.4e-05 | 9e-06 | 9.533E-06 | 4.225E-06 | 5.600E-06 | 1.400E-05 |
| 7399 | 370 | L370Y | A | 8.9e-06 | 4.8e-06 | 1.2e-05 | 8.567E-06 | 3.612E-06 | 4.800E-06 | 1.200E-05 |
| 7400 | 370 | L370V | A | 1.2e-05 | 3.5e-06 | 8.9e-06 | 8.133E-06 | 4.302E-06 | 3.500E-06 | 1.200E-05 |
| 7401 | 371 | Q371A | A | 6.4e-06 | 7.3e-06 | 6e-06 | 6.567E-06 | 6.658E-07 | 6.000E-06 | 7.300E-06 |
| 7402 | 371 | Q371R | A | 1.3e-05 | 1.1e-05 | 7.3e-06 | 1.043E-05 | 2.892E-06 | 7.300E-06 | 1.300E-05 |
| 7403 | 371 | Q371N | A | 8.4e-06 | 6.2e-06 | 9.6e-06 | 8.067E-06 | 1.724E-06 | 6.200E-06 | 9.600E-06 |
| 7404 | 371 | Q371D | A | 7.6e-06 | 6.1e-06 | 9e-06 | 7.567E-06 | 1.450E-06 | 6.100E-06 | 9.000E-06 |
| 7405 | 371 | Q371C | A | 6.7e-06 | 7.5e-06 | 6.5e-06 | 6.900E-06 | 5.292E-07 | 6.500E-06 | 7.500E-06 |
| 7406 | 371 | Q371E | A | 9.4e-06 | 9.2e-06 | 1e-05 | 9.533E-06 | 4.163E-07 | 9.200E-06 | 1.000E-05 |
| 7407 | 371 | Q371Q | A | 1e-05 | 5.4e-06 | 8e-06 | 7.800E-06 | 2.307E-06 | 5.400E-06 | 1.000E-05 |
| 7408 | 371 | Q371G | A | 6.9e-06 | 5.7e-06 | 9.9e-06 | 7.500E-06 | 2.163E-06 | 5.700E-06 | 9.900E-06 |
| 7409 | 371 | Q371H | A | 5.2e-06 | 6.3e-06 | 6.9e-06 | 6.133E-06 | 8.622E-07 | 5.200E-06 | 6.900E-06 |
| 7410 | 371 | Q371I | A | 3.9e-06 | 5.6e-06 | 5e-06 | 4.833E-06 | 8.622E-07 | 3.900E-06 | 5.600E-06 |
| 7411 | 371 | Q371L | A | 7.6e-06 | 9.6e-06 | 9.6e-06 | 8.933E-06 | 1.155E-06 | 7.600E-06 | 9.600E-06 |
| 7412 | 371 | Q371K | A | 9.8e-06 | 4.5e-06 | 7.9e-06 | 7.400E-06 | 2.685E-06 | 4.500E-06 | 9.800E-06 |
| 7413 | 371 | Q371M | A | 8.3e-06 | 9.9e-06 | 1e-05 | 9.400E-06 | 9.539E-07 | 8.300E-06 | 1.000E-05 |
| 7414 | 371 | Q371F | A | 6.3e-06 | 7.7e-06 | 1.4e-05 | 9.333E-06 | 4.102E-06 | 6.300E-06 | 1.400E-05 |
| 7415 | 371 | Q371P | A | 9.3e-06 | 7.8e-06 | 5.3e-06 | 7.467E-06 | 2.021E-06 | 5.300E-06 | 9.300E-06 |
| 7416 | 371 | Q371S | A | 1.1e-05 | 9.7e-06 | 7.4e-06 | 9.367E-06 | 1.823E-06 | 7.400E-06 | 1.100E-05 |
| 7417 | 371 | Q371T | A | 1e-05 | 7.4e-06 | 6.4e-06 | 7.933E-06 | 1.858E-06 | 6.400E-06 | 1.000E-05 |
| 7418 | 371 | Q371W | A | 9.8e-06 | 6.7e-06 | 6.8e-06 | 7.767E-06 | 1.762E-06 | 6.700E-06 | 9.800E-06 |
| 7419 | 371 | Q371Y | A | 1e-05 | 6.9e-06 | 8.1e-06 | 8.333E-06 | 1.563E-06 | 6.900E-06 | 1.000E-05 |
| 7420 | 371 | Q371V | A | 6.3e-06 | 1.3e-05 | 1e-05 | 9.767E-06 | 3.356E-06 | 6.300E-06 | 1.300E-05 |
| 7421 | 372 | P372A | A | 1.5e-05 | 7e-06 | 7.9e-06 | 9.967E-06 | 4.382E-06 | 7.000E-06 | 1.500E-05 |
| 7422 | 372 | P372R | A | 9.9e-06 | 8e-06 | 4.6e-06 | 7.500E-06 | 2.685E-06 | 4.600E-06 | 9.900E-06 |
| 7423 | 372 | P372N | A | 4.9e-06 | 1.4e-05 | 5.4e-06 | 8.100E-06 | 5.116E-06 | 4.900E-06 | 1.400E-05 |
| 7424 | 372 | P372D | A | 4.5e-06 | 1.1e-05 | 6.2e-06 | 7.233E-06 | 3.371E-06 | 4.500E-06 | 1.100E-05 |
| 7425 | 372 | P372C | A | 4.6e-06 | 7.4e-06 | 1.2e-05 | 8.000E-06 | 3.736E-06 | 4.600E-06 | 1.200E-05 |
| 7426 | 372 | P372E | A | 6.4e-06 | 5.9e-06 | 6.5e-06 | 6.267E-06 | 3.215E-07 | 5.900E-06 | 6.500E-06 |
| 7427 | 372 | P372Q | A | 7.6e-06 | 5.7e-06 | 1.2e-05 | 8.433E-06 | 3.232E-06 | 5.700E-06 | 1.200E-05 |
| 7428 | 372 | P372G | A | 9.3e-06 | 8.8e-06 | 1.1e-05 | 9.700E-06 | 1.153E-06 | 8.800E-06 | 1.100E-05 |
| 7429 | 372 | P372H | A | 9.8e-06 | 1e-05 | 9.2e-06 | 9.667E-06 | 4.163E-07 | 9.200E-06 | 1.000E-05 |
| 7430 | 372 | P372I | A | 5.6e-06 | 1.2e-05 | 1e-05 | 9.200E-06 | 3.274E-06 | 5.600E-06 | 1.200E-05 |
| 7431 | 372 | P372L | A | 1e-05 | 1e-05 | 6.9e-06 | 8.967E-06 | 1.790E-06 | 6.900E-06 | 1.000E-05 |
| 7432 | 372 | P372K | A | 9.8e-06 | 8.2e-06 | 8.8e-06 | 8.933E-06 | 8.083E-07 | 8.200E-06 | 9.800E-06 |
| 7433 | 372 | P372M | A | 9.4e-06 | 5.5e-06 | 9e-06 | 7.967E-06 | 2.146E-06 | 5.500E-06 | 9.400E-06 |
| 7434 | 372 | P372F | A | 8.3e-06 | 6.3e-06 | 9e-06 | 7.867E-06 | 1.401E-06 | 6.300E-06 | 9.000E-06 |
| 7435 | 372 | P372P | A | 8.3e-06 | 1.2e-05 | 5.2e-06 | 8.500E-06 | 3.404E-06 | 5.200E-06 | 1.200E-05 |
| 7436 | 372 | P372S | A | 9.8e-06 | 5.5e-06 | 7.5e-06 | 7.600E-06 | 2.152E-06 | 5.500E-06 | 9.800E-06 |
| 7437 | 372 | P372T | A | 1.2e-05 | 9e-06 | 7.5e-06 | 9.500E-06 | 2.291E-06 | 7.500E-06 | 1.200E-05 |
| 7438 | 372 | P372W | A | 4.3e-06 | 1.3e-05 | 7.2e-06 | 8.167E-06 | 4.430E-06 | 4.300E-06 | 1.300E-05 |
| 7439 | 372 | P372Y | A | 6.1e-06 | 8.7e-06 | 5.3e-06 | 6.700E-06 | 1.778E-06 | 5.300E-06 | 8.700E-06 |
| 7440 | 372 | P372V | A | 6.6e-06 | 8.6e-06 | 9.8e-06 | 8.333E-06 | 1.617E-06 | 6.600E-06 | 9.800E-06 |
| 7441 | 373 | E373A | A | 8.1e-06 | 9.9e-06 | 8.9e-06 | 8.967E-06 | 9.018E-07 | 8.100E-06 | 9.900E-06 |
| 7442 | 373 | E373R | A | 7.4e-06 | 1e-05 | 5.6e-06 | 7.667E-06 | 2.212E-06 | 5.600E-06 | 1.000E-05 |
| 7443 | 373 | E373N | A | 1.4e-05 | 8.8e-06 | 5.6e-06 | 9.467E-06 | 4.239E-06 | 5.600E-06 | 1.400E-05 |
| 7444 | 373 | E373D | A | 1.1e-05 | 6.8e-06 | 1.1e-05 | 9.600E-06 | 2.425E-06 | 6.800E-06 | 1.100E-05 |
| 7445 | 373 | E373C | A | 7.3e-06 | 1e-05 | 6.8e-06 | 8.033E-06 | 1.721E-06 | 6.800E-06 | 1.000E-05 |
| 7446 | 373 | E373E | A | 5.3e-06 | 1.4e-05 | 8.8e-06 | 9.367E-06 | 4.378E-06 | 5.300E-06 | 1.400E-05 |
| 7447 | 373 | E373Q | A | 1.4e-05 | 1.2e-05 | 8.6e-06 | 1.153E-05 | 2.730E-06 | 8.600E-06 | 1.400E-05 |
| 7448 | 373 | E373G | A | 6.3e-06 | 9.5e-06 | 9.4e-06 | 8.400E-06 | 1.819E-06 | 6.300E-06 | 9.500E-06 |
| 7449 | 373 | E373H | A | 6.2e-06 | 7.6e-06 | 6.5e-06 | 6.767E-06 | 7.371E-07 | 6.200E-06 | 7.600E-06 |
| 7450 | 373 | E373I | A | 7.7e-06 | 1.4e-05 | 1.3e-05 | 1.157E-05 | 3.386E-06 | 7.700E-06 | 1.400E-05 |
| 7451 | 373 | E373L | A | 9e-06 | 1.1e-05 | 1.2e-05 | 1.067E-05 | 1.528E-06 | 9.000E-06 | 1.200E-05 |
| 7452 | 373 | E373K | A | 6.4e-06 | 7.2e-06 | 1.1e-05 | 8.200E-06 | 2.458E-06 | 6.400E-06 | 1.100E-05 |
| 7453 | 373 | E373M | A | 6.4e-06 | 8.3e-06 | 5.2e-06 | 6.633E-06 | 1.563E-06 | 5.200E-06 | 8.300E-06 |
| 7454 | 373 | E373F | A | 1e-05 | 9.1e-06 | 9.7e-06 | 9.600E-06 | 4.583E-07 | 9.100E-06 | 1.000E-05 |
| 7455 | 373 | E373P | A | 5.6e-06 | 5.4e-06 | 5.8e-06 | 5.600E-06 | 2.000E-07 | 5.400E-06 | 5.800E-06 |
| 7456 | 373 | E373S | A | 6.5e-06 | 1e-05 | 5.8e-06 | 7.433E-06 | 2.250E-06 | 5.800E-06 | 1.000E-05 |
| 7457 | 373 | E373T | A | 6.7e-06 | 9.9e-06 | 6.3e-06 | 7.633E-06 | 1.973E-06 | 6.300E-06 | 9.900E-06 |
| 7458 | 373 | E373W | A | 7.5e-06 | 8.4e-06 | 6.2e-06 | 7.367E-06 | 1.106E-06 | 6.200E-06 | 8.400E-06 |
| 7459 | 373 | E373Y | A | 6.2e-06 | 1.2e-05 | 1.2e-05 | 1.007E-05 | 3.349E-06 | 6.200E-06 | 1.200E-05 |
| 7460 | 373 | E373V | A | 5.6e-06 | 5e-06 | 6.5e-06 | 5.700E-06 | 7.550E-07 | 5.000E-06 | 6.500E-06 |
| 7461 | 374 | Q374A | A | 1.1e-05 | 9.1e-06 | 9.3e-06 | 9.800E-06 | 1.044E-06 | 9.100E-06 | 1.100E-05 |
| 7462 | 374 | Q374R | A | 1.3e-05 | 9.7e-06 | 1.1e-05 | 1.123E-05 | 1.662E-06 | 9.700E-06 | 1.300E-05 |
| 7463 | 374 | Q374N | A | 5.7e-06 | 7.8e-06 | 6.6e-06 | 6.700E-06 | 1.054E-06 | 5.700E-06 | 7.800E-06 |
| 7464 | 374 | Q374D | A | 7.9e-06 | 7.4e-06 | 8.8e-06 | 8.033E-06 | 7.095E-07 | 7.400E-06 | 8.800E-06 |
| 7465 | 374 | Q374C | A | 6.5e-06 | 7.9e-06 | 1e-05 | 8.133E-06 | 1.762E-06 | 6.500E-06 | 1.000E-05 |
| 7466 | 374 | Q374E | A | 6.4e-06 | 7e-06 | 6.8e-06 | 6.733E-06 | 3.055E-07 | 6.400E-06 | 7.000E-06 |
| 7467 | 374 | Q374Q | A | 8.3e-06 | 4.8e-06 | 8.5e-06 | 7.200E-06 | 2.081E-06 | 4.800E-06 | 8.500E-06 |
| 7468 | 374 | Q374G | A | 6.9e-06 | 1.3e-05 | 6.6e-06 | 8.833E-06 | 3.612E-06 | 6.600E-06 | 1.300E-05 |
| 7469 | 374 | Q374H | A | 9.6e-06 | 3.6e-06 | 6.7e-06 | 6.633E-06 | 3.001E-06 | 3.600E-06 | 9.600E-06 |
| 7470 | 374 | Q374I | A | 1.3e-05 | 7.3e-06 | 4.9e-06 | 8.400E-06 | 4.161E-06 | 4.900E-06 | 1.300E-05 |
| 7471 | 374 | Q374L | A | 1.2e-05 | 7.2e-06 | 4e-06 | 7.733E-06 | 4.027E-06 | 4.000E-06 | 1.200E-05 |
| 7472 | 374 | Q374K | A | 8.5e-06 | 9.1e-06 | 8.9e-06 | 8.833E-06 | 3.055E-07 | 8.500E-06 | 9.100E-06 |
| 7473 | 374 | Q374M | A | 7.2e-06 | 7.3e-06 | 7.7e-06 | 7.400E-06 | 2.646E-07 | 7.200E-06 | 7.700E-06 |
| 7474 | 374 | Q374F | A | 6.1e-06 | 6.1e-06 | 1e-05 | 7.400E-06 | 2.252E-06 | 6.100E-06 | 1.000E-05 |
| 7475 | 374 | Q374P | A | 1.1e-05 | 8.3e-06 | 1.3e-05 | 1.077E-05 | 2.359E-06 | 8.300E-06 | 1.300E-05 |
| 7476 | 374 | Q374S | A | 9.2e-06 | 1.4e-05 | 1e-05 | 1.107E-05 | 2.572E-06 | 9.200E-06 | 1.400E-05 |
| 7477 | 374 | Q374T | A | 8.5e-06 | 1.6e-05 | 6.6e-06 | 1.037E-05 | 4.970E-06 | 6.600E-06 | 1.600E-05 |
| 7478 | 374 | Q374W | A | 4.3e-06 | 8.1e-06 | 7.4e-06 | 6.600E-06 | 2.022E-06 | 4.300E-06 | 8.100E-06 |
| 7479 | 374 | Q374Y | A | 8.1e-06 | 9.9e-06 | 8.3e-06 | 8.767E-06 | 9.866E-07 | 8.100E-06 | 9.900E-06 |

| | | | | | | | | | | |
|---|---|---|---|---|---|---|---|---|---|---|
| 7480 | 374 | Q374V | A | 1.1e-05 | 5.2e-06 | 6e-06 | 7.400E-06 | 3.143E-06 | 5.200E-06 | 1.100E-05 |
| 7481 | 375 | L375A | A | 1.4e-05 | 6.5e-06 | 8.5e-06 | 9.667E-06 | 3.884E-06 | 6.500E-06 | 1.400E-05 |
| 7482 | 375 | L375R | A | 1.8e-05 | 3.7e-06 | 9.6e-06 | 1.043E-05 | 7.186E-06 | 3.700E-06 | 1.800E-05 |
| 7483 | 375 | L375N | A | 1.2e-05 | 8.1e-06 | 6.7e-06 | 8.933E-06 | 2.747E-06 | 6.700E-06 | 1.200E-05 |
| 7484 | 375 | L375D | A | 1e-05 | 4.7e-06 | 9.1e-06 | 7.933E-06 | 2.836E-06 | 4.700E-06 | 1.000E-05 |
| 7485 | 375 | L375C | A | 9.2e-06 | 8.6e-06 | 6.5e-06 | 8.100E-06 | 1.418E-06 | 6.500E-06 | 9.200E-06 |
| 7486 | 375 | L375E | A | 6.4e-06 | 8.9e-06 | 1.1e-05 | 8.767E-06 | 2.303E-06 | 6.400E-06 | 1.100E-05 |
| 7487 | 375 | L375Q | A | 7.6e-06 | 9.5e-06 | 4.5e-06 | 7.200E-06 | 2.524E-06 | 4.500E-06 | 9.500E-06 |
| 7488 | 375 | L375G | A | 4.4e-06 | 1e-05 | 5.8e-06 | 6.733E-06 | 2.914E-06 | 4.400E-06 | 1.000E-05 |
| 7489 | 375 | L375H | A | 5.5e-06 | 1e-05 | 9e-06 | 8.167E-06 | 2.363E-06 | 5.500E-06 | 1.000E-05 |
| 7490 | 375 | L375I | A | 6.6e-06 | 5.7e-06 | 1.2e-05 | 8.100E-06 | 3.407E-06 | 5.700E-06 | 1.200E-05 |
| 7491 | 375 | L375L | A | 9.3e-06 | 6e-06 | 1.1e-05 | 8.767E-06 | 2.542E-06 | 6.000E-06 | 1.100E-05 |
| 7492 | 375 | L375K | A | 9.5e-06 | 9.9e-06 | 7.5e-06 | 8.967E-06 | 1.286E-06 | 7.500E-06 | 9.900E-06 |
| 7493 | 375 | L375M | A | 6.8e-06 | 7.3e-06 | 6.7e-06 | 6.933E-06 | 3.215E-07 | 6.700E-06 | 7.300E-06 |
| 7494 | 375 | L375F | A | 4.3e-06 | 6.4e-06 | 8.8e-06 | 6.500E-06 | 2.252E-06 | 4.300E-06 | 8.800E-06 |
| 7495 | 375 | L375P | A | 6e-06 | 7.9e-06 | 1.1e-05 | 8.300E-06 | 2.524E-06 | 6.000E-06 | 1.100E-05 |
| 7496 | 375 | L375S | A | 7.3e-06 | 8.1e-06 | 1.4e-05 | 9.800E-06 | 3.659E-06 | 7.300E-06 | 1.400E-05 |
| 7497 | 375 | L375T | A | 7.5e-06 | 8.4e-06 | 8e-06 | 7.967E-06 | 4.509E-07 | 7.500E-06 | 8.400E-06 |
| 7498 | 375 | L375W | A | 9.1e-06 | 1.4e-05 | 1.2e-05 | 1.170E-05 | 2.464E-06 | 9.100E-06 | 1.400E-05 |
| 7499 | 375 | L375Y | A | 1.1e-05 | 8.1e-06 | 9.3e-06 | 9.467E-06 | 1.457E-06 | 8.100E-06 | 1.100E-05 |
| 7500 | 375 | L375V | A | 6.4e-06 | 3.4e-06 | 1.1e-05 | 6.933E-06 | 3.828E-06 | 3.400E-06 | 1.100E-05 |
| 7501 | 376 | Q376A | A | 6.3e-06 | 6.1e-06 | 8.2e-06 | 6.867E-06 | 1.159E-06 | 6.100E-06 | 8.200E-06 |
| 7502 | 376 | Q376R | A | 6.9e-06 | 5.7e-06 | 1.4e-05 | 8.867E-06 | 4.486E-06 | 5.700E-06 | 1.400E-05 |
| 7503 | 376 | Q376N | A | 6e-06 | 9.3e-06 | 1.2e-05 | 9.100E-06 | 3.005E-06 | 6.000E-06 | 1.200E-05 |
| 7504 | 376 | Q376D | A | 8.2e-06 | 7.5e-06 | 6.1e-06 | 7.267E-06 | 1.069E-06 | 6.100E-06 | 8.200E-06 |
| 7505 | 376 | Q376C | A | 6.8e-06 | 9.6e-06 | 8.3e-06 | 8.233E-06 | 1.401E-06 | 6.800E-06 | 9.600E-06 |
| 7506 | 376 | Q376E | A | 6.7e-06 | 9.5e-06 | 1.2e-05 | 9.400E-06 | 2.651E-06 | 6.700E-06 | 1.200E-05 |
| 7507 | 376 | Q376Q | A | 9.7e-06 | 1.4e-05 | 6.3e-06 | 1.000E-05 | 3.859E-06 | 6.300E-06 | 1.400E-05 |
| 7508 | 376 | Q376G | A | 1e-05 | 6.7e-06 | 7.7e-06 | 8.133E-06 | 1.692E-06 | 6.700E-06 | 1.000E-05 |
| 7509 | 376 | Q376H | A | 6.2e-06 | 1.5e-05 | 9.8e-06 | 1.033E-05 | 4.424E-06 | 6.200E-06 | 1.500E-05 |
| 7510 | 376 | Q376I | A | 1.2e-05 | 6.4e-06 | 1.3e-05 | 1.047E-05 | 3.557E-06 | 6.400E-06 | 1.300E-05 |
| 7511 | 376 | Q376L | A | 1.1e-05 | 8.7e-06 | 5.5e-06 | 8.400E-06 | 2.762E-06 | 5.500E-06 | 1.100E-05 |
| 7512 | 376 | Q376K | A | 7.8e-06 | 8.3e-06 | 7.5e-06 | 7.867E-06 | 4.041E-07 | 7.500E-06 | 8.300E-06 |
| 7513 | 376 | Q376M | A | 7.9e-06 | 9.9e-06 | 8.3e-06 | 8.700E-06 | 1.058E-06 | 7.900E-06 | 9.900E-06 |
| 7514 | 376 | Q376F | A | 4.4e-06 | 1e-05 | 5.6e-06 | 6.667E-06 | 2.948E-06 | 4.400E-06 | 1.000E-05 |
| 7515 | 376 | Q376P | A | 4.7e-06 | 8e-06 | 6.7e-06 | 6.467E-06 | 1.662E-06 | 4.700E-06 | 8.000E-06 |
| 7516 | 376 | Q376S | A | 6.1e-06 | 5.5e-06 | 8e-06 | 6.533E-06 | 1.305E-06 | 5.500E-06 | 8.000E-06 |
| 7517 | 376 | Q376T | A | 8.5e-06 | 1.3e-05 | 7.5e-06 | 9.667E-06 | 2.930E-06 | 7.500E-06 | 1.300E-05 |
| 7518 | 376 | Q376W | A | 6.3e-06 | 6.4e-06 | 7.9e-06 | 6.867E-06 | 8.963E-07 | 6.300E-06 | 7.900E-06 |
| 7519 | 376 | Q376Y | A | 6.8e-06 | 7.5e-06 | 7.3e-06 | 7.200E-06 | 3.606E-07 | 6.800E-06 | 7.500E-06 |
| 7520 | 376 | Q376V | A | 4.8e-06 | 9.1e-06 | 1.1e-05 | 8.300E-06 | 3.176E-06 | 4.800E-06 | 1.100E-05 |
| 7521 | 377 | V377A | A | 1.9e-05 | 1.2e-05 | 9e-06 | 1.333E-05 | 5.132E-06 | 9.000E-06 | 1.900E-05 |
| 7522 | 377 | V377R | A | 1.1e-05 | 8.8e-06 | 9.7e-06 | 9.833E-06 | 1.106E-06 | 8.800E-06 | 1.100E-05 |
| 7523 | 377 | V377N | A | 5.9e-06 | 9.7e-06 | 6.7e-06 | 7.433E-06 | 2.003E-06 | 5.900E-06 | 9.700E-06 |
| 7524 | 377 | V377D | A | 1.1e-05 | 9.5e-06 | 7.4e-06 | 9.300E-06 | 1.808E-06 | 7.400E-06 | 1.100E-05 |
| 7525 | 377 | V377C | A | 6e-06 | 6.9e-06 | 6.2e-06 | 6.367E-06 | 4.726E-07 | 6.000E-06 | 6.900E-06 |
| 7526 | 377 | V377E | A | 7e-06 | 8.9e-06 | 1.2e-05 | 9.300E-06 | 2.524E-06 | 7.000E-06 | 1.200E-05 |
| 7527 | 377 | V377Q | A | 1.3e-05 | 1.3e-05 | 8.4e-06 | 1.147E-05 | 2.656E-06 | 8.400E-06 | 1.300E-05 |
| 7528 | 377 | V377G | A | 9.5e-06 | 1.2e-05 | 6.3e-06 | 9.267E-06 | 2.857E-06 | 6.300E-06 | 1.200E-05 |
| 7529 | 377 | V377H | A | 9.4e-06 | 9.6e-06 | 1.4e-05 | 1.100E-05 | 2.600E-06 | 9.400E-06 | 1.400E-05 |
| 7530 | 377 | V377I | A | 5e-06 | 7e-06 | 6.3e-06 | 6.100E-06 | 1.015E-06 | 5.000E-06 | 7.000E-06 |
| 7531 | 377 | V377L | A | 6.4e-06 | 6.4e-06 | 5.8e-06 | 6.200E-06 | 3.464E-07 | 5.800E-06 | 6.400E-06 |
| 7532 | 377 | V377K | A | 9.8e-06 | 6.6e-06 | 4.4e-06 | 6.933E-06 | 2.715E-06 | 4.400E-06 | 9.800E-06 |
| 7533 | 377 | V377M | A | 9.3e-06 | 1.1e-05 | 1.2e-05 | 1.077E-05 | 1.365E-06 | 9.300E-06 | 1.200E-05 |
| 7534 | 377 | V377F | A | 5.3e-06 | 8.6e-06 | 1.4e-05 | 9.300E-06 | 4.392E-06 | 5.300E-06 | 1.400E-05 |
| 7535 | 377 | V377P | A | 6.9e-06 | 7.7e-06 | 7.5e-06 | 7.367E-06 | 4.163E-07 | 6.900E-06 | 7.700E-06 |
| 7536 | 377 | V377S | A | 4.4e-06 | 1e-05 | 6.3e-06 | 6.900E-06 | 2.848E-06 | 4.400E-06 | 1.000E-05 |
| 7537 | 377 | V377T | A | 1.1e-05 | 8.7e-06 | 1.1e-05 | 1.023E-05 | 1.328E-06 | 8.700E-06 | 1.100E-05 |
| 7538 | 377 | V377W | A | 6.5e-06 | 1.2e-05 | 1.3e-05 | 1.050E-05 | 3.500E-06 | 6.500E-06 | 1.300E-05 |
| 7539 | 377 | V377Y | A | 6.3e-06 | 9e-06 | 1.3e-05 | 9.433E-06 | 3.371E-06 | 6.300E-06 | 1.300E-05 |
| 7540 | 377 | V377V | A | 9.7e-06 | 1e-05 | 5.3e-06 | 8.333E-06 | 2.631E-06 | 5.300E-06 | 1.000E-05 |
| 7541 | 378 | F378A | A | 7.1e-06 | 1.2e-05 | 6.4e-06 | 8.500E-06 | 3.051E-06 | 6.400E-06 | 1.200E-05 |
| 7542 | 378 | F378R | A | 1.5e-05 | 3.8e-06 | 1.1e-05 | 9.933E-06 | 5.676E-06 | 3.800E-06 | 1.500E-05 |
| 7543 | 378 | F378N | A | 1e-05 | 8.8e-06 | 8.5e-06 | 9.100E-06 | 7.937E-07 | 8.500E-06 | 1.000E-05 |
| 7544 | 378 | F378D | A | 5.7e-06 | 9.2e-06 | 8.5e-06 | 7.800E-06 | 1.852E-06 | 5.700E-06 | 9.200E-06 |
| 7545 | 378 | F378C | A | 4.5e-06 | 1e-05 | 9.3e-06 | 7.933E-06 | 2.994E-06 | 4.500E-06 | 1.000E-05 |
| 7546 | 378 | F378E | A | 5.5e-06 | 5.4e-06 | 1e-05 | 6.967E-06 | 2.627E-06 | 5.400E-06 | 1.000E-05 |
| 7547 | 378 | F378Q | A | 4.9e-06 | 6.1e-06 | 2.4e-06 | 4.467E-06 | 1.888E-06 | 2.400E-06 | 6.100E-06 |
| 7548 | 378 | F378G | A | 1.1e-05 | 8.8e-06 | 4.7e-06 | 8.167E-06 | 3.197E-06 | 4.700E-06 | 1.100E-05 |
| 7549 | 378 | F378H | A | 7.1e-06 | 5.4e-06 | 9.3e-06 | 7.267E-06 | 1.955E-06 | 5.400E-06 | 9.300E-06 |
| 7550 | 378 | F378I | A | 7.6e-06 | 7.6e-06 | 7.6e-06 | 7.600E-06 | 1.037E-21 | 7.600E-06 | 7.600E-06 |
| 7551 | 378 | F378L | A | 5.8e-06 | 4.6e-06 | 1.1e-05 | 7.133E-06 | 3.402E-06 | 4.600E-06 | 1.100E-05 |
| 7552 | 378 | F378K | A | 7.8e-06 | 9.9e-06 | 1.3e-05 | 1.023E-05 | 2.616E-06 | 7.800E-06 | 1.300E-05 |
| 7553 | 378 | F378M | A | 1.2e-05 | 5e-06 | 7.1e-06 | 8.033E-06 | 3.592E-06 | 5.000E-06 | 1.200E-05 |
| 7554 | 378 | F378F | A | 1.1e-05 | 6.7e-06 | 8.2e-06 | 8.633E-06 | 2.183E-06 | 6.700E-06 | 1.100E-05 |
| 7555 | 378 | F378P | A | 7.7e-06 | 6.6e-06 | 9.1e-06 | 7.800E-06 | 1.253E-06 | 6.600E-06 | 9.100E-06 |
| 7556 | 378 | F378S | A | 7.7e-06 | 7.9e-06 | 9.1e-06 | 8.233E-06 | 7.572E-07 | 7.700E-06 | 9.100E-06 |
| 7557 | 378 | F378T | A | 6.3e-06 | 8.6e-06 | 8.4e-06 | 7.767E-06 | 1.274E-06 | 6.300E-06 | 8.600E-06 |
| 7558 | 378 | F378W | A | 6.5e-06 | 7.2e-06 | 1.3e-05 | 8.900E-06 | 3.568E-06 | 6.500E-06 | 1.300E-05 |
| 7559 | 378 | F378Y | A | 6e-06 | 4.3e-06 | 6.4e-06 | 5.567E-06 | 1.115E-06 | 4.300E-06 | 6.400E-06 |
| 7560 | 378 | F378V | A | 5.6e-06 | 8.4e-06 | 1.1e-05 | 8.333E-06 | 2.701E-06 | 5.600E-06 | 1.100E-05 |
| 7561 | 379 | E379A | A | 6.8e-06 | 4.8e-06 | 1.3e-05 | 8.200E-06 | 4.276E-06 | 4.800E-06 | 1.300E-05 |
| 7562 | 379 | E379R | A | 5.6e-06 | 9.6e-06 | 4.4e-06 | 6.533E-06 | 2.723E-06 | 4.400E-06 | 9.600E-06 |
| 7563 | 379 | E379N | A | 5.7e-06 | 1.1e-05 | 1.7e-05 | 1.123E-05 | 5.654E-06 | 5.700E-06 | 1.700E-05 |
| 7564 | 379 | E379D | A | 8.4e-06 | 7.2e-06 | 9.6e-06 | 8.400E-06 | 1.200E-06 | 7.200E-06 | 9.600E-06 |

| | | | | | | | | | | |
|---|---|---|---|---|---|---|---|---|---|---|
| 7565 | 379 | E379C | A | 1.3e-05 | 7.4e-06 | 1e-05 | 1.013E-05 | 2.802E-06 | 7.400E-06 | 1.300E-05 |
| 7566 | 379 | E379E | A | 6.8e-06 | 1.4e-05 | 7.2e-06 | 9.333E-06 | 4.046E-06 | 6.800E-06 | 1.400E-05 |
| 7567 | 379 | E379Q | A | 9.8e-06 | 6.4e-06 | 8.8e-06 | 8.333E-06 | 1.747E-06 | 6.400E-06 | 9.800E-06 |
| 7568 | 379 | E379G | A | 9.9e-06 | 7.3e-06 | 8.5e-06 | 8.567E-06 | 1.301E-06 | 7.300E-06 | 9.900E-06 |
| 7569 | 379 | E379H | A | 1.2e-05 | 5.2e-06 | 8.7e-06 | 8.633E-06 | 3.400E-06 | 5.200E-06 | 1.200E-05 |
| 7570 | 379 | E379I | A | 9e-06 | 8.8e-06 | 1e-05 | 9.267E-06 | 6.429E-07 | 8.800E-06 | 1.000E-05 |
| 7571 | 379 | E379L | A | 8.2e-06 | 7.1e-06 | 5.2e-06 | 6.833E-06 | 1.518E-06 | 5.200E-06 | 8.200E-06 |
| 7572 | 379 | E379K | A | 9.2e-06 | 6.9e-06 | 1.4e-05 | 1.003E-05 | 3.623E-06 | 6.900E-06 | 1.400E-05 |
| 7573 | 379 | E379M | A | 9.3e-06 | 6.4e-06 | 1.2e-05 | 9.233E-06 | 2.801E-06 | 6.400E-06 | 1.200E-05 |
| 7574 | 379 | E379F | A | 8.5e-06 | 9.7e-06 | 9.1e-06 | 9.100E-06 | 6.000E-07 | 8.500E-06 | 9.700E-06 |
| 7575 | 379 | E379P | A | 7.8e-06 | 1e-05 | 7.2e-06 | 8.333E-06 | 1.474E-06 | 7.200E-06 | 1.000E-05 |
| 7576 | 379 | E379S | A | 8.2e-06 | 8.7e-06 | 6.6e-06 | 7.833E-06 | 1.097E-06 | 6.600E-06 | 8.700E-06 |
| 7577 | 379 | E379T | A | 9.2e-06 | 9.5e-06 | 4.3e-06 | 7.667E-06 | 2.919E-06 | 4.300E-06 | 9.500E-06 |
| 7578 | 379 | E379W | A | 8.4e-06 | 9e-06 | 9.5e-06 | 8.967E-06 | 5.508E-07 | 8.400E-06 | 9.500E-06 |
| 7579 | 379 | E379Y | A | 8.5e-06 | 9e-06 | 6.8e-06 | 8.100E-06 | 1.153E-06 | 6.800E-06 | 9.000E-06 |
| 7580 | 379 | E379V | A | 5.4e-06 | 1e-05 | 1.3e-05 | 9.467E-06 | 3.828E-06 | 5.400E-06 | 1.300E-05 |
| 7581 | 380 | T380A | A | 1.4e-05 | 6.8e-06 | 9.3e-06 | 1.003E-05 | 3.656E-06 | 6.800E-06 | 1.400E-05 |
| 7582 | 380 | T380R | A | 8.1e-06 | 6.2e-06 | 6.4e-06 | 6.900E-06 | 1.044E-06 | 6.200E-06 | 8.100E-06 |
| 7583 | 380 | T380N | A | 1.1e-05 | 6.3e-06 | 1.1e-05 | 9.433E-06 | 2.714E-06 | 6.300E-06 | 1.100E-05 |
| 7584 | 380 | T380D | A | 9.3e-06 | 7.1e-06 | 1.1e-05 | 9.133E-06 | 1.955E-06 | 7.100E-06 | 1.100E-05 |
| 7585 | 380 | T380C | A | 9.7e-06 | 7.5e-06 | 8.6e-06 | 8.600E-06 | 1.100E-06 | 7.500E-06 | 9.700E-06 |
| 7586 | 380 | T380E | A | 9.7e-06 | 8.8e-06 | 4.7e-06 | 7.733E-06 | 2.665E-06 | 4.700E-06 | 9.700E-06 |
| 7587 | 380 | T380Q | A | 4.6e-06 | 8.6e-06 | 5e-06 | 6.067E-06 | 2.203E-06 | 4.600E-06 | 8.600E-06 |
| 7588 | 380 | T380G | A | 9.3e-06 | 7.9e-06 | 8.2e-06 | 8.467E-06 | 7.371E-07 | 7.900E-06 | 9.300E-06 |
| 7589 | 380 | T380H | A | 5.2e-06 | 6.8e-06 | 1.3e-05 | 8.333E-06 | 4.120E-06 | 5.200E-06 | 1.300E-05 |
| 7590 | 380 | T380I | A | 8.8e-06 | 9.6e-06 | 5.4e-06 | 7.933E-06 | 2.230E-06 | 5.400E-06 | 9.600E-06 |
| 7591 | 380 | T380L | A | 5.5e-06 | 3.9e-06 | 8.9e-06 | 6.100E-06 | 2.553E-06 | 3.900E-06 | 8.900E-06 |
| 7592 | 380 | T380K | A | 7.2e-06 | 5.5e-06 | 7.2e-06 | 6.633E-06 | 9.815E-07 | 5.500E-06 | 7.200E-06 |
| 7593 | 380 | T380M | A | 1.5e-05 | 9.4e-06 | 7.3e-06 | 1.057E-05 | 3.980E-06 | 7.300E-06 | 1.500E-05 |
| 7594 | 380 | T380F | A | 6.8e-06 | 8e-06 | 4.2e-06 | 6.333E-06 | 1.943E-06 | 4.200E-06 | 8.000E-06 |
| 7595 | 380 | T380P | A | 9.3e-06 | 1.4e-05 | 7.5e-06 | 1.027E-05 | 3.356E-06 | 7.500E-06 | 1.400E-05 |
| 7596 | 380 | T380S | A | 1.1e-05 | 6.4e-06 | 8.7e-06 | 8.700E-06 | 2.300E-06 | 6.400E-06 | 1.100E-05 |
| 7597 | 380 | T380T | A | 6.7e-06 | 6.3e-06 | 6e-06 | 6.333E-06 | 3.512E-07 | 6.000E-06 | 6.700E-06 |
| 7598 | 380 | T380W | A | 7.4e-06 | 9.1e-06 | 7.2e-06 | 7.900E-06 | 1.044E-06 | 7.200E-06 | 9.100E-06 |
| 7599 | 380 | T380Y | A | 9.4e-06 | 5.8e-06 | 6.5e-06 | 7.233E-06 | 1.909E-06 | 5.800E-06 | 9.400E-06 |
| 7600 | 380 | T380V | A | 1.1e-05 | 6.9e-06 | 7.1e-06 | 8.333E-06 | 2.312E-06 | 6.900E-06 | 1.100E-05 |
| 7601 | 381 | L381A | A | 5.7e-06 | 9.3e-06 | 1.4e-05 | 9.667E-06 | 4.162E-06 | 5.700E-06 | 1.400E-05 |
| 7602 | 381 | L381R | A | 5.8e-06 | 1.1e-05 | 7.7e-06 | 8.167E-06 | 2.631E-06 | 5.800E-06 | 1.100E-05 |
| 7603 | 381 | L381N | A | 8.6e-06 | 5.3e-06 | 6.7e-06 | 6.867E-06 | 1.656E-06 | 5.300E-06 | 8.600E-06 |
| 7604 | 381 | L381D | A | 8.6e-06 | 1.3e-05 | 7e-06 | 9.533E-06 | 3.107E-06 | 7.000E-06 | 1.300E-05 |
| 7605 | 381 | L381C | A | 1.1e-05 | 9.6e-06 | 1.2e-05 | 1.087E-05 | 1.206E-06 | 9.600E-06 | 1.200E-05 |
| 7606 | 381 | L381E | A | 8.1e-06 | 1.4e-05 | 5.5e-06 | 9.200E-06 | 4.355E-06 | 5.500E-06 | 1.400E-05 |
| 7607 | 381 | L381Q | A | 4.8e-06 | 8.1e-06 | 6.5e-06 | 6.467E-06 | 1.650E-06 | 4.800E-06 | 8.100E-06 |
| 7608 | 381 | L381G | A | 9.2e-06 | 7.9e-06 | 9.7e-06 | 8.933E-06 | 9.292E-07 | 7.900E-06 | 9.700E-06 |
| 7609 | 381 | L381H | A | 5.2e-06 | 6.7e-06 | 1e-05 | 7.300E-06 | 2.456E-06 | 5.200E-06 | 1.000E-05 |
| 7610 | 381 | L381I | A | 8.7e-06 | 7.9e-06 | 1.3e-05 | 9.867E-06 | 2.743E-06 | 7.900E-06 | 1.300E-05 |
| 7611 | 381 | L381L | A | 7.3e-06 | 6.8e-06 | 8.4e-06 | 7.500E-06 | 8.185E-07 | 6.800E-06 | 8.400E-06 |
| 7612 | 381 | L381K | A | 8.5e-06 | 7.8e-06 | 6.5e-06 | 7.600E-06 | 1.015E-06 | 6.500E-06 | 8.500E-06 |
| 7613 | 381 | L381M | A | 9.4e-06 | 4.3e-06 | 8.6e-06 | 7.433E-06 | 2.743E-06 | 4.300E-06 | 9.400E-06 |
| 7614 | 381 | L381F | A | 4e-06 | 5.1e-06 | 6.8e-06 | 5.300E-06 | 1.411E-06 | 4.000E-06 | 6.800E-06 |
| 7615 | 381 | L381P | A | 7.3e-06 | 4.9e-06 | 7e-06 | 6.400E-06 | 1.308E-06 | 4.900E-06 | 7.300E-06 |
| 7616 | 381 | L381S | A | 6e-06 | 6.2e-06 | 4.2e-06 | 5.467E-06 | 1.102E-06 | 4.200E-06 | 6.200E-06 |
| 7617 | 381 | L381T | A | 7.5e-06 | 4.8e-06 | 1e-05 | 7.433E-06 | 2.601E-06 | 4.800E-06 | 1.000E-05 |
| 7618 | 381 | L381W | A | 7.1e-06 | 8.8e-06 | 6.1e-06 | 7.333E-06 | 1.365E-06 | 6.100E-06 | 8.800E-06 |
| 7619 | 381 | L381Y | A | 8.8e-06 | 1.2e-05 | 5.6e-06 | 8.800E-06 | 3.200E-06 | 5.600E-06 | 1.200E-05 |
| 7620 | 381 | L381V | A | 1e-05 | 6.4e-06 | 8.9e-06 | 8.433E-06 | 1.845E-06 | 6.400E-06 | 1.000E-05 |
| 7621 | 382 | E382A | A | 9.7e-06 | 5.6e-06 | 8.2e-06 | 7.833E-06 | 2.074E-06 | 5.600E-06 | 9.700E-06 |
| 7622 | 382 | E382R | A | 1.2e-05 | 8e-06 | 5.6e-06 | 8.533E-06 | 3.233E-06 | 5.600E-06 | 1.200E-05 |
| 7623 | 382 | E382N | A | 9.7e-06 | 7.4e-06 | 7.4e-06 | 8.167E-06 | 1.328E-06 | 7.400E-06 | 9.700E-06 |
| 7624 | 382 | E382D | A | 8.8e-06 | 5.6e-06 | 6.6e-06 | 7.000E-06 | 1.637E-06 | 5.600E-06 | 8.800E-06 |
| 7625 | 382 | E382C | A | 6.2e-06 | 1e-05 | 7.7e-06 | 7.967E-06 | 1.914E-06 | 6.200E-06 | 1.000E-05 |
| 7626 | 382 | E382E | A | 6e-06 | 1e-05 | 4e-06 | 6.667E-06 | 3.055E-06 | 4.000E-06 | 1.000E-05 |
| 7627 | 382 | E382Q | A | 1.1e-05 | 1.4e-05 | 8.3e-06 | 1.110E-05 | 2.851E-06 | 8.300E-06 | 1.400E-05 |
| 7628 | 382 | E382G | A | 7.8e-06 | 9.4e-06 | 9.4e-06 | 8.867E-06 | 9.238E-07 | 7.800E-06 | 9.400E-06 |
| 7629 | 382 | E382H | A | 1.7e-05 | 5.2e-06 | 5.2e-06 | 9.133E-06 | 6.813E-06 | 5.200E-06 | 1.700E-05 |
| 7630 | 382 | E382I | A | 9.5e-06 | 4.4e-06 | 8e-06 | 7.300E-06 | 2.621E-06 | 4.400E-06 | 9.500E-06 |
| 7631 | 382 | E382L | A | 6.3e-06 | 1.3e-05 | 4.3e-06 | 7.867E-06 | 4.557E-06 | 4.300E-06 | 1.300E-05 |
| 7632 | 382 | E382K | A | 5.3e-06 | 8.7e-06 | 6.3e-06 | 6.767E-06 | 1.747E-06 | 5.300E-06 | 8.700E-06 |
| 7633 | 382 | E382M | A | 9.5e-06 | 6.2e-06 | 9.7e-06 | 8.467E-06 | 1.966E-06 | 6.200E-06 | 9.700E-06 |
| 7634 | 382 | E382F | A | 7.9e-06 | 1.3e-05 | 7.1e-06 | 9.333E-06 | 3.201E-06 | 7.100E-06 | 1.300E-05 |
| 7635 | 382 | E382P | A | 1e-05 | 1.3e-05 | 8.4e-06 | 1.047E-05 | 2.335E-06 | 8.400E-06 | 1.300E-05 |
| 7636 | 382 | E382S | A | 5.2e-06 | 1.3e-05 | 2.7e-06 | 6.967E-06 | 5.372E-06 | 2.700E-06 | 1.300E-05 |
| 7637 | 382 | E382T | A | 7e-06 | 7.6e-06 | 8.6e-06 | 7.733E-06 | 8.083E-07 | 7.000E-06 | 8.600E-06 |
| 7638 | 382 | E382W | A | 6.8e-06 | 8.8e-06 | 6e-06 | 7.200E-06 | 1.442E-06 | 6.000E-06 | 8.800E-06 |
| 7639 | 382 | E382Y | A | 7.9e-06 | 6.9e-06 | 6.7e-06 | 7.167E-06 | 6.429E-07 | 6.700E-06 | 7.900E-06 |
| 7640 | 382 | E382V | A | 9.9e-06 | 6.9e-06 | 9e-06 | 8.600E-06 | 1.539E-06 | 6.900E-06 | 9.900E-06 |
| 7641 | 383 | E383A | A | 1.4e-05 | 5.7e-06 | 6.7e-06 | 8.800E-06 | 4.531E-06 | 5.700E-06 | 1.400E-05 |
| 7642 | 383 | E383R | A | 3.8e-06 | 9e-06 | 5.8e-06 | 6.200E-06 | 2.623E-06 | 3.800E-06 | 9.000E-06 |
| 7643 | 383 | E383N | A | 9.3e-06 | 7.8e-06 | 5.3e-06 | 7.467E-06 | 2.021E-06 | 5.300E-06 | 9.300E-06 |
| 7644 | 383 | E383D | A | 8.7e-06 | 1e-05 | 7.3e-06 | 8.667E-06 | 1.350E-06 | 7.300E-06 | 1.000E-05 |
| 7645 | 383 | E383C | A | 1.1e-05 | 7.3e-06 | 5.9e-06 | 8.067E-06 | 2.635E-06 | 5.900E-06 | 1.100E-05 |
| 7646 | 383 | E383E | A | 7.4e-06 | 7.3e-06 | 6.4e-06 | 7.033E-06 | 5.508E-07 | 6.400E-06 | 7.400E-06 |
| 7647 | 383 | E383Q | A | 9.3e-06 | 8.5e-06 | 1.3e-05 | 1.027E-05 | 2.401E-06 | 8.500E-06 | 1.300E-05 |
| 7648 | 383 | E383G | A | 7.2e-06 | 1.2e-05 | 8.6e-06 | 9.267E-06 | 2.468E-06 | 7.200E-06 | 1.200E-05 |
| 7649 | 383 | E383H | A | 9.8e-06 | 7.4e-06 | 6.9e-06 | 8.033E-06 | 1.550E-06 | 6.900E-06 | 9.800E-06 |

| | | | | | | | | | | |
|---|---|---|---|---|---|---|---|---|---|---|
| 7650 | 383 | E383I | A | 4.4e-06 | 6e-06 | 9.8e-06 | 6.733E-06 | 2.774E-06 | 4.400E-06 | 9.800E-06 |
| 7651 | 383 | E383L | A | 1.1e-05 | 1e-05 | 9.8e-06 | 1.027E-05 | 6.429E-07 | 9.800E-06 | 1.100E-05 |
| 7652 | 383 | E383K | A | 7.3e-06 | 1.4e-05 | 7.8e-06 | 9.700E-06 | 3.732E-06 | 7.300E-06 | 1.400E-05 |
| 7653 | 383 | E383M | A | 6.1e-06 | 7.4e-06 | 7.9e-06 | 7.133E-06 | 9.292E-07 | 6.100E-06 | 7.900E-06 |
| 7654 | 383 | E383F | A | 9.6e-06 | 6.5e-06 | 5.4e-06 | 7.167E-06 | 2.178E-06 | 5.400E-06 | 9.600E-06 |
| 7655 | 383 | E383P | A | 8.2e-06 | 1.1e-05 | 1.1e-05 | 1.007E-05 | 1.617E-06 | 8.200E-06 | 1.100E-05 |
| 7656 | 383 | E383S | A | 1.6e-05 | 4.9e-06 | 8.5e-06 | 9.800E-06 | 5.663E-06 | 4.900E-06 | 1.600E-05 |
| 7657 | 383 | E383T | A | 5e-06 | 1.1e-05 | 8.5e-06 | 8.167E-06 | 3.014E-06 | 5.000E-06 | 1.100E-05 |
| 7658 | 383 | E383W | A | 8.8e-06 | 9.6e-06 | 6.9e-06 | 8.433E-06 | 1.387E-06 | 6.900E-06 | 9.600E-06 |
| 7659 | 383 | E383Y | A | 5.1e-06 | 1.2e-05 | 7.4e-06 | 8.167E-06 | 3.513E-06 | 5.100E-06 | 1.200E-05 |
| 7660 | 383 | E383V | A | 9.5e-06 | 7.5e-06 | 9.1e-06 | 8.700E-06 | 1.058E-06 | 7.500E-06 | 9.500E-06 |
| 7661 | 384 | I384A | A | 1.8e-05 | 8.7e-06 | 8.7e-06 | 1.180E-05 | 5.369E-06 | 8.700E-06 | 1.800E-05 |
| 7662 | 384 | I384R | A | 8.9e-06 | 6.6e-06 | 6.2e-06 | 7.233E-06 | 1.457E-06 | 6.200E-06 | 8.900E-06 |
| 7663 | 384 | I384N | A | 1.4e-05 | 8e-06 | 1.1e-05 | 1.100E-05 | 3.000E-06 | 8.000E-06 | 1.400E-05 |
| 7664 | 384 | I384D | A | 7.2e-06 | 7.8e-06 | 8.5e-06 | 7.833E-06 | 6.506E-07 | 7.200E-06 | 8.500E-06 |
| 7665 | 384 | I384C | A | 1.4e-05 | 1.1e-05 | 6.3e-06 | 1.043E-05 | 3.881E-06 | 6.300E-06 | 1.400E-05 |
| 7666 | 384 | I384E | A | 9e-06 | 7.5e-06 | 7.5e-06 | 8.000E-06 | 8.660E-07 | 7.500E-06 | 9.000E-06 |
| 7667 | 384 | I384Q | A | 6.2e-06 | 1.1e-05 | 9.7e-06 | 8.967E-06 | 2.483E-06 | 6.200E-06 | 1.100E-05 |
| 7668 | 384 | I384G | A | 9.6e-06 | 5.7e-06 | 4.7e-06 | 6.667E-06 | 2.589E-06 | 4.700E-06 | 9.600E-06 |
| 7669 | 384 | I384H | A | 7.4e-06 | 8.7e-06 | 5.7e-06 | 7.267E-06 | 1.504E-06 | 5.700E-06 | 8.700E-06 |
| 7670 | 384 | I384I | A | 7e-06 | 8.5e-06 | 6.7e-06 | 7.400E-06 | 9.644E-07 | 6.700E-06 | 8.500E-06 |
| 7671 | 384 | I384L | A | 7.1e-06 | 1e-05 | 9.6e-06 | 8.900E-06 | 1.572E-06 | 7.100E-06 | 1.000E-05 |
| 7672 | 384 | I384K | A | 1.5e-05 | 9.1e-06 | 8.6e-06 | 1.090E-05 | 3.559E-06 | 8.600E-06 | 1.500E-05 |
| 7673 | 384 | I384M | A | 5.5e-06 | 1e-05 | 1.2e-05 | 9.167E-06 | 3.329E-06 | 5.500E-06 | 1.200E-05 |
| 7674 | 384 | I384F | A | 9.7e-06 | 7.3e-06 | 1.4e-05 | 1.033E-05 | 3.395E-06 | 7.300E-06 | 1.400E-05 |
| 7675 | 384 | I384P | A | 5.3e-06 | 9.1e-06 | 9.2e-06 | 7.867E-06 | 2.223E-06 | 5.300E-06 | 9.200E-06 |
| 7676 | 384 | I384S | A | 6.3e-06 | 7.7e-06 | 8.8e-06 | 7.600E-06 | 1.253E-06 | 6.300E-06 | 8.800E-06 |
| 7677 | 384 | I384T | A | 7.1e-06 | 9.6e-06 | 1.1e-05 | 9.233E-06 | 1.976E-06 | 7.100E-06 | 1.100E-05 |
| 7678 | 384 | I384W | A | 8.9e-06 | 1e-05 | 8.8e-06 | 9.233E-06 | 6.658E-07 | 8.800E-06 | 1.000E-05 |
| 7679 | 384 | I384Y | A | 8.2e-06 | 6.6e-06 | 1.1e-05 | 8.600E-06 | 2.227E-06 | 6.600E-06 | 1.100E-05 |
| 7680 | 384 | I384V | A | 8.5e-06 | 7.9e-06 | 1.3e-05 | 9.800E-06 | 2.787E-06 | 7.900E-06 | 1.300E-05 |
| 7681 | 385 | T385A | A | 9.5e-06 | 9.6e-06 | 7.8e-06 | 8.967E-06 | 1.012E-06 | 7.800E-06 | 9.600E-06 |
| 7682 | 385 | T385R | A | 6.6e-06 | 7.6e-06 | 9.4e-06 | 7.867E-06 | 1.419E-06 | 6.600E-06 | 9.400E-06 |
| 7683 | 385 | T385N | A | 4.5e-06 | 9.2e-06 | 8.3e-06 | 7.333E-06 | 2.495E-06 | 4.500E-06 | 9.200E-06 |
| 7684 | 385 | T385D | A | 6.2e-06 | 1.2e-05 | 7.2e-06 | 8.467E-06 | 3.101E-06 | 6.200E-06 | 1.200E-05 |
| 7685 | 385 | T385C | A | 8.5e-06 | 4.8e-06 | 9.2e-06 | 7.500E-06 | 2.364E-06 | 4.800E-06 | 9.200E-06 |
| 7686 | 385 | T385E | A | 6.8e-06 | 1e-05 | 8.3e-06 | 8.367E-06 | 1.601E-06 | 6.800E-06 | 1.000E-05 |
| 7687 | 385 | T385Q | A | 9.7e-06 | 6.4e-06 | 1.2e-05 | 9.367E-06 | 2.815E-06 | 6.400E-06 | 1.200E-05 |
| 7688 | 385 | T385G | A | 1.4e-05 | 6e-06 | 4.3e-06 | 8.100E-06 | 5.180E-06 | 4.300E-06 | 1.400E-05 |
| 7689 | 385 | T385H | A | 8.7e-06 | 5.3e-06 | 6.1e-06 | 6.700E-06 | 1.778E-06 | 5.300E-06 | 8.700E-06 |
| 7690 | 385 | T385I | A | 1.3e-05 | 6.4e-06 | 1e-05 | 9.800E-06 | 3.305E-06 | 6.400E-06 | 1.300E-05 |
| 7691 | 385 | T385L | A | 9.8e-06 | 1.1e-05 | 4.7e-06 | 8.500E-06 | 3.345E-06 | 4.700E-06 | 1.100E-05 |
| 7692 | 385 | T385K | A | 1.1e-05 | 1.3e-05 | 7.9e-06 | 1.063E-05 | 2.570E-06 | 7.900E-06 | 1.300E-05 |
| 7693 | 385 | T385M | A | 7.2e-06 | 7.6e-06 | 1.2e-05 | 8.933E-06 | 2.663E-06 | 7.200E-06 | 1.200E-05 |
| 7694 | 385 | T385F | A | 4.6e-06 | 1e-05 | 7e-06 | 7.200E-06 | 2.706E-06 | 4.600E-06 | 1.000E-05 |
| 7695 | 385 | T385P | A | 1.2e-05 | 9.9e-06 | 1.2e-05 | 1.130E-05 | 1.212E-06 | 9.900E-06 | 1.200E-05 |
| 7696 | 385 | T385S | A | 8.6e-06 | 7.9e-06 | 6.6e-06 | 7.700E-06 | 1.015E-06 | 6.600E-06 | 8.600E-06 |
| 7697 | 385 | T385T | A | 1.2e-05 | 5e-06 | 7e-06 | 8.000E-06 | 3.606E-06 | 5.000E-06 | 1.200E-05 |
| 7698 | 385 | T385W | A | 5.8e-06 | 1.3e-05 | 6.9e-06 | 8.567E-06 | 3.879E-06 | 5.800E-06 | 1.300E-05 |
| 7699 | 385 | T385Y | A | 6.2e-06 | 9.3e-06 | 7.9e-06 | 7.800E-06 | 1.552E-06 | 6.200E-06 | 9.300E-06 |
| 7700 | 385 | T385V | A | 6.7e-06 | 5.5e-06 | 6.8e-06 | 6.333E-06 | 7.234E-07 | 5.500E-06 | 6.800E-06 |
| 7701 | 386 | G386A | A | 9.9e-06 | 4.4e-06 | 1e-05 | 8.100E-06 | 3.205E-06 | 4.400E-06 | 1.000E-05 |
| 7702 | 386 | G386R | A | 8.9e-06 | 1.5e-05 | 4.8e-06 | 9.567E-06 | 5.133E-06 | 4.800E-06 | 1.500E-05 |
| 7703 | 386 | G386N | A | 8.9e-06 | 8.3e-06 | 1.1e-05 | 9.400E-06 | 1.418E-06 | 8.300E-06 | 1.100E-05 |
| 7704 | 386 | G386D | A | 6.6e-06 | 6.1e-06 | 1e-05 | 7.567E-06 | 2.122E-06 | 6.100E-06 | 1.000E-05 |
| 7705 | 386 | G386C | A | 6.3e-06 | 8.6e-06 | 8.8e-06 | 7.900E-06 | 1.389E-06 | 6.300E-06 | 8.800E-06 |
| 7706 | 386 | G386E | A | 1e-05 | 7.7e-06 | 6.8e-06 | 8.167E-06 | 1.650E-06 | 6.800E-06 | 1.000E-05 |
| 7707 | 386 | G386Q | A | 6.7e-06 | 7.9e-06 | 4.2e-06 | 6.267E-06 | 1.888E-06 | 4.200E-06 | 7.900E-06 |
| 7708 | 386 | G386G | A | 3.4e-06 | 7.2e-06 | 1e-05 | 6.867E-06 | 3.313E-06 | 3.400E-06 | 1.000E-05 |
| 7709 | 386 | G386H | A | 6.3e-06 | 7.1e-06 | 6.4e-06 | 6.600E-06 | 4.359E-07 | 6.300E-06 | 7.100E-06 |
| 7710 | 386 | G386I | A | 5.1e-06 | 7.5e-06 | 7.7e-06 | 6.767E-06 | 1.447E-06 | 5.100E-06 | 7.700E-06 |
| 7711 | 386 | G386L | A | 1.3e-05 | 5.4e-06 | 8.9e-06 | 9.100E-06 | 3.804E-06 | 5.400E-06 | 1.300E-05 |
| 7712 | 386 | G386K | A | 9e-06 | 7.6e-06 | 4.5e-06 | 7.033E-06 | 2.303E-06 | 4.500E-06 | 9.000E-06 |
| 7713 | 386 | G386M | A | 4.7e-06 | 1e-05 | 8.3e-06 | 7.667E-06 | 2.706E-06 | 4.700E-06 | 1.000E-05 |
| 7714 | 386 | G386F | A | 4.9e-06 | 6.4e-06 | 5.6e-06 | 5.633E-06 | 7.506E-07 | 4.900E-06 | 6.400E-06 |
| 7715 | 386 | G386P | A | 6.8e-06 | 1.2e-05 | 4.6e-06 | 7.800E-06 | 3.800E-06 | 4.600E-06 | 1.200E-05 |
| 7716 | 386 | G386S | A | 1.4e-05 | 7.9e-06 | 8.7e-06 | 1.020E-05 | 3.315E-06 | 7.900E-06 | 1.400E-05 |
| 7717 | 386 | G386T | A | 8.8e-06 | 6.7e-06 | 6.3e-06 | 7.267E-06 | 1.343E-06 | 6.300E-06 | 8.800E-06 |
| 7718 | 386 | G386W | A | 4.5e-06 | 1.2e-05 | 7.8e-06 | 8.100E-06 | 3.759E-06 | 4.500E-06 | 1.200E-05 |
| 7719 | 386 | G386Y | A | 8.8e-06 | 1e-05 | 5.8e-06 | 8.200E-06 | 2.163E-06 | 5.800E-06 | 1.000E-05 |
| 7720 | 386 | G386V | A | 8.9e-06 | 7.1e-06 | 4.8e-06 | 6.933E-06 | 2.055E-06 | 4.800E-06 | 8.900E-06 |
| 7721 | 387 | Y387A | A | 1.3e-05 | 6e-06 | 9.6e-06 | 9.533E-06 | 3.500E-06 | 6.000E-06 | 1.300E-05 |
| 7722 | 387 | Y387R | A | 9.6e-06 | 6.3e-06 | 8.8e-06 | 8.233E-06 | 1.721E-06 | 6.300E-06 | 9.600E-06 |
| 7723 | 387 | Y387N | A | 3.4e-06 | 9.2e-06 | 9.2e-06 | 7.267E-06 | 3.349E-06 | 3.400E-06 | 9.200E-06 |
| 7724 | 387 | Y387D | A | 7.6e-06 | 8.8e-06 | 9.7e-06 | 8.700E-06 | 1.054E-06 | 7.600E-06 | 9.700E-06 |
| 7725 | 387 | Y387C | A | 7.9e-06 | 9.8e-06 | 5.4e-06 | 7.700E-06 | 2.207E-06 | 5.400E-06 | 9.800E-06 |
| 7726 | 387 | Y387E | A | 5.5e-06 | 6.9e-06 | 1.4e-05 | 8.800E-06 | 4.557E-06 | 5.500E-06 | 1.400E-05 |
| 7727 | 387 | Y387Q | A | 1.2e-05 | 1.3e-05 | 5.7e-06 | 1.023E-05 | 3.958E-06 | 5.700E-06 | 1.300E-05 |
| 7728 | 387 | Y387G | A | 9e-06 | 6.3e-06 | 1.3e-05 | 9.433E-06 | 3.371E-06 | 6.300E-06 | 1.300E-05 |
| 7729 | 387 | Y387H | A | 1.1e-05 | 9.6e-06 | 1e-05 | 1.020E-05 | 7.211E-07 | 9.600E-06 | 1.100E-05 |
| 7730 | 387 | Y387I | A | 1.2e-05 | 6.6e-06 | 8.9e-06 | 9.167E-06 | 2.710E-06 | 6.600E-06 | 1.200E-05 |
| 7731 | 387 | Y387L | A | 1e-05 | 5.4e-06 | 9.8e-06 | 8.400E-06 | 2.600E-06 | 5.400E-06 | 1.000E-05 |
| 7732 | 387 | Y387K | A | 1.1e-05 | 1.1e-05 | 1.5e-05 | 1.233E-05 | 2.309E-06 | 1.100E-05 | 1.500E-05 |
| 7733 | 387 | Y387M | A | 9.2e-06 | 6.1e-06 | 1.4e-05 | 9.767E-06 | 3.980E-06 | 6.100E-06 | 1.400E-05 |
| 7734 | 387 | Y387F | A | 1.4e-05 | 7.8e-06 | 1e-05 | 1.060E-05 | 3.143E-06 | 7.800E-06 | 1.400E-05 |

| | | | | | | | | | | |
|---|---|---|---|---|---|---|---|---|---|---|
| 7735 | 387 | Y387P | A | 7.6e-06 | 1.2e-05 | 1.2e-05 | 1.053E-05 | 2.540E-06 | 7.600E-06 | 1.200E-05 |
| 7736 | 387 | Y387S | A | 7.1e-06 | 9.5e-06 | 5e-06 | 7.200E-06 | 2.252E-06 | 5.000E-06 | 9.500E-06 |
| 7737 | 387 | Y387T | A | 9.5e-06 | 1.4e-05 | 8.2e-06 | 1.057E-05 | 3.044E-06 | 8.200E-06 | 1.400E-05 |
| 7738 | 387 | Y387W | A | 8.8e-06 | 7.3e-06 | 7.1e-06 | 7.733E-06 | 9.292E-07 | 7.100E-06 | 8.800E-06 |
| 7739 | 387 | Y387Y | A | 4.8e-06 | 7.5e-06 | 5e-06 | 5.767E-06 | 1.504E-06 | 4.800E-06 | 7.500E-06 |
| 7740 | 387 | Y387V | A | 6.6e-06 | 7.1e-06 | 4.3e-06 | 6.000E-06 | 1.493E-06 | 4.300E-06 | 7.100E-06 |
| 7741 | 388 | L388A | A | 8.3e-06 | 1e-05 | 1e-05 | 9.433E-06 | 9.815E-07 | 8.300E-06 | 1.000E-05 |
| 7742 | 388 | L388R | A | 7e-06 | 9.2e-06 | 1e-05 | 8.733E-06 | 1.553E-06 | 7.000E-06 | 1.000E-05 |
| 7743 | 388 | L388N | A | 8.8e-06 | 8.2e-06 | 5.4e-06 | 7.467E-06 | 1.815E-06 | 5.400E-06 | 8.800E-06 |
| 7744 | 388 | L388D | A | 9.1e-06 | 1.3e-05 | 8.2e-06 | 1.010E-05 | 2.551E-06 | 8.200E-06 | 1.300E-05 |
| 7745 | 388 | L388C | A | 5.8e-06 | 6.9e-06 | 8e-06 | 6.900E-06 | 1.100E-06 | 5.800E-06 | 8.000E-06 |
| 7746 | 388 | L388E | A | 1.3e-05 | 5.3e-06 | 7.4e-06 | 8.567E-06 | 3.980E-06 | 5.300E-06 | 1.300E-05 |
| 7747 | 388 | L388Q | A | 6.4e-06 | 1e-05 | 6.5e-06 | 7.633E-06 | 2.050E-06 | 6.400E-06 | 1.000E-05 |
| 7748 | 388 | L388G | A | 9.6e-06 | 6.9e-06 | 1.1e-05 | 9.167E-06 | 2.084E-06 | 6.900E-06 | 1.100E-05 |
| 7749 | 388 | L388H | A | 8.5e-06 | 9e-06 | 5e-06 | 7.500E-06 | 2.179E-06 | 5.000E-06 | 9.000E-06 |
| 7750 | 388 | L388I | A | 6.3e-06 | 3.2e-06 | 1.1e-05 | 6.833E-06 | 3.927E-06 | 3.200E-06 | 1.100E-05 |
| 7751 | 388 | L388L | A | 1.3e-05 | 8.3e-06 | 7.1e-06 | 9.467E-06 | 3.118E-06 | 7.100E-06 | 1.300E-05 |
| 7752 | 388 | L388K | A | 6.9e-06 | 7.2e-06 | 5.4e-06 | 6.500E-06 | 9.644E-07 | 5.400E-06 | 7.200E-06 |
| 7753 | 388 | L388M | A | 9.5e-06 | 9.5e-06 | 1.1e-05 | 1.000E-05 | 8.660E-07 | 9.500E-06 | 1.100E-05 |
| 7754 | 388 | L388F | A | 7e-06 | 6.3e-06 | 9.6e-06 | 7.633E-06 | 1.739E-06 | 6.300E-06 | 9.600E-06 |
| 7755 | 388 | L388P | A | 9.7e-06 | 9e-06 | 1.6e-05 | 1.157E-05 | 3.855E-06 | 9.000E-06 | 1.600E-05 |
| 7756 | 388 | L388S | A | 5e-06 | 7.4e-06 | 4.5e-06 | 5.633E-06 | 1.550E-06 | 4.500E-06 | 7.400E-06 |
| 7757 | 388 | L388T | A | 8.2e-06 | 5.9e-06 | 8.1e-06 | 7.400E-06 | 1.300E-06 | 5.900E-06 | 8.200E-06 |
| 7758 | 388 | L388W | A | 5.4e-06 | 5.3e-06 | 5.4e-06 | 5.367E-06 | 5.774E-08 | 5.300E-06 | 5.400E-06 |
| 7759 | 388 | L388Y | A | 9.7e-06 | 4.6e-06 | 4.9e-06 | 6.400E-06 | 2.862E-06 | 4.600E-06 | 9.700E-06 |
| 7760 | 388 | L388V | A | 8.5e-06 | 5.6e-06 | 6.5e-06 | 6.867E-06 | 1.484E-06 | 5.600E-06 | 8.500E-06 |
| 7761 | 389 | Y389A | A | 7.8e-06 | 5.1e-06 | 5.6e-06 | 6.167E-06 | 1.436E-06 | 5.100E-06 | 7.800E-06 |
| 7762 | 389 | Y389R | A | 8.5e-06 | 7.7e-06 | 1.2e-05 | 9.400E-06 | 2.287E-06 | 7.700E-06 | 1.200E-05 |
| 7763 | 389 | Y389N | A | 7.4e-06 | 6.3e-06 | 7e-06 | 6.900E-06 | 5.568E-07 | 6.300E-06 | 7.400E-06 |
| 7764 | 389 | Y389D | A | 1.3e-05 | 4.7e-06 | 8.5e-06 | 8.733E-06 | 4.155E-06 | 4.700E-06 | 1.300E-05 |
| 7765 | 389 | Y389C | A | 1.2e-05 | 9.1e-06 | 6.3e-06 | 9.133E-06 | 2.850E-06 | 6.300E-06 | 1.200E-05 |
| 7766 | 389 | Y389E | A | 6.6e-06 | 6.3e-06 | 9.3e-06 | 7.400E-06 | 1.652E-06 | 6.300E-06 | 9.300E-06 |
| 7767 | 389 | Y389Q | A | 6.6e-06 | 8.6e-06 | 1.4e-05 | 9.733E-06 | 3.828E-06 | 6.600E-06 | 1.400E-05 |
| 7768 | 389 | Y389G | A | 7.9e-06 | 1.1e-05 | 8.9e-06 | 9.267E-06 | 1.582E-06 | 7.900E-06 | 1.100E-05 |
| 7769 | 389 | Y389H | A | 7.6e-06 | 5.7e-06 | 9.5e-06 | 7.600E-06 | 1.900E-06 | 5.700E-06 | 9.500E-06 |
| 7770 | 389 | Y389I | A | 3.9e-06 | 8.7e-06 | 1.1e-05 | 7.867E-06 | 3.623E-06 | 3.900E-06 | 1.100E-05 |
| 7771 | 389 | Y389L | A | 7.7e-06 | 1.4e-05 | 6.9e-06 | 9.533E-06 | 3.889E-06 | 6.900E-06 | 1.400E-05 |
| 7772 | 389 | Y389K | A | 7.6e-06 | 7.5e-06 | 1.1e-05 | 8.700E-06 | 1.992E-06 | 7.500E-06 | 1.100E-05 |
| 7773 | 389 | Y389M | A | 1.1e-05 | 9.3e-06 | 5e-06 | 8.433E-06 | 3.092E-06 | 5.000E-06 | 1.100E-05 |
| 7774 | 389 | Y389F | A | 9.5e-06 | 9.1e-06 | 5.2e-06 | 7.933E-06 | 2.376E-06 | 5.200E-06 | 9.500E-06 |
| 7775 | 389 | Y389P | A | 1e-05 | 9.5e-06 | 9.4e-06 | 9.633E-06 | 3.215E-07 | 9.400E-06 | 1.000E-05 |
| 7776 | 389 | Y389S | A | 6.4e-06 | 8.2e-06 | 6.1e-06 | 6.900E-06 | 1.136E-06 | 6.100E-06 | 8.200E-06 |
| 7777 | 389 | Y389T | A | 1e-05 | 5e-06 | 5.4e-06 | 6.800E-06 | 2.778E-06 | 5.000E-06 | 1.000E-05 |
| 7778 | 389 | Y389W | A | 9.2e-06 | 6.2e-06 | 9.3e-06 | 8.233E-06 | 1.762E-06 | 6.200E-06 | 9.300E-06 |
| 7779 | 389 | Y389Y | A | 6.7e-06 | 9.1e-06 | 7.5e-06 | 7.767E-06 | 1.222E-06 | 6.700E-06 | 9.100E-06 |
| 7780 | 389 | Y389V | A | 9.1e-06 | 9.9e-06 | 9.3e-06 | 9.433E-06 | 4.163E-07 | 9.100E-06 | 9.900E-06 |
| 7781 | 390 | I390A | A | 8.7e-06 | 9.9e-06 | 6.5e-06 | 8.367E-06 | 1.724E-06 | 6.500E-06 | 9.900E-06 |
| 7782 | 390 | I390R | A | 6.4e-06 | 1.1e-05 | 1.3e-05 | 1.013E-05 | 3.384E-06 | 6.400E-06 | 1.300E-05 |
| 7783 | 390 | I390N | A | 7.8e-06 | 4.6e-06 | 5e-06 | 5.800E-06 | 1.744E-06 | 4.600E-06 | 7.800E-06 |
| 7784 | 390 | I390D | A | 8.1e-06 | 9e-06 | 1e-05 | 9.033E-06 | 9.504E-07 | 8.100E-06 | 1.000E-05 |
| 7785 | 390 | I390C | A | 9.4e-06 | 9.2e-06 | 1.1e-05 | 9.867E-06 | 9.866E-07 | 9.200E-06 | 1.100E-05 |
| 7786 | 390 | I390E | A | 1e-05 | 6.2e-06 | 1e-05 | 8.733E-06 | 2.194E-06 | 6.200E-06 | 1.000E-05 |
| 7787 | 390 | I390Q | A | 1.2e-05 | 6.9e-06 | 1.1e-05 | 9.967E-06 | 2.702E-06 | 6.900E-06 | 1.200E-05 |
| 7788 | 390 | I390G | A | 6.3e-06 | 9.6e-06 | 8.1e-06 | 8.000E-06 | 1.652E-06 | 6.300E-06 | 9.600E-06 |
| 7789 | 390 | I390H | A | 8e-06 | 8.8e-06 | 9.2e-06 | 8.667E-06 | 6.110E-07 | 8.000E-06 | 9.200E-06 |
| 7790 | 390 | I390I | A | 1.2e-05 | 1.1e-05 | 6.8e-06 | 9.933E-06 | 2.759E-06 | 6.800E-06 | 1.200E-05 |
| 7791 | 390 | I390L | A | 8.7e-06 | 8.7e-06 | 7.6e-06 | 8.333E-06 | 6.351E-07 | 7.600E-06 | 8.700E-06 |
| 7792 | 390 | I390K | A | 9e-06 | 6.8e-06 | 4.7e-06 | 6.833E-06 | 2.150E-06 | 4.700E-06 | 9.000E-06 |
| 7793 | 390 | I390M | A | 9.1e-06 | 5.1e-06 | 9.9e-06 | 8.033E-06 | 2.572E-06 | 5.100E-06 | 9.900E-06 |
| 7794 | 390 | I390F | A | 6.8e-06 | 5.2e-06 | 9.4e-06 | 7.133E-06 | 2.120E-06 | 5.200E-06 | 9.400E-06 |
| 7795 | 390 | I390P | A | 7.8e-06 | 7.9e-06 | 4.8e-06 | 6.833E-06 | 1.762E-06 | 4.800E-06 | 7.900E-06 |
| 7796 | 390 | I390S | A | 8.7e-06 | 7.2e-06 | 8.8e-06 | 8.233E-06 | 8.963E-07 | 7.200E-06 | 8.800E-06 |
| 7797 | 390 | I390T | A | 9.5e-06 | 5.3e-06 | 9.8e-06 | 8.200E-06 | 2.516E-06 | 5.300E-06 | 9.800E-06 |
| 7798 | 390 | I390W | A | 6.2e-06 | 1e-05 | 1.3e-05 | 9.733E-06 | 3.408E-06 | 6.200E-06 | 1.300E-05 |
| 7799 | 390 | I390Y | A | 6.7e-06 | 1.5e-05 | 8.8e-06 | 1.017E-05 | 4.315E-06 | 6.700E-06 | 1.500E-05 |
| 7800 | 390 | I390V | A | 5.8e-06 | 1e-05 | 5.2e-06 | 7.000E-06 | 2.615E-06 | 5.200E-06 | 1.000E-05 |
| 7801 | 391 | S391A | A | 1e-05 | 9.2e-06 | 1.5e-05 | 1.140E-05 | 3.143E-06 | 9.200E-06 | 1.500E-05 |
| 7802 | 391 | S391R | A | 7.4e-06 | 6.3e-06 | 8.2e-06 | 7.300E-06 | 9.539E-07 | 6.300E-06 | 8.200E-06 |
| 7803 | 391 | S391N | A | 5.9e-06 | 6.6e-06 | 4.8e-06 | 5.767E-06 | 9.074E-07 | 4.800E-06 | 6.600E-06 |
| 7804 | 391 | S391D | A | 9.9e-06 | 7.9e-06 | 4.8e-06 | 7.533E-06 | 2.570E-06 | 4.800E-06 | 9.900E-06 |
| 7805 | 391 | S391C | A | 7.2e-06 | 8e-06 | 8.5e-06 | 7.900E-06 | 6.557E-07 | 7.200E-06 | 8.500E-06 |
| 7806 | 391 | S391E | A | 5.4e-06 | 3.8e-06 | 1.1e-05 | 6.733E-06 | 3.781E-06 | 3.800E-06 | 1.100E-05 |
| 7807 | 391 | S391Q | A | 4.9e-06 | 5.6e-06 | 6e-06 | 5.500E-06 | 5.568E-07 | 4.900E-06 | 6.000E-06 |
| 7808 | 391 | S391G | A | 5e-06 | 9.9e-06 | 6.8e-06 | 7.233E-06 | 2.479E-06 | 5.000E-06 | 9.900E-06 |
| 7809 | 391 | S391H | A | 1.5e-05 | 8.2e-06 | 8.7e-06 | 1.063E-05 | 3.790E-06 | 8.200E-06 | 1.500E-05 |
| 7810 | 391 | S391I | A | 4.8e-06 | 8e-06 | 1.3e-05 | 8.600E-06 | 4.133E-06 | 4.800E-06 | 1.300E-05 |
| 7811 | 391 | S391L | A | 9e-06 | 7.8e-06 | 6.8e-06 | 7.867E-06 | 1.102E-06 | 6.800E-06 | 9.000E-06 |
| 7812 | 391 | S391K | A | 5.9e-06 | 1.2e-05 | 5.7e-06 | 7.867E-06 | 3.581E-06 | 5.700E-06 | 1.200E-05 |
| 7813 | 391 | S391M | A | 1e-05 | 5.6e-06 | 5.2e-06 | 6.933E-06 | 2.663E-06 | 5.200E-06 | 1.000E-05 |
| 7814 | 391 | S391F | A | 5.2e-06 | 5.7e-06 | 9.6e-06 | 6.833E-06 | 2.409E-06 | 5.200E-06 | 9.600E-06 |
| 7815 | 391 | S391P | A | 9.5e-06 | 6.6e-06 | 1.2e-05 | 9.367E-06 | 2.702E-06 | 6.600E-06 | 1.200E-05 |
| 7816 | 391 | S391S | A | 8.7e-06 | 7.6e-06 | 6.2e-06 | 7.500E-06 | 1.253E-06 | 6.200E-06 | 8.700E-06 |
| 7817 | 391 | S391T | A | 8.2e-06 | 6.4e-06 | 5.7e-06 | 6.767E-06 | 1.290E-06 | 5.700E-06 | 8.200E-06 |
| 7818 | 391 | S391W | A | 9.7e-06 | 6.7e-06 | 4.7e-06 | 7.033E-06 | 2.517E-06 | 4.700E-06 | 9.700E-06 |
| 7819 | 391 | S391Y | A | 6.8e-06 | 7.9e-06 | 5.4e-06 | 6.700E-06 | 1.253E-06 | 5.400E-06 | 7.900E-06 |

| | | | | | | | | | | |
|---|---|---|---|---|---|---|---|---|---|---|
| 7820 | 391 | S391V | A | 6.4e-06 | 6.7e-06 | 6.6e-06 | 6.567E-06 | 1.528E-07 | 6.400E-06 | 6.700E-06 |
| 7821 | 392 | A392A | A | 9.3e-06 | 6.6e-06 | 1.2e-05 | 9.300E-06 | 2.700E-06 | 6.600E-06 | 1.200E-05 |
| 7822 | 392 | A392R | A | 3.4e-06 | 6.6e-06 | 6.4e-06 | 5.467E-06 | 1.793E-06 | 3.400E-06 | 6.600E-06 |
| 7823 | 392 | A392N | A | 8.3e-06 | 5e-06 | 9.2e-06 | 7.500E-06 | 2.211E-06 | 5.000E-06 | 9.200E-06 |
| 7824 | 392 | A392D | A | 6.8e-06 | 7e-06 | 5.6e-06 | 6.467E-06 | 7.572E-07 | 5.600E-06 | 7.000E-06 |
| 7825 | 392 | A392C | A | 1.3e-05 | 1.1e-05 | 1.3e-05 | 1.233E-05 | 1.155E-06 | 1.100E-05 | 1.300E-05 |
| 7826 | 392 | A392E | A | 9.7e-06 | 1.3e-05 | 7.8e-06 | 1.017E-05 | 2.631E-06 | 7.800E-06 | 1.300E-05 |
| 7827 | 392 | A392Q | A | 1.3e-05 | 9.8e-06 | 6.5e-06 | 9.767E-06 | 3.250E-06 | 6.500E-06 | 1.300E-05 |
| 7828 | 392 | A392G | A | 6.7e-06 | 5.4e-06 | 7.3e-06 | 6.467E-06 | 9.713E-07 | 5.400E-06 | 7.300E-06 |
| 7829 | 392 | A392H | A | 6.1e-06 | 4.8e-06 | 9.3e-06 | 6.733E-06 | 2.316E-06 | 4.800E-06 | 9.300E-06 |
| 7830 | 392 | A392I | A | 4.7e-06 | 7.5e-06 | 6.2e-06 | 6.133E-06 | 1.401E-06 | 4.700E-06 | 7.500E-06 |
| 7831 | 392 | A392L | A | 9.7e-06 | 5.8e-06 | 8.5e-06 | 8.000E-06 | 1.997E-06 | 5.800E-06 | 9.700E-06 |
| 7832 | 392 | A392K | A | 8.6e-06 | 6e-06 | 6e-06 | 6.867E-06 | 1.501E-06 | 6.000E-06 | 8.600E-06 |
| 7833 | 392 | A392M | A | 6.1e-06 | 5.5e-06 | 9.5e-06 | 7.033E-06 | 2.157E-06 | 5.500E-06 | 9.500E-06 |
| 7834 | 392 | A392F | A | 4e-06 | 9.3e-06 | 8.1e-06 | 7.133E-06 | 2.779E-06 | 4.000E-06 | 9.300E-06 |
| 7835 | 392 | A392P | A | 1e-05 | 7e-06 | 6.9e-06 | 7.967E-06 | 1.762E-06 | 6.900E-06 | 1.000E-05 |
| 7836 | 392 | A392S | A | 5.7e-06 | 7.9e-06 | 1.3e-05 | 8.867E-06 | 3.745E-06 | 5.700E-06 | 1.300E-05 |
| 7837 | 392 | A392T | A | 8.9e-06 | 7.2e-06 | 4.5e-06 | 6.867E-06 | 2.219E-06 | 4.500E-06 | 8.900E-06 |
| 7838 | 392 | A392W | A | 9.4e-06 | 6.5e-06 | 5.3e-06 | 7.067E-06 | 2.108E-06 | 5.300E-06 | 9.400E-06 |
| 7839 | 392 | A392Y | A | 6.5e-06 | 8e-06 | 8.5e-06 | 7.667E-06 | 1.041E-06 | 6.500E-06 | 8.500E-06 |
| 7840 | 392 | A392V | A | 8.1e-06 | 4.2e-06 | 1e-05 | 7.433E-06 | 2.957E-06 | 4.200E-06 | 1.000E-05 |
| 7841 | 393 | W393A | A | 8.3e-06 | 8.4e-06 | 9.6e-06 | 8.767E-06 | 7.234E-07 | 8.300E-06 | 9.600E-06 |
| 7842 | 393 | W393R | A | 1.1e-05 | 9.7e-06 | 7.2e-06 | 9.300E-06 | 1.931E-06 | 7.200E-06 | 1.100E-05 |
| 7843 | 393 | W393N | A | 9.5e-06 | 6e-06 | 1.5e-05 | 1.017E-05 | 4.537E-06 | 6.000E-06 | 1.500E-05 |
| 7844 | 393 | W393D | A | 4.8e-06 | 5.2e-06 | 7.4e-06 | 5.800E-06 | 1.400E-06 | 4.800E-06 | 7.400E-06 |
| 7845 | 393 | W393C | A | 5.9e-06 | 1e-05 | 6.5e-06 | 7.467E-06 | 2.214E-06 | 5.900E-06 | 1.000E-05 |
| 7846 | 393 | W393E | A | 7e-06 | 9.4e-06 | 4.7e-06 | 7.033E-06 | 2.350E-06 | 4.700E-06 | 9.400E-06 |
| 7847 | 393 | W393Q | A | 6.1e-06 | 1.1e-05 | 6.5e-06 | 7.867E-06 | 2.721E-06 | 6.100E-06 | 1.100E-05 |
| 7848 | 393 | W393G | A | 8.6e-06 | 7.2e-06 | 9.5e-06 | 8.433E-06 | 1.159E-06 | 7.200E-06 | 9.500E-06 |
| 7849 | 393 | W393H | A | 1.3e-05 | 8e-06 | 1.1e-05 | 1.067E-05 | 2.517E-06 | 8.000E-06 | 1.300E-05 |
| 7850 | 393 | W393I | A | 7.2e-06 | 1e-05 | 9.9e-06 | 9.033E-06 | 1.589E-06 | 7.200E-06 | 1.000E-05 |
| 7851 | 393 | W393L | A | 9.3e-06 | 9.9e-06 | 9.4e-06 | 9.533E-06 | 3.215E-07 | 9.300E-06 | 9.900E-06 |
| 7852 | 393 | W393K | A | 7.1e-06 | 1.2e-05 | 5.2e-06 | 8.100E-06 | 3.509E-06 | 5.200E-06 | 1.200E-05 |
| 7853 | 393 | W393M | A | 1.1e-05 | 5.6e-06 | 7e-06 | 7.867E-06 | 2.802E-06 | 5.600E-06 | 1.100E-05 |
| 7854 | 393 | W393F | A | 4.7e-06 | 6.6e-06 | 1.2e-05 | 7.767E-06 | 3.787E-06 | 4.700E-06 | 1.200E-05 |
| 7855 | 393 | W393P | A | 8.8e-06 | 8.8e-06 | 6.8e-06 | 8.133E-06 | 1.155E-06 | 6.800E-06 | 8.800E-06 |
| 7856 | 393 | W393S | A | 1.2e-05 | 1.5e-05 | 3e-06 | 1.000E-05 | 6.245E-06 | 3.000E-06 | 1.500E-05 |
| 7857 | 393 | W393T | A | 6.2e-06 | 7.8e-06 | 9.9e-06 | 7.967E-06 | 1.856E-06 | 6.200E-06 | 9.900E-06 |
| 7858 | 393 | W393W | A | 1.1e-05 | 1e-05 | 7e-06 | 9.333E-06 | 2.082E-06 | 7.000E-06 | 1.100E-05 |
| 7859 | 393 | W393Y | A | 3.6e-06 | 9.5e-06 | 8.5e-06 | 7.200E-06 | 3.158E-06 | 3.600E-06 | 9.500E-06 |
| 7860 | 393 | W393V | A | 8.3e-06 | 5.5e-06 | 8.7e-06 | 7.500E-06 | 1.744E-06 | 5.500E-06 | 8.700E-06 |
| 7861 | 394 | P394A | A | 1.2e-05 | 8.9e-06 | 1.3e-05 | 1.130E-05 | 2.138E-06 | 8.900E-06 | 1.300E-05 |
| 7862 | 394 | P394R | A | 4.3e-06 | 1.3e-05 | 8.5e-06 | 8.600E-06 | 4.351E-06 | 4.300E-06 | 1.300E-05 |
| 7863 | 394 | P394N | A | 1.2e-05 | 7.4e-06 | 6.3e-06 | 8.567E-06 | 3.024E-06 | 6.300E-06 | 1.200E-05 |
| 7864 | 394 | P394D | A | 7.1e-06 | 3.7e-06 | 9.2e-06 | 6.667E-06 | 2.775E-06 | 3.700E-06 | 9.200E-06 |
| 7865 | 394 | P394C | A | 1.1e-05 | 6.7e-06 | 7.1e-06 | 8.267E-06 | 2.376E-06 | 6.700E-06 | 1.100E-05 |
| 7866 | 394 | P394E | A | 1.5e-05 | 4.6e-06 | 1.2e-05 | 1.053E-05 | 5.353E-06 | 4.600E-06 | 1.500E-05 |
| 7867 | 394 | P394Q | A | 8.1e-06 | 6.6e-06 | 9.4e-06 | 8.033E-06 | 1.401E-06 | 6.600E-06 | 9.400E-06 |
| 7868 | 394 | P394G | A | 9.3e-06 | 7.3e-06 | 1e-05 | 8.867E-06 | 1.401E-06 | 7.300E-06 | 1.000E-05 |
| 7869 | 394 | P394H | A | 9.3e-06 | 9.5e-06 | 1.1e-05 | 9.933E-06 | 9.292E-07 | 9.300E-06 | 1.100E-05 |
| 7870 | 394 | P394I | A | 7.9e-06 | 7.4e-06 | 9.4e-06 | 8.233E-06 | 1.041E-06 | 7.400E-06 | 9.400E-06 |
| 7871 | 394 | P394L | A | 1e-05 | 6.6e-06 | 6.9e-06 | 7.833E-06 | 1.882E-06 | 6.600E-06 | 1.000E-05 |
| 7872 | 394 | P394K | A | 5.5e-06 | 7.9e-06 | 5.2e-06 | 6.200E-06 | 1.480E-06 | 5.200E-06 | 7.900E-06 |
| 7873 | 394 | P394M | A | 5.3e-06 | 1e-05 | 5.8e-06 | 7.033E-06 | 2.581E-06 | 5.300E-06 | 1.000E-05 |
| 7874 | 394 | P394F | A | 1.1e-05 | 7e-06 | 4.5e-06 | 7.500E-06 | 3.279E-06 | 4.500E-06 | 1.100E-05 |
| 7875 | 394 | P394P | A | 4.7e-06 | 6.5e-06 | 6.4e-06 | 5.867E-06 | 1.012E-06 | 4.700E-06 | 6.500E-06 |
| 7876 | 394 | P394S | A | 6.2e-06 | 6.2e-06 | 9.4e-06 | 7.267E-06 | 1.848E-06 | 6.200E-06 | 9.400E-06 |
| 7877 | 394 | P394T | A | 9.6e-06 | 1e-05 | 5.6e-06 | 8.400E-06 | 2.433E-06 | 5.600E-06 | 1.000E-05 |
| 7878 | 394 | P394W | A | 6.7e-06 | 6.5e-06 | 1e-05 | 7.733E-06 | 1.966E-06 | 6.500E-06 | 1.000E-05 |
| 7879 | 394 | P394Y | A | 5.6e-06 | 7e-06 | 7.5e-06 | 6.700E-06 | 9.849E-07 | 5.600E-06 | 7.500E-06 |
| 7880 | 394 | P394V | A | 1.5e-05 | 6.3e-06 | 5.6e-06 | 8.967E-06 | 5.237E-06 | 5.600E-06 | 1.500E-05 |
| 7881 | 395 | D395A | A | 4.8e-06 | 8.9e-06 | 9.4e-06 | 7.700E-06 | 2.524E-06 | 4.800E-06 | 9.400E-06 |
| 7882 | 395 | D395R | A | 6.8e-06 | 7.2e-06 | 7.1e-06 | 7.033E-06 | 2.082E-07 | 6.800E-06 | 7.200E-06 |
| 7883 | 395 | D395N | A | 5.6e-06 | 1.1e-05 | 6.2e-06 | 7.600E-06 | 2.960E-06 | 5.600E-06 | 1.100E-05 |
| 7884 | 395 | D395D | A | 6.6e-06 | 1.2e-05 | 8.3e-06 | 8.967E-06 | 2.761E-06 | 6.600E-06 | 1.200E-05 |
| 7885 | 395 | D395C | A | 7.1e-06 | 8.7e-06 | 9e-06 | 8.267E-06 | 1.021E-06 | 7.100E-06 | 9.000E-06 |
| 7886 | 395 | D395E | A | 4.4e-06 | 5.5e-06 | 7.8e-06 | 5.900E-06 | 1.735E-06 | 4.400E-06 | 7.800E-06 |
| 7887 | 395 | D395Q | A | 5.5e-06 | 1.3e-05 | 1.1e-05 | 9.833E-06 | 3.884E-06 | 5.500E-06 | 1.300E-05 |
| 7888 | 395 | D395G | A | 4.6e-06 | 1e-05 | 9.6e-06 | 8.067E-06 | 3.009E-06 | 4.600E-06 | 1.000E-05 |
| 7889 | 395 | D395H | A | 4.9e-06 | 9.1e-06 | 9.1e-06 | 7.700E-06 | 2.425E-06 | 4.900E-06 | 9.100E-06 |
| 7890 | 395 | D395I | A | 6.8e-06 | 9e-06 | 8.6e-06 | 8.133E-06 | 1.172E-06 | 6.800E-06 | 9.000E-06 |
| 7891 | 395 | D395L | A | 4e-06 | 5.8e-06 | 8.8e-06 | 6.200E-06 | 2.425E-06 | 4.000E-06 | 8.800E-06 |
| 7892 | 395 | D395K | A | 6.3e-06 | 6e-06 | 8e-06 | 6.767E-06 | 1.079E-06 | 6.000E-06 | 8.000E-06 |
| 7893 | 395 | D395M | A | 1.4e-05 | 9.5e-06 | 7.7e-06 | 1.040E-05 | 3.245E-06 | 7.700E-06 | 1.400E-05 |
| 7894 | 395 | D395F | A | 5.7e-06 | 8.2e-06 | 6e-06 | 6.633E-06 | 1.365E-06 | 5.700E-06 | 8.200E-06 |
| 7895 | 395 | D395P | A | 7.7e-06 | 4.7e-06 | 1.2e-05 | 8.133E-06 | 3.669E-06 | 4.700E-06 | 1.200E-05 |
| 7896 | 395 | D395S | A | 6.7e-06 | 7.6e-06 | 4.5e-06 | 6.267E-06 | 1.595E-06 | 4.500E-06 | 7.600E-06 |
| 7897 | 395 | D395T | A | 8.8e-06 | 5.8e-06 | 7.4e-06 | 7.333E-06 | 1.501E-06 | 5.800E-06 | 8.800E-06 |
| 7898 | 395 | D395W | A | 7.6e-06 | 5.2e-06 | 9.1e-06 | 7.300E-06 | 1.967E-06 | 5.200E-06 | 9.100E-06 |
| 7899 | 395 | D395Y | A | 1.1e-05 | 1e-05 | 8.5e-06 | 9.833E-06 | 1.258E-06 | 8.500E-06 | 1.100E-05 |
| 7900 | 395 | D395V | A | 6.4e-06 | 7.3e-06 | 1e-05 | 7.900E-06 | 1.873E-06 | 6.400E-06 | 1.000E-05 |
| 7901 | 396 | S396A | A | 1e-05 | 9.4e-06 | 5.7e-06 | 8.367E-06 | 2.329E-06 | 5.700E-06 | 1.000E-05 |
| 7902 | 396 | S396R | A | 6.6e-06 | 1e-05 | 9.5e-06 | 8.700E-06 | 1.836E-06 | 6.600E-06 | 1.000E-05 |
| 7903 | 396 | S396N | A | 6.8e-06 | 7.6e-06 | 1.4e-05 | 9.467E-06 | 3.946E-06 | 6.800E-06 | 1.400E-05 |
| 7904 | 396 | S396D | A | 9.2e-06 | 6.6e-06 | 9.1e-06 | 8.300E-06 | 1.473E-06 | 6.600E-06 | 9.200E-06 |

| | | | | | | | | | | |
|---|---|---|---|---|---|---|---|---|---|---|
| 7905 | 396 | S396C | A | 9.1e-06 | 3.3e-06 | 9.5e-06 | 7.300E-06 | 3.470E-06 | 3.300E-06 | 9.500E-06 |
| 7906 | 396 | S396E | A | 1.9e-05 | 8.3e-06 | 4.3e-06 | 1.053E-05 | 7.600E-06 | 4.300E-06 | 1.900E-05 |
| 7907 | 396 | S396Q | A | 8.8e-06 | 5e-06 | 8.6e-06 | 7.467E-06 | 2.139E-06 | 5.000E-06 | 8.800E-06 |
| 7908 | 396 | S396G | A | 6.4e-06 | 6.8e-06 | 5.1e-06 | 6.100E-06 | 8.888E-07 | 5.100E-06 | 6.800E-06 |
| 7909 | 396 | S396H | A | 6.5e-06 | 8e-06 | 5.9e-06 | 6.800E-06 | 1.082E-06 | 5.900E-06 | 8.000E-06 |
| 7910 | 396 | S396I | A | 1.1e-05 | 9e-06 | 9.4e-06 | 9.800E-06 | 1.058E-06 | 9.000E-06 | 1.100E-05 |
| 7911 | 396 | S396L | A | 6.1e-06 | 7.3e-06 | 7.4e-06 | 6.933E-06 | 7.234E-07 | 6.100E-06 | 7.400E-06 |
| 7912 | 396 | S396K | A | 8.3e-06 | 8.7e-06 | 6.6e-06 | 7.867E-06 | 1.115E-06 | 6.600E-06 | 8.700E-06 |
| 7913 | 396 | S396M | A | 1.2e-05 | 1.2e-05 | 8.7e-06 | 1.090E-05 | 1.905E-06 | 8.700E-06 | 1.200E-05 |
| 7914 | 396 | S396F | A | 8.8e-06 | 9.1e-06 | 6.7e-06 | 8.200E-06 | 1.308E-06 | 6.700E-06 | 9.100E-06 |
| 7915 | 396 | S396P | A | 1.3e-05 | 6e-06 | 4.7e-06 | 7.900E-06 | 4.464E-06 | 4.700E-06 | 1.300E-05 |
| 7916 | 396 | S396S | A | 9.6e-06 | 1.3e-05 | 6.7e-06 | 9.767E-06 | 3.153E-06 | 6.700E-06 | 1.300E-05 |
| 7917 | 396 | S396T | A | 6.4e-06 | 9e-06 | 1.1e-05 | 8.800E-06 | 2.307E-06 | 6.400E-06 | 1.100E-05 |
| 7918 | 396 | S396W | A | 7.5e-06 | 8.4e-06 | 1.2e-05 | 9.300E-06 | 2.381E-06 | 7.500E-06 | 1.200E-05 |
| 7919 | 396 | S396Y | A | 6.5e-06 | 1e-05 | 9.9e-06 | 8.800E-06 | 1.992E-06 | 6.500E-06 | 1.000E-05 |
| 7920 | 396 | S396V | A | 5e-06 | 9.5e-06 | 9.8e-06 | 8.100E-06 | 2.689E-06 | 5.000E-06 | 9.800E-06 |
| 7921 | 397 | L397A | A | 4.5e-06 | 4.9e-06 | 6.2e-06 | 5.200E-06 | 8.888E-07 | 4.500E-06 | 6.200E-06 |
| 7922 | 397 | L397R | A | 6.3e-06 | 6.6e-06 | 1.1e-05 | 7.967E-06 | 2.631E-06 | 6.300E-06 | 1.100E-05 |
| 7923 | 397 | L397N | A | 1.3e-05 | 9.8e-06 | 4.4e-06 | 9.067E-06 | 4.347E-06 | 4.400E-06 | 1.300E-05 |
| 7924 | 397 | L397D | A | 1.1e-05 | 9.5e-06 | 4.4e-06 | 8.300E-06 | 3.460E-06 | 4.400E-06 | 1.100E-05 |
| 7925 | 397 | L397C | A | 5.8e-06 | 7.6e-06 | 7.3e-06 | 6.900E-06 | 9.644E-07 | 5.800E-06 | 7.600E-06 |
| 7926 | 397 | L397E | A | 5.3e-06 | 5.6e-06 | 7.9e-06 | 6.267E-06 | 1.422E-06 | 5.300E-06 | 7.900E-06 |
| 7927 | 397 | L397Q | A | 8.5e-06 | 4.7e-06 | 7.1e-06 | 6.767E-06 | 1.922E-06 | 4.700E-06 | 8.500E-06 |
| 7928 | 397 | L397G | A | 1.3e-05 | 1e-05 | 4.2e-06 | 9.067E-06 | 4.474E-06 | 4.200E-06 | 1.300E-05 |
| 7929 | 397 | L397H | A | 7.8e-06 | 4.5e-06 | 6.3e-06 | 6.200E-06 | 1.652E-06 | 4.500E-06 | 7.800E-06 |
| 7930 | 397 | L397I | A | 6.5e-06 | 5e-06 | 6.5e-06 | 6.000E-06 | 8.660E-07 | 5.000E-06 | 6.500E-06 |
| 7931 | 397 | L397L | A | 7.8e-06 | 9.4e-06 | 7.9e-06 | 8.367E-06 | 8.963E-07 | 7.800E-06 | 9.400E-06 |
| 7932 | 397 | L397K | A | 7.4e-06 | 8.4e-06 | 9.3e-06 | 8.367E-06 | 9.504E-07 | 7.400E-06 | 9.300E-06 |
| 7933 | 397 | L397M | A | 9.3e-06 | 7.1e-06 | 7.5e-06 | 7.967E-06 | 1.172E-06 | 7.100E-06 | 9.300E-06 |
| 7934 | 397 | L397F | A | 9.5e-06 | 8.9e-06 | 6.6e-06 | 8.333E-06 | 1.531E-06 | 6.600E-06 | 9.500E-06 |
| 7935 | 397 | L397P | A | 9e-06 | 6.5e-06 | 8.6e-06 | 8.033E-06 | 1.343E-06 | 6.500E-06 | 9.000E-06 |
| 7936 | 397 | L397S | A | 3.9e-06 | 9.1e-06 | 8.8e-06 | 7.267E-06 | 2.919E-06 | 3.900E-06 | 9.100E-06 |
| 7937 | 397 | L397T | A | 5.5e-06 | 1.1e-05 | 4.7e-06 | 7.067E-06 | 3.430E-06 | 4.700E-06 | 1.100E-05 |
| 7938 | 397 | L397W | A | 1.1e-05 | 6.2e-06 | 3.9e-06 | 7.033E-06 | 3.623E-06 | 3.900E-06 | 1.100E-05 |
| 7939 | 397 | L397Y | A | 1.2e-05 | 8.9e-06 | 7.4e-06 | 9.433E-06 | 2.346E-06 | 7.400E-06 | 1.200E-05 |
| 7940 | 397 | L397V | A | 4.7e-06 | 5.5e-06 | 7e-06 | 5.733E-06 | 1.168E-06 | 4.700E-06 | 7.000E-06 |
| 7941 | 398 | P398A | A | 8.9e-06 | 5.5e-06 | 8.9e-06 | 7.767E-06 | 1.963E-06 | 5.500E-06 | 8.900E-06 |
| 7942 | 398 | P398R | A | 9.5e-06 | 6.5e-06 | 5.9e-06 | 7.300E-06 | 1.929E-06 | 5.900E-06 | 9.500E-06 |
| 7943 | 398 | P398N | A | 4.7e-06 | 1e-05 | 7e-06 | 7.233E-06 | 2.658E-06 | 4.700E-06 | 1.000E-05 |
| 7944 | 398 | P398D | A | 7.8e-06 | 7e-06 | 9.9e-06 | 8.233E-06 | 1.498E-06 | 7.000E-06 | 9.900E-06 |
| 7945 | 398 | P398C | A | 1.3e-05 | 1.1e-05 | 9.2e-06 | 1.107E-05 | 1.901E-06 | 9.200E-06 | 1.300E-05 |
| 7946 | 398 | P398E | A | 1.3e-05 | 1.3e-05 | 9e-06 | 1.167E-05 | 2.309E-06 | 9.000E-06 | 1.300E-05 |
| 7947 | 398 | P398Q | A | 6.3e-06 | 6.5e-06 | 6.3e-06 | 6.367E-06 | 1.155E-07 | 6.300E-06 | 6.500E-06 |
| 7948 | 398 | P398G | A | 1.2e-05 | 7.2e-06 | 6.8e-06 | 8.667E-06 | 2.894E-06 | 6.800E-06 | 1.200E-05 |
| 7949 | 398 | P398H | A | 4.8e-06 | 5e-06 | 6.6e-06 | 5.467E-06 | 9.866E-07 | 4.800E-06 | 6.600E-06 |
| 7950 | 398 | P398I | A | 3.9e-06 | 7.8e-06 | 9.2e-06 | 6.967E-06 | 2.747E-06 | 3.900E-06 | 9.200E-06 |
| 7951 | 398 | P398L | A | 9.7e-06 | 6.2e-06 | 7.4e-06 | 7.767E-06 | 1.779E-06 | 6.200E-06 | 9.700E-06 |
| 7952 | 398 | P398K | A | 9.5e-06 | 7.4e-06 | 7e-06 | 7.967E-06 | 1.343E-06 | 7.000E-06 | 9.500E-06 |
| 7953 | 398 | P398M | A | 7e-06 | 9e-06 | 4.3e-06 | 6.767E-06 | 2.359E-06 | 4.300E-06 | 9.000E-06 |
| 7954 | 398 | P398F | A | 5.3e-06 | 6.8e-06 | 6.9e-06 | 6.333E-06 | 8.963E-07 | 5.300E-06 | 6.900E-06 |
| 7955 | 398 | P398P | A | 9e-06 | 8e-06 | 7.2e-06 | 8.067E-06 | 9.018E-07 | 7.200E-06 | 9.000E-06 |
| 7956 | 398 | P398S | A | 1.3e-05 | 7.6e-06 | 6.5e-06 | 9.033E-06 | 3.479E-06 | 6.500E-06 | 1.300E-05 |
| 7957 | 398 | P398T | A | 6.1e-06 | 1.2e-05 | 6.4e-06 | 8.167E-06 | 3.323E-06 | 6.100E-06 | 1.200E-05 |
| 7958 | 398 | P398W | A | 1e-05 | 5.3e-06 | 1.3e-05 | 9.433E-06 | 3.881E-06 | 5.300E-06 | 1.300E-05 |
| 7959 | 398 | P398Y | A | 7.9e-06 | 6.7e-06 | 1.1e-05 | 8.533E-06 | 2.219E-06 | 6.700E-06 | 1.100E-05 |
| 7960 | 398 | P398V | A | 6.4e-06 | 6.3e-06 | 7.4e-06 | 6.700E-06 | 6.083E-07 | 6.300E-06 | 7.400E-06 |
| 7961 | 399 | D399A | A | 7.1e-06 | 1e-05 | 7.9e-06 | 8.333E-06 | 1.498E-06 | 7.100E-06 | 1.000E-05 |
| 7962 | 399 | D399R | A | 8.5e-06 | 5.6e-06 | 4.2e-06 | 6.100E-06 | 2.193E-06 | 4.200E-06 | 8.500E-06 |
| 7963 | 399 | D399N | A | 6.9e-06 | 9.4e-06 | 5e-06 | 7.100E-06 | 2.207E-06 | 5.000E-06 | 9.400E-06 |
| 7964 | 399 | D399D | A | 1.2e-05 | 6.2e-06 | 1.2e-05 | 1.007E-05 | 3.349E-06 | 6.200E-06 | 1.200E-05 |
| 7965 | 399 | D399C | A | 6.7e-06 | 4.5e-06 | 7.5e-06 | 6.233E-06 | 1.553E-06 | 4.500E-06 | 7.500E-06 |
| 7966 | 399 | D399E | A | 8.6e-06 | 8.3e-06 | 6.3e-06 | 7.733E-06 | 1.250E-06 | 6.300E-06 | 8.600E-06 |
| 7967 | 399 | D399Q | A | 9.3e-06 | 9.4e-06 | 8.4e-06 | 9.033E-06 | 5.508E-07 | 8.400E-06 | 9.400E-06 |
| 7968 | 399 | D399G | A | 5e-06 | 1.1e-05 | 7.1e-06 | 7.700E-06 | 3.045E-06 | 5.000E-06 | 1.100E-05 |
| 7969 | 399 | D399H | A | 8.8e-06 | 5.7e-06 | 1.3e-05 | 9.167E-06 | 3.664E-06 | 5.700E-06 | 1.300E-05 |
| 7970 | 399 | D399I | A | 1.2e-05 | 7.8e-06 | 6.8e-06 | 8.867E-06 | 2.759E-06 | 6.800E-06 | 1.200E-05 |
| 7971 | 399 | D399L | A | 8.4e-06 | 5.8e-06 | 1.6e-05 | 1.007E-05 | 5.300E-06 | 5.800E-06 | 1.600E-05 |
| 7972 | 399 | D399K | A | 1.3e-05 | 5.9e-06 | 8.3e-06 | 9.067E-06 | 3.612E-06 | 5.900E-06 | 1.300E-05 |
| 7973 | 399 | D399M | A | 1.1e-05 | 8.9e-06 | 1.1e-05 | 1.030E-05 | 1.212E-06 | 8.900E-06 | 1.100E-05 |
| 7974 | 399 | D399F | A | 6.1e-06 | 6.7e-06 | 6.7e-06 | 6.500E-06 | 3.464E-07 | 6.100E-06 | 6.700E-06 |
| 7975 | 399 | D399P | A | 6.6e-06 | 8.4e-06 | 9.7e-06 | 8.233E-06 | 1.557E-06 | 6.600E-06 | 9.700E-06 |
| 7976 | 399 | D399S | A | 9.3e-06 | 9.2e-06 | 6.2e-06 | 8.233E-06 | 1.762E-06 | 6.200E-06 | 9.300E-06 |
| 7977 | 399 | D399T | A | 7.3e-06 | 5.9e-06 | 8.8e-06 | 7.333E-06 | 1.450E-06 | 5.900E-06 | 8.800E-06 |
| 7978 | 399 | D399W | A | 3.7e-06 | 4.9e-06 | 6.8e-06 | 5.133E-06 | 1.563E-06 | 3.700E-06 | 6.800E-06 |
| 7979 | 399 | D399Y | A | 1.3e-05 | 5.4e-06 | 6.7e-06 | 8.367E-06 | 4.065E-06 | 5.400E-06 | 1.300E-05 |
| 7980 | 399 | D399V | A | 6.4e-06 | 1.2e-05 | 9.3e-06 | 9.233E-06 | 2.801E-06 | 6.400E-06 | 1.200E-05 |
| 7981 | 400 | L400A | A | 7.4e-06 | 7.5e-06 | 6.8e-06 | 7.233E-06 | 3.786E-07 | 6.800E-06 | 7.500E-06 |
| 7982 | 400 | L400R | A | 7.7e-06 | 7.2e-06 | 6.5e-06 | 7.133E-06 | 6.028E-07 | 6.500E-06 | 7.700E-06 |
| 7983 | 400 | L400N | A | 6.6e-06 | 8.5e-06 | 8.7e-06 | 7.933E-06 | 1.159E-06 | 6.600E-06 | 8.700E-06 |
| 7984 | 400 | L400D | A | 7.5e-06 | 9.8e-06 | 6.8e-06 | 8.033E-06 | 1.570E-06 | 6.800E-06 | 9.800E-06 |
| 7985 | 400 | L400C | A | 4.2e-06 | 7.8e-06 | 1e-05 | 7.333E-06 | 2.928E-06 | 4.200E-06 | 1.000E-05 |
| 7986 | 400 | L400E | A | 8.7e-06 | 8e-06 | 7.7e-06 | 8.133E-06 | 5.132E-07 | 7.700E-06 | 8.700E-06 |
| 7987 | 400 | L400Q | A | 8.2e-06 | 4.7e-06 | 7.8e-06 | 6.900E-06 | 1.916E-06 | 4.700E-06 | 8.200E-06 |
| 7988 | 400 | L400G | A | 1.1e-05 | 5.6e-06 | 1.2e-05 | 9.533E-06 | 3.443E-06 | 5.600E-06 | 1.200E-05 |
| 7989 | 400 | L400H | A | 1.1e-05 | 7.5e-06 | 9e-06 | 9.167E-06 | 1.756E-06 | 7.500E-06 | 1.100E-05 |

| | | | | | | | | | | |
|---|---|---|---|---|---|---|---|---|---|---|
| 7990 | 400 | L400I | A | 1.2e-05 | 1.2e-05 | 1e-05 | 1.133E-05 | 1.155E-06 | 1.000E-05 | 1.200E-05 |
| 7991 | 400 | L400L | A | 5.7e-06 | 7.3e-06 | 1e-05 | 7.667E-06 | 2.173E-06 | 5.700E-06 | 1.000E-05 |
| 7992 | 400 | L400K | A | 5.5e-06 | 9e-06 | 1.3e-05 | 9.167E-06 | 3.753E-06 | 5.500E-06 | 1.300E-05 |
| 7993 | 400 | L400M | A | 6.8e-06 | 7.7e-06 | 6.5e-06 | 7.000E-06 | 6.245E-07 | 6.500E-06 | 7.700E-06 |
| 7994 | 400 | L400F | A | 9.3e-06 | 7.8e-06 | 7.4e-06 | 8.167E-06 | 1.002E-06 | 7.400E-06 | 9.300E-06 |
| 7995 | 400 | L400P | A | 8e-06 | 1e-05 | 8e-06 | 8.667E-06 | 1.155E-06 | 8.000E-06 | 1.000E-05 |
| 7996 | 400 | L400S | A | 9.1e-06 | 8e-06 | 7.6e-06 | 8.233E-06 | 7.767E-07 | 7.600E-06 | 9.100E-06 |
| 7997 | 400 | L400T | A | 9.9e-06 | 1.2e-05 | 6.2e-06 | 9.367E-06 | 2.937E-06 | 6.200E-06 | 1.200E-05 |
| 7998 | 400 | L400W | A | 8.6e-06 | 5.1e-06 | 6.9e-06 | 6.867E-06 | 1.750E-06 | 5.100E-06 | 8.600E-06 |
| 7999 | 400 | L400Y | A | 8.2e-06 | 7.8e-06 | 9.3e-06 | 8.433E-06 | 7.767E-07 | 7.800E-06 | 9.300E-06 |
| 8000 | 400 | L400V | A | 6.1e-06 | 1.3e-05 | 6.7e-06 | 8.600E-06 | 3.822E-06 | 6.100E-06 | 1.300E-05 |
| 8001 | 401 | S401A | A | 6.6e-06 | 1e-05 | 7.1e-06 | 7.900E-06 | 1.836E-06 | 6.600E-06 | 1.000E-05 |
| 8002 | 401 | S401R | A | 9.5e-06 | 5.6e-06 | 6.9e-06 | 7.333E-06 | 1.986E-06 | 5.600E-06 | 9.500E-06 |
| 8003 | 401 | S401N | A | 6.9e-06 | 8.7e-06 | 9.8e-06 | 8.467E-06 | 1.464E-06 | 6.900E-06 | 9.800E-06 |
| 8004 | 401 | S401D | A | 9.1e-06 | 4.7e-06 | 1.2e-05 | 8.600E-06 | 3.676E-06 | 4.700E-06 | 1.200E-05 |
| 8005 | 401 | S401C | A | 8.2e-06 | 6.8e-06 | 6.8e-06 | 7.267E-06 | 8.083E-07 | 6.800E-06 | 8.200E-06 |
| 8006 | 401 | S401E | A | 9.1e-06 | 8.4e-06 | 6.9e-06 | 8.133E-06 | 1.124E-06 | 6.900E-06 | 9.100E-06 |
| 8007 | 401 | S401Q | A | 5.1e-06 | 6e-06 | 7.1e-06 | 6.067E-06 | 1.002E-06 | 5.100E-06 | 7.100E-06 |
| 8008 | 401 | S401G | A | 8.9e-06 | 5.7e-06 | 1.9e-05 | 1.120E-05 | 6.942E-06 | 5.700E-06 | 1.900E-05 |
| 8009 | 401 | S401H | A | 1.1e-05 | 9.7e-06 | 6.2e-06 | 8.967E-06 | 2.483E-06 | 6.200E-06 | 1.100E-05 |
| 8010 | 401 | S401I | A | 1.1e-05 | 9.3e-06 | 6.9e-06 | 9.067E-06 | 2.060E-06 | 6.900E-06 | 1.100E-05 |
| 8011 | 401 | S401L | A | 5.2e-06 | 6.9e-06 | 7.2e-06 | 6.433E-06 | 1.079E-06 | 5.200E-06 | 7.200E-06 |
| 8012 | 401 | S401K | A | 7.4e-06 | 7.2e-06 | 1.5e-05 | 9.867E-06 | 4.447E-06 | 7.200E-06 | 1.500E-05 |
| 8013 | 401 | S401M | A | 7e-06 | 4.8e-06 | 8.9e-06 | 6.900E-06 | 2.052E-06 | 4.800E-06 | 8.900E-06 |
| 8014 | 401 | S401F | A | 9.4e-06 | 7.4e-06 | 5.6e-06 | 7.467E-06 | 1.901E-06 | 5.600E-06 | 9.400E-06 |
| 8015 | 401 | S401P | A | 7.1e-06 | 7.7e-06 | 4.2e-06 | 6.333E-06 | 1.872E-06 | 4.200E-06 | 7.700E-06 |
| 8016 | 401 | S401S | A | 8e-06 | 8.3e-06 | 9.3e-06 | 8.533E-06 | 6.807E-07 | 8.000E-06 | 9.300E-06 |
| 8017 | 401 | S401T | A | 1e-05 | 4.1e-06 | 8.6e-06 | 7.567E-06 | 3.083E-06 | 4.100E-06 | 1.000E-05 |
| 8018 | 401 | S401W | A | 7.2e-06 | 9e-06 | 6.8e-06 | 7.667E-06 | 1.172E-06 | 6.800E-06 | 9.000E-06 |
| 8019 | 401 | S401Y | A | 5.5e-06 | 6.8e-06 | 7.2e-06 | 6.500E-06 | 8.888E-07 | 5.500E-06 | 7.200E-06 |
| 8020 | 401 | S401V | A | 6.7e-06 | 6.5e-06 | 1.2e-05 | 8.400E-06 | 3.119E-06 | 6.500E-06 | 1.200E-05 |
| 8021 | 402 | V402A | A | 1.4e-05 | 5.4e-06 | 8.4e-06 | 9.267E-06 | 4.365E-06 | 5.400E-06 | 1.400E-05 |
| 8022 | 402 | V402R | A | 1e-05 | 4.9e-06 | 1e-05 | 8.300E-06 | 2.944E-06 | 4.900E-06 | 1.000E-05 |
| 8023 | 402 | V402N | A | 5.8e-06 | 8e-06 | 1.2e-05 | 8.600E-06 | 3.143E-06 | 5.800E-06 | 1.200E-05 |
| 8024 | 402 | V402D | A | 1e-05 | 6.2e-06 | 7.7e-06 | 7.967E-06 | 1.914E-06 | 6.200E-06 | 1.000E-05 |
| 8025 | 402 | V402C | A | 1e-05 | 5.4e-06 | 5.8e-06 | 7.067E-06 | 2.548E-06 | 5.400E-06 | 1.000E-05 |
| 8026 | 402 | V402E | A | 6.3e-06 | 9.6e-06 | 6.5e-06 | 7.467E-06 | 1.850E-06 | 6.300E-06 | 9.600E-06 |
| 8027 | 402 | V402Q | A | 7.2e-06 | 9.9e-06 | 7.1e-06 | 8.067E-06 | 1.589E-06 | 7.100E-06 | 9.900E-06 |
| 8028 | 402 | V402G | A | 1.2e-05 | 7.1e-06 | 9.9e-06 | 9.667E-06 | 2.458E-06 | 7.100E-06 | 1.200E-05 |
| 8029 | 402 | V402H | A | 1.4e-05 | 5.7e-06 | 9.7e-06 | 9.800E-06 | 4.151E-06 | 5.700E-06 | 1.400E-05 |
| 8030 | 402 | V402I | A | 1.6e-05 | 6.1e-06 | 7e-06 | 9.700E-06 | 5.474E-06 | 6.100E-06 | 1.600E-05 |
| 8031 | 402 | V402L | A | 9.8e-06 | 1.1e-05 | 5.5e-06 | 8.767E-06 | 2.892E-06 | 5.500E-06 | 1.100E-05 |
| 8032 | 402 | V402K | A | 5e-06 | 8.8e-06 | 6.2e-06 | 6.667E-06 | 1.943E-06 | 5.000E-06 | 8.800E-06 |
| 8033 | 402 | V402M | A | 9.6e-06 | 7.9e-06 | 9.5e-06 | 9.000E-06 | 9.539E-07 | 7.900E-06 | 9.600E-06 |
| 8034 | 402 | V402F | A | 6.1e-06 | 7.7e-06 | 7.1e-06 | 6.967E-06 | 8.083E-07 | 6.100E-06 | 7.700E-06 |
| 8035 | 402 | V402P | A | 8.6e-06 | 1.2e-05 | 8.9e-06 | 9.833E-06 | 1.882E-06 | 8.600E-06 | 1.200E-05 |
| 8036 | 402 | V402S | A | 4.4e-06 | 6.2e-06 | 1.2e-05 | 7.533E-06 | 3.972E-06 | 4.400E-06 | 1.200E-05 |
| 8037 | 402 | V402T | A | 8.4e-06 | 6.6e-06 | 6.3e-06 | 7.100E-06 | 1.136E-06 | 6.300E-06 | 8.400E-06 |
| 8038 | 402 | V402W | A | 1.7e-05 | 6.2e-06 | 1e-05 | 1.107E-05 | 5.478E-06 | 6.200E-06 | 1.700E-05 |
| 8039 | 402 | V402Y | A | 6.4e-06 | 9.8e-06 | 1.4e-05 | 1.007E-05 | 3.807E-06 | 6.400E-06 | 1.400E-05 |
| 8040 | 402 | V402V | A | 9.7e-06 | 1.1e-05 | 1.3e-05 | 1.123E-05 | 1.662E-06 | 9.700E-06 | 1.300E-05 |
| 8041 | 403 | F403A | A | 9.4e-06 | 7.8e-06 | 1.3e-05 | 1.007E-05 | 2.663E-06 | 7.800E-06 | 1.300E-05 |
| 8042 | 403 | F403R | A | 6.7e-06 | 7.4e-06 | 6.1e-06 | 6.733E-06 | 6.506E-07 | 6.100E-06 | 7.400E-06 |
| 8043 | 403 | F403N | A | 6.2e-06 | 4.4e-06 | 9.7e-06 | 6.767E-06 | 2.695E-06 | 4.400E-06 | 9.700E-06 |
| 8044 | 403 | F403D | A | 8.7e-06 | 6.1e-06 | 5.9e-06 | 6.900E-06 | 1.562E-06 | 5.900E-06 | 8.700E-06 |
| 8045 | 403 | F403C | A | 9.8e-06 | 4.8e-06 | 9.9e-06 | 8.167E-06 | 2.916E-06 | 4.800E-06 | 9.900E-06 |
| 8046 | 403 | F403E | A | 9.9e-06 | 1.3e-05 | 8.7e-06 | 1.053E-05 | 2.219E-06 | 8.700E-06 | 1.300E-05 |
| 8047 | 403 | F403Q | A | 8.9e-06 | 3.9e-06 | 8.6e-06 | 7.133E-06 | 2.804E-06 | 3.900E-06 | 8.900E-06 |
| 8048 | 403 | F403G | A | 7.3e-06 | 8.8e-06 | 1.1e-05 | 9.033E-06 | 1.861E-06 | 7.300E-06 | 1.100E-05 |
| 8049 | 403 | F403H | A | 6.1e-06 | 5e-06 | 4.2e-06 | 5.100E-06 | 9.539E-07 | 4.200E-06 | 6.100E-06 |
| 8050 | 403 | F403I | A | 8.8e-06 | 8.9e-06 | 9.1e-06 | 8.933E-06 | 1.528E-07 | 8.800E-06 | 9.100E-06 |
| 8051 | 403 | F403L | A | 1.3e-05 | 4.8e-06 | 8.4e-06 | 8.733E-06 | 4.110E-06 | 4.800E-06 | 1.300E-05 |
| 8052 | 403 | F403K | A | 6.5e-06 | 8.6e-06 | 5.8e-06 | 6.967E-06 | 1.457E-06 | 5.800E-06 | 8.600E-06 |
| 8053 | 403 | F403M | A | 7.6e-06 | 1.2e-05 | 6.6e-06 | 8.733E-06 | 2.873E-06 | 6.600E-06 | 1.200E-05 |
| 8054 | 403 | F403F | A | 6e-06 | 8e-06 | 5.4e-06 | 6.467E-06 | 1.361E-06 | 5.400E-06 | 8.000E-06 |
| 8055 | 403 | F403P | A | 1.1e-05 | 5.1e-06 | 8.8e-06 | 8.300E-06 | 2.982E-06 | 5.100E-06 | 1.100E-05 |
| 8056 | 403 | F403S | A | 9.4e-06 | 8.8e-06 | 8.2e-06 | 8.800E-06 | 6.000E-07 | 8.200E-06 | 9.400E-06 |
| 8057 | 403 | F403T | A | 5.1e-06 | 7e-06 | 9.3e-06 | 7.133E-06 | 2.103E-06 | 5.100E-06 | 9.300E-06 |
| 8058 | 403 | F403W | A | 9.2e-06 | 5.7e-06 | 1.1e-05 | 8.633E-06 | 2.695E-06 | 5.700E-06 | 1.100E-05 |
| 8059 | 403 | F403Y | A | 5.3e-06 | 8.3e-06 | 9.8e-06 | 7.800E-06 | 2.291E-06 | 5.300E-06 | 9.800E-06 |
| 8060 | 403 | F403V | A | 4.5e-06 | 8.7e-06 | 8.3e-06 | 7.167E-06 | 2.318E-06 | 4.500E-06 | 8.700E-06 |
| 8061 | 404 | Q404A | A | 6.8e-06 | 1.2e-05 | 1e-05 | 9.600E-06 | 2.623E-06 | 6.800E-06 | 1.200E-05 |
| 8062 | 404 | Q404R | A | 8.5e-06 | 1.4e-05 | 1.3e-05 | 1.183E-05 | 2.930E-06 | 8.500E-06 | 1.400E-05 |
| 8063 | 404 | Q404N | A | 7.5e-06 | 1.4e-05 | 5.8e-06 | 9.100E-06 | 4.328E-06 | 5.800E-06 | 1.400E-05 |
| 8064 | 404 | Q404D | A | 5.5e-06 | 1.3e-05 | 1e-05 | 9.500E-06 | 3.775E-06 | 5.500E-06 | 1.300E-05 |
| 8065 | 404 | Q404C | A | 4.7e-06 | 5.8e-06 | 4.7e-06 | 5.067E-06 | 6.351E-07 | 4.700E-06 | 5.800E-06 |
| 8066 | 404 | Q404E | A | 6.8e-06 | 8.8e-06 | 1.4e-05 | 9.867E-06 | 3.717E-06 | 6.800E-06 | 1.400E-05 |
| 8067 | 404 | Q404Q | A | 6.3e-06 | 1.2e-05 | 8.5e-06 | 8.933E-06 | 2.875E-06 | 6.300E-06 | 1.200E-05 |
| 8068 | 404 | Q404G | A | 8e-06 | 8e-06 | 9.9e-06 | 8.633E-06 | 1.097E-06 | 8.000E-06 | 9.900E-06 |
| 8069 | 404 | Q404H | A | 7.2e-06 | 1.8e-05 | 1.6e-05 | 1.373E-05 | 5.746E-06 | 7.200E-06 | 1.800E-05 |
| 8070 | 404 | Q404I | A | 8.4e-06 | 5.3e-06 | 4.2e-06 | 5.967E-06 | 2.178E-06 | 4.200E-06 | 8.400E-06 |
| 8071 | 404 | Q404L | A | 6.5e-06 | 7.2e-06 | 6.4e-06 | 6.700E-06 | 4.359E-07 | 6.400E-06 | 7.200E-06 |
| 8072 | 404 | Q404K | A | 1.2e-05 | 7e-06 | 6.8e-06 | 8.600E-06 | 2.946E-06 | 6.800E-06 | 1.200E-05 |
| 8073 | 404 | Q404M | A | 1.1e-05 | 1.1e-05 | 6e-06 | 9.333E-06 | 2.887E-06 | 6.000E-06 | 1.100E-05 |
| 8074 | 404 | Q404F | A | 1.3e-05 | 9.6e-06 | 9.7e-06 | 1.077E-05 | 1.935E-06 | 9.600E-06 | 1.300E-05 |

| | | | | | | | | | | |
|---|---|---|---|---|---|---|---|---|---|---|
| 8075 | 404 | Q404P | A | 8.3e-06 | 7e-06 | 6.4e-06 | 7.233E-06 | 9.713E-07 | 6.400E-06 | 8.300E-06 |
| 8076 | 404 | Q404S | A | 8.4e-06 | 9.8e-06 | 7e-06 | 8.400E-06 | 1.400E-06 | 7.000E-06 | 9.800E-06 |
| 8077 | 404 | Q404T | A | 7.4e-06 | 8.3e-06 | 5.2e-06 | 6.967E-06 | 1.595E-06 | 5.200E-06 | 8.300E-06 |
| 8078 | 404 | Q404W | A | 7.4e-06 | 6.2e-06 | 5.5e-06 | 6.367E-06 | 9.609E-07 | 5.500E-06 | 7.400E-06 |
| 8079 | 404 | Q404Y | A | 6.2e-06 | 1e-05 | 4.5e-06 | 6.900E-06 | 2.816E-06 | 4.500E-06 | 1.000E-05 |
| 8080 | 404 | Q404V | A | 4.7e-06 | 7.6e-06 | 7.2e-06 | 6.500E-06 | 1.572E-06 | 4.700E-06 | 7.600E-06 |
| 8081 | 405 | N405A | A | 9.7e-06 | 7.5e-06 | 5e-06 | 7.400E-06 | 2.352E-06 | 5.000E-06 | 9.700E-06 |
| 8082 | 405 | N405R | A | 6.4e-06 | 4.9e-06 | 1.3e-05 | 8.100E-06 | 4.309E-06 | 4.900E-06 | 1.300E-05 |
| 8083 | 405 | N405N | A | 4.2e-06 | 6.5e-06 | 9.1e-06 | 6.600E-06 | 2.452E-06 | 4.200E-06 | 9.100E-06 |
| 8084 | 405 | N405D | A | 7.7e-06 | 6.6e-06 | 1.2e-05 | 8.767E-06 | 2.854E-06 | 6.600E-06 | 1.200E-05 |
| 8085 | 405 | N405C | A | 1e-05 | 7.3e-06 | 8.4e-06 | 8.567E-06 | 1.358E-06 | 7.300E-06 | 1.000E-05 |
| 8086 | 405 | N405E | A | 7.7e-06 | 5.8e-06 | 5.9e-06 | 6.467E-06 | 1.069E-06 | 5.800E-06 | 7.700E-06 |
| 8087 | 405 | N405Q | A | 8.3e-06 | 7.5e-06 | 1.5e-05 | 1.027E-05 | 4.119E-06 | 7.500E-06 | 1.500E-05 |
| 8088 | 405 | N405G | A | 4.4e-06 | 7.5e-06 | 7.2e-06 | 6.367E-06 | 1.710E-06 | 4.400E-06 | 7.500E-06 |
| 8089 | 405 | N405H | A | 8.7e-06 | 9.6e-06 | 1.2e-05 | 1.010E-05 | 1.706E-06 | 8.700E-06 | 1.200E-05 |
| 8090 | 405 | N405I | A | 9.3e-06 | 1.2e-05 | 4.9e-06 | 8.733E-06 | 3.584E-06 | 4.900E-06 | 1.200E-05 |
| 8091 | 405 | N405L | A | 8.3e-06 | 1.4e-05 | 7.6e-06 | 9.967E-06 | 3.510E-06 | 7.600E-06 | 1.400E-05 |
| 8092 | 405 | N405K | A | 7.2e-06 | 5.1e-06 | 9.2e-06 | 7.167E-06 | 2.050E-06 | 5.100E-06 | 9.200E-06 |
| 8093 | 405 | N405M | A | 1.2e-05 | 4.5e-06 | 6.6e-06 | 7.700E-06 | 3.869E-06 | 4.500E-06 | 1.200E-05 |
| 8094 | 405 | N405F | A | 5.2e-06 | 1.3e-05 | 5.9e-06 | 8.033E-06 | 4.315E-06 | 5.200E-06 | 1.300E-05 |
| 8095 | 405 | N405P | A | 6.1e-06 | 9.8e-06 | 8.1e-06 | 8.000E-06 | 1.852E-06 | 6.100E-06 | 9.800E-06 |
| 8096 | 405 | N405S | A | 4.9e-06 | 7e-06 | 1.1e-05 | 7.633E-06 | 3.099E-06 | 4.900E-06 | 1.100E-05 |
| 8097 | 405 | N405T | A | 7.4e-06 | 6.9e-06 | 9.7e-06 | 8.000E-06 | 1.493E-06 | 6.900E-06 | 9.700E-06 |
| 8098 | 405 | N405W | A | 1.1e-05 | 1.1e-05 | 9.3e-06 | 1.043E-05 | 9.815E-07 | 9.300E-06 | 1.100E-05 |
| 8099 | 405 | N405Y | A | 6.5e-06 | 9.2e-06 | 1.1e-05 | 8.900E-06 | 2.265E-06 | 6.500E-06 | 1.100E-05 |
| 8100 | 405 | N405V | A | 9.1e-06 | 1.1e-05 | 8.5e-06 | 9.533E-06 | 1.305E-06 | 8.500E-06 | 1.100E-05 |
| 8101 | 406 | L406A | A | 8.2e-06 | 6.4e-06 | 5.6e-06 | 6.733E-06 | 1.332E-06 | 5.600E-06 | 8.200E-06 |
| 8102 | 406 | L406R | A | 5.4e-06 | 4e-06 | 1.1e-05 | 6.800E-06 | 3.704E-06 | 4.000E-06 | 1.100E-05 |
| 8103 | 406 | L406N | A | 9.7e-06 | 8.4e-06 | 6.6e-06 | 8.233E-06 | 1.557E-06 | 6.600E-06 | 9.700E-06 |
| 8104 | 406 | L406D | A | 9e-06 | 1.2e-05 | 1e-05 | 1.033E-05 | 1.528E-06 | 9.000E-06 | 1.200E-05 |
| 8105 | 406 | L406C | A | 9e-06 | 2.7e-06 | 1.8e-05 | 9.900E-06 | 7.690E-06 | 2.700E-06 | 1.800E-05 |
| 8106 | 406 | L406E | A | 5.9e-06 | 4.2e-06 | 4.7e-06 | 4.933E-06 | 8.737E-07 | 4.200E-06 | 5.900E-06 |
| 8107 | 406 | L406Q | A | 1.1e-05 | 1.4e-05 | 1.1e-05 | 1.200E-05 | 1.732E-06 | 1.100E-05 | 1.400E-05 |
| 8108 | 406 | L406G | A | 5.4e-06 | 1.1e-05 | 8.4e-06 | 8.267E-06 | 2.802E-06 | 5.400E-06 | 1.100E-05 |
| 8109 | 406 | L406H | A | 1e-05 | 6.1e-06 | 1.2e-05 | 9.367E-06 | 3.001E-06 | 6.100E-06 | 1.200E-05 |
| 8110 | 406 | L406I | A | 1.3e-05 | 8.5e-06 | 1.4e-05 | 1.183E-05 | 2.930E-06 | 8.500E-06 | 1.400E-05 |
| 8111 | 406 | L406L | A | 7.4e-06 | 1.3e-05 | 4.3e-06 | 8.233E-06 | 4.409E-06 | 4.300E-06 | 1.300E-05 |
| 8112 | 406 | L406K | A | 3.3e-06 | 8.1e-06 | 6.2e-06 | 5.867E-06 | 2.417E-06 | 3.300E-06 | 8.100E-06 |
| 8113 | 406 | L406M | A | 9.8e-06 | 8.5e-06 | 9.8e-06 | 9.367E-06 | 7.506E-07 | 8.500E-06 | 9.800E-06 |
| 8114 | 406 | L406F | A | 7e-06 | 6.6e-06 | 9.9e-06 | 7.833E-06 | 1.801E-06 | 6.600E-06 | 9.900E-06 |
| 8115 | 406 | L406P | A | 9.4e-06 | 5.3e-06 | 6.7e-06 | 7.133E-06 | 2.084E-06 | 5.300E-06 | 9.400E-06 |
| 8116 | 406 | L406S | A | 8.3e-06 | 8.7e-06 | 9.1e-06 | 8.700E-06 | 4.000E-07 | 8.300E-06 | 9.100E-06 |
| 8117 | 406 | L406T | A | 5.3e-06 | 1.3e-05 | 1e-05 | 9.433E-06 | 3.881E-06 | 5.300E-06 | 1.300E-05 |
| 8118 | 406 | L406W | A | 7.8e-06 | 5.1e-06 | 5.6e-06 | 6.167E-06 | 1.436E-06 | 5.100E-06 | 7.800E-06 |
| 8119 | 406 | L406Y | A | 3.6e-06 | 3.9e-06 | 8e-06 | 5.167E-06 | 2.458E-06 | 3.600E-06 | 8.000E-06 |
| 8120 | 406 | L406V | A | 5.2e-06 | 8.5e-06 | 9.4e-06 | 7.700E-06 | 2.211E-06 | 5.200E-06 | 9.400E-06 |
| 8121 | 407 | Q407A | A | 1.1e-05 | 1.2e-05 | 1.3e-05 | 1.200E-05 | 1.000E-06 | 1.100E-05 | 1.300E-05 |
| 8122 | 407 | Q407R | A | 6.6e-06 | 8.7e-06 | 7.3e-06 | 7.533E-06 | 1.069E-06 | 6.600E-06 | 8.700E-06 |
| 8123 | 407 | Q407N | A | 9.2e-06 | 7.8e-06 | 5.8e-06 | 7.600E-06 | 1.709E-06 | 5.800E-06 | 9.200E-06 |
| 8124 | 407 | Q407D | A | 1e-05 | 1.2e-05 | 6.2e-06 | 9.400E-06 | 2.946E-06 | 6.200E-06 | 1.200E-05 |
| 8125 | 407 | Q407C | A | 5.5e-06 | 1.1e-05 | 6.5e-06 | 7.667E-06 | 2.930E-06 | 5.500E-06 | 1.100E-05 |
| 8126 | 407 | Q407E | A | 8.8e-06 | 6.1e-06 | 5.9e-06 | 6.933E-06 | 1.620E-06 | 5.900E-06 | 8.800E-06 |
| 8127 | 407 | Q407Q | A | 1.2e-05 | 9.4e-06 | 4.9e-06 | 8.767E-06 | 3.592E-06 | 4.900E-06 | 1.200E-05 |
| 8128 | 407 | Q407G | A | 6.4e-06 | 6.4e-06 | 9.7e-06 | 7.500E-06 | 1.905E-06 | 6.400E-06 | 9.700E-06 |
| 8129 | 407 | Q407H | A | 4.7e-06 | 6.3e-06 | 5.5e-06 | 5.500E-06 | 8.000E-07 | 4.700E-06 | 6.300E-06 |
| 8130 | 407 | Q407I | A | 5.6e-06 | 8.7e-06 | 9.8e-06 | 8.033E-06 | 2.178E-06 | 5.600E-06 | 9.800E-06 |
| 8131 | 407 | Q407L | A | 1.2e-05 | 6.9e-06 | 9.8e-06 | 9.567E-06 | 2.558E-06 | 6.900E-06 | 1.200E-05 |
| 8132 | 407 | Q407K | A | 5.4e-06 | 5.6e-06 | 6.2e-06 | 5.733E-06 | 4.163E-07 | 5.400E-06 | 6.200E-06 |
| 8133 | 407 | Q407M | A | 8.2e-06 | 8.5e-06 | 9.8e-06 | 8.833E-06 | 8.505E-07 | 8.200E-06 | 9.800E-06 |
| 8134 | 407 | Q407F | A | 5.1e-06 | 9.1e-06 | 1e-05 | 8.067E-06 | 2.608E-06 | 5.100E-06 | 1.000E-05 |
| 8135 | 407 | Q407P | A | 8.7e-06 | 9.5e-06 | 7.4e-06 | 8.533E-06 | 1.060E-06 | 7.400E-06 | 9.500E-06 |
| 8136 | 407 | Q407S | A | 8.6e-06 | 5.5e-06 | 1.3e-05 | 9.033E-06 | 3.769E-06 | 5.500E-06 | 1.300E-05 |
| 8137 | 407 | Q407T | A | 9.5e-06 | 9.4e-06 | 7.5e-06 | 8.800E-06 | 1.127E-06 | 7.500E-06 | 9.500E-06 |
| 8138 | 407 | Q407W | A | 8.8e-06 | 5.5e-06 | 7.4e-06 | 7.233E-06 | 1.656E-06 | 5.500E-06 | 8.800E-06 |
| 8139 | 407 | Q407Y | A | 6.6e-06 | 1.2e-05 | 7.8e-06 | 8.800E-06 | 2.835E-06 | 6.600E-06 | 1.200E-05 |
| 8140 | 407 | Q407V | A | 9e-06 | 1.1e-05 | 3.6e-06 | 7.867E-06 | 3.828E-06 | 3.600E-06 | 1.100E-05 |
| 8141 | 408 | V408A | A | 8.4e-06 | 1e-05 | 5.3e-06 | 7.900E-06 | 2.390E-06 | 5.300E-06 | 1.000E-05 |
| 8142 | 408 | V408R | A | 6.2e-06 | 8.8e-06 | 7.7e-06 | 7.567E-06 | 1.305E-06 | 6.200E-06 | 8.800E-06 |
| 8143 | 408 | V408N | A | 9e-06 | 9.6e-06 | 6.1e-06 | 8.233E-06 | 1.872E-06 | 6.100E-06 | 9.600E-06 |
| 8144 | 408 | V408D | A | 9.5e-06 | 7.5e-06 | 3.7e-06 | 6.900E-06 | 2.946E-06 | 3.700E-06 | 9.500E-06 |
| 8145 | 408 | V408C | A | 7.5e-06 | 4.9e-06 | 6.8e-06 | 6.400E-06 | 1.345E-06 | 4.900E-06 | 7.500E-06 |
| 8146 | 408 | V408E | A | 6.3e-06 | 6.6e-06 | 9.1e-06 | 7.333E-06 | 1.537E-06 | 6.300E-06 | 9.100E-06 |
| 8147 | 408 | V408Q | A | 6.1e-06 | 1.3e-05 | 1.3e-05 | 1.070E-05 | 3.984E-06 | 6.100E-06 | 1.300E-05 |
| 8148 | 408 | V408G | A | 1.2e-05 | 5.7e-06 | 7e-06 | 8.233E-06 | 3.326E-06 | 5.700E-06 | 1.200E-05 |
| 8149 | 408 | V408H | A | 7.3e-06 | 7.7e-06 | 6.7e-06 | 7.233E-06 | 5.033E-07 | 6.700E-06 | 7.700E-06 |
| 8150 | 408 | V408I | A | 6.7e-06 | 1.3e-05 | 9.5e-06 | 9.733E-06 | 3.156E-06 | 6.700E-06 | 1.300E-05 |
| 8151 | 408 | V408L | A | 8.8e-06 | 7.9e-06 | 1e-05 | 8.900E-06 | 1.054E-06 | 7.900E-06 | 1.000E-05 |
| 8152 | 408 | V408K | A | 1.1e-05 | 1e-05 | 4.8e-06 | 8.600E-06 | 3.329E-06 | 4.800E-06 | 1.100E-05 |
| 8153 | 408 | V408M | A | 6e-06 | 6.9e-06 | 6.6e-06 | 6.500E-06 | 4.583E-07 | 6.000E-06 | 6.900E-06 |
| 8154 | 408 | V408F | A | 8.5e-06 | 4.6e-06 | 7.7e-06 | 6.933E-06 | 2.060E-06 | 4.600E-06 | 8.500E-06 |
| 8155 | 408 | V408P | A | 6e-06 | 1.4e-05 | 1.3e-05 | 1.100E-05 | 4.359E-06 | 6.000E-06 | 1.400E-05 |
| 8156 | 408 | V408S | A | 9.2e-06 | 5.9e-06 | 6.5e-06 | 7.200E-06 | 1.758E-06 | 5.900E-06 | 9.200E-06 |
| 8157 | 408 | V408T | A | 1.2e-05 | 8.3e-06 | 9.7e-06 | 1.000E-05 | 1.868E-06 | 8.300E-06 | 1.200E-05 |
| 8158 | 408 | V408W | A | 1.1e-05 | 9.5e-06 | 1.2e-05 | 1.083E-05 | 1.258E-06 | 9.500E-06 | 1.200E-05 |
| 8159 | 408 | V408Y | A | 6.4e-06 | 8.5e-06 | 9.6e-06 | 8.167E-06 | 1.626E-06 | 6.400E-06 | 9.600E-06 |

| | | | | | | | | | | |
|---|---|---|---|---|---|---|---|---|---|---|
| 8160 | 408 | V408V | A | 1.2e-05 | 6.7e-06 | 6.2e-06 | 8.300E-06 | 3.214E-06 | 6.200E-06 | 1.200E-05 |
| 8161 | 409 | I409A | A | 1.2e-05 | 1.3e-05 | 9.7e-06 | 1.157E-05 | 1.692E-06 | 9.700E-06 | 1.300E-05 |
| 8162 | 409 | I409R | A | 9.9e-06 | 6.4e-06 | 1.4e-05 | 1.010E-05 | 3.804E-06 | 6.400E-06 | 1.400E-05 |
| 8163 | 409 | I409N | A | 1.1e-05 | 9.2e-06 | 8.1e-06 | 9.433E-06 | 1.464E-06 | 8.100E-06 | 1.100E-05 |
| 8164 | 409 | I409D | A | 6.7e-06 | 6.9e-06 | 9.8e-06 | 7.800E-06 | 1.735E-06 | 6.700E-06 | 9.800E-06 |
| 8165 | 409 | I409C | A | 4.5e-06 | 5.1e-06 | 6.4e-06 | 5.333E-06 | 9.713E-07 | 4.500E-06 | 6.400E-06 |
| 8166 | 409 | I409E | A | 7.3e-06 | 7.5e-06 | 5.7e-06 | 6.833E-06 | 9.866E-07 | 5.700E-06 | 7.500E-06 |
| 8167 | 409 | I409Q | A | 7.8e-06 | 9.8e-06 | 9.7e-06 | 9.100E-06 | 1.127E-06 | 7.800E-06 | 9.800E-06 |
| 8168 | 409 | I409G | A | 1.1e-05 | 6.3e-06 | 6e-06 | 7.767E-06 | 2.804E-06 | 6.000E-06 | 1.100E-05 |
| 8169 | 409 | I409H | A | 6e-06 | 7.4e-06 | 9e-06 | 7.467E-06 | 1.501E-06 | 6.000E-06 | 9.000E-06 |
| 8170 | 409 | I409I | A | 5.3e-06 | 5.2e-06 | 5.1e-06 | 5.200E-06 | 1.000E-07 | 5.100E-06 | 5.300E-06 |
| 8171 | 409 | I409L | A | 6.9e-06 | 6.1e-06 | 6.8e-06 | 6.600E-06 | 4.359E-07 | 6.100E-06 | 6.900E-06 |
| 8172 | 409 | I409K | A | 6.7e-06 | 7.2e-06 | 8.9e-06 | 7.600E-06 | 1.153E-06 | 6.700E-06 | 8.900E-06 |
| 8173 | 409 | I409M | A | 1e-05 | 4.4e-06 | 8.7e-06 | 7.700E-06 | 2.931E-06 | 4.400E-06 | 1.000E-05 |
| 8174 | 409 | I409F | A | 8.6e-06 | 3.5e-06 | 1.4e-05 | 8.700E-06 | 5.251E-06 | 3.500E-06 | 1.400E-05 |
| 8175 | 409 | I409P | A | 1e-05 | 4.1e-06 | 8e-06 | 7.367E-06 | 3.001E-06 | 4.100E-06 | 1.000E-05 |
| 8176 | 409 | I409S | A | 8.8e-06 | 1.4e-05 | 9.2e-06 | 1.067E-05 | 2.894E-06 | 8.800E-06 | 1.400E-05 |
| 8177 | 409 | I409T | A | 1.3e-05 | 9e-06 | 9.6e-06 | 1.053E-05 | 2.157E-06 | 9.000E-06 | 1.300E-05 |
| 8178 | 409 | I409W | A | 3.7e-06 | 8.9e-06 | 6.7e-06 | 6.433E-06 | 2.610E-06 | 3.700E-06 | 8.900E-06 |
| 8179 | 409 | I409Y | A | 5.3e-06 | 6.8e-06 | 1.4e-05 | 8.700E-06 | 4.651E-06 | 5.300E-06 | 1.400E-05 |
| 8180 | 409 | I409V | A | 1.1e-05 | 5.8e-06 | 1.3e-05 | 9.933E-06 | 3.717E-06 | 5.800E-06 | 1.300E-05 |
| 8181 | 410 | R410A | A | 9.7e-06 | 9e-06 | 7e-06 | 8.567E-06 | 1.401E-06 | 7.000E-06 | 9.700E-06 |
| 8182 | 410 | R410R | A | 1.2e-05 | 5.6e-06 | 8.7e-06 | 8.767E-06 | 3.201E-06 | 5.600E-06 | 1.200E-05 |
| 8183 | 410 | R410N | A | 9.1e-06 | 6e-06 | 6e-06 | 7.033E-06 | 1.790E-06 | 6.000E-06 | 9.100E-06 |
| 8184 | 410 | R410D | A | 6.1e-06 | 7.5e-06 | 1.4e-05 | 9.200E-06 | 4.215E-06 | 6.100E-06 | 1.400E-05 |
| 8185 | 410 | R410C | A | 5.2e-06 | 1.1e-05 | 6.7e-06 | 7.633E-06 | 3.011E-06 | 5.200E-06 | 1.100E-05 |
| 8186 | 410 | R410E | A | 3.9e-06 | 1.3e-05 | 1.1e-05 | 9.300E-06 | 4.782E-06 | 3.900E-06 | 1.300E-05 |
| 8187 | 410 | R410Q | A | 8.5e-06 | 9.7e-06 | 6.6e-06 | 8.267E-06 | 1.563E-06 | 6.600E-06 | 9.700E-06 |
| 8188 | 410 | R410G | A | 5.1e-06 | 9.3e-06 | 1.1e-05 | 8.467E-06 | 3.037E-06 | 5.100E-06 | 1.100E-05 |
| 8189 | 410 | R410H | A | 9.3e-06 | 7.1e-06 | 1e-05 | 8.800E-06 | 1.513E-06 | 7.100E-06 | 1.000E-05 |
| 8190 | 410 | R410I | A | 7.3e-06 | 1e-05 | 6.1e-06 | 7.800E-06 | 1.997E-06 | 6.100E-06 | 1.000E-05 |
| 8191 | 410 | R410L | A | 7.1e-06 | 5.5e-06 | 9e-06 | 7.200E-06 | 1.752E-06 | 5.500E-06 | 9.000E-06 |
| 8192 | 410 | R410K | A | 7.3e-06 | 7.7e-06 | 8.5e-06 | 7.833E-06 | 6.110E-07 | 7.300E-06 | 8.500E-06 |
| 8193 | 410 | R410M | A | 7.6e-06 | 6.6e-06 | 6.5e-06 | 6.900E-06 | 6.083E-07 | 6.500E-06 | 7.600E-06 |
| 8194 | 410 | R410F | A | 8.2e-06 | 1.3e-05 | 1.1e-05 | 1.073E-05 | 2.411E-06 | 8.200E-06 | 1.300E-05 |
| 8195 | 410 | R410P | A | 6.3e-06 | 5.6e-06 | 6.3e-06 | 6.067E-06 | 4.041E-07 | 5.600E-06 | 6.300E-06 |
| 8196 | 410 | R410S | A | 9.3e-06 | 7.9e-06 | 1e-05 | 9.067E-06 | 1.069E-06 | 7.900E-06 | 1.000E-05 |
| 8197 | 410 | R410T | A | 6.7e-06 | 1e-05 | 5.9e-06 | 7.533E-06 | 2.173E-06 | 5.900E-06 | 1.000E-05 |
| 8198 | 410 | R410W | A | 6.3e-06 | 6.7e-06 | 5.3e-06 | 6.100E-06 | 7.211E-07 | 5.300E-06 | 6.700E-06 |
| 8199 | 410 | R410Y | A | 1.2e-05 | 9.2e-06 | 5.6e-06 | 8.933E-06 | 3.208E-06 | 5.600E-06 | 1.200E-05 |
| 8200 | 410 | R410V | A | 6.6e-06 | 4.4e-06 | 7.6e-06 | 6.200E-06 | 1.637E-06 | 4.400E-06 | 7.600E-06 |
| 8201 | 411 | G411A | A | 8e-06 | 8.4e-06 | 1.2e-05 | 9.467E-06 | 2.203E-06 | 8.000E-06 | 1.200E-05 |
| 8202 | 411 | G411R | A | 1.5e-05 | 7.7e-06 | 5e-06 | 9.233E-06 | 5.173E-06 | 5.000E-06 | 1.500E-05 |
| 8203 | 411 | G411N | A | 7.2e-06 | 1.2e-05 | 3.9e-06 | 7.700E-06 | 4.073E-06 | 3.900E-06 | 1.200E-05 |
| 8204 | 411 | G411D | A | 9.8e-06 | 9e-06 | 3.5e-06 | 7.433E-06 | 3.430E-06 | 3.500E-06 | 9.800E-06 |
| 8205 | 411 | G411C | A | 8.6e-06 | 8.6e-06 | 8.6e-06 | 8.600E-06 | 0.000E+00 | 8.600E-06 | 8.600E-06 |
| 8206 | 411 | G411E | A | 9.3e-06 | 1.4e-05 | 6.2e-06 | 9.833E-06 | 3.927E-06 | 6.200E-06 | 1.400E-05 |
| 8207 | 411 | G411Q | A | 8.8e-06 | 8.7e-06 | 5.1e-06 | 7.533E-06 | 2.108E-06 | 5.100E-06 | 8.800E-06 |
| 8208 | 411 | G411G | A | 8.4e-06 | 5.9e-06 | 9.7e-06 | 8.000E-06 | 1.931E-06 | 5.900E-06 | 9.700E-06 |
| 8209 | 411 | G411H | A | 1e-05 | 9.4e-06 | 1e-05 | 9.800E-06 | 3.464E-07 | 9.400E-06 | 1.000E-05 |
| 8210 | 411 | G411I | A | 4.2e-06 | 4.8e-06 | 7.4e-06 | 5.467E-06 | 1.701E-06 | 4.200E-06 | 7.400E-06 |
| 8211 | 411 | G411L | A | 1.1e-05 | 1.2e-05 | 7.5e-06 | 1.017E-05 | 2.363E-06 | 7.500E-06 | 1.200E-05 |
| 8212 | 411 | G411K | A | 9.6e-06 | 5.1e-06 | 9.8e-06 | 8.167E-06 | 2.658E-06 | 5.100E-06 | 9.800E-06 |
| 8213 | 411 | G411M | A | 1.3e-05 | 1.9e-05 | 8.4e-06 | 1.347E-05 | 5.315E-06 | 8.400E-06 | 1.900E-05 |
| 8214 | 411 | G411F | A | 8.1e-06 | 7.7e-06 | 4.3e-06 | 6.700E-06 | 2.088E-06 | 4.300E-06 | 8.100E-06 |
| 8215 | 411 | G411P | A | 1.2e-05 | 8.4e-06 | 1.9e-05 | 1.313E-05 | 5.390E-06 | 8.400E-06 | 1.900E-05 |
| 8216 | 411 | G411S | A | 1.2e-05 | 9.1e-06 | 5.9e-06 | 9.000E-06 | 3.051E-06 | 5.900E-06 | 1.200E-05 |
| 8217 | 411 | G411T | A | 1.3e-05 | 5.9e-06 | 1.4e-05 | 1.097E-05 | 4.416E-06 | 5.900E-06 | 1.400E-05 |
| 8218 | 411 | G411W | A | 1e-05 | 1.4e-05 | 1e-05 | 1.133E-05 | 2.309E-06 | 1.000E-05 | 1.400E-05 |
| 8219 | 411 | G411Y | A | 5.2e-06 | 6.5e-06 | 8.2e-06 | 6.633E-06 | 1.504E-06 | 5.200E-06 | 8.200E-06 |
| 8220 | 411 | G411V | A | 9.1e-06 | 9.3e-06 | 8.2e-06 | 8.867E-06 | 5.859E-07 | 8.200E-06 | 9.300E-06 |
| 8221 | 412 | R412A | A | 5.6e-06 | 1.1e-05 | 6.5e-06 | 7.700E-06 | 2.893E-06 | 5.600E-06 | 1.100E-05 |
| 8222 | 412 | R412R | A | 1.1e-05 | 8.6e-06 | 7.6e-06 | 9.067E-06 | 1.747E-06 | 7.600E-06 | 1.100E-05 |
| 8223 | 412 | R412N | A | 4.7e-06 | 8.9e-06 | 9.7e-06 | 7.767E-06 | 2.686E-06 | 4.700E-06 | 9.700E-06 |
| 8224 | 412 | R412D | A | 9.9e-06 | 7.6e-06 | 8.1e-06 | 8.533E-06 | 1.210E-06 | 7.600E-06 | 9.900E-06 |
| 8225 | 412 | R412C | A | 1.2e-05 | 5.7e-06 | 6.1e-06 | 7.933E-06 | 3.528E-06 | 5.700E-06 | 1.200E-05 |
| 8226 | 412 | R412E | A | 9.8e-06 | 6.6e-06 | 6.9e-06 | 7.767E-06 | 1.767E-06 | 6.600E-06 | 9.800E-06 |
| 8227 | 412 | R412Q | A | 8.2e-06 | 7.4e-06 | 9.4e-06 | 8.333E-06 | 1.007E-06 | 7.400E-06 | 9.400E-06 |
| 8228 | 412 | R412G | A | 7.8e-06 | 7.8e-06 | 8.5e-06 | 8.033E-06 | 4.041E-07 | 7.800E-06 | 8.500E-06 |
| 8229 | 412 | R412H | A | 1.1e-05 | 3.8e-06 | 1.7e-05 | 1.060E-05 | 6.609E-06 | 3.800E-06 | 1.700E-05 |
| 8230 | 412 | R412I | A | 7.5e-06 | 5.5e-06 | 1.3e-05 | 8.667E-06 | 3.884E-06 | 5.500E-06 | 1.300E-05 |
| 8231 | 412 | R412L | A | 5.9e-06 | 4e-06 | 1.2e-05 | 7.300E-06 | 4.180E-06 | 4.000E-06 | 1.200E-05 |
| 8232 | 412 | R412K | A | 9.3e-06 | 4.5e-06 | 6.7e-06 | 6.833E-06 | 2.403E-06 | 4.500E-06 | 9.300E-06 |
| 8233 | 412 | R412M | A | 8e-06 | 9.7e-06 | 5.7e-06 | 7.800E-06 | 2.007E-06 | 5.700E-06 | 9.700E-06 |
| 8234 | 412 | R412F | A | 9.3e-06 | 1e-05 | 5.7e-06 | 8.333E-06 | 2.307E-06 | 5.700E-06 | 1.000E-05 |
| 8235 | 412 | R412P | A | 6.8e-06 | 9.7e-06 | 4.4e-06 | 6.967E-06 | 2.654E-06 | 4.400E-06 | 9.700E-06 |
| 8236 | 412 | R412S | A | 6.6e-06 | 1.1e-05 | 8.2e-06 | 8.600E-06 | 2.227E-06 | 6.600E-06 | 1.100E-05 |
| 8237 | 412 | R412T | A | 8.2e-06 | 9.6e-06 | 8.9e-06 | 8.900E-06 | 7.000E-07 | 8.200E-06 | 9.600E-06 |
| 8238 | 412 | R412W | A | 6.9e-06 | 6.8e-06 | 5.7e-06 | 6.467E-06 | 6.658E-07 | 5.700E-06 | 6.900E-06 |
| 8239 | 412 | R412Y | A | 5.9e-06 | 9.2e-06 | 6.3e-06 | 7.133E-06 | 1.801E-06 | 5.900E-06 | 9.200E-06 |
| 8240 | 412 | R412V | A | 5.1e-06 | 1.8e-05 | 1.3e-05 | 1.203E-05 | 6.504E-06 | 5.100E-06 | 1.800E-05 |
| 8241 | 413 | I413A | A | 8.9e-06 | 6.8e-06 | 8e-06 | 7.900E-06 | 1.054E-06 | 6.800E-06 | 8.900E-06 |
| 8242 | 413 | I413R | A | 8.5e-06 | 1.1e-05 | 6.3e-06 | 8.600E-06 | 2.352E-06 | 6.300E-06 | 1.100E-05 |
| 8243 | 413 | I413N | A | 6.7e-06 | 8.5e-06 | 1.5e-05 | 1.007E-05 | 4.366E-06 | 6.700E-06 | 1.500E-05 |
| 8244 | 413 | I413D | A | 9.1e-06 | 1.2e-05 | 7.4e-06 | 9.500E-06 | 2.326E-06 | 7.400E-06 | 1.200E-05 |

| | | | | | | | | | | |
|---|---|---|---|---|---|---|---|---|---|---|
| 8245 | 413 | I413C | A | 9.1e-06 | 7.5e-06 | 1.1e-05 | 9.200E-06 | 1.752E-06 | 7.500E-06 | 1.100E-05 |
| 8246 | 413 | I413E | A | 9.4e-06 | 1.3e-05 | 1.4e-05 | 1.213E-05 | 2.419E-06 | 9.400E-06 | 1.400E-05 |
| 8247 | 413 | I413Q | A | 8.7e-06 | 1.4e-05 | 8.7e-06 | 1.047E-05 | 3.060E-06 | 8.700E-06 | 1.400E-05 |
| 8248 | 413 | I413G | A | 1.3e-05 | 6.3e-06 | 1.4e-05 | 1.110E-05 | 4.187E-06 | 6.300E-06 | 1.400E-05 |
| 8249 | 413 | I413H | A | 5.4e-06 | 7.5e-06 | 6.3e-06 | 6.400E-06 | 1.054E-06 | 5.400E-06 | 7.500E-06 |
| 8250 | 413 | I413I | A | 6.5e-06 | 7.6e-06 | 7.9e-06 | 7.333E-06 | 7.371E-07 | 6.500E-06 | 7.900E-06 |
| 8251 | 413 | I413L | A | 6.2e-06 | 9.6e-06 | 4.5e-06 | 6.767E-06 | 2.597E-06 | 4.500E-06 | 9.600E-06 |
| 8252 | 413 | I413K | A | 6.2e-06 | 7.5e-06 | 1.1e-05 | 8.233E-06 | 2.483E-06 | 6.200E-06 | 1.100E-05 |
| 8253 | 413 | I413M | A | 7.6e-06 | 8.5e-06 | 1.1e-05 | 9.033E-06 | 1.762E-06 | 7.600E-06 | 1.100E-05 |
| 8254 | 413 | I413F | A | 7e-06 | 6e-06 | 5e-06 | 6.000E-06 | 1.000E-06 | 5.000E-06 | 7.000E-06 |
| 8255 | 413 | I413P | A | 1e-05 | 5.4e-06 | 8.6e-06 | 8.000E-06 | 2.358E-06 | 5.400E-06 | 1.000E-05 |
| 8256 | 413 | I413S | A | 1.1e-05 | 8.9e-06 | 1.1e-05 | 1.030E-05 | 1.212E-06 | 8.900E-06 | 1.100E-05 |
| 8257 | 413 | I413T | A | 5.5e-06 | 9.7e-06 | 6.5e-06 | 7.233E-06 | 2.194E-06 | 5.500E-06 | 9.700E-06 |
| 8258 | 413 | I413W | A | 1e-05 | 5e-06 | 6.6e-06 | 7.200E-06 | 2.553E-06 | 5.000E-06 | 1.000E-05 |
| 8259 | 413 | I413Y | A | 7e-06 | 8.7e-06 | 4.9e-06 | 6.867E-06 | 1.904E-06 | 4.900E-06 | 8.700E-06 |
| 8260 | 413 | I413V | A | 6.5e-06 | 8.3e-06 | 1.2e-05 | 8.933E-06 | 2.804E-06 | 6.500E-06 | 1.200E-05 |
| 8261 | 414 | L414A | A | 1.3e-05 | 6.4e-06 | 8.7e-06 | 9.367E-06 | 3.350E-06 | 6.400E-06 | 1.300E-05 |
| 8262 | 414 | L414R | A | 6e-06 | 5.6e-06 | 6.5e-06 | 6.033E-06 | 4.509E-07 | 5.600E-06 | 6.500E-06 |
| 8263 | 414 | L414N | A | 4.8e-06 | 6.8e-06 | 6.6e-06 | 6.067E-06 | 1.102E-06 | 4.800E-06 | 6.800E-06 |
| 8264 | 414 | L414D | A | 6.4e-06 | 8.3e-06 | 1.3e-05 | 9.233E-06 | 3.398E-06 | 6.400E-06 | 1.300E-05 |
| 8265 | 414 | L414C | A | 1.2e-05 | 9.3e-06 | 1.2e-05 | 1.110E-05 | 1.559E-06 | 9.300E-06 | 1.200E-05 |
| 8266 | 414 | L414E | A | 5.9e-06 | 7.2e-06 | 1.3e-05 | 8.700E-06 | 3.780E-06 | 5.900E-06 | 1.300E-05 |
| 8267 | 414 | L414Q | A | 5e-06 | 7.6e-06 | 4.3e-06 | 5.633E-06 | 1.739E-06 | 4.300E-06 | 7.600E-06 |
| 8268 | 414 | L414G | A | 6.5e-06 | 5.4e-06 | 5.1e-06 | 5.667E-06 | 7.371E-07 | 5.100E-06 | 6.500E-06 |
| 8269 | 414 | L414H | A | 4.2e-06 | 9.1e-06 | 9.6e-06 | 7.633E-06 | 2.984E-06 | 4.200E-06 | 9.600E-06 |
| 8270 | 414 | L414I | A | 2.9e-06 | 7.8e-06 | 7.8e-06 | 6.167E-06 | 2.829E-06 | 2.900E-06 | 7.800E-06 |
| 8271 | 414 | L414L | A | 8.7e-06 | 9e-06 | 1.1e-05 | 9.567E-06 | 1.250E-06 | 8.700E-06 | 1.100E-05 |
| 8272 | 414 | L414K | A | 8.3e-06 | 6e-06 | 9.8e-06 | 8.033E-06 | 1.914E-06 | 6.000E-06 | 9.800E-06 |
| 8273 | 414 | L414M | A | 1e-05 | 1.1e-05 | 4.5e-06 | 8.500E-06 | 3.500E-06 | 4.500E-06 | 1.100E-05 |
| 8274 | 414 | L414F | A | 7.5e-06 | 8.4e-06 | 1e-05 | 8.633E-06 | 1.266E-06 | 7.500E-06 | 1.000E-05 |
| 8275 | 414 | L414P | A | 9.9e-06 | 5.2e-06 | 7.3e-06 | 7.467E-06 | 2.354E-06 | 5.200E-06 | 9.900E-06 |
| 8276 | 414 | L414S | A | 8.1e-06 | 1.3e-05 | 9.5e-06 | 1.020E-05 | 2.524E-06 | 8.100E-06 | 1.300E-05 |
| 8277 | 414 | L414T | A | 5.3e-06 | 5.7e-06 | 9.1e-06 | 6.700E-06 | 2.088E-06 | 5.300E-06 | 9.100E-06 |
| 8278 | 414 | L414W | A | 8.6e-06 | 6.1e-06 | 1e-05 | 8.233E-06 | 1.976E-06 | 6.100E-06 | 1.000E-05 |
| 8279 | 414 | L414Y | A | 6.2e-06 | 1.7e-05 | 7.5e-06 | 1.023E-05 | 5.896E-06 | 6.200E-06 | 1.700E-05 |
| 8280 | 414 | L414V | A | 8e-06 | 1.4e-05 | 1e-05 | 1.067E-05 | 3.055E-06 | 8.000E-06 | 1.400E-05 |
| 8281 | 415 | H415A | A | 9.4e-06 | 4.3e-06 | 7.1e-06 | 6.933E-06 | 2.554E-06 | 4.300E-06 | 9.400E-06 |
| 8282 | 415 | H415R | A | 8.6e-06 | 6.3e-06 | 7.9e-06 | 7.600E-06 | 1.179E-06 | 6.300E-06 | 8.600E-06 |
| 8283 | 415 | H415N | A | 4.9e-06 | 1.1e-05 | 5.5e-06 | 7.133E-06 | 3.362E-06 | 4.900E-06 | 1.100E-05 |
| 8284 | 415 | H415D | A | 5e-06 | 1.1e-05 | 5e-06 | 7.000E-06 | 3.464E-06 | 5.000E-06 | 1.100E-05 |
| 8285 | 415 | H415C | A | 6.5e-06 | 9.4e-06 | 7.7e-06 | 7.867E-06 | 1.457E-06 | 6.500E-06 | 9.400E-06 |
| 8286 | 415 | H415E | A | 9.4e-06 | 9.5e-06 | 1.1e-05 | 9.967E-06 | 8.963E-07 | 9.400E-06 | 1.100E-05 |
| 8287 | 415 | H415Q | A | 6.8e-06 | 5.8e-06 | 7e-06 | 6.533E-06 | 6.429E-07 | 5.800E-06 | 7.000E-06 |
| 8288 | 415 | H415G | A | 7.2e-06 | 5.3e-06 | 4.1e-06 | 5.533E-06 | 1.563E-06 | 4.100E-06 | 7.200E-06 |
| 8289 | 415 | H415H | A | 5.4e-06 | 6.7e-06 | 1.1e-05 | 7.700E-06 | 2.931E-06 | 5.400E-06 | 1.100E-05 |
| 8290 | 415 | H415I | A | 1.5e-05 | 7e-06 | 7.3e-06 | 9.767E-06 | 4.535E-06 | 7.000E-06 | 1.500E-05 |
| 8291 | 415 | H415L | A | 7.5e-06 | 8.3e-06 | 8.1e-06 | 7.967E-06 | 4.163E-07 | 7.500E-06 | 8.300E-06 |
| 8292 | 415 | H415K | A | 1.4e-05 | 6.7e-06 | 4.9e-06 | 8.533E-06 | 4.819E-06 | 4.900E-06 | 1.400E-05 |
| 8293 | 415 | H415M | A | 1.2e-05 | 8e-06 | 1e-05 | 1.000E-05 | 2.000E-06 | 8.000E-06 | 1.200E-05 |
| 8294 | 415 | H415F | A | 1.5e-05 | 1.1e-05 | 8.1e-06 | 1.137E-05 | 3.465E-06 | 8.100E-06 | 1.500E-05 |
| 8295 | 415 | H415P | A | 1.1e-05 | 9.8e-06 | 9.8e-06 | 1.020E-05 | 6.928E-07 | 9.800E-06 | 1.100E-05 |
| 8296 | 415 | H415S | A | 1.2e-05 | 6.5e-06 | 5.4e-06 | 7.967E-06 | 3.536E-06 | 5.400E-06 | 1.200E-05 |
| 8297 | 415 | H415T | A | 1.4e-05 | 4.1e-06 | 8.5e-06 | 8.867E-06 | 4.960E-06 | 4.100E-06 | 1.400E-05 |
| 8298 | 415 | H415W | A | 5e-06 | 1e-05 | 1.1e-05 | 8.667E-06 | 3.215E-06 | 5.000E-06 | 1.100E-05 |
| 8299 | 415 | H415Y | A | 6.6e-06 | 9.5e-06 | 4.6e-06 | 6.900E-06 | 2.464E-06 | 4.600E-06 | 9.500E-06 |
| 8300 | 415 | H415V | A | 7e-06 | 8.4e-06 | 7.5e-06 | 7.633E-06 | 7.095E-07 | 7.000E-06 | 8.400E-06 |
| 8301 | 416 | N416A | A | 1e-05 | 6.2e-06 | 3.8e-06 | 6.667E-06 | 3.126E-06 | 3.800E-06 | 1.000E-05 |
| 8302 | 416 | N416R | A | 5.2e-06 | 1.1e-05 | 1e-05 | 8.733E-06 | 3.101E-06 | 5.200E-06 | 1.100E-05 |
| 8303 | 416 | N416N | A | 8.4e-06 | 9.8e-06 | 9.1e-06 | 9.100E-06 | 7.000E-07 | 8.400E-06 | 9.800E-06 |
| 8304 | 416 | N416D | A | 8.3e-06 | 5.6e-06 | 1.3e-05 | 8.967E-06 | 3.745E-06 | 5.600E-06 | 1.300E-05 |
| 8305 | 416 | N416C | A | 7e-06 | 9.9e-06 | 4.3e-06 | 7.067E-06 | 2.801E-06 | 4.300E-06 | 9.900E-06 |
| 8306 | 416 | N416E | A | 1.1e-05 | 1.3e-05 | 6.9e-06 | 1.030E-05 | 3.110E-06 | 6.900E-06 | 1.300E-05 |
| 8307 | 416 | N416Q | A | 1.3e-05 | 1e-05 | 7.3e-06 | 1.010E-05 | 2.851E-06 | 7.300E-06 | 1.300E-05 |
| 8308 | 416 | N416G | A | 4.1e-06 | 1.1e-05 | 6.6e-06 | 7.233E-06 | 3.493E-06 | 4.100E-06 | 1.100E-05 |
| 8309 | 416 | N416H | A | 8.5e-06 | 5.2e-06 | 9.9e-06 | 7.867E-06 | 2.413E-06 | 5.200E-06 | 9.900E-06 |
| 8310 | 416 | N416I | A | 7.5e-06 | 7.7e-06 | 9.5e-06 | 8.233E-06 | 1.102E-06 | 7.500E-06 | 9.500E-06 |
| 8311 | 416 | N416L | A | 7.1e-06 | 1.1e-05 | 8.8e-06 | 8.967E-06 | 1.955E-06 | 7.100E-06 | 1.100E-05 |
| 8312 | 416 | N416K | A | 8.5e-06 | 1.4e-05 | 3.3e-06 | 8.600E-06 | 5.351E-06 | 3.300E-06 | 1.400E-05 |
| 8313 | 416 | N416M | A | 9.9e-06 | 8e-06 | 7.1e-06 | 8.333E-06 | 1.429E-06 | 7.100E-06 | 9.900E-06 |
| 8314 | 416 | N416F | A | 4.9e-06 | 4.6e-06 | 5.7e-06 | 5.067E-06 | 5.686E-07 | 4.600E-06 | 5.700E-06 |
| 8315 | 416 | N416P | A | 7.3e-06 | 6.8e-06 | 7.9e-06 | 7.333E-06 | 5.508E-07 | 6.800E-06 | 7.900E-06 |
| 8316 | 416 | N416S | A | 4.5e-06 | 9.1e-06 | 6.3e-06 | 6.633E-06 | 2.318E-06 | 4.500E-06 | 9.100E-06 |
| 8317 | 416 | N416T | A | 8.7e-06 | 3.9e-06 | 9.1e-06 | 7.233E-06 | 2.894E-06 | 3.900E-06 | 9.100E-06 |
| 8318 | 416 | N416W | A | 9.7e-06 | 1.1e-05 | 6.6e-06 | 9.100E-06 | 2.261E-06 | 6.600E-06 | 1.100E-05 |
| 8319 | 416 | N416Y | A | 7.7e-06 | 6.6e-06 | 6e-06 | 6.767E-06 | 8.622E-07 | 6.000E-06 | 7.700E-06 |
| 8320 | 416 | N416V | A | 7e-06 | 1e-05 | 6.7e-06 | 7.900E-06 | 1.825E-06 | 6.700E-06 | 1.000E-05 |
| 8321 | 417 | G417A | A | 8e-06 | 9.3e-06 | 7.9e-06 | 8.400E-06 | 7.810E-07 | 7.900E-06 | 9.300E-06 |
| 8322 | 417 | G417R | A | 6.8e-06 | 7.8e-06 | 5.1e-06 | 6.567E-06 | 1.365E-06 | 5.100E-06 | 7.800E-06 |
| 8323 | 417 | G417N | A | 7.8e-06 | 6.4e-06 | 7.7e-06 | 7.300E-06 | 7.810E-07 | 6.400E-06 | 7.800E-06 |
| 8324 | 417 | G417D | A | 6e-06 | 1.3e-05 | 6.3e-06 | 8.433E-06 | 3.958E-06 | 6.000E-06 | 1.300E-05 |
| 8325 | 417 | G417C | A | 6.8e-06 | 1.2e-05 | 7e-06 | 8.600E-06 | 2.946E-06 | 6.800E-06 | 1.200E-05 |
| 8326 | 417 | G417E | A | 5.2e-06 | 1.3e-05 | 1.1e-05 | 9.733E-06 | 4.051E-06 | 5.200E-06 | 1.300E-05 |
| 8327 | 417 | G417Q | A | 3e-06 | 9.3e-06 | 1.4e-05 | 8.767E-06 | 5.519E-06 | 3.000E-06 | 1.400E-05 |
| 8328 | 417 | G417G | A | 8.6e-06 | 5.1e-06 | 1.4e-05 | 9.233E-06 | 4.484E-06 | 5.100E-06 | 1.400E-05 |
| 8329 | 417 | G417H | A | 9.3e-06 | 8.4e-06 | 5.1e-06 | 7.600E-06 | 2.211E-06 | 5.100E-06 | 9.300E-06 |

| | | | | | | | | | | |
|---|---|---|---|---|---|---|---|---|---|---|
| 8330 | 417 | G417I | A | 6.2e-06 | 7.5e-06 | 9.8e-06 | 7.833E-06 | 1.823E-06 | 6.200E-06 | 9.800E-06 |
| 8331 | 417 | G417L | A | 9.8e-06 | 8.5e-06 | 9.7e-06 | 9.333E-06 | 7.234E-07 | 8.500E-06 | 9.800E-06 |
| 8332 | 417 | G417K | A | 6.1e-06 | 8.4e-06 | 9.6e-06 | 8.033E-06 | 1.779E-06 | 6.100E-06 | 9.600E-06 |
| 8333 | 417 | G417M | A | 6e-06 | 7.9e-06 | 3.9e-06 | 5.933E-06 | 2.001E-06 | 3.900E-06 | 7.900E-06 |
| 8334 | 417 | G417F | A | 5.8e-06 | 1.1e-05 | 1.1e-05 | 9.267E-06 | 3.002E-06 | 5.800E-06 | 1.100E-05 |
| 8335 | 417 | G417P | A | 8.4e-06 | 3.3e-06 | 1.2e-05 | 7.900E-06 | 4.371E-06 | 3.300E-06 | 1.200E-05 |
| 8336 | 417 | G417S | A | 8.4e-06 | 5.2e-06 | 7.3e-06 | 6.967E-06 | 1.626E-06 | 5.200E-06 | 8.400E-06 |
| 8337 | 417 | G417T | A | 7.1e-06 | 6.5e-06 | 8.2e-06 | 7.267E-06 | 8.622E-07 | 6.500E-06 | 8.200E-06 |
| 8338 | 417 | G417W | A | 5.3e-06 | 4.5e-06 | 9e-06 | 6.267E-06 | 2.401E-06 | 4.500E-06 | 9.000E-06 |
| 8339 | 417 | G417Y | A | 1.5e-05 | 1.1e-05 | 9.3e-06 | 1.177E-05 | 2.926E-06 | 9.300E-06 | 1.500E-05 |
| 8340 | 417 | G417V | A | 9e-06 | 4.5e-06 | 9.4e-06 | 7.633E-06 | 2.721E-06 | 4.500E-06 | 9.400E-06 |
| 8341 | 418 | A418A | A | 7e-06 | 3.3e-06 | 5.8e-06 | 5.367E-06 | 1.888E-06 | 3.300E-06 | 7.000E-06 |
| 8342 | 418 | A418R | A | 6.6e-06 | 1.7e-05 | 9e-06 | 1.087E-05 | 5.445E-06 | 6.600E-06 | 1.700E-05 |
| 8343 | 418 | A418N | A | 5.6e-06 | 8.5e-06 | 8.7e-06 | 7.600E-06 | 1.735E-06 | 5.600E-06 | 8.700E-06 |
| 8344 | 418 | A418D | A | 7e-06 | 4.7e-06 | 6.6e-06 | 6.100E-06 | 1.229E-06 | 4.700E-06 | 7.000E-06 |
| 8345 | 418 | A418C | A | 5.2e-06 | 8.5e-06 | 1.9e-05 | 1.090E-05 | 7.206E-06 | 5.200E-06 | 1.900E-05 |
| 8346 | 418 | A418E | A | 6.2e-06 | 9.5e-06 | 1.1e-05 | 8.900E-06 | 2.456E-06 | 6.200E-06 | 1.100E-05 |
| 8347 | 418 | A418Q | A | 6.4e-06 | 1.6e-05 | 9.6e-06 | 1.067E-05 | 4.888E-06 | 6.400E-06 | 1.600E-05 |
| 8348 | 418 | A418G | A | 1.1e-05 | 2.1e-05 | 9.5e-06 | 1.383E-05 | 6.252E-06 | 9.500E-06 | 2.100E-05 |
| 8349 | 418 | A418H | A | 4.2e-06 | 6.3e-06 | 7.2e-06 | 5.900E-06 | 1.539E-06 | 4.200E-06 | 7.200E-06 |
| 8350 | 418 | A418I | A | 8.8e-06 | 7.5e-06 | 1.3e-05 | 9.767E-06 | 2.875E-06 | 7.500E-06 | 1.300E-05 |
| 8351 | 418 | A418L | A | 5.7e-06 | 5.5e-06 | 6.8e-06 | 6.000E-06 | 7.000E-07 | 5.500E-06 | 6.800E-06 |
| 8352 | 418 | A418K | A | 7.7e-06 | 1.4e-05 | 4.7e-06 | 8.800E-06 | 4.747E-06 | 4.700E-06 | 1.400E-05 |
| 8353 | 418 | A418M | A | 4.2e-06 | 8.2e-06 | 4.4e-06 | 5.600E-06 | 2.254E-06 | 4.200E-06 | 8.200E-06 |
| 8354 | 418 | A418F | A | 8.8e-06 | 5.1e-06 | 4.7e-06 | 6.200E-06 | 2.261E-06 | 4.700E-06 | 8.800E-06 |
| 8355 | 418 | A418P | A | 6.5e-06 | 7.3e-06 | 5.1e-06 | 6.300E-06 | 1.114E-06 | 5.100E-06 | 7.300E-06 |
| 8356 | 418 | A418S | A | 9.7e-06 | 7.4e-06 | 8.1e-06 | 8.400E-06 | 1.179E-06 | 7.400E-06 | 9.700E-06 |
| 8357 | 418 | A418T | A | 8.5e-06 | 7.9e-06 | 9.7e-06 | 8.700E-06 | 9.165E-07 | 7.900E-06 | 9.700E-06 |
| 8358 | 418 | A418W | A | 8.7e-06 | 8.6e-06 | 7.2e-06 | 8.167E-06 | 8.386E-07 | 7.200E-06 | 8.700E-06 |
| 8359 | 418 | A418Y | A | 1.3e-05 | 6.3e-06 | 6.1e-06 | 8.467E-06 | 3.927E-06 | 6.100E-06 | 1.300E-05 |
| 8360 | 418 | A418V | A | 6.9e-06 | 5.1e-06 | 8e-06 | 6.667E-06 | 1.464E-06 | 5.100E-06 | 8.000E-06 |
| 8361 | 419 | Y419A | A | 5.9e-06 | 9.7e-06 | 6e-06 | 7.200E-06 | 2.166E-06 | 5.900E-06 | 9.700E-06 |
| 8362 | 419 | Y419R | A | 1.2e-05 | 9e-06 | 1.3e-05 | 1.133E-05 | 2.082E-06 | 9.000E-06 | 1.300E-05 |
| 8363 | 419 | Y419N | A | 1.3e-05 | 5.8e-06 | 7.7e-06 | 8.833E-06 | 3.731E-06 | 5.800E-06 | 1.300E-05 |
| 8364 | 419 | Y419D | A | 5.8e-06 | 1.2e-05 | 1.6e-05 | 1.127E-05 | 5.139E-06 | 5.800E-06 | 1.600E-05 |
| 8365 | 419 | Y419C | A | 1.2e-05 | 8.5e-06 | 6.1e-06 | 8.867E-06 | 2.967E-06 | 6.100E-06 | 1.200E-05 |
| 8366 | 419 | Y419E | A | 1e-05 | 1.1e-05 | 7.4e-06 | 9.467E-06 | 1.858E-06 | 7.400E-06 | 1.100E-05 |
| 8367 | 419 | Y419Q | A | 8.5e-06 | 4.7e-06 | 8.6e-06 | 7.267E-06 | 2.223E-06 | 4.700E-06 | 8.600E-06 |
| 8368 | 419 | Y419G | A | 7.4e-06 | 6.7e-06 | 8.8e-06 | 7.633E-06 | 1.069E-06 | 6.700E-06 | 8.800E-06 |
| 8369 | 419 | Y419H | A | 8.3e-06 | 7.2e-06 | 9.1e-06 | 8.200E-06 | 9.539E-07 | 7.200E-06 | 9.100E-06 |
| 8370 | 419 | Y419I | A | 8.1e-06 | 9.5e-06 | 1e-05 | 9.200E-06 | 9.849E-07 | 8.100E-06 | 1.000E-05 |
| 8371 | 419 | Y419L | A | 5.6e-06 | 7.3e-06 | 1.2e-05 | 8.300E-06 | 3.315E-06 | 5.600E-06 | 1.200E-05 |
| 8372 | 419 | Y419K | A | 1.1e-05 | 8.1e-06 | 6.8e-06 | 8.633E-06 | 2.150E-06 | 6.800E-06 | 1.100E-05 |
| 8373 | 419 | Y419M | A | 6.1e-06 | 1e-05 | 6.4e-06 | 7.500E-06 | 2.170E-06 | 6.100E-06 | 1.000E-05 |
| 8374 | 419 | Y419F | A | 1e-05 | 8.7e-06 | 8.8e-06 | 9.167E-06 | 7.234E-07 | 8.700E-06 | 1.000E-05 |
| 8375 | 419 | Y419P | A | 5.7e-06 | 1.4e-05 | 5.8e-06 | 8.500E-06 | 4.763E-06 | 5.700E-06 | 1.400E-05 |
| 8376 | 419 | Y419S | A | 8.5e-06 | 8.1e-06 | 6.7e-06 | 7.767E-06 | 9.452E-07 | 6.700E-06 | 8.500E-06 |
| 8377 | 419 | Y419T | A | 1.4e-05 | 8.7e-06 | 6.3e-06 | 9.667E-06 | 3.940E-06 | 6.300E-06 | 1.400E-05 |
| 8378 | 419 | Y419W | A | 6.7e-06 | 8.1e-06 | 9.1e-06 | 7.967E-06 | 1.206E-06 | 6.700E-06 | 9.100E-06 |
| 8379 | 419 | Y419Y | A | 9.2e-06 | 8.5e-06 | 8.3e-06 | 8.667E-06 | 4.726E-07 | 8.300E-06 | 9.200E-06 |
| 8380 | 419 | Y419V | A | 8.4e-06 | 6.7e-06 | 5.6e-06 | 6.900E-06 | 1.411E-06 | 5.600E-06 | 8.400E-06 |
| 8381 | 420 | S420A | A | 7.3e-06 | 8.4e-06 | 1.3e-05 | 9.567E-06 | 3.024E-06 | 7.300E-06 | 1.300E-05 |
| 8382 | 420 | S420R | A | 9.4e-06 | 6e-06 | 4.7e-06 | 6.700E-06 | 2.427E-06 | 4.700E-06 | 9.400E-06 |
| 8383 | 420 | S420N | A | 4.4e-06 | 9.8e-06 | 6.6e-06 | 6.933E-06 | 2.715E-06 | 4.400E-06 | 9.800E-06 |
| 8384 | 420 | S420D | A | 5.1e-06 | 4.5e-06 | 9.9e-06 | 6.500E-06 | 2.960E-06 | 4.500E-06 | 9.900E-06 |
| 8385 | 420 | S420C | A | 1.2e-05 | 9.3e-06 | 8e-06 | 9.767E-06 | 2.040E-06 | 8.000E-06 | 1.200E-05 |
| 8386 | 420 | S420E | A | 5.1e-06 | 8.7e-06 | 1.2e-05 | 8.600E-06 | 3.451E-06 | 5.100E-06 | 1.200E-05 |
| 8387 | 420 | S420Q | A | 5.6e-06 | 9.2e-06 | 8.4e-06 | 7.733E-06 | 1.890E-06 | 5.600E-06 | 9.200E-06 |
| 8388 | 420 | S420G | A | 6.1e-06 | 1.2e-05 | 8.4e-06 | 8.833E-06 | 2.974E-06 | 6.100E-06 | 1.200E-05 |
| 8389 | 420 | S420H | A | 9.7e-06 | 9e-06 | 1.1e-05 | 9.900E-06 | 1.015E-06 | 9.000E-06 | 1.100E-05 |
| 8390 | 420 | S420I | A | 4.7e-06 | 7.6e-06 | 7e-06 | 6.433E-06 | 1.531E-06 | 4.700E-06 | 7.600E-06 |
| 8391 | 420 | S420L | A | 9.6e-06 | 7.9e-06 | 6.1e-06 | 7.867E-06 | 1.750E-06 | 6.100E-06 | 9.600E-06 |
| 8392 | 420 | S420K | A | 1e-05 | 8.1e-06 | 6.7e-06 | 8.267E-06 | 1.656E-06 | 6.700E-06 | 1.000E-05 |
| 8393 | 420 | S420M | A | 5.3e-06 | 9.7e-06 | 7.6e-06 | 7.533E-06 | 2.201E-06 | 5.300E-06 | 9.700E-06 |
| 8394 | 420 | S420F | A | 9.6e-06 | 9.9e-06 | 9.1e-06 | 9.533E-06 | 4.041E-07 | 9.100E-06 | 9.900E-06 |
| 8395 | 420 | S420P | A | 7.8e-06 | 5.6e-06 | 8.3e-06 | 7.233E-06 | 1.436E-06 | 5.600E-06 | 8.300E-06 |
| 8396 | 420 | S420S | A | 8.3e-06 | 8.2e-06 | 3.7e-06 | 6.733E-06 | 2.627E-06 | 3.700E-06 | 8.300E-06 |
| 8397 | 420 | S420T | A | 6.2e-06 | 7.6e-06 | 1e-05 | 7.933E-06 | 1.922E-06 | 6.200E-06 | 1.000E-05 |
| 8398 | 420 | S420W | A | 7.1e-06 | 5.3e-06 | 9e-06 | 7.133E-06 | 1.850E-06 | 5.300E-06 | 9.000E-06 |
| 8399 | 420 | S420Y | A | 1.2e-05 | 8.1e-06 | 6.7e-06 | 8.933E-06 | 2.747E-06 | 6.700E-06 | 1.200E-05 |
| 8400 | 420 | S420V | A | 1e-05 | 7.4e-06 | 6.3e-06 | 7.900E-06 | 1.900E-06 | 6.300E-06 | 1.000E-05 |
| 8401 | 421 | L421A | A | 6.8e-06 | 5.4e-06 | 1.1e-05 | 7.733E-06 | 2.914E-06 | 5.400E-06 | 1.100E-05 |
| 8402 | 421 | L421R | A | 1.1e-05 | 4.5e-06 | 1.4e-05 | 9.833E-06 | 4.856E-06 | 4.500E-06 | 1.400E-05 |
| 8403 | 421 | L421N | A | 5.8e-06 | 8.2e-06 | 1.1e-05 | 8.333E-06 | 2.603E-06 | 5.800E-06 | 1.100E-05 |
| 8404 | 421 | L421D | A | 7.5e-06 | 7e-06 | 9.5e-06 | 8.000E-06 | 1.323E-06 | 7.000E-06 | 9.500E-06 |
| 8405 | 421 | L421C | A | 8.1e-06 | 5.5e-06 | 9.2e-06 | 7.600E-06 | 1.900E-06 | 5.500E-06 | 9.200E-06 |
| 8406 | 421 | L421E | A | 5.4e-06 | 3.4e-06 | 4.6e-06 | 4.467E-06 | 1.007E-06 | 3.400E-06 | 5.400E-06 |
| 8407 | 421 | L421Q | A | 9.3e-06 | 7.9e-06 | 6.8e-06 | 8.000E-06 | 1.253E-06 | 6.800E-06 | 9.300E-06 |
| 8408 | 421 | L421G | A | 8.1e-06 | 1.2e-05 | 1.3e-05 | 1.103E-05 | 2.589E-06 | 8.100E-06 | 1.300E-05 |
| 8409 | 421 | L421H | A | 1.2e-05 | 6.3e-06 | 8.5e-06 | 8.933E-06 | 2.875E-06 | 6.300E-06 | 1.200E-05 |
| 8410 | 421 | L421I | A | 7.4e-06 | 2.5e-06 | 8.9e-06 | 6.267E-06 | 3.347E-06 | 2.500E-06 | 8.900E-06 |
| 8411 | 421 | L421L | A | 7.1e-06 | 8.9e-06 | 9.3e-06 | 8.433E-06 | 1.172E-06 | 7.100E-06 | 9.300E-06 |
| 8412 | 421 | L421K | A | 1e-05 | 7.2e-06 | 1.2e-05 | 9.733E-06 | 2.411E-06 | 7.200E-06 | 1.200E-05 |
| 8413 | 421 | L421M | A | 5e-06 | 6.4e-06 | 6.4e-06 | 5.933E-06 | 8.083E-07 | 5.000E-06 | 6.400E-06 |
| 8414 | 421 | L421F | A | 9.4e-06 | 5.5e-06 | 5.4e-06 | 6.767E-06 | 2.281E-06 | 5.400E-06 | 9.400E-06 |

| | | | | | | | | | | |
|---|---|---|---|---|---|---|---|---|---|---|
| 8415 | 421 | L421P | A | 1.2e-05 | 1.1e-05 | 1.4e-05 | 1.233E-05 | 1.528E-06 | 1.100E-05 | 1.400E-05 |
| 8416 | 421 | L421S | A | 5.1e-06 | 7e-06 | 9.8e-06 | 7.300E-06 | 2.364E-06 | 5.100E-06 | 9.800E-06 |
| 8417 | 421 | L421T | A | 7.3e-06 | 9.1e-06 | 9.4e-06 | 8.600E-06 | 1.136E-06 | 7.300E-06 | 9.400E-06 |
| 8418 | 421 | L421W | A | 5.2e-06 | 1.2e-05 | 7.4e-06 | 8.200E-06 | 3.470E-06 | 5.200E-06 | 1.200E-05 |
| 8419 | 421 | L421Y | A | 4.5e-06 | 9.6e-06 | 7.7e-06 | 7.267E-06 | 2.577E-06 | 4.500E-06 | 9.600E-06 |
| 8420 | 421 | L421V | A | 8.7e-06 | 8.7e-06 | 7.7e-06 | 8.367E-06 | 5.774E-07 | 7.700E-06 | 8.700E-06 |
| 8421 | 422 | T422A | A | 9.4e-06 | 9.1e-06 | 9e-06 | 9.167E-06 | 2.082E-07 | 9.000E-06 | 9.400E-06 |
| 8422 | 422 | T422R | A | 3.5e-06 | 8e-06 | 1e-05 | 7.167E-06 | 3.329E-06 | 3.500E-06 | 1.000E-05 |
| 8423 | 422 | T422N | A | 6.2e-06 | 4.6e-06 | 1.1e-05 | 7.267E-06 | 3.331E-06 | 4.600E-06 | 1.100E-05 |
| 8424 | 422 | T422D | A | 7.6e-06 | 1.2e-05 | 6.7e-06 | 8.767E-06 | 2.836E-06 | 6.700E-06 | 1.200E-05 |
| 8425 | 422 | T422C | A | 9.3e-06 | 8.9e-06 | 9.4e-06 | 9.200E-06 | 2.646E-07 | 8.900E-06 | 9.400E-06 |
| 8426 | 422 | T422E | A | 6.5e-06 | 8.4e-06 | 5.7e-06 | 6.867E-06 | 1.387E-06 | 5.700E-06 | 8.400E-06 |
| 8427 | 422 | T422Q | A | 6.2e-06 | 9.7e-06 | 5.7e-06 | 7.200E-06 | 2.179E-06 | 5.700E-06 | 9.700E-06 |
| 8428 | 422 | T422G | A | 5e-06 | 4.6e-06 | 6.9e-06 | 5.500E-06 | 1.229E-06 | 4.600E-06 | 6.900E-06 |
| 8429 | 422 | T422H | A | 5.8e-06 | 1.1e-05 | 1e-05 | 8.933E-06 | 2.759E-06 | 5.800E-06 | 1.100E-05 |
| 8430 | 422 | T422I | A | 1.3e-05 | 5.7e-06 | 6.9e-06 | 8.533E-06 | 3.915E-06 | 5.700E-06 | 1.300E-05 |
| 8431 | 422 | T422L | A | 8.7e-06 | 7.9e-06 | 1.4e-05 | 1.020E-05 | 3.315E-06 | 7.900E-06 | 1.400E-05 |
| 8432 | 422 | T422K | A | 8.5e-06 | 1e-05 | 8.8e-06 | 9.100E-06 | 7.937E-07 | 8.500E-06 | 1.000E-05 |
| 8433 | 422 | T422M | A | 1.3e-05 | 6.7e-06 | 8.6e-06 | 9.433E-06 | 3.232E-06 | 6.700E-06 | 1.300E-05 |
| 8434 | 422 | T422F | A | 5.2e-06 | 7.2e-06 | 1.4e-05 | 8.800E-06 | 4.613E-06 | 5.200E-06 | 1.400E-05 |
| 8435 | 422 | T422P | A | 1.4e-05 | 6.8e-06 | 1.1e-05 | 1.060E-05 | 3.617E-06 | 6.800E-06 | 1.400E-05 |
| 8436 | 422 | T422S | A | 1.1e-05 | 7.5e-06 | 5.3e-06 | 7.933E-06 | 2.875E-06 | 5.300E-06 | 1.100E-05 |
| 8437 | 422 | T422T | A | 8.6e-06 | 6.2e-06 | 8e-06 | 7.600E-06 | 1.249E-06 | 6.200E-06 | 8.600E-06 |
| 8438 | 422 | T422W | A | 7e-06 | 1.1e-05 | 1e-05 | 9.333E-06 | 2.082E-06 | 7.000E-06 | 1.100E-05 |
| 8439 | 422 | T422Y | A | 1e-05 | 8e-06 | 6.4e-06 | 8.133E-06 | 1.804E-06 | 6.400E-06 | 1.000E-05 |
| 8440 | 422 | T422V | A | 9.6e-06 | 7.4e-06 | 7e-06 | 8.000E-06 | 1.400E-06 | 7.000E-06 | 9.600E-06 |
| 8441 | 423 | L423A | A | 1.1e-05 | 4.1e-06 | 9.1e-06 | 8.067E-06 | 3.564E-06 | 4.100E-06 | 1.100E-05 |
| 8442 | 423 | L423R | A | 4.5e-06 | 1e-05 | 9.2e-06 | 7.900E-06 | 2.972E-06 | 4.500E-06 | 1.000E-05 |
| 8443 | 423 | L423N | A | 6.3e-06 | 7e-06 | 1e-05 | 7.767E-06 | 1.966E-06 | 6.300E-06 | 1.000E-05 |
| 8444 | 423 | L423D | A | 6.8e-06 | 7.3e-06 | 1.6e-05 | 1.003E-05 | 5.173E-06 | 6.800E-06 | 1.600E-05 |
| 8445 | 423 | L423C | A | 9.1e-06 | 5.4e-06 | 1.2e-05 | 8.833E-06 | 3.308E-06 | 5.400E-06 | 1.200E-05 |
| 8446 | 423 | L423E | A | 1e-05 | 6.3e-06 | 7.7e-06 | 8.000E-06 | 1.868E-06 | 6.300E-06 | 1.000E-05 |
| 8447 | 423 | L423Q | A | 9.3e-06 | 1.3e-05 | 9.7e-06 | 1.067E-05 | 2.031E-06 | 9.300E-06 | 1.300E-05 |
| 8448 | 423 | L423G | A | 9.4e-06 | 8.2e-06 | 7.5e-06 | 8.367E-06 | 9.609E-07 | 7.500E-06 | 9.400E-06 |
| 8449 | 423 | L423H | A | 4.9e-06 | 5.3e-06 | 8.1e-06 | 6.100E-06 | 1.744E-06 | 4.900E-06 | 8.100E-06 |
| 8450 | 423 | L423I | A | 1.4e-05 | 8.4e-06 | 7e-06 | 9.800E-06 | 3.704E-06 | 7.000E-06 | 1.400E-05 |
| 8451 | 423 | L423L | A | 9.9e-06 | 1.4e-05 | 8.4e-06 | 1.077E-05 | 2.899E-06 | 8.400E-06 | 1.400E-05 |
| 8452 | 423 | L423K | A | 9.7e-06 | 1.1e-05 | 6.8e-06 | 9.167E-06 | 2.150E-06 | 6.800E-06 | 1.100E-05 |
| 8453 | 423 | L423M | A | 9e-06 | 1.4e-05 | 5.4e-06 | 9.467E-06 | 4.319E-06 | 5.400E-06 | 1.400E-05 |
| 8454 | 423 | L423F | A | 9.2e-06 | 7.2e-06 | 9.5e-06 | 8.633E-06 | 1.250E-06 | 7.200E-06 | 9.500E-06 |
| 8455 | 423 | L423P | A | 1e-05 | 6.9e-06 | 8.4e-06 | 8.433E-06 | 1.550E-06 | 6.900E-06 | 1.000E-05 |
| 8456 | 423 | L423S | A | 7.5e-06 | 5.9e-06 | 9.3e-06 | 7.567E-06 | 1.701E-06 | 5.900E-06 | 9.300E-06 |
| 8457 | 423 | L423T | A | 9.5e-06 | 6.4e-06 | 9.2e-06 | 8.367E-06 | 1.710E-06 | 6.400E-06 | 9.500E-06 |
| 8458 | 423 | L423W | A | 1.3e-05 | 4.7e-06 | 1.2e-05 | 9.900E-06 | 4.531E-06 | 4.700E-06 | 1.300E-05 |
| 8459 | 423 | L423Y | A | 9.6e-06 | 1e-05 | 4.6e-06 | 8.067E-06 | 3.009E-06 | 4.600E-06 | 1.000E-05 |
| 8460 | 423 | L423V | A | 1.3e-05 | 7.3e-06 | 6e-06 | 8.767E-06 | 3.723E-06 | 6.000E-06 | 1.300E-05 |
| 8461 | 424 | Q424A | A | 1e-05 | 5e-06 | 1.5e-05 | 1.000E-05 | 5.000E-06 | 5.000E-06 | 1.500E-05 |
| 8462 | 424 | Q424R | A | 5.3e-06 | 6.1e-06 | 1.6e-05 | 9.133E-06 | 5.960E-06 | 5.300E-06 | 1.600E-05 |
| 8463 | 424 | Q424N | A | 4.6e-06 | 7.9e-06 | 6.4e-06 | 6.300E-06 | 1.652E-06 | 4.600E-06 | 7.900E-06 |
| 8464 | 424 | Q424D | A | 1.3e-05 | 5.6e-06 | 6.7e-06 | 8.433E-06 | 3.993E-06 | 5.600E-06 | 1.300E-05 |
| 8465 | 424 | Q424C | A | 8.7e-06 | 1.3e-05 | 8.2e-06 | 9.967E-06 | 2.639E-06 | 8.200E-06 | 1.300E-05 |
| 8466 | 424 | Q424E | A | 4.7e-06 | 1e-05 | 4e-06 | 6.233E-06 | 3.281E-06 | 4.000E-06 | 1.000E-05 |
| 8467 | 424 | Q424Q | A | 9.2e-06 | 4.4e-06 | 8.8e-06 | 7.467E-06 | 2.663E-06 | 4.400E-06 | 9.200E-06 |
| 8468 | 424 | Q424G | A | 6.7e-06 | 7.3e-06 | 6.9e-06 | 6.967E-06 | 3.055E-07 | 6.700E-06 | 7.300E-06 |
| 8469 | 424 | Q424H | A | 5.9e-06 | 6.4e-06 | 8.8e-06 | 7.033E-06 | 1.550E-06 | 5.900E-06 | 8.800E-06 |
| 8470 | 424 | Q424I | A | 8e-06 | 3.3e-06 | 4.5e-06 | 5.267E-06 | 2.442E-06 | 3.300E-06 | 8.000E-06 |
| 8471 | 424 | Q424L | A | 8.5e-06 | 1.3e-05 | 9.2e-06 | 1.023E-05 | 2.421E-06 | 8.500E-06 | 1.300E-05 |
| 8472 | 424 | Q424K | A | 8.8e-06 | 1.8e-05 | 1e-05 | 1.227E-05 | 5.001E-06 | 8.800E-06 | 1.800E-05 |
| 8473 | 424 | Q424M | A | 7.2e-06 | 4.7e-06 | 1.9e-05 | 1.030E-05 | 7.637E-06 | 4.700E-06 | 1.900E-05 |
| 8474 | 424 | Q424F | A | 5.4e-06 | 9.9e-06 | 6.2e-06 | 7.167E-06 | 2.401E-06 | 5.400E-06 | 9.900E-06 |
| 8475 | 424 | Q424P | A | 9.6e-06 | 7.9e-06 | 8.9e-06 | 8.800E-06 | 8.544E-07 | 7.900E-06 | 9.600E-06 |
| 8476 | 424 | Q424S | A | 8.8e-06 | 4.4e-06 | 8.7e-06 | 7.300E-06 | 2.512E-06 | 4.400E-06 | 8.800E-06 |
| 8477 | 424 | Q424T | A | 7.4e-06 | 4.8e-06 | 5.2e-06 | 5.800E-06 | 1.400E-06 | 4.800E-06 | 7.400E-06 |
| 8478 | 424 | Q424W | A | 1.4e-05 | 1.2e-05 | 1.1e-05 | 1.233E-05 | 1.528E-06 | 1.100E-05 | 1.400E-05 |
| 8479 | 424 | Q424Y | A | 7.2e-06 | 5.1e-06 | 7.3e-06 | 6.533E-06 | 1.242E-06 | 5.100E-06 | 7.300E-06 |
| 8480 | 424 | Q424V | A | 4.9e-06 | 7.2e-06 | 1.1e-05 | 7.700E-06 | 3.081E-06 | 4.900E-06 | 1.100E-05 |
| 8481 | 425 | G425A | A | 6.4e-06 | 7e-06 | 8.4e-06 | 7.267E-06 | 1.026E-06 | 6.400E-06 | 8.400E-06 |
| 8482 | 425 | G425R | A | 5.3e-06 | 1e-05 | 1.2e-05 | 9.100E-06 | 3.439E-06 | 5.300E-06 | 1.200E-05 |
| 8483 | 425 | G425N | A | 7.7e-06 | 5.2e-06 | 9.9e-06 | 7.600E-06 | 2.352E-06 | 5.200E-06 | 9.900E-06 |
| 8484 | 425 | G425D | A | 4.6e-06 | 8.9e-06 | 8.4e-06 | 7.300E-06 | 2.352E-06 | 4.600E-06 | 8.900E-06 |
| 8485 | 425 | G425C | A | 4.4e-06 | 9.5e-06 | 9.1e-06 | 7.667E-06 | 2.836E-06 | 4.400E-06 | 9.500E-06 |
| 8486 | 425 | G425E | A | 9.4e-06 | 4.9e-06 | 8.1e-06 | 7.467E-06 | 2.316E-06 | 4.900E-06 | 9.400E-06 |
| 8487 | 425 | G425Q | A | 7.1e-06 | 6.5e-06 | 9.8e-06 | 7.800E-06 | 1.758E-06 | 6.500E-06 | 9.800E-06 |
| 8488 | 425 | G425G | A | 1.3e-05 | 9.6e-06 | 8.7e-06 | 1.043E-05 | 2.268E-06 | 8.700E-06 | 1.300E-05 |
| 8489 | 425 | G425H | A | 6.1e-06 | 4e-06 | 4.1e-06 | 4.733E-06 | 1.185E-06 | 4.000E-06 | 6.100E-06 |
| 8490 | 425 | G425I | A | 7.9e-06 | 8.2e-06 | 8.1e-06 | 8.067E-06 | 1.528E-07 | 7.900E-06 | 8.200E-06 |
| 8491 | 425 | G425L | A | 7.3e-06 | 7.2e-06 | 1e-05 | 8.167E-06 | 1.589E-06 | 7.200E-06 | 1.000E-05 |
| 8492 | 425 | G425K | A | 8.2e-06 | 8.2e-06 | 3.8e-06 | 6.733E-06 | 2.540E-06 | 3.800E-06 | 8.200E-06 |
| 8493 | 425 | G425M | A | 1.1e-05 | 6.3e-06 | 7.2e-06 | 8.167E-06 | 2.495E-06 | 6.300E-06 | 1.100E-05 |
| 8494 | 425 | G425F | A | 9.9e-06 | 1e-05 | 1.1e-05 | 1.030E-05 | 6.083E-07 | 9.900E-06 | 1.100E-05 |
| 8495 | 425 | G425P | A | 4.8e-06 | 9e-06 | 6.5e-06 | 6.767E-06 | 2.113E-06 | 4.800E-06 | 9.000E-06 |
| 8496 | 425 | G425S | A | 6e-06 | 8.1e-06 | 7.3e-06 | 7.133E-06 | 1.060E-06 | 6.000E-06 | 8.100E-06 |
| 8497 | 425 | G425T | A | 7.5e-06 | 4.3e-06 | 4.6e-06 | 5.467E-06 | 1.767E-06 | 4.300E-06 | 7.500E-06 |
| 8498 | 425 | G425W | A | 6.9e-06 | 6.4e-06 | 5.6e-06 | 6.300E-06 | 6.557E-07 | 5.600E-06 | 6.900E-06 |
| 8499 | 425 | G425Y | A | 6.1e-06 | 5.7e-06 | 7.8e-06 | 6.533E-06 | 1.115E-06 | 5.700E-06 | 7.800E-06 |

| | | | | | | | | | | |
|---|---|---|---|---|---|---|---|---|---|---|
| 8500 | 425 | G425V | A | 8.8e-06 | 1.1e-05 | 6.9e-06 | 8.900E-06 | 2.052E-06 | 6.900E-06 | 1.100E-05 |
| 8501 | 426 | L426A | A | 4.5e-06 | 6.7e-06 | 5.6e-06 | 5.600E-06 | 1.100E-06 | 4.500E-06 | 6.700E-06 |
| 8502 | 426 | L426R | A | 9.2e-06 | 8.9e-06 | 9.2e-06 | 9.100E-06 | 1.732E-07 | 8.900E-06 | 9.200E-06 |
| 8503 | 426 | L426N | A | 6.9e-06 | 1.1e-05 | 5.6e-06 | 7.833E-06 | 2.818E-06 | 5.600E-06 | 1.100E-05 |
| 8504 | 426 | L426D | A | 5.8e-06 | 9.5e-06 | 6.2e-06 | 7.167E-06 | 2.031E-06 | 5.800E-06 | 9.500E-06 |
| 8505 | 426 | L426C | A | 5.8e-06 | 1.3e-05 | 8.6e-06 | 9.133E-06 | 3.630E-06 | 5.800E-06 | 1.300E-05 |
| 8506 | 426 | L426E | A | 7.5e-06 | 7.5e-06 | 6.7e-06 | 7.233E-06 | 4.619E-07 | 6.700E-06 | 7.500E-06 |
| 8507 | 426 | L426Q | A | 6.4e-06 | 4.6e-06 | 6.8e-06 | 5.933E-06 | 1.172E-06 | 4.600E-06 | 6.800E-06 |
| 8508 | 426 | L426G | A | 8.9e-06 | 8e-06 | 7.9e-06 | 8.267E-06 | 5.508E-07 | 7.900E-06 | 8.900E-06 |
| 8509 | 426 | L426H | A | 9e-06 | 5.2e-06 | 7.2e-06 | 7.133E-06 | 1.901E-06 | 5.200E-06 | 9.000E-06 |
| 8510 | 426 | L426I | A | 6e-06 | 7.2e-06 | 1e-05 | 7.733E-06 | 2.053E-06 | 6.000E-06 | 1.000E-05 |
| 8511 | 426 | L426L | A | 4.4e-06 | 8.2e-06 | 4.2e-06 | 5.600E-06 | 2.254E-06 | 4.200E-06 | 8.200E-06 |
| 8512 | 426 | L426K | A | 4.9e-06 | 5e-06 | 4.8e-06 | 4.900E-06 | 1.000E-07 | 4.800E-06 | 5.000E-06 |
| 8513 | 426 | L426M | A | 7.2e-06 | 2e-05 | 5.1e-06 | 1.077E-05 | 8.065E-06 | 5.100E-06 | 2.000E-05 |
| 8514 | 426 | L426F | A | 4.5e-06 | 7e-06 | 9e-06 | 6.833E-06 | 2.255E-06 | 4.500E-06 | 9.000E-06 |
| 8515 | 426 | L426P | A | 9.4e-06 | 9.4e-06 | 6.4e-06 | 8.400E-06 | 1.732E-06 | 6.400E-06 | 9.400E-06 |
| 8516 | 426 | L426S | A | 7.4e-06 | 7.7e-06 | 4.1e-06 | 6.400E-06 | 1.997E-06 | 4.100E-06 | 7.700E-06 |
| 8517 | 426 | L426T | A | 2e-05 | 9e-06 | 5.3e-06 | 1.143E-05 | 7.646E-06 | 5.300E-06 | 2.000E-05 |
| 8518 | 426 | L426W | A | 7.1e-06 | 1e-05 | 1e-05 | 9.033E-06 | 1.674E-06 | 7.100E-06 | 1.000E-05 |
| 8519 | 426 | L426Y | A | 9.3e-06 | 3.5e-06 | 1.1e-05 | 7.933E-06 | 3.932E-06 | 3.500E-06 | 1.100E-05 |
| 8520 | 426 | L426V | A | 9e-06 | 5.2e-06 | 9.1e-06 | 7.767E-06 | 2.223E-06 | 5.200E-06 | 9.100E-06 |
| 8521 | 427 | G427A | A | 1.3e-05 | 1.3e-05 | 8.4e-06 | 1.147E-05 | 2.656E-06 | 8.400E-06 | 1.300E-05 |
| 8522 | 427 | G427R | A | 8.1e-06 | 7e-06 | 5.2e-06 | 6.767E-06 | 1.464E-06 | 5.200E-06 | 8.100E-06 |
| 8523 | 427 | G427N | A | 9e-06 | 4.9e-06 | 6.3e-06 | 6.733E-06 | 2.084E-06 | 4.900E-06 | 9.000E-06 |
| 8524 | 427 | G427D | A | 6.4e-06 | 5.7e-06 | 6.5e-06 | 6.200E-06 | 4.359E-07 | 5.700E-06 | 6.500E-06 |
| 8525 | 427 | G427C | A | 9.7e-06 | 4.5e-06 | 9e-06 | 7.733E-06 | 2.822E-06 | 4.500E-06 | 9.700E-06 |
| 8526 | 427 | G427E | A | 1e-05 | 1.1e-05 | 6.7e-06 | 9.233E-06 | 2.250E-06 | 6.700E-06 | 1.100E-05 |
| 8527 | 427 | G427Q | A | 3.9e-06 | 7.5e-06 | 5.9e-06 | 5.767E-06 | 1.804E-06 | 3.900E-06 | 7.500E-06 |
| 8528 | 427 | G427G | A | 1.1e-05 | 1e-05 | 6.6e-06 | 9.200E-06 | 2.307E-06 | 6.600E-06 | 1.100E-05 |
| 8529 | 427 | G427H | A | 6.1e-06 | 7.9e-06 | 4.9e-06 | 6.300E-06 | 1.510E-06 | 4.900E-06 | 7.900E-06 |
| 8530 | 427 | G427I | A | 6.9e-06 | 9.4e-06 | 7.3e-06 | 7.867E-06 | 1.343E-06 | 6.900E-06 | 9.400E-06 |
| 8531 | 427 | G427L | A | 8.9e-06 | 9.4e-06 | 5.6e-06 | 7.967E-06 | 2.065E-06 | 5.600E-06 | 9.400E-06 |
| 8532 | 427 | G427K | A | 5.4e-06 | 7.1e-06 | 9.4e-06 | 7.300E-06 | 2.007E-06 | 5.400E-06 | 9.400E-06 |
| 8533 | 427 | G427M | A | 1.1e-05 | 5.4e-06 | 5.9e-06 | 7.433E-06 | 3.099E-06 | 5.400E-06 | 1.100E-05 |
| 8534 | 427 | G427F | A | 7.5e-06 | 7e-06 | 1.3e-05 | 9.167E-06 | 3.329E-06 | 7.000E-06 | 1.300E-05 |
| 8535 | 427 | G427P | A | 4.4e-06 | 1.3e-05 | 8.8e-06 | 8.733E-06 | 4.300E-06 | 4.400E-06 | 1.300E-05 |
| 8536 | 427 | G427S | A | 6.3e-06 | 9.6e-06 | 6.2e-06 | 7.367E-06 | 1.935E-06 | 6.200E-06 | 9.600E-06 |
| 8537 | 427 | G427T | A | 1.2e-05 | 4.3e-06 | 7.8e-06 | 8.033E-06 | 3.855E-06 | 4.300E-06 | 1.200E-05 |
| 8538 | 427 | G427W | A | 6.6e-06 | 1.3e-05 | 8.9e-06 | 9.500E-06 | 3.242E-06 | 6.600E-06 | 1.300E-05 |
| 8539 | 427 | G427Y | A | 5.3e-06 | 7.3e-06 | 7.1e-06 | 6.567E-06 | 1.102E-06 | 5.300E-06 | 7.300E-06 |
| 8540 | 427 | G427V | A | 5.6e-06 | 9.9e-06 | 5.3e-06 | 6.933E-06 | 2.574E-06 | 5.300E-06 | 9.900E-06 |
| 8541 | 428 | I428A | A | 1.1e-05 | 9.6e-06 | 5.4e-06 | 8.667E-06 | 2.914E-06 | 5.400E-06 | 1.100E-05 |
| 8542 | 428 | I428R | A | 6.3e-06 | 4.2e-06 | 7.9e-06 | 6.133E-06 | 1.856E-06 | 4.200E-06 | 7.900E-06 |
| 8543 | 428 | I428N | A | 9.7e-06 | 7.3e-06 | 7.9e-06 | 8.300E-06 | 1.249E-06 | 7.300E-06 | 9.700E-06 |
| 8544 | 428 | I428D | A | 4.8e-06 | 1e-05 | 7.5e-06 | 7.433E-06 | 2.601E-06 | 4.800E-06 | 1.000E-05 |
| 8545 | 428 | I428C | A | 7.1e-06 | 6.4e-06 | 1.1e-05 | 8.167E-06 | 2.479E-06 | 6.400E-06 | 1.100E-05 |
| 8546 | 428 | I428E | A | 7.8e-06 | 1e-05 | 9.3e-06 | 9.033E-06 | 1.124E-06 | 7.800E-06 | 1.000E-05 |
| 8547 | 428 | I428Q | A | 9.1e-06 | 6.7e-06 | 6.8e-06 | 7.533E-06 | 1.358E-06 | 6.700E-06 | 9.100E-06 |
| 8548 | 428 | I428G | A | 7e-06 | 6e-06 | 1.3e-05 | 8.667E-06 | 3.786E-06 | 6.000E-06 | 1.300E-05 |
| 8549 | 428 | I428H | A | 4.6e-06 | 4.3e-06 | 4.2e-06 | 4.367E-06 | 2.082E-07 | 4.200E-06 | 4.600E-06 |
| 8550 | 428 | I428I | A | 9.1e-06 | 8.1e-06 | 1e-05 | 9.067E-06 | 9.504E-07 | 8.100E-06 | 1.000E-05 |
| 8551 | 428 | I428L | A | 5e-06 | 6.6e-06 | 1e-05 | 7.200E-06 | 2.553E-06 | 5.000E-06 | 1.000E-05 |
| 8552 | 428 | I428K | A | 6.8e-06 | 8e-06 | 7.7e-06 | 7.500E-06 | 6.245E-07 | 6.800E-06 | 8.000E-06 |
| 8553 | 428 | I428M | A | 9e-06 | 9e-06 | 1e-05 | 9.333E-06 | 5.774E-07 | 9.000E-06 | 1.000E-05 |
| 8554 | 428 | I428F | A | 1.5e-05 | 6e-06 | 9.1e-06 | 1.003E-05 | 4.572E-06 | 6.000E-06 | 1.500E-05 |
| 8555 | 428 | I428P | A | 8.2e-06 | 6.4e-06 | 8.6e-06 | 7.733E-06 | 1.172E-06 | 6.400E-06 | 8.600E-06 |
| 8556 | 428 | I428S | A | 1e-05 | 7.1e-06 | 5.9e-06 | 7.667E-06 | 2.108E-06 | 5.900E-06 | 1.000E-05 |
| 8557 | 428 | I428T | A | 4.4e-06 | 1.2e-05 | 6e-06 | 7.467E-06 | 4.007E-06 | 4.400E-06 | 1.200E-05 |
| 8558 | 428 | I428W | A | 6.2e-06 | 1.2e-05 | 5.8e-06 | 8.000E-06 | 3.470E-06 | 5.800E-06 | 1.200E-05 |
| 8559 | 428 | I428Y | A | 7.4e-06 | 4.6e-06 | 1e-05 | 7.333E-06 | 2.701E-06 | 4.600E-06 | 1.000E-05 |
| 8560 | 428 | I428V | A | 5.1e-06 | 4.7e-06 | 9.7e-06 | 6.500E-06 | 2.778E-06 | 4.700E-06 | 9.700E-06 |
| 8561 | 429 | S429A | A | 9.7e-06 | 8.8e-06 | 6.3e-06 | 8.267E-06 | 1.762E-06 | 6.300E-06 | 9.700E-06 |
| 8562 | 429 | S429R | A | 1e-05 | 6.3e-06 | 4.7e-06 | 7.000E-06 | 2.718E-06 | 4.700E-06 | 1.000E-05 |
| 8563 | 429 | S429N | A | 5.9e-06 | 6.5e-06 | 4.7e-06 | 5.700E-06 | 9.165E-07 | 4.700E-06 | 6.500E-06 |
| 8564 | 429 | S429D | A | 1.4e-05 | 7e-06 | 5e-06 | 8.667E-06 | 4.726E-06 | 5.000E-06 | 1.400E-05 |
| 8565 | 429 | S429C | A | 4.7e-06 | 7.9e-06 | 7.7e-06 | 6.767E-06 | 1.793E-06 | 4.700E-06 | 7.900E-06 |
| 8566 | 429 | S429E | A | 9.7e-06 | 8.9e-06 | 5.1e-06 | 7.900E-06 | 2.458E-06 | 5.100E-06 | 9.700E-06 |
| 8567 | 429 | S429Q | A | 1.2e-05 | 8e-06 | 6.6e-06 | 8.867E-06 | 2.802E-06 | 6.600E-06 | 1.200E-05 |
| 8568 | 429 | S429G | A | 1.2e-05 | 1.1e-05 | 7.3e-06 | 1.010E-05 | 2.476E-06 | 7.300E-06 | 1.200E-05 |
| 8569 | 429 | S429H | A | 1e-05 | 9.8e-06 | 1.5e-05 | 1.160E-05 | 2.946E-06 | 9.800E-06 | 1.500E-05 |
| 8570 | 429 | S429I | A | 7.9e-06 | 8.5e-06 | 1.1e-05 | 9.133E-06 | 1.644E-06 | 7.900E-06 | 1.100E-05 |
| 8571 | 429 | S429L | A | 1e-05 | 5e-06 | 8e-06 | 7.667E-06 | 2.517E-06 | 5.000E-06 | 1.000E-05 |
| 8572 | 429 | S429K | A | 6.9e-06 | 9.3e-06 | 4.4e-06 | 6.867E-06 | 2.450E-06 | 4.400E-06 | 9.300E-06 |
| 8573 | 429 | S429M | A | 5.9e-06 | 5.5e-06 | 7.7e-06 | 6.367E-06 | 1.172E-06 | 5.500E-06 | 7.700E-06 |
| 8574 | 429 | S429F | A | 7.1e-06 | 1e-05 | 9.6e-06 | 8.900E-06 | 1.572E-06 | 7.100E-06 | 1.000E-05 |
| 8575 | 429 | S429P | A | 1.1e-05 | 4.6e-06 | 8.2e-06 | 7.933E-06 | 3.208E-06 | 4.600E-06 | 1.100E-05 |
| 8576 | 429 | S429S | A | 7.4e-06 | 6.9e-06 | 4.6e-06 | 6.300E-06 | 1.493E-06 | 4.600E-06 | 7.400E-06 |
| 8577 | 429 | S429T | A | 8.7e-06 | 4.4e-06 | 7.4e-06 | 6.833E-06 | 2.205E-06 | 4.400E-06 | 8.700E-06 |
| 8578 | 429 | S429W | A | 1.2e-05 | 7.3e-06 | 8e-06 | 9.100E-06 | 2.536E-06 | 7.300E-06 | 1.200E-05 |
| 8579 | 429 | S429Y | A | 1.1e-05 | 6.9e-06 | 8.4e-06 | 8.767E-06 | 2.074E-06 | 6.900E-06 | 1.100E-05 |
| 8580 | 429 | S429V | A | 7.8e-06 | 3.5e-06 | 6.7e-06 | 6.000E-06 | 2.234E-06 | 3.500E-06 | 7.800E-06 |
| 8581 | 430 | W430A | A | 8.3e-06 | 8.3e-06 | 9.7e-06 | 8.767E-06 | 8.083E-07 | 8.300E-06 | 9.700E-06 |
| 8582 | 430 | W430R | A | 4.8e-06 | 4.9e-06 | 9.2e-06 | 6.300E-06 | 2.512E-06 | 4.800E-06 | 9.200E-06 |
| 8583 | 430 | W430N | A | 3.3e-06 | 8.9e-06 | 7.9e-06 | 6.700E-06 | 2.987E-06 | 3.300E-06 | 8.900E-06 |
| 8584 | 430 | W430D | A | 8.6e-06 | 7.9e-06 | 1.3e-05 | 9.833E-06 | 2.765E-06 | 7.900E-06 | 1.300E-05 |

| | | | | | | | | | | |
|---|---|---|---|---|---|---|---|---|---|---|
| 8585 | 430 | W430C | A | 6.7e-06 | 7.4e-06 | 8.3e-06 | 7.467E-06 | 8.021E-07 | 6.700E-06 | 8.300E-06 |
| 8586 | 430 | W430E | A | 1.4e-05 | 1.3e-05 | 8.5e-06 | 1.183E-05 | 2.930E-06 | 8.500E-06 | 1.400E-05 |
| 8587 | 430 | W430Q | A | 5.3e-06 | 1.1e-05 | 6.3e-06 | 7.533E-06 | 3.044E-06 | 5.300E-06 | 1.100E-05 |
| 8588 | 430 | W430G | A | 1.3e-05 | 1.1e-05 | 9.3e-06 | 1.110E-05 | 1.852E-06 | 9.300E-06 | 1.300E-05 |
| 8589 | 430 | W430H | A | 9.1e-06 | 1.4e-05 | 4.6e-06 | 9.233E-06 | 4.701E-06 | 4.600E-06 | 1.400E-05 |
| 8590 | 430 | W430I | A | 9.1e-06 | 9.5e-06 | 9.4e-06 | 9.333E-06 | 2.082E-07 | 9.100E-06 | 9.500E-06 |
| 8591 | 430 | W430L | A | 9.6e-06 | 7.6e-06 | 5.4e-06 | 7.533E-06 | 2.101E-06 | 5.400E-06 | 9.600E-06 |
| 8592 | 430 | W430K | A | 7.4e-06 | 1e-05 | 7.2e-06 | 8.200E-06 | 1.562E-06 | 7.200E-06 | 1.000E-05 |
| 8593 | 430 | W430M | A | 6e-06 | 8e-06 | 5.3e-06 | 6.433E-06 | 1.401E-06 | 5.300E-06 | 8.000E-06 |
| 8594 | 430 | W430F | A | 5.2e-06 | 4.3e-06 | 7.4e-06 | 5.633E-06 | 1.595E-06 | 4.300E-06 | 7.400E-06 |
| 8595 | 430 | W430P | A | 8e-06 | 6.4e-06 | 5.2e-06 | 6.533E-06 | 1.405E-06 | 5.200E-06 | 8.000E-06 |
| 8596 | 430 | W430S | A | 8.3e-06 | 8.9e-06 | 7.5e-06 | 8.233E-06 | 7.024E-07 | 7.500E-06 | 8.900E-06 |
| 8597 | 430 | W430T | A | 8.1e-06 | 9.3e-06 | 5.4e-06 | 7.600E-06 | 1.997E-06 | 5.400E-06 | 9.300E-06 |
| 8598 | 430 | W430W | A | 7.4e-06 | 6.5e-06 | 7.7e-06 | 7.200E-06 | 6.245E-07 | 6.500E-06 | 7.700E-06 |
| 8599 | 430 | W430Y | A | 1.2e-05 | 1.1e-05 | 5.2e-06 | 9.400E-06 | 3.672E-06 | 5.200E-06 | 1.200E-05 |
| 8600 | 430 | W430V | A | 1e-05 | 1.4e-05 | 1e-05 | 1.133E-05 | 2.309E-06 | 1.000E-05 | 1.400E-05 |
| 8601 | 431 | L431A | A | 1.4e-05 | 1.1e-05 | 8.5e-06 | 1.117E-05 | 2.754E-06 | 8.500E-06 | 1.400E-05 |
| 8602 | 431 | L431R | A | 9.2e-06 | 8.8e-06 | 8.7e-06 | 8.900E-06 | 2.646E-07 | 8.700E-06 | 9.200E-06 |
| 8603 | 431 | L431N | A | 7.1e-06 | 5.8e-06 | 1.2e-05 | 8.300E-06 | 3.270E-06 | 5.800E-06 | 1.200E-05 |
| 8604 | 431 | L431D | A | 7.6e-06 | 8.9e-06 | 7.9e-06 | 8.133E-06 | 6.807E-07 | 7.600E-06 | 8.900E-06 |
| 8605 | 431 | L431C | A | 1.7e-05 | 4.8e-06 | 6.2e-06 | 9.333E-06 | 6.676E-06 | 4.800E-06 | 1.700E-05 |
| 8606 | 431 | L431E | A | 7.3e-06 | 7.2e-06 | 4.4e-06 | 6.300E-06 | 1.646E-06 | 4.400E-06 | 7.300E-06 |
| 8607 | 431 | L431Q | A | 1.3e-05 | 8.4e-06 | 5.1e-06 | 8.833E-06 | 3.968E-06 | 5.100E-06 | 1.300E-05 |
| 8608 | 431 | L431G | A | 8.9e-06 | 1e-05 | 1e-05 | 9.633E-06 | 6.351E-07 | 8.900E-06 | 1.000E-05 |
| 8609 | 431 | L431H | A | 5.2e-06 | 8.6e-06 | 1e-05 | 7.933E-06 | 2.468E-06 | 5.200E-06 | 1.000E-05 |
| 8610 | 431 | L431I | A | 8.6e-06 | 7.6e-06 | 9.7e-06 | 8.633E-06 | 1.050E-06 | 7.600E-06 | 9.700E-06 |
| 8611 | 431 | L431L | A | 6.8e-06 | 9.7e-06 | 6.1e-06 | 7.533E-06 | 1.909E-06 | 6.100E-06 | 9.700E-06 |
| 8612 | 431 | L431K | A | 1e-05 | 1.3e-05 | 6.6e-06 | 9.867E-06 | 3.202E-06 | 6.600E-06 | 1.300E-05 |
| 8613 | 431 | L431M | A | 9.3e-06 | 7.7e-06 | 1e-05 | 9.000E-06 | 1.179E-06 | 7.700E-06 | 1.000E-05 |
| 8614 | 431 | L431F | A | 9.6e-06 | 1.3e-05 | 7.8e-06 | 1.013E-05 | 2.641E-06 | 7.800E-06 | 1.300E-05 |
| 8615 | 431 | L431P | A | 1.1e-05 | 8.1e-06 | 8.2e-06 | 9.100E-06 | 1.646E-06 | 8.100E-06 | 1.100E-05 |
| 8616 | 431 | L431S | A | 6.6e-06 | 7.5e-06 | 1.1e-05 | 8.367E-06 | 2.325E-06 | 6.600E-06 | 1.100E-05 |
| 8617 | 431 | L431T | A | 7.7e-06 | 9.7e-06 | 7.6e-06 | 8.333E-06 | 1.185E-06 | 7.600E-06 | 9.700E-06 |
| 8618 | 431 | L431W | A | 9.1e-06 | 9.2e-06 | 6.7e-06 | 8.333E-06 | 1.415E-06 | 6.700E-06 | 9.200E-06 |
| 8619 | 431 | L431Y | A | 6.5e-06 | 6.2e-06 | 6.8e-06 | 6.500E-06 | 3.000E-07 | 6.200E-06 | 6.800E-06 |
| 8620 | 431 | L431V | A | 1.1e-05 | 9.2e-06 | 7.1e-06 | 9.100E-06 | 1.952E-06 | 7.100E-06 | 1.100E-05 |
| 8621 | 432 | G432A | A | 1.1e-05 | 1.3e-05 | 4.6e-06 | 9.533E-06 | 4.388E-06 | 4.600E-06 | 1.300E-05 |
| 8622 | 432 | G432R | A | 1e-05 | 6.8e-06 | 1e-05 | 8.933E-06 | 1.848E-06 | 6.800E-06 | 1.000E-05 |
| 8623 | 432 | G432N | A | 6.8e-06 | 3.8e-06 | 1.2e-05 | 7.533E-06 | 4.149E-06 | 3.800E-06 | 1.200E-05 |
| 8624 | 432 | G432D | A | 2.6e-06 | 1.1e-05 | 8.3e-06 | 7.300E-06 | 4.288E-06 | 2.600E-06 | 1.100E-05 |
| 8625 | 432 | G432C | A | 8.6e-06 | 1e-05 | 1.1e-05 | 9.867E-06 | 1.206E-06 | 8.600E-06 | 1.100E-05 |
| 8626 | 432 | G432E | A | 3.6e-06 | 7.4e-06 | 5.1e-06 | 5.367E-06 | 1.914E-06 | 3.600E-06 | 7.400E-06 |
| 8627 | 432 | G432Q | A | 8.2e-06 | 6e-06 | 5.7e-06 | 6.633E-06 | 1.365E-06 | 5.700E-06 | 8.200E-06 |
| 8628 | 432 | G432G | A | 1e-05 | 9.5e-06 | 9.5e-06 | 9.667E-06 | 2.887E-07 | 9.500E-06 | 1.000E-05 |
| 8629 | 432 | G432H | A | 1e-05 | 7.6e-06 | 6.7e-06 | 8.100E-06 | 1.706E-06 | 6.700E-06 | 1.000E-05 |
| 8630 | 432 | G432I | A | 7.8e-06 | 1.4e-05 | 9.5e-06 | 1.043E-05 | 3.204E-06 | 7.800E-06 | 1.400E-05 |
| 8631 | 432 | G432L | A | 8.8e-06 | 8.1e-06 | 9.4e-06 | 8.767E-06 | 6.506E-07 | 8.100E-06 | 9.400E-06 |
| 8632 | 432 | G432K | A | 5.7e-06 | 1.3e-05 | 1.2e-05 | 1.023E-05 | 3.958E-06 | 5.700E-06 | 1.300E-05 |
| 8633 | 432 | G432M | A | 7.6e-06 | 1.3e-05 | 6.5e-06 | 9.033E-06 | 3.479E-06 | 6.500E-06 | 1.300E-05 |
| 8634 | 432 | G432F | A | 4.7e-06 | 1.5e-05 | 7.1e-06 | 8.933E-06 | 5.389E-06 | 4.700E-06 | 1.500E-05 |
| 8635 | 432 | G432P | A | 5e-06 | 7.5e-06 | 1.2e-05 | 8.167E-06 | 3.547E-06 | 5.000E-06 | 1.200E-05 |
| 8636 | 432 | G432S | A | 7.7e-06 | 8.8e-06 | 6.4e-06 | 7.633E-06 | 1.201E-06 | 6.400E-06 | 8.800E-06 |
| 8637 | 432 | G432T | A | 4.5e-06 | 8.4e-06 | 6.7e-06 | 6.533E-06 | 1.955E-06 | 4.500E-06 | 8.400E-06 |
| 8638 | 432 | G432W | A | 5.4e-06 | 7.4e-06 | 1.1e-05 | 7.933E-06 | 2.838E-06 | 5.400E-06 | 1.100E-05 |
| 8639 | 432 | G432Y | A | 1.2e-05 | 8.8e-06 | 6.4e-06 | 9.067E-06 | 2.810E-06 | 6.400E-06 | 1.200E-05 |
| 8640 | 432 | G432V | A | 7.2e-06 | 8.4e-06 | 6.7e-06 | 7.433E-06 | 8.737E-07 | 6.700E-06 | 8.400E-06 |
| 8641 | 433 | L433A | A | 1.1e-05 | 1.3e-05 | 6e-06 | 1.000E-05 | 3.606E-06 | 6.000E-06 | 1.300E-05 |
| 8642 | 433 | L433R | A | 7.4e-06 | 9.5e-06 | 1.4e-05 | 1.030E-05 | 3.372E-06 | 7.400E-06 | 1.400E-05 |
| 8643 | 433 | L433N | A | 4.5e-06 | 8.8e-06 | 8.5e-06 | 7.267E-06 | 2.401E-06 | 4.500E-06 | 8.800E-06 |
| 8644 | 433 | L433D | A | 7.1e-06 | 7.5e-06 | 9.8e-06 | 8.133E-06 | 1.457E-06 | 7.100E-06 | 9.800E-06 |
| 8645 | 433 | L433C | A | 8.6e-06 | 9.5e-06 | 8.5e-06 | 8.867E-06 | 5.508E-07 | 8.500E-06 | 9.500E-06 |
| 8646 | 433 | L433E | A | 8e-06 | 5.4e-06 | 5e-06 | 6.133E-06 | 1.629E-06 | 5.000E-06 | 8.000E-06 |
| 8647 | 433 | L433Q | A | 9e-06 | 6e-06 | 5e-06 | 6.667E-06 | 2.082E-06 | 5.000E-06 | 9.000E-06 |
| 8648 | 433 | L433G | A | 8.4e-06 | 4.7e-06 | 7.4e-06 | 6.833E-06 | 1.914E-06 | 4.700E-06 | 8.400E-06 |
| 8649 | 433 | L433H | A | 9.3e-06 | 8.5e-06 | 1.4e-05 | 1.060E-05 | 2.972E-06 | 8.500E-06 | 1.400E-05 |
| 8650 | 433 | L433I | A | 9.7e-06 | 1.3e-05 | 9.2e-06 | 1.063E-05 | 2.065E-06 | 9.200E-06 | 1.300E-05 |
| 8651 | 433 | L433L | A | 6.6e-06 | 6.4e-06 | 7e-06 | 6.667E-06 | 3.055E-07 | 6.400E-06 | 7.000E-06 |
| 8652 | 433 | L433K | A | 1.2e-05 | 1.3e-05 | 6.5e-06 | 1.050E-05 | 3.500E-06 | 6.500E-06 | 1.300E-05 |
| 8653 | 433 | L433M | A | 1e-05 | 6.4e-06 | 8.4e-06 | 8.267E-06 | 1.804E-06 | 6.400E-06 | 1.000E-05 |
| 8654 | 433 | L433F | A | 1.3e-05 | 1.1e-05 | 1e-05 | 1.133E-05 | 1.528E-06 | 1.000E-05 | 1.300E-05 |
| 8655 | 433 | L433P | A | 1.3e-05 | 1.3e-05 | 5.6e-06 | 1.053E-05 | 4.272E-06 | 5.600E-06 | 1.300E-05 |
| 8656 | 433 | L433S | A | 1.3e-05 | 6.9e-06 | 6.3e-06 | 8.733E-06 | 3.707E-06 | 6.300E-06 | 1.300E-05 |
| 8657 | 433 | L433T | A | 6.5e-06 | 7.8e-06 | 9.5e-06 | 7.933E-06 | 1.504E-06 | 6.500E-06 | 9.500E-06 |
| 8658 | 433 | L433W | A | 5.5e-06 | 4.7e-06 | 1.3e-05 | 7.733E-06 | 4.579E-06 | 4.700E-06 | 1.300E-05 |
| 8659 | 433 | L433Y | A | 9.7e-06 | 1e-05 | 8.6e-06 | 9.433E-06 | 7.371E-07 | 8.600E-06 | 1.000E-05 |
| 8660 | 433 | L433V | A | 7.7e-06 | 4.4e-06 | 6.5e-06 | 6.200E-06 | 1.670E-06 | 4.400E-06 | 7.700E-06 |
| 8661 | 434 | R434A | A | 5.9e-06 | 8.6e-06 | 5e-06 | 6.500E-06 | 1.873E-06 | 5.000E-06 | 8.600E-06 |
| 8662 | 434 | R434R | A | 8.8e-06 | 6.7e-06 | 5.6e-06 | 7.033E-06 | 1.626E-06 | 5.600E-06 | 8.800E-06 |
| 8663 | 434 | R434N | A | 9.2e-06 | 6.6e-06 | 9.9e-06 | 8.567E-06 | 1.739E-06 | 6.600E-06 | 9.900E-06 |
| 8664 | 434 | R434D | A | 1.3e-05 | 6.7e-06 | 7.1e-06 | 8.933E-06 | 3.528E-06 | 6.700E-06 | 1.300E-05 |
| 8665 | 434 | R434C | A | 1.5e-05 | 6.1e-06 | 8.5e-06 | 9.867E-06 | 4.605E-06 | 6.100E-06 | 1.500E-05 |
| 8666 | 434 | R434E | A | 1.2e-05 | 7e-06 | 1.3e-05 | 1.067E-05 | 3.215E-06 | 7.000E-06 | 1.300E-05 |
| 8667 | 434 | R434Q | A | 8.7e-06 | 8.2e-06 | 9e-06 | 8.633E-06 | 4.041E-07 | 8.200E-06 | 9.000E-06 |
| 8668 | 434 | R434G | A | 8.5e-06 | 1.4e-05 | 8.2e-06 | 1.023E-05 | 3.265E-06 | 8.200E-06 | 1.400E-05 |
| 8669 | 434 | R434H | A | 1e-05 | 1.3e-05 | 9e-06 | 1.067E-05 | 2.082E-06 | 9.000E-06 | 1.300E-05 |

| | | | | | | | | | | |
|---|---|---|---|---|---|---|---|---|---|---|
| 8670 | 434 | R434I | A | 5.7e-06 | 6e-06 | 4.7e-06 | 5.467E-06 | 6.807E-07 | 4.700E-06 | 6.000E-06 |
| 8671 | 434 | R434L | A | 7.1e-06 | 8.3e-06 | 8.7e-06 | 8.033E-06 | 8.327E-07 | 7.100E-06 | 8.700E-06 |
| 8672 | 434 | R434K | A | 1.4e-05 | 9.1e-06 | 6.7e-06 | 9.933E-06 | 3.721E-06 | 6.700E-06 | 1.400E-05 |
| 8673 | 434 | R434M | A | 1.3e-05 | 6.5e-06 | 4e-06 | 7.833E-06 | 4.646E-06 | 4.000E-06 | 1.300E-05 |
| 8674 | 434 | R434F | A | 9e-06 | 8.6e-06 | 4.3e-06 | 7.300E-06 | 2.606E-06 | 4.300E-06 | 9.000E-06 |
| 8675 | 434 | R434P | A | 5.8e-06 | 7.1e-06 | 5.5e-06 | 6.133E-06 | 8.505E-07 | 5.500E-06 | 7.100E-06 |
| 8676 | 434 | R434S | A | 5.7e-06 | 6.1e-06 | 5.2e-06 | 5.667E-06 | 4.509E-07 | 5.200E-06 | 6.100E-06 |
| 8677 | 434 | R434T | A | 4.8e-06 | 1.3e-05 | 5.6e-06 | 7.800E-06 | 4.521E-06 | 4.800E-06 | 1.300E-05 |
| 8678 | 434 | R434W | A | 6.5e-06 | 1.2e-05 | 1.1e-05 | 9.833E-06 | 2.930E-06 | 6.500E-06 | 1.200E-05 |
| 8679 | 434 | R434Y | A | 4.2e-06 | 7.9e-06 | 1.9e-05 | 1.037E-05 | 7.702E-06 | 4.200E-06 | 1.900E-05 |
| 8680 | 434 | R434V | A | 6.1e-06 | 1e-05 | 9.4e-06 | 8.500E-06 | 2.100E-06 | 6.100E-06 | 1.000E-05 |
| 8681 | 435 | S435A | A | 7e-06 | 6.9e-06 | 8.9e-06 | 7.600E-06 | 1.127E-06 | 6.900E-06 | 8.900E-06 |
| 8682 | 435 | S435R | A | 4.2e-06 | 1.3e-05 | 1.1e-05 | 9.400E-06 | 4.613E-06 | 4.200E-06 | 1.300E-05 |
| 8683 | 435 | S435N | A | 6.1e-06 | 6.9e-06 | 1.3e-05 | 8.667E-06 | 3.774E-06 | 6.100E-06 | 1.300E-05 |
| 8684 | 435 | S435D | A | 4.7e-06 | 6.2e-06 | 1e-05 | 6.967E-06 | 2.732E-06 | 4.700E-06 | 1.000E-05 |
| 8685 | 435 | S435C | A | 9.6e-06 | 5.9e-06 | 1.1e-05 | 8.833E-06 | 2.635E-06 | 5.900E-06 | 1.100E-05 |
| 8686 | 435 | S435E | A | 5.2e-06 | 6.8e-06 | 6.4e-06 | 6.133E-06 | 8.327E-07 | 5.200E-06 | 6.800E-06 |
| 8687 | 435 | S435Q | A | 1.1e-05 | 5.5e-06 | 8.6e-06 | 8.367E-06 | 2.757E-06 | 5.500E-06 | 1.100E-05 |
| 8688 | 435 | S435G | A | 3.8e-06 | 7.3e-06 | 6.7e-06 | 5.933E-06 | 1.872E-06 | 3.800E-06 | 7.300E-06 |
| 8689 | 435 | S435H | A | 6.1e-06 | 9e-06 | 6.7e-06 | 7.267E-06 | 1.531E-06 | 6.100E-06 | 9.000E-06 |
| 8690 | 435 | S435I | A | 6e-06 | 7e-06 | 7e-06 | 6.667E-06 | 5.774E-07 | 6.000E-06 | 7.000E-06 |
| 8691 | 435 | S435L | A | 9.9e-06 | 9.9e-06 | 1e-05 | 9.933E-06 | 5.774E-08 | 9.900E-06 | 1.000E-05 |
| 8692 | 435 | S435K | A | 6.5e-06 | 1.3e-05 | 4.7e-06 | 8.067E-06 | 4.366E-06 | 4.700E-06 | 1.300E-05 |
| 8693 | 435 | S435M | A | 1.1e-05 | 1.5e-05 | 6.5e-06 | 1.083E-05 | 4.252E-06 | 6.500E-06 | 1.500E-05 |
| 8694 | 435 | S435F | A | 5.7e-06 | 1.2e-05 | 1.3e-05 | 1.023E-05 | 3.958E-06 | 5.700E-06 | 1.300E-05 |
| 8695 | 435 | S435P | A | 6.3e-06 | 1.5e-05 | 4.7e-06 | 8.667E-06 | 5.543E-06 | 4.700E-06 | 1.500E-05 |
| 8696 | 435 | S435S | A | 5.4e-06 | 6.4e-06 | 4.4e-06 | 5.400E-06 | 1.000E-06 | 4.400E-06 | 6.400E-06 |
| 8697 | 435 | S435T | A | 1.3e-05 | 1e-05 | 7.5e-06 | 1.017E-05 | 2.754E-06 | 7.500E-06 | 1.300E-05 |
| 8698 | 435 | S435W | A | 7.4e-06 | 9.4e-06 | 5.7e-06 | 7.500E-06 | 1.852E-06 | 5.700E-06 | 9.400E-06 |
| 8699 | 435 | S435Y | A | 8.9e-06 | 1.4e-05 | 8.6e-06 | 1.050E-05 | 3.035E-06 | 8.600E-06 | 1.400E-05 |
| 8700 | 435 | S435V | A | 6.5e-06 | 8.3e-06 | 9.5e-06 | 8.100E-06 | 1.510E-06 | 6.500E-06 | 9.500E-06 |
| 8701 | 436 | L436A | A | 1.2e-05 | 1.4e-05 | 6.6e-06 | 1.087E-05 | 3.828E-06 | 6.600E-06 | 1.400E-05 |
| 8702 | 436 | L436R | A | 5.2e-06 | 6.7e-06 | 6.5e-06 | 6.133E-06 | 8.145E-07 | 5.200E-06 | 6.700E-06 |
| 8703 | 436 | L436N | A | 5.1e-06 | 7.9e-06 | 7.6e-06 | 6.867E-06 | 1.537E-06 | 5.100E-06 | 7.900E-06 |
| 8704 | 436 | L436D | A | 6.3e-06 | 9.7e-06 | 1.1e-05 | 9.000E-06 | 2.427E-06 | 6.300E-06 | 1.100E-05 |
| 8705 | 436 | L436C | A | 6.5e-06 | 9.3e-06 | 7e-06 | 7.600E-06 | 1.493E-06 | 6.500E-06 | 9.300E-06 |
| 8706 | 436 | L436E | A | 8.8e-06 | 1.3e-05 | 1e-05 | 1.060E-05 | 2.163E-06 | 8.800E-06 | 1.300E-05 |
| 8707 | 436 | L436Q | A | 1.3e-05 | 6.7e-06 | 1.3e-05 | 1.090E-05 | 3.637E-06 | 6.700E-06 | 1.300E-05 |
| 8708 | 436 | L436G | A | 6.2e-06 | 7.9e-06 | 7.1e-06 | 7.067E-06 | 8.505E-07 | 6.200E-06 | 7.900E-06 |
| 8709 | 436 | L436H | A | 1e-05 | 1.2e-05 | 4.5e-06 | 8.833E-06 | 3.884E-06 | 4.500E-06 | 1.200E-05 |
| 8710 | 436 | L436I | A | 9.2e-06 | 5e-06 | 5.6e-06 | 6.600E-06 | 2.272E-06 | 5.000E-06 | 9.200E-06 |
| 8711 | 436 | L436L | A | 9e-06 | 9.5e-06 | 9e-06 | 9.167E-06 | 2.887E-07 | 9.000E-06 | 9.500E-06 |
| 8712 | 436 | L436K | A | 9.6e-06 | 7.4e-06 | 9.6e-06 | 8.867E-06 | 1.270E-06 | 7.400E-06 | 9.600E-06 |
| 8713 | 436 | L436M | A | 1.1e-05 | 6.6e-06 | 5.4e-06 | 7.667E-06 | 2.948E-06 | 5.400E-06 | 1.100E-05 |
| 8714 | 436 | L436F | A | 4.6e-06 | 5.6e-06 | 6.3e-06 | 5.500E-06 | 8.544E-07 | 4.600E-06 | 6.300E-06 |
| 8715 | 436 | L436P | A | 9.1e-06 | 9.9e-06 | 6.6e-06 | 8.533E-06 | 1.721E-06 | 6.600E-06 | 9.900E-06 |
| 8716 | 436 | L436S | A | 1.2e-05 | 9.8e-06 | 1.4e-05 | 1.193E-05 | 2.101E-06 | 9.800E-06 | 1.400E-05 |
| 8717 | 436 | L436T | A | 1.1e-05 | 4.9e-06 | 7e-06 | 7.633E-06 | 3.099E-06 | 4.900E-06 | 1.100E-05 |
| 8718 | 436 | L436W | A | 8.2e-06 | 9.5e-06 | 7.5e-06 | 8.400E-06 | 1.015E-06 | 7.500E-06 | 9.500E-06 |
| 8719 | 436 | L436Y | A | 7.6e-06 | 5.7e-06 | 7.6e-06 | 6.967E-06 | 1.097E-06 | 5.700E-06 | 7.600E-06 |
| 8720 | 436 | L436V | A | 5.8e-06 | 9.5e-06 | 1.4e-05 | 9.767E-06 | 4.106E-06 | 5.800E-06 | 1.400E-05 |
| 8721 | 437 | R437A | A | 7.6e-06 | 1.1e-05 | 5.2e-06 | 7.933E-06 | 2.914E-06 | 5.200E-06 | 1.100E-05 |
| 8722 | 437 | R437R | A | 1e-05 | 4.7e-06 | 6.8e-06 | 7.167E-06 | 2.669E-06 | 4.700E-06 | 1.000E-05 |
| 8723 | 437 | R437N | A | 1.2e-05 | 4.1e-06 | 1e-05 | 8.700E-06 | 4.107E-06 | 4.100E-06 | 1.200E-05 |
| 8724 | 437 | R437D | A | 5.8e-06 | 1.1e-05 | 9.5e-06 | 8.767E-06 | 2.676E-06 | 5.800E-06 | 1.100E-05 |
| 8725 | 437 | R437C | A | 9.9e-06 | 7.5e-06 | 5.9e-06 | 7.767E-06 | 2.013E-06 | 5.900E-06 | 9.900E-06 |
| 8726 | 437 | R437E | A | 1e-05 | 7.8e-06 | 8.6e-06 | 8.800E-06 | 1.114E-06 | 7.800E-06 | 1.000E-05 |
| 8727 | 437 | R437Q | A | 6.1e-06 | 8.6e-06 | 1.2e-05 | 8.900E-06 | 2.961E-06 | 6.100E-06 | 1.200E-05 |
| 8728 | 437 | R437G | A | 5.9e-06 | 8.4e-06 | 6.8e-06 | 7.033E-06 | 1.266E-06 | 5.900E-06 | 8.400E-06 |
| 8729 | 437 | R437H | A | 1.3e-05 | 4.6e-06 | 4.7e-06 | 7.433E-06 | 4.821E-06 | 4.600E-06 | 1.300E-05 |
| 8730 | 437 | R437I | A | 6.8e-06 | 5.6e-06 | 5e-06 | 5.800E-06 | 9.165E-07 | 5.000E-06 | 6.800E-06 |
| 8731 | 437 | R437L | A | 4.8e-06 | 1.3e-05 | 1.1e-05 | 9.600E-06 | 4.276E-06 | 4.800E-06 | 1.300E-05 |
| 8732 | 437 | R437K | A | 6.3e-06 | 6.4e-06 | 6.3e-06 | 6.333E-06 | 5.774E-08 | 6.300E-06 | 6.400E-06 |
| 8733 | 437 | R437M | A | 8.7e-06 | 6.4e-06 | 8e-06 | 7.700E-06 | 1.179E-06 | 6.400E-06 | 8.700E-06 |
| 8734 | 437 | R437F | A | 6.6e-06 | 8e-06 | 8e-06 | 7.533E-06 | 8.083E-07 | 6.600E-06 | 8.000E-06 |
| 8735 | 437 | R437P | A | 5.8e-06 | 9e-06 | 6.5e-06 | 7.100E-06 | 1.682E-06 | 5.800E-06 | 9.000E-06 |
| 8736 | 437 | R437S | A | 6.3e-06 | 1.3e-05 | 6.7e-06 | 8.667E-06 | 3.758E-06 | 6.300E-06 | 1.300E-05 |
| 8737 | 437 | R437T | A | 9.8e-06 | 8.2e-06 | 8.8e-06 | 8.933E-06 | 8.083E-07 | 8.200E-06 | 9.800E-06 |
| 8738 | 437 | R437W | A | 4.5e-06 | 1e-05 | 8.5e-06 | 7.667E-06 | 2.843E-06 | 4.500E-06 | 1.000E-05 |
| 8739 | 437 | R437Y | A | 5.6e-06 | 7e-06 | 6.7e-06 | 6.433E-06 | 7.371E-07 | 5.600E-06 | 7.000E-06 |
| 8740 | 437 | R437V | A | 8.4e-06 | 8.3e-06 | 9.5e-06 | 8.733E-06 | 6.658E-07 | 8.300E-06 | 9.500E-06 |
| 8741 | 438 | E438A | A | 9.1e-06 | 9.3e-06 | 6.4e-06 | 8.267E-06 | 1.620E-06 | 6.400E-06 | 9.300E-06 |
| 8742 | 438 | E438R | A | 9.4e-06 | 6.8e-06 | 5.7e-06 | 7.300E-06 | 1.900E-06 | 5.700E-06 | 9.400E-06 |
| 8743 | 438 | E438N | A | 8.6e-06 | 7.7e-06 | 1.3e-05 | 9.767E-06 | 2.836E-06 | 7.700E-06 | 1.300E-05 |
| 8744 | 438 | E438D | A | 1.3e-05 | 8.2e-06 | 1.3e-05 | 1.140E-05 | 2.771E-06 | 8.200E-06 | 1.300E-05 |
| 8745 | 438 | E438C | A | 1.4e-05 | 6e-06 | 6.9e-06 | 8.967E-06 | 4.382E-06 | 6.000E-06 | 1.400E-05 |
| 8746 | 438 | E438E | A | 7.3e-06 | 9.8e-06 | 1.1e-05 | 9.367E-06 | 1.888E-06 | 7.300E-06 | 1.100E-05 |
| 8747 | 438 | E438Q | A | 9.5e-06 | 1.2e-05 | 9.1e-06 | 1.020E-05 | 1.572E-06 | 9.100E-06 | 1.200E-05 |
| 8748 | 438 | E438G | A | 6.1e-06 | 8.8e-06 | 1.1e-05 | 8.633E-06 | 2.454E-06 | 6.100E-06 | 1.100E-05 |
| 8749 | 438 | E438H | A | 8.2e-06 | 4.9e-06 | 5.1e-06 | 6.067E-06 | 1.850E-06 | 4.900E-06 | 8.200E-06 |
| 8750 | 438 | E438I | A | 9.3e-06 | 8.9e-06 | 1.1e-05 | 9.733E-06 | 1.115E-06 | 8.900E-06 | 1.100E-05 |
| 8751 | 438 | E438L | A | 9.7e-06 | 7.6e-06 | 1.1e-05 | 9.433E-06 | 1.716E-06 | 7.600E-06 | 1.100E-05 |
| 8752 | 438 | E438K | A | 6.7e-06 | 1.3e-05 | 1.3e-05 | 1.090E-05 | 3.637E-06 | 6.700E-06 | 1.300E-05 |
| 8753 | 438 | E438M | A | 7.6e-06 | 5.8e-06 | 8.1e-06 | 7.167E-06 | 1.210E-06 | 5.800E-06 | 8.100E-06 |
| 8754 | 438 | E438F | A | 6.8e-06 | 5e-06 | 1e-05 | 7.267E-06 | 2.532E-06 | 5.000E-06 | 1.000E-05 |

| | | | | | | | | | | |
|---|---|---|---|---|---|---|---|---|---|---|
| 8755 | 438 | E438P | A | 9.2e-06 | 7.1e-06 | 9.2e-06 | 8.500E-06 | 1.212E-06 | 7.100E-06 | 9.200E-06 |
| 8756 | 438 | E438S | A | 8.2e-06 | 6.5e-06 | 9.3e-06 | 8.000E-06 | 1.411E-06 | 6.500E-06 | 9.300E-06 |
| 8757 | 438 | E438T | A | 5.2e-06 | 7.1e-06 | 5.1e-06 | 5.800E-06 | 1.127E-06 | 5.100E-06 | 7.100E-06 |
| 8758 | 438 | E438W | A | 9.4e-06 | 3.4e-06 | 8.4e-06 | 7.067E-06 | 3.215E-06 | 3.400E-06 | 9.400E-06 |
| 8759 | 438 | E438Y | A | 9.5e-06 | 5.5e-06 | 8.4e-06 | 7.800E-06 | 2.066E-06 | 5.500E-06 | 9.500E-06 |
| 8760 | 438 | E438V | A | 1e-05 | 7.5e-06 | 9.6e-06 | 9.033E-06 | 1.343E-06 | 7.500E-06 | 1.000E-05 |
| 8761 | 439 | L439A | A | 9.8e-06 | 5.1e-06 | 7.6e-06 | 7.500E-06 | 2.352E-06 | 5.100E-06 | 9.800E-06 |
| 8762 | 439 | L439R | A | 4.5e-06 | 4.8e-06 | 3.9e-06 | 4.400E-06 | 4.583E-07 | 3.900E-06 | 4.800E-06 |
| 8763 | 439 | L439N | A | 8.4e-06 | 1.4e-05 | 8.5e-06 | 1.030E-05 | 3.205E-06 | 8.400E-06 | 1.400E-05 |
| 8764 | 439 | L439D | A | 5.6e-06 | 1.4e-05 | 9.7e-06 | 9.767E-06 | 4.200E-06 | 5.600E-06 | 1.400E-05 |
| 8765 | 439 | L439C | A | 1.2e-05 | 1.3e-05 | 6.5e-06 | 1.050E-05 | 3.500E-06 | 6.500E-06 | 1.300E-05 |
| 8766 | 439 | L439E | A | 1.2e-05 | 9.5e-06 | 6.1e-06 | 9.200E-06 | 2.961E-06 | 6.100E-06 | 1.200E-05 |
| 8767 | 439 | L439Q | A | 8.8e-06 | 6.2e-06 | 6.7e-06 | 7.233E-06 | 1.380E-06 | 6.200E-06 | 8.800E-06 |
| 8768 | 439 | L439G | A | 4.9e-06 | 4.6e-06 | 1.1e-05 | 6.833E-06 | 3.612E-06 | 4.600E-06 | 1.100E-05 |
| 8769 | 439 | L439H | A | 7.1e-06 | 7.4e-06 | 8.4e-06 | 7.633E-06 | 6.807E-07 | 7.100E-06 | 8.400E-06 |
| 8770 | 439 | L439I | A | 7.2e-06 | 8.7e-06 | 6.9e-06 | 7.600E-06 | 9.644E-07 | 6.900E-06 | 8.700E-06 |
| 8771 | 439 | L439L | A | 5.1e-06 | 5.4e-06 | 8.4e-06 | 6.300E-06 | 1.825E-06 | 5.100E-06 | 8.400E-06 |
| 8772 | 439 | L439K | A | 6e-06 | 7.7e-06 | 5.2e-06 | 6.300E-06 | 1.277E-06 | 5.200E-06 | 7.700E-06 |
| 8773 | 439 | L439M | A | 9.8e-06 | 5.1e-06 | 6.6e-06 | 7.167E-06 | 2.401E-06 | 5.100E-06 | 9.800E-06 |
| 8774 | 439 | L439F | A | 1.1e-05 | 5.1e-06 | 7.3e-06 | 7.800E-06 | 2.982E-06 | 5.100E-06 | 1.100E-05 |
| 8775 | 439 | L439P | A | 7.7e-06 | 7.7e-06 | 9.5e-06 | 8.300E-06 | 1.039E-06 | 7.700E-06 | 9.500E-06 |
| 8776 | 439 | L439S | A | 6e-06 | 7.1e-06 | 7.9e-06 | 7.000E-06 | 9.539E-07 | 6.000E-06 | 7.900E-06 |
| 8777 | 439 | L439T | A | 1e-05 | 5e-06 | 6.1e-06 | 7.033E-06 | 2.627E-06 | 5.000E-06 | 1.000E-05 |
| 8778 | 439 | L439W | A | 6.3e-06 | 9.9e-06 | 7e-06 | 7.733E-06 | 1.909E-06 | 6.300E-06 | 9.900E-06 |
| 8779 | 439 | L439Y | A | 6.4e-06 | 6.8e-06 | 6.6e-06 | 6.600E-06 | 2.000E-07 | 6.400E-06 | 6.800E-06 |
| 8780 | 439 | L439V | A | 5.2e-06 | 7e-06 | 9.8e-06 | 7.333E-06 | 2.318E-06 | 5.200E-06 | 9.800E-06 |
| 8781 | 440 | G440A | A | 6.7e-06 | 6.3e-06 | 1.6e-05 | 9.667E-06 | 5.488E-06 | 6.300E-06 | 1.600E-05 |
| 8782 | 440 | G440R | A | 4.9e-06 | 1.1e-05 | 3.8e-06 | 6.567E-06 | 3.879E-06 | 3.800E-06 | 1.100E-05 |
| 8783 | 440 | G440N | A | 1.1e-05 | 9.1e-06 | 9.8e-06 | 9.967E-06 | 9.609E-07 | 9.100E-06 | 1.100E-05 |
| 8784 | 440 | G440D | A | 7.5e-06 | 1e-05 | 1e-05 | 9.167E-06 | 1.443E-06 | 7.500E-06 | 1.000E-05 |
| 8785 | 440 | G440C | A | 1.1e-05 | 1.5e-05 | 6.6e-06 | 1.087E-05 | 4.202E-06 | 6.600E-06 | 1.500E-05 |
| 8786 | 440 | G440E | A | 9.4e-06 | 1.2e-05 | 4.4e-06 | 8.600E-06 | 3.863E-06 | 4.400E-06 | 1.200E-05 |
| 8787 | 440 | G440Q | A | 8.1e-06 | 5.6e-06 | 6.9e-06 | 6.867E-06 | 1.250E-06 | 5.600E-06 | 8.100E-06 |
| 8788 | 440 | G440G | A | 9.8e-06 | 8.6e-06 | 9.4e-06 | 9.267E-06 | 6.110E-07 | 8.600E-06 | 9.800E-06 |
| 8789 | 440 | G440H | A | 7.6e-06 | 6.5e-06 | 7.9e-06 | 7.333E-06 | 7.371E-07 | 6.500E-06 | 7.900E-06 |
| 8790 | 440 | G440I | A | 7.5e-06 | 1.3e-05 | 5.2e-06 | 8.567E-06 | 4.008E-06 | 5.200E-06 | 1.300E-05 |
| 8791 | 440 | G440L | A | 7e-06 | 5.4e-06 | 1.3e-05 | 8.467E-06 | 4.007E-06 | 5.400E-06 | 1.300E-05 |
| 8792 | 440 | G440K | A | 1.3e-05 | 1.2e-05 | 6.9e-06 | 1.063E-05 | 3.272E-06 | 6.900E-06 | 1.300E-05 |
| 8793 | 440 | G440M | A | 4.8e-06 | 9.7e-06 | 9.5e-06 | 8.000E-06 | 2.773E-06 | 4.800E-06 | 9.700E-06 |
| 8794 | 440 | G440F | A | 9.1e-06 | 6.3e-06 | 6e-06 | 7.133E-06 | 1.710E-06 | 6.000E-06 | 9.100E-06 |
| 8795 | 440 | G440P | A | 8.2e-06 | 5.4e-06 | 6.7e-06 | 6.767E-06 | 1.401E-06 | 5.400E-06 | 8.200E-06 |
| 8796 | 440 | G440S | A | 8.4e-06 | 1e-05 | 6e-06 | 8.133E-06 | 2.013E-06 | 6.000E-06 | 1.000E-05 |
| 8797 | 440 | G440T | A | 4.9e-06 | 6.7e-06 | 7e-06 | 6.200E-06 | 1.136E-06 | 4.900E-06 | 7.000E-06 |
| 8798 | 440 | G440W | A | 6e-06 | 1.2e-05 | 5.8e-06 | 7.933E-06 | 3.523E-06 | 5.800E-06 | 1.200E-05 |
| 8799 | 440 | G440Y | A | 7.5e-06 | 1.3e-05 | 6.5e-06 | 9.000E-06 | 3.500E-06 | 6.500E-06 | 1.300E-05 |
| 8800 | 440 | G440V | A | 7.3e-06 | 5.5e-06 | 6.5e-06 | 6.433E-06 | 9.018E-07 | 5.500E-06 | 7.300E-06 |
| 8801 | 441 | S441A | A | 1.3e-05 | 8.2e-06 | 5.9e-06 | 9.033E-06 | 3.623E-06 | 5.900E-06 | 1.300E-05 |
| 8802 | 441 | S441R | A | 1.6e-05 | 9.4e-06 | 1.2e-05 | 1.247E-05 | 3.325E-06 | 9.400E-06 | 1.600E-05 |
| 8803 | 441 | S441N | A | 9.2e-06 | 5.8e-06 | 6.5e-06 | 7.167E-06 | 1.795E-06 | 5.800E-06 | 9.200E-06 |
| 8804 | 441 | S441D | A | 6.8e-06 | 9.4e-06 | 5.1e-06 | 7.100E-06 | 2.166E-06 | 5.100E-06 | 9.400E-06 |
| 8805 | 441 | S441C | A | 8.9e-06 | 1.1e-05 | 6.6e-06 | 8.833E-06 | 2.201E-06 | 6.600E-06 | 1.100E-05 |
| 8806 | 441 | S441E | A | 8.1e-06 | 8.4e-06 | 8.8e-06 | 8.433E-06 | 3.512E-07 | 8.100E-06 | 8.800E-06 |
| 8807 | 441 | S441Q | A | 7.4e-06 | 9.7e-06 | 1.3e-05 | 1.003E-05 | 2.815E-06 | 7.400E-06 | 1.300E-05 |
| 8808 | 441 | S441G | A | 9.4e-06 | 6.5e-06 | 6.4e-06 | 7.433E-06 | 1.704E-06 | 6.400E-06 | 9.400E-06 |
| 8809 | 441 | S441H | A | 4.2e-06 | 6.2e-06 | 6.7e-06 | 5.700E-06 | 1.323E-06 | 4.200E-06 | 6.700E-06 |
| 8810 | 441 | S441I | A | 1e-05 | 6.7e-06 | 1.2e-05 | 9.567E-06 | 2.676E-06 | 6.700E-06 | 1.200E-05 |
| 8811 | 441 | S441L | A | 8.1e-06 | 1e-05 | 6.9e-06 | 8.333E-06 | 1.563E-06 | 6.900E-06 | 1.000E-05 |
| 8812 | 441 | S441K | A | 7.1e-06 | 5.9e-06 | 1.2e-05 | 8.333E-06 | 3.232E-06 | 5.900E-06 | 1.200E-05 |
| 8813 | 441 | S441M | A | 8.1e-06 | 5.1e-06 | 6.5e-06 | 6.567E-06 | 1.501E-06 | 5.100E-06 | 8.100E-06 |
| 8814 | 441 | S441F | A | 9.3e-06 | 1.2e-05 | 7.5e-06 | 9.600E-06 | 2.265E-06 | 7.500E-06 | 1.200E-05 |
| 8815 | 441 | S441P | A | 7.2e-06 | 1.3e-05 | 9.9e-06 | 1.003E-05 | 2.902E-06 | 7.200E-06 | 1.300E-05 |
| 8816 | 441 | S441S | A | 5.7e-06 | 9.6e-06 | 5.6e-06 | 6.967E-06 | 2.281E-06 | 5.600E-06 | 9.600E-06 |
| 8817 | 441 | S441T | A | 5.8e-06 | 1.2e-05 | 8.1e-06 | 8.633E-06 | 3.134E-06 | 5.800E-06 | 1.200E-05 |
| 8818 | 441 | S441W | A | 9.2e-06 | 8.8e-06 | 6.7e-06 | 8.233E-06 | 1.343E-06 | 6.700E-06 | 9.200E-06 |
| 8819 | 441 | S441Y | A | 7.9e-06 | 1.2e-05 | 9.2e-06 | 9.700E-06 | 2.095E-06 | 7.900E-06 | 1.200E-05 |
| 8820 | 441 | S441V | A | 1.4e-05 | 4.5e-06 | 1.3e-05 | 1.050E-05 | 5.220E-06 | 4.500E-06 | 1.400E-05 |
| 8821 | 442 | G442A | A | 8.8e-06 | 5.4e-06 | 4.7e-06 | 6.300E-06 | 2.193E-06 | 4.700E-06 | 8.800E-06 |
| 8822 | 442 | G442R | A | 8e-06 | 8.9e-06 | 1.1e-05 | 9.300E-06 | 1.539E-06 | 8.000E-06 | 1.100E-05 |
| 8823 | 442 | G442N | A | 1.1e-05 | 9.2e-06 | 6e-06 | 8.733E-06 | 2.532E-06 | 6.000E-06 | 1.100E-05 |
| 8824 | 442 | G442D | A | 7.1e-06 | 5e-06 | 1.2e-05 | 8.033E-06 | 3.592E-06 | 5.000E-06 | 1.200E-05 |
| 8825 | 442 | G442C | A | 1.2e-05 | 7e-06 | 5.7e-06 | 8.233E-06 | 3.326E-06 | 5.700E-06 | 1.200E-05 |
| 8826 | 442 | G442E | A | 6.7e-06 | 5e-06 | 5.8e-06 | 5.833E-06 | 8.505E-07 | 5.000E-06 | 6.700E-06 |
| 8827 | 442 | G442Q | A | 7.2e-06 | 7.5e-06 | 1.7e-05 | 1.057E-05 | 5.573E-06 | 7.200E-06 | 1.700E-05 |
| 8828 | 442 | G442G | A | 6.8e-06 | 8.5e-06 | 7e-06 | 7.433E-06 | 9.292E-07 | 6.800E-06 | 8.500E-06 |
| 8829 | 442 | G442H | A | 6.6e-06 | 8.8e-06 | 7.7e-06 | 7.700E-06 | 1.100E-06 | 6.600E-06 | 8.800E-06 |
| 8830 | 442 | G442I | A | 4.4e-06 | 4.5e-06 | 9.3e-06 | 6.067E-06 | 2.801E-06 | 4.400E-06 | 9.300E-06 |
| 8831 | 442 | G442L | A | 9.7e-06 | 1.1e-05 | 1e-05 | 1.023E-05 | 6.807E-07 | 9.700E-06 | 1.100E-05 |
| 8832 | 442 | G442K | A | 5.8e-06 | 1.2e-05 | 9.3e-06 | 9.033E-06 | 3.109E-06 | 5.800E-06 | 1.200E-05 |
| 8833 | 442 | G442M | A | 6.4e-06 | 4.6e-06 | 1.3e-05 | 8.000E-06 | 4.423E-06 | 4.600E-06 | 1.300E-05 |
| 8834 | 442 | G442F | A | 7.5e-06 | 1.4e-05 | 6.3e-06 | 9.267E-06 | 4.143E-06 | 6.300E-06 | 1.400E-05 |
| 8835 | 442 | G442P | A | 1.2e-05 | 9e-06 | 8.6e-06 | 9.867E-06 | 1.858E-06 | 8.600E-06 | 1.200E-05 |
| 8836 | 442 | G442S | A | 4.8e-06 | 9.4e-06 | 1.4e-05 | 9.400E-06 | 4.600E-06 | 4.800E-06 | 1.400E-05 |
| 8837 | 442 | G442T | A | 1.1e-05 | 9e-06 | 6.6e-06 | 8.867E-06 | 2.203E-06 | 6.600E-06 | 1.100E-05 |
| 8838 | 442 | G442W | A | 6.3e-06 | 7e-06 | 8.3e-06 | 7.200E-06 | 1.015E-06 | 6.300E-06 | 8.300E-06 |
| 8839 | 442 | G442Y | A | 1.1e-05 | 6.1e-06 | 4.6e-06 | 7.233E-06 | 3.347E-06 | 4.600E-06 | 1.100E-05 |

| | | | | | | | | | | |
|---|---|---|---|---|---|---|---|---|---|---|
| 8840 | 442 | G442V | A | 9.2e-06 | 3.9e-06 | 1.1e-05 | 8.033E-06 | 3.691E-06 | 3.900E-06 | 1.100E-05 |
| 8841 | 443 | L443A | A | 1.5e-05 | 9.6e-06 | 6e-06 | 1.020E-05 | 4.530E-06 | 6.000E-06 | 1.500E-05 |
| 8842 | 443 | L443R | A | 7.7e-06 | 4.5e-06 | 6.9e-06 | 6.367E-06 | 1.665E-06 | 4.500E-06 | 7.700E-06 |
| 8843 | 443 | L443N | A | 7.1e-06 | 1.3e-05 | 5.1e-06 | 8.400E-06 | 4.107E-06 | 5.100E-06 | 1.300E-05 |
| 8844 | 443 | L443D | A | 1.5e-05 | 9.6e-06 | 1e-05 | 1.153E-05 | 3.009E-06 | 9.600E-06 | 1.500E-05 |
| 8845 | 443 | L443C | A | 1.1e-05 | 5.2e-06 | 1.1e-05 | 9.067E-06 | 3.349E-06 | 5.200E-06 | 1.100E-05 |
| 8846 | 443 | L443E | A | 6.7e-06 | 7e-06 | 1.2e-05 | 8.567E-06 | 2.977E-06 | 6.700E-06 | 1.200E-05 |
| 8847 | 443 | L443Q | A | 5.4e-06 | 6.8e-06 | 7.1e-06 | 6.433E-06 | 9.074E-07 | 5.400E-06 | 7.100E-06 |
| 8848 | 443 | L443G | A | 7.1e-06 | 6.4e-06 | 1.1e-05 | 8.167E-06 | 2.479E-06 | 6.400E-06 | 1.100E-05 |
| 8849 | 443 | L443H | A | 1.2e-05 | 8.5e-06 | 1.1e-05 | 1.050E-05 | 1.803E-06 | 8.500E-06 | 1.200E-05 |
| 8850 | 443 | L443I | A | 8.4e-06 | 6.7e-06 | 5.7e-06 | 6.933E-06 | 1.365E-06 | 5.700E-06 | 8.400E-06 |
| 8851 | 443 | L443L | A | 5.8e-06 | 9.6e-06 | 1.3e-05 | 9.467E-06 | 3.602E-06 | 5.800E-06 | 1.300E-05 |
| 8852 | 443 | L443K | A | 9.5e-06 | 1.3e-05 | 4.2e-06 | 8.900E-06 | 4.431E-06 | 4.200E-06 | 1.300E-05 |
| 8853 | 443 | L443M | A | 4.9e-06 | 8.7e-06 | 7.3e-06 | 6.967E-06 | 1.922E-06 | 4.900E-06 | 8.700E-06 |
| 8854 | 443 | L443F | A | 6.6e-06 | 5.9e-06 | 1.4e-05 | 8.833E-06 | 4.488E-06 | 5.900E-06 | 1.400E-05 |
| 8855 | 443 | L443P | A | 6.9e-06 | 7.3e-06 | 1.1e-05 | 8.400E-06 | 2.261E-06 | 6.900E-06 | 1.100E-05 |
| 8856 | 443 | L443S | A | 5.8e-06 | 1.4e-05 | 1.1e-05 | 1.027E-05 | 4.149E-06 | 5.800E-06 | 1.400E-05 |
| 8857 | 443 | L443T | A | 1e-05 | 9.6e-06 | 1.2e-05 | 1.053E-05 | 1.286E-06 | 9.600E-06 | 1.200E-05 |
| 8858 | 443 | L443W | A | 1e-05 | 6.6e-06 | 7.7e-06 | 8.100E-06 | 1.735E-06 | 6.600E-06 | 1.000E-05 |
| 8859 | 443 | L443Y | A | 5.4e-06 | 8.8e-06 | 6.6e-06 | 6.933E-06 | 1.724E-06 | 5.400E-06 | 8.800E-06 |
| 8860 | 443 | L443V | A | 9.8e-06 | 5e-06 | 8.5e-06 | 7.767E-06 | 2.483E-06 | 5.000E-06 | 9.800E-06 |
| 8861 | 444 | A444A | A | 1.1e-05 | 8.9e-06 | 6.4e-06 | 8.767E-06 | 2.303E-06 | 6.400E-06 | 1.100E-05 |
| 8862 | 444 | A444R | A | 6.8e-06 | 8.2e-06 | 1.1e-05 | 8.667E-06 | 2.139E-06 | 6.800E-06 | 1.100E-05 |
| 8863 | 444 | A444N | A | 1.3e-05 | 1.5e-05 | 8.4e-06 | 1.213E-05 | 3.384E-06 | 8.400E-06 | 1.500E-05 |
| 8864 | 444 | A444D | A | 9.4e-06 | 1.3e-05 | 9.9e-06 | 1.077E-05 | 1.950E-06 | 9.400E-06 | 1.300E-05 |
| 8865 | 444 | A444C | A | 6.3e-06 | 5.8e-06 | 7.1e-06 | 6.400E-06 | 6.557E-07 | 5.800E-06 | 7.100E-06 |
| 8866 | 444 | A444E | A | 8e-06 | 5.5e-06 | 8.4e-06 | 7.300E-06 | 1.572E-06 | 5.500E-06 | 8.400E-06 |
| 8867 | 444 | A444Q | A | 7.6e-06 | 9.9e-06 | 9.5e-06 | 9.000E-06 | 1.229E-06 | 7.600E-06 | 9.900E-06 |
| 8868 | 444 | A444G | A | 1.2e-05 | 5.4e-06 | 7.3e-06 | 8.233E-06 | 3.398E-06 | 5.400E-06 | 1.200E-05 |
| 8869 | 444 | A444H | A | 8.6e-06 | 8.3e-06 | 9.9e-06 | 8.933E-06 | 8.505E-07 | 8.300E-06 | 9.900E-06 |
| 8870 | 444 | A444I | A | 7.7e-06 | 5.9e-06 | 5.1e-06 | 6.233E-06 | 1.332E-06 | 5.100E-06 | 7.700E-06 |
| 8871 | 444 | A444L | A | 5.9e-06 | 7.4e-06 | 7.3e-06 | 6.867E-06 | 8.386E-07 | 5.900E-06 | 7.400E-06 |
| 8872 | 444 | A444K | A | 9.1e-06 | 8e-06 | 5.6e-06 | 7.567E-06 | 1.790E-06 | 5.600E-06 | 9.100E-06 |
| 8873 | 444 | A444M | A | 9.8e-06 | 1.1e-05 | 5.7e-06 | 8.833E-06 | 2.779E-06 | 5.700E-06 | 1.100E-05 |
| 8874 | 444 | A444F | A | 6.3e-06 | 6.6e-06 | 5.1e-06 | 6.000E-06 | 7.937E-07 | 5.100E-06 | 6.600E-06 |
| 8875 | 444 | A444P | A | 1.1e-05 | 7.9e-06 | 8.5e-06 | 9.133E-06 | 1.644E-06 | 7.900E-06 | 1.100E-05 |
| 8876 | 444 | A444S | A | 5.8e-06 | 4.8e-06 | 1e-05 | 6.867E-06 | 2.759E-06 | 4.800E-06 | 1.000E-05 |
| 8877 | 444 | A444T | A | 1.1e-05 | 9.3e-06 | 1.2e-05 | 1.077E-05 | 1.365E-06 | 9.300E-06 | 1.200E-05 |
| 8878 | 444 | A444W | A | 9.8e-06 | 6.6e-06 | 8.2e-06 | 8.200E-06 | 1.600E-06 | 6.600E-06 | 9.800E-06 |
| 8879 | 444 | A444Y | A | 7.1e-06 | 5.6e-06 | 9.7e-06 | 7.467E-06 | 2.074E-06 | 5.600E-06 | 9.700E-06 |
| 8880 | 444 | A444V | A | 8.2e-06 | 4.8e-06 | 8.9e-06 | 7.300E-06 | 2.193E-06 | 4.800E-06 | 8.900E-06 |
| 8881 | 445 | L445A | A | 9.1e-06 | 6.6e-06 | 8.6e-06 | 8.100E-06 | 1.323E-06 | 6.600E-06 | 9.100E-06 |
| 8882 | 445 | L445R | A | 1.2e-05 | 5.7e-06 | 6.3e-06 | 8.000E-06 | 3.477E-06 | 5.700E-06 | 1.200E-05 |
| 8883 | 445 | L445N | A | 6.3e-06 | 9.6e-06 | 9.6e-06 | 8.500E-06 | 1.905E-06 | 6.300E-06 | 9.600E-06 |
| 8884 | 445 | L445D | A | 5.7e-06 | 9.1e-06 | 4.4e-06 | 6.400E-06 | 2.427E-06 | 4.400E-06 | 9.100E-06 |
| 8885 | 445 | L445C | A | 7.5e-06 | 8e-06 | 5.8e-06 | 7.100E-06 | 1.153E-06 | 5.800E-06 | 8.000E-06 |
| 8886 | 445 | L445E | A | 9.4e-06 | 7.3e-06 | 4.9e-06 | 7.200E-06 | 2.252E-06 | 4.900E-06 | 9.400E-06 |
| 8887 | 445 | L445Q | A | 6.7e-06 | 1.3e-05 | 1.1e-05 | 1.023E-05 | 3.219E-06 | 6.700E-06 | 1.300E-05 |
| 8888 | 445 | L445G | A | 6.8e-06 | 6.2e-06 | 6.7e-06 | 6.567E-06 | 3.215E-07 | 6.200E-06 | 6.800E-06 |
| 8889 | 445 | L445H | A | 7.7e-06 | 9.2e-06 | 5.6e-06 | 7.500E-06 | 1.808E-06 | 5.600E-06 | 9.200E-06 |
| 8890 | 445 | L445I | A | 5.8e-06 | 9.3e-06 | 8.5e-06 | 7.867E-06 | 1.834E-06 | 5.800E-06 | 9.300E-06 |
| 8891 | 445 | L445L | A | 9.9e-06 | 6.4e-06 | 9.1e-06 | 8.467E-06 | 1.834E-06 | 6.400E-06 | 9.900E-06 |
| 8892 | 445 | L445K | A | 1.3e-05 | 6.6e-06 | 6.7e-06 | 8.767E-06 | 3.667E-06 | 6.600E-06 | 1.300E-05 |
| 8893 | 445 | L445M | A | 8.4e-06 | 8.6e-06 | 7.6e-06 | 8.200E-06 | 5.292E-07 | 7.600E-06 | 8.600E-06 |
| 8894 | 445 | L445F | A | 1e-05 | 6.1e-06 | 9.7e-06 | 8.600E-06 | 2.170E-06 | 6.100E-06 | 1.000E-05 |
| 8895 | 445 | L445P | A | 5.7e-06 | 5.5e-06 | 7.1e-06 | 6.100E-06 | 8.718E-07 | 5.500E-06 | 7.100E-06 |
| 8896 | 445 | L445S | A | 5.9e-06 | 1.2e-05 | 1.1e-05 | 9.633E-06 | 3.272E-06 | 5.900E-06 | 1.200E-05 |
| 8897 | 445 | L445T | A | 4.7e-06 | 5.3e-06 | 9.7e-06 | 6.567E-06 | 2.730E-06 | 4.700E-06 | 9.700E-06 |
| 8898 | 445 | L445W | A | 5.8e-06 | 6.8e-06 | 8.5e-06 | 7.033E-06 | 1.365E-06 | 5.800E-06 | 8.500E-06 |
| 8899 | 445 | L445Y | A | 8.7e-06 | 5.2e-06 | 1.1e-05 | 8.300E-06 | 2.921E-06 | 5.200E-06 | 1.100E-05 |
| 8900 | 445 | L445V | A | 9e-06 | 5.6e-06 | 8.4e-06 | 7.667E-06 | 1.815E-06 | 5.600E-06 | 9.000E-06 |
| 8901 | 446 | I446A | A | 1.1e-05 | 1.8e-05 | 7.1e-06 | 1.203E-05 | 5.523E-06 | 7.100E-06 | 1.800E-05 |
| 8902 | 446 | I446R | A | 7.2e-06 | 5.1e-06 | 7.1e-06 | 6.467E-06 | 1.185E-06 | 5.100E-06 | 7.200E-06 |
| 8903 | 446 | I446N | A | 8.1e-06 | 5.5e-06 | 5.8e-06 | 6.467E-06 | 1.422E-06 | 5.500E-06 | 8.100E-06 |
| 8904 | 446 | I446D | A | 8.3e-06 | 7.8e-06 | 1.7e-05 | 1.103E-05 | 5.173E-06 | 7.800E-06 | 1.700E-05 |
| 8905 | 446 | I446C | A | 5.7e-06 | 6.1e-06 | 1.2e-05 | 7.933E-06 | 3.528E-06 | 5.700E-06 | 1.200E-05 |
| 8906 | 446 | I446E | A | 3.4e-06 | 6.8e-06 | 9.4e-06 | 6.533E-06 | 3.009E-06 | 3.400E-06 | 9.400E-06 |
| 8907 | 446 | I446Q | A | 8.7e-06 | 6.8e-06 | 7.8e-06 | 7.767E-06 | 9.504E-07 | 6.800E-06 | 8.700E-06 |
| 8908 | 446 | I446G | A | 9.1e-06 | 5.7e-06 | 5.1e-06 | 6.633E-06 | 2.157E-06 | 5.100E-06 | 9.100E-06 |
| 8909 | 446 | I446H | A | 9.2e-06 | 8.2e-06 | 7.9e-06 | 8.433E-06 | 6.807E-07 | 7.900E-06 | 9.200E-06 |
| 8910 | 446 | I446I | A | 1.4e-05 | 7.9e-06 | 9.5e-06 | 1.047E-05 | 3.163E-06 | 7.900E-06 | 1.400E-05 |
| 8911 | 446 | I446L | A | 5.6e-06 | 6.5e-06 | 8.2e-06 | 6.767E-06 | 1.320E-06 | 5.600E-06 | 8.200E-06 |
| 8912 | 446 | I446K | A | 5.1e-06 | 1e-05 | 6.1e-06 | 7.067E-06 | 2.589E-06 | 5.100E-06 | 1.000E-05 |
| 8913 | 446 | I446M | A | 1.1e-05 | 7.2e-06 | 7.5e-06 | 8.567E-06 | 2.113E-06 | 7.200E-06 | 1.100E-05 |
| 8914 | 446 | I446F | A | 5e-06 | 1.3e-05 | 6.9e-06 | 8.300E-06 | 4.180E-06 | 5.000E-06 | 1.300E-05 |
| 8915 | 446 | I446P | A | 1.4e-05 | 5.2e-06 | 5.9e-06 | 8.367E-06 | 4.891E-06 | 5.200E-06 | 1.400E-05 |
| 8916 | 446 | I446S | A | 7.2e-06 | 1e-05 | 1e-05 | 9.067E-06 | 1.617E-06 | 7.200E-06 | 1.000E-05 |
| 8917 | 446 | I446T | A | 1.1e-05 | 1.3e-05 | 8.4e-06 | 1.080E-05 | 2.307E-06 | 8.400E-06 | 1.300E-05 |
| 8918 | 446 | I446W | A | 9.2e-06 | 5.2e-06 | 6.5e-06 | 6.967E-06 | 2.040E-06 | 5.200E-06 | 9.200E-06 |
| 8919 | 446 | I446Y | A | 5.8e-06 | 9e-06 | 9.5e-06 | 8.100E-06 | 2.007E-06 | 5.800E-06 | 9.500E-06 |
| 8920 | 446 | I446V | A | 9.4e-06 | 5.8e-06 | 9.3e-06 | 8.167E-06 | 2.050E-06 | 5.800E-06 | 9.400E-06 |
| 8921 | 447 | H447A | A | 6.6e-06 | 8.4e-06 | 1e-05 | 8.333E-06 | 1.701E-06 | 6.600E-06 | 1.000E-05 |
| 8922 | 447 | H447R | A | 6.6e-06 | 5.1e-06 | 7.1e-06 | 6.267E-06 | 1.041E-06 | 5.100E-06 | 7.100E-06 |
| 8923 | 447 | H447N | A | 8.1e-06 | 9.7e-06 | 7.1e-06 | 8.300E-06 | 1.311E-06 | 7.100E-06 | 9.700E-06 |
| 8924 | 447 | H447D | A | 8.7e-06 | 4.9e-06 | 7.5e-06 | 7.033E-06 | 1.943E-06 | 4.900E-06 | 8.700E-06 |

| | | | | | | | | | | |
|---|---|---|---|---|---|---|---|---|---|---|
| 8925 | 447 | H447C | A | 6e-06 | 7.3e-06 | 1e-05 | 7.767E-06 | 2.040E-06 | 6.000E-06 | 1.000E-05 |
| 8926 | 447 | H447E | A | 8.9e-06 | 7.9e-06 | 7.8e-06 | 8.200E-06 | 6.083E-07 | 7.800E-06 | 8.900E-06 |
| 8927 | 447 | H447Q | A | 5.6e-06 | 7e-06 | 7e-06 | 6.533E-06 | 8.083E-07 | 5.600E-06 | 7.000E-06 |
| 8928 | 447 | H447G | A | 7.8e-06 | 8e-06 | 7e-06 | 7.600E-06 | 5.292E-07 | 7.000E-06 | 8.000E-06 |
| 8929 | 447 | H447H | A | 6.6e-06 | 6.3e-06 | 1.3e-05 | 8.633E-06 | 3.785E-06 | 6.300E-06 | 1.300E-05 |
| 8930 | 447 | H447I | A | 5.2e-06 | 5e-06 | 6.6e-06 | 5.600E-06 | 8.718E-07 | 5.000E-06 | 6.600E-06 |
| 8931 | 447 | H447L | A | 7.3e-06 | 6.4e-06 | 9.8e-06 | 7.833E-06 | 1.762E-06 | 6.400E-06 | 9.800E-06 |
| 8932 | 447 | H447K | A | 8.5e-06 | 8.4e-06 | 6.4e-06 | 7.767E-06 | 1.185E-06 | 6.400E-06 | 8.500E-06 |
| 8933 | 447 | H447M | A | 8.9e-06 | 8.6e-06 | 8e-06 | 8.500E-06 | 4.583E-07 | 8.000E-06 | 8.900E-06 |
| 8934 | 447 | H447F | A | 7.5e-06 | 1.3e-05 | 6.6e-06 | 9.033E-06 | 3.465E-06 | 6.600E-06 | 1.300E-05 |
| 8935 | 447 | H447P | A | 1.2e-05 | 1e-05 | 5.7e-06 | 9.233E-06 | 3.219E-06 | 5.700E-06 | 1.200E-05 |
| 8936 | 447 | H447S | A | 8.4e-06 | 8.4e-06 | 1e-05 | 8.933E-06 | 9.238E-07 | 8.400E-06 | 1.000E-05 |
| 8937 | 447 | H447T | A | 7.2e-06 | 4.2e-06 | 1.3e-05 | 8.133E-06 | 4.474E-06 | 4.200E-06 | 1.300E-05 |
| 8938 | 447 | H447W | A | 6.8e-06 | 3.3e-06 | 9.4e-06 | 6.500E-06 | 3.061E-06 | 3.300E-06 | 9.400E-06 |
| 8939 | 447 | H447Y | A | 6.6e-06 | 5.5e-06 | 7.3e-06 | 6.467E-06 | 9.074E-07 | 5.500E-06 | 7.300E-06 |
| 8940 | 447 | H447V | A | 1e-05 | 8.9e-06 | 1.2e-05 | 1.030E-05 | 1.572E-06 | 8.900E-06 | 1.200E-05 |
| 8941 | 448 | H448A | A | 4.6e-06 | 1.1e-05 | 4.2e-06 | 6.600E-06 | 3.816E-06 | 4.200E-06 | 1.100E-05 |
| 8942 | 448 | H448R | A | 4.3e-06 | 8e-06 | 8.8e-06 | 7.033E-06 | 2.401E-06 | 4.300E-06 | 8.800E-06 |
| 8943 | 448 | H448N | A | 5.9e-06 | 7.5e-06 | 7.4e-06 | 6.933E-06 | 8.963E-07 | 5.900E-06 | 7.500E-06 |
| 8944 | 448 | H448D | A | 6.8e-06 | 8.8e-06 | 1e-05 | 8.533E-06 | 1.617E-06 | 6.800E-06 | 1.000E-05 |
| 8945 | 448 | H448C | A | 4.9e-06 | 7.1e-06 | 5.4e-06 | 5.800E-06 | 1.153E-06 | 4.900E-06 | 7.100E-06 |
| 8946 | 448 | H448E | A | 1.8e-05 | 1.3e-05 | 1.1e-05 | 1.400E-05 | 3.606E-06 | 1.100E-05 | 1.800E-05 |
| 8947 | 448 | H448Q | A | 9.1e-06 | 6.6e-06 | 1.1e-05 | 8.900E-06 | 2.207E-06 | 6.600E-06 | 1.100E-05 |
| 8948 | 448 | H448G | A | 6e-06 | 6.4e-06 | 1.1e-05 | 7.800E-06 | 2.778E-06 | 6.000E-06 | 1.100E-05 |
| 8949 | 448 | H448H | A | 4e-06 | 8e-06 | 6e-06 | 6.000E-06 | 2.000E-06 | 4.000E-06 | 8.000E-06 |
| 8950 | 448 | H448I | A | 1e-05 | 4.3e-06 | 5.1e-06 | 6.467E-06 | 3.086E-06 | 4.300E-06 | 1.000E-05 |
| 8951 | 448 | H448L | A | 8.8e-06 | 9.4e-06 | 8.8e-06 | 9.000E-06 | 3.464E-07 | 8.800E-06 | 9.400E-06 |
| 8952 | 448 | H448K | A | 7.6e-06 | 7.5e-06 | 4.2e-06 | 6.433E-06 | 1.935E-06 | 4.200E-06 | 7.600E-06 |
| 8953 | 448 | H448M | A | 7.7e-06 | 1.5e-05 | 7.3e-06 | 1.000E-05 | 4.335E-06 | 7.300E-06 | 1.500E-05 |
| 8954 | 448 | H448F | A | 9.9e-06 | 6.5e-06 | 1.3e-05 | 9.800E-06 | 3.251E-06 | 6.500E-06 | 1.300E-05 |
| 8955 | 448 | H448P | A | 8e-06 | 7.4e-06 | 1.2e-05 | 9.133E-06 | 2.501E-06 | 7.400E-06 | 1.200E-05 |
| 8956 | 448 | H448S | A | 1.6e-05 | 9.7e-06 | 9.1e-06 | 1.160E-05 | 3.822E-06 | 9.100E-06 | 1.600E-05 |
| 8957 | 448 | H448T | A | 9.2e-06 | 7.8e-06 | 5.1e-06 | 7.367E-06 | 2.084E-06 | 5.100E-06 | 9.200E-06 |
| 8958 | 448 | H448W | A | 8e-06 | 9.9e-06 | 1.3e-05 | 1.030E-05 | 2.524E-06 | 8.000E-06 | 1.300E-05 |
| 8959 | 448 | H448Y | A | 9.7e-06 | 5e-06 | 8.6e-06 | 7.767E-06 | 2.458E-06 | 5.000E-06 | 9.700E-06 |
| 8960 | 448 | H448V | A | 9.3e-06 | 7.8e-06 | 7e-06 | 8.033E-06 | 1.168E-06 | 7.000E-06 | 9.300E-06 |
| 8961 | 449 | N449A | A | 1e-05 | 9.1e-06 | 5.6e-06 | 8.233E-06 | 2.325E-06 | 5.600E-06 | 1.000E-05 |
| 8962 | 449 | N449R | A | 4.4e-06 | 6.4e-06 | 6.9e-06 | 5.900E-06 | 1.323E-06 | 4.400E-06 | 6.900E-06 |
| 8963 | 449 | N449N | A | 6e-06 | 6.4e-06 | 1.1e-05 | 7.800E-06 | 2.778E-06 | 6.000E-06 | 1.100E-05 |
| 8964 | 449 | N449D | A | 7.8e-06 | 1e-05 | 9e-06 | 8.933E-06 | 1.102E-06 | 7.800E-06 | 1.000E-05 |
| 8965 | 449 | N449C | A | 5.4e-06 | 7e-06 | 6.4e-06 | 6.267E-06 | 8.083E-07 | 5.400E-06 | 7.000E-06 |
| 8966 | 449 | N449E | A | 4.9e-06 | 7.8e-06 | 7.8e-06 | 6.833E-06 | 1.674E-06 | 4.900E-06 | 7.800E-06 |
| 8967 | 449 | N449Q | A | 9.4e-06 | 9.7e-06 | 6.8e-06 | 8.633E-06 | 1.595E-06 | 6.800E-06 | 9.700E-06 |
| 8968 | 449 | N449G | A | 9e-06 | 4.7e-06 | 4.9e-06 | 6.200E-06 | 2.427E-06 | 4.700E-06 | 9.000E-06 |
| 8969 | 449 | N449H | A | 9.6e-06 | 4.9e-06 | 1.1e-05 | 8.500E-06 | 3.195E-06 | 4.900E-06 | 1.100E-05 |
| 8970 | 449 | N449I | A | 5.3e-06 | 4e-06 | 6.1e-06 | 5.133E-06 | 1.060E-06 | 4.000E-06 | 6.100E-06 |
| 8971 | 449 | N449L | A | 5.6e-06 | 7.9e-06 | 1.1e-05 | 8.167E-06 | 2.710E-06 | 5.600E-06 | 1.100E-05 |
| 8972 | 449 | N449K | A | 9e-06 | 1.3e-05 | 5.2e-06 | 9.067E-06 | 3.900E-06 | 5.200E-06 | 1.300E-05 |
| 8973 | 449 | N449M | A | 4.9e-06 | 6.5e-06 | 7.6e-06 | 6.333E-06 | 1.358E-06 | 4.900E-06 | 7.600E-06 |
| 8974 | 449 | N449F | A | 9.7e-06 | 7.9e-06 | 5.4e-06 | 7.667E-06 | 2.159E-06 | 5.400E-06 | 9.700E-06 |
| 8975 | 449 | N449P | A | 4.4e-06 | 5.5e-06 | 9.5e-06 | 6.467E-06 | 2.684E-06 | 4.400E-06 | 9.500E-06 |
| 8976 | 449 | N449S | A | 1.1e-05 | 9e-06 | 7.5e-06 | 9.167E-06 | 1.756E-06 | 7.500E-06 | 1.100E-05 |
| 8977 | 449 | N449T | A | 1.4e-05 | 7.5e-06 | 7.2e-06 | 9.567E-06 | 3.842E-06 | 7.200E-06 | 1.400E-05 |
| 8978 | 449 | N449W | A | 4.3e-06 | 1.1e-05 | 6.6e-06 | 7.300E-06 | 3.404E-06 | 4.300E-06 | 1.100E-05 |
| 8979 | 449 | N449Y | A | 4.8e-06 | 8.9e-06 | 4.7e-06 | 6.133E-06 | 2.397E-06 | 4.700E-06 | 8.900E-06 |
| 8980 | 449 | N449V | A | 1.1e-05 | 1.2e-05 | 5.3e-06 | 9.433E-06 | 3.614E-06 | 5.300E-06 | 1.200E-05 |
| 8981 | 450 | T450A | A | 5.8e-06 | 8.4e-06 | 7.6e-06 | 7.267E-06 | 1.332E-06 | 5.800E-06 | 8.400E-06 |
| 8982 | 450 | T450R | A | 9.5e-06 | 6.1e-06 | 1.2e-05 | 9.200E-06 | 2.961E-06 | 6.100E-06 | 1.200E-05 |
| 8983 | 450 | T450N | A | 7.8e-06 | 7.2e-06 | 5.5e-06 | 6.833E-06 | 1.193E-06 | 5.500E-06 | 7.800E-06 |
| 8984 | 450 | T450D | A | 4.9e-06 | 7.4e-06 | 8.1e-06 | 6.800E-06 | 1.682E-06 | 4.900E-06 | 8.100E-06 |
| 8985 | 450 | T450C | A | 9.4e-06 | 6.5e-06 | 8.7e-06 | 8.200E-06 | 1.513E-06 | 6.500E-06 | 9.400E-06 |
| 8986 | 450 | T450E | A | 7.8e-06 | 8.2e-06 | 4.8e-06 | 6.933E-06 | 1.858E-06 | 4.800E-06 | 8.200E-06 |
| 8987 | 450 | T450Q | A | 9.1e-06 | 1.1e-05 | 9.8e-06 | 9.967E-06 | 9.609E-07 | 9.100E-06 | 1.100E-05 |
| 8988 | 450 | T450G | A | 8.8e-06 | 7.3e-06 | 4.4e-06 | 6.833E-06 | 2.237E-06 | 4.400E-06 | 8.800E-06 |
| 8989 | 450 | T450H | A | 4.2e-06 | 9.3e-06 | 7.3e-06 | 6.933E-06 | 2.570E-06 | 4.200E-06 | 9.300E-06 |
| 8990 | 450 | T450I | A | 6.8e-06 | 1.3e-05 | 7.4e-06 | 9.067E-06 | 3.420E-06 | 6.800E-06 | 1.300E-05 |
| 8991 | 450 | T450L | A | 1.2e-05 | 1.1e-05 | 8.3e-06 | 1.043E-05 | 1.914E-06 | 8.300E-06 | 1.200E-05 |
| 8992 | 450 | T450K | A | 8.2e-06 | 5e-06 | 9e-06 | 7.400E-06 | 2.117E-06 | 5.000E-06 | 9.000E-06 |
| 8993 | 450 | T450M | A | 5.8e-06 | 7.1e-06 | 1.4e-05 | 8.967E-06 | 4.407E-06 | 5.800E-06 | 1.400E-05 |
| 8994 | 450 | T450F | A | 8.4e-06 | 9.9e-06 | 8.4e-06 | 8.900E-06 | 8.660E-07 | 8.400E-06 | 9.900E-06 |
| 8995 | 450 | T450P | A | 1e-05 | 8e-06 | 1.1e-05 | 9.667E-06 | 1.528E-06 | 8.000E-06 | 1.100E-05 |
| 8996 | 450 | T450S | A | 7.4e-06 | 8.8e-06 | 7.3e-06 | 7.833E-06 | 8.386E-07 | 7.300E-06 | 8.800E-06 |
| 8997 | 450 | T450T | A | 7e-06 | 1.2e-05 | 6.8e-06 | 8.600E-06 | 2.946E-06 | 6.800E-06 | 1.200E-05 |
| 8998 | 450 | T450W | A | 7.4e-06 | 9.9e-06 | 7.1e-06 | 8.133E-06 | 1.537E-06 | 7.100E-06 | 9.900E-06 |
| 8999 | 450 | T450Y | A | 9.6e-06 | 1.4e-05 | 1e-05 | 1.120E-05 | 2.433E-06 | 9.600E-06 | 1.400E-05 |
| 9000 | 450 | T450V | A | 7.2e-06 | 9.2e-06 | 1.3e-05 | 9.800E-06 | 2.946E-06 | 7.200E-06 | 1.300E-05 |
| 9001 | 451 | H451A | A | 5.6e-06 | 1.1e-05 | 7.6e-06 | 8.067E-06 | 2.730E-06 | 5.600E-06 | 1.100E-05 |
| 9002 | 451 | H451R | A | 7e-06 | 9.4e-06 | 9.9e-06 | 8.767E-06 | 1.550E-06 | 7.000E-06 | 9.900E-06 |
| 9003 | 451 | H451N | A | 4.7e-06 | 1.9e-05 | 9.6e-06 | 1.110E-05 | 7.267E-06 | 4.700E-06 | 1.900E-05 |
| 9004 | 451 | H451D | A | 1.1e-05 | 4.8e-06 | 9.1e-06 | 8.300E-06 | 3.176E-06 | 4.800E-06 | 1.100E-05 |
| 9005 | 451 | H451C | A | 5.9e-06 | 3.7e-06 | 6.7e-06 | 5.433E-06 | 1.553E-06 | 3.700E-06 | 6.700E-06 |
| 9006 | 451 | H451E | A | 9.4e-06 | 7.8e-06 | 1.2e-05 | 9.733E-06 | 2.120E-06 | 7.800E-06 | 1.200E-05 |
| 9007 | 451 | H451Q | A | 6.6e-06 | 4.7e-06 | 7.4e-06 | 6.233E-06 | 1.387E-06 | 4.700E-06 | 7.400E-06 |
| 9008 | 451 | H451G | A | 1e-05 | 5.4e-06 | 1.2e-05 | 9.133E-06 | 3.384E-06 | 5.400E-06 | 1.200E-05 |
| 9009 | 451 | H451H | A | 6.3e-06 | 1e-05 | 1.3e-05 | 9.767E-06 | 3.356E-06 | 6.300E-06 | 1.300E-05 |

| | | | | | | | | | | |
|---|---|---|---|---|---|---|---|---|---|---|
| 9010 | 451 | H451I | A | 1.4e-05 | 7.7e-06 | 1.1e-05 | 1.090E-05 | 3.151E-06 | 7.700E-06 | 1.400E-05 |
| 9011 | 451 | H451L | A | 1e-05 | 6.4e-06 | 8e-06 | 8.133E-06 | 1.804E-06 | 6.400E-06 | 1.000E-05 |
| 9012 | 451 | H451K | A | 7.4e-06 | 6.6e-06 | 5.8e-06 | 6.600E-06 | 8.000E-07 | 5.800E-06 | 7.400E-06 |
| 9013 | 451 | H451M | A | 8.5e-06 | 6.7e-06 | 5.1e-06 | 6.767E-06 | 1.701E-06 | 5.100E-06 | 8.500E-06 |
| 9014 | 451 | H451F | A | 1.3e-05 | 7e-06 | 6.7e-06 | 8.900E-06 | 3.554E-06 | 6.700E-06 | 1.300E-05 |
| 9015 | 451 | H451P | A | 5.7e-06 | 8.4e-06 | 8.8e-06 | 7.633E-06 | 1.686E-06 | 5.700E-06 | 8.800E-06 |
| 9016 | 451 | H451S | A | 1e-05 | 9.1e-06 | 8.5e-06 | 9.200E-06 | 7.550E-07 | 8.500E-06 | 1.000E-05 |
| 9017 | 451 | H451T | A | 4.5e-06 | 7.7e-06 | 8e-06 | 6.733E-06 | 1.940E-06 | 4.500E-06 | 8.000E-06 |
| 9018 | 451 | H451W | A | 5e-06 | 8.4e-06 | 6.2e-06 | 6.533E-06 | 1.724E-06 | 5.000E-06 | 8.400E-06 |
| 9019 | 451 | H451Y | A | 7.9e-06 | 1.1e-05 | 5.9e-06 | 8.267E-06 | 2.570E-06 | 5.900E-06 | 1.100E-05 |
| 9020 | 451 | H451V | A | 6.6e-06 | 9.8e-06 | 1.4e-05 | 1.013E-05 | 3.711E-06 | 6.600E-06 | 1.400E-05 |
| 9021 | 452 | L452A | A | 6e-06 | 1.4e-05 | 9.3e-06 | 9.767E-06 | 4.020E-06 | 6.000E-06 | 1.400E-05 |
| 9022 | 452 | L452R | A | 6.6e-06 | 1e-05 | 9.3e-06 | 8.633E-06 | 1.795E-06 | 6.600E-06 | 1.000E-05 |
| 9023 | 452 | L452N | A | 7.5e-06 | 9.3e-06 | 4.7e-06 | 7.167E-06 | 2.318E-06 | 4.700E-06 | 9.300E-06 |
| 9024 | 452 | L452D | A | 8.9e-06 | 1.1e-05 | 1.4e-05 | 1.130E-05 | 2.563E-06 | 8.900E-06 | 1.400E-05 |
| 9025 | 452 | L452C | A | 1.1e-05 | 6.4e-06 | 8.2e-06 | 8.533E-06 | 2.318E-06 | 6.400E-06 | 1.100E-05 |
| 9026 | 452 | L452E | A | 9.1e-06 | 1.2e-05 | 5e-06 | 8.700E-06 | 3.517E-06 | 5.000E-06 | 1.200E-05 |
| 9027 | 452 | L452Q | A | 5.8e-06 | 6.6e-06 | 8.7e-06 | 7.033E-06 | 1.498E-06 | 5.800E-06 | 8.700E-06 |
| 9028 | 452 | L452G | A | 7.6e-06 | 9.9e-06 | 6.4e-06 | 7.967E-06 | 1.779E-06 | 6.400E-06 | 9.900E-06 |
| 9029 | 452 | L452H | A | 9.3e-06 | 9e-06 | 1.1e-05 | 9.767E-06 | 1.079E-06 | 9.000E-06 | 1.100E-05 |
| 9030 | 452 | L452I | A | 5.1e-06 | 6.9e-06 | 9.1e-06 | 7.033E-06 | 2.003E-06 | 5.100E-06 | 9.100E-06 |
| 9031 | 452 | L452L | A | 9e-06 | 8.2e-06 | 9.6e-06 | 8.933E-06 | 7.024E-07 | 8.200E-06 | 9.600E-06 |
| 9032 | 452 | L452K | A | 4.7e-06 | 4.3e-06 | 9.1e-06 | 6.033E-06 | 2.663E-06 | 4.300E-06 | 9.100E-06 |
| 9033 | 452 | L452M | A | 5.9e-06 | 9.1e-06 | 9.2e-06 | 8.067E-06 | 1.877E-06 | 5.900E-06 | 9.200E-06 |
| 9034 | 452 | L452F | A | 1.4e-05 | 4.9e-06 | 6.5e-06 | 8.467E-06 | 4.858E-06 | 4.900E-06 | 1.400E-05 |
| 9035 | 452 | L452P | A | 5.6e-06 | 1e-05 | 1.3e-05 | 9.533E-06 | 3.722E-06 | 5.600E-06 | 1.300E-05 |
| 9036 | 452 | L452S | A | 6.8e-06 | 7.7e-06 | 7.7e-06 | 7.400E-06 | 5.196E-07 | 6.800E-06 | 7.700E-06 |
| 9037 | 452 | L452T | A | 8.9e-06 | 8.5e-06 | 8.8e-06 | 8.733E-06 | 2.082E-07 | 8.500E-06 | 8.900E-06 |
| 9038 | 452 | L452W | A | 9.6e-06 | 9.1e-06 | 8e-06 | 8.900E-06 | 8.185E-07 | 8.000E-06 | 9.600E-06 |
| 9039 | 452 | L452Y | A | 7.3e-06 | 7e-06 | 6.7e-06 | 7.000E-06 | 3.000E-07 | 6.700E-06 | 7.300E-06 |
| 9040 | 452 | L452V | A | 5.2e-06 | 1.1e-05 | 9.7e-06 | 8.633E-06 | 3.044E-06 | 5.200E-06 | 1.100E-05 |
| 9041 | 453 | C453A | A | 1.3e-05 | 7e-06 | 8.8e-06 | 9.600E-06 | 3.079E-06 | 7.000E-06 | 1.300E-05 |
| 9042 | 453 | C453R | A | 5.3e-06 | 6.7e-06 | 5.5e-06 | 5.833E-06 | 7.572E-07 | 5.300E-06 | 6.700E-06 |
| 9043 | 453 | C453N | A | 6.8e-06 | 4.2e-06 | 5.9e-06 | 5.633E-06 | 1.320E-06 | 4.200E-06 | 6.800E-06 |
| 9044 | 453 | C453D | A | 1e-05 | 9.5e-06 | 6.2e-06 | 8.567E-06 | 2.065E-06 | 6.200E-06 | 1.000E-05 |
| 9045 | 453 | C453C | A | 5.6e-06 | 3.2e-06 | 5.6e-06 | 4.800E-06 | 1.386E-06 | 3.200E-06 | 5.600E-06 |
| 9046 | 453 | C453E | A | 1.2e-05 | 1.1e-05 | 9.6e-06 | 1.087E-05 | 1.206E-06 | 9.600E-06 | 1.200E-05 |
| 9047 | 453 | C453Q | A | 1.9e-05 | 8.5e-06 | 9.3e-06 | 1.227E-05 | 5.845E-06 | 8.500E-06 | 1.900E-05 |
| 9048 | 453 | C453G | A | 8.2e-06 | 7e-06 | 9.1e-06 | 8.100E-06 | 1.054E-06 | 7.000E-06 | 9.100E-06 |
| 9049 | 453 | C453H | A | 9.2e-06 | 5.9e-06 | 8.2e-06 | 7.767E-06 | 1.692E-06 | 5.900E-06 | 9.200E-06 |
| 9050 | 453 | C453I | A | 9.3e-06 | 8.5e-06 | 9.5e-06 | 9.100E-06 | 5.292E-07 | 8.500E-06 | 9.500E-06 |
| 9051 | 453 | C453L | A | 4.6e-06 | 8.5e-06 | 5.9e-06 | 6.333E-06 | 1.986E-06 | 4.600E-06 | 8.500E-06 |
| 9052 | 453 | C453K | A | 8.2e-06 | 1.3e-05 | 6.5e-06 | 9.233E-06 | 3.371E-06 | 6.500E-06 | 1.300E-05 |
| 9053 | 453 | C453M | A | 9.1e-06 | 8e-06 | 9.1e-06 | 8.733E-06 | 6.351E-07 | 8.000E-06 | 9.100E-06 |
| 9054 | 453 | C453F | A | 8.2e-06 | 8e-06 | 9.6e-06 | 8.600E-06 | 8.718E-07 | 8.000E-06 | 9.600E-06 |
| 9055 | 453 | C453P | A | 7.9e-06 | 8.2e-06 | 4.9e-06 | 7.000E-06 | 1.825E-06 | 4.900E-06 | 8.200E-06 |
| 9056 | 453 | C453S | A | 1e-05 | 8.7e-06 | 6.4e-06 | 8.367E-06 | 1.823E-06 | 6.400E-06 | 1.000E-05 |
| 9057 | 453 | C453T | A | 6.7e-06 | 7.8e-06 | 7.3e-06 | 7.267E-06 | 5.508E-07 | 6.700E-06 | 7.800E-06 |
| 9058 | 453 | C453W | A | 4.9e-06 | 9.7e-06 | 7.1e-06 | 7.233E-06 | 2.403E-06 | 4.900E-06 | 9.700E-06 |
| 9059 | 453 | C453Y | A | 6.9e-06 | 4.6e-06 | 7.5e-06 | 6.333E-06 | 1.531E-06 | 4.600E-06 | 7.500E-06 |
| 9060 | 453 | C453V | A | 1.1e-05 | 7.6e-06 | 5.6e-06 | 8.067E-06 | 2.730E-06 | 5.600E-06 | 1.100E-05 |
| 9061 | 454 | F454A | A | 6.2e-06 | 7.4e-06 | 9.9e-06 | 7.833E-06 | 1.888E-06 | 6.200E-06 | 9.900E-06 |
| 9062 | 454 | F454R | A | 6.6e-06 | 9.4e-06 | 1.4e-05 | 1.000E-05 | 3.736E-06 | 6.600E-06 | 1.400E-05 |
| 9063 | 454 | F454N | A | 4e-06 | 8.6e-06 | 9.8e-06 | 7.467E-06 | 3.062E-06 | 4.000E-06 | 9.800E-06 |
| 9064 | 454 | F454D | A | 1.5e-05 | 6.6e-06 | 4.8e-06 | 8.800E-06 | 5.444E-06 | 4.800E-06 | 1.500E-05 |
| 9065 | 454 | F454C | A | 6.9e-06 | 7.7e-06 | 1.2e-05 | 8.867E-06 | 2.743E-06 | 6.900E-06 | 1.200E-05 |
| 9066 | 454 | F454E | A | 1e-05 | 8.1e-06 | 8e-06 | 8.700E-06 | 1.127E-06 | 8.000E-06 | 1.000E-05 |
| 9067 | 454 | F454Q | A | 5.4e-06 | 6.5e-06 | 6.1e-06 | 6.000E-06 | 5.568E-07 | 5.400E-06 | 6.500E-06 |
| 9068 | 454 | F454G | A | 5.5e-06 | 9.7e-06 | 4.8e-06 | 6.667E-06 | 2.650E-06 | 4.800E-06 | 9.700E-06 |
| 9069 | 454 | F454H | A | 7.2e-06 | 8.7e-06 | 7.1e-06 | 7.667E-06 | 8.963E-07 | 7.100E-06 | 8.700E-06 |
| 9070 | 454 | F454I | A | 5.2e-06 | 8.8e-06 | 7.4e-06 | 7.133E-06 | 1.815E-06 | 5.200E-06 | 8.800E-06 |
| 9071 | 454 | F454L | A | 8.5e-06 | 8.1e-06 | 9.6e-06 | 8.733E-06 | 7.767E-07 | 8.100E-06 | 9.600E-06 |
| 9072 | 454 | F454K | A | 5.9e-06 | 8e-06 | 4.6e-06 | 6.167E-06 | 1.716E-06 | 4.600E-06 | 8.000E-06 |
| 9073 | 454 | F454M | A | 7.7e-06 | 9.6e-06 | 5.7e-06 | 7.667E-06 | 1.950E-06 | 5.700E-06 | 9.600E-06 |
| 9074 | 454 | F454F | A | 1e-05 | 8.3e-06 | 1e-05 | 9.433E-06 | 9.815E-07 | 8.300E-06 | 1.000E-05 |
| 9075 | 454 | F454P | A | 6e-06 | 6.2e-06 | 9.3e-06 | 7.167E-06 | 1.850E-06 | 6.000E-06 | 9.300E-06 |
| 9076 | 454 | F454S | A | 4.9e-06 | 7.7e-06 | 1.1e-05 | 7.867E-06 | 3.053E-06 | 4.900E-06 | 1.100E-05 |
| 9077 | 454 | F454T | A | 4.3e-06 | 9.5e-06 | 1e-05 | 7.933E-06 | 3.156E-06 | 4.300E-06 | 1.000E-05 |
| 9078 | 454 | F454W | A | 9.1e-06 | 6.2e-06 | 5.8e-06 | 7.033E-06 | 1.801E-06 | 5.800E-06 | 9.100E-06 |
| 9079 | 454 | F454Y | A | 1.1e-05 | 7.3e-06 | 4.3e-06 | 7.533E-06 | 3.356E-06 | 4.300E-06 | 1.100E-05 |
| 9080 | 454 | F454V | A | 1.6e-05 | 1.4e-05 | 1.2e-05 | 1.400E-05 | 2.000E-06 | 1.200E-05 | 1.600E-05 |
| 9081 | 455 | V455A | A | 5.2e-06 | 1.3e-05 | 6e-06 | 8.067E-06 | 4.291E-06 | 5.200E-06 | 1.300E-05 |
| 9082 | 455 | V455R | A | 9.7e-06 | 8.9e-06 | 8.8e-06 | 9.133E-06 | 4.933E-07 | 8.800E-06 | 9.700E-06 |
| 9083 | 455 | V455N | A | 1e-05 | 9.7e-06 | 6.4e-06 | 8.700E-06 | 1.997E-06 | 6.400E-06 | 1.000E-05 |
| 9084 | 455 | V455D | A | 6.5e-06 | 5.1e-06 | 6.5e-06 | 6.033E-06 | 8.083E-07 | 5.100E-06 | 6.500E-06 |
| 9085 | 455 | V455C | A | 8.3e-06 | 4.9e-06 | 9.4e-06 | 7.533E-06 | 2.346E-06 | 4.900E-06 | 9.400E-06 |
| 9086 | 455 | V455E | A | 4.7e-06 | 8.6e-06 | 6.7e-06 | 6.667E-06 | 1.950E-06 | 4.700E-06 | 8.600E-06 |
| 9087 | 455 | V455Q | A | 1.2e-05 | 1.2e-05 | 7.1e-06 | 1.037E-05 | 2.829E-06 | 7.100E-06 | 1.200E-05 |
| 9088 | 455 | V455G | A | 9.2e-06 | 6.1e-06 | 6.7e-06 | 7.333E-06 | 1.644E-06 | 6.100E-06 | 9.200E-06 |
| 9089 | 455 | V455H | A | 4.9e-06 | 8.5e-06 | 7e-06 | 6.800E-06 | 1.808E-06 | 4.900E-06 | 8.500E-06 |
| 9090 | 455 | V455I | A | 1.1e-05 | 9.2e-06 | 6e-06 | 8.733E-06 | 2.532E-06 | 6.000E-06 | 1.100E-05 |
| 9091 | 455 | V455L | A | 1.1e-05 | 9.4e-06 | 6.7e-06 | 9.033E-06 | 2.173E-06 | 6.700E-06 | 1.100E-05 |
| 9092 | 455 | V455K | A | 1.3e-05 | 8.6e-06 | 6.1e-06 | 9.233E-06 | 3.493E-06 | 6.100E-06 | 1.300E-05 |
| 9093 | 455 | V455M | A | 1.5e-05 | 7.3e-06 | 5.5e-06 | 9.267E-06 | 5.046E-06 | 5.500E-06 | 1.500E-05 |
| 9094 | 455 | V455F | A | 1.3e-05 | 5.4e-06 | 9.7e-06 | 9.367E-06 | 3.811E-06 | 5.400E-06 | 1.300E-05 |

| | | | | | | | | | | |
|---|---|---|---|---|---|---|---|---|---|---|
| 9095 | 455 | V455P | A | 5.7e-06 | 9.8e-06 | 6.5e-06 | 7.333E-06 | 2.173E-06 | 5.700E-06 | 9.800E-06 |
| 9096 | 455 | V455S | A | 7.4e-06 | 6.4e-06 | 8.1e-06 | 7.300E-06 | 8.544E-07 | 6.400E-06 | 8.100E-06 |
| 9097 | 455 | V455T | A | 9.4e-06 | 1.1e-05 | 4.4e-06 | 8.267E-06 | 3.443E-06 | 4.400E-06 | 1.100E-05 |
| 9098 | 455 | V455W | A | 1.3e-05 | 8.5e-06 | 7.7e-06 | 9.733E-06 | 2.857E-06 | 7.700E-06 | 1.300E-05 |
| 9099 | 455 | V455Y | A | 1.1e-05 | 4.2e-06 | 1.2e-05 | 9.067E-06 | 4.244E-06 | 4.200E-06 | 1.200E-05 |
| 9100 | 455 | V455V | A | 4.6e-06 | 2e-05 | 8.8e-06 | 1.113E-05 | 7.961E-06 | 4.600E-06 | 2.000E-05 |
| 9101 | 456 | H456A | A | 8.1e-06 | 4.6e-06 | 8.3e-06 | 7.000E-06 | 2.081E-06 | 4.600E-06 | 8.300E-06 |
| 9102 | 456 | H456R | A | 7.2e-06 | 8.5e-06 | 3.6e-06 | 6.433E-06 | 2.538E-06 | 3.600E-06 | 8.500E-06 |
| 9103 | 456 | H456N | A | 8.2e-06 | 5.8e-06 | 6.3e-06 | 6.767E-06 | 1.266E-06 | 5.800E-06 | 8.200E-06 |
| 9104 | 456 | H456D | A | 6.8e-06 | 6.5e-06 | 7.9e-06 | 7.067E-06 | 7.371E-07 | 6.500E-06 | 7.900E-06 |
| 9105 | 456 | H456C | A | 6.3e-06 | 9.3e-06 | 4.5e-06 | 6.700E-06 | 2.425E-06 | 4.500E-06 | 9.300E-06 |
| 9106 | 456 | H456E | A | 9.5e-06 | 4.3e-06 | 7.3e-06 | 7.033E-06 | 2.610E-06 | 4.300E-06 | 9.500E-06 |
| 9107 | 456 | H456Q | A | 9.7e-06 | 7.9e-06 | 8e-06 | 8.533E-06 | 1.012E-06 | 7.900E-06 | 9.700E-06 |
| 9108 | 456 | H456G | A | 8.2e-06 | 7.5e-06 | 7.9e-06 | 7.867E-06 | 3.512E-07 | 7.500E-06 | 8.200E-06 |
| 9109 | 456 | H456H | A | 4.8e-06 | 9e-06 | 5.3e-06 | 6.367E-06 | 2.294E-06 | 4.800E-06 | 9.000E-06 |
| 9110 | 456 | H456I | A | 1.2e-05 | 1e-05 | 8.5e-06 | 1.017E-05 | 1.756E-06 | 8.500E-06 | 1.200E-05 |
| 9111 | 456 | H456L | A | 6.1e-06 | 9.9e-06 | 9.3e-06 | 8.433E-06 | 2.043E-06 | 6.100E-06 | 9.900E-06 |
| 9112 | 456 | H456K | A | 1.3e-05 | 9.8e-06 | 6.1e-06 | 9.633E-06 | 3.453E-06 | 6.100E-06 | 1.300E-05 |
| 9113 | 456 | H456M | A | 8.2e-06 | 9.1e-06 | 8.9e-06 | 8.733E-06 | 4.726E-07 | 8.200E-06 | 9.100E-06 |
| 9114 | 456 | H456F | A | 1.5e-05 | 9.2e-06 | 9e-06 | 1.107E-05 | 3.408E-06 | 9.000E-06 | 1.500E-05 |
| 9115 | 456 | H456P | A | 5.3e-06 | 1.2e-05 | 1e-05 | 9.100E-06 | 3.439E-06 | 5.300E-06 | 1.200E-05 |
| 9116 | 456 | H456S | A | 6.4e-06 | 5.2e-06 | 4.3e-06 | 5.300E-06 | 1.054E-06 | 4.300E-06 | 6.400E-06 |
| 9117 | 456 | H456T | A | 4e-06 | 6.3e-06 | 1e-05 | 6.767E-06 | 3.027E-06 | 4.000E-06 | 1.000E-05 |
| 9118 | 456 | H456W | A | 9.9e-06 | 5.4e-06 | 4.1e-06 | 6.467E-06 | 3.044E-06 | 4.100E-06 | 9.900E-06 |
| 9119 | 456 | H456Y | A | 8.1e-06 | 1.2e-05 | 6.8e-06 | 8.967E-06 | 2.706E-06 | 6.800E-06 | 1.200E-05 |
| 9120 | 456 | H456V | A | 7.5e-06 | 6.7e-06 | 8.5e-06 | 7.567E-06 | 9.018E-07 | 6.700E-06 | 8.500E-06 |
| 9121 | 457 | T457A | A | 5e-06 | 8.1e-06 | 7.4e-06 | 6.833E-06 | 1.626E-06 | 5.000E-06 | 8.100E-06 |
| 9122 | 457 | T457R | A | 6.5e-06 | 9.8e-06 | 7.2e-06 | 7.833E-06 | 1.739E-06 | 6.500E-06 | 9.800E-06 |
| 9123 | 457 | T457N | A | 4.9e-06 | 6.5e-06 | 1.2e-05 | 7.800E-06 | 3.724E-06 | 4.900E-06 | 1.200E-05 |
| 9124 | 457 | T457D | A | 8.6e-06 | 7.1e-06 | 5.1e-06 | 6.933E-06 | 1.756E-06 | 5.100E-06 | 8.600E-06 |
| 9125 | 457 | T457C | A | 8.9e-06 | 1e-05 | 8.6e-06 | 9.167E-06 | 7.371E-07 | 8.600E-06 | 1.000E-05 |
| 9126 | 457 | T457E | A | 1.3e-05 | 9.6e-06 | 6.7e-06 | 9.767E-06 | 3.153E-06 | 6.700E-06 | 1.300E-05 |
| 9127 | 457 | T457Q | A | 6.2e-06 | 6.6e-06 | 8.1e-06 | 6.967E-06 | 1.002E-06 | 6.200E-06 | 8.100E-06 |
| 9128 | 457 | T457G | A | 9.2e-06 | 8.3e-06 | 6.8e-06 | 8.100E-06 | 1.212E-06 | 6.800E-06 | 9.200E-06 |
| 9129 | 457 | T457H | A | 9.6e-06 | 6.8e-06 | 8.2e-06 | 8.200E-06 | 1.400E-06 | 6.800E-06 | 9.600E-06 |
| 9130 | 457 | T457I | A | 7.6e-06 | 7e-06 | 1.4e-05 | 9.533E-06 | 3.880E-06 | 7.000E-06 | 1.400E-05 |
| 9131 | 457 | T457L | A | 6.9e-06 | 8.2e-06 | 1.1e-05 | 8.700E-06 | 2.095E-06 | 6.900E-06 | 1.100E-05 |
| 9132 | 457 | T457K | A | 6.4e-06 | 7.5e-06 | 5.7e-06 | 6.533E-06 | 9.074E-07 | 5.700E-06 | 7.500E-06 |
| 9133 | 457 | T457M | A | 1.1e-05 | 5.7e-06 | 6.5e-06 | 7.733E-06 | 2.857E-06 | 5.700E-06 | 1.100E-05 |
| 9134 | 457 | T457F | A | 5.4e-06 | 1.1e-05 | 7.3e-06 | 7.900E-06 | 2.848E-06 | 5.400E-06 | 1.100E-05 |
| 9135 | 457 | T457P | A | 5.8e-06 | 1.1e-05 | 6.4e-06 | 7.733E-06 | 2.845E-06 | 5.800E-06 | 1.100E-05 |
| 9136 | 457 | T457S | A | 9.5e-06 | 8.8e-06 | 7.3e-06 | 8.533E-06 | 1.124E-06 | 7.300E-06 | 9.500E-06 |
| 9137 | 457 | T457T | A | 6.1e-06 | 5e-06 | 7.6e-06 | 6.233E-06 | 1.305E-06 | 5.000E-06 | 7.600E-06 |
| 9138 | 457 | T457W | A | 8.4e-06 | 7.3e-06 | 5.9e-06 | 7.200E-06 | 1.253E-06 | 5.900E-06 | 8.400E-06 |
| 9139 | 457 | T457Y | A | 8e-06 | 6.7e-06 | 5.4e-06 | 6.700E-06 | 1.300E-06 | 5.400E-06 | 8.000E-06 |
| 9140 | 457 | T457V | A | 5.6e-06 | 8.2e-06 | 9.8e-06 | 7.867E-06 | 2.120E-06 | 5.600E-06 | 9.800E-06 |
| 9141 | 458 | V458A | A | 6.3e-06 | 5.9e-06 | 8.7e-06 | 6.967E-06 | 1.514E-06 | 5.900E-06 | 8.700E-06 |
| 9142 | 458 | V458R | A | 8.2e-06 | 8.3e-06 | 6.4e-06 | 7.633E-06 | 1.069E-06 | 6.400E-06 | 8.300E-06 |
| 9143 | 458 | V458N | A | 9.4e-06 | 1.2e-05 | 1e-05 | 1.047E-05 | 1.361E-06 | 9.400E-06 | 1.200E-05 |
| 9144 | 458 | V458D | A | 8e-06 | 1.3e-05 | 7.8e-06 | 9.600E-06 | 2.946E-06 | 7.800E-06 | 1.300E-05 |
| 9145 | 458 | V458C | A | 7.6e-06 | 1.1e-05 | 6.6e-06 | 8.400E-06 | 2.307E-06 | 6.600E-06 | 1.100E-05 |
| 9146 | 458 | V458E | A | 1.4e-05 | 1.3e-05 | 3.8e-06 | 1.027E-05 | 5.623E-06 | 3.800E-06 | 1.400E-05 |
| 9147 | 458 | V458Q | A | 6.7e-06 | 8.2e-06 | 1e-05 | 8.300E-06 | 1.652E-06 | 6.700E-06 | 1.000E-05 |
| 9148 | 458 | V458G | A | 1.3e-05 | 5.2e-06 | 9.4e-06 | 9.200E-06 | 3.904E-06 | 5.200E-06 | 1.300E-05 |
| 9149 | 458 | V458H | A | 9.1e-06 | 7.9e-06 | 3.7e-06 | 6.900E-06 | 2.835E-06 | 3.700E-06 | 9.100E-06 |
| 9150 | 458 | V458I | A | 6.6e-06 | 9.4e-06 | 8.4e-06 | 8.133E-06 | 1.419E-06 | 6.600E-06 | 9.400E-06 |
| 9151 | 458 | V458L | A | 1.3e-05 | 6.8e-06 | 9.8e-06 | 9.867E-06 | 3.101E-06 | 6.800E-06 | 1.300E-05 |
| 9152 | 458 | V458K | A | 1.6e-05 | 6.8e-06 | 6.4e-06 | 9.733E-06 | 5.431E-06 | 6.400E-06 | 1.600E-05 |
| 9153 | 458 | V458M | A | 1.2e-05 | 7.2e-06 | 6.9e-06 | 8.700E-06 | 2.862E-06 | 6.900E-06 | 1.200E-05 |
| 9154 | 458 | V458F | A | 6.8e-06 | 9e-06 | 7.2e-06 | 7.667E-06 | 1.172E-06 | 6.800E-06 | 9.000E-06 |
| 9155 | 458 | V458P | A | 1.1e-05 | 1e-05 | 7.5e-06 | 9.500E-06 | 1.803E-06 | 7.500E-06 | 1.100E-05 |
| 9156 | 458 | V458S | A | 7.8e-06 | 9.9e-06 | 6.9e-06 | 8.200E-06 | 1.539E-06 | 6.900E-06 | 9.900E-06 |
| 9157 | 458 | V458T | A | 8.7e-06 | 4.6e-06 | 1.1e-05 | 8.100E-06 | 3.242E-06 | 4.600E-06 | 1.100E-05 |
| 9158 | 458 | V458W | A | 1e-05 | 6.5e-06 | 1.3e-05 | 9.833E-06 | 3.253E-06 | 6.500E-06 | 1.300E-05 |
| 9159 | 458 | V458Y | A | 6e-06 | 1.2e-05 | 1.3e-05 | 1.033E-05 | 3.786E-06 | 6.000E-06 | 1.300E-05 |
| 9160 | 458 | V458V | A | 9.4e-06 | 7.8e-06 | 7.9e-06 | 8.367E-06 | 8.963E-07 | 7.800E-06 | 9.400E-06 |
| 9161 | 459 | P459A | A | 6e-06 | 6.4e-06 | 3.3e-06 | 5.233E-06 | 1.686E-06 | 3.300E-06 | 6.400E-06 |
| 9162 | 459 | P459R | A | 8.3e-06 | 7.1e-06 | 4.1e-06 | 6.500E-06 | 2.163E-06 | 4.100E-06 | 8.300E-06 |
| 9163 | 459 | P459N | A | 7e-06 | 3.2e-06 | 9.1e-06 | 6.433E-06 | 2.991E-06 | 3.200E-06 | 9.100E-06 |
| 9164 | 459 | P459D | A | 1.1e-05 | 1.2e-05 | 4.5e-06 | 9.167E-06 | 4.072E-06 | 4.500E-06 | 1.200E-05 |
| 9165 | 459 | P459C | A | 1.4e-05 | 5.1e-06 | 4.2e-06 | 7.767E-06 | 5.417E-06 | 4.200E-06 | 1.400E-05 |
| 9166 | 459 | P459E | A | 5.6e-06 | 6.7e-06 | 5.6e-06 | 5.967E-06 | 6.351E-07 | 5.600E-06 | 6.700E-06 |
| 9167 | 459 | P459Q | A | 7.8e-06 | 8.3e-06 | 9.4e-06 | 8.500E-06 | 8.185E-07 | 7.800E-06 | 9.400E-06 |
| 9168 | 459 | P459G | A | 1.3e-05 | 5.8e-06 | 1.1e-05 | 9.933E-06 | 3.717E-06 | 5.800E-06 | 1.300E-05 |
| 9169 | 459 | P459H | A | 7.3e-06 | 1e-05 | 5.6e-06 | 7.633E-06 | 2.219E-06 | 5.600E-06 | 1.000E-05 |
| 9170 | 459 | P459I | A | 5.8e-06 | 1.1e-05 | 7e-06 | 7.933E-06 | 2.723E-06 | 5.800E-06 | 1.100E-05 |
| 9171 | 459 | P459L | A | 9.2e-06 | 6.9e-06 | 6.8e-06 | 7.633E-06 | 1.358E-06 | 6.800E-06 | 9.200E-06 |
| 9172 | 459 | P459K | A | 1.3e-05 | 3.7e-06 | 1.1e-05 | 9.233E-06 | 4.895E-06 | 3.700E-06 | 1.300E-05 |
| 9173 | 459 | P459M | A | 7.6e-06 | 5.2e-06 | 7.5e-06 | 6.767E-06 | 1.358E-06 | 5.200E-06 | 7.600E-06 |
| 9174 | 459 | P459F | A | 8.9e-06 | 3.4e-06 | 4.3e-06 | 5.533E-06 | 2.950E-06 | 3.400E-06 | 8.900E-06 |
| 9175 | 459 | P459P | A | 6.6e-06 | 5.5e-06 | 9.8e-06 | 7.300E-06 | 2.234E-06 | 5.500E-06 | 9.800E-06 |
| 9176 | 459 | P459S | A | 7.9e-06 | 4.1e-06 | 9.1e-06 | 7.033E-06 | 2.610E-06 | 4.100E-06 | 9.100E-06 |
| 9177 | 459 | P459T | A | 1.3e-05 | 9.8e-06 | 9.6e-06 | 1.080E-05 | 1.908E-06 | 9.600E-06 | 1.300E-05 |
| 9178 | 459 | P459W | A | 4.7e-06 | 8.1e-06 | 7.9e-06 | 6.900E-06 | 1.908E-06 | 4.700E-06 | 8.100E-06 |
| 9179 | 459 | P459Y | A | 5.8e-06 | 5.5e-06 | 1e-05 | 7.100E-06 | 2.516E-06 | 5.500E-06 | 1.000E-05 |

| | | | | | | | | | | |
|---|---|---|---|---|---|---|---|---|---|---|
| 9180 | 459 | P459V | A | 7.3e-06 | 1.1e-05 | 6.2e-06 | 8.167E-06 | 2.515E-06 | 6.200E-06 | 1.100E-05 |
| 9181 | 460 | W460A | A | 8.7e-06 | 8e-06 | 5.8e-06 | 7.500E-06 | 1.513E-06 | 5.800E-06 | 8.700E-06 |
| 9182 | 460 | W460R | A | 1.4e-05 | 8.3e-06 | 4.6e-06 | 8.967E-06 | 4.735E-06 | 4.600E-06 | 1.400E-05 |
| 9183 | 460 | W460N | A | 7.8e-06 | 9.4e-06 | 8.4e-06 | 8.533E-06 | 8.083E-07 | 7.800E-06 | 9.400E-06 |
| 9184 | 460 | W460D | A | 7.8e-06 | 8.6e-06 | 8.1e-06 | 8.167E-06 | 4.041E-07 | 7.800E-06 | 8.600E-06 |
| 9185 | 460 | W460C | A | 7e-06 | 1e-05 | 9.4e-06 | 8.800E-06 | 1.587E-06 | 7.000E-06 | 1.000E-05 |
| 9186 | 460 | W460E | A | 7.2e-06 | 9.2e-06 | 4.4e-06 | 6.933E-06 | 2.411E-06 | 4.400E-06 | 9.200E-06 |
| 9187 | 460 | W460Q | A | 1.5e-05 | 5.4e-06 | 7.4e-06 | 9.267E-06 | 5.065E-06 | 5.400E-06 | 1.500E-05 |
| 9188 | 460 | W460G | A | 6.5e-06 | 6.8e-06 | 8.6e-06 | 7.300E-06 | 1.136E-06 | 6.500E-06 | 8.600E-06 |
| 9189 | 460 | W460H | A | 7.7e-06 | 8e-06 | 6.2e-06 | 7.300E-06 | 9.644E-07 | 6.200E-06 | 8.000E-06 |
| 9190 | 460 | W460I | A | 6.1e-06 | 6.1e-06 | 5.7e-06 | 5.967E-06 | 2.309E-07 | 5.700E-06 | 6.100E-06 |
| 9191 | 460 | W460L | A | 7.3e-06 | 1e-05 | 5.8e-06 | 7.700E-06 | 2.128E-06 | 5.800E-06 | 1.000E-05 |
| 9192 | 460 | W460K | A | 1.2e-05 | 7.4e-06 | 9.9e-06 | 9.767E-06 | 2.303E-06 | 7.400E-06 | 1.200E-05 |
| 9193 | 460 | W460M | A | 9.5e-06 | 6.3e-06 | 7.7e-06 | 7.833E-06 | 1.604E-06 | 6.300E-06 | 9.500E-06 |
| 9194 | 460 | W460F | A | 1.2e-05 | 5.2e-06 | 6.1e-06 | 7.767E-06 | 3.694E-06 | 5.200E-06 | 1.200E-05 |
| 9195 | 460 | W460P | A | 8.5e-06 | 1e-05 | 8.1e-06 | 8.867E-06 | 1.002E-06 | 8.100E-06 | 1.000E-05 |
| 9196 | 460 | W460S | A | 1.4e-05 | 1.6e-05 | 8.6e-06 | 1.287E-05 | 3.828E-06 | 8.600E-06 | 1.600E-05 |
| 9197 | 460 | W460T | A | 1.4e-05 | 6.4e-06 | 7e-06 | 9.133E-06 | 4.225E-06 | 6.400E-06 | 1.400E-05 |
| 9198 | 460 | W460W | A | 5.4e-06 | 4.7e-06 | 6.5e-06 | 5.533E-06 | 9.074E-07 | 4.700E-06 | 6.500E-06 |
| 9199 | 460 | W460Y | A | 1.1e-05 | 6.2e-06 | 1.3e-05 | 1.007E-05 | 3.495E-06 | 6.200E-06 | 1.300E-05 |
| 9200 | 460 | W460V | A | 5.3e-06 | 1.1e-05 | 7e-06 | 7.767E-06 | 2.926E-06 | 5.300E-06 | 1.100E-05 |
| 9201 | 461 | D461A | A | 7.7e-06 | 5.5e-06 | 1e-05 | 7.733E-06 | 2.250E-06 | 5.500E-06 | 1.000E-05 |
| 9202 | 461 | D461R | A | 8.1e-06 | 7.3e-06 | 5.3e-06 | 6.900E-06 | 1.442E-06 | 5.300E-06 | 8.100E-06 |
| 9203 | 461 | D461N | A | 9.1e-06 | 5.9e-06 | 8.5e-06 | 7.833E-06 | 1.701E-06 | 5.900E-06 | 9.100E-06 |
| 9204 | 461 | D461D | A | 4.9e-06 | 9e-06 | 5.1e-06 | 6.333E-06 | 2.312E-06 | 4.900E-06 | 9.000E-06 |
| 9205 | 461 | D461C | A | 6e-06 | 4.9e-06 | 9.5e-06 | 6.800E-06 | 2.402E-06 | 4.900E-06 | 9.500E-06 |
| 9206 | 461 | D461E | A | 7.7e-06 | 7.5e-06 | 1.1e-05 | 8.733E-06 | 1.966E-06 | 7.500E-06 | 1.100E-05 |
| 9207 | 461 | D461Q | A | 7.9e-06 | 1.2e-05 | 7.3e-06 | 9.067E-06 | 2.558E-06 | 7.300E-06 | 1.200E-05 |
| 9208 | 461 | D461G | A | 6.2e-06 | 6.9e-06 | 7.6e-06 | 6.900E-06 | 7.000E-07 | 6.200E-06 | 7.600E-06 |
| 9209 | 461 | D461H | A | 1.2e-05 | 5.4e-06 | 8e-06 | 8.467E-06 | 3.325E-06 | 5.400E-06 | 1.200E-05 |
| 9210 | 461 | D461I | A | 1.1e-05 | 8.2e-06 | 5.6e-06 | 8.267E-06 | 2.701E-06 | 5.600E-06 | 1.100E-05 |
| 9211 | 461 | D461L | A | 6.8e-06 | 8.3e-06 | 9.3e-06 | 8.133E-06 | 1.258E-06 | 6.800E-06 | 9.300E-06 |
| 9212 | 461 | D461K | A | 5.5e-06 | 7.5e-06 | 1.2e-05 | 8.333E-06 | 3.329E-06 | 5.500E-06 | 1.200E-05 |
| 9213 | 461 | D461M | A | 7.9e-06 | 1.2e-05 | 1.1e-05 | 1.030E-05 | 2.138E-06 | 7.900E-06 | 1.200E-05 |
| 9214 | 461 | D461F | A | 5e-06 | 1.1e-05 | 8.5e-06 | 8.167E-06 | 3.014E-06 | 5.000E-06 | 1.100E-05 |
| 9215 | 461 | D461P | A | 9.5e-06 | 5.8e-06 | 8.6e-06 | 7.967E-06 | 1.930E-06 | 5.800E-06 | 9.500E-06 |
| 9216 | 461 | D461S | A | 8.3e-06 | 6.8e-06 | 8.5e-06 | 7.867E-06 | 9.292E-07 | 6.800E-06 | 8.500E-06 |
| 9217 | 461 | D461T | A | 6.3e-06 | 1.2e-05 | 9.9e-06 | 9.400E-06 | 2.883E-06 | 6.300E-06 | 1.200E-05 |
| 9218 | 461 | D461W | A | 4.8e-06 | 6.3e-06 | 1e-05 | 7.033E-06 | 2.676E-06 | 4.800E-06 | 1.000E-05 |
| 9219 | 461 | D461Y | A | 9.6e-06 | 9.1e-06 | 5.6e-06 | 8.100E-06 | 2.179E-06 | 5.600E-06 | 9.600E-06 |
| 9220 | 461 | D461V | A | 6.4e-06 | 3.8e-06 | 5.3e-06 | 5.167E-06 | 1.305E-06 | 3.800E-06 | 6.400E-06 |
| 9221 | 462 | Q462A | A | 6.2e-06 | 6.5e-06 | 8e-06 | 6.900E-06 | 9.644E-07 | 6.200E-06 | 8.000E-06 |
| 9222 | 462 | Q462R | A | 1e-05 | 1.1e-05 | 6.3e-06 | 9.100E-06 | 2.476E-06 | 6.300E-06 | 1.100E-05 |
| 9223 | 462 | Q462N | A | 1.1e-05 | 5.6e-06 | 9.3e-06 | 8.633E-06 | 2.761E-06 | 5.600E-06 | 1.100E-05 |
| 9224 | 462 | Q462D | A | 7.6e-06 | 8.6e-06 | 9.6e-06 | 8.600E-06 | 1.000E-06 | 7.600E-06 | 9.600E-06 |
| 9225 | 462 | Q462C | A | 9e-06 | 6.7e-06 | 7.1e-06 | 7.600E-06 | 1.229E-06 | 6.700E-06 | 9.000E-06 |
| 9226 | 462 | Q462E | A | 9.6e-06 | 7.7e-06 | 1e-05 | 9.100E-06 | 1.229E-06 | 7.700E-06 | 1.000E-05 |
| 9227 | 462 | Q462Q | A | 1.3e-05 | 1.3e-05 | 8.1e-06 | 1.137E-05 | 2.829E-06 | 8.100E-06 | 1.300E-05 |
| 9228 | 462 | Q462G | A | 9.2e-06 | 6.7e-06 | 1.1e-05 | 8.967E-06 | 2.159E-06 | 6.700E-06 | 1.100E-05 |
| 9229 | 462 | Q462H | A | 9.4e-06 | 9.6e-06 | 1.1e-05 | 1.000E-05 | 8.718E-07 | 9.400E-06 | 1.100E-05 |
| 9230 | 462 | Q462I | A | 9.3e-06 | 9.7e-06 | 8.8e-06 | 9.267E-06 | 4.509E-07 | 8.800E-06 | 9.700E-06 |
| 9231 | 462 | Q462L | A | 5.7e-06 | 7.9e-06 | 7.7e-06 | 7.100E-06 | 1.217E-06 | 5.700E-06 | 7.900E-06 |
| 9232 | 462 | Q462K | A | 1.1e-05 | 4.7e-06 | 4.9e-06 | 6.867E-06 | 3.581E-06 | 4.700E-06 | 1.100E-05 |
| 9233 | 462 | Q462M | A | 3.9e-06 | 5e-06 | 1e-05 | 6.300E-06 | 3.251E-06 | 3.900E-06 | 1.000E-05 |
| 9234 | 462 | Q462F | A | 6.6e-06 | 6e-06 | 8.9e-06 | 7.167E-06 | 1.531E-06 | 6.000E-06 | 8.900E-06 |
| 9235 | 462 | Q462P | A | 1.1e-05 | 9.1e-06 | 6.1e-06 | 8.733E-06 | 2.470E-06 | 6.100E-06 | 1.100E-05 |
| 9236 | 462 | Q462S | A | 8.8e-06 | 1e-05 | 7e-06 | 8.600E-06 | 1.510E-06 | 7.000E-06 | 1.000E-05 |
| 9237 | 462 | Q462T | A | 7.2e-06 | 7.1e-06 | 9.2e-06 | 7.833E-06 | 1.185E-06 | 7.100E-06 | 9.200E-06 |
| 9238 | 462 | Q462W | A | 1.1e-05 | 5.3e-06 | 1.4e-05 | 1.010E-05 | 4.419E-06 | 5.300E-06 | 1.400E-05 |
| 9239 | 462 | Q462Y | A | 8.6e-06 | 1.7e-05 | 5.3e-06 | 1.030E-05 | 6.032E-06 | 5.300E-06 | 1.700E-05 |
| 9240 | 462 | Q462V | A | 9.1e-06 | 9.4e-06 | 7.3e-06 | 8.600E-06 | 1.136E-06 | 7.300E-06 | 9.400E-06 |
| 9241 | 463 | L463A | A | 1.2e-05 | 1.4e-05 | 1.3e-05 | 1.300E-05 | 1.000E-06 | 1.200E-05 | 1.400E-05 |
| 9242 | 463 | L463R | A | 4.5e-06 | 1e-05 | 1e-05 | 8.167E-06 | 3.175E-06 | 4.500E-06 | 1.000E-05 |
| 9243 | 463 | L463N | A | 1.3e-05 | 1e-05 | 6.7e-06 | 9.900E-06 | 3.151E-06 | 6.700E-06 | 1.300E-05 |
| 9244 | 463 | L463D | A | 6.8e-06 | 6.7e-06 | 9.7e-06 | 7.733E-06 | 1.704E-06 | 6.700E-06 | 9.700E-06 |
| 9245 | 463 | L463C | A | 1.2e-05 | 7.6e-06 | 5.6e-06 | 8.400E-06 | 3.274E-06 | 5.600E-06 | 1.200E-05 |
| 9246 | 463 | L463E | A | 7.5e-06 | 5.4e-06 | 6.6e-06 | 6.500E-06 | 1.054E-06 | 5.400E-06 | 7.500E-06 |
| 9247 | 463 | L463Q | A | 1.1e-05 | 5.1e-06 | 9.6e-06 | 8.567E-06 | 3.083E-06 | 5.100E-06 | 1.100E-05 |
| 9248 | 463 | L463G | A | 5.9e-06 | 4.3e-06 | 6.9e-06 | 5.700E-06 | 1.311E-06 | 4.300E-06 | 6.900E-06 |
| 9249 | 463 | L463H | A | 4.1e-06 | 4.4e-06 | 7.1e-06 | 5.200E-06 | 1.652E-06 | 4.100E-06 | 7.100E-06 |
| 9250 | 463 | L463I | A | 6.3e-06 | 5.2e-06 | 8.6e-06 | 6.700E-06 | 1.735E-06 | 5.200E-06 | 8.600E-06 |
| 9251 | 463 | L463L | A | 7e-06 | 8.4e-06 | 5.1e-06 | 6.833E-06 | 1.656E-06 | 5.100E-06 | 8.400E-06 |
| 9252 | 463 | L463K | A | 3.9e-06 | 4.6e-06 | 7.2e-06 | 5.233E-06 | 1.739E-06 | 3.900E-06 | 7.200E-06 |
| 9253 | 463 | L463M | A | 1.1e-05 | 4.9e-06 | 5.4e-06 | 7.100E-06 | 3.387E-06 | 4.900E-06 | 1.100E-05 |
| 9254 | 463 | L463F | A | 6.4e-06 | 9e-06 | 8.7e-06 | 8.033E-06 | 1.422E-06 | 6.400E-06 | 9.000E-06 |
| 9255 | 463 | L463P | A | 8.9e-06 | 5.9e-06 | 8e-06 | 7.600E-06 | 1.539E-06 | 5.900E-06 | 8.900E-06 |
| 9256 | 463 | L463S | A | 4.3e-06 | 5e-06 | 5.8e-06 | 5.033E-06 | 7.506E-07 | 4.300E-06 | 5.800E-06 |
| 9257 | 463 | L463T | A | 8.3e-06 | 4.7e-06 | 1e-05 | 7.667E-06 | 2.706E-06 | 4.700E-06 | 1.000E-05 |
| 9258 | 463 | L463W | A | 9.9e-06 | 7.2e-06 | 1.1e-05 | 9.367E-06 | 1.955E-06 | 7.200E-06 | 1.100E-05 |
| 9259 | 463 | L463Y | A | 5.9e-06 | 8.6e-06 | 6.6e-06 | 7.033E-06 | 1.401E-06 | 5.900E-06 | 8.600E-06 |
| 9260 | 463 | L463V | A | 1.5e-05 | 9.8e-06 | 5e-06 | 9.933E-06 | 5.001E-06 | 5.000E-06 | 1.500E-05 |
| 9261 | 464 | F464A | A | 1e-05 | 7.5e-06 | 9.6e-06 | 9.033E-06 | 1.343E-06 | 7.500E-06 | 1.000E-05 |
| 9262 | 464 | F464R | A | 8.5e-06 | 1.3e-05 | 7.9e-06 | 9.800E-06 | 2.787E-06 | 7.900E-06 | 1.300E-05 |
| 9263 | 464 | F464N | A | 9.8e-06 | 5.8e-06 | 8.1e-06 | 7.900E-06 | 2.007E-06 | 5.800E-06 | 9.800E-06 |
| 9264 | 464 | F464D | A | 6.6e-06 | 1.4e-05 | 9e-06 | 9.867E-06 | 3.775E-06 | 6.600E-06 | 1.400E-05 |

| | | | | | | | | | | |
|---|---|---|---|---|---|---|---|---|---|---|
| 9265 | 464 | F464C | A | 7.3e-06 | 6.2e-06 | 6.1e-06 | 6.533E-06 | 6.658E-07 | 6.100E-06 | 7.300E-06 |
| 9266 | 464 | F464E | A | 8.7e-06 | 1.2e-05 | 6.9e-06 | 9.200E-06 | 2.587E-06 | 6.900E-06 | 1.200E-05 |
| 9267 | 464 | F464Q | A | 8.5e-06 | 4.4e-06 | 8.6e-06 | 7.167E-06 | 2.397E-06 | 4.400E-06 | 8.600E-06 |
| 9268 | 464 | F464G | A | 8.4e-06 | 7.8e-06 | 1e-05 | 8.733E-06 | 1.137E-06 | 7.800E-06 | 1.000E-05 |
| 9269 | 464 | F464H | A | 9.9e-06 | 8.7e-06 | 8e-06 | 8.867E-06 | 9.609E-07 | 8.000E-06 | 9.900E-06 |
| 9270 | 464 | F464I | A | 8.3e-06 | 8e-06 | 9.8e-06 | 8.700E-06 | 9.644E-07 | 8.000E-06 | 9.800E-06 |
| 9271 | 464 | F464L | A | 7.8e-06 | 1.1e-05 | 5e-06 | 7.933E-06 | 3.002E-06 | 5.000E-06 | 1.100E-05 |
| 9272 | 464 | F464K | A | 6.6e-06 | 3.9e-06 | 1.4e-05 | 8.167E-06 | 5.229E-06 | 3.900E-06 | 1.400E-05 |
| 9273 | 464 | F464M | A | 4.7e-06 | 1e-05 | 8.1e-06 | 7.600E-06 | 2.685E-06 | 4.700E-06 | 1.000E-05 |
| 9274 | 464 | F464F | A | 5.8e-06 | 7.6e-06 | 8.1e-06 | 7.167E-06 | 1.210E-06 | 5.800E-06 | 8.100E-06 |
| 9275 | 464 | F464P | A | 7.6e-06 | 7.4e-06 | 9.4e-06 | 8.133E-06 | 1.102E-06 | 7.400E-06 | 9.400E-06 |
| 9276 | 464 | F464S | A | 6.9e-06 | 7.9e-06 | 1e-05 | 8.267E-06 | 1.582E-06 | 6.900E-06 | 1.000E-05 |
| 9277 | 464 | F464T | A | 8.7e-06 | 1e-05 | 6.6e-06 | 8.433E-06 | 1.716E-06 | 6.600E-06 | 1.000E-05 |
| 9278 | 464 | F464W | A | 6.9e-06 | 1.4e-05 | 5.2e-06 | 8.700E-06 | 4.668E-06 | 5.200E-06 | 1.400E-05 |
| 9279 | 464 | F464Y | A | 6.7e-06 | 7.7e-06 | 1.1e-05 | 8.467E-06 | 2.250E-06 | 6.700E-06 | 1.100E-05 |
| 9280 | 464 | F464V | A | 5.9e-06 | 8.1e-06 | 5.5e-06 | 6.500E-06 | 1.400E-06 | 5.500E-06 | 8.100E-06 |
| 9281 | 465 | R465A | A | 7.9e-06 | 5.8e-06 | 1.2e-05 | 8.567E-06 | 3.153E-06 | 5.800E-06 | 1.200E-05 |
| 9282 | 465 | R465R | A | 6.1e-06 | 7.8e-06 | 1.2e-05 | 8.633E-06 | 3.037E-06 | 6.100E-06 | 1.200E-05 |
| 9283 | 465 | R465N | A | 5.7e-06 | 5.6e-06 | 1e-05 | 7.100E-06 | 2.512E-06 | 5.600E-06 | 1.000E-05 |
| 9284 | 465 | R465D | A | 8.5e-06 | 8e-06 | 8.6e-06 | 8.367E-06 | 3.215E-07 | 8.000E-06 | 8.600E-06 |
| 9285 | 465 | R465C | A | 7.5e-06 | 8e-06 | 9.3e-06 | 8.267E-06 | 9.292E-07 | 7.500E-06 | 9.300E-06 |
| 9286 | 465 | R465E | A | 7e-06 | 8.5e-06 | 9.5e-06 | 8.333E-06 | 1.258E-06 | 7.000E-06 | 9.500E-06 |
| 9287 | 465 | R465Q | A | 7.3e-06 | 5e-06 | 7.5e-06 | 6.600E-06 | 1.389E-06 | 5.000E-06 | 7.500E-06 |
| 9288 | 465 | R465G | A | 1.2e-05 | 9.4e-06 | 6.2e-06 | 9.200E-06 | 2.905E-06 | 6.200E-06 | 1.200E-05 |
| 9289 | 465 | R465H | A | 7.9e-06 | 5.9e-06 | 1e-05 | 7.933E-06 | 2.050E-06 | 5.900E-06 | 1.000E-05 |
| 9290 | 465 | R465I | A | 5.5e-06 | 7.9e-06 | 8.4e-06 | 7.267E-06 | 1.550E-06 | 5.500E-06 | 8.400E-06 |
| 9291 | 465 | R465L | A | 9.4e-06 | 5.5e-06 | 9.1e-06 | 8.000E-06 | 2.170E-06 | 5.500E-06 | 9.400E-06 |
| 9292 | 465 | R465K | A | 8.5e-06 | 1.1e-05 | 7.7e-06 | 9.067E-06 | 1.721E-06 | 7.700E-06 | 1.100E-05 |
| 9293 | 465 | R465M | A | 7.3e-06 | 6.7e-06 | 8e-06 | 7.333E-06 | 6.506E-07 | 6.700E-06 | 8.000E-06 |
| 9294 | 465 | R465F | A | 8.2e-06 | 5.2e-06 | 1.2e-05 | 8.467E-06 | 3.408E-06 | 5.200E-06 | 1.200E-05 |
| 9295 | 465 | R465P | A | 1.1e-05 | 1.2e-05 | 7.8e-06 | 1.027E-05 | 2.194E-06 | 7.800E-06 | 1.200E-05 |
| 9296 | 465 | R465S | A | 5.4e-06 | 9.5e-06 | 9.1e-06 | 8.000E-06 | 2.261E-06 | 5.400E-06 | 9.500E-06 |
| 9297 | 465 | R465T | A | 9.5e-06 | 7.4e-06 | 1.4e-05 | 1.030E-05 | 3.372E-06 | 7.400E-06 | 1.400E-05 |
| 9298 | 465 | R465W | A | 1.3e-05 | 8.8e-06 | 6.6e-06 | 9.467E-06 | 3.252E-06 | 6.600E-06 | 1.300E-05 |
| 9299 | 465 | R465Y | A | 9.7e-06 | 8.1e-06 | 3.5e-06 | 7.100E-06 | 3.219E-06 | 3.500E-06 | 9.700E-06 |
| 9300 | 465 | R465V | A | 1.3e-05 | 1.9e-05 | 5.9e-06 | 1.263E-05 | 6.558E-06 | 5.900E-06 | 1.900E-05 |
| 9301 | 466 | N466A | A | 7.5e-06 | 9.9e-06 | 1e-05 | 9.133E-06 | 1.415E-06 | 7.500E-06 | 1.000E-05 |
| 9302 | 466 | N466R | A | 1.2e-05 | 9.4e-06 | 5.2e-06 | 8.867E-06 | 3.431E-06 | 5.200E-06 | 1.200E-05 |
| 9303 | 466 | N466N | A | 6.2e-06 | 1.4e-05 | 6.7e-06 | 8.967E-06 | 4.366E-06 | 6.200E-06 | 1.400E-05 |
| 9304 | 466 | N466D | A | 6.7e-06 | 8.1e-06 | 5.1e-06 | 6.633E-06 | 1.501E-06 | 5.100E-06 | 8.100E-06 |
| 9305 | 466 | N466C | A | 7.4e-06 | 5.6e-06 | 8e-06 | 7.000E-06 | 1.249E-06 | 5.600E-06 | 8.000E-06 |
| 9306 | 466 | N466E | A | 9.2e-06 | 8.2e-06 | 6.5e-06 | 7.967E-06 | 1.365E-06 | 6.500E-06 | 9.200E-06 |
| 9307 | 466 | N466Q | A | 9.2e-06 | 9.2e-06 | 6.5e-06 | 8.300E-06 | 1.559E-06 | 6.500E-06 | 9.200E-06 |
| 9308 | 466 | N466G | A | 1.6e-05 | 7.2e-06 | 8.1e-06 | 1.043E-05 | 4.842E-06 | 7.200E-06 | 1.600E-05 |
| 9309 | 466 | N466H | A | 1.3e-05 | 6.7e-06 | 9.1e-06 | 9.600E-06 | 3.180E-06 | 6.700E-06 | 1.300E-05 |
| 9310 | 466 | N466I | A | 9.9e-06 | 6.7e-06 | 5.2e-06 | 7.267E-06 | 2.401E-06 | 5.200E-06 | 9.900E-06 |
| 9311 | 466 | N466L | A | 8.1e-06 | 1.1e-05 | 4.8e-06 | 7.967E-06 | 3.102E-06 | 4.800E-06 | 1.100E-05 |
| 9312 | 466 | N466K | A | 7.4e-06 | 6.4e-06 | 6.4e-06 | 6.733E-06 | 5.774E-07 | 6.400E-06 | 7.400E-06 |
| 9313 | 466 | N466M | A | 1.2e-05 | 9.8e-06 | 7.8e-06 | 9.867E-06 | 2.101E-06 | 7.800E-06 | 1.200E-05 |
| 9314 | 466 | N466F | A | 7.8e-06 | 5.8e-06 | 7.1e-06 | 6.900E-06 | 1.015E-06 | 5.800E-06 | 7.800E-06 |
| 9315 | 466 | N466P | A | 4.8e-06 | 8.4e-06 | 7.7e-06 | 6.967E-06 | 1.909E-06 | 4.800E-06 | 8.400E-06 |
| 9316 | 466 | N466S | A | 3.9e-06 | 1.3e-05 | 6.9e-06 | 7.933E-06 | 4.637E-06 | 3.900E-06 | 1.300E-05 |
| 9317 | 466 | N466T | A | 6.2e-06 | 6.2e-06 | 7.8e-06 | 6.733E-06 | 9.238E-07 | 6.200E-06 | 7.800E-06 |
| 9318 | 466 | N466W | A | 9.3e-06 | 5.8e-06 | 5.4e-06 | 6.833E-06 | 2.146E-06 | 5.400E-06 | 9.300E-06 |
| 9319 | 466 | N466Y | A | 4e-06 | 5.4e-06 | 5.2e-06 | 4.867E-06 | 7.572E-07 | 4.000E-06 | 5.400E-06 |
| 9320 | 466 | N466V | A | 9.9e-06 | 4.1e-06 | 7.3e-06 | 7.100E-06 | 2.905E-06 | 4.100E-06 | 9.900E-06 |
| 9321 | 467 | P467A | A | 1.1e-05 | 1e-05 | 4.8e-06 | 8.600E-06 | 3.329E-06 | 4.800E-06 | 1.100E-05 |
| 9322 | 467 | P467R | A | 6.8e-06 | 1.3e-05 | 7e-06 | 8.933E-06 | 3.523E-06 | 6.800E-06 | 1.300E-05 |
| 9323 | 467 | P467N | A | 6.6e-06 | 1.3e-05 | 1.6e-05 | 1.187E-05 | 4.801E-06 | 6.600E-06 | 1.600E-05 |
| 9324 | 467 | P467D | A | 6.3e-06 | 8.3e-06 | 1e-05 | 8.200E-06 | 1.852E-06 | 6.300E-06 | 1.000E-05 |
| 9325 | 467 | P467C | A | 8.3e-06 | 9.6e-06 | 7.2e-06 | 8.367E-06 | 1.201E-06 | 7.200E-06 | 9.600E-06 |
| 9326 | 467 | P467E | A | 9e-06 | 6.2e-06 | 8.6e-06 | 7.933E-06 | 1.514E-06 | 6.200E-06 | 9.000E-06 |
| 9327 | 467 | P467Q | A | 4.3e-06 | 6.8e-06 | 7.3e-06 | 6.133E-06 | 1.607E-06 | 4.300E-06 | 7.300E-06 |
| 9328 | 467 | P467G | A | 5.5e-06 | 1.2e-05 | 9.6e-06 | 9.033E-06 | 3.287E-06 | 5.500E-06 | 1.200E-05 |
| 9329 | 467 | P467H | A | 9.3e-06 | 6.4e-06 | 8.6e-06 | 8.100E-06 | 1.513E-06 | 6.400E-06 | 9.300E-06 |
| 9330 | 467 | P467I | A | 6.3e-06 | 6.8e-06 | 8e-06 | 7.033E-06 | 8.737E-07 | 6.300E-06 | 8.000E-06 |
| 9331 | 467 | P467L | A | 9e-06 | 6.7e-06 | 6.3e-06 | 7.333E-06 | 1.457E-06 | 6.300E-06 | 9.000E-06 |
| 9332 | 467 | P467K | A | 4e-06 | 7.4e-06 | 1e-05 | 7.133E-06 | 3.009E-06 | 4.000E-06 | 1.000E-05 |
| 9333 | 467 | P467M | A | 7.6e-06 | 9.7e-06 | 8.1e-06 | 8.467E-06 | 1.097E-06 | 7.600E-06 | 9.700E-06 |
| 9334 | 467 | P467F | A | 1.2e-05 | 7.4e-06 | 5.5e-06 | 8.300E-06 | 3.342E-06 | 5.500E-06 | 1.200E-05 |
| 9335 | 467 | P467P | A | 5.8e-06 | 7.1e-06 | 8.7e-06 | 7.200E-06 | 1.453E-06 | 5.800E-06 | 8.700E-06 |
| 9336 | 467 | P467S | A | 4.4e-06 | 1e-05 | 4.4e-06 | 6.267E-06 | 3.233E-06 | 4.400E-06 | 1.000E-05 |
| 9337 | 467 | P467T | A | 3.6e-06 | 9.8e-06 | 8.1e-06 | 7.167E-06 | 3.204E-06 | 3.600E-06 | 9.800E-06 |
| 9338 | 467 | P467W | A | 1.2e-05 | 1.4e-05 | 8.1e-06 | 1.137E-05 | 3.001E-06 | 8.100E-06 | 1.400E-05 |
| 9339 | 467 | P467Y | A | 5e-06 | 7e-06 | 1e-05 | 7.333E-06 | 2.517E-06 | 5.000E-06 | 1.000E-05 |
| 9340 | 467 | P467V | A | 1e-05 | 8.1e-06 | 7.5e-06 | 8.533E-06 | 1.305E-06 | 7.500E-06 | 1.000E-05 |
| 9341 | 468 | H468A | A | 5.9e-06 | 8.1e-06 | 1.3e-05 | 9.000E-06 | 3.635E-06 | 5.900E-06 | 1.300E-05 |
| 9342 | 468 | H468R | A | 6.4e-06 | 1.4e-05 | 1.5e-05 | 1.180E-05 | 4.703E-06 | 6.400E-06 | 1.500E-05 |
| 9343 | 468 | H468N | A | 7.5e-06 | 8.5e-06 | 6.6e-06 | 7.533E-06 | 9.504E-07 | 6.600E-06 | 8.500E-06 |
| 9344 | 468 | H468D | A | 1.4e-05 | 6e-06 | 5.8e-06 | 8.600E-06 | 4.678E-06 | 5.800E-06 | 1.400E-05 |
| 9345 | 468 | H468C | A | 5.7e-06 | 1.4e-05 | 1.1e-05 | 1.023E-05 | 4.203E-06 | 5.700E-06 | 1.400E-05 |
| 9346 | 468 | H468E | A | 9.3e-06 | 7.2e-06 | 5.4e-06 | 7.300E-06 | 1.952E-06 | 5.400E-06 | 9.300E-06 |
| 9347 | 468 | H468Q | A | 5.7e-06 | 8.6e-06 | 5.5e-06 | 6.600E-06 | 1.735E-06 | 5.500E-06 | 8.600E-06 |
| 9348 | 468 | H468G | A | 9.2e-06 | 7.5e-06 | 1.3e-05 | 9.900E-06 | 2.816E-06 | 7.500E-06 | 1.300E-05 |
| 9349 | 468 | H468H | A | 8.6e-06 | 9.5e-06 | 9.7e-06 | 9.267E-06 | 5.859E-07 | 8.600E-06 | 9.700E-06 |

| | | | | | | | | | | |
|---|---|---|---|---|---|---|---|---|---|---|
| 9350 | 468 | H468I | A | 1.4e-05 | 8.6e-06 | 8e-06 | 1.020E-05 | 3.305E-06 | 8.000E-06 | 1.400E-05 |
| 9351 | 468 | H468L | A | 8.1e-06 | 9e-06 | 5.9e-06 | 7.667E-06 | 1.595E-06 | 5.900E-06 | 9.000E-06 |
| 9352 | 468 | H468K | A | 7.9e-06 | 9.4e-06 | 7.1e-06 | 8.133E-06 | 1.168E-06 | 7.100E-06 | 9.400E-06 |
| 9353 | 468 | H468M | A | 1e-05 | 9.1e-06 | 1.3e-05 | 1.070E-05 | 2.042E-06 | 9.100E-06 | 1.300E-05 |
| 9354 | 468 | H468F | A | 6.6e-06 | 4.4e-06 | 7.4e-06 | 6.133E-06 | 1.553E-06 | 4.400E-06 | 7.400E-06 |
| 9355 | 468 | H468P | A | 1.3e-05 | 8.1e-06 | 3.3e-06 | 8.133E-06 | 4.850E-06 | 3.300E-06 | 1.300E-05 |
| 9356 | 468 | H468S | A | 1e-05 | 6.6e-06 | 4.2e-06 | 6.933E-06 | 2.914E-06 | 4.200E-06 | 1.000E-05 |
| 9357 | 468 | H468T | A | 7.7e-06 | 3.5e-06 | 5.4e-06 | 5.533E-06 | 2.103E-06 | 3.500E-06 | 7.700E-06 |
| 9358 | 468 | H468W | A | 6.1e-06 | 6.8e-06 | 8.3e-06 | 7.067E-06 | 1.124E-06 | 6.100E-06 | 8.300E-06 |
| 9359 | 468 | H468Y | A | 1.2e-05 | 8e-06 | 7.7e-06 | 9.233E-06 | 2.401E-06 | 7.700E-06 | 1.200E-05 |
| 9360 | 468 | H468V | A | 4.9e-06 | 6.9e-06 | 6.8e-06 | 6.200E-06 | 1.127E-06 | 4.900E-06 | 6.900E-06 |
| 9361 | 469 | Q469A | A | 4.6e-06 | 8.8e-06 | 3.2e-06 | 5.533E-06 | 2.914E-06 | 3.200E-06 | 8.800E-06 |
| 9362 | 469 | Q469R | A | 1e-05 | 1.1e-05 | 9.1e-06 | 1.003E-05 | 9.504E-07 | 9.100E-06 | 1.100E-05 |
| 9363 | 469 | Q469N | A | 1e-05 | 7.4e-06 | 6.6e-06 | 8.000E-06 | 1.778E-06 | 6.600E-06 | 1.000E-05 |
| 9364 | 469 | Q469D | A | 7.3e-06 | 9.6e-06 | 9e-06 | 8.633E-06 | 1.193E-06 | 7.300E-06 | 9.600E-06 |
| 9365 | 469 | Q469C | A | 1.3e-05 | 7.7e-06 | 1.5e-05 | 1.190E-05 | 3.772E-06 | 7.700E-06 | 1.500E-05 |
| 9366 | 469 | Q469E | A | 4.2e-06 | 6.7e-06 | 1.1e-05 | 7.300E-06 | 3.439E-06 | 4.200E-06 | 1.100E-05 |
| 9367 | 469 | Q469Q | A | 1e-05 | 1e-05 | 1e-05 | 1.000E-05 | 0.000E+00 | 1.000E-05 | 1.000E-05 |
| 9368 | 469 | Q469G | A | 9.5e-06 | 6.8e-06 | 6.7e-06 | 7.667E-06 | 1.589E-06 | 6.700E-06 | 9.500E-06 |
| 9369 | 469 | Q469H | A | 7.7e-06 | 1.1e-05 | 8.9e-06 | 9.200E-06 | 1.670E-06 | 7.700E-06 | 1.100E-05 |
| 9370 | 469 | Q469I | A | 1.1e-05 | 7.4e-06 | 7.1e-06 | 8.500E-06 | 2.170E-06 | 7.100E-06 | 1.100E-05 |
| 9371 | 469 | Q469L | A | 9e-06 | 7.9e-06 | 8e-06 | 8.300E-06 | 6.083E-07 | 7.900E-06 | 9.000E-06 |
| 9372 | 469 | Q469K | A | 1e-05 | 6.2e-06 | 4.6e-06 | 6.933E-06 | 2.774E-06 | 4.600E-06 | 1.000E-05 |
| 9373 | 469 | Q469M | A | 7.5e-06 | 1.9e-05 | 5.4e-06 | 1.063E-05 | 7.321E-06 | 5.400E-06 | 1.900E-05 |
| 9374 | 469 | Q469F | A | 7.9e-06 | 8.9e-06 | 9.1e-06 | 8.633E-06 | 6.429E-07 | 7.900E-06 | 9.100E-06 |
| 9375 | 469 | Q469P | A | 3e-06 | 1.5e-05 | 4.1e-06 | 7.367E-06 | 6.634E-06 | 3.000E-06 | 1.500E-05 |
| 9376 | 469 | Q469S | A | 1.3e-05 | 1.4e-05 | 9.8e-06 | 1.227E-05 | 2.194E-06 | 9.800E-06 | 1.400E-05 |
| 9377 | 469 | Q469T | A | 5.2e-06 | 1.3e-05 | 7.2e-06 | 8.467E-06 | 4.051E-06 | 5.200E-06 | 1.300E-05 |
| 9378 | 469 | Q469W | A | 1.2e-05 | 9.5e-06 | 6.5e-06 | 9.333E-06 | 2.754E-06 | 6.500E-06 | 1.200E-05 |
| 9379 | 469 | Q469Y | A | 1.3e-05 | 4.5e-06 | 9.5e-06 | 9.000E-06 | 4.272E-06 | 4.500E-06 | 1.300E-05 |
| 9380 | 469 | Q469V | A | 6.8e-06 | 7.4e-06 | 9e-06 | 7.733E-06 | 1.137E-06 | 6.800E-06 | 9.000E-06 |
| 9381 | 470 | A470A | A | 3.8e-06 | 6.7e-06 | 9.7e-06 | 6.733E-06 | 2.950E-06 | 3.800E-06 | 9.700E-06 |
| 9382 | 470 | A470R | A | 4.1e-06 | 4.7e-06 | 9.3e-06 | 6.033E-06 | 2.845E-06 | 4.100E-06 | 9.300E-06 |
| 9383 | 470 | A470N | A | 8.7e-06 | 1e-05 | 5.9e-06 | 8.200E-06 | 2.095E-06 | 5.900E-06 | 1.000E-05 |
| 9384 | 470 | A470D | A | 7.7e-06 | 6.1e-06 | 9.8e-06 | 7.867E-06 | 1.856E-06 | 6.100E-06 | 9.800E-06 |
| 9385 | 470 | A470C | A | 6.2e-06 | 7.4e-06 | 5.5e-06 | 6.367E-06 | 9.609E-07 | 5.500E-06 | 7.400E-06 |
| 9386 | 470 | A470E | A | 1.4e-05 | 4.9e-06 | 7.1e-06 | 8.667E-06 | 4.748E-06 | 4.900E-06 | 1.400E-05 |
| 9387 | 470 | A470Q | A | 1.2e-05 | 8.4e-06 | 7e-06 | 9.133E-06 | 2.579E-06 | 7.000E-06 | 1.200E-05 |
| 9388 | 470 | A470G | A | 8.7e-06 | 5.5e-06 | 1e-05 | 8.067E-06 | 2.316E-06 | 5.500E-06 | 1.000E-05 |
| 9389 | 470 | A470H | A | 5.7e-06 | 9.6e-06 | 1.1e-05 | 8.767E-06 | 2.747E-06 | 5.700E-06 | 1.100E-05 |
| 9390 | 470 | A470I | A | 6.4e-06 | 6.6e-06 | 7.8e-06 | 6.933E-06 | 7.572E-07 | 6.400E-06 | 7.800E-06 |
| 9391 | 470 | A470L | A | 9.8e-06 | 4.6e-06 | 5.9e-06 | 6.767E-06 | 2.706E-06 | 4.600E-06 | 9.800E-06 |
| 9392 | 470 | A470K | A | 8.1e-06 | 4.9e-06 | 1.1e-05 | 8.000E-06 | 3.051E-06 | 4.900E-06 | 1.100E-05 |
| 9393 | 470 | A470M | A | 5.8e-06 | 1e-05 | 1.4e-05 | 9.933E-06 | 4.100E-06 | 5.800E-06 | 1.400E-05 |
| 9394 | 470 | A470F | A | 9.3e-06 | 8e-06 | 1.1e-05 | 9.433E-06 | 1.504E-06 | 8.000E-06 | 1.100E-05 |
| 9395 | 470 | A470P | A | 7e-06 | 7.6e-06 | 7.4e-06 | 7.333E-06 | 3.055E-07 | 7.000E-06 | 7.600E-06 |
| 9396 | 470 | A470S | A | 1.2e-05 | 8.1e-06 | 1.1e-05 | 1.037E-05 | 2.026E-06 | 8.100E-06 | 1.200E-05 |
| 9397 | 470 | A470T | A | 9.4e-06 | 9.2e-06 | 6.2e-06 | 8.267E-06 | 1.793E-06 | 6.200E-06 | 9.400E-06 |
| 9398 | 470 | A470W | A | 1e-05 | 7.7e-06 | 5.2e-06 | 7.633E-06 | 2.401E-06 | 5.200E-06 | 1.000E-05 |
| 9399 | 470 | A470Y | A | 4e-06 | 1e-05 | 6.4e-06 | 6.800E-06 | 3.020E-06 | 4.000E-06 | 1.000E-05 |
| 9400 | 470 | A470V | A | 9.5e-06 | 8.8e-06 | 5.6e-06 | 7.967E-06 | 2.079E-06 | 5.600E-06 | 9.500E-06 |
| 9401 | 471 | L471A | A | 7.2e-06 | 7.5e-06 | 4.1e-06 | 6.267E-06 | 1.882E-06 | 4.100E-06 | 7.500E-06 |
| 9402 | 471 | L471R | A | 9e-06 | 4.3e-06 | 9.6e-06 | 7.633E-06 | 2.902E-06 | 4.300E-06 | 9.600E-06 |
| 9403 | 471 | L471N | A | 6.4e-06 | 5.4e-06 | 6.3e-06 | 6.033E-06 | 5.508E-07 | 5.400E-06 | 6.400E-06 |
| 9404 | 471 | L471D | A | 5.3e-06 | 6.4e-06 | 1.7e-05 | 9.567E-06 | 6.461E-06 | 5.300E-06 | 1.700E-05 |
| 9405 | 471 | L471C | A | 7.1e-06 | 5.6e-06 | 9.3e-06 | 7.333E-06 | 1.861E-06 | 5.600E-06 | 9.300E-06 |
| 9406 | 471 | L471E | A | 7.3e-06 | 7.6e-06 | 9.3e-06 | 8.067E-06 | 1.079E-06 | 7.300E-06 | 9.300E-06 |
| 9407 | 471 | L471Q | A | 8.9e-06 | 7.4e-06 | 1.3e-05 | 9.767E-06 | 2.899E-06 | 7.400E-06 | 1.300E-05 |
| 9408 | 471 | L471G | A | 6.6e-06 | 7.5e-06 | 9.6e-06 | 7.900E-06 | 1.539E-06 | 6.600E-06 | 9.600E-06 |
| 9409 | 471 | L471H | A | 5e-06 | 6.7e-06 | 4.9e-06 | 5.533E-06 | 1.012E-06 | 4.900E-06 | 6.700E-06 |
| 9410 | 471 | L471I | A | 1.3e-05 | 9.4e-06 | 7.7e-06 | 1.003E-05 | 2.706E-06 | 7.700E-06 | 1.300E-05 |
| 9411 | 471 | L471L | A | 6.6e-06 | 9.9e-06 | 8.7e-06 | 8.400E-06 | 1.670E-06 | 6.600E-06 | 9.900E-06 |
| 9412 | 471 | L471K | A | 7.4e-06 | 9.7e-06 | 1.3e-05 | 1.003E-05 | 2.815E-06 | 7.400E-06 | 1.300E-05 |
| 9413 | 471 | L471M | A | 6.9e-06 | 1.4e-05 | 1e-05 | 1.030E-05 | 3.559E-06 | 6.900E-06 | 1.400E-05 |
| 9414 | 471 | L471F | A | 1e-05 | 6.6e-06 | 1.4e-05 | 1.020E-05 | 3.704E-06 | 6.600E-06 | 1.400E-05 |
| 9415 | 471 | L471P | A | 9.2e-06 | 8e-06 | 1e-05 | 9.067E-06 | 1.007E-06 | 8.000E-06 | 1.000E-05 |
| 9416 | 471 | L471S | A | 1.5e-05 | 6.6e-06 | 7.1e-06 | 9.567E-06 | 4.712E-06 | 6.600E-06 | 1.500E-05 |
| 9417 | 471 | L471T | A | 6e-06 | 4.5e-06 | 9.7e-06 | 6.733E-06 | 2.676E-06 | 4.500E-06 | 9.700E-06 |
| 9418 | 471 | L471W | A | 8.5e-06 | 6.5e-06 | 9.7e-06 | 8.233E-06 | 1.617E-06 | 6.500E-06 | 9.700E-06 |
| 9419 | 471 | L471Y | A | 1.2e-05 | 9.4e-06 | 6.4e-06 | 9.267E-06 | 2.802E-06 | 6.400E-06 | 1.200E-05 |
| 9420 | 471 | L471V | A | 7.9e-06 | 5.3e-06 | 6.8e-06 | 6.667E-06 | 1.305E-06 | 5.300E-06 | 7.900E-06 |
| 9421 | 472 | L472A | A | 6.5e-06 | 8.2e-06 | 9.7e-06 | 8.133E-06 | 1.601E-06 | 6.500E-06 | 9.700E-06 |
| 9422 | 472 | L472R | A | 9.6e-06 | 8.4e-06 | 4.6e-06 | 7.533E-06 | 2.610E-06 | 4.600E-06 | 9.600E-06 |
| 9423 | 472 | L472N | A | 7.8e-06 | 1.2e-05 | 6.9e-06 | 8.900E-06 | 2.722E-06 | 6.900E-06 | 1.200E-05 |
| 9424 | 472 | L472D | A | 5.9e-06 | 8.9e-06 | 3.2e-06 | 6.000E-06 | 2.851E-06 | 3.200E-06 | 8.900E-06 |
| 9425 | 472 | L472C | A | 6.5e-06 | 1.1e-05 | 9.2e-06 | 8.900E-06 | 2.265E-06 | 6.500E-06 | 1.100E-05 |
| 9426 | 472 | L472E | A | 7.8e-06 | 6.9e-06 | 7.1e-06 | 7.267E-06 | 4.726E-07 | 6.900E-06 | 7.800E-06 |
| 9427 | 472 | L472Q | A | 4e-06 | 6.5e-06 | 9.1e-06 | 6.533E-06 | 2.550E-06 | 4.000E-06 | 9.100E-06 |
| 9428 | 472 | L472G | A | 1.1e-05 | 9.2e-06 | 5.8e-06 | 8.667E-06 | 2.641E-06 | 5.800E-06 | 1.100E-05 |
| 9429 | 472 | L472H | A | 6.4e-06 | 7.7e-06 | 4.8e-06 | 6.300E-06 | 1.453E-06 | 4.800E-06 | 7.700E-06 |
| 9430 | 472 | L472I | A | 8e-06 | 8.5e-06 | 8.2e-06 | 8.233E-06 | 2.517E-07 | 8.000E-06 | 8.500E-06 |
| 9431 | 472 | L472L | A | 9.4e-06 | 1.1e-05 | 5.9e-06 | 8.767E-06 | 2.608E-06 | 5.900E-06 | 1.100E-05 |
| 9432 | 472 | L472K | A | 7.1e-06 | 6.4e-06 | 6.9e-06 | 6.800E-06 | 3.606E-07 | 6.400E-06 | 7.100E-06 |
| 9433 | 472 | L472M | A | 6.9e-06 | 5.2e-06 | 4.7e-06 | 5.600E-06 | 1.153E-06 | 4.700E-06 | 6.900E-06 |
| 9434 | 472 | L472F | A | 7.5e-06 | 8.3e-06 | 6.2e-06 | 7.333E-06 | 1.060E-06 | 6.200E-06 | 8.300E-06 |

| | | | | | | | | | | |
|---|---|---|---|---|---|---|---|---|---|---|
| 9435 | 472 | L472P | A | 7.9e-06 | 8.4e-06 | 1e-05 | 8.767E-06 | 1.097E-06 | 7.900E-06 | 1.000E-05 |
| 9436 | 472 | L472S | A | 8.9e-06 | 4.2e-06 | 5.6e-06 | 6.233E-06 | 2.413E-06 | 4.200E-06 | 8.900E-06 |
| 9437 | 472 | L472T | A | 6.7e-06 | 8.2e-06 | 9.9e-06 | 8.267E-06 | 1.601E-06 | 6.700E-06 | 9.900E-06 |
| 9438 | 472 | L472W | A | 9.5e-06 | 8.3e-06 | 6.6e-06 | 8.133E-06 | 1.457E-06 | 6.600E-06 | 9.500E-06 |
| 9439 | 472 | L472Y | A | 5.3e-06 | 1.1e-05 | 5.3e-06 | 7.200E-06 | 3.291E-06 | 5.300E-06 | 1.100E-05 |
| 9440 | 472 | L472V | A | 1.2e-05 | 1.2e-05 | 8.7e-06 | 1.090E-05 | 1.905E-06 | 8.700E-06 | 1.200E-05 |
| 9441 | 473 | H473A | A | 9.4e-06 | 6.9e-06 | 7.6e-06 | 7.967E-06 | 1.290E-06 | 6.900E-06 | 9.400E-06 |
| 9442 | 473 | H473R | A | 1.1e-05 | 9.7e-06 | 6.3e-06 | 9.000E-06 | 2.427E-06 | 6.300E-06 | 1.100E-05 |
| 9443 | 473 | H473N | A | 4.2e-06 | 7.2e-06 | 7.4e-06 | 6.267E-06 | 1.793E-06 | 4.200E-06 | 7.400E-06 |
| 9444 | 473 | H473D | A | 6.4e-06 | 8.5e-06 | 8.5e-06 | 7.800E-06 | 1.212E-06 | 6.400E-06 | 8.500E-06 |
| 9445 | 473 | H473C | A | 9.3e-06 | 3.7e-06 | 5.5e-06 | 6.167E-06 | 2.859E-06 | 3.700E-06 | 9.300E-06 |
| 9446 | 473 | H473E | A | 1.3e-05 | 3.2e-06 | 1.1e-05 | 9.067E-06 | 5.178E-06 | 3.200E-06 | 1.300E-05 |
| 9447 | 473 | H473Q | A | 9.2e-06 | 1.1e-05 | 6e-06 | 8.733E-06 | 2.532E-06 | 6.000E-06 | 1.100E-05 |
| 9448 | 473 | H473G | A | 1.1e-05 | 4.6e-06 | 5.8e-06 | 7.133E-06 | 3.402E-06 | 4.600E-06 | 1.100E-05 |
| 9449 | 473 | H473H | A | 6e-06 | 8.7e-06 | 5.8e-06 | 6.833E-06 | 1.620E-06 | 5.800E-06 | 8.700E-06 |
| 9450 | 473 | H473I | A | 1.2e-05 | 1e-05 | 7.4e-06 | 9.800E-06 | 2.307E-06 | 7.400E-06 | 1.200E-05 |
| 9451 | 473 | H473L | A | 7.7e-06 | 7.9e-06 | 6.5e-06 | 7.367E-06 | 7.572E-07 | 6.500E-06 | 7.900E-06 |
| 9452 | 473 | H473K | A | 6.9e-06 | 6.7e-06 | 1e-05 | 7.867E-06 | 1.850E-06 | 6.700E-06 | 1.000E-05 |
| 9453 | 473 | H473M | A | 6.2e-06 | 1.4e-05 | 5.8e-06 | 8.667E-06 | 4.623E-06 | 5.800E-06 | 1.400E-05 |
| 9454 | 473 | H473F | A | 8.9e-06 | 6.2e-06 | 6.3e-06 | 7.133E-06 | 1.531E-06 | 6.200E-06 | 8.900E-06 |
| 9455 | 473 | H473P | A | 1.4e-05 | 7.9e-06 | 9e-06 | 1.030E-05 | 3.251E-06 | 7.900E-06 | 1.400E-05 |
| 9456 | 473 | H473S | A | 5.9e-06 | 1.1e-05 | 7e-06 | 7.967E-06 | 2.684E-06 | 5.900E-06 | 1.100E-05 |
| 9457 | 473 | H473T | A | 6.3e-06 | 1.4e-05 | 6.2e-06 | 8.833E-06 | 4.475E-06 | 6.200E-06 | 1.400E-05 |
| 9458 | 473 | H473W | A | 8.6e-06 | 6.5e-06 | 7e-06 | 7.367E-06 | 1.097E-06 | 6.500E-06 | 8.600E-06 |
| 9459 | 473 | H473Y | A | 1.2e-05 | 9.1e-06 | 7.5e-06 | 9.533E-06 | 2.281E-06 | 7.500E-06 | 1.200E-05 |
| 9460 | 473 | H473V | A | 6.8e-06 | 8.5e-06 | 8.2e-06 | 7.833E-06 | 9.074E-07 | 6.800E-06 | 8.500E-06 |
| 9461 | 474 | T474A | A | 9.6e-06 | 1e-05 | 6e-06 | 8.533E-06 | 2.203E-06 | 6.000E-06 | 1.000E-05 |
| 9462 | 474 | T474R | A | 5.7e-06 | 9.8e-06 | 6.1e-06 | 7.200E-06 | 2.261E-06 | 5.700E-06 | 9.800E-06 |
| 9463 | 474 | T474N | A | 1.3e-05 | 6.4e-06 | 6.7e-06 | 8.700E-06 | 3.727E-06 | 6.400E-06 | 1.300E-05 |
| 9464 | 474 | T474D | A | 9.5e-06 | 1e-05 | 1.3e-05 | 1.083E-05 | 1.893E-06 | 9.500E-06 | 1.300E-05 |
| 9465 | 474 | T474C | A | 1.3e-05 | 6.7e-06 | 5.7e-06 | 8.467E-06 | 3.958E-06 | 5.700E-06 | 1.300E-05 |
| 9466 | 474 | T474E | A | 9e-06 | 9.7e-06 | 1.3e-05 | 1.057E-05 | 2.136E-06 | 9.000E-06 | 1.300E-05 |
| 9467 | 474 | T474Q | A | 7e-06 | 1.2e-05 | 7.2e-06 | 8.733E-06 | 2.831E-06 | 7.000E-06 | 1.200E-05 |
| 9468 | 474 | T474G | A | 1.1e-05 | 8.5e-06 | 1.1e-05 | 1.017E-05 | 1.443E-06 | 8.500E-06 | 1.100E-05 |
| 9469 | 474 | T474H | A | 9e-06 | 6.9e-06 | 1e-05 | 8.633E-06 | 1.582E-06 | 6.900E-06 | 1.000E-05 |
| 9470 | 474 | T474I | A | 6.5e-06 | 9.3e-06 | 9.5e-06 | 8.433E-06 | 1.677E-06 | 6.500E-06 | 9.500E-06 |
| 9471 | 474 | T474L | A | 9.7e-06 | 1.2e-05 | 9.3e-06 | 1.033E-05 | 1.457E-06 | 9.300E-06 | 1.200E-05 |
| 9472 | 474 | T474K | A | 7.4e-06 | 1e-05 | 8.3e-06 | 8.567E-06 | 1.320E-06 | 7.400E-06 | 1.000E-05 |
| 9473 | 474 | T474M | A | 1.4e-05 | 1.1e-05 | 3.2e-06 | 9.400E-06 | 5.575E-06 | 3.200E-06 | 1.400E-05 |
| 9474 | 474 | T474F | A | 8.8e-06 | 1e-05 | 7.4e-06 | 8.733E-06 | 1.301E-06 | 7.400E-06 | 1.000E-05 |
| 9475 | 474 | T474P | A | 6.9e-06 | 1.4e-05 | 1.3e-05 | 1.130E-05 | 3.843E-06 | 6.900E-06 | 1.400E-05 |
| 9476 | 474 | T474S | A | 7.2e-06 | 7.5e-06 | 5.2e-06 | 6.633E-06 | 1.250E-06 | 5.200E-06 | 7.500E-06 |
| 9477 | 474 | T474T | A | 1e-05 | 6.2e-06 | 1.1e-05 | 9.067E-06 | 2.532E-06 | 6.200E-06 | 1.100E-05 |
| 9478 | 474 | T474W | A | 6.9e-06 | 5.4e-06 | 5.5e-06 | 5.933E-06 | 8.386E-07 | 5.400E-06 | 6.900E-06 |
| 9479 | 474 | T474Y | A | 8.9e-06 | 9.2e-06 | 1.1e-05 | 9.700E-06 | 1.136E-06 | 8.900E-06 | 1.100E-05 |
| 9480 | 474 | T474V | A | 1.1e-05 | 6.9e-06 | 1.3e-05 | 1.030E-05 | 3.110E-06 | 6.900E-06 | 1.300E-05 |
| 9481 | 475 | A475A | A | 9.6e-06 | 9.8e-06 | 1.1e-05 | 1.013E-05 | 7.572E-07 | 9.600E-06 | 1.100E-05 |
| 9482 | 475 | A475R | A | 5.8e-06 | 8.3e-06 | 5.8e-06 | 6.633E-06 | 1.443E-06 | 5.800E-06 | 8.300E-06 |
| 9483 | 475 | A475N | A | 1e-05 | 6.4e-06 | 1.3e-05 | 9.800E-06 | 3.305E-06 | 6.400E-06 | 1.300E-05 |
| 9484 | 475 | A475D | A | 4.7e-06 | 7e-06 | 9.4e-06 | 7.033E-06 | 2.350E-06 | 4.700E-06 | 9.400E-06 |
| 9485 | 475 | A475C | A | 9.2e-06 | 1.3e-05 | 9.9e-06 | 1.070E-05 | 2.022E-06 | 9.200E-06 | 1.300E-05 |
| 9486 | 475 | A475E | A | 1.2e-05 | 7e-06 | 9.5e-06 | 9.500E-06 | 2.500E-06 | 7.000E-06 | 1.200E-05 |
| 9487 | 475 | A475Q | A | 9e-06 | 9.7e-06 | 5e-06 | 7.900E-06 | 2.536E-06 | 5.000E-06 | 9.700E-06 |
| 9488 | 475 | A475G | A | 5.3e-06 | 8.4e-06 | 9.8e-06 | 7.833E-06 | 2.303E-06 | 5.300E-06 | 9.800E-06 |
| 9489 | 475 | A475H | A | 3.3e-06 | 6.8e-06 | 6.5e-06 | 5.533E-06 | 1.940E-06 | 3.300E-06 | 6.800E-06 |
| 9490 | 475 | A475I | A | 8.6e-06 | 7.5e-06 | 9.9e-06 | 8.667E-06 | 1.201E-06 | 7.500E-06 | 9.900E-06 |
| 9491 | 475 | A475L | A | 7.3e-06 | 4e-06 | 7.7e-06 | 6.333E-06 | 2.031E-06 | 4.000E-06 | 7.700E-06 |
| 9492 | 475 | A475K | A | 8.4e-06 | 9.7e-06 | 7.6e-06 | 8.567E-06 | 1.060E-06 | 7.600E-06 | 9.700E-06 |
| 9493 | 475 | A475M | A | 6.6e-06 | 8.9e-06 | 7.4e-06 | 7.633E-06 | 1.168E-06 | 6.600E-06 | 8.900E-06 |
| 9494 | 475 | A475F | A | 1.3e-05 | 5.7e-06 | 5.2e-06 | 7.967E-06 | 4.366E-06 | 5.200E-06 | 1.300E-05 |
| 9495 | 475 | A475P | A | 1.3e-05 | 7.7e-06 | 5.5e-06 | 8.733E-06 | 3.855E-06 | 5.500E-06 | 1.300E-05 |
| 9496 | 475 | A475S | A | 4.1e-06 | 9.6e-06 | 6.6e-06 | 6.767E-06 | 2.754E-06 | 4.100E-06 | 9.600E-06 |
| 9497 | 475 | A475T | A | 1.8e-05 | 4.7e-06 | 9.8e-06 | 1.083E-05 | 6.710E-06 | 4.700E-06 | 1.800E-05 |
| 9498 | 475 | A475W | A | 6.2e-06 | 9.2e-06 | 8.4e-06 | 7.933E-06 | 1.553E-06 | 6.200E-06 | 9.200E-06 |
| 9499 | 475 | A475Y | A | 9.4e-06 | 9.9e-06 | 4.1e-06 | 7.800E-06 | 3.214E-06 | 4.100E-06 | 9.900E-06 |
| 9500 | 475 | A475V | A | 6.5e-06 | 1.3e-05 | 1e-05 | 9.833E-06 | 3.253E-06 | 6.500E-06 | 1.300E-05 |
| 9501 | 476 | N476A | A | 6.4e-06 | 8.2e-06 | 9.4e-06 | 8.000E-06 | 1.510E-06 | 6.400E-06 | 9.400E-06 |
| 9502 | 476 | N476R | A | 9.3e-06 | 1.3e-05 | 7.4e-06 | 9.900E-06 | 2.848E-06 | 7.400E-06 | 1.300E-05 |
| 9503 | 476 | N476N | A | 8.3e-06 | 7e-06 | 8.4e-06 | 7.900E-06 | 7.810E-07 | 7.000E-06 | 8.400E-06 |
| 9504 | 476 | N476D | A | 1.1e-05 | 4.8e-06 | 1.3e-05 | 9.600E-06 | 4.276E-06 | 4.800E-06 | 1.300E-05 |
| 9505 | 476 | N476C | A | 7.6e-06 | 8.7e-06 | 6.9e-06 | 7.733E-06 | 9.074E-07 | 6.900E-06 | 8.700E-06 |
| 9506 | 476 | N476E | A | 1e-05 | 1.3e-05 | 7.2e-06 | 1.007E-05 | 2.901E-06 | 7.200E-06 | 1.300E-05 |
| 9507 | 476 | N476Q | A | 1.1e-05 | 6e-06 | 4e-06 | 7.000E-06 | 3.606E-06 | 4.000E-06 | 1.100E-05 |
| 9508 | 476 | N476G | A | 9.7e-06 | 8.5e-06 | 4.6e-06 | 7.600E-06 | 2.666E-06 | 4.600E-06 | 9.700E-06 |
| 9509 | 476 | N476H | A | 7.3e-06 | 6.9e-06 | 7.5e-06 | 7.233E-06 | 3.055E-07 | 6.900E-06 | 7.500E-06 |
| 9510 | 476 | N476I | A | 6.3e-06 | 1.3e-05 | 8.9e-06 | 9.400E-06 | 3.378E-06 | 6.300E-06 | 1.300E-05 |
| 9511 | 476 | N476L | A | 4.7e-06 | 1.1e-05 | 6.6e-06 | 7.433E-06 | 3.232E-06 | 4.700E-06 | 1.100E-05 |
| 9512 | 476 | N476K | A | 7.1e-06 | 8.6e-06 | 1.2e-05 | 9.233E-06 | 2.511E-06 | 7.100E-06 | 1.200E-05 |
| 9513 | 476 | N476M | A | 5.5e-06 | 7.5e-06 | 1.1e-05 | 8.000E-06 | 2.784E-06 | 5.500E-06 | 1.100E-05 |
| 9514 | 476 | N476F | A | 8e-06 | 5.4e-06 | 1.4e-05 | 9.133E-06 | 4.411E-06 | 5.400E-06 | 1.400E-05 |
| 9515 | 476 | N476P | A | 8.9e-06 | 7.5e-06 | 9.7e-06 | 8.700E-06 | 1.114E-06 | 7.500E-06 | 9.700E-06 |
| 9516 | 476 | N476S | A | 9.1e-06 | 5.6e-06 | 6e-06 | 6.900E-06 | 1.916E-06 | 5.600E-06 | 9.100E-06 |
| 9517 | 476 | N476T | A | 8e-06 | 7.5e-06 | 1e-05 | 8.500E-06 | 1.323E-06 | 7.500E-06 | 1.000E-05 |
| 9518 | 476 | N476W | A | 1.3e-05 | 5.1e-06 | 7.4e-06 | 8.500E-06 | 4.063E-06 | 5.100E-06 | 1.300E-05 |
| 9519 | 476 | N476Y | A | 1.5e-05 | 5.1e-06 | 5.1e-06 | 8.400E-06 | 5.716E-06 | 5.100E-06 | 1.500E-05 |

| | | | | | | | | | | |
|---|---|---|---|---|---|---|---|---|---|---|
| 9520 | 476 | N476V | A | 5.4e-06 | 1.1e-05 | 6.6e-06 | 7.667E-06 | 2.948E-06 | 5.400E-06 | 1.100E-05 |
| 9521 | 477 | R477A | A | 1.1e-05 | 1.3e-05 | 4.6e-06 | 9.533E-06 | 4.388E-06 | 4.600E-06 | 1.300E-05 |
| 9522 | 477 | R477R | A | 5e-06 | 6e-06 | 5.3e-06 | 5.433E-06 | 5.132E-07 | 5.000E-06 | 6.000E-06 |
| 9523 | 477 | R477N | A | 6.5e-06 | 6.5e-06 | 6.8e-06 | 6.600E-06 | 1.732E-07 | 6.500E-06 | 6.800E-06 |
| 9524 | 477 | R477D | A | 6.2e-06 | 6.5e-06 | 9.1e-06 | 7.267E-06 | 1.595E-06 | 6.200E-06 | 9.100E-06 |
| 9525 | 477 | R477C | A | 6.9e-06 | 9.8e-06 | 9.6e-06 | 8.767E-06 | 1.620E-06 | 6.900E-06 | 9.800E-06 |
| 9526 | 477 | R477E | A | 6.5e-06 | 9.9e-06 | 5.9e-06 | 7.433E-06 | 2.157E-06 | 5.900E-06 | 9.900E-06 |
| 9527 | 477 | R477Q | A | 1.1e-05 | 8.9e-06 | 8.3e-06 | 9.400E-06 | 1.418E-06 | 8.300E-06 | 1.100E-05 |
| 9528 | 477 | R477G | A | 8.2e-06 | 1.1e-05 | 6.7e-06 | 8.633E-06 | 2.183E-06 | 6.700E-06 | 1.100E-05 |
| 9529 | 477 | R477H | A | 1.2e-05 | 1.3e-05 | 8.2e-06 | 1.107E-05 | 2.532E-06 | 8.200E-06 | 1.300E-05 |
| 9530 | 477 | R477I | A | 6.4e-06 | 7.7e-06 | 4.3e-06 | 6.133E-06 | 1.716E-06 | 4.300E-06 | 7.700E-06 |
| 9531 | 477 | R477L | A | 6.3e-06 | 7.4e-06 | 8.3e-06 | 7.333E-06 | 1.002E-06 | 6.300E-06 | 8.300E-06 |
| 9532 | 477 | R477K | A | 4.3e-06 | 5.6e-06 | 9e-06 | 6.300E-06 | 2.427E-06 | 4.300E-06 | 9.000E-06 |
| 9533 | 477 | R477M | A | 9.4e-06 | 6.6e-06 | 5.7e-06 | 7.233E-06 | 1.930E-06 | 5.700E-06 | 9.400E-06 |
| 9534 | 477 | R477F | A | 1e-05 | 9.7e-06 | 8.9e-06 | 9.533E-06 | 5.686E-07 | 8.900E-06 | 1.000E-05 |
| 9535 | 477 | R477P | A | 1.1e-05 | 6.8e-06 | 7.1e-06 | 8.300E-06 | 2.343E-06 | 6.800E-06 | 1.100E-05 |
| 9536 | 477 | R477S | A | 5.5e-06 | 9.9e-06 | 5.3e-06 | 6.900E-06 | 2.600E-06 | 5.300E-06 | 9.900E-06 |
| 9537 | 477 | R477T | A | 6.5e-06 | 6.5e-06 | 7.5e-06 | 6.833E-06 | 5.774E-07 | 6.500E-06 | 7.500E-06 |
| 9538 | 477 | R477W | A | 7e-06 | 5.5e-06 | 1.2e-05 | 8.167E-06 | 3.403E-06 | 5.500E-06 | 1.200E-05 |
| 9539 | 477 | R477Y | A | 1e-05 | 6.6e-06 | 9.8e-06 | 8.800E-06 | 1.908E-06 | 6.600E-06 | 1.000E-05 |
| 9540 | 477 | R477V | A | 6.8e-06 | 6.9e-06 | 1.3e-05 | 8.900E-06 | 3.551E-06 | 6.800E-06 | 1.300E-05 |
| 9541 | 478 | P478A | A | 6.6e-06 | 7.6e-06 | 1.3e-05 | 9.067E-06 | 3.443E-06 | 6.600E-06 | 1.300E-05 |
| 9542 | 478 | P478R | A | 7e-06 | 8e-06 | 4.9e-06 | 6.633E-06 | 1.582E-06 | 4.900E-06 | 8.000E-06 |
| 9543 | 478 | P478N | A | 5.7e-06 | 8.9e-06 | 9.9e-06 | 8.167E-06 | 2.194E-06 | 5.700E-06 | 9.900E-06 |
| 9544 | 478 | P478D | A | 5.5e-06 | 7.7e-06 | 8.6e-06 | 7.267E-06 | 1.595E-06 | 5.500E-06 | 8.600E-06 |
| 9545 | 478 | P478C | A | 7.6e-06 | 6.3e-06 | 8.2e-06 | 7.367E-06 | 9.713E-07 | 6.300E-06 | 8.200E-06 |
| 9546 | 478 | P478E | A | 5.8e-06 | 9.8e-06 | 9.9e-06 | 8.500E-06 | 2.339E-06 | 5.800E-06 | 9.900E-06 |
| 9547 | 478 | P478Q | A | 6.8e-06 | 7.3e-06 | 8.6e-06 | 7.567E-06 | 9.292E-07 | 6.800E-06 | 8.600E-06 |
| 9548 | 478 | P478G | A | 8.1e-06 | 8.6e-06 | 1.2e-05 | 9.567E-06 | 2.122E-06 | 8.100E-06 | 1.200E-05 |
| 9549 | 478 | P478H | A | 6.4e-06 | 4.9e-06 | 1e-05 | 7.100E-06 | 2.621E-06 | 4.900E-06 | 1.000E-05 |
| 9550 | 478 | P478I | A | 7.8e-06 | 9e-06 | 8.7e-06 | 8.500E-06 | 6.245E-07 | 7.800E-06 | 9.000E-06 |
| 9551 | 478 | P478L | A | 9.1e-06 | 6.6e-06 | 4.6e-06 | 6.767E-06 | 2.255E-06 | 4.600E-06 | 9.100E-06 |
| 9552 | 478 | P478K | A | 9.4e-06 | 1.2e-05 | 8.7e-06 | 1.003E-05 | 1.739E-06 | 8.700E-06 | 1.200E-05 |
| 9553 | 478 | P478M | A | 9.8e-06 | 8.7e-06 | 5.5e-06 | 8.000E-06 | 2.234E-06 | 5.500E-06 | 9.800E-06 |
| 9554 | 478 | P478F | A | 1.2e-05 | 1.3e-05 | 8.8e-06 | 1.127E-05 | 2.194E-06 | 8.800E-06 | 1.300E-05 |
| 9555 | 478 | P478P | A | 8.7e-06 | 8.9e-06 | 5e-06 | 7.533E-06 | 2.196E-06 | 5.000E-06 | 8.900E-06 |
| 9556 | 478 | P478S | A | 6.3e-06 | 6.6e-06 | 6.5e-06 | 6.467E-06 | 1.528E-07 | 6.300E-06 | 6.600E-06 |
| 9557 | 478 | P478T | A | 1.3e-05 | 7.8e-06 | 6.6e-06 | 9.133E-06 | 3.402E-06 | 6.600E-06 | 1.300E-05 |
| 9558 | 478 | P478W | A | 8.8e-06 | 4.3e-06 | 1.2e-05 | 8.367E-06 | 3.868E-06 | 4.300E-06 | 1.200E-05 |
| 9559 | 478 | P478Y | A | 6.5e-06 | 9.2e-06 | 8.3e-06 | 8.000E-06 | 1.375E-06 | 6.500E-06 | 9.200E-06 |
| 9560 | 478 | P478V | A | 9.2e-06 | 9.3e-06 | 1e-05 | 9.500E-06 | 4.359E-07 | 9.200E-06 | 1.000E-05 |
| 9561 | 479 | E479A | A | 6.8e-06 | 7e-06 | 7.9e-06 | 7.233E-06 | 5.859E-07 | 6.800E-06 | 7.900E-06 |
| 9562 | 479 | E479R | A | 5.2e-06 | 7.3e-06 | 6.3e-06 | 6.267E-06 | 1.050E-06 | 5.200E-06 | 7.300E-06 |
| 9563 | 479 | E479N | A | 1.1e-05 | 9.5e-06 | 7.2e-06 | 9.233E-06 | 1.914E-06 | 7.200E-06 | 1.100E-05 |
| 9564 | 479 | E479D | A | 8.8e-06 | 8.6e-06 | 1.3e-05 | 1.013E-05 | 2.485E-06 | 8.600E-06 | 1.300E-05 |
| 9565 | 479 | E479C | A | 8.6e-06 | 6.7e-06 | 1e-05 | 8.433E-06 | 1.656E-06 | 6.700E-06 | 1.000E-05 |
| 9566 | 479 | E479E | A | 2.9e-06 | 9.6e-06 | 5.8e-06 | 6.100E-06 | 3.360E-06 | 2.900E-06 | 9.600E-06 |
| 9567 | 479 | E479Q | A | 5.3e-06 | 6.8e-06 | 1.1e-05 | 7.700E-06 | 2.955E-06 | 5.300E-06 | 1.100E-05 |
| 9568 | 479 | E479G | A | 5.1e-06 | 1e-05 | 5.2e-06 | 6.767E-06 | 2.801E-06 | 5.100E-06 | 1.000E-05 |
| 9569 | 479 | E479H | A | 1e-05 | 1.2e-05 | 4.2e-06 | 8.733E-06 | 4.051E-06 | 4.200E-06 | 1.200E-05 |
| 9570 | 479 | E479I | A | 5.3e-06 | 9.6e-06 | 9.4e-06 | 8.100E-06 | 2.427E-06 | 5.300E-06 | 9.600E-06 |
| 9571 | 479 | E479L | A | 6.9e-06 | 9e-06 | 6.4e-06 | 7.433E-06 | 1.380E-06 | 6.400E-06 | 9.000E-06 |
| 9572 | 479 | E479K | A | 6.6e-06 | 1.2e-05 | 8.5e-06 | 9.033E-06 | 2.739E-06 | 6.600E-06 | 1.200E-05 |
| 9573 | 479 | E479M | A | 6.9e-06 | 8.8e-06 | 4.9e-06 | 6.867E-06 | 1.950E-06 | 4.900E-06 | 8.800E-06 |
| 9574 | 479 | E479F | A | 9.5e-06 | 8e-06 | 8.7e-06 | 8.733E-06 | 7.506E-07 | 8.000E-06 | 9.500E-06 |
| 9575 | 479 | E479P | A | 1.2e-05 | 9.1e-06 | 1.4e-05 | 1.170E-05 | 2.464E-06 | 9.100E-06 | 1.400E-05 |
| 9576 | 479 | E479S | A | 5.3e-06 | 7.6e-06 | 9.1e-06 | 7.333E-06 | 1.914E-06 | 5.300E-06 | 9.100E-06 |
| 9577 | 479 | E479T | A | 6.3e-06 | 3.1e-06 | 8.8e-06 | 6.067E-06 | 2.857E-06 | 3.100E-06 | 8.800E-06 |
| 9578 | 479 | E479W | A | 7.3e-06 | 1.3e-05 | 4.8e-06 | 8.367E-06 | 4.203E-06 | 4.800E-06 | 1.300E-05 |
| 9579 | 479 | E479Y | A | 5.9e-06 | 6.6e-06 | 6.8e-06 | 6.433E-06 | 4.726E-07 | 5.900E-06 | 6.800E-06 |
| 9580 | 479 | E479V | A | 7.5e-06 | 9.6e-06 | 1e-05 | 9.033E-06 | 1.343E-06 | 7.500E-06 | 1.000E-05 |
| 9581 | 480 | D480A | A | 1e-05 | 1.1e-05 | 5.3e-06 | 8.767E-06 | 3.044E-06 | 5.300E-06 | 1.100E-05 |
| 9582 | 480 | D480R | A | 1.3e-05 | 1.2e-05 | 4.5e-06 | 9.833E-06 | 4.646E-06 | 4.500E-06 | 1.300E-05 |
| 9583 | 480 | D480N | A | 1e-05 | 1.1e-05 | 1e-05 | 1.033E-05 | 5.774E-07 | 1.000E-05 | 1.100E-05 |
| 9584 | 480 | D480D | A | 8.5e-06 | 9.6e-06 | 1e-05 | 9.367E-06 | 7.767E-07 | 8.500E-06 | 1.000E-05 |
| 9585 | 480 | D480C | A | 8.6e-06 | 6.1e-06 | 7e-06 | 7.233E-06 | 1.266E-06 | 6.100E-06 | 8.600E-06 |
| 9586 | 480 | D480E | A | 9.8e-06 | 9.3e-06 | 1e-05 | 9.700E-06 | 3.606E-07 | 9.300E-06 | 1.000E-05 |
| 9587 | 480 | D480Q | A | 1.2e-05 | 8e-06 | 5.8e-06 | 8.600E-06 | 3.143E-06 | 5.800E-06 | 1.200E-05 |
| 9588 | 480 | D480G | A | 9.6e-06 | 8.2e-06 | 8.5e-06 | 8.767E-06 | 7.371E-07 | 8.200E-06 | 9.600E-06 |
| 9589 | 480 | D480H | A | 1.1e-05 | 1.6e-05 | 8.4e-06 | 1.180E-05 | 3.863E-06 | 8.400E-06 | 1.600E-05 |
| 9590 | 480 | D480I | A | 7.1e-06 | 9.9e-06 | 4.1e-06 | 7.033E-06 | 2.901E-06 | 4.100E-06 | 9.900E-06 |
| 9591 | 480 | D480L | A | 7.9e-06 | 6.8e-06 | 4.8e-06 | 6.500E-06 | 1.572E-06 | 4.800E-06 | 7.900E-06 |
| 9592 | 480 | D480K | A | 6.2e-06 | 1.1e-05 | 1.2e-05 | 9.733E-06 | 3.101E-06 | 6.200E-06 | 1.200E-05 |
| 9593 | 480 | D480M | A | 5.8e-06 | 7.1e-06 | 7.3e-06 | 6.733E-06 | 8.145E-07 | 5.800E-06 | 7.300E-06 |
| 9594 | 480 | D480F | A | 6.7e-06 | 9.2e-06 | 1.4e-05 | 9.967E-06 | 3.710E-06 | 6.700E-06 | 1.400E-05 |
| 9595 | 480 | D480P | A | 9.3e-06 | 1.5e-05 | 6.2e-06 | 1.017E-05 | 4.464E-06 | 6.200E-06 | 1.500E-05 |
| 9596 | 480 | D480S | A | 7.4e-06 | 1e-05 | 7e-06 | 8.133E-06 | 1.629E-06 | 7.000E-06 | 1.000E-05 |
| 9597 | 480 | D480T | A | 7.8e-06 | 6.1e-06 | 9.2e-06 | 7.700E-06 | 1.552E-06 | 6.100E-06 | 9.200E-06 |
| 9598 | 480 | D480W | A | 6.4e-06 | 8.6e-06 | 6.8e-06 | 7.267E-06 | 1.172E-06 | 6.400E-06 | 8.600E-06 |
| 9599 | 480 | D480Y | A | 5.6e-06 | 7.5e-06 | 6e-06 | 6.367E-06 | 1.002E-06 | 5.600E-06 | 7.500E-06 |
| 9600 | 480 | D480V | A | 1.2e-05 | 8.1e-06 | 8.5e-06 | 9.533E-06 | 2.146E-06 | 8.100E-06 | 1.200E-05 |
| 9601 | 481 | E481A | A | 9.2e-06 | 8.8e-06 | 1.1e-05 | 9.667E-06 | 1.172E-06 | 8.800E-06 | 1.100E-05 |
| 9602 | 481 | E481R | A | 4.9e-06 | 6e-06 | 7.6e-06 | 6.167E-06 | 1.358E-06 | 4.900E-06 | 7.600E-06 |
| 9603 | 481 | E481N | A | 1.4e-05 | 1.4e-05 | 8.3e-06 | 1.210E-05 | 3.291E-06 | 8.300E-06 | 1.400E-05 |
| 9604 | 481 | E481D | A | 1e-05 | 7.2e-06 | 1.1e-05 | 9.400E-06 | 1.970E-06 | 7.200E-06 | 1.100E-05 |

| | | | | | | | | | | |
|---|---|---|---|---|---|---|---|---|---|---|
| 9605 | 481 | E481C | A | 5.7e-06 | 1e-05 | 1.4e-05 | 9.900E-06 | 4.151E-06 | 5.700E-06 | 1.400E-05 |
| 9606 | 481 | E481E | A | 4.8e-06 | 5.3e-06 | 1e-05 | 6.700E-06 | 2.869E-06 | 4.800E-06 | 1.000E-05 |
| 9607 | 481 | E481Q | A | 8.3e-06 | 7.5e-06 | 4.1e-06 | 6.633E-06 | 2.230E-06 | 4.100E-06 | 8.300E-06 |
| 9608 | 481 | E481G | A | 6e-06 | 6.9e-06 | 9.1e-06 | 7.333E-06 | 1.595E-06 | 6.000E-06 | 9.100E-06 |
| 9609 | 481 | E481H | A | 1.1e-05 | 8.4e-06 | 7.4e-06 | 8.933E-06 | 1.858E-06 | 7.400E-06 | 1.100E-05 |
| 9610 | 481 | E481I | A | 8.4e-06 | 7.1e-06 | 6.4e-06 | 7.300E-06 | 1.015E-06 | 6.400E-06 | 8.400E-06 |
| 9611 | 481 | E481L | A | 1.2e-05 | 4.8e-06 | 7.9e-06 | 8.233E-06 | 3.612E-06 | 4.800E-06 | 1.200E-05 |
| 9612 | 481 | E481K | A | 6.9e-06 | 7.3e-06 | 1.1e-05 | 8.400E-06 | 2.261E-06 | 6.900E-06 | 1.100E-05 |
| 9613 | 481 | E481M | A | 8.5e-06 | 6.6e-06 | 9.1e-06 | 8.067E-06 | 1.305E-06 | 6.600E-06 | 9.100E-06 |
| 9614 | 481 | E481F | A | 6.5e-06 | 9.3e-06 | 6.6e-06 | 7.467E-06 | 1.589E-06 | 6.500E-06 | 9.300E-06 |
| 9615 | 481 | E481P | A | 8.2e-06 | 1.1e-05 | 6.7e-06 | 8.633E-06 | 2.183E-06 | 6.700E-06 | 1.100E-05 |
| 9616 | 481 | E481S | A | 9e-06 | 6.6e-06 | 3.1e-06 | 6.233E-06 | 2.967E-06 | 3.100E-06 | 9.000E-06 |
| 9617 | 481 | E481T | A | 8.7e-06 | 4.9e-06 | 1e-05 | 7.867E-06 | 2.650E-06 | 4.900E-06 | 1.000E-05 |
| 9618 | 481 | E481W | A | 6.1e-06 | 1e-05 | 8.3e-06 | 8.133E-06 | 1.955E-06 | 6.100E-06 | 1.000E-05 |
| 9619 | 481 | E481Y | A | 9.4e-06 | 1.6e-05 | 7.7e-06 | 1.103E-05 | 4.384E-06 | 7.700E-06 | 1.600E-05 |
| 9620 | 481 | E481V | A | 5.4e-06 | 6.5e-06 | 1.4e-05 | 8.633E-06 | 4.680E-06 | 5.400E-06 | 1.400E-05 |
| 9621 | 482 | C482A | A | 9.5e-06 | 5e-06 | 9.2e-06 | 7.900E-06 | 2.516E-06 | 5.000E-06 | 9.500E-06 |
| 9622 | 482 | C482R | A | 5.2e-06 | 3.1e-06 | 1e-05 | 6.100E-06 | 3.537E-06 | 3.100E-06 | 1.000E-05 |
| 9623 | 482 | C482N | A | 1.1e-05 | 1.4e-05 | 5e-06 | 1.000E-05 | 4.583E-06 | 5.000E-06 | 1.400E-05 |
| 9624 | 482 | C482D | A | 8.1e-06 | 7.8e-06 | 6.2e-06 | 7.367E-06 | 1.021E-06 | 6.200E-06 | 8.100E-06 |
| 9625 | 482 | C482C | A | 9.5e-06 | 8.6e-06 | 4.9e-06 | 7.667E-06 | 2.438E-06 | 4.900E-06 | 9.500E-06 |
| 9626 | 482 | C482E | A | 8.4e-06 | 9e-06 | 5.1e-06 | 7.500E-06 | 2.100E-06 | 5.100E-06 | 9.000E-06 |
| 9627 | 482 | C482Q | A | 1e-05 | 1.2e-05 | 8.2e-06 | 1.007E-05 | 1.901E-06 | 8.200E-06 | 1.200E-05 |
| 9628 | 482 | C482G | A | 8.2e-06 | 1.6e-05 | 6.6e-06 | 1.027E-05 | 5.029E-06 | 6.600E-06 | 1.600E-05 |
| 9629 | 482 | C482H | A | 5e-06 | 5.5e-06 | 1.2e-05 | 7.500E-06 | 3.905E-06 | 5.000E-06 | 1.200E-05 |
| 9630 | 482 | C482I | A | 9.5e-06 | 8.9e-06 | 1.5e-05 | 1.113E-05 | 3.362E-06 | 8.900E-06 | 1.500E-05 |
| 9631 | 482 | C482L | A | 6.9e-06 | 6.7e-06 | 6e-06 | 6.533E-06 | 4.726E-07 | 6.000E-06 | 6.900E-06 |
| 9632 | 482 | C482K | A | 5.6e-06 | 1.1e-05 | 1.1e-05 | 9.200E-06 | 3.118E-06 | 5.600E-06 | 1.100E-05 |
| 9633 | 482 | C482M | A | 7.4e-06 | 8.9e-06 | 5.6e-06 | 7.300E-06 | 1.652E-06 | 5.600E-06 | 8.900E-06 |
| 9634 | 482 | C482F | A | 9.7e-06 | 6.6e-06 | 1.6e-05 | 1.077E-05 | 4.790E-06 | 6.600E-06 | 1.600E-05 |
| 9635 | 482 | C482P | A | 8.2e-06 | 8.8e-06 | 9.6e-06 | 8.867E-06 | 7.024E-07 | 8.200E-06 | 9.600E-06 |
| 9636 | 482 | C482S | A | 1.5e-05 | 6.6e-06 | 6.9e-06 | 9.500E-06 | 4.766E-06 | 6.600E-06 | 1.500E-05 |
| 9637 | 482 | C482T | A | 6e-06 | 5.2e-06 | 7e-06 | 6.067E-06 | 9.018E-07 | 5.200E-06 | 7.000E-06 |
| 9638 | 482 | C482W | A | 1.3e-05 | 7.6e-06 | 5.7e-06 | 8.767E-06 | 3.787E-06 | 5.700E-06 | 1.300E-05 |
| 9639 | 482 | C482Y | A | 4.2e-06 | 7.1e-06 | 6.5e-06 | 5.933E-06 | 1.531E-06 | 4.200E-06 | 7.100E-06 |
| 9640 | 482 | C482V | A | 1.2e-05 | 9.9e-06 | 7.4e-06 | 9.767E-06 | 2.303E-06 | 7.400E-06 | 1.200E-05 |
| 9641 | 483 | V483A | A | 5.2e-06 | 5.3e-06 | 1.3e-05 | 7.833E-06 | 4.475E-06 | 5.200E-06 | 1.300E-05 |
| 9642 | 483 | V483R | A | 1e-05 | 1.3e-05 | 7.3e-06 | 1.010E-05 | 2.851E-06 | 7.300E-06 | 1.300E-05 |
| 9643 | 483 | V483N | A | 6.1e-06 | 5e-06 | 1.2e-05 | 7.700E-06 | 3.764E-06 | 5.000E-06 | 1.200E-05 |
| 9644 | 483 | V483D | A | 6.5e-06 | 6.7e-06 | 6.2e-06 | 6.467E-06 | 2.517E-07 | 6.200E-06 | 6.700E-06 |
| 9645 | 483 | V483C | A | 9.8e-06 | 5.8e-06 | 8.1e-06 | 7.900E-06 | 2.007E-06 | 5.800E-06 | 9.800E-06 |
| 9646 | 483 | V483E | A | 7.9e-06 | 9.5e-06 | 1e-05 | 9.133E-06 | 1.097E-06 | 7.900E-06 | 1.000E-05 |
| 9647 | 483 | V483Q | A | 9.4e-06 | 7.4e-06 | 9.5e-06 | 8.767E-06 | 1.185E-06 | 7.400E-06 | 9.500E-06 |
| 9648 | 483 | V483G | A | 9.2e-06 | 9.8e-06 | 9e-06 | 9.333E-06 | 4.163E-07 | 9.000E-06 | 9.800E-06 |
| 9649 | 483 | V483H | A | 1.4e-05 | 1e-05 | 5.8e-06 | 9.933E-06 | 4.100E-06 | 5.800E-06 | 1.400E-05 |
| 9650 | 483 | V483I | A | 1e-05 | 1.4e-05 | 1e-05 | 1.133E-05 | 2.309E-06 | 1.000E-05 | 1.400E-05 |
| 9651 | 483 | V483L | A | 8.3e-06 | 8.4e-06 | 1.3e-05 | 9.900E-06 | 2.685E-06 | 8.300E-06 | 1.300E-05 |
| 9652 | 483 | V483K | A | 1e-05 | 7.6e-06 | 8.3e-06 | 8.633E-06 | 1.234E-06 | 7.600E-06 | 1.000E-05 |
| 9653 | 483 | V483M | A | 9.7e-06 | 9.7e-06 | 1.4e-05 | 1.113E-05 | 2.483E-06 | 9.700E-06 | 1.400E-05 |
| 9654 | 483 | V483F | A | 7.2e-06 | 6.9e-06 | 6.6e-06 | 6.900E-06 | 3.000E-07 | 6.600E-06 | 7.200E-06 |
| 9655 | 483 | V483P | A | 6.3e-06 | 1.1e-05 | 5.3e-06 | 7.533E-06 | 3.044E-06 | 5.300E-06 | 1.100E-05 |
| 9656 | 483 | V483S | A | 1.1e-05 | 6.9e-06 | 1.1e-05 | 9.633E-06 | 2.367E-06 | 6.900E-06 | 1.100E-05 |
| 9657 | 483 | V483T | A | 5.9e-06 | 1.4e-05 | 1.3e-05 | 1.097E-05 | 4.416E-06 | 5.900E-06 | 1.400E-05 |
| 9658 | 483 | V483W | A | 6.7e-06 | 4.7e-06 | 9.8e-06 | 7.067E-06 | 2.570E-06 | 4.700E-06 | 9.800E-06 |
| 9659 | 483 | V483Y | A | 8.2e-06 | 5e-06 | 9.5e-06 | 7.567E-06 | 2.316E-06 | 5.000E-06 | 9.500E-06 |
| 9660 | 483 | V483V | A | 5e-06 | 6.5e-06 | 5.9e-06 | 5.800E-06 | 7.550E-07 | 5.000E-06 | 6.500E-06 |
| 9661 | 484 | G484A | A | 9.1e-06 | 9.9e-06 | 8.7e-06 | 9.233E-06 | 6.110E-07 | 8.700E-06 | 9.900E-06 |
| 9662 | 484 | G484R | A | 1.1e-05 | 1.2e-05 | 1.1e-05 | 1.133E-05 | 5.774E-07 | 1.100E-05 | 1.200E-05 |
| 9663 | 484 | G484N | A | 6.1e-06 | 9.4e-06 | 6.2e-06 | 7.233E-06 | 1.877E-06 | 6.100E-06 | 9.400E-06 |
| 9664 | 484 | G484D | A | 8.1e-06 | 7.4e-06 | 6.5e-06 | 7.333E-06 | 8.021E-07 | 6.500E-06 | 8.100E-06 |
| 9665 | 484 | G484C | A | 8.9e-06 | 5.5e-06 | 7.5e-06 | 7.300E-06 | 1.709E-06 | 5.500E-06 | 8.900E-06 |
| 9666 | 484 | G484E | A | 1e-05 | 8.7e-06 | 1.2e-05 | 1.023E-05 | 1.662E-06 | 8.700E-06 | 1.200E-05 |
| 9667 | 484 | G484Q | A | 7.2e-06 | 5.8e-06 | 1.1e-05 | 8.000E-06 | 2.691E-06 | 5.800E-06 | 1.100E-05 |
| 9668 | 484 | G484G | A | 1.2e-05 | 6.5e-06 | 9.2e-06 | 9.233E-06 | 2.750E-06 | 6.500E-06 | 1.200E-05 |
| 9669 | 484 | G484H | A | 1.1e-05 | 5.6e-06 | 6.9e-06 | 7.833E-06 | 2.818E-06 | 5.600E-06 | 1.100E-05 |
| 9670 | 484 | G484I | A | 6.9e-06 | 9.7e-06 | 2.1e-05 | 1.253E-05 | 7.465E-06 | 6.900E-06 | 2.100E-05 |
| 9671 | 484 | G484L | A | 4.9e-06 | 6.4e-06 | 7.3e-06 | 6.200E-06 | 1.212E-06 | 4.900E-06 | 7.300E-06 |
| 9672 | 484 | G484K | A | 1.2e-05 | 1.1e-05 | 6.9e-06 | 9.967E-06 | 2.702E-06 | 6.900E-06 | 1.200E-05 |
| 9673 | 484 | G484M | A | 4.8e-06 | 1.3e-05 | 4.3e-06 | 7.367E-06 | 4.885E-06 | 4.300E-06 | 1.300E-05 |
| 9674 | 484 | G484F | A | 1e-05 | 9.3e-06 | 6.9e-06 | 8.733E-06 | 1.626E-06 | 6.900E-06 | 1.000E-05 |
| 9675 | 484 | G484P | A | 7.1e-06 | 7e-06 | 7.6e-06 | 7.233E-06 | 3.215E-07 | 7.000E-06 | 7.600E-06 |
| 9676 | 484 | G484S | A | 1.3e-05 | 4.6e-06 | 8.4e-06 | 8.667E-06 | 4.206E-06 | 4.600E-06 | 1.300E-05 |
| 9677 | 484 | G484T | A | 4.7e-06 | 7.7e-06 | 1e-05 | 7.467E-06 | 2.658E-06 | 4.700E-06 | 1.000E-05 |
| 9678 | 484 | G484W | A | 5.9e-06 | 8.6e-06 | 1e-05 | 8.167E-06 | 2.084E-06 | 5.900E-06 | 1.000E-05 |
| 9679 | 484 | G484Y | A | 1e-05 | 1.3e-05 | 6.5e-06 | 9.833E-06 | 3.253E-06 | 6.500E-06 | 1.300E-05 |
| 9680 | 484 | G484V | A | 9.3e-06 | 4.8e-06 | 6.9e-06 | 7.000E-06 | 2.252E-06 | 4.800E-06 | 9.300E-06 |
| 9681 | 485 | E485A | A | 6.2e-06 | 1.2e-05 | 8.1e-06 | 8.767E-06 | 2.957E-06 | 6.200E-06 | 1.200E-05 |
| 9682 | 485 | E485R | A | 1e-05 | 1.1e-05 | 8.8e-06 | 9.933E-06 | 1.102E-06 | 8.800E-06 | 1.100E-05 |
| 9683 | 485 | E485N | A | 5.5e-06 | 1.4e-05 | 9.5e-06 | 9.667E-06 | 4.252E-06 | 5.500E-06 | 1.400E-05 |
| 9684 | 485 | E485D | A | 6.8e-06 | 1.2e-05 | 9.7e-06 | 9.500E-06 | 2.606E-06 | 6.800E-06 | 1.200E-05 |
| 9685 | 485 | E485C | A | 1.3e-05 | 1.5e-05 | 9.3e-06 | 1.243E-05 | 2.892E-06 | 9.300E-06 | 1.500E-05 |
| 9686 | 485 | E485E | A | 1.4e-05 | 4.1e-06 | 9.4e-06 | 9.167E-06 | 4.954E-06 | 4.100E-06 | 1.400E-05 |
| 9687 | 485 | E485Q | A | 1.1e-05 | 6.1e-06 | 4.8e-06 | 7.300E-06 | 3.270E-06 | 4.800E-06 | 1.100E-05 |
| 9688 | 485 | E485G | A | 9.4e-06 | 8.5e-06 | 9.3e-06 | 9.067E-06 | 4.933E-07 | 8.500E-06 | 9.400E-06 |
| 9689 | 485 | E485H | A | 9.3e-06 | 7.4e-06 | 6.5e-06 | 7.733E-06 | 1.429E-06 | 6.500E-06 | 9.300E-06 |

| | | | | | | | | | | |
|---|---|---|---|---|---|---|---|---|---|---|
| 9690 | 485 | E485I | A | 1.8e-05 | 8.5e-06 | 9.3e-06 | 1.193E-05 | 5.269E-06 | 8.500E-06 | 1.800E-05 |
| 9691 | 485 | E485L | A | 4.6e-06 | 1.3e-05 | 9.2e-06 | 8.933E-06 | 4.206E-06 | 4.600E-06 | 1.300E-05 |
| 9692 | 485 | E485K | A | 8.4e-06 | 3.8e-06 | 8.8e-06 | 7.000E-06 | 2.778E-06 | 3.800E-06 | 8.800E-06 |
| 9693 | 485 | E485M | A | 7.6e-06 | 8.7e-06 | 5.2e-06 | 7.167E-06 | 1.790E-06 | 5.200E-06 | 8.700E-06 |
| 9694 | 485 | E485F | A | 1e-05 | 6.4e-06 | 7.2e-06 | 7.867E-06 | 1.890E-06 | 6.400E-06 | 1.000E-05 |
| 9695 | 485 | E485P | A | 9.8e-06 | 9e-06 | 1.3e-05 | 1.060E-05 | 2.117E-06 | 9.000E-06 | 1.300E-05 |
| 9696 | 485 | E485S | A | 6.8e-06 | 6.6e-06 | 8.4e-06 | 7.267E-06 | 9.866E-07 | 6.600E-06 | 8.400E-06 |
| 9697 | 485 | E485T | A | 6.1e-06 | 9.9e-06 | 1.1e-05 | 9.000E-06 | 2.571E-06 | 6.100E-06 | 1.100E-05 |
| 9698 | 485 | E485W | A | 4.8e-06 | 8.3e-06 | 9.6e-06 | 7.567E-06 | 2.483E-06 | 4.800E-06 | 9.600E-06 |
| 9699 | 485 | E485Y | A | 6.6e-06 | 7.4e-06 | 7.7e-06 | 7.233E-06 | 5.686E-07 | 6.600E-06 | 7.700E-06 |
| 9700 | 485 | E485V | A | 8.9e-06 | 7.5e-06 | 7.8e-06 | 8.067E-06 | 7.371E-07 | 7.500E-06 | 8.900E-06 |
| 9701 | 486 | G486A | A | 6.9e-06 | 6.5e-06 | 9.7e-06 | 7.700E-06 | 1.744E-06 | 6.500E-06 | 9.700E-06 |
| 9702 | 486 | G486R | A | 6.5e-06 | 7.8e-06 | 1.1e-05 | 8.433E-06 | 2.316E-06 | 6.500E-06 | 1.100E-05 |
| 9703 | 486 | G486N | A | 8.4e-06 | 6.7e-06 | 6.2e-06 | 7.100E-06 | 1.153E-06 | 6.200E-06 | 8.400E-06 |
| 9704 | 486 | G486D | A | 1.3e-05 | 5.7e-06 | 1.3e-05 | 1.057E-05 | 4.215E-06 | 5.700E-06 | 1.300E-05 |
| 9705 | 486 | G486C | A | 9.3e-06 | 7.4e-06 | 7.8e-06 | 8.167E-06 | 1.002E-06 | 7.400E-06 | 9.300E-06 |
| 9706 | 486 | G486E | A | 7.3e-06 | 7.7e-06 | 8.6e-06 | 7.867E-06 | 6.658E-07 | 7.300E-06 | 8.600E-06 |
| 9707 | 486 | G486Q | A | 6.2e-06 | 7.2e-06 | 9.1e-06 | 7.500E-06 | 1.473E-06 | 6.200E-06 | 9.100E-06 |
| 9708 | 486 | G486G | A | 1.3e-05 | 4.6e-06 | 1.3e-05 | 1.020E-05 | 4.850E-06 | 4.600E-06 | 1.300E-05 |
| 9709 | 486 | G486H | A | 6.5e-06 | 8e-06 | 4.2e-06 | 6.233E-06 | 1.914E-06 | 4.200E-06 | 8.000E-06 |
| 9710 | 486 | G486I | A | 5e-06 | 9e-06 | 1.3e-05 | 9.000E-06 | 4.000E-06 | 5.000E-06 | 1.300E-05 |
| 9711 | 486 | G486L | A | 5.7e-06 | 6e-06 | 7.3e-06 | 6.333E-06 | 8.505E-07 | 5.700E-06 | 7.300E-06 |
| 9712 | 486 | G486K | A | 5.8e-06 | 3.2e-06 | 6.4e-06 | 5.133E-06 | 1.701E-06 | 3.200E-06 | 6.400E-06 |
| 9713 | 486 | G486M | A | 7.9e-06 | 9.7e-06 | 1.8e-05 | 1.187E-05 | 5.387E-06 | 7.900E-06 | 1.800E-05 |
| 9714 | 486 | G486F | A | 6.2e-06 | 1e-05 | 8.1e-06 | 8.100E-06 | 1.900E-06 | 6.200E-06 | 1.000E-05 |
| 9715 | 486 | G486P | A | 5.1e-06 | 6.5e-06 | 9.6e-06 | 7.067E-06 | 2.303E-06 | 5.100E-06 | 9.600E-06 |
| 9716 | 486 | G486S | A | 7.6e-06 | 8.4e-06 | 6.2e-06 | 7.400E-06 | 1.114E-06 | 6.200E-06 | 8.400E-06 |
| 9717 | 486 | G486T | A | 8.6e-06 | 6.5e-06 | 5.7e-06 | 6.933E-06 | 1.498E-06 | 5.700E-06 | 8.600E-06 |
| 9718 | 486 | G486W | A | 5.8e-06 | 1.3e-05 | 1.1e-05 | 9.933E-06 | 3.717E-06 | 5.800E-06 | 1.300E-05 |
| 9719 | 486 | G486Y | A | 8.8e-06 | 1.2e-05 | 7.6e-06 | 9.467E-06 | 2.274E-06 | 7.600E-06 | 1.200E-05 |
| 9720 | 486 | G486V | A | 5.5e-06 | 8.4e-06 | 7e-06 | 6.967E-06 | 1.450E-06 | 5.500E-06 | 8.400E-06 |
| 9721 | 487 | L487A | A | 1.3e-05 | 7.5e-06 | 2.7e-06 | 7.733E-06 | 5.154E-06 | 2.700E-06 | 1.300E-05 |
| 9722 | 487 | L487R | A | 7.1e-06 | 4.9e-06 | 4.3e-06 | 5.433E-06 | 1.474E-06 | 4.300E-06 | 7.100E-06 |
| 9723 | 487 | L487N | A | 1.1e-05 | 1e-05 | 9.6e-06 | 1.020E-05 | 7.211E-07 | 9.600E-06 | 1.100E-05 |
| 9724 | 487 | L487D | A | 5.9e-06 | 7.6e-06 | 8.7e-06 | 7.400E-06 | 1.411E-06 | 5.900E-06 | 8.700E-06 |
| 9725 | 487 | L487C | A | 7e-06 | 1.2e-05 | 8.7e-06 | 9.233E-06 | 2.542E-06 | 7.000E-06 | 1.200E-05 |
| 9726 | 487 | L487E | A | 9.2e-06 | 8.5e-06 | 9.9e-06 | 9.200E-06 | 7.000E-07 | 8.500E-06 | 9.900E-06 |
| 9727 | 487 | L487Q | A | 1e-05 | 6.9e-06 | 9.3e-06 | 8.733E-06 | 1.626E-06 | 6.900E-06 | 1.000E-05 |
| 9728 | 487 | L487G | A | 9.8e-06 | 7.3e-06 | 1e-05 | 9.033E-06 | 1.504E-06 | 7.300E-06 | 1.000E-05 |
| 9729 | 487 | L487H | A | 4.3e-06 | 1.3e-05 | 8.1e-06 | 8.467E-06 | 4.362E-06 | 4.300E-06 | 1.300E-05 |
| 9730 | 487 | L487I | A | 7.4e-06 | 8e-06 | 1.1e-05 | 8.800E-06 | 1.929E-06 | 7.400E-06 | 1.100E-05 |
| 9731 | 487 | L487L | A | 6.3e-06 | 6.9e-06 | 4.3e-06 | 5.833E-06 | 1.361E-06 | 4.300E-06 | 6.900E-06 |
| 9732 | 487 | L487K | A | 6.4e-06 | 9.1e-06 | 6.4e-06 | 7.300E-06 | 1.559E-06 | 6.400E-06 | 9.100E-06 |
| 9733 | 487 | L487M | A | 9.3e-06 | 7e-06 | 7.1e-06 | 7.800E-06 | 1.300E-06 | 7.000E-06 | 9.300E-06 |
| 9734 | 487 | L487F | A | 6.3e-06 | 7.3e-06 | 6e-06 | 6.533E-06 | 6.807E-07 | 6.000E-06 | 7.300E-06 |
| 9735 | 487 | L487P | A | 9e-06 | 9.7e-06 | 9.4e-06 | 9.367E-06 | 3.512E-07 | 9.000E-06 | 9.700E-06 |
| 9736 | 487 | L487S | A | 9.7e-06 | 1.5e-05 | 5.6e-06 | 1.010E-05 | 4.713E-06 | 5.600E-06 | 1.500E-05 |
| 9737 | 487 | L487T | A | 8.8e-06 | 9.4e-06 | 9.4e-06 | 9.200E-06 | 3.464E-07 | 8.800E-06 | 9.400E-06 |
| 9738 | 487 | L487W | A | 9.2e-06 | 5e-06 | 1e-05 | 8.067E-06 | 2.686E-06 | 5.000E-06 | 1.000E-05 |
| 9739 | 487 | L487Y | A | 6.9e-06 | 8.4e-06 | 1.2e-05 | 9.100E-06 | 2.621E-06 | 6.900E-06 | 1.200E-05 |
| 9740 | 487 | L487V | A | 9.6e-06 | 1.1e-05 | 1.2e-05 | 1.087E-05 | 1.206E-06 | 9.600E-06 | 1.200E-05 |
| 9741 | 488 | A488A | A | 1.4e-05 | 9.8e-06 | 7.7e-06 | 1.050E-05 | 3.208E-06 | 7.700E-06 | 1.400E-05 |
| 9742 | 488 | A488R | A | 1e-05 | 7.4e-06 | 1e-05 | 9.133E-06 | 1.501E-06 | 7.400E-06 | 1.000E-05 |
| 9743 | 488 | A488N | A | 4.9e-06 | 1.1e-05 | 1.1e-05 | 8.967E-06 | 3.522E-06 | 4.900E-06 | 1.100E-05 |
| 9744 | 488 | A488D | A | 7.5e-06 | 6e-06 | 5.6e-06 | 6.367E-06 | 1.002E-06 | 5.600E-06 | 7.500E-06 |
| 9745 | 488 | A488C | A | 6.5e-06 | 7.3e-06 | 4.5e-06 | 6.100E-06 | 1.442E-06 | 4.500E-06 | 7.300E-06 |
| 9746 | 488 | A488E | A | 9.6e-06 | 8.2e-06 | 8.9e-06 | 8.900E-06 | 7.000E-07 | 8.200E-06 | 9.600E-06 |
| 9747 | 488 | A488Q | A | 5.6e-06 | 1e-05 | 7.1e-06 | 7.567E-06 | 2.237E-06 | 5.600E-06 | 1.000E-05 |
| 9748 | 488 | A488G | A | 9.2e-06 | 9.3e-06 | 4.6e-06 | 7.700E-06 | 2.685E-06 | 4.600E-06 | 9.300E-06 |
| 9749 | 488 | A488H | A | 7.5e-06 | 4.1e-06 | 9.8e-06 | 7.133E-06 | 2.868E-06 | 4.100E-06 | 9.800E-06 |
| 9750 | 488 | A488I | A | 9e-06 | 8e-06 | 8.4e-06 | 8.467E-06 | 5.033E-07 | 8.000E-06 | 9.000E-06 |
| 9751 | 488 | A488L | A | 9e-06 | 7e-06 | 1e-05 | 8.667E-06 | 1.528E-06 | 7.000E-06 | 1.000E-05 |
| 9752 | 488 | A488K | A | 8.2e-06 | 1.1e-05 | 6e-06 | 8.400E-06 | 2.506E-06 | 6.000E-06 | 1.100E-05 |
| 9753 | 488 | A488M | A | 9.3e-06 | 8.1e-06 | 4.8e-06 | 7.400E-06 | 2.330E-06 | 4.800E-06 | 9.300E-06 |
| 9754 | 488 | A488F | A | 5.7e-06 | 8.5e-06 | 6.5e-06 | 6.900E-06 | 1.442E-06 | 5.700E-06 | 8.500E-06 |
| 9755 | 488 | A488P | A | 3.5e-06 | 7.4e-06 | 7.3e-06 | 6.067E-06 | 2.223E-06 | 3.500E-06 | 7.400E-06 |
| 9756 | 488 | A488S | A | 1e-05 | 8.3e-06 | 1.1e-05 | 9.767E-06 | 1.365E-06 | 8.300E-06 | 1.100E-05 |
| 9757 | 488 | A488T | A | 8.3e-06 | 1.2e-05 | 8.6e-06 | 9.633E-06 | 2.055E-06 | 8.300E-06 | 1.200E-05 |
| 9758 | 488 | A488W | A | 8.7e-06 | 1.3e-05 | 6.2e-06 | 9.300E-06 | 3.439E-06 | 6.200E-06 | 1.300E-05 |
| 9759 | 488 | A488Y | A | 5.9e-06 | 5.3e-06 | 6.1e-06 | 5.767E-06 | 4.163E-07 | 5.300E-06 | 6.100E-06 |
| 9760 | 488 | A488V | A | 6.5e-06 | 7.1e-06 | 4.8e-06 | 6.133E-06 | 1.193E-06 | 4.800E-06 | 7.100E-06 |
| 9761 | 489 | C489A | A | 8e-06 | 4.3e-06 | 5.5e-06 | 5.933E-06 | 1.888E-06 | 4.300E-06 | 8.000E-06 |
| 9762 | 489 | C489R | A | 1.1e-05 | 5.1e-06 | 7.5e-06 | 7.867E-06 | 2.967E-06 | 5.100E-06 | 1.100E-05 |
| 9763 | 489 | C489N | A | 9.5e-06 | 8e-06 | 1e-05 | 9.167E-06 | 1.041E-06 | 8.000E-06 | 1.000E-05 |
| 9764 | 489 | C489D | A | 6.6e-06 | 1.1e-05 | 1.6e-05 | 1.120E-05 | 4.703E-06 | 6.600E-06 | 1.600E-05 |
| 9765 | 489 | C489C | A | 1.4e-05 | 5.9e-06 | 5.6e-06 | 8.500E-06 | 4.766E-06 | 5.600E-06 | 1.400E-05 |
| 9766 | 489 | C489E | A | 6e-06 | 1.4e-05 | 9.6e-06 | 9.867E-06 | 4.007E-06 | 6.000E-06 | 1.400E-05 |
| 9767 | 489 | C489Q | A | 9.8e-06 | 6.3e-06 | 8.2e-06 | 8.100E-06 | 1.752E-06 | 6.300E-06 | 9.800E-06 |
| 9768 | 489 | C489G | A | 1e-05 | 1.8e-05 | 1.3e-05 | 1.367E-05 | 4.041E-06 | 1.000E-05 | 1.800E-05 |
| 9769 | 489 | C489H | A | 9.8e-06 | 6.7e-06 | 6.2e-06 | 7.567E-06 | 1.950E-06 | 6.200E-06 | 9.800E-06 |
| 9770 | 489 | C489I | A | 1.4e-05 | 9.9e-06 | 1e-05 | 1.130E-05 | 2.339E-06 | 9.900E-06 | 1.400E-05 |
| 9771 | 489 | C489L | A | 8.2e-06 | 9.2e-06 | 9e-06 | 8.800E-06 | 5.292E-07 | 8.200E-06 | 9.200E-06 |
| 9772 | 489 | C489K | A | 6.6e-06 | 6.3e-06 | 9.9e-06 | 7.600E-06 | 1.997E-06 | 6.300E-06 | 9.900E-06 |
| 9773 | 489 | C489M | A | 9e-06 | 6.8e-06 | 6.8e-06 | 7.533E-06 | 1.270E-06 | 6.800E-06 | 9.000E-06 |
| 9774 | 489 | C489F | A | 1e-05 | 6.2e-06 | 6.2e-06 | 7.467E-06 | 2.194E-06 | 6.200E-06 | 1.000E-05 |

| | | | | | | | | | | |
|---|---|---|---|---|---|---|---|---|---|---|
| 9775 | 489 | C489P | A | 5.6e-06 | 6.3e-06 | 4.9e-06 | 5.600E-06 | 7.000E-07 | 4.900E-06 | 6.300E-06 |
| 9776 | 489 | C489S | A | 5.7e-06 | 6.5e-06 | 7.2e-06 | 6.467E-06 | 7.506E-07 | 5.700E-06 | 7.200E-06 |
| 9777 | 489 | C489T | A | 1e-05 | 8.7e-06 | 7.1e-06 | 8.600E-06 | 1.453E-06 | 7.100E-06 | 1.000E-05 |
| 9778 | 489 | C489W | A | 6.2e-06 | 9e-06 | 6.7e-06 | 7.300E-06 | 1.493E-06 | 6.200E-06 | 9.000E-06 |
| 9779 | 489 | C489Y | A | 9.9e-06 | 4.9e-06 | 5.8e-06 | 6.867E-06 | 2.665E-06 | 4.900E-06 | 9.900E-06 |
| 9780 | 489 | C489V | A | 9.3e-06 | 1e-05 | 8.3e-06 | 9.200E-06 | 8.544E-07 | 8.300E-06 | 1.000E-05 |
| 9781 | 490 | H490A | A | 7.1e-06 | 9e-06 | 1.3e-05 | 9.700E-06 | 3.012E-06 | 7.100E-06 | 1.300E-05 |
| 9782 | 490 | H490R | A | 1e-05 | 7.9e-06 | 5.5e-06 | 7.800E-06 | 2.252E-06 | 5.500E-06 | 1.000E-05 |
| 9783 | 490 | H490N | A | 9.2e-06 | 5.6e-06 | 7.4e-06 | 7.400E-06 | 1.800E-06 | 5.600E-06 | 9.200E-06 |
| 9784 | 490 | H490D | A | 1e-05 | 6.2e-06 | 8.4e-06 | 8.200E-06 | 1.908E-06 | 6.200E-06 | 1.000E-05 |
| 9785 | 490 | H490C | A | 7.3e-06 | 8.3e-06 | 8.1e-06 | 7.900E-06 | 5.292E-07 | 7.300E-06 | 8.300E-06 |
| 9786 | 490 | H490E | A | 9.1e-06 | 8.4e-06 | 5.2e-06 | 7.567E-06 | 2.079E-06 | 5.200E-06 | 9.100E-06 |
| 9787 | 490 | H490Q | A | 6.6e-06 | 4.6e-06 | 1.3e-05 | 8.067E-06 | 4.388E-06 | 4.600E-06 | 1.300E-05 |
| 9788 | 490 | H490G | A | 7.7e-06 | 1e-05 | 6e-06 | 7.900E-06 | 2.007E-06 | 6.000E-06 | 1.000E-05 |
| 9789 | 490 | H490H | A | 4.7e-06 | 1.1e-05 | 6.6e-06 | 7.433E-06 | 3.232E-06 | 4.700E-06 | 1.100E-05 |
| 9790 | 490 | H490I | A | 1.3e-05 | 7.2e-06 | 8.3e-06 | 9.500E-06 | 3.081E-06 | 7.200E-06 | 1.300E-05 |
| 9791 | 490 | H490L | A | 6.4e-06 | 7.9e-06 | 1.4e-05 | 9.433E-06 | 4.025E-06 | 6.400E-06 | 1.400E-05 |
| 9792 | 490 | H490K | A | 1.5e-05 | 7e-06 | 8.3e-06 | 1.010E-05 | 4.293E-06 | 7.000E-06 | 1.500E-05 |
| 9793 | 490 | H490M | A | 1e-05 | 8.5e-06 | 6.5e-06 | 8.333E-06 | 1.756E-06 | 6.500E-06 | 1.000E-05 |
| 9794 | 490 | H490F | A | 4.1e-06 | 1.1e-05 | 1e-05 | 8.367E-06 | 3.729E-06 | 4.100E-06 | 1.100E-05 |
| 9795 | 490 | H490P | A | 7e-06 | 1.1e-05 | 8.9e-06 | 8.967E-06 | 2.001E-06 | 7.000E-06 | 1.100E-05 |
| 9796 | 490 | H490S | A | 9e-06 | 5.4e-06 | 1e-05 | 8.133E-06 | 2.419E-06 | 5.400E-06 | 1.000E-05 |
| 9797 | 490 | H490T | A | 1.1e-05 | 7.9e-06 | 6.9e-06 | 8.600E-06 | 2.138E-06 | 6.900E-06 | 1.100E-05 |
| 9798 | 490 | H490W | A | 9.9e-06 | 9.4e-06 | 7.9e-06 | 9.067E-06 | 1.041E-06 | 7.900E-06 | 9.900E-06 |
| 9799 | 490 | H490Y | A | 6.8e-06 | 5.3e-06 | 6.1e-06 | 6.067E-06 | 7.506E-07 | 5.300E-06 | 6.800E-06 |
| 9800 | 490 | H490V | A | 4e-06 | 6.5e-06 | 1.4e-05 | 8.167E-06 | 5.204E-06 | 4.000E-06 | 1.400E-05 |
| 9801 | 491 | Q491A | A | 1.2e-05 | 4.7e-06 | 1.3e-05 | 9.900E-06 | 4.531E-06 | 4.700E-06 | 1.300E-05 |
| 9802 | 491 | Q491R | A | 5.2e-06 | 7e-06 | 7.2e-06 | 6.467E-06 | 1.102E-06 | 5.200E-06 | 7.200E-06 |
| 9803 | 491 | Q491N | A | 7.9e-06 | 5.3e-06 | 6.7e-06 | 6.633E-06 | 1.301E-06 | 5.300E-06 | 7.900E-06 |
| 9804 | 491 | Q491D | A | 1.2e-05 | 9.8e-06 | 5.3e-06 | 9.033E-06 | 3.415E-06 | 5.300E-06 | 1.200E-05 |
| 9805 | 491 | Q491C | A | 9.2e-06 | 1.2e-05 | 8.7e-06 | 9.967E-06 | 1.779E-06 | 8.700E-06 | 1.200E-05 |
| 9806 | 491 | Q491E | A | 1.4e-05 | 4.8e-06 | 1.3e-05 | 1.060E-05 | 5.048E-06 | 4.800E-06 | 1.400E-05 |
| 9807 | 491 | Q491Q | A | 9.8e-06 | 5.8e-06 | 9.4e-06 | 8.333E-06 | 2.203E-06 | 5.800E-06 | 9.800E-06 |
| 9808 | 491 | Q491G | A | 7.2e-06 | 5.7e-06 | 1.4e-05 | 8.967E-06 | 4.423E-06 | 5.700E-06 | 1.400E-05 |
| 9809 | 491 | Q491H | A | 9.7e-06 | 9.6e-06 | 8.6e-06 | 9.300E-06 | 6.083E-07 | 8.600E-06 | 9.700E-06 |
| 9810 | 491 | Q491I | A | 9.9e-06 | 8.3e-06 | 1.4e-05 | 1.073E-05 | 2.940E-06 | 8.300E-06 | 1.400E-05 |
| 9811 | 491 | Q491L | A | 1.2e-05 | 1.3e-05 | 1.6e-05 | 1.367E-05 | 2.082E-06 | 1.200E-05 | 1.600E-05 |
| 9812 | 491 | Q491K | A | 8.5e-06 | 6.7e-06 | 6.6e-06 | 7.267E-06 | 1.069E-06 | 6.600E-06 | 8.500E-06 |
| 9813 | 491 | Q491M | A | 9.8e-06 | 5.6e-06 | 9.4e-06 | 8.267E-06 | 2.318E-06 | 5.600E-06 | 9.800E-06 |
| 9814 | 491 | Q491F | A | 6.8e-06 | 9.5e-06 | 5.2e-06 | 7.167E-06 | 2.173E-06 | 5.200E-06 | 9.500E-06 |
| 9815 | 491 | Q491P | A | 1.1e-05 | 1.1e-05 | 8.4e-06 | 1.013E-05 | 1.501E-06 | 8.400E-06 | 1.100E-05 |
| 9816 | 491 | Q491S | A | 1e-05 | 6.1e-06 | 7e-06 | 7.700E-06 | 2.042E-06 | 6.100E-06 | 1.000E-05 |
| 9817 | 491 | Q491T | A | 9.9e-06 | 7.5e-06 | 8.7e-06 | 8.700E-06 | 1.200E-06 | 7.500E-06 | 9.900E-06 |
| 9818 | 491 | Q491W | A | 5.5e-06 | 7.6e-06 | 6.3e-06 | 6.467E-06 | 1.060E-06 | 5.500E-06 | 7.600E-06 |
| 9819 | 491 | Q491Y | A | 9.7e-06 | 2.8e-06 | 4.2e-06 | 5.567E-06 | 3.647E-06 | 2.800E-06 | 9.700E-06 |
| 9820 | 491 | Q491V | A | 8.2e-06 | 4.5e-06 | 9.5e-06 | 7.400E-06 | 2.594E-06 | 4.500E-06 | 9.500E-06 |
| 9821 | 492 | L492A | A | 6.8e-06 | 8.4e-06 | 7.4e-06 | 7.533E-06 | 8.083E-07 | 6.800E-06 | 8.400E-06 |
| 9822 | 492 | L492R | A | 1.7e-05 | 6.9e-06 | 5.4e-06 | 9.767E-06 | 6.309E-06 | 5.400E-06 | 1.700E-05 |
| 9823 | 492 | L492N | A | 1.2e-05 | 8.5e-06 | 8.3e-06 | 9.600E-06 | 2.081E-06 | 8.300E-06 | 1.200E-05 |
| 9824 | 492 | L492D | A | 6.2e-06 | 1.1e-05 | 6.2e-06 | 7.800E-06 | 2.771E-06 | 6.200E-06 | 1.100E-05 |
| 9825 | 492 | L492C | A | 9e-06 | 8.4e-06 | 6.7e-06 | 8.033E-06 | 1.193E-06 | 6.700E-06 | 9.000E-06 |
| 9826 | 492 | L492E | A | 5.6e-06 | 6.9e-06 | 7.1e-06 | 6.533E-06 | 8.145E-07 | 5.600E-06 | 7.100E-06 |
| 9827 | 492 | L492Q | A | 7.6e-06 | 1.1e-05 | 3.2e-06 | 7.267E-06 | 3.911E-06 | 3.200E-06 | 1.100E-05 |
| 9828 | 492 | L492G | A | 7.8e-06 | 1.1e-05 | 1.3e-05 | 1.060E-05 | 2.623E-06 | 7.800E-06 | 1.300E-05 |
| 9829 | 492 | L492H | A | 1.1e-05 | 6.6e-06 | 6.6e-06 | 8.067E-06 | 2.540E-06 | 6.600E-06 | 1.100E-05 |
| 9830 | 492 | L492I | A | 6e-06 | 1.3e-05 | 1e-05 | 9.667E-06 | 3.512E-06 | 6.000E-06 | 1.300E-05 |
| 9831 | 492 | L492L | A | 5.5e-06 | 1e-05 | 7.9e-06 | 7.800E-06 | 2.252E-06 | 5.500E-06 | 1.000E-05 |
| 9832 | 492 | L492K | A | 1.3e-05 | 8.5e-06 | 6.4e-06 | 9.300E-06 | 3.372E-06 | 6.400E-06 | 1.300E-05 |
| 9833 | 492 | L492M | A | 8.5e-06 | 1e-05 | 9.8e-06 | 9.433E-06 | 8.145E-07 | 8.500E-06 | 1.000E-05 |
| 9834 | 492 | L492F | A | 8.2e-06 | 7.9e-06 | 1.1e-05 | 9.033E-06 | 1.710E-06 | 7.900E-06 | 1.100E-05 |
| 9835 | 492 | L492P | A | 8.7e-06 | 8.2e-06 | 6.3e-06 | 7.733E-06 | 1.266E-06 | 6.300E-06 | 8.700E-06 |
| 9836 | 492 | L492S | A | 4.6e-06 | 7.3e-06 | 6.1e-06 | 6.000E-06 | 1.353E-06 | 4.600E-06 | 7.300E-06 |
| 9837 | 492 | L492T | A | 9.4e-06 | 1.8e-05 | 8.3e-06 | 1.190E-05 | 5.311E-06 | 8.300E-06 | 1.800E-05 |
| 9838 | 492 | L492W | A | 5.6e-06 | 5.2e-06 | 5.3e-06 | 5.367E-06 | 2.082E-07 | 5.200E-06 | 5.600E-06 |
| 9839 | 492 | L492Y | A | 1.1e-05 | 1.2e-05 | 8.3e-06 | 1.043E-05 | 1.914E-06 | 8.300E-06 | 1.200E-05 |
| 9840 | 492 | L492V | A | 9.8e-06 | 6.8e-06 | 8.6e-06 | 8.400E-06 | 1.510E-06 | 6.800E-06 | 9.800E-06 |
| 9841 | 493 | C493A | A | 8.7e-06 | 6.6e-06 | 9.3e-06 | 8.200E-06 | 1.418E-06 | 6.600E-06 | 9.300E-06 |
| 9842 | 493 | C493R | A | 5.7e-06 | 7.1e-06 | 5.6e-06 | 6.133E-06 | 8.386E-07 | 5.600E-06 | 7.100E-06 |
| 9843 | 493 | C493N | A | 1.3e-05 | 1.4e-05 | 6.8e-06 | 1.127E-05 | 3.900E-06 | 6.800E-06 | 1.400E-05 |
| 9844 | 493 | C493D | A | 9.4e-06 | 5.9e-06 | 9.5e-06 | 8.267E-06 | 2.050E-06 | 5.900E-06 | 9.500E-06 |
| 9845 | 493 | C493C | A | 1.3e-05 | 8.5e-06 | 1.1e-05 | 1.083E-05 | 2.255E-06 | 8.500E-06 | 1.300E-05 |
| 9846 | 493 | C493E | A | 1.4e-05 | 1.3e-05 | 6.6e-06 | 1.120E-05 | 4.015E-06 | 6.600E-06 | 1.400E-05 |
| 9847 | 493 | C493Q | A | 9.8e-06 | 8.7e-06 | 9.3e-06 | 9.267E-06 | 5.508E-07 | 8.700E-06 | 9.800E-06 |
| 9848 | 493 | C493G | A | 5.9e-06 | 1.2e-05 | 1.4e-05 | 1.063E-05 | 4.219E-06 | 5.900E-06 | 1.400E-05 |
| 9849 | 493 | C493H | A | 6.4e-06 | 1.1e-05 | 6.5e-06 | 7.967E-06 | 2.627E-06 | 6.400E-06 | 1.100E-05 |
| 9850 | 493 | C493I | A | 9.7e-06 | 5.9e-06 | 5.8e-06 | 7.133E-06 | 2.223E-06 | 5.800E-06 | 9.700E-06 |
| 9851 | 493 | C493L | A | 7.6e-06 | 1.3e-05 | 8.7e-06 | 9.767E-06 | 2.854E-06 | 7.600E-06 | 1.300E-05 |
| 9852 | 493 | C493K | A | 1e-05 | 6.1e-06 | 5.6e-06 | 7.233E-06 | 2.409E-06 | 5.600E-06 | 1.000E-05 |
| 9853 | 493 | C493M | A | 1.3e-05 | 7.6e-06 | 6.6e-06 | 9.067E-06 | 3.443E-06 | 6.600E-06 | 1.300E-05 |
| 9854 | 493 | C493F | A | 7.1e-06 | 5.3e-06 | 9.5e-06 | 7.300E-06 | 2.107E-06 | 5.300E-06 | 9.500E-06 |
| 9855 | 493 | C493P | A | 9.9e-06 | 6.4e-06 | 6.6e-06 | 7.633E-06 | 1.966E-06 | 6.400E-06 | 9.900E-06 |
| 9856 | 493 | C493S | A | 9.7e-06 | 1.9e-05 | 9.5e-06 | 1.273E-05 | 5.428E-06 | 9.500E-06 | 1.900E-05 |
| 9857 | 493 | C493T | A | 7e-06 | 9.3e-06 | 8.2e-06 | 8.167E-06 | 1.150E-06 | 7.000E-06 | 9.300E-06 |
| 9858 | 493 | C493W | A | 4.6e-06 | 7.4e-06 | 6.5e-06 | 6.167E-06 | 1.429E-06 | 4.600E-06 | 7.400E-06 |
| 9859 | 493 | C493Y | A | 7.7e-06 | 7.4e-06 | 9.2e-06 | 8.100E-06 | 9.644E-07 | 7.400E-06 | 9.200E-06 |

| | | | | | | | | | | |
|---|---|---|---|---|---|---|---|---|---|---|
| 9860 | 493 | C493V | A | 1e-05 | 6e-06 | 1.5e-05 | 1.033E-05 | 4.509E-06 | 6.000E-06 | 1.500E-05 |
| 9861 | 494 | A494A | A | 1.3e-05 | 6.4e-06 | 6.9e-06 | 8.767E-06 | 3.675E-06 | 6.400E-06 | 1.300E-05 |
| 9862 | 494 | A494R | A | 4.7e-06 | 1.1e-05 | 7e-06 | 7.567E-06 | 3.188E-06 | 4.700E-06 | 1.100E-05 |
| 9863 | 494 | A494N | A | 1.4e-05 | 1.3e-05 | 6.7e-06 | 1.123E-05 | 3.958E-06 | 6.700E-06 | 1.400E-05 |
| 9864 | 494 | A494D | A | 5.8e-06 | 7.3e-06 | 8.5e-06 | 7.200E-06 | 1.353E-06 | 5.800E-06 | 8.500E-06 |
| 9865 | 494 | A494C | A | 5.7e-06 | 1.5e-05 | 5.2e-06 | 8.633E-06 | 5.519E-06 | 5.200E-06 | 1.500E-05 |
| 9866 | 494 | A494E | A | 1e-05 | 6.2e-06 | 8e-06 | 8.067E-06 | 1.901E-06 | 6.200E-06 | 1.000E-05 |
| 9867 | 494 | A494Q | A | 6.2e-06 | 8.2e-06 | 4.5e-06 | 6.300E-06 | 1.852E-06 | 4.500E-06 | 8.200E-06 |
| 9868 | 494 | A494G | A | 8.4e-06 | 1.2e-05 | 9.3e-06 | 9.900E-06 | 1.873E-06 | 8.400E-06 | 1.200E-05 |
| 9869 | 494 | A494H | A | 1e-05 | 9.7e-06 | 9.9e-06 | 9.867E-06 | 1.528E-07 | 9.700E-06 | 1.000E-05 |
| 9870 | 494 | A494I | A | 1.4e-05 | 1.3e-05 | 8.9e-06 | 1.197E-05 | 2.702E-06 | 8.900E-06 | 1.400E-05 |
| 9871 | 494 | A494L | A | 9.8e-06 | 1.3e-05 | 1.4e-05 | 1.227E-05 | 2.194E-06 | 9.800E-06 | 1.400E-05 |
| 9872 | 494 | A494K | A | 6.6e-06 | 6.6e-06 | 1.1e-05 | 8.067E-06 | 2.540E-06 | 6.600E-06 | 1.100E-05 |
| 9873 | 494 | A494M | A | 8.8e-06 | 9.9e-06 | 5.1e-06 | 7.933E-06 | 2.515E-06 | 5.100E-06 | 9.900E-06 |
| 9874 | 494 | A494F | A | 7.7e-06 | 7e-06 | 5.6e-06 | 6.767E-06 | 1.069E-06 | 5.600E-06 | 7.700E-06 |
| 9875 | 494 | A494P | A | 7.5e-06 | 1.1e-05 | 7.8e-06 | 8.767E-06 | 1.940E-06 | 7.500E-06 | 1.100E-05 |
| 9876 | 494 | A494S | A | 1e-05 | 9.6e-06 | 7e-06 | 8.867E-06 | 1.629E-06 | 7.000E-06 | 1.000E-05 |
| 9877 | 494 | A494T | A | 8.8e-06 | 1e-05 | 1.1e-05 | 9.933E-06 | 1.102E-06 | 8.800E-06 | 1.100E-05 |
| 9878 | 494 | A494W | A | 9.9e-06 | 7.1e-06 | 8.6e-06 | 8.533E-06 | 1.401E-06 | 7.100E-06 | 9.900E-06 |
| 9879 | 494 | A494Y | A | 8.9e-06 | 8.8e-06 | 1.3e-05 | 1.023E-05 | 2.397E-06 | 8.800E-06 | 1.300E-05 |
| 9880 | 494 | A494V | A | 6.9e-06 | 6.6e-06 | 7.1e-06 | 6.867E-06 | 2.517E-07 | 6.600E-06 | 7.100E-06 |
| 9881 | 495 | R495A | A | 4e-06 | 9.6e-06 | 9.1e-06 | 7.567E-06 | 3.099E-06 | 4.000E-06 | 9.600E-06 |
| 9882 | 495 | R495R | A | 9.2e-06 | 9.7e-06 | 5.9e-06 | 8.267E-06 | 2.065E-06 | 5.900E-06 | 9.700E-06 |
| 9883 | 495 | R495N | A | 1.4e-05 | 1e-05 | 9.6e-06 | 1.120E-05 | 2.433E-06 | 9.600E-06 | 1.400E-05 |
| 9884 | 495 | R495D | A | 8.3e-06 | 6.8e-06 | 5.9e-06 | 7.000E-06 | 1.212E-06 | 5.900E-06 | 8.300E-06 |
| 9885 | 495 | R495C | A | 7.2e-06 | 4.3e-06 | 9.6e-06 | 7.033E-06 | 2.654E-06 | 4.300E-06 | 9.600E-06 |
| 9886 | 495 | R495E | A | 8.3e-06 | 1.5e-05 | 8.8e-06 | 1.070E-05 | 3.732E-06 | 8.300E-06 | 1.500E-05 |
| 9887 | 495 | R495Q | A | 7.3e-06 | 6.5e-06 | 4.3e-06 | 6.033E-06 | 1.553E-06 | 4.300E-06 | 7.300E-06 |
| 9888 | 495 | R495G | A | 9.3e-06 | 4.2e-06 | 8.2e-06 | 7.233E-06 | 2.684E-06 | 4.200E-06 | 9.300E-06 |
| 9889 | 495 | R495H | A | 8e-06 | 9.4e-06 | 1.2e-05 | 9.800E-06 | 2.030E-06 | 8.000E-06 | 1.200E-05 |
| 9890 | 495 | R495I | A | 9.3e-06 | 8.8e-06 | 6.7e-06 | 8.267E-06 | 1.380E-06 | 6.700E-06 | 9.300E-06 |
| 9891 | 495 | R495L | A | 6.8e-06 | 1.1e-05 | 6.3e-06 | 8.033E-06 | 2.581E-06 | 6.300E-06 | 1.100E-05 |
| 9892 | 495 | R495K | A | 9e-06 | 1.1e-05 | 6.2e-06 | 8.733E-06 | 2.411E-06 | 6.200E-06 | 1.100E-05 |
| 9893 | 495 | R495M | A | 9e-06 | 9.8e-06 | 9.3e-06 | 9.367E-06 | 4.041E-07 | 9.000E-06 | 9.800E-06 |
| 9894 | 495 | R495F | A | 7.5e-06 | 9.9e-06 | 1.1e-05 | 9.467E-06 | 1.790E-06 | 7.500E-06 | 1.100E-05 |
| 9895 | 495 | R495P | A | 5e-06 | 9.9e-06 | 9e-06 | 7.967E-06 | 2.608E-06 | 5.000E-06 | 9.900E-06 |
| 9896 | 495 | R495S | A | 5.8e-06 | 8.6e-06 | 5.1e-06 | 6.500E-06 | 1.852E-06 | 5.100E-06 | 8.600E-06 |
| 9897 | 495 | R495T | A | 6.5e-06 | 6e-06 | 1.1e-05 | 7.833E-06 | 2.754E-06 | 6.000E-06 | 1.100E-05 |
| 9898 | 495 | R495W | A | 1.2e-05 | 1.2e-05 | 7e-06 | 1.033E-05 | 2.887E-06 | 7.000E-06 | 1.200E-05 |
| 9899 | 495 | R495Y | A | 9.5e-06 | 5.7e-06 | 7.5e-06 | 7.567E-06 | 1.901E-06 | 5.700E-06 | 9.500E-06 |
| 9900 | 495 | R495V | A | 5.1e-06 | 8e-06 | 6e-06 | 6.367E-06 | 1.484E-06 | 5.100E-06 | 8.000E-06 |
| 9901 | 496 | G496A | A | 6.6e-06 | 9.5e-06 | 6.7e-06 | 7.600E-06 | 1.646E-06 | 6.600E-06 | 9.500E-06 |
| 9902 | 496 | G496R | A | 6.7e-06 | 5.2e-06 | 8.9e-06 | 6.933E-06 | 1.861E-06 | 5.200E-06 | 8.900E-06 |
| 9903 | 496 | G496N | A | 4.9e-06 | 1e-05 | 7.5e-06 | 7.467E-06 | 2.550E-06 | 4.900E-06 | 1.000E-05 |
| 9904 | 496 | G496D | A | 6.4e-06 | 5.8e-06 | 6.2e-06 | 6.133E-06 | 3.055E-07 | 5.800E-06 | 6.400E-06 |
| 9905 | 496 | G496C | A | 1.3e-05 | 7.8e-06 | 8.1e-06 | 9.633E-06 | 2.919E-06 | 7.800E-06 | 1.300E-05 |
| 9906 | 496 | G496E | A | 7.6e-06 | 5.2e-06 | 8.5e-06 | 7.100E-06 | 1.706E-06 | 5.200E-06 | 8.500E-06 |
| 9907 | 496 | G496Q | A | 8.4e-06 | 3.7e-06 | 8.9e-06 | 7.000E-06 | 2.869E-06 | 3.700E-06 | 8.900E-06 |
| 9908 | 496 | G496G | A | 5.3e-06 | 8.2e-06 | 1e-05 | 7.833E-06 | 2.371E-06 | 5.300E-06 | 1.000E-05 |
| 9909 | 496 | G496H | A | 7.9e-06 | 5.8e-06 | 5.8e-06 | 6.500E-06 | 1.212E-06 | 5.800E-06 | 7.900E-06 |
| 9910 | 496 | G496I | A | 1.2e-05 | 8e-06 | 6.6e-06 | 8.867E-06 | 2.802E-06 | 6.600E-06 | 1.200E-05 |
| 9911 | 496 | G496L | A | 9e-06 | 9e-06 | 4.5e-06 | 7.500E-06 | 2.598E-06 | 4.500E-06 | 9.000E-06 |
| 9912 | 496 | G496K | A | 8.7e-06 | 5.2e-06 | 7.5e-06 | 7.133E-06 | 1.779E-06 | 5.200E-06 | 8.700E-06 |
| 9913 | 496 | G496M | A | 9.4e-06 | 1e-05 | 9.9e-06 | 9.767E-06 | 3.215E-07 | 9.400E-06 | 1.000E-05 |
| 9914 | 496 | G496F | A | 9.9e-06 | 1.4e-05 | 9.9e-06 | 1.127E-05 | 2.367E-06 | 9.900E-06 | 1.400E-05 |
| 9915 | 496 | G496P | A | 9.8e-06 | 9e-06 | 1.2e-05 | 1.027E-05 | 1.553E-06 | 9.000E-06 | 1.200E-05 |
| 9916 | 496 | G496S | A | 9.1e-06 | 1.7e-05 | 5.7e-06 | 1.060E-05 | 5.797E-06 | 5.700E-06 | 1.700E-05 |
| 9917 | 496 | G496T | A | 7.3e-06 | 7.2e-06 | 1.1e-05 | 8.500E-06 | 2.166E-06 | 7.200E-06 | 1.100E-05 |
| 9918 | 496 | G496W | A | 8.8e-06 | 1.2e-05 | 5.1e-06 | 8.633E-06 | 3.453E-06 | 5.100E-06 | 1.200E-05 |
| 9919 | 496 | G496Y | A | 1.4e-05 | 4.9e-06 | 9.4e-06 | 9.433E-06 | 4.550E-06 | 4.900E-06 | 1.400E-05 |
| 9920 | 496 | G496V | A | 8.1e-06 | 9.2e-06 | 7.3e-06 | 8.200E-06 | 9.539E-07 | 7.300E-06 | 9.200E-06 |
| 9921 | 497 | H497A | A | 8.9e-06 | 1e-05 | 3.3e-06 | 7.400E-06 | 3.593E-06 | 3.300E-06 | 1.000E-05 |
| 9922 | 497 | H497R | A | 8.9e-06 | 9.1e-06 | 1e-05 | 9.333E-06 | 5.859E-07 | 8.900E-06 | 1.000E-05 |
| 9923 | 497 | H497N | A | 5.8e-06 | 7.8e-06 | 7e-06 | 6.867E-06 | 1.007E-06 | 5.800E-06 | 7.800E-06 |
| 9924 | 497 | H497D | A | 1.1e-05 | 6.6e-06 | 7.3e-06 | 8.300E-06 | 2.364E-06 | 6.600E-06 | 1.100E-05 |
| 9925 | 497 | H497C | A | 9.3e-06 | 1.3e-05 | 5.7e-06 | 9.333E-06 | 3.650E-06 | 5.700E-06 | 1.300E-05 |
| 9926 | 497 | H497E | A | 5.4e-06 | 8.7e-06 | 1e-05 | 8.033E-06 | 2.371E-06 | 5.400E-06 | 1.000E-05 |
| 9927 | 497 | H497Q | A | 9.6e-06 | 9.4e-06 | 8.6e-06 | 9.200E-06 | 5.292E-07 | 8.600E-06 | 9.600E-06 |
| 9928 | 497 | H497G | A | 6.5e-06 | 1.2e-05 | 1.6e-05 | 1.150E-05 | 4.770E-06 | 6.500E-06 | 1.600E-05 |
| 9929 | 497 | H497H | A | 1e-05 | 1.3e-05 | 8.9e-06 | 1.063E-05 | 2.122E-06 | 8.900E-06 | 1.300E-05 |
| 9930 | 497 | H497I | A | 6.2e-06 | 5.2e-06 | 8.9e-06 | 6.767E-06 | 1.914E-06 | 5.200E-06 | 8.900E-06 |
| 9931 | 497 | H497L | A | 4.3e-06 | 8.3e-06 | 4.6e-06 | 5.733E-06 | 2.228E-06 | 4.300E-06 | 8.300E-06 |
| 9932 | 497 | H497K | A | 1e-05 | 6.6e-06 | 1e-05 | 8.867E-06 | 1.963E-06 | 6.600E-06 | 1.000E-05 |
| 9933 | 497 | H497M | A | 8.1e-06 | 6.4e-06 | 6.9e-06 | 7.133E-06 | 8.737E-07 | 6.400E-06 | 8.100E-06 |
| 9934 | 497 | H497F | A | 4.9e-06 | 4.8e-06 | 1e-05 | 6.567E-06 | 2.974E-06 | 4.800E-06 | 1.000E-05 |
| 9935 | 497 | H497P | A | 8.1e-06 | 1e-05 | 7.2e-06 | 8.433E-06 | 1.429E-06 | 7.200E-06 | 1.000E-05 |
| 9936 | 497 | H497S | A | 3.9e-06 | 1.2e-05 | 1e-05 | 8.633E-06 | 4.219E-06 | 3.900E-06 | 1.200E-05 |
| 9937 | 497 | H497T | A | 6.5e-06 | 6.6e-06 | 1e-05 | 7.700E-06 | 1.992E-06 | 6.500E-06 | 1.000E-05 |
| 9938 | 497 | H497W | A | 7.8e-06 | 9.3e-06 | 7.9e-06 | 8.333E-06 | 8.386E-07 | 7.800E-06 | 9.300E-06 |
| 9939 | 497 | H497Y | A | 8.8e-06 | 6.6e-06 | 7.8e-06 | 7.733E-06 | 1.102E-06 | 6.600E-06 | 8.800E-06 |
| 9940 | 497 | H497V | A | 7.5e-06 | 9.3e-06 | 5.7e-06 | 7.500E-06 | 1.800E-06 | 5.700E-06 | 9.300E-06 |
| 9941 | 498 | C498A | A | 1.3e-05 | 9.6e-06 | 5.9e-06 | 9.500E-06 | 3.551E-06 | 5.900E-06 | 1.300E-05 |
| 9942 | 498 | C498R | A | 6.7e-06 | 6.6e-06 | 7.3e-06 | 6.867E-06 | 3.786E-07 | 6.600E-06 | 7.300E-06 |
| 9943 | 498 | C498N | A | 5.1e-06 | 7.4e-06 | 1.3e-05 | 8.500E-06 | 4.063E-06 | 5.100E-06 | 1.300E-05 |
| 9944 | 498 | C498D | A | 6.4e-06 | 8.7e-06 | 5.4e-06 | 6.833E-06 | 1.692E-06 | 5.400E-06 | 8.700E-06 |

| | | | | | | | | | | |
|---|---|---|---|---|---|---|---|---|---|---|
| 9945 | 498 | C498C | A | 1e-05 | 1e-05 | 4.9e-06 | 8.300E-06 | 2.944E-06 | 4.900E-06 | 1.000E-05 |
| 9946 | 498 | C498E | A | 5e-06 | 5.6e-06 | 6.9e-06 | 5.833E-06 | 9.713E-07 | 5.000E-06 | 6.900E-06 |
| 9947 | 498 | C498Q | A | 1e-05 | 6.3e-06 | 6.7e-06 | 7.667E-06 | 2.031E-06 | 6.300E-06 | 1.000E-05 |
| 9948 | 498 | C498G | A | 7.6e-06 | 9.4e-06 | 1.2e-05 | 9.667E-06 | 2.212E-06 | 7.600E-06 | 1.200E-05 |
| 9949 | 498 | C498H | A | 1.1e-05 | 6.3e-06 | 8.4e-06 | 8.567E-06 | 2.354E-06 | 6.300E-06 | 1.100E-05 |
| 9950 | 498 | C498I | A | 5.5e-06 | 1.1e-05 | 1.3e-05 | 9.833E-06 | 3.884E-06 | 5.500E-06 | 1.300E-05 |
| 9951 | 498 | C498L | A | 6.4e-06 | 4.3e-06 | 4e-06 | 4.900E-06 | 1.308E-06 | 4.000E-06 | 6.400E-06 |
| 9952 | 498 | C498K | A | 1.1e-05 | 7.8e-06 | 5.4e-06 | 8.067E-06 | 2.810E-06 | 5.400E-06 | 1.100E-05 |
| 9953 | 498 | C498M | A | 1.3e-05 | 9.2e-06 | 5.1e-06 | 9.100E-06 | 3.951E-06 | 5.100E-06 | 1.300E-05 |
| 9954 | 498 | C498F | A | 1.9e-05 | 5.1e-06 | 7.1e-06 | 1.040E-05 | 7.515E-06 | 5.100E-06 | 1.900E-05 |
| 9955 | 498 | C498P | A | 9.1e-06 | 8.3e-06 | 7.4e-06 | 8.267E-06 | 8.505E-07 | 7.400E-06 | 9.100E-06 |
| 9956 | 498 | C498S | A | 4.6e-06 | 1.2e-05 | 7.9e-06 | 8.167E-06 | 3.707E-06 | 4.600E-06 | 1.200E-05 |
| 9957 | 498 | C498T | A | 1.2e-05 | 4.8e-06 | 1.3e-05 | 9.933E-06 | 4.474E-06 | 4.800E-06 | 1.300E-05 |
| 9958 | 498 | C498W | A | 9.9e-06 | 4.3e-06 | 4.7e-06 | 6.300E-06 | 3.124E-06 | 4.300E-06 | 9.900E-06 |
| 9959 | 498 | C498Y | A | 8e-06 | 7.9e-06 | 4e-06 | 6.633E-06 | 2.281E-06 | 4.000E-06 | 8.000E-06 |
| 9960 | 498 | C498V | A | 5.6e-06 | 7e-06 | 9.8e-06 | 7.467E-06 | 2.139E-06 | 5.600E-06 | 9.800E-06 |
| 9961 | 499 | W499A | A | 8.1e-06 | 4.9e-06 | 1e-05 | 7.667E-06 | 2.577E-06 | 4.900E-06 | 1.000E-05 |
| 9962 | 499 | W499R | A | 6.3e-06 | 9.8e-06 | 8.7e-06 | 8.267E-06 | 1.790E-06 | 6.300E-06 | 9.800E-06 |
| 9963 | 499 | W499N | A | 9.5e-06 | 1e-05 | 8.4e-06 | 9.300E-06 | 8.185E-07 | 8.400E-06 | 1.000E-05 |
| 9964 | 499 | W499D | A | 1.3e-05 | 5.1e-06 | 5.1e-06 | 7.733E-06 | 4.561E-06 | 5.100E-06 | 1.300E-05 |
| 9965 | 499 | W499C | A | 9.6e-06 | 7.9e-06 | 7.1e-06 | 8.200E-06 | 1.277E-06 | 7.100E-06 | 9.600E-06 |
| 9966 | 499 | W499E | A | 6.3e-06 | 8.5e-06 | 7e-06 | 7.267E-06 | 1.124E-06 | 6.300E-06 | 8.500E-06 |
| 9967 | 499 | W499Q | A | 5.6e-06 | 1e-05 | 7.2e-06 | 7.600E-06 | 2.227E-06 | 5.600E-06 | 1.000E-05 |
| 9968 | 499 | W499G | A | 1.3e-05 | 6.5e-06 | 8.5e-06 | 9.333E-06 | 3.329E-06 | 6.500E-06 | 1.300E-05 |
| 9969 | 499 | W499H | A | 5.7e-06 | 9e-06 | 1.2e-05 | 8.900E-06 | 3.151E-06 | 5.700E-06 | 1.200E-05 |
| 9970 | 499 | W499I | A | 6.2e-06 | 1e-05 | 9.5e-06 | 8.567E-06 | 2.065E-06 | 6.200E-06 | 1.000E-05 |
| 9971 | 499 | W499L | A | 9e-06 | 9.1e-06 | 6.9e-06 | 8.333E-06 | 1.242E-06 | 6.900E-06 | 9.100E-06 |
| 9972 | 499 | W499K | A | 1.1e-05 | 9.8e-06 | 1.4e-05 | 1.160E-05 | 2.163E-06 | 9.800E-06 | 1.400E-05 |
| 9973 | 499 | W499M | A | 6.7e-06 | 5.3e-06 | 8.1e-06 | 6.700E-06 | 1.400E-06 | 5.300E-06 | 8.100E-06 |
| 9974 | 499 | W499F | A | 5.2e-06 | 8e-06 | 7.4e-06 | 6.867E-06 | 1.474E-06 | 5.200E-06 | 8.000E-06 |
| 9975 | 499 | W499P | A | 8.8e-06 | 1.5e-05 | 6.5e-06 | 1.010E-05 | 4.397E-06 | 6.500E-06 | 1.500E-05 |
| 9976 | 499 | W499S | A | 7.2e-06 | 5e-06 | 8.2e-06 | 6.800E-06 | 1.637E-06 | 5.000E-06 | 8.200E-06 |
| 9977 | 499 | W499T | A | 8.2e-06 | 8.1e-06 | 1.5e-05 | 1.043E-05 | 3.955E-06 | 8.100E-06 | 1.500E-05 |
| 9978 | 499 | W499W | A | 1.9e-05 | 9.3e-06 | 6.5e-06 | 1.160E-05 | 6.560E-06 | 6.500E-06 | 1.900E-05 |
| 9979 | 499 | W499Y | A | 6.5e-06 | 6.2e-06 | 8.5e-06 | 7.067E-06 | 1.250E-06 | 6.200E-06 | 8.500E-06 |
| 9980 | 499 | W499V | A | 4.8e-06 | 5.9e-06 | 1.6e-05 | 8.900E-06 | 6.173E-06 | 4.800E-06 | 1.600E-05 |
| 9981 | 500 | G500A | A | 4.6e-06 | 9e-06 | 6.2e-06 | 6.600E-06 | 2.227E-06 | 4.600E-06 | 9.000E-06 |
| 9982 | 500 | G500R | A | 7.7e-06 | 4.6e-06 | 6.7e-06 | 6.333E-06 | 1.582E-06 | 4.600E-06 | 7.700E-06 |
| 9983 | 500 | G500N | A | 6.8e-06 | 8.1e-06 | 8.8e-06 | 7.900E-06 | 1.015E-06 | 6.800E-06 | 8.800E-06 |
| 9984 | 500 | G500D | A | 1.2e-05 | 8.9e-06 | 4.6e-06 | 8.500E-06 | 3.716E-06 | 4.600E-06 | 1.200E-05 |
| 9985 | 500 | G500C | A | 1.4e-05 | 8.7e-06 | 9.7e-06 | 1.080E-05 | 2.816E-06 | 8.700E-06 | 1.400E-05 |
| 9986 | 500 | G500E | A | 1.2e-05 | 7.8e-06 | 7.6e-06 | 9.133E-06 | 2.485E-06 | 7.600E-06 | 1.200E-05 |
| 9987 | 500 | G500Q | A | 4.2e-06 | 5.3e-06 | 8.7e-06 | 6.067E-06 | 2.346E-06 | 4.200E-06 | 8.700E-06 |
| 9988 | 500 | G500G | A | 7.9e-06 | 6.1e-06 | 7.9e-06 | 7.300E-06 | 1.039E-06 | 6.100E-06 | 7.900E-06 |
| 9989 | 500 | G500H | A | 7.5e-06 | 9.2e-06 | 6.4e-06 | 7.700E-06 | 1.411E-06 | 6.400E-06 | 9.200E-06 |
| 9990 | 500 | G500I | A | 7.6e-06 | 6.6e-06 | 7.2e-06 | 7.133E-06 | 5.033E-07 | 6.600E-06 | 7.600E-06 |
| 9991 | 500 | G500L | A | 8.4e-06 | 7.7e-06 | 6.2e-06 | 7.433E-06 | 1.124E-06 | 6.200E-06 | 8.400E-06 |
| 9992 | 500 | G500K | A | 6.2e-06 | 5.7e-06 | 8.5e-06 | 6.800E-06 | 1.493E-06 | 5.700E-06 | 8.500E-06 |
| 9993 | 500 | G500M | A | 3.1e-06 | 9.9e-06 | 9.3e-06 | 7.433E-06 | 3.765E-06 | 3.100E-06 | 9.900E-06 |
| 9994 | 500 | G500F | A | 9.3e-06 | 8.3e-06 | 1.1e-05 | 9.533E-06 | 1.365E-06 | 8.300E-06 | 1.100E-05 |
| 9995 | 500 | G500P | A | 9.5e-06 | 6.7e-06 | 4.8e-06 | 7.000E-06 | 2.364E-06 | 4.800E-06 | 9.500E-06 |
| 9996 | 500 | G500S | A | 7.2e-06 | 9.9e-06 | 4.7e-06 | 7.267E-06 | 2.601E-06 | 4.700E-06 | 9.900E-06 |
| 9997 | 500 | G500T | A | 9.9e-06 | 8.6e-06 | 5e-06 | 7.833E-06 | 2.538E-06 | 5.000E-06 | 9.900E-06 |
| 9998 | 500 | G500W | A | 1.3e-05 | 9.9e-06 | 5.3e-06 | 9.400E-06 | 3.874E-06 | 5.300E-06 | 1.300E-05 |
| 9999 | 500 | G500Y | A | 7.6e-06 | 4.9e-06 | 8.3e-06 | 6.933E-06 | 1.795E-06 | 4.900E-06 | 8.300E-06 |
| 10000 | 500 | G500V | A | 6.1e-06 | 6.2e-06 | 1e-05 | 7.433E-06 | 2.223E-06 | 6.100E-06 | 1.000E-05 |
| 10001 | 501 | P501A | A | 6.8e-06 | 1.1e-05 | 5.4e-06 | 7.733E-06 | 2.914E-06 | 5.400E-06 | 1.100E-05 |
| 10002 | 501 | P501R | A | 1.4e-05 | 8.2e-06 | 8.9e-06 | 1.037E-05 | 3.166E-06 | 8.200E-06 | 1.400E-05 |
| 10003 | 501 | P501N | A | 4.6e-06 | 6e-06 | 1.3e-05 | 7.867E-06 | 4.500E-06 | 4.600E-06 | 1.300E-05 |
| 10004 | 501 | P501D | A | 6.9e-06 | 9.8e-06 | 1.1e-05 | 9.233E-06 | 2.108E-06 | 6.900E-06 | 1.100E-05 |
| 10005 | 501 | P501C | A | 1.3e-05 | 8.8e-06 | 6.3e-06 | 9.367E-06 | 3.386E-06 | 6.300E-06 | 1.300E-05 |
| 10006 | 501 | P501E | A | 1e-05 | 6.5e-06 | 4.7e-06 | 7.067E-06 | 2.695E-06 | 4.700E-06 | 1.000E-05 |
| 10007 | 501 | P501Q | A | 1.4e-05 | 9.1e-06 | 9.7e-06 | 1.093E-05 | 2.673E-06 | 9.100E-06 | 1.400E-05 |
| 10008 | 501 | P501G | A | 7.4e-06 | 9.4e-06 | 8.4e-06 | 8.400E-06 | 1.000E-06 | 7.400E-06 | 9.400E-06 |
| 10009 | 501 | P501H | A | 5.6e-06 | 5.4e-06 | 4.9e-06 | 5.300E-06 | 3.606E-07 | 4.900E-06 | 5.600E-06 |
| 10010 | 501 | P501I | A | 4.8e-06 | 8.7e-06 | 9.8e-06 | 7.767E-06 | 2.627E-06 | 4.800E-06 | 9.800E-06 |
| 10011 | 501 | P501L | A | 9e-06 | 6.1e-06 | 6.7e-06 | 7.267E-06 | 1.531E-06 | 6.100E-06 | 9.000E-06 |
| 10012 | 501 | P501K | A | 8e-06 | 5.8e-06 | 4.7e-06 | 6.167E-06 | 1.680E-06 | 4.700E-06 | 8.000E-06 |
| 10013 | 501 | P501M | A | 7.6e-06 | 6.2e-06 | 1.2e-05 | 8.600E-06 | 3.027E-06 | 6.200E-06 | 1.200E-05 |
| 10014 | 501 | P501F | A | 7.4e-06 | 7.4e-06 | 1e-05 | 8.267E-06 | 1.501E-06 | 7.400E-06 | 1.000E-05 |
| 10015 | 501 | P501P | A | 1.3e-05 | 1.1e-05 | 8.8e-06 | 1.093E-05 | 2.101E-06 | 8.800E-06 | 1.300E-05 |
| 10016 | 501 | P501S | A | 9.5e-06 | 1e-05 | 8e-06 | 9.167E-06 | 1.041E-06 | 8.000E-06 | 1.000E-05 |
| 10017 | 501 | P501T | A | 5.3e-06 | 1.1e-05 | 7.1e-06 | 7.800E-06 | 2.914E-06 | 5.300E-06 | 1.100E-05 |
| 10018 | 501 | P501W | A | 1.1e-05 | 9.4e-06 | 6e-06 | 8.800E-06 | 2.553E-06 | 6.000E-06 | 1.100E-05 |
| 10019 | 501 | P501Y | A | 6.1e-06 | 4.6e-06 | 9.6e-06 | 6.767E-06 | 2.566E-06 | 4.600E-06 | 9.600E-06 |
| 10020 | 501 | P501V | A | 9.9e-06 | 7.1e-06 | 5e-06 | 7.333E-06 | 2.458E-06 | 5.000E-06 | 9.900E-06 |
| 10021 | 502 | G502A | A | 4.6e-06 | 6e-06 | 1.8e-05 | 9.533E-06 | 7.366E-06 | 4.600E-06 | 1.800E-05 |
| 10022 | 502 | G502R | A | 9.5e-06 | 5.2e-06 | 8.7e-06 | 7.800E-06 | 2.287E-06 | 5.200E-06 | 9.500E-06 |
| 10023 | 502 | G502N | A | 8.6e-06 | 8.9e-06 | 1.4e-05 | 1.050E-05 | 3.035E-06 | 8.600E-06 | 1.400E-05 |
| 10024 | 502 | G502D | A | 3.9e-06 | 8.3e-06 | 1.3e-05 | 8.400E-06 | 4.551E-06 | 3.900E-06 | 1.300E-05 |
| 10025 | 502 | G502C | A | 8.7e-06 | 7.5e-06 | 7e-06 | 7.733E-06 | 8.737E-07 | 7.000E-06 | 8.700E-06 |
| 10026 | 502 | G502E | A | 1.1e-05 | 4.6e-06 | 9.4e-06 | 8.333E-06 | 3.331E-06 | 4.600E-06 | 1.100E-05 |
| 10027 | 502 | G502Q | A | 7.5e-06 | 8.6e-06 | 6.5e-06 | 7.533E-06 | 1.050E-06 | 6.500E-06 | 8.600E-06 |
| 10028 | 502 | G502G | A | 1.2e-05 | 6.5e-06 | 5.9e-06 | 8.133E-06 | 3.362E-06 | 5.900E-06 | 1.200E-05 |
| 10029 | 502 | G502H | A | 4.5e-06 | 5.6e-06 | 7.6e-06 | 5.900E-06 | 1.572E-06 | 4.500E-06 | 7.600E-06 |

| | | | | | | | | | | |
|---|---|---|---|---|---|---|---|---|---|---|
| 10030 | 502 | G502I | A | 8.7e-06 | 6.5e-06 | 6.2e-06 | 7.133E-06 | 1.365E-06 | 6.200E-06 | 8.700E-06 |
| 10031 | 502 | G502L | A | 1e-05 | 1.4e-05 | 9.6e-06 | 1.120E-05 | 2.433E-06 | 9.600E-06 | 1.400E-05 |
| 10032 | 502 | G502K | A | 4.4e-06 | 1.2e-05 | 1e-05 | 8.800E-06 | 3.940E-06 | 4.400E-06 | 1.200E-05 |
| 10033 | 502 | G502M | A | 1.4e-05 | 1.2e-05 | 8.1e-06 | 1.137E-05 | 3.001E-06 | 8.100E-06 | 1.400E-05 |
| 10034 | 502 | G502F | A | 1.9e-05 | 4.7e-06 | 5.2e-06 | 9.633E-06 | 8.116E-06 | 4.700E-06 | 1.900E-05 |
| 10035 | 502 | G502P | A | 8.8e-06 | 7.4e-06 | 6.4e-06 | 7.533E-06 | 1.206E-06 | 6.400E-06 | 8.800E-06 |
| 10036 | 502 | G502S | A | 4.9e-06 | 5.2e-06 | 6.8e-06 | 5.633E-06 | 1.021E-06 | 4.900E-06 | 6.800E-06 |
| 10037 | 502 | G502T | A | 1.4e-05 | 1e-05 | 5.4e-06 | 9.800E-06 | 4.303E-06 | 5.400E-06 | 1.400E-05 |
| 10038 | 502 | G502W | A | 1.4e-05 | 9.2e-06 | 6.4e-06 | 9.867E-06 | 3.844E-06 | 6.400E-06 | 1.400E-05 |
| 10039 | 502 | G502Y | A | 1.4e-05 | 9.5e-06 | 1.1e-05 | 1.150E-05 | 2.291E-06 | 9.500E-06 | 1.400E-05 |
| 10040 | 502 | G502V | A | 6.8e-06 | 7.8e-06 | 5.8e-06 | 6.800E-06 | 1.000E-06 | 5.800E-06 | 7.800E-06 |
| 10041 | 503 | P503A | A | 1.7e-05 | 1.4e-05 | 9.3e-06 | 1.343E-05 | 3.881E-06 | 9.300E-06 | 1.700E-05 |
| 10042 | 503 | P503R | A | 4.6e-06 | 1.1e-05 | 1.4e-05 | 9.867E-06 | 4.801E-06 | 4.600E-06 | 1.400E-05 |
| 10043 | 503 | P503N | A | 4.4e-06 | 1.9e-05 | 1e-05 | 1.113E-05 | 7.366E-06 | 4.400E-06 | 1.900E-05 |
| 10044 | 503 | P503D | A | 5.3e-06 | 6.7e-06 | 8.2e-06 | 6.733E-06 | 1.450E-06 | 5.300E-06 | 8.200E-06 |
| 10045 | 503 | P503C | A | 6e-06 | 6.1e-06 | 1.6e-05 | 9.367E-06 | 5.745E-06 | 6.000E-06 | 1.600E-05 |
| 10046 | 503 | P503E | A | 7.2e-06 | 9.3e-06 | 8.1e-06 | 8.200E-06 | 1.054E-06 | 7.200E-06 | 9.300E-06 |
| 10047 | 503 | P503Q | A | 7.4e-06 | 6e-06 | 7.6e-06 | 7.000E-06 | 8.718E-07 | 6.000E-06 | 7.600E-06 |
| 10048 | 503 | P503G | A | 9.9e-06 | 1.4e-05 | 6.6e-06 | 1.017E-05 | 3.707E-06 | 6.600E-06 | 1.400E-05 |
| 10049 | 503 | P503H | A | 8.8e-06 | 7.3e-06 | 1.6e-05 | 1.070E-05 | 4.651E-06 | 7.300E-06 | 1.600E-05 |
| 10050 | 503 | P503I | A | 5.9e-06 | 6.8e-06 | 1.2e-05 | 8.233E-06 | 3.293E-06 | 5.900E-06 | 1.200E-05 |
| 10051 | 503 | P503L | A | 9.2e-06 | 9.7e-06 | 1.2e-05 | 1.030E-05 | 1.493E-06 | 9.200E-06 | 1.200E-05 |
| 10052 | 503 | P503K | A | 6.8e-06 | 9.9e-06 | 9.5e-06 | 8.733E-06 | 1.686E-06 | 6.800E-06 | 9.900E-06 |
| 10053 | 503 | P503M | A | 9.4e-06 | 6.1e-06 | 8.5e-06 | 8.000E-06 | 1.706E-06 | 6.100E-06 | 9.400E-06 |
| 10054 | 503 | P503F | A | 8.6e-06 | 4.5e-06 | 1.2e-05 | 8.367E-06 | 3.755E-06 | 4.500E-06 | 1.200E-05 |
| 10055 | 503 | P503P | A | 1.2e-05 | 5.7e-06 | 8.8e-06 | 8.833E-06 | 3.150E-06 | 5.700E-06 | 1.200E-05 |
| 10056 | 503 | P503S | A | 5.8e-06 | 7.2e-06 | 6.9e-06 | 6.633E-06 | 7.371E-07 | 5.800E-06 | 7.200E-06 |
| 10057 | 503 | P503T | A | 6.3e-06 | 6.8e-06 | 8.6e-06 | 7.233E-06 | 1.210E-06 | 6.300E-06 | 8.600E-06 |
| 10058 | 503 | P503W | A | 1.3e-05 | 1.1e-05 | 7.3e-06 | 1.043E-05 | 2.892E-06 | 7.300E-06 | 1.300E-05 |
| 10059 | 503 | P503Y | A | 7.6e-06 | 6.5e-06 | 8.9e-06 | 7.667E-06 | 1.201E-06 | 6.500E-06 | 8.900E-06 |
| 10060 | 503 | P503V | A | 7.5e-06 | 8.3e-06 | 8.2e-06 | 8.000E-06 | 4.359E-07 | 7.500E-06 | 8.300E-06 |
| 10061 | 504 | T504A | A | 4.8e-06 | 8.7e-06 | 5.3e-06 | 6.267E-06 | 2.122E-06 | 4.800E-06 | 8.700E-06 |
| 10062 | 504 | T504R | A | 7.6e-06 | 7.8e-06 | 9.9e-06 | 8.433E-06 | 1.274E-06 | 7.600E-06 | 9.900E-06 |
| 10063 | 504 | T504N | A | 4.9e-06 | 8e-06 | 4.5e-06 | 5.800E-06 | 1.916E-06 | 4.500E-06 | 8.000E-06 |
| 10064 | 504 | T504D | A | 7.6e-06 | 7.7e-06 | 7.7e-06 | 7.667E-06 | 5.774E-08 | 7.600E-06 | 7.700E-06 |
| 10065 | 504 | T504C | A | 5e-06 | 9.3e-06 | 8.1e-06 | 7.467E-06 | 2.219E-06 | 5.000E-06 | 9.300E-06 |
| 10066 | 504 | T504E | A | 6.4e-06 | 6.7e-06 | 1e-05 | 7.700E-06 | 1.997E-06 | 6.400E-06 | 1.000E-05 |
| 10067 | 504 | T504Q | A | 6.5e-06 | 8.1e-06 | 9.8e-06 | 8.133E-06 | 1.650E-06 | 6.500E-06 | 9.800E-06 |
| 10068 | 504 | T504G | A | 4.8e-06 | 1e-05 | 5e-06 | 6.600E-06 | 2.946E-06 | 4.800E-06 | 1.000E-05 |
| 10069 | 504 | T504H | A | 9.6e-06 | 9.6e-06 | 6.1e-06 | 8.433E-06 | 2.021E-06 | 6.100E-06 | 9.600E-06 |
| 10070 | 504 | T504I | A | 6.7e-06 | 5.9e-06 | 8.8e-06 | 7.133E-06 | 1.498E-06 | 5.900E-06 | 8.800E-06 |
| 10071 | 504 | T504L | A | 3e-06 | 8e-06 | 6.8e-06 | 5.933E-06 | 2.610E-06 | 3.000E-06 | 8.000E-06 |
| 10072 | 504 | T504K | A | 6.2e-06 | 9.1e-06 | 7.9e-06 | 7.733E-06 | 1.457E-06 | 6.200E-06 | 9.100E-06 |
| 10073 | 504 | T504M | A | 7.8e-06 | 1e-05 | 1.1e-05 | 9.600E-06 | 1.637E-06 | 7.800E-06 | 1.100E-05 |
| 10074 | 504 | T504F | A | 1e-05 | 4e-06 | 5.8e-06 | 6.600E-06 | 3.079E-06 | 4.000E-06 | 1.000E-05 |
| 10075 | 504 | T504P | A | 1e-05 | 6.3e-06 | 6.7e-06 | 7.667E-06 | 2.031E-06 | 6.300E-06 | 1.000E-05 |
| 10076 | 504 | T504S | A | 6.5e-06 | 1.2e-05 | 7.4e-06 | 8.633E-06 | 2.950E-06 | 6.500E-06 | 1.200E-05 |
| 10077 | 504 | T504T | A | 5.6e-06 | 7.6e-06 | 8.9e-06 | 7.367E-06 | 1.662E-06 | 5.600E-06 | 8.900E-06 |
| 10078 | 504 | T504W | A | 4.4e-06 | 1.4e-05 | 5.5e-06 | 7.967E-06 | 5.254E-06 | 4.400E-06 | 1.400E-05 |
| 10079 | 504 | T504Y | A | 6.8e-06 | 6.4e-06 | 1.3e-05 | 8.733E-06 | 3.700E-06 | 6.400E-06 | 1.300E-05 |
| 10080 | 504 | T504V | A | 6.3e-06 | 7.5e-06 | 4.9e-06 | 6.233E-06 | 1.301E-06 | 4.900E-06 | 7.500E-06 |
| 10081 | 505 | Q505A | A | 8e-06 | 5.2e-06 | 6.2e-06 | 6.467E-06 | 1.419E-06 | 5.200E-06 | 8.000E-06 |
| 10082 | 505 | Q505R | A | 9.1e-06 | 7.4e-06 | 8.5e-06 | 8.333E-06 | 8.622E-07 | 7.400E-06 | 9.100E-06 |
| 10083 | 505 | Q505N | A | 1.3e-05 | 9.2e-06 | 5.6e-06 | 9.267E-06 | 3.700E-06 | 5.600E-06 | 1.300E-05 |
| 10084 | 505 | Q505D | A | 4.6e-06 | 6.9e-06 | 1.2e-05 | 7.833E-06 | 3.787E-06 | 4.600E-06 | 1.200E-05 |
| 10085 | 505 | Q505C | A | 8.4e-06 | 4.4e-06 | 6.9e-06 | 6.567E-06 | 2.021E-06 | 4.400E-06 | 8.400E-06 |
| 10086 | 505 | Q505E | A | 1.2e-05 | 6.7e-06 | 1.3e-05 | 1.057E-05 | 3.386E-06 | 6.700E-06 | 1.300E-05 |
| 10087 | 505 | Q505Q | A | 8.2e-06 | 6.1e-06 | 8.3e-06 | 7.533E-06 | 1.242E-06 | 6.100E-06 | 8.300E-06 |
| 10088 | 505 | Q505G | A | 6.6e-06 | 8.9e-06 | 4.6e-06 | 6.700E-06 | 2.152E-06 | 4.600E-06 | 8.900E-06 |
| 10089 | 505 | Q505H | A | 3.5e-06 | 6.8e-06 | 5.9e-06 | 5.400E-06 | 1.706E-06 | 3.500E-06 | 6.800E-06 |
| 10090 | 505 | Q505I | A | 9.8e-06 | 5.4e-06 | 7.4e-06 | 7.533E-06 | 2.203E-06 | 5.400E-06 | 9.800E-06 |
| 10091 | 505 | Q505L | A | 7.1e-06 | 6.7e-06 | 1e-05 | 7.933E-06 | 1.801E-06 | 6.700E-06 | 1.000E-05 |
| 10092 | 505 | Q505K | A | 6.7e-06 | 7.2e-06 | 9.9e-06 | 7.933E-06 | 1.721E-06 | 6.700E-06 | 9.900E-06 |
| 10093 | 505 | Q505M | A | 6e-06 | 6.4e-06 | 8.6e-06 | 7.000E-06 | 1.400E-06 | 6.000E-06 | 8.600E-06 |
| 10094 | 505 | Q505F | A | 1.2e-05 | 5e-06 | 8.6e-06 | 8.533E-06 | 3.500E-06 | 5.000E-06 | 1.200E-05 |
| 10095 | 505 | Q505P | A | 1.1e-05 | 9e-06 | 3.2e-06 | 7.733E-06 | 4.051E-06 | 3.200E-06 | 1.100E-05 |
| 10096 | 505 | Q505S | A | 5.2e-06 | 1.3e-05 | 7.9e-06 | 8.700E-06 | 3.961E-06 | 5.200E-06 | 1.300E-05 |
| 10097 | 505 | Q505T | A | 7.5e-06 | 9.1e-06 | 9.1e-06 | 8.567E-06 | 9.238E-07 | 7.500E-06 | 9.100E-06 |
| 10098 | 505 | Q505W | A | 6.9e-06 | 3.1e-06 | 1.3e-05 | 7.667E-06 | 4.994E-06 | 3.100E-06 | 1.300E-05 |
| 10099 | 505 | Q505Y | A | 7.9e-06 | 1e-05 | 9.1e-06 | 9.000E-06 | 1.054E-06 | 7.900E-06 | 1.000E-05 |
| 10100 | 505 | Q505V | A | 7.3e-06 | 6.4e-06 | 8.8e-06 | 7.500E-06 | 1.212E-06 | 6.400E-06 | 8.800E-06 |
| 10101 | 506 | C506A | A | 1.2e-05 | 8.5e-06 | 1.2e-05 | 1.083E-05 | 2.021E-06 | 8.500E-06 | 1.200E-05 |
| 10102 | 506 | C506R | A | 5.1e-06 | 1.1e-05 | 9.3e-06 | 8.467E-06 | 3.037E-06 | 5.100E-06 | 1.100E-05 |
| 10103 | 506 | C506N | A | 8.3e-06 | 1.1e-05 | 7.8e-06 | 9.033E-06 | 1.721E-06 | 7.800E-06 | 1.100E-05 |
| 10104 | 506 | C506D | A | 5.6e-06 | 8.4e-06 | 1.2e-05 | 8.667E-06 | 3.208E-06 | 5.600E-06 | 1.200E-05 |
| 10105 | 506 | C506C | A | 1.2e-05 | 4.2e-06 | 6.9e-06 | 7.700E-06 | 3.961E-06 | 4.200E-06 | 1.200E-05 |
| 10106 | 506 | C506E | A | 1.7e-05 | 9.3e-06 | 7.2e-06 | 1.117E-05 | 5.160E-06 | 7.200E-06 | 1.700E-05 |
| 10107 | 506 | C506Q | A | 7e-06 | 9e-06 | 9.8e-06 | 8.600E-06 | 1.442E-06 | 7.000E-06 | 9.800E-06 |
| 10108 | 506 | C506G | A | 6.6e-06 | 1.1e-05 | 7.8e-06 | 8.467E-06 | 2.274E-06 | 6.600E-06 | 1.100E-05 |
| 10109 | 506 | C506H | A | 5.9e-06 | 5e-06 | 5.1e-06 | 5.333E-06 | 4.933E-07 | 5.000E-06 | 5.900E-06 |
| 10110 | 506 | C506I | A | 4.2e-06 | 7.5e-06 | 4.9e-06 | 5.533E-06 | 1.739E-06 | 4.200E-06 | 7.500E-06 |
| 10111 | 506 | C506L | A | 8.5e-06 | 6.5e-06 | 1.3e-05 | 9.333E-06 | 3.329E-06 | 6.500E-06 | 1.300E-05 |
| 10112 | 506 | C506K | A | 5.4e-06 | 1e-05 | 1e-05 | 8.467E-06 | 2.656E-06 | 5.400E-06 | 1.000E-05 |
| 10113 | 506 | C506M | A | 8.5e-06 | 1.2e-05 | 1.4e-05 | 1.150E-05 | 2.784E-06 | 8.500E-06 | 1.400E-05 |
| 10114 | 506 | C506F | A | 1.1e-05 | 8.5e-06 | 1.1e-05 | 1.017E-05 | 1.443E-06 | 8.500E-06 | 1.100E-05 |

| | | | | | | | | | | |
|---|---|---|---|---|---|---|---|---|---|---|
| 10115 | 506 | C506P | A | 8.2e-06 | 4.1e-06 | 5.2e-06 | 5.833E-06 | 2.122E-06 | 4.100E-06 | 8.200E-06 |
| 10116 | 506 | C506S | A | 8.1e-06 | 1.2e-05 | 7.9e-06 | 9.333E-06 | 2.312E-06 | 7.900E-06 | 1.200E-05 |
| 10117 | 506 | C506T | A | 1.3e-05 | 6.2e-06 | 7.7e-06 | 8.967E-06 | 3.573E-06 | 6.200E-06 | 1.300E-05 |
| 10118 | 506 | C506W | A | 8.6e-06 | 6.3e-06 | 1e-05 | 8.300E-06 | 1.868E-06 | 6.300E-06 | 1.000E-05 |
| 10119 | 506 | C506Y | A | 8e-06 | 9e-06 | 8.2e-06 | 8.400E-06 | 5.292E-07 | 8.000E-06 | 9.000E-06 |
| 10120 | 506 | C506V | A | 8.8e-06 | 1.4e-05 | 4.4e-06 | 9.067E-06 | 4.806E-06 | 4.400E-06 | 1.400E-05 |
| 10121 | 507 | V507A | A | 1.1e-05 | 4.2e-06 | 6.6e-06 | 7.267E-06 | 3.449E-06 | 4.200E-06 | 1.100E-05 |
| 10122 | 507 | V507R | A | 4.8e-06 | 4.6e-06 | 6.2e-06 | 5.200E-06 | 8.718E-07 | 4.600E-06 | 6.200E-06 |
| 10123 | 507 | V507N | A | 6.8e-06 | 1.3e-05 | 5.9e-06 | 8.567E-06 | 3.866E-06 | 5.900E-06 | 1.300E-05 |
| 10124 | 507 | V507D | A | 9.6e-06 | 7.4e-06 | 5.4e-06 | 7.467E-06 | 2.101E-06 | 5.400E-06 | 9.600E-06 |
| 10125 | 507 | V507C | A | 6.6e-06 | 8.8e-06 | 8.5e-06 | 7.967E-06 | 1.193E-06 | 6.600E-06 | 8.800E-06 |
| 10126 | 507 | V507E | A | 7.1e-06 | 7.1e-06 | 5.7e-06 | 6.633E-06 | 8.083E-07 | 5.700E-06 | 7.100E-06 |
| 10127 | 507 | V507Q | A | 6.4e-06 | 7.1e-06 | 6.5e-06 | 6.667E-06 | 3.786E-07 | 6.400E-06 | 7.100E-06 |
| 10128 | 507 | V507G | A | 9.5e-06 | 8.9e-06 | 9.6e-06 | 9.333E-06 | 3.786E-07 | 8.900E-06 | 9.600E-06 |
| 10129 | 507 | V507H | A | 4.7e-06 | 9.1e-06 | 6.9e-06 | 6.900E-06 | 2.200E-06 | 4.700E-06 | 9.100E-06 |
| 10130 | 507 | V507I | A | 1e-05 | 6.4e-06 | 6.2e-06 | 7.533E-06 | 2.139E-06 | 6.200E-06 | 1.000E-05 |
| 10131 | 507 | V507L | A | 5.1e-06 | 1.4e-05 | 7.4e-06 | 8.833E-06 | 4.620E-06 | 5.100E-06 | 1.400E-05 |
| 10132 | 507 | V507K | A | 1e-05 | 9.5e-06 | 1.3e-05 | 1.083E-05 | 1.893E-06 | 9.500E-06 | 1.300E-05 |
| 10133 | 507 | V507M | A | 9.4e-06 | 7.1e-06 | 1.1e-05 | 9.167E-06 | 1.960E-06 | 7.100E-06 | 1.100E-05 |
| 10134 | 507 | V507F | A | 1.2e-05 | 1.3e-05 | 8.6e-06 | 1.120E-05 | 2.307E-06 | 8.600E-06 | 1.300E-05 |
| 10135 | 507 | V507P | A | 1.3e-05 | 6.9e-06 | 5.5e-06 | 8.467E-06 | 3.988E-06 | 5.500E-06 | 1.300E-05 |
| 10136 | 507 | V507S | A | 9.6e-06 | 1e-05 | 5.9e-06 | 8.500E-06 | 2.261E-06 | 5.900E-06 | 1.000E-05 |
| 10137 | 507 | V507T | A | 7.9e-06 | 7.3e-06 | 8.5e-06 | 7.900E-06 | 6.000E-07 | 7.300E-06 | 8.500E-06 |
| 10138 | 507 | V507W | A | 7.2e-06 | 1.1e-05 | 6.6e-06 | 8.267E-06 | 2.386E-06 | 6.600E-06 | 1.100E-05 |
| 10139 | 507 | V507Y | A | 6.2e-06 | 6.2e-06 | 4.2e-06 | 5.533E-06 | 1.155E-06 | 4.200E-06 | 6.200E-06 |
| 10140 | 507 | V507V | A | 6.5e-06 | 8.3e-06 | 9.6e-06 | 8.133E-06 | 1.557E-06 | 6.500E-06 | 9.600E-06 |
| 10141 | 508 | N508A | A | 9.8e-06 | 8.8e-06 | 1.3e-05 | 1.053E-05 | 2.194E-06 | 8.800E-06 | 1.300E-05 |
| 10142 | 508 | N508R | A | 6.5e-06 | 6.3e-06 | 4e-06 | 5.600E-06 | 1.389E-06 | 4.000E-06 | 6.500E-06 |
| 10143 | 508 | N508N | A | 8.9e-06 | 5.8e-06 | 1.3e-05 | 9.233E-06 | 3.612E-06 | 5.800E-06 | 1.300E-05 |
| 10144 | 508 | N508D | A | 3.5e-06 | 1.1e-05 | 9.8e-06 | 8.100E-06 | 4.029E-06 | 3.500E-06 | 1.100E-05 |
| 10145 | 508 | N508C | A | 6.3e-06 | 3.8e-06 | 4.6e-06 | 4.900E-06 | 1.277E-06 | 3.800E-06 | 6.300E-06 |
| 10146 | 508 | N508E | A | 3.3e-06 | 7.2e-06 | 9.7e-06 | 6.733E-06 | 3.225E-06 | 3.300E-06 | 9.700E-06 |
| 10147 | 508 | N508Q | A | 6.8e-06 | 7.8e-06 | 4.9e-06 | 6.500E-06 | 1.473E-06 | 4.900E-06 | 7.800E-06 |
| 10148 | 508 | N508G | A | 1.3e-05 | 1e-05 | 9.7e-06 | 1.090E-05 | 1.825E-06 | 9.700E-06 | 1.300E-05 |
| 10149 | 508 | N508H | A | 1.1e-05 | 1.2e-05 | 7.9e-06 | 1.030E-05 | 2.138E-06 | 7.900E-06 | 1.200E-05 |
| 10150 | 508 | N508I | A | 7.1e-06 | 6.9e-06 | 7.6e-06 | 7.200E-06 | 3.606E-07 | 6.900E-06 | 7.600E-06 |
| 10151 | 508 | N508L | A | 6.6e-06 | 4.8e-06 | 7e-06 | 6.133E-06 | 1.172E-06 | 4.800E-06 | 7.000E-06 |
| 10152 | 508 | N508K | A | 5e-06 | 2e-05 | 1e-05 | 1.167E-05 | 7.638E-06 | 5.000E-06 | 2.000E-05 |
| 10153 | 508 | N508M | A | 8.5e-06 | 5.3e-06 | 1.3e-05 | 8.933E-06 | 3.868E-06 | 5.300E-06 | 1.300E-05 |
| 10154 | 508 | N508F | A | 9e-06 | 1.1e-05 | 1e-05 | 1.000E-05 | 1.000E-06 | 9.000E-06 | 1.100E-05 |
| 10155 | 508 | N508P | A | 6.5e-06 | 1.3e-05 | 6.3e-06 | 8.600E-06 | 3.812E-06 | 6.300E-06 | 1.300E-05 |
| 10156 | 508 | N508S | A | 1e-05 | 4.5e-06 | 8.1e-06 | 7.533E-06 | 2.793E-06 | 4.500E-06 | 1.000E-05 |
| 10157 | 508 | N508T | A | 4.4e-06 | 1.3e-05 | 6.9e-06 | 8.100E-06 | 4.424E-06 | 4.400E-06 | 1.300E-05 |
| 10158 | 508 | N508W | A | 7.5e-06 | 9.7e-06 | 3.8e-06 | 7.000E-06 | 2.982E-06 | 3.800E-06 | 9.700E-06 |
| 10159 | 508 | N508Y | A | 7.8e-06 | 8.6e-06 | 1.5e-05 | 1.047E-05 | 3.946E-06 | 7.800E-06 | 1.500E-05 |
| 10160 | 508 | N508V | A | 9.3e-06 | 1.1e-05 | 9.2e-06 | 9.833E-06 | 1.012E-06 | 9.200E-06 | 1.100E-05 |
| 10161 | 509 | C509A | A | 6e-06 | 7.2e-06 | 6.9e-06 | 6.700E-06 | 6.245E-07 | 6.000E-06 | 7.200E-06 |
| 10162 | 509 | C509R | A | 1e-05 | 5.5e-06 | 1.1e-05 | 8.833E-06 | 2.930E-06 | 5.500E-06 | 1.100E-05 |
| 10163 | 509 | C509N | A | 6.7e-06 | 4.3e-06 | 9.2e-06 | 6.733E-06 | 2.450E-06 | 4.300E-06 | 9.200E-06 |
| 10164 | 509 | C509D | A | 1.3e-05 | 7.9e-06 | 6.9e-06 | 9.267E-06 | 3.272E-06 | 6.900E-06 | 1.300E-05 |
| 10165 | 509 | C509C | A | 7.7e-06 | 7.4e-06 | 5.3e-06 | 6.800E-06 | 1.308E-06 | 5.300E-06 | 7.700E-06 |
| 10166 | 509 | C509E | A | 1.2e-05 | 7e-06 | 1.4e-05 | 1.100E-05 | 3.606E-06 | 7.000E-06 | 1.400E-05 |
| 10167 | 509 | C509Q | A | 6e-06 | 8.1e-06 | 9.7e-06 | 7.933E-06 | 1.856E-06 | 6.000E-06 | 9.700E-06 |
| 10168 | 509 | C509G | A | 1.1e-05 | 1.1e-05 | 1.4e-05 | 1.200E-05 | 1.732E-06 | 1.100E-05 | 1.400E-05 |
| 10169 | 509 | C509H | A | 1.3e-05 | 5.6e-06 | 6.3e-06 | 8.300E-06 | 4.085E-06 | 5.600E-06 | 1.300E-05 |
| 10170 | 509 | C509I | A | 7.8e-06 | 5.5e-06 | 1e-05 | 7.767E-06 | 2.250E-06 | 5.500E-06 | 1.000E-05 |
| 10171 | 509 | C509L | A | 5.3e-06 | 1.4e-05 | 7.4e-06 | 8.900E-06 | 4.540E-06 | 5.300E-06 | 1.400E-05 |
| 10172 | 509 | C509K | A | 1.3e-05 | 7e-06 | 7.1e-06 | 9.033E-06 | 3.436E-06 | 7.000E-06 | 1.300E-05 |
| 10173 | 509 | C509M | A | 5e-06 | 6.9e-06 | 1e-05 | 7.300E-06 | 2.524E-06 | 5.000E-06 | 1.000E-05 |
| 10174 | 509 | C509F | A | 9.3e-06 | 5.4e-06 | 6.4e-06 | 7.033E-06 | 2.026E-06 | 5.400E-06 | 9.300E-06 |
| 10175 | 509 | C509P | A | 1.2e-05 | 5.6e-06 | 1e-05 | 9.200E-06 | 3.274E-06 | 5.600E-06 | 1.200E-05 |
| 10176 | 509 | C509S | A | 5.5e-06 | 8.2e-06 | 4.8e-06 | 6.167E-06 | 1.795E-06 | 4.800E-06 | 8.200E-06 |
| 10177 | 509 | C509T | A | 7.1e-06 | 5.3e-06 | 3.5e-06 | 5.300E-06 | 1.800E-06 | 3.500E-06 | 7.100E-06 |
| 10178 | 509 | C509W | A | 5.4e-06 | 9.8e-06 | 8.5e-06 | 7.900E-06 | 2.261E-06 | 5.400E-06 | 9.800E-06 |
| 10179 | 509 | C509Y | A | 7.3e-06 | 5.3e-06 | 1.1e-05 | 7.867E-06 | 2.892E-06 | 5.300E-06 | 1.100E-05 |
| 10180 | 509 | C509V | A | 7e-06 | 1.2e-05 | 7.4e-06 | 8.800E-06 | 2.778E-06 | 7.000E-06 | 1.200E-05 |
| 10181 | 510 | S510A | A | 6.9e-06 | 6.6e-06 | 1e-05 | 7.833E-06 | 1.882E-06 | 6.600E-06 | 1.000E-05 |
| 10182 | 510 | S510R | A | 8.1e-06 | 1.3e-05 | 1.1e-05 | 1.070E-05 | 2.464E-06 | 8.100E-06 | 1.300E-05 |
| 10183 | 510 | S510N | A | 9.3e-06 | 8.1e-06 | 8.5e-06 | 8.633E-06 | 6.110E-07 | 8.100E-06 | 9.300E-06 |
| 10184 | 510 | S510D | A | 8.9e-06 | 6.8e-06 | 1.8e-05 | 1.123E-05 | 5.953E-06 | 6.800E-06 | 1.800E-05 |
| 10185 | 510 | S510C | A | 6.5e-06 | 6e-06 | 4.6e-06 | 5.700E-06 | 9.849E-07 | 4.600E-06 | 6.500E-06 |
| 10186 | 510 | S510E | A | 5.9e-06 | 6.6e-06 | 8.3e-06 | 6.933E-06 | 1.234E-06 | 5.900E-06 | 8.300E-06 |
| 10187 | 510 | S510Q | A | 7.6e-06 | 6.7e-06 | 9.9e-06 | 8.067E-06 | 1.650E-06 | 6.700E-06 | 9.900E-06 |
| 10188 | 510 | S510G | A | 1.8e-05 | 1.2e-05 | 6.7e-06 | 1.223E-05 | 5.654E-06 | 6.700E-06 | 1.800E-05 |
| 10189 | 510 | S510H | A | 1.3e-05 | 7.8e-06 | 5.2e-06 | 8.667E-06 | 3.972E-06 | 5.200E-06 | 1.300E-05 |
| 10190 | 510 | S510I | A | 1.3e-05 | 1.3e-05 | 5.1e-06 | 1.037E-05 | 4.561E-06 | 5.100E-06 | 1.300E-05 |
| 10191 | 510 | S510L | A | 1.2e-05 | 1.3e-05 | 5.6e-06 | 1.020E-05 | 4.015E-06 | 5.600E-06 | 1.300E-05 |
| 10192 | 510 | S510K | A | 5.8e-06 | 8.4e-06 | 2e-05 | 1.140E-05 | 7.560E-06 | 5.800E-06 | 2.000E-05 |
| 10193 | 510 | S510M | A | 5.8e-06 | 6.5e-06 | 8.7e-06 | 7.000E-06 | 1.513E-06 | 5.800E-06 | 8.700E-06 |
| 10194 | 510 | S510F | A | 6.9e-06 | 6.8e-06 | 1.1e-05 | 8.233E-06 | 2.397E-06 | 6.800E-06 | 1.100E-05 |
| 10195 | 510 | S510P | A | 7.1e-06 | 7.7e-06 | 9.1e-06 | 7.967E-06 | 1.026E-06 | 7.100E-06 | 9.100E-06 |
| 10196 | 510 | S510S | A | 9.8e-06 | 1e-05 | 9.7e-06 | 9.833E-06 | 1.528E-07 | 9.700E-06 | 1.000E-05 |
| 10197 | 510 | S510T | A | 5.6e-06 | 6.4e-06 | 7.8e-06 | 6.600E-06 | 1.114E-06 | 5.600E-06 | 7.800E-06 |
| 10198 | 510 | S510W | A | 6.5e-06 | 8.2e-06 | 9e-06 | 7.900E-06 | 1.277E-06 | 6.500E-06 | 9.000E-06 |
| 10199 | 510 | S510Y | A | 3.7e-06 | 5.4e-06 | 6e-06 | 5.033E-06 | 1.193E-06 | 3.700E-06 | 6.000E-06 |

| | | | | | | | | | | |
|---|---|---|---|---|---|---|---|---|---|---|
| 10200 | 510 | S510V | A | 5.4e-06 | 6.7e-06 | 8.1e-06 | 6.733E-06 | 1.350E-06 | 5.400E-06 | 8.100E-06 |
| 10201 | 511 | Q511A | A | 4.6e-06 | 1.1e-05 | 6.4e-06 | 7.333E-06 | 3.301E-06 | 4.600E-06 | 1.100E-05 |
| 10202 | 511 | Q511R | A | 7.6e-06 | 4.7e-06 | 6e-06 | 6.100E-06 | 1.453E-06 | 4.700E-06 | 7.600E-06 |
| 10203 | 511 | Q511N | A | 8.6e-06 | 6.8e-06 | 7.7e-06 | 7.700E-06 | 9.000E-07 | 6.800E-06 | 8.600E-06 |
| 10204 | 511 | Q511D | A | 1.2e-05 | 1.4e-05 | 9.3e-06 | 1.177E-05 | 2.359E-06 | 9.300E-06 | 1.400E-05 |
| 10205 | 511 | Q511C | A | 6.9e-06 | 8.6e-06 | 8.7e-06 | 8.067E-06 | 1.012E-06 | 6.900E-06 | 8.700E-06 |
| 10206 | 511 | Q511E | A | 5.4e-06 | 7.8e-06 | 1.1e-05 | 8.067E-06 | 2.810E-06 | 5.400E-06 | 1.100E-05 |
| 10207 | 511 | Q511Q | A | 7.7e-06 | 9.4e-06 | 1.1e-05 | 9.367E-06 | 1.650E-06 | 7.700E-06 | 1.100E-05 |
| 10208 | 511 | Q511G | A | 5.1e-06 | 9.8e-06 | 1.3e-05 | 9.300E-06 | 3.974E-06 | 5.100E-06 | 1.300E-05 |
| 10209 | 511 | Q511H | A | 6.5e-06 | 9.1e-06 | 5.4e-06 | 7.000E-06 | 1.900E-06 | 5.400E-06 | 9.100E-06 |
| 10210 | 511 | Q511I | A | 6.4e-06 | 4.7e-06 | 1.2e-05 | 7.700E-06 | 3.820E-06 | 4.700E-06 | 1.200E-05 |
| 10211 | 511 | Q511L | A | 8.5e-06 | 7.1e-06 | 5e-06 | 6.867E-06 | 1.762E-06 | 5.000E-06 | 8.500E-06 |
| 10212 | 511 | Q511K | A | 9.8e-06 | 1.2e-05 | 1e-05 | 1.060E-05 | 1.217E-06 | 9.800E-06 | 1.200E-05 |
| 10213 | 511 | Q511M | A | 1.2e-05 | 1e-05 | 1.4e-05 | 1.200E-05 | 2.000E-06 | 1.000E-05 | 1.400E-05 |
| 10214 | 511 | Q511F | A | 8.1e-06 | 9e-06 | 5.5e-06 | 7.533E-06 | 1.818E-06 | 5.500E-06 | 9.000E-06 |
| 10215 | 511 | Q511P | A | 7.6e-06 | 7.2e-06 | 6.9e-06 | 7.233E-06 | 3.512E-07 | 6.900E-06 | 7.600E-06 |
| 10216 | 511 | Q511S | A | 5.4e-06 | 9.4e-06 | 6.4e-06 | 7.067E-06 | 2.082E-06 | 5.400E-06 | 9.400E-06 |
| 10217 | 511 | Q511T | A | 9.4e-06 | 1.2e-05 | 4.5e-06 | 8.633E-06 | 3.808E-06 | 4.500E-06 | 1.200E-05 |
| 10218 | 511 | Q511W | A | 4.9e-06 | 5.7e-06 | 6.9e-06 | 5.833E-06 | 1.007E-06 | 4.900E-06 | 6.900E-06 |
| 10219 | 511 | Q511Y | A | 8.6e-06 | 6.3e-06 | 1.2e-05 | 8.967E-06 | 2.868E-06 | 6.300E-06 | 1.200E-05 |
| 10220 | 511 | Q511V | A | 9.6e-06 | 6.2e-06 | 8.2e-06 | 8.000E-06 | 1.709E-06 | 6.200E-06 | 9.600E-06 |
| 10221 | 512 | F512A | A | 4.4e-06 | 7.6e-06 | 5.8e-06 | 5.933E-06 | 1.604E-06 | 4.400E-06 | 7.600E-06 |
| 10222 | 512 | F512R | A | 6.5e-06 | 8.6e-06 | 1.3e-05 | 9.367E-06 | 3.317E-06 | 6.500E-06 | 1.300E-05 |
| 10223 | 512 | F512N | A | 5.3e-06 | 4e-06 | 5.5e-06 | 4.933E-06 | 8.145E-07 | 4.000E-06 | 5.500E-06 |
| 10224 | 512 | F512D | A | 8.8e-06 | 1e-05 | 7.9e-06 | 8.900E-06 | 1.054E-06 | 7.900E-06 | 1.000E-05 |
| 10225 | 512 | F512C | A | 6.5e-06 | 1.1e-05 | 6.5e-06 | 8.000E-06 | 2.598E-06 | 6.500E-06 | 1.100E-05 |
| 10226 | 512 | F512E | A | 1.2e-05 | 8.8e-06 | 6.3e-06 | 9.033E-06 | 2.857E-06 | 6.300E-06 | 1.200E-05 |
| 10227 | 512 | F512Q | A | 1e-05 | 6.5e-06 | 4.4e-06 | 6.967E-06 | 2.829E-06 | 4.400E-06 | 1.000E-05 |
| 10228 | 512 | F512G | A | 1.1e-05 | 7e-06 | 9.6e-06 | 9.200E-06 | 2.030E-06 | 7.000E-06 | 1.100E-05 |
| 10229 | 512 | F512H | A | 8.5e-06 | 7.9e-06 | 9.7e-06 | 8.700E-06 | 9.165E-07 | 7.900E-06 | 9.700E-06 |
| 10230 | 512 | F512I | A | 1.1e-05 | 6.7e-06 | 4.7e-06 | 7.467E-06 | 3.219E-06 | 4.700E-06 | 1.100E-05 |
| 10231 | 512 | F512L | A | 9.8e-06 | 6.8e-06 | 9.6e-06 | 8.733E-06 | 1.677E-06 | 6.800E-06 | 9.800E-06 |
| 10232 | 512 | F512K | A | 8.2e-06 | 1.3e-05 | 1.4e-05 | 1.173E-05 | 3.101E-06 | 8.200E-06 | 1.400E-05 |
| 10233 | 512 | F512M | A | 6.9e-06 | 9.7e-06 | 7.1e-06 | 7.900E-06 | 1.562E-06 | 6.900E-06 | 9.700E-06 |
| 10234 | 512 | F512F | A | 1e-05 | 5.3e-06 | 8.7e-06 | 8.000E-06 | 2.427E-06 | 5.300E-06 | 1.000E-05 |
| 10235 | 512 | F512P | A | 9.8e-06 | 1e-05 | 3.3e-06 | 7.700E-06 | 3.812E-06 | 3.300E-06 | 1.000E-05 |
| 10236 | 512 | F512S | A | 4.4e-06 | 1e-05 | 6.7e-06 | 7.033E-06 | 2.815E-06 | 4.400E-06 | 1.000E-05 |
| 10237 | 512 | F512T | A | 8.6e-06 | 8.1e-06 | 5.9e-06 | 7.533E-06 | 1.436E-06 | 5.900E-06 | 8.600E-06 |
| 10238 | 512 | F512W | A | 6.6e-06 | 7.1e-06 | 6e-06 | 6.567E-06 | 5.508E-07 | 6.000E-06 | 7.100E-06 |
| 10239 | 512 | F512Y | A | 9.5e-06 | 9e-06 | 8e-06 | 8.833E-06 | 7.638E-07 | 8.000E-06 | 9.500E-06 |
| 10240 | 512 | F512V | A | 8.5e-06 | 9.1e-06 | 4.9e-06 | 7.500E-06 | 2.272E-06 | 4.900E-06 | 9.100E-06 |
| 10241 | 513 | L513A | A | 5.4e-06 | 8.9e-06 | 1e-05 | 8.100E-06 | 2.402E-06 | 5.400E-06 | 1.000E-05 |
| 10242 | 513 | L513R | A | 9.3e-06 | 6.4e-06 | 7e-06 | 7.567E-06 | 1.531E-06 | 6.400E-06 | 9.300E-06 |
| 10243 | 513 | L513N | A | 8.8e-06 | 8.9e-06 | 1.1e-05 | 9.567E-06 | 1.242E-06 | 8.800E-06 | 1.100E-05 |
| 10244 | 513 | L513D | A | 1.2e-05 | 9.6e-06 | 1e-05 | 1.053E-05 | 1.286E-06 | 9.600E-06 | 1.200E-05 |
| 10245 | 513 | L513C | A | 7.5e-06 | 4.3e-06 | 1.1e-05 | 7.600E-06 | 3.351E-06 | 4.300E-06 | 1.100E-05 |
| 10246 | 513 | L513E | A | 9.3e-06 | 6e-06 | 6.5e-06 | 7.267E-06 | 1.779E-06 | 6.000E-06 | 9.300E-06 |
| 10247 | 513 | L513Q | A | 8.7e-06 | 6.2e-06 | 4.5e-06 | 6.467E-06 | 2.113E-06 | 4.500E-06 | 8.700E-06 |
| 10248 | 513 | L513G | A | 6.1e-06 | 1.2e-05 | 8.2e-06 | 8.767E-06 | 2.991E-06 | 6.100E-06 | 1.200E-05 |
| 10249 | 513 | L513H | A | 7.4e-06 | 4.9e-06 | 1e-05 | 7.433E-06 | 2.550E-06 | 4.900E-06 | 1.000E-05 |
| 10250 | 513 | L513I | A | 7e-06 | 8.8e-06 | 8.4e-06 | 8.067E-06 | 9.452E-07 | 7.000E-06 | 8.800E-06 |
| 10251 | 513 | L513L | A | 6.3e-06 | 4e-06 | 4.7e-06 | 5.000E-06 | 1.179E-06 | 4.000E-06 | 6.300E-06 |
| 10252 | 513 | L513K | A | 8.3e-06 | 4.8e-06 | 6e-06 | 6.367E-06 | 1.779E-06 | 4.800E-06 | 8.300E-06 |
| 10253 | 513 | L513M | A | 6.9e-06 | 7.6e-06 | 8.3e-06 | 7.600E-06 | 7.000E-07 | 6.900E-06 | 8.300E-06 |
| 10254 | 513 | L513F | A | 9.1e-06 | 5.6e-06 | 8.5e-06 | 7.733E-06 | 1.872E-06 | 5.600E-06 | 9.100E-06 |
| 10255 | 513 | L513P | A | 7.7e-06 | 6.8e-06 | 9.2e-06 | 7.900E-06 | 1.212E-06 | 6.800E-06 | 9.200E-06 |
| 10256 | 513 | L513S | A | 5.5e-06 | 7.1e-06 | 7e-06 | 6.533E-06 | 8.963E-07 | 5.500E-06 | 7.100E-06 |
| 10257 | 513 | L513T | A | 8.7e-06 | 6.5e-06 | 7.8e-06 | 7.667E-06 | 1.106E-06 | 6.500E-06 | 8.700E-06 |
| 10258 | 513 | L513W | A | 8.5e-06 | 8.1e-06 | 1.1e-05 | 9.200E-06 | 1.572E-06 | 8.100E-06 | 1.100E-05 |
| 10259 | 513 | L513Y | A | 1.2e-05 | 1.3e-05 | 5.2e-06 | 1.007E-05 | 4.244E-06 | 5.200E-06 | 1.300E-05 |
| 10260 | 513 | L513V | A | 1e-05 | 5.8e-06 | 1e-05 | 8.600E-06 | 2.425E-06 | 5.800E-06 | 1.000E-05 |
| 10261 | 514 | R514A | A | 5.6e-06 | 1.1e-05 | 9.1e-06 | 8.567E-06 | 2.739E-06 | 5.600E-06 | 1.100E-05 |
| 10262 | 514 | R514R | A | 7.1e-06 | 9.4e-06 | 5.9e-06 | 7.467E-06 | 1.779E-06 | 5.900E-06 | 9.400E-06 |
| 10263 | 514 | R514N | A | 9.2e-06 | 4.2e-06 | 5.3e-06 | 6.233E-06 | 2.627E-06 | 4.200E-06 | 9.200E-06 |
| 10264 | 514 | R514D | A | 9.9e-06 | 7.3e-06 | 7.5e-06 | 8.233E-06 | 1.447E-06 | 7.300E-06 | 9.900E-06 |
| 10265 | 514 | R514C | A | 4.8e-06 | 8.1e-06 | 7.3e-06 | 6.733E-06 | 1.721E-06 | 4.800E-06 | 8.100E-06 |
| 10266 | 514 | R514E | A | 9.7e-06 | 5.5e-06 | 7.4e-06 | 7.533E-06 | 2.103E-06 | 5.500E-06 | 9.700E-06 |
| 10267 | 514 | R514Q | A | 9.7e-06 | 5.5e-06 | 9.4e-06 | 8.200E-06 | 2.343E-06 | 5.500E-06 | 9.700E-06 |
| 10268 | 514 | R514G | A | 5.5e-06 | 9.4e-06 | 6.9e-06 | 7.267E-06 | 1.976E-06 | 5.500E-06 | 9.400E-06 |
| 10269 | 514 | R514H | A | 1.1e-05 | 5.2e-06 | 6.1e-06 | 7.433E-06 | 3.121E-06 | 5.200E-06 | 1.100E-05 |
| 10270 | 514 | R514I | A | 6.8e-06 | 6.7e-06 | 1.3e-05 | 8.833E-06 | 3.609E-06 | 6.700E-06 | 1.300E-05 |
| 10271 | 514 | R514L | A | 1.3e-05 | 5.8e-06 | 6.8e-06 | 8.533E-06 | 3.900E-06 | 5.800E-06 | 1.300E-05 |
| 10272 | 514 | R514K | A | 1.3e-05 | 6.5e-06 | 5.3e-06 | 8.267E-06 | 4.143E-06 | 5.300E-06 | 1.300E-05 |
| 10273 | 514 | R514M | A | 9.2e-06 | 6.3e-06 | 4.8e-06 | 6.767E-06 | 2.237E-06 | 4.800E-06 | 9.200E-06 |
| 10274 | 514 | R514F | A | 1.1e-05 | 5.1e-06 | 5.5e-06 | 7.200E-06 | 3.297E-06 | 5.100E-06 | 1.100E-05 |
| 10275 | 514 | R514P | A | 1.3e-05 | 1.1e-05 | 1.1e-05 | 1.167E-05 | 1.155E-06 | 1.100E-05 | 1.300E-05 |
| 10276 | 514 | R514S | A | 9.5e-06 | 7.2e-06 | 5.7e-06 | 7.467E-06 | 1.914E-06 | 5.700E-06 | 9.500E-06 |
| 10277 | 514 | R514T | A | 1.2e-05 | 1.1e-05 | 6.7e-06 | 9.900E-06 | 2.816E-06 | 6.700E-06 | 1.200E-05 |
| 10278 | 514 | R514W | A | 9.5e-06 | 1.3e-05 | 5.3e-06 | 9.267E-06 | 3.855E-06 | 5.300E-06 | 1.300E-05 |
| 10279 | 514 | R514Y | A | 6e-06 | 8.3e-06 | 1e-05 | 8.100E-06 | 2.007E-06 | 6.000E-06 | 1.000E-05 |
| 10280 | 514 | R514V | A | 4.2e-06 | 8.2e-06 | 1e-05 | 7.467E-06 | 2.969E-06 | 4.200E-06 | 1.000E-05 |
| 10281 | 515 | G515A | A | 4.8e-06 | 6.8e-06 | 6.9e-06 | 6.167E-06 | 1.185E-06 | 4.800E-06 | 6.900E-06 |
| 10282 | 515 | G515R | A | 9.4e-06 | 9.4e-06 | 8.3e-06 | 9.033E-06 | 6.351E-07 | 8.300E-06 | 9.400E-06 |
| 10283 | 515 | G515N | A | 7.8e-06 | 9.1e-06 | 6e-06 | 7.633E-06 | 1.557E-06 | 6.000E-06 | 9.100E-06 |
| 10284 | 515 | G515D | A | 6.2e-06 | 8.3e-06 | 6.5e-06 | 7.000E-06 | 1.136E-06 | 6.200E-06 | 8.300E-06 |

| | | | | | | | | | | |
|---|---|---|---|---|---|---|---|---|---|---|
| 10285 | 515 | G515C | A | 3.2e-06 | 8.4e-06 | 4.3e-06 | 5.300E-06 | 2.740E-06 | 3.200E-06 | 8.400E-06 |
| 10286 | 515 | G515E | A | 7.6e-06 | 5.4e-06 | 5e-06 | 6.000E-06 | 1.400E-06 | 5.000E-06 | 7.600E-06 |
| 10287 | 515 | G515Q | A | 7.9e-06 | 1.5e-05 | 9.4e-06 | 1.077E-05 | 3.742E-06 | 7.900E-06 | 1.500E-05 |
| 10288 | 515 | G515G | A | 9.7e-06 | 9.8e-06 | 4.3e-06 | 7.933E-06 | 3.147E-06 | 4.300E-06 | 9.800E-06 |
| 10289 | 515 | G515H | A | 1e-05 | 9.6e-06 | 5.5e-06 | 8.367E-06 | 2.491E-06 | 5.500E-06 | 1.000E-05 |
| 10290 | 515 | G515I | A | 7e-06 | 8.7e-06 | 5.8e-06 | 7.167E-06 | 1.457E-06 | 5.800E-06 | 8.700E-06 |
| 10291 | 515 | G515L | A | 7.5e-06 | 7e-06 | 1.2e-05 | 8.833E-06 | 2.754E-06 | 7.000E-06 | 1.200E-05 |
| 10292 | 515 | G515K | A | 1e-05 | 8.5e-06 | 9.8e-06 | 9.433E-06 | 8.145E-07 | 8.500E-06 | 1.000E-05 |
| 10293 | 515 | G515M | A | 8.7e-06 | 1.1e-05 | 1.1e-05 | 1.023E-05 | 1.328E-06 | 8.700E-06 | 1.100E-05 |
| 10294 | 515 | G515F | A | 1e-05 | 1.7e-05 | 1.1e-05 | 1.267E-05 | 3.786E-06 | 1.000E-05 | 1.700E-05 |
| 10295 | 515 | G515P | A | 6.8e-06 | 9.6e-06 | 4.7e-06 | 7.033E-06 | 2.458E-06 | 4.700E-06 | 9.600E-06 |
| 10296 | 515 | G515S | A | 1.1e-05 | 6.8e-06 | 5.7e-06 | 7.833E-06 | 2.797E-06 | 5.700E-06 | 1.100E-05 |
| 10297 | 515 | G515T | A | 8.7e-06 | 9.2e-06 | 6.8e-06 | 8.233E-06 | 1.266E-06 | 6.800E-06 | 9.200E-06 |
| 10298 | 515 | G515W | A | 9.9e-06 | 1.3e-05 | 5.7e-06 | 9.533E-06 | 3.664E-06 | 5.700E-06 | 1.300E-05 |
| 10299 | 515 | G515Y | A | 9.7e-06 | 9.3e-06 | 7.8e-06 | 8.933E-06 | 1.002E-06 | 7.800E-06 | 9.700E-06 |
| 10300 | 515 | G515V | A | 5.1e-06 | 4.3e-06 | 1.6e-05 | 8.467E-06 | 6.536E-06 | 4.300E-06 | 1.600E-05 |
| 10301 | 516 | Q516A | A | 6.7e-06 | 1.4e-05 | 5.9e-06 | 8.867E-06 | 4.464E-06 | 5.900E-06 | 1.400E-05 |
| 10302 | 516 | Q516R | A | 1.5e-05 | 7.2e-06 | 6e-06 | 9.400E-06 | 4.887E-06 | 6.000E-06 | 1.500E-05 |
| 10303 | 516 | Q516N | A | 1e-05 | 8e-06 | 1.3e-05 | 1.033E-05 | 2.517E-06 | 8.000E-06 | 1.300E-05 |
| 10304 | 516 | Q516D | A | 7.3e-06 | 5.5e-06 | 9.9e-06 | 7.567E-06 | 2.212E-06 | 5.500E-06 | 9.900E-06 |
| 10305 | 516 | Q516C | A | 9.9e-06 | 8.3e-06 | 4.1e-06 | 7.433E-06 | 2.996E-06 | 4.100E-06 | 9.900E-06 |
| 10306 | 516 | Q516E | A | 8.2e-06 | 9.6e-06 | 1e-05 | 9.267E-06 | 9.452E-07 | 8.200E-06 | 1.000E-05 |
| 10307 | 516 | Q516Q | A | 9e-06 | 9.6e-06 | 8.7e-06 | 9.100E-06 | 4.583E-07 | 8.700E-06 | 9.600E-06 |
| 10308 | 516 | Q516G | A | 9.4e-06 | 8.2e-06 | 5.3e-06 | 7.633E-06 | 2.108E-06 | 5.300E-06 | 9.400E-06 |
| 10309 | 516 | Q516H | A | 7.1e-06 | 1.2e-05 | 1e-05 | 9.700E-06 | 2.464E-06 | 7.100E-06 | 1.200E-05 |
| 10310 | 516 | Q516I | A | 6.4e-06 | 1.1e-05 | 9.8e-06 | 9.067E-06 | 2.386E-06 | 6.400E-06 | 1.100E-05 |
| 10311 | 516 | Q516L | A | 9.8e-06 | 8.5e-06 | 4.9e-06 | 7.733E-06 | 2.538E-06 | 4.900E-06 | 9.800E-06 |
| 10312 | 516 | Q516K | A | 6.7e-06 | 4e-06 | 5.8e-06 | 5.500E-06 | 1.375E-06 | 4.000E-06 | 6.700E-06 |
| 10313 | 516 | Q516M | A | 9.7e-06 | 9.2e-06 | 9.1e-06 | 9.333E-06 | 3.215E-07 | 9.100E-06 | 9.700E-06 |
| 10314 | 516 | Q516F | A | 6.5e-06 | 1.2e-05 | 8.5e-06 | 9.000E-06 | 2.784E-06 | 6.500E-06 | 1.200E-05 |
| 10315 | 516 | Q516P | A | 8.1e-06 | 7.4e-06 | 7.3e-06 | 7.600E-06 | 4.359E-07 | 7.300E-06 | 8.100E-06 |
| 10316 | 516 | Q516S | A | 6.4e-06 | 5.3e-06 | 9.5e-06 | 7.067E-06 | 2.178E-06 | 5.300E-06 | 9.500E-06 |
| 10317 | 516 | Q516T | A | 9.3e-06 | 3.4e-06 | 6.6e-06 | 6.433E-06 | 2.954E-06 | 3.400E-06 | 9.300E-06 |
| 10318 | 516 | Q516W | A | 5.9e-06 | 8.8e-06 | 1e-05 | 8.233E-06 | 2.108E-06 | 5.900E-06 | 1.000E-05 |
| 10319 | 516 | Q516Y | A | 1e-05 | 1e-05 | 7.2e-06 | 9.067E-06 | 1.617E-06 | 7.200E-06 | 1.000E-05 |
| 10320 | 516 | Q516V | A | 5e-06 | 7.5e-06 | 7.1e-06 | 6.533E-06 | 1.343E-06 | 5.000E-06 | 7.500E-06 |
| 10321 | 517 | E517A | A | 8.8e-06 | 9e-06 | 8.8e-06 | 8.867E-06 | 1.155E-07 | 8.800E-06 | 9.000E-06 |
| 10322 | 517 | E517R | A | 8e-06 | 8.1e-06 | 9.6e-06 | 8.567E-06 | 8.963E-07 | 8.000E-06 | 9.600E-06 |
| 10323 | 517 | E517N | A | 1e-05 | 8.5e-06 | 9.2e-06 | 9.233E-06 | 7.506E-07 | 8.500E-06 | 1.000E-05 |
| 10324 | 517 | E517D | A | 6.8e-06 | 6.7e-06 | 7.6e-06 | 7.033E-06 | 4.933E-07 | 6.700E-06 | 7.600E-06 |
| 10325 | 517 | E517C | A | 8.4e-06 | 1e-05 | 8.9e-06 | 9.100E-06 | 8.185E-07 | 8.400E-06 | 1.000E-05 |
| 10326 | 517 | E517E | A | 5.8e-06 | 1.5e-05 | 9.9e-06 | 1.023E-05 | 4.609E-06 | 5.800E-06 | 1.500E-05 |
| 10327 | 517 | E517Q | A | 8.2e-06 | 6.9e-06 | 1.3e-05 | 9.367E-06 | 3.213E-06 | 6.900E-06 | 1.300E-05 |
| 10328 | 517 | E517G | A | 7e-06 | 4.5e-06 | 9.2e-06 | 6.900E-06 | 2.352E-06 | 4.500E-06 | 9.200E-06 |
| 10329 | 517 | E517H | A | 9.6e-06 | 8e-06 | 8.7e-06 | 8.767E-06 | 8.021E-07 | 8.000E-06 | 9.600E-06 |
| 10330 | 517 | E517I | A | 7e-06 | 9.6e-06 | 7.4e-06 | 8.000E-06 | 1.400E-06 | 7.000E-06 | 9.600E-06 |
| 10331 | 517 | E517L | A | 2.5e-05 | 1.3e-05 | 9.5e-06 | 1.583E-05 | 8.129E-06 | 9.500E-06 | 2.500E-05 |
| 10332 | 517 | E517K | A | 8e-06 | 4.4e-06 | 7e-06 | 6.467E-06 | 1.858E-06 | 4.400E-06 | 8.000E-06 |
| 10333 | 517 | E517M | A | 6.6e-06 | 8.2e-06 | 8.2e-06 | 7.667E-06 | 9.238E-07 | 6.600E-06 | 8.200E-06 |
| 10334 | 517 | E517F | A | 6.6e-06 | 7.6e-06 | 3.4e-06 | 5.867E-06 | 2.194E-06 | 3.400E-06 | 7.600E-06 |
| 10335 | 517 | E517P | A | 6.4e-06 | 8e-06 | 1.2e-05 | 8.800E-06 | 2.884E-06 | 6.400E-06 | 1.200E-05 |
| 10336 | 517 | E517S | A | 1.2e-05 | 8e-06 | 8.5e-06 | 9.500E-06 | 2.179E-06 | 8.000E-06 | 1.200E-05 |
| 10337 | 517 | E517T | A | 6.5e-06 | 1.1e-05 | 7.1e-06 | 8.200E-06 | 2.443E-06 | 6.500E-06 | 1.100E-05 |
| 10338 | 517 | E517W | A | 1e-05 | 7.2e-06 | 1.1e-05 | 9.400E-06 | 1.970E-06 | 7.200E-06 | 1.100E-05 |
| 10339 | 517 | E517Y | A | 6.3e-06 | 1.3e-05 | 8.3e-06 | 9.200E-06 | 3.439E-06 | 6.300E-06 | 1.300E-05 |
| 10340 | 517 | E517V | A | 5e-06 | 6.4e-06 | 8.1e-06 | 6.500E-06 | 1.552E-06 | 5.000E-06 | 8.100E-06 |
| 10341 | 518 | C518A | A | 7e-06 | 1.4e-05 | 9.5e-06 | 1.017E-05 | 3.547E-06 | 7.000E-06 | 1.400E-05 |
| 10342 | 518 | C518R | A | 9.5e-06 | 8.3e-06 | 1.2e-05 | 9.933E-06 | 1.888E-06 | 8.300E-06 | 1.200E-05 |
| 10343 | 518 | C518N | A | 4.4e-06 | 6.6e-06 | 8e-06 | 6.333E-06 | 1.815E-06 | 4.400E-06 | 8.000E-06 |
| 10344 | 518 | C518D | A | 9.8e-06 | 1.1e-05 | 4.7e-06 | 8.500E-06 | 3.345E-06 | 4.700E-06 | 1.100E-05 |
| 10345 | 518 | C518C | A | 8.2e-06 | 6.1e-06 | 1e-05 | 8.100E-06 | 1.952E-06 | 6.100E-06 | 1.000E-05 |
| 10346 | 518 | C518E | A | 7.9e-06 | 5.9e-06 | 6.5e-06 | 6.767E-06 | 1.026E-06 | 5.900E-06 | 7.900E-06 |
| 10347 | 518 | C518Q | A | 4.6e-06 | 5.4e-06 | 7.7e-06 | 5.900E-06 | 1.609E-06 | 4.600E-06 | 7.700E-06 |
| 10348 | 518 | C518G | A | 7.3e-06 | 7.1e-06 | 7.3e-06 | 7.233E-06 | 1.155E-07 | 7.100E-06 | 7.300E-06 |
| 10349 | 518 | C518H | A | 9.9e-06 | 6.2e-06 | 6.4e-06 | 7.500E-06 | 2.081E-06 | 6.200E-06 | 9.900E-06 |
| 10350 | 518 | C518I | A | 1.1e-05 | 7.3e-06 | 7.5e-06 | 8.600E-06 | 2.081E-06 | 7.300E-06 | 1.100E-05 |
| 10351 | 518 | C518L | A | 5.3e-06 | 1.1e-05 | 9e-06 | 8.433E-06 | 2.892E-06 | 5.300E-06 | 1.100E-05 |
| 10352 | 518 | C518K | A | 7.4e-06 | 1.9e-05 | 4.8e-06 | 1.040E-05 | 7.560E-06 | 4.800E-06 | 1.900E-05 |
| 10353 | 518 | C518M | A | 9.9e-06 | 8.5e-06 | 8.3e-06 | 8.900E-06 | 8.718E-07 | 8.300E-06 | 9.900E-06 |
| 10354 | 518 | C518F | A | 8.7e-06 | 1.3e-05 | 1e-05 | 1.057E-05 | 2.205E-06 | 8.700E-06 | 1.300E-05 |
| 10355 | 518 | C518P | A | 9.2e-06 | 7.4e-06 | 5.7e-06 | 7.433E-06 | 1.750E-06 | 5.700E-06 | 9.200E-06 |
| 10356 | 518 | C518S | A | 7.2e-06 | 8.7e-06 | 7e-06 | 7.633E-06 | 9.292E-07 | 7.000E-06 | 8.700E-06 |
| 10357 | 518 | C518T | A | 6.7e-06 | 6.6e-06 | 8.5e-06 | 7.267E-06 | 1.069E-06 | 6.600E-06 | 8.500E-06 |
| 10358 | 518 | C518W | A | 7.3e-06 | 1.3e-05 | 8.6e-06 | 9.633E-06 | 2.987E-06 | 7.300E-06 | 1.300E-05 |
| 10359 | 518 | C518Y | A | 9.3e-06 | 9.1e-06 | 1.6e-05 | 1.147E-05 | 3.927E-06 | 9.100E-06 | 1.600E-05 |
| 10360 | 518 | C518V | A | 6.5e-06 | 3.4e-06 | 1.4e-05 | 7.967E-06 | 5.450E-06 | 3.400E-06 | 1.400E-05 |
| 10361 | 519 | V519A | A | 5.2e-06 | 5.4e-06 | 8.5e-06 | 6.367E-06 | 1.850E-06 | 5.200E-06 | 8.500E-06 |
| 10362 | 519 | V519R | A | 1.8e-05 | 7.7e-06 | 5.9e-06 | 1.053E-05 | 6.529E-06 | 5.900E-06 | 1.800E-05 |
| 10363 | 519 | V519N | A | 6.7e-06 | 6.3e-06 | 4.9e-06 | 5.967E-06 | 9.452E-07 | 4.900E-06 | 6.700E-06 |
| 10364 | 519 | V519D | A | 7.8e-06 | 8.8e-06 | 1.4e-05 | 1.020E-05 | 3.329E-06 | 7.800E-06 | 1.400E-05 |
| 10365 | 519 | V519C | A | 8.3e-06 | 6.5e-06 | 9.9e-06 | 8.233E-06 | 1.701E-06 | 6.500E-06 | 9.900E-06 |
| 10366 | 519 | V519E | A | 6.3e-06 | 5.6e-06 | 1.3e-05 | 8.300E-06 | 4.085E-06 | 5.600E-06 | 1.300E-05 |
| 10367 | 519 | V519Q | A | 6.1e-06 | 7.2e-06 | 8e-06 | 7.100E-06 | 9.539E-07 | 6.100E-06 | 8.000E-06 |
| 10368 | 519 | V519G | A | 4.9e-06 | 6.8e-06 | 7.9e-06 | 6.533E-06 | 1.518E-06 | 4.900E-06 | 7.900E-06 |
| 10369 | 519 | V519H | A | 1.2e-05 | 4.3e-06 | 5.8e-06 | 7.367E-06 | 4.082E-06 | 4.300E-06 | 1.200E-05 |

| | | | | | | | | | | |
|---|---|---|---|---|---|---|---|---|---|---|
| 10370 | 519 | V519I | A | 1.4e-05 | 4.4e-06 | 9.9e-06 | 9.433E-06 | 4.817E-06 | 4.400E-06 | 1.400E-05 |
| 10371 | 519 | V519L | A | 7e-06 | 5.3e-06 | 1.1e-05 | 7.767E-06 | 2.926E-06 | 5.300E-06 | 1.100E-05 |
| 10372 | 519 | V519K | A | 1.4e-05 | 4.7e-06 | 9.3e-06 | 9.333E-06 | 4.650E-06 | 4.700E-06 | 1.400E-05 |
| 10373 | 519 | V519M | A | 5.9e-06 | 4.4e-06 | 7.7e-06 | 6.000E-06 | 1.652E-06 | 4.400E-06 | 7.700E-06 |
| 10374 | 519 | V519F | A | 6.9e-06 | 8.5e-06 | 7.2e-06 | 7.533E-06 | 8.505E-07 | 6.900E-06 | 8.500E-06 |
| 10375 | 519 | V519P | A | 7e-06 | 9.6e-06 | 6e-06 | 7.533E-06 | 1.858E-06 | 6.000E-06 | 9.600E-06 |
| 10376 | 519 | V519S | A | 8.6e-06 | 6e-06 | 9.7e-06 | 8.100E-06 | 1.900E-06 | 6.000E-06 | 9.700E-06 |
| 10377 | 519 | V519T | A | 7.9e-06 | 7.8e-06 | 7.4e-06 | 7.700E-06 | 2.646E-07 | 7.400E-06 | 7.900E-06 |
| 10378 | 519 | V519W | A | 8.5e-06 | 9.2e-06 | 8.7e-06 | 8.800E-06 | 3.606E-07 | 8.500E-06 | 9.200E-06 |
| 10379 | 519 | V519Y | A | 6.9e-06 | 6.1e-06 | 1.4e-05 | 9.000E-06 | 4.349E-06 | 6.100E-06 | 1.400E-05 |
| 10380 | 519 | V519V | A | 7.6e-06 | 6.6e-06 | 9.8e-06 | 8.000E-06 | 1.637E-06 | 6.600E-06 | 9.800E-06 |
| 10381 | 520 | E520A | A | 5.9e-06 | 6.9e-06 | 1.4e-05 | 8.933E-06 | 4.416E-06 | 5.900E-06 | 1.400E-05 |
| 10382 | 520 | E520R | A | 1.1e-05 | 1.1e-05 | 5.7e-06 | 9.233E-06 | 3.060E-06 | 5.700E-06 | 1.100E-05 |
| 10383 | 520 | E520N | A | 5e-06 | 8.3e-06 | 9.1e-06 | 7.467E-06 | 2.173E-06 | 5.000E-06 | 9.100E-06 |
| 10384 | 520 | E520D | A | 1.2e-05 | 6.3e-06 | 6.7e-06 | 8.333E-06 | 3.182E-06 | 6.300E-06 | 1.200E-05 |
| 10385 | 520 | E520C | A | 9.2e-06 | 5.3e-06 | 1e-05 | 8.167E-06 | 2.515E-06 | 5.300E-06 | 1.000E-05 |
| 10386 | 520 | E520E | A | 4.4e-06 | 6.2e-06 | 8.3e-06 | 6.300E-06 | 1.952E-06 | 4.400E-06 | 8.300E-06 |
| 10387 | 520 | E520Q | A | 5.9e-06 | 4.9e-06 | 6.1e-06 | 5.633E-06 | 6.429E-07 | 4.900E-06 | 6.100E-06 |
| 10388 | 520 | E520G | A | 6.3e-06 | 5.1e-06 | 9.7e-06 | 7.033E-06 | 2.386E-06 | 5.100E-06 | 9.700E-06 |
| 10389 | 520 | E520H | A | 1.1e-05 | 7.8e-06 | 9.5e-06 | 9.433E-06 | 1.601E-06 | 7.800E-06 | 1.100E-05 |
| 10390 | 520 | E520I | A | 7.1e-06 | 9.2e-06 | 1.2e-05 | 9.433E-06 | 2.458E-06 | 7.100E-06 | 1.200E-05 |
| 10391 | 520 | E520L | A | 8.7e-06 | 1e-05 | 6.9e-06 | 8.533E-06 | 1.557E-06 | 6.900E-06 | 1.000E-05 |
| 10392 | 520 | E520K | A | 3.3e-06 | 5.4e-06 | 5.1e-06 | 4.600E-06 | 1.136E-06 | 3.300E-06 | 5.400E-06 |
| 10393 | 520 | E520M | A | 6.2e-06 | 6.8e-06 | 1e-05 | 7.667E-06 | 2.043E-06 | 6.200E-06 | 1.000E-05 |
| 10394 | 520 | E520F | A | 1e-05 | 6.5e-06 | 7.7e-06 | 8.067E-06 | 1.779E-06 | 6.500E-06 | 1.000E-05 |
| 10395 | 520 | E520P | A | 9.7e-06 | 8.9e-06 | 6.7e-06 | 8.433E-06 | 1.553E-06 | 6.700E-06 | 9.700E-06 |
| 10396 | 520 | E520S | A | 9.3e-06 | 1.4e-05 | 1.1e-05 | 1.143E-05 | 2.380E-06 | 9.300E-06 | 1.400E-05 |
| 10397 | 520 | E520T | A | 7.9e-06 | 4.7e-06 | 9.8e-06 | 7.467E-06 | 2.577E-06 | 4.700E-06 | 9.800E-06 |
| 10398 | 520 | E520W | A | 8.6e-06 | 8.6e-06 | 1.3e-05 | 1.007E-05 | 2.540E-06 | 8.600E-06 | 1.300E-05 |
| 10399 | 520 | E520Y | A | 9.1e-06 | 8.2e-06 | 1.1e-05 | 9.433E-06 | 1.429E-06 | 8.200E-06 | 1.100E-05 |
| 10400 | 520 | E520V | A | 9.8e-06 | 8.9e-06 | 9.6e-06 | 9.433E-06 | 4.726E-07 | 8.900E-06 | 9.800E-06 |
| 10401 | 521 | E521A | A | 8.3e-06 | 1.2e-05 | 6.1e-06 | 8.800E-06 | 2.982E-06 | 6.100E-06 | 1.200E-05 |
| 10402 | 521 | E521R | A | 8.8e-06 | 6.7e-06 | 1e-05 | 8.500E-06 | 1.670E-06 | 6.700E-06 | 1.000E-05 |
| 10403 | 521 | E521N | A | 6.8e-06 | 1.4e-05 | 7.4e-06 | 9.400E-06 | 3.995E-06 | 6.800E-06 | 1.400E-05 |
| 10404 | 521 | E521D | A | 9.1e-06 | 9.8e-06 | 8.9e-06 | 9.267E-06 | 4.726E-07 | 8.900E-06 | 9.800E-06 |
| 10405 | 521 | E521C | A | 1.1e-05 | 9.2e-06 | 9.4e-06 | 9.867E-06 | 9.866E-07 | 9.200E-06 | 1.100E-05 |
| 10406 | 521 | E521E | A | 8.2e-06 | 6.7e-06 | 9.1e-06 | 8.000E-06 | 1.212E-06 | 6.700E-06 | 9.100E-06 |
| 10407 | 521 | E521Q | A | 8.8e-06 | 6.7e-06 | 8.9e-06 | 8.133E-06 | 1.242E-06 | 6.700E-06 | 8.900E-06 |
| 10408 | 521 | E521G | A | 5.3e-06 | 1.5e-05 | 4.6e-06 | 8.300E-06 | 5.813E-06 | 4.600E-06 | 1.500E-05 |
| 10409 | 521 | E521H | A | 9.7e-06 | 1.2e-05 | 5.8e-06 | 9.167E-06 | 3.134E-06 | 5.800E-06 | 1.200E-05 |
| 10410 | 521 | E521I | A | 6.9e-06 | 8.4e-06 | 1.5e-05 | 1.010E-05 | 4.309E-06 | 6.900E-06 | 1.500E-05 |
| 10411 | 521 | E521L | A | 7.4e-06 | 6.8e-06 | 1.1e-05 | 8.400E-06 | 2.272E-06 | 6.800E-06 | 1.100E-05 |
| 10412 | 521 | E521K | A | 9.4e-06 | 1e-05 | 7.4e-06 | 8.933E-06 | 1.361E-06 | 7.400E-06 | 1.000E-05 |
| 10413 | 521 | E521M | A | 9.3e-06 | 5.4e-06 | 1.2e-05 | 8.900E-06 | 3.318E-06 | 5.400E-06 | 1.200E-05 |
| 10414 | 521 | E521F | A | 6.8e-06 | 9.6e-06 | 3.3e-06 | 6.567E-06 | 3.156E-06 | 3.300E-06 | 9.600E-06 |
| 10415 | 521 | E521P | A | 6.3e-06 | 8.8e-06 | 5.9e-06 | 7.000E-06 | 1.572E-06 | 5.900E-06 | 8.800E-06 |
| 10416 | 521 | E521S | A | 9.6e-06 | 1.3e-05 | 6.7e-06 | 9.767E-06 | 3.153E-06 | 6.700E-06 | 1.300E-05 |
| 10417 | 521 | E521T | A | 7.4e-06 | 7.4e-06 | 7.6e-06 | 7.467E-06 | 1.155E-07 | 7.400E-06 | 7.600E-06 |
| 10418 | 521 | E521W | A | 6.9e-06 | 1.1e-05 | 7e-06 | 8.300E-06 | 2.339E-06 | 6.900E-06 | 1.100E-05 |
| 10419 | 521 | E521Y | A | 8.6e-06 | 9.1e-06 | 6.2e-06 | 7.967E-06 | 1.550E-06 | 6.200E-06 | 9.100E-06 |
| 10420 | 521 | E521V | A | 9.7e-06 | 1.3e-05 | 1.1e-05 | 1.123E-05 | 1.662E-06 | 9.700E-06 | 1.300E-05 |
| 10421 | 522 | C522A | A | 7.2e-06 | 7.1e-06 | 1e-05 | 8.100E-06 | 1.646E-06 | 7.100E-06 | 1.000E-05 |
| 10422 | 522 | C522R | A | 9.5e-06 | 7.7e-06 | 7.7e-06 | 8.300E-06 | 1.039E-06 | 7.700E-06 | 9.500E-06 |
| 10423 | 522 | C522N | A | 4.5e-06 | 4.8e-06 | 8.1e-06 | 5.800E-06 | 1.997E-06 | 4.500E-06 | 8.100E-06 |
| 10424 | 522 | C522D | A | 1.2e-05 | 1e-05 | 6.2e-06 | 9.400E-06 | 2.946E-06 | 6.200E-06 | 1.200E-05 |
| 10425 | 522 | C522C | A | 6.1e-06 | 1.1e-05 | 6e-06 | 7.700E-06 | 2.858E-06 | 6.000E-06 | 1.100E-05 |
| 10426 | 522 | C522E | A | 6.1e-06 | 9.3e-06 | 1.1e-05 | 8.800E-06 | 2.488E-06 | 6.100E-06 | 1.100E-05 |
| 10427 | 522 | C522Q | A | 7e-06 | 1.1e-05 | 8.9e-06 | 8.967E-06 | 2.001E-06 | 7.000E-06 | 1.100E-05 |
| 10428 | 522 | C522G | A | 8.4e-06 | 9.6e-06 | 9.5e-06 | 9.167E-06 | 6.658E-07 | 8.400E-06 | 9.600E-06 |
| 10429 | 522 | C522H | A | 7.3e-06 | 5.1e-06 | 6.9e-06 | 6.433E-06 | 1.172E-06 | 5.100E-06 | 7.300E-06 |
| 10430 | 522 | C522I | A | 8.8e-06 | 9.5e-06 | 1e-05 | 9.433E-06 | 6.028E-07 | 8.800E-06 | 1.000E-05 |
| 10431 | 522 | C522L | A | 7.5e-06 | 8.2e-06 | 9.1e-06 | 8.267E-06 | 8.021E-07 | 7.500E-06 | 9.100E-06 |
| 10432 | 522 | C522K | A | 1.1e-05 | 8.9e-06 | 7.8e-06 | 9.233E-06 | 1.626E-06 | 7.800E-06 | 1.100E-05 |
| 10433 | 522 | C522M | A | 9.8e-06 | 1.3e-05 | 6.1e-06 | 9.633E-06 | 3.453E-06 | 6.100E-06 | 1.300E-05 |
| 10434 | 522 | C522F | A | 5.9e-06 | 1.3e-05 | 1e-05 | 9.633E-06 | 3.564E-06 | 5.900E-06 | 1.300E-05 |
| 10435 | 522 | C522P | A | 6.8e-06 | 7.9e-06 | 9.8e-06 | 8.167E-06 | 1.518E-06 | 6.800E-06 | 9.800E-06 |
| 10436 | 522 | C522S | A | 5.8e-06 | 8.2e-06 | 6.6e-06 | 6.867E-06 | 1.222E-06 | 5.800E-06 | 8.200E-06 |
| 10437 | 522 | C522T | A | 1.3e-05 | 5.5e-06 | 6.7e-06 | 8.400E-06 | 4.029E-06 | 5.500E-06 | 1.300E-05 |
| 10438 | 522 | C522W | A | 7.3e-06 | 9.7e-06 | 1e-05 | 9.000E-06 | 1.480E-06 | 7.300E-06 | 1.000E-05 |
| 10439 | 522 | C522Y | A | 8.8e-06 | 5.3e-06 | 4.8e-06 | 6.300E-06 | 2.179E-06 | 4.800E-06 | 8.800E-06 |
| 10440 | 522 | C522V | A | 1.3e-05 | 6.1e-06 | 6.9e-06 | 8.667E-06 | 3.774E-06 | 6.100E-06 | 1.300E-05 |
| 10441 | 523 | R523A | A | 5.5e-06 | 6.6e-06 | 1.3e-05 | 8.367E-06 | 4.050E-06 | 5.500E-06 | 1.300E-05 |
| 10442 | 523 | R523R | A | 6.7e-06 | 1.4e-05 | 4.8e-06 | 8.500E-06 | 4.857E-06 | 4.800E-06 | 1.400E-05 |
| 10443 | 523 | R523N | A | 6.8e-06 | 9.9e-06 | 6.4e-06 | 7.700E-06 | 1.916E-06 | 6.400E-06 | 9.900E-06 |
| 10444 | 523 | R523D | A | 6.4e-06 | 5.5e-06 | 9.6e-06 | 7.167E-06 | 2.155E-06 | 5.500E-06 | 9.600E-06 |
| 10445 | 523 | R523C | A | 9.3e-06 | 1e-05 | 5.7e-06 | 8.333E-06 | 2.307E-06 | 5.700E-06 | 1.000E-05 |
| 10446 | 523 | R523E | A | 7e-06 | 1.2e-05 | 6.9e-06 | 8.633E-06 | 2.916E-06 | 6.900E-06 | 1.200E-05 |
| 10447 | 523 | R523Q | A | 6.7e-06 | 1.4e-05 | 7e-06 | 9.233E-06 | 4.131E-06 | 6.700E-06 | 1.400E-05 |
| 10448 | 523 | R523G | A | 9.6e-06 | 7.3e-06 | 8.3e-06 | 8.400E-06 | 1.153E-06 | 7.300E-06 | 9.600E-06 |
| 10449 | 523 | R523H | A | 5.6e-06 | 9.7e-06 | 8e-06 | 7.767E-06 | 2.060E-06 | 5.600E-06 | 9.700E-06 |
| 10450 | 523 | R523I | A | 1.2e-05 | 6.2e-06 | 1.2e-05 | 1.007E-05 | 3.349E-06 | 6.200E-06 | 1.200E-05 |
| 10451 | 523 | R523L | A | 6.7e-06 | 9.3e-06 | 7.1e-06 | 7.700E-06 | 1.400E-06 | 6.700E-06 | 9.300E-06 |
| 10452 | 523 | R523K | A | 1e-05 | 9e-06 | 6.3e-06 | 8.433E-06 | 1.914E-06 | 6.300E-06 | 1.000E-05 |
| 10453 | 523 | R523M | A | 8.8e-06 | 4.7e-06 | 4.6e-06 | 6.033E-06 | 2.397E-06 | 4.600E-06 | 8.800E-06 |
| 10454 | 523 | R523F | A | 5.6e-06 | 5.6e-06 | 7.3e-06 | 6.167E-06 | 9.815E-07 | 5.600E-06 | 7.300E-06 |

| | | | | | | | | | | |
|---|---|---|---|---|---|---|---|---|---|---|
| 10455 | 523 | R523P | A | 6.9e-06 | 5.2e-06 | 6.7e-06 | 6.267E-06 | 9.292E-07 | 5.200E-06 | 6.900E-06 |
| 10456 | 523 | R523S | A | 6.4e-06 | 9.4e-06 | 9.4e-06 | 8.400E-06 | 1.732E-06 | 6.400E-06 | 9.400E-06 |
| 10457 | 523 | R523T | A | 4.4e-06 | 9.7e-06 | 6.8e-06 | 6.967E-06 | 2.654E-06 | 4.400E-06 | 9.700E-06 |
| 10458 | 523 | R523W | A | 1.1e-05 | 8.5e-06 | 8.8e-06 | 9.433E-06 | 1.365E-06 | 8.500E-06 | 1.100E-05 |
| 10459 | 523 | R523Y | A | 7.7e-06 | 6.2e-06 | 6.8e-06 | 6.900E-06 | 7.550E-07 | 6.200E-06 | 7.700E-06 |
| 10460 | 523 | R523V | A | 9.8e-06 | 9.4e-06 | 1.3e-05 | 1.073E-05 | 1.973E-06 | 9.400E-06 | 1.300E-05 |
| 10461 | 524 | V524A | A | 8.9e-06 | 5.3e-06 | 4.4e-06 | 6.200E-06 | 2.381E-06 | 4.400E-06 | 8.900E-06 |
| 10462 | 524 | V524R | A | 1.1e-05 | 3.2e-06 | 5e-06 | 6.400E-06 | 4.084E-06 | 3.200E-06 | 1.100E-05 |
| 10463 | 524 | V524N | A | 8.5e-06 | 5.9e-06 | 5.5e-06 | 6.633E-06 | 1.629E-06 | 5.500E-06 | 8.500E-06 |
| 10464 | 524 | V524D | A | 4.7e-06 | 6.6e-06 | 8.2e-06 | 6.500E-06 | 1.752E-06 | 4.700E-06 | 8.200E-06 |
| 10465 | 524 | V524C | A | 6.4e-06 | 1.7e-05 | 8.5e-06 | 1.063E-05 | 5.613E-06 | 6.400E-06 | 1.700E-05 |
| 10466 | 524 | V524E | A | 9.4e-06 | 9.3e-06 | 7.7e-06 | 8.800E-06 | 9.539E-07 | 7.700E-06 | 9.400E-06 |
| 10467 | 524 | V524Q | A | 8.9e-06 | 9.1e-06 | 9.1e-06 | 9.033E-06 | 1.155E-07 | 8.900E-06 | 9.100E-06 |
| 10468 | 524 | V524G | A | 9.1e-06 | 1.4e-05 | 8.3e-06 | 1.047E-05 | 3.086E-06 | 8.300E-06 | 1.400E-05 |
| 10469 | 524 | V524H | A | 7.5e-06 | 6.7e-06 | 8.5e-06 | 7.567E-06 | 9.018E-07 | 6.700E-06 | 8.500E-06 |
| 10470 | 524 | V524I | A | 1.3e-05 | 1.9e-05 | 7.1e-06 | 1.303E-05 | 5.950E-06 | 7.100E-06 | 1.900E-05 |
| 10471 | 524 | V524L | A | 7.1e-06 | 9e-06 | 9.2e-06 | 8.433E-06 | 1.159E-06 | 7.100E-06 | 9.200E-06 |
| 10472 | 524 | V524K | A | 1.2e-05 | 6.2e-06 | 5.3e-06 | 7.833E-06 | 3.636E-06 | 5.300E-06 | 1.200E-05 |
| 10473 | 524 | V524M | A | 8e-06 | 7.2e-06 | 5.5e-06 | 6.900E-06 | 1.277E-06 | 5.500E-06 | 8.000E-06 |
| 10474 | 524 | V524F | A | 3.8e-06 | 4.9e-06 | 1.1e-05 | 6.567E-06 | 3.879E-06 | 3.800E-06 | 1.100E-05 |
| 10475 | 524 | V524P | A | 8.3e-06 | 1.1e-05 | 5.2e-06 | 8.167E-06 | 2.902E-06 | 5.200E-06 | 1.100E-05 |
| 10476 | 524 | V524S | A | 1.2e-05 | 1e-05 | 8e-06 | 1.000E-05 | 2.000E-06 | 8.000E-06 | 1.200E-05 |
| 10477 | 524 | V524T | A | 6.7e-06 | 9.9e-06 | 9.4e-06 | 8.667E-06 | 1.721E-06 | 6.700E-06 | 9.900E-06 |
| 10478 | 524 | V524W | A | 7e-06 | 9.8e-06 | 7.4e-06 | 8.067E-06 | 1.514E-06 | 7.000E-06 | 9.800E-06 |
| 10479 | 524 | V524Y | A | 8.1e-06 | 5.9e-06 | 5.2e-06 | 6.400E-06 | 1.513E-06 | 5.200E-06 | 8.100E-06 |
| 10480 | 524 | V524V | A | 6.2e-06 | 1.1e-05 | 8.9e-06 | 8.700E-06 | 2.406E-06 | 6.200E-06 | 1.100E-05 |
| 10481 | 525 | L525A | A | 9.7e-06 | 6.4e-06 | 9.2e-06 | 8.433E-06 | 1.779E-06 | 6.400E-06 | 9.700E-06 |
| 10482 | 525 | L525R | A | 6.6e-06 | 9.2e-06 | 6.3e-06 | 7.367E-06 | 1.595E-06 | 6.300E-06 | 9.200E-06 |
| 10483 | 525 | L525N | A | 7.3e-06 | 6.9e-06 | 6.8e-06 | 7.000E-06 | 2.646E-07 | 6.800E-06 | 7.300E-06 |
| 10484 | 525 | L525D | A | 4.6e-06 | 8.8e-06 | 1.1e-05 | 8.133E-06 | 3.252E-06 | 4.600E-06 | 1.100E-05 |
| 10485 | 525 | L525C | A | 9.8e-06 | 6.1e-06 | 1.1e-05 | 8.967E-06 | 2.554E-06 | 6.100E-06 | 1.100E-05 |
| 10486 | 525 | L525E | A | 7.6e-06 | 5.2e-06 | 7.1e-06 | 6.633E-06 | 1.266E-06 | 5.200E-06 | 7.600E-06 |
| 10487 | 525 | L525Q | A | 6.4e-06 | 7e-06 | 9.6e-06 | 7.667E-06 | 1.701E-06 | 6.400E-06 | 9.600E-06 |
| 10488 | 525 | L525G | A | 7.9e-06 | 4.9e-06 | 5.8e-06 | 6.200E-06 | 1.539E-06 | 4.900E-06 | 7.900E-06 |
| 10489 | 525 | L525H | A | 4.4e-06 | 5.7e-06 | 9.8e-06 | 6.633E-06 | 2.818E-06 | 4.400E-06 | 9.800E-06 |
| 10490 | 525 | L525I | A | 9.4e-06 | 9.8e-06 | 9.2e-06 | 9.467E-06 | 3.055E-07 | 9.200E-06 | 9.800E-06 |
| 10491 | 525 | L525L | A | 9.2e-06 | 6.8e-06 | 1e-05 | 8.667E-06 | 1.665E-06 | 6.800E-06 | 1.000E-05 |
| 10492 | 525 | L525K | A | 4.2e-06 | 9.3e-06 | 1.3e-05 | 8.833E-06 | 4.419E-06 | 4.200E-06 | 1.300E-05 |
| 10493 | 525 | L525M | A | 7.6e-06 | 5.2e-06 | 6.5e-06 | 6.433E-06 | 1.201E-06 | 5.200E-06 | 7.600E-06 |
| 10494 | 525 | L525F | A | 6.2e-06 | 1.1e-05 | 9e-06 | 8.733E-06 | 2.411E-06 | 6.200E-06 | 1.100E-05 |
| 10495 | 525 | L525P | A | 6.6e-06 | 8e-06 | 1.1e-05 | 8.533E-06 | 2.248E-06 | 6.600E-06 | 1.100E-05 |
| 10496 | 525 | L525S | A | 6e-06 | 8.3e-06 | 1.3e-05 | 9.100E-06 | 3.568E-06 | 6.000E-06 | 1.300E-05 |
| 10497 | 525 | L525T | A | 3.4e-06 | 1e-05 | 6.6e-06 | 6.667E-06 | 3.301E-06 | 3.400E-06 | 1.000E-05 |
| 10498 | 525 | L525W | A | 1.1e-05 | 1.1e-05 | 1.3e-05 | 1.167E-05 | 1.155E-06 | 1.100E-05 | 1.300E-05 |
| 10499 | 525 | L525Y | A | 7.4e-06 | 7.7e-06 | 9.3e-06 | 8.133E-06 | 1.021E-06 | 7.400E-06 | 9.300E-06 |
| 10500 | 525 | L525V | A | 7.2e-06 | 5.8e-06 | 6.5e-06 | 6.500E-06 | 7.000E-07 | 5.800E-06 | 7.200E-06 |
| 10501 | 526 | Q526A | A | 9.4e-06 | 6.6e-06 | 1.1e-05 | 9.000E-06 | 2.227E-06 | 6.600E-06 | 1.100E-05 |
| 10502 | 526 | Q526R | A | 7.8e-06 | 6.9e-06 | 8e-06 | 7.567E-06 | 5.859E-07 | 6.900E-06 | 8.000E-06 |
| 10503 | 526 | Q526N | A | 8.7e-06 | 7.5e-06 | 1e-05 | 8.733E-06 | 1.250E-06 | 7.500E-06 | 1.000E-05 |
| 10504 | 526 | Q526D | A | 8.1e-06 | 1.4e-05 | 6.9e-06 | 9.667E-06 | 3.800E-06 | 6.900E-06 | 1.400E-05 |
| 10505 | 526 | Q526C | A | 8.1e-06 | 8.9e-06 | 9.5e-06 | 8.833E-06 | 7.024E-07 | 8.100E-06 | 9.500E-06 |
| 10506 | 526 | Q526E | A | 6.3e-06 | 7.1e-06 | 6.9e-06 | 6.767E-06 | 4.163E-07 | 6.300E-06 | 7.100E-06 |
| 10507 | 526 | Q526Q | A | 6.4e-06 | 7.1e-06 | 9.2e-06 | 7.567E-06 | 1.457E-06 | 6.400E-06 | 9.200E-06 |
| 10508 | 526 | Q526G | A | 6.8e-06 | 1.2e-05 | 1.1e-05 | 9.933E-06 | 2.759E-06 | 6.800E-06 | 1.200E-05 |
| 10509 | 526 | Q526H | A | 1e-05 | 1e-05 | 8.2e-06 | 9.400E-06 | 1.039E-06 | 8.200E-06 | 1.000E-05 |
| 10510 | 526 | Q526I | A | 8.6e-06 | 7.7e-06 | 7.4e-06 | 7.900E-06 | 6.245E-07 | 7.400E-06 | 8.600E-06 |
| 10511 | 526 | Q526L | A | 6.5e-06 | 5.5e-06 | 8.9e-06 | 6.967E-06 | 1.747E-06 | 5.500E-06 | 8.900E-06 |
| 10512 | 526 | Q526K | A | 7.8e-06 | 9.5e-06 | 5e-06 | 7.433E-06 | 2.272E-06 | 5.000E-06 | 9.500E-06 |
| 10513 | 526 | Q526M | A | 6.8e-06 | 8.4e-06 | 5.4e-06 | 6.867E-06 | 1.501E-06 | 5.400E-06 | 8.400E-06 |
| 10514 | 526 | Q526F | A | 9.5e-06 | 1.1e-05 | 5.4e-06 | 8.633E-06 | 2.899E-06 | 5.400E-06 | 1.100E-05 |
| 10515 | 526 | Q526P | A | 1.1e-05 | 9.9e-06 | 7.2e-06 | 9.367E-06 | 1.955E-06 | 7.200E-06 | 1.100E-05 |
| 10516 | 526 | Q526S | A | 1.4e-05 | 9.8e-06 | 4.1e-06 | 9.300E-06 | 4.969E-06 | 4.100E-06 | 1.400E-05 |
| 10517 | 526 | Q526T | A | 5.7e-06 | 4.8e-06 | 1.3e-05 | 7.833E-06 | 4.497E-06 | 4.800E-06 | 1.300E-05 |
| 10518 | 526 | Q526W | A | 5.8e-06 | 6.5e-06 | 3.1e-06 | 5.133E-06 | 1.795E-06 | 3.100E-06 | 6.500E-06 |
| 10519 | 526 | Q526Y | A | 6.8e-06 | 4.6e-06 | 9.2e-06 | 6.867E-06 | 2.301E-06 | 4.600E-06 | 9.200E-06 |
| 10520 | 526 | Q526V | A | 9.8e-06 | 6.7e-06 | 1.1e-05 | 9.167E-06 | 2.219E-06 | 6.700E-06 | 1.100E-05 |
| 10521 | 527 | G527A | A | 4.5e-06 | 9.4e-06 | 1.1e-05 | 8.300E-06 | 3.387E-06 | 4.500E-06 | 1.100E-05 |
| 10522 | 527 | G527R | A | 5.6e-06 | 7.5e-06 | 1.1e-05 | 8.033E-06 | 2.739E-06 | 5.600E-06 | 1.100E-05 |
| 10523 | 527 | G527N | A | 8.1e-06 | 1.5e-05 | 6.3e-06 | 9.800E-06 | 4.592E-06 | 6.300E-06 | 1.500E-05 |
| 10524 | 527 | G527D | A | 6.1e-06 | 1.3e-05 | 5.5e-06 | 8.200E-06 | 4.168E-06 | 5.500E-06 | 1.300E-05 |
| 10525 | 527 | G527C | A | 1.2e-05 | 6.7e-06 | 8.2e-06 | 8.967E-06 | 2.732E-06 | 6.700E-06 | 1.200E-05 |
| 10526 | 527 | G527E | A | 7.2e-06 | 1.4e-05 | 6.8e-06 | 9.333E-06 | 4.046E-06 | 6.800E-06 | 1.400E-05 |
| 10527 | 527 | G527Q | A | 1e-05 | 4.8e-06 | 6.1e-06 | 6.967E-06 | 2.706E-06 | 4.800E-06 | 1.000E-05 |
| 10528 | 527 | G527G | A | 7.4e-06 | 1.1e-05 | 5e-06 | 7.800E-06 | 3.020E-06 | 5.000E-06 | 1.100E-05 |
| 10529 | 527 | G527H | A | 9.2e-06 | 9.3e-06 | 7.5e-06 | 8.667E-06 | 1.012E-06 | 7.500E-06 | 9.300E-06 |
| 10530 | 527 | G527I | A | 6.7e-06 | 7.3e-06 | 5.1e-06 | 6.367E-06 | 1.137E-06 | 5.100E-06 | 7.300E-06 |
| 10531 | 527 | G527L | A | 9.4e-06 | 1.2e-05 | 1e-05 | 1.047E-05 | 1.361E-06 | 9.400E-06 | 1.200E-05 |
| 10532 | 527 | G527K | A | 9.6e-06 | 1.1e-05 | 5.1e-06 | 8.567E-06 | 3.083E-06 | 5.100E-06 | 1.100E-05 |
| 10533 | 527 | G527M | A | 9.2e-06 | 5.9e-06 | 1e-05 | 8.367E-06 | 2.173E-06 | 5.900E-06 | 1.000E-05 |
| 10534 | 527 | G527F | A | 1.2e-05 | 8.3e-06 | 1.1e-05 | 1.043E-05 | 1.914E-06 | 8.300E-06 | 1.200E-05 |
| 10535 | 527 | G527P | A | 1.1e-05 | 4.9e-06 | 6.5e-06 | 7.467E-06 | 3.163E-06 | 4.900E-06 | 1.100E-05 |
| 10536 | 527 | G527S | A | 1.2e-05 | 7.4e-06 | 1.1e-05 | 1.013E-05 | 2.419E-06 | 7.400E-06 | 1.200E-05 |
| 10537 | 527 | G527T | A | 1.2e-05 | 6.1e-06 | 1.1e-05 | 9.700E-06 | 3.158E-06 | 6.100E-06 | 1.200E-05 |
| 10538 | 527 | G527W | A | 9.8e-06 | 5e-06 | 9.1e-06 | 7.967E-06 | 2.593E-06 | 5.000E-06 | 9.800E-06 |
| 10539 | 527 | G527Y | A | 8.9e-06 | 3.2e-06 | 7.6e-06 | 6.567E-06 | 2.987E-06 | 3.200E-06 | 8.900E-06 |

| | | | | | | | | | | |
|---|---|---|---|---|---|---|---|---|---|---|
| 10540 | 527 | G527V | A | 6.5e-06 | 8.7e-06 | 9.9e-06 | 8.367E-06 | 1.724E-06 | 6.500E-06 | 9.900E-06 |
| 10541 | 528 | L528A | A | 4.4e-06 | 6.4e-06 | 5.6e-06 | 5.467E-06 | 1.007E-06 | 4.400E-06 | 6.400E-06 |
| 10542 | 528 | L528R | A | 7.3e-06 | 5.7e-06 | 5.2e-06 | 6.067E-06 | 1.097E-06 | 5.200E-06 | 7.300E-06 |
| 10543 | 528 | L528N | A | 7.8e-06 | 5.7e-06 | 7.9e-06 | 7.133E-06 | 1.242E-06 | 5.700E-06 | 7.900E-06 |
| 10544 | 528 | L528D | A | 1.3e-05 | 1.2e-05 | 6.4e-06 | 1.047E-05 | 3.557E-06 | 6.400E-06 | 1.300E-05 |
| 10545 | 528 | L528C | A | 8.8e-06 | 4.8e-06 | 6.2e-06 | 6.600E-06 | 2.030E-06 | 4.800E-06 | 8.800E-06 |
| 10546 | 528 | L528E | A | 9.2e-06 | 9.5e-06 | 6.5e-06 | 8.400E-06 | 1.652E-06 | 6.500E-06 | 9.500E-06 |
| 10547 | 528 | L528Q | A | 9.9e-06 | 9.2e-06 | 1.2e-05 | 1.037E-05 | 1.457E-06 | 9.200E-06 | 1.200E-05 |
| 10548 | 528 | L528G | A | 7e-06 | 4.1e-06 | 5e-06 | 5.367E-06 | 1.484E-06 | 4.100E-06 | 7.000E-06 |
| 10549 | 528 | L528H | A | 9.7e-06 | 7.6e-06 | 7.3e-06 | 8.200E-06 | 1.308E-06 | 7.300E-06 | 9.700E-06 |
| 10550 | 528 | L528I | A | 6.1e-06 | 7.4e-06 | 7.7e-06 | 7.067E-06 | 8.505E-07 | 6.100E-06 | 7.700E-06 |
| 10551 | 528 | L528L | A | 1e-05 | 1.1e-05 | 1.3e-05 | 1.133E-05 | 1.528E-06 | 1.000E-05 | 1.300E-05 |
| 10552 | 528 | L528K | A | 1.3e-05 | 7.2e-06 | 1.7e-05 | 1.240E-05 | 4.927E-06 | 7.200E-06 | 1.700E-05 |
| 10553 | 528 | L528M | A | 1.3e-05 | 6.7e-06 | 1.4e-05 | 1.123E-05 | 3.958E-06 | 6.700E-06 | 1.400E-05 |
| 10554 | 528 | L528F | A | 7.2e-06 | 1.1e-05 | 9.2e-06 | 9.133E-06 | 1.901E-06 | 7.200E-06 | 1.100E-05 |
| 10555 | 528 | L528P | A | 8.1e-06 | 6e-06 | 4.2e-06 | 6.100E-06 | 1.952E-06 | 4.200E-06 | 8.100E-06 |
| 10556 | 528 | L528S | A | 9.9e-06 | 6.8e-06 | 6.2e-06 | 7.633E-06 | 1.986E-06 | 6.200E-06 | 9.900E-06 |
| 10557 | 528 | L528T | A | 5.8e-06 | 6.3e-06 | 1e-05 | 7.367E-06 | 2.294E-06 | 5.800E-06 | 1.000E-05 |
| 10558 | 528 | L528W | A | 9.1e-06 | 1.2e-05 | 1.3e-05 | 1.137E-05 | 2.026E-06 | 9.100E-06 | 1.300E-05 |
| 10559 | 528 | L528Y | A | 8.9e-06 | 4.7e-06 | 9.6e-06 | 7.733E-06 | 2.650E-06 | 4.700E-06 | 9.600E-06 |
| 10560 | 528 | L528V | A | 9.1e-06 | 7.8e-06 | 8e-06 | 8.300E-06 | 7.000E-07 | 7.800E-06 | 9.100E-06 |
| 10561 | 529 | P529A | A | 7.7e-06 | 5.8e-06 | 6.2e-06 | 6.567E-06 | 1.002E-06 | 5.800E-06 | 7.700E-06 |
| 10562 | 529 | P529R | A | 7.7e-06 | 9.9e-06 | 7.4e-06 | 8.333E-06 | 1.365E-06 | 7.400E-06 | 9.900E-06 |
| 10563 | 529 | P529N | A | 7e-06 | 1e-05 | 5.8e-06 | 7.600E-06 | 2.163E-06 | 5.800E-06 | 1.000E-05 |
| 10564 | 529 | P529D | A | 8.8e-06 | 5.2e-06 | 5.4e-06 | 6.467E-06 | 2.023E-06 | 5.200E-06 | 8.800E-06 |
| 10565 | 529 | P529C | A | 6.1e-06 | 6e-06 | 8.3e-06 | 6.800E-06 | 1.300E-06 | 6.000E-06 | 8.300E-06 |
| 10566 | 529 | P529E | A | 7.6e-06 | 4.2e-06 | 7.4e-06 | 6.400E-06 | 1.908E-06 | 4.200E-06 | 7.600E-06 |
| 10567 | 529 | P529Q | A | 5.8e-06 | 7.2e-06 | 9.3e-06 | 7.433E-06 | 1.762E-06 | 5.800E-06 | 9.300E-06 |
| 10568 | 529 | P529G | A | 7.7e-06 | 8.1e-06 | 1.1e-05 | 8.933E-06 | 1.801E-06 | 7.700E-06 | 1.100E-05 |
| 10569 | 529 | P529H | A | 6.8e-06 | 1.2e-05 | 4.7e-06 | 7.833E-06 | 3.758E-06 | 4.700E-06 | 1.200E-05 |
| 10570 | 529 | P529I | A | 1.2e-05 | 8.7e-06 | 9.9e-06 | 1.020E-05 | 1.670E-06 | 8.700E-06 | 1.200E-05 |
| 10571 | 529 | P529L | A | 1.4e-05 | 9e-06 | 8.4e-06 | 1.047E-05 | 3.075E-06 | 8.400E-06 | 1.400E-05 |
| 10572 | 529 | P529K | A | 4.6e-06 | 6.8e-06 | 5.4e-06 | 5.600E-06 | 1.114E-06 | 4.600E-06 | 6.800E-06 |
| 10573 | 529 | P529M | A | 1e-05 | 4.3e-06 | 1e-05 | 8.100E-06 | 3.291E-06 | 4.300E-06 | 1.000E-05 |
| 10574 | 529 | P529F | A | 5.5e-06 | 4.5e-06 | 6.9e-06 | 5.633E-06 | 1.206E-06 | 4.500E-06 | 6.900E-06 |
| 10575 | 529 | P529P | A | 9.2e-06 | 9.8e-06 | 6.4e-06 | 8.467E-06 | 1.815E-06 | 6.400E-06 | 9.800E-06 |
| 10576 | 529 | P529S | A | 6.9e-06 | 1e-05 | 8.6e-06 | 8.500E-06 | 1.552E-06 | 6.900E-06 | 1.000E-05 |
| 10577 | 529 | P529T | A | 8.1e-06 | 9.1e-06 | 6e-06 | 7.733E-06 | 1.582E-06 | 6.000E-06 | 9.100E-06 |
| 10578 | 529 | P529W | A | 6.3e-06 | 9.2e-06 | 6.5e-06 | 7.333E-06 | 1.620E-06 | 6.300E-06 | 9.200E-06 |
| 10579 | 529 | P529Y | A | 6e-06 | 4.8e-06 | 5.6e-06 | 5.467E-06 | 6.110E-07 | 4.800E-06 | 6.000E-06 |
| 10580 | 529 | P529V | A | 6.6e-06 | 9.4e-06 | 4.8e-06 | 6.933E-06 | 2.318E-06 | 4.800E-06 | 9.400E-06 |
| 10581 | 530 | R530A | A | 6.6e-06 | 3.9e-06 | 1.3e-05 | 7.833E-06 | 4.674E-06 | 3.900E-06 | 1.300E-05 |
| 10582 | 530 | R530R | A | 1.2e-05 | 1.3e-05 | 6.2e-06 | 1.040E-05 | 3.672E-06 | 6.200E-06 | 1.300E-05 |
| 10583 | 530 | R530N | A | 1.1e-05 | 5.9e-06 | 8.2e-06 | 8.367E-06 | 2.554E-06 | 5.900E-06 | 1.100E-05 |
| 10584 | 530 | R530D | A | 1.1e-05 | 1.1e-05 | 7.4e-06 | 9.800E-06 | 2.078E-06 | 7.400E-06 | 1.100E-05 |
| 10585 | 530 | R530C | A | 6.9e-06 | 7.4e-06 | 1.1e-05 | 8.433E-06 | 2.237E-06 | 6.900E-06 | 1.100E-05 |
| 10586 | 530 | R530E | A | 6.7e-06 | 9.1e-06 | 6.7e-06 | 7.500E-06 | 1.386E-06 | 6.700E-06 | 9.100E-06 |
| 10587 | 530 | R530Q | A | 8.5e-06 | 6.1e-06 | 9.4e-06 | 8.000E-06 | 1.706E-06 | 6.100E-06 | 9.400E-06 |
| 10588 | 530 | R530G | A | 7.3e-06 | 7.7e-06 | 1.2e-05 | 9.000E-06 | 2.606E-06 | 7.300E-06 | 1.200E-05 |
| 10589 | 530 | R530H | A | 8.5e-06 | 6.7e-06 | 7.6e-06 | 7.600E-06 | 9.000E-07 | 6.700E-06 | 8.500E-06 |
| 10590 | 530 | R530I | A | 8.6e-06 | 9.1e-06 | 8.2e-06 | 8.633E-06 | 4.509E-07 | 8.200E-06 | 9.100E-06 |
| 10591 | 530 | R530L | A | 8.1e-06 | 3.8e-06 | 5.5e-06 | 5.800E-06 | 2.166E-06 | 3.800E-06 | 8.100E-06 |
| 10592 | 530 | R530K | A | 5.6e-06 | 9.4e-06 | 1.2e-05 | 9.000E-06 | 3.219E-06 | 5.600E-06 | 1.200E-05 |
| 10593 | 530 | R530M | A | 1.4e-05 | 1.1e-05 | 9.1e-06 | 1.137E-05 | 2.470E-06 | 9.100E-06 | 1.400E-05 |
| 10594 | 530 | R530F | A | 8.3e-06 | 9.7e-06 | 7.2e-06 | 8.400E-06 | 1.253E-06 | 7.200E-06 | 9.700E-06 |
| 10595 | 530 | R530P | A | 1.3e-05 | 1.4e-05 | 1.4e-05 | 1.367E-05 | 5.774E-07 | 1.300E-05 | 1.400E-05 |
| 10596 | 530 | R530S | A | 4.6e-06 | 4.5e-06 | 6.7e-06 | 5.267E-06 | 1.242E-06 | 4.500E-06 | 6.700E-06 |
| 10597 | 530 | R530T | A | 9.2e-06 | 1.1e-05 | 6.9e-06 | 9.033E-06 | 2.055E-06 | 6.900E-06 | 1.100E-05 |
| 10598 | 530 | R530W | A | 5.1e-06 | 7.7e-06 | 1.2e-05 | 8.267E-06 | 3.485E-06 | 5.100E-06 | 1.200E-05 |
| 10599 | 530 | R530Y | A | 8.5e-06 | 5.1e-06 | 1.5e-05 | 9.533E-06 | 5.030E-06 | 5.100E-06 | 1.500E-05 |
| 10600 | 530 | R530V | A | 9.1e-06 | 3e-06 | 7.9e-06 | 6.667E-06 | 3.232E-06 | 3.000E-06 | 9.100E-06 |
| 10601 | 531 | E531A | A | 1.2e-05 | 6.8e-06 | 9.4e-06 | 9.400E-06 | 2.600E-06 | 6.800E-06 | 1.200E-05 |
| 10602 | 531 | E531R | A | 9.2e-06 | 6.7e-06 | 8e-06 | 7.967E-06 | 1.250E-06 | 6.700E-06 | 9.200E-06 |
| 10603 | 531 | E531N | A | 1.2e-05 | 7.9e-06 | 9.1e-06 | 9.667E-06 | 2.108E-06 | 7.900E-06 | 1.200E-05 |
| 10604 | 531 | E531D | A | 7.8e-06 | 1e-05 | 5.3e-06 | 7.700E-06 | 2.352E-06 | 5.300E-06 | 1.000E-05 |
| 10605 | 531 | E531C | A | 7.5e-06 | 1.4e-05 | 1.3e-05 | 1.150E-05 | 3.500E-06 | 7.500E-06 | 1.400E-05 |
| 10606 | 531 | E531E | A | 4.6e-06 | 6.7e-06 | 9.4e-06 | 6.900E-06 | 2.406E-06 | 4.600E-06 | 9.400E-06 |
| 10607 | 531 | E531Q | A | 6.4e-06 | 8.8e-06 | 8.4e-06 | 7.867E-06 | 1.286E-06 | 6.400E-06 | 8.800E-06 |
| 10608 | 531 | E531G | A | 9.6e-06 | 1.4e-05 | 1.1e-05 | 1.153E-05 | 2.248E-06 | 9.600E-06 | 1.400E-05 |
| 10609 | 531 | E531H | A | 9.5e-06 | 5.6e-06 | 6.1e-06 | 7.067E-06 | 2.122E-06 | 5.600E-06 | 9.500E-06 |
| 10610 | 531 | E531I | A | 1.2e-05 | 1e-05 | 5.5e-06 | 9.167E-06 | 3.329E-06 | 5.500E-06 | 1.200E-05 |
| 10611 | 531 | E531L | A | 1.1e-05 | 1e-05 | 6.9e-06 | 9.300E-06 | 2.138E-06 | 6.900E-06 | 1.100E-05 |
| 10612 | 531 | E531K | A | 1e-05 | 6.4e-06 | 6.5e-06 | 7.633E-06 | 2.050E-06 | 6.400E-06 | 1.000E-05 |
| 10613 | 531 | E531M | A | 7.3e-06 | 6.9e-06 | 8.2e-06 | 7.467E-06 | 6.658E-07 | 6.900E-06 | 8.200E-06 |
| 10614 | 531 | E531F | A | 8.6e-06 | 9.1e-06 | 9.7e-06 | 9.133E-06 | 5.508E-07 | 8.600E-06 | 9.700E-06 |
| 10615 | 531 | E531P | A | 1.5e-05 | 8e-06 | 5.7e-06 | 9.567E-06 | 4.844E-06 | 5.700E-06 | 1.500E-05 |
| 10616 | 531 | E531S | A | 7.6e-06 | 5.5e-06 | 1.4e-05 | 9.033E-06 | 4.428E-06 | 5.500E-06 | 1.400E-05 |
| 10617 | 531 | E531T | A | 8.2e-06 | 9.8e-06 | 8.8e-06 | 8.933E-06 | 8.083E-07 | 8.200E-06 | 9.800E-06 |
| 10618 | 531 | E531W | A | 7.4e-06 | 6.2e-06 | 5e-06 | 6.200E-06 | 1.200E-06 | 5.000E-06 | 7.400E-06 |
| 10619 | 531 | E531Y | A | 8.4e-06 | 8.2e-06 | 1.2e-05 | 9.533E-06 | 2.139E-06 | 8.200E-06 | 1.200E-05 |
| 10620 | 531 | E531V | A | 1.3e-05 | 1.6e-05 | 6.1e-06 | 1.170E-05 | 5.076E-06 | 6.100E-06 | 1.600E-05 |
| 10621 | 532 | Y532A | A | 6.5e-06 | 1.1e-05 | 1.1e-05 | 9.500E-06 | 2.598E-06 | 6.500E-06 | 1.100E-05 |
| 10622 | 532 | Y532R | A | 5e-06 | 3.7e-06 | 9.4e-06 | 6.033E-06 | 2.987E-06 | 3.700E-06 | 9.400E-06 |
| 10623 | 532 | Y532N | A | 8.7e-06 | 4.9e-06 | 1.3e-05 | 8.867E-06 | 4.053E-06 | 4.900E-06 | 1.300E-05 |
| 10624 | 532 | Y532D | A | 9.8e-06 | 1e-05 | 7.7e-06 | 9.167E-06 | 1.274E-06 | 7.700E-06 | 1.000E-05 |

| | | | | | | | | | | |
|---|---|---|---|---|---|---|---|---|---|---|
| 10625 | 532 | Y532C | A | 4.3e-06 | 6.5e-06 | 6.5e-06 | 5.767E-06 | 1.270E-06 | 4.300E-06 | 6.500E-06 |
| 10626 | 532 | Y532E | A | 5.8e-06 | 1.5e-05 | 6.1e-06 | 8.967E-06 | 5.227E-06 | 5.800E-06 | 1.500E-05 |
| 10627 | 532 | Y532Q | A | 1.4e-05 | 4e-06 | 8.7e-06 | 8.900E-06 | 5.003E-06 | 4.000E-06 | 1.400E-05 |
| 10628 | 532 | Y532G | A | 8.2e-06 | 7.1e-06 | 4.2e-06 | 6.500E-06 | 2.066E-06 | 4.200E-06 | 8.200E-06 |
| 10629 | 532 | Y532H | A | 5.7e-06 | 5.4e-06 | 8.7e-06 | 6.600E-06 | 1.825E-06 | 5.400E-06 | 8.700E-06 |
| 10630 | 532 | Y532I | A | 6.7e-06 | 6.8e-06 | 1.1e-05 | 8.167E-06 | 2.454E-06 | 6.700E-06 | 1.100E-05 |
| 10631 | 532 | Y532L | A | 9e-06 | 8.8e-06 | 6.5e-06 | 8.100E-06 | 1.389E-06 | 6.500E-06 | 9.000E-06 |
| 10632 | 532 | Y532K | A | 5.1e-06 | 8.1e-06 | 6.1e-06 | 6.433E-06 | 1.528E-06 | 5.100E-06 | 8.100E-06 |
| 10633 | 532 | Y532M | A | 1.1e-05 | 8e-06 | 1e-05 | 9.667E-06 | 1.528E-06 | 8.000E-06 | 1.100E-05 |
| 10634 | 532 | Y532F | A | 1e-05 | 1.3e-05 | 7.2e-06 | 1.007E-05 | 2.901E-06 | 7.200E-06 | 1.300E-05 |
| 10635 | 532 | Y532P | A | 8.6e-06 | 1.1e-05 | 1.1e-05 | 1.020E-05 | 1.386E-06 | 8.600E-06 | 1.100E-05 |
| 10636 | 532 | Y532S | A | 9.8e-06 | 8.4e-06 | 1.1e-05 | 9.733E-06 | 1.301E-06 | 8.400E-06 | 1.100E-05 |
| 10637 | 532 | Y532T | A | 9.1e-06 | 7.1e-06 | 4.4e-06 | 6.867E-06 | 2.359E-06 | 4.400E-06 | 9.100E-06 |
| 10638 | 532 | Y532W | A | 1.1e-05 | 4.8e-06 | 8.3e-06 | 8.033E-06 | 3.109E-06 | 4.800E-06 | 1.100E-05 |
| 10639 | 532 | Y532Y | A | 1.2e-05 | 9e-06 | 6.8e-06 | 9.267E-06 | 2.610E-06 | 6.800E-06 | 1.200E-05 |
| 10640 | 532 | Y532V | A | 1.1e-05 | 1e-05 | 9.3e-06 | 1.010E-05 | 8.544E-07 | 9.300E-06 | 1.100E-05 |
| 10641 | 533 | V533A | A | 7.2e-06 | 6.2e-06 | 7.9e-06 | 7.100E-06 | 8.544E-07 | 6.200E-06 | 7.900E-06 |
| 10642 | 533 | V533R | A | 9.8e-06 | 8.6e-06 | 8.2e-06 | 8.867E-06 | 8.327E-07 | 8.200E-06 | 9.800E-06 |
| 10643 | 533 | V533N | A | 7.8e-06 | 8.1e-06 | 5.9e-06 | 7.267E-06 | 1.193E-06 | 5.900E-06 | 8.100E-06 |
| 10644 | 533 | V533D | A | 5.4e-06 | 1.3e-05 | 4.7e-06 | 7.700E-06 | 4.603E-06 | 4.700E-06 | 1.300E-05 |
| 10645 | 533 | V533C | A | 5.6e-06 | 1e-05 | 9.8e-06 | 8.467E-06 | 2.485E-06 | 5.600E-06 | 1.000E-05 |
| 10646 | 533 | V533E | A | 6.8e-06 | 8.2e-06 | 8.7e-06 | 7.900E-06 | 9.849E-07 | 6.800E-06 | 8.700E-06 |
| 10647 | 533 | V533Q | A | 1.1e-05 | 7.1e-06 | 7.2e-06 | 8.433E-06 | 2.223E-06 | 7.100E-06 | 1.100E-05 |
| 10648 | 533 | V533G | A | 4.2e-06 | 1e-05 | 8.6e-06 | 7.600E-06 | 3.027E-06 | 4.200E-06 | 1.000E-05 |
| 10649 | 533 | V533H | A | 7.5e-06 | 1.3e-05 | 9.4e-06 | 9.967E-06 | 2.793E-06 | 7.500E-06 | 1.300E-05 |
| 10650 | 533 | V533I | A | 7.2e-06 | 9.6e-06 | 1.2e-05 | 9.600E-06 | 2.400E-06 | 7.200E-06 | 1.200E-05 |
| 10651 | 533 | V533L | A | 8.3e-06 | 8.8e-06 | 8.8e-06 | 8.633E-06 | 2.887E-07 | 8.300E-06 | 8.800E-06 |
| 10652 | 533 | V533K | A | 6.2e-06 | 8.5e-06 | 6.6e-06 | 7.100E-06 | 1.229E-06 | 6.200E-06 | 8.500E-06 |
| 10653 | 533 | V533M | A | 6.5e-06 | 5.6e-06 | 9.5e-06 | 7.200E-06 | 2.042E-06 | 5.600E-06 | 9.500E-06 |
| 10654 | 533 | V533F | A | 6.6e-06 | 1.9e-05 | 1.1e-05 | 1.220E-05 | 6.286E-06 | 6.600E-06 | 1.900E-05 |
| 10655 | 533 | V533P | A | 1.1e-05 | 7.6e-06 | 9.7e-06 | 9.433E-06 | 1.716E-06 | 7.600E-06 | 1.100E-05 |
| 10656 | 533 | V533S | A | 5.8e-06 | 3.8e-06 | 5.9e-06 | 5.167E-06 | 1.185E-06 | 3.800E-06 | 5.900E-06 |
| 10657 | 533 | V533T | A | 9.6e-06 | 7.7e-06 | 8.1e-06 | 8.467E-06 | 1.002E-06 | 7.700E-06 | 9.600E-06 |
| 10658 | 533 | V533W | A | 5.5e-06 | 1.2e-05 | 6.5e-06 | 8.000E-06 | 3.500E-06 | 5.500E-06 | 1.200E-05 |
| 10659 | 533 | V533Y | A | 4.7e-06 | 1.1e-05 | 8.5e-06 | 8.067E-06 | 3.172E-06 | 4.700E-06 | 1.100E-05 |
| 10660 | 533 | V533V | A | 6.6e-06 | 6.2e-06 | 6.8e-06 | 6.533E-06 | 3.055E-07 | 6.200E-06 | 6.800E-06 |
| 10661 | 534 | N534A | A | 5.7e-06 | 7.7e-06 | 9e-06 | 7.467E-06 | 1.662E-06 | 5.700E-06 | 9.000E-06 |
| 10662 | 534 | N534R | A | 1.2e-05 | 1e-05 | 1.4e-05 | 1.200E-05 | 2.000E-06 | 1.000E-05 | 1.400E-05 |
| 10663 | 534 | N534N | A | 6.7e-06 | 1.2e-05 | 5.2e-06 | 7.967E-06 | 3.573E-06 | 5.200E-06 | 1.200E-05 |
| 10664 | 534 | N534D | A | 7.3e-06 | 1.3e-05 | 6.9e-06 | 9.067E-06 | 3.412E-06 | 6.900E-06 | 1.300E-05 |
| 10665 | 534 | N534C | A | 5.8e-06 | 8.1e-06 | 8.3e-06 | 7.400E-06 | 1.389E-06 | 5.800E-06 | 8.300E-06 |
| 10666 | 534 | N534E | A | 1.2e-05 | 6.6e-06 | 1.2e-05 | 1.020E-05 | 3.118E-06 | 6.600E-06 | 1.200E-05 |
| 10667 | 534 | N534Q | A | 9e-06 | 6.9e-06 | 5.5e-06 | 7.133E-06 | 1.762E-06 | 5.500E-06 | 9.000E-06 |
| 10668 | 534 | N534G | A | 6.6e-06 | 8.4e-06 | 7.4e-06 | 7.467E-06 | 9.018E-07 | 6.600E-06 | 8.400E-06 |
| 10669 | 534 | N534H | A | 8.2e-06 | 7.4e-06 | 7.5e-06 | 7.700E-06 | 4.359E-07 | 7.400E-06 | 8.200E-06 |
| 10670 | 534 | N534I | A | 7.9e-06 | 9.1e-06 | 8.5e-06 | 8.500E-06 | 6.000E-07 | 7.900E-06 | 9.100E-06 |
| 10671 | 534 | N534L | A | 4.4e-06 | 8.9e-06 | 8.7e-06 | 7.333E-06 | 2.542E-06 | 4.400E-06 | 8.900E-06 |
| 10672 | 534 | N534K | A | 8.3e-06 | 1.3e-05 | 6.5e-06 | 9.267E-06 | 3.356E-06 | 6.500E-06 | 1.300E-05 |
| 10673 | 534 | N534M | A | 6e-06 | 7.5e-06 | 1.3e-05 | 8.833E-06 | 3.686E-06 | 6.000E-06 | 1.300E-05 |
| 10674 | 534 | N534F | A | 5.5e-06 | 9.1e-06 | 1.3e-05 | 9.200E-06 | 3.751E-06 | 5.500E-06 | 1.300E-05 |
| 10675 | 534 | N534P | A | 1.3e-05 | 6.4e-06 | 5.1e-06 | 8.167E-06 | 4.236E-06 | 5.100E-06 | 1.300E-05 |
| 10676 | 534 | N534S | A | 5.6e-06 | 6e-06 | 1.8e-05 | 9.867E-06 | 7.047E-06 | 5.600E-06 | 1.800E-05 |
| 10677 | 534 | N534T | A | 1.4e-05 | 1.4e-05 | 1.2e-05 | 1.333E-05 | 1.155E-06 | 1.200E-05 | 1.400E-05 |
| 10678 | 534 | N534W | A | 1.2e-05 | 6.9e-06 | 9.8e-06 | 9.567E-06 | 2.558E-06 | 6.900E-06 | 1.200E-05 |
| 10679 | 534 | N534Y | A | 8.9e-06 | 8.5e-06 | 7.2e-06 | 8.200E-06 | 8.888E-07 | 7.200E-06 | 8.900E-06 |
| 10680 | 534 | N534V | A | 4.6e-06 | 6.4e-06 | 9.8e-06 | 6.933E-06 | 2.641E-06 | 4.600E-06 | 9.800E-06 |
| 10681 | 535 | A535A | A | 1.1e-05 | 6.1e-06 | 4.7e-06 | 7.267E-06 | 3.308E-06 | 4.700E-06 | 1.100E-05 |
| 10682 | 535 | A535R | A | 5.2e-06 | 9.5e-06 | 8.5e-06 | 7.733E-06 | 2.250E-06 | 5.200E-06 | 9.500E-06 |
| 10683 | 535 | A535N | A | 6.6e-06 | 9.8e-06 | 9e-06 | 8.467E-06 | 1.665E-06 | 6.600E-06 | 9.800E-06 |
| 10684 | 535 | A535D | A | 1.3e-05 | 5.7e-06 | 5.3e-06 | 8.000E-06 | 4.335E-06 | 5.300E-06 | 1.300E-05 |
| 10685 | 535 | A535C | A | 8.2e-06 | 7.4e-06 | 1e-05 | 8.533E-06 | 1.332E-06 | 7.400E-06 | 1.000E-05 |
| 10686 | 535 | A535E | A | 1.1e-05 | 7.3e-06 | 1e-05 | 9.433E-06 | 1.914E-06 | 7.300E-06 | 1.100E-05 |
| 10687 | 535 | A535Q | A | 4.3e-06 | 7.6e-06 | 6.3e-06 | 6.067E-06 | 1.662E-06 | 4.300E-06 | 7.600E-06 |
| 10688 | 535 | A535G | A | 1.3e-05 | 6e-06 | 7.3e-06 | 8.767E-06 | 3.723E-06 | 6.000E-06 | 1.300E-05 |
| 10689 | 535 | A535H | A | 1.1e-05 | 9.7e-06 | 5.3e-06 | 8.667E-06 | 2.987E-06 | 5.300E-06 | 1.100E-05 |
| 10690 | 535 | A535I | A | 1.1e-05 | 7.1e-06 | 7e-06 | 8.367E-06 | 2.281E-06 | 7.000E-06 | 1.100E-05 |
| 10691 | 535 | A535L | A | 7.2e-06 | 2.3e-06 | 1.4e-05 | 7.833E-06 | 5.876E-06 | 2.300E-06 | 1.400E-05 |
| 10692 | 535 | A535K | A | 7.3e-06 | 8.9e-06 | 8.9e-06 | 8.367E-06 | 9.238E-07 | 7.300E-06 | 8.900E-06 |
| 10693 | 535 | A535M | A | 8.6e-06 | 9.5e-06 | 4.9e-06 | 7.667E-06 | 2.438E-06 | 4.900E-06 | 9.500E-06 |
| 10694 | 535 | A535F | A | 5.9e-06 | 6.9e-06 | 8.3e-06 | 7.033E-06 | 1.206E-06 | 5.900E-06 | 8.300E-06 |
| 10695 | 535 | A535P | A | 5e-06 | 7.6e-06 | 7e-06 | 6.533E-06 | 1.361E-06 | 5.000E-06 | 7.600E-06 |
| 10696 | 535 | A535S | A | 8e-06 | 5e-06 | 8.4e-06 | 7.133E-06 | 1.858E-06 | 5.000E-06 | 8.400E-06 |
| 10697 | 535 | A535T | A | 6.9e-06 | 7.1e-06 | 5.9e-06 | 6.633E-06 | 6.429E-07 | 5.900E-06 | 7.100E-06 |
| 10698 | 535 | A535W | A | 5.5e-06 | 6.7e-06 | 6.6e-06 | 6.267E-06 | 6.658E-07 | 5.500E-06 | 6.700E-06 |
| 10699 | 535 | A535Y | A | 1e-05 | 9.8e-06 | 4.5e-06 | 8.100E-06 | 3.119E-06 | 4.500E-06 | 1.000E-05 |
| 10700 | 535 | A535V | A | 8.4e-06 | 1.1e-05 | 1.2e-05 | 1.047E-05 | 1.858E-06 | 8.400E-06 | 1.200E-05 |
| 10701 | 536 | R536A | A | 6.8e-06 | 9.2e-06 | 1.4e-05 | 1.000E-05 | 3.666E-06 | 6.800E-06 | 1.400E-05 |
| 10702 | 536 | R536R | A | 6.6e-06 | 1.4e-05 | 8.3e-06 | 9.633E-06 | 3.876E-06 | 6.600E-06 | 1.400E-05 |
| 10703 | 536 | R536N | A | 9.8e-06 | 1.3e-05 | 8.6e-06 | 1.047E-05 | 2.274E-06 | 8.600E-06 | 1.300E-05 |
| 10704 | 536 | R536D | A | 5.4e-06 | 9.6e-06 | 6.6e-06 | 7.200E-06 | 2.163E-06 | 5.400E-06 | 9.600E-06 |
| 10705 | 536 | R536C | A | 9.5e-06 | 6.8e-06 | 6.7e-06 | 7.667E-06 | 1.589E-06 | 6.700E-06 | 9.500E-06 |
| 10706 | 536 | R536E | A | 1e-05 | 7.6e-06 | 7.9e-06 | 8.500E-06 | 1.308E-06 | 7.600E-06 | 1.000E-05 |
| 10707 | 536 | R536Q | A | 6.4e-06 | 1.3e-05 | 6e-06 | 8.467E-06 | 3.931E-06 | 6.000E-06 | 1.300E-05 |
| 10708 | 536 | R536G | A | 1.1e-05 | 9.9e-06 | 5.8e-06 | 8.900E-06 | 2.740E-06 | 5.800E-06 | 1.100E-05 |
| 10709 | 536 | R536H | A | 4.7e-06 | 4.8e-06 | 7.4e-06 | 5.633E-06 | 1.531E-06 | 4.700E-06 | 7.400E-06 |

| | | | | | | | | | | |
|---|---|---|---|---|---|---|---|---|---|---|
| 10710 | 536 | R536I | A | 1.1e-05 | 5.7e-06 | 6.6e-06 | 7.767E-06 | 2.836E-06 | 5.700E-06 | 1.100E-05 |
| 10711 | 536 | R536L | A | 8.9e-06 | 6.2e-06 | 6.9e-06 | 7.333E-06 | 1.401E-06 | 6.200E-06 | 8.900E-06 |
| 10712 | 536 | R536K | A | 6.6e-06 | 1.4e-05 | 7.8e-06 | 9.467E-06 | 3.972E-06 | 6.600E-06 | 1.400E-05 |
| 10713 | 536 | R536M | A | 9.8e-06 | 7.2e-06 | 1.3e-05 | 1.000E-05 | 2.905E-06 | 7.200E-06 | 1.300E-05 |
| 10714 | 536 | R536F | A | 5.7e-06 | 6.4e-06 | 9.7e-06 | 7.267E-06 | 2.136E-06 | 5.700E-06 | 9.700E-06 |
| 10715 | 536 | R536P | A | 6e-06 | 5e-06 | 7.6e-06 | 6.200E-06 | 1.311E-06 | 5.000E-06 | 7.600E-06 |
| 10716 | 536 | R536S | A | 5.9e-06 | 9.8e-06 | 5.2e-06 | 6.967E-06 | 2.479E-06 | 5.200E-06 | 9.800E-06 |
| 10717 | 536 | R536T | A | 7.2e-06 | 6.7e-06 | 5.6e-06 | 6.500E-06 | 8.185E-07 | 5.600E-06 | 7.200E-06 |
| 10718 | 536 | R536W | A | 5.8e-06 | 6.4e-06 | 6.5e-06 | 6.233E-06 | 3.786E-07 | 5.800E-06 | 6.500E-06 |
| 10719 | 536 | R536Y | A | 7.5e-06 | 4.8e-06 | 6.8e-06 | 6.367E-06 | 1.401E-06 | 4.800E-06 | 7.500E-06 |
| 10720 | 536 | R536V | A | 8.1e-06 | 1.2e-05 | 1.4e-05 | 1.137E-05 | 3.001E-06 | 8.100E-06 | 1.400E-05 |
| 10721 | 537 | H537A | A | 4.9e-06 | 6.6e-06 | 5.9e-06 | 5.800E-06 | 8.544E-07 | 4.900E-06 | 6.600E-06 |
| 10722 | 537 | H537R | A | 8.6e-06 | 1.6e-05 | 4.6e-06 | 9.733E-06 | 5.784E-06 | 4.600E-06 | 1.600E-05 |
| 10723 | 537 | H537N | A | 7.8e-06 | 9.2e-06 | 1.2e-05 | 9.667E-06 | 2.139E-06 | 7.800E-06 | 1.200E-05 |
| 10724 | 537 | H537D | A | 8.4e-06 | 9.1e-06 | 8.7e-06 | 8.733E-06 | 3.512E-07 | 8.400E-06 | 9.100E-06 |
| 10725 | 537 | H537C | A | 6.8e-06 | 1.1e-05 | 7.1e-06 | 8.300E-06 | 2.343E-06 | 6.800E-06 | 1.100E-05 |
| 10726 | 537 | H537E | A | 1.2e-05 | 5.1e-06 | 3.6e-06 | 6.900E-06 | 4.480E-06 | 3.600E-06 | 1.200E-05 |
| 10727 | 537 | H537Q | A | 1.3e-05 | 1.2e-05 | 1e-05 | 1.167E-05 | 1.528E-06 | 1.000E-05 | 1.300E-05 |
| 10728 | 537 | H537G | A | 4.6e-06 | 4.4e-06 | 6.8e-06 | 5.267E-06 | 1.332E-06 | 4.400E-06 | 6.800E-06 |
| 10729 | 537 | H537H | A | 9e-06 | 6.9e-06 | 1e-05 | 8.633E-06 | 1.582E-06 | 6.900E-06 | 1.000E-05 |
| 10730 | 537 | H537I | A | 7.4e-06 | 4.7e-06 | 8.8e-06 | 6.967E-06 | 2.084E-06 | 4.700E-06 | 8.800E-06 |
| 10731 | 537 | H537L | A | 1.1e-05 | 7.1e-06 | 1.2e-05 | 1.003E-05 | 2.589E-06 | 7.100E-06 | 1.200E-05 |
| 10732 | 537 | H537K | A | 5.4e-06 | 9.4e-06 | 7e-06 | 7.267E-06 | 2.013E-06 | 5.400E-06 | 9.400E-06 |
| 10733 | 537 | H537M | A | 6.8e-06 | 4.5e-06 | 8.5e-06 | 6.600E-06 | 2.007E-06 | 4.500E-06 | 8.500E-06 |
| 10734 | 537 | H537F | A | 1.7e-05 | 8.4e-06 | 6.4e-06 | 1.060E-05 | 5.632E-06 | 6.400E-06 | 1.700E-05 |
| 10735 | 537 | H537P | A | 6.9e-06 | 7.6e-06 | 6.7e-06 | 7.067E-06 | 4.726E-07 | 6.700E-06 | 7.600E-06 |
| 10736 | 537 | H537S | A | 6.4e-06 | 9.9e-06 | 6.2e-06 | 7.500E-06 | 2.081E-06 | 6.200E-06 | 9.900E-06 |
| 10737 | 537 | H537T | A | 9e-06 | 9e-06 | 7.8e-06 | 8.600E-06 | 6.928E-07 | 7.800E-06 | 9.000E-06 |
| 10738 | 537 | H537W | A | 6.8e-06 | 1.9e-05 | 6.4e-06 | 1.073E-05 | 7.162E-06 | 6.400E-06 | 1.900E-05 |
| 10739 | 537 | H537Y | A | 8.1e-06 | 7.3e-06 | 5.4e-06 | 6.933E-06 | 1.387E-06 | 5.400E-06 | 8.100E-06 |
| 10740 | 537 | H537V | A | 1.4e-05 | 6e-06 | 9.1e-06 | 9.700E-06 | 4.034E-06 | 6.000E-06 | 1.400E-05 |
| 10741 | 538 | C538A | A | 9e-06 | 7.6e-06 | 9e-06 | 8.533E-06 | 8.083E-07 | 7.600E-06 | 9.000E-06 |
| 10742 | 538 | C538R | A | 6.5e-06 | 9e-06 | 1.3e-05 | 9.500E-06 | 3.279E-06 | 6.500E-06 | 1.300E-05 |
| 10743 | 538 | C538N | A | 4.7e-06 | 5.2e-06 | 5.4e-06 | 5.100E-06 | 3.606E-07 | 4.700E-06 | 5.400E-06 |
| 10744 | 538 | C538D | A | 9.8e-06 | 1e-05 | 8e-06 | 9.267E-06 | 1.102E-06 | 8.000E-06 | 1.000E-05 |
| 10745 | 538 | C538C | A | 9.1e-06 | 1e-05 | 6.3e-06 | 8.467E-06 | 1.930E-06 | 6.300E-06 | 1.000E-05 |
| 10746 | 538 | C538E | A | 7.8e-06 | 1.1e-05 | 1.2e-05 | 1.027E-05 | 2.194E-06 | 7.800E-06 | 1.200E-05 |
| 10747 | 538 | C538Q | A | 9.9e-06 | 9.6e-06 | 4.5e-06 | 8.000E-06 | 3.035E-06 | 4.500E-06 | 9.900E-06 |
| 10748 | 538 | C538G | A | 7.2e-06 | 6.5e-06 | 6.2e-06 | 6.633E-06 | 5.132E-07 | 6.200E-06 | 7.200E-06 |
| 10749 | 538 | C538H | A | 6.1e-06 | 5.7e-06 | 6.8e-06 | 6.200E-06 | 5.568E-07 | 5.700E-06 | 6.800E-06 |
| 10750 | 538 | C538I | A | 8.4e-06 | 6.6e-06 | 4.7e-06 | 6.567E-06 | 1.850E-06 | 4.700E-06 | 8.400E-06 |
| 10751 | 538 | C538L | A | 4.9e-06 | 5.5e-06 | 7e-06 | 5.800E-06 | 1.082E-06 | 4.900E-06 | 7.000E-06 |
| 10752 | 538 | C538K | A | 8e-06 | 9.7e-06 | 7e-06 | 8.233E-06 | 1.365E-06 | 7.000E-06 | 9.700E-06 |
| 10753 | 538 | C538M | A | 9.7e-06 | 7.1e-06 | 6.5e-06 | 7.767E-06 | 1.701E-06 | 6.500E-06 | 9.700E-06 |
| 10754 | 538 | C538F | A | 7.5e-06 | 5.9e-06 | 9.4e-06 | 7.600E-06 | 1.752E-06 | 5.900E-06 | 9.400E-06 |
| 10755 | 538 | C538P | A | 7e-06 | 9.2e-06 | 6.3e-06 | 7.500E-06 | 1.513E-06 | 6.300E-06 | 9.200E-06 |
| 10756 | 538 | C538S | A | 1.1e-05 | 8.5e-06 | 8.2e-06 | 9.233E-06 | 1.537E-06 | 8.200E-06 | 1.100E-05 |
| 10757 | 538 | C538T | A | 1.1e-05 | 6.5e-06 | 6.9e-06 | 8.133E-06 | 2.491E-06 | 6.500E-06 | 1.100E-05 |
| 10758 | 538 | C538W | A | 1.1e-05 | 7.6e-06 | 1.2e-05 | 1.020E-05 | 2.307E-06 | 7.600E-06 | 1.200E-05 |
| 10759 | 538 | C538Y | A | 1e-05 | 6.2e-06 | 7.3e-06 | 7.833E-06 | 1.955E-06 | 6.200E-06 | 1.000E-05 |
| 10760 | 538 | C538V | A | 5.8e-06 | 4.8e-06 | 8.4e-06 | 6.333E-06 | 1.858E-06 | 4.800E-06 | 8.400E-06 |
| 10761 | 539 | L539A | A | 7e-06 | 5.8e-06 | 5.5e-06 | 6.100E-06 | 7.937E-07 | 5.500E-06 | 7.000E-06 |
| 10762 | 539 | L539R | A | 6.1e-06 | 4.8e-06 | 6.4e-06 | 5.767E-06 | 8.505E-07 | 4.800E-06 | 6.400E-06 |
| 10763 | 539 | L539N | A | 5.9e-06 | 1.2e-05 | 9.9e-06 | 9.267E-06 | 3.099E-06 | 5.900E-06 | 1.200E-05 |
| 10764 | 539 | L539D | A | 6.7e-06 | 7.9e-06 | 1.2e-05 | 8.867E-06 | 2.779E-06 | 6.700E-06 | 1.200E-05 |
| 10765 | 539 | L539C | A | 4.5e-06 | 5.5e-06 | 7.6e-06 | 5.867E-06 | 1.582E-06 | 4.500E-06 | 7.600E-06 |
| 10766 | 539 | L539E | A | 9.9e-06 | 5.9e-06 | 9.4e-06 | 8.400E-06 | 2.179E-06 | 5.900E-06 | 9.900E-06 |
| 10767 | 539 | L539Q | A | 1.2e-05 | 1e-05 | 7.4e-06 | 9.800E-06 | 2.307E-06 | 7.400E-06 | 1.200E-05 |
| 10768 | 539 | L539G | A | 1.5e-05 | 1.8e-05 | 5.8e-06 | 1.293E-05 | 6.357E-06 | 5.800E-06 | 1.800E-05 |
| 10769 | 539 | L539H | A | 9.1e-06 | 6.7e-06 | 8.1e-06 | 7.967E-06 | 1.206E-06 | 6.700E-06 | 9.100E-06 |
| 10770 | 539 | L539I | A | 8.1e-06 | 8.4e-06 | 6.5e-06 | 7.667E-06 | 1.021E-06 | 6.500E-06 | 8.400E-06 |
| 10771 | 539 | L539L | A | 3.7e-06 | 8.3e-06 | 1e-05 | 7.333E-06 | 3.259E-06 | 3.700E-06 | 1.000E-05 |
| 10772 | 539 | L539K | A | 1.2e-05 | 5.8e-06 | 6.4e-06 | 8.067E-06 | 3.420E-06 | 5.800E-06 | 1.200E-05 |
| 10773 | 539 | L539M | A | 8.9e-06 | 1.5e-05 | 9e-06 | 1.097E-05 | 3.493E-06 | 8.900E-06 | 1.500E-05 |
| 10774 | 539 | L539F | A | 1.3e-05 | 1.5e-05 | 6.4e-06 | 1.147E-05 | 4.500E-06 | 6.400E-06 | 1.500E-05 |
| 10775 | 539 | L539P | A | 1.1e-05 | 9.5e-06 | 5.3e-06 | 8.600E-06 | 2.955E-06 | 5.300E-06 | 1.100E-05 |
| 10776 | 539 | L539S | A | 6.2e-06 | 6.9e-06 | 5.9e-06 | 6.333E-06 | 5.132E-07 | 5.900E-06 | 6.900E-06 |
| 10777 | 539 | L539T | A | 6e-06 | 1.1e-05 | 1.2e-05 | 9.667E-06 | 3.215E-06 | 6.000E-06 | 1.200E-05 |
| 10778 | 539 | L539W | A | 7.8e-06 | 5.2e-06 | 5.9e-06 | 6.300E-06 | 1.345E-06 | 5.200E-06 | 7.800E-06 |
| 10779 | 539 | L539Y | A | 7.9e-06 | 6.1e-06 | 5.5e-06 | 6.500E-06 | 1.249E-06 | 5.500E-06 | 7.900E-06 |
| 10780 | 539 | L539V | A | 4.6e-06 | 1.5e-05 | 5.4e-06 | 8.333E-06 | 5.787E-06 | 4.600E-06 | 1.500E-05 |
| 10781 | 540 | P540A | A | 5.1e-06 | 6.7e-06 | 1e-05 | 7.267E-06 | 2.499E-06 | 5.100E-06 | 1.000E-05 |
| 10782 | 540 | P540R | A | 1e-05 | 2.1e-05 | 8.6e-06 | 1.320E-05 | 6.791E-06 | 8.600E-06 | 2.100E-05 |
| 10783 | 540 | P540N | A | 4.4e-06 | 7.6e-06 | 1.1e-05 | 7.667E-06 | 3.301E-06 | 4.400E-06 | 1.100E-05 |
| 10784 | 540 | P540D | A | 7.1e-06 | 8.3e-06 | 8.7e-06 | 8.033E-06 | 8.327E-07 | 7.100E-06 | 8.700E-06 |
| 10785 | 540 | P540C | A | 9.3e-06 | 8.4e-06 | 4.3e-06 | 7.333E-06 | 2.665E-06 | 4.300E-06 | 9.300E-06 |
| 10786 | 540 | P540E | A | 1e-05 | 5.8e-06 | 6.7e-06 | 7.500E-06 | 2.211E-06 | 5.800E-06 | 1.000E-05 |
| 10787 | 540 | P540Q | A | 1.4e-05 | 5.8e-06 | 1.1e-05 | 1.027E-05 | 4.149E-06 | 5.800E-06 | 1.400E-05 |
| 10788 | 540 | P540G | A | 9.7e-06 | 1.1e-05 | 1.3e-05 | 1.123E-05 | 1.662E-06 | 9.700E-06 | 1.300E-05 |
| 10789 | 540 | P540H | A | 7e-06 | 6.8e-06 | 4.6e-06 | 6.133E-06 | 1.332E-06 | 4.600E-06 | 7.000E-06 |
| 10790 | 540 | P540I | A | 9.9e-06 | 1e-05 | 7.9e-06 | 9.267E-06 | 1.185E-06 | 7.900E-06 | 1.000E-05 |
| 10791 | 540 | P540L | A | 7.4e-06 | 7.8e-06 | 1.1e-05 | 8.733E-06 | 1.973E-06 | 7.400E-06 | 1.100E-05 |
| 10792 | 540 | P540K | A | 1.3e-05 | 4.5e-06 | 8e-06 | 8.500E-06 | 4.272E-06 | 4.500E-06 | 1.300E-05 |
| 10793 | 540 | P540M | A | 1.3e-05 | 8.7e-06 | 6.7e-06 | 9.467E-06 | 3.219E-06 | 6.700E-06 | 1.300E-05 |
| 10794 | 540 | P540F | A | 2.1e-05 | 1.3e-05 | 5.2e-06 | 1.307E-05 | 7.900E-06 | 5.200E-06 | 2.100E-05 |

| | | | | | | | | | | |
|---|---|---|---|---|---|---|---|---|---|---|
| 10795 | 540 | P540P | A | 4.9e-06 | 9.8e-06 | 1.4e-05 | 9.567E-06 | 4.554E-06 | 4.900E-06 | 1.400E-05 |
| 10796 | 540 | P540S | A | 6.5e-06 | 7.6e-06 | 1.4e-05 | 9.367E-06 | 4.050E-06 | 6.500E-06 | 1.400E-05 |
| 10797 | 540 | P540T | A | 6.1e-06 | 4.5e-06 | 1.4e-05 | 8.200E-06 | 5.086E-06 | 4.500E-06 | 1.400E-05 |
| 10798 | 540 | P540W | A | 1.3e-05 | 1.1e-05 | 6.8e-06 | 1.027E-05 | 3.164E-06 | 6.800E-06 | 1.300E-05 |
| 10799 | 540 | P540Y | A | 6.3e-06 | 8.1e-06 | 8.3e-06 | 7.567E-06 | 1.102E-06 | 6.300E-06 | 8.300E-06 |
| 10800 | 540 | P540V | A | 8e-06 | 8.9e-06 | 6.1e-06 | 7.667E-06 | 1.429E-06 | 6.100E-06 | 8.900E-06 |
| 10801 | 541 | C541A | A | 8e-06 | 9.2e-06 | 7.7e-06 | 8.300E-06 | 7.937E-07 | 7.700E-06 | 9.200E-06 |
| 10802 | 541 | C541R | A | 5.7e-06 | 1.1e-05 | 1.4e-05 | 1.023E-05 | 4.203E-06 | 5.700E-06 | 1.400E-05 |
| 10803 | 541 | C541N | A | 8.2e-06 | 1e-05 | 7.5e-06 | 8.567E-06 | 1.290E-06 | 7.500E-06 | 1.000E-05 |
| 10804 | 541 | C541D | A | 1.5e-05 | 5.2e-06 | 9.2e-06 | 9.800E-06 | 4.927E-06 | 5.200E-06 | 1.500E-05 |
| 10805 | 541 | C541C | A | 6.2e-06 | 7e-06 | 1.1e-05 | 8.067E-06 | 2.572E-06 | 6.200E-06 | 1.100E-05 |
| 10806 | 541 | C541E | A | 9.2e-06 | 6.6e-06 | 4.8e-06 | 6.867E-06 | 2.212E-06 | 4.800E-06 | 9.200E-06 |
| 10807 | 541 | C541Q | A | 6.7e-06 | 6.9e-06 | 8.2e-06 | 7.267E-06 | 8.145E-07 | 6.700E-06 | 8.200E-06 |
| 10808 | 541 | C541G | A | 9.8e-06 | 9.6e-06 | 5.1e-06 | 8.167E-06 | 2.658E-06 | 5.100E-06 | 9.800E-06 |
| 10809 | 541 | C541H | A | 8.2e-06 | 7.3e-06 | 5.9e-06 | 7.133E-06 | 1.159E-06 | 5.900E-06 | 8.200E-06 |
| 10810 | 541 | C541I | A | 6.6e-06 | 7.5e-06 | 6.7e-06 | 6.933E-06 | 4.933E-07 | 6.600E-06 | 7.500E-06 |
| 10811 | 541 | C541L | A | 5.9e-06 | 6.3e-06 | 8.5e-06 | 6.900E-06 | 1.400E-06 | 5.900E-06 | 8.500E-06 |
| 10812 | 541 | C541K | A | 5.9e-06 | 6.5e-06 | 7.8e-06 | 6.733E-06 | 9.713E-07 | 5.900E-06 | 7.800E-06 |
| 10813 | 541 | C541M | A | 1.4e-05 | 9.3e-06 | 5.7e-06 | 9.667E-06 | 4.162E-06 | 5.700E-06 | 1.400E-05 |
| 10814 | 541 | C541F | A | 7.1e-06 | 1e-05 | 8.2e-06 | 8.433E-06 | 1.464E-06 | 7.100E-06 | 1.000E-05 |
| 10815 | 541 | C541P | A | 9e-06 | 6.6e-06 | 8.1e-06 | 7.900E-06 | 1.212E-06 | 6.600E-06 | 9.000E-06 |
| 10816 | 541 | C541S | A | 6.3e-06 | 5.5e-06 | 9.8e-06 | 7.200E-06 | 2.287E-06 | 5.500E-06 | 9.800E-06 |
| 10817 | 541 | C541T | A | 9.3e-06 | 1e-05 | 8.2e-06 | 9.167E-06 | 9.074E-07 | 8.200E-06 | 1.000E-05 |
| 10818 | 541 | C541W | A | 8.4e-06 | 5.5e-06 | 7.7e-06 | 7.200E-06 | 1.513E-06 | 5.500E-06 | 8.400E-06 |
| 10819 | 541 | C541Y | A | 9.4e-06 | 1.2e-05 | 1e-05 | 1.047E-05 | 1.361E-06 | 9.400E-06 | 1.200E-05 |
| 10820 | 541 | C541V | A | 5.2e-06 | 9.9e-06 | 9.5e-06 | 8.200E-06 | 2.606E-06 | 5.200E-06 | 9.900E-06 |
| 10821 | 542 | H542A | A | 5e-06 | 5.9e-06 | 6.3e-06 | 5.733E-06 | 6.658E-07 | 5.000E-06 | 6.300E-06 |
| 10822 | 542 | H542R | A | 5.1e-06 | 9.6e-06 | 7.6e-06 | 7.433E-06 | 2.255E-06 | 5.100E-06 | 9.600E-06 |
| 10823 | 542 | H542N | A | 1.1e-05 | 8.7e-06 | 9.7e-06 | 9.800E-06 | 1.153E-06 | 8.700E-06 | 1.100E-05 |
| 10824 | 542 | H542D | A | 1e-05 | 1.6e-05 | 8.1e-06 | 1.137E-05 | 4.124E-06 | 8.100E-06 | 1.600E-05 |
| 10825 | 542 | H542C | A | 4.2e-06 | 4.1e-06 | 6.6e-06 | 4.967E-06 | 1.415E-06 | 4.100E-06 | 6.600E-06 |
| 10826 | 542 | H542E | A | 6.4e-06 | 8.9e-06 | 6.8e-06 | 7.367E-06 | 1.343E-06 | 6.400E-06 | 8.900E-06 |
| 10827 | 542 | H542Q | A | 8.4e-06 | 7.1e-06 | 1.4e-05 | 9.833E-06 | 3.667E-06 | 7.100E-06 | 1.400E-05 |
| 10828 | 542 | H542G | A | 4.5e-06 | 6.3e-06 | 7.6e-06 | 6.133E-06 | 1.557E-06 | 4.500E-06 | 7.600E-06 |
| 10829 | 542 | H542H | A | 1e-05 | 1.3e-05 | 9.7e-06 | 1.090E-05 | 1.825E-06 | 9.700E-06 | 1.300E-05 |
| 10830 | 542 | H542I | A | 6.6e-06 | 9.6e-06 | 4.4e-06 | 6.867E-06 | 2.610E-06 | 4.400E-06 | 9.600E-06 |
| 10831 | 542 | H542L | A | 7.2e-06 | 1.3e-05 | 6.6e-06 | 8.933E-06 | 3.535E-06 | 6.600E-06 | 1.300E-05 |
| 10832 | 542 | H542K | A | 5.4e-06 | 7e-06 | 4.4e-06 | 5.600E-06 | 1.311E-06 | 4.400E-06 | 7.000E-06 |
| 10833 | 542 | H542M | A | 1.2e-05 | 8.9e-06 | 1.5e-05 | 1.197E-05 | 3.050E-06 | 8.900E-06 | 1.500E-05 |
| 10834 | 542 | H542F | A | 6.9e-06 | 9.9e-06 | 6.3e-06 | 7.700E-06 | 1.929E-06 | 6.300E-06 | 9.900E-06 |
| 10835 | 542 | H542P | A | 6.8e-06 | 4.3e-06 | 9.9e-06 | 7.000E-06 | 2.805E-06 | 4.300E-06 | 9.900E-06 |
| 10836 | 542 | H542S | A | 8.2e-06 | 9.9e-06 | 9.8e-06 | 9.300E-06 | 9.539E-07 | 8.200E-06 | 9.900E-06 |
| 10837 | 542 | H542T | A | 5.7e-06 | 1.4e-05 | 1e-05 | 9.900E-06 | 4.151E-06 | 5.700E-06 | 1.400E-05 |
| 10838 | 542 | H542W | A | 1.2e-05 | 9.8e-06 | 8e-06 | 9.933E-06 | 2.003E-06 | 8.000E-06 | 1.200E-05 |
| 10839 | 542 | H542Y | A | 8.4e-06 | 6.4e-06 | 1.3e-05 | 9.267E-06 | 3.384E-06 | 6.400E-06 | 1.300E-05 |
| 10840 | 542 | H542V | A | 6.8e-06 | 9e-06 | 5.8e-06 | 7.200E-06 | 1.637E-06 | 5.800E-06 | 9.000E-06 |
| 10841 | 543 | P543A | A | 3.2e-06 | 7.5e-06 | 4.7e-06 | 5.133E-06 | 2.183E-06 | 3.200E-06 | 7.500E-06 |
| 10842 | 543 | P543R | A | 8.4e-06 | 1.8e-05 | 6.4e-06 | 1.093E-05 | 6.201E-06 | 6.400E-06 | 1.800E-05 |
| 10843 | 543 | P543N | A | 9.3e-06 | 7.3e-06 | 7.3e-06 | 7.967E-06 | 1.155E-06 | 7.300E-06 | 9.300E-06 |
| 10844 | 543 | P543D | A | 7.3e-06 | 9.4e-06 | 8.1e-06 | 8.267E-06 | 1.060E-06 | 7.300E-06 | 9.400E-06 |
| 10845 | 543 | P543C | A | 1.7e-05 | 5.7e-06 | 4.2e-06 | 8.967E-06 | 6.997E-06 | 4.200E-06 | 1.700E-05 |
| 10846 | 543 | P543E | A | 6.1e-06 | 4.6e-06 | 5.3e-06 | 5.333E-06 | 7.506E-07 | 4.600E-06 | 6.100E-06 |
| 10847 | 543 | P543Q | A | 6.3e-06 | 1.3e-05 | 6.9e-06 | 8.733E-06 | 3.707E-06 | 6.300E-06 | 1.300E-05 |
| 10848 | 543 | P543G | A | 8.1e-06 | 7.3e-06 | 8.5e-06 | 7.967E-06 | 6.110E-07 | 7.300E-06 | 8.500E-06 |
| 10849 | 543 | P543H | A | 9.1e-06 | 9.5e-06 | 8.3e-06 | 8.967E-06 | 6.110E-07 | 8.300E-06 | 9.500E-06 |
| 10850 | 543 | P543I | A | 9.6e-06 | 7.2e-06 | 1e-05 | 8.933E-06 | 1.514E-06 | 7.200E-06 | 1.000E-05 |
| 10851 | 543 | P543L | A | 1.1e-05 | 7.5e-06 | 6.7e-06 | 8.400E-06 | 2.287E-06 | 6.700E-06 | 1.100E-05 |
| 10852 | 543 | P543K | A | 1.5e-05 | 9.8e-06 | 9.3e-06 | 1.137E-05 | 3.156E-06 | 9.300E-06 | 1.500E-05 |
| 10853 | 543 | P543M | A | 6.9e-06 | 5e-06 | 1.2e-05 | 7.967E-06 | 3.620E-06 | 5.000E-06 | 1.200E-05 |
| 10854 | 543 | P543F | A | 8.4e-06 | 1.3e-05 | 6.6e-06 | 9.333E-06 | 3.301E-06 | 6.600E-06 | 1.300E-05 |
| 10855 | 543 | P543P | A | 8.4e-06 | 8.5e-06 | 6e-06 | 7.633E-06 | 1.415E-06 | 6.000E-06 | 8.500E-06 |
| 10856 | 543 | P543S | A | 6e-06 | 8.1e-06 | 6.8e-06 | 6.967E-06 | 1.060E-06 | 6.000E-06 | 8.100E-06 |
| 10857 | 543 | P543T | A | 5.6e-06 | 5.1e-06 | 5e-06 | 5.233E-06 | 3.215E-07 | 5.000E-06 | 5.600E-06 |
| 10858 | 543 | P543W | A | 7e-06 | 4.6e-06 | 6.2e-06 | 5.933E-06 | 1.222E-06 | 4.600E-06 | 7.000E-06 |
| 10859 | 543 | P543Y | A | 8.6e-06 | 6.6e-06 | 9.5e-06 | 8.233E-06 | 1.484E-06 | 6.600E-06 | 9.500E-06 |
| 10860 | 543 | P543V | A | 1e-05 | 7.2e-06 | 6.9e-06 | 8.033E-06 | 1.710E-06 | 6.900E-06 | 1.000E-05 |
| 10861 | 544 | E544A | A | 9.5e-06 | 6.5e-06 | 9.1e-06 | 8.367E-06 | 1.629E-06 | 6.500E-06 | 9.500E-06 |
| 10862 | 544 | E544R | A | 4.8e-06 | 8.9e-06 | 4.4e-06 | 6.033E-06 | 2.491E-06 | 4.400E-06 | 8.900E-06 |
| 10863 | 544 | E544N | A | 6.7e-06 | 8.4e-06 | 6.7e-06 | 7.267E-06 | 9.815E-07 | 6.700E-06 | 8.400E-06 |
| 10864 | 544 | E544D | A | 7.3e-06 | 5.8e-06 | 8.6e-06 | 7.233E-06 | 1.401E-06 | 5.800E-06 | 8.600E-06 |
| 10865 | 544 | E544C | A | 7.2e-06 | 1.3e-05 | 1.2e-05 | 1.073E-05 | 3.101E-06 | 7.200E-06 | 1.300E-05 |
| 10866 | 544 | E544E | A | 4.9e-06 | 1e-05 | 4.5e-06 | 6.467E-06 | 3.066E-06 | 4.500E-06 | 1.000E-05 |
| 10867 | 544 | E544Q | A | 9.3e-06 | 6.8e-06 | 9.6e-06 | 8.567E-06 | 1.537E-06 | 6.800E-06 | 9.600E-06 |
| 10868 | 544 | E544G | A | 6.6e-06 | 6.8e-06 | 6.7e-06 | 6.700E-06 | 1.000E-07 | 6.600E-06 | 6.800E-06 |
| 10869 | 544 | E544H | A | 7.8e-06 | 1.3e-05 | 7.7e-06 | 9.500E-06 | 3.032E-06 | 7.700E-06 | 1.300E-05 |
| 10870 | 544 | E544I | A | 1e-05 | 4.2e-06 | 9.1e-06 | 7.767E-06 | 3.121E-06 | 4.200E-06 | 1.000E-05 |
| 10871 | 544 | E544L | A | 6.9e-06 | 6.8e-06 | 1e-05 | 7.900E-06 | 1.819E-06 | 6.800E-06 | 1.000E-05 |
| 10872 | 544 | E544K | A | 9.1e-06 | 6.2e-06 | 1.2e-05 | 9.100E-06 | 2.900E-06 | 6.200E-06 | 1.200E-05 |
| 10873 | 544 | E544M | A | 8.7e-06 | 8.6e-06 | 5.7e-06 | 7.667E-06 | 1.704E-06 | 5.700E-06 | 8.700E-06 |
| 10874 | 544 | E544F | A | 7.4e-06 | 5.1e-06 | 5.6e-06 | 6.033E-06 | 1.210E-06 | 5.100E-06 | 7.400E-06 |
| 10875 | 544 | E544P | A | 9.9e-06 | 9e-06 | 9.8e-06 | 9.567E-06 | 4.933E-07 | 9.000E-06 | 9.900E-06 |
| 10876 | 544 | E544S | A | 6.4e-06 | 4.5e-06 | 7e-06 | 5.967E-06 | 1.305E-06 | 4.500E-06 | 7.000E-06 |
| 10877 | 544 | E544T | A | 6.5e-06 | 9e-06 | 5.8e-06 | 7.100E-06 | 1.682E-06 | 5.800E-06 | 9.000E-06 |
| 10878 | 544 | E544W | A | 1e-05 | 5.9e-06 | 6.4e-06 | 7.433E-06 | 2.237E-06 | 5.900E-06 | 1.000E-05 |
| 10879 | 544 | E544Y | A | 7e-06 | 1.3e-05 | 1.1e-05 | 1.033E-05 | 3.055E-06 | 7.000E-06 | 1.300E-05 |

| | | | | | | | | | | |
|---|---|---|---|---|---|---|---|---|---|---|
| 10880 | 544 | E544V | A | 1.3e-05 | 6.9e-06 | 7.8e-06 | 9.233E-06 | 3.293E-06 | 6.900E-06 | 1.300E-05 |
| 10881 | 545 | C545A | A | 8.5e-06 | 9.6e-06 | 6.6e-06 | 8.233E-06 | 1.518E-06 | 6.600E-06 | 9.600E-06 |
| 10882 | 545 | C545R | A | 6.5e-06 | 9.7e-06 | 7.6e-06 | 7.933E-06 | 1.626E-06 | 6.500E-06 | 9.700E-06 |
| 10883 | 545 | C545N | A | 6e-06 | 9.1e-06 | 9.5e-06 | 8.200E-06 | 1.916E-06 | 6.000E-06 | 9.500E-06 |
| 10884 | 545 | C545D | A | 8.3e-06 | 1.2e-05 | 6.8e-06 | 9.033E-06 | 2.676E-06 | 6.800E-06 | 1.200E-05 |
| 10885 | 545 | C545C | A | 7.9e-06 | 8.4e-06 | 5.3e-06 | 7.200E-06 | 1.664E-06 | 5.300E-06 | 8.400E-06 |
| 10886 | 545 | C545E | A | 6.9e-06 | 3.6e-06 | 8.5e-06 | 6.333E-06 | 2.499E-06 | 3.600E-06 | 8.500E-06 |
| 10887 | 545 | C545Q | A | 1.3e-05 | 6.9e-06 | 7.2e-06 | 9.033E-06 | 3.439E-06 | 6.900E-06 | 1.300E-05 |
| 10888 | 545 | C545G | A | 7.1e-06 | 7.2e-06 | 4.7e-06 | 6.333E-06 | 1.415E-06 | 4.700E-06 | 7.200E-06 |
| 10889 | 545 | C545H | A | 9.1e-06 | 6.7e-06 | 1.5e-05 | 1.027E-05 | 4.271E-06 | 6.700E-06 | 1.500E-05 |
| 10890 | 545 | C545I | A | 4.7e-06 | 6.2e-06 | 8e-06 | 6.300E-06 | 1.652E-06 | 4.700E-06 | 8.000E-06 |
| 10891 | 545 | C545L | A | 5.8e-06 | 8.7e-06 | 9.7e-06 | 8.067E-06 | 2.026E-06 | 5.800E-06 | 9.700E-06 |
| 10892 | 545 | C545K | A | 3.3e-06 | 6.6e-06 | 5.6e-06 | 5.167E-06 | 1.692E-06 | 3.300E-06 | 6.600E-06 |
| 10893 | 545 | C545M | A | 9.6e-06 | 6.2e-06 | 1.1e-05 | 8.933E-06 | 2.468E-06 | 6.200E-06 | 1.100E-05 |
| 10894 | 545 | C545F | A | 1.3e-05 | 7.4e-06 | 4.7e-06 | 8.367E-06 | 4.234E-06 | 4.700E-06 | 1.300E-05 |
| 10895 | 545 | C545P | A | 7.3e-06 | 7.6e-06 | 6.4e-06 | 7.100E-06 | 6.245E-07 | 6.400E-06 | 7.600E-06 |
| 10896 | 545 | C545S | A | 6.5e-06 | 7e-06 | 6.9e-06 | 6.800E-06 | 2.646E-07 | 6.500E-06 | 7.000E-06 |
| 10897 | 545 | C545T | A | 1.2e-05 | 1.1e-05 | 1.1e-05 | 1.133E-05 | 5.774E-07 | 1.100E-05 | 1.200E-05 |
| 10898 | 545 | C545W | A | 8e-06 | 9.2e-06 | 6.7e-06 | 7.967E-06 | 1.250E-06 | 6.700E-06 | 9.200E-06 |
| 10899 | 545 | C545Y | A | 8.4e-06 | 6.4e-06 | 8.9e-06 | 7.900E-06 | 1.323E-06 | 6.400E-06 | 8.900E-06 |
| 10900 | 545 | C545V | A | 1.4e-05 | 5.3e-06 | 7.8e-06 | 9.033E-06 | 4.479E-06 | 5.300E-06 | 1.400E-05 |
| 10901 | 546 | Q546A | A | 1.4e-05 | 1.1e-05 | 3.7e-06 | 9.567E-06 | 5.297E-06 | 3.700E-06 | 1.400E-05 |
| 10902 | 546 | Q546R | A | 9.7e-06 | 6.2e-06 | 8.9e-06 | 8.267E-06 | 1.834E-06 | 6.200E-06 | 9.700E-06 |
| 10903 | 546 | Q546N | A | 4.7e-06 | 7.4e-06 | 1.5e-05 | 9.033E-06 | 5.341E-06 | 4.700E-06 | 1.500E-05 |
| 10904 | 546 | Q546D | A | 1.1e-05 | 5.1e-06 | 6.2e-06 | 7.433E-06 | 3.137E-06 | 5.100E-06 | 1.100E-05 |
| 10905 | 546 | Q546C | A | 4.8e-06 | 4.3e-06 | 3e-06 | 4.033E-06 | 9.292E-07 | 3.000E-06 | 4.800E-06 |
| 10906 | 546 | Q546E | A | 6.7e-06 | 6.9e-06 | 6.1e-06 | 6.567E-06 | 4.163E-07 | 6.100E-06 | 6.900E-06 |
| 10907 | 546 | Q546Q | A | 5.4e-06 | 1.1e-05 | 5.9e-06 | 7.433E-06 | 3.099E-06 | 5.400E-06 | 1.100E-05 |
| 10908 | 546 | Q546G | A | 6.1e-06 | 6.2e-06 | 1.2e-05 | 8.100E-06 | 3.378E-06 | 6.100E-06 | 1.200E-05 |
| 10909 | 546 | Q546H | A | 1.4e-05 | 9.9e-06 | 5.7e-06 | 9.867E-06 | 4.150E-06 | 5.700E-06 | 1.400E-05 |
| 10910 | 546 | Q546I | A | 9.6e-06 | 4.8e-06 | 9.2e-06 | 7.867E-06 | 2.663E-06 | 4.800E-06 | 9.600E-06 |
| 10911 | 546 | Q546L | A | 5e-06 | 1.5e-05 | 7.3e-06 | 9.100E-06 | 5.237E-06 | 5.000E-06 | 1.500E-05 |
| 10912 | 546 | Q546K | A | 6.2e-06 | 4.6e-06 | 9.4e-06 | 6.733E-06 | 2.444E-06 | 4.600E-06 | 9.400E-06 |
| 10913 | 546 | Q546M | A | 5.9e-06 | 7e-06 | 8e-06 | 6.967E-06 | 1.050E-06 | 5.900E-06 | 8.000E-06 |
| 10914 | 546 | Q546F | A | 6.5e-06 | 6.7e-06 | 6.2e-06 | 6.467E-06 | 2.517E-07 | 6.200E-06 | 6.700E-06 |
| 10915 | 546 | Q546P | A | 4.9e-06 | 9.8e-06 | 6.5e-06 | 7.067E-06 | 2.499E-06 | 4.900E-06 | 9.800E-06 |
| 10916 | 546 | Q546S | A | 6.6e-06 | 6.4e-06 | 8.3e-06 | 7.100E-06 | 1.044E-06 | 6.400E-06 | 8.300E-06 |
| 10917 | 546 | Q546T | A | 9.1e-06 | 7.2e-06 | 1.4e-05 | 1.010E-05 | 3.509E-06 | 7.200E-06 | 1.400E-05 |
| 10918 | 546 | Q546W | A | 4.5e-06 | 6.3e-06 | 7.6e-06 | 6.133E-06 | 1.557E-06 | 4.500E-06 | 7.600E-06 |
| 10919 | 546 | Q546Y | A | 1.3e-05 | 9.6e-06 | 1e-05 | 1.087E-05 | 1.858E-06 | 9.600E-06 | 1.300E-05 |
| 10920 | 546 | Q546V | A | 6.2e-06 | 5e-06 | 7.1e-06 | 6.100E-06 | 1.054E-06 | 5.000E-06 | 7.100E-06 |
| 10921 | 547 | P547A | A | 5.5e-06 | 8.1e-06 | 6e-06 | 6.533E-06 | 1.380E-06 | 5.500E-06 | 8.100E-06 |
| 10922 | 547 | P547R | A | 1.4e-05 | 3.9e-06 | 8.3e-06 | 8.733E-06 | 5.064E-06 | 3.900E-06 | 1.400E-05 |
| 10923 | 547 | P547N | A | 6e-06 | 4.7e-06 | 6.7e-06 | 5.800E-06 | 1.015E-06 | 4.700E-06 | 6.700E-06 |
| 10924 | 547 | P547D | A | 5.4e-06 | 6.4e-06 | 9.3e-06 | 7.033E-06 | 2.026E-06 | 5.400E-06 | 9.300E-06 |
| 10925 | 547 | P547C | A | 6e-06 | 9.6e-06 | 7.2e-06 | 7.600E-06 | 1.833E-06 | 6.000E-06 | 9.600E-06 |
| 10926 | 547 | P547E | A | 6.5e-06 | 6.7e-06 | 6.9e-06 | 6.700E-06 | 2.000E-07 | 6.500E-06 | 6.900E-06 |
| 10927 | 547 | P547Q | A | 8e-06 | 1.1e-05 | 9.8e-06 | 9.600E-06 | 1.510E-06 | 8.000E-06 | 1.100E-05 |
| 10928 | 547 | P547G | A | 6.1e-06 | 6.7e-06 | 8.3e-06 | 7.033E-06 | 1.137E-06 | 6.100E-06 | 8.300E-06 |
| 10929 | 547 | P547H | A | 5.5e-06 | 1.2e-05 | 4.3e-06 | 7.267E-06 | 4.143E-06 | 4.300E-06 | 1.200E-05 |
| 10930 | 547 | P547I | A | 5.3e-06 | 1e-05 | 7.3e-06 | 7.533E-06 | 2.359E-06 | 5.300E-06 | 1.000E-05 |
| 10931 | 547 | P547L | A | 1.1e-05 | 1.2e-05 | 5.7e-06 | 9.567E-06 | 3.386E-06 | 5.700E-06 | 1.200E-05 |
| 10932 | 547 | P547K | A | 8.1e-06 | 6.2e-06 | 8.7e-06 | 7.667E-06 | 1.305E-06 | 6.200E-06 | 8.700E-06 |
| 10933 | 547 | P547M | A | 9.9e-06 | 1.2e-05 | 7.3e-06 | 9.733E-06 | 2.354E-06 | 7.300E-06 | 1.200E-05 |
| 10934 | 547 | P547F | A | 8.7e-06 | 9.3e-06 | 7e-06 | 8.333E-06 | 1.193E-06 | 7.000E-06 | 9.300E-06 |
| 10935 | 547 | P547P | A | 1.1e-05 | 4.1e-06 | 8.9e-06 | 8.000E-06 | 3.537E-06 | 4.100E-06 | 1.100E-05 |
| 10936 | 547 | P547S | A | 1.2e-05 | 7.3e-06 | 7.2e-06 | 8.833E-06 | 2.743E-06 | 7.200E-06 | 1.200E-05 |
| 10937 | 547 | P547T | A | 1.4e-05 | 5.3e-06 | 5e-06 | 8.100E-06 | 5.112E-06 | 5.000E-06 | 1.400E-05 |
| 10938 | 547 | P547W | A | 7e-06 | 4.7e-06 | 7.6e-06 | 6.433E-06 | 1.531E-06 | 4.700E-06 | 7.600E-06 |
| 10939 | 547 | P547Y | A | 5.2e-06 | 1.1e-05 | 1.2e-05 | 9.400E-06 | 3.672E-06 | 5.200E-06 | 1.200E-05 |
| 10940 | 547 | P547V | A | 9e-06 | 1.3e-05 | 6.7e-06 | 9.567E-06 | 3.188E-06 | 6.700E-06 | 1.300E-05 |
| 10941 | 548 | Q548A | A | 9.9e-06 | 9.8e-06 | 1.8e-05 | 1.257E-05 | 4.706E-06 | 9.800E-06 | 1.800E-05 |
| 10942 | 548 | Q548R | A | 1.6e-05 | 9.1e-06 | 8.3e-06 | 1.113E-05 | 4.234E-06 | 8.300E-06 | 1.600E-05 |
| 10943 | 548 | Q548N | A | 7.2e-06 | 5.2e-06 | 9.2e-06 | 7.200E-06 | 2.000E-06 | 5.200E-06 | 9.200E-06 |
| 10944 | 548 | Q548D | A | 9.3e-06 | 4.8e-06 | 8.4e-06 | 7.500E-06 | 2.381E-06 | 4.800E-06 | 9.300E-06 |
| 10945 | 548 | Q548C | A | 6.4e-06 | 1.1e-05 | 9.5e-06 | 8.967E-06 | 2.346E-06 | 6.400E-06 | 1.100E-05 |
| 10946 | 548 | Q548E | A | 1.6e-05 | 1e-05 | 6.6e-06 | 1.087E-05 | 4.760E-06 | 6.600E-06 | 1.600E-05 |
| 10947 | 548 | Q548Q | A | 6.5e-06 | 7.2e-06 | 7.7e-06 | 7.133E-06 | 6.028E-07 | 6.500E-06 | 7.700E-06 |
| 10948 | 548 | Q548G | A | 1.2e-05 | 7.8e-06 | 7.4e-06 | 9.067E-06 | 2.548E-06 | 7.400E-06 | 1.200E-05 |
| 10949 | 548 | Q548H | A | 7.3e-06 | 1.3e-05 | 1.2e-05 | 1.077E-05 | 3.044E-06 | 7.300E-06 | 1.300E-05 |
| 10950 | 548 | Q548I | A | 7.5e-06 | 6.7e-06 | 6.9e-06 | 7.033E-06 | 4.163E-07 | 6.700E-06 | 7.500E-06 |
| 10951 | 548 | Q548L | A | 5.8e-06 | 4.7e-06 | 1.1e-05 | 7.167E-06 | 3.365E-06 | 4.700E-06 | 1.100E-05 |
| 10952 | 548 | Q548K | A | 6.8e-06 | 3.9e-06 | 1.1e-05 | 7.233E-06 | 3.570E-06 | 3.900E-06 | 1.100E-05 |
| 10953 | 548 | Q548M | A | 1.2e-05 | 1.4e-05 | 5.3e-06 | 1.043E-05 | 4.557E-06 | 5.300E-06 | 1.400E-05 |
| 10954 | 548 | Q548F | A | 1e-05 | 6.8e-06 | 1e-05 | 8.933E-06 | 1.848E-06 | 6.800E-06 | 1.000E-05 |
| 10955 | 548 | Q548P | A | 4.9e-06 | 9.7e-06 | 1.1e-05 | 8.533E-06 | 3.213E-06 | 4.900E-06 | 1.100E-05 |
| 10956 | 548 | Q548S | A | 6.8e-06 | 7.7e-06 | 5.7e-06 | 6.733E-06 | 1.002E-06 | 5.700E-06 | 7.700E-06 |
| 10957 | 548 | Q548T | A | 1.1e-05 | 7.3e-06 | 6.6e-06 | 8.300E-06 | 2.364E-06 | 6.600E-06 | 1.100E-05 |
| 10958 | 548 | Q548W | A | 1.2e-05 | 9.6e-06 | 6.3e-06 | 9.300E-06 | 2.862E-06 | 6.300E-06 | 1.200E-05 |
| 10959 | 548 | Q548Y | A | 6.5e-06 | 9.6e-06 | 9.4e-06 | 8.500E-06 | 1.735E-06 | 6.500E-06 | 9.600E-06 |
| 10960 | 548 | Q548V | A | 1.3e-05 | 3.3e-06 | 1e-05 | 8.767E-06 | 4.966E-06 | 3.300E-06 | 1.300E-05 |
| 10961 | 549 | N549A | A | 5.8e-06 | 5.4e-06 | 6e-06 | 5.733E-06 | 3.055E-07 | 5.400E-06 | 6.000E-06 |
| 10962 | 549 | N549R | A | 6.3e-06 | 6.7e-06 | 1e-05 | 7.667E-06 | 2.031E-06 | 6.300E-06 | 1.000E-05 |
| 10963 | 549 | N549N | A | 8.1e-06 | 5.8e-06 | 7.6e-06 | 7.167E-06 | 1.210E-06 | 5.800E-06 | 8.100E-06 |
| 10964 | 549 | N549D | A | 1.3e-05 | 5.3e-06 | 7.5e-06 | 8.600E-06 | 3.966E-06 | 5.300E-06 | 1.300E-05 |

| | | | | | | | | | | |
|---|---|---|---|---|---|---|---|---|---|---|
| 10965 | 549 | N549C | A | 9.4e-06 | 8.3e-06 | 8.6e-06 | 8.767E-06 | 5.686E-07 | 8.300E-06 | 9.400E-06 |
| 10966 | 549 | N549E | A | 1.2e-05 | 3.3e-06 | 9.6e-06 | 8.300E-06 | 4.493E-06 | 3.300E-06 | 1.200E-05 |
| 10967 | 549 | N549Q | A | 9.2e-06 | 4.8e-06 | 5.9e-06 | 6.633E-06 | 2.290E-06 | 4.800E-06 | 9.200E-06 |
| 10968 | 549 | N549G | A | 6.1e-06 | 6.9e-06 | 9.4e-06 | 7.467E-06 | 1.721E-06 | 6.100E-06 | 9.400E-06 |
| 10969 | 549 | N549H | A | 9.5e-06 | 6.5e-06 | 6.6e-06 | 7.533E-06 | 1.704E-06 | 6.500E-06 | 9.500E-06 |
| 10970 | 549 | N549I | A | 2e-05 | 5.5e-06 | 1e-05 | 1.183E-05 | 7.422E-06 | 5.500E-06 | 2.000E-05 |
| 10971 | 549 | N549L | A | 6.7e-06 | 9.5e-06 | 1.8e-05 | 1.140E-05 | 5.885E-06 | 6.700E-06 | 1.800E-05 |
| 10972 | 549 | N549K | A | 4.7e-06 | 6.5e-06 | 1.8e-05 | 9.733E-06 | 7.215E-06 | 4.700E-06 | 1.800E-05 |
| 10973 | 549 | N549M | A | 6.5e-06 | 6.7e-06 | 8.8e-06 | 7.333E-06 | 1.274E-06 | 6.500E-06 | 8.800E-06 |
| 10974 | 549 | N549F | A | 1.2e-05 | 5.5e-06 | 9.5e-06 | 9.000E-06 | 3.279E-06 | 5.500E-06 | 1.200E-05 |
| 10975 | 549 | N549P | A | 7.7e-06 | 1.1e-05 | 7.5e-06 | 8.733E-06 | 1.966E-06 | 7.500E-06 | 1.100E-05 |
| 10976 | 549 | N549S | A | 1e-05 | 6.4e-06 | 7.4e-06 | 7.933E-06 | 1.858E-06 | 6.400E-06 | 1.000E-05 |
| 10977 | 549 | N549T | A | 1.2e-05 | 5.4e-06 | 9.2e-06 | 8.867E-06 | 3.313E-06 | 5.400E-06 | 1.200E-05 |
| 10978 | 549 | N549W | A | 4.6e-06 | 6.3e-06 | 6.5e-06 | 5.800E-06 | 1.044E-06 | 4.600E-06 | 6.500E-06 |
| 10979 | 549 | N549Y | A | 6.1e-06 | 6.9e-06 | 1.2e-05 | 8.333E-06 | 3.201E-06 | 6.100E-06 | 1.200E-05 |
| 10980 | 549 | N549V | A | 6.8e-06 | 1.2e-05 | 6.8e-06 | 8.533E-06 | 3.002E-06 | 6.800E-06 | 1.200E-05 |
| 10981 | 550 | G550A | A | 8.6e-06 | 6.5e-06 | 1e-05 | 8.367E-06 | 1.762E-06 | 6.500E-06 | 1.000E-05 |
| 10982 | 550 | G550R | A | 1e-05 | 1.3e-05 | 8.2e-06 | 1.040E-05 | 2.425E-06 | 8.200E-06 | 1.300E-05 |
| 10983 | 550 | G550N | A | 1.3e-05 | 7e-06 | 1e-05 | 1.000E-05 | 3.000E-06 | 7.000E-06 | 1.300E-05 |
| 10984 | 550 | G550D | A | 1.2e-05 | 8.5e-06 | 8.1e-06 | 9.533E-06 | 2.146E-06 | 8.100E-06 | 1.200E-05 |
| 10985 | 550 | G550C | A | 9.7e-06 | 6.7e-06 | 6e-06 | 7.467E-06 | 1.966E-06 | 6.000E-06 | 9.700E-06 |
| 10986 | 550 | G550E | A | 6.8e-06 | 7.4e-06 | 7.6e-06 | 7.267E-06 | 4.163E-07 | 6.800E-06 | 7.600E-06 |
| 10987 | 550 | G550Q | A | 7.8e-06 | 1.1e-05 | 9.4e-06 | 9.400E-06 | 1.600E-06 | 7.800E-06 | 1.100E-05 |
| 10988 | 550 | G550G | A | 9.6e-06 | 6.7e-06 | 5.1e-06 | 7.133E-06 | 2.281E-06 | 5.100E-06 | 9.600E-06 |
| 10989 | 550 | G550H | A | 8.5e-06 | 4.3e-06 | 7e-06 | 6.600E-06 | 2.128E-06 | 4.300E-06 | 8.500E-06 |
| 10990 | 550 | G550I | A | 4.6e-06 | 4.3e-06 | 7.9e-06 | 5.600E-06 | 1.997E-06 | 4.300E-06 | 7.900E-06 |
| 10991 | 550 | G550L | A | 6.7e-06 | 1.1e-05 | 8.4e-06 | 8.700E-06 | 2.166E-06 | 6.700E-06 | 1.100E-05 |
| 10992 | 550 | G550K | A | 2.4e-05 | 7.3e-06 | 1.1e-05 | 1.410E-05 | 8.771E-06 | 7.300E-06 | 2.400E-05 |
| 10993 | 550 | G550M | A | 6.5e-06 | 1.4e-05 | 5.5e-06 | 8.667E-06 | 4.646E-06 | 5.500E-06 | 1.400E-05 |
| 10994 | 550 | G550F | A | 5.8e-06 | 8.3e-06 | 5.4e-06 | 6.500E-06 | 1.572E-06 | 5.400E-06 | 8.300E-06 |
| 10995 | 550 | G550P | A | 7.4e-06 | 6.6e-06 | 1.1e-05 | 8.333E-06 | 2.344E-06 | 6.600E-06 | 1.100E-05 |
| 10996 | 550 | G550S | A | 6.9e-06 | 9.1e-06 | 6.7e-06 | 7.567E-06 | 1.332E-06 | 6.700E-06 | 9.100E-06 |
| 10997 | 550 | G550T | A | 4.8e-06 | 7.7e-06 | 1e-05 | 7.500E-06 | 2.606E-06 | 4.800E-06 | 1.000E-05 |
| 10998 | 550 | G550W | A | 5.8e-06 | 7.1e-06 | 5.9e-06 | 6.267E-06 | 7.234E-07 | 5.800E-06 | 7.100E-06 |
| 10999 | 550 | G550Y | A | 9.4e-06 | 4.1e-06 | 7.8e-06 | 7.100E-06 | 2.718E-06 | 4.100E-06 | 9.400E-06 |
| 11000 | 550 | G550V | A | 8.8e-06 | 8.9e-06 | 8.9e-06 | 8.867E-06 | 5.774E-08 | 8.800E-06 | 8.900E-06 |
| 11001 | 551 | S551A | A | 8.8e-06 | 1.1e-05 | 6.5e-06 | 8.767E-06 | 2.250E-06 | 6.500E-06 | 1.100E-05 |
| 11002 | 551 | S551R | A | 5.6e-06 | 7.5e-06 | 9.2e-06 | 7.433E-06 | 1.801E-06 | 5.600E-06 | 9.200E-06 |
| 11003 | 551 | S551N | A | 9.2e-06 | 8.9e-06 | 9.6e-06 | 9.233E-06 | 3.512E-07 | 8.900E-06 | 9.600E-06 |
| 11004 | 551 | S551D | A | 6.6e-06 | 9.5e-06 | 6.9e-06 | 7.667E-06 | 1.595E-06 | 6.600E-06 | 9.500E-06 |
| 11005 | 551 | S551C | A | 9.2e-06 | 6.7e-06 | 6.9e-06 | 7.600E-06 | 1.389E-06 | 6.700E-06 | 9.200E-06 |
| 11006 | 551 | S551E | A | 1.3e-05 | 1.4e-05 | 5.5e-06 | 1.083E-05 | 4.646E-06 | 5.500E-06 | 1.400E-05 |
| 11007 | 551 | S551Q | A | 1.3e-05 | 6.9e-06 | 7e-06 | 8.967E-06 | 3.493E-06 | 6.900E-06 | 1.300E-05 |
| 11008 | 551 | S551G | A | 6.5e-06 | 6.3e-06 | 1e-05 | 7.600E-06 | 2.081E-06 | 6.300E-06 | 1.000E-05 |
| 11009 | 551 | S551H | A | 1.7e-05 | 7.6e-06 | 8.8e-06 | 1.113E-05 | 5.116E-06 | 7.600E-06 | 1.700E-05 |
| 11010 | 551 | S551I | A | 7.9e-06 | 4.2e-06 | 6.9e-06 | 6.333E-06 | 1.914E-06 | 4.200E-06 | 7.900E-06 |
| 11011 | 551 | S551L | A | 7.1e-06 | 1.3e-05 | 8.9e-06 | 9.667E-06 | 3.024E-06 | 7.100E-06 | 1.300E-05 |
| 11012 | 551 | S551K | A | 7e-06 | 9.8e-06 | 8.9e-06 | 8.567E-06 | 1.429E-06 | 7.000E-06 | 9.800E-06 |
| 11013 | 551 | S551M | A | 8.4e-06 | 7.3e-06 | 7.2e-06 | 7.633E-06 | 6.658E-07 | 7.200E-06 | 8.400E-06 |
| 11014 | 551 | S551F | A | 8.3e-06 | 1.2e-05 | 7.7e-06 | 9.333E-06 | 2.329E-06 | 7.700E-06 | 1.200E-05 |
| 11015 | 551 | S551P | A | 1.1e-05 | 9.3e-06 | 4.9e-06 | 8.400E-06 | 3.148E-06 | 4.900E-06 | 1.100E-05 |
| 11016 | 551 | S551S | A | 1e-05 | 8.2e-06 | 7.6e-06 | 8.600E-06 | 1.249E-06 | 7.600E-06 | 1.000E-05 |
| 11017 | 551 | S551T | A | 1e-05 | 7.8e-06 | 8.8e-06 | 8.867E-06 | 1.102E-06 | 7.800E-06 | 1.000E-05 |
| 11018 | 551 | S551W | A | 8.5e-06 | 7.1e-06 | 8.6e-06 | 8.067E-06 | 8.386E-07 | 7.100E-06 | 8.600E-06 |
| 11019 | 551 | S551Y | A | 7.6e-06 | 8.8e-06 | 7.7e-06 | 8.033E-06 | 6.658E-07 | 7.600E-06 | 8.800E-06 |
| 11020 | 551 | S551V | A | 7.2e-06 | 1.1e-05 | 1.7e-05 | 1.173E-05 | 4.941E-06 | 7.200E-06 | 1.700E-05 |
| 11021 | 552 | V552A | A | 1e-05 | 7.7e-06 | 9.5e-06 | 9.067E-06 | 1.210E-06 | 7.700E-06 | 1.000E-05 |
| 11022 | 552 | V552R | A | 9.6e-06 | 9.7e-06 | 8.2e-06 | 9.167E-06 | 8.386E-07 | 8.200E-06 | 9.700E-06 |
| 11023 | 552 | V552N | A | 6.5e-06 | 1.3e-05 | 7.8e-06 | 9.100E-06 | 3.439E-06 | 6.500E-06 | 1.300E-05 |
| 11024 | 552 | V552D | A | 6.1e-06 | 5.6e-06 | 9.3e-06 | 7.000E-06 | 2.007E-06 | 5.600E-06 | 9.300E-06 |
| 11025 | 552 | V552C | A | 6.6e-06 | 9.1e-06 | 1.3e-05 | 9.567E-06 | 3.225E-06 | 6.600E-06 | 1.300E-05 |
| 11026 | 552 | V552E | A | 8e-06 | 9.4e-06 | 7.8e-06 | 8.400E-06 | 8.718E-07 | 7.800E-06 | 9.400E-06 |
| 11027 | 552 | V552Q | A | 9.7e-06 | 9.9e-06 | 7.9e-06 | 9.167E-06 | 1.102E-06 | 7.900E-06 | 9.900E-06 |
| 11028 | 552 | V552G | A | 8.1e-06 | 8.2e-06 | 5.8e-06 | 7.367E-06 | 1.358E-06 | 5.800E-06 | 8.200E-06 |
| 11029 | 552 | V552H | A | 6.6e-06 | 8.2e-06 | 6.8e-06 | 7.200E-06 | 8.718E-07 | 6.600E-06 | 8.200E-06 |
| 11030 | 552 | V552I | A | 9.6e-06 | 1.1e-05 | 1e-05 | 1.020E-05 | 7.211E-07 | 9.600E-06 | 1.100E-05 |
| 11031 | 552 | V552L | A | 1.2e-05 | 1e-05 | 1.2e-05 | 1.133E-05 | 1.155E-06 | 1.000E-05 | 1.200E-05 |
| 11032 | 552 | V552K | A | 8.1e-06 | 7e-06 | 6.6e-06 | 7.233E-06 | 7.767E-07 | 6.600E-06 | 8.100E-06 |
| 11033 | 552 | V552M | A | 6.2e-06 | 7.2e-06 | 9.7e-06 | 7.700E-06 | 1.803E-06 | 6.200E-06 | 9.700E-06 |
| 11034 | 552 | V552F | A | 5.8e-06 | 6.3e-06 | 7.8e-06 | 6.633E-06 | 1.041E-06 | 5.800E-06 | 7.800E-06 |
| 11035 | 552 | V552P | A | 6.8e-06 | 5.4e-06 | 7.5e-06 | 6.567E-06 | 1.069E-06 | 5.400E-06 | 7.500E-06 |
| 11036 | 552 | V552S | A | 7.3e-06 | 1.2e-05 | 9.7e-06 | 9.667E-06 | 2.350E-06 | 7.300E-06 | 1.200E-05 |
| 11037 | 552 | V552T | A | 9.9e-06 | 9e-06 | 5.1e-06 | 8.000E-06 | 2.551E-06 | 5.100E-06 | 9.900E-06 |
| 11038 | 552 | V552W | A | 1.1e-05 | 9.7e-06 | 5.5e-06 | 8.733E-06 | 2.875E-06 | 5.500E-06 | 1.100E-05 |
| 11039 | 552 | V552Y | A | 1.2e-05 | 1.4e-05 | 1.2e-05 | 1.267E-05 | 1.155E-06 | 1.200E-05 | 1.400E-05 |
| 11040 | 552 | V552V | A | 7.5e-06 | 1.4e-05 | 1.2e-05 | 1.117E-05 | 3.329E-06 | 7.500E-06 | 1.400E-05 |
| 11041 | 553 | T553A | A | 7.7e-06 | 9.5e-06 | 6.4e-06 | 7.867E-06 | 1.557E-06 | 6.400E-06 | 9.500E-06 |
| 11042 | 553 | T553R | A | 1.1e-05 | 8.1e-06 | 6e-06 | 8.367E-06 | 2.511E-06 | 6.000E-06 | 1.100E-05 |
| 11043 | 553 | T553N | A | 9.7e-06 | 7e-06 | 8.1e-06 | 8.267E-06 | 1.358E-06 | 7.000E-06 | 9.700E-06 |
| 11044 | 553 | T553D | A | 4.8e-06 | 7e-06 | 9.4e-06 | 7.067E-06 | 2.301E-06 | 4.800E-06 | 9.400E-06 |
| 11045 | 553 | T553C | A | 5.7e-06 | 8.4e-06 | 9.9e-06 | 8.000E-06 | 2.128E-06 | 5.700E-06 | 9.900E-06 |
| 11046 | 553 | T553E | A | 6.3e-06 | 6.7e-06 | 5.3e-06 | 6.100E-06 | 7.211E-07 | 5.300E-06 | 6.700E-06 |
| 11047 | 553 | T553Q | A | 1e-05 | 1.2e-05 | 6.6e-06 | 9.533E-06 | 2.730E-06 | 6.600E-06 | 1.200E-05 |
| 11048 | 553 | T553G | A | 8.8e-06 | 6.4e-06 | 7.1e-06 | 7.433E-06 | 1.234E-06 | 6.400E-06 | 8.800E-06 |
| 11049 | 553 | T553H | A | 7.1e-06 | 6.5e-06 | 9.8e-06 | 7.800E-06 | 1.758E-06 | 6.500E-06 | 9.800E-06 |

| | | | | | | | | | | |
|---|---|---|---|---|---|---|---|---|---|---|
| 11050 | 553 | T553I | A | 1.3e-05 | 9.1e-06 | 9.4e-06 | 1.050E-05 | 2.170E-06 | 9.100E-06 | 1.300E-05 |
| 11051 | 553 | T553L | A | 4.6e-06 | 6.4e-06 | 6.3e-06 | 5.767E-06 | 1.012E-06 | 4.600E-06 | 6.400E-06 |
| 11052 | 553 | T553K | A | 8.6e-06 | 1.5e-05 | 6.2e-06 | 9.933E-06 | 4.549E-06 | 6.200E-06 | 1.500E-05 |
| 11053 | 553 | T553M | A | 5.8e-06 | 6.2e-06 | 1e-05 | 7.333E-06 | 2.318E-06 | 5.800E-06 | 1.000E-05 |
| 11054 | 553 | T553F | A | 5.7e-06 | 6e-06 | 1e-05 | 7.233E-06 | 2.401E-06 | 5.700E-06 | 1.000E-05 |
| 11055 | 553 | T553P | A | 6.7e-06 | 8e-06 | 5.2e-06 | 6.633E-06 | 1.401E-06 | 5.200E-06 | 8.000E-06 |
| 11056 | 553 | T553S | A | 8.7e-06 | 8.2e-06 | 8e-06 | 8.300E-06 | 3.606E-07 | 8.000E-06 | 8.700E-06 |
| 11057 | 553 | T553T | A | 5.5e-06 | 7.1e-06 | 7.5e-06 | 6.700E-06 | 1.058E-06 | 5.500E-06 | 7.500E-06 |
| 11058 | 553 | T553W | A | 1.2e-05 | 4.2e-06 | 6.1e-06 | 7.433E-06 | 4.067E-06 | 4.200E-06 | 1.200E-05 |
| 11059 | 553 | T553Y | A | 7.3e-06 | 9e-06 | 1.3e-05 | 9.767E-06 | 2.926E-06 | 7.300E-06 | 1.300E-05 |
| 11060 | 553 | T553V | A | 1.4e-05 | 4.7e-06 | 6.2e-06 | 8.300E-06 | 4.993E-06 | 4.700E-06 | 1.400E-05 |
| 11061 | 554 | C554A | A | 6.3e-06 | 4.5e-06 | 1e-05 | 6.933E-06 | 2.804E-06 | 4.500E-06 | 1.000E-05 |
| 11062 | 554 | C554R | A | 8.7e-06 | 6e-06 | 9.8e-06 | 8.167E-06 | 1.955E-06 | 6.000E-06 | 9.800E-06 |
| 11063 | 554 | C554N | A | 7.9e-06 | 3.9e-06 | 1.8e-05 | 9.933E-06 | 7.267E-06 | 3.900E-06 | 1.800E-05 |
| 11064 | 554 | C554D | A | 6.2e-06 | 6.9e-06 | 5.1e-06 | 6.067E-06 | 9.074E-07 | 5.100E-06 | 6.900E-06 |
| 11065 | 554 | C554C | A | 8.6e-06 | 1e-05 | 9.3e-06 | 9.300E-06 | 7.000E-07 | 8.600E-06 | 1.000E-05 |
| 11066 | 554 | C554E | A | 5.7e-06 | 5.8e-06 | 8.5e-06 | 6.667E-06 | 1.589E-06 | 5.700E-06 | 8.500E-06 |
| 11067 | 554 | C554Q | A | 7.1e-06 | 6.4e-06 | 7.7e-06 | 7.067E-06 | 6.506E-07 | 6.400E-06 | 7.700E-06 |
| 11068 | 554 | C554G | A | 6.8e-06 | 5.1e-06 | 6.3e-06 | 6.067E-06 | 8.737E-07 | 5.100E-06 | 6.800E-06 |
| 11069 | 554 | C554H | A | 9.4e-06 | 6.6e-06 | 1.3e-05 | 9.667E-06 | 3.208E-06 | 6.600E-06 | 1.300E-05 |
| 11070 | 554 | C554I | A | 8.8e-06 | 6e-06 | 4.8e-06 | 6.533E-06 | 2.053E-06 | 4.800E-06 | 8.800E-06 |
| 11071 | 554 | C554L | A | 4.5e-06 | 7.7e-06 | 9.4e-06 | 7.200E-06 | 2.488E-06 | 4.500E-06 | 9.400E-06 |
| 11072 | 554 | C554K | A | 7.7e-06 | 6.9e-06 | 1.2e-05 | 8.867E-06 | 2.743E-06 | 6.900E-06 | 1.200E-05 |
| 11073 | 554 | C554M | A | 5.6e-06 | 8.9e-06 | 1.5e-05 | 9.833E-06 | 4.769E-06 | 5.600E-06 | 1.500E-05 |
| 11074 | 554 | C554F | A | 1.3e-05 | 3.2e-06 | 4.7e-06 | 6.967E-06 | 5.279E-06 | 3.200E-06 | 1.300E-05 |
| 11075 | 554 | C554P | A | 7.1e-06 | 5.9e-06 | 9.4e-06 | 7.467E-06 | 1.779E-06 | 5.900E-06 | 9.400E-06 |
| 11076 | 554 | C554S | A | 6e-06 | 6.8e-06 | 1.3e-05 | 8.600E-06 | 3.831E-06 | 6.000E-06 | 1.300E-05 |
| 11077 | 554 | C554T | A | 6e-06 | 7.8e-06 | 6.1e-06 | 6.633E-06 | 1.012E-06 | 6.000E-06 | 7.800E-06 |
| 11078 | 554 | C554W | A | 4.8e-06 | 4.1e-06 | 4.8e-06 | 4.567E-06 | 4.041E-07 | 4.100E-06 | 4.800E-06 |
| 11079 | 554 | C554Y | A | 8.8e-06 | 1e-05 | 8.2e-06 | 9.000E-06 | 9.165E-07 | 8.200E-06 | 1.000E-05 |
| 11080 | 554 | C554V | A | 7.5e-06 | 5.4e-06 | 1.1e-05 | 7.967E-06 | 2.829E-06 | 5.400E-06 | 1.100E-05 |
| 11081 | 555 | F555A | A | 1.2e-05 | 6.8e-06 | 6.5e-06 | 8.433E-06 | 3.092E-06 | 6.500E-06 | 1.200E-05 |
| 11082 | 555 | F555R | A | 8e-06 | 5.3e-06 | 6.9e-06 | 6.733E-06 | 1.358E-06 | 5.300E-06 | 8.000E-06 |
| 11083 | 555 | F555N | A | 9.1e-06 | 7.9e-06 | 4.1e-06 | 7.033E-06 | 2.610E-06 | 4.100E-06 | 9.100E-06 |
| 11084 | 555 | F555D | A | 1.3e-05 | 9.6e-06 | 6.9e-06 | 9.833E-06 | 3.057E-06 | 6.900E-06 | 1.300E-05 |
| 11085 | 555 | F555C | A | 7.9e-06 | 4.6e-06 | 1.1e-05 | 7.833E-06 | 3.201E-06 | 4.600E-06 | 1.100E-05 |
| 11086 | 555 | F555E | A | 9.8e-06 | 9.9e-06 | 1.4e-05 | 1.123E-05 | 2.397E-06 | 9.800E-06 | 1.400E-05 |
| 11087 | 555 | F555Q | A | 5.6e-06 | 5.8e-06 | 5.6e-06 | 5.667E-06 | 1.155E-07 | 5.600E-06 | 5.800E-06 |
| 11088 | 555 | F555G | A | 6.2e-06 | 8.1e-06 | 6.8e-06 | 7.033E-06 | 9.713E-07 | 6.200E-06 | 8.100E-06 |
| 11089 | 555 | F555H | A | 5e-06 | 1.1e-05 | 6.7e-06 | 7.567E-06 | 3.092E-06 | 5.000E-06 | 1.100E-05 |
| 11090 | 555 | F555I | A | 5.2e-06 | 1.1e-05 | 7.8e-06 | 8.000E-06 | 2.905E-06 | 5.200E-06 | 1.100E-05 |
| 11091 | 555 | F555L | A | 1.2e-05 | 1.2e-05 | 5.9e-06 | 9.967E-06 | 3.522E-06 | 5.900E-06 | 1.200E-05 |
| 11092 | 555 | F555K | A | 9.7e-06 | 4.6e-06 | 7e-06 | 7.100E-06 | 2.551E-06 | 4.600E-06 | 9.700E-06 |
| 11093 | 555 | F555M | A | 5.7e-06 | 8.7e-06 | 6.5e-06 | 6.967E-06 | 1.553E-06 | 5.700E-06 | 8.700E-06 |
| 11094 | 555 | F555F | A | 9.2e-06 | 9.5e-06 | 4.5e-06 | 7.733E-06 | 2.804E-06 | 4.500E-06 | 9.500E-06 |
| 11095 | 555 | F555P | A | 3.7e-06 | 7.4e-06 | 9.2e-06 | 6.767E-06 | 2.804E-06 | 3.700E-06 | 9.200E-06 |
| 11096 | 555 | F555S | A | 1.2e-05 | 7.8e-06 | 6.2e-06 | 8.667E-06 | 2.996E-06 | 6.200E-06 | 1.200E-05 |
| 11097 | 555 | F555T | A | 6.5e-06 | 9.5e-06 | 3.8e-06 | 6.600E-06 | 2.851E-06 | 3.800E-06 | 9.500E-06 |
| 11098 | 555 | F555W | A | 7.5e-06 | 7.3e-06 | 7.2e-06 | 7.333E-06 | 1.528E-07 | 7.200E-06 | 7.500E-06 |
| 11099 | 555 | F555Y | A | 8.8e-06 | 4.4e-06 | 9.3e-06 | 7.500E-06 | 2.696E-06 | 4.400E-06 | 9.300E-06 |
| 11100 | 555 | F555V | A | 4e-06 | 1e-05 | 1e-05 | 8.000E-06 | 3.464E-06 | 4.000E-06 | 1.000E-05 |
| 11101 | 556 | G556A | A | 6.7e-06 | 4.2e-06 | 6.4e-06 | 5.767E-06 | 1.365E-06 | 4.200E-06 | 6.700E-06 |
| 11102 | 556 | G556R | A | 7.3e-06 | 8e-06 | 7e-06 | 7.433E-06 | 5.132E-07 | 7.000E-06 | 8.000E-06 |
| 11103 | 556 | G556N | A | 1.1e-05 | 7.2e-06 | 7e-06 | 8.400E-06 | 2.254E-06 | 7.000E-06 | 1.100E-05 |
| 11104 | 556 | G556D | A | 4e-06 | 4.8e-06 | 8.3e-06 | 5.700E-06 | 2.287E-06 | 4.000E-06 | 8.300E-06 |
| 11105 | 556 | G556C | A | 6.9e-06 | 8.1e-06 | 8.6e-06 | 7.867E-06 | 8.737E-07 | 6.900E-06 | 8.600E-06 |
| 11106 | 556 | G556E | A | 6.7e-06 | 9.7e-06 | 7.3e-06 | 7.900E-06 | 1.587E-06 | 6.700E-06 | 9.700E-06 |
| 11107 | 556 | G556Q | A | 1.2e-05 | 4.5e-06 | 8.9e-06 | 8.467E-06 | 3.769E-06 | 4.500E-06 | 1.200E-05 |
| 11108 | 556 | G556G | A | 8.1e-06 | 4.5e-06 | 9.3e-06 | 7.300E-06 | 2.498E-06 | 4.500E-06 | 9.300E-06 |
| 11109 | 556 | G556H | A | 8e-06 | 8.1e-06 | 7.6e-06 | 7.900E-06 | 2.646E-07 | 7.600E-06 | 8.100E-06 |
| 11110 | 556 | G556I | A | 7.4e-06 | 6.5e-06 | 7.2e-06 | 7.033E-06 | 4.726E-07 | 6.500E-06 | 7.400E-06 |
| 11111 | 556 | G556L | A | 9.1e-06 | 7.8e-06 | 9.5e-06 | 8.800E-06 | 8.888E-07 | 7.800E-06 | 9.500E-06 |
| 11112 | 556 | G556K | A | 8.4e-06 | 9.5e-06 | 9.2e-06 | 9.033E-06 | 5.686E-07 | 8.400E-06 | 9.500E-06 |
| 11113 | 556 | G556M | A | 1.2e-05 | 8.9e-06 | 5.2e-06 | 8.700E-06 | 3.404E-06 | 5.200E-06 | 1.200E-05 |
| 11114 | 556 | G556F | A | 1.2e-05 | 9.3e-06 | 9.4e-06 | 1.023E-05 | 1.531E-06 | 9.300E-06 | 1.200E-05 |
| 11115 | 556 | G556P | A | 2.1e-05 | 1.9e-05 | 6.7e-06 | 1.557E-05 | 7.744E-06 | 6.700E-06 | 2.100E-05 |
| 11116 | 556 | G556S | A | 5.5e-06 | 9.4e-06 | 1.3e-05 | 9.300E-06 | 3.751E-06 | 5.500E-06 | 1.300E-05 |
| 11117 | 556 | G556T | A | 6.7e-06 | 7.7e-06 | 6.6e-06 | 7.000E-06 | 6.083E-07 | 6.600E-06 | 7.700E-06 |
| 11118 | 556 | G556W | A | 9.8e-06 | 9.7e-06 | 4.6e-06 | 8.033E-06 | 2.974E-06 | 4.600E-06 | 9.800E-06 |
| 11119 | 556 | G556Y | A | 2.7e-06 | 8.6e-06 | 8.6e-06 | 6.633E-06 | 3.406E-06 | 2.700E-06 | 8.600E-06 |
| 11120 | 556 | G556V | A | 9.3e-06 | 9.5e-06 | 7.8e-06 | 8.867E-06 | 9.292E-07 | 7.800E-06 | 9.500E-06 |
| 11121 | 557 | P557A | A | 7.2e-06 | 4.7e-06 | 8.7e-06 | 6.867E-06 | 2.021E-06 | 4.700E-06 | 8.700E-06 |
| 11122 | 557 | P557R | A | 5e-06 | 9.8e-06 | 4.5e-06 | 6.433E-06 | 2.926E-06 | 4.500E-06 | 9.800E-06 |
| 11123 | 557 | P557N | A | 1.2e-05 | 1.1e-05 | 8.1e-06 | 1.037E-05 | 2.026E-06 | 8.100E-06 | 1.200E-05 |
| 11124 | 557 | P557D | A | 6.4e-06 | 5.9e-06 | 3.6e-06 | 5.300E-06 | 1.493E-06 | 3.600E-06 | 6.400E-06 |
| 11125 | 557 | P557C | A | 1.2e-05 | 1.2e-05 | 9.1e-06 | 1.103E-05 | 1.674E-06 | 9.100E-06 | 1.200E-05 |
| 11126 | 557 | P557E | A | 1e-05 | 8.6e-06 | 6e-06 | 8.200E-06 | 2.030E-06 | 6.000E-06 | 1.000E-05 |
| 11127 | 557 | P557Q | A | 7.7e-06 | 7.4e-06 | 6.3e-06 | 7.133E-06 | 7.371E-07 | 6.300E-06 | 7.700E-06 |
| 11128 | 557 | P557G | A | 9e-06 | 5.1e-06 | 3.8e-06 | 5.967E-06 | 2.706E-06 | 3.800E-06 | 9.000E-06 |
| 11129 | 557 | P557H | A | 9.5e-06 | 6.5e-06 | 7.8e-06 | 7.933E-06 | 1.504E-06 | 6.500E-06 | 9.500E-06 |
| 11130 | 557 | P557I | A | 7.1e-06 | 5.7e-06 | 8.6e-06 | 7.133E-06 | 1.450E-06 | 5.700E-06 | 8.600E-06 |
| 11131 | 557 | P557L | A | 8.7e-06 | 1.4e-05 | 8.9e-06 | 1.053E-05 | 3.004E-06 | 8.700E-06 | 1.400E-05 |
| 11132 | 557 | P557K | A | 1.2e-05 | 1.1e-05 | 9.8e-06 | 1.093E-05 | 1.102E-06 | 9.800E-06 | 1.200E-05 |
| 11133 | 557 | P557M | A | 6.6e-06 | 8.9e-06 | 7.8e-06 | 7.767E-06 | 1.150E-06 | 6.600E-06 | 8.900E-06 |
| 11134 | 557 | P557F | A | 6.9e-06 | 5.8e-06 | 7.5e-06 | 6.733E-06 | 8.622E-07 | 5.800E-06 | 7.500E-06 |

| | | | | | | | | | | |
|---|---|---|---|---|---|---|---|---|---|---|
| 11135 | 557 | P557P | A | 7.2e-06 | 4.9e-06 | 6.7e-06 | 6.267E-06 | 1.210E-06 | 4.900E-06 | 7.200E-06 |
| 11136 | 557 | P557S | A | 7.4e-06 | 7.1e-06 | 1e-05 | 8.167E-06 | 1.595E-06 | 7.100E-06 | 1.000E-05 |
| 11137 | 557 | P557T | A | 4.3e-06 | 9.3e-06 | 5.9e-06 | 6.500E-06 | 2.553E-06 | 4.300E-06 | 9.300E-06 |
| 11138 | 557 | P557W | A | 4.8e-06 | 7.3e-06 | 5.1e-06 | 5.733E-06 | 1.365E-06 | 4.800E-06 | 7.300E-06 |
| 11139 | 557 | P557Y | A | 1e-05 | 6.3e-06 | 1e-05 | 8.767E-06 | 2.136E-06 | 6.300E-06 | 1.000E-05 |
| 11140 | 557 | P557V | A | 1.1e-05 | 7e-06 | 1.1e-05 | 9.667E-06 | 2.309E-06 | 7.000E-06 | 1.100E-05 |
| 11141 | 558 | E558A | A | 5.3e-06 | 6.9e-06 | 9.5e-06 | 7.233E-06 | 2.120E-06 | 5.300E-06 | 9.500E-06 |
| 11142 | 558 | E558R | A | 9.6e-06 | 9.4e-06 | 7e-06 | 8.667E-06 | 1.447E-06 | 7.000E-06 | 9.600E-06 |
| 11143 | 558 | E558N | A | 9.8e-06 | 1e-05 | 6.5e-06 | 8.767E-06 | 1.966E-06 | 6.500E-06 | 1.000E-05 |
| 11144 | 558 | E558D | A | 9.9e-06 | 1.2e-05 | 1.2e-05 | 1.130E-05 | 1.212E-06 | 9.900E-06 | 1.200E-05 |
| 11145 | 558 | E558C | A | 9.1e-06 | 5.4e-06 | 8.1e-06 | 7.533E-06 | 1.914E-06 | 5.400E-06 | 9.100E-06 |
| 11146 | 558 | E558E | A | 1.4e-05 | 7.2e-06 | 1.3e-05 | 1.140E-05 | 3.672E-06 | 7.200E-06 | 1.400E-05 |
| 11147 | 558 | E558Q | A | 1.1e-05 | 4.4e-06 | 1.6e-05 | 1.047E-05 | 5.818E-06 | 4.400E-06 | 1.600E-05 |
| 11148 | 558 | E558G | A | 4.6e-06 | 9.4e-06 | 9.3e-06 | 7.767E-06 | 2.743E-06 | 4.600E-06 | 9.400E-06 |
| 11149 | 558 | E558H | A | 4.6e-06 | 8.9e-06 | 7.3e-06 | 6.933E-06 | 2.173E-06 | 4.600E-06 | 8.900E-06 |
| 11150 | 558 | E558I | A | 9.2e-06 | 9.8e-06 | 1.4e-05 | 1.100E-05 | 2.615E-06 | 9.200E-06 | 1.400E-05 |
| 11151 | 558 | E558L | A | 8.9e-06 | 6.2e-06 | 1.1e-05 | 8.700E-06 | 2.406E-06 | 6.200E-06 | 1.100E-05 |
| 11152 | 558 | E558K | A | 6.8e-06 | 1.2e-05 | 4.7e-06 | 7.833E-06 | 3.758E-06 | 4.700E-06 | 1.200E-05 |
| 11153 | 558 | E558M | A | 6e-06 | 9.8e-06 | 8.3e-06 | 8.033E-06 | 1.914E-06 | 6.000E-06 | 9.800E-06 |
| 11154 | 558 | E558F | A | 1.2e-05 | 7.2e-06 | 7.5e-06 | 8.900E-06 | 2.689E-06 | 7.200E-06 | 1.200E-05 |
| 11155 | 558 | E558P | A | 8.2e-06 | 5.8e-06 | 9.7e-06 | 7.900E-06 | 1.967E-06 | 5.800E-06 | 9.700E-06 |
| 11156 | 558 | E558S | A | 1.1e-05 | 9.9e-06 | 9.1e-06 | 1.000E-05 | 9.539E-07 | 9.100E-06 | 1.100E-05 |
| 11157 | 558 | E558T | A | 9.1e-06 | 5.9e-06 | 8.2e-06 | 7.733E-06 | 1.650E-06 | 5.900E-06 | 9.100E-06 |
| 11158 | 558 | E558W | A | 1.2e-05 | 1.5e-05 | 1.2e-05 | 1.300E-05 | 1.732E-06 | 1.200E-05 | 1.500E-05 |
| 11159 | 558 | E558Y | A | 6.6e-06 | 7.9e-06 | 7.1e-06 | 7.200E-06 | 6.557E-07 | 6.600E-06 | 7.900E-06 |
| 11160 | 558 | E558V | A | 9.1e-06 | 1.3e-05 | 7.9e-06 | 1.000E-05 | 2.666E-06 | 7.900E-06 | 1.300E-05 |
| 11161 | 559 | A559A | A | 1.5e-05 | 3.8e-06 | 1.4e-05 | 1.093E-05 | 6.198E-06 | 3.800E-06 | 1.500E-05 |
| 11162 | 559 | A559R | A | 1.1e-05 | 7.9e-06 | 1e-05 | 9.633E-06 | 1.582E-06 | 7.900E-06 | 1.100E-05 |
| 11163 | 559 | A559N | A | 9.6e-06 | 8.1e-06 | 8.7e-06 | 8.800E-06 | 7.550E-07 | 8.100E-06 | 9.600E-06 |
| 11164 | 559 | A559D | A | 6.7e-06 | 9.6e-06 | 1e-05 | 8.767E-06 | 1.801E-06 | 6.700E-06 | 1.000E-05 |
| 11165 | 559 | A559C | A | 8.3e-06 | 9.1e-06 | 1e-05 | 9.133E-06 | 8.505E-07 | 8.300E-06 | 1.000E-05 |
| 11166 | 559 | A559E | A | 8.9e-06 | 9.8e-06 | 1.2e-05 | 1.023E-05 | 1.595E-06 | 8.900E-06 | 1.200E-05 |
| 11167 | 559 | A559Q | A | 1.1e-05 | 7.9e-06 | 1.3e-05 | 1.063E-05 | 2.570E-06 | 7.900E-06 | 1.300E-05 |
| 11168 | 559 | A559G | A | 4.7e-06 | 8.9e-06 | 5.6e-06 | 6.400E-06 | 2.211E-06 | 4.700E-06 | 8.900E-06 |
| 11169 | 559 | A559H | A | 4.5e-06 | 9.9e-06 | 9e-06 | 7.800E-06 | 2.893E-06 | 4.500E-06 | 9.900E-06 |
| 11170 | 559 | A559I | A | 6.7e-06 | 8.7e-06 | 1.3e-05 | 9.467E-06 | 3.219E-06 | 6.700E-06 | 1.300E-05 |
| 11171 | 559 | A559L | A | 6.9e-06 | 8.1e-06 | 4.2e-06 | 6.400E-06 | 1.997E-06 | 4.200E-06 | 8.100E-06 |
| 11172 | 559 | A559K | A | 1.9e-05 | 8.2e-06 | 8.4e-06 | 1.187E-05 | 6.178E-06 | 8.200E-06 | 1.900E-05 |
| 11173 | 559 | A559M | A | 9.2e-06 | 7.6e-06 | 4.8e-06 | 7.200E-06 | 2.227E-06 | 4.800E-06 | 9.200E-06 |
| 11174 | 559 | A559F | A | 8.7e-06 | 8e-06 | 9.6e-06 | 8.767E-06 | 8.021E-07 | 8.000E-06 | 9.600E-06 |
| 11175 | 559 | A559P | A | 9.3e-06 | 9.6e-06 | 8.6e-06 | 9.167E-06 | 5.132E-07 | 8.600E-06 | 9.600E-06 |
| 11176 | 559 | A559S | A | 1e-05 | 8.5e-06 | 4.3e-06 | 7.600E-06 | 2.955E-06 | 4.300E-06 | 1.000E-05 |
| 11177 | 559 | A559T | A | 7.3e-06 | 1.3e-05 | 9.7e-06 | 1.000E-05 | 2.862E-06 | 7.300E-06 | 1.300E-05 |
| 11178 | 559 | A559W | A | 1e-05 | 8.3e-06 | 1.5e-05 | 1.110E-05 | 3.483E-06 | 8.300E-06 | 1.500E-05 |
| 11179 | 559 | A559Y | A | 1.3e-05 | 9.4e-06 | 8.3e-06 | 1.023E-05 | 2.458E-06 | 8.300E-06 | 1.300E-05 |
| 11180 | 559 | A559V | A | 6.2e-06 | 1.3e-05 | 5.4e-06 | 8.200E-06 | 4.176E-06 | 5.400E-06 | 1.300E-05 |
| 11181 | 560 | D560A | A | 5.4e-06 | 8.3e-06 | 5.2e-06 | 6.300E-06 | 1.735E-06 | 5.200E-06 | 8.300E-06 |
| 11182 | 560 | D560R | A | 6.3e-06 | 9.1e-06 | 7.7e-06 | 7.700E-06 | 1.400E-06 | 6.300E-06 | 9.100E-06 |
| 11183 | 560 | D560N | A | 1.2e-05 | 1.5e-05 | 1.5e-05 | 1.400E-05 | 1.732E-06 | 1.200E-05 | 1.500E-05 |
| 11184 | 560 | D560D | A | 5.8e-06 | 6.6e-06 | 1.3e-05 | 8.467E-06 | 3.946E-06 | 5.800E-06 | 1.300E-05 |
| 11185 | 560 | D560C | A | 5.6e-06 | 4.2e-06 | 4.4e-06 | 4.733E-06 | 7.572E-07 | 4.200E-06 | 5.600E-06 |
| 11186 | 560 | D560E | A | 4.7e-06 | 1.1e-05 | 5e-06 | 6.900E-06 | 3.554E-06 | 4.700E-06 | 1.100E-05 |
| 11187 | 560 | D560Q | A | 1.2e-05 | 1.4e-05 | 1.4e-05 | 1.333E-05 | 1.155E-06 | 1.200E-05 | 1.400E-05 |
| 11188 | 560 | D560G | A | 6.2e-06 | 8e-06 | 6.6e-06 | 6.933E-06 | 9.452E-07 | 6.200E-06 | 8.000E-06 |
| 11189 | 560 | D560H | A | 5.1e-06 | 9.4e-06 | 8e-06 | 7.500E-06 | 2.193E-06 | 5.100E-06 | 9.400E-06 |
| 11190 | 560 | D560I | A | 5e-06 | 2.7e-06 | 3.4e-06 | 3.700E-06 | 1.179E-06 | 2.700E-06 | 5.000E-06 |
| 11191 | 560 | D560L | A | 2.1e-06 | 4.2e-06 | 3.3e-06 | 3.200E-06 | 1.054E-06 | 2.100E-06 | 4.200E-06 |
| 11192 | 560 | D560K | A | 1.2e-05 | 7.7e-06 | 8.7e-06 | 9.467E-06 | 2.250E-06 | 7.700E-06 | 1.200E-05 |
| 11193 | 560 | D560M | A | 3e-06 | 5.9e-06 | 3.9e-06 | 4.267E-06 | 1.484E-06 | 3.000E-06 | 5.900E-06 |
| 11194 | 560 | D560F | A | 2.3e-06 | 3.9e-06 | 6.5e-06 | 4.233E-06 | 2.120E-06 | 2.300E-06 | 6.500E-06 |
| 11195 | 560 | D560P | A | 4.4e-06 | 9.3e-06 | 4.3e-06 | 6.000E-06 | 2.858E-06 | 4.300E-06 | 9.300E-06 |
| 11196 | 560 | D560S | A | 1.5e-05 | 1.1e-05 | 1.1e-05 | 1.233E-05 | 2.309E-06 | 1.100E-05 | 1.500E-05 |
| 11197 | 560 | D560T | A | 1.5e-05 | 1.1e-05 | 7.2e-06 | 1.107E-05 | 3.900E-06 | 7.200E-06 | 1.500E-05 |
| 11198 | 560 | D560W | A | 2.9e-06 | 3.1e-06 | 2.6e-06 | 2.867E-06 | 2.517E-07 | 2.600E-06 | 3.100E-06 |
| 11199 | 560 | D560Y | A | 4.5e-06 | 4.8e-06 | 4.3e-06 | 4.533E-06 | 2.517E-07 | 4.300E-06 | 4.800E-06 |
| 11200 | 560 | D560V | A | 4.5e-06 | 6.3e-06 | 4.6e-06 | 5.133E-06 | 1.012E-06 | 4.500E-06 | 6.300E-06 |
| 11201 | 561 | Q561A | A | 6.7e-06 | 1e-05 | 7.3e-06 | 8.000E-06 | 1.758E-06 | 6.700E-06 | 1.000E-05 |
| 11202 | 561 | Q561R | A | 8.9e-06 | 5e-06 | 6.9e-06 | 6.933E-06 | 1.950E-06 | 5.000E-06 | 8.900E-06 |
| 11203 | 561 | Q561N | A | 1e-05 | 7.2e-06 | 1.1e-05 | 9.400E-06 | 1.970E-06 | 7.200E-06 | 1.100E-05 |
| 11204 | 561 | Q561D | A | 8.4e-06 | 4.6e-06 | 9.7e-06 | 7.567E-06 | 2.650E-06 | 4.600E-06 | 9.700E-06 |
| 11205 | 561 | Q561C | A | 2.9e-06 | 1e-05 | 6.3e-06 | 6.400E-06 | 3.551E-06 | 2.900E-06 | 1.000E-05 |
| 11206 | 561 | Q561E | A | 6.5e-06 | 5e-06 | 4.9e-06 | 5.467E-06 | 8.963E-07 | 4.900E-06 | 6.500E-06 |
| 11207 | 561 | Q561Q | A | 1.1e-05 | 9.7e-06 | 1.3e-05 | 1.123E-05 | 1.662E-06 | 9.700E-06 | 1.300E-05 |
| 11208 | 561 | Q561G | A | 4.4e-06 | 1e-05 | 8e-06 | 7.467E-06 | 2.838E-06 | 4.400E-06 | 1.000E-05 |
| 11209 | 561 | Q561H | A | 4.2e-06 | 9.9e-06 | 5e-06 | 6.367E-06 | 3.086E-06 | 4.200E-06 | 9.900E-06 |
| 11210 | 561 | Q561I | A | 6.3e-06 | 1.2e-05 | 5.7e-06 | 8.000E-06 | 3.477E-06 | 5.700E-06 | 1.200E-05 |
| 11211 | 561 | Q561L | A | 4.7e-06 | 4.5e-06 | 3.8e-06 | 4.333E-06 | 4.726E-07 | 3.800E-06 | 4.700E-06 |
| 11212 | 561 | Q561K | A | 9.6e-06 | 5.8e-06 | 3.9e-06 | 6.433E-06 | 2.902E-06 | 3.900E-06 | 9.600E-06 |
| 11213 | 561 | Q561M | A | 5.2e-06 | 5.8e-06 | 4e-06 | 5.000E-06 | 9.165E-07 | 4.000E-06 | 5.800E-06 |
| 11214 | 561 | Q561F | A | 3.8e-06 | 6.7e-06 | 6.9e-06 | 5.800E-06 | 1.735E-06 | 3.800E-06 | 6.900E-06 |
| 11215 | 561 | Q561P | A | 7.4e-06 | 9.4e-06 | 8.8e-06 | 8.533E-06 | 1.026E-06 | 7.400E-06 | 9.400E-06 |
| 11216 | 561 | Q561S | A | 9.1e-06 | 6.4e-06 | 8.6e-06 | 8.033E-06 | 1.436E-06 | 6.400E-06 | 9.100E-06 |
| 11217 | 561 | Q561T | A | 5.1e-06 | 8.9e-06 | 7e-06 | 7.000E-06 | 1.900E-06 | 5.100E-06 | 8.900E-06 |
| 11218 | 561 | Q561W | A | 6.8e-06 | 2.9e-06 | 6.5e-06 | 5.400E-06 | 2.170E-06 | 2.900E-06 | 6.800E-06 |
| 11219 | 561 | Q561Y | A | 6.4e-06 | 5.6e-06 | 4.6e-06 | 5.533E-06 | 9.018E-07 | 4.600E-06 | 6.400E-06 |

| | | | | | | | | | | |
|---|---|---|---|---|---|---|---|---|---|---|
| 11220 | 561 | Q561V | A | 4.6e-06 | 4.3e-06 | 7.1e-06 | 5.333E-06 | 1.537E-06 | 4.300E-06 | 7.100E-06 |
| 11221 | 562 | C562A | A | 6.4e-06 | 6.2e-06 | 1.5e-05 | 9.200E-06 | 5.024E-06 | 6.200E-06 | 1.500E-05 |
| 11222 | 562 | C562R | A | 6e-06 | 1.2e-05 | 5.2e-06 | 7.733E-06 | 3.717E-06 | 5.200E-06 | 1.200E-05 |
| 11223 | 562 | C562N | A | 7.5e-06 | 1.2e-05 | 7.2e-06 | 8.900E-06 | 2.689E-06 | 7.200E-06 | 1.200E-05 |
| 11224 | 562 | C562D | A | 5.1e-06 | 6.1e-06 | 9.5e-06 | 6.900E-06 | 2.307E-06 | 5.100E-06 | 9.500E-06 |
| 11225 | 562 | C562C | A | 6.4e-06 | 1.1e-05 | 9.8e-06 | 9.067E-06 | 2.386E-06 | 6.400E-06 | 1.100E-05 |
| 11226 | 562 | C562E | A | 1.2e-05 | 1.3e-05 | 9.2e-06 | 1.140E-05 | 1.970E-06 | 9.200E-06 | 1.300E-05 |
| 11227 | 562 | C562Q | A | 5.2e-06 | 1.2e-05 | 7.5e-06 | 8.233E-06 | 3.459E-06 | 5.200E-06 | 1.200E-05 |
| 11228 | 562 | C562G | A | 6.3e-06 | 6e-06 | 1.3e-05 | 8.433E-06 | 3.958E-06 | 6.000E-06 | 1.300E-05 |
| 11229 | 562 | C562H | A | 6.2e-06 | 9.1e-06 | 1.2e-05 | 9.100E-06 | 2.900E-06 | 6.200E-06 | 1.200E-05 |
| 11230 | 562 | C562I | A | 1.3e-05 | 8.3e-06 | 1.3e-05 | 1.143E-05 | 2.714E-06 | 8.300E-06 | 1.300E-05 |
| 11231 | 562 | C562L | A | 8.2e-06 | 7.3e-06 | 6.6e-06 | 7.367E-06 | 8.021E-07 | 6.600E-06 | 8.200E-06 |
| 11232 | 562 | C562K | A | 1.2e-05 | 7.2e-06 | 1.1e-05 | 1.007E-05 | 2.532E-06 | 7.200E-06 | 1.200E-05 |
| 11233 | 562 | C562M | A | 8.6e-06 | 9.2e-06 | 7.7e-06 | 8.500E-06 | 7.550E-07 | 7.700E-06 | 9.200E-06 |
| 11234 | 562 | C562F | A | 5.6e-06 | 7.5e-06 | 4.9e-06 | 6.000E-06 | 1.345E-06 | 4.900E-06 | 7.500E-06 |
| 11235 | 562 | C562P | A | 6e-06 | 6.8e-06 | 8.2e-06 | 7.000E-06 | 1.114E-06 | 6.000E-06 | 8.200E-06 |
| 11236 | 562 | C562S | A | 7.7e-06 | 8e-06 | 1.4e-05 | 9.900E-06 | 3.554E-06 | 7.700E-06 | 1.400E-05 |
| 11237 | 562 | C562T | A | 5.4e-06 | 9.2e-06 | 8.7e-06 | 7.767E-06 | 2.065E-06 | 5.400E-06 | 9.200E-06 |
| 11238 | 562 | C562W | A | 5.1e-06 | 1.3e-05 | 4e-06 | 7.367E-06 | 4.910E-06 | 4.000E-06 | 1.300E-05 |
| 11239 | 562 | C562Y | A | 4e-06 | 9.9e-06 | 9.2e-06 | 7.700E-06 | 3.223E-06 | 4.000E-06 | 9.900E-06 |
| 11240 | 562 | C562V | A | 4.3e-06 | 1.2e-05 | 9.3e-06 | 8.533E-06 | 3.907E-06 | 4.300E-06 | 1.200E-05 |
| 11241 | 563 | V563A | A | 8.6e-06 | 6.7e-06 | 1.2e-05 | 9.100E-06 | 2.685E-06 | 6.700E-06 | 1.200E-05 |
| 11242 | 563 | V563R | A | 1.1e-05 | 5.6e-06 | 6.2e-06 | 7.600E-06 | 2.960E-06 | 5.600E-06 | 1.100E-05 |
| 11243 | 563 | V563N | A | 9.6e-06 | 6.8e-06 | 8.5e-06 | 8.300E-06 | 1.411E-06 | 6.800E-06 | 9.600E-06 |
| 11244 | 563 | V563D | A | 7.3e-06 | 6.6e-06 | 9.2e-06 | 7.700E-06 | 1.345E-06 | 6.600E-06 | 9.200E-06 |
| 11245 | 563 | V563C | A | 7.1e-06 | 6.6e-06 | 4.3e-06 | 6.000E-06 | 1.493E-06 | 4.300E-06 | 7.100E-06 |
| 11246 | 563 | V563E | A | 5.4e-06 | 1.1e-05 | 1.2e-05 | 9.467E-06 | 3.557E-06 | 5.400E-06 | 1.200E-05 |
| 11247 | 563 | V563Q | A | 1e-05 | 1.1e-05 | 1.4e-05 | 1.167E-05 | 2.082E-06 | 1.000E-05 | 1.400E-05 |
| 11248 | 563 | V563G | A | 1e-05 | 9.7e-06 | 8.9e-06 | 9.533E-06 | 5.686E-07 | 8.900E-06 | 1.000E-05 |
| 11249 | 563 | V563H | A | 1.2e-05 | 8.3e-06 | 8.5e-06 | 9.600E-06 | 2.081E-06 | 8.300E-06 | 1.200E-05 |
| 11250 | 563 | V563I | A | 9.4e-06 | 9.8e-06 | 1.6e-05 | 1.173E-05 | 3.700E-06 | 9.400E-06 | 1.600E-05 |
| 11251 | 563 | V563L | A | 1.3e-05 | 6e-06 | 6.2e-06 | 8.400E-06 | 3.985E-06 | 6.000E-06 | 1.300E-05 |
| 11252 | 563 | V563K | A | 8.7e-06 | 6.6e-06 | 7.9e-06 | 7.733E-06 | 1.060E-06 | 6.600E-06 | 8.700E-06 |
| 11253 | 563 | V563M | A | 6.3e-06 | 1.3e-05 | 1e-05 | 9.767E-06 | 3.356E-06 | 6.300E-06 | 1.300E-05 |
| 11254 | 563 | V563F | A | 9.2e-06 | 7.6e-06 | 9.2e-06 | 8.667E-06 | 9.238E-07 | 7.600E-06 | 9.200E-06 |
| 11255 | 563 | V563P | A | 9e-06 | 5.2e-06 | 1.1e-05 | 8.400E-06 | 2.946E-06 | 5.200E-06 | 1.100E-05 |
| 11256 | 563 | V563S | A | 1e-05 | 7.1e-06 | 9.9e-06 | 9.000E-06 | 1.646E-06 | 7.100E-06 | 1.000E-05 |
| 11257 | 563 | V563T | A | 6.7e-06 | 9.1e-06 | 5.8e-06 | 7.200E-06 | 1.706E-06 | 5.800E-06 | 9.100E-06 |
| 11258 | 563 | V563W | A | 5.6e-06 | 5.7e-06 | 4.4e-06 | 5.233E-06 | 7.234E-07 | 4.400E-06 | 5.700E-06 |
| 11259 | 563 | V563Y | A | 9e-06 | 5.3e-06 | 7.8e-06 | 7.367E-06 | 1.888E-06 | 5.300E-06 | 9.000E-06 |
| 11260 | 563 | V563V | A | 6.7e-06 | 6e-06 | 1.2e-05 | 8.233E-06 | 3.281E-06 | 6.000E-06 | 1.200E-05 |
| 11261 | 564 | A564A | A | 7.9e-06 | 7.4e-06 | 1.2e-05 | 9.100E-06 | 2.524E-06 | 7.400E-06 | 1.200E-05 |
| 11262 | 564 | A564R | A | 6.9e-06 | 1.3e-05 | 7e-06 | 8.967E-06 | 3.493E-06 | 6.900E-06 | 1.300E-05 |
| 11263 | 564 | A564N | A | 3.6e-06 | 5.2e-06 | 4.8e-06 | 4.533E-06 | 8.327E-07 | 3.600E-06 | 5.200E-06 |
| 11264 | 564 | A564D | A | 5.6e-06 | 5.9e-06 | 5.5e-06 | 5.667E-06 | 2.082E-07 | 5.500E-06 | 5.900E-06 |
| 11265 | 564 | A564C | A | 8.3e-06 | 6e-06 | 3.2e-06 | 5.833E-06 | 2.554E-06 | 3.200E-06 | 8.300E-06 |
| 11266 | 564 | A564E | A | 7.1e-06 | 7.1e-06 | 8.2e-06 | 7.467E-06 | 6.351E-07 | 7.100E-06 | 8.200E-06 |
| 11267 | 564 | A564Q | A | 5.6e-06 | 7.8e-06 | 4.7e-06 | 6.033E-06 | 1.595E-06 | 4.700E-06 | 7.800E-06 |
| 11268 | 564 | A564G | A | 7.4e-06 | 7.6e-06 | 4.4e-06 | 6.467E-06 | 1.793E-06 | 4.400E-06 | 7.600E-06 |
| 11269 | 564 | A564H | A | 8.4e-06 | 5.5e-06 | 9.3e-06 | 7.733E-06 | 1.986E-06 | 5.500E-06 | 9.300E-06 |
| 11270 | 564 | A564I | A | 4.9e-06 | 4.8e-06 | 9.9e-06 | 6.533E-06 | 2.916E-06 | 4.800E-06 | 9.900E-06 |
| 11271 | 564 | A564L | A | 1.2e-05 | 6e-06 | 5.9e-06 | 7.967E-06 | 3.493E-06 | 5.900E-06 | 1.200E-05 |
| 11272 | 564 | A564K | A | 5.7e-06 | 6.3e-06 | 4.2e-06 | 5.400E-06 | 1.082E-06 | 4.200E-06 | 6.300E-06 |
| 11273 | 564 | A564M | A | 6.7e-06 | 4.8e-06 | 5.7e-06 | 5.733E-06 | 9.504E-07 | 4.800E-06 | 6.700E-06 |
| 11274 | 564 | A564F | A | 9.6e-06 | 5e-06 | 5.4e-06 | 6.667E-06 | 2.548E-06 | 5.000E-06 | 9.600E-06 |
| 11275 | 564 | A564P | A | 9.3e-06 | 4.5e-06 | 8.8e-06 | 7.533E-06 | 2.639E-06 | 4.500E-06 | 9.300E-06 |
| 11276 | 564 | A564S | A | 9.9e-06 | 6.2e-06 | 1e-05 | 8.700E-06 | 2.166E-06 | 6.200E-06 | 1.000E-05 |
| 11277 | 564 | A564T | A | 5.9e-06 | 5.3e-06 | 6.1e-06 | 5.767E-06 | 4.163E-07 | 5.300E-06 | 6.100E-06 |
| 11278 | 564 | A564W | A | 1.2e-05 | 8.1e-06 | 9e-06 | 9.700E-06 | 2.042E-06 | 8.100E-06 | 1.200E-05 |
| 11279 | 564 | A564Y | A | 4.8e-06 | 1.1e-05 | 5.5e-06 | 7.100E-06 | 3.396E-06 | 4.800E-06 | 1.100E-05 |
| 11280 | 564 | A564V | A | 1.1e-05 | 5.5e-06 | 6.6e-06 | 7.700E-06 | 2.910E-06 | 5.500E-06 | 1.100E-05 |
| 11281 | 565 | C565A | A | 8.8e-06 | 8.5e-06 | 6.8e-06 | 8.033E-06 | 1.079E-06 | 6.800E-06 | 8.800E-06 |
| 11282 | 565 | C565R | A | 6.5e-06 | 6.9e-06 | 8.9e-06 | 7.433E-06 | 1.286E-06 | 6.500E-06 | 8.900E-06 |
| 11283 | 565 | C565N | A | 1.7e-05 | 3.1e-06 | 4.4e-06 | 8.167E-06 | 7.677E-06 | 3.100E-06 | 1.700E-05 |
| 11284 | 565 | C565D | A | 8.8e-06 | 7.4e-06 | 5e-06 | 7.067E-06 | 1.922E-06 | 5.000E-06 | 8.800E-06 |
| 11285 | 565 | C565C | A | 6.4e-06 | 4.7e-06 | 9.1e-06 | 6.733E-06 | 2.219E-06 | 4.700E-06 | 9.100E-06 |
| 11286 | 565 | C565E | A | 6.5e-06 | 1.5e-05 | 7.5e-06 | 9.667E-06 | 4.646E-06 | 6.500E-06 | 1.500E-05 |
| 11287 | 565 | C565Q | A | 9.9e-06 | 1.1e-05 | 9.3e-06 | 1.007E-05 | 8.622E-07 | 9.300E-06 | 1.100E-05 |
| 11288 | 565 | C565G | A | 1e-05 | 5.5e-06 | 7.4e-06 | 7.633E-06 | 2.259E-06 | 5.500E-06 | 1.000E-05 |
| 11289 | 565 | C565H | A | 7.2e-06 | 8e-06 | 8e-06 | 7.733E-06 | 4.619E-07 | 7.200E-06 | 8.000E-06 |
| 11290 | 565 | C565I | A | 8.4e-06 | 1.5e-05 | 6.7e-06 | 1.003E-05 | 4.384E-06 | 6.700E-06 | 1.500E-05 |
| 11291 | 565 | C565L | A | 7.5e-06 | 6.7e-06 | 8.2e-06 | 7.467E-06 | 7.506E-07 | 6.700E-06 | 8.200E-06 |
| 11292 | 565 | C565K | A | 1.1e-05 | 5e-06 | 1.1e-05 | 9.000E-06 | 3.464E-06 | 5.000E-06 | 1.100E-05 |
| 11293 | 565 | C565M | A | 8.5e-06 | 3.8e-06 | 6.3e-06 | 6.200E-06 | 2.352E-06 | 3.800E-06 | 8.500E-06 |
| 11294 | 565 | C565F | A | 1.2e-05 | 5.7e-06 | 6.4e-06 | 8.033E-06 | 3.453E-06 | 5.700E-06 | 1.200E-05 |
| 11295 | 565 | C565P | A | 5e-06 | 8.9e-06 | 6.2e-06 | 6.700E-06 | 1.997E-06 | 5.000E-06 | 8.900E-06 |
| 11296 | 565 | C565S | A | 8.6e-06 | 7.3e-06 | 1.4e-05 | 9.967E-06 | 3.553E-06 | 7.300E-06 | 1.400E-05 |
| 11297 | 565 | C565T | A | 4.8e-06 | 1.2e-05 | 5.5e-06 | 7.433E-06 | 3.970E-06 | 4.800E-06 | 1.200E-05 |
| 11298 | 565 | C565W | A | 1.2e-05 | 1e-05 | 1e-05 | 1.067E-05 | 1.155E-06 | 1.000E-05 | 1.200E-05 |
| 11299 | 565 | C565Y | A | 6.5e-06 | 8.8e-06 | 1e-05 | 8.433E-06 | 1.779E-06 | 6.500E-06 | 1.000E-05 |
| 11300 | 565 | C565V | A | 1.1e-05 | 7.3e-06 | 7.9e-06 | 8.733E-06 | 1.986E-06 | 7.300E-06 | 1.100E-05 |
| 11301 | 566 | A566A | A | 1e-05 | 6.7e-06 | 4.1e-06 | 6.933E-06 | 2.957E-06 | 4.100E-06 | 1.000E-05 |
| 11302 | 566 | A566R | A | 5.5e-06 | 7.9e-06 | 1.1e-05 | 8.133E-06 | 2.757E-06 | 5.500E-06 | 1.100E-05 |
| 11303 | 566 | A566N | A | 1.1e-05 | 7.6e-06 | 4.7e-06 | 7.767E-06 | 3.153E-06 | 4.700E-06 | 1.100E-05 |
| 11304 | 566 | A566D | A | 6.9e-06 | 4.3e-06 | 8.9e-06 | 6.700E-06 | 2.307E-06 | 4.300E-06 | 8.900E-06 |

| | | | | | | | | | | |
|---|---|---|---|---|---|---|---|---|---|---|
| 11305 | 566 | A566C | A | 3e-06 | 6.9e-06 | 1.1e-05 | 6.967E-06 | 4.000E-06 | 3.000E-06 | 1.100E-05 |
| 11306 | 566 | A566E | A | 9.1e-06 | 8.8e-06 | 9.5e-06 | 9.133E-06 | 3.512E-07 | 8.800E-06 | 9.500E-06 |
| 11307 | 566 | A566Q | A | 6.5e-06 | 6.5e-06 | 9.9e-06 | 7.633E-06 | 1.963E-06 | 6.500E-06 | 9.900E-06 |
| 11308 | 566 | A566G | A | 7.4e-06 | 1e-05 | 1.2e-05 | 9.800E-06 | 2.307E-06 | 7.400E-06 | 1.200E-05 |
| 11309 | 566 | A566H | A | 6.7e-06 | 1.2e-05 | 7.2e-06 | 8.633E-06 | 2.926E-06 | 6.700E-06 | 1.200E-05 |
| 11310 | 566 | A566I | A | 5e-06 | 9.6e-06 | 1.1e-05 | 8.533E-06 | 3.139E-06 | 5.000E-06 | 1.100E-05 |
| 11311 | 566 | A566L | A | 6.6e-06 | 8e-06 | 3.5e-06 | 6.033E-06 | 2.303E-06 | 3.500E-06 | 8.000E-06 |
| 11312 | 566 | A566K | A | 7.1e-06 | 1e-05 | 1.3e-05 | 1.003E-05 | 2.950E-06 | 7.100E-06 | 1.300E-05 |
| 11313 | 566 | A566M | A | 7.2e-06 | 5.1e-06 | 6.5e-06 | 6.267E-06 | 1.069E-06 | 5.100E-06 | 7.200E-06 |
| 11314 | 566 | A566F | A | 8.8e-06 | 1e-05 | 9.3e-06 | 9.367E-06 | 6.028E-07 | 8.800E-06 | 1.000E-05 |
| 11315 | 566 | A566P | A | 5.9e-06 | 6.6e-06 | 1.4e-05 | 8.833E-06 | 4.488E-06 | 5.900E-06 | 1.400E-05 |
| 11316 | 566 | A566S | A | 9.9e-06 | 8e-06 | 7.4e-06 | 8.433E-06 | 1.305E-06 | 7.400E-06 | 9.900E-06 |
| 11317 | 566 | A566T | A | 4.4e-06 | 3.6e-06 | 5.8e-06 | 4.600E-06 | 1.114E-06 | 3.600E-06 | 5.800E-06 |
| 11318 | 566 | A566W | A | 5.6e-06 | 8.3e-06 | 6.8e-06 | 6.900E-06 | 1.353E-06 | 5.600E-06 | 8.300E-06 |
| 11319 | 566 | A566Y | A | 5.9e-06 | 6.8e-06 | 9.5e-06 | 7.400E-06 | 1.873E-06 | 5.900E-06 | 9.500E-06 |
| 11320 | 566 | A566V | A | 7e-06 | 6.4e-06 | 5.4e-06 | 6.267E-06 | 8.083E-07 | 5.400E-06 | 7.000E-06 |
| 11321 | 567 | H567A | A | 1.1e-05 | 6.6e-06 | 1.1e-05 | 9.533E-06 | 2.540E-06 | 6.600E-06 | 1.100E-05 |
| 11322 | 567 | H567R | A | 6.9e-06 | 7.3e-06 | 1.1e-05 | 8.400E-06 | 2.261E-06 | 6.900E-06 | 1.100E-05 |
| 11323 | 567 | H567N | A | 7.8e-06 | 8.6e-06 | 8.8e-06 | 8.400E-06 | 5.292E-07 | 7.800E-06 | 8.800E-06 |
| 11324 | 567 | H567D | A | 4.9e-06 | 4.9e-06 | 1.4e-05 | 7.933E-06 | 5.254E-06 | 4.900E-06 | 1.400E-05 |
| 11325 | 567 | H567C | A | 1e-05 | 7.4e-06 | 1.5e-05 | 1.080E-05 | 3.863E-06 | 7.400E-06 | 1.500E-05 |
| 11326 | 567 | H567E | A | 1.1e-05 | 1.5e-05 | 6.4e-06 | 1.080E-05 | 4.303E-06 | 6.400E-06 | 1.500E-05 |
| 11327 | 567 | H567Q | A | 9.6e-06 | 1.4e-05 | 1e-05 | 1.120E-05 | 2.433E-06 | 9.600E-06 | 1.400E-05 |
| 11328 | 567 | H567G | A | 6.4e-06 | 6e-06 | 1.2e-05 | 8.133E-06 | 3.355E-06 | 6.000E-06 | 1.200E-05 |
| 11329 | 567 | H567H | A | 7.5e-06 | 1.6e-05 | 7.6e-06 | 1.037E-05 | 4.879E-06 | 7.500E-06 | 1.600E-05 |
| 11330 | 567 | H567I | A | 1.3e-05 | 6.6e-06 | 1.3e-05 | 1.087E-05 | 3.695E-06 | 6.600E-06 | 1.300E-05 |
| 11331 | 567 | H567L | A | 9.7e-06 | 6.9e-06 | 1.4e-05 | 1.020E-05 | 3.576E-06 | 6.900E-06 | 1.400E-05 |
| 11332 | 567 | H567K | A | 1.1e-05 | 7.9e-06 | 6.7e-06 | 8.533E-06 | 2.219E-06 | 6.700E-06 | 1.100E-05 |
| 11333 | 567 | H567M | A | 9e-06 | 6.8e-06 | 5.3e-06 | 7.033E-06 | 1.861E-06 | 5.300E-06 | 9.000E-06 |
| 11334 | 567 | H567F | A | 5.4e-06 | 1.1e-05 | 1e-05 | 8.800E-06 | 2.987E-06 | 5.400E-06 | 1.100E-05 |
| 11335 | 567 | H567P | A | 8.9e-06 | 6.4e-06 | 8.8e-06 | 8.033E-06 | 1.415E-06 | 6.400E-06 | 8.900E-06 |
| 11336 | 567 | H567S | A | 7.7e-06 | 9.8e-06 | 9.6e-06 | 9.033E-06 | 1.159E-06 | 7.700E-06 | 9.800E-06 |
| 11337 | 567 | H567T | A | 7.3e-06 | 1.2e-05 | 4.4e-06 | 7.900E-06 | 3.835E-06 | 4.400E-06 | 1.200E-05 |
| 11338 | 567 | H567W | A | 7.6e-06 | 6.5e-06 | 1e-05 | 8.033E-06 | 1.790E-06 | 6.500E-06 | 1.000E-05 |
| 11339 | 567 | H567Y | A | 6.3e-06 | 5e-06 | 7.1e-06 | 6.133E-06 | 1.060E-06 | 5.000E-06 | 7.100E-06 |
| 11340 | 567 | H567V | A | 5.6e-06 | 5.7e-06 | 8.6e-06 | 6.633E-06 | 1.704E-06 | 5.600E-06 | 8.600E-06 |
| 11341 | 568 | Y568A | A | 8.8e-06 | 7.4e-06 | 6.5e-06 | 7.567E-06 | 1.159E-06 | 6.500E-06 | 8.800E-06 |
| 11342 | 568 | Y568R | A | 4.6e-06 | 4.9e-06 | 6.3e-06 | 5.267E-06 | 9.074E-07 | 4.600E-06 | 6.300E-06 |
| 11343 | 568 | Y568N | A | 1.3e-05 | 6.1e-06 | 4.8e-06 | 7.967E-06 | 4.407E-06 | 4.800E-06 | 1.300E-05 |
| 11344 | 568 | Y568D | A | 5e-06 | 5.2e-06 | 6.7e-06 | 5.633E-06 | 9.292E-07 | 5.000E-06 | 6.700E-06 |
| 11345 | 568 | Y568C | A | 5e-06 | 6.1e-06 | 7.7e-06 | 6.267E-06 | 1.358E-06 | 5.000E-06 | 7.700E-06 |
| 11346 | 568 | Y568E | A | 6.5e-06 | 5e-06 | 7.9e-06 | 6.467E-06 | 1.450E-06 | 5.000E-06 | 7.900E-06 |
| 11347 | 568 | Y568Q | A | 6.1e-06 | 6.5e-06 | 9e-06 | 7.200E-06 | 1.572E-06 | 6.100E-06 | 9.000E-06 |
| 11348 | 568 | Y568G | A | 5.2e-06 | 7.3e-06 | 1.4e-05 | 8.833E-06 | 4.596E-06 | 5.200E-06 | 1.400E-05 |
| 11349 | 568 | Y568H | A | 7.5e-06 | 5e-06 | 1.4e-05 | 8.833E-06 | 4.646E-06 | 5.000E-06 | 1.400E-05 |
| 11350 | 568 | Y568I | A | 6.9e-06 | 6.7e-06 | 7.3e-06 | 6.967E-06 | 3.055E-07 | 6.700E-06 | 7.300E-06 |
| 11351 | 568 | Y568L | A | 1.3e-05 | 1e-05 | 4.6e-06 | 9.200E-06 | 4.257E-06 | 4.600E-06 | 1.300E-05 |
| 11352 | 568 | Y568K | A | 6.8e-06 | 6.4e-06 | 9.8e-06 | 7.667E-06 | 1.858E-06 | 6.400E-06 | 9.800E-06 |
| 11353 | 568 | Y568M | A | 8.3e-06 | 7.6e-06 | 8.2e-06 | 8.033E-06 | 3.786E-07 | 7.600E-06 | 8.300E-06 |
| 11354 | 568 | Y568F | A | 9.8e-06 | 5.4e-06 | 1e-05 | 8.400E-06 | 2.600E-06 | 5.400E-06 | 1.000E-05 |
| 11355 | 568 | Y568P | A | 6.9e-06 | 7.2e-06 | 8e-06 | 7.367E-06 | 5.686E-07 | 6.900E-06 | 8.000E-06 |
| 11356 | 568 | Y568S | A | 6.8e-06 | 6e-06 | 1.1e-05 | 7.933E-06 | 2.686E-06 | 6.000E-06 | 1.100E-05 |
| 11357 | 568 | Y568T | A | 4.4e-06 | 6e-06 | 6.3e-06 | 5.567E-06 | 1.021E-06 | 4.400E-06 | 6.300E-06 |
| 11358 | 568 | Y568W | A | 1.1e-05 | 4.6e-06 | 1.3e-05 | 9.533E-06 | 4.388E-06 | 4.600E-06 | 1.300E-05 |
| 11359 | 568 | Y568Y | A | 5.2e-06 | 7.1e-06 | 6.9e-06 | 6.400E-06 | 1.044E-06 | 5.200E-06 | 7.100E-06 |
| 11360 | 568 | Y568V | A | 7.1e-06 | 8.5e-06 | 9.2e-06 | 8.267E-06 | 1.069E-06 | 7.100E-06 | 9.200E-06 |
| 11361 | 569 | K569A | A | 9.2e-06 | 8.3e-06 | 1.6e-05 | 1.117E-05 | 4.210E-06 | 8.300E-06 | 1.600E-05 |
| 11362 | 569 | K569R | A | 8.1e-06 | 3.6e-06 | 6.5e-06 | 6.067E-06 | 2.281E-06 | 3.600E-06 | 8.100E-06 |
| 11363 | 569 | K569N | A | 1.1e-05 | 5.8e-06 | 6.3e-06 | 7.700E-06 | 2.869E-06 | 5.800E-06 | 1.100E-05 |
| 11364 | 569 | K569D | A | 5.9e-06 | 6.2e-06 | 8.7e-06 | 6.933E-06 | 1.537E-06 | 5.900E-06 | 8.700E-06 |
| 11365 | 569 | K569C | A | 5.5e-06 | 9.1e-06 | 6.4e-06 | 7.000E-06 | 1.873E-06 | 5.500E-06 | 9.100E-06 |
| 11366 | 569 | K569E | A | 6e-06 | 7.7e-06 | 1.3e-05 | 8.900E-06 | 3.651E-06 | 6.000E-06 | 1.300E-05 |
| 11367 | 569 | K569Q | A | 7e-06 | 6e-06 | 5.9e-06 | 6.300E-06 | 6.083E-07 | 5.900E-06 | 7.000E-06 |
| 11368 | 569 | K569G | A | 9.6e-06 | 1e-05 | 1.2e-05 | 1.053E-05 | 1.286E-06 | 9.600E-06 | 1.200E-05 |
| 11369 | 569 | K569H | A | 9.3e-06 | 1e-05 | 1.4e-05 | 1.110E-05 | 2.536E-06 | 9.300E-06 | 1.400E-05 |
| 11370 | 569 | K569I | A | 9.9e-06 | 1.1e-05 | 9.1e-06 | 1.000E-05 | 9.539E-07 | 9.100E-06 | 1.100E-05 |
| 11371 | 569 | K569L | A | 1.2e-05 | 6.6e-06 | 1.2e-05 | 1.020E-05 | 3.118E-06 | 6.600E-06 | 1.200E-05 |
| 11372 | 569 | K569K | A | 6.7e-06 | 9e-06 | 1.4e-05 | 9.900E-06 | 3.732E-06 | 6.700E-06 | 1.400E-05 |
| 11373 | 569 | K569M | A | 8.8e-06 | 4.7e-06 | 8.2e-06 | 7.233E-06 | 2.214E-06 | 4.700E-06 | 8.800E-06 |
| 11374 | 569 | K569F | A | 1.1e-05 | 3.1e-06 | 7.8e-06 | 7.300E-06 | 3.974E-06 | 3.100E-06 | 1.100E-05 |
| 11375 | 569 | K569P | A | 7.2e-06 | 5.8e-06 | 9.5e-06 | 7.500E-06 | 1.868E-06 | 5.800E-06 | 9.500E-06 |
| 11376 | 569 | K569S | A | 7.4e-06 | 5.1e-06 | 1e-05 | 7.500E-06 | 2.452E-06 | 5.100E-06 | 1.000E-05 |
| 11377 | 569 | K569T | A | 7.1e-06 | 5.5e-06 | 5.5e-06 | 6.033E-06 | 9.238E-07 | 5.500E-06 | 7.100E-06 |
| 11378 | 569 | K569W | A | 8.1e-06 | 6.8e-06 | 5.5e-06 | 6.800E-06 | 1.300E-06 | 5.500E-06 | 8.100E-06 |
| 11379 | 569 | K569Y | A | 7.6e-06 | 4.4e-06 | 1.1e-05 | 7.667E-06 | 3.301E-06 | 4.400E-06 | 1.100E-05 |
| 11380 | 569 | K569V | A | 8.1e-06 | 1.1e-05 | 1e-05 | 9.700E-06 | 1.473E-06 | 8.100E-06 | 1.100E-05 |
| 11381 | 570 | D570A | A | 6.3e-06 | 9.2e-06 | 9.2e-06 | 8.233E-06 | 1.674E-06 | 6.300E-06 | 9.200E-06 |
| 11382 | 570 | D570R | A | 8.2e-06 | 9.2e-06 | 9.3e-06 | 8.900E-06 | 6.083E-07 | 8.200E-06 | 9.300E-06 |
| 11383 | 570 | D570N | A | 7.3e-06 | 6.4e-06 | 1.3e-05 | 8.900E-06 | 3.579E-06 | 6.400E-06 | 1.300E-05 |
| 11384 | 570 | D570D | A | 8.2e-06 | 4.8e-06 | 8.6e-06 | 7.200E-06 | 2.088E-06 | 4.800E-06 | 8.600E-06 |
| 11385 | 570 | D570C | A | 4.5e-06 | 4.5e-06 | 6.2e-06 | 5.067E-06 | 9.815E-07 | 4.500E-06 | 6.200E-06 |
| 11386 | 570 | D570E | A | 8.7e-06 | 8.7e-06 | 6.2e-06 | 7.867E-06 | 1.443E-06 | 6.200E-06 | 8.700E-06 |
| 11387 | 570 | D570Q | A | 1.2e-05 | 8.2e-06 | 6.6e-06 | 8.933E-06 | 2.774E-06 | 6.600E-06 | 1.200E-05 |
| 11388 | 570 | D570G | A | 5.7e-06 | 8.3e-06 | 2e-05 | 1.133E-05 | 7.617E-06 | 5.700E-06 | 2.000E-05 |
| 11389 | 570 | D570H | A | 1.1e-05 | 5e-06 | 1.1e-05 | 9.000E-06 | 3.464E-06 | 5.000E-06 | 1.100E-05 |

| | | | | | | | | | | |
|---|---|---|---|---|---|---|---|---|---|---|
| 11390 | 570 | D570I | A | 1.3e-05 | 4.5e-06 | 1.2e-05 | 9.833E-06 | 4.646E-06 | 4.500E-06 | 1.300E-05 |
| 11391 | 570 | D570L | A | 9.3e-06 | 8.8e-06 | 9.4e-06 | 9.167E-06 | 3.215E-07 | 8.800E-06 | 9.400E-06 |
| 11392 | 570 | D570K | A | 9e-06 | 9.1e-06 | 7e-06 | 8.367E-06 | 1.185E-06 | 7.000E-06 | 9.100E-06 |
| 11393 | 570 | D570M | A | 5.7e-06 | 6.4e-06 | 7.8e-06 | 6.633E-06 | 1.069E-06 | 5.700E-06 | 7.800E-06 |
| 11394 | 570 | D570F | A | 7.8e-06 | 4.5e-06 | 8.9e-06 | 7.067E-06 | 2.290E-06 | 4.500E-06 | 8.900E-06 |
| 11395 | 570 | D570P | A | 7.2e-06 | 6.7e-06 | 4.5e-06 | 6.133E-06 | 1.436E-06 | 4.500E-06 | 7.200E-06 |
| 11396 | 570 | D570S | A | 4.7e-06 | 7.6e-06 | 5.4e-06 | 5.900E-06 | 1.513E-06 | 4.700E-06 | 7.600E-06 |
| 11397 | 570 | D570T | A | 4.8e-06 | 6.9e-06 | 5.9e-06 | 5.867E-06 | 1.050E-06 | 4.800E-06 | 6.900E-06 |
| 11398 | 570 | D570W | A | 1.3e-05 | 3.7e-06 | 5.9e-06 | 7.533E-06 | 4.860E-06 | 3.700E-06 | 1.300E-05 |
| 11399 | 570 | D570Y | A | 3.5e-06 | 6e-06 | 1e-05 | 6.500E-06 | 3.279E-06 | 3.500E-06 | 1.000E-05 |
| 11400 | 570 | D570V | A | 5.9e-06 | 6.2e-06 | 1.2e-05 | 8.033E-06 | 3.439E-06 | 5.900E-06 | 1.200E-05 |
| 11401 | 571 | P571A | A | 6.5e-06 | 1e-05 | 7.8e-06 | 8.100E-06 | 1.769E-06 | 6.500E-06 | 1.000E-05 |
| 11402 | 571 | P571R | A | 1.1e-05 | 4.4e-06 | 4.3e-06 | 6.567E-06 | 3.840E-06 | 4.300E-06 | 1.100E-05 |
| 11403 | 571 | P571N | A | 1.3e-05 | 1.1e-05 | 9.8e-06 | 1.127E-05 | 1.617E-06 | 9.800E-06 | 1.300E-05 |
| 11404 | 571 | P571D | A | 5.6e-06 | 6.8e-06 | 8.4e-06 | 6.933E-06 | 1.405E-06 | 5.600E-06 | 8.400E-06 |
| 11405 | 571 | P571C | A | 8.7e-06 | 4.1e-06 | 6.1e-06 | 6.300E-06 | 2.307E-06 | 4.100E-06 | 8.700E-06 |
| 11406 | 571 | P571E | A | 8.2e-06 | 7.8e-06 | 1e-05 | 8.667E-06 | 1.172E-06 | 7.800E-06 | 1.000E-05 |
| 11407 | 571 | P571Q | A | 1.3e-05 | 1e-05 | 1.7e-05 | 1.333E-05 | 3.512E-06 | 1.000E-05 | 1.700E-05 |
| 11408 | 571 | P571G | A | 1.3e-05 | 1e-05 | 1.3e-05 | 1.200E-05 | 1.732E-06 | 1.000E-05 | 1.300E-05 |
| 11409 | 571 | P571H | A | 6.1e-06 | 6.8e-06 | 9.1e-06 | 7.333E-06 | 1.570E-06 | 6.100E-06 | 9.100E-06 |
| 11410 | 571 | P571I | A | 5.1e-06 | 8.1e-06 | 3.6e-06 | 5.600E-06 | 2.291E-06 | 3.600E-06 | 8.100E-06 |
| 11411 | 571 | P571L | A | 3.4e-06 | 3.5e-06 | 3.5e-06 | 3.467E-06 | 5.774E-08 | 3.400E-06 | 3.500E-06 |
| 11412 | 571 | P571K | A | 5.8e-06 | 6.8e-06 | 1e-05 | 7.533E-06 | 2.194E-06 | 5.800E-06 | 1.000E-05 |
| 11413 | 571 | P571M | A | 3.7e-06 | 3.1e-06 | 4.1e-06 | 3.633E-06 | 5.033E-07 | 3.100E-06 | 4.100E-06 |
| 11414 | 571 | P571F | A | 3.8e-06 | 6.9e-06 | 3e-06 | 4.567E-06 | 2.060E-06 | 3.000E-06 | 6.900E-06 |
| 11415 | 571 | P571P | A | 7e-06 | 1.2e-05 | 9.5e-06 | 9.500E-06 | 2.500E-06 | 7.000E-06 | 1.200E-05 |
| 11416 | 571 | P571S | A | 7.1e-06 | 5.7e-06 | 6.8e-06 | 6.533E-06 | 7.371E-07 | 5.700E-06 | 7.100E-06 |
| 11417 | 571 | P571T | A | 7.6e-06 | 5.1e-06 | 6.4e-06 | 6.367E-06 | 1.250E-06 | 5.100E-06 | 7.600E-06 |
| 11418 | 571 | P571W | A | 2.9e-06 | 6.6e-06 | 2.7e-06 | 4.067E-06 | 2.196E-06 | 2.700E-06 | 6.600E-06 |
| 11419 | 571 | P571Y | A | 4e-06 | 3.7e-06 | 4.5e-06 | 4.067E-06 | 4.041E-07 | 3.700E-06 | 4.500E-06 |
| 11420 | 571 | P571V | A | 4.6e-06 | 3e-06 | 4.5e-06 | 4.033E-06 | 8.963E-07 | 3.000E-06 | 4.600E-06 |
| 11421 | 572 | P572A | A | 1.3e-05 | 3.4e-06 | 6.1e-06 | 7.500E-06 | 4.951E-06 | 3.400E-06 | 1.300E-05 |
| 11422 | 572 | P572R | A | 1.2e-05 | 2.6e-05 | 1.8e-05 | 1.867E-05 | 7.024E-06 | 1.200E-05 | 2.600E-05 |
| 11423 | 572 | P572N | A | 2.5e-05 | 2.9e-05 | 1.6e-05 | 2.333E-05 | 6.658E-06 | 1.600E-05 | 2.900E-05 |
| 11424 | 572 | P572D | A | 1.1e-05 | 1.7e-05 | 1.3e-05 | 1.367E-05 | 3.055E-06 | 1.100E-05 | 1.700E-05 |
| 11425 | 572 | P572C | A | 6.9e-06 | 7.6e-06 | 8.5e-06 | 7.667E-06 | 8.021E-07 | 6.900E-06 | 8.500E-06 |
| 11426 | 572 | P572E | A | 2.7e-05 | 1.1e-05 | 6.8e-06 | 1.493E-05 | 1.066E-05 | 6.800E-06 | 2.700E-05 |
| 11427 | 572 | P572Q | A | 2.9e-05 | 5.7e-06 | 5.5e-05 | 2.990E-05 | 2.466E-05 | 5.700E-06 | 5.500E-05 |
| 11428 | 572 | P572G | A | 7.8e-06 | 1.1e-05 | 7.7e-06 | 8.833E-06 | 1.877E-06 | 7.700E-06 | 1.100E-05 |
| 11429 | 572 | P572H | A | 1.7e-05 | 1.6e-05 | 9.2e-06 | 1.407E-05 | 4.244E-06 | 9.200E-06 | 1.700E-05 |
| 11430 | 572 | P572I | A | 5.8e-06 | 1.1e-05 | 7.3e-06 | 8.033E-06 | 2.676E-06 | 5.800E-06 | 1.100E-05 |
| 11431 | 572 | P572L | A | 3e-06 | 6.4e-06 | 4.9e-06 | 4.767E-06 | 1.704E-06 | 3.000E-06 | 6.400E-06 |
| 11432 | 572 | P572K | A | 9.1e-06 | 1.6e-05 | 1.1e-05 | 1.203E-05 | 3.564E-06 | 9.100E-06 | 1.600E-05 |
| 11433 | 572 | P572M | A | 1.6e-05 | 8e-06 | 4.4e-06 | 9.467E-06 | 5.937E-06 | 4.400E-06 | 1.600E-05 |
| 11434 | 572 | P572F | A | 7.1e-06 | 5.2e-06 | 7.2e-06 | 6.500E-06 | 1.127E-06 | 5.200E-06 | 7.200E-06 |
| 11435 | 572 | P572P | A | 9.7e-06 | 6.5e-06 | 9.7e-06 | 8.633E-06 | 1.848E-06 | 6.500E-06 | 9.700E-06 |
| 11436 | 572 | P572S | A | 3.5e-05 | 2.6e-05 | 1.7e-05 | 2.600E-05 | 9.000E-06 | 1.700E-05 | 3.500E-05 |
| 11437 | 572 | P572T | A | 2.2e-05 | 2.6e-05 | 1.9e-05 | 2.233E-05 | 3.512E-06 | 1.900E-05 | 2.600E-05 |
| 11438 | 572 | P572W | A | 6.9e-06 | 8.7e-06 | 6.2e-06 | 7.267E-06 | 1.290E-06 | 6.200E-06 | 8.700E-06 |
| 11439 | 572 | P572Y | A | 7.2e-06 | 9e-06 | 4.8e-06 | 7.000E-06 | 2.107E-06 | 4.800E-06 | 9.000E-06 |
| 11440 | 572 | P572V | A | 5.9e-06 | 9.3e-06 | 8.5e-06 | 7.900E-06 | 1.778E-06 | 5.900E-06 | 9.300E-06 |
| 11441 | 573 | F573A | A | 1.2e-05 | 9.4e-06 | 7.1e-06 | 9.500E-06 | 2.452E-06 | 7.100E-06 | 1.200E-05 |
| 11442 | 573 | F573R | A | 5.6e-06 | 5.8e-06 | 6.6e-06 | 6.000E-06 | 5.292E-07 | 5.600E-06 | 6.600E-06 |
| 11443 | 573 | F573N | A | 1.4e-05 | 9.8e-06 | 6.8e-06 | 1.020E-05 | 3.617E-06 | 6.800E-06 | 1.400E-05 |
| 11444 | 573 | F573D | A | 9.3e-06 | 6.9e-06 | 5.7e-06 | 7.300E-06 | 1.833E-06 | 5.700E-06 | 9.300E-06 |
| 11445 | 573 | F573C | A | 4.9e-06 | 7.4e-06 | 1.3e-05 | 8.433E-06 | 4.148E-06 | 4.900E-06 | 1.300E-05 |
| 11446 | 573 | F573E | A | 5.6e-06 | 6.4e-06 | 9.2e-06 | 7.067E-06 | 1.890E-06 | 5.600E-06 | 9.200E-06 |
| 11447 | 573 | F573Q | A | 5.7e-06 | 6.2e-06 | 6.7e-06 | 6.200E-06 | 5.000E-07 | 5.700E-06 | 6.700E-06 |
| 11448 | 573 | F573G | A | 6.5e-06 | 9.3e-06 | 5.2e-06 | 7.000E-06 | 2.095E-06 | 5.200E-06 | 9.300E-06 |
| 11449 | 573 | F573H | A | 9.3e-06 | 7.8e-06 | 1.1e-05 | 9.367E-06 | 1.601E-06 | 7.800E-06 | 1.100E-05 |
| 11450 | 573 | F573I | A | 7.6e-06 | 9.7e-06 | 7.7e-06 | 8.333E-06 | 1.185E-06 | 7.600E-06 | 9.700E-06 |
| 11451 | 573 | F573L | A | 8.9e-06 | 6.8e-06 | 9.4e-06 | 8.367E-06 | 1.380E-06 | 6.800E-06 | 9.400E-06 |
| 11452 | 573 | F573K | A | 7.7e-06 | 1.5e-05 | 1.2e-05 | 1.157E-05 | 3.669E-06 | 7.700E-06 | 1.500E-05 |
| 11453 | 573 | F573M | A | 1e-05 | 5.9e-06 | 6e-06 | 7.300E-06 | 2.339E-06 | 5.900E-06 | 1.000E-05 |
| 11454 | 573 | F573F | A | 1e-05 | 6.7e-06 | 8e-06 | 8.233E-06 | 1.662E-06 | 6.700E-06 | 1.000E-05 |
| 11455 | 573 | F573P | A | 2e-05 | 1.3e-05 | 1e-05 | 1.433E-05 | 5.132E-06 | 1.000E-05 | 2.000E-05 |
| 11456 | 573 | F573S | A | 1.4e-05 | 5.7e-06 | 1e-05 | 9.900E-06 | 4.151E-06 | 5.700E-06 | 1.400E-05 |
| 11457 | 573 | F573T | A | 4.8e-06 | 4.6e-06 | 6.5e-06 | 5.300E-06 | 1.044E-06 | 4.600E-06 | 6.500E-06 |
| 11458 | 573 | F573W | A | 1.1e-05 | 6.2e-06 | 7e-06 | 8.067E-06 | 2.572E-06 | 6.200E-06 | 1.100E-05 |
| 11459 | 573 | F573Y | A | 9.1e-06 | 8.8e-06 | 8.5e-06 | 8.800E-06 | 3.000E-07 | 8.500E-06 | 9.100E-06 |
| 11460 | 573 | F573V | A | 1e-05 | 8.9e-06 | 4.3e-06 | 7.733E-06 | 3.024E-06 | 4.300E-06 | 1.000E-05 |
| 11461 | 574 | C574A | A | 7.1e-06 | 7.4e-06 | 9.8e-06 | 8.100E-06 | 1.480E-06 | 7.100E-06 | 9.800E-06 |
| 11462 | 574 | C574R | A | 5.5e-06 | 9.5e-06 | 1.4e-05 | 9.667E-06 | 4.252E-06 | 5.500E-06 | 1.400E-05 |
| 11463 | 574 | C574N | A | 4.7e-06 | 6e-06 | 9.2e-06 | 6.633E-06 | 2.316E-06 | 4.700E-06 | 9.200E-06 |
| 11464 | 574 | C574D | A | 9.9e-06 | 5.1e-06 | 1.1e-05 | 8.667E-06 | 3.137E-06 | 5.100E-06 | 1.100E-05 |
| 11465 | 574 | C574C | A | 7e-06 | 8.6e-06 | 6.7e-06 | 7.433E-06 | 1.021E-06 | 6.700E-06 | 8.600E-06 |
| 11466 | 574 | C574E | A | 5.3e-06 | 9.8e-06 | 9.5e-06 | 8.200E-06 | 2.516E-06 | 5.300E-06 | 9.800E-06 |
| 11467 | 574 | C574Q | A | 9.5e-06 | 6.8e-06 | 8.6e-06 | 8.300E-06 | 1.375E-06 | 6.800E-06 | 9.500E-06 |
| 11468 | 574 | C574G | A | 3.8e-06 | 5.5e-06 | 6.5e-06 | 5.267E-06 | 1.365E-06 | 3.800E-06 | 6.500E-06 |
| 11469 | 574 | C574H | A | 8.8e-06 | 6.7e-06 | 1.2e-05 | 9.167E-06 | 2.669E-06 | 6.700E-06 | 1.200E-05 |
| 11470 | 574 | C574I | A | 7e-06 | 5.8e-06 | 4.4e-06 | 5.733E-06 | 1.301E-06 | 4.400E-06 | 7.000E-06 |
| 11471 | 574 | C574L | A | 6.8e-06 | 6.4e-06 | 1.1e-05 | 8.067E-06 | 2.548E-06 | 6.400E-06 | 1.100E-05 |
| 11472 | 574 | C574K | A | 1.1e-05 | 6.2e-06 | 8.8e-06 | 8.667E-06 | 2.403E-06 | 6.200E-06 | 1.100E-05 |
| 11473 | 574 | C574M | A | 1.1e-05 | 6.6e-06 | 6.9e-06 | 8.167E-06 | 2.458E-06 | 6.600E-06 | 1.100E-05 |
| 11474 | 574 | C574F | A | 4.3e-06 | 1.1e-05 | 1.3e-05 | 9.433E-06 | 4.557E-06 | 4.300E-06 | 1.300E-05 |

| | | | | | | | | | | |
|---|---|---|---|---|---|---|---|---|---|---|
| 11475 | 574 | C574P | A | 8.7e-06 | 8e-06 | 8.8e-06 | 8.500E-06 | 4.359E-07 | 8.000E-06 | 8.800E-06 |
| 11476 | 574 | C574S | A | 9.7e-06 | 7.1e-06 | 3.5e-06 | 6.767E-06 | 3.113E-06 | 3.500E-06 | 9.700E-06 |
| 11477 | 574 | C574T | A | 8.8e-06 | 5.1e-06 | 1.2e-05 | 8.633E-06 | 3.453E-06 | 5.100E-06 | 1.200E-05 |
| 11478 | 574 | C574W | A | 4.4e-06 | 7.9e-06 | 6.7e-06 | 6.333E-06 | 1.779E-06 | 4.400E-06 | 7.900E-06 |
| 11479 | 574 | C574Y | A | 7.8e-06 | 6.6e-06 | 8.9e-06 | 7.767E-06 | 1.150E-06 | 6.600E-06 | 8.900E-06 |
| 11480 | 574 | C574V | A | 9.1e-06 | 1.1e-05 | 8.4e-06 | 9.500E-06 | 1.345E-06 | 8.400E-06 | 1.100E-05 |
| 11481 | 575 | V575A | A | 9.9e-06 | 6.9e-06 | 7.4e-06 | 8.067E-06 | 1.607E-06 | 6.900E-06 | 9.900E-06 |
| 11482 | 575 | V575R | A | 5.7e-06 | 6.7e-06 | 7.9e-06 | 6.767E-06 | 1.102E-06 | 5.700E-06 | 7.900E-06 |
| 11483 | 575 | V575N | A | 6.1e-06 | 4.7e-06 | 4.8e-06 | 5.200E-06 | 7.810E-07 | 4.700E-06 | 6.100E-06 |
| 11484 | 575 | V575D | A | 9.3e-06 | 9.8e-06 | 6.3e-06 | 8.467E-06 | 1.893E-06 | 6.300E-06 | 9.800E-06 |
| 11485 | 575 | V575C | A | 4.9e-06 | 6.8e-06 | 4.8e-06 | 5.500E-06 | 1.127E-06 | 4.800E-06 | 6.800E-06 |
| 11486 | 575 | V575E | A | 4.5e-06 | 8.2e-06 | 4e-06 | 5.567E-06 | 2.294E-06 | 4.000E-06 | 8.200E-06 |
| 11487 | 575 | V575Q | A | 1.2e-05 | 6.5e-06 | 6.3e-06 | 8.267E-06 | 3.235E-06 | 6.300E-06 | 1.200E-05 |
| 11488 | 575 | V575G | A | 8.4e-06 | 6.4e-06 | 6.1e-06 | 6.967E-06 | 1.250E-06 | 6.100E-06 | 8.400E-06 |
| 11489 | 575 | V575H | A | 5.6e-06 | 3.9e-06 | 6.8e-06 | 5.433E-06 | 1.457E-06 | 3.900E-06 | 6.800E-06 |
| 11490 | 575 | V575I | A | 6.8e-06 | 9.9e-06 | 7.4e-06 | 8.033E-06 | 1.644E-06 | 6.800E-06 | 9.900E-06 |
| 11491 | 575 | V575L | A | 1.2e-05 | 1.2e-05 | 4.4e-06 | 9.467E-06 | 4.388E-06 | 4.400E-06 | 1.200E-05 |
| 11492 | 575 | V575K | A | 1.4e-05 | 6.2e-06 | 7.1e-06 | 9.100E-06 | 4.267E-06 | 6.200E-06 | 1.400E-05 |
| 11493 | 575 | V575M | A | 1e-05 | 7.7e-06 | 5.6e-06 | 7.767E-06 | 2.201E-06 | 5.600E-06 | 1.000E-05 |
| 11494 | 575 | V575F | A | 8.3e-06 | 7.8e-06 | 6.5e-06 | 7.533E-06 | 9.292E-07 | 6.500E-06 | 8.300E-06 |
| 11495 | 575 | V575P | A | 4.9e-06 | 1.5e-05 | 9.9e-06 | 9.933E-06 | 5.050E-06 | 4.900E-06 | 1.500E-05 |
| 11496 | 575 | V575S | A | 8.4e-06 | 8.3e-06 | 9.3e-06 | 8.667E-06 | 5.508E-07 | 8.300E-06 | 9.300E-06 |
| 11497 | 575 | V575T | A | 4.7e-06 | 9.4e-06 | 5.7e-06 | 6.600E-06 | 2.476E-06 | 4.700E-06 | 9.400E-06 |
| 11498 | 575 | V575W | A | 4.9e-06 | 6.2e-06 | 5.7e-06 | 5.600E-06 | 6.557E-07 | 4.900E-06 | 6.200E-06 |
| 11499 | 575 | V575Y | A | 7.3e-06 | 9e-06 | 5.7e-06 | 7.333E-06 | 1.650E-06 | 5.700E-06 | 9.000E-06 |
| 11500 | 575 | V575V | A | 4.6e-06 | 1.2e-05 | 6.2e-06 | 7.600E-06 | 3.894E-06 | 4.600E-06 | 1.200E-05 |
| 11501 | 576 | A576A | A | 6.2e-06 | 5.5e-06 | 6.7e-06 | 6.133E-06 | 6.028E-07 | 5.500E-06 | 6.700E-06 |
| 11502 | 576 | A576R | A | 6.6e-06 | 9.1e-06 | 9.3e-06 | 8.333E-06 | 1.504E-06 | 6.600E-06 | 9.300E-06 |
| 11503 | 576 | A576N | A | 1e-05 | 1.1e-05 | 8.6e-06 | 9.867E-06 | 1.206E-06 | 8.600E-06 | 1.100E-05 |
| 11504 | 576 | A576D | A | 8.7e-06 | 9e-06 | 1.1e-05 | 9.567E-06 | 1.250E-06 | 8.700E-06 | 1.100E-05 |
| 11505 | 576 | A576C | A | 8.1e-06 | 1.4e-05 | 5.1e-06 | 9.067E-06 | 4.528E-06 | 5.100E-06 | 1.400E-05 |
| 11506 | 576 | A576E | A | 7.7e-06 | 7e-06 | 5e-06 | 6.567E-06 | 1.401E-06 | 5.000E-06 | 7.700E-06 |
| 11507 | 576 | A576Q | A | 9.6e-06 | 6.7e-06 | 7.7e-06 | 8.000E-06 | 1.473E-06 | 6.700E-06 | 9.600E-06 |
| 11508 | 576 | A576G | A | 9.2e-06 | 6.5e-06 | 5.5e-06 | 7.067E-06 | 1.914E-06 | 5.500E-06 | 9.200E-06 |
| 11509 | 576 | A576H | A | 8.9e-06 | 9.4e-06 | 6.4e-06 | 8.233E-06 | 1.607E-06 | 6.400E-06 | 9.400E-06 |
| 11510 | 576 | A576I | A | 4.4e-06 | 4.6e-06 | 7.6e-06 | 5.533E-06 | 1.793E-06 | 4.400E-06 | 7.600E-06 |
| 11511 | 576 | A576L | A | 5.7e-06 | 6e-06 | 9.8e-06 | 7.167E-06 | 2.285E-06 | 5.700E-06 | 9.800E-06 |
| 11512 | 576 | A576K | A | 8.7e-06 | 6.3e-06 | 5.6e-06 | 6.867E-06 | 1.626E-06 | 5.600E-06 | 8.700E-06 |
| 11513 | 576 | A576M | A | 3.9e-06 | 4.9e-06 | 9.2e-06 | 6.000E-06 | 2.816E-06 | 3.900E-06 | 9.200E-06 |
| 11514 | 576 | A576F | A | 6.5e-06 | 8.9e-06 | 1e-05 | 8.467E-06 | 1.790E-06 | 6.500E-06 | 1.000E-05 |
| 11515 | 576 | A576P | A | 8.5e-06 | 6.3e-06 | 6.9e-06 | 7.233E-06 | 1.137E-06 | 6.300E-06 | 8.500E-06 |
| 11516 | 576 | A576S | A | 1.3e-05 | 1e-05 | 4.6e-06 | 9.200E-06 | 4.257E-06 | 4.600E-06 | 1.300E-05 |
| 11517 | 576 | A576T | A | 1.1e-05 | 6.7e-06 | 8.7e-06 | 8.800E-06 | 2.152E-06 | 6.700E-06 | 1.100E-05 |
| 11518 | 576 | A576W | A | 6.5e-06 | 8e-06 | 4.5e-06 | 6.333E-06 | 1.756E-06 | 4.500E-06 | 8.000E-06 |
| 11519 | 576 | A576Y | A | 8e-06 | 6.9e-06 | 1.2e-05 | 8.967E-06 | 2.684E-06 | 6.900E-06 | 1.200E-05 |
| 11520 | 576 | A576V | A | 6.4e-06 | 9.6e-06 | 1.6e-05 | 1.067E-05 | 4.888E-06 | 6.400E-06 | 1.600E-05 |
| 11521 | 577 | R577A | A | 6.8e-06 | 7.2e-06 | 1.2e-05 | 8.667E-06 | 2.894E-06 | 6.800E-06 | 1.200E-05 |
| 11522 | 577 | R577R | A | 1.5e-05 | 4.6e-06 | 4.5e-06 | 8.033E-06 | 6.034E-06 | 4.500E-06 | 1.500E-05 |
| 11523 | 577 | R577N | A | 9.2e-06 | 4.4e-06 | 6.2e-06 | 6.600E-06 | 2.425E-06 | 4.400E-06 | 9.200E-06 |
| 11524 | 577 | R577D | A | 6.5e-06 | 9.9e-06 | 1e-05 | 8.800E-06 | 1.992E-06 | 6.500E-06 | 1.000E-05 |
| 11525 | 577 | R577C | A | 8.8e-06 | 9.7e-06 | 1e-05 | 9.500E-06 | 6.245E-07 | 8.800E-06 | 1.000E-05 |
| 11526 | 577 | R577E | A | 8.4e-06 | 9.2e-06 | 6.2e-06 | 7.933E-06 | 1.553E-06 | 6.200E-06 | 9.200E-06 |
| 11527 | 577 | R577Q | A | 4.5e-06 | 7.4e-06 | 7.9e-06 | 6.600E-06 | 1.836E-06 | 4.500E-06 | 7.900E-06 |
| 11528 | 577 | R577G | A | 8.5e-06 | 5.5e-06 | 5.1e-06 | 6.367E-06 | 1.858E-06 | 5.100E-06 | 8.500E-06 |
| 11529 | 577 | R577H | A | 5.9e-06 | 1e-05 | 1.4e-05 | 9.967E-06 | 4.050E-06 | 5.900E-06 | 1.400E-05 |
| 11530 | 577 | R577I | A | 7.5e-06 | 5.3e-06 | 1.3e-05 | 8.600E-06 | 3.966E-06 | 5.300E-06 | 1.300E-05 |
| 11531 | 577 | R577L | A | 8.6e-06 | 8.2e-06 | 8.6e-06 | 8.467E-06 | 2.309E-07 | 8.200E-06 | 8.600E-06 |
| 11532 | 577 | R577K | A | 1e-05 | 8.8e-06 | 5.3e-06 | 8.033E-06 | 2.442E-06 | 5.300E-06 | 1.000E-05 |
| 11533 | 577 | R577M | A | 1.2e-05 | 4.4e-06 | 6.7e-06 | 7.700E-06 | 3.897E-06 | 4.400E-06 | 1.200E-05 |
| 11534 | 577 | R577F | A | 8.6e-06 | 5.5e-06 | 1.1e-05 | 8.367E-06 | 2.757E-06 | 5.500E-06 | 1.100E-05 |
| 11535 | 577 | R577P | A | 1.1e-05 | 6.7e-06 | 4.2e-06 | 7.300E-06 | 3.439E-06 | 4.200E-06 | 1.100E-05 |
| 11536 | 577 | R577S | A | 7.1e-06 | 1.3e-05 | 4.6e-06 | 8.233E-06 | 4.313E-06 | 4.600E-06 | 1.300E-05 |
| 11537 | 577 | R577T | A | 9.2e-06 | 9.7e-06 | 5.7e-06 | 8.200E-06 | 2.179E-06 | 5.700E-06 | 9.700E-06 |
| 11538 | 577 | R577W | A | 7.7e-06 | 7.6e-06 | 6e-06 | 7.100E-06 | 9.539E-07 | 6.000E-06 | 7.700E-06 |
| 11539 | 577 | R577Y | A | 6.3e-06 | 8.5e-06 | 7.4e-06 | 7.400E-06 | 1.100E-06 | 6.300E-06 | 8.500E-06 |
| 11540 | 577 | R577V | A | 7.1e-06 | 8.6e-06 | 5.9e-06 | 7.200E-06 | 1.353E-06 | 5.900E-06 | 8.600E-06 |
| 11541 | 578 | C578A | A | 1.4e-05 | 1.1e-05 | 1.2e-05 | 1.233E-05 | 1.528E-06 | 1.100E-05 | 1.400E-05 |
| 11542 | 578 | C578R | A | 7.1e-06 | 9.7e-06 | 1e-05 | 8.933E-06 | 1.595E-06 | 7.100E-06 | 1.000E-05 |
| 11543 | 578 | C578N | A | 6.1e-06 | 4.9e-06 | 6.3e-06 | 5.767E-06 | 7.572E-07 | 4.900E-06 | 6.300E-06 |
| 11544 | 578 | C578D | A | 6.5e-06 | 5.7e-06 | 7.7e-06 | 6.633E-06 | 1.007E-06 | 5.700E-06 | 7.700E-06 |
| 11545 | 578 | C578C | A | 1.2e-05 | 8.2e-06 | 6.5e-06 | 8.900E-06 | 2.816E-06 | 6.500E-06 | 1.200E-05 |
| 11546 | 578 | C578E | A | 4.5e-06 | 5.6e-06 | 6.9e-06 | 5.667E-06 | 1.201E-06 | 4.500E-06 | 6.900E-06 |
| 11547 | 578 | C578Q | A | 9.7e-06 | 7.2e-06 | 1.2e-05 | 9.633E-06 | 2.401E-06 | 7.200E-06 | 1.200E-05 |
| 11548 | 578 | C578G | A | 1.2e-05 | 1e-05 | 4.3e-06 | 8.767E-06 | 3.995E-06 | 4.300E-06 | 1.200E-05 |
| 11549 | 578 | C578H | A | 6.2e-06 | 1.7e-05 | 5.6e-06 | 9.600E-06 | 6.416E-06 | 5.600E-06 | 1.700E-05 |
| 11550 | 578 | C578I | A | 9.2e-06 | 8.9e-06 | 1e-05 | 9.367E-06 | 5.686E-07 | 8.900E-06 | 1.000E-05 |
| 11551 | 578 | C578L | A | 8.3e-06 | 9.3e-06 | 7e-06 | 8.200E-06 | 1.153E-06 | 7.000E-06 | 9.300E-06 |
| 11552 | 578 | C578K | A | 1.3e-05 | 6.4e-06 | 5.8e-06 | 8.400E-06 | 3.995E-06 | 5.800E-06 | 1.300E-05 |
| 11553 | 578 | C578M | A | 7.3e-06 | 5.7e-06 | 1e-05 | 7.667E-06 | 2.173E-06 | 5.700E-06 | 1.000E-05 |
| 11554 | 578 | C578F | A | 1.2e-05 | 4.7e-06 | 8.7e-06 | 8.467E-06 | 3.656E-06 | 4.700E-06 | 1.200E-05 |
| 11555 | 578 | C578P | A | 9.7e-06 | 6.3e-06 | 1.8e-05 | 1.133E-05 | 6.019E-06 | 6.300E-06 | 1.800E-05 |
| 11556 | 578 | C578S | A | 6.1e-06 | 1.1e-05 | 5.5e-06 | 7.533E-06 | 3.017E-06 | 5.500E-06 | 1.100E-05 |
| 11557 | 578 | C578T | A | 1.2e-05 | 1.2e-05 | 6.7e-06 | 1.023E-05 | 3.060E-06 | 6.700E-06 | 1.200E-05 |
| 11558 | 578 | C578W | A | 5.5e-06 | 7.3e-06 | 5e-06 | 5.933E-06 | 1.210E-06 | 5.000E-06 | 7.300E-06 |
| 11559 | 578 | C578Y | A | 6.8e-06 | 5.3e-06 | 6e-06 | 6.033E-06 | 7.506E-07 | 5.300E-06 | 6.800E-06 |

| | | | | | | | | | | |
|---|---|---|---|---|---|---|---|---|---|---|
| 11560 | 578 | C578V | A | 1.2e-05 | 6.6e-06 | 5.9e-06 | 8.167E-06 | 3.338E-06 | 5.900E-06 | 1.200E-05 |
| 11561 | 579 | P579A | A | 7e-06 | 1.8e-05 | 1.3e-05 | 1.267E-05 | 5.508E-06 | 7.000E-06 | 1.800E-05 |
| 11562 | 579 | P579R | A | 6.9e-06 | 7.7e-06 | 9e-06 | 7.867E-06 | 1.060E-06 | 6.900E-06 | 9.000E-06 |
| 11563 | 579 | P579N | A | 1.1e-05 | 6.2e-06 | 1.4e-05 | 1.040E-05 | 3.934E-06 | 6.200E-06 | 1.400E-05 |
| 11564 | 579 | P579D | A | 8.7e-06 | 7.5e-06 | 4.2e-06 | 6.800E-06 | 2.330E-06 | 4.200E-06 | 8.700E-06 |
| 11565 | 579 | P579C | A | 5.7e-06 | 6.4e-06 | 4.8e-06 | 5.633E-06 | 8.021E-07 | 4.800E-06 | 6.400E-06 |
| 11566 | 579 | P579E | A | 7.7e-06 | 8.9e-06 | 1.2e-05 | 9.533E-06 | 2.219E-06 | 7.700E-06 | 1.200E-05 |
| 11567 | 579 | P579Q | A | 9.1e-06 | 6.8e-06 | 6e-06 | 7.300E-06 | 1.609E-06 | 6.000E-06 | 9.100E-06 |
| 11568 | 579 | P579G | A | 4.9e-06 | 9e-06 | 8.5e-06 | 7.467E-06 | 2.237E-06 | 4.900E-06 | 9.000E-06 |
| 11569 | 579 | P579H | A | 9.3e-06 | 8.8e-06 | 5.3e-06 | 7.800E-06 | 2.179E-06 | 5.300E-06 | 9.300E-06 |
| 11570 | 579 | P579I | A | 3.8e-06 | 1e-05 | 1e-05 | 7.933E-06 | 3.580E-06 | 3.800E-06 | 1.000E-05 |
| 11571 | 579 | P579L | A | 1e-05 | 7.9e-06 | 7.3e-06 | 8.400E-06 | 1.418E-06 | 7.300E-06 | 1.000E-05 |
| 11572 | 579 | P579K | A | 1.5e-05 | 9e-06 | 1.3e-05 | 1.233E-05 | 3.055E-06 | 9.000E-06 | 1.500E-05 |
| 11573 | 579 | P579M | A | 1.2e-05 | 2e-05 | 8.3e-06 | 1.343E-05 | 5.980E-06 | 8.300E-06 | 2.000E-05 |
| 11574 | 579 | P579F | A | 4.5e-06 | 9.2e-06 | 1e-05 | 7.900E-06 | 2.972E-06 | 4.500E-06 | 1.000E-05 |
| 11575 | 579 | P579P | A | 9.3e-06 | 5.4e-06 | 7.9e-06 | 7.533E-06 | 1.976E-06 | 5.400E-06 | 9.300E-06 |
| 11576 | 579 | P579S | A | 8.1e-06 | 5e-06 | 8.7e-06 | 7.267E-06 | 1.986E-06 | 5.000E-06 | 8.700E-06 |
| 11577 | 579 | P579T | A | 4.4e-06 | 7.8e-06 | 6.2e-06 | 6.133E-06 | 1.701E-06 | 4.400E-06 | 7.800E-06 |
| 11578 | 579 | P579W | A | 9e-06 | 8e-06 | 1e-05 | 9.000E-06 | 1.000E-06 | 8.000E-06 | 1.000E-05 |
| 11579 | 579 | P579Y | A | 6.8e-06 | 5.3e-06 | 4.4e-06 | 5.500E-06 | 1.212E-06 | 4.400E-06 | 6.800E-06 |
| 11580 | 579 | P579V | A | 5.8e-06 | 6.1e-06 | 6.7e-06 | 6.200E-06 | 4.583E-07 | 5.800E-06 | 6.700E-06 |
| 11581 | 580 | S580A | A | 6.5e-06 | 8.9e-06 | 6.2e-06 | 7.200E-06 | 1.480E-06 | 6.200E-06 | 8.900E-06 |
| 11582 | 580 | S580R | A | 7.5e-06 | 8.8e-06 | 8.5e-06 | 8.267E-06 | 6.807E-07 | 7.500E-06 | 8.800E-06 |
| 11583 | 580 | S580N | A | 1.8e-05 | 1e-05 | 6.8e-06 | 1.160E-05 | 5.769E-06 | 6.800E-06 | 1.800E-05 |
| 11584 | 580 | S580D | A | 7.6e-06 | 8.4e-06 | 1.1e-05 | 9.000E-06 | 1.778E-06 | 7.600E-06 | 1.100E-05 |
| 11585 | 580 | S580C | A | 6.3e-06 | 4.5e-06 | 9.9e-06 | 6.900E-06 | 2.750E-06 | 4.500E-06 | 9.900E-06 |
| 11586 | 580 | S580E | A | 9.4e-06 | 8.5e-06 | 8.8e-06 | 8.900E-06 | 4.583E-07 | 8.500E-06 | 9.400E-06 |
| 11587 | 580 | S580Q | A | 7e-06 | 5.6e-06 | 5.9e-06 | 6.167E-06 | 7.371E-07 | 5.600E-06 | 7.000E-06 |
| 11588 | 580 | S580G | A | 5.5e-06 | 8.1e-06 | 1e-05 | 7.867E-06 | 2.259E-06 | 5.500E-06 | 1.000E-05 |
| 11589 | 580 | S580H | A | 7.7e-06 | 1.3e-05 | 8.9e-06 | 9.867E-06 | 2.779E-06 | 7.700E-06 | 1.300E-05 |
| 11590 | 580 | S580I | A | 5.7e-06 | 9.8e-06 | 9e-06 | 8.167E-06 | 2.173E-06 | 5.700E-06 | 9.800E-06 |
| 11591 | 580 | S580L | A | 9.7e-06 | 7e-06 | 7.9e-06 | 8.200E-06 | 1.375E-06 | 7.000E-06 | 9.700E-06 |
| 11592 | 580 | S580K | A | 6.3e-06 | 6.5e-06 | 7e-06 | 6.600E-06 | 3.606E-07 | 6.300E-06 | 7.000E-06 |
| 11593 | 580 | S580M | A | 8e-06 | 8.7e-06 | 9.6e-06 | 8.767E-06 | 8.021E-07 | 8.000E-06 | 9.600E-06 |
| 11594 | 580 | S580F | A | 9.5e-06 | 1.3e-05 | 9.5e-06 | 1.067E-05 | 2.021E-06 | 9.500E-06 | 1.300E-05 |
| 11595 | 580 | S580P | A | 7.6e-06 | 7.4e-06 | 1.1e-05 | 8.667E-06 | 2.023E-06 | 7.400E-06 | 1.100E-05 |
| 11596 | 580 | S580S | A | 6.8e-06 | 4.3e-06 | 4.3e-06 | 5.133E-06 | 1.443E-06 | 4.300E-06 | 6.800E-06 |
| 11597 | 580 | S580T | A | 8.5e-06 | 1.1e-05 | 9.8e-06 | 9.767E-06 | 1.250E-06 | 8.500E-06 | 1.100E-05 |
| 11598 | 580 | S580W | A | 6.5e-06 | 1.9e-05 | 9.1e-06 | 1.153E-05 | 6.596E-06 | 6.500E-06 | 1.900E-05 |
| 11599 | 580 | S580Y | A | 6.5e-06 | 5.3e-06 | 5.9e-06 | 5.900E-06 | 6.000E-07 | 5.300E-06 | 6.500E-06 |
| 11600 | 580 | S580V | A | 9.8e-06 | 1e-05 | 6e-06 | 8.600E-06 | 2.254E-06 | 6.000E-06 | 1.000E-05 |
| 11601 | 581 | G581A | A | 9.9e-06 | 7.6e-06 | 1.4e-05 | 1.050E-05 | 3.242E-06 | 7.600E-06 | 1.400E-05 |
| 11602 | 581 | G581R | A | 9.4e-06 | 6.7e-06 | 7.9e-06 | 8.000E-06 | 1.353E-06 | 6.700E-06 | 9.400E-06 |
| 11603 | 581 | G581N | A | 5.5e-06 | 9.1e-06 | 4.8e-06 | 6.467E-06 | 2.307E-06 | 4.800E-06 | 9.100E-06 |
| 11604 | 581 | G581D | A | 8.1e-06 | 7.3e-06 | 9.2e-06 | 8.200E-06 | 9.539E-07 | 7.300E-06 | 9.200E-06 |
| 11605 | 581 | G581C | A | 1.2e-05 | 6.4e-06 | 6.5e-06 | 8.300E-06 | 3.205E-06 | 6.400E-06 | 1.200E-05 |
| 11606 | 581 | G581E | A | 6.4e-06 | 7e-06 | 7.6e-06 | 7.000E-06 | 6.000E-07 | 6.400E-06 | 7.600E-06 |
| 11607 | 581 | G581Q | A | 7.6e-06 | 1e-05 | 1e-05 | 9.200E-06 | 1.386E-06 | 7.600E-06 | 1.000E-05 |
| 11608 | 581 | G581G | A | 7e-06 | 6.6e-06 | 8.5e-06 | 7.367E-06 | 1.002E-06 | 6.600E-06 | 8.500E-06 |
| 11609 | 581 | G581H | A | 6.2e-06 | 9.2e-06 | 6.2e-06 | 7.200E-06 | 1.732E-06 | 6.200E-06 | 9.200E-06 |
| 11610 | 581 | G581I | A | 9.9e-06 | 9.2e-06 | 6.4e-06 | 8.500E-06 | 1.852E-06 | 6.400E-06 | 9.900E-06 |
| 11611 | 581 | G581L | A | 6.6e-06 | 9.7e-06 | 7.7e-06 | 8.000E-06 | 1.572E-06 | 6.600E-06 | 9.700E-06 |
| 11612 | 581 | G581K | A | 6.6e-06 | 6.4e-06 | 1.4e-05 | 9.000E-06 | 4.331E-06 | 6.400E-06 | 1.400E-05 |
| 11613 | 581 | G581M | A | 7.4e-06 | 9.9e-06 | 8.5e-06 | 8.600E-06 | 1.253E-06 | 7.400E-06 | 9.900E-06 |
| 11614 | 581 | G581F | A | 6.7e-06 | 1.3e-05 | 6.5e-06 | 8.733E-06 | 3.696E-06 | 6.500E-06 | 1.300E-05 |
| 11615 | 581 | G581P | A | 9.4e-06 | 9.2e-06 | 8.5e-06 | 9.033E-06 | 4.726E-07 | 8.500E-06 | 9.400E-06 |
| 11616 | 581 | G581S | A | 5.3e-06 | 1.3e-05 | 7.9e-06 | 8.733E-06 | 3.917E-06 | 5.300E-06 | 1.300E-05 |
| 11617 | 581 | G581T | A | 5.4e-06 | 7.3e-06 | 5e-06 | 5.900E-06 | 1.229E-06 | 5.000E-06 | 7.300E-06 |
| 11618 | 581 | G581W | A | 1e-05 | 4.9e-06 | 5.3e-06 | 6.733E-06 | 2.836E-06 | 4.900E-06 | 1.000E-05 |
| 11619 | 581 | G581Y | A | 5.8e-06 | 1.4e-05 | 6.8e-06 | 8.867E-06 | 4.474E-06 | 5.800E-06 | 1.400E-05 |
| 11620 | 581 | G581V | A | 8e-06 | 7.3e-06 | 1.2e-05 | 9.100E-06 | 2.536E-06 | 7.300E-06 | 1.200E-05 |
| 11621 | 582 | V582A | A | 1.3e-05 | 8.7e-06 | 6.1e-06 | 9.267E-06 | 3.485E-06 | 6.100E-06 | 1.300E-05 |
| 11622 | 582 | V582R | A | 9.2e-06 | 8.1e-06 | 5.7e-06 | 7.667E-06 | 1.790E-06 | 5.700E-06 | 9.200E-06 |
| 11623 | 582 | V582N | A | 6.2e-06 | 1.1e-05 | 7e-06 | 8.067E-06 | 2.572E-06 | 6.200E-06 | 1.100E-05 |
| 11624 | 582 | V582D | A | 8.1e-06 | 7.8e-06 | 6.6e-06 | 7.500E-06 | 7.937E-07 | 6.600E-06 | 8.100E-06 |
| 11625 | 582 | V582C | A | 5.9e-06 | 9.5e-06 | 5.5e-06 | 6.967E-06 | 2.203E-06 | 5.500E-06 | 9.500E-06 |
| 11626 | 582 | V582E | A | 9.9e-06 | 6.4e-06 | 9.6e-06 | 8.633E-06 | 1.940E-06 | 6.400E-06 | 9.900E-06 |
| 11627 | 582 | V582Q | A | 7.5e-06 | 8.2e-06 | 4.1e-06 | 6.600E-06 | 2.193E-06 | 4.100E-06 | 8.200E-06 |
| 11628 | 582 | V582G | A | 7.1e-06 | 8.1e-06 | 9e-06 | 8.067E-06 | 9.504E-07 | 7.100E-06 | 9.000E-06 |
| 11629 | 582 | V582H | A | 6.7e-06 | 9.7e-06 | 6.1e-06 | 7.500E-06 | 1.929E-06 | 6.100E-06 | 9.700E-06 |
| 11630 | 582 | V582I | A | 5.6e-06 | 4.6e-06 | 9.8e-06 | 6.667E-06 | 2.759E-06 | 4.600E-06 | 9.800E-06 |
| 11631 | 582 | V582L | A | 5.5e-06 | 6.8e-06 | 8.2e-06 | 6.833E-06 | 1.350E-06 | 5.500E-06 | 8.200E-06 |
| 11632 | 582 | V582K | A | 6.5e-06 | 5.5e-06 | 7.1e-06 | 6.367E-06 | 8.083E-07 | 5.500E-06 | 7.100E-06 |
| 11633 | 582 | V582M | A | 1.4e-05 | 9.7e-06 | 8.3e-06 | 1.067E-05 | 2.970E-06 | 8.300E-06 | 1.400E-05 |
| 11634 | 582 | V582F | A | 9.7e-06 | 9e-06 | 8.4e-06 | 9.033E-06 | 6.506E-07 | 8.400E-06 | 9.700E-06 |
| 11635 | 582 | V582P | A | 4.2e-06 | 8e-06 | 4.1e-06 | 5.433E-06 | 2.223E-06 | 4.100E-06 | 8.000E-06 |
| 11636 | 582 | V582S | A | 7.6e-06 | 6.6e-06 | 5.7e-06 | 6.633E-06 | 9.504E-07 | 5.700E-06 | 7.600E-06 |
| 11637 | 582 | V582T | A | 4.9e-06 | 5.7e-06 | 7.4e-06 | 6.000E-06 | 1.277E-06 | 4.900E-06 | 7.400E-06 |
| 11638 | 582 | V582W | A | 4.1e-06 | 8.9e-06 | 1.1e-05 | 8.000E-06 | 3.537E-06 | 4.100E-06 | 1.100E-05 |
| 11639 | 582 | V582Y | A | 7.1e-06 | 9.8e-06 | 6.9e-06 | 7.933E-06 | 1.620E-06 | 6.900E-06 | 9.800E-06 |
| 11640 | 582 | V582V | A | 6.2e-06 | 1.1e-05 | 9.8e-06 | 9.000E-06 | 2.498E-06 | 6.200E-06 | 1.100E-05 |
| 11641 | 583 | K583A | A | 6.5e-06 | 9.7e-06 | 8e-06 | 8.067E-06 | 1.601E-06 | 6.500E-06 | 9.700E-06 |
| 11642 | 583 | K583R | A | 5.8e-06 | 1.3e-05 | 5e-06 | 7.933E-06 | 4.406E-06 | 5.000E-06 | 1.300E-05 |
| 11643 | 583 | K583N | A | 1.1e-05 | 1.3e-05 | 5.5e-06 | 9.833E-06 | 3.884E-06 | 5.500E-06 | 1.300E-05 |
| 11644 | 583 | K583D | A | 1.1e-05 | 8.2e-06 | 9.5e-06 | 9.567E-06 | 1.401E-06 | 8.200E-06 | 1.100E-05 |

| | | | | | | | | | | |
|---|---|---|---|---|---|---|---|---|---|---|
| 11645 | 583 | K583C | A | 7.1e-06 | 1.3e-05 | 8.1e-06 | 9.400E-06 | 3.158E-06 | 7.100E-06 | 1.300E-05 |
| 11646 | 583 | K583E | A | 6.9e-06 | 9.9e-06 | 8e-06 | 8.267E-06 | 1.518E-06 | 6.900E-06 | 9.900E-06 |
| 11647 | 583 | K583Q | A | 1.3e-05 | 6.6e-06 | 5.8e-06 | 8.467E-06 | 3.946E-06 | 5.800E-06 | 1.300E-05 |
| 11648 | 583 | K583G | A | 6.4e-06 | 6.7e-06 | 5.7e-06 | 6.267E-06 | 5.132E-07 | 5.700E-06 | 6.700E-06 |
| 11649 | 583 | K583H | A | 1e-05 | 3.6e-06 | 8.4e-06 | 7.333E-06 | 3.331E-06 | 3.600E-06 | 1.000E-05 |
| 11650 | 583 | K583I | A | 9e-06 | 8.2e-06 | 1e-05 | 9.067E-06 | 9.018E-07 | 8.200E-06 | 1.000E-05 |
| 11651 | 583 | K583L | A | 7.6e-06 | 5.9e-06 | 1.8e-05 | 1.050E-05 | 6.551E-06 | 5.900E-06 | 1.800E-05 |
| 11652 | 583 | K583K | A | 4.4e-06 | 8e-06 | 6.6e-06 | 6.333E-06 | 1.815E-06 | 4.400E-06 | 8.000E-06 |
| 11653 | 583 | K583M | A | 9.7e-06 | 5.6e-06 | 7.1e-06 | 7.467E-06 | 2.074E-06 | 5.600E-06 | 9.700E-06 |
| 11654 | 583 | K583F | A | 1e-05 | 4.8e-06 | 1.4e-05 | 9.600E-06 | 4.613E-06 | 4.800E-06 | 1.400E-05 |
| 11655 | 583 | K583P | A | 9.8e-06 | 6.6e-06 | 4.5e-06 | 6.967E-06 | 2.669E-06 | 4.500E-06 | 9.800E-06 |
| 11656 | 583 | K583S | A | 4.5e-06 | 5.2e-06 | 6.5e-06 | 5.400E-06 | 1.015E-06 | 4.500E-06 | 6.500E-06 |
| 11657 | 583 | K583T | A | 6.1e-06 | 9.4e-06 | 8.6e-06 | 8.033E-06 | 1.721E-06 | 6.100E-06 | 9.400E-06 |
| 11658 | 583 | K583W | A | 6.7e-06 | 7.7e-06 | 9.8e-06 | 8.067E-06 | 1.582E-06 | 6.700E-06 | 9.800E-06 |
| 11659 | 583 | K583Y | A | 5.3e-06 | 7.6e-06 | 6.4e-06 | 6.433E-06 | 1.150E-06 | 5.300E-06 | 7.600E-06 |
| 11660 | 583 | K583V | A | 4.7e-06 | 8.1e-06 | 5e-06 | 5.933E-06 | 1.882E-06 | 4.700E-06 | 8.100E-06 |
| 11661 | 584 | P584A | A | 7.3e-06 | 1.3e-05 | 7.5e-06 | 9.267E-06 | 3.235E-06 | 7.300E-06 | 1.300E-05 |
| 11662 | 584 | P584R | A | 7.6e-06 | 5.2e-06 | 1.1e-05 | 7.933E-06 | 2.914E-06 | 5.200E-06 | 1.100E-05 |
| 11663 | 584 | P584N | A | 9.1e-06 | 8.1e-06 | 5e-06 | 7.400E-06 | 2.138E-06 | 5.000E-06 | 9.100E-06 |
| 11664 | 584 | P584D | A | 6.6e-06 | 1.1e-05 | 7.5e-06 | 8.367E-06 | 2.325E-06 | 6.600E-06 | 1.100E-05 |
| 11665 | 584 | P584C | A | 5.9e-06 | 7.5e-06 | 5.3e-06 | 6.233E-06 | 1.137E-06 | 5.300E-06 | 7.500E-06 |
| 11666 | 584 | P584E | A | 8.7e-06 | 9.3e-06 | 6.8e-06 | 8.267E-06 | 1.305E-06 | 6.800E-06 | 9.300E-06 |
| 11667 | 584 | P584Q | A | 6.2e-06 | 4.6e-06 | 9.6e-06 | 6.800E-06 | 2.553E-06 | 4.600E-06 | 9.600E-06 |
| 11668 | 584 | P584G | A | 5.4e-06 | 5.5e-06 | 5.4e-06 | 5.433E-06 | 5.774E-08 | 5.400E-06 | 5.500E-06 |
| 11669 | 584 | P584H | A | 7.9e-06 | 9.3e-06 | 4.3e-06 | 7.167E-06 | 2.579E-06 | 4.300E-06 | 9.300E-06 |
| 11670 | 584 | P584I | A | 1.3e-05 | 1.1e-05 | 5.3e-06 | 9.767E-06 | 3.995E-06 | 5.300E-06 | 1.300E-05 |
| 11671 | 584 | P584L | A | 6.7e-06 | 1.4e-05 | 5.1e-06 | 8.600E-06 | 4.744E-06 | 5.100E-06 | 1.400E-05 |
| 11672 | 584 | P584K | A | 6.7e-06 | 7.1e-06 | 1.3e-05 | 8.933E-06 | 3.528E-06 | 6.700E-06 | 1.300E-05 |
| 11673 | 584 | P584M | A | 1.2e-05 | 4.3e-06 | 1.3e-05 | 9.767E-06 | 4.761E-06 | 4.300E-06 | 1.300E-05 |
| 11674 | 584 | P584F | A | 3.2e-06 | 6.6e-06 | 8e-06 | 5.933E-06 | 2.468E-06 | 3.200E-06 | 8.000E-06 |
| 11675 | 584 | P584P | A | 5.6e-06 | 5.4e-06 | 5.1e-06 | 5.367E-06 | 2.517E-07 | 5.100E-06 | 5.600E-06 |
| 11676 | 584 | P584S | A | 8.3e-06 | 6.5e-06 | 5.1e-06 | 6.633E-06 | 1.604E-06 | 5.100E-06 | 8.300E-06 |
| 11677 | 584 | P584T | A | 1.3e-05 | 4.6e-06 | 1.8e-05 | 1.187E-05 | 6.772E-06 | 4.600E-06 | 1.800E-05 |
| 11678 | 584 | P584W | A | 8.9e-06 | 4.2e-06 | 6.4e-06 | 6.500E-06 | 2.352E-06 | 4.200E-06 | 8.900E-06 |
| 11679 | 584 | P584Y | A | 8.7e-06 | 6.6e-06 | 5.3e-06 | 6.867E-06 | 1.716E-06 | 5.300E-06 | 8.700E-06 |
| 11680 | 584 | P584V | A | 6.3e-06 | 1.2e-05 | 1.2e-05 | 1.010E-05 | 3.291E-06 | 6.300E-06 | 1.200E-05 |
| 11681 | 585 | D585A | A | 4.6e-06 | 6.4e-06 | 4.9e-06 | 5.300E-06 | 9.644E-07 | 4.600E-06 | 6.400E-06 |
| 11682 | 585 | D585R | A | 4.7e-06 | 1.2e-05 | 6.3e-06 | 7.667E-06 | 3.837E-06 | 4.700E-06 | 1.200E-05 |
| 11683 | 585 | D585N | A | 1.2e-05 | 1e-05 | 9.5e-06 | 1.050E-05 | 1.323E-06 | 9.500E-06 | 1.200E-05 |
| 11684 | 585 | D585D | A | 7.1e-06 | 6.3e-06 | 9.9e-06 | 7.767E-06 | 1.890E-06 | 6.300E-06 | 9.900E-06 |
| 11685 | 585 | D585C | A | 1.3e-05 | 9.1e-06 | 6.3e-06 | 9.467E-06 | 3.365E-06 | 6.300E-06 | 1.300E-05 |
| 11686 | 585 | D585E | A | 3.9e-06 | 9.7e-06 | 6.8e-06 | 6.800E-06 | 2.900E-06 | 3.900E-06 | 9.700E-06 |
| 11687 | 585 | D585Q | A | 6.1e-06 | 5.1e-06 | 6.4e-06 | 5.867E-06 | 6.807E-07 | 5.100E-06 | 6.400E-06 |
| 11688 | 585 | D585G | A | 6.5e-06 | 4.4e-06 | 6.4e-06 | 5.767E-06 | 1.185E-06 | 4.400E-06 | 6.500E-06 |
| 11689 | 585 | D585H | A | 5.8e-06 | 6.2e-06 | 9.3e-06 | 7.100E-06 | 1.916E-06 | 5.800E-06 | 9.300E-06 |
| 11690 | 585 | D585I | A | 7.5e-06 | 7.9e-06 | 6.2e-06 | 7.200E-06 | 8.888E-07 | 6.200E-06 | 7.900E-06 |
| 11691 | 585 | D585L | A | 5e-06 | 9.2e-06 | 1e-05 | 8.067E-06 | 2.686E-06 | 5.000E-06 | 1.000E-05 |
| 11692 | 585 | D585K | A | 6.7e-06 | 1.4e-05 | 1.2e-05 | 1.090E-05 | 3.772E-06 | 6.700E-06 | 1.400E-05 |
| 11693 | 585 | D585M | A | 6e-06 | 5.8e-06 | 6.7e-06 | 6.167E-06 | 4.726E-07 | 5.800E-06 | 6.700E-06 |
| 11694 | 585 | D585F | A | 7.6e-06 | 3.5e-06 | 5.1e-06 | 5.400E-06 | 2.066E-06 | 3.500E-06 | 7.600E-06 |
| 11695 | 585 | D585P | A | 7.8e-06 | 7.1e-06 | 1e-05 | 8.300E-06 | 1.513E-06 | 7.100E-06 | 1.000E-05 |
| 11696 | 585 | D585S | A | 7.2e-06 | 6.1e-06 | 8.2e-06 | 7.167E-06 | 1.050E-06 | 6.100E-06 | 8.200E-06 |
| 11697 | 585 | D585T | A | 1.2e-05 | 6.4e-06 | 8.3e-06 | 8.900E-06 | 2.848E-06 | 6.400E-06 | 1.200E-05 |
| 11698 | 585 | D585W | A | 5.4e-06 | 6.9e-06 | 8.4e-06 | 6.900E-06 | 1.500E-06 | 5.400E-06 | 8.400E-06 |
| 11699 | 585 | D585Y | A | 9.8e-06 | 1.4e-05 | 6.2e-06 | 1.000E-05 | 3.904E-06 | 6.200E-06 | 1.400E-05 |
| 11700 | 585 | D585V | A | 4.8e-06 | 1.3e-05 | 6.3e-06 | 8.033E-06 | 4.366E-06 | 4.800E-06 | 1.300E-05 |
| 11701 | 586 | L586A | A | 8.6e-06 | 1.1e-05 | 4.3e-06 | 7.967E-06 | 3.395E-06 | 4.300E-06 | 1.100E-05 |
| 11702 | 586 | L586R | A | 5.8e-06 | 7.6e-06 | 7.9e-06 | 7.100E-06 | 1.136E-06 | 5.800E-06 | 7.900E-06 |
| 11703 | 586 | L586N | A | 8.6e-06 | 8.4e-06 | 4.3e-06 | 7.100E-06 | 2.427E-06 | 4.300E-06 | 8.600E-06 |
| 11704 | 586 | L586D | A | 4.1e-06 | 5.8e-06 | 5.5e-06 | 5.133E-06 | 9.074E-07 | 4.100E-06 | 5.800E-06 |
| 11705 | 586 | L586C | A | 1.2e-05 | 6.2e-06 | 9.3e-06 | 9.167E-06 | 2.902E-06 | 6.200E-06 | 1.200E-05 |
| 11706 | 586 | L586E | A | 8.8e-06 | 1e-05 | 6.5e-06 | 8.433E-06 | 1.779E-06 | 6.500E-06 | 1.000E-05 |
| 11707 | 586 | L586Q | A | 8.3e-06 | 9.4e-06 | 8.5e-06 | 8.733E-06 | 5.859E-07 | 8.300E-06 | 9.400E-06 |
| 11708 | 586 | L586G | A | 1.3e-05 | 6.1e-06 | 8.7e-06 | 9.267E-06 | 3.485E-06 | 6.100E-06 | 1.300E-05 |
| 11709 | 586 | L586H | A | 1.1e-05 | 5.7e-06 | 8.1e-06 | 8.267E-06 | 2.654E-06 | 5.700E-06 | 1.100E-05 |
| 11710 | 586 | L586I | A | 7.5e-06 | 6.3e-06 | 6.2e-06 | 6.667E-06 | 7.234E-07 | 6.200E-06 | 7.500E-06 |
| 11711 | 586 | L586L | A | 9.2e-06 | 1.2e-05 | 1.4e-05 | 1.173E-05 | 2.411E-06 | 9.200E-06 | 1.400E-05 |
| 11712 | 586 | L586K | A | 7.8e-06 | 5.2e-06 | 7.6e-06 | 6.867E-06 | 1.447E-06 | 5.200E-06 | 7.800E-06 |
| 11713 | 586 | L586M | A | 6.4e-06 | 1e-05 | 8.3e-06 | 8.233E-06 | 1.801E-06 | 6.400E-06 | 1.000E-05 |
| 11714 | 586 | L586F | A | 9.6e-06 | 8.6e-06 | 8.7e-06 | 8.967E-06 | 5.508E-07 | 8.600E-06 | 9.600E-06 |
| 11715 | 586 | L586P | A | 9.6e-06 | 1.4e-05 | 1.2e-05 | 1.187E-05 | 2.203E-06 | 9.600E-06 | 1.400E-05 |
| 11716 | 586 | L586S | A | 9e-06 | 6.4e-06 | 5.1e-06 | 6.833E-06 | 1.986E-06 | 5.100E-06 | 9.000E-06 |
| 11717 | 586 | L586T | A | 4.3e-06 | 6.9e-06 | 8.3e-06 | 6.500E-06 | 2.030E-06 | 4.300E-06 | 8.300E-06 |
| 11718 | 586 | L586W | A | 5e-06 | 7.2e-06 | 1.3e-05 | 8.400E-06 | 4.133E-06 | 5.000E-06 | 1.300E-05 |
| 11719 | 586 | L586Y | A | 4.7e-06 | 7e-06 | 6.9e-06 | 6.200E-06 | 1.300E-06 | 4.700E-06 | 7.000E-06 |
| 11720 | 586 | L586V | A | 6.5e-06 | 9.1e-06 | 7e-06 | 7.533E-06 | 1.380E-06 | 6.500E-06 | 9.100E-06 |
| 11721 | 587 | S587A | A | 1.1e-05 | 8.5e-06 | 5.8e-06 | 8.433E-06 | 2.601E-06 | 5.800E-06 | 1.100E-05 |
| 11722 | 587 | S587R | A | 9.6e-06 | 7.9e-06 | 6.6e-06 | 8.033E-06 | 1.504E-06 | 6.600E-06 | 9.600E-06 |
| 11723 | 587 | S587N | A | 1.1e-05 | 9.7e-06 | 4.9e-06 | 8.533E-06 | 3.213E-06 | 4.900E-06 | 1.100E-05 |
| 11724 | 587 | S587D | A | 1.3e-05 | 9.9e-06 | 6.6e-06 | 9.833E-06 | 3.201E-06 | 6.600E-06 | 1.300E-05 |
| 11725 | 587 | S587C | A | 1e-05 | 7.1e-06 | 6e-06 | 7.700E-06 | 2.066E-06 | 6.000E-06 | 1.000E-05 |
| 11726 | 587 | S587E | A | 5.6e-06 | 7.3e-06 | 6.9e-06 | 6.600E-06 | 8.888E-07 | 5.600E-06 | 7.300E-06 |
| 11727 | 587 | S587Q | A | 9.3e-06 | 1.1e-05 | 6.1e-06 | 8.800E-06 | 2.488E-06 | 6.100E-06 | 1.100E-05 |
| 11728 | 587 | S587G | A | 9.9e-06 | 7.3e-06 | 1e-05 | 9.067E-06 | 1.531E-06 | 7.300E-06 | 1.000E-05 |
| 11729 | 587 | S587H | A | 6.4e-06 | 9.2e-06 | 1.3e-05 | 9.533E-06 | 3.313E-06 | 6.400E-06 | 1.300E-05 |

| | | | | | | | | | | |
|---|---|---|---|---|---|---|---|---|---|---|
| 11730 | 587 | S587I | A | 5.2e-06 | 7e-06 | 1.1e-05 | 7.733E-06 | 2.969E-06 | 5.200E-06 | 1.100E-05 |
| 11731 | 587 | S587L | A | 5.7e-06 | 6.9e-06 | 1e-05 | 7.533E-06 | 2.219E-06 | 5.700E-06 | 1.000E-05 |
| 11732 | 587 | S587K | A | 9.9e-06 | 8.5e-06 | 7.6e-06 | 8.667E-06 | 1.159E-06 | 7.600E-06 | 9.900E-06 |
| 11733 | 587 | S587M | A | 9.5e-06 | 9.7e-06 | 1e-05 | 9.733E-06 | 2.517E-07 | 9.500E-06 | 1.000E-05 |
| 11734 | 587 | S587F | A | 5.9e-06 | 6.2e-06 | 9.5e-06 | 7.200E-06 | 1.997E-06 | 5.900E-06 | 9.500E-06 |
| 11735 | 587 | S587P | A | 9.4e-06 | 1.2e-05 | 1.4e-05 | 1.180E-05 | 2.307E-06 | 9.400E-06 | 1.400E-05 |
| 11736 | 587 | S587S | A | 5.5e-06 | 8.9e-06 | 5.9e-06 | 6.767E-06 | 1.858E-06 | 5.500E-06 | 8.900E-06 |
| 11737 | 587 | S587T | A | 6.8e-06 | 1e-05 | 8.2e-06 | 8.333E-06 | 1.604E-06 | 6.800E-06 | 1.000E-05 |
| 11738 | 587 | S587W | A | 6.3e-06 | 3.4e-06 | 7.6e-06 | 5.767E-06 | 2.150E-06 | 3.400E-06 | 7.600E-06 |
| 11739 | 587 | S587Y | A | 6.5e-06 | 5.5e-06 | 5.7e-06 | 5.900E-06 | 5.292E-07 | 5.500E-06 | 6.500E-06 |
| 11740 | 587 | S587V | A | 9e-06 | 6.7e-06 | 9.6e-06 | 8.433E-06 | 1.531E-06 | 6.700E-06 | 9.600E-06 |
| 11741 | 588 | Y588A | A | 1.4e-05 | 1.3e-05 | 1.4e-05 | 1.367E-05 | 5.774E-07 | 1.300E-05 | 1.400E-05 |
| 11742 | 588 | Y588R | A | 9e-06 | 1e-05 | 7.7e-06 | 8.900E-06 | 1.153E-06 | 7.700E-06 | 1.000E-05 |
| 11743 | 588 | Y588N | A | 2.9e-05 | 7e-06 | 1.4e-05 | 1.667E-05 | 1.124E-05 | 7.000E-06 | 2.900E-05 |
| 11744 | 588 | Y588D | A | 1.3e-05 | 1.3e-05 | 6.7e-06 | 1.090E-05 | 3.637E-06 | 6.700E-06 | 1.300E-05 |
| 11745 | 588 | Y588C | A | 1.1e-05 | 8.4e-06 | 9.6e-06 | 9.667E-06 | 1.301E-06 | 8.400E-06 | 1.100E-05 |
| 11746 | 588 | Y588E | A | 4.1e-06 | 7.1e-06 | 1.2e-05 | 7.733E-06 | 3.988E-06 | 4.100E-06 | 1.200E-05 |
| 11747 | 588 | Y588Q | A | 1e-05 | 7.3e-06 | 7.2e-06 | 8.167E-06 | 1.589E-06 | 7.200E-06 | 1.000E-05 |
| 11748 | 588 | Y588G | A | 1.9e-05 | 8.4e-06 | 1.8e-05 | 1.513E-05 | 5.853E-06 | 8.400E-06 | 1.900E-05 |
| 11749 | 588 | Y588H | A | 1.2e-05 | 1.1e-05 | 1.1e-05 | 1.133E-05 | 5.774E-07 | 1.100E-05 | 1.200E-05 |
| 11750 | 588 | Y588I | A | 1.3e-05 | 1.3e-05 | 1.9e-05 | 1.500E-05 | 3.464E-06 | 1.300E-05 | 1.900E-05 |
| 11751 | 588 | Y588L | A | 1.1e-05 | 1.2e-05 | 1.3e-05 | 1.200E-05 | 1.000E-06 | 1.100E-05 | 1.300E-05 |
| 11752 | 588 | Y588K | A | 5.3e-06 | 8.6e-06 | 5.6e-06 | 6.500E-06 | 1.825E-06 | 5.300E-06 | 8.600E-06 |
| 11753 | 588 | Y588M | A | 1.1e-05 | 1.3e-05 | 6.7e-06 | 1.023E-05 | 3.219E-06 | 6.700E-06 | 1.300E-05 |
| 11754 | 588 | Y588F | A | 9.6e-06 | 4.8e-06 | 1e-05 | 8.133E-06 | 2.894E-06 | 4.800E-06 | 1.000E-05 |
| 11755 | 588 | Y588P | A | 1e-05 | 7.9e-06 | 8.2e-06 | 8.700E-06 | 1.136E-06 | 7.900E-06 | 1.000E-05 |
| 11756 | 588 | Y588S | A | 1.3e-05 | 8.2e-06 | 7.3e-06 | 9.500E-06 | 3.064E-06 | 7.300E-06 | 1.300E-05 |
| 11757 | 588 | Y588T | A | 9.5e-06 | 2e-05 | 1.5e-05 | 1.483E-05 | 5.252E-06 | 9.500E-06 | 2.000E-05 |
| 11758 | 588 | Y588W | A | 8.7e-06 | 1.4e-05 | 1.1e-05 | 1.123E-05 | 2.658E-06 | 8.700E-06 | 1.400E-05 |
| 11759 | 588 | Y588Y | A | 1e-05 | 1.1e-05 | 1.1e-05 | 1.067E-05 | 5.774E-07 | 1.000E-05 | 1.100E-05 |
| 11760 | 588 | Y588V | A | 9.6e-06 | 1.7e-05 | 1.3e-05 | 1.320E-05 | 3.704E-06 | 9.600E-06 | 1.700E-05 |
| 11761 | 589 | M589A | A | 9.7e-06 | 4.7e-06 | 6.4e-06 | 6.933E-06 | 2.542E-06 | 4.700E-06 | 9.700E-06 |
| 11762 | 589 | M589R | A | 8e-06 | 5e-06 | 7.1e-06 | 6.700E-06 | 1.539E-06 | 5.000E-06 | 8.000E-06 |
| 11763 | 589 | M589N | A | 7.5e-06 | 1.2e-05 | 5e-06 | 8.167E-06 | 3.547E-06 | 5.000E-06 | 1.200E-05 |
| 11764 | 589 | M589D | A | 1e-05 | 9.5e-06 | 5.8e-06 | 8.433E-06 | 2.294E-06 | 5.800E-06 | 1.000E-05 |
| 11765 | 589 | M589C | A | 1.3e-05 | 9.9e-06 | 7.5e-06 | 1.013E-05 | 2.757E-06 | 7.500E-06 | 1.300E-05 |
| 11766 | 589 | M589E | A | 6.9e-06 | 1.1e-05 | 6.7e-06 | 8.200E-06 | 2.427E-06 | 6.700E-06 | 1.100E-05 |
| 11767 | 589 | M589Q | A | 6.4e-06 | 4.5e-06 | 6.6e-06 | 5.833E-06 | 1.159E-06 | 4.500E-06 | 6.600E-06 |
| 11768 | 589 | M589G | A | 1.7e-05 | 1.5e-05 | 1e-05 | 1.400E-05 | 3.606E-06 | 1.000E-05 | 1.700E-05 |
| 11769 | 589 | M589H | A | 8.6e-06 | 6.3e-06 | 7.4e-06 | 7.433E-06 | 1.150E-06 | 6.300E-06 | 8.600E-06 |
| 11770 | 589 | M589I | A | 6.1e-06 | 5.4e-06 | 5.1e-06 | 5.533E-06 | 5.132E-07 | 5.100E-06 | 6.100E-06 |
| 11771 | 589 | M589L | A | 6.6e-06 | 6.3e-06 | 4.6e-06 | 5.833E-06 | 1.079E-06 | 4.600E-06 | 6.600E-06 |
| 11772 | 589 | M589K | A | 7.6e-06 | 7.2e-06 | 7.3e-06 | 7.367E-06 | 2.082E-07 | 7.200E-06 | 7.600E-06 |
| 11773 | 589 | M589M | A | 8.9e-06 | 1.3e-05 | 1e-05 | 1.063E-05 | 2.122E-06 | 8.900E-06 | 1.300E-05 |
| 11774 | 589 | M589F | A | 9.4e-06 | 6.6e-06 | 5.2e-06 | 7.067E-06 | 2.139E-06 | 5.200E-06 | 9.400E-06 |
| 11775 | 589 | M589P | A | 5.6e-06 | 1.3e-05 | 6.5e-06 | 8.367E-06 | 4.038E-06 | 5.600E-06 | 1.300E-05 |
| 11776 | 589 | M589S | A | 5.6e-06 | 1.1e-05 | 1.2e-05 | 9.533E-06 | 3.443E-06 | 5.600E-06 | 1.200E-05 |
| 11777 | 589 | M589T | A | 9.2e-06 | 7.6e-06 | 8.9e-06 | 8.567E-06 | 8.505E-07 | 7.600E-06 | 9.200E-06 |
| 11778 | 589 | M589W | A | 1.4e-05 | 6.9e-06 | 5.8e-06 | 8.900E-06 | 4.451E-06 | 5.800E-06 | 1.400E-05 |
| 11779 | 589 | M589Y | A | 8.8e-06 | 4.4e-06 | 6.5e-06 | 6.567E-06 | 2.201E-06 | 4.400E-06 | 8.800E-06 |
| 11780 | 589 | M589V | A | 6.3e-06 | 1.2e-05 | 9.3e-06 | 9.200E-06 | 2.851E-06 | 6.300E-06 | 1.200E-05 |
| 11781 | 590 | P590A | A | 6.5e-06 | 1.3e-05 | 6.5e-06 | 8.667E-06 | 3.753E-06 | 6.500E-06 | 1.300E-05 |
| 11782 | 590 | P590R | A | 9.4e-06 | 5.5e-06 | 9e-06 | 7.967E-06 | 2.146E-06 | 5.500E-06 | 9.400E-06 |
| 11783 | 590 | P590N | A | 1.3e-05 | 9.1e-06 | 8.2e-06 | 1.010E-05 | 2.551E-06 | 8.200E-06 | 1.300E-05 |
| 11784 | 590 | P590D | A | 5.4e-06 | 7.3e-06 | 1.1e-05 | 7.900E-06 | 2.848E-06 | 5.400E-06 | 1.100E-05 |
| 11785 | 590 | P590C | A | 8.3e-06 | 5.5e-06 | 4.2e-06 | 6.000E-06 | 2.095E-06 | 4.200E-06 | 8.300E-06 |
| 11786 | 590 | P590E | A | 5.8e-06 | 8e-06 | 9.9e-06 | 7.900E-06 | 2.052E-06 | 5.800E-06 | 9.900E-06 |
| 11787 | 590 | P590Q | A | 1e-05 | 9.2e-06 | 8.2e-06 | 9.133E-06 | 9.018E-07 | 8.200E-06 | 1.000E-05 |
| 11788 | 590 | P590G | A | 9.2e-06 | 1.1e-05 | 7.5e-06 | 9.233E-06 | 1.750E-06 | 7.500E-06 | 1.100E-05 |
| 11789 | 590 | P590H | A | 5.6e-06 | 6.5e-06 | 6.9e-06 | 6.333E-06 | 6.658E-07 | 5.600E-06 | 6.900E-06 |
| 11790 | 590 | P590I | A | 6.4e-06 | 8.9e-06 | 8.5e-06 | 7.933E-06 | 1.343E-06 | 6.400E-06 | 8.900E-06 |
| 11791 | 590 | P590L | A | 6.3e-06 | 8.7e-06 | 8.2e-06 | 7.733E-06 | 1.266E-06 | 6.300E-06 | 8.700E-06 |
| 11792 | 590 | P590K | A | 9.2e-06 | 4.9e-06 | 1.4e-05 | 9.367E-06 | 4.552E-06 | 4.900E-06 | 1.400E-05 |
| 11793 | 590 | P590M | A | 8.1e-06 | 8.9e-06 | 9.2e-06 | 8.733E-06 | 5.686E-07 | 8.100E-06 | 9.200E-06 |
| 11794 | 590 | P590F | A | 1.3e-05 | 7.1e-06 | 4.4e-06 | 8.167E-06 | 4.398E-06 | 4.400E-06 | 1.300E-05 |
| 11795 | 590 | P590P | A | 8.4e-06 | 7.3e-06 | 8.4e-06 | 8.033E-06 | 6.351E-07 | 7.300E-06 | 8.400E-06 |
| 11796 | 590 | P590S | A | 7.8e-06 | 7.1e-06 | 7.3e-06 | 7.400E-06 | 3.606E-07 | 7.100E-06 | 7.800E-06 |
| 11797 | 590 | P590T | A | 4.7e-06 | 1.1e-05 | 8.4e-06 | 8.033E-06 | 3.166E-06 | 4.700E-06 | 1.100E-05 |
| 11798 | 590 | P590W | A | 7.3e-06 | 6.9e-06 | 7.9e-06 | 7.367E-06 | 5.033E-07 | 6.900E-06 | 7.900E-06 |
| 11799 | 590 | P590Y | A | 9.2e-06 | 1e-05 | 8.7e-06 | 9.300E-06 | 6.557E-07 | 8.700E-06 | 1.000E-05 |
| 11800 | 590 | P590V | A | 6.3e-06 | 6.8e-06 | 5.2e-06 | 6.100E-06 | 8.185E-07 | 5.200E-06 | 6.800E-06 |
| 11801 | 591 | I591A | A | 7.3e-06 | 6.4e-06 | 1.1e-05 | 8.233E-06 | 2.438E-06 | 6.400E-06 | 1.100E-05 |
| 11802 | 591 | I591R | A | 7.8e-06 | 8.6e-06 | 7.9e-06 | 8.100E-06 | 4.359E-07 | 7.800E-06 | 8.600E-06 |
| 11803 | 591 | I591N | A | 1.5e-05 | 1.2e-05 | 1e-05 | 1.233E-05 | 2.517E-06 | 1.000E-05 | 1.500E-05 |
| 11804 | 591 | I591D | A | 8.2e-06 | 4.7e-06 | 6.7e-06 | 6.533E-06 | 1.756E-06 | 4.700E-06 | 8.200E-06 |
| 11805 | 591 | I591C | A | 4.6e-06 | 8.1e-06 | 6e-06 | 6.233E-06 | 1.762E-06 | 4.600E-06 | 8.100E-06 |
| 11806 | 591 | I591E | A | 1.1e-05 | 1.6e-05 | 8.4e-06 | 1.180E-05 | 3.863E-06 | 8.400E-06 | 1.600E-05 |
| 11807 | 591 | I591Q | A | 9.1e-06 | 6.3e-06 | 8e-06 | 7.800E-06 | 1.411E-06 | 6.300E-06 | 9.100E-06 |
| 11808 | 591 | I591G | A | 4.2e-06 | 1e-05 | 1e-05 | 8.067E-06 | 3.349E-06 | 4.200E-06 | 1.000E-05 |
| 11809 | 591 | I591H | A | 6.6e-06 | 1.2e-05 | 1.2e-05 | 1.020E-05 | 3.118E-06 | 6.600E-06 | 1.200E-05 |
| 11810 | 591 | I591I | A | 1e-05 | 1.5e-05 | 9e-06 | 1.133E-05 | 3.215E-06 | 9.000E-06 | 1.500E-05 |
| 11811 | 591 | I591L | A | 7.5e-06 | 4.6e-06 | 9.3e-06 | 7.133E-06 | 2.371E-06 | 4.600E-06 | 9.300E-06 |
| 11812 | 591 | I591K | A | 8.5e-06 | 9.2e-06 | 7.7e-06 | 8.467E-06 | 7.506E-07 | 7.700E-06 | 9.200E-06 |
| 11813 | 591 | I591M | A | 8.5e-06 | 5.6e-06 | 5.8e-06 | 6.633E-06 | 1.620E-06 | 5.600E-06 | 8.500E-06 |
| 11814 | 591 | I591F | A | 1.1e-05 | 4.9e-06 | 1.7e-05 | 1.097E-05 | 6.050E-06 | 4.900E-06 | 1.700E-05 |

| | | | | | | | | | | |
|---|---|---|---|---|---|---|---|---|---|---|
| 11815 | 591 | I591P | A | 1.3e-05 | 1.4e-05 | 4.2e-06 | 1.040E-05 | 5.393E-06 | 4.200E-06 | 1.400E-05 |
| 11816 | 591 | I591S | A | 6.7e-06 | 6.6e-06 | 5.2e-06 | 6.167E-06 | 8.386E-07 | 5.200E-06 | 6.700E-06 |
| 11817 | 591 | I591T | A | 4.9e-06 | 6.3e-06 | 8.6e-06 | 6.600E-06 | 1.868E-06 | 4.900E-06 | 8.600E-06 |
| 11818 | 591 | I591W | A | 8.3e-06 | 8.1e-06 | 6.7e-06 | 7.700E-06 | 8.718E-07 | 6.700E-06 | 8.300E-06 |
| 11819 | 591 | I591Y | A | 1.3e-05 | 1.2e-05 | 5.6e-06 | 1.020E-05 | 4.015E-06 | 5.600E-06 | 1.300E-05 |
| 11820 | 591 | I591V | A | 6.3e-06 | 7.7e-06 | 5.9e-06 | 6.633E-06 | 9.452E-07 | 5.900E-06 | 7.700E-06 |
| 11821 | 592 | W592A | A | 8.2e-06 | 6.8e-06 | 8.3e-06 | 7.767E-06 | 8.386E-07 | 6.800E-06 | 8.300E-06 |
| 11822 | 592 | W592R | A | 6.7e-06 | 8.7e-06 | 8e-06 | 7.800E-06 | 1.015E-06 | 6.700E-06 | 8.700E-06 |
| 11823 | 592 | W592N | A | 8.7e-06 | 6.5e-06 | 1.8e-05 | 1.107E-05 | 6.104E-06 | 6.500E-06 | 1.800E-05 |
| 11824 | 592 | W592D | A | 5.8e-06 | 4.8e-06 | 4.3e-06 | 4.967E-06 | 7.638E-07 | 4.300E-06 | 5.800E-06 |
| 11825 | 592 | W592C | A | 1.4e-05 | 7.4e-06 | 1.5e-05 | 1.213E-05 | 4.130E-06 | 7.400E-06 | 1.500E-05 |
| 11826 | 592 | W592E | A | 9.3e-06 | 7.8e-06 | 8.7e-06 | 8.600E-06 | 7.550E-07 | 7.800E-06 | 9.300E-06 |
| 11827 | 592 | W592Q | A | 1.2e-05 | 9.1e-06 | 6e-06 | 9.033E-06 | 3.001E-06 | 6.000E-06 | 1.200E-05 |
| 11828 | 592 | W592G | A | 1.4e-05 | 9.4e-06 | 1.2e-05 | 1.180E-05 | 2.307E-06 | 9.400E-06 | 1.400E-05 |
| 11829 | 592 | W592H | A | 6.3e-06 | 9.9e-06 | 1e-05 | 8.733E-06 | 2.108E-06 | 6.300E-06 | 1.000E-05 |
| 11830 | 592 | W592I | A | 7.4e-06 | 1.3e-05 | 7.3e-06 | 9.233E-06 | 3.262E-06 | 7.300E-06 | 1.300E-05 |
| 11831 | 592 | W592L | A | 8.2e-06 | 7e-06 | 9.5e-06 | 8.233E-06 | 1.250E-06 | 7.000E-06 | 9.500E-06 |
| 11832 | 592 | W592K | A | 5.5e-06 | 6e-06 | 6.2e-06 | 5.900E-06 | 3.606E-07 | 5.500E-06 | 6.200E-06 |
| 11833 | 592 | W592M | A | 5.2e-06 | 4.6e-06 | 1e-05 | 6.600E-06 | 2.960E-06 | 4.600E-06 | 1.000E-05 |
| 11834 | 592 | W592F | A | 8.9e-06 | 1.1e-05 | 9.6e-06 | 9.833E-06 | 1.069E-06 | 8.900E-06 | 1.100E-05 |
| 11835 | 592 | W592P | A | 8.7e-06 | 1e-05 | 7.6e-06 | 8.767E-06 | 1.201E-06 | 7.600E-06 | 1.000E-05 |
| 11836 | 592 | W592S | A | 9.4e-06 | 7.4e-06 | 6.1e-06 | 7.633E-06 | 1.662E-06 | 6.100E-06 | 9.400E-06 |
| 11837 | 592 | W592T | A | 8.7e-06 | 8.7e-06 | 1.9e-05 | 1.213E-05 | 5.947E-06 | 8.700E-06 | 1.900E-05 |
| 11838 | 592 | W592W | A | 5.7e-06 | 5.1e-06 | 1.4e-05 | 8.267E-06 | 4.974E-06 | 5.100E-06 | 1.400E-05 |
| 11839 | 592 | W592Y | A | 9.7e-06 | 9.8e-06 | 1.1e-05 | 1.017E-05 | 7.234E-07 | 9.700E-06 | 1.100E-05 |
| 11840 | 592 | W592V | A | 5.3e-06 | 6.7e-06 | 7.5e-06 | 6.500E-06 | 1.114E-06 | 5.300E-06 | 7.500E-06 |
| 11841 | 593 | K593A | A | 1e-05 | 8.3e-06 | 1.1e-05 | 9.767E-06 | 1.365E-06 | 8.300E-06 | 1.100E-05 |
| 11842 | 593 | K593R | A | 9.9e-06 | 9.7e-06 | 7.9e-06 | 9.167E-06 | 1.102E-06 | 7.900E-06 | 9.900E-06 |
| 11843 | 593 | K593N | A | 4.9e-06 | 1.5e-05 | 9.8e-06 | 9.900E-06 | 5.051E-06 | 4.900E-06 | 1.500E-05 |
| 11844 | 593 | K593D | A | 6e-06 | 5e-06 | 6.8e-06 | 5.933E-06 | 9.018E-07 | 5.000E-06 | 6.800E-06 |
| 11845 | 593 | K593C | A | 4.8e-06 | 5.7e-06 | 1.2e-05 | 7.500E-06 | 3.923E-06 | 4.800E-06 | 1.200E-05 |
| 11846 | 593 | K593E | A | 1e-05 | 7e-06 | 5.9e-06 | 7.633E-06 | 2.122E-06 | 5.900E-06 | 1.000E-05 |
| 11847 | 593 | K593Q | A | 7.4e-06 | 6.8e-06 | 1.1e-05 | 8.400E-06 | 2.272E-06 | 6.800E-06 | 1.100E-05 |
| 11848 | 593 | K593G | A | 7.1e-06 | 7.3e-06 | 9.8e-06 | 8.067E-06 | 1.504E-06 | 7.100E-06 | 9.800E-06 |
| 11849 | 593 | K593H | A | 9.9e-06 | 9.7e-06 | 7.9e-06 | 9.167E-06 | 1.102E-06 | 7.900E-06 | 9.900E-06 |
| 11850 | 593 | K593I | A | 1.2e-05 | 1.2e-05 | 8.3e-06 | 1.077E-05 | 2.136E-06 | 8.300E-06 | 1.200E-05 |
| 11851 | 593 | K593L | A | 6.4e-06 | 9.9e-06 | 8.9e-06 | 8.400E-06 | 1.803E-06 | 6.400E-06 | 9.900E-06 |
| 11852 | 593 | K593K | A | 7.2e-06 | 1.2e-05 | 7.8e-06 | 9.000E-06 | 2.615E-06 | 7.200E-06 | 1.200E-05 |
| 11853 | 593 | K593M | A | 4.2e-06 | 6.4e-06 | 6.5e-06 | 5.700E-06 | 1.300E-06 | 4.200E-06 | 6.500E-06 |
| 11854 | 593 | K593F | A | 8.8e-06 | 6.8e-06 | 6.7e-06 | 7.433E-06 | 1.185E-06 | 6.700E-06 | 8.800E-06 |
| 11855 | 593 | K593P | A | 5.7e-06 | 1.1e-05 | 9.6e-06 | 8.767E-06 | 2.747E-06 | 5.700E-06 | 1.100E-05 |
| 11856 | 593 | K593S | A | 4.7e-06 | 6.3e-06 | 7.6e-06 | 6.200E-06 | 1.453E-06 | 4.700E-06 | 7.600E-06 |
| 11857 | 593 | K593T | A | 1e-05 | 1.2e-05 | 9.8e-06 | 1.060E-05 | 1.217E-06 | 9.800E-06 | 1.200E-05 |
| 11858 | 593 | K593W | A | 7.7e-06 | 9.3e-06 | 7.3e-06 | 8.100E-06 | 1.058E-06 | 7.300E-06 | 9.300E-06 |
| 11859 | 593 | K593Y | A | 1.4e-05 | 5.5e-06 | 6.6e-06 | 8.700E-06 | 4.623E-06 | 5.500E-06 | 1.400E-05 |
| 11860 | 593 | K593V | A | 1.3e-05 | 5.4e-06 | 6e-06 | 8.133E-06 | 4.225E-06 | 5.400E-06 | 1.300E-05 |
| 11861 | 594 | F594A | A | 6.7e-06 | 1.1e-05 | 5e-06 | 7.567E-06 | 3.092E-06 | 5.000E-06 | 1.100E-05 |
| 11862 | 594 | F594R | A | 9.8e-06 | 6.5e-06 | 1.2e-05 | 9.433E-06 | 2.768E-06 | 6.500E-06 | 1.200E-05 |
| 11863 | 594 | F594N | A | 8.3e-06 | 7.5e-06 | 7.1e-06 | 7.633E-06 | 6.110E-07 | 7.100E-06 | 8.300E-06 |
| 11864 | 594 | F594D | A | 1.2e-05 | 1.2e-05 | 1.1e-05 | 1.167E-05 | 5.774E-07 | 1.100E-05 | 1.200E-05 |
| 11865 | 594 | F594C | A | 1.2e-05 | 5.7e-06 | 8.8e-06 | 8.833E-06 | 3.150E-06 | 5.700E-06 | 1.200E-05 |
| 11866 | 594 | F594E | A | 6.4e-06 | 5.8e-06 | 1.2e-05 | 8.067E-06 | 3.420E-06 | 5.800E-06 | 1.200E-05 |
| 11867 | 594 | F594Q | A | 6.2e-06 | 9e-06 | 6e-06 | 7.067E-06 | 1.677E-06 | 6.000E-06 | 9.000E-06 |
| 11868 | 594 | F594G | A | 8.3e-06 | 5.3e-06 | 9.1e-06 | 7.567E-06 | 2.003E-06 | 5.300E-06 | 9.100E-06 |
| 11869 | 594 | F594H | A | 1.3e-05 | 1.3e-05 | 5.2e-06 | 1.040E-05 | 4.503E-06 | 5.200E-06 | 1.300E-05 |
| 11870 | 594 | F594I | A | 8.3e-06 | 5.7e-06 | 6.1e-06 | 6.700E-06 | 1.400E-06 | 5.700E-06 | 8.300E-06 |
| 11871 | 594 | F594L | A | 9.8e-06 | 8e-06 | 5.8e-06 | 7.867E-06 | 2.003E-06 | 5.800E-06 | 9.800E-06 |
| 11872 | 594 | F594K | A | 1.3e-05 | 1.2e-05 | 7.3e-06 | 1.077E-05 | 3.044E-06 | 7.300E-06 | 1.300E-05 |
| 11873 | 594 | F594M | A | 9.7e-06 | 7e-06 | 6.4e-06 | 7.700E-06 | 1.758E-06 | 6.400E-06 | 9.700E-06 |
| 11874 | 594 | F594F | A | 9.5e-06 | 7.2e-06 | 6.5e-06 | 7.733E-06 | 1.570E-06 | 6.500E-06 | 9.500E-06 |
| 11875 | 594 | F594P | A | 7.5e-06 | 4.1e-06 | 1.3e-05 | 8.200E-06 | 4.491E-06 | 4.100E-06 | 1.300E-05 |
| 11876 | 594 | F594S | A | 6.1e-06 | 4.5e-06 | 6.6e-06 | 5.733E-06 | 1.097E-06 | 4.500E-06 | 6.600E-06 |
| 11877 | 594 | F594T | A | 6.5e-06 | 5.1e-06 | 6.8e-06 | 6.133E-06 | 9.074E-07 | 5.100E-06 | 6.800E-06 |
| 11878 | 594 | F594W | A | 8.4e-06 | 1.1e-05 | 1.2e-05 | 1.047E-05 | 1.858E-06 | 8.400E-06 | 1.200E-05 |
| 11879 | 594 | F594Y | A | 5.7e-06 | 7.4e-06 | 7.1e-06 | 6.733E-06 | 9.074E-07 | 5.700E-06 | 7.400E-06 |
| 11880 | 594 | F594V | A | 8.7e-06 | 5.2e-06 | 7.5e-06 | 7.133E-06 | 1.779E-06 | 5.200E-06 | 8.700E-06 |
| 11881 | 595 | P595A | A | 7.4e-06 | 8e-06 | 1.4e-05 | 9.800E-06 | 3.650E-06 | 7.400E-06 | 1.400E-05 |
| 11882 | 595 | P595R | A | 8.7e-06 | 8.6e-06 | 4.9e-06 | 7.400E-06 | 2.166E-06 | 4.900E-06 | 8.700E-06 |
| 11883 | 595 | P595N | A | 8.5e-06 | 6e-06 | 4.9e-06 | 6.467E-06 | 1.845E-06 | 4.900E-06 | 8.500E-06 |
| 11884 | 595 | P595D | A | 7.8e-06 | 1.2e-05 | 5.5e-06 | 8.433E-06 | 3.296E-06 | 5.500E-06 | 1.200E-05 |
| 11885 | 595 | P595C | A | 1e-05 | 1.1e-05 | 7.4e-06 | 9.467E-06 | 1.858E-06 | 7.400E-06 | 1.100E-05 |
| 11886 | 595 | P595E | A | 5.1e-06 | 7.8e-06 | 8.5e-06 | 7.133E-06 | 1.795E-06 | 5.100E-06 | 8.500E-06 |
| 11887 | 595 | P595Q | A | 6.1e-06 | 6.8e-06 | 6.3e-06 | 6.400E-06 | 3.606E-07 | 6.100E-06 | 6.800E-06 |
| 11888 | 595 | P595G | A | 1.3e-05 | 9.3e-06 | 9.3e-06 | 1.053E-05 | 2.136E-06 | 9.300E-06 | 1.300E-05 |
| 11889 | 595 | P595H | A | 7.1e-06 | 5.6e-06 | 6.3e-06 | 6.333E-06 | 7.506E-07 | 5.600E-06 | 7.100E-06 |
| 11890 | 595 | P595I | A | 5.2e-06 | 7.9e-06 | 8.7e-06 | 7.267E-06 | 1.834E-06 | 5.200E-06 | 8.700E-06 |
| 11891 | 595 | P595L | A | 6.1e-06 | 4.5e-06 | 6.3e-06 | 5.633E-06 | 9.866E-07 | 4.500E-06 | 6.300E-06 |
| 11892 | 595 | P595K | A | 6.4e-06 | 9.1e-06 | 5.4e-06 | 6.967E-06 | 1.914E-06 | 5.400E-06 | 9.100E-06 |
| 11893 | 595 | P595M | A | 6.5e-06 | 9.1e-06 | 1.1e-05 | 8.867E-06 | 2.259E-06 | 6.500E-06 | 1.100E-05 |
| 11894 | 595 | P595F | A | 8.4e-06 | 9.5e-06 | 8e-06 | 8.633E-06 | 7.767E-07 | 8.000E-06 | 9.500E-06 |
| 11895 | 595 | P595P | A | 4.7e-06 | 1e-05 | 5.2e-06 | 6.633E-06 | 2.926E-06 | 4.700E-06 | 1.000E-05 |
| 11896 | 595 | P595S | A | 1.2e-05 | 8.8e-06 | 7.1e-06 | 9.300E-06 | 2.488E-06 | 7.100E-06 | 1.200E-05 |
| 11897 | 595 | P595T | A | 3.9e-06 | 8.4e-06 | 9.3e-06 | 7.200E-06 | 2.893E-06 | 3.900E-06 | 9.300E-06 |
| 11898 | 595 | P595W | A | 8e-06 | 9.5e-06 | 6.7e-06 | 8.067E-06 | 1.401E-06 | 6.700E-06 | 9.500E-06 |
| 11899 | 595 | P595Y | A | 5.9e-06 | 1.2e-05 | 4.5e-06 | 7.467E-06 | 3.988E-06 | 4.500E-06 | 1.200E-05 |

| | | | | | | | | | | |
|---|---|---|---|---|---|---|---|---|---|---|
| 11900 | 595 | P595V | A | 4.9e-06 | 9.1e-06 | 9.1e-06 | 7.700E-06 | 2.425E-06 | 4.900E-06 | 9.100E-06 |
| 11901 | 596 | D596A | A | 5.5e-06 | 4.9e-06 | 7e-06 | 5.800E-06 | 1.082E-06 | 4.900E-06 | 7.000E-06 |
| 11902 | 596 | D596R | A | 5.8e-06 | 6.9e-06 | 6.6e-06 | 6.433E-06 | 5.686E-07 | 5.800E-06 | 6.900E-06 |
| 11903 | 596 | D596N | A | 7.8e-06 | 9.3e-06 | 1.9e-05 | 1.203E-05 | 6.080E-06 | 7.800E-06 | 1.900E-05 |
| 11904 | 596 | D596D | A | 4.5e-06 | 1e-05 | 9.2e-06 | 7.900E-06 | 2.972E-06 | 4.500E-06 | 1.000E-05 |
| 11905 | 596 | D596C | A | 2.2e-06 | 6.8e-06 | 1.8e-05 | 9.000E-06 | 8.126E-06 | 2.200E-06 | 1.800E-05 |
| 11906 | 596 | D596E | A | 8.6e-06 | 9e-06 | 9.9e-06 | 9.167E-06 | 6.658E-07 | 8.600E-06 | 9.900E-06 |
| 11907 | 596 | D596Q | A | 1.9e-05 | 9.3e-06 | 1.5e-05 | 1.443E-05 | 4.875E-06 | 9.300E-06 | 1.900E-05 |
| 11908 | 596 | D596G | A | 1.2e-05 | 6.7e-06 | 5.3e-06 | 8.000E-06 | 3.534E-06 | 5.300E-06 | 1.200E-05 |
| 11909 | 596 | D596H | A | 6.6e-06 | 2.2e-05 | 7.7e-06 | 1.210E-05 | 8.591E-06 | 6.600E-06 | 2.200E-05 |
| 11910 | 596 | D596I | A | 6.5e-06 | 4.1e-06 | 2.3e-06 | 4.300E-06 | 2.107E-06 | 2.300E-06 | 6.500E-06 |
| 11911 | 596 | D596L | A | 4e-06 | 2.5e-06 | 3.3e-06 | 3.267E-06 | 7.506E-07 | 2.500E-06 | 4.000E-06 |
| 11912 | 596 | D596K | A | 9e-06 | 6.3e-06 | 9.3e-06 | 8.200E-06 | 1.652E-06 | 6.300E-06 | 9.300E-06 |
| 11913 | 596 | D596M | A | 6.5e-06 | 3e-06 | 9.1e-06 | 6.200E-06 | 3.061E-06 | 3.000E-06 | 9.100E-06 |
| 11914 | 596 | D596F | A | 4.2e-06 | 5.3e-06 | 2.6e-06 | 4.033E-06 | 1.358E-06 | 2.600E-06 | 5.300E-06 |
| 11915 | 596 | D596P | A | 1.8e-05 | 1.6e-05 | 4.9e-06 | 1.297E-05 | 7.057E-06 | 4.900E-06 | 1.800E-05 |
| 11916 | 596 | D596S | A | 1.3e-05 | 9.4e-06 | 6.9e-06 | 9.767E-06 | 3.066E-06 | 6.900E-06 | 1.300E-05 |
| 11917 | 596 | D596T | A | 7.6e-06 | 1.1e-05 | 5.8e-06 | 8.133E-06 | 2.641E-06 | 5.800E-06 | 1.100E-05 |
| 11918 | 596 | D596W | A | 5.5e-06 | 5.8e-06 | 4.8e-06 | 5.367E-06 | 5.132E-07 | 4.800E-06 | 5.800E-06 |
| 11919 | 596 | D596Y | A | 4.1e-06 | 6.3e-06 | 8.2e-06 | 6.200E-06 | 2.052E-06 | 4.100E-06 | 8.200E-06 |
| 11920 | 596 | D596V | A | 1.1e-05 | 1.8e-05 | 4.5e-06 | 1.117E-05 | 6.752E-06 | 4.500E-06 | 1.800E-05 |
| 11921 | 597 | E597A | A | 7.4e-06 | 7.7e-06 | 8.4e-06 | 7.833E-06 | 5.132E-07 | 7.400E-06 | 8.400E-06 |
| 11922 | 597 | E597R | A | 5.6e-06 | 9e-06 | 1.2e-05 | 8.867E-06 | 3.202E-06 | 5.600E-06 | 1.200E-05 |
| 11923 | 597 | E597N | A | 7.7e-06 | 8e-06 | 1.1e-05 | 8.900E-06 | 1.825E-06 | 7.700E-06 | 1.100E-05 |
| 11924 | 597 | E597D | A | 9.7e-06 | 7e-06 | 6.9e-06 | 7.867E-06 | 1.589E-06 | 6.900E-06 | 9.700E-06 |
| 11925 | 597 | E597C | A | 8.6e-06 | 6.2e-06 | 1.3e-05 | 9.267E-06 | 3.449E-06 | 6.200E-06 | 1.300E-05 |
| 11926 | 597 | E597E | A | 6e-06 | 6.4e-06 | 4.8e-06 | 5.733E-06 | 8.327E-07 | 4.800E-06 | 6.400E-06 |
| 11927 | 597 | E597Q | A | 6.5e-06 | 1e-05 | 7.2e-06 | 7.900E-06 | 1.852E-06 | 6.500E-06 | 1.000E-05 |
| 11928 | 597 | E597G | A | 1.2e-05 | 6.5e-06 | 9.7e-06 | 9.400E-06 | 2.762E-06 | 6.500E-06 | 1.200E-05 |
| 11929 | 597 | E597H | A | 9.5e-06 | 6.6e-06 | 9.6e-06 | 8.567E-06 | 1.704E-06 | 6.600E-06 | 9.600E-06 |
| 11930 | 597 | E597I | A | 7.2e-06 | 7.3e-06 | 9.7e-06 | 8.067E-06 | 1.415E-06 | 7.200E-06 | 9.700E-06 |
| 11931 | 597 | E597L | A | 6.7e-06 | 5.3e-06 | 9.4e-06 | 7.133E-06 | 2.084E-06 | 5.300E-06 | 9.400E-06 |
| 11932 | 597 | E597K | A | 1.3e-05 | 1.3e-05 | 6.4e-06 | 1.080E-05 | 3.811E-06 | 6.400E-06 | 1.300E-05 |
| 11933 | 597 | E597M | A | 8.5e-06 | 9.7e-06 | 8.1e-06 | 8.767E-06 | 8.327E-07 | 8.100E-06 | 9.700E-06 |
| 11934 | 597 | E597F | A | 7.3e-06 | 7.8e-06 | 7.2e-06 | 7.433E-06 | 3.215E-07 | 7.200E-06 | 7.800E-06 |
| 11935 | 597 | E597P | A | 9.8e-06 | 1.4e-05 | 7.4e-06 | 1.040E-05 | 3.341E-06 | 7.400E-06 | 1.400E-05 |
| 11936 | 597 | E597S | A | 6.8e-06 | 7.3e-06 | 8.7e-06 | 7.600E-06 | 9.849E-07 | 6.800E-06 | 8.700E-06 |
| 11937 | 597 | E597T | A | 1e-05 | 8e-06 | 7.3e-06 | 8.433E-06 | 1.401E-06 | 7.300E-06 | 1.000E-05 |
| 11938 | 597 | E597W | A | 8.5e-06 | 3.7e-06 | 6.7e-06 | 6.300E-06 | 2.425E-06 | 3.700E-06 | 8.500E-06 |
| 11939 | 597 | E597Y | A | 6e-06 | 4.8e-06 | 7.7e-06 | 6.167E-06 | 1.457E-06 | 4.800E-06 | 7.700E-06 |
| 11940 | 597 | E597V | A | 8.1e-06 | 1.4e-05 | 7.8e-06 | 9.967E-06 | 3.496E-06 | 7.800E-06 | 1.400E-05 |
| 11941 | 598 | E598A | A | 6.7e-06 | 4.8e-06 | 6.2e-06 | 5.900E-06 | 9.849E-07 | 4.800E-06 | 6.700E-06 |
| 11942 | 598 | E598R | A | 7.9e-06 | 4.2e-06 | 6.4e-06 | 6.167E-06 | 1.861E-06 | 4.200E-06 | 7.900E-06 |
| 11943 | 598 | E598N | A | 1.8e-05 | 5.7e-06 | 6e-06 | 9.900E-06 | 7.016E-06 | 5.700E-06 | 1.800E-05 |
| 11944 | 598 | E598D | A | 1.2e-05 | 7.4e-06 | 6.9e-06 | 8.767E-06 | 2.811E-06 | 6.900E-06 | 1.200E-05 |
| 11945 | 598 | E598C | A | 4.2e-06 | 5.9e-06 | 6.8e-06 | 5.633E-06 | 1.320E-06 | 4.200E-06 | 6.800E-06 |
| 11946 | 598 | E598E | A | 6.6e-06 | 7.1e-06 | 1e-05 | 7.900E-06 | 1.836E-06 | 6.600E-06 | 1.000E-05 |
| 11947 | 598 | E598Q | A | 1.2e-05 | 1.1e-05 | 7e-06 | 1.000E-05 | 2.646E-06 | 7.000E-06 | 1.200E-05 |
| 11948 | 598 | E598G | A | 1e-05 | 9e-06 | 6.6e-06 | 8.533E-06 | 1.747E-06 | 6.600E-06 | 1.000E-05 |
| 11949 | 598 | E598H | A | 1e-05 | 9.7e-06 | 2.6e-05 | 1.523E-05 | 9.325E-06 | 9.700E-06 | 2.600E-05 |
| 11950 | 598 | E598I | A | 4.6e-06 | 4.5e-06 | 4.9e-06 | 4.667E-06 | 2.082E-07 | 4.500E-06 | 4.900E-06 |
| 11951 | 598 | E598L | A | 6e-06 | 4.5e-06 | 5.4e-06 | 5.300E-06 | 7.550E-07 | 4.500E-06 | 6.000E-06 |
| 11952 | 598 | E598K | A | 6.6e-06 | 5.1e-06 | 9.5e-06 | 7.067E-06 | 2.237E-06 | 5.100E-06 | 9.500E-06 |
| 11953 | 598 | E598M | A | 8.9e-06 | 8e-06 | 7.2e-06 | 8.033E-06 | 8.505E-07 | 7.200E-06 | 8.900E-06 |
| 11954 | 598 | E598F | A | 6.1e-06 | 4.4e-06 | 6.6e-06 | 5.700E-06 | 1.153E-06 | 4.400E-06 | 6.600E-06 |
| 11955 | 598 | E598P | A | 1.4e-05 | 6.5e-06 | 6e-06 | 8.833E-06 | 4.481E-06 | 6.000E-06 | 1.400E-05 |
| 11956 | 598 | E598S | A | 7.9e-06 | 1.2e-05 | 1.7e-05 | 1.230E-05 | 4.557E-06 | 7.900E-06 | 1.700E-05 |
| 11957 | 598 | E598T | A | 1.2e-05 | 1.2e-05 | 9e-06 | 1.100E-05 | 1.732E-06 | 9.000E-06 | 1.200E-05 |
| 11958 | 598 | E598W | A | 7.8e-06 | 3.9e-06 | 3.5e-06 | 5.067E-06 | 2.376E-06 | 3.500E-06 | 7.800E-06 |
| 11959 | 598 | E598Y | A | 9.3e-06 | 6.3e-06 | 3.7e-06 | 6.433E-06 | 2.802E-06 | 3.700E-06 | 9.300E-06 |
| 11960 | 598 | E598V | A | 7.7e-06 | 7.4e-06 | 6.8e-06 | 7.300E-06 | 4.583E-07 | 6.800E-06 | 7.700E-06 |
| 11961 | 599 | G599A | A | 5e-06 | 1.5e-05 | 5.6e-06 | 8.533E-06 | 5.608E-06 | 5.000E-06 | 1.500E-05 |
| 11962 | 599 | G599R | A | 6e-06 | 5.6e-06 | 6.4e-06 | 6.000E-06 | 4.000E-07 | 5.600E-06 | 6.400E-06 |
| 11963 | 599 | G599N | A | 5.4e-06 | 8.6e-06 | 9e-06 | 7.667E-06 | 1.973E-06 | 5.400E-06 | 9.000E-06 |
| 11964 | 599 | G599D | A | 6.9e-06 | 7.2e-06 | 4.8e-06 | 6.300E-06 | 1.308E-06 | 4.800E-06 | 7.200E-06 |
| 11965 | 599 | G599C | A | 9.1e-06 | 1e-05 | 1.3e-05 | 1.070E-05 | 2.042E-06 | 9.100E-06 | 1.300E-05 |
| 11966 | 599 | G599E | A | 9.3e-06 | 7.7e-06 | 3.9e-06 | 6.967E-06 | 2.774E-06 | 3.900E-06 | 9.300E-06 |
| 11967 | 599 | G599Q | A | 7.4e-06 | 9.8e-06 | 5.9e-06 | 7.700E-06 | 1.967E-06 | 5.900E-06 | 9.800E-06 |
| 11968 | 599 | G599G | A | 5.1e-06 | 8.2e-06 | 9e-06 | 7.433E-06 | 2.060E-06 | 5.100E-06 | 9.000E-06 |
| 11969 | 599 | G599H | A | 6.1e-06 | 9.6e-06 | 7.6e-06 | 7.767E-06 | 1.756E-06 | 6.100E-06 | 9.600E-06 |
| 11970 | 599 | G599I | A | 5.9e-06 | 8.2e-06 | 6e-06 | 6.700E-06 | 1.300E-06 | 5.900E-06 | 8.200E-06 |
| 11971 | 599 | G599L | A | 6.6e-06 | 8.1e-06 | 7.9e-06 | 7.533E-06 | 8.145E-07 | 6.600E-06 | 8.100E-06 |
| 11972 | 599 | G599K | A | 5.2e-06 | 6.4e-06 | 7e-06 | 6.200E-06 | 9.165E-07 | 5.200E-06 | 7.000E-06 |
| 11973 | 599 | G599M | A | 4.1e-06 | 5.9e-06 | 7.5e-06 | 5.833E-06 | 1.701E-06 | 4.100E-06 | 7.500E-06 |
| 11974 | 599 | G599F | A | 1.2e-05 | 7.1e-06 | 7.4e-06 | 8.833E-06 | 2.747E-06 | 7.100E-06 | 1.200E-05 |
| 11975 | 599 | G599P | A | 5.2e-06 | 3.7e-06 | 9.2e-06 | 6.033E-06 | 2.843E-06 | 3.700E-06 | 9.200E-06 |
| 11976 | 599 | G599S | A | 6.3e-06 | 6.6e-06 | 8e-06 | 6.967E-06 | 9.074E-07 | 6.300E-06 | 8.000E-06 |
| 11977 | 599 | G599T | A | 6e-06 | 1e-05 | 1.2e-05 | 9.333E-06 | 3.055E-06 | 6.000E-06 | 1.200E-05 |
| 11978 | 599 | G599W | A | 1.3e-05 | 7e-06 | 8.1e-06 | 9.367E-06 | 3.194E-06 | 7.000E-06 | 1.300E-05 |
| 11979 | 599 | G599Y | A | 9.6e-06 | 9.4e-06 | 7.4e-06 | 8.800E-06 | 1.217E-06 | 7.400E-06 | 9.600E-06 |
| 11980 | 599 | G599V | A | 9.9e-06 | 2.2e-05 | 9.2e-06 | 1.370E-05 | 7.197E-06 | 9.200E-06 | 2.200E-05 |
| 11981 | 600 | A600A | A | 6.9e-06 | 8.9e-06 | 6.8e-06 | 7.533E-06 | 1.185E-06 | 6.800E-06 | 8.900E-06 |
| 11982 | 600 | A600R | A | 8.7e-06 | 8.2e-06 | 9.6e-06 | 8.833E-06 | 7.095E-07 | 8.200E-06 | 9.600E-06 |
| 11983 | 600 | A600N | A | 2.7e-05 | 1.5e-05 | 1.6e-05 | 1.933E-05 | 6.658E-06 | 1.500E-05 | 2.700E-05 |
| 11984 | 600 | A600D | A | 1.1e-05 | 1.7e-05 | 8.2e-06 | 1.207E-05 | 4.496E-06 | 8.200E-06 | 1.700E-05 |

| | | | | | | | | | | |
|---|---|---|---|---|---|---|---|---|---|---|
| 11985 | 600 | A600C | A | 8.5e-06 | 5.1e-06 | 1.2e-05 | 8.533E-06 | 3.450E-06 | 5.100E-06 | 1.200E-05 |
| 11986 | 600 | A600E | A | 7.3e-06 | 1.3e-05 | 1.3e-05 | 1.110E-05 | 3.291E-06 | 7.300E-06 | 1.300E-05 |
| 11987 | 600 | A600Q | A | 3.8e-05 | 1.7e-05 | 2.1e-05 | 2.533E-05 | 1.115E-05 | 1.700E-05 | 3.800E-05 |
| 11988 | 600 | A600G | A | 1.2e-05 | 2.9e-05 | 1.5e-05 | 1.867E-05 | 9.074E-06 | 1.200E-05 | 2.900E-05 |
| 11989 | 600 | A600H | A | 1e-05 | 9.1e-06 | 2.7e-05 | 1.537E-05 | 1.008E-05 | 9.100E-06 | 2.700E-05 |
| 11990 | 600 | A600I | A | 5e-06 | 9e-06 | 4.9e-06 | 6.300E-06 | 2.339E-06 | 4.900E-06 | 9.000E-06 |
| 11991 | 600 | A600L | A | 7.9e-06 | 9.2e-06 | 4.7e-06 | 7.267E-06 | 2.316E-06 | 4.700E-06 | 9.200E-06 |
| 11992 | 600 | A600K | A | 2.5e-05 | 1e-05 | 1.4e-05 | 1.633E-05 | 7.767E-06 | 1.000E-05 | 2.500E-05 |
| 11993 | 600 | A600M | A | 9.7e-06 | 9.1e-06 | 1.2e-05 | 1.027E-05 | 1.531E-06 | 9.100E-06 | 1.200E-05 |
| 11994 | 600 | A600F | A | 4.7e-06 | 5.5e-06 | 8.5e-06 | 6.233E-06 | 2.003E-06 | 4.700E-06 | 8.500E-06 |
| 11995 | 600 | A600P | A | 6.9e-06 | 5e-06 | 3.2e-06 | 5.033E-06 | 1.850E-06 | 3.200E-06 | 6.900E-06 |
| 11996 | 600 | A600S | A | 4.7e-05 | 2.7e-05 | 3.8e-05 | 3.733E-05 | 1.002E-05 | 2.700E-05 | 4.700E-05 |
| 11997 | 600 | A600T | A | 2.8e-05 | 1.8e-05 | 2.2e-05 | 2.267E-05 | 5.033E-06 | 1.800E-05 | 2.800E-05 |
| 11998 | 600 | A600W | A | 6.5e-06 | 1.3e-05 | 7.8e-06 | 9.100E-06 | 3.439E-06 | 6.500E-06 | 1.300E-05 |
| 11999 | 600 | A600Y | A | 1.1e-05 | 8.2e-06 | 1.6e-05 | 1.173E-05 | 3.951E-06 | 8.200E-06 | 1.600E-05 |
| 12000 | 600 | A600V | A | 1.2e-05 | 6.7e-06 | 8.3e-06 | 9.000E-06 | 2.718E-06 | 6.700E-06 | 1.200E-05 |
| 12001 | 601 | C601A | A | 9.4e-06 | 1.1e-05 | 7.4e-06 | 9.267E-06 | 1.804E-06 | 7.400E-06 | 1.100E-05 |
| 12002 | 601 | C601R | A | 3.9e-06 | 9.3e-06 | 4.4e-06 | 5.867E-06 | 2.984E-06 | 3.900E-06 | 9.300E-06 |
| 12003 | 601 | C601N | A | 6.7e-06 | 6.1e-06 | 9.4e-06 | 7.400E-06 | 1.758E-06 | 6.100E-06 | 9.400E-06 |
| 12004 | 601 | C601D | A | 9.1e-06 | 1.3e-05 | 6.2e-06 | 9.433E-06 | 3.412E-06 | 6.200E-06 | 1.300E-05 |
| 12005 | 601 | C601C | A | 5.1e-06 | 4.4e-06 | 8.8e-06 | 6.100E-06 | 2.364E-06 | 4.400E-06 | 8.800E-06 |
| 12006 | 601 | C601E | A | 7.6e-06 | 5.6e-06 | 8.8e-06 | 7.333E-06 | 1.617E-06 | 5.600E-06 | 8.800E-06 |
| 12007 | 601 | C601Q | A | 8e-06 | 5.1e-06 | 3.4e-06 | 5.500E-06 | 2.326E-06 | 3.400E-06 | 8.000E-06 |
| 12008 | 601 | C601G | A | 8.9e-06 | 7.3e-06 | 7.1e-06 | 7.767E-06 | 9.866E-07 | 7.100E-06 | 8.900E-06 |
| 12009 | 601 | C601H | A | 5.1e-06 | 4.6e-06 | 6.4e-06 | 5.367E-06 | 9.292E-07 | 4.600E-06 | 6.400E-06 |
| 12010 | 601 | C601I | A | 7.3e-06 | 1.4e-05 | 1.2e-05 | 1.110E-05 | 3.439E-06 | 7.300E-06 | 1.400E-05 |
| 12011 | 601 | C601L | A | 1e-05 | 9.6e-06 | 9.8e-06 | 9.800E-06 | 2.000E-07 | 9.600E-06 | 1.000E-05 |
| 12012 | 601 | C601K | A | 8.3e-06 | 1.3e-05 | 7.2e-06 | 9.500E-06 | 3.081E-06 | 7.200E-06 | 1.300E-05 |
| 12013 | 601 | C601M | A | 9.7e-06 | 5.6e-06 | 8.7e-06 | 8.000E-06 | 2.138E-06 | 5.600E-06 | 9.700E-06 |
| 12014 | 601 | C601F | A | 6.5e-06 | 6.6e-06 | 9e-06 | 7.367E-06 | 1.415E-06 | 6.500E-06 | 9.000E-06 |
| 12015 | 601 | C601P | A | 6.1e-06 | 1.3e-05 | 7.1e-06 | 8.733E-06 | 3.729E-06 | 6.100E-06 | 1.300E-05 |
| 12016 | 601 | C601S | A | 8.2e-06 | 6.8e-06 | 7.2e-06 | 7.400E-06 | 7.211E-07 | 6.800E-06 | 8.200E-06 |
| 12017 | 601 | C601T | A | 1e-05 | 1e-05 | 9.3e-06 | 9.767E-06 | 4.041E-07 | 9.300E-06 | 1.000E-05 |
| 12018 | 601 | C601W | A | 4.4e-06 | 8.1e-06 | 6.6e-06 | 6.367E-06 | 1.861E-06 | 4.400E-06 | 8.100E-06 |
| 12019 | 601 | C601Y | A | 5e-06 | 5.5e-06 | 5e-06 | 5.167E-06 | 2.887E-07 | 5.000E-06 | 5.500E-06 |
| 12020 | 601 | C601V | A | 6.7e-06 | 1.3e-05 | 5.5e-06 | 8.400E-06 | 4.029E-06 | 5.500E-06 | 1.300E-05 |
| 12021 | 602 | Q602A | A | 4.2e-06 | 3.1e-06 | 2.4e-06 | 3.233E-06 | 9.074E-07 | 2.400E-06 | 4.200E-06 |
| 12022 | 602 | Q602R | A | 2e-06 | 3.7e-06 | 3.5e-06 | 3.067E-06 | 9.292E-07 | 2.000E-06 | 3.700E-06 |
| 12023 | 602 | Q602N | A | 8.1e-06 | 7.1e-06 | 6.9e-06 | 7.367E-06 | 6.429E-07 | 6.900E-06 | 8.100E-06 |
| 12024 | 602 | Q602D | A | 3.9e-06 | 3.3e-06 | 2.4e-06 | 3.200E-06 | 7.550E-07 | 2.400E-06 | 3.900E-06 |
| 12025 | 602 | Q602C | A | 1.9e-06 | 2.1e-06 | 4e-06 | 2.667E-06 | 1.159E-06 | 1.900E-06 | 4.000E-06 |
| 12026 | 602 | Q602E | A | 4.3e-06 | 3.8e-06 | 4.6e-06 | 4.233E-06 | 4.041E-07 | 3.800E-06 | 4.600E-06 |
| 12027 | 602 | Q602Q | A | 1e-05 | 6.1e-06 | 1.5e-05 | 1.037E-05 | 4.461E-06 | 6.100E-06 | 1.500E-05 |
| 12028 | 602 | Q602G | A | 4.2e-06 | 2.6e-06 | 4.6e-06 | 3.800E-06 | 1.058E-06 | 2.600E-06 | 4.600E-06 |
| 12029 | 602 | Q602H | A | 4e-06 | 3.1e-06 | 5.2e-06 | 4.100E-06 | 1.054E-06 | 3.100E-06 | 5.200E-06 |
| 12030 | 602 | Q602I | A | 2.9e-06 | 2.1e-06 | 1.9e-06 | 2.300E-06 | 5.292E-07 | 1.900E-06 | 2.900E-06 |
| 12031 | 602 | Q602L | A | 4.8e-06 | 3.1e-06 | 3.1e-06 | 3.667E-06 | 9.815E-07 | 3.100E-06 | 4.800E-06 |
| 12032 | 602 | Q602K | A | 2.3e-06 | 3.3e-06 | 4.7e-06 | 3.433E-06 | 1.206E-06 | 2.300E-06 | 4.700E-06 |
| 12033 | 602 | Q602M | A | 4.4e-06 | 2.4e-06 | 2.9e-06 | 3.233E-06 | 1.041E-06 | 2.400E-06 | 4.400E-06 |
| 12034 | 602 | Q602F | A | 1.9e-06 | 9.6e-07 | 2e-06 | 1.620E-06 | 5.738E-07 | 9.600E-07 | 2.000E-06 |
| 12035 | 602 | Q602P | A | 4.9e-06 | 2.6e-06 | 3.2e-06 | 3.567E-06 | 1.193E-06 | 2.600E-06 | 4.900E-06 |
| 12036 | 602 | Q602S | A | 4.8e-06 | 3.9e-06 | 3.7e-06 | 4.133E-06 | 5.859E-07 | 3.700E-06 | 4.800E-06 |
| 12037 | 602 | Q602T | A | 5.2e-06 | 3.7e-06 | 2.7e-06 | 3.867E-06 | 1.258E-06 | 2.700E-06 | 5.200E-06 |
| 12038 | 602 | Q602W | A | 2.3e-06 | 3.1e-06 | 1.1e-06 | 2.167E-06 | 1.007E-06 | 1.100E-06 | 3.100E-06 |
| 12039 | 602 | Q602Y | A | 1.5e-06 | 2.3e-06 | 1.4e-06 | 1.733E-06 | 4.933E-07 | 1.400E-06 | 2.300E-06 |
| 12040 | 602 | Q602V | A | 2e-06 | 3.4e-06 | 2.8e-06 | 2.733E-06 | 7.024E-07 | 2.000E-06 | 3.400E-06 |
| 12041 | 603 | P603A | A | 1.3e-05 | 1.3e-05 | 6.5e-06 | 1.083E-05 | 3.753E-06 | 6.500E-06 | 1.300E-05 |
| 12042 | 603 | P603R | A | 1.1e-05 | 8.1e-06 | 9.4e-06 | 9.500E-06 | 1.453E-06 | 8.100E-06 | 1.100E-05 |
| 12043 | 603 | P603N | A | 6.8e-06 | 4.6e-06 | 8.4e-06 | 6.600E-06 | 1.908E-06 | 4.600E-06 | 8.400E-06 |
| 12044 | 603 | P603D | A | 8.3e-06 | 1.1e-05 | 9.1e-06 | 9.467E-06 | 1.387E-06 | 8.300E-06 | 1.100E-05 |
| 12045 | 603 | P603C | A | 5.6e-06 | 5.7e-06 | 4.7e-06 | 5.333E-06 | 5.508E-07 | 4.700E-06 | 5.700E-06 |
| 12046 | 603 | P603E | A | 6.6e-06 | 9.5e-06 | 6e-06 | 7.367E-06 | 1.872E-06 | 6.000E-06 | 9.500E-06 |
| 12047 | 603 | P603Q | A | 8.4e-06 | 5.4e-06 | 1.2e-05 | 8.600E-06 | 3.305E-06 | 5.400E-06 | 1.200E-05 |
| 12048 | 603 | P603G | A | 1.1e-05 | 1.5e-05 | 1e-05 | 1.200E-05 | 2.646E-06 | 1.000E-05 | 1.500E-05 |
| 12049 | 603 | P603H | A | 1.1e-05 | 1e-05 | 1.6e-05 | 1.233E-05 | 3.215E-06 | 1.000E-05 | 1.600E-05 |
| 12050 | 603 | P603I | A | 6.9e-06 | 6.9e-06 | 9.5e-06 | 7.767E-06 | 1.501E-06 | 6.900E-06 | 9.500E-06 |
| 12051 | 603 | P603L | A | 9.9e-06 | 6.5e-06 | 1.3e-05 | 9.800E-06 | 3.251E-06 | 6.500E-06 | 1.300E-05 |
| 12052 | 603 | P603K | A | 1.2e-05 | 6.6e-06 | 9.1e-06 | 9.233E-06 | 2.702E-06 | 6.600E-06 | 1.200E-05 |
| 12053 | 603 | P603M | A | 3.5e-06 | 1.2e-05 | 4.4e-06 | 6.633E-06 | 4.669E-06 | 3.500E-06 | 1.200E-05 |
| 12054 | 603 | P603F | A | 7.6e-06 | 6e-06 | 1.4e-05 | 9.200E-06 | 4.233E-06 | 6.000E-06 | 1.400E-05 |
| 12055 | 603 | P603P | A | 3.3e-06 | 4.6e-06 | 5e-06 | 4.300E-06 | 8.888E-07 | 3.300E-06 | 5.000E-06 |
| 12056 | 603 | P603S | A | 8.8e-06 | 1.1e-05 | 8.1e-06 | 9.300E-06 | 1.513E-06 | 8.100E-06 | 1.100E-05 |
| 12057 | 603 | P603T | A | 8e-06 | 6.8e-06 | 6.4e-06 | 7.067E-06 | 8.327E-07 | 6.400E-06 | 8.000E-06 |
| 12058 | 603 | P603W | A | 6.5e-06 | 6.2e-06 | 1e-05 | 7.567E-06 | 2.113E-06 | 6.200E-06 | 1.000E-05 |
| 12059 | 603 | P603Y | A | 4.3e-06 | 5.2e-06 | 1e-05 | 6.500E-06 | 3.064E-06 | 4.300E-06 | 1.000E-05 |
| 12060 | 603 | P603V | A | 6.3e-06 | 8.2e-06 | 6.7e-06 | 7.067E-06 | 1.002E-06 | 6.300E-06 | 8.200E-06 |
| 12061 | 604 | C604A | A | 1.5e-05 | 4.6e-06 | 5.7e-06 | 8.433E-06 | 5.713E-06 | 4.600E-06 | 1.500E-05 |
| 12062 | 604 | C604R | A | 1.1e-05 | 7.1e-06 | 7.4e-06 | 8.500E-06 | 2.170E-06 | 7.100E-06 | 1.100E-05 |
| 12063 | 604 | C604N | A | 6e-06 | 6.3e-06 | 6e-06 | 6.100E-06 | 1.732E-07 | 6.000E-06 | 6.300E-06 |
| 12064 | 604 | C604D | A | 6.2e-06 | 6.4e-06 | 6.1e-06 | 6.233E-06 | 1.528E-07 | 6.100E-06 | 6.400E-06 |
| 12065 | 604 | C604C | A | 5.2e-06 | 4.9e-06 | 1.4e-05 | 8.033E-06 | 5.169E-06 | 4.900E-06 | 1.400E-05 |
| 12066 | 604 | C604E | A | 1.2e-05 | 1.1e-05 | 1.1e-05 | 1.133E-05 | 5.774E-07 | 1.100E-05 | 1.200E-05 |
| 12067 | 604 | C604Q | A | 4.3e-06 | 4.2e-06 | 5.1e-06 | 4.533E-06 | 4.933E-07 | 4.200E-06 | 5.100E-06 |
| 12068 | 604 | C604G | A | 9.8e-06 | 8.1e-06 | 6.8e-06 | 8.233E-06 | 1.504E-06 | 6.800E-06 | 9.800E-06 |
| 12069 | 604 | C604H | A | 7.9e-06 | 8.1e-06 | 5.9e-06 | 7.300E-06 | 1.217E-06 | 5.900E-06 | 8.100E-06 |

| | | | | | | | | | | |
|---|---|---|---|---|---|---|---|---|---|---|
| 12070 | 604 | C604I | A | 7.5e-06 | 5.9e-06 | 8.1e-06 | 7.167E-06 | 1.137E-06 | 5.900E-06 | 8.100E-06 |
| 12071 | 604 | C604L | A | 1.2e-05 | 1.1e-05 | 8.4e-06 | 1.047E-05 | 1.858E-06 | 8.400E-06 | 1.200E-05 |
| 12072 | 604 | C604K | A | 6.8e-06 | 8.4e-06 | 1.1e-05 | 8.733E-06 | 2.120E-06 | 6.800E-06 | 1.100E-05 |
| 12073 | 604 | C604M | A | 6.6e-06 | 7.2e-06 | 4.7e-06 | 6.167E-06 | 1.305E-06 | 4.700E-06 | 7.200E-06 |
| 12074 | 604 | C604F | A | 9.9e-06 | 5.3e-06 | 7.5e-06 | 7.567E-06 | 2.301E-06 | 5.300E-06 | 9.900E-06 |
| 12075 | 604 | C604P | A | 8.3e-06 | 1e-05 | 7.2e-06 | 8.500E-06 | 1.411E-06 | 7.200E-06 | 1.000E-05 |
| 12076 | 604 | C604S | A | 4.6e-06 | 4.7e-06 | 8.2e-06 | 5.833E-06 | 2.050E-06 | 4.600E-06 | 8.200E-06 |
| 12077 | 604 | C604T | A | 9.4e-06 | 7e-06 | 6.4e-06 | 7.600E-06 | 1.587E-06 | 6.400E-06 | 9.400E-06 |
| 12078 | 604 | C604W | A | 1.4e-05 | 1e-05 | 1.1e-05 | 1.167E-05 | 2.082E-06 | 1.000E-05 | 1.400E-05 |
| 12079 | 604 | C604Y | A | 1e-05 | 4.3e-06 | 7.3e-06 | 7.200E-06 | 2.851E-06 | 4.300E-06 | 1.000E-05 |
| 12080 | 604 | C604V | A | 8.4e-06 | 4.9e-06 | 8.1e-06 | 7.133E-06 | 1.940E-06 | 4.900E-06 | 8.400E-06 |
| 12081 | 605 | P605A | A | 9.5e-06 | 1e-05 | 4.2e-06 | 7.900E-06 | 3.214E-06 | 4.200E-06 | 1.000E-05 |
| 12082 | 605 | P605R | A | 5.7e-06 | 7.7e-06 | 9.1e-06 | 7.500E-06 | 1.709E-06 | 5.700E-06 | 9.100E-06 |
| 12083 | 605 | P605N | A | 6.6e-06 | 9.9e-06 | 5.7e-06 | 7.400E-06 | 2.211E-06 | 5.700E-06 | 9.900E-06 |
| 12084 | 605 | P605D | A | 8.1e-06 | 1e-05 | 5.4e-06 | 7.833E-06 | 2.312E-06 | 5.400E-06 | 1.000E-05 |
| 12085 | 605 | P605C | A | 5.9e-06 | 9.5e-06 | 7.5e-06 | 7.633E-06 | 1.804E-06 | 5.900E-06 | 9.500E-06 |
| 12086 | 605 | P605E | A | 9.2e-06 | 1.5e-05 | 9.9e-06 | 1.137E-05 | 3.166E-06 | 9.200E-06 | 1.500E-05 |
| 12087 | 605 | P605Q | A | 5.6e-06 | 6.9e-06 | 4.7e-06 | 5.733E-06 | 1.106E-06 | 4.700E-06 | 6.900E-06 |
| 12088 | 605 | P605G | A | 1.2e-05 | 6.9e-06 | 8.3e-06 | 9.067E-06 | 2.635E-06 | 6.900E-06 | 1.200E-05 |
| 12089 | 605 | P605H | A | 6.3e-06 | 6.8e-06 | 4e-06 | 5.700E-06 | 1.493E-06 | 4.000E-06 | 6.800E-06 |
| 12090 | 605 | P605I | A | 6.8e-06 | 1.3e-05 | 6.6e-06 | 8.800E-06 | 3.639E-06 | 6.600E-06 | 1.300E-05 |
| 12091 | 605 | P605L | A | 8e-06 | 1.1e-05 | 6.3e-06 | 8.433E-06 | 2.380E-06 | 6.300E-06 | 1.100E-05 |
| 12092 | 605 | P605K | A | 9.9e-06 | 1.1e-05 | 6e-06 | 8.967E-06 | 2.627E-06 | 6.000E-06 | 1.100E-05 |
| 12093 | 605 | P605M | A | 5.8e-06 | 7.1e-06 | 5.7e-06 | 6.200E-06 | 7.810E-07 | 5.700E-06 | 7.100E-06 |
| 12094 | 605 | P605F | A | 7.1e-06 | 4e-06 | 5.2e-06 | 5.433E-06 | 1.563E-06 | 4.000E-06 | 7.100E-06 |
| 12095 | 605 | P605P | A | 6.6e-06 | 6.3e-06 | 9.9e-06 | 7.600E-06 | 1.997E-06 | 6.300E-06 | 9.900E-06 |
| 12096 | 605 | P605S | A | 4.4e-06 | 6.5e-06 | 5.1e-06 | 5.333E-06 | 1.069E-06 | 4.400E-06 | 6.500E-06 |
| 12097 | 605 | P605T | A | 4.3e-06 | 6.7e-06 | 3.5e-06 | 4.833E-06 | 1.665E-06 | 3.500E-06 | 6.700E-06 |
| 12098 | 605 | P605W | A | 9e-06 | 6.1e-06 | 1.1e-05 | 8.700E-06 | 2.464E-06 | 6.100E-06 | 1.100E-05 |
| 12099 | 605 | P605Y | A | 6.2e-06 | 7.6e-06 | 5.5e-06 | 6.433E-06 | 1.069E-06 | 5.500E-06 | 7.600E-06 |
| 12100 | 605 | P605V | A | 6e-06 | 1.3e-05 | 8.7e-06 | 9.233E-06 | 3.530E-06 | 6.000E-06 | 1.300E-05 |
| 12101 | 606 | I606A | A | 8.8e-06 | 4.3e-06 | 9.4e-06 | 7.500E-06 | 2.787E-06 | 4.300E-06 | 9.400E-06 |
| 12102 | 606 | I606R | A | 6.1e-06 | 1.2e-05 | 7.2e-06 | 8.433E-06 | 3.137E-06 | 6.100E-06 | 1.200E-05 |
| 12103 | 606 | I606N | A | 3.2e-06 | 3.3e-06 | 4.5e-06 | 3.667E-06 | 7.234E-07 | 3.200E-06 | 4.500E-06 |
| 12104 | 606 | I606D | A | 6.5e-06 | 7.1e-06 | 6.2e-06 | 6.600E-06 | 4.583E-07 | 6.200E-06 | 7.100E-06 |
| 12105 | 606 | I606C | A | 6.6e-06 | 5.4e-06 | 8e-06 | 6.667E-06 | 1.301E-06 | 5.400E-06 | 8.000E-06 |
| 12106 | 606 | I606E | A | 6.5e-06 | 5.4e-06 | 7.1e-06 | 6.333E-06 | 8.622E-07 | 5.400E-06 | 7.100E-06 |
| 12107 | 606 | I606Q | A | 9.3e-06 | 5.8e-06 | 4.9e-06 | 6.667E-06 | 2.325E-06 | 4.900E-06 | 9.300E-06 |
| 12108 | 606 | I606G | A | 6.4e-06 | 8e-06 | 7.1e-06 | 7.167E-06 | 8.021E-07 | 6.400E-06 | 8.000E-06 |
| 12109 | 606 | I606H | A | 7.1e-06 | 5.4e-06 | 5.9e-06 | 6.133E-06 | 8.737E-07 | 5.400E-06 | 7.100E-06 |
| 12110 | 606 | I606I | A | 4.9e-06 | 1.8e-05 | 8.4e-06 | 1.043E-05 | 6.783E-06 | 4.900E-06 | 1.800E-05 |
| 12111 | 606 | I606L | A | 6e-06 | 1.1e-05 | 3.6e-06 | 6.867E-06 | 3.775E-06 | 3.600E-06 | 1.100E-05 |
| 12112 | 606 | I606K | A | 8.3e-06 | 8.2e-06 | 7.3e-06 | 7.933E-06 | 5.508E-07 | 7.300E-06 | 8.300E-06 |
| 12113 | 606 | I606M | A | 7.4e-06 | 1.3e-05 | 8.6e-06 | 9.667E-06 | 2.948E-06 | 7.400E-06 | 1.300E-05 |
| 12114 | 606 | I606F | A | 8.6e-06 | 9.9e-06 | 5e-06 | 7.833E-06 | 2.538E-06 | 5.000E-06 | 9.900E-06 |
| 12115 | 606 | I606P | A | 1e-05 | 5.5e-06 | 6.4e-06 | 7.300E-06 | 2.381E-06 | 5.500E-06 | 1.000E-05 |
| 12116 | 606 | I606S | A | 5.1e-06 | 1.5e-05 | 5.2e-06 | 8.433E-06 | 5.687E-06 | 5.100E-06 | 1.500E-05 |
| 12117 | 606 | I606T | A | 8.6e-06 | 6.6e-06 | 5e-06 | 6.733E-06 | 1.804E-06 | 5.000E-06 | 8.600E-06 |
| 12118 | 606 | I606W | A | 9.9e-06 | 8.2e-06 | 4.6e-06 | 7.567E-06 | 2.706E-06 | 4.600E-06 | 9.900E-06 |
| 12119 | 606 | I606Y | A | 5e-06 | 8.6e-06 | 1.3e-05 | 8.867E-06 | 4.007E-06 | 5.000E-06 | 1.300E-05 |
| 12120 | 606 | I606V | A | 8e-06 | 6.2e-06 | 6.7e-06 | 6.967E-06 | 9.292E-07 | 6.200E-06 | 8.000E-06 |
| 12121 | 607 | N607A | A | 6.8e-06 | 7.6e-06 | 1e-05 | 8.133E-06 | 1.665E-06 | 6.800E-06 | 1.000E-05 |
| 12122 | 607 | N607R | A | 8.4e-06 | 6.9e-06 | 8.1e-06 | 7.800E-06 | 7.937E-07 | 6.900E-06 | 8.400E-06 |
| 12123 | 607 | N607N | A | 1.3e-05 | 5.6e-06 | 7.2e-06 | 8.600E-06 | 3.894E-06 | 5.600E-06 | 1.300E-05 |
| 12124 | 607 | N607D | A | 9.9e-06 | 8.5e-06 | 6.7e-06 | 8.367E-06 | 1.604E-06 | 6.700E-06 | 9.900E-06 |
| 12125 | 607 | N607C | A | 1.4e-05 | 6.1e-06 | 8.7e-06 | 9.600E-06 | 4.026E-06 | 6.100E-06 | 1.400E-05 |
| 12126 | 607 | N607E | A | 5.5e-06 | 5.5e-06 | 7.3e-06 | 6.100E-06 | 1.039E-06 | 5.500E-06 | 7.300E-06 |
| 12127 | 607 | N607Q | A | 1.4e-05 | 7.9e-06 | 6e-06 | 9.300E-06 | 4.180E-06 | 6.000E-06 | 1.400E-05 |
| 12128 | 607 | N607G | A | 9.7e-06 | 1.2e-05 | 1.3e-05 | 1.157E-05 | 1.692E-06 | 9.700E-06 | 1.300E-05 |
| 12129 | 607 | N607H | A | 4.6e-06 | 7.6e-06 | 1.3e-05 | 8.400E-06 | 4.257E-06 | 4.600E-06 | 1.300E-05 |
| 12130 | 607 | N607I | A | 1.6e-05 | 9.4e-06 | 7.2e-06 | 1.087E-05 | 4.580E-06 | 7.200E-06 | 1.600E-05 |
| 12131 | 607 | N607L | A | 6.4e-06 | 1.1e-05 | 6.9e-06 | 8.100E-06 | 2.524E-06 | 6.400E-06 | 1.100E-05 |
| 12132 | 607 | N607K | A | 1.2e-05 | 6.5e-06 | 1.1e-05 | 9.833E-06 | 2.930E-06 | 6.500E-06 | 1.200E-05 |
| 12133 | 607 | N607M | A | 9.1e-06 | 1e-05 | 9.2e-06 | 9.433E-06 | 4.933E-07 | 9.100E-06 | 1.000E-05 |
| 12134 | 607 | N607F | A | 9.1e-06 | 6.9e-06 | 5.3e-06 | 7.100E-06 | 1.908E-06 | 5.300E-06 | 9.100E-06 |
| 12135 | 607 | N607P | A | 7e-06 | 5.2e-06 | 5.2e-06 | 5.800E-06 | 1.039E-06 | 5.200E-06 | 7.000E-06 |
| 12136 | 607 | N607S | A | 6.7e-06 | 9.8e-06 | 1.1e-05 | 9.167E-06 | 2.219E-06 | 6.700E-06 | 1.100E-05 |
| 12137 | 607 | N607T | A | 7.1e-06 | 9.9e-06 | 1.1e-05 | 9.333E-06 | 2.011E-06 | 7.100E-06 | 1.100E-05 |
| 12138 | 607 | N607W | A | 2.8e-06 | 6.5e-06 | 7.3e-06 | 5.533E-06 | 2.401E-06 | 2.800E-06 | 7.300E-06 |
| 12139 | 607 | N607Y | A | 8.8e-06 | 1.1e-05 | 8e-06 | 9.267E-06 | 1.553E-06 | 8.000E-06 | 1.100E-05 |
| 12140 | 607 | N607V | A | 1.4e-05 | 7.5e-06 | 1.3e-05 | 1.150E-05 | 3.500E-06 | 7.500E-06 | 1.400E-05 |
| 12141 | 608 | C608A | A | 9.7e-06 | 6.7e-06 | 7.2e-06 | 7.867E-06 | 1.607E-06 | 6.700E-06 | 9.700E-06 |
| 12142 | 608 | C608R | A | 6.5e-06 | 9.9e-06 | 9.6e-06 | 8.667E-06 | 1.882E-06 | 6.500E-06 | 9.900E-06 |
| 12143 | 608 | C608N | A | 5.6e-06 | 7.2e-06 | 7.5e-06 | 6.767E-06 | 1.021E-06 | 5.600E-06 | 7.500E-06 |
| 12144 | 608 | C608D | A | 9.7e-06 | 7.5e-06 | 7.6e-06 | 8.267E-06 | 1.242E-06 | 7.500E-06 | 9.700E-06 |
| 12145 | 608 | C608C | A | 7.2e-06 | 5.6e-06 | 1e-05 | 7.600E-06 | 2.227E-06 | 5.600E-06 | 1.000E-05 |
| 12146 | 608 | C608E | A | 7.7e-06 | 1e-05 | 9.4e-06 | 9.033E-06 | 1.193E-06 | 7.700E-06 | 1.000E-05 |
| 12147 | 608 | C608Q | A | 1.5e-05 | 1.4e-05 | 9.3e-06 | 1.277E-05 | 3.044E-06 | 9.300E-06 | 1.500E-05 |
| 12148 | 608 | C608G | A | 8.5e-06 | 1.2e-05 | 5.8e-06 | 8.767E-06 | 3.109E-06 | 5.800E-06 | 1.200E-05 |
| 12149 | 608 | C608H | A | 6.5e-06 | 5.2e-06 | 9.8e-06 | 7.167E-06 | 2.371E-06 | 5.200E-06 | 9.800E-06 |
| 12150 | 608 | C608I | A | 8.2e-06 | 8.7e-06 | 7.6e-06 | 8.167E-06 | 5.508E-07 | 7.600E-06 | 8.700E-06 |
| 12151 | 608 | C608L | A | 1e-05 | 1.1e-05 | 1.2e-05 | 1.100E-05 | 1.000E-06 | 1.000E-05 | 1.200E-05 |
| 12152 | 608 | C608K | A | 8e-06 | 9.9e-06 | 1e-05 | 9.300E-06 | 1.127E-06 | 8.000E-06 | 1.000E-05 |
| 12153 | 608 | C608M | A | 1e-05 | 7.1e-06 | 8.8e-06 | 8.633E-06 | 1.457E-06 | 7.100E-06 | 1.000E-05 |
| 12154 | 608 | C608F | A | 8.9e-06 | 7.5e-06 | 9.7e-06 | 8.700E-06 | 1.114E-06 | 7.500E-06 | 9.700E-06 |

| | | | | | | | | | | |
|---|---|---|---|---|---|---|---|---|---|---|
| 12155 | 608 | C608P | A | 3.7e-06 | 6.3e-06 | 9.8e-06 | 6.600E-06 | 3.061E-06 | 3.700E-06 | 9.800E-06 |
| 12156 | 608 | C608S | A | 9.1e-06 | 8.2e-06 | 5.9e-06 | 7.733E-06 | 1.650E-06 | 5.900E-06 | 9.100E-06 |
| 12157 | 608 | C608T | A | 6.8e-06 | 1.5e-05 | 1.3e-05 | 1.160E-05 | 4.276E-06 | 6.800E-06 | 1.500E-05 |
| 12158 | 608 | C608W | A | 1.3e-05 | 5e-06 | 6.1e-06 | 8.033E-06 | 4.336E-06 | 5.000E-06 | 1.300E-05 |
| 12159 | 608 | C608Y | A | 9.8e-06 | 9.3e-06 | 8.6e-06 | 9.233E-06 | 6.028E-07 | 8.600E-06 | 9.800E-06 |
| 12160 | 608 | C608V | A | 1e-05 | 5e-06 | 9.6e-06 | 8.200E-06 | 2.778E-06 | 5.000E-06 | 1.000E-05 |
| 12161 | 609 | T609A | A | 5e-06 | 6.1e-06 | 7.1e-06 | 6.067E-06 | 1.050E-06 | 5.000E-06 | 7.100E-06 |
| 12162 | 609 | T609R | A | 9.3e-06 | 1.3e-05 | 6.1e-06 | 9.467E-06 | 3.453E-06 | 6.100E-06 | 1.300E-05 |
| 12163 | 609 | T609N | A | 9.4e-06 | 1.1e-05 | 7.2e-06 | 9.200E-06 | 1.908E-06 | 7.200E-06 | 1.100E-05 |
| 12164 | 609 | T609D | A | 3.6e-06 | 5.3e-06 | 3.3e-06 | 4.067E-06 | 1.079E-06 | 3.300E-06 | 5.300E-06 |
| 12165 | 609 | T609C | A | 7.1e-06 | 1.2e-05 | 5e-06 | 8.033E-06 | 3.592E-06 | 5.000E-06 | 1.200E-05 |
| 12166 | 609 | T609E | A | 1.7e-05 | 8.5e-06 | 9.3e-06 | 1.160E-05 | 4.694E-06 | 8.500E-06 | 1.700E-05 |
| 12167 | 609 | T609Q | A | 1e-05 | 1.2e-05 | 9e-06 | 1.033E-05 | 1.528E-06 | 9.000E-06 | 1.200E-05 |
| 12168 | 609 | T609G | A | 1.2e-05 | 8.5e-06 | 1e-05 | 1.017E-05 | 1.756E-06 | 8.500E-06 | 1.200E-05 |
| 12169 | 609 | T609H | A | 1.2e-05 | 1.1e-05 | 1.1e-05 | 1.133E-05 | 5.774E-07 | 1.100E-05 | 1.200E-05 |
| 12170 | 609 | T609I | A | 1.5e-05 | 8.8e-06 | 1.4e-05 | 1.260E-05 | 3.329E-06 | 8.800E-06 | 1.500E-05 |
| 12171 | 609 | T609L | A | 4.8e-06 | 3.5e-06 | 7.8e-06 | 5.367E-06 | 2.205E-06 | 3.500E-06 | 7.800E-06 |
| 12172 | 609 | T609K | A | 1.1e-05 | 6.8e-06 | 9.5e-06 | 9.100E-06 | 2.128E-06 | 6.800E-06 | 1.100E-05 |
| 12173 | 609 | T609M | A | 6.5e-06 | 5.6e-06 | 7.1e-06 | 6.400E-06 | 7.550E-07 | 5.600E-06 | 7.100E-06 |
| 12174 | 609 | T609F | A | 7.1e-06 | 7.5e-06 | 9.5e-06 | 8.033E-06 | 1.286E-06 | 7.100E-06 | 9.500E-06 |
| 12175 | 609 | T609P | A | 9.4e-06 | 8.5e-06 | 8.3e-06 | 8.733E-06 | 5.859E-07 | 8.300E-06 | 9.400E-06 |
| 12176 | 609 | T609S | A | 1.3e-05 | 5.5e-06 | 4.7e-06 | 7.733E-06 | 4.579E-06 | 4.700E-06 | 1.300E-05 |
| 12177 | 609 | T609T | A | 8e-06 | 7e-06 | 1e-05 | 8.333E-06 | 1.528E-06 | 7.000E-06 | 1.000E-05 |
| 12178 | 609 | T609W | A | 9.5e-06 | 4e-06 | 7.4e-06 | 6.967E-06 | 2.775E-06 | 4.000E-06 | 9.500E-06 |
| 12179 | 609 | T609Y | A | 6e-06 | 1.1e-05 | 7.8e-06 | 8.267E-06 | 2.532E-06 | 6.000E-06 | 1.100E-05 |
| 12180 | 609 | T609V | A | 5.5e-06 | 7.3e-06 | 4.8e-06 | 5.867E-06 | 1.290E-06 | 4.800E-06 | 7.300E-06 |
| 12181 | 610 | H610A | A | 4.9e-06 | 1e-05 | 5.6e-06 | 6.833E-06 | 2.765E-06 | 4.900E-06 | 1.000E-05 |
| 12182 | 610 | H610R | A | 5.3e-06 | 6.2e-06 | 6.6e-06 | 6.033E-06 | 6.658E-07 | 5.300E-06 | 6.600E-06 |
| 12183 | 610 | H610N | A | 8e-06 | 1e-05 | 5.2e-06 | 7.733E-06 | 2.411E-06 | 5.200E-06 | 1.000E-05 |
| 12184 | 610 | H610D | A | 1.9e-05 | 5.6e-06 | 8.2e-06 | 1.093E-05 | 7.106E-06 | 5.600E-06 | 1.900E-05 |
| 12185 | 610 | H610C | A | 8.6e-06 | 1e-05 | 7.2e-06 | 8.600E-06 | 1.400E-06 | 7.200E-06 | 1.000E-05 |
| 12186 | 610 | H610E | A | 6.3e-06 | 1.5e-05 | 1.4e-05 | 1.177E-05 | 4.761E-06 | 6.300E-06 | 1.500E-05 |
| 12187 | 610 | H610Q | A | 7.3e-06 | 9.1e-06 | 9.4e-06 | 8.600E-06 | 1.136E-06 | 7.300E-06 | 9.400E-06 |
| 12188 | 610 | H610G | A | 7.4e-06 | 6.1e-06 | 6.8e-06 | 6.767E-06 | 6.506E-07 | 6.100E-06 | 7.400E-06 |
| 12189 | 610 | H610H | A | 1e-05 | 9.5e-06 | 6.5e-06 | 8.667E-06 | 1.893E-06 | 6.500E-06 | 1.000E-05 |
| 12190 | 610 | H610I | A | 1.3e-05 | 1.3e-05 | 1e-05 | 1.200E-05 | 1.732E-06 | 1.000E-05 | 1.300E-05 |
| 12191 | 610 | H610L | A | 1.1e-05 | 4.6e-06 | 1.1e-05 | 8.867E-06 | 3.695E-06 | 4.600E-06 | 1.100E-05 |
| 12192 | 610 | H610K | A | 9.9e-06 | 7.4e-06 | 7.3e-06 | 8.200E-06 | 1.473E-06 | 7.300E-06 | 9.900E-06 |
| 12193 | 610 | H610M | A | 8.3e-06 | 9.7e-06 | 6.3e-06 | 8.100E-06 | 1.709E-06 | 6.300E-06 | 9.700E-06 |
| 12194 | 610 | H610F | A | 7.4e-06 | 9.1e-06 | 7.3e-06 | 7.933E-06 | 1.012E-06 | 7.300E-06 | 9.100E-06 |
| 12195 | 610 | H610P | A | 1.2e-05 | 5.4e-06 | 1e-05 | 9.133E-06 | 3.384E-06 | 5.400E-06 | 1.200E-05 |
| 12196 | 610 | H610S | A | 6.6e-06 | 8.2e-06 | 1e-05 | 8.267E-06 | 1.701E-06 | 6.600E-06 | 1.000E-05 |
| 12197 | 610 | H610T | A | 1.1e-05 | 8.2e-06 | 1.1e-05 | 1.007E-05 | 1.617E-06 | 8.200E-06 | 1.100E-05 |
| 12198 | 610 | H610W | A | 9.1e-06 | 4.4e-06 | 5.2e-06 | 6.233E-06 | 2.515E-06 | 4.400E-06 | 9.100E-06 |
| 12199 | 610 | H610Y | A | 9.7e-06 | 4.5e-06 | 5.4e-06 | 6.533E-06 | 2.779E-06 | 4.500E-06 | 9.700E-06 |
| 12200 | 610 | H610V | A | 1.3e-05 | 6.5e-06 | 8.9e-06 | 9.467E-06 | 3.287E-06 | 6.500E-06 | 1.300E-05 |
| 12201 | 611 | S611A | A | 9.7e-06 | 6.8e-06 | 1e-05 | 8.833E-06 | 1.767E-06 | 6.800E-06 | 1.000E-05 |
| 12202 | 611 | S611R | A | 4.8e-06 | 4.9e-06 | 1.1e-05 | 6.900E-06 | 3.551E-06 | 4.800E-06 | 1.100E-05 |
| 12203 | 611 | S611N | A | 4.8e-06 | 7e-06 | 9.4e-06 | 7.067E-06 | 2.301E-06 | 4.800E-06 | 9.400E-06 |
| 12204 | 611 | S611D | A | 6.5e-06 | 5.6e-06 | 8.3e-06 | 6.800E-06 | 1.375E-06 | 5.600E-06 | 8.300E-06 |
| 12205 | 611 | S611C | A | 1.1e-05 | 7.4e-06 | 1.2e-05 | 1.013E-05 | 2.419E-06 | 7.400E-06 | 1.200E-05 |
| 12206 | 611 | S611E | A | 8.8e-06 | 8.3e-06 | 9.1e-06 | 8.733E-06 | 4.041E-07 | 8.300E-06 | 9.100E-06 |
| 12207 | 611 | S611Q | A | 3.3e-06 | 6.3e-06 | 8.5e-06 | 6.033E-06 | 2.610E-06 | 3.300E-06 | 8.500E-06 |
| 12208 | 611 | S611G | A | 9.5e-06 | 1.6e-05 | 1e-05 | 1.183E-05 | 3.617E-06 | 9.500E-06 | 1.600E-05 |
| 12209 | 611 | S611H | A | 9.5e-06 | 6.8e-06 | 9.9e-06 | 8.733E-06 | 1.686E-06 | 6.800E-06 | 9.900E-06 |
| 12210 | 611 | S611I | A | 1.3e-05 | 1.4e-05 | 7.8e-06 | 1.160E-05 | 3.329E-06 | 7.800E-06 | 1.400E-05 |
| 12211 | 611 | S611L | A | 9.2e-06 | 2.2e-05 | 9.1e-06 | 1.343E-05 | 7.419E-06 | 9.100E-06 | 2.200E-05 |
| 12212 | 611 | S611K | A | 6.5e-06 | 9.6e-06 | 4.2e-06 | 6.767E-06 | 2.710E-06 | 4.200E-06 | 9.600E-06 |
| 12213 | 611 | S611M | A | 6e-06 | 6.1e-06 | 1.4e-05 | 8.700E-06 | 4.590E-06 | 6.000E-06 | 1.400E-05 |
| 12214 | 611 | S611F | A | 6.8e-06 | 5e-06 | 8.2e-06 | 6.667E-06 | 1.604E-06 | 5.000E-06 | 8.200E-06 |
| 12215 | 611 | S611P | A | 9.1e-06 | 1.1e-05 | 4.6e-06 | 8.233E-06 | 3.287E-06 | 4.600E-06 | 1.100E-05 |
| 12216 | 611 | S611S | A | 4.9e-06 | 5.8e-06 | 9.4e-06 | 6.700E-06 | 2.381E-06 | 4.900E-06 | 9.400E-06 |
| 12217 | 611 | S611T | A | 6.2e-06 | 1.2e-05 | 9.5e-06 | 9.233E-06 | 2.909E-06 | 6.200E-06 | 1.200E-05 |
| 12218 | 611 | S611W | A | 6.5e-06 | 4.3e-06 | 6e-06 | 5.600E-06 | 1.153E-06 | 4.300E-06 | 6.500E-06 |
| 12219 | 611 | S611Y | A | 8.5e-06 | 1.4e-05 | 1.5e-05 | 1.250E-05 | 3.500E-06 | 8.500E-06 | 1.500E-05 |
| 12220 | 611 | S611V | A | 6.2e-06 | 4.9e-06 | 7.2e-06 | 6.100E-06 | 1.153E-06 | 4.900E-06 | 7.200E-06 |
| 12221 | 612 | C612A | A | 5e-06 | 1e-05 | 1e-05 | 8.333E-06 | 2.887E-06 | 5.000E-06 | 1.000E-05 |
| 12222 | 612 | C612R | A | 8.9e-06 | 9.4e-06 | 6.1e-06 | 8.133E-06 | 1.779E-06 | 6.100E-06 | 9.400E-06 |
| 12223 | 612 | C612N | A | 4.6e-06 | 1.4e-05 | 9.4e-06 | 9.333E-06 | 4.700E-06 | 4.600E-06 | 1.400E-05 |
| 12224 | 612 | C612D | A | 7.6e-06 | 9.2e-06 | 8.8e-06 | 8.533E-06 | 8.327E-07 | 7.600E-06 | 9.200E-06 |
| 12225 | 612 | C612C | A | 1.2e-05 | 1.1e-05 | 1.2e-05 | 1.167E-05 | 5.774E-07 | 1.100E-05 | 1.200E-05 |
| 12226 | 612 | C612E | A | 1.2e-05 | 9.1e-06 | 6.6e-06 | 9.233E-06 | 2.702E-06 | 6.600E-06 | 1.200E-05 |
| 12227 | 612 | C612Q | A | 5e-06 | 9.2e-06 | 4.8e-06 | 6.333E-06 | 2.485E-06 | 4.800E-06 | 9.200E-06 |
| 12228 | 612 | C612G | A | 5.8e-06 | 7.8e-06 | 7.5e-06 | 7.033E-06 | 1.079E-06 | 5.800E-06 | 7.800E-06 |
| 12229 | 612 | C612H | A | 8.4e-06 | 7.5e-06 | 6.8e-06 | 7.567E-06 | 8.021E-07 | 6.800E-06 | 8.400E-06 |
| 12230 | 612 | C612I | A | 5.1e-06 | 6.8e-06 | 7.4e-06 | 6.433E-06 | 1.193E-06 | 5.100E-06 | 7.400E-06 |
| 12231 | 612 | C612L | A | 7.4e-06 | 1.2e-05 | 8.9e-06 | 9.433E-06 | 2.346E-06 | 7.400E-06 | 1.200E-05 |
| 12232 | 612 | C612K | A | 9.9e-06 | 5.8e-06 | 1.1e-05 | 8.900E-06 | 2.740E-06 | 5.800E-06 | 1.100E-05 |
| 12233 | 612 | C612M | A | 7.3e-06 | 5.3e-06 | 8.5e-06 | 7.033E-06 | 1.617E-06 | 5.300E-06 | 8.500E-06 |
| 12234 | 612 | C612F | A | 8.3e-06 | 9e-06 | 8.4e-06 | 8.567E-06 | 3.786E-07 | 8.300E-06 | 9.000E-06 |
| 12235 | 612 | C612P | A | 5.3e-06 | 9.2e-06 | 7.6e-06 | 7.367E-06 | 1.960E-06 | 5.300E-06 | 9.200E-06 |
| 12236 | 612 | C612S | A | 4.9e-06 | 5.5e-06 | 1e-05 | 6.800E-06 | 2.787E-06 | 4.900E-06 | 1.000E-05 |
| 12237 | 612 | C612T | A | 6e-06 | 6.3e-06 | 9.2e-06 | 7.167E-06 | 1.767E-06 | 6.000E-06 | 9.200E-06 |
| 12238 | 612 | C612W | A | 6.3e-06 | 7.7e-06 | 4.5e-06 | 6.167E-06 | 1.604E-06 | 4.500E-06 | 7.700E-06 |
| 12239 | 612 | C612Y | A | 8.1e-06 | 4.6e-06 | 1.3e-05 | 8.567E-06 | 4.219E-06 | 4.600E-06 | 1.300E-05 |

| | | | | | | | | | | |
|---|---|---|---|---|---|---|---|---|---|---|
| 12240 | 612 | C612V | A | 5.9e-06 | 4.9e-06 | 9.4e-06 | 6.733E-06 | 2.363E-06 | 4.900E-06 | 9.400E-06 |
| 12241 | 613 | V613A | A | 8.8e-06 | 6e-06 | 2.1e-05 | 1.193E-05 | 7.976E-06 | 6.000E-06 | 2.100E-05 |
| 12242 | 613 | V613R | A | 6.9e-06 | 6.8e-06 | 1e-05 | 7.900E-06 | 1.819E-06 | 6.800E-06 | 1.000E-05 |
| 12243 | 613 | V613N | A | 9.1e-06 | 5.5e-06 | 6.2e-06 | 6.933E-06 | 1.909E-06 | 5.500E-06 | 9.100E-06 |
| 12244 | 613 | V613D | A | 6.7e-06 | 8.5e-06 | 4.2e-06 | 6.467E-06 | 2.159E-06 | 4.200E-06 | 8.500E-06 |
| 12245 | 613 | V613C | A | 8e-06 | 1.1e-05 | 8.3e-06 | 9.100E-06 | 1.652E-06 | 8.000E-06 | 1.100E-05 |
| 12246 | 613 | V613E | A | 8.8e-06 | 6.6e-06 | 6.3e-06 | 7.233E-06 | 1.365E-06 | 6.300E-06 | 8.800E-06 |
| 12247 | 613 | V613Q | A | 4.5e-06 | 8.7e-06 | 9.7e-06 | 7.633E-06 | 2.759E-06 | 4.500E-06 | 9.700E-06 |
| 12248 | 613 | V613G | A | 5.5e-06 | 6.5e-06 | 8.1e-06 | 6.700E-06 | 1.311E-06 | 5.500E-06 | 8.100E-06 |
| 12249 | 613 | V613H | A | 9.1e-06 | 6.9e-06 | 1.1e-05 | 9.000E-06 | 2.052E-06 | 6.900E-06 | 1.100E-05 |
| 12250 | 613 | V613I | A | 7.3e-06 | 7.6e-06 | 1.4e-05 | 9.633E-06 | 3.785E-06 | 7.300E-06 | 1.400E-05 |
| 12251 | 613 | V613L | A | 6.4e-06 | 8.9e-06 | 8.2e-06 | 7.833E-06 | 1.290E-06 | 6.400E-06 | 8.900E-06 |
| 12252 | 613 | V613K | A | 4.5e-06 | 1.3e-05 | 5.4e-06 | 7.633E-06 | 4.669E-06 | 4.500E-06 | 1.300E-05 |
| 12253 | 613 | V613M | A | 9.8e-06 | 9.5e-06 | 7.3e-06 | 8.867E-06 | 1.365E-06 | 7.300E-06 | 9.800E-06 |
| 12254 | 613 | V613F | A | 8.9e-06 | 5.6e-06 | 1.4e-05 | 9.500E-06 | 4.232E-06 | 5.600E-06 | 1.400E-05 |
| 12255 | 613 | V613P | A | 9.7e-06 | 1.1e-05 | 4.5e-06 | 8.400E-06 | 3.439E-06 | 4.500E-06 | 1.100E-05 |
| 12256 | 613 | V613S | A | 8.7e-06 | 7e-06 | 9.7e-06 | 8.467E-06 | 1.365E-06 | 7.000E-06 | 9.700E-06 |
| 12257 | 613 | V613T | A | 9.9e-06 | 8.7e-06 | 6.7e-06 | 8.433E-06 | 1.617E-06 | 6.700E-06 | 9.900E-06 |
| 12258 | 613 | V613W | A | 6.6e-06 | 6.6e-06 | 1.4e-05 | 9.067E-06 | 4.272E-06 | 6.600E-06 | 1.400E-05 |
| 12259 | 613 | V613Y | A | 9e-06 | 1.4e-05 | 1.2e-05 | 1.167E-05 | 2.517E-06 | 9.000E-06 | 1.400E-05 |
| 12260 | 613 | V613V | A | 8.1e-06 | 4.2e-06 | 5.4e-06 | 5.900E-06 | 1.997E-06 | 4.200E-06 | 8.100E-06 |
| 12261 | 614 | D614A | A | 5.2e-06 | 8e-06 | 7.1e-06 | 6.767E-06 | 1.429E-06 | 5.200E-06 | 8.000E-06 |
| 12262 | 614 | D614R | A | 1e-05 | 9.3e-06 | 8.9e-06 | 9.400E-06 | 5.568E-07 | 8.900E-06 | 1.000E-05 |
| 12263 | 614 | D614N | A | 6.6e-06 | 8.6e-06 | 5.7e-06 | 6.967E-06 | 1.484E-06 | 5.700E-06 | 8.600E-06 |
| 12264 | 614 | D614D | A | 8.2e-06 | 8.7e-06 | 6.4e-06 | 7.767E-06 | 1.210E-06 | 6.400E-06 | 8.700E-06 |
| 12265 | 614 | D614C | A | 5.5e-06 | 8.3e-06 | 9.8e-06 | 7.867E-06 | 2.183E-06 | 5.500E-06 | 9.800E-06 |
| 12266 | 614 | D614E | A | 1.3e-05 | 8.7e-06 | 8.3e-06 | 1.000E-05 | 2.606E-06 | 8.300E-06 | 1.300E-05 |
| 12267 | 614 | D614Q | A | 8.8e-06 | 9.5e-06 | 7.7e-06 | 8.667E-06 | 9.074E-07 | 7.700E-06 | 9.500E-06 |
| 12268 | 614 | D614G | A | 7.6e-06 | 7.1e-06 | 5.9e-06 | 6.867E-06 | 8.737E-07 | 5.900E-06 | 7.600E-06 |
| 12269 | 614 | D614H | A | 6.9e-06 | 1.2e-05 | 9.5e-06 | 9.467E-06 | 2.550E-06 | 6.900E-06 | 1.200E-05 |
| 12270 | 614 | D614I | A | 1.6e-05 | 1.4e-05 | 7e-06 | 1.233E-05 | 4.726E-06 | 7.000E-06 | 1.600E-05 |
| 12271 | 614 | D614L | A | 6.9e-06 | 1.4e-05 | 8.8e-06 | 9.900E-06 | 3.676E-06 | 6.900E-06 | 1.400E-05 |
| 12272 | 614 | D614K | A | 9.8e-06 | 1.1e-05 | 4.6e-06 | 8.467E-06 | 3.402E-06 | 4.600E-06 | 1.100E-05 |
| 12273 | 614 | D614M | A | 7.8e-06 | 6.2e-06 | 6.6e-06 | 6.867E-06 | 8.327E-07 | 6.200E-06 | 7.800E-06 |
| 12274 | 614 | D614F | A | 5.8e-06 | 9.2e-06 | 9.3e-06 | 8.100E-06 | 1.992E-06 | 5.800E-06 | 9.300E-06 |
| 12275 | 614 | D614P | A | 6.9e-06 | 7.9e-06 | 7.8e-06 | 7.533E-06 | 5.508E-07 | 6.900E-06 | 7.900E-06 |
| 12276 | 614 | D614S | A | 1e-05 | 4.8e-06 | 8.4e-06 | 7.733E-06 | 2.663E-06 | 4.800E-06 | 1.000E-05 |
| 12277 | 614 | D614T | A | 6.8e-06 | 1.3e-05 | 8.4e-06 | 9.400E-06 | 3.219E-06 | 6.800E-06 | 1.300E-05 |
| 12278 | 614 | D614W | A | 1.4e-05 | 7.9e-06 | 5.2e-06 | 9.033E-06 | 4.508E-06 | 5.200E-06 | 1.400E-05 |
| 12279 | 614 | D614Y | A | 9.6e-06 | 5.7e-06 | 9.3e-06 | 8.200E-06 | 2.170E-06 | 5.700E-06 | 9.600E-06 |
| 12280 | 614 | D614V | A | 9.8e-06 | 5.2e-06 | 5.5e-06 | 6.833E-06 | 2.574E-06 | 5.200E-06 | 9.800E-06 |
| 12281 | 615 | L615A | A | 8.1e-06 | 1.3e-05 | 6.7e-06 | 9.267E-06 | 3.308E-06 | 6.700E-06 | 1.300E-05 |
| 12282 | 615 | L615R | A | 6.1e-06 | 6e-06 | 7.3e-06 | 6.467E-06 | 7.234E-07 | 6.000E-06 | 7.300E-06 |
| 12283 | 615 | L615N | A | 1.3e-05 | 8.5e-06 | 5.2e-06 | 8.900E-06 | 3.915E-06 | 5.200E-06 | 1.300E-05 |
| 12284 | 615 | L615D | A | 1.1e-05 | 6e-06 | 7.3e-06 | 8.100E-06 | 2.594E-06 | 6.000E-06 | 1.100E-05 |
| 12285 | 615 | L615C | A | 8e-06 | 6.4e-06 | 1.2e-05 | 8.800E-06 | 2.884E-06 | 6.400E-06 | 1.200E-05 |
| 12286 | 615 | L615E | A | 1.1e-05 | 8.9e-06 | 1.1e-05 | 1.030E-05 | 1.212E-06 | 8.900E-06 | 1.100E-05 |
| 12287 | 615 | L615Q | A | 8.2e-06 | 7.3e-06 | 7.1e-06 | 7.533E-06 | 5.859E-07 | 7.100E-06 | 8.200E-06 |
| 12288 | 615 | L615G | A | 1.6e-05 | 9.6e-06 | 5.3e-06 | 1.030E-05 | 5.384E-06 | 5.300E-06 | 1.600E-05 |
| 12289 | 615 | L615H | A | 9.4e-06 | 9e-06 | 6.5e-06 | 8.300E-06 | 1.572E-06 | 6.500E-06 | 9.400E-06 |
| 12290 | 615 | L615I | A | 1e-05 | 5e-06 | 5.4e-06 | 6.800E-06 | 2.778E-06 | 5.000E-06 | 1.000E-05 |
| 12291 | 615 | L615L | A | 4.9e-06 | 8.6e-06 | 4.4e-06 | 5.967E-06 | 2.294E-06 | 4.400E-06 | 8.600E-06 |
| 12292 | 615 | L615K | A | 6.7e-06 | 7e-06 | 4.1e-06 | 5.933E-06 | 1.595E-06 | 4.100E-06 | 7.000E-06 |
| 12293 | 615 | L615M | A | 5.6e-06 | 4e-06 | 9.7e-06 | 6.433E-06 | 2.940E-06 | 4.000E-06 | 9.700E-06 |
| 12294 | 615 | L615F | A | 5.4e-06 | 1.1e-05 | 3.8e-06 | 6.733E-06 | 3.781E-06 | 3.800E-06 | 1.100E-05 |
| 12295 | 615 | L615P | A | 9.9e-06 | 4.9e-06 | 8.8e-06 | 7.867E-06 | 2.627E-06 | 4.900E-06 | 9.900E-06 |
| 12296 | 615 | L615S | A | 1.3e-05 | 9e-06 | 6.6e-06 | 9.533E-06 | 3.233E-06 | 6.600E-06 | 1.300E-05 |
| 12297 | 615 | L615T | A | 1.3e-05 | 5.6e-06 | 6.3e-06 | 8.300E-06 | 4.085E-06 | 5.600E-06 | 1.300E-05 |
| 12298 | 615 | L615W | A | 8.9e-06 | 9.6e-06 | 1.2e-05 | 1.017E-05 | 1.626E-06 | 8.900E-06 | 1.200E-05 |
| 12299 | 615 | L615Y | A | 1.2e-05 | 6.1e-06 | 7.2e-06 | 8.433E-06 | 3.137E-06 | 6.100E-06 | 1.200E-05 |
| 12300 | 615 | L615V | A | 8.6e-06 | 6.4e-06 | 1.1e-05 | 8.667E-06 | 2.301E-06 | 6.400E-06 | 1.100E-05 |
| 12301 | 616 | D616A | A | 5.9e-06 | 9.2e-06 | 1.1e-05 | 8.700E-06 | 2.587E-06 | 5.900E-06 | 1.100E-05 |
| 12302 | 616 | D616R | A | 1.1e-05 | 1.2e-05 | 8.3e-06 | 1.043E-05 | 1.914E-06 | 8.300E-06 | 1.200E-05 |
| 12303 | 616 | D616N | A | 1.5e-05 | 6.8e-06 | 7e-06 | 9.600E-06 | 4.678E-06 | 6.800E-06 | 1.500E-05 |
| 12304 | 616 | D616D | A | 5.6e-06 | 1.2e-05 | 6.7e-06 | 8.100E-06 | 3.422E-06 | 5.600E-06 | 1.200E-05 |
| 12305 | 616 | D616C | A | 8.3e-06 | 1.2e-05 | 8.6e-06 | 9.633E-06 | 2.055E-06 | 8.300E-06 | 1.200E-05 |
| 12306 | 616 | D616E | A | 7.9e-06 | 1e-05 | 7.7e-06 | 8.533E-06 | 1.274E-06 | 7.700E-06 | 1.000E-05 |
| 12307 | 616 | D616Q | A | 3.8e-06 | 9.4e-06 | 8.7e-06 | 7.300E-06 | 3.051E-06 | 3.800E-06 | 9.400E-06 |
| 12308 | 616 | D616G | A | 9.8e-06 | 5.4e-06 | 6.5e-06 | 7.233E-06 | 2.290E-06 | 5.400E-06 | 9.800E-06 |
| 12309 | 616 | D616H | A | 1.2e-05 | 8.8e-06 | 4.2e-06 | 8.333E-06 | 3.921E-06 | 4.200E-06 | 1.200E-05 |
| 12310 | 616 | D616I | A | 6.6e-06 | 5.7e-06 | 1.2e-05 | 8.100E-06 | 3.407E-06 | 5.700E-06 | 1.200E-05 |
| 12311 | 616 | D616L | A | 9.3e-06 | 9.8e-06 | 9e-06 | 9.367E-06 | 4.041E-07 | 9.000E-06 | 9.800E-06 |
| 12312 | 616 | D616K | A | 5.5e-06 | 8.2e-06 | 5.9e-06 | 6.533E-06 | 1.457E-06 | 5.500E-06 | 8.200E-06 |
| 12313 | 616 | D616M | A | 7.9e-06 | 1.2e-05 | 9.8e-06 | 9.900E-06 | 2.052E-06 | 7.900E-06 | 1.200E-05 |
| 12314 | 616 | D616F | A | 8.9e-06 | 5e-06 | 6.9e-06 | 6.933E-06 | 1.950E-06 | 5.000E-06 | 8.900E-06 |
| 12315 | 616 | D616P | A | 8.6e-06 | 6.6e-06 | 5e-06 | 6.733E-06 | 1.804E-06 | 5.000E-06 | 8.600E-06 |
| 12316 | 616 | D616S | A | 5.8e-06 | 6.3e-06 | 1e-05 | 7.367E-06 | 2.294E-06 | 5.800E-06 | 1.000E-05 |
| 12317 | 616 | D616T | A | 9e-06 | 9.4e-06 | 8.2e-06 | 8.867E-06 | 6.110E-07 | 8.200E-06 | 9.400E-06 |
| 12318 | 616 | D616W | A | 5.9e-06 | 9.5e-06 | 4.4e-06 | 6.600E-06 | 2.621E-06 | 4.400E-06 | 9.500E-06 |
| 12319 | 616 | D616Y | A | 5.5e-06 | 6.7e-06 | 6.3e-06 | 6.167E-06 | 6.110E-07 | 5.500E-06 | 6.700E-06 |
| 12320 | 616 | D616V | A | 8.8e-06 | 1e-05 | 5e-06 | 7.933E-06 | 2.610E-06 | 5.000E-06 | 1.000E-05 |
| 12321 | 617 | D617A | A | 7.8e-06 | 5.5e-06 | 7.3e-06 | 6.867E-06 | 1.210E-06 | 5.500E-06 | 7.800E-06 |
| 12322 | 617 | D617R | A | 7.3e-06 | 4.8e-06 | 1.1e-05 | 7.700E-06 | 3.119E-06 | 4.800E-06 | 1.100E-05 |
| 12323 | 617 | D617N | A | 6e-06 | 5.4e-06 | 6.6e-06 | 6.000E-06 | 6.000E-07 | 5.400E-06 | 6.600E-06 |
| 12324 | 617 | D617D | A | 7.8e-06 | 6.6e-06 | 9e-06 | 7.800E-06 | 1.200E-06 | 6.600E-06 | 9.000E-06 |

| | | | | | | | | | | |
|---|---|---|---|---|---|---|---|---|---|---|
| 12325 | 617 | D617C | A | 1.2e-05 | 4.6e-06 | 6.8e-06 | 7.800E-06 | 3.800E-06 | 4.600E-06 | 1.200E-05 |
| 12326 | 617 | D617E | A | 8.6e-06 | 5.2e-06 | 5.1e-06 | 6.300E-06 | 1.992E-06 | 5.100E-06 | 8.600E-06 |
| 12327 | 617 | D617Q | A | 1e-05 | 7.6e-06 | 9.4e-06 | 9.000E-06 | 1.249E-06 | 7.600E-06 | 1.000E-05 |
| 12328 | 617 | D617G | A | 9.9e-06 | 9.3e-06 | 6.2e-06 | 8.467E-06 | 1.986E-06 | 6.200E-06 | 9.900E-06 |
| 12329 | 617 | D617H | A | 1.2e-05 | 1.3e-05 | 5.1e-06 | 1.003E-05 | 4.302E-06 | 5.100E-06 | 1.300E-05 |
| 12330 | 617 | D617I | A | 5.1e-06 | 1e-05 | 8.6e-06 | 7.900E-06 | 2.524E-06 | 5.100E-06 | 1.000E-05 |
| 12331 | 617 | D617L | A | 7.5e-06 | 1.3e-05 | 5.7e-06 | 8.733E-06 | 3.803E-06 | 5.700E-06 | 1.300E-05 |
| 12332 | 617 | D617K | A | 5.3e-06 | 5.7e-06 | 9.6e-06 | 6.867E-06 | 2.376E-06 | 5.300E-06 | 9.600E-06 |
| 12333 | 617 | D617M | A | 1.2e-05 | 4.9e-06 | 9.4e-06 | 8.767E-06 | 3.592E-06 | 4.900E-06 | 1.200E-05 |
| 12334 | 617 | D617F | A | 6.8e-06 | 1.4e-05 | 9.5e-06 | 1.010E-05 | 3.637E-06 | 6.800E-06 | 1.400E-05 |
| 12335 | 617 | D617P | A | 1.3e-05 | 6.1e-06 | 4.7e-06 | 7.933E-06 | 4.443E-06 | 4.700E-06 | 1.300E-05 |
| 12336 | 617 | D617S | A | 8.1e-06 | 1.2e-05 | 1.3e-05 | 1.103E-05 | 2.589E-06 | 8.100E-06 | 1.300E-05 |
| 12337 | 617 | D617T | A | 7.7e-06 | 5.2e-06 | 9.4e-06 | 7.433E-06 | 2.113E-06 | 5.200E-06 | 9.400E-06 |
| 12338 | 617 | D617W | A | 6e-06 | 5.5e-06 | 6.3e-06 | 5.933E-06 | 4.041E-07 | 5.500E-06 | 6.300E-06 |
| 12339 | 617 | D617Y | A | 7.2e-06 | 1.3e-05 | 8.4e-06 | 9.533E-06 | 3.062E-06 | 7.200E-06 | 1.300E-05 |
| 12340 | 617 | D617V | A | 7.1e-06 | 1.3e-05 | 1.2e-05 | 1.070E-05 | 3.158E-06 | 7.100E-06 | 1.300E-05 |
| 12341 | 618 | K618A | A | 6.3e-06 | 6.4e-06 | 9.8e-06 | 7.500E-06 | 1.992E-06 | 6.300E-06 | 9.800E-06 |
| 12342 | 618 | K618R | A | 6.5e-06 | 5.7e-06 | 5.2e-06 | 5.800E-06 | 6.557E-07 | 5.200E-06 | 6.500E-06 |
| 12343 | 618 | K618N | A | 8.9e-06 | 9.4e-06 | 7.6e-06 | 8.633E-06 | 9.292E-07 | 7.600E-06 | 9.400E-06 |
| 12344 | 618 | K618D | A | 9.1e-06 | 1.1e-05 | 6e-06 | 8.700E-06 | 2.524E-06 | 6.000E-06 | 1.100E-05 |
| 12345 | 618 | K618C | A | 2.2e-05 | 6.4e-06 | 9e-06 | 1.247E-05 | 8.358E-06 | 6.400E-06 | 2.200E-05 |
| 12346 | 618 | K618E | A | 6.1e-06 | 5.6e-06 | 7.6e-06 | 6.433E-06 | 1.041E-06 | 5.600E-06 | 7.600E-06 |
| 12347 | 618 | K618Q | A | 1e-05 | 1e-05 | 7.2e-06 | 9.067E-06 | 1.617E-06 | 7.200E-06 | 1.000E-05 |
| 12348 | 618 | K618G | A | 6.8e-06 | 9.1e-06 | 6e-06 | 7.300E-06 | 1.609E-06 | 6.000E-06 | 9.100E-06 |
| 12349 | 618 | K618H | A | 1.4e-05 | 9.6e-06 | 9.3e-06 | 1.097E-05 | 2.631E-06 | 9.300E-06 | 1.400E-05 |
| 12350 | 618 | K618I | A | 6.6e-06 | 1.1e-05 | 1.3e-05 | 1.020E-05 | 3.274E-06 | 6.600E-06 | 1.300E-05 |
| 12351 | 618 | K618L | A | 6.5e-06 | 8.6e-06 | 7.5e-06 | 7.533E-06 | 1.050E-06 | 6.500E-06 | 8.600E-06 |
| 12352 | 618 | K618K | A | 9.6e-06 | 9.2e-06 | 8e-06 | 8.933E-06 | 8.327E-07 | 8.000E-06 | 9.600E-06 |
| 12353 | 618 | K618M | A | 7.3e-06 | 8.8e-06 | 6.7e-06 | 7.600E-06 | 1.082E-06 | 6.700E-06 | 8.800E-06 |
| 12354 | 618 | K618F | A | 9.1e-06 | 7.5e-06 | 4.6e-06 | 7.067E-06 | 2.281E-06 | 4.600E-06 | 9.100E-06 |
| 12355 | 618 | K618P | A | 1.1e-05 | 9.8e-06 | 1.1e-05 | 1.060E-05 | 6.928E-07 | 9.800E-06 | 1.100E-05 |
| 12356 | 618 | K618S | A | 4.2e-06 | 4.8e-06 | 6.7e-06 | 5.233E-06 | 1.305E-06 | 4.200E-06 | 6.700E-06 |
| 12357 | 618 | K618T | A | 9.1e-06 | 3.5e-06 | 1e-05 | 7.533E-06 | 3.522E-06 | 3.500E-06 | 1.000E-05 |
| 12358 | 618 | K618W | A | 6.3e-06 | 4.9e-06 | 6.7e-06 | 5.967E-06 | 9.452E-07 | 4.900E-06 | 6.700E-06 |
| 12359 | 618 | K618Y | A | 1.2e-05 | 7.4e-06 | 8e-06 | 9.133E-06 | 2.501E-06 | 7.400E-06 | 1.200E-05 |
| 12360 | 618 | K618V | A | 9e-06 | 8.1e-06 | 7.2e-06 | 8.100E-06 | 9.000E-07 | 7.200E-06 | 9.000E-06 |
| 12361 | 619 | G619A | A | 9.1e-06 | 7e-06 | 1.3e-05 | 9.700E-06 | 3.045E-06 | 7.000E-06 | 1.300E-05 |
| 12362 | 619 | G619R | A | 1.2e-05 | 8.5e-06 | 4.8e-06 | 8.433E-06 | 3.600E-06 | 4.800E-06 | 1.200E-05 |
| 12363 | 619 | G619N | A | 6.6e-06 | 8.3e-06 | 4.6e-06 | 6.500E-06 | 1.852E-06 | 4.600E-06 | 8.300E-06 |
| 12364 | 619 | G619D | A | 1.3e-05 | 4.8e-06 | 9.2e-06 | 9.000E-06 | 4.104E-06 | 4.800E-06 | 1.300E-05 |
| 12365 | 619 | G619C | A | 6e-06 | 7e-06 | 4.4e-06 | 5.800E-06 | 1.311E-06 | 4.400E-06 | 7.000E-06 |
| 12366 | 619 | G619E | A | 1.3e-05 | 1.2e-05 | 4.3e-06 | 9.767E-06 | 4.761E-06 | 4.300E-06 | 1.300E-05 |
| 12367 | 619 | G619Q | A | 1.2e-05 | 7.8e-06 | 7.2e-06 | 9.000E-06 | 2.615E-06 | 7.200E-06 | 1.200E-05 |
| 12368 | 619 | G619G | A | 4.2e-06 | 1.1e-05 | 7.1e-06 | 7.433E-06 | 3.412E-06 | 4.200E-06 | 1.100E-05 |
| 12369 | 619 | G619H | A | 5.3e-06 | 6.2e-06 | 6.6e-06 | 6.033E-06 | 6.658E-07 | 5.300E-06 | 6.600E-06 |
| 12370 | 619 | G619I | A | 1.1e-05 | 9.6e-06 | 1.4e-05 | 1.153E-05 | 2.248E-06 | 9.600E-06 | 1.400E-05 |
| 12371 | 619 | G619L | A | 1.3e-05 | 1e-05 | 9.1e-06 | 1.070E-05 | 2.042E-06 | 9.100E-06 | 1.300E-05 |
| 12372 | 619 | G619K | A | 1e-05 | 6.1e-06 | 4.6e-06 | 6.900E-06 | 2.787E-06 | 4.600E-06 | 1.000E-05 |
| 12373 | 619 | G619M | A | 8.7e-06 | 6.7e-06 | 5.8e-06 | 7.067E-06 | 1.484E-06 | 5.800E-06 | 8.700E-06 |
| 12374 | 619 | G619F | A | 6.3e-06 | 6.2e-06 | 1e-05 | 7.500E-06 | 2.166E-06 | 6.200E-06 | 1.000E-05 |
| 12375 | 619 | G619P | A | 6.9e-06 | 6.9e-06 | 4.6e-06 | 6.133E-06 | 1.328E-06 | 4.600E-06 | 6.900E-06 |
| 12376 | 619 | G619S | A | 1.3e-05 | 5.6e-06 | 9.1e-06 | 9.233E-06 | 3.702E-06 | 5.600E-06 | 1.300E-05 |
| 12377 | 619 | G619T | A | 6e-06 | 6.9e-06 | 8.1e-06 | 7.000E-06 | 1.054E-06 | 6.000E-06 | 8.100E-06 |
| 12378 | 619 | G619W | A | 8e-06 | 5.1e-06 | 9.6e-06 | 7.567E-06 | 2.281E-06 | 5.100E-06 | 9.600E-06 |
| 12379 | 619 | G619Y | A | 6.4e-06 | 5.3e-06 | 6.2e-06 | 5.967E-06 | 5.859E-07 | 5.300E-06 | 6.400E-06 |
| 12380 | 619 | G619V | A | 5.3e-06 | 9.8e-06 | 8.3e-06 | 7.800E-06 | 2.291E-06 | 5.300E-06 | 9.800E-06 |
| 12381 | 620 | C620A | A | 9.1e-06 | 6.8e-06 | 8e-06 | 7.967E-06 | 1.150E-06 | 6.800E-06 | 9.100E-06 |
| 12382 | 620 | C620R | A | 5.2e-06 | 1e-05 | 7.5e-06 | 7.567E-06 | 2.401E-06 | 5.200E-06 | 1.000E-05 |
| 12383 | 620 | C620N | A | 7.2e-06 | 1.2e-05 | 5.6e-06 | 8.267E-06 | 3.331E-06 | 5.600E-06 | 1.200E-05 |
| 12384 | 620 | C620D | A | 8.1e-06 | 4.8e-06 | 9.2e-06 | 7.367E-06 | 2.290E-06 | 4.800E-06 | 9.200E-06 |
| 12385 | 620 | C620C | A | 5.7e-06 | 1.1e-05 | 8.7e-06 | 8.467E-06 | 2.658E-06 | 5.700E-06 | 1.100E-05 |
| 12386 | 620 | C620E | A | 7.9e-06 | 5.7e-06 | 7.6e-06 | 7.067E-06 | 1.193E-06 | 5.700E-06 | 7.900E-06 |
| 12387 | 620 | C620Q | A | 1e-05 | 6.9e-06 | 6.3e-06 | 7.733E-06 | 1.986E-06 | 6.300E-06 | 1.000E-05 |
| 12388 | 620 | C620G | A | 7.2e-06 | 8.9e-06 | 5.7e-06 | 7.267E-06 | 1.601E-06 | 5.700E-06 | 8.900E-06 |
| 12389 | 620 | C620H | A | 6.8e-06 | 6.9e-06 | 9.4e-06 | 7.700E-06 | 1.473E-06 | 6.800E-06 | 9.400E-06 |
| 12390 | 620 | C620I | A | 7.2e-06 | 7.2e-06 | 8.8e-06 | 7.733E-06 | 9.238E-07 | 7.200E-06 | 8.800E-06 |
| 12391 | 620 | C620L | A | 1e-05 | 9.4e-06 | 7.9e-06 | 9.100E-06 | 1.082E-06 | 7.900E-06 | 1.000E-05 |
| 12392 | 620 | C620K | A | 8.2e-06 | 1.1e-05 | 9.3e-06 | 9.500E-06 | 1.411E-06 | 8.200E-06 | 1.100E-05 |
| 12393 | 620 | C620M | A | 5.1e-06 | 1e-05 | 6.6e-06 | 7.233E-06 | 2.511E-06 | 5.100E-06 | 1.000E-05 |
| 12394 | 620 | C620F | A | 5.3e-06 | 1.5e-05 | 7.9e-06 | 9.400E-06 | 5.021E-06 | 5.300E-06 | 1.500E-05 |
| 12395 | 620 | C620P | A | 1.4e-05 | 6.9e-06 | 7.4e-06 | 9.433E-06 | 3.963E-06 | 6.900E-06 | 1.400E-05 |
| 12396 | 620 | C620S | A | 6.8e-06 | 6.2e-06 | 9.9e-06 | 7.633E-06 | 1.986E-06 | 6.200E-06 | 9.900E-06 |
| 12397 | 620 | C620T | A | 9.2e-06 | 1.2e-05 | 1e-05 | 1.040E-05 | 1.442E-06 | 9.200E-06 | 1.200E-05 |
| 12398 | 620 | C620W | A | 7.4e-06 | 8.8e-06 | 7e-06 | 7.733E-06 | 9.452E-07 | 7.000E-06 | 8.800E-06 |
| 12399 | 620 | C620Y | A | 8.5e-06 | 9.6e-06 | 9.5e-06 | 9.200E-06 | 6.083E-07 | 8.500E-06 | 9.600E-06 |
| 12400 | 620 | C620V | A | 7.4e-06 | 7.4e-06 | 9.4e-06 | 8.067E-06 | 1.155E-06 | 7.400E-06 | 9.400E-06 |
| 12401 | 621 | P621A | A | 9.6e-06 | 7e-06 | 9.8e-06 | 8.800E-06 | 1.562E-06 | 7.000E-06 | 9.800E-06 |
| 12402 | 621 | P621R | A | 8.1e-06 | 6.3e-06 | 1.3e-05 | 9.133E-06 | 3.467E-06 | 6.300E-06 | 1.300E-05 |
| 12403 | 621 | P621N | A | 7.5e-06 | 4.4e-06 | 9.4e-06 | 7.100E-06 | 2.524E-06 | 4.400E-06 | 9.400E-06 |
| 12404 | 621 | P621D | A | 8.5e-06 | 9.5e-06 | 1e-05 | 9.333E-06 | 7.638E-07 | 8.500E-06 | 1.000E-05 |
| 12405 | 621 | P621C | A | 7.3e-06 | 1.1e-05 | 4.2e-06 | 7.500E-06 | 3.404E-06 | 4.200E-06 | 1.100E-05 |
| 12406 | 621 | P621E | A | 5.5e-06 | 1.2e-05 | 1.3e-05 | 1.017E-05 | 4.072E-06 | 5.500E-06 | 1.300E-05 |
| 12407 | 621 | P621Q | A | 8.4e-06 | 7.3e-06 | 5.1e-06 | 6.933E-06 | 1.680E-06 | 5.100E-06 | 8.400E-06 |
| 12408 | 621 | P621G | A | 5e-06 | 7.2e-06 | 8.8e-06 | 7.000E-06 | 1.908E-06 | 5.000E-06 | 8.800E-06 |
| 12409 | 621 | P621H | A | 4.9e-06 | 6.9e-06 | 6.8e-06 | 6.200E-06 | 1.127E-06 | 4.900E-06 | 6.900E-06 |

| | | | | | | | | | | |
|---|---|---|---|---|---|---|---|---|---|---|
| 12410 | 621 | P621I | A | 8.2e-06 | 7.7e-06 | 6.6e-06 | 7.500E-06 | 8.185E-07 | 6.600E-06 | 8.200E-06 |
| 12411 | 621 | P621L | A | 8e-06 | 1.2e-05 | 8.5e-06 | 9.500E-06 | 2.179E-06 | 8.000E-06 | 1.200E-05 |
| 12412 | 621 | P621K | A | 8.4e-06 | 5.3e-06 | 1.3e-05 | 8.900E-06 | 3.874E-06 | 5.300E-06 | 1.300E-05 |
| 12413 | 621 | P621M | A | 6.6e-06 | 6.8e-06 | 6.6e-06 | 6.667E-06 | 1.155E-07 | 6.600E-06 | 6.800E-06 |
| 12414 | 621 | P621F | A | 9.5e-06 | 8e-06 | 1.1e-05 | 9.500E-06 | 1.500E-06 | 8.000E-06 | 1.100E-05 |
| 12415 | 621 | P621P | A | 4.2e-06 | 5.9e-06 | 1.1e-05 | 7.033E-06 | 3.539E-06 | 4.200E-06 | 1.100E-05 |
| 12416 | 621 | P621S | A | 9e-06 | 8.7e-06 | 7.5e-06 | 8.400E-06 | 7.937E-07 | 7.500E-06 | 9.000E-06 |
| 12417 | 621 | P621T | A | 8.1e-06 | 7e-06 | 4.7e-06 | 6.600E-06 | 1.735E-06 | 4.700E-06 | 8.100E-06 |
| 12418 | 621 | P621W | A | 1e-05 | 5.5e-06 | 6.2e-06 | 7.233E-06 | 2.421E-06 | 5.500E-06 | 1.000E-05 |
| 12419 | 621 | P621Y | A | 5.6e-06 | 1.3e-05 | 6.3e-06 | 8.300E-06 | 4.085E-06 | 5.600E-06 | 1.300E-05 |
| 12420 | 621 | P621V | A | 9.6e-06 | 6.8e-06 | 1.1e-05 | 9.133E-06 | 2.139E-06 | 6.800E-06 | 1.100E-05 |
| 12421 | 622 | A622A | A | 9.6e-06 | 1.1e-05 | 7.3e-06 | 9.300E-06 | 1.868E-06 | 7.300E-06 | 1.100E-05 |
| 12422 | 622 | A622R | A | 5.6e-06 | 7.4e-06 | 5.1e-06 | 6.033E-06 | 1.210E-06 | 5.100E-06 | 7.400E-06 |
| 12423 | 622 | A622N | A | 1.2e-05 | 5.4e-06 | 8.8e-06 | 8.733E-06 | 3.301E-06 | 5.400E-06 | 1.200E-05 |
| 12424 | 622 | A622D | A | 1.3e-05 | 8.9e-06 | 6e-06 | 9.300E-06 | 3.517E-06 | 6.000E-06 | 1.300E-05 |
| 12425 | 622 | A622C | A | 8e-06 | 8.9e-06 | 4.4e-06 | 7.100E-06 | 2.381E-06 | 4.400E-06 | 8.900E-06 |
| 12426 | 622 | A622E | A | 6.2e-06 | 9.5e-06 | 1.3e-05 | 9.567E-06 | 3.400E-06 | 6.200E-06 | 1.300E-05 |
| 12427 | 622 | A622Q | A | 8.2e-06 | 1.1e-05 | 8.8e-06 | 9.333E-06 | 1.474E-06 | 8.200E-06 | 1.100E-05 |
| 12428 | 622 | A622G | A | 7.1e-06 | 4.5e-06 | 7.6e-06 | 6.400E-06 | 1.664E-06 | 4.500E-06 | 7.600E-06 |
| 12429 | 622 | A622H | A | 6.1e-06 | 6.2e-06 | 6.3e-06 | 6.200E-06 | 1.000E-07 | 6.100E-06 | 6.300E-06 |
| 12430 | 622 | A622I | A | 8.1e-06 | 4.6e-06 | 8.9e-06 | 7.200E-06 | 2.287E-06 | 4.600E-06 | 8.900E-06 |
| 12431 | 622 | A622L | A | 1e-05 | 1.4e-05 | 8.3e-06 | 1.077E-05 | 2.926E-06 | 8.300E-06 | 1.400E-05 |
| 12432 | 622 | A622K | A | 6.9e-06 | 8.5e-06 | 1.1e-05 | 8.800E-06 | 2.066E-06 | 6.900E-06 | 1.100E-05 |
| 12433 | 622 | A622M | A | 4.6e-06 | 6.3e-06 | 9.9e-06 | 6.933E-06 | 2.706E-06 | 4.600E-06 | 9.900E-06 |
| 12434 | 622 | A622F | A | 1.2e-05 | 4.7e-06 | 8.2e-06 | 8.300E-06 | 3.651E-06 | 4.700E-06 | 1.200E-05 |
| 12435 | 622 | A622P | A | 7e-06 | 6.4e-06 | 1.1e-05 | 8.133E-06 | 2.501E-06 | 6.400E-06 | 1.100E-05 |
| 12436 | 622 | A622S | A | 1e-05 | 1.2e-05 | 1.3e-05 | 1.167E-05 | 1.528E-06 | 1.000E-05 | 1.300E-05 |
| 12437 | 622 | A622T | A | 9.8e-06 | 1.1e-05 | 7e-06 | 9.267E-06 | 2.053E-06 | 7.000E-06 | 1.100E-05 |
| 12438 | 622 | A622W | A | 7e-06 | 8.5e-06 | 5.9e-06 | 7.133E-06 | 1.305E-06 | 5.900E-06 | 8.500E-06 |
| 12439 | 622 | A622Y | A | 9.3e-06 | 7.3e-06 | 9.1e-06 | 8.567E-06 | 1.102E-06 | 7.300E-06 | 9.300E-06 |
| 12440 | 622 | A622V | A | 6.7e-06 | 6.8e-06 | 9.2e-06 | 7.567E-06 | 1.415E-06 | 6.700E-06 | 9.200E-06 |
| 12441 | 623 | D623A | B | 1e-05 | 1e-05 | 1.2e-05 | 1.067E-05 | 1.155E-06 | 1.000E-05 | 1.200E-05 |
| 12442 | 623 | D623R | B | 1.2e-05 | 6.2e-06 | 7.7e-06 | 8.633E-06 | 3.011E-06 | 6.200E-06 | 1.200E-05 |
| 12443 | 623 | D623N | B | 4.8e-06 | 9.2e-06 | 1.1e-05 | 8.333E-06 | 3.190E-06 | 4.800E-06 | 1.100E-05 |
| 12444 | 623 | D623D | B | 5.8e-06 | 1.3e-05 | 7.3e-06 | 8.700E-06 | 3.799E-06 | 5.800E-06 | 1.300E-05 |
| 12445 | 623 | D623C | B | 9.5e-06 | 1.1e-05 | 8.3e-06 | 9.600E-06 | 1.353E-06 | 8.300E-06 | 1.100E-05 |
| 12446 | 623 | D623E | B | 6.6e-06 | 1.1e-05 | 9.2e-06 | 8.933E-06 | 2.212E-06 | 6.600E-06 | 1.100E-05 |
| 12447 | 623 | D623Q | B | 5.1e-06 | 8.1e-06 | 9.4e-06 | 7.533E-06 | 2.205E-06 | 5.100E-06 | 9.400E-06 |
| 12448 | 623 | D623G | B | 9.5e-06 | 8.3e-06 | 1.2e-05 | 9.933E-06 | 1.888E-06 | 8.300E-06 | 1.200E-05 |
| 12449 | 623 | D623H | B | 6.6e-06 | 9e-06 | 6.7e-06 | 7.433E-06 | 1.358E-06 | 6.600E-06 | 9.000E-06 |
| 12450 | 623 | D623I | B | 9.3e-06 | 5.1e-06 | 4.7e-06 | 6.367E-06 | 2.548E-06 | 4.700E-06 | 9.300E-06 |
| 12451 | 623 | D623L | B | 8.7e-06 | 6.7e-06 | 8.2e-06 | 7.867E-06 | 1.041E-06 | 6.700E-06 | 8.700E-06 |
| 12452 | 623 | D623K | B | 6.3e-06 | 5.6e-06 | 6.2e-06 | 6.033E-06 | 3.786E-07 | 5.600E-06 | 6.300E-06 |
| 12453 | 623 | D623M | B | 8e-06 | 4.1e-06 | 9.5e-06 | 7.200E-06 | 2.787E-06 | 4.100E-06 | 9.500E-06 |
| 12454 | 623 | D623F | B | 1.4e-05 | 1.3e-05 | 9.3e-06 | 1.210E-05 | 2.476E-06 | 9.300E-06 | 1.400E-05 |
| 12455 | 623 | D623P | B | 1.3e-05 | 6.8e-06 | 4.6e-06 | 8.133E-06 | 4.356E-06 | 4.600E-06 | 1.300E-05 |
| 12456 | 623 | D623S | B | 1.3e-05 | 6.3e-06 | 7.7e-06 | 9.000E-06 | 3.534E-06 | 6.300E-06 | 1.300E-05 |
| 12457 | 623 | D623T | B | 8.3e-06 | 7.6e-06 | 6e-06 | 7.300E-06 | 1.179E-06 | 6.000E-06 | 8.300E-06 |
| 12458 | 623 | D623W | B | 8.5e-06 | 9.1e-06 | 9.4e-06 | 9.000E-06 | 4.583E-07 | 8.500E-06 | 9.400E-06 |
| 12459 | 623 | D623Y | B | 6.5e-06 | 9.5e-06 | 9.3e-06 | 8.433E-06 | 1.677E-06 | 6.500E-06 | 9.500E-06 |
| 12460 | 623 | D623V | B | 1.3e-05 | 1e-05 | 1e-05 | 1.100E-05 | 1.732E-06 | 1.000E-05 | 1.300E-05 |
| 12461 | 624 | I624A | B | 5e-06 | 1.1e-05 | 1.4e-05 | 1.000E-05 | 4.583E-06 | 5.000E-06 | 1.400E-05 |
| 12462 | 624 | I624R | B | 1e-05 | 8.7e-06 | 1.3e-05 | 1.057E-05 | 2.205E-06 | 8.700E-06 | 1.300E-05 |
| 12463 | 624 | I624N | B | 7e-06 | 1e-05 | 7.1e-06 | 8.033E-06 | 1.704E-06 | 7.000E-06 | 1.000E-05 |
| 12464 | 624 | I624D | B | 1e-05 | 8.5e-06 | 6.7e-06 | 8.400E-06 | 1.652E-06 | 6.700E-06 | 1.000E-05 |
| 12465 | 624 | I624C | B | 9.1e-06 | 1.5e-05 | 1.1e-05 | 1.170E-05 | 3.012E-06 | 9.100E-06 | 1.500E-05 |
| 12466 | 624 | I624E | B | 9.6e-06 | 1.2e-05 | 9e-06 | 1.020E-05 | 1.587E-06 | 9.000E-06 | 1.200E-05 |
| 12467 | 624 | I624Q | B | 6.8e-06 | 6.7e-06 | 8e-06 | 7.167E-06 | 7.234E-07 | 6.700E-06 | 8.000E-06 |
| 12468 | 624 | I624G | B | 1.3e-05 | 4.8e-06 | 7.2e-06 | 8.333E-06 | 4.216E-06 | 4.800E-06 | 1.300E-05 |
| 12469 | 624 | I624H | B | 9.8e-06 | 5.1e-06 | 1.3e-05 | 9.300E-06 | 3.974E-06 | 5.100E-06 | 1.300E-05 |
| 12470 | 624 | I624I | B | 6.1e-06 | 6.3e-06 | 7.1e-06 | 6.500E-06 | 5.292E-07 | 6.100E-06 | 7.100E-06 |
| 12471 | 624 | I624L | B | 5.1e-06 | 8.6e-06 | 5.6e-06 | 6.433E-06 | 1.893E-06 | 5.100E-06 | 8.600E-06 |
| 12472 | 624 | I624K | B | 7.7e-06 | 9.1e-06 | 6.2e-06 | 7.667E-06 | 1.450E-06 | 6.200E-06 | 9.100E-06 |
| 12473 | 624 | I624M | B | 9.1e-06 | 1.3e-05 | 9.5e-06 | 1.053E-05 | 2.146E-06 | 9.100E-06 | 1.300E-05 |
| 12474 | 624 | I624F | B | 9.6e-06 | 7.6e-06 | 9.8e-06 | 9.000E-06 | 1.217E-06 | 7.600E-06 | 9.800E-06 |
| 12475 | 624 | I624P | B | 9.4e-06 | 1.1e-05 | 8.1e-06 | 9.500E-06 | 1.453E-06 | 8.100E-06 | 1.100E-05 |
| 12476 | 624 | I624S | B | 6.9e-06 | 8.3e-06 | 8.3e-06 | 7.833E-06 | 8.083E-07 | 6.900E-06 | 8.300E-06 |
| 12477 | 624 | I624T | B | 5.6e-06 | 7e-06 | 9.5e-06 | 7.367E-06 | 1.976E-06 | 5.600E-06 | 9.500E-06 |
| 12478 | 624 | I624W | B | 6.5e-06 | 6.5e-06 | 6.8e-06 | 6.600E-06 | 1.732E-07 | 6.500E-06 | 6.800E-06 |
| 12479 | 624 | I624Y | B | 8.8e-06 | 9.5e-06 | 4.7e-06 | 7.667E-06 | 2.593E-06 | 4.700E-06 | 9.500E-06 |
| 12480 | 624 | I624V | B | 7.5e-06 | 7.3e-06 | 8.1e-06 | 7.633E-06 | 4.163E-07 | 7.300E-06 | 8.100E-06 |
| 12481 | 625 | Q625A | B | 7.1e-06 | 9.1e-06 | 8.9e-06 | 8.367E-06 | 1.102E-06 | 7.100E-06 | 9.100E-06 |
| 12482 | 625 | Q625R | B | 5.2e-06 | 9.8e-06 | 7.9e-06 | 7.633E-06 | 2.312E-06 | 5.200E-06 | 9.800E-06 |
| 12483 | 625 | Q625N | B | 7.2e-06 | 7.2e-06 | 9e-06 | 7.800E-06 | 1.039E-06 | 7.200E-06 | 9.000E-06 |
| 12484 | 625 | Q625D | B | 7.2e-06 | 1.1e-05 | 4.4e-06 | 7.533E-06 | 3.313E-06 | 4.400E-06 | 1.100E-05 |
| 12485 | 625 | Q625C | B | 9.5e-06 | 5.6e-06 | 6.3e-06 | 7.133E-06 | 2.079E-06 | 5.600E-06 | 9.500E-06 |
| 12486 | 625 | Q625E | B | 1.5e-05 | 1.3e-05 | 1.5e-05 | 1.433E-05 | 1.155E-06 | 1.300E-05 | 1.500E-05 |
| 12487 | 625 | Q625Q | B | 9.3e-06 | 6.9e-06 | 7.4e-06 | 7.867E-06 | 1.266E-06 | 6.900E-06 | 9.300E-06 |
| 12488 | 625 | Q625G | B | 1.4e-05 | 6.7e-06 | 9.2e-06 | 9.967E-06 | 3.710E-06 | 6.700E-06 | 1.400E-05 |
| 12489 | 625 | Q625H | B | 1.2e-05 | 1.1e-05 | 1e-05 | 1.100E-05 | 1.000E-06 | 1.000E-05 | 1.200E-05 |
| 12490 | 625 | Q625I | B | 8.9e-06 | 1.1e-05 | 6.5e-06 | 8.800E-06 | 2.252E-06 | 6.500E-06 | 1.100E-05 |
| 12491 | 625 | Q625L | B | 9.7e-06 | 1.1e-05 | 1.3e-05 | 1.123E-05 | 1.662E-06 | 9.700E-06 | 1.300E-05 |
| 12492 | 625 | Q625K | B | 7.5e-06 | 5.1e-06 | 1.2e-05 | 8.200E-06 | 3.503E-06 | 5.100E-06 | 1.200E-05 |
| 12493 | 625 | Q625M | B | 5.8e-06 | 6e-06 | 6.1e-06 | 5.967E-06 | 1.528E-07 | 5.800E-06 | 6.100E-06 |
| 12494 | 625 | Q625F | B | 5.5e-06 | 9.6e-06 | 6.3e-06 | 7.133E-06 | 2.173E-06 | 5.500E-06 | 9.600E-06 |

| | | | | | | | | | | |
|---|---|---|---|---|---|---|---|---|---|---|
| 12495 | 625 | Q625P | B | 1e-05 | 7.2e-06 | 3.3e-06 | 6.833E-06 | 3.365E-06 | 3.300E-06 | 1.000E-05 |
| 12496 | 625 | Q625S | B | 6.9e-06 | 8.2e-06 | 4.4e-06 | 6.500E-06 | 1.931E-06 | 4.400E-06 | 8.200E-06 |
| 12497 | 625 | Q625T | B | 9.2e-06 | 9.1e-06 | 1e-05 | 9.433E-06 | 4.933E-07 | 9.100E-06 | 1.000E-05 |
| 12498 | 625 | Q625W | B | 8.9e-06 | 4.8e-06 | 1.5e-05 | 9.567E-06 | 5.133E-06 | 4.800E-06 | 1.500E-05 |
| 12499 | 625 | Q625Y | B | 1.2e-05 | 1.5e-05 | 8.8e-06 | 1.193E-05 | 3.101E-06 | 8.800E-06 | 1.500E-05 |
| 12500 | 625 | Q625V | B | 5.6e-06 | 6.4e-06 | 1.1e-05 | 7.667E-06 | 2.914E-06 | 5.600E-06 | 1.100E-05 |
| 12501 | 626 | M626A | B | 9.8e-06 | 9.6e-06 | 8.9e-06 | 9.433E-06 | 4.726E-07 | 8.900E-06 | 9.800E-06 |
| 12502 | 626 | M626R | B | 8.9e-06 | 9.5e-06 | 6.9e-06 | 8.433E-06 | 1.361E-06 | 6.900E-06 | 9.500E-06 |
| 12503 | 626 | M626N | B | 6.1e-06 | 5e-06 | 1.2e-05 | 7.700E-06 | 3.764E-06 | 5.000E-06 | 1.200E-05 |
| 12504 | 626 | M626D | B | 1e-05 | 6.4e-06 | 1.3e-05 | 9.800E-06 | 3.305E-06 | 6.400E-06 | 1.300E-05 |
| 12505 | 626 | M626C | B | 6.2e-06 | 9.6e-06 | 2.1e-05 | 1.227E-05 | 7.752E-06 | 6.200E-06 | 2.100E-05 |
| 12506 | 626 | M626E | B | 1.4e-05 | 1.2e-05 | 7.7e-06 | 1.123E-05 | 3.219E-06 | 7.700E-06 | 1.400E-05 |
| 12507 | 626 | M626Q | B | 9e-06 | 1e-05 | 1.1e-05 | 1.000E-05 | 1.000E-06 | 9.000E-06 | 1.100E-05 |
| 12508 | 626 | M626G | B | 7.9e-06 | 9.7e-06 | 1.3e-05 | 1.020E-05 | 2.587E-06 | 7.900E-06 | 1.300E-05 |
| 12509 | 626 | M626H | B | 4.8e-06 | 6.3e-06 | 7e-06 | 6.033E-06 | 1.124E-06 | 4.800E-06 | 7.000E-06 |
| 12510 | 626 | M626I | B | 8.1e-06 | 9.8e-06 | 9.2e-06 | 9.033E-06 | 8.622E-07 | 8.100E-06 | 9.800E-06 |
| 12511 | 626 | M626L | B | 6.8e-06 | 7.3e-06 | 5.9e-06 | 6.667E-06 | 7.095E-07 | 5.900E-06 | 7.300E-06 |
| 12512 | 626 | M626K | B | 5.7e-06 | 4.9e-06 | 1.2e-05 | 7.533E-06 | 3.889E-06 | 4.900E-06 | 1.200E-05 |
| 12513 | 626 | M626M | B | 9e-06 | 4.5e-06 | 4.5e-06 | 6.000E-06 | 2.598E-06 | 4.500E-06 | 9.000E-06 |
| 12514 | 626 | M626F | B | 4.1e-06 | 1e-05 | 6e-06 | 6.700E-06 | 3.012E-06 | 4.100E-06 | 1.000E-05 |
| 12515 | 626 | M626P | B | 1.4e-05 | 6.7e-06 | 1e-05 | 1.023E-05 | 3.656E-06 | 6.700E-06 | 1.400E-05 |
| 12516 | 626 | M626S | B | 1.2e-05 | 6.3e-06 | 9.2e-06 | 9.167E-06 | 2.850E-06 | 6.300E-06 | 1.200E-05 |
| 12517 | 626 | M626T | B | 8.3e-06 | 8.2e-06 | 6.4e-06 | 7.633E-06 | 1.069E-06 | 6.400E-06 | 8.300E-06 |
| 12518 | 626 | M626W | B | 3.5e-06 | 1.4e-05 | 1.2e-05 | 9.833E-06 | 5.575E-06 | 3.500E-06 | 1.400E-05 |
| 12519 | 626 | M626Y | B | 5.5e-06 | 5.4e-06 | 9e-06 | 6.633E-06 | 2.050E-06 | 5.400E-06 | 9.000E-06 |
| 12520 | 626 | M626V | B | 8.9e-06 | 4.5e-06 | 6.9e-06 | 6.767E-06 | 2.203E-06 | 4.500E-06 | 8.900E-06 |
| 12521 | 627 | T627A | B | 6.3e-06 | 9.8e-06 | 8.1e-06 | 8.067E-06 | 1.750E-06 | 6.300E-06 | 9.800E-06 |
| 12522 | 627 | T627R | B | 7.7e-06 | 7.9e-06 | 7.1e-06 | 7.567E-06 | 4.163E-07 | 7.100E-06 | 7.900E-06 |
| 12523 | 627 | T627N | B | 9.4e-06 | 1e-05 | 1.4e-05 | 1.113E-05 | 2.501E-06 | 9.400E-06 | 1.400E-05 |
| 12524 | 627 | T627D | B | 9.5e-06 | 7.7e-06 | 1.2e-05 | 9.733E-06 | 2.159E-06 | 7.700E-06 | 1.200E-05 |
| 12525 | 627 | T627C | B | 9.7e-06 | 7.1e-06 | 1.1e-05 | 9.267E-06 | 1.986E-06 | 7.100E-06 | 1.100E-05 |
| 12526 | 627 | T627E | B | 1.2e-05 | 5.1e-06 | 9.7e-06 | 8.933E-06 | 3.513E-06 | 5.100E-06 | 1.200E-05 |
| 12527 | 627 | T627Q | B | 7.4e-06 | 4.6e-06 | 6e-06 | 6.000E-06 | 1.400E-06 | 4.600E-06 | 7.400E-06 |
| 12528 | 627 | T627G | B | 6.4e-06 | 5.2e-06 | 1.1e-05 | 7.533E-06 | 3.062E-06 | 5.200E-06 | 1.100E-05 |
| 12529 | 627 | T627H | B | 9.2e-06 | 1.2e-05 | 6.7e-06 | 9.300E-06 | 2.651E-06 | 6.700E-06 | 1.200E-05 |
| 12530 | 627 | T627I | B | 7.9e-06 | 8.9e-06 | 6.2e-06 | 7.667E-06 | 1.365E-06 | 6.200E-06 | 8.900E-06 |
| 12531 | 627 | T627L | B | 6e-06 | 6.5e-06 | 5.4e-06 | 5.967E-06 | 5.508E-07 | 5.400E-06 | 6.500E-06 |
| 12532 | 627 | T627K | B | 1.2e-05 | 5.3e-06 | 7.7e-06 | 8.333E-06 | 3.395E-06 | 5.300E-06 | 1.200E-05 |
| 12533 | 627 | T627M | B | 9.2e-06 | 1e-05 | 8.9e-06 | 9.367E-06 | 5.686E-07 | 8.900E-06 | 1.000E-05 |
| 12534 | 627 | T627F | B | 5.3e-06 | 9.5e-06 | 4.1e-06 | 6.300E-06 | 2.835E-06 | 4.100E-06 | 9.500E-06 |
| 12535 | 627 | T627P | B | 7.2e-06 | 1.2e-05 | 7.4e-06 | 8.867E-06 | 2.715E-06 | 7.200E-06 | 1.200E-05 |
| 12536 | 627 | T627S | B | 6.2e-06 | 5.5e-06 | 5.2e-06 | 5.633E-06 | 5.132E-07 | 5.200E-06 | 6.200E-06 |
| 12537 | 627 | T627T | B | 8e-06 | 7.7e-06 | 9.5e-06 | 8.400E-06 | 9.644E-07 | 7.700E-06 | 9.500E-06 |
| 12538 | 627 | T627W | B | 9.5e-06 | 1e-05 | 6.1e-06 | 8.533E-06 | 2.122E-06 | 6.100E-06 | 1.000E-05 |
| 12539 | 627 | T627Y | B | 8.5e-06 | 8.4e-06 | 6.2e-06 | 7.700E-06 | 1.300E-06 | 6.200E-06 | 8.500E-06 |
| 12540 | 627 | T627V | B | 1e-05 | 1.1e-05 | 1.1e-05 | 1.067E-05 | 5.774E-07 | 1.000E-05 | 1.100E-05 |
| 12541 | 628 | Q628A | B | 1.1e-05 | 1.1e-05 | 6.8e-06 | 9.600E-06 | 2.425E-06 | 6.800E-06 | 1.100E-05 |
| 12542 | 628 | Q628R | B | 9.2e-06 | 3.9e-06 | 7e-06 | 6.700E-06 | 2.663E-06 | 3.900E-06 | 9.200E-06 |
| 12543 | 628 | Q628N | B | 7.7e-06 | 5.1e-06 | 5.6e-06 | 6.133E-06 | 1.380E-06 | 5.100E-06 | 7.700E-06 |
| 12544 | 628 | Q628D | B | 8.1e-06 | 8.1e-06 | 6.3e-06 | 7.500E-06 | 1.039E-06 | 6.300E-06 | 8.100E-06 |
| 12545 | 628 | Q628C | B | 9.7e-06 | 6.4e-06 | 4.5e-06 | 6.867E-06 | 2.631E-06 | 4.500E-06 | 9.700E-06 |
| 12546 | 628 | Q628E | B | 5.7e-06 | 6.6e-06 | 5.1e-06 | 5.800E-06 | 7.550E-07 | 5.100E-06 | 6.600E-06 |
| 12547 | 628 | Q628Q | B | 1.4e-05 | 1.4e-05 | 5.3e-06 | 1.110E-05 | 5.023E-06 | 5.300E-06 | 1.400E-05 |
| 12548 | 628 | Q628G | B | 7.5e-06 | 7.7e-06 | 6.7e-06 | 7.300E-06 | 5.292E-07 | 6.700E-06 | 7.700E-06 |
| 12549 | 628 | Q628H | B | 6.5e-06 | 7.5e-06 | 9.6e-06 | 7.867E-06 | 1.582E-06 | 6.500E-06 | 9.600E-06 |
| 12550 | 628 | Q628I | B | 1.5e-05 | 1.2e-05 | 6.9e-06 | 1.130E-05 | 4.095E-06 | 6.900E-06 | 1.500E-05 |
| 12551 | 628 | Q628L | B | 9.7e-06 | 7.2e-06 | 1.2e-05 | 9.633E-06 | 2.401E-06 | 7.200E-06 | 1.200E-05 |
| 12552 | 628 | Q628K | B | 5.5e-06 | 1e-05 | 6.1e-06 | 7.200E-06 | 2.443E-06 | 5.500E-06 | 1.000E-05 |
| 12553 | 628 | Q628M | B | 6.3e-06 | 8.9e-06 | 1.3e-05 | 9.400E-06 | 3.378E-06 | 6.300E-06 | 1.300E-05 |
| 12554 | 628 | Q628F | B | 5.9e-06 | 8.6e-06 | 1.5e-05 | 9.833E-06 | 4.674E-06 | 5.900E-06 | 1.500E-05 |
| 12555 | 628 | Q628P | B | 6.7e-06 | 1.1e-05 | 1e-05 | 9.233E-06 | 2.250E-06 | 6.700E-06 | 1.100E-05 |
| 12556 | 628 | Q628S | B | 7.6e-06 | 9.5e-06 | 7.3e-06 | 8.133E-06 | 1.193E-06 | 7.300E-06 | 9.500E-06 |
| 12557 | 628 | Q628T | B | 8.4e-06 | 9.6e-06 | 6.9e-06 | 8.300E-06 | 1.353E-06 | 6.900E-06 | 9.600E-06 |
| 12558 | 628 | Q628W | B | 8.7e-06 | 7e-06 | 5.7e-06 | 7.133E-06 | 1.504E-06 | 5.700E-06 | 8.700E-06 |
| 12559 | 628 | Q628Y | B | 8.1e-06 | 1.1e-05 | 6.8e-06 | 8.633E-06 | 2.150E-06 | 6.800E-06 | 1.100E-05 |
| 12560 | 628 | Q628V | B | 6.4e-06 | 8.5e-06 | 1.4e-05 | 9.633E-06 | 3.925E-06 | 6.400E-06 | 1.400E-05 |
| 12561 | 629 | S629A | B | 1.1e-05 | 6e-06 | 8.1e-06 | 8.367E-06 | 2.511E-06 | 6.000E-06 | 1.100E-05 |
| 12562 | 629 | S629R | B | 8e-06 | 7.6e-06 | 9.2e-06 | 8.267E-06 | 8.327E-07 | 7.600E-06 | 9.200E-06 |
| 12563 | 629 | S629N | B | 1.4e-05 | 8.1e-06 | 4.6e-06 | 8.900E-06 | 4.751E-06 | 4.600E-06 | 1.400E-05 |
| 12564 | 629 | S629D | B | 6.3e-06 | 1e-05 | 9e-06 | 8.433E-06 | 1.914E-06 | 6.300E-06 | 1.000E-05 |
| 12565 | 629 | S629C | B | 8.5e-06 | 8.8e-06 | 1e-05 | 9.100E-06 | 7.937E-07 | 8.500E-06 | 1.000E-05 |
| 12566 | 629 | S629E | B | 5.6e-06 | 6.7e-06 | 9.1e-06 | 7.133E-06 | 1.790E-06 | 5.600E-06 | 9.100E-06 |
| 12567 | 629 | S629Q | B | 7.4e-06 | 1e-05 | 9e-06 | 8.800E-06 | 1.311E-06 | 7.400E-06 | 1.000E-05 |
| 12568 | 629 | S629G | B | 7.5e-06 | 1.1e-05 | 5.7e-06 | 8.067E-06 | 2.695E-06 | 5.700E-06 | 1.100E-05 |
| 12569 | 629 | S629H | B | 6.8e-06 | 4.7e-06 | 1.2e-05 | 7.833E-06 | 3.758E-06 | 4.700E-06 | 1.200E-05 |
| 12570 | 629 | S629I | B | 6.5e-06 | 1.4e-05 | 5.2e-06 | 8.567E-06 | 4.750E-06 | 5.200E-06 | 1.400E-05 |
| 12571 | 629 | S629L | B | 6.9e-06 | 1.1e-05 | 1.1e-05 | 9.633E-06 | 2.367E-06 | 6.900E-06 | 1.100E-05 |
| 12572 | 629 | S629K | B | 1.1e-05 | 7.9e-06 | 6.4e-06 | 8.433E-06 | 2.346E-06 | 6.400E-06 | 1.100E-05 |
| 12573 | 629 | S629M | B | 9.7e-06 | 6.4e-06 | 6.8e-06 | 7.633E-06 | 1.801E-06 | 6.400E-06 | 9.700E-06 |
| 12574 | 629 | S629F | B | 8.4e-06 | 1.1e-05 | 1.2e-05 | 1.047E-05 | 1.858E-06 | 8.400E-06 | 1.200E-05 |
| 12575 | 629 | S629P | B | 4.8e-06 | 1.3e-05 | 1.2e-05 | 9.933E-06 | 4.474E-06 | 4.800E-06 | 1.300E-05 |
| 12576 | 629 | S629S | B | 5.9e-06 | 9.6e-06 | 7.5e-06 | 7.667E-06 | 1.856E-06 | 5.900E-06 | 9.600E-06 |
| 12577 | 629 | S629T | B | 1.2e-05 | 8.2e-06 | 9.3e-06 | 9.833E-06 | 1.955E-06 | 8.200E-06 | 1.200E-05 |
| 12578 | 629 | S629W | B | 6.6e-06 | 4.6e-06 | 7.6e-06 | 6.267E-06 | 1.528E-06 | 4.600E-06 | 7.600E-06 |
| 12579 | 629 | S629Y | B | 9.4e-06 | 5.1e-06 | 1e-05 | 8.167E-06 | 2.673E-06 | 5.100E-06 | 1.000E-05 |

| | | | | | | | | | | |
|---|---|---|---|---|---|---|---|---|---|---|
| 12580 | 629 | S629V | B | 1.1e-05 | 9.2e-06 | 6.4e-06 | 8.867E-06 | 2.318E-06 | 6.400E-06 | 1.100E-05 |
| 12581 | 630 | P630A | B | 9.2e-06 | 8.4e-06 | 6.7e-06 | 8.100E-06 | 1.277E-06 | 6.700E-06 | 9.200E-06 |
| 12582 | 630 | P630R | B | 9.8e-06 | 5.6e-06 | 6.1e-06 | 7.167E-06 | 2.294E-06 | 5.600E-06 | 9.800E-06 |
| 12583 | 630 | P630N | B | 7.7e-06 | 6.4e-06 | 6.8e-06 | 6.967E-06 | 6.658E-07 | 6.400E-06 | 7.700E-06 |
| 12584 | 630 | P630D | B | 6.6e-06 | 7.5e-06 | 7.7e-06 | 7.267E-06 | 5.859E-07 | 6.600E-06 | 7.700E-06 |
| 12585 | 630 | P630C | B | 5.9e-06 | 9.1e-06 | 9.6e-06 | 8.200E-06 | 2.007E-06 | 5.900E-06 | 9.600E-06 |
| 12586 | 630 | P630E | B | 8.4e-06 | 7.4e-06 | 8.8e-06 | 8.200E-06 | 7.211E-07 | 7.400E-06 | 8.800E-06 |
| 12587 | 630 | P630Q | B | 6.5e-06 | 1.1e-05 | 8.5e-06 | 8.667E-06 | 2.255E-06 | 6.500E-06 | 1.100E-05 |
| 12588 | 630 | P630G | B | 8.7e-06 | 7.8e-06 | 1.1e-05 | 9.167E-06 | 1.650E-06 | 7.800E-06 | 1.100E-05 |
| 12589 | 630 | P630H | B | 1.3e-05 | 7.4e-06 | 1.4e-05 | 1.147E-05 | 3.557E-06 | 7.400E-06 | 1.400E-05 |
| 12590 | 630 | P630I | B | 5.2e-06 | 8.3e-06 | 1.1e-05 | 8.167E-06 | 2.902E-06 | 5.200E-06 | 1.100E-05 |
| 12591 | 630 | P630L | B | 8.3e-06 | 9.8e-06 | 1.1e-05 | 9.700E-06 | 1.353E-06 | 8.300E-06 | 1.100E-05 |
| 12592 | 630 | P630K | B | 1.1e-05 | 1.2e-05 | 1.1e-05 | 1.133E-05 | 5.774E-07 | 1.100E-05 | 1.200E-05 |
| 12593 | 630 | P630M | B | 8.3e-06 | 6.9e-06 | 8.6e-06 | 7.933E-06 | 9.074E-07 | 6.900E-06 | 8.600E-06 |
| 12594 | 630 | P630F | B | 8.1e-06 | 9.3e-06 | 1.3e-05 | 1.013E-05 | 2.554E-06 | 8.100E-06 | 1.300E-05 |
| 12595 | 630 | P630P | B | 1.4e-05 | 9.9e-06 | 1.6e-05 | 1.330E-05 | 3.110E-06 | 9.900E-06 | 1.600E-05 |
| 12596 | 630 | P630S | B | 8.2e-06 | 9.3e-06 | 7.6e-06 | 8.367E-06 | 8.622E-07 | 7.600E-06 | 9.300E-06 |
| 12597 | 630 | P630T | B | 9.8e-06 | 6.1e-06 | 5.9e-06 | 7.267E-06 | 2.196E-06 | 5.900E-06 | 9.800E-06 |
| 12598 | 630 | P630W | B | 1.3e-05 | 1.1e-05 | 8.7e-06 | 1.090E-05 | 2.152E-06 | 8.700E-06 | 1.300E-05 |
| 12599 | 630 | P630Y | B | 5.2e-06 | 5.9e-06 | 6.8e-06 | 5.967E-06 | 8.021E-07 | 5.200E-06 | 6.800E-06 |
| 12600 | 630 | P630V | B | 1.1e-05 | 7.5e-06 | 9.5e-06 | 9.333E-06 | 1.756E-06 | 7.500E-06 | 1.100E-05 |
| 12601 | 631 | S631A | B | 6.5e-06 | 4.3e-06 | 1.3e-05 | 7.933E-06 | 4.524E-06 | 4.300E-06 | 1.300E-05 |
| 12602 | 631 | S631R | B | 6.7e-06 | 9.8e-06 | 5e-06 | 7.167E-06 | 2.434E-06 | 5.000E-06 | 9.800E-06 |
| 12603 | 631 | S631N | B | 1e-05 | 1.1e-05 | 6.9e-06 | 9.300E-06 | 2.138E-06 | 6.900E-06 | 1.100E-05 |
| 12604 | 631 | S631D | B | 5.4e-06 | 9.7e-06 | 1.1e-05 | 8.700E-06 | 2.931E-06 | 5.400E-06 | 1.100E-05 |
| 12605 | 631 | S631C | B | 4.7e-06 | 9.2e-06 | 7.1e-06 | 7.000E-06 | 2.252E-06 | 4.700E-06 | 9.200E-06 |
| 12606 | 631 | S631E | B | 5.7e-06 | 7.3e-06 | 6.1e-06 | 6.367E-06 | 8.327E-07 | 5.700E-06 | 7.300E-06 |
| 12607 | 631 | S631Q | B | 6.6e-06 | 9.7e-06 | 1.1e-05 | 9.100E-06 | 2.261E-06 | 6.600E-06 | 1.100E-05 |
| 12608 | 631 | S631G | B | 1.3e-05 | 6e-06 | 1.4e-05 | 1.100E-05 | 4.359E-06 | 6.000E-06 | 1.400E-05 |
| 12609 | 631 | S631H | B | 5.6e-06 | 7.2e-06 | 8.9e-06 | 7.233E-06 | 1.650E-06 | 5.600E-06 | 8.900E-06 |
| 12610 | 631 | S631I | B | 6.7e-06 | 1.1e-05 | 1.1e-05 | 9.567E-06 | 2.483E-06 | 6.700E-06 | 1.100E-05 |
| 12611 | 631 | S631L | B | 4.6e-06 | 5.7e-06 | 1.3e-05 | 7.767E-06 | 4.565E-06 | 4.600E-06 | 1.300E-05 |
| 12612 | 631 | S631K | B | 5.9e-06 | 8.7e-06 | 7.7e-06 | 7.433E-06 | 1.419E-06 | 5.900E-06 | 8.700E-06 |
| 12613 | 631 | S631M | B | 5.8e-06 | 6.6e-06 | 5.5e-06 | 5.967E-06 | 5.686E-07 | 5.500E-06 | 6.600E-06 |
| 12614 | 631 | S631F | B | 7.3e-06 | 9.6e-06 | 6.7e-06 | 7.867E-06 | 1.531E-06 | 6.700E-06 | 9.600E-06 |
| 12615 | 631 | S631P | B | 7.9e-06 | 8.3e-06 | 8.9e-06 | 8.367E-06 | 5.033E-07 | 7.900E-06 | 8.900E-06 |
| 12616 | 631 | S631S | B | 9.6e-06 | 6.6e-06 | 6.1e-06 | 7.433E-06 | 1.893E-06 | 6.100E-06 | 9.600E-06 |
| 12617 | 631 | S631T | B | 5.9e-06 | 5.1e-06 | 9.4e-06 | 6.800E-06 | 2.287E-06 | 5.100E-06 | 9.400E-06 |
| 12618 | 631 | S631W | B | 4.9e-06 | 4.4e-06 | 8e-06 | 5.767E-06 | 1.950E-06 | 4.400E-06 | 8.000E-06 |
| 12619 | 631 | S631Y | B | 5.1e-06 | 1.3e-05 | 9.4e-06 | 9.167E-06 | 3.955E-06 | 5.100E-06 | 1.300E-05 |
| 12620 | 631 | S631V | B | 1.4e-05 | 9.8e-06 | 9e-06 | 1.093E-05 | 2.686E-06 | 9.000E-06 | 1.400E-05 |
| 12621 | 632 | S632A | B | 9.4e-06 | 5.4e-06 | 1e-05 | 8.267E-06 | 2.501E-06 | 5.400E-06 | 1.000E-05 |
| 12622 | 632 | S632R | B | 6.1e-06 | 8.7e-06 | 9e-06 | 7.933E-06 | 1.595E-06 | 6.100E-06 | 9.000E-06 |
| 12623 | 632 | S632N | B | 1.1e-05 | 6.9e-06 | 1.4e-05 | 1.063E-05 | 3.564E-06 | 6.900E-06 | 1.400E-05 |
| 12624 | 632 | S632D | B | 8.8e-06 | 9.9e-06 | 9.7e-06 | 9.467E-06 | 5.859E-07 | 8.800E-06 | 9.900E-06 |
| 12625 | 632 | S632C | B | 6.9e-06 | 1.4e-05 | 6e-06 | 8.967E-06 | 4.382E-06 | 6.000E-06 | 1.400E-05 |
| 12626 | 632 | S632E | B | 9.6e-06 | 7.4e-06 | 7.2e-06 | 8.067E-06 | 1.332E-06 | 7.200E-06 | 9.600E-06 |
| 12627 | 632 | S632Q | B | 6.3e-06 | 7e-06 | 9.2e-06 | 7.500E-06 | 1.513E-06 | 6.300E-06 | 9.200E-06 |
| 12628 | 632 | S632G | B | 6.1e-06 | 1.1e-05 | 1.1e-05 | 9.367E-06 | 2.829E-06 | 6.100E-06 | 1.100E-05 |
| 12629 | 632 | S632H | B | 7.6e-06 | 7.9e-06 | 8.4e-06 | 7.967E-06 | 4.041E-07 | 7.600E-06 | 8.400E-06 |
| 12630 | 632 | S632I | B | 1.2e-05 | 7.7e-06 | 6.7e-06 | 8.800E-06 | 2.816E-06 | 6.700E-06 | 1.200E-05 |
| 12631 | 632 | S632L | B | 9.6e-06 | 9e-06 | 5.4e-06 | 8.000E-06 | 2.272E-06 | 5.400E-06 | 9.600E-06 |
| 12632 | 632 | S632K | B | 9.7e-06 | 4.7e-06 | 6.7e-06 | 7.033E-06 | 2.517E-06 | 4.700E-06 | 9.700E-06 |
| 12633 | 632 | S632M | B | 1.5e-05 | 9.2e-06 | 9.5e-06 | 1.123E-05 | 3.265E-06 | 9.200E-06 | 1.500E-05 |
| 12634 | 632 | S632F | B | 6.4e-06 | 4.8e-06 | 9.9e-06 | 7.033E-06 | 2.608E-06 | 4.800E-06 | 9.900E-06 |
| 12635 | 632 | S632P | B | 7.1e-06 | 8.6e-06 | 7.3e-06 | 7.667E-06 | 8.145E-07 | 7.100E-06 | 8.600E-06 |
| 12636 | 632 | S632S | B | 9.8e-06 | 1.2e-05 | 7e-06 | 9.600E-06 | 2.506E-06 | 7.000E-06 | 1.200E-05 |
| 12637 | 632 | S632T | B | 1.4e-05 | 5.4e-06 | 5e-06 | 8.133E-06 | 5.085E-06 | 5.000E-06 | 1.400E-05 |
| 12638 | 632 | S632W | B | 5.7e-06 | 6.3e-06 | 1.3e-05 | 8.333E-06 | 4.053E-06 | 5.700E-06 | 1.300E-05 |
| 12639 | 632 | S632Y | B | 9.6e-06 | 6.5e-06 | 5.7e-06 | 7.267E-06 | 2.060E-06 | 5.700E-06 | 9.600E-06 |
| 12640 | 632 | S632V | B | 7.2e-06 | 1e-05 | 1.3e-05 | 1.007E-05 | 2.901E-06 | 7.200E-06 | 1.300E-05 |
| 12641 | 633 | L633A | B | 9.1e-06 | 1.3e-05 | 6.8e-06 | 9.633E-06 | 3.134E-06 | 6.800E-06 | 1.300E-05 |
| 12642 | 633 | L633R | B | 6.8e-06 | 9.9e-06 | 7.1e-06 | 7.933E-06 | 1.710E-06 | 6.800E-06 | 9.900E-06 |
| 12643 | 633 | L633N | B | 8e-06 | 9.7e-06 | 9e-06 | 8.900E-06 | 8.544E-07 | 8.000E-06 | 9.700E-06 |
| 12644 | 633 | L633D | B | 6.1e-06 | 7.5e-06 | 5.2e-06 | 6.267E-06 | 1.159E-06 | 5.200E-06 | 7.500E-06 |
| 12645 | 633 | L633C | B | 8.4e-06 | 7.6e-06 | 8e-06 | 8.000E-06 | 4.000E-07 | 7.600E-06 | 8.400E-06 |
| 12646 | 633 | L633E | B | 7.4e-06 | 1.3e-05 | 6.7e-06 | 9.033E-06 | 3.453E-06 | 6.700E-06 | 1.300E-05 |
| 12647 | 633 | L633Q | B | 6.1e-06 | 9.6e-06 | 9.2e-06 | 8.300E-06 | 1.916E-06 | 6.100E-06 | 9.600E-06 |
| 12648 | 633 | L633G | B | 6.2e-06 | 1e-05 | 8.8e-06 | 8.333E-06 | 1.943E-06 | 6.200E-06 | 1.000E-05 |
| 12649 | 633 | L633H | B | 6.4e-06 | 6.8e-06 | 7.4e-06 | 6.867E-06 | 5.033E-07 | 6.400E-06 | 7.400E-06 |
| 12650 | 633 | L633I | B | 9.1e-06 | 4.6e-06 | 1e-05 | 7.900E-06 | 2.893E-06 | 4.600E-06 | 1.000E-05 |
| 12651 | 633 | L633L | B | 8.8e-06 | 7.7e-06 | 1.8e-05 | 1.150E-05 | 5.656E-06 | 7.700E-06 | 1.800E-05 |
| 12652 | 633 | L633K | B | 1.3e-05 | 8.9e-06 | 9.1e-06 | 1.033E-05 | 2.312E-06 | 8.900E-06 | 1.300E-05 |
| 12653 | 633 | L633M | B | 8.7e-06 | 9.3e-06 | 6.6e-06 | 8.200E-06 | 1.418E-06 | 6.600E-06 | 9.300E-06 |
| 12654 | 633 | L633F | B | 6.5e-06 | 7.4e-06 | 4.1e-06 | 6.000E-06 | 1.706E-06 | 4.100E-06 | 7.400E-06 |
| 12655 | 633 | L633P | B | 1.1e-05 | 4.4e-06 | 1.5e-05 | 1.013E-05 | 5.353E-06 | 4.400E-06 | 1.500E-05 |
| 12656 | 633 | L633S | B | 1.3e-05 | 9.3e-06 | 8.9e-06 | 1.040E-05 | 2.261E-06 | 8.900E-06 | 1.300E-05 |
| 12657 | 633 | L633T | B | 9.3e-06 | 8.6e-06 | 4.4e-06 | 7.433E-06 | 2.650E-06 | 4.400E-06 | 9.300E-06 |
| 12658 | 633 | L633W | B | 8e-06 | 1e-05 | 9.2e-06 | 9.067E-06 | 1.007E-06 | 8.000E-06 | 1.000E-05 |
| 12659 | 633 | L633Y | B | 8e-06 | 7.5e-06 | 7e-06 | 7.500E-06 | 5.000E-07 | 7.000E-06 | 8.000E-06 |
| 12660 | 633 | L633V | B | 7.1e-06 | 7.1e-06 | 7.8e-06 | 7.333E-06 | 4.041E-07 | 7.100E-06 | 7.800E-06 |
| 12661 | 634 | S634A | B | 6.6e-06 | 6.7e-06 | 1.1e-05 | 8.100E-06 | 2.512E-06 | 6.600E-06 | 1.100E-05 |
| 12662 | 634 | S634R | B | 8.5e-06 | 6.6e-06 | 9.7e-06 | 8.267E-06 | 1.563E-06 | 6.600E-06 | 9.700E-06 |
| 12663 | 634 | S634N | B | 1.2e-05 | 9.1e-06 | 5.7e-06 | 8.933E-06 | 3.153E-06 | 5.700E-06 | 1.200E-05 |
| 12664 | 634 | S634D | B | 6.5e-06 | 9.6e-06 | 1.8e-05 | 1.137E-05 | 5.950E-06 | 6.500E-06 | 1.800E-05 |

| | | | | | | | | | | |
|---|---|---|---|---|---|---|---|---|---|---|
| 12665 | 634 | S634C | B | 8.9e-06 | 9.2e-06 | 6.9e-06 | 8.333E-06 | 1.250E-06 | 6.900E-06 | 9.200E-06 |
| 12666 | 634 | S634E | B | 9.8e-06 | 6.1e-06 | 7e-06 | 7.633E-06 | 1.930E-06 | 6.100E-06 | 9.800E-06 |
| 12667 | 634 | S634Q | B | 7e-06 | 1.2e-05 | 1.1e-05 | 1.000E-05 | 2.646E-06 | 7.000E-06 | 1.200E-05 |
| 12668 | 634 | S634G | B | 1e-05 | 1e-05 | 6.8e-06 | 8.933E-06 | 1.848E-06 | 6.800E-06 | 1.000E-05 |
| 12669 | 634 | S634H | B | 3.3e-06 | 7.4e-06 | 1.2e-05 | 7.567E-06 | 4.352E-06 | 3.300E-06 | 1.200E-05 |
| 12670 | 634 | S634I | B | 5.6e-06 | 7.7e-06 | 1.3e-05 | 8.767E-06 | 3.814E-06 | 5.600E-06 | 1.300E-05 |
| 12671 | 634 | S634L | B | 1.3e-05 | 7.8e-06 | 7e-06 | 9.267E-06 | 3.258E-06 | 7.000E-06 | 1.300E-05 |
| 12672 | 634 | S634K | B | 1.5e-05 | 1.1e-05 | 8.2e-06 | 1.140E-05 | 3.418E-06 | 8.200E-06 | 1.500E-05 |
| 12673 | 634 | S634M | B | 7.8e-06 | 8.7e-06 | 6.2e-06 | 7.567E-06 | 1.266E-06 | 6.200E-06 | 8.700E-06 |
| 12674 | 634 | S634F | B | 6.8e-06 | 8.9e-06 | 6.5e-06 | 7.400E-06 | 1.308E-06 | 6.500E-06 | 8.900E-06 |
| 12675 | 634 | S634P | B | 3.2e-06 | 8.1e-06 | 5.7e-06 | 5.667E-06 | 2.450E-06 | 3.200E-06 | 8.100E-06 |
| 12676 | 634 | S634S | B | 1.2e-05 | 1.2e-05 | 1.2e-05 | 1.200E-05 | 0.000E+00 | 1.200E-05 | 1.200E-05 |
| 12677 | 634 | S634T | B | 8.6e-06 | 6.9e-06 | 7.9e-06 | 7.800E-06 | 8.544E-07 | 6.900E-06 | 8.600E-06 |
| 12678 | 634 | S634W | B | 6.4e-06 | 9.9e-06 | 6.5e-06 | 7.600E-06 | 1.992E-06 | 6.400E-06 | 9.900E-06 |
| 12679 | 634 | S634Y | B | 8.2e-06 | 7e-06 | 7.3e-06 | 7.500E-06 | 6.245E-07 | 7.000E-06 | 8.200E-06 |
| 12680 | 634 | S634V | B | 8.5e-06 | 5.4e-06 | 8.9e-06 | 7.600E-06 | 1.916E-06 | 5.400E-06 | 8.900E-06 |
| 12681 | 635 | A635A | B | 5.8e-06 | 7.6e-06 | 5.1e-06 | 6.167E-06 | 1.290E-06 | 5.100E-06 | 7.600E-06 |
| 12682 | 635 | A635R | B | 1.2e-05 | 7.1e-06 | 8.5e-06 | 9.200E-06 | 2.524E-06 | 7.100E-06 | 1.200E-05 |
| 12683 | 635 | A635N | B | 5.7e-06 | 7.4e-06 | 6.7e-06 | 6.600E-06 | 8.544E-07 | 5.700E-06 | 7.400E-06 |
| 12684 | 635 | A635D | B | 1.1e-05 | 8.4e-06 | 6.2e-06 | 8.533E-06 | 2.403E-06 | 6.200E-06 | 1.100E-05 |
| 12685 | 635 | A635C | B | 1.3e-05 | 6.4e-06 | 7.5e-06 | 8.967E-06 | 3.536E-06 | 6.400E-06 | 1.300E-05 |
| 12686 | 635 | A635E | B | 1e-05 | 7.1e-06 | 5.5e-06 | 7.533E-06 | 2.281E-06 | 5.500E-06 | 1.000E-05 |
| 12687 | 635 | A635Q | B | 1.1e-05 | 3.8e-06 | 6.2e-06 | 7.000E-06 | 3.666E-06 | 3.800E-06 | 1.100E-05 |
| 12688 | 635 | A635G | B | 6.4e-06 | 1e-05 | 8.4e-06 | 8.267E-06 | 1.804E-06 | 6.400E-06 | 1.000E-05 |
| 12689 | 635 | A635H | B | 4.7e-06 | 8.6e-06 | 5e-06 | 6.100E-06 | 2.170E-06 | 4.700E-06 | 8.600E-06 |
| 12690 | 635 | A635I | B | 6.8e-06 | 4.1e-06 | 9.5e-06 | 6.800E-06 | 2.700E-06 | 4.100E-06 | 9.500E-06 |
| 12691 | 635 | A635L | B | 6.2e-06 | 6.4e-06 | 7.9e-06 | 6.833E-06 | 9.292E-07 | 6.200E-06 | 7.900E-06 |
| 12692 | 635 | A635K | B | 1.3e-05 | 1.4e-05 | 8.3e-06 | 1.177E-05 | 3.044E-06 | 8.300E-06 | 1.400E-05 |
| 12693 | 635 | A635M | B | 7.5e-06 | 8.6e-06 | 1.3e-05 | 9.700E-06 | 2.910E-06 | 7.500E-06 | 1.300E-05 |
| 12694 | 635 | A635F | B | 9.1e-06 | 1.3e-05 | 8.7e-06 | 1.027E-05 | 2.376E-06 | 8.700E-06 | 1.300E-05 |
| 12695 | 635 | A635P | B | 5e-06 | 1.2e-05 | 5.3e-06 | 7.433E-06 | 3.958E-06 | 5.000E-06 | 1.200E-05 |
| 12696 | 635 | A635S | B | 5e-06 | 8.2e-06 | 8.6e-06 | 7.267E-06 | 1.973E-06 | 5.000E-06 | 8.600E-06 |
| 12697 | 635 | A635T | B | 8e-06 | 9.3e-06 | 6.5e-06 | 7.933E-06 | 1.401E-06 | 6.500E-06 | 9.300E-06 |
| 12698 | 635 | A635W | B | 5.9e-06 | 6.2e-06 | 1.1e-05 | 7.700E-06 | 2.862E-06 | 5.900E-06 | 1.100E-05 |
| 12699 | 635 | A635Y | B | 4.4e-06 | 5.6e-06 | 5.7e-06 | 5.233E-06 | 7.234E-07 | 4.400E-06 | 5.700E-06 |
| 12700 | 635 | A635V | B | 6.8e-06 | 9.7e-06 | 6.7e-06 | 7.733E-06 | 1.704E-06 | 6.700E-06 | 9.700E-06 |
| 12701 | 636 | S636A | B | 4.5e-06 | 7.1e-06 | 6.1e-06 | 5.900E-06 | 1.311E-06 | 4.500E-06 | 7.100E-06 |
| 12702 | 636 | S636R | B | 1.3e-05 | 6.2e-06 | 8e-06 | 9.067E-06 | 3.523E-06 | 6.200E-06 | 1.300E-05 |
| 12703 | 636 | S636N | B | 8.7e-06 | 4.3e-06 | 4.7e-06 | 5.900E-06 | 2.433E-06 | 4.300E-06 | 8.700E-06 |
| 12704 | 636 | S636D | B | 5.3e-06 | 1.2e-05 | 1.4e-05 | 1.043E-05 | 4.557E-06 | 5.300E-06 | 1.400E-05 |
| 12705 | 636 | S636C | B | 7.9e-06 | 9.6e-06 | 9.2e-06 | 8.900E-06 | 8.888E-07 | 7.900E-06 | 9.600E-06 |
| 12706 | 636 | S636E | B | 6.1e-06 | 9e-06 | 1.2e-05 | 9.033E-06 | 2.950E-06 | 6.100E-06 | 1.200E-05 |
| 12707 | 636 | S636Q | B | 6.4e-06 | 1e-05 | 1.1e-05 | 9.133E-06 | 2.419E-06 | 6.400E-06 | 1.100E-05 |
| 12708 | 636 | S636G | B | 7.8e-06 | 5.1e-06 | 9.2e-06 | 7.367E-06 | 2.084E-06 | 5.100E-06 | 9.200E-06 |
| 12709 | 636 | S636H | B | 7.5e-06 | 6.1e-06 | 7.6e-06 | 7.067E-06 | 8.386E-07 | 6.100E-06 | 7.600E-06 |
| 12710 | 636 | S636I | B | 4.7e-06 | 1.4e-05 | 7.8e-06 | 8.833E-06 | 4.735E-06 | 4.700E-06 | 1.400E-05 |
| 12711 | 636 | S636L | B | 1e-05 | 6.8e-06 | 6.7e-06 | 7.833E-06 | 1.877E-06 | 6.700E-06 | 1.000E-05 |
| 12712 | 636 | S636K | B | 8.9e-06 | 8.5e-06 | 1.2e-05 | 9.800E-06 | 1.916E-06 | 8.500E-06 | 1.200E-05 |
| 12713 | 636 | S636M | B | 9.6e-06 | 7.3e-06 | 1.8e-05 | 1.163E-05 | 5.632E-06 | 7.300E-06 | 1.800E-05 |
| 12714 | 636 | S636F | B | 1.3e-05 | 9.7e-06 | 8.7e-06 | 1.047E-05 | 2.250E-06 | 8.700E-06 | 1.300E-05 |
| 12715 | 636 | S636P | B | 1.1e-05 | 6e-06 | 1.3e-05 | 1.000E-05 | 3.606E-06 | 6.000E-06 | 1.300E-05 |
| 12716 | 636 | S636S | B | 9.2e-06 | 4.8e-06 | 6.1e-06 | 6.700E-06 | 2.261E-06 | 4.800E-06 | 9.200E-06 |
| 12717 | 636 | S636T | B | 6.2e-06 | 1.1e-05 | 9.7e-06 | 8.967E-06 | 2.483E-06 | 6.200E-06 | 1.100E-05 |
| 12718 | 636 | S636W | B | 4.9e-06 | 8.5e-06 | 5.9e-06 | 6.433E-06 | 1.858E-06 | 4.900E-06 | 8.500E-06 |
| 12719 | 636 | S636Y | B | 8.1e-06 | 1.3e-05 | 9.9e-06 | 1.033E-05 | 2.479E-06 | 8.100E-06 | 1.300E-05 |
| 12720 | 636 | S636V | B | 6.2e-06 | 9.2e-06 | 4.6e-06 | 6.667E-06 | 2.335E-06 | 4.600E-06 | 9.200E-06 |
| 12721 | 637 | V637A | B | 1.2e-05 | 8.7e-06 | 8.7e-06 | 9.800E-06 | 1.905E-06 | 8.700E-06 | 1.200E-05 |
| 12722 | 637 | V637R | B | 9e-06 | 8.4e-06 | 1e-05 | 9.133E-06 | 8.083E-07 | 8.400E-06 | 1.000E-05 |
| 12723 | 637 | V637N | B | 1.1e-05 | 1.1e-05 | 9.4e-06 | 1.047E-05 | 9.238E-07 | 9.400E-06 | 1.100E-05 |
| 12724 | 637 | V637D | B | 1.2e-05 | 7.8e-06 | 1.1e-05 | 1.027E-05 | 2.194E-06 | 7.800E-06 | 1.200E-05 |
| 12725 | 637 | V637C | B | 8.9e-06 | 1.1e-05 | 1.2e-05 | 1.063E-05 | 1.582E-06 | 8.900E-06 | 1.200E-05 |
| 12726 | 637 | V637E | B | 5.7e-06 | 5e-06 | 7.1e-06 | 5.933E-06 | 1.069E-06 | 5.000E-06 | 7.100E-06 |
| 12727 | 637 | V637Q | B | 7.5e-06 | 5.1e-06 | 1.2e-05 | 8.200E-06 | 3.503E-06 | 5.100E-06 | 1.200E-05 |
| 12728 | 637 | V637G | B | 5.5e-06 | 1e-05 | 1.3e-05 | 9.500E-06 | 3.775E-06 | 5.500E-06 | 1.300E-05 |
| 12729 | 637 | V637H | B | 1.1e-05 | 1.1e-05 | 7.2e-06 | 9.733E-06 | 2.194E-06 | 7.200E-06 | 1.100E-05 |
| 12730 | 637 | V637I | B | 8.1e-06 | 1.2e-05 | 6.9e-06 | 9.000E-06 | 2.666E-06 | 6.900E-06 | 1.200E-05 |
| 12731 | 637 | V637L | B | 6.8e-06 | 6.7e-06 | 8.4e-06 | 7.300E-06 | 9.539E-07 | 6.700E-06 | 8.400E-06 |
| 12732 | 637 | V637K | B | 5.2e-06 | 4.4e-06 | 7.2e-06 | 5.600E-06 | 1.442E-06 | 4.400E-06 | 7.200E-06 |
| 12733 | 637 | V637M | B | 5.2e-06 | 5.7e-06 | 8.5e-06 | 6.467E-06 | 1.779E-06 | 5.200E-06 | 8.500E-06 |
| 12734 | 637 | V637F | B | 5e-06 | 1.2e-05 | 6.6e-06 | 7.867E-06 | 3.668E-06 | 5.000E-06 | 1.200E-05 |
| 12735 | 637 | V637P | B | 7.3e-06 | 7.1e-06 | 6.8e-06 | 7.067E-06 | 2.517E-07 | 6.800E-06 | 7.300E-06 |
| 12736 | 637 | V637S | B | 6.5e-06 | 4.4e-06 | 6.2e-06 | 5.700E-06 | 1.136E-06 | 4.400E-06 | 6.500E-06 |
| 12737 | 637 | V637T | B | 6.6e-06 | 9.2e-06 | 7.6e-06 | 7.800E-06 | 1.311E-06 | 6.600E-06 | 9.200E-06 |
| 12738 | 637 | V637W | B | 6.6e-06 | 5.8e-06 | 8.3e-06 | 6.900E-06 | 1.277E-06 | 5.800E-06 | 8.300E-06 |
| 12739 | 637 | V637Y | B | 9.1e-06 | 5.8e-06 | 9.2e-06 | 8.033E-06 | 1.935E-06 | 5.800E-06 | 9.200E-06 |
| 12740 | 637 | V637V | B | 9e-06 | 7e-06 | 1.1e-05 | 9.000E-06 | 2.000E-06 | 7.000E-06 | 1.100E-05 |
| 12741 | 638 | G638A | B | 5.9e-06 | 8e-06 | 6.7e-06 | 6.867E-06 | 1.060E-06 | 5.900E-06 | 8.000E-06 |
| 12742 | 638 | G638R | B | 6.2e-06 | 6.3e-06 | 9.2e-06 | 7.233E-06 | 1.704E-06 | 6.200E-06 | 9.200E-06 |
| 12743 | 638 | G638N | B | 6.5e-06 | 1.3e-05 | 5.4e-06 | 8.300E-06 | 4.107E-06 | 5.400E-06 | 1.300E-05 |
| 12744 | 638 | G638D | B | 7.3e-06 | 6.3e-06 | 9e-06 | 7.533E-06 | 1.365E-06 | 6.300E-06 | 9.000E-06 |
| 12745 | 638 | G638C | B | 8.5e-06 | 4.4e-06 | 7.9e-06 | 6.933E-06 | 2.214E-06 | 4.400E-06 | 8.500E-06 |
| 12746 | 638 | G638E | B | 7.6e-06 | 5.4e-06 | 9.2e-06 | 7.400E-06 | 1.908E-06 | 5.400E-06 | 9.200E-06 |
| 12747 | 638 | G638Q | B | 8.5e-06 | 8.8e-06 | 8.6e-06 | 8.633E-06 | 1.528E-07 | 8.500E-06 | 8.800E-06 |
| 12748 | 638 | G638G | B | 1.1e-05 | 1.4e-05 | 1.2e-05 | 1.233E-05 | 1.528E-06 | 1.100E-05 | 1.400E-05 |
| 12749 | 638 | G638H | B | 6.1e-06 | 4.5e-06 | 9.2e-06 | 6.600E-06 | 2.390E-06 | 4.500E-06 | 9.200E-06 |

| | | | | | | | | | | |
|---|---|---|---|---|---|---|---|---|---|---|
| 12750 | 638 | G638I | B | 9.5e-06 | 6.2e-06 | 5.1e-06 | 6.933E-06 | 2.290E-06 | 5.100E-06 | 9.500E-06 |
| 12751 | 638 | G638L | B | 7.5e-06 | 6.5e-06 | 4.5e-06 | 6.167E-06 | 1.528E-06 | 4.500E-06 | 7.500E-06 |
| 12752 | 638 | G638K | B | 6.4e-06 | 5.1e-06 | 8.2e-06 | 6.567E-06 | 1.557E-06 | 5.100E-06 | 8.200E-06 |
| 12753 | 638 | G638M | B | 6.7e-06 | 1e-05 | 9e-06 | 8.567E-06 | 1.692E-06 | 6.700E-06 | 1.000E-05 |
| 12754 | 638 | G638F | B | 9.5e-06 | 7.8e-06 | 8.9e-06 | 8.733E-06 | 8.622E-07 | 7.800E-06 | 9.500E-06 |
| 12755 | 638 | G638P | B | 7.5e-06 | 8.7e-06 | 1.1e-05 | 9.067E-06 | 1.779E-06 | 7.500E-06 | 1.100E-05 |
| 12756 | 638 | G638S | B | 4.4e-06 | 6.4e-06 | 6e-06 | 5.600E-06 | 1.058E-06 | 4.400E-06 | 6.400E-06 |
| 12757 | 638 | G638T | B | 1e-05 | 5.8e-06 | 5.1e-06 | 6.967E-06 | 2.650E-06 | 5.100E-06 | 1.000E-05 |
| 12758 | 638 | G638W | B | 7e-06 | 5e-06 | 7e-06 | 6.333E-06 | 1.155E-06 | 5.000E-06 | 7.000E-06 |
| 12759 | 638 | G638Y | B | 6.9e-06 | 6.9e-06 | 8.7e-06 | 7.500E-06 | 1.039E-06 | 6.900E-06 | 8.700E-06 |
| 12760 | 638 | G638V | B | 5.3e-06 | 8e-06 | 6.8e-06 | 6.700E-06 | 1.353E-06 | 5.300E-06 | 8.000E-06 |
| 12761 | 639 | D639A | B | 8.4e-06 | 7.3e-06 | 1.4e-05 | 9.900E-06 | 3.593E-06 | 7.300E-06 | 1.400E-05 |
| 12762 | 639 | D639R | B | 9.5e-06 | 4.7e-06 | 6.4e-06 | 6.867E-06 | 2.434E-06 | 4.700E-06 | 9.500E-06 |
| 12763 | 639 | D639N | B | 6.9e-06 | 6e-06 | 7.8e-06 | 6.900E-06 | 9.000E-07 | 6.000E-06 | 7.800E-06 |
| 12764 | 639 | D639D | B | 5e-06 | 1e-05 | 8e-06 | 7.667E-06 | 2.517E-06 | 5.000E-06 | 1.000E-05 |
| 12765 | 639 | D639C | B | 1.5e-05 | 6.9e-06 | 8.1e-06 | 1.000E-05 | 4.371E-06 | 6.900E-06 | 1.500E-05 |
| 12766 | 639 | D639E | B | 8.2e-06 | 6.2e-06 | 6.7e-06 | 7.033E-06 | 1.041E-06 | 6.200E-06 | 8.200E-06 |
| 12767 | 639 | D639Q | B | 9.2e-06 | 6.4e-06 | 7.1e-06 | 7.567E-06 | 1.457E-06 | 6.400E-06 | 9.200E-06 |
| 12768 | 639 | D639G | B | 6.4e-06 | 5.7e-06 | 7.9e-06 | 6.667E-06 | 1.124E-06 | 5.700E-06 | 7.900E-06 |
| 12769 | 639 | D639H | B | 8.9e-06 | 1.3e-05 | 4.9e-06 | 8.933E-06 | 4.050E-06 | 4.900E-06 | 1.300E-05 |
| 12770 | 639 | D639I | B | 4.3e-06 | 6.8e-06 | 1e-05 | 7.033E-06 | 2.857E-06 | 4.300E-06 | 1.000E-05 |
| 12771 | 639 | D639L | B | 9.6e-06 | 5e-06 | 9e-06 | 7.867E-06 | 2.501E-06 | 5.000E-06 | 9.600E-06 |
| 12772 | 639 | D639K | B | 1.3e-05 | 1.1e-05 | 9.4e-06 | 1.113E-05 | 1.804E-06 | 9.400E-06 | 1.300E-05 |
| 12773 | 639 | D639M | B | 8.9e-06 | 9.2e-06 | 6e-06 | 8.033E-06 | 1.767E-06 | 6.000E-06 | 9.200E-06 |
| 12774 | 639 | D639F | B | 7.5e-06 | 7.8e-06 | 4.5e-06 | 6.600E-06 | 1.825E-06 | 4.500E-06 | 7.800E-06 |
| 12775 | 639 | D639P | B | 8.1e-06 | 6.3e-06 | 8.5e-06 | 7.633E-06 | 1.172E-06 | 6.300E-06 | 8.500E-06 |
| 12776 | 639 | D639S | B | 8.4e-06 | 5.2e-06 | 7.1e-06 | 6.900E-06 | 1.609E-06 | 5.200E-06 | 8.400E-06 |
| 12777 | 639 | D639T | B | 1.2e-05 | 6.4e-06 | 6.8e-06 | 8.400E-06 | 3.124E-06 | 6.400E-06 | 1.200E-05 |
| 12778 | 639 | D639W | B | 9.4e-06 | 5.8e-06 | 5e-06 | 6.733E-06 | 2.344E-06 | 5.000E-06 | 9.400E-06 |
| 12779 | 639 | D639Y | B | 1.3e-05 | 6.9e-06 | 9.3e-06 | 9.733E-06 | 3.073E-06 | 6.900E-06 | 1.300E-05 |
| 12780 | 639 | D639V | B | 6.7e-06 | 7.7e-06 | 6.8e-06 | 7.067E-06 | 5.508E-07 | 6.700E-06 | 7.700E-06 |
| 12781 | 640 | R640A | B | 9.9e-06 | 8.2e-06 | 7.7e-06 | 8.600E-06 | 1.153E-06 | 7.700E-06 | 9.900E-06 |
| 12782 | 640 | R640R | B | 1.1e-05 | 1.1e-05 | 8.5e-06 | 1.017E-05 | 1.443E-06 | 8.500E-06 | 1.100E-05 |
| 12783 | 640 | R640N | B | 7.3e-06 | 7.8e-06 | 8.3e-06 | 7.800E-06 | 5.000E-07 | 7.300E-06 | 8.300E-06 |
| 12784 | 640 | R640D | B | 8.2e-06 | 6.4e-06 | 5.9e-06 | 6.833E-06 | 1.210E-06 | 5.900E-06 | 8.200E-06 |
| 12785 | 640 | R640C | B | 9.5e-06 | 9.7e-06 | 6e-06 | 8.400E-06 | 2.081E-06 | 6.000E-06 | 9.700E-06 |
| 12786 | 640 | R640E | B | 8e-06 | 1.2e-05 | 7.4e-06 | 9.133E-06 | 2.501E-06 | 7.400E-06 | 1.200E-05 |
| 12787 | 640 | R640Q | B | 2.1e-05 | 4.9e-06 | 4.5e-06 | 1.013E-05 | 9.413E-06 | 4.500E-06 | 2.100E-05 |
| 12788 | 640 | R640G | B | 9.5e-06 | 5.3e-06 | 4.3e-06 | 6.367E-06 | 2.759E-06 | 4.300E-06 | 9.500E-06 |
| 12789 | 640 | R640H | B | 9.6e-06 | 6.7e-06 | 8.5e-06 | 8.267E-06 | 1.464E-06 | 6.700E-06 | 9.600E-06 |
| 12790 | 640 | R640I | B | 8.4e-06 | 4.7e-06 | 7.4e-06 | 6.833E-06 | 1.914E-06 | 4.700E-06 | 8.400E-06 |
| 12791 | 640 | R640L | B | 7.4e-06 | 8e-06 | 8.2e-06 | 7.867E-06 | 4.163E-07 | 7.400E-06 | 8.200E-06 |
| 12792 | 640 | R640K | B | 5.4e-06 | 6.2e-06 | 1.2e-05 | 7.867E-06 | 3.602E-06 | 5.400E-06 | 1.200E-05 |
| 12793 | 640 | R640M | B | 8.5e-06 | 9.2e-06 | 9.3e-06 | 9.000E-06 | 4.359E-07 | 8.500E-06 | 9.300E-06 |
| 12794 | 640 | R640F | B | 9.4e-06 | 9.8e-06 | 1e-05 | 9.733E-06 | 3.055E-07 | 9.400E-06 | 1.000E-05 |
| 12795 | 640 | R640P | B | 8.8e-06 | 8.2e-06 | 9.1e-06 | 8.700E-06 | 4.583E-07 | 8.200E-06 | 9.100E-06 |
| 12796 | 640 | R640S | B | 8.9e-06 | 9e-06 | 6.4e-06 | 8.100E-06 | 1.473E-06 | 6.400E-06 | 9.000E-06 |
| 12797 | 640 | R640T | B | 1.2e-05 | 5.8e-06 | 6.8e-06 | 8.200E-06 | 3.329E-06 | 5.800E-06 | 1.200E-05 |
| 12798 | 640 | R640W | B | 6.5e-06 | 1.1e-05 | 1.3e-05 | 1.017E-05 | 3.329E-06 | 6.500E-06 | 1.300E-05 |
| 12799 | 640 | R640Y | B | 7.5e-06 | 9.8e-06 | 5.9e-06 | 7.733E-06 | 1.960E-06 | 5.900E-06 | 9.800E-06 |
| 12800 | 640 | R640V | B | 5.7e-06 | 6.6e-06 | 8.9e-06 | 7.067E-06 | 1.650E-06 | 5.700E-06 | 8.900E-06 |
| 12801 | 641 | V641A | B | 6.9e-06 | 3.7e-06 | 7.2e-06 | 5.933E-06 | 1.940E-06 | 3.700E-06 | 7.200E-06 |
| 12802 | 641 | V641R | B | 7.1e-06 | 7e-06 | 8.6e-06 | 7.567E-06 | 8.963E-07 | 7.000E-06 | 8.600E-06 |
| 12803 | 641 | V641N | B | 1.2e-05 | 8.5e-06 | 7.5e-06 | 9.333E-06 | 2.363E-06 | 7.500E-06 | 1.200E-05 |
| 12804 | 641 | V641D | B | 1.1e-05 | 4.7e-06 | 6.4e-06 | 7.367E-06 | 3.259E-06 | 4.700E-06 | 1.100E-05 |
| 12805 | 641 | V641C | B | 5.1e-06 | 8.8e-06 | 1e-05 | 7.967E-06 | 2.554E-06 | 5.100E-06 | 1.000E-05 |
| 12806 | 641 | V641E | B | 6.1e-06 | 6.7e-06 | 6.9e-06 | 6.567E-06 | 4.163E-07 | 6.100E-06 | 6.900E-06 |
| 12807 | 641 | V641Q | B | 6.5e-06 | 1.3e-05 | 6.3e-06 | 8.600E-06 | 3.812E-06 | 6.300E-06 | 1.300E-05 |
| 12808 | 641 | V641G | B | 6.5e-06 | 6.7e-06 | 1e-05 | 7.733E-06 | 1.966E-06 | 6.500E-06 | 1.000E-05 |
| 12809 | 641 | V641H | B | 1.2e-05 | 9.9e-06 | 1.4e-05 | 1.197E-05 | 2.050E-06 | 9.900E-06 | 1.400E-05 |
| 12810 | 641 | V641I | B | 7.8e-06 | 4.8e-06 | 1e-05 | 7.533E-06 | 2.610E-06 | 4.800E-06 | 1.000E-05 |
| 12811 | 641 | V641L | B | 8e-06 | 8.8e-06 | 6.5e-06 | 7.767E-06 | 1.168E-06 | 6.500E-06 | 8.800E-06 |
| 12812 | 641 | V641K | B | 7.6e-06 | 6.5e-06 | 5.1e-06 | 6.400E-06 | 1.253E-06 | 5.100E-06 | 7.600E-06 |
| 12813 | 641 | V641M | B | 5.3e-06 | 8.6e-06 | 1e-05 | 7.967E-06 | 2.413E-06 | 5.300E-06 | 1.000E-05 |
| 12814 | 641 | V641F | B | 6.6e-06 | 6.7e-06 | 8.2e-06 | 7.167E-06 | 8.963E-07 | 6.600E-06 | 8.200E-06 |
| 12815 | 641 | V641P | B | 9.5e-06 | 1.3e-05 | 2e-05 | 1.417E-05 | 5.346E-06 | 9.500E-06 | 2.000E-05 |
| 12816 | 641 | V641S | B | 5.1e-06 | 5.4e-06 | 7.2e-06 | 5.900E-06 | 1.136E-06 | 5.100E-06 | 7.200E-06 |
| 12817 | 641 | V641T | B | 5e-06 | 3.2e-06 | 1e-05 | 6.067E-06 | 3.523E-06 | 3.200E-06 | 1.000E-05 |
| 12818 | 641 | V641W | B | 8.5e-06 | 8e-06 | 7.1e-06 | 7.867E-06 | 7.095E-07 | 7.100E-06 | 8.500E-06 |
| 12819 | 641 | V641Y | B | 4.4e-06 | 6.9e-06 | 5.6e-06 | 5.633E-06 | 1.250E-06 | 4.400E-06 | 6.900E-06 |
| 12820 | 641 | V641V | B | 8.3e-06 | 9.1e-06 | 8.6e-06 | 8.667E-06 | 4.041E-07 | 8.300E-06 | 9.100E-06 |
| 12821 | 642 | T642A | B | 7.7e-06 | 7e-06 | 6.7e-06 | 7.133E-06 | 5.132E-07 | 6.700E-06 | 7.700E-06 |
| 12822 | 642 | T642R | B | 9.2e-06 | 9e-06 | 9.3e-06 | 9.167E-06 | 1.528E-07 | 9.000E-06 | 9.300E-06 |
| 12823 | 642 | T642N | B | 6.9e-06 | 8.9e-06 | 8e-06 | 7.933E-06 | 1.002E-06 | 6.900E-06 | 8.900E-06 |
| 12824 | 642 | T642D | B | 5.7e-06 | 8.7e-06 | 8.2e-06 | 7.533E-06 | 1.607E-06 | 5.700E-06 | 8.700E-06 |
| 12825 | 642 | T642C | B | 1.1e-05 | 9.9e-06 | 6.8e-06 | 9.233E-06 | 2.178E-06 | 6.800E-06 | 1.100E-05 |
| 12826 | 642 | T642E | B | 1e-05 | 4.5e-06 | 1.2e-05 | 8.833E-06 | 3.884E-06 | 4.500E-06 | 1.200E-05 |
| 12827 | 642 | T642Q | B | 9.5e-06 | 7.7e-06 | 7.9e-06 | 8.367E-06 | 9.866E-07 | 7.700E-06 | 9.500E-06 |
| 12828 | 642 | T642G | B | 9.8e-06 | 5.8e-06 | 9.5e-06 | 8.367E-06 | 2.228E-06 | 5.800E-06 | 9.800E-06 |
| 12829 | 642 | T642H | B | 1.3e-05 | 7.4e-06 | 9.4e-06 | 9.933E-06 | 2.838E-06 | 7.400E-06 | 1.300E-05 |
| 12830 | 642 | T642I | B | 9.5e-06 | 8.6e-06 | 1.1e-05 | 9.700E-06 | 1.212E-06 | 8.600E-06 | 1.100E-05 |
| 12831 | 642 | T642L | B | 1.4e-05 | 6.4e-06 | 4.8e-06 | 8.400E-06 | 4.915E-06 | 4.800E-06 | 1.400E-05 |
| 12832 | 642 | T642K | B | 6.9e-06 | 8.1e-06 | 6.7e-06 | 7.233E-06 | 7.572E-07 | 6.700E-06 | 8.100E-06 |
| 12833 | 642 | T642M | B | 9.7e-06 | 7e-06 | 9.1e-06 | 8.600E-06 | 1.418E-06 | 7.000E-06 | 9.700E-06 |
| 12834 | 642 | T642F | B | 6.8e-06 | 9.8e-06 | 6.6e-06 | 7.733E-06 | 1.793E-06 | 6.600E-06 | 9.800E-06 |

| | | | | | | | | | | |
|---|---|---|---|---|---|---|---|---|---|---|
| 12835 | 642 | T642P | B | 1.7e-05 | 1.3e-05 | 4.5e-06 | 1.150E-05 | 6.384E-06 | 4.500E-06 | 1.700E-05 |
| 12836 | 642 | T642S | B | 6.6e-06 | 8.3e-06 | 5.5e-06 | 6.800E-06 | 1.411E-06 | 5.500E-06 | 8.300E-06 |
| 12837 | 642 | T642T | B | 5.3e-06 | 9e-06 | 1e-05 | 8.100E-06 | 2.476E-06 | 5.300E-06 | 1.000E-05 |
| 12838 | 642 | T642W | B | 4.4e-06 | 1.3e-05 | 8e-06 | 8.467E-06 | 4.319E-06 | 4.400E-06 | 1.300E-05 |
| 12839 | 642 | T642Y | B | 9e-06 | 9.4e-06 | 7.5e-06 | 8.633E-06 | 1.002E-06 | 7.500E-06 | 9.400E-06 |
| 12840 | 642 | T642V | B | 7.1e-06 | 1.1e-05 | 5.4e-06 | 7.833E-06 | 2.871E-06 | 5.400E-06 | 1.100E-05 |
| 12841 | 643 | I643A | B | 8.8e-06 | 7.4e-06 | 8.9e-06 | 8.367E-06 | 8.386E-07 | 7.400E-06 | 8.900E-06 |
| 12842 | 643 | I643R | B | 6e-06 | 9e-06 | 1.2e-05 | 9.000E-06 | 3.000E-06 | 6.000E-06 | 1.200E-05 |
| 12843 | 643 | I643N | B | 5.6e-06 | 1.1e-05 | 5.1e-06 | 7.233E-06 | 3.272E-06 | 5.100E-06 | 1.100E-05 |
| 12844 | 643 | I643D | B | 8.2e-06 | 8.5e-06 | 7e-06 | 7.900E-06 | 7.937E-07 | 7.000E-06 | 8.500E-06 |
| 12845 | 643 | I643C | B | 1.2e-05 | 1.1e-05 | 3.9e-06 | 8.967E-06 | 4.416E-06 | 3.900E-06 | 1.200E-05 |
| 12846 | 643 | I643E | B | 1e-05 | 6.3e-06 | 5.7e-06 | 7.333E-06 | 2.329E-06 | 5.700E-06 | 1.000E-05 |
| 12847 | 643 | I643Q | B | 7.3e-06 | 5.6e-06 | 4.5e-06 | 5.800E-06 | 1.411E-06 | 4.500E-06 | 7.300E-06 |
| 12848 | 643 | I643G | B | 6.4e-06 | 1.5e-05 | 9e-06 | 1.013E-05 | 4.411E-06 | 6.400E-06 | 1.500E-05 |
| 12849 | 643 | I643H | B | 9e-06 | 1.3e-05 | 6.2e-06 | 9.400E-06 | 3.418E-06 | 6.200E-06 | 1.300E-05 |
| 12850 | 643 | I643I | B | 9.1e-06 | 6.9e-06 | 4.8e-06 | 6.933E-06 | 2.150E-06 | 4.800E-06 | 9.100E-06 |
| 12851 | 643 | I643L | B | 6.6e-06 | 7.6e-06 | 1e-05 | 8.067E-06 | 1.747E-06 | 6.600E-06 | 1.000E-05 |
| 12852 | 643 | I643K | B | 8.2e-06 | 9.6e-06 | 7.3e-06 | 8.367E-06 | 1.159E-06 | 7.300E-06 | 9.600E-06 |
| 12853 | 643 | I643M | B | 9.5e-06 | 7.2e-06 | 4.3e-06 | 7.000E-06 | 2.606E-06 | 4.300E-06 | 9.500E-06 |
| 12854 | 643 | I643F | B | 9.2e-06 | 7.2e-06 | 8.1e-06 | 8.167E-06 | 1.002E-06 | 7.200E-06 | 9.200E-06 |
| 12855 | 643 | I643P | B | 9.7e-06 | 7.7e-06 | 1e-05 | 9.133E-06 | 1.250E-06 | 7.700E-06 | 1.000E-05 |
| 12856 | 643 | I643S | B | 1.3e-05 | 6.8e-06 | 1e-05 | 9.933E-06 | 3.101E-06 | 6.800E-06 | 1.300E-05 |
| 12857 | 643 | I643T | B | 1.3e-05 | 1.3e-05 | 7.1e-06 | 1.103E-05 | 3.406E-06 | 7.100E-06 | 1.300E-05 |
| 12858 | 643 | I643W | B | 1.3e-05 | 6.2e-06 | 6.6e-06 | 8.600E-06 | 3.816E-06 | 6.200E-06 | 1.300E-05 |
| 12859 | 643 | I643Y | B | 9.5e-06 | 1.3e-05 | 6.5e-06 | 9.667E-06 | 3.253E-06 | 6.500E-06 | 1.300E-05 |
| 12860 | 643 | I643V | B | 6.1e-06 | 1.6e-05 | 3.9e-06 | 8.667E-06 | 6.445E-06 | 3.900E-06 | 1.600E-05 |
| 12861 | 644 | T644A | B | 6.2e-06 | 5.4e-06 | 9.1e-06 | 6.900E-06 | 1.947E-06 | 5.400E-06 | 9.100E-06 |
| 12862 | 644 | T644R | B | 6.6e-06 | 6.7e-06 | 8.8e-06 | 7.367E-06 | 1.242E-06 | 6.600E-06 | 8.800E-06 |
| 12863 | 644 | T644N | B | 7.6e-06 | 7.4e-06 | 8.3e-06 | 7.767E-06 | 4.726E-07 | 7.400E-06 | 8.300E-06 |
| 12864 | 644 | T644D | B | 9.9e-06 | 3.9e-06 | 9.7e-06 | 7.833E-06 | 3.408E-06 | 3.900E-06 | 9.900E-06 |
| 12865 | 644 | T644C | B | 5.9e-06 | 6.4e-06 | 4.7e-06 | 5.667E-06 | 8.737E-07 | 4.700E-06 | 6.400E-06 |
| 12866 | 644 | T644E | B | 1.1e-05 | 9.8e-06 | 7.6e-06 | 9.467E-06 | 1.724E-06 | 7.600E-06 | 1.100E-05 |
| 12867 | 644 | T644Q | B | 1.3e-05 | 1.5e-05 | 2.5e-05 | 1.767E-05 | 6.429E-06 | 1.300E-05 | 2.500E-05 |
| 12868 | 644 | T644G | B | 4.6e-06 | 4.8e-06 | 1.4e-05 | 7.800E-06 | 5.370E-06 | 4.600E-06 | 1.400E-05 |
| 12869 | 644 | T644H | B | 6.3e-06 | 6.8e-06 | 6.4e-06 | 6.500E-06 | 2.646E-07 | 6.300E-06 | 6.800E-06 |
| 12870 | 644 | T644I | B | 9.3e-06 | 1.4e-05 | 9.2e-06 | 1.083E-05 | 2.743E-06 | 9.200E-06 | 1.400E-05 |
| 12871 | 644 | T644L | B | 1.1e-05 | 6.5e-06 | 5e-06 | 7.500E-06 | 3.122E-06 | 5.000E-06 | 1.100E-05 |
| 12872 | 644 | T644K | B | 7e-06 | 9e-06 | 9.1e-06 | 8.367E-06 | 1.185E-06 | 7.000E-06 | 9.100E-06 |
| 12873 | 644 | T644M | B | 5.6e-06 | 6.2e-06 | 1.3e-05 | 8.267E-06 | 4.110E-06 | 5.600E-06 | 1.300E-05 |
| 12874 | 644 | T644F | B | 6.7e-06 | 9.8e-06 | 4.6e-06 | 7.033E-06 | 2.616E-06 | 4.600E-06 | 9.800E-06 |
| 12875 | 644 | T644P | B | 9.8e-06 | 1e-05 | 1.3e-05 | 1.093E-05 | 1.793E-06 | 9.800E-06 | 1.300E-05 |
| 12876 | 644 | T644S | B | 5.5e-06 | 3.6e-06 | 5.8e-06 | 4.967E-06 | 1.193E-06 | 3.600E-06 | 5.800E-06 |
| 12877 | 644 | T644T | B | 6.9e-06 | 9.6e-06 | 9.3e-06 | 8.600E-06 | 1.480E-06 | 6.900E-06 | 9.600E-06 |
| 12878 | 644 | T644W | B | 9.6e-06 | 9.7e-06 | 1.8e-05 | 1.243E-05 | 4.821E-06 | 9.600E-06 | 1.800E-05 |
| 12879 | 644 | T644Y | B | 7.3e-06 | 9.1e-06 | 1.2e-05 | 9.467E-06 | 2.371E-06 | 7.300E-06 | 1.200E-05 |
| 12880 | 644 | T644V | B | 1.1e-05 | 5.2e-06 | 1.1e-05 | 9.067E-06 | 3.349E-06 | 5.200E-06 | 1.100E-05 |
| 12881 | 645 | C645A | B | 1.1e-05 | 1.2e-05 | 8e-06 | 1.033E-05 | 2.082E-06 | 8.000E-06 | 1.200E-05 |
| 12882 | 645 | C645R | B | 1e-05 | 5.7e-06 | 1.2e-05 | 9.233E-06 | 3.219E-06 | 5.700E-06 | 1.200E-05 |
| 12883 | 645 | C645N | B | 7.1e-06 | 7.5e-06 | 7.1e-06 | 7.233E-06 | 2.309E-07 | 7.100E-06 | 7.500E-06 |
| 12884 | 645 | C645D | B | 7.6e-06 | 1.2e-05 | 8.6e-06 | 9.400E-06 | 2.307E-06 | 7.600E-06 | 1.200E-05 |
| 12885 | 645 | C645C | B | 5.3e-06 | 6.6e-06 | 8.5e-06 | 6.800E-06 | 1.609E-06 | 5.300E-06 | 8.500E-06 |
| 12886 | 645 | C645E | B | 9.2e-06 | 1.4e-05 | 8.2e-06 | 1.047E-05 | 3.101E-06 | 8.200E-06 | 1.400E-05 |
| 12887 | 645 | C645Q | B | 8.2e-06 | 5e-06 | 9.1e-06 | 7.433E-06 | 2.155E-06 | 5.000E-06 | 9.100E-06 |
| 12888 | 645 | C645G | B | 4.7e-06 | 1e-05 | 6.4e-06 | 7.033E-06 | 2.706E-06 | 4.700E-06 | 1.000E-05 |
| 12889 | 645 | C645H | B | 4.5e-06 | 1.2e-05 | 6.4e-06 | 7.633E-06 | 3.899E-06 | 4.500E-06 | 1.200E-05 |
| 12890 | 645 | C645I | B | 9.2e-06 | 6.7e-06 | 6.4e-06 | 7.433E-06 | 1.537E-06 | 6.400E-06 | 9.200E-06 |
| 12891 | 645 | C645L | B | 6.2e-06 | 6.4e-06 | 6.1e-06 | 6.233E-06 | 1.528E-07 | 6.100E-06 | 6.400E-06 |
| 12892 | 645 | C645K | B | 1.3e-05 | 9.6e-06 | 9.2e-06 | 1.060E-05 | 2.088E-06 | 9.200E-06 | 1.300E-05 |
| 12893 | 645 | C645M | B | 6.5e-06 | 5.2e-06 | 6.3e-06 | 6.000E-06 | 7.000E-07 | 5.200E-06 | 6.500E-06 |
| 12894 | 645 | C645F | B | 6.2e-06 | 1e-05 | 1e-05 | 8.733E-06 | 2.194E-06 | 6.200E-06 | 1.000E-05 |
| 12895 | 645 | C645P | B | 8.2e-06 | 7e-06 | 7.9e-06 | 7.700E-06 | 6.245E-07 | 7.000E-06 | 8.200E-06 |
| 12896 | 645 | C645S | B | 7.2e-06 | 8e-06 | 6.7e-06 | 7.300E-06 | 6.557E-07 | 6.700E-06 | 8.000E-06 |
| 12897 | 645 | C645T | B | 5.8e-06 | 1.1e-05 | 6.6e-06 | 7.800E-06 | 2.800E-06 | 5.800E-06 | 1.100E-05 |
| 12898 | 645 | C645W | B | 5.6e-06 | 1.4e-05 | 5.1e-06 | 8.233E-06 | 5.000E-06 | 5.100E-06 | 1.400E-05 |
| 12899 | 645 | C645Y | B | 4.7e-06 | 6.4e-06 | 5.2e-06 | 5.433E-06 | 8.737E-07 | 4.700E-06 | 6.400E-06 |
| 12900 | 645 | C645V | B | 8.5e-06 | 9.5e-06 | 5.4e-06 | 7.800E-06 | 2.138E-06 | 5.400E-06 | 9.500E-06 |
| 12901 | 646 | K646A | B | 8.2e-06 | 8.4e-06 | 9.4e-06 | 8.667E-06 | 6.429E-07 | 8.200E-06 | 9.400E-06 |
| 12902 | 646 | K646R | B | 1.8e-05 | 5e-06 | 9.2e-06 | 1.073E-05 | 6.634E-06 | 5.000E-06 | 1.800E-05 |
| 12903 | 646 | K646N | B | 9.5e-06 | 7e-06 | 8.9e-06 | 8.467E-06 | 1.305E-06 | 7.000E-06 | 9.500E-06 |
| 12904 | 646 | K646D | B | 9.8e-06 | 7.1e-06 | 5.3e-06 | 7.400E-06 | 2.265E-06 | 5.300E-06 | 9.800E-06 |
| 12905 | 646 | K646C | B | 7.3e-06 | 9.2e-06 | 9e-06 | 8.500E-06 | 1.044E-06 | 7.300E-06 | 9.200E-06 |
| 12906 | 646 | K646E | B | 9.1e-06 | 5.6e-06 | 3.8e-06 | 6.167E-06 | 2.695E-06 | 3.800E-06 | 9.100E-06 |
| 12907 | 646 | K646Q | B | 4.6e-06 | 7.2e-06 | 7.7e-06 | 6.500E-06 | 1.664E-06 | 4.600E-06 | 7.700E-06 |
| 12908 | 646 | K646G | B | 9e-06 | 1.1e-05 | 4.8e-06 | 8.267E-06 | 3.164E-06 | 4.800E-06 | 1.100E-05 |
| 12909 | 646 | K646H | B | 5.8e-06 | 9e-06 | 1.1e-05 | 8.600E-06 | 2.623E-06 | 5.800E-06 | 1.100E-05 |
| 12910 | 646 | K646I | B | 4.8e-06 | 7e-06 | 1.1e-05 | 7.600E-06 | 3.143E-06 | 4.800E-06 | 1.100E-05 |
| 12911 | 646 | K646L | B | 7.1e-06 | 6.3e-06 | 5.3e-06 | 6.233E-06 | 9.018E-07 | 5.300E-06 | 7.100E-06 |
| 12912 | 646 | K646K | B | 9.2e-06 | 6.9e-06 | 9.7e-06 | 8.600E-06 | 1.493E-06 | 6.900E-06 | 9.700E-06 |
| 12913 | 646 | K646M | B | 7.5e-06 | 1.2e-05 | 8.5e-06 | 9.333E-06 | 2.363E-06 | 7.500E-06 | 1.200E-05 |
| 12914 | 646 | K646F | B | 4.7e-06 | 1.1e-05 | 7.7e-06 | 7.800E-06 | 3.151E-06 | 4.700E-06 | 1.100E-05 |
| 12915 | 646 | K646P | B | 9e-06 | 7.5e-06 | 5.5e-06 | 7.333E-06 | 1.756E-06 | 5.500E-06 | 9.000E-06 |
| 12916 | 646 | K646S | B | 9e-06 | 9.7e-06 | 7.2e-06 | 8.633E-06 | 1.290E-06 | 7.200E-06 | 9.700E-06 |
| 12917 | 646 | K646T | B | 9.2e-06 | 1.3e-05 | 6.9e-06 | 9.700E-06 | 3.081E-06 | 6.900E-06 | 1.300E-05 |
| 12918 | 646 | K646W | B | 4.7e-06 | 5.9e-06 | 8.7e-06 | 6.433E-06 | 2.053E-06 | 4.700E-06 | 8.700E-06 |
| 12919 | 646 | K646Y | B | 8.9e-06 | 7e-06 | 6.5e-06 | 7.467E-06 | 1.266E-06 | 6.500E-06 | 8.900E-06 |

| | | | | | | | | | | |
|---|---|---|---|---|---|---|---|---|---|---|
| 12920 | 646 | K646V | B | 8.5e-06 | 8.1e-06 | 1.1e-05 | 9.200E-06 | 1.572E-06 | 8.100E-06 | 1.100E-05 |
| 12921 | 647 | A647A | B | 7.7e-06 | 9.1e-06 | 1.1e-05 | 9.267E-06 | 1.656E-06 | 7.700E-06 | 1.100E-05 |
| 12922 | 647 | A647R | B | 1.5e-05 | 1e-05 | 8.2e-06 | 1.107E-05 | 3.523E-06 | 8.200E-06 | 1.500E-05 |
| 12923 | 647 | A647N | B | 1e-05 | 9.3e-06 | 6.6e-06 | 8.633E-06 | 1.795E-06 | 6.600E-06 | 1.000E-05 |
| 12924 | 647 | A647D | B | 6.7e-06 | 8.9e-06 | 8.7e-06 | 8.100E-06 | 1.217E-06 | 6.700E-06 | 8.900E-06 |
| 12925 | 647 | A647C | B | 5.4e-06 | 7e-06 | 1e-05 | 7.467E-06 | 2.335E-06 | 5.400E-06 | 1.000E-05 |
| 12926 | 647 | A647E | B | 5.6e-06 | 8.2e-06 | 1.2e-05 | 8.600E-06 | 3.219E-06 | 5.600E-06 | 1.200E-05 |
| 12927 | 647 | A647Q | B | 6.6e-06 | 9.1e-06 | 1.1e-05 | 8.900E-06 | 2.207E-06 | 6.600E-06 | 1.100E-05 |
| 12928 | 647 | A647G | B | 5.9e-06 | 7.2e-06 | 7.8e-06 | 6.967E-06 | 9.713E-07 | 5.900E-06 | 7.800E-06 |
| 12929 | 647 | A647H | B | 6.3e-06 | 4.5e-06 | 6.4e-06 | 5.733E-06 | 1.069E-06 | 4.500E-06 | 6.400E-06 |
| 12930 | 647 | A647I | B | 1.1e-05 | 1.1e-05 | 7.1e-06 | 9.700E-06 | 2.252E-06 | 7.100E-06 | 1.100E-05 |
| 12931 | 647 | A647L | B | 4.8e-06 | 1.1e-05 | 1.2e-05 | 9.267E-06 | 3.900E-06 | 4.800E-06 | 1.200E-05 |
| 12932 | 647 | A647K | B | 6.3e-06 | 9.3e-06 | 7.1e-06 | 7.567E-06 | 1.553E-06 | 6.300E-06 | 9.300E-06 |
| 12933 | 647 | A647M | B | 1.2e-05 | 9.2e-06 | 8.5e-06 | 9.900E-06 | 1.852E-06 | 8.500E-06 | 1.200E-05 |
| 12934 | 647 | A647F | B | 9.8e-06 | 9.9e-06 | 5.5e-06 | 8.400E-06 | 2.512E-06 | 5.500E-06 | 9.900E-06 |
| 12935 | 647 | A647P | B | 6.6e-06 | 9.4e-06 | 1e-05 | 8.667E-06 | 1.815E-06 | 6.600E-06 | 1.000E-05 |
| 12936 | 647 | A647S | B | 9.2e-06 | 9.4e-06 | 5.5e-06 | 8.033E-06 | 2.196E-06 | 5.500E-06 | 9.400E-06 |
| 12937 | 647 | A647T | B | 4.5e-06 | 9.3e-06 | 3.7e-06 | 5.833E-06 | 3.029E-06 | 3.700E-06 | 9.300E-06 |
| 12938 | 647 | A647W | B | 7.9e-06 | 9.9e-06 | 5.1e-06 | 7.633E-06 | 2.411E-06 | 5.100E-06 | 9.900E-06 |
| 12939 | 647 | A647Y | B | 6.2e-06 | 8.3e-06 | 6e-06 | 6.833E-06 | 1.274E-06 | 6.000E-06 | 8.300E-06 |
| 12940 | 647 | A647V | B | 6.9e-06 | 5.2e-06 | 1.2e-05 | 8.033E-06 | 3.539E-06 | 5.200E-06 | 1.200E-05 |
| 12941 | 648 | S648A | B | 6.1e-06 | 8.4e-06 | 8.2e-06 | 7.567E-06 | 1.274E-06 | 6.100E-06 | 8.400E-06 |
| 12942 | 648 | S648R | B | 6.3e-06 | 8.1e-06 | 6e-06 | 6.800E-06 | 1.136E-06 | 6.000E-06 | 8.100E-06 |
| 12943 | 648 | S648N | B | 9.7e-06 | 5.2e-06 | 4.5e-06 | 6.467E-06 | 2.822E-06 | 4.500E-06 | 9.700E-06 |
| 12944 | 648 | S648D | B | 6.4e-06 | 1.2e-05 | 6.7e-06 | 8.367E-06 | 3.150E-06 | 6.400E-06 | 1.200E-05 |
| 12945 | 648 | S648C | B | 8.6e-06 | 7e-06 | 1e-05 | 8.533E-06 | 1.501E-06 | 7.000E-06 | 1.000E-05 |
| 12946 | 648 | S648E | B | 6.2e-06 | 6.5e-06 | 6.9e-06 | 6.533E-06 | 3.512E-07 | 6.200E-06 | 6.900E-06 |
| 12947 | 648 | S648Q | B | 4.6e-06 | 5.1e-06 | 1.1e-05 | 6.900E-06 | 3.559E-06 | 4.600E-06 | 1.100E-05 |
| 12948 | 648 | S648G | B | 9e-06 | 8.3e-06 | 7e-06 | 8.100E-06 | 1.015E-06 | 7.000E-06 | 9.000E-06 |
| 12949 | 648 | S648H | B | 7e-06 | 7.1e-06 | 7.7e-06 | 7.267E-06 | 3.786E-07 | 7.000E-06 | 7.700E-06 |
| 12950 | 648 | S648I | B | 6.4e-06 | 7.2e-06 | 4.5e-06 | 6.033E-06 | 1.387E-06 | 4.500E-06 | 7.200E-06 |
| 12951 | 648 | S648L | B | 9.6e-06 | 5.3e-06 | 1.4e-05 | 9.633E-06 | 4.350E-06 | 5.300E-06 | 1.400E-05 |
| 12952 | 648 | S648K | B | 8.3e-06 | 7.5e-06 | 5.2e-06 | 7.000E-06 | 1.609E-06 | 5.200E-06 | 8.300E-06 |
| 12953 | 648 | S648M | B | 5.2e-06 | 8.1e-06 | 5.2e-06 | 6.167E-06 | 1.674E-06 | 5.200E-06 | 8.100E-06 |
| 12954 | 648 | S648F | B | 1e-05 | 9.9e-06 | 4.6e-06 | 8.167E-06 | 3.089E-06 | 4.600E-06 | 1.000E-05 |
| 12955 | 648 | S648P | B | 9.7e-06 | 8.2e-06 | 9.6e-06 | 9.167E-06 | 8.386E-07 | 8.200E-06 | 9.700E-06 |
| 12956 | 648 | S648S | B | 8.6e-06 | 7.5e-06 | 7.1e-06 | 7.733E-06 | 7.767E-07 | 7.100E-06 | 8.600E-06 |
| 12957 | 648 | S648T | B | 5.5e-06 | 8.6e-06 | 6.4e-06 | 6.833E-06 | 1.595E-06 | 5.500E-06 | 8.600E-06 |
| 12958 | 648 | S648W | B | 7.9e-06 | 1e-05 | 5.9e-06 | 7.933E-06 | 2.050E-06 | 5.900E-06 | 1.000E-05 |
| 12959 | 648 | S648Y | B | 7.3e-06 | 8.5e-06 | 8.5e-06 | 8.100E-06 | 6.928E-07 | 7.300E-06 | 8.500E-06 |
| 12960 | 648 | S648V | B | 7e-06 | 1.1e-05 | 1e-05 | 9.333E-06 | 2.082E-06 | 7.000E-06 | 1.100E-05 |
| 12961 | 649 | Q649A | B | 1.1e-05 | 1.3e-05 | 8.5e-06 | 1.083E-05 | 2.255E-06 | 8.500E-06 | 1.300E-05 |
| 12962 | 649 | Q649R | B | 1e-05 | 1e-05 | 1.3e-05 | 1.100E-05 | 1.732E-06 | 1.000E-05 | 1.300E-05 |
| 12963 | 649 | Q649N | B | 1.3e-05 | 8.9e-06 | 1.3e-05 | 1.163E-05 | 2.367E-06 | 8.900E-06 | 1.300E-05 |
| 12964 | 649 | Q649D | B | 6.2e-06 | 1.1e-05 | 6.4e-06 | 7.867E-06 | 2.715E-06 | 6.200E-06 | 1.100E-05 |
| 12965 | 649 | Q649C | B | 8e-06 | 9.1e-06 | 8.2e-06 | 8.433E-06 | 5.859E-07 | 8.000E-06 | 9.100E-06 |
| 12966 | 649 | Q649E | B | 8.6e-06 | 9.7e-06 | 9.9e-06 | 9.400E-06 | 7.000E-07 | 8.600E-06 | 9.900E-06 |
| 12967 | 649 | Q649Q | B | 1.4e-05 | 9.1e-06 | 7.5e-06 | 1.020E-05 | 3.387E-06 | 7.500E-06 | 1.400E-05 |
| 12968 | 649 | Q649G | B | 6.5e-06 | 1.2e-05 | 6.5e-06 | 8.333E-06 | 3.175E-06 | 6.500E-06 | 1.200E-05 |
| 12969 | 649 | Q649H | B | 6.7e-06 | 7.1e-06 | 6.8e-06 | 6.867E-06 | 2.082E-07 | 6.700E-06 | 7.100E-06 |
| 12970 | 649 | Q649I | B | 4.3e-06 | 7.5e-06 | 9.7e-06 | 7.167E-06 | 2.715E-06 | 4.300E-06 | 9.700E-06 |
| 12971 | 649 | Q649L | B | 7e-06 | 8.1e-06 | 9.9e-06 | 8.333E-06 | 1.464E-06 | 7.000E-06 | 9.900E-06 |
| 12972 | 649 | Q649K | B | 1.2e-05 | 5.9e-06 | 5e-06 | 7.633E-06 | 3.808E-06 | 5.000E-06 | 1.200E-05 |
| 12973 | 649 | Q649M | B | 5.9e-06 | 9.5e-06 | 6.2e-06 | 7.200E-06 | 1.997E-06 | 5.900E-06 | 9.500E-06 |
| 12974 | 649 | Q649F | B | 7.4e-06 | 4.7e-06 | 6e-06 | 6.033E-06 | 1.350E-06 | 4.700E-06 | 7.400E-06 |
| 12975 | 649 | Q649P | B | 1.3e-05 | 7e-06 | 1.3e-05 | 1.100E-05 | 3.464E-06 | 7.000E-06 | 1.300E-05 |
| 12976 | 649 | Q649S | B | 1.1e-05 | 1.3e-05 | 1e-05 | 1.133E-05 | 1.528E-06 | 1.000E-05 | 1.300E-05 |
| 12977 | 649 | Q649T | B | 7.4e-06 | 7.6e-06 | 1.4e-05 | 9.667E-06 | 3.754E-06 | 7.400E-06 | 1.400E-05 |
| 12978 | 649 | Q649W | B | 7.5e-06 | 9.4e-06 | 6e-06 | 7.633E-06 | 1.704E-06 | 6.000E-06 | 9.400E-06 |
| 12979 | 649 | Q649Y | B | 1.3e-05 | 7.4e-06 | 1e-05 | 1.013E-05 | 2.802E-06 | 7.400E-06 | 1.300E-05 |
| 12980 | 649 | Q649V | B | 1.5e-05 | 5.7e-06 | 4.7e-06 | 8.467E-06 | 5.680E-06 | 4.700E-06 | 1.500E-05 |
| 12981 | 650 | D650A | B | 6.6e-06 | 8.3e-06 | 7.3e-06 | 7.400E-06 | 8.544E-07 | 6.600E-06 | 8.300E-06 |
| 12982 | 650 | D650R | B | 6.4e-06 | 7.1e-06 | 1e-05 | 7.833E-06 | 1.909E-06 | 6.400E-06 | 1.000E-05 |
| 12983 | 650 | D650N | B | 8.4e-06 | 3.4e-06 | 1.2e-05 | 7.933E-06 | 4.319E-06 | 3.400E-06 | 1.200E-05 |
| 12984 | 650 | D650D | B | 1e-05 | 6.1e-06 | 4.5e-06 | 6.867E-06 | 2.829E-06 | 4.500E-06 | 1.000E-05 |
| 12985 | 650 | D650C | B | 6.4e-06 | 6.2e-06 | 5.7e-06 | 6.100E-06 | 3.606E-07 | 5.700E-06 | 6.400E-06 |
| 12986 | 650 | D650E | B | 5.4e-06 | 9.2e-06 | 6.1e-06 | 6.900E-06 | 2.022E-06 | 5.400E-06 | 9.200E-06 |
| 12987 | 650 | D650Q | B | 7.6e-06 | 5.5e-06 | 8.5e-06 | 7.200E-06 | 1.539E-06 | 5.500E-06 | 8.500E-06 |
| 12988 | 650 | D650G | B | 1.3e-05 | 5.8e-06 | 8.2e-06 | 9.000E-06 | 3.666E-06 | 5.800E-06 | 1.300E-05 |
| 12989 | 650 | D650H | B | 4.6e-06 | 8.2e-06 | 9.3e-06 | 7.367E-06 | 2.458E-06 | 4.600E-06 | 9.300E-06 |
| 12990 | 650 | D650I | B | 7.2e-06 | 3.8e-06 | 9.7e-06 | 6.900E-06 | 2.961E-06 | 3.800E-06 | 9.700E-06 |
| 12991 | 650 | D650L | B | 7.4e-06 | 7.7e-06 | 4.6e-06 | 6.567E-06 | 1.710E-06 | 4.600E-06 | 7.700E-06 |
| 12992 | 650 | D650K | B | 5.1e-06 | 8.9e-06 | 6.7e-06 | 6.900E-06 | 1.908E-06 | 5.100E-06 | 8.900E-06 |
| 12993 | 650 | D650M | B | 1.4e-05 | 8e-06 | 6.2e-06 | 9.400E-06 | 4.084E-06 | 6.200E-06 | 1.400E-05 |
| 12994 | 650 | D650F | B | 7.8e-06 | 5.7e-06 | 3.6e-06 | 5.700E-06 | 2.100E-06 | 3.600E-06 | 7.800E-06 |
| 12995 | 650 | D650P | B | 8.1e-06 | 7.6e-06 | 6.1e-06 | 7.267E-06 | 1.041E-06 | 6.100E-06 | 8.100E-06 |
| 12996 | 650 | D650S | B | 1.2e-05 | 7.1e-06 | 4.8e-06 | 7.967E-06 | 3.677E-06 | 4.800E-06 | 1.200E-05 |
| 12997 | 650 | D650T | B | 6.9e-06 | 6.7e-06 | 7e-06 | 6.867E-06 | 1.528E-07 | 6.700E-06 | 7.000E-06 |
| 12998 | 650 | D650W | B | 3.3e-06 | 7.9e-06 | 2.1e-05 | 1.073E-05 | 9.184E-06 | 3.300E-06 | 2.100E-05 |
| 12999 | 650 | D650Y | B | 5.1e-06 | 6.7e-06 | 5.8e-06 | 5.867E-06 | 8.021E-07 | 5.100E-06 | 6.700E-06 |
| 13000 | 650 | D650V | B | 5.3e-06 | 1.3e-05 | 5.2e-06 | 7.833E-06 | 4.475E-06 | 5.200E-06 | 1.300E-05 |
| 13001 | 651 | V651A | B | 8.1e-06 | 3.7e-06 | 8.3e-06 | 6.700E-06 | 2.600E-06 | 3.700E-06 | 8.300E-06 |
| 13002 | 651 | V651R | B | 8.8e-06 | 4.8e-06 | 7.3e-06 | 6.967E-06 | 2.021E-06 | 4.800E-06 | 8.800E-06 |
| 13003 | 651 | V651N | B | 5.2e-06 | 7.9e-06 | 8.9e-06 | 7.333E-06 | 1.914E-06 | 5.200E-06 | 8.900E-06 |
| 13004 | 651 | V651D | B | 1.3e-05 | 1.2e-05 | 1.3e-05 | 1.267E-05 | 5.774E-07 | 1.200E-05 | 1.300E-05 |

| | | | | | | | | | | |
|---|---|---|---|---|---|---|---|---|---|---|
| 13005 | 651 | V651C | B | 8.9e-06 | 1e-05 | 8.7e-06 | 9.200E-06 | 7.000E-07 | 8.700E-06 | 1.000E-05 |
| 13006 | 651 | V651E | B | 7.1e-06 | 6.5e-06 | 1.2e-05 | 8.533E-06 | 3.017E-06 | 6.500E-06 | 1.200E-05 |
| 13007 | 651 | V651Q | B | 4.7e-06 | 1.1e-05 | 8.1e-06 | 7.933E-06 | 3.153E-06 | 4.700E-06 | 1.100E-05 |
| 13008 | 651 | V651G | B | 5.5e-06 | 8.9e-06 | 1.1e-05 | 8.467E-06 | 2.775E-06 | 5.500E-06 | 1.100E-05 |
| 13009 | 651 | V651H | B | 6e-06 | 6.9e-06 | 9e-06 | 7.300E-06 | 1.539E-06 | 6.000E-06 | 9.000E-06 |
| 13010 | 651 | V651I | B | 7e-06 | 8.4e-06 | 6.1e-06 | 7.167E-06 | 1.159E-06 | 6.100E-06 | 8.400E-06 |
| 13011 | 651 | V651L | B | 9.3e-06 | 1e-05 | 1.2e-05 | 1.043E-05 | 1.401E-06 | 9.300E-06 | 1.200E-05 |
| 13012 | 651 | V651K | B | 7.1e-06 | 6.3e-06 | 4.9e-06 | 6.100E-06 | 1.114E-06 | 4.900E-06 | 7.100E-06 |
| 13013 | 651 | V651M | B | 4.4e-06 | 1e-05 | 6.4e-06 | 6.933E-06 | 2.838E-06 | 4.400E-06 | 1.000E-05 |
| 13014 | 651 | V651F | B | 7e-06 | 8.5e-06 | 7.7e-06 | 7.733E-06 | 7.506E-07 | 7.000E-06 | 8.500E-06 |
| 13015 | 651 | V651P | B | 1.1e-05 | 7.7e-06 | 7e-06 | 8.567E-06 | 2.136E-06 | 7.000E-06 | 1.100E-05 |
| 13016 | 651 | V651S | B | 9.3e-06 | 1.1e-05 | 7.6e-06 | 9.300E-06 | 1.700E-06 | 7.600E-06 | 1.100E-05 |
| 13017 | 651 | V651T | B | 8e-06 | 6.6e-06 | 5.6e-06 | 6.733E-06 | 1.206E-06 | 5.600E-06 | 8.000E-06 |
| 13018 | 651 | V651W | B | 9.4e-06 | 9.4e-06 | 4.8e-06 | 7.867E-06 | 2.656E-06 | 4.800E-06 | 9.400E-06 |
| 13019 | 651 | V651Y | B | 6.1e-06 | 5.2e-06 | 6.7e-06 | 6.000E-06 | 7.550E-07 | 5.200E-06 | 6.700E-06 |
| 13020 | 651 | V651V | B | 1.1e-05 | 1.4e-05 | 7e-06 | 1.067E-05 | 3.512E-06 | 7.000E-06 | 1.400E-05 |
| 13021 | 652 | S652A | B | 6.8e-06 | 9.8e-06 | 8.8e-06 | 8.467E-06 | 1.528E-06 | 6.800E-06 | 9.800E-06 |
| 13022 | 652 | S652R | B | 7e-06 | 7e-06 | 1.3e-05 | 9.000E-06 | 3.464E-06 | 7.000E-06 | 1.300E-05 |
| 13023 | 652 | S652N | B | 4.9e-06 | 6.4e-06 | 9.3e-06 | 6.867E-06 | 2.237E-06 | 4.900E-06 | 9.300E-06 |
| 13024 | 652 | S652D | B | 9.5e-06 | 9.5e-06 | 7.3e-06 | 8.767E-06 | 1.270E-06 | 7.300E-06 | 9.500E-06 |
| 13025 | 652 | S652C | B | 8.8e-06 | 9.2e-06 | 5.3e-06 | 7.767E-06 | 2.146E-06 | 5.300E-06 | 9.200E-06 |
| 13026 | 652 | S652E | B | 7.9e-06 | 5.9e-06 | 8.9e-06 | 7.567E-06 | 1.528E-06 | 5.900E-06 | 8.900E-06 |
| 13027 | 652 | S652Q | B | 1e-05 | 5.8e-06 | 6.1e-06 | 7.300E-06 | 2.343E-06 | 5.800E-06 | 1.000E-05 |
| 13028 | 652 | S652G | B | 7.8e-06 | 1e-05 | 6.8e-06 | 8.200E-06 | 1.637E-06 | 6.800E-06 | 1.000E-05 |
| 13029 | 652 | S652H | B | 8.9e-06 | 9.1e-06 | 4.7e-06 | 7.567E-06 | 2.485E-06 | 4.700E-06 | 9.100E-06 |
| 13030 | 652 | S652I | B | 9.3e-06 | 7.8e-06 | 9.4e-06 | 8.833E-06 | 8.963E-07 | 7.800E-06 | 9.400E-06 |
| 13031 | 652 | S652L | B | 2.9e-06 | 6.2e-06 | 1.2e-05 | 7.033E-06 | 4.607E-06 | 2.900E-06 | 1.200E-05 |
| 13032 | 652 | S652K | B | 1.3e-05 | 1e-05 | 1.2e-05 | 1.167E-05 | 1.528E-06 | 1.000E-05 | 1.300E-05 |
| 13033 | 652 | S652M | B | 5.6e-06 | 7.8e-06 | 1.5e-05 | 9.467E-06 | 4.917E-06 | 5.600E-06 | 1.500E-05 |
| 13034 | 652 | S652F | B | 1.8e-05 | 5.4e-06 | 8.3e-06 | 1.057E-05 | 6.599E-06 | 5.400E-06 | 1.800E-05 |
| 13035 | 652 | S652P | B | 1.2e-05 | 7.9e-06 | 9.1e-06 | 9.667E-06 | 2.108E-06 | 7.900E-06 | 1.200E-05 |
| 13036 | 652 | S652S | B | 4.3e-06 | 1.1e-05 | 1.3e-05 | 9.433E-06 | 4.557E-06 | 4.300E-06 | 1.300E-05 |
| 13037 | 652 | S652T | B | 1e-05 | 8.6e-06 | 3.4e-06 | 7.333E-06 | 3.478E-06 | 3.400E-06 | 1.000E-05 |
| 13038 | 652 | S652W | B | 4.6e-06 | 6.8e-06 | 7.3e-06 | 6.233E-06 | 1.436E-06 | 4.600E-06 | 7.300E-06 |
| 13039 | 652 | S652Y | B | 6.5e-06 | 5.2e-06 | 8e-06 | 6.567E-06 | 1.401E-06 | 5.200E-06 | 8.000E-06 |
| 13040 | 652 | S652V | B | 6.1e-06 | 7.6e-06 | 6.2e-06 | 6.633E-06 | 8.386E-07 | 6.100E-06 | 7.600E-06 |
| 13041 | 653 | I653A | B | 9.4e-06 | 2e-05 | 3.7e-06 | 1.103E-05 | 8.272E-06 | 3.700E-06 | 2.000E-05 |
| 13042 | 653 | I653R | B | 5.2e-06 | 1.2e-05 | 8.3e-06 | 8.500E-06 | 3.404E-06 | 5.200E-06 | 1.200E-05 |
| 13043 | 653 | I653N | B | 8.7e-06 | 4.5e-06 | 9.6e-06 | 7.600E-06 | 2.722E-06 | 4.500E-06 | 9.600E-06 |
| 13044 | 653 | I653D | B | 8.1e-06 | 1.3e-05 | 6.4e-06 | 9.167E-06 | 3.427E-06 | 6.400E-06 | 1.300E-05 |
| 13045 | 653 | I653C | B | 1.1e-05 | 3.8e-06 | 1.3e-05 | 9.267E-06 | 4.839E-06 | 3.800E-06 | 1.300E-05 |
| 13046 | 653 | I653E | B | 1.1e-05 | 4.2e-06 | 8.2e-06 | 7.800E-06 | 3.418E-06 | 4.200E-06 | 1.100E-05 |
| 13047 | 653 | I653Q | B | 7.5e-06 | 9.3e-06 | 6.9e-06 | 7.900E-06 | 1.249E-06 | 6.900E-06 | 9.300E-06 |
| 13048 | 653 | I653G | B | 1.1e-05 | 7.8e-06 | 5.3e-06 | 8.033E-06 | 2.857E-06 | 5.300E-06 | 1.100E-05 |
| 13049 | 653 | I653H | B | 5.7e-06 | 6.3e-06 | 6.5e-06 | 6.167E-06 | 4.163E-07 | 5.700E-06 | 6.500E-06 |
| 13050 | 653 | I653I | B | 6.7e-06 | 7.2e-06 | 1.2e-05 | 8.633E-06 | 2.926E-06 | 6.700E-06 | 1.200E-05 |
| 13051 | 653 | I653L | B | 1e-05 | 9e-06 | 6.5e-06 | 8.500E-06 | 1.803E-06 | 6.500E-06 | 1.000E-05 |
| 13052 | 653 | I653K | B | 5.8e-06 | 5e-06 | 5.6e-06 | 5.467E-06 | 4.163E-07 | 5.000E-06 | 5.800E-06 |
| 13053 | 653 | I653M | B | 1.1e-05 | 6.8e-06 | 7.9e-06 | 8.567E-06 | 2.178E-06 | 6.800E-06 | 1.100E-05 |
| 13054 | 653 | I653F | B | 8.8e-06 | 6.9e-06 | 9e-06 | 8.233E-06 | 1.159E-06 | 6.900E-06 | 9.000E-06 |
| 13055 | 653 | I653P | B | 1e-05 | 6.6e-06 | 8.5e-06 | 8.367E-06 | 1.704E-06 | 6.600E-06 | 1.000E-05 |
| 13056 | 653 | I653S | B | 5.9e-06 | 1.4e-05 | 7.8e-06 | 9.233E-06 | 4.236E-06 | 5.900E-06 | 1.400E-05 |
| 13057 | 653 | I653T | B | 7.5e-06 | 6.7e-06 | 1e-05 | 8.067E-06 | 1.721E-06 | 6.700E-06 | 1.000E-05 |
| 13058 | 653 | I653W | B | 6.6e-06 | 6.1e-06 | 9.3e-06 | 7.333E-06 | 1.721E-06 | 6.100E-06 | 9.300E-06 |
| 13059 | 653 | I653Y | B | 1.1e-05 | 6.1e-06 | 7.7e-06 | 8.267E-06 | 2.499E-06 | 6.100E-06 | 1.100E-05 |
| 13060 | 653 | I653V | B | 4.8e-06 | 8.6e-06 | 9.3e-06 | 7.567E-06 | 2.421E-06 | 4.800E-06 | 9.300E-06 |
| 13061 | 654 | G654A | B | 7.9e-06 | 6.6e-06 | 1.1e-05 | 8.500E-06 | 2.261E-06 | 6.600E-06 | 1.100E-05 |
| 13062 | 654 | G654R | B | 7.2e-06 | 5.5e-06 | 1e-05 | 7.567E-06 | 2.272E-06 | 5.500E-06 | 1.000E-05 |
| 13063 | 654 | G654N | B | 1.1e-05 | 4.3e-06 | 6.6e-06 | 7.300E-06 | 3.404E-06 | 4.300E-06 | 1.100E-05 |
| 13064 | 654 | G654D | B | 1.3e-05 | 9.1e-06 | 6.2e-06 | 9.433E-06 | 3.412E-06 | 6.200E-06 | 1.300E-05 |
| 13065 | 654 | G654C | B | 8.7e-06 | 4.7e-06 | 4.6e-06 | 6.000E-06 | 2.339E-06 | 4.600E-06 | 8.700E-06 |
| 13066 | 654 | G654E | B | 5.5e-06 | 7e-06 | 9e-06 | 7.167E-06 | 1.756E-06 | 5.500E-06 | 9.000E-06 |
| 13067 | 654 | G654Q | B | 8e-06 | 1.3e-05 | 1.1e-05 | 1.067E-05 | 2.517E-06 | 8.000E-06 | 1.300E-05 |
| 13068 | 654 | G654G | B | 1.1e-05 | 1.1e-05 | 6.3e-06 | 9.433E-06 | 2.714E-06 | 6.300E-06 | 1.100E-05 |
| 13069 | 654 | G654H | B | 4.6e-06 | 5.3e-06 | 1e-05 | 6.633E-06 | 2.937E-06 | 4.600E-06 | 1.000E-05 |
| 13070 | 654 | G654I | B | 8.1e-06 | 1e-05 | 1.5e-05 | 1.103E-05 | 3.564E-06 | 8.100E-06 | 1.500E-05 |
| 13071 | 654 | G654L | B | 1e-05 | 4.4e-06 | 9.8e-06 | 8.067E-06 | 3.177E-06 | 4.400E-06 | 1.000E-05 |
| 13072 | 654 | G654K | B | 1.2e-05 | 7e-06 | 5.1e-06 | 8.033E-06 | 3.564E-06 | 5.100E-06 | 1.200E-05 |
| 13073 | 654 | G654M | B | 1.4e-05 | 8.1e-06 | 6.1e-06 | 9.400E-06 | 4.107E-06 | 6.100E-06 | 1.400E-05 |
| 13074 | 654 | G654F | B | 1e-05 | 8.5e-06 | 8.5e-06 | 9.000E-06 | 8.660E-07 | 8.500E-06 | 1.000E-05 |
| 13075 | 654 | G654P | B | 8.6e-06 | 9.1e-06 | 7.9e-06 | 8.533E-06 | 6.028E-07 | 7.900E-06 | 9.100E-06 |
| 13076 | 654 | G654S | B | 6.5e-06 | 5.9e-06 | 5.7e-06 | 6.033E-06 | 4.163E-07 | 5.700E-06 | 6.500E-06 |
| 13077 | 654 | G654T | B | 8.2e-06 | 7.2e-06 | 6.6e-06 | 7.333E-06 | 8.083E-07 | 6.600E-06 | 8.200E-06 |
| 13078 | 654 | G654W | B | 7.8e-06 | 1.7e-05 | 9.5e-06 | 1.143E-05 | 4.895E-06 | 7.800E-06 | 1.700E-05 |
| 13079 | 654 | G654Y | B | 1.2e-05 | 8.5e-06 | 9e-06 | 9.833E-06 | 1.893E-06 | 8.500E-06 | 1.200E-05 |
| 13080 | 654 | G654V | B | 9.2e-06 | 6.1e-06 | 8.4e-06 | 7.900E-06 | 1.609E-06 | 6.100E-06 | 9.200E-06 |
| 13081 | 655 | V655A | B | 4.7e-06 | 5.5e-06 | 6.4e-06 | 5.533E-06 | 8.505E-07 | 4.700E-06 | 6.400E-06 |
| 13082 | 655 | V655R | B | 1.7e-05 | 6.3e-06 | 7e-06 | 1.010E-05 | 5.986E-06 | 6.300E-06 | 1.700E-05 |
| 13083 | 655 | V655N | B | 9.8e-06 | 1.2e-05 | 8.1e-06 | 9.967E-06 | 1.955E-06 | 8.100E-06 | 1.200E-05 |
| 13084 | 655 | V655D | B | 9.5e-06 | 9.9e-06 | 4.3e-06 | 7.900E-06 | 3.124E-06 | 4.300E-06 | 9.900E-06 |
| 13085 | 655 | V655C | B | 4e-06 | 1.5e-05 | 8.4e-06 | 9.133E-06 | 5.537E-06 | 4.000E-06 | 1.500E-05 |
| 13086 | 655 | V655E | B | 1.3e-05 | 1.3e-05 | 1e-05 | 1.200E-05 | 1.732E-06 | 1.000E-05 | 1.300E-05 |
| 13087 | 655 | V655Q | B | 6.3e-06 | 1.3e-05 | 8.6e-06 | 9.300E-06 | 3.404E-06 | 6.300E-06 | 1.300E-05 |
| 13088 | 655 | V655G | B | 1e-05 | 9.4e-06 | 8.2e-06 | 9.200E-06 | 9.165E-07 | 8.200E-06 | 1.000E-05 |
| 13089 | 655 | V655H | B | 6.9e-06 | 7.2e-06 | 1e-05 | 8.033E-06 | 1.710E-06 | 6.900E-06 | 1.000E-05 |

| | | | | | | | | | | |
|---|---|---|---|---|---|---|---|---|---|---|
| 13090 | 655 | V655I | B | 8.8e-06 | 4.8e-06 | 1.4e-05 | 9.200E-06 | 4.613E-06 | 4.800E-06 | 1.400E-05 |
| 13091 | 655 | V655L | B | 7.2e-06 | 8.2e-06 | 7.3e-06 | 7.567E-06 | 5.508E-07 | 7.200E-06 | 8.200E-06 |
| 13092 | 655 | V655K | B | 1e-05 | 1.4e-05 | 8.4e-06 | 1.080E-05 | 2.884E-06 | 8.400E-06 | 1.400E-05 |
| 13093 | 655 | V655M | B | 1.8e-05 | 8.5e-06 | 4.3e-06 | 1.027E-05 | 7.019E-06 | 4.300E-06 | 1.800E-05 |
| 13094 | 655 | V655F | B | 9e-06 | 1.1e-05 | 9.1e-06 | 9.700E-06 | 1.127E-06 | 9.000E-06 | 1.100E-05 |
| 13095 | 655 | V655P | B | 5.2e-06 | 9.5e-06 | 1.1e-05 | 8.567E-06 | 3.011E-06 | 5.200E-06 | 1.100E-05 |
| 13096 | 655 | V655S | B | 9.1e-06 | 1.4e-05 | 4.7e-06 | 9.267E-06 | 4.652E-06 | 4.700E-06 | 1.400E-05 |
| 13097 | 655 | V655T | B | 5.7e-06 | 4.6e-06 | 1e-05 | 6.767E-06 | 2.854E-06 | 4.600E-06 | 1.000E-05 |
| 13098 | 655 | V655W | B | 1.1e-05 | 8.4e-06 | 1.1e-05 | 1.013E-05 | 1.501E-06 | 8.400E-06 | 1.100E-05 |
| 13099 | 655 | V655Y | B | 1.2e-05 | 1e-05 | 9.7e-06 | 1.057E-05 | 1.250E-06 | 9.700E-06 | 1.200E-05 |
| 13100 | 655 | V655V | B | 9.1e-06 | 1e-05 | 1.3e-05 | 1.070E-05 | 2.042E-06 | 9.100E-06 | 1.300E-05 |
| 13101 | 656 | A656A | B | 9.9e-06 | 1.2e-05 | 9e-06 | 1.030E-05 | 1.539E-06 | 9.000E-06 | 1.200E-05 |
| 13102 | 656 | A656R | B | 5e-06 | 5.6e-06 | 5.6e-06 | 5.400E-06 | 3.464E-07 | 5.000E-06 | 5.600E-06 |
| 13103 | 656 | A656N | B | 8.6e-06 | 8.9e-06 | 9.8e-06 | 9.100E-06 | 6.245E-07 | 8.600E-06 | 9.800E-06 |
| 13104 | 656 | A656D | B | 5.8e-06 | 5.6e-06 | 8.2e-06 | 6.533E-06 | 1.447E-06 | 5.600E-06 | 8.200E-06 |
| 13105 | 656 | A656C | B | 8.2e-06 | 6.3e-06 | 6.1e-06 | 6.867E-06 | 1.159E-06 | 6.100E-06 | 8.200E-06 |
| 13106 | 656 | A656E | B | 1.1e-05 | 1.2e-05 | 6e-06 | 9.667E-06 | 3.215E-06 | 6.000E-06 | 1.200E-05 |
| 13107 | 656 | A656Q | B | 6.1e-06 | 7.1e-06 | 8.5e-06 | 7.233E-06 | 1.206E-06 | 6.100E-06 | 8.500E-06 |
| 13108 | 656 | A656G | B | 1.9e-05 | 7.7e-06 | 7.9e-06 | 1.153E-05 | 6.467E-06 | 7.700E-06 | 1.900E-05 |
| 13109 | 656 | A656H | B | 8.3e-06 | 1.4e-05 | 7e-06 | 9.767E-06 | 3.723E-06 | 7.000E-06 | 1.400E-05 |
| 13110 | 656 | A656I | B | 7.3e-06 | 4.4e-06 | 9.3e-06 | 7.000E-06 | 2.464E-06 | 4.400E-06 | 9.300E-06 |
| 13111 | 656 | A656L | B | 1.3e-05 | 5e-06 | 6.1e-06 | 8.033E-06 | 4.336E-06 | 5.000E-06 | 1.300E-05 |
| 13112 | 656 | A656K | B | 7.5e-06 | 6.2e-06 | 1.2e-05 | 8.567E-06 | 3.044E-06 | 6.200E-06 | 1.200E-05 |
| 13113 | 656 | A656M | B | 1e-05 | 6.3e-06 | 7.7e-06 | 8.000E-06 | 1.868E-06 | 6.300E-06 | 1.000E-05 |
| 13114 | 656 | A656F | B | 1.1e-05 | 8.6e-06 | 9.1e-06 | 9.567E-06 | 1.266E-06 | 8.600E-06 | 1.100E-05 |
| 13115 | 656 | A656P | B | 1.3e-05 | 1.2e-05 | 4.5e-06 | 9.833E-06 | 4.646E-06 | 4.500E-06 | 1.300E-05 |
| 13116 | 656 | A656S | B | 1.3e-05 | 1e-05 | 8.2e-06 | 1.040E-05 | 2.425E-06 | 8.200E-06 | 1.300E-05 |
| 13117 | 656 | A656T | B | 4.6e-06 | 1.4e-05 | 7.1e-06 | 8.567E-06 | 4.869E-06 | 4.600E-06 | 1.400E-05 |
| 13118 | 656 | A656W | B | 5.6e-06 | 6.6e-06 | 4.8e-06 | 5.667E-06 | 9.018E-07 | 4.800E-06 | 6.600E-06 |
| 13119 | 656 | A656Y | B | 5.9e-06 | 6.7e-06 | 1.4e-05 | 8.867E-06 | 4.464E-06 | 5.900E-06 | 1.400E-05 |
| 13120 | 656 | A656V | B | 9.1e-06 | 8.1e-06 | 9.1e-06 | 8.767E-06 | 5.774E-07 | 8.100E-06 | 9.100E-06 |
| 13121 | 657 | W657A | B | 1e-05 | 4.4e-06 | 1e-05 | 8.133E-06 | 3.233E-06 | 4.400E-06 | 1.000E-05 |
| 13122 | 657 | W657R | B | 8e-06 | 6.8e-06 | 1.1e-05 | 8.600E-06 | 2.163E-06 | 6.800E-06 | 1.100E-05 |
| 13123 | 657 | W657N | B | 5.9e-06 | 8.7e-06 | 6.6e-06 | 7.067E-06 | 1.457E-06 | 5.900E-06 | 8.700E-06 |
| 13124 | 657 | W657D | B | 6.6e-06 | 9.3e-06 | 6.3e-06 | 7.400E-06 | 1.652E-06 | 6.300E-06 | 9.300E-06 |
| 13125 | 657 | W657C | B | 9e-06 | 9.4e-06 | 5.8e-06 | 8.067E-06 | 1.973E-06 | 5.800E-06 | 9.400E-06 |
| 13126 | 657 | W657E | B | 9.4e-06 | 7.9e-06 | 6.4e-06 | 7.900E-06 | 1.500E-06 | 6.400E-06 | 9.400E-06 |
| 13127 | 657 | W657Q | B | 7.8e-06 | 1.6e-05 | 1.5e-05 | 1.293E-05 | 4.474E-06 | 7.800E-06 | 1.600E-05 |
| 13128 | 657 | W657G | B | 8.3e-06 | 5.8e-06 | 6.5e-06 | 6.867E-06 | 1.290E-06 | 5.800E-06 | 8.300E-06 |
| 13129 | 657 | W657H | B | 1.1e-05 | 5.8e-06 | 1.2e-05 | 9.600E-06 | 3.329E-06 | 5.800E-06 | 1.200E-05 |
| 13130 | 657 | W657I | B | 1e-05 | 8.7e-06 | 8.6e-06 | 9.100E-06 | 7.810E-07 | 8.600E-06 | 1.000E-05 |
| 13131 | 657 | W657L | B | 1e-05 | 5.1e-06 | 1.2e-05 | 9.033E-06 | 3.550E-06 | 5.100E-06 | 1.200E-05 |
| 13132 | 657 | W657K | B | 8.2e-06 | 1.3e-05 | 1.1e-05 | 1.073E-05 | 2.411E-06 | 8.200E-06 | 1.300E-05 |
| 13133 | 657 | W657M | B | 7e-06 | 1.4e-05 | 1e-05 | 1.033E-05 | 3.512E-06 | 7.000E-06 | 1.400E-05 |
| 13134 | 657 | W657F | B | 7.3e-06 | 6.7e-06 | 6.4e-06 | 6.800E-06 | 4.583E-07 | 6.400E-06 | 7.300E-06 |
| 13135 | 657 | W657P | B | 6.4e-06 | 1.1e-05 | 1.5e-05 | 1.080E-05 | 4.303E-06 | 6.400E-06 | 1.500E-05 |
| 13136 | 657 | W657S | B | 5.3e-06 | 1.5e-05 | 1.3e-05 | 1.110E-05 | 5.122E-06 | 5.300E-06 | 1.500E-05 |
| 13137 | 657 | W657T | B | 1.3e-05 | 9.2e-06 | 1.3e-05 | 1.173E-05 | 2.194E-06 | 9.200E-06 | 1.300E-05 |
| 13138 | 657 | W657W | B | 4.6e-06 | 5.4e-06 | 7.8e-06 | 5.933E-06 | 1.665E-06 | 4.600E-06 | 7.800E-06 |
| 13139 | 657 | W657Y | B | 8.8e-06 | 1.6e-05 | 8.9e-06 | 1.123E-05 | 4.128E-06 | 8.800E-06 | 1.600E-05 |
| 13140 | 657 | W657V | B | 1e-05 | 7e-06 | 7e-06 | 8.000E-06 | 1.732E-06 | 7.000E-06 | 1.000E-05 |
| 13141 | 658 | Y658A | B | 7.3e-06 | 6.4e-06 | 5.6e-06 | 6.433E-06 | 8.505E-07 | 5.600E-06 | 7.300E-06 |
| 13142 | 658 | Y658R | B | 1.1e-05 | 1.1e-05 | 7.8e-06 | 9.933E-06 | 1.848E-06 | 7.800E-06 | 1.100E-05 |
| 13143 | 658 | Y658N | B | 8.9e-06 | 5.6e-06 | 5.5e-06 | 6.667E-06 | 1.935E-06 | 5.500E-06 | 8.900E-06 |
| 13144 | 658 | Y658D | B | 6.3e-06 | 8.1e-06 | 9.2e-06 | 7.867E-06 | 1.464E-06 | 6.300E-06 | 9.200E-06 |
| 13145 | 658 | Y658C | B | 1.2e-05 | 5.3e-06 | 1.1e-05 | 9.433E-06 | 3.614E-06 | 5.300E-06 | 1.200E-05 |
| 13146 | 658 | Y658E | B | 1.5e-05 | 1.7e-05 | 5.4e-06 | 1.247E-05 | 6.201E-06 | 5.400E-06 | 1.700E-05 |
| 13147 | 658 | Y658Q | B | 8.6e-06 | 1.2e-05 | 7e-06 | 9.200E-06 | 2.553E-06 | 7.000E-06 | 1.200E-05 |
| 13148 | 658 | Y658G | B | 7.1e-06 | 1e-05 | 1.3e-05 | 1.003E-05 | 2.950E-06 | 7.100E-06 | 1.300E-05 |
| 13149 | 658 | Y658H | B | 5.1e-06 | 6.4e-06 | 5.2e-06 | 5.567E-06 | 7.234E-07 | 5.100E-06 | 6.400E-06 |
| 13150 | 658 | Y658I | B | 8.1e-06 | 1e-05 | 9.1e-06 | 9.067E-06 | 9.504E-07 | 8.100E-06 | 1.000E-05 |
| 13151 | 658 | Y658L | B | 7.8e-06 | 6.8e-06 | 6.2e-06 | 6.933E-06 | 8.083E-07 | 6.200E-06 | 7.800E-06 |
| 13152 | 658 | Y658K | B | 9.3e-06 | 8.6e-06 | 1.3e-05 | 1.030E-05 | 2.364E-06 | 8.600E-06 | 1.300E-05 |
| 13153 | 658 | Y658M | B | 5e-06 | 8.4e-06 | 5.8e-06 | 6.400E-06 | 1.778E-06 | 5.000E-06 | 8.400E-06 |
| 13154 | 658 | Y658F | B | 6.2e-06 | 1.1e-05 | 1.2e-05 | 9.733E-06 | 3.101E-06 | 6.200E-06 | 1.200E-05 |
| 13155 | 658 | Y658P | B | 1.4e-05 | 8.1e-06 | 9.3e-06 | 1.047E-05 | 3.118E-06 | 8.100E-06 | 1.400E-05 |
| 13156 | 658 | Y658S | B | 7e-06 | 9.8e-06 | 7.2e-06 | 8.000E-06 | 1.562E-06 | 7.000E-06 | 9.800E-06 |
| 13157 | 658 | Y658T | B | 8.6e-06 | 7.8e-06 | 5.4e-06 | 7.267E-06 | 1.665E-06 | 5.400E-06 | 8.600E-06 |
| 13158 | 658 | Y658W | B | 4.8e-06 | 9.8e-06 | 1e-05 | 8.200E-06 | 2.946E-06 | 4.800E-06 | 1.000E-05 |
| 13159 | 658 | Y658Y | B | 9.7e-06 | 6.1e-06 | 8.6e-06 | 8.133E-06 | 1.845E-06 | 6.100E-06 | 9.700E-06 |
| 13160 | 658 | Y658V | B | 5.3e-06 | 1.4e-05 | 7.7e-06 | 9.000E-06 | 4.493E-06 | 5.300E-06 | 1.400E-05 |
| 13161 | 659 | Q659A | B | 9.3e-06 | 9e-06 | 6.5e-06 | 8.267E-06 | 1.537E-06 | 6.500E-06 | 9.300E-06 |
| 13162 | 659 | Q659R | B | 6.6e-06 | 9.4e-06 | 5.6e-06 | 7.200E-06 | 1.970E-06 | 5.600E-06 | 9.400E-06 |
| 13163 | 659 | Q659N | B | 4.4e-06 | 6.6e-06 | 8.1e-06 | 6.367E-06 | 1.861E-06 | 4.400E-06 | 8.100E-06 |
| 13164 | 659 | Q659D | B | 1.5e-05 | 6.2e-06 | 5e-06 | 8.733E-06 | 5.460E-06 | 5.000E-06 | 1.500E-05 |
| 13165 | 659 | Q659C | B | 9.5e-06 | 5.3e-06 | 1.2e-05 | 8.933E-06 | 3.386E-06 | 5.300E-06 | 1.200E-05 |
| 13166 | 659 | Q659E | B | 4.7e-06 | 6.4e-06 | 4.2e-06 | 5.100E-06 | 1.153E-06 | 4.200E-06 | 6.400E-06 |
| 13167 | 659 | Q659Q | B | 8.8e-06 | 8.4e-06 | 1.2e-05 | 9.733E-06 | 1.973E-06 | 8.400E-06 | 1.200E-05 |
| 13168 | 659 | Q659G | B | 8.7e-06 | 1.3e-05 | 6.1e-06 | 9.267E-06 | 3.485E-06 | 6.100E-06 | 1.300E-05 |
| 13169 | 659 | Q659H | B | 5.1e-06 | 1.4e-05 | 6.1e-06 | 8.400E-06 | 4.875E-06 | 5.100E-06 | 1.400E-05 |
| 13170 | 659 | Q659I | B | 9.6e-06 | 1e-05 | 8.8e-06 | 9.467E-06 | 6.110E-07 | 8.800E-06 | 1.000E-05 |
| 13171 | 659 | Q659L | B | 1.1e-05 | 4.8e-06 | 1.5e-05 | 1.027E-05 | 5.139E-06 | 4.800E-06 | 1.500E-05 |
| 13172 | 659 | Q659K | B | 5.9e-06 | 7.8e-06 | 1.3e-05 | 8.900E-06 | 3.676E-06 | 5.900E-06 | 1.300E-05 |
| 13173 | 659 | Q659M | B | 9.2e-06 | 6.5e-06 | 5.7e-06 | 7.133E-06 | 1.834E-06 | 5.700E-06 | 9.200E-06 |
| 13174 | 659 | Q659F | B | 1.2e-05 | 1.7e-05 | 1.5e-05 | 1.467E-05 | 2.517E-06 | 1.200E-05 | 1.700E-05 |

| | | | | | | | | | | |
|---|---|---|---|---|---|---|---|---|---|---|
| 13175 | 659 | Q659P | B | 6.4e-06 | 8.1e-06 | 4.1e-06 | 6.200E-06 | 2.007E-06 | 4.100E-06 | 8.100E-06 |
| 13176 | 659 | Q659S | B | 5.2e-06 | 9.5e-06 | 1e-05 | 8.233E-06 | 2.639E-06 | 5.200E-06 | 1.000E-05 |
| 13177 | 659 | Q659T | B | 4.3e-06 | 6.6e-06 | 4.1e-06 | 5.000E-06 | 1.389E-06 | 4.100E-06 | 6.600E-06 |
| 13178 | 659 | Q659W | B | 6.5e-06 | 9.5e-06 | 9.8e-06 | 8.600E-06 | 1.825E-06 | 6.500E-06 | 9.800E-06 |
| 13179 | 659 | Q659Y | B | 6.6e-06 | 6.2e-06 | 6.4e-06 | 6.400E-06 | 2.000E-07 | 6.200E-06 | 6.600E-06 |
| 13180 | 659 | Q659V | B | 6.1e-06 | 8.2e-06 | 3.9e-06 | 6.067E-06 | 2.150E-06 | 3.900E-06 | 8.200E-06 |
| 13181 | 660 | Q660A | B | 9.7e-06 | 1e-05 | 6.6e-06 | 8.767E-06 | 1.882E-06 | 6.600E-06 | 1.000E-05 |
| 13182 | 660 | Q660R | B | 9.1e-06 | 9.2e-06 | 7.3e-06 | 8.533E-06 | 1.069E-06 | 7.300E-06 | 9.200E-06 |
| 13183 | 660 | Q660N | B | 7.2e-06 | 1.2e-05 | 4.9e-06 | 8.033E-06 | 3.623E-06 | 4.900E-06 | 1.200E-05 |
| 13184 | 660 | Q660D | B | 8.2e-06 | 8.8e-06 | 6.6e-06 | 7.867E-06 | 1.137E-06 | 6.600E-06 | 8.800E-06 |
| 13185 | 660 | Q660C | B | 1.1e-05 | 5.9e-06 | 5.3e-06 | 7.400E-06 | 3.132E-06 | 5.300E-06 | 1.100E-05 |
| 13186 | 660 | Q660E | B | 9.6e-06 | 4.8e-06 | 4.1e-06 | 6.167E-06 | 2.994E-06 | 4.100E-06 | 9.600E-06 |
| 13187 | 660 | Q660Q | B | 1.4e-05 | 7.6e-06 | 6.5e-06 | 9.367E-06 | 4.050E-06 | 6.500E-06 | 1.400E-05 |
| 13188 | 660 | Q660G | B | 7.1e-06 | 7.5e-06 | 1e-05 | 8.200E-06 | 1.572E-06 | 7.100E-06 | 1.000E-05 |
| 13189 | 660 | Q660H | B | 6.5e-06 | 7.5e-06 | 6e-06 | 6.667E-06 | 7.638E-07 | 6.000E-06 | 7.500E-06 |
| 13190 | 660 | Q660I | B | 8.7e-06 | 7.5e-06 | 7.7e-06 | 7.967E-06 | 6.429E-07 | 7.500E-06 | 8.700E-06 |
| 13191 | 660 | Q660L | B | 9.9e-06 | 1.5e-05 | 5.8e-06 | 1.023E-05 | 4.609E-06 | 5.800E-06 | 1.500E-05 |
| 13192 | 660 | Q660K | B | 5.7e-06 | 9.7e-06 | 9e-06 | 8.133E-06 | 2.136E-06 | 5.700E-06 | 9.700E-06 |
| 13193 | 660 | Q660M | B | 7.4e-06 | 6.8e-06 | 6.1e-06 | 6.767E-06 | 6.506E-07 | 6.100E-06 | 7.400E-06 |
| 13194 | 660 | Q660F | B | 5.6e-06 | 8.2e-06 | 9.2e-06 | 7.667E-06 | 1.858E-06 | 5.600E-06 | 9.200E-06 |
| 13195 | 660 | Q660P | B | 4.2e-06 | 9.5e-06 | 8.4e-06 | 7.367E-06 | 2.797E-06 | 4.200E-06 | 9.500E-06 |
| 13196 | 660 | Q660S | B | 1e-05 | 9e-06 | 1.2e-05 | 1.033E-05 | 1.528E-06 | 9.000E-06 | 1.200E-05 |
| 13197 | 660 | Q660T | B | 6.5e-06 | 5.2e-06 | 7.5e-06 | 6.400E-06 | 1.153E-06 | 5.200E-06 | 7.500E-06 |
| 13198 | 660 | Q660W | B | 1e-05 | 9.5e-06 | 5.9e-06 | 8.467E-06 | 2.237E-06 | 5.900E-06 | 1.000E-05 |
| 13199 | 660 | Q660Y | B | 6.3e-06 | 1e-05 | 8.5e-06 | 8.267E-06 | 1.861E-06 | 6.300E-06 | 1.000E-05 |
| 13200 | 660 | Q660V | B | 9.4e-06 | 1.3e-05 | 1.1e-05 | 1.113E-05 | 1.804E-06 | 9.400E-06 | 1.300E-05 |
| 13201 | 661 | K661A | B | 8.1e-06 | 6.3e-06 | 4.3e-06 | 6.233E-06 | 1.901E-06 | 4.300E-06 | 8.100E-06 |
| 13202 | 661 | K661R | B | 6e-06 | 1.3e-05 | 7.7e-06 | 8.900E-06 | 3.651E-06 | 6.000E-06 | 1.300E-05 |
| 13203 | 661 | K661N | B | 6.5e-06 | 8.3e-06 | 4.8e-06 | 6.533E-06 | 1.750E-06 | 4.800E-06 | 8.300E-06 |
| 13204 | 661 | K661D | B | 1.8e-05 | 4.8e-06 | 6.7e-06 | 9.833E-06 | 7.136E-06 | 4.800E-06 | 1.800E-05 |
| 13205 | 661 | K661C | B | 5.1e-06 | 6.8e-06 | 9.3e-06 | 7.067E-06 | 2.113E-06 | 5.100E-06 | 9.300E-06 |
| 13206 | 661 | K661E | B | 1.3e-05 | 9.7e-06 | 8.2e-06 | 1.030E-05 | 2.456E-06 | 8.200E-06 | 1.300E-05 |
| 13207 | 661 | K661Q | B | 9.4e-06 | 6.2e-06 | 9.7e-06 | 8.433E-06 | 1.940E-06 | 6.200E-06 | 9.700E-06 |
| 13208 | 661 | K661G | B | 9.3e-06 | 8.2e-06 | 1.4e-05 | 1.050E-05 | 3.081E-06 | 8.200E-06 | 1.400E-05 |
| 13209 | 661 | K661H | B | 4.7e-06 | 1e-05 | 1.3e-05 | 9.233E-06 | 4.203E-06 | 4.700E-06 | 1.300E-05 |
| 13210 | 661 | K661I | B | 9.7e-06 | 4.8e-06 | 7.5e-06 | 7.333E-06 | 2.454E-06 | 4.800E-06 | 9.700E-06 |
| 13211 | 661 | K661L | B | 9.9e-06 | 1.2e-05 | 5e-06 | 8.967E-06 | 3.592E-06 | 5.000E-06 | 1.200E-05 |
| 13212 | 661 | K661K | B | 9.8e-06 | 6.4e-06 | 9.4e-06 | 8.533E-06 | 1.858E-06 | 6.400E-06 | 9.800E-06 |
| 13213 | 661 | K661M | B | 5.7e-06 | 1.3e-05 | 8.2e-06 | 8.967E-06 | 3.710E-06 | 5.700E-06 | 1.300E-05 |
| 13214 | 661 | K661F | B | 4.4e-06 | 4.3e-06 | 1.1e-05 | 6.567E-06 | 3.840E-06 | 4.300E-06 | 1.100E-05 |
| 13215 | 661 | K661P | B | 1.3e-05 | 4.5e-06 | 5.3e-06 | 7.600E-06 | 4.694E-06 | 4.500E-06 | 1.300E-05 |
| 13216 | 661 | K661S | B | 9.5e-06 | 7.6e-06 | 3.8e-06 | 6.967E-06 | 2.902E-06 | 3.800E-06 | 9.500E-06 |
| 13217 | 661 | K661T | B | 6.9e-06 | 5.8e-06 | 4.3e-06 | 5.667E-06 | 1.305E-06 | 4.300E-06 | 6.900E-06 |
| 13218 | 661 | K661W | B | 9.5e-06 | 7e-06 | 1.5e-05 | 1.050E-05 | 4.093E-06 | 7.000E-06 | 1.500E-05 |
| 13219 | 661 | K661Y | B | 9e-06 | 6.1e-06 | 6.9e-06 | 7.333E-06 | 1.498E-06 | 6.100E-06 | 9.000E-06 |
| 13220 | 661 | K661V | B | 7e-06 | 1.3e-05 | 7.7e-06 | 9.233E-06 | 3.281E-06 | 7.000E-06 | 1.300E-05 |
| 13221 | 662 | P662A | B | 1e-05 | 4.6e-06 | 8.9e-06 | 7.833E-06 | 2.854E-06 | 4.600E-06 | 1.000E-05 |
| 13222 | 662 | P662R | B | 6.1e-06 | 7e-06 | 8.9e-06 | 7.333E-06 | 1.429E-06 | 6.100E-06 | 8.900E-06 |
| 13223 | 662 | P662N | B | 7e-06 | 9.2e-06 | 6.5e-06 | 7.567E-06 | 1.436E-06 | 6.500E-06 | 9.200E-06 |
| 13224 | 662 | P662D | B | 1.4e-05 | 8.5e-06 | 9.6e-06 | 1.070E-05 | 2.910E-06 | 8.500E-06 | 1.400E-05 |
| 13225 | 662 | P662C | B | 8.4e-06 | 7.6e-06 | 1.1e-05 | 9.000E-06 | 1.778E-06 | 7.600E-06 | 1.100E-05 |
| 13226 | 662 | P662E | B | 6.4e-06 | 1.5e-05 | 9.2e-06 | 1.020E-05 | 4.386E-06 | 6.400E-06 | 1.500E-05 |
| 13227 | 662 | P662Q | B | 1.4e-05 | 9.6e-06 | 9.4e-06 | 1.100E-05 | 2.600E-06 | 9.400E-06 | 1.400E-05 |
| 13228 | 662 | P662G | B | 9.6e-06 | 4.6e-06 | 1.6e-05 | 1.007E-05 | 5.714E-06 | 4.600E-06 | 1.600E-05 |
| 13229 | 662 | P662H | B | 1e-05 | 7e-06 | 6.1e-06 | 7.700E-06 | 2.042E-06 | 6.100E-06 | 1.000E-05 |
| 13230 | 662 | P662I | B | 7.3e-06 | 8.9e-06 | 1.5e-05 | 1.040E-05 | 4.063E-06 | 7.300E-06 | 1.500E-05 |
| 13231 | 662 | P662L | B | 1.3e-05 | 8.1e-06 | 4.2e-06 | 8.433E-06 | 4.409E-06 | 4.200E-06 | 1.300E-05 |
| 13232 | 662 | P662K | B | 8.8e-06 | 6.9e-06 | 7.8e-06 | 7.833E-06 | 9.504E-07 | 6.900E-06 | 8.800E-06 |
| 13233 | 662 | P662M | B | 8.3e-06 | 9.4e-06 | 7.5e-06 | 8.400E-06 | 9.539E-07 | 7.500E-06 | 9.400E-06 |
| 13234 | 662 | P662F | B | 5.2e-06 | 8.9e-06 | 9.7e-06 | 7.933E-06 | 2.401E-06 | 5.200E-06 | 9.700E-06 |
| 13235 | 662 | P662P | B | 7.5e-06 | 1.4e-05 | 1.4e-05 | 1.183E-05 | 3.753E-06 | 7.500E-06 | 1.400E-05 |
| 13236 | 662 | P662S | B | 8e-06 | 5.5e-06 | 5.7e-06 | 6.400E-06 | 1.389E-06 | 5.500E-06 | 8.000E-06 |
| 13237 | 662 | P662T | B | 9.2e-06 | 9.1e-06 | 5.7e-06 | 8.000E-06 | 1.992E-06 | 5.700E-06 | 9.200E-06 |
| 13238 | 662 | P662W | B | 1.8e-05 | 9.5e-06 | 6.1e-06 | 1.120E-05 | 6.129E-06 | 6.100E-06 | 1.800E-05 |
| 13239 | 662 | P662Y | B | 1.1e-05 | 8.8e-06 | 1e-05 | 9.933E-06 | 1.102E-06 | 8.800E-06 | 1.100E-05 |
| 13240 | 662 | P662V | B | 6.3e-06 | 1e-05 | 7e-06 | 7.767E-06 | 1.966E-06 | 6.300E-06 | 1.000E-05 |
| 13241 | 663 | G663A | B | 7.1e-06 | 9.7e-06 | 5.3e-06 | 7.367E-06 | 2.212E-06 | 5.300E-06 | 9.700E-06 |
| 13242 | 663 | G663R | B | 7.1e-06 | 8.7e-06 | 1.1e-05 | 8.933E-06 | 1.960E-06 | 7.100E-06 | 1.100E-05 |
| 13243 | 663 | G663N | B | 5e-06 | 1.2e-05 | 1.1e-05 | 9.333E-06 | 3.786E-06 | 5.000E-06 | 1.200E-05 |
| 13244 | 663 | G663D | B | 6.5e-06 | 9.7e-06 | 6.9e-06 | 7.700E-06 | 1.744E-06 | 6.500E-06 | 9.700E-06 |
| 13245 | 663 | G663C | B | 6.1e-06 | 5.9e-06 | 9.2e-06 | 7.067E-06 | 1.850E-06 | 5.900E-06 | 9.200E-06 |
| 13246 | 663 | G663E | B | 1.1e-05 | 1.1e-05 | 9.1e-06 | 1.037E-05 | 1.097E-06 | 9.100E-06 | 1.100E-05 |
| 13247 | 663 | G663Q | B | 6.3e-06 | 7.1e-06 | 6.1e-06 | 6.500E-06 | 5.292E-07 | 6.100E-06 | 7.100E-06 |
| 13248 | 663 | G663G | B | 9.4e-06 | 7e-06 | 7e-06 | 7.800E-06 | 1.386E-06 | 7.000E-06 | 9.400E-06 |
| 13249 | 663 | G663H | B | 5.4e-06 | 8.1e-06 | 7.8e-06 | 7.100E-06 | 1.480E-06 | 5.400E-06 | 8.100E-06 |
| 13250 | 663 | G663I | B | 7.4e-06 | 1.1e-05 | 6.1e-06 | 8.167E-06 | 2.538E-06 | 6.100E-06 | 1.100E-05 |
| 13251 | 663 | G663L | B | 8.6e-06 | 9.2e-06 | 5.8e-06 | 7.867E-06 | 1.815E-06 | 5.800E-06 | 9.200E-06 |
| 13252 | 663 | G663K | B | 7.2e-06 | 9.5e-06 | 1.3e-05 | 9.900E-06 | 2.921E-06 | 7.200E-06 | 1.300E-05 |
| 13253 | 663 | G663M | B | 7.3e-06 | 8.9e-06 | 9.1e-06 | 8.433E-06 | 9.866E-07 | 7.300E-06 | 9.100E-06 |
| 13254 | 663 | G663F | B | 9.4e-06 | 4.2e-06 | 6.8e-06 | 6.800E-06 | 2.600E-06 | 4.200E-06 | 9.400E-06 |
| 13255 | 663 | G663P | B | 6.3e-06 | 9.4e-06 | 5.4e-06 | 7.033E-06 | 2.098E-06 | 5.400E-06 | 9.400E-06 |
| 13256 | 663 | G663S | B | 6.1e-06 | 7e-06 | 4e-06 | 5.700E-06 | 1.539E-06 | 4.000E-06 | 7.000E-06 |
| 13257 | 663 | G663T | B | 8.9e-06 | 1.4e-05 | 1.1e-05 | 1.130E-05 | 2.563E-06 | 8.900E-06 | 1.400E-05 |
| 13258 | 663 | G663W | B | 6.6e-06 | 7.1e-06 | 8.6e-06 | 7.433E-06 | 1.041E-06 | 6.600E-06 | 8.600E-06 |
| 13259 | 663 | G663Y | B | 6.3e-06 | 7e-06 | 7.6e-06 | 6.967E-06 | 6.506E-07 | 6.300E-06 | 7.600E-06 |

| | | | | | | | | | | |
|---|---|---|---|---|---|---|---|---|---|---|
| 13260 | 663 | G663V | B | 5e-06 | 1.1e-05 | 7e-06 | 7.667E-06 | 3.055E-06 | 5.000E-06 | 1.100E-05 |
| 13261 | 664 | K664A | B | 7.4e-06 | 5.9e-06 | 5e-06 | 6.100E-06 | 1.212E-06 | 5.000E-06 | 7.400E-06 |
| 13262 | 664 | K664R | B | 8.1e-06 | 4.9e-06 | 7.4e-06 | 6.800E-06 | 1.682E-06 | 4.900E-06 | 8.100E-06 |
| 13263 | 664 | K664N | B | 5.9e-06 | 9.7e-06 | 5.3e-06 | 6.967E-06 | 2.386E-06 | 5.300E-06 | 9.700E-06 |
| 13264 | 664 | K664D | B | 8.9e-06 | 8.4e-06 | 9.5e-06 | 8.933E-06 | 5.508E-07 | 8.400E-06 | 9.500E-06 |
| 13265 | 664 | K664C | B | 7.1e-06 | 1.1e-05 | 7.7e-06 | 8.600E-06 | 2.100E-06 | 7.100E-06 | 1.100E-05 |
| 13266 | 664 | K664E | B | 1.5e-05 | 1e-05 | 5.5e-06 | 1.017E-05 | 4.752E-06 | 5.500E-06 | 1.500E-05 |
| 13267 | 664 | K664Q | B | 1.1e-05 | 1.3e-05 | 8.1e-06 | 1.070E-05 | 2.464E-06 | 8.100E-06 | 1.300E-05 |
| 13268 | 664 | K664G | B | 6.7e-06 | 6.4e-06 | 4e-06 | 5.700E-06 | 1.480E-06 | 4.000E-06 | 6.700E-06 |
| 13269 | 664 | K664H | B | 9e-06 | 6.8e-06 | 6.8e-06 | 7.533E-06 | 1.270E-06 | 6.800E-06 | 9.000E-06 |
| 13270 | 664 | K664I | B | 9e-06 | 4.4e-06 | 4e-06 | 5.800E-06 | 2.778E-06 | 4.000E-06 | 9.000E-06 |
| 13271 | 664 | K664L | B | 4.9e-06 | 1.3e-05 | 9.3e-06 | 9.067E-06 | 4.055E-06 | 4.900E-06 | 1.300E-05 |
| 13272 | 664 | K664K | B | 1e-05 | 1.1e-05 | 4.1e-06 | 8.367E-06 | 3.729E-06 | 4.100E-06 | 1.100E-05 |
| 13273 | 664 | K664M | B | 7.9e-06 | 7e-06 | 9.2e-06 | 8.033E-06 | 1.106E-06 | 7.000E-06 | 9.200E-06 |
| 13274 | 664 | K664F | B | 1.3e-05 | 9.8e-06 | 6.4e-06 | 9.733E-06 | 3.301E-06 | 6.400E-06 | 1.300E-05 |
| 13275 | 664 | K664P | B | 6.8e-06 | 1e-05 | 9.5e-06 | 8.767E-06 | 1.721E-06 | 6.800E-06 | 1.000E-05 |
| 13276 | 664 | K664S | B | 8.6e-06 | 7e-06 | 1.4e-05 | 9.867E-06 | 3.668E-06 | 7.000E-06 | 1.400E-05 |
| 13277 | 664 | K664T | B | 5.5e-06 | 1.3e-05 | 4.8e-06 | 7.767E-06 | 4.546E-06 | 4.800E-06 | 1.300E-05 |
| 13278 | 664 | K664W | B | 3.4e-06 | 8.5e-06 | 6.1e-06 | 6.000E-06 | 2.551E-06 | 3.400E-06 | 8.500E-06 |
| 13279 | 664 | K664Y | B | 6.6e-06 | 1.3e-05 | 5.7e-06 | 8.433E-06 | 3.980E-06 | 5.700E-06 | 1.300E-05 |
| 13280 | 664 | K664V | B | 7.9e-06 | 8.1e-06 | 9.2e-06 | 8.400E-06 | 7.000E-07 | 7.900E-06 | 9.200E-06 |
| 13281 | 665 | A665A | B | 6.4e-06 | 8.1e-06 | 8.7e-06 | 7.733E-06 | 1.193E-06 | 6.400E-06 | 8.700E-06 |
| 13282 | 665 | A665R | B | 5.3e-06 | 1.2e-05 | 4e-06 | 7.100E-06 | 4.293E-06 | 4.000E-06 | 1.200E-05 |
| 13283 | 665 | A665N | B | 9.3e-06 | 9e-06 | 6.4e-06 | 8.233E-06 | 1.595E-06 | 6.400E-06 | 9.300E-06 |
| 13284 | 665 | A665D | B | 6.7e-06 | 1e-05 | 1.3e-05 | 9.900E-06 | 3.151E-06 | 6.700E-06 | 1.300E-05 |
| 13285 | 665 | A665C | B | 7.9e-06 | 9.8e-06 | 1.1e-05 | 9.567E-06 | 1.563E-06 | 7.900E-06 | 1.100E-05 |
| 13286 | 665 | A665E | B | 7.8e-06 | 7.2e-06 | 1.2e-05 | 9.000E-06 | 2.615E-06 | 7.200E-06 | 1.200E-05 |
| 13287 | 665 | A665Q | B | 1.4e-05 | 7.7e-06 | 9.2e-06 | 1.030E-05 | 3.291E-06 | 7.700E-06 | 1.400E-05 |
| 13288 | 665 | A665G | B | 8e-06 | 1.2e-05 | 1.3e-05 | 1.100E-05 | 2.646E-06 | 8.000E-06 | 1.300E-05 |
| 13289 | 665 | A665H | B | 6.2e-06 | 9.1e-06 | 7.5e-06 | 7.600E-06 | 1.453E-06 | 6.200E-06 | 9.100E-06 |
| 13290 | 665 | A665I | B | 9e-06 | 7.5e-06 | 6.2e-06 | 7.567E-06 | 1.401E-06 | 6.200E-06 | 9.000E-06 |
| 13291 | 665 | A665L | B | 9.4e-06 | 7.9e-06 | 5e-06 | 7.433E-06 | 2.237E-06 | 5.000E-06 | 9.400E-06 |
| 13292 | 665 | A665K | B | 6.8e-06 | 8.7e-06 | 4.4e-06 | 6.633E-06 | 2.155E-06 | 4.400E-06 | 8.700E-06 |
| 13293 | 665 | A665M | B | 1e-05 | 8.1e-06 | 8.2e-06 | 8.767E-06 | 1.069E-06 | 8.100E-06 | 1.000E-05 |
| 13294 | 665 | A665F | B | 1e-05 | 1.1e-05 | 6.4e-06 | 9.133E-06 | 2.419E-06 | 6.400E-06 | 1.100E-05 |
| 13295 | 665 | A665P | B | 7.2e-06 | 5.2e-06 | 7.4e-06 | 6.600E-06 | 1.217E-06 | 5.200E-06 | 7.400E-06 |
| 13296 | 665 | A665S | B | 8.5e-06 | 9.4e-06 | 7.3e-06 | 8.400E-06 | 1.054E-06 | 7.300E-06 | 9.400E-06 |
| 13297 | 665 | A665T | B | 4.9e-06 | 7.8e-06 | 1.1e-05 | 7.900E-06 | 3.051E-06 | 4.900E-06 | 1.100E-05 |
| 13298 | 665 | A665W | B | 1.2e-05 | 4.5e-06 | 8.3e-06 | 8.267E-06 | 3.750E-06 | 4.500E-06 | 1.200E-05 |
| 13299 | 665 | A665Y | B | 9.7e-06 | 8.8e-06 | 8.1e-06 | 8.867E-06 | 8.021E-07 | 8.100E-06 | 9.700E-06 |
| 13300 | 665 | A665V | B | 6.6e-06 | 8.2e-06 | 8.6e-06 | 7.800E-06 | 1.058E-06 | 6.600E-06 | 8.600E-06 |
| 13301 | 666 | P666A | B | 7.1e-06 | 7.3e-06 | 8.5e-06 | 7.633E-06 | 7.572E-07 | 7.100E-06 | 8.500E-06 |
| 13302 | 666 | P666R | B | 1.1e-05 | 7.8e-06 | 8.7e-06 | 9.167E-06 | 1.650E-06 | 7.800E-06 | 1.100E-05 |
| 13303 | 666 | P666N | B | 1.2e-05 | 7.6e-06 | 7.2e-06 | 8.933E-06 | 2.663E-06 | 7.200E-06 | 1.200E-05 |
| 13304 | 666 | P666D | B | 1e-05 | 9.3e-06 | 7.6e-06 | 8.967E-06 | 1.234E-06 | 7.600E-06 | 1.000E-05 |
| 13305 | 666 | P666C | B | 1.4e-05 | 5.8e-06 | 5.6e-06 | 8.467E-06 | 4.793E-06 | 5.600E-06 | 1.400E-05 |
| 13306 | 666 | P666E | B | 6.7e-06 | 9.7e-06 | 7.9e-06 | 8.100E-06 | 1.510E-06 | 6.700E-06 | 9.700E-06 |
| 13307 | 666 | P666Q | B | 4.8e-06 | 1e-05 | 7.1e-06 | 7.300E-06 | 2.606E-06 | 4.800E-06 | 1.000E-05 |
| 13308 | 666 | P666G | B | 7.3e-06 | 4.5e-06 | 9.1e-06 | 6.967E-06 | 2.318E-06 | 4.500E-06 | 9.100E-06 |
| 13309 | 666 | P666H | B | 6.7e-06 | 7.3e-06 | 4.9e-06 | 6.300E-06 | 1.249E-06 | 4.900E-06 | 7.300E-06 |
| 13310 | 666 | P666I | B | 7.5e-06 | 1.1e-05 | 6.9e-06 | 8.467E-06 | 2.214E-06 | 6.900E-06 | 1.100E-05 |
| 13311 | 666 | P666L | B | 9.6e-06 | 3e-06 | 6.7e-06 | 6.433E-06 | 3.308E-06 | 3.000E-06 | 9.600E-06 |
| 13312 | 666 | P666K | B | 6.4e-06 | 4.5e-06 | 7e-06 | 5.967E-06 | 1.305E-06 | 4.500E-06 | 7.000E-06 |
| 13313 | 666 | P666M | B | 6.1e-06 | 7e-06 | 5.8e-06 | 6.300E-06 | 6.245E-07 | 5.800E-06 | 7.000E-06 |
| 13314 | 666 | P666F | B | 6.7e-06 | 6.2e-06 | 1.2e-05 | 8.300E-06 | 3.214E-06 | 6.200E-06 | 1.200E-05 |
| 13315 | 666 | P666P | B | 9.2e-06 | 4.9e-06 | 8.1e-06 | 7.400E-06 | 2.234E-06 | 4.900E-06 | 9.200E-06 |
| 13316 | 666 | P666S | B | 7.4e-06 | 3.9e-06 | 7e-06 | 6.100E-06 | 1.916E-06 | 3.900E-06 | 7.400E-06 |
| 13317 | 666 | P666T | B | 4.8e-06 | 7.6e-06 | 8.4e-06 | 6.933E-06 | 1.890E-06 | 4.800E-06 | 8.400E-06 |
| 13318 | 666 | P666W | B | 6.1e-06 | 7.6e-06 | 1.2e-05 | 8.567E-06 | 3.066E-06 | 6.100E-06 | 1.200E-05 |
| 13319 | 666 | P666Y | B | 5.6e-06 | 5.9e-06 | 1.2e-05 | 7.833E-06 | 3.612E-06 | 5.600E-06 | 1.200E-05 |
| 13320 | 666 | P666V | B | 5.5e-06 | 5.9e-06 | 6.1e-06 | 5.833E-06 | 3.055E-07 | 5.500E-06 | 6.100E-06 |
| 13321 | 667 | K667A | B | 8.3e-06 | 7e-06 | 9.6e-06 | 8.300E-06 | 1.300E-06 | 7.000E-06 | 9.600E-06 |
| 13322 | 667 | K667R | B | 5.8e-06 | 8.7e-06 | 6.6e-06 | 7.033E-06 | 1.498E-06 | 5.800E-06 | 8.700E-06 |
| 13323 | 667 | K667N | B | 4.5e-06 | 6.3e-06 | 1.2e-05 | 7.600E-06 | 3.915E-06 | 4.500E-06 | 1.200E-05 |
| 13324 | 667 | K667D | B | 4.6e-06 | 1e-05 | 7.5e-06 | 7.367E-06 | 2.702E-06 | 4.600E-06 | 1.000E-05 |
| 13325 | 667 | K667C | B | 3.1e-06 | 6.9e-06 | 7.3e-06 | 5.767E-06 | 2.318E-06 | 3.100E-06 | 7.300E-06 |
| 13326 | 667 | K667E | B | 6.3e-06 | 1e-05 | 9e-06 | 8.433E-06 | 1.914E-06 | 6.300E-06 | 1.000E-05 |
| 13327 | 667 | K667Q | B | 6.1e-06 | 5.9e-06 | 1e-05 | 7.333E-06 | 2.312E-06 | 5.900E-06 | 1.000E-05 |
| 13328 | 667 | K667G | B | 7.1e-06 | 6.1e-06 | 9.2e-06 | 7.467E-06 | 1.582E-06 | 6.100E-06 | 9.200E-06 |
| 13329 | 667 | K667H | B | 1.4e-05 | 9e-06 | 1.3e-05 | 1.200E-05 | 2.646E-06 | 9.000E-06 | 1.400E-05 |
| 13330 | 667 | K667I | B | 9.4e-06 | 8.3e-06 | 1e-05 | 9.233E-06 | 8.622E-07 | 8.300E-06 | 1.000E-05 |
| 13331 | 667 | K667L | B | 5e-06 | 5.8e-06 | 7e-06 | 5.933E-06 | 1.007E-06 | 5.000E-06 | 7.000E-06 |
| 13332 | 667 | K667K | B | 6.5e-06 | 7.6e-06 | 7e-06 | 7.033E-06 | 5.508E-07 | 6.500E-06 | 7.600E-06 |
| 13333 | 667 | K667M | B | 1e-05 | 9e-06 | 6.6e-06 | 8.533E-06 | 1.747E-06 | 6.600E-06 | 1.000E-05 |
| 13334 | 667 | K667F | B | 1.3e-05 | 6.2e-06 | 6.6e-06 | 8.600E-06 | 3.816E-06 | 6.200E-06 | 1.300E-05 |
| 13335 | 667 | K667P | B | 9.7e-06 | 9.9e-06 | 9.5e-06 | 9.700E-06 | 2.000E-07 | 9.500E-06 | 9.900E-06 |
| 13336 | 667 | K667S | B | 5.8e-06 | 5e-06 | 6.7e-06 | 5.833E-06 | 8.505E-07 | 5.000E-06 | 6.700E-06 |
| 13337 | 667 | K667T | B | 5.2e-06 | 6.6e-06 | 4.7e-06 | 5.500E-06 | 9.849E-07 | 4.700E-06 | 6.600E-06 |
| 13338 | 667 | K667W | B | 6.2e-06 | 1.4e-05 | 9.6e-06 | 9.933E-06 | 3.911E-06 | 6.200E-06 | 1.400E-05 |
| 13339 | 667 | K667Y | B | 1.2e-05 | 1.3e-05 | 6.5e-06 | 1.050E-05 | 3.500E-06 | 6.500E-06 | 1.300E-05 |
| 13340 | 667 | K667V | B | 5.8e-06 | 8.8e-06 | 9.5e-06 | 8.033E-06 | 1.966E-06 | 5.800E-06 | 9.500E-06 |
| 13341 | 668 | L668A | B | 8.7e-06 | 7.7e-06 | 1.3e-05 | 9.800E-06 | 2.816E-06 | 7.700E-06 | 1.300E-05 |
| 13342 | 668 | L668R | B | 5.5e-06 | 6.5e-06 | 7.9e-06 | 6.633E-06 | 1.206E-06 | 5.500E-06 | 7.900E-06 |
| 13343 | 668 | L668N | B | 5.7e-06 | 6.5e-06 | 3e-06 | 5.067E-06 | 1.834E-06 | 3.000E-06 | 6.500E-06 |
| 13344 | 668 | L668D | B | 5.2e-06 | 7.1e-06 | 8.7e-06 | 7.000E-06 | 1.752E-06 | 5.200E-06 | 8.700E-06 |

| | | | | | | | | | | |
|---|---|---|---|---|---|---|---|---|---|---|
| 13345 | 668 | L668C | B | 8.6e-06 | 8e-06 | 8.4e-06 | 8.333E-06 | 3.055E-07 | 8.000E-06 | 8.600E-06 |
| 13346 | 668 | L668E | B | 1.1e-05 | 6.2e-06 | 1.2e-05 | 9.733E-06 | 3.101E-06 | 6.200E-06 | 1.200E-05 |
| 13347 | 668 | L668Q | B | 9.1e-06 | 6.7e-06 | 4.3e-06 | 6.700E-06 | 2.400E-06 | 4.300E-06 | 9.100E-06 |
| 13348 | 668 | L668G | B | 8.3e-06 | 6.7e-06 | 6.1e-06 | 7.033E-06 | 1.137E-06 | 6.100E-06 | 8.300E-06 |
| 13349 | 668 | L668H | B | 1.1e-05 | 1.4e-05 | 1.2e-05 | 1.233E-05 | 1.528E-06 | 1.100E-05 | 1.400E-05 |
| 13350 | 668 | L668I | B | 9.2e-06 | 8e-06 | 6.3e-06 | 7.833E-06 | 1.457E-06 | 6.300E-06 | 9.200E-06 |
| 13351 | 668 | L668L | B | 8.3e-06 | 1.2e-05 | 7.7e-06 | 9.333E-06 | 2.329E-06 | 7.700E-06 | 1.200E-05 |
| 13352 | 668 | L668K | B | 7.6e-06 | 8.8e-06 | 8.6e-06 | 8.333E-06 | 6.429E-07 | 7.600E-06 | 8.800E-06 |
| 13353 | 668 | L668M | B | 1.1e-05 | 6.9e-06 | 1.1e-05 | 9.633E-06 | 2.367E-06 | 6.900E-06 | 1.100E-05 |
| 13354 | 668 | L668F | B | 1e-05 | 8.9e-06 | 9.3e-06 | 9.400E-06 | 5.568E-07 | 8.900E-06 | 1.000E-05 |
| 13355 | 668 | L668P | B | 6.7e-06 | 1e-05 | 7.6e-06 | 8.100E-06 | 1.706E-06 | 6.700E-06 | 1.000E-05 |
| 13356 | 668 | L668S | B | 7.7e-06 | 1.1e-05 | 9.2e-06 | 9.300E-06 | 1.652E-06 | 7.700E-06 | 1.100E-05 |
| 13357 | 668 | L668T | B | 8e-06 | 6.1e-06 | 8.2e-06 | 7.433E-06 | 1.159E-06 | 6.100E-06 | 8.200E-06 |
| 13358 | 668 | L668W | B | 9.6e-06 | 8.5e-06 | 1.5e-05 | 1.103E-05 | 3.479E-06 | 8.500E-06 | 1.500E-05 |
| 13359 | 668 | L668Y | B | 9.6e-06 | 7.8e-06 | 6.6e-06 | 8.000E-06 | 1.510E-06 | 6.600E-06 | 9.600E-06 |
| 13360 | 668 | L668V | B | 9.6e-06 | 7.9e-06 | 9.9e-06 | 9.133E-06 | 1.079E-06 | 7.900E-06 | 9.900E-06 |
| 13361 | 669 | L669A | B | 6.7e-06 | 8.1e-06 | 6.4e-06 | 7.067E-06 | 9.074E-07 | 6.400E-06 | 8.100E-06 |
| 13362 | 669 | L669R | B | 4.2e-06 | 6.4e-06 | 9.8e-06 | 6.800E-06 | 2.821E-06 | 4.200E-06 | 9.800E-06 |
| 13363 | 669 | L669N | B | 9.6e-06 | 7.1e-06 | 9.5e-06 | 8.733E-06 | 1.415E-06 | 7.100E-06 | 9.600E-06 |
| 13364 | 669 | L669D | B | 4.8e-06 | 1.3e-05 | 8e-06 | 8.600E-06 | 4.133E-06 | 4.800E-06 | 1.300E-05 |
| 13365 | 669 | L669C | B | 7.2e-06 | 6.1e-06 | 1.3e-05 | 8.767E-06 | 3.707E-06 | 6.100E-06 | 1.300E-05 |
| 13366 | 669 | L669E | B | 4.7e-06 | 6.5e-06 | 6.8e-06 | 6.000E-06 | 1.136E-06 | 4.700E-06 | 6.800E-06 |
| 13367 | 669 | L669Q | B | 7.7e-06 | 1.2e-05 | 6.8e-06 | 8.833E-06 | 2.779E-06 | 6.800E-06 | 1.200E-05 |
| 13368 | 669 | L669G | B | 7.3e-06 | 8e-06 | 2e-05 | 1.177E-05 | 7.139E-06 | 7.300E-06 | 2.000E-05 |
| 13369 | 669 | L669H | B | 7e-06 | 5.6e-06 | 8.8e-06 | 7.133E-06 | 1.604E-06 | 5.600E-06 | 8.800E-06 |
| 13370 | 669 | L669I | B | 7.2e-06 | 9.8e-06 | 8.7e-06 | 8.567E-06 | 1.305E-06 | 7.200E-06 | 9.800E-06 |
| 13371 | 669 | L669L | B | 8.7e-06 | 1.1e-05 | 5.8e-06 | 8.500E-06 | 2.606E-06 | 5.800E-06 | 1.100E-05 |
| 13372 | 669 | L669K | B | 4.8e-06 | 6.8e-06 | 1.2e-05 | 7.867E-06 | 3.717E-06 | 4.800E-06 | 1.200E-05 |
| 13373 | 669 | L669M | B | 1.1e-05 | 6.2e-06 | 7.1e-06 | 8.100E-06 | 2.551E-06 | 6.200E-06 | 1.100E-05 |
| 13374 | 669 | L669F | B | 6.3e-06 | 7.7e-06 | 5.3e-06 | 6.433E-06 | 1.206E-06 | 5.300E-06 | 7.700E-06 |
| 13375 | 669 | L669P | B | 9.5e-06 | 5.1e-06 | 7.1e-06 | 7.233E-06 | 2.203E-06 | 5.100E-06 | 9.500E-06 |
| 13376 | 669 | L669S | B | 6.7e-06 | 7.7e-06 | 6.9e-06 | 7.100E-06 | 5.292E-07 | 6.700E-06 | 7.700E-06 |
| 13377 | 669 | L669T | B | 6.8e-06 | 5e-06 | 8.3e-06 | 6.700E-06 | 1.652E-06 | 5.000E-06 | 8.300E-06 |
| 13378 | 669 | L669W | B | 9.9e-06 | 9.8e-06 | 5.6e-06 | 8.433E-06 | 2.454E-06 | 5.600E-06 | 9.900E-06 |
| 13379 | 669 | L669Y | B | 6.9e-06 | 7.9e-06 | 7.6e-06 | 7.467E-06 | 5.132E-07 | 6.900E-06 | 7.900E-06 |
| 13380 | 669 | L669V | B | 5.8e-06 | 5.3e-06 | 5.2e-06 | 5.433E-06 | 3.215E-07 | 5.200E-06 | 5.800E-06 |
| 13381 | 670 | I670A | B | 1.2e-05 | 1.1e-05 | 4.4e-06 | 9.133E-06 | 4.130E-06 | 4.400E-06 | 1.200E-05 |
| 13382 | 670 | I670R | B | 1.1e-05 | 8.3e-06 | 5.3e-06 | 8.200E-06 | 2.851E-06 | 5.300E-06 | 1.100E-05 |
| 13383 | 670 | I670N | B | 6.2e-06 | 7.6e-06 | 6.4e-06 | 6.733E-06 | 7.572E-07 | 6.200E-06 | 7.600E-06 |
| 13384 | 670 | I670D | B | 7.8e-06 | 7.9e-06 | 8.9e-06 | 8.200E-06 | 6.083E-07 | 7.800E-06 | 8.900E-06 |
| 13385 | 670 | I670C | B | 4.2e-06 | 8.8e-06 | 1e-05 | 7.667E-06 | 3.062E-06 | 4.200E-06 | 1.000E-05 |
| 13386 | 670 | I670E | B | 1e-05 | 9e-06 | 9e-06 | 9.333E-06 | 5.774E-07 | 9.000E-06 | 1.000E-05 |
| 13387 | 670 | I670Q | B | 9.1e-06 | 5.4e-06 | 5.2e-06 | 6.567E-06 | 2.196E-06 | 5.200E-06 | 9.100E-06 |
| 13388 | 670 | I670G | B | 5.7e-06 | 1e-05 | 4.2e-06 | 6.633E-06 | 3.011E-06 | 4.200E-06 | 1.000E-05 |
| 13389 | 670 | I670H | B | 5.9e-06 | 1.8e-05 | 9.8e-06 | 1.123E-05 | 6.176E-06 | 5.900E-06 | 1.800E-05 |
| 13390 | 670 | I670I | B | 6.3e-06 | 1e-05 | 6.7e-06 | 7.667E-06 | 2.031E-06 | 6.300E-06 | 1.000E-05 |
| 13391 | 670 | I670L | B | 5.2e-06 | 6.6e-06 | 8.9e-06 | 6.900E-06 | 1.868E-06 | 5.200E-06 | 8.900E-06 |
| 13392 | 670 | I670K | B | 8e-06 | 6.9e-06 | 1.1e-05 | 8.633E-06 | 2.122E-06 | 6.900E-06 | 1.100E-05 |
| 13393 | 670 | I670M | B | 6.4e-06 | 9.2e-06 | 9.4e-06 | 8.333E-06 | 1.677E-06 | 6.400E-06 | 9.400E-06 |
| 13394 | 670 | I670F | B | 7.8e-06 | 6.8e-06 | 9.7e-06 | 8.100E-06 | 1.473E-06 | 6.800E-06 | 9.700E-06 |
| 13395 | 670 | I670P | B | 4.6e-06 | 1.4e-05 | 1.1e-05 | 9.867E-06 | 4.801E-06 | 4.600E-06 | 1.400E-05 |
| 13396 | 670 | I670S | B | 7.2e-06 | 1.1e-05 | 7.6e-06 | 8.600E-06 | 2.088E-06 | 7.200E-06 | 1.100E-05 |
| 13397 | 670 | I670T | B | 6.5e-06 | 8.6e-06 | 4.7e-06 | 6.600E-06 | 1.952E-06 | 4.700E-06 | 8.600E-06 |
| 13398 | 670 | I670W | B | 6.9e-06 | 6.2e-06 | 9e-06 | 7.367E-06 | 1.457E-06 | 6.200E-06 | 9.000E-06 |
| 13399 | 670 | I670Y | B | 9.7e-06 | 6.5e-06 | 7.1e-06 | 7.767E-06 | 1.701E-06 | 6.500E-06 | 9.700E-06 |
| 13400 | 670 | I670V | B | 7.7e-06 | 7.3e-06 | 9.4e-06 | 8.133E-06 | 1.115E-06 | 7.300E-06 | 9.400E-06 |
| 13401 | 671 | Y671A | B | 4.9e-06 | 9.4e-06 | 8.2e-06 | 7.500E-06 | 2.330E-06 | 4.900E-06 | 9.400E-06 |
| 13402 | 671 | Y671R | B | 9e-06 | 4.7e-06 | 9.3e-06 | 7.667E-06 | 2.574E-06 | 4.700E-06 | 9.300E-06 |
| 13403 | 671 | Y671N | B | 1.3e-05 | 7.3e-06 | 8.8e-06 | 9.700E-06 | 2.955E-06 | 7.300E-06 | 1.300E-05 |
| 13404 | 671 | Y671D | B | 1e-05 | 5.6e-06 | 1.5e-05 | 1.020E-05 | 4.703E-06 | 5.600E-06 | 1.500E-05 |
| 13405 | 671 | Y671C | B | 5.1e-06 | 6.6e-06 | 7.8e-06 | 6.500E-06 | 1.353E-06 | 5.100E-06 | 7.800E-06 |
| 13406 | 671 | Y671E | B | 9.6e-06 | 6.8e-06 | 1.5e-05 | 1.047E-05 | 4.168E-06 | 6.800E-06 | 1.500E-05 |
| 13407 | 671 | Y671Q | B | 5.5e-06 | 7.5e-06 | 6.7e-06 | 6.567E-06 | 1.007E-06 | 5.500E-06 | 7.500E-06 |
| 13408 | 671 | Y671G | B | 9.7e-06 | 9.3e-06 | 7.6e-06 | 8.867E-06 | 1.115E-06 | 7.600E-06 | 9.700E-06 |
| 13409 | 671 | Y671H | B | 1.3e-05 | 5.9e-06 | 1.7e-05 | 1.197E-05 | 5.622E-06 | 5.900E-06 | 1.700E-05 |
| 13410 | 671 | Y671I | B | 7.4e-06 | 8.7e-06 | 9e-06 | 8.367E-06 | 8.505E-07 | 7.400E-06 | 9.000E-06 |
| 13411 | 671 | Y671L | B | 1.1e-05 | 1.3e-05 | 5.8e-06 | 9.933E-06 | 3.717E-06 | 5.800E-06 | 1.300E-05 |
| 13412 | 671 | Y671K | B | 9.6e-06 | 7.1e-06 | 6e-06 | 7.567E-06 | 1.845E-06 | 6.000E-06 | 9.600E-06 |
| 13413 | 671 | Y671M | B | 5.7e-06 | 1e-05 | 9.4e-06 | 8.367E-06 | 2.329E-06 | 5.700E-06 | 1.000E-05 |
| 13414 | 671 | Y671F | B | 9.2e-06 | 6.9e-06 | 6.4e-06 | 7.500E-06 | 1.493E-06 | 6.400E-06 | 9.200E-06 |
| 13415 | 671 | Y671P | B | 8e-06 | 1.7e-05 | 6.6e-06 | 1.053E-05 | 5.644E-06 | 6.600E-06 | 1.700E-05 |
| 13416 | 671 | Y671S | B | 8.3e-06 | 1e-05 | 4.9e-06 | 7.733E-06 | 2.597E-06 | 4.900E-06 | 1.000E-05 |
| 13417 | 671 | Y671T | B | 7.3e-06 | 7.5e-06 | 5e-06 | 6.600E-06 | 1.389E-06 | 5.000E-06 | 7.500E-06 |
| 13418 | 671 | Y671W | B | 8.6e-06 | 5.9e-06 | 6.8e-06 | 7.100E-06 | 1.375E-06 | 5.900E-06 | 8.600E-06 |
| 13419 | 671 | Y671Y | B | 1e-05 | 1.1e-05 | 8.8e-06 | 9.933E-06 | 1.102E-06 | 8.800E-06 | 1.100E-05 |
| 13420 | 671 | Y671V | B | 1.3e-05 | 9e-06 | 6.7e-06 | 9.567E-06 | 3.188E-06 | 6.700E-06 | 1.300E-05 |
| 13421 | 672 | S672A | B | 4.5e-06 | 9.3e-06 | 6.8e-06 | 6.867E-06 | 2.401E-06 | 4.500E-06 | 9.300E-06 |
| 13422 | 672 | S672R | B | 7.8e-06 | 9.4e-06 | 1e-05 | 9.067E-06 | 1.137E-06 | 7.800E-06 | 1.000E-05 |
| 13423 | 672 | S672N | B | 1e-05 | 1.2e-05 | 9.1e-06 | 1.037E-05 | 1.484E-06 | 9.100E-06 | 1.200E-05 |
| 13424 | 672 | S672D | B | 5.9e-06 | 6.6e-06 | 6e-06 | 6.167E-06 | 3.786E-07 | 5.900E-06 | 6.600E-06 |
| 13425 | 672 | S672C | B | 1.2e-05 | 8.5e-06 | 5.9e-06 | 8.800E-06 | 3.061E-06 | 5.900E-06 | 1.200E-05 |
| 13426 | 672 | S672E | B | 8.3e-06 | 6.8e-06 | 8.8e-06 | 7.967E-06 | 1.041E-06 | 6.800E-06 | 8.800E-06 |
| 13427 | 672 | S672Q | B | 1.1e-05 | 7.6e-06 | 6.1e-06 | 8.233E-06 | 2.511E-06 | 6.100E-06 | 1.100E-05 |
| 13428 | 672 | S672G | B | 5.8e-06 | 6.7e-06 | 1.4e-05 | 8.833E-06 | 4.497E-06 | 5.800E-06 | 1.400E-05 |
| 13429 | 672 | S672H | B | 6.7e-06 | 1.9e-05 | 7.3e-06 | 1.100E-05 | 6.935E-06 | 6.700E-06 | 1.900E-05 |

| | | | | | | | | | | |
|---|---|---|---|---|---|---|---|---|---|---|
| 13430 | 672 | S672I | B | 1.1e-05 | 4.8e-06 | 6.8e-06 | 7.533E-06 | 3.164E-06 | 4.800E-06 | 1.100E-05 |
| 13431 | 672 | S672L | B | 8.8e-06 | 9.4e-06 | 1.5e-05 | 1.107E-05 | 3.420E-06 | 8.800E-06 | 1.500E-05 |
| 13432 | 672 | S672K | B | 6.1e-06 | 1e-05 | 5.8e-06 | 7.300E-06 | 2.343E-06 | 5.800E-06 | 1.000E-05 |
| 13433 | 672 | S672M | B | 6.5e-06 | 6.1e-06 | 6.8e-06 | 6.467E-06 | 3.512E-07 | 6.100E-06 | 6.800E-06 |
| 13434 | 672 | S672F | B | 6.9e-06 | 1e-05 | 6.7e-06 | 7.867E-06 | 1.850E-06 | 6.700E-06 | 1.000E-05 |
| 13435 | 672 | S672P | B | 7.3e-06 | 5.6e-06 | 6.3e-06 | 6.400E-06 | 8.544E-07 | 5.600E-06 | 7.300E-06 |
| 13436 | 672 | S672S | B | 3.2e-06 | 6.7e-06 | 5.1e-06 | 5.000E-06 | 1.752E-06 | 3.200E-06 | 6.700E-06 |
| 13437 | 672 | S672T | B | 1.2e-05 | 9.8e-06 | 4.6e-06 | 8.800E-06 | 3.800E-06 | 4.600E-06 | 1.200E-05 |
| 13438 | 672 | S672W | B | 3.2e-06 | 1.3e-05 | 6.6e-06 | 7.600E-06 | 4.976E-06 | 3.200E-06 | 1.300E-05 |
| 13439 | 672 | S672Y | B | 1.1e-05 | 1.1e-05 | 1e-05 | 1.067E-05 | 5.774E-07 | 1.000E-05 | 1.100E-05 |
| 13440 | 672 | S672V | B | 5.4e-06 | 8.4e-06 | 7.8e-06 | 7.200E-06 | 1.587E-06 | 5.400E-06 | 8.400E-06 |
| 13441 | 673 | A673A | B | 1e-05 | 6.2e-06 | 9.9e-06 | 8.700E-06 | 2.166E-06 | 6.200E-06 | 1.000E-05 |
| 13442 | 673 | A673R | B | 5.7e-06 | 1.1e-05 | 1.5e-05 | 1.057E-05 | 4.665E-06 | 5.700E-06 | 1.500E-05 |
| 13443 | 673 | A673N | B | 6.1e-06 | 8.3e-06 | 1e-05 | 8.133E-06 | 1.955E-06 | 6.100E-06 | 1.000E-05 |
| 13444 | 673 | A673D | B | 7.9e-06 | 6.3e-06 | 7.3e-06 | 7.167E-06 | 8.083E-07 | 6.300E-06 | 7.900E-06 |
| 13445 | 673 | A673C | B | 5.9e-06 | 7.6e-06 | 5.7e-06 | 6.400E-06 | 1.044E-06 | 5.700E-06 | 7.600E-06 |
| 13446 | 673 | A673E | B | 9e-06 | 1.3e-05 | 9.6e-06 | 1.053E-05 | 2.157E-06 | 9.000E-06 | 1.300E-05 |
| 13447 | 673 | A673Q | B | 6.7e-06 | 1.1e-05 | 7.8e-06 | 8.500E-06 | 2.234E-06 | 6.700E-06 | 1.100E-05 |
| 13448 | 673 | A673G | B | 7.3e-06 | 1.3e-05 | 1.3e-05 | 1.110E-05 | 3.291E-06 | 7.300E-06 | 1.300E-05 |
| 13449 | 673 | A673H | B | 5.6e-06 | 6.8e-06 | 9.3e-06 | 7.233E-06 | 1.888E-06 | 5.600E-06 | 9.300E-06 |
| 13450 | 673 | A673I | B | 1.3e-05 | 4.7e-06 | 7.9e-06 | 8.533E-06 | 4.186E-06 | 4.700E-06 | 1.300E-05 |
| 13451 | 673 | A673L | B | 1e-05 | 1.4e-05 | 9.4e-06 | 1.113E-05 | 2.501E-06 | 9.400E-06 | 1.400E-05 |
| 13452 | 673 | A673K | B | 1e-05 | 1.1e-05 | 8.6e-06 | 9.867E-06 | 1.206E-06 | 8.600E-06 | 1.100E-05 |
| 13453 | 673 | A673M | B | 9.4e-06 | 5e-06 | 6.5e-06 | 6.967E-06 | 2.237E-06 | 5.000E-06 | 9.400E-06 |
| 13454 | 673 | A673F | B | 1.3e-05 | 1.1e-05 | 6.2e-06 | 1.007E-05 | 3.495E-06 | 6.200E-06 | 1.300E-05 |
| 13455 | 673 | A673P | B | 4.3e-06 | 5.3e-06 | 6.7e-06 | 5.433E-06 | 1.206E-06 | 4.300E-06 | 6.700E-06 |
| 13456 | 673 | A673S | B | 9e-06 | 5.8e-06 | 5.6e-06 | 6.800E-06 | 1.908E-06 | 5.600E-06 | 9.000E-06 |
| 13457 | 673 | A673T | B | 4.7e-06 | 7e-06 | 5.3e-06 | 5.667E-06 | 1.193E-06 | 4.700E-06 | 7.000E-06 |
| 13458 | 673 | A673W | B | 6.7e-06 | 1.1e-05 | 7.6e-06 | 8.433E-06 | 2.268E-06 | 6.700E-06 | 1.100E-05 |
| 13459 | 673 | A673Y | B | 7.7e-06 | 3.8e-06 | 8.9e-06 | 6.800E-06 | 2.666E-06 | 3.800E-06 | 8.900E-06 |
| 13460 | 673 | A673V | B | 7.6e-06 | 1.2e-05 | 9.7e-06 | 9.767E-06 | 2.201E-06 | 7.600E-06 | 1.200E-05 |
| 13461 | 674 | S674A | B | 6.7e-06 | 9.9e-06 | 6.3e-06 | 7.633E-06 | 1.973E-06 | 6.300E-06 | 9.900E-06 |
| 13462 | 674 | S674R | B | 6.1e-06 | 8.7e-06 | 8.9e-06 | 7.900E-06 | 1.562E-06 | 6.100E-06 | 8.900E-06 |
| 13463 | 674 | S674N | B | 8e-06 | 6.5e-06 | 9.3e-06 | 7.933E-06 | 1.401E-06 | 6.500E-06 | 9.300E-06 |
| 13464 | 674 | S674D | B | 1.4e-05 | 6.9e-06 | 4.4e-06 | 8.433E-06 | 4.980E-06 | 4.400E-06 | 1.400E-05 |
| 13465 | 674 | S674C | B | 8.7e-06 | 1e-05 | 9.1e-06 | 9.267E-06 | 6.658E-07 | 8.700E-06 | 1.000E-05 |
| 13466 | 674 | S674E | B | 7.5e-06 | 5.8e-06 | 6.4e-06 | 6.567E-06 | 8.622E-07 | 5.800E-06 | 7.500E-06 |
| 13467 | 674 | S674Q | B | 6.1e-06 | 4.6e-06 | 1e-05 | 6.900E-06 | 2.787E-06 | 4.600E-06 | 1.000E-05 |
| 13468 | 674 | S674G | B | 4.7e-06 | 8.8e-06 | 7e-06 | 6.833E-06 | 2.055E-06 | 4.700E-06 | 8.800E-06 |
| 13469 | 674 | S674H | B | 4.7e-06 | 6.6e-06 | 8.2e-06 | 6.500E-06 | 1.752E-06 | 4.700E-06 | 8.200E-06 |
| 13470 | 674 | S674I | B | 1.2e-05 | 9.5e-06 | 6.5e-06 | 9.333E-06 | 2.754E-06 | 6.500E-06 | 1.200E-05 |
| 13471 | 674 | S674L | B | 7.7e-06 | 6.6e-06 | 1.1e-05 | 8.433E-06 | 2.290E-06 | 6.600E-06 | 1.100E-05 |
| 13472 | 674 | S674K | B | 7.9e-06 | 8.4e-06 | 7e-06 | 7.767E-06 | 7.095E-07 | 7.000E-06 | 8.400E-06 |
| 13473 | 674 | S674M | B | 6.4e-06 | 7.1e-06 | 6.3e-06 | 6.600E-06 | 4.359E-07 | 6.300E-06 | 7.100E-06 |
| 13474 | 674 | S674F | B | 9.9e-06 | 5.7e-06 | 8.1e-06 | 7.900E-06 | 2.107E-06 | 5.700E-06 | 9.900E-06 |
| 13475 | 674 | S674P | B | 7.7e-06 | 1.2e-05 | 1.3e-05 | 1.090E-05 | 2.816E-06 | 7.700E-06 | 1.300E-05 |
| 13476 | 674 | S674S | B | 8e-06 | 6.5e-06 | 7.3e-06 | 7.267E-06 | 7.506E-07 | 6.500E-06 | 8.000E-06 |
| 13477 | 674 | S674T | B | 8.9e-06 | 9.6e-06 | 5e-06 | 7.833E-06 | 2.479E-06 | 5.000E-06 | 9.600E-06 |
| 13478 | 674 | S674W | B | 2.1e-05 | 1.2e-05 | 5.9e-06 | 1.297E-05 | 7.596E-06 | 5.900E-06 | 2.100E-05 |
| 13479 | 674 | S674Y | B | 9.2e-06 | 7.6e-06 | 9e-06 | 8.600E-06 | 8.718E-07 | 7.600E-06 | 9.200E-06 |
| 13480 | 674 | S674V | B | 1e-05 | 9e-06 | 6.9e-06 | 8.633E-06 | 1.582E-06 | 6.900E-06 | 1.000E-05 |
| 13481 | 675 | Y675A | B | 9.3e-06 | 4.4e-06 | 6.7e-06 | 6.800E-06 | 2.452E-06 | 4.400E-06 | 9.300E-06 |
| 13482 | 675 | Y675R | B | 1.2e-05 | 8.7e-06 | 8.9e-06 | 9.867E-06 | 1.850E-06 | 8.700E-06 | 1.200E-05 |
| 13483 | 675 | Y675N | B | 8.5e-06 | 6.6e-06 | 6.7e-06 | 7.267E-06 | 1.069E-06 | 6.600E-06 | 8.500E-06 |
| 13484 | 675 | Y675D | B | 9.7e-06 | 5.1e-06 | 8e-06 | 7.600E-06 | 2.326E-06 | 5.100E-06 | 9.700E-06 |
| 13485 | 675 | Y675C | B | 8.7e-06 | 6.1e-06 | 9.2e-06 | 8.000E-06 | 1.664E-06 | 6.100E-06 | 9.200E-06 |
| 13486 | 675 | Y675E | B | 4.5e-06 | 9.8e-06 | 8.6e-06 | 7.633E-06 | 2.779E-06 | 4.500E-06 | 9.800E-06 |
| 13487 | 675 | Y675Q | B | 1.1e-05 | 8.3e-06 | 6.6e-06 | 8.633E-06 | 2.219E-06 | 6.600E-06 | 1.100E-05 |
| 13488 | 675 | Y675G | B | 7.2e-06 | 6.9e-06 | 6.5e-06 | 6.867E-06 | 3.512E-07 | 6.500E-06 | 7.200E-06 |
| 13489 | 675 | Y675H | B | 1.1e-05 | 1.4e-05 | 9.7e-06 | 1.157E-05 | 2.205E-06 | 9.700E-06 | 1.400E-05 |
| 13490 | 675 | Y675I | B | 8.5e-06 | 9.5e-06 | 1.1e-05 | 9.667E-06 | 1.258E-06 | 8.500E-06 | 1.100E-05 |
| 13491 | 675 | Y675L | B | 1.1e-05 | 9.5e-06 | 8.9e-06 | 9.800E-06 | 1.082E-06 | 8.900E-06 | 1.100E-05 |
| 13492 | 675 | Y675K | B | 6.8e-06 | 6.6e-06 | 9.1e-06 | 7.500E-06 | 1.389E-06 | 6.600E-06 | 9.100E-06 |
| 13493 | 675 | Y675M | B | 9.3e-06 | 1.4e-05 | 7.7e-06 | 1.033E-05 | 3.275E-06 | 7.700E-06 | 1.400E-05 |
| 13494 | 675 | Y675F | B | 7.1e-06 | 7.7e-06 | 1e-05 | 8.267E-06 | 1.531E-06 | 7.100E-06 | 1.000E-05 |
| 13495 | 675 | Y675P | B | 9.3e-06 | 5.1e-06 | 1.1e-05 | 8.467E-06 | 3.037E-06 | 5.100E-06 | 1.100E-05 |
| 13496 | 675 | Y675S | B | 7e-06 | 6.4e-06 | 4.8e-06 | 6.067E-06 | 1.137E-06 | 4.800E-06 | 7.000E-06 |
| 13497 | 675 | Y675T | B | 7.7e-06 | 1.1e-05 | 6.7e-06 | 8.467E-06 | 2.250E-06 | 6.700E-06 | 1.100E-05 |
| 13498 | 675 | Y675W | B | 9.6e-06 | 8.6e-06 | 9e-06 | 9.067E-06 | 5.033E-07 | 8.600E-06 | 9.600E-06 |
| 13499 | 675 | Y675Y | B | 8.8e-06 | 1.1e-05 | 8e-06 | 9.267E-06 | 1.553E-06 | 8.000E-06 | 1.100E-05 |
| 13500 | 675 | Y675V | B | 7.6e-06 | 1.2e-05 | 1.3e-05 | 1.087E-05 | 2.873E-06 | 7.600E-06 | 1.300E-05 |
| 13501 | 676 | R676A | B | 8.3e-06 | 5.5e-06 | 4.5e-06 | 6.100E-06 | 1.970E-06 | 4.500E-06 | 8.300E-06 |
| 13502 | 676 | R676R | B | 7.3e-06 | 8e-06 | 1.1e-05 | 8.767E-06 | 1.966E-06 | 7.300E-06 | 1.100E-05 |
| 13503 | 676 | R676N | B | 6.7e-06 | 5.9e-06 | 8.9e-06 | 7.167E-06 | 1.553E-06 | 5.900E-06 | 8.900E-06 |
| 13504 | 676 | R676D | B | 1e-05 | 6.8e-06 | 9.5e-06 | 8.767E-06 | 1.721E-06 | 6.800E-06 | 1.000E-05 |
| 13505 | 676 | R676C | B | 1.1e-05 | 9.7e-06 | 3.7e-06 | 8.133E-06 | 3.894E-06 | 3.700E-06 | 1.100E-05 |
| 13506 | 676 | R676E | B | 5.7e-06 | 9e-06 | 9.9e-06 | 8.200E-06 | 2.211E-06 | 5.700E-06 | 9.900E-06 |
| 13507 | 676 | R676Q | B | 6.9e-06 | 5.5e-06 | 1e-05 | 7.467E-06 | 2.303E-06 | 5.500E-06 | 1.000E-05 |
| 13508 | 676 | R676G | B | 1.2e-05 | 9.2e-06 | 6.3e-06 | 9.167E-06 | 2.850E-06 | 6.300E-06 | 1.200E-05 |
| 13509 | 676 | R676H | B | 8.4e-06 | 7.2e-06 | 7.9e-06 | 7.833E-06 | 6.028E-07 | 7.200E-06 | 8.400E-06 |
| 13510 | 676 | R676I | B | 1.3e-05 | 7.9e-06 | 5.3e-06 | 8.733E-06 | 3.917E-06 | 5.300E-06 | 1.300E-05 |
| 13511 | 676 | R676L | B | 6.4e-06 | 1.1e-05 | 1.2e-05 | 9.800E-06 | 2.987E-06 | 6.400E-06 | 1.200E-05 |
| 13512 | 676 | R676K | B | 5.3e-06 | 8.8e-06 | 8.5e-06 | 7.533E-06 | 1.940E-06 | 5.300E-06 | 8.800E-06 |
| 13513 | 676 | R676M | B | 1.3e-05 | 7.6e-06 | 7.7e-06 | 9.433E-06 | 3.089E-06 | 7.600E-06 | 1.300E-05 |
| 13514 | 676 | R676F | B | 1e-05 | 1.2e-05 | 5e-06 | 9.000E-06 | 3.606E-06 | 5.000E-06 | 1.200E-05 |

| | | | | | | | | | | |
|---|---|---|---|---|---|---|---|---|---|---|
| 13515 | 676 | R676P | B | 7.8e-06 | 6.2e-06 | 9.2e-06 | 7.733E-06 | 1.501E-06 | 6.200E-06 | 9.200E-06 |
| 13516 | 676 | R676S | B | 9.2e-06 | 7.7e-06 | 8.1e-06 | 8.333E-06 | 7.767E-07 | 7.700E-06 | 9.200E-06 |
| 13517 | 676 | R676T | B | 6.5e-06 | 7.9e-06 | 6.5e-06 | 6.967E-06 | 8.083E-07 | 6.500E-06 | 7.900E-06 |
| 13518 | 676 | R676W | B | 6.1e-06 | 9.3e-06 | 9e-06 | 8.133E-06 | 1.767E-06 | 6.100E-06 | 9.300E-06 |
| 13519 | 676 | R676Y | B | 7.1e-06 | 6.1e-06 | 7.3e-06 | 6.833E-06 | 6.429E-07 | 6.100E-06 | 7.300E-06 |
| 13520 | 676 | R676V | B | 5.4e-06 | 5.9e-06 | 1.2e-05 | 7.767E-06 | 3.675E-06 | 5.400E-06 | 1.200E-05 |
| 13521 | 677 | Y677A | B | 8.5e-06 | 4.1e-06 | 6.6e-06 | 6.400E-06 | 2.207E-06 | 4.100E-06 | 8.500E-06 |
| 13522 | 677 | Y677R | B | 8.5e-06 | 8.3e-06 | 6.6e-06 | 7.800E-06 | 1.044E-06 | 6.600E-06 | 8.500E-06 |
| 13523 | 677 | Y677N | B | 7.4e-06 | 1.4e-05 | 7.7e-06 | 9.700E-06 | 3.727E-06 | 7.400E-06 | 1.400E-05 |
| 13524 | 677 | Y677D | B | 4.4e-06 | 6.4e-06 | 4.6e-06 | 5.133E-06 | 1.102E-06 | 4.400E-06 | 6.400E-06 |
| 13525 | 677 | Y677C | B | 1.1e-05 | 9.8e-06 | 1.9e-05 | 1.327E-05 | 5.001E-06 | 9.800E-06 | 1.900E-05 |
| 13526 | 677 | Y677E | B | 9.7e-06 | 9e-06 | 6.6e-06 | 8.433E-06 | 1.626E-06 | 6.600E-06 | 9.700E-06 |
| 13527 | 677 | Y677Q | B | 6.3e-06 | 6.1e-06 | 8.7e-06 | 7.033E-06 | 1.447E-06 | 6.100E-06 | 8.700E-06 |
| 13528 | 677 | Y677G | B | 7e-06 | 6e-06 | 1e-05 | 7.667E-06 | 2.082E-06 | 6.000E-06 | 1.000E-05 |
| 13529 | 677 | Y677H | B | 6e-06 | 6.8e-06 | 8.4e-06 | 7.067E-06 | 1.222E-06 | 6.000E-06 | 8.400E-06 |
| 13530 | 677 | Y677I | B | 1.4e-05 | 6.7e-06 | 9.9e-06 | 1.020E-05 | 3.659E-06 | 6.700E-06 | 1.400E-05 |
| 13531 | 677 | Y677L | B | 1e-05 | 7.1e-06 | 6.5e-06 | 7.867E-06 | 1.872E-06 | 6.500E-06 | 1.000E-05 |
| 13532 | 677 | Y677K | B | 5e-06 | 8.2e-06 | 7.2e-06 | 6.800E-06 | 1.637E-06 | 5.000E-06 | 8.200E-06 |
| 13533 | 677 | Y677M | B | 5e-06 | 1.9e-05 | 8.9e-06 | 1.097E-05 | 7.225E-06 | 5.000E-06 | 1.900E-05 |
| 13534 | 677 | Y677F | B | 6.9e-06 | 5.9e-06 | 9.2e-06 | 7.333E-06 | 1.692E-06 | 5.900E-06 | 9.200E-06 |
| 13535 | 677 | Y677P | B | 8.3e-06 | 8e-06 | 6.8e-06 | 7.700E-06 | 7.937E-07 | 6.800E-06 | 8.300E-06 |
| 13536 | 677 | Y677S | B | 6.6e-06 | 1.8e-05 | 1.3e-05 | 1.253E-05 | 5.714E-06 | 6.600E-06 | 1.800E-05 |
| 13537 | 677 | Y677T | B | 5.5e-06 | 6.7e-06 | 1.1e-05 | 7.733E-06 | 2.892E-06 | 5.500E-06 | 1.100E-05 |
| 13538 | 677 | Y677W | B | 7.9e-06 | 1.2e-05 | 9e-06 | 9.633E-06 | 2.122E-06 | 7.900E-06 | 1.200E-05 |
| 13539 | 677 | Y677Y | B | 7.6e-06 | 7.8e-06 | 6.3e-06 | 7.233E-06 | 8.145E-07 | 6.300E-06 | 7.800E-06 |
| 13540 | 677 | Y677V | B | 7.7e-06 | 6.5e-06 | 1.2e-05 | 8.733E-06 | 2.892E-06 | 6.500E-06 | 1.200E-05 |
| 13541 | 678 | T678A | B | 8.8e-06 | 5.3e-06 | 1.4e-05 | 9.367E-06 | 4.378E-06 | 5.300E-06 | 1.400E-05 |
| 13542 | 678 | T678R | B | 1e-05 | 6.4e-06 | 6.6e-06 | 7.667E-06 | 2.023E-06 | 6.400E-06 | 1.000E-05 |
| 13543 | 678 | T678N | B | 1.4e-05 | 4.5e-06 | 4.6e-06 | 7.700E-06 | 5.456E-06 | 4.500E-06 | 1.400E-05 |
| 13544 | 678 | T678D | B | 7e-06 | 1.7e-05 | 8.1e-06 | 1.070E-05 | 5.484E-06 | 7.000E-06 | 1.700E-05 |
| 13545 | 678 | T678C | B | 5.6e-06 | 7.7e-06 | 9.7e-06 | 7.667E-06 | 2.050E-06 | 5.600E-06 | 9.700E-06 |
| 13546 | 678 | T678E | B | 9.1e-06 | 9.1e-06 | 1e-05 | 9.400E-06 | 5.196E-07 | 9.100E-06 | 1.000E-05 |
| 13547 | 678 | T678Q | B | 6e-06 | 5.7e-06 | 1e-05 | 7.233E-06 | 2.401E-06 | 5.700E-06 | 1.000E-05 |
| 13548 | 678 | T678G | B | 1e-05 | 1.5e-05 | 1.2e-05 | 1.233E-05 | 2.517E-06 | 1.000E-05 | 1.500E-05 |
| 13549 | 678 | T678H | B | 9.6e-06 | 9.2e-06 | 1e-05 | 9.600E-06 | 4.000E-07 | 9.200E-06 | 1.000E-05 |
| 13550 | 678 | T678I | B | 1e-05 | 6.7e-06 | 6.8e-06 | 7.833E-06 | 1.877E-06 | 6.700E-06 | 1.000E-05 |
| 13551 | 678 | T678L | B | 6e-06 | 8.2e-06 | 7.5e-06 | 7.233E-06 | 1.124E-06 | 6.000E-06 | 8.200E-06 |
| 13552 | 678 | T678K | B | 9.7e-06 | 7.1e-06 | 7.8e-06 | 8.200E-06 | 1.345E-06 | 7.100E-06 | 9.700E-06 |
| 13553 | 678 | T678M | B | 1.2e-05 | 7.4e-06 | 7e-06 | 8.800E-06 | 2.778E-06 | 7.000E-06 | 1.200E-05 |
| 13554 | 678 | T678F | B | 9.5e-06 | 1.4e-05 | 1.5e-05 | 1.283E-05 | 2.930E-06 | 9.500E-06 | 1.500E-05 |
| 13555 | 678 | T678P | B | 5.8e-06 | 4.8e-06 | 9.8e-06 | 6.800E-06 | 2.646E-06 | 4.800E-06 | 9.800E-06 |
| 13556 | 678 | T678S | B | 4.4e-06 | 1e-05 | 9.7e-06 | 8.033E-06 | 3.150E-06 | 4.400E-06 | 1.000E-05 |
| 13557 | 678 | T678T | B | 4.7e-06 | 5.6e-06 | 6.2e-06 | 5.500E-06 | 7.550E-07 | 4.700E-06 | 6.200E-06 |
| 13558 | 678 | T678W | B | 8.7e-06 | 1e-05 | 9e-06 | 9.233E-06 | 6.807E-07 | 8.700E-06 | 1.000E-05 |
| 13559 | 678 | T678Y | B | 1.1e-05 | 5e-06 | 1e-05 | 8.667E-06 | 3.215E-06 | 5.000E-06 | 1.100E-05 |
| 13560 | 678 | T678V | B | 9.8e-06 | 1e-05 | 4.1e-06 | 7.967E-06 | 3.350E-06 | 4.100E-06 | 1.000E-05 |
| 13561 | 679 | G679A | B | 9.8e-06 | 8.8e-06 | 7.5e-06 | 8.700E-06 | 1.153E-06 | 7.500E-06 | 9.800E-06 |
| 13562 | 679 | G679R | B | 9.7e-06 | 5.5e-06 | 9.7e-06 | 8.300E-06 | 2.425E-06 | 5.500E-06 | 9.700E-06 |
| 13563 | 679 | G679N | B | 9.7e-06 | 6.9e-06 | 4.6e-06 | 7.067E-06 | 2.554E-06 | 4.600E-06 | 9.700E-06 |
| 13564 | 679 | G679D | B | 8.9e-06 | 7.4e-06 | 4.4e-06 | 6.900E-06 | 2.291E-06 | 4.400E-06 | 8.900E-06 |
| 13565 | 679 | G679C | B | 1.2e-05 | 9.8e-06 | 7.9e-06 | 9.900E-06 | 2.052E-06 | 7.900E-06 | 1.200E-05 |
| 13566 | 679 | G679E | B | 1.4e-05 | 8.2e-06 | 7.5e-06 | 9.900E-06 | 3.568E-06 | 7.500E-06 | 1.400E-05 |
| 13567 | 679 | G679Q | B | 8.3e-06 | 7e-06 | 6.3e-06 | 7.200E-06 | 1.015E-06 | 6.300E-06 | 8.300E-06 |
| 13568 | 679 | G679G | B | 1.4e-05 | 9.4e-06 | 1e-05 | 1.113E-05 | 2.501E-06 | 9.400E-06 | 1.400E-05 |
| 13569 | 679 | G679H | B | 6.2e-06 | 5.9e-06 | 9.2e-06 | 7.100E-06 | 1.825E-06 | 5.900E-06 | 9.200E-06 |
| 13570 | 679 | G679I | B | 6.7e-06 | 9.3e-06 | 8.1e-06 | 8.033E-06 | 1.301E-06 | 6.700E-06 | 9.300E-06 |
| 13571 | 679 | G679L | B | 1.3e-05 | 4.9e-06 | 7.7e-06 | 8.533E-06 | 4.114E-06 | 4.900E-06 | 1.300E-05 |
| 13572 | 679 | G679K | B | 1.9e-05 | 1e-05 | 6.7e-06 | 1.190E-05 | 6.366E-06 | 6.700E-06 | 1.900E-05 |
| 13573 | 679 | G679M | B | 6.2e-06 | 6.6e-06 | 4.7e-06 | 5.833E-06 | 1.002E-06 | 4.700E-06 | 6.600E-06 |
| 13574 | 679 | G679F | B | 3.3e-06 | 1.2e-05 | 8.2e-06 | 7.833E-06 | 4.362E-06 | 3.300E-06 | 1.200E-05 |
| 13575 | 679 | G679P | B | 4.7e-06 | 8e-06 | 8.5e-06 | 7.067E-06 | 2.065E-06 | 4.700E-06 | 8.500E-06 |
| 13576 | 679 | G679S | B | 8.1e-06 | 7.2e-06 | 9.5e-06 | 8.267E-06 | 1.159E-06 | 7.200E-06 | 9.500E-06 |
| 13577 | 679 | G679T | B | 7.2e-06 | 6.9e-06 | 4.9e-06 | 6.333E-06 | 1.250E-06 | 4.900E-06 | 7.200E-06 |
| 13578 | 679 | G679W | B | 6e-06 | 1.1e-05 | 8.8e-06 | 8.600E-06 | 2.506E-06 | 6.000E-06 | 1.100E-05 |
| 13579 | 679 | G679Y | B | 8.1e-06 | 5.9e-06 | 8e-06 | 7.333E-06 | 1.242E-06 | 5.900E-06 | 8.100E-06 |
| 13580 | 679 | G679V | B | 5.2e-06 | 7.5e-06 | 5.8e-06 | 6.167E-06 | 1.193E-06 | 5.200E-06 | 7.500E-06 |
| 13581 | 680 | V680A | B | 9.2e-06 | 1.2e-05 | 7.4e-06 | 9.533E-06 | 2.318E-06 | 7.400E-06 | 1.200E-05 |
| 13582 | 680 | V680R | B | 4.4e-06 | 1e-05 | 5.7e-06 | 6.700E-06 | 2.931E-06 | 4.400E-06 | 1.000E-05 |
| 13583 | 680 | V680N | B | 6.9e-06 | 6e-06 | 9.2e-06 | 7.367E-06 | 1.650E-06 | 6.000E-06 | 9.200E-06 |
| 13584 | 680 | V680D | B | 4e-06 | 1.3e-05 | 9e-06 | 8.667E-06 | 4.509E-06 | 4.000E-06 | 1.300E-05 |
| 13585 | 680 | V680C | B | 6e-06 | 8.5e-06 | 6.7e-06 | 7.067E-06 | 1.290E-06 | 6.000E-06 | 8.500E-06 |
| 13586 | 680 | V680E | B | 6.6e-06 | 6e-06 | 6e-06 | 6.200E-06 | 3.464E-07 | 6.000E-06 | 6.600E-06 |
| 13587 | 680 | V680Q | B | 5.9e-06 | 1.2e-05 | 6.8e-06 | 8.233E-06 | 3.293E-06 | 5.900E-06 | 1.200E-05 |
| 13588 | 680 | V680G | B | 6.5e-06 | 9.9e-06 | 7e-06 | 7.800E-06 | 1.836E-06 | 6.500E-06 | 9.900E-06 |
| 13589 | 680 | V680H | B | 6.7e-06 | 9.1e-06 | 6.7e-06 | 7.500E-06 | 1.386E-06 | 6.700E-06 | 9.100E-06 |
| 13590 | 680 | V680I | B | 9.6e-06 | 8.1e-06 | 7.7e-06 | 8.467E-06 | 1.002E-06 | 7.700E-06 | 9.600E-06 |
| 13591 | 680 | V680L | B | 1.2e-05 | 9.3e-06 | 8.6e-06 | 9.967E-06 | 1.795E-06 | 8.600E-06 | 1.200E-05 |
| 13592 | 680 | V680K | B | 7.6e-06 | 6e-06 | 7.3e-06 | 6.967E-06 | 8.505E-07 | 6.000E-06 | 7.600E-06 |
| 13593 | 680 | V680M | B | 8.2e-06 | 7.4e-06 | 7.2e-06 | 7.600E-06 | 5.292E-07 | 7.200E-06 | 8.200E-06 |
| 13594 | 680 | V680F | B | 6.1e-06 | 9.7e-06 | 6.4e-06 | 7.400E-06 | 1.997E-06 | 6.100E-06 | 9.700E-06 |
| 13595 | 680 | V680P | B | 7.6e-06 | 6e-06 | 9.3e-06 | 7.633E-06 | 1.650E-06 | 6.000E-06 | 9.300E-06 |
| 13596 | 680 | V680S | B | 8.4e-06 | 4.4e-06 | 8.3e-06 | 7.033E-06 | 2.281E-06 | 4.400E-06 | 8.400E-06 |
| 13597 | 680 | V680T | B | 5.7e-06 | 6.2e-06 | 7.7e-06 | 6.533E-06 | 1.041E-06 | 5.700E-06 | 7.700E-06 |
| 13598 | 680 | V680W | B | 8.4e-06 | 6.2e-06 | 6.5e-06 | 7.033E-06 | 1.193E-06 | 6.200E-06 | 8.400E-06 |
| 13599 | 680 | V680Y | B | 9.3e-06 | 8.4e-06 | 6.2e-06 | 7.967E-06 | 1.595E-06 | 6.200E-06 | 9.300E-06 |

| | | | | | | | | | | |
|---|---|---|---|---|---|---|---|---|---|---|
| 13600 | 680 | V680V | B | 9.9e-06 | 6.5e-06 | 1.1e-05 | 9.133E-06 | 2.346E-06 | 6.500E-06 | 1.100E-05 |
| 13601 | 681 | P681A | B | 5.8e-06 | 1e-05 | 7.2e-06 | 7.667E-06 | 2.139E-06 | 5.800E-06 | 1.000E-05 |
| 13602 | 681 | P681R | B | 1.1e-05 | 7.2e-06 | 9.7e-06 | 9.300E-06 | 1.931E-06 | 7.200E-06 | 1.100E-05 |
| 13603 | 681 | P681N | B | 8.5e-06 | 8.1e-06 | 6.6e-06 | 7.733E-06 | 1.002E-06 | 6.600E-06 | 8.500E-06 |
| 13604 | 681 | P681D | B | 1e-05 | 5.6e-06 | 1.6e-05 | 1.053E-05 | 5.220E-06 | 5.600E-06 | 1.600E-05 |
| 13605 | 681 | P681C | B | 9.5e-06 | 6.6e-06 | 5.4e-06 | 7.167E-06 | 2.108E-06 | 5.400E-06 | 9.500E-06 |
| 13606 | 681 | P681E | B | 8.2e-06 | 6.5e-06 | 6.4e-06 | 7.033E-06 | 1.012E-06 | 6.400E-06 | 8.200E-06 |
| 13607 | 681 | P681Q | B | 1.1e-05 | 8.4e-06 | 7.9e-06 | 9.100E-06 | 1.664E-06 | 7.900E-06 | 1.100E-05 |
| 13608 | 681 | P681G | B | 9e-06 | 1.8e-05 | 9.4e-06 | 1.213E-05 | 5.085E-06 | 9.000E-06 | 1.800E-05 |
| 13609 | 681 | P681H | B | 6.5e-06 | 7e-06 | 6.4e-06 | 6.633E-06 | 3.215E-07 | 6.400E-06 | 7.000E-06 |
| 13610 | 681 | P681I | B | 7.4e-06 | 6.2e-06 | 9.2e-06 | 7.600E-06 | 1.510E-06 | 6.200E-06 | 9.200E-06 |
| 13611 | 681 | P681L | B | 3.9e-06 | 7.6e-06 | 1.1e-05 | 7.500E-06 | 3.551E-06 | 3.900E-06 | 1.100E-05 |
| 13612 | 681 | P681K | B | 1.3e-05 | 7.4e-06 | 6.5e-06 | 8.967E-06 | 3.522E-06 | 6.500E-06 | 1.300E-05 |
| 13613 | 681 | P681M | B | 7.5e-06 | 9.1e-06 | 5.1e-06 | 7.233E-06 | 2.013E-06 | 5.100E-06 | 9.100E-06 |
| 13614 | 681 | P681F | B | 1.3e-05 | 5.8e-06 | 9e-06 | 9.267E-06 | 3.607E-06 | 5.800E-06 | 1.300E-05 |
| 13615 | 681 | P681P | B | 9e-06 | 8.1e-06 | 1e-05 | 9.033E-06 | 9.504E-07 | 8.100E-06 | 1.000E-05 |
| 13616 | 681 | P681S | B | 1e-05 | 9.4e-06 | 6.9e-06 | 8.767E-06 | 1.644E-06 | 6.900E-06 | 1.000E-05 |
| 13617 | 681 | P681T | B | 1.1e-05 | 6.8e-06 | 9.9e-06 | 9.233E-06 | 2.178E-06 | 6.800E-06 | 1.100E-05 |
| 13618 | 681 | P681W | B | 1.3e-05 | 8.6e-06 | 8e-06 | 9.867E-06 | 2.730E-06 | 8.000E-06 | 1.300E-05 |
| 13619 | 681 | P681Y | B | 8e-06 | 6.8e-06 | 9e-06 | 7.933E-06 | 1.102E-06 | 6.800E-06 | 9.000E-06 |
| 13620 | 681 | P681V | B | 8.4e-06 | 8.3e-06 | 1.2e-05 | 9.567E-06 | 2.108E-06 | 8.300E-06 | 1.200E-05 |
| 13621 | 682 | S682A | B | 5.1e-06 | 6.4e-06 | 6.8e-06 | 6.100E-06 | 8.888E-07 | 5.100E-06 | 6.800E-06 |
| 13622 | 682 | S682R | B | 9.2e-06 | 7.2e-06 | 9.2e-06 | 8.533E-06 | 1.155E-06 | 7.200E-06 | 9.200E-06 |
| 13623 | 682 | S682N | B | 8.5e-06 | 8.3e-06 | 8.8e-06 | 8.533E-06 | 2.517E-07 | 8.300E-06 | 8.800E-06 |
| 13624 | 682 | S682D | B | 5.7e-06 | 5e-06 | 6.6e-06 | 5.767E-06 | 8.021E-07 | 5.000E-06 | 6.600E-06 |
| 13625 | 682 | S682C | B | 6.2e-06 | 1.1e-05 | 9.1e-06 | 8.767E-06 | 2.417E-06 | 6.200E-06 | 1.100E-05 |
| 13626 | 682 | S682E | B | 5.1e-06 | 6.7e-06 | 4.8e-06 | 5.533E-06 | 1.021E-06 | 4.800E-06 | 6.700E-06 |
| 13627 | 682 | S682Q | B | 1.1e-05 | 9.1e-06 | 4.5e-06 | 8.200E-06 | 3.342E-06 | 4.500E-06 | 1.100E-05 |
| 13628 | 682 | S682G | B | 5.9e-06 | 1.2e-05 | 8.7e-06 | 8.867E-06 | 3.053E-06 | 5.900E-06 | 1.200E-05 |
| 13629 | 682 | S682H | B | 6.6e-06 | 6.8e-06 | 8.7e-06 | 7.367E-06 | 1.159E-06 | 6.600E-06 | 8.700E-06 |
| 13630 | 682 | S682I | B | 6.3e-06 | 1.2e-05 | 3.4e-06 | 7.233E-06 | 4.375E-06 | 3.400E-06 | 1.200E-05 |
| 13631 | 682 | S682L | B | 7.5e-06 | 7.9e-06 | 6.9e-06 | 7.433E-06 | 5.033E-07 | 6.900E-06 | 7.900E-06 |
| 13632 | 682 | S682K | B | 7e-06 | 1e-05 | 6.1e-06 | 7.700E-06 | 2.042E-06 | 6.100E-06 | 1.000E-05 |
| 13633 | 682 | S682M | B | 6.5e-06 | 1.1e-05 | 1.1e-05 | 9.500E-06 | 2.598E-06 | 6.500E-06 | 1.100E-05 |
| 13634 | 682 | S682F | B | 9.5e-06 | 9.6e-06 | 6.9e-06 | 8.667E-06 | 1.531E-06 | 6.900E-06 | 9.600E-06 |
| 13635 | 682 | S682P | B | 6.9e-06 | 5.1e-06 | 1e-05 | 7.333E-06 | 2.479E-06 | 5.100E-06 | 1.000E-05 |
| 13636 | 682 | S682S | B | 5.4e-06 | 7.8e-06 | 1.3e-05 | 8.733E-06 | 3.885E-06 | 5.400E-06 | 1.300E-05 |
| 13637 | 682 | S682T | B | 8.3e-06 | 7.5e-06 | 6.3e-06 | 7.367E-06 | 1.007E-06 | 6.300E-06 | 8.300E-06 |
| 13638 | 682 | S682W | B | 5.5e-06 | 1.2e-05 | 1e-05 | 9.167E-06 | 3.329E-06 | 5.500E-06 | 1.200E-05 |
| 13639 | 682 | S682Y | B | 8.2e-06 | 7.6e-06 | 6.3e-06 | 7.367E-06 | 9.713E-07 | 6.300E-06 | 8.200E-06 |
| 13640 | 682 | S682V | B | 1.1e-05 | 7.3e-06 | 6.5e-06 | 8.267E-06 | 2.401E-06 | 6.500E-06 | 1.100E-05 |
| 13641 | 683 | R683A | B | 9e-06 | 5.9e-06 | 6.7e-06 | 7.200E-06 | 1.609E-06 | 5.900E-06 | 9.000E-06 |
| 13642 | 683 | R683R | B | 6.6e-06 | 5.9e-06 | 8.1e-06 | 6.867E-06 | 1.124E-06 | 5.900E-06 | 8.100E-06 |
| 13643 | 683 | R683N | B | 9.3e-06 | 8.8e-06 | 2.9e-06 | 7.000E-06 | 3.559E-06 | 2.900E-06 | 9.300E-06 |
| 13644 | 683 | R683D | B | 5.6e-06 | 7.2e-06 | 9.4e-06 | 7.400E-06 | 1.908E-06 | 5.600E-06 | 9.400E-06 |
| 13645 | 683 | R683C | B | 6e-06 | 4.2e-06 | 6.7e-06 | 5.633E-06 | 1.290E-06 | 4.200E-06 | 6.700E-06 |
| 13646 | 683 | R683E | B | 1.2e-05 | 7.2e-06 | 6.9e-06 | 8.700E-06 | 2.862E-06 | 6.900E-06 | 1.200E-05 |
| 13647 | 683 | R683Q | B | 1.2e-05 | 4.5e-06 | 4.6e-06 | 7.033E-06 | 4.302E-06 | 4.500E-06 | 1.200E-05 |
| 13648 | 683 | R683G | B | 8.6e-06 | 9.8e-06 | 1e-05 | 9.467E-06 | 7.572E-07 | 8.600E-06 | 1.000E-05 |
| 13649 | 683 | R683H | B | 9.7e-06 | 1.2e-05 | 5.8e-06 | 9.167E-06 | 3.134E-06 | 5.800E-06 | 1.200E-05 |
| 13650 | 683 | R683I | B | 9.2e-06 | 8.6e-06 | 8.2e-06 | 8.667E-06 | 5.033E-07 | 8.200E-06 | 9.200E-06 |
| 13651 | 683 | R683L | B | 7.6e-06 | 8.4e-06 | 9.6e-06 | 8.533E-06 | 1.007E-06 | 7.600E-06 | 9.600E-06 |
| 13652 | 683 | R683K | B | 5.3e-06 | 7.5e-06 | 8.2e-06 | 7.000E-06 | 1.513E-06 | 5.300E-06 | 8.200E-06 |
| 13653 | 683 | R683M | B | 1.1e-05 | 1.5e-05 | 1.1e-05 | 1.233E-05 | 2.309E-06 | 1.100E-05 | 1.500E-05 |
| 13654 | 683 | R683F | B | 5.8e-06 | 1.3e-05 | 5.2e-06 | 8.000E-06 | 4.341E-06 | 5.200E-06 | 1.300E-05 |
| 13655 | 683 | R683P | B | 6.7e-06 | 6.1e-06 | 7.1e-06 | 6.633E-06 | 5.033E-07 | 6.100E-06 | 7.100E-06 |
| 13656 | 683 | R683S | B | 8e-06 | 5e-06 | 6.5e-06 | 6.500E-06 | 1.500E-06 | 5.000E-06 | 8.000E-06 |
| 13657 | 683 | R683T | B | 8.1e-06 | 8.8e-06 | 1.2e-05 | 9.633E-06 | 2.079E-06 | 8.100E-06 | 1.200E-05 |
| 13658 | 683 | R683W | B | 8.4e-06 | 8.6e-06 | 9.3e-06 | 8.767E-06 | 4.726E-07 | 8.400E-06 | 9.300E-06 |
| 13659 | 683 | R683Y | B | 8.2e-06 | 1e-05 | 9.7e-06 | 9.300E-06 | 9.644E-07 | 8.200E-06 | 1.000E-05 |
| 13660 | 683 | R683V | B | 7.4e-06 | 6.5e-06 | 1e-05 | 7.967E-06 | 1.818E-06 | 6.500E-06 | 1.000E-05 |
| 13661 | 684 | F684A | B | 5.4e-06 | 1e-05 | 5.4e-06 | 6.933E-06 | 2.656E-06 | 5.400E-06 | 1.000E-05 |
| 13662 | 684 | F684R | B | 6.5e-06 | 1.3e-05 | 7.2e-06 | 8.900E-06 | 3.568E-06 | 6.500E-06 | 1.300E-05 |
| 13663 | 684 | F684N | B | 1.1e-05 | 7.7e-06 | 8.8e-06 | 9.167E-06 | 1.680E-06 | 7.700E-06 | 1.100E-05 |
| 13664 | 684 | F684D | B | 7.5e-06 | 1e-05 | 7.3e-06 | 8.267E-06 | 1.504E-06 | 7.300E-06 | 1.000E-05 |
| 13665 | 684 | F684C | B | 8.1e-06 | 5e-06 | 6.7e-06 | 6.600E-06 | 1.552E-06 | 5.000E-06 | 8.100E-06 |
| 13666 | 684 | F684E | B | 8.1e-06 | 5.8e-06 | 9.1e-06 | 7.667E-06 | 1.692E-06 | 5.800E-06 | 9.100E-06 |
| 13667 | 684 | F684Q | B | 8.3e-06 | 4.7e-06 | 5.9e-06 | 6.300E-06 | 1.833E-06 | 4.700E-06 | 8.300E-06 |
| 13668 | 684 | F684G | B | 5.8e-06 | 9.4e-06 | 9.2e-06 | 8.133E-06 | 2.023E-06 | 5.800E-06 | 9.400E-06 |
| 13669 | 684 | F684H | B | 1.1e-05 | 1e-05 | 7.7e-06 | 9.567E-06 | 1.692E-06 | 7.700E-06 | 1.100E-05 |
| 13670 | 684 | F684I | B | 8.2e-06 | 6.1e-06 | 1.2e-05 | 8.767E-06 | 2.991E-06 | 6.100E-06 | 1.200E-05 |
| 13671 | 684 | F684L | B | 4.9e-06 | 7.4e-06 | 6.8e-06 | 6.367E-06 | 1.305E-06 | 4.900E-06 | 7.400E-06 |
| 13672 | 684 | F684K | B | 9.7e-06 | 5.4e-06 | 5.7e-06 | 6.933E-06 | 2.401E-06 | 5.400E-06 | 9.700E-06 |
| 13673 | 684 | F684M | B | 8e-06 | 7.1e-06 | 1.2e-05 | 9.033E-06 | 2.608E-06 | 7.100E-06 | 1.200E-05 |
| 13674 | 684 | F684F | B | 9.5e-06 | 9.2e-06 | 6.5e-06 | 8.400E-06 | 1.652E-06 | 6.500E-06 | 9.500E-06 |
| 13675 | 684 | F684P | B | 5e-06 | 1e-05 | 9.8e-06 | 8.267E-06 | 2.831E-06 | 5.000E-06 | 1.000E-05 |
| 13676 | 684 | F684S | B | 1.2e-05 | 6.6e-06 | 5.2e-06 | 7.933E-06 | 3.591E-06 | 5.200E-06 | 1.200E-05 |
| 13677 | 684 | F684T | B | 6.6e-06 | 7.2e-06 | 6.9e-06 | 6.900E-06 | 3.000E-07 | 6.600E-06 | 7.200E-06 |
| 13678 | 684 | F684W | B | 8.5e-06 | 5.4e-06 | 6e-06 | 6.633E-06 | 1.644E-06 | 5.400E-06 | 8.500E-06 |
| 13679 | 684 | F684Y | B | 4.5e-06 | 1e-05 | 4.8e-06 | 6.433E-06 | 3.092E-06 | 4.500E-06 | 1.000E-05 |
| 13680 | 684 | F684V | B | 4.6e-06 | 9.2e-06 | 5.2e-06 | 6.333E-06 | 2.501E-06 | 4.600E-06 | 9.200E-06 |
| 13681 | 685 | S685A | B | 8.7e-06 | 7e-06 | 6e-06 | 7.233E-06 | 1.365E-06 | 6.000E-06 | 8.700E-06 |
| 13682 | 685 | S685R | B | 5.2e-06 | 7.3e-06 | 7e-06 | 6.500E-06 | 1.136E-06 | 5.200E-06 | 7.300E-06 |
| 13683 | 685 | S685N | B | 6.6e-06 | 1e-05 | 6.5e-06 | 7.700E-06 | 1.992E-06 | 6.500E-06 | 1.000E-05 |
| 13684 | 685 | S685D | B | 6.2e-06 | 9.4e-06 | 9.4e-06 | 8.333E-06 | 1.848E-06 | 6.200E-06 | 9.400E-06 |

| | | | | | | | | | | |
|---|---|---|---|---|---|---|---|---|---|---|
| 13685 | 685 | S685C | B | 6.1e-06 | 5.6e-06 | 6e-06 | 5.900E-06 | 2.646E-07 | 5.600E-06 | 6.100E-06 |
| 13686 | 685 | S685E | B | 7.3e-06 | 9.6e-06 | 7.1e-06 | 8.000E-06 | 1.389E-06 | 7.100E-06 | 9.600E-06 |
| 13687 | 685 | S685Q | B | 7.2e-06 | 1e-05 | 4.5e-06 | 7.233E-06 | 2.750E-06 | 4.500E-06 | 1.000E-05 |
| 13688 | 685 | S685G | B | 7.5e-06 | 9e-06 | 1.2e-05 | 9.500E-06 | 2.291E-06 | 7.500E-06 | 1.200E-05 |
| 13689 | 685 | S685H | B | 1e-05 | 6.7e-06 | 8.4e-06 | 8.367E-06 | 1.650E-06 | 6.700E-06 | 1.000E-05 |
| 13690 | 685 | S685I | B | 6e-06 | 6.6e-06 | 1.2e-05 | 8.200E-06 | 3.305E-06 | 6.000E-06 | 1.200E-05 |
| 13691 | 685 | S685L | B | 6.5e-06 | 6.6e-06 | 9.1e-06 | 7.400E-06 | 1.473E-06 | 6.500E-06 | 9.100E-06 |
| 13692 | 685 | S685K | B | 9.1e-06 | 1.3e-05 | 1.3e-05 | 1.170E-05 | 2.252E-06 | 9.100E-06 | 1.300E-05 |
| 13693 | 685 | S685M | B | 5.6e-06 | 5.3e-06 | 4.7e-06 | 5.200E-06 | 4.583E-07 | 4.700E-06 | 5.600E-06 |
| 13694 | 685 | S685F | B | 9.8e-06 | 6.8e-06 | 9.7e-06 | 8.767E-06 | 1.704E-06 | 6.800E-06 | 9.800E-06 |
| 13695 | 685 | S685P | B | 6e-06 | 7.9e-06 | 6.4e-06 | 6.767E-06 | 1.002E-06 | 6.000E-06 | 7.900E-06 |
| 13696 | 685 | S685S | B | 1.3e-05 | 6.9e-06 | 1e-05 | 9.967E-06 | 3.050E-06 | 6.900E-06 | 1.300E-05 |
| 13697 | 685 | S685T | B | 7.9e-06 | 9.7e-06 | 5.4e-06 | 7.667E-06 | 2.159E-06 | 5.400E-06 | 9.700E-06 |
| 13698 | 685 | S685W | B | 6.2e-06 | 6.6e-06 | 6.2e-06 | 6.333E-06 | 2.309E-07 | 6.200E-06 | 6.600E-06 |
| 13699 | 685 | S685Y | B | 9.1e-06 | 1.4e-05 | 8.4e-06 | 1.050E-05 | 3.051E-06 | 8.400E-06 | 1.400E-05 |
| 13700 | 685 | S685V | B | 9.8e-06 | 1e-05 | 9.3e-06 | 9.700E-06 | 3.606E-07 | 9.300E-06 | 1.000E-05 |
| 13701 | 686 | G686A | B | 1.4e-05 | 6.1e-06 | 7.2e-06 | 9.100E-06 | 4.279E-06 | 6.100E-06 | 1.400E-05 |
| 13702 | 686 | G686R | B | 5.8e-06 | 5.9e-06 | 9.9e-06 | 7.200E-06 | 2.339E-06 | 5.800E-06 | 9.900E-06 |
| 13703 | 686 | G686N | B | 6.9e-06 | 4.8e-06 | 1.3e-05 | 8.233E-06 | 4.259E-06 | 4.800E-06 | 1.300E-05 |
| 13704 | 686 | G686D | B | 8.7e-06 | 5.3e-06 | 8.5e-06 | 7.500E-06 | 1.908E-06 | 5.300E-06 | 8.700E-06 |
| 13705 | 686 | G686C | B | 5.8e-06 | 4.7e-06 | 7e-06 | 5.833E-06 | 1.150E-06 | 4.700E-06 | 7.000E-06 |
| 13706 | 686 | G686E | B | 5.5e-06 | 9.9e-06 | 8.3e-06 | 7.900E-06 | 2.227E-06 | 5.500E-06 | 9.900E-06 |
| 13707 | 686 | G686Q | B | 5.7e-06 | 7e-06 | 5.1e-06 | 5.933E-06 | 9.713E-07 | 5.100E-06 | 7.000E-06 |
| 13708 | 686 | G686G | B | 7e-06 | 5.6e-06 | 6.2e-06 | 6.267E-06 | 7.024E-07 | 5.600E-06 | 7.000E-06 |
| 13709 | 686 | G686H | B | 8.6e-06 | 4.1e-06 | 8e-06 | 6.900E-06 | 2.443E-06 | 4.100E-06 | 8.600E-06 |
| 13710 | 686 | G686I | B | 7e-06 | 8.4e-06 | 6e-06 | 7.133E-06 | 1.206E-06 | 6.000E-06 | 8.400E-06 |
| 13711 | 686 | G686L | B | 6.4e-06 | 8e-06 | 1.2e-05 | 8.800E-06 | 2.884E-06 | 6.400E-06 | 1.200E-05 |
| 13712 | 686 | G686K | B | 6.7e-06 | 8.6e-06 | 1.1e-05 | 8.767E-06 | 2.155E-06 | 6.700E-06 | 1.100E-05 |
| 13713 | 686 | G686M | B | 1.2e-05 | 1.2e-05 | 8.5e-06 | 1.083E-05 | 2.021E-06 | 8.500E-06 | 1.200E-05 |
| 13714 | 686 | G686F | B | 7.7e-06 | 4.7e-06 | 7e-06 | 6.467E-06 | 1.570E-06 | 4.700E-06 | 7.700E-06 |
| 13715 | 686 | G686P | B | 7.4e-06 | 1.2e-05 | 7.1e-06 | 8.833E-06 | 2.747E-06 | 7.100E-06 | 1.200E-05 |
| 13716 | 686 | G686S | B | 6.2e-06 | 5.2e-06 | 1.3e-05 | 8.133E-06 | 4.244E-06 | 5.200E-06 | 1.300E-05 |
| 13717 | 686 | G686T | B | 1.2e-05 | 5e-06 | 7.3e-06 | 8.100E-06 | 3.568E-06 | 5.000E-06 | 1.200E-05 |
| 13718 | 686 | G686W | B | 9.3e-06 | 9.1e-06 | 3.4e-06 | 7.267E-06 | 3.350E-06 | 3.400E-06 | 9.300E-06 |
| 13719 | 686 | G686Y | B | 1.1e-05 | 6.6e-06 | 1e-05 | 9.200E-06 | 2.307E-06 | 6.600E-06 | 1.100E-05 |
| 13720 | 686 | G686V | B | 4.1e-06 | 6.8e-06 | 8.4e-06 | 6.433E-06 | 2.173E-06 | 4.100E-06 | 8.400E-06 |
| 13721 | 687 | S687A | B | 1.3e-05 | 4.8e-06 | 8.9e-06 | 8.900E-06 | 4.100E-06 | 4.800E-06 | 1.300E-05 |
| 13722 | 687 | S687R | B | 6.7e-06 | 9.1e-06 | 4.9e-06 | 6.900E-06 | 2.107E-06 | 4.900E-06 | 9.100E-06 |
| 13723 | 687 | S687N | B | 1e-05 | 8.3e-06 | 6.1e-06 | 8.133E-06 | 1.955E-06 | 6.100E-06 | 1.000E-05 |
| 13724 | 687 | S687D | B | 7.7e-06 | 8.8e-06 | 9.1e-06 | 8.533E-06 | 7.371E-07 | 7.700E-06 | 9.100E-06 |
| 13725 | 687 | S687C | B | 8.8e-06 | 5.6e-06 | 8e-06 | 7.467E-06 | 1.665E-06 | 5.600E-06 | 8.800E-06 |
| 13726 | 687 | S687E | B | 9.9e-06 | 6.4e-06 | 4.8e-06 | 7.033E-06 | 2.608E-06 | 4.800E-06 | 9.900E-06 |
| 13727 | 687 | S687Q | B | 9.1e-06 | 4.8e-06 | 1e-05 | 7.967E-06 | 2.779E-06 | 4.800E-06 | 1.000E-05 |
| 13728 | 687 | S687G | B | 9.5e-06 | 7.2e-06 | 1.4e-05 | 1.023E-05 | 3.459E-06 | 7.200E-06 | 1.400E-05 |
| 13729 | 687 | S687H | B | 8.9e-06 | 7e-06 | 1.4e-05 | 9.967E-06 | 3.620E-06 | 7.000E-06 | 1.400E-05 |
| 13730 | 687 | S687I | B | 7.9e-06 | 5.4e-06 | 1e-05 | 7.767E-06 | 2.303E-06 | 5.400E-06 | 1.000E-05 |
| 13731 | 687 | S687L | B | 8e-06 | 9.1e-06 | 8.4e-06 | 8.500E-06 | 5.568E-07 | 8.000E-06 | 9.100E-06 |
| 13732 | 687 | S687K | B | 8.1e-06 | 3.6e-06 | 7.7e-06 | 6.467E-06 | 2.491E-06 | 3.600E-06 | 8.100E-06 |
| 13733 | 687 | S687M | B | 1.3e-05 | 8.9e-06 | 8.4e-06 | 1.010E-05 | 2.524E-06 | 8.400E-06 | 1.300E-05 |
| 13734 | 687 | S687F | B | 1.1e-05 | 8.1e-06 | 7.2e-06 | 8.767E-06 | 1.986E-06 | 7.200E-06 | 1.100E-05 |
| 13735 | 687 | S687P | B | 1.6e-05 | 7e-06 | 5.9e-06 | 9.633E-06 | 5.541E-06 | 5.900E-06 | 1.600E-05 |
| 13736 | 687 | S687S | B | 1.1e-05 | 8.9e-06 | 1.3e-05 | 1.097E-05 | 2.050E-06 | 8.900E-06 | 1.300E-05 |
| 13737 | 687 | S687T | B | 6.4e-06 | 1.2e-05 | 6.9e-06 | 8.433E-06 | 3.099E-06 | 6.400E-06 | 1.200E-05 |
| 13738 | 687 | S687W | B | 8.6e-06 | 9e-06 | 4.6e-06 | 7.400E-06 | 2.433E-06 | 4.600E-06 | 9.000E-06 |
| 13739 | 687 | S687Y | B | 1e-05 | 4.3e-06 | 7.1e-06 | 7.133E-06 | 2.850E-06 | 4.300E-06 | 1.000E-05 |
| 13740 | 687 | S687V | B | 9.6e-06 | 8.1e-06 | 9.4e-06 | 9.033E-06 | 8.145E-07 | 8.100E-06 | 9.600E-06 |
| 13741 | 688 | G688A | B | 8.6e-06 | 5e-06 | 9.3e-06 | 7.633E-06 | 2.307E-06 | 5.000E-06 | 9.300E-06 |
| 13742 | 688 | G688R | B | 5.9e-06 | 8.2e-06 | 6.3e-06 | 6.800E-06 | 1.229E-06 | 5.900E-06 | 8.200E-06 |
| 13743 | 688 | G688N | B | 5.6e-06 | 1.2e-05 | 7.1e-06 | 8.233E-06 | 3.347E-06 | 5.600E-06 | 1.200E-05 |
| 13744 | 688 | G688D | B | 7.6e-06 | 1.2e-05 | 8.7e-06 | 9.433E-06 | 2.290E-06 | 7.600E-06 | 1.200E-05 |
| 13745 | 688 | G688C | B | 6.3e-06 | 8.6e-06 | 5.5e-06 | 6.800E-06 | 1.609E-06 | 5.500E-06 | 8.600E-06 |
| 13746 | 688 | G688E | B | 1.2e-05 | 8e-06 | 1e-05 | 1.000E-05 | 2.000E-06 | 8.000E-06 | 1.200E-05 |
| 13747 | 688 | G688Q | B | 1.3e-05 | 5.7e-06 | 1.5e-05 | 1.123E-05 | 4.895E-06 | 5.700E-06 | 1.500E-05 |
| 13748 | 688 | G688G | B | 1.1e-05 | 1.3e-05 | 6.7e-06 | 1.023E-05 | 3.219E-06 | 6.700E-06 | 1.300E-05 |
| 13749 | 688 | G688H | B | 1e-05 | 4.8e-06 | 6.6e-06 | 7.133E-06 | 2.641E-06 | 4.800E-06 | 1.000E-05 |
| 13750 | 688 | G688I | B | 6.7e-06 | 6.7e-06 | 1.1e-05 | 8.133E-06 | 2.483E-06 | 6.700E-06 | 1.100E-05 |
| 13751 | 688 | G688L | B | 1.3e-05 | 6.6e-06 | 9.9e-06 | 9.833E-06 | 3.201E-06 | 6.600E-06 | 1.300E-05 |
| 13752 | 688 | G688K | B | 7.2e-06 | 9.8e-06 | 7.1e-06 | 8.033E-06 | 1.531E-06 | 7.100E-06 | 9.800E-06 |
| 13753 | 688 | G688M | B | 8.6e-06 | 6.5e-06 | 6.7e-06 | 7.267E-06 | 1.159E-06 | 6.500E-06 | 8.600E-06 |
| 13754 | 688 | G688F | B | 1.5e-05 | 3.5e-06 | 1.1e-05 | 9.833E-06 | 5.838E-06 | 3.500E-06 | 1.500E-05 |
| 13755 | 688 | G688P | B | 7.3e-06 | 5.1e-06 | 8e-06 | 6.800E-06 | 1.513E-06 | 5.100E-06 | 8.000E-06 |
| 13756 | 688 | G688S | B | 4.5e-06 | 7e-06 | 8.7e-06 | 6.733E-06 | 2.113E-06 | 4.500E-06 | 8.700E-06 |
| 13757 | 688 | G688T | B | 1.2e-05 | 1.2e-05 | 7e-06 | 1.033E-05 | 2.887E-06 | 7.000E-06 | 1.200E-05 |
| 13758 | 688 | G688W | B | 8.5e-06 | 8.8e-06 | 1.6e-05 | 1.110E-05 | 4.246E-06 | 8.500E-06 | 1.600E-05 |
| 13759 | 688 | G688Y | B | 8.6e-06 | 7.5e-06 | 9.1e-06 | 8.400E-06 | 8.185E-07 | 7.500E-06 | 9.100E-06 |
| 13760 | 688 | G688V | B | 1.2e-05 | 9.2e-06 | 7.4e-06 | 9.533E-06 | 2.318E-06 | 7.400E-06 | 1.200E-05 |
| 13761 | 689 | S689A | B | 6.6e-06 | 6.9e-06 | 8.3e-06 | 7.267E-06 | 9.074E-07 | 6.600E-06 | 8.300E-06 |
| 13762 | 689 | S689R | B | 1.1e-05 | 4.5e-06 | 1e-05 | 8.500E-06 | 3.500E-06 | 4.500E-06 | 1.100E-05 |
| 13763 | 689 | S689N | B | 1e-05 | 7.4e-06 | 7.7e-06 | 8.367E-06 | 1.422E-06 | 7.400E-06 | 1.000E-05 |
| 13764 | 689 | S689D | B | 9.3e-06 | 8.4e-06 | 8.2e-06 | 8.633E-06 | 5.859E-07 | 8.200E-06 | 9.300E-06 |
| 13765 | 689 | S689C | B | 1.6e-05 | 6.9e-06 | 6.2e-06 | 9.700E-06 | 5.467E-06 | 6.200E-06 | 1.600E-05 |
| 13766 | 689 | S689E | B | 9.2e-06 | 1.3e-05 | 1.1e-05 | 1.107E-05 | 1.901E-06 | 9.200E-06 | 1.300E-05 |
| 13767 | 689 | S689Q | B | 1.1e-05 | 5e-06 | 9.9e-06 | 8.633E-06 | 3.194E-06 | 5.000E-06 | 1.100E-05 |
| 13768 | 689 | S689G | B | 7.9e-06 | 9.1e-06 | 1.2e-05 | 9.667E-06 | 2.108E-06 | 7.900E-06 | 1.200E-05 |
| 13769 | 689 | S689H | B | 1e-05 | 5.6e-06 | 1.2e-05 | 9.200E-06 | 3.274E-06 | 5.600E-06 | 1.200E-05 |

| | | | | | | | | | | |
|---|---|---|---|---|---|---|---|---|---|---|
| 13770 | 689 | S689I | B | 6.7e-06 | 9e-06 | 6.6e-06 | 7.433E-06 | 1.358E-06 | 6.600E-06 | 9.000E-06 |
| 13771 | 689 | S689L | B | 1e-05 | 5.7e-06 | 9.2e-06 | 8.300E-06 | 2.287E-06 | 5.700E-06 | 1.000E-05 |
| 13772 | 689 | S689K | B | 4.2e-06 | 5.3e-06 | 5.4e-06 | 4.967E-06 | 6.658E-07 | 4.200E-06 | 5.400E-06 |
| 13773 | 689 | S689M | B | 1.5e-05 | 7.8e-06 | 3.7e-06 | 8.833E-06 | 5.720E-06 | 3.700E-06 | 1.500E-05 |
| 13774 | 689 | S689F | B | 4.7e-06 | 1.3e-05 | 7e-06 | 8.233E-06 | 4.285E-06 | 4.700E-06 | 1.300E-05 |
| 13775 | 689 | S689P | B | 5.2e-06 | 6.1e-06 | 7.5e-06 | 6.267E-06 | 1.159E-06 | 5.200E-06 | 7.500E-06 |
| 13776 | 689 | S689S | B | 7.2e-06 | 9.9e-06 | 8.1e-06 | 8.400E-06 | 1.375E-06 | 7.200E-06 | 9.900E-06 |
| 13777 | 689 | S689T | B | 1.9e-05 | 8.3e-06 | 6.5e-06 | 1.127E-05 | 6.757E-06 | 6.500E-06 | 1.900E-05 |
| 13778 | 689 | S689W | B | 1.2e-05 | 5.9e-06 | 7.7e-06 | 8.533E-06 | 3.134E-06 | 5.900E-06 | 1.200E-05 |
| 13779 | 689 | S689Y | B | 1e-05 | 5.8e-06 | 7.4e-06 | 7.733E-06 | 2.120E-06 | 5.800E-06 | 1.000E-05 |
| 13780 | 689 | S689V | B | 8.8e-06 | 1.9e-05 | 3.9e-06 | 1.057E-05 | 7.703E-06 | 3.900E-06 | 1.900E-05 |
| 13781 | 690 | G690A | B | 1.8e-05 | 9.4e-06 | 1.1e-05 | 1.280E-05 | 4.574E-06 | 9.400E-06 | 1.800E-05 |
| 13782 | 690 | G690R | B | 8.8e-06 | 6.4e-06 | 8.1e-06 | 7.767E-06 | 1.234E-06 | 6.400E-06 | 8.800E-06 |
| 13783 | 690 | G690N | B | 1.2e-05 | 7.3e-06 | 8.6e-06 | 9.300E-06 | 2.427E-06 | 7.300E-06 | 1.200E-05 |
| 13784 | 690 | G690D | B | 1.4e-05 | 8.8e-06 | 7.6e-06 | 1.013E-05 | 3.402E-06 | 7.600E-06 | 1.400E-05 |
| 13785 | 690 | G690C | B | 4.1e-06 | 7e-06 | 9e-06 | 6.700E-06 | 2.464E-06 | 4.100E-06 | 9.000E-06 |
| 13786 | 690 | G690E | B | 9.3e-06 | 7.7e-06 | 8.1e-06 | 8.367E-06 | 8.327E-07 | 7.700E-06 | 9.300E-06 |
| 13787 | 690 | G690Q | B | 8.7e-06 | 6.7e-06 | 8.8e-06 | 8.067E-06 | 1.185E-06 | 6.700E-06 | 8.800E-06 |
| 13788 | 690 | G690G | B | 1.4e-05 | 6.3e-06 | 7e-06 | 9.100E-06 | 4.258E-06 | 6.300E-06 | 1.400E-05 |
| 13789 | 690 | G690H | B | 6.9e-06 | 4.5e-06 | 9.4e-06 | 6.933E-06 | 2.450E-06 | 4.500E-06 | 9.400E-06 |
| 13790 | 690 | G690I | B | 1e-05 | 9.9e-06 | 6.7e-06 | 8.867E-06 | 1.877E-06 | 6.700E-06 | 1.000E-05 |
| 13791 | 690 | G690L | B | 6.9e-06 | 9.5e-06 | 1.3e-05 | 9.800E-06 | 3.061E-06 | 6.900E-06 | 1.300E-05 |
| 13792 | 690 | G690K | B | 6.1e-06 | 1.2e-05 | 6.1e-06 | 8.067E-06 | 3.406E-06 | 6.100E-06 | 1.200E-05 |
| 13793 | 690 | G690M | B | 1.4e-05 | 8.7e-06 | 8.8e-06 | 1.050E-05 | 3.032E-06 | 8.700E-06 | 1.400E-05 |
| 13794 | 690 | G690F | B | 7e-06 | 7.6e-06 | 8.7e-06 | 7.767E-06 | 8.622E-07 | 7.000E-06 | 8.700E-06 |
| 13795 | 690 | G690P | B | 5.5e-06 | 7.5e-06 | 5.5e-06 | 6.167E-06 | 1.155E-06 | 5.500E-06 | 7.500E-06 |
| 13796 | 690 | G690S | B | 6.3e-06 | 1e-05 | 4.6e-06 | 6.967E-06 | 2.761E-06 | 4.600E-06 | 1.000E-05 |
| 13797 | 690 | G690T | B | 5.8e-06 | 5.6e-06 | 9.6e-06 | 7.000E-06 | 2.254E-06 | 5.600E-06 | 9.600E-06 |
| 13798 | 690 | G690W | B | 9.8e-06 | 7.7e-06 | 9.1e-06 | 8.867E-06 | 1.069E-06 | 7.700E-06 | 9.800E-06 |
| 13799 | 690 | G690Y | B | 7.4e-06 | 6.9e-06 | 6.7e-06 | 7.000E-06 | 3.606E-07 | 6.700E-06 | 7.400E-06 |
| 13800 | 690 | G690V | B | 1.3e-05 | 7.2e-06 | 9e-06 | 9.733E-06 | 2.969E-06 | 7.200E-06 | 1.300E-05 |
| 13801 | 691 | T691A | B | 1.1e-05 | 7.2e-06 | 1.1e-05 | 9.733E-06 | 2.194E-06 | 7.200E-06 | 1.100E-05 |
| 13802 | 691 | T691R | B | 7.8e-06 | 1e-05 | 1.1e-05 | 9.600E-06 | 1.637E-06 | 7.800E-06 | 1.100E-05 |
| 13803 | 691 | T691N | B | 1.3e-05 | 7.6e-06 | 5e-06 | 8.533E-06 | 4.081E-06 | 5.000E-06 | 1.300E-05 |
| 13804 | 691 | T691D | B | 8e-06 | 1.3e-05 | 8.8e-06 | 9.933E-06 | 2.686E-06 | 8.000E-06 | 1.300E-05 |
| 13805 | 691 | T691C | B | 8.6e-06 | 7.4e-06 | 5.7e-06 | 7.233E-06 | 1.457E-06 | 5.700E-06 | 8.600E-06 |
| 13806 | 691 | T691E | B | 6.7e-06 | 8.3e-06 | 5.2e-06 | 6.733E-06 | 1.550E-06 | 5.200E-06 | 8.300E-06 |
| 13807 | 691 | T691Q | B | 8.8e-06 | 7.2e-06 | 5.9e-06 | 7.300E-06 | 1.453E-06 | 5.900E-06 | 8.800E-06 |
| 13808 | 691 | T691G | B | 1e-05 | 1.2e-05 | 7.1e-06 | 9.700E-06 | 2.464E-06 | 7.100E-06 | 1.200E-05 |
| 13809 | 691 | T691H | B | 1.8e-05 | 6.4e-06 | 1.1e-05 | 1.180E-05 | 5.841E-06 | 6.400E-06 | 1.800E-05 |
| 13810 | 691 | T691I | B | 6.9e-06 | 8.3e-06 | 6.1e-06 | 7.100E-06 | 1.114E-06 | 6.100E-06 | 8.300E-06 |
| 13811 | 691 | T691L | B | 6.3e-06 | 8e-06 | 9.1e-06 | 7.800E-06 | 1.411E-06 | 6.300E-06 | 9.100E-06 |
| 13812 | 691 | T691K | B | 8.7e-06 | 5.7e-06 | 5e-06 | 6.467E-06 | 1.966E-06 | 5.000E-06 | 8.700E-06 |
| 13813 | 691 | T691M | B | 8.9e-06 | 1.3e-05 | 4.9e-06 | 8.933E-06 | 4.050E-06 | 4.900E-06 | 1.300E-05 |
| 13814 | 691 | T691F | B | 1.2e-05 | 8.3e-06 | 8.8e-06 | 9.700E-06 | 2.007E-06 | 8.300E-06 | 1.200E-05 |
| 13815 | 691 | T691P | B | 8.5e-06 | 1e-05 | 6.4e-06 | 8.300E-06 | 1.808E-06 | 6.400E-06 | 1.000E-05 |
| 13816 | 691 | T691S | B | 6.6e-06 | 7.1e-06 | 5.4e-06 | 6.367E-06 | 8.737E-07 | 5.400E-06 | 7.100E-06 |
| 13817 | 691 | T691T | B | 5.1e-06 | 8.5e-06 | 7.3e-06 | 6.967E-06 | 1.724E-06 | 5.100E-06 | 8.500E-06 |
| 13818 | 691 | T691W | B | 7.1e-06 | 9.6e-06 | 1e-05 | 8.900E-06 | 1.572E-06 | 7.100E-06 | 1.000E-05 |
| 13819 | 691 | T691Y | B | 5e-06 | 1e-05 | 6.3e-06 | 7.100E-06 | 2.594E-06 | 5.000E-06 | 1.000E-05 |
| 13820 | 691 | T691V | B | 1e-05 | 9.9e-06 | 9.6e-06 | 9.833E-06 | 2.082E-07 | 9.600E-06 | 1.000E-05 |
| 13821 | 692 | D692A | B | 1.5e-05 | 5.5e-06 | 5.3e-06 | 8.600E-06 | 5.543E-06 | 5.300E-06 | 1.500E-05 |
| 13822 | 692 | D692R | B | 9.8e-06 | 2.1e-05 | 5.5e-06 | 1.210E-05 | 8.002E-06 | 5.500E-06 | 2.100E-05 |
| 13823 | 692 | D692N | B | 9.2e-06 | 7.5e-06 | 4.8e-06 | 7.167E-06 | 2.219E-06 | 4.800E-06 | 9.200E-06 |
| 13824 | 692 | D692D | B | 9.3e-06 | 7.7e-06 | 1e-05 | 9.000E-06 | 1.179E-06 | 7.700E-06 | 1.000E-05 |
| 13825 | 692 | D692C | B | 9.8e-06 | 4.7e-06 | 9.1e-06 | 7.867E-06 | 2.765E-06 | 4.700E-06 | 9.800E-06 |
| 13826 | 692 | D692E | B | 1.3e-05 | 6.1e-06 | 8.7e-06 | 9.267E-06 | 3.485E-06 | 6.100E-06 | 1.300E-05 |
| 13827 | 692 | D692Q | B | 8.6e-06 | 5e-06 | 1.3e-05 | 8.867E-06 | 4.007E-06 | 5.000E-06 | 1.300E-05 |
| 13828 | 692 | D692G | B | 9.3e-06 | 9.2e-06 | 1.1e-05 | 9.833E-06 | 1.012E-06 | 9.200E-06 | 1.100E-05 |
| 13829 | 692 | D692H | B | 9.8e-06 | 7.9e-06 | 8.2e-06 | 8.633E-06 | 1.021E-06 | 7.900E-06 | 9.800E-06 |
| 13830 | 692 | D692I | B | 1.8e-05 | 6.8e-06 | 9.9e-06 | 1.157E-05 | 5.783E-06 | 6.800E-06 | 1.800E-05 |
| 13831 | 692 | D692L | B | 8.9e-06 | 6.1e-06 | 7e-06 | 7.333E-06 | 1.429E-06 | 6.100E-06 | 8.900E-06 |
| 13832 | 692 | D692K | B | 7.4e-06 | 1e-05 | 8.5e-06 | 8.633E-06 | 1.305E-06 | 7.400E-06 | 1.000E-05 |
| 13833 | 692 | D692M | B | 1.3e-05 | 1.2e-05 | 5.9e-06 | 1.030E-05 | 3.843E-06 | 5.900E-06 | 1.300E-05 |
| 13834 | 692 | D692F | B | 5.4e-06 | 6.7e-06 | 7.9e-06 | 6.667E-06 | 1.250E-06 | 5.400E-06 | 7.900E-06 |
| 13835 | 692 | D692P | B | 6.6e-06 | 8.8e-06 | 7e-06 | 7.467E-06 | 1.172E-06 | 6.600E-06 | 8.800E-06 |
| 13836 | 692 | D692S | B | 7e-06 | 9.6e-06 | 8.5e-06 | 8.367E-06 | 1.305E-06 | 7.000E-06 | 9.600E-06 |
| 13837 | 692 | D692T | B | 8.5e-06 | 1.3e-05 | 1.1e-05 | 1.083E-05 | 2.255E-06 | 8.500E-06 | 1.300E-05 |
| 13838 | 692 | D692W | B | 9.6e-06 | 8.1e-06 | 8.2e-06 | 8.633E-06 | 8.386E-07 | 8.100E-06 | 9.600E-06 |
| 13839 | 692 | D692Y | B | 7e-06 | 7.6e-06 | 5.5e-06 | 6.700E-06 | 1.082E-06 | 5.500E-06 | 7.600E-06 |
| 13840 | 692 | D692V | B | 6.8e-06 | 8.6e-06 | 1.2e-05 | 9.133E-06 | 2.641E-06 | 6.800E-06 | 1.200E-05 |
| 13841 | 693 | F693A | B | 9.5e-06 | 8.8e-06 | 7.9e-06 | 8.733E-06 | 8.021E-07 | 7.900E-06 | 9.500E-06 |
| 13842 | 693 | F693R | B | 6e-06 | 4.2e-06 | 8.1e-06 | 6.100E-06 | 1.952E-06 | 4.200E-06 | 8.100E-06 |
| 13843 | 693 | F693N | B | 5.2e-06 | 7.3e-06 | 1.2e-05 | 8.167E-06 | 3.482E-06 | 5.200E-06 | 1.200E-05 |
| 13844 | 693 | F693D | B | 9.8e-06 | 5.6e-06 | 1.1e-05 | 8.800E-06 | 2.835E-06 | 5.600E-06 | 1.100E-05 |
| 13845 | 693 | F693C | B | 6.6e-06 | 4.8e-06 | 9.7e-06 | 7.033E-06 | 2.479E-06 | 4.800E-06 | 9.700E-06 |
| 13846 | 693 | F693E | B | 1.1e-05 | 1.1e-05 | 9.3e-06 | 1.043E-05 | 9.815E-07 | 9.300E-06 | 1.100E-05 |
| 13847 | 693 | F693Q | B | 4.6e-06 | 1.1e-05 | 4.9e-06 | 6.833E-06 | 3.612E-06 | 4.600E-06 | 1.100E-05 |
| 13848 | 693 | F693G | B | 1.1e-05 | 1.1e-05 | 7.5e-06 | 9.833E-06 | 2.021E-06 | 7.500E-06 | 1.100E-05 |
| 13849 | 693 | F693H | B | 6.2e-06 | 3.9e-06 | 7.2e-06 | 5.767E-06 | 1.692E-06 | 3.900E-06 | 7.200E-06 |
| 13850 | 693 | F693I | B | 7.4e-06 | 5e-06 | 8.2e-06 | 6.867E-06 | 1.665E-06 | 5.000E-06 | 8.200E-06 |
| 13851 | 693 | F693L | B | 1.2e-05 | 3.4e-06 | 1.2e-05 | 9.133E-06 | 4.965E-06 | 3.400E-06 | 1.200E-05 |
| 13852 | 693 | F693K | B | 9.3e-06 | 6.8e-06 | 7.2e-06 | 7.767E-06 | 1.343E-06 | 6.800E-06 | 9.300E-06 |
| 13853 | 693 | F693M | B | 4.5e-06 | 8.6e-06 | 9.6e-06 | 7.567E-06 | 2.702E-06 | 4.500E-06 | 9.600E-06 |
| 13854 | 693 | F693F | B | 9.2e-06 | 6.8e-06 | 7.8e-06 | 7.933E-06 | 1.206E-06 | 6.800E-06 | 9.200E-06 |

| | | | | | | | | | | |
|---|---|---|---|---|---|---|---|---|---|---|
| 13855 | 693 | F693P | B | 6.3e-06 | 9.6e-06 | 6.7e-06 | 7.533E-06 | 1.801E-06 | 6.300E-06 | 9.600E-06 |
| 13856 | 693 | F693S | B | 7.1e-06 | 4.8e-06 | 8.6e-06 | 6.833E-06 | 1.914E-06 | 4.800E-06 | 8.600E-06 |
| 13857 | 693 | F693T | B | 8.5e-06 | 9e-06 | 1.3e-05 | 1.017E-05 | 2.466E-06 | 8.500E-06 | 1.300E-05 |
| 13858 | 693 | F693W | B | 7.6e-06 | 6.1e-06 | 5.1e-06 | 6.267E-06 | 1.258E-06 | 5.100E-06 | 7.600E-06 |
| 13859 | 693 | F693Y | B | 4.7e-06 | 5.4e-06 | 6.3e-06 | 5.467E-06 | 8.021E-07 | 4.700E-06 | 6.300E-06 |
| 13860 | 693 | F693V | B | 9e-06 | 8.4e-06 | 7.7e-06 | 8.367E-06 | 6.506E-07 | 7.700E-06 | 9.000E-06 |
| 13861 | 694 | T694A | B | 8.7e-06 | 1.4e-05 | 2e-05 | 1.423E-05 | 5.654E-06 | 8.700E-06 | 2.000E-05 |
| 13862 | 694 | T694R | B | 5.2e-06 | 3.2e-06 | 1.2e-05 | 6.800E-06 | 4.613E-06 | 3.200E-06 | 1.200E-05 |
| 13863 | 694 | T694N | B | 1.1e-05 | 3.5e-06 | 5.9e-06 | 6.800E-06 | 3.830E-06 | 3.500E-06 | 1.100E-05 |
| 13864 | 694 | T694D | B | 6.1e-06 | 6.9e-06 | 1.2e-05 | 8.333E-06 | 3.201E-06 | 6.100E-06 | 1.200E-05 |
| 13865 | 694 | T694C | B | 7.5e-06 | 9e-06 | 3.2e-06 | 6.567E-06 | 3.011E-06 | 3.200E-06 | 9.000E-06 |
| 13866 | 694 | T694E | B | 1e-05 | 9.1e-06 | 6e-06 | 8.367E-06 | 2.098E-06 | 6.000E-06 | 1.000E-05 |
| 13867 | 694 | T694Q | B | 4.9e-06 | 9.6e-06 | 6.6e-06 | 7.033E-06 | 2.380E-06 | 4.900E-06 | 9.600E-06 |
| 13868 | 694 | T694G | B | 6.8e-06 | 6.5e-06 | 1.3e-05 | 8.767E-06 | 3.669E-06 | 6.500E-06 | 1.300E-05 |
| 13869 | 694 | T694H | B | 4.7e-06 | 8.7e-06 | 4.2e-06 | 5.867E-06 | 2.466E-06 | 4.200E-06 | 8.700E-06 |
| 13870 | 694 | T694I | B | 9.1e-06 | 1.1e-05 | 5.1e-06 | 8.400E-06 | 3.012E-06 | 5.100E-06 | 1.100E-05 |
| 13871 | 694 | T694L | B | 2.5e-06 | 8.3e-06 | 1.1e-05 | 7.267E-06 | 4.343E-06 | 2.500E-06 | 1.100E-05 |
| 13872 | 694 | T694K | B | 1.1e-05 | 5.2e-06 | 7.8e-06 | 8.000E-06 | 2.905E-06 | 5.200E-06 | 1.100E-05 |
| 13873 | 694 | T694M | B | 1.3e-05 | 6.9e-06 | 6.2e-06 | 8.700E-06 | 3.740E-06 | 6.200E-06 | 1.300E-05 |
| 13874 | 694 | T694F | B | 5.7e-06 | 6.8e-06 | 8.4e-06 | 6.967E-06 | 1.358E-06 | 5.700E-06 | 8.400E-06 |
| 13875 | 694 | T694P | B | 5.9e-06 | 7.1e-06 | 1.5e-05 | 9.333E-06 | 4.944E-06 | 5.900E-06 | 1.500E-05 |
| 13876 | 694 | T694S | B | 6.5e-06 | 1.2e-05 | 1.3e-05 | 1.050E-05 | 3.500E-06 | 6.500E-06 | 1.300E-05 |
| 13877 | 694 | T694T | B | 6.2e-06 | 7.6e-06 | 2.2e-05 | 1.193E-05 | 8.746E-06 | 6.200E-06 | 2.200E-05 |
| 13878 | 694 | T694W | B | 4.7e-06 | 4.4e-06 | 9.6e-06 | 6.233E-06 | 2.919E-06 | 4.400E-06 | 9.600E-06 |
| 13879 | 694 | T694Y | B | 8.7e-06 | 9.8e-06 | 7.4e-06 | 8.633E-06 | 1.201E-06 | 7.400E-06 | 9.800E-06 |
| 13880 | 694 | T694V | B | 5.5e-06 | 1.4e-05 | 4.6e-06 | 8.033E-06 | 5.187E-06 | 4.600E-06 | 1.400E-05 |
| 13881 | 695 | L695A | B | 9.1e-06 | 1.1e-05 | 6.6e-06 | 8.900E-06 | 2.207E-06 | 6.600E-06 | 1.100E-05 |
| 13882 | 695 | L695R | B | 5e-06 | 5.4e-06 | 6.4e-06 | 5.600E-06 | 7.211E-07 | 5.000E-06 | 6.400E-06 |
| 13883 | 695 | L695N | B | 6.8e-06 | 9.5e-06 | 5.6e-06 | 7.300E-06 | 1.997E-06 | 5.600E-06 | 9.500E-06 |
| 13884 | 695 | L695D | B | 6.8e-06 | 1.2e-05 | 1.1e-05 | 9.933E-06 | 2.759E-06 | 6.800E-06 | 1.200E-05 |
| 13885 | 695 | L695C | B | 6.8e-06 | 6.6e-06 | 9.3e-06 | 7.567E-06 | 1.504E-06 | 6.600E-06 | 9.300E-06 |
| 13886 | 695 | L695E | B | 7.3e-06 | 5.6e-06 | 5.7e-06 | 6.200E-06 | 9.539E-07 | 5.600E-06 | 7.300E-06 |
| 13887 | 695 | L695Q | B | 6.3e-06 | 4e-06 | 8.9e-06 | 6.400E-06 | 2.452E-06 | 4.000E-06 | 8.900E-06 |
| 13888 | 695 | L695G | B | 6.9e-06 | 4.6e-06 | 6e-06 | 5.833E-06 | 1.159E-06 | 4.600E-06 | 6.900E-06 |
| 13889 | 695 | L695H | B | 8.8e-06 | 8.5e-06 | 4.9e-06 | 7.400E-06 | 2.170E-06 | 4.900E-06 | 8.800E-06 |
| 13890 | 695 | L695I | B | 9.5e-06 | 1.3e-05 | 8.5e-06 | 1.033E-05 | 2.363E-06 | 8.500E-06 | 1.300E-05 |
| 13891 | 695 | L695L | B | 6.4e-06 | 5.9e-06 | 7.4e-06 | 6.567E-06 | 7.638E-07 | 5.900E-06 | 7.400E-06 |
| 13892 | 695 | L695K | B | 9.8e-06 | 3.7e-06 | 4.3e-06 | 5.933E-06 | 3.362E-06 | 3.700E-06 | 9.800E-06 |
| 13893 | 695 | L695M | B | 8.9e-06 | 6.1e-06 | 5.4e-06 | 6.800E-06 | 1.852E-06 | 5.400E-06 | 8.900E-06 |
| 13894 | 695 | L695F | B | 1.3e-05 | 5.1e-06 | 9.2e-06 | 9.100E-06 | 3.951E-06 | 5.100E-06 | 1.300E-05 |
| 13895 | 695 | L695P | B | 1.1e-05 | 1.3e-05 | 1.4e-05 | 1.267E-05 | 1.528E-06 | 1.100E-05 | 1.400E-05 |
| 13896 | 695 | L695S | B | 7.3e-06 | 9.6e-06 | 7.4e-06 | 8.100E-06 | 1.300E-06 | 7.300E-06 | 9.600E-06 |
| 13897 | 695 | L695T | B | 7.5e-06 | 9.4e-06 | 7.1e-06 | 8.000E-06 | 1.229E-06 | 7.100E-06 | 9.400E-06 |
| 13898 | 695 | L695W | B | 8.9e-06 | 3.8e-06 | 5.5e-06 | 6.067E-06 | 2.597E-06 | 3.800E-06 | 8.900E-06 |
| 13899 | 695 | L695Y | B | 1.3e-05 | 1.3e-05 | 8.9e-06 | 1.163E-05 | 2.367E-06 | 8.900E-06 | 1.300E-05 |
| 13900 | 695 | L695V | B | 9.6e-06 | 6.3e-06 | 1.3e-05 | 9.633E-06 | 3.350E-06 | 6.300E-06 | 1.300E-05 |
| 13901 | 696 | T696A | B | 1.3e-05 | 4.9e-06 | 1.4e-05 | 1.063E-05 | 4.990E-06 | 4.900E-06 | 1.400E-05 |
| 13902 | 696 | T696R | B | 5.5e-06 | 7.7e-06 | 4.6e-06 | 5.933E-06 | 1.595E-06 | 4.600E-06 | 7.700E-06 |
| 13903 | 696 | T696N | B | 8.6e-06 | 8.2e-06 | 5.9e-06 | 7.567E-06 | 1.457E-06 | 5.900E-06 | 8.600E-06 |
| 13904 | 696 | T696D | B | 7e-06 | 1.4e-05 | 9.9e-06 | 1.030E-05 | 3.517E-06 | 7.000E-06 | 1.400E-05 |
| 13905 | 696 | T696C | B | 1e-05 | 6.6e-06 | 6.5e-06 | 7.700E-06 | 1.992E-06 | 6.500E-06 | 1.000E-05 |
| 13906 | 696 | T696E | B | 1e-05 | 5.3e-06 | 1.3e-05 | 9.433E-06 | 3.881E-06 | 5.300E-06 | 1.300E-05 |
| 13907 | 696 | T696Q | B | 6.9e-06 | 5.6e-06 | 6.1e-06 | 6.200E-06 | 6.557E-07 | 5.600E-06 | 6.900E-06 |
| 13908 | 696 | T696G | B | 5.7e-06 | 7.2e-06 | 9.5e-06 | 7.467E-06 | 1.914E-06 | 5.700E-06 | 9.500E-06 |
| 13909 | 696 | T696H | B | 7.3e-06 | 5e-06 | 1.2e-05 | 8.100E-06 | 3.568E-06 | 5.000E-06 | 1.200E-05 |
| 13910 | 696 | T696I | B | 1.1e-05 | 9.5e-06 | 1.3e-05 | 1.117E-05 | 1.756E-06 | 9.500E-06 | 1.300E-05 |
| 13911 | 696 | T696L | B | 1.2e-05 | 7.5e-06 | 6.6e-06 | 8.700E-06 | 2.893E-06 | 6.600E-06 | 1.200E-05 |
| 13912 | 696 | T696K | B | 9.8e-06 | 1.2e-05 | 5.5e-06 | 9.100E-06 | 3.306E-06 | 5.500E-06 | 1.200E-05 |
| 13913 | 696 | T696M | B | 1.3e-05 | 1.3e-05 | 6.7e-06 | 1.090E-05 | 3.637E-06 | 6.700E-06 | 1.300E-05 |
| 13914 | 696 | T696F | B | 9.4e-06 | 7.8e-06 | 7.5e-06 | 8.233E-06 | 1.021E-06 | 7.500E-06 | 9.400E-06 |
| 13915 | 696 | T696P | B | 9.8e-06 | 7.3e-06 | 6.2e-06 | 7.767E-06 | 1.845E-06 | 6.200E-06 | 9.800E-06 |
| 13916 | 696 | T696S | B | 7.4e-06 | 8e-06 | 5.5e-06 | 6.967E-06 | 1.305E-06 | 5.500E-06 | 8.000E-06 |
| 13917 | 696 | T696T | B | 9.3e-06 | 8.9e-06 | 7.2e-06 | 8.467E-06 | 1.115E-06 | 7.200E-06 | 9.300E-06 |
| 13918 | 696 | T696W | B | 4.7e-06 | 1.5e-05 | 6.6e-06 | 8.767E-06 | 5.481E-06 | 4.700E-06 | 1.500E-05 |
| 13919 | 696 | T696Y | B | 1.3e-05 | 4e-06 | 9e-06 | 8.667E-06 | 4.509E-06 | 4.000E-06 | 1.300E-05 |
| 13920 | 696 | T696V | B | 6.2e-06 | 6.1e-06 | 7.1e-06 | 6.467E-06 | 5.508E-07 | 6.100E-06 | 7.100E-06 |
| 13921 | 697 | I697A | B | 8.5e-06 | 6.2e-06 | 1.2e-05 | 8.900E-06 | 2.921E-06 | 6.200E-06 | 1.200E-05 |
| 13922 | 697 | I697R | B | 9.7e-06 | 9.6e-06 | 7.9e-06 | 9.067E-06 | 1.012E-06 | 7.900E-06 | 9.700E-06 |
| 13923 | 697 | I697N | B | 8.3e-06 | 9.7e-06 | 9.6e-06 | 9.200E-06 | 7.810E-07 | 8.300E-06 | 9.700E-06 |
| 13924 | 697 | I697D | B | 9.5e-06 | 7.3e-06 | 6.8e-06 | 7.867E-06 | 1.436E-06 | 6.800E-06 | 9.500E-06 |
| 13925 | 697 | I697C | B | 5.9e-06 | 1e-05 | 7.8e-06 | 7.900E-06 | 2.052E-06 | 5.900E-06 | 1.000E-05 |
| 13926 | 697 | I697E | B | 8.9e-06 | 8.2e-06 | 9.5e-06 | 8.867E-06 | 6.506E-07 | 8.200E-06 | 9.500E-06 |
| 13927 | 697 | I697Q | B | 6.3e-06 | 7.7e-06 | 6.3e-06 | 6.767E-06 | 8.083E-07 | 6.300E-06 | 7.700E-06 |
| 13928 | 697 | I697G | B | 5.3e-06 | 6.7e-06 | 1e-05 | 7.333E-06 | 2.413E-06 | 5.300E-06 | 1.000E-05 |
| 13929 | 697 | I697H | B | 6.8e-06 | 6.7e-06 | 7.6e-06 | 7.033E-06 | 4.933E-07 | 6.700E-06 | 7.600E-06 |
| 13930 | 697 | I697I | B | 9.2e-06 | 9e-06 | 7e-06 | 8.400E-06 | 1.217E-06 | 7.000E-06 | 9.200E-06 |
| 13931 | 697 | I697L | B | 5.3e-06 | 7.2e-06 | 1.4e-05 | 8.833E-06 | 4.574E-06 | 5.300E-06 | 1.400E-05 |
| 13932 | 697 | I697K | B | 1.1e-05 | 8.7e-06 | 8.6e-06 | 9.433E-06 | 1.358E-06 | 8.600E-06 | 1.100E-05 |
| 13933 | 697 | I697M | B | 8.1e-06 | 8.1e-06 | 5.5e-06 | 7.233E-06 | 1.501E-06 | 5.500E-06 | 8.100E-06 |
| 13934 | 697 | I697F | B | 1.3e-05 | 5.1e-06 | 9.6e-06 | 9.233E-06 | 3.963E-06 | 5.100E-06 | 1.300E-05 |
| 13935 | 697 | I697P | B | 1e-05 | 9.9e-06 | 6.3e-06 | 8.733E-06 | 2.108E-06 | 6.300E-06 | 1.000E-05 |
| 13936 | 697 | I697S | B | 5.4e-06 | 9e-06 | 4.4e-06 | 6.267E-06 | 2.419E-06 | 4.400E-06 | 9.000E-06 |
| 13937 | 697 | I697T | B | 9.6e-06 | 8e-06 | 5.9e-06 | 7.833E-06 | 1.856E-06 | 5.900E-06 | 9.600E-06 |
| 13938 | 697 | I697W | B | 9.7e-06 | 1.3e-05 | 6.6e-06 | 9.767E-06 | 3.201E-06 | 6.600E-06 | 1.300E-05 |
| 13939 | 697 | I697Y | B | 1.1e-05 | 8.9e-06 | 4.5e-06 | 8.133E-06 | 3.317E-06 | 4.500E-06 | 1.100E-05 |

| | | | | | | | | | | |
|---|---|---|---|---|---|---|---|---|---|---|
| 13940 | 697 | I697V | B | 7.7e-06 | 5.9e-06 | 4.6e-06 | 6.067E-06 | 1.557E-06 | 4.600E-06 | 7.700E-06 |
| 13941 | 698 | S698A | B | 4.2e-06 | 8.7e-06 | 4.3e-06 | 5.733E-06 | 2.570E-06 | 4.200E-06 | 8.700E-06 |
| 13942 | 698 | S698R | B | 7.5e-06 | 1.1e-05 | 7.7e-06 | 8.733E-06 | 1.966E-06 | 7.500E-06 | 1.100E-05 |
| 13943 | 698 | S698N | B | 8.6e-06 | 9.3e-06 | 9.8e-06 | 9.233E-06 | 6.028E-07 | 8.600E-06 | 9.800E-06 |
| 13944 | 698 | S698D | B | 8e-06 | 9.8e-06 | 9e-06 | 8.933E-06 | 9.018E-07 | 8.000E-06 | 9.800E-06 |
| 13945 | 698 | S698C | B | 1.1e-05 | 8.1e-06 | 5.3e-06 | 8.133E-06 | 2.850E-06 | 5.300E-06 | 1.100E-05 |
| 13946 | 698 | S698E | B | 6.5e-06 | 5.5e-06 | 5.5e-06 | 5.833E-06 | 5.774E-07 | 5.500E-06 | 6.500E-06 |
| 13947 | 698 | S698Q | B | 6.5e-06 | 7.6e-06 | 1.2e-05 | 8.700E-06 | 2.910E-06 | 6.500E-06 | 1.200E-05 |
| 13948 | 698 | S698G | B | 8.7e-06 | 1.7e-05 | 9.1e-06 | 1.160E-05 | 4.681E-06 | 8.700E-06 | 1.700E-05 |
| 13949 | 698 | S698H | B | 4.3e-06 | 6.4e-06 | 9.8e-06 | 6.833E-06 | 2.775E-06 | 4.300E-06 | 9.800E-06 |
| 13950 | 698 | S698I | B | 1.3e-05 | 1.3e-05 | 1.3e-05 | 1.300E-05 | 0.000E+00 | 1.300E-05 | 1.300E-05 |
| 13951 | 698 | S698L | B | 7.1e-06 | 1.1e-05 | 9e-06 | 9.033E-06 | 1.950E-06 | 7.100E-06 | 1.100E-05 |
| 13952 | 698 | S698K | B | 1e-05 | 1.4e-05 | 6.9e-06 | 1.030E-05 | 3.559E-06 | 6.900E-06 | 1.400E-05 |
| 13953 | 698 | S698M | B | 8.1e-06 | 1.1e-05 | 8.2e-06 | 9.100E-06 | 1.646E-06 | 8.100E-06 | 1.100E-05 |
| 13954 | 698 | S698F | B | 4.1e-06 | 6.1e-06 | 5.9e-06 | 5.367E-06 | 1.102E-06 | 4.100E-06 | 6.100E-06 |
| 13955 | 698 | S698P | B | 5.4e-06 | 7.1e-06 | 1.1e-05 | 7.833E-06 | 2.871E-06 | 5.400E-06 | 1.100E-05 |
| 13956 | 698 | S698S | B | 7.8e-06 | 5.4e-06 | 1e-05 | 7.733E-06 | 2.301E-06 | 5.400E-06 | 1.000E-05 |
| 13957 | 698 | S698T | B | 1.2e-05 | 1.3e-05 | 9.4e-06 | 1.147E-05 | 1.858E-06 | 9.400E-06 | 1.300E-05 |
| 13958 | 698 | S698W | B | 4.9e-06 | 6.9e-06 | 9.7e-06 | 7.167E-06 | 2.411E-06 | 4.900E-06 | 9.700E-06 |
| 13959 | 698 | S698Y | B | 5.6e-06 | 1.1e-05 | 1.3e-05 | 9.867E-06 | 3.828E-06 | 5.600E-06 | 1.300E-05 |
| 13960 | 698 | S698V | B | 6.4e-06 | 6.4e-06 | 1.9e-05 | 1.060E-05 | 7.275E-06 | 6.400E-06 | 1.900E-05 |
| 13961 | 699 | S699A | B | 3.8e-06 | 6.5e-06 | 4.3e-06 | 4.867E-06 | 1.436E-06 | 3.800E-06 | 6.500E-06 |
| 13962 | 699 | S699R | B | 5e-06 | 1.3e-05 | 9.5e-06 | 9.167E-06 | 4.010E-06 | 5.000E-06 | 1.300E-05 |
| 13963 | 699 | S699N | B | 3e-06 | 1.3e-05 | 9.5e-06 | 8.500E-06 | 5.074E-06 | 3.000E-06 | 1.300E-05 |
| 13964 | 699 | S699D | B | 1.5e-05 | 6.2e-06 | 1.3e-05 | 1.140E-05 | 4.613E-06 | 6.200E-06 | 1.500E-05 |
| 13965 | 699 | S699C | B | 4.6e-06 | 9.8e-06 | 6.9e-06 | 7.100E-06 | 2.606E-06 | 4.600E-06 | 9.800E-06 |
| 13966 | 699 | S699E | B | 6.4e-06 | 8.8e-06 | 1.2e-05 | 9.067E-06 | 2.810E-06 | 6.400E-06 | 1.200E-05 |
| 13967 | 699 | S699Q | B | 6.5e-06 | 6.3e-06 | 6.5e-06 | 6.433E-06 | 1.155E-07 | 6.300E-06 | 6.500E-06 |
| 13968 | 699 | S699G | B | 1e-05 | 7.6e-06 | 8.1e-06 | 8.567E-06 | 1.266E-06 | 7.600E-06 | 1.000E-05 |
| 13969 | 699 | S699H | B | 4.9e-06 | 6.1e-06 | 6.1e-06 | 5.700E-06 | 6.928E-07 | 4.900E-06 | 6.100E-06 |
| 13970 | 699 | S699I | B | 7.3e-06 | 1e-05 | 6.5e-06 | 7.933E-06 | 1.834E-06 | 6.500E-06 | 1.000E-05 |
| 13971 | 699 | S699L | B | 1e-05 | 7e-06 | 6.9e-06 | 7.967E-06 | 1.762E-06 | 6.900E-06 | 1.000E-05 |
| 13972 | 699 | S699K | B | 5.5e-06 | 7.1e-06 | 6.7e-06 | 6.433E-06 | 8.327E-07 | 5.500E-06 | 7.100E-06 |
| 13973 | 699 | S699M | B | 9.5e-06 | 7.5e-06 | 9e-06 | 8.667E-06 | 1.041E-06 | 7.500E-06 | 9.500E-06 |
| 13974 | 699 | S699F | B | 1e-05 | 9.7e-06 | 1.4e-05 | 1.123E-05 | 2.401E-06 | 9.700E-06 | 1.400E-05 |
| 13975 | 699 | S699P | B | 5.2e-06 | 1.1e-05 | 4.4e-06 | 6.867E-06 | 3.602E-06 | 4.400E-06 | 1.100E-05 |
| 13976 | 699 | S699S | B | 5.7e-06 | 8.8e-06 | 8.4e-06 | 7.633E-06 | 1.686E-06 | 5.700E-06 | 8.800E-06 |
| 13977 | 699 | S699T | B | 1.6e-05 | 4.5e-06 | 9.9e-06 | 1.013E-05 | 5.754E-06 | 4.500E-06 | 1.600E-05 |
| 13978 | 699 | S699W | B | 8.3e-06 | 5.1e-06 | 9.4e-06 | 7.600E-06 | 2.234E-06 | 5.100E-06 | 9.400E-06 |
| 13979 | 699 | S699Y | B | 9.9e-06 | 8.8e-06 | 9.1e-06 | 9.267E-06 | 5.686E-07 | 8.800E-06 | 9.900E-06 |
| 13980 | 699 | S699V | B | 4.8e-06 | 7.6e-06 | 9.8e-06 | 7.400E-06 | 2.506E-06 | 4.800E-06 | 9.800E-06 |
| 13981 | 700 | L700A | B | 7.4e-06 | 7.7e-06 | 4.5e-06 | 6.533E-06 | 1.767E-06 | 4.500E-06 | 7.700E-06 |
| 13982 | 700 | L700R | B | 1.4e-05 | 6.7e-06 | 8.1e-06 | 9.600E-06 | 3.874E-06 | 6.700E-06 | 1.400E-05 |
| 13983 | 700 | L700N | B | 3.9e-06 | 8.6e-06 | 5.5e-06 | 6.000E-06 | 2.390E-06 | 3.900E-06 | 8.600E-06 |
| 13984 | 700 | L700D | B | 3.8e-06 | 1e-05 | 6.7e-06 | 6.833E-06 | 3.102E-06 | 3.800E-06 | 1.000E-05 |
| 13985 | 700 | L700C | B | 9.8e-06 | 9.4e-06 | 4e-06 | 7.733E-06 | 3.239E-06 | 4.000E-06 | 9.800E-06 |
| 13986 | 700 | L700E | B | 9e-06 | 1.1e-05 | 8.2e-06 | 9.400E-06 | 1.442E-06 | 8.200E-06 | 1.100E-05 |
| 13987 | 700 | L700Q | B | 1.1e-05 | 6.9e-06 | 9.3e-06 | 9.067E-06 | 2.060E-06 | 6.900E-06 | 1.100E-05 |
| 13988 | 700 | L700G | B | 1.2e-05 | 7e-06 | 6e-06 | 8.333E-06 | 3.215E-06 | 6.000E-06 | 1.200E-05 |
| 13989 | 700 | L700H | B | 6.4e-06 | 9.7e-06 | 9.3e-06 | 8.467E-06 | 1.801E-06 | 6.400E-06 | 9.700E-06 |
| 13990 | 700 | L700I | B | 1.1e-05 | 6.6e-06 | 4.9e-06 | 7.500E-06 | 3.148E-06 | 4.900E-06 | 1.100E-05 |
| 13991 | 700 | L700L | B | 9e-06 | 5.2e-06 | 6.7e-06 | 6.967E-06 | 1.914E-06 | 5.200E-06 | 9.000E-06 |
| 13992 | 700 | L700K | B | 9.6e-06 | 8.9e-06 | 1.1e-05 | 9.833E-06 | 1.069E-06 | 8.900E-06 | 1.100E-05 |
| 13993 | 700 | L700M | B | 7.2e-06 | 9.6e-06 | 8.5e-06 | 8.433E-06 | 1.201E-06 | 7.200E-06 | 9.600E-06 |
| 13994 | 700 | L700F | B | 1e-05 | 6.8e-06 | 9.2e-06 | 8.667E-06 | 1.665E-06 | 6.800E-06 | 1.000E-05 |
| 13995 | 700 | L700P | B | 4.4e-06 | 6.2e-06 | 5.2e-06 | 5.267E-06 | 9.018E-07 | 4.400E-06 | 6.200E-06 |
| 13996 | 700 | L700S | B | 9.6e-06 | 6.4e-06 | 4.7e-06 | 6.900E-06 | 2.488E-06 | 4.700E-06 | 9.600E-06 |
| 13997 | 700 | L700T | B | 6.8e-06 | 7.1e-06 | 6.1e-06 | 6.667E-06 | 5.132E-07 | 6.100E-06 | 7.100E-06 |
| 13998 | 700 | L700W | B | 8.4e-06 | 8.2e-06 | 1.2e-05 | 9.533E-06 | 2.139E-06 | 8.200E-06 | 1.200E-05 |
| 13999 | 700 | L700Y | B | 6.9e-06 | 6.8e-06 | 7.5e-06 | 7.067E-06 | 3.786E-07 | 6.800E-06 | 7.500E-06 |
| 14000 | 700 | L700V | B | 7.6e-06 | 1.2e-05 | 7.2e-06 | 8.933E-06 | 2.663E-06 | 7.200E-06 | 1.200E-05 |
| 14001 | 701 | Q701A | B | 9.2e-06 | 6.8e-06 | 7.5e-06 | 7.833E-06 | 1.234E-06 | 6.800E-06 | 9.200E-06 |
| 14002 | 701 | Q701R | B | 7.7e-06 | 8.2e-06 | 1.5e-05 | 1.030E-05 | 4.078E-06 | 7.700E-06 | 1.500E-05 |
| 14003 | 701 | Q701N | B | 6.1e-06 | 6.2e-06 | 4.8e-06 | 5.700E-06 | 7.810E-07 | 4.800E-06 | 6.200E-06 |
| 14004 | 701 | Q701D | B | 1.1e-05 | 8.7e-06 | 6.5e-06 | 8.733E-06 | 2.250E-06 | 6.500E-06 | 1.100E-05 |
| 14005 | 701 | Q701C | B | 7e-06 | 8.1e-06 | 6.6e-06 | 7.233E-06 | 7.767E-07 | 6.600E-06 | 8.100E-06 |
| 14006 | 701 | Q701E | B | 5.8e-06 | 7.1e-06 | 1.1e-05 | 7.967E-06 | 2.706E-06 | 5.800E-06 | 1.100E-05 |
| 14007 | 701 | Q701Q | B | 7.5e-06 | 6.2e-06 | 4.6e-06 | 6.100E-06 | 1.453E-06 | 4.600E-06 | 7.500E-06 |
| 14008 | 701 | Q701G | B | 9.6e-06 | 7.7e-06 | 8.9e-06 | 8.733E-06 | 9.609E-07 | 7.700E-06 | 9.600E-06 |
| 14009 | 701 | Q701H | B | 6e-06 | 1.3e-05 | 9.3e-06 | 9.433E-06 | 3.502E-06 | 6.000E-06 | 1.300E-05 |
| 14010 | 701 | Q701I | B | 6.5e-06 | 9.2e-06 | 7.9e-06 | 7.867E-06 | 1.350E-06 | 6.500E-06 | 9.200E-06 |
| 14011 | 701 | Q701L | B | 5e-06 | 1.1e-05 | 6.4e-06 | 7.467E-06 | 3.139E-06 | 5.000E-06 | 1.100E-05 |
| 14012 | 701 | Q701K | B | 7.1e-06 | 1.1e-05 | 7.6e-06 | 8.567E-06 | 2.122E-06 | 7.100E-06 | 1.100E-05 |
| 14013 | 701 | Q701M | B | 4e-06 | 8.7e-06 | 6.2e-06 | 6.300E-06 | 2.352E-06 | 4.000E-06 | 8.700E-06 |
| 14014 | 701 | Q701F | B | 9.8e-06 | 1e-05 | 6.4e-06 | 8.733E-06 | 2.023E-06 | 6.400E-06 | 1.000E-05 |
| 14015 | 701 | Q701P | B | 8.6e-06 | 6e-06 | 9.2e-06 | 7.933E-06 | 1.701E-06 | 6.000E-06 | 9.200E-06 |
| 14016 | 701 | Q701S | B | 4.3e-06 | 3.9e-06 | 6.8e-06 | 5.000E-06 | 1.572E-06 | 3.900E-06 | 6.800E-06 |
| 14017 | 701 | Q701T | B | 5.2e-06 | 1.3e-05 | 9.3e-06 | 9.167E-06 | 3.902E-06 | 5.200E-06 | 1.300E-05 |
| 14018 | 701 | Q701W | B | 4.7e-06 | 6.6e-06 | 6.3e-06 | 5.867E-06 | 1.021E-06 | 4.700E-06 | 6.600E-06 |
| 14019 | 701 | Q701Y | B | 4.5e-06 | 1e-05 | 6.7e-06 | 7.067E-06 | 2.768E-06 | 4.500E-06 | 1.000E-05 |
| 14020 | 701 | Q701V | B | 7.1e-06 | 5.4e-06 | 7.9e-06 | 6.800E-06 | 1.277E-06 | 5.400E-06 | 7.900E-06 |
| 14021 | 702 | P702A | B | 5.4e-06 | 8.2e-06 | 7.4e-06 | 7.000E-06 | 1.442E-06 | 5.400E-06 | 8.200E-06 |
| 14022 | 702 | P702R | B | 9.4e-06 | 4.8e-06 | 8.7e-06 | 7.633E-06 | 2.479E-06 | 4.800E-06 | 9.400E-06 |
| 14023 | 702 | P702N | B | 4.9e-06 | 1.2e-05 | 1e-05 | 8.967E-06 | 3.661E-06 | 4.900E-06 | 1.200E-05 |
| 14024 | 702 | P702D | B | 7e-06 | 6.9e-06 | 3.3e-06 | 5.733E-06 | 2.108E-06 | 3.300E-06 | 7.000E-06 |

| | | | | | | | | | | |
|---|---|---|---|---|---|---|---|---|---|---|
| 14025 | 702 | P702C | B | 6.6e-06 | 8.2e-06 | 9.3e-06 | 8.033E-06 | 1.358E-06 | 6.600E-06 | 9.300E-06 |
| 14026 | 702 | P702E | B | 6.5e-06 | 6.4e-06 | 4.6e-06 | 5.833E-06 | 1.069E-06 | 4.600E-06 | 6.500E-06 |
| 14027 | 702 | P702Q | B | 5.6e-06 | 1.4e-05 | 1.3e-05 | 1.087E-05 | 4.588E-06 | 5.600E-06 | 1.400E-05 |
| 14028 | 702 | P702G | B | 6.5e-06 | 1e-05 | 6.4e-06 | 7.633E-06 | 2.050E-06 | 6.400E-06 | 1.000E-05 |
| 14029 | 702 | P702H | B | 5.3e-06 | 5.4e-06 | 8e-06 | 6.233E-06 | 1.531E-06 | 5.300E-06 | 8.000E-06 |
| 14030 | 702 | P702I | B | 5.9e-06 | 9.1e-06 | 7.5e-06 | 7.500E-06 | 1.600E-06 | 5.900E-06 | 9.100E-06 |
| 14031 | 702 | P702L | B | 9.9e-06 | 8.1e-06 | 5.7e-06 | 7.900E-06 | 2.107E-06 | 5.700E-06 | 9.900E-06 |
| 14032 | 702 | P702K | B | 1e-05 | 9.7e-06 | 8.3e-06 | 9.333E-06 | 9.074E-07 | 8.300E-06 | 1.000E-05 |
| 14033 | 702 | P702M | B | 1.2e-05 | 5.5e-06 | 7.1e-06 | 8.200E-06 | 3.387E-06 | 5.500E-06 | 1.200E-05 |
| 14034 | 702 | P702F | B | 8.1e-06 | 8.1e-06 | 6.8e-06 | 7.667E-06 | 7.506E-07 | 6.800E-06 | 8.100E-06 |
| 14035 | 702 | P702P | B | 8.5e-06 | 8.8e-06 | 7.7e-06 | 8.333E-06 | 5.686E-07 | 7.700E-06 | 8.800E-06 |
| 14036 | 702 | P702S | B | 9.2e-06 | 1e-05 | 8.4e-06 | 9.200E-06 | 8.000E-07 | 8.400E-06 | 1.000E-05 |
| 14037 | 702 | P702T | B | 4.6e-06 | 1e-05 | 6.7e-06 | 7.100E-06 | 2.722E-06 | 4.600E-06 | 1.000E-05 |
| 14038 | 702 | P702W | B | 8.4e-06 | 9.8e-06 | 1.4e-05 | 1.073E-05 | 2.914E-06 | 8.400E-06 | 1.400E-05 |
| 14039 | 702 | P702Y | B | 9.3e-06 | 6.3e-06 | 6.4e-06 | 7.333E-06 | 1.704E-06 | 6.300E-06 | 9.300E-06 |
| 14040 | 702 | P702V | B | 6.9e-06 | 8.6e-06 | 7.7e-06 | 7.733E-06 | 8.505E-07 | 6.900E-06 | 8.600E-06 |
| 14041 | 703 | E703A | B | 9.3e-06 | 1e-05 | 8.8e-06 | 9.367E-06 | 6.028E-07 | 8.800E-06 | 1.000E-05 |
| 14042 | 703 | E703R | B | 7.1e-06 | 6.8e-06 | 4.8e-06 | 6.233E-06 | 1.250E-06 | 4.800E-06 | 7.100E-06 |
| 14043 | 703 | E703N | B | 6e-06 | 8.9e-06 | 6.3e-06 | 7.067E-06 | 1.595E-06 | 6.000E-06 | 8.900E-06 |
| 14044 | 703 | E703D | B | 6.3e-06 | 8.4e-06 | 7.6e-06 | 7.433E-06 | 1.060E-06 | 6.300E-06 | 8.400E-06 |
| 14045 | 703 | E703C | B | 6.5e-06 | 8.7e-06 | 9.7e-06 | 8.300E-06 | 1.637E-06 | 6.500E-06 | 9.700E-06 |
| 14046 | 703 | E703E | B | 1.2e-05 | 7.8e-06 | 8.9e-06 | 9.567E-06 | 2.178E-06 | 7.800E-06 | 1.200E-05 |
| 14047 | 703 | E703Q | B | 8.7e-06 | 3.9e-06 | 6.5e-06 | 6.367E-06 | 2.403E-06 | 3.900E-06 | 8.700E-06 |
| 14048 | 703 | E703G | B | 1.3e-05 | 4.7e-06 | 4.6e-06 | 7.433E-06 | 4.821E-06 | 4.600E-06 | 1.300E-05 |
| 14049 | 703 | E703H | B | 5.9e-06 | 5.1e-06 | 3.3e-06 | 4.767E-06 | 1.332E-06 | 3.300E-06 | 5.900E-06 |
| 14050 | 703 | E703I | B | 8.7e-06 | 9.9e-06 | 7.8e-06 | 8.800E-06 | 1.054E-06 | 7.800E-06 | 9.900E-06 |
| 14051 | 703 | E703L | B | 5.7e-06 | 6.5e-06 | 8.9e-06 | 7.033E-06 | 1.665E-06 | 5.700E-06 | 8.900E-06 |
| 14052 | 703 | E703K | B | 9.2e-06 | 6.7e-06 | 8e-06 | 7.967E-06 | 1.250E-06 | 6.700E-06 | 9.200E-06 |
| 14053 | 703 | E703M | B | 5.1e-06 | 5.8e-06 | 7.6e-06 | 6.167E-06 | 1.290E-06 | 5.100E-06 | 7.600E-06 |
| 14054 | 703 | E703F | B | 7.6e-06 | 6.9e-06 | 8e-06 | 7.500E-06 | 5.568E-07 | 6.900E-06 | 8.000E-06 |
| 14055 | 703 | E703P | B | 8e-06 | 5.8e-06 | 6.8e-06 | 6.867E-06 | 1.102E-06 | 5.800E-06 | 8.000E-06 |
| 14056 | 703 | E703S | B | 1.4e-05 | 6.7e-06 | 9.8e-06 | 1.017E-05 | 3.664E-06 | 6.700E-06 | 1.400E-05 |
| 14057 | 703 | E703T | B | 5.6e-06 | 6.2e-06 | 5.7e-06 | 5.833E-06 | 3.215E-07 | 5.600E-06 | 6.200E-06 |
| 14058 | 703 | E703W | B | 6e-06 | 6.6e-06 | 8e-06 | 6.867E-06 | 1.026E-06 | 6.000E-06 | 8.000E-06 |
| 14059 | 703 | E703Y | B | 1.1e-05 | 8.8e-06 | 1.2e-05 | 1.060E-05 | 1.637E-06 | 8.800E-06 | 1.200E-05 |
| 14060 | 703 | E703V | B | 8.9e-06 | 1.1e-05 | 8.7e-06 | 9.533E-06 | 1.274E-06 | 8.700E-06 | 1.100E-05 |
| 14061 | 704 | D704A | B | 4.9e-06 | 6.6e-06 | 7.3e-06 | 6.267E-06 | 1.234E-06 | 4.900E-06 | 7.300E-06 |
| 14062 | 704 | D704R | B | 6.3e-06 | 6.6e-06 | 9e-06 | 7.300E-06 | 1.480E-06 | 6.300E-06 | 9.000E-06 |
| 14063 | 704 | D704N | B | 4.9e-06 | 6e-06 | 9.1e-06 | 6.667E-06 | 2.178E-06 | 4.900E-06 | 9.100E-06 |
| 14064 | 704 | D704D | B | 8.3e-06 | 4.6e-06 | 6.2e-06 | 6.367E-06 | 1.856E-06 | 4.600E-06 | 8.300E-06 |
| 14065 | 704 | D704C | B | 1.2e-05 | 5.1e-06 | 4.4e-06 | 7.167E-06 | 4.200E-06 | 4.400E-06 | 1.200E-05 |
| 14066 | 704 | D704E | B | 6.3e-06 | 1.2e-05 | 1.3e-05 | 1.043E-05 | 3.614E-06 | 6.300E-06 | 1.300E-05 |
| 14067 | 704 | D704Q | B | 5.8e-06 | 6.6e-06 | 4.8e-06 | 5.733E-06 | 9.018E-07 | 4.800E-06 | 6.600E-06 |
| 14068 | 704 | D704G | B | 5.4e-06 | 9.8e-06 | 1e-05 | 8.400E-06 | 2.600E-06 | 5.400E-06 | 1.000E-05 |
| 14069 | 704 | D704H | B | 9.5e-06 | 1.1e-05 | 8.3e-06 | 9.600E-06 | 1.353E-06 | 8.300E-06 | 1.100E-05 |
| 14070 | 704 | D704I | B | 8.2e-06 | 1.1e-05 | 4.8e-06 | 8.000E-06 | 3.105E-06 | 4.800E-06 | 1.100E-05 |
| 14071 | 704 | D704L | B | 9e-06 | 1e-05 | 9.2e-06 | 9.400E-06 | 5.292E-07 | 9.000E-06 | 1.000E-05 |
| 14072 | 704 | D704K | B | 9.8e-06 | 4.5e-06 | 9.9e-06 | 8.067E-06 | 3.089E-06 | 4.500E-06 | 9.900E-06 |
| 14073 | 704 | D704M | B | 6.9e-06 | 8.7e-06 | 6.5e-06 | 7.367E-06 | 1.172E-06 | 6.500E-06 | 8.700E-06 |
| 14074 | 704 | D704F | B | 9.2e-06 | 8.6e-06 | 1.1e-05 | 9.600E-06 | 1.249E-06 | 8.600E-06 | 1.100E-05 |
| 14075 | 704 | D704P | B | 6.7e-06 | 8.3e-06 | 7.5e-06 | 7.500E-06 | 8.000E-07 | 6.700E-06 | 8.300E-06 |
| 14076 | 704 | D704S | B | 8.1e-06 | 6.1e-06 | 7.3e-06 | 7.167E-06 | 1.007E-06 | 6.100E-06 | 8.100E-06 |
| 14077 | 704 | D704T | B | 1.3e-05 | 6.5e-06 | 7.4e-06 | 8.967E-06 | 3.522E-06 | 6.500E-06 | 1.300E-05 |
| 14078 | 704 | D704W | B | 1.5e-05 | 8.7e-06 | 6.2e-06 | 9.967E-06 | 4.535E-06 | 6.200E-06 | 1.500E-05 |
| 14079 | 704 | D704Y | B | 6.3e-06 | 8e-06 | 9.6e-06 | 7.967E-06 | 1.650E-06 | 6.300E-06 | 9.600E-06 |
| 14080 | 704 | D704V | B | 7.7e-06 | 5.4e-06 | 6.7e-06 | 6.600E-06 | 1.153E-06 | 5.400E-06 | 7.700E-06 |
| 14081 | 705 | F705A | B | 1e-05 | 8e-06 | 6.3e-06 | 8.100E-06 | 1.852E-06 | 6.300E-06 | 1.000E-05 |
| 14082 | 705 | F705R | B | 5.8e-06 | 4.7e-06 | 6.8e-06 | 5.767E-06 | 1.050E-06 | 4.700E-06 | 6.800E-06 |
| 14083 | 705 | F705N | B | 6.6e-06 | 9.4e-06 | 1.1e-05 | 9.000E-06 | 2.227E-06 | 6.600E-06 | 1.100E-05 |
| 14084 | 705 | F705D | B | 6.8e-06 | 1e-05 | 8e-06 | 8.267E-06 | 1.617E-06 | 6.800E-06 | 1.000E-05 |
| 14085 | 705 | F705C | B | 7.1e-06 | 5.7e-06 | 1e-05 | 7.600E-06 | 2.193E-06 | 5.700E-06 | 1.000E-05 |
| 14086 | 705 | F705E | B | 8.4e-06 | 8.6e-06 | 4.9e-06 | 7.300E-06 | 2.081E-06 | 4.900E-06 | 8.600E-06 |
| 14087 | 705 | F705Q | B | 7.7e-06 | 8.9e-06 | 7.4e-06 | 8.000E-06 | 7.937E-07 | 7.400E-06 | 8.900E-06 |
| 14088 | 705 | F705G | B | 8.9e-06 | 1.5e-05 | 8.8e-06 | 1.090E-05 | 3.551E-06 | 8.800E-06 | 1.500E-05 |
| 14089 | 705 | F705H | B | 5.6e-06 | 1.3e-05 | 6.6e-06 | 8.400E-06 | 4.015E-06 | 5.600E-06 | 1.300E-05 |
| 14090 | 705 | F705I | B | 1.1e-05 | 1.1e-05 | 6.2e-06 | 9.400E-06 | 2.771E-06 | 6.200E-06 | 1.100E-05 |
| 14091 | 705 | F705L | B | 7.2e-06 | 1e-05 | 4.9e-06 | 7.367E-06 | 2.554E-06 | 4.900E-06 | 1.000E-05 |
| 14092 | 705 | F705K | B | 5.7e-06 | 4.6e-06 | 5.4e-06 | 5.233E-06 | 5.686E-07 | 4.600E-06 | 5.700E-06 |
| 14093 | 705 | F705M | B | 8.2e-06 | 8e-06 | 9e-06 | 8.400E-06 | 5.292E-07 | 8.000E-06 | 9.000E-06 |
| 14094 | 705 | F705F | B | 8.2e-06 | 9.9e-06 | 8.9e-06 | 9.000E-06 | 8.544E-07 | 8.200E-06 | 9.900E-06 |
| 14095 | 705 | F705P | B | 1.1e-05 | 7.5e-06 | 9e-06 | 9.167E-06 | 1.756E-06 | 7.500E-06 | 1.100E-05 |
| 14096 | 705 | F705S | B | 9.6e-06 | 7e-06 | 8e-06 | 8.200E-06 | 1.311E-06 | 7.000E-06 | 9.600E-06 |
| 14097 | 705 | F705T | B | 8e-06 | 8.2e-06 | 7.3e-06 | 7.833E-06 | 4.726E-07 | 7.300E-06 | 8.200E-06 |
| 14098 | 705 | F705W | B | 8.2e-06 | 1.1e-05 | 9.7e-06 | 9.633E-06 | 1.401E-06 | 8.200E-06 | 1.100E-05 |
| 14099 | 705 | F705Y | B | 1.4e-05 | 6.7e-06 | 8.5e-06 | 9.733E-06 | 3.803E-06 | 6.700E-06 | 1.400E-05 |
| 14100 | 705 | F705V | B | 1e-05 | 7.1e-06 | 7.5e-06 | 8.200E-06 | 1.572E-06 | 7.100E-06 | 1.000E-05 |
| 14101 | 706 | A706A | B | 6.3e-06 | 6.4e-06 | 9.4e-06 | 7.367E-06 | 1.762E-06 | 6.300E-06 | 9.400E-06 |
| 14102 | 706 | A706R | B | 5e-06 | 1e-05 | 6.5e-06 | 7.167E-06 | 2.566E-06 | 5.000E-06 | 1.000E-05 |
| 14103 | 706 | A706N | B | 3.7e-06 | 5.3e-06 | 4.7e-06 | 4.567E-06 | 8.083E-07 | 3.700E-06 | 5.300E-06 |
| 14104 | 706 | A706D | B | 8.4e-06 | 7.4e-06 | 6.3e-06 | 7.367E-06 | 1.050E-06 | 6.300E-06 | 8.400E-06 |
| 14105 | 706 | A706C | B | 6.8e-06 | 8.1e-06 | 4.1e-06 | 6.333E-06 | 2.040E-06 | 4.100E-06 | 8.100E-06 |
| 14106 | 706 | A706E | B | 9e-06 | 6.9e-06 | 9e-06 | 8.300E-06 | 1.212E-06 | 6.900E-06 | 9.000E-06 |
| 14107 | 706 | A706Q | B | 7.8e-06 | 9.3e-06 | 5.8e-06 | 7.633E-06 | 1.756E-06 | 5.800E-06 | 9.300E-06 |
| 14108 | 706 | A706G | B | 6.6e-06 | 5.2e-06 | 4.7e-06 | 5.500E-06 | 9.849E-07 | 4.700E-06 | 6.600E-06 |
| 14109 | 706 | A706H | B | 8.9e-06 | 5.7e-06 | 7e-06 | 7.200E-06 | 1.609E-06 | 5.700E-06 | 8.900E-06 |

| | | | | | | | | | | |
|---|---|---|---|---|---|---|---|---|---|---|
| 14110 | 706 | A706I | B | 7.9e-06 | 1.2e-05 | 4.7e-06 | 8.200E-06 | 3.659E-06 | 4.700E-06 | 1.200E-05 |
| 14111 | 706 | A706L | B | 7.1e-06 | 9.1e-06 | 5e-06 | 7.067E-06 | 2.050E-06 | 5.000E-06 | 9.100E-06 |
| 14112 | 706 | A706K | B | 1.2e-05 | 6.1e-06 | 5.8e-06 | 7.967E-06 | 3.496E-06 | 5.800E-06 | 1.200E-05 |
| 14113 | 706 | A706M | B | 9.7e-06 | 8e-06 | 9.1e-06 | 8.933E-06 | 8.622E-07 | 8.000E-06 | 9.700E-06 |
| 14114 | 706 | A706F | B | 1.4e-05 | 5.8e-06 | 5.6e-06 | 8.467E-06 | 4.793E-06 | 5.600E-06 | 1.400E-05 |
| 14115 | 706 | A706P | B | 7.1e-06 | 4.7e-06 | 7.2e-06 | 6.333E-06 | 1.415E-06 | 4.700E-06 | 7.200E-06 |
| 14116 | 706 | A706S | B | 1.3e-05 | 8.4e-06 | 6e-06 | 9.133E-06 | 3.557E-06 | 6.000E-06 | 1.300E-05 |
| 14117 | 706 | A706T | B | 9.9e-06 | 9.5e-06 | 5.4e-06 | 8.267E-06 | 2.491E-06 | 5.400E-06 | 9.900E-06 |
| 14118 | 706 | A706W | B | 7.1e-06 | 4.4e-06 | 6.4e-06 | 5.967E-06 | 1.401E-06 | 4.400E-06 | 7.100E-06 |
| 14119 | 706 | A706Y | B | 7.6e-06 | 8.8e-06 | 9.3e-06 | 8.567E-06 | 8.737E-07 | 7.600E-06 | 9.300E-06 |
| 14120 | 706 | A706V | B | 5.2e-06 | 5.7e-06 | 9.8e-06 | 6.900E-06 | 2.524E-06 | 5.200E-06 | 9.800E-06 |
| 14121 | 707 | T707A | B | 5.1e-06 | 5.6e-06 | 7.6e-06 | 6.100E-06 | 1.323E-06 | 5.100E-06 | 7.600E-06 |
| 14122 | 707 | T707R | B | 8.2e-06 | 7.5e-06 | 5.9e-06 | 7.200E-06 | 1.179E-06 | 5.900E-06 | 8.200E-06 |
| 14123 | 707 | T707N | B | 1.3e-05 | 7.1e-06 | 8.5e-06 | 9.533E-06 | 3.083E-06 | 7.100E-06 | 1.300E-05 |
| 14124 | 707 | T707D | B | 6.8e-06 | 7.2e-06 | 4.7e-06 | 6.233E-06 | 1.343E-06 | 4.700E-06 | 7.200E-06 |
| 14125 | 707 | T707C | B | 9.3e-06 | 9.2e-06 | 1e-05 | 9.500E-06 | 4.359E-07 | 9.200E-06 | 1.000E-05 |
| 14126 | 707 | T707E | B | 4.6e-06 | 5.6e-06 | 9.5e-06 | 6.567E-06 | 2.589E-06 | 4.600E-06 | 9.500E-06 |
| 14127 | 707 | T707Q | B | 1.3e-05 | 6.1e-06 | 1.1e-05 | 1.003E-05 | 3.550E-06 | 6.100E-06 | 1.300E-05 |
| 14128 | 707 | T707G | B | 7.7e-06 | 8.9e-06 | 7.2e-06 | 7.933E-06 | 8.737E-07 | 7.200E-06 | 8.900E-06 |
| 14129 | 707 | T707H | B | 5.1e-06 | 9.7e-06 | 6.8e-06 | 7.200E-06 | 2.326E-06 | 5.100E-06 | 9.700E-06 |
| 14130 | 707 | T707I | B | 1.4e-05 | 6.9e-06 | 9e-06 | 9.967E-06 | 3.647E-06 | 6.900E-06 | 1.400E-05 |
| 14131 | 707 | T707L | B | 8.1e-06 | 5.8e-06 | 5.3e-06 | 6.400E-06 | 1.493E-06 | 5.300E-06 | 8.100E-06 |
| 14132 | 707 | T707K | B | 7.3e-06 | 9.3e-06 | 8.7e-06 | 8.433E-06 | 1.026E-06 | 7.300E-06 | 9.300E-06 |
| 14133 | 707 | T707M | B | 1.4e-05 | 5.5e-06 | 6.5e-06 | 8.667E-06 | 4.646E-06 | 5.500E-06 | 1.400E-05 |
| 14134 | 707 | T707F | B | 9e-06 | 6.5e-06 | 7.2e-06 | 7.567E-06 | 1.290E-06 | 6.500E-06 | 9.000E-06 |
| 14135 | 707 | T707P | B | 8.1e-06 | 2.9e-06 | 5.8e-06 | 5.600E-06 | 2.606E-06 | 2.900E-06 | 8.100E-06 |
| 14136 | 707 | T707S | B | 9.6e-06 | 9.3e-06 | 1.2e-05 | 1.030E-05 | 1.480E-06 | 9.300E-06 | 1.200E-05 |
| 14137 | 707 | T707T | B | 6.3e-06 | 5.5e-06 | 7.3e-06 | 6.367E-06 | 9.018E-07 | 5.500E-06 | 7.300E-06 |
| 14138 | 707 | T707W | B | 8.2e-06 | 9.2e-06 | 1e-05 | 9.133E-06 | 9.018E-07 | 8.200E-06 | 1.000E-05 |
| 14139 | 707 | T707Y | B | 6.2e-06 | 1.1e-05 | 6.6e-06 | 7.933E-06 | 2.663E-06 | 6.200E-06 | 1.100E-05 |
| 14140 | 707 | T707V | B | 7.4e-06 | 1.2e-05 | 5.7e-06 | 8.367E-06 | 3.259E-06 | 5.700E-06 | 1.200E-05 |
| 14141 | 708 | Y708A | B | 1e-05 | 7.4e-06 | 9.8e-06 | 9.067E-06 | 1.447E-06 | 7.400E-06 | 1.000E-05 |
| 14142 | 708 | Y708R | B | 5.5e-06 | 1.3e-05 | 1.1e-05 | 9.833E-06 | 3.884E-06 | 5.500E-06 | 1.300E-05 |
| 14143 | 708 | Y708N | B | 1.6e-05 | 7.4e-06 | 7e-06 | 1.013E-05 | 5.085E-06 | 7.000E-06 | 1.600E-05 |
| 14144 | 708 | Y708D | B | 6.1e-06 | 8.4e-06 | 1.5e-05 | 9.833E-06 | 4.620E-06 | 6.100E-06 | 1.500E-05 |
| 14145 | 708 | Y708C | B | 6.6e-06 | 7.5e-06 | 6.5e-06 | 6.867E-06 | 5.508E-07 | 6.500E-06 | 7.500E-06 |
| 14146 | 708 | Y708E | B | 9.2e-06 | 8.6e-06 | 4.8e-06 | 7.533E-06 | 2.386E-06 | 4.800E-06 | 9.200E-06 |
| 14147 | 708 | Y708Q | B | 8.4e-06 | 1.3e-05 | 1.3e-05 | 1.147E-05 | 2.656E-06 | 8.400E-06 | 1.300E-05 |
| 14148 | 708 | Y708G | B | 6.6e-06 | 4.7e-06 | 6e-06 | 5.767E-06 | 9.713E-07 | 4.700E-06 | 6.600E-06 |
| 14149 | 708 | Y708H | B | 6.8e-06 | 6.6e-06 | 5.9e-06 | 6.433E-06 | 4.726E-07 | 5.900E-06 | 6.800E-06 |
| 14150 | 708 | Y708I | B | 1e-05 | 9.7e-06 | 5.3e-06 | 8.333E-06 | 2.631E-06 | 5.300E-06 | 1.000E-05 |
| 14151 | 708 | Y708L | B | 4.7e-06 | 1.6e-05 | 9.7e-06 | 1.013E-05 | 5.662E-06 | 4.700E-06 | 1.600E-05 |
| 14152 | 708 | Y708K | B | 5.5e-06 | 9.1e-06 | 7e-06 | 7.200E-06 | 1.808E-06 | 5.500E-06 | 9.100E-06 |
| 14153 | 708 | Y708M | B | 4.6e-06 | 6.4e-06 | 8.4e-06 | 6.467E-06 | 1.901E-06 | 4.600E-06 | 8.400E-06 |
| 14154 | 708 | Y708F | B | 1.3e-05 | 8.6e-06 | 9.9e-06 | 1.050E-05 | 2.261E-06 | 8.600E-06 | 1.300E-05 |
| 14155 | 708 | Y708P | B | 8e-06 | 4.3e-06 | 5e-06 | 5.767E-06 | 1.966E-06 | 4.300E-06 | 8.000E-06 |
| 14156 | 708 | Y708S | B | 1.9e-05 | 6.3e-06 | 1e-05 | 1.177E-05 | 6.532E-06 | 6.300E-06 | 1.900E-05 |
| 14157 | 708 | Y708T | B | 1.1e-05 | 9.6e-06 | 6.3e-06 | 8.967E-06 | 2.413E-06 | 6.300E-06 | 1.100E-05 |
| 14158 | 708 | Y708W | B | 8.5e-06 | 9.1e-06 | 1.4e-05 | 1.053E-05 | 3.017E-06 | 8.500E-06 | 1.400E-05 |
| 14159 | 708 | Y708Y | B | 7.1e-06 | 6.7e-06 | 6.5e-06 | 6.767E-06 | 3.055E-07 | 6.500E-06 | 7.100E-06 |
| 14160 | 708 | Y708V | B | 7.4e-06 | 6.9e-06 | 6.4e-06 | 6.900E-06 | 5.000E-07 | 6.400E-06 | 7.400E-06 |
| 14161 | 709 | Y709A | B | 8.7e-06 | 9.3e-06 | 6.5e-06 | 8.167E-06 | 1.474E-06 | 6.500E-06 | 9.300E-06 |
| 14162 | 709 | Y709R | B | 7.3e-06 | 7.5e-06 | 1.1e-05 | 8.600E-06 | 2.081E-06 | 7.300E-06 | 1.100E-05 |
| 14163 | 709 | Y709N | B | 6.6e-06 | 5.7e-06 | 9.7e-06 | 7.333E-06 | 2.098E-06 | 5.700E-06 | 9.700E-06 |
| 14164 | 709 | Y709D | B | 1.4e-05 | 1.1e-05 | 7.7e-06 | 1.090E-05 | 3.151E-06 | 7.700E-06 | 1.400E-05 |
| 14165 | 709 | Y709C | B | 6.8e-06 | 5e-06 | 6.8e-06 | 6.200E-06 | 1.039E-06 | 5.000E-06 | 6.800E-06 |
| 14166 | 709 | Y709E | B | 6.5e-06 | 6.9e-06 | 6.5e-06 | 6.633E-06 | 2.309E-07 | 6.500E-06 | 6.900E-06 |
| 14167 | 709 | Y709Q | B | 8.8e-06 | 1.4e-05 | 9.4e-06 | 1.073E-05 | 2.845E-06 | 8.800E-06 | 1.400E-05 |
| 14168 | 709 | Y709G | B | 1e-05 | 1e-05 | 9.6e-06 | 9.867E-06 | 2.309E-07 | 9.600E-06 | 1.000E-05 |
| 14169 | 709 | Y709H | B | 1.2e-05 | 6.6e-06 | 8.9e-06 | 9.167E-06 | 2.710E-06 | 6.600E-06 | 1.200E-05 |
| 14170 | 709 | Y709I | B | 7.6e-06 | 5e-06 | 9.1e-06 | 7.233E-06 | 2.074E-06 | 5.000E-06 | 9.100E-06 |
| 14171 | 709 | Y709L | B | 1.4e-05 | 4.8e-06 | 6.4e-06 | 8.400E-06 | 4.915E-06 | 4.800E-06 | 1.400E-05 |
| 14172 | 709 | Y709K | B | 1.2e-05 | 8.6e-06 | 7e-06 | 9.200E-06 | 2.553E-06 | 7.000E-06 | 1.200E-05 |
| 14173 | 709 | Y709M | B | 9.4e-06 | 7.2e-06 | 5.2e-06 | 7.267E-06 | 2.101E-06 | 5.200E-06 | 9.400E-06 |
| 14174 | 709 | Y709F | B | 6.5e-06 | 5.6e-06 | 8.9e-06 | 7.000E-06 | 1.706E-06 | 5.600E-06 | 8.900E-06 |
| 14175 | 709 | Y709P | B | 1.2e-05 | 6.9e-06 | 9.8e-06 | 9.567E-06 | 2.558E-06 | 6.900E-06 | 1.200E-05 |
| 14176 | 709 | Y709S | B | 8.4e-06 | 6e-06 | 6.7e-06 | 7.033E-06 | 1.234E-06 | 6.000E-06 | 8.400E-06 |
| 14177 | 709 | Y709T | B | 9.1e-06 | 6.8e-06 | 9.4e-06 | 8.433E-06 | 1.422E-06 | 6.800E-06 | 9.400E-06 |
| 14178 | 709 | Y709W | B | 9.1e-06 | 9.6e-06 | 6.2e-06 | 8.300E-06 | 1.836E-06 | 6.200E-06 | 9.600E-06 |
| 14179 | 709 | Y709Y | B | 8.6e-06 | 8.5e-06 | 6.5e-06 | 7.867E-06 | 1.185E-06 | 6.500E-06 | 8.600E-06 |
| 14180 | 709 | Y709V | B | 8.8e-06 | 1.4e-05 | 4.6e-06 | 9.133E-06 | 4.709E-06 | 4.600E-06 | 1.400E-05 |
| 14181 | 710 | C710A | B | 7.9e-06 | 5e-06 | 1.3e-05 | 8.633E-06 | 4.050E-06 | 5.000E-06 | 1.300E-05 |
| 14182 | 710 | C710R | B | 9.4e-06 | 8.5e-06 | 9.6e-06 | 9.167E-06 | 5.859E-07 | 8.500E-06 | 9.600E-06 |
| 14183 | 710 | C710N | B | 7.2e-06 | 7.7e-06 | 8.8e-06 | 7.900E-06 | 8.185E-07 | 7.200E-06 | 8.800E-06 |
| 14184 | 710 | C710D | B | 6.5e-06 | 5.8e-06 | 1.9e-05 | 1.043E-05 | 7.427E-06 | 5.800E-06 | 1.900E-05 |
| 14185 | 710 | C710C | B | 8.5e-06 | 6.3e-06 | 1.3e-05 | 9.267E-06 | 3.415E-06 | 6.300E-06 | 1.300E-05 |
| 14186 | 710 | C710E | B | 9.5e-06 | 5.2e-06 | 1.3e-05 | 9.233E-06 | 3.907E-06 | 5.200E-06 | 1.300E-05 |
| 14187 | 710 | C710Q | B | 1.1e-05 | 5.6e-06 | 5.4e-06 | 7.333E-06 | 3.177E-06 | 5.400E-06 | 1.100E-05 |
| 14188 | 710 | C710G | B | 6.8e-06 | 1.1e-05 | 8.4e-06 | 8.733E-06 | 2.120E-06 | 6.800E-06 | 1.100E-05 |
| 14189 | 710 | C710H | B | 3.6e-06 | 8.4e-06 | 7.2e-06 | 6.400E-06 | 2.498E-06 | 3.600E-06 | 8.400E-06 |
| 14190 | 710 | C710I | B | 5.9e-06 | 7.2e-06 | 6.6e-06 | 6.567E-06 | 6.506E-07 | 5.900E-06 | 7.200E-06 |
| 14191 | 710 | C710L | B | 9.3e-06 | 5.9e-06 | 1.4e-05 | 9.733E-06 | 4.067E-06 | 5.900E-06 | 1.400E-05 |
| 14192 | 710 | C710K | B | 9e-06 | 8.8e-06 | 6.6e-06 | 8.133E-06 | 1.332E-06 | 6.600E-06 | 9.000E-06 |
| 14193 | 710 | C710M | B | 6.7e-06 | 1.1e-05 | 7.4e-06 | 8.367E-06 | 2.307E-06 | 6.700E-06 | 1.100E-05 |
| 14194 | 710 | C710F | B | 5.3e-06 | 7.1e-06 | 1.2e-05 | 8.133E-06 | 3.467E-06 | 5.300E-06 | 1.200E-05 |

| | | | | | | | | | | |
|---|---|---|---|---|---|---|---|---|---|---|
| 14195 | 710 | C710P | B | 9.3e-06 | 8e-06 | 8.3e-06 | 8.533E-06 | 6.807E-07 | 8.000E-06 | 9.300E-06 |
| 14196 | 710 | C710S | B | 9.6e-06 | 6.4e-06 | 8e-06 | 8.000E-06 | 1.600E-06 | 6.400E-06 | 9.600E-06 |
| 14197 | 710 | C710T | B | 5.8e-06 | 6.8e-06 | 6.5e-06 | 6.367E-06 | 5.132E-07 | 5.800E-06 | 6.800E-06 |
| 14198 | 710 | C710W | B | 1.3e-05 | 8.1e-06 | 6.9e-06 | 9.333E-06 | 3.232E-06 | 6.900E-06 | 1.300E-05 |
| 14199 | 710 | C710Y | B | 6.9e-06 | 7.7e-06 | 9.6e-06 | 8.067E-06 | 1.387E-06 | 6.900E-06 | 9.600E-06 |
| 14200 | 710 | C710V | B | 7.1e-06 | 6.1e-06 | 9.3e-06 | 7.500E-06 | 1.637E-06 | 6.100E-06 | 9.300E-06 |
| 14201 | 711 | Q711A | B | 8.5e-06 | 8.7e-06 | 5.2e-06 | 7.467E-06 | 1.966E-06 | 5.200E-06 | 8.700E-06 |
| 14202 | 711 | Q711R | B | 9.4e-06 | 6.5e-06 | 8.6e-06 | 8.167E-06 | 1.498E-06 | 6.500E-06 | 9.400E-06 |
| 14203 | 711 | Q711N | B | 1.2e-05 | 1e-05 | 4.7e-06 | 8.900E-06 | 3.772E-06 | 4.700E-06 | 1.200E-05 |
| 14204 | 711 | Q711D | B | 6.1e-06 | 5.1e-06 | 5.3e-06 | 5.500E-06 | 5.292E-07 | 5.100E-06 | 6.100E-06 |
| 14205 | 711 | Q711C | B | 6e-06 | 5.4e-06 | 7.4e-06 | 6.267E-06 | 1.026E-06 | 5.400E-06 | 7.400E-06 |
| 14206 | 711 | Q711E | B | 6.9e-06 | 7.8e-06 | 1.1e-05 | 8.567E-06 | 2.155E-06 | 6.900E-06 | 1.100E-05 |
| 14207 | 711 | Q711Q | B | 6.1e-06 | 5.9e-06 | 8.5e-06 | 6.833E-06 | 1.447E-06 | 5.900E-06 | 8.500E-06 |
| 14208 | 711 | Q711G | B | 1.1e-05 | 1.2e-05 | 1.2e-05 | 1.167E-05 | 5.774E-07 | 1.100E-05 | 1.200E-05 |
| 14209 | 711 | Q711H | B | 6.9e-06 | 7.9e-06 | 6.9e-06 | 7.233E-06 | 5.774E-07 | 6.900E-06 | 7.900E-06 |
| 14210 | 711 | Q711I | B | 5.2e-06 | 4.3e-06 | 6.7e-06 | 5.400E-06 | 1.212E-06 | 4.300E-06 | 6.700E-06 |
| 14211 | 711 | Q711L | B | 6.6e-06 | 1.1e-05 | 6.8e-06 | 8.133E-06 | 2.485E-06 | 6.600E-06 | 1.100E-05 |
| 14212 | 711 | Q711K | B | 1.3e-05 | 1e-05 | 1.1e-05 | 1.133E-05 | 1.528E-06 | 1.000E-05 | 1.300E-05 |
| 14213 | 711 | Q711M | B | 4.6e-06 | 1e-05 | 9.2e-06 | 7.933E-06 | 2.914E-06 | 4.600E-06 | 1.000E-05 |
| 14214 | 711 | Q711F | B | 5.3e-06 | 9.1e-06 | 9.5e-06 | 7.967E-06 | 2.318E-06 | 5.300E-06 | 9.500E-06 |
| 14215 | 711 | Q711P | B | 1.2e-05 | 8.8e-06 | 4.7e-06 | 8.500E-06 | 3.659E-06 | 4.700E-06 | 1.200E-05 |
| 14216 | 711 | Q711S | B | 9.2e-06 | 4.8e-06 | 9.4e-06 | 7.800E-06 | 2.600E-06 | 4.800E-06 | 9.400E-06 |
| 14217 | 711 | Q711T | B | 7.9e-06 | 6.4e-06 | 9.3e-06 | 7.867E-06 | 1.450E-06 | 6.400E-06 | 9.300E-06 |
| 14218 | 711 | Q711W | B | 1e-05 | 7.8e-06 | 5.8e-06 | 7.867E-06 | 2.101E-06 | 5.800E-06 | 1.000E-05 |
| 14219 | 711 | Q711Y | B | 4e-06 | 1e-05 | 1.1e-05 | 8.333E-06 | 3.786E-06 | 4.000E-06 | 1.100E-05 |
| 14220 | 711 | Q711V | B | 6.7e-06 | 1e-05 | 1.4e-05 | 1.023E-05 | 3.656E-06 | 6.700E-06 | 1.400E-05 |
| 14221 | 712 | Q712A | B | 9.4e-06 | 4.8e-06 | 9.7e-06 | 7.967E-06 | 2.747E-06 | 4.800E-06 | 9.700E-06 |
| 14222 | 712 | Q712R | B | 4.6e-06 | 5.8e-06 | 7.6e-06 | 6.000E-06 | 1.510E-06 | 4.600E-06 | 7.600E-06 |
| 14223 | 712 | Q712N | B | 1.3e-05 | 7e-06 | 7.1e-06 | 9.033E-06 | 3.436E-06 | 7.000E-06 | 1.300E-05 |
| 14224 | 712 | Q712D | B | 4.6e-06 | 4.3e-06 | 7e-06 | 5.300E-06 | 1.480E-06 | 4.300E-06 | 7.000E-06 |
| 14225 | 712 | Q712C | B | 5.6e-06 | 6.3e-06 | 5e-06 | 5.633E-06 | 6.506E-07 | 5.000E-06 | 6.300E-06 |
| 14226 | 712 | Q712E | B | 5.5e-06 | 7.3e-06 | 6.6e-06 | 6.467E-06 | 9.074E-07 | 5.500E-06 | 7.300E-06 |
| 14227 | 712 | Q712Q | B | 6.7e-06 | 9.6e-06 | 9.1e-06 | 8.467E-06 | 1.550E-06 | 6.700E-06 | 9.600E-06 |
| 14228 | 712 | Q712G | B | 8e-06 | 6.5e-06 | 6.2e-06 | 6.900E-06 | 9.644E-07 | 6.200E-06 | 8.000E-06 |
| 14229 | 712 | Q712H | B | 8.1e-06 | 9.3e-06 | 9.4e-06 | 8.933E-06 | 7.234E-07 | 8.100E-06 | 9.400E-06 |
| 14230 | 712 | Q712I | B | 1.4e-05 | 1.3e-05 | 6.5e-06 | 1.117E-05 | 4.072E-06 | 6.500E-06 | 1.400E-05 |
| 14231 | 712 | Q712L | B | 1.2e-05 | 1e-05 | 6.3e-06 | 9.433E-06 | 2.892E-06 | 6.300E-06 | 1.200E-05 |
| 14232 | 712 | Q712K | B | 1.1e-05 | 6.1e-06 | 7e-06 | 8.033E-06 | 2.608E-06 | 6.100E-06 | 1.100E-05 |
| 14233 | 712 | Q712M | B | 9.1e-06 | 4.9e-06 | 1.2e-05 | 8.667E-06 | 3.570E-06 | 4.900E-06 | 1.200E-05 |
| 14234 | 712 | Q712F | B | 5.4e-06 | 9.6e-06 | 1e-05 | 8.333E-06 | 2.548E-06 | 5.400E-06 | 1.000E-05 |
| 14235 | 712 | Q712P | B | 6.5e-06 | 9e-06 | 1.3e-05 | 9.500E-06 | 3.279E-06 | 6.500E-06 | 1.300E-05 |
| 14236 | 712 | Q712S | B | 1.4e-05 | 1.3e-05 | 8.6e-06 | 1.187E-05 | 2.873E-06 | 8.600E-06 | 1.400E-05 |
| 14237 | 712 | Q712T | B | 7.1e-06 | 1e-05 | 6.9e-06 | 8.000E-06 | 1.735E-06 | 6.900E-06 | 1.000E-05 |
| 14238 | 712 | Q712W | B | 3.8e-06 | 6.6e-06 | 6.2e-06 | 5.533E-06 | 1.514E-06 | 3.800E-06 | 6.600E-06 |
| 14239 | 712 | Q712Y | B | 6.7e-06 | 7.4e-06 | 1e-05 | 8.033E-06 | 1.739E-06 | 6.700E-06 | 1.000E-05 |
| 14240 | 712 | Q712V | B | 6.8e-06 | 1.2e-05 | 5.5e-06 | 8.100E-06 | 3.439E-06 | 5.500E-06 | 1.200E-05 |
| 14241 | 713 | Y713A | B | 8.4e-06 | 9e-06 | 1.1e-05 | 9.467E-06 | 1.361E-06 | 8.400E-06 | 1.100E-05 |
| 14242 | 713 | Y713R | B | 6.1e-06 | 7.5e-06 | 8.2e-06 | 7.267E-06 | 1.069E-06 | 6.100E-06 | 8.200E-06 |
| 14243 | 713 | Y713N | B | 9.5e-06 | 1.7e-05 | 6.6e-06 | 1.103E-05 | 5.367E-06 | 6.600E-06 | 1.700E-05 |
| 14244 | 713 | Y713D | B | 1.1e-05 | 1.2e-05 | 1.2e-05 | 1.167E-05 | 5.774E-07 | 1.100E-05 | 1.200E-05 |
| 14245 | 713 | Y713C | B | 1e-05 | 1.2e-05 | 9.7e-06 | 1.057E-05 | 1.250E-06 | 9.700E-06 | 1.200E-05 |
| 14246 | 713 | Y713E | B | 6.7e-06 | 6.1e-06 | 8.6e-06 | 7.133E-06 | 1.305E-06 | 6.100E-06 | 8.600E-06 |
| 14247 | 713 | Y713Q | B | 8.8e-06 | 5.6e-06 | 9.1e-06 | 7.833E-06 | 1.940E-06 | 5.600E-06 | 9.100E-06 |
| 14248 | 713 | Y713G | B | 1.1e-05 | 1.6e-05 | 3.2e-06 | 1.007E-05 | 6.451E-06 | 3.200E-06 | 1.600E-05 |
| 14249 | 713 | Y713H | B | 4.3e-06 | 9.2e-06 | 9.4e-06 | 7.633E-06 | 2.888E-06 | 4.300E-06 | 9.400E-06 |
| 14250 | 713 | Y713I | B | 1e-05 | 1e-05 | 6.4e-06 | 8.800E-06 | 2.078E-06 | 6.400E-06 | 1.000E-05 |
| 14251 | 713 | Y713L | B | 9.5e-06 | 6.6e-06 | 9.4e-06 | 8.500E-06 | 1.646E-06 | 6.600E-06 | 9.500E-06 |
| 14252 | 713 | Y713K | B | 1.3e-05 | 6.6e-06 | 7.7e-06 | 9.100E-06 | 3.422E-06 | 6.600E-06 | 1.300E-05 |
| 14253 | 713 | Y713M | B | 9.8e-06 | 8.8e-06 | 9.7e-06 | 9.433E-06 | 5.508E-07 | 8.800E-06 | 9.800E-06 |
| 14254 | 713 | Y713F | B | 1.1e-05 | 6.9e-06 | 1e-05 | 9.300E-06 | 2.138E-06 | 6.900E-06 | 1.100E-05 |
| 14255 | 713 | Y713P | B | 7.9e-06 | 5.3e-06 | 7.2e-06 | 6.800E-06 | 1.345E-06 | 5.300E-06 | 7.900E-06 |
| 14256 | 713 | Y713S | B | 6.7e-06 | 4.7e-06 | 5.4e-06 | 5.600E-06 | 1.015E-06 | 4.700E-06 | 6.700E-06 |
| 14257 | 713 | Y713T | B | 9e-06 | 4.9e-06 | 9.9e-06 | 7.933E-06 | 2.665E-06 | 4.900E-06 | 9.900E-06 |
| 14258 | 713 | Y713W | B | 1e-05 | 6.9e-06 | 5.8e-06 | 7.567E-06 | 2.178E-06 | 5.800E-06 | 1.000E-05 |
| 14259 | 713 | Y713Y | B | 1e-05 | 9.5e-06 | 6.4e-06 | 8.633E-06 | 1.950E-06 | 6.400E-06 | 1.000E-05 |
| 14260 | 713 | Y713V | B | 6.1e-06 | 7.1e-06 | 1.2e-05 | 8.400E-06 | 3.158E-06 | 6.100E-06 | 1.200E-05 |
| 14261 | 714 | Y714A | B | 5.5e-06 | 8.5e-06 | 7.2e-06 | 7.067E-06 | 1.504E-06 | 5.500E-06 | 8.500E-06 |
| 14262 | 714 | Y714R | B | 9.2e-06 | 1e-05 | 1e-05 | 9.733E-06 | 4.619E-07 | 9.200E-06 | 1.000E-05 |
| 14263 | 714 | Y714N | B | 4.6e-06 | 1.3e-05 | 4.7e-06 | 7.433E-06 | 4.821E-06 | 4.600E-06 | 1.300E-05 |
| 14264 | 714 | Y714D | B | 1.1e-05 | 7.9e-06 | 5.6e-06 | 8.167E-06 | 2.710E-06 | 5.600E-06 | 1.100E-05 |
| 14265 | 714 | Y714C | B | 5.3e-06 | 9.6e-06 | 1e-05 | 8.300E-06 | 2.606E-06 | 5.300E-06 | 1.000E-05 |
| 14266 | 714 | Y714E | B | 9.5e-06 | 9.5e-06 | 6.1e-06 | 8.367E-06 | 1.963E-06 | 6.100E-06 | 9.500E-06 |
| 14267 | 714 | Y714Q | B | 1.3e-05 | 6.7e-06 | 9.1e-06 | 9.600E-06 | 3.180E-06 | 6.700E-06 | 1.300E-05 |
| 14268 | 714 | Y714G | B | 6.1e-06 | 1.2e-05 | 1.1e-05 | 9.700E-06 | 3.158E-06 | 6.100E-06 | 1.200E-05 |
| 14269 | 714 | Y714H | B | 6.4e-06 | 7.7e-06 | 8.1e-06 | 7.400E-06 | 8.888E-07 | 6.400E-06 | 8.100E-06 |
| 14270 | 714 | Y714I | B | 6.9e-06 | 5.5e-06 | 6e-06 | 6.133E-06 | 7.095E-07 | 5.500E-06 | 6.900E-06 |
| 14271 | 714 | Y714L | B | 6.8e-06 | 5.6e-06 | 8.1e-06 | 6.833E-06 | 1.250E-06 | 5.600E-06 | 8.100E-06 |
| 14272 | 714 | Y714K | B | 6.9e-06 | 7.4e-06 | 5.6e-06 | 6.633E-06 | 9.292E-07 | 5.600E-06 | 7.400E-06 |
| 14273 | 714 | Y714M | B | 6.7e-06 | 8e-06 | 9e-06 | 7.900E-06 | 1.153E-06 | 6.700E-06 | 9.000E-06 |
| 14274 | 714 | Y714F | B | 6.4e-06 | 7.4e-06 | 4.7e-06 | 6.167E-06 | 1.365E-06 | 4.700E-06 | 7.400E-06 |
| 14275 | 714 | Y714P | B | 7e-06 | 9.1e-06 | 8.7e-06 | 8.267E-06 | 1.115E-06 | 7.000E-06 | 9.100E-06 |
| 14276 | 714 | Y714S | B | 8.3e-06 | 8.9e-06 | 9.4e-06 | 8.867E-06 | 5.508E-07 | 8.300E-06 | 9.400E-06 |
| 14277 | 714 | Y714T | B | 8.8e-06 | 5.7e-06 | 6.9e-06 | 7.133E-06 | 1.563E-06 | 5.700E-06 | 8.800E-06 |
| 14278 | 714 | Y714W | B | 1.3e-05 | 8.8e-06 | 6.2e-06 | 9.333E-06 | 3.431E-06 | 6.200E-06 | 1.300E-05 |
| 14279 | 714 | Y714Y | B | 6.7e-06 | 7.3e-06 | 6.7e-06 | 6.900E-06 | 3.464E-07 | 6.700E-06 | 7.300E-06 |

| | | | | | | | | | | |
|---|---|---|---|---|---|---|---|---|---|---|
| 14280 | 714 | Y714V | B | 4.5e-06 | 6.6e-06 | 9.5e-06 | 6.867E-06 | 2.511E-06 | 4.500E-06 | 9.500E-06 |
| 14281 | 715 | I715A | B | 8.9e-06 | 3.5e-06 | 6.9e-06 | 6.433E-06 | 2.730E-06 | 3.500E-06 | 8.900E-06 |
| 14282 | 715 | I715R | B | 9e-06 | 9.8e-06 | 5.5e-06 | 8.100E-06 | 2.287E-06 | 5.500E-06 | 9.800E-06 |
| 14283 | 715 | I715N | B | 6.9e-06 | 8.1e-06 | 9.8e-06 | 8.267E-06 | 1.457E-06 | 6.900E-06 | 9.800E-06 |
| 14284 | 715 | I715D | B | 6.8e-06 | 5.8e-06 | 7e-06 | 6.533E-06 | 6.429E-07 | 5.800E-06 | 7.000E-06 |
| 14285 | 715 | I715C | B | 7.6e-06 | 1e-05 | 8.1e-06 | 8.567E-06 | 1.266E-06 | 7.600E-06 | 1.000E-05 |
| 14286 | 715 | I715E | B | 9.6e-06 | 5.8e-06 | 8.6e-06 | 8.000E-06 | 1.970E-06 | 5.800E-06 | 9.600E-06 |
| 14287 | 715 | I715Q | B | 9.1e-06 | 8.7e-06 | 1e-05 | 9.267E-06 | 6.658E-07 | 8.700E-06 | 1.000E-05 |
| 14288 | 715 | I715G | B | 5.6e-06 | 8.7e-06 | 9.7e-06 | 8.000E-06 | 2.138E-06 | 5.600E-06 | 9.700E-06 |
| 14289 | 715 | I715H | B | 6.4e-06 | 8.5e-06 | 1.2e-05 | 8.967E-06 | 2.829E-06 | 6.400E-06 | 1.200E-05 |
| 14290 | 715 | I715I | B | 6.8e-06 | 6.3e-06 | 1.2e-05 | 8.367E-06 | 3.156E-06 | 6.300E-06 | 1.200E-05 |
| 14291 | 715 | I715L | B | 6.7e-06 | 4.9e-06 | 8.4e-06 | 6.667E-06 | 1.750E-06 | 4.900E-06 | 8.400E-06 |
| 14292 | 715 | I715K | B | 1.9e-05 | 8.9e-06 | 1.3e-05 | 1.363E-05 | 5.080E-06 | 8.900E-06 | 1.900E-05 |
| 14293 | 715 | I715M | B | 8.7e-06 | 1.1e-05 | 9.9e-06 | 9.867E-06 | 1.150E-06 | 8.700E-06 | 1.100E-05 |
| 14294 | 715 | I715F | B | 6.4e-06 | 7.6e-06 | 7.2e-06 | 7.067E-06 | 6.110E-07 | 6.400E-06 | 7.600E-06 |
| 14295 | 715 | I715P | B | 5.4e-06 | 8.9e-06 | 1e-05 | 8.100E-06 | 2.402E-06 | 5.400E-06 | 1.000E-05 |
| 14296 | 715 | I715S | B | 8.8e-06 | 6.1e-06 | 5.6e-06 | 6.833E-06 | 1.721E-06 | 5.600E-06 | 8.800E-06 |
| 14297 | 715 | I715T | B | 9.2e-06 | 6.6e-06 | 5.4e-06 | 7.067E-06 | 1.943E-06 | 5.400E-06 | 9.200E-06 |
| 14298 | 715 | I715W | B | 9.2e-06 | 1.2e-05 | 6.4e-06 | 9.200E-06 | 2.800E-06 | 6.400E-06 | 1.200E-05 |
| 14299 | 715 | I715Y | B | 6.5e-06 | 5.7e-06 | 8.8e-06 | 7.000E-06 | 1.609E-06 | 5.700E-06 | 8.800E-06 |
| 14300 | 715 | I715V | B | 5.3e-06 | 7.3e-06 | 4.3e-06 | 5.633E-06 | 1.528E-06 | 4.300E-06 | 7.300E-06 |
| 14301 | 716 | Y716A | B | 7.8e-06 | 1.7e-05 | 4.4e-06 | 9.733E-06 | 6.519E-06 | 4.400E-06 | 1.700E-05 |
| 14302 | 716 | Y716R | B | 9.4e-06 | 5.4e-06 | 6.2e-06 | 7.000E-06 | 2.117E-06 | 5.400E-06 | 9.400E-06 |
| 14303 | 716 | Y716N | B | 8e-06 | 8.8e-06 | 6.7e-06 | 7.833E-06 | 1.060E-06 | 6.700E-06 | 8.800E-06 |
| 14304 | 716 | Y716D | B | 5e-06 | 7.1e-06 | 1.2e-05 | 8.033E-06 | 3.592E-06 | 5.000E-06 | 1.200E-05 |
| 14305 | 716 | Y716C | B | 7.5e-06 | 9.4e-06 | 1.1e-05 | 9.300E-06 | 1.752E-06 | 7.500E-06 | 1.100E-05 |
| 14306 | 716 | Y716E | B | 6.6e-06 | 9.7e-06 | 1e-05 | 8.767E-06 | 1.882E-06 | 6.600E-06 | 1.000E-05 |
| 14307 | 716 | Y716Q | B | 5.5e-06 | 6.7e-06 | 6.8e-06 | 6.333E-06 | 7.234E-07 | 5.500E-06 | 6.800E-06 |
| 14308 | 716 | Y716G | B | 6.6e-06 | 6.5e-06 | 7.9e-06 | 7.000E-06 | 7.810E-07 | 6.500E-06 | 7.900E-06 |
| 14309 | 716 | Y716H | B | 7.8e-06 | 8.2e-06 | 1.8e-05 | 1.133E-05 | 5.777E-06 | 7.800E-06 | 1.800E-05 |
| 14310 | 716 | Y716I | B | 9.4e-06 | 5.2e-06 | 9.8e-06 | 8.133E-06 | 2.548E-06 | 5.200E-06 | 9.800E-06 |
| 14311 | 716 | Y716L | B | 9.2e-06 | 5.2e-06 | 1.5e-05 | 9.800E-06 | 4.927E-06 | 5.200E-06 | 1.500E-05 |
| 14312 | 716 | Y716K | B | 8.5e-06 | 1.3e-05 | 1.5e-05 | 1.217E-05 | 3.329E-06 | 8.500E-06 | 1.500E-05 |
| 14313 | 716 | Y716M | B | 9.3e-06 | 6.6e-06 | 6.9e-06 | 7.600E-06 | 1.480E-06 | 6.600E-06 | 9.300E-06 |
| 14314 | 716 | Y716F | B | 5.1e-06 | 5.1e-06 | 9.3e-06 | 6.500E-06 | 2.425E-06 | 5.100E-06 | 9.300E-06 |
| 14315 | 716 | Y716P | B | 3.5e-06 | 7.2e-06 | 4.9e-06 | 5.200E-06 | 1.868E-06 | 3.500E-06 | 7.200E-06 |
| 14316 | 716 | Y716S | B | 1.1e-05 | 9.3e-06 | 6.8e-06 | 9.033E-06 | 2.113E-06 | 6.800E-06 | 1.100E-05 |
| 14317 | 716 | Y716T | B | 8.4e-06 | 7.1e-06 | 5.3e-06 | 6.933E-06 | 1.557E-06 | 5.300E-06 | 8.400E-06 |
| 14318 | 716 | Y716W | B | 5.1e-06 | 7.2e-06 | 7.2e-06 | 6.500E-06 | 1.212E-06 | 5.100E-06 | 7.200E-06 |
| 14319 | 716 | Y716Y | B | 4.8e-06 | 8.7e-06 | 9.7e-06 | 7.733E-06 | 2.589E-06 | 4.800E-06 | 9.700E-06 |
| 14320 | 716 | Y716V | B | 6.4e-06 | 4.7e-06 | 9.9e-06 | 7.000E-06 | 2.651E-06 | 4.700E-06 | 9.900E-06 |
| 14321 | 717 | P717A | B | 6.4e-06 | 5.2e-06 | 1.5e-05 | 8.867E-06 | 5.345E-06 | 5.200E-06 | 1.500E-05 |
| 14322 | 717 | P717R | B | 1e-05 | 4.6e-06 | 4.4e-06 | 6.333E-06 | 3.177E-06 | 4.400E-06 | 1.000E-05 |
| 14323 | 717 | P717N | B | 4.9e-06 | 7.2e-06 | 1.4e-05 | 8.700E-06 | 4.732E-06 | 4.900E-06 | 1.400E-05 |
| 14324 | 717 | P717D | B | 8.8e-06 | 8.9e-06 | 5.6e-06 | 7.767E-06 | 1.877E-06 | 5.600E-06 | 8.900E-06 |
| 14325 | 717 | P717C | B | 7.3e-06 | 7.4e-06 | 7.1e-06 | 7.267E-06 | 1.528E-07 | 7.100E-06 | 7.400E-06 |
| 14326 | 717 | P717E | B | 9.3e-06 | 5.6e-06 | 6.9e-06 | 7.267E-06 | 1.877E-06 | 5.600E-06 | 9.300E-06 |
| 14327 | 717 | P717Q | B | 6.3e-06 | 6.9e-06 | 6.9e-06 | 6.700E-06 | 3.464E-07 | 6.300E-06 | 6.900E-06 |
| 14328 | 717 | P717G | B | 1e-05 | 5.3e-06 | 6.6e-06 | 7.300E-06 | 2.427E-06 | 5.300E-06 | 1.000E-05 |
| 14329 | 717 | P717H | B | 5.7e-06 | 8.7e-06 | 5.7e-06 | 6.700E-06 | 1.732E-06 | 5.700E-06 | 8.700E-06 |
| 14330 | 717 | P717I | B | 1.7e-05 | 8.9e-06 | 9.5e-06 | 1.180E-05 | 4.513E-06 | 8.900E-06 | 1.700E-05 |
| 14331 | 717 | P717L | B | 9.4e-06 | 6.5e-06 | 7.5e-06 | 7.800E-06 | 1.473E-06 | 6.500E-06 | 9.400E-06 |
| 14332 | 717 | P717K | B | 1e-05 | 5.5e-06 | 7.3e-06 | 7.600E-06 | 2.265E-06 | 5.500E-06 | 1.000E-05 |
| 14333 | 717 | P717M | B | 5e-06 | 1.4e-05 | 9.7e-06 | 9.567E-06 | 4.501E-06 | 5.000E-06 | 1.400E-05 |
| 14334 | 717 | P717F | B | 4.9e-06 | 6.6e-06 | 6.2e-06 | 5.900E-06 | 8.888E-07 | 4.900E-06 | 6.600E-06 |
| 14335 | 717 | P717P | B | 8.9e-06 | 6.7e-06 | 4.9e-06 | 6.833E-06 | 2.003E-06 | 4.900E-06 | 8.900E-06 |
| 14336 | 717 | P717S | B | 5.5e-06 | 7.1e-06 | 9.5e-06 | 7.367E-06 | 2.013E-06 | 5.500E-06 | 9.500E-06 |
| 14337 | 717 | P717T | B | 4.2e-06 | 6.8e-06 | 1.3e-05 | 8.000E-06 | 4.521E-06 | 4.200E-06 | 1.300E-05 |
| 14338 | 717 | P717W | B | 1.3e-05 | 9.4e-06 | 8.6e-06 | 1.033E-05 | 2.344E-06 | 8.600E-06 | 1.300E-05 |
| 14339 | 717 | P717Y | B | 6.5e-06 | 7e-06 | 9.9e-06 | 7.800E-06 | 1.836E-06 | 6.500E-06 | 9.900E-06 |
| 14340 | 717 | P717V | B | 8.3e-06 | 4.2e-06 | 6.5e-06 | 6.333E-06 | 2.055E-06 | 4.200E-06 | 8.300E-06 |
| 14341 | 718 | Y718A | B | 5.8e-06 | 8.2e-06 | 5.7e-06 | 6.567E-06 | 1.415E-06 | 5.700E-06 | 8.200E-06 |
| 14342 | 718 | Y718R | B | 8.7e-06 | 9e-06 | 8.9e-06 | 8.867E-06 | 1.528E-07 | 8.700E-06 | 9.000E-06 |
| 14343 | 718 | Y718N | B | 7.6e-06 | 5.4e-06 | 7.4e-06 | 6.800E-06 | 1.217E-06 | 5.400E-06 | 7.600E-06 |
| 14344 | 718 | Y718D | B | 1.5e-05 | 7e-06 | 8.2e-06 | 1.007E-05 | 4.314E-06 | 7.000E-06 | 1.500E-05 |
| 14345 | 718 | Y718C | B | 1.1e-05 | 1.3e-05 | 9.1e-06 | 1.103E-05 | 1.950E-06 | 9.100E-06 | 1.300E-05 |
| 14346 | 718 | Y718E | B | 6e-06 | 7.4e-06 | 1.3e-05 | 8.800E-06 | 3.704E-06 | 6.000E-06 | 1.300E-05 |
| 14347 | 718 | Y718Q | B | 8.3e-06 | 7.3e-06 | 6.3e-06 | 7.300E-06 | 1.000E-06 | 6.300E-06 | 8.300E-06 |
| 14348 | 718 | Y718G | B | 5.5e-06 | 7.5e-06 | 1.3e-05 | 8.667E-06 | 3.884E-06 | 5.500E-06 | 1.300E-05 |
| 14349 | 718 | Y718H | B | 6.9e-06 | 9.7e-06 | 9e-06 | 8.533E-06 | 1.457E-06 | 6.900E-06 | 9.700E-06 |
| 14350 | 718 | Y718I | B | 8.4e-06 | 4.8e-06 | 9.8e-06 | 7.667E-06 | 2.579E-06 | 4.800E-06 | 9.800E-06 |
| 14351 | 718 | Y718L | B | 7.8e-06 | 3.6e-06 | 9.7e-06 | 7.033E-06 | 3.121E-06 | 3.600E-06 | 9.700E-06 |
| 14352 | 718 | Y718K | B | 1.1e-05 | 1.2e-05 | 6.3e-06 | 9.767E-06 | 3.044E-06 | 6.300E-06 | 1.200E-05 |
| 14353 | 718 | Y718M | B | 4.7e-06 | 1.2e-05 | 7.8e-06 | 8.167E-06 | 3.664E-06 | 4.700E-06 | 1.200E-05 |
| 14354 | 718 | Y718F | B | 1e-05 | 1.3e-05 | 1.2e-05 | 1.167E-05 | 1.528E-06 | 1.000E-05 | 1.300E-05 |
| 14355 | 718 | Y718P | B | 1e-05 | 7.3e-06 | 9.1e-06 | 8.800E-06 | 1.375E-06 | 7.300E-06 | 1.000E-05 |
| 14356 | 718 | Y718S | B | 6.7e-06 | 1e-05 | 6.4e-06 | 7.700E-06 | 1.997E-06 | 6.400E-06 | 1.000E-05 |
| 14357 | 718 | Y718T | B | 5e-06 | 8.7e-06 | 6e-06 | 6.567E-06 | 1.914E-06 | 5.000E-06 | 8.700E-06 |
| 14358 | 718 | Y718W | B | 4.4e-06 | 7.1e-06 | 1.3e-05 | 8.167E-06 | 4.398E-06 | 4.400E-06 | 1.300E-05 |
| 14359 | 718 | Y718Y | B | 5.3e-06 | 5.3e-06 | 1.3e-05 | 7.867E-06 | 4.446E-06 | 5.300E-06 | 1.300E-05 |
| 14360 | 718 | Y718V | B | 9.4e-06 | 6.6e-06 | 7.5e-06 | 7.833E-06 | 1.429E-06 | 6.600E-06 | 9.400E-06 |
| 14361 | 719 | T719A | B | 9.7e-06 | 9.8e-06 | 9e-06 | 9.500E-06 | 4.359E-07 | 9.000E-06 | 9.800E-06 |
| 14362 | 719 | T719R | B | 8.3e-06 | 6.4e-06 | 1.4e-05 | 9.567E-06 | 3.955E-06 | 6.400E-06 | 1.400E-05 |
| 14363 | 719 | T719N | B | 1.8e-05 | 6.6e-06 | 8.5e-06 | 1.103E-05 | 6.108E-06 | 6.600E-06 | 1.800E-05 |
| 14364 | 719 | T719D | B | 6.4e-06 | 7.5e-06 | 7.4e-06 | 7.100E-06 | 6.083E-07 | 6.400E-06 | 7.500E-06 |

| | | | | | | | | | | |
|---|---|---|---|---|---|---|---|---|---|---|
| 14365 | 719 | T719C | B | 7.4e-06 | 9.8e-06 | 1.3e-05 | 1.007E-05 | 2.810E-06 | 7.400E-06 | 1.300E-05 |
| 14366 | 719 | T719E | B | 1.4e-05 | 7.6e-06 | 1.3e-05 | 1.153E-05 | 3.443E-06 | 7.600E-06 | 1.400E-05 |
| 14367 | 719 | T719Q | B | 7.8e-06 | 3.1e-06 | 1.1e-05 | 7.300E-06 | 3.974E-06 | 3.100E-06 | 1.100E-05 |
| 14368 | 719 | T719G | B | 5.1e-06 | 7e-06 | 8.4e-06 | 6.833E-06 | 1.656E-06 | 5.100E-06 | 8.400E-06 |
| 14369 | 719 | T719H | B | 9.3e-06 | 1.2e-05 | 8.2e-06 | 9.833E-06 | 1.955E-06 | 8.200E-06 | 1.200E-05 |
| 14370 | 719 | T719I | B | 8.9e-06 | 4e-06 | 7e-06 | 6.633E-06 | 2.470E-06 | 4.000E-06 | 8.900E-06 |
| 14371 | 719 | T719L | B | 1.1e-05 | 6.9e-06 | 9.9e-06 | 9.267E-06 | 2.122E-06 | 6.900E-06 | 1.100E-05 |
| 14372 | 719 | T719K | B | 6.1e-06 | 1.1e-05 | 7.2e-06 | 8.100E-06 | 2.571E-06 | 6.100E-06 | 1.100E-05 |
| 14373 | 719 | T719M | B | 1e-05 | 5.7e-06 | 6e-06 | 7.233E-06 | 2.401E-06 | 5.700E-06 | 1.000E-05 |
| 14374 | 719 | T719F | B | 8.7e-06 | 5.7e-06 | 4.5e-06 | 6.300E-06 | 2.163E-06 | 4.500E-06 | 8.700E-06 |
| 14375 | 719 | T719P | B | 7.3e-06 | 1.4e-05 | 1.4e-05 | 1.177E-05 | 3.868E-06 | 7.300E-06 | 1.400E-05 |
| 14376 | 719 | T719S | B | 6.5e-06 | 1.2e-05 | 7.6e-06 | 8.700E-06 | 2.910E-06 | 6.500E-06 | 1.200E-05 |
| 14377 | 719 | T719T | B | 1e-05 | 7.2e-06 | 6.7e-06 | 7.967E-06 | 1.779E-06 | 6.700E-06 | 1.000E-05 |
| 14378 | 719 | T719W | B | 6.6e-06 | 1.3e-05 | 4.6e-06 | 8.067E-06 | 4.388E-06 | 4.600E-06 | 1.300E-05 |
| 14379 | 719 | T719Y | B | 5.9e-06 | 1.3e-05 | 1.4e-05 | 1.097E-05 | 4.416E-06 | 5.900E-06 | 1.400E-05 |
| 14380 | 719 | T719V | B | 1e-05 | 6.8e-06 | 9.4e-06 | 8.733E-06 | 1.701E-06 | 6.800E-06 | 1.000E-05 |
| 14381 | 720 | F720A | B | 1.3e-05 | 6.9e-06 | 5.5e-06 | 8.467E-06 | 3.988E-06 | 5.500E-06 | 1.300E-05 |
| 14382 | 720 | F720R | B | 8.7e-06 | 5.6e-06 | 6.5e-06 | 6.933E-06 | 1.595E-06 | 5.600E-06 | 8.700E-06 |
| 14383 | 720 | F720N | B | 5.2e-06 | 8.1e-06 | 1.3e-05 | 8.767E-06 | 3.943E-06 | 5.200E-06 | 1.300E-05 |
| 14384 | 720 | F720D | B | 8.2e-06 | 5.8e-06 | 8.7e-06 | 7.567E-06 | 1.550E-06 | 5.800E-06 | 8.700E-06 |
| 14385 | 720 | F720C | B | 5e-06 | 8.7e-06 | 7.5e-06 | 7.067E-06 | 1.888E-06 | 5.000E-06 | 8.700E-06 |
| 14386 | 720 | F720E | B | 9.9e-06 | 6.8e-06 | 9.1e-06 | 8.600E-06 | 1.609E-06 | 6.800E-06 | 9.900E-06 |
| 14387 | 720 | F720Q | B | 1.3e-05 | 1.4e-05 | 4.3e-06 | 1.043E-05 | 5.335E-06 | 4.300E-06 | 1.400E-05 |
| 14388 | 720 | F720G | B | 6.6e-06 | 1.7e-05 | 1.2e-05 | 1.187E-05 | 5.201E-06 | 6.600E-06 | 1.700E-05 |
| 14389 | 720 | F720H | B | 1.2e-05 | 9.3e-06 | 6.1e-06 | 9.133E-06 | 2.954E-06 | 6.100E-06 | 1.200E-05 |
| 14390 | 720 | F720I | B | 8.9e-06 | 8.6e-06 | 4.6e-06 | 7.367E-06 | 2.401E-06 | 4.600E-06 | 8.900E-06 |
| 14391 | 720 | F720L | B | 9.9e-06 | 1.2e-05 | 8.9e-06 | 1.027E-05 | 1.582E-06 | 8.900E-06 | 1.200E-05 |
| 14392 | 720 | F720K | B | 9.8e-06 | 5.7e-06 | 5.1e-06 | 6.867E-06 | 2.558E-06 | 5.100E-06 | 9.800E-06 |
| 14393 | 720 | F720M | B | 4.1e-06 | 7.7e-06 | 8.3e-06 | 6.700E-06 | 2.272E-06 | 4.100E-06 | 8.300E-06 |
| 14394 | 720 | F720F | B | 1.1e-05 | 8e-06 | 8.6e-06 | 9.200E-06 | 1.587E-06 | 8.000E-06 | 1.100E-05 |
| 14395 | 720 | F720P | B | 5.4e-06 | 6.8e-06 | 9.1e-06 | 7.100E-06 | 1.868E-06 | 5.400E-06 | 9.100E-06 |
| 14396 | 720 | F720S | B | 9e-06 | 7.5e-06 | 8.8e-06 | 8.433E-06 | 8.145E-07 | 7.500E-06 | 9.000E-06 |
| 14397 | 720 | F720T | B | 9.6e-06 | 9.5e-06 | 9e-06 | 9.367E-06 | 3.215E-07 | 9.000E-06 | 9.600E-06 |
| 14398 | 720 | F720W | B | 1.3e-05 | 5.5e-06 | 8.9e-06 | 9.133E-06 | 3.755E-06 | 5.500E-06 | 1.300E-05 |
| 14399 | 720 | F720Y | B | 8.4e-06 | 6.4e-06 | 5.4e-06 | 6.733E-06 | 1.528E-06 | 5.400E-06 | 8.400E-06 |
| 14400 | 720 | F720V | B | 5e-06 | 5.8e-06 | 5.8e-06 | 5.533E-06 | 4.619E-07 | 5.000E-06 | 5.800E-06 |
| 14401 | 721 | G721A | B | 1.1e-05 | 5.6e-06 | 7.9e-06 | 8.167E-06 | 2.710E-06 | 5.600E-06 | 1.100E-05 |
| 14402 | 721 | G721R | B | 6e-06 | 7.2e-06 | 6.9e-06 | 6.700E-06 | 6.245E-07 | 6.000E-06 | 7.200E-06 |
| 14403 | 721 | G721N | B | 8.5e-06 | 7.3e-06 | 8.1e-06 | 7.967E-06 | 6.110E-07 | 7.300E-06 | 8.500E-06 |
| 14404 | 721 | G721D | B | 1.3e-05 | 4.5e-06 | 5.5e-06 | 7.667E-06 | 4.646E-06 | 4.500E-06 | 1.300E-05 |
| 14405 | 721 | G721C | B | 8.1e-06 | 8.2e-06 | 9.6e-06 | 8.633E-06 | 8.386E-07 | 8.100E-06 | 9.600E-06 |
| 14406 | 721 | G721E | B | 6.4e-06 | 4.6e-06 | 6.8e-06 | 5.933E-06 | 1.172E-06 | 4.600E-06 | 6.800E-06 |
| 14407 | 721 | G721Q | B | 9.9e-06 | 9.7e-06 | 6e-06 | 8.533E-06 | 2.196E-06 | 6.000E-06 | 9.900E-06 |
| 14408 | 721 | G721G | B | 7e-06 | 1.3e-05 | 7.6e-06 | 9.200E-06 | 3.305E-06 | 7.000E-06 | 1.300E-05 |
| 14409 | 721 | G721H | B | 9.5e-06 | 1.1e-05 | 5.3e-06 | 8.600E-06 | 2.955E-06 | 5.300E-06 | 1.100E-05 |
| 14410 | 721 | G721I | B | 8.5e-06 | 5.5e-06 | 9.1e-06 | 7.700E-06 | 1.929E-06 | 5.500E-06 | 9.100E-06 |
| 14411 | 721 | G721L | B | 9.3e-06 | 6.3e-06 | 7.3e-06 | 7.633E-06 | 1.528E-06 | 6.300E-06 | 9.300E-06 |
| 14412 | 721 | G721K | B | 7.1e-06 | 5e-06 | 1.3e-05 | 8.367E-06 | 4.148E-06 | 5.000E-06 | 1.300E-05 |
| 14413 | 721 | G721M | B | 3.8e-06 | 9e-06 | 1.1e-05 | 7.933E-06 | 3.717E-06 | 3.800E-06 | 1.100E-05 |
| 14414 | 721 | G721F | B | 6.1e-06 | 1e-05 | 3.4e-06 | 6.500E-06 | 3.318E-06 | 3.400E-06 | 1.000E-05 |
| 14415 | 721 | G721P | B | 7.3e-06 | 1e-05 | 5.5e-06 | 7.600E-06 | 2.265E-06 | 5.500E-06 | 1.000E-05 |
| 14416 | 721 | G721S | B | 1.3e-05 | 4.5e-06 | 5e-06 | 7.500E-06 | 4.770E-06 | 4.500E-06 | 1.300E-05 |
| 14417 | 721 | G721T | B | 6.4e-06 | 6.5e-06 | 4.5e-06 | 5.800E-06 | 1.127E-06 | 4.500E-06 | 6.500E-06 |
| 14418 | 721 | G721W | B | 6.2e-06 | 6.4e-06 | 6.1e-06 | 6.233E-06 | 1.528E-07 | 6.100E-06 | 6.400E-06 |
| 14419 | 721 | G721Y | B | 1.1e-05 | 6e-06 | 7.8e-06 | 8.267E-06 | 2.532E-06 | 6.000E-06 | 1.100E-05 |
| 14420 | 721 | G721V | B | 8.5e-06 | 1e-05 | 1e-05 | 9.500E-06 | 8.660E-07 | 8.500E-06 | 1.000E-05 |
| 14421 | 722 | Q722A | B | 9.8e-06 | 6.9e-06 | 1.4e-05 | 1.023E-05 | 3.570E-06 | 6.900E-06 | 1.400E-05 |
| 14422 | 722 | Q722R | B | 7.6e-06 | 9.7e-06 | 8.4e-06 | 8.567E-06 | 1.060E-06 | 7.600E-06 | 9.700E-06 |
| 14423 | 722 | Q722N | B | 1.1e-05 | 7.6e-06 | 5.8e-06 | 8.133E-06 | 2.641E-06 | 5.800E-06 | 1.100E-05 |
| 14424 | 722 | Q722D | B | 1e-05 | 1.1e-05 | 8.8e-06 | 9.933E-06 | 1.102E-06 | 8.800E-06 | 1.100E-05 |
| 14425 | 722 | Q722C | B | 9.3e-06 | 1e-05 | 1.3e-05 | 1.077E-05 | 1.966E-06 | 9.300E-06 | 1.300E-05 |
| 14426 | 722 | Q722E | B | 9.6e-06 | 7.5e-06 | 6.1e-06 | 7.733E-06 | 1.762E-06 | 6.100E-06 | 9.600E-06 |
| 14427 | 722 | Q722Q | B | 1.1e-05 | 1e-05 | 7.7e-06 | 9.567E-06 | 1.692E-06 | 7.700E-06 | 1.100E-05 |
| 14428 | 722 | Q722G | B | 8e-06 | 8.4e-06 | 8.4e-06 | 8.267E-06 | 2.309E-07 | 8.000E-06 | 8.400E-06 |
| 14429 | 722 | Q722H | B | 8.5e-06 | 7.5e-06 | 1.1e-05 | 9.000E-06 | 1.803E-06 | 7.500E-06 | 1.100E-05 |
| 14430 | 722 | Q722I | B | 6.4e-06 | 6.2e-06 | 6.1e-06 | 6.233E-06 | 1.528E-07 | 6.100E-06 | 6.400E-06 |
| 14431 | 722 | Q722L | B | 7.8e-06 | 8.2e-06 | 6.3e-06 | 7.433E-06 | 1.002E-06 | 6.300E-06 | 8.200E-06 |
| 14432 | 722 | Q722K | B | 8.1e-06 | 6.7e-06 | 6.9e-06 | 7.233E-06 | 7.572E-07 | 6.700E-06 | 8.100E-06 |
| 14433 | 722 | Q722M | B | 4.7e-06 | 9.5e-06 | 8.5e-06 | 7.567E-06 | 2.532E-06 | 4.700E-06 | 9.500E-06 |
| 14434 | 722 | Q722F | B | 5.9e-06 | 6.2e-06 | 8.3e-06 | 6.800E-06 | 1.308E-06 | 5.900E-06 | 8.300E-06 |
| 14435 | 722 | Q722P | B | 8.4e-06 | 1.3e-05 | 5.3e-06 | 8.900E-06 | 3.874E-06 | 5.300E-06 | 1.300E-05 |
| 14436 | 722 | Q722S | B | 8.9e-06 | 9.7e-06 | 6.5e-06 | 8.367E-06 | 1.665E-06 | 6.500E-06 | 9.700E-06 |
| 14437 | 722 | Q722T | B | 9.2e-06 | 5.8e-06 | 8.5e-06 | 7.833E-06 | 1.795E-06 | 5.800E-06 | 9.200E-06 |
| 14438 | 722 | Q722W | B | 4.7e-06 | 7.8e-06 | 1.4e-05 | 8.833E-06 | 4.735E-06 | 4.700E-06 | 1.400E-05 |
| 14439 | 722 | Q722Y | B | 8.5e-06 | 4.6e-06 | 1.8e-05 | 1.037E-05 | 6.892E-06 | 4.600E-06 | 1.800E-05 |
| 14440 | 722 | Q722V | B | 9.7e-06 | 7.3e-06 | 1e-05 | 9.000E-06 | 1.480E-06 | 7.300E-06 | 1.000E-05 |
| 14441 | 723 | G723A | B | 4.3e-06 | 8.1e-06 | 7.4e-06 | 6.600E-06 | 2.022E-06 | 4.300E-06 | 8.100E-06 |
| 14442 | 723 | G723R | B | 4.9e-06 | 7.4e-06 | 6.3e-06 | 6.200E-06 | 1.253E-06 | 4.900E-06 | 7.400E-06 |
| 14443 | 723 | G723N | B | 4.6e-06 | 8.3e-06 | 6e-06 | 6.300E-06 | 1.868E-06 | 4.600E-06 | 8.300E-06 |
| 14444 | 723 | G723D | B | 4.9e-06 | 1.1e-05 | 1.4e-05 | 9.967E-06 | 4.637E-06 | 4.900E-06 | 1.400E-05 |
| 14445 | 723 | G723C | B | 9.4e-06 | 8.8e-06 | 7.3e-06 | 8.500E-06 | 1.082E-06 | 7.300E-06 | 9.400E-06 |
| 14446 | 723 | G723E | B | 1.2e-05 | 8e-06 | 6.6e-06 | 8.867E-06 | 2.802E-06 | 6.600E-06 | 1.200E-05 |
| 14447 | 723 | G723Q | B | 6.4e-06 | 1.4e-05 | 6.7e-06 | 9.033E-06 | 4.304E-06 | 6.400E-06 | 1.400E-05 |
| 14448 | 723 | G723G | B | 5e-06 | 9e-06 | 9.8e-06 | 7.933E-06 | 2.572E-06 | 5.000E-06 | 9.800E-06 |
| 14449 | 723 | G723H | B | 9.2e-06 | 8.1e-06 | 6.6e-06 | 7.967E-06 | 1.305E-06 | 6.600E-06 | 9.200E-06 |

| | | | | | | | | | | |
|---|---|---|---|---|---|---|---|---|---|---|
| 14450 | 723 | G723I | B | 7.2e-06 | 9.6e-06 | 1.3e-05 | 9.933E-06 | 2.914E-06 | 7.200E-06 | 1.300E-05 |
| 14451 | 723 | G723L | B | 7.3e-06 | 5.8e-06 | 6.2e-06 | 6.433E-06 | 7.767E-07 | 5.800E-06 | 7.300E-06 |
| 14452 | 723 | G723K | B | 9.7e-06 | 1.3e-05 | 8e-06 | 1.023E-05 | 2.542E-06 | 8.000E-06 | 1.300E-05 |
| 14453 | 723 | G723M | B | 8e-06 | 5.9e-06 | 5e-06 | 6.300E-06 | 1.539E-06 | 5.000E-06 | 8.000E-06 |
| 14454 | 723 | G723F | B | 8.3e-06 | 7.1e-06 | 1.2e-05 | 9.133E-06 | 2.554E-06 | 7.100E-06 | 1.200E-05 |
| 14455 | 723 | G723P | B | 7.1e-06 | 7.8e-06 | 6.8e-06 | 7.233E-06 | 5.132E-07 | 6.800E-06 | 7.800E-06 |
| 14456 | 723 | G723S | B | 1.3e-05 | 9.9e-06 | 6.3e-06 | 9.733E-06 | 3.353E-06 | 6.300E-06 | 1.300E-05 |
| 14457 | 723 | G723T | B | 9.4e-06 | 9.3e-06 | 1.1e-05 | 9.900E-06 | 9.539E-07 | 9.300E-06 | 1.100E-05 |
| 14458 | 723 | G723W | B | 9e-06 | 9.9e-06 | 8.4e-06 | 9.100E-06 | 7.550E-07 | 8.400E-06 | 9.900E-06 |
| 14459 | 723 | G723Y | B | 8.5e-06 | 5.6e-06 | 9.5e-06 | 7.867E-06 | 2.026E-06 | 5.600E-06 | 9.500E-06 |
| 14460 | 723 | G723V | B | 8.5e-06 | 6.3e-06 | 6.9e-06 | 7.233E-06 | 1.137E-06 | 6.300E-06 | 8.500E-06 |
| 14461 | 724 | T724A | B | 1.1e-05 | 7.1e-06 | 6.4e-06 | 8.167E-06 | 2.479E-06 | 6.400E-06 | 1.100E-05 |
| 14462 | 724 | T724R | B | 4.9e-06 | 8.5e-06 | 1.2e-05 | 8.467E-06 | 3.550E-06 | 4.900E-06 | 1.200E-05 |
| 14463 | 724 | T724N | B | 4.7e-06 | 7.2e-06 | 9.4e-06 | 7.100E-06 | 2.352E-06 | 4.700E-06 | 9.400E-06 |
| 14464 | 724 | T724D | B | 9.2e-06 | 8.8e-06 | 7.3e-06 | 8.433E-06 | 1.002E-06 | 7.300E-06 | 9.200E-06 |
| 14465 | 724 | T724C | B | 6.6e-06 | 6.9e-06 | 6.8e-06 | 6.767E-06 | 1.528E-07 | 6.600E-06 | 6.900E-06 |
| 14466 | 724 | T724E | B | 6.9e-06 | 6.8e-06 | 6.5e-06 | 6.733E-06 | 2.082E-07 | 6.500E-06 | 6.900E-06 |
| 14467 | 724 | T724Q | B | 6.5e-06 | 8.7e-06 | 6.4e-06 | 7.200E-06 | 1.300E-06 | 6.400E-06 | 8.700E-06 |
| 14468 | 724 | T724G | B | 7.2e-06 | 1.4e-05 | 3.2e-06 | 8.133E-06 | 5.460E-06 | 3.200E-06 | 1.400E-05 |
| 14469 | 724 | T724H | B | 5.7e-06 | 1e-05 | 9.9e-06 | 8.533E-06 | 2.454E-06 | 5.700E-06 | 1.000E-05 |
| 14470 | 724 | T724I | B | 8.9e-06 | 7.6e-06 | 4.4e-06 | 6.967E-06 | 2.316E-06 | 4.400E-06 | 8.900E-06 |
| 14471 | 724 | T724L | B | 8.9e-06 | 6.8e-06 | 9.1e-06 | 8.267E-06 | 1.274E-06 | 6.800E-06 | 9.100E-06 |
| 14472 | 724 | T724K | B | 5.2e-06 | 6.4e-06 | 9.4e-06 | 7.000E-06 | 2.163E-06 | 5.200E-06 | 9.400E-06 |
| 14473 | 724 | T724M | B | 8.8e-06 | 4.9e-06 | 6.7e-06 | 6.800E-06 | 1.952E-06 | 4.900E-06 | 8.800E-06 |
| 14474 | 724 | T724F | B | 5.5e-06 | 5.5e-06 | 3.6e-06 | 4.867E-06 | 1.097E-06 | 3.600E-06 | 5.500E-06 |
| 14475 | 724 | T724P | B | 4.6e-06 | 8.5e-06 | 8.4e-06 | 7.167E-06 | 2.223E-06 | 4.600E-06 | 8.500E-06 |
| 14476 | 724 | T724S | B | 5.9e-06 | 6e-06 | 1e-05 | 7.300E-06 | 2.339E-06 | 5.900E-06 | 1.000E-05 |
| 14477 | 724 | T724T | B | 8.8e-06 | 6.1e-06 | 1.3e-05 | 9.300E-06 | 3.477E-06 | 6.100E-06 | 1.300E-05 |
| 14478 | 724 | T724W | B | 1.1e-05 | 1.3e-05 | 1.5e-05 | 1.300E-05 | 2.000E-06 | 1.100E-05 | 1.500E-05 |
| 14479 | 724 | T724Y | B | 1.6e-05 | 6.3e-06 | 8.2e-06 | 1.017E-05 | 5.140E-06 | 6.300E-06 | 1.600E-05 |
| 14480 | 724 | T724V | B | 6.3e-06 | 5.8e-06 | 9.6e-06 | 7.233E-06 | 2.065E-06 | 5.800E-06 | 9.600E-06 |
| 14481 | 725 | K725A | B | 1.2e-05 | 8.1e-06 | 6.8e-06 | 8.967E-06 | 2.706E-06 | 6.800E-06 | 1.200E-05 |
| 14482 | 725 | K725R | B | 6.9e-06 | 5.4e-06 | 7e-06 | 6.433E-06 | 8.963E-07 | 5.400E-06 | 7.000E-06 |
| 14483 | 725 | K725N | B | 9.2e-06 | 6.1e-06 | 7.6e-06 | 7.633E-06 | 1.550E-06 | 6.100E-06 | 9.200E-06 |
| 14484 | 725 | K725D | B | 4.2e-06 | 1e-05 | 3.9e-06 | 6.033E-06 | 3.439E-06 | 3.900E-06 | 1.000E-05 |
| 14485 | 725 | K725C | B | 8e-06 | 6.9e-06 | 8.1e-06 | 7.667E-06 | 6.658E-07 | 6.900E-06 | 8.100E-06 |
| 14486 | 725 | K725E | B | 8.4e-06 | 5.6e-06 | 6.6e-06 | 6.867E-06 | 1.419E-06 | 5.600E-06 | 8.400E-06 |
| 14487 | 725 | K725Q | B | 5.8e-06 | 3.3e-06 | 1.4e-05 | 7.700E-06 | 5.597E-06 | 3.300E-06 | 1.400E-05 |
| 14488 | 725 | K725G | B | 1.1e-05 | 4.6e-06 | 4.6e-06 | 6.733E-06 | 3.695E-06 | 4.600E-06 | 1.100E-05 |
| 14489 | 725 | K725H | B | 7.3e-06 | 1.2e-05 | 9.7e-06 | 9.667E-06 | 2.350E-06 | 7.300E-06 | 1.200E-05 |
| 14490 | 725 | K725I | B | 7.6e-06 | 6.6e-06 | 1e-05 | 8.067E-06 | 1.747E-06 | 6.600E-06 | 1.000E-05 |
| 14491 | 725 | K725L | B | 7.6e-06 | 4.2e-06 | 5.8e-06 | 5.867E-06 | 1.701E-06 | 4.200E-06 | 7.600E-06 |
| 14492 | 725 | K725K | B | 4.3e-06 | 6.9e-06 | 5.4e-06 | 5.533E-06 | 1.305E-06 | 4.300E-06 | 6.900E-06 |
| 14493 | 725 | K725M | B | 9.1e-06 | 1.1e-05 | 7.4e-06 | 9.167E-06 | 1.801E-06 | 7.400E-06 | 1.100E-05 |
| 14494 | 725 | K725F | B | 6.6e-06 | 6.7e-06 | 1e-05 | 7.767E-06 | 1.935E-06 | 6.600E-06 | 1.000E-05 |
| 14495 | 725 | K725P | B | 5.9e-06 | 4.8e-06 | 5.6e-06 | 5.433E-06 | 5.686E-07 | 4.800E-06 | 5.900E-06 |
| 14496 | 725 | K725S | B | 7.2e-06 | 8.9e-06 | 8.4e-06 | 8.167E-06 | 8.737E-07 | 7.200E-06 | 8.900E-06 |
| 14497 | 725 | K725T | B | 6.2e-06 | 7.7e-06 | 9.3e-06 | 7.733E-06 | 1.550E-06 | 6.200E-06 | 9.300E-06 |
| 14498 | 725 | K725W | B | 8.6e-06 | 4.8e-06 | 6.2e-06 | 6.533E-06 | 1.922E-06 | 4.800E-06 | 8.600E-06 |
| 14499 | 725 | K725Y | B | 6.8e-06 | 1.2e-05 | 1.1e-05 | 9.933E-06 | 2.759E-06 | 6.800E-06 | 1.200E-05 |
| 14500 | 725 | K725V | B | 7.5e-06 | 7.5e-06 | 8e-06 | 7.667E-06 | 2.887E-07 | 7.500E-06 | 8.000E-06 |
| 14501 | 726 | V726A | B | 5.2e-06 | 5.4e-06 | 7.7e-06 | 6.100E-06 | 1.389E-06 | 5.200E-06 | 7.700E-06 |
| 14502 | 726 | V726R | B | 8.6e-06 | 6.1e-06 | 8.8e-06 | 7.833E-06 | 1.504E-06 | 6.100E-06 | 8.800E-06 |
| 14503 | 726 | V726N | B | 1.2e-05 | 9.7e-06 | 5.6e-06 | 9.100E-06 | 3.242E-06 | 5.600E-06 | 1.200E-05 |
| 14504 | 726 | V726D | B | 9.8e-06 | 1.3e-05 | 9.1e-06 | 1.063E-05 | 2.079E-06 | 9.100E-06 | 1.300E-05 |
| 14505 | 726 | V726C | B | 9.6e-06 | 6.9e-06 | 7.4e-06 | 7.967E-06 | 1.436E-06 | 6.900E-06 | 9.600E-06 |
| 14506 | 726 | V726E | B | 8.9e-06 | 1.1e-05 | 5.8e-06 | 8.567E-06 | 2.616E-06 | 5.800E-06 | 1.100E-05 |
| 14507 | 726 | V726Q | B | 8.8e-06 | 6.3e-06 | 9e-06 | 8.033E-06 | 1.504E-06 | 6.300E-06 | 9.000E-06 |
| 14508 | 726 | V726G | B | 9.2e-06 | 7e-06 | 8.3e-06 | 8.167E-06 | 1.106E-06 | 7.000E-06 | 9.200E-06 |
| 14509 | 726 | V726H | B | 7.3e-06 | 7.5e-06 | 3.8e-06 | 6.200E-06 | 2.081E-06 | 3.800E-06 | 7.500E-06 |
| 14510 | 726 | V726I | B | 1e-05 | 1.3e-05 | 8.5e-06 | 1.050E-05 | 2.291E-06 | 8.500E-06 | 1.300E-05 |
| 14511 | 726 | V726L | B | 8.2e-06 | 8.5e-06 | 9e-06 | 8.567E-06 | 4.041E-07 | 8.200E-06 | 9.000E-06 |
| 14512 | 726 | V726K | B | 7.6e-06 | 1.1e-05 | 1.4e-05 | 1.087E-05 | 3.202E-06 | 7.600E-06 | 1.400E-05 |
| 14513 | 726 | V726M | B | 1e-05 | 7.5e-06 | 6.9e-06 | 8.133E-06 | 1.644E-06 | 6.900E-06 | 1.000E-05 |
| 14514 | 726 | V726F | B | 9.3e-06 | 4.3e-06 | 6.5e-06 | 6.700E-06 | 2.506E-06 | 4.300E-06 | 9.300E-06 |
| 14515 | 726 | V726P | B | 8.1e-06 | 4.6e-06 | 4.9e-06 | 5.867E-06 | 1.940E-06 | 4.600E-06 | 8.100E-06 |
| 14516 | 726 | V726S | B | 1.4e-05 | 8.8e-06 | 8.6e-06 | 1.047E-05 | 3.062E-06 | 8.600E-06 | 1.400E-05 |
| 14517 | 726 | V726T | B | 9.4e-06 | 7.3e-06 | 5.4e-06 | 7.367E-06 | 2.001E-06 | 5.400E-06 | 9.400E-06 |
| 14518 | 726 | V726W | B | 7e-06 | 8.3e-06 | 4.8e-06 | 6.700E-06 | 1.769E-06 | 4.800E-06 | 8.300E-06 |
| 14519 | 726 | V726Y | B | 5e-06 | 7.7e-06 | 9e-06 | 7.233E-06 | 2.040E-06 | 5.000E-06 | 9.000E-06 |
| 14520 | 726 | V726V | B | 4.1e-06 | 6.3e-06 | 1.1e-05 | 7.133E-06 | 3.525E-06 | 4.100E-06 | 1.100E-05 |
| 14521 | 727 | E727A | B | 5.4e-06 | 6.6e-06 | 6.3e-06 | 6.100E-06 | 6.245E-07 | 5.400E-06 | 6.600E-06 |
| 14522 | 727 | E727R | B | 8.9e-06 | 4.9e-06 | 1.3e-05 | 8.933E-06 | 4.050E-06 | 4.900E-06 | 1.300E-05 |
| 14523 | 727 | E727N | B | 8.3e-06 | 7.8e-06 | 1.5e-05 | 1.037E-05 | 4.020E-06 | 7.800E-06 | 1.500E-05 |
| 14524 | 727 | E727D | B | 6.5e-06 | 7.9e-06 | 7.6e-06 | 7.333E-06 | 7.371E-07 | 6.500E-06 | 7.900E-06 |
| 14525 | 727 | E727C | B | 6.4e-06 | 6.2e-06 | 7.6e-06 | 6.733E-06 | 7.572E-07 | 6.200E-06 | 7.600E-06 |
| 14526 | 727 | E727E | B | 4.3e-06 | 5.5e-06 | 8e-06 | 5.933E-06 | 1.888E-06 | 4.300E-06 | 8.000E-06 |
| 14527 | 727 | E727Q | B | 1.2e-05 | 1e-05 | 6.7e-06 | 9.567E-06 | 2.676E-06 | 6.700E-06 | 1.200E-05 |
| 14528 | 727 | E727G | B | 9.3e-06 | 9.7e-06 | 7.2e-06 | 8.733E-06 | 1.343E-06 | 7.200E-06 | 9.700E-06 |
| 14529 | 727 | E727H | B | 8.3e-06 | 4.9e-06 | 5.4e-06 | 6.200E-06 | 1.836E-06 | 4.900E-06 | 8.300E-06 |
| 14530 | 727 | E727I | B | 1.3e-05 | 8.2e-06 | 9.9e-06 | 1.037E-05 | 2.434E-06 | 8.200E-06 | 1.300E-05 |
| 14531 | 727 | E727L | B | 7.6e-06 | 5.6e-06 | 9.8e-06 | 7.667E-06 | 2.101E-06 | 5.600E-06 | 9.800E-06 |
| 14532 | 727 | E727K | B | 1.1e-05 | 6.1e-06 | 1.4e-05 | 1.037E-05 | 3.988E-06 | 6.100E-06 | 1.400E-05 |
| 14533 | 727 | E727M | B | 9.2e-06 | 6.4e-06 | 8e-06 | 7.867E-06 | 1.405E-06 | 6.400E-06 | 9.200E-06 |
| 14534 | 727 | E727F | B | 7.2e-06 | 6.3e-06 | 7.4e-06 | 6.967E-06 | 5.859E-07 | 6.300E-06 | 7.400E-06 |

| | | | | | | | | | | |
|---|---|---|---|---|---|---|---|---|---|---|
| 14535 | 727 | E727P | B | 5.8e-06 | 9e-06 | 6.6e-06 | 7.133E-06 | 1.665E-06 | 5.800E-06 | 9.000E-06 |
| 14536 | 727 | E727S | B | 5.3e-06 | 9.3e-06 | 8e-06 | 7.533E-06 | 2.040E-06 | 5.300E-06 | 9.300E-06 |
| 14537 | 727 | E727T | B | 7.7e-06 | 3.8e-06 | 9.1e-06 | 6.867E-06 | 2.747E-06 | 3.800E-06 | 9.100E-06 |
| 14538 | 727 | E727W | B | 1.1e-05 | 9.3e-06 | 6.3e-06 | 8.867E-06 | 2.380E-06 | 6.300E-06 | 1.100E-05 |
| 14539 | 727 | E727Y | B | 6.6e-06 | 4.8e-06 | 8.5e-06 | 6.633E-06 | 1.850E-06 | 4.800E-06 | 8.500E-06 |
| 14540 | 727 | E727V | B | 9.3e-06 | 6.4e-06 | 9.3e-06 | 8.333E-06 | 1.674E-06 | 6.400E-06 | 9.300E-06 |
| 14541 | 728 | I728A | B | 6.1e-06 | 9.5e-06 | 4.6e-06 | 6.733E-06 | 2.511E-06 | 4.600E-06 | 9.500E-06 |
| 14542 | 728 | I728R | B | 6.5e-06 | 1.1e-05 | 5.1e-06 | 7.533E-06 | 3.083E-06 | 5.100E-06 | 1.100E-05 |
| 14543 | 728 | I728N | B | 6.4e-06 | 4.2e-06 | 1.2e-05 | 7.533E-06 | 4.022E-06 | 4.200E-06 | 1.200E-05 |
| 14544 | 728 | I728D | B | 6.9e-06 | 7.2e-06 | 1.2e-05 | 8.700E-06 | 2.862E-06 | 6.900E-06 | 1.200E-05 |
| 14545 | 728 | I728C | B | 6.9e-06 | 7.8e-06 | 5.8e-06 | 6.833E-06 | 1.002E-06 | 5.800E-06 | 7.800E-06 |
| 14546 | 728 | I728E | B | 8e-06 | 9.6e-06 | 4.9e-06 | 7.500E-06 | 2.390E-06 | 4.900E-06 | 9.600E-06 |
| 14547 | 728 | I728Q | B | 6.7e-06 | 9.9e-06 | 8.8e-06 | 8.467E-06 | 1.626E-06 | 6.700E-06 | 9.900E-06 |
| 14548 | 728 | I728G | B | 7.9e-06 | 1e-05 | 8.7e-06 | 8.867E-06 | 1.060E-06 | 7.900E-06 | 1.000E-05 |
| 14549 | 728 | I728H | B | 9.2e-06 | 1.2e-05 | 6e-06 | 9.067E-06 | 3.002E-06 | 6.000E-06 | 1.200E-05 |
| 14550 | 728 | I728I | B | 8.1e-06 | 8.8e-06 | 8.4e-06 | 8.433E-06 | 3.512E-07 | 8.100E-06 | 8.800E-06 |
| 14551 | 728 | I728L | B | 7.1e-06 | 1.1e-05 | 5.9e-06 | 8.000E-06 | 2.666E-06 | 5.900E-06 | 1.100E-05 |
| 14552 | 728 | I728K | B | 4.6e-06 | 6.8e-06 | 5.3e-06 | 5.567E-06 | 1.124E-06 | 4.600E-06 | 6.800E-06 |
| 14553 | 728 | I728M | B | 1.1e-05 | 5e-06 | 1.3e-05 | 9.667E-06 | 4.163E-06 | 5.000E-06 | 1.300E-05 |
| 14554 | 728 | I728F | B | 6.8e-06 | 7.7e-06 | 1.3e-05 | 9.167E-06 | 3.350E-06 | 6.800E-06 | 1.300E-05 |
| 14555 | 728 | I728P | B | 6e-06 | 8.5e-06 | 1.2e-05 | 8.833E-06 | 3.014E-06 | 6.000E-06 | 1.200E-05 |
| 14556 | 728 | I728S | B | 1e-05 | 5.1e-06 | 4.9e-06 | 6.667E-06 | 2.888E-06 | 4.900E-06 | 1.000E-05 |
| 14557 | 728 | I728T | B | 4.3e-06 | 8.5e-06 | 1.1e-05 | 7.933E-06 | 3.386E-06 | 4.300E-06 | 1.100E-05 |
| 14558 | 728 | I728W | B | 4.4e-06 | 8.4e-06 | 7.6e-06 | 6.800E-06 | 2.117E-06 | 4.400E-06 | 8.400E-06 |
| 14559 | 728 | I728Y | B | 6.8e-06 | 1e-05 | 8.7e-06 | 8.500E-06 | 1.609E-06 | 6.800E-06 | 1.000E-05 |
| 14560 | 728 | I728V | B | 6.3e-06 | 8.9e-06 | 4.4e-06 | 6.533E-06 | 2.259E-06 | 4.400E-06 | 8.900E-06 |
| 14561 | 729 | K729A | B | 6.9e-06 | 8.5e-06 | 9.7e-06 | 8.367E-06 | 1.405E-06 | 6.900E-06 | 9.700E-06 |
| 14562 | 729 | K729R | B | 8.5e-06 | 1.4e-05 | 7.5e-06 | 1.000E-05 | 3.500E-06 | 7.500E-06 | 1.400E-05 |
| 14563 | 729 | K729N | B | 6.2e-06 | 8.7e-06 | 1.1e-05 | 8.633E-06 | 2.401E-06 | 6.200E-06 | 1.100E-05 |
| 14564 | 729 | K729D | B | 1.3e-05 | 5.2e-06 | 6.6e-06 | 8.267E-06 | 4.159E-06 | 5.200E-06 | 1.300E-05 |
| 14565 | 729 | K729C | B | 1.1e-05 | 8.1e-06 | 4.4e-06 | 7.833E-06 | 3.308E-06 | 4.400E-06 | 1.100E-05 |
| 14566 | 729 | K729E | B | 4.9e-06 | 8.8e-06 | 7e-06 | 6.900E-06 | 1.952E-06 | 4.900E-06 | 8.800E-06 |
| 14567 | 729 | K729Q | B | 6.4e-06 | 9e-06 | 1.3e-05 | 9.467E-06 | 3.325E-06 | 6.400E-06 | 1.300E-05 |
| 14568 | 729 | K729G | B | 7.6e-06 | 4.7e-06 | 6.6e-06 | 6.300E-06 | 1.473E-06 | 4.700E-06 | 7.600E-06 |
| 14569 | 729 | K729H | B | 1.2e-05 | 8.4e-06 | 8.8e-06 | 9.733E-06 | 1.973E-06 | 8.400E-06 | 1.200E-05 |
| 14570 | 729 | K729I | B | 1.1e-05 | 9.3e-06 | 6e-06 | 8.767E-06 | 2.542E-06 | 6.000E-06 | 1.100E-05 |
| 14571 | 729 | K729L | B | 9.2e-06 | 6.9e-06 | 9.2e-06 | 8.433E-06 | 1.328E-06 | 6.900E-06 | 9.200E-06 |
| 14572 | 729 | K729K | B | 7.3e-06 | 6.4e-06 | 1.2e-05 | 8.567E-06 | 3.007E-06 | 6.400E-06 | 1.200E-05 |
| 14573 | 729 | K729M | B | 9.3e-06 | 4.5e-06 | 8.8e-06 | 7.533E-06 | 2.639E-06 | 4.500E-06 | 9.300E-06 |
| 14574 | 729 | K729F | B | 5.6e-06 | 5.7e-06 | 1.2e-05 | 7.767E-06 | 3.667E-06 | 5.600E-06 | 1.200E-05 |
| 14575 | 729 | K729P | B | 6.5e-06 | 5.8e-06 | 5.2e-06 | 5.833E-06 | 6.506E-07 | 5.200E-06 | 6.500E-06 |
| 14576 | 729 | K729S | B | 9.9e-06 | 4.2e-06 | 9.3e-06 | 7.800E-06 | 3.132E-06 | 4.200E-06 | 9.900E-06 |
| 14577 | 729 | K729T | B | 5.3e-06 | 9.8e-06 | 6.8e-06 | 7.300E-06 | 2.291E-06 | 5.300E-06 | 9.800E-06 |
| 14578 | 729 | K729W | B | 1.2e-05 | 6.1e-06 | 5.6e-06 | 7.900E-06 | 3.559E-06 | 5.600E-06 | 1.200E-05 |
| 14579 | 729 | K729Y | B | 8e-06 | 4.4e-06 | 8.6e-06 | 7.000E-06 | 2.272E-06 | 4.400E-06 | 8.600E-06 |
| 14580 | 729 | K729V | B | 6.3e-06 | 7.6e-06 | 6e-06 | 6.633E-06 | 8.505E-07 | 6.000E-06 | 7.600E-06 |
| 14581 | 730 | R730A | B | 7.3e-06 | 9.9e-06 | 1e-05 | 9.067E-06 | 1.531E-06 | 7.300E-06 | 1.000E-05 |
| 14582 | 730 | R730R | B | 6.7e-06 | 5e-06 | 4.8e-06 | 5.500E-06 | 1.044E-06 | 4.800E-06 | 6.700E-06 |
| 14583 | 730 | R730N | B | 9.7e-06 | 7e-06 | 1e-05 | 8.900E-06 | 1.652E-06 | 7.000E-06 | 1.000E-05 |
| 14584 | 730 | R730D | B | 9.2e-06 | 9.9e-06 | 6.7e-06 | 8.600E-06 | 1.682E-06 | 6.700E-06 | 9.900E-06 |
| 14585 | 730 | R730C | B | 7.7e-06 | 9.9e-06 | 6e-06 | 7.867E-06 | 1.955E-06 | 6.000E-06 | 9.900E-06 |
| 14586 | 730 | R730E | B | 1.1e-05 | 5.5e-06 | 9.5e-06 | 8.667E-06 | 2.843E-06 | 5.500E-06 | 1.100E-05 |
| 14587 | 730 | R730Q | B | 1.2e-05 | 9e-06 | 9.2e-06 | 1.007E-05 | 1.677E-06 | 9.000E-06 | 1.200E-05 |
| 14588 | 730 | R730G | B | 8.5e-06 | 5.6e-06 | 6.5e-06 | 6.867E-06 | 1.484E-06 | 5.600E-06 | 8.500E-06 |
| 14589 | 730 | R730H | B | 1.8e-05 | 1.4e-05 | 8.7e-06 | 1.357E-05 | 4.665E-06 | 8.700E-06 | 1.800E-05 |
| 14590 | 730 | R730I | B | 1.4e-05 | 4.9e-06 | 1e-05 | 9.633E-06 | 4.561E-06 | 4.900E-06 | 1.400E-05 |
| 14591 | 730 | R730L | B | 8.1e-06 | 9.6e-06 | 6.5e-06 | 8.067E-06 | 1.550E-06 | 6.500E-06 | 9.600E-06 |
| 14592 | 730 | R730K | B | 6.3e-06 | 8.6e-06 | 1.2e-05 | 8.967E-06 | 2.868E-06 | 6.300E-06 | 1.200E-05 |
| 14593 | 730 | R730M | B | 6.8e-06 | 8.5e-06 | 8.8e-06 | 8.033E-06 | 1.079E-06 | 6.800E-06 | 8.800E-06 |
| 14594 | 730 | R730F | B | 6.5e-06 | 4.8e-06 | 1.1e-05 | 7.433E-06 | 3.204E-06 | 4.800E-06 | 1.100E-05 |
| 14595 | 730 | R730P | B | 7.1e-06 | 4.4e-06 | 6.3e-06 | 5.933E-06 | 1.387E-06 | 4.400E-06 | 7.100E-06 |
| 14596 | 730 | R730S | B | 1.2e-05 | 5.8e-06 | 9.8e-06 | 9.200E-06 | 3.143E-06 | 5.800E-06 | 1.200E-05 |
| 14597 | 730 | R730T | B | 8.9e-06 | 8.8e-06 | 1.1e-05 | 9.567E-06 | 1.242E-06 | 8.800E-06 | 1.100E-05 |
| 14598 | 730 | R730W | B | 6.7e-06 | 8.5e-06 | 8.6e-06 | 7.933E-06 | 1.069E-06 | 6.700E-06 | 8.600E-06 |
| 14599 | 730 | R730Y | B | 2.1e-05 | 9.7e-06 | 1.1e-05 | 1.390E-05 | 6.183E-06 | 9.700E-06 | 2.100E-05 |
| 14600 | 730 | R730V | B | 8.4e-06 | 5.3e-06 | 1.5e-05 | 9.567E-06 | 4.954E-06 | 5.300E-06 | 1.500E-05 |
| 14601 | 731 | T731A | B | 1.3e-05 | 6.4e-06 | 1.2e-05 | 1.047E-05 | 3.557E-06 | 6.400E-06 | 1.300E-05 |
| 14602 | 731 | T731R | B | 1e-05 | 9.1e-06 | 7.9e-06 | 9.000E-06 | 1.054E-06 | 7.900E-06 | 1.000E-05 |
| 14603 | 731 | T731N | B | 5.9e-06 | 9.5e-06 | 6.5e-06 | 7.300E-06 | 1.929E-06 | 5.900E-06 | 9.500E-06 |
| 14604 | 731 | T731D | B | 7.9e-06 | 8.1e-06 | 6.7e-06 | 7.567E-06 | 7.572E-07 | 6.700E-06 | 8.100E-06 |
| 14605 | 731 | T731C | B | 4.7e-06 | 1.2e-05 | 6.2e-06 | 7.633E-06 | 3.855E-06 | 4.700E-06 | 1.200E-05 |
| 14606 | 731 | T731E | B | 6.8e-06 | 6.7e-06 | 7.4e-06 | 6.967E-06 | 3.786E-07 | 6.700E-06 | 7.400E-06 |
| 14607 | 731 | T731Q | B | 4.6e-06 | 8e-06 | 7.1e-06 | 6.567E-06 | 1.762E-06 | 4.600E-06 | 8.000E-06 |
| 14608 | 731 | T731G | B | 6.3e-06 | 6.5e-06 | 8.2e-06 | 7.000E-06 | 1.044E-06 | 6.300E-06 | 8.200E-06 |
| 14609 | 731 | T731H | B | 5.5e-06 | 8.5e-06 | 4.3e-06 | 6.100E-06 | 2.163E-06 | 4.300E-06 | 8.500E-06 |
| 14610 | 731 | T731I | B | 8.2e-06 | 6e-06 | 1.1e-05 | 8.400E-06 | 2.506E-06 | 6.000E-06 | 1.100E-05 |
| 14611 | 731 | T731L | B | 1.7e-05 | 8.8e-06 | 6.3e-06 | 1.070E-05 | 5.597E-06 | 6.300E-06 | 1.700E-05 |
| 14612 | 731 | T731K | B | 9e-06 | 9.3e-06 | 5.9e-06 | 8.067E-06 | 1.882E-06 | 5.900E-06 | 9.300E-06 |
| 14613 | 731 | T731M | B | 1.1e-05 | 9.5e-06 | 6.5e-06 | 9.000E-06 | 2.291E-06 | 6.500E-06 | 1.100E-05 |
| 14614 | 731 | T731F | B | 5.1e-06 | 7.2e-06 | 1e-05 | 7.433E-06 | 2.458E-06 | 5.100E-06 | 1.000E-05 |
| 14615 | 731 | T731P | B | 1.5e-05 | 9.3e-06 | 8.6e-06 | 1.097E-05 | 3.510E-06 | 8.600E-06 | 1.500E-05 |
| 14616 | 731 | T731S | B | 5.9e-06 | 7.8e-06 | 7.9e-06 | 7.200E-06 | 1.127E-06 | 5.900E-06 | 7.900E-06 |
| 14617 | 731 | T731T | B | 8.7e-06 | 7.1e-06 | 1.1e-05 | 8.933E-06 | 1.960E-06 | 7.100E-06 | 1.100E-05 |
| 14618 | 731 | T731W | B | 1.3e-05 | 1.1e-05 | 9.1e-06 | 1.103E-05 | 1.950E-06 | 9.100E-06 | 1.300E-05 |
| 14619 | 731 | T731Y | B | 9.2e-06 | 1e-05 | 1e-05 | 9.733E-06 | 4.619E-07 | 9.200E-06 | 1.000E-05 |

| | | | | | | | | | | |
|---|---|---|---|---|---|---|---|---|---|---|
| 14620 | 731 | T731V | B | 5.5e-06 | 6.9e-06 | 6.9e-06 | 6.433E-06 | 8.083E-07 | 5.500E-06 | 6.900E-06 |
| 14621 | 732 | V732A | B | 4e-06 | 1e-05 | 1.1e-05 | 8.333E-06 | 3.786E-06 | 4.000E-06 | 1.100E-05 |
| 14622 | 732 | V732R | B | 9.1e-06 | 9.6e-06 | 9.5e-06 | 9.400E-06 | 2.646E-07 | 9.100E-06 | 9.600E-06 |
| 14623 | 732 | V732N | B | 6.5e-06 | 1.2e-05 | 1e-05 | 9.500E-06 | 2.784E-06 | 6.500E-06 | 1.200E-05 |
| 14624 | 732 | V732D | B | 1.4e-05 | 3.4e-06 | 1.5e-05 | 1.080E-05 | 6.428E-06 | 3.400E-06 | 1.500E-05 |
| 14625 | 732 | V732C | B | 9.7e-06 | 6.9e-06 | 1.3e-05 | 9.867E-06 | 3.053E-06 | 6.900E-06 | 1.300E-05 |
| 14626 | 732 | V732E | B | 5.3e-06 | 6.6e-06 | 5.6e-06 | 5.833E-06 | 6.807E-07 | 5.300E-06 | 6.600E-06 |
| 14627 | 732 | V732Q | B | 7.7e-06 | 1.5e-05 | 6.5e-06 | 9.733E-06 | 4.600E-06 | 6.500E-06 | 1.500E-05 |
| 14628 | 732 | V732G | B | 7.5e-06 | 4.6e-06 | 6.7e-06 | 6.267E-06 | 1.498E-06 | 4.600E-06 | 7.500E-06 |
| 14629 | 732 | V732H | B | 4.7e-06 | 6.7e-06 | 9.1e-06 | 6.833E-06 | 2.203E-06 | 4.700E-06 | 9.100E-06 |
| 14630 | 732 | V732I | B | 6.7e-06 | 1.2e-05 | 1.2e-05 | 1.023E-05 | 3.060E-06 | 6.700E-06 | 1.200E-05 |
| 14631 | 732 | V732L | B | 9.9e-06 | 5.7e-06 | 6.4e-06 | 7.333E-06 | 2.250E-06 | 5.700E-06 | 9.900E-06 |
| 14632 | 732 | V732K | B | 6.3e-06 | 6.5e-06 | 9.1e-06 | 7.300E-06 | 1.562E-06 | 6.300E-06 | 9.100E-06 |
| 14633 | 732 | V732M | B | 4.7e-06 | 6.5e-06 | 6.9e-06 | 6.033E-06 | 1.172E-06 | 4.700E-06 | 6.900E-06 |
| 14634 | 732 | V732F | B | 6.7e-06 | 1.3e-05 | 7e-06 | 8.900E-06 | 3.554E-06 | 6.700E-06 | 1.300E-05 |
| 14635 | 732 | V732P | B | 7.6e-06 | 9.5e-06 | 8.4e-06 | 8.500E-06 | 9.539E-07 | 7.600E-06 | 9.500E-06 |
| 14636 | 732 | V732S | B | 8.1e-06 | 6e-06 | 8.5e-06 | 7.533E-06 | 1.343E-06 | 6.000E-06 | 8.500E-06 |
| 14637 | 732 | V732T | B | 6e-06 | 5.5e-06 | 6.7e-06 | 6.067E-06 | 6.028E-07 | 5.500E-06 | 6.700E-06 |
| 14638 | 732 | V732W | B | 5.1e-06 | 8.3e-06 | 5.7e-06 | 6.367E-06 | 1.701E-06 | 5.100E-06 | 8.300E-06 |
| 14639 | 732 | V732Y | B | 1e-05 | 8.9e-06 | 7.5e-06 | 8.800E-06 | 1.253E-06 | 7.500E-06 | 1.000E-05 |
| 14640 | 732 | V732V | B | 1.1e-05 | 5.4e-06 | 1e-05 | 8.800E-06 | 2.987E-06 | 5.400E-06 | 1.100E-05 |
| 14641 | 733 | A733A | B | 1e-05 | 6.5e-06 | 1.5e-05 | 1.050E-05 | 4.272E-06 | 6.500E-06 | 1.500E-05 |
| 14642 | 733 | A733R | B | 7.2e-06 | 6.9e-06 | 5e-06 | 6.367E-06 | 1.193E-06 | 5.000E-06 | 7.200E-06 |
| 14643 | 733 | A733N | B | 8.1e-06 | 6.7e-06 | 1e-05 | 8.267E-06 | 1.656E-06 | 6.700E-06 | 1.000E-05 |
| 14644 | 733 | A733D | B | 9.7e-06 | 8.4e-06 | 9e-06 | 9.033E-06 | 6.506E-07 | 8.400E-06 | 9.700E-06 |
| 14645 | 733 | A733C | B | 7.6e-06 | 6.4e-06 | 4.6e-06 | 6.200E-06 | 1.510E-06 | 4.600E-06 | 7.600E-06 |
| 14646 | 733 | A733E | B | 9.4e-06 | 4.6e-06 | 7.6e-06 | 7.200E-06 | 2.425E-06 | 4.600E-06 | 9.400E-06 |
| 14647 | 733 | A733Q | B | 5.8e-06 | 7.9e-06 | 1.2e-05 | 8.567E-06 | 3.153E-06 | 5.800E-06 | 1.200E-05 |
| 14648 | 733 | A733G | B | 6.5e-06 | 1e-05 | 7.3e-06 | 7.933E-06 | 1.834E-06 | 6.500E-06 | 1.000E-05 |
| 14649 | 733 | A733H | B | 6.9e-06 | 1.2e-05 | 6.1e-06 | 8.333E-06 | 3.201E-06 | 6.100E-06 | 1.200E-05 |
| 14650 | 733 | A733I | B | 8.2e-06 | 8.8e-06 | 5.8e-06 | 7.600E-06 | 1.587E-06 | 5.800E-06 | 8.800E-06 |
| 14651 | 733 | A733L | B | 6.8e-06 | 9.1e-06 | 5.8e-06 | 7.233E-06 | 1.692E-06 | 5.800E-06 | 9.100E-06 |
| 14652 | 733 | A733K | B | 6.1e-06 | 9.1e-06 | 1.3e-05 | 9.400E-06 | 3.460E-06 | 6.100E-06 | 1.300E-05 |
| 14653 | 733 | A733M | B | 6.7e-06 | 9.2e-06 | 6.1e-06 | 7.333E-06 | 1.644E-06 | 6.100E-06 | 9.200E-06 |
| 14654 | 733 | A733F | B | 9.3e-06 | 6.7e-06 | 8.2e-06 | 8.067E-06 | 1.305E-06 | 6.700E-06 | 9.300E-06 |
| 14655 | 733 | A733P | B | 1.3e-05 | 1.5e-05 | 7.4e-06 | 1.180E-05 | 3.940E-06 | 7.400E-06 | 1.500E-05 |
| 14656 | 733 | A733S | B | 7.9e-06 | 1.6e-05 | 8.6e-06 | 1.083E-05 | 4.488E-06 | 7.900E-06 | 1.600E-05 |
| 14657 | 733 | A733T | B | 9.2e-06 | 4.3e-06 | 8e-06 | 7.167E-06 | 2.554E-06 | 4.300E-06 | 9.200E-06 |
| 14658 | 733 | A733W | B | 8.8e-06 | 5.9e-06 | 1.3e-05 | 9.233E-06 | 3.570E-06 | 5.900E-06 | 1.300E-05 |
| 14659 | 733 | A733Y | B | 9.9e-06 | 5.6e-06 | 8e-06 | 7.833E-06 | 2.155E-06 | 5.600E-06 | 9.900E-06 |
| 14660 | 733 | A733V | B | 1.1e-05 | 7.5e-06 | 5.6e-06 | 8.033E-06 | 2.739E-06 | 5.600E-06 | 1.100E-05 |
| 14661 | 734 | A734A | B | 5.6e-06 | 6.4e-06 | 1.2e-05 | 8.000E-06 | 3.487E-06 | 5.600E-06 | 1.200E-05 |
| 14662 | 734 | A734R | B | 6.6e-06 | 7.4e-06 | 7.2e-06 | 7.067E-06 | 4.163E-07 | 6.600E-06 | 7.400E-06 |
| 14663 | 734 | A734N | B | 6.4e-06 | 9.8e-06 | 1.2e-05 | 9.400E-06 | 2.821E-06 | 6.400E-06 | 1.200E-05 |
| 14664 | 734 | A734D | B | 6.1e-06 | 6.6e-06 | 5.4e-06 | 6.033E-06 | 6.028E-07 | 5.400E-06 | 6.600E-06 |
| 14665 | 734 | A734C | B | 5.2e-06 | 8.3e-06 | 6.7e-06 | 6.733E-06 | 1.550E-06 | 5.200E-06 | 8.300E-06 |
| 14666 | 734 | A734E | B | 1e-05 | 8.1e-06 | 8.5e-06 | 8.867E-06 | 1.002E-06 | 8.100E-06 | 1.000E-05 |
| 14667 | 734 | A734Q | B | 1.2e-05 | 5e-06 | 5.4e-06 | 7.467E-06 | 3.931E-06 | 5.000E-06 | 1.200E-05 |
| 14668 | 734 | A734G | B | 1e-05 | 6.3e-06 | 8.7e-06 | 8.333E-06 | 1.877E-06 | 6.300E-06 | 1.000E-05 |
| 14669 | 734 | A734H | B | 8.2e-06 | 7.8e-06 | 6.2e-06 | 7.400E-06 | 1.058E-06 | 6.200E-06 | 8.200E-06 |
| 14670 | 734 | A734I | B | 1e-05 | 7e-06 | 7.1e-06 | 8.033E-06 | 1.704E-06 | 7.000E-06 | 1.000E-05 |
| 14671 | 734 | A734L | B | 7e-06 | 9.5e-06 | 8.7e-06 | 8.400E-06 | 1.277E-06 | 7.000E-06 | 9.500E-06 |
| 14672 | 734 | A734K | B | 8e-06 | 9.7e-06 | 1.3e-05 | 1.023E-05 | 2.542E-06 | 8.000E-06 | 1.300E-05 |
| 14673 | 734 | A734M | B | 6.7e-06 | 8.4e-06 | 1e-05 | 8.367E-06 | 1.650E-06 | 6.700E-06 | 1.000E-05 |
| 14674 | 734 | A734F | B | 6.9e-06 | 5.4e-06 | 3.6e-06 | 5.300E-06 | 1.652E-06 | 3.600E-06 | 6.900E-06 |
| 14675 | 734 | A734P | B | 7.6e-06 | 8e-06 | 5.6e-06 | 7.067E-06 | 1.286E-06 | 5.600E-06 | 8.000E-06 |
| 14676 | 734 | A734S | B | 1e-05 | 9.2e-06 | 8.9e-06 | 9.367E-06 | 5.686E-07 | 8.900E-06 | 1.000E-05 |
| 14677 | 734 | A734T | B | 7.7e-06 | 1.5e-05 | 8.6e-06 | 1.043E-05 | 3.980E-06 | 7.700E-06 | 1.500E-05 |
| 14678 | 734 | A734W | B | 7.5e-06 | 1.2e-05 | 6.9e-06 | 8.800E-06 | 2.787E-06 | 6.900E-06 | 1.200E-05 |
| 14679 | 734 | A734Y | B | 7.1e-06 | 7e-06 | 5e-06 | 6.367E-06 | 1.185E-06 | 5.000E-06 | 7.100E-06 |
| 14680 | 734 | A734V | B | 7.9e-06 | 5.3e-06 | 5.7e-06 | 6.300E-06 | 1.400E-06 | 5.300E-06 | 7.900E-06 |
| 14681 | 735 | P735A | B | 2e-05 | 1e-05 | 1e-05 | 1.333E-05 | 5.774E-06 | 1.000E-05 | 2.000E-05 |
| 14682 | 735 | P735R | B | 6.3e-06 | 1e-05 | 6e-06 | 7.433E-06 | 2.228E-06 | 6.000E-06 | 1.000E-05 |
| 14683 | 735 | P735N | B | 6.6e-06 | 8e-06 | 5e-06 | 6.533E-06 | 1.501E-06 | 5.000E-06 | 8.000E-06 |
| 14684 | 735 | P735D | B | 9.3e-06 | 4.4e-06 | 9.6e-06 | 7.767E-06 | 2.919E-06 | 4.400E-06 | 9.600E-06 |
| 14685 | 735 | P735C | B | 1.7e-05 | 1.3e-05 | 8.9e-06 | 1.297E-05 | 4.050E-06 | 8.900E-06 | 1.700E-05 |
| 14686 | 735 | P735E | B | 1e-05 | 6.2e-06 | 9.4e-06 | 8.533E-06 | 2.043E-06 | 6.200E-06 | 1.000E-05 |
| 14687 | 735 | P735Q | B | 7.5e-06 | 8.4e-06 | 8.5e-06 | 8.133E-06 | 5.508E-07 | 7.500E-06 | 8.500E-06 |
| 14688 | 735 | P735G | B | 5.9e-06 | 5.3e-06 | 8.8e-06 | 6.667E-06 | 1.872E-06 | 5.300E-06 | 8.800E-06 |
| 14689 | 735 | P735H | B | 8.3e-06 | 1.2e-05 | 9.9e-06 | 1.007E-05 | 1.856E-06 | 8.300E-06 | 1.200E-05 |
| 14690 | 735 | P735I | B | 8.6e-06 | 1.3e-05 | 6.5e-06 | 9.367E-06 | 3.317E-06 | 6.500E-06 | 1.300E-05 |
| 14691 | 735 | P735L | B | 5.6e-06 | 8.4e-06 | 6.1e-06 | 6.700E-06 | 1.493E-06 | 5.600E-06 | 8.400E-06 |
| 14692 | 735 | P735K | B | 7.6e-06 | 6.9e-06 | 4.2e-06 | 6.233E-06 | 1.795E-06 | 4.200E-06 | 7.600E-06 |
| 14693 | 735 | P735M | B | 5.8e-06 | 8.6e-06 | 3.9e-06 | 6.100E-06 | 2.364E-06 | 3.900E-06 | 8.600E-06 |
| 14694 | 735 | P735F | B | 8.6e-06 | 9.4e-06 | 5.1e-06 | 7.700E-06 | 2.287E-06 | 5.100E-06 | 9.400E-06 |
| 14695 | 735 | P735P | B | 5.6e-06 | 7.7e-06 | 9.4e-06 | 7.567E-06 | 1.904E-06 | 5.600E-06 | 9.400E-06 |
| 14696 | 735 | P735S | B | 9e-06 | 4.6e-06 | 1.1e-05 | 8.200E-06 | 3.274E-06 | 4.600E-06 | 1.100E-05 |
| 14697 | 735 | P735T | B | 1.1e-05 | 1.1e-05 | 9.3e-06 | 1.043E-05 | 9.815E-07 | 9.300E-06 | 1.100E-05 |
| 14698 | 735 | P735W | B | 7.4e-06 | 6.8e-06 | 9.3e-06 | 7.833E-06 | 1.305E-06 | 6.800E-06 | 9.300E-06 |
| 14699 | 735 | P735Y | B | 5.3e-06 | 8.7e-06 | 9.8e-06 | 7.933E-06 | 2.346E-06 | 5.300E-06 | 9.800E-06 |
| 14700 | 735 | P735V | B | 6.6e-06 | 7.9e-06 | 8.5e-06 | 7.667E-06 | 9.713E-07 | 6.600E-06 | 8.500E-06 |
| 14701 | 736 | S736A | B | 1.1e-05 | 7e-06 | 1.1e-05 | 9.667E-06 | 2.309E-06 | 7.000E-06 | 1.100E-05 |
| 14702 | 736 | S736R | B | 1e-05 | 6.6e-06 | 8.4e-06 | 8.333E-06 | 1.701E-06 | 6.600E-06 | 1.000E-05 |
| 14703 | 736 | S736N | B | 1e-05 | 6.7e-06 | 9.9e-06 | 8.867E-06 | 1.877E-06 | 6.700E-06 | 1.000E-05 |
| 14704 | 736 | S736D | B | 4.8e-06 | 1e-05 | 7.7e-06 | 7.500E-06 | 2.606E-06 | 4.800E-06 | 1.000E-05 |

| | | | | | | | | | | |
|---|---|---|---|---|---|---|---|---|---|---|
| 14705 | 736 | S736C | B | 1.1e-05 | 7.8e-06 | 5.6e-06 | 8.133E-06 | 2.715E-06 | 5.600E-06 | 1.100E-05 |
| 14706 | 736 | S736E | B | 6e-06 | 6.1e-06 | 1.4e-05 | 8.700E-06 | 4.590E-06 | 6.000E-06 | 1.400E-05 |
| 14707 | 736 | S736Q | B | 6.3e-06 | 9.9e-06 | 8.9e-06 | 8.367E-06 | 1.858E-06 | 6.300E-06 | 9.900E-06 |
| 14708 | 736 | S736G | B | 9.8e-06 | 1.1e-05 | 1.2e-05 | 1.093E-05 | 1.102E-06 | 9.800E-06 | 1.200E-05 |
| 14709 | 736 | S736H | B | 1.4e-05 | 6.9e-06 | 1.4e-05 | 1.163E-05 | 4.099E-06 | 6.900E-06 | 1.400E-05 |
| 14710 | 736 | S736I | B | 7.5e-06 | 7e-06 | 5.6e-06 | 6.700E-06 | 9.849E-07 | 5.600E-06 | 7.500E-06 |
| 14711 | 736 | S736L | B | 5.2e-06 | 8.6e-06 | 6.5e-06 | 6.767E-06 | 1.716E-06 | 5.200E-06 | 8.600E-06 |
| 14712 | 736 | S736K | B | 8.3e-06 | 8.3e-06 | 5.5e-06 | 7.367E-06 | 1.617E-06 | 5.500E-06 | 8.300E-06 |
| 14713 | 736 | S736M | B | 4.8e-06 | 1.2e-05 | 6.5e-06 | 7.767E-06 | 3.763E-06 | 4.800E-06 | 1.200E-05 |
| 14714 | 736 | S736F | B | 9.9e-06 | 6.2e-06 | 4.9e-06 | 7.000E-06 | 2.594E-06 | 4.900E-06 | 9.900E-06 |
| 14715 | 736 | S736P | B | 8.6e-06 | 9.7e-06 | 1.1e-05 | 9.767E-06 | 1.201E-06 | 8.600E-06 | 1.100E-05 |
| 14716 | 736 | S736S | B | 5.1e-06 | 8.6e-06 | 7.1e-06 | 6.933E-06 | 1.756E-06 | 5.100E-06 | 8.600E-06 |
| 14717 | 736 | S736T | B | 1e-05 | 8.3e-06 | 6.3e-06 | 8.200E-06 | 1.852E-06 | 6.300E-06 | 1.000E-05 |
| 14718 | 736 | S736W | B | 7.5e-06 | 9.5e-06 | 6.1e-06 | 7.700E-06 | 1.709E-06 | 6.100E-06 | 9.500E-06 |
| 14719 | 736 | S736Y | B | 8.5e-06 | 7.6e-06 | 6.6e-06 | 7.567E-06 | 9.504E-07 | 6.600E-06 | 8.500E-06 |
| 14720 | 736 | S736V | B | 7.9e-06 | 8.9e-06 | 1.2e-05 | 9.600E-06 | 2.138E-06 | 7.900E-06 | 1.200E-05 |
| 14721 | 737 | V737A | B | 9.5e-06 | 8.7e-06 | 9.5e-06 | 9.233E-06 | 4.619E-07 | 8.700E-06 | 9.500E-06 |
| 14722 | 737 | V737R | B | 5e-06 | 5.9e-06 | 1e-05 | 6.967E-06 | 2.665E-06 | 5.000E-06 | 1.000E-05 |
| 14723 | 737 | V737N | B | 8e-06 | 1.1e-05 | 7.7e-06 | 8.900E-06 | 1.825E-06 | 7.700E-06 | 1.100E-05 |
| 14724 | 737 | V737D | B | 7.5e-06 | 1e-05 | 4.3e-06 | 7.267E-06 | 2.857E-06 | 4.300E-06 | 1.000E-05 |
| 14725 | 737 | V737C | B | 6.9e-06 | 7e-06 | 4.5e-06 | 6.133E-06 | 1.415E-06 | 4.500E-06 | 7.000E-06 |
| 14726 | 737 | V737E | B | 4.5e-06 | 6.5e-06 | 5.9e-06 | 5.633E-06 | 1.026E-06 | 4.500E-06 | 6.500E-06 |
| 14727 | 737 | V737Q | B | 1.4e-05 | 1.3e-05 | 9e-06 | 1.200E-05 | 2.646E-06 | 9.000E-06 | 1.400E-05 |
| 14728 | 737 | V737G | B | 1.3e-05 | 8.2e-06 | 1.1e-05 | 1.073E-05 | 2.411E-06 | 8.200E-06 | 1.300E-05 |
| 14729 | 737 | V737H | B | 7.5e-06 | 5.3e-06 | 8.4e-06 | 7.067E-06 | 1.595E-06 | 5.300E-06 | 8.400E-06 |
| 14730 | 737 | V737I | B | 9e-06 | 7.8e-06 | 4.5e-06 | 7.100E-06 | 2.330E-06 | 4.500E-06 | 9.000E-06 |
| 14731 | 737 | V737L | B | 1e-05 | 9.4e-06 | 8.2e-06 | 9.200E-06 | 9.165E-07 | 8.200E-06 | 1.000E-05 |
| 14732 | 737 | V737K | B | 9.7e-06 | 9.3e-06 | 8.5e-06 | 9.167E-06 | 6.110E-07 | 8.500E-06 | 9.700E-06 |
| 14733 | 737 | V737M | B | 4.8e-06 | 1.5e-05 | 9.4e-06 | 9.733E-06 | 5.108E-06 | 4.800E-06 | 1.500E-05 |
| 14734 | 737 | V737F | B | 9.8e-06 | 9.7e-06 | 6.3e-06 | 8.600E-06 | 1.992E-06 | 6.300E-06 | 9.800E-06 |
| 14735 | 737 | V737P | B | 8.9e-06 | 6.7e-06 | 1.1e-05 | 8.867E-06 | 2.150E-06 | 6.700E-06 | 1.100E-05 |
| 14736 | 737 | V737S | B | 9.3e-06 | 8.1e-06 | 1.3e-05 | 1.013E-05 | 2.554E-06 | 8.100E-06 | 1.300E-05 |
| 14737 | 737 | V737T | B | 9.4e-06 | 6.5e-06 | 7.1e-06 | 7.667E-06 | 1.531E-06 | 6.500E-06 | 9.400E-06 |
| 14738 | 737 | V737W | B | 5.5e-06 | 5.2e-06 | 5.1e-06 | 5.267E-06 | 2.082E-07 | 5.100E-06 | 5.500E-06 |
| 14739 | 737 | V737Y | B | 7.7e-06 | 5.8e-06 | 7.6e-06 | 7.033E-06 | 1.069E-06 | 5.800E-06 | 7.700E-06 |
| 14740 | 737 | V737V | B | 9.3e-06 | 9.4e-06 | 1.1e-05 | 9.900E-06 | 9.539E-07 | 9.300E-06 | 1.100E-05 |
| 14741 | 738 | F738A | B | 6.5e-06 | 8.6e-06 | 7.2e-06 | 7.433E-06 | 1.069E-06 | 6.500E-06 | 8.600E-06 |
| 14742 | 738 | F738R | B | 8.4e-06 | 1.2e-05 | 1e-05 | 1.013E-05 | 1.804E-06 | 8.400E-06 | 1.200E-05 |
| 14743 | 738 | F738N | B | 7.8e-06 | 8.9e-06 | 1.3e-05 | 9.900E-06 | 2.740E-06 | 7.800E-06 | 1.300E-05 |
| 14744 | 738 | F738D | B | 9.1e-06 | 4.9e-06 | 1.1e-05 | 8.333E-06 | 3.121E-06 | 4.900E-06 | 1.100E-05 |
| 14745 | 738 | F738C | B | 8.7e-06 | 9.7e-06 | 9.3e-06 | 9.233E-06 | 5.033E-07 | 8.700E-06 | 9.700E-06 |
| 14746 | 738 | F738E | B | 9.5e-06 | 7.2e-06 | 9.7e-06 | 8.800E-06 | 1.389E-06 | 7.200E-06 | 9.700E-06 |
| 14747 | 738 | F738Q | B | 7.9e-06 | 9e-06 | 1.2e-05 | 9.633E-06 | 2.122E-06 | 7.900E-06 | 1.200E-05 |
| 14748 | 738 | F738G | B | 6.5e-06 | 5.3e-06 | 9.6e-06 | 7.133E-06 | 2.219E-06 | 5.300E-06 | 9.600E-06 |
| 14749 | 738 | F738H | B | 8.9e-06 | 1.2e-05 | 5.3e-06 | 8.733E-06 | 3.353E-06 | 5.300E-06 | 1.200E-05 |
| 14750 | 738 | F738I | B | 9.3e-06 | 1.1e-05 | 6.4e-06 | 8.900E-06 | 2.326E-06 | 6.400E-06 | 1.100E-05 |
| 14751 | 738 | F738L | B | 5.3e-06 | 5.3e-06 | 1.4e-05 | 8.200E-06 | 5.023E-06 | 5.300E-06 | 1.400E-05 |
| 14752 | 738 | F738K | B | 6.2e-06 | 9.3e-06 | 8.6e-06 | 8.033E-06 | 1.626E-06 | 6.200E-06 | 9.300E-06 |
| 14753 | 738 | F738M | B | 4.5e-06 | 6.9e-06 | 5.6e-06 | 5.667E-06 | 1.201E-06 | 4.500E-06 | 6.900E-06 |
| 14754 | 738 | F738F | B | 5.8e-06 | 5e-06 | 8.5e-06 | 6.433E-06 | 1.834E-06 | 5.000E-06 | 8.500E-06 |
| 14755 | 738 | F738P | B | 9e-06 | 5.7e-06 | 9.3e-06 | 8.000E-06 | 1.997E-06 | 5.700E-06 | 9.300E-06 |
| 14756 | 738 | F738S | B | 1.2e-05 | 8.5e-06 | 9.3e-06 | 9.933E-06 | 1.834E-06 | 8.500E-06 | 1.200E-05 |
| 14757 | 738 | F738T | B | 3.9e-06 | 1.3e-05 | 5.7e-06 | 7.533E-06 | 4.819E-06 | 3.900E-06 | 1.300E-05 |
| 14758 | 738 | F738W | B | 8.5e-06 | 4.6e-06 | 6.8e-06 | 6.633E-06 | 1.955E-06 | 4.600E-06 | 8.500E-06 |
| 14759 | 738 | F738Y | B | 4.7e-06 | 9.5e-06 | 5.5e-06 | 6.567E-06 | 2.572E-06 | 4.700E-06 | 9.500E-06 |
| 14760 | 738 | F738V | B | 6.1e-06 | 4.7e-06 | 6.9e-06 | 5.900E-06 | 1.114E-06 | 4.700E-06 | 6.900E-06 |
| 14761 | 739 | I739A | B | 2e-05 | 6e-06 | 4.7e-06 | 1.023E-05 | 8.483E-06 | 4.700E-06 | 2.000E-05 |
| 14762 | 739 | I739R | B | 1.2e-05 | 1.4e-05 | 7.9e-06 | 1.130E-05 | 3.110E-06 | 7.900E-06 | 1.400E-05 |
| 14763 | 739 | I739N | B | 1.4e-05 | 3.7e-06 | 7.1e-06 | 8.267E-06 | 5.248E-06 | 3.700E-06 | 1.400E-05 |
| 14764 | 739 | I739D | B | 9.9e-06 | 6.8e-06 | 1e-05 | 8.900E-06 | 1.819E-06 | 6.800E-06 | 1.000E-05 |
| 14765 | 739 | I739C | B | 9.5e-06 | 6.6e-06 | 6.1e-06 | 7.400E-06 | 1.836E-06 | 6.100E-06 | 9.500E-06 |
| 14766 | 739 | I739E | B | 9.1e-06 | 6.9e-06 | 6.9e-06 | 7.633E-06 | 1.270E-06 | 6.900E-06 | 9.100E-06 |
| 14767 | 739 | I739Q | B | 7.2e-06 | 6.2e-06 | 8.2e-06 | 7.200E-06 | 1.000E-06 | 6.200E-06 | 8.200E-06 |
| 14768 | 739 | I739G | B | 8.5e-06 | 6.7e-06 | 1.3e-05 | 9.400E-06 | 3.245E-06 | 6.700E-06 | 1.300E-05 |
| 14769 | 739 | I739H | B | 8.1e-06 | 1e-05 | 4.6e-06 | 7.567E-06 | 2.739E-06 | 4.600E-06 | 1.000E-05 |
| 14770 | 739 | I739I | B | 1.2e-05 | 4.3e-06 | 7.9e-06 | 8.067E-06 | 3.853E-06 | 4.300E-06 | 1.200E-05 |
| 14771 | 739 | I739L | B | 4.6e-06 | 7.2e-06 | 9.9e-06 | 7.233E-06 | 2.650E-06 | 4.600E-06 | 9.900E-06 |
| 14772 | 739 | I739K | B | 1.2e-05 | 1.3e-05 | 5.6e-06 | 1.020E-05 | 4.015E-06 | 5.600E-06 | 1.300E-05 |
| 14773 | 739 | I739M | B | 7.2e-06 | 1.2e-05 | 9.6e-06 | 9.600E-06 | 2.400E-06 | 7.200E-06 | 1.200E-05 |
| 14774 | 739 | I739F | B | 8.2e-06 | 7.6e-06 | 4.8e-06 | 6.867E-06 | 1.815E-06 | 4.800E-06 | 8.200E-06 |
| 14775 | 739 | I739P | B | 1.1e-05 | 1.2e-05 | 1.3e-05 | 1.200E-05 | 1.000E-06 | 1.100E-05 | 1.300E-05 |
| 14776 | 739 | I739S | B | 9.9e-06 | 9.5e-06 | 8.6e-06 | 9.333E-06 | 6.658E-07 | 8.600E-06 | 9.900E-06 |
| 14777 | 739 | I739T | B | 6e-06 | 9.4e-06 | 9.7e-06 | 8.367E-06 | 2.055E-06 | 6.000E-06 | 9.700E-06 |
| 14778 | 739 | I739W | B | 5.6e-06 | 5.2e-06 | 1.2e-05 | 7.600E-06 | 3.816E-06 | 5.200E-06 | 1.200E-05 |
| 14779 | 739 | I739Y | B | 6.4e-06 | 4.3e-06 | 7.5e-06 | 6.067E-06 | 1.626E-06 | 4.300E-06 | 7.500E-06 |
| 14780 | 739 | I739V | B | 1.3e-05 | 1.3e-05 | 8.1e-06 | 1.137E-05 | 2.829E-06 | 8.100E-06 | 1.300E-05 |
| 14781 | 740 | F740A | B | 5.8e-06 | 9.5e-06 | 7.5e-06 | 7.600E-06 | 1.852E-06 | 5.800E-06 | 9.500E-06 |
| 14782 | 740 | F740R | B | 8.8e-06 | 6.5e-06 | 8.4e-06 | 7.900E-06 | 1.229E-06 | 6.500E-06 | 8.800E-06 |
| 14783 | 740 | F740N | B | 9.7e-06 | 6.3e-06 | 8.6e-06 | 8.200E-06 | 1.735E-06 | 6.300E-06 | 9.700E-06 |
| 14784 | 740 | F740D | B | 7.3e-06 | 8e-06 | 5.2e-06 | 6.833E-06 | 1.457E-06 | 5.200E-06 | 8.000E-06 |
| 14785 | 740 | F740C | B | 4.4e-06 | 9.4e-06 | 7.4e-06 | 7.067E-06 | 2.517E-06 | 4.400E-06 | 9.400E-06 |
| 14786 | 740 | F740E | B | 6.6e-06 | 8.3e-06 | 1.5e-05 | 9.967E-06 | 4.441E-06 | 6.600E-06 | 1.500E-05 |
| 14787 | 740 | F740Q | B | 6.8e-06 | 1.2e-05 | 6e-06 | 8.267E-06 | 3.258E-06 | 6.000E-06 | 1.200E-05 |
| 14788 | 740 | F740G | B | 1.4e-05 | 9.4e-06 | 6.8e-06 | 1.007E-05 | 3.646E-06 | 6.800E-06 | 1.400E-05 |
| 14789 | 740 | F740H | B | 9e-06 | 8.3e-06 | 7.7e-06 | 8.333E-06 | 6.506E-07 | 7.700E-06 | 9.000E-06 |

| | | | | | | | | | | |
|---|---|---|---|---|---|---|---|---|---|---|
| 14790 | 740 | F740I | B | 1.2e-05 | 8.5e-06 | 6.9e-06 | 9.133E-06 | 2.608E-06 | 6.900E-06 | 1.200E-05 |
| 14791 | 740 | F740L | B | 6.8e-06 | 3.4e-06 | 9.3e-06 | 6.500E-06 | 2.961E-06 | 3.400E-06 | 9.300E-06 |
| 14792 | 740 | F740K | B | 7.5e-06 | 5.6e-06 | 6.2e-06 | 6.433E-06 | 9.713E-07 | 5.600E-06 | 7.500E-06 |
| 14793 | 740 | F740M | B | 6.6e-06 | 5.3e-06 | 4.5e-06 | 5.467E-06 | 1.060E-06 | 4.500E-06 | 6.600E-06 |
| 14794 | 740 | F740F | B | 9.6e-06 | 1e-05 | 8.9e-06 | 9.500E-06 | 5.568E-07 | 8.900E-06 | 1.000E-05 |
| 14795 | 740 | F740P | B | 9.2e-06 | 5.4e-06 | 9.5e-06 | 8.033E-06 | 2.285E-06 | 5.400E-06 | 9.500E-06 |
| 14796 | 740 | F740S | B | 8.8e-06 | 9.5e-06 | 7e-06 | 8.433E-06 | 1.290E-06 | 7.000E-06 | 9.500E-06 |
| 14797 | 740 | F740T | B | 1.5e-05 | 8.4e-06 | 9.9e-06 | 1.110E-05 | 3.460E-06 | 8.400E-06 | 1.500E-05 |
| 14798 | 740 | F740W | B | 5.3e-06 | 6e-06 | 1.3e-05 | 8.100E-06 | 4.258E-06 | 5.300E-06 | 1.300E-05 |
| 14799 | 740 | F740Y | B | 4.7e-06 | 9.4e-06 | 5e-06 | 6.367E-06 | 2.631E-06 | 4.700E-06 | 9.400E-06 |
| 14800 | 740 | F740V | B | 1e-05 | 5e-06 | 9.2e-06 | 8.067E-06 | 2.686E-06 | 5.000E-06 | 1.000E-05 |
| 14801 | 741 | P741A | B | 9.6e-06 | 6.1e-06 | 6.4e-06 | 7.367E-06 | 1.940E-06 | 6.100E-06 | 9.600E-06 |
| 14802 | 741 | P741R | B | 7e-06 | 5.1e-06 | 8.3e-06 | 6.800E-06 | 1.609E-06 | 5.100E-06 | 8.300E-06 |
| 14803 | 741 | P741N | B | 6.8e-06 | 8.5e-06 | 7.1e-06 | 7.467E-06 | 9.074E-07 | 6.800E-06 | 8.500E-06 |
| 14804 | 741 | P741D | B | 6.6e-06 | 7e-06 | 1.2e-05 | 8.533E-06 | 3.009E-06 | 6.600E-06 | 1.200E-05 |
| 14805 | 741 | P741C | B | 1e-05 | 5.2e-06 | 5.5e-06 | 6.900E-06 | 2.689E-06 | 5.200E-06 | 1.000E-05 |
| 14806 | 741 | P741E | B | 8.6e-06 | 1.2e-05 | 9.4e-06 | 1.000E-05 | 1.778E-06 | 8.600E-06 | 1.200E-05 |
| 14807 | 741 | P741Q | B | 5.9e-06 | 1.8e-05 | 7e-06 | 1.030E-05 | 6.691E-06 | 5.900E-06 | 1.800E-05 |
| 14808 | 741 | P741G | B | 9.9e-06 | 7e-06 | 3.6e-06 | 6.833E-06 | 3.153E-06 | 3.600E-06 | 9.900E-06 |
| 14809 | 741 | P741H | B | 6.2e-06 | 7.3e-06 | 8.8e-06 | 7.433E-06 | 1.305E-06 | 6.200E-06 | 8.800E-06 |
| 14810 | 741 | P741I | B | 8.4e-06 | 5.6e-06 | 1.1e-05 | 8.333E-06 | 2.701E-06 | 5.600E-06 | 1.100E-05 |
| 14811 | 741 | P741L | B | 6.2e-06 | 6.1e-06 | 4.4e-06 | 5.567E-06 | 1.012E-06 | 4.400E-06 | 6.200E-06 |
| 14812 | 741 | P741K | B | 7.6e-06 | 1.2e-05 | 8.4e-06 | 9.333E-06 | 2.344E-06 | 7.600E-06 | 1.200E-05 |
| 14813 | 741 | P741M | B | 7.4e-06 | 5.3e-06 | 8.9e-06 | 7.200E-06 | 1.808E-06 | 5.300E-06 | 8.900E-06 |
| 14814 | 741 | P741F | B | 6e-06 | 1.4e-05 | 4.7e-06 | 8.233E-06 | 5.036E-06 | 4.700E-06 | 1.400E-05 |
| 14815 | 741 | P741P | B | 1.2e-05 | 5.8e-06 | 5.8e-06 | 7.867E-06 | 3.580E-06 | 5.800E-06 | 1.200E-05 |
| 14816 | 741 | P741S | B | 9.2e-06 | 7.4e-06 | 1.9e-05 | 1.187E-05 | 6.243E-06 | 7.400E-06 | 1.900E-05 |
| 14817 | 741 | P741T | B | 9.6e-06 | 6.5e-06 | 6.3e-06 | 7.467E-06 | 1.850E-06 | 6.300E-06 | 9.600E-06 |
| 14818 | 741 | P741W | B | 1.2e-05 | 8e-06 | 6.5e-06 | 8.833E-06 | 2.843E-06 | 6.500E-06 | 1.200E-05 |
| 14819 | 741 | P741Y | B | 6.5e-06 | 8.6e-06 | 9.3e-06 | 8.133E-06 | 1.457E-06 | 6.500E-06 | 9.300E-06 |
| 14820 | 741 | P741V | B | 6.6e-06 | 6.2e-06 | 5.5e-06 | 6.100E-06 | 5.568E-07 | 5.500E-06 | 6.600E-06 |
| 14821 | 742 | P742A | B | 1.1e-05 | 1.4e-05 | 5.8e-06 | 1.027E-05 | 4.149E-06 | 5.800E-06 | 1.400E-05 |
| 14822 | 742 | P742R | B | 8.5e-06 | 8.6e-06 | 1.1e-05 | 9.367E-06 | 1.415E-06 | 8.500E-06 | 1.100E-05 |
| 14823 | 742 | P742N | B | 5.1e-06 | 9e-06 | 5.5e-06 | 6.533E-06 | 2.146E-06 | 5.100E-06 | 9.000E-06 |
| 14824 | 742 | P742D | B | 4.5e-06 | 5e-06 | 7.6e-06 | 5.700E-06 | 1.664E-06 | 4.500E-06 | 7.600E-06 |
| 14825 | 742 | P742C | B | 9.1e-06 | 7e-06 | 7.3e-06 | 7.800E-06 | 1.136E-06 | 7.000E-06 | 9.100E-06 |
| 14826 | 742 | P742E | B | 9.5e-06 | 1.3e-05 | 9.7e-06 | 1.073E-05 | 1.966E-06 | 9.500E-06 | 1.300E-05 |
| 14827 | 742 | P742Q | B | 7e-06 | 7.7e-06 | 9e-06 | 7.900E-06 | 1.015E-06 | 7.000E-06 | 9.000E-06 |
| 14828 | 742 | P742G | B | 9.3e-06 | 1.3e-05 | 1.4e-05 | 1.210E-05 | 2.476E-06 | 9.300E-06 | 1.400E-05 |
| 14829 | 742 | P742H | B | 3.8e-06 | 6.4e-06 | 4.5e-06 | 4.900E-06 | 1.345E-06 | 3.800E-06 | 6.400E-06 |
| 14830 | 742 | P742I | B | 9.9e-06 | 7.3e-06 | 1e-05 | 9.067E-06 | 1.531E-06 | 7.300E-06 | 1.000E-05 |
| 14831 | 742 | P742L | B | 8.5e-06 | 1.3e-05 | 6.1e-06 | 9.200E-06 | 3.503E-06 | 6.100E-06 | 1.300E-05 |
| 14832 | 742 | P742K | B | 9.9e-06 | 8.4e-06 | 1.1e-05 | 9.767E-06 | 1.305E-06 | 8.400E-06 | 1.100E-05 |
| 14833 | 742 | P742M | B | 6.5e-06 | 1.2e-05 | 9.3e-06 | 9.267E-06 | 2.750E-06 | 6.500E-06 | 1.200E-05 |
| 14834 | 742 | P742F | B | 7.6e-06 | 9.3e-06 | 8.8e-06 | 8.567E-06 | 8.737E-07 | 7.600E-06 | 9.300E-06 |
| 14835 | 742 | P742P | B | 1e-05 | 6.8e-06 | 7.7e-06 | 8.167E-06 | 1.650E-06 | 6.800E-06 | 1.000E-05 |
| 14836 | 742 | P742S | B | 4.5e-06 | 8.9e-06 | 7.4e-06 | 6.933E-06 | 2.237E-06 | 4.500E-06 | 8.900E-06 |
| 14837 | 742 | P742T | B | 6.2e-06 | 9.4e-06 | 8.5e-06 | 8.033E-06 | 1.650E-06 | 6.200E-06 | 9.400E-06 |
| 14838 | 742 | P742W | B | 8.7e-06 | 9.1e-06 | 5.7e-06 | 7.833E-06 | 1.858E-06 | 5.700E-06 | 9.100E-06 |
| 14839 | 742 | P742Y | B | 7.9e-06 | 1.4e-05 | 1.3e-05 | 1.163E-05 | 3.272E-06 | 7.900E-06 | 1.400E-05 |
| 14840 | 742 | P742V | B | 7.3e-06 | 5.9e-06 | 8e-06 | 7.067E-06 | 1.069E-06 | 5.900E-06 | 8.000E-06 |
| 14841 | 743 | S743A | B | 7.1e-06 | 6.7e-06 | 1e-05 | 7.933E-06 | 1.801E-06 | 6.700E-06 | 1.000E-05 |
| 14842 | 743 | S743R | B | 5.4e-06 | 6.6e-06 | 6.1e-06 | 6.033E-06 | 6.028E-07 | 5.400E-06 | 6.600E-06 |
| 14843 | 743 | S743N | B | 8.9e-06 | 1e-05 | 4.4e-06 | 7.767E-06 | 2.967E-06 | 4.400E-06 | 1.000E-05 |
| 14844 | 743 | S743D | B | 6.7e-06 | 4.4e-06 | 6.5e-06 | 5.867E-06 | 1.274E-06 | 4.400E-06 | 6.700E-06 |
| 14845 | 743 | S743C | B | 8.5e-06 | 9.2e-06 | 1.5e-05 | 1.090E-05 | 3.568E-06 | 8.500E-06 | 1.500E-05 |
| 14846 | 743 | S743E | B | 8.3e-06 | 1.5e-05 | 1e-05 | 1.110E-05 | 3.483E-06 | 8.300E-06 | 1.500E-05 |
| 14847 | 743 | S743Q | B | 9.6e-06 | 6.3e-06 | 5.1e-06 | 7.000E-06 | 2.330E-06 | 5.100E-06 | 9.600E-06 |
| 14848 | 743 | S743G | B | 4.9e-06 | 9.8e-06 | 1.4e-05 | 9.567E-06 | 4.554E-06 | 4.900E-06 | 1.400E-05 |
| 14849 | 743 | S743H | B | 5.3e-06 | 4.5e-06 | 6.7e-06 | 5.500E-06 | 1.114E-06 | 4.500E-06 | 6.700E-06 |
| 14850 | 743 | S743I | B | 1.1e-05 | 8.8e-06 | 5.3e-06 | 8.367E-06 | 2.875E-06 | 5.300E-06 | 1.100E-05 |
| 14851 | 743 | S743L | B | 3.4e-06 | 6.3e-06 | 3.3e-06 | 4.333E-06 | 1.704E-06 | 3.300E-06 | 6.300E-06 |
| 14852 | 743 | S743K | B | 1.2e-05 | 8.7e-06 | 7.1e-06 | 9.267E-06 | 2.499E-06 | 7.100E-06 | 1.200E-05 |
| 14853 | 743 | S743M | B | 7.3e-06 | 8.6e-06 | 4.8e-06 | 6.900E-06 | 1.931E-06 | 4.800E-06 | 8.600E-06 |
| 14854 | 743 | S743F | B | 4.6e-06 | 5.1e-06 | 1.3e-05 | 7.567E-06 | 4.712E-06 | 4.600E-06 | 1.300E-05 |
| 14855 | 743 | S743P | B | 8.5e-06 | 6.1e-06 | 1.2e-05 | 8.867E-06 | 2.967E-06 | 6.100E-06 | 1.200E-05 |
| 14856 | 743 | S743S | B | 7.2e-06 | 6.2e-06 | 6.6e-06 | 6.667E-06 | 5.033E-07 | 6.200E-06 | 7.200E-06 |
| 14857 | 743 | S743T | B | 1.1e-05 | 4.5e-06 | 7.7e-06 | 7.733E-06 | 3.250E-06 | 4.500E-06 | 1.100E-05 |
| 14858 | 743 | S743W | B | 4.3e-06 | 5e-06 | 6.6e-06 | 5.300E-06 | 1.179E-06 | 4.300E-06 | 6.600E-06 |
| 14859 | 743 | S743Y | B | 9.4e-06 | 1.1e-05 | 6.6e-06 | 9.000E-06 | 2.227E-06 | 6.600E-06 | 1.100E-05 |
| 14860 | 743 | S743V | B | 1e-05 | 6.7e-06 | 5.6e-06 | 7.433E-06 | 2.290E-06 | 5.600E-06 | 1.000E-05 |
| 14861 | 744 | D744A | B | 6.6e-06 | 7.4e-06 | 1.2e-05 | 8.667E-06 | 2.914E-06 | 6.600E-06 | 1.200E-05 |
| 14862 | 744 | D744R | B | 8.6e-06 | 1.1e-05 | 6.3e-06 | 8.633E-06 | 2.350E-06 | 6.300E-06 | 1.100E-05 |
| 14863 | 744 | D744N | B | 5.9e-06 | 8.7e-06 | 5.4e-06 | 6.667E-06 | 1.779E-06 | 5.400E-06 | 8.700E-06 |
| 14864 | 744 | D744D | B | 4.1e-06 | 8.8e-06 | 4.7e-06 | 5.867E-06 | 2.558E-06 | 4.100E-06 | 8.800E-06 |
| 14865 | 744 | D744C | B | 9.2e-06 | 8.3e-06 | 7.9e-06 | 8.467E-06 | 6.658E-07 | 7.900E-06 | 9.200E-06 |
| 14866 | 744 | D744E | B | 5.9e-06 | 7.3e-06 | 5.7e-06 | 6.300E-06 | 8.718E-07 | 5.700E-06 | 7.300E-06 |
| 14867 | 744 | D744Q | B | 7.6e-06 | 6.5e-06 | 8.5e-06 | 7.533E-06 | 1.002E-06 | 6.500E-06 | 8.500E-06 |
| 14868 | 744 | D744G | B | 6.3e-06 | 6.3e-06 | 8.6e-06 | 7.067E-06 | 1.328E-06 | 6.300E-06 | 8.600E-06 |
| 14869 | 744 | D744H | B | 8.1e-06 | 9.3e-06 | 8e-06 | 8.467E-06 | 7.234E-07 | 8.000E-06 | 9.300E-06 |
| 14870 | 744 | D744I | B | 7.9e-06 | 7.2e-06 | 9e-06 | 8.033E-06 | 9.074E-07 | 7.200E-06 | 9.000E-06 |
| 14871 | 744 | D744L | B | 9.8e-06 | 7.4e-06 | 5.3e-06 | 7.500E-06 | 2.252E-06 | 5.300E-06 | 9.800E-06 |
| 14872 | 744 | D744K | B | 6.2e-06 | 1.3e-05 | 9.6e-06 | 9.600E-06 | 3.400E-06 | 6.200E-06 | 1.300E-05 |
| 14873 | 744 | D744M | B | 5.9e-06 | 6.4e-06 | 4.8e-06 | 5.700E-06 | 8.185E-07 | 4.800E-06 | 6.400E-06 |
| 14874 | 744 | D744F | B | 9.5e-06 | 7.9e-06 | 8.6e-06 | 8.667E-06 | 8.021E-07 | 7.900E-06 | 9.500E-06 |

| | | | | | | | | | | |
|---|---|---|---|---|---|---|---|---|---|---|
| 14875 | 744 | D744P | B | 6.2e-06 | 1.2e-05 | 1.4e-05 | 1.073E-05 | 4.051E-06 | 6.200E-06 | 1.400E-05 |
| 14876 | 744 | D744S | B | 9.8e-06 | 6.8e-06 | 5.4e-06 | 7.333E-06 | 2.248E-06 | 5.400E-06 | 9.800E-06 |
| 14877 | 744 | D744T | B | 1e-05 | 5.9e-06 | 7.3e-06 | 7.733E-06 | 2.084E-06 | 5.900E-06 | 1.000E-05 |
| 14878 | 744 | D744W | B | 1.2e-05 | 9.9e-06 | 6.6e-06 | 9.500E-06 | 2.722E-06 | 6.600E-06 | 1.200E-05 |
| 14879 | 744 | D744Y | B | 5.6e-06 | 8.2e-06 | 6.3e-06 | 6.700E-06 | 1.345E-06 | 5.600E-06 | 8.200E-06 |
| 14880 | 744 | D744V | B | 7.1e-06 | 1.1e-05 | 6.7e-06 | 8.267E-06 | 2.376E-06 | 6.700E-06 | 1.100E-05 |
| 14881 | 745 | E745A | B | 6.4e-06 | 5.9e-06 | 7.7e-06 | 6.667E-06 | 9.292E-07 | 5.900E-06 | 7.700E-06 |
| 14882 | 745 | E745R | B | 4.5e-06 | 1.3e-05 | 8.3e-06 | 8.600E-06 | 4.258E-06 | 4.500E-06 | 1.300E-05 |
| 14883 | 745 | E745N | B | 9.4e-06 | 4.9e-06 | 1.3e-05 | 9.100E-06 | 4.058E-06 | 4.900E-06 | 1.300E-05 |
| 14884 | 745 | E745D | B | 5.4e-06 | 6.8e-06 | 1e-05 | 7.400E-06 | 2.358E-06 | 5.400E-06 | 1.000E-05 |
| 14885 | 745 | E745C | B | 7.1e-06 | 6.8e-06 | 5.2e-06 | 6.367E-06 | 1.021E-06 | 5.200E-06 | 7.100E-06 |
| 14886 | 745 | E745E | B | 5.5e-06 | 6.4e-06 | 5.4e-06 | 5.767E-06 | 5.508E-07 | 5.400E-06 | 6.400E-06 |
| 14887 | 745 | E745Q | B | 9.4e-06 | 7.9e-06 | 7.1e-06 | 8.133E-06 | 1.168E-06 | 7.100E-06 | 9.400E-06 |
| 14888 | 745 | E745G | B | 4.4e-06 | 1.6e-05 | 6.5e-06 | 8.967E-06 | 6.181E-06 | 4.400E-06 | 1.600E-05 |
| 14889 | 745 | E745H | B | 1.3e-05 | 6.3e-06 | 6e-06 | 8.433E-06 | 3.958E-06 | 6.000E-06 | 1.300E-05 |
| 14890 | 745 | E745I | B | 1.2e-05 | 9e-06 | 7e-06 | 9.333E-06 | 2.517E-06 | 7.000E-06 | 1.200E-05 |
| 14891 | 745 | E745L | B | 7.7e-06 | 8.5e-06 | 1.3e-05 | 9.733E-06 | 2.857E-06 | 7.700E-06 | 1.300E-05 |
| 14892 | 745 | E745K | B | 1.3e-05 | 7.6e-06 | 6e-06 | 8.867E-06 | 3.668E-06 | 6.000E-06 | 1.300E-05 |
| 14893 | 745 | E745M | B | 8.4e-06 | 8.5e-06 | 6.2e-06 | 7.700E-06 | 1.300E-06 | 6.200E-06 | 8.500E-06 |
| 14894 | 745 | E745F | B | 1.5e-05 | 5e-06 | 1e-05 | 1.000E-05 | 5.000E-06 | 5.000E-06 | 1.500E-05 |
| 14895 | 745 | E745P | B | 1.3e-05 | 6e-06 | 1.3e-05 | 1.067E-05 | 4.041E-06 | 6.000E-06 | 1.300E-05 |
| 14896 | 745 | E745S | B | 8.3e-06 | 5.9e-06 | 1.1e-05 | 8.400E-06 | 2.551E-06 | 5.900E-06 | 1.100E-05 |
| 14897 | 745 | E745T | B | 6.3e-06 | 1.5e-05 | 5e-06 | 8.767E-06 | 5.437E-06 | 5.000E-06 | 1.500E-05 |
| 14898 | 745 | E745W | B | 9.6e-06 | 1.2e-05 | 8.4e-06 | 1.000E-05 | 1.833E-06 | 8.400E-06 | 1.200E-05 |
| 14899 | 745 | E745Y | B | 8.9e-06 | 6.9e-06 | 1.2e-05 | 9.267E-06 | 2.570E-06 | 6.900E-06 | 1.200E-05 |
| 14900 | 745 | E745V | B | 8e-06 | 9.6e-06 | 6.7e-06 | 8.100E-06 | 1.453E-06 | 6.700E-06 | 9.600E-06 |
| 14901 | 746 | Q746A | B | 1e-05 | 6.7e-06 | 3.8e-06 | 6.833E-06 | 3.102E-06 | 3.800E-06 | 1.000E-05 |
| 14902 | 746 | Q746R | B | 6.4e-06 | 9.1e-06 | 9.3e-06 | 8.267E-06 | 1.620E-06 | 6.400E-06 | 9.300E-06 |
| 14903 | 746 | Q746N | B | 8.1e-06 | 6.7e-06 | 9.4e-06 | 8.067E-06 | 1.350E-06 | 6.700E-06 | 9.400E-06 |
| 14904 | 746 | Q746D | B | 7.8e-06 | 6.9e-06 | 1.1e-05 | 8.567E-06 | 2.155E-06 | 6.900E-06 | 1.100E-05 |
| 14905 | 746 | Q746C | B | 6.1e-06 | 9.3e-06 | 8.9e-06 | 8.100E-06 | 1.744E-06 | 6.100E-06 | 9.300E-06 |
| 14906 | 746 | Q746E | B | 6e-06 | 1.2e-05 | 9.9e-06 | 9.300E-06 | 3.045E-06 | 6.000E-06 | 1.200E-05 |
| 14907 | 746 | Q746Q | B | 1.3e-05 | 1.2e-05 | 8.8e-06 | 1.127E-05 | 2.194E-06 | 8.800E-06 | 1.300E-05 |
| 14908 | 746 | Q746G | B | 5.6e-06 | 7.7e-06 | 1.1e-05 | 8.100E-06 | 2.722E-06 | 5.600E-06 | 1.100E-05 |
| 14909 | 746 | Q746H | B | 6.5e-06 | 5.6e-06 | 4.9e-06 | 5.667E-06 | 8.021E-07 | 4.900E-06 | 6.500E-06 |
| 14910 | 746 | Q746I | B | 7.6e-06 | 6.9e-06 | 9.3e-06 | 7.933E-06 | 1.234E-06 | 6.900E-06 | 9.300E-06 |
| 14911 | 746 | Q746L | B | 7.1e-06 | 9.7e-06 | 5.4e-06 | 7.400E-06 | 2.166E-06 | 5.400E-06 | 9.700E-06 |
| 14912 | 746 | Q746K | B | 9.3e-06 | 9e-06 | 8.5e-06 | 8.933E-06 | 4.041E-07 | 8.500E-06 | 9.300E-06 |
| 14913 | 746 | Q746M | B | 8.2e-06 | 8.8e-06 | 9.4e-06 | 8.800E-06 | 6.000E-07 | 8.200E-06 | 9.400E-06 |
| 14914 | 746 | Q746F | B | 9.5e-06 | 9.4e-06 | 6.3e-06 | 8.400E-06 | 1.819E-06 | 6.300E-06 | 9.500E-06 |
| 14915 | 746 | Q746P | B | 6.4e-06 | 7.3e-06 | 9.4e-06 | 7.700E-06 | 1.539E-06 | 6.400E-06 | 9.400E-06 |
| 14916 | 746 | Q746S | B | 5.4e-06 | 8.9e-06 | 2.9e-06 | 5.733E-06 | 3.014E-06 | 2.900E-06 | 8.900E-06 |
| 14917 | 746 | Q746T | B | 7.4e-06 | 4.5e-06 | 1.3e-05 | 8.300E-06 | 4.321E-06 | 4.500E-06 | 1.300E-05 |
| 14918 | 746 | Q746W | B | 8.4e-06 | 5.2e-06 | 1.4e-05 | 9.200E-06 | 4.454E-06 | 5.200E-06 | 1.400E-05 |
| 14919 | 746 | Q746Y | B | 7.6e-06 | 1.1e-05 | 7.1e-06 | 8.567E-06 | 2.122E-06 | 7.100E-06 | 1.100E-05 |
| 14920 | 746 | Q746V | B | 7.5e-06 | 4.8e-06 | 6.8e-06 | 6.367E-06 | 1.401E-06 | 4.800E-06 | 7.500E-06 |
| 14921 | 747 | L747A | B | 8.8e-06 | 4.6e-06 | 4.5e-06 | 5.967E-06 | 2.454E-06 | 4.500E-06 | 8.800E-06 |
| 14922 | 747 | L747R | B | 9.8e-06 | 6.2e-06 | 8.3e-06 | 8.100E-06 | 1.808E-06 | 6.200E-06 | 9.800E-06 |
| 14923 | 747 | L747N | B | 1e-05 | 7.2e-06 | 1.3e-05 | 1.007E-05 | 2.901E-06 | 7.200E-06 | 1.300E-05 |
| 14924 | 747 | L747D | B | 7.6e-06 | 1e-05 | 6.4e-06 | 8.000E-06 | 1.833E-06 | 6.400E-06 | 1.000E-05 |
| 14925 | 747 | L747C | B | 5e-06 | 6.8e-06 | 9.4e-06 | 7.067E-06 | 2.212E-06 | 5.000E-06 | 9.400E-06 |
| 14926 | 747 | L747E | B | 1e-05 | 7.5e-06 | 7.3e-06 | 8.267E-06 | 1.504E-06 | 7.300E-06 | 1.000E-05 |
| 14927 | 747 | L747Q | B | 4.5e-06 | 6.3e-06 | 4.3e-06 | 5.033E-06 | 1.102E-06 | 4.300E-06 | 6.300E-06 |
| 14928 | 747 | L747G | B | 8.8e-06 | 1.2e-05 | 1.3e-05 | 1.127E-05 | 2.194E-06 | 8.800E-06 | 1.300E-05 |
| 14929 | 747 | L747H | B | 6.5e-06 | 7.9e-06 | 6e-06 | 6.800E-06 | 9.849E-07 | 6.000E-06 | 7.900E-06 |
| 14930 | 747 | L747I | B | 1.4e-05 | 7.8e-06 | 8.9e-06 | 1.023E-05 | 3.308E-06 | 7.800E-06 | 1.400E-05 |
| 14931 | 747 | L747L | B | 8.2e-06 | 8.9e-06 | 9.1e-06 | 8.733E-06 | 4.726E-07 | 8.200E-06 | 9.100E-06 |
| 14932 | 747 | L747K | B | 4.8e-06 | 9.6e-06 | 6.9e-06 | 7.100E-06 | 2.406E-06 | 4.800E-06 | 9.600E-06 |
| 14933 | 747 | L747M | B | 1.4e-05 | 9.1e-06 | 4.6e-06 | 9.233E-06 | 4.701E-06 | 4.600E-06 | 1.400E-05 |
| 14934 | 747 | L747F | B | 6.9e-06 | 6.2e-06 | 8.8e-06 | 7.300E-06 | 1.345E-06 | 6.200E-06 | 8.800E-06 |
| 14935 | 747 | L747P | B | 4.6e-06 | 1.1e-05 | 4.6e-06 | 6.733E-06 | 3.695E-06 | 4.600E-06 | 1.100E-05 |
| 14936 | 747 | L747S | B | 5.6e-06 | 7.3e-06 | 5.4e-06 | 6.100E-06 | 1.044E-06 | 5.400E-06 | 7.300E-06 |
| 14937 | 747 | L747T | B | 5.3e-06 | 6.8e-06 | 6.9e-06 | 6.333E-06 | 8.963E-07 | 5.300E-06 | 6.900E-06 |
| 14938 | 747 | L747W | B | 6.4e-06 | 9.2e-06 | 8.1e-06 | 7.900E-06 | 1.411E-06 | 6.400E-06 | 9.200E-06 |
| 14939 | 747 | L747Y | B | 8e-06 | 9.3e-06 | 8.3e-06 | 8.533E-06 | 6.807E-07 | 8.000E-06 | 9.300E-06 |
| 14940 | 747 | L747V | B | 8.9e-06 | 4.7e-06 | 6.6e-06 | 6.733E-06 | 2.103E-06 | 4.700E-06 | 8.900E-06 |
| 14941 | 748 | K748A | B | 6.6e-06 | 8.6e-06 | 6e-06 | 7.067E-06 | 1.361E-06 | 6.000E-06 | 8.600E-06 |
| 14942 | 748 | K748R | B | 1.4e-05 | 5.4e-06 | 1.1e-05 | 1.013E-05 | 4.365E-06 | 5.400E-06 | 1.400E-05 |
| 14943 | 748 | K748N | B | 9.4e-06 | 5.9e-06 | 8.9e-06 | 8.067E-06 | 1.893E-06 | 5.900E-06 | 9.400E-06 |
| 14944 | 748 | K748D | B | 6.8e-06 | 6.4e-06 | 1.1e-05 | 8.067E-06 | 2.548E-06 | 6.400E-06 | 1.100E-05 |
| 14945 | 748 | K748C | B | 6.3e-06 | 4.7e-06 | 7e-06 | 6.000E-06 | 1.179E-06 | 4.700E-06 | 7.000E-06 |
| 14946 | 748 | K748E | B | 6.5e-06 | 5.6e-06 | 6.8e-06 | 6.300E-06 | 6.245E-07 | 5.600E-06 | 6.800E-06 |
| 14947 | 748 | K748Q | B | 2e-05 | 5.2e-06 | 7.1e-06 | 1.077E-05 | 8.053E-06 | 5.200E-06 | 2.000E-05 |
| 14948 | 748 | K748G | B | 7.8e-06 | 9.8e-06 | 7.1e-06 | 8.233E-06 | 1.401E-06 | 7.100E-06 | 9.800E-06 |
| 14949 | 748 | K748H | B | 1.3e-05 | 9.4e-06 | 6.2e-06 | 9.533E-06 | 3.402E-06 | 6.200E-06 | 1.300E-05 |
| 14950 | 748 | K748I | B | 5.8e-06 | 5.8e-06 | 8.8e-06 | 6.800E-06 | 1.732E-06 | 5.800E-06 | 8.800E-06 |
| 14951 | 748 | K748L | B | 4.9e-06 | 6.3e-06 | 9.1e-06 | 6.767E-06 | 2.139E-06 | 4.900E-06 | 9.100E-06 |
| 14952 | 748 | K748K | B | 7.8e-06 | 7.9e-06 | 5.4e-06 | 7.033E-06 | 1.415E-06 | 5.400E-06 | 7.900E-06 |
| 14953 | 748 | K748M | B | 8.4e-06 | 7.7e-06 | 7.1e-06 | 7.733E-06 | 6.506E-07 | 7.100E-06 | 8.400E-06 |
| 14954 | 748 | K748F | B | 1.4e-05 | 1.2e-05 | 9.2e-06 | 1.173E-05 | 2.411E-06 | 9.200E-06 | 1.400E-05 |
| 14955 | 748 | K748P | B | 9.1e-06 | 4.2e-06 | 5.9e-06 | 6.400E-06 | 2.488E-06 | 4.200E-06 | 9.100E-06 |
| 14956 | 748 | K748S | B | 6.3e-06 | 7.3e-06 | 7.9e-06 | 7.167E-06 | 8.083E-07 | 6.300E-06 | 7.900E-06 |
| 14957 | 748 | K748T | B | 1.2e-05 | 7.7e-06 | 9.3e-06 | 9.667E-06 | 2.173E-06 | 7.700E-06 | 1.200E-05 |
| 14958 | 748 | K748W | B | 8.6e-06 | 5.3e-06 | 7e-06 | 6.967E-06 | 1.650E-06 | 5.300E-06 | 8.600E-06 |
| 14959 | 748 | K748Y | B | 6.2e-06 | 9.5e-06 | 5.8e-06 | 7.167E-06 | 2.031E-06 | 5.800E-06 | 9.500E-06 |

| | | | | | | | | | | |
|---|---|---|---|---|---|---|---|---|---|---|
| 14960 | 748 | K748V | B | 5e-06 | 7.8e-06 | 8.6e-06 | 7.133E-06 | 1.890E-06 | 5.000E-06 | 8.600E-06 |
| 14961 | 749 | S749A | B | 6.8e-06 | 6.6e-06 | 9.6e-06 | 7.667E-06 | 1.677E-06 | 6.600E-06 | 9.600E-06 |
| 14962 | 749 | S749R | B | 1.4e-05 | 5e-06 | 9.8e-06 | 9.600E-06 | 4.503E-06 | 5.000E-06 | 1.400E-05 |
| 14963 | 749 | S749N | B | 6.7e-06 | 6.7e-06 | 4.1e-06 | 5.833E-06 | 1.501E-06 | 4.100E-06 | 6.700E-06 |
| 14964 | 749 | S749D | B | 8.7e-06 | 9.9e-06 | 5.2e-06 | 7.933E-06 | 2.442E-06 | 5.200E-06 | 9.900E-06 |
| 14965 | 749 | S749C | B | 7.6e-06 | 1.3e-05 | 1.1e-05 | 1.053E-05 | 2.730E-06 | 7.600E-06 | 1.300E-05 |
| 14966 | 749 | S749E | B | 6.8e-06 | 6.1e-06 | 6.6e-06 | 6.500E-06 | 3.606E-07 | 6.100E-06 | 6.800E-06 |
| 14967 | 749 | S749Q | B | 4.5e-06 | 9.1e-06 | 8.7e-06 | 7.433E-06 | 2.548E-06 | 4.500E-06 | 9.100E-06 |
| 14968 | 749 | S749G | B | 9.3e-06 | 1.4e-05 | 7.8e-06 | 1.037E-05 | 3.235E-06 | 7.800E-06 | 1.400E-05 |
| 14969 | 749 | S749H | B | 8.7e-06 | 8.6e-06 | 1e-05 | 9.100E-06 | 7.810E-07 | 8.600E-06 | 1.000E-05 |
| 14970 | 749 | S749I | B | 4.5e-06 | 7.6e-06 | 2.1e-05 | 1.103E-05 | 8.769E-06 | 4.500E-06 | 2.100E-05 |
| 14971 | 749 | S749L | B | 7.1e-06 | 5.8e-06 | 7.2e-06 | 6.700E-06 | 7.810E-07 | 5.800E-06 | 7.200E-06 |
| 14972 | 749 | S749K | B | 4.2e-06 | 6.3e-06 | 7.7e-06 | 6.067E-06 | 1.762E-06 | 4.200E-06 | 7.700E-06 |
| 14973 | 749 | S749M | B | 6.7e-06 | 6.8e-06 | 6.7e-06 | 6.733E-06 | 5.774E-08 | 6.700E-06 | 6.800E-06 |
| 14974 | 749 | S749F | B | 8.1e-06 | 2e-05 | 1.2e-05 | 1.337E-05 | 6.067E-06 | 8.100E-06 | 2.000E-05 |
| 14975 | 749 | S749P | B | 8.9e-06 | 1.4e-05 | 7e-06 | 9.967E-06 | 3.620E-06 | 7.000E-06 | 1.400E-05 |
| 14976 | 749 | S749S | B | 8.1e-06 | 6e-06 | 7e-06 | 7.033E-06 | 1.050E-06 | 6.000E-06 | 8.100E-06 |
| 14977 | 749 | S749T | B | 5.1e-06 | 6.6e-06 | 4.9e-06 | 5.533E-06 | 9.292E-07 | 4.900E-06 | 6.600E-06 |
| 14978 | 749 | S749W | B | 1.8e-05 | 6.6e-06 | 5.8e-06 | 1.013E-05 | 6.824E-06 | 5.800E-06 | 1.800E-05 |
| 14979 | 749 | S749Y | B | 1.2e-05 | 6.8e-06 | 1.4e-05 | 1.093E-05 | 3.717E-06 | 6.800E-06 | 1.400E-05 |
| 14980 | 749 | S749V | B | 8.6e-06 | 9.7e-06 | 7.2e-06 | 8.500E-06 | 1.253E-06 | 7.200E-06 | 9.700E-06 |
| 14981 | 750 | G750A | B | 6.8e-06 | 4.6e-06 | 9.4e-06 | 6.933E-06 | 2.403E-06 | 4.600E-06 | 9.400E-06 |
| 14982 | 750 | G750R | B | 5.8e-06 | 9e-06 | 4.6e-06 | 6.467E-06 | 2.274E-06 | 4.600E-06 | 9.000E-06 |
| 14983 | 750 | G750N | B | 5.6e-06 | 1.1e-05 | 1e-05 | 8.867E-06 | 2.873E-06 | 5.600E-06 | 1.100E-05 |
| 14984 | 750 | G750D | B | 6.9e-06 | 1.1e-05 | 7.4e-06 | 8.433E-06 | 2.237E-06 | 6.900E-06 | 1.100E-05 |
| 14985 | 750 | G750C | B | 9.5e-06 | 6.5e-06 | 4.7e-06 | 6.900E-06 | 2.425E-06 | 4.700E-06 | 9.500E-06 |
| 14986 | 750 | G750E | B | 8.8e-06 | 7.4e-06 | 1e-05 | 8.733E-06 | 1.301E-06 | 7.400E-06 | 1.000E-05 |
| 14987 | 750 | G750Q | B | 8.4e-06 | 1.1e-05 | 5.9e-06 | 8.433E-06 | 2.550E-06 | 5.900E-06 | 1.100E-05 |
| 14988 | 750 | G750G | B | 5.5e-06 | 8.6e-06 | 1e-05 | 8.033E-06 | 2.303E-06 | 5.500E-06 | 1.000E-05 |
| 14989 | 750 | G750H | B | 5.1e-06 | 1.3e-05 | 7.2e-06 | 8.433E-06 | 4.092E-06 | 5.100E-06 | 1.300E-05 |
| 14990 | 750 | G750I | B | 9.1e-06 | 1e-05 | 1.4e-05 | 1.103E-05 | 2.608E-06 | 9.100E-06 | 1.400E-05 |
| 14991 | 750 | G750L | B | 3.7e-06 | 6e-06 | 9.3e-06 | 6.333E-06 | 2.815E-06 | 3.700E-06 | 9.300E-06 |
| 14992 | 750 | G750K | B | 1.5e-05 | 1.5e-05 | 8.7e-06 | 1.290E-05 | 3.637E-06 | 8.700E-06 | 1.500E-05 |
| 14993 | 750 | G750M | B | 4.3e-06 | 9.2e-06 | 8.6e-06 | 7.367E-06 | 2.673E-06 | 4.300E-06 | 9.200E-06 |
| 14994 | 750 | G750F | B | 7.5e-06 | 4.6e-06 | 1.4e-05 | 8.700E-06 | 4.814E-06 | 4.600E-06 | 1.400E-05 |
| 14995 | 750 | G750P | B | 9.4e-06 | 7e-06 | 4.2e-06 | 6.867E-06 | 2.603E-06 | 4.200E-06 | 9.400E-06 |
| 14996 | 750 | G750S | B | 7.8e-06 | 1e-05 | 7.5e-06 | 8.433E-06 | 1.365E-06 | 7.500E-06 | 1.000E-05 |
| 14997 | 750 | G750T | B | 9.8e-06 | 6.6e-06 | 7.7e-06 | 8.033E-06 | 1.626E-06 | 6.600E-06 | 9.800E-06 |
| 14998 | 750 | G750W | B | 6.9e-06 | 4e-06 | 1.2e-05 | 7.633E-06 | 4.050E-06 | 4.000E-06 | 1.200E-05 |
| 14999 | 750 | G750Y | B | 5.9e-06 | 6.9e-06 | 5.1e-06 | 5.967E-06 | 9.018E-07 | 5.100E-06 | 6.900E-06 |
| 15000 | 750 | G750V | B | 6.5e-06 | 9e-06 | 5.1e-06 | 6.867E-06 | 1.976E-06 | 5.100E-06 | 9.000E-06 |
| 15001 | 751 | T751A | B | 9.1e-06 | 8.9e-06 | 5.6e-06 | 7.867E-06 | 1.966E-06 | 5.600E-06 | 9.100E-06 |
| 15002 | 751 | T751R | B | 1e-05 | 8.6e-06 | 6.2e-06 | 8.267E-06 | 1.922E-06 | 6.200E-06 | 1.000E-05 |
| 15003 | 751 | T751N | B | 1.3e-05 | 6.2e-06 | 1e-05 | 9.733E-06 | 3.408E-06 | 6.200E-06 | 1.300E-05 |
| 15004 | 751 | T751D | B | 4.9e-06 | 9.6e-06 | 5.1e-06 | 6.533E-06 | 2.658E-06 | 4.900E-06 | 9.600E-06 |
| 15005 | 751 | T751C | B | 9.6e-06 | 6.6e-06 | 7.4e-06 | 7.867E-06 | 1.553E-06 | 6.600E-06 | 9.600E-06 |
| 15006 | 751 | T751E | B | 8.7e-06 | 6.3e-06 | 9.3e-06 | 8.100E-06 | 1.587E-06 | 6.300E-06 | 9.300E-06 |
| 15007 | 751 | T751Q | B | 9.3e-06 | 1.1e-05 | 9e-06 | 9.767E-06 | 1.079E-06 | 9.000E-06 | 1.100E-05 |
| 15008 | 751 | T751G | B | 7.3e-06 | 8.9e-06 | 1.1e-05 | 9.067E-06 | 1.856E-06 | 7.300E-06 | 1.100E-05 |
| 15009 | 751 | T751H | B | 1.4e-05 | 6.4e-06 | 6.4e-06 | 8.933E-06 | 4.388E-06 | 6.400E-06 | 1.400E-05 |
| 15010 | 751 | T751I | B | 8.6e-06 | 8.2e-06 | 7.8e-06 | 8.200E-06 | 4.000E-07 | 7.800E-06 | 8.600E-06 |
| 15011 | 751 | T751L | B | 7.4e-06 | 1.1e-05 | 6.4e-06 | 8.267E-06 | 2.419E-06 | 6.400E-06 | 1.100E-05 |
| 15012 | 751 | T751K | B | 7.3e-06 | 7.7e-06 | 1.3e-05 | 9.333E-06 | 3.182E-06 | 7.300E-06 | 1.300E-05 |
| 15013 | 751 | T751M | B | 6.7e-06 | 5.9e-06 | 7.5e-06 | 6.700E-06 | 8.000E-07 | 5.900E-06 | 7.500E-06 |
| 15014 | 751 | T751F | B | 6.5e-06 | 7.6e-06 | 5.6e-06 | 6.567E-06 | 1.002E-06 | 5.600E-06 | 7.600E-06 |
| 15015 | 751 | T751P | B | 9.2e-06 | 6.6e-06 | 9.6e-06 | 8.467E-06 | 1.629E-06 | 6.600E-06 | 9.600E-06 |
| 15016 | 751 | T751S | B | 1e-05 | 7.7e-06 | 1.4e-05 | 1.057E-05 | 3.188E-06 | 7.700E-06 | 1.400E-05 |
| 15017 | 751 | T751T | B | 6.9e-06 | 6.8e-06 | 4.7e-06 | 6.133E-06 | 1.242E-06 | 4.700E-06 | 6.900E-06 |
| 15018 | 751 | T751W | B | 7.2e-06 | 8.8e-06 | 9.4e-06 | 8.467E-06 | 1.137E-06 | 7.200E-06 | 9.400E-06 |
| 15019 | 751 | T751Y | B | 4.9e-06 | 1.4e-05 | 5.9e-06 | 8.267E-06 | 4.990E-06 | 4.900E-06 | 1.400E-05 |
| 15020 | 751 | T751V | B | 6e-06 | 5.9e-06 | 9.1e-06 | 7.000E-06 | 1.819E-06 | 5.900E-06 | 9.100E-06 |
| 15021 | 752 | A752A | B | 5e-06 | 7.4e-06 | 4.8e-06 | 5.733E-06 | 1.447E-06 | 4.800E-06 | 7.400E-06 |
| 15022 | 752 | A752R | B | 6.3e-06 | 5.9e-06 | 6.5e-06 | 6.233E-06 | 3.055E-07 | 5.900E-06 | 6.500E-06 |
| 15023 | 752 | A752N | B | 7.8e-06 | 8e-06 | 1.4e-05 | 9.933E-06 | 3.523E-06 | 7.800E-06 | 1.400E-05 |
| 15024 | 752 | A752D | B | 1.3e-05 | 6.8e-06 | 4.8e-06 | 8.200E-06 | 4.276E-06 | 4.800E-06 | 1.300E-05 |
| 15025 | 752 | A752C | B | 7.3e-06 | 1.4e-05 | 4.2e-06 | 8.500E-06 | 5.009E-06 | 4.200E-06 | 1.400E-05 |
| 15026 | 752 | A752E | B | 4.5e-06 | 1.4e-05 | 9.7e-06 | 9.400E-06 | 4.757E-06 | 4.500E-06 | 1.400E-05 |
| 15027 | 752 | A752Q | B | 1.1e-05 | 8.4e-06 | 1.8e-05 | 1.247E-05 | 4.965E-06 | 8.400E-06 | 1.800E-05 |
| 15028 | 752 | A752G | B | 1.1e-05 | 9.1e-06 | 9.1e-06 | 9.733E-06 | 1.097E-06 | 9.100E-06 | 1.100E-05 |
| 15029 | 752 | A752H | B | 6.9e-06 | 9.1e-06 | 1.4e-05 | 1.000E-05 | 3.635E-06 | 6.900E-06 | 1.400E-05 |
| 15030 | 752 | A752I | B | 7.2e-06 | 9.3e-06 | 9.3e-06 | 8.600E-06 | 1.212E-06 | 7.200E-06 | 9.300E-06 |
| 15031 | 752 | A752L | B | 7.1e-06 | 9.2e-06 | 6.7e-06 | 7.667E-06 | 1.343E-06 | 6.700E-06 | 9.200E-06 |
| 15032 | 752 | A752K | B | 5.1e-06 | 7.2e-06 | 1e-05 | 7.433E-06 | 2.458E-06 | 5.100E-06 | 1.000E-05 |
| 15033 | 752 | A752M | B | 9.1e-06 | 6.3e-06 | 1.8e-05 | 1.113E-05 | 6.109E-06 | 6.300E-06 | 1.800E-05 |
| 15034 | 752 | A752F | B | 7.4e-06 | 7e-06 | 4.3e-06 | 6.233E-06 | 1.686E-06 | 4.300E-06 | 7.400E-06 |
| 15035 | 752 | A752P | B | 4.9e-06 | 8.5e-06 | 6.8e-06 | 6.733E-06 | 1.801E-06 | 4.900E-06 | 8.500E-06 |
| 15036 | 752 | A752S | B | 6.8e-06 | 1.1e-05 | 6.7e-06 | 8.167E-06 | 2.454E-06 | 6.700E-06 | 1.100E-05 |
| 15037 | 752 | A752T | B | 6.2e-06 | 1.3e-05 | 7.4e-06 | 8.867E-06 | 3.630E-06 | 6.200E-06 | 1.300E-05 |
| 15038 | 752 | A752W | B | 6.3e-06 | 9.2e-06 | 6.8e-06 | 7.433E-06 | 1.550E-06 | 6.300E-06 | 9.200E-06 |
| 15039 | 752 | A752Y | B | 2e-05 | 7.1e-06 | 6.6e-06 | 1.123E-05 | 7.596E-06 | 6.600E-06 | 2.000E-05 |
| 15040 | 752 | A752V | B | 3.7e-06 | 5.7e-06 | 8.6e-06 | 6.000E-06 | 2.464E-06 | 3.700E-06 | 8.600E-06 |
| 15041 | 753 | S753A | B | 8.8e-06 | 6.5e-06 | 5.2e-06 | 6.833E-06 | 1.823E-06 | 5.200E-06 | 8.800E-06 |
| 15042 | 753 | S753R | B | 9.4e-06 | 5.4e-06 | 7.8e-06 | 7.533E-06 | 2.013E-06 | 5.400E-06 | 9.400E-06 |
| 15043 | 753 | S753N | B | 6.5e-06 | 1.3e-05 | 4.8e-06 | 8.100E-06 | 4.328E-06 | 4.800E-06 | 1.300E-05 |
| 15044 | 753 | S753D | B | 7.6e-06 | 3e-06 | 1.1e-05 | 7.200E-06 | 4.015E-06 | 3.000E-06 | 1.100E-05 |

| | | | | | | | | | | |
|---|---|---|---|---|---|---|---|---|---|---|
| 15045 | 753 | S753C | B | 1e-05 | 6e-06 | 7.2e-06 | 7.733E-06 | 2.053E-06 | 6.000E-06 | 1.000E-05 |
| 15046 | 753 | S753E | B | 9.2e-06 | 5.9e-06 | 5.1e-06 | 6.733E-06 | 2.173E-06 | 5.100E-06 | 9.200E-06 |
| 15047 | 753 | S753Q | B | 6.9e-06 | 6.8e-06 | 1e-05 | 7.900E-06 | 1.819E-06 | 6.800E-06 | 1.000E-05 |
| 15048 | 753 | S753G | B | 4.8e-06 | 5.6e-06 | 8.1e-06 | 6.167E-06 | 1.721E-06 | 4.800E-06 | 8.100E-06 |
| 15049 | 753 | S753H | B | 6.7e-06 | 5.1e-06 | 5.1e-06 | 5.633E-06 | 9.238E-07 | 5.100E-06 | 6.700E-06 |
| 15050 | 753 | S753I | B | 6.8e-06 | 7.2e-06 | 9.7e-06 | 7.900E-06 | 1.572E-06 | 6.800E-06 | 9.700E-06 |
| 15051 | 753 | S753L | B | 5.4e-06 | 1.3e-05 | 5.5e-06 | 7.967E-06 | 4.359E-06 | 5.400E-06 | 1.300E-05 |
| 15052 | 753 | S753K | B | 8.9e-06 | 1e-05 | 1.4e-05 | 1.097E-05 | 2.684E-06 | 8.900E-06 | 1.400E-05 |
| 15053 | 753 | S753M | B | 2.7e-06 | 5.5e-06 | 1.2e-05 | 6.733E-06 | 4.771E-06 | 2.700E-06 | 1.200E-05 |
| 15054 | 753 | S753F | B | 9.1e-06 | 6.4e-06 | 7.9e-06 | 7.800E-06 | 1.353E-06 | 6.400E-06 | 9.100E-06 |
| 15055 | 753 | S753P | B | 6.7e-06 | 4.5e-06 | 4.8e-06 | 5.333E-06 | 1.193E-06 | 4.500E-06 | 6.700E-06 |
| 15056 | 753 | S753S | B | 8.2e-06 | 5.7e-06 | 4.6e-06 | 6.167E-06 | 1.845E-06 | 4.600E-06 | 8.200E-06 |
| 15057 | 753 | S753T | B | 6.9e-06 | 6.8e-06 | 9.7e-06 | 7.800E-06 | 1.646E-06 | 6.800E-06 | 9.700E-06 |
| 15058 | 753 | S753W | B | 6e-06 | 7.2e-06 | 1.2e-05 | 8.400E-06 | 3.175E-06 | 6.000E-06 | 1.200E-05 |
| 15059 | 753 | S753Y | B | 6.5e-06 | 8.7e-06 | 6.6e-06 | 7.267E-06 | 1.242E-06 | 6.500E-06 | 8.700E-06 |
| 15060 | 753 | S753V | B | 8.6e-06 | 8.4e-06 | 1.4e-05 | 1.033E-05 | 3.177E-06 | 8.400E-06 | 1.400E-05 |
| 15061 | 754 | V754A | B | 1.1e-05 | 5.9e-06 | 9.9e-06 | 8.933E-06 | 2.684E-06 | 5.900E-06 | 1.100E-05 |
| 15062 | 754 | V754R | B | 5.6e-06 | 6.8e-06 | 8e-06 | 6.800E-06 | 1.200E-06 | 5.600E-06 | 8.000E-06 |
| 15063 | 754 | V754N | B | 5.6e-06 | 6.9e-06 | 8e-06 | 6.833E-06 | 1.201E-06 | 5.600E-06 | 8.000E-06 |
| 15064 | 754 | V754D | B | 7.3e-06 | 8.8e-06 | 5.2e-06 | 7.100E-06 | 1.808E-06 | 5.200E-06 | 8.800E-06 |
| 15065 | 754 | V754C | B | 7.3e-06 | 3.8e-06 | 7.6e-06 | 6.233E-06 | 2.113E-06 | 3.800E-06 | 7.600E-06 |
| 15066 | 754 | V754E | B | 5.1e-06 | 6.8e-06 | 7.5e-06 | 6.467E-06 | 1.234E-06 | 5.100E-06 | 7.500E-06 |
| 15067 | 754 | V754Q | B | 9.8e-06 | 7.7e-06 | 1e-05 | 9.167E-06 | 1.274E-06 | 7.700E-06 | 1.000E-05 |
| 15068 | 754 | V754G | B | 8.7e-06 | 6.3e-06 | 3.3e-06 | 6.100E-06 | 2.706E-06 | 3.300E-06 | 8.700E-06 |
| 15069 | 754 | V754H | B | 5.1e-06 | 6.6e-06 | 7e-06 | 6.233E-06 | 1.002E-06 | 5.100E-06 | 7.000E-06 |
| 15070 | 754 | V754I | B | 7.5e-06 | 8.4e-06 | 5.6e-06 | 7.167E-06 | 1.429E-06 | 5.600E-06 | 8.400E-06 |
| 15071 | 754 | V754L | B | 5.3e-06 | 5.4e-06 | 8.8e-06 | 6.500E-06 | 1.992E-06 | 5.300E-06 | 8.800E-06 |
| 15072 | 754 | V754K | B | 4.3e-06 | 8.7e-06 | 6e-06 | 6.333E-06 | 2.219E-06 | 4.300E-06 | 8.700E-06 |
| 15073 | 754 | V754M | B | 5.8e-06 | 1e-05 | 6e-06 | 7.267E-06 | 2.369E-06 | 5.800E-06 | 1.000E-05 |
| 15074 | 754 | V754F | B | 9.6e-06 | 8.3e-06 | 1.3e-05 | 1.030E-05 | 2.427E-06 | 8.300E-06 | 1.300E-05 |
| 15075 | 754 | V754P | B | 6.7e-06 | 1.2e-05 | 8.2e-06 | 8.967E-06 | 2.732E-06 | 6.700E-06 | 1.200E-05 |
| 15076 | 754 | V754S | B | 1.3e-05 | 7.7e-06 | 9.7e-06 | 1.013E-05 | 2.676E-06 | 7.700E-06 | 1.300E-05 |
| 15077 | 754 | V754T | B | 9.6e-06 | 6.7e-06 | 3.9e-06 | 6.733E-06 | 2.850E-06 | 3.900E-06 | 9.600E-06 |
| 15078 | 754 | V754W | B | 5.2e-06 | 9.3e-06 | 8e-06 | 7.500E-06 | 2.095E-06 | 5.200E-06 | 9.300E-06 |
| 15079 | 754 | V754Y | B | 8.5e-06 | 1.2e-05 | 4.8e-06 | 8.433E-06 | 3.600E-06 | 4.800E-06 | 1.200E-05 |
| 15080 | 754 | V754V | B | 6.6e-06 | 1.4e-05 | 5.3e-06 | 8.633E-06 | 4.693E-06 | 5.300E-06 | 1.400E-05 |
| 15081 | 755 | V755A | B | 1.3e-05 | 4.2e-06 | 8.5e-06 | 8.567E-06 | 4.400E-06 | 4.200E-06 | 1.300E-05 |
| 15082 | 755 | V755R | B | 6.1e-06 | 8e-06 | 9.3e-06 | 7.800E-06 | 1.609E-06 | 6.100E-06 | 9.300E-06 |
| 15083 | 755 | V755N | B | 5.6e-06 | 1e-05 | 6.1e-06 | 7.233E-06 | 2.409E-06 | 5.600E-06 | 1.000E-05 |
| 15084 | 755 | V755D | B | 1.4e-05 | 6.8e-06 | 7.3e-06 | 9.367E-06 | 4.020E-06 | 6.800E-06 | 1.400E-05 |
| 15085 | 755 | V755C | B | 6.4e-06 | 6.1e-06 | 8.5e-06 | 7.000E-06 | 1.308E-06 | 6.100E-06 | 8.500E-06 |
| 15086 | 755 | V755E | B | 6.1e-06 | 5.2e-06 | 9.7e-06 | 7.000E-06 | 2.381E-06 | 5.200E-06 | 9.700E-06 |
| 15087 | 755 | V755Q | B | 4.9e-06 | 6.9e-06 | 9.4e-06 | 7.067E-06 | 2.255E-06 | 4.900E-06 | 9.400E-06 |
| 15088 | 755 | V755G | B | 1e-05 | 8.5e-06 | 7.1e-06 | 8.533E-06 | 1.450E-06 | 7.100E-06 | 1.000E-05 |
| 15089 | 755 | V755H | B | 9.5e-06 | 6.4e-06 | 6.5e-06 | 7.467E-06 | 1.762E-06 | 6.400E-06 | 9.500E-06 |
| 15090 | 755 | V755I | B | 9.3e-06 | 5.6e-06 | 6.6e-06 | 7.167E-06 | 1.914E-06 | 5.600E-06 | 9.300E-06 |
| 15091 | 755 | V755L | B | 9.6e-06 | 9.7e-06 | 6.7e-06 | 8.667E-06 | 1.704E-06 | 6.700E-06 | 9.700E-06 |
| 15092 | 755 | V755K | B | 7.1e-06 | 9.9e-06 | 6.4e-06 | 7.800E-06 | 1.852E-06 | 6.400E-06 | 9.900E-06 |
| 15093 | 755 | V755M | B | 7.8e-06 | 1.2e-05 | 6.6e-06 | 8.800E-06 | 2.835E-06 | 6.600E-06 | 1.200E-05 |
| 15094 | 755 | V755F | B | 9.5e-06 | 6.9e-06 | 7.8e-06 | 8.067E-06 | 1.320E-06 | 6.900E-06 | 9.500E-06 |
| 15095 | 755 | V755P | B | 1e-05 | 3.4e-06 | 9.5e-06 | 7.633E-06 | 3.675E-06 | 3.400E-06 | 1.000E-05 |
| 15096 | 755 | V755S | B | 4.5e-06 | 5.7e-06 | 8e-06 | 6.067E-06 | 1.779E-06 | 4.500E-06 | 8.000E-06 |
| 15097 | 755 | V755T | B | 6.5e-06 | 6.2e-06 | 6.8e-06 | 6.500E-06 | 3.000E-07 | 6.200E-06 | 6.800E-06 |
| 15098 | 755 | V755W | B | 7.5e-06 | 7.1e-06 | 4.4e-06 | 6.333E-06 | 1.686E-06 | 4.400E-06 | 7.500E-06 |
| 15099 | 755 | V755Y | B | 8.5e-06 | 7.5e-06 | 1.2e-05 | 9.333E-06 | 2.363E-06 | 7.500E-06 | 1.200E-05 |
| 15100 | 755 | V755V | B | 1.3e-05 | 9.4e-06 | 7e-06 | 9.800E-06 | 3.020E-06 | 7.000E-06 | 1.300E-05 |
| 15101 | 756 | C756A | B | 6.4e-06 | 8.4e-06 | 8.7e-06 | 7.833E-06 | 1.250E-06 | 6.400E-06 | 8.700E-06 |
| 15102 | 756 | C756R | B | 1.1e-05 | 5.8e-06 | 7.2e-06 | 8.000E-06 | 2.691E-06 | 5.800E-06 | 1.100E-05 |
| 15103 | 756 | C756N | B | 6.3e-06 | 8.5e-06 | 5.6e-06 | 6.800E-06 | 1.513E-06 | 5.600E-06 | 8.500E-06 |
| 15104 | 756 | C756D | B | 7.2e-06 | 5.2e-06 | 9.7e-06 | 7.367E-06 | 2.255E-06 | 5.200E-06 | 9.700E-06 |
| 15105 | 756 | C756C | B | 8.2e-06 | 1.1e-05 | 9.1e-06 | 9.433E-06 | 1.429E-06 | 8.200E-06 | 1.100E-05 |
| 15106 | 756 | C756E | B | 1e-05 | 6.7e-06 | 7.3e-06 | 8.000E-06 | 1.758E-06 | 6.700E-06 | 1.000E-05 |
| 15107 | 756 | C756Q | B | 9.3e-06 | 4.9e-06 | 1.2e-05 | 8.733E-06 | 3.584E-06 | 4.900E-06 | 1.200E-05 |
| 15108 | 756 | C756G | B | 7.3e-06 | 6.4e-06 | 6.7e-06 | 6.800E-06 | 4.583E-07 | 6.400E-06 | 7.300E-06 |
| 15109 | 756 | C756H | B | 7e-06 | 6.6e-06 | 4.6e-06 | 6.067E-06 | 1.286E-06 | 4.600E-06 | 7.000E-06 |
| 15110 | 756 | C756I | B | 1.2e-05 | 9.6e-06 | 7.8e-06 | 9.800E-06 | 2.107E-06 | 7.800E-06 | 1.200E-05 |
| 15111 | 756 | C756L | B | 8.1e-06 | 7.4e-06 | 6.2e-06 | 7.233E-06 | 9.609E-07 | 6.200E-06 | 8.100E-06 |
| 15112 | 756 | C756K | B | 8e-06 | 1.3e-05 | 7.6e-06 | 9.533E-06 | 3.009E-06 | 7.600E-06 | 1.300E-05 |
| 15113 | 756 | C756M | B | 6.2e-06 | 4.4e-06 | 5.8e-06 | 5.467E-06 | 9.452E-07 | 4.400E-06 | 6.200E-06 |
| 15114 | 756 | C756F | B | 7.9e-06 | 1.1e-05 | 1.4e-05 | 1.097E-05 | 3.050E-06 | 7.900E-06 | 1.400E-05 |
| 15115 | 756 | C756P | B | 5.8e-06 | 9.5e-06 | 9.6e-06 | 8.300E-06 | 2.166E-06 | 5.800E-06 | 9.600E-06 |
| 15116 | 756 | C756S | B | 9.9e-06 | 5.2e-06 | 7.2e-06 | 7.433E-06 | 2.359E-06 | 5.200E-06 | 9.900E-06 |
| 15117 | 756 | C756T | B | 4.9e-06 | 5.7e-06 | 7.9e-06 | 6.167E-06 | 1.553E-06 | 4.900E-06 | 7.900E-06 |
| 15118 | 756 | C756W | B | 6.6e-06 | 4.8e-06 | 7.9e-06 | 6.433E-06 | 1.557E-06 | 4.800E-06 | 7.900E-06 |
| 15119 | 756 | C756Y | B | 5.8e-06 | 1.3e-05 | 1.1e-05 | 9.933E-06 | 3.717E-06 | 5.800E-06 | 1.300E-05 |
| 15120 | 756 | C756V | B | 9.3e-06 | 1.3e-05 | 6.7e-06 | 9.667E-06 | 3.166E-06 | 6.700E-06 | 1.300E-05 |
| 15121 | 757 | L757A | B | 7.3e-06 | 6.2e-06 | 9e-06 | 7.500E-06 | 1.411E-06 | 6.200E-06 | 9.000E-06 |
| 15122 | 757 | L757R | B | 1e-05 | 4.1e-06 | 4.5e-06 | 6.200E-06 | 3.297E-06 | 4.100E-06 | 1.000E-05 |
| 15123 | 757 | L757N | B | 6.5e-06 | 8.9e-06 | 9.3e-06 | 8.233E-06 | 1.514E-06 | 6.500E-06 | 9.300E-06 |
| 15124 | 757 | L757D | B | 7e-06 | 4.6e-06 | 8.3e-06 | 6.633E-06 | 1.877E-06 | 4.600E-06 | 8.300E-06 |
| 15125 | 757 | L757C | B | 8.6e-06 | 6.1e-06 | 1e-05 | 8.233E-06 | 1.976E-06 | 6.100E-06 | 1.000E-05 |
| 15126 | 757 | L757E | B | 9.3e-06 | 1.2e-05 | 5.7e-06 | 9.000E-06 | 3.161E-06 | 5.700E-06 | 1.200E-05 |
| 15127 | 757 | L757Q | B | 8.6e-06 | 1.1e-05 | 8.4e-06 | 9.333E-06 | 1.447E-06 | 8.400E-06 | 1.100E-05 |
| 15128 | 757 | L757G | B | 9.2e-06 | 6.3e-06 | 6.7e-06 | 7.400E-06 | 1.572E-06 | 6.300E-06 | 9.200E-06 |
| 15129 | 757 | L757H | B | 6.7e-06 | 6.1e-06 | 1.3e-05 | 8.600E-06 | 3.822E-06 | 6.100E-06 | 1.300E-05 |

| | | | | | | | | | | |
|---|---|---|---|---|---|---|---|---|---|---|
| 15130 | 757 | L757I | B | 9.1e-06 | 6.6e-06 | 4.3e-06 | 6.667E-06 | 2.401E-06 | 4.300E-06 | 9.100E-06 |
| 15131 | 757 | L757L | B | 5.2e-06 | 1.4e-05 | 8.3e-06 | 9.167E-06 | 4.464E-06 | 5.200E-06 | 1.400E-05 |
| 15132 | 757 | L757K | B | 9.1e-06 | 7.4e-06 | 5.3e-06 | 7.267E-06 | 1.904E-06 | 5.300E-06 | 9.100E-06 |
| 15133 | 757 | L757M | B | 1.1e-05 | 8.4e-06 | 8.4e-06 | 9.267E-06 | 1.501E-06 | 8.400E-06 | 1.100E-05 |
| 15134 | 757 | L757F | B | 5.9e-06 | 7e-06 | 6.4e-06 | 6.433E-06 | 5.508E-07 | 5.900E-06 | 7.000E-06 |
| 15135 | 757 | L757P | B | 5.8e-06 | 6.5e-06 | 1.6e-05 | 9.433E-06 | 5.698E-06 | 5.800E-06 | 1.600E-05 |
| 15136 | 757 | L757S | B | 7.4e-06 | 7e-06 | 6.8e-06 | 7.067E-06 | 3.055E-07 | 6.800E-06 | 7.400E-06 |
| 15137 | 757 | L757T | B | 5.8e-06 | 5.2e-06 | 1.2e-05 | 7.667E-06 | 3.765E-06 | 5.200E-06 | 1.200E-05 |
| 15138 | 757 | L757W | B | 3.8e-06 | 8.8e-06 | 1.2e-05 | 8.200E-06 | 4.133E-06 | 3.800E-06 | 1.200E-05 |
| 15139 | 757 | L757Y | B | 6e-06 | 8.6e-06 | 6.4e-06 | 7.000E-06 | 1.400E-06 | 6.000E-06 | 8.600E-06 |
| 15140 | 757 | L757V | B | 8.6e-06 | 8.4e-06 | 1.1e-05 | 9.333E-06 | 1.447E-06 | 8.400E-06 | 1.100E-05 |
| 15141 | 758 | L758A | B | 1.2e-05 | 6.2e-06 | 8.2e-06 | 8.800E-06 | 2.946E-06 | 6.200E-06 | 1.200E-05 |
| 15142 | 758 | L758R | B | 4.9e-06 | 9.3e-06 | 8.6e-06 | 7.600E-06 | 2.364E-06 | 4.900E-06 | 9.300E-06 |
| 15143 | 758 | L758N | B | 8.3e-06 | 1.2e-05 | 1e-05 | 1.010E-05 | 1.852E-06 | 8.300E-06 | 1.200E-05 |
| 15144 | 758 | L758D | B | 1.1e-05 | 1.5e-05 | 1e-05 | 1.200E-05 | 2.646E-06 | 1.000E-05 | 1.500E-05 |
| 15145 | 758 | L758C | B | 8.8e-06 | 7.8e-06 | 6.5e-06 | 7.700E-06 | 1.153E-06 | 6.500E-06 | 8.800E-06 |
| 15146 | 758 | L758E | B | 7.7e-06 | 6.3e-06 | 1e-05 | 8.000E-06 | 1.868E-06 | 6.300E-06 | 1.000E-05 |
| 15147 | 758 | L758Q | B | 9.1e-06 | 4.6e-06 | 4.6e-06 | 6.100E-06 | 2.598E-06 | 4.600E-06 | 9.100E-06 |
| 15148 | 758 | L758G | B | 7.1e-06 | 6.4e-06 | 6.3e-06 | 6.600E-06 | 4.359E-07 | 6.300E-06 | 7.100E-06 |
| 15149 | 758 | L758H | B | 8.6e-06 | 9.3e-06 | 1.2e-05 | 9.967E-06 | 1.795E-06 | 8.600E-06 | 1.200E-05 |
| 15150 | 758 | L758I | B | 9.6e-06 | 1e-05 | 9.8e-06 | 9.800E-06 | 2.000E-07 | 9.600E-06 | 1.000E-05 |
| 15151 | 758 | L758L | B | 5.5e-06 | 1e-05 | 4.6e-06 | 6.700E-06 | 2.893E-06 | 4.600E-06 | 1.000E-05 |
| 15152 | 758 | L758K | B | 5.8e-06 | 1.1e-05 | 9.1e-06 | 8.633E-06 | 2.631E-06 | 5.800E-06 | 1.100E-05 |
| 15153 | 758 | L758M | B | 6.3e-06 | 8.3e-06 | 9.6e-06 | 8.067E-06 | 1.662E-06 | 6.300E-06 | 9.600E-06 |
| 15154 | 758 | L758F | B | 8.9e-06 | 7e-06 | 4.9e-06 | 6.933E-06 | 2.001E-06 | 4.900E-06 | 8.900E-06 |
| 15155 | 758 | L758P | B | 1.3e-05 | 1.5e-05 | 1e-05 | 1.267E-05 | 2.517E-06 | 1.000E-05 | 1.500E-05 |
| 15156 | 758 | L758S | B | 8.5e-06 | 1.2e-05 | 1e-05 | 1.017E-05 | 1.756E-06 | 8.500E-06 | 1.200E-05 |
| 15157 | 758 | L758T | B | 1e-05 | 7.1e-06 | 5.5e-06 | 7.533E-06 | 2.281E-06 | 5.500E-06 | 1.000E-05 |
| 15158 | 758 | L758W | B | 9.7e-06 | 1.4e-05 | 7.4e-06 | 1.037E-05 | 3.350E-06 | 7.400E-06 | 1.400E-05 |
| 15159 | 758 | L758Y | B | 3.4e-06 | 7.6e-06 | 6.6e-06 | 5.867E-06 | 2.194E-06 | 3.400E-06 | 7.600E-06 |
| 15160 | 758 | L758V | B | 5.4e-06 | 5.9e-06 | 5e-06 | 5.433E-06 | 4.509E-07 | 5.000E-06 | 5.900E-06 |
| 15161 | 759 | N759A | B | 9.4e-06 | 5.1e-06 | 9.2e-06 | 7.900E-06 | 2.427E-06 | 5.100E-06 | 9.400E-06 |
| 15162 | 759 | N759R | B | 5.9e-06 | 8.9e-06 | 1.3e-05 | 9.267E-06 | 3.564E-06 | 5.900E-06 | 1.300E-05 |
| 15163 | 759 | N759N | B | 9.2e-06 | 8.4e-06 | 6.6e-06 | 8.067E-06 | 1.332E-06 | 6.600E-06 | 9.200E-06 |
| 15164 | 759 | N759D | B | 1.1e-05 | 6.6e-06 | 1.1e-05 | 9.533E-06 | 2.540E-06 | 6.600E-06 | 1.100E-05 |
| 15165 | 759 | N759C | B | 9.1e-06 | 6.4e-06 | 7.2e-06 | 7.567E-06 | 1.387E-06 | 6.400E-06 | 9.100E-06 |
| 15166 | 759 | N759E | B | 1e-05 | 9.7e-06 | 5.9e-06 | 8.533E-06 | 2.285E-06 | 5.900E-06 | 1.000E-05 |
| 15167 | 759 | N759Q | B | 5.1e-06 | 8.7e-06 | 1.2e-05 | 8.600E-06 | 3.451E-06 | 5.100E-06 | 1.200E-05 |
| 15168 | 759 | N759G | B | 6.4e-06 | 1.1e-05 | 9.3e-06 | 8.900E-06 | 2.326E-06 | 6.400E-06 | 1.100E-05 |
| 15169 | 759 | N759H | B | 3.6e-06 | 7.2e-06 | 6.4e-06 | 5.733E-06 | 1.890E-06 | 3.600E-06 | 7.200E-06 |
| 15170 | 759 | N759I | B | 7.1e-06 | 1.1e-05 | 5.7e-06 | 7.933E-06 | 2.747E-06 | 5.700E-06 | 1.100E-05 |
| 15171 | 759 | N759L | B | 9.1e-06 | 8.4e-06 | 9.8e-06 | 9.100E-06 | 7.000E-07 | 8.400E-06 | 9.800E-06 |
| 15172 | 759 | N759K | B | 7.5e-06 | 7.7e-06 | 9.4e-06 | 8.200E-06 | 1.044E-06 | 7.500E-06 | 9.400E-06 |
| 15173 | 759 | N759M | B | 1.3e-05 | 4e-06 | 3.6e-06 | 6.867E-06 | 5.315E-06 | 3.600E-06 | 1.300E-05 |
| 15174 | 759 | N759F | B | 1.1e-05 | 1.2e-05 | 8.6e-06 | 1.053E-05 | 1.747E-06 | 8.600E-06 | 1.200E-05 |
| 15175 | 759 | N759P | B | 1.4e-05 | 1.1e-05 | 1.1e-05 | 1.200E-05 | 1.732E-06 | 1.100E-05 | 1.400E-05 |
| 15176 | 759 | N759S | B | 8.8e-06 | 5.1e-06 | 1.3e-05 | 8.967E-06 | 3.953E-06 | 5.100E-06 | 1.300E-05 |
| 15177 | 759 | N759T | B | 3.4e-06 | 8.5e-06 | 6.4e-06 | 6.100E-06 | 2.563E-06 | 3.400E-06 | 8.500E-06 |
| 15178 | 759 | N759W | B | 5.6e-06 | 8.1e-06 | 9.9e-06 | 7.867E-06 | 2.159E-06 | 5.600E-06 | 9.900E-06 |
| 15179 | 759 | N759Y | B | 6e-06 | 6.8e-06 | 5.9e-06 | 6.233E-06 | 4.933E-07 | 5.900E-06 | 6.800E-06 |
| 15180 | 759 | N759V | B | 7.2e-06 | 6.4e-06 | 6e-06 | 6.533E-06 | 6.110E-07 | 6.000E-06 | 7.200E-06 |
| 15181 | 760 | N760A | B | 6.8e-06 | 7e-06 | 1.4e-05 | 9.267E-06 | 4.100E-06 | 6.800E-06 | 1.400E-05 |
| 15182 | 760 | N760R | B | 7e-06 | 1.5e-05 | 7.1e-06 | 9.700E-06 | 4.590E-06 | 7.000E-06 | 1.500E-05 |
| 15183 | 760 | N760N | B | 7.5e-06 | 7.4e-06 | 6.5e-06 | 7.133E-06 | 5.508E-07 | 6.500E-06 | 7.500E-06 |
| 15184 | 760 | N760D | B | 7.8e-06 | 1.6e-05 | 6.1e-06 | 9.967E-06 | 5.294E-06 | 6.100E-06 | 1.600E-05 |
| 15185 | 760 | N760C | B | 7.7e-06 | 7.3e-06 | 6.8e-06 | 7.267E-06 | 4.509E-07 | 6.800E-06 | 7.700E-06 |
| 15186 | 760 | N760E | B | 8e-06 | 1e-05 | 6.3e-06 | 8.100E-06 | 1.852E-06 | 6.300E-06 | 1.000E-05 |
| 15187 | 760 | N760Q | B | 8e-06 | 7.9e-06 | 7.6e-06 | 7.833E-06 | 2.082E-07 | 7.600E-06 | 8.000E-06 |
| 15188 | 760 | N760G | B | 4.8e-06 | 7.4e-06 | 8.1e-06 | 6.767E-06 | 1.739E-06 | 4.800E-06 | 8.100E-06 |
| 15189 | 760 | N760H | B | 3.5e-06 | 7e-06 | 6.4e-06 | 5.633E-06 | 1.872E-06 | 3.500E-06 | 7.000E-06 |
| 15190 | 760 | N760I | B | 9.7e-06 | 7.7e-06 | 1.3e-05 | 1.013E-05 | 2.676E-06 | 7.700E-06 | 1.300E-05 |
| 15191 | 760 | N760L | B | 1.3e-05 | 8.8e-06 | 6.7e-06 | 9.500E-06 | 3.208E-06 | 6.700E-06 | 1.300E-05 |
| 15192 | 760 | N760K | B | 3.9e-06 | 1e-05 | 1.9e-05 | 1.097E-05 | 7.596E-06 | 3.900E-06 | 1.900E-05 |
| 15193 | 760 | N760M | B | 6e-06 | 7.1e-06 | 7.7e-06 | 6.933E-06 | 8.622E-07 | 6.000E-06 | 7.700E-06 |
| 15194 | 760 | N760F | B | 6.3e-06 | 8.9e-06 | 6e-06 | 7.067E-06 | 1.595E-06 | 6.000E-06 | 8.900E-06 |
| 15195 | 760 | N760P | B | 5.8e-06 | 6.2e-06 | 1e-05 | 7.333E-06 | 2.318E-06 | 5.800E-06 | 1.000E-05 |
| 15196 | 760 | N760S | B | 9.3e-06 | 1.1e-05 | 8.4e-06 | 9.567E-06 | 1.320E-06 | 8.400E-06 | 1.100E-05 |
| 15197 | 760 | N760T | B | 7e-06 | 8.7e-06 | 7.6e-06 | 7.767E-06 | 8.622E-07 | 7.000E-06 | 8.700E-06 |
| 15198 | 760 | N760W | B | 9.3e-06 | 8e-06 | 6.6e-06 | 7.967E-06 | 1.350E-06 | 6.600E-06 | 9.300E-06 |
| 15199 | 760 | N760Y | B | 1.2e-05 | 1.1e-05 | 1.1e-05 | 1.133E-05 | 5.774E-07 | 1.100E-05 | 1.200E-05 |
| 15200 | 760 | N760V | B | 8.7e-06 | 9.2e-06 | 8.8e-06 | 8.900E-06 | 2.646E-07 | 8.700E-06 | 9.200E-06 |
| 15201 | 761 | F761A | B | 4.3e-06 | 5.9e-06 | 8.4e-06 | 6.200E-06 | 2.066E-06 | 4.300E-06 | 8.400E-06 |
| 15202 | 761 | F761R | B | 7.9e-06 | 8.9e-06 | 4.9e-06 | 7.233E-06 | 2.082E-06 | 4.900E-06 | 8.900E-06 |
| 15203 | 761 | F761N | B | 9e-06 | 6.2e-06 | 6.5e-06 | 7.233E-06 | 1.537E-06 | 6.200E-06 | 9.000E-06 |
| 15204 | 761 | F761D | B | 6e-06 | 8.1e-06 | 6.2e-06 | 6.767E-06 | 1.159E-06 | 6.000E-06 | 8.100E-06 |
| 15205 | 761 | F761C | B | 5.6e-06 | 1.3e-05 | 8.7e-06 | 9.100E-06 | 3.716E-06 | 5.600E-06 | 1.300E-05 |
| 15206 | 761 | F761E | B | 4.9e-06 | 7.4e-06 | 8.2e-06 | 6.833E-06 | 1.721E-06 | 4.900E-06 | 8.200E-06 |
| 15207 | 761 | F761Q | B | 4.7e-06 | 8.8e-06 | 1.1e-05 | 8.167E-06 | 3.197E-06 | 4.700E-06 | 1.100E-05 |
| 15208 | 761 | F761G | B | 8.1e-06 | 4.9e-06 | 6.4e-06 | 6.467E-06 | 1.601E-06 | 4.900E-06 | 8.100E-06 |
| 15209 | 761 | F761H | B | 6e-06 | 8.7e-06 | 4.5e-06 | 6.400E-06 | 2.128E-06 | 4.500E-06 | 8.700E-06 |
| 15210 | 761 | F761I | B | 1.7e-05 | 1.4e-05 | 1.1e-05 | 1.400E-05 | 3.000E-06 | 1.100E-05 | 1.700E-05 |
| 15211 | 761 | F761L | B | 8.8e-06 | 4.7e-06 | 3.3e-06 | 5.600E-06 | 2.858E-06 | 3.300E-06 | 8.800E-06 |
| 15212 | 761 | F761K | B | 8e-06 | 8.2e-06 | 1.4e-05 | 1.007E-05 | 3.408E-06 | 8.000E-06 | 1.400E-05 |
| 15213 | 761 | F761M | B | 1.1e-05 | 8.1e-06 | 6.7e-06 | 8.600E-06 | 2.193E-06 | 6.700E-06 | 1.100E-05 |
| 15214 | 761 | F761F | B | 7e-06 | 5.6e-06 | 6.2e-06 | 6.267E-06 | 7.024E-07 | 5.600E-06 | 7.000E-06 |

| | | | | | | | | | | |
|---|---|---|---|---|---|---|---|---|---|---|
| 15215 | 761 | F761P | B | 1.2e-05 | 6.1e-06 | 8e-06 | 8.700E-06 | 3.012E-06 | 6.100E-06 | 1.200E-05 |
| 15216 | 761 | F761S | B | 7e-06 | 4.6e-06 | 4.7e-06 | 5.433E-06 | 1.358E-06 | 4.600E-06 | 7.000E-06 |
| 15217 | 761 | F761T | B | 8.4e-06 | 6.3e-06 | 5.8e-06 | 6.833E-06 | 1.380E-06 | 5.800E-06 | 8.400E-06 |
| 15218 | 761 | F761W | B | 9.1e-06 | 7.5e-06 | 8.5e-06 | 8.367E-06 | 8.083E-07 | 7.500E-06 | 9.100E-06 |
| 15219 | 761 | F761Y | B | 7.3e-06 | 9.6e-06 | 1.1e-05 | 9.300E-06 | 1.868E-06 | 7.300E-06 | 1.100E-05 |
| 15220 | 761 | F761V | B | 4.8e-06 | 7.3e-06 | 3.5e-06 | 5.200E-06 | 1.931E-06 | 3.500E-06 | 7.300E-06 |
| 15221 | 762 | Y762A | B | 7.7e-06 | 8.3e-06 | 5.3e-06 | 7.100E-06 | 1.587E-06 | 5.300E-06 | 8.300E-06 |
| 15222 | 762 | Y762R | B | 8.4e-06 | 8.3e-06 | 6.5e-06 | 7.733E-06 | 1.069E-06 | 6.500E-06 | 8.400E-06 |
| 15223 | 762 | Y762N | B | 4.8e-06 | 9.2e-06 | 1.4e-05 | 9.333E-06 | 4.601E-06 | 4.800E-06 | 1.400E-05 |
| 15224 | 762 | Y762D | B | 6.3e-06 | 1e-05 | 1.3e-05 | 9.767E-06 | 3.356E-06 | 6.300E-06 | 1.300E-05 |
| 15225 | 762 | Y762C | B | 1.4e-05 | 6.9e-06 | 6.7e-06 | 9.200E-06 | 4.158E-06 | 6.700E-06 | 1.400E-05 |
| 15226 | 762 | Y762E | B | 8.8e-06 | 1.1e-05 | 7.5e-06 | 9.100E-06 | 1.769E-06 | 7.500E-06 | 1.100E-05 |
| 15227 | 762 | Y762Q | B | 1.3e-05 | 5.2e-06 | 4.5e-06 | 7.567E-06 | 4.718E-06 | 4.500E-06 | 1.300E-05 |
| 15228 | 762 | Y762G | B | 9.2e-06 | 9.6e-06 | 4.5e-06 | 7.767E-06 | 2.836E-06 | 4.500E-06 | 9.600E-06 |
| 15229 | 762 | Y762H | B | 8.1e-06 | 5.5e-06 | 5.7e-06 | 6.433E-06 | 1.447E-06 | 5.500E-06 | 8.100E-06 |
| 15230 | 762 | Y762I | B | 6.5e-06 | 8.3e-06 | 8e-06 | 7.600E-06 | 9.644E-07 | 6.500E-06 | 8.300E-06 |
| 15231 | 762 | Y762L | B | 7e-06 | 8.2e-06 | 1.1e-05 | 8.733E-06 | 2.053E-06 | 7.000E-06 | 1.100E-05 |
| 15232 | 762 | Y762K | B | 9.2e-06 | 1.1e-05 | 1.1e-05 | 1.040E-05 | 1.039E-06 | 9.200E-06 | 1.100E-05 |
| 15233 | 762 | Y762M | B | 1.3e-05 | 1.4e-05 | 8.4e-06 | 1.180E-05 | 2.987E-06 | 8.400E-06 | 1.400E-05 |
| 15234 | 762 | Y762F | B | 1e-05 | 7e-06 | 1.1e-05 | 9.333E-06 | 2.082E-06 | 7.000E-06 | 1.100E-05 |
| 15235 | 762 | Y762P | B | 7.9e-06 | 5.9e-06 | 8e-06 | 7.267E-06 | 1.185E-06 | 5.900E-06 | 8.000E-06 |
| 15236 | 762 | Y762S | B | 8.2e-06 | 1.3e-05 | 5.6e-06 | 8.933E-06 | 3.754E-06 | 5.600E-06 | 1.300E-05 |
| 15237 | 762 | Y762T | B | 5e-06 | 1.4e-05 | 1.4e-05 | 1.100E-05 | 5.196E-06 | 5.000E-06 | 1.400E-05 |
| 15238 | 762 | Y762W | B | 1.2e-05 | 8.7e-06 | 6.2e-06 | 8.967E-06 | 2.909E-06 | 6.200E-06 | 1.200E-05 |
| 15239 | 762 | Y762Y | B | 1.3e-05 | 5.6e-06 | 5.2e-06 | 7.933E-06 | 4.392E-06 | 5.200E-06 | 1.300E-05 |
| 15240 | 762 | Y762V | B | 9.6e-06 | 6.9e-06 | 9.7e-06 | 8.733E-06 | 1.589E-06 | 6.900E-06 | 9.700E-06 |
| 15241 | 763 | P763A | B | 9.5e-06 | 9.6e-06 | 6.5e-06 | 8.533E-06 | 1.762E-06 | 6.500E-06 | 9.600E-06 |
| 15242 | 763 | P763R | B | 1.2e-05 | 1.4e-05 | 7.2e-06 | 1.107E-05 | 3.495E-06 | 7.200E-06 | 1.400E-05 |
| 15243 | 763 | P763N | B | 7.8e-06 | 1.4e-05 | 1.4e-05 | 1.193E-05 | 3.580E-06 | 7.800E-06 | 1.400E-05 |
| 15244 | 763 | P763D | B | 7.9e-06 | 4.3e-06 | 1.1e-05 | 7.733E-06 | 3.353E-06 | 4.300E-06 | 1.100E-05 |
| 15245 | 763 | P763C | B | 6.2e-06 | 9.4e-06 | 9.5e-06 | 8.367E-06 | 1.877E-06 | 6.200E-06 | 9.500E-06 |
| 15246 | 763 | P763E | B | 9.2e-06 | 9.8e-06 | 8.6e-06 | 9.200E-06 | 6.000E-07 | 8.600E-06 | 9.800E-06 |
| 15247 | 763 | P763Q | B | 8.3e-06 | 5.7e-06 | 1.1e-05 | 8.333E-06 | 2.650E-06 | 5.700E-06 | 1.100E-05 |
| 15248 | 763 | P763G | B | 4.3e-06 | 1.2e-05 | 6.7e-06 | 7.667E-06 | 3.940E-06 | 4.300E-06 | 1.200E-05 |
| 15249 | 763 | P763H | B | 5e-06 | 1.1e-05 | 9.4e-06 | 8.467E-06 | 3.107E-06 | 5.000E-06 | 1.100E-05 |
| 15250 | 763 | P763I | B | 8.2e-06 | 6.5e-06 | 8.3e-06 | 7.667E-06 | 1.012E-06 | 6.500E-06 | 8.300E-06 |
| 15251 | 763 | P763L | B | 9.1e-06 | 6.5e-06 | 9.7e-06 | 8.433E-06 | 1.701E-06 | 6.500E-06 | 9.700E-06 |
| 15252 | 763 | P763K | B | 7.6e-06 | 4.6e-06 | 4.3e-06 | 5.500E-06 | 1.825E-06 | 4.300E-06 | 7.600E-06 |
| 15253 | 763 | P763M | B | 7.3e-06 | 9.7e-06 | 9.3e-06 | 8.767E-06 | 1.286E-06 | 7.300E-06 | 9.700E-06 |
| 15254 | 763 | P763F | B | 6.9e-06 | 8.3e-06 | 9.4e-06 | 8.200E-06 | 1.253E-06 | 6.900E-06 | 9.400E-06 |
| 15255 | 763 | P763P | B | 8.6e-06 | 5.4e-06 | 8.1e-06 | 7.367E-06 | 1.721E-06 | 5.400E-06 | 8.600E-06 |
| 15256 | 763 | P763S | B | 6.4e-06 | 5.1e-06 | 8.6e-06 | 6.700E-06 | 1.769E-06 | 5.100E-06 | 8.600E-06 |
| 15257 | 763 | P763T | B | 7.7e-06 | 1.3e-05 | 9.6e-06 | 1.010E-05 | 2.685E-06 | 7.700E-06 | 1.300E-05 |
| 15258 | 763 | P763W | B | 4.4e-06 | 7.5e-06 | 1.1e-05 | 7.633E-06 | 3.302E-06 | 4.400E-06 | 1.100E-05 |
| 15259 | 763 | P763Y | B | 7e-06 | 6.5e-06 | 9e-06 | 7.500E-06 | 1.323E-06 | 6.500E-06 | 9.000E-06 |
| 15260 | 763 | P763V | B | 8.7e-06 | 3.3e-06 | 8.7e-06 | 6.900E-06 | 3.118E-06 | 3.300E-06 | 8.700E-06 |
| 15261 | 764 | R764A | B | 9.1e-06 | 5.3e-06 | 1.3e-05 | 9.133E-06 | 3.850E-06 | 5.300E-06 | 1.300E-05 |
| 15262 | 764 | R764R | B | 7.7e-06 | 4.8e-06 | 8.4e-06 | 6.967E-06 | 1.909E-06 | 4.800E-06 | 8.400E-06 |
| 15263 | 764 | R764N | B | 6.6e-06 | 6.2e-06 | 7.7e-06 | 6.833E-06 | 7.767E-07 | 6.200E-06 | 7.700E-06 |
| 15264 | 764 | R764D | B | 5.5e-06 | 7.4e-06 | 4.2e-06 | 5.700E-06 | 1.609E-06 | 4.200E-06 | 7.400E-06 |
| 15265 | 764 | R764C | B | 7.9e-06 | 5.6e-06 | 6.6e-06 | 6.700E-06 | 1.153E-06 | 5.600E-06 | 7.900E-06 |
| 15266 | 764 | R764E | B | 6.2e-06 | 9e-06 | 5.6e-06 | 6.933E-06 | 1.815E-06 | 5.600E-06 | 9.000E-06 |
| 15267 | 764 | R764Q | B | 7.8e-06 | 9.3e-06 | 6.2e-06 | 7.767E-06 | 1.550E-06 | 6.200E-06 | 9.300E-06 |
| 15268 | 764 | R764G | B | 5.8e-06 | 8.1e-06 | 5.5e-06 | 6.467E-06 | 1.422E-06 | 5.500E-06 | 8.100E-06 |
| 15269 | 764 | R764H | B | 7.6e-06 | 9.3e-06 | 8.8e-06 | 8.567E-06 | 8.737E-07 | 7.600E-06 | 9.300E-06 |
| 15270 | 764 | R764I | B | 6.8e-06 | 9.5e-06 | 6.3e-06 | 7.533E-06 | 1.721E-06 | 6.300E-06 | 9.500E-06 |
| 15271 | 764 | R764L | B | 1.4e-05 | 8.1e-06 | 1.4e-05 | 1.203E-05 | 3.406E-06 | 8.100E-06 | 1.400E-05 |
| 15272 | 764 | R764K | B | 6.3e-06 | 8.7e-06 | 1.2e-05 | 9.000E-06 | 2.862E-06 | 6.300E-06 | 1.200E-05 |
| 15273 | 764 | R764M | B | 9.2e-06 | 4.4e-06 | 8.7e-06 | 7.433E-06 | 2.639E-06 | 4.400E-06 | 9.200E-06 |
| 15274 | 764 | R764F | B | 6.7e-06 | 7.9e-06 | 6.8e-06 | 7.133E-06 | 6.658E-07 | 6.700E-06 | 7.900E-06 |
| 15275 | 764 | R764P | B | 1.1e-05 | 6.9e-06 | 8.2e-06 | 8.700E-06 | 2.095E-06 | 6.900E-06 | 1.100E-05 |
| 15276 | 764 | R764S | B | 6.2e-06 | 6.5e-06 | 7e-06 | 6.567E-06 | 4.041E-07 | 6.200E-06 | 7.000E-06 |
| 15277 | 764 | R764T | B | 4.6e-06 | 6.8e-06 | 7.2e-06 | 6.200E-06 | 1.400E-06 | 4.600E-06 | 7.200E-06 |
| 15278 | 764 | R764W | B | 6.7e-06 | 9.6e-06 | 8.6e-06 | 8.300E-06 | 1.473E-06 | 6.700E-06 | 9.600E-06 |
| 15279 | 764 | R764Y | B | 6.1e-06 | 1.2e-05 | 1.2e-05 | 1.003E-05 | 3.406E-06 | 6.100E-06 | 1.200E-05 |
| 15280 | 764 | R764V | B | 1.4e-05 | 6e-06 | 7.2e-06 | 9.067E-06 | 4.314E-06 | 6.000E-06 | 1.400E-05 |
| 15281 | 765 | E765A | B | 9.1e-06 | 4.8e-06 | 8.8e-06 | 7.567E-06 | 2.401E-06 | 4.800E-06 | 9.100E-06 |
| 15282 | 765 | E765R | B | 6.4e-06 | 9.3e-06 | 7.2e-06 | 7.633E-06 | 1.498E-06 | 6.400E-06 | 9.300E-06 |
| 15283 | 765 | E765N | B | 6.6e-06 | 7.1e-06 | 6.7e-06 | 6.800E-06 | 2.646E-07 | 6.600E-06 | 7.100E-06 |
| 15284 | 765 | E765D | B | 9.3e-06 | 1.3e-05 | 9.1e-06 | 1.047E-05 | 2.196E-06 | 9.100E-06 | 1.300E-05 |
| 15285 | 765 | E765C | B | 6.8e-06 | 9.6e-06 | 1.2e-05 | 9.467E-06 | 2.603E-06 | 6.800E-06 | 1.200E-05 |
| 15286 | 765 | E765E | B | 4.9e-06 | 1.2e-05 | 6.6e-06 | 7.833E-06 | 3.707E-06 | 4.900E-06 | 1.200E-05 |
| 15287 | 765 | E765Q | B | 5e-06 | 7.1e-06 | 7.4e-06 | 6.500E-06 | 1.308E-06 | 5.000E-06 | 7.400E-06 |
| 15288 | 765 | E765G | B | 1.1e-05 | 7e-06 | 1e-05 | 9.333E-06 | 2.082E-06 | 7.000E-06 | 1.100E-05 |
| 15289 | 765 | E765H | B | 9.6e-06 | 9.3e-06 | 5.4e-06 | 8.100E-06 | 2.343E-06 | 5.400E-06 | 9.600E-06 |
| 15290 | 765 | E765I | B | 1.3e-05 | 8.2e-06 | 6.1e-06 | 9.100E-06 | 3.537E-06 | 6.100E-06 | 1.300E-05 |
| 15291 | 765 | E765L | B | 1.5e-05 | 1.3e-05 | 1.1e-05 | 1.300E-05 | 2.000E-06 | 1.100E-05 | 1.500E-05 |
| 15292 | 765 | E765K | B | 7.7e-06 | 1.6e-05 | 9e-06 | 1.090E-05 | 4.464E-06 | 7.700E-06 | 1.600E-05 |
| 15293 | 765 | E765M | B | 8.1e-06 | 1.2e-05 | 6.2e-06 | 8.767E-06 | 2.957E-06 | 6.200E-06 | 1.200E-05 |
| 15294 | 765 | E765F | B | 8.1e-06 | 7.8e-06 | 7.8e-06 | 7.900E-06 | 1.732E-07 | 7.800E-06 | 8.100E-06 |
| 15295 | 765 | E765P | B | 8.6e-06 | 3.8e-06 | 7.8e-06 | 6.733E-06 | 2.572E-06 | 3.800E-06 | 8.600E-06 |
| 15296 | 765 | E765S | B | 9.6e-06 | 7.2e-06 | 1.1e-05 | 9.267E-06 | 1.922E-06 | 7.200E-06 | 1.100E-05 |
| 15297 | 765 | E765T | B | 8.7e-06 | 1e-05 | 6.4e-06 | 8.367E-06 | 1.823E-06 | 6.400E-06 | 1.000E-05 |
| 15298 | 765 | E765W | B | 7.8e-06 | 9.3e-06 | 4.9e-06 | 7.333E-06 | 2.237E-06 | 4.900E-06 | 9.300E-06 |
| 15299 | 765 | E765Y | B | 6.4e-06 | 1.3e-05 | 9.3e-06 | 9.567E-06 | 3.308E-06 | 6.400E-06 | 1.300E-05 |

| | | | | | | | | | | |
|---|---|---|---|---|---|---|---|---|---|---|
| 15300 | 765 | E765V | B | 1.3e-05 | 8e-06 | 6.5e-06 | 9.167E-06 | 3.403E-06 | 6.500E-06 | 1.300E-05 |
| 15301 | 766 | A766A | B | 9.5e-06 | 7.2e-06 | 9e-06 | 8.567E-06 | 1.210E-06 | 7.200E-06 | 9.500E-06 |
| 15302 | 766 | A766R | B | 1.5e-05 | 8.3e-06 | 7e-06 | 1.010E-05 | 4.293E-06 | 7.000E-06 | 1.500E-05 |
| 15303 | 766 | A766N | B | 5e-06 | 5.6e-06 | 1.2e-05 | 7.533E-06 | 3.880E-06 | 5.000E-06 | 1.200E-05 |
| 15304 | 766 | A766D | B | 9.2e-06 | 1e-05 | 1e-05 | 9.733E-06 | 4.619E-07 | 9.200E-06 | 1.000E-05 |
| 15305 | 766 | A766C | B | 6.1e-06 | 5.1e-06 | 6.7e-06 | 5.967E-06 | 8.083E-07 | 5.100E-06 | 6.700E-06 |
| 15306 | 766 | A766E | B | 5.8e-06 | 9.4e-06 | 6.1e-06 | 7.100E-06 | 1.997E-06 | 5.800E-06 | 9.400E-06 |
| 15307 | 766 | A766Q | B | 9.7e-06 | 7.8e-06 | 9.8e-06 | 9.100E-06 | 1.127E-06 | 7.800E-06 | 9.800E-06 |
| 15308 | 766 | A766G | B | 1.3e-05 | 7.9e-06 | 8.5e-06 | 9.800E-06 | 2.787E-06 | 7.900E-06 | 1.300E-05 |
| 15309 | 766 | A766H | B | 5.4e-06 | 1.2e-05 | 5.7e-06 | 7.700E-06 | 3.727E-06 | 5.400E-06 | 1.200E-05 |
| 15310 | 766 | A766I | B | 8.9e-06 | 9.9e-06 | 4.2e-06 | 7.667E-06 | 3.044E-06 | 4.200E-06 | 9.900E-06 |
| 15311 | 766 | A766L | B | 8.5e-06 | 8.9e-06 | 9.5e-06 | 8.967E-06 | 5.033E-07 | 8.500E-06 | 9.500E-06 |
| 15312 | 766 | A766K | B | 9.2e-06 | 6.6e-06 | 8.5e-06 | 8.100E-06 | 1.345E-06 | 6.600E-06 | 9.200E-06 |
| 15313 | 766 | A766M | B | 6.9e-06 | 7e-06 | 1.4e-05 | 9.300E-06 | 4.071E-06 | 6.900E-06 | 1.400E-05 |
| 15314 | 766 | A766F | B | 1e-05 | 5.1e-06 | 1e-05 | 8.367E-06 | 2.829E-06 | 5.100E-06 | 1.000E-05 |
| 15315 | 766 | A766P | B | 7.9e-06 | 8.8e-06 | 1.8e-05 | 1.157E-05 | 5.590E-06 | 7.900E-06 | 1.800E-05 |
| 15316 | 766 | A766S | B | 7.3e-06 | 1e-05 | 1e-05 | 9.100E-06 | 1.559E-06 | 7.300E-06 | 1.000E-05 |
| 15317 | 766 | A766T | B | 9e-06 | 6.7e-06 | 6.2e-06 | 7.300E-06 | 1.493E-06 | 6.200E-06 | 9.000E-06 |
| 15318 | 766 | A766W | B | 8.8e-06 | 2.3e-06 | 6.7e-06 | 5.933E-06 | 3.317E-06 | 2.300E-06 | 8.800E-06 |
| 15319 | 766 | A766Y | B | 6.4e-06 | 6.4e-06 | 1.3e-05 | 8.600E-06 | 3.811E-06 | 6.400E-06 | 1.300E-05 |
| 15320 | 766 | A766V | B | 8.4e-06 | 6.1e-06 | 6e-06 | 6.833E-06 | 1.358E-06 | 6.000E-06 | 8.400E-06 |
| 15321 | 767 | K767A | B | 1.3e-05 | 1.1e-05 | 1.1e-05 | 1.167E-05 | 1.155E-06 | 1.100E-05 | 1.300E-05 |
| 15322 | 767 | K767R | B | 6e-06 | 9.3e-06 | 5.5e-06 | 6.933E-06 | 2.065E-06 | 5.500E-06 | 9.300E-06 |
| 15323 | 767 | K767N | B | 9e-06 | 7.2e-06 | 7.9e-06 | 8.033E-06 | 9.074E-07 | 7.200E-06 | 9.000E-06 |
| 15324 | 767 | K767D | B | 2e-05 | 5.8e-06 | 6.6e-06 | 1.080E-05 | 7.977E-06 | 5.800E-06 | 2.000E-05 |
| 15325 | 767 | K767C | B | 1e-05 | 8.5e-06 | 6.2e-06 | 8.233E-06 | 1.914E-06 | 6.200E-06 | 1.000E-05 |
| 15326 | 767 | K767E | B | 5.6e-06 | 6.3e-06 | 7.4e-06 | 6.433E-06 | 9.074E-07 | 5.600E-06 | 7.400E-06 |
| 15327 | 767 | K767Q | B | 5.8e-06 | 5.8e-06 | 6.7e-06 | 6.100E-06 | 5.196E-07 | 5.800E-06 | 6.700E-06 |
| 15328 | 767 | K767G | B | 5.9e-06 | 4.6e-06 | 7.8e-06 | 6.100E-06 | 1.609E-06 | 4.600E-06 | 7.800E-06 |
| 15329 | 767 | K767H | B | 8.8e-06 | 1.3e-05 | 6.5e-06 | 9.433E-06 | 3.296E-06 | 6.500E-06 | 1.300E-05 |
| 15330 | 767 | K767I | B | 8.6e-06 | 7.9e-06 | 8.9e-06 | 8.467E-06 | 5.132E-07 | 7.900E-06 | 8.900E-06 |
| 15331 | 767 | K767L | B | 9.3e-06 | 9.8e-06 | 5.8e-06 | 8.300E-06 | 2.179E-06 | 5.800E-06 | 9.800E-06 |
| 15332 | 767 | K767K | B | 1e-05 | 8.3e-06 | 5.9e-06 | 8.067E-06 | 2.060E-06 | 5.900E-06 | 1.000E-05 |
| 15333 | 767 | K767M | B | 5.6e-06 | 8.6e-06 | 7.1e-06 | 7.100E-06 | 1.500E-06 | 5.600E-06 | 8.600E-06 |
| 15334 | 767 | K767F | B | 6.4e-06 | 1.3e-05 | 1e-05 | 9.800E-06 | 3.305E-06 | 6.400E-06 | 1.300E-05 |
| 15335 | 767 | K767P | B | 1.1e-05 | 6.5e-06 | 5.2e-06 | 7.567E-06 | 3.044E-06 | 5.200E-06 | 1.100E-05 |
| 15336 | 767 | K767S | B | 5.8e-06 | 6.3e-06 | 6.1e-06 | 6.067E-06 | 2.517E-07 | 5.800E-06 | 6.300E-06 |
| 15337 | 767 | K767T | B | 1e-05 | 4.4e-06 | 8.4e-06 | 7.600E-06 | 2.884E-06 | 4.400E-06 | 1.000E-05 |
| 15338 | 767 | K767W | B | 6.8e-06 | 1.3e-05 | 6.5e-06 | 8.767E-06 | 3.669E-06 | 6.500E-06 | 1.300E-05 |
| 15339 | 767 | K767Y | B | 4.5e-06 | 1.1e-05 | 6.5e-06 | 7.333E-06 | 3.329E-06 | 4.500E-06 | 1.100E-05 |
| 15340 | 767 | K767V | B | 1.1e-05 | 6.4e-06 | 9.7e-06 | 9.033E-06 | 2.371E-06 | 6.400E-06 | 1.100E-05 |
| 15341 | 768 | V768A | B | 9.9e-06 | 9.1e-06 | 1.3e-05 | 1.067E-05 | 2.060E-06 | 9.100E-06 | 1.300E-05 |
| 15342 | 768 | V768R | B | 1e-05 | 5e-06 | 9.6e-06 | 8.200E-06 | 2.778E-06 | 5.000E-06 | 1.000E-05 |
| 15343 | 768 | V768N | B | 6.2e-06 | 9.3e-06 | 7.3e-06 | 7.600E-06 | 1.572E-06 | 6.200E-06 | 9.300E-06 |
| 15344 | 768 | V768D | B | 5.1e-06 | 1.1e-05 | 6.7e-06 | 7.600E-06 | 3.051E-06 | 5.100E-06 | 1.100E-05 |
| 15345 | 768 | V768C | B | 6.7e-06 | 7.7e-06 | 8.8e-06 | 7.733E-06 | 1.050E-06 | 6.700E-06 | 8.800E-06 |
| 15346 | 768 | V768E | B | 1.3e-05 | 7.3e-06 | 6.6e-06 | 8.967E-06 | 3.510E-06 | 6.600E-06 | 1.300E-05 |
| 15347 | 768 | V768Q | B | 8.5e-06 | 1.2e-05 | 1.2e-05 | 1.083E-05 | 2.021E-06 | 8.500E-06 | 1.200E-05 |
| 15348 | 768 | V768G | B | 6.3e-06 | 9.8e-06 | 1e-05 | 8.700E-06 | 2.081E-06 | 6.300E-06 | 1.000E-05 |
| 15349 | 768 | V768H | B | 5e-06 | 1e-05 | 8.3e-06 | 7.767E-06 | 2.542E-06 | 5.000E-06 | 1.000E-05 |
| 15350 | 768 | V768I | B | 6.5e-06 | 4.7e-06 | 5.3e-06 | 5.500E-06 | 9.165E-07 | 4.700E-06 | 6.500E-06 |
| 15351 | 768 | V768L | B | 1.1e-05 | 9.6e-06 | 9.3e-06 | 9.967E-06 | 9.074E-07 | 9.300E-06 | 1.100E-05 |
| 15352 | 768 | V768K | B | 6.1e-06 | 5.4e-06 | 8.8e-06 | 6.767E-06 | 1.795E-06 | 5.400E-06 | 8.800E-06 |
| 15353 | 768 | V768M | B | 1.2e-05 | 8e-06 | 6.8e-06 | 8.933E-06 | 2.723E-06 | 6.800E-06 | 1.200E-05 |
| 15354 | 768 | V768F | B | 8.4e-06 | 8.6e-06 | 1.5e-05 | 1.067E-05 | 3.754E-06 | 8.400E-06 | 1.500E-05 |
| 15355 | 768 | V768P | B | 1.2e-05 | 1.3e-05 | 1.3e-05 | 1.267E-05 | 5.774E-07 | 1.200E-05 | 1.300E-05 |
| 15356 | 768 | V768S | B | 8.5e-06 | 1e-05 | 9.8e-06 | 9.433E-06 | 8.145E-07 | 8.500E-06 | 1.000E-05 |
| 15357 | 768 | V768T | B | 7.5e-06 | 5.5e-06 | 9.5e-06 | 7.500E-06 | 2.000E-06 | 5.500E-06 | 9.500E-06 |
| 15358 | 768 | V768W | B | 8.9e-06 | 9e-06 | 9.5e-06 | 9.133E-06 | 3.215E-07 | 8.900E-06 | 9.500E-06 |
| 15359 | 768 | V768Y | B | 4.8e-06 | 7.4e-06 | 3.9e-06 | 5.367E-06 | 1.818E-06 | 3.900E-06 | 7.400E-06 |
| 15360 | 768 | V768V | B | 6.5e-06 | 6.6e-06 | 5.2e-06 | 6.100E-06 | 7.810E-07 | 5.200E-06 | 6.600E-06 |
| 15361 | 769 | Q769A | B | 4.1e-06 | 4.9e-06 | 8.4e-06 | 5.800E-06 | 2.287E-06 | 4.100E-06 | 8.400E-06 |
| 15362 | 769 | Q769R | B | 3.5e-06 | 7.7e-06 | 8.5e-06 | 6.567E-06 | 2.686E-06 | 3.500E-06 | 8.500E-06 |
| 15363 | 769 | Q769N | B | 7.1e-06 | 7.2e-06 | 1e-05 | 8.100E-06 | 1.646E-06 | 7.100E-06 | 1.000E-05 |
| 15364 | 769 | Q769D | B | 5.2e-06 | 7.6e-06 | 7.1e-06 | 6.633E-06 | 1.266E-06 | 5.200E-06 | 7.600E-06 |
| 15365 | 769 | Q769C | B | 7.1e-06 | 4.8e-06 | 8.1e-06 | 6.667E-06 | 1.692E-06 | 4.800E-06 | 8.100E-06 |
| 15366 | 769 | Q769E | B | 4.7e-06 | 5e-06 | 4.4e-06 | 4.700E-06 | 3.000E-07 | 4.400E-06 | 5.000E-06 |
| 15367 | 769 | Q769Q | B | 7.9e-06 | 6.9e-06 | 7.6e-06 | 7.467E-06 | 5.132E-07 | 6.900E-06 | 7.900E-06 |
| 15368 | 769 | Q769G | B | 6.6e-06 | 5.5e-06 | 5.9e-06 | 6.000E-06 | 5.568E-07 | 5.500E-06 | 6.600E-06 |
| 15369 | 769 | Q769H | B | 9.9e-06 | 4.6e-06 | 1.4e-05 | 9.500E-06 | 4.713E-06 | 4.600E-06 | 1.400E-05 |
| 15370 | 769 | Q769I | B | 5.6e-06 | 7.6e-06 | 1e-05 | 7.733E-06 | 2.203E-06 | 5.600E-06 | 1.000E-05 |
| 15371 | 769 | Q769L | B | 4.8e-06 | 6.5e-06 | 8.1e-06 | 6.467E-06 | 1.650E-06 | 4.800E-06 | 8.100E-06 |
| 15372 | 769 | Q769K | B | 5.2e-06 | 4.4e-06 | 3.7e-06 | 4.433E-06 | 7.506E-07 | 3.700E-06 | 5.200E-06 |
| 15373 | 769 | Q769M | B | 6.6e-06 | 9.7e-06 | 4.9e-06 | 7.067E-06 | 2.434E-06 | 4.900E-06 | 9.700E-06 |
| 15374 | 769 | Q769F | B | 6.7e-06 | 1.9e-05 | 6.1e-06 | 1.060E-05 | 7.281E-06 | 6.100E-06 | 1.900E-05 |
| 15375 | 769 | Q769P | B | 7.1e-06 | 7.2e-06 | 6.9e-06 | 7.067E-06 | 1.528E-07 | 6.900E-06 | 7.200E-06 |
| 15376 | 769 | Q769S | B | 1.7e-05 | 8.7e-06 | 9.9e-06 | 1.187E-05 | 4.486E-06 | 8.700E-06 | 1.700E-05 |
| 15377 | 769 | Q769T | B | 5.5e-06 | 9.2e-06 | 7.8e-06 | 7.500E-06 | 1.868E-06 | 5.500E-06 | 9.200E-06 |
| 15378 | 769 | Q769W | B | 8.2e-06 | 6.1e-06 | 6.9e-06 | 7.067E-06 | 1.060E-06 | 6.100E-06 | 8.200E-06 |
| 15379 | 769 | Q769Y | B | 8.9e-06 | 8.6e-06 | 8.4e-06 | 8.633E-06 | 2.517E-07 | 8.400E-06 | 8.900E-06 |
| 15380 | 769 | Q769V | B | 1.5e-05 | 7.5e-06 | 1.1e-05 | 1.117E-05 | 3.753E-06 | 7.500E-06 | 1.500E-05 |
| 15381 | 770 | W770A | B | 1.3e-05 | 4.7e-06 | 6.7e-06 | 8.133E-06 | 4.332E-06 | 4.700E-06 | 1.300E-05 |
| 15382 | 770 | W770R | B | 9.6e-06 | 6.6e-06 | 9.5e-06 | 8.567E-06 | 1.704E-06 | 6.600E-06 | 9.600E-06 |
| 15383 | 770 | W770N | B | 9.6e-06 | 5e-06 | 6.9e-06 | 7.167E-06 | 2.312E-06 | 5.000E-06 | 9.600E-06 |
| 15384 | 770 | W770D | B | 9.5e-06 | 6.4e-06 | 7.3e-06 | 7.733E-06 | 1.595E-06 | 6.400E-06 | 9.500E-06 |

| | | | | | | | | | | |
|---|---|---|---|---|---|---|---|---|---|---|
| 15385 | 770 | W770C | B | 9.4e-06 | 1.1e-05 | 6.7e-06 | 9.033E-06 | 2.173E-06 | 6.700E-06 | 1.100E-05 |
| 15386 | 770 | W770E | B | 3.9e-06 | 1.4e-05 | 6.2e-06 | 8.033E-06 | 5.294E-06 | 3.900E-06 | 1.400E-05 |
| 15387 | 770 | W770Q | B | 1.7e-05 | 6.9e-06 | 7.7e-06 | 1.053E-05 | 5.615E-06 | 6.900E-06 | 1.700E-05 |
| 15388 | 770 | W770G | B | 9.5e-06 | 8.7e-06 | 8.5e-06 | 8.900E-06 | 5.292E-07 | 8.500E-06 | 9.500E-06 |
| 15389 | 770 | W770H | B | 7.3e-06 | 1.4e-05 | 4.5e-06 | 8.600E-06 | 4.882E-06 | 4.500E-06 | 1.400E-05 |
| 15390 | 770 | W770I | B | 6.9e-06 | 7.1e-06 | 5.1e-06 | 6.367E-06 | 1.102E-06 | 5.100E-06 | 7.100E-06 |
| 15391 | 770 | W770L | B | 9.7e-06 | 1e-05 | 1e-05 | 9.900E-06 | 1.732E-07 | 9.700E-06 | 1.000E-05 |
| 15392 | 770 | W770K | B | 6.8e-06 | 9.3e-06 | 1.3e-05 | 9.700E-06 | 3.119E-06 | 6.800E-06 | 1.300E-05 |
| 15393 | 770 | W770M | B | 1.1e-05 | 7.6e-06 | 1.3e-05 | 1.053E-05 | 2.730E-06 | 7.600E-06 | 1.300E-05 |
| 15394 | 770 | W770F | B | 1e-05 | 1.1e-05 | 6.9e-06 | 9.300E-06 | 2.138E-06 | 6.900E-06 | 1.100E-05 |
| 15395 | 770 | W770P | B | 8.2e-06 | 7.9e-06 | 5.8e-06 | 7.300E-06 | 1.308E-06 | 5.800E-06 | 8.200E-06 |
| 15396 | 770 | W770S | B | 8.6e-06 | 8.9e-06 | 7.7e-06 | 8.400E-06 | 6.245E-07 | 7.700E-06 | 8.900E-06 |
| 15397 | 770 | W770T | B | 6.2e-06 | 1.1e-05 | 5.2e-06 | 7.467E-06 | 3.101E-06 | 5.200E-06 | 1.100E-05 |
| 15398 | 770 | W770W | B | 1.1e-05 | 9.4e-06 | 5.4e-06 | 8.600E-06 | 2.884E-06 | 5.400E-06 | 1.100E-05 |
| 15399 | 770 | W770Y | B | 1.1e-05 | 6.8e-06 | 5.2e-06 | 7.667E-06 | 2.996E-06 | 5.200E-06 | 1.100E-05 |
| 15400 | 770 | W770V | B | 6.1e-06 | 1.3e-05 | 6.6e-06 | 8.567E-06 | 3.848E-06 | 6.100E-06 | 1.300E-05 |
| 15401 | 771 | K771A | B | 7.3e-06 | 9.8e-06 | 1.1e-05 | 9.367E-06 | 1.888E-06 | 7.300E-06 | 1.100E-05 |
| 15402 | 771 | K771R | B | 8.7e-06 | 9.6e-06 | 9.4e-06 | 9.233E-06 | 4.726E-07 | 8.700E-06 | 9.600E-06 |
| 15403 | 771 | K771N | B | 8.3e-06 | 4.8e-06 | 8.6e-06 | 7.233E-06 | 2.113E-06 | 4.800E-06 | 8.600E-06 |
| 15404 | 771 | K771D | B | 9.7e-06 | 4.6e-06 | 7e-06 | 7.100E-06 | 2.551E-06 | 4.600E-06 | 9.700E-06 |
| 15405 | 771 | K771C | B | 7.2e-06 | 8.3e-06 | 1.1e-05 | 8.833E-06 | 1.955E-06 | 7.200E-06 | 1.100E-05 |
| 15406 | 771 | K771E | B | 7e-06 | 8.4e-06 | 1.5e-05 | 1.013E-05 | 4.272E-06 | 7.000E-06 | 1.500E-05 |
| 15407 | 771 | K771Q | B | 6.3e-06 | 8.7e-06 | 6.5e-06 | 7.167E-06 | 1.332E-06 | 6.300E-06 | 8.700E-06 |
| 15408 | 771 | K771G | B | 1.2e-05 | 5e-06 | 8.5e-06 | 8.500E-06 | 3.500E-06 | 5.000E-06 | 1.200E-05 |
| 15409 | 771 | K771H | B | 5.6e-06 | 6.5e-06 | 9.9e-06 | 7.333E-06 | 2.268E-06 | 5.600E-06 | 9.900E-06 |
| 15410 | 771 | K771I | B | 4.5e-06 | 7.2e-06 | 5.7e-06 | 5.800E-06 | 1.353E-06 | 4.500E-06 | 7.200E-06 |
| 15411 | 771 | K771L | B | 1.3e-05 | 7e-06 | 6.5e-06 | 8.833E-06 | 3.617E-06 | 6.500E-06 | 1.300E-05 |
| 15412 | 771 | K771K | B | 8.4e-06 | 1.8e-05 | 7e-06 | 1.113E-05 | 5.988E-06 | 7.000E-06 | 1.800E-05 |
| 15413 | 771 | K771M | B | 1e-05 | 6.8e-06 | 1.1e-05 | 9.267E-06 | 2.194E-06 | 6.800E-06 | 1.100E-05 |
| 15414 | 771 | K771F | B | 7.3e-06 | 5.6e-06 | 6.4e-06 | 6.433E-06 | 8.505E-07 | 5.600E-06 | 7.300E-06 |
| 15415 | 771 | K771P | B | 8.8e-06 | 7.2e-06 | 7.7e-06 | 7.900E-06 | 8.185E-07 | 7.200E-06 | 8.800E-06 |
| 15416 | 771 | K771S | B | 5.6e-06 | 4.1e-06 | 7.3e-06 | 5.667E-06 | 1.601E-06 | 4.100E-06 | 7.300E-06 |
| 15417 | 771 | K771T | B | 5.2e-06 | 8.2e-06 | 7.8e-06 | 7.067E-06 | 1.629E-06 | 5.200E-06 | 8.200E-06 |
| 15418 | 771 | K771W | B | 1.2e-05 | 9.2e-06 | 7.7e-06 | 9.633E-06 | 2.183E-06 | 7.700E-06 | 1.200E-05 |
| 15419 | 771 | K771Y | B | 9.5e-06 | 8.7e-06 | 8.6e-06 | 8.933E-06 | 4.933E-07 | 8.600E-06 | 9.500E-06 |
| 15420 | 771 | K771V | B | 9.1e-06 | 9.6e-06 | 6.7e-06 | 8.467E-06 | 1.550E-06 | 6.700E-06 | 9.600E-06 |
| 15421 | 772 | V772A | B | 7.2e-06 | 4.5e-06 | 6.8e-06 | 6.167E-06 | 1.457E-06 | 4.500E-06 | 7.200E-06 |
| 15422 | 772 | V772R | B | 5.5e-06 | 7.7e-06 | 9.4e-06 | 7.533E-06 | 1.955E-06 | 5.500E-06 | 9.400E-06 |
| 15423 | 772 | V772N | B | 5.9e-06 | 1e-05 | 1e-05 | 8.633E-06 | 2.367E-06 | 5.900E-06 | 1.000E-05 |
| 15424 | 772 | V772D | B | 7.7e-06 | 1e-05 | 6.4e-06 | 8.033E-06 | 1.823E-06 | 6.400E-06 | 1.000E-05 |
| 15425 | 772 | V772C | B | 1.4e-05 | 9.7e-06 | 1.6e-05 | 1.323E-05 | 3.219E-06 | 9.700E-06 | 1.600E-05 |
| 15426 | 772 | V772E | B | 6.8e-06 | 7e-06 | 8.4e-06 | 7.400E-06 | 8.718E-07 | 6.800E-06 | 8.400E-06 |
| 15427 | 772 | V772Q | B | 9e-06 | 8.9e-06 | 6.6e-06 | 8.167E-06 | 1.358E-06 | 6.600E-06 | 9.000E-06 |
| 15428 | 772 | V772G | B | 9.1e-06 | 8.5e-06 | 6.9e-06 | 8.167E-06 | 1.137E-06 | 6.900E-06 | 9.100E-06 |
| 15429 | 772 | V772H | B | 7e-06 | 7e-06 | 5.6e-06 | 6.533E-06 | 8.083E-07 | 5.600E-06 | 7.000E-06 |
| 15430 | 772 | V772I | B | 7e-06 | 1.2e-05 | 7.9e-06 | 8.967E-06 | 2.665E-06 | 7.000E-06 | 1.200E-05 |
| 15431 | 772 | V772L | B | 8.2e-06 | 1.2e-05 | 8.8e-06 | 9.667E-06 | 2.043E-06 | 8.200E-06 | 1.200E-05 |
| 15432 | 772 | V772K | B | 5.4e-06 | 6.6e-06 | 5.9e-06 | 5.967E-06 | 6.028E-07 | 5.400E-06 | 6.600E-06 |
| 15433 | 772 | V772M | B | 5.4e-06 | 6.6e-06 | 6.5e-06 | 6.167E-06 | 6.658E-07 | 5.400E-06 | 6.600E-06 |
| 15434 | 772 | V772F | B | 5.8e-06 | 4.1e-06 | 8.2e-06 | 6.033E-06 | 2.060E-06 | 4.100E-06 | 8.200E-06 |
| 15435 | 772 | V772P | B | 7.8e-06 | 1e-05 | 9.3e-06 | 9.033E-06 | 1.124E-06 | 7.800E-06 | 1.000E-05 |
| 15436 | 772 | V772S | B | 1.2e-05 | 6.7e-06 | 9e-06 | 9.233E-06 | 2.658E-06 | 6.700E-06 | 1.200E-05 |
| 15437 | 772 | V772T | B | 1.2e-05 | 6e-06 | 5.4e-06 | 7.800E-06 | 3.650E-06 | 5.400E-06 | 1.200E-05 |
| 15438 | 772 | V772W | B | 6.6e-06 | 6.1e-06 | 6.4e-06 | 6.367E-06 | 2.517E-07 | 6.100E-06 | 6.600E-06 |
| 15439 | 772 | V772Y | B | 9e-06 | 7.5e-06 | 3.5e-06 | 6.667E-06 | 2.843E-06 | 3.500E-06 | 9.000E-06 |
| 15440 | 772 | V772V | B | 5.8e-06 | 6.5e-06 | 6.9e-06 | 6.400E-06 | 5.568E-07 | 5.800E-06 | 6.900E-06 |
| 15441 | 773 | D773A | B | 7.6e-06 | 1.4e-05 | 5e-06 | 8.867E-06 | 4.632E-06 | 5.000E-06 | 1.400E-05 |
| 15442 | 773 | D773R | B | 6.2e-06 | 1.3e-05 | 9.9e-06 | 9.700E-06 | 3.404E-06 | 6.200E-06 | 1.300E-05 |
| 15443 | 773 | D773N | B | 9.2e-06 | 1.1e-05 | 9.4e-06 | 9.867E-06 | 9.866E-07 | 9.200E-06 | 1.100E-05 |
| 15444 | 773 | D773D | B | 8.3e-06 | 9.4e-06 | 8.2e-06 | 8.633E-06 | 6.658E-07 | 8.200E-06 | 9.400E-06 |
| 15445 | 773 | D773C | B | 1e-05 | 8.7e-06 | 7.9e-06 | 8.867E-06 | 1.060E-06 | 7.900E-06 | 1.000E-05 |
| 15446 | 773 | D773E | B | 1.5e-05 | 1.1e-05 | 6.2e-06 | 1.073E-05 | 4.406E-06 | 6.200E-06 | 1.500E-05 |
| 15447 | 773 | D773Q | B | 7.3e-06 | 1.3e-05 | 4.7e-06 | 8.333E-06 | 4.245E-06 | 4.700E-06 | 1.300E-05 |
| 15448 | 773 | D773G | B | 1e-05 | 7.1e-06 | 7e-06 | 8.033E-06 | 1.704E-06 | 7.000E-06 | 1.000E-05 |
| 15449 | 773 | D773H | B | 7.1e-06 | 9e-06 | 7.6e-06 | 7.900E-06 | 9.849E-07 | 7.100E-06 | 9.000E-06 |
| 15450 | 773 | D773I | B | 1e-05 | 8e-06 | 1e-05 | 9.333E-06 | 1.155E-06 | 8.000E-06 | 1.000E-05 |
| 15451 | 773 | D773L | B | 5.6e-06 | 5.2e-06 | 6.9e-06 | 5.900E-06 | 8.888E-07 | 5.200E-06 | 6.900E-06 |
| 15452 | 773 | D773K | B | 8.1e-06 | 5.5e-06 | 9.7e-06 | 7.767E-06 | 2.120E-06 | 5.500E-06 | 9.700E-06 |
| 15453 | 773 | D773M | B | 8.2e-06 | 9.4e-06 | 9.3e-06 | 8.967E-06 | 6.658E-07 | 8.200E-06 | 9.400E-06 |
| 15454 | 773 | D773F | B | 1e-05 | 6.7e-06 | 5.2e-06 | 7.300E-06 | 2.456E-06 | 5.200E-06 | 1.000E-05 |
| 15455 | 773 | D773P | B | 9.9e-06 | 9.1e-06 | 8.4e-06 | 9.133E-06 | 7.506E-07 | 8.400E-06 | 9.900E-06 |
| 15456 | 773 | D773S | B | 7.5e-06 | 1e-05 | 7.6e-06 | 8.367E-06 | 1.415E-06 | 7.500E-06 | 1.000E-05 |
| 15457 | 773 | D773T | B | 6.5e-06 | 6.4e-06 | 1.4e-05 | 8.967E-06 | 4.359E-06 | 6.400E-06 | 1.400E-05 |
| 15458 | 773 | D773W | B | 1.3e-05 | 6.4e-06 | 8.4e-06 | 9.267E-06 | 3.384E-06 | 6.400E-06 | 1.300E-05 |
| 15459 | 773 | D773Y | B | 9.8e-06 | 7.1e-06 | 7.4e-06 | 8.100E-06 | 1.480E-06 | 7.100E-06 | 9.800E-06 |
| 15460 | 773 | D773V | B | 9.6e-06 | 6.5e-06 | 1.2e-05 | 9.367E-06 | 2.757E-06 | 6.500E-06 | 1.200E-05 |
| 15461 | 774 | N774A | B | 6.9e-06 | 4.9e-06 | 5.5e-06 | 5.767E-06 | 1.026E-06 | 4.900E-06 | 6.900E-06 |
| 15462 | 774 | N774R | B | 8.9e-06 | 7.2e-06 | 6.8e-06 | 7.633E-06 | 1.115E-06 | 6.800E-06 | 8.900E-06 |
| 15463 | 774 | N774N | B | 1e-05 | 3.1e-06 | 5.7e-06 | 6.267E-06 | 3.485E-06 | 3.100E-06 | 1.000E-05 |
| 15464 | 774 | N774D | B | 7.6e-06 | 7.6e-06 | 9.5e-06 | 8.233E-06 | 1.097E-06 | 7.600E-06 | 9.500E-06 |
| 15465 | 774 | N774C | B | 9e-06 | 1.3e-05 | 4.8e-06 | 8.933E-06 | 4.100E-06 | 4.800E-06 | 1.300E-05 |
| 15466 | 774 | N774E | B | 1.3e-05 | 8.1e-06 | 1.4e-05 | 1.170E-05 | 3.158E-06 | 8.100E-06 | 1.400E-05 |
| 15467 | 774 | N774Q | B | 8.7e-06 | 4.8e-06 | 7.5e-06 | 7.000E-06 | 1.997E-06 | 4.800E-06 | 8.700E-06 |
| 15468 | 774 | N774G | B | 1.4e-05 | 5.2e-06 | 5.4e-06 | 8.200E-06 | 5.024E-06 | 5.200E-06 | 1.400E-05 |
| 15469 | 774 | N774H | B | 6.5e-06 | 4.8e-06 | 1e-05 | 7.100E-06 | 2.651E-06 | 4.800E-06 | 1.000E-05 |

| | | | | | | | | | | |
|---|---|---|---|---|---|---|---|---|---|---|
| 15470 | 774 | N774I | B | 1e-05 | 3.6e-06 | 6.4e-06 | 6.667E-06 | 3.208E-06 | 3.600E-06 | 1.000E-05 |
| 15471 | 774 | N774L | B | 9.7e-06 | 1.4e-05 | 4.6e-06 | 9.433E-06 | 4.706E-06 | 4.600E-06 | 1.400E-05 |
| 15472 | 774 | N774K | B | 9.1e-06 | 4.8e-06 | 9.2e-06 | 7.700E-06 | 2.512E-06 | 4.800E-06 | 9.200E-06 |
| 15473 | 774 | N774M | B | 9.1e-06 | 9e-06 | 6.1e-06 | 8.067E-06 | 1.704E-06 | 6.100E-06 | 9.100E-06 |
| 15474 | 774 | N774F | B | 1.1e-05 | 8.3e-06 | 9.8e-06 | 9.700E-06 | 1.353E-06 | 8.300E-06 | 1.100E-05 |
| 15475 | 774 | N774P | B | 7.9e-06 | 5.6e-06 | 1.1e-05 | 8.167E-06 | 2.710E-06 | 5.600E-06 | 1.100E-05 |
| 15476 | 774 | N774S | B | 4.9e-06 | 6.2e-06 | 8e-06 | 6.367E-06 | 1.557E-06 | 4.900E-06 | 8.000E-06 |
| 15477 | 774 | N774T | B | 8.4e-06 | 7.8e-06 | 9.5e-06 | 8.567E-06 | 8.622E-07 | 7.800E-06 | 9.500E-06 |
| 15478 | 774 | N774W | B | 1.3e-05 | 5.6e-06 | 9.9e-06 | 9.500E-06 | 3.716E-06 | 5.600E-06 | 1.300E-05 |
| 15479 | 774 | N774Y | B | 1.3e-05 | 5.9e-06 | 8.4e-06 | 9.100E-06 | 3.601E-06 | 5.900E-06 | 1.300E-05 |
| 15480 | 774 | N774V | B | 6.4e-06 | 6.4e-06 | 6.8e-06 | 6.533E-06 | 2.309E-07 | 6.400E-06 | 6.800E-06 |
| 15481 | 775 | A775A | B | 5.1e-06 | 4.7e-06 | 8.3e-06 | 6.033E-06 | 1.973E-06 | 4.700E-06 | 8.300E-06 |
| 15482 | 775 | A775R | B | 8.6e-06 | 5.5e-06 | 9.9e-06 | 8.000E-06 | 2.261E-06 | 5.500E-06 | 9.900E-06 |
| 15483 | 775 | A775N | B | 4.5e-06 | 8.1e-06 | 5.7e-06 | 6.100E-06 | 1.833E-06 | 4.500E-06 | 8.100E-06 |
| 15484 | 775 | A775D | B | 7.7e-06 | 6.6e-06 | 9.8e-06 | 8.033E-06 | 1.626E-06 | 6.600E-06 | 9.800E-06 |
| 15485 | 775 | A775C | B | 8.5e-06 | 6.3e-06 | 1e-05 | 8.267E-06 | 1.861E-06 | 6.300E-06 | 1.000E-05 |
| 15486 | 775 | A775E | B | 7e-06 | 1.5e-05 | 1.1e-05 | 1.100E-05 | 4.000E-06 | 7.000E-06 | 1.500E-05 |
| 15487 | 775 | A775Q | B | 7.1e-06 | 6.6e-06 | 8.7e-06 | 7.467E-06 | 1.097E-06 | 6.600E-06 | 8.700E-06 |
| 15488 | 775 | A775G | B | 6.3e-06 | 1e-05 | 7.1e-06 | 7.800E-06 | 1.947E-06 | 6.300E-06 | 1.000E-05 |
| 15489 | 775 | A775H | B | 4e-06 | 8.8e-06 | 6.1e-06 | 6.300E-06 | 2.406E-06 | 4.000E-06 | 8.800E-06 |
| 15490 | 775 | A775I | B | 8.8e-06 | 9.2e-06 | 8.7e-06 | 8.900E-06 | 2.646E-07 | 8.700E-06 | 9.200E-06 |
| 15491 | 775 | A775L | B | 1e-05 | 5.4e-06 | 1e-05 | 8.467E-06 | 2.656E-06 | 5.400E-06 | 1.000E-05 |
| 15492 | 775 | A775K | B | 3.5e-06 | 5.3e-06 | 7.9e-06 | 5.567E-06 | 2.212E-06 | 3.500E-06 | 7.900E-06 |
| 15493 | 775 | A775M | B | 9.1e-06 | 1e-05 | 6.7e-06 | 8.600E-06 | 1.706E-06 | 6.700E-06 | 1.000E-05 |
| 15494 | 775 | A775F | B | 4.9e-06 | 7.9e-06 | 9e-06 | 7.267E-06 | 2.122E-06 | 4.900E-06 | 9.000E-06 |
| 15495 | 775 | A775P | B | 1.3e-05 | 6.4e-06 | 1.5e-05 | 1.147E-05 | 4.500E-06 | 6.400E-06 | 1.500E-05 |
| 15496 | 775 | A775S | B | 5.5e-06 | 8.5e-06 | 9.6e-06 | 7.867E-06 | 2.122E-06 | 5.500E-06 | 9.600E-06 |
| 15497 | 775 | A775T | B | 1.4e-05 | 3.2e-06 | 4.2e-06 | 7.133E-06 | 5.968E-06 | 3.200E-06 | 1.400E-05 |
| 15498 | 775 | A775W | B | 6.8e-06 | 7.3e-06 | 5e-06 | 6.367E-06 | 1.210E-06 | 5.000E-06 | 7.300E-06 |
| 15499 | 775 | A775Y | B | 1.4e-05 | 5.6e-06 | 8.1e-06 | 9.233E-06 | 4.313E-06 | 5.600E-06 | 1.400E-05 |
| 15500 | 775 | A775V | B | 1.1e-05 | 8.4e-06 | 5.1e-06 | 8.167E-06 | 2.957E-06 | 5.100E-06 | 1.100E-05 |
| 15501 | 776 | L776A | B | 1.2e-05 | 5.3e-06 | 7.4e-06 | 8.233E-06 | 3.427E-06 | 5.300E-06 | 1.200E-05 |
| 15502 | 776 | L776R | B | 6.7e-06 | 7.5e-06 | 9.5e-06 | 7.900E-06 | 1.442E-06 | 6.700E-06 | 9.500E-06 |
| 15503 | 776 | L776N | B | 6.7e-06 | 8.6e-06 | 5.3e-06 | 6.867E-06 | 1.656E-06 | 5.300E-06 | 8.600E-06 |
| 15504 | 776 | L776D | B | 3.7e-06 | 8.9e-06 | 5e-06 | 5.867E-06 | 2.706E-06 | 3.700E-06 | 8.900E-06 |
| 15505 | 776 | L776C | B | 6.8e-06 | 5e-06 | 8e-06 | 6.600E-06 | 1.510E-06 | 5.000E-06 | 8.000E-06 |
| 15506 | 776 | L776E | B | 8.8e-06 | 6.5e-06 | 9.6e-06 | 8.300E-06 | 1.609E-06 | 6.500E-06 | 9.600E-06 |
| 15507 | 776 | L776Q | B | 9.7e-06 | 1.6e-05 | 7.9e-06 | 1.120E-05 | 4.253E-06 | 7.900E-06 | 1.600E-05 |
| 15508 | 776 | L776G | B | 6.6e-06 | 7.3e-06 | 5.4e-06 | 6.433E-06 | 9.609E-07 | 5.400E-06 | 7.300E-06 |
| 15509 | 776 | L776H | B | 1e-05 | 1.6e-05 | 5.3e-06 | 1.043E-05 | 5.363E-06 | 5.300E-06 | 1.600E-05 |
| 15510 | 776 | L776I | B | 6.7e-06 | 1e-05 | 3.6e-06 | 6.767E-06 | 3.201E-06 | 3.600E-06 | 1.000E-05 |
| 15511 | 776 | L776L | B | 6.6e-06 | 7e-06 | 1.2e-05 | 8.533E-06 | 3.009E-06 | 6.600E-06 | 1.200E-05 |
| 15512 | 776 | L776K | B | 1.4e-05 | 6.1e-06 | 8.7e-06 | 9.600E-06 | 4.026E-06 | 6.100E-06 | 1.400E-05 |
| 15513 | 776 | L776M | B | 6.9e-06 | 7.2e-06 | 7.6e-06 | 7.233E-06 | 3.512E-07 | 6.900E-06 | 7.600E-06 |
| 15514 | 776 | L776F | B | 9.9e-06 | 9.4e-06 | 1.2e-05 | 1.043E-05 | 1.380E-06 | 9.400E-06 | 1.200E-05 |
| 15515 | 776 | L776P | B | 7.8e-06 | 6.7e-06 | 5.1e-06 | 6.533E-06 | 1.358E-06 | 5.100E-06 | 7.800E-06 |
| 15516 | 776 | L776S | B | 9.2e-06 | 6.3e-06 | 9.7e-06 | 8.400E-06 | 1.836E-06 | 6.300E-06 | 9.700E-06 |
| 15517 | 776 | L776T | B | 8.7e-06 | 9.7e-06 | 9.2e-06 | 9.200E-06 | 5.000E-07 | 8.700E-06 | 9.700E-06 |
| 15518 | 776 | L776W | B | 7e-06 | 1.3e-05 | 1e-05 | 1.000E-05 | 3.000E-06 | 7.000E-06 | 1.300E-05 |
| 15519 | 776 | L776Y | B | 9.6e-06 | 9.1e-06 | 1e-05 | 9.567E-06 | 4.509E-07 | 9.100E-06 | 1.000E-05 |
| 15520 | 776 | L776V | B | 7.4e-06 | 5.8e-06 | 6.6e-06 | 6.600E-06 | 8.000E-07 | 5.800E-06 | 7.400E-06 |
| 15521 | 777 | Q777A | B | 7.2e-06 | 9.2e-06 | 4.9e-06 | 7.100E-06 | 2.152E-06 | 4.900E-06 | 9.200E-06 |
| 15522 | 777 | Q777R | B | 1.2e-05 | 5.1e-06 | 5.6e-06 | 7.567E-06 | 3.848E-06 | 5.100E-06 | 1.200E-05 |
| 15523 | 777 | Q777N | B | 4.3e-06 | 5.8e-06 | 6.6e-06 | 5.567E-06 | 1.168E-06 | 4.300E-06 | 6.600E-06 |
| 15524 | 777 | Q777D | B | 5.6e-06 | 6.4e-06 | 6.6e-06 | 6.200E-06 | 5.292E-07 | 5.600E-06 | 6.600E-06 |
| 15525 | 777 | Q777C | B | 5e-06 | 7.4e-06 | 9.3e-06 | 7.233E-06 | 2.155E-06 | 5.000E-06 | 9.300E-06 |
| 15526 | 777 | Q777E | B | 5.6e-06 | 1.3e-05 | 1e-05 | 9.533E-06 | 3.722E-06 | 5.600E-06 | 1.300E-05 |
| 15527 | 777 | Q777Q | B | 4.6e-06 | 1e-05 | 1.2e-05 | 8.867E-06 | 3.828E-06 | 4.600E-06 | 1.200E-05 |
| 15528 | 777 | Q777G | B | 4.6e-06 | 1e-05 | 9.4e-06 | 8.000E-06 | 2.960E-06 | 4.600E-06 | 1.000E-05 |
| 15529 | 777 | Q777H | B | 6.3e-06 | 5.2e-06 | 1.2e-05 | 7.833E-06 | 3.650E-06 | 5.200E-06 | 1.200E-05 |
| 15530 | 777 | Q777I | B | 7.4e-06 | 6.5e-06 | 1e-05 | 7.967E-06 | 1.818E-06 | 6.500E-06 | 1.000E-05 |
| 15531 | 777 | Q777L | B | 5.3e-06 | 6.6e-06 | 5.4e-06 | 5.767E-06 | 7.234E-07 | 5.300E-06 | 6.600E-06 |
| 15532 | 777 | Q777K | B | 9.4e-06 | 9.1e-06 | 1e-05 | 9.500E-06 | 4.583E-07 | 9.100E-06 | 1.000E-05 |
| 15533 | 777 | Q777M | B | 7e-06 | 9.6e-06 | 7e-06 | 7.867E-06 | 1.501E-06 | 7.000E-06 | 9.600E-06 |
| 15534 | 777 | Q777F | B | 1e-05 | 4.6e-06 | 1.1e-05 | 8.533E-06 | 3.443E-06 | 4.600E-06 | 1.100E-05 |
| 15535 | 777 | Q777P | B | 7.2e-06 | 1.1e-05 | 6.1e-06 | 8.100E-06 | 2.571E-06 | 6.100E-06 | 1.100E-05 |
| 15536 | 777 | Q777S | B | 7.8e-06 | 5.9e-06 | 6.5e-06 | 6.733E-06 | 9.713E-07 | 5.900E-06 | 7.800E-06 |
| 15537 | 777 | Q777T | B | 1e-05 | 5.7e-06 | 1.2e-05 | 9.233E-06 | 3.219E-06 | 5.700E-06 | 1.200E-05 |
| 15538 | 777 | Q777W | B | 8.6e-06 | 5.1e-06 | 9.3e-06 | 7.667E-06 | 2.250E-06 | 5.100E-06 | 9.300E-06 |
| 15539 | 777 | Q777Y | B | 1.3e-05 | 5.8e-06 | 6.1e-06 | 8.300E-06 | 4.073E-06 | 5.800E-06 | 1.300E-05 |
| 15540 | 777 | Q777V | B | 8.8e-06 | 1e-05 | 7.9e-06 | 8.900E-06 | 1.054E-06 | 7.900E-06 | 1.000E-05 |
| 15541 | 778 | S778A | B | 6.4e-06 | 1.1e-05 | 1.2e-05 | 9.800E-06 | 2.987E-06 | 6.400E-06 | 1.200E-05 |
| 15542 | 778 | S778R | B | 9.1e-06 | 8.8e-06 | 1.3e-05 | 1.030E-05 | 2.343E-06 | 8.800E-06 | 1.300E-05 |
| 15543 | 778 | S778N | B | 6.8e-06 | 6.6e-06 | 7e-06 | 6.800E-06 | 2.000E-07 | 6.600E-06 | 7.000E-06 |
| 15544 | 778 | S778D | B | 1e-05 | 7.4e-06 | 1.1e-05 | 9.467E-06 | 1.858E-06 | 7.400E-06 | 1.100E-05 |
| 15545 | 778 | S778C | B | 6.6e-06 | 5.5e-06 | 5e-06 | 5.700E-06 | 8.185E-07 | 5.000E-06 | 6.600E-06 |
| 15546 | 778 | S778E | B | 7.5e-06 | 8.6e-06 | 7.3e-06 | 7.800E-06 | 7.000E-07 | 7.300E-06 | 8.600E-06 |
| 15547 | 778 | S778Q | B | 7.1e-06 | 6e-06 | 1.5e-05 | 9.367E-06 | 4.910E-06 | 6.000E-06 | 1.500E-05 |
| 15548 | 778 | S778G | B | 1e-05 | 6e-06 | 4.2e-06 | 6.733E-06 | 2.969E-06 | 4.200E-06 | 1.000E-05 |
| 15549 | 778 | S778H | B | 1.3e-05 | 9.3e-06 | 7.1e-06 | 9.800E-06 | 2.982E-06 | 7.100E-06 | 1.300E-05 |
| 15550 | 778 | S778I | B | 6.6e-06 | 6.3e-06 | 1e-05 | 7.633E-06 | 2.055E-06 | 6.300E-06 | 1.000E-05 |
| 15551 | 778 | S778L | B | 6.5e-06 | 6.1e-06 | 9.3e-06 | 7.300E-06 | 1.744E-06 | 6.100E-06 | 9.300E-06 |
| 15552 | 778 | S778K | B | 9.8e-06 | 7e-06 | 5.5e-06 | 7.433E-06 | 2.183E-06 | 5.500E-06 | 9.800E-06 |
| 15553 | 778 | S778M | B | 5e-06 | 5.5e-06 | 7.1e-06 | 5.867E-06 | 1.097E-06 | 5.000E-06 | 7.100E-06 |
| 15554 | 778 | S778F | B | 1.1e-05 | 9.2e-06 | 1.1e-05 | 1.040E-05 | 1.039E-06 | 9.200E-06 | 1.100E-05 |

| | | | | | | | | | | |
|---|---|---|---|---|---|---|---|---|---|---|
| 15555 | 778 | S778P | B | 6.3e-06 | 1.3e-05 | 8.6e-06 | 9.300E-06 | 3.404E-06 | 6.300E-06 | 1.300E-05 |
| 15556 | 778 | S778S | B | 6.2e-06 | 4.7e-06 | 9.1e-06 | 6.667E-06 | 2.237E-06 | 4.700E-06 | 9.100E-06 |
| 15557 | 778 | S778T | B | 8.4e-06 | 1.2e-05 | 9.1e-06 | 9.833E-06 | 1.909E-06 | 8.400E-06 | 1.200E-05 |
| 15558 | 778 | S778W | B | 9.3e-06 | 9.8e-06 | 6.2e-06 | 8.433E-06 | 1.950E-06 | 6.200E-06 | 9.800E-06 |
| 15559 | 778 | S778Y | B | 9.4e-06 | 9.2e-06 | 5.4e-06 | 8.000E-06 | 2.254E-06 | 5.400E-06 | 9.400E-06 |
| 15560 | 778 | S778V | B | 1.5e-05 | 5.6e-06 | 1.1e-05 | 1.053E-05 | 4.717E-06 | 5.600E-06 | 1.500E-05 |
| 15561 | 779 | G779A | B | 9.4e-06 | 1e-05 | 8e-06 | 9.133E-06 | 1.026E-06 | 8.000E-06 | 1.000E-05 |
| 15562 | 779 | G779R | B | 1.3e-05 | 6.4e-06 | 6.7e-06 | 8.700E-06 | 3.727E-06 | 6.400E-06 | 1.300E-05 |
| 15563 | 779 | G779N | B | 1.3e-05 | 7.9e-06 | 8.3e-06 | 9.733E-06 | 2.836E-06 | 7.900E-06 | 1.300E-05 |
| 15564 | 779 | G779D | B | 8.1e-06 | 5.2e-06 | 5.5e-06 | 6.267E-06 | 1.595E-06 | 5.200E-06 | 8.100E-06 |
| 15565 | 779 | G779C | B | 1.3e-05 | 6.1e-06 | 1e-05 | 9.700E-06 | 3.460E-06 | 6.100E-06 | 1.300E-05 |
| 15566 | 779 | G779E | B | 9.4e-06 | 9.3e-06 | 5e-06 | 7.900E-06 | 2.512E-06 | 5.000E-06 | 9.400E-06 |
| 15567 | 779 | G779Q | B | 6.8e-06 | 8.8e-06 | 7.5e-06 | 7.700E-06 | 1.015E-06 | 6.800E-06 | 8.800E-06 |
| 15568 | 779 | G779G | B | 4.8e-06 | 1e-05 | 6e-06 | 6.933E-06 | 2.723E-06 | 4.800E-06 | 1.000E-05 |
| 15569 | 779 | G779H | B | 9e-06 | 1e-05 | 9.2e-06 | 9.400E-06 | 5.292E-07 | 9.000E-06 | 1.000E-05 |
| 15570 | 779 | G779I | B | 1e-05 | 8.1e-06 | 1.2e-05 | 1.003E-05 | 1.950E-06 | 8.100E-06 | 1.200E-05 |
| 15571 | 779 | G779L | B | 5.4e-06 | 9e-06 | 7.5e-06 | 7.300E-06 | 1.808E-06 | 5.400E-06 | 9.000E-06 |
| 15572 | 779 | G779K | B | 8.7e-06 | 8e-06 | 9.2e-06 | 8.633E-06 | 6.028E-07 | 8.000E-06 | 9.200E-06 |
| 15573 | 779 | G779M | B | 5.3e-06 | 7.3e-06 | 1.2e-05 | 8.200E-06 | 3.439E-06 | 5.300E-06 | 1.200E-05 |
| 15574 | 779 | G779F | B | 8.5e-06 | 8.2e-06 | 5.7e-06 | 7.467E-06 | 1.537E-06 | 5.700E-06 | 8.500E-06 |
| 15575 | 779 | G779P | B | 9.3e-06 | 1.2e-05 | 4.8e-06 | 8.700E-06 | 3.637E-06 | 4.800E-06 | 1.200E-05 |
| 15576 | 779 | G779S | B | 4.7e-06 | 6.4e-06 | 8.9e-06 | 6.667E-06 | 2.113E-06 | 4.700E-06 | 8.900E-06 |
| 15577 | 779 | G779T | B | 7.2e-06 | 9.2e-06 | 4.1e-06 | 6.833E-06 | 2.570E-06 | 4.100E-06 | 9.200E-06 |
| 15578 | 779 | G779W | B | 7.5e-06 | 6.7e-06 | 8.9e-06 | 7.700E-06 | 1.114E-06 | 6.700E-06 | 8.900E-06 |
| 15579 | 779 | G779Y | B | 8.9e-06 | 9e-06 | 8.7e-06 | 8.867E-06 | 1.528E-07 | 8.700E-06 | 9.000E-06 |
| 15580 | 779 | G779V | B | 5.5e-06 | 9.2e-06 | 6.3e-06 | 7.000E-06 | 1.947E-06 | 5.500E-06 | 9.200E-06 |
| 15581 | 780 | N780A | B | 8.5e-06 | 5.2e-06 | 8.8e-06 | 7.500E-06 | 1.997E-06 | 5.200E-06 | 8.800E-06 |
| 15582 | 780 | N780R | B | 9.4e-06 | 9e-06 | 9e-06 | 9.133E-06 | 2.309E-07 | 9.000E-06 | 9.400E-06 |
| 15583 | 780 | N780N | B | 8.6e-06 | 4.9e-06 | 6.7e-06 | 6.733E-06 | 1.850E-06 | 4.900E-06 | 8.600E-06 |
| 15584 | 780 | N780D | B | 1e-05 | 6.1e-06 | 1.5e-05 | 1.037E-05 | 4.461E-06 | 6.100E-06 | 1.500E-05 |
| 15585 | 780 | N780C | B | 5.2e-06 | 6.9e-06 | 5.3e-06 | 5.800E-06 | 9.539E-07 | 5.200E-06 | 6.900E-06 |
| 15586 | 780 | N780E | B | 6.5e-06 | 9.4e-06 | 8.9e-06 | 8.267E-06 | 1.550E-06 | 6.500E-06 | 9.400E-06 |
| 15587 | 780 | N780Q | B | 5.3e-06 | 5.8e-06 | 1.3e-05 | 8.033E-06 | 4.309E-06 | 5.300E-06 | 1.300E-05 |
| 15588 | 780 | N780G | B | 1.1e-05 | 1e-05 | 8.5e-06 | 9.833E-06 | 1.258E-06 | 8.500E-06 | 1.100E-05 |
| 15589 | 780 | N780H | B | 7.4e-06 | 6.8e-06 | 5.5e-06 | 6.567E-06 | 9.713E-07 | 5.500E-06 | 7.400E-06 |
| 15590 | 780 | N780I | B | 7.9e-06 | 9.8e-06 | 9e-06 | 8.900E-06 | 9.539E-07 | 7.900E-06 | 9.800E-06 |
| 15591 | 780 | N780L | B | 1.3e-05 | 1.1e-05 | 6.4e-06 | 1.013E-05 | 3.384E-06 | 6.400E-06 | 1.300E-05 |
| 15592 | 780 | N780K | B | 1.1e-05 | 4.3e-06 | 8.5e-06 | 7.933E-06 | 3.386E-06 | 4.300E-06 | 1.100E-05 |
| 15593 | 780 | N780M | B | 1.6e-05 | 7.6e-06 | 1.1e-05 | 1.153E-05 | 4.225E-06 | 7.600E-06 | 1.600E-05 |
| 15594 | 780 | N780F | B | 1e-05 | 8.9e-06 | 6.8e-06 | 8.567E-06 | 1.626E-06 | 6.800E-06 | 1.000E-05 |
| 15595 | 780 | N780P | B | 1.1e-05 | 4.8e-06 | 7.4e-06 | 7.733E-06 | 3.113E-06 | 4.800E-06 | 1.100E-05 |
| 15596 | 780 | N780S | B | 9.1e-06 | 8e-06 | 5.7e-06 | 7.600E-06 | 1.735E-06 | 5.700E-06 | 9.100E-06 |
| 15597 | 780 | N780T | B | 1.1e-05 | 6e-06 | 9.7e-06 | 8.900E-06 | 2.594E-06 | 6.000E-06 | 1.100E-05 |
| 15598 | 780 | N780W | B | 5.9e-06 | 9.5e-06 | 6.5e-06 | 7.300E-06 | 1.929E-06 | 5.900E-06 | 9.500E-06 |
| 15599 | 780 | N780Y | B | 6.8e-06 | 7.9e-06 | 5.7e-06 | 6.800E-06 | 1.100E-06 | 5.700E-06 | 7.900E-06 |
| 15600 | 780 | N780V | B | 1.1e-05 | 1.3e-05 | 1.1e-05 | 1.167E-05 | 1.155E-06 | 1.100E-05 | 1.300E-05 |
| 15601 | 781 | S781A | B | 8e-06 | 5.8e-06 | 8.3e-06 | 7.367E-06 | 1.365E-06 | 5.800E-06 | 8.300E-06 |
| 15602 | 781 | S781R | B | 1.2e-05 | 6.6e-06 | 1.1e-05 | 9.867E-06 | 2.873E-06 | 6.600E-06 | 1.200E-05 |
| 15603 | 781 | S781N | B | 6.7e-06 | 8.9e-06 | 6.9e-06 | 7.500E-06 | 1.217E-06 | 6.700E-06 | 8.900E-06 |
| 15604 | 781 | S781D | B | 9e-06 | 1e-05 | 1.3e-05 | 1.067E-05 | 2.082E-06 | 9.000E-06 | 1.300E-05 |
| 15605 | 781 | S781C | B | 9e-06 | 1e-05 | 6.1e-06 | 8.367E-06 | 2.026E-06 | 6.100E-06 | 1.000E-05 |
| 15606 | 781 | S781E | B | 9.8e-06 | 8.5e-06 | 9.8e-06 | 9.367E-06 | 7.506E-07 | 8.500E-06 | 9.800E-06 |
| 15607 | 781 | S781Q | B | 4.3e-06 | 7.1e-06 | 9e-06 | 6.800E-06 | 2.364E-06 | 4.300E-06 | 9.000E-06 |
| 15608 | 781 | S781G | B | 1.3e-05 | 5.5e-06 | 6.5e-06 | 8.333E-06 | 4.072E-06 | 5.500E-06 | 1.300E-05 |
| 15609 | 781 | S781H | B | 8.8e-06 | 1.4e-05 | 9.7e-06 | 1.083E-05 | 2.779E-06 | 8.800E-06 | 1.400E-05 |
| 15610 | 781 | S781I | B | 9.7e-06 | 7.8e-06 | 5.3e-06 | 7.600E-06 | 2.207E-06 | 5.300E-06 | 9.700E-06 |
| 15611 | 781 | S781L | B | 9.6e-06 | 8.8e-06 | 4.4e-06 | 7.600E-06 | 2.800E-06 | 4.400E-06 | 9.600E-06 |
| 15612 | 781 | S781K | B | 1.1e-05 | 4.9e-06 | 1.1e-05 | 8.967E-06 | 3.522E-06 | 4.900E-06 | 1.100E-05 |
| 15613 | 781 | S781M | B | 1e-05 | 8.9e-06 | 8.8e-06 | 9.233E-06 | 6.658E-07 | 8.800E-06 | 1.000E-05 |
| 15614 | 781 | S781F | B | 1.5e-05 | 1.2e-05 | 9.1e-06 | 1.203E-05 | 2.950E-06 | 9.100E-06 | 1.500E-05 |
| 15615 | 781 | S781P | B | 1.1e-05 | 5e-06 | 8.9e-06 | 8.300E-06 | 3.045E-06 | 5.000E-06 | 1.100E-05 |
| 15616 | 781 | S781S | B | 4.5e-06 | 5.3e-06 | 5.8e-06 | 5.200E-06 | 6.557E-07 | 4.500E-06 | 5.800E-06 |
| 15617 | 781 | S781T | B | 1e-05 | 1.4e-05 | 7.2e-06 | 1.040E-05 | 3.418E-06 | 7.200E-06 | 1.400E-05 |
| 15618 | 781 | S781W | B | 6.6e-06 | 8.2e-06 | 3.1e-06 | 5.967E-06 | 2.608E-06 | 3.100E-06 | 8.200E-06 |
| 15619 | 781 | S781Y | B | 8.8e-06 | 7.6e-06 | 1.3e-05 | 9.800E-06 | 2.835E-06 | 7.600E-06 | 1.300E-05 |
| 15620 | 781 | S781V | B | 9.7e-06 | 7e-06 | 5.2e-06 | 7.300E-06 | 2.265E-06 | 5.200E-06 | 9.700E-06 |
| 15621 | 782 | Q782A | B | 1.4e-05 | 6.4e-06 | 5.3e-06 | 8.567E-06 | 4.737E-06 | 5.300E-06 | 1.400E-05 |
| 15622 | 782 | Q782R | B | 1.1e-05 | 5.8e-06 | 9.7e-06 | 8.833E-06 | 2.706E-06 | 5.800E-06 | 1.100E-05 |
| 15623 | 782 | Q782N | B | 5.7e-06 | 1.2e-05 | 9.3e-06 | 9.000E-06 | 3.161E-06 | 5.700E-06 | 1.200E-05 |
| 15624 | 782 | Q782D | B | 8e-06 | 9.6e-06 | 5.9e-06 | 7.833E-06 | 1.856E-06 | 5.900E-06 | 9.600E-06 |
| 15625 | 782 | Q782C | B | 8.4e-06 | 8.3e-06 | 6.8e-06 | 7.833E-06 | 8.963E-07 | 6.800E-06 | 8.400E-06 |
| 15626 | 782 | Q782E | B | 5.1e-06 | 9.4e-06 | 5.3e-06 | 6.600E-06 | 2.427E-06 | 5.100E-06 | 9.400E-06 |
| 15627 | 782 | Q782Q | B | 8.2e-06 | 1.3e-05 | 1.4e-05 | 1.173E-05 | 3.101E-06 | 8.200E-06 | 1.400E-05 |
| 15628 | 782 | Q782G | B | 6.3e-06 | 5.2e-06 | 7.7e-06 | 6.400E-06 | 1.253E-06 | 5.200E-06 | 7.700E-06 |
| 15629 | 782 | Q782H | B | 4.8e-06 | 4.6e-06 | 8.1e-06 | 5.833E-06 | 1.966E-06 | 4.600E-06 | 8.100E-06 |
| 15630 | 782 | Q782I | B | 1e-05 | 1.4e-05 | 1.4e-05 | 1.267E-05 | 2.309E-06 | 1.000E-05 | 1.400E-05 |
| 15631 | 782 | Q782L | B | 7.4e-06 | 6e-06 | 7.6e-06 | 7.000E-06 | 8.718E-07 | 6.000E-06 | 7.600E-06 |
| 15632 | 782 | Q782K | B | 4.9e-06 | 6.7e-06 | 9.5e-06 | 7.033E-06 | 2.318E-06 | 4.900E-06 | 9.500E-06 |
| 15633 | 782 | Q782M | B | 9e-06 | 8.1e-06 | 9e-06 | 8.700E-06 | 5.196E-07 | 8.100E-06 | 9.000E-06 |
| 15634 | 782 | Q782F | B | 9e-06 | 8.7e-06 | 9.5e-06 | 9.067E-06 | 4.041E-07 | 8.700E-06 | 9.500E-06 |
| 15635 | 782 | Q782P | B | 9.5e-06 | 6.9e-06 | 9e-06 | 8.467E-06 | 1.380E-06 | 6.900E-06 | 9.500E-06 |
| 15636 | 782 | Q782S | B | 7.3e-06 | 6.6e-06 | 1e-05 | 7.967E-06 | 1.795E-06 | 6.600E-06 | 1.000E-05 |
| 15637 | 782 | Q782T | B | 9.8e-06 | 4.2e-06 | 6.7e-06 | 6.900E-06 | 2.805E-06 | 4.200E-06 | 9.800E-06 |
| 15638 | 782 | Q782W | B | 8.6e-06 | 6.8e-06 | 5.6e-06 | 7.000E-06 | 1.510E-06 | 5.600E-06 | 8.600E-06 |
| 15639 | 782 | Q782Y | B | 1.4e-05 | 5.5e-06 | 7.9e-06 | 9.133E-06 | 4.382E-06 | 5.500E-06 | 1.400E-05 |

| | | | | | | | | | | |
|---|---|---|---|---|---|---|---|---|---|---|
| 15640 | 782 | Q782V | B | 5.3e-06 | 6.1e-06 | 6.9e-06 | 6.100E-06 | 8.000E-07 | 5.300E-06 | 6.900E-06 |
| 15641 | 783 | E783A | B | 9.2e-06 | 5.9e-06 | 5.5e-06 | 6.867E-06 | 2.031E-06 | 5.500E-06 | 9.200E-06 |
| 15642 | 783 | E783R | B | 6.1e-06 | 1e-05 | 9e-06 | 8.367E-06 | 2.026E-06 | 6.100E-06 | 1.000E-05 |
| 15643 | 783 | E783N | B | 1.3e-05 | 1.1e-05 | 8.6e-06 | 1.087E-05 | 2.203E-06 | 8.600E-06 | 1.300E-05 |
| 15644 | 783 | E783D | B | 1.2e-05 | 1.3e-05 | 7.4e-06 | 1.080E-05 | 2.987E-06 | 7.400E-06 | 1.300E-05 |
| 15645 | 783 | E783C | B | 6.3e-06 | 6.8e-06 | 9.4e-06 | 7.500E-06 | 1.664E-06 | 6.300E-06 | 9.400E-06 |
| 15646 | 783 | E783E | B | 6.2e-06 | 1e-05 | 6.3e-06 | 7.500E-06 | 2.166E-06 | 6.200E-06 | 1.000E-05 |
| 15647 | 783 | E783Q | B | 8.3e-06 | 9e-06 | 5e-06 | 7.433E-06 | 2.136E-06 | 5.000E-06 | 9.000E-06 |
| 15648 | 783 | E783G | B | 5.1e-06 | 6e-06 | 1.1e-05 | 7.367E-06 | 3.179E-06 | 5.100E-06 | 1.100E-05 |
| 15649 | 783 | E783H | B | 1.1e-05 | 1.3e-05 | 8.9e-06 | 1.097E-05 | 2.050E-06 | 8.900E-06 | 1.300E-05 |
| 15650 | 783 | E783I | B | 9.6e-06 | 7.8e-06 | 9.9e-06 | 9.100E-06 | 1.136E-06 | 7.800E-06 | 9.900E-06 |
| 15651 | 783 | E783L | B | 6.6e-06 | 5.1e-06 | 6.9e-06 | 6.200E-06 | 9.644E-07 | 5.100E-06 | 6.900E-06 |
| 15652 | 783 | E783K | B | 4.5e-06 | 8.2e-06 | 5.6e-06 | 6.100E-06 | 1.900E-06 | 4.500E-06 | 8.200E-06 |
| 15653 | 783 | E783M | B | 7.7e-06 | 7.8e-06 | 1.1e-05 | 8.833E-06 | 1.877E-06 | 7.700E-06 | 1.100E-05 |
| 15654 | 783 | E783F | B | 8.6e-06 | 4.1e-06 | 5.2e-06 | 5.967E-06 | 2.346E-06 | 4.100E-06 | 8.600E-06 |
| 15655 | 783 | E783P | B | 1e-05 | 4.6e-06 | 1.2e-05 | 8.867E-06 | 3.828E-06 | 4.600E-06 | 1.200E-05 |
| 15656 | 783 | E783S | B | 7.8e-06 | 6.8e-06 | 6.7e-06 | 7.100E-06 | 6.083E-07 | 6.700E-06 | 7.800E-06 |
| 15657 | 783 | E783T | B | 4.2e-06 | 8.6e-06 | 5.8e-06 | 6.200E-06 | 2.227E-06 | 4.200E-06 | 8.600E-06 |
| 15658 | 783 | E783W | B | 9.1e-06 | 5.3e-06 | 5.7e-06 | 6.700E-06 | 2.088E-06 | 5.300E-06 | 9.100E-06 |
| 15659 | 783 | E783Y | B | 5e-06 | 4.7e-06 | 8.2e-06 | 5.967E-06 | 1.940E-06 | 4.700E-06 | 8.200E-06 |
| 15660 | 783 | E783V | B | 6.2e-06 | 1.1e-05 | 6.6e-06 | 7.933E-06 | 2.663E-06 | 6.200E-06 | 1.100E-05 |
| 15661 | 784 | S784A | B | 5.6e-06 | 9.9e-06 | 6.6e-06 | 7.367E-06 | 2.250E-06 | 5.600E-06 | 9.900E-06 |
| 15662 | 784 | S784R | B | 1.4e-05 | 9.8e-06 | 8.2e-06 | 1.067E-05 | 2.996E-06 | 8.200E-06 | 1.400E-05 |
| 15663 | 784 | S784N | B | 1.4e-05 | 1.3e-05 | 6.1e-06 | 1.103E-05 | 4.302E-06 | 6.100E-06 | 1.400E-05 |
| 15664 | 784 | S784D | B | 8.6e-06 | 1e-05 | 1.4e-05 | 1.087E-05 | 2.802E-06 | 8.600E-06 | 1.400E-05 |
| 15665 | 784 | S784C | B | 6.2e-06 | 6.5e-06 | 8.9e-06 | 7.200E-06 | 1.480E-06 | 6.200E-06 | 8.900E-06 |
| 15666 | 784 | S784E | B | 9.2e-06 | 4.8e-06 | 7.3e-06 | 7.100E-06 | 2.207E-06 | 4.800E-06 | 9.200E-06 |
| 15667 | 784 | S784Q | B | 1.1e-05 | 6.4e-06 | 8.3e-06 | 8.567E-06 | 2.312E-06 | 6.400E-06 | 1.100E-05 |
| 15668 | 784 | S784G | B | 9e-06 | 1e-05 | 9.8e-06 | 9.600E-06 | 5.292E-07 | 9.000E-06 | 1.000E-05 |
| 15669 | 784 | S784H | B | 7.1e-06 | 8.1e-06 | 6.5e-06 | 7.233E-06 | 8.083E-07 | 6.500E-06 | 8.100E-06 |
| 15670 | 784 | S784I | B | 9.3e-06 | 1.3e-05 | 5.4e-06 | 9.233E-06 | 3.800E-06 | 5.400E-06 | 1.300E-05 |
| 15671 | 784 | S784L | B | 1.5e-05 | 1.1e-05 | 1.5e-05 | 1.367E-05 | 2.309E-06 | 1.100E-05 | 1.500E-05 |
| 15672 | 784 | S784K | B | 7.5e-06 | 8.8e-06 | 3.4e-06 | 6.567E-06 | 2.818E-06 | 3.400E-06 | 8.800E-06 |
| 15673 | 784 | S784M | B | 7.2e-06 | 8.6e-06 | 7.1e-06 | 7.633E-06 | 8.386E-07 | 7.100E-06 | 8.600E-06 |
| 15674 | 784 | S784F | B | 9.7e-06 | 6.1e-06 | 8.4e-06 | 8.067E-06 | 1.823E-06 | 6.100E-06 | 9.700E-06 |
| 15675 | 784 | S784P | B | 6.9e-06 | 1e-05 | 9.5e-06 | 8.800E-06 | 1.664E-06 | 6.900E-06 | 1.000E-05 |
| 15676 | 784 | S784S | B | 7.5e-06 | 8.8e-06 | 6.6e-06 | 7.633E-06 | 1.106E-06 | 6.600E-06 | 8.800E-06 |
| 15677 | 784 | S784T | B | 5.3e-06 | 1.5e-05 | 9.5e-06 | 9.933E-06 | 4.864E-06 | 5.300E-06 | 1.500E-05 |
| 15678 | 784 | S784W | B | 4.7e-06 | 1.4e-05 | 9e-06 | 9.233E-06 | 4.654E-06 | 4.700E-06 | 1.400E-05 |
| 15679 | 784 | S784Y | B | 4.3e-06 | 9.3e-06 | 4.9e-06 | 6.167E-06 | 2.730E-06 | 4.300E-06 | 9.300E-06 |
| 15680 | 784 | S784V | B | 8.8e-06 | 6.4e-06 | 8.7e-06 | 7.967E-06 | 1.358E-06 | 6.400E-06 | 8.800E-06 |
| 15681 | 785 | V785A | B | 4.2e-06 | 1.4e-05 | 6.7e-06 | 8.300E-06 | 5.092E-06 | 4.200E-06 | 1.400E-05 |
| 15682 | 785 | V785R | B | 1e-05 | 4.3e-06 | 1.3e-05 | 9.100E-06 | 4.419E-06 | 4.300E-06 | 1.300E-05 |
| 15683 | 785 | V785N | B | 6.1e-06 | 7.7e-06 | 1.4e-05 | 9.267E-06 | 4.177E-06 | 6.100E-06 | 1.400E-05 |
| 15684 | 785 | V785D | B | 8.9e-06 | 1.5e-05 | 1.1e-05 | 1.163E-05 | 3.099E-06 | 8.900E-06 | 1.500E-05 |
| 15685 | 785 | V785C | B | 5.6e-06 | 5.7e-06 | 6e-06 | 5.767E-06 | 2.082E-07 | 5.600E-06 | 6.000E-06 |
| 15686 | 785 | V785E | B | 6.6e-06 | 4.8e-06 | 9.2e-06 | 6.867E-06 | 2.212E-06 | 4.800E-06 | 9.200E-06 |
| 15687 | 785 | V785Q | B | 6.1e-06 | 7.5e-06 | 8.8e-06 | 7.467E-06 | 1.350E-06 | 6.100E-06 | 8.800E-06 |
| 15688 | 785 | V785G | B | 5.6e-06 | 8.8e-06 | 9.9e-06 | 8.100E-06 | 2.234E-06 | 5.600E-06 | 9.900E-06 |
| 15689 | 785 | V785H | B | 5.3e-06 | 6.4e-06 | 7e-06 | 6.233E-06 | 8.622E-07 | 5.300E-06 | 7.000E-06 |
| 15690 | 785 | V785I | B | 4.2e-06 | 5.1e-06 | 8.2e-06 | 5.833E-06 | 2.098E-06 | 4.200E-06 | 8.200E-06 |
| 15691 | 785 | V785L | B | 6.6e-06 | 6.6e-06 | 7.6e-06 | 6.933E-06 | 5.774E-07 | 6.600E-06 | 7.600E-06 |
| 15692 | 785 | V785K | B | 9.2e-06 | 6.5e-06 | 6.8e-06 | 7.500E-06 | 1.480E-06 | 6.500E-06 | 9.200E-06 |
| 15693 | 785 | V785M | B | 1.2e-05 | 9.4e-06 | 9.2e-06 | 1.020E-05 | 1.562E-06 | 9.200E-06 | 1.200E-05 |
| 15694 | 785 | V785F | B | 4.7e-06 | 6.9e-06 | 6.9e-06 | 6.167E-06 | 1.270E-06 | 4.700E-06 | 6.900E-06 |
| 15695 | 785 | V785P | B | 1e-05 | 8e-06 | 5.7e-06 | 7.900E-06 | 2.152E-06 | 5.700E-06 | 1.000E-05 |
| 15696 | 785 | V785S | B | 6.6e-06 | 9.9e-06 | 5.1e-06 | 7.200E-06 | 2.456E-06 | 5.100E-06 | 9.900E-06 |
| 15697 | 785 | V785T | B | 8.9e-06 | 1.2e-05 | 5.2e-06 | 8.700E-06 | 3.404E-06 | 5.200E-06 | 1.200E-05 |
| 15698 | 785 | V785W | B | 5.3e-06 | 4.3e-06 | 9.2e-06 | 6.267E-06 | 2.589E-06 | 4.300E-06 | 9.200E-06 |
| 15699 | 785 | V785Y | B | 6.1e-06 | 4.3e-06 | 8.3e-06 | 6.233E-06 | 2.003E-06 | 4.300E-06 | 8.300E-06 |
| 15700 | 785 | V785V | B | 4.7e-06 | 1.2e-05 | 5.6e-06 | 7.433E-06 | 3.980E-06 | 4.700E-06 | 1.200E-05 |
| 15701 | 786 | T786A | B | 7.5e-06 | 1.3e-05 | 7.9e-06 | 9.467E-06 | 3.066E-06 | 7.500E-06 | 1.300E-05 |
| 15702 | 786 | T786R | B | 5.9e-06 | 5.3e-06 | 9.4e-06 | 6.867E-06 | 2.214E-06 | 5.300E-06 | 9.400E-06 |
| 15703 | 786 | T786N | B | 5.8e-06 | 4.8e-06 | 6.8e-06 | 5.800E-06 | 1.000E-06 | 4.800E-06 | 6.800E-06 |
| 15704 | 786 | T786D | B | 7.2e-06 | 8.9e-06 | 9.3e-06 | 8.467E-06 | 1.115E-06 | 7.200E-06 | 9.300E-06 |
| 15705 | 786 | T786C | B | 7.7e-06 | 5.4e-06 | 1.4e-05 | 9.033E-06 | 4.452E-06 | 5.400E-06 | 1.400E-05 |
| 15706 | 786 | T786E | B | 6e-06 | 6e-06 | 6.8e-06 | 6.267E-06 | 4.619E-07 | 6.000E-06 | 6.800E-06 |
| 15707 | 786 | T786Q | B | 1.3e-05 | 1.1e-05 | 9e-06 | 1.100E-05 | 2.000E-06 | 9.000E-06 | 1.300E-05 |
| 15708 | 786 | T786G | B | 8.7e-06 | 6.1e-06 | 8.8e-06 | 7.867E-06 | 1.531E-06 | 6.100E-06 | 8.800E-06 |
| 15709 | 786 | T786H | B | 6.4e-06 | 9.4e-06 | 4.5e-06 | 6.767E-06 | 2.470E-06 | 4.500E-06 | 9.400E-06 |
| 15710 | 786 | T786I | B | 1.3e-05 | 7.3e-06 | 1.1e-05 | 1.043E-05 | 2.892E-06 | 7.300E-06 | 1.300E-05 |
| 15711 | 786 | T786L | B | 8.5e-06 | 1.9e-05 | 6.7e-06 | 1.140E-05 | 6.643E-06 | 6.700E-06 | 1.900E-05 |
| 15712 | 786 | T786K | B | 7.5e-06 | 9.3e-06 | 9.3e-06 | 8.700E-06 | 1.039E-06 | 7.500E-06 | 9.300E-06 |
| 15713 | 786 | T786M | B | 6e-06 | 9.1e-06 | 1.1e-05 | 8.700E-06 | 2.524E-06 | 6.000E-06 | 1.100E-05 |
| 15714 | 786 | T786F | B | 1.4e-05 | 4.4e-06 | 9.4e-06 | 9.267E-06 | 4.801E-06 | 4.400E-06 | 1.400E-05 |
| 15715 | 786 | T786P | B | 1.3e-05 | 6.2e-06 | 1e-05 | 9.733E-06 | 3.408E-06 | 6.200E-06 | 1.300E-05 |
| 15716 | 786 | T786S | B | 4.8e-06 | 9.2e-06 | 9.4e-06 | 7.800E-06 | 2.600E-06 | 4.800E-06 | 9.400E-06 |
| 15717 | 786 | T786T | B | 9.2e-06 | 7.9e-06 | 9.1e-06 | 8.733E-06 | 7.234E-07 | 7.900E-06 | 9.200E-06 |
| 15718 | 786 | T786W | B | 6.4e-06 | 1.4e-05 | 9.4e-06 | 9.933E-06 | 3.828E-06 | 6.400E-06 | 1.400E-05 |
| 15719 | 786 | T786Y | B | 5.3e-06 | 1.3e-05 | 5.1e-06 | 7.800E-06 | 4.504E-06 | 5.100E-06 | 1.300E-05 |
| 15720 | 786 | T786V | B | 9.4e-06 | 6.9e-06 | 9.2e-06 | 8.500E-06 | 1.389E-06 | 6.900E-06 | 9.400E-06 |
| 15721 | 787 | E787A | B | 8.2e-06 | 7.1e-06 | 7.1e-06 | 7.467E-06 | 6.351E-07 | 7.100E-06 | 8.200E-06 |
| 15722 | 787 | E787R | B | 6.7e-06 | 6.4e-06 | 6.8e-06 | 6.633E-06 | 2.082E-07 | 6.400E-06 | 6.800E-06 |
| 15723 | 787 | E787N | B | 9.9e-06 | 9.4e-06 | 3.7e-06 | 7.667E-06 | 3.444E-06 | 3.700E-06 | 9.900E-06 |
| 15724 | 787 | E787D | B | 1e-05 | 1.4e-05 | 7.1e-06 | 1.037E-05 | 3.465E-06 | 7.100E-06 | 1.400E-05 |

| | | | | | | | | | | |
|---|---|---|---|---|---|---|---|---|---|---|
| 15725 | 787 | E787C | B | 1e-05 | 6e-06 | 1.1e-05 | 9.000E-06 | 2.646E-06 | 6.000E-06 | 1.100E-05 |
| 15726 | 787 | E787E | B | 5.3e-06 | 1.2e-05 | 8.6e-06 | 8.633E-06 | 3.350E-06 | 5.300E-06 | 1.200E-05 |
| 15727 | 787 | E787Q | B | 6.7e-06 | 8.9e-06 | 6.2e-06 | 7.267E-06 | 1.436E-06 | 6.200E-06 | 8.900E-06 |
| 15728 | 787 | E787G | B | 4.9e-06 | 9.6e-06 | 9.5e-06 | 8.000E-06 | 2.685E-06 | 4.900E-06 | 9.600E-06 |
| 15729 | 787 | E787H | B | 9.1e-06 | 1e-05 | 1.1e-05 | 1.003E-05 | 9.504E-07 | 9.100E-06 | 1.100E-05 |
| 15730 | 787 | E787I | B | 5.5e-06 | 9.4e-06 | 6.7e-06 | 7.200E-06 | 1.997E-06 | 5.500E-06 | 9.400E-06 |
| 15731 | 787 | E787L | B | 6.9e-06 | 7.8e-06 | 9.8e-06 | 8.167E-06 | 1.484E-06 | 6.900E-06 | 9.800E-06 |
| 15732 | 787 | E787K | B | 4.8e-06 | 5.1e-06 | 6.5e-06 | 5.467E-06 | 9.074E-07 | 4.800E-06 | 6.500E-06 |
| 15733 | 787 | E787M | B | 6.7e-06 | 5.4e-06 | 8.3e-06 | 6.800E-06 | 1.453E-06 | 5.400E-06 | 8.300E-06 |
| 15734 | 787 | E787F | B | 5e-06 | 6.8e-06 | 9.9e-06 | 7.233E-06 | 2.479E-06 | 5.000E-06 | 9.900E-06 |
| 15735 | 787 | E787P | B | 7.2e-06 | 1.2e-05 | 5.9e-06 | 8.367E-06 | 3.213E-06 | 5.900E-06 | 1.200E-05 |
| 15736 | 787 | E787S | B | 7.7e-06 | 8.7e-06 | 5.7e-06 | 7.367E-06 | 1.528E-06 | 5.700E-06 | 8.700E-06 |
| 15737 | 787 | E787T | B | 6.2e-06 | 9.1e-06 | 7.1e-06 | 7.467E-06 | 1.484E-06 | 6.200E-06 | 9.100E-06 |
| 15738 | 787 | E787W | B | 7.7e-06 | 6.7e-06 | 5e-06 | 6.467E-06 | 1.365E-06 | 5.000E-06 | 7.700E-06 |
| 15739 | 787 | E787Y | B | 5.2e-06 | 1.2e-05 | 8.9e-06 | 8.700E-06 | 3.404E-06 | 5.200E-06 | 1.200E-05 |
| 15740 | 787 | E787V | B | 8.7e-06 | 8.3e-06 | 7.9e-06 | 8.300E-06 | 4.000E-07 | 7.900E-06 | 8.700E-06 |
| 15741 | 788 | Q788A | B | 6.6e-06 | 1e-05 | 4.7e-06 | 7.100E-06 | 2.685E-06 | 4.700E-06 | 1.000E-05 |
| 15742 | 788 | Q788R | B | 7.1e-06 | 1.4e-05 | 4.5e-06 | 8.533E-06 | 4.910E-06 | 4.500E-06 | 1.400E-05 |
| 15743 | 788 | Q788N | B | 7.7e-06 | 4.5e-06 | 7.8e-06 | 6.667E-06 | 1.877E-06 | 4.500E-06 | 7.800E-06 |
| 15744 | 788 | Q788D | B | 6.6e-06 | 1.3e-05 | 9.2e-06 | 9.600E-06 | 3.219E-06 | 6.600E-06 | 1.300E-05 |
| 15745 | 788 | Q788C | B | 8e-06 | 7.9e-06 | 8.1e-06 | 8.000E-06 | 1.000E-07 | 7.900E-06 | 8.100E-06 |
| 15746 | 788 | Q788E | B | 4.4e-06 | 9.8e-06 | 9.3e-06 | 7.833E-06 | 2.984E-06 | 4.400E-06 | 9.800E-06 |
| 15747 | 788 | Q788Q | B | 9.7e-06 | 7.6e-06 | 1.6e-05 | 1.110E-05 | 4.371E-06 | 7.600E-06 | 1.600E-05 |
| 15748 | 788 | Q788G | B | 4.6e-06 | 7.7e-06 | 7.6e-06 | 6.633E-06 | 1.762E-06 | 4.600E-06 | 7.700E-06 |
| 15749 | 788 | Q788H | B | 6.1e-06 | 6.7e-06 | 5.1e-06 | 5.967E-06 | 8.083E-07 | 5.100E-06 | 6.700E-06 |
| 15750 | 788 | Q788I | B | 1e-05 | 1.4e-05 | 7.3e-06 | 1.043E-05 | 3.371E-06 | 7.300E-06 | 1.400E-05 |
| 15751 | 788 | Q788L | B | 7.2e-06 | 1.1e-05 | 6.3e-06 | 8.167E-06 | 2.495E-06 | 6.300E-06 | 1.100E-05 |
| 15752 | 788 | Q788K | B | 1.4e-05 | 5.8e-06 | 5.6e-06 | 8.467E-06 | 4.793E-06 | 5.600E-06 | 1.400E-05 |
| 15753 | 788 | Q788M | B | 6e-06 | 7e-06 | 9.5e-06 | 7.500E-06 | 1.803E-06 | 6.000E-06 | 9.500E-06 |
| 15754 | 788 | Q788F | B | 8.5e-06 | 5.9e-06 | 5e-06 | 6.467E-06 | 1.818E-06 | 5.000E-06 | 8.500E-06 |
| 15755 | 788 | Q788P | B | 5.2e-06 | 1e-05 | 7e-06 | 7.400E-06 | 2.425E-06 | 5.200E-06 | 1.000E-05 |
| 15756 | 788 | Q788S | B | 9.4e-06 | 4.6e-06 | 1.2e-05 | 8.667E-06 | 3.754E-06 | 4.600E-06 | 1.200E-05 |
| 15757 | 788 | Q788T | B | 9.3e-06 | 6.7e-06 | 1e-05 | 8.667E-06 | 1.739E-06 | 6.700E-06 | 1.000E-05 |
| 15758 | 788 | Q788W | B | 7.6e-06 | 7.5e-06 | 8.6e-06 | 7.900E-06 | 6.083E-07 | 7.500E-06 | 8.600E-06 |
| 15759 | 788 | Q788Y | B | 1e-05 | 9.3e-06 | 9.6e-06 | 9.633E-06 | 3.512E-07 | 9.300E-06 | 1.000E-05 |
| 15760 | 788 | Q788V | B | 7.6e-06 | 9.2e-06 | 7.2e-06 | 8.000E-06 | 1.058E-06 | 7.200E-06 | 9.200E-06 |
| 15761 | 789 | D789A | B | 7.4e-06 | 4.4e-06 | 9e-06 | 6.933E-06 | 2.335E-06 | 4.400E-06 | 9.000E-06 |
| 15762 | 789 | D789R | B | 8.6e-06 | 7.1e-06 | 7.6e-06 | 7.767E-06 | 7.638E-07 | 7.100E-06 | 8.600E-06 |
| 15763 | 789 | D789N | B | 1.2e-05 | 1.3e-05 | 1e-05 | 1.167E-05 | 1.528E-06 | 1.000E-05 | 1.300E-05 |
| 15764 | 789 | D789D | B | 6e-06 | 8.3e-06 | 7e-06 | 7.100E-06 | 1.153E-06 | 6.000E-06 | 8.300E-06 |
| 15765 | 789 | D789C | B | 9.9e-06 | 1.1e-05 | 4.7e-06 | 8.533E-06 | 3.365E-06 | 4.700E-06 | 1.100E-05 |
| 15766 | 789 | D789E | B | 8.5e-06 | 5e-06 | 1.1e-05 | 8.167E-06 | 3.014E-06 | 5.000E-06 | 1.100E-05 |
| 15767 | 789 | D789Q | B | 7.8e-06 | 1e-05 | 1.2e-05 | 9.933E-06 | 2.101E-06 | 7.800E-06 | 1.200E-05 |
| 15768 | 789 | D789G | B | 6.9e-06 | 1e-05 | 7.4e-06 | 8.100E-06 | 1.664E-06 | 6.900E-06 | 1.000E-05 |
| 15769 | 789 | D789H | B | 9.9e-06 | 1.3e-05 | 4.8e-06 | 9.233E-06 | 4.140E-06 | 4.800E-06 | 1.300E-05 |
| 15770 | 789 | D789I | B | 5.3e-06 | 7.7e-06 | 5.9e-06 | 6.300E-06 | 1.249E-06 | 5.300E-06 | 7.700E-06 |
| 15771 | 789 | D789L | B | 1.5e-05 | 9.6e-06 | 1.4e-05 | 1.287E-05 | 2.873E-06 | 9.600E-06 | 1.500E-05 |
| 15772 | 789 | D789K | B | 8.9e-06 | 9.9e-06 | 9.1e-06 | 9.300E-06 | 5.292E-07 | 8.900E-06 | 9.900E-06 |
| 15773 | 789 | D789M | B | 8e-06 | 1e-05 | 6.2e-06 | 8.067E-06 | 1.901E-06 | 6.200E-06 | 1.000E-05 |
| 15774 | 789 | D789F | B | 8.5e-06 | 1.3e-05 | 7.8e-06 | 9.767E-06 | 2.822E-06 | 7.800E-06 | 1.300E-05 |
| 15775 | 789 | D789P | B | 8.8e-06 | 1.5e-05 | 1.4e-05 | 1.260E-05 | 3.329E-06 | 8.800E-06 | 1.500E-05 |
| 15776 | 789 | D789S | B | 1e-05 | 8.5e-06 | 9.3e-06 | 9.267E-06 | 7.506E-07 | 8.500E-06 | 1.000E-05 |
| 15777 | 789 | D789T | B | 6.5e-06 | 7.1e-06 | 7.3e-06 | 6.967E-06 | 4.163E-07 | 6.500E-06 | 7.300E-06 |
| 15778 | 789 | D789W | B | 7.7e-06 | 1.1e-05 | 6.5e-06 | 8.400E-06 | 2.330E-06 | 6.500E-06 | 1.100E-05 |
| 15779 | 789 | D789Y | B | 9.2e-06 | 8.5e-06 | 7.9e-06 | 8.533E-06 | 6.506E-07 | 7.900E-06 | 9.200E-06 |
| 15780 | 789 | D789V | B | 1.1e-05 | 1.1e-05 | 7.8e-06 | 9.933E-06 | 1.848E-06 | 7.800E-06 | 1.100E-05 |
| 15781 | 790 | S790A | B | 1.4e-05 | 1.3e-05 | 5.3e-06 | 1.077E-05 | 4.761E-06 | 5.300E-06 | 1.400E-05 |
| 15782 | 790 | S790R | B | 1.3e-05 | 7.4e-06 | 7.1e-06 | 9.167E-06 | 3.323E-06 | 7.100E-06 | 1.300E-05 |
| 15783 | 790 | S790N | B | 5.7e-06 | 6.6e-06 | 4.9e-06 | 5.733E-06 | 8.505E-07 | 4.900E-06 | 6.600E-06 |
| 15784 | 790 | S790D | B | 3.8e-06 | 6.8e-06 | 1.1e-05 | 7.200E-06 | 3.617E-06 | 3.800E-06 | 1.100E-05 |
| 15785 | 790 | S790C | B | 8.3e-06 | 1.2e-05 | 4.3e-06 | 8.200E-06 | 3.851E-06 | 4.300E-06 | 1.200E-05 |
| 15786 | 790 | S790E | B | 1.5e-05 | 1.5e-05 | 5.6e-06 | 1.187E-05 | 5.427E-06 | 5.600E-06 | 1.500E-05 |
| 15787 | 790 | S790Q | B | 5.9e-06 | 1.1e-05 | 9.1e-06 | 8.667E-06 | 2.577E-06 | 5.900E-06 | 1.100E-05 |
| 15788 | 790 | S790G | B | 9.5e-06 | 8.1e-06 | 7.2e-06 | 8.267E-06 | 1.159E-06 | 7.200E-06 | 9.500E-06 |
| 15789 | 790 | S790H | B | 1.1e-05 | 4.7e-06 | 4e-06 | 6.567E-06 | 3.855E-06 | 4.000E-06 | 1.100E-05 |
| 15790 | 790 | S790I | B | 1.9e-05 | 5.1e-06 | 6.5e-06 | 1.020E-05 | 7.653E-06 | 5.100E-06 | 1.900E-05 |
| 15791 | 790 | S790L | B | 6.8e-06 | 9.8e-06 | 6.5e-06 | 7.700E-06 | 1.825E-06 | 6.500E-06 | 9.800E-06 |
| 15792 | 790 | S790K | B | 1.2e-05 | 8.8e-06 | 6.3e-06 | 9.033E-06 | 2.857E-06 | 6.300E-06 | 1.200E-05 |
| 15793 | 790 | S790M | B | 7.6e-06 | 7.9e-06 | 7.6e-06 | 7.700E-06 | 1.732E-07 | 7.600E-06 | 7.900E-06 |
| 15794 | 790 | S790F | B | 1.5e-05 | 1.5e-05 | 4.8e-06 | 1.160E-05 | 5.889E-06 | 4.800E-06 | 1.500E-05 |
| 15795 | 790 | S790P | B | 6.8e-06 | 1.1e-05 | 9.8e-06 | 9.200E-06 | 2.163E-06 | 6.800E-06 | 1.100E-05 |
| 15796 | 790 | S790S | B | 1.5e-05 | 7.1e-06 | 6.7e-06 | 9.600E-06 | 4.681E-06 | 6.700E-06 | 1.500E-05 |
| 15797 | 790 | S790T | B | 6.6e-06 | 6.9e-06 | 7.4e-06 | 6.967E-06 | 4.041E-07 | 6.600E-06 | 7.400E-06 |
| 15798 | 790 | S790W | B | 5.7e-06 | 9.4e-06 | 5.8e-06 | 6.967E-06 | 2.108E-06 | 5.700E-06 | 9.400E-06 |
| 15799 | 790 | S790Y | B | 5.7e-06 | 7.5e-06 | 9.1e-06 | 7.433E-06 | 1.701E-06 | 5.700E-06 | 9.100E-06 |
| 15800 | 790 | S790V | B | 5.7e-06 | 3.7e-06 | 1.1e-05 | 6.800E-06 | 3.772E-06 | 3.700E-06 | 1.100E-05 |
| 15801 | 791 | K791A | B | 1e-05 | 5.4e-06 | 7e-06 | 7.467E-06 | 2.335E-06 | 5.400E-06 | 1.000E-05 |
| 15802 | 791 | K791R | B | 5.8e-06 | 9.7e-06 | 1.1e-05 | 8.833E-06 | 2.706E-06 | 5.800E-06 | 1.100E-05 |
| 15803 | 791 | K791N | B | 9.2e-06 | 9.7e-06 | 1.1e-05 | 9.967E-06 | 9.292E-07 | 9.200E-06 | 1.100E-05 |
| 15804 | 791 | K791D | B | 5.8e-06 | 7.4e-06 | 5.7e-06 | 6.300E-06 | 9.539E-07 | 5.700E-06 | 7.400E-06 |
| 15805 | 791 | K791C | B | 4.8e-06 | 6.4e-06 | 6.4e-06 | 5.867E-06 | 9.238E-07 | 4.800E-06 | 6.400E-06 |
| 15806 | 791 | K791E | B | 6.5e-06 | 7.2e-06 | 9.6e-06 | 7.767E-06 | 1.626E-06 | 6.500E-06 | 9.600E-06 |
| 15807 | 791 | K791Q | B | 6.1e-06 | 4.6e-06 | 4.6e-06 | 5.100E-06 | 8.660E-07 | 4.600E-06 | 6.100E-06 |
| 15808 | 791 | K791G | B | 1.2e-05 | 6.8e-06 | 9.8e-06 | 9.533E-06 | 2.610E-06 | 6.800E-06 | 1.200E-05 |
| 15809 | 791 | K791H | B | 8e-06 | 6.3e-06 | 9e-06 | 7.767E-06 | 1.365E-06 | 6.300E-06 | 9.000E-06 |

| | | | | | | | | | | |
|---|---|---|---|---|---|---|---|---|---|---|
| 15810 | 791 | K791I | B | 8.8e-06 | 4.3e-06 | 8.5e-06 | 7.200E-06 | 2.516E-06 | 4.300E-06 | 8.800E-06 |
| 15811 | 791 | K791L | B | 6.4e-06 | 9.1e-06 | 6.5e-06 | 7.333E-06 | 1.531E-06 | 6.400E-06 | 9.100E-06 |
| 15812 | 791 | K791K | B | 4.2e-06 | 6.9e-06 | 8.8e-06 | 6.633E-06 | 2.312E-06 | 4.200E-06 | 8.800E-06 |
| 15813 | 791 | K791M | B | 5.2e-06 | 1.3e-05 | 7.7e-06 | 8.633E-06 | 3.983E-06 | 5.200E-06 | 1.300E-05 |
| 15814 | 791 | K791F | B | 9e-06 | 6.6e-06 | 5.6e-06 | 7.067E-06 | 1.747E-06 | 5.600E-06 | 9.000E-06 |
| 15815 | 791 | K791P | B | 5.6e-06 | 8.5e-06 | 6.7e-06 | 6.933E-06 | 1.464E-06 | 5.600E-06 | 8.500E-06 |
| 15816 | 791 | K791S | B | 6e-06 | 1.3e-05 | 7.7e-06 | 8.900E-06 | 3.651E-06 | 6.000E-06 | 1.300E-05 |
| 15817 | 791 | K791T | B | 5.4e-06 | 1e-05 | 6.1e-06 | 7.167E-06 | 2.479E-06 | 5.400E-06 | 1.000E-05 |
| 15818 | 791 | K791W | B | 9.6e-06 | 7.3e-06 | 8.1e-06 | 8.333E-06 | 1.168E-06 | 7.300E-06 | 9.600E-06 |
| 15819 | 791 | K791Y | B | 5.8e-06 | 6.7e-06 | 7e-06 | 6.500E-06 | 6.245E-07 | 5.800E-06 | 7.000E-06 |
| 15820 | 791 | K791V | B | 6.4e-06 | 1.1e-05 | 9.2e-06 | 8.867E-06 | 2.318E-06 | 6.400E-06 | 1.100E-05 |
| 15821 | 792 | D792A | B | 3.3e-06 | 6.8e-06 | 8.8e-06 | 6.300E-06 | 2.784E-06 | 3.300E-06 | 8.800E-06 |
| 15822 | 792 | D792R | B | 4.9e-06 | 1.2e-05 | 7.8e-06 | 8.233E-06 | 3.570E-06 | 4.900E-06 | 1.200E-05 |
| 15823 | 792 | D792N | B | 8.1e-06 | 6.1e-06 | 9.5e-06 | 7.900E-06 | 1.709E-06 | 6.100E-06 | 9.500E-06 |
| 15824 | 792 | D792D | B | 1.2e-05 | 1e-05 | 7.9e-06 | 9.967E-06 | 2.050E-06 | 7.900E-06 | 1.200E-05 |
| 15825 | 792 | D792C | B | 6.7e-06 | 7.4e-06 | 1.2e-05 | 8.700E-06 | 2.879E-06 | 6.700E-06 | 1.200E-05 |
| 15826 | 792 | D792E | B | 4.7e-06 | 5e-06 | 9.1e-06 | 6.267E-06 | 2.458E-06 | 4.700E-06 | 9.100E-06 |
| 15827 | 792 | D792Q | B | 9.3e-06 | 9e-06 | 7.5e-06 | 8.600E-06 | 9.644E-07 | 7.500E-06 | 9.300E-06 |
| 15828 | 792 | D792G | B | 4.9e-06 | 6e-06 | 7e-06 | 5.967E-06 | 1.050E-06 | 4.900E-06 | 7.000E-06 |
| 15829 | 792 | D792H | B | 8.7e-06 | 4.9e-06 | 8.9e-06 | 7.500E-06 | 2.254E-06 | 4.900E-06 | 8.900E-06 |
| 15830 | 792 | D792I | B | 7.9e-06 | 5.5e-06 | 6.1e-06 | 6.500E-06 | 1.249E-06 | 5.500E-06 | 7.900E-06 |
| 15831 | 792 | D792L | B | 6.4e-06 | 7.3e-06 | 9.1e-06 | 7.600E-06 | 1.375E-06 | 6.400E-06 | 9.100E-06 |
| 15832 | 792 | D792K | B | 1.3e-05 | 1.1e-05 | 6.7e-06 | 1.023E-05 | 3.219E-06 | 6.700E-06 | 1.300E-05 |
| 15833 | 792 | D792M | B | 9.3e-06 | 5.7e-06 | 1.4e-05 | 9.667E-06 | 4.162E-06 | 5.700E-06 | 1.400E-05 |
| 15834 | 792 | D792F | B | 7.3e-06 | 4.2e-06 | 7.4e-06 | 6.300E-06 | 1.819E-06 | 4.200E-06 | 7.400E-06 |
| 15835 | 792 | D792P | B | 8.7e-06 | 1.5e-05 | 9e-06 | 1.090E-05 | 3.554E-06 | 8.700E-06 | 1.500E-05 |
| 15836 | 792 | D792S | B | 6.3e-06 | 8.5e-06 | 7.3e-06 | 7.367E-06 | 1.102E-06 | 6.300E-06 | 8.500E-06 |
| 15837 | 792 | D792T | B | 8.9e-06 | 5.8e-06 | 7.4e-06 | 7.367E-06 | 1.550E-06 | 5.800E-06 | 8.900E-06 |
| 15838 | 792 | D792W | B | 1.4e-05 | 7.8e-06 | 8.6e-06 | 1.013E-05 | 3.372E-06 | 7.800E-06 | 1.400E-05 |
| 15839 | 792 | D792Y | B | 9.2e-06 | 8.3e-06 | 8e-06 | 8.500E-06 | 6.245E-07 | 8.000E-06 | 9.200E-06 |
| 15840 | 792 | D792V | B | 1.4e-05 | 8.1e-06 | 1.1e-05 | 1.103E-05 | 2.950E-06 | 8.100E-06 | 1.400E-05 |
| 15841 | 793 | S793A | B | 5.6e-06 | 7e-06 | 8.6e-06 | 7.067E-06 | 1.501E-06 | 5.600E-06 | 8.600E-06 |
| 15842 | 793 | S793R | B | 7.7e-06 | 8.4e-06 | 5.4e-06 | 7.167E-06 | 1.570E-06 | 5.400E-06 | 8.400E-06 |
| 15843 | 793 | S793N | B | 6.1e-06 | 9.2e-06 | 8.1e-06 | 7.800E-06 | 1.572E-06 | 6.100E-06 | 9.200E-06 |
| 15844 | 793 | S793D | B | 6.7e-06 | 6.7e-06 | 4.4e-06 | 5.933E-06 | 1.328E-06 | 4.400E-06 | 6.700E-06 |
| 15845 | 793 | S793C | B | 1.3e-05 | 4.3e-06 | 4.8e-06 | 7.367E-06 | 4.885E-06 | 4.300E-06 | 1.300E-05 |
| 15846 | 793 | S793E | B | 9.2e-06 | 1e-05 | 7.4e-06 | 8.867E-06 | 1.332E-06 | 7.400E-06 | 1.000E-05 |
| 15847 | 793 | S793Q | B | 1.2e-05 | 3e-06 | 1.1e-05 | 8.667E-06 | 4.933E-06 | 3.000E-06 | 1.200E-05 |
| 15848 | 793 | S793G | B | 8.4e-06 | 9.9e-06 | 4.4e-06 | 7.567E-06 | 2.843E-06 | 4.400E-06 | 9.900E-06 |
| 15849 | 793 | S793H | B | 1.1e-05 | 4.7e-06 | 1.2e-05 | 9.233E-06 | 3.958E-06 | 4.700E-06 | 1.200E-05 |
| 15850 | 793 | S793I | B | 6.8e-06 | 6.2e-06 | 4.7e-06 | 5.900E-06 | 1.082E-06 | 4.700E-06 | 6.800E-06 |
| 15851 | 793 | S793L | B | 7.1e-06 | 6.3e-06 | 1.4e-05 | 9.133E-06 | 4.234E-06 | 6.300E-06 | 1.400E-05 |
| 15852 | 793 | S793K | B | 1e-05 | 5.4e-06 | 5.5e-06 | 6.967E-06 | 2.627E-06 | 5.400E-06 | 1.000E-05 |
| 15853 | 793 | S793M | B | 4.7e-06 | 6.6e-06 | 9.2e-06 | 6.833E-06 | 2.259E-06 | 4.700E-06 | 9.200E-06 |
| 15854 | 793 | S793F | B | 1.4e-05 | 1.1e-05 | 9.8e-06 | 1.160E-05 | 2.163E-06 | 9.800E-06 | 1.400E-05 |
| 15855 | 793 | S793P | B | 1.6e-05 | 1.3e-05 | 9.2e-06 | 1.273E-05 | 3.408E-06 | 9.200E-06 | 1.600E-05 |
| 15856 | 793 | S793S | B | 6.7e-06 | 5e-06 | 1.4e-05 | 8.567E-06 | 4.782E-06 | 5.000E-06 | 1.400E-05 |
| 15857 | 793 | S793T | B | 9.6e-06 | 7.1e-06 | 1e-05 | 8.900E-06 | 1.572E-06 | 7.100E-06 | 1.000E-05 |
| 15858 | 793 | S793W | B | 9e-06 | 6.6e-06 | 1.1e-05 | 8.867E-06 | 2.203E-06 | 6.600E-06 | 1.100E-05 |
| 15859 | 793 | S793Y | B | 7.5e-06 | 8.9e-06 | 6.6e-06 | 7.667E-06 | 1.159E-06 | 6.600E-06 | 8.900E-06 |
| 15860 | 793 | S793V | B | 5.4e-06 | 9.6e-06 | 9.5e-06 | 8.167E-06 | 2.397E-06 | 5.400E-06 | 9.600E-06 |
| 15861 | 794 | T794A | B | 4.9e-06 | 1e-05 | 7.7e-06 | 7.533E-06 | 2.554E-06 | 4.900E-06 | 1.000E-05 |
| 15862 | 794 | T794R | B | 1.4e-05 | 8.7e-06 | 9.8e-06 | 1.083E-05 | 2.797E-06 | 8.700E-06 | 1.400E-05 |
| 15863 | 794 | T794N | B | 7.2e-06 | 1e-05 | 6.7e-06 | 7.967E-06 | 1.779E-06 | 6.700E-06 | 1.000E-05 |
| 15864 | 794 | T794D | B | 1.1e-05 | 1.3e-05 | 6e-06 | 1.000E-05 | 3.606E-06 | 6.000E-06 | 1.300E-05 |
| 15865 | 794 | T794C | B | 1.2e-05 | 1.5e-05 | 7.4e-06 | 1.147E-05 | 3.828E-06 | 7.400E-06 | 1.500E-05 |
| 15866 | 794 | T794E | B | 7.3e-06 | 6.2e-06 | 6.4e-06 | 6.633E-06 | 5.859E-07 | 6.200E-06 | 7.300E-06 |
| 15867 | 794 | T794Q | B | 9.3e-06 | 6.7e-06 | 9e-06 | 8.333E-06 | 1.422E-06 | 6.700E-06 | 9.300E-06 |
| 15868 | 794 | T794G | B | 8.7e-06 | 1.1e-05 | 4.9e-06 | 8.200E-06 | 3.081E-06 | 4.900E-06 | 1.100E-05 |
| 15869 | 794 | T794H | B | 1.1e-05 | 9.5e-06 | 7.7e-06 | 9.400E-06 | 1.652E-06 | 7.700E-06 | 1.100E-05 |
| 15870 | 794 | T794I | B | 6.2e-06 | 1.1e-05 | 8.9e-06 | 8.700E-06 | 2.406E-06 | 6.200E-06 | 1.100E-05 |
| 15871 | 794 | T794L | B | 1.1e-05 | 7.4e-06 | 7e-06 | 8.467E-06 | 2.203E-06 | 7.000E-06 | 1.100E-05 |
| 15872 | 794 | T794K | B | 8.2e-06 | 1.5e-05 | 8.9e-06 | 1.070E-05 | 3.740E-06 | 8.200E-06 | 1.500E-05 |
| 15873 | 794 | T794M | B | 7e-06 | 6.9e-06 | 8.4e-06 | 7.433E-06 | 8.386E-07 | 6.900E-06 | 8.400E-06 |
| 15874 | 794 | T794F | B | 7.8e-06 | 1.2e-05 | 9.8e-06 | 9.867E-06 | 2.101E-06 | 7.800E-06 | 1.200E-05 |
| 15875 | 794 | T794P | B | 6.6e-06 | 6.3e-06 | 5.9e-06 | 6.267E-06 | 3.512E-07 | 5.900E-06 | 6.600E-06 |
| 15876 | 794 | T794S | B | 9.3e-06 | 9.5e-06 | 1.4e-05 | 1.093E-05 | 2.658E-06 | 9.300E-06 | 1.400E-05 |
| 15877 | 794 | T794T | B | 5.9e-06 | 7.4e-06 | 5e-06 | 6.100E-06 | 1.212E-06 | 5.000E-06 | 7.400E-06 |
| 15878 | 794 | T794W | B | 8.1e-06 | 9.2e-06 | 1.1e-05 | 9.433E-06 | 1.464E-06 | 8.100E-06 | 1.100E-05 |
| 15879 | 794 | T794Y | B | 5e-06 | 1.2e-05 | 5.4e-06 | 7.467E-06 | 3.931E-06 | 5.000E-06 | 1.200E-05 |
| 15880 | 794 | T794V | B | 9.1e-06 | 8.8e-06 | 1.2e-05 | 9.967E-06 | 1.767E-06 | 8.800E-06 | 1.200E-05 |
| 15881 | 795 | Y795A | B | 7.3e-06 | 7.7e-06 | 8.6e-06 | 7.867E-06 | 6.658E-07 | 7.300E-06 | 8.600E-06 |
| 15882 | 795 | Y795R | B | 1.4e-05 | 1.4e-05 | 5.5e-06 | 1.117E-05 | 4.907E-06 | 5.500E-06 | 1.400E-05 |
| 15883 | 795 | Y795N | B | 6.5e-06 | 7.5e-06 | 6.8e-06 | 6.933E-06 | 5.132E-07 | 6.500E-06 | 7.500E-06 |
| 15884 | 795 | Y795D | B | 6.5e-06 | 6.7e-06 | 4.9e-06 | 6.033E-06 | 9.866E-07 | 4.900E-06 | 6.700E-06 |
| 15885 | 795 | Y795C | B | 6.7e-06 | 8.7e-06 | 1.2e-05 | 9.133E-06 | 2.676E-06 | 6.700E-06 | 1.200E-05 |
| 15886 | 795 | Y795E | B | 9.8e-06 | 9.9e-06 | 5.1e-06 | 8.267E-06 | 2.743E-06 | 5.100E-06 | 9.900E-06 |
| 15887 | 795 | Y795Q | B | 8.2e-06 | 5.9e-06 | 6.5e-06 | 6.867E-06 | 1.193E-06 | 5.900E-06 | 8.200E-06 |
| 15888 | 795 | Y795G | B | 6.5e-06 | 4.4e-06 | 6.4e-06 | 5.767E-06 | 1.185E-06 | 4.400E-06 | 6.500E-06 |
| 15889 | 795 | Y795H | B | 5.4e-06 | 1e-05 | 3.8e-06 | 6.400E-06 | 3.219E-06 | 3.800E-06 | 1.000E-05 |
| 15890 | 795 | Y795I | B | 7e-06 | 5.3e-06 | 1e-05 | 7.433E-06 | 2.380E-06 | 5.300E-06 | 1.000E-05 |
| 15891 | 795 | Y795L | B | 7.4e-06 | 7.4e-06 | 1.1e-05 | 8.600E-06 | 2.078E-06 | 7.400E-06 | 1.100E-05 |
| 15892 | 795 | Y795K | B | 7.3e-06 | 7.9e-06 | 9.7e-06 | 8.300E-06 | 1.249E-06 | 7.300E-06 | 9.700E-06 |
| 15893 | 795 | Y795M | B | 1.2e-05 | 6.8e-06 | 9e-06 | 9.267E-06 | 2.610E-06 | 6.800E-06 | 1.200E-05 |
| 15894 | 795 | Y795F | B | 8.9e-06 | 8.8e-06 | 9.5e-06 | 9.067E-06 | 3.786E-07 | 8.800E-06 | 9.500E-06 |

| | | | | | | | | | | |
|---|---|---|---|---|---|---|---|---|---|---|
| 15895 | 795 | Y795P | B | 9.6e-06 | 1.5e-05 | 1.1e-05 | 1.187E-05 | 2.802E-06 | 9.600E-06 | 1.500E-05 |
| 15896 | 795 | Y795S | B | 7.9e-06 | 8.2e-06 | 6.6e-06 | 7.567E-06 | 8.505E-07 | 6.600E-06 | 8.200E-06 |
| 15897 | 795 | Y795T | B | 7.2e-06 | 7.1e-06 | 7e-06 | 7.100E-06 | 1.000E-07 | 7.000E-06 | 7.200E-06 |
| 15898 | 795 | Y795W | B | 1.2e-05 | 6.2e-06 | 9.6e-06 | 9.267E-06 | 2.914E-06 | 6.200E-06 | 1.200E-05 |
| 15899 | 795 | Y795Y | B | 1.3e-05 | 7.2e-06 | 8.5e-06 | 9.567E-06 | 3.044E-06 | 7.200E-06 | 1.300E-05 |
| 15900 | 795 | Y795V | B | 9.8e-06 | 8.3e-06 | 9.8e-06 | 9.300E-06 | 8.660E-07 | 8.300E-06 | 9.800E-06 |
| 15901 | 796 | S796A | B | 4.8e-06 | 8.9e-06 | 9.3e-06 | 7.667E-06 | 2.491E-06 | 4.800E-06 | 9.300E-06 |
| 15902 | 796 | S796R | B | 6.6e-06 | 9.1e-06 | 9e-06 | 8.233E-06 | 1.415E-06 | 6.600E-06 | 9.100E-06 |
| 15903 | 796 | S796N | B | 8.9e-06 | 3.3e-06 | 2.9e-06 | 5.033E-06 | 3.355E-06 | 2.900E-06 | 8.900E-06 |
| 15904 | 796 | S796D | B | 7.6e-06 | 6.7e-06 | 1.9e-05 | 1.110E-05 | 6.856E-06 | 6.700E-06 | 1.900E-05 |
| 15905 | 796 | S796C | B | 8.8e-06 | 1e-05 | 5.4e-06 | 8.067E-06 | 2.386E-06 | 5.400E-06 | 1.000E-05 |
| 15906 | 796 | S796E | B | 7.1e-06 | 9.2e-06 | 5.5e-06 | 7.267E-06 | 1.856E-06 | 5.500E-06 | 9.200E-06 |
| 15907 | 796 | S796Q | B | 6.5e-06 | 8.7e-06 | 1e-05 | 8.400E-06 | 1.769E-06 | 6.500E-06 | 1.000E-05 |
| 15908 | 796 | S796G | B | 9e-06 | 1.2e-05 | 1e-05 | 1.033E-05 | 1.528E-06 | 9.000E-06 | 1.200E-05 |
| 15909 | 796 | S796H | B | 7.4e-06 | 5.2e-06 | 1e-05 | 7.533E-06 | 2.403E-06 | 5.200E-06 | 1.000E-05 |
| 15910 | 796 | S796I | B | 5.6e-06 | 7.9e-06 | 1.1e-05 | 8.167E-06 | 2.710E-06 | 5.600E-06 | 1.100E-05 |
| 15911 | 796 | S796L | B | 9.6e-06 | 1.2e-05 | 8e-06 | 9.867E-06 | 2.013E-06 | 8.000E-06 | 1.200E-05 |
| 15912 | 796 | S796K | B | 3.1e-06 | 6.4e-06 | 7.7e-06 | 5.733E-06 | 2.371E-06 | 3.100E-06 | 7.700E-06 |
| 15913 | 796 | S796M | B | 7.2e-06 | 1.1e-05 | 9.5e-06 | 9.233E-06 | 1.914E-06 | 7.200E-06 | 1.100E-05 |
| 15914 | 796 | S796F | B | 6.2e-06 | 8.5e-06 | 4.5e-06 | 6.400E-06 | 2.007E-06 | 4.500E-06 | 8.500E-06 |
| 15915 | 796 | S796P | B | 4.3e-06 | 1e-05 | 9.1e-06 | 7.800E-06 | 3.064E-06 | 4.300E-06 | 1.000E-05 |
| 15916 | 796 | S796S | B | 8.1e-06 | 7.2e-06 | 5.1e-06 | 6.800E-06 | 1.539E-06 | 5.100E-06 | 8.100E-06 |
| 15917 | 796 | S796T | B | 9.6e-06 | 6.5e-06 | 5.6e-06 | 7.233E-06 | 2.098E-06 | 5.600E-06 | 9.600E-06 |
| 15918 | 796 | S796W | B | 5.9e-06 | 1.4e-05 | 6.9e-06 | 8.933E-06 | 4.416E-06 | 5.900E-06 | 1.400E-05 |
| 15919 | 796 | S796Y | B | 1.2e-05 | 9.7e-06 | 4.6e-06 | 8.767E-06 | 3.787E-06 | 4.600E-06 | 1.200E-05 |
| 15920 | 796 | S796V | B | 7.6e-06 | 1.1e-05 | 8.9e-06 | 9.167E-06 | 1.716E-06 | 7.600E-06 | 1.100E-05 |
| 15921 | 797 | L797A | B | 7.3e-06 | 5.2e-06 | 5.4e-06 | 5.967E-06 | 1.159E-06 | 5.200E-06 | 7.300E-06 |
| 15922 | 797 | L797R | B | 4.7e-06 | 6.6e-06 | 8.9e-06 | 6.733E-06 | 2.103E-06 | 4.700E-06 | 8.900E-06 |
| 15923 | 797 | L797N | B | 1.6e-05 | 8.2e-06 | 9e-06 | 1.107E-05 | 4.291E-06 | 8.200E-06 | 1.600E-05 |
| 15924 | 797 | L797D | B | 6.9e-06 | 6.6e-06 | 1.4e-05 | 9.167E-06 | 4.188E-06 | 6.600E-06 | 1.400E-05 |
| 15925 | 797 | L797C | B | 7.8e-06 | 6.4e-06 | 1.2e-05 | 8.733E-06 | 2.914E-06 | 6.400E-06 | 1.200E-05 |
| 15926 | 797 | L797E | B | 4e-06 | 6.6e-06 | 5.1e-06 | 5.233E-06 | 1.305E-06 | 4.000E-06 | 6.600E-06 |
| 15927 | 797 | L797Q | B | 5.2e-06 | 7.5e-06 | 5.3e-06 | 6.000E-06 | 1.300E-06 | 5.200E-06 | 7.500E-06 |
| 15928 | 797 | L797G | B | 6.5e-06 | 6.2e-06 | 8.9e-06 | 7.200E-06 | 1.480E-06 | 6.200E-06 | 8.900E-06 |
| 15929 | 797 | L797H | B | 4.1e-06 | 5.7e-06 | 8.7e-06 | 6.167E-06 | 2.335E-06 | 4.100E-06 | 8.700E-06 |
| 15930 | 797 | L797I | B | 1.3e-05 | 7.5e-06 | 6.9e-06 | 9.133E-06 | 3.362E-06 | 6.900E-06 | 1.300E-05 |
| 15931 | 797 | L797L | B | 5.9e-06 | 8.9e-06 | 1.1e-05 | 8.600E-06 | 2.563E-06 | 5.900E-06 | 1.100E-05 |
| 15932 | 797 | L797K | B | 1e-05 | 1.3e-05 | 3.5e-06 | 8.833E-06 | 4.856E-06 | 3.500E-06 | 1.300E-05 |
| 15933 | 797 | L797M | B | 8.5e-06 | 4.4e-06 | 9.8e-06 | 7.567E-06 | 2.818E-06 | 4.400E-06 | 9.800E-06 |
| 15934 | 797 | L797F | B | 6.2e-06 | 4.3e-06 | 5.3e-06 | 5.267E-06 | 9.504E-07 | 4.300E-06 | 6.200E-06 |
| 15935 | 797 | L797P | B | 6.1e-06 | 8.4e-06 | 7.8e-06 | 7.433E-06 | 1.193E-06 | 6.100E-06 | 8.400E-06 |
| 15936 | 797 | L797S | B | 9.1e-06 | 1.2e-05 | 7.1e-06 | 9.400E-06 | 2.464E-06 | 7.100E-06 | 1.200E-05 |
| 15937 | 797 | L797T | B | 7.3e-06 | 7.5e-06 | 5.9e-06 | 6.900E-06 | 8.718E-07 | 5.900E-06 | 7.500E-06 |
| 15938 | 797 | L797W | B | 5.5e-06 | 4e-06 | 9e-06 | 6.167E-06 | 2.566E-06 | 4.000E-06 | 9.000E-06 |
| 15939 | 797 | L797Y | B | 6.4e-06 | 5.5e-06 | 6.7e-06 | 6.200E-06 | 6.245E-07 | 5.500E-06 | 6.700E-06 |
| 15940 | 797 | L797V | B | 7e-06 | 9.1e-06 | 7e-06 | 7.700E-06 | 1.212E-06 | 7.000E-06 | 9.100E-06 |
| 15941 | 798 | S798A | B | 1e-05 | 5.9e-06 | 1.1e-05 | 8.967E-06 | 2.702E-06 | 5.900E-06 | 1.100E-05 |
| 15942 | 798 | S798R | B | 9.3e-06 | 7.2e-06 | 4.7e-06 | 7.067E-06 | 2.303E-06 | 4.700E-06 | 9.300E-06 |
| 15943 | 798 | S798N | B | 6.3e-06 | 8.2e-06 | 9.8e-06 | 8.100E-06 | 1.752E-06 | 6.300E-06 | 9.800E-06 |
| 15944 | 798 | S798D | B | 7.6e-06 | 9.1e-06 | 5e-06 | 7.233E-06 | 2.074E-06 | 5.000E-06 | 9.100E-06 |
| 15945 | 798 | S798C | B | 7.4e-06 | 1.1e-05 | 8.7e-06 | 9.033E-06 | 1.823E-06 | 7.400E-06 | 1.100E-05 |
| 15946 | 798 | S798E | B | 9.9e-06 | 6.6e-06 | 6.4e-06 | 7.633E-06 | 1.966E-06 | 6.400E-06 | 9.900E-06 |
| 15947 | 798 | S798Q | B | 1.4e-05 | 6.7e-06 | 4.3e-06 | 8.333E-06 | 5.052E-06 | 4.300E-06 | 1.400E-05 |
| 15948 | 798 | S798G | B | 6.6e-06 | 7.2e-06 | 5.7e-06 | 6.500E-06 | 7.550E-07 | 5.700E-06 | 7.200E-06 |
| 15949 | 798 | S798H | B | 3.9e-06 | 9.6e-06 | 1.2e-05 | 8.500E-06 | 4.161E-06 | 3.900E-06 | 1.200E-05 |
| 15950 | 798 | S798I | B | 7.7e-06 | 1.1e-05 | 5.2e-06 | 7.967E-06 | 2.909E-06 | 5.200E-06 | 1.100E-05 |
| 15951 | 798 | S798L | B | 7e-06 | 5.3e-06 | 8.2e-06 | 6.833E-06 | 1.457E-06 | 5.300E-06 | 8.200E-06 |
| 15952 | 798 | S798K | B | 8.2e-06 | 6.3e-06 | 4.4e-06 | 6.300E-06 | 1.900E-06 | 4.400E-06 | 8.200E-06 |
| 15953 | 798 | S798M | B | 6.7e-06 | 1.1e-05 | 6.8e-06 | 8.167E-06 | 2.454E-06 | 6.700E-06 | 1.100E-05 |
| 15954 | 798 | S798F | B | 5e-06 | 1.1e-05 | 9.4e-06 | 8.467E-06 | 3.107E-06 | 5.000E-06 | 1.100E-05 |
| 15955 | 798 | S798P | B | 7.9e-06 | 6.3e-06 | 7.2e-06 | 7.133E-06 | 8.021E-07 | 6.300E-06 | 7.900E-06 |
| 15956 | 798 | S798S | B | 9.6e-06 | 7.4e-06 | 9.4e-06 | 8.800E-06 | 1.217E-06 | 7.400E-06 | 9.600E-06 |
| 15957 | 798 | S798T | B | 4.6e-06 | 9.4e-06 | 6.8e-06 | 6.933E-06 | 2.403E-06 | 4.600E-06 | 9.400E-06 |
| 15958 | 798 | S798W | B | 1.5e-05 | 6.7e-06 | 6.6e-06 | 9.433E-06 | 4.821E-06 | 6.600E-06 | 1.500E-05 |
| 15959 | 798 | S798Y | B | 8.5e-06 | 1e-05 | 1.1e-05 | 9.833E-06 | 1.258E-06 | 8.500E-06 | 1.100E-05 |
| 15960 | 798 | S798V | B | 6.6e-06 | 8.3e-06 | 9.9e-06 | 8.267E-06 | 1.650E-06 | 6.600E-06 | 9.900E-06 |
| 15961 | 799 | S799A | B | 1e-05 | 9.4e-06 | 5.4e-06 | 8.267E-06 | 2.501E-06 | 5.400E-06 | 1.000E-05 |
| 15962 | 799 | S799R | B | 8.2e-06 | 9.4e-06 | 1.1e-05 | 9.533E-06 | 1.405E-06 | 8.200E-06 | 1.100E-05 |
| 15963 | 799 | S799N | B | 9.6e-06 | 7.9e-06 | 1.4e-05 | 1.050E-05 | 3.148E-06 | 7.900E-06 | 1.400E-05 |
| 15964 | 799 | S799D | B | 8.8e-06 | 3.7e-06 | 8.9e-06 | 7.133E-06 | 2.974E-06 | 3.700E-06 | 8.900E-06 |
| 15965 | 799 | S799C | B | 4.6e-06 | 8e-06 | 1.4e-05 | 8.867E-06 | 4.760E-06 | 4.600E-06 | 1.400E-05 |
| 15966 | 799 | S799E | B | 5.6e-06 | 9.9e-06 | 1e-05 | 8.500E-06 | 2.512E-06 | 5.600E-06 | 1.000E-05 |
| 15967 | 799 | S799Q | B | 5.3e-06 | 5.2e-06 | 5.3e-06 | 5.267E-06 | 5.774E-08 | 5.200E-06 | 5.300E-06 |
| 15968 | 799 | S799G | B | 5.9e-06 | 9.3e-06 | 8.3e-06 | 7.833E-06 | 1.747E-06 | 5.900E-06 | 9.300E-06 |
| 15969 | 799 | S799H | B | 5.4e-06 | 9.5e-06 | 5.8e-06 | 6.900E-06 | 2.261E-06 | 5.400E-06 | 9.500E-06 |
| 15970 | 799 | S799I | B | 1.2e-05 | 9.6e-06 | 9.1e-06 | 1.023E-05 | 1.550E-06 | 9.100E-06 | 1.200E-05 |
| 15971 | 799 | S799L | B | 6.9e-06 | 7.4e-06 | 7.3e-06 | 7.200E-06 | 2.646E-07 | 6.900E-06 | 7.400E-06 |
| 15972 | 799 | S799K | B | 1.9e-05 | 7.1e-06 | 7.9e-06 | 1.133E-05 | 6.652E-06 | 7.100E-06 | 1.900E-05 |
| 15973 | 799 | S799M | B | 7.3e-06 | 9.3e-06 | 1.3e-05 | 9.867E-06 | 2.892E-06 | 7.300E-06 | 1.300E-05 |
| 15974 | 799 | S799F | B | 9.3e-06 | 7.3e-06 | 5.8e-06 | 7.467E-06 | 1.756E-06 | 5.800E-06 | 9.300E-06 |
| 15975 | 799 | S799P | B | 9.2e-06 | 1.2e-05 | 6.8e-06 | 9.333E-06 | 2.603E-06 | 6.800E-06 | 1.200E-05 |
| 15976 | 799 | S799S | B | 1.1e-05 | 1.3e-05 | 5.8e-06 | 9.933E-06 | 3.717E-06 | 5.800E-06 | 1.300E-05 |
| 15977 | 799 | S799T | B | 7.9e-06 | 1.2e-05 | 6.9e-06 | 8.933E-06 | 2.702E-06 | 6.900E-06 | 1.200E-05 |
| 15978 | 799 | S799W | B | 9e-06 | 7.5e-06 | 9.5e-06 | 8.667E-06 | 1.041E-06 | 7.500E-06 | 9.500E-06 |
| 15979 | 799 | S799Y | B | 1.4e-05 | 9.7e-06 | 9.2e-06 | 1.097E-05 | 2.639E-06 | 9.200E-06 | 1.400E-05 |

| | | | | | | | | | | |
|---|---|---|---|---|---|---|---|---|---|---|
| 15980 | 799 | S799V | B | 1.7e-05 | 7e-06 | 4.7e-06 | 9.567E-06 | 6.539E-06 | 4.700E-06 | 1.700E-05 |
| 15981 | 800 | T800A | B | 8e-06 | 6.7e-06 | 7.3e-06 | 7.333E-06 | 6.506E-07 | 6.700E-06 | 8.000E-06 |
| 15982 | 800 | T800R | B | 9.5e-06 | 6.8e-06 | 7.6e-06 | 7.967E-06 | 1.387E-06 | 6.800E-06 | 9.500E-06 |
| 15983 | 800 | T800N | B | 5.5e-06 | 8.6e-06 | 1e-05 | 8.033E-06 | 2.303E-06 | 5.500E-06 | 1.000E-05 |
| 15984 | 800 | T800D | B | 9.4e-06 | 7.1e-06 | 1.3e-05 | 9.833E-06 | 2.974E-06 | 7.100E-06 | 1.300E-05 |
| 15985 | 800 | T800C | B | 1.3e-05 | 8.4e-06 | 6.7e-06 | 9.367E-06 | 3.259E-06 | 6.700E-06 | 1.300E-05 |
| 15986 | 800 | T800E | B | 4.7e-06 | 9.6e-06 | 1e-05 | 8.100E-06 | 2.951E-06 | 4.700E-06 | 1.000E-05 |
| 15987 | 800 | T800Q | B | 8e-06 | 7.9e-06 | 4.5e-06 | 6.800E-06 | 1.992E-06 | 4.500E-06 | 8.000E-06 |
| 15988 | 800 | T800G | B | 1.3e-05 | 8.5e-06 | 1.3e-05 | 1.150E-05 | 2.598E-06 | 8.500E-06 | 1.300E-05 |
| 15989 | 800 | T800H | B | 1.2e-05 | 1e-05 | 1.3e-05 | 1.167E-05 | 1.528E-06 | 1.000E-05 | 1.300E-05 |
| 15990 | 800 | T800I | B | 5.9e-06 | 6.3e-06 | 1.4e-05 | 8.733E-06 | 4.565E-06 | 5.900E-06 | 1.400E-05 |
| 15991 | 800 | T800L | B | 8.3e-06 | 4.6e-06 | 1.3e-05 | 8.633E-06 | 4.210E-06 | 4.600E-06 | 1.300E-05 |
| 15992 | 800 | T800K | B | 1.1e-05 | 5.7e-06 | 1.3e-05 | 9.900E-06 | 3.772E-06 | 5.700E-06 | 1.300E-05 |
| 15993 | 800 | T800M | B | 6e-06 | 9.8e-06 | 7.8e-06 | 7.867E-06 | 1.901E-06 | 6.000E-06 | 9.800E-06 |
| 15994 | 800 | T800F | B | 6.9e-06 | 1.1e-05 | 1e-05 | 9.300E-06 | 2.138E-06 | 6.900E-06 | 1.100E-05 |
| 15995 | 800 | T800P | B | 1.4e-05 | 5.7e-06 | 9.4e-06 | 9.700E-06 | 4.158E-06 | 5.700E-06 | 1.400E-05 |
| 15996 | 800 | T800S | B | 9.1e-06 | 9.8e-06 | 7.1e-06 | 8.667E-06 | 1.401E-06 | 7.100E-06 | 9.800E-06 |
| 15997 | 800 | T800T | B | 8.4e-06 | 9.2e-06 | 4.6e-06 | 7.400E-06 | 2.458E-06 | 4.600E-06 | 9.200E-06 |
| 15998 | 800 | T800W | B | 7.3e-06 | 7.8e-06 | 7.7e-06 | 7.600E-06 | 2.646E-07 | 7.300E-06 | 7.800E-06 |
| 15999 | 800 | T800Y | B | 7.2e-06 | 1.1e-05 | 8.8e-06 | 9.000E-06 | 1.908E-06 | 7.200E-06 | 1.100E-05 |
| 16000 | 800 | T800V | B | 9.9e-06 | 1.1e-05 | 6e-06 | 8.967E-06 | 2.627E-06 | 6.000E-06 | 1.100E-05 |
| 16001 | 801 | L801A | B | 6.1e-06 | 8.3e-06 | 8.7e-06 | 7.700E-06 | 1.400E-06 | 6.100E-06 | 8.700E-06 |
| 16002 | 801 | L801R | B | 4.7e-06 | 4.4e-06 | 5.7e-06 | 4.933E-06 | 6.807E-07 | 4.400E-06 | 5.700E-06 |
| 16003 | 801 | L801N | B | 6.9e-06 | 8.4e-06 | 9.4e-06 | 8.233E-06 | 1.258E-06 | 6.900E-06 | 9.400E-06 |
| 16004 | 801 | L801D | B | 4.4e-06 | 5.8e-06 | 8.7e-06 | 6.300E-06 | 2.193E-06 | 4.400E-06 | 8.700E-06 |
| 16005 | 801 | L801C | B | 6.9e-06 | 6e-06 | 5.7e-06 | 6.200E-06 | 6.245E-07 | 5.700E-06 | 6.900E-06 |
| 16006 | 801 | L801E | B | 1.7e-05 | 9.8e-06 | 6.6e-06 | 1.113E-05 | 5.327E-06 | 6.600E-06 | 1.700E-05 |
| 16007 | 801 | L801Q | B | 7.6e-06 | 4.9e-06 | 7.5e-06 | 6.667E-06 | 1.531E-06 | 4.900E-06 | 7.600E-06 |
| 16008 | 801 | L801G | B | 6.9e-06 | 6.1e-06 | 6.7e-06 | 6.567E-06 | 4.163E-07 | 6.100E-06 | 6.900E-06 |
| 16009 | 801 | L801H | B | 1e-05 | 7.3e-06 | 8.6e-06 | 8.633E-06 | 1.350E-06 | 7.300E-06 | 1.000E-05 |
| 16010 | 801 | L801I | B | 1.1e-05 | 1.1e-05 | 8.5e-06 | 1.017E-05 | 1.443E-06 | 8.500E-06 | 1.100E-05 |
| 16011 | 801 | L801L | B | 1.7e-05 | 9.6e-06 | 3.6e-06 | 1.007E-05 | 6.712E-06 | 3.600E-06 | 1.700E-05 |
| 16012 | 801 | L801K | B | 9.6e-06 | 8.8e-06 | 1.1e-05 | 9.800E-06 | 1.114E-06 | 8.800E-06 | 1.100E-05 |
| 16013 | 801 | L801M | B | 1e-05 | 5.8e-06 | 8.8e-06 | 8.200E-06 | 2.163E-06 | 5.800E-06 | 1.000E-05 |
| 16014 | 801 | L801F | B | 9.2e-06 | 7e-06 | 6.1e-06 | 7.433E-06 | 1.595E-06 | 6.100E-06 | 9.200E-06 |
| 16015 | 801 | L801P | B | 1.1e-05 | 5.5e-06 | 9.6e-06 | 8.700E-06 | 2.858E-06 | 5.500E-06 | 1.100E-05 |
| 16016 | 801 | L801S | B | 9.8e-06 | 1e-05 | 9.3e-06 | 9.700E-06 | 3.606E-07 | 9.300E-06 | 1.000E-05 |
| 16017 | 801 | L801T | B | 1e-05 | 7.4e-06 | 9.2e-06 | 8.867E-06 | 1.332E-06 | 7.400E-06 | 1.000E-05 |
| 16018 | 801 | L801W | B | 7.8e-06 | 1.4e-05 | 1.9e-05 | 1.360E-05 | 5.611E-06 | 7.800E-06 | 1.900E-05 |
| 16019 | 801 | L801Y | B | 7.1e-06 | 4.6e-06 | 1.1e-05 | 7.567E-06 | 3.225E-06 | 4.600E-06 | 1.100E-05 |
| 16020 | 801 | L801V | B | 9.1e-06 | 7.5e-06 | 8.1e-06 | 8.233E-06 | 8.083E-07 | 7.500E-06 | 9.100E-06 |
| 16021 | 802 | T802A | B | 1.2e-05 | 9.4e-06 | 1.1e-05 | 1.080E-05 | 1.311E-06 | 9.400E-06 | 1.200E-05 |
| 16022 | 802 | T802R | B | 6.6e-06 | 1e-05 | 1.2e-05 | 9.533E-06 | 2.730E-06 | 6.600E-06 | 1.200E-05 |
| 16023 | 802 | T802N | B | 6.4e-06 | 9.5e-06 | 4.1e-06 | 6.667E-06 | 2.710E-06 | 4.100E-06 | 9.500E-06 |
| 16024 | 802 | T802D | B | 8.5e-06 | 7.1e-06 | 6.7e-06 | 7.433E-06 | 9.452E-07 | 6.700E-06 | 8.500E-06 |
| 16025 | 802 | T802C | B | 8.3e-06 | 8.9e-06 | 6.9e-06 | 8.033E-06 | 1.026E-06 | 6.900E-06 | 8.900E-06 |
| 16026 | 802 | T802E | B | 4.3e-06 | 1.3e-05 | 7.1e-06 | 8.133E-06 | 4.441E-06 | 4.300E-06 | 1.300E-05 |
| 16027 | 802 | T802Q | B | 1e-05 | 6.6e-06 | 9.3e-06 | 8.633E-06 | 1.795E-06 | 6.600E-06 | 1.000E-05 |
| 16028 | 802 | T802G | B | 7.6e-06 | 9e-06 | 7.7e-06 | 8.100E-06 | 7.810E-07 | 7.600E-06 | 9.000E-06 |
| 16029 | 802 | T802H | B | 9.8e-06 | 9e-06 | 6.7e-06 | 8.500E-06 | 1.609E-06 | 6.700E-06 | 9.800E-06 |
| 16030 | 802 | T802I | B | 6.8e-06 | 9.3e-06 | 9.6e-06 | 8.567E-06 | 1.537E-06 | 6.800E-06 | 9.600E-06 |
| 16031 | 802 | T802L | B | 1e-05 | 1e-05 | 1e-05 | 1.000E-05 | 0.000E+00 | 1.000E-05 | 1.000E-05 |
| 16032 | 802 | T802K | B | 5.9e-06 | 5.6e-06 | 7.6e-06 | 6.367E-06 | 1.079E-06 | 5.600E-06 | 7.600E-06 |
| 16033 | 802 | T802M | B | 7.1e-06 | 1.5e-05 | 9.8e-06 | 1.063E-05 | 4.015E-06 | 7.100E-06 | 1.500E-05 |
| 16034 | 802 | T802F | B | 9.8e-06 | 7.2e-06 | 7.8e-06 | 8.267E-06 | 1.361E-06 | 7.200E-06 | 9.800E-06 |
| 16035 | 802 | T802P | B | 9.9e-06 | 6.9e-06 | 1.4e-05 | 1.027E-05 | 3.564E-06 | 6.900E-06 | 1.400E-05 |
| 16036 | 802 | T802S | B | 9e-06 | 6.6e-06 | 6.2e-06 | 7.267E-06 | 1.514E-06 | 6.200E-06 | 9.000E-06 |
| 16037 | 802 | T802T | B | 7.4e-06 | 1.2e-05 | 5e-06 | 8.133E-06 | 3.557E-06 | 5.000E-06 | 1.200E-05 |
| 16038 | 802 | T802W | B | 1.3e-05 | 9.2e-06 | 6.5e-06 | 9.567E-06 | 3.265E-06 | 6.500E-06 | 1.300E-05 |
| 16039 | 802 | T802Y | B | 5.4e-06 | 6.5e-06 | 1e-05 | 7.300E-06 | 2.402E-06 | 5.400E-06 | 1.000E-05 |
| 16040 | 802 | T802V | B | 6.2e-06 | 8.9e-06 | 8.5e-06 | 7.867E-06 | 1.457E-06 | 6.200E-06 | 8.900E-06 |
| 16041 | 803 | L803A | B | 9.8e-06 | 1e-05 | 3.8e-06 | 7.867E-06 | 3.523E-06 | 3.800E-06 | 1.000E-05 |
| 16042 | 803 | L803R | B | 1e-05 | 5.7e-06 | 9.6e-06 | 8.433E-06 | 2.376E-06 | 5.700E-06 | 1.000E-05 |
| 16043 | 803 | L803N | B | 6.5e-06 | 8.4e-06 | 5.3e-06 | 6.733E-06 | 1.563E-06 | 5.300E-06 | 8.400E-06 |
| 16044 | 803 | L803D | B | 7e-06 | 8.6e-06 | 6e-06 | 7.200E-06 | 1.311E-06 | 6.000E-06 | 8.600E-06 |
| 16045 | 803 | L803C | B | 9.6e-06 | 3.9e-06 | 1.2e-05 | 8.500E-06 | 4.161E-06 | 3.900E-06 | 1.200E-05 |
| 16046 | 803 | L803E | B | 1e-05 | 1.1e-05 | 1.4e-05 | 1.167E-05 | 2.082E-06 | 1.000E-05 | 1.400E-05 |
| 16047 | 803 | L803Q | B | 8.8e-06 | 6.5e-06 | 8.1e-06 | 7.800E-06 | 1.179E-06 | 6.500E-06 | 8.800E-06 |
| 16048 | 803 | L803G | B | 5.2e-06 | 1.3e-05 | 1.4e-05 | 1.073E-05 | 4.818E-06 | 5.200E-06 | 1.400E-05 |
| 16049 | 803 | L803H | B | 7.1e-06 | 7.2e-06 | 9.4e-06 | 7.900E-06 | 1.300E-06 | 7.100E-06 | 9.400E-06 |
| 16050 | 803 | L803I | B | 7.2e-06 | 1.2e-05 | 6.2e-06 | 8.467E-06 | 3.101E-06 | 6.200E-06 | 1.200E-05 |
| 16051 | 803 | L803L | B | 5.7e-06 | 7e-06 | 9.9e-06 | 7.533E-06 | 2.150E-06 | 5.700E-06 | 9.900E-06 |
| 16052 | 803 | L803K | B | 6.7e-06 | 1.2e-05 | 6.6e-06 | 8.433E-06 | 3.089E-06 | 6.600E-06 | 1.200E-05 |
| 16053 | 803 | L803M | B | 5.5e-06 | 1e-05 | 7.3e-06 | 7.600E-06 | 2.265E-06 | 5.500E-06 | 1.000E-05 |
| 16054 | 803 | L803F | B | 4.5e-06 | 6.1e-06 | 6.9e-06 | 5.833E-06 | 1.222E-06 | 4.500E-06 | 6.900E-06 |
| 16055 | 803 | L803P | B | 1.3e-05 | 8e-06 | 9.9e-06 | 1.030E-05 | 2.524E-06 | 8.000E-06 | 1.300E-05 |
| 16056 | 803 | L803S | B | 7e-06 | 7.1e-06 | 9.6e-06 | 7.900E-06 | 1.473E-06 | 7.000E-06 | 9.600E-06 |
| 16057 | 803 | L803T | B | 8e-06 | 1.2e-05 | 8.6e-06 | 9.533E-06 | 2.157E-06 | 8.000E-06 | 1.200E-05 |
| 16058 | 803 | L803W | B | 9.5e-06 | 6.8e-06 | 4.9e-06 | 7.067E-06 | 2.312E-06 | 4.900E-06 | 9.500E-06 |
| 16059 | 803 | L803Y | B | 5e-06 | 5.6e-06 | 5.7e-06 | 5.433E-06 | 3.786E-07 | 5.000E-06 | 5.700E-06 |
| 16060 | 803 | L803V | B | 1e-05 | 6e-06 | 7.6e-06 | 7.867E-06 | 2.013E-06 | 6.000E-06 | 1.000E-05 |
| 16061 | 804 | S804A | B | 6.8e-06 | 7.6e-06 | 1e-05 | 8.133E-06 | 1.665E-06 | 6.800E-06 | 1.000E-05 |
| 16062 | 804 | S804R | B | 4.9e-06 | 6.5e-06 | 1.3e-05 | 8.133E-06 | 4.290E-06 | 4.900E-06 | 1.300E-05 |
| 16063 | 804 | S804N | B | 8.5e-06 | 8.4e-06 | 8.1e-06 | 8.333E-06 | 2.082E-07 | 8.100E-06 | 8.500E-06 |
| 16064 | 804 | S804D | B | 9.3e-06 | 1.5e-05 | 1.1e-05 | 1.177E-05 | 2.926E-06 | 9.300E-06 | 1.500E-05 |

| | | | | | | | | | | |
|---|---|---|---|---|---|---|---|---|---|---|
| 16065 | 804 | S804C | B | 1e-05 | 1e-05 | 9.3e-06 | 9.767E-06 | 4.041E-07 | 9.300E-06 | 1.000E-05 |
| 16066 | 804 | S804E | B | 4.9e-06 | 9.8e-06 | 4e-06 | 6.233E-06 | 3.121E-06 | 4.000E-06 | 9.800E-06 |
| 16067 | 804 | S804Q | B | 1.8e-05 | 4.5e-06 | 6.8e-06 | 9.767E-06 | 7.222E-06 | 4.500E-06 | 1.800E-05 |
| 16068 | 804 | S804G | B | 7.5e-06 | 7.4e-06 | 4.8e-06 | 6.567E-06 | 1.531E-06 | 4.800E-06 | 7.500E-06 |
| 16069 | 804 | S804H | B | 6.6e-06 | 1.1e-05 | 6.5e-06 | 8.033E-06 | 2.570E-06 | 6.500E-06 | 1.100E-05 |
| 16070 | 804 | S804I | B | 1e-05 | 4.8e-06 | 1e-05 | 8.267E-06 | 3.002E-06 | 4.800E-06 | 1.000E-05 |
| 16071 | 804 | S804L | B | 6.5e-06 | 1.4e-05 | 7.7e-06 | 9.400E-06 | 4.029E-06 | 6.500E-06 | 1.400E-05 |
| 16072 | 804 | S804K | B | 8.2e-06 | 5.8e-06 | 1.2e-05 | 8.667E-06 | 3.126E-06 | 5.800E-06 | 1.200E-05 |
| 16073 | 804 | S804M | B | 8e-06 | 8.9e-06 | 7.7e-06 | 8.200E-06 | 6.245E-07 | 7.700E-06 | 8.900E-06 |
| 16074 | 804 | S804F | B | 7e-06 | 7.7e-06 | 9.7e-06 | 8.133E-06 | 1.401E-06 | 7.000E-06 | 9.700E-06 |
| 16075 | 804 | S804P | B | 8.2e-06 | 8.1e-06 | 5.1e-06 | 7.133E-06 | 1.762E-06 | 5.100E-06 | 8.200E-06 |
| 16076 | 804 | S804S | B | 9.4e-06 | 5.9e-06 | 9.4e-06 | 8.233E-06 | 2.021E-06 | 5.900E-06 | 9.400E-06 |
| 16077 | 804 | S804T | B | 7e-06 | 1.9e-05 | 1.2e-05 | 1.267E-05 | 6.028E-06 | 7.000E-06 | 1.900E-05 |
| 16078 | 804 | S804W | B | 9.3e-06 | 6.2e-06 | 8.3e-06 | 7.933E-06 | 1.582E-06 | 6.200E-06 | 9.300E-06 |
| 16079 | 804 | S804Y | B | 6.2e-06 | 8.4e-06 | 6.8e-06 | 7.133E-06 | 1.137E-06 | 6.200E-06 | 8.400E-06 |
| 16080 | 804 | S804V | B | 9.6e-06 | 7.5e-06 | 6.2e-06 | 7.767E-06 | 1.716E-06 | 6.200E-06 | 9.600E-06 |
| 16081 | 805 | K805A | B | 6.5e-06 | 6.3e-06 | 1e-05 | 7.600E-06 | 2.081E-06 | 6.300E-06 | 1.000E-05 |
| 16082 | 805 | K805R | B | 9.8e-06 | 7.9e-06 | 1.1e-05 | 9.567E-06 | 1.563E-06 | 7.900E-06 | 1.100E-05 |
| 16083 | 805 | K805N | B | 8.6e-06 | 6.4e-06 | 7e-06 | 7.333E-06 | 1.137E-06 | 6.400E-06 | 8.600E-06 |
| 16084 | 805 | K805D | B | 4.7e-06 | 1.2e-05 | 7.1e-06 | 7.933E-06 | 3.721E-06 | 4.700E-06 | 1.200E-05 |
| 16085 | 805 | K805C | B | 9.3e-06 | 1.3e-05 | 7.3e-06 | 9.867E-06 | 2.892E-06 | 7.300E-06 | 1.300E-05 |
| 16086 | 805 | K805E | B | 8.4e-06 | 1.1e-05 | 1.3e-05 | 1.080E-05 | 2.307E-06 | 8.400E-06 | 1.300E-05 |
| 16087 | 805 | K805Q | B | 7.8e-06 | 6.5e-06 | 7.6e-06 | 7.300E-06 | 7.000E-07 | 6.500E-06 | 7.800E-06 |
| 16088 | 805 | K805G | B | 7e-06 | 8.5e-06 | 6.9e-06 | 7.467E-06 | 8.963E-07 | 6.900E-06 | 8.500E-06 |
| 16089 | 805 | K805H | B | 1.3e-05 | 1.3e-05 | 6.5e-06 | 1.083E-05 | 3.753E-06 | 6.500E-06 | 1.300E-05 |
| 16090 | 805 | K805I | B | 6.7e-06 | 6.8e-06 | 1.4e-05 | 9.167E-06 | 4.186E-06 | 6.700E-06 | 1.400E-05 |
| 16091 | 805 | K805L | B | 4.7e-06 | 5.5e-06 | 5.9e-06 | 5.367E-06 | 6.110E-07 | 4.700E-06 | 5.900E-06 |
| 16092 | 805 | K805K | B | 3.7e-06 | 9.7e-06 | 1.2e-05 | 8.467E-06 | 4.285E-06 | 3.700E-06 | 1.200E-05 |
| 16093 | 805 | K805M | B | 7e-06 | 5.9e-06 | 1.1e-05 | 7.967E-06 | 2.684E-06 | 5.900E-06 | 1.100E-05 |
| 16094 | 805 | K805F | B | 9.6e-06 | 4.7e-06 | 7.2e-06 | 7.167E-06 | 2.450E-06 | 4.700E-06 | 9.600E-06 |
| 16095 | 805 | K805P | B | 8.9e-06 | 1.1e-05 | 6.7e-06 | 8.867E-06 | 2.150E-06 | 6.700E-06 | 1.100E-05 |
| 16096 | 805 | K805S | B | 6e-06 | 9.5e-06 | 9.4e-06 | 8.300E-06 | 1.992E-06 | 6.000E-06 | 9.500E-06 |
| 16097 | 805 | K805T | B | 6e-06 | 8.7e-06 | 5.4e-06 | 6.700E-06 | 1.758E-06 | 5.400E-06 | 8.700E-06 |
| 16098 | 805 | K805W | B | 7.8e-06 | 1.3e-05 | 9.4e-06 | 1.007E-05 | 2.663E-06 | 7.800E-06 | 1.300E-05 |
| 16099 | 805 | K805Y | B | 8.3e-06 | 1.2e-05 | 8.1e-06 | 9.467E-06 | 2.196E-06 | 8.100E-06 | 1.200E-05 |
| 16100 | 805 | K805V | B | 4.3e-06 | 8.8e-06 | 4.5e-06 | 5.867E-06 | 2.542E-06 | 4.300E-06 | 8.800E-06 |
| 16101 | 806 | A806A | B | 8.5e-06 | 6.3e-06 | 8.7e-06 | 7.833E-06 | 1.332E-06 | 6.300E-06 | 8.700E-06 |
| 16102 | 806 | A806R | B | 4.6e-06 | 7.7e-06 | 7.2e-06 | 6.500E-06 | 1.664E-06 | 4.600E-06 | 7.700E-06 |
| 16103 | 806 | A806N | B | 9e-06 | 3.8e-06 | 4.5e-06 | 5.767E-06 | 2.822E-06 | 3.800E-06 | 9.000E-06 |
| 16104 | 806 | A806D | B | 6.5e-06 | 8.8e-06 | 6.7e-06 | 7.333E-06 | 1.274E-06 | 6.500E-06 | 8.800E-06 |
| 16105 | 806 | A806C | B | 1.9e-05 | 5.4e-06 | 7e-06 | 1.047E-05 | 7.433E-06 | 5.400E-06 | 1.900E-05 |
| 16106 | 806 | A806E | B | 7.9e-06 | 1.3e-05 | 8.5e-06 | 9.800E-06 | 2.787E-06 | 7.900E-06 | 1.300E-05 |
| 16107 | 806 | A806Q | B | 6.3e-06 | 5.3e-06 | 8.5e-06 | 6.700E-06 | 1.637E-06 | 5.300E-06 | 8.500E-06 |
| 16108 | 806 | A806G | B | 6.8e-06 | 5.7e-06 | 1.1e-05 | 7.833E-06 | 2.797E-06 | 5.700E-06 | 1.100E-05 |
| 16109 | 806 | A806H | B | 6.5e-06 | 1.1e-05 | 7.7e-06 | 8.400E-06 | 2.330E-06 | 6.500E-06 | 1.100E-05 |
| 16110 | 806 | A806I | B | 9.9e-06 | 1.1e-05 | 5.6e-06 | 8.833E-06 | 2.854E-06 | 5.600E-06 | 1.100E-05 |
| 16111 | 806 | A806L | B | 7.7e-06 | 8.2e-06 | 6.4e-06 | 7.433E-06 | 9.292E-07 | 6.400E-06 | 8.200E-06 |
| 16112 | 806 | A806K | B | 1.1e-05 | 1.4e-05 | 1e-05 | 1.167E-05 | 2.082E-06 | 1.000E-05 | 1.400E-05 |
| 16113 | 806 | A806M | B | 4.1e-06 | 9.9e-06 | 6.6e-06 | 6.867E-06 | 2.909E-06 | 4.100E-06 | 9.900E-06 |
| 16114 | 806 | A806F | B | 1e-05 | 6.4e-06 | 4.3e-06 | 6.900E-06 | 2.883E-06 | 4.300E-06 | 1.000E-05 |
| 16115 | 806 | A806P | B | 4.9e-06 | 8.8e-06 | 1e-05 | 7.900E-06 | 2.666E-06 | 4.900E-06 | 1.000E-05 |
| 16116 | 806 | A806S | B | 8.7e-06 | 1.2e-05 | 4.1e-06 | 8.267E-06 | 3.968E-06 | 4.100E-06 | 1.200E-05 |
| 16117 | 806 | A806T | B | 5.3e-06 | 7.3e-06 | 6.2e-06 | 6.267E-06 | 1.002E-06 | 5.300E-06 | 7.300E-06 |
| 16118 | 806 | A806W | B | 6.7e-06 | 6.4e-06 | 4.4e-06 | 5.833E-06 | 1.250E-06 | 4.400E-06 | 6.700E-06 |
| 16119 | 806 | A806Y | B | 6.5e-06 | 6e-06 | 8.5e-06 | 7.000E-06 | 1.323E-06 | 6.000E-06 | 8.500E-06 |
| 16120 | 806 | A806V | B | 9.5e-06 | 6.6e-06 | 5.4e-06 | 7.167E-06 | 2.108E-06 | 5.400E-06 | 9.500E-06 |
| 16121 | 807 | D807A | B | 5.2e-06 | 7.6e-06 | 6.3e-06 | 6.367E-06 | 1.201E-06 | 5.200E-06 | 7.600E-06 |
| 16122 | 807 | D807R | B | 1e-05 | 1e-05 | 6.1e-06 | 8.700E-06 | 2.252E-06 | 6.100E-06 | 1.000E-05 |
| 16123 | 807 | D807N | B | 5.7e-06 | 1e-05 | 9.9e-06 | 8.533E-06 | 2.454E-06 | 5.700E-06 | 1.000E-05 |
| 16124 | 807 | D807D | B | 1.4e-05 | 6.2e-06 | 9.6e-06 | 9.933E-06 | 3.911E-06 | 6.200E-06 | 1.400E-05 |
| 16125 | 807 | D807C | B | 1.2e-05 | 1.1e-05 | 6.4e-06 | 9.800E-06 | 2.987E-06 | 6.400E-06 | 1.200E-05 |
| 16126 | 807 | D807E | B | 5.2e-06 | 7e-06 | 5.8e-06 | 6.000E-06 | 9.165E-07 | 5.200E-06 | 7.000E-06 |
| 16127 | 807 | D807Q | B | 6.7e-06 | 8.3e-06 | 7.2e-06 | 7.400E-06 | 8.185E-07 | 6.700E-06 | 8.300E-06 |
| 16128 | 807 | D807G | B | 9.4e-06 | 9e-06 | 4.9e-06 | 7.767E-06 | 2.491E-06 | 4.900E-06 | 9.400E-06 |
| 16129 | 807 | D807H | B | 1.3e-05 | 1.1e-05 | 6.7e-06 | 1.023E-05 | 3.219E-06 | 6.700E-06 | 1.300E-05 |
| 16130 | 807 | D807I | B | 7.6e-06 | 6.9e-06 | 6.5e-06 | 7.000E-06 | 5.568E-07 | 6.500E-06 | 7.600E-06 |
| 16131 | 807 | D807L | B | 6.6e-06 | 7.5e-06 | 6.3e-06 | 6.800E-06 | 6.245E-07 | 6.300E-06 | 7.500E-06 |
| 16132 | 807 | D807K | B | 8.2e-06 | 5.1e-06 | 8.2e-06 | 7.167E-06 | 1.790E-06 | 5.100E-06 | 8.200E-06 |
| 16133 | 807 | D807M | B | 4.9e-06 | 7.4e-06 | 8.4e-06 | 6.900E-06 | 1.803E-06 | 4.900E-06 | 8.400E-06 |
| 16134 | 807 | D807F | B | 5.7e-06 | 9.1e-06 | 6.2e-06 | 7.000E-06 | 1.836E-06 | 5.700E-06 | 9.100E-06 |
| 16135 | 807 | D807P | B | 8.7e-06 | 6.2e-06 | 9.6e-06 | 8.167E-06 | 1.762E-06 | 6.200E-06 | 9.600E-06 |
| 16136 | 807 | D807S | B | 7.3e-06 | 5.7e-06 | 9.7e-06 | 7.567E-06 | 2.013E-06 | 5.700E-06 | 9.700E-06 |
| 16137 | 807 | D807T | B | 1.3e-05 | 5.8e-06 | 7.9e-06 | 8.900E-06 | 3.703E-06 | 5.800E-06 | 1.300E-05 |
| 16138 | 807 | D807W | B | 5.3e-06 | 3.4e-06 | 5.3e-06 | 4.667E-06 | 1.097E-06 | 3.400E-06 | 5.300E-06 |
| 16139 | 807 | D807Y | B | 5.5e-06 | 9.1e-06 | 8.9e-06 | 7.833E-06 | 2.023E-06 | 5.500E-06 | 9.100E-06 |
| 16140 | 807 | D807V | B | 8.4e-06 | 7.5e-06 | 5e-06 | 6.967E-06 | 1.762E-06 | 5.000E-06 | 8.400E-06 |
| 16141 | 808 | Y808A | B | 9.8e-06 | 5.1e-06 | 9e-06 | 7.967E-06 | 2.515E-06 | 5.100E-06 | 9.800E-06 |
| 16142 | 808 | Y808R | B | 5.9e-06 | 9.9e-06 | 9.8e-06 | 8.533E-06 | 2.281E-06 | 5.900E-06 | 9.900E-06 |
| 16143 | 808 | Y808N | B | 5.9e-06 | 8.3e-06 | 5.1e-06 | 6.433E-06 | 1.665E-06 | 5.100E-06 | 8.300E-06 |
| 16144 | 808 | Y808D | B | 5.6e-06 | 1.1e-05 | 8.1e-06 | 8.233E-06 | 2.702E-06 | 5.600E-06 | 1.100E-05 |
| 16145 | 808 | Y808C | B | 8.4e-06 | 8.4e-06 | 6.5e-06 | 7.767E-06 | 1.097E-06 | 6.500E-06 | 8.400E-06 |
| 16146 | 808 | Y808E | B | 7.9e-06 | 6.2e-06 | 7.4e-06 | 7.167E-06 | 8.737E-07 | 6.200E-06 | 7.900E-06 |
| 16147 | 808 | Y808Q | B | 6.2e-06 | 9.2e-06 | 5.9e-06 | 7.100E-06 | 1.825E-06 | 5.900E-06 | 9.200E-06 |
| 16148 | 808 | Y808G | B | 1.3e-05 | 6.5e-06 | 9.6e-06 | 9.700E-06 | 3.251E-06 | 6.500E-06 | 1.300E-05 |
| 16149 | 808 | Y808H | B | 1.1e-05 | 4.4e-06 | 8e-06 | 7.800E-06 | 3.305E-06 | 4.400E-06 | 1.100E-05 |

| | | | | | | | | | | |
|---|---|---|---|---|---|---|---|---|---|---|
| 16150 | 808 | Y808I | B | 9.2e-06 | 8.2e-06 | 1.1e-05 | 9.467E-06 | 1.419E-06 | 8.200E-06 | 1.100E-05 |
| 16151 | 808 | Y808L | B | 1.5e-05 | 9.4e-06 | 1e-05 | 1.147E-05 | 3.075E-06 | 9.400E-06 | 1.500E-05 |
| 16152 | 808 | Y808K | B | 9.5e-06 | 4.5e-06 | 7.7e-06 | 7.233E-06 | 2.532E-06 | 4.500E-06 | 9.500E-06 |
| 16153 | 808 | Y808M | B | 6.5e-06 | 7e-06 | 9.4e-06 | 7.633E-06 | 1.550E-06 | 6.500E-06 | 9.400E-06 |
| 16154 | 808 | Y808F | B | 5.4e-06 | 1.3e-05 | 8.4e-06 | 8.933E-06 | 3.828E-06 | 5.400E-06 | 1.300E-05 |
| 16155 | 808 | Y808P | B | 1e-05 | 1.4e-05 | 8.5e-06 | 1.083E-05 | 2.843E-06 | 8.500E-06 | 1.400E-05 |
| 16156 | 808 | Y808S | B | 5.9e-06 | 1e-05 | 8.9e-06 | 8.267E-06 | 2.122E-06 | 5.900E-06 | 1.000E-05 |
| 16157 | 808 | Y808T | B | 9.9e-06 | 7.8e-06 | 9.4e-06 | 9.033E-06 | 1.097E-06 | 7.800E-06 | 9.900E-06 |
| 16158 | 808 | Y808W | B | 9.5e-06 | 5.5e-06 | 6.2e-06 | 7.067E-06 | 2.136E-06 | 5.500E-06 | 9.500E-06 |
| 16159 | 808 | Y808Y | B | 1.2e-05 | 9.3e-06 | 7.2e-06 | 9.500E-06 | 2.406E-06 | 7.200E-06 | 1.200E-05 |
| 16160 | 808 | Y808V | B | 9.2e-06 | 4.5e-06 | 6.6e-06 | 6.767E-06 | 2.354E-06 | 4.500E-06 | 9.200E-06 |
| 16161 | 809 | E809A | B | 5.4e-06 | 8.4e-06 | 8.9e-06 | 7.567E-06 | 1.893E-06 | 5.400E-06 | 8.900E-06 |
| 16162 | 809 | E809R | B | 4.3e-06 | 9.2e-06 | 5.1e-06 | 6.200E-06 | 2.629E-06 | 4.300E-06 | 9.200E-06 |
| 16163 | 809 | E809N | B | 8.2e-06 | 9.1e-06 | 9.9e-06 | 9.067E-06 | 8.505E-07 | 8.200E-06 | 9.900E-06 |
| 16164 | 809 | E809D | B | 9.1e-06 | 1.1e-05 | 1e-05 | 1.003E-05 | 9.504E-07 | 9.100E-06 | 1.100E-05 |
| 16165 | 809 | E809C | B | 6.3e-06 | 4.8e-06 | 7.1e-06 | 6.067E-06 | 1.168E-06 | 4.800E-06 | 7.100E-06 |
| 16166 | 809 | E809E | B | 9.4e-06 | 1.2e-05 | 8e-06 | 9.800E-06 | 2.030E-06 | 8.000E-06 | 1.200E-05 |
| 16167 | 809 | E809Q | B | 5.7e-06 | 4.6e-06 | 6.7e-06 | 5.667E-06 | 1.050E-06 | 4.600E-06 | 6.700E-06 |
| 16168 | 809 | E809G | B | 5.1e-06 | 7.1e-06 | 6.7e-06 | 6.300E-06 | 1.058E-06 | 5.100E-06 | 7.100E-06 |
| 16169 | 809 | E809H | B | 1.2e-05 | 8.3e-06 | 8.5e-06 | 9.600E-06 | 2.081E-06 | 8.300E-06 | 1.200E-05 |
| 16170 | 809 | E809I | B | 4.8e-06 | 9.5e-06 | 9.7e-06 | 8.000E-06 | 2.773E-06 | 4.800E-06 | 9.700E-06 |
| 16171 | 809 | E809L | B | 7.9e-06 | 1e-05 | 5.9e-06 | 7.933E-06 | 2.050E-06 | 5.900E-06 | 1.000E-05 |
| 16172 | 809 | E809K | B | 8e-06 | 7.1e-06 | 1e-05 | 8.367E-06 | 1.484E-06 | 7.100E-06 | 1.000E-05 |
| 16173 | 809 | E809M | B | 1.2e-05 | 6.5e-06 | 9.7e-06 | 9.400E-06 | 2.762E-06 | 6.500E-06 | 1.200E-05 |
| 16174 | 809 | E809F | B | 7.7e-06 | 8.7e-06 | 9.9e-06 | 8.767E-06 | 1.102E-06 | 7.700E-06 | 9.900E-06 |
| 16175 | 809 | E809P | B | 9.9e-06 | 8e-06 | 7.3e-06 | 8.400E-06 | 1.345E-06 | 7.300E-06 | 9.900E-06 |
| 16176 | 809 | E809S | B | 9.7e-06 | 9.4e-06 | 1.3e-05 | 1.070E-05 | 1.997E-06 | 9.400E-06 | 1.300E-05 |
| 16177 | 809 | E809T | B | 7.6e-06 | 1.1e-05 | 5.3e-06 | 7.967E-06 | 2.868E-06 | 5.300E-06 | 1.100E-05 |
| 16178 | 809 | E809W | B | 6.2e-06 | 9.6e-06 | 7.3e-06 | 7.700E-06 | 1.735E-06 | 6.200E-06 | 9.600E-06 |
| 16179 | 809 | E809Y | B | 5.7e-06 | 1e-05 | 7e-06 | 7.567E-06 | 2.205E-06 | 5.700E-06 | 1.000E-05 |
| 16180 | 809 | E809V | B | 6.9e-06 | 8.5e-06 | 5.9e-06 | 7.100E-06 | 1.311E-06 | 5.900E-06 | 8.500E-06 |
| 16181 | 810 | K810A | B | 1.1e-05 | 8.8e-06 | 5.2e-06 | 8.333E-06 | 2.928E-06 | 5.200E-06 | 1.100E-05 |
| 16182 | 810 | K810R | B | 9e-06 | 8.3e-06 | 8.1e-06 | 8.467E-06 | 4.726E-07 | 8.100E-06 | 9.000E-06 |
| 16183 | 810 | K810N | B | 8.6e-06 | 9.6e-06 | 7.1e-06 | 8.433E-06 | 1.258E-06 | 7.100E-06 | 9.600E-06 |
| 16184 | 810 | K810D | B | 4.9e-06 | 8.8e-06 | 1.1e-05 | 8.233E-06 | 3.089E-06 | 4.900E-06 | 1.100E-05 |
| 16185 | 810 | K810C | B | 6.6e-06 | 6.8e-06 | 6.6e-06 | 6.667E-06 | 1.155E-07 | 6.600E-06 | 6.800E-06 |
| 16186 | 810 | K810E | B | 5.3e-06 | 5.4e-06 | 9.2e-06 | 6.633E-06 | 2.223E-06 | 5.300E-06 | 9.200E-06 |
| 16187 | 810 | K810Q | B | 1.3e-05 | 3.5e-06 | 1.5e-05 | 1.050E-05 | 6.144E-06 | 3.500E-06 | 1.500E-05 |
| 16188 | 810 | K810G | B | 7.8e-06 | 1.2e-05 | 1.3e-05 | 1.093E-05 | 2.759E-06 | 7.800E-06 | 1.300E-05 |
| 16189 | 810 | K810H | B | 1.2e-05 | 6e-06 | 1.1e-05 | 9.667E-06 | 3.215E-06 | 6.000E-06 | 1.200E-05 |
| 16190 | 810 | K810I | B | 1.1e-05 | 5.6e-06 | 9.9e-06 | 8.833E-06 | 2.854E-06 | 5.600E-06 | 1.100E-05 |
| 16191 | 810 | K810L | B | 1.2e-05 | 9.3e-06 | 4.9e-06 | 8.733E-06 | 3.584E-06 | 4.900E-06 | 1.200E-05 |
| 16192 | 810 | K810K | B | 5.6e-06 | 1.1e-05 | 8.2e-06 | 8.267E-06 | 2.701E-06 | 5.600E-06 | 1.100E-05 |
| 16193 | 810 | K810M | B | 7.7e-06 | 5.5e-06 | 8.1e-06 | 7.100E-06 | 1.400E-06 | 5.500E-06 | 8.100E-06 |
| 16194 | 810 | K810F | B | 1.5e-05 | 8.4e-06 | 6.7e-06 | 1.003E-05 | 4.384E-06 | 6.700E-06 | 1.500E-05 |
| 16195 | 810 | K810P | B | 1e-05 | 7.1e-06 | 5.5e-06 | 7.533E-06 | 2.281E-06 | 5.500E-06 | 1.000E-05 |
| 16196 | 810 | K810S | B | 1e-05 | 7e-06 | 6.2e-06 | 7.733E-06 | 2.003E-06 | 6.200E-06 | 1.000E-05 |
| 16197 | 810 | K810T | B | 1e-05 | 9.5e-06 | 9.4e-06 | 9.633E-06 | 3.215E-07 | 9.400E-06 | 1.000E-05 |
| 16198 | 810 | K810W | B | 5e-06 | 1.2e-05 | 6.5e-06 | 7.833E-06 | 3.686E-06 | 5.000E-06 | 1.200E-05 |
| 16199 | 810 | K810Y | B | 8.5e-06 | 9.8e-06 | 8.2e-06 | 8.833E-06 | 8.505E-07 | 8.200E-06 | 9.800E-06 |
| 16200 | 810 | K810V | B | 7e-06 | 6.1e-06 | 1e-05 | 7.700E-06 | 2.042E-06 | 6.100E-06 | 1.000E-05 |
| 16201 | 811 | H811A | B | 7.4e-06 | 7.6e-06 | 8e-06 | 7.667E-06 | 3.055E-07 | 7.400E-06 | 8.000E-06 |
| 16202 | 811 | H811R | B | 7.3e-06 | 7.5e-06 | 9.5e-06 | 8.100E-06 | 1.217E-06 | 7.300E-06 | 9.500E-06 |
| 16203 | 811 | H811N | B | 1.9e-05 | 5.2e-06 | 6.9e-06 | 1.037E-05 | 7.525E-06 | 5.200E-06 | 1.900E-05 |
| 16204 | 811 | H811D | B | 7.6e-06 | 6.6e-06 | 1.4e-05 | 9.400E-06 | 4.015E-06 | 6.600E-06 | 1.400E-05 |
| 16205 | 811 | H811C | B | 7.7e-06 | 6.9e-06 | 7.4e-06 | 7.333E-06 | 4.041E-07 | 6.900E-06 | 7.700E-06 |
| 16206 | 811 | H811E | B | 4.6e-06 | 5.9e-06 | 7.3e-06 | 5.933E-06 | 1.350E-06 | 4.600E-06 | 7.300E-06 |
| 16207 | 811 | H811Q | B | 1.3e-05 | 5.6e-06 | 6.3e-06 | 8.300E-06 | 4.085E-06 | 5.600E-06 | 1.300E-05 |
| 16208 | 811 | H811G | B | 7.5e-06 | 6.2e-06 | 6e-06 | 6.567E-06 | 8.145E-07 | 6.000E-06 | 7.500E-06 |
| 16209 | 811 | H811H | B | 9.2e-06 | 1.3e-05 | 6.7e-06 | 9.633E-06 | 3.172E-06 | 6.700E-06 | 1.300E-05 |
| 16210 | 811 | H811I | B | 9.3e-06 | 6.8e-06 | 1.1e-05 | 9.033E-06 | 2.113E-06 | 6.800E-06 | 1.100E-05 |
| 16211 | 811 | H811L | B | 1.5e-05 | 4.6e-06 | 5.2e-06 | 8.267E-06 | 5.839E-06 | 4.600E-06 | 1.500E-05 |
| 16212 | 811 | H811K | B | 1.2e-05 | 7.5e-06 | 6.2e-06 | 8.567E-06 | 3.044E-06 | 6.200E-06 | 1.200E-05 |
| 16213 | 811 | H811M | B | 1.1e-05 | 4.7e-06 | 4.6e-06 | 6.767E-06 | 3.667E-06 | 4.600E-06 | 1.100E-05 |
| 16214 | 811 | H811F | B | 9.2e-06 | 1.5e-05 | 7e-06 | 1.040E-05 | 4.133E-06 | 7.000E-06 | 1.500E-05 |
| 16215 | 811 | H811P | B | 1.1e-05 | 9.5e-06 | 8.6e-06 | 9.700E-06 | 1.212E-06 | 8.600E-06 | 1.100E-05 |
| 16216 | 811 | H811S | B | 8.9e-06 | 1e-05 | 8.3e-06 | 9.067E-06 | 8.622E-07 | 8.300E-06 | 1.000E-05 |
| 16217 | 811 | H811T | B | 7.3e-06 | 8e-06 | 6.1e-06 | 7.133E-06 | 9.609E-07 | 6.100E-06 | 8.000E-06 |
| 16218 | 811 | H811W | B | 8.6e-06 | 5.6e-06 | 9.6e-06 | 7.933E-06 | 2.082E-06 | 5.600E-06 | 9.600E-06 |
| 16219 | 811 | H811Y | B | 1e-05 | 8.9e-06 | 7.1e-06 | 8.667E-06 | 1.464E-06 | 7.100E-06 | 1.000E-05 |
| 16220 | 811 | H811V | B | 5.9e-06 | 9.9e-06 | 1.5e-05 | 1.027E-05 | 4.561E-06 | 5.900E-06 | 1.500E-05 |
| 16221 | 812 | K812A | B | 6.3e-06 | 1.2e-05 | 1e-05 | 9.433E-06 | 2.892E-06 | 6.300E-06 | 1.200E-05 |
| 16222 | 812 | K812R | B | 9.1e-06 | 1.2e-05 | 6.9e-06 | 9.333E-06 | 2.558E-06 | 6.900E-06 | 1.200E-05 |
| 16223 | 812 | K812N | B | 9.6e-06 | 1.1e-05 | 8.7e-06 | 9.767E-06 | 1.159E-06 | 8.700E-06 | 1.100E-05 |
| 16224 | 812 | K812D | B | 8.1e-06 | 5.1e-06 | 4.4e-06 | 5.867E-06 | 1.966E-06 | 4.400E-06 | 8.100E-06 |
| 16225 | 812 | K812C | B | 7.8e-06 | 9.2e-06 | 1.4e-05 | 1.033E-05 | 3.252E-06 | 7.800E-06 | 1.400E-05 |
| 16226 | 812 | K812E | B | 3.8e-06 | 1.1e-05 | 1e-05 | 8.267E-06 | 3.900E-06 | 3.800E-06 | 1.100E-05 |
| 16227 | 812 | K812Q | B | 5e-06 | 1.3e-05 | 5.7e-06 | 7.900E-06 | 4.431E-06 | 5.000E-06 | 1.300E-05 |
| 16228 | 812 | K812G | B | 1e-05 | 9.4e-06 | 8.1e-06 | 9.167E-06 | 9.713E-07 | 8.100E-06 | 1.000E-05 |
| 16229 | 812 | K812H | B | 8.8e-06 | 7.8e-06 | 5.7e-06 | 7.433E-06 | 1.582E-06 | 5.700E-06 | 8.800E-06 |
| 16230 | 812 | K812I | B | 1.3e-05 | 9.1e-06 | 8.8e-06 | 1.030E-05 | 2.343E-06 | 8.800E-06 | 1.300E-05 |
| 16231 | 812 | K812L | B | 6.7e-06 | 5.6e-06 | 6.6e-06 | 6.300E-06 | 6.083E-07 | 5.600E-06 | 6.700E-06 |
| 16232 | 812 | K812K | B | 7.5e-06 | 7.4e-06 | 7.5e-06 | 7.467E-06 | 5.774E-08 | 7.400E-06 | 7.500E-06 |
| 16233 | 812 | K812M | B | 6e-06 | 7.6e-06 | 1.1e-05 | 8.200E-06 | 2.553E-06 | 6.000E-06 | 1.100E-05 |
| 16234 | 812 | K812F | B | 9.5e-06 | 7.7e-06 | 5.9e-06 | 7.700E-06 | 1.800E-06 | 5.900E-06 | 9.500E-06 |

| | | | | | | | | | | |
|---|---|---|---|---|---|---|---|---|---|---|
| 16235 | 812 | K812P | B | 7.6e-06 | 7.1e-06 | 6.7e-06 | 7.133E-06 | 4.509E-07 | 6.700E-06 | 7.600E-06 |
| 16236 | 812 | K812S | B | 7.3e-06 | 7.9e-06 | 6e-06 | 7.067E-06 | 9.713E-07 | 6.000E-06 | 7.900E-06 |
| 16237 | 812 | K812T | B | 1e-05 | 9.2e-06 | 6.3e-06 | 8.500E-06 | 1.947E-06 | 6.300E-06 | 1.000E-05 |
| 16238 | 812 | K812W | B | 1.4e-05 | 9.3e-06 | 7.3e-06 | 1.020E-05 | 3.439E-06 | 7.300E-06 | 1.400E-05 |
| 16239 | 812 | K812Y | B | 2.1e-05 | 1e-05 | 9.6e-06 | 1.353E-05 | 6.469E-06 | 9.600E-06 | 2.100E-05 |
| 16240 | 812 | K812V | B | 7.9e-06 | 6.9e-06 | 3.1e-06 | 5.967E-06 | 2.532E-06 | 3.100E-06 | 7.900E-06 |
| 16241 | 813 | V813A | B | 5.2e-06 | 1e-05 | 6.9e-06 | 7.367E-06 | 2.434E-06 | 5.200E-06 | 1.000E-05 |
| 16242 | 813 | V813R | B | 8.7e-06 | 7.2e-06 | 7.1e-06 | 7.667E-06 | 8.963E-07 | 7.100E-06 | 8.700E-06 |
| 16243 | 813 | V813N | B | 6.5e-06 | 8.7e-06 | 7.8e-06 | 7.667E-06 | 1.106E-06 | 6.500E-06 | 8.700E-06 |
| 16244 | 813 | V813D | B | 5.9e-06 | 6.4e-06 | 9.1e-06 | 7.133E-06 | 1.721E-06 | 5.900E-06 | 9.100E-06 |
| 16245 | 813 | V813C | B | 9.3e-06 | 9.2e-06 | 1.4e-05 | 1.083E-05 | 2.743E-06 | 9.200E-06 | 1.400E-05 |
| 16246 | 813 | V813E | B | 1e-05 | 8e-06 | 6.3e-06 | 8.100E-06 | 1.852E-06 | 6.300E-06 | 1.000E-05 |
| 16247 | 813 | V813Q | B | 9.6e-06 | 7.9e-06 | 8.5e-06 | 8.667E-06 | 8.622E-07 | 7.900E-06 | 9.600E-06 |
| 16248 | 813 | V813G | B | 6.2e-06 | 9.5e-06 | 7.6e-06 | 7.767E-06 | 1.656E-06 | 6.200E-06 | 9.500E-06 |
| 16249 | 813 | V813H | B | 5e-06 | 6.4e-06 | 6.8e-06 | 6.067E-06 | 9.452E-07 | 5.000E-06 | 6.800E-06 |
| 16250 | 813 | V813I | B | 6.8e-06 | 1.4e-05 | 5.5e-06 | 8.767E-06 | 4.579E-06 | 5.500E-06 | 1.400E-05 |
| 16251 | 813 | V813L | B | 6.2e-06 | 8.7e-06 | 8e-06 | 7.633E-06 | 1.290E-06 | 6.200E-06 | 8.700E-06 |
| 16252 | 813 | V813K | B | 7.3e-06 | 8.8e-06 | 8e-06 | 8.033E-06 | 7.506E-07 | 7.300E-06 | 8.800E-06 |
| 16253 | 813 | V813M | B | 1e-05 | 1.1e-05 | 7.3e-06 | 9.433E-06 | 1.914E-06 | 7.300E-06 | 1.100E-05 |
| 16254 | 813 | V813F | B | 1.4e-05 | 6.5e-06 | 5.7e-06 | 8.733E-06 | 4.579E-06 | 5.700E-06 | 1.400E-05 |
| 16255 | 813 | V813P | B | 7.6e-06 | 6.3e-06 | 7.1e-06 | 7.000E-06 | 6.557E-07 | 6.300E-06 | 7.600E-06 |
| 16256 | 813 | V813S | B | 6.5e-06 | 9.2e-06 | 9.1e-06 | 8.267E-06 | 1.531E-06 | 6.500E-06 | 9.200E-06 |
| 16257 | 813 | V813T | B | 8.5e-06 | 8.7e-06 | 9.3e-06 | 8.833E-06 | 4.163E-07 | 8.500E-06 | 9.300E-06 |
| 16258 | 813 | V813W | B | 1.1e-05 | 6.3e-06 | 8.6e-06 | 8.633E-06 | 2.350E-06 | 6.300E-06 | 1.100E-05 |
| 16259 | 813 | V813Y | B | 6.7e-06 | 9e-06 | 9.7e-06 | 8.467E-06 | 1.570E-06 | 6.700E-06 | 9.700E-06 |
| 16260 | 813 | V813V | B | 2.8e-06 | 5.7e-06 | 1.1e-05 | 6.500E-06 | 4.158E-06 | 2.800E-06 | 1.100E-05 |
| 16261 | 814 | Y814A | B | 1.1e-05 | 6.7e-06 | 9.2e-06 | 8.967E-06 | 2.159E-06 | 6.700E-06 | 1.100E-05 |
| 16262 | 814 | Y814R | B | 1.1e-05 | 8.4e-06 | 6.7e-06 | 8.700E-06 | 2.166E-06 | 6.700E-06 | 1.100E-05 |
| 16263 | 814 | Y814N | B | 8.5e-06 | 3.7e-06 | 1.2e-05 | 8.067E-06 | 4.167E-06 | 3.700E-06 | 1.200E-05 |
| 16264 | 814 | Y814D | B | 1.3e-05 | 9e-06 | 5.2e-06 | 9.067E-06 | 3.900E-06 | 5.200E-06 | 1.300E-05 |
| 16265 | 814 | Y814C | B | 5.1e-06 | 7.2e-06 | 5.1e-06 | 5.800E-06 | 1.212E-06 | 5.100E-06 | 7.200E-06 |
| 16266 | 814 | Y814E | B | 5.5e-06 | 1.1e-05 | 6.7e-06 | 7.733E-06 | 2.892E-06 | 5.500E-06 | 1.100E-05 |
| 16267 | 814 | Y814Q | B | 1e-05 | 4.4e-06 | 5.1e-06 | 6.500E-06 | 3.051E-06 | 4.400E-06 | 1.000E-05 |
| 16268 | 814 | Y814G | B | 8.1e-06 | 1.6e-05 | 5.5e-06 | 9.867E-06 | 5.468E-06 | 5.500E-06 | 1.600E-05 |
| 16269 | 814 | Y814H | B | 6.1e-06 | 4.8e-06 | 7.2e-06 | 6.033E-06 | 1.201E-06 | 4.800E-06 | 7.200E-06 |
| 16270 | 814 | Y814I | B | 1.1e-05 | 7.1e-06 | 6.4e-06 | 8.167E-06 | 2.479E-06 | 6.400E-06 | 1.100E-05 |
| 16271 | 814 | Y814L | B | 8.1e-06 | 1.2e-05 | 6.8e-06 | 8.967E-06 | 2.706E-06 | 6.800E-06 | 1.200E-05 |
| 16272 | 814 | Y814K | B | 6.6e-06 | 1.3e-05 | 4.5e-06 | 8.033E-06 | 4.428E-06 | 4.500E-06 | 1.300E-05 |
| 16273 | 814 | Y814M | B | 7.2e-06 | 1e-05 | 9.6e-06 | 8.933E-06 | 1.514E-06 | 7.200E-06 | 1.000E-05 |
| 16274 | 814 | Y814F | B | 1.5e-05 | 6.9e-06 | 9e-06 | 1.030E-05 | 4.204E-06 | 6.900E-06 | 1.500E-05 |
| 16275 | 814 | Y814P | B | 6e-06 | 2e-05 | 6e-06 | 1.067E-05 | 8.083E-06 | 6.000E-06 | 2.000E-05 |
| 16276 | 814 | Y814S | B | 5.7e-06 | 6.7e-06 | 6e-06 | 6.133E-06 | 5.132E-07 | 5.700E-06 | 6.700E-06 |
| 16277 | 814 | Y814T | B | 8.3e-06 | 5.7e-06 | 7.5e-06 | 7.167E-06 | 1.332E-06 | 5.700E-06 | 8.300E-06 |
| 16278 | 814 | Y814W | B | 6.1e-06 | 1.5e-05 | 1e-05 | 1.037E-05 | 4.461E-06 | 6.100E-06 | 1.500E-05 |
| 16279 | 814 | Y814Y | B | 6.8e-06 | 5.3e-06 | 5.8e-06 | 5.967E-06 | 7.638E-07 | 5.300E-06 | 6.800E-06 |
| 16280 | 814 | Y814V | B | 5.9e-06 | 1.3e-05 | 1.3e-05 | 1.063E-05 | 4.099E-06 | 5.900E-06 | 1.300E-05 |
| 16281 | 815 | A815A | B | 7.8e-06 | 6.6e-06 | 5.8e-06 | 6.733E-06 | 1.007E-06 | 5.800E-06 | 7.800E-06 |
| 16282 | 815 | A815R | B | 1.5e-05 | 6.2e-06 | 8.8e-06 | 1.000E-05 | 4.521E-06 | 6.200E-06 | 1.500E-05 |
| 16283 | 815 | A815N | B | 6.2e-06 | 9.3e-06 | 6.7e-06 | 7.400E-06 | 1.664E-06 | 6.200E-06 | 9.300E-06 |
| 16284 | 815 | A815D | B | 9.6e-06 | 5.6e-06 | 1e-05 | 8.400E-06 | 2.433E-06 | 5.600E-06 | 1.000E-05 |
| 16285 | 815 | A815C | B | 6.2e-06 | 1e-05 | 7.2e-06 | 7.800E-06 | 1.970E-06 | 6.200E-06 | 1.000E-05 |
| 16286 | 815 | A815E | B | 1.3e-05 | 4.6e-06 | 4.8e-06 | 7.467E-06 | 4.793E-06 | 4.600E-06 | 1.300E-05 |
| 16287 | 815 | A815Q | B | 6.8e-06 | 9.4e-06 | 1.2e-05 | 9.400E-06 | 2.600E-06 | 6.800E-06 | 1.200E-05 |
| 16288 | 815 | A815G | B | 5.7e-06 | 4.9e-06 | 8.7e-06 | 6.433E-06 | 2.003E-06 | 4.900E-06 | 8.700E-06 |
| 16289 | 815 | A815H | B | 9.6e-06 | 7.2e-06 | 6.6e-06 | 7.800E-06 | 1.587E-06 | 6.600E-06 | 9.600E-06 |
| 16290 | 815 | A815I | B | 7.8e-06 | 7.5e-06 | 1.3e-05 | 9.433E-06 | 3.092E-06 | 7.500E-06 | 1.300E-05 |
| 16291 | 815 | A815L | B | 5.7e-06 | 5.3e-06 | 6.8e-06 | 5.933E-06 | 7.767E-07 | 5.300E-06 | 6.800E-06 |
| 16292 | 815 | A815K | B | 1.3e-05 | 1.1e-05 | 3.9e-06 | 9.300E-06 | 4.782E-06 | 3.900E-06 | 1.300E-05 |
| 16293 | 815 | A815M | B | 8.1e-06 | 9.5e-06 | 9.4e-06 | 9.000E-06 | 7.810E-07 | 8.100E-06 | 9.500E-06 |
| 16294 | 815 | A815F | B | 1.3e-05 | 8.1e-06 | 6.3e-06 | 9.133E-06 | 3.467E-06 | 6.300E-06 | 1.300E-05 |
| 16295 | 815 | A815P | B | 9.5e-06 | 8.6e-06 | 6.7e-06 | 8.267E-06 | 1.429E-06 | 6.700E-06 | 9.500E-06 |
| 16296 | 815 | A815S | B | 5.9e-06 | 7.8e-06 | 7.4e-06 | 7.033E-06 | 1.002E-06 | 5.900E-06 | 7.800E-06 |
| 16297 | 815 | A815T | B | 9.6e-06 | 7.2e-06 | 8.4e-06 | 8.400E-06 | 1.200E-06 | 7.200E-06 | 9.600E-06 |
| 16298 | 815 | A815W | B | 8.4e-06 | 5.7e-06 | 5.4e-06 | 6.500E-06 | 1.652E-06 | 5.400E-06 | 8.400E-06 |
| 16299 | 815 | A815Y | B | 8.6e-06 | 6.3e-06 | 7.1e-06 | 7.333E-06 | 1.168E-06 | 6.300E-06 | 8.600E-06 |
| 16300 | 815 | A815V | B | 1.4e-05 | 5.6e-06 | 8.7e-06 | 9.433E-06 | 4.248E-06 | 5.600E-06 | 1.400E-05 |
| 16301 | 816 | C816A | B | 1e-05 | 5.8e-06 | 4.9e-06 | 6.900E-06 | 2.722E-06 | 4.900E-06 | 1.000E-05 |
| 16302 | 816 | C816R | B | 1.3e-05 | 1.2e-05 | 4.4e-06 | 9.800E-06 | 4.703E-06 | 4.400E-06 | 1.300E-05 |
| 16303 | 816 | C816N | B | 9.5e-06 | 6.7e-06 | 1.1e-05 | 9.067E-06 | 2.183E-06 | 6.700E-06 | 1.100E-05 |
| 16304 | 816 | C816D | B | 6.4e-06 | 8.9e-06 | 7.7e-06 | 7.667E-06 | 1.250E-06 | 6.400E-06 | 8.900E-06 |
| 16305 | 816 | C816C | B | 6.3e-06 | 1e-05 | 7.4e-06 | 7.900E-06 | 1.900E-06 | 6.300E-06 | 1.000E-05 |
| 16306 | 816 | C816E | B | 7.9e-06 | 7.6e-06 | 4.7e-06 | 6.733E-06 | 1.767E-06 | 4.700E-06 | 7.900E-06 |
| 16307 | 816 | C816Q | B | 6.5e-06 | 6.5e-06 | 8e-06 | 7.000E-06 | 8.660E-07 | 6.500E-06 | 8.000E-06 |
| 16308 | 816 | C816G | B | 6.6e-06 | 1.3e-05 | 6e-06 | 8.533E-06 | 3.880E-06 | 6.000E-06 | 1.300E-05 |
| 16309 | 816 | C816H | B | 6.9e-06 | 8.6e-06 | 7.6e-06 | 7.700E-06 | 8.544E-07 | 6.900E-06 | 8.600E-06 |
| 16310 | 816 | C816I | B | 9.5e-06 | 8.7e-06 | 6.8e-06 | 8.333E-06 | 1.387E-06 | 6.800E-06 | 9.500E-06 |
| 16311 | 816 | C816L | B | 6.9e-06 | 6.4e-06 | 9.9e-06 | 7.733E-06 | 1.893E-06 | 6.400E-06 | 9.900E-06 |
| 16312 | 816 | C816K | B | 9.2e-06 | 7.2e-06 | 7.8e-06 | 8.067E-06 | 1.026E-06 | 7.200E-06 | 9.200E-06 |
| 16313 | 816 | C816M | B | 6.8e-06 | 1.4e-05 | 7.3e-06 | 9.367E-06 | 4.020E-06 | 6.800E-06 | 1.400E-05 |
| 16314 | 816 | C816F | B | 7.5e-06 | 1.1e-05 | 9.2e-06 | 9.233E-06 | 1.750E-06 | 7.500E-06 | 1.100E-05 |
| 16315 | 816 | C816P | B | 9.9e-06 | 1e-05 | 1e-05 | 9.967E-06 | 5.774E-08 | 9.900E-06 | 1.000E-05 |
| 16316 | 816 | C816S | B | 4.6e-06 | 8.5e-06 | 1.3e-05 | 8.700E-06 | 4.204E-06 | 4.600E-06 | 1.300E-05 |
| 16317 | 816 | C816T | B | 8.1e-06 | 7.7e-06 | 1.3e-05 | 9.600E-06 | 2.951E-06 | 7.700E-06 | 1.300E-05 |
| 16318 | 816 | C816W | B | 1e-05 | 8.5e-06 | 7.1e-06 | 8.533E-06 | 1.450E-06 | 7.100E-06 | 1.000E-05 |
| 16319 | 816 | C816Y | B | 7.9e-06 | 7.1e-06 | 6.8e-06 | 7.267E-06 | 5.686E-07 | 6.800E-06 | 7.900E-06 |

| | | | | | | | | | | |
|---|---|---|---|---|---|---|---|---|---|---|
| 16320 | 816 | C816V | B | 9.9e-06 | 1e-05 | 4.1e-06 | 8.000E-06 | 3.378E-06 | 4.100E-06 | 1.000E-05 |
| 16321 | 817 | E817A | B | 6.2e-06 | 8.8e-06 | 9e-06 | 8.000E-06 | 1.562E-06 | 6.200E-06 | 9.000E-06 |
| 16322 | 817 | E817R | B | 1.8e-05 | 8.1e-06 | 6.4e-06 | 1.083E-05 | 6.264E-06 | 6.400E-06 | 1.800E-05 |
| 16323 | 817 | E817N | B | 8.9e-06 | 9.6e-06 | 9.3e-06 | 9.267E-06 | 3.512E-07 | 8.900E-06 | 9.600E-06 |
| 16324 | 817 | E817D | B | 9.3e-06 | 1.1e-05 | 6.9e-06 | 9.067E-06 | 2.060E-06 | 6.900E-06 | 1.100E-05 |
| 16325 | 817 | E817C | B | 5.7e-06 | 1.2e-05 | 5.7e-06 | 7.800E-06 | 3.637E-06 | 5.700E-06 | 1.200E-05 |
| 16326 | 817 | E817E | B | 5.2e-06 | 9.3e-06 | 4.5e-06 | 6.333E-06 | 2.593E-06 | 4.500E-06 | 9.300E-06 |
| 16327 | 817 | E817Q | B | 9.5e-06 | 6.7e-06 | 1.1e-05 | 9.067E-06 | 2.183E-06 | 6.700E-06 | 1.100E-05 |
| 16328 | 817 | E817G | B | 4.6e-06 | 7e-06 | 9.3e-06 | 6.967E-06 | 2.350E-06 | 4.600E-06 | 9.300E-06 |
| 16329 | 817 | E817H | B | 1.4e-05 | 8.6e-06 | 9.2e-06 | 1.060E-05 | 2.960E-06 | 8.600E-06 | 1.400E-05 |
| 16330 | 817 | E817I | B | 1.5e-05 | 6.1e-06 | 7e-06 | 9.367E-06 | 4.899E-06 | 6.100E-06 | 1.500E-05 |
| 16331 | 817 | E817L | B | 4.7e-06 | 1.2e-05 | 9.7e-06 | 8.800E-06 | 3.732E-06 | 4.700E-06 | 1.200E-05 |
| 16332 | 817 | E817K | B | 1.2e-05 | 1.5e-05 | 9.3e-06 | 1.210E-05 | 2.851E-06 | 9.300E-06 | 1.500E-05 |
| 16333 | 817 | E817M | B | 1e-05 | 8.4e-06 | 7.8e-06 | 8.733E-06 | 1.137E-06 | 7.800E-06 | 1.000E-05 |
| 16334 | 817 | E817F | B | 5e-06 | 8.7e-06 | 7.2e-06 | 6.967E-06 | 1.861E-06 | 5.000E-06 | 8.700E-06 |
| 16335 | 817 | E817P | B | 2e-05 | 6.8e-06 | 1.1e-05 | 1.260E-05 | 6.744E-06 | 6.800E-06 | 2.000E-05 |
| 16336 | 817 | E817S | B | 7.8e-06 | 7.6e-06 | 7.3e-06 | 7.567E-06 | 2.517E-07 | 7.300E-06 | 7.800E-06 |
| 16337 | 817 | E817T | B | 6e-06 | 1.1e-05 | 8.4e-06 | 8.467E-06 | 2.501E-06 | 6.000E-06 | 1.100E-05 |
| 16338 | 817 | E817W | B | 8.9e-06 | 7.7e-06 | 7.7e-06 | 8.100E-06 | 6.928E-07 | 7.700E-06 | 8.900E-06 |
| 16339 | 817 | E817Y | B | 7.6e-06 | 7.6e-06 | 1.3e-05 | 9.400E-06 | 3.118E-06 | 7.600E-06 | 1.300E-05 |
| 16340 | 817 | E817V | B | 6e-06 | 8.8e-06 | 9.4e-06 | 8.067E-06 | 1.815E-06 | 6.000E-06 | 9.400E-06 |
| 16341 | 818 | V818A | B | 8.3e-06 | 9.2e-06 | 9.5e-06 | 9.000E-06 | 6.245E-07 | 8.300E-06 | 9.500E-06 |
| 16342 | 818 | V818R | B | 8.4e-06 | 8.9e-06 | 9.7e-06 | 9.000E-06 | 6.557E-07 | 8.400E-06 | 9.700E-06 |
| 16343 | 818 | V818N | B | 6.9e-06 | 8.7e-06 | 8.4e-06 | 8.000E-06 | 9.644E-07 | 6.900E-06 | 8.700E-06 |
| 16344 | 818 | V818D | B | 6.3e-06 | 1.2e-05 | 7e-06 | 8.433E-06 | 3.109E-06 | 6.300E-06 | 1.200E-05 |
| 16345 | 818 | V818C | B | 9.8e-06 | 5.3e-06 | 6.2e-06 | 7.100E-06 | 2.381E-06 | 5.300E-06 | 9.800E-06 |
| 16346 | 818 | V818E | B | 1.2e-05 | 3.5e-06 | 8.9e-06 | 8.133E-06 | 4.302E-06 | 3.500E-06 | 1.200E-05 |
| 16347 | 818 | V818Q | B | 9.1e-06 | 8.5e-06 | 8e-06 | 8.533E-06 | 5.508E-07 | 8.000E-06 | 9.100E-06 |
| 16348 | 818 | V818G | B | 4.4e-06 | 1.2e-05 | 5e-06 | 7.133E-06 | 4.225E-06 | 4.400E-06 | 1.200E-05 |
| 16349 | 818 | V818H | B | 8.8e-06 | 4.6e-06 | 4.9e-06 | 6.100E-06 | 2.343E-06 | 4.600E-06 | 8.800E-06 |
| 16350 | 818 | V818I | B | 6.8e-06 | 7.1e-06 | 7.2e-06 | 7.033E-06 | 2.082E-07 | 6.800E-06 | 7.200E-06 |
| 16351 | 818 | V818L | B | 1.1e-05 | 1.2e-05 | 4.5e-06 | 9.167E-06 | 4.072E-06 | 4.500E-06 | 1.200E-05 |
| 16352 | 818 | V818K | B | 6.7e-06 | 5e-06 | 8.6e-06 | 6.767E-06 | 1.801E-06 | 5.000E-06 | 8.600E-06 |
| 16353 | 818 | V818M | B | 1.5e-05 | 1.2e-05 | 8.5e-06 | 1.183E-05 | 3.253E-06 | 8.500E-06 | 1.500E-05 |
| 16354 | 818 | V818F | B | 3.6e-06 | 4.9e-06 | 4.5e-06 | 4.333E-06 | 6.658E-07 | 3.600E-06 | 4.900E-06 |
| 16355 | 818 | V818P | B | 7e-06 | 6.3e-06 | 4.8e-06 | 6.033E-06 | 1.124E-06 | 4.800E-06 | 7.000E-06 |
| 16356 | 818 | V818S | B | 9.3e-06 | 9.6e-06 | 4.8e-06 | 7.900E-06 | 2.689E-06 | 4.800E-06 | 9.600E-06 |
| 16357 | 818 | V818T | B | 1.2e-05 | 5.1e-06 | 9.9e-06 | 9.000E-06 | 3.537E-06 | 5.100E-06 | 1.200E-05 |
| 16358 | 818 | V818W | B | 1e-05 | 6.4e-06 | 9.8e-06 | 8.733E-06 | 2.023E-06 | 6.400E-06 | 1.000E-05 |
| 16359 | 818 | V818Y | B | 6.1e-06 | 7.2e-06 | 7.9e-06 | 7.067E-06 | 9.074E-07 | 6.100E-06 | 7.900E-06 |
| 16360 | 818 | V818V | B | 8.1e-06 | 5.6e-06 | 1.1e-05 | 8.233E-06 | 2.702E-06 | 5.600E-06 | 1.100E-05 |
| 16361 | 819 | T819A | B | 8.5e-06 | 8.9e-06 | 8.5e-06 | 8.633E-06 | 2.309E-07 | 8.500E-06 | 8.900E-06 |
| 16362 | 819 | T819R | B | 4.5e-06 | 9.3e-06 | 8.1e-06 | 7.300E-06 | 2.498E-06 | 4.500E-06 | 9.300E-06 |
| 16363 | 819 | T819N | B | 1.3e-05 | 6.7e-06 | 1.3e-05 | 1.090E-05 | 3.637E-06 | 6.700E-06 | 1.300E-05 |
| 16364 | 819 | T819D | B | 1e-05 | 8.9e-06 | 6.7e-06 | 8.533E-06 | 1.680E-06 | 6.700E-06 | 1.000E-05 |
| 16365 | 819 | T819C | B | 8.7e-06 | 6.2e-06 | 8.1e-06 | 7.667E-06 | 1.305E-06 | 6.200E-06 | 8.700E-06 |
| 16366 | 819 | T819E | B | 9e-06 | 5.9e-06 | 6e-06 | 6.967E-06 | 1.762E-06 | 5.900E-06 | 9.000E-06 |
| 16367 | 819 | T819Q | B | 6.6e-06 | 1e-05 | 5.4e-06 | 7.333E-06 | 2.386E-06 | 5.400E-06 | 1.000E-05 |
| 16368 | 819 | T819G | B | 7.8e-06 | 9.7e-06 | 9.5e-06 | 9.000E-06 | 1.044E-06 | 7.800E-06 | 9.700E-06 |
| 16369 | 819 | T819H | B | 9.9e-06 | 6.3e-06 | 8.8e-06 | 8.333E-06 | 1.845E-06 | 6.300E-06 | 9.900E-06 |
| 16370 | 819 | T819I | B | 8.5e-06 | 5.3e-06 | 7.7e-06 | 7.167E-06 | 1.665E-06 | 5.300E-06 | 8.500E-06 |
| 16371 | 819 | T819L | B | 8.2e-06 | 5.8e-06 | 7.1e-06 | 7.033E-06 | 1.201E-06 | 5.800E-06 | 8.200E-06 |
| 16372 | 819 | T819K | B | 1.1e-05 | 7.8e-06 | 1.1e-05 | 9.933E-06 | 1.848E-06 | 7.800E-06 | 1.100E-05 |
| 16373 | 819 | T819M | B | 1e-05 | 6.6e-06 | 1.1e-05 | 9.200E-06 | 2.307E-06 | 6.600E-06 | 1.100E-05 |
| 16374 | 819 | T819F | B | 1e-05 | 6.3e-06 | 9.7e-06 | 8.667E-06 | 2.055E-06 | 6.300E-06 | 1.000E-05 |
| 16375 | 819 | T819P | B | 1.2e-05 | 6.6e-06 | 9.2e-06 | 9.267E-06 | 2.701E-06 | 6.600E-06 | 1.200E-05 |
| 16376 | 819 | T819S | B | 5.3e-06 | 7.4e-06 | 7.8e-06 | 6.833E-06 | 1.343E-06 | 5.300E-06 | 7.800E-06 |
| 16377 | 819 | T819T | B | 9.3e-06 | 3.4e-06 | 1e-05 | 7.567E-06 | 3.625E-06 | 3.400E-06 | 1.000E-05 |
| 16378 | 819 | T819W | B | 1.2e-05 | 6.7e-06 | 8.6e-06 | 9.100E-06 | 2.685E-06 | 6.700E-06 | 1.200E-05 |
| 16379 | 819 | T819Y | B | 5.6e-06 | 8.8e-06 | 4e-06 | 6.133E-06 | 2.444E-06 | 4.000E-06 | 8.800E-06 |
| 16380 | 819 | T819V | B | 1e-05 | 1.4e-05 | 2e-05 | 1.467E-05 | 5.033E-06 | 1.000E-05 | 2.000E-05 |
| 16381 | 820 | H820A | B | 1.4e-05 | 7.5e-06 | 5.9e-06 | 9.133E-06 | 4.290E-06 | 5.900E-06 | 1.400E-05 |
| 16382 | 820 | H820R | B | 1.3e-05 | 8.6e-06 | 7.9e-06 | 9.833E-06 | 2.765E-06 | 7.900E-06 | 1.300E-05 |
| 16383 | 820 | H820N | B | 8.9e-06 | 1.1e-05 | 7.3e-06 | 9.067E-06 | 1.856E-06 | 7.300E-06 | 1.100E-05 |
| 16384 | 820 | H820D | B | 5.3e-06 | 1e-05 | 5.8e-06 | 7.033E-06 | 2.581E-06 | 5.300E-06 | 1.000E-05 |
| 16385 | 820 | H820C | B | 6.5e-06 | 1.1e-05 | 1.2e-05 | 9.833E-06 | 2.930E-06 | 6.500E-06 | 1.200E-05 |
| 16386 | 820 | H820E | B | 9e-06 | 7e-06 | 4.6e-06 | 6.867E-06 | 2.203E-06 | 4.600E-06 | 9.000E-06 |
| 16387 | 820 | H820Q | B | 9.9e-06 | 5.5e-06 | 2e-05 | 1.180E-05 | 7.434E-06 | 5.500E-06 | 2.000E-05 |
| 16388 | 820 | H820G | B | 4.6e-06 | 4.5e-06 | 9.5e-06 | 6.200E-06 | 2.858E-06 | 4.500E-06 | 9.500E-06 |
| 16389 | 820 | H820H | B | 6.3e-06 | 9.6e-06 | 7.4e-06 | 7.767E-06 | 1.680E-06 | 6.300E-06 | 9.600E-06 |
| 16390 | 820 | H820I | B | 4.8e-06 | 1.1e-05 | 1e-05 | 8.600E-06 | 3.329E-06 | 4.800E-06 | 1.100E-05 |
| 16391 | 820 | H820L | B | 1.2e-05 | 1.3e-05 | 1.1e-05 | 1.200E-05 | 1.000E-06 | 1.100E-05 | 1.300E-05 |
| 16392 | 820 | H820K | B | 7e-06 | 5.4e-06 | 1.5e-05 | 9.133E-06 | 5.143E-06 | 5.400E-06 | 1.500E-05 |
| 16393 | 820 | H820M | B | 9.8e-06 | 9.1e-06 | 1e-05 | 9.633E-06 | 4.726E-07 | 9.100E-06 | 1.000E-05 |
| 16394 | 820 | H820F | B | 9.1e-06 | 9.9e-06 | 9.8e-06 | 9.600E-06 | 4.359E-07 | 9.100E-06 | 9.900E-06 |
| 16395 | 820 | H820P | B | 4.5e-06 | 9.2e-06 | 5.4e-06 | 6.367E-06 | 2.495E-06 | 4.500E-06 | 9.200E-06 |
| 16396 | 820 | H820S | B | 8e-06 | 9.3e-06 | 7e-06 | 8.100E-06 | 1.153E-06 | 7.000E-06 | 9.300E-06 |
| 16397 | 820 | H820T | B | 8.4e-06 | 6.9e-06 | 5.5e-06 | 6.933E-06 | 1.450E-06 | 5.500E-06 | 8.400E-06 |
| 16398 | 820 | H820W | B | 9.3e-06 | 1.2e-05 | 1.3e-05 | 1.143E-05 | 1.914E-06 | 9.300E-06 | 1.300E-05 |
| 16399 | 820 | H820Y | B | 9.5e-06 | 8.7e-06 | 5.9e-06 | 8.033E-06 | 1.890E-06 | 5.900E-06 | 9.500E-06 |
| 16400 | 820 | H820V | B | 7.9e-06 | 5.7e-06 | 1.2e-05 | 8.533E-06 | 3.197E-06 | 5.700E-06 | 1.200E-05 |
| 16401 | 821 | Q821A | B | 6.3e-06 | 1.1e-05 | 1e-05 | 9.100E-06 | 2.476E-06 | 6.300E-06 | 1.100E-05 |
| 16402 | 821 | Q821R | B | 3.8e-06 | 8.5e-06 | 6.5e-06 | 6.267E-06 | 2.359E-06 | 3.800E-06 | 8.500E-06 |
| 16403 | 821 | Q821N | B | 6.8e-06 | 7.3e-06 | 7.1e-06 | 7.067E-06 | 2.517E-07 | 6.800E-06 | 7.300E-06 |
| 16404 | 821 | Q821D | B | 5.4e-06 | 4.3e-06 | 6.8e-06 | 5.500E-06 | 1.253E-06 | 4.300E-06 | 6.800E-06 |

| | | | | | | | | | | |
|---|---|---|---|---|---|---|---|---|---|---|
| 16405 | 821 | Q821C | B | 6.6e-06 | 5e-06 | 5e-06 | 5.533E-06 | 9.238E-07 | 5.000E-06 | 6.600E-06 |
| 16406 | 821 | Q821E | B | 7.2e-06 | 9.5e-06 | 7.8e-06 | 8.167E-06 | 1.193E-06 | 7.200E-06 | 9.500E-06 |
| 16407 | 821 | Q821Q | B | 7.4e-06 | 6.5e-06 | 4.9e-06 | 6.267E-06 | 1.266E-06 | 4.900E-06 | 7.400E-06 |
| 16408 | 821 | Q821G | B | 5.9e-06 | 1.5e-05 | 1.3e-05 | 1.130E-05 | 4.782E-06 | 5.900E-06 | 1.500E-05 |
| 16409 | 821 | Q821H | B | 6e-06 | 5.9e-06 | 5.7e-06 | 5.867E-06 | 1.528E-07 | 5.700E-06 | 6.000E-06 |
| 16410 | 821 | Q821I | B | 9.8e-06 | 8.9e-06 | 5e-06 | 7.900E-06 | 2.551E-06 | 5.000E-06 | 9.800E-06 |
| 16411 | 821 | Q821L | B | 7.3e-06 | 9.8e-06 | 6.7e-06 | 7.933E-06 | 1.644E-06 | 6.700E-06 | 9.800E-06 |
| 16412 | 821 | Q821K | B | 6.5e-06 | 5.9e-06 | 9.1e-06 | 7.167E-06 | 1.701E-06 | 5.900E-06 | 9.100E-06 |
| 16413 | 821 | Q821M | B | 5.2e-06 | 1e-05 | 6.3e-06 | 7.167E-06 | 2.515E-06 | 5.200E-06 | 1.000E-05 |
| 16414 | 821 | Q821F | B | 1.1e-05 | 5.3e-06 | 6.8e-06 | 7.700E-06 | 2.955E-06 | 5.300E-06 | 1.100E-05 |
| 16415 | 821 | Q821P | B | 1.4e-05 | 8.9e-06 | 1e-05 | 1.097E-05 | 2.684E-06 | 8.900E-06 | 1.400E-05 |
| 16416 | 821 | Q821S | B | 6.8e-06 | 5.3e-06 | 1.2e-05 | 8.033E-06 | 3.516E-06 | 5.300E-06 | 1.200E-05 |
| 16417 | 821 | Q821T | B | 1.2e-05 | 6.4e-06 | 5.9e-06 | 8.100E-06 | 3.387E-06 | 5.900E-06 | 1.200E-05 |
| 16418 | 821 | Q821W | B | 8.9e-06 | 9.1e-06 | 6.8e-06 | 8.267E-06 | 1.274E-06 | 6.800E-06 | 9.100E-06 |
| 16419 | 821 | Q821Y | B | 1.1e-05 | 7.4e-06 | 6.8e-06 | 8.400E-06 | 2.272E-06 | 6.800E-06 | 1.100E-05 |
| 16420 | 821 | Q821V | B | 4.8e-06 | 9e-06 | 5.9e-06 | 6.567E-06 | 2.178E-06 | 4.800E-06 | 9.000E-06 |
| 16421 | 822 | G822A | B | 5.4e-06 | 1e-05 | 1.3e-05 | 9.467E-06 | 3.828E-06 | 5.400E-06 | 1.300E-05 |
| 16422 | 822 | G822R | B | 7.4e-06 | 8.4e-06 | 1.2e-05 | 9.267E-06 | 2.419E-06 | 7.400E-06 | 1.200E-05 |
| 16423 | 822 | G822N | B | 7.7e-06 | 8.9e-06 | 4.3e-06 | 6.967E-06 | 2.386E-06 | 4.300E-06 | 8.900E-06 |
| 16424 | 822 | G822D | B | 7.7e-06 | 7.5e-06 | 1e-05 | 8.400E-06 | 1.389E-06 | 7.500E-06 | 1.000E-05 |
| 16425 | 822 | G822C | B | 1.5e-05 | 1.9e-05 | 7.2e-06 | 1.373E-05 | 6.001E-06 | 7.200E-06 | 1.900E-05 |
| 16426 | 822 | G822E | B | 5.5e-06 | 4.8e-06 | 1.2e-05 | 7.433E-06 | 3.970E-06 | 4.800E-06 | 1.200E-05 |
| 16427 | 822 | G822Q | B | 1.4e-05 | 9.3e-06 | 5.8e-06 | 9.700E-06 | 4.115E-06 | 5.800E-06 | 1.400E-05 |
| 16428 | 822 | G822G | B | 1.8e-05 | 5e-06 | 1.1e-05 | 1.133E-05 | 6.506E-06 | 5.000E-06 | 1.800E-05 |
| 16429 | 822 | G822H | B | 7e-06 | 6.1e-06 | 9.7e-06 | 7.600E-06 | 1.873E-06 | 6.100E-06 | 9.700E-06 |
| 16430 | 822 | G822I | B | 1.3e-05 | 9.9e-06 | 8e-06 | 1.030E-05 | 2.524E-06 | 8.000E-06 | 1.300E-05 |
| 16431 | 822 | G822L | B | 7.6e-06 | 8.4e-06 | 6.3e-06 | 7.433E-06 | 1.060E-06 | 6.300E-06 | 8.400E-06 |
| 16432 | 822 | G822K | B | 8.5e-06 | 5.9e-06 | 7e-06 | 7.133E-06 | 1.305E-06 | 5.900E-06 | 8.500E-06 |
| 16433 | 822 | G822M | B | 1.4e-05 | 7.8e-06 | 7.9e-06 | 9.900E-06 | 3.551E-06 | 7.800E-06 | 1.400E-05 |
| 16434 | 822 | G822F | B | 1.1e-05 | 7.6e-06 | 8.4e-06 | 9.000E-06 | 1.778E-06 | 7.600E-06 | 1.100E-05 |
| 16435 | 822 | G822P | B | 1.1e-05 | 6.8e-06 | 7.1e-06 | 8.300E-06 | 2.343E-06 | 6.800E-06 | 1.100E-05 |
| 16436 | 822 | G822S | B | 7.7e-06 | 9.3e-06 | 1.1e-05 | 9.333E-06 | 1.650E-06 | 7.700E-06 | 1.100E-05 |
| 16437 | 822 | G822T | B | 9.8e-06 | 1.2e-05 | 5e-06 | 8.933E-06 | 3.580E-06 | 5.000E-06 | 1.200E-05 |
| 16438 | 822 | G822W | B | 1.2e-05 | 4.8e-06 | 1.2e-05 | 9.600E-06 | 4.157E-06 | 4.800E-06 | 1.200E-05 |
| 16439 | 822 | G822Y | B | 8e-06 | 8.3e-06 | 8.9e-06 | 8.400E-06 | 4.583E-07 | 8.000E-06 | 8.900E-06 |
| 16440 | 822 | G822V | B | 6.2e-06 | 9.6e-06 | 6.3e-06 | 7.367E-06 | 1.935E-06 | 6.200E-06 | 9.600E-06 |
| 16441 | 823 | L823A | B | 7e-06 | 1.6e-05 | 7.7e-06 | 1.023E-05 | 5.006E-06 | 7.000E-06 | 1.600E-05 |
| 16442 | 823 | L823R | B | 7.5e-06 | 6.7e-06 | 7.5e-06 | 7.233E-06 | 4.619E-07 | 6.700E-06 | 7.500E-06 |
| 16443 | 823 | L823N | B | 7.5e-06 | 1.2e-05 | 6.3e-06 | 8.600E-06 | 3.005E-06 | 6.300E-06 | 1.200E-05 |
| 16444 | 823 | L823D | B | 5e-06 | 1.2e-05 | 9.2e-06 | 8.733E-06 | 3.523E-06 | 5.000E-06 | 1.200E-05 |
| 16445 | 823 | L823C | B | 3.1e-06 | 9.6e-06 | 1.3e-05 | 8.567E-06 | 5.030E-06 | 3.100E-06 | 1.300E-05 |
| 16446 | 823 | L823E | B | 6.4e-06 | 7e-06 | 5.7e-06 | 6.367E-06 | 6.506E-07 | 5.700E-06 | 7.000E-06 |
| 16447 | 823 | L823Q | B | 5.2e-06 | 8.3e-06 | 7.5e-06 | 7.000E-06 | 1.609E-06 | 5.200E-06 | 8.300E-06 |
| 16448 | 823 | L823G | B | 6e-06 | 9.7e-06 | 5.5e-06 | 7.067E-06 | 2.294E-06 | 5.500E-06 | 9.700E-06 |
| 16449 | 823 | L823H | B | 1.3e-05 | 6.3e-06 | 8.9e-06 | 9.400E-06 | 3.378E-06 | 6.300E-06 | 1.300E-05 |
| 16450 | 823 | L823I | B | 7.9e-06 | 8.7e-06 | 8.3e-06 | 8.300E-06 | 4.000E-07 | 7.900E-06 | 8.700E-06 |
| 16451 | 823 | L823L | B | 5.7e-06 | 1.3e-05 | 7.6e-06 | 8.767E-06 | 3.787E-06 | 5.700E-06 | 1.300E-05 |
| 16452 | 823 | L823K | B | 9.5e-06 | 5.7e-06 | 1.3e-05 | 9.400E-06 | 3.651E-06 | 5.700E-06 | 1.300E-05 |
| 16453 | 823 | L823M | B | 1.2e-05 | 8.9e-06 | 1e-05 | 1.030E-05 | 1.572E-06 | 8.900E-06 | 1.200E-05 |
| 16454 | 823 | L823F | B | 1.1e-05 | 7.3e-06 | 4.2e-06 | 7.500E-06 | 3.404E-06 | 4.200E-06 | 1.100E-05 |
| 16455 | 823 | L823P | B | 1.2e-05 | 5.2e-06 | 4.4e-06 | 7.200E-06 | 4.176E-06 | 4.400E-06 | 1.200E-05 |
| 16456 | 823 | L823S | B | 1.3e-05 | 7.7e-06 | 5.3e-06 | 8.667E-06 | 3.940E-06 | 5.300E-06 | 1.300E-05 |
| 16457 | 823 | L823T | B | 1.3e-05 | 8e-06 | 6e-06 | 9.000E-06 | 3.606E-06 | 6.000E-06 | 1.300E-05 |
| 16458 | 823 | L823W | B | 6e-06 | 9.9e-06 | 7e-06 | 7.633E-06 | 2.026E-06 | 6.000E-06 | 9.900E-06 |
| 16459 | 823 | L823Y | B | 6.5e-06 | 1.1e-05 | 8.6e-06 | 8.700E-06 | 2.252E-06 | 6.500E-06 | 1.100E-05 |
| 16460 | 823 | L823V | B | 1.5e-05 | 1.6e-05 | 9.2e-06 | 1.340E-05 | 3.672E-06 | 9.200E-06 | 1.600E-05 |
| 16461 | 824 | S824A | B | 6.4e-06 | 9.6e-06 | 5.9e-06 | 7.300E-06 | 2.007E-06 | 5.900E-06 | 9.600E-06 |
| 16462 | 824 | S824R | B | 6.8e-06 | 3.4e-06 | 7e-06 | 5.733E-06 | 2.023E-06 | 3.400E-06 | 7.000E-06 |
| 16463 | 824 | S824N | B | 8.3e-06 | 9.9e-06 | 4.6e-06 | 7.600E-06 | 2.718E-06 | 4.600E-06 | 9.900E-06 |
| 16464 | 824 | S824D | B | 4.3e-06 | 4.3e-06 | 1.3e-05 | 7.200E-06 | 5.023E-06 | 4.300E-06 | 1.300E-05 |
| 16465 | 824 | S824C | B | 9.4e-06 | 6.2e-06 | 9.8e-06 | 8.467E-06 | 1.973E-06 | 6.200E-06 | 9.800E-06 |
| 16466 | 824 | S824E | B | 7.2e-06 | 1.1e-05 | 6.8e-06 | 8.333E-06 | 2.318E-06 | 6.800E-06 | 1.100E-05 |
| 16467 | 824 | S824Q | B | 7.6e-06 | 1.2e-05 | 4.6e-06 | 8.067E-06 | 3.722E-06 | 4.600E-06 | 1.200E-05 |
| 16468 | 824 | S824G | B | 5.2e-06 | 9e-06 | 7.4e-06 | 7.200E-06 | 1.908E-06 | 5.200E-06 | 9.000E-06 |
| 16469 | 824 | S824H | B | 8.7e-06 | 1.1e-05 | 1.3e-05 | 1.090E-05 | 2.152E-06 | 8.700E-06 | 1.300E-05 |
| 16470 | 824 | S824I | B | 8.3e-06 | 5.4e-06 | 7.2e-06 | 6.967E-06 | 1.464E-06 | 5.400E-06 | 8.300E-06 |
| 16471 | 824 | S824L | B | 1.4e-05 | 1.9e-05 | 1.1e-05 | 1.467E-05 | 4.041E-06 | 1.100E-05 | 1.900E-05 |
| 16472 | 824 | S824K | B | 6.9e-06 | 4.8e-06 | 5.8e-06 | 5.833E-06 | 1.050E-06 | 4.800E-06 | 6.900E-06 |
| 16473 | 824 | S824M | B | 5.5e-06 | 7.6e-06 | 9.6e-06 | 7.567E-06 | 2.050E-06 | 5.500E-06 | 9.600E-06 |
| 16474 | 824 | S824F | B | 5.9e-06 | 6.2e-06 | 7.7e-06 | 6.600E-06 | 9.644E-07 | 5.900E-06 | 7.700E-06 |
| 16475 | 824 | S824P | B | 4.9e-06 | 1.2e-05 | 7.7e-06 | 8.200E-06 | 3.576E-06 | 4.900E-06 | 1.200E-05 |
| 16476 | 824 | S824S | B | 6.2e-06 | 8.8e-06 | 1.1e-05 | 8.667E-06 | 2.403E-06 | 6.200E-06 | 1.100E-05 |
| 16477 | 824 | S824T | B | 6.8e-06 | 4.3e-06 | 5.7e-06 | 5.600E-06 | 1.253E-06 | 4.300E-06 | 6.800E-06 |
| 16478 | 824 | S824W | B | 5.2e-06 | 1.2e-05 | 6.9e-06 | 8.033E-06 | 3.539E-06 | 5.200E-06 | 1.200E-05 |
| 16479 | 824 | S824Y | B | 9.6e-06 | 6.6e-06 | 6.3e-06 | 7.500E-06 | 1.825E-06 | 6.300E-06 | 9.600E-06 |
| 16480 | 824 | S824V | B | 9.3e-06 | 7.4e-06 | 8e-06 | 8.233E-06 | 9.713E-07 | 7.400E-06 | 9.300E-06 |
| 16481 | 825 | S825A | B | 4.7e-06 | 1.1e-05 | 5.2e-06 | 6.967E-06 | 3.502E-06 | 4.700E-06 | 1.100E-05 |
| 16482 | 825 | S825R | B | 9e-06 | 8.4e-06 | 1.3e-05 | 1.013E-05 | 2.501E-06 | 8.400E-06 | 1.300E-05 |
| 16483 | 825 | S825N | B | 9.1e-06 | 4.7e-06 | 1.7e-05 | 1.027E-05 | 6.232E-06 | 4.700E-06 | 1.700E-05 |
| 16484 | 825 | S825D | B | 8.9e-06 | 1e-05 | 9.4e-06 | 9.433E-06 | 5.508E-07 | 8.900E-06 | 1.000E-05 |
| 16485 | 825 | S825C | B | 5.6e-06 | 6.8e-06 | 1e-05 | 7.467E-06 | 2.274E-06 | 5.600E-06 | 1.000E-05 |
| 16486 | 825 | S825E | B | 7.8e-06 | 8.9e-06 | 1.1e-05 | 9.233E-06 | 1.626E-06 | 7.800E-06 | 1.100E-05 |
| 16487 | 825 | S825Q | B | 8.2e-06 | 6e-06 | 1.1e-05 | 8.400E-06 | 2.506E-06 | 6.000E-06 | 1.100E-05 |
| 16488 | 825 | S825G | B | 6.3e-06 | 9.9e-06 | 8.6e-06 | 8.267E-06 | 1.823E-06 | 6.300E-06 | 9.900E-06 |
| 16489 | 825 | S825H | B | 4.5e-06 | 7e-06 | 6.2e-06 | 5.900E-06 | 1.277E-06 | 4.500E-06 | 7.000E-06 |

| | | | | | | | | | | |
|---|---|---|---|---|---|---|---|---|---|---|
| 16490 | 825 | S825I | B | 9.3e-06 | 6.3e-06 | 9.4e-06 | 8.333E-06 | 1.762E-06 | 6.300E-06 | 9.400E-06 |
| 16491 | 825 | S825L | B | 5.8e-06 | 1.2e-05 | 9.2e-06 | 9.000E-06 | 3.105E-06 | 5.800E-06 | 1.200E-05 |
| 16492 | 825 | S825K | B | 7.1e-06 | 1e-05 | 9.4e-06 | 8.833E-06 | 1.531E-06 | 7.100E-06 | 1.000E-05 |
| 16493 | 825 | S825M | B | 1.9e-05 | 4.5e-06 | 5.8e-06 | 9.767E-06 | 8.023E-06 | 4.500E-06 | 1.900E-05 |
| 16494 | 825 | S825F | B | 9.3e-06 | 5.1e-06 | 5.2e-06 | 6.533E-06 | 2.397E-06 | 5.100E-06 | 9.300E-06 |
| 16495 | 825 | S825P | B | 9.5e-06 | 9.7e-06 | 5.5e-06 | 8.233E-06 | 2.369E-06 | 5.500E-06 | 9.700E-06 |
| 16496 | 825 | S825S | B | 9.4e-06 | 1e-05 | 5.2e-06 | 8.200E-06 | 2.615E-06 | 5.200E-06 | 1.000E-05 |
| 16497 | 825 | S825T | B | 9.9e-06 | 6.4e-06 | 8.1e-06 | 8.133E-06 | 1.750E-06 | 6.400E-06 | 9.900E-06 |
| 16498 | 825 | S825W | B | 7.4e-06 | 1e-05 | 6.1e-06 | 7.833E-06 | 1.986E-06 | 6.100E-06 | 1.000E-05 |
| 16499 | 825 | S825Y | B | 7.4e-06 | 6.9e-06 | 1.2e-05 | 8.767E-06 | 2.811E-06 | 6.900E-06 | 1.200E-05 |
| 16500 | 825 | S825V | B | 7.3e-06 | 7.6e-06 | 7.5e-06 | 7.467E-06 | 1.528E-07 | 7.300E-06 | 7.600E-06 |
| 16501 | 826 | P826A | B | 9e-06 | 9.4e-06 | 9.1e-06 | 9.167E-06 | 2.082E-07 | 9.000E-06 | 9.400E-06 |
| 16502 | 826 | P826R | B | 9e-06 | 1.1e-05 | 8.6e-06 | 9.533E-06 | 1.286E-06 | 8.600E-06 | 1.100E-05 |
| 16503 | 826 | P826N | B | 4.2e-06 | 8.7e-06 | 5.1e-06 | 6.000E-06 | 2.381E-06 | 4.200E-06 | 8.700E-06 |
| 16504 | 826 | P826D | B | 7.1e-06 | 9.6e-06 | 6.6e-06 | 7.767E-06 | 1.607E-06 | 6.600E-06 | 9.600E-06 |
| 16505 | 826 | P826C | B | 9.6e-06 | 7.4e-06 | 8.6e-06 | 8.533E-06 | 1.102E-06 | 7.400E-06 | 9.600E-06 |
| 16506 | 826 | P826E | B | 4.8e-06 | 1e-05 | 7.2e-06 | 7.333E-06 | 2.603E-06 | 4.800E-06 | 1.000E-05 |
| 16507 | 826 | P826Q | B | 1.1e-05 | 6.8e-06 | 1.2e-05 | 9.933E-06 | 2.759E-06 | 6.800E-06 | 1.200E-05 |
| 16508 | 826 | P826G | B | 9.7e-06 | 4.9e-06 | 1.1e-05 | 8.533E-06 | 3.213E-06 | 4.900E-06 | 1.100E-05 |
| 16509 | 826 | P826H | B | 6.2e-06 | 8.4e-06 | 8.7e-06 | 7.767E-06 | 1.365E-06 | 6.200E-06 | 8.700E-06 |
| 16510 | 826 | P826I | B | 6.7e-06 | 1.3e-05 | 9.8e-06 | 9.833E-06 | 3.150E-06 | 6.700E-06 | 1.300E-05 |
| 16511 | 826 | P826L | B | 6.4e-06 | 4.3e-06 | 9.8e-06 | 6.833E-06 | 2.775E-06 | 4.300E-06 | 9.800E-06 |
| 16512 | 826 | P826K | B | 7.7e-06 | 5.9e-06 | 4.4e-06 | 6.000E-06 | 1.652E-06 | 4.400E-06 | 7.700E-06 |
| 16513 | 826 | P826M | B | 1.4e-05 | 7.3e-06 | 9.7e-06 | 1.033E-05 | 3.395E-06 | 7.300E-06 | 1.400E-05 |
| 16514 | 826 | P826F | B | 6.3e-06 | 1.4e-05 | 8.7e-06 | 9.667E-06 | 3.940E-06 | 6.300E-06 | 1.400E-05 |
| 16515 | 826 | P826P | B | 1e-05 | 5.3e-06 | 1e-05 | 8.433E-06 | 2.714E-06 | 5.300E-06 | 1.000E-05 |
| 16516 | 826 | P826S | B | 6.7e-06 | 5.5e-06 | 6.3e-06 | 6.167E-06 | 6.110E-07 | 5.500E-06 | 6.700E-06 |
| 16517 | 826 | P826T | B | 6.5e-06 | 8.4e-06 | 3.1e-06 | 6.000E-06 | 2.685E-06 | 3.100E-06 | 8.400E-06 |
| 16518 | 826 | P826W | B | 7e-06 | 9.6e-06 | 7.2e-06 | 7.933E-06 | 1.447E-06 | 7.000E-06 | 9.600E-06 |
| 16519 | 826 | P826Y | B | 6.4e-06 | 4e-06 | 6.6e-06 | 5.667E-06 | 1.447E-06 | 4.000E-06 | 6.600E-06 |
| 16520 | 826 | P826V | B | 1.1e-05 | 6.3e-06 | 8.8e-06 | 8.700E-06 | 2.352E-06 | 6.300E-06 | 1.100E-05 |
| 16521 | 827 | V827A | B | 7.4e-06 | 6.9e-06 | 7.2e-06 | 7.167E-06 | 2.517E-07 | 6.900E-06 | 7.400E-06 |
| 16522 | 827 | V827R | B | 1.1e-05 | 4.5e-06 | 7.4e-06 | 7.633E-06 | 3.256E-06 | 4.500E-06 | 1.100E-05 |
| 16523 | 827 | V827N | B | 4.5e-06 | 1.2e-05 | 8.3e-06 | 8.267E-06 | 3.750E-06 | 4.500E-06 | 1.200E-05 |
| 16524 | 827 | V827D | B | 9e-06 | 7.1e-06 | 6.9e-06 | 7.667E-06 | 1.159E-06 | 6.900E-06 | 9.000E-06 |
| 16525 | 827 | V827C | B | 7.1e-06 | 1.4e-05 | 4.4e-06 | 8.500E-06 | 4.951E-06 | 4.400E-06 | 1.400E-05 |
| 16526 | 827 | V827E | B | 9.5e-06 | 8.2e-06 | 1.1e-05 | 9.567E-06 | 1.401E-06 | 8.200E-06 | 1.100E-05 |
| 16527 | 827 | V827Q | B | 8.3e-06 | 9.3e-06 | 7.4e-06 | 8.333E-06 | 9.504E-07 | 7.400E-06 | 9.300E-06 |
| 16528 | 827 | V827G | B | 8.5e-06 | 9.9e-06 | 7.8e-06 | 8.733E-06 | 1.069E-06 | 7.800E-06 | 9.900E-06 |
| 16529 | 827 | V827H | B | 9.2e-06 | 4.4e-06 | 8.1e-06 | 7.233E-06 | 2.515E-06 | 4.400E-06 | 9.200E-06 |
| 16530 | 827 | V827I | B | 5.5e-06 | 8.4e-06 | 7.2e-06 | 7.033E-06 | 1.457E-06 | 5.500E-06 | 8.400E-06 |
| 16531 | 827 | V827L | B | 1.1e-05 | 9.6e-06 | 6.3e-06 | 8.967E-06 | 2.413E-06 | 6.300E-06 | 1.100E-05 |
| 16532 | 827 | V827K | B | 5.1e-06 | 8.2e-06 | 9.4e-06 | 7.567E-06 | 2.219E-06 | 5.100E-06 | 9.400E-06 |
| 16533 | 827 | V827M | B | 6.3e-06 | 5.7e-06 | 9.3e-06 | 7.100E-06 | 1.929E-06 | 5.700E-06 | 9.300E-06 |
| 16534 | 827 | V827F | B | 5.3e-06 | 8.8e-06 | 5.8e-06 | 6.633E-06 | 1.893E-06 | 5.300E-06 | 8.800E-06 |
| 16535 | 827 | V827P | B | 1e-05 | 4.9e-06 | 9e-06 | 7.967E-06 | 2.702E-06 | 4.900E-06 | 1.000E-05 |
| 16536 | 827 | V827S | B | 6e-06 | 7.9e-06 | 1e-05 | 7.967E-06 | 2.001E-06 | 6.000E-06 | 1.000E-05 |
| 16537 | 827 | V827T | B | 3.5e-06 | 7.1e-06 | 7.5e-06 | 6.033E-06 | 2.203E-06 | 3.500E-06 | 7.500E-06 |
| 16538 | 827 | V827W | B | 8.2e-06 | 5.5e-06 | 9.7e-06 | 7.800E-06 | 2.128E-06 | 5.500E-06 | 9.700E-06 |
| 16539 | 827 | V827Y | B | 1.1e-05 | 7e-06 | 6.4e-06 | 8.133E-06 | 2.501E-06 | 6.400E-06 | 1.100E-05 |
| 16540 | 827 | V827V | B | 6.1e-06 | 7.6e-06 | 1.4e-05 | 9.233E-06 | 4.196E-06 | 6.100E-06 | 1.400E-05 |
| 16541 | 828 | T828A | B | 7.2e-06 | 1.4e-05 | 7.9e-06 | 9.700E-06 | 3.740E-06 | 7.200E-06 | 1.400E-05 |
| 16542 | 828 | T828R | B | 5.4e-06 | 7.5e-06 | 6.4e-06 | 6.433E-06 | 1.050E-06 | 5.400E-06 | 7.500E-06 |
| 16543 | 828 | T828N | B | 7.6e-06 | 8.8e-06 | 6.4e-06 | 7.600E-06 | 1.200E-06 | 6.400E-06 | 8.800E-06 |
| 16544 | 828 | T828D | B | 7.4e-06 | 6.3e-06 | 5.9e-06 | 6.533E-06 | 7.767E-07 | 5.900E-06 | 7.400E-06 |
| 16545 | 828 | T828C | B | 6.7e-06 | 7.3e-06 | 8.1e-06 | 7.367E-06 | 7.024E-07 | 6.700E-06 | 8.100E-06 |
| 16546 | 828 | T828E | B | 6.7e-06 | 7e-06 | 9.4e-06 | 7.700E-06 | 1.480E-06 | 6.700E-06 | 9.400E-06 |
| 16547 | 828 | T828Q | B | 7.9e-06 | 6.2e-06 | 5e-06 | 6.367E-06 | 1.457E-06 | 5.000E-06 | 7.900E-06 |
| 16548 | 828 | T828G | B | 6.8e-06 | 5.7e-06 | 9.8e-06 | 7.433E-06 | 2.122E-06 | 5.700E-06 | 9.800E-06 |
| 16549 | 828 | T828H | B | 9.8e-06 | 8.3e-06 | 7.5e-06 | 8.533E-06 | 1.168E-06 | 7.500E-06 | 9.800E-06 |
| 16550 | 828 | T828I | B | 1.2e-05 | 5.8e-06 | 1.4e-05 | 1.060E-05 | 4.276E-06 | 5.800E-06 | 1.400E-05 |
| 16551 | 828 | T828L | B | 6.4e-06 | 6.8e-06 | 9.3e-06 | 7.500E-06 | 1.572E-06 | 6.400E-06 | 9.300E-06 |
| 16552 | 828 | T828K | B | 9.4e-06 | 1.3e-05 | 4.5e-06 | 8.967E-06 | 4.267E-06 | 4.500E-06 | 1.300E-05 |
| 16553 | 828 | T828M | B | 1.3e-05 | 5.5e-06 | 1e-05 | 9.500E-06 | 3.775E-06 | 5.500E-06 | 1.300E-05 |
| 16554 | 828 | T828F | B | 6.6e-06 | 9.5e-06 | 1.1e-05 | 9.033E-06 | 2.237E-06 | 6.600E-06 | 1.100E-05 |
| 16555 | 828 | T828P | B | 6.1e-06 | 9.4e-06 | 5.1e-06 | 6.867E-06 | 2.250E-06 | 5.100E-06 | 9.400E-06 |
| 16556 | 828 | T828S | B | 5.9e-06 | 7.9e-06 | 6.9e-06 | 6.900E-06 | 1.000E-06 | 5.900E-06 | 7.900E-06 |
| 16557 | 828 | T828T | B | 8.5e-06 | 4.9e-06 | 1.1e-05 | 8.133E-06 | 3.066E-06 | 4.900E-06 | 1.100E-05 |
| 16558 | 828 | T828W | B | 9.9e-06 | 5.6e-06 | 7.8e-06 | 7.767E-06 | 2.150E-06 | 5.600E-06 | 9.900E-06 |
| 16559 | 828 | T828Y | B | 1e-05 | 9.3e-06 | 3.3e-06 | 7.533E-06 | 3.683E-06 | 3.300E-06 | 1.000E-05 |
| 16560 | 828 | T828V | B | 4.6e-06 | 9.4e-06 | 9.5e-06 | 7.833E-06 | 2.801E-06 | 4.600E-06 | 9.500E-06 |
| 16561 | 829 | K829A | B | 1e-05 | 5.8e-06 | 7e-06 | 7.600E-06 | 2.163E-06 | 5.800E-06 | 1.000E-05 |
| 16562 | 829 | K829R | B | 6.5e-06 | 8.8e-06 | 9.6e-06 | 8.300E-06 | 1.609E-06 | 6.500E-06 | 9.600E-06 |
| 16563 | 829 | K829N | B | 1.1e-05 | 6e-06 | 8.7e-06 | 8.567E-06 | 2.503E-06 | 6.000E-06 | 1.100E-05 |
| 16564 | 829 | K829D | B | 5.7e-06 | 9.6e-06 | 1.6e-05 | 1.043E-05 | 5.200E-06 | 5.700E-06 | 1.600E-05 |
| 16565 | 829 | K829C | B | 7.7e-06 | 8.2e-06 | 5.1e-06 | 7.000E-06 | 1.664E-06 | 5.100E-06 | 8.200E-06 |
| 16566 | 829 | K829E | B | 3.6e-06 | 8.4e-06 | 6.1e-06 | 6.033E-06 | 2.401E-06 | 3.600E-06 | 8.400E-06 |
| 16567 | 829 | K829Q | B | 1.1e-05 | 4.4e-06 | 7.8e-06 | 7.733E-06 | 3.301E-06 | 4.400E-06 | 1.100E-05 |
| 16568 | 829 | K829G | B | 1e-05 | 7.6e-06 | 1.1e-05 | 9.533E-06 | 1.747E-06 | 7.600E-06 | 1.100E-05 |
| 16569 | 829 | K829H | B | 9e-06 | 7.3e-06 | 1.1e-05 | 9.100E-06 | 1.852E-06 | 7.300E-06 | 1.100E-05 |
| 16570 | 829 | K829I | B | 6.1e-06 | 7e-06 | 1.3e-05 | 8.700E-06 | 3.751E-06 | 6.100E-06 | 1.300E-05 |
| 16571 | 829 | K829L | B | 9.5e-06 | 4.4e-06 | 9.5e-06 | 7.800E-06 | 2.944E-06 | 4.400E-06 | 9.500E-06 |
| 16572 | 829 | K829K | B | 4.5e-06 | 5.5e-06 | 9e-06 | 6.333E-06 | 2.363E-06 | 4.500E-06 | 9.000E-06 |
| 16573 | 829 | K829M | B | 5.8e-06 | 7.1e-06 | 7.7e-06 | 6.867E-06 | 9.713E-07 | 5.800E-06 | 7.700E-06 |
| 16574 | 829 | K829F | B | 8.9e-06 | 5e-06 | 7.8e-06 | 7.233E-06 | 2.011E-06 | 5.000E-06 | 8.900E-06 |

| | | | | | | | | | | |
|---|---|---|---|---|---|---|---|---|---|---|
| 16575 | 829 | K829P | B | 5.1e-06 | 6.2e-06 | 9.4e-06 | 6.900E-06 | 2.234E-06 | 5.100E-06 | 9.400E-06 |
| 16576 | 829 | K829S | B | 9e-06 | 8.2e-06 | 8.9e-06 | 8.700E-06 | 4.359E-07 | 8.200E-06 | 9.000E-06 |
| 16577 | 829 | K829T | B | 8.5e-06 | 6.7e-06 | 1.1e-05 | 8.733E-06 | 2.159E-06 | 6.700E-06 | 1.100E-05 |
| 16578 | 829 | K829W | B | 9.8e-06 | 6.1e-06 | 1.1e-05 | 8.967E-06 | 2.554E-06 | 6.100E-06 | 1.100E-05 |
| 16579 | 829 | K829Y | B | 3.7e-06 | 4.1e-06 | 8.1e-06 | 5.300E-06 | 2.433E-06 | 3.700E-06 | 8.100E-06 |
| 16580 | 829 | K829V | B | 8.6e-06 | 7.5e-06 | 9.7e-06 | 8.600E-06 | 1.100E-06 | 7.500E-06 | 9.700E-06 |
| 16581 | 830 | S830A | B | 1.3e-05 | 6.8e-06 | 9.1e-06 | 9.633E-06 | 3.134E-06 | 6.800E-06 | 1.300E-05 |
| 16582 | 830 | S830R | B | 6.1e-06 | 7.9e-06 | 5.3e-06 | 6.433E-06 | 1.332E-06 | 5.300E-06 | 7.900E-06 |
| 16583 | 830 | S830N | B | 7.3e-06 | 1e-05 | 5.9e-06 | 7.733E-06 | 2.084E-06 | 5.900E-06 | 1.000E-05 |
| 16584 | 830 | S830D | B | 6.7e-06 | 7.7e-06 | 6.7e-06 | 7.033E-06 | 5.774E-07 | 6.700E-06 | 7.700E-06 |
| 16585 | 830 | S830C | B | 7.3e-06 | 9.7e-06 | 6e-06 | 7.667E-06 | 1.877E-06 | 6.000E-06 | 9.700E-06 |
| 16586 | 830 | S830E | B | 7.7e-06 | 1e-05 | 6.1e-06 | 7.933E-06 | 1.960E-06 | 6.100E-06 | 1.000E-05 |
| 16587 | 830 | S830Q | B | 1.1e-05 | 1.1e-05 | 5.8e-06 | 9.267E-06 | 3.002E-06 | 5.800E-06 | 1.100E-05 |
| 16588 | 830 | S830G | B | 1.2e-05 | 4.3e-06 | 8.9e-06 | 8.400E-06 | 3.874E-06 | 4.300E-06 | 1.200E-05 |
| 16589 | 830 | S830H | B | 9.7e-06 | 7.9e-06 | 7e-06 | 8.200E-06 | 1.375E-06 | 7.000E-06 | 9.700E-06 |
| 16590 | 830 | S830I | B | 1.1e-05 | 7.9e-06 | 8.4e-06 | 9.100E-06 | 1.664E-06 | 7.900E-06 | 1.100E-05 |
| 16591 | 830 | S830L | B | 1.3e-05 | 6.4e-06 | 1.2e-05 | 1.047E-05 | 3.557E-06 | 6.400E-06 | 1.300E-05 |
| 16592 | 830 | S830K | B | 8.8e-06 | 8.2e-06 | 7.2e-06 | 8.067E-06 | 8.083E-07 | 7.200E-06 | 8.800E-06 |
| 16593 | 830 | S830M | B | 1.4e-05 | 1.5e-05 | 1.2e-05 | 1.367E-05 | 1.528E-06 | 1.200E-05 | 1.500E-05 |
| 16594 | 830 | S830F | B | 6.3e-06 | 1e-05 | 9.4e-06 | 8.567E-06 | 1.986E-06 | 6.300E-06 | 1.000E-05 |
| 16595 | 830 | S830P | B | 5.3e-06 | 1e-05 | 7e-06 | 7.433E-06 | 2.380E-06 | 5.300E-06 | 1.000E-05 |
| 16596 | 830 | S830S | B | 8.1e-06 | 6e-06 | 6.8e-06 | 6.967E-06 | 1.060E-06 | 6.000E-06 | 8.100E-06 |
| 16597 | 830 | S830T | B | 7.5e-06 | 1.6e-05 | 1.2e-05 | 1.183E-05 | 4.252E-06 | 7.500E-06 | 1.600E-05 |
| 16598 | 830 | S830W | B | 7.1e-06 | 1.2e-05 | 7.7e-06 | 8.933E-06 | 2.673E-06 | 7.100E-06 | 1.200E-05 |
| 16599 | 830 | S830Y | B | 1.3e-05 | 7.5e-06 | 1.1e-05 | 1.050E-05 | 2.784E-06 | 7.500E-06 | 1.300E-05 |
| 16600 | 830 | S830V | B | 8.6e-06 | 5.5e-06 | 7.3e-06 | 7.133E-06 | 1.557E-06 | 5.500E-06 | 8.600E-06 |
| 16601 | 831 | F831A | B | 9.2e-06 | 1.3e-05 | 6.4e-06 | 9.533E-06 | 3.313E-06 | 6.400E-06 | 1.300E-05 |
| 16602 | 831 | F831R | B | 6.7e-06 | 9e-06 | 5.5e-06 | 7.067E-06 | 1.779E-06 | 5.500E-06 | 9.000E-06 |
| 16603 | 831 | F831N | B | 7.8e-06 | 9.2e-06 | 8e-06 | 8.333E-06 | 7.572E-07 | 7.800E-06 | 9.200E-06 |
| 16604 | 831 | F831D | B | 9.9e-06 | 8.6e-06 | 9.3e-06 | 9.267E-06 | 6.506E-07 | 8.600E-06 | 9.900E-06 |
| 16605 | 831 | F831C | B | 4.7e-06 | 8.8e-06 | 2.1e-05 | 1.150E-05 | 8.479E-06 | 4.700E-06 | 2.100E-05 |
| 16606 | 831 | F831E | B | 5.3e-06 | 8e-06 | 1e-05 | 7.767E-06 | 2.359E-06 | 5.300E-06 | 1.000E-05 |
| 16607 | 831 | F831Q | B | 7.7e-06 | 1.3e-05 | 7.7e-06 | 9.467E-06 | 3.060E-06 | 7.700E-06 | 1.300E-05 |
| 16608 | 831 | F831G | B | 7.3e-06 | 8.5e-06 | 9e-06 | 8.267E-06 | 8.737E-07 | 7.300E-06 | 9.000E-06 |
| 16609 | 831 | F831H | B | 8.4e-06 | 7.3e-06 | 1.3e-05 | 9.567E-06 | 3.024E-06 | 7.300E-06 | 1.300E-05 |
| 16610 | 831 | F831I | B | 5.6e-06 | 4e-06 | 1.1e-05 | 6.867E-06 | 3.668E-06 | 4.000E-06 | 1.100E-05 |
| 16611 | 831 | F831L | B | 9.9e-06 | 9.2e-06 | 5.7e-06 | 8.267E-06 | 2.250E-06 | 5.700E-06 | 9.900E-06 |
| 16612 | 831 | F831K | B | 6.9e-06 | 9.3e-06 | 3.9e-06 | 6.700E-06 | 2.706E-06 | 3.900E-06 | 9.300E-06 |
| 16613 | 831 | F831M | B | 9.2e-06 | 7.1e-06 | 4.1e-06 | 6.800E-06 | 2.563E-06 | 4.100E-06 | 9.200E-06 |
| 16614 | 831 | F831F | B | 7.3e-06 | 6.7e-06 | 1e-05 | 8.000E-06 | 1.758E-06 | 6.700E-06 | 1.000E-05 |
| 16615 | 831 | F831P | B | 4.9e-06 | 8.1e-06 | 7.4e-06 | 6.800E-06 | 1.682E-06 | 4.900E-06 | 8.100E-06 |
| 16616 | 831 | F831S | B | 1.8e-05 | 5.4e-06 | 8.4e-06 | 1.060E-05 | 6.582E-06 | 5.400E-06 | 1.800E-05 |
| 16617 | 831 | F831T | B | 6.6e-06 | 8.2e-06 | 6.4e-06 | 7.067E-06 | 9.866E-07 | 6.400E-06 | 8.200E-06 |
| 16618 | 831 | F831W | B | 5.8e-06 | 9.1e-06 | 4.3e-06 | 6.400E-06 | 2.456E-06 | 4.300E-06 | 9.100E-06 |
| 16619 | 831 | F831Y | B | 6.6e-06 | 1e-05 | 1.3e-05 | 9.867E-06 | 3.202E-06 | 6.600E-06 | 1.300E-05 |
| 16620 | 831 | F831V | B | 5.6e-06 | 8.5e-06 | 9.9e-06 | 8.000E-06 | 2.193E-06 | 5.600E-06 | 9.900E-06 |
| 16621 | 832 | N832A | B | 7.1e-06 | 1.1e-05 | 4.3e-06 | 7.467E-06 | 3.365E-06 | 4.300E-06 | 1.100E-05 |
| 16622 | 832 | N832R | B | 4.9e-06 | 1.4e-05 | 4.4e-06 | 7.767E-06 | 5.404E-06 | 4.400E-06 | 1.400E-05 |
| 16623 | 832 | N832N | B | 4.9e-06 | 9.7e-06 | 9.7e-06 | 8.100E-06 | 2.771E-06 | 4.900E-06 | 9.700E-06 |
| 16624 | 832 | N832D | B | 7.1e-06 | 8.4e-06 | 8.7e-06 | 8.067E-06 | 8.505E-07 | 7.100E-06 | 8.700E-06 |
| 16625 | 832 | N832C | B | 7.3e-06 | 4.3e-06 | 1.1e-05 | 7.533E-06 | 3.356E-06 | 4.300E-06 | 1.100E-05 |
| 16626 | 832 | N832E | B | 1.1e-05 | 7.4e-06 | 8.4e-06 | 8.933E-06 | 1.858E-06 | 7.400E-06 | 1.100E-05 |
| 16627 | 832 | N832Q | B | 1.4e-05 | 9e-06 | 7.7e-06 | 1.023E-05 | 3.326E-06 | 7.700E-06 | 1.400E-05 |
| 16628 | 832 | N832G | B | 7.1e-06 | 1e-05 | 1e-05 | 9.033E-06 | 1.674E-06 | 7.100E-06 | 1.000E-05 |
| 16629 | 832 | N832H | B | 1.3e-05 | 5.6e-06 | 8.5e-06 | 9.033E-06 | 3.729E-06 | 5.600E-06 | 1.300E-05 |
| 16630 | 832 | N832I | B | 6.8e-06 | 8.8e-06 | 7.9e-06 | 7.833E-06 | 1.002E-06 | 6.800E-06 | 8.800E-06 |
| 16631 | 832 | N832L | B | 9.4e-06 | 8.6e-06 | 8.4e-06 | 8.800E-06 | 5.292E-07 | 8.400E-06 | 9.400E-06 |
| 16632 | 832 | N832K | B | 1.3e-05 | 1e-05 | 5.3e-06 | 9.433E-06 | 3.881E-06 | 5.300E-06 | 1.300E-05 |
| 16633 | 832 | N832M | B | 7.4e-06 | 9.9e-06 | 6.3e-06 | 7.867E-06 | 1.845E-06 | 6.300E-06 | 9.900E-06 |
| 16634 | 832 | N832F | B | 6.1e-06 | 5.7e-06 | 9.3e-06 | 7.033E-06 | 1.973E-06 | 5.700E-06 | 9.300E-06 |
| 16635 | 832 | N832P | B | 7.8e-06 | 7.1e-06 | 4.8e-06 | 6.567E-06 | 1.570E-06 | 4.800E-06 | 7.800E-06 |
| 16636 | 832 | N832S | B | 5.5e-06 | 8.6e-06 | 1.2e-05 | 8.700E-06 | 3.251E-06 | 5.500E-06 | 1.200E-05 |
| 16637 | 832 | N832T | B | 7.5e-06 | 5.5e-06 | 1.5e-05 | 9.333E-06 | 5.008E-06 | 5.500E-06 | 1.500E-05 |
| 16638 | 832 | N832W | B | 8.9e-06 | 9.5e-06 | 5.6e-06 | 8.000E-06 | 2.100E-06 | 5.600E-06 | 9.500E-06 |
| 16639 | 832 | N832Y | B | 9.8e-06 | 5.6e-06 | 4e-06 | 6.467E-06 | 2.996E-06 | 4.000E-06 | 9.800E-06 |
| 16640 | 832 | N832V | B | 8.4e-06 | 6.7e-06 | 1.1e-05 | 8.700E-06 | 2.166E-06 | 6.700E-06 | 1.100E-05 |
| 16641 | 833 | R833A | B | 1e-05 | 6.8e-06 | 7e-06 | 7.933E-06 | 1.793E-06 | 6.800E-06 | 1.000E-05 |
| 16642 | 833 | R833R | B | 5.2e-06 | 8.8e-06 | 1e-05 | 8.000E-06 | 2.498E-06 | 5.200E-06 | 1.000E-05 |
| 16643 | 833 | R833N | B | 9.7e-06 | 1.4e-05 | 8e-06 | 1.057E-05 | 3.092E-06 | 8.000E-06 | 1.400E-05 |
| 16644 | 833 | R833D | B | 1e-05 | 5.5e-06 | 1.3e-05 | 9.500E-06 | 3.775E-06 | 5.500E-06 | 1.300E-05 |
| 16645 | 833 | R833C | B | 1.5e-05 | 8.8e-06 | 5.8e-06 | 9.867E-06 | 4.692E-06 | 5.800E-06 | 1.500E-05 |
| 16646 | 833 | R833E | B | 8.9e-06 | 8.3e-06 | 6.6e-06 | 7.933E-06 | 1.193E-06 | 6.600E-06 | 8.900E-06 |
| 16647 | 833 | R833Q | B | 6.2e-06 | 4.6e-06 | 7.7e-06 | 6.167E-06 | 1.550E-06 | 4.600E-06 | 7.700E-06 |
| 16648 | 833 | R833G | B | 6.5e-06 | 9.2e-06 | 8.1e-06 | 7.933E-06 | 1.358E-06 | 6.500E-06 | 9.200E-06 |
| 16649 | 833 | R833H | B | 7.1e-06 | 7.1e-06 | 4.6e-06 | 6.267E-06 | 1.443E-06 | 4.600E-06 | 7.100E-06 |
| 16650 | 833 | R833I | B | 9.1e-06 | 5.4e-06 | 8.4e-06 | 7.633E-06 | 1.966E-06 | 5.400E-06 | 9.100E-06 |
| 16651 | 833 | R833L | B | 6e-06 | 7.8e-06 | 1e-05 | 7.933E-06 | 2.003E-06 | 6.000E-06 | 1.000E-05 |
| 16652 | 833 | R833K | B | 7.1e-06 | 6.9e-06 | 4.5e-06 | 6.167E-06 | 1.447E-06 | 4.500E-06 | 7.100E-06 |
| 16653 | 833 | R833M | B | 1.4e-05 | 1.3e-05 | 7.9e-06 | 1.163E-05 | 3.272E-06 | 7.900E-06 | 1.400E-05 |
| 16654 | 833 | R833F | B | 8.1e-06 | 1.3e-05 | 7.6e-06 | 9.567E-06 | 2.984E-06 | 7.600E-06 | 1.300E-05 |
| 16655 | 833 | R833P | B | 1.1e-05 | 8.9e-06 | 1.1e-05 | 1.030E-05 | 1.212E-06 | 8.900E-06 | 1.100E-05 |
| 16656 | 833 | R833S | B | 6.3e-06 | 4.8e-06 | 5.9e-06 | 5.667E-06 | 7.767E-07 | 4.800E-06 | 6.300E-06 |
| 16657 | 833 | R833T | B | 4.2e-06 | 4.7e-06 | 8.3e-06 | 5.733E-06 | 2.237E-06 | 4.200E-06 | 8.300E-06 |
| 16658 | 833 | R833W | B | 7e-06 | 4.4e-06 | 6.4e-06 | 5.933E-06 | 1.361E-06 | 4.400E-06 | 7.000E-06 |
| 16659 | 833 | R833Y | B | 5.7e-06 | 5.4e-06 | 8.2e-06 | 6.433E-06 | 1.537E-06 | 5.400E-06 | 8.200E-06 |

| | | | | | | | | | | |
|---|---|---|---|---|---|---|---|---|---|---|
| 16660 | 833 | R833V | B | 8.3e-06 | 8.8e-06 | 6.9e-06 | 8.000E-06 | 9.849E-07 | 6.900E-06 | 8.800E-06 |
| 16661 | 834 | G834A | B | 1.1e-05 | 5.7e-06 | 9.7e-06 | 8.800E-06 | 2.762E-06 | 5.700E-06 | 1.100E-05 |
| 16662 | 834 | G834R | B | 4.3e-06 | 4.9e-06 | 9.2e-06 | 6.133E-06 | 2.673E-06 | 4.300E-06 | 9.200E-06 |
| 16663 | 834 | G834N | B | 5.8e-06 | 5.1e-06 | 7.4e-06 | 6.100E-06 | 1.179E-06 | 5.100E-06 | 7.400E-06 |
| 16664 | 834 | G834D | B | 6.3e-06 | 8.1e-06 | 9.8e-06 | 8.067E-06 | 1.750E-06 | 6.300E-06 | 9.800E-06 |
| 16665 | 834 | G834C | B | 9.8e-06 | 8.6e-06 | 1.3e-05 | 1.047E-05 | 2.274E-06 | 8.600E-06 | 1.300E-05 |
| 16666 | 834 | G834E | B | 6.9e-06 | 1e-05 | 6.9e-06 | 7.933E-06 | 1.790E-06 | 6.900E-06 | 1.000E-05 |
| 16667 | 834 | G834Q | B | 6.2e-06 | 6.3e-06 | 4.6e-06 | 5.700E-06 | 9.539E-07 | 4.600E-06 | 6.300E-06 |
| 16668 | 834 | G834G | B | 1e-05 | 1e-05 | 8.3e-06 | 9.433E-06 | 9.815E-07 | 8.300E-06 | 1.000E-05 |
| 16669 | 834 | G834H | B | 6.5e-06 | 8.9e-06 | 8.9e-06 | 8.100E-06 | 1.386E-06 | 6.500E-06 | 8.900E-06 |
| 16670 | 834 | G834I | B | 5.1e-06 | 4.7e-06 | 6.6e-06 | 5.467E-06 | 1.002E-06 | 4.700E-06 | 6.600E-06 |
| 16671 | 834 | G834L | B | 9.2e-06 | 6.1e-06 | 3.4e-06 | 6.233E-06 | 2.902E-06 | 3.400E-06 | 9.200E-06 |
| 16672 | 834 | G834K | B | 7e-06 | 9.9e-06 | 7.5e-06 | 8.133E-06 | 1.550E-06 | 7.000E-06 | 9.900E-06 |
| 16673 | 834 | G834M | B | 9.7e-06 | 7.1e-06 | 1.2e-05 | 9.600E-06 | 2.452E-06 | 7.100E-06 | 1.200E-05 |
| 16674 | 834 | G834F | B | 1e-05 | 6.9e-06 | 8.4e-06 | 8.433E-06 | 1.550E-06 | 6.900E-06 | 1.000E-05 |
| 16675 | 834 | G834P | B | 1.1e-05 | 9.7e-06 | 8.6e-06 | 9.767E-06 | 1.201E-06 | 8.600E-06 | 1.100E-05 |
| 16676 | 834 | G834S | B | 1.3e-05 | 1.2e-05 | 9.5e-06 | 1.150E-05 | 1.803E-06 | 9.500E-06 | 1.300E-05 |
| 16677 | 834 | G834T | B | 1.7e-05 | 1e-05 | 5.9e-06 | 1.097E-05 | 5.613E-06 | 5.900E-06 | 1.700E-05 |
| 16678 | 834 | G834W | B | 9.7e-06 | 6.8e-06 | 4.9e-06 | 7.133E-06 | 2.417E-06 | 4.900E-06 | 9.700E-06 |
| 16679 | 834 | G834Y | B | 9.8e-06 | 8.1e-06 | 8.3e-06 | 8.733E-06 | 9.292E-07 | 8.100E-06 | 9.800E-06 |
| 16680 | 834 | G834V | B | 5.7e-06 | 6.5e-06 | 4e-06 | 5.400E-06 | 1.277E-06 | 4.000E-06 | 6.500E-06 |
| 16681 | 835 | E835A | B | 4.5e-06 | 6.6e-06 | 6.3e-06 | 5.800E-06 | 1.136E-06 | 4.500E-06 | 6.600E-06 |
| 16682 | 835 | E835R | B | 5e-06 | 8.9e-06 | 9e-06 | 7.633E-06 | 2.281E-06 | 5.000E-06 | 9.000E-06 |
| 16683 | 835 | E835N | B | 7.1e-06 | 1.3e-05 | 2.1e-05 | 1.370E-05 | 6.976E-06 | 7.100E-06 | 2.100E-05 |
| 16684 | 835 | E835D | B | 7.2e-06 | 7.3e-06 | 9.1e-06 | 7.867E-06 | 1.069E-06 | 7.200E-06 | 9.100E-06 |
| 16685 | 835 | E835C | B | 9e-06 | 6.1e-06 | 6.9e-06 | 7.333E-06 | 1.498E-06 | 6.100E-06 | 9.000E-06 |
| 16686 | 835 | E835E | B | 5.6e-06 | 5.8e-06 | 5.8e-06 | 5.733E-06 | 1.155E-07 | 5.600E-06 | 5.800E-06 |
| 16687 | 835 | E835Q | B | 9.1e-06 | 8.6e-06 | 9.8e-06 | 9.167E-06 | 6.028E-07 | 8.600E-06 | 9.800E-06 |
| 16688 | 835 | E835G | B | 8.4e-06 | 8.1e-06 | 8.2e-06 | 8.233E-06 | 1.528E-07 | 8.100E-06 | 8.400E-06 |
| 16689 | 835 | E835H | B | 4.1e-06 | 7.2e-06 | 6.4e-06 | 5.900E-06 | 1.609E-06 | 4.100E-06 | 7.200E-06 |
| 16690 | 835 | E835I | B | 1e-05 | 8.4e-06 | 1.4e-05 | 1.080E-05 | 2.884E-06 | 8.400E-06 | 1.400E-05 |
| 16691 | 835 | E835L | B | 6.7e-06 | 8.9e-06 | 8.3e-06 | 7.967E-06 | 1.137E-06 | 6.700E-06 | 8.900E-06 |
| 16692 | 835 | E835K | B | 5.2e-06 | 4.5e-06 | 1.3e-05 | 7.567E-06 | 4.718E-06 | 4.500E-06 | 1.300E-05 |
| 16693 | 835 | E835M | B | 4.7e-06 | 9e-06 | 9.1e-06 | 7.600E-06 | 2.512E-06 | 4.700E-06 | 9.100E-06 |
| 16694 | 835 | E835F | B | 4.4e-06 | 1.4e-05 | 9.6e-06 | 9.333E-06 | 4.806E-06 | 4.400E-06 | 1.400E-05 |
| 16695 | 835 | E835P | B | 7.5e-06 | 5.7e-06 | 6.9e-06 | 6.700E-06 | 9.165E-07 | 5.700E-06 | 7.500E-06 |
| 16696 | 835 | E835S | B | 1e-05 | 3.8e-06 | 5.5e-06 | 6.433E-06 | 3.204E-06 | 3.800E-06 | 1.000E-05 |
| 16697 | 835 | E835T | B | 5.1e-06 | 7.3e-06 | 7.3e-06 | 6.567E-06 | 1.270E-06 | 5.100E-06 | 7.300E-06 |
| 16698 | 835 | E835W | B | 6.4e-06 | 5.9e-06 | 6.3e-06 | 6.200E-06 | 2.646E-07 | 5.900E-06 | 6.400E-06 |
| 16699 | 835 | E835Y | B | 6.2e-06 | 5e-06 | 8.9e-06 | 6.700E-06 | 1.997E-06 | 5.000E-06 | 8.900E-06 |
| 16700 | 835 | E835V | B | 4.5e-06 | 9.9e-06 | 6.6e-06 | 7.000E-06 | 2.722E-06 | 4.500E-06 | 9.900E-06 |
| 16701 | 836 | C836A | B | 6.9e-06 | 1.7e-05 | 4.6e-06 | 9.500E-06 | 6.596E-06 | 4.600E-06 | 1.700E-05 |
| 16702 | 836 | C836R | B | 9.6e-06 | 7.1e-06 | 5.2e-06 | 7.300E-06 | 2.207E-06 | 5.200E-06 | 9.600E-06 |
| 16703 | 836 | C836N | B | 6.4e-06 | 6.2e-06 | 6.1e-06 | 6.233E-06 | 1.528E-07 | 6.100E-06 | 6.400E-06 |
| 16704 | 836 | C836D | B | 1.2e-05 | 7.4e-06 | 1.1e-05 | 1.013E-05 | 2.419E-06 | 7.400E-06 | 1.200E-05 |
| 16705 | 836 | C836C | B | 4.9e-06 | 6.9e-06 | 7.8e-06 | 6.533E-06 | 1.484E-06 | 4.900E-06 | 7.800E-06 |
| 16706 | 836 | C836E | B | 8.3e-06 | 6.7e-06 | 7.8e-06 | 7.600E-06 | 8.185E-07 | 6.700E-06 | 8.300E-06 |
| 16707 | 836 | C836Q | B | 1.2e-05 | 6.7e-06 | 1.6e-05 | 1.157E-05 | 4.665E-06 | 6.700E-06 | 1.600E-05 |
| 16708 | 836 | C836G | B | 1.2e-05 | 6.6e-06 | 1e-05 | 9.533E-06 | 2.730E-06 | 6.600E-06 | 1.200E-05 |
| 16709 | 836 | C836H | B | 8.4e-06 | 8.6e-06 | 8e-06 | 8.333E-06 | 3.055E-07 | 8.000E-06 | 8.600E-06 |
| 16710 | 836 | C836I | B | 8.9e-06 | 1.1e-05 | 1.4e-05 | 1.130E-05 | 2.563E-06 | 8.900E-06 | 1.400E-05 |
| 16711 | 836 | C836L | B | 8.4e-06 | 1e-05 | 7.9e-06 | 8.767E-06 | 1.097E-06 | 7.900E-06 | 1.000E-05 |
| 16712 | 836 | C836K | B | 9.2e-06 | 1.4e-05 | 1.5e-05 | 1.273E-05 | 3.101E-06 | 9.200E-06 | 1.500E-05 |
| 16713 | 836 | C836M | B | 6.7e-06 | 4.4e-06 | 6.9e-06 | 6.000E-06 | 1.389E-06 | 4.400E-06 | 6.900E-06 |
| 16714 | 836 | C836F | B | 2.5e-05 | 1e-05 | 1.2e-05 | 1.567E-05 | 8.145E-06 | 1.000E-05 | 2.500E-05 |
| 16715 | 836 | C836P | B | 7.2e-06 | 7e-06 | 6.5e-06 | 6.900E-06 | 3.606E-07 | 6.500E-06 | 7.200E-06 |
| 16716 | 836 | C836S | B | 7.3e-06 | 7.6e-06 | 8.8e-06 | 7.900E-06 | 7.937E-07 | 7.300E-06 | 8.800E-06 |
| 16717 | 836 | C836T | B | 8.5e-06 | 7e-06 | 1.2e-05 | 9.167E-06 | 2.566E-06 | 7.000E-06 | 1.200E-05 |
| 16718 | 836 | C836W | B | 1.1e-05 | 7e-06 | 1e-05 | 9.333E-06 | 2.082E-06 | 7.000E-06 | 1.100E-05 |
| 16719 | 836 | C836Y | B | 5.4e-06 | 6e-06 | 1.1e-05 | 7.467E-06 | 3.075E-06 | 5.400E-06 | 1.100E-05 |
| 16720 | 836 | C836V | B | 7.1e-06 | 1.1e-05 | 5e-06 | 7.700E-06 | 3.045E-06 | 5.000E-06 | 1.100E-05 |
| 16721 | 837 | E837A | C | 9.9e-06 | 1e-05 | 6.5e-06 | 8.800E-06 | 1.992E-06 | 6.500E-06 | 1.000E-05 |
| 16722 | 837 | E837R | C | 1.1e-05 | 8.8e-06 | 6e-06 | 8.600E-06 | 2.506E-06 | 6.000E-06 | 1.100E-05 |
| 16723 | 837 | E837N | C | 8.7e-06 | 6.4e-06 | 8.5e-06 | 7.867E-06 | 1.274E-06 | 6.400E-06 | 8.700E-06 |
| 16724 | 837 | E837D | C | 8.9e-06 | 9.9e-06 | 1e-05 | 9.600E-06 | 6.083E-07 | 8.900E-06 | 1.000E-05 |
| 16725 | 837 | E837C | C | 1.1e-05 | 7.3e-06 | 5e-06 | 7.767E-06 | 3.027E-06 | 5.000E-06 | 1.100E-05 |
| 16726 | 837 | E837E | C | 7.1e-06 | 1e-05 | 8.3e-06 | 8.467E-06 | 1.457E-06 | 7.100E-06 | 1.000E-05 |
| 16727 | 837 | E837Q | C | 4.5e-06 | 9.7e-06 | 8.2e-06 | 7.467E-06 | 2.676E-06 | 4.500E-06 | 9.700E-06 |
| 16728 | 837 | E837G | C | 7.6e-06 | 8.6e-06 | 5.7e-06 | 7.300E-06 | 1.473E-06 | 5.700E-06 | 8.600E-06 |
| 16729 | 837 | E837H | C | 6.2e-06 | 1.2e-05 | 9.8e-06 | 9.333E-06 | 2.928E-06 | 6.200E-06 | 1.200E-05 |
| 16730 | 837 | E837I | C | 9.6e-06 | 7e-06 | 7.2e-06 | 7.933E-06 | 1.447E-06 | 7.000E-06 | 9.600E-06 |
| 16731 | 837 | E837L | C | 7.5e-06 | 1.4e-05 | 5.3e-06 | 8.933E-06 | 4.524E-06 | 5.300E-06 | 1.400E-05 |
| 16732 | 837 | E837K | C | 6.4e-06 | 9e-06 | 1.2e-05 | 9.133E-06 | 2.802E-06 | 6.400E-06 | 1.200E-05 |
| 16733 | 837 | E837M | C | 9.9e-06 | 8.7e-06 | 9.9e-06 | 9.500E-06 | 6.928E-07 | 8.700E-06 | 9.900E-06 |
| 16734 | 837 | E837F | C | 9.7e-06 | 9e-06 | 1.3e-05 | 1.057E-05 | 2.136E-06 | 9.000E-06 | 1.300E-05 |
| 16735 | 837 | E837P | C | 6.9e-06 | 5.6e-06 | 6.3e-06 | 6.267E-06 | 6.506E-07 | 5.600E-06 | 6.900E-06 |
| 16736 | 837 | E837S | C | 6.3e-06 | 5.3e-06 | 1.3e-05 | 8.200E-06 | 4.187E-06 | 5.300E-06 | 1.300E-05 |
| 16737 | 837 | E837T | C | 6.7e-06 | 4.9e-06 | 5.6e-06 | 5.733E-06 | 9.074E-07 | 4.900E-06 | 6.700E-06 |
| 16738 | 837 | E837W | C | 3.1e-06 | 7.6e-06 | 9.4e-06 | 6.700E-06 | 3.245E-06 | 3.100E-06 | 9.400E-06 |
| 16739 | 837 | E837Y | C | 4.6e-06 | 5.6e-06 | 8.6e-06 | 6.267E-06 | 2.082E-06 | 4.600E-06 | 8.600E-06 |
| 16740 | 837 | E837V | C | 9.8e-06 | 1.3e-05 | 8.1e-06 | 1.030E-05 | 2.488E-06 | 8.100E-06 | 1.300E-05 |
| 16741 | 838 | V838A | C | 9.3e-06 | 5.1e-06 | 7.1e-06 | 7.167E-06 | 2.101E-06 | 5.100E-06 | 9.300E-06 |
| 16742 | 838 | V838R | C | 7.4e-06 | 6.2e-06 | 1e-05 | 7.867E-06 | 1.943E-06 | 6.200E-06 | 1.000E-05 |
| 16743 | 838 | V838N | C | 9.8e-06 | 9.5e-06 | 3.3e-06 | 7.533E-06 | 3.669E-06 | 3.300E-06 | 9.800E-06 |
| 16744 | 838 | V838D | C | 9.3e-06 | 9e-06 | 9.6e-06 | 9.300E-06 | 3.000E-07 | 9.000E-06 | 9.600E-06 |

| | | | | | | | | | | |
|---|---|---|---|---|---|---|---|---|---|---|
| 16745 | 838 | V838C | C | 9.7e-06 | 9.6e-06 | 1.3e-05 | 1.077E-05 | 1.935E-06 | 9.600E-06 | 1.300E-05 |
| 16746 | 838 | V838E | C | 8.4e-06 | 1.2e-05 | 1.4e-05 | 1.147E-05 | 2.838E-06 | 8.400E-06 | 1.400E-05 |
| 16747 | 838 | V838Q | C | 1.4e-05 | 1.1e-05 | 9.2e-06 | 1.140E-05 | 2.425E-06 | 9.200E-06 | 1.400E-05 |
| 16748 | 838 | V838G | C | 9.3e-06 | 1.4e-05 | 6.1e-06 | 9.800E-06 | 3.974E-06 | 6.100E-06 | 1.400E-05 |
| 16749 | 838 | V838H | C | 6.3e-06 | 5.5e-06 | 6.1e-06 | 5.967E-06 | 4.163E-07 | 5.500E-06 | 6.300E-06 |
| 16750 | 838 | V838I | C | 1.2e-05 | 9.4e-06 | 5.5e-06 | 8.967E-06 | 3.272E-06 | 5.500E-06 | 1.200E-05 |
| 16751 | 838 | V838L | C | 1e-05 | 1e-05 | 4e-06 | 8.000E-06 | 3.464E-06 | 4.000E-06 | 1.000E-05 |
| 16752 | 838 | V838K | C | 9.8e-06 | 4.2e-06 | 1e-05 | 8.000E-06 | 3.292E-06 | 4.200E-06 | 1.000E-05 |
| 16753 | 838 | V838M | C | 6e-06 | 8.4e-06 | 8.3e-06 | 7.567E-06 | 1.358E-06 | 6.000E-06 | 8.400E-06 |
| 16754 | 838 | V838F | C | 7.1e-06 | 6.6e-06 | 8.9e-06 | 7.533E-06 | 1.210E-06 | 6.600E-06 | 8.900E-06 |
| 16755 | 838 | V838P | C | 1e-05 | 6.5e-06 | 6.2e-06 | 7.567E-06 | 2.113E-06 | 6.200E-06 | 1.000E-05 |
| 16756 | 838 | V838S | C | 9.8e-06 | 8.8e-06 | 4.5e-06 | 7.700E-06 | 2.816E-06 | 4.500E-06 | 9.800E-06 |
| 16757 | 838 | V838T | C | 6.8e-06 | 9.6e-06 | 8.2e-06 | 8.200E-06 | 1.400E-06 | 6.800E-06 | 9.600E-06 |
| 16758 | 838 | V838W | C | 1.2e-05 | 9.5e-06 | 3.9e-06 | 8.467E-06 | 4.148E-06 | 3.900E-06 | 1.200E-05 |
| 16759 | 838 | V838Y | C | 6.6e-06 | 5.6e-06 | 1.1e-05 | 7.733E-06 | 2.873E-06 | 5.600E-06 | 1.100E-05 |
| 16760 | 838 | V838V | C | 7.8e-06 | 1.3e-05 | 1.1e-05 | 1.060E-05 | 2.623E-06 | 7.800E-06 | 1.300E-05 |
| 16761 | 839 | Q839A | C | 9.6e-06 | 9.6e-06 | 6.4e-06 | 8.533E-06 | 1.848E-06 | 6.400E-06 | 9.600E-06 |
| 16762 | 839 | Q839R | C | 8.5e-06 | 6.3e-06 | 4.9e-06 | 6.567E-06 | 1.815E-06 | 4.900E-06 | 8.500E-06 |
| 16763 | 839 | Q839N | C | 6e-06 | 5.6e-06 | 4.9e-06 | 5.500E-06 | 5.568E-07 | 4.900E-06 | 6.000E-06 |
| 16764 | 839 | Q839D | C | 1.2e-05 | 1.1e-05 | 7.4e-06 | 1.013E-05 | 2.419E-06 | 7.400E-06 | 1.200E-05 |
| 16765 | 839 | Q839C | C | 6.5e-06 | 8.2e-06 | 6.1e-06 | 6.933E-06 | 1.115E-06 | 6.100E-06 | 8.200E-06 |
| 16766 | 839 | Q839E | C | 8e-06 | 6.1e-06 | 5.5e-06 | 6.533E-06 | 1.305E-06 | 5.500E-06 | 8.000E-06 |
| 16767 | 839 | Q839Q | C | 4.7e-06 | 7.1e-06 | 7e-06 | 6.267E-06 | 1.358E-06 | 4.700E-06 | 7.100E-06 |
| 16768 | 839 | Q839G | C | 7.4e-06 | 5.3e-06 | 4.9e-06 | 5.867E-06 | 1.343E-06 | 4.900E-06 | 7.400E-06 |
| 16769 | 839 | Q839H | C | 9.4e-06 | 9.9e-06 | 6.3e-06 | 8.533E-06 | 1.950E-06 | 6.300E-06 | 9.900E-06 |
| 16770 | 839 | Q839I | C | 7.1e-06 | 4.8e-06 | 7e-06 | 6.300E-06 | 1.300E-06 | 4.800E-06 | 7.100E-06 |
| 16771 | 839 | Q839L | C | 1.1e-05 | 9.6e-06 | 9.6e-06 | 1.007E-05 | 8.083E-07 | 9.600E-06 | 1.100E-05 |
| 16772 | 839 | Q839K | C | 7e-06 | 7.7e-06 | 8.9e-06 | 7.867E-06 | 9.609E-07 | 7.000E-06 | 8.900E-06 |
| 16773 | 839 | Q839M | C | 5.4e-06 | 5.2e-06 | 1.4e-05 | 8.200E-06 | 5.024E-06 | 5.200E-06 | 1.400E-05 |
| 16774 | 839 | Q839F | C | 6.1e-06 | 6.2e-06 | 8.4e-06 | 6.900E-06 | 1.300E-06 | 6.100E-06 | 8.400E-06 |
| 16775 | 839 | Q839P | C | 5.9e-06 | 5.4e-06 | 7.1e-06 | 6.133E-06 | 8.737E-07 | 5.400E-06 | 7.100E-06 |
| 16776 | 839 | Q839S | C | 5.1e-06 | 8e-06 | 4.9e-06 | 6.000E-06 | 1.735E-06 | 4.900E-06 | 8.000E-06 |
| 16777 | 839 | Q839T | C | 1e-05 | 5.1e-06 | 9.2e-06 | 8.100E-06 | 2.629E-06 | 5.100E-06 | 1.000E-05 |
| 16778 | 839 | Q839W | C | 3.6e-06 | 4.3e-06 | 6.3e-06 | 4.733E-06 | 1.401E-06 | 3.600E-06 | 6.300E-06 |
| 16779 | 839 | Q839Y | C | 6.5e-06 | 7.4e-06 | 8.4e-06 | 7.433E-06 | 9.504E-07 | 6.500E-06 | 8.400E-06 |
| 16780 | 839 | Q839V | C | 1e-05 | 1.2e-05 | 1e-05 | 1.067E-05 | 1.155E-06 | 1.000E-05 | 1.200E-05 |
| 16781 | 840 | L840A | C | 1.4e-05 | 4.3e-06 | 1e-05 | 9.433E-06 | 4.875E-06 | 4.300E-06 | 1.400E-05 |
| 16782 | 840 | L840R | C | 8.7e-06 | 6.3e-06 | 5.9e-06 | 6.967E-06 | 1.514E-06 | 5.900E-06 | 8.700E-06 |
| 16783 | 840 | L840N | C | 4.6e-06 | 6.7e-06 | 6.6e-06 | 5.967E-06 | 1.185E-06 | 4.600E-06 | 6.700E-06 |
| 16784 | 840 | L840D | C | 4.4e-06 | 6.5e-06 | 9.4e-06 | 6.767E-06 | 2.511E-06 | 4.400E-06 | 9.400E-06 |
| 16785 | 840 | L840C | C | 1.6e-05 | 1.1e-05 | 7.4e-06 | 1.147E-05 | 4.319E-06 | 7.400E-06 | 1.600E-05 |
| 16786 | 840 | L840E | C | 6.9e-06 | 8.4e-06 | 4.7e-06 | 6.667E-06 | 1.861E-06 | 4.700E-06 | 8.400E-06 |
| 16787 | 840 | L840Q | C | 1e-05 | 4.7e-06 | 4.6e-06 | 6.433E-06 | 3.089E-06 | 4.600E-06 | 1.000E-05 |
| 16788 | 840 | L840G | C | 1e-05 | 7.8e-06 | 5.4e-06 | 7.733E-06 | 2.301E-06 | 5.400E-06 | 1.000E-05 |
| 16789 | 840 | L840H | C | 1e-05 | 6.2e-06 | 6.6e-06 | 7.600E-06 | 2.088E-06 | 6.200E-06 | 1.000E-05 |
| 16790 | 840 | L840I | C | 7.7e-06 | 5.8e-06 | 1.1e-05 | 8.167E-06 | 2.631E-06 | 5.800E-06 | 1.100E-05 |
| 16791 | 840 | L840L | C | 7.4e-06 | 7.2e-06 | 8.6e-06 | 7.733E-06 | 7.572E-07 | 7.200E-06 | 8.600E-06 |
| 16792 | 840 | L840K | C | 4.5e-06 | 5.6e-06 | 7.1e-06 | 5.733E-06 | 1.305E-06 | 4.500E-06 | 7.100E-06 |
| 16793 | 840 | L840M | C | 1.4e-05 | 8e-06 | 9e-06 | 1.033E-05 | 3.215E-06 | 8.000E-06 | 1.400E-05 |
| 16794 | 840 | L840F | C | 7.6e-06 | 4.4e-06 | 1e-05 | 7.333E-06 | 2.810E-06 | 4.400E-06 | 1.000E-05 |
| 16795 | 840 | L840P | C | 7.9e-06 | 6.7e-06 | 4.8e-06 | 6.467E-06 | 1.563E-06 | 4.800E-06 | 7.900E-06 |
| 16796 | 840 | L840S | C | 5.7e-06 | 7.4e-06 | 6.6e-06 | 6.567E-06 | 8.505E-07 | 5.700E-06 | 7.400E-06 |
| 16797 | 840 | L840T | C | 9e-06 | 5.9e-06 | 9.7e-06 | 8.200E-06 | 2.022E-06 | 5.900E-06 | 9.700E-06 |
| 16798 | 840 | L840W | C | 7.5e-06 | 8.4e-06 | 1.3e-05 | 9.633E-06 | 2.950E-06 | 7.500E-06 | 1.300E-05 |
| 16799 | 840 | L840Y | C | 9.5e-06 | 1e-05 | 6e-06 | 8.500E-06 | 2.179E-06 | 6.000E-06 | 1.000E-05 |
| 16800 | 840 | L840V | C | 5.1e-06 | 8.9e-06 | 1.1e-05 | 8.333E-06 | 2.991E-06 | 5.100E-06 | 1.100E-05 |
| 16801 | 841 | V841A | C | 1e-05 | 1e-05 | 7.1e-06 | 9.033E-06 | 1.674E-06 | 7.100E-06 | 1.000E-05 |
| 16802 | 841 | V841R | C | 9.7e-06 | 1.1e-05 | 8.2e-06 | 9.633E-06 | 1.401E-06 | 8.200E-06 | 1.100E-05 |
| 16803 | 841 | V841N | C | 6.5e-06 | 8.3e-06 | 6.7e-06 | 7.167E-06 | 9.866E-07 | 6.500E-06 | 8.300E-06 |
| 16804 | 841 | V841D | C | 7.2e-06 | 7.9e-06 | 4.3e-06 | 6.467E-06 | 1.909E-06 | 4.300E-06 | 7.900E-06 |
| 16805 | 841 | V841C | C | 6e-06 | 6.3e-06 | 1.4e-05 | 8.767E-06 | 4.535E-06 | 6.000E-06 | 1.400E-05 |
| 16806 | 841 | V841E | C | 9e-06 | 5.2e-06 | 1.3e-05 | 9.067E-06 | 3.900E-06 | 5.200E-06 | 1.300E-05 |
| 16807 | 841 | V841Q | C | 9.8e-06 | 9.1e-06 | 7e-06 | 8.633E-06 | 1.457E-06 | 7.000E-06 | 9.800E-06 |
| 16808 | 841 | V841G | C | 7.4e-06 | 9.9e-06 | 7.1e-06 | 8.133E-06 | 1.537E-06 | 7.100E-06 | 9.900E-06 |
| 16809 | 841 | V841H | C | 2.6e-06 | 6.6e-06 | 9.1e-06 | 6.100E-06 | 3.279E-06 | 2.600E-06 | 9.100E-06 |
| 16810 | 841 | V841I | C | 3.4e-06 | 8.6e-06 | 6.8e-06 | 6.267E-06 | 2.641E-06 | 3.400E-06 | 8.600E-06 |
| 16811 | 841 | V841L | C | 4.8e-06 | 1e-05 | 6.8e-06 | 7.200E-06 | 2.623E-06 | 4.800E-06 | 1.000E-05 |
| 16812 | 841 | V841K | C | 5.2e-06 | 8.3e-06 | 9.4e-06 | 7.633E-06 | 2.178E-06 | 5.200E-06 | 9.400E-06 |
| 16813 | 841 | V841M | C | 6.4e-06 | 8.3e-06 | 4.4e-06 | 6.367E-06 | 1.950E-06 | 4.400E-06 | 8.300E-06 |
| 16814 | 841 | V841F | C | 7.9e-06 | 9.6e-06 | 9.5e-06 | 9.000E-06 | 9.539E-07 | 7.900E-06 | 9.600E-06 |
| 16815 | 841 | V841P | C | 1.5e-05 | 1e-05 | 9.4e-06 | 1.147E-05 | 3.075E-06 | 9.400E-06 | 1.500E-05 |
| 16816 | 841 | V841S | C | 9.6e-06 | 5e-06 | 8.3e-06 | 7.633E-06 | 2.371E-06 | 5.000E-06 | 9.600E-06 |
| 16817 | 841 | V841T | C | 9.1e-06 | 7.7e-06 | 6.6e-06 | 7.800E-06 | 1.253E-06 | 6.600E-06 | 9.100E-06 |
| 16818 | 841 | V841W | C | 7.9e-06 | 5.4e-06 | 6.3e-06 | 6.533E-06 | 1.266E-06 | 5.400E-06 | 7.900E-06 |
| 16819 | 841 | V841Y | C | 6.8e-06 | 5e-06 | 9.2e-06 | 7.000E-06 | 2.107E-06 | 5.000E-06 | 9.200E-06 |
| 16820 | 841 | V841V | C | 9.3e-06 | 4.9e-06 | 1.2e-05 | 8.733E-06 | 3.584E-06 | 4.900E-06 | 1.200E-05 |
| 16821 | 842 | E842A | C | 9.4e-06 | 9.6e-06 | 7.3e-06 | 8.767E-06 | 1.274E-06 | 7.300E-06 | 9.600E-06 |
| 16822 | 842 | E842R | C | 1.1e-05 | 9.4e-06 | 4.8e-06 | 8.400E-06 | 3.219E-06 | 4.800E-06 | 1.100E-05 |
| 16823 | 842 | E842N | C | 4.5e-06 | 7e-06 | 6e-06 | 5.833E-06 | 1.258E-06 | 4.500E-06 | 7.000E-06 |
| 16824 | 842 | E842D | C | 1.1e-05 | 7.7e-06 | 6e-06 | 8.233E-06 | 2.542E-06 | 6.000E-06 | 1.100E-05 |
| 16825 | 842 | E842C | C | 8.6e-06 | 5.9e-06 | 1.1e-05 | 8.500E-06 | 2.551E-06 | 5.900E-06 | 1.100E-05 |
| 16826 | 842 | E842E | C | 7e-06 | 6.8e-06 | 1.4e-05 | 9.267E-06 | 4.100E-06 | 6.800E-06 | 1.400E-05 |
| 16827 | 842 | E842Q | C | 9.9e-06 | 1.2e-05 | 6.6e-06 | 9.500E-06 | 2.722E-06 | 6.600E-06 | 1.200E-05 |
| 16828 | 842 | E842G | C | 1.5e-05 | 7.8e-06 | 5.1e-06 | 9.300E-06 | 5.118E-06 | 5.100E-06 | 1.500E-05 |
| 16829 | 842 | E842H | C | 1e-05 | 5.9e-06 | 6.5e-06 | 7.467E-06 | 2.214E-06 | 5.900E-06 | 1.000E-05 |

| | | | | | | | | | | |
|---|---|---|---|---|---|---|---|---|---|---|
| 16830 | 842 | E842I | C | 3.9e-06 | 8.5e-06 | 1.3e-05 | 8.467E-06 | 4.550E-06 | 3.900E-06 | 1.300E-05 |
| 16831 | 842 | E842L | C | 5.6e-06 | 7.2e-06 | 9.3e-06 | 7.367E-06 | 1.856E-06 | 5.600E-06 | 9.300E-06 |
| 16832 | 842 | E842K | C | 8.1e-06 | 7.8e-06 | 8.6e-06 | 8.167E-06 | 4.041E-07 | 7.800E-06 | 8.600E-06 |
| 16833 | 842 | E842M | C | 1.1e-05 | 1.4e-05 | 6.9e-06 | 1.063E-05 | 3.564E-06 | 6.900E-06 | 1.400E-05 |
| 16834 | 842 | E842F | C | 7.3e-06 | 1e-05 | 1.3e-05 | 1.010E-05 | 2.851E-06 | 7.300E-06 | 1.300E-05 |
| 16835 | 842 | E842P | C | 7.2e-06 | 1e-05 | 7.2e-06 | 8.133E-06 | 1.617E-06 | 7.200E-06 | 1.000E-05 |
| 16836 | 842 | E842S | C | 4.8e-06 | 1.3e-05 | 5.2e-06 | 7.667E-06 | 4.623E-06 | 4.800E-06 | 1.300E-05 |
| 16837 | 842 | E842T | C | 5.4e-06 | 6.2e-06 | 5.8e-06 | 5.800E-06 | 4.000E-07 | 5.400E-06 | 6.200E-06 |
| 16838 | 842 | E842W | C | 6.8e-06 | 4.4e-06 | 8.5e-06 | 6.567E-06 | 2.060E-06 | 4.400E-06 | 8.500E-06 |
| 16839 | 842 | E842Y | C | 6.6e-06 | 3.5e-06 | 6.9e-06 | 5.667E-06 | 1.882E-06 | 3.500E-06 | 6.900E-06 |
| 16840 | 842 | E842V | C | 3.9e-06 | 1e-05 | 9e-06 | 7.633E-06 | 3.272E-06 | 3.900E-06 | 1.000E-05 |
| 16841 | 843 | S843A | C | 8.1e-06 | 9.7e-06 | 9.6e-06 | 9.133E-06 | 8.963E-07 | 8.100E-06 | 9.700E-06 |
| 16842 | 843 | S843R | C | 9.8e-06 | 7.3e-06 | 7.6e-06 | 8.233E-06 | 1.365E-06 | 7.300E-06 | 9.800E-06 |
| 16843 | 843 | S843N | C | 6.1e-06 | 7.4e-06 | 9.2e-06 | 7.567E-06 | 1.557E-06 | 6.100E-06 | 9.200E-06 |
| 16844 | 843 | S843D | C | 8.7e-06 | 8.7e-06 | 5.3e-06 | 7.567E-06 | 1.963E-06 | 5.300E-06 | 8.700E-06 |
| 16845 | 843 | S843C | C | 6.5e-06 | 4.9e-06 | 6.1e-06 | 5.833E-06 | 8.327E-07 | 4.900E-06 | 6.500E-06 |
| 16846 | 843 | S843E | C | 8.7e-06 | 8.5e-06 | 9.9e-06 | 9.033E-06 | 7.572E-07 | 8.500E-06 | 9.900E-06 |
| 16847 | 843 | S843Q | C | 6.8e-06 | 9.4e-06 | 6.9e-06 | 7.700E-06 | 1.473E-06 | 6.800E-06 | 9.400E-06 |
| 16848 | 843 | S843G | C | 7.3e-06 | 4.8e-06 | 9.2e-06 | 7.100E-06 | 2.207E-06 | 4.800E-06 | 9.200E-06 |
| 16849 | 843 | S843H | C | 9.2e-06 | 4.5e-06 | 7e-06 | 6.900E-06 | 2.352E-06 | 4.500E-06 | 9.200E-06 |
| 16850 | 843 | S843I | C | 1e-05 | 7.2e-06 | 9.4e-06 | 8.867E-06 | 1.474E-06 | 7.200E-06 | 1.000E-05 |
| 16851 | 843 | S843L | C | 7.6e-06 | 6.7e-06 | 5.7e-06 | 6.667E-06 | 9.504E-07 | 5.700E-06 | 7.600E-06 |
| 16852 | 843 | S843K | C | 5.6e-06 | 7.9e-06 | 1.1e-05 | 8.167E-06 | 2.710E-06 | 5.600E-06 | 1.100E-05 |
| 16853 | 843 | S843M | C | 1.2e-05 | 1e-05 | 6.7e-06 | 9.567E-06 | 2.676E-06 | 6.700E-06 | 1.200E-05 |
| 16854 | 843 | S843F | C | 1.3e-05 | 5.6e-06 | 4.6e-06 | 7.733E-06 | 4.588E-06 | 4.600E-06 | 1.300E-05 |
| 16855 | 843 | S843P | C | 6.6e-06 | 8.1e-06 | 5.7e-06 | 6.800E-06 | 1.212E-06 | 5.700E-06 | 8.100E-06 |
| 16856 | 843 | S843S | C | 1.1e-05 | 1.3e-05 | 5.1e-06 | 9.700E-06 | 4.107E-06 | 5.100E-06 | 1.300E-05 |
| 16857 | 843 | S843T | C | 6.2e-06 | 6.5e-06 | 8.3e-06 | 7.000E-06 | 1.136E-06 | 6.200E-06 | 8.300E-06 |
| 16858 | 843 | S843W | C | 1.3e-05 | 7e-06 | 8.2e-06 | 9.400E-06 | 3.175E-06 | 7.000E-06 | 1.300E-05 |
| 16859 | 843 | S843Y | C | 9.6e-06 | 8e-06 | 8.3e-06 | 8.633E-06 | 8.505E-07 | 8.000E-06 | 9.600E-06 |
| 16860 | 843 | S843V | C | 1.1e-05 | 9.3e-06 | 9.4e-06 | 9.900E-06 | 9.539E-07 | 9.300E-06 | 1.100E-05 |
| 16861 | 844 | G844A | C | 6.5e-06 | 8.7e-06 | 8.6e-06 | 7.933E-06 | 1.242E-06 | 6.500E-06 | 8.700E-06 |
| 16862 | 844 | G844R | C | 1.1e-05 | 6.6e-06 | 6.6e-06 | 8.067E-06 | 2.540E-06 | 6.600E-06 | 1.100E-05 |
| 16863 | 844 | G844N | C | 4.7e-06 | 1.2e-05 | 5.3e-06 | 7.333E-06 | 4.053E-06 | 4.700E-06 | 1.200E-05 |
| 16864 | 844 | G844D | C | 7.3e-06 | 5.1e-06 | 9.3e-06 | 7.233E-06 | 2.101E-06 | 5.100E-06 | 9.300E-06 |
| 16865 | 844 | G844C | C | 9.1e-06 | 7.4e-06 | 9.8e-06 | 8.767E-06 | 1.234E-06 | 7.400E-06 | 9.800E-06 |
| 16866 | 844 | G844E | C | 8.3e-06 | 4.6e-06 | 5.9e-06 | 6.267E-06 | 1.877E-06 | 4.600E-06 | 8.300E-06 |
| 16867 | 844 | G844Q | C | 5.6e-06 | 9.5e-06 | 4.2e-06 | 6.433E-06 | 2.747E-06 | 4.200E-06 | 9.500E-06 |
| 16868 | 844 | G844G | C | 8.3e-06 | 8.4e-06 | 5.4e-06 | 7.367E-06 | 1.704E-06 | 5.400E-06 | 8.400E-06 |
| 16869 | 844 | G844H | C | 6.3e-06 | 9.8e-06 | 9.4e-06 | 8.500E-06 | 1.916E-06 | 6.300E-06 | 9.800E-06 |
| 16870 | 844 | G844I | C | 8.7e-06 | 5.4e-06 | 6.7e-06 | 6.933E-06 | 1.662E-06 | 5.400E-06 | 8.700E-06 |
| 16871 | 844 | G844L | C | 7.2e-06 | 6.3e-06 | 1.2e-05 | 8.500E-06 | 3.064E-06 | 6.300E-06 | 1.200E-05 |
| 16872 | 844 | G844K | C | 6.1e-06 | 6.6e-06 | 9.2e-06 | 7.300E-06 | 1.664E-06 | 6.100E-06 | 9.200E-06 |
| 16873 | 844 | G844M | C | 1.2e-05 | 4.9e-06 | 6e-06 | 7.633E-06 | 3.821E-06 | 4.900E-06 | 1.200E-05 |
| 16874 | 844 | G844F | C | 6.9e-06 | 6.1e-06 | 9.7e-06 | 7.567E-06 | 1.890E-06 | 6.100E-06 | 9.700E-06 |
| 16875 | 844 | G844P | C | 8.2e-06 | 7.5e-06 | 6.4e-06 | 7.367E-06 | 9.074E-07 | 6.400E-06 | 8.200E-06 |
| 16876 | 844 | G844S | C | 1.1e-05 | 7.8e-06 | 9.9e-06 | 9.567E-06 | 1.626E-06 | 7.800E-06 | 1.100E-05 |
| 16877 | 844 | G844T | C | 7.4e-06 | 7.5e-06 | 7.7e-06 | 7.533E-06 | 1.528E-07 | 7.400E-06 | 7.700E-06 |
| 16878 | 844 | G844W | C | 9.2e-06 | 1.4e-05 | 7.6e-06 | 1.027E-05 | 3.331E-06 | 7.600E-06 | 1.400E-05 |
| 16879 | 844 | G844Y | C | 9.3e-06 | 1.1e-05 | 1.2e-05 | 1.077E-05 | 1.365E-06 | 9.300E-06 | 1.200E-05 |
| 16880 | 844 | G844V | C | 1e-05 | 7.8e-06 | 1.1e-05 | 9.600E-06 | 1.637E-06 | 7.800E-06 | 1.100E-05 |
| 16881 | 845 | G845A | C | 6e-06 | 1.2e-05 | 7.3e-06 | 8.433E-06 | 3.156E-06 | 6.000E-06 | 1.200E-05 |
| 16882 | 845 | G845R | C | 6.1e-06 | 8.4e-06 | 6e-06 | 6.833E-06 | 1.358E-06 | 6.000E-06 | 8.400E-06 |
| 16883 | 845 | G845N | C | 6.5e-06 | 8.3e-06 | 4.5e-06 | 6.433E-06 | 1.901E-06 | 4.500E-06 | 8.300E-06 |
| 16884 | 845 | G845D | C | 1e-05 | 6.3e-06 | 7.1e-06 | 7.800E-06 | 1.947E-06 | 6.300E-06 | 1.000E-05 |
| 16885 | 845 | G845C | C | 6.1e-06 | 6.7e-06 | 1.3e-05 | 8.600E-06 | 3.822E-06 | 6.100E-06 | 1.300E-05 |
| 16886 | 845 | G845E | C | 1.3e-05 | 8.1e-06 | 5.9e-06 | 9.000E-06 | 3.635E-06 | 5.900E-06 | 1.300E-05 |
| 16887 | 845 | G845Q | C | 8.2e-06 | 8.1e-06 | 3.9e-06 | 6.733E-06 | 2.454E-06 | 3.900E-06 | 8.200E-06 |
| 16888 | 845 | G845G | C | 4.5e-06 | 1e-05 | 5.4e-06 | 6.633E-06 | 2.950E-06 | 4.500E-06 | 1.000E-05 |
| 16889 | 845 | G845H | C | 8.1e-06 | 5.6e-06 | 5.5e-06 | 6.400E-06 | 1.473E-06 | 5.500E-06 | 8.100E-06 |
| 16890 | 845 | G845I | C | 1.2e-05 | 4.7e-06 | 9.6e-06 | 8.767E-06 | 3.721E-06 | 4.700E-06 | 1.200E-05 |
| 16891 | 845 | G845L | C | 8.9e-06 | 5.6e-06 | 1.1e-05 | 8.500E-06 | 2.722E-06 | 5.600E-06 | 1.100E-05 |
| 16892 | 845 | G845K | C | 7.3e-06 | 6.1e-06 | 5.9e-06 | 6.433E-06 | 7.572E-07 | 5.900E-06 | 7.300E-06 |
| 16893 | 845 | G845M | C | 9.1e-06 | 6.5e-06 | 6.5e-06 | 7.367E-06 | 1.501E-06 | 6.500E-06 | 9.100E-06 |
| 16894 | 845 | G845F | C | 7.8e-06 | 4e-06 | 9.9e-06 | 7.233E-06 | 2.991E-06 | 4.000E-06 | 9.900E-06 |
| 16895 | 845 | G845P | C | 7.4e-06 | 5.1e-06 | 9.7e-06 | 7.400E-06 | 2.300E-06 | 5.100E-06 | 9.700E-06 |
| 16896 | 845 | G845S | C | 1.1e-05 | 1e-05 | 1.1e-05 | 1.067E-05 | 5.774E-07 | 1.000E-05 | 1.100E-05 |
| 16897 | 845 | G845T | C | 1.3e-05 | 1.1e-05 | 1e-05 | 1.133E-05 | 1.528E-06 | 1.000E-05 | 1.300E-05 |
| 16898 | 845 | G845W | C | 1.1e-05 | 8.6e-06 | 8.4e-06 | 9.333E-06 | 1.447E-06 | 8.400E-06 | 1.100E-05 |
| 16899 | 845 | G845Y | C | 9.8e-06 | 8.2e-06 | 8.7e-06 | 8.900E-06 | 8.185E-07 | 8.200E-06 | 9.800E-06 |
| 16900 | 845 | G845V | C | 6.4e-06 | 9.9e-06 | 8.5e-06 | 8.267E-06 | 1.762E-06 | 6.400E-06 | 9.900E-06 |
| 16901 | 846 | G846A | C | 7e-06 | 6.8e-06 | 1.4e-05 | 9.267E-06 | 4.100E-06 | 6.800E-06 | 1.400E-05 |
| 16902 | 846 | G846R | C | 6.7e-06 | 8.9e-06 | 9.6e-06 | 8.400E-06 | 1.513E-06 | 6.700E-06 | 9.600E-06 |
| 16903 | 846 | G846N | C | 8.8e-06 | 6.4e-06 | 5.5e-06 | 6.900E-06 | 1.706E-06 | 5.500E-06 | 8.800E-06 |
| 16904 | 846 | G846D | C | 4.9e-06 | 7.6e-06 | 9e-06 | 7.167E-06 | 2.084E-06 | 4.900E-06 | 9.000E-06 |
| 16905 | 846 | G846C | C | 9e-06 | 1.1e-05 | 6.3e-06 | 8.767E-06 | 2.359E-06 | 6.300E-06 | 1.100E-05 |
| 16906 | 846 | G846E | C | 5.5e-06 | 7.8e-06 | 1.4e-05 | 9.100E-06 | 4.397E-06 | 5.500E-06 | 1.400E-05 |
| 16907 | 846 | G846Q | C | 8.3e-06 | 7.6e-06 | 6.6e-06 | 7.500E-06 | 8.544E-07 | 6.600E-06 | 8.300E-06 |
| 16908 | 846 | G846G | C | 7.6e-06 | 8.5e-06 | 6.3e-06 | 7.467E-06 | 1.106E-06 | 6.300E-06 | 8.500E-06 |
| 16909 | 846 | G846H | C | 7.4e-06 | 9e-06 | 6.2e-06 | 7.533E-06 | 1.405E-06 | 6.200E-06 | 9.000E-06 |
| 16910 | 846 | G846I | C | 1.2e-05 | 6.4e-06 | 6.1e-06 | 8.167E-06 | 3.323E-06 | 6.100E-06 | 1.200E-05 |
| 16911 | 846 | G846L | C | 6.7e-06 | 1.3e-05 | 6.8e-06 | 8.833E-06 | 3.609E-06 | 6.700E-06 | 1.300E-05 |
| 16912 | 846 | G846K | C | 1.1e-05 | 8.1e-06 | 8.5e-06 | 9.200E-06 | 1.572E-06 | 8.100E-06 | 1.100E-05 |
| 16913 | 846 | G846M | C | 9.1e-06 | 6.3e-06 | 7.3e-06 | 7.567E-06 | 1.419E-06 | 6.300E-06 | 9.100E-06 |
| 16914 | 846 | G846F | C | 1.3e-05 | 4.3e-06 | 1.1e-05 | 9.433E-06 | 4.557E-06 | 4.300E-06 | 1.300E-05 |

| | | | | | | | | | | |
|---|---|---|---|---|---|---|---|---|---|---|
| 16915 | 846 | G846P | C | 7.5e-06 | 6.8e-06 | 6.6e-06 | 6.967E-06 | 4.726E-07 | 6.600E-06 | 7.500E-06 |
| 16916 | 846 | G846S | C | 6.3e-06 | 1.2e-05 | 8.7e-06 | 9.000E-06 | 2.862E-06 | 6.300E-06 | 1.200E-05 |
| 16917 | 846 | G846T | C | 4.1e-06 | 7.2e-06 | 8.8e-06 | 6.700E-06 | 2.390E-06 | 4.100E-06 | 8.800E-06 |
| 16918 | 846 | G846W | C | 9.7e-06 | 9.4e-06 | 9.1e-06 | 9.400E-06 | 3.000E-07 | 9.100E-06 | 9.700E-06 |
| 16919 | 846 | G846Y | C | 1.3e-05 | 5.9e-06 | 7e-06 | 8.633E-06 | 3.821E-06 | 5.900E-06 | 1.300E-05 |
| 16920 | 846 | G846V | C | 1e-05 | 4.2e-06 | 8.9e-06 | 7.700E-06 | 3.081E-06 | 4.200E-06 | 1.000E-05 |
| 16921 | 847 | L847A | C | 5.5e-06 | 7.5e-06 | 8.1e-06 | 7.033E-06 | 1.361E-06 | 5.500E-06 | 8.100E-06 |
| 16922 | 847 | L847R | C | 1e-05 | 1.2e-05 | 1e-05 | 1.067E-05 | 1.155E-06 | 1.000E-05 | 1.200E-05 |
| 16923 | 847 | L847N | C | 7e-06 | 1.3e-05 | 4.5e-06 | 8.167E-06 | 4.368E-06 | 4.500E-06 | 1.300E-05 |
| 16924 | 847 | L847D | C | 4.7e-06 | 6.3e-06 | 6.4e-06 | 5.800E-06 | 9.539E-07 | 4.700E-06 | 6.400E-06 |
| 16925 | 847 | L847C | C | 1.5e-05 | 5.3e-06 | 8.5e-06 | 9.600E-06 | 4.943E-06 | 5.300E-06 | 1.500E-05 |
| 16926 | 847 | L847E | C | 5.4e-06 | 9.2e-06 | 6.2e-06 | 6.933E-06 | 2.003E-06 | 5.400E-06 | 9.200E-06 |
| 16927 | 847 | L847Q | C | 5.9e-06 | 1.2e-05 | 6.3e-06 | 8.067E-06 | 3.412E-06 | 5.900E-06 | 1.200E-05 |
| 16928 | 847 | L847G | C | 8.9e-06 | 6.3e-06 | 2.1e-05 | 1.207E-05 | 7.845E-06 | 6.300E-06 | 2.100E-05 |
| 16929 | 847 | L847H | C | 5.9e-06 | 8.9e-06 | 1e-05 | 8.267E-06 | 2.122E-06 | 5.900E-06 | 1.000E-05 |
| 16930 | 847 | L847I | C | 9.9e-06 | 5e-06 | 7e-06 | 7.300E-06 | 2.464E-06 | 5.000E-06 | 9.900E-06 |
| 16931 | 847 | L847L | C | 4.3e-06 | 1.2e-05 | 1.3e-05 | 9.767E-06 | 4.761E-06 | 4.300E-06 | 1.300E-05 |
| 16932 | 847 | L847K | C | 9.6e-06 | 1e-05 | 7.4e-06 | 9.000E-06 | 1.400E-06 | 7.400E-06 | 1.000E-05 |
| 16933 | 847 | L847M | C | 1.2e-05 | 1.1e-05 | 1.1e-05 | 1.133E-05 | 5.774E-07 | 1.100E-05 | 1.200E-05 |
| 16934 | 847 | L847F | C | 8.5e-06 | 1.1e-05 | 6.7e-06 | 8.733E-06 | 2.159E-06 | 6.700E-06 | 1.100E-05 |
| 16935 | 847 | L847P | C | 6.5e-06 | 9.4e-06 | 6.3e-06 | 7.400E-06 | 1.735E-06 | 6.300E-06 | 9.400E-06 |
| 16936 | 847 | L847S | C | 7.3e-06 | 7.8e-06 | 1.2e-05 | 9.033E-06 | 2.581E-06 | 7.300E-06 | 1.200E-05 |
| 16937 | 847 | L847T | C | 1.1e-05 | 9.3e-06 | 9.3e-06 | 9.867E-06 | 9.815E-07 | 9.300E-06 | 1.100E-05 |
| 16938 | 847 | L847W | C | 6.4e-06 | 9.2e-06 | 7.4e-06 | 7.667E-06 | 1.419E-06 | 6.400E-06 | 9.200E-06 |
| 16939 | 847 | L847Y | C | 5.1e-06 | 6.7e-06 | 1e-05 | 7.267E-06 | 2.499E-06 | 5.100E-06 | 1.000E-05 |
| 16940 | 847 | L847V | C | 1.3e-05 | 4.4e-06 | 1.3e-05 | 1.013E-05 | 4.965E-06 | 4.400E-06 | 1.300E-05 |
| 16941 | 848 | V848A | C | 5.1e-06 | 1.4e-05 | 8.7e-06 | 9.267E-06 | 4.477E-06 | 5.100E-06 | 1.400E-05 |
| 16942 | 848 | V848R | C | 6.7e-06 | 5.2e-06 | 7e-06 | 6.300E-06 | 9.644E-07 | 5.200E-06 | 7.000E-06 |
| 16943 | 848 | V848N | C | 1.1e-05 | 4.6e-06 | 9.1e-06 | 8.233E-06 | 3.287E-06 | 4.600E-06 | 1.100E-05 |
| 16944 | 848 | V848D | C | 1.4e-05 | 8.9e-06 | 1.3e-05 | 1.197E-05 | 2.702E-06 | 8.900E-06 | 1.400E-05 |
| 16945 | 848 | V848C | C | 8.1e-06 | 9.9e-06 | 8.8e-06 | 8.933E-06 | 9.074E-07 | 8.100E-06 | 9.900E-06 |
| 16946 | 848 | V848E | C | 8.7e-06 | 8.9e-06 | 8.8e-06 | 8.800E-06 | 1.000E-07 | 8.700E-06 | 8.900E-06 |
| 16947 | 848 | V848Q | C | 8.3e-06 | 1.1e-05 | 1e-05 | 9.767E-06 | 1.365E-06 | 8.300E-06 | 1.100E-05 |
| 16948 | 848 | V848G | C | 4.8e-06 | 1.1e-05 | 7.8e-06 | 7.867E-06 | 3.101E-06 | 4.800E-06 | 1.100E-05 |
| 16949 | 848 | V848H | C | 5.4e-06 | 1.1e-05 | 8.5e-06 | 8.300E-06 | 2.805E-06 | 5.400E-06 | 1.100E-05 |
| 16950 | 848 | V848I | C | 8.2e-06 | 5.7e-06 | 1.2e-05 | 8.633E-06 | 3.172E-06 | 5.700E-06 | 1.200E-05 |
| 16951 | 848 | V848L | C | 9.2e-06 | 7.6e-06 | 6.6e-06 | 7.800E-06 | 1.311E-06 | 6.600E-06 | 9.200E-06 |
| 16952 | 848 | V848K | C | 4.8e-06 | 6.5e-06 | 5.3e-06 | 5.533E-06 | 8.737E-07 | 4.800E-06 | 6.500E-06 |
| 16953 | 848 | V848M | C | 8e-06 | 1.2e-05 | 1.9e-05 | 1.300E-05 | 5.568E-06 | 8.000E-06 | 1.900E-05 |
| 16954 | 848 | V848F | C | 7e-06 | 1.2e-05 | 1.2e-05 | 1.033E-05 | 2.887E-06 | 7.000E-06 | 1.200E-05 |
| 16955 | 848 | V848P | C | 1.4e-05 | 6e-06 | 7e-06 | 9.000E-06 | 4.359E-06 | 6.000E-06 | 1.400E-05 |
| 16956 | 848 | V848S | C | 6.4e-06 | 6.2e-06 | 7.4e-06 | 6.667E-06 | 6.429E-07 | 6.200E-06 | 7.400E-06 |
| 16957 | 848 | V848T | C | 1.3e-05 | 7.6e-06 | 1.2e-05 | 1.087E-05 | 2.873E-06 | 7.600E-06 | 1.300E-05 |
| 16958 | 848 | V848W | C | 6.7e-06 | 1.4e-05 | 6.7e-06 | 9.133E-06 | 4.215E-06 | 6.700E-06 | 1.400E-05 |
| 16959 | 848 | V848Y | C | 9.6e-06 | 9e-06 | 1.4e-05 | 1.087E-05 | 2.730E-06 | 9.000E-06 | 1.400E-05 |
| 16960 | 848 | V848V | C | 6.3e-06 | 6.2e-06 | 9.4e-06 | 7.300E-06 | 1.819E-06 | 6.200E-06 | 9.400E-06 |
| 16961 | 849 | Q849A | C | 9.7e-06 | 1.2e-05 | 6.7e-06 | 9.467E-06 | 2.658E-06 | 6.700E-06 | 1.200E-05 |
| 16962 | 849 | Q849R | C | 7.1e-06 | 1.2e-05 | 7.6e-06 | 8.900E-06 | 2.696E-06 | 7.100E-06 | 1.200E-05 |
| 16963 | 849 | Q849N | C | 4e-06 | 7.6e-06 | 4.1e-06 | 5.233E-06 | 2.050E-06 | 4.000E-06 | 7.600E-06 |
| 16964 | 849 | Q849D | C | 7.9e-06 | 9.5e-06 | 1.6e-05 | 1.113E-05 | 4.290E-06 | 7.900E-06 | 1.600E-05 |
| 16965 | 849 | Q849C | C | 6.1e-06 | 9.8e-06 | 5.3e-06 | 7.067E-06 | 2.401E-06 | 5.300E-06 | 9.800E-06 |
| 16966 | 849 | Q849E | C | 8.6e-06 | 1.3e-05 | 6.4e-06 | 9.333E-06 | 3.361E-06 | 6.400E-06 | 1.300E-05 |
| 16967 | 849 | Q849Q | C | 4.6e-06 | 6.4e-06 | 8.8e-06 | 6.600E-06 | 2.107E-06 | 4.600E-06 | 8.800E-06 |
| 16968 | 849 | Q849G | C | 6.1e-06 | 9.6e-06 | 1.5e-05 | 1.023E-05 | 4.484E-06 | 6.100E-06 | 1.500E-05 |
| 16969 | 849 | Q849H | C | 5.4e-06 | 8.5e-06 | 4.9e-06 | 6.267E-06 | 1.950E-06 | 4.900E-06 | 8.500E-06 |
| 16970 | 849 | Q849I | C | 2e-05 | 7e-06 | 7.1e-06 | 1.137E-05 | 7.477E-06 | 7.000E-06 | 2.000E-05 |
| 16971 | 849 | Q849L | C | 6.5e-06 | 1.6e-05 | 6.7e-06 | 9.733E-06 | 5.428E-06 | 6.500E-06 | 1.600E-05 |
| 16972 | 849 | Q849K | C | 8.5e-06 | 7.6e-06 | 1.4e-05 | 1.003E-05 | 3.465E-06 | 7.600E-06 | 1.400E-05 |
| 16973 | 849 | Q849M | C | 5.9e-06 | 9.1e-06 | 1.4e-05 | 9.667E-06 | 4.080E-06 | 5.900E-06 | 1.400E-05 |
| 16974 | 849 | Q849F | C | 6.9e-06 | 5.4e-06 | 1e-05 | 7.433E-06 | 2.346E-06 | 5.400E-06 | 1.000E-05 |
| 16975 | 849 | Q849P | C | 5.5e-06 | 5.3e-06 | 5.8e-06 | 5.533E-06 | 2.517E-07 | 5.300E-06 | 5.800E-06 |
| 16976 | 849 | Q849S | C | 7.8e-06 | 7.5e-06 | 7.7e-06 | 7.667E-06 | 1.528E-07 | 7.500E-06 | 7.800E-06 |
| 16977 | 849 | Q849T | C | 7.7e-06 | 7.4e-06 | 8.4e-06 | 7.833E-06 | 5.132E-07 | 7.400E-06 | 8.400E-06 |
| 16978 | 849 | Q849W | C | 6.6e-06 | 1e-05 | 7.4e-06 | 8.000E-06 | 1.778E-06 | 6.600E-06 | 1.000E-05 |
| 16979 | 849 | Q849Y | C | 8e-06 | 9.6e-06 | 7.7e-06 | 8.433E-06 | 1.021E-06 | 7.700E-06 | 9.600E-06 |
| 16980 | 849 | Q849V | C | 8e-06 | 8.5e-06 | 7.2e-06 | 7.900E-06 | 6.557E-07 | 7.200E-06 | 8.500E-06 |
| 16981 | 850 | P850A | C | 5.1e-06 | 5.1e-06 | 7e-06 | 5.733E-06 | 1.097E-06 | 5.100E-06 | 7.000E-06 |
| 16982 | 850 | P850R | C | 7.5e-06 | 6.8e-06 | 1.4e-05 | 9.433E-06 | 3.970E-06 | 6.800E-06 | 1.400E-05 |
| 16983 | 850 | P850N | C | 8.9e-06 | 6e-06 | 1.6e-05 | 1.030E-05 | 5.145E-06 | 6.000E-06 | 1.600E-05 |
| 16984 | 850 | P850D | C | 1.1e-05 | 6.9e-06 | 1.1e-05 | 9.633E-06 | 2.367E-06 | 6.900E-06 | 1.100E-05 |
| 16985 | 850 | P850C | C | 5.9e-06 | 5.5e-06 | 4.4e-06 | 5.267E-06 | 7.767E-07 | 4.400E-06 | 5.900E-06 |
| 16986 | 850 | P850E | C | 3.8e-06 | 9.9e-06 | 8.8e-06 | 7.500E-06 | 3.251E-06 | 3.800E-06 | 9.900E-06 |
| 16987 | 850 | P850Q | C | 7.5e-06 | 4.8e-06 | 6.7e-06 | 6.333E-06 | 1.387E-06 | 4.800E-06 | 7.500E-06 |
| 16988 | 850 | P850G | C | 7e-06 | 8.5e-06 | 8.2e-06 | 7.900E-06 | 7.937E-07 | 7.000E-06 | 8.500E-06 |
| 16989 | 850 | P850H | C | 1.3e-05 | 6.4e-06 | 7.3e-06 | 8.900E-06 | 3.579E-06 | 6.400E-06 | 1.300E-05 |
| 16990 | 850 | P850I | C | 7.7e-06 | 7.6e-06 | 5.9e-06 | 7.067E-06 | 1.012E-06 | 5.900E-06 | 7.700E-06 |
| 16991 | 850 | P850L | C | 1.1e-05 | 6.7e-06 | 1.4e-05 | 1.057E-05 | 3.669E-06 | 6.700E-06 | 1.400E-05 |
| 16992 | 850 | P850K | C | 7.4e-06 | 7.2e-06 | 9.6e-06 | 8.067E-06 | 1.332E-06 | 7.200E-06 | 9.600E-06 |
| 16993 | 850 | P850M | C | 1.1e-05 | 6.8e-06 | 9.2e-06 | 9.000E-06 | 2.107E-06 | 6.800E-06 | 1.100E-05 |
| 16994 | 850 | P850F | C | 1.1e-05 | 1e-05 | 9.5e-06 | 1.017E-05 | 7.638E-07 | 9.500E-06 | 1.100E-05 |
| 16995 | 850 | P850P | C | 8.6e-06 | 1.3e-05 | 7.5e-06 | 9.700E-06 | 2.910E-06 | 7.500E-06 | 1.300E-05 |
| 16996 | 850 | P850S | C | 7.2e-06 | 1.9e-05 | 5e-06 | 1.040E-05 | 7.529E-06 | 5.000E-06 | 1.900E-05 |
| 16997 | 850 | P850T | C | 7.1e-06 | 5.3e-06 | 9.7e-06 | 7.367E-06 | 2.212E-06 | 5.300E-06 | 9.700E-06 |
| 16998 | 850 | P850W | C | 7.1e-06 | 8.3e-06 | 4.7e-06 | 6.700E-06 | 1.833E-06 | 4.700E-06 | 8.300E-06 |
| 16999 | 850 | P850Y | C | 1.3e-05 | 7.5e-06 | 7e-06 | 9.167E-06 | 3.329E-06 | 7.000E-06 | 1.300E-05 |

| | | | | | | | | | | |
|---|---|---|---|---|---|---|---|---|---|---|
| 17000 | 850 | P850V | C | 8.6e-06 | 1.1e-05 | 1e-05 | 9.867E-06 | 1.206E-06 | 8.600E-06 | 1.100E-05 |
| 17001 | 851 | G851A | C | 5.8e-06 | 9.1e-06 | 7.4e-06 | 7.433E-06 | 1.650E-06 | 5.800E-06 | 9.100E-06 |
| 17002 | 851 | G851R | C | 1e-05 | 6.1e-06 | 6.5e-06 | 7.533E-06 | 2.146E-06 | 6.100E-06 | 1.000E-05 |
| 17003 | 851 | G851N | C | 9.4e-06 | 6.7e-06 | 5e-06 | 7.033E-06 | 2.219E-06 | 5.000E-06 | 9.400E-06 |
| 17004 | 851 | G851D | C | 6.3e-06 | 1.2e-05 | 9.5e-06 | 9.267E-06 | 2.857E-06 | 6.300E-06 | 1.200E-05 |
| 17005 | 851 | G851C | C | 5.6e-06 | 1.1e-05 | 5.6e-06 | 7.400E-06 | 3.118E-06 | 5.600E-06 | 1.100E-05 |
| 17006 | 851 | G851E | C | 9.4e-06 | 1.1e-05 | 8.5e-06 | 9.633E-06 | 1.266E-06 | 8.500E-06 | 1.100E-05 |
| 17007 | 851 | G851Q | C | 6.7e-06 | 6.8e-06 | 9.9e-06 | 7.800E-06 | 1.819E-06 | 6.700E-06 | 9.900E-06 |
| 17008 | 851 | G851G | C | 9.3e-06 | 7e-06 | 7.9e-06 | 8.067E-06 | 1.159E-06 | 7.000E-06 | 9.300E-06 |
| 17009 | 851 | G851H | C | 4.5e-06 | 8.9e-06 | 1e-05 | 7.800E-06 | 2.910E-06 | 4.500E-06 | 1.000E-05 |
| 17010 | 851 | G851I | C | 9.6e-06 | 1.2e-05 | 6.5e-06 | 9.367E-06 | 2.757E-06 | 6.500E-06 | 1.200E-05 |
| 17011 | 851 | G851L | C | 1.2e-05 | 6.8e-06 | 1.1e-05 | 9.933E-06 | 2.759E-06 | 6.800E-06 | 1.200E-05 |
| 17012 | 851 | G851K | C | 9.4e-06 | 8.2e-06 | 6.4e-06 | 8.000E-06 | 1.510E-06 | 6.400E-06 | 9.400E-06 |
| 17013 | 851 | G851M | C | 9.9e-06 | 9.6e-06 | 9.7e-06 | 9.733E-06 | 1.528E-07 | 9.600E-06 | 9.900E-06 |
| 17014 | 851 | G851F | C | 7e-06 | 6.4e-06 | 9.7e-06 | 7.700E-06 | 1.758E-06 | 6.400E-06 | 9.700E-06 |
| 17015 | 851 | G851P | C | 6.3e-06 | 8.4e-06 | 7e-06 | 7.233E-06 | 1.069E-06 | 6.300E-06 | 8.400E-06 |
| 17016 | 851 | G851S | C | 1e-05 | 6.2e-06 | 7e-06 | 7.733E-06 | 2.003E-06 | 6.200E-06 | 1.000E-05 |
| 17017 | 851 | G851T | C | 6.5e-06 | 9.8e-06 | 6.4e-06 | 7.567E-06 | 1.935E-06 | 6.400E-06 | 9.800E-06 |
| 17018 | 851 | G851W | C | 5.1e-06 | 7.9e-06 | 1.3e-05 | 8.667E-06 | 4.005E-06 | 5.100E-06 | 1.300E-05 |
| 17019 | 851 | G851Y | C | 4.7e-06 | 9.3e-06 | 8.2e-06 | 7.400E-06 | 2.402E-06 | 4.700E-06 | 9.300E-06 |
| 17020 | 851 | G851V | C | 5.3e-06 | 9.8e-06 | 1.2e-05 | 9.033E-06 | 3.415E-06 | 5.300E-06 | 1.200E-05 |
| 17021 | 852 | G852A | C | 1.1e-05 | 9.5e-06 | 1.1e-05 | 1.050E-05 | 8.660E-07 | 9.500E-06 | 1.100E-05 |
| 17022 | 852 | G852R | C | 6.8e-06 | 7.9e-06 | 6.2e-06 | 6.967E-06 | 8.622E-07 | 6.200E-06 | 7.900E-06 |
| 17023 | 852 | G852N | C | 6.1e-06 | 1e-05 | 4.8e-06 | 6.967E-06 | 2.706E-06 | 4.800E-06 | 1.000E-05 |
| 17024 | 852 | G852D | C | 7.1e-06 | 5.6e-06 | 6.9e-06 | 6.533E-06 | 8.145E-07 | 5.600E-06 | 7.100E-06 |
| 17025 | 852 | G852C | C | 1.1e-05 | 7.3e-06 | 8.9e-06 | 9.067E-06 | 1.856E-06 | 7.300E-06 | 1.100E-05 |
| 17026 | 852 | G852E | C | 9.6e-06 | 9.2e-06 | 8.4e-06 | 9.067E-06 | 6.110E-07 | 8.400E-06 | 9.600E-06 |
| 17027 | 852 | G852Q | C | 7.6e-06 | 7.8e-06 | 1.3e-05 | 9.467E-06 | 3.062E-06 | 7.600E-06 | 1.300E-05 |
| 17028 | 852 | G852G | C | 6.4e-06 | 7.9e-06 | 8.4e-06 | 7.567E-06 | 1.041E-06 | 6.400E-06 | 8.400E-06 |
| 17029 | 852 | G852H | C | 1.2e-05 | 9.1e-06 | 7.1e-06 | 9.400E-06 | 2.464E-06 | 7.100E-06 | 1.200E-05 |
| 17030 | 852 | G852I | C | 4.6e-06 | 8.9e-06 | 7e-06 | 6.833E-06 | 2.155E-06 | 4.600E-06 | 8.900E-06 |
| 17031 | 852 | G852L | C | 8.1e-06 | 1e-05 | 7.6e-06 | 8.567E-06 | 1.266E-06 | 7.600E-06 | 1.000E-05 |
| 17032 | 852 | G852K | C | 7.8e-06 | 1.1e-05 | 7e-06 | 8.600E-06 | 2.117E-06 | 7.000E-06 | 1.100E-05 |
| 17033 | 852 | G852M | C | 7.8e-06 | 1.1e-05 | 5.9e-06 | 8.233E-06 | 2.577E-06 | 5.900E-06 | 1.100E-05 |
| 17034 | 852 | G852F | C | 6.9e-06 | 7.4e-06 | 5.9e-06 | 6.733E-06 | 7.638E-07 | 5.900E-06 | 7.400E-06 |
| 17035 | 852 | G852P | C | 5.4e-06 | 8.3e-06 | 9.1e-06 | 7.600E-06 | 1.947E-06 | 5.400E-06 | 9.100E-06 |
| 17036 | 852 | G852S | C | 6.7e-06 | 7.2e-06 | 1e-05 | 7.967E-06 | 1.779E-06 | 6.700E-06 | 1.000E-05 |
| 17037 | 852 | G852T | C | 1.2e-05 | 9.6e-06 | 7.8e-06 | 9.800E-06 | 2.107E-06 | 7.800E-06 | 1.200E-05 |
| 17038 | 852 | G852W | C | 4.6e-06 | 7e-06 | 5.3e-06 | 5.633E-06 | 1.234E-06 | 4.600E-06 | 7.000E-06 |
| 17039 | 852 | G852Y | C | 6.2e-06 | 6e-06 | 3.5e-06 | 5.233E-06 | 1.504E-06 | 3.500E-06 | 6.200E-06 |
| 17040 | 852 | G852V | C | 1.3e-05 | 9.3e-06 | 5.1e-06 | 9.133E-06 | 3.953E-06 | 5.100E-06 | 1.300E-05 |
| 17041 | 853 | S853A | C | 1.2e-05 | 9.6e-06 | 9.2e-06 | 1.027E-05 | 1.514E-06 | 9.200E-06 | 1.200E-05 |
| 17042 | 853 | S853R | C | 9.5e-06 | 8.5e-06 | 1.1e-05 | 9.667E-06 | 1.258E-06 | 8.500E-06 | 1.100E-05 |
| 17043 | 853 | S853N | C | 1e-05 | 5.6e-06 | 1.2e-05 | 9.200E-06 | 3.274E-06 | 5.600E-06 | 1.200E-05 |
| 17044 | 853 | S853D | C | 7.2e-06 | 8.8e-06 | 6.8e-06 | 7.600E-06 | 1.058E-06 | 6.800E-06 | 8.800E-06 |
| 17045 | 853 | S853C | C | 9e-06 | 9.8e-06 | 6.7e-06 | 8.500E-06 | 1.609E-06 | 6.700E-06 | 9.800E-06 |
| 17046 | 853 | S853E | C | 1.3e-05 | 6.5e-06 | 7.1e-06 | 8.867E-06 | 3.592E-06 | 6.500E-06 | 1.300E-05 |
| 17047 | 853 | S853Q | C | 8.7e-06 | 1.3e-05 | 9.3e-06 | 1.033E-05 | 2.329E-06 | 8.700E-06 | 1.300E-05 |
| 17048 | 853 | S853G | C | 4.4e-06 | 1e-05 | 7.4e-06 | 7.267E-06 | 2.802E-06 | 4.400E-06 | 1.000E-05 |
| 17049 | 853 | S853H | C | 7.1e-06 | 5.3e-06 | 7.9e-06 | 6.767E-06 | 1.332E-06 | 5.300E-06 | 7.900E-06 |
| 17050 | 853 | S853I | C | 7e-06 | 6.6e-06 | 1.1e-05 | 8.200E-06 | 2.433E-06 | 6.600E-06 | 1.100E-05 |
| 17051 | 853 | S853L | C | 1.4e-05 | 7.6e-06 | 5.5e-06 | 9.033E-06 | 4.428E-06 | 5.500E-06 | 1.400E-05 |
| 17052 | 853 | S853K | C | 6.2e-06 | 6.9e-06 | 1.5e-05 | 9.367E-06 | 4.891E-06 | 6.200E-06 | 1.500E-05 |
| 17053 | 853 | S853M | C | 1.3e-05 | 7.3e-06 | 7.3e-06 | 9.200E-06 | 3.291E-06 | 7.300E-06 | 1.300E-05 |
| 17054 | 853 | S853F | C | 6.5e-06 | 5.9e-06 | 9.3e-06 | 7.233E-06 | 1.815E-06 | 5.900E-06 | 9.300E-06 |
| 17055 | 853 | S853P | C | 9.8e-06 | 8.9e-06 | 1.2e-05 | 1.023E-05 | 1.595E-06 | 8.900E-06 | 1.200E-05 |
| 17056 | 853 | S853S | C | 9.3e-06 | 8.7e-06 | 8.2e-06 | 8.733E-06 | 5.508E-07 | 8.200E-06 | 9.300E-06 |
| 17057 | 853 | S853T | C | 8.9e-06 | 7e-06 | 8e-06 | 7.967E-06 | 9.504E-07 | 7.000E-06 | 8.900E-06 |
| 17058 | 853 | S853W | C | 8.7e-06 | 6.5e-06 | 8e-06 | 7.733E-06 | 1.124E-06 | 6.500E-06 | 8.700E-06 |
| 17059 | 853 | S853Y | C | 7.2e-06 | 8.7e-06 | 6.7e-06 | 7.533E-06 | 1.041E-06 | 6.700E-06 | 8.700E-06 |
| 17060 | 853 | S853V | C | 7.7e-06 | 6.1e-06 | 8.1e-06 | 7.300E-06 | 1.058E-06 | 6.100E-06 | 8.100E-06 |
| 17061 | 854 | L854A | C | 4.3e-06 | 3e-06 | 3.6e-06 | 3.633E-06 | 6.506E-07 | 3.000E-06 | 4.300E-06 |
| 17062 | 854 | L854R | C | 1.3e-05 | 1e-05 | 5.6e-06 | 9.533E-06 | 3.722E-06 | 5.600E-06 | 1.300E-05 |
| 17063 | 854 | L854N | C | 1e-05 | 6.7e-06 | 9.2e-06 | 8.633E-06 | 1.721E-06 | 6.700E-06 | 1.000E-05 |
| 17064 | 854 | L854D | C | 9.9e-06 | 7.9e-06 | 9.3e-06 | 9.033E-06 | 1.026E-06 | 7.900E-06 | 9.900E-06 |
| 17065 | 854 | L854C | C | 9e-06 | 9.8e-06 | 1.1e-05 | 9.933E-06 | 1.007E-06 | 9.000E-06 | 1.100E-05 |
| 17066 | 854 | L854E | C | 8.8e-06 | 9e-06 | 9.6e-06 | 9.133E-06 | 4.163E-07 | 8.800E-06 | 9.600E-06 |
| 17067 | 854 | L854Q | C | 8.9e-06 | 3.9e-06 | 8.1e-06 | 6.967E-06 | 2.686E-06 | 3.900E-06 | 8.900E-06 |
| 17068 | 854 | L854G | C | 8.4e-06 | 5.2e-06 | 6.9e-06 | 6.833E-06 | 1.601E-06 | 5.200E-06 | 8.400E-06 |
| 17069 | 854 | L854H | C | 9.7e-06 | 7.3e-06 | 5.1e-06 | 7.367E-06 | 2.301E-06 | 5.100E-06 | 9.700E-06 |
| 17070 | 854 | L854I | C | 5.3e-06 | 7.2e-06 | 1.3e-05 | 8.500E-06 | 4.011E-06 | 5.300E-06 | 1.300E-05 |
| 17071 | 854 | L854L | C | 1.1e-05 | 1.4e-05 | 1.1e-05 | 1.200E-05 | 1.732E-06 | 1.100E-05 | 1.400E-05 |
| 17072 | 854 | L854K | C | 5.3e-06 | 6.6e-06 | 8e-06 | 6.633E-06 | 1.350E-06 | 5.300E-06 | 8.000E-06 |
| 17073 | 854 | L854M | C | 1.1e-05 | 7.3e-06 | 3.8e-06 | 7.367E-06 | 3.600E-06 | 3.800E-06 | 1.100E-05 |
| 17074 | 854 | L854F | C | 7.2e-06 | 7.7e-06 | 7.2e-06 | 7.367E-06 | 2.887E-07 | 7.200E-06 | 7.700E-06 |
| 17075 | 854 | L854P | C | 4.8e-06 | 9.1e-06 | 6.1e-06 | 6.667E-06 | 2.205E-06 | 4.800E-06 | 9.100E-06 |
| 17076 | 854 | L854S | C | 7.5e-06 | 1e-05 | 9.9e-06 | 9.133E-06 | 1.415E-06 | 7.500E-06 | 1.000E-05 |
| 17077 | 854 | L854T | C | 1.3e-05 | 1e-05 | 7.5e-06 | 1.017E-05 | 2.754E-06 | 7.500E-06 | 1.300E-05 |
| 17078 | 854 | L854W | C | 4.1e-06 | 1.2e-05 | 1.4e-05 | 1.003E-05 | 5.235E-06 | 4.100E-06 | 1.400E-05 |
| 17079 | 854 | L854Y | C | 9.5e-06 | 9.8e-06 | 5.1e-06 | 8.133E-06 | 2.631E-06 | 5.100E-06 | 9.800E-06 |
| 17080 | 854 | L854V | C | 8.4e-06 | 8.6e-06 | 1.2e-05 | 9.667E-06 | 2.023E-06 | 8.400E-06 | 1.200E-05 |
| 17081 | 855 | R855A | C | 7e-06 | 1.4e-05 | 9.2e-06 | 1.007E-05 | 3.580E-06 | 7.000E-06 | 1.400E-05 |
| 17082 | 855 | R855R | C | 8.7e-06 | 6.8e-06 | 8.9e-06 | 8.133E-06 | 1.159E-06 | 6.800E-06 | 8.900E-06 |
| 17083 | 855 | R855N | C | 6e-06 | 6.8e-06 | 9.3e-06 | 7.367E-06 | 1.721E-06 | 6.000E-06 | 9.300E-06 |
| 17084 | 855 | R855D | C | 4.8e-06 | 8.9e-06 | 9.9e-06 | 7.867E-06 | 2.702E-06 | 4.800E-06 | 9.900E-06 |

| | | | | | | | | | | |
|---|---|---|---|---|---|---|---|---|---|---|
| 17085 | 855 | R855C | C | 8.4e-06 | 6.8e-06 | 4.6e-06 | 6.600E-06 | 1.908E-06 | 4.600E-06 | 8.400E-06 |
| 17086 | 855 | R855E | C | 6.8e-06 | 9.1e-06 | 9.5e-06 | 8.467E-06 | 1.457E-06 | 6.800E-06 | 9.500E-06 |
| 17087 | 855 | R855Q | C | 6.8e-06 | 1.3e-05 | 6e-06 | 8.600E-06 | 3.831E-06 | 6.000E-06 | 1.300E-05 |
| 17088 | 855 | R855G | C | 7.5e-06 | 7.3e-06 | 7.7e-06 | 7.500E-06 | 2.000E-07 | 7.300E-06 | 7.700E-06 |
| 17089 | 855 | R855H | C | 8.6e-06 | 7.1e-06 | 8.3e-06 | 8.000E-06 | 7.937E-07 | 7.100E-06 | 8.600E-06 |
| 17090 | 855 | R855I | C | 6.8e-06 | 8.7e-06 | 7.2e-06 | 7.567E-06 | 1.002E-06 | 6.800E-06 | 8.700E-06 |
| 17091 | 855 | R855L | C | 1e-05 | 8.5e-06 | 6.7e-06 | 8.400E-06 | 1.652E-06 | 6.700E-06 | 1.000E-05 |
| 17092 | 855 | R855K | C | 1.7e-05 | 7.8e-06 | 8.7e-06 | 1.117E-05 | 5.072E-06 | 7.800E-06 | 1.700E-05 |
| 17093 | 855 | R855M | C | 1.3e-05 | 1.4e-05 | 1.1e-05 | 1.267E-05 | 1.528E-06 | 1.100E-05 | 1.400E-05 |
| 17094 | 855 | R855F | C | 1e-05 | 4.9e-06 | 5e-06 | 6.633E-06 | 2.916E-06 | 4.900E-06 | 1.000E-05 |
| 17095 | 855 | R855P | C | 6.4e-06 | 8.9e-06 | 6.2e-06 | 7.167E-06 | 1.504E-06 | 6.200E-06 | 8.900E-06 |
| 17096 | 855 | R855S | C | 6.5e-06 | 5e-06 | 9e-06 | 6.833E-06 | 2.021E-06 | 5.000E-06 | 9.000E-06 |
| 17097 | 855 | R855T | C | 2.9e-06 | 7.2e-06 | 6.4e-06 | 5.500E-06 | 2.287E-06 | 2.900E-06 | 7.200E-06 |
| 17098 | 855 | R855W | C | 7.9e-06 | 5.3e-06 | 9.9e-06 | 7.700E-06 | 2.307E-06 | 5.300E-06 | 9.900E-06 |
| 17099 | 855 | R855Y | C | 5.3e-06 | 5.2e-06 | 4.4e-06 | 4.967E-06 | 4.933E-07 | 4.400E-06 | 5.300E-06 |
| 17100 | 855 | R855V | C | 1e-05 | 1.6e-05 | 7.3e-06 | 1.110E-05 | 4.453E-06 | 7.300E-06 | 1.600E-05 |
| 17101 | 856 | L856A | C | 6.7e-06 | 1.5e-05 | 1.1e-05 | 1.090E-05 | 4.151E-06 | 6.700E-06 | 1.500E-05 |
| 17102 | 856 | L856R | C | 9.5e-06 | 5.4e-06 | 6.9e-06 | 7.267E-06 | 2.074E-06 | 5.400E-06 | 9.500E-06 |
| 17103 | 856 | L856N | C | 6.8e-06 | 8.6e-06 | 5.8e-06 | 7.067E-06 | 1.419E-06 | 5.800E-06 | 8.600E-06 |
| 17104 | 856 | L856D | C | 1.2e-05 | 5.7e-06 | 5.5e-06 | 7.733E-06 | 3.696E-06 | 5.500E-06 | 1.200E-05 |
| 17105 | 856 | L856C | C | 5.2e-06 | 4.3e-06 | 6.7e-06 | 5.400E-06 | 1.212E-06 | 4.300E-06 | 6.700E-06 |
| 17106 | 856 | L856E | C | 7.1e-06 | 1.1e-05 | 8e-06 | 8.700E-06 | 2.042E-06 | 7.100E-06 | 1.100E-05 |
| 17107 | 856 | L856Q | C | 7.2e-06 | 1.2e-05 | 9e-06 | 9.400E-06 | 2.425E-06 | 7.200E-06 | 1.200E-05 |
| 17108 | 856 | L856G | C | 8e-06 | 6.7e-06 | 1.3e-05 | 9.233E-06 | 3.326E-06 | 6.700E-06 | 1.300E-05 |
| 17109 | 856 | L856H | C | 6.5e-06 | 6.1e-06 | 8.5e-06 | 7.033E-06 | 1.286E-06 | 6.100E-06 | 8.500E-06 |
| 17110 | 856 | L856I | C | 4.8e-06 | 5.8e-06 | 9.4e-06 | 6.667E-06 | 2.419E-06 | 4.800E-06 | 9.400E-06 |
| 17111 | 856 | L856L | C | 8.3e-06 | 8.1e-06 | 9.4e-06 | 8.600E-06 | 7.000E-07 | 8.100E-06 | 9.400E-06 |
| 17112 | 856 | L856K | C | 1.1e-05 | 7.9e-06 | 8.9e-06 | 9.267E-06 | 1.582E-06 | 7.900E-06 | 1.100E-05 |
| 17113 | 856 | L856M | C | 8.7e-06 | 1.7e-05 | 7.4e-06 | 1.103E-05 | 5.208E-06 | 7.400E-06 | 1.700E-05 |
| 17114 | 856 | L856F | C | 5.9e-06 | 3.8e-06 | 8.3e-06 | 6.000E-06 | 2.252E-06 | 3.800E-06 | 8.300E-06 |
| 17115 | 856 | L856P | C | 8.5e-06 | 9.7e-06 | 8.7e-06 | 8.967E-06 | 6.429E-07 | 8.500E-06 | 9.700E-06 |
| 17116 | 856 | L856S | C | 4e-06 | 9.5e-06 | 1.1e-05 | 8.167E-06 | 3.686E-06 | 4.000E-06 | 1.100E-05 |
| 17117 | 856 | L856T | C | 8.6e-06 | 1.1e-05 | 8e-06 | 9.200E-06 | 1.587E-06 | 8.000E-06 | 1.100E-05 |
| 17118 | 856 | L856W | C | 6.7e-06 | 5.5e-06 | 5.7e-06 | 5.967E-06 | 6.429E-07 | 5.500E-06 | 6.700E-06 |
| 17119 | 856 | L856Y | C | 6.6e-06 | 8.1e-06 | 1.4e-05 | 9.567E-06 | 3.912E-06 | 6.600E-06 | 1.400E-05 |
| 17120 | 856 | L856V | C | 7.6e-06 | 1e-05 | 5.5e-06 | 7.700E-06 | 2.252E-06 | 5.500E-06 | 1.000E-05 |
| 17121 | 857 | S857A | C | 6.7e-06 | 5.8e-06 | 1e-05 | 7.500E-06 | 2.211E-06 | 5.800E-06 | 1.000E-05 |
| 17122 | 857 | S857R | C | 6.4e-06 | 7.5e-06 | 7e-06 | 6.967E-06 | 5.508E-07 | 6.400E-06 | 7.500E-06 |
| 17123 | 857 | S857N | C | 8.8e-06 | 8.3e-06 | 1.3e-05 | 1.003E-05 | 2.581E-06 | 8.300E-06 | 1.300E-05 |
| 17124 | 857 | S857D | C | 6.7e-06 | 6e-06 | 7.7e-06 | 6.800E-06 | 8.544E-07 | 6.000E-06 | 7.700E-06 |
| 17125 | 857 | S857C | C | 9.7e-06 | 8e-06 | 6.7e-06 | 8.133E-06 | 1.504E-06 | 6.700E-06 | 9.700E-06 |
| 17126 | 857 | S857E | C | 7.6e-06 | 9.5e-06 | 6.2e-06 | 7.767E-06 | 1.656E-06 | 6.200E-06 | 9.500E-06 |
| 17127 | 857 | S857Q | C | 9.2e-06 | 5.8e-06 | 6e-06 | 7.000E-06 | 1.908E-06 | 5.800E-06 | 9.200E-06 |
| 17128 | 857 | S857G | C | 6.5e-06 | 6.8e-06 | 1.2e-05 | 8.433E-06 | 3.092E-06 | 6.500E-06 | 1.200E-05 |
| 17129 | 857 | S857H | C | 6.6e-06 | 4.8e-06 | 9.2e-06 | 6.867E-06 | 2.212E-06 | 4.800E-06 | 9.200E-06 |
| 17130 | 857 | S857I | C | 6.6e-06 | 7e-06 | 3.8e-06 | 5.800E-06 | 1.744E-06 | 3.800E-06 | 7.000E-06 |
| 17131 | 857 | S857L | C | 1.1e-05 | 7.2e-06 | 1.4e-05 | 1.073E-05 | 3.408E-06 | 7.200E-06 | 1.400E-05 |
| 17132 | 857 | S857K | C | 1e-05 | 1e-05 | 7.5e-06 | 9.167E-06 | 1.443E-06 | 7.500E-06 | 1.000E-05 |
| 17133 | 857 | S857M | C | 1.1e-05 | 3.8e-06 | 6.7e-06 | 7.167E-06 | 3.623E-06 | 3.800E-06 | 1.100E-05 |
| 17134 | 857 | S857F | C | 7.3e-06 | 4.6e-06 | 6.9e-06 | 6.267E-06 | 1.457E-06 | 4.600E-06 | 7.300E-06 |
| 17135 | 857 | S857P | C | 8.4e-06 | 9.5e-06 | 4.7e-06 | 7.533E-06 | 2.515E-06 | 4.700E-06 | 9.500E-06 |
| 17136 | 857 | S857S | C | 8.2e-06 | 7.6e-06 | 6.2e-06 | 7.333E-06 | 1.026E-06 | 6.200E-06 | 8.200E-06 |
| 17137 | 857 | S857T | C | 3.3e-06 | 4.6e-06 | 9.8e-06 | 5.900E-06 | 3.439E-06 | 3.300E-06 | 9.800E-06 |
| 17138 | 857 | S857W | C | 8.6e-06 | 6.7e-06 | 7.8e-06 | 7.700E-06 | 9.539E-07 | 6.700E-06 | 8.600E-06 |
| 17139 | 857 | S857Y | C | 4.8e-06 | 5.1e-06 | 1.8e-05 | 9.300E-06 | 7.536E-06 | 4.800E-06 | 1.800E-05 |
| 17140 | 857 | S857V | C | 4.7e-06 | 7.2e-06 | 5.9e-06 | 5.933E-06 | 1.250E-06 | 4.700E-06 | 7.200E-06 |
| 17141 | 858 | C858A | C | 4.4e-06 | 8.6e-06 | 6.5e-06 | 6.500E-06 | 2.100E-06 | 4.400E-06 | 8.600E-06 |
| 17142 | 858 | C858R | C | 1.3e-05 | 7e-06 | 9.2e-06 | 9.733E-06 | 3.035E-06 | 7.000E-06 | 1.300E-05 |
| 17143 | 858 | C858N | C | 1.3e-05 | 4.9e-06 | 5.7e-06 | 7.867E-06 | 4.464E-06 | 4.900E-06 | 1.300E-05 |
| 17144 | 858 | C858D | C | 8.8e-06 | 1.4e-05 | 5e-06 | 9.267E-06 | 4.518E-06 | 5.000E-06 | 1.400E-05 |
| 17145 | 858 | C858C | C | 5e-06 | 8.9e-06 | 6.9e-06 | 6.933E-06 | 1.950E-06 | 5.000E-06 | 8.900E-06 |
| 17146 | 858 | C858E | C | 6.2e-06 | 1.2e-05 | 8.5e-06 | 8.900E-06 | 2.921E-06 | 6.200E-06 | 1.200E-05 |
| 17147 | 858 | C858Q | C | 4.9e-06 | 4.6e-06 | 6.5e-06 | 5.333E-06 | 1.021E-06 | 4.600E-06 | 6.500E-06 |
| 17148 | 858 | C858G | C | 5e-06 | 6.3e-06 | 8.1e-06 | 6.467E-06 | 1.557E-06 | 5.000E-06 | 8.100E-06 |
| 17149 | 858 | C858H | C | 1.4e-05 | 8.7e-06 | 7.6e-06 | 1.010E-05 | 3.422E-06 | 7.600E-06 | 1.400E-05 |
| 17150 | 858 | C858I | C | 1e-05 | 8.2e-06 | 3e-06 | 7.067E-06 | 3.635E-06 | 3.000E-06 | 1.000E-05 |
| 17151 | 858 | C858L | C | 9.1e-06 | 7.2e-06 | 3.2e-06 | 6.500E-06 | 3.012E-06 | 3.200E-06 | 9.100E-06 |
| 17152 | 858 | C858K | C | 9.7e-06 | 5.7e-06 | 7.1e-06 | 7.500E-06 | 2.030E-06 | 5.700E-06 | 9.700E-06 |
| 17153 | 858 | C858M | C | 6.9e-06 | 1.1e-05 | 8.5e-06 | 8.800E-06 | 2.066E-06 | 6.900E-06 | 1.100E-05 |
| 17154 | 858 | C858F | C | 1.2e-05 | 8.4e-06 | 9.6e-06 | 1.000E-05 | 1.833E-06 | 8.400E-06 | 1.200E-05 |
| 17155 | 858 | C858P | C | 5.8e-06 | 8.1e-06 | 1.1e-05 | 8.300E-06 | 2.606E-06 | 5.800E-06 | 1.100E-05 |
| 17156 | 858 | C858S | C | 1e-05 | 1e-05 | 8.3e-06 | 9.433E-06 | 9.815E-07 | 8.300E-06 | 1.000E-05 |
| 17157 | 858 | C858T | C | 9.6e-06 | 8.4e-06 | 9.4e-06 | 9.133E-06 | 6.429E-07 | 8.400E-06 | 9.600E-06 |
| 17158 | 858 | C858W | C | 5.9e-06 | 8.9e-06 | 6.9e-06 | 7.233E-06 | 1.528E-06 | 5.900E-06 | 8.900E-06 |
| 17159 | 858 | C858Y | C | 7.1e-06 | 7.2e-06 | 6.9e-06 | 7.067E-06 | 1.528E-07 | 6.900E-06 | 7.200E-06 |
| 17160 | 858 | C858V | C | 9.3e-06 | 9e-06 | 1.4e-05 | 1.077E-05 | 2.804E-06 | 9.000E-06 | 1.400E-05 |
| 17161 | 859 | A859A | C | 6.5e-06 | 6.2e-06 | 5.6e-06 | 6.100E-06 | 4.583E-07 | 5.600E-06 | 6.500E-06 |
| 17162 | 859 | A859R | C | 3.1e-06 | 8.3e-06 | 5.6e-06 | 5.667E-06 | 2.601E-06 | 3.100E-06 | 8.300E-06 |
| 17163 | 859 | A859N | C | 6.7e-06 | 1.5e-05 | 9e-06 | 1.023E-05 | 4.285E-06 | 6.700E-06 | 1.500E-05 |
| 17164 | 859 | A859D | C | 1.1e-05 | 9.8e-06 | 1.3e-05 | 1.127E-05 | 1.617E-06 | 9.800E-06 | 1.300E-05 |
| 17165 | 859 | A859C | C | 6.3e-06 | 7.8e-06 | 1.1e-05 | 8.367E-06 | 2.401E-06 | 6.300E-06 | 1.100E-05 |
| 17166 | 859 | A859E | C | 4.7e-06 | 9.5e-06 | 7e-06 | 7.067E-06 | 2.401E-06 | 4.700E-06 | 9.500E-06 |
| 17167 | 859 | A859Q | C | 6.2e-06 | 6.6e-06 | 8.3e-06 | 7.033E-06 | 1.115E-06 | 6.200E-06 | 8.300E-06 |
| 17168 | 859 | A859G | C | 6.8e-06 | 1.1e-05 | 8.4e-06 | 8.733E-06 | 2.120E-06 | 6.800E-06 | 1.100E-05 |
| 17169 | 859 | A859H | C | 1.3e-05 | 8.6e-06 | 9.2e-06 | 1.027E-05 | 2.386E-06 | 8.600E-06 | 1.300E-05 |

| | | | | | | | | | | |
|---|---|---|---|---|---|---|---|---|---|---|
| 17170 | 859 | A859I | C | 9.7e-06 | 5.6e-06 | 6.4e-06 | 7.233E-06 | 2.173E-06 | 5.600E-06 | 9.700E-06 |
| 17171 | 859 | A859L | C | 6.2e-06 | 6.5e-06 | 1.1e-05 | 7.900E-06 | 2.689E-06 | 6.200E-06 | 1.100E-05 |
| 17172 | 859 | A859K | C | 1.2e-05 | 1.4e-05 | 5.7e-06 | 1.057E-05 | 4.332E-06 | 5.700E-06 | 1.400E-05 |
| 17173 | 859 | A859M | C | 9e-06 | 4e-06 | 5.1e-06 | 6.033E-06 | 2.627E-06 | 4.000E-06 | 9.000E-06 |
| 17174 | 859 | A859F | C | 7.3e-06 | 7.1e-06 | 7.8e-06 | 7.400E-06 | 3.606E-07 | 7.100E-06 | 7.800E-06 |
| 17175 | 859 | A859P | C | 6.8e-06 | 7.5e-06 | 1.1e-05 | 8.433E-06 | 2.250E-06 | 6.800E-06 | 1.100E-05 |
| 17176 | 859 | A859S | C | 4.7e-06 | 6.2e-06 | 4.2e-06 | 5.033E-06 | 1.041E-06 | 4.200E-06 | 6.200E-06 |
| 17177 | 859 | A859T | C | 6e-06 | 9.5e-06 | 6.6e-06 | 7.367E-06 | 1.872E-06 | 6.000E-06 | 9.500E-06 |
| 17178 | 859 | A859W | C | 6.3e-06 | 1.2e-05 | 9.2e-06 | 9.167E-06 | 2.850E-06 | 6.300E-06 | 1.200E-05 |
| 17179 | 859 | A859Y | C | 8.9e-06 | 4.6e-06 | 6.8e-06 | 6.767E-06 | 2.150E-06 | 4.600E-06 | 8.900E-06 |
| 17180 | 859 | A859V | C | 1.3e-05 | 9.8e-06 | 6.5e-06 | 9.767E-06 | 3.250E-06 | 6.500E-06 | 1.300E-05 |
| 17181 | 860 | A860A | C | 6.6e-06 | 9.2e-06 | 1.2e-05 | 9.267E-06 | 2.701E-06 | 6.600E-06 | 1.200E-05 |
| 17182 | 860 | A860R | C | 6.8e-06 | 9.9e-06 | 8.4e-06 | 8.367E-06 | 1.550E-06 | 6.800E-06 | 9.900E-06 |
| 17183 | 860 | A860N | C | 1.1e-05 | 9.1e-06 | 6.9e-06 | 9.000E-06 | 2.052E-06 | 6.900E-06 | 1.100E-05 |
| 17184 | 860 | A860D | C | 6.5e-06 | 4.6e-06 | 7.7e-06 | 6.267E-06 | 1.563E-06 | 4.600E-06 | 7.700E-06 |
| 17185 | 860 | A860C | C | 7e-06 | 6.9e-06 | 1.1e-05 | 8.300E-06 | 2.339E-06 | 6.900E-06 | 1.100E-05 |
| 17186 | 860 | A860E | C | 5.8e-06 | 8.2e-06 | 9.3e-06 | 7.767E-06 | 1.790E-06 | 5.800E-06 | 9.300E-06 |
| 17187 | 860 | A860Q | C | 9.5e-06 | 8.6e-06 | 8.7e-06 | 8.933E-06 | 4.933E-07 | 8.600E-06 | 9.500E-06 |
| 17188 | 860 | A860G | C | 9.9e-06 | 5.3e-06 | 1.2e-05 | 9.067E-06 | 3.427E-06 | 5.300E-06 | 1.200E-05 |
| 17189 | 860 | A860H | C | 6.1e-06 | 7.8e-06 | 8.1e-06 | 7.333E-06 | 1.079E-06 | 6.100E-06 | 8.100E-06 |
| 17190 | 860 | A860I | C | 1.4e-05 | 6.5e-06 | 4.4e-06 | 8.300E-06 | 5.047E-06 | 4.400E-06 | 1.400E-05 |
| 17191 | 860 | A860L | C | 8.8e-06 | 1.4e-05 | 1.3e-05 | 1.193E-05 | 2.759E-06 | 8.800E-06 | 1.400E-05 |
| 17192 | 860 | A860K | C | 9.3e-06 | 1e-05 | 7e-06 | 8.767E-06 | 1.570E-06 | 7.000E-06 | 1.000E-05 |
| 17193 | 860 | A860M | C | 9.5e-06 | 7.3e-06 | 6.7e-06 | 7.833E-06 | 1.474E-06 | 6.700E-06 | 9.500E-06 |
| 17194 | 860 | A860F | C | 8e-06 | 4.8e-06 | 1.3e-05 | 8.600E-06 | 4.133E-06 | 4.800E-06 | 1.300E-05 |
| 17195 | 860 | A860P | C | 5.6e-06 | 6.7e-06 | 5.6e-06 | 5.967E-06 | 6.351E-07 | 5.600E-06 | 6.700E-06 |
| 17196 | 860 | A860S | C | 6.2e-06 | 3.9e-06 | 9.7e-06 | 6.600E-06 | 2.921E-06 | 3.900E-06 | 9.700E-06 |
| 17197 | 860 | A860T | C | 4.2e-06 | 6e-06 | 7.7e-06 | 5.967E-06 | 1.750E-06 | 4.200E-06 | 7.700E-06 |
| 17198 | 860 | A860W | C | 1.1e-05 | 6e-06 | 4.4e-06 | 7.133E-06 | 3.443E-06 | 4.400E-06 | 1.100E-05 |
| 17199 | 860 | A860Y | C | 8.5e-06 | 1.1e-05 | 1.1e-05 | 1.017E-05 | 1.443E-06 | 8.500E-06 | 1.100E-05 |
| 17200 | 860 | A860V | C | 9.7e-06 | 6.5e-06 | 5.8e-06 | 7.333E-06 | 2.079E-06 | 5.800E-06 | 9.700E-06 |
| 17201 | 861 | S861A | C | 1.1e-05 | 7.8e-06 | 3.2e-06 | 7.333E-06 | 3.921E-06 | 3.200E-06 | 1.100E-05 |
| 17202 | 861 | S861R | C | 6.6e-06 | 6e-06 | 8.9e-06 | 7.167E-06 | 1.531E-06 | 6.000E-06 | 8.900E-06 |
| 17203 | 861 | S861N | C | 1.3e-05 | 6.5e-06 | 5.5e-06 | 8.333E-06 | 4.072E-06 | 5.500E-06 | 1.300E-05 |
| 17204 | 861 | S861D | C | 8.7e-06 | 4.5e-06 | 5.5e-06 | 6.233E-06 | 2.194E-06 | 4.500E-06 | 8.700E-06 |
| 17205 | 861 | S861C | C | 5.6e-06 | 7.9e-06 | 1.2e-05 | 8.500E-06 | 3.242E-06 | 5.600E-06 | 1.200E-05 |
| 17206 | 861 | S861E | C | 5.8e-06 | 6.5e-06 | 7.7e-06 | 6.667E-06 | 9.609E-07 | 5.800E-06 | 7.700E-06 |
| 17207 | 861 | S861Q | C | 6.1e-06 | 1.3e-05 | 7.8e-06 | 8.967E-06 | 3.595E-06 | 6.100E-06 | 1.300E-05 |
| 17208 | 861 | S861G | C | 9.6e-06 | 4.7e-06 | 1e-05 | 8.100E-06 | 2.951E-06 | 4.700E-06 | 1.000E-05 |
| 17209 | 861 | S861H | C | 7.6e-06 | 1.7e-05 | 5.2e-06 | 9.933E-06 | 6.236E-06 | 5.200E-06 | 1.700E-05 |
| 17210 | 861 | S861I | C | 8.6e-06 | 4.7e-06 | 4.2e-06 | 5.833E-06 | 2.409E-06 | 4.200E-06 | 8.600E-06 |
| 17211 | 861 | S861L | C | 7.9e-06 | 8.1e-06 | 8.8e-06 | 8.267E-06 | 4.726E-07 | 7.900E-06 | 8.800E-06 |
| 17212 | 861 | S861K | C | 8.3e-06 | 6.3e-06 | 1.2e-05 | 8.867E-06 | 2.892E-06 | 6.300E-06 | 1.200E-05 |
| 17213 | 861 | S861M | C | 4.7e-06 | 7.7e-06 | 8.4e-06 | 6.933E-06 | 1.966E-06 | 4.700E-06 | 8.400E-06 |
| 17214 | 861 | S861F | C | 8.7e-06 | 4.7e-06 | 6.9e-06 | 6.767E-06 | 2.003E-06 | 4.700E-06 | 8.700E-06 |
| 17215 | 861 | S861P | C | 1.3e-05 | 1.3e-05 | 1.4e-05 | 1.333E-05 | 5.774E-07 | 1.300E-05 | 1.400E-05 |
| 17216 | 861 | S861S | C | 6.4e-06 | 4e-06 | 6.1e-06 | 5.500E-06 | 1.308E-06 | 4.000E-06 | 6.400E-06 |
| 17217 | 861 | S861T | C | 1e-05 | 4.9e-06 | 7.3e-06 | 7.400E-06 | 2.551E-06 | 4.900E-06 | 1.000E-05 |
| 17218 | 861 | S861W | C | 8.6e-06 | 6e-06 | 6.4e-06 | 7.000E-06 | 1.400E-06 | 6.000E-06 | 8.600E-06 |
| 17219 | 861 | S861Y | C | 7.3e-06 | 6.9e-06 | 4.4e-06 | 6.200E-06 | 1.572E-06 | 4.400E-06 | 7.300E-06 |
| 17220 | 861 | S861V | C | 7.8e-06 | 5e-06 | 5.5e-06 | 6.100E-06 | 1.493E-06 | 5.000E-06 | 7.800E-06 |
| 17221 | 862 | G862A | C | 5.6e-06 | 3.8e-06 | 6.7e-06 | 5.367E-06 | 1.464E-06 | 3.800E-06 | 6.700E-06 |
| 17222 | 862 | G862R | C | 1.2e-05 | 9.4e-06 | 6.6e-06 | 9.333E-06 | 2.701E-06 | 6.600E-06 | 1.200E-05 |
| 17223 | 862 | G862N | C | 5.4e-06 | 5.3e-06 | 9.2e-06 | 6.633E-06 | 2.223E-06 | 5.300E-06 | 9.200E-06 |
| 17224 | 862 | G862D | C | 7.8e-06 | 5.5e-06 | 6.2e-06 | 6.500E-06 | 1.179E-06 | 5.500E-06 | 7.800E-06 |
| 17225 | 862 | G862C | C | 9.5e-06 | 7.8e-06 | 6.1e-06 | 7.800E-06 | 1.700E-06 | 6.100E-06 | 9.500E-06 |
| 17226 | 862 | G862E | C | 1.3e-05 | 9e-06 | 5.7e-06 | 9.233E-06 | 3.656E-06 | 5.700E-06 | 1.300E-05 |
| 17227 | 862 | G862Q | C | 6.7e-06 | 9.3e-06 | 6.6e-06 | 7.533E-06 | 1.531E-06 | 6.600E-06 | 9.300E-06 |
| 17228 | 862 | G862G | C | 8.2e-06 | 9.9e-06 | 7.8e-06 | 8.633E-06 | 1.115E-06 | 7.800E-06 | 9.900E-06 |
| 17229 | 862 | G862H | C | 6.8e-06 | 1.7e-05 | 7.8e-06 | 1.053E-05 | 5.623E-06 | 6.800E-06 | 1.700E-05 |
| 17230 | 862 | G862I | C | 4.2e-06 | 1.2e-05 | 8.6e-06 | 8.267E-06 | 3.911E-06 | 4.200E-06 | 1.200E-05 |
| 17231 | 862 | G862L | C | 5.9e-06 | 7.2e-06 | 1.1e-05 | 8.033E-06 | 2.650E-06 | 5.900E-06 | 1.100E-05 |
| 17232 | 862 | G862K | C | 1.1e-05 | 5.4e-06 | 7.4e-06 | 7.933E-06 | 2.838E-06 | 5.400E-06 | 1.100E-05 |
| 17233 | 862 | G862M | C | 9.5e-06 | 7.8e-06 | 8.9e-06 | 8.733E-06 | 8.622E-07 | 7.800E-06 | 9.500E-06 |
| 17234 | 862 | G862F | C | 8.6e-06 | 1.1e-05 | 1e-05 | 9.867E-06 | 1.206E-06 | 8.600E-06 | 1.100E-05 |
| 17235 | 862 | G862P | C | 5.5e-06 | 6.9e-06 | 9.2e-06 | 7.200E-06 | 1.868E-06 | 5.500E-06 | 9.200E-06 |
| 17236 | 862 | G862S | C | 1e-05 | 4.2e-06 | 7.5e-06 | 7.233E-06 | 2.909E-06 | 4.200E-06 | 1.000E-05 |
| 17237 | 862 | G862T | C | 6e-06 | 8.6e-06 | 6.6e-06 | 7.067E-06 | 1.361E-06 | 6.000E-06 | 8.600E-06 |
| 17238 | 862 | G862W | C | 6e-06 | 8.6e-06 | 4.9e-06 | 6.500E-06 | 1.900E-06 | 4.900E-06 | 8.600E-06 |
| 17239 | 862 | G862Y | C | 9e-06 | 4.2e-06 | 6.3e-06 | 6.500E-06 | 2.406E-06 | 4.200E-06 | 9.000E-06 |
| 17240 | 862 | G862V | C | 6.6e-06 | 5.9e-06 | 8.1e-06 | 6.867E-06 | 1.124E-06 | 5.900E-06 | 8.100E-06 |
| 17241 | 863 | F863A | C | 1.4e-05 | 9.3e-06 | 4.9e-06 | 9.400E-06 | 4.551E-06 | 4.900E-06 | 1.400E-05 |
| 17242 | 863 | F863R | C | 6.5e-06 | 1.2e-05 | 6.2e-06 | 8.233E-06 | 3.265E-06 | 6.200E-06 | 1.200E-05 |
| 17243 | 863 | F863N | C | 9.5e-06 | 1e-05 | 1.3e-05 | 1.083E-05 | 1.893E-06 | 9.500E-06 | 1.300E-05 |
| 17244 | 863 | F863D | C | 9.9e-06 | 7.3e-06 | 6.7e-06 | 7.967E-06 | 1.701E-06 | 6.700E-06 | 9.900E-06 |
| 17245 | 863 | F863C | C | 5.7e-06 | 8.3e-06 | 5.6e-06 | 6.533E-06 | 1.531E-06 | 5.600E-06 | 8.300E-06 |
| 17246 | 863 | F863E | C | 8.1e-06 | 7.1e-06 | 9.9e-06 | 8.367E-06 | 1.419E-06 | 7.100E-06 | 9.900E-06 |
| 17247 | 863 | F863Q | C | 6.2e-06 | 1.1e-05 | 6.8e-06 | 8.000E-06 | 2.615E-06 | 6.200E-06 | 1.100E-05 |
| 17248 | 863 | F863G | C | 4e-06 | 4.9e-06 | 3.8e-06 | 4.233E-06 | 5.859E-07 | 3.800E-06 | 4.900E-06 |
| 17249 | 863 | F863H | C | 1.1e-05 | 5e-06 | 4.9e-06 | 6.967E-06 | 3.493E-06 | 4.900E-06 | 1.100E-05 |
| 17250 | 863 | F863I | C | 9.3e-06 | 9.8e-06 | 7e-06 | 8.700E-06 | 1.493E-06 | 7.000E-06 | 9.800E-06 |
| 17251 | 863 | F863L | C | 7.4e-06 | 6.1e-06 | 1e-05 | 7.833E-06 | 1.986E-06 | 6.100E-06 | 1.000E-05 |
| 17252 | 863 | F863K | C | 8.4e-06 | 5.4e-06 | 5.7e-06 | 6.500E-06 | 1.652E-06 | 5.400E-06 | 8.400E-06 |
| 17253 | 863 | F863M | C | 8.3e-06 | 4.7e-06 | 5.8e-06 | 6.267E-06 | 1.845E-06 | 4.700E-06 | 8.300E-06 |
| 17254 | 863 | F863F | C | 9.3e-06 | 9.8e-06 | 5.3e-06 | 8.133E-06 | 2.466E-06 | 5.300E-06 | 9.800E-06 |

| | | | | | | | | | | |
|---|---|---|---|---|---|---|---|---|---|---|
| 17255 | 863 | F863P | C | 7.1e-06 | 8.9e-06 | 7.1e-06 | 7.700E-06 | 1.039E-06 | 7.100E-06 | 8.900E-06 |
| 17256 | 863 | F863S | C | 4.7e-06 | 5.8e-06 | 6.1e-06 | 5.533E-06 | 7.371E-07 | 4.700E-06 | 6.100E-06 |
| 17257 | 863 | F863T | C | 6.1e-06 | 1e-05 | 7e-06 | 7.700E-06 | 2.042E-06 | 6.100E-06 | 1.000E-05 |
| 17258 | 863 | F863W | C | 7.7e-06 | 1.3e-05 | 4.3e-06 | 8.333E-06 | 4.384E-06 | 4.300E-06 | 1.300E-05 |
| 17259 | 863 | F863Y | C | 7.2e-06 | 5.1e-06 | 5.9e-06 | 6.067E-06 | 1.060E-06 | 5.100E-06 | 7.200E-06 |
| 17260 | 863 | F863V | C | 4.7e-06 | 7.1e-06 | 1.6e-05 | 9.267E-06 | 5.953E-06 | 4.700E-06 | 1.600E-05 |
| 17261 | 864 | T864A | C | 1e-05 | 4.8e-06 | 1.2e-05 | 8.933E-06 | 3.717E-06 | 4.800E-06 | 1.200E-05 |
| 17262 | 864 | T864R | C | 6.9e-06 | 9.4e-06 | 8.9e-06 | 8.400E-06 | 1.323E-06 | 6.900E-06 | 9.400E-06 |
| 17263 | 864 | T864N | C | 4.6e-06 | 6.2e-06 | 6.2e-06 | 5.667E-06 | 9.238E-07 | 4.600E-06 | 6.200E-06 |
| 17264 | 864 | T864D | C | 1.2e-05 | 1.4e-05 | 5.5e-06 | 1.050E-05 | 4.444E-06 | 5.500E-06 | 1.400E-05 |
| 17265 | 864 | T864C | C | 1.2e-05 | 1e-05 | 6.8e-06 | 9.600E-06 | 2.623E-06 | 6.800E-06 | 1.200E-05 |
| 17266 | 864 | T864E | C | 9.1e-06 | 7.1e-06 | 9.2e-06 | 8.467E-06 | 1.185E-06 | 7.100E-06 | 9.200E-06 |
| 17267 | 864 | T864Q | C | 6e-06 | 5.1e-06 | 1.8e-05 | 9.700E-06 | 7.202E-06 | 5.100E-06 | 1.800E-05 |
| 17268 | 864 | T864G | C | 6.9e-06 | 9.7e-06 | 6.1e-06 | 7.567E-06 | 1.890E-06 | 6.100E-06 | 9.700E-06 |
| 17269 | 864 | T864H | C | 6.2e-06 | 1e-05 | 9.7e-06 | 8.633E-06 | 2.113E-06 | 6.200E-06 | 1.000E-05 |
| 17270 | 864 | T864I | C | 6.8e-06 | 5.8e-06 | 8.4e-06 | 7.000E-06 | 1.311E-06 | 5.800E-06 | 8.400E-06 |
| 17271 | 864 | T864L | C | 9.2e-06 | 5.3e-06 | 1e-05 | 8.167E-06 | 2.515E-06 | 5.300E-06 | 1.000E-05 |
| 17272 | 864 | T864K | C | 5.7e-06 | 5.3e-06 | 1e-05 | 7.000E-06 | 2.606E-06 | 5.300E-06 | 1.000E-05 |
| 17273 | 864 | T864M | C | 4.8e-06 | 8.3e-06 | 1.3e-05 | 8.700E-06 | 4.115E-06 | 4.800E-06 | 1.300E-05 |
| 17274 | 864 | T864F | C | 5.7e-06 | 6.1e-06 | 7.3e-06 | 6.367E-06 | 8.327E-07 | 5.700E-06 | 7.300E-06 |
| 17275 | 864 | T864P | C | 7.9e-06 | 6.8e-06 | 9.9e-06 | 8.200E-06 | 1.572E-06 | 6.800E-06 | 9.900E-06 |
| 17276 | 864 | T864S | C | 8.2e-06 | 5.1e-06 | 1e-05 | 7.767E-06 | 2.479E-06 | 5.100E-06 | 1.000E-05 |
| 17277 | 864 | T864T | C | 8.7e-06 | 1.3e-05 | 1.4e-05 | 1.190E-05 | 2.816E-06 | 8.700E-06 | 1.400E-05 |
| 17278 | 864 | T864W | C | 7.5e-06 | 4.3e-06 | 5.2e-06 | 5.667E-06 | 1.650E-06 | 4.300E-06 | 7.500E-06 |
| 17279 | 864 | T864Y | C | 8.2e-06 | 9e-06 | 8.9e-06 | 8.700E-06 | 4.359E-07 | 8.200E-06 | 9.000E-06 |
| 17280 | 864 | T864V | C | 1e-05 | 6.4e-06 | 7.4e-06 | 7.933E-06 | 1.858E-06 | 6.400E-06 | 1.000E-05 |
| 17281 | 865 | F865A | C | 8.4e-06 | 9.1e-06 | 7.3e-06 | 8.267E-06 | 9.074E-07 | 7.300E-06 | 9.100E-06 |
| 17282 | 865 | F865R | C | 1.4e-05 | 9.2e-06 | 6.6e-06 | 9.933E-06 | 3.754E-06 | 6.600E-06 | 1.400E-05 |
| 17283 | 865 | F865N | C | 7.4e-06 | 8.7e-06 | 9.8e-06 | 8.633E-06 | 1.201E-06 | 7.400E-06 | 9.800E-06 |
| 17284 | 865 | F865D | C | 5e-06 | 7.1e-06 | 9.5e-06 | 7.200E-06 | 2.252E-06 | 5.000E-06 | 9.500E-06 |
| 17285 | 865 | F865C | C | 1e-05 | 7.2e-06 | 1.3e-05 | 1.007E-05 | 2.901E-06 | 7.200E-06 | 1.300E-05 |
| 17286 | 865 | F865E | C | 4.4e-06 | 6.9e-06 | 8.5e-06 | 6.600E-06 | 2.066E-06 | 4.400E-06 | 8.500E-06 |
| 17287 | 865 | F865Q | C | 1.1e-05 | 7.1e-06 | 9.7e-06 | 9.267E-06 | 1.986E-06 | 7.100E-06 | 1.100E-05 |
| 17288 | 865 | F865G | C | 9.4e-06 | 8.3e-06 | 9.5e-06 | 9.067E-06 | 6.658E-07 | 8.300E-06 | 9.500E-06 |
| 17289 | 865 | F865H | C | 1.4e-05 | 4.8e-06 | 7.2e-06 | 8.667E-06 | 4.772E-06 | 4.800E-06 | 1.400E-05 |
| 17290 | 865 | F865I | C | 8.1e-06 | 9.1e-06 | 5.4e-06 | 7.533E-06 | 1.914E-06 | 5.400E-06 | 9.100E-06 |
| 17291 | 865 | F865L | C | 9.3e-06 | 8.1e-06 | 1e-05 | 9.133E-06 | 9.609E-07 | 8.100E-06 | 1.000E-05 |
| 17292 | 865 | F865K | C | 6.8e-06 | 1.4e-05 | 1.4e-05 | 1.160E-05 | 4.157E-06 | 6.800E-06 | 1.400E-05 |
| 17293 | 865 | F865M | C | 5.8e-06 | 1.2e-05 | 8.6e-06 | 8.800E-06 | 3.105E-06 | 5.800E-06 | 1.200E-05 |
| 17294 | 865 | F865F | C | 4.7e-06 | 6.2e-06 | 1e-05 | 6.967E-06 | 2.732E-06 | 4.700E-06 | 1.000E-05 |
| 17295 | 865 | F865P | C | 6.7e-06 | 8.6e-06 | 1.2e-05 | 9.100E-06 | 2.685E-06 | 6.700E-06 | 1.200E-05 |
| 17296 | 865 | F865S | C | 9.4e-06 | 2e-05 | 7.8e-06 | 1.240E-05 | 6.630E-06 | 7.800E-06 | 2.000E-05 |
| 17297 | 865 | F865T | C | 5.7e-06 | 7.7e-06 | 7.9e-06 | 7.100E-06 | 1.217E-06 | 5.700E-06 | 7.900E-06 |
| 17298 | 865 | F865W | C | 7.8e-06 | 7.6e-06 | 8e-06 | 7.800E-06 | 2.000E-07 | 7.600E-06 | 8.000E-06 |
| 17299 | 865 | F865Y | C | 4.9e-06 | 7.8e-06 | 8.3e-06 | 7.000E-06 | 1.836E-06 | 4.900E-06 | 8.300E-06 |
| 17300 | 865 | F865V | C | 9.3e-06 | 6.5e-06 | 1e-05 | 8.600E-06 | 1.852E-06 | 6.500E-06 | 1.000E-05 |
| 17301 | 866 | T866A | C | 7e-06 | 8.8e-06 | 6.7e-06 | 7.500E-06 | 1.136E-06 | 6.700E-06 | 8.800E-06 |
| 17302 | 866 | T866R | C | 6.5e-06 | 6.8e-06 | 9.5e-06 | 7.600E-06 | 1.652E-06 | 6.500E-06 | 9.500E-06 |
| 17303 | 866 | T866N | C | 8.4e-06 | 1e-05 | 4.6e-06 | 7.667E-06 | 2.774E-06 | 4.600E-06 | 1.000E-05 |
| 17304 | 866 | T866D | C | 6e-06 | 1e-05 | 1.3e-05 | 9.667E-06 | 3.512E-06 | 6.000E-06 | 1.300E-05 |
| 17305 | 866 | T866C | C | 6.1e-06 | 6.7e-06 | 4.8e-06 | 5.867E-06 | 9.713E-07 | 4.800E-06 | 6.700E-06 |
| 17306 | 866 | T866E | C | 5.9e-06 | 4.8e-06 | 5.8e-06 | 5.500E-06 | 6.083E-07 | 4.800E-06 | 5.900E-06 |
| 17307 | 866 | T866Q | C | 1.1e-05 | 9.5e-06 | 7.2e-06 | 9.233E-06 | 1.914E-06 | 7.200E-06 | 1.100E-05 |
| 17308 | 866 | T866G | C | 1e-05 | 7.3e-06 | 8e-06 | 8.433E-06 | 1.401E-06 | 7.300E-06 | 1.000E-05 |
| 17309 | 866 | T866H | C | 5e-06 | 6e-06 | 9.7e-06 | 6.900E-06 | 2.476E-06 | 5.000E-06 | 9.700E-06 |
| 17310 | 866 | T866I | C | 6.7e-06 | 8.1e-06 | 6.8e-06 | 7.200E-06 | 7.810E-07 | 6.700E-06 | 8.100E-06 |
| 17311 | 866 | T866L | C | 9.4e-06 | 4.3e-06 | 1e-05 | 7.900E-06 | 3.132E-06 | 4.300E-06 | 1.000E-05 |
| 17312 | 866 | T866K | C | 6.5e-06 | 6.6e-06 | 5.3e-06 | 6.133E-06 | 7.234E-07 | 5.300E-06 | 6.600E-06 |
| 17313 | 866 | T866M | C | 8.4e-06 | 7.2e-06 | 1.1e-05 | 8.867E-06 | 1.943E-06 | 7.200E-06 | 1.100E-05 |
| 17314 | 866 | T866F | C | 4.9e-06 | 7.8e-06 | 6.5e-06 | 6.400E-06 | 1.453E-06 | 4.900E-06 | 7.800E-06 |
| 17315 | 866 | T866P | C | 8.3e-06 | 4.1e-06 | 1e-05 | 7.467E-06 | 3.037E-06 | 4.100E-06 | 1.000E-05 |
| 17316 | 866 | T866S | C | 9.6e-06 | 7e-06 | 7.4e-06 | 8.000E-06 | 1.400E-06 | 7.000E-06 | 9.600E-06 |
| 17317 | 866 | T866T | C | 1.5e-05 | 8.5e-06 | 9.1e-06 | 1.087E-05 | 3.592E-06 | 8.500E-06 | 1.500E-05 |
| 17318 | 866 | T866W | C | 5e-06 | 6.2e-06 | 6.5e-06 | 5.900E-06 | 7.937E-07 | 5.000E-06 | 6.500E-06 |
| 17319 | 866 | T866Y | C | 8.1e-06 | 7.3e-06 | 1.2e-05 | 9.133E-06 | 2.515E-06 | 7.300E-06 | 1.200E-05 |
| 17320 | 866 | T866V | C | 7e-06 | 8.8e-06 | 6.4e-06 | 7.400E-06 | 1.249E-06 | 6.400E-06 | 8.800E-06 |
| 17321 | 867 | D867A | C | 1e-05 | 5.7e-06 | 9.1e-06 | 8.267E-06 | 2.268E-06 | 5.700E-06 | 1.000E-05 |
| 17322 | 867 | D867R | C | 9.9e-06 | 6.6e-06 | 6.9e-06 | 7.800E-06 | 1.825E-06 | 6.600E-06 | 9.900E-06 |
| 17323 | 867 | D867N | C | 7.4e-06 | 1e-05 | 4.9e-06 | 7.433E-06 | 2.550E-06 | 4.900E-06 | 1.000E-05 |
| 17324 | 867 | D867D | C | 5.7e-06 | 1e-05 | 9.4e-06 | 8.367E-06 | 2.329E-06 | 5.700E-06 | 1.000E-05 |
| 17325 | 867 | D867C | C | 1.2e-05 | 9.3e-06 | 7e-06 | 9.433E-06 | 2.503E-06 | 7.000E-06 | 1.200E-05 |
| 17326 | 867 | D867E | C | 8.2e-06 | 1.3e-05 | 8.1e-06 | 9.767E-06 | 2.801E-06 | 8.100E-06 | 1.300E-05 |
| 17327 | 867 | D867Q | C | 7.4e-06 | 1e-05 | 6.8e-06 | 8.067E-06 | 1.701E-06 | 6.800E-06 | 1.000E-05 |
| 17328 | 867 | D867G | C | 5.9e-06 | 7.3e-06 | 1.1e-05 | 8.067E-06 | 2.635E-06 | 5.900E-06 | 1.100E-05 |
| 17329 | 867 | D867H | C | 1.3e-05 | 8e-06 | 7.7e-06 | 9.567E-06 | 2.977E-06 | 7.700E-06 | 1.300E-05 |
| 17330 | 867 | D867I | C | 5.7e-06 | 9.9e-06 | 9e-06 | 8.200E-06 | 2.211E-06 | 5.700E-06 | 9.900E-06 |
| 17331 | 867 | D867L | C | 5.1e-06 | 9e-06 | 8e-06 | 7.367E-06 | 2.026E-06 | 5.100E-06 | 9.000E-06 |
| 17332 | 867 | D867K | C | 5.9e-06 | 6.6e-06 | 1e-05 | 7.500E-06 | 2.193E-06 | 5.900E-06 | 1.000E-05 |
| 17333 | 867 | D867M | C | 5.9e-06 | 9.2e-06 | 7.3e-06 | 7.467E-06 | 1.656E-06 | 5.900E-06 | 9.200E-06 |
| 17334 | 867 | D867F | C | 5.1e-06 | 1.4e-05 | 6.1e-06 | 8.400E-06 | 4.875E-06 | 5.100E-06 | 1.400E-05 |
| 17335 | 867 | D867P | C | 1.1e-05 | 1.3e-05 | 7.2e-06 | 1.040E-05 | 2.946E-06 | 7.200E-06 | 1.300E-05 |
| 17336 | 867 | D867S | C | 8.7e-06 | 5.9e-06 | 6.6e-06 | 7.067E-06 | 1.457E-06 | 5.900E-06 | 8.700E-06 |
| 17337 | 867 | D867T | C | 8.6e-06 | 8.3e-06 | 9.6e-06 | 8.833E-06 | 6.807E-07 | 8.300E-06 | 9.600E-06 |
| 17338 | 867 | D867W | C | 9.9e-06 | 6.5e-06 | 1.5e-05 | 1.047E-05 | 4.278E-06 | 6.500E-06 | 1.500E-05 |
| 17339 | 867 | D867Y | C | 9.1e-06 | 4.4e-06 | 4.9e-06 | 6.133E-06 | 2.581E-06 | 4.400E-06 | 9.100E-06 |

| | | | | | | | | | | |
|---|---|---|---|---|---|---|---|---|---|---|
| 17340 | 867 | D867V | C | 7.7e-06 | 6.1e-06 | 5.6e-06 | 6.467E-06 | 1.097E-06 | 5.600E-06 | 7.700E-06 |
| 17341 | 868 | Y868A | C | 6.5e-06 | 6e-06 | 4.7e-06 | 5.733E-06 | 9.292E-07 | 4.700E-06 | 6.500E-06 |
| 17342 | 868 | Y868R | C | 1.3e-05 | 9.1e-06 | 9.4e-06 | 1.050E-05 | 2.170E-06 | 9.100E-06 | 1.300E-05 |
| 17343 | 868 | Y868N | C | 1.3e-05 | 1.1e-05 | 5.9e-06 | 9.967E-06 | 3.661E-06 | 5.900E-06 | 1.300E-05 |
| 17344 | 868 | Y868D | C | 8.8e-06 | 5.9e-06 | 1.1e-05 | 8.567E-06 | 2.558E-06 | 5.900E-06 | 1.100E-05 |
| 17345 | 868 | Y868C | C | 8.7e-06 | 1.2e-05 | 8.7e-06 | 9.800E-06 | 1.905E-06 | 8.700E-06 | 1.200E-05 |
| 17346 | 868 | Y868E | C | 7e-06 | 6.3e-06 | 9.3e-06 | 7.533E-06 | 1.570E-06 | 6.300E-06 | 9.300E-06 |
| 17347 | 868 | Y868Q | C | 8.6e-06 | 1.1e-05 | 5.2e-06 | 8.267E-06 | 2.914E-06 | 5.200E-06 | 1.100E-05 |
| 17348 | 868 | Y868G | C | 7.5e-06 | 1.7e-05 | 8.8e-06 | 1.110E-05 | 5.151E-06 | 7.500E-06 | 1.700E-05 |
| 17349 | 868 | Y868H | C | 7.2e-06 | 6.5e-06 | 9.5e-06 | 7.733E-06 | 1.570E-06 | 6.500E-06 | 9.500E-06 |
| 17350 | 868 | Y868I | C | 6.3e-06 | 1.3e-05 | 8.4e-06 | 9.233E-06 | 3.427E-06 | 6.300E-06 | 1.300E-05 |
| 17351 | 868 | Y868L | C | 1.3e-05 | 7.6e-06 | 5.2e-06 | 8.600E-06 | 3.995E-06 | 5.200E-06 | 1.300E-05 |
| 17352 | 868 | Y868K | C | 7e-06 | 5.6e-06 | 1e-05 | 7.533E-06 | 2.248E-06 | 5.600E-06 | 1.000E-05 |
| 17353 | 868 | Y868M | C | 7.2e-06 | 1.1e-05 | 5.5e-06 | 7.900E-06 | 2.816E-06 | 5.500E-06 | 1.100E-05 |
| 17354 | 868 | Y868F | C | 5.6e-06 | 9.7e-06 | 7e-06 | 7.433E-06 | 2.084E-06 | 5.600E-06 | 9.700E-06 |
| 17355 | 868 | Y868P | C | 9.1e-06 | 9.2e-06 | 1.1e-05 | 9.767E-06 | 1.069E-06 | 9.100E-06 | 1.100E-05 |
| 17356 | 868 | Y868S | C | 9e-06 | 6.4e-06 | 6.5e-06 | 7.300E-06 | 1.473E-06 | 6.400E-06 | 9.000E-06 |
| 17357 | 868 | Y868T | C | 1.2e-05 | 4.8e-06 | 1.4e-05 | 1.027E-05 | 4.839E-06 | 4.800E-06 | 1.400E-05 |
| 17358 | 868 | Y868W | C | 1.1e-05 | 7.2e-06 | 6.9e-06 | 8.367E-06 | 2.285E-06 | 6.900E-06 | 1.100E-05 |
| 17359 | 868 | Y868Y | C | 1.1e-05 | 5e-06 | 4.3e-06 | 6.767E-06 | 3.683E-06 | 4.300E-06 | 1.100E-05 |
| 17360 | 868 | Y868V | C | 6e-06 | 8.8e-06 | 6.7e-06 | 7.167E-06 | 1.457E-06 | 6.000E-06 | 8.800E-06 |
| 17361 | 869 | T869A | C | 4.7e-06 | 5.6e-06 | 6.2e-06 | 5.500E-06 | 7.550E-07 | 4.700E-06 | 6.200E-06 |
| 17362 | 869 | T869R | C | 8.3e-06 | 8.9e-06 | 1e-05 | 9.067E-06 | 8.622E-07 | 8.300E-06 | 1.000E-05 |
| 17363 | 869 | T869N | C | 8.8e-06 | 6.9e-06 | 1.2e-05 | 9.233E-06 | 2.577E-06 | 6.900E-06 | 1.200E-05 |
| 17364 | 869 | T869D | C | 8.8e-06 | 8.4e-06 | 1.3e-05 | 1.007E-05 | 2.548E-06 | 8.400E-06 | 1.300E-05 |
| 17365 | 869 | T869C | C | 9e-06 | 6.2e-06 | 6.9e-06 | 7.367E-06 | 1.457E-06 | 6.200E-06 | 9.000E-06 |
| 17366 | 869 | T869E | C | 1.2e-05 | 7.4e-06 | 8.9e-06 | 9.433E-06 | 2.346E-06 | 7.400E-06 | 1.200E-05 |
| 17367 | 869 | T869Q | C | 9.3e-06 | 9.7e-06 | 8.3e-06 | 9.100E-06 | 7.211E-07 | 8.300E-06 | 9.700E-06 |
| 17368 | 869 | T869G | C | 7.2e-06 | 9.6e-06 | 4.8e-06 | 7.200E-06 | 2.400E-06 | 4.800E-06 | 9.600E-06 |
| 17369 | 869 | T869H | C | 5.9e-06 | 9.9e-06 | 5.5e-06 | 7.100E-06 | 2.433E-06 | 5.500E-06 | 9.900E-06 |
| 17370 | 869 | T869I | C | 7.5e-06 | 7.3e-06 | 1.2e-05 | 8.933E-06 | 2.658E-06 | 7.300E-06 | 1.200E-05 |
| 17371 | 869 | T869L | C | 7.7e-06 | 6.5e-06 | 8.7e-06 | 7.633E-06 | 1.102E-06 | 6.500E-06 | 8.700E-06 |
| 17372 | 869 | T869K | C | 1.3e-05 | 9.8e-06 | 6.9e-06 | 9.900E-06 | 3.051E-06 | 6.900E-06 | 1.300E-05 |
| 17373 | 869 | T869M | C | 1.5e-05 | 7.6e-06 | 8.7e-06 | 1.043E-05 | 3.993E-06 | 7.600E-06 | 1.500E-05 |
| 17374 | 869 | T869F | C | 5.1e-06 | 1.3e-05 | 7.5e-06 | 8.533E-06 | 4.050E-06 | 5.100E-06 | 1.300E-05 |
| 17375 | 869 | T869P | C | 9.9e-06 | 1.1e-05 | 1e-05 | 1.030E-05 | 6.083E-07 | 9.900E-06 | 1.100E-05 |
| 17376 | 869 | T869S | C | 1.2e-05 | 1.2e-05 | 6.9e-06 | 1.030E-05 | 2.944E-06 | 6.900E-06 | 1.200E-05 |
| 17377 | 869 | T869T | C | 1.2e-05 | 6.7e-06 | 1.1e-05 | 9.900E-06 | 2.816E-06 | 6.700E-06 | 1.200E-05 |
| 17378 | 869 | T869W | C | 6.9e-06 | 1.1e-05 | 1.4e-05 | 1.063E-05 | 3.564E-06 | 6.900E-06 | 1.400E-05 |
| 17379 | 869 | T869Y | C | 1.1e-05 | 9.5e-06 | 8.9e-06 | 9.800E-06 | 1.082E-06 | 8.900E-06 | 1.100E-05 |
| 17380 | 869 | T869V | C | 1e-05 | 3.7e-06 | 9.4e-06 | 7.700E-06 | 3.477E-06 | 3.700E-06 | 1.000E-05 |
| 17381 | 870 | M870A | C | 4.6e-06 | 4.4e-06 | 7.4e-06 | 5.467E-06 | 1.677E-06 | 4.400E-06 | 7.400E-06 |
| 17382 | 870 | M870R | C | 7.2e-06 | 8.3e-06 | 8.1e-06 | 7.867E-06 | 5.859E-07 | 7.200E-06 | 8.300E-06 |
| 17383 | 870 | M870N | C | 5.8e-06 | 7.1e-06 | 1e-05 | 7.633E-06 | 2.150E-06 | 5.800E-06 | 1.000E-05 |
| 17384 | 870 | M870D | C | 8.7e-06 | 1.7e-05 | 7.9e-06 | 1.120E-05 | 5.039E-06 | 7.900E-06 | 1.700E-05 |
| 17385 | 870 | M870C | C | 9.4e-06 | 4.8e-06 | 6.6e-06 | 6.933E-06 | 2.318E-06 | 4.800E-06 | 9.400E-06 |
| 17386 | 870 | M870E | C | 1.1e-05 | 4.8e-06 | 7.5e-06 | 7.767E-06 | 3.109E-06 | 4.800E-06 | 1.100E-05 |
| 17387 | 870 | M870Q | C | 7.3e-06 | 4.8e-06 | 3.3e-06 | 5.133E-06 | 2.021E-06 | 3.300E-06 | 7.300E-06 |
| 17388 | 870 | M870G | C | 7.6e-06 | 9.4e-06 | 1.5e-05 | 1.067E-05 | 3.859E-06 | 7.600E-06 | 1.500E-05 |
| 17389 | 870 | M870H | C | 9.4e-06 | 6.3e-06 | 3.1e-06 | 6.267E-06 | 3.150E-06 | 3.100E-06 | 9.400E-06 |
| 17390 | 870 | M870I | C | 8.4e-06 | 6.2e-06 | 1.1e-05 | 8.533E-06 | 2.403E-06 | 6.200E-06 | 1.100E-05 |
| 17391 | 870 | M870L | C | 5.3e-06 | 1.2e-05 | 9.1e-06 | 8.800E-06 | 3.360E-06 | 5.300E-06 | 1.200E-05 |
| 17392 | 870 | M870K | C | 1.1e-05 | 1.1e-05 | 6.9e-06 | 9.633E-06 | 2.367E-06 | 6.900E-06 | 1.100E-05 |
| 17393 | 870 | M870M | C | 1.3e-05 | 4.3e-06 | 6.5e-06 | 7.933E-06 | 4.524E-06 | 4.300E-06 | 1.300E-05 |
| 17394 | 870 | M870F | C | 6e-06 | 1.1e-05 | 7.9e-06 | 8.300E-06 | 2.524E-06 | 6.000E-06 | 1.100E-05 |
| 17395 | 870 | M870P | C | 5.1e-06 | 6.5e-06 | 5.8e-06 | 5.800E-06 | 7.000E-07 | 5.100E-06 | 6.500E-06 |
| 17396 | 870 | M870S | C | 9.9e-06 | 6.2e-06 | 5.8e-06 | 7.300E-06 | 2.261E-06 | 5.800E-06 | 9.900E-06 |
| 17397 | 870 | M870T | C | 3.5e-06 | 8.2e-06 | 5.6e-06 | 5.767E-06 | 2.354E-06 | 3.500E-06 | 8.200E-06 |
| 17398 | 870 | M870W | C | 6.5e-06 | 1.3e-05 | 7.4e-06 | 8.967E-06 | 3.522E-06 | 6.500E-06 | 1.300E-05 |
| 17399 | 870 | M870Y | C | 1.1e-05 | 1.1e-05 | 8.4e-06 | 1.013E-05 | 1.501E-06 | 8.400E-06 | 1.100E-05 |
| 17400 | 870 | M870V | C | 8.7e-06 | 6.3e-06 | 1.1e-05 | 8.667E-06 | 2.350E-06 | 6.300E-06 | 1.100E-05 |
| 17401 | 871 | D871A | C | 5.4e-06 | 6.4e-06 | 5.6e-06 | 5.800E-06 | 5.292E-07 | 5.400E-06 | 6.400E-06 |
| 17402 | 871 | D871R | C | 6.8e-06 | 6.6e-06 | 1.4e-05 | 9.133E-06 | 4.216E-06 | 6.600E-06 | 1.400E-05 |
| 17403 | 871 | D871N | C | 6.3e-06 | 6.4e-06 | 7.9e-06 | 6.867E-06 | 8.963E-07 | 6.300E-06 | 7.900E-06 |
| 17404 | 871 | D871D | C | 5.4e-06 | 9.5e-06 | 5.5e-06 | 6.800E-06 | 2.339E-06 | 5.400E-06 | 9.500E-06 |
| 17405 | 871 | D871C | C | 1.2e-05 | 6.7e-06 | 6.6e-06 | 8.433E-06 | 3.089E-06 | 6.600E-06 | 1.200E-05 |
| 17406 | 871 | D871E | C | 7.3e-06 | 5.3e-06 | 4.7e-06 | 5.767E-06 | 1.361E-06 | 4.700E-06 | 7.300E-06 |
| 17407 | 871 | D871Q | C | 9.7e-06 | 1.5e-05 | 6.3e-06 | 1.033E-05 | 4.384E-06 | 6.300E-06 | 1.500E-05 |
| 17408 | 871 | D871G | C | 1.2e-05 | 8.3e-06 | 1.2e-05 | 1.077E-05 | 2.136E-06 | 8.300E-06 | 1.200E-05 |
| 17409 | 871 | D871H | C | 7.3e-06 | 5.5e-06 | 9.2e-06 | 7.333E-06 | 1.850E-06 | 5.500E-06 | 9.200E-06 |
| 17410 | 871 | D871I | C | 4.8e-06 | 1.2e-05 | 7.3e-06 | 8.033E-06 | 3.656E-06 | 4.800E-06 | 1.200E-05 |
| 17411 | 871 | D871L | C | 4.4e-06 | 4.8e-06 | 9.1e-06 | 6.100E-06 | 2.606E-06 | 4.400E-06 | 9.100E-06 |
| 17412 | 871 | D871K | C | 9.5e-06 | 1.3e-05 | 5.1e-06 | 9.200E-06 | 3.959E-06 | 5.100E-06 | 1.300E-05 |
| 17413 | 871 | D871M | C | 6e-06 | 5.2e-06 | 4.4e-06 | 5.200E-06 | 8.000E-07 | 4.400E-06 | 6.000E-06 |
| 17414 | 871 | D871F | C | 1e-05 | 6.4e-06 | 8e-06 | 8.133E-06 | 1.804E-06 | 6.400E-06 | 1.000E-05 |
| 17415 | 871 | D871P | C | 9.5e-06 | 1.1e-05 | 8.7e-06 | 9.733E-06 | 1.168E-06 | 8.700E-06 | 1.100E-05 |
| 17416 | 871 | D871S | C | 7.2e-06 | 3.8e-06 | 1.3e-05 | 8.000E-06 | 4.652E-06 | 3.800E-06 | 1.300E-05 |
| 17417 | 871 | D871T | C | 9.1e-06 | 8.9e-06 | 7.6e-06 | 8.533E-06 | 8.145E-07 | 7.600E-06 | 9.100E-06 |
| 17418 | 871 | D871W | C | 8.7e-06 | 8.1e-06 | 1.3e-05 | 9.933E-06 | 2.673E-06 | 8.100E-06 | 1.300E-05 |
| 17419 | 871 | D871Y | C | 7.8e-06 | 1.2e-05 | 9.7e-06 | 9.833E-06 | 2.103E-06 | 7.800E-06 | 1.200E-05 |
| 17420 | 871 | D871V | C | 7.5e-06 | 9.2e-06 | 5.9e-06 | 7.533E-06 | 1.650E-06 | 5.900E-06 | 9.200E-06 |
| 17421 | 872 | W872A | C | 4.6e-06 | 8.6e-06 | 6.9e-06 | 6.700E-06 | 2.007E-06 | 4.600E-06 | 8.600E-06 |
| 17422 | 872 | W872R | C | 1.1e-05 | 4.8e-06 | 4.5e-06 | 6.767E-06 | 3.669E-06 | 4.500E-06 | 1.100E-05 |
| 17423 | 872 | W872N | C | 7e-06 | 1.5e-05 | 7.7e-06 | 9.900E-06 | 4.431E-06 | 7.000E-06 | 1.500E-05 |
| 17424 | 872 | W872D | C | 8.6e-06 | 1e-05 | 7e-06 | 8.533E-06 | 1.501E-06 | 7.000E-06 | 1.000E-05 |

| | | | | | | | | | | |
|---|---|---|---|---|---|---|---|---|---|---|
| 17425 | 872 | W872C | C | 8e-06 | 6.1e-06 | 7e-06 | 7.033E-06 | 9.504E-07 | 6.100E-06 | 8.000E-06 |
| 17426 | 872 | W872E | C | 6.9e-06 | 7.2e-06 | 8.6e-06 | 7.567E-06 | 9.074E-07 | 6.900E-06 | 8.600E-06 |
| 17427 | 872 | W872Q | C | 4.9e-06 | 7e-06 | 6.2e-06 | 6.033E-06 | 1.060E-06 | 4.900E-06 | 7.000E-06 |
| 17428 | 872 | W872G | C | 6.4e-06 | 6.4e-06 | 9.3e-06 | 7.367E-06 | 1.674E-06 | 6.400E-06 | 9.300E-06 |
| 17429 | 872 | W872H | C | 7.4e-06 | 8.9e-06 | 9.3e-06 | 8.533E-06 | 1.002E-06 | 7.400E-06 | 9.300E-06 |
| 17430 | 872 | W872I | C | 8.3e-06 | 1.5e-05 | 9.7e-06 | 1.100E-05 | 3.534E-06 | 8.300E-06 | 1.500E-05 |
| 17431 | 872 | W872L | C | 3e-06 | 7.1e-06 | 1.3e-05 | 7.700E-06 | 5.027E-06 | 3.000E-06 | 1.300E-05 |
| 17432 | 872 | W872K | C | 8.7e-06 | 6.7e-06 | 7.6e-06 | 7.667E-06 | 1.002E-06 | 6.700E-06 | 8.700E-06 |
| 17433 | 872 | W872M | C | 8.9e-06 | 6.7e-06 | 7.5e-06 | 7.700E-06 | 1.114E-06 | 6.700E-06 | 8.900E-06 |
| 17434 | 872 | W872F | C | 6.1e-06 | 4.4e-06 | 1.4e-05 | 8.167E-06 | 5.123E-06 | 4.400E-06 | 1.400E-05 |
| 17435 | 872 | W872P | C | 1.3e-05 | 6.4e-06 | 9.8e-06 | 9.733E-06 | 3.301E-06 | 6.400E-06 | 1.300E-05 |
| 17436 | 872 | W872S | C | 2.8e-05 | 5.1e-06 | 4.9e-06 | 1.267E-05 | 1.328E-05 | 4.900E-06 | 2.800E-05 |
| 17437 | 872 | W872T | C | 8.6e-06 | 7.4e-06 | 9.3e-06 | 8.433E-06 | 9.609E-07 | 7.400E-06 | 9.300E-06 |
| 17438 | 872 | W872W | C | 1e-05 | 5.2e-06 | 8.8e-06 | 8.000E-06 | 2.498E-06 | 5.200E-06 | 1.000E-05 |
| 17439 | 872 | W872Y | C | 4.9e-06 | 9.5e-06 | 4.7e-06 | 6.367E-06 | 2.715E-06 | 4.700E-06 | 9.500E-06 |
| 17440 | 872 | W872V | C | 5e-06 | 6.3e-06 | 1.9e-05 | 1.010E-05 | 7.735E-06 | 5.000E-06 | 1.900E-05 |
| 17441 | 873 | V873A | C | 5.1e-06 | 9.4e-06 | 8.5e-06 | 7.667E-06 | 2.268E-06 | 5.100E-06 | 9.400E-06 |
| 17442 | 873 | V873R | C | 6.4e-06 | 9.9e-06 | 4.7e-06 | 7.000E-06 | 2.651E-06 | 4.700E-06 | 9.900E-06 |
| 17443 | 873 | V873N | C | 1.1e-05 | 8e-06 | 8.3e-06 | 9.100E-06 | 1.652E-06 | 8.000E-06 | 1.100E-05 |
| 17444 | 873 | V873D | C | 6.5e-06 | 7.1e-06 | 1.2e-05 | 8.533E-06 | 3.017E-06 | 6.500E-06 | 1.200E-05 |
| 17445 | 873 | V873C | C | 6.6e-06 | 6.8e-06 | 6.1e-06 | 6.500E-06 | 3.606E-07 | 6.100E-06 | 6.800E-06 |
| 17446 | 873 | V873E | C | 1.2e-05 | 9.2e-06 | 7.2e-06 | 9.467E-06 | 2.411E-06 | 7.200E-06 | 1.200E-05 |
| 17447 | 873 | V873Q | C | 8.4e-06 | 7.6e-06 | 1e-05 | 8.667E-06 | 1.222E-06 | 7.600E-06 | 1.000E-05 |
| 17448 | 873 | V873G | C | 9.5e-06 | 5.8e-06 | 6.7e-06 | 7.333E-06 | 1.930E-06 | 5.800E-06 | 9.500E-06 |
| 17449 | 873 | V873H | C | 8e-06 | 4.6e-06 | 7.5e-06 | 6.700E-06 | 1.836E-06 | 4.600E-06 | 8.000E-06 |
| 17450 | 873 | V873I | C | 1.2e-05 | 9.6e-06 | 7e-06 | 9.533E-06 | 2.501E-06 | 7.000E-06 | 1.200E-05 |
| 17451 | 873 | V873L | C | 1.2e-05 | 8.5e-06 | 9e-06 | 9.833E-06 | 1.893E-06 | 8.500E-06 | 1.200E-05 |
| 17452 | 873 | V873K | C | 7.5e-06 | 6.5e-06 | 9.2e-06 | 7.733E-06 | 1.365E-06 | 6.500E-06 | 9.200E-06 |
| 17453 | 873 | V873M | C | 6.5e-06 | 6.6e-06 | 1e-05 | 7.700E-06 | 1.992E-06 | 6.500E-06 | 1.000E-05 |
| 17454 | 873 | V873F | C | 8.5e-06 | 6.4e-06 | 7.9e-06 | 7.600E-06 | 1.082E-06 | 6.400E-06 | 8.500E-06 |
| 17455 | 873 | V873P | C | 9.6e-06 | 6.5e-06 | 3.5e-06 | 6.533E-06 | 3.050E-06 | 3.500E-06 | 9.600E-06 |
| 17456 | 873 | V873S | C | 4.7e-06 | 5.8e-06 | 8.4e-06 | 6.300E-06 | 1.900E-06 | 4.700E-06 | 8.400E-06 |
| 17457 | 873 | V873T | C | 7.6e-06 | 6.9e-06 | 7.7e-06 | 7.400E-06 | 4.359E-07 | 6.900E-06 | 7.700E-06 |
| 17458 | 873 | V873W | C | 5.9e-06 | 8.5e-06 | 1.1e-05 | 8.467E-06 | 2.550E-06 | 5.900E-06 | 1.100E-05 |
| 17459 | 873 | V873Y | C | 8.2e-06 | 7e-06 | 7.2e-06 | 7.467E-06 | 6.429E-07 | 7.000E-06 | 8.200E-06 |
| 17460 | 873 | V873V | C | 8.7e-06 | 1.1e-05 | 7.5e-06 | 9.067E-06 | 1.779E-06 | 7.500E-06 | 1.100E-05 |
| 17461 | 874 | R874A | C | 7.2e-06 | 8.7e-06 | 1.3e-05 | 9.633E-06 | 3.011E-06 | 7.200E-06 | 1.300E-05 |
| 17462 | 874 | R874R | C | 6.2e-06 | 1.5e-05 | 4.9e-06 | 8.700E-06 | 5.495E-06 | 4.900E-06 | 1.500E-05 |
| 17463 | 874 | R874N | C | 9e-06 | 7.9e-06 | 4.7e-06 | 7.200E-06 | 2.234E-06 | 4.700E-06 | 9.000E-06 |
| 17464 | 874 | R874D | C | 7.6e-06 | 5.3e-06 | 1.1e-05 | 7.967E-06 | 2.868E-06 | 5.300E-06 | 1.100E-05 |
| 17465 | 874 | R874C | C | 8.5e-06 | 9.9e-06 | 5.2e-06 | 7.867E-06 | 2.413E-06 | 5.200E-06 | 9.900E-06 |
| 17466 | 874 | R874E | C | 6.7e-06 | 1.2e-05 | 6e-06 | 8.233E-06 | 3.281E-06 | 6.000E-06 | 1.200E-05 |
| 17467 | 874 | R874Q | C | 6.2e-06 | 8e-06 | 6.1e-06 | 6.767E-06 | 1.069E-06 | 6.100E-06 | 8.000E-06 |
| 17468 | 874 | R874G | C | 6.1e-06 | 9.5e-06 | 7.8e-06 | 7.800E-06 | 1.700E-06 | 6.100E-06 | 9.500E-06 |
| 17469 | 874 | R874H | C | 6.7e-06 | 7.7e-06 | 1.5e-05 | 9.800E-06 | 4.531E-06 | 6.700E-06 | 1.500E-05 |
| 17470 | 874 | R874I | C | 9.1e-06 | 7.9e-06 | 9.6e-06 | 8.867E-06 | 8.737E-07 | 7.900E-06 | 9.600E-06 |
| 17471 | 874 | R874L | C | 9.8e-06 | 5.9e-06 | 6.2e-06 | 7.300E-06 | 2.170E-06 | 5.900E-06 | 9.800E-06 |
| 17472 | 874 | R874K | C | 7.1e-06 | 9e-06 | 5.4e-06 | 7.167E-06 | 1.801E-06 | 5.400E-06 | 9.000E-06 |
| 17473 | 874 | R874M | C | 6.7e-06 | 4.6e-06 | 6.3e-06 | 5.867E-06 | 1.115E-06 | 4.600E-06 | 6.700E-06 |
| 17474 | 874 | R874F | C | 1e-05 | 7.8e-06 | 1.2e-05 | 9.933E-06 | 2.101E-06 | 7.800E-06 | 1.200E-05 |
| 17475 | 874 | R874P | C | 9.8e-06 | 7.8e-06 | 9.2e-06 | 8.933E-06 | 1.026E-06 | 7.800E-06 | 9.800E-06 |
| 17476 | 874 | R874S | C | 9.1e-06 | 8.5e-06 | 1.3e-05 | 1.020E-05 | 2.443E-06 | 8.500E-06 | 1.300E-05 |
| 17477 | 874 | R874T | C | 1.3e-05 | 1.1e-05 | 6.2e-06 | 1.007E-05 | 3.495E-06 | 6.200E-06 | 1.300E-05 |
| 17478 | 874 | R874W | C | 6.1e-06 | 6.6e-06 | 7.3e-06 | 6.667E-06 | 6.028E-07 | 6.100E-06 | 7.300E-06 |
| 17479 | 874 | R874Y | C | 8.8e-06 | 3.6e-06 | 8.4e-06 | 6.933E-06 | 2.894E-06 | 3.600E-06 | 8.800E-06 |
| 17480 | 874 | R874V | C | 9.2e-06 | 6.8e-06 | 1.1e-05 | 9.000E-06 | 2.107E-06 | 6.800E-06 | 1.100E-05 |
| 17481 | 875 | Q875A | C | 9.5e-06 | 1e-05 | 4.8e-06 | 8.100E-06 | 2.869E-06 | 4.800E-06 | 1.000E-05 |
| 17482 | 875 | Q875R | C | 1.4e-05 | 6.6e-06 | 8.9e-06 | 9.833E-06 | 3.787E-06 | 6.600E-06 | 1.400E-05 |
| 17483 | 875 | Q875N | C | 7e-06 | 5.5e-06 | 8.1e-06 | 6.867E-06 | 1.305E-06 | 5.500E-06 | 8.100E-06 |
| 17484 | 875 | Q875D | C | 6.7e-06 | 8.4e-06 | 1e-05 | 8.367E-06 | 1.650E-06 | 6.700E-06 | 1.000E-05 |
| 17485 | 875 | Q875C | C | 6.2e-06 | 4.7e-06 | 1e-05 | 6.967E-06 | 2.732E-06 | 4.700E-06 | 1.000E-05 |
| 17486 | 875 | Q875E | C | 9.2e-06 | 8e-06 | 1.1e-05 | 9.400E-06 | 1.510E-06 | 8.000E-06 | 1.100E-05 |
| 17487 | 875 | Q875Q | C | 5.1e-06 | 5.8e-06 | 9.5e-06 | 6.800E-06 | 2.364E-06 | 5.100E-06 | 9.500E-06 |
| 17488 | 875 | Q875G | C | 1.5e-05 | 9e-06 | 1e-05 | 1.133E-05 | 3.215E-06 | 9.000E-06 | 1.500E-05 |
| 17489 | 875 | Q875H | C | 5.9e-06 | 5.9e-06 | 1.2e-05 | 7.933E-06 | 3.522E-06 | 5.900E-06 | 1.200E-05 |
| 17490 | 875 | Q875I | C | 7.3e-06 | 1e-05 | 4.8e-06 | 7.367E-06 | 2.601E-06 | 4.800E-06 | 1.000E-05 |
| 17491 | 875 | Q875L | C | 9.6e-06 | 9e-06 | 1.4e-05 | 1.087E-05 | 2.730E-06 | 9.000E-06 | 1.400E-05 |
| 17492 | 875 | Q875K | C | 4.7e-06 | 1.1e-05 | 4.3e-06 | 6.667E-06 | 3.758E-06 | 4.300E-06 | 1.100E-05 |
| 17493 | 875 | Q875M | C | 6.3e-06 | 9.8e-06 | 9.4e-06 | 8.500E-06 | 1.916E-06 | 6.300E-06 | 9.800E-06 |
| 17494 | 875 | Q875F | C | 6.7e-06 | 8.1e-06 | 4.4e-06 | 6.400E-06 | 1.868E-06 | 4.400E-06 | 8.100E-06 |
| 17495 | 875 | Q875P | C | 1.4e-05 | 6.6e-06 | 7e-06 | 9.200E-06 | 4.162E-06 | 6.600E-06 | 1.400E-05 |
| 17496 | 875 | Q875S | C | 4.6e-06 | 9.5e-06 | 4e-06 | 6.033E-06 | 3.017E-06 | 4.000E-06 | 9.500E-06 |
| 17497 | 875 | Q875T | C | 6.8e-06 | 9e-06 | 9.5e-06 | 8.433E-06 | 1.436E-06 | 6.800E-06 | 9.500E-06 |
| 17498 | 875 | Q875W | C | 9.6e-06 | 7.1e-06 | 6e-06 | 7.567E-06 | 1.845E-06 | 6.000E-06 | 9.600E-06 |
| 17499 | 875 | Q875Y | C | 1.3e-05 | 1.2e-05 | 4.2e-06 | 9.733E-06 | 4.818E-06 | 4.200E-06 | 1.300E-05 |
| 17500 | 875 | Q875V | C | 6.1e-06 | 1e-05 | 9.8e-06 | 8.633E-06 | 2.196E-06 | 6.100E-06 | 1.000E-05 |
| 17501 | 876 | A876A | C | 1.8e-05 | 6.5e-06 | 7.5e-06 | 1.067E-05 | 6.371E-06 | 6.500E-06 | 1.800E-05 |
| 17502 | 876 | A876R | C | 8.9e-06 | 7.7e-06 | 6.9e-06 | 7.833E-06 | 1.007E-06 | 6.900E-06 | 8.900E-06 |
| 17503 | 876 | A876N | C | 5.5e-06 | 3.7e-06 | 9.8e-06 | 6.333E-06 | 3.134E-06 | 3.700E-06 | 9.800E-06 |
| 17504 | 876 | A876D | C | 7.7e-06 | 1.2e-05 | 5.2e-06 | 8.300E-06 | 3.439E-06 | 5.200E-06 | 1.200E-05 |
| 17505 | 876 | A876C | C | 8.3e-06 | 5.6e-06 | 7.6e-06 | 7.167E-06 | 1.401E-06 | 5.600E-06 | 8.300E-06 |
| 17506 | 876 | A876E | C | 8e-06 | 6e-06 | 1.1e-05 | 8.333E-06 | 2.517E-06 | 6.000E-06 | 1.100E-05 |
| 17507 | 876 | A876Q | C | 8.7e-06 | 9.5e-06 | 6.7e-06 | 8.300E-06 | 1.442E-06 | 6.700E-06 | 9.500E-06 |
| 17508 | 876 | A876G | C | 5.2e-06 | 1.3e-05 | 7.5e-06 | 8.567E-06 | 4.008E-06 | 5.200E-06 | 1.300E-05 |
| 17509 | 876 | A876H | C | 8.5e-06 | 7.9e-06 | 9.5e-06 | 8.633E-06 | 8.083E-07 | 7.900E-06 | 9.500E-06 |

| | | | | | | | | | | |
|---|---|---|---|---|---|---|---|---|---|---|
| 17510 | 876 | A876I | C | 9.6e-06 | 1.2e-05 | 6.9e-06 | 9.500E-06 | 2.551E-06 | 6.900E-06 | 1.200E-05 |
| 17511 | 876 | A876L | C | 1.1e-05 | 1.5e-05 | 9.6e-06 | 1.187E-05 | 2.802E-06 | 9.600E-06 | 1.500E-05 |
| 17512 | 876 | A876K | C | 1.4e-05 | 1e-05 | 1.3e-05 | 1.233E-05 | 2.082E-06 | 1.000E-05 | 1.400E-05 |
| 17513 | 876 | A876M | C | 9.8e-06 | 9.9e-06 | 9.4e-06 | 9.700E-06 | 2.646E-07 | 9.400E-06 | 9.900E-06 |
| 17514 | 876 | A876F | C | 3.6e-06 | 9e-06 | 9.3e-06 | 7.300E-06 | 3.208E-06 | 3.600E-06 | 9.300E-06 |
| 17515 | 876 | A876P | C | 9.4e-06 | 5.5e-06 | 7.9e-06 | 7.600E-06 | 1.967E-06 | 5.500E-06 | 9.400E-06 |
| 17516 | 876 | A876S | C | 5.2e-06 | 8e-06 | 5.9e-06 | 6.367E-06 | 1.457E-06 | 5.200E-06 | 8.000E-06 |
| 17517 | 876 | A876T | C | 4.7e-06 | 1.2e-05 | 5.3e-06 | 7.333E-06 | 4.053E-06 | 4.700E-06 | 1.200E-05 |
| 17518 | 876 | A876W | C | 7.7e-06 | 7.5e-06 | 6.8e-06 | 7.333E-06 | 4.726E-07 | 6.800E-06 | 7.700E-06 |
| 17519 | 876 | A876Y | C | 6.6e-06 | 4.8e-06 | 8.2e-06 | 6.533E-06 | 1.701E-06 | 4.800E-06 | 8.200E-06 |
| 17520 | 876 | A876V | C | 6.6e-06 | 6.9e-06 | 9.3e-06 | 7.600E-06 | 1.480E-06 | 6.600E-06 | 9.300E-06 |
| 17521 | 877 | P877A | C | 1.4e-05 | 1.1e-05 | 3.5e-06 | 9.500E-06 | 5.408E-06 | 3.500E-06 | 1.400E-05 |
| 17522 | 877 | P877R | C | 2e-05 | 9.3e-06 | 9.2e-06 | 1.283E-05 | 6.207E-06 | 9.200E-06 | 2.000E-05 |
| 17523 | 877 | P877N | C | 5.8e-06 | 1.5e-05 | 6.8e-06 | 9.200E-06 | 5.048E-06 | 5.800E-06 | 1.500E-05 |
| 17524 | 877 | P877D | C | 9.1e-06 | 8.3e-06 | 1.4e-05 | 1.047E-05 | 3.086E-06 | 8.300E-06 | 1.400E-05 |
| 17525 | 877 | P877C | C | 9.1e-06 | 1e-05 | 9e-06 | 9.367E-06 | 5.508E-07 | 9.000E-06 | 1.000E-05 |
| 17526 | 877 | P877E | C | 1e-05 | 9.7e-06 | 8.2e-06 | 9.300E-06 | 9.644E-07 | 8.200E-06 | 1.000E-05 |
| 17527 | 877 | P877Q | C | 8.1e-06 | 6e-06 | 8.8e-06 | 7.633E-06 | 1.457E-06 | 6.000E-06 | 8.800E-06 |
| 17528 | 877 | P877G | C | 5.4e-06 | 5.1e-06 | 1e-05 | 6.833E-06 | 2.747E-06 | 5.100E-06 | 1.000E-05 |
| 17529 | 877 | P877H | C | 7.2e-06 | 9.2e-06 | 9.5e-06 | 8.633E-06 | 1.250E-06 | 7.200E-06 | 9.500E-06 |
| 17530 | 877 | P877I | C | 3.1e-06 | 4.3e-06 | 9.2e-06 | 5.533E-06 | 3.232E-06 | 3.100E-06 | 9.200E-06 |
| 17531 | 877 | P877L | C | 8.6e-06 | 1.1e-05 | 8.9e-06 | 9.500E-06 | 1.308E-06 | 8.600E-06 | 1.100E-05 |
| 17532 | 877 | P877K | C | 1.1e-05 | 4.4e-06 | 7.3e-06 | 7.567E-06 | 3.308E-06 | 4.400E-06 | 1.100E-05 |
| 17533 | 877 | P877M | C | 1e-05 | 1e-05 | 5.9e-06 | 8.633E-06 | 2.367E-06 | 5.900E-06 | 1.000E-05 |
| 17534 | 877 | P877F | C | 9.5e-06 | 9.4e-06 | 5.9e-06 | 8.267E-06 | 2.050E-06 | 5.900E-06 | 9.500E-06 |
| 17535 | 877 | P877P | C | 8.2e-06 | 4.6e-06 | 4.5e-06 | 5.767E-06 | 2.108E-06 | 4.500E-06 | 8.200E-06 |
| 17536 | 877 | P877S | C | 9.2e-06 | 8.1e-06 | 6e-06 | 7.767E-06 | 1.626E-06 | 6.000E-06 | 9.200E-06 |
| 17537 | 877 | P877T | C | 1.5e-05 | 9.7e-06 | 7.4e-06 | 1.070E-05 | 3.897E-06 | 7.400E-06 | 1.500E-05 |
| 17538 | 877 | P877W | C | 4.9e-06 | 6e-06 | 8e-06 | 6.300E-06 | 1.572E-06 | 4.900E-06 | 8.000E-06 |
| 17539 | 877 | P877Y | C | 1.5e-05 | 7.5e-06 | 5.8e-06 | 9.433E-06 | 4.895E-06 | 5.800E-06 | 1.500E-05 |
| 17540 | 877 | P877V | C | 1.2e-05 | 8.2e-06 | 6.4e-06 | 8.867E-06 | 2.859E-06 | 6.400E-06 | 1.200E-05 |
| 17541 | 878 | G878A | C | 8.3e-06 | 9.4e-06 | 1.5e-05 | 1.090E-05 | 3.593E-06 | 8.300E-06 | 1.500E-05 |
| 17542 | 878 | G878R | C | 1.2e-05 | 5.6e-06 | 1.2e-05 | 9.867E-06 | 3.695E-06 | 5.600E-06 | 1.200E-05 |
| 17543 | 878 | G878N | C | 6.1e-06 | 7.6e-06 | 8.9e-06 | 7.533E-06 | 1.401E-06 | 6.100E-06 | 8.900E-06 |
| 17544 | 878 | G878D | C | 7e-06 | 7.9e-06 | 6.5e-06 | 7.133E-06 | 7.095E-07 | 6.500E-06 | 7.900E-06 |
| 17545 | 878 | G878C | C | 6.7e-06 | 5.7e-06 | 7.9e-06 | 6.767E-06 | 1.102E-06 | 5.700E-06 | 7.900E-06 |
| 17546 | 878 | G878E | C | 4.1e-06 | 5.8e-06 | 9.8e-06 | 6.567E-06 | 2.926E-06 | 4.100E-06 | 9.800E-06 |
| 17547 | 878 | G878Q | C | 1e-05 | 5.8e-06 | 8.3e-06 | 8.033E-06 | 2.113E-06 | 5.800E-06 | 1.000E-05 |
| 17548 | 878 | G878G | C | 7.2e-06 | 9.1e-06 | 8.7e-06 | 8.333E-06 | 1.002E-06 | 7.200E-06 | 9.100E-06 |
| 17549 | 878 | G878H | C | 6.2e-06 | 6.3e-06 | 6.2e-06 | 6.233E-06 | 5.774E-08 | 6.200E-06 | 6.300E-06 |
| 17550 | 878 | G878I | C | 6e-06 | 7.4e-06 | 5.7e-06 | 6.367E-06 | 9.074E-07 | 5.700E-06 | 7.400E-06 |
| 17551 | 878 | G878L | C | 7.5e-06 | 8.2e-06 | 1.1e-05 | 8.900E-06 | 1.852E-06 | 7.500E-06 | 1.100E-05 |
| 17552 | 878 | G878K | C | 6e-06 | 7.2e-06 | 7.4e-06 | 6.867E-06 | 7.572E-07 | 6.000E-06 | 7.400E-06 |
| 17553 | 878 | G878M | C | 9.7e-06 | 9e-06 | 1.4e-05 | 1.090E-05 | 2.707E-06 | 9.000E-06 | 1.400E-05 |
| 17554 | 878 | G878F | C | 5.3e-06 | 6e-06 | 6.3e-06 | 5.867E-06 | 5.132E-07 | 5.300E-06 | 6.300E-06 |
| 17555 | 878 | G878P | C | 5.8e-06 | 8.7e-06 | 6.7e-06 | 7.067E-06 | 1.484E-06 | 5.800E-06 | 8.700E-06 |
| 17556 | 878 | G878S | C | 5.7e-06 | 4.2e-06 | 1.3e-05 | 7.633E-06 | 4.708E-06 | 4.200E-06 | 1.300E-05 |
| 17557 | 878 | G878T | C | 8e-06 | 1.3e-05 | 6.8e-06 | 9.267E-06 | 3.288E-06 | 6.800E-06 | 1.300E-05 |
| 17558 | 878 | G878W | C | 5.8e-06 | 8.8e-06 | 8e-06 | 7.533E-06 | 1.553E-06 | 5.800E-06 | 8.800E-06 |
| 17559 | 878 | G878Y | C | 6.2e-06 | 1.7e-05 | 9e-06 | 1.073E-05 | 5.605E-06 | 6.200E-06 | 1.700E-05 |
| 17560 | 878 | G878V | C | 7.3e-06 | 9.2e-06 | 1.4e-05 | 1.017E-05 | 3.453E-06 | 7.300E-06 | 1.400E-05 |
| 17561 | 879 | K879A | C | 7.4e-06 | 1.4e-05 | 7e-06 | 9.467E-06 | 3.931E-06 | 7.000E-06 | 1.400E-05 |
| 17562 | 879 | K879R | C | 1.2e-05 | 1e-05 | 6.4e-06 | 9.467E-06 | 2.838E-06 | 6.400E-06 | 1.200E-05 |
| 17563 | 879 | K879N | C | 7.1e-06 | 1.1e-05 | 7.1e-06 | 8.400E-06 | 2.252E-06 | 7.100E-06 | 1.100E-05 |
| 17564 | 879 | K879D | C | 4.8e-06 | 1.4e-05 | 6.1e-06 | 8.300E-06 | 4.979E-06 | 4.800E-06 | 1.400E-05 |
| 17565 | 879 | K879C | C | 5.5e-06 | 9.5e-06 | 6.5e-06 | 7.167E-06 | 2.082E-06 | 5.500E-06 | 9.500E-06 |
| 17566 | 879 | K879E | C | 6e-06 | 6.6e-06 | 9.4e-06 | 7.333E-06 | 1.815E-06 | 6.000E-06 | 9.400E-06 |
| 17567 | 879 | K879Q | C | 8.5e-06 | 1.4e-05 | 6.4e-06 | 9.633E-06 | 3.925E-06 | 6.400E-06 | 1.400E-05 |
| 17568 | 879 | K879G | C | 8.4e-06 | 1.2e-05 | 6.6e-06 | 9.000E-06 | 2.750E-06 | 6.600E-06 | 1.200E-05 |
| 17569 | 879 | K879H | C | 6.4e-06 | 7e-06 | 8.1e-06 | 7.167E-06 | 8.622E-07 | 6.400E-06 | 8.100E-06 |
| 17570 | 879 | K879I | C | 1.3e-05 | 1e-05 | 9.7e-06 | 1.090E-05 | 1.825E-06 | 9.700E-06 | 1.300E-05 |
| 17571 | 879 | K879L | C | 8.7e-06 | 8e-06 | 8.3e-06 | 8.333E-06 | 3.512E-07 | 8.000E-06 | 8.700E-06 |
| 17572 | 879 | K879K | C | 8.3e-06 | 8e-06 | 1.5e-05 | 1.043E-05 | 3.958E-06 | 8.000E-06 | 1.500E-05 |
| 17573 | 879 | K879M | C | 5.5e-06 | 1.1e-05 | 4.9e-06 | 7.133E-06 | 3.362E-06 | 4.900E-06 | 1.100E-05 |
| 17574 | 879 | K879F | C | 6.7e-06 | 6.3e-06 | 7.2e-06 | 6.733E-06 | 4.509E-07 | 6.300E-06 | 7.200E-06 |
| 17575 | 879 | K879P | C | 9.7e-06 | 6.7e-06 | 6.2e-06 | 7.533E-06 | 1.893E-06 | 6.200E-06 | 9.700E-06 |
| 17576 | 879 | K879S | C | 7.1e-06 | 8.1e-06 | 7.2e-06 | 7.467E-06 | 5.508E-07 | 7.100E-06 | 8.100E-06 |
| 17577 | 879 | K879T | C | 1.2e-05 | 7e-06 | 9.5e-06 | 9.500E-06 | 2.500E-06 | 7.000E-06 | 1.200E-05 |
| 17578 | 879 | K879W | C | 8.5e-06 | 5.4e-06 | 5.7e-06 | 6.533E-06 | 1.710E-06 | 5.400E-06 | 8.500E-06 |
| 17579 | 879 | K879Y | C | 9.4e-06 | 7.9e-06 | 6.2e-06 | 7.833E-06 | 1.601E-06 | 6.200E-06 | 9.400E-06 |
| 17580 | 879 | K879V | C | 1e-05 | 6.8e-06 | 6.2e-06 | 7.667E-06 | 2.043E-06 | 6.200E-06 | 1.000E-05 |
| 17581 | 880 | G880A | C | 8.5e-06 | 6.9e-06 | 9.5e-06 | 8.300E-06 | 1.311E-06 | 6.900E-06 | 9.500E-06 |
| 17582 | 880 | G880R | C | 5.7e-06 | 7.1e-06 | 8.7e-06 | 7.167E-06 | 1.501E-06 | 5.700E-06 | 8.700E-06 |
| 17583 | 880 | G880N | C | 4.4e-06 | 9.7e-06 | 7.2e-06 | 7.100E-06 | 2.651E-06 | 4.400E-06 | 9.700E-06 |
| 17584 | 880 | G880D | C | 9e-06 | 9.9e-06 | 8.4e-06 | 9.100E-06 | 7.550E-07 | 8.400E-06 | 9.900E-06 |
| 17585 | 880 | G880C | C | 5.8e-06 | 8.3e-06 | 8.7e-06 | 7.600E-06 | 1.572E-06 | 5.800E-06 | 8.700E-06 |
| 17586 | 880 | G880E | C | 4.6e-06 | 7.4e-06 | 7.9e-06 | 6.633E-06 | 1.779E-06 | 4.600E-06 | 7.900E-06 |
| 17587 | 880 | G880Q | C | 1.2e-05 | 8.5e-06 | 1.3e-05 | 1.117E-05 | 2.363E-06 | 8.500E-06 | 1.300E-05 |
| 17588 | 880 | G880G | C | 6.2e-06 | 7.3e-06 | 8.6e-06 | 7.367E-06 | 1.201E-06 | 6.200E-06 | 8.600E-06 |
| 17589 | 880 | G880H | C | 8.8e-06 | 8.8e-06 | 4.4e-06 | 7.333E-06 | 2.540E-06 | 4.400E-06 | 8.800E-06 |
| 17590 | 880 | G880I | C | 1e-05 | 7.5e-06 | 5.6e-06 | 7.700E-06 | 2.207E-06 | 5.600E-06 | 1.000E-05 |
| 17591 | 880 | G880L | C | 4.4e-06 | 6.8e-06 | 1.3e-05 | 8.067E-06 | 4.438E-06 | 4.400E-06 | 1.300E-05 |
| 17592 | 880 | G880K | C | 4.5e-06 | 7.3e-06 | 8.8e-06 | 6.867E-06 | 2.183E-06 | 4.500E-06 | 8.800E-06 |
| 17593 | 880 | G880M | C | 7.5e-06 | 8.3e-06 | 7.8e-06 | 7.867E-06 | 4.041E-07 | 7.500E-06 | 8.300E-06 |
| 17594 | 880 | G880F | C | 7e-06 | 5.3e-06 | 9.9e-06 | 7.400E-06 | 2.326E-06 | 5.300E-06 | 9.900E-06 |

| | | | | | | | | | | |
|---|---|---|---|---|---|---|---|---|---|---|
| 17595 | 880 | G880P | C | 1e-05 | 1.1e-05 | 8.5e-06 | 9.833E-06 | 1.258E-06 | 8.500E-06 | 1.100E-05 |
| 17596 | 880 | G880S | C | 6.6e-06 | 8.7e-06 | 6.9e-06 | 7.400E-06 | 1.136E-06 | 6.600E-06 | 8.700E-06 |
| 17597 | 880 | G880T | C | 9.3e-06 | 4.6e-06 | 6.7e-06 | 6.867E-06 | 2.354E-06 | 4.600E-06 | 9.300E-06 |
| 17598 | 880 | G880W | C | 8.8e-06 | 8.4e-06 | 7.4e-06 | 8.200E-06 | 7.211E-07 | 7.400E-06 | 8.800E-06 |
| 17599 | 880 | G880Y | C | 6.7e-06 | 6.5e-06 | 8.8e-06 | 7.333E-06 | 1.274E-06 | 6.500E-06 | 8.800E-06 |
| 17600 | 880 | G880V | C | 7.2e-06 | 5.3e-06 | 7.8e-06 | 6.767E-06 | 1.305E-06 | 5.300E-06 | 7.800E-06 |
| 17601 | 881 | L881A | C | 9.2e-06 | 1.2e-05 | 1.2e-05 | 1.107E-05 | 1.617E-06 | 9.200E-06 | 1.200E-05 |
| 17602 | 881 | L881R | C | 5.8e-06 | 1.3e-05 | 6.5e-06 | 8.433E-06 | 3.970E-06 | 5.800E-06 | 1.300E-05 |
| 17603 | 881 | L881N | C | 9.7e-06 | 6.3e-06 | 7.7e-06 | 7.900E-06 | 1.709E-06 | 6.300E-06 | 9.700E-06 |
| 17604 | 881 | L881D | C | 8.5e-06 | 8e-06 | 8.7e-06 | 8.400E-06 | 3.606E-07 | 8.000E-06 | 8.700E-06 |
| 17605 | 881 | L881C | C | 8e-06 | 9.3e-06 | 6.6e-06 | 7.967E-06 | 1.350E-06 | 6.600E-06 | 9.300E-06 |
| 17606 | 881 | L881E | C | 7.4e-06 | 7.7e-06 | 1.1e-05 | 8.700E-06 | 1.997E-06 | 7.400E-06 | 1.100E-05 |
| 17607 | 881 | L881Q | C | 6.4e-06 | 9.5e-06 | 6.1e-06 | 7.333E-06 | 1.882E-06 | 6.100E-06 | 9.500E-06 |
| 17608 | 881 | L881G | C | 4.9e-06 | 1.1e-05 | 9.9e-06 | 8.600E-06 | 3.251E-06 | 4.900E-06 | 1.100E-05 |
| 17609 | 881 | L881H | C | 7.4e-06 | 3.1e-06 | 1.2e-05 | 7.500E-06 | 4.451E-06 | 3.100E-06 | 1.200E-05 |
| 17610 | 881 | L881I | C | 1.2e-05 | 9e-06 | 1e-05 | 1.033E-05 | 1.528E-06 | 9.000E-06 | 1.200E-05 |
| 17611 | 881 | L881L | C | 9.4e-06 | 8.8e-06 | 4.7e-06 | 7.633E-06 | 2.558E-06 | 4.700E-06 | 9.400E-06 |
| 17612 | 881 | L881K | C | 7e-06 | 8.6e-06 | 8.2e-06 | 7.933E-06 | 8.327E-07 | 7.000E-06 | 8.600E-06 |
| 17613 | 881 | L881M | C | 6.1e-06 | 8e-06 | 9.4e-06 | 7.833E-06 | 1.656E-06 | 6.100E-06 | 9.400E-06 |
| 17614 | 881 | L881F | C | 8.8e-06 | 6.2e-06 | 9.6e-06 | 8.200E-06 | 1.778E-06 | 6.200E-06 | 9.600E-06 |
| 17615 | 881 | L881P | C | 7e-06 | 6.1e-06 | 7e-06 | 6.700E-06 | 5.196E-07 | 6.100E-06 | 7.000E-06 |
| 17616 | 881 | L881S | C | 6.5e-06 | 7.2e-06 | 1.2e-05 | 8.567E-06 | 2.994E-06 | 6.500E-06 | 1.200E-05 |
| 17617 | 881 | L881T | C | 6.5e-06 | 9.9e-06 | 6.8e-06 | 7.733E-06 | 1.882E-06 | 6.500E-06 | 9.900E-06 |
| 17618 | 881 | L881W | C | 1.4e-05 | 6.7e-06 | 5.6e-06 | 8.767E-06 | 4.565E-06 | 5.600E-06 | 1.400E-05 |
| 17619 | 881 | L881Y | C | 6.7e-06 | 1.3e-05 | 9.3e-06 | 9.667E-06 | 3.166E-06 | 6.700E-06 | 1.300E-05 |
| 17620 | 881 | L881V | C | 8.9e-06 | 8.3e-06 | 6.7e-06 | 7.967E-06 | 1.137E-06 | 6.700E-06 | 8.900E-06 |
| 17621 | 882 | E882A | C | 7.4e-06 | 6e-06 | 5.5e-06 | 6.300E-06 | 9.849E-07 | 5.500E-06 | 7.400E-06 |
| 17622 | 882 | E882R | C | 6.1e-06 | 6.8e-06 | 1.3e-05 | 8.633E-06 | 3.798E-06 | 6.100E-06 | 1.300E-05 |
| 17623 | 882 | E882N | C | 9.4e-06 | 6.5e-06 | 6.4e-06 | 7.433E-06 | 1.704E-06 | 6.400E-06 | 9.400E-06 |
| 17624 | 882 | E882D | C | 6.6e-06 | 8.9e-06 | 9.2e-06 | 8.233E-06 | 1.422E-06 | 6.600E-06 | 9.200E-06 |
| 17625 | 882 | E882C | C | 3.1e-06 | 4.8e-06 | 7.3e-06 | 5.067E-06 | 2.113E-06 | 3.100E-06 | 7.300E-06 |
| 17626 | 882 | E882E | C | 5.6e-06 | 8.3e-06 | 1e-05 | 7.967E-06 | 2.219E-06 | 5.600E-06 | 1.000E-05 |
| 17627 | 882 | E882Q | C | 8.8e-06 | 7.4e-06 | 5.3e-06 | 7.167E-06 | 1.762E-06 | 5.300E-06 | 8.800E-06 |
| 17628 | 882 | E882G | C | 9.2e-06 | 7.1e-06 | 7e-06 | 7.767E-06 | 1.242E-06 | 7.000E-06 | 9.200E-06 |
| 17629 | 882 | E882H | C | 7.6e-06 | 1.6e-05 | 7.9e-06 | 1.050E-05 | 4.766E-06 | 7.600E-06 | 1.600E-05 |
| 17630 | 882 | E882I | C | 8.7e-06 | 1.3e-05 | 7.2e-06 | 9.633E-06 | 3.011E-06 | 7.200E-06 | 1.300E-05 |
| 17631 | 882 | E882L | C | 1e-05 | 5.3e-06 | 8e-06 | 7.767E-06 | 2.359E-06 | 5.300E-06 | 1.000E-05 |
| 17632 | 882 | E882K | C | 5.4e-06 | 6.9e-06 | 9.9e-06 | 7.400E-06 | 2.291E-06 | 5.400E-06 | 9.900E-06 |
| 17633 | 882 | E882M | C | 6.5e-06 | 1.9e-05 | 5.8e-06 | 1.043E-05 | 7.427E-06 | 5.800E-06 | 1.900E-05 |
| 17634 | 882 | E882F | C | 8.4e-06 | 6.4e-06 | 1.3e-05 | 9.267E-06 | 3.384E-06 | 6.400E-06 | 1.300E-05 |
| 17635 | 882 | E882P | C | 8.5e-06 | 8e-06 | 1e-05 | 8.833E-06 | 1.041E-06 | 8.000E-06 | 1.000E-05 |
| 17636 | 882 | E882S | C | 7.6e-06 | 7.1e-06 | 6.4e-06 | 7.033E-06 | 6.028E-07 | 6.400E-06 | 7.600E-06 |
| 17637 | 882 | E882T | C | 5.9e-06 | 8.4e-06 | 1.1e-05 | 8.433E-06 | 2.550E-06 | 5.900E-06 | 1.100E-05 |
| 17638 | 882 | E882W | C | 9.3e-06 | 8.7e-06 | 5.8e-06 | 7.933E-06 | 1.872E-06 | 5.800E-06 | 9.300E-06 |
| 17639 | 882 | E882Y | C | 9.6e-06 | 7.1e-06 | 1.1e-05 | 9.233E-06 | 1.976E-06 | 7.100E-06 | 1.100E-05 |
| 17640 | 882 | E882V | C | 9.5e-06 | 1.3e-05 | 1.8e-05 | 1.350E-05 | 4.272E-06 | 9.500E-06 | 1.800E-05 |
| 17641 | 883 | W883A | C | 7.2e-06 | 6.9e-06 | 7e-06 | 7.033E-06 | 1.528E-07 | 6.900E-06 | 7.200E-06 |
| 17642 | 883 | W883R | C | 9.1e-06 | 4e-06 | 8.5e-06 | 7.200E-06 | 2.787E-06 | 4.000E-06 | 9.100E-06 |
| 17643 | 883 | W883N | C | 9.7e-06 | 6.4e-06 | 1.1e-05 | 9.033E-06 | 2.371E-06 | 6.400E-06 | 1.100E-05 |
| 17644 | 883 | W883D | C | 1.1e-05 | 5.8e-06 | 5e-06 | 7.267E-06 | 3.258E-06 | 5.000E-06 | 1.100E-05 |
| 17645 | 883 | W883C | C | 5.6e-06 | 1.3e-05 | 1.2e-05 | 1.020E-05 | 4.015E-06 | 5.600E-06 | 1.300E-05 |
| 17646 | 883 | W883E | C | 6.7e-06 | 8.5e-06 | 1.4e-05 | 9.733E-06 | 3.803E-06 | 6.700E-06 | 1.400E-05 |
| 17647 | 883 | W883Q | C | 6.7e-06 | 6.6e-06 | 6.6e-06 | 6.633E-06 | 5.774E-08 | 6.600E-06 | 6.700E-06 |
| 17648 | 883 | W883G | C | 7.6e-06 | 4.8e-06 | 9.5e-06 | 7.300E-06 | 2.364E-06 | 4.800E-06 | 9.500E-06 |
| 17649 | 883 | W883H | C | 7.3e-06 | 5.6e-06 | 9.3e-06 | 7.400E-06 | 1.852E-06 | 5.600E-06 | 9.300E-06 |
| 17650 | 883 | W883I | C | 6e-06 | 6.4e-06 | 6.7e-06 | 6.367E-06 | 3.512E-07 | 6.000E-06 | 6.700E-06 |
| 17651 | 883 | W883L | C | 1.3e-05 | 6e-06 | 7.6e-06 | 8.867E-06 | 3.668E-06 | 6.000E-06 | 1.300E-05 |
| 17652 | 883 | W883K | C | 7.2e-06 | 8.2e-06 | 7.9e-06 | 7.767E-06 | 5.132E-07 | 7.200E-06 | 8.200E-06 |
| 17653 | 883 | W883M | C | 4.6e-06 | 7.4e-06 | 7e-06 | 6.333E-06 | 1.514E-06 | 4.600E-06 | 7.400E-06 |
| 17654 | 883 | W883F | C | 6.6e-06 | 5.2e-06 | 6.7e-06 | 6.167E-06 | 8.386E-07 | 5.200E-06 | 6.700E-06 |
| 17655 | 883 | W883P | C | 7e-06 | 5.4e-06 | 1.3e-05 | 8.467E-06 | 4.007E-06 | 5.400E-06 | 1.300E-05 |
| 17656 | 883 | W883S | C | 1.2e-05 | 6e-06 | 6e-06 | 8.000E-06 | 3.464E-06 | 6.000E-06 | 1.200E-05 |
| 17657 | 883 | W883T | C | 7.4e-06 | 8.6e-06 | 6.7e-06 | 7.567E-06 | 9.609E-07 | 6.700E-06 | 8.600E-06 |
| 17658 | 883 | W883W | C | 8.6e-06 | 8.8e-06 | 6.7e-06 | 8.033E-06 | 1.159E-06 | 6.700E-06 | 8.800E-06 |
| 17659 | 883 | W883Y | C | 1.3e-05 | 9.6e-06 | 8.6e-06 | 1.040E-05 | 2.307E-06 | 8.600E-06 | 1.300E-05 |
| 17660 | 883 | W883V | C | 1.1e-05 | 7.2e-06 | 1.5e-05 | 1.107E-05 | 3.900E-06 | 7.200E-06 | 1.500E-05 |
| 17661 | 884 | V884A | C | 3.2e-06 | 8.6e-06 | 6.4e-06 | 6.067E-06 | 2.715E-06 | 3.200E-06 | 8.600E-06 |
| 17662 | 884 | V884R | C | 4.3e-06 | 7.2e-06 | 9.3e-06 | 6.933E-06 | 2.511E-06 | 4.300E-06 | 9.300E-06 |
| 17663 | 884 | V884N | C | 9.9e-06 | 8.1e-06 | 8.9e-06 | 8.967E-06 | 9.018E-07 | 8.100E-06 | 9.900E-06 |
| 17664 | 884 | V884D | C | 7.3e-06 | 9e-06 | 1.3e-05 | 9.767E-06 | 2.926E-06 | 7.300E-06 | 1.300E-05 |
| 17665 | 884 | V884C | C | 5.9e-06 | 4.3e-06 | 1.4e-05 | 8.067E-06 | 5.200E-06 | 4.300E-06 | 1.400E-05 |
| 17666 | 884 | V884E | C | 9.6e-06 | 8.8e-06 | 7.8e-06 | 8.733E-06 | 9.018E-07 | 7.800E-06 | 9.600E-06 |
| 17667 | 884 | V884Q | C | 8.9e-06 | 7.3e-06 | 8.5e-06 | 8.233E-06 | 8.327E-07 | 7.300E-06 | 8.900E-06 |
| 17668 | 884 | V884G | C | 7.9e-06 | 5.7e-06 | 9.9e-06 | 7.833E-06 | 2.101E-06 | 5.700E-06 | 9.900E-06 |
| 17669 | 884 | V884H | C | 4.8e-06 | 5.8e-06 | 6.1e-06 | 5.567E-06 | 6.807E-07 | 4.800E-06 | 6.100E-06 |
| 17670 | 884 | V884I | C | 6.7e-06 | 7.6e-06 | 9.2e-06 | 7.833E-06 | 1.266E-06 | 6.700E-06 | 9.200E-06 |
| 17671 | 884 | V884L | C | 9.4e-06 | 5.5e-06 | 1.6e-05 | 1.030E-05 | 5.308E-06 | 5.500E-06 | 1.600E-05 |
| 17672 | 884 | V884K | C | 1.1e-05 | 1.3e-05 | 7.1e-06 | 1.037E-05 | 3.001E-06 | 7.100E-06 | 1.300E-05 |
| 17673 | 884 | V884M | C | 6.7e-06 | 5.7e-06 | 7.5e-06 | 6.633E-06 | 9.018E-07 | 5.700E-06 | 7.500E-06 |
| 17674 | 884 | V884F | C | 6.4e-06 | 1.3e-05 | 5.7e-06 | 8.367E-06 | 4.028E-06 | 5.700E-06 | 1.300E-05 |
| 17675 | 884 | V884P | C | 8.5e-06 | 9.8e-06 | 6.3e-06 | 8.200E-06 | 1.769E-06 | 6.300E-06 | 9.800E-06 |
| 17676 | 884 | V884S | C | 1.3e-05 | 8.3e-06 | 7.6e-06 | 9.633E-06 | 2.937E-06 | 7.600E-06 | 1.300E-05 |
| 17677 | 884 | V884T | C | 9.1e-06 | 8e-06 | 5.1e-06 | 7.400E-06 | 2.066E-06 | 5.100E-06 | 9.100E-06 |
| 17678 | 884 | V884W | C | 1.3e-05 | 1.3e-05 | 6.8e-06 | 1.093E-05 | 3.580E-06 | 6.800E-06 | 1.300E-05 |
| 17679 | 884 | V884Y | C | 8.3e-06 | 8.1e-06 | 1.4e-05 | 1.013E-05 | 3.350E-06 | 8.100E-06 | 1.400E-05 |

| | | | | | | | | | | |
|---|---|---|---|---|---|---|---|---|---|---|
| 17680 | 884 | V884V | C | 7.3e-06 | 9.1e-06 | 5.2e-06 | 7.200E-06 | 1.952E-06 | 5.200E-06 | 9.100E-06 |
| 17681 | 885 | A885A | C | 1e-05 | 4.9e-06 | 1e-05 | 8.300E-06 | 2.944E-06 | 4.900E-06 | 1.000E-05 |
| 17682 | 885 | A885R | C | 4.7e-06 | 1.2e-05 | 4.7e-06 | 7.133E-06 | 4.215E-06 | 4.700E-06 | 1.200E-05 |
| 17683 | 885 | A885N | C | 1.3e-05 | 1.5e-05 | 1.3e-05 | 1.367E-05 | 1.155E-06 | 1.300E-05 | 1.500E-05 |
| 17684 | 885 | A885D | C | 6e-06 | 8.1e-06 | 6.7e-06 | 6.933E-06 | 1.069E-06 | 6.000E-06 | 8.100E-06 |
| 17685 | 885 | A885C | C | 9.1e-06 | 6.8e-06 | 1e-05 | 8.633E-06 | 1.650E-06 | 6.800E-06 | 1.000E-05 |
| 17686 | 885 | A885E | C | 9.2e-06 | 7.1e-06 | 6.3e-06 | 7.533E-06 | 1.498E-06 | 6.300E-06 | 9.200E-06 |
| 17687 | 885 | A885Q | C | 1.2e-05 | 8.8e-06 | 8e-06 | 9.600E-06 | 2.117E-06 | 8.000E-06 | 1.200E-05 |
| 17688 | 885 | A885G | C | 5.9e-06 | 8.3e-06 | 6.6e-06 | 6.933E-06 | 1.234E-06 | 5.900E-06 | 8.300E-06 |
| 17689 | 885 | A885H | C | 9.2e-06 | 6.7e-06 | 6.1e-06 | 7.333E-06 | 1.644E-06 | 6.100E-06 | 9.200E-06 |
| 17690 | 885 | A885I | C | 8.6e-06 | 1.5e-05 | 1.1e-05 | 1.153E-05 | 3.233E-06 | 8.600E-06 | 1.500E-05 |
| 17691 | 885 | A885L | C | 7.1e-06 | 9.5e-06 | 5.8e-06 | 7.467E-06 | 1.877E-06 | 5.800E-06 | 9.500E-06 |
| 17692 | 885 | A885K | C | 6.4e-06 | 5.8e-06 | 8.2e-06 | 6.800E-06 | 1.249E-06 | 5.800E-06 | 8.200E-06 |
| 17693 | 885 | A885M | C | 5.1e-06 | 4.8e-06 | 6.3e-06 | 5.400E-06 | 7.937E-07 | 4.800E-06 | 6.300E-06 |
| 17694 | 885 | A885F | C | 9.5e-06 | 6.7e-06 | 7.3e-06 | 7.833E-06 | 1.474E-06 | 6.700E-06 | 9.500E-06 |
| 17695 | 885 | A885P | C | 4.7e-06 | 2.2e-05 | 5.8e-06 | 1.083E-05 | 9.686E-06 | 4.700E-06 | 2.200E-05 |
| 17696 | 885 | A885S | C | 8e-06 | 6.7e-06 | 1.3e-05 | 9.233E-06 | 3.326E-06 | 6.700E-06 | 1.300E-05 |
| 17697 | 885 | A885T | C | 6e-06 | 6.6e-06 | 1e-05 | 7.533E-06 | 2.157E-06 | 6.000E-06 | 1.000E-05 |
| 17698 | 885 | A885W | C | 1.3e-05 | 8e-06 | 1.1e-05 | 1.067E-05 | 2.517E-06 | 8.000E-06 | 1.300E-05 |
| 17699 | 885 | A885Y | C | 1.1e-05 | 2.8e-06 | 5.5e-06 | 6.433E-06 | 4.179E-06 | 2.800E-06 | 1.100E-05 |
| 17700 | 885 | A885V | C | 6e-06 | 2.9e-06 | 5.4e-06 | 4.767E-06 | 1.644E-06 | 2.900E-06 | 6.000E-06 |
| 17701 | 886 | D886A | C | 6.5e-06 | 1.1e-05 | 7.5e-06 | 8.333E-06 | 2.363E-06 | 6.500E-06 | 1.100E-05 |
| 17702 | 886 | D886R | C | 9.4e-06 | 1.1e-05 | 1.2e-05 | 1.080E-05 | 1.311E-06 | 9.400E-06 | 1.200E-05 |
| 17703 | 886 | D886N | C | 1e-05 | 6.4e-06 | 7.6e-06 | 8.000E-06 | 1.833E-06 | 6.400E-06 | 1.000E-05 |
| 17704 | 886 | D886D | C | 1.2e-05 | 5.7e-06 | 1e-05 | 9.233E-06 | 3.219E-06 | 5.700E-06 | 1.200E-05 |
| 17705 | 886 | D886C | C | 9.3e-06 | 6.4e-06 | 9.8e-06 | 8.500E-06 | 1.836E-06 | 6.400E-06 | 9.800E-06 |
| 17706 | 886 | D886E | C | 1.1e-05 | 7e-06 | 1.1e-05 | 9.667E-06 | 2.309E-06 | 7.000E-06 | 1.100E-05 |
| 17707 | 886 | D886Q | C | 6.9e-06 | 1.2e-05 | 8e-06 | 8.967E-06 | 2.684E-06 | 6.900E-06 | 1.200E-05 |
| 17708 | 886 | D886G | C | 7.3e-06 | 9.6e-06 | 9.6e-06 | 8.833E-06 | 1.328E-06 | 7.300E-06 | 9.600E-06 |
| 17709 | 886 | D886H | C | 1.3e-05 | 7.1e-06 | 5.1e-06 | 8.400E-06 | 4.107E-06 | 5.100E-06 | 1.300E-05 |
| 17710 | 886 | D886I | C | 5.6e-06 | 1.1e-05 | 6.5e-06 | 7.700E-06 | 2.893E-06 | 5.600E-06 | 1.100E-05 |
| 17711 | 886 | D886L | C | 5.6e-06 | 7.6e-06 | 6.7e-06 | 6.633E-06 | 1.002E-06 | 5.600E-06 | 7.600E-06 |
| 17712 | 886 | D886K | C | 6.6e-06 | 7.1e-06 | 8.1e-06 | 7.267E-06 | 7.638E-07 | 6.600E-06 | 8.100E-06 |
| 17713 | 886 | D886M | C | 7.8e-06 | 1.4e-05 | 5.2e-06 | 9.000E-06 | 4.521E-06 | 5.200E-06 | 1.400E-05 |
| 17714 | 886 | D886F | C | 6.2e-06 | 7.9e-06 | 7.3e-06 | 7.133E-06 | 8.622E-07 | 6.200E-06 | 7.900E-06 |
| 17715 | 886 | D886P | C | 4.1e-06 | 8.2e-06 | 6.4e-06 | 6.233E-06 | 2.055E-06 | 4.100E-06 | 8.200E-06 |
| 17716 | 886 | D886S | C | 4.9e-06 | 6.6e-06 | 1.1e-05 | 7.500E-06 | 3.148E-06 | 4.900E-06 | 1.100E-05 |
| 17717 | 886 | D886T | C | 1.1e-05 | 5.1e-06 | 9.9e-06 | 8.667E-06 | 3.137E-06 | 5.100E-06 | 1.100E-05 |
| 17718 | 886 | D886W | C | 6.7e-06 | 1.2e-05 | 8e-06 | 8.900E-06 | 2.762E-06 | 6.700E-06 | 1.200E-05 |
| 17719 | 886 | D886Y | C | 1.7e-05 | 7.3e-06 | 7e-06 | 1.043E-05 | 5.689E-06 | 7.000E-06 | 1.700E-05 |
| 17720 | 886 | D886V | C | 6.9e-06 | 5.9e-06 | 9.9e-06 | 7.567E-06 | 2.082E-06 | 5.900E-06 | 9.900E-06 |
| 17721 | 887 | V887A | C | 7.2e-06 | 1.2e-05 | 1e-05 | 9.733E-06 | 2.411E-06 | 7.200E-06 | 1.200E-05 |
| 17722 | 887 | V887R | C | 7.3e-06 | 1.3e-05 | 4.7e-06 | 8.333E-06 | 4.245E-06 | 4.700E-06 | 1.300E-05 |
| 17723 | 887 | V887N | C | 8.8e-06 | 8.9e-06 | 8.9e-06 | 8.867E-06 | 5.774E-08 | 8.800E-06 | 8.900E-06 |
| 17724 | 887 | V887D | C | 6.1e-06 | 7e-06 | 5.8e-06 | 6.300E-06 | 6.245E-07 | 5.800E-06 | 7.000E-06 |
| 17725 | 887 | V887C | C | 8.2e-06 | 5.7e-06 | 3.8e-06 | 5.900E-06 | 2.207E-06 | 3.800E-06 | 8.200E-06 |
| 17726 | 887 | V887E | C | 8.2e-06 | 5.5e-06 | 4.9e-06 | 6.200E-06 | 1.758E-06 | 4.900E-06 | 8.200E-06 |
| 17727 | 887 | V887Q | C | 9e-06 | 5.8e-06 | 6.6e-06 | 7.133E-06 | 1.665E-06 | 5.800E-06 | 9.000E-06 |
| 17728 | 887 | V887G | C | 5.6e-06 | 1e-05 | 4.8e-06 | 6.800E-06 | 2.800E-06 | 4.800E-06 | 1.000E-05 |
| 17729 | 887 | V887H | C | 3.3e-06 | 7.2e-06 | 1.2e-05 | 7.500E-06 | 4.358E-06 | 3.300E-06 | 1.200E-05 |
| 17730 | 887 | V887I | C | 7.5e-06 | 1e-05 | 8e-06 | 8.500E-06 | 1.323E-06 | 7.500E-06 | 1.000E-05 |
| 17731 | 887 | V887L | C | 5.7e-06 | 8.9e-06 | 8.5e-06 | 7.700E-06 | 1.744E-06 | 5.700E-06 | 8.900E-06 |
| 17732 | 887 | V887K | C | 8.1e-06 | 1e-05 | 1.3e-05 | 1.037E-05 | 2.470E-06 | 8.100E-06 | 1.300E-05 |
| 17733 | 887 | V887M | C | 1.1e-05 | 5.4e-06 | 1.3e-05 | 9.800E-06 | 3.940E-06 | 5.400E-06 | 1.300E-05 |
| 17734 | 887 | V887F | C | 1.3e-05 | 7.8e-06 | 6.7e-06 | 9.167E-06 | 3.365E-06 | 6.700E-06 | 1.300E-05 |
| 17735 | 887 | V887P | C | 1.4e-05 | 9.6e-06 | 7.5e-06 | 1.037E-05 | 3.317E-06 | 7.500E-06 | 1.400E-05 |
| 17736 | 887 | V887S | C | 7.2e-06 | 7e-06 | 8.9e-06 | 7.700E-06 | 1.044E-06 | 7.000E-06 | 8.900E-06 |
| 17737 | 887 | V887T | C | 5.8e-06 | 1.3e-05 | 7.7e-06 | 8.833E-06 | 3.731E-06 | 5.800E-06 | 1.300E-05 |
| 17738 | 887 | V887W | C | 7.5e-06 | 6.2e-06 | 8.3e-06 | 7.333E-06 | 1.060E-06 | 6.200E-06 | 8.300E-06 |
| 17739 | 887 | V887Y | C | 5.9e-06 | 9.9e-06 | 9.5e-06 | 8.433E-06 | 2.203E-06 | 5.900E-06 | 9.900E-06 |
| 17740 | 887 | V887V | C | 6.3e-06 | 1.2e-05 | 7.9e-06 | 8.733E-06 | 2.940E-06 | 6.300E-06 | 1.200E-05 |
| 17741 | 888 | N888A | C | 7.1e-06 | 1.3e-05 | 8.9e-06 | 9.667E-06 | 3.024E-06 | 7.100E-06 | 1.300E-05 |
| 17742 | 888 | N888R | C | 8.7e-06 | 9.2e-06 | 6.3e-06 | 8.067E-06 | 1.550E-06 | 6.300E-06 | 9.200E-06 |
| 17743 | 888 | N888N | C | 9.2e-06 | 9.5e-06 | 6e-06 | 8.233E-06 | 1.940E-06 | 6.000E-06 | 9.500E-06 |
| 17744 | 888 | N888D | C | 4.4e-06 | 5.9e-06 | 3.2e-06 | 4.500E-06 | 1.353E-06 | 3.200E-06 | 5.900E-06 |
| 17745 | 888 | N888C | C | 8.4e-06 | 5.6e-06 | 9.8e-06 | 7.933E-06 | 2.139E-06 | 5.600E-06 | 9.800E-06 |
| 17746 | 888 | N888E | C | 6.8e-06 | 6.6e-06 | 8e-06 | 7.133E-06 | 7.572E-07 | 6.600E-06 | 8.000E-06 |
| 17747 | 888 | N888Q | C | 6.5e-06 | 1.3e-05 | 5e-06 | 8.167E-06 | 4.252E-06 | 5.000E-06 | 1.300E-05 |
| 17748 | 888 | N888G | C | 7e-06 | 1.4e-05 | 9.5e-06 | 1.017E-05 | 3.547E-06 | 7.000E-06 | 1.400E-05 |
| 17749 | 888 | N888H | C | 9.3e-06 | 8.7e-06 | 9.2e-06 | 9.067E-06 | 3.215E-07 | 8.700E-06 | 9.300E-06 |
| 17750 | 888 | N888I | C | 5e-06 | 9.9e-06 | 7.1e-06 | 7.333E-06 | 2.458E-06 | 5.000E-06 | 9.900E-06 |
| 17751 | 888 | N888L | C | 1e-05 | 1e-05 | 8.4e-06 | 9.467E-06 | 9.238E-07 | 8.400E-06 | 1.000E-05 |
| 17752 | 888 | N888K | C | 9.7e-06 | 1.4e-05 | 6.2e-06 | 9.967E-06 | 3.907E-06 | 6.200E-06 | 1.400E-05 |
| 17753 | 888 | N888M | C | 8.6e-06 | 1.1e-05 | 1.5e-05 | 1.153E-05 | 3.233E-06 | 8.600E-06 | 1.500E-05 |
| 17754 | 888 | N888F | C | 1e-05 | 6.9e-06 | 6.4e-06 | 7.767E-06 | 1.950E-06 | 6.400E-06 | 1.000E-05 |
| 17755 | 888 | N888P | C | 6.5e-06 | 6.3e-06 | 1.1e-05 | 7.933E-06 | 2.658E-06 | 6.300E-06 | 1.100E-05 |
| 17756 | 888 | N888S | C | 1.1e-05 | 8.5e-06 | 9.5e-06 | 9.667E-06 | 1.258E-06 | 8.500E-06 | 1.100E-05 |
| 17757 | 888 | N888T | C | 7.1e-06 | 8.1e-06 | 1.5e-05 | 1.007E-05 | 4.302E-06 | 7.100E-06 | 1.500E-05 |
| 17758 | 888 | N888W | C | 6.4e-06 | 1.5e-05 | 9.5e-06 | 1.030E-05 | 4.355E-06 | 6.400E-06 | 1.500E-05 |
| 17759 | 888 | N888Y | C | 1.1e-05 | 1e-05 | 3.8e-06 | 8.267E-06 | 3.900E-06 | 3.800E-06 | 1.100E-05 |
| 17760 | 888 | N888V | C | 9.6e-06 | 7.4e-06 | 8.4e-06 | 8.467E-06 | 1.102E-06 | 7.400E-06 | 9.600E-06 |
| 17761 | 889 | P889A | C | 6.3e-06 | 9.8e-06 | 8.1e-06 | 8.067E-06 | 1.750E-06 | 6.300E-06 | 9.800E-06 |
| 17762 | 889 | P889R | C | 5e-06 | 8.6e-06 | 1.5e-05 | 9.533E-06 | 5.065E-06 | 5.000E-06 | 1.500E-05 |
| 17763 | 889 | P889N | C | 8.9e-06 | 7.8e-06 | 5.7e-06 | 7.467E-06 | 1.626E-06 | 5.700E-06 | 8.900E-06 |
| 17764 | 889 | P889D | C | 8.9e-06 | 8.2e-06 | 9.8e-06 | 8.967E-06 | 8.021E-07 | 8.200E-06 | 9.800E-06 |

| | | | | | | | | | | |
|---|---|---|---|---|---|---|---|---|---|---|
| 17765 | 889 | P889C | C | 3.8e-06 | 4.6e-06 | 1.1e-05 | 6.467E-06 | 3.946E-06 | 3.800E-06 | 1.100E-05 |
| 17766 | 889 | P889E | C | 7.3e-06 | 6.3e-06 | 1.1e-05 | 8.200E-06 | 2.476E-06 | 6.300E-06 | 1.100E-05 |
| 17767 | 889 | P889Q | C | 8.4e-06 | 9.4e-06 | 7.7e-06 | 8.500E-06 | 8.544E-07 | 7.700E-06 | 9.400E-06 |
| 17768 | 889 | P889G | C | 7.1e-06 | 5.1e-06 | 9e-06 | 7.067E-06 | 1.950E-06 | 5.100E-06 | 9.000E-06 |
| 17769 | 889 | P889H | C | 6.4e-06 | 9.7e-06 | 6.1e-06 | 7.400E-06 | 1.997E-06 | 6.100E-06 | 9.700E-06 |
| 17770 | 889 | P889I | C | 9.6e-06 | 9.5e-06 | 6.1e-06 | 8.400E-06 | 1.992E-06 | 6.100E-06 | 9.600E-06 |
| 17771 | 889 | P889L | C | 7.7e-06 | 9.6e-06 | 1.1e-05 | 9.433E-06 | 1.656E-06 | 7.700E-06 | 1.100E-05 |
| 17772 | 889 | P889K | C | 1.9e-05 | 7.3e-06 | 6.7e-06 | 1.100E-05 | 6.935E-06 | 6.700E-06 | 1.900E-05 |
| 17773 | 889 | P889M | C | 6.6e-06 | 4.3e-06 | 8.1e-06 | 6.333E-06 | 1.914E-06 | 4.300E-06 | 8.100E-06 |
| 17774 | 889 | P889F | C | 1.3e-05 | 8.1e-06 | 7.4e-06 | 9.500E-06 | 3.051E-06 | 7.400E-06 | 1.300E-05 |
| 17775 | 889 | P889P | C | 5e-06 | 6.6e-06 | 7.9e-06 | 6.500E-06 | 1.453E-06 | 5.000E-06 | 7.900E-06 |
| 17776 | 889 | P889S | C | 9.2e-06 | 9e-06 | 3.3e-06 | 7.167E-06 | 3.350E-06 | 3.300E-06 | 9.200E-06 |
| 17777 | 889 | P889T | C | 1.2e-05 | 8.2e-06 | 7e-06 | 9.067E-06 | 2.610E-06 | 7.000E-06 | 1.200E-05 |
| 17778 | 889 | P889W | C | 5.6e-06 | 6.4e-06 | 2e-05 | 1.067E-05 | 8.093E-06 | 5.600E-06 | 2.000E-05 |
| 17779 | 889 | P889Y | C | 1.2e-05 | 6.4e-06 | 8.2e-06 | 8.867E-06 | 2.859E-06 | 6.400E-06 | 1.200E-05 |
| 17780 | 889 | P889V | C | 1.1e-05 | 6.4e-06 | 9.6e-06 | 9.000E-06 | 2.358E-06 | 6.400E-06 | 1.100E-05 |
| 17781 | 890 | N890A | C | 7.2e-06 | 6.9e-06 | 6.5e-06 | 6.867E-06 | 3.512E-07 | 6.500E-06 | 7.200E-06 |
| 17782 | 890 | N890R | C | 7.1e-06 | 7.2e-06 | 2e-05 | 1.143E-05 | 7.419E-06 | 7.100E-06 | 2.000E-05 |
| 17783 | 890 | N890N | C | 8.7e-06 | 8.9e-06 | 8.7e-06 | 8.767E-06 | 1.155E-07 | 8.700E-06 | 8.900E-06 |
| 17784 | 890 | N890D | C | 8e-06 | 1.4e-05 | 7.6e-06 | 9.867E-06 | 3.585E-06 | 7.600E-06 | 1.400E-05 |
| 17785 | 890 | N890C | C | 5.4e-06 | 5.6e-06 | 9.1e-06 | 6.700E-06 | 2.081E-06 | 5.400E-06 | 9.100E-06 |
| 17786 | 890 | N890E | C | 8.9e-06 | 5.3e-06 | 6.6e-06 | 6.933E-06 | 1.823E-06 | 5.300E-06 | 8.900E-06 |
| 17787 | 890 | N890Q | C | 8.2e-06 | 9.7e-06 | 5e-06 | 7.633E-06 | 2.401E-06 | 5.000E-06 | 9.700E-06 |
| 17788 | 890 | N890G | C | 8.2e-06 | 7e-06 | 1e-05 | 8.400E-06 | 1.510E-06 | 7.000E-06 | 1.000E-05 |
| 17789 | 890 | N890H | C | 1.2e-05 | 7.9e-06 | 9.2e-06 | 9.700E-06 | 2.095E-06 | 7.900E-06 | 1.200E-05 |
| 17790 | 890 | N890I | C | 6.3e-06 | 9.2e-06 | 8.6e-06 | 8.033E-06 | 1.531E-06 | 6.300E-06 | 9.200E-06 |
| 17791 | 890 | N890L | C | 4.3e-06 | 9.2e-06 | 6.5e-06 | 6.667E-06 | 2.454E-06 | 4.300E-06 | 9.200E-06 |
| 17792 | 890 | N890K | C | 4.4e-06 | 9.8e-06 | 6.7e-06 | 6.967E-06 | 2.710E-06 | 4.400E-06 | 9.800E-06 |
| 17793 | 890 | N890M | C | 7.8e-06 | 7.3e-06 | 1e-05 | 8.367E-06 | 1.436E-06 | 7.300E-06 | 1.000E-05 |
| 17794 | 890 | N890F | C | 7.7e-06 | 6.9e-06 | 7.6e-06 | 7.400E-06 | 4.359E-07 | 6.900E-06 | 7.700E-06 |
| 17795 | 890 | N890P | C | 1.3e-05 | 8e-06 | 7.6e-06 | 9.533E-06 | 3.009E-06 | 7.600E-06 | 1.300E-05 |
| 17796 | 890 | N890S | C | 4.8e-06 | 6.3e-06 | 7.8e-06 | 6.300E-06 | 1.500E-06 | 4.800E-06 | 7.800E-06 |
| 17797 | 890 | N890T | C | 6.9e-06 | 6.2e-06 | 1.3e-05 | 8.700E-06 | 3.740E-06 | 6.200E-06 | 1.300E-05 |
| 17798 | 890 | N890W | C | 4.1e-06 | 6.9e-06 | 7.9e-06 | 6.300E-06 | 1.970E-06 | 4.100E-06 | 7.900E-06 |
| 17799 | 890 | N890Y | C | 1.2e-05 | 6.4e-06 | 1e-05 | 9.467E-06 | 2.838E-06 | 6.400E-06 | 1.200E-05 |
| 17800 | 890 | N890V | C | 6.5e-06 | 9.1e-06 | 6.2e-06 | 7.267E-06 | 1.595E-06 | 6.200E-06 | 9.100E-06 |
| 17801 | 891 | S891A | C | 1.9e-05 | 5.6e-06 | 7.9e-06 | 1.083E-05 | 7.165E-06 | 5.600E-06 | 1.900E-05 |
| 17802 | 891 | S891R | C | 1.4e-05 | 1e-05 | 7.8e-06 | 1.060E-05 | 3.143E-06 | 7.800E-06 | 1.400E-05 |
| 17803 | 891 | S891N | C | 9.2e-06 | 5e-06 | 7.1e-06 | 7.100E-06 | 2.100E-06 | 5.000E-06 | 9.200E-06 |
| 17804 | 891 | S891D | C | 1.3e-05 | 9.5e-06 | 8.8e-06 | 1.043E-05 | 2.250E-06 | 8.800E-06 | 1.300E-05 |
| 17805 | 891 | S891C | C | 6.7e-06 | 9.5e-06 | 8e-06 | 8.067E-06 | 1.401E-06 | 6.700E-06 | 9.500E-06 |
| 17806 | 891 | S891E | C | 8.8e-06 | 8.2e-06 | 1.2e-05 | 9.667E-06 | 2.043E-06 | 8.200E-06 | 1.200E-05 |
| 17807 | 891 | S891Q | C | 5.3e-06 | 1.2e-05 | 4.4e-06 | 7.233E-06 | 4.153E-06 | 4.400E-06 | 1.200E-05 |
| 17808 | 891 | S891G | C | 1e-05 | 5.9e-06 | 1.1e-05 | 8.967E-06 | 2.702E-06 | 5.900E-06 | 1.100E-05 |
| 17809 | 891 | S891H | C | 7.5e-06 | 1e-05 | 8e-06 | 8.500E-06 | 1.323E-06 | 7.500E-06 | 1.000E-05 |
| 17810 | 891 | S891I | C | 9e-06 | 8e-06 | 7.7e-06 | 8.233E-06 | 6.807E-07 | 7.700E-06 | 9.000E-06 |
| 17811 | 891 | S891L | C | 4.4e-06 | 4.8e-06 | 1.4e-05 | 7.733E-06 | 5.431E-06 | 4.400E-06 | 1.400E-05 |
| 17812 | 891 | S891K | C | 6.7e-06 | 8.4e-06 | 8.6e-06 | 7.900E-06 | 1.044E-06 | 6.700E-06 | 8.600E-06 |
| 17813 | 891 | S891M | C | 8.7e-06 | 1.1e-05 | 9.3e-06 | 9.667E-06 | 1.193E-06 | 8.700E-06 | 1.100E-05 |
| 17814 | 891 | S891F | C | 9.2e-06 | 4.2e-06 | 5e-06 | 6.133E-06 | 2.686E-06 | 4.200E-06 | 9.200E-06 |
| 17815 | 891 | S891P | C | 1.1e-05 | 4.5e-06 | 1.1e-05 | 8.833E-06 | 3.753E-06 | 4.500E-06 | 1.100E-05 |
| 17816 | 891 | S891S | C | 6.9e-06 | 4.9e-06 | 8.1e-06 | 6.633E-06 | 1.617E-06 | 4.900E-06 | 8.100E-06 |
| 17817 | 891 | S891T | C | 8.2e-06 | 7.3e-06 | 1e-05 | 8.500E-06 | 1.375E-06 | 7.300E-06 | 1.000E-05 |
| 17818 | 891 | S891W | C | 1.1e-05 | 9.8e-06 | 7e-06 | 9.267E-06 | 2.053E-06 | 7.000E-06 | 1.100E-05 |
| 17819 | 891 | S891Y | C | 8.7e-06 | 5.4e-06 | 8.5e-06 | 7.533E-06 | 1.850E-06 | 5.400E-06 | 8.700E-06 |
| 17820 | 891 | S891V | C | 6.5e-06 | 6e-06 | 9.4e-06 | 7.300E-06 | 1.836E-06 | 6.000E-06 | 9.400E-06 |
| 17821 | 892 | G892A | C | 9.7e-06 | 1.3e-05 | 6.4e-06 | 9.700E-06 | 3.300E-06 | 6.400E-06 | 1.300E-05 |
| 17822 | 892 | G892R | C | 1e-05 | 1.4e-05 | 1.2e-05 | 1.200E-05 | 2.000E-06 | 1.000E-05 | 1.400E-05 |
| 17823 | 892 | G892N | C | 4.6e-06 | 1.4e-05 | 5.1e-06 | 7.900E-06 | 5.289E-06 | 4.600E-06 | 1.400E-05 |
| 17824 | 892 | G892D | C | 1.3e-05 | 8.9e-06 | 8.3e-06 | 1.007E-05 | 2.558E-06 | 8.300E-06 | 1.300E-05 |
| 17825 | 892 | G892C | C | 5.8e-06 | 5.2e-06 | 7.9e-06 | 6.300E-06 | 1.418E-06 | 5.200E-06 | 7.900E-06 |
| 17826 | 892 | G892E | C | 9e-06 | 1e-05 | 8.4e-06 | 9.133E-06 | 8.083E-07 | 8.400E-06 | 1.000E-05 |
| 17827 | 892 | G892Q | C | 7.8e-06 | 6.2e-06 | 1e-05 | 8.000E-06 | 1.908E-06 | 6.200E-06 | 1.000E-05 |
| 17828 | 892 | G892G | C | 1.2e-05 | 1.2e-05 | 1e-05 | 1.133E-05 | 1.155E-06 | 1.000E-05 | 1.200E-05 |
| 17829 | 892 | G892H | C | 7.5e-06 | 1e-05 | 4.2e-06 | 7.233E-06 | 2.909E-06 | 4.200E-06 | 1.000E-05 |
| 17830 | 892 | G892I | C | 8.6e-06 | 7e-06 | 7e-06 | 7.533E-06 | 9.238E-07 | 7.000E-06 | 8.600E-06 |
| 17831 | 892 | G892L | C | 8.7e-06 | 7.5e-06 | 6.9e-06 | 7.700E-06 | 9.165E-07 | 6.900E-06 | 8.700E-06 |
| 17832 | 892 | G892K | C | 1.3e-05 | 5.7e-06 | 9.8e-06 | 9.500E-06 | 3.659E-06 | 5.700E-06 | 1.300E-05 |
| 17833 | 892 | G892M | C | 8.9e-06 | 8e-06 | 1.6e-05 | 1.097E-05 | 4.382E-06 | 8.000E-06 | 1.600E-05 |
| 17834 | 892 | G892F | C | 6.3e-06 | 1.2e-05 | 5.8e-06 | 8.033E-06 | 3.444E-06 | 5.800E-06 | 1.200E-05 |
| 17835 | 892 | G892P | C | 6.7e-06 | 6.9e-06 | 8.9e-06 | 7.500E-06 | 1.217E-06 | 6.700E-06 | 8.900E-06 |
| 17836 | 892 | G892S | C | 8.8e-06 | 5.3e-06 | 5.5e-06 | 6.533E-06 | 1.966E-06 | 5.300E-06 | 8.800E-06 |
| 17837 | 892 | G892T | C | 7.4e-06 | 7.9e-06 | 1.1e-05 | 8.767E-06 | 1.950E-06 | 7.400E-06 | 1.100E-05 |
| 17838 | 892 | G892W | C | 5.8e-06 | 5.2e-06 | 6.4e-06 | 5.800E-06 | 6.000E-07 | 5.200E-06 | 6.400E-06 |
| 17839 | 892 | G892Y | C | 8e-06 | 7.8e-06 | 6.7e-06 | 7.500E-06 | 7.000E-07 | 6.700E-06 | 8.000E-06 |
| 17840 | 892 | G892V | C | 6.2e-06 | 5.6e-06 | 4.9e-06 | 5.567E-06 | 6.506E-07 | 4.900E-06 | 6.200E-06 |
| 17841 | 893 | G893A | C | 8.8e-06 | 7.2e-06 | 8.5e-06 | 8.167E-06 | 8.505E-07 | 7.200E-06 | 8.800E-06 |
| 17842 | 893 | G893R | C | 1.3e-05 | 1.1e-05 | 8.2e-06 | 1.073E-05 | 2.411E-06 | 8.200E-06 | 1.300E-05 |
| 17843 | 893 | G893N | C | 9e-06 | 6.1e-06 | 8.9e-06 | 8.000E-06 | 1.646E-06 | 6.100E-06 | 9.000E-06 |
| 17844 | 893 | G893D | C | 3.5e-06 | 8.5e-06 | 7.7e-06 | 6.567E-06 | 2.686E-06 | 3.500E-06 | 8.500E-06 |
| 17845 | 893 | G893C | C | 7.3e-06 | 1.2e-05 | 6.6e-06 | 8.633E-06 | 2.937E-06 | 6.600E-06 | 1.200E-05 |
| 17846 | 893 | G893E | C | 1e-05 | 7.2e-06 | 5.9e-06 | 7.700E-06 | 2.095E-06 | 5.900E-06 | 1.000E-05 |
| 17847 | 893 | G893Q | C | 6.1e-06 | 9.6e-06 | 5e-06 | 6.900E-06 | 2.402E-06 | 5.000E-06 | 9.600E-06 |
| 17848 | 893 | G893G | C | 1.5e-05 | 3e-06 | 7.2e-06 | 8.400E-06 | 6.089E-06 | 3.000E-06 | 1.500E-05 |
| 17849 | 893 | G893H | C | 6.2e-06 | 5.9e-06 | 5.1e-06 | 5.733E-06 | 5.686E-07 | 5.100E-06 | 6.200E-06 |

| | | | | | | | | | | |
|---|---|---|---|---|---|---|---|---|---|---|
| 17850 | 893 | G893I | C | 7.2e-06 | 1e-05 | 1.1e-05 | 9.400E-06 | 1.970E-06 | 7.200E-06 | 1.100E-05 |
| 17851 | 893 | G893L | C | 6.3e-06 | 1.3e-05 | 1.4e-05 | 1.110E-05 | 4.187E-06 | 6.300E-06 | 1.400E-05 |
| 17852 | 893 | G893K | C | 9.8e-06 | 7e-06 | 6.8e-06 | 7.867E-06 | 1.677E-06 | 6.800E-06 | 9.800E-06 |
| 17853 | 893 | G893M | C | 1.1e-05 | 1.3e-05 | 9.2e-06 | 1.107E-05 | 1.901E-06 | 9.200E-06 | 1.300E-05 |
| 17854 | 893 | G893F | C | 7e-06 | 1.2e-05 | 7.5e-06 | 8.833E-06 | 2.754E-06 | 7.000E-06 | 1.200E-05 |
| 17855 | 893 | G893P | C | 8.2e-06 | 1.1e-05 | 4.7e-06 | 7.967E-06 | 3.156E-06 | 4.700E-06 | 1.100E-05 |
| 17856 | 893 | G893S | C | 4.7e-06 | 5.8e-06 | 7.7e-06 | 6.067E-06 | 1.518E-06 | 4.700E-06 | 7.700E-06 |
| 17857 | 893 | G893T | C | 1e-05 | 8e-06 | 1e-05 | 9.333E-06 | 1.155E-06 | 8.000E-06 | 1.000E-05 |
| 17858 | 893 | G893W | C | 4.5e-06 | 5.4e-06 | 1.3e-05 | 7.633E-06 | 4.669E-06 | 4.500E-06 | 1.300E-05 |
| 17859 | 893 | G893Y | C | 9.1e-06 | 4.3e-06 | 7.9e-06 | 7.100E-06 | 2.498E-06 | 4.300E-06 | 9.100E-06 |
| 17860 | 893 | G893V | C | 9.3e-06 | 1.4e-05 | 1.7e-05 | 1.343E-05 | 3.881E-06 | 9.300E-06 | 1.700E-05 |
| 17861 | 894 | S894A | C | 1.4e-05 | 8.1e-06 | 1.4e-05 | 1.203E-05 | 3.406E-06 | 8.100E-06 | 1.400E-05 |
| 17862 | 894 | S894R | C | 6.1e-06 | 5e-06 | 6.3e-06 | 5.800E-06 | 7.000E-07 | 5.000E-06 | 6.300E-06 |
| 17863 | 894 | S894N | C | 8.3e-06 | 1.2e-05 | 6.2e-06 | 8.833E-06 | 2.937E-06 | 6.200E-06 | 1.200E-05 |
| 17864 | 894 | S894D | C | 9.6e-06 | 5.9e-06 | 7.8e-06 | 7.767E-06 | 1.850E-06 | 5.900E-06 | 9.600E-06 |
| 17865 | 894 | S894C | C | 6.2e-06 | 7.7e-06 | 6.5e-06 | 6.800E-06 | 7.937E-07 | 6.200E-06 | 7.700E-06 |
| 17866 | 894 | S894E | C | 4.6e-06 | 8.1e-06 | 5.5e-06 | 6.067E-06 | 1.818E-06 | 4.600E-06 | 8.100E-06 |
| 17867 | 894 | S894Q | C | 6.7e-06 | 1.3e-05 | 1.3e-05 | 1.090E-05 | 3.637E-06 | 6.700E-06 | 1.300E-05 |
| 17868 | 894 | S894G | C | 9.7e-06 | 1.2e-05 | 8e-06 | 9.900E-06 | 2.007E-06 | 8.000E-06 | 1.200E-05 |
| 17869 | 894 | S894H | C | 5.6e-06 | 8.4e-06 | 5.4e-06 | 6.467E-06 | 1.677E-06 | 5.400E-06 | 8.400E-06 |
| 17870 | 894 | S894I | C | 1.3e-05 | 7e-06 | 8.1e-06 | 9.367E-06 | 3.194E-06 | 7.000E-06 | 1.300E-05 |
| 17871 | 894 | S894L | C | 1.2e-05 | 5e-06 | 9.4e-06 | 8.800E-06 | 3.538E-06 | 5.000E-06 | 1.200E-05 |
| 17872 | 894 | S894K | C | 7.2e-06 | 1.1e-05 | 7.4e-06 | 8.533E-06 | 2.139E-06 | 7.200E-06 | 1.100E-05 |
| 17873 | 894 | S894M | C | 9.4e-06 | 6e-06 | 4.9e-06 | 6.767E-06 | 2.346E-06 | 4.900E-06 | 9.400E-06 |
| 17874 | 894 | S894F | C | 7.6e-06 | 1.5e-05 | 5.4e-06 | 9.333E-06 | 5.029E-06 | 5.400E-06 | 1.500E-05 |
| 17875 | 894 | S894P | C | 8.1e-06 | 7.6e-06 | 9.2e-06 | 8.300E-06 | 8.185E-07 | 7.600E-06 | 9.200E-06 |
| 17876 | 894 | S894S | C | 4.3e-06 | 1.2e-05 | 6.5e-06 | 7.600E-06 | 3.966E-06 | 4.300E-06 | 1.200E-05 |
| 17877 | 894 | S894T | C | 1.1e-05 | 6.7e-06 | 1.3e-05 | 1.023E-05 | 3.219E-06 | 6.700E-06 | 1.300E-05 |
| 17878 | 894 | S894W | C | 5.2e-06 | 1.1e-05 | 4.2e-06 | 6.800E-06 | 3.672E-06 | 4.200E-06 | 1.100E-05 |
| 17879 | 894 | S894Y | C | 4.8e-06 | 5e-06 | 8.9e-06 | 6.233E-06 | 2.312E-06 | 4.800E-06 | 8.900E-06 |
| 17880 | 894 | S894V | C | 7.5e-06 | 1e-05 | 9.3e-06 | 8.933E-06 | 1.290E-06 | 7.500E-06 | 1.000E-05 |
| 17881 | 895 | I895A | C | 5.7e-06 | 8.2e-06 | 7.3e-06 | 7.067E-06 | 1.266E-06 | 5.700E-06 | 8.200E-06 |
| 17882 | 895 | I895R | C | 5.5e-06 | 9.4e-06 | 8.8e-06 | 7.900E-06 | 2.100E-06 | 5.500E-06 | 9.400E-06 |
| 17883 | 895 | I895N | C | 9.6e-06 | 8.6e-06 | 6.9e-06 | 8.367E-06 | 1.365E-06 | 6.900E-06 | 9.600E-06 |
| 17884 | 895 | I895D | C | 7.6e-06 | 8.3e-06 | 9.5e-06 | 8.467E-06 | 9.609E-07 | 7.600E-06 | 9.500E-06 |
| 17885 | 895 | I895C | C | 6.9e-06 | 6.3e-06 | 9.6e-06 | 7.600E-06 | 1.758E-06 | 6.300E-06 | 9.600E-06 |
| 17886 | 895 | I895E | C | 1.4e-05 | 1.2e-05 | 1.1e-05 | 1.233E-05 | 1.528E-06 | 1.100E-05 | 1.400E-05 |
| 17887 | 895 | I895Q | C | 1.3e-05 | 3.3e-06 | 6.8e-06 | 7.700E-06 | 4.912E-06 | 3.300E-06 | 1.300E-05 |
| 17888 | 895 | I895G | C | 6.1e-06 | 7.6e-06 | 8e-06 | 7.233E-06 | 1.002E-06 | 6.100E-06 | 8.000E-06 |
| 17889 | 895 | I895H | C | 8.7e-06 | 4.2e-06 | 6.5e-06 | 6.467E-06 | 2.250E-06 | 4.200E-06 | 8.700E-06 |
| 17890 | 895 | I895I | C | 1e-05 | 7.9e-06 | 1e-05 | 9.300E-06 | 1.212E-06 | 7.900E-06 | 1.000E-05 |
| 17891 | 895 | I895L | C | 9.7e-06 | 7.1e-06 | 1.1e-05 | 9.267E-06 | 1.986E-06 | 7.100E-06 | 1.100E-05 |
| 17892 | 895 | I895K | C | 8.2e-06 | 9.3e-06 | 1.5e-05 | 1.083E-05 | 3.650E-06 | 8.200E-06 | 1.500E-05 |
| 17893 | 895 | I895M | C | 7.6e-06 | 1.1e-05 | 4.9e-06 | 7.833E-06 | 3.057E-06 | 4.900E-06 | 1.100E-05 |
| 17894 | 895 | I895F | C | 8.5e-06 | 1.5e-05 | 7.2e-06 | 1.023E-05 | 4.179E-06 | 7.200E-06 | 1.500E-05 |
| 17895 | 895 | I895P | C | 1.1e-05 | 1.4e-05 | 5.8e-06 | 1.027E-05 | 4.149E-06 | 5.800E-06 | 1.400E-05 |
| 17896 | 895 | I895S | C | 3.7e-06 | 6e-06 | 1.3e-05 | 7.567E-06 | 4.844E-06 | 3.700E-06 | 1.300E-05 |
| 17897 | 895 | I895T | C | 8.8e-06 | 9.5e-06 | 6.1e-06 | 8.133E-06 | 1.795E-06 | 6.100E-06 | 9.500E-06 |
| 17898 | 895 | I895W | C | 6e-06 | 1e-05 | 6.3e-06 | 7.433E-06 | 2.228E-06 | 6.000E-06 | 1.000E-05 |
| 17899 | 895 | I895Y | C | 1.1e-05 | 1e-05 | 8.6e-06 | 9.867E-06 | 1.206E-06 | 8.600E-06 | 1.100E-05 |
| 17900 | 895 | I895V | C | 4.1e-06 | 1e-05 | 1.1e-05 | 8.367E-06 | 3.729E-06 | 4.100E-06 | 1.100E-05 |
| 17901 | 896 | Y896A | C | 4.1e-06 | 9.9e-06 | 9.9e-06 | 7.967E-06 | 3.349E-06 | 4.100E-06 | 9.900E-06 |
| 17902 | 896 | Y896R | C | 7.7e-06 | 7.3e-06 | 1.2e-05 | 9.000E-06 | 2.606E-06 | 7.300E-06 | 1.200E-05 |
| 17903 | 896 | Y896N | C | 5.7e-06 | 1.1e-05 | 9.1e-06 | 8.600E-06 | 2.685E-06 | 5.700E-06 | 1.100E-05 |
| 17904 | 896 | Y896D | C | 7.3e-06 | 6.7e-06 | 6.4e-06 | 6.800E-06 | 4.583E-07 | 6.400E-06 | 7.300E-06 |
| 17905 | 896 | Y896C | C | 7.5e-06 | 8.5e-06 | 1.2e-05 | 9.333E-06 | 2.363E-06 | 7.500E-06 | 1.200E-05 |
| 17906 | 896 | Y896E | C | 6.5e-06 | 1.2e-05 | 7e-06 | 8.500E-06 | 3.041E-06 | 6.500E-06 | 1.200E-05 |
| 17907 | 896 | Y896Q | C | 6e-06 | 8.6e-06 | 4.6e-06 | 6.400E-06 | 2.030E-06 | 4.600E-06 | 8.600E-06 |
| 17908 | 896 | Y896G | C | 7.1e-06 | 1.1e-05 | 1.3e-05 | 1.037E-05 | 3.001E-06 | 7.100E-06 | 1.300E-05 |
| 17909 | 896 | Y896H | C | 5.9e-06 | 5.1e-06 | 9.4e-06 | 6.800E-06 | 2.287E-06 | 5.100E-06 | 9.400E-06 |
| 17910 | 896 | Y896I | C | 5.3e-06 | 1.1e-05 | 6.6e-06 | 7.633E-06 | 2.987E-06 | 5.300E-06 | 1.100E-05 |
| 17911 | 896 | Y896L | C | 1.3e-05 | 9.1e-06 | 6.4e-06 | 9.500E-06 | 3.318E-06 | 6.400E-06 | 1.300E-05 |
| 17912 | 896 | Y896K | C | 5.9e-06 | 6.7e-06 | 6.8e-06 | 6.467E-06 | 4.933E-07 | 5.900E-06 | 6.800E-06 |
| 17913 | 896 | Y896M | C | 8.6e-06 | 9.3e-06 | 6.3e-06 | 8.067E-06 | 1.570E-06 | 6.300E-06 | 9.300E-06 |
| 17914 | 896 | Y896F | C | 1.3e-05 | 8.3e-06 | 7.2e-06 | 9.500E-06 | 3.081E-06 | 7.200E-06 | 1.300E-05 |
| 17915 | 896 | Y896P | C | 1.4e-05 | 6.8e-06 | 9.1e-06 | 9.967E-06 | 3.677E-06 | 6.800E-06 | 1.400E-05 |
| 17916 | 896 | Y896S | C | 1.6e-05 | 1.1e-05 | 7.6e-06 | 1.153E-05 | 4.225E-06 | 7.600E-06 | 1.600E-05 |
| 17917 | 896 | Y896T | C | 7.3e-06 | 7.4e-06 | 6.6e-06 | 7.100E-06 | 4.359E-07 | 6.600E-06 | 7.400E-06 |
| 17918 | 896 | Y896W | C | 6.8e-06 | 9.1e-06 | 7.3e-06 | 7.733E-06 | 1.210E-06 | 6.800E-06 | 9.100E-06 |
| 17919 | 896 | Y896Y | C | 1.2e-05 | 6.9e-06 | 9.2e-06 | 9.367E-06 | 2.554E-06 | 6.900E-06 | 1.200E-05 |
| 17920 | 896 | Y896V | C | 4.8e-06 | 6.6e-06 | 5.5e-06 | 5.633E-06 | 9.074E-07 | 4.800E-06 | 6.600E-06 |
| 17921 | 897 | N897A | C | 9.9e-06 | 1.3e-05 | 7.8e-06 | 1.023E-05 | 2.616E-06 | 7.800E-06 | 1.300E-05 |
| 17922 | 897 | N897R | C | 5.6e-06 | 6.2e-06 | 7.8e-06 | 6.533E-06 | 1.137E-06 | 5.600E-06 | 7.800E-06 |
| 17923 | 897 | N897N | C | 8.6e-06 | 5.3e-06 | 1e-05 | 7.967E-06 | 2.413E-06 | 5.300E-06 | 1.000E-05 |
| 17924 | 897 | N897D | C | 9.2e-06 | 7.7e-06 | 4.7e-06 | 7.200E-06 | 2.291E-06 | 4.700E-06 | 9.200E-06 |
| 17925 | 897 | N897C | C | 9.2e-06 | 1e-05 | 4.2e-06 | 7.800E-06 | 3.143E-06 | 4.200E-06 | 1.000E-05 |
| 17926 | 897 | N897E | C | 4.5e-06 | 4.7e-06 | 6.9e-06 | 5.367E-06 | 1.332E-06 | 4.500E-06 | 6.900E-06 |
| 17927 | 897 | N897Q | C | 7.3e-06 | 9.4e-06 | 9.3e-06 | 8.667E-06 | 1.185E-06 | 7.300E-06 | 9.400E-06 |
| 17928 | 897 | N897G | C | 7.7e-06 | 8e-06 | 1.4e-05 | 9.900E-06 | 3.554E-06 | 7.700E-06 | 1.400E-05 |
| 17929 | 897 | N897H | C | 5.5e-06 | 4.3e-06 | 5e-06 | 4.933E-06 | 6.028E-07 | 4.300E-06 | 5.500E-06 |
| 17930 | 897 | N897I | C | 8.6e-06 | 5.8e-06 | 8.8e-06 | 7.733E-06 | 1.677E-06 | 5.800E-06 | 8.800E-06 |
| 17931 | 897 | N897L | C | 6.1e-06 | 7.3e-06 | 7.1e-06 | 6.833E-06 | 6.429E-07 | 6.100E-06 | 7.300E-06 |
| 17932 | 897 | N897K | C | 9e-06 | 6.1e-06 | 1e-05 | 8.367E-06 | 2.026E-06 | 6.100E-06 | 1.000E-05 |
| 17933 | 897 | N897M | C | 9.1e-06 | 9.5e-06 | 6.7e-06 | 8.433E-06 | 1.514E-06 | 6.700E-06 | 9.500E-06 |
| 17934 | 897 | N897F | C | 1.5e-05 | 6.8e-06 | 1.4e-05 | 1.193E-05 | 4.474E-06 | 6.800E-06 | 1.500E-05 |

| | | | | | | | | | | |
|---|---|---|---|---|---|---|---|---|---|---|
| 17935 | 897 | N897P | C | 6.3e-06 | 1.3e-05 | 1.4e-05 | 1.110E-05 | 4.187E-06 | 6.300E-06 | 1.400E-05 |
| 17936 | 897 | N897S | C | 1e-05 | 9.8e-06 | 6e-06 | 8.600E-06 | 2.254E-06 | 6.000E-06 | 1.000E-05 |
| 17937 | 897 | N897T | C | 1.2e-05 | 3.2e-06 | 6.7e-06 | 7.300E-06 | 4.431E-06 | 3.200E-06 | 1.200E-05 |
| 17938 | 897 | N897W | C | 8.4e-06 | 6.6e-06 | 9.4e-06 | 8.133E-06 | 1.419E-06 | 6.600E-06 | 9.400E-06 |
| 17939 | 897 | N897Y | C | 7.1e-06 | 8.4e-06 | 9.3e-06 | 8.267E-06 | 1.106E-06 | 7.100E-06 | 9.300E-06 |
| 17940 | 897 | N897V | C | 9.2e-06 | 6.2e-06 | 7.3e-06 | 7.567E-06 | 1.518E-06 | 6.200E-06 | 9.200E-06 |
| 17941 | 898 | Q898A | C | 5.4e-06 | 6.4e-06 | 7.8e-06 | 6.533E-06 | 1.206E-06 | 5.400E-06 | 7.800E-06 |
| 17942 | 898 | Q898R | C | 4.9e-06 | 7.9e-06 | 8.7e-06 | 7.167E-06 | 2.003E-06 | 4.900E-06 | 8.700E-06 |
| 17943 | 898 | Q898N | C | 6.8e-06 | 7.2e-06 | 9.9e-06 | 7.967E-06 | 1.686E-06 | 6.800E-06 | 9.900E-06 |
| 17944 | 898 | Q898D | C | 5.9e-06 | 8.2e-06 | 5.9e-06 | 6.667E-06 | 1.328E-06 | 5.900E-06 | 8.200E-06 |
| 17945 | 898 | Q898C | C | 6.2e-06 | 7e-06 | 6.8e-06 | 6.667E-06 | 4.163E-07 | 6.200E-06 | 7.000E-06 |
| 17946 | 898 | Q898E | C | 9.2e-06 | 6.6e-06 | 8e-06 | 7.933E-06 | 1.301E-06 | 6.600E-06 | 9.200E-06 |
| 17947 | 898 | Q898Q | C | 8e-06 | 1.1e-05 | 5.1e-06 | 8.033E-06 | 2.950E-06 | 5.100E-06 | 1.100E-05 |
| 17948 | 898 | Q898G | C | 8.4e-06 | 5.4e-06 | 8.8e-06 | 7.533E-06 | 1.858E-06 | 5.400E-06 | 8.800E-06 |
| 17949 | 898 | Q898H | C | 8e-06 | 7.7e-06 | 6.8e-06 | 7.500E-06 | 6.245E-07 | 6.800E-06 | 8.000E-06 |
| 17950 | 898 | Q898I | C | 9.7e-06 | 4.8e-06 | 9.7e-06 | 8.067E-06 | 2.829E-06 | 4.800E-06 | 9.700E-06 |
| 17951 | 898 | Q898L | C | 9.6e-06 | 1.3e-05 | 7.9e-06 | 1.017E-05 | 2.597E-06 | 7.900E-06 | 1.300E-05 |
| 17952 | 898 | Q898K | C | 9.5e-06 | 9.3e-06 | 6.7e-06 | 8.500E-06 | 1.562E-06 | 6.700E-06 | 9.500E-06 |
| 17953 | 898 | Q898M | C | 9.5e-06 | 9e-06 | 8.8e-06 | 9.100E-06 | 3.606E-07 | 8.800E-06 | 9.500E-06 |
| 17954 | 898 | Q898F | C | 1.3e-05 | 7.2e-06 | 6.3e-06 | 8.833E-06 | 3.636E-06 | 6.300E-06 | 1.300E-05 |
| 17955 | 898 | Q898P | C | 1.4e-05 | 8.9e-06 | 8.9e-06 | 1.060E-05 | 2.944E-06 | 8.900E-06 | 1.400E-05 |
| 17956 | 898 | Q898S | C | 7.8e-06 | 7.9e-06 | 5.6e-06 | 7.100E-06 | 1.300E-06 | 5.600E-06 | 7.900E-06 |
| 17957 | 898 | Q898T | C | 1.5e-05 | 8.3e-06 | 6.4e-06 | 9.900E-06 | 4.518E-06 | 6.400E-06 | 1.500E-05 |
| 17958 | 898 | Q898W | C | 5.2e-06 | 7e-06 | 8.4e-06 | 6.867E-06 | 1.604E-06 | 5.200E-06 | 8.400E-06 |
| 17959 | 898 | Q898Y | C | 1.5e-05 | 6.4e-06 | 1.3e-05 | 1.147E-05 | 4.500E-06 | 6.400E-06 | 1.500E-05 |
| 17960 | 898 | Q898V | C | 8.9e-06 | 8.3e-06 | 5.4e-06 | 7.533E-06 | 1.872E-06 | 5.400E-06 | 8.900E-06 |
| 17961 | 899 | R899A | C | 6.6e-06 | 1e-05 | 1.1e-05 | 9.200E-06 | 2.307E-06 | 6.600E-06 | 1.100E-05 |
| 17962 | 899 | R899R | C | 4.8e-06 | 4.6e-06 | 5.8e-06 | 5.067E-06 | 6.429E-07 | 4.600E-06 | 5.800E-06 |
| 17963 | 899 | R899N | C | 1.5e-05 | 7e-06 | 9.4e-06 | 1.047E-05 | 4.105E-06 | 7.000E-06 | 1.500E-05 |
| 17964 | 899 | R899D | C | 8.4e-06 | 9.1e-06 | 4.7e-06 | 7.400E-06 | 2.364E-06 | 4.700E-06 | 9.100E-06 |
| 17965 | 899 | R899C | C | 5.6e-06 | 9e-06 | 6.3e-06 | 6.967E-06 | 1.795E-06 | 5.600E-06 | 9.000E-06 |
| 17966 | 899 | R899E | C | 1.3e-05 | 6.4e-06 | 9.5e-06 | 9.633E-06 | 3.302E-06 | 6.400E-06 | 1.300E-05 |
| 17967 | 899 | R899Q | C | 6.8e-06 | 1.2e-05 | 6.8e-06 | 8.533E-06 | 3.002E-06 | 6.800E-06 | 1.200E-05 |
| 17968 | 899 | R899G | C | 9e-06 | 9.8e-06 | 5.1e-06 | 7.967E-06 | 2.515E-06 | 5.100E-06 | 9.800E-06 |
| 17969 | 899 | R899H | C | 1.9e-05 | 5.7e-06 | 7.6e-06 | 1.077E-05 | 7.193E-06 | 5.700E-06 | 1.900E-05 |
| 17970 | 899 | R899I | C | 1.2e-05 | 4.7e-06 | 7.3e-06 | 8.000E-06 | 3.700E-06 | 4.700E-06 | 1.200E-05 |
| 17971 | 899 | R899L | C | 9.6e-06 | 7.9e-06 | 4e-06 | 7.167E-06 | 2.871E-06 | 4.000E-06 | 9.600E-06 |
| 17972 | 899 | R899K | C | 6.3e-06 | 6.7e-06 | 6.3e-06 | 6.433E-06 | 2.309E-07 | 6.300E-06 | 6.700E-06 |
| 17973 | 899 | R899M | C | 9.6e-06 | 4.2e-06 | 6.4e-06 | 6.733E-06 | 2.715E-06 | 4.200E-06 | 9.600E-06 |
| 17974 | 899 | R899F | C | 8.7e-06 | 9.4e-06 | 1.2e-05 | 1.003E-05 | 1.739E-06 | 8.700E-06 | 1.200E-05 |
| 17975 | 899 | R899P | C | 8.7e-06 | 6.7e-06 | 7.5e-06 | 7.633E-06 | 1.007E-06 | 6.700E-06 | 8.700E-06 |
| 17976 | 899 | R899S | C | 8.2e-06 | 6.1e-06 | 6.3e-06 | 6.867E-06 | 1.159E-06 | 6.100E-06 | 8.200E-06 |
| 17977 | 899 | R899T | C | 6.4e-06 | 7.1e-06 | 6.1e-06 | 6.533E-06 | 5.132E-07 | 6.100E-06 | 7.100E-06 |
| 17978 | 899 | R899W | C | 8.8e-06 | 7.6e-06 | 4.3e-06 | 6.900E-06 | 2.330E-06 | 4.300E-06 | 8.800E-06 |
| 17979 | 899 | R899Y | C | 7.5e-06 | 5.9e-06 | 9.3e-06 | 7.567E-06 | 1.701E-06 | 5.900E-06 | 9.300E-06 |
| 17980 | 899 | R899V | C | 7.3e-06 | 4.7e-06 | 8e-06 | 6.667E-06 | 1.739E-06 | 4.700E-06 | 8.000E-06 |
| 17981 | 900 | F900A | C | 1.2e-05 | 1.2e-05 | 1e-05 | 1.133E-05 | 1.155E-06 | 1.000E-05 | 1.200E-05 |
| 17982 | 900 | F900R | C | 8.7e-06 | 7.1e-06 | 6.7e-06 | 7.500E-06 | 1.058E-06 | 6.700E-06 | 8.700E-06 |
| 17983 | 900 | F900N | C | 7.6e-06 | 6.4e-06 | 7.5e-06 | 7.167E-06 | 6.658E-07 | 6.400E-06 | 7.600E-06 |
| 17984 | 900 | F900D | C | 1.4e-05 | 9.1e-06 | 8.2e-06 | 1.043E-05 | 3.121E-06 | 8.200E-06 | 1.400E-05 |
| 17985 | 900 | F900C | C | 5.8e-06 | 7e-06 | 5.3e-06 | 6.033E-06 | 8.737E-07 | 5.300E-06 | 7.000E-06 |
| 17986 | 900 | F900E | C | 7e-06 | 1e-05 | 4.4e-06 | 7.133E-06 | 2.802E-06 | 4.400E-06 | 1.000E-05 |
| 17987 | 900 | F900Q | C | 1.3e-05 | 8.1e-06 | 8.7e-06 | 9.933E-06 | 2.673E-06 | 8.100E-06 | 1.300E-05 |
| 17988 | 900 | F900G | C | 6.6e-06 | 1.1e-05 | 1.3e-05 | 1.020E-05 | 3.274E-06 | 6.600E-06 | 1.300E-05 |
| 17989 | 900 | F900H | C | 8.2e-06 | 9.7e-06 | 4.5e-06 | 7.467E-06 | 2.676E-06 | 4.500E-06 | 9.700E-06 |
| 17990 | 900 | F900I | C | 6.7e-06 | 9.1e-06 | 8.3e-06 | 8.033E-06 | 1.222E-06 | 6.700E-06 | 9.100E-06 |
| 17991 | 900 | F900L | C | 1.7e-05 | 6.3e-06 | 1.2e-05 | 1.177E-05 | 5.354E-06 | 6.300E-06 | 1.700E-05 |
| 17992 | 900 | F900K | C | 8e-06 | 8.9e-06 | 6.9e-06 | 7.933E-06 | 1.002E-06 | 6.900E-06 | 8.900E-06 |
| 17993 | 900 | F900M | C | 5e-06 | 6.6e-06 | 6.8e-06 | 6.133E-06 | 9.866E-07 | 5.000E-06 | 6.800E-06 |
| 17994 | 900 | F900F | C | 6.7e-06 | 9.6e-06 | 6.9e-06 | 7.733E-06 | 1.620E-06 | 6.700E-06 | 9.600E-06 |
| 17995 | 900 | F900P | C | 6.7e-06 | 7.6e-06 | 1.1e-05 | 8.433E-06 | 2.268E-06 | 6.700E-06 | 1.100E-05 |
| 17996 | 900 | F900S | C | 1.2e-05 | 9.8e-06 | 6.1e-06 | 9.300E-06 | 2.982E-06 | 6.100E-06 | 1.200E-05 |
| 17997 | 900 | F900T | C | 4.6e-06 | 1.3e-05 | 6.3e-06 | 7.967E-06 | 4.441E-06 | 4.600E-06 | 1.300E-05 |
| 17998 | 900 | F900W | C | 7.9e-06 | 5.7e-06 | 5.4e-06 | 6.333E-06 | 1.365E-06 | 5.400E-06 | 7.900E-06 |
| 17999 | 900 | F900Y | C | 1.8e-05 | 9.1e-06 | 4.8e-06 | 1.063E-05 | 6.732E-06 | 4.800E-06 | 1.800E-05 |
| 18000 | 900 | F900V | C | 4.6e-06 | 1.2e-05 | 1e-05 | 8.867E-06 | 3.828E-06 | 4.600E-06 | 1.200E-05 |
| 18001 | 901 | K901A | C | 8.3e-06 | 9.7e-06 | 1.1e-05 | 9.667E-06 | 1.350E-06 | 8.300E-06 | 1.100E-05 |
| 18002 | 901 | K901R | C | 4.1e-06 | 1.4e-05 | 6.9e-06 | 8.333E-06 | 5.103E-06 | 4.100E-06 | 1.400E-05 |
| 18003 | 901 | K901N | C | 6.1e-06 | 6.8e-06 | 8.3e-06 | 7.067E-06 | 1.124E-06 | 6.100E-06 | 8.300E-06 |
| 18004 | 901 | K901D | C | 1.1e-05 | 6.2e-06 | 5.3e-06 | 7.500E-06 | 3.064E-06 | 5.300E-06 | 1.100E-05 |
| 18005 | 901 | K901C | C | 9.4e-06 | 1.2e-05 | 3.9e-06 | 8.433E-06 | 4.136E-06 | 3.900E-06 | 1.200E-05 |
| 18006 | 901 | K901E | C | 5.8e-06 | 6.3e-06 | 6.7e-06 | 6.267E-06 | 4.509E-07 | 5.800E-06 | 6.700E-06 |
| 18007 | 901 | K901Q | C | 8.8e-06 | 9.8e-06 | 6.4e-06 | 8.333E-06 | 1.747E-06 | 6.400E-06 | 9.800E-06 |
| 18008 | 901 | K901G | C | 7.1e-06 | 1.1e-05 | 8.1e-06 | 8.733E-06 | 2.026E-06 | 7.100E-06 | 1.100E-05 |
| 18009 | 901 | K901H | C | 7.1e-06 | 9.2e-06 | 8.5e-06 | 8.267E-06 | 1.069E-06 | 7.100E-06 | 9.200E-06 |
| 18010 | 901 | K901I | C | 9.6e-06 | 7e-06 | 1e-05 | 8.867E-06 | 1.629E-06 | 7.000E-06 | 1.000E-05 |
| 18011 | 901 | K901L | C | 6.3e-06 | 8.6e-06 | 6.6e-06 | 7.167E-06 | 1.250E-06 | 6.300E-06 | 8.600E-06 |
| 18012 | 901 | K901K | C | 5.5e-06 | 1.6e-05 | 9.5e-06 | 1.033E-05 | 5.299E-06 | 5.500E-06 | 1.600E-05 |
| 18013 | 901 | K901M | C | 1.2e-05 | 9.2e-06 | 1.2e-05 | 1.107E-05 | 1.617E-06 | 9.200E-06 | 1.200E-05 |
| 18014 | 901 | K901F | C | 8.9e-06 | 8.7e-06 | 8.5e-06 | 8.700E-06 | 2.000E-07 | 8.500E-06 | 8.900E-06 |
| 18015 | 901 | K901P | C | 7e-06 | 8e-06 | 6.8e-06 | 7.267E-06 | 6.429E-07 | 6.800E-06 | 8.000E-06 |
| 18016 | 901 | K901S | C | 6.3e-06 | 7e-06 | 1.1e-05 | 8.100E-06 | 2.536E-06 | 6.300E-06 | 1.100E-05 |
| 18017 | 901 | K901T | C | 5.9e-06 | 6.5e-06 | 9.2e-06 | 7.200E-06 | 1.758E-06 | 5.900E-06 | 9.200E-06 |
| 18018 | 901 | K901W | C | 6.7e-06 | 7.8e-06 | 9.6e-06 | 8.033E-06 | 1.464E-06 | 6.700E-06 | 9.600E-06 |
| 18019 | 901 | K901Y | C | 1.1e-05 | 5.4e-06 | 1e-05 | 8.800E-06 | 2.987E-06 | 5.400E-06 | 1.100E-05 |

| | | | | | | | | | | |
|---|---|---|---|---|---|---|---|---|---|---|
| 18020 | 901 | K901V | C | 1.1e-05 | 1.2e-05 | 7.1e-06 | 1.003E-05 | 2.589E-06 | 7.100E-06 | 1.200E-05 |
| 18021 | 902 | G902A | C | 1.1e-05 | 1.2e-05 | 1.4e-05 | 1.233E-05 | 1.528E-06 | 1.100E-05 | 1.400E-05 |
| 18022 | 902 | G902R | C | 9.7e-06 | 1e-05 | 6.2e-06 | 8.633E-06 | 2.113E-06 | 6.200E-06 | 1.000E-05 |
| 18023 | 902 | G902N | C | 8.2e-06 | 1.3e-05 | 9e-06 | 1.007E-05 | 2.572E-06 | 8.200E-06 | 1.300E-05 |
| 18024 | 902 | G902D | C | 8.2e-06 | 5.7e-06 | 6.7e-06 | 6.867E-06 | 1.258E-06 | 5.700E-06 | 8.200E-06 |
| 18025 | 902 | G902C | C | 9.6e-06 | 4.9e-06 | 8.9e-06 | 7.800E-06 | 2.536E-06 | 4.900E-06 | 9.600E-06 |
| 18026 | 902 | G902E | C | 1e-05 | 6.4e-06 | 1.2e-05 | 9.467E-06 | 2.838E-06 | 6.400E-06 | 1.200E-05 |
| 18027 | 902 | G902Q | C | 7.2e-06 | 1.9e-05 | 4.7e-06 | 1.030E-05 | 7.637E-06 | 4.700E-06 | 1.900E-05 |
| 18028 | 902 | G902G | C | 6.9e-06 | 7.3e-06 | 1.2e-05 | 8.733E-06 | 2.836E-06 | 6.900E-06 | 1.200E-05 |
| 18029 | 902 | G902H | C | 5.3e-06 | 1.2e-05 | 5.6e-06 | 7.633E-06 | 3.785E-06 | 5.300E-06 | 1.200E-05 |
| 18030 | 902 | G902I | C | 9.1e-06 | 7.9e-06 | 6.5e-06 | 7.833E-06 | 1.301E-06 | 6.500E-06 | 9.100E-06 |
| 18031 | 902 | G902L | C | 1.4e-05 | 6.6e-06 | 4.9e-06 | 8.500E-06 | 4.838E-06 | 4.900E-06 | 1.400E-05 |
| 18032 | 902 | G902K | C | 8.5e-06 | 9.2e-06 | 6.8e-06 | 8.167E-06 | 1.234E-06 | 6.800E-06 | 9.200E-06 |
| 18033 | 902 | G902M | C | 7.4e-06 | 6.9e-06 | 6.4e-06 | 6.900E-06 | 5.000E-07 | 6.400E-06 | 7.400E-06 |
| 18034 | 902 | G902F | C | 9.5e-06 | 5e-06 | 3.5e-06 | 6.000E-06 | 3.122E-06 | 3.500E-06 | 9.500E-06 |
| 18035 | 902 | G902P | C | 6.4e-06 | 5e-06 | 5e-06 | 5.467E-06 | 8.083E-07 | 5.000E-06 | 6.400E-06 |
| 18036 | 902 | G902S | C | 9.2e-06 | 3.3e-06 | 7.1e-06 | 6.533E-06 | 2.991E-06 | 3.300E-06 | 9.200E-06 |
| 18037 | 902 | G902T | C | 9.9e-06 | 1.2e-05 | 9e-06 | 1.030E-05 | 1.539E-06 | 9.000E-06 | 1.200E-05 |
| 18038 | 902 | G902W | C | 9.6e-06 | 1.1e-05 | 8.8e-06 | 9.800E-06 | 1.114E-06 | 8.800E-06 | 1.100E-05 |
| 18039 | 902 | G902Y | C | 7.1e-06 | 7e-06 | 5.4e-06 | 6.500E-06 | 9.539E-07 | 5.400E-06 | 7.100E-06 |
| 18040 | 902 | G902V | C | 6.3e-06 | 4.7e-06 | 8.8e-06 | 6.600E-06 | 2.066E-06 | 4.700E-06 | 8.800E-06 |
| 18041 | 903 | R903A | C | 9.8e-06 | 7.8e-06 | 6.5e-06 | 8.033E-06 | 1.662E-06 | 6.500E-06 | 9.800E-06 |
| 18042 | 903 | R903R | C | 7.7e-06 | 4.5e-06 | 5.9e-06 | 6.033E-06 | 1.604E-06 | 4.500E-06 | 7.700E-06 |
| 18043 | 903 | R903N | C | 1.3e-05 | 9.9e-06 | 5.4e-06 | 9.433E-06 | 3.821E-06 | 5.400E-06 | 1.300E-05 |
| 18044 | 903 | R903D | C | 9.2e-06 | 8.6e-06 | 4.4e-06 | 7.400E-06 | 2.615E-06 | 4.400E-06 | 9.200E-06 |
| 18045 | 903 | R903C | C | 6.5e-06 | 1e-05 | 6.5e-06 | 7.667E-06 | 2.021E-06 | 6.500E-06 | 1.000E-05 |
| 18046 | 903 | R903E | C | 9.7e-06 | 7.5e-06 | 9e-06 | 8.733E-06 | 1.124E-06 | 7.500E-06 | 9.700E-06 |
| 18047 | 903 | R903Q | C | 6.6e-06 | 1.2e-05 | 8.1e-06 | 8.900E-06 | 2.787E-06 | 6.600E-06 | 1.200E-05 |
| 18048 | 903 | R903G | C | 9.1e-06 | 9.1e-06 | 8.1e-06 | 8.767E-06 | 5.774E-07 | 8.100E-06 | 9.100E-06 |
| 18049 | 903 | R903H | C | 7.4e-06 | 6.8e-06 | 7.6e-06 | 7.267E-06 | 4.163E-07 | 6.800E-06 | 7.600E-06 |
| 18050 | 903 | R903I | C | 5.8e-06 | 8.4e-06 | 6.7e-06 | 6.967E-06 | 1.320E-06 | 5.800E-06 | 8.400E-06 |
| 18051 | 903 | R903L | C | 1.2e-05 | 8.9e-06 | 1e-05 | 1.030E-05 | 1.572E-06 | 8.900E-06 | 1.200E-05 |
| 18052 | 903 | R903K | C | 1.1e-05 | 8.6e-06 | 1.4e-05 | 1.120E-05 | 2.706E-06 | 8.600E-06 | 1.400E-05 |
| 18053 | 903 | R903M | C | 1e-05 | 6.4e-06 | 6.7e-06 | 7.700E-06 | 1.997E-06 | 6.400E-06 | 1.000E-05 |
| 18054 | 903 | R903F | C | 6e-06 | 1.3e-05 | 5.8e-06 | 8.267E-06 | 4.100E-06 | 5.800E-06 | 1.300E-05 |
| 18055 | 903 | R903P | C | 9.6e-06 | 1.4e-05 | 1.2e-05 | 1.187E-05 | 2.203E-06 | 9.600E-06 | 1.400E-05 |
| 18056 | 903 | R903S | C | 8.5e-06 | 5.7e-06 | 9.3e-06 | 7.833E-06 | 1.890E-06 | 5.700E-06 | 9.300E-06 |
| 18057 | 903 | R903T | C | 9.3e-06 | 8.7e-06 | 7.3e-06 | 8.433E-06 | 1.026E-06 | 7.300E-06 | 9.300E-06 |
| 18058 | 903 | R903W | C | 5.2e-06 | 7e-06 | 8.3e-06 | 6.833E-06 | 1.557E-06 | 5.200E-06 | 8.300E-06 |
| 18059 | 903 | R903Y | C | 5.5e-06 | 1.3e-05 | 8.1e-06 | 8.867E-06 | 3.808E-06 | 5.500E-06 | 1.300E-05 |
| 18060 | 903 | R903V | C | 6.1e-06 | 1.1e-05 | 7.4e-06 | 8.167E-06 | 2.538E-06 | 6.100E-06 | 1.100E-05 |
| 18061 | 904 | F904A | C | 8.9e-06 | 1.5e-05 | 1.1e-05 | 1.163E-05 | 3.099E-06 | 8.900E-06 | 1.500E-05 |
| 18062 | 904 | F904R | C | 5.9e-06 | 6.9e-06 | 9.7e-06 | 7.500E-06 | 1.970E-06 | 5.900E-06 | 9.700E-06 |
| 18063 | 904 | F904N | C | 1.2e-05 | 7.3e-06 | 6.3e-06 | 8.533E-06 | 3.044E-06 | 6.300E-06 | 1.200E-05 |
| 18064 | 904 | F904D | C | 7.7e-06 | 9.6e-06 | 4.9e-06 | 7.400E-06 | 2.364E-06 | 4.900E-06 | 9.600E-06 |
| 18065 | 904 | F904C | C | 9.9e-06 | 9.1e-06 | 6.7e-06 | 8.567E-06 | 1.665E-06 | 6.700E-06 | 9.900E-06 |
| 18066 | 904 | F904E | C | 8.8e-06 | 6.1e-06 | 9.8e-06 | 8.233E-06 | 1.914E-06 | 6.100E-06 | 9.800E-06 |
| 18067 | 904 | F904Q | C | 6.3e-06 | 1.2e-05 | 9.9e-06 | 9.400E-06 | 2.883E-06 | 6.300E-06 | 1.200E-05 |
| 18068 | 904 | F904G | C | 9.3e-06 | 8.4e-06 | 5.3e-06 | 7.667E-06 | 2.098E-06 | 5.300E-06 | 9.300E-06 |
| 18069 | 904 | F904H | C | 1e-05 | 4.6e-06 | 5.3e-06 | 6.633E-06 | 2.937E-06 | 4.600E-06 | 1.000E-05 |
| 18070 | 904 | F904I | C | 1e-05 | 4.6e-06 | 7.3e-06 | 7.300E-06 | 2.700E-06 | 4.600E-06 | 1.000E-05 |
| 18071 | 904 | F904L | C | 1e-05 | 8.5e-06 | 9.5e-06 | 9.333E-06 | 7.638E-07 | 8.500E-06 | 1.000E-05 |
| 18072 | 904 | F904K | C | 9e-06 | 1.2e-05 | 1.3e-05 | 1.133E-05 | 2.082E-06 | 9.000E-06 | 1.300E-05 |
| 18073 | 904 | F904M | C | 8.3e-06 | 6.8e-06 | 6.8e-06 | 7.300E-06 | 8.660E-07 | 6.800E-06 | 8.300E-06 |
| 18074 | 904 | F904F | C | 7.1e-06 | 5.4e-06 | 8.1e-06 | 6.867E-06 | 1.365E-06 | 5.400E-06 | 8.100E-06 |
| 18075 | 904 | F904P | C | 7.9e-06 | 6.3e-06 | 5.1e-06 | 6.433E-06 | 1.405E-06 | 5.100E-06 | 7.900E-06 |
| 18076 | 904 | F904S | C | 1.4e-05 | 8.8e-06 | 6.1e-06 | 9.633E-06 | 4.015E-06 | 6.100E-06 | 1.400E-05 |
| 18077 | 904 | F904T | C | 5e-06 | 9.2e-06 | 1.1e-05 | 8.400E-06 | 3.079E-06 | 5.000E-06 | 1.100E-05 |
| 18078 | 904 | F904W | C | 7.7e-06 | 1.1e-05 | 7.6e-06 | 8.767E-06 | 1.935E-06 | 7.600E-06 | 1.100E-05 |
| 18079 | 904 | F904Y | C | 8.1e-06 | 8.8e-06 | 1e-05 | 8.967E-06 | 9.609E-07 | 8.100E-06 | 1.000E-05 |
| 18080 | 904 | F904V | C | 6.5e-06 | 8.5e-06 | 1.6e-05 | 1.033E-05 | 5.008E-06 | 6.500E-06 | 1.600E-05 |
| 18081 | 905 | T905A | C | 6.2e-06 | 4.7e-06 | 9.7e-06 | 6.867E-06 | 2.566E-06 | 4.700E-06 | 9.700E-06 |
| 18082 | 905 | T905R | C | 9.4e-06 | 1e-05 | 7.2e-06 | 8.867E-06 | 1.474E-06 | 7.200E-06 | 1.000E-05 |
| 18083 | 905 | T905N | C | 5.1e-06 | 4.3e-06 | 7e-06 | 5.467E-06 | 1.387E-06 | 4.300E-06 | 7.000E-06 |
| 18084 | 905 | T905D | C | 1.4e-05 | 9.9e-06 | 7.4e-06 | 1.043E-05 | 3.332E-06 | 7.400E-06 | 1.400E-05 |
| 18085 | 905 | T905C | C | 8.2e-06 | 8.4e-06 | 6.5e-06 | 7.700E-06 | 1.044E-06 | 6.500E-06 | 8.400E-06 |
| 18086 | 905 | T905E | C | 1.1e-05 | 1.2e-05 | 4.4e-06 | 9.133E-06 | 4.130E-06 | 4.400E-06 | 1.200E-05 |
| 18087 | 905 | T905Q | C | 6.7e-06 | 1.2e-05 | 6e-06 | 8.233E-06 | 3.281E-06 | 6.000E-06 | 1.200E-05 |
| 18088 | 905 | T905G | C | 5.4e-06 | 3.4e-06 | 9e-06 | 5.933E-06 | 2.838E-06 | 3.400E-06 | 9.000E-06 |
| 18089 | 905 | T905H | C | 7e-06 | 6.5e-06 | 9.8e-06 | 7.767E-06 | 1.779E-06 | 6.500E-06 | 9.800E-06 |
| 18090 | 905 | T905I | C | 6.9e-06 | 1.2e-05 | 6.4e-06 | 8.433E-06 | 3.099E-06 | 6.400E-06 | 1.200E-05 |
| 18091 | 905 | T905L | C | 8.5e-06 | 1e-05 | 8.9e-06 | 9.133E-06 | 7.767E-07 | 8.500E-06 | 1.000E-05 |
| 18092 | 905 | T905K | C | 1e-05 | 6.7e-06 | 8.4e-06 | 8.367E-06 | 1.650E-06 | 6.700E-06 | 1.000E-05 |
| 18093 | 905 | T905M | C | 1e-05 | 5.7e-06 | 8.6e-06 | 8.100E-06 | 2.193E-06 | 5.700E-06 | 1.000E-05 |
| 18094 | 905 | T905F | C | 5.8e-06 | 9.5e-06 | 8.5e-06 | 7.933E-06 | 1.914E-06 | 5.800E-06 | 9.500E-06 |
| 18095 | 905 | T905P | C | 8.6e-06 | 4.7e-06 | 1.3e-05 | 8.767E-06 | 4.153E-06 | 4.700E-06 | 1.300E-05 |
| 18096 | 905 | T905S | C | 1.6e-05 | 7.2e-06 | 1e-05 | 1.107E-05 | 4.496E-06 | 7.200E-06 | 1.600E-05 |
| 18097 | 905 | T905T | C | 1.3e-05 | 7.1e-06 | 8.5e-06 | 9.533E-06 | 3.083E-06 | 7.100E-06 | 1.300E-05 |
| 18098 | 905 | T905W | C | 6.5e-06 | 7.5e-06 | 1.8e-05 | 1.067E-05 | 6.371E-06 | 6.500E-06 | 1.800E-05 |
| 18099 | 905 | T905Y | C | 5.6e-06 | 1e-05 | 7.6e-06 | 7.733E-06 | 2.203E-06 | 5.600E-06 | 1.000E-05 |
| 18100 | 905 | T905V | C | 1.3e-05 | 9.6e-06 | 5.3e-06 | 9.300E-06 | 3.859E-06 | 5.300E-06 | 1.300E-05 |
| 18101 | 906 | L906A | C | 1e-05 | 9.6e-06 | 1e-05 | 9.867E-06 | 2.309E-07 | 9.600E-06 | 1.000E-05 |
| 18102 | 906 | L906R | C | 7.6e-06 | 1.3e-05 | 7.8e-06 | 9.467E-06 | 3.062E-06 | 7.600E-06 | 1.300E-05 |
| 18103 | 906 | L906N | C | 7.1e-06 | 4.7e-06 | 5.3e-06 | 5.700E-06 | 1.249E-06 | 4.700E-06 | 7.100E-06 |
| 18104 | 906 | L906D | C | 8.8e-06 | 1.1e-05 | 1.2e-05 | 1.060E-05 | 1.637E-06 | 8.800E-06 | 1.200E-05 |

| | | | | | | | | | | |
|---|---|---|---|---|---|---|---|---|---|---|
| 18105 | 906 | L906C | C | 4.3e-06 | 7.7e-06 | 6.8e-06 | 6.267E-06 | 1.762E-06 | 4.300E-06 | 7.700E-06 |
| 18106 | 906 | L906E | C | 5e-06 | 6e-06 | 8e-06 | 6.333E-06 | 1.528E-06 | 5.000E-06 | 8.000E-06 |
| 18107 | 906 | L906Q | C | 6.5e-06 | 1.5e-05 | 5.2e-06 | 8.900E-06 | 5.323E-06 | 5.200E-06 | 1.500E-05 |
| 18108 | 906 | L906G | C | 1.2e-05 | 5.7e-06 | 7.1e-06 | 8.267E-06 | 3.308E-06 | 5.700E-06 | 1.200E-05 |
| 18109 | 906 | L906H | C | 5.8e-06 | 6.7e-06 | 7.4e-06 | 6.633E-06 | 8.021E-07 | 5.800E-06 | 7.400E-06 |
| 18110 | 906 | L906I | C | 1.1e-05 | 6.8e-06 | 7.5e-06 | 8.433E-06 | 2.250E-06 | 6.800E-06 | 1.100E-05 |
| 18111 | 906 | L906L | C | 5.9e-06 | 6e-06 | 6.4e-06 | 6.100E-06 | 2.646E-07 | 5.900E-06 | 6.400E-06 |
| 18112 | 906 | L906K | C | 8.4e-06 | 6e-06 | 1.2e-05 | 8.800E-06 | 3.020E-06 | 6.000E-06 | 1.200E-05 |
| 18113 | 906 | L906M | C | 8e-06 | 1.3e-05 | 7.2e-06 | 9.400E-06 | 3.143E-06 | 7.200E-06 | 1.300E-05 |
| 18114 | 906 | L906F | C | 8.8e-06 | 4.3e-06 | 9.2e-06 | 7.433E-06 | 2.721E-06 | 4.300E-06 | 9.200E-06 |
| 18115 | 906 | L906P | C | 8.5e-06 | 1e-05 | 6.3e-06 | 8.267E-06 | 1.861E-06 | 6.300E-06 | 1.000E-05 |
| 18116 | 906 | L906S | C | 8.4e-06 | 1e-05 | 6.7e-06 | 8.367E-06 | 1.650E-06 | 6.700E-06 | 1.000E-05 |
| 18117 | 906 | L906T | C | 6.4e-06 | 1e-05 | 9.9e-06 | 8.767E-06 | 2.050E-06 | 6.400E-06 | 1.000E-05 |
| 18118 | 906 | L906W | C | 9.6e-06 | 5.8e-06 | 4.5e-06 | 6.633E-06 | 2.650E-06 | 4.500E-06 | 9.600E-06 |
| 18119 | 906 | L906Y | C | 6.5e-06 | 1.4e-05 | 1e-05 | 1.017E-05 | 3.753E-06 | 6.500E-06 | 1.400E-05 |
| 18120 | 906 | L906V | C | 9.6e-06 | 1.4e-05 | 8.1e-06 | 1.057E-05 | 3.066E-06 | 8.100E-06 | 1.400E-05 |
| 18121 | 907 | S907A | C | 9.5e-06 | 9.8e-06 | 7.2e-06 | 8.833E-06 | 1.422E-06 | 7.200E-06 | 9.800E-06 |
| 18122 | 907 | S907R | C | 1.3e-05 | 3.6e-06 | 9.5e-06 | 8.700E-06 | 4.751E-06 | 3.600E-06 | 1.300E-05 |
| 18123 | 907 | S907N | C | 7.4e-06 | 1.2e-05 | 6e-06 | 8.467E-06 | 3.139E-06 | 6.000E-06 | 1.200E-05 |
| 18124 | 907 | S907D | C | 7.8e-06 | 7.3e-06 | 9.2e-06 | 8.100E-06 | 9.849E-07 | 7.300E-06 | 9.200E-06 |
| 18125 | 907 | S907C | C | 9.9e-06 | 8.2e-06 | 7.2e-06 | 8.433E-06 | 1.365E-06 | 7.200E-06 | 9.900E-06 |
| 18126 | 907 | S907E | C | 9.7e-06 | 5.8e-06 | 7.3e-06 | 7.600E-06 | 1.967E-06 | 5.800E-06 | 9.700E-06 |
| 18127 | 907 | S907Q | C | 8.4e-06 | 7.6e-06 | 8.8e-06 | 8.267E-06 | 6.110E-07 | 7.600E-06 | 8.800E-06 |
| 18128 | 907 | S907G | C | 1e-05 | 1.3e-05 | 8.5e-06 | 1.050E-05 | 2.291E-06 | 8.500E-06 | 1.300E-05 |
| 18129 | 907 | S907H | C | 8.3e-06 | 7.3e-06 | 1.3e-05 | 9.533E-06 | 3.044E-06 | 7.300E-06 | 1.300E-05 |
| 18130 | 907 | S907I | C | 4.1e-06 | 6.6e-06 | 7.4e-06 | 6.033E-06 | 1.721E-06 | 4.100E-06 | 7.400E-06 |
| 18131 | 907 | S907L | C | 7.7e-06 | 9.3e-06 | 9.9e-06 | 8.967E-06 | 1.137E-06 | 7.700E-06 | 9.900E-06 |
| 18132 | 907 | S907K | C | 9.4e-06 | 6.8e-06 | 7.3e-06 | 7.833E-06 | 1.380E-06 | 6.800E-06 | 9.400E-06 |
| 18133 | 907 | S907M | C | 1.2e-05 | 6.3e-06 | 6.1e-06 | 8.133E-06 | 3.350E-06 | 6.100E-06 | 1.200E-05 |
| 18134 | 907 | S907F | C | 1e-05 | 6.8e-06 | 6.2e-06 | 7.667E-06 | 2.043E-06 | 6.200E-06 | 1.000E-05 |
| 18135 | 907 | S907P | C | 9.2e-06 | 5.7e-06 | 1.5e-05 | 9.967E-06 | 4.697E-06 | 5.700E-06 | 1.500E-05 |
| 18136 | 907 | S907S | C | 5.7e-06 | 8e-06 | 6.8e-06 | 6.833E-06 | 1.150E-06 | 5.700E-06 | 8.000E-06 |
| 18137 | 907 | S907T | C | 1.1e-05 | 1.1e-05 | 7.5e-06 | 9.833E-06 | 2.021E-06 | 7.500E-06 | 1.100E-05 |
| 18138 | 907 | S907W | C | 1.1e-05 | 9.1e-06 | 5.7e-06 | 8.600E-06 | 2.685E-06 | 5.700E-06 | 1.100E-05 |
| 18139 | 907 | S907Y | C | 6.3e-06 | 3.9e-06 | 7.2e-06 | 5.800E-06 | 1.706E-06 | 3.900E-06 | 7.200E-06 |
| 18140 | 907 | S907V | C | 1.1e-05 | 1.4e-05 | 1.3e-05 | 1.267E-05 | 1.528E-06 | 1.100E-05 | 1.400E-05 |
| 18141 | 908 | V908A | C | 7.2e-06 | 6.4e-06 | 7.2e-06 | 6.933E-06 | 4.619E-07 | 6.400E-06 | 7.200E-06 |
| 18142 | 908 | V908R | C | 8.6e-06 | 5.7e-06 | 6.5e-06 | 6.933E-06 | 1.498E-06 | 5.700E-06 | 8.600E-06 |
| 18143 | 908 | V908N | C | 8.6e-06 | 9.4e-06 | 8.1e-06 | 8.700E-06 | 6.557E-07 | 8.100E-06 | 9.400E-06 |
| 18144 | 908 | V908D | C | 9.9e-06 | 8.9e-06 | 9.2e-06 | 9.333E-06 | 5.132E-07 | 8.900E-06 | 9.900E-06 |
| 18145 | 908 | V908C | C | 6.8e-06 | 6.7e-06 | 4.4e-06 | 5.967E-06 | 1.358E-06 | 4.400E-06 | 6.800E-06 |
| 18146 | 908 | V908E | C | 8.3e-06 | 6.8e-06 | 8.9e-06 | 8.000E-06 | 1.082E-06 | 6.800E-06 | 8.900E-06 |
| 18147 | 908 | V908Q | C | 9.6e-06 | 8.5e-06 | 5.3e-06 | 7.800E-06 | 2.234E-06 | 5.300E-06 | 9.600E-06 |
| 18148 | 908 | V908G | C | 1.1e-05 | 1e-05 | 6.4e-06 | 9.133E-06 | 2.419E-06 | 6.400E-06 | 1.100E-05 |
| 18149 | 908 | V908H | C | 8.9e-06 | 1.2e-05 | 1.3e-05 | 1.130E-05 | 2.138E-06 | 8.900E-06 | 1.300E-05 |
| 18150 | 908 | V908I | C | 5.6e-06 | 9.4e-06 | 8.8e-06 | 7.933E-06 | 2.043E-06 | 5.600E-06 | 9.400E-06 |
| 18151 | 908 | V908L | C | 6.8e-06 | 6.1e-06 | 9.5e-06 | 7.467E-06 | 1.795E-06 | 6.100E-06 | 9.500E-06 |
| 18152 | 908 | V908K | C | 6.4e-06 | 6.9e-06 | 6e-06 | 6.433E-06 | 4.509E-07 | 6.000E-06 | 6.900E-06 |
| 18153 | 908 | V908M | C | 1.2e-05 | 8.7e-06 | 1.3e-05 | 1.123E-05 | 2.250E-06 | 8.700E-06 | 1.300E-05 |
| 18154 | 908 | V908F | C | 3.1e-06 | 8.1e-06 | 7.7e-06 | 6.300E-06 | 2.778E-06 | 3.100E-06 | 8.100E-06 |
| 18155 | 908 | V908P | C | 1.3e-05 | 7.9e-06 | 7.8e-06 | 9.567E-06 | 2.974E-06 | 7.800E-06 | 1.300E-05 |
| 18156 | 908 | V908S | C | 9.2e-06 | 1.2e-05 | 5.6e-06 | 8.933E-06 | 3.208E-06 | 5.600E-06 | 1.200E-05 |
| 18157 | 908 | V908T | C | 1.3e-05 | 4.2e-06 | 6.8e-06 | 8.000E-06 | 4.521E-06 | 4.200E-06 | 1.300E-05 |
| 18158 | 908 | V908W | C | 6.5e-06 | 6.5e-06 | 5.7e-06 | 6.233E-06 | 4.619E-07 | 5.700E-06 | 6.500E-06 |
| 18159 | 908 | V908Y | C | 8.4e-06 | 7.5e-06 | 1.1e-05 | 8.967E-06 | 1.818E-06 | 7.500E-06 | 1.100E-05 |
| 18160 | 908 | V908V | C | 4.6e-06 | 1.2e-05 | 5.1e-06 | 7.233E-06 | 4.136E-06 | 4.600E-06 | 1.200E-05 |
| 18161 | 909 | D909A | C | 8.4e-06 | 9.9e-06 | 6.5e-06 | 8.267E-06 | 1.704E-06 | 6.500E-06 | 9.900E-06 |
| 18162 | 909 | D909R | C | 1.8e-05 | 7.3e-06 | 8.2e-06 | 1.117E-05 | 5.935E-06 | 7.300E-06 | 1.800E-05 |
| 18163 | 909 | D909N | C | 7.7e-06 | 5.4e-06 | 9.6e-06 | 7.567E-06 | 2.103E-06 | 5.400E-06 | 9.600E-06 |
| 18164 | 909 | D909D | C | 4.4e-06 | 6.3e-06 | 1.2e-05 | 7.567E-06 | 3.955E-06 | 4.400E-06 | 1.200E-05 |
| 18165 | 909 | D909C | C | 7.3e-06 | 6.1e-06 | 7.8e-06 | 7.067E-06 | 8.737E-07 | 6.100E-06 | 7.800E-06 |
| 18166 | 909 | D909E | C | 9e-06 | 6.6e-06 | 8.1e-06 | 7.900E-06 | 1.212E-06 | 6.600E-06 | 9.000E-06 |
| 18167 | 909 | D909Q | C | 7.8e-06 | 8.4e-06 | 5.6e-06 | 7.267E-06 | 1.474E-06 | 5.600E-06 | 8.400E-06 |
| 18168 | 909 | D909G | C | 7.9e-06 | 9.3e-06 | 1.2e-05 | 9.733E-06 | 2.084E-06 | 7.900E-06 | 1.200E-05 |
| 18169 | 909 | D909H | C | 8.3e-06 | 7.9e-06 | 8.9e-06 | 8.367E-06 | 5.033E-07 | 7.900E-06 | 8.900E-06 |
| 18170 | 909 | D909I | C | 1.1e-05 | 7.1e-06 | 1e-05 | 9.367E-06 | 2.026E-06 | 7.100E-06 | 1.100E-05 |
| 18171 | 909 | D909L | C | 7.4e-06 | 9.3e-06 | 5e-06 | 7.233E-06 | 2.155E-06 | 5.000E-06 | 9.300E-06 |
| 18172 | 909 | D909K | C | 3.2e-06 | 8e-06 | 9.3e-06 | 6.833E-06 | 3.213E-06 | 3.200E-06 | 9.300E-06 |
| 18173 | 909 | D909M | C | 7e-06 | 9.7e-06 | 9.8e-06 | 8.833E-06 | 1.589E-06 | 7.000E-06 | 9.800E-06 |
| 18174 | 909 | D909F | C | 6.1e-06 | 6.8e-06 | 8.8e-06 | 7.233E-06 | 1.401E-06 | 6.100E-06 | 8.800E-06 |
| 18175 | 909 | D909P | C | 8.6e-06 | 9.8e-06 | 5e-06 | 7.800E-06 | 2.498E-06 | 5.000E-06 | 9.800E-06 |
| 18176 | 909 | D909S | C | 6.2e-06 | 1.7e-05 | 6.6e-06 | 9.933E-06 | 6.123E-06 | 6.200E-06 | 1.700E-05 |
| 18177 | 909 | D909T | C | 7e-06 | 8.9e-06 | 9.4e-06 | 8.433E-06 | 1.266E-06 | 7.000E-06 | 9.400E-06 |
| 18178 | 909 | D909W | C | 1.2e-05 | 9.1e-06 | 8.5e-06 | 9.867E-06 | 1.872E-06 | 8.500E-06 | 1.200E-05 |
| 18179 | 909 | D909Y | C | 6.2e-06 | 9.7e-06 | 8.5e-06 | 8.133E-06 | 1.779E-06 | 6.200E-06 | 9.700E-06 |
| 18180 | 909 | D909V | C | 1.4e-05 | 1.1e-05 | 1.1e-05 | 1.200E-05 | 1.732E-06 | 1.100E-05 | 1.400E-05 |
| 18181 | 910 | R910A | C | 8.8e-06 | 9e-06 | 5.9e-06 | 7.900E-06 | 1.735E-06 | 5.900E-06 | 9.000E-06 |
| 18182 | 910 | R910R | C | 6.4e-06 | 4.2e-06 | 8.6e-06 | 6.400E-06 | 2.200E-06 | 4.200E-06 | 8.600E-06 |
| 18183 | 910 | R910N | C | 8.9e-06 | 5.6e-06 | 8.4e-06 | 7.633E-06 | 1.779E-06 | 5.600E-06 | 8.900E-06 |
| 18184 | 910 | R910D | C | 5.6e-06 | 6.5e-06 | 9e-06 | 7.033E-06 | 1.762E-06 | 5.600E-06 | 9.000E-06 |
| 18185 | 910 | R910C | C | 6.9e-06 | 3.7e-06 | 8e-06 | 6.200E-06 | 2.234E-06 | 3.700E-06 | 8.000E-06 |
| 18186 | 910 | R910E | C | 6.4e-06 | 6.4e-06 | 1.4e-05 | 8.933E-06 | 4.388E-06 | 6.400E-06 | 1.400E-05 |
| 18187 | 910 | R910Q | C | 9.2e-06 | 1.1e-05 | 6.6e-06 | 8.933E-06 | 2.212E-06 | 6.600E-06 | 1.100E-05 |
| 18188 | 910 | R910G | C | 1.1e-05 | 9.7e-06 | 7.7e-06 | 9.467E-06 | 1.662E-06 | 7.700E-06 | 1.100E-05 |
| 18189 | 910 | R910H | C | 6.3e-06 | 8e-06 | 6.5e-06 | 6.933E-06 | 9.292E-07 | 6.300E-06 | 8.000E-06 |

| | | | | | | | | | | |
|---|---|---|---|---|---|---|---|---|---|---|
| 18190 | 910 | R910I | C | 4.3e-06 | 7.1e-06 | 9.7e-06 | 7.033E-06 | 2.701E-06 | 4.300E-06 | 9.700E-06 |
| 18191 | 910 | R910L | C | 9.6e-06 | 9.9e-06 | 7.2e-06 | 8.900E-06 | 1.480E-06 | 7.200E-06 | 9.900E-06 |
| 18192 | 910 | R910K | C | 1.7e-05 | 8.7e-06 | 6.2e-06 | 1.063E-05 | 5.654E-06 | 6.200E-06 | 1.700E-05 |
| 18193 | 910 | R910M | C | 8.9e-06 | 1.4e-05 | 1.3e-05 | 1.197E-05 | 2.702E-06 | 8.900E-06 | 1.400E-05 |
| 18194 | 910 | R910F | C | 6.4e-06 | 1.4e-05 | 5.8e-06 | 8.733E-06 | 4.571E-06 | 5.800E-06 | 1.400E-05 |
| 18195 | 910 | R910P | C | 6.8e-06 | 5.7e-06 | 6.5e-06 | 6.333E-06 | 5.686E-07 | 5.700E-06 | 6.800E-06 |
| 18196 | 910 | R910S | C | 3.7e-06 | 5.5e-06 | 9.9e-06 | 6.367E-06 | 3.190E-06 | 3.700E-06 | 9.900E-06 |
| 18197 | 910 | R910T | C | 9.7e-06 | 8.8e-06 | 6.6e-06 | 8.367E-06 | 1.595E-06 | 6.600E-06 | 9.700E-06 |
| 18198 | 910 | R910W | C | 4.6e-06 | 8.6e-06 | 4.6e-06 | 5.933E-06 | 2.309E-06 | 4.600E-06 | 8.600E-06 |
| 18199 | 910 | R910Y | C | 1.1e-05 | 6.7e-06 | 1.1e-05 | 9.567E-06 | 2.483E-06 | 6.700E-06 | 1.100E-05 |
| 18200 | 910 | R910V | C | 7.8e-06 | 7.3e-06 | 1.1e-05 | 8.700E-06 | 2.007E-06 | 7.300E-06 | 1.100E-05 |
| 18201 | 911 | S911A | C | 6.9e-06 | 9.3e-06 | 6.2e-06 | 7.467E-06 | 1.626E-06 | 6.200E-06 | 9.300E-06 |
| 18202 | 911 | S911R | C | 6.4e-06 | 8.5e-06 | 7.3e-06 | 7.400E-06 | 1.054E-06 | 6.400E-06 | 8.500E-06 |
| 18203 | 911 | S911N | C | 9.2e-06 | 1e-05 | 6.6e-06 | 8.600E-06 | 1.778E-06 | 6.600E-06 | 1.000E-05 |
| 18204 | 911 | S911D | C | 8.9e-06 | 8.7e-06 | 9.6e-06 | 9.067E-06 | 4.726E-07 | 8.700E-06 | 9.600E-06 |
| 18205 | 911 | S911C | C | 9.2e-06 | 1.4e-05 | 9.8e-06 | 1.100E-05 | 2.615E-06 | 9.200E-06 | 1.400E-05 |
| 18206 | 911 | S911E | C | 9.6e-06 | 9.2e-06 | 5.6e-06 | 8.133E-06 | 2.203E-06 | 5.600E-06 | 9.600E-06 |
| 18207 | 911 | S911Q | C | 5.7e-06 | 5.7e-06 | 7e-06 | 6.133E-06 | 7.506E-07 | 5.700E-06 | 7.000E-06 |
| 18208 | 911 | S911G | C | 8.9e-06 | 7.3e-06 | 6.4e-06 | 7.533E-06 | 1.266E-06 | 6.400E-06 | 8.900E-06 |
| 18209 | 911 | S911H | C | 4.5e-06 | 8.1e-06 | 4.6e-06 | 5.733E-06 | 2.050E-06 | 4.500E-06 | 8.100E-06 |
| 18210 | 911 | S911I | C | 1.2e-05 | 1e-05 | 1.5e-05 | 1.233E-05 | 2.517E-06 | 1.000E-05 | 1.500E-05 |
| 18211 | 911 | S911L | C | 1e-05 | 1.2e-05 | 5.8e-06 | 9.267E-06 | 3.164E-06 | 5.800E-06 | 1.200E-05 |
| 18212 | 911 | S911K | C | 1e-05 | 9.8e-06 | 6.6e-06 | 8.800E-06 | 1.908E-06 | 6.600E-06 | 1.000E-05 |
| 18213 | 911 | S911M | C | 5.3e-06 | 9.1e-06 | 3.8e-06 | 6.067E-06 | 2.732E-06 | 3.800E-06 | 9.100E-06 |
| 18214 | 911 | S911F | C | 7.7e-06 | 1e-05 | 1.1e-05 | 9.567E-06 | 1.692E-06 | 7.700E-06 | 1.100E-05 |
| 18215 | 911 | S911P | C | 8.8e-06 | 5.1e-06 | 7.3e-06 | 7.067E-06 | 1.861E-06 | 5.100E-06 | 8.800E-06 |
| 18216 | 911 | S911S | C | 6.9e-06 | 1.3e-05 | 8.1e-06 | 9.333E-06 | 3.232E-06 | 6.900E-06 | 1.300E-05 |
| 18217 | 911 | S911T | C | 7e-06 | 9.8e-06 | 1.1e-05 | 9.267E-06 | 2.053E-06 | 7.000E-06 | 1.100E-05 |
| 18218 | 911 | S911W | C | 1.1e-05 | 7e-06 | 5.9e-06 | 7.967E-06 | 2.684E-06 | 5.900E-06 | 1.100E-05 |
| 18219 | 911 | S911Y | C | 7.7e-06 | 5.3e-06 | 9.2e-06 | 7.400E-06 | 1.967E-06 | 5.300E-06 | 9.200E-06 |
| 18220 | 911 | S911V | C | 1.4e-05 | 9.6e-06 | 1.2e-05 | 1.187E-05 | 2.203E-06 | 9.600E-06 | 1.400E-05 |
| 18221 | 912 | K912A | C | 1e-05 | 7.8e-06 | 6.6e-06 | 8.133E-06 | 1.724E-06 | 6.600E-06 | 1.000E-05 |
| 18222 | 912 | K912R | C | 1.1e-05 | 9.9e-06 | 9.7e-06 | 1.020E-05 | 7.000E-07 | 9.700E-06 | 1.100E-05 |
| 18223 | 912 | K912N | C | 6.7e-06 | 1.3e-05 | 4.6e-06 | 8.100E-06 | 4.371E-06 | 4.600E-06 | 1.300E-05 |
| 18224 | 912 | K912D | C | 6.6e-06 | 9.3e-06 | 1.4e-05 | 9.967E-06 | 3.745E-06 | 6.600E-06 | 1.400E-05 |
| 18225 | 912 | K912C | C | 6e-06 | 7.3e-06 | 4.9e-06 | 6.067E-06 | 1.201E-06 | 4.900E-06 | 7.300E-06 |
| 18226 | 912 | K912E | C | 4e-06 | 1.2e-05 | 7e-06 | 7.667E-06 | 4.041E-06 | 4.000E-06 | 1.200E-05 |
| 18227 | 912 | K912Q | C | 8.3e-06 | 9.8e-06 | 8.4e-06 | 8.833E-06 | 8.386E-07 | 8.300E-06 | 9.800E-06 |
| 18228 | 912 | K912G | C | 4.6e-06 | 5.9e-06 | 9.1e-06 | 6.533E-06 | 2.316E-06 | 4.600E-06 | 9.100E-06 |
| 18229 | 912 | K912H | C | 6.8e-06 | 6e-06 | 9.7e-06 | 7.500E-06 | 1.947E-06 | 6.000E-06 | 9.700E-06 |
| 18230 | 912 | K912I | C | 6.2e-06 | 1.2e-05 | 7.4e-06 | 8.533E-06 | 3.062E-06 | 6.200E-06 | 1.200E-05 |
| 18231 | 912 | K912L | C | 4.2e-06 | 8.4e-06 | 8e-06 | 6.867E-06 | 2.318E-06 | 4.200E-06 | 8.400E-06 |
| 18232 | 912 | K912K | C | 1e-05 | 1.2e-05 | 8.8e-06 | 1.027E-05 | 1.617E-06 | 8.800E-06 | 1.200E-05 |
| 18233 | 912 | K912M | C | 9e-06 | 9.2e-06 | 9.3e-06 | 9.167E-06 | 1.528E-07 | 9.000E-06 | 9.300E-06 |
| 18234 | 912 | K912F | C | 6.4e-06 | 8.5e-06 | 5.7e-06 | 6.867E-06 | 1.457E-06 | 5.700E-06 | 8.500E-06 |
| 18235 | 912 | K912P | C | 9e-06 | 6.4e-06 | 9.4e-06 | 8.267E-06 | 1.629E-06 | 6.400E-06 | 9.400E-06 |
| 18236 | 912 | K912S | C | 7.9e-06 | 5.8e-06 | 7.4e-06 | 7.033E-06 | 1.097E-06 | 5.800E-06 | 7.900E-06 |
| 18237 | 912 | K912T | C | 1.1e-05 | 1e-05 | 5.8e-06 | 8.933E-06 | 2.759E-06 | 5.800E-06 | 1.100E-05 |
| 18238 | 912 | K912W | C | 8.8e-06 | 1.1e-05 | 1.1e-05 | 1.027E-05 | 1.270E-06 | 8.800E-06 | 1.100E-05 |
| 18239 | 912 | K912Y | C | 7e-06 | 1.3e-05 | 9.9e-06 | 9.967E-06 | 3.001E-06 | 7.000E-06 | 1.300E-05 |
| 18240 | 912 | K912V | C | 8.7e-06 | 6.4e-06 | 6.9e-06 | 7.333E-06 | 1.210E-06 | 6.400E-06 | 8.700E-06 |
| 18241 | 913 | N913A | C | 6.9e-06 | 9.8e-06 | 9.2e-06 | 8.633E-06 | 1.531E-06 | 6.900E-06 | 9.800E-06 |
| 18242 | 913 | N913R | C | 5.9e-06 | 7.7e-06 | 1.3e-05 | 8.867E-06 | 3.691E-06 | 5.900E-06 | 1.300E-05 |
| 18243 | 913 | N913N | C | 1.1e-05 | 8.5e-06 | 3.9e-06 | 7.800E-06 | 3.601E-06 | 3.900E-06 | 1.100E-05 |
| 18244 | 913 | N913D | C | 8e-06 | 6.2e-06 | 7.7e-06 | 7.300E-06 | 9.644E-07 | 6.200E-06 | 8.000E-06 |
| 18245 | 913 | N913C | C | 1.4e-05 | 7.4e-06 | 5.6e-06 | 9.000E-06 | 4.423E-06 | 5.600E-06 | 1.400E-05 |
| 18246 | 913 | N913E | C | 7.2e-06 | 1e-05 | 1.4e-05 | 1.040E-05 | 3.418E-06 | 7.200E-06 | 1.400E-05 |
| 18247 | 913 | N913Q | C | 1e-05 | 6.6e-06 | 6.8e-06 | 7.800E-06 | 1.908E-06 | 6.600E-06 | 1.000E-05 |
| 18248 | 913 | N913G | C | 9e-06 | 9.9e-06 | 6.1e-06 | 8.333E-06 | 1.986E-06 | 6.100E-06 | 9.900E-06 |
| 18249 | 913 | N913H | C | 9.1e-06 | 6.6e-06 | 9.3e-06 | 8.333E-06 | 1.504E-06 | 6.600E-06 | 9.300E-06 |
| 18250 | 913 | N913I | C | 5.1e-06 | 6.4e-06 | 7.1e-06 | 6.200E-06 | 1.015E-06 | 5.100E-06 | 7.100E-06 |
| 18251 | 913 | N913L | C | 1.5e-05 | 6.9e-06 | 9.8e-06 | 1.057E-05 | 4.104E-06 | 6.900E-06 | 1.500E-05 |
| 18252 | 913 | N913K | C | 6.8e-06 | 9.4e-06 | 8.2e-06 | 8.133E-06 | 1.301E-06 | 6.800E-06 | 9.400E-06 |
| 18253 | 913 | N913M | C | 8.3e-06 | 8.9e-06 | 6.7e-06 | 7.967E-06 | 1.137E-06 | 6.700E-06 | 8.900E-06 |
| 18254 | 913 | N913F | C | 5e-06 | 6e-06 | 6.7e-06 | 5.900E-06 | 8.544E-07 | 5.000E-06 | 6.700E-06 |
| 18255 | 913 | N913P | C | 5.6e-06 | 5.4e-06 | 8.4e-06 | 6.467E-06 | 1.677E-06 | 5.400E-06 | 8.400E-06 |
| 18256 | 913 | N913S | C | 7.2e-06 | 9.9e-06 | 6.1e-06 | 7.733E-06 | 1.955E-06 | 6.100E-06 | 9.900E-06 |
| 18257 | 913 | N913T | C | 6.1e-06 | 9.3e-06 | 6.2e-06 | 7.200E-06 | 1.819E-06 | 6.100E-06 | 9.300E-06 |
| 18258 | 913 | N913W | C | 1.2e-05 | 1e-05 | 9.1e-06 | 1.037E-05 | 1.484E-06 | 9.100E-06 | 1.200E-05 |
| 18259 | 913 | N913Y | C | 1.1e-05 | 7.2e-06 | 8.3e-06 | 8.833E-06 | 1.955E-06 | 7.200E-06 | 1.100E-05 |
| 18260 | 913 | N913V | C | 8.7e-06 | 6.6e-06 | 7.2e-06 | 7.500E-06 | 1.082E-06 | 6.600E-06 | 8.700E-06 |
| 18261 | 914 | T914A | C | 1.2e-05 | 1.2e-05 | 7.6e-06 | 1.053E-05 | 2.540E-06 | 7.600E-06 | 1.200E-05 |
| 18262 | 914 | T914R | C | 1.4e-05 | 6.7e-06 | 9.7e-06 | 1.013E-05 | 3.669E-06 | 6.700E-06 | 1.400E-05 |
| 18263 | 914 | T914N | C | 5.5e-06 | 4.9e-06 | 8.5e-06 | 6.300E-06 | 1.929E-06 | 4.900E-06 | 8.500E-06 |
| 18264 | 914 | T914D | C | 7.3e-06 | 5.1e-06 | 1.1e-05 | 7.800E-06 | 2.982E-06 | 5.100E-06 | 1.100E-05 |
| 18265 | 914 | T914C | C | 7.6e-06 | 6.3e-06 | 1.5e-05 | 9.633E-06 | 4.693E-06 | 6.300E-06 | 1.500E-05 |
| 18266 | 914 | T914E | C | 5.8e-06 | 6.8e-06 | 5.5e-06 | 6.033E-06 | 6.807E-07 | 5.500E-06 | 6.800E-06 |
| 18267 | 914 | T914Q | C | 9.7e-06 | 9.6e-06 | 9e-06 | 9.433E-06 | 3.786E-07 | 9.000E-06 | 9.700E-06 |
| 18268 | 914 | T914G | C | 7.6e-06 | 6.9e-06 | 5.5e-06 | 6.667E-06 | 1.069E-06 | 5.500E-06 | 7.600E-06 |
| 18269 | 914 | T914H | C | 1e-05 | 6.7e-06 | 7.3e-06 | 8.000E-06 | 1.758E-06 | 6.700E-06 | 1.000E-05 |
| 18270 | 914 | T914I | C | 9.6e-06 | 5.5e-06 | 5.9e-06 | 7.000E-06 | 2.261E-06 | 5.500E-06 | 9.600E-06 |
| 18271 | 914 | T914L | C | 1.8e-05 | 1.5e-05 | 7.2e-06 | 1.340E-05 | 5.575E-06 | 7.200E-06 | 1.800E-05 |
| 18272 | 914 | T914K | C | 5.2e-06 | 6.6e-06 | 6.3e-06 | 6.033E-06 | 7.371E-07 | 5.200E-06 | 6.600E-06 |
| 18273 | 914 | T914M | C | 1.2e-05 | 9.1e-06 | 5e-06 | 8.700E-06 | 3.517E-06 | 5.000E-06 | 1.200E-05 |
| 18274 | 914 | T914F | C | 4.3e-06 | 1e-05 | 4.6e-06 | 6.300E-06 | 3.208E-06 | 4.300E-06 | 1.000E-05 |

| | | | | | | | | | | |
|---|---|---|---|---|---|---|---|---|---|---|
| 18275 | 914 | T914P | C | 8.7e-06 | 6.8e-06 | 5.5e-06 | 7.000E-06 | 1.609E-06 | 5.500E-06 | 8.700E-06 |
| 18276 | 914 | T914S | C | 7.2e-06 | 1e-05 | 5.9e-06 | 7.700E-06 | 2.095E-06 | 5.900E-06 | 1.000E-05 |
| 18277 | 914 | T914T | C | 8.3e-06 | 1.4e-05 | 7.1e-06 | 9.800E-06 | 3.686E-06 | 7.100E-06 | 1.400E-05 |
| 18278 | 914 | T914W | C | 1.1e-05 | 5.4e-06 | 7.4e-06 | 7.933E-06 | 2.838E-06 | 5.400E-06 | 1.100E-05 |
| 18279 | 914 | T914Y | C | 1.1e-05 | 7.8e-06 | 8.6e-06 | 9.133E-06 | 1.665E-06 | 7.800E-06 | 1.100E-05 |
| 18280 | 914 | T914V | C | 8e-06 | 9.5e-06 | 7.5e-06 | 8.333E-06 | 1.041E-06 | 7.500E-06 | 9.500E-06 |
| 18281 | 915 | L915A | C | 7.9e-06 | 1.2e-05 | 1.4e-05 | 1.130E-05 | 3.110E-06 | 7.900E-06 | 1.400E-05 |
| 18282 | 915 | L915R | C | 9.4e-06 | 7.4e-06 | 1.2e-05 | 9.600E-06 | 2.307E-06 | 7.400E-06 | 1.200E-05 |
| 18283 | 915 | L915N | C | 5.4e-06 | 9.1e-06 | 1.1e-05 | 8.500E-06 | 2.848E-06 | 5.400E-06 | 1.100E-05 |
| 18284 | 915 | L915D | C | 7.5e-06 | 1.3e-05 | 6.1e-06 | 8.867E-06 | 3.647E-06 | 6.100E-06 | 1.300E-05 |
| 18285 | 915 | L915C | C | 7.1e-06 | 1.1e-05 | 6.7e-06 | 8.267E-06 | 2.376E-06 | 6.700E-06 | 1.100E-05 |
| 18286 | 915 | L915E | C | 1.3e-05 | 6.2e-06 | 6.4e-06 | 8.533E-06 | 3.870E-06 | 6.200E-06 | 1.300E-05 |
| 18287 | 915 | L915Q | C | 1.9e-05 | 4.4e-06 | 7e-06 | 1.013E-05 | 7.788E-06 | 4.400E-06 | 1.900E-05 |
| 18288 | 915 | L915G | C | 5e-06 | 7e-06 | 1e-05 | 7.333E-06 | 2.517E-06 | 5.000E-06 | 1.000E-05 |
| 18289 | 915 | L915H | C | 5.2e-06 | 8.7e-06 | 4.9e-06 | 6.267E-06 | 2.113E-06 | 4.900E-06 | 8.700E-06 |
| 18290 | 915 | L915I | C | 8.5e-06 | 1e-05 | 1.4e-05 | 1.083E-05 | 2.843E-06 | 8.500E-06 | 1.400E-05 |
| 18291 | 915 | L915L | C | 6.2e-06 | 7.9e-06 | 7.8e-06 | 7.300E-06 | 9.539E-07 | 6.200E-06 | 7.900E-06 |
| 18292 | 915 | L915K | C | 5.9e-06 | 4.6e-06 | 8e-06 | 6.167E-06 | 1.716E-06 | 4.600E-06 | 8.000E-06 |
| 18293 | 915 | L915M | C | 1.3e-05 | 7.8e-06 | 7.2e-06 | 9.333E-06 | 3.190E-06 | 7.200E-06 | 1.300E-05 |
| 18294 | 915 | L915F | C | 9.2e-06 | 8.6e-06 | 5.3e-06 | 7.700E-06 | 2.100E-06 | 5.300E-06 | 9.200E-06 |
| 18295 | 915 | L915P | C | 7.2e-06 | 6.4e-06 | 9.6e-06 | 7.733E-06 | 1.665E-06 | 6.400E-06 | 9.600E-06 |
| 18296 | 915 | L915S | C | 4.7e-06 | 8.2e-06 | 1.3e-05 | 8.633E-06 | 4.167E-06 | 4.700E-06 | 1.300E-05 |
| 18297 | 915 | L915T | C | 6.1e-06 | 6.7e-06 | 6.2e-06 | 6.333E-06 | 3.215E-07 | 6.100E-06 | 6.700E-06 |
| 18298 | 915 | L915W | C | 7.1e-06 | 8.3e-06 | 1.2e-05 | 9.133E-06 | 2.554E-06 | 7.100E-06 | 1.200E-05 |
| 18299 | 915 | L915Y | C | 1.1e-05 | 5.7e-06 | 6.9e-06 | 7.867E-06 | 2.779E-06 | 5.700E-06 | 1.100E-05 |
| 18300 | 915 | L915V | C | 1.3e-05 | 6e-06 | 8.6e-06 | 9.200E-06 | 3.538E-06 | 6.000E-06 | 1.300E-05 |
| 18301 | 916 | Y916A | C | 8.7e-06 | 5e-06 | 5.4e-06 | 6.367E-06 | 2.031E-06 | 5.000E-06 | 8.700E-06 |
| 18302 | 916 | Y916R | C | 6e-06 | 6.9e-06 | 5.2e-06 | 6.033E-06 | 8.505E-07 | 5.200E-06 | 6.900E-06 |
| 18303 | 916 | Y916N | C | 1.2e-05 | 8.1e-06 | 4.5e-06 | 8.200E-06 | 3.751E-06 | 4.500E-06 | 1.200E-05 |
| 18304 | 916 | Y916D | C | 9.3e-06 | 7.8e-06 | 6.6e-06 | 7.900E-06 | 1.353E-06 | 6.600E-06 | 9.300E-06 |
| 18305 | 916 | Y916C | C | 6.3e-06 | 1.2e-05 | 8.6e-06 | 8.967E-06 | 2.868E-06 | 6.300E-06 | 1.200E-05 |
| 18306 | 916 | Y916E | C | 9e-06 | 6.4e-06 | 7.8e-06 | 7.733E-06 | 1.301E-06 | 6.400E-06 | 9.000E-06 |
| 18307 | 916 | Y916Q | C | 7.3e-06 | 3.6e-06 | 1.1e-05 | 7.300E-06 | 3.700E-06 | 3.600E-06 | 1.100E-05 |
| 18308 | 916 | Y916G | C | 1.6e-05 | 9.4e-06 | 8.1e-06 | 1.117E-05 | 4.236E-06 | 8.100E-06 | 1.600E-05 |
| 18309 | 916 | Y916H | C | 5.4e-06 | 9e-06 | 9.7e-06 | 8.033E-06 | 2.307E-06 | 5.400E-06 | 9.700E-06 |
| 18310 | 916 | Y916I | C | 8.1e-06 | 8.5e-06 | 7.4e-06 | 8.000E-06 | 5.568E-07 | 7.400E-06 | 8.500E-06 |
| 18311 | 916 | Y916L | C | 1.1e-05 | 8.3e-06 | 4e-06 | 7.767E-06 | 3.530E-06 | 4.000E-06 | 1.100E-05 |
| 18312 | 916 | Y916K | C | 5.2e-06 | 7.9e-06 | 9.6e-06 | 7.567E-06 | 2.219E-06 | 5.200E-06 | 9.600E-06 |
| 18313 | 916 | Y916M | C | 5.5e-06 | 6.4e-06 | 6.3e-06 | 6.067E-06 | 4.933E-07 | 5.500E-06 | 6.400E-06 |
| 18314 | 916 | Y916F | C | 1.2e-05 | 7.2e-06 | 1e-05 | 9.733E-06 | 2.411E-06 | 7.200E-06 | 1.200E-05 |
| 18315 | 916 | Y916P | C | 8.6e-06 | 7.7e-06 | 1e-05 | 8.767E-06 | 1.159E-06 | 7.700E-06 | 1.000E-05 |
| 18316 | 916 | Y916S | C | 6e-06 | 7.8e-06 | 7.6e-06 | 7.133E-06 | 9.866E-07 | 6.000E-06 | 7.800E-06 |
| 18317 | 916 | Y916T | C | 6.5e-06 | 9.2e-06 | 9.3e-06 | 8.333E-06 | 1.589E-06 | 6.500E-06 | 9.300E-06 |
| 18318 | 916 | Y916W | C | 7.6e-06 | 7.1e-06 | 7.9e-06 | 7.533E-06 | 4.041E-07 | 7.100E-06 | 7.900E-06 |
| 18319 | 916 | Y916Y | C | 1.1e-05 | 9.7e-06 | 6.5e-06 | 9.067E-06 | 2.316E-06 | 6.500E-06 | 1.100E-05 |
| 18320 | 916 | Y916V | C | 7.5e-06 | 9.7e-06 | 4.8e-06 | 7.333E-06 | 2.454E-06 | 4.800E-06 | 9.700E-06 |
| 18321 | 917 | L917A | C | 9.8e-06 | 4.8e-06 | 7.2e-06 | 7.267E-06 | 2.501E-06 | 4.800E-06 | 9.800E-06 |
| 18322 | 917 | L917R | C | 5.9e-06 | 5.8e-06 | 6.2e-06 | 5.967E-06 | 2.082E-07 | 5.800E-06 | 6.200E-06 |
| 18323 | 917 | L917N | C | 4.8e-06 | 9e-06 | 6.6e-06 | 6.800E-06 | 2.107E-06 | 4.800E-06 | 9.000E-06 |
| 18324 | 917 | L917D | C | 7.8e-06 | 6.6e-06 | 9.3e-06 | 7.900E-06 | 1.353E-06 | 6.600E-06 | 9.300E-06 |
| 18325 | 917 | L917C | C | 1.2e-05 | 1.3e-05 | 1.3e-05 | 1.267E-05 | 5.774E-07 | 1.200E-05 | 1.300E-05 |
| 18326 | 917 | L917E | C | 9.1e-06 | 5.9e-06 | 7.2e-06 | 7.400E-06 | 1.609E-06 | 5.900E-06 | 9.100E-06 |
| 18327 | 917 | L917Q | C | 6.5e-06 | 8.6e-06 | 7.7e-06 | 7.600E-06 | 1.054E-06 | 6.500E-06 | 8.600E-06 |
| 18328 | 917 | L917G | C | 4.7e-06 | 1.4e-05 | 5.3e-06 | 8.000E-06 | 5.205E-06 | 4.700E-06 | 1.400E-05 |
| 18329 | 917 | L917H | C | 8.5e-06 | 8.4e-06 | 5.4e-06 | 7.433E-06 | 1.762E-06 | 5.400E-06 | 8.500E-06 |
| 18330 | 917 | L917I | C | 7.9e-06 | 4.6e-06 | 7.6e-06 | 6.700E-06 | 1.825E-06 | 4.600E-06 | 7.900E-06 |
| 18331 | 917 | L917L | C | 5.7e-06 | 8.1e-06 | 5.5e-06 | 6.433E-06 | 1.447E-06 | 5.500E-06 | 8.100E-06 |
| 18332 | 917 | L917K | C | 6e-06 | 9.4e-06 | 8.7e-06 | 8.033E-06 | 1.795E-06 | 6.000E-06 | 9.400E-06 |
| 18333 | 917 | L917M | C | 6.4e-06 | 6.8e-06 | 8e-06 | 7.067E-06 | 8.327E-07 | 6.400E-06 | 8.000E-06 |
| 18334 | 917 | L917F | C | 6.8e-06 | 7.5e-06 | 5e-06 | 6.433E-06 | 1.290E-06 | 5.000E-06 | 7.500E-06 |
| 18335 | 917 | L917P | C | 1.1e-05 | 8.7e-06 | 9e-06 | 9.567E-06 | 1.250E-06 | 8.700E-06 | 1.100E-05 |
| 18336 | 917 | L917S | C | 1.4e-05 | 1.2e-05 | 8.7e-06 | 1.157E-05 | 2.676E-06 | 8.700E-06 | 1.400E-05 |
| 18337 | 917 | L917T | C | 7.7e-06 | 1.4e-05 | 8.4e-06 | 1.003E-05 | 3.453E-06 | 7.700E-06 | 1.400E-05 |
| 18338 | 917 | L917W | C | 1.9e-05 | 7.3e-06 | 4.4e-06 | 1.023E-05 | 7.729E-06 | 4.400E-06 | 1.900E-05 |
| 18339 | 917 | L917Y | C | 5.7e-06 | 4.2e-06 | 1.1e-05 | 6.967E-06 | 3.573E-06 | 4.200E-06 | 1.100E-05 |
| 18340 | 917 | L917V | C | 1.4e-05 | 6.2e-06 | 9.1e-06 | 9.767E-06 | 3.943E-06 | 6.200E-06 | 1.400E-05 |
| 18341 | 918 | Q918A | C | 6.4e-06 | 7e-06 | 9.1e-06 | 7.500E-06 | 1.418E-06 | 6.400E-06 | 9.100E-06 |
| 18342 | 918 | Q918R | C | 7.4e-06 | 9.1e-06 | 7.1e-06 | 7.867E-06 | 1.079E-06 | 7.100E-06 | 9.100E-06 |
| 18343 | 918 | Q918N | C | 9.8e-06 | 3.8e-06 | 5.4e-06 | 6.333E-06 | 3.107E-06 | 3.800E-06 | 9.800E-06 |
| 18344 | 918 | Q918D | C | 7.3e-06 | 4.9e-06 | 3.5e-06 | 5.233E-06 | 1.922E-06 | 3.500E-06 | 7.300E-06 |
| 18345 | 918 | Q918C | C | 6.9e-06 | 6.4e-06 | 7.8e-06 | 7.033E-06 | 7.095E-07 | 6.400E-06 | 7.800E-06 |
| 18346 | 918 | Q918E | C | 5.2e-06 | 9.5e-06 | 1.4e-05 | 9.567E-06 | 4.400E-06 | 5.200E-06 | 1.400E-05 |
| 18347 | 918 | Q918Q | C | 3.8e-06 | 8e-06 | 8e-06 | 6.600E-06 | 2.425E-06 | 3.800E-06 | 8.000E-06 |
| 18348 | 918 | Q918G | C | 6.3e-06 | 5.4e-06 | 9.5e-06 | 7.067E-06 | 2.155E-06 | 5.400E-06 | 9.500E-06 |
| 18349 | 918 | Q918H | C | 7.5e-06 | 6.7e-06 | 8.4e-06 | 7.533E-06 | 8.505E-07 | 6.700E-06 | 8.400E-06 |
| 18350 | 918 | Q918I | C | 1.1e-05 | 6.4e-06 | 7.9e-06 | 8.433E-06 | 2.346E-06 | 6.400E-06 | 1.100E-05 |
| 18351 | 918 | Q918L | C | 6.2e-06 | 4.5e-06 | 5.9e-06 | 5.533E-06 | 9.074E-07 | 4.500E-06 | 6.200E-06 |
| 18352 | 918 | Q918K | C | 9.3e-06 | 6.5e-06 | 5.6e-06 | 7.133E-06 | 1.930E-06 | 5.600E-06 | 9.300E-06 |
| 18353 | 918 | Q918M | C | 7.3e-06 | 6.1e-06 | 5.4e-06 | 6.267E-06 | 9.609E-07 | 5.400E-06 | 7.300E-06 |
| 18354 | 918 | Q918F | C | 9.4e-06 | 6.7e-06 | 1.8e-05 | 1.137E-05 | 5.901E-06 | 6.700E-06 | 1.800E-05 |
| 18355 | 918 | Q918P | C | 1.1e-05 | 6.9e-06 | 9.8e-06 | 9.233E-06 | 2.108E-06 | 6.900E-06 | 1.100E-05 |
| 18356 | 918 | Q918S | C | 6e-06 | 6.3e-06 | 8.8e-06 | 7.033E-06 | 1.537E-06 | 6.000E-06 | 8.800E-06 |
| 18357 | 918 | Q918T | C | 1.2e-05 | 7.9e-06 | 9.2e-06 | 9.700E-06 | 2.095E-06 | 7.900E-06 | 1.200E-05 |
| 18358 | 918 | Q918W | C | 9.9e-06 | 9.1e-06 | 6.2e-06 | 8.400E-06 | 1.947E-06 | 6.200E-06 | 9.900E-06 |
| 18359 | 918 | Q918Y | C | 4.3e-06 | 1.3e-05 | 6.5e-06 | 7.933E-06 | 4.524E-06 | 4.300E-06 | 1.300E-05 |

| | | | | | | | | | | |
|---|---|---|---|---|---|---|---|---|---|---|
| 18360 | 918 | Q918V | C | 1.3e-05 | 7.5e-06 | 9e-06 | 9.833E-06 | 2.843E-06 | 7.500E-06 | 1.300E-05 |
| 18361 | 919 | M919A | C | 6.4e-06 | 7.3e-06 | 9.9e-06 | 7.867E-06 | 1.818E-06 | 6.400E-06 | 9.900E-06 |
| 18362 | 919 | M919R | C | 8.4e-06 | 5.7e-06 | 7.4e-06 | 7.167E-06 | 1.365E-06 | 5.700E-06 | 8.400E-06 |
| 18363 | 919 | M919N | C | 1.4e-05 | 6.1e-06 | 7e-06 | 9.033E-06 | 4.325E-06 | 6.100E-06 | 1.400E-05 |
| 18364 | 919 | M919D | C | 6.4e-06 | 6.6e-06 | 5.6e-06 | 6.200E-06 | 5.292E-07 | 5.600E-06 | 6.600E-06 |
| 18365 | 919 | M919C | C | 1.3e-05 | 6.5e-06 | 7.1e-06 | 8.867E-06 | 3.592E-06 | 6.500E-06 | 1.300E-05 |
| 18366 | 919 | M919E | C | 6.7e-06 | 1.1e-05 | 1.3e-05 | 1.023E-05 | 3.219E-06 | 6.700E-06 | 1.300E-05 |
| 18367 | 919 | M919Q | C | 8.5e-06 | 7.8e-06 | 5.2e-06 | 7.167E-06 | 1.739E-06 | 5.200E-06 | 8.500E-06 |
| 18368 | 919 | M919G | C | 1.1e-05 | 8.5e-06 | 8.9e-06 | 9.467E-06 | 1.343E-06 | 8.500E-06 | 1.100E-05 |
| 18369 | 919 | M919H | C | 9.5e-06 | 5.8e-06 | 1.2e-05 | 9.100E-06 | 3.119E-06 | 5.800E-06 | 1.200E-05 |
| 18370 | 919 | M919I | C | 3.4e-06 | 1.4e-05 | 1.4e-05 | 1.047E-05 | 6.120E-06 | 3.400E-06 | 1.400E-05 |
| 18371 | 919 | M919L | C | 8.2e-06 | 6.1e-06 | 1.4e-05 | 9.433E-06 | 4.092E-06 | 6.100E-06 | 1.400E-05 |
| 18372 | 919 | M919K | C | 1.5e-05 | 9.5e-06 | 9.5e-06 | 1.133E-05 | 3.175E-06 | 9.500E-06 | 1.500E-05 |
| 18373 | 919 | M919M | C | 1.1e-05 | 5e-06 | 9.5e-06 | 8.500E-06 | 3.122E-06 | 5.000E-06 | 1.100E-05 |
| 18374 | 919 | M919F | C | 7.6e-06 | 9.8e-06 | 6.6e-06 | 8.000E-06 | 1.637E-06 | 6.600E-06 | 9.800E-06 |
| 18375 | 919 | M919P | C | 9.4e-06 | 1.1e-05 | 8.6e-06 | 9.667E-06 | 1.222E-06 | 8.600E-06 | 1.100E-05 |
| 18376 | 919 | M919S | C | 6e-06 | 1.1e-05 | 6.4e-06 | 7.800E-06 | 2.778E-06 | 6.000E-06 | 1.100E-05 |
| 18377 | 919 | M919T | C | 8.2e-06 | 6.9e-06 | 6e-06 | 7.033E-06 | 1.106E-06 | 6.000E-06 | 8.200E-06 |
| 18378 | 919 | M919W | C | 8.7e-06 | 7.7e-06 | 8.4e-06 | 8.267E-06 | 5.132E-07 | 7.700E-06 | 8.700E-06 |
| 18379 | 919 | M919Y | C | 6.2e-06 | 4.3e-06 | 6.2e-06 | 5.567E-06 | 1.097E-06 | 4.300E-06 | 6.200E-06 |
| 18380 | 919 | M919V | C | 4.7e-06 | 1.2e-05 | 5.3e-06 | 7.333E-06 | 4.053E-06 | 4.700E-06 | 1.200E-05 |
| 18381 | 920 | N920A | C | 7.6e-06 | 1.2e-05 | 4.5e-06 | 8.033E-06 | 3.769E-06 | 4.500E-06 | 1.200E-05 |
| 18382 | 920 | N920R | C | 5.1e-06 | 7.9e-06 | 9.1e-06 | 7.367E-06 | 2.053E-06 | 5.100E-06 | 9.100E-06 |
| 18383 | 920 | N920N | C | 5.2e-06 | 7.4e-06 | 1.1e-05 | 7.867E-06 | 2.928E-06 | 5.200E-06 | 1.100E-05 |
| 18384 | 920 | N920D | C | 6.9e-06 | 5.4e-06 | 1.2e-05 | 8.100E-06 | 3.460E-06 | 5.400E-06 | 1.200E-05 |
| 18385 | 920 | N920C | C | 9.9e-06 | 1.3e-05 | 9.5e-06 | 1.080E-05 | 1.916E-06 | 9.500E-06 | 1.300E-05 |
| 18386 | 920 | N920E | C | 7.4e-06 | 8.8e-06 | 8.8e-06 | 8.333E-06 | 8.083E-07 | 7.400E-06 | 8.800E-06 |
| 18387 | 920 | N920Q | C | 1.4e-05 | 9.2e-06 | 7.9e-06 | 1.037E-05 | 3.213E-06 | 7.900E-06 | 1.400E-05 |
| 18388 | 920 | N920G | C | 5.8e-06 | 9.6e-06 | 7.1e-06 | 7.500E-06 | 1.931E-06 | 5.800E-06 | 9.600E-06 |
| 18389 | 920 | N920H | C | 7e-06 | 4.7e-06 | 4.7e-06 | 5.467E-06 | 1.328E-06 | 4.700E-06 | 7.000E-06 |
| 18390 | 920 | N920I | C | 6e-06 | 9.9e-06 | 1.5e-05 | 1.030E-05 | 4.513E-06 | 6.000E-06 | 1.500E-05 |
| 18391 | 920 | N920L | C | 7.6e-06 | 4.7e-06 | 7.7e-06 | 6.667E-06 | 1.704E-06 | 4.700E-06 | 7.700E-06 |
| 18392 | 920 | N920K | C | 1.7e-05 | 7.9e-06 | 6.6e-06 | 1.050E-05 | 5.667E-06 | 6.600E-06 | 1.700E-05 |
| 18393 | 920 | N920M | C | 7.5e-06 | 1e-05 | 7.3e-06 | 8.267E-06 | 1.504E-06 | 7.300E-06 | 1.000E-05 |
| 18394 | 920 | N920F | C | 8e-06 | 1.1e-05 | 5.7e-06 | 8.233E-06 | 2.658E-06 | 5.700E-06 | 1.100E-05 |
| 18395 | 920 | N920P | C | 8.4e-06 | 1.1e-05 | 8e-06 | 9.133E-06 | 1.629E-06 | 8.000E-06 | 1.100E-05 |
| 18396 | 920 | N920S | C | 8.2e-06 | 7.8e-06 | 1e-05 | 8.667E-06 | 1.172E-06 | 7.800E-06 | 1.000E-05 |
| 18397 | 920 | N920T | C | 9.5e-06 | 1.2e-05 | 8.6e-06 | 1.003E-05 | 1.762E-06 | 8.600E-06 | 1.200E-05 |
| 18398 | 920 | N920W | C | 9.3e-06 | 6e-06 | 8.3e-06 | 7.867E-06 | 1.692E-06 | 6.000E-06 | 9.300E-06 |
| 18399 | 920 | N920Y | C | 8.9e-06 | 1.2e-05 | 8.7e-06 | 9.867E-06 | 1.850E-06 | 8.700E-06 | 1.200E-05 |
| 18400 | 920 | N920V | C | 4.9e-06 | 9.6e-06 | 1.1e-05 | 8.500E-06 | 3.195E-06 | 4.900E-06 | 1.100E-05 |
| 18401 | 921 | S921A | C | 8.4e-06 | 9.6e-06 | 7.4e-06 | 8.467E-06 | 1.102E-06 | 7.400E-06 | 9.600E-06 |
| 18402 | 921 | S921R | C | 9.4e-06 | 8.8e-06 | 8.3e-06 | 8.833E-06 | 5.508E-07 | 8.300E-06 | 9.400E-06 |
| 18403 | 921 | S921N | C | 6.9e-06 | 9.1e-06 | 6.1e-06 | 7.367E-06 | 1.553E-06 | 6.100E-06 | 9.100E-06 |
| 18404 | 921 | S921D | C | 4.7e-06 | 7e-06 | 8.8e-06 | 6.833E-06 | 2.055E-06 | 4.700E-06 | 8.800E-06 |
| 18405 | 921 | S921C | C | 6.6e-06 | 7.6e-06 | 5.4e-06 | 6.533E-06 | 1.102E-06 | 5.400E-06 | 7.600E-06 |
| 18406 | 921 | S921E | C | 9.5e-06 | 5.1e-06 | 6.3e-06 | 6.967E-06 | 2.274E-06 | 5.100E-06 | 9.500E-06 |
| 18407 | 921 | S921Q | C | 9.2e-06 | 9e-06 | 7.6e-06 | 8.600E-06 | 8.718E-07 | 7.600E-06 | 9.200E-06 |
| 18408 | 921 | S921G | C | 5.2e-06 | 8.3e-06 | 7e-06 | 6.833E-06 | 1.557E-06 | 5.200E-06 | 8.300E-06 |
| 18409 | 921 | S921H | C | 6e-06 | 4e-06 | 1.2e-05 | 7.333E-06 | 4.163E-06 | 4.000E-06 | 1.200E-05 |
| 18410 | 921 | S921I | C | 1.1e-05 | 5.8e-06 | 1e-05 | 8.933E-06 | 2.759E-06 | 5.800E-06 | 1.100E-05 |
| 18411 | 921 | S921L | C | 8.2e-06 | 7e-06 | 7.6e-06 | 7.600E-06 | 6.000E-07 | 7.000E-06 | 8.200E-06 |
| 18412 | 921 | S921K | C | 6.8e-06 | 9.8e-06 | 4.6e-06 | 7.067E-06 | 2.610E-06 | 4.600E-06 | 9.800E-06 |
| 18413 | 921 | S921M | C | 7.3e-06 | 1.4e-05 | 4.6e-06 | 8.633E-06 | 4.840E-06 | 4.600E-06 | 1.400E-05 |
| 18414 | 921 | S921F | C | 6.2e-06 | 6.7e-06 | 8.6e-06 | 7.167E-06 | 1.266E-06 | 6.200E-06 | 8.600E-06 |
| 18415 | 921 | S921P | C | 8.8e-06 | 1e-05 | 6.8e-06 | 8.533E-06 | 1.617E-06 | 6.800E-06 | 1.000E-05 |
| 18416 | 921 | S921S | C | 5.5e-06 | 9.2e-06 | 1.2e-05 | 8.900E-06 | 3.260E-06 | 5.500E-06 | 1.200E-05 |
| 18417 | 921 | S921T | C | 6.9e-06 | 9.2e-06 | 1.3e-05 | 9.700E-06 | 3.081E-06 | 6.900E-06 | 1.300E-05 |
| 18418 | 921 | S921W | C | 1e-05 | 4.4e-06 | 9.8e-06 | 8.067E-06 | 3.177E-06 | 4.400E-06 | 1.000E-05 |
| 18419 | 921 | S921Y | C | 1.3e-05 | 6.2e-06 | 1.2e-05 | 1.040E-05 | 3.672E-06 | 6.200E-06 | 1.300E-05 |
| 18420 | 921 | S921V | C | 1.2e-05 | 8.9e-06 | 1e-05 | 1.030E-05 | 1.572E-06 | 8.900E-06 | 1.200E-05 |
| 18421 | 922 | L922A | C | 7e-06 | 6.7e-06 | 5.1e-06 | 6.267E-06 | 1.021E-06 | 5.100E-06 | 7.000E-06 |
| 18422 | 922 | L922R | C | 5e-06 | 6.1e-06 | 5.3e-06 | 5.467E-06 | 5.686E-07 | 5.000E-06 | 6.100E-06 |
| 18423 | 922 | L922N | C | 8.6e-06 | 7.5e-06 | 6.5e-06 | 7.533E-06 | 1.050E-06 | 6.500E-06 | 8.600E-06 |
| 18424 | 922 | L922D | C | 1.5e-05 | 6e-06 | 6.3e-06 | 9.100E-06 | 5.112E-06 | 6.000E-06 | 1.500E-05 |
| 18425 | 922 | L922C | C | 8.5e-06 | 1e-05 | 7.7e-06 | 8.733E-06 | 1.168E-06 | 7.700E-06 | 1.000E-05 |
| 18426 | 922 | L922E | C | 3.7e-06 | 5.9e-06 | 7.5e-06 | 5.700E-06 | 1.908E-06 | 3.700E-06 | 7.500E-06 |
| 18427 | 922 | L922Q | C | 7.4e-06 | 4.2e-06 | 5.9e-06 | 5.833E-06 | 1.601E-06 | 4.200E-06 | 7.400E-06 |
| 18428 | 922 | L922G | C | 4.9e-06 | 8.2e-06 | 7e-06 | 6.700E-06 | 1.670E-06 | 4.900E-06 | 8.200E-06 |
| 18429 | 922 | L922H | C | 8.7e-06 | 8.8e-06 | 1e-05 | 9.167E-06 | 7.234E-07 | 8.700E-06 | 1.000E-05 |
| 18430 | 922 | L922I | C | 1.1e-05 | 1e-05 | 5e-06 | 8.667E-06 | 3.215E-06 | 5.000E-06 | 1.100E-05 |
| 18431 | 922 | L922L | C | 6.3e-06 | 4.9e-06 | 1.3e-05 | 8.067E-06 | 4.329E-06 | 4.900E-06 | 1.300E-05 |
| 18432 | 922 | L922K | C | 5.5e-06 | 6.4e-06 | 7.3e-06 | 6.400E-06 | 9.000E-07 | 5.500E-06 | 7.300E-06 |
| 18433 | 922 | L922M | C | 4.7e-06 | 1.3e-05 | 7.2e-06 | 8.300E-06 | 4.258E-06 | 4.700E-06 | 1.300E-05 |
| 18434 | 922 | L922F | C | 7.7e-06 | 7.2e-06 | 9e-06 | 7.967E-06 | 9.292E-07 | 7.200E-06 | 9.000E-06 |
| 18435 | 922 | L922P | C | 9.4e-06 | 9.2e-06 | 9.5e-06 | 9.367E-06 | 1.528E-07 | 9.200E-06 | 9.500E-06 |
| 18436 | 922 | L922S | C | 7.2e-06 | 9.9e-06 | 9.6e-06 | 8.900E-06 | 1.480E-06 | 7.200E-06 | 9.900E-06 |
| 18437 | 922 | L922T | C | 5.1e-06 | 5.6e-06 | 9.5e-06 | 6.733E-06 | 2.409E-06 | 5.100E-06 | 9.500E-06 |
| 18438 | 922 | L922W | C | 1e-05 | 7e-06 | 1.1e-05 | 9.333E-06 | 2.082E-06 | 7.000E-06 | 1.100E-05 |
| 18439 | 922 | L922Y | C | 6.6e-06 | 8.5e-06 | 9.3e-06 | 8.133E-06 | 1.387E-06 | 6.600E-06 | 9.300E-06 |
| 18440 | 922 | L922V | C | 7.1e-06 | 6.8e-06 | 7.4e-06 | 7.100E-06 | 3.000E-07 | 6.800E-06 | 7.400E-06 |
| 18441 | 923 | R923A | C | 1e-05 | 7e-06 | 8e-06 | 8.333E-06 | 1.528E-06 | 7.000E-06 | 1.000E-05 |
| 18442 | 923 | R923R | C | 1.3e-05 | 9.8e-06 | 4.6e-06 | 9.133E-06 | 4.239E-06 | 4.600E-06 | 1.300E-05 |
| 18443 | 923 | R923N | C | 9.7e-06 | 8.9e-06 | 4.5e-06 | 7.700E-06 | 2.800E-06 | 4.500E-06 | 9.700E-06 |
| 18444 | 923 | R923D | C | 1.5e-05 | 5.5e-06 | 8.6e-06 | 9.700E-06 | 4.845E-06 | 5.500E-06 | 1.500E-05 |

| | | | | | | | | | | |
|---|---|---|---|---|---|---|---|---|---|---|
| 18445 | 923 | R923C | C | 9.7e-06 | 9.8e-06 | 3.9e-06 | 7.800E-06 | 3.378E-06 | 3.900E-06 | 9.800E-06 |
| 18446 | 923 | R923E | C | 6e-06 | 9.3e-06 | 9.2e-06 | 8.167E-06 | 1.877E-06 | 6.000E-06 | 9.300E-06 |
| 18447 | 923 | R923Q | C | 8.6e-06 | 9e-06 | 1e-05 | 9.200E-06 | 7.211E-07 | 8.600E-06 | 1.000E-05 |
| 18448 | 923 | R923G | C | 9.7e-06 | 5.6e-06 | 9.8e-06 | 8.367E-06 | 2.397E-06 | 5.600E-06 | 9.800E-06 |
| 18449 | 923 | R923H | C | 5.7e-06 | 1.4e-05 | 6.9e-06 | 8.867E-06 | 4.486E-06 | 5.700E-06 | 1.400E-05 |
| 18450 | 923 | R923I | C | 8e-06 | 6.4e-06 | 4.7e-06 | 6.367E-06 | 1.650E-06 | 4.700E-06 | 8.000E-06 |
| 18451 | 923 | R923L | C | 6.4e-06 | 9.4e-06 | 5.6e-06 | 7.133E-06 | 2.003E-06 | 5.600E-06 | 9.400E-06 |
| 18452 | 923 | R923K | C | 9e-06 | 6.8e-06 | 6.5e-06 | 7.433E-06 | 1.365E-06 | 6.500E-06 | 9.000E-06 |
| 18453 | 923 | R923M | C | 7.6e-06 | 6.5e-06 | 6.7e-06 | 6.933E-06 | 5.859E-07 | 6.500E-06 | 7.600E-06 |
| 18454 | 923 | R923F | C | 7.3e-06 | 1.4e-05 | 1.1e-05 | 1.077E-05 | 3.356E-06 | 7.300E-06 | 1.400E-05 |
| 18455 | 923 | R923P | C | 7.4e-06 | 8.9e-06 | 6.2e-06 | 7.500E-06 | 1.353E-06 | 6.200E-06 | 8.900E-06 |
| 18456 | 923 | R923S | C | 9.2e-06 | 4.4e-06 | 4.8e-06 | 6.133E-06 | 2.663E-06 | 4.400E-06 | 9.200E-06 |
| 18457 | 923 | R923T | C | 8.5e-06 | 8.6e-06 | 8.8e-06 | 8.633E-06 | 1.528E-07 | 8.500E-06 | 8.800E-06 |
| 18458 | 923 | R923W | C | 8.1e-06 | 6.4e-06 | 4.9e-06 | 6.467E-06 | 1.601E-06 | 4.900E-06 | 8.100E-06 |
| 18459 | 923 | R923Y | C | 9.2e-06 | 7.6e-06 | 8.1e-06 | 8.300E-06 | 8.185E-07 | 7.600E-06 | 9.200E-06 |
| 18460 | 923 | R923V | C | 1.2e-05 | 1.2e-05 | 6.9e-06 | 1.030E-05 | 2.944E-06 | 6.900E-06 | 1.200E-05 |
| 18461 | 924 | A924A | C | 8.4e-06 | 4.5e-06 | 1.2e-05 | 8.300E-06 | 3.751E-06 | 4.500E-06 | 1.200E-05 |
| 18462 | 924 | A924R | C | 5.2e-06 | 7.7e-06 | 5.2e-06 | 6.033E-06 | 1.443E-06 | 5.200E-06 | 7.700E-06 |
| 18463 | 924 | A924N | C | 3.8e-06 | 6e-06 | 1.2e-05 | 7.267E-06 | 4.244E-06 | 3.800E-06 | 1.200E-05 |
| 18464 | 924 | A924D | C | 1.5e-05 | 6.1e-06 | 6.5e-06 | 9.200E-06 | 5.027E-06 | 6.100E-06 | 1.500E-05 |
| 18465 | 924 | A924C | C | 1.3e-05 | 9.1e-06 | 1.4e-05 | 1.203E-05 | 2.589E-06 | 9.100E-06 | 1.400E-05 |
| 18466 | 924 | A924E | C | 9.7e-06 | 8.3e-06 | 6.2e-06 | 8.067E-06 | 1.762E-06 | 6.200E-06 | 9.700E-06 |
| 18467 | 924 | A924Q | C | 7.8e-06 | 3.1e-06 | 6.5e-06 | 5.800E-06 | 2.427E-06 | 3.100E-06 | 7.800E-06 |
| 18468 | 924 | A924G | C | 9.4e-06 | 9.9e-06 | 9.9e-06 | 9.733E-06 | 2.887E-07 | 9.400E-06 | 9.900E-06 |
| 18469 | 924 | A924H | C | 6.5e-06 | 8.6e-06 | 6.9e-06 | 7.333E-06 | 1.115E-06 | 6.500E-06 | 8.600E-06 |
| 18470 | 924 | A924I | C | 1.3e-05 | 4.7e-06 | 1.2e-05 | 9.900E-06 | 4.531E-06 | 4.700E-06 | 1.300E-05 |
| 18471 | 924 | A924L | C | 6.8e-06 | 8.8e-06 | 9.1e-06 | 8.233E-06 | 1.250E-06 | 6.800E-06 | 9.100E-06 |
| 18472 | 924 | A924K | C | 1e-05 | 4.2e-06 | 1.2e-05 | 8.733E-06 | 4.051E-06 | 4.200E-06 | 1.200E-05 |
| 18473 | 924 | A924M | C | 2.1e-06 | 1.2e-05 | 9.6e-06 | 7.900E-06 | 5.164E-06 | 2.100E-06 | 1.200E-05 |
| 18474 | 924 | A924F | C | 1.1e-05 | 9.6e-06 | 1e-05 | 1.020E-05 | 7.211E-07 | 9.600E-06 | 1.100E-05 |
| 18475 | 924 | A924P | C | 1.1e-05 | 9.9e-06 | 7.7e-06 | 9.533E-06 | 1.680E-06 | 7.700E-06 | 1.100E-05 |
| 18476 | 924 | A924S | C | 5.2e-06 | 6.7e-06 | 4.3e-06 | 5.400E-06 | 1.212E-06 | 4.300E-06 | 6.700E-06 |
| 18477 | 924 | A924T | C | 6.8e-06 | 7.9e-06 | 8.7e-06 | 7.800E-06 | 9.539E-07 | 6.800E-06 | 8.700E-06 |
| 18478 | 924 | A924W | C | 6.7e-06 | 1e-05 | 9.1e-06 | 8.600E-06 | 1.706E-06 | 6.700E-06 | 1.000E-05 |
| 18479 | 924 | A924Y | C | 6e-06 | 7.8e-06 | 1.1e-05 | 8.267E-06 | 2.532E-06 | 6.000E-06 | 1.100E-05 |
| 18480 | 924 | A924V | C | 1.4e-05 | 7.4e-06 | 5.6e-06 | 9.000E-06 | 4.423E-06 | 5.600E-06 | 1.400E-05 |
| 18481 | 925 | E925A | C | 8.3e-06 | 1.3e-05 | 6.6e-06 | 9.300E-06 | 3.315E-06 | 6.600E-06 | 1.300E-05 |
| 18482 | 925 | E925R | C | 6e-06 | 1e-05 | 6.9e-06 | 7.633E-06 | 2.098E-06 | 6.000E-06 | 1.000E-05 |
| 18483 | 925 | E925N | C | 9.9e-06 | 8.1e-06 | 7.3e-06 | 8.433E-06 | 1.332E-06 | 7.300E-06 | 9.900E-06 |
| 18484 | 925 | E925D | C | 1.4e-05 | 1.4e-05 | 6.4e-06 | 1.147E-05 | 4.388E-06 | 6.400E-06 | 1.400E-05 |
| 18485 | 925 | E925C | C | 8.7e-06 | 6.6e-06 | 1.3e-05 | 9.433E-06 | 3.262E-06 | 6.600E-06 | 1.300E-05 |
| 18486 | 925 | E925E | C | 6.1e-06 | 8.4e-06 | 1.2e-05 | 8.833E-06 | 2.974E-06 | 6.100E-06 | 1.200E-05 |
| 18487 | 925 | E925Q | C | 6.9e-06 | 5.7e-06 | 8.2e-06 | 6.933E-06 | 1.250E-06 | 5.700E-06 | 8.200E-06 |
| 18488 | 925 | E925G | C | 6e-06 | 1.3e-05 | 1e-05 | 9.667E-06 | 3.512E-06 | 6.000E-06 | 1.300E-05 |
| 18489 | 925 | E925H | C | 8e-06 | 6.8e-06 | 5.9e-06 | 6.900E-06 | 1.054E-06 | 5.900E-06 | 8.000E-06 |
| 18490 | 925 | E925I | C | 7.8e-06 | 5.5e-06 | 7.6e-06 | 6.967E-06 | 1.274E-06 | 5.500E-06 | 7.800E-06 |
| 18491 | 925 | E925L | C | 8.4e-06 | 1.6e-05 | 5.3e-06 | 9.900E-06 | 5.505E-06 | 5.300E-06 | 1.600E-05 |
| 18492 | 925 | E925K | C | 1.4e-05 | 9.1e-06 | 8.6e-06 | 1.057E-05 | 2.984E-06 | 8.600E-06 | 1.400E-05 |
| 18493 | 925 | E925M | C | 4.3e-06 | 4.7e-06 | 9.5e-06 | 6.167E-06 | 2.894E-06 | 4.300E-06 | 9.500E-06 |
| 18494 | 925 | E925F | C | 6.9e-06 | 9.4e-06 | 8.3e-06 | 8.200E-06 | 1.253E-06 | 6.900E-06 | 9.400E-06 |
| 18495 | 925 | E925P | C | 4.1e-06 | 8.9e-06 | 1.1e-05 | 8.000E-06 | 3.537E-06 | 4.100E-06 | 1.100E-05 |
| 18496 | 925 | E925S | C | 5.5e-06 | 5.9e-06 | 7.1e-06 | 6.167E-06 | 8.327E-07 | 5.500E-06 | 7.100E-06 |
| 18497 | 925 | E925T | C | 8.5e-06 | 6.3e-06 | 8.8e-06 | 7.867E-06 | 1.365E-06 | 6.300E-06 | 8.800E-06 |
| 18498 | 925 | E925W | C | 6.1e-06 | 6.2e-06 | 6e-06 | 6.100E-06 | 1.000E-07 | 6.000E-06 | 6.200E-06 |
| 18499 | 925 | E925Y | C | 9.4e-06 | 9.4e-06 | 1.3e-05 | 1.060E-05 | 2.078E-06 | 9.400E-06 | 1.300E-05 |
| 18500 | 925 | E925V | C | 9.9e-06 | 9.8e-06 | 6.2e-06 | 8.633E-06 | 2.108E-06 | 6.200E-06 | 9.900E-06 |
| 18501 | 926 | D926A | C | 8.4e-06 | 1.1e-05 | 5.9e-06 | 8.433E-06 | 2.550E-06 | 5.900E-06 | 1.100E-05 |
| 18502 | 926 | D926R | C | 8.2e-06 | 7.6e-06 | 7e-06 | 7.600E-06 | 6.000E-07 | 7.000E-06 | 8.200E-06 |
| 18503 | 926 | D926N | C | 6.1e-06 | 9.9e-06 | 8.9e-06 | 8.300E-06 | 1.970E-06 | 6.100E-06 | 9.900E-06 |
| 18504 | 926 | D926D | C | 8.5e-06 | 6.5e-06 | 8.3e-06 | 7.767E-06 | 1.102E-06 | 6.500E-06 | 8.500E-06 |
| 18505 | 926 | D926C | C | 8.4e-06 | 1.3e-05 | 1.2e-05 | 1.113E-05 | 2.419E-06 | 8.400E-06 | 1.300E-05 |
| 18506 | 926 | D926E | C | 7.4e-06 | 1.3e-05 | 7.6e-06 | 9.333E-06 | 3.177E-06 | 7.400E-06 | 1.300E-05 |
| 18507 | 926 | D926Q | C | 1.3e-05 | 8.4e-06 | 9.6e-06 | 1.033E-05 | 2.386E-06 | 8.400E-06 | 1.300E-05 |
| 18508 | 926 | D926G | C | 1e-05 | 1.1e-05 | 1.2e-05 | 1.100E-05 | 1.000E-06 | 1.000E-05 | 1.200E-05 |
| 18509 | 926 | D926H | C | 7e-06 | 7.3e-06 | 8.3e-06 | 7.533E-06 | 6.807E-07 | 7.000E-06 | 8.300E-06 |
| 18510 | 926 | D926I | C | 6.4e-06 | 5.1e-06 | 9.5e-06 | 7.000E-06 | 2.261E-06 | 5.100E-06 | 9.500E-06 |
| 18511 | 926 | D926L | C | 8.9e-06 | 6.3e-06 | 8.2e-06 | 7.800E-06 | 1.345E-06 | 6.300E-06 | 8.900E-06 |
| 18512 | 926 | D926K | C | 8.1e-06 | 7.2e-06 | 4.5e-06 | 6.600E-06 | 1.873E-06 | 4.500E-06 | 8.100E-06 |
| 18513 | 926 | D926M | C | 6.9e-06 | 9e-06 | 8e-06 | 7.967E-06 | 1.050E-06 | 6.900E-06 | 9.000E-06 |
| 18514 | 926 | D926F | C | 6.3e-06 | 9.3e-06 | 1e-05 | 8.533E-06 | 1.966E-06 | 6.300E-06 | 1.000E-05 |
| 18515 | 926 | D926P | C | 5.8e-06 | 8e-06 | 5.6e-06 | 6.467E-06 | 1.332E-06 | 5.600E-06 | 8.000E-06 |
| 18516 | 926 | D926S | C | 8e-06 | 1.9e-05 | 5.9e-06 | 1.097E-05 | 7.036E-06 | 5.900E-06 | 1.900E-05 |
| 18517 | 926 | D926T | C | 9.9e-06 | 1.1e-05 | 8.3e-06 | 9.733E-06 | 1.358E-06 | 8.300E-06 | 1.100E-05 |
| 18518 | 926 | D926W | C | 8.2e-06 | 4.9e-06 | 4e-06 | 5.700E-06 | 2.211E-06 | 4.000E-06 | 8.200E-06 |
| 18519 | 926 | D926Y | C | 6.6e-06 | 1.8e-05 | 8.3e-06 | 1.097E-05 | 6.150E-06 | 6.600E-06 | 1.800E-05 |
| 18520 | 926 | D926V | C | 1.4e-05 | 8.2e-06 | 6.8e-06 | 9.667E-06 | 3.818E-06 | 6.800E-06 | 1.400E-05 |
| 18521 | 927 | T927A | C | 9.2e-06 | 1.2e-05 | 7.9e-06 | 9.700E-06 | 2.095E-06 | 7.900E-06 | 1.200E-05 |
| 18522 | 927 | T927R | C | 8e-06 | 1e-05 | 5.2e-06 | 7.733E-06 | 2.411E-06 | 5.200E-06 | 1.000E-05 |
| 18523 | 927 | T927N | C | 5.4e-06 | 8.7e-06 | 1.1e-05 | 8.367E-06 | 2.815E-06 | 5.400E-06 | 1.100E-05 |
| 18524 | 927 | T927D | C | 9.7e-06 | 9.2e-06 | 9.8e-06 | 9.567E-06 | 3.215E-07 | 9.200E-06 | 9.800E-06 |
| 18525 | 927 | T927C | C | 1.2e-05 | 1.5e-05 | 6.6e-06 | 1.120E-05 | 4.257E-06 | 6.600E-06 | 1.500E-05 |
| 18526 | 927 | T927E | C | 4.5e-06 | 9.6e-06 | 5.3e-06 | 6.467E-06 | 2.743E-06 | 4.500E-06 | 9.600E-06 |
| 18527 | 927 | T927Q | C | 1.2e-05 | 9.1e-06 | 4.4e-06 | 8.500E-06 | 3.835E-06 | 4.400E-06 | 1.200E-05 |
| 18528 | 927 | T927G | C | 6.8e-06 | 9.6e-06 | 1.4e-05 | 1.013E-05 | 3.630E-06 | 6.800E-06 | 1.400E-05 |
| 18529 | 927 | T927H | C | 6.5e-06 | 7.8e-06 | 4.6e-06 | 6.300E-06 | 1.609E-06 | 4.600E-06 | 7.800E-06 |

| | | | | | | | | | | |
|---|---|---|---|---|---|---|---|---|---|---|
| 18530 | 927 | T927I | C | 8.4e-06 | 1e-05 | 1.3e-05 | 1.047E-05 | 2.335E-06 | 8.400E-06 | 1.300E-05 |
| 18531 | 927 | T927L | C | 9.1e-06 | 6.3e-06 | 9.5e-06 | 8.300E-06 | 1.744E-06 | 6.300E-06 | 9.500E-06 |
| 18532 | 927 | T927K | C | 6.6e-06 | 4.7e-06 | 6.4e-06 | 5.900E-06 | 1.044E-06 | 4.700E-06 | 6.600E-06 |
| 18533 | 927 | T927M | C | 6.6e-06 | 1.1e-05 | 1.4e-05 | 1.053E-05 | 3.722E-06 | 6.600E-06 | 1.400E-05 |
| 18534 | 927 | T927F | C | 9.9e-06 | 7.9e-06 | 6.5e-06 | 8.100E-06 | 1.709E-06 | 6.500E-06 | 9.900E-06 |
| 18535 | 927 | T927P | C | 6.6e-06 | 6.8e-06 | 6.7e-06 | 6.700E-06 | 1.000E-07 | 6.600E-06 | 6.800E-06 |
| 18536 | 927 | T927S | C | 9e-06 | 6.1e-06 | 1.2e-05 | 9.033E-06 | 2.950E-06 | 6.100E-06 | 1.200E-05 |
| 18537 | 927 | T927T | C | 8.9e-06 | 4.1e-06 | 5.2e-06 | 6.067E-06 | 2.515E-06 | 4.100E-06 | 8.900E-06 |
| 18538 | 927 | T927W | C | 9.3e-06 | 5.9e-06 | 8.7e-06 | 7.967E-06 | 1.815E-06 | 5.900E-06 | 9.300E-06 |
| 18539 | 927 | T927Y | C | 1.4e-05 | 7.7e-06 | 1e-05 | 1.057E-05 | 3.188E-06 | 7.700E-06 | 1.400E-05 |
| 18540 | 927 | T927V | C | 7.1e-06 | 6.4e-06 | 5.8e-06 | 6.433E-06 | 6.506E-07 | 5.800E-06 | 7.100E-06 |
| 18541 | 928 | A928A | C | 4.5e-06 | 6.1e-06 | 9.4e-06 | 6.667E-06 | 2.499E-06 | 4.500E-06 | 9.400E-06 |
| 18542 | 928 | A928R | C | 8.8e-06 | 1.2e-05 | 6.4e-06 | 9.067E-06 | 2.810E-06 | 6.400E-06 | 1.200E-05 |
| 18543 | 928 | A928N | C | 7.4e-06 | 5.9e-06 | 9.7e-06 | 7.667E-06 | 1.914E-06 | 5.900E-06 | 9.700E-06 |
| 18544 | 928 | A928D | C | 8.2e-06 | 5.2e-06 | 9.7e-06 | 7.700E-06 | 2.291E-06 | 5.200E-06 | 9.700E-06 |
| 18545 | 928 | A928C | C | 8e-06 | 7.1e-06 | 9.2e-06 | 8.100E-06 | 1.054E-06 | 7.100E-06 | 9.200E-06 |
| 18546 | 928 | A928E | C | 8.8e-06 | 7.9e-06 | 5.6e-06 | 7.433E-06 | 1.650E-06 | 5.600E-06 | 8.800E-06 |
| 18547 | 928 | A928Q | C | 5.3e-06 | 4.7e-06 | 5.2e-06 | 5.067E-06 | 3.215E-07 | 4.700E-06 | 5.300E-06 |
| 18548 | 928 | A928G | C | 5.9e-06 | 9.1e-06 | 9.3e-06 | 8.100E-06 | 1.908E-06 | 5.900E-06 | 9.300E-06 |
| 18549 | 928 | A928H | C | 1.2e-05 | 6.8e-06 | 1.2e-05 | 1.027E-05 | 3.002E-06 | 6.800E-06 | 1.200E-05 |
| 18550 | 928 | A928I | C | 7.2e-06 | 9.3e-06 | 8.2e-06 | 8.233E-06 | 1.050E-06 | 7.200E-06 | 9.300E-06 |
| 18551 | 928 | A928L | C | 6.6e-06 | 6.7e-06 | 8.6e-06 | 7.300E-06 | 1.127E-06 | 6.600E-06 | 8.600E-06 |
| 18552 | 928 | A928K | C | 1e-05 | 4.7e-06 | 5.5e-06 | 6.733E-06 | 2.857E-06 | 4.700E-06 | 1.000E-05 |
| 18553 | 928 | A928M | C | 1e-05 | 8.1e-06 | 8e-06 | 8.700E-06 | 1.127E-06 | 8.000E-06 | 1.000E-05 |
| 18554 | 928 | A928F | C | 9.7e-06 | 1.3e-05 | 8.5e-06 | 1.040E-05 | 2.330E-06 | 8.500E-06 | 1.300E-05 |
| 18555 | 928 | A928P | C | 6.1e-06 | 8.2e-06 | 9.2e-06 | 7.833E-06 | 1.582E-06 | 6.100E-06 | 9.200E-06 |
| 18556 | 928 | A928S | C | 8.9e-06 | 1e-05 | 3.6e-06 | 7.500E-06 | 3.422E-06 | 3.600E-06 | 1.000E-05 |
| 18557 | 928 | A928T | C | 6e-06 | 9.7e-06 | 9.4e-06 | 8.367E-06 | 2.055E-06 | 6.000E-06 | 9.700E-06 |
| 18558 | 928 | A928W | C | 6e-06 | 1.4e-05 | 5.9e-06 | 8.633E-06 | 4.648E-06 | 5.900E-06 | 1.400E-05 |
| 18559 | 928 | A928Y | C | 1.3e-05 | 8.8e-06 | 1.4e-05 | 1.193E-05 | 2.759E-06 | 8.800E-06 | 1.400E-05 |
| 18560 | 928 | A928V | C | 6e-06 | 4.4e-06 | 6.3e-06 | 5.567E-06 | 1.021E-06 | 4.400E-06 | 6.300E-06 |
| 18561 | 929 | V929A | C | 9.5e-06 | 6.9e-06 | 4.6e-06 | 7.000E-06 | 2.452E-06 | 4.600E-06 | 9.500E-06 |
| 18562 | 929 | V929R | C | 8.5e-06 | 9.8e-06 | 5.2e-06 | 7.833E-06 | 2.371E-06 | 5.200E-06 | 9.800E-06 |
| 18563 | 929 | V929N | C | 6.8e-06 | 5.7e-06 | 9.5e-06 | 7.333E-06 | 1.955E-06 | 5.700E-06 | 9.500E-06 |
| 18564 | 929 | V929D | C | 6.6e-06 | 1.3e-05 | 7.9e-06 | 9.167E-06 | 3.383E-06 | 6.600E-06 | 1.300E-05 |
| 18565 | 929 | V929C | C | 1.1e-05 | 8.3e-06 | 1.2e-05 | 1.043E-05 | 1.914E-06 | 8.300E-06 | 1.200E-05 |
| 18566 | 929 | V929E | C | 6.6e-06 | 1.6e-05 | 1.3e-05 | 1.187E-05 | 4.801E-06 | 6.600E-06 | 1.600E-05 |
| 18567 | 929 | V929Q | C | 6.8e-06 | 7.4e-06 | 6.6e-06 | 6.933E-06 | 4.163E-07 | 6.600E-06 | 7.400E-06 |
| 18568 | 929 | V929G | C | 6.2e-06 | 6.5e-06 | 1e-05 | 7.567E-06 | 2.113E-06 | 6.200E-06 | 1.000E-05 |
| 18569 | 929 | V929H | C | 8.4e-06 | 5.7e-06 | 1.3e-05 | 9.033E-06 | 3.691E-06 | 5.700E-06 | 1.300E-05 |
| 18570 | 929 | V929I | C | 1e-05 | 4.7e-06 | 4.8e-06 | 6.500E-06 | 3.032E-06 | 4.700E-06 | 1.000E-05 |
| 18571 | 929 | V929L | C | 7.7e-06 | 5.5e-06 | 1.6e-05 | 9.733E-06 | 5.537E-06 | 5.500E-06 | 1.600E-05 |
| 18572 | 929 | V929K | C | 3.9e-06 | 1.2e-05 | 1.3e-05 | 9.633E-06 | 4.990E-06 | 3.900E-06 | 1.300E-05 |
| 18573 | 929 | V929M | C | 1.1e-05 | 8.6e-06 | 1.2e-05 | 1.053E-05 | 1.747E-06 | 8.600E-06 | 1.200E-05 |
| 18574 | 929 | V929F | C | 9.3e-06 | 9.7e-06 | 9.4e-06 | 9.467E-06 | 2.082E-07 | 9.300E-06 | 9.700E-06 |
| 18575 | 929 | V929P | C | 8e-06 | 5.6e-06 | 9.4e-06 | 7.667E-06 | 1.922E-06 | 5.600E-06 | 9.400E-06 |
| 18576 | 929 | V929S | C | 5e-06 | 6.2e-06 | 8.9e-06 | 6.700E-06 | 1.997E-06 | 5.000E-06 | 8.900E-06 |
| 18577 | 929 | V929T | C | 9.5e-06 | 5e-06 | 8.4e-06 | 7.633E-06 | 2.346E-06 | 5.000E-06 | 9.500E-06 |
| 18578 | 929 | V929W | C | 8.9e-06 | 4e-06 | 4.7e-06 | 5.867E-06 | 2.650E-06 | 4.000E-06 | 8.900E-06 |
| 18579 | 929 | V929Y | C | 9e-06 | 9.5e-06 | 8.7e-06 | 9.067E-06 | 4.041E-07 | 8.700E-06 | 9.500E-06 |
| 18580 | 929 | V929V | C | 9.3e-06 | 6.4e-06 | 1.3e-05 | 9.567E-06 | 3.308E-06 | 6.400E-06 | 1.300E-05 |
| 18581 | 930 | Y930A | C | 5.3e-06 | 5.4e-06 | 8.8e-06 | 6.500E-06 | 1.992E-06 | 5.300E-06 | 8.800E-06 |
| 18582 | 930 | Y930R | C | 1.3e-05 | 9.2e-06 | 1e-05 | 1.073E-05 | 2.003E-06 | 9.200E-06 | 1.300E-05 |
| 18583 | 930 | Y930N | C | 8.2e-06 | 1.9e-05 | 5.7e-06 | 1.097E-05 | 7.068E-06 | 5.700E-06 | 1.900E-05 |
| 18584 | 930 | Y930D | C | 1.4e-05 | 5.4e-06 | 1.1e-05 | 1.013E-05 | 4.365E-06 | 5.400E-06 | 1.400E-05 |
| 18585 | 930 | Y930C | C | 8e-06 | 5.6e-06 | 1.2e-05 | 8.533E-06 | 3.233E-06 | 5.600E-06 | 1.200E-05 |
| 18586 | 930 | Y930E | C | 9.9e-06 | 9.3e-06 | 1.7e-05 | 1.207E-05 | 4.283E-06 | 9.300E-06 | 1.700E-05 |
| 18587 | 930 | Y930Q | C | 1.5e-05 | 7.4e-06 | 9e-06 | 1.047E-05 | 4.007E-06 | 7.400E-06 | 1.500E-05 |
| 18588 | 930 | Y930G | C | 7.4e-06 | 8.7e-06 | 1e-05 | 8.700E-06 | 1.300E-06 | 7.400E-06 | 1.000E-05 |
| 18589 | 930 | Y930H | C | 6.7e-06 | 6.4e-06 | 7.8e-06 | 6.967E-06 | 7.371E-07 | 6.400E-06 | 7.800E-06 |
| 18590 | 930 | Y930I | C | 6.3e-06 | 5.8e-06 | 1.5e-05 | 9.033E-06 | 5.173E-06 | 5.800E-06 | 1.500E-05 |
| 18591 | 930 | Y930L | C | 4.7e-06 | 8.3e-06 | 7.8e-06 | 6.933E-06 | 1.950E-06 | 4.700E-06 | 8.300E-06 |
| 18592 | 930 | Y930K | C | 1e-05 | 6.4e-06 | 9.8e-06 | 8.733E-06 | 2.023E-06 | 6.400E-06 | 1.000E-05 |
| 18593 | 930 | Y930M | C | 9.1e-06 | 1.1e-05 | 1.1e-05 | 1.037E-05 | 1.097E-06 | 9.100E-06 | 1.100E-05 |
| 18594 | 930 | Y930F | C | 1.3e-05 | 6.8e-06 | 7e-06 | 8.933E-06 | 3.523E-06 | 6.800E-06 | 1.300E-05 |
| 18595 | 930 | Y930P | C | 8.8e-06 | 5.6e-06 | 1.1e-05 | 8.467E-06 | 2.715E-06 | 5.600E-06 | 1.100E-05 |
| 18596 | 930 | Y930S | C | 4.9e-06 | 4.7e-06 | 1e-05 | 6.533E-06 | 3.004E-06 | 4.700E-06 | 1.000E-05 |
| 18597 | 930 | Y930T | C | 6.5e-06 | 5.5e-06 | 4.8e-06 | 5.600E-06 | 8.544E-07 | 4.800E-06 | 6.500E-06 |
| 18598 | 930 | Y930W | C | 1e-05 | 9.8e-06 | 1e-05 | 9.933E-06 | 1.155E-07 | 9.800E-06 | 1.000E-05 |
| 18599 | 930 | Y930Y | C | 9.7e-06 | 8.8e-06 | 8e-06 | 8.833E-06 | 8.505E-07 | 8.000E-06 | 9.700E-06 |
| 18600 | 930 | Y930V | C | 9.3e-06 | 1.4e-05 | 8.6e-06 | 1.063E-05 | 2.937E-06 | 8.600E-06 | 1.400E-05 |
| 18601 | 931 | Y931A | C | 1.3e-05 | 9.5e-06 | 7.5e-06 | 1.000E-05 | 2.784E-06 | 7.500E-06 | 1.300E-05 |
| 18602 | 931 | Y931R | C | 6.5e-06 | 8.3e-06 | 1.2e-05 | 8.933E-06 | 2.804E-06 | 6.500E-06 | 1.200E-05 |
| 18603 | 931 | Y931N | C | 9.5e-06 | 7.3e-06 | 1.3e-05 | 9.933E-06 | 2.875E-06 | 7.300E-06 | 1.300E-05 |
| 18604 | 931 | Y931D | C | 9.8e-06 | 9.6e-06 | 1e-05 | 9.800E-06 | 2.000E-07 | 9.600E-06 | 1.000E-05 |
| 18605 | 931 | Y931C | C | 1.1e-05 | 4e-06 | 1.1e-05 | 8.667E-06 | 4.041E-06 | 4.000E-06 | 1.100E-05 |
| 18606 | 931 | Y931E | C | 1.5e-05 | 9.3e-06 | 8.4e-06 | 1.090E-05 | 3.579E-06 | 8.400E-06 | 1.500E-05 |
| 18607 | 931 | Y931Q | C | 1e-05 | 6.9e-06 | 6.8e-06 | 7.900E-06 | 1.819E-06 | 6.800E-06 | 1.000E-05 |
| 18608 | 931 | Y931G | C | 1.2e-05 | 8.7e-06 | 9.5e-06 | 1.007E-05 | 1.721E-06 | 8.700E-06 | 1.200E-05 |
| 18609 | 931 | Y931H | C | 1.4e-05 | 8.5e-06 | 1e-05 | 1.083E-05 | 2.843E-06 | 8.500E-06 | 1.400E-05 |
| 18610 | 931 | Y931I | C | 7e-06 | 1.1e-05 | 6.5e-06 | 8.167E-06 | 2.466E-06 | 6.500E-06 | 1.100E-05 |
| 18611 | 931 | Y931L | C | 7e-06 | 1.4e-05 | 6.6e-06 | 9.200E-06 | 4.162E-06 | 6.600E-06 | 1.400E-05 |
| 18612 | 931 | Y931K | C | 1e-05 | 1.1e-05 | 9.5e-06 | 1.017E-05 | 7.638E-07 | 9.500E-06 | 1.100E-05 |
| 18613 | 931 | Y931M | C | 5.4e-06 | 1.3e-05 | 6.5e-06 | 8.300E-06 | 4.107E-06 | 5.400E-06 | 1.300E-05 |
| 18614 | 931 | Y931F | C | 6.5e-06 | 7.4e-06 | 1.4e-05 | 9.300E-06 | 4.095E-06 | 6.500E-06 | 1.400E-05 |

| | | | | | | | | | | |
|---|---|---|---|---|---|---|---|---|---|---|
| 18615 | 931 | Y931P | C | 6.7e-06 | 6.1e-06 | 7e-06 | 6.600E-06 | 4.583E-07 | 6.100E-06 | 7.000E-06 |
| 18616 | 931 | Y931S | C | 6e-06 | 8.3e-06 | 4.9e-06 | 6.400E-06 | 1.735E-06 | 4.900E-06 | 8.300E-06 |
| 18617 | 931 | Y931T | C | 1.4e-05 | 5.8e-06 | 9e-06 | 9.600E-06 | 4.133E-06 | 5.800E-06 | 1.400E-05 |
| 18618 | 931 | Y931W | C | 9e-06 | 7.1e-06 | 8.7e-06 | 8.267E-06 | 1.021E-06 | 7.100E-06 | 9.000E-06 |
| 18619 | 931 | Y931Y | C | 1e-05 | 4.4e-06 | 1.3e-05 | 9.133E-06 | 4.365E-06 | 4.400E-06 | 1.300E-05 |
| 18620 | 931 | Y931V | C | 4.9e-06 | 6.6e-06 | 7.7e-06 | 6.400E-06 | 1.411E-06 | 4.900E-06 | 7.700E-06 |
| 18621 | 932 | C932A | C | 1e-05 | 1.1e-05 | 1.1e-05 | 1.067E-05 | 5.774E-07 | 1.000E-05 | 1.100E-05 |
| 18622 | 932 | C932R | C | 1.2e-05 | 6e-06 | 7e-06 | 8.333E-06 | 3.215E-06 | 6.000E-06 | 1.200E-05 |
| 18623 | 932 | C932N | C | 5.4e-06 | 6.6e-06 | 9.8e-06 | 7.267E-06 | 2.274E-06 | 5.400E-06 | 9.800E-06 |
| 18624 | 932 | C932D | C | 6.7e-06 | 8.6e-06 | 5.9e-06 | 7.067E-06 | 1.387E-06 | 5.900E-06 | 8.600E-06 |
| 18625 | 932 | C932C | C | 7.2e-06 | 8.2e-06 | 1.1e-05 | 8.800E-06 | 1.970E-06 | 7.200E-06 | 1.100E-05 |
| 18626 | 932 | C932E | C | 7e-06 | 6.8e-06 | 5.2e-06 | 6.333E-06 | 9.866E-07 | 5.200E-06 | 7.000E-06 |
| 18627 | 932 | C932Q | C | 3.6e-06 | 5.1e-06 | 5.9e-06 | 4.867E-06 | 1.168E-06 | 3.600E-06 | 5.900E-06 |
| 18628 | 932 | C932G | C | 6.6e-06 | 6.4e-06 | 1.3e-05 | 8.667E-06 | 3.754E-06 | 6.400E-06 | 1.300E-05 |
| 18629 | 932 | C932H | C | 1e-05 | 5.6e-06 | 9.4e-06 | 8.333E-06 | 2.386E-06 | 5.600E-06 | 1.000E-05 |
| 18630 | 932 | C932I | C | 1.4e-05 | 1.6e-05 | 1.2e-05 | 1.400E-05 | 2.000E-06 | 1.200E-05 | 1.600E-05 |
| 18631 | 932 | C932L | C | 6.1e-06 | 4.8e-06 | 1e-05 | 6.967E-06 | 2.706E-06 | 4.800E-06 | 1.000E-05 |
| 18632 | 932 | C932K | C | 8.7e-06 | 1.2e-05 | 8.6e-06 | 9.767E-06 | 1.935E-06 | 8.600E-06 | 1.200E-05 |
| 18633 | 932 | C932M | C | 9.5e-06 | 7.6e-06 | 8.3e-06 | 8.467E-06 | 9.609E-07 | 7.600E-06 | 9.500E-06 |
| 18634 | 932 | C932F | C | 1.2e-05 | 4.1e-06 | 1.1e-05 | 9.033E-06 | 4.302E-06 | 4.100E-06 | 1.200E-05 |
| 18635 | 932 | C932P | C | 5.5e-06 | 9.4e-06 | 9.3e-06 | 8.067E-06 | 2.223E-06 | 5.500E-06 | 9.400E-06 |
| 18636 | 932 | C932S | C | 5.7e-06 | 9.2e-06 | 1.3e-05 | 9.300E-06 | 3.651E-06 | 5.700E-06 | 1.300E-05 |
| 18637 | 932 | C932T | C | 9e-06 | 2.3e-06 | 6.1e-06 | 5.800E-06 | 3.360E-06 | 2.300E-06 | 9.000E-06 |
| 18638 | 932 | C932W | C | 7e-06 | 9.5e-06 | 8.1e-06 | 8.200E-06 | 1.253E-06 | 7.000E-06 | 9.500E-06 |
| 18639 | 932 | C932Y | C | 8e-06 | 5.7e-06 | 5.1e-06 | 6.267E-06 | 1.531E-06 | 5.100E-06 | 8.000E-06 |
| 18640 | 932 | C932V | C | 1.1e-05 | 1.4e-05 | 7e-06 | 1.067E-05 | 3.512E-06 | 7.000E-06 | 1.400E-05 |
| 18641 | 933 | A933A | C | 8e-06 | 7.8e-06 | 1e-05 | 8.600E-06 | 1.217E-06 | 7.800E-06 | 1.000E-05 |
| 18642 | 933 | A933R | C | 8.4e-06 | 1.1e-05 | 1.1e-05 | 1.013E-05 | 1.501E-06 | 8.400E-06 | 1.100E-05 |
| 18643 | 933 | A933N | C | 7.3e-06 | 8.7e-06 | 1.1e-05 | 9.000E-06 | 1.868E-06 | 7.300E-06 | 1.100E-05 |
| 18644 | 933 | A933D | C | 9.7e-06 | 9.6e-06 | 8e-06 | 9.100E-06 | 9.539E-07 | 8.000E-06 | 9.700E-06 |
| 18645 | 933 | A933C | C | 5.8e-06 | 6.2e-06 | 9e-06 | 7.000E-06 | 1.744E-06 | 5.800E-06 | 9.000E-06 |
| 18646 | 933 | A933E | C | 4.9e-06 | 1.3e-05 | 1.4e-05 | 1.063E-05 | 4.990E-06 | 4.900E-06 | 1.400E-05 |
| 18647 | 933 | A933Q | C | 5.5e-06 | 9.5e-06 | 8.7e-06 | 7.900E-06 | 2.117E-06 | 5.500E-06 | 9.500E-06 |
| 18648 | 933 | A933G | C | 7e-06 | 6.2e-06 | 1.2e-05 | 8.400E-06 | 3.143E-06 | 6.200E-06 | 1.200E-05 |
| 18649 | 933 | A933H | C | 5.5e-06 | 6.5e-06 | 9.9e-06 | 7.300E-06 | 2.307E-06 | 5.500E-06 | 9.900E-06 |
| 18650 | 933 | A933I | C | 6.7e-06 | 9.6e-06 | 8.4e-06 | 8.233E-06 | 1.457E-06 | 6.700E-06 | 9.600E-06 |
| 18651 | 933 | A933L | C | 9.4e-06 | 5.9e-06 | 6.6e-06 | 7.300E-06 | 1.852E-06 | 5.900E-06 | 9.400E-06 |
| 18652 | 933 | A933K | C | 8.6e-06 | 5.9e-06 | 8e-06 | 7.500E-06 | 1.418E-06 | 5.900E-06 | 8.600E-06 |
| 18653 | 933 | A933M | C | 8.9e-06 | 1.3e-05 | 1.1e-05 | 1.097E-05 | 2.050E-06 | 8.900E-06 | 1.300E-05 |
| 18654 | 933 | A933F | C | 1e-05 | 4.7e-06 | 4.9e-06 | 6.533E-06 | 3.004E-06 | 4.700E-06 | 1.000E-05 |
| 18655 | 933 | A933P | C | 9.5e-06 | 5.2e-06 | 8.2e-06 | 7.633E-06 | 2.205E-06 | 5.200E-06 | 9.500E-06 |
| 18656 | 933 | A933S | C | 8.3e-06 | 4.7e-06 | 1.3e-05 | 8.667E-06 | 4.162E-06 | 4.700E-06 | 1.300E-05 |
| 18657 | 933 | A933T | C | 1e-05 | 1.2e-05 | 6.5e-06 | 9.500E-06 | 2.784E-06 | 6.500E-06 | 1.200E-05 |
| 18658 | 933 | A933W | C | 1.1e-05 | 6e-06 | 1e-05 | 9.000E-06 | 2.646E-06 | 6.000E-06 | 1.100E-05 |
| 18659 | 933 | A933Y | C | 4.8e-06 | 6.3e-06 | 7e-06 | 6.033E-06 | 1.124E-06 | 4.800E-06 | 7.000E-06 |
| 18660 | 933 | A933V | C | 8.7e-06 | 9.7e-06 | 7.5e-06 | 8.633E-06 | 1.102E-06 | 7.500E-06 | 9.700E-06 |
| 18661 | 934 | R934A | C | 6.7e-06 | 9.5e-06 | 1.1e-05 | 9.067E-06 | 2.183E-06 | 6.700E-06 | 1.100E-05 |
| 18662 | 934 | R934R | C | 7.6e-06 | 1.1e-05 | 1e-05 | 9.533E-06 | 1.747E-06 | 7.600E-06 | 1.100E-05 |
| 18663 | 934 | R934N | C | 6.2e-06 | 1e-05 | 9.4e-06 | 8.533E-06 | 2.043E-06 | 6.200E-06 | 1.000E-05 |
| 18664 | 934 | R934D | C | 6.9e-06 | 9.7e-06 | 9.6e-06 | 8.733E-06 | 1.589E-06 | 6.900E-06 | 9.700E-06 |
| 18665 | 934 | R934C | C | 1.6e-05 | 5.6e-06 | 1.3e-05 | 1.153E-05 | 5.353E-06 | 5.600E-06 | 1.600E-05 |
| 18666 | 934 | R934E | C | 8.8e-06 | 9.4e-06 | 9.8e-06 | 9.333E-06 | 5.033E-07 | 8.800E-06 | 9.800E-06 |
| 18667 | 934 | R934Q | C | 1.3e-05 | 1.4e-05 | 5.3e-06 | 1.077E-05 | 4.761E-06 | 5.300E-06 | 1.400E-05 |
| 18668 | 934 | R934G | C | 6.7e-06 | 9.7e-06 | 6.7e-06 | 7.700E-06 | 1.732E-06 | 6.700E-06 | 9.700E-06 |
| 18669 | 934 | R934H | C | 3.3e-06 | 1.1e-05 | 9.6e-06 | 7.967E-06 | 4.102E-06 | 3.300E-06 | 1.100E-05 |
| 18670 | 934 | R934I | C | 1.3e-05 | 9.7e-06 | 1e-05 | 1.090E-05 | 1.825E-06 | 9.700E-06 | 1.300E-05 |
| 18671 | 934 | R934L | C | 9.4e-06 | 1e-05 | 1.3e-05 | 1.080E-05 | 1.929E-06 | 9.400E-06 | 1.300E-05 |
| 18672 | 934 | R934K | C | 8.5e-06 | 3.4e-06 | 7.1e-06 | 6.333E-06 | 2.635E-06 | 3.400E-06 | 8.500E-06 |
| 18673 | 934 | R934M | C | 6.1e-06 | 1.2e-05 | 1.1e-05 | 9.700E-06 | 3.158E-06 | 6.100E-06 | 1.200E-05 |
| 18674 | 934 | R934F | C | 8.3e-06 | 8.2e-06 | 7.7e-06 | 8.067E-06 | 3.215E-07 | 7.700E-06 | 8.300E-06 |
| 18675 | 934 | R934P | C | 7.9e-06 | 6.4e-06 | 9e-06 | 7.767E-06 | 1.305E-06 | 6.400E-06 | 9.000E-06 |
| 18676 | 934 | R934S | C | 1.1e-05 | 1e-05 | 3.4e-06 | 8.133E-06 | 4.130E-06 | 3.400E-06 | 1.100E-05 |
| 18677 | 934 | R934T | C | 8.8e-06 | 6.5e-06 | 7.1e-06 | 7.467E-06 | 1.193E-06 | 6.500E-06 | 8.800E-06 |
| 18678 | 934 | R934W | C | 7.2e-06 | 1.1e-05 | 1.3e-05 | 1.040E-05 | 2.946E-06 | 7.200E-06 | 1.300E-05 |
| 18679 | 934 | R934Y | C | 8.8e-06 | 1e-05 | 5.4e-06 | 8.067E-06 | 2.386E-06 | 5.400E-06 | 1.000E-05 |
| 18680 | 934 | R934V | C | 1.3e-05 | 6.8e-06 | 9.2e-06 | 9.667E-06 | 3.126E-06 | 6.800E-06 | 1.300E-05 |
| 18681 | 935 | N935A | C | 6.3e-06 | 6.1e-06 | 7.3e-06 | 6.567E-06 | 6.429E-07 | 6.100E-06 | 7.300E-06 |
| 18682 | 935 | N935R | C | 8.8e-06 | 6.5e-06 | 7.3e-06 | 7.533E-06 | 1.168E-06 | 6.500E-06 | 8.800E-06 |
| 18683 | 935 | N935N | C | 1.4e-05 | 4.6e-06 | 7.8e-06 | 8.800E-06 | 4.779E-06 | 4.600E-06 | 1.400E-05 |
| 18684 | 935 | N935D | C | 5.6e-06 | 1.4e-05 | 5.3e-06 | 8.300E-06 | 4.939E-06 | 5.300E-06 | 1.400E-05 |
| 18685 | 935 | N935C | C | 1.8e-05 | 7.5e-06 | 9.1e-06 | 1.153E-05 | 5.657E-06 | 7.500E-06 | 1.800E-05 |
| 18686 | 935 | N935E | C | 4.5e-06 | 1e-05 | 1e-05 | 8.167E-06 | 3.175E-06 | 4.500E-06 | 1.000E-05 |
| 18687 | 935 | N935Q | C | 5.2e-06 | 1.5e-05 | 1.2e-05 | 1.073E-05 | 5.021E-06 | 5.200E-06 | 1.500E-05 |
| 18688 | 935 | N935G | C | 9.3e-06 | 5.5e-06 | 1.5e-05 | 9.933E-06 | 4.782E-06 | 5.500E-06 | 1.500E-05 |
| 18689 | 935 | N935H | C | 4e-06 | 8.8e-06 | 4.6e-06 | 5.800E-06 | 2.615E-06 | 4.000E-06 | 8.800E-06 |
| 18690 | 935 | N935I | C | 1e-05 | 6.1e-06 | 5.6e-06 | 7.233E-06 | 2.409E-06 | 5.600E-06 | 1.000E-05 |
| 18691 | 935 | N935L | C | 5.9e-06 | 7e-06 | 1.2e-05 | 8.300E-06 | 3.251E-06 | 5.900E-06 | 1.200E-05 |
| 18692 | 935 | N935K | C | 7e-06 | 6.7e-06 | 9.5e-06 | 7.733E-06 | 1.537E-06 | 6.700E-06 | 9.500E-06 |
| 18693 | 935 | N935M | C | 5.8e-06 | 6.9e-06 | 1.4e-05 | 8.900E-06 | 4.451E-06 | 5.800E-06 | 1.400E-05 |
| 18694 | 935 | N935F | C | 6.3e-06 | 8.5e-06 | 8.5e-06 | 7.767E-06 | 1.270E-06 | 6.300E-06 | 8.500E-06 |
| 18695 | 935 | N935P | C | 9.2e-06 | 7e-06 | 7.5e-06 | 7.900E-06 | 1.153E-06 | 7.000E-06 | 9.200E-06 |
| 18696 | 935 | N935S | C | 7.7e-06 | 4.6e-06 | 7.2e-06 | 6.500E-06 | 1.664E-06 | 4.600E-06 | 7.700E-06 |
| 18697 | 935 | N935T | C | 9.5e-06 | 1.2e-05 | 6.9e-06 | 9.467E-06 | 2.550E-06 | 6.900E-06 | 1.200E-05 |
| 18698 | 935 | N935W | C | 7e-06 | 7.9e-06 | 9.8e-06 | 8.233E-06 | 1.429E-06 | 7.000E-06 | 9.800E-06 |
| 18699 | 935 | N935Y | C | 9e-06 | 8.3e-06 | 1.1e-05 | 9.433E-06 | 1.401E-06 | 8.300E-06 | 1.100E-05 |

| | | | | | | | | | | |
|---|---|---|---|---|---|---|---|---|---|---|
| 18700 | 935 | N935V | C | 1.7e-05 | 6.5e-06 | 7.8e-06 | 1.043E-05 | 5.724E-06 | 6.500E-06 | 1.700E-05 |
| 18701 | 936 | L936A | C | 7.4e-06 | 9.5e-06 | 7.8e-06 | 8.233E-06 | 1.115E-06 | 7.400E-06 | 9.500E-06 |
| 18702 | 936 | L936R | C | 7.7e-06 | 8.7e-06 | 6.7e-06 | 7.700E-06 | 1.000E-06 | 6.700E-06 | 8.700E-06 |
| 18703 | 936 | L936N | C | 3.7e-06 | 5.9e-06 | 4.9e-06 | 4.833E-06 | 1.102E-06 | 3.700E-06 | 5.900E-06 |
| 18704 | 936 | L936D | C | 7.9e-06 | 1.2e-05 | 9.2e-06 | 9.700E-06 | 2.095E-06 | 7.900E-06 | 1.200E-05 |
| 18705 | 936 | L936C | C | 9.3e-06 | 1.1e-05 | 6.1e-06 | 8.800E-06 | 2.488E-06 | 6.100E-06 | 1.100E-05 |
| 18706 | 936 | L936E | C | 1.2e-05 | 9.2e-06 | 4.6e-06 | 8.600E-06 | 3.736E-06 | 4.600E-06 | 1.200E-05 |
| 18707 | 936 | L936Q | C | 7.9e-06 | 9.2e-06 | 9.1e-06 | 8.733E-06 | 7.234E-07 | 7.900E-06 | 9.200E-06 |
| 18708 | 936 | L936G | C | 6.2e-06 | 8.2e-06 | 6.9e-06 | 7.100E-06 | 1.015E-06 | 6.200E-06 | 8.200E-06 |
| 18709 | 936 | L936H | C | 6.4e-06 | 6.9e-06 | 6.7e-06 | 6.667E-06 | 2.517E-07 | 6.400E-06 | 6.900E-06 |
| 18710 | 936 | L936I | C | 9.6e-06 | 9.5e-06 | 6.9e-06 | 8.667E-06 | 1.531E-06 | 6.900E-06 | 9.600E-06 |
| 18711 | 936 | L936L | C | 1e-05 | 9.3e-06 | 5.7e-06 | 8.333E-06 | 2.307E-06 | 5.700E-06 | 1.000E-05 |
| 18712 | 936 | L936K | C | 1.2e-05 | 1.1e-05 | 8.5e-06 | 1.050E-05 | 1.803E-06 | 8.500E-06 | 1.200E-05 |
| 18713 | 936 | L936M | C | 9.2e-06 | 9.1e-06 | 7.1e-06 | 8.467E-06 | 1.185E-06 | 7.100E-06 | 9.200E-06 |
| 18714 | 936 | L936F | C | 4.5e-06 | 8.7e-06 | 9.3e-06 | 7.500E-06 | 2.615E-06 | 4.500E-06 | 9.300E-06 |
| 18715 | 936 | L936P | C | 1e-05 | 6.9e-06 | 5.9e-06 | 7.600E-06 | 2.138E-06 | 5.900E-06 | 1.000E-05 |
| 18716 | 936 | L936S | C | 6.1e-06 | 1e-05 | 5.8e-06 | 7.300E-06 | 2.343E-06 | 5.800E-06 | 1.000E-05 |
| 18717 | 936 | L936T | C | 5.7e-06 | 6.7e-06 | 9.9e-06 | 7.433E-06 | 2.194E-06 | 5.700E-06 | 9.900E-06 |
| 18718 | 936 | L936W | C | 1e-05 | 1.1e-05 | 8.3e-06 | 9.767E-06 | 1.365E-06 | 8.300E-06 | 1.100E-05 |
| 18719 | 936 | L936Y | C | 6.3e-06 | 7e-06 | 7e-06 | 6.767E-06 | 4.041E-07 | 6.300E-06 | 7.000E-06 |
| 18720 | 936 | L936V | C | 9e-06 | 6.6e-06 | 1e-05 | 8.533E-06 | 1.747E-06 | 6.600E-06 | 1.000E-05 |
| 18721 | 937 | G937A | C | 1e-05 | 8e-06 | 7.2e-06 | 8.400E-06 | 1.442E-06 | 7.200E-06 | 1.000E-05 |
| 18722 | 937 | G937R | C | 7.5e-06 | 1.2e-05 | 6.8e-06 | 8.767E-06 | 2.822E-06 | 6.800E-06 | 1.200E-05 |
| 18723 | 937 | G937N | C | 6.9e-06 | 1.1e-05 | 6.7e-06 | 8.200E-06 | 2.427E-06 | 6.700E-06 | 1.100E-05 |
| 18724 | 937 | G937D | C | 4.7e-06 | 5.1e-06 | 9.5e-06 | 6.433E-06 | 2.663E-06 | 4.700E-06 | 9.500E-06 |
| 18725 | 937 | G937C | C | 5.6e-06 | 4.7e-06 | 6.6e-06 | 5.633E-06 | 9.504E-07 | 4.700E-06 | 6.600E-06 |
| 18726 | 937 | G937E | C | 4.8e-06 | 9e-06 | 4.9e-06 | 6.233E-06 | 2.397E-06 | 4.800E-06 | 9.000E-06 |
| 18727 | 937 | G937Q | C | 7e-06 | 7e-06 | 9.6e-06 | 7.867E-06 | 1.501E-06 | 7.000E-06 | 9.600E-06 |
| 18728 | 937 | G937G | C | 5.7e-06 | 6e-06 | 1e-05 | 7.233E-06 | 2.401E-06 | 5.700E-06 | 1.000E-05 |
| 18729 | 937 | G937H | C | 5.3e-06 | 4.4e-06 | 7e-06 | 5.567E-06 | 1.320E-06 | 4.400E-06 | 7.000E-06 |
| 18730 | 937 | G937I | C | 5.7e-06 | 7.4e-06 | 7.6e-06 | 6.900E-06 | 1.044E-06 | 5.700E-06 | 7.600E-06 |
| 18731 | 937 | G937L | C | 9.8e-06 | 1e-05 | 6.3e-06 | 8.700E-06 | 2.081E-06 | 6.300E-06 | 1.000E-05 |
| 18732 | 937 | G937K | C | 1.4e-05 | 9.7e-06 | 7.1e-06 | 1.027E-05 | 3.485E-06 | 7.100E-06 | 1.400E-05 |
| 18733 | 937 | G937M | C | 1.2e-05 | 8.3e-06 | 5.3e-06 | 8.533E-06 | 3.356E-06 | 5.300E-06 | 1.200E-05 |
| 18734 | 937 | G937F | C | 1.6e-05 | 1e-05 | 1.1e-05 | 1.233E-05 | 3.215E-06 | 1.000E-05 | 1.600E-05 |
| 18735 | 937 | G937P | C | 7.5e-06 | 7.4e-06 | 1.3e-05 | 9.300E-06 | 3.205E-06 | 7.400E-06 | 1.300E-05 |
| 18736 | 937 | G937S | C | 4.8e-06 | 6.6e-06 | 4.7e-06 | 5.367E-06 | 1.069E-06 | 4.700E-06 | 6.600E-06 |
| 18737 | 937 | G937T | C | 9.1e-06 | 3.2e-06 | 8.6e-06 | 6.967E-06 | 3.272E-06 | 3.200E-06 | 9.100E-06 |
| 18738 | 937 | G937W | C | 6.7e-06 | 8e-06 | 5.8e-06 | 6.833E-06 | 1.106E-06 | 5.800E-06 | 8.000E-06 |
| 18739 | 937 | G937Y | C | 7e-06 | 9.5e-06 | 6.9e-06 | 7.800E-06 | 1.473E-06 | 6.900E-06 | 9.500E-06 |
| 18740 | 937 | G937V | C | 1.2e-05 | 1.1e-05 | 6.9e-06 | 9.967E-06 | 2.702E-06 | 6.900E-06 | 1.200E-05 |
| 18741 | 938 | P938A | C | 1.3e-05 | 1.7e-05 | 5.1e-06 | 1.170E-05 | 6.056E-06 | 5.100E-06 | 1.700E-05 |
| 18742 | 938 | P938R | C | 9.8e-06 | 6.6e-06 | 1.2e-05 | 9.467E-06 | 2.715E-06 | 6.600E-06 | 1.200E-05 |
| 18743 | 938 | P938N | C | 8.2e-06 | 6e-06 | 4.6e-06 | 6.267E-06 | 1.815E-06 | 4.600E-06 | 8.200E-06 |
| 18744 | 938 | P938D | C | 1e-05 | 1.1e-05 | 6.1e-06 | 9.033E-06 | 2.589E-06 | 6.100E-06 | 1.100E-05 |
| 18745 | 938 | P938C | C | 7.8e-06 | 1e-05 | 5.5e-06 | 7.767E-06 | 2.250E-06 | 5.500E-06 | 1.000E-05 |
| 18746 | 938 | P938E | C | 4.9e-06 | 7e-06 | 4.5e-06 | 5.467E-06 | 1.343E-06 | 4.500E-06 | 7.000E-06 |
| 18747 | 938 | P938Q | C | 3.1e-06 | 5.1e-06 | 1e-05 | 6.067E-06 | 3.550E-06 | 3.100E-06 | 1.000E-05 |
| 18748 | 938 | P938G | C | 1e-05 | 9.2e-06 | 1.2e-05 | 1.040E-05 | 1.442E-06 | 9.200E-06 | 1.200E-05 |
| 18749 | 938 | P938H | C | 6.2e-06 | 5.2e-06 | 6.7e-06 | 6.033E-06 | 7.638E-07 | 5.200E-06 | 6.700E-06 |
| 18750 | 938 | P938I | C | 8.3e-06 | 5.9e-06 | 8.8e-06 | 7.667E-06 | 1.550E-06 | 5.900E-06 | 8.800E-06 |
| 18751 | 938 | P938L | C | 6.9e-06 | 9.6e-06 | 6.3e-06 | 7.600E-06 | 1.758E-06 | 6.300E-06 | 9.600E-06 |
| 18752 | 938 | P938K | C | 9.3e-06 | 6.2e-06 | 6.1e-06 | 7.200E-06 | 1.819E-06 | 6.100E-06 | 9.300E-06 |
| 18753 | 938 | P938M | C | 4.7e-06 | 9.1e-06 | 6.6e-06 | 6.800E-06 | 2.207E-06 | 4.700E-06 | 9.100E-06 |
| 18754 | 938 | P938F | C | 6.2e-06 | 8.6e-06 | 7.1e-06 | 7.300E-06 | 1.212E-06 | 6.200E-06 | 8.600E-06 |
| 18755 | 938 | P938P | C | 5.8e-06 | 8.4e-06 | 5.8e-06 | 6.667E-06 | 1.501E-06 | 5.800E-06 | 8.400E-06 |
| 18756 | 938 | P938S | C | 6.4e-06 | 4.7e-06 | 7.1e-06 | 6.067E-06 | 1.234E-06 | 4.700E-06 | 7.100E-06 |
| 18757 | 938 | P938T | C | 9.8e-06 | 6.6e-06 | 5.2e-06 | 7.200E-06 | 2.358E-06 | 5.200E-06 | 9.800E-06 |
| 18758 | 938 | P938W | C | 6.5e-06 | 5.9e-06 | 8.2e-06 | 6.867E-06 | 1.193E-06 | 5.900E-06 | 8.200E-06 |
| 18759 | 938 | P938Y | C | 6.5e-06 | 9.4e-06 | 5.2e-06 | 7.033E-06 | 2.150E-06 | 5.200E-06 | 9.400E-06 |
| 18760 | 938 | P938V | C | 1e-05 | 9.7e-06 | 9.5e-06 | 9.733E-06 | 2.517E-07 | 9.500E-06 | 1.000E-05 |
| 18761 | 939 | S939A | C | 1.4e-05 | 5.5e-06 | 6e-06 | 8.500E-06 | 4.770E-06 | 5.500E-06 | 1.400E-05 |
| 18762 | 939 | S939R | C | 8.2e-06 | 1.3e-05 | 1.5e-05 | 1.207E-05 | 3.495E-06 | 8.200E-06 | 1.500E-05 |
| 18763 | 939 | S939N | C | 4.5e-06 | 5.8e-06 | 5.1e-06 | 5.133E-06 | 6.506E-07 | 4.500E-06 | 5.800E-06 |
| 18764 | 939 | S939D | C | 5.8e-06 | 7.6e-06 | 6.7e-06 | 6.700E-06 | 9.000E-07 | 5.800E-06 | 7.600E-06 |
| 18765 | 939 | S939C | C | 9.9e-06 | 1.2e-05 | 1e-05 | 1.063E-05 | 1.185E-06 | 9.900E-06 | 1.200E-05 |
| 18766 | 939 | S939E | C | 4.9e-06 | 8.2e-06 | 4.6e-06 | 5.900E-06 | 1.997E-06 | 4.600E-06 | 8.200E-06 |
| 18767 | 939 | S939Q | C | 1e-05 | 6.8e-06 | 8.6e-06 | 8.467E-06 | 1.604E-06 | 6.800E-06 | 1.000E-05 |
| 18768 | 939 | S939G | C | 9.8e-06 | 1.1e-05 | 6.1e-06 | 8.967E-06 | 2.554E-06 | 6.100E-06 | 1.100E-05 |
| 18769 | 939 | S939H | C | 1e-05 | 7.5e-06 | 7.6e-06 | 8.367E-06 | 1.415E-06 | 7.500E-06 | 1.000E-05 |
| 18770 | 939 | S939I | C | 7.8e-06 | 5.7e-06 | 4.7e-06 | 6.067E-06 | 1.582E-06 | 4.700E-06 | 7.800E-06 |
| 18771 | 939 | S939L | C | 1.2e-05 | 7.7e-06 | 3.9e-06 | 7.867E-06 | 4.053E-06 | 3.900E-06 | 1.200E-05 |
| 18772 | 939 | S939K | C | 6.9e-06 | 7.9e-06 | 8.8e-06 | 7.867E-06 | 9.504E-07 | 6.900E-06 | 8.800E-06 |
| 18773 | 939 | S939M | C | 1.1e-05 | 1.2e-05 | 9.9e-06 | 1.097E-05 | 1.050E-06 | 9.900E-06 | 1.200E-05 |
| 18774 | 939 | S939F | C | 1.3e-05 | 6.8e-06 | 9.8e-06 | 9.867E-06 | 3.101E-06 | 6.800E-06 | 1.300E-05 |
| 18775 | 939 | S939P | C | 7.7e-06 | 5.5e-06 | 5.7e-06 | 6.300E-06 | 1.217E-06 | 5.500E-06 | 7.700E-06 |
| 18776 | 939 | S939S | C | 9.4e-06 | 7.7e-06 | 1.1e-05 | 9.367E-06 | 1.650E-06 | 7.700E-06 | 1.100E-05 |
| 18777 | 939 | S939T | C | 1.3e-05 | 8.6e-06 | 8.3e-06 | 9.967E-06 | 2.631E-06 | 8.300E-06 | 1.300E-05 |
| 18778 | 939 | S939W | C | 6.2e-06 | 9e-06 | 1.6e-05 | 1.040E-05 | 5.048E-06 | 6.200E-06 | 1.600E-05 |
| 18779 | 939 | S939Y | C | 8.5e-06 | 4.9e-06 | 1.3e-05 | 8.800E-06 | 4.058E-06 | 4.900E-06 | 1.300E-05 |
| 18780 | 939 | S939V | C | 7.9e-06 | 1.2e-05 | 1.1e-05 | 1.030E-05 | 2.138E-06 | 7.900E-06 | 1.200E-05 |
| 18781 | 940 | F940A | C | 8.6e-06 | 6.1e-06 | 4.4e-06 | 6.367E-06 | 2.113E-06 | 4.400E-06 | 8.600E-06 |
| 18782 | 940 | F940R | C | 4.6e-06 | 6.3e-06 | 1.3e-05 | 7.967E-06 | 4.441E-06 | 4.600E-06 | 1.300E-05 |
| 18783 | 940 | F940N | C | 9.6e-06 | 7.3e-06 | 3.3e-06 | 6.733E-06 | 3.188E-06 | 3.300E-06 | 9.600E-06 |
| 18784 | 940 | F940D | C | 8.8e-06 | 1e-05 | 8.5e-06 | 9.100E-06 | 7.937E-07 | 8.500E-06 | 1.000E-05 |

| | | | | | | | | | | |
|---|---|---|---|---|---|---|---|---|---|---|
| 18785 | 940 | F940C | C | 7.6e-06 | 6.1e-06 | 5.1e-06 | 6.267E-06 | 1.258E-06 | 5.100E-06 | 7.600E-06 |
| 18786 | 940 | F940E | C | 9e-06 | 9.1e-06 | 7.1e-06 | 8.400E-06 | 1.127E-06 | 7.100E-06 | 9.100E-06 |
| 18787 | 940 | F940Q | C | 8.7e-06 | 6.5e-06 | 6e-06 | 7.067E-06 | 1.436E-06 | 6.000E-06 | 8.700E-06 |
| 18788 | 940 | F940G | C | 7.8e-06 | 1e-05 | 7.8e-06 | 8.533E-06 | 1.270E-06 | 7.800E-06 | 1.000E-05 |
| 18789 | 940 | F940H | C | 1.3e-05 | 8.2e-06 | 9.7e-06 | 1.030E-05 | 2.456E-06 | 8.200E-06 | 1.300E-05 |
| 18790 | 940 | F940I | C | 1e-05 | 1.2e-05 | 6.4e-06 | 9.467E-06 | 2.838E-06 | 6.400E-06 | 1.200E-05 |
| 18791 | 940 | F940L | C | 1.1e-05 | 8.9e-06 | 1.2e-05 | 1.063E-05 | 1.582E-06 | 8.900E-06 | 1.200E-05 |
| 18792 | 940 | F940K | C | 9.4e-06 | 9.5e-06 | 6.8e-06 | 8.567E-06 | 1.531E-06 | 6.800E-06 | 9.500E-06 |
| 18793 | 940 | F940M | C | 9.6e-06 | 8e-06 | 1.2e-05 | 9.867E-06 | 2.013E-06 | 8.000E-06 | 1.200E-05 |
| 18794 | 940 | F940F | C | 8.1e-06 | 7.3e-06 | 8.8e-06 | 8.067E-06 | 7.506E-07 | 7.300E-06 | 8.800E-06 |
| 18795 | 940 | F940P | C | 4.4e-06 | 9.8e-06 | 7.1e-06 | 7.100E-06 | 2.700E-06 | 4.400E-06 | 9.800E-06 |
| 18796 | 940 | F940S | C | 7e-06 | 5.2e-06 | 8.6e-06 | 6.933E-06 | 1.701E-06 | 5.200E-06 | 8.600E-06 |
| 18797 | 940 | F940T | C | 7e-06 | 1e-05 | 5.8e-06 | 7.600E-06 | 2.163E-06 | 5.800E-06 | 1.000E-05 |
| 18798 | 940 | F940W | C | 4.3e-06 | 8.8e-06 | 8.4e-06 | 7.167E-06 | 2.491E-06 | 4.300E-06 | 8.800E-06 |
| 18799 | 940 | F940Y | C | 5.2e-06 | 8e-06 | 5.5e-06 | 6.233E-06 | 1.537E-06 | 5.200E-06 | 8.000E-06 |
| 18800 | 940 | F940V | C | 5.8e-06 | 6.9e-06 | 6.5e-06 | 6.400E-06 | 5.568E-07 | 5.800E-06 | 6.900E-06 |
| 18801 | 941 | Y941A | C | 1.1e-05 | 1.3e-05 | 8.6e-06 | 1.087E-05 | 2.203E-06 | 8.600E-06 | 1.300E-05 |
| 18802 | 941 | Y941R | C | 1.5e-05 | 1.1e-05 | 4.3e-06 | 1.010E-05 | 5.406E-06 | 4.300E-06 | 1.500E-05 |
| 18803 | 941 | Y941N | C | 9.6e-06 | 6.8e-06 | 5e-06 | 7.133E-06 | 2.318E-06 | 5.000E-06 | 9.600E-06 |
| 18804 | 941 | Y941D | C | 6.6e-06 | 7.4e-06 | 6.1e-06 | 6.700E-06 | 6.557E-07 | 6.100E-06 | 7.400E-06 |
| 18805 | 941 | Y941C | C | 1.4e-05 | 4.7e-06 | 9.8e-06 | 9.500E-06 | 4.657E-06 | 4.700E-06 | 1.400E-05 |
| 18806 | 941 | Y941E | C | 8.6e-06 | 5.2e-06 | 1.1e-05 | 8.267E-06 | 2.914E-06 | 5.200E-06 | 1.100E-05 |
| 18807 | 941 | Y941Q | C | 6.9e-06 | 1.5e-05 | 3.1e-06 | 8.333E-06 | 6.078E-06 | 3.100E-06 | 1.500E-05 |
| 18808 | 941 | Y941G | C | 5.8e-06 | 4.1e-06 | 9e-06 | 6.300E-06 | 2.488E-06 | 4.100E-06 | 9.000E-06 |
| 18809 | 941 | Y941H | C | 9.1e-06 | 4.6e-06 | 6.6e-06 | 6.767E-06 | 2.255E-06 | 4.600E-06 | 9.100E-06 |
| 18810 | 941 | Y941I | C | 1.2e-05 | 8.1e-06 | 1e-05 | 1.003E-05 | 1.950E-06 | 8.100E-06 | 1.200E-05 |
| 18811 | 941 | Y941L | C | 9.1e-06 | 1.5e-05 | 8e-06 | 1.070E-05 | 3.764E-06 | 8.000E-06 | 1.500E-05 |
| 18812 | 941 | Y941K | C | 1.4e-05 | 8.6e-06 | 5.8e-06 | 9.467E-06 | 4.168E-06 | 5.800E-06 | 1.400E-05 |
| 18813 | 941 | Y941M | C | 5.4e-06 | 9.5e-06 | 5.1e-06 | 6.667E-06 | 2.458E-06 | 5.100E-06 | 9.500E-06 |
| 18814 | 941 | Y941F | C | 1.2e-05 | 1.3e-05 | 5.5e-06 | 1.017E-05 | 4.072E-06 | 5.500E-06 | 1.300E-05 |
| 18815 | 941 | Y941P | C | 6.2e-06 | 4.4e-06 | 1.7e-05 | 9.200E-06 | 6.815E-06 | 4.400E-06 | 1.700E-05 |
| 18816 | 941 | Y941S | C | 6.3e-06 | 7.1e-06 | 9e-06 | 7.467E-06 | 1.387E-06 | 6.300E-06 | 9.000E-06 |
| 18817 | 941 | Y941T | C | 6.6e-06 | 7.3e-06 | 4.8e-06 | 6.233E-06 | 1.290E-06 | 4.800E-06 | 7.300E-06 |
| 18818 | 941 | Y941W | C | 9.2e-06 | 1e-05 | 1.2e-05 | 1.040E-05 | 1.442E-06 | 9.200E-06 | 1.200E-05 |
| 18819 | 941 | Y941Y | C | 6.2e-06 | 8.3e-06 | 4.6e-06 | 6.367E-06 | 1.856E-06 | 4.600E-06 | 8.300E-06 |
| 18820 | 941 | Y941V | C | 5e-06 | 4.9e-06 | 1e-05 | 6.633E-06 | 2.916E-06 | 4.900E-06 | 1.000E-05 |
| 18821 | 942 | F942A | C | 6.8e-06 | 4.7e-06 | 8.7e-06 | 6.733E-06 | 2.001E-06 | 4.700E-06 | 8.700E-06 |
| 18822 | 942 | F942R | C | 7.5e-06 | 8.6e-06 | 1.1e-05 | 9.033E-06 | 1.790E-06 | 7.500E-06 | 1.100E-05 |
| 18823 | 942 | F942N | C | 7.5e-06 | 7.2e-06 | 8.3e-06 | 7.667E-06 | 5.686E-07 | 7.200E-06 | 8.300E-06 |
| 18824 | 942 | F942D | C | 5.8e-06 | 1e-05 | 1.2e-05 | 9.267E-06 | 3.164E-06 | 5.800E-06 | 1.200E-05 |
| 18825 | 942 | F942C | C | 1.2e-05 | 6.5e-06 | 8.6e-06 | 9.033E-06 | 2.775E-06 | 6.500E-06 | 1.200E-05 |
| 18826 | 942 | F942E | C | 1.2e-05 | 5.6e-06 | 1e-05 | 9.200E-06 | 3.274E-06 | 5.600E-06 | 1.200E-05 |
| 18827 | 942 | F942Q | C | 8.4e-06 | 1e-05 | 9e-06 | 9.133E-06 | 8.083E-07 | 8.400E-06 | 1.000E-05 |
| 18828 | 942 | F942G | C | 8.4e-06 | 5.3e-06 | 1.3e-05 | 8.900E-06 | 3.874E-06 | 5.300E-06 | 1.300E-05 |
| 18829 | 942 | F942H | C | 1.1e-05 | 7.3e-06 | 5e-06 | 7.767E-06 | 3.027E-06 | 5.000E-06 | 1.100E-05 |
| 18830 | 942 | F942I | C | 8.4e-06 | 8.7e-06 | 6.2e-06 | 7.767E-06 | 1.365E-06 | 6.200E-06 | 8.700E-06 |
| 18831 | 942 | F942L | C | 9.4e-06 | 8.8e-06 | 8.5e-06 | 8.900E-06 | 4.583E-07 | 8.500E-06 | 9.400E-06 |
| 18832 | 942 | F942K | C | 7.8e-06 | 7e-06 | 7e-06 | 7.267E-06 | 4.619E-07 | 7.000E-06 | 7.800E-06 |
| 18833 | 942 | F942M | C | 9.9e-06 | 4.6e-06 | 1.4e-05 | 9.500E-06 | 4.713E-06 | 4.600E-06 | 1.400E-05 |
| 18834 | 942 | F942F | C | 1.3e-05 | 6e-06 | 9.3e-06 | 9.433E-06 | 3.502E-06 | 6.000E-06 | 1.300E-05 |
| 18835 | 942 | F942P | C | 1e-05 | 6.5e-06 | 6.4e-06 | 7.633E-06 | 2.050E-06 | 6.400E-06 | 1.000E-05 |
| 18836 | 942 | F942S | C | 5.4e-06 | 6.6e-06 | 1.3e-05 | 8.333E-06 | 4.086E-06 | 5.400E-06 | 1.300E-05 |
| 18837 | 942 | F942T | C | 8.3e-06 | 5.7e-06 | 8.5e-06 | 7.500E-06 | 1.562E-06 | 5.700E-06 | 8.500E-06 |
| 18838 | 942 | F942W | C | 7.5e-06 | 3.5e-06 | 5.8e-06 | 5.600E-06 | 2.007E-06 | 3.500E-06 | 7.500E-06 |
| 18839 | 942 | F942Y | C | 5.6e-06 | 9e-06 | 6.4e-06 | 7.000E-06 | 1.778E-06 | 5.600E-06 | 9.000E-06 |
| 18840 | 942 | F942V | C | 9.5e-06 | 9.8e-06 | 1.3e-05 | 1.077E-05 | 1.940E-06 | 9.500E-06 | 1.300E-05 |
| 18841 | 943 | D943A | C | 9e-06 | 8.9e-06 | 1.4e-05 | 1.063E-05 | 2.916E-06 | 8.900E-06 | 1.400E-05 |
| 18842 | 943 | D943R | C | 5.6e-06 | 1.3e-05 | 6.7e-06 | 8.433E-06 | 3.993E-06 | 5.600E-06 | 1.300E-05 |
| 18843 | 943 | D943N | C | 1.3e-05 | 7.8e-06 | 6e-06 | 8.933E-06 | 3.635E-06 | 6.000E-06 | 1.300E-05 |
| 18844 | 943 | D943D | C | 6.6e-06 | 6.5e-06 | 6.3e-06 | 6.467E-06 | 1.528E-07 | 6.300E-06 | 6.600E-06 |
| 18845 | 943 | D943C | C | 9e-06 | 7.8e-06 | 1.2e-05 | 9.600E-06 | 2.163E-06 | 7.800E-06 | 1.200E-05 |
| 18846 | 943 | D943E | C | 1.1e-05 | 7.2e-06 | 1e-05 | 9.400E-06 | 1.970E-06 | 7.200E-06 | 1.100E-05 |
| 18847 | 943 | D943Q | C | 7.1e-06 | 7.5e-06 | 4.6e-06 | 6.400E-06 | 1.572E-06 | 4.600E-06 | 7.500E-06 |
| 18848 | 943 | D943G | C | 9.1e-06 | 1.3e-05 | 9.4e-06 | 1.050E-05 | 2.170E-06 | 9.100E-06 | 1.300E-05 |
| 18849 | 943 | D943H | C | 7.5e-06 | 9.1e-06 | 6.4e-06 | 7.667E-06 | 1.358E-06 | 6.400E-06 | 9.100E-06 |
| 18850 | 943 | D943I | C | 1.3e-05 | 8.1e-06 | 1e-05 | 1.037E-05 | 2.470E-06 | 8.100E-06 | 1.300E-05 |
| 18851 | 943 | D943L | C | 9.2e-06 | 5.2e-06 | 5.7e-06 | 6.700E-06 | 2.179E-06 | 5.200E-06 | 9.200E-06 |
| 18852 | 943 | D943K | C | 8.3e-06 | 7e-06 | 3.9e-06 | 6.400E-06 | 2.261E-06 | 3.900E-06 | 8.300E-06 |
| 18853 | 943 | D943M | C | 6.9e-06 | 1.2e-05 | 7.7e-06 | 8.867E-06 | 2.743E-06 | 6.900E-06 | 1.200E-05 |
| 18854 | 943 | D943F | C | 9.9e-06 | 6.2e-06 | 3.4e-06 | 6.500E-06 | 3.260E-06 | 3.400E-06 | 9.900E-06 |
| 18855 | 943 | D943P | C | 5.1e-06 | 6.2e-06 | 9e-06 | 6.767E-06 | 2.011E-06 | 5.100E-06 | 9.000E-06 |
| 18856 | 943 | D943S | C | 6.3e-06 | 9e-06 | 6.1e-06 | 7.133E-06 | 1.620E-06 | 6.100E-06 | 9.000E-06 |
| 18857 | 943 | D943T | C | 6.2e-06 | 8.5e-06 | 6e-06 | 6.900E-06 | 1.389E-06 | 6.000E-06 | 8.500E-06 |
| 18858 | 943 | D943W | C | 9e-06 | 4.2e-06 | 5.6e-06 | 6.267E-06 | 2.468E-06 | 4.200E-06 | 9.000E-06 |
| 18859 | 943 | D943Y | C | 9.7e-06 | 5.2e-06 | 6.2e-06 | 7.033E-06 | 2.363E-06 | 5.200E-06 | 9.700E-06 |
| 18860 | 943 | D943V | C | 5.1e-06 | 8.5e-06 | 8.2e-06 | 7.267E-06 | 1.882E-06 | 5.100E-06 | 8.500E-06 |
| 18861 | 944 | Y944A | C | 7.4e-06 | 8.6e-06 | 6.6e-06 | 7.533E-06 | 1.007E-06 | 6.600E-06 | 8.600E-06 |
| 18862 | 944 | Y944R | C | 5.3e-06 | 1.3e-05 | 9.4e-06 | 9.233E-06 | 3.853E-06 | 5.300E-06 | 1.300E-05 |
| 18863 | 944 | Y944N | C | 6.6e-06 | 1e-05 | 6.1e-06 | 7.567E-06 | 2.122E-06 | 6.100E-06 | 1.000E-05 |
| 18864 | 944 | Y944D | C | 1e-05 | 8.3e-06 | 7.7e-06 | 8.667E-06 | 1.193E-06 | 7.700E-06 | 1.000E-05 |
| 18865 | 944 | Y944C | C | 5.8e-06 | 4.2e-06 | 1.3e-05 | 7.667E-06 | 4.688E-06 | 4.200E-06 | 1.300E-05 |
| 18866 | 944 | Y944E | C | 4.7e-06 | 9.4e-06 | 7.2e-06 | 7.100E-06 | 2.352E-06 | 4.700E-06 | 9.400E-06 |
| 18867 | 944 | Y944Q | C | 1.1e-05 | 1.3e-05 | 8.2e-06 | 1.073E-05 | 2.411E-06 | 8.200E-06 | 1.300E-05 |
| 18868 | 944 | Y944G | C | 4.8e-06 | 7.3e-06 | 8.6e-06 | 6.900E-06 | 1.931E-06 | 4.800E-06 | 8.600E-06 |
| 18869 | 944 | Y944H | C | 6.8e-06 | 1.2e-05 | 8.7e-06 | 9.167E-06 | 2.631E-06 | 6.800E-06 | 1.200E-05 |

| | | | | | | | | | | |
|---|---|---|---|---|---|---|---|---|---|---|
| 18870 | 944 | Y944I | C | 9.7e-06 | 4.9e-06 | 1.2e-05 | 8.867E-06 | 3.623E-06 | 4.900E-06 | 1.200E-05 |
| 18871 | 944 | Y944L | C | 6.8e-06 | 1.5e-05 | 1.2e-05 | 1.127E-05 | 4.149E-06 | 6.800E-06 | 1.500E-05 |
| 18872 | 944 | Y944K | C | 7.9e-06 | 1.5e-05 | 6.4e-06 | 9.767E-06 | 4.594E-06 | 6.400E-06 | 1.500E-05 |
| 18873 | 944 | Y944M | C | 6.2e-06 | 7.8e-06 | 6.9e-06 | 6.967E-06 | 8.021E-07 | 6.200E-06 | 7.800E-06 |
| 18874 | 944 | Y944F | C | 9.5e-06 | 5.2e-06 | 9.3e-06 | 8.000E-06 | 2.427E-06 | 5.200E-06 | 9.500E-06 |
| 18875 | 944 | Y944P | C | 6.6e-06 | 7.7e-06 | 6.8e-06 | 7.033E-06 | 5.859E-07 | 6.600E-06 | 7.700E-06 |
| 18876 | 944 | Y944S | C | 9.2e-06 | 5.6e-06 | 8.6e-06 | 7.800E-06 | 1.929E-06 | 5.600E-06 | 9.200E-06 |
| 18877 | 944 | Y944T | C | 1.1e-05 | 5.7e-06 | 9e-06 | 8.567E-06 | 2.676E-06 | 5.700E-06 | 1.100E-05 |
| 18878 | 944 | Y944W | C | 9.7e-06 | 7.1e-06 | 4e-06 | 6.933E-06 | 2.854E-06 | 4.000E-06 | 9.700E-06 |
| 18879 | 944 | Y944Y | C | 8.8e-06 | 9.1e-06 | 1e-05 | 9.300E-06 | 6.245E-07 | 8.800E-06 | 1.000E-05 |
| 18880 | 944 | Y944V | C | 1e-05 | 1.5e-05 | 9.7e-06 | 1.157E-05 | 2.977E-06 | 9.700E-06 | 1.500E-05 |
| 18881 | 945 | W945A | C | 6.4e-06 | 1e-05 | 1.2e-05 | 9.467E-06 | 2.838E-06 | 6.400E-06 | 1.200E-05 |
| 18882 | 945 | W945R | C | 6.8e-06 | 1.1e-05 | 1.4e-05 | 1.060E-05 | 3.617E-06 | 6.800E-06 | 1.400E-05 |
| 18883 | 945 | W945N | C | 6.5e-06 | 9e-06 | 6.8e-06 | 7.433E-06 | 1.365E-06 | 6.500E-06 | 9.000E-06 |
| 18884 | 945 | W945D | C | 7e-06 | 8e-06 | 1.2e-05 | 9.000E-06 | 2.646E-06 | 7.000E-06 | 1.200E-05 |
| 18885 | 945 | W945C | C | 8.7e-06 | 1.2e-05 | 1.3e-05 | 1.123E-05 | 2.250E-06 | 8.700E-06 | 1.300E-05 |
| 18886 | 945 | W945E | C | 1.1e-05 | 1e-05 | 5.8e-06 | 8.933E-06 | 2.759E-06 | 5.800E-06 | 1.100E-05 |
| 18887 | 945 | W945Q | C | 1.2e-05 | 1e-05 | 6.2e-06 | 9.400E-06 | 2.946E-06 | 6.200E-06 | 1.200E-05 |
| 18888 | 945 | W945G | C | 6.2e-06 | 6.3e-06 | 7.6e-06 | 6.700E-06 | 7.810E-07 | 6.200E-06 | 7.600E-06 |
| 18889 | 945 | W945H | C | 7.6e-06 | 6.8e-06 | 9.2e-06 | 7.867E-06 | 1.222E-06 | 6.800E-06 | 9.200E-06 |
| 18890 | 945 | W945I | C | 6.3e-06 | 8.2e-06 | 9e-06 | 7.833E-06 | 1.387E-06 | 6.300E-06 | 9.000E-06 |
| 18891 | 945 | W945L | C | 6.3e-06 | 1.3e-05 | 7.2e-06 | 8.833E-06 | 3.636E-06 | 6.300E-06 | 1.300E-05 |
| 18892 | 945 | W945K | C | 1e-05 | 4.8e-06 | 6.5e-06 | 7.100E-06 | 2.651E-06 | 4.800E-06 | 1.000E-05 |
| 18893 | 945 | W945M | C | 6.8e-06 | 9.4e-06 | 6.5e-06 | 7.567E-06 | 1.595E-06 | 6.500E-06 | 9.400E-06 |
| 18894 | 945 | W945F | C | 1.2e-05 | 8.6e-06 | 9e-06 | 9.867E-06 | 1.858E-06 | 8.600E-06 | 1.200E-05 |
| 18895 | 945 | W945P | C | 1e-05 | 1e-05 | 1.6e-05 | 1.200E-05 | 3.464E-06 | 1.000E-05 | 1.600E-05 |
| 18896 | 945 | W945S | C | 8.8e-06 | 1e-05 | 9.8e-06 | 9.533E-06 | 6.429E-07 | 8.800E-06 | 1.000E-05 |
| 18897 | 945 | W945T | C | 9.7e-06 | 6.4e-06 | 1.4e-05 | 1.003E-05 | 3.811E-06 | 6.400E-06 | 1.400E-05 |
| 18898 | 945 | W945W | C | 5.9e-06 | 1e-05 | 8.9e-06 | 8.267E-06 | 2.122E-06 | 5.900E-06 | 1.000E-05 |
| 18899 | 945 | W945Y | C | 1.1e-05 | 9.2e-06 | 1.3e-05 | 1.107E-05 | 1.901E-06 | 9.200E-06 | 1.300E-05 |
| 18900 | 945 | W945V | C | 8.1e-06 | 7e-06 | 1.3e-05 | 9.367E-06 | 3.194E-06 | 7.000E-06 | 1.300E-05 |
| 18901 | 946 | G946A | C | 8.8e-06 | 7.6e-06 | 4.9e-06 | 7.100E-06 | 1.997E-06 | 4.900E-06 | 8.800E-06 |
| 18902 | 946 | G946R | C | 4.7e-06 | 5.7e-06 | 5.2e-06 | 5.200E-06 | 5.000E-07 | 4.700E-06 | 5.700E-06 |
| 18903 | 946 | G946N | C | 1.1e-05 | 9.2e-06 | 5.9e-06 | 8.700E-06 | 2.587E-06 | 5.900E-06 | 1.100E-05 |
| 18904 | 946 | G946D | C | 8.9e-06 | 6.8e-06 | 4.3e-06 | 6.667E-06 | 2.303E-06 | 4.300E-06 | 8.900E-06 |
| 18905 | 946 | G946C | C | 5e-06 | 9.6e-06 | 6e-06 | 6.867E-06 | 2.419E-06 | 5.000E-06 | 9.600E-06 |
| 18906 | 946 | G946E | C | 1.3e-05 | 8.2e-06 | 6.5e-06 | 9.233E-06 | 3.371E-06 | 6.500E-06 | 1.300E-05 |
| 18907 | 946 | G946Q | C | 4.9e-06 | 8.5e-06 | 1.4e-05 | 9.133E-06 | 4.583E-06 | 4.900E-06 | 1.400E-05 |
| 18908 | 946 | G946G | C | 5e-06 | 8.9e-06 | 6.1e-06 | 6.667E-06 | 2.011E-06 | 5.000E-06 | 8.900E-06 |
| 18909 | 946 | G946H | C | 8.2e-06 | 1.2e-05 | 8.6e-06 | 9.600E-06 | 2.088E-06 | 8.200E-06 | 1.200E-05 |
| 18910 | 946 | G946I | C | 9.7e-06 | 8.2e-06 | 9.7e-06 | 9.200E-06 | 8.660E-07 | 8.200E-06 | 9.700E-06 |
| 18911 | 946 | G946L | C | 5.5e-06 | 8.6e-06 | 1e-05 | 8.033E-06 | 2.303E-06 | 5.500E-06 | 1.000E-05 |
| 18912 | 946 | G946K | C | 6.9e-06 | 6.5e-06 | 7.8e-06 | 7.067E-06 | 6.658E-07 | 6.500E-06 | 7.800E-06 |
| 18913 | 946 | G946M | C | 8.6e-06 | 1.2e-05 | 7.4e-06 | 9.333E-06 | 2.386E-06 | 7.400E-06 | 1.200E-05 |
| 18914 | 946 | G946F | C | 9.6e-06 | 9e-06 | 4.5e-06 | 7.700E-06 | 2.787E-06 | 4.500E-06 | 9.600E-06 |
| 18915 | 946 | G946P | C | 7.8e-06 | 6.1e-06 | 5.4e-06 | 6.433E-06 | 1.234E-06 | 5.400E-06 | 7.800E-06 |
| 18916 | 946 | G946S | C | 1e-05 | 6.5e-06 | 5.3e-06 | 7.267E-06 | 2.442E-06 | 5.300E-06 | 1.000E-05 |
| 18917 | 946 | G946T | C | 9.7e-06 | 8.4e-06 | 8.1e-06 | 8.733E-06 | 8.505E-07 | 8.100E-06 | 9.700E-06 |
| 18918 | 946 | G946W | C | 5.5e-06 | 7.2e-06 | 4.8e-06 | 5.833E-06 | 1.234E-06 | 4.800E-06 | 7.200E-06 |
| 18919 | 946 | G946Y | C | 7.6e-06 | 8.7e-06 | 6.9e-06 | 7.733E-06 | 9.074E-07 | 6.900E-06 | 8.700E-06 |
| 18920 | 946 | G946V | C | 5.7e-06 | 9.9e-06 | 1.1e-05 | 8.867E-06 | 2.797E-06 | 5.700E-06 | 1.100E-05 |
| 18921 | 947 | Q947A | C | 1e-05 | 7.3e-06 | 5.3e-06 | 7.533E-06 | 2.359E-06 | 5.300E-06 | 1.000E-05 |
| 18922 | 947 | Q947R | C | 1e-05 | 6.2e-06 | 7.7e-06 | 7.967E-06 | 1.914E-06 | 6.200E-06 | 1.000E-05 |
| 18923 | 947 | Q947N | C | 6.6e-06 | 9.6e-06 | 1.4e-05 | 1.007E-05 | 3.722E-06 | 6.600E-06 | 1.400E-05 |
| 18924 | 947 | Q947D | C | 8.1e-06 | 6.9e-06 | 5e-06 | 6.667E-06 | 1.563E-06 | 5.000E-06 | 8.100E-06 |
| 18925 | 947 | Q947C | C | 8.4e-06 | 1.3e-05 | 1e-05 | 1.047E-05 | 2.335E-06 | 8.400E-06 | 1.300E-05 |
| 18926 | 947 | Q947E | C | 8.2e-06 | 4.9e-06 | 1.5e-05 | 9.367E-06 | 5.150E-06 | 4.900E-06 | 1.500E-05 |
| 18927 | 947 | Q947Q | C | 6.8e-06 | 7.4e-06 | 9.6e-06 | 7.933E-06 | 1.474E-06 | 6.800E-06 | 9.600E-06 |
| 18928 | 947 | Q947G | C | 6.7e-06 | 7.6e-06 | 1.3e-05 | 9.100E-06 | 3.407E-06 | 6.700E-06 | 1.300E-05 |
| 18929 | 947 | Q947H | C | 8.6e-06 | 7e-06 | 1e-05 | 8.533E-06 | 1.501E-06 | 7.000E-06 | 1.000E-05 |
| 18930 | 947 | Q947I | C | 1e-05 | 7.3e-06 | 5.4e-06 | 7.567E-06 | 2.312E-06 | 5.400E-06 | 1.000E-05 |
| 18931 | 947 | Q947L | C | 1e-05 | 1.5e-05 | 1.4e-05 | 1.300E-05 | 2.646E-06 | 1.000E-05 | 1.500E-05 |
| 18932 | 947 | Q947K | C | 6.4e-06 | 6.9e-06 | 9.4e-06 | 7.567E-06 | 1.607E-06 | 6.400E-06 | 9.400E-06 |
| 18933 | 947 | Q947M | C | 8.9e-06 | 9.6e-06 | 1.6e-05 | 1.150E-05 | 3.913E-06 | 8.900E-06 | 1.600E-05 |
| 18934 | 947 | Q947F | C | 1e-05 | 7.4e-06 | 1.3e-05 | 1.013E-05 | 2.802E-06 | 7.400E-06 | 1.300E-05 |
| 18935 | 947 | Q947P | C | 5.9e-06 | 8.2e-06 | 7.5e-06 | 7.200E-06 | 1.179E-06 | 5.900E-06 | 8.200E-06 |
| 18936 | 947 | Q947S | C | 9.4e-06 | 7.5e-06 | 6.9e-06 | 7.933E-06 | 1.305E-06 | 6.900E-06 | 9.400E-06 |
| 18937 | 947 | Q947T | C | 8.2e-06 | 6.3e-06 | 7.4e-06 | 7.300E-06 | 9.539E-07 | 6.300E-06 | 8.200E-06 |
| 18938 | 947 | Q947W | C | 9.3e-06 | 7.8e-06 | 7.2e-06 | 8.100E-06 | 1.082E-06 | 7.200E-06 | 9.300E-06 |
| 18939 | 947 | Q947Y | C | 7e-06 | 5.8e-06 | 4.8e-06 | 5.867E-06 | 1.102E-06 | 4.800E-06 | 7.000E-06 |
| 18940 | 947 | Q947V | C | 1e-05 | 7.9e-06 | 9.7e-06 | 9.200E-06 | 1.136E-06 | 7.900E-06 | 1.000E-05 |
| 18941 | 948 | G948A | C | 4.8e-06 | 6.4e-06 | 1.2e-05 | 7.733E-06 | 3.781E-06 | 4.800E-06 | 1.200E-05 |
| 18942 | 948 | G948R | C | 1.2e-05 | 8.4e-06 | 7.4e-06 | 9.267E-06 | 2.419E-06 | 7.400E-06 | 1.200E-05 |
| 18943 | 948 | G948N | C | 6.8e-06 | 6.1e-06 | 4.6e-06 | 5.833E-06 | 1.124E-06 | 4.600E-06 | 6.800E-06 |
| 18944 | 948 | G948D | C | 9.1e-06 | 8.3e-06 | 6.5e-06 | 7.967E-06 | 1.332E-06 | 6.500E-06 | 9.100E-06 |
| 18945 | 948 | G948C | C | 6.6e-06 | 7.1e-06 | 5.2e-06 | 6.300E-06 | 9.849E-07 | 5.200E-06 | 7.100E-06 |
| 18946 | 948 | G948E | C | 7.5e-06 | 8.9e-06 | 8.4e-06 | 8.267E-06 | 7.095E-07 | 7.500E-06 | 8.900E-06 |
| 18947 | 948 | G948Q | C | 8.2e-06 | 1.2e-05 | 9.1e-06 | 9.767E-06 | 1.986E-06 | 8.200E-06 | 1.200E-05 |
| 18948 | 948 | G948G | C | 1.3e-05 | 6.1e-06 | 7e-06 | 8.700E-06 | 3.751E-06 | 6.100E-06 | 1.300E-05 |
| 18949 | 948 | G948H | C | 8.2e-06 | 9.3e-06 | 6.6e-06 | 8.033E-06 | 1.358E-06 | 6.600E-06 | 9.300E-06 |
| 18950 | 948 | G948I | C | 6.1e-06 | 6.1e-06 | 5.1e-06 | 5.767E-06 | 5.774E-07 | 5.100E-06 | 6.100E-06 |
| 18951 | 948 | G948L | C | 6.8e-06 | 7.2e-06 | 6.9e-06 | 6.967E-06 | 2.082E-07 | 6.800E-06 | 7.200E-06 |
| 18952 | 948 | G948K | C | 1.1e-05 | 8.4e-06 | 5e-06 | 8.133E-06 | 3.009E-06 | 5.000E-06 | 1.100E-05 |
| 18953 | 948 | G948M | C | 5.9e-06 | 6.4e-06 | 1.2e-05 | 8.100E-06 | 3.387E-06 | 5.900E-06 | 1.200E-05 |
| 18954 | 948 | G948F | C | 8.5e-06 | 1.4e-05 | 6.7e-06 | 9.733E-06 | 3.803E-06 | 6.700E-06 | 1.400E-05 |

| | | | | | | | | | | |
|---|---|---|---|---|---|---|---|---|---|---|
| 18955 | 948 | G948P | C | 9.3e-06 | 8.1e-06 | 1.1e-05 | 9.467E-06 | 1.457E-06 | 8.100E-06 | 1.100E-05 |
| 18956 | 948 | G948S | C | 6.4e-06 | 5.2e-06 | 1.2e-05 | 7.867E-06 | 3.630E-06 | 5.200E-06 | 1.200E-05 |
| 18957 | 948 | G948T | C | 5.9e-06 | 5.9e-06 | 1e-05 | 7.267E-06 | 2.367E-06 | 5.900E-06 | 1.000E-05 |
| 18958 | 948 | G948W | C | 1.2e-05 | 6.4e-06 | 7.6e-06 | 8.667E-06 | 2.948E-06 | 6.400E-06 | 1.200E-05 |
| 18959 | 948 | G948Y | C | 1.8e-05 | 1.3e-05 | 1.1e-05 | 1.400E-05 | 3.606E-06 | 1.100E-05 | 1.800E-05 |
| 18960 | 948 | G948V | C | 6.1e-06 | 7e-06 | 1.1e-05 | 8.033E-06 | 2.608E-06 | 6.100E-06 | 1.100E-05 |
| 18961 | 949 | T949A | C | 8.5e-06 | 6.6e-06 | 9.2e-06 | 8.100E-06 | 1.345E-06 | 6.600E-06 | 9.200E-06 |
| 18962 | 949 | T949R | C | 7e-06 | 6.9e-06 | 7.7e-06 | 7.200E-06 | 4.359E-07 | 6.900E-06 | 7.700E-06 |
| 18963 | 949 | T949N | C | 5.3e-06 | 5.7e-06 | 6.4e-06 | 5.800E-06 | 5.568E-07 | 5.300E-06 | 6.400E-06 |
| 18964 | 949 | T949D | C | 6.5e-06 | 7.5e-06 | 1.1e-05 | 8.333E-06 | 2.363E-06 | 6.500E-06 | 1.100E-05 |
| 18965 | 949 | T949C | C | 7e-06 | 5e-06 | 6.8e-06 | 6.267E-06 | 1.102E-06 | 5.000E-06 | 7.000E-06 |
| 18966 | 949 | T949E | C | 7.5e-06 | 4.6e-06 | 9.6e-06 | 7.233E-06 | 2.511E-06 | 4.600E-06 | 9.600E-06 |
| 18967 | 949 | T949Q | C | 7.9e-06 | 8.1e-06 | 4e-06 | 6.667E-06 | 2.312E-06 | 4.000E-06 | 8.100E-06 |
| 18968 | 949 | T949G | C | 9.3e-06 | 5.2e-06 | 8.2e-06 | 7.567E-06 | 2.122E-06 | 5.200E-06 | 9.300E-06 |
| 18969 | 949 | T949H | C | 5.6e-06 | 9e-06 | 5.7e-06 | 6.767E-06 | 1.935E-06 | 5.600E-06 | 9.000E-06 |
| 18970 | 949 | T949I | C | 8.8e-06 | 9.9e-06 | 1e-05 | 9.567E-06 | 6.658E-07 | 8.800E-06 | 1.000E-05 |
| 18971 | 949 | T949L | C | 1e-05 | 9.9e-06 | 5.3e-06 | 8.400E-06 | 2.685E-06 | 5.300E-06 | 1.000E-05 |
| 18972 | 949 | T949K | C | 6.4e-06 | 5.5e-06 | 7.8e-06 | 6.567E-06 | 1.159E-06 | 5.500E-06 | 7.800E-06 |
| 18973 | 949 | T949M | C | 1e-05 | 1e-05 | 8.4e-06 | 9.467E-06 | 9.238E-07 | 8.400E-06 | 1.000E-05 |
| 18974 | 949 | T949F | C | 9.2e-06 | 9.4e-06 | 1.3e-05 | 1.053E-05 | 2.139E-06 | 9.200E-06 | 1.300E-05 |
| 18975 | 949 | T949P | C | 2e-05 | 1e-05 | 8.1e-06 | 1.270E-05 | 6.393E-06 | 8.100E-06 | 2.000E-05 |
| 18976 | 949 | T949S | C | 7.4e-06 | 7.4e-06 | 7.9e-06 | 7.567E-06 | 2.887E-07 | 7.400E-06 | 7.900E-06 |
| 18977 | 949 | T949T | C | 8.1e-06 | 6.3e-06 | 1e-05 | 8.133E-06 | 1.850E-06 | 6.300E-06 | 1.000E-05 |
| 18978 | 949 | T949W | C | 8e-06 | 1.1e-05 | 5e-06 | 8.000E-06 | 3.000E-06 | 5.000E-06 | 1.100E-05 |
| 18979 | 949 | T949Y | C | 1e-05 | 6.6e-06 | 8.6e-06 | 8.400E-06 | 1.709E-06 | 6.600E-06 | 1.000E-05 |
| 18980 | 949 | T949V | C | 8.9e-06 | 9e-06 | 1.8e-05 | 1.197E-05 | 5.225E-06 | 8.900E-06 | 1.800E-05 |
| 18981 | 950 | L950A | C | 6.9e-06 | 1e-05 | 8.5e-06 | 8.467E-06 | 1.550E-06 | 6.900E-06 | 1.000E-05 |
| 18982 | 950 | L950R | C | 6.5e-06 | 1.2e-05 | 5e-06 | 7.833E-06 | 3.686E-06 | 5.000E-06 | 1.200E-05 |
| 18983 | 950 | L950N | C | 6.3e-06 | 9.5e-06 | 7.7e-06 | 7.833E-06 | 1.604E-06 | 6.300E-06 | 9.500E-06 |
| 18984 | 950 | L950D | C | 8.4e-06 | 8.2e-06 | 1.2e-05 | 9.533E-06 | 2.139E-06 | 8.200E-06 | 1.200E-05 |
| 18985 | 950 | L950C | C | 8.6e-06 | 9.6e-06 | 1.3e-05 | 1.040E-05 | 2.307E-06 | 8.600E-06 | 1.300E-05 |
| 18986 | 950 | L950E | C | 1.3e-05 | 1.2e-05 | 8.6e-06 | 1.120E-05 | 2.307E-06 | 8.600E-06 | 1.300E-05 |
| 18987 | 950 | L950Q | C | 7.4e-06 | 1e-05 | 6.8e-06 | 8.067E-06 | 1.701E-06 | 6.800E-06 | 1.000E-05 |
| 18988 | 950 | L950G | C | 1e-05 | 4.7e-06 | 6.5e-06 | 7.067E-06 | 2.695E-06 | 4.700E-06 | 1.000E-05 |
| 18989 | 950 | L950H | C | 6.8e-06 | 1.2e-05 | 4e-06 | 7.600E-06 | 4.060E-06 | 4.000E-06 | 1.200E-05 |
| 18990 | 950 | L950I | C | 5.4e-06 | 5.3e-06 | 4.4e-06 | 5.033E-06 | 5.508E-07 | 4.400E-06 | 5.400E-06 |
| 18991 | 950 | L950L | C | 1e-05 | 6.7e-06 | 7.3e-06 | 8.000E-06 | 1.758E-06 | 6.700E-06 | 1.000E-05 |
| 18992 | 950 | L950K | C | 7.1e-06 | 6.4e-06 | 9.9e-06 | 7.800E-06 | 1.852E-06 | 6.400E-06 | 9.900E-06 |
| 18993 | 950 | L950M | C | 7.5e-06 | 8.5e-06 | 8.9e-06 | 8.300E-06 | 7.211E-07 | 7.500E-06 | 8.900E-06 |
| 18994 | 950 | L950F | C | 9.1e-06 | 1.4e-05 | 5.8e-06 | 9.633E-06 | 4.126E-06 | 5.800E-06 | 1.400E-05 |
| 18995 | 950 | L950P | C | 1.1e-05 | 4.6e-06 | 9e-06 | 8.200E-06 | 3.274E-06 | 4.600E-06 | 1.100E-05 |
| 18996 | 950 | L950S | C | 7.1e-06 | 1.7e-05 | 9.7e-06 | 1.127E-05 | 5.133E-06 | 7.100E-06 | 1.700E-05 |
| 18997 | 950 | L950T | C | 7e-06 | 5.8e-06 | 4.9e-06 | 5.900E-06 | 1.054E-06 | 4.900E-06 | 7.000E-06 |
| 18998 | 950 | L950W | C | 6.2e-06 | 6.9e-06 | 5.9e-06 | 6.333E-06 | 5.132E-07 | 5.900E-06 | 6.900E-06 |
| 18999 | 950 | L950Y | C | 9.8e-06 | 4e-06 | 1.1e-05 | 8.267E-06 | 3.743E-06 | 4.000E-06 | 1.100E-05 |
| 19000 | 950 | L950V | C | 1.3e-05 | 1.2e-05 | 9.1e-06 | 1.137E-05 | 2.026E-06 | 9.100E-06 | 1.300E-05 |
| 19001 | 951 | V951A | C | 5.1e-06 | 1.3e-05 | 1.1e-05 | 9.700E-06 | 4.107E-06 | 5.100E-06 | 1.300E-05 |
| 19002 | 951 | V951R | C | 6.9e-06 | 1.4e-05 | 7.1e-06 | 9.333E-06 | 4.043E-06 | 6.900E-06 | 1.400E-05 |
| 19003 | 951 | V951N | C | 8.2e-06 | 8.2e-06 | 9e-06 | 8.467E-06 | 4.619E-07 | 8.200E-06 | 9.000E-06 |
| 19004 | 951 | V951D | C | 1.1e-05 | 1.1e-05 | 7.9e-06 | 9.967E-06 | 1.790E-06 | 7.900E-06 | 1.100E-05 |
| 19005 | 951 | V951C | C | 1.1e-05 | 9.5e-06 | 4.2e-06 | 8.233E-06 | 3.573E-06 | 4.200E-06 | 1.100E-05 |
| 19006 | 951 | V951E | C | 1.2e-05 | 3.8e-06 | 9.7e-06 | 8.500E-06 | 4.230E-06 | 3.800E-06 | 1.200E-05 |
| 19007 | 951 | V951Q | C | 6.5e-06 | 4.7e-06 | 9.3e-06 | 6.833E-06 | 2.318E-06 | 4.700E-06 | 9.300E-06 |
| 19008 | 951 | V951G | C | 6.9e-06 | 8.9e-06 | 5.4e-06 | 7.067E-06 | 1.756E-06 | 5.400E-06 | 8.900E-06 |
| 19009 | 951 | V951H | C | 8.3e-06 | 1.1e-05 | 8.8e-06 | 9.367E-06 | 1.436E-06 | 8.300E-06 | 1.100E-05 |
| 19010 | 951 | V951I | C | 1e-05 | 1.1e-05 | 6.9e-06 | 9.300E-06 | 2.138E-06 | 6.900E-06 | 1.100E-05 |
| 19011 | 951 | V951L | C | 1.3e-05 | 1e-05 | 7.8e-06 | 1.027E-05 | 2.610E-06 | 7.800E-06 | 1.300E-05 |
| 19012 | 951 | V951K | C | 6.7e-06 | 1.4e-05 | 4.4e-06 | 8.367E-06 | 5.012E-06 | 4.400E-06 | 1.400E-05 |
| 19013 | 951 | V951M | C | 9.3e-06 | 1.6e-05 | 9.6e-06 | 1.163E-05 | 3.785E-06 | 9.300E-06 | 1.600E-05 |
| 19014 | 951 | V951F | C | 6.7e-06 | 5.3e-06 | 9.7e-06 | 7.233E-06 | 2.248E-06 | 5.300E-06 | 9.700E-06 |
| 19015 | 951 | V951P | C | 1.1e-05 | 8.9e-06 | 1.4e-05 | 1.130E-05 | 2.563E-06 | 8.900E-06 | 1.400E-05 |
| 19016 | 951 | V951S | C | 5.2e-06 | 8.1e-06 | 1.1e-05 | 8.100E-06 | 2.900E-06 | 5.200E-06 | 1.100E-05 |
| 19017 | 951 | V951T | C | 1.1e-05 | 6.8e-06 | 4.8e-06 | 7.533E-06 | 3.164E-06 | 4.800E-06 | 1.100E-05 |
| 19018 | 951 | V951W | C | 9.8e-06 | 1e-05 | 7e-06 | 8.933E-06 | 1.677E-06 | 7.000E-06 | 1.000E-05 |
| 19019 | 951 | V951Y | C | 7.6e-06 | 5.1e-06 | 6.2e-06 | 6.300E-06 | 1.253E-06 | 5.100E-06 | 7.600E-06 |
| 19020 | 951 | V951V | C | 8.2e-06 | 5.9e-06 | 6.1e-06 | 6.733E-06 | 1.274E-06 | 5.900E-06 | 8.200E-06 |
| 19021 | 952 | T952A | C | 8.3e-06 | 1.2e-05 | 1.3e-05 | 1.110E-05 | 2.476E-06 | 8.300E-06 | 1.300E-05 |
| 19022 | 952 | T952R | C | 1.2e-05 | 1.3e-05 | 5.1e-06 | 1.003E-05 | 4.302E-06 | 5.100E-06 | 1.300E-05 |
| 19023 | 952 | T952N | C | 8.1e-06 | 8.5e-06 | 9e-06 | 8.533E-06 | 4.509E-07 | 8.100E-06 | 9.000E-06 |
| 19024 | 952 | T952D | C | 7e-06 | 7.8e-06 | 9.8e-06 | 8.200E-06 | 1.442E-06 | 7.000E-06 | 9.800E-06 |
| 19025 | 952 | T952C | C | 9.6e-06 | 1.1e-05 | 8.2e-06 | 9.600E-06 | 1.400E-06 | 8.200E-06 | 1.100E-05 |
| 19026 | 952 | T952E | C | 9.1e-06 | 1.4e-05 | 5.3e-06 | 9.467E-06 | 4.362E-06 | 5.300E-06 | 1.400E-05 |
| 19027 | 952 | T952Q | C | 5.8e-06 | 1e-05 | 6.1e-06 | 7.300E-06 | 2.343E-06 | 5.800E-06 | 1.000E-05 |
| 19028 | 952 | T952G | C | 1e-05 | 6e-06 | 7e-06 | 7.667E-06 | 2.082E-06 | 6.000E-06 | 1.000E-05 |
| 19029 | 952 | T952H | C | 6.9e-06 | 1.3e-05 | 6.6e-06 | 8.833E-06 | 3.612E-06 | 6.600E-06 | 1.300E-05 |
| 19030 | 952 | T952I | C | 6.3e-06 | 9.4e-06 | 7e-06 | 7.567E-06 | 1.626E-06 | 6.300E-06 | 9.400E-06 |
| 19031 | 952 | T952L | C | 6.8e-06 | 1.1e-05 | 1.3e-05 | 1.027E-05 | 3.164E-06 | 6.800E-06 | 1.300E-05 |
| 19032 | 952 | T952K | C | 7.7e-06 | 7.1e-06 | 1.4e-05 | 9.600E-06 | 3.822E-06 | 7.100E-06 | 1.400E-05 |
| 19033 | 952 | T952M | C | 5.4e-06 | 5.1e-06 | 6.9e-06 | 5.800E-06 | 9.644E-07 | 5.100E-06 | 6.900E-06 |
| 19034 | 952 | T952F | C | 9e-06 | 5.1e-06 | 7.2e-06 | 7.100E-06 | 1.952E-06 | 5.100E-06 | 9.000E-06 |
| 19035 | 952 | T952P | C | 7.8e-06 | 9.6e-06 | 7.5e-06 | 8.300E-06 | 1.136E-06 | 7.500E-06 | 9.600E-06 |
| 19036 | 952 | T952S | C | 2e-05 | 1e-05 | 9.3e-06 | 1.310E-05 | 5.986E-06 | 9.300E-06 | 2.000E-05 |
| 19037 | 952 | T952T | C | 6.6e-06 | 1.2e-05 | 1.4e-05 | 1.087E-05 | 3.828E-06 | 6.600E-06 | 1.400E-05 |
| 19038 | 952 | T952W | C | 6e-06 | 1.3e-05 | 7.4e-06 | 8.800E-06 | 3.704E-06 | 6.000E-06 | 1.300E-05 |
| 19039 | 952 | T952Y | C | 8.6e-06 | 6.8e-06 | 7.6e-06 | 7.667E-06 | 9.018E-07 | 6.800E-06 | 8.600E-06 |

| | | | | | | | | | | |
|---|---|---|---|---|---|---|---|---|---|---|
| 19040 | 952 | T952V | C | 5.5e-06 | 4.1e-06 | 1.3e-05 | 7.533E-06 | 4.786E-06 | 4.100E-06 | 1.300E-05 |
| 19041 | 953 | V953A | C | 1.3e-05 | 1.1e-05 | 5.5e-06 | 9.833E-06 | 3.884E-06 | 5.500E-06 | 1.300E-05 |
| 19042 | 953 | V953R | C | 8.7e-06 | 6.5e-06 | 7.9e-06 | 7.700E-06 | 1.114E-06 | 6.500E-06 | 8.700E-06 |
| 19043 | 953 | V953N | C | 6.9e-06 | 5.4e-06 | 9.5e-06 | 7.267E-06 | 2.074E-06 | 5.400E-06 | 9.500E-06 |
| 19044 | 953 | V953D | C | 6.7e-06 | 1.2e-05 | 9.9e-06 | 9.533E-06 | 2.669E-06 | 6.700E-06 | 1.200E-05 |
| 19045 | 953 | V953C | C | 8.4e-06 | 7.8e-06 | 6.8e-06 | 7.667E-06 | 8.083E-07 | 6.800E-06 | 8.400E-06 |
| 19046 | 953 | V953E | C | 8.8e-06 | 9.9e-06 | 6.2e-06 | 8.300E-06 | 1.900E-06 | 6.200E-06 | 9.900E-06 |
| 19047 | 953 | V953Q | C | 7.6e-06 | 9.7e-06 | 7.6e-06 | 8.300E-06 | 1.212E-06 | 7.600E-06 | 9.700E-06 |
| 19048 | 953 | V953G | C | 9.5e-06 | 1.1e-05 | 1e-05 | 1.017E-05 | 7.638E-07 | 9.500E-06 | 1.100E-05 |
| 19049 | 953 | V953H | C | 1.1e-05 | 7.7e-06 | 6.5e-06 | 8.400E-06 | 2.330E-06 | 6.500E-06 | 1.100E-05 |
| 19050 | 953 | V953I | C | 5.2e-06 | 1.2e-05 | 7.3e-06 | 8.167E-06 | 3.482E-06 | 5.200E-06 | 1.200E-05 |
| 19051 | 953 | V953L | C | 7e-06 | 1.1e-05 | 1.4e-05 | 1.067E-05 | 3.512E-06 | 7.000E-06 | 1.400E-05 |
| 19052 | 953 | V953K | C | 8.7e-06 | 6.3e-06 | 5.2e-06 | 6.733E-06 | 1.790E-06 | 5.200E-06 | 8.700E-06 |
| 19053 | 953 | V953M | C | 6.6e-06 | 9.3e-06 | 9.1e-06 | 8.333E-06 | 1.504E-06 | 6.600E-06 | 9.300E-06 |
| 19054 | 953 | V953F | C | 6.4e-06 | 1.2e-05 | 6.7e-06 | 8.367E-06 | 3.150E-06 | 6.400E-06 | 1.200E-05 |
| 19055 | 953 | V953P | C | 1.4e-05 | 4.1e-06 | 6.6e-06 | 8.233E-06 | 5.148E-06 | 4.100E-06 | 1.400E-05 |
| 19056 | 953 | V953S | C | 8.1e-06 | 4.2e-06 | 7.9e-06 | 6.733E-06 | 2.196E-06 | 4.200E-06 | 8.100E-06 |
| 19057 | 953 | V953T | C | 1.2e-05 | 8.1e-06 | 6.8e-06 | 8.967E-06 | 2.706E-06 | 6.800E-06 | 1.200E-05 |
| 19058 | 953 | V953W | C | 9.2e-06 | 1e-05 | 7.9e-06 | 9.033E-06 | 1.060E-06 | 7.900E-06 | 1.000E-05 |
| 19059 | 953 | V953Y | C | 8.9e-06 | 4.6e-06 | 1.2e-05 | 8.500E-06 | 3.716E-06 | 4.600E-06 | 1.200E-05 |
| 19060 | 953 | V953V | C | 9.2e-06 | 7.8e-06 | 9.7e-06 | 8.900E-06 | 9.849E-07 | 7.800E-06 | 9.700E-06 |
| 19061 | 954 | S954A | C | 8.6e-06 | 9e-06 | 1.2e-05 | 9.867E-06 | 1.858E-06 | 8.600E-06 | 1.200E-05 |
| 19062 | 954 | S954R | C | 5.9e-06 | 7.2e-06 | 1.3e-05 | 8.700E-06 | 3.780E-06 | 5.900E-06 | 1.300E-05 |
| 19063 | 954 | S954N | C | 9.7e-06 | 5.6e-06 | 7.2e-06 | 7.500E-06 | 2.066E-06 | 5.600E-06 | 9.700E-06 |
| 19064 | 954 | S954D | C | 1.2e-05 | 9.9e-06 | 7.9e-06 | 9.933E-06 | 2.050E-06 | 7.900E-06 | 1.200E-05 |
| 19065 | 954 | S954C | C | 1.1e-05 | 7.4e-06 | 9.3e-06 | 9.233E-06 | 1.801E-06 | 7.400E-06 | 1.100E-05 |
| 19066 | 954 | S954E | C | 1.1e-05 | 6e-06 | 5.8e-06 | 7.600E-06 | 2.946E-06 | 5.800E-06 | 1.100E-05 |
| 19067 | 954 | S954Q | C | 1e-05 | 9.7e-06 | 1e-05 | 9.900E-06 | 1.732E-07 | 9.700E-06 | 1.000E-05 |
| 19068 | 954 | S954G | C | 1.2e-05 | 8.8e-06 | 1.4e-05 | 1.160E-05 | 2.623E-06 | 8.800E-06 | 1.400E-05 |
| 19069 | 954 | S954H | C | 5.7e-06 | 9e-06 | 7.7e-06 | 7.467E-06 | 1.662E-06 | 5.700E-06 | 9.000E-06 |
| 19070 | 954 | S954I | C | 7.1e-06 | 6.8e-06 | 2e-05 | 1.130E-05 | 7.536E-06 | 6.800E-06 | 2.000E-05 |
| 19071 | 954 | S954L | C | 1.2e-05 | 9.2e-06 | 4.8e-06 | 8.667E-06 | 3.630E-06 | 4.800E-06 | 1.200E-05 |
| 19072 | 954 | S954K | C | 1.4e-05 | 6.1e-06 | 5.9e-06 | 8.667E-06 | 4.620E-06 | 5.900E-06 | 1.400E-05 |
| 19073 | 954 | S954M | C | 1.1e-05 | 1.1e-05 | 9.3e-06 | 1.043E-05 | 9.815E-07 | 9.300E-06 | 1.100E-05 |
| 19074 | 954 | S954F | C | 6.5e-06 | 6.4e-06 | 6.8e-06 | 6.567E-06 | 2.082E-07 | 6.400E-06 | 6.800E-06 |
| 19075 | 954 | S954P | C | 7.3e-06 | 8.1e-06 | 9.4e-06 | 8.267E-06 | 1.060E-06 | 7.300E-06 | 9.400E-06 |
| 19076 | 954 | S954S | C | 9.9e-06 | 6.1e-06 | 1.1e-05 | 9.000E-06 | 2.571E-06 | 6.100E-06 | 1.100E-05 |
| 19077 | 954 | S954T | C | 4.5e-06 | 5.4e-06 | 1.3e-05 | 7.633E-06 | 4.669E-06 | 4.500E-06 | 1.300E-05 |
| 19078 | 954 | S954W | C | 8.4e-06 | 9.4e-06 | 1.1e-05 | 9.600E-06 | 1.311E-06 | 8.400E-06 | 1.100E-05 |
| 19079 | 954 | S954Y | C | 6.2e-06 | 9.9e-06 | 1e-05 | 8.700E-06 | 2.166E-06 | 6.200E-06 | 1.000E-05 |
| 19080 | 954 | S954V | C | 6e-06 | 1.5e-05 | 9.4e-06 | 1.013E-05 | 4.545E-06 | 6.000E-06 | 1.500E-05 |
| 19081 | 955 | S955A | C | 5.6e-06 | 8.1e-06 | 6.5e-06 | 6.733E-06 | 1.266E-06 | 5.600E-06 | 8.100E-06 |
| 19082 | 955 | S955R | C | 8.8e-06 | 1.4e-05 | 7.8e-06 | 1.020E-05 | 3.329E-06 | 7.800E-06 | 1.400E-05 |
| 19083 | 955 | S955N | C | 1.8e-05 | 9.1e-06 | 7.3e-06 | 1.147E-05 | 5.729E-06 | 7.300E-06 | 1.800E-05 |
| 19084 | 955 | S955D | C | 8e-06 | 9.5e-06 | 6.1e-06 | 7.867E-06 | 1.704E-06 | 6.100E-06 | 9.500E-06 |
| 19085 | 955 | S955C | C | 7.4e-06 | 5.2e-06 | 7.9e-06 | 6.833E-06 | 1.436E-06 | 5.200E-06 | 7.900E-06 |
| 19086 | 955 | S955E | C | 4.7e-06 | 8.5e-06 | 7.7e-06 | 6.967E-06 | 2.003E-06 | 4.700E-06 | 8.500E-06 |
| 19087 | 955 | S955Q | C | 8.5e-06 | 5e-06 | 1.2e-05 | 8.500E-06 | 3.500E-06 | 5.000E-06 | 1.200E-05 |
| 19088 | 955 | S955G | C | 7.9e-06 | 8.6e-06 | 9.8e-06 | 8.767E-06 | 9.609E-07 | 7.900E-06 | 9.800E-06 |
| 19089 | 955 | S955H | C | 9.2e-06 | 1.1e-05 | 6.9e-06 | 9.033E-06 | 2.055E-06 | 6.900E-06 | 1.100E-05 |
| 19090 | 955 | S955I | C | 6.3e-06 | 7.2e-06 | 1.5e-05 | 9.500E-06 | 4.784E-06 | 6.300E-06 | 1.500E-05 |
| 19091 | 955 | S955L | C | 7.6e-06 | 1.2e-05 | 8.4e-06 | 9.333E-06 | 2.344E-06 | 7.600E-06 | 1.200E-05 |
| 19092 | 955 | S955K | C | 7e-06 | 9.7e-06 | 6.3e-06 | 7.667E-06 | 1.795E-06 | 6.300E-06 | 9.700E-06 |
| 19093 | 955 | S955M | C | 6.5e-06 | 7.5e-06 | 5.7e-06 | 6.567E-06 | 9.018E-07 | 5.700E-06 | 7.500E-06 |
| 19094 | 955 | S955F | C | 4.9e-06 | 1.1e-05 | 1.3e-05 | 9.633E-06 | 4.219E-06 | 4.900E-06 | 1.300E-05 |
| 19095 | 955 | S955P | C | 7.9e-06 | 7.1e-06 | 1e-05 | 8.333E-06 | 1.498E-06 | 7.100E-06 | 1.000E-05 |
| 19096 | 955 | S955S | C | 1.3e-05 | 5.7e-06 | 7.6e-06 | 8.767E-06 | 3.787E-06 | 5.700E-06 | 1.300E-05 |
| 19097 | 955 | S955T | C | 5.4e-06 | 9.1e-06 | 1.3e-05 | 9.167E-06 | 3.800E-06 | 5.400E-06 | 1.300E-05 |
| 19098 | 955 | S955W | C | 7.2e-06 | 8.2e-06 | 9.7e-06 | 8.367E-06 | 1.258E-06 | 7.200E-06 | 9.700E-06 |
| 19099 | 955 | S955Y | C | 7.1e-06 | 9.3e-06 | 6.3e-06 | 7.567E-06 | 1.553E-06 | 6.300E-06 | 9.300E-06 |
| 19100 | 955 | S955V | C | 8.9e-06 | 1.4e-05 | 1.3e-05 | 1.197E-05 | 2.702E-06 | 8.900E-06 | 1.400E-05 |
| 19101 | 956 | A956A | C | 5.3e-06 | 6.6e-06 | 6.6e-06 | 6.167E-06 | 7.506E-07 | 5.300E-06 | 6.600E-06 |
| 19102 | 956 | A956R | C | 6.7e-06 | 6.1e-06 | 6.8e-06 | 6.533E-06 | 3.786E-07 | 6.100E-06 | 6.800E-06 |
| 19103 | 956 | A956N | C | 3.5e-06 | 5.6e-06 | 5.8e-06 | 4.967E-06 | 1.274E-06 | 3.500E-06 | 5.800E-06 |
| 19104 | 956 | A956D | C | 6.7e-06 | 7.7e-06 | 4.7e-06 | 6.367E-06 | 1.528E-06 | 4.700E-06 | 7.700E-06 |
| 19105 | 956 | A956C | C | 1.2e-05 | 7.1e-06 | 8.8e-06 | 9.300E-06 | 2.488E-06 | 7.100E-06 | 1.200E-05 |
| 19106 | 956 | A956E | C | 5.6e-06 | 5.5e-06 | 3.8e-06 | 4.967E-06 | 1.012E-06 | 3.800E-06 | 5.600E-06 |
| 19107 | 956 | A956Q | C | 1.2e-05 | 5.9e-06 | 1.5e-05 | 1.097E-05 | 4.637E-06 | 5.900E-06 | 1.500E-05 |
| 19108 | 956 | A956G | C | 5.6e-06 | 9.1e-06 | 8.2e-06 | 7.633E-06 | 1.818E-06 | 5.600E-06 | 9.100E-06 |
| 19109 | 956 | A956H | C | 7.8e-06 | 3.3e-06 | 4.6e-06 | 5.233E-06 | 2.316E-06 | 3.300E-06 | 7.800E-06 |
| 19110 | 956 | A956I | C | 1.1e-05 | 1.3e-05 | 7.1e-06 | 1.037E-05 | 3.001E-06 | 7.100E-06 | 1.300E-05 |
| 19111 | 956 | A956L | C | 1e-05 | 9.3e-06 | 8.2e-06 | 9.167E-06 | 9.074E-07 | 8.200E-06 | 1.000E-05 |
| 19112 | 956 | A956K | C | 1.3e-05 | 7.8e-06 | 8.5e-06 | 9.767E-06 | 2.822E-06 | 7.800E-06 | 1.300E-05 |
| 19113 | 956 | A956M | C | 4.6e-06 | 5.6e-06 | 9.4e-06 | 6.533E-06 | 2.532E-06 | 4.600E-06 | 9.400E-06 |
| 19114 | 956 | A956F | C | 6.7e-06 | 6.5e-06 | 7e-06 | 6.733E-06 | 2.517E-07 | 6.500E-06 | 7.000E-06 |
| 19115 | 956 | A956P | C | 5.1e-06 | 6e-06 | 1.1e-05 | 7.367E-06 | 3.179E-06 | 5.100E-06 | 1.100E-05 |
| 19116 | 956 | A956S | C | 7.4e-06 | 8.6e-06 | 9.9e-06 | 8.633E-06 | 1.250E-06 | 7.400E-06 | 9.900E-06 |
| 19117 | 956 | A956T | C | 8e-06 | 4.8e-06 | 6.2e-06 | 6.333E-06 | 1.604E-06 | 4.800E-06 | 8.000E-06 |
| 19118 | 956 | A956W | C | 8.7e-06 | 8.4e-06 | 1e-05 | 9.033E-06 | 8.505E-07 | 8.400E-06 | 1.000E-05 |
| 19119 | 956 | A956Y | C | 1.5e-05 | 6.7e-06 | 5.8e-06 | 9.167E-06 | 5.072E-06 | 5.800E-06 | 1.500E-05 |
| 19120 | 956 | A956V | C | 6.3e-06 | 8.2e-06 | 4.8e-06 | 6.433E-06 | 1.704E-06 | 4.800E-06 | 8.200E-06 |
| 19121 | 957 | S957A | C | 7.6e-06 | 1.1e-05 | 1.5e-05 | 1.120E-05 | 3.704E-06 | 7.600E-06 | 1.500E-05 |
| 19122 | 957 | S957R | C | 5.2e-06 | 7.8e-06 | 6.8e-06 | 6.600E-06 | 1.311E-06 | 5.200E-06 | 7.800E-06 |
| 19123 | 957 | S957N | C | 9.4e-06 | 1e-05 | 6.7e-06 | 8.700E-06 | 1.758E-06 | 6.700E-06 | 1.000E-05 |
| 19124 | 957 | S957D | C | 8.2e-06 | 1e-05 | 1.3e-05 | 1.040E-05 | 2.425E-06 | 8.200E-06 | 1.300E-05 |

| | | | | | | | | | | |
|---|---|---|---|---|---|---|---|---|---|---|
| 19125 | 957 | S957C | C | 5.7e-06 | 9e-06 | 5.1e-06 | 6.600E-06 | 2.100E-06 | 5.100E-06 | 9.000E-06 |
| 19126 | 957 | S957E | C | 6.6e-06 | 1.4e-05 | 8e-06 | 9.533E-06 | 3.931E-06 | 6.600E-06 | 1.400E-05 |
| 19127 | 957 | S957Q | C | 1.2e-05 | 7.2e-06 | 1.1e-05 | 1.007E-05 | 2.532E-06 | 7.200E-06 | 1.200E-05 |
| 19128 | 957 | S957G | C | 7e-06 | 5.3e-06 | 5.3e-06 | 5.867E-06 | 9.815E-07 | 5.300E-06 | 7.000E-06 |
| 19129 | 957 | S957H | C | 8.1e-06 | 1.6e-05 | 6.1e-06 | 1.007E-05 | 5.235E-06 | 6.100E-06 | 1.600E-05 |
| 19130 | 957 | S957I | C | 9.2e-06 | 6e-06 | 9.2e-06 | 8.133E-06 | 1.848E-06 | 6.000E-06 | 9.200E-06 |
| 19131 | 957 | S957L | C | 7.5e-06 | 8.4e-06 | 6.5e-06 | 7.467E-06 | 9.504E-07 | 6.500E-06 | 8.400E-06 |
| 19132 | 957 | S957K | C | 8.6e-06 | 7e-06 | 1e-05 | 8.533E-06 | 1.501E-06 | 7.000E-06 | 1.000E-05 |
| 19133 | 957 | S957M | C | 6.2e-06 | 2.3e-05 | 7.2e-06 | 1.213E-05 | 9.424E-06 | 6.200E-06 | 2.300E-05 |
| 19134 | 957 | S957F | C | 1.3e-05 | 1.1e-05 | 5.5e-06 | 9.833E-06 | 3.884E-06 | 5.500E-06 | 1.300E-05 |
| 19135 | 957 | S957P | C | 6.9e-06 | 6.8e-06 | 8.1e-06 | 7.267E-06 | 7.234E-07 | 6.800E-06 | 8.100E-06 |
| 19136 | 957 | S957S | C | 1e-05 | 1.2e-05 | 8.7e-06 | 1.023E-05 | 1.662E-06 | 8.700E-06 | 1.200E-05 |
| 19137 | 957 | S957T | C | 1.3e-05 | 4.9e-06 | 8.7e-06 | 8.867E-06 | 4.053E-06 | 4.900E-06 | 1.300E-05 |
| 19138 | 957 | S957W | C | 6e-06 | 4.4e-06 | 1e-05 | 6.800E-06 | 2.884E-06 | 4.400E-06 | 1.000E-05 |
| 19139 | 957 | S957Y | C | 1.1e-05 | 8.2e-06 | 9.2e-06 | 9.467E-06 | 1.419E-06 | 8.200E-06 | 1.100E-05 |
| 19140 | 957 | S957V | C | 8.2e-06 | 1e-05 | 7.3e-06 | 8.500E-06 | 1.375E-06 | 7.300E-06 | 1.000E-05 |
| 19141 | 958 | T958A | C | 5.8e-06 | 6.4e-06 | 1.1e-05 | 7.733E-06 | 2.845E-06 | 5.800E-06 | 1.100E-05 |
| 19142 | 958 | T958R | C | 7.4e-06 | 1e-05 | 9e-06 | 8.800E-06 | 1.311E-06 | 7.400E-06 | 1.000E-05 |
| 19143 | 958 | T958N | C | 7.4e-06 | 1.1e-05 | 4.6e-06 | 7.667E-06 | 3.208E-06 | 4.600E-06 | 1.100E-05 |
| 19144 | 958 | T958D | C | 9.3e-06 | 6.5e-06 | 8.7e-06 | 8.167E-06 | 1.474E-06 | 6.500E-06 | 9.300E-06 |
| 19145 | 958 | T958C | C | 9.3e-06 | 1.1e-05 | 1.2e-05 | 1.077E-05 | 1.365E-06 | 9.300E-06 | 1.200E-05 |
| 19146 | 958 | T958E | C | 1.1e-05 | 7.3e-06 | 7.9e-06 | 8.733E-06 | 1.986E-06 | 7.300E-06 | 1.100E-05 |
| 19147 | 958 | T958Q | C | 9e-06 | 1.3e-05 | 6.2e-06 | 9.400E-06 | 3.418E-06 | 6.200E-06 | 1.300E-05 |
| 19148 | 958 | T958G | C | 5.4e-06 | 9e-06 | 5.2e-06 | 6.533E-06 | 2.139E-06 | 5.200E-06 | 9.000E-06 |
| 19149 | 958 | T958H | C | 1e-05 | 1.4e-05 | 1.1e-05 | 1.167E-05 | 2.082E-06 | 1.000E-05 | 1.400E-05 |
| 19150 | 958 | T958I | C | 1e-05 | 1e-05 | 5.8e-06 | 8.600E-06 | 2.425E-06 | 5.800E-06 | 1.000E-05 |
| 19151 | 958 | T958L | C | 1.7e-05 | 1.1e-05 | 4.9e-06 | 1.097E-05 | 6.050E-06 | 4.900E-06 | 1.700E-05 |
| 19152 | 958 | T958K | C | 1.1e-05 | 1.4e-05 | 6.5e-06 | 1.050E-05 | 3.775E-06 | 6.500E-06 | 1.400E-05 |
| 19153 | 958 | T958M | C | 1.3e-05 | 6.5e-06 | 1.3e-05 | 1.083E-05 | 3.753E-06 | 6.500E-06 | 1.300E-05 |
| 19154 | 958 | T958F | C | 7.9e-06 | 9.9e-06 | 7.1e-06 | 8.300E-06 | 1.442E-06 | 7.100E-06 | 9.900E-06 |
| 19155 | 958 | T958P | C | 8.3e-06 | 6.4e-06 | 4.8e-06 | 6.500E-06 | 1.752E-06 | 4.800E-06 | 8.300E-06 |
| 19156 | 958 | T958S | C | 5.4e-06 | 9.1e-06 | 4.1e-06 | 6.200E-06 | 2.594E-06 | 4.100E-06 | 9.100E-06 |
| 19157 | 958 | T958T | C | 9.6e-06 | 4.8e-06 | 6e-06 | 6.800E-06 | 2.498E-06 | 4.800E-06 | 9.600E-06 |
| 19158 | 958 | T958W | C | 5.5e-06 | 6.8e-06 | 6.7e-06 | 6.333E-06 | 7.234E-07 | 5.500E-06 | 6.800E-06 |
| 19159 | 958 | T958Y | C | 1e-05 | 7.1e-06 | 8.5e-06 | 8.533E-06 | 1.450E-06 | 7.100E-06 | 1.000E-05 |
| 19160 | 958 | T958V | C | 7.9e-06 | 8.2e-06 | 3.5e-06 | 6.533E-06 | 2.631E-06 | 3.500E-06 | 8.200E-06 |
| 19161 | 959 | K959A | C | 7.1e-06 | 8.2e-06 | 6.5e-06 | 7.267E-06 | 8.622E-07 | 6.500E-06 | 8.200E-06 |
| 19162 | 959 | K959R | C | 1.2e-05 | 7e-06 | 6.2e-06 | 8.400E-06 | 3.143E-06 | 6.200E-06 | 1.200E-05 |
| 19163 | 959 | K959N | C | 9.2e-06 | 6.7e-06 | 7.7e-06 | 7.867E-06 | 1.258E-06 | 6.700E-06 | 9.200E-06 |
| 19164 | 959 | K959D | C | 1.1e-05 | 6.6e-06 | 6.8e-06 | 8.133E-06 | 2.485E-06 | 6.600E-06 | 1.100E-05 |
| 19165 | 959 | K959C | C | 8.3e-06 | 4.9e-06 | 1.1e-05 | 8.067E-06 | 3.057E-06 | 4.900E-06 | 1.100E-05 |
| 19166 | 959 | K959E | C | 1.4e-05 | 7.1e-06 | 6.1e-06 | 9.067E-06 | 4.302E-06 | 6.100E-06 | 1.400E-05 |
| 19167 | 959 | K959Q | C | 8.6e-06 | 7.5e-06 | 4.5e-06 | 6.867E-06 | 2.122E-06 | 4.500E-06 | 8.600E-06 |
| 19168 | 959 | K959G | C | 6.2e-06 | 6.3e-06 | 7.8e-06 | 6.767E-06 | 8.963E-07 | 6.200E-06 | 7.800E-06 |
| 19169 | 959 | K959H | C | 4e-06 | 4.6e-06 | 8.9e-06 | 5.833E-06 | 2.673E-06 | 4.000E-06 | 8.900E-06 |
| 19170 | 959 | K959I | C | 6.7e-06 | 4.6e-06 | 7.3e-06 | 6.200E-06 | 1.418E-06 | 4.600E-06 | 7.300E-06 |
| 19171 | 959 | K959L | C | 9.8e-06 | 4.9e-06 | 5.3e-06 | 6.667E-06 | 2.721E-06 | 4.900E-06 | 9.800E-06 |
| 19172 | 959 | K959K | C | 8.8e-06 | 7.4e-06 | 1.1e-05 | 9.067E-06 | 1.815E-06 | 7.400E-06 | 1.100E-05 |
| 19173 | 959 | K959M | C | 5.3e-06 | 1.2e-05 | 9.8e-06 | 9.033E-06 | 3.415E-06 | 5.300E-06 | 1.200E-05 |
| 19174 | 959 | K959F | C | 6.4e-06 | 4.3e-06 | 9.1e-06 | 6.600E-06 | 2.406E-06 | 4.300E-06 | 9.100E-06 |
| 19175 | 959 | K959P | C | 7.4e-06 | 9.6e-06 | 7.5e-06 | 8.167E-06 | 1.242E-06 | 7.400E-06 | 9.600E-06 |
| 19176 | 959 | K959S | C | 6.2e-06 | 8.7e-06 | 1.3e-05 | 9.300E-06 | 3.439E-06 | 6.200E-06 | 1.300E-05 |
| 19177 | 959 | K959T | C | 6.5e-06 | 9.9e-06 | 5.2e-06 | 7.200E-06 | 2.427E-06 | 5.200E-06 | 9.900E-06 |
| 19178 | 959 | K959W | C | 1e-05 | 5.9e-06 | 6.3e-06 | 7.400E-06 | 2.261E-06 | 5.900E-06 | 1.000E-05 |
| 19179 | 959 | K959Y | C | 1.2e-05 | 9.3e-06 | 5.5e-06 | 8.933E-06 | 3.265E-06 | 5.500E-06 | 1.200E-05 |
| 19180 | 959 | K959V | C | 7.6e-06 | 6.4e-06 | 1e-05 | 8.000E-06 | 1.833E-06 | 6.400E-06 | 1.000E-05 |
| 19181 | 960 | G960A | C | 9.2e-06 | 9.5e-06 | 1.2e-05 | 1.023E-05 | 1.537E-06 | 9.200E-06 | 1.200E-05 |
| 19182 | 960 | G960R | C | 6.7e-06 | 1.1e-05 | 7.7e-06 | 8.467E-06 | 2.250E-06 | 6.700E-06 | 1.100E-05 |
| 19183 | 960 | G960N | C | 8.7e-06 | 6.1e-06 | 8.5e-06 | 7.767E-06 | 1.447E-06 | 6.100E-06 | 8.700E-06 |
| 19184 | 960 | G960D | C | 6.5e-06 | 5.7e-06 | 1.3e-05 | 8.400E-06 | 4.004E-06 | 5.700E-06 | 1.300E-05 |
| 19185 | 960 | G960C | C | 1.3e-05 | 9.2e-06 | 7.2e-06 | 9.800E-06 | 2.946E-06 | 7.200E-06 | 1.300E-05 |
| 19186 | 960 | G960E | C | 1.4e-05 | 7.3e-06 | 8.3e-06 | 9.867E-06 | 3.614E-06 | 7.300E-06 | 1.400E-05 |
| 19187 | 960 | G960Q | C | 1.3e-05 | 1.2e-05 | 9.6e-06 | 1.153E-05 | 1.747E-06 | 9.600E-06 | 1.300E-05 |
| 19188 | 960 | G960G | C | 3.4e-06 | 5.8e-06 | 4.5e-06 | 4.567E-06 | 1.201E-06 | 3.400E-06 | 5.800E-06 |
| 19189 | 960 | G960H | C | 5.7e-06 | 8.8e-06 | 1.2e-05 | 8.833E-06 | 3.150E-06 | 5.700E-06 | 1.200E-05 |
| 19190 | 960 | G960I | C | 4.4e-06 | 6.1e-06 | 8.5e-06 | 6.333E-06 | 2.060E-06 | 4.400E-06 | 8.500E-06 |
| 19191 | 960 | G960L | C | 1.3e-05 | 7e-06 | 7.1e-06 | 9.033E-06 | 3.436E-06 | 7.000E-06 | 1.300E-05 |
| 19192 | 960 | G960K | C | 1.4e-05 | 5.7e-06 | 7e-06 | 8.900E-06 | 4.464E-06 | 5.700E-06 | 1.400E-05 |
| 19193 | 960 | G960M | C | 6.7e-06 | 5.1e-06 | 7.1e-06 | 6.300E-06 | 1.058E-06 | 5.100E-06 | 7.100E-06 |
| 19194 | 960 | G960F | C | 1.1e-05 | 4.5e-06 | 1.9e-05 | 1.150E-05 | 7.263E-06 | 4.500E-06 | 1.900E-05 |
| 19195 | 960 | G960P | C | 6.4e-06 | 7.5e-06 | 6.1e-06 | 6.667E-06 | 7.371E-07 | 6.100E-06 | 7.500E-06 |
| 19196 | 960 | G960S | C | 5.5e-06 | 5.7e-06 | 9e-06 | 6.733E-06 | 1.966E-06 | 5.500E-06 | 9.000E-06 |
| 19197 | 960 | G960T | C | 8.6e-06 | 6.2e-06 | 7.6e-06 | 7.467E-06 | 1.206E-06 | 6.200E-06 | 8.600E-06 |
| 19198 | 960 | G960W | C | 8e-06 | 7.4e-06 | 1.3e-05 | 9.467E-06 | 3.075E-06 | 7.400E-06 | 1.300E-05 |
| 19199 | 960 | G960Y | C | 6.7e-06 | 9e-06 | 1e-05 | 8.567E-06 | 1.692E-06 | 6.700E-06 | 1.000E-05 |
| 19200 | 960 | G960V | C | 6.6e-06 | 9.1e-06 | 8.8e-06 | 8.167E-06 | 1.365E-06 | 6.600E-06 | 9.100E-06 |
| 19201 | 961 | P961A | C | 1.3e-05 | 5e-06 | 7.3e-06 | 8.433E-06 | 4.119E-06 | 5.000E-06 | 1.300E-05 |
| 19202 | 961 | P961R | C | 4e-06 | 9.5e-06 | 5.5e-06 | 6.333E-06 | 2.843E-06 | 4.000E-06 | 9.500E-06 |
| 19203 | 961 | P961N | C | 9.6e-06 | 8.5e-06 | 9.4e-06 | 9.167E-06 | 5.859E-07 | 8.500E-06 | 9.600E-06 |
| 19204 | 961 | P961D | C | 6.7e-06 | 1.2e-05 | 8.3e-06 | 9.000E-06 | 2.718E-06 | 6.700E-06 | 1.200E-05 |
| 19205 | 961 | P961C | C | 1e-05 | 1.1e-05 | 2.5e-05 | 1.533E-05 | 8.386E-06 | 1.000E-05 | 2.500E-05 |
| 19206 | 961 | P961E | C | 9.3e-06 | 7.2e-06 | 5.2e-06 | 7.233E-06 | 2.050E-06 | 5.200E-06 | 9.300E-06 |
| 19207 | 961 | P961Q | C | 1.1e-05 | 8.8e-06 | 9.7e-06 | 9.833E-06 | 1.106E-06 | 8.800E-06 | 1.100E-05 |
| 19208 | 961 | P961G | C | 8.7e-06 | 1.2e-05 | 4.3e-06 | 8.333E-06 | 3.863E-06 | 4.300E-06 | 1.200E-05 |
| 19209 | 961 | P961H | C | 8.5e-06 | 1.3e-05 | 6e-06 | 9.167E-06 | 3.547E-06 | 6.000E-06 | 1.300E-05 |

| | | | | | | | | | | |
|---|---|---|---|---|---|---|---|---|---|---|
| 19210 | 961 | P961I | C | 1e-05 | 8.1e-06 | 6.1e-06 | 8.067E-06 | 1.950E-06 | 6.100E-06 | 1.000E-05 |
| 19211 | 961 | P961L | C | 6.8e-06 | 1.2e-05 | 9.1e-06 | 9.300E-06 | 2.606E-06 | 6.800E-06 | 1.200E-05 |
| 19212 | 961 | P961K | C | 8.8e-06 | 8.7e-06 | 5.9e-06 | 7.800E-06 | 1.646E-06 | 5.900E-06 | 8.800E-06 |
| 19213 | 961 | P961M | C | 8.5e-06 | 1.8e-05 | 6.6e-06 | 1.103E-05 | 6.108E-06 | 6.600E-06 | 1.800E-05 |
| 19214 | 961 | P961F | C | 7.5e-06 | 8.6e-06 | 6.9e-06 | 7.667E-06 | 8.622E-07 | 6.900E-06 | 8.600E-06 |
| 19215 | 961 | P961P | C | 8.5e-06 | 9.9e-06 | 8.5e-06 | 8.967E-06 | 8.083E-07 | 8.500E-06 | 9.900E-06 |
| 19216 | 961 | P961S | C | 1.7e-05 | 6.5e-06 | 6.7e-06 | 1.007E-05 | 6.005E-06 | 6.500E-06 | 1.700E-05 |
| 19217 | 961 | P961T | C | 9.5e-06 | 5.6e-06 | 7.1e-06 | 7.400E-06 | 1.967E-06 | 5.600E-06 | 9.500E-06 |
| 19218 | 961 | P961W | C | 1.1e-05 | 1.1e-05 | 9.2e-06 | 1.040E-05 | 1.039E-06 | 9.200E-06 | 1.100E-05 |
| 19219 | 961 | P961Y | C | 5.5e-06 | 4.4e-06 | 5.6e-06 | 5.167E-06 | 6.658E-07 | 4.400E-06 | 5.600E-06 |
| 19220 | 961 | P961V | C | 5.5e-06 | 1e-05 | 1.2e-05 | 9.167E-06 | 3.329E-06 | 5.500E-06 | 1.200E-05 |
| 19221 | 962 | S962A | C | 9.7e-06 | 6e-06 | 1e-05 | 8.567E-06 | 2.228E-06 | 6.000E-06 | 1.000E-05 |
| 19222 | 962 | S962R | C | 9.7e-06 | 5.4e-06 | 5.3e-06 | 6.800E-06 | 2.512E-06 | 5.300E-06 | 9.700E-06 |
| 19223 | 962 | S962N | C | 8e-06 | 9.2e-06 | 5.1e-06 | 7.433E-06 | 2.108E-06 | 5.100E-06 | 9.200E-06 |
| 19224 | 962 | S962D | C | 1.4e-05 | 7e-06 | 1.3e-05 | 1.133E-05 | 3.786E-06 | 7.000E-06 | 1.400E-05 |
| 19225 | 962 | S962C | C | 5.3e-06 | 8.1e-06 | 5.4e-06 | 6.267E-06 | 1.589E-06 | 5.300E-06 | 8.100E-06 |
| 19226 | 962 | S962E | C | 1.2e-05 | 8.5e-06 | 9e-06 | 9.833E-06 | 1.893E-06 | 8.500E-06 | 1.200E-05 |
| 19227 | 962 | S962Q | C | 9.8e-06 | 7.8e-06 | 8.2e-06 | 8.600E-06 | 1.058E-06 | 7.800E-06 | 9.800E-06 |
| 19228 | 962 | S962G | C | 1.5e-05 | 7.7e-06 | 7.2e-06 | 9.967E-06 | 4.366E-06 | 7.200E-06 | 1.500E-05 |
| 19229 | 962 | S962H | C | 7.6e-06 | 6.5e-06 | 6.5e-06 | 6.867E-06 | 6.351E-07 | 6.500E-06 | 7.600E-06 |
| 19230 | 962 | S962I | C | 9.4e-06 | 1.3e-05 | 4.3e-06 | 8.900E-06 | 4.371E-06 | 4.300E-06 | 1.300E-05 |
| 19231 | 962 | S962L | C | 9.4e-06 | 6.6e-06 | 1e-05 | 8.667E-06 | 1.815E-06 | 6.600E-06 | 1.000E-05 |
| 19232 | 962 | S962K | C | 5e-06 | 7.4e-06 | 6.4e-06 | 6.267E-06 | 1.206E-06 | 5.000E-06 | 7.400E-06 |
| 19233 | 962 | S962M | C | 1.1e-05 | 3.8e-06 | 8.2e-06 | 7.667E-06 | 3.630E-06 | 3.800E-06 | 1.100E-05 |
| 19234 | 962 | S962F | C | 8.2e-06 | 5.3e-06 | 9.9e-06 | 7.800E-06 | 2.326E-06 | 5.300E-06 | 9.900E-06 |
| 19235 | 962 | S962P | C | 1.9e-05 | 5.4e-06 | 9.3e-06 | 1.123E-05 | 7.003E-06 | 5.400E-06 | 1.900E-05 |
| 19236 | 962 | S962S | C | 8.3e-06 | 1e-05 | 7.5e-06 | 8.600E-06 | 1.277E-06 | 7.500E-06 | 1.000E-05 |
| 19237 | 962 | S962T | C | 7.9e-06 | 8.5e-06 | 7.9e-06 | 8.100E-06 | 3.464E-07 | 7.900E-06 | 8.500E-06 |
| 19238 | 962 | S962W | C | 7.1e-06 | 5.5e-06 | 1.1e-05 | 7.867E-06 | 2.829E-06 | 5.500E-06 | 1.100E-05 |
| 19239 | 962 | S962Y | C | 2.7e-06 | 4e-06 | 9.4e-06 | 5.367E-06 | 3.553E-06 | 2.700E-06 | 9.400E-06 |
| 19240 | 962 | S962V | C | 4.9e-06 | 7.9e-06 | 8.1e-06 | 6.967E-06 | 1.793E-06 | 4.900E-06 | 8.100E-06 |
| 19241 | 963 | V963A | C | 7e-06 | 5.2e-06 | 1.1e-05 | 7.733E-06 | 2.969E-06 | 5.200E-06 | 1.100E-05 |
| 19242 | 963 | V963R | C | 7.6e-06 | 7.1e-06 | 4.1e-06 | 6.267E-06 | 1.893E-06 | 4.100E-06 | 7.600E-06 |
| 19243 | 963 | V963N | C | 9e-06 | 8.2e-06 | 1.1e-05 | 9.400E-06 | 1.442E-06 | 8.200E-06 | 1.100E-05 |
| 19244 | 963 | V963D | C | 6.9e-06 | 1e-05 | 1.3e-05 | 9.967E-06 | 3.050E-06 | 6.900E-06 | 1.300E-05 |
| 19245 | 963 | V963C | C | 5.9e-06 | 1e-05 | 1.8e-05 | 1.130E-05 | 6.154E-06 | 5.900E-06 | 1.800E-05 |
| 19246 | 963 | V963E | C | 5e-06 | 6.4e-06 | 5.8e-06 | 5.733E-06 | 7.024E-07 | 5.000E-06 | 6.400E-06 |
| 19247 | 963 | V963Q | C | 6.5e-06 | 7e-06 | 7.4e-06 | 6.967E-06 | 4.509E-07 | 6.500E-06 | 7.400E-06 |
| 19248 | 963 | V963G | C | 6.9e-06 | 9.2e-06 | 7.2e-06 | 7.767E-06 | 1.250E-06 | 6.900E-06 | 9.200E-06 |
| 19249 | 963 | V963H | C | 1e-05 | 6.9e-06 | 5.3e-06 | 7.400E-06 | 2.390E-06 | 5.300E-06 | 1.000E-05 |
| 19250 | 963 | V963I | C | 8.6e-06 | 1.1e-05 | 7.7e-06 | 9.100E-06 | 1.706E-06 | 7.700E-06 | 1.100E-05 |
| 19251 | 963 | V963L | C | 7.1e-06 | 7.2e-06 | 9.1e-06 | 7.800E-06 | 1.127E-06 | 7.100E-06 | 9.100E-06 |
| 19252 | 963 | V963K | C | 8.9e-06 | 1.1e-05 | 5e-06 | 8.300E-06 | 3.045E-06 | 5.000E-06 | 1.100E-05 |
| 19253 | 963 | V963M | C | 7.4e-06 | 6.9e-06 | 9.3e-06 | 7.867E-06 | 1.266E-06 | 6.900E-06 | 9.300E-06 |
| 19254 | 963 | V963F | C | 1.3e-05 | 9.5e-06 | 6.4e-06 | 9.633E-06 | 3.302E-06 | 6.400E-06 | 1.300E-05 |
| 19255 | 963 | V963P | C | 9.5e-06 | 8.5e-06 | 6.9e-06 | 8.300E-06 | 1.311E-06 | 6.900E-06 | 9.500E-06 |
| 19256 | 963 | V963S | C | 8.6e-06 | 1.1e-05 | 1.3e-05 | 1.087E-05 | 2.203E-06 | 8.600E-06 | 1.300E-05 |
| 19257 | 963 | V963T | C | 9.2e-06 | 7.2e-06 | 8.4e-06 | 8.267E-06 | 1.007E-06 | 7.200E-06 | 9.200E-06 |
| 19258 | 963 | V963W | C | 1.2e-05 | 4.9e-06 | 9.3e-06 | 8.733E-06 | 3.584E-06 | 4.900E-06 | 1.200E-05 |
| 19259 | 963 | V963Y | C | 9.5e-06 | 5.8e-06 | 8.3e-06 | 7.867E-06 | 1.888E-06 | 5.800E-06 | 9.500E-06 |
| 19260 | 963 | V963V | C | 1.3e-05 | 9e-06 | 5.8e-06 | 9.267E-06 | 3.607E-06 | 5.800E-06 | 1.300E-05 |
| 19261 | 964 | F964A | C | 1e-05 | 9.7e-06 | 9.9e-06 | 9.867E-06 | 1.528E-07 | 9.700E-06 | 1.000E-05 |
| 19262 | 964 | F964R | C | 8.1e-06 | 4.1e-06 | 7.6e-06 | 6.600E-06 | 2.179E-06 | 4.100E-06 | 8.100E-06 |
| 19263 | 964 | F964N | C | 6.2e-06 | 4.9e-06 | 9.7e-06 | 6.933E-06 | 2.483E-06 | 4.900E-06 | 9.700E-06 |
| 19264 | 964 | F964D | C | 5.6e-06 | 8.8e-06 | 9.4e-06 | 7.933E-06 | 2.043E-06 | 5.600E-06 | 9.400E-06 |
| 19265 | 964 | F964C | C | 9.1e-06 | 8.8e-06 | 1.3e-05 | 1.030E-05 | 2.343E-06 | 8.800E-06 | 1.300E-05 |
| 19266 | 964 | F964E | C | 1.3e-05 | 8.3e-06 | 9.4e-06 | 1.023E-05 | 2.458E-06 | 8.300E-06 | 1.300E-05 |
| 19267 | 964 | F964Q | C | 3.2e-06 | 9.1e-06 | 7.4e-06 | 6.567E-06 | 3.037E-06 | 3.200E-06 | 9.100E-06 |
| 19268 | 964 | F964G | C | 1.3e-05 | 6.4e-06 | 8.3e-06 | 9.233E-06 | 3.398E-06 | 6.400E-06 | 1.300E-05 |
| 19269 | 964 | F964H | C | 8e-06 | 7.6e-06 | 3.3e-06 | 6.300E-06 | 2.606E-06 | 3.300E-06 | 8.000E-06 |
| 19270 | 964 | F964I | C | 6.9e-06 | 9.4e-06 | 1e-05 | 8.767E-06 | 1.644E-06 | 6.900E-06 | 1.000E-05 |
| 19271 | 964 | F964L | C | 8.6e-06 | 9.4e-06 | 8.7e-06 | 8.900E-06 | 4.359E-07 | 8.600E-06 | 9.400E-06 |
| 19272 | 964 | F964K | C | 6.8e-06 | 9.8e-06 | 1.3e-05 | 9.867E-06 | 3.101E-06 | 6.800E-06 | 1.300E-05 |
| 19273 | 964 | F964M | C | 6.8e-06 | 9.5e-06 | 1.5e-05 | 1.043E-05 | 4.179E-06 | 6.800E-06 | 1.500E-05 |
| 19274 | 964 | F964F | C | 8e-06 | 1.4e-05 | 9.9e-06 | 1.063E-05 | 3.066E-06 | 8.000E-06 | 1.400E-05 |
| 19275 | 964 | F964P | C | 1.2e-05 | 9.2e-06 | 6e-06 | 9.067E-06 | 3.002E-06 | 6.000E-06 | 1.200E-05 |
| 19276 | 964 | F964S | C | 8.3e-06 | 1e-05 | 6.9e-06 | 8.400E-06 | 1.552E-06 | 6.900E-06 | 1.000E-05 |
| 19277 | 964 | F964T | C | 9.1e-06 | 8.8e-06 | 9.5e-06 | 9.133E-06 | 3.512E-07 | 8.800E-06 | 9.500E-06 |
| 19278 | 964 | F964W | C | 1.1e-05 | 5.9e-06 | 1.5e-05 | 1.063E-05 | 4.561E-06 | 5.900E-06 | 1.500E-05 |
| 19279 | 964 | F964Y | C | 9.4e-06 | 8.4e-06 | 6.3e-06 | 8.033E-06 | 1.582E-06 | 6.300E-06 | 9.400E-06 |
| 19280 | 964 | F964V | C | 8.9e-06 | 1.3e-05 | 8.9e-06 | 1.027E-05 | 2.367E-06 | 8.900E-06 | 1.300E-05 |
| 19281 | 965 | P965A | C | 8.8e-06 | 9.7e-06 | 5.2e-06 | 7.900E-06 | 2.381E-06 | 5.200E-06 | 9.700E-06 |
| 19282 | 965 | P965R | C | 8.8e-06 | 4.9e-06 | 6.7e-06 | 6.800E-06 | 1.952E-06 | 4.900E-06 | 8.800E-06 |
| 19283 | 965 | P965N | C | 9.1e-06 | 4.4e-06 | 6.8e-06 | 6.767E-06 | 2.350E-06 | 4.400E-06 | 9.100E-06 |
| 19284 | 965 | P965D | C | 5.4e-06 | 6.7e-06 | 3.3e-06 | 5.133E-06 | 1.716E-06 | 3.300E-06 | 6.700E-06 |
| 19285 | 965 | P965C | C | 9.4e-06 | 9.7e-06 | 7.9e-06 | 9.000E-06 | 9.644E-07 | 7.900E-06 | 9.700E-06 |
| 19286 | 965 | P965E | C | 1.1e-05 | 7.8e-06 | 9e-06 | 9.267E-06 | 1.617E-06 | 7.800E-06 | 1.100E-05 |
| 19287 | 965 | P965Q | C | 6.2e-06 | 6.8e-06 | 7.9e-06 | 6.967E-06 | 8.622E-07 | 6.200E-06 | 7.900E-06 |
| 19288 | 965 | P965G | C | 4.1e-06 | 5e-06 | 8.3e-06 | 5.800E-06 | 2.211E-06 | 4.100E-06 | 8.300E-06 |
| 19289 | 965 | P965H | C | 1.4e-05 | 6.5e-06 | 8.9e-06 | 9.800E-06 | 3.830E-06 | 6.500E-06 | 1.400E-05 |
| 19290 | 965 | P965I | C | 8.3e-06 | 9e-06 | 4.3e-06 | 7.200E-06 | 2.536E-06 | 4.300E-06 | 9.000E-06 |
| 19291 | 965 | P965L | C | 1.1e-05 | 8.1e-06 | 7e-06 | 8.700E-06 | 2.066E-06 | 7.000E-06 | 1.100E-05 |
| 19292 | 965 | P965K | C | 7e-06 | 7.3e-06 | 7.2e-06 | 7.167E-06 | 1.528E-07 | 7.000E-06 | 7.300E-06 |
| 19293 | 965 | P965M | C | 1.3e-05 | 5.7e-06 | 5.2e-06 | 7.967E-06 | 4.366E-06 | 5.200E-06 | 1.300E-05 |
| 19294 | 965 | P965F | C | 1.4e-05 | 4.1e-06 | 6.8e-06 | 8.300E-06 | 5.118E-06 | 4.100E-06 | 1.400E-05 |

| | | | | | | | | | | |
|---|---|---|---|---|---|---|---|---|---|---|
| 19295 | 965 | P965P | C | 9.1e-06 | 7.2e-06 | 6e-06 | 7.433E-06 | 1.563E-06 | 6.000E-06 | 9.100E-06 |
| 19296 | 965 | P965S | C | 1.3e-05 | 8.4e-06 | 4.7e-06 | 8.700E-06 | 4.158E-06 | 4.700E-06 | 1.300E-05 |
| 19297 | 965 | P965T | C | 7.5e-06 | 7.2e-06 | 5.8e-06 | 6.833E-06 | 9.074E-07 | 5.800E-06 | 7.500E-06 |
| 19298 | 965 | P965W | C | 8.2e-06 | 8.4e-06 | 9.7e-06 | 8.767E-06 | 8.145E-07 | 8.200E-06 | 9.700E-06 |
| 19299 | 965 | P965Y | C | 1.3e-05 | 6.5e-06 | 9.7e-06 | 9.733E-06 | 3.250E-06 | 6.500E-06 | 1.300E-05 |
| 19300 | 965 | P965V | C | 7.8e-06 | 6.4e-06 | 1e-05 | 8.067E-06 | 1.815E-06 | 6.400E-06 | 1.000E-05 |
| 19301 | 966 | L966A | C | 8.5e-06 | 6.6e-06 | 1e-05 | 8.367E-06 | 1.704E-06 | 6.600E-06 | 1.000E-05 |
| 19302 | 966 | L966R | C | 1.8e-05 | 7e-06 | 3.9e-06 | 9.633E-06 | 7.410E-06 | 3.900E-06 | 1.800E-05 |
| 19303 | 966 | L966N | C | 1.1e-05 | 1.1e-05 | 8.5e-06 | 1.017E-05 | 1.443E-06 | 8.500E-06 | 1.100E-05 |
| 19304 | 966 | L966D | C | 5.4e-06 | 4.6e-06 | 6.7e-06 | 5.567E-06 | 1.060E-06 | 4.600E-06 | 6.700E-06 |
| 19305 | 966 | L966C | C | 5.5e-06 | 5.3e-06 | 8e-06 | 6.267E-06 | 1.504E-06 | 5.300E-06 | 8.000E-06 |
| 19306 | 966 | L966E | C | 6.6e-06 | 1.2e-05 | 3.7e-06 | 7.433E-06 | 4.212E-06 | 3.700E-06 | 1.200E-05 |
| 19307 | 966 | L966Q | C | 6.4e-06 | 8.2e-06 | 8.8e-06 | 7.800E-06 | 1.249E-06 | 6.400E-06 | 8.800E-06 |
| 19308 | 966 | L966G | C | 1.1e-05 | 8.3e-06 | 1.3e-05 | 1.077E-05 | 2.359E-06 | 8.300E-06 | 1.300E-05 |
| 19309 | 966 | L966H | C | 6.6e-06 | 8.8e-06 | 8.1e-06 | 7.833E-06 | 1.124E-06 | 6.600E-06 | 8.800E-06 |
| 19310 | 966 | L966I | C | 9.8e-06 | 9.5e-06 | 5.8e-06 | 8.367E-06 | 2.228E-06 | 5.800E-06 | 9.800E-06 |
| 19311 | 966 | L966L | C | 4.1e-06 | 8.5e-06 | 6.8e-06 | 6.467E-06 | 2.219E-06 | 4.100E-06 | 8.500E-06 |
| 19312 | 966 | L966K | C | 6.5e-06 | 5.3e-06 | 4.7e-06 | 5.500E-06 | 9.165E-07 | 4.700E-06 | 6.500E-06 |
| 19313 | 966 | L966M | C | 8.3e-06 | 8.9e-06 | 4.7e-06 | 7.300E-06 | 2.272E-06 | 4.700E-06 | 8.900E-06 |
| 19314 | 966 | L966F | C | 8.8e-06 | 6.3e-06 | 6.2e-06 | 7.100E-06 | 1.473E-06 | 6.200E-06 | 8.800E-06 |
| 19315 | 966 | L966P | C | 5.6e-06 | 5.4e-06 | 9e-06 | 6.667E-06 | 2.023E-06 | 5.400E-06 | 9.000E-06 |
| 19316 | 966 | L966S | C | 8.8e-06 | 1.1e-05 | 5.1e-06 | 8.300E-06 | 2.982E-06 | 5.100E-06 | 1.100E-05 |
| 19317 | 966 | L966T | C | 1.3e-05 | 4.7e-06 | 8.8e-06 | 8.833E-06 | 4.150E-06 | 4.700E-06 | 1.300E-05 |
| 19318 | 966 | L966W | C | 9.2e-06 | 7.3e-06 | 1.2e-05 | 9.500E-06 | 2.364E-06 | 7.300E-06 | 1.200E-05 |
| 19319 | 966 | L966Y | C | 1.1e-05 | 9.9e-06 | 9.3e-06 | 1.007E-05 | 8.622E-07 | 9.300E-06 | 1.100E-05 |
| 19320 | 966 | L966V | C | 3.1e-06 | 5.1e-06 | 8.8e-06 | 5.667E-06 | 2.892E-06 | 3.100E-06 | 8.800E-06 |
| 19321 | 967 | A967A | C | 1.3e-05 | 9.4e-06 | 6.9e-06 | 9.767E-06 | 3.066E-06 | 6.900E-06 | 1.300E-05 |
| 19322 | 967 | A967R | C | 7.4e-06 | 8.5e-06 | 1e-05 | 8.633E-06 | 1.305E-06 | 7.400E-06 | 1.000E-05 |
| 19323 | 967 | A967N | C | 8.1e-06 | 1.5e-05 | 1.3e-05 | 1.203E-05 | 3.550E-06 | 8.100E-06 | 1.500E-05 |
| 19324 | 967 | A967D | C | 6.2e-06 | 8.2e-06 | 7.7e-06 | 7.367E-06 | 1.041E-06 | 6.200E-06 | 8.200E-06 |
| 19325 | 967 | A967C | C | 9e-06 | 4.4e-06 | 8.1e-06 | 7.167E-06 | 2.438E-06 | 4.400E-06 | 9.000E-06 |
| 19326 | 967 | A967E | C | 9.2e-06 | 8.2e-06 | 5.9e-06 | 7.767E-06 | 1.692E-06 | 5.900E-06 | 9.200E-06 |
| 19327 | 967 | A967Q | C | 6.5e-06 | 1.2e-05 | 1e-05 | 9.500E-06 | 2.784E-06 | 6.500E-06 | 1.200E-05 |
| 19328 | 967 | A967G | C | 9.2e-06 | 7.5e-06 | 6.6e-06 | 7.767E-06 | 1.320E-06 | 6.600E-06 | 9.200E-06 |
| 19329 | 967 | A967H | C | 1.1e-05 | 7.8e-06 | 6.2e-06 | 8.333E-06 | 2.444E-06 | 6.200E-06 | 1.100E-05 |
| 19330 | 967 | A967I | C | 9.9e-06 | 8.8e-06 | 4.7e-06 | 7.800E-06 | 2.740E-06 | 4.700E-06 | 9.900E-06 |
| 19331 | 967 | A967L | C | 1.2e-05 | 1.2e-05 | 1.1e-05 | 1.167E-05 | 5.774E-07 | 1.100E-05 | 1.200E-05 |
| 19332 | 967 | A967K | C | 6e-06 | 1.1e-05 | 6.2e-06 | 7.733E-06 | 2.831E-06 | 6.000E-06 | 1.100E-05 |
| 19333 | 967 | A967M | C | 6.7e-06 | 7.7e-06 | 6.7e-06 | 7.033E-06 | 5.774E-07 | 6.700E-06 | 7.700E-06 |
| 19334 | 967 | A967F | C | 4.8e-06 | 1.1e-05 | 8.6e-06 | 8.133E-06 | 3.126E-06 | 4.800E-06 | 1.100E-05 |
| 19335 | 967 | A967P | C | 4.8e-06 | 5.7e-06 | 7.5e-06 | 6.000E-06 | 1.375E-06 | 4.800E-06 | 7.500E-06 |
| 19336 | 967 | A967S | C | 6.7e-06 | 1.5e-05 | 9.1e-06 | 1.027E-05 | 4.271E-06 | 6.700E-06 | 1.500E-05 |
| 19337 | 967 | A967T | C | 4.8e-06 | 7.9e-06 | 5.8e-06 | 6.167E-06 | 1.582E-06 | 4.800E-06 | 7.900E-06 |
| 19338 | 967 | A967W | C | 1e-05 | 1e-05 | 6.1e-06 | 8.700E-06 | 2.252E-06 | 6.100E-06 | 1.000E-05 |
| 19339 | 967 | A967Y | C | 6.8e-06 | 5.3e-06 | 7.9e-06 | 6.667E-06 | 1.305E-06 | 5.300E-06 | 7.900E-06 |
| 19340 | 967 | A967V | C | 7.8e-06 | 4.5e-06 | 1.2e-05 | 8.100E-06 | 3.759E-06 | 4.500E-06 | 1.200E-05 |
| 19341 | 968 | P968A | C | 1.1e-05 | 9.7e-06 | 6.5e-06 | 9.067E-06 | 2.316E-06 | 6.500E-06 | 1.100E-05 |
| 19342 | 968 | P968R | C | 1.3e-05 | 4.5e-06 | 5.2e-06 | 7.567E-06 | 4.718E-06 | 4.500E-06 | 1.300E-05 |
| 19343 | 968 | P968N | C | 9.4e-06 | 1.2e-05 | 5.4e-06 | 8.933E-06 | 3.325E-06 | 5.400E-06 | 1.200E-05 |
| 19344 | 968 | P968D | C | 1e-05 | 1e-05 | 1.1e-05 | 1.033E-05 | 5.774E-07 | 1.000E-05 | 1.100E-05 |
| 19345 | 968 | P968C | C | 8.9e-06 | 5.8e-06 | 6.9e-06 | 7.200E-06 | 1.572E-06 | 5.800E-06 | 8.900E-06 |
| 19346 | 968 | P968E | C | 6.9e-06 | 1.4e-05 | 1.2e-05 | 1.097E-05 | 3.661E-06 | 6.900E-06 | 1.400E-05 |
| 19347 | 968 | P968Q | C | 6.3e-06 | 1.3e-05 | 1.1e-05 | 1.010E-05 | 3.439E-06 | 6.300E-06 | 1.300E-05 |
| 19348 | 968 | P968G | C | 8e-06 | 8.1e-06 | 9.4e-06 | 8.500E-06 | 7.810E-07 | 8.000E-06 | 9.400E-06 |
| 19349 | 968 | P968H | C | 9e-06 | 8.3e-06 | 1.2e-05 | 9.767E-06 | 1.966E-06 | 8.300E-06 | 1.200E-05 |
| 19350 | 968 | P968I | C | 6e-06 | 1.1e-05 | 4.7e-06 | 7.233E-06 | 3.326E-06 | 4.700E-06 | 1.100E-05 |
| 19351 | 968 | P968L | C | 8.8e-06 | 1.1e-05 | 5.2e-06 | 8.333E-06 | 2.928E-06 | 5.200E-06 | 1.100E-05 |
| 19352 | 968 | P968K | C | 6.9e-06 | 5.1e-06 | 6.7e-06 | 6.233E-06 | 9.866E-07 | 5.100E-06 | 6.900E-06 |
| 19353 | 968 | P968M | C | 6.3e-06 | 1.1e-05 | 7.7e-06 | 8.333E-06 | 2.413E-06 | 6.300E-06 | 1.100E-05 |
| 19354 | 968 | P968F | C | 9.7e-06 | 4.9e-06 | 1e-05 | 8.200E-06 | 2.862E-06 | 4.900E-06 | 1.000E-05 |
| 19355 | 968 | P968P | C | 7.9e-06 | 1.4e-05 | 8.5e-06 | 1.013E-05 | 3.362E-06 | 7.900E-06 | 1.400E-05 |
| 19356 | 968 | P968S | C | 9e-06 | 1e-05 | 8.6e-06 | 9.200E-06 | 7.211E-07 | 8.600E-06 | 1.000E-05 |
| 19357 | 968 | P968T | C | 4.4e-06 | 4.3e-06 | 5.3e-06 | 4.667E-06 | 5.508E-07 | 4.300E-06 | 5.300E-06 |
| 19358 | 968 | P968W | C | 5.9e-06 | 1.4e-05 | 9.6e-06 | 9.833E-06 | 4.055E-06 | 5.900E-06 | 1.400E-05 |
| 19359 | 968 | P968Y | C | 4.6e-06 | 9.7e-06 | 9.6e-06 | 7.967E-06 | 2.916E-06 | 4.600E-06 | 9.700E-06 |
| 19360 | 968 | P968V | C | 1.2e-05 | 9.8e-06 | 4.6e-06 | 8.800E-06 | 3.800E-06 | 4.600E-06 | 1.200E-05 |
| 19361 | 969 | S969A | C | 8.5e-06 | 1e-05 | 8.5e-06 | 9.000E-06 | 8.660E-07 | 8.500E-06 | 1.000E-05 |
| 19362 | 969 | S969R | C | 1.2e-05 | 7e-06 | 9.8e-06 | 9.600E-06 | 2.506E-06 | 7.000E-06 | 1.200E-05 |
| 19363 | 969 | S969N | C | 9.2e-06 | 4.9e-06 | 7e-06 | 7.033E-06 | 2.150E-06 | 4.900E-06 | 9.200E-06 |
| 19364 | 969 | S969D | C | 1.2e-05 | 6.9e-06 | 1.2e-05 | 1.030E-05 | 2.944E-06 | 6.900E-06 | 1.200E-05 |
| 19365 | 969 | S969C | C | 6.1e-06 | 5e-06 | 1.3e-05 | 8.033E-06 | 4.336E-06 | 5.000E-06 | 1.300E-05 |
| 19366 | 969 | S969E | C | 9.8e-06 | 6.5e-06 | 6.1e-06 | 7.467E-06 | 2.031E-06 | 6.100E-06 | 9.800E-06 |
| 19367 | 969 | S969Q | C | 4.2e-06 | 5.3e-06 | 5.4e-06 | 4.967E-06 | 6.658E-07 | 4.200E-06 | 5.400E-06 |
| 19368 | 969 | S969G | C | 8.1e-06 | 9.8e-06 | 6.3e-06 | 8.067E-06 | 1.750E-06 | 6.300E-06 | 9.800E-06 |
| 19369 | 969 | S969H | C | 9.1e-06 | 4.2e-06 | 8.7e-06 | 7.333E-06 | 2.721E-06 | 4.200E-06 | 9.100E-06 |
| 19370 | 969 | S969I | C | 6.8e-06 | 8.5e-06 | 7.4e-06 | 7.567E-06 | 8.622E-07 | 6.800E-06 | 8.500E-06 |
| 19371 | 969 | S969L | C | 6.3e-06 | 5e-06 | 7.5e-06 | 6.267E-06 | 1.250E-06 | 5.000E-06 | 7.500E-06 |
| 19372 | 969 | S969K | C | 1.5e-05 | 8.8e-06 | 6.4e-06 | 1.007E-05 | 4.438E-06 | 6.400E-06 | 1.500E-05 |
| 19373 | 969 | S969M | C | 1e-05 | 6.8e-06 | 1.2e-05 | 9.600E-06 | 2.623E-06 | 6.800E-06 | 1.200E-05 |
| 19374 | 969 | S969F | C | 1.4e-05 | 1e-05 | 6.2e-06 | 1.007E-05 | 3.900E-06 | 6.200E-06 | 1.400E-05 |
| 19375 | 969 | S969P | C | 6.6e-06 | 5.8e-06 | 6.9e-06 | 6.433E-06 | 5.686E-07 | 5.800E-06 | 6.900E-06 |
| 19376 | 969 | S969S | C | 1.1e-05 | 6.7e-06 | 9.7e-06 | 9.133E-06 | 2.205E-06 | 6.700E-06 | 1.100E-05 |
| 19377 | 969 | S969T | C | 1.4e-05 | 1.3e-05 | 5.8e-06 | 1.093E-05 | 4.474E-06 | 5.800E-06 | 1.400E-05 |
| 19378 | 969 | S969W | C | 1.1e-05 | 6.3e-06 | 8.7e-06 | 8.667E-06 | 2.350E-06 | 6.300E-06 | 1.100E-05 |
| 19379 | 969 | S969Y | C | 8.5e-06 | 7.3e-06 | 9.8e-06 | 8.533E-06 | 1.250E-06 | 7.300E-06 | 9.800E-06 |

| | | | | | | | | | | |
|---|---|---|---|---|---|---|---|---|---|---|
| 19380 | 969 | S969V | C | 9.8e-06 | 3.8e-06 | 8.8e-06 | 7.467E-06 | 3.215E-06 | 3.800E-06 | 9.800E-06 |
| 19381 | 970 | S970A | C | 1.1e-05 | 6.5e-06 | 9.6e-06 | 9.033E-06 | 2.303E-06 | 6.500E-06 | 1.100E-05 |
| 19382 | 970 | S970R | C | 1.3e-05 | 5.6e-06 | 6.5e-06 | 8.367E-06 | 4.038E-06 | 5.600E-06 | 1.300E-05 |
| 19383 | 970 | S970N | C | 9.6e-06 | 6.2e-06 | 7e-06 | 7.600E-06 | 1.778E-06 | 6.200E-06 | 9.600E-06 |
| 19384 | 970 | S970D | C | 7.7e-06 | 4.8e-06 | 7.2e-06 | 6.567E-06 | 1.550E-06 | 4.800E-06 | 7.700E-06 |
| 19385 | 970 | S970C | C | 8.6e-06 | 8.7e-06 | 5.4e-06 | 7.567E-06 | 1.877E-06 | 5.400E-06 | 8.700E-06 |
| 19386 | 970 | S970E | C | 6.4e-06 | 5.4e-06 | 5.6e-06 | 5.800E-06 | 5.292E-07 | 5.400E-06 | 6.400E-06 |
| 19387 | 970 | S970Q | C | 1.4e-05 | 1e-05 | 1e-05 | 1.133E-05 | 2.309E-06 | 1.000E-05 | 1.400E-05 |
| 19388 | 970 | S970G | C | 5.5e-06 | 6.6e-06 | 8.3e-06 | 6.800E-06 | 1.411E-06 | 5.500E-06 | 8.300E-06 |
| 19389 | 970 | S970H | C | 6.2e-06 | 4.9e-06 | 8.9e-06 | 6.667E-06 | 2.040E-06 | 4.900E-06 | 8.900E-06 |
| 19390 | 970 | S970I | C | 1.2e-05 | 1e-05 | 6.1e-06 | 9.367E-06 | 3.001E-06 | 6.100E-06 | 1.200E-05 |
| 19391 | 970 | S970L | C | 1e-05 | 1e-05 | 6.6e-06 | 8.867E-06 | 1.963E-06 | 6.600E-06 | 1.000E-05 |
| 19392 | 970 | S970K | C | 5e-06 | 8.3e-06 | 5.8e-06 | 6.367E-06 | 1.721E-06 | 5.000E-06 | 8.300E-06 |
| 19393 | 970 | S970M | C | 1e-05 | 5.2e-06 | 1.1e-05 | 8.733E-06 | 3.101E-06 | 5.200E-06 | 1.100E-05 |
| 19394 | 970 | S970F | C | 6.9e-06 | 9.5e-06 | 7.7e-06 | 8.033E-06 | 1.332E-06 | 6.900E-06 | 9.500E-06 |
| 19395 | 970 | S970P | C | 5.2e-06 | 6.8e-06 | 7.3e-06 | 6.433E-06 | 1.097E-06 | 5.200E-06 | 7.300E-06 |
| 19396 | 970 | S970S | C | 8.2e-06 | 5.3e-06 | 8.4e-06 | 7.300E-06 | 1.735E-06 | 5.300E-06 | 8.400E-06 |
| 19397 | 970 | S970T | C | 4e-06 | 4.8e-06 | 6.2e-06 | 5.000E-06 | 1.114E-06 | 4.000E-06 | 6.200E-06 |
| 19398 | 970 | S970W | C | 1.3e-05 | 6.1e-06 | 9.8e-06 | 9.633E-06 | 3.453E-06 | 6.100E-06 | 1.300E-05 |
| 19399 | 970 | S970Y | C | 5.8e-06 | 5e-06 | 6.5e-06 | 5.767E-06 | 7.506E-07 | 5.000E-06 | 6.500E-06 |
| 19400 | 970 | S970V | C | 5.6e-06 | 7.6e-06 | 7.1e-06 | 6.767E-06 | 1.041E-06 | 5.600E-06 | 7.600E-06 |
| 19401 | 971 | K971A | C | 7.6e-06 | 7.8e-06 | 6.4e-06 | 7.267E-06 | 7.572E-07 | 6.400E-06 | 7.800E-06 |
| 19402 | 971 | K971R | C | 6.9e-06 | 7.3e-06 | 9.5e-06 | 7.900E-06 | 1.400E-06 | 6.900E-06 | 9.500E-06 |
| 19403 | 971 | K971N | C | 9e-06 | 7.4e-06 | 6.8e-06 | 7.733E-06 | 1.137E-06 | 6.800E-06 | 9.000E-06 |
| 19404 | 971 | K971D | C | 5.1e-06 | 9.1e-06 | 5.5e-06 | 6.567E-06 | 2.203E-06 | 5.100E-06 | 9.100E-06 |
| 19405 | 971 | K971C | C | 1.1e-05 | 4.6e-06 | 6.9e-06 | 7.500E-06 | 3.242E-06 | 4.600E-06 | 1.100E-05 |
| 19406 | 971 | K971E | C | 6e-06 | 4.7e-06 | 8.5e-06 | 6.400E-06 | 1.931E-06 | 4.700E-06 | 8.500E-06 |
| 19407 | 971 | K971Q | C | 8e-06 | 6.3e-06 | 6.6e-06 | 6.967E-06 | 9.074E-07 | 6.300E-06 | 8.000E-06 |
| 19408 | 971 | K971G | C | 1.2e-05 | 1.3e-05 | 8.9e-06 | 1.130E-05 | 2.138E-06 | 8.900E-06 | 1.300E-05 |
| 19409 | 971 | K971H | C | 5.9e-06 | 5.3e-06 | 9.8e-06 | 7.000E-06 | 2.443E-06 | 5.300E-06 | 9.800E-06 |
| 19410 | 971 | K971I | C | 6.3e-06 | 5.8e-06 | 7e-06 | 6.367E-06 | 6.028E-07 | 5.800E-06 | 7.000E-06 |
| 19411 | 971 | K971L | C | 9.5e-06 | 9.8e-06 | 1.1e-05 | 1.010E-05 | 7.937E-07 | 9.500E-06 | 1.100E-05 |
| 19412 | 971 | K971K | C | 8.3e-06 | 6.7e-06 | 5.3e-06 | 6.767E-06 | 1.501E-06 | 5.300E-06 | 8.300E-06 |
| 19413 | 971 | K971M | C | 6.7e-06 | 1.2e-05 | 4.5e-06 | 7.733E-06 | 3.855E-06 | 4.500E-06 | 1.200E-05 |
| 19414 | 971 | K971F | C | 6.8e-06 | 7.3e-06 | 6e-06 | 6.700E-06 | 6.557E-07 | 6.000E-06 | 7.300E-06 |
| 19415 | 971 | K971P | C | 5.7e-06 | 6.5e-06 | 4.6e-06 | 5.600E-06 | 9.539E-07 | 4.600E-06 | 6.500E-06 |
| 19416 | 971 | K971S | C | 1.2e-05 | 8.2e-06 | 3.5e-06 | 7.900E-06 | 4.258E-06 | 3.500E-06 | 1.200E-05 |
| 19417 | 971 | K971T | C | 1.1e-05 | 9.2e-06 | 1e-05 | 1.007E-05 | 9.018E-07 | 9.200E-06 | 1.100E-05 |
| 19418 | 971 | K971W | C | 8.2e-06 | 9.6e-06 | 8.3e-06 | 8.700E-06 | 7.810E-07 | 8.200E-06 | 9.600E-06 |
| 19419 | 971 | K971Y | C | 1.1e-05 | 5.4e-06 | 6.7e-06 | 7.700E-06 | 2.931E-06 | 5.400E-06 | 1.100E-05 |
| 19420 | 971 | K971V | C | 1.3e-05 | 7.3e-06 | 6.5e-06 | 8.933E-06 | 3.544E-06 | 6.500E-06 | 1.300E-05 |
| 19421 | 972 | S972A | C | 1e-05 | 5.3e-06 | 8e-06 | 7.767E-06 | 2.359E-06 | 5.300E-06 | 1.000E-05 |
| 19422 | 972 | S972R | C | 9.9e-06 | 4.3e-06 | 7.8e-06 | 7.333E-06 | 2.829E-06 | 4.300E-06 | 9.900E-06 |
| 19423 | 972 | S972N | C | 5.3e-06 | 1.3e-05 | 6.3e-06 | 8.200E-06 | 4.187E-06 | 5.300E-06 | 1.300E-05 |
| 19424 | 972 | S972D | C | 8.7e-06 | 7.6e-06 | 7.7e-06 | 8.000E-06 | 6.083E-07 | 7.600E-06 | 8.700E-06 |
| 19425 | 972 | S972C | C | 1.3e-05 | 1.9e-05 | 1.2e-05 | 1.467E-05 | 3.786E-06 | 1.200E-05 | 1.900E-05 |
| 19426 | 972 | S972E | C | 6.7e-06 | 7.2e-06 | 8.3e-06 | 7.400E-06 | 8.185E-07 | 6.700E-06 | 8.300E-06 |
| 19427 | 972 | S972Q | C | 6.7e-06 | 7.5e-06 | 6.5e-06 | 6.900E-06 | 5.292E-07 | 6.500E-06 | 7.500E-06 |
| 19428 | 972 | S972G | C | 5.7e-06 | 1.4e-05 | 7.9e-06 | 9.200E-06 | 4.300E-06 | 5.700E-06 | 1.400E-05 |
| 19429 | 972 | S972H | C | 6.9e-06 | 6e-06 | 9e-06 | 7.300E-06 | 1.539E-06 | 6.000E-06 | 9.000E-06 |
| 19430 | 972 | S972I | C | 5.2e-06 | 5.1e-06 | 6.8e-06 | 5.700E-06 | 9.539E-07 | 5.100E-06 | 6.800E-06 |
| 19431 | 972 | S972L | C | 4.7e-06 | 8.4e-06 | 4.6e-06 | 5.900E-06 | 2.166E-06 | 4.600E-06 | 8.400E-06 |
| 19432 | 972 | S972K | C | 9.1e-06 | 7.4e-06 | 1e-05 | 8.833E-06 | 1.320E-06 | 7.400E-06 | 1.000E-05 |
| 19433 | 972 | S972M | C | 9.9e-06 | 7.5e-06 | 1.4e-05 | 1.047E-05 | 3.287E-06 | 7.500E-06 | 1.400E-05 |
| 19434 | 972 | S972F | C | 5.5e-06 | 8.6e-06 | 1.2e-05 | 8.700E-06 | 3.251E-06 | 5.500E-06 | 1.200E-05 |
| 19435 | 972 | S972P | C | 6.5e-06 | 9.5e-06 | 1.3e-05 | 9.667E-06 | 3.253E-06 | 6.500E-06 | 1.300E-05 |
| 19436 | 972 | S972S | C | 1.3e-05 | 1.1e-05 | 9.6e-06 | 1.120E-05 | 1.709E-06 | 9.600E-06 | 1.300E-05 |
| 19437 | 972 | S972T | C | 6.6e-06 | 1.4e-05 | 5.8e-06 | 8.800E-06 | 4.521E-06 | 5.800E-06 | 1.400E-05 |
| 19438 | 972 | S972W | C | 7.7e-06 | 7.5e-06 | 8.2e-06 | 7.800E-06 | 3.606E-07 | 7.500E-06 | 8.200E-06 |
| 19439 | 972 | S972Y | C | 8.3e-06 | 6.5e-06 | 6.6e-06 | 7.133E-06 | 1.012E-06 | 6.500E-06 | 8.300E-06 |
| 19440 | 972 | S972V | C | 1.1e-05 | 1e-05 | 6.2e-06 | 9.067E-06 | 2.532E-06 | 6.200E-06 | 1.100E-05 |
| 19441 | 973 | T973A | C | 5e-06 | 1.3e-05 | 1e-05 | 9.333E-06 | 4.041E-06 | 5.000E-06 | 1.300E-05 |
| 19442 | 973 | T973R | C | 5.8e-06 | 6.9e-06 | 1.2e-05 | 8.233E-06 | 3.308E-06 | 5.800E-06 | 1.200E-05 |
| 19443 | 973 | T973N | C | 5.5e-06 | 6.3e-06 | 4.9e-06 | 5.567E-06 | 7.024E-07 | 4.900E-06 | 6.300E-06 |
| 19444 | 973 | T973D | C | 9.4e-06 | 1e-05 | 5.3e-06 | 8.233E-06 | 2.558E-06 | 5.300E-06 | 1.000E-05 |
| 19445 | 973 | T973C | C | 1.6e-05 | 6.3e-06 | 7.8e-06 | 1.003E-05 | 5.221E-06 | 6.300E-06 | 1.600E-05 |
| 19446 | 973 | T973E | C | 8.6e-06 | 1e-05 | 5.4e-06 | 8.000E-06 | 2.358E-06 | 5.400E-06 | 1.000E-05 |
| 19447 | 973 | T973Q | C | 1.1e-05 | 9.5e-06 | 1e-05 | 1.017E-05 | 7.638E-07 | 9.500E-06 | 1.100E-05 |
| 19448 | 973 | T973G | C | 6.4e-06 | 7.3e-06 | 6.9e-06 | 6.867E-06 | 4.509E-07 | 6.400E-06 | 7.300E-06 |
| 19449 | 973 | T973H | C | 9.1e-06 | 4.7e-06 | 5.7e-06 | 6.500E-06 | 2.307E-06 | 4.700E-06 | 9.100E-06 |
| 19450 | 973 | T973I | C | 5.1e-06 | 4.4e-06 | 6.7e-06 | 5.400E-06 | 1.179E-06 | 4.400E-06 | 6.700E-06 |
| 19451 | 973 | T973L | C | 9.3e-06 | 1e-05 | 9.8e-06 | 9.700E-06 | 3.606E-07 | 9.300E-06 | 1.000E-05 |
| 19452 | 973 | T973K | C | 9.1e-06 | 1.1e-05 | 8.9e-06 | 9.667E-06 | 1.159E-06 | 8.900E-06 | 1.100E-05 |
| 19453 | 973 | T973M | C | 1e-05 | 9.1e-06 | 5.1e-06 | 8.067E-06 | 2.608E-06 | 5.100E-06 | 1.000E-05 |
| 19454 | 973 | T973F | C | 7.5e-06 | 5.8e-06 | 6.4e-06 | 6.567E-06 | 8.622E-07 | 5.800E-06 | 7.500E-06 |
| 19455 | 973 | T973P | C | 8e-06 | 4.8e-06 | 7.3e-06 | 6.700E-06 | 1.682E-06 | 4.800E-06 | 8.000E-06 |
| 19456 | 973 | T973S | C | 1.1e-05 | 5.7e-06 | 1.3e-05 | 9.900E-06 | 3.772E-06 | 5.700E-06 | 1.300E-05 |
| 19457 | 973 | T973T | C | 8.5e-06 | 5.1e-06 | 6e-06 | 6.533E-06 | 1.762E-06 | 5.100E-06 | 8.500E-06 |
| 19458 | 973 | T973W | C | 9.7e-06 | 6.5e-06 | 7e-06 | 7.733E-06 | 1.721E-06 | 6.500E-06 | 9.700E-06 |
| 19459 | 973 | T973Y | C | 6.5e-06 | 9.3e-06 | 6e-06 | 7.267E-06 | 1.779E-06 | 6.000E-06 | 9.300E-06 |
| 19460 | 973 | T973V | C | 1.6e-05 | 8.1e-06 | 9.8e-06 | 1.130E-05 | 4.158E-06 | 8.100E-06 | 1.600E-05 |
| 19461 | 974 | S974A | C | 1e-05 | 1.1e-05 | 5.3e-06 | 8.767E-06 | 3.044E-06 | 5.300E-06 | 1.100E-05 |
| 19462 | 974 | S974R | C | 8.6e-06 | 6.4e-06 | 1e-05 | 8.333E-06 | 1.815E-06 | 6.400E-06 | 1.000E-05 |
| 19463 | 974 | S974N | C | 8.1e-06 | 9.4e-06 | 8.4e-06 | 8.633E-06 | 6.807E-07 | 8.100E-06 | 9.400E-06 |
| 19464 | 974 | S974D | C | 9.1e-06 | 2.1e-05 | 1.2e-05 | 1.403E-05 | 6.205E-06 | 9.100E-06 | 2.100E-05 |

| | | | | | | | | | | |
|---|---|---|---|---|---|---|---|---|---|---|
| 19465 | 974 | S974C | C | 9.5e-06 | 4.5e-06 | 9.7e-06 | 7.900E-06 | 2.946E-06 | 4.500E-06 | 9.700E-06 |
| 19466 | 974 | S974E | C | 6.3e-06 | 9.9e-06 | 1.1e-05 | 9.067E-06 | 2.458E-06 | 6.300E-06 | 1.100E-05 |
| 19467 | 974 | S974Q | C | 6.6e-06 | 6.8e-06 | 9e-06 | 7.467E-06 | 1.332E-06 | 6.600E-06 | 9.000E-06 |
| 19468 | 974 | S974G | C | 9.8e-06 | 6.8e-06 | 1.3e-05 | 9.867E-06 | 3.101E-06 | 6.800E-06 | 1.300E-05 |
| 19469 | 974 | S974H | C | 8e-06 | 5.7e-06 | 1.1e-05 | 8.233E-06 | 2.658E-06 | 5.700E-06 | 1.100E-05 |
| 19470 | 974 | S974I | C | 6.8e-06 | 6.7e-06 | 8.5e-06 | 7.333E-06 | 1.012E-06 | 6.700E-06 | 8.500E-06 |
| 19471 | 974 | S974L | C | 8.4e-06 | 4.3e-06 | 6.9e-06 | 6.533E-06 | 2.074E-06 | 4.300E-06 | 8.400E-06 |
| 19472 | 974 | S974K | C | 9.9e-06 | 6.4e-06 | 9.2e-06 | 8.500E-06 | 1.852E-06 | 6.400E-06 | 9.900E-06 |
| 19473 | 974 | S974M | C | 5.1e-06 | 1.2e-05 | 6.8e-06 | 7.967E-06 | 3.595E-06 | 5.100E-06 | 1.200E-05 |
| 19474 | 974 | S974F | C | 1.9e-05 | 1.1e-05 | 8.8e-06 | 1.293E-05 | 5.368E-06 | 8.800E-06 | 1.900E-05 |
| 19475 | 974 | S974P | C | 1e-05 | 4.6e-06 | 9.2e-06 | 7.933E-06 | 2.914E-06 | 4.600E-06 | 1.000E-05 |
| 19476 | 974 | S974S | C | 8.8e-06 | 1.2e-05 | 7e-06 | 9.267E-06 | 2.532E-06 | 7.000E-06 | 1.200E-05 |
| 19477 | 974 | S974T | C | 6e-06 | 1e-05 | 4.7e-06 | 6.900E-06 | 2.762E-06 | 4.700E-06 | 1.000E-05 |
| 19478 | 974 | S974W | C | 1.9e-05 | 8.9e-06 | 6.4e-06 | 1.143E-05 | 6.671E-06 | 6.400E-06 | 1.900E-05 |
| 19479 | 974 | S974Y | C | 1.1e-05 | 6.3e-06 | 6.9e-06 | 8.067E-06 | 2.558E-06 | 6.300E-06 | 1.100E-05 |
| 19480 | 974 | S974V | C | 5.5e-06 | 8e-06 | 9.1e-06 | 7.533E-06 | 1.845E-06 | 5.500E-06 | 9.100E-06 |
| 19481 | 975 | G975A | C | 9.1e-06 | 9.5e-06 | 6.1e-06 | 8.233E-06 | 1.858E-06 | 6.100E-06 | 9.500E-06 |
| 19482 | 975 | G975R | C | 8.3e-06 | 6.4e-06 | 1.9e-05 | 1.123E-05 | 6.793E-06 | 6.400E-06 | 1.900E-05 |
| 19483 | 975 | G975N | C | 4.7e-06 | 6.4e-06 | 6.9e-06 | 6.000E-06 | 1.153E-06 | 4.700E-06 | 6.900E-06 |
| 19484 | 975 | G975D | C | 1.3e-05 | 5.9e-06 | 1.3e-05 | 1.063E-05 | 4.099E-06 | 5.900E-06 | 1.300E-05 |
| 19485 | 975 | G975C | C | 9.2e-06 | 4.5e-06 | 1.2e-05 | 8.567E-06 | 3.790E-06 | 4.500E-06 | 1.200E-05 |
| 19486 | 975 | G975E | C | 6.9e-06 | 4.8e-06 | 1.3e-05 | 8.233E-06 | 4.259E-06 | 4.800E-06 | 1.300E-05 |
| 19487 | 975 | G975Q | C | 1e-05 | 9.3e-06 | 9.6e-06 | 9.633E-06 | 3.512E-07 | 9.300E-06 | 1.000E-05 |
| 19488 | 975 | G975G | C | 1.2e-05 | 4.8e-06 | 9.7e-06 | 8.833E-06 | 3.677E-06 | 4.800E-06 | 1.200E-05 |
| 19489 | 975 | G975H | C | 6.7e-06 | 4.5e-06 | 8.5e-06 | 6.567E-06 | 2.003E-06 | 4.500E-06 | 8.500E-06 |
| 19490 | 975 | G975I | C | 7.1e-06 | 7.7e-06 | 7.8e-06 | 7.533E-06 | 3.786E-07 | 7.100E-06 | 7.800E-06 |
| 19491 | 975 | G975L | C | 5.9e-06 | 8.5e-06 | 1.4e-05 | 9.467E-06 | 4.136E-06 | 5.900E-06 | 1.400E-05 |
| 19492 | 975 | G975K | C | 1.3e-05 | 7.4e-06 | 6.6e-06 | 9.000E-06 | 3.487E-06 | 6.600E-06 | 1.300E-05 |
| 19493 | 975 | G975M | C | 4.5e-06 | 7.4e-06 | 9.2e-06 | 7.033E-06 | 2.371E-06 | 4.500E-06 | 9.200E-06 |
| 19494 | 975 | G975F | C | 8.7e-06 | 5.5e-06 | 5.9e-06 | 6.700E-06 | 1.744E-06 | 5.500E-06 | 8.700E-06 |
| 19495 | 975 | G975P | C | 8.8e-06 | 1e-05 | 9.2e-06 | 9.333E-06 | 6.110E-07 | 8.800E-06 | 1.000E-05 |
| 19496 | 975 | G975S | C | 7.3e-06 | 5.3e-06 | 5.9e-06 | 6.167E-06 | 1.026E-06 | 5.300E-06 | 7.300E-06 |
| 19497 | 975 | G975T | C | 9.3e-06 | 9.4e-06 | 4.8e-06 | 7.833E-06 | 2.627E-06 | 4.800E-06 | 9.400E-06 |
| 19498 | 975 | G975W | C | 1.1e-05 | 8.6e-06 | 1.2e-05 | 1.053E-05 | 1.747E-06 | 8.600E-06 | 1.200E-05 |
| 19499 | 975 | G975Y | C | 1.1e-05 | 1.3e-05 | 7.1e-06 | 1.037E-05 | 3.001E-06 | 7.100E-06 | 1.300E-05 |
| 19500 | 975 | G975V | C | 8.2e-06 | 6.7e-06 | 1.4e-05 | 9.633E-06 | 3.855E-06 | 6.700E-06 | 1.400E-05 |
| 19501 | 976 | G976A | C | 6.4e-06 | 6.1e-06 | 9.6e-06 | 7.367E-06 | 1.940E-06 | 6.100E-06 | 9.600E-06 |
| 19502 | 976 | G976R | C | 5.3e-06 | 9.5e-06 | 6.9e-06 | 7.233E-06 | 2.120E-06 | 5.300E-06 | 9.500E-06 |
| 19503 | 976 | G976N | C | 8.1e-06 | 7.2e-06 | 7.7e-06 | 7.667E-06 | 4.509E-07 | 7.200E-06 | 8.100E-06 |
| 19504 | 976 | G976D | C | 8.3e-06 | 9.8e-06 | 1.2e-05 | 1.003E-05 | 1.861E-06 | 8.300E-06 | 1.200E-05 |
| 19505 | 976 | G976C | C | 7.6e-06 | 1.1e-05 | 5.9e-06 | 8.167E-06 | 2.597E-06 | 5.900E-06 | 1.100E-05 |
| 19506 | 976 | G976E | C | 1.2e-05 | 8.1e-06 | 1.3e-05 | 1.103E-05 | 2.589E-06 | 8.100E-06 | 1.300E-05 |
| 19507 | 976 | G976Q | C | 9.3e-06 | 6e-06 | 1.2e-05 | 9.100E-06 | 3.005E-06 | 6.000E-06 | 1.200E-05 |
| 19508 | 976 | G976G | C | 8.8e-06 | 5.5e-06 | 7.7e-06 | 7.333E-06 | 1.680E-06 | 5.500E-06 | 8.800E-06 |
| 19509 | 976 | G976H | C | 1.2e-05 | 7.2e-06 | 6e-06 | 8.400E-06 | 3.175E-06 | 6.000E-06 | 1.200E-05 |
| 19510 | 976 | G976I | C | 1.3e-05 | 8.3e-06 | 6.8e-06 | 9.367E-06 | 3.235E-06 | 6.800E-06 | 1.300E-05 |
| 19511 | 976 | G976L | C | 7e-06 | 6.3e-06 | 1e-05 | 7.767E-06 | 1.966E-06 | 6.300E-06 | 1.000E-05 |
| 19512 | 976 | G976K | C | 7.4e-06 | 8.9e-06 | 8.7e-06 | 8.333E-06 | 8.145E-07 | 7.400E-06 | 8.900E-06 |
| 19513 | 976 | G976M | C | 7.8e-06 | 5.4e-06 | 4.4e-06 | 5.867E-06 | 1.747E-06 | 4.400E-06 | 7.800E-06 |
| 19514 | 976 | G976F | C | 3.9e-06 | 1.3e-05 | 8.7e-06 | 8.533E-06 | 4.552E-06 | 3.900E-06 | 1.300E-05 |
| 19515 | 976 | G976P | C | 7.2e-06 | 1.1e-05 | 9.5e-06 | 9.233E-06 | 1.914E-06 | 7.200E-06 | 1.100E-05 |
| 19516 | 976 | G976S | C | 5.8e-06 | 1.4e-05 | 8.2e-06 | 9.333E-06 | 4.216E-06 | 5.800E-06 | 1.400E-05 |
| 19517 | 976 | G976T | C | 8.6e-06 | 6.4e-06 | 7.8e-06 | 7.600E-06 | 1.114E-06 | 6.400E-06 | 8.600E-06 |
| 19518 | 976 | G976W | C | 7.4e-06 | 4.9e-06 | 6.2e-06 | 6.167E-06 | 1.250E-06 | 4.900E-06 | 7.400E-06 |
| 19519 | 976 | G976Y | C | 7.4e-06 | 7e-06 | 7.1e-06 | 7.167E-06 | 2.082E-07 | 7.000E-06 | 7.400E-06 |
| 19520 | 976 | G976V | C | 5.3e-06 | 1e-05 | 5.6e-06 | 6.967E-06 | 2.631E-06 | 5.300E-06 | 1.000E-05 |
| 19521 | 977 | T977A | C | 6.7e-06 | 1.3e-05 | 7.6e-06 | 9.100E-06 | 3.407E-06 | 6.700E-06 | 1.300E-05 |
| 19522 | 977 | T977R | C | 6.8e-06 | 9.7e-06 | 8.3e-06 | 8.267E-06 | 1.450E-06 | 6.800E-06 | 9.700E-06 |
| 19523 | 977 | T977N | C | 9.6e-06 | 7.5e-06 | 9.6e-06 | 8.900E-06 | 1.212E-06 | 7.500E-06 | 9.600E-06 |
| 19524 | 977 | T977D | C | 6.8e-06 | 1.4e-05 | 8e-06 | 9.600E-06 | 3.857E-06 | 6.800E-06 | 1.400E-05 |
| 19525 | 977 | T977C | C | 1.3e-05 | 9e-06 | 8.6e-06 | 1.020E-05 | 2.433E-06 | 8.600E-06 | 1.300E-05 |
| 19526 | 977 | T977E | C | 8.7e-06 | 9.2e-06 | 1.3e-05 | 1.030E-05 | 2.352E-06 | 8.700E-06 | 1.300E-05 |
| 19527 | 977 | T977Q | C | 7.2e-06 | 4.9e-06 | 5.3e-06 | 5.800E-06 | 1.229E-06 | 4.900E-06 | 7.200E-06 |
| 19528 | 977 | T977G | C | 9.6e-06 | 9.5e-06 | 6e-06 | 8.367E-06 | 2.050E-06 | 6.000E-06 | 9.600E-06 |
| 19529 | 977 | T977H | C | 9e-06 | 4.5e-06 | 8.5e-06 | 7.333E-06 | 2.466E-06 | 4.500E-06 | 9.000E-06 |
| 19530 | 977 | T977I | C | 5.9e-06 | 8.9e-06 | 1.6e-05 | 1.027E-05 | 5.187E-06 | 5.900E-06 | 1.600E-05 |
| 19531 | 977 | T977L | C | 7.1e-06 | 8.2e-06 | 6.9e-06 | 7.400E-06 | 7.000E-07 | 6.900E-06 | 8.200E-06 |
| 19532 | 977 | T977K | C | 1.2e-05 | 6.8e-06 | 7.2e-06 | 8.667E-06 | 2.894E-06 | 6.800E-06 | 1.200E-05 |
| 19533 | 977 | T977M | C | 9.4e-06 | 8.4e-06 | 3.3e-06 | 7.033E-06 | 3.272E-06 | 3.300E-06 | 9.400E-06 |
| 19534 | 977 | T977F | C | 7e-06 | 5e-06 | 4.8e-06 | 5.600E-06 | 1.217E-06 | 4.800E-06 | 7.000E-06 |
| 19535 | 977 | T977P | C | 7.5e-06 | 8.3e-06 | 8.5e-06 | 8.100E-06 | 5.292E-07 | 7.500E-06 | 8.500E-06 |
| 19536 | 977 | T977S | C | 1e-05 | 6.5e-06 | 1.4e-05 | 1.017E-05 | 3.753E-06 | 6.500E-06 | 1.400E-05 |
| 19537 | 977 | T977T | C | 1.1e-05 | 1e-05 | 6.8e-06 | 9.267E-06 | 2.194E-06 | 6.800E-06 | 1.100E-05 |
| 19538 | 977 | T977W | C | 9.9e-06 | 3.1e-06 | 9.1e-06 | 7.367E-06 | 3.717E-06 | 3.100E-06 | 9.900E-06 |
| 19539 | 977 | T977Y | C | 8.9e-06 | 6e-06 | 9.6e-06 | 8.167E-06 | 1.909E-06 | 6.000E-06 | 9.600E-06 |
| 19540 | 977 | T977V | C | 8.1e-06 | 6.3e-06 | 1.2e-05 | 8.800E-06 | 2.914E-06 | 6.300E-06 | 1.200E-05 |
| 19541 | 978 | A978A | C | 7.3e-06 | 7e-06 | 8.3e-06 | 7.533E-06 | 6.807E-07 | 7.000E-06 | 8.300E-06 |
| 19542 | 978 | A978R | C | 8.7e-06 | 6.5e-06 | 8.9e-06 | 8.033E-06 | 1.332E-06 | 6.500E-06 | 8.900E-06 |
| 19543 | 978 | A978N | C | 5e-06 | 1.3e-05 | 6e-06 | 8.000E-06 | 4.359E-06 | 5.000E-06 | 1.300E-05 |
| 19544 | 978 | A978D | C | 9.8e-06 | 5.9e-06 | 7.5e-06 | 7.733E-06 | 1.960E-06 | 5.900E-06 | 9.800E-06 |
| 19545 | 978 | A978C | C | 6.5e-06 | 6.8e-06 | 6.2e-06 | 6.500E-06 | 3.000E-07 | 6.200E-06 | 6.800E-06 |
| 19546 | 978 | A978E | C | 8.3e-06 | 7.5e-06 | 8.9e-06 | 8.233E-06 | 7.024E-07 | 7.500E-06 | 8.900E-06 |
| 19547 | 978 | A978Q | C | 4.3e-06 | 1.4e-05 | 8.5e-06 | 8.933E-06 | 4.864E-06 | 4.300E-06 | 1.400E-05 |
| 19548 | 978 | A978G | C | 9.2e-06 | 7.4e-06 | 8.5e-06 | 8.367E-06 | 9.074E-07 | 7.400E-06 | 9.200E-06 |
| 19549 | 978 | A978H | C | 7e-06 | 8.4e-06 | 9.8e-06 | 8.400E-06 | 1.400E-06 | 7.000E-06 | 9.800E-06 |

| | | | | | | | | | | |
|---|---|---|---|---|---|---|---|---|---|---|
| 19550 | 978 | A978I | C | 7.3e-06 | 8.1e-06 | 9.5e-06 | 8.300E-06 | 1.114E-06 | 7.300E-06 | 9.500E-06 |
| 19551 | 978 | A978L | C | 9.5e-06 | 7e-06 | 6.4e-06 | 7.633E-06 | 1.644E-06 | 6.400E-06 | 9.500E-06 |
| 19552 | 978 | A978K | C | 9.2e-06 | 8.3e-06 | 9.5e-06 | 9.000E-06 | 6.245E-07 | 8.300E-06 | 9.500E-06 |
| 19553 | 978 | A978M | C | 5.1e-06 | 1.6e-05 | 6.5e-06 | 9.200E-06 | 5.930E-06 | 5.100E-06 | 1.600E-05 |
| 19554 | 978 | A978F | C | 1.3e-05 | 7.4e-06 | 9.8e-06 | 1.007E-05 | 2.810E-06 | 7.400E-06 | 1.300E-05 |
| 19555 | 978 | A978P | C | 4.8e-06 | 1e-05 | 8.1e-06 | 7.633E-06 | 2.631E-06 | 4.800E-06 | 1.000E-05 |
| 19556 | 978 | A978S | C | 1.1e-05 | 1e-05 | 7.5e-06 | 9.500E-06 | 1.803E-06 | 7.500E-06 | 1.100E-05 |
| 19557 | 978 | A978T | C | 4.8e-06 | 5.2e-06 | 3.6e-06 | 4.533E-06 | 8.327E-07 | 3.600E-06 | 5.200E-06 |
| 19558 | 978 | A978W | C | 1.1e-05 | 6.5e-06 | 8e-06 | 8.500E-06 | 2.291E-06 | 6.500E-06 | 1.100E-05 |
| 19559 | 978 | A978Y | C | 4.4e-06 | 7.4e-06 | 1e-05 | 7.267E-06 | 2.802E-06 | 4.400E-06 | 1.000E-05 |
| 19560 | 978 | A978V | C | 4.7e-06 | 1e-05 | 1.2e-05 | 8.900E-06 | 3.772E-06 | 4.700E-06 | 1.200E-05 |
| 19561 | 979 | A979A | C | 1.4e-05 | 9.5e-06 | 1.1e-05 | 1.150E-05 | 2.291E-06 | 9.500E-06 | 1.400E-05 |
| 19562 | 979 | A979R | C | 9.3e-06 | 3.7e-06 | 8.6e-06 | 7.200E-06 | 3.051E-06 | 3.700E-06 | 9.300E-06 |
| 19563 | 979 | A979N | C | 4.8e-06 | 9.5e-06 | 8.9e-06 | 7.733E-06 | 2.558E-06 | 4.800E-06 | 9.500E-06 |
| 19564 | 979 | A979D | C | 7.9e-06 | 9.4e-06 | 7.6e-06 | 8.300E-06 | 9.644E-07 | 7.600E-06 | 9.400E-06 |
| 19565 | 979 | A979C | C | 1.3e-05 | 6e-06 | 9.2e-06 | 9.400E-06 | 3.504E-06 | 6.000E-06 | 1.300E-05 |
| 19566 | 979 | A979E | C | 1.1e-05 | 5.6e-06 | 1.2e-05 | 9.533E-06 | 3.443E-06 | 5.600E-06 | 1.200E-05 |
| 19567 | 979 | A979Q | C | 1e-05 | 5.5e-06 | 1.2e-05 | 9.167E-06 | 3.329E-06 | 5.500E-06 | 1.200E-05 |
| 19568 | 979 | A979G | C | 8.4e-06 | 6.6e-06 | 4.6e-06 | 6.533E-06 | 1.901E-06 | 4.600E-06 | 8.400E-06 |
| 19569 | 979 | A979H | C | 9.8e-06 | 8.5e-06 | 7.8e-06 | 8.700E-06 | 1.015E-06 | 7.800E-06 | 9.800E-06 |
| 19570 | 979 | A979I | C | 9.8e-06 | 4.5e-06 | 5.1e-06 | 6.467E-06 | 2.902E-06 | 4.500E-06 | 9.800E-06 |
| 19571 | 979 | A979L | C | 8.7e-06 | 7.6e-06 | 6.1e-06 | 7.467E-06 | 1.305E-06 | 6.100E-06 | 8.700E-06 |
| 19572 | 979 | A979K | C | 3.9e-06 | 6.3e-06 | 7.8e-06 | 6.000E-06 | 1.967E-06 | 3.900E-06 | 7.800E-06 |
| 19573 | 979 | A979M | C | 1.1e-05 | 6.1e-06 | 6.5e-06 | 7.867E-06 | 2.721E-06 | 6.100E-06 | 1.100E-05 |
| 19574 | 979 | A979F | C | 1.6e-05 | 1.3e-05 | 8e-06 | 1.233E-05 | 4.041E-06 | 8.000E-06 | 1.600E-05 |
| 19575 | 979 | A979P | C | 1.8e-05 | 9e-06 | 7.9e-06 | 1.163E-05 | 5.541E-06 | 7.900E-06 | 1.800E-05 |
| 19576 | 979 | A979S | C | 7.8e-06 | 4.3e-06 | 7.7e-06 | 6.600E-06 | 1.992E-06 | 4.300E-06 | 7.800E-06 |
| 19577 | 979 | A979T | C | 1.2e-05 | 6.9e-06 | 6.2e-06 | 8.367E-06 | 3.166E-06 | 6.200E-06 | 1.200E-05 |
| 19578 | 979 | A979W | C | 6.3e-06 | 6.4e-06 | 7.8e-06 | 6.833E-06 | 8.386E-07 | 6.300E-06 | 7.800E-06 |
| 19579 | 979 | A979Y | C | 4.8e-06 | 6.7e-06 | 8.1e-06 | 6.533E-06 | 1.656E-06 | 4.800E-06 | 8.100E-06 |
| 19580 | 979 | A979V | C | 8e-06 | 7.8e-06 | 7.3e-06 | 7.700E-06 | 3.606E-07 | 7.300E-06 | 8.000E-06 |
| 19581 | 980 | L980A | C | 7.3e-06 | 8.3e-06 | 7.6e-06 | 7.733E-06 | 5.132E-07 | 7.300E-06 | 8.300E-06 |
| 19582 | 980 | L980R | C | 7.8e-06 | 7e-06 | 7.5e-06 | 7.433E-06 | 4.041E-07 | 7.000E-06 | 7.800E-06 |
| 19583 | 980 | L980N | C | 8.5e-06 | 1.2e-05 | 1.1e-05 | 1.050E-05 | 1.803E-06 | 8.500E-06 | 1.200E-05 |
| 19584 | 980 | L980D | C | 9.6e-06 | 6.6e-06 | 9.4e-06 | 8.533E-06 | 1.677E-06 | 6.600E-06 | 9.600E-06 |
| 19585 | 980 | L980C | C | 1e-05 | 1.1e-05 | 8.1e-06 | 9.700E-06 | 1.473E-06 | 8.100E-06 | 1.100E-05 |
| 19586 | 980 | L980E | C | 5e-06 | 9.4e-06 | 7.9e-06 | 7.433E-06 | 2.237E-06 | 5.000E-06 | 9.400E-06 |
| 19587 | 980 | L980Q | C | 1.1e-05 | 1.2e-05 | 8.8e-06 | 1.060E-05 | 1.637E-06 | 8.800E-06 | 1.200E-05 |
| 19588 | 980 | L980G | C | 8.4e-06 | 8.6e-06 | 6.8e-06 | 7.933E-06 | 9.866E-07 | 6.800E-06 | 8.600E-06 |
| 19589 | 980 | L980H | C | 7e-06 | 9.1e-06 | 7.6e-06 | 7.900E-06 | 1.082E-06 | 7.000E-06 | 9.100E-06 |
| 19590 | 980 | L980I | C | 6.5e-06 | 7.8e-06 | 7.2e-06 | 7.167E-06 | 6.506E-07 | 6.500E-06 | 7.800E-06 |
| 19591 | 980 | L980L | C | 1.1e-05 | 1.1e-05 | 1.4e-05 | 1.200E-05 | 1.732E-06 | 1.100E-05 | 1.400E-05 |
| 19592 | 980 | L980K | C | 8.2e-06 | 9.8e-06 | 7.7e-06 | 8.567E-06 | 1.097E-06 | 7.700E-06 | 9.800E-06 |
| 19593 | 980 | L980M | C | 6e-06 | 8.7e-06 | 7.7e-06 | 7.467E-06 | 1.365E-06 | 6.000E-06 | 8.700E-06 |
| 19594 | 980 | L980F | C | 5.4e-06 | 6.9e-06 | 8.8e-06 | 7.033E-06 | 1.704E-06 | 5.400E-06 | 8.800E-06 |
| 19595 | 980 | L980P | C | 7.2e-06 | 8.7e-06 | 7.4e-06 | 7.767E-06 | 8.145E-07 | 7.200E-06 | 8.700E-06 |
| 19596 | 980 | L980S | C | 7.8e-06 | 7.7e-06 | 9.5e-06 | 8.333E-06 | 1.012E-06 | 7.700E-06 | 9.500E-06 |
| 19597 | 980 | L980T | C | 1.4e-05 | 7.2e-06 | 5.7e-06 | 8.967E-06 | 4.423E-06 | 5.700E-06 | 1.400E-05 |
| 19598 | 980 | L980W | C | 6e-06 | 9.3e-06 | 8.3e-06 | 7.867E-06 | 1.692E-06 | 6.000E-06 | 9.300E-06 |
| 19599 | 980 | L980Y | C | 4.1e-06 | 8.4e-06 | 1.1e-05 | 7.833E-06 | 3.485E-06 | 4.100E-06 | 1.100E-05 |
| 19600 | 980 | L980V | C | 1.4e-05 | 5e-06 | 7.3e-06 | 8.767E-06 | 4.676E-06 | 5.000E-06 | 1.400E-05 |
| 19601 | 981 | G981A | C | 7e-06 | 8e-06 | 6.7e-06 | 7.233E-06 | 6.807E-07 | 6.700E-06 | 8.000E-06 |
| 19602 | 981 | G981R | C | 9.1e-06 | 7.1e-06 | 7.5e-06 | 7.900E-06 | 1.058E-06 | 7.100E-06 | 9.100E-06 |
| 19603 | 981 | G981N | C | 6.5e-06 | 7.5e-06 | 5.9e-06 | 6.633E-06 | 8.083E-07 | 5.900E-06 | 7.500E-06 |
| 19604 | 981 | G981D | C | 5.4e-06 | 5.9e-06 | 1.1e-05 | 7.433E-06 | 3.099E-06 | 5.400E-06 | 1.100E-05 |
| 19605 | 981 | G981C | C | 6.9e-06 | 8.2e-06 | 9.2e-06 | 8.100E-06 | 1.153E-06 | 6.900E-06 | 9.200E-06 |
| 19606 | 981 | G981E | C | 9.1e-06 | 3e-06 | 6.6e-06 | 6.233E-06 | 3.066E-06 | 3.000E-06 | 9.100E-06 |
| 19607 | 981 | G981Q | C | 9.9e-06 | 1e-05 | 1.1e-05 | 1.030E-05 | 6.083E-07 | 9.900E-06 | 1.100E-05 |
| 19608 | 981 | G981G | C | 8.4e-06 | 8.4e-06 | 6.6e-06 | 7.800E-06 | 1.039E-06 | 6.600E-06 | 8.400E-06 |
| 19609 | 981 | G981H | C | 4.1e-06 | 9.5e-06 | 1.5e-05 | 9.533E-06 | 5.450E-06 | 4.100E-06 | 1.500E-05 |
| 19610 | 981 | G981I | C | 1.3e-05 | 8.3e-06 | 1e-05 | 1.043E-05 | 2.380E-06 | 8.300E-06 | 1.300E-05 |
| 19611 | 981 | G981L | C | 7.2e-06 | 9.5e-06 | 9.6e-06 | 8.767E-06 | 1.358E-06 | 7.200E-06 | 9.600E-06 |
| 19612 | 981 | G981K | C | 4.3e-06 | 1.3e-05 | 5.9e-06 | 7.733E-06 | 4.631E-06 | 4.300E-06 | 1.300E-05 |
| 19613 | 981 | G981M | C | 6.9e-06 | 7.2e-06 | 8.2e-06 | 7.433E-06 | 6.807E-07 | 6.900E-06 | 8.200E-06 |
| 19614 | 981 | G981F | C | 7.1e-06 | 1.1e-05 | 8.7e-06 | 8.933E-06 | 1.960E-06 | 7.100E-06 | 1.100E-05 |
| 19615 | 981 | G981P | C | 9.4e-06 | 8.6e-06 | 1.3e-05 | 1.033E-05 | 2.344E-06 | 8.600E-06 | 1.300E-05 |
| 19616 | 981 | G981S | C | 5.5e-06 | 6.6e-06 | 5.7e-06 | 5.933E-06 | 5.859E-07 | 5.500E-06 | 6.600E-06 |
| 19617 | 981 | G981T | C | 6e-06 | 7.1e-06 | 7.2e-06 | 6.767E-06 | 6.658E-07 | 6.000E-06 | 7.200E-06 |
| 19618 | 981 | G981W | C | 6.3e-06 | 8.8e-06 | 1.8e-05 | 1.103E-05 | 6.161E-06 | 6.300E-06 | 1.800E-05 |
| 19619 | 981 | G981Y | C | 7.2e-06 | 9.8e-06 | 9.7e-06 | 8.900E-06 | 1.473E-06 | 7.200E-06 | 9.800E-06 |
| 19620 | 981 | G981V | C | 6.8e-06 | 1e-05 | 6.7e-06 | 7.833E-06 | 1.877E-06 | 6.700E-06 | 1.000E-05 |
| 19621 | 982 | C982A | C | 6.9e-06 | 1.3e-05 | 5.1e-06 | 8.333E-06 | 4.140E-06 | 5.100E-06 | 1.300E-05 |
| 19622 | 982 | C982R | C | 9.3e-06 | 6e-06 | 9.1e-06 | 8.133E-06 | 1.850E-06 | 6.000E-06 | 9.300E-06 |
| 19623 | 982 | C982N | C | 7.1e-06 | 8.7e-06 | 8.6e-06 | 8.133E-06 | 8.963E-07 | 7.100E-06 | 8.700E-06 |
| 19624 | 982 | C982D | C | 1.9e-05 | 3.2e-06 | 7.1e-06 | 9.767E-06 | 8.231E-06 | 3.200E-06 | 1.900E-05 |
| 19625 | 982 | C982C | C | 9.2e-06 | 6.9e-06 | 8.8e-06 | 8.300E-06 | 1.229E-06 | 6.900E-06 | 9.200E-06 |
| 19626 | 982 | C982E | C | 7.7e-06 | 1e-05 | 6.8e-06 | 8.167E-06 | 1.650E-06 | 6.800E-06 | 1.000E-05 |
| 19627 | 982 | C982Q | C | 8.3e-06 | 5.9e-06 | 6.6e-06 | 6.933E-06 | 1.234E-06 | 5.900E-06 | 8.300E-06 |
| 19628 | 982 | C982G | C | 5.9e-06 | 9.7e-06 | 1.1e-05 | 8.867E-06 | 2.650E-06 | 5.900E-06 | 1.100E-05 |
| 19629 | 982 | C982H | C | 5.8e-06 | 7.8e-06 | 8.8e-06 | 7.467E-06 | 1.528E-06 | 5.800E-06 | 8.800E-06 |
| 19630 | 982 | C982I | C | 7.6e-06 | 8.3e-06 | 8.8e-06 | 8.233E-06 | 6.028E-07 | 7.600E-06 | 8.800E-06 |
| 19631 | 982 | C982L | C | 4.9e-06 | 5.5e-06 | 4.7e-06 | 5.033E-06 | 4.163E-07 | 4.700E-06 | 5.500E-06 |
| 19632 | 982 | C982K | C | 5.9e-06 | 5.6e-06 | 1e-05 | 7.167E-06 | 2.458E-06 | 5.600E-06 | 1.000E-05 |
| 19633 | 982 | C982M | C | 1.4e-05 | 4.8e-06 | 6.9e-06 | 8.567E-06 | 4.821E-06 | 4.800E-06 | 1.400E-05 |
| 19634 | 982 | C982F | C | 5.4e-06 | 7.6e-06 | 6.7e-06 | 6.567E-06 | 1.106E-06 | 5.400E-06 | 7.600E-06 |

| | | | | | | | | | | |
|---|---|---|---|---|---|---|---|---|---|---|
| 19635 | 982 | C982P | C | 9.6e-06 | 1.4e-05 | 1.1e-05 | 1.153E-05 | 2.248E-06 | 9.600E-06 | 1.400E-05 |
| 19636 | 982 | C982S | C | 7.9e-06 | 7.3e-06 | 4.1e-06 | 6.433E-06 | 2.043E-06 | 4.100E-06 | 7.900E-06 |
| 19637 | 982 | C982T | C | 6.6e-06 | 1e-05 | 6.8e-06 | 7.800E-06 | 1.908E-06 | 6.600E-06 | 1.000E-05 |
| 19638 | 982 | C982W | C | 6.9e-06 | 6e-06 | 6.8e-06 | 6.567E-06 | 4.933E-07 | 6.000E-06 | 6.900E-06 |
| 19639 | 982 | C982Y | C | 9.1e-06 | 9.2e-06 | 5.8e-06 | 8.033E-06 | 1.935E-06 | 5.800E-06 | 9.200E-06 |
| 19640 | 982 | C982V | C | 9.7e-06 | 8.9e-06 | 1.2e-05 | 1.020E-05 | 1.609E-06 | 8.900E-06 | 1.200E-05 |
| 19641 | 983 | L983A | C | 6.7e-06 | 1e-05 | 6.7e-06 | 7.800E-06 | 1.905E-06 | 6.700E-06 | 1.000E-05 |
| 19642 | 983 | L983R | C | 1.2e-05 | 9e-06 | 8.9e-06 | 9.967E-06 | 1.762E-06 | 8.900E-06 | 1.200E-05 |
| 19643 | 983 | L983N | C | 6.5e-06 | 7.4e-06 | 8.1e-06 | 7.333E-06 | 8.021E-07 | 6.500E-06 | 8.100E-06 |
| 19644 | 983 | L983D | C | 1.1e-05 | 7.2e-06 | 8.9e-06 | 9.033E-06 | 1.904E-06 | 7.200E-06 | 1.100E-05 |
| 19645 | 983 | L983C | C | 6.3e-06 | 9.6e-06 | 1.4e-05 | 9.967E-06 | 3.863E-06 | 6.300E-06 | 1.400E-05 |
| 19646 | 983 | L983E | C | 9.5e-06 | 8.9e-06 | 3.7e-06 | 7.367E-06 | 3.190E-06 | 3.700E-06 | 9.500E-06 |
| 19647 | 983 | L983Q | C | 8.2e-06 | 5.5e-06 | 8.4e-06 | 7.367E-06 | 1.620E-06 | 5.500E-06 | 8.400E-06 |
| 19648 | 983 | L983G | C | 6.6e-06 | 8.1e-06 | 7.2e-06 | 7.300E-06 | 7.550E-07 | 6.600E-06 | 8.100E-06 |
| 19649 | 983 | L983H | C | 7.5e-06 | 7.1e-06 | 6.4e-06 | 7.000E-06 | 5.568E-07 | 6.400E-06 | 7.500E-06 |
| 19650 | 983 | L983I | C | 7.2e-06 | 7.8e-06 | 1.2e-05 | 9.000E-06 | 2.615E-06 | 7.200E-06 | 1.200E-05 |
| 19651 | 983 | L983L | C | 6.5e-06 | 1.4e-05 | 5.9e-06 | 8.800E-06 | 4.513E-06 | 5.900E-06 | 1.400E-05 |
| 19652 | 983 | L983K | C | 1.1e-05 | 1.2e-05 | 6.4e-06 | 9.800E-06 | 2.987E-06 | 6.400E-06 | 1.200E-05 |
| 19653 | 983 | L983M | C | 1.2e-05 | 5e-06 | 1.5e-05 | 1.067E-05 | 5.132E-06 | 5.000E-06 | 1.500E-05 |
| 19654 | 983 | L983F | C | 8.4e-06 | 7.7e-06 | 5.6e-06 | 7.233E-06 | 1.457E-06 | 5.600E-06 | 8.400E-06 |
| 19655 | 983 | L983P | C | 7.4e-06 | 8.6e-06 | 6.2e-06 | 7.400E-06 | 1.200E-06 | 6.200E-06 | 8.600E-06 |
| 19656 | 983 | L983S | C | 1e-05 | 5.1e-06 | 1.2e-05 | 9.033E-06 | 3.550E-06 | 5.100E-06 | 1.200E-05 |
| 19657 | 983 | L983T | C | 6.9e-06 | 6.4e-06 | 7.5e-06 | 6.933E-06 | 5.508E-07 | 6.400E-06 | 7.500E-06 |
| 19658 | 983 | L983W | C | 6.5e-06 | 1.6e-05 | 8.7e-06 | 1.040E-05 | 4.973E-06 | 6.500E-06 | 1.600E-05 |
| 19659 | 983 | L983Y | C | 7.7e-06 | 6.2e-06 | 6.9e-06 | 6.933E-06 | 7.506E-07 | 6.200E-06 | 7.700E-06 |
| 19660 | 983 | L983V | C | 5e-06 | 8.6e-06 | 9.7e-06 | 7.767E-06 | 2.458E-06 | 5.000E-06 | 9.700E-06 |
| 19661 | 984 | V984A | C | 1e-05 | 9.2e-06 | 6.9e-06 | 8.700E-06 | 1.609E-06 | 6.900E-06 | 1.000E-05 |
| 19662 | 984 | V984R | C | 8.6e-06 | 6.1e-06 | 1.1e-05 | 8.567E-06 | 2.450E-06 | 6.100E-06 | 1.100E-05 |
| 19663 | 984 | V984N | C | 9.3e-06 | 1e-05 | 6.6e-06 | 8.633E-06 | 1.795E-06 | 6.600E-06 | 1.000E-05 |
| 19664 | 984 | V984D | C | 5.4e-06 | 5e-06 | 7.6e-06 | 6.000E-06 | 1.400E-06 | 5.000E-06 | 7.600E-06 |
| 19665 | 984 | V984C | C | 7.7e-06 | 6.4e-06 | 9.8e-06 | 7.967E-06 | 1.716E-06 | 6.400E-06 | 9.800E-06 |
| 19666 | 984 | V984E | C | 5.1e-06 | 1.3e-05 | 8.8e-06 | 8.967E-06 | 3.953E-06 | 5.100E-06 | 1.300E-05 |
| 19667 | 984 | V984Q | C | 1.2e-05 | 8.6e-06 | 9.9e-06 | 1.017E-05 | 1.716E-06 | 8.600E-06 | 1.200E-05 |
| 19668 | 984 | V984G | C | 9e-06 | 5.1e-06 | 9.6e-06 | 7.900E-06 | 2.443E-06 | 5.100E-06 | 9.600E-06 |
| 19669 | 984 | V984H | C | 6.5e-06 | 6.4e-06 | 7.5e-06 | 6.800E-06 | 6.083E-07 | 6.400E-06 | 7.500E-06 |
| 19670 | 984 | V984I | C | 6.1e-06 | 5.1e-06 | 7.5e-06 | 6.233E-06 | 1.206E-06 | 5.100E-06 | 7.500E-06 |
| 19671 | 984 | V984L | C | 1e-05 | 6.1e-06 | 6.4e-06 | 7.500E-06 | 2.170E-06 | 6.100E-06 | 1.000E-05 |
| 19672 | 984 | V984K | C | 6.8e-06 | 7.7e-06 | 1.3e-05 | 9.167E-06 | 3.350E-06 | 6.800E-06 | 1.300E-05 |
| 19673 | 984 | V984M | C | 6.9e-06 | 8.4e-06 | 9.8e-06 | 8.367E-06 | 1.450E-06 | 6.900E-06 | 9.800E-06 |
| 19674 | 984 | V984F | C | 8.3e-06 | 4e-06 | 1.3e-05 | 8.433E-06 | 4.501E-06 | 4.000E-06 | 1.300E-05 |
| 19675 | 984 | V984P | C | 1.1e-05 | 6.9e-06 | 7.7e-06 | 8.533E-06 | 2.173E-06 | 6.900E-06 | 1.100E-05 |
| 19676 | 984 | V984S | C | 8.5e-06 | 7.9e-06 | 6.5e-06 | 7.633E-06 | 1.026E-06 | 6.500E-06 | 8.500E-06 |
| 19677 | 984 | V984T | C | 1.3e-05 | 1.3e-05 | 6.4e-06 | 1.080E-05 | 3.811E-06 | 6.400E-06 | 1.300E-05 |
| 19678 | 984 | V984W | C | 7.5e-06 | 6.6e-06 | 9.4e-06 | 7.833E-06 | 1.429E-06 | 6.600E-06 | 9.400E-06 |
| 19679 | 984 | V984Y | C | 8.3e-06 | 8.9e-06 | 9.1e-06 | 8.767E-06 | 4.163E-07 | 8.300E-06 | 9.100E-06 |
| 19680 | 984 | V984V | C | 8.7e-06 | 1.3e-05 | 4.7e-06 | 8.800E-06 | 4.151E-06 | 4.700E-06 | 1.300E-05 |
| 19681 | 985 | K985A | C | 8.1e-06 | 6.6e-06 | 5.5e-06 | 6.733E-06 | 1.305E-06 | 5.500E-06 | 8.100E-06 |
| 19682 | 985 | K985R | C | 7.3e-06 | 4.5e-06 | 6.8e-06 | 6.200E-06 | 1.493E-06 | 4.500E-06 | 7.300E-06 |
| 19683 | 985 | K985N | C | 7.3e-06 | 8.8e-06 | 9.4e-06 | 8.500E-06 | 1.082E-06 | 7.300E-06 | 9.400E-06 |
| 19684 | 985 | K985D | C | 8.1e-06 | 1.2e-05 | 8.2e-06 | 9.433E-06 | 2.223E-06 | 8.100E-06 | 1.200E-05 |
| 19685 | 985 | K985C | C | 4.3e-06 | 7.6e-06 | 1e-05 | 7.300E-06 | 2.862E-06 | 4.300E-06 | 1.000E-05 |
| 19686 | 985 | K985E | C | 9.2e-06 | 1.1e-05 | 1.2e-05 | 1.073E-05 | 1.419E-06 | 9.200E-06 | 1.200E-05 |
| 19687 | 985 | K985Q | C | 6.2e-06 | 4.8e-06 | 6.3e-06 | 5.767E-06 | 8.386E-07 | 4.800E-06 | 6.300E-06 |
| 19688 | 985 | K985G | C | 1.2e-05 | 8.5e-06 | 1e-05 | 1.017E-05 | 1.756E-06 | 8.500E-06 | 1.200E-05 |
| 19689 | 985 | K985H | C | 6.5e-06 | 6.3e-06 | 9.4e-06 | 7.400E-06 | 1.735E-06 | 6.300E-06 | 9.400E-06 |
| 19690 | 985 | K985I | C | 9.1e-06 | 1.4e-05 | 6.8e-06 | 9.967E-06 | 3.677E-06 | 6.800E-06 | 1.400E-05 |
| 19691 | 985 | K985L | C | 1e-05 | 5.2e-06 | 6e-06 | 7.067E-06 | 2.572E-06 | 5.200E-06 | 1.000E-05 |
| 19692 | 985 | K985K | C | 1.4e-05 | 6e-06 | 4.3e-06 | 8.100E-06 | 5.180E-06 | 4.300E-06 | 1.400E-05 |
| 19693 | 985 | K985M | C | 6.2e-06 | 9.5e-06 | 1.1e-05 | 8.900E-06 | 2.456E-06 | 6.200E-06 | 1.100E-05 |
| 19694 | 985 | K985F | C | 1.3e-05 | 1.3e-05 | 6.8e-06 | 1.093E-05 | 3.580E-06 | 6.800E-06 | 1.300E-05 |
| 19695 | 985 | K985P | C | 9.8e-06 | 8.2e-06 | 6.9e-06 | 8.300E-06 | 1.453E-06 | 6.900E-06 | 9.800E-06 |
| 19696 | 985 | K985S | C | 6.1e-06 | 8.6e-06 | 9.6e-06 | 8.100E-06 | 1.803E-06 | 6.100E-06 | 9.600E-06 |
| 19697 | 985 | K985T | C | 5.8e-06 | 1.3e-05 | 7.5e-06 | 8.767E-06 | 3.763E-06 | 5.800E-06 | 1.300E-05 |
| 19698 | 985 | K985W | C | 1.3e-05 | 5.7e-06 | 5.9e-06 | 8.200E-06 | 4.158E-06 | 5.700E-06 | 1.300E-05 |
| 19699 | 985 | K985Y | C | 7.8e-06 | 9.1e-06 | 9.4e-06 | 8.767E-06 | 8.505E-07 | 7.800E-06 | 9.400E-06 |
| 19700 | 985 | K985V | C | 9.2e-06 | 6.7e-06 | 5.5e-06 | 7.133E-06 | 1.888E-06 | 5.500E-06 | 9.200E-06 |
| 19701 | 986 | D986A | C | 1.1e-05 | 8.5e-06 | 9.2e-06 | 9.567E-06 | 1.290E-06 | 8.500E-06 | 1.100E-05 |
| 19702 | 986 | D986R | C | 6.3e-06 | 8.7e-06 | 5.6e-06 | 6.867E-06 | 1.626E-06 | 5.600E-06 | 8.700E-06 |
| 19703 | 986 | D986N | C | 6.2e-06 | 9.3e-06 | 9.7e-06 | 8.400E-06 | 1.916E-06 | 6.200E-06 | 9.700E-06 |
| 19704 | 986 | D986D | C | 6.6e-06 | 9e-06 | 9.9e-06 | 8.500E-06 | 1.706E-06 | 6.600E-06 | 9.900E-06 |
| 19705 | 986 | D986C | C | 6.7e-06 | 9.4e-06 | 9.2e-06 | 8.433E-06 | 1.504E-06 | 6.700E-06 | 9.400E-06 |
| 19706 | 986 | D986E | C | 5.5e-06 | 7.1e-06 | 1.2e-05 | 8.200E-06 | 3.387E-06 | 5.500E-06 | 1.200E-05 |
| 19707 | 986 | D986Q | C | 5.6e-06 | 1.5e-05 | 8.4e-06 | 9.667E-06 | 4.826E-06 | 5.600E-06 | 1.500E-05 |
| 19708 | 986 | D986G | C | 5.3e-06 | 6.3e-06 | 6.4e-06 | 6.000E-06 | 6.083E-07 | 5.300E-06 | 6.400E-06 |
| 19709 | 986 | D986H | C | 5.6e-06 | 6.6e-06 | 6.4e-06 | 6.200E-06 | 5.292E-07 | 5.600E-06 | 6.600E-06 |
| 19710 | 986 | D986I | C | 8.3e-06 | 4.2e-06 | 9.7e-06 | 7.400E-06 | 2.858E-06 | 4.200E-06 | 9.700E-06 |
| 19711 | 986 | D986L | C | 7.6e-06 | 9.5e-06 | 1.3e-05 | 1.003E-05 | 2.739E-06 | 7.600E-06 | 1.300E-05 |
| 19712 | 986 | D986K | C | 6.4e-06 | 9.5e-06 | 7.2e-06 | 7.700E-06 | 1.609E-06 | 6.400E-06 | 9.500E-06 |
| 19713 | 986 | D986M | C | 1.3e-05 | 4.6e-06 | 8.8e-06 | 8.800E-06 | 4.200E-06 | 4.600E-06 | 1.300E-05 |
| 19714 | 986 | D986F | C | 9.7e-06 | 9e-06 | 1.3e-05 | 1.057E-05 | 2.136E-06 | 9.000E-06 | 1.300E-05 |
| 19715 | 986 | D986P | C | 6.6e-06 | 5.2e-06 | 9e-06 | 6.933E-06 | 1.922E-06 | 5.200E-06 | 9.000E-06 |
| 19716 | 986 | D986S | C | 7.4e-06 | 7.6e-06 | 8.5e-06 | 7.833E-06 | 5.859E-07 | 7.400E-06 | 8.500E-06 |
| 19717 | 986 | D986T | C | 7.1e-06 | 1.7e-05 | 1.4e-05 | 1.270E-05 | 5.076E-06 | 7.100E-06 | 1.700E-05 |
| 19718 | 986 | D986W | C | 1e-05 | 8.8e-06 | 1.2e-05 | 1.027E-05 | 1.617E-06 | 8.800E-06 | 1.200E-05 |
| 19719 | 986 | D986Y | C | 5.9e-06 | 6.3e-06 | 1e-05 | 7.400E-06 | 2.261E-06 | 5.900E-06 | 1.000E-05 |

| | | | | | | | | | | |
|---|---|---|---|---|---|---|---|---|---|---|
| 19720 | 986 | D986V | C | 9.4e-06 | 6.7e-06 | 4.1e-06 | 6.733E-06 | 2.650E-06 | 4.100E-06 | 9.400E-06 |
| 19721 | 987 | Y987A | C | 4.6e-06 | 9.8e-06 | 8.5e-06 | 7.633E-06 | 2.706E-06 | 4.600E-06 | 9.800E-06 |
| 19722 | 987 | Y987R | C | 6.9e-06 | 5e-06 | 5.2e-06 | 5.700E-06 | 1.044E-06 | 5.000E-06 | 6.900E-06 |
| 19723 | 987 | Y987N | C | 6.9e-06 | 1.3e-05 | 8.9e-06 | 9.600E-06 | 3.110E-06 | 6.900E-06 | 1.300E-05 |
| 19724 | 987 | Y987D | C | 7e-06 | 6.3e-06 | 1.2e-05 | 8.433E-06 | 3.109E-06 | 6.300E-06 | 1.200E-05 |
| 19725 | 987 | Y987C | C | 7.9e-06 | 5.9e-06 | 8.5e-06 | 7.433E-06 | 1.361E-06 | 5.900E-06 | 8.500E-06 |
| 19726 | 987 | Y987E | C | 8e-06 | 9.6e-06 | 1.1e-05 | 9.533E-06 | 1.501E-06 | 8.000E-06 | 1.100E-05 |
| 19727 | 987 | Y987Q | C | 9.8e-06 | 1.4e-05 | 1.1e-05 | 1.160E-05 | 2.163E-06 | 9.800E-06 | 1.400E-05 |
| 19728 | 987 | Y987G | C | 5e-06 | 6.2e-06 | 9.7e-06 | 6.967E-06 | 2.442E-06 | 5.000E-06 | 9.700E-06 |
| 19729 | 987 | Y987H | C | 7.7e-06 | 8.1e-06 | 7.2e-06 | 7.667E-06 | 4.509E-07 | 7.200E-06 | 8.100E-06 |
| 19730 | 987 | Y987I | C | 6.5e-06 | 1.1e-05 | 5.4e-06 | 7.633E-06 | 2.967E-06 | 5.400E-06 | 1.100E-05 |
| 19731 | 987 | Y987L | C | 1.3e-05 | 6.4e-06 | 6.4e-06 | 8.600E-06 | 3.811E-06 | 6.400E-06 | 1.300E-05 |
| 19732 | 987 | Y987K | C | 8.5e-06 | 1.1e-05 | 7.4e-06 | 8.967E-06 | 1.845E-06 | 7.400E-06 | 1.100E-05 |
| 19733 | 987 | Y987M | C | 7.9e-06 | 1.6e-05 | 6.8e-06 | 1.023E-05 | 5.024E-06 | 6.800E-06 | 1.600E-05 |
| 19734 | 987 | Y987F | C | 3.6e-06 | 1.1e-05 | 5.7e-06 | 6.767E-06 | 3.814E-06 | 3.600E-06 | 1.100E-05 |
| 19735 | 987 | Y987P | C | 9.2e-06 | 6.1e-06 | 1e-05 | 8.433E-06 | 2.060E-06 | 6.100E-06 | 1.000E-05 |
| 19736 | 987 | Y987S | C | 8e-06 | 7.4e-06 | 8.9e-06 | 8.100E-06 | 7.550E-07 | 7.400E-06 | 8.900E-06 |
| 19737 | 987 | Y987T | C | 5.9e-06 | 9.8e-06 | 6.6e-06 | 7.433E-06 | 2.079E-06 | 5.900E-06 | 9.800E-06 |
| 19738 | 987 | Y987W | C | 5.2e-06 | 6.6e-06 | 8.2e-06 | 6.667E-06 | 1.501E-06 | 5.200E-06 | 8.200E-06 |
| 19739 | 987 | Y987Y | C | 8.5e-06 | 7.5e-06 | 4.9e-06 | 6.967E-06 | 1.858E-06 | 4.900E-06 | 8.500E-06 |
| 19740 | 987 | Y987V | C | 1.1e-05 | 9.3e-06 | 4.9e-06 | 8.400E-06 | 3.148E-06 | 4.900E-06 | 1.100E-05 |
| 19741 | 988 | F988A | C | 6.8e-06 | 7.6e-06 | 6.7e-06 | 7.033E-06 | 4.933E-07 | 6.700E-06 | 7.600E-06 |
| 19742 | 988 | F988R | C | 5.4e-06 | 5e-06 | 6.3e-06 | 5.567E-06 | 6.658E-07 | 5.000E-06 | 6.300E-06 |
| 19743 | 988 | F988N | C | 1.1e-05 | 7.6e-06 | 9.1e-06 | 9.233E-06 | 1.704E-06 | 7.600E-06 | 1.100E-05 |
| 19744 | 988 | F988D | C | 4.5e-06 | 9.1e-06 | 6.8e-06 | 6.800E-06 | 2.300E-06 | 4.500E-06 | 9.100E-06 |
| 19745 | 988 | F988C | C | 1.1e-05 | 7.7e-06 | 1.4e-05 | 1.090E-05 | 3.151E-06 | 7.700E-06 | 1.400E-05 |
| 19746 | 988 | F988E | C | 7e-06 | 6.6e-06 | 4.6e-06 | 6.067E-06 | 1.286E-06 | 4.600E-06 | 7.000E-06 |
| 19747 | 988 | F988Q | C | 4.6e-06 | 8.9e-06 | 5.4e-06 | 6.300E-06 | 2.287E-06 | 4.600E-06 | 8.900E-06 |
| 19748 | 988 | F988G | C | 9.8e-06 | 5.6e-06 | 9.3e-06 | 8.233E-06 | 2.294E-06 | 5.600E-06 | 9.800E-06 |
| 19749 | 988 | F988H | C | 8.6e-06 | 9e-06 | 5e-06 | 7.533E-06 | 2.203E-06 | 5.000E-06 | 9.000E-06 |
| 19750 | 988 | F988I | C | 6.4e-06 | 1.1e-05 | 9.9e-06 | 9.100E-06 | 2.402E-06 | 6.400E-06 | 1.100E-05 |
| 19751 | 988 | F988L | C | 1e-05 | 6e-06 | 6.5e-06 | 7.500E-06 | 2.179E-06 | 6.000E-06 | 1.000E-05 |
| 19752 | 988 | F988K | C | 8.7e-06 | 7.2e-06 | 4.6e-06 | 6.833E-06 | 2.074E-06 | 4.600E-06 | 8.700E-06 |
| 19753 | 988 | F988M | C | 1.3e-05 | 9.3e-06 | 4.8e-06 | 9.033E-06 | 4.106E-06 | 4.800E-06 | 1.300E-05 |
| 19754 | 988 | F988F | C | 6.5e-06 | 6.2e-06 | 6e-06 | 6.233E-06 | 2.517E-07 | 6.000E-06 | 6.500E-06 |
| 19755 | 988 | F988P | C | 8.9e-06 | 6.3e-06 | 8.1e-06 | 7.767E-06 | 1.332E-06 | 6.300E-06 | 8.900E-06 |
| 19756 | 988 | F988S | C | 5.9e-06 | 1e-05 | 9.8e-06 | 8.567E-06 | 2.312E-06 | 5.900E-06 | 1.000E-05 |
| 19757 | 988 | F988T | C | 5.3e-06 | 8.2e-06 | 5.1e-06 | 6.200E-06 | 1.735E-06 | 5.100E-06 | 8.200E-06 |
| 19758 | 988 | F988W | C | 6.2e-06 | 1.2e-05 | 6.8e-06 | 8.333E-06 | 3.190E-06 | 6.200E-06 | 1.200E-05 |
| 19759 | 988 | F988Y | C | 8.8e-06 | 4.5e-06 | 4.3e-06 | 5.867E-06 | 2.542E-06 | 4.300E-06 | 8.800E-06 |
| 19760 | 988 | F988V | C | 6.2e-06 | 4.9e-06 | 1e-05 | 7.033E-06 | 2.650E-06 | 4.900E-06 | 1.000E-05 |
| 19761 | 989 | P989A | C | 5.5e-06 | 4.9e-06 | 6.1e-06 | 5.500E-06 | 6.000E-07 | 4.900E-06 | 6.100E-06 |
| 19762 | 989 | P989R | C | 1e-05 | 1e-05 | 6.9e-06 | 8.967E-06 | 1.790E-06 | 6.900E-06 | 1.000E-05 |
| 19763 | 989 | P989N | C | 8.5e-06 | 9.1e-06 | 5.5e-06 | 7.700E-06 | 1.929E-06 | 5.500E-06 | 9.100E-06 |
| 19764 | 989 | P989D | C | 8.9e-06 | 6.7e-06 | 6.5e-06 | 7.367E-06 | 1.332E-06 | 6.500E-06 | 8.900E-06 |
| 19765 | 989 | P989C | C | 7.6e-06 | 1.1e-05 | 9.5e-06 | 9.367E-06 | 1.704E-06 | 7.600E-06 | 1.100E-05 |
| 19766 | 989 | P989E | C | 8.3e-06 | 1.5e-05 | 1.4e-05 | 1.243E-05 | 3.614E-06 | 8.300E-06 | 1.500E-05 |
| 19767 | 989 | P989Q | C | 6.8e-06 | 6.3e-06 | 1.3e-05 | 8.700E-06 | 3.732E-06 | 6.300E-06 | 1.300E-05 |
| 19768 | 989 | P989G | C | 1.6e-05 | 9.6e-06 | 6.5e-06 | 1.070E-05 | 4.845E-06 | 6.500E-06 | 1.600E-05 |
| 19769 | 989 | P989H | C | 4.5e-06 | 5.2e-06 | 8.7e-06 | 6.133E-06 | 2.250E-06 | 4.500E-06 | 8.700E-06 |
| 19770 | 989 | P989I | C | 9.6e-06 | 8.8e-06 | 6e-06 | 8.133E-06 | 1.890E-06 | 6.000E-06 | 9.600E-06 |
| 19771 | 989 | P989L | C | 6.9e-06 | 1e-05 | 1.1e-05 | 9.300E-06 | 2.138E-06 | 6.900E-06 | 1.100E-05 |
| 19772 | 989 | P989K | C | 9.1e-06 | 1e-05 | 5.8e-06 | 8.300E-06 | 2.211E-06 | 5.800E-06 | 1.000E-05 |
| 19773 | 989 | P989M | C | 3.8e-06 | 7.6e-06 | 6.8e-06 | 6.067E-06 | 2.003E-06 | 3.800E-06 | 7.600E-06 |
| 19774 | 989 | P989F | C | 8.3e-06 | 4.5e-06 | 1.3e-05 | 8.600E-06 | 4.258E-06 | 4.500E-06 | 1.300E-05 |
| 19775 | 989 | P989P | C | 1e-05 | 6.4e-06 | 9.9e-06 | 8.767E-06 | 2.050E-06 | 6.400E-06 | 1.000E-05 |
| 19776 | 989 | P989S | C | 4.8e-06 | 9.4e-06 | 9.4e-06 | 7.867E-06 | 2.656E-06 | 4.800E-06 | 9.400E-06 |
| 19777 | 989 | P989T | C | 4.3e-06 | 1.3e-05 | 4.8e-06 | 7.367E-06 | 4.885E-06 | 4.300E-06 | 1.300E-05 |
| 19778 | 989 | P989W | C | 6.8e-06 | 7.6e-06 | 1.2e-05 | 8.800E-06 | 2.800E-06 | 6.800E-06 | 1.200E-05 |
| 19779 | 989 | P989Y | C | 9.6e-06 | 5.2e-06 | 1.2e-05 | 8.933E-06 | 3.449E-06 | 5.200E-06 | 1.200E-05 |
| 19780 | 989 | P989V | C | 4.8e-06 | 6.8e-06 | 1.3e-05 | 8.200E-06 | 4.276E-06 | 4.800E-06 | 1.300E-05 |
| 19781 | 990 | E990A | C | 4e-06 | 7.9e-06 | 6.1e-06 | 6.000E-06 | 1.952E-06 | 4.000E-06 | 7.900E-06 |
| 19782 | 990 | E990R | C | 9.2e-06 | 8.6e-06 | 8.1e-06 | 8.633E-06 | 5.508E-07 | 8.100E-06 | 9.200E-06 |
| 19783 | 990 | E990N | C | 4.7e-06 | 1.1e-05 | 6e-06 | 7.233E-06 | 3.326E-06 | 4.700E-06 | 1.100E-05 |
| 19784 | 990 | E990D | C | 5.3e-06 | 8.4e-06 | 9.7e-06 | 7.800E-06 | 2.261E-06 | 5.300E-06 | 9.700E-06 |
| 19785 | 990 | E990C | C | 9.8e-06 | 6.7e-06 | 8.5e-06 | 8.333E-06 | 1.557E-06 | 6.700E-06 | 9.800E-06 |
| 19786 | 990 | E990E | C | 8e-06 | 6.4e-06 | 1e-05 | 8.133E-06 | 1.804E-06 | 6.400E-06 | 1.000E-05 |
| 19787 | 990 | E990Q | C | 1.4e-05 | 1.2e-05 | 7e-06 | 1.100E-05 | 3.606E-06 | 7.000E-06 | 1.400E-05 |
| 19788 | 990 | E990G | C | 4.2e-06 | 8.3e-06 | 5.6e-06 | 6.033E-06 | 2.084E-06 | 4.200E-06 | 8.300E-06 |
| 19789 | 990 | E990H | C | 1.3e-05 | 4.7e-06 | 1.3e-05 | 1.023E-05 | 4.792E-06 | 4.700E-06 | 1.300E-05 |
| 19790 | 990 | E990I | C | 6.3e-06 | 7.3e-06 | 9.8e-06 | 7.800E-06 | 1.803E-06 | 6.300E-06 | 9.800E-06 |
| 19791 | 990 | E990L | C | 9.2e-06 | 7.7e-06 | 6.6e-06 | 7.833E-06 | 1.305E-06 | 6.600E-06 | 9.200E-06 |
| 19792 | 990 | E990K | C | 6.7e-06 | 7.1e-06 | 9.6e-06 | 7.800E-06 | 1.572E-06 | 6.700E-06 | 9.600E-06 |
| 19793 | 990 | E990M | C | 7e-06 | 4.7e-06 | 9.6e-06 | 7.100E-06 | 2.452E-06 | 4.700E-06 | 9.600E-06 |
| 19794 | 990 | E990F | C | 1.2e-05 | 8.8e-06 | 1e-05 | 1.027E-05 | 1.617E-06 | 8.800E-06 | 1.200E-05 |
| 19795 | 990 | E990P | C | 7.8e-06 | 8.9e-06 | 5.3e-06 | 7.333E-06 | 1.845E-06 | 5.300E-06 | 8.900E-06 |
| 19796 | 990 | E990S | C | 6.3e-06 | 8.4e-06 | 1e-05 | 8.233E-06 | 1.856E-06 | 6.300E-06 | 1.000E-05 |
| 19797 | 990 | E990T | C | 9e-06 | 7.4e-06 | 5.1e-06 | 7.167E-06 | 1.960E-06 | 5.100E-06 | 9.000E-06 |
| 19798 | 990 | E990W | C | 9.3e-06 | 9e-06 | 6.3e-06 | 8.200E-06 | 1.652E-06 | 6.300E-06 | 9.300E-06 |
| 19799 | 990 | E990Y | C | 4.9e-06 | 8e-06 | 1e-05 | 7.633E-06 | 2.570E-06 | 4.900E-06 | 1.000E-05 |
| 19800 | 990 | E990V | C | 1.3e-05 | 9.1e-06 | 6.3e-06 | 9.467E-06 | 3.365E-06 | 6.300E-06 | 1.300E-05 |
| 19801 | 991 | P991A | C | 6.2e-06 | 1.1e-05 | 5.9e-06 | 7.700E-06 | 2.862E-06 | 5.900E-06 | 1.100E-05 |
| 19802 | 991 | P991R | C | 8.5e-06 | 8.8e-06 | 5e-06 | 7.433E-06 | 2.113E-06 | 5.000E-06 | 8.800E-06 |
| 19803 | 991 | P991N | C | 5.1e-06 | 8.7e-06 | 4.6e-06 | 6.133E-06 | 2.237E-06 | 4.600E-06 | 8.700E-06 |
| 19804 | 991 | P991D | C | 7.4e-06 | 8.8e-06 | 1.4e-05 | 1.007E-05 | 3.478E-06 | 7.400E-06 | 1.400E-05 |

| | | | | | | | | | | |
|---|---|---|---|---|---|---|---|---|---|---|
| 19805 | 991 | P991C | C | 8.7e-06 | 9.2e-06 | 5.7e-06 | 7.867E-06 | 1.893E-06 | 5.700E-06 | 9.200E-06 |
| 19806 | 991 | P991E | C | 8.4e-06 | 8.5e-06 | 1.2e-05 | 9.633E-06 | 2.050E-06 | 8.400E-06 | 1.200E-05 |
| 19807 | 991 | P991Q | C | 6e-06 | 6.7e-06 | 6.9e-06 | 6.533E-06 | 4.726E-07 | 6.000E-06 | 6.900E-06 |
| 19808 | 991 | P991G | C | 8.4e-06 | 8.4e-06 | 9.2e-06 | 8.667E-06 | 4.619E-07 | 8.400E-06 | 9.200E-06 |
| 19809 | 991 | P991H | C | 9.3e-06 | 6e-06 | 8.8e-06 | 8.033E-06 | 1.779E-06 | 6.000E-06 | 9.300E-06 |
| 19810 | 991 | P991I | C | 7.1e-06 | 8.2e-06 | 7.8e-06 | 7.700E-06 | 5.568E-07 | 7.100E-06 | 8.200E-06 |
| 19811 | 991 | P991L | C | 8.2e-06 | 1.4e-05 | 5.4e-06 | 9.200E-06 | 4.386E-06 | 5.400E-06 | 1.400E-05 |
| 19812 | 991 | P991K | C | 1.3e-05 | 5.5e-06 | 6.2e-06 | 8.233E-06 | 4.143E-06 | 5.500E-06 | 1.300E-05 |
| 19813 | 991 | P991M | C | 1e-05 | 5.4e-06 | 1e-05 | 8.467E-06 | 2.656E-06 | 5.400E-06 | 1.000E-05 |
| 19814 | 991 | P991F | C | 7.4e-06 | 9.5e-06 | 3.8e-06 | 6.900E-06 | 2.883E-06 | 3.800E-06 | 9.500E-06 |
| 19815 | 991 | P991P | C | 8.7e-06 | 7.5e-06 | 7.4e-06 | 7.867E-06 | 7.234E-07 | 7.400E-06 | 8.700E-06 |
| 19816 | 991 | P991S | C | 6.5e-06 | 5.7e-06 | 7.2e-06 | 6.467E-06 | 7.506E-07 | 5.700E-06 | 7.200E-06 |
| 19817 | 991 | P991T | C | 4.6e-06 | 6.5e-06 | 1e-05 | 7.033E-06 | 2.739E-06 | 4.600E-06 | 1.000E-05 |
| 19818 | 991 | P991W | C | 1e-05 | 5.2e-06 | 6.8e-06 | 7.333E-06 | 2.444E-06 | 5.200E-06 | 1.000E-05 |
| 19819 | 991 | P991Y | C | 5.9e-06 | 1.1e-05 | 7.9e-06 | 8.267E-06 | 2.570E-06 | 5.900E-06 | 1.100E-05 |
| 19820 | 991 | P991V | C | 8.6e-06 | 9.5e-06 | 4.6e-06 | 7.567E-06 | 2.608E-06 | 4.600E-06 | 9.500E-06 |
| 19821 | 992 | V992A | C | 1e-05 | 1.3e-05 | 1.2e-05 | 1.167E-05 | 1.528E-06 | 1.000E-05 | 1.300E-05 |
| 19822 | 992 | V992R | C | 6.1e-06 | 1.3e-05 | 4.6e-06 | 7.900E-06 | 4.480E-06 | 4.600E-06 | 1.300E-05 |
| 19823 | 992 | V992N | C | 5.8e-06 | 4.8e-06 | 4.4e-06 | 5.000E-06 | 7.211E-07 | 4.400E-06 | 5.800E-06 |
| 19824 | 992 | V992D | C | 5.5e-06 | 9.2e-06 | 5.5e-06 | 6.733E-06 | 2.136E-06 | 5.500E-06 | 9.200E-06 |
| 19825 | 992 | V992C | C | 9.6e-06 | 6.6e-06 | 5.9e-06 | 7.367E-06 | 1.966E-06 | 5.900E-06 | 9.600E-06 |
| 19826 | 992 | V992E | C | 1.4e-05 | 8.4e-06 | 4.3e-06 | 8.900E-06 | 4.869E-06 | 4.300E-06 | 1.400E-05 |
| 19827 | 992 | V992Q | C | 4.7e-06 | 1.1e-05 | 7.4e-06 | 7.700E-06 | 3.161E-06 | 4.700E-06 | 1.100E-05 |
| 19828 | 992 | V992G | C | 9.3e-06 | 9.4e-06 | 7.7e-06 | 8.800E-06 | 9.539E-07 | 7.700E-06 | 9.400E-06 |
| 19829 | 992 | V992H | C | 9.1e-06 | 4.8e-06 | 8.4e-06 | 7.433E-06 | 2.307E-06 | 4.800E-06 | 9.100E-06 |
| 19830 | 992 | V992I | C | 1.2e-05 | 9.4e-06 | 9.4e-06 | 1.027E-05 | 1.501E-06 | 9.400E-06 | 1.200E-05 |
| 19831 | 992 | V992L | C | 1.1e-05 | 8.7e-06 | 1e-05 | 9.900E-06 | 1.153E-06 | 8.700E-06 | 1.100E-05 |
| 19832 | 992 | V992K | C | 8.1e-06 | 4.8e-06 | 6.2e-06 | 6.367E-06 | 1.656E-06 | 4.800E-06 | 8.100E-06 |
| 19833 | 992 | V992M | C | 9.2e-06 | 7e-06 | 1e-05 | 8.733E-06 | 1.553E-06 | 7.000E-06 | 1.000E-05 |
| 19834 | 992 | V992F | C | 8.9e-06 | 8.5e-06 | 1e-05 | 9.133E-06 | 7.767E-07 | 8.500E-06 | 1.000E-05 |
| 19835 | 992 | V992P | C | 1.2e-05 | 1.2e-05 | 7.2e-06 | 1.040E-05 | 2.771E-06 | 7.200E-06 | 1.200E-05 |
| 19836 | 992 | V992S | C | 7e-06 | 8.6e-06 | 8.9e-06 | 8.167E-06 | 1.021E-06 | 7.000E-06 | 8.900E-06 |
| 19837 | 992 | V992T | C | 8.2e-06 | 7.7e-06 | 5.9e-06 | 7.267E-06 | 1.210E-06 | 5.900E-06 | 8.200E-06 |
| 19838 | 992 | V992W | C | 1.8e-05 | 7.2e-06 | 7.7e-06 | 1.097E-05 | 6.096E-06 | 7.200E-06 | 1.800E-05 |
| 19839 | 992 | V992Y | C | 7.3e-06 | 5.4e-06 | 5.5e-06 | 6.067E-06 | 1.069E-06 | 5.400E-06 | 7.300E-06 |
| 19840 | 992 | V992V | C | 9.1e-06 | 4.8e-06 | 6.6e-06 | 6.833E-06 | 2.159E-06 | 4.800E-06 | 9.100E-06 |
| 19841 | 993 | T993A | C | 4.7e-06 | 8.5e-06 | 7e-06 | 6.733E-06 | 1.914E-06 | 4.700E-06 | 8.500E-06 |
| 19842 | 993 | T993R | C | 1.2e-05 | 7e-06 | 4.5e-06 | 7.833E-06 | 3.819E-06 | 4.500E-06 | 1.200E-05 |
| 19843 | 993 | T993N | C | 6.9e-06 | 6.9e-06 | 7.1e-06 | 6.967E-06 | 1.155E-07 | 6.900E-06 | 7.100E-06 |
| 19844 | 993 | T993D | C | 5.2e-06 | 9.1e-06 | 4.4e-06 | 6.233E-06 | 2.515E-06 | 4.400E-06 | 9.100E-06 |
| 19845 | 993 | T993C | C | 6e-06 | 6.7e-06 | 5.7e-06 | 6.133E-06 | 5.132E-07 | 5.700E-06 | 6.700E-06 |
| 19846 | 993 | T993E | C | 1.3e-05 | 9.2e-06 | 9.7e-06 | 1.063E-05 | 2.065E-06 | 9.200E-06 | 1.300E-05 |
| 19847 | 993 | T993Q | C | 7.6e-06 | 1e-05 | 7.6e-06 | 8.400E-06 | 1.386E-06 | 7.600E-06 | 1.000E-05 |
| 19848 | 993 | T993G | C | 7.7e-06 | 9.3e-06 | 1.2e-05 | 9.667E-06 | 2.173E-06 | 7.700E-06 | 1.200E-05 |
| 19849 | 993 | T993H | C | 4.6e-06 | 1.3e-05 | 7.9e-06 | 8.500E-06 | 4.232E-06 | 4.600E-06 | 1.300E-05 |
| 19850 | 993 | T993I | C | 9.5e-06 | 5.4e-06 | 7.3e-06 | 7.400E-06 | 2.052E-06 | 5.400E-06 | 9.500E-06 |
| 19851 | 993 | T993L | C | 1.2e-05 | 6.2e-06 | 1.4e-05 | 1.073E-05 | 4.051E-06 | 6.200E-06 | 1.400E-05 |
| 19852 | 993 | T993K | C | 1.1e-05 | 6.7e-06 | 5.1e-06 | 7.600E-06 | 3.051E-06 | 5.100E-06 | 1.100E-05 |
| 19853 | 993 | T993M | C | 8.1e-06 | 5.9e-06 | 4.8e-06 | 6.267E-06 | 1.680E-06 | 4.800E-06 | 8.100E-06 |
| 19854 | 993 | T993F | C | 8.7e-06 | 6.9e-06 | 9.8e-06 | 8.467E-06 | 1.464E-06 | 6.900E-06 | 9.800E-06 |
| 19855 | 993 | T993P | C | 6.3e-06 | 7.5e-06 | 1.1e-05 | 8.267E-06 | 2.442E-06 | 6.300E-06 | 1.100E-05 |
| 19856 | 993 | T993S | C | 3.8e-06 | 9.6e-06 | 5.1e-06 | 6.167E-06 | 3.044E-06 | 3.800E-06 | 9.600E-06 |
| 19857 | 993 | T993T | C | 7.2e-06 | 7e-06 | 6.6e-06 | 6.933E-06 | 3.055E-07 | 6.600E-06 | 7.200E-06 |
| 19858 | 993 | T993W | C | 8.9e-06 | 1.5e-05 | 5.5e-06 | 9.800E-06 | 4.814E-06 | 5.500E-06 | 1.500E-05 |
| 19859 | 993 | T993Y | C | 1e-05 | 9.8e-06 | 8.1e-06 | 9.300E-06 | 1.044E-06 | 8.100E-06 | 1.000E-05 |
| 19860 | 993 | T993V | C | 8.8e-06 | 8e-06 | 7.5e-06 | 8.100E-06 | 6.557E-07 | 7.500E-06 | 8.800E-06 |
| 19861 | 994 | V994A | C | 8e-06 | 7.1e-06 | 5.5e-06 | 6.867E-06 | 1.266E-06 | 5.500E-06 | 8.000E-06 |
| 19862 | 994 | V994R | C | 1.4e-05 | 7e-06 | 9.3e-06 | 1.010E-05 | 3.568E-06 | 7.000E-06 | 1.400E-05 |
| 19863 | 994 | V994N | C | 8.8e-06 | 9.2e-06 | 6.9e-06 | 8.300E-06 | 1.229E-06 | 6.900E-06 | 9.200E-06 |
| 19864 | 994 | V994D | C | 9.6e-06 | 8.5e-06 | 4.3e-06 | 7.467E-06 | 2.797E-06 | 4.300E-06 | 9.600E-06 |
| 19865 | 994 | V994C | C | 4.3e-06 | 8.9e-06 | 1.3e-05 | 8.733E-06 | 4.352E-06 | 4.300E-06 | 1.300E-05 |
| 19866 | 994 | V994E | C | 4.7e-06 | 6.8e-06 | 7.4e-06 | 6.300E-06 | 1.418E-06 | 4.700E-06 | 7.400E-06 |
| 19867 | 994 | V994Q | C | 1.3e-05 | 6.2e-06 | 8.6e-06 | 9.267E-06 | 3.449E-06 | 6.200E-06 | 1.300E-05 |
| 19868 | 994 | V994G | C | 8.2e-06 | 1e-05 | 7e-06 | 8.400E-06 | 1.510E-06 | 7.000E-06 | 1.000E-05 |
| 19869 | 994 | V994H | C | 9.8e-06 | 9.3e-06 | 1.2e-05 | 1.037E-05 | 1.436E-06 | 9.300E-06 | 1.200E-05 |
| 19870 | 994 | V994I | C | 9e-06 | 8.3e-06 | 5.1e-06 | 7.467E-06 | 2.079E-06 | 5.100E-06 | 9.000E-06 |
| 19871 | 994 | V994L | C | 6.2e-06 | 7.1e-06 | 7.6e-06 | 6.967E-06 | 7.095E-07 | 6.200E-06 | 7.600E-06 |
| 19872 | 994 | V994K | C | 6.7e-06 | 6.7e-06 | 1.1e-05 | 8.133E-06 | 2.483E-06 | 6.700E-06 | 1.100E-05 |
| 19873 | 994 | V994M | C | 7.1e-06 | 7.6e-06 | 9.1e-06 | 7.933E-06 | 1.041E-06 | 7.100E-06 | 9.100E-06 |
| 19874 | 994 | V994F | C | 5.1e-06 | 8.1e-06 | 6.8e-06 | 6.667E-06 | 1.504E-06 | 5.100E-06 | 8.100E-06 |
| 19875 | 994 | V994P | C | 7.1e-06 | 6.5e-06 | 8.5e-06 | 7.367E-06 | 1.026E-06 | 6.500E-06 | 8.500E-06 |
| 19876 | 994 | V994S | C | 6.2e-06 | 1.3e-05 | 1.2e-05 | 1.040E-05 | 3.672E-06 | 6.200E-06 | 1.300E-05 |
| 19877 | 994 | V994T | C | 1.1e-05 | 4.9e-06 | 5.6e-06 | 7.167E-06 | 3.338E-06 | 4.900E-06 | 1.100E-05 |
| 19878 | 994 | V994W | C | 9.1e-06 | 7e-06 | 6.6e-06 | 7.567E-06 | 1.343E-06 | 6.600E-06 | 9.100E-06 |
| 19879 | 994 | V994Y | C | 9.8e-06 | 7.7e-06 | 1e-05 | 9.167E-06 | 1.274E-06 | 7.700E-06 | 1.000E-05 |
| 19880 | 994 | V994V | C | 1.2e-05 | 7.1e-06 | 1e-05 | 9.700E-06 | 2.464E-06 | 7.100E-06 | 1.200E-05 |
| 19881 | 995 | S995A | C | 8.4e-06 | 1.1e-05 | 6.6e-06 | 8.667E-06 | 2.212E-06 | 6.600E-06 | 1.100E-05 |
| 19882 | 995 | S995R | C | 4.4e-06 | 7.7e-06 | 4.6e-06 | 5.567E-06 | 1.850E-06 | 4.400E-06 | 7.700E-06 |
| 19883 | 995 | S995N | C | 1e-05 | 5.3e-06 | 1e-05 | 8.433E-06 | 2.714E-06 | 5.300E-06 | 1.000E-05 |
| 19884 | 995 | S995D | C | 5.6e-06 | 6.5e-06 | 7.5e-06 | 6.533E-06 | 9.504E-07 | 5.600E-06 | 7.500E-06 |
| 19885 | 995 | S995C | C | 6.8e-06 | 9.2e-06 | 1.3e-05 | 9.667E-06 | 3.126E-06 | 6.800E-06 | 1.300E-05 |
| 19886 | 995 | S995E | C | 6.3e-06 | 7.5e-06 | 5.5e-06 | 6.433E-06 | 1.007E-06 | 5.500E-06 | 7.500E-06 |
| 19887 | 995 | S995Q | C | 5.4e-06 | 5.8e-06 | 9.5e-06 | 6.900E-06 | 2.261E-06 | 5.400E-06 | 9.500E-06 |
| 19888 | 995 | S995G | C | 9.8e-06 | 1.2e-05 | 9.3e-06 | 1.037E-05 | 1.436E-06 | 9.300E-06 | 1.200E-05 |
| 19889 | 995 | S995H | C | 3.7e-06 | 5.3e-06 | 8.1e-06 | 5.700E-06 | 2.227E-06 | 3.700E-06 | 8.100E-06 |

| | | | | | | | | | | |
|---|---|---|---|---|---|---|---|---|---|---|
| 19890 | 995 | S995I | C | 6.8e-06 | 6.5e-06 | 5.3e-06 | 6.200E-06 | 7.937E-07 | 5.300E-06 | 6.800E-06 |
| 19891 | 995 | S995L | C | 8e-06 | 7.7e-06 | 1.3e-05 | 9.567E-06 | 2.977E-06 | 7.700E-06 | 1.300E-05 |
| 19892 | 995 | S995K | C | 8.8e-06 | 5.7e-06 | 7.9e-06 | 7.467E-06 | 1.595E-06 | 5.700E-06 | 8.800E-06 |
| 19893 | 995 | S995M | C | 4.8e-06 | 8.1e-06 | 7.2e-06 | 6.700E-06 | 1.706E-06 | 4.800E-06 | 8.100E-06 |
| 19894 | 995 | S995F | C | 9.6e-06 | 8.7e-06 | 1.3e-05 | 1.043E-05 | 2.268E-06 | 8.700E-06 | 1.300E-05 |
| 19895 | 995 | S995P | C | 4.7e-06 | 1.2e-05 | 7.8e-06 | 8.167E-06 | 3.664E-06 | 4.700E-06 | 1.200E-05 |
| 19896 | 995 | S995S | C | 9.9e-06 | 1.1e-05 | 7e-06 | 9.300E-06 | 2.066E-06 | 7.000E-06 | 1.100E-05 |
| 19897 | 995 | S995T | C | 5.8e-06 | 6.7e-06 | 1e-05 | 7.500E-06 | 2.211E-06 | 5.800E-06 | 1.000E-05 |
| 19898 | 995 | S995W | C | 4e-06 | 1e-05 | 6.8e-06 | 6.933E-06 | 3.002E-06 | 4.000E-06 | 1.000E-05 |
| 19899 | 995 | S995Y | C | 1e-05 | 5.3e-06 | 8.5e-06 | 7.933E-06 | 2.401E-06 | 5.300E-06 | 1.000E-05 |
| 19900 | 995 | S995V | C | 7.4e-06 | 5.1e-06 | 8.3e-06 | 6.933E-06 | 1.650E-06 | 5.100E-06 | 8.300E-06 |
| 19901 | 996 | W996A | C | 6.9e-06 | 6.7e-06 | 1.1e-05 | 8.200E-06 | 2.427E-06 | 6.700E-06 | 1.100E-05 |
| 19902 | 996 | W996R | C | 6.9e-06 | 9.2e-06 | 1.2e-05 | 9.367E-06 | 2.554E-06 | 6.900E-06 | 1.200E-05 |
| 19903 | 996 | W996N | C | 6.9e-06 | 4.5e-06 | 1.2e-05 | 7.800E-06 | 3.830E-06 | 4.500E-06 | 1.200E-05 |
| 19904 | 996 | W996D | C | 1.2e-05 | 6.4e-06 | 1.2e-05 | 1.013E-05 | 3.233E-06 | 6.400E-06 | 1.200E-05 |
| 19905 | 996 | W996C | C | 1e-05 | 9.4e-06 | 1.1e-05 | 1.013E-05 | 8.083E-07 | 9.400E-06 | 1.100E-05 |
| 19906 | 996 | W996E | C | 5.6e-06 | 1e-05 | 1.2e-05 | 9.200E-06 | 3.274E-06 | 5.600E-06 | 1.200E-05 |
| 19907 | 996 | W996Q | C | 7.9e-06 | 9.4e-06 | 7.7e-06 | 8.333E-06 | 9.292E-07 | 7.700E-06 | 9.400E-06 |
| 19908 | 996 | W996G | C | 7.5e-06 | 7.4e-06 | 5.2e-06 | 6.700E-06 | 1.300E-06 | 5.200E-06 | 7.500E-06 |
| 19909 | 996 | W996H | C | 1.2e-05 | 1.6e-05 | 1.5e-05 | 1.433E-05 | 2.082E-06 | 1.200E-05 | 1.600E-05 |
| 19910 | 996 | W996I | C | 4.1e-06 | 8.2e-06 | 5e-06 | 5.767E-06 | 2.155E-06 | 4.100E-06 | 8.200E-06 |
| 19911 | 996 | W996L | C | 6.9e-06 | 4.1e-06 | 9.9e-06 | 6.967E-06 | 2.901E-06 | 4.100E-06 | 9.900E-06 |
| 19912 | 996 | W996K | C | 9.4e-06 | 6.7e-06 | 7.6e-06 | 7.900E-06 | 1.375E-06 | 6.700E-06 | 9.400E-06 |
| 19913 | 996 | W996M | C | 1e-05 | 8.1e-06 | 1.1e-05 | 9.700E-06 | 1.473E-06 | 8.100E-06 | 1.100E-05 |
| 19914 | 996 | W996F | C | 9.8e-06 | 1.1e-05 | 9.7e-06 | 1.017E-05 | 7.234E-07 | 9.700E-06 | 1.100E-05 |
| 19915 | 996 | W996P | C | 4.9e-06 | 7.1e-06 | 6.2e-06 | 6.067E-06 | 1.106E-06 | 4.900E-06 | 7.100E-06 |
| 19916 | 996 | W996S | C | 9.3e-06 | 7.8e-06 | 5.9e-06 | 7.667E-06 | 1.704E-06 | 5.900E-06 | 9.300E-06 |
| 19917 | 996 | W996T | C | 9.6e-06 | 6.1e-06 | 6.6e-06 | 7.433E-06 | 1.893E-06 | 6.100E-06 | 9.600E-06 |
| 19918 | 996 | W996W | C | 8.3e-06 | 7e-06 | 6.8e-06 | 7.367E-06 | 8.145E-07 | 6.800E-06 | 8.300E-06 |
| 19919 | 996 | W996Y | C | 7.6e-06 | 1.4e-05 | 7.3e-06 | 9.633E-06 | 3.785E-06 | 7.300E-06 | 1.400E-05 |
| 19920 | 996 | W996V | C | 7.4e-06 | 1.5e-05 | 1.1e-05 | 1.113E-05 | 3.802E-06 | 7.400E-06 | 1.500E-05 |
| 19921 | 997 | N997A | C | 1.4e-05 | 1.3e-05 | 6.5e-06 | 1.117E-05 | 4.072E-06 | 6.500E-06 | 1.400E-05 |
| 19922 | 997 | N997R | C | 6e-06 | 6.8e-06 | 8.6e-06 | 7.133E-06 | 1.332E-06 | 6.000E-06 | 8.600E-06 |
| 19923 | 997 | N997N | C | 5.8e-06 | 9.7e-06 | 5.6e-06 | 7.033E-06 | 2.312E-06 | 5.600E-06 | 9.700E-06 |
| 19924 | 997 | N997D | C | 4e-06 | 8.7e-06 | 9.1e-06 | 7.267E-06 | 2.836E-06 | 4.000E-06 | 9.100E-06 |
| 19925 | 997 | N997C | C | 1.2e-05 | 9.1e-06 | 7.3e-06 | 9.467E-06 | 2.371E-06 | 7.300E-06 | 1.200E-05 |
| 19926 | 997 | N997E | C | 6.3e-06 | 8.8e-06 | 6.1e-06 | 7.067E-06 | 1.504E-06 | 6.100E-06 | 8.800E-06 |
| 19927 | 997 | N997Q | C | 7.7e-06 | 7.1e-06 | 8e-06 | 7.600E-06 | 4.583E-07 | 7.100E-06 | 8.000E-06 |
| 19928 | 997 | N997G | C | 7.9e-06 | 7.9e-06 | 5.7e-06 | 7.167E-06 | 1.270E-06 | 5.700E-06 | 7.900E-06 |
| 19929 | 997 | N997H | C | 7.3e-06 | 9.3e-06 | 8.9e-06 | 8.500E-06 | 1.058E-06 | 7.300E-06 | 9.300E-06 |
| 19930 | 997 | N997I | C | 7.5e-06 | 1.2e-05 | 6.5e-06 | 8.667E-06 | 2.930E-06 | 6.500E-06 | 1.200E-05 |
| 19931 | 997 | N997L | C | 4.5e-06 | 7.7e-06 | 1.3e-05 | 8.400E-06 | 4.293E-06 | 4.500E-06 | 1.300E-05 |
| 19932 | 997 | N997K | C | 5.9e-06 | 6.8e-06 | 6.6e-06 | 6.433E-06 | 4.726E-07 | 5.900E-06 | 6.800E-06 |
| 19933 | 997 | N997M | C | 1.4e-05 | 6.1e-06 | 1.2e-05 | 1.070E-05 | 4.107E-06 | 6.100E-06 | 1.400E-05 |
| 19934 | 997 | N997F | C | 8.5e-06 | 7.2e-06 | 1.3e-05 | 9.567E-06 | 3.044E-06 | 7.200E-06 | 1.300E-05 |
| 19935 | 997 | N997P | C | 1e-05 | 7.6e-06 | 7.3e-06 | 8.300E-06 | 1.480E-06 | 7.300E-06 | 1.000E-05 |
| 19936 | 997 | N997S | C | 6.7e-06 | 5.8e-06 | 1e-05 | 7.500E-06 | 2.211E-06 | 5.800E-06 | 1.000E-05 |
| 19937 | 997 | N997T | C | 5.6e-06 | 5.9e-06 | 8.8e-06 | 6.767E-06 | 1.767E-06 | 5.600E-06 | 8.800E-06 |
| 19938 | 997 | N997W | C | 1e-05 | 1e-05 | 6.6e-06 | 8.867E-06 | 1.963E-06 | 6.600E-06 | 1.000E-05 |
| 19939 | 997 | N997Y | C | 5.6e-06 | 5.1e-06 | 1.3e-05 | 7.900E-06 | 4.424E-06 | 5.100E-06 | 1.300E-05 |
| 19940 | 997 | N997V | C | 5.6e-06 | 9.7e-06 | 1e-05 | 8.433E-06 | 2.458E-06 | 5.600E-06 | 1.000E-05 |
| 19941 | 998 | S998A | C | 6.7e-06 | 6.1e-06 | 7.9e-06 | 6.900E-06 | 9.165E-07 | 6.100E-06 | 7.900E-06 |
| 19942 | 998 | S998R | C | 8.4e-06 | 6.1e-06 | 9.9e-06 | 8.133E-06 | 1.914E-06 | 6.100E-06 | 9.900E-06 |
| 19943 | 998 | S998N | C | 6.2e-06 | 3.5e-06 | 7.4e-06 | 5.700E-06 | 1.997E-06 | 3.500E-06 | 7.400E-06 |
| 19944 | 998 | S998D | C | 1.6e-05 | 6.8e-06 | 9.8e-06 | 1.087E-05 | 4.692E-06 | 6.800E-06 | 1.600E-05 |
| 19945 | 998 | S998C | C | 8.3e-06 | 3.7e-06 | 9.4e-06 | 7.133E-06 | 3.024E-06 | 3.700E-06 | 9.400E-06 |
| 19946 | 998 | S998E | C | 4.7e-06 | 7.6e-06 | 1e-05 | 7.433E-06 | 2.654E-06 | 4.700E-06 | 1.000E-05 |
| 19947 | 998 | S998Q | C | 6.2e-06 | 6.6e-06 | 8.1e-06 | 6.967E-06 | 1.002E-06 | 6.200E-06 | 8.100E-06 |
| 19948 | 998 | S998G | C | 1.1e-05 | 7.5e-06 | 6.4e-06 | 8.300E-06 | 2.402E-06 | 6.400E-06 | 1.100E-05 |
| 19949 | 998 | S998H | C | 1.2e-05 | 6.2e-06 | 9.2e-06 | 9.133E-06 | 2.901E-06 | 6.200E-06 | 1.200E-05 |
| 19950 | 998 | S998I | C | 9.8e-06 | 8.8e-06 | 7.3e-06 | 8.633E-06 | 1.258E-06 | 7.300E-06 | 9.800E-06 |
| 19951 | 998 | S998L | C | 7.6e-06 | 9.9e-06 | 5.5e-06 | 7.667E-06 | 2.201E-06 | 5.500E-06 | 9.900E-06 |
| 19952 | 998 | S998K | C | 4.5e-06 | 6.8e-06 | 4.6e-06 | 5.300E-06 | 1.300E-06 | 4.500E-06 | 6.800E-06 |
| 19953 | 998 | S998M | C | 7e-06 | 8.5e-06 | 8.9e-06 | 8.133E-06 | 1.002E-06 | 7.000E-06 | 8.900E-06 |
| 19954 | 998 | S998F | C | 9.5e-06 | 6.4e-06 | 5.9e-06 | 7.267E-06 | 1.950E-06 | 5.900E-06 | 9.500E-06 |
| 19955 | 998 | S998P | C | 4.3e-06 | 9e-06 | 8.1e-06 | 7.133E-06 | 2.495E-06 | 4.300E-06 | 9.000E-06 |
| 19956 | 998 | S998S | C | 6.3e-06 | 5.1e-06 | 7.3e-06 | 6.233E-06 | 1.102E-06 | 5.100E-06 | 7.300E-06 |
| 19957 | 998 | S998T | C | 8.4e-06 | 9e-06 | 1.5e-05 | 1.080E-05 | 3.650E-06 | 8.400E-06 | 1.500E-05 |
| 19958 | 998 | S998W | C | 5.3e-06 | 8.9e-06 | 7.1e-06 | 7.100E-06 | 1.800E-06 | 5.300E-06 | 8.900E-06 |
| 19959 | 998 | S998Y | C | 7e-06 | 1.1e-05 | 1.2e-05 | 1.000E-05 | 2.646E-06 | 7.000E-06 | 1.200E-05 |
| 19960 | 998 | S998V | C | 1.1e-05 | 6.4e-06 | 1.4e-05 | 1.047E-05 | 3.828E-06 | 6.400E-06 | 1.400E-05 |
| 19961 | 999 | G999A | C | 7.3e-06 | 8.5e-06 | 5.9e-06 | 7.233E-06 | 1.301E-06 | 5.900E-06 | 8.500E-06 |
| 19962 | 999 | G999R | C | 9.8e-06 | 5.9e-06 | 7.9e-06 | 7.867E-06 | 1.950E-06 | 5.900E-06 | 9.800E-06 |
| 19963 | 999 | G999N | C | 8.9e-06 | 8.5e-06 | 8.5e-06 | 8.633E-06 | 2.309E-07 | 8.500E-06 | 8.900E-06 |
| 19964 | 999 | G999D | C | 5.9e-06 | 9.2e-06 | 6.6e-06 | 7.233E-06 | 1.739E-06 | 5.900E-06 | 9.200E-06 |
| 19965 | 999 | G999C | C | 1e-05 | 6.1e-06 | 4.8e-06 | 6.967E-06 | 2.706E-06 | 4.800E-06 | 1.000E-05 |
| 19966 | 999 | G999E | C | 8.4e-06 | 6.9e-06 | 1.8e-05 | 1.110E-05 | 6.022E-06 | 6.900E-06 | 1.800E-05 |
| 19967 | 999 | G999Q | C | 5.9e-06 | 7e-06 | 1.1e-05 | 7.967E-06 | 2.684E-06 | 5.900E-06 | 1.100E-05 |
| 19968 | 999 | G999G | C | 9.6e-06 | 1.2e-05 | 9.5e-06 | 1.037E-05 | 1.415E-06 | 9.500E-06 | 1.200E-05 |
| 19969 | 999 | G999H | C | 5e-06 | 8.4e-06 | 8.1e-06 | 7.167E-06 | 1.882E-06 | 5.000E-06 | 8.400E-06 |
| 19970 | 999 | G999I | C | 5.7e-06 | 8.1e-06 | 9.8e-06 | 7.867E-06 | 2.060E-06 | 5.700E-06 | 9.800E-06 |
| 19971 | 999 | G999L | C | 9.1e-06 | 6.6e-06 | 6e-06 | 7.233E-06 | 1.644E-06 | 6.000E-06 | 9.100E-06 |
| 19972 | 999 | G999K | C | 1e-05 | 4.4e-06 | 6.1e-06 | 6.833E-06 | 2.871E-06 | 4.400E-06 | 1.000E-05 |
| 19973 | 999 | G999M | C | 7.9e-06 | 8.7e-06 | 4.8e-06 | 7.133E-06 | 2.060E-06 | 4.800E-06 | 8.700E-06 |
| 19974 | 999 | G999F | C | 7.4e-06 | 5e-06 | 5.2e-06 | 5.867E-06 | 1.332E-06 | 5.000E-06 | 7.400E-06 |

| | | | | | | | | | | |
|---|---|---|---|---|---|---|---|---|---|---|
| 19975 | 999 | G999P | C | 8e-06 | 6.8e-06 | 9.9e-06 | 8.233E-06 | 1.563E-06 | 6.800E-06 | 9.900E-06 |
| 19976 | 999 | G999S | C | 9.6e-06 | 6.3e-06 | 4.7e-06 | 6.867E-06 | 2.499E-06 | 4.700E-06 | 9.600E-06 |
| 19977 | 999 | G999T | C | 1e-05 | 5.2e-06 | 1.2e-05 | 9.067E-06 | 3.495E-06 | 5.200E-06 | 1.200E-05 |
| 19978 | 999 | G999W | C | 1e-05 | 5.8e-06 | 7e-06 | 7.600E-06 | 2.163E-06 | 5.800E-06 | 1.000E-05 |
| 19979 | 999 | G999Y | C | 8.9e-06 | 1.2e-05 | 7.6e-06 | 9.500E-06 | 2.261E-06 | 7.600E-06 | 1.200E-05 |
| 19980 | 999 | G999V | C | 9.1e-06 | 1e-05 | 1.7e-05 | 1.203E-05 | 4.325E-06 | 9.100E-06 | 1.700E-05 |
| 19981 | 1000 | A1000A | C | 1.1e-05 | 7.7e-06 | 6.6e-06 | 8.433E-06 | 2.290E-06 | 6.600E-06 | 1.100E-05 |
| 19982 | 1000 | A1000R | C | 1e-05 | 8.7e-06 | 1e-05 | 9.567E-06 | 7.506E-07 | 8.700E-06 | 1.000E-05 |
| 19983 | 1000 | A1000N | C | 6.5e-06 | 6e-06 | 1.3e-05 | 8.500E-06 | 3.905E-06 | 6.000E-06 | 1.300E-05 |
| 19984 | 1000 | A1000D | C | 7.2e-06 | 1.2e-05 | 8.9e-06 | 9.367E-06 | 2.434E-06 | 7.200E-06 | 1.200E-05 |
| 19985 | 1000 | A1000C | C | 6.4e-06 | 8.9e-06 | 1e-05 | 8.433E-06 | 1.845E-06 | 6.400E-06 | 1.000E-05 |
| 19986 | 1000 | A1000E | C | 9.9e-06 | 4.8e-06 | 8.3e-06 | 7.667E-06 | 2.608E-06 | 4.800E-06 | 9.900E-06 |
| 19987 | 1000 | A1000Q | C | 1e-05 | 6.4e-06 | 9.6e-06 | 8.667E-06 | 1.973E-06 | 6.400E-06 | 1.000E-05 |
| 19988 | 1000 | A1000G | C | 1.8e-05 | 4.9e-06 | 9.7e-06 | 1.087E-05 | 6.627E-06 | 4.900E-06 | 1.800E-05 |
| 19989 | 1000 | A1000H | C | 1.3e-05 | 1.3e-05 | 9.4e-06 | 1.180E-05 | 2.078E-06 | 9.400E-06 | 1.300E-05 |
| 19990 | 1000 | A1000I | C | 8.6e-06 | 5e-06 | 5.7e-06 | 6.433E-06 | 1.909E-06 | 5.000E-06 | 8.600E-06 |
| 19991 | 1000 | A1000L | C | 1.1e-05 | 1.7e-05 | 7.7e-06 | 1.190E-05 | 4.715E-06 | 7.700E-06 | 1.700E-05 |
| 19992 | 1000 | A1000K | C | 7.4e-06 | 1e-05 | 6.5e-06 | 7.967E-06 | 1.818E-06 | 6.500E-06 | 1.000E-05 |
| 19993 | 1000 | A1000M | C | 4.6e-06 | 9e-06 | 7e-06 | 6.867E-06 | 2.203E-06 | 4.600E-06 | 9.000E-06 |
| 19994 | 1000 | A1000F | C | 1.5e-05 | 8.2e-06 | 1.3e-05 | 1.207E-05 | 3.495E-06 | 8.200E-06 | 1.500E-05 |
| 19995 | 1000 | A1000P | C | 8.5e-06 | 1.4e-05 | 1.3e-05 | 1.183E-05 | 2.930E-06 | 8.500E-06 | 1.400E-05 |
| 19996 | 1000 | A1000S | C | 7.4e-06 | 7.5e-06 | 6.7e-06 | 7.200E-06 | 4.359E-07 | 6.700E-06 | 7.500E-06 |
| 19997 | 1000 | A1000T | C | 1.4e-05 | 9.6e-06 | 6.6e-06 | 1.007E-05 | 3.722E-06 | 6.600E-06 | 1.400E-05 |
| 19998 | 1000 | A1000W | C | 1e-05 | 7.3e-06 | 1e-05 | 9.100E-06 | 1.559E-06 | 7.300E-06 | 1.000E-05 |
| 19999 | 1000 | A1000Y | C | 8.4e-06 | 9.7e-06 | 6.9e-06 | 8.333E-06 | 1.401E-06 | 6.900E-06 | 9.700E-06 |
| 20000 | 1000 | A1000V | C | 7.1e-06 | 5.1e-06 | 4.9e-06 | 5.700E-06 | 1.217E-06 | 4.900E-06 | 7.100E-06 |
| 20001 | 1001 | L1001A | C | 5.2e-06 | 6.1e-06 | 1.4e-05 | 8.433E-06 | 4.842E-06 | 5.200E-06 | 1.400E-05 |
| 20002 | 1001 | L1001R | C | 9.6e-06 | 7.5e-06 | 8.6e-06 | 8.567E-06 | 1.050E-06 | 7.500E-06 | 9.600E-06 |
| 20003 | 1001 | L1001N | C | 6.5e-06 | 6.7e-06 | 8.6e-06 | 7.267E-06 | 1.159E-06 | 6.500E-06 | 8.600E-06 |
| 20004 | 1001 | L1001D | C | 5.8e-06 | 6.2e-06 | 6.9e-06 | 6.300E-06 | 5.568E-07 | 5.800E-06 | 6.900E-06 |
| 20005 | 1001 | L1001C | C | 1.2e-05 | 6.2e-06 | 5.6e-06 | 7.933E-06 | 3.535E-06 | 5.600E-06 | 1.200E-05 |
| 20006 | 1001 | L1001E | C | 4.6e-06 | 7.4e-06 | 1e-05 | 7.333E-06 | 2.701E-06 | 4.600E-06 | 1.000E-05 |
| 20007 | 1001 | L1001Q | C | 8.4e-06 | 9.8e-06 | 6.1e-06 | 8.100E-06 | 1.868E-06 | 6.100E-06 | 9.800E-06 |
| 20008 | 1001 | L1001G | C | 1.4e-05 | 4.8e-06 | 1.3e-05 | 1.060E-05 | 5.048E-06 | 4.800E-06 | 1.400E-05 |
| 20009 | 1001 | L1001H | C | 1e-05 | 6e-06 | 1e-05 | 8.667E-06 | 2.309E-06 | 6.000E-06 | 1.000E-05 |
| 20010 | 1001 | L1001I | C | 1.1e-05 | 1.3e-05 | 5.1e-06 | 9.700E-06 | 4.107E-06 | 5.100E-06 | 1.300E-05 |
| 20011 | 1001 | L1001L | C | 1.2e-05 | 4.2e-06 | 6.6e-06 | 7.600E-06 | 3.995E-06 | 4.200E-06 | 1.200E-05 |
| 20012 | 1001 | L1001K | C | 6.6e-06 | 5.3e-06 | 5.9e-06 | 5.933E-06 | 6.506E-07 | 5.300E-06 | 6.600E-06 |
| 20013 | 1001 | L1001M | C | 1e-05 | 1e-05 | 9.5e-06 | 9.833E-06 | 2.887E-07 | 9.500E-06 | 1.000E-05 |
| 20014 | 1001 | L1001F | C | 8.8e-06 | 7.1e-06 | 5.6e-06 | 7.167E-06 | 1.601E-06 | 5.600E-06 | 8.800E-06 |
| 20015 | 1001 | L1001P | C | 1.5e-05 | 1.4e-05 | 6.9e-06 | 1.197E-05 | 4.416E-06 | 6.900E-06 | 1.500E-05 |
| 20016 | 1001 | L1001S | C | 1.3e-05 | 7.2e-06 | 5.7e-06 | 8.633E-06 | 3.855E-06 | 5.700E-06 | 1.300E-05 |
| 20017 | 1001 | L1001T | C | 4.8e-06 | 8.7e-06 | 8.5e-06 | 7.333E-06 | 2.196E-06 | 4.800E-06 | 8.700E-06 |
| 20018 | 1001 | L1001W | C | 6.7e-06 | 9.6e-06 | 4.9e-06 | 7.067E-06 | 2.371E-06 | 4.900E-06 | 9.600E-06 |
| 20019 | 1001 | L1001Y | C | 1.3e-05 | 8.7e-06 | 6.4e-06 | 9.367E-06 | 3.350E-06 | 6.400E-06 | 1.300E-05 |
| 20020 | 1001 | L1001V | C | 8.4e-06 | 9.8e-06 | 5.8e-06 | 8.000E-06 | 2.030E-06 | 5.800E-06 | 9.800E-06 |
| 20021 | 1002 | T1002A | C | 9.6e-06 | 6e-06 | 9.3e-06 | 8.300E-06 | 1.997E-06 | 6.000E-06 | 9.600E-06 |
| 20022 | 1002 | T1002R | C | 6.5e-06 | 7.4e-06 | 5.5e-06 | 6.467E-06 | 9.504E-07 | 5.500E-06 | 7.400E-06 |
| 20023 | 1002 | T1002N | C | 9.3e-06 | 9.8e-06 | 1.2e-05 | 1.037E-05 | 1.436E-06 | 9.300E-06 | 1.200E-05 |
| 20024 | 1002 | T1002D | C | 6.8e-06 | 1e-05 | 1.4e-05 | 1.027E-05 | 3.607E-06 | 6.800E-06 | 1.400E-05 |
| 20025 | 1002 | T1002C | C | 6e-06 | 9.4e-06 | 1e-05 | 8.467E-06 | 2.157E-06 | 6.000E-06 | 1.000E-05 |
| 20026 | 1002 | T1002E | C | 7.3e-06 | 6.2e-06 | 7.3e-06 | 6.933E-06 | 6.351E-07 | 6.200E-06 | 7.300E-06 |
| 20027 | 1002 | T1002Q | C | 1.4e-05 | 5.4e-06 | 1.1e-05 | 1.013E-05 | 4.365E-06 | 5.400E-06 | 1.400E-05 |
| 20028 | 1002 | T1002G | C | 4.9e-06 | 7.6e-06 | 5.2e-06 | 5.900E-06 | 1.480E-06 | 4.900E-06 | 7.600E-06 |
| 20029 | 1002 | T1002H | C | 6.9e-06 | 6.8e-06 | 1.1e-05 | 8.233E-06 | 2.397E-06 | 6.800E-06 | 1.100E-05 |
| 20030 | 1002 | T1002I | C | 5.7e-06 | 1.1e-05 | 1.3e-05 | 9.900E-06 | 3.772E-06 | 5.700E-06 | 1.300E-05 |
| 20031 | 1002 | T1002L | C | 7.7e-06 | 7.2e-06 | 1.1e-05 | 8.633E-06 | 2.065E-06 | 7.200E-06 | 1.100E-05 |
| 20032 | 1002 | T1002K | C | 5.8e-06 | 7.2e-06 | 6.7e-06 | 6.567E-06 | 7.095E-07 | 5.800E-06 | 7.200E-06 |
| 20033 | 1002 | T1002M | C | 9.6e-06 | 6.7e-06 | 7e-06 | 7.767E-06 | 1.595E-06 | 6.700E-06 | 9.600E-06 |
| 20034 | 1002 | T1002F | C | 8.5e-06 | 6.8e-06 | 8.9e-06 | 8.067E-06 | 1.115E-06 | 6.800E-06 | 8.900E-06 |
| 20035 | 1002 | T1002P | C | 1.1e-05 | 1.1e-05 | 9.3e-06 | 1.043E-05 | 9.815E-07 | 9.300E-06 | 1.100E-05 |
| 20036 | 1002 | T1002S | C | 6e-06 | 1e-05 | 6.2e-06 | 7.400E-06 | 2.254E-06 | 6.000E-06 | 1.000E-05 |
| 20037 | 1002 | T1002T | C | 6.6e-06 | 8.6e-06 | 8.5e-06 | 7.900E-06 | 1.127E-06 | 6.600E-06 | 8.600E-06 |
| 20038 | 1002 | T1002W | C | 7.1e-06 | 5.8e-06 | 6.5e-06 | 6.467E-06 | 6.506E-07 | 5.800E-06 | 7.100E-06 |
| 20039 | 1002 | T1002Y | C | 1.9e-05 | 7.3e-06 | 9e-06 | 1.177E-05 | 6.322E-06 | 7.300E-06 | 1.900E-05 |
| 20040 | 1002 | T1002V | C | 7.6e-06 | 5.8e-06 | 6.6e-06 | 6.667E-06 | 9.018E-07 | 5.800E-06 | 7.600E-06 |
| 20041 | 1003 | S1003A | C | 7.5e-06 | 6.6e-06 | 5.9e-06 | 6.667E-06 | 8.021E-07 | 5.900E-06 | 7.500E-06 |
| 20042 | 1003 | S1003R | C | 1.9e-05 | 6.8e-06 | 9.1e-06 | 1.163E-05 | 6.483E-06 | 6.800E-06 | 1.900E-05 |
| 20043 | 1003 | S1003N | C | 6.5e-06 | 5.4e-06 | 6.8e-06 | 6.233E-06 | 7.371E-07 | 5.400E-06 | 6.800E-06 |
| 20044 | 1003 | S1003D | C | 1e-05 | 7.2e-06 | 5.1e-06 | 7.433E-06 | 2.458E-06 | 5.100E-06 | 1.000E-05 |
| 20045 | 1003 | S1003C | C | 1e-05 | 9.5e-06 | 7.6e-06 | 9.033E-06 | 1.266E-06 | 7.600E-06 | 1.000E-05 |
| 20046 | 1003 | S1003E | C | 1e-05 | 4.8e-06 | 1.1e-05 | 8.600E-06 | 3.329E-06 | 4.800E-06 | 1.100E-05 |
| 20047 | 1003 | S1003Q | C | 8.3e-06 | 1.5e-05 | 6e-06 | 9.767E-06 | 4.676E-06 | 6.000E-06 | 1.500E-05 |
| 20048 | 1003 | S1003G | C | 1.3e-05 | 5.3e-06 | 1.2e-05 | 1.010E-05 | 4.187E-06 | 5.300E-06 | 1.300E-05 |
| 20049 | 1003 | S1003H | C | 7.8e-06 | 6.2e-06 | 5.2e-06 | 6.400E-06 | 1.311E-06 | 5.200E-06 | 7.800E-06 |
| 20050 | 1003 | S1003I | C | 1e-05 | 6.5e-06 | 7.3e-06 | 7.933E-06 | 1.834E-06 | 6.500E-06 | 1.000E-05 |
| 20051 | 1003 | S1003L | C | 9.9e-06 | 8.7e-06 | 9.4e-06 | 9.333E-06 | 6.028E-07 | 8.700E-06 | 9.900E-06 |
| 20052 | 1003 | S1003K | C | 5.5e-06 | 1.4e-05 | 5.8e-06 | 8.433E-06 | 4.823E-06 | 5.500E-06 | 1.400E-05 |
| 20053 | 1003 | S1003M | C | 7.3e-06 | 1.3e-05 | 1.3e-05 | 1.110E-05 | 3.291E-06 | 7.300E-06 | 1.300E-05 |
| 20054 | 1003 | S1003F | C | 9.4e-06 | 7e-06 | 8.3e-06 | 8.233E-06 | 1.201E-06 | 7.000E-06 | 9.400E-06 |
| 20055 | 1003 | S1003P | C | 1.1e-05 | 7e-06 | 9.7e-06 | 9.233E-06 | 2.040E-06 | 7.000E-06 | 1.100E-05 |
| 20056 | 1003 | S1003S | C | 6.3e-06 | 8.2e-06 | 8.7e-06 | 7.733E-06 | 1.266E-06 | 6.300E-06 | 8.700E-06 |
| 20057 | 1003 | S1003T | C | 5.5e-06 | 8.3e-06 | 3.7e-06 | 5.833E-06 | 2.318E-06 | 3.700E-06 | 8.300E-06 |
| 20058 | 1003 | S1003W | C | 9.2e-06 | 9.9e-06 | 1.2e-05 | 1.037E-05 | 1.457E-06 | 9.200E-06 | 1.200E-05 |
| 20059 | 1003 | S1003Y | C | 6.9e-06 | 7e-06 | 6.9e-06 | 6.933E-06 | 5.774E-08 | 6.900E-06 | 7.000E-06 |

| | | | | | | | | | | |
|---|---|---|---|---|---|---|---|---|---|---|
| 20060 | 1003 | S1003V | C | 8.9e-06 | 1e-05 | 1.2e-05 | 1.030E-05 | 1.572E-06 | 8.900E-06 | 1.200E-05 |
| 20061 | 1004 | G1004A | C | 1.3e-05 | 5.7e-06 | 4.8e-06 | 7.833E-06 | 4.497E-06 | 4.800E-06 | 1.300E-05 |
| 20062 | 1004 | G1004R | C | 5.6e-06 | 8.2e-06 | 4.9e-06 | 6.233E-06 | 1.739E-06 | 4.900E-06 | 8.200E-06 |
| 20063 | 1004 | G1004N | C | 5.1e-06 | 8.7e-06 | 4.6e-06 | 6.133E-06 | 2.237E-06 | 4.600E-06 | 8.700E-06 |
| 20064 | 1004 | G1004D | C | 9.3e-06 | 4.5e-06 | 7.6e-06 | 7.133E-06 | 2.434E-06 | 4.500E-06 | 9.300E-06 |
| 20065 | 1004 | G1004C | C | 9.4e-06 | 1e-05 | 7.8e-06 | 9.067E-06 | 1.137E-06 | 7.800E-06 | 1.000E-05 |
| 20066 | 1004 | G1004E | C | 1.1e-05 | 6.3e-06 | 9.5e-06 | 8.933E-06 | 2.401E-06 | 6.300E-06 | 1.100E-05 |
| 20067 | 1004 | G1004Q | C | 6.3e-06 | 6.3e-06 | 6.4e-06 | 6.333E-06 | 5.774E-08 | 6.300E-06 | 6.400E-06 |
| 20068 | 1004 | G1004G | C | 8.9e-06 | 4.6e-06 | 5.3e-06 | 6.267E-06 | 2.307E-06 | 4.600E-06 | 8.900E-06 |
| 20069 | 1004 | G1004H | C | 7.7e-06 | 6.4e-06 | 6.3e-06 | 6.800E-06 | 7.810E-07 | 6.300E-06 | 7.700E-06 |
| 20070 | 1004 | G1004I | C | 7.7e-06 | 9.9e-06 | 1.6e-05 | 1.120E-05 | 4.300E-06 | 7.700E-06 | 1.600E-05 |
| 20071 | 1004 | G1004L | C | 6.7e-06 | 7.7e-06 | 5.7e-06 | 6.700E-06 | 1.000E-06 | 5.700E-06 | 7.700E-06 |
| 20072 | 1004 | G1004K | C | 1.3e-05 | 4.8e-06 | 1.1e-05 | 9.600E-06 | 4.276E-06 | 4.800E-06 | 1.300E-05 |
| 20073 | 1004 | G1004M | C | 4.9e-06 | 1e-05 | 8.3e-06 | 7.733E-06 | 2.597E-06 | 4.900E-06 | 1.000E-05 |
| 20074 | 1004 | G1004F | C | 5e-06 | 8.1e-06 | 6.6e-06 | 6.567E-06 | 1.550E-06 | 5.000E-06 | 8.100E-06 |
| 20075 | 1004 | G1004P | C | 1e-05 | 7.2e-06 | 1.4e-05 | 1.040E-05 | 3.418E-06 | 7.200E-06 | 1.400E-05 |
| 20076 | 1004 | G1004S | C | 4.6e-06 | 5.2e-06 | 9.4e-06 | 6.400E-06 | 2.615E-06 | 4.600E-06 | 9.400E-06 |
| 20077 | 1004 | G1004T | C | 7.7e-06 | 6.9e-06 | 5.5e-06 | 6.700E-06 | 1.114E-06 | 5.500E-06 | 7.700E-06 |
| 20078 | 1004 | G1004W | C | 6.8e-06 | 9.7e-06 | 6.5e-06 | 7.667E-06 | 1.767E-06 | 6.500E-06 | 9.700E-06 |
| 20079 | 1004 | G1004Y | C | 1.3e-05 | 1.3e-05 | 6.1e-06 | 1.070E-05 | 3.984E-06 | 6.100E-06 | 1.300E-05 |
| 20080 | 1004 | G1004V | C | 4.2e-06 | 8.3e-06 | 8.8e-06 | 7.100E-06 | 2.524E-06 | 4.200E-06 | 8.800E-06 |
| 20081 | 1005 | V1005A | C | 7.6e-06 | 6.7e-06 | 1e-05 | 8.100E-06 | 1.706E-06 | 6.700E-06 | 1.000E-05 |
| 20082 | 1005 | V1005R | C | 5.6e-06 | 7.6e-06 | 1.3e-05 | 8.733E-06 | 3.828E-06 | 5.600E-06 | 1.300E-05 |
| 20083 | 1005 | V1005N | C | 8.3e-06 | 5e-06 | 9.8e-06 | 7.700E-06 | 2.456E-06 | 5.000E-06 | 9.800E-06 |
| 20084 | 1005 | V1005D | C | 1.1e-05 | 6.9e-06 | 6.8e-06 | 8.233E-06 | 2.397E-06 | 6.800E-06 | 1.100E-05 |
| 20085 | 1005 | V1005C | C | 7.2e-06 | 6.6e-06 | 7.2e-06 | 7.000E-06 | 3.464E-07 | 6.600E-06 | 7.200E-06 |
| 20086 | 1005 | V1005E | C | 1.2e-05 | 7.4e-06 | 9.4e-06 | 9.600E-06 | 2.307E-06 | 7.400E-06 | 1.200E-05 |
| 20087 | 1005 | V1005Q | C | 1.1e-05 | 6.8e-06 | 5.2e-06 | 7.667E-06 | 2.996E-06 | 5.200E-06 | 1.100E-05 |
| 20088 | 1005 | V1005G | C | 4.8e-06 | 6.9e-06 | 7.9e-06 | 6.533E-06 | 1.582E-06 | 4.800E-06 | 7.900E-06 |
| 20089 | 1005 | V1005H | C | 7.1e-06 | 5e-06 | 6.7e-06 | 6.267E-06 | 1.115E-06 | 5.000E-06 | 7.100E-06 |
| 20090 | 1005 | V1005I | C | 1e-05 | 8.7e-06 | 7.8e-06 | 8.833E-06 | 1.106E-06 | 7.800E-06 | 1.000E-05 |
| 20091 | 1005 | V1005L | C | 8.2e-06 | 3.7e-06 | 9.4e-06 | 7.100E-06 | 3.005E-06 | 3.700E-06 | 9.400E-06 |
| 20092 | 1005 | V1005K | C | 7.4e-06 | 1.1e-05 | 1.9e-05 | 1.247E-05 | 5.937E-06 | 7.400E-06 | 1.900E-05 |
| 20093 | 1005 | V1005M | C | 8.3e-06 | 9.1e-06 | 8.7e-06 | 8.700E-06 | 4.000E-07 | 8.300E-06 | 9.100E-06 |
| 20094 | 1005 | V1005F | C | 6e-06 | 8.6e-06 | 9.1e-06 | 7.900E-06 | 1.664E-06 | 6.000E-06 | 9.100E-06 |
| 20095 | 1005 | V1005P | C | 8.5e-06 | 7.1e-06 | 5.9e-06 | 7.167E-06 | 1.301E-06 | 5.900E-06 | 8.500E-06 |
| 20096 | 1005 | V1005S | C | 9.5e-06 | 1.1e-05 | 8.8e-06 | 9.767E-06 | 1.124E-06 | 8.800E-06 | 1.100E-05 |
| 20097 | 1005 | V1005T | C | 9.1e-06 | 5.9e-06 | 5.9e-06 | 6.967E-06 | 1.848E-06 | 5.900E-06 | 9.100E-06 |
| 20098 | 1005 | V1005W | C | 9.4e-06 | 5.9e-06 | 5.6e-06 | 6.967E-06 | 2.113E-06 | 5.600E-06 | 9.400E-06 |
| 20099 | 1005 | V1005Y | C | 8.3e-06 | 1e-05 | 7.4e-06 | 8.567E-06 | 1.320E-06 | 7.400E-06 | 1.000E-05 |
| 20100 | 1005 | V1005V | C | 1.1e-05 | 1.3e-05 | 6e-06 | 1.000E-05 | 3.606E-06 | 6.000E-06 | 1.300E-05 |
| 20101 | 1006 | H1006A | C | 1e-05 | 1.5e-05 | 9.6e-06 | 1.153E-05 | 3.009E-06 | 9.600E-06 | 1.500E-05 |
| 20102 | 1006 | H1006R | C | 5.3e-06 | 8.4e-06 | 1.2e-05 | 8.567E-06 | 3.353E-06 | 5.300E-06 | 1.200E-05 |
| 20103 | 1006 | H1006N | C | 4.2e-06 | 7.8e-06 | 6.3e-06 | 6.100E-06 | 1.808E-06 | 4.200E-06 | 7.800E-06 |
| 20104 | 1006 | H1006D | C | 9.2e-06 | 5e-06 | 6.8e-06 | 7.000E-06 | 2.107E-06 | 5.000E-06 | 9.200E-06 |
| 20105 | 1006 | H1006C | C | 7e-06 | 1.2e-05 | 5e-06 | 8.000E-06 | 3.606E-06 | 5.000E-06 | 1.200E-05 |
| 20106 | 1006 | H1006E | C | 1.3e-05 | 7.3e-06 | 5.9e-06 | 8.733E-06 | 3.761E-06 | 5.900E-06 | 1.300E-05 |
| 20107 | 1006 | H1006Q | C | 9.2e-06 | 4.8e-06 | 1.1e-05 | 8.333E-06 | 3.190E-06 | 4.800E-06 | 1.100E-05 |
| 20108 | 1006 | H1006G | C | 8e-06 | 1.1e-05 | 8e-06 | 9.000E-06 | 1.732E-06 | 8.000E-06 | 1.100E-05 |
| 20109 | 1006 | H1006H | C | 1.1e-05 | 3.6e-06 | 7e-06 | 7.200E-06 | 3.704E-06 | 3.600E-06 | 1.100E-05 |
| 20110 | 1006 | H1006I | C | 1e-05 | 6.3e-06 | 1.2e-05 | 9.433E-06 | 2.892E-06 | 6.300E-06 | 1.200E-05 |
| 20111 | 1006 | H1006L | C | 1e-05 | 5.2e-06 | 6e-06 | 7.067E-06 | 2.572E-06 | 5.200E-06 | 1.000E-05 |
| 20112 | 1006 | H1006K | C | 7e-06 | 7.1e-06 | 9.5e-06 | 7.867E-06 | 1.415E-06 | 7.000E-06 | 9.500E-06 |
| 20113 | 1006 | H1006M | C | 9.7e-06 | 8.7e-06 | 4.8e-06 | 7.733E-06 | 2.589E-06 | 4.800E-06 | 9.700E-06 |
| 20114 | 1006 | H1006F | C | 8e-06 | 9.2e-06 | 9.5e-06 | 8.900E-06 | 7.937E-07 | 8.000E-06 | 9.500E-06 |
| 20115 | 1006 | H1006P | C | 5.8e-06 | 8.1e-06 | 1e-05 | 7.967E-06 | 2.103E-06 | 5.800E-06 | 1.000E-05 |
| 20116 | 1006 | H1006S | C | 1e-05 | 9.2e-06 | 1.2e-05 | 1.040E-05 | 1.442E-06 | 9.200E-06 | 1.200E-05 |
| 20117 | 1006 | H1006T | C | 9e-06 | 7.4e-06 | 6.3e-06 | 7.567E-06 | 1.358E-06 | 6.300E-06 | 9.000E-06 |
| 20118 | 1006 | H1006W | C | 7.2e-06 | 9.1e-06 | 8.4e-06 | 8.233E-06 | 9.609E-07 | 7.200E-06 | 9.100E-06 |
| 20119 | 1006 | H1006Y | C | 8e-06 | 4.7e-06 | 7e-06 | 6.567E-06 | 1.692E-06 | 4.700E-06 | 8.000E-06 |
| 20120 | 1006 | H1006V | C | 6e-06 | 8.5e-06 | 8.2e-06 | 7.567E-06 | 1.365E-06 | 6.000E-06 | 8.500E-06 |
| 20121 | 1007 | T1007A | C | 5.7e-06 | 4.3e-06 | 5e-06 | 5.000E-06 | 7.000E-07 | 4.300E-06 | 5.700E-06 |
| 20122 | 1007 | T1007R | C | 6.8e-06 | 6.6e-06 | 9.4e-06 | 7.600E-06 | 1.562E-06 | 6.600E-06 | 9.400E-06 |
| 20123 | 1007 | T1007N | C | 7.5e-06 | 7.2e-06 | 9.1e-06 | 7.933E-06 | 1.021E-06 | 7.200E-06 | 9.100E-06 |
| 20124 | 1007 | T1007D | C | 1.2e-05 | 8.3e-06 | 1e-05 | 1.010E-05 | 1.852E-06 | 8.300E-06 | 1.200E-05 |
| 20125 | 1007 | T1007C | C | 5.6e-06 | 8.9e-06 | 9.4e-06 | 7.967E-06 | 2.065E-06 | 5.600E-06 | 9.400E-06 |
| 20126 | 1007 | T1007E | C | 9.6e-06 | 1.5e-05 | 8.9e-06 | 1.117E-05 | 3.338E-06 | 8.900E-06 | 1.500E-05 |
| 20127 | 1007 | T1007Q | C | 1.2e-05 | 5.9e-06 | 5.5e-06 | 7.800E-06 | 3.643E-06 | 5.500E-06 | 1.200E-05 |
| 20128 | 1007 | T1007G | C | 8e-06 | 3.8e-06 | 1.5e-05 | 8.933E-06 | 5.658E-06 | 3.800E-06 | 1.500E-05 |
| 20129 | 1007 | T1007H | C | 6e-06 | 7.3e-06 | 1.2e-05 | 8.433E-06 | 3.156E-06 | 6.000E-06 | 1.200E-05 |
| 20130 | 1007 | T1007I | C | 8.4e-06 | 9.6e-06 | 6.4e-06 | 8.133E-06 | 1.617E-06 | 6.400E-06 | 9.600E-06 |
| 20131 | 1007 | T1007L | C | 1e-05 | 4.6e-06 | 7.8e-06 | 7.467E-06 | 2.715E-06 | 4.600E-06 | 1.000E-05 |
| 20132 | 1007 | T1007K | C | 1.1e-05 | 7.8e-06 | 1.5e-05 | 1.127E-05 | 3.607E-06 | 7.800E-06 | 1.500E-05 |
| 20133 | 1007 | T1007M | C | 1.3e-05 | 1.2e-05 | 1e-05 | 1.167E-05 | 1.528E-06 | 1.000E-05 | 1.300E-05 |
| 20134 | 1007 | T1007F | C | 6.7e-06 | 6.8e-06 | 5.7e-06 | 6.400E-06 | 6.083E-07 | 5.700E-06 | 6.800E-06 |
| 20135 | 1007 | T1007P | C | 9.6e-06 | 8.9e-06 | 1.3e-05 | 1.050E-05 | 2.193E-06 | 8.900E-06 | 1.300E-05 |
| 20136 | 1007 | T1007S | C | 6.6e-06 | 5.7e-06 | 6.6e-06 | 6.300E-06 | 5.196E-07 | 5.700E-06 | 6.600E-06 |
| 20137 | 1007 | T1007T | C | 9.6e-06 | 7.8e-06 | 5.1e-06 | 7.500E-06 | 2.265E-06 | 5.100E-06 | 9.600E-06 |
| 20138 | 1007 | T1007W | C | 1e-05 | 7.1e-06 | 9.5e-06 | 8.867E-06 | 1.550E-06 | 7.100E-06 | 1.000E-05 |
| 20139 | 1007 | T1007Y | C | 8.8e-06 | 8.9e-06 | 1.3e-05 | 1.023E-05 | 2.397E-06 | 8.800E-06 | 1.300E-05 |
| 20140 | 1007 | T1007V | C | 6.2e-06 | 1.3e-05 | 9.7e-06 | 9.633E-06 | 3.400E-06 | 6.200E-06 | 1.300E-05 |
| 20141 | 1008 | F1008A | C | 8e-06 | 8.7e-06 | 5.4e-06 | 7.367E-06 | 1.739E-06 | 5.400E-06 | 8.700E-06 |
| 20142 | 1008 | F1008R | C | 1.3e-05 | 6.8e-06 | 1.1e-05 | 1.027E-05 | 3.164E-06 | 6.800E-06 | 1.300E-05 |
| 20143 | 1008 | F1008N | C | 7.1e-06 | 9.6e-06 | 4.9e-06 | 7.200E-06 | 2.352E-06 | 4.900E-06 | 9.600E-06 |
| 20144 | 1008 | F1008D | C | 9.4e-06 | 4.9e-06 | 8.1e-06 | 7.467E-06 | 2.316E-06 | 4.900E-06 | 9.400E-06 |

| | | | | | | | | | | |
|---|---|---|---|---|---|---|---|---|---|---|
| 20145 | 1008 | F1008C | C | 8.1e-06 | 1.4e-05 | 9e-06 | 1.037E-05 | 3.179E-06 | 8.100E-06 | 1.400E-05 |
| 20146 | 1008 | F1008E | C | 9.3e-06 | 6.7e-06 | 6.5e-06 | 7.500E-06 | 1.562E-06 | 6.500E-06 | 9.300E-06 |
| 20147 | 1008 | F1008Q | C | 5.4e-06 | 7.1e-06 | 9.8e-06 | 7.433E-06 | 2.219E-06 | 5.400E-06 | 9.800E-06 |
| 20148 | 1008 | F1008G | C | 6.6e-06 | 9.9e-06 | 6.5e-06 | 7.667E-06 | 1.935E-06 | 6.500E-06 | 9.900E-06 |
| 20149 | 1008 | F1008H | C | 4.6e-06 | 4.8e-06 | 6.8e-06 | 5.400E-06 | 1.217E-06 | 4.600E-06 | 6.800E-06 |
| 20150 | 1008 | F1008I | C | 9.8e-06 | 5.4e-06 | 7e-06 | 7.400E-06 | 2.227E-06 | 5.400E-06 | 9.800E-06 |
| 20151 | 1008 | F1008L | C | 8.8e-06 | 8.8e-06 | 6.2e-06 | 7.933E-06 | 1.501E-06 | 6.200E-06 | 8.800E-06 |
| 20152 | 1008 | F1008K | C | 5.8e-06 | 8e-06 | 9.5e-06 | 7.767E-06 | 1.861E-06 | 5.800E-06 | 9.500E-06 |
| 20153 | 1008 | F1008M | C | 1e-05 | 1e-05 | 6.6e-06 | 8.867E-06 | 1.963E-06 | 6.600E-06 | 1.000E-05 |
| 20154 | 1008 | F1008F | C | 4.7e-06 | 6.7e-06 | 5.1e-06 | 5.500E-06 | 1.058E-06 | 4.700E-06 | 6.700E-06 |
| 20155 | 1008 | F1008P | C | 7.5e-06 | 7.9e-06 | 7.2e-06 | 7.533E-06 | 3.512E-07 | 7.200E-06 | 7.900E-06 |
| 20156 | 1008 | F1008S | C | 1.1e-05 | 8e-06 | 8.8e-06 | 9.267E-06 | 1.553E-06 | 8.000E-06 | 1.100E-05 |
| 20157 | 1008 | F1008T | C | 7.4e-06 | 1.4e-05 | 8.6e-06 | 1.000E-05 | 3.516E-06 | 7.400E-06 | 1.400E-05 |
| 20158 | 1008 | F1008W | C | 1.1e-05 | 6.1e-06 | 1.2e-05 | 9.700E-06 | 3.158E-06 | 6.100E-06 | 1.200E-05 |
| 20159 | 1008 | F1008Y | C | 4.3e-06 | 9.5e-06 | 7.2e-06 | 7.000E-06 | 2.606E-06 | 4.300E-06 | 9.500E-06 |
| 20160 | 1008 | F1008V | C | 1.1e-05 | 7.8e-06 | 6.4e-06 | 8.400E-06 | 2.358E-06 | 6.400E-06 | 1.100E-05 |
| 20161 | 1009 | P1009A | C | 1e-05 | 7.7e-06 | 1.3e-05 | 1.023E-05 | 2.658E-06 | 7.700E-06 | 1.300E-05 |
| 20162 | 1009 | P1009R | C | 6.8e-06 | 1.3e-05 | 1.3e-05 | 1.093E-05 | 3.580E-06 | 6.800E-06 | 1.300E-05 |
| 20163 | 1009 | P1009N | C | 6.7e-06 | 8.1e-06 | 1.4e-05 | 9.600E-06 | 3.874E-06 | 6.700E-06 | 1.400E-05 |
| 20164 | 1009 | P1009D | C | 9.5e-06 | 7.7e-06 | 8.8e-06 | 8.667E-06 | 9.074E-07 | 7.700E-06 | 9.500E-06 |
| 20165 | 1009 | P1009C | C | 4.7e-06 | 8.1e-06 | 8.5e-06 | 7.100E-06 | 2.088E-06 | 4.700E-06 | 8.500E-06 |
| 20166 | 1009 | P1009E | C | 5.9e-06 | 9.4e-06 | 9.1e-06 | 8.133E-06 | 1.940E-06 | 5.900E-06 | 9.400E-06 |
| 20167 | 1009 | P1009Q | C | 6.2e-06 | 7.4e-06 | 9.9e-06 | 7.833E-06 | 1.888E-06 | 6.200E-06 | 9.900E-06 |
| 20168 | 1009 | P1009G | C | 6.6e-06 | 8e-06 | 6.9e-06 | 7.167E-06 | 7.371E-07 | 6.600E-06 | 8.000E-06 |
| 20169 | 1009 | P1009H | C | 9.2e-06 | 5.3e-06 | 7.2e-06 | 7.233E-06 | 1.950E-06 | 5.300E-06 | 9.200E-06 |
| 20170 | 1009 | P1009I | C | 4.1e-06 | 9.2e-06 | 5.8e-06 | 6.367E-06 | 2.597E-06 | 4.100E-06 | 9.200E-06 |
| 20171 | 1009 | P1009L | C | 8.1e-06 | 5.4e-06 | 1e-05 | 7.833E-06 | 2.312E-06 | 5.400E-06 | 1.000E-05 |
| 20172 | 1009 | P1009K | C | 6.8e-06 | 6.9e-06 | 6e-06 | 6.567E-06 | 4.933E-07 | 6.000E-06 | 6.900E-06 |
| 20173 | 1009 | P1009M | C | 5.8e-06 | 8.6e-06 | 5.8e-06 | 6.733E-06 | 1.617E-06 | 5.800E-06 | 8.600E-06 |
| 20174 | 1009 | P1009F | C | 1.3e-05 | 9.9e-06 | 1.3e-05 | 1.197E-05 | 1.790E-06 | 9.900E-06 | 1.300E-05 |
| 20175 | 1009 | P1009P | C | 7.5e-06 | 7.2e-06 | 1.6e-05 | 1.023E-05 | 4.996E-06 | 7.200E-06 | 1.600E-05 |
| 20176 | 1009 | P1009S | C | 8.3e-06 | 9.3e-06 | 1.2e-05 | 9.867E-06 | 1.914E-06 | 8.300E-06 | 1.200E-05 |
| 20177 | 1009 | P1009T | C | 1.2e-05 | 7e-06 | 9e-06 | 9.333E-06 | 2.517E-06 | 7.000E-06 | 1.200E-05 |
| 20178 | 1009 | P1009W | C | 8.9e-06 | 9.4e-06 | 7.7e-06 | 8.667E-06 | 8.737E-07 | 7.700E-06 | 9.400E-06 |
| 20179 | 1009 | P1009Y | C | 7.7e-06 | 1.4e-05 | 9.3e-06 | 1.033E-05 | 3.275E-06 | 7.700E-06 | 1.400E-05 |
| 20180 | 1009 | P1009V | C | 8.7e-06 | 1.2e-05 | 6.7e-06 | 9.133E-06 | 2.676E-06 | 6.700E-06 | 1.200E-05 |
| 20181 | 1010 | A1010A | C | 1e-05 | 8.1e-06 | 7.3e-06 | 8.467E-06 | 1.387E-06 | 7.300E-06 | 1.000E-05 |
| 20182 | 1010 | A1010R | C | 8.8e-06 | 1.3e-05 | 9.2e-06 | 1.033E-05 | 2.318E-06 | 8.800E-06 | 1.300E-05 |
| 20183 | 1010 | A1010N | C | 5.1e-06 | 7.1e-06 | 1.1e-05 | 7.733E-06 | 3.001E-06 | 5.100E-06 | 1.100E-05 |
| 20184 | 1010 | A1010D | C | 1.3e-05 | 1e-05 | 9e-06 | 1.067E-05 | 2.082E-06 | 9.000E-06 | 1.300E-05 |
| 20185 | 1010 | A1010C | C | 1.4e-05 | 9.1e-06 | 6.8e-06 | 9.967E-06 | 3.677E-06 | 6.800E-06 | 1.400E-05 |
| 20186 | 1010 | A1010E | C | 1.4e-05 | 9e-06 | 5.8e-06 | 9.600E-06 | 4.133E-06 | 5.800E-06 | 1.400E-05 |
| 20187 | 1010 | A1010Q | C | 1.3e-05 | 6.1e-06 | 8.6e-06 | 9.233E-06 | 3.493E-06 | 6.100E-06 | 1.300E-05 |
| 20188 | 1010 | A1010G | C | 6.8e-06 | 5.9e-06 | 8.7e-06 | 7.133E-06 | 1.429E-06 | 5.900E-06 | 8.700E-06 |
| 20189 | 1010 | A1010H | C | 8e-06 | 7e-06 | 1.2e-05 | 9.000E-06 | 2.646E-06 | 7.000E-06 | 1.200E-05 |
| 20190 | 1010 | A1010I | C | 9.1e-06 | 5.6e-06 | 7e-06 | 7.233E-06 | 1.762E-06 | 5.600E-06 | 9.100E-06 |
| 20191 | 1010 | A1010L | C | 1.3e-05 | 6.2e-06 | 1e-05 | 9.733E-06 | 3.408E-06 | 6.200E-06 | 1.300E-05 |
| 20192 | 1010 | A1010K | C | 6.5e-06 | 7.8e-06 | 8.9e-06 | 7.733E-06 | 1.201E-06 | 6.500E-06 | 8.900E-06 |
| 20193 | 1010 | A1010M | C | 6.8e-06 | 8e-06 | 1.5e-05 | 9.933E-06 | 4.429E-06 | 6.800E-06 | 1.500E-05 |
| 20194 | 1010 | A1010F | C | 9.7e-06 | 5e-06 | 6.2e-06 | 6.967E-06 | 2.442E-06 | 5.000E-06 | 9.700E-06 |
| 20195 | 1010 | A1010P | C | 1e-05 | 1.4e-05 | 6.6e-06 | 1.020E-05 | 3.704E-06 | 6.600E-06 | 1.400E-05 |
| 20196 | 1010 | A1010S | C | 5.2e-06 | 8.1e-06 | 1.5e-05 | 9.433E-06 | 5.034E-06 | 5.200E-06 | 1.500E-05 |
| 20197 | 1010 | A1010T | C | 6.9e-06 | 7.3e-06 | 1.2e-05 | 8.733E-06 | 2.836E-06 | 6.900E-06 | 1.200E-05 |
| 20198 | 1010 | A1010W | C | 5.6e-06 | 6.5e-06 | 8e-06 | 6.700E-06 | 1.212E-06 | 5.600E-06 | 8.000E-06 |
| 20199 | 1010 | A1010Y | C | 4.7e-06 | 4.3e-06 | 6.5e-06 | 5.167E-06 | 1.172E-06 | 4.300E-06 | 6.500E-06 |
| 20200 | 1010 | A1010V | C | 8.8e-06 | 1e-05 | 6.3e-06 | 8.367E-06 | 1.888E-06 | 6.300E-06 | 1.000E-05 |
| 20201 | 1011 | V1011A | C | 6.5e-06 | 5.6e-06 | 8.9e-06 | 7.000E-06 | 1.706E-06 | 5.600E-06 | 8.900E-06 |
| 20202 | 1011 | V1011R | C | 6.2e-06 | 9.8e-06 | 1.4e-05 | 1.000E-05 | 3.904E-06 | 6.200E-06 | 1.400E-05 |
| 20203 | 1011 | V1011N | C | 6.2e-06 | 1.1e-05 | 8.4e-06 | 8.533E-06 | 2.403E-06 | 6.200E-06 | 1.100E-05 |
| 20204 | 1011 | V1011D | C | 5e-06 | 5.1e-06 | 9.7e-06 | 6.600E-06 | 2.685E-06 | 5.000E-06 | 9.700E-06 |
| 20205 | 1011 | V1011C | C | 1.2e-05 | 1.2e-05 | 5.2e-06 | 9.733E-06 | 3.926E-06 | 5.200E-06 | 1.200E-05 |
| 20206 | 1011 | V1011E | C | 8e-06 | 5.6e-06 | 1.3e-05 | 8.867E-06 | 3.775E-06 | 5.600E-06 | 1.300E-05 |
| 20207 | 1011 | V1011Q | C | 7.7e-06 | 1.2e-05 | 1.3e-05 | 1.090E-05 | 2.816E-06 | 7.700E-06 | 1.300E-05 |
| 20208 | 1011 | V1011G | C | 7.1e-06 | 1.2e-05 | 8.5e-06 | 9.200E-06 | 2.524E-06 | 7.100E-06 | 1.200E-05 |
| 20209 | 1011 | V1011H | C | 1.3e-05 | 7.6e-06 | 3.5e-06 | 8.033E-06 | 4.765E-06 | 3.500E-06 | 1.300E-05 |
| 20210 | 1011 | V1011I | C | 8.6e-06 | 1.1e-05 | 4.6e-06 | 8.067E-06 | 3.233E-06 | 4.600E-06 | 1.100E-05 |
| 20211 | 1011 | V1011L | C | 4.1e-06 | 1.3e-05 | 1.2e-05 | 9.700E-06 | 4.875E-06 | 4.100E-06 | 1.300E-05 |
| 20212 | 1011 | V1011K | C | 8.2e-06 | 7.2e-06 | 4.5e-06 | 6.633E-06 | 1.914E-06 | 4.500E-06 | 8.200E-06 |
| 20213 | 1011 | V1011M | C | 4.8e-06 | 8.6e-06 | 5.3e-06 | 6.233E-06 | 2.065E-06 | 4.800E-06 | 8.600E-06 |
| 20214 | 1011 | V1011F | C | 7e-06 | 1.4e-05 | 1.1e-05 | 1.067E-05 | 3.512E-06 | 7.000E-06 | 1.400E-05 |
| 20215 | 1011 | V1011P | C | 7e-06 | 6.6e-06 | 8.6e-06 | 7.400E-06 | 1.058E-06 | 6.600E-06 | 8.600E-06 |
| 20216 | 1011 | V1011S | C | 4.4e-06 | 8.3e-06 | 7.4e-06 | 6.700E-06 | 2.042E-06 | 4.400E-06 | 8.300E-06 |
| 20217 | 1011 | V1011T | C | 4.3e-06 | 6.2e-06 | 6.3e-06 | 5.600E-06 | 1.127E-06 | 4.300E-06 | 6.300E-06 |
| 20218 | 1011 | V1011W | C | 8.6e-06 | 7.6e-06 | 4e-06 | 6.733E-06 | 2.419E-06 | 4.000E-06 | 8.600E-06 |
| 20219 | 1011 | V1011Y | C | 4.1e-06 | 9.8e-06 | 1.4e-05 | 9.300E-06 | 4.969E-06 | 4.100E-06 | 1.400E-05 |
| 20220 | 1011 | V1011V | C | 6.7e-06 | 4.9e-06 | 8e-06 | 6.533E-06 | 1.557E-06 | 4.900E-06 | 8.000E-06 |
| 20221 | 1012 | L1012A | C | 9.5e-06 | 1e-05 | 8.8e-06 | 9.433E-06 | 6.028E-07 | 8.800E-06 | 1.000E-05 |
| 20222 | 1012 | L1012R | C | 7.4e-06 | 1.1e-05 | 5.9e-06 | 8.100E-06 | 2.621E-06 | 5.900E-06 | 1.100E-05 |
| 20223 | 1012 | L1012N | C | 7e-06 | 7.9e-06 | 1.2e-05 | 8.967E-06 | 2.665E-06 | 7.000E-06 | 1.200E-05 |
| 20224 | 1012 | L1012D | C | 1e-05 | 8.7e-06 | 9.1e-06 | 9.267E-06 | 6.658E-07 | 8.700E-06 | 1.000E-05 |
| 20225 | 1012 | L1012C | C | 1.1e-05 | 8.1e-06 | 4.8e-06 | 7.967E-06 | 3.102E-06 | 4.800E-06 | 1.100E-05 |
| 20226 | 1012 | L1012E | C | 9.5e-06 | 9.7e-06 | 8.3e-06 | 9.167E-06 | 7.572E-07 | 8.300E-06 | 9.700E-06 |
| 20227 | 1012 | L1012Q | C | 3.3e-06 | 1.3e-05 | 6.9e-06 | 7.733E-06 | 4.903E-06 | 3.300E-06 | 1.300E-05 |
| 20228 | 1012 | L1012G | C | 6.5e-06 | 1e-05 | 9.6e-06 | 8.700E-06 | 1.916E-06 | 6.500E-06 | 1.000E-05 |
| 20229 | 1012 | L1012H | C | 9.5e-06 | 9.3e-06 | 1e-05 | 9.600E-06 | 3.606E-07 | 9.300E-06 | 1.000E-05 |

| | | | | | | | | | | |
|---|---|---|---|---|---|---|---|---|---|---|
| 20230 | 1012 | L1012I | C | 7.7e-06 | 8.1e-06 | 4.5e-06 | 6.767E-06 | 1.973E-06 | 4.500E-06 | 8.100E-06 |
| 20231 | 1012 | L1012L | C | 1.5e-05 | 7.6e-06 | 9.7e-06 | 1.077E-05 | 3.814E-06 | 7.600E-06 | 1.500E-05 |
| 20232 | 1012 | L1012K | C | 6e-06 | 7.3e-06 | 6.2e-06 | 6.500E-06 | 7.000E-07 | 6.000E-06 | 7.300E-06 |
| 20233 | 1012 | L1012M | C | 9e-06 | 8.9e-06 | 1.1e-05 | 9.633E-06 | 1.185E-06 | 8.900E-06 | 1.100E-05 |
| 20234 | 1012 | L1012F | C | 7.1e-06 | 4.8e-06 | 6e-06 | 5.967E-06 | 1.150E-06 | 4.800E-06 | 7.100E-06 |
| 20235 | 1012 | L1012P | C | 8.1e-06 | 5.3e-06 | 8.2e-06 | 7.200E-06 | 1.646E-06 | 5.300E-06 | 8.200E-06 |
| 20236 | 1012 | L1012S | C | 8.5e-06 | 1.1e-05 | 7.1e-06 | 8.867E-06 | 1.976E-06 | 7.100E-06 | 1.100E-05 |
| 20237 | 1012 | L1012T | C | 1.1e-05 | 1e-05 | 5.7e-06 | 8.900E-06 | 2.816E-06 | 5.700E-06 | 1.100E-05 |
| 20238 | 1012 | L1012W | C | 5.1e-06 | 6.2e-06 | 9.1e-06 | 6.800E-06 | 2.066E-06 | 5.100E-06 | 9.100E-06 |
| 20239 | 1012 | L1012Y | C | 1.1e-05 | 4.3e-06 | 6.7e-06 | 7.333E-06 | 3.395E-06 | 4.300E-06 | 1.100E-05 |
| 20240 | 1012 | L1012V | C | 9.8e-06 | 9.5e-06 | 6.6e-06 | 8.633E-06 | 1.767E-06 | 6.600E-06 | 9.800E-06 |
| 20241 | 1013 | Q1013A | C | 5.5e-06 | 1.3e-05 | 6.9e-06 | 8.467E-06 | 3.988E-06 | 5.500E-06 | 1.300E-05 |
| 20242 | 1013 | Q1013R | C | 5.4e-06 | 7.1e-06 | 6.9e-06 | 6.467E-06 | 9.292E-07 | 5.400E-06 | 7.100E-06 |
| 20243 | 1013 | Q1013N | C | 6.9e-06 | 1.2e-05 | 8.5e-06 | 9.133E-06 | 2.608E-06 | 6.900E-06 | 1.200E-05 |
| 20244 | 1013 | Q1013D | C | 9.6e-06 | 9.5e-06 | 8.8e-06 | 9.300E-06 | 4.359E-07 | 8.800E-06 | 9.600E-06 |
| 20245 | 1013 | Q1013C | C | 1.1e-05 | 5.7e-06 | 5.6e-06 | 7.433E-06 | 3.089E-06 | 5.600E-06 | 1.100E-05 |
| 20246 | 1013 | Q1013E | C | 6.6e-06 | 9.7e-06 | 6.7e-06 | 7.667E-06 | 1.762E-06 | 6.600E-06 | 9.700E-06 |
| 20247 | 1013 | Q1013Q | C | 4.9e-06 | 7.5e-06 | 6.2e-06 | 6.200E-06 | 1.300E-06 | 4.900E-06 | 7.500E-06 |
| 20248 | 1013 | Q1013G | C | 1.3e-05 | 1e-05 | 7.9e-06 | 1.030E-05 | 2.563E-06 | 7.900E-06 | 1.300E-05 |
| 20249 | 1013 | Q1013H | C | 1.2e-05 | 4.8e-06 | 7.3e-06 | 8.033E-06 | 3.656E-06 | 4.800E-06 | 1.200E-05 |
| 20250 | 1013 | Q1013I | C | 1.4e-05 | 6.4e-06 | 7.3e-06 | 9.233E-06 | 4.153E-06 | 6.400E-06 | 1.400E-05 |
| 20251 | 1013 | Q1013L | C | 6.1e-06 | 5.1e-06 | 7.4e-06 | 6.200E-06 | 1.153E-06 | 5.100E-06 | 7.400E-06 |
| 20252 | 1013 | Q1013K | C | 8.1e-06 | 7.2e-06 | 1.2e-05 | 9.100E-06 | 2.551E-06 | 7.200E-06 | 1.200E-05 |
| 20253 | 1013 | Q1013M | C | 7.2e-06 | 9e-06 | 7.6e-06 | 7.933E-06 | 9.452E-07 | 7.200E-06 | 9.000E-06 |
| 20254 | 1013 | Q1013F | C | 1.6e-05 | 4.7e-06 | 1e-05 | 1.023E-05 | 5.654E-06 | 4.700E-06 | 1.600E-05 |
| 20255 | 1013 | Q1013P | C | 8.3e-06 | 1.5e-05 | 1e-05 | 1.110E-05 | 3.483E-06 | 8.300E-06 | 1.500E-05 |
| 20256 | 1013 | Q1013S | C | 9e-06 | 6.6e-06 | 1.3e-05 | 9.533E-06 | 3.233E-06 | 6.600E-06 | 1.300E-05 |
| 20257 | 1013 | Q1013T | C | 7.6e-06 | 6.7e-06 | 6.5e-06 | 6.933E-06 | 5.859E-07 | 6.500E-06 | 7.600E-06 |
| 20258 | 1013 | Q1013W | C | 1.1e-05 | 7.2e-06 | 6.6e-06 | 8.267E-06 | 2.386E-06 | 6.600E-06 | 1.100E-05 |
| 20259 | 1013 | Q1013Y | C | 8.6e-06 | 1.8e-05 | 3e-06 | 9.867E-06 | 7.580E-06 | 3.000E-06 | 1.800E-05 |
| 20260 | 1013 | Q1013V | C | 8e-06 | 6.8e-06 | 1.3e-05 | 9.267E-06 | 3.288E-06 | 6.800E-06 | 1.300E-05 |
| 20261 | 1014 | S1014A | C | 9e-06 | 7.4e-06 | 9.6e-06 | 8.667E-06 | 1.137E-06 | 7.400E-06 | 9.600E-06 |
| 20262 | 1014 | S1014R | C | 6.7e-06 | 6.1e-06 | 7.2e-06 | 6.667E-06 | 5.508E-07 | 6.100E-06 | 7.200E-06 |
| 20263 | 1014 | S1014N | C | 2.6e-05 | 8e-06 | 1e-05 | 1.467E-05 | 9.866E-06 | 8.000E-06 | 2.600E-05 |
| 20264 | 1014 | S1014D | C | 8.1e-06 | 7.1e-06 | 1e-05 | 8.400E-06 | 1.473E-06 | 7.100E-06 | 1.000E-05 |
| 20265 | 1014 | S1014C | C | 1.4e-05 | 5.7e-06 | 7e-06 | 8.900E-06 | 4.464E-06 | 5.700E-06 | 1.400E-05 |
| 20266 | 1014 | S1014E | C | 6.9e-06 | 5.1e-06 | 8.1e-06 | 6.700E-06 | 1.510E-06 | 5.100E-06 | 8.100E-06 |
| 20267 | 1014 | S1014Q | C | 4.2e-06 | 5.8e-06 | 5.8e-06 | 5.267E-06 | 9.238E-07 | 4.200E-06 | 5.800E-06 |
| 20268 | 1014 | S1014G | C | 7.1e-06 | 4.2e-06 | 8.5e-06 | 6.600E-06 | 2.193E-06 | 4.200E-06 | 8.500E-06 |
| 20269 | 1014 | S1014H | C | 7e-06 | 9.2e-06 | 1.4e-05 | 1.007E-05 | 3.580E-06 | 7.000E-06 | 1.400E-05 |
| 20270 | 1014 | S1014I | C | 7.6e-06 | 5.1e-06 | 7.5e-06 | 6.733E-06 | 1.415E-06 | 5.100E-06 | 7.600E-06 |
| 20271 | 1014 | S1014L | C | 8.5e-06 | 1.1e-05 | 6.5e-06 | 8.667E-06 | 2.255E-06 | 6.500E-06 | 1.100E-05 |
| 20272 | 1014 | S1014K | C | 8.8e-06 | 9e-06 | 5.5e-06 | 7.767E-06 | 1.966E-06 | 5.500E-06 | 9.000E-06 |
| 20273 | 1014 | S1014M | C | 8.1e-06 | 6.1e-06 | 1e-05 | 8.067E-06 | 1.950E-06 | 6.100E-06 | 1.000E-05 |
| 20274 | 1014 | S1014F | C | 1.5e-05 | 1.2e-05 | 5.6e-06 | 1.087E-05 | 4.801E-06 | 5.600E-06 | 1.500E-05 |
| 20275 | 1014 | S1014P | C | 9.7e-06 | 7.1e-06 | 1.3e-05 | 9.933E-06 | 2.957E-06 | 7.100E-06 | 1.300E-05 |
| 20276 | 1014 | S1014S | C | 9.8e-06 | 7.3e-06 | 5e-06 | 7.367E-06 | 2.401E-06 | 5.000E-06 | 9.800E-06 |
| 20277 | 1014 | S1014T | C | 5.6e-06 | 8.4e-06 | 8.5e-06 | 7.500E-06 | 1.646E-06 | 5.600E-06 | 8.500E-06 |
| 20278 | 1014 | S1014W | C | 9.8e-06 | 7.8e-06 | 5.2e-06 | 7.600E-06 | 2.307E-06 | 5.200E-06 | 9.800E-06 |
| 20279 | 1014 | S1014Y | C | 7.6e-06 | 8.2e-06 | 6.2e-06 | 7.333E-06 | 1.026E-06 | 6.200E-06 | 8.200E-06 |
| 20280 | 1014 | S1014V | C | 8.3e-06 | 6.4e-06 | 1.4e-05 | 9.567E-06 | 3.955E-06 | 6.400E-06 | 1.400E-05 |
| 20281 | 1015 | S1015A | C | 7.3e-06 | 1e-05 | 6.6e-06 | 7.967E-06 | 1.795E-06 | 6.600E-06 | 1.000E-05 |
| 20282 | 1015 | S1015R | C | 5e-06 | 8.1e-06 | 9.3e-06 | 7.467E-06 | 2.219E-06 | 5.000E-06 | 9.300E-06 |
| 20283 | 1015 | S1015N | C | 8.3e-06 | 1.3e-05 | 1.2e-05 | 1.110E-05 | 2.476E-06 | 8.300E-06 | 1.300E-05 |
| 20284 | 1015 | S1015D | C | 9.3e-06 | 4.8e-06 | 6e-06 | 6.700E-06 | 2.330E-06 | 4.800E-06 | 9.300E-06 |
| 20285 | 1015 | S1015C | C | 1.3e-05 | 5.9e-06 | 9.8e-06 | 9.567E-06 | 3.556E-06 | 5.900E-06 | 1.300E-05 |
| 20286 | 1015 | S1015E | C | 8.5e-06 | 7.4e-06 | 8.8e-06 | 8.233E-06 | 7.371E-07 | 7.400E-06 | 8.800E-06 |
| 20287 | 1015 | S1015Q | C | 5.6e-06 | 1.1e-05 | 1.1e-05 | 9.200E-06 | 3.118E-06 | 5.600E-06 | 1.100E-05 |
| 20288 | 1015 | S1015G | C | 7.5e-06 | 9.8e-06 | 1.1e-05 | 9.433E-06 | 1.779E-06 | 7.500E-06 | 1.100E-05 |
| 20289 | 1015 | S1015H | C | 5.9e-06 | 7e-06 | 5.8e-06 | 6.233E-06 | 6.658E-07 | 5.800E-06 | 7.000E-06 |
| 20290 | 1015 | S1015I | C | 5.1e-06 | 6.9e-06 | 1.1e-05 | 7.667E-06 | 3.024E-06 | 5.100E-06 | 1.100E-05 |
| 20291 | 1015 | S1015L | C | 5.7e-06 | 4.9e-06 | 7.8e-06 | 6.133E-06 | 1.498E-06 | 4.900E-06 | 7.800E-06 |
| 20292 | 1015 | S1015K | C | 6.2e-06 | 5.2e-06 | 6.9e-06 | 6.100E-06 | 8.544E-07 | 5.200E-06 | 6.900E-06 |
| 20293 | 1015 | S1015M | C | 1.3e-05 | 9.7e-06 | 6.6e-06 | 9.767E-06 | 3.201E-06 | 6.600E-06 | 1.300E-05 |
| 20294 | 1015 | S1015F | C | 4.9e-06 | 8.4e-06 | 9.4e-06 | 7.567E-06 | 2.363E-06 | 4.900E-06 | 9.400E-06 |
| 20295 | 1015 | S1015P | C | 7e-06 | 4.6e-06 | 1.1e-05 | 7.533E-06 | 3.233E-06 | 4.600E-06 | 1.100E-05 |
| 20296 | 1015 | S1015S | C | 4.8e-06 | 6.9e-06 | 7.7e-06 | 6.467E-06 | 1.498E-06 | 4.800E-06 | 7.700E-06 |
| 20297 | 1015 | S1015T | C | 7.9e-06 | 4.8e-06 | 5.2e-06 | 5.967E-06 | 1.686E-06 | 4.800E-06 | 7.900E-06 |
| 20298 | 1015 | S1015W | C | 8.1e-06 | 1e-05 | 1.8e-05 | 1.203E-05 | 5.254E-06 | 8.100E-06 | 1.800E-05 |
| 20299 | 1015 | S1015Y | C | 5e-06 | 9.3e-06 | 1e-05 | 8.100E-06 | 2.707E-06 | 5.000E-06 | 1.000E-05 |
| 20300 | 1015 | S1015V | C | 7e-06 | 5.3e-06 | 9.7e-06 | 7.333E-06 | 2.219E-06 | 5.300E-06 | 9.700E-06 |
| 20301 | 1016 | G1016A | C | 6.7e-06 | 6.4e-06 | 7.5e-06 | 6.867E-06 | 5.686E-07 | 6.400E-06 | 7.500E-06 |
| 20302 | 1016 | G1016R | C | 9.4e-06 | 6.7e-06 | 9.6e-06 | 8.567E-06 | 1.620E-06 | 6.700E-06 | 9.600E-06 |
| 20303 | 1016 | G1016N | C | 1.6e-05 | 5.3e-06 | 5.7e-06 | 9.000E-06 | 6.065E-06 | 5.300E-06 | 1.600E-05 |
| 20304 | 1016 | G1016D | C | 5.6e-06 | 7.9e-06 | 1.1e-05 | 8.167E-06 | 2.710E-06 | 5.600E-06 | 1.100E-05 |
| 20305 | 1016 | G1016C | C | 8.2e-06 | 9.8e-06 | 5.8e-06 | 7.933E-06 | 2.013E-06 | 5.800E-06 | 9.800E-06 |
| 20306 | 1016 | G1016E | C | 9.1e-06 | 6.7e-06 | 1.1e-05 | 8.933E-06 | 2.155E-06 | 6.700E-06 | 1.100E-05 |
| 20307 | 1016 | G1016Q | C | 1.1e-05 | 6.2e-06 | 5.5e-06 | 7.567E-06 | 2.994E-06 | 5.500E-06 | 1.100E-05 |
| 20308 | 1016 | G1016G | C | 1.2e-05 | 5.6e-06 | 6.5e-06 | 8.033E-06 | 3.465E-06 | 5.600E-06 | 1.200E-05 |
| 20309 | 1016 | G1016H | C | 8.7e-06 | 1.3e-05 | 7e-06 | 9.567E-06 | 3.092E-06 | 7.000E-06 | 1.300E-05 |
| 20310 | 1016 | G1016I | C | 7.8e-06 | 4.2e-06 | 7.9e-06 | 6.633E-06 | 2.108E-06 | 4.200E-06 | 7.900E-06 |
| 20311 | 1016 | G1016L | C | 7.9e-06 | 9.5e-06 | 5.8e-06 | 7.733E-06 | 1.856E-06 | 5.800E-06 | 9.500E-06 |
| 20312 | 1016 | G1016K | C | 1.3e-05 | 5.8e-06 | 9.1e-06 | 9.300E-06 | 3.604E-06 | 5.800E-06 | 1.300E-05 |
| 20313 | 1016 | G1016M | C | 7.5e-06 | 1.5e-05 | 6.2e-06 | 9.567E-06 | 4.750E-06 | 6.200E-06 | 1.500E-05 |
| 20314 | 1016 | G1016F | C | 4.6e-06 | 1e-05 | 1.4e-05 | 9.533E-06 | 4.717E-06 | 4.600E-06 | 1.400E-05 |

| | | | | | | | | | | |
|---|---|---|---|---|---|---|---|---|---|---|
| 20315 | 1016 | G1016P | C | 7.7e-06 | 5.1e-06 | 9.9e-06 | 7.567E-06 | 2.403E-06 | 5.100E-06 | 9.900E-06 |
| 20316 | 1016 | G1016S | C | 9.1e-06 | 1e-05 | 6.1e-06 | 8.400E-06 | 2.042E-06 | 6.100E-06 | 1.000E-05 |
| 20317 | 1016 | G1016T | C | 1e-05 | 9e-06 | 4.8e-06 | 7.933E-06 | 2.759E-06 | 4.800E-06 | 1.000E-05 |
| 20318 | 1016 | G1016W | C | 9.7e-06 | 7.7e-06 | 8.5e-06 | 8.633E-06 | 1.007E-06 | 7.700E-06 | 9.700E-06 |
| 20319 | 1016 | G1016Y | C | 5.8e-06 | 8.9e-06 | 5.7e-06 | 6.800E-06 | 1.819E-06 | 5.700E-06 | 8.900E-06 |
| 20320 | 1016 | G1016V | C | 6.4e-06 | 4.7e-06 | 7.6e-06 | 6.233E-06 | 1.457E-06 | 4.700E-06 | 7.600E-06 |
| 20321 | 1017 | L1017A | C | 7.6e-06 | 8.3e-06 | 1e-05 | 8.633E-06 | 1.234E-06 | 7.600E-06 | 1.000E-05 |
| 20322 | 1017 | L1017R | C | 7.4e-06 | 1.3e-05 | 1.6e-05 | 1.213E-05 | 4.365E-06 | 7.400E-06 | 1.600E-05 |
| 20323 | 1017 | L1017N | C | 6.2e-06 | 4.4e-06 | 6.1e-06 | 5.567E-06 | 1.012E-06 | 4.400E-06 | 6.200E-06 |
| 20324 | 1017 | L1017D | C | 6.6e-06 | 5.7e-06 | 9.2e-06 | 7.167E-06 | 1.818E-06 | 5.700E-06 | 9.200E-06 |
| 20325 | 1017 | L1017C | C | 5.6e-06 | 5.7e-06 | 8.2e-06 | 6.500E-06 | 1.473E-06 | 5.600E-06 | 8.200E-06 |
| 20326 | 1017 | L1017E | C | 9.2e-06 | 1.2e-05 | 1.4e-05 | 1.173E-05 | 2.411E-06 | 9.200E-06 | 1.400E-05 |
| 20327 | 1017 | L1017Q | C | 6.1e-06 | 4.7e-06 | 5.8e-06 | 5.533E-06 | 7.371E-07 | 4.700E-06 | 6.100E-06 |
| 20328 | 1017 | L1017G | C | 8.1e-06 | 6.4e-06 | 5.6e-06 | 6.700E-06 | 1.277E-06 | 5.600E-06 | 8.100E-06 |
| 20329 | 1017 | L1017H | C | 6.3e-06 | 6.2e-06 | 9.3e-06 | 7.267E-06 | 1.762E-06 | 6.200E-06 | 9.300E-06 |
| 20330 | 1017 | L1017I | C | 9.2e-06 | 8e-06 | 1e-05 | 9.067E-06 | 1.007E-06 | 8.000E-06 | 1.000E-05 |
| 20331 | 1017 | L1017L | C | 8e-06 | 6e-06 | 9.8e-06 | 7.933E-06 | 1.901E-06 | 6.000E-06 | 9.800E-06 |
| 20332 | 1017 | L1017K | C | 6.6e-06 | 7e-06 | 5.1e-06 | 6.233E-06 | 1.002E-06 | 5.100E-06 | 7.000E-06 |
| 20333 | 1017 | L1017M | C | 9e-06 | 6.3e-06 | 7.8e-06 | 7.700E-06 | 1.353E-06 | 6.300E-06 | 9.000E-06 |
| 20334 | 1017 | L1017F | C | 1.5e-05 | 9.5e-06 | 7e-06 | 1.050E-05 | 4.093E-06 | 7.000E-06 | 1.500E-05 |
| 20335 | 1017 | L1017P | C | 1e-05 | 5.9e-06 | 7.4e-06 | 7.767E-06 | 2.074E-06 | 5.900E-06 | 1.000E-05 |
| 20336 | 1017 | L1017S | C | 7.1e-06 | 7.1e-06 | 7.7e-06 | 7.300E-06 | 3.464E-07 | 7.100E-06 | 7.700E-06 |
| 20337 | 1017 | L1017T | C | 9.3e-06 | 1e-05 | 9.4e-06 | 9.567E-06 | 3.786E-07 | 9.300E-06 | 1.000E-05 |
| 20338 | 1017 | L1017W | C | 5.8e-06 | 1e-05 | 4.4e-06 | 6.733E-06 | 2.914E-06 | 4.400E-06 | 1.000E-05 |
| 20339 | 1017 | L1017Y | C | 7e-06 | 9.3e-06 | 9.3e-06 | 8.533E-06 | 1.328E-06 | 7.000E-06 | 9.300E-06 |
| 20340 | 1017 | L1017V | C | 4.7e-06 | 7.4e-06 | 9.8e-06 | 7.300E-06 | 2.551E-06 | 4.700E-06 | 9.800E-06 |
| 20341 | 1018 | Y1018A | C | 6.7e-06 | 1.4e-05 | 8.1e-06 | 9.600E-06 | 3.874E-06 | 6.700E-06 | 1.400E-05 |
| 20342 | 1018 | Y1018R | C | 1e-05 | 1.3e-05 | 7.7e-06 | 1.023E-05 | 2.658E-06 | 7.700E-06 | 1.300E-05 |
| 20343 | 1018 | Y1018N | C | 7.2e-06 | 4.5e-06 | 1.1e-05 | 7.567E-06 | 3.265E-06 | 4.500E-06 | 1.100E-05 |
| 20344 | 1018 | Y1018D | C | 1e-05 | 1.6e-05 | 5.1e-06 | 1.037E-05 | 5.459E-06 | 5.100E-06 | 1.600E-05 |
| 20345 | 1018 | Y1018C | C | 7.8e-06 | 9.9e-06 | 5.9e-06 | 7.867E-06 | 2.001E-06 | 5.900E-06 | 9.900E-06 |
| 20346 | 1018 | Y1018E | C | 1.4e-05 | 1e-05 | 6.1e-06 | 1.003E-05 | 3.950E-06 | 6.100E-06 | 1.400E-05 |
| 20347 | 1018 | Y1018Q | C | 7.6e-06 | 9.4e-06 | 6.4e-06 | 7.800E-06 | 1.510E-06 | 6.400E-06 | 9.400E-06 |
| 20348 | 1018 | Y1018G | C | 6.8e-06 | 6.6e-06 | 5e-06 | 6.133E-06 | 9.866E-07 | 5.000E-06 | 6.800E-06 |
| 20349 | 1018 | Y1018H | C | 7.3e-06 | 6.2e-06 | 9.1e-06 | 7.533E-06 | 1.464E-06 | 6.200E-06 | 9.100E-06 |
| 20350 | 1018 | Y1018I | C | 7.8e-06 | 1.2e-05 | 8.6e-06 | 9.467E-06 | 2.230E-06 | 7.800E-06 | 1.200E-05 |
| 20351 | 1018 | Y1018L | C | 1.2e-05 | 4.8e-06 | 8.6e-06 | 8.467E-06 | 3.602E-06 | 4.800E-06 | 1.200E-05 |
| 20352 | 1018 | Y1018K | C | 9.7e-06 | 4.9e-06 | 1e-05 | 8.200E-06 | 2.862E-06 | 4.900E-06 | 1.000E-05 |
| 20353 | 1018 | Y1018M | C | 6.1e-06 | 4.8e-06 | 8.9e-06 | 6.600E-06 | 2.095E-06 | 4.800E-06 | 8.900E-06 |
| 20354 | 1018 | Y1018F | C | 6.6e-06 | 7.1e-06 | 7.5e-06 | 7.067E-06 | 4.509E-07 | 6.600E-06 | 7.500E-06 |
| 20355 | 1018 | Y1018P | C | 5.4e-06 | 8.4e-06 | 9.8e-06 | 7.867E-06 | 2.248E-06 | 5.400E-06 | 9.800E-06 |
| 20356 | 1018 | Y1018S | C | 1.4e-05 | 8.6e-06 | 9.4e-06 | 1.067E-05 | 2.914E-06 | 8.600E-06 | 1.400E-05 |
| 20357 | 1018 | Y1018T | C | 8.1e-06 | 7.5e-06 | 8.8e-06 | 8.133E-06 | 6.506E-07 | 7.500E-06 | 8.800E-06 |
| 20358 | 1018 | Y1018W | C | 1.8e-05 | 5.6e-06 | 7.3e-06 | 1.030E-05 | 6.722E-06 | 5.600E-06 | 1.800E-05 |
| 20359 | 1018 | Y1018Y | C | 6.9e-06 | 9.5e-06 | 6.2e-06 | 7.533E-06 | 1.739E-06 | 6.200E-06 | 9.500E-06 |
| 20360 | 1018 | Y1018V | C | 1.3e-05 | 9.5e-06 | 4.8e-06 | 9.100E-06 | 4.115E-06 | 4.800E-06 | 1.300E-05 |
| 20361 | 1019 | S1019A | C | 1.1e-05 | 7.7e-06 | 1.3e-05 | 1.057E-05 | 2.676E-06 | 7.700E-06 | 1.300E-05 |
| 20362 | 1019 | S1019R | C | 9e-06 | 1.2e-05 | 9.8e-06 | 1.027E-05 | 1.553E-06 | 9.000E-06 | 1.200E-05 |
| 20363 | 1019 | S1019N | C | 1.1e-05 | 9.7e-06 | 9.4e-06 | 1.003E-05 | 8.505E-07 | 9.400E-06 | 1.100E-05 |
| 20364 | 1019 | S1019D | C | 7.5e-06 | 4.7e-06 | 1e-05 | 7.400E-06 | 2.651E-06 | 4.700E-06 | 1.000E-05 |
| 20365 | 1019 | S1019C | C | 8.4e-06 | 4.7e-06 | 1e-05 | 7.700E-06 | 2.718E-06 | 4.700E-06 | 1.000E-05 |
| 20366 | 1019 | S1019E | C | 5.9e-06 | 5.1e-06 | 9.3e-06 | 6.767E-06 | 2.230E-06 | 5.100E-06 | 9.300E-06 |
| 20367 | 1019 | S1019Q | C | 9.6e-06 | 8.9e-06 | 1.3e-05 | 1.050E-05 | 2.193E-06 | 8.900E-06 | 1.300E-05 |
| 20368 | 1019 | S1019G | C | 8.1e-06 | 9.2e-06 | 8.5e-06 | 8.600E-06 | 5.568E-07 | 8.100E-06 | 9.200E-06 |
| 20369 | 1019 | S1019H | C | 5e-06 | 1.2e-05 | 4.6e-06 | 7.200E-06 | 4.162E-06 | 4.600E-06 | 1.200E-05 |
| 20370 | 1019 | S1019I | C | 6.5e-06 | 8.7e-06 | 1.2e-05 | 9.067E-06 | 2.768E-06 | 6.500E-06 | 1.200E-05 |
| 20371 | 1019 | S1019L | C | 1.4e-05 | 8.3e-06 | 1e-05 | 1.077E-05 | 2.926E-06 | 8.300E-06 | 1.400E-05 |
| 20372 | 1019 | S1019K | C | 7e-06 | 9e-06 | 7.1e-06 | 7.700E-06 | 1.127E-06 | 7.000E-06 | 9.000E-06 |
| 20373 | 1019 | S1019M | C | 1.2e-05 | 7e-06 | 9.7e-06 | 9.567E-06 | 2.503E-06 | 7.000E-06 | 1.200E-05 |
| 20374 | 1019 | S1019F | C | 6.9e-06 | 1.1e-05 | 1.2e-05 | 9.967E-06 | 2.702E-06 | 6.900E-06 | 1.200E-05 |
| 20375 | 1019 | S1019P | C | 4.5e-06 | 6.5e-06 | 1.6e-05 | 9.000E-06 | 6.144E-06 | 4.500E-06 | 1.600E-05 |
| 20376 | 1019 | S1019S | C | 5.1e-06 | 8.5e-06 | 4.5e-06 | 6.033E-06 | 2.157E-06 | 4.500E-06 | 8.500E-06 |
| 20377 | 1019 | S1019T | C | 1.1e-05 | 3.6e-06 | 4.4e-06 | 6.333E-06 | 4.061E-06 | 3.600E-06 | 1.100E-05 |
| 20378 | 1019 | S1019W | C | 5.3e-06 | 7.3e-06 | 9.2e-06 | 7.267E-06 | 1.950E-06 | 5.300E-06 | 9.200E-06 |
| 20379 | 1019 | S1019Y | C | 4.6e-06 | 6.9e-06 | 1e-05 | 7.167E-06 | 2.710E-06 | 4.600E-06 | 1.000E-05 |
| 20380 | 1019 | S1019V | C | 6.4e-06 | 1e-05 | 5.4e-06 | 7.267E-06 | 2.419E-06 | 5.400E-06 | 1.000E-05 |
| 20381 | 1020 | L1020A | C | 8.6e-06 | 5.6e-06 | 1.5e-05 | 9.733E-06 | 4.801E-06 | 5.600E-06 | 1.500E-05 |
| 20382 | 1020 | L1020R | C | 5.9e-06 | 8.6e-06 | 1.1e-05 | 8.500E-06 | 2.551E-06 | 5.900E-06 | 1.100E-05 |
| 20383 | 1020 | L1020N | C | 1.5e-05 | 1.6e-05 | 5.1e-06 | 1.203E-05 | 6.025E-06 | 5.100E-06 | 1.600E-05 |
| 20384 | 1020 | L1020D | C | 1e-05 | 5.8e-06 | 1e-05 | 8.600E-06 | 2.425E-06 | 5.800E-06 | 1.000E-05 |
| 20385 | 1020 | L1020C | C | 1.3e-05 | 1e-05 | 7.4e-06 | 1.013E-05 | 2.802E-06 | 7.400E-06 | 1.300E-05 |
| 20386 | 1020 | L1020E | C | 6.9e-06 | 6.9e-06 | 9.5e-06 | 7.767E-06 | 1.501E-06 | 6.900E-06 | 9.500E-06 |
| 20387 | 1020 | L1020Q | C | 1.1e-05 | 4.7e-06 | 7.5e-06 | 7.733E-06 | 3.156E-06 | 4.700E-06 | 1.100E-05 |
| 20388 | 1020 | L1020G | C | 5.3e-06 | 1e-05 | 6.9e-06 | 7.400E-06 | 2.390E-06 | 5.300E-06 | 1.000E-05 |
| 20389 | 1020 | L1020H | C | 5.7e-06 | 5.6e-06 | 6.9e-06 | 6.067E-06 | 7.234E-07 | 5.600E-06 | 6.900E-06 |
| 20390 | 1020 | L1020I | C | 5.8e-06 | 5.8e-06 | 7.5e-06 | 6.367E-06 | 9.815E-07 | 5.800E-06 | 7.500E-06 |
| 20391 | 1020 | L1020L | C | 1.1e-05 | 4.8e-06 | 6.4e-06 | 7.400E-06 | 3.219E-06 | 4.800E-06 | 1.100E-05 |
| 20392 | 1020 | L1020K | C | 5.3e-06 | 6.3e-06 | 1.2e-05 | 7.867E-06 | 3.614E-06 | 5.300E-06 | 1.200E-05 |
| 20393 | 1020 | L1020M | C | 9.6e-06 | 6.4e-06 | 8.6e-06 | 8.200E-06 | 1.637E-06 | 6.400E-06 | 9.600E-06 |
| 20394 | 1020 | L1020F | C | 8.7e-06 | 8.6e-06 | 1.3e-05 | 1.010E-05 | 2.512E-06 | 8.600E-06 | 1.300E-05 |
| 20395 | 1020 | L1020P | C | 8.2e-06 | 7.2e-06 | 4.3e-06 | 6.567E-06 | 2.026E-06 | 4.300E-06 | 8.200E-06 |
| 20396 | 1020 | L1020S | C | 4.4e-06 | 6.4e-06 | 9.5e-06 | 6.767E-06 | 2.570E-06 | 4.400E-06 | 9.500E-06 |
| 20397 | 1020 | L1020T | C | 9.9e-06 | 9e-06 | 6.2e-06 | 8.367E-06 | 1.930E-06 | 6.200E-06 | 9.900E-06 |
| 20398 | 1020 | L1020W | C | 6.6e-06 | 6.5e-06 | 5.4e-06 | 6.167E-06 | 6.658E-07 | 5.400E-06 | 6.600E-06 |
| 20399 | 1020 | L1020Y | C | 6.6e-06 | 9.4e-06 | 4.5e-06 | 6.833E-06 | 2.458E-06 | 4.500E-06 | 9.400E-06 |

| | | | | | | | | | | |
|---|---|---|---|---|---|---|---|---|---|---|
| 20400 | 1020 | L1020V | C | 8.7e-06 | 5.9e-06 | 8.9e-06 | 7.833E-06 | 1.677E-06 | 5.900E-06 | 8.900E-06 |
| 20401 | 1021 | S1021A | C | 1.9e-05 | 9.3e-06 | 7e-06 | 1.177E-05 | 6.369E-06 | 7.000E-06 | 1.900E-05 |
| 20402 | 1021 | S1021R | C | 6.1e-06 | 1.3e-05 | 6.6e-06 | 8.567E-06 | 3.848E-06 | 6.100E-06 | 1.300E-05 |
| 20403 | 1021 | S1021N | C | 5.5e-06 | 5.6e-06 | 1.3e-05 | 8.033E-06 | 4.302E-06 | 5.500E-06 | 1.300E-05 |
| 20404 | 1021 | S1021D | C | 6.4e-06 | 5.5e-06 | 7.7e-06 | 6.533E-06 | 1.106E-06 | 5.500E-06 | 7.700E-06 |
| 20405 | 1021 | S1021C | C | 8.9e-06 | 9.2e-06 | 7.2e-06 | 8.433E-06 | 1.079E-06 | 7.200E-06 | 9.200E-06 |
| 20406 | 1021 | S1021E | C | 5.3e-06 | 1.3e-05 | 8.7e-06 | 9.000E-06 | 3.859E-06 | 5.300E-06 | 1.300E-05 |
| 20407 | 1021 | S1021Q | C | 7.3e-06 | 5.3e-06 | 1.9e-05 | 1.053E-05 | 7.400E-06 | 5.300E-06 | 1.900E-05 |
| 20408 | 1021 | S1021G | C | 7.2e-06 | 6.2e-06 | 8.2e-06 | 7.200E-06 | 1.000E-06 | 6.200E-06 | 8.200E-06 |
| 20409 | 1021 | S1021H | C | 6.8e-06 | 7.6e-06 | 6.8e-06 | 7.067E-06 | 4.619E-07 | 6.800E-06 | 7.600E-06 |
| 20410 | 1021 | S1021I | C | 5.7e-06 | 1e-05 | 1e-05 | 8.567E-06 | 2.483E-06 | 5.700E-06 | 1.000E-05 |
| 20411 | 1021 | S1021L | C | 4.9e-06 | 1.4e-05 | 5.9e-06 | 8.267E-06 | 4.990E-06 | 4.900E-06 | 1.400E-05 |
| 20412 | 1021 | S1021K | C | 1.3e-05 | 5.4e-06 | 1e-05 | 9.467E-06 | 3.828E-06 | 5.400E-06 | 1.300E-05 |
| 20413 | 1021 | S1021M | C | 1.4e-05 | 6.4e-06 | 7.7e-06 | 9.367E-06 | 4.065E-06 | 6.400E-06 | 1.400E-05 |
| 20414 | 1021 | S1021F | C | 6.5e-06 | 8e-06 | 8.4e-06 | 7.633E-06 | 1.002E-06 | 6.500E-06 | 8.400E-06 |
| 20415 | 1021 | S1021P | C | 5.2e-06 | 6.2e-06 | 9.1e-06 | 6.833E-06 | 2.026E-06 | 5.200E-06 | 9.100E-06 |
| 20416 | 1021 | S1021S | C | 9.4e-06 | 4.5e-06 | 5.8e-06 | 6.567E-06 | 2.538E-06 | 4.500E-06 | 9.400E-06 |
| 20417 | 1021 | S1021T | C | 8.5e-06 | 7e-06 | 7.1e-06 | 7.533E-06 | 8.386E-07 | 7.000E-06 | 8.500E-06 |
| 20418 | 1021 | S1021W | C | 5.7e-06 | 6.1e-06 | 8.6e-06 | 6.800E-06 | 1.572E-06 | 5.700E-06 | 8.600E-06 |
| 20419 | 1021 | S1021Y | C | 6.1e-06 | 6.3e-06 | 9.3e-06 | 7.233E-06 | 1.793E-06 | 6.100E-06 | 9.300E-06 |
| 20420 | 1021 | S1021V | C | 5.6e-06 | 8.6e-06 | 1.4e-05 | 9.400E-06 | 4.257E-06 | 5.600E-06 | 1.400E-05 |
| 20421 | 1022 | S1022A | C | 9e-06 | 8.1e-06 | 7.2e-06 | 8.100E-06 | 9.000E-07 | 7.200E-06 | 9.000E-06 |
| 20422 | 1022 | S1022R | C | 6.4e-06 | 7.7e-06 | 7.3e-06 | 7.133E-06 | 6.658E-07 | 6.400E-06 | 7.700E-06 |
| 20423 | 1022 | S1022N | C | 7.4e-06 | 4.9e-06 | 4.9e-06 | 5.733E-06 | 1.443E-06 | 4.900E-06 | 7.400E-06 |
| 20424 | 1022 | S1022D | C | 1e-05 | 5.6e-06 | 1.1e-05 | 8.867E-06 | 2.873E-06 | 5.600E-06 | 1.100E-05 |
| 20425 | 1022 | S1022C | C | 9.4e-06 | 7e-06 | 6e-06 | 7.467E-06 | 1.747E-06 | 6.000E-06 | 9.400E-06 |
| 20426 | 1022 | S1022E | C | 1.3e-05 | 7.6e-06 | 6.5e-06 | 9.033E-06 | 3.479E-06 | 6.500E-06 | 1.300E-05 |
| 20427 | 1022 | S1022Q | C | 6.4e-06 | 9.7e-06 | 9.4e-06 | 8.500E-06 | 1.825E-06 | 6.400E-06 | 9.700E-06 |
| 20428 | 1022 | S1022G | C | 6.9e-06 | 9.7e-06 | 6.5e-06 | 7.700E-06 | 1.744E-06 | 6.500E-06 | 9.700E-06 |
| 20429 | 1022 | S1022H | C | 4.7e-06 | 5.5e-06 | 6.2e-06 | 5.467E-06 | 7.506E-07 | 4.700E-06 | 6.200E-06 |
| 20430 | 1022 | S1022I | C | 7.1e-06 | 5.5e-06 | 6.2e-06 | 6.267E-06 | 8.021E-07 | 5.500E-06 | 7.100E-06 |
| 20431 | 1022 | S1022L | C | 5.4e-06 | 6.3e-06 | 8.8e-06 | 6.833E-06 | 1.762E-06 | 5.400E-06 | 8.800E-06 |
| 20432 | 1022 | S1022K | C | 7e-06 | 2.9e-06 | 1.3e-05 | 7.633E-06 | 5.080E-06 | 2.900E-06 | 1.300E-05 |
| 20433 | 1022 | S1022M | C | 1.5e-05 | 8.9e-06 | 6.2e-06 | 1.003E-05 | 4.508E-06 | 6.200E-06 | 1.500E-05 |
| 20434 | 1022 | S1022F | C | 1.5e-05 | 5e-06 | 8.1e-06 | 9.367E-06 | 5.119E-06 | 5.000E-06 | 1.500E-05 |
| 20435 | 1022 | S1022P | C | 4.6e-06 | 7.8e-06 | 7.7e-06 | 6.700E-06 | 1.819E-06 | 4.600E-06 | 7.800E-06 |
| 20436 | 1022 | S1022S | C | 1.3e-05 | 6.5e-06 | 5.9e-06 | 8.467E-06 | 3.937E-06 | 5.900E-06 | 1.300E-05 |
| 20437 | 1022 | S1022T | C | 1.1e-05 | 8.5e-06 | 9.7e-06 | 9.733E-06 | 1.250E-06 | 8.500E-06 | 1.100E-05 |
| 20438 | 1022 | S1022W | C | 1.1e-05 | 4.9e-06 | 9.1e-06 | 8.333E-06 | 3.121E-06 | 4.900E-06 | 1.100E-05 |
| 20439 | 1022 | S1022Y | C | 5.7e-06 | 5.3e-06 | 1.2e-05 | 7.667E-06 | 3.758E-06 | 5.300E-06 | 1.200E-05 |
| 20440 | 1022 | S1022V | C | 9e-06 | 5.7e-06 | 1.4e-05 | 9.567E-06 | 4.179E-06 | 5.700E-06 | 1.400E-05 |
| 20441 | 1023 | V1023A | C | 6.6e-06 | 7e-06 | 7.2e-06 | 6.933E-06 | 3.055E-07 | 6.600E-06 | 7.200E-06 |
| 20442 | 1023 | V1023R | C | 9.3e-06 | 9.3e-06 | 9.7e-06 | 9.433E-06 | 2.309E-07 | 9.300E-06 | 9.700E-06 |
| 20443 | 1023 | V1023N | C | 8.8e-06 | 1.4e-05 | 8.4e-06 | 1.040E-05 | 3.124E-06 | 8.400E-06 | 1.400E-05 |
| 20444 | 1023 | V1023D | C | 7.9e-06 | 7.6e-06 | 9.3e-06 | 8.267E-06 | 9.074E-07 | 7.600E-06 | 9.300E-06 |
| 20445 | 1023 | V1023C | C | 9.8e-06 | 8.1e-06 | 5.5e-06 | 7.800E-06 | 2.166E-06 | 5.500E-06 | 9.800E-06 |
| 20446 | 1023 | V1023E | C | 1.3e-05 | 5.3e-06 | 1.3e-05 | 1.043E-05 | 4.446E-06 | 5.300E-06 | 1.300E-05 |
| 20447 | 1023 | V1023Q | C | 8.4e-06 | 7.8e-06 | 5.8e-06 | 7.333E-06 | 1.361E-06 | 5.800E-06 | 8.400E-06 |
| 20448 | 1023 | V1023G | C | 5e-06 | 8.7e-06 | 8.6e-06 | 7.433E-06 | 2.108E-06 | 5.000E-06 | 8.700E-06 |
| 20449 | 1023 | V1023H | C | 9.7e-06 | 9.2e-06 | 6.7e-06 | 8.533E-06 | 1.607E-06 | 6.700E-06 | 9.700E-06 |
| 20450 | 1023 | V1023I | C | 9.3e-06 | 7.3e-06 | 9.6e-06 | 8.733E-06 | 1.250E-06 | 7.300E-06 | 9.600E-06 |
| 20451 | 1023 | V1023L | C | 4.5e-06 | 4.7e-06 | 1.2e-05 | 7.067E-06 | 4.274E-06 | 4.500E-06 | 1.200E-05 |
| 20452 | 1023 | V1023K | C | 8.7e-06 | 8e-06 | 6.6e-06 | 7.767E-06 | 1.069E-06 | 6.600E-06 | 8.700E-06 |
| 20453 | 1023 | V1023M | C | 1.5e-05 | 1.5e-05 | 7.2e-06 | 1.240E-05 | 4.503E-06 | 7.200E-06 | 1.500E-05 |
| 20454 | 1023 | V1023F | C | 5.8e-06 | 7.2e-06 | 6.9e-06 | 6.633E-06 | 7.371E-07 | 5.800E-06 | 7.200E-06 |
| 20455 | 1023 | V1023P | C | 6.6e-06 | 7.3e-06 | 7e-06 | 6.967E-06 | 3.512E-07 | 6.600E-06 | 7.300E-06 |
| 20456 | 1023 | V1023S | C | 7.5e-06 | 5.9e-06 | 6.7e-06 | 6.700E-06 | 8.000E-07 | 5.900E-06 | 7.500E-06 |
| 20457 | 1023 | V1023T | C | 8.6e-06 | 1.4e-05 | 5.6e-06 | 9.400E-06 | 4.257E-06 | 5.600E-06 | 1.400E-05 |
| 20458 | 1023 | V1023W | C | 8.8e-06 | 5.5e-06 | 8.7e-06 | 7.667E-06 | 1.877E-06 | 5.500E-06 | 8.800E-06 |
| 20459 | 1023 | V1023Y | C | 5.7e-06 | 8.9e-06 | 8.5e-06 | 7.700E-06 | 1.744E-06 | 5.700E-06 | 8.900E-06 |
| 20460 | 1023 | V1023V | C | 1.4e-05 | 5.7e-06 | 7.6e-06 | 9.100E-06 | 4.349E-06 | 5.700E-06 | 1.400E-05 |
| 20461 | 1024 | V1024A | C | 5.1e-06 | 5.6e-06 | 1.1e-05 | 7.233E-06 | 3.272E-06 | 5.100E-06 | 1.100E-05 |
| 20462 | 1024 | V1024R | C | 9.5e-06 | 1e-05 | 5.3e-06 | 8.267E-06 | 2.581E-06 | 5.300E-06 | 1.000E-05 |
| 20463 | 1024 | V1024N | C | 9.8e-06 | 6.5e-06 | 8.2e-06 | 8.167E-06 | 1.650E-06 | 6.500E-06 | 9.800E-06 |
| 20464 | 1024 | V1024D | C | 3.7e-06 | 9.6e-06 | 6.8e-06 | 6.700E-06 | 2.951E-06 | 3.700E-06 | 9.600E-06 |
| 20465 | 1024 | V1024C | C | 9.6e-06 | 5.2e-06 | 8e-06 | 7.600E-06 | 2.227E-06 | 5.200E-06 | 9.600E-06 |
| 20466 | 1024 | V1024E | C | 6.6e-06 | 7.2e-06 | 8.3e-06 | 7.367E-06 | 8.622E-07 | 6.600E-06 | 8.300E-06 |
| 20467 | 1024 | V1024Q | C | 1e-05 | 4.6e-06 | 6.7e-06 | 7.100E-06 | 2.722E-06 | 4.600E-06 | 1.000E-05 |
| 20468 | 1024 | V1024G | C | 7.1e-06 | 1e-05 | 1.3e-05 | 1.003E-05 | 2.950E-06 | 7.100E-06 | 1.300E-05 |
| 20469 | 1024 | V1024H | C | 1e-05 | 8.8e-06 | 8.5e-06 | 9.100E-06 | 7.937E-07 | 8.500E-06 | 1.000E-05 |
| 20470 | 1024 | V1024I | C | 5.8e-06 | 7.4e-06 | 8.3e-06 | 7.167E-06 | 1.266E-06 | 5.800E-06 | 8.300E-06 |
| 20471 | 1024 | V1024L | C | 6.6e-06 | 8.3e-06 | 8e-06 | 7.633E-06 | 9.074E-07 | 6.600E-06 | 8.300E-06 |
| 20472 | 1024 | V1024K | C | 8.9e-06 | 7.6e-06 | 1e-05 | 8.833E-06 | 1.201E-06 | 7.600E-06 | 1.000E-05 |
| 20473 | 1024 | V1024M | C | 6.8e-06 | 8.1e-06 | 7.6e-06 | 7.500E-06 | 6.557E-07 | 6.800E-06 | 8.100E-06 |
| 20474 | 1024 | V1024F | C | 6.9e-06 | 4.4e-06 | 6.4e-06 | 5.900E-06 | 1.323E-06 | 4.400E-06 | 6.900E-06 |
| 20475 | 1024 | V1024P | C | 5.8e-06 | 7.1e-06 | 9e-06 | 7.300E-06 | 1.609E-06 | 5.800E-06 | 9.000E-06 |
| 20476 | 1024 | V1024S | C | 1.4e-05 | 1.2e-05 | 1.1e-05 | 1.233E-05 | 1.528E-06 | 1.100E-05 | 1.400E-05 |
| 20477 | 1024 | V1024T | C | 6.8e-06 | 9.4e-06 | 8.7e-06 | 8.300E-06 | 1.345E-06 | 6.800E-06 | 9.400E-06 |
| 20478 | 1024 | V1024W | C | 8e-06 | 6.3e-06 | 7e-06 | 7.100E-06 | 8.544E-07 | 6.300E-06 | 8.000E-06 |
| 20479 | 1024 | V1024Y | C | 9.4e-06 | 9.4e-06 | 3.4e-06 | 7.400E-06 | 3.464E-06 | 3.400E-06 | 9.400E-06 |
| 20480 | 1024 | V1024V | C | 7.2e-06 | 7.6e-06 | 1.1e-05 | 8.600E-06 | 2.088E-06 | 7.200E-06 | 1.100E-05 |
| 20481 | 1025 | T1025A | C | 9e-06 | 9.8e-06 | 4.9e-06 | 7.900E-06 | 2.629E-06 | 4.900E-06 | 9.800E-06 |
| 20482 | 1025 | T1025R | C | 1.3e-05 | 1e-05 | 9e-06 | 1.067E-05 | 2.082E-06 | 9.000E-06 | 1.300E-05 |
| 20483 | 1025 | T1025N | C | 7.6e-06 | 1e-05 | 8.2e-06 | 8.600E-06 | 1.249E-06 | 7.600E-06 | 1.000E-05 |
| 20484 | 1025 | T1025D | C | 3.9e-06 | 8.2e-06 | 9.9e-06 | 7.333E-06 | 3.092E-06 | 3.900E-06 | 9.900E-06 |

| | | | | | | | | | | |
|---|---|---|---|---|---|---|---|---|---|---|
| 20485 | 1025 | T1025C | C | 5.1e-06 | 6.9e-06 | 1.4e-05 | 8.667E-06 | 4.706E-06 | 5.100E-06 | 1.400E-05 |
| 20486 | 1025 | T1025E | C | 8.4e-06 | 4.8e-06 | 7.1e-06 | 6.767E-06 | 1.823E-06 | 4.800E-06 | 8.400E-06 |
| 20487 | 1025 | T1025Q | C | 6e-06 | 1.2e-05 | 8.7e-06 | 8.900E-06 | 3.005E-06 | 6.000E-06 | 1.200E-05 |
| 20488 | 1025 | T1025G | C | 7.5e-06 | 1.1e-05 | 7.3e-06 | 8.600E-06 | 2.081E-06 | 7.300E-06 | 1.100E-05 |
| 20489 | 1025 | T1025H | C | 7.3e-06 | 1.4e-05 | 8.5e-06 | 9.933E-06 | 3.573E-06 | 7.300E-06 | 1.400E-05 |
| 20490 | 1025 | T1025I | C | 1.1e-05 | 9.4e-06 | 7.2e-06 | 9.200E-06 | 1.908E-06 | 7.200E-06 | 1.100E-05 |
| 20491 | 1025 | T1025L | C | 1.2e-05 | 6.7e-06 | 6.4e-06 | 8.367E-06 | 3.150E-06 | 6.400E-06 | 1.200E-05 |
| 20492 | 1025 | T1025K | C | 4.6e-06 | 2.1e-05 | 5.6e-06 | 1.040E-05 | 9.193E-06 | 4.600E-06 | 2.100E-05 |
| 20493 | 1025 | T1025M | C | 1e-05 | 6.2e-06 | 1.1e-05 | 9.067E-06 | 2.532E-06 | 6.200E-06 | 1.100E-05 |
| 20494 | 1025 | T1025F | C | 6.6e-06 | 8e-06 | 7e-06 | 7.200E-06 | 7.211E-07 | 6.600E-06 | 8.000E-06 |
| 20495 | 1025 | T1025P | C | 5.7e-06 | 7.7e-06 | 9.6e-06 | 7.667E-06 | 1.950E-06 | 5.700E-06 | 9.600E-06 |
| 20496 | 1025 | T1025S | C | 7.7e-06 | 5.7e-06 | 8.8e-06 | 7.400E-06 | 1.572E-06 | 5.700E-06 | 8.800E-06 |
| 20497 | 1025 | T1025T | C | 9.6e-06 | 7.6e-06 | 9e-06 | 8.733E-06 | 1.026E-06 | 7.600E-06 | 9.600E-06 |
| 20498 | 1025 | T1025W | C | 9.6e-06 | 6.3e-06 | 5.6e-06 | 7.167E-06 | 2.136E-06 | 5.600E-06 | 9.600E-06 |
| 20499 | 1025 | T1025Y | C | 9.3e-06 | 6.6e-06 | 8.8e-06 | 8.233E-06 | 1.436E-06 | 6.600E-06 | 9.300E-06 |
| 20500 | 1025 | T1025V | C | 6.3e-06 | 6.2e-06 | 7.6e-06 | 6.700E-06 | 7.810E-07 | 6.200E-06 | 7.600E-06 |
| 20501 | 1026 | V1026A | C | 6.9e-06 | 1.3e-05 | 8.6e-06 | 9.500E-06 | 3.148E-06 | 6.900E-06 | 1.300E-05 |
| 20502 | 1026 | V1026R | C | 1.3e-05 | 1.4e-05 | 8e-06 | 1.167E-05 | 3.215E-06 | 8.000E-06 | 1.400E-05 |
| 20503 | 1026 | V1026N | C | 6.8e-06 | 4.4e-06 | 6.5e-06 | 5.900E-06 | 1.308E-06 | 4.400E-06 | 6.800E-06 |
| 20504 | 1026 | V1026D | C | 1.3e-05 | 1.3e-05 | 1.4e-05 | 1.333E-05 | 5.774E-07 | 1.300E-05 | 1.400E-05 |
| 20505 | 1026 | V1026C | C | 5.4e-06 | 7.3e-06 | 8.9e-06 | 7.200E-06 | 1.752E-06 | 5.400E-06 | 8.900E-06 |
| 20506 | 1026 | V1026E | C | 9.2e-06 | 6.9e-06 | 8.5e-06 | 8.200E-06 | 1.179E-06 | 6.900E-06 | 9.200E-06 |
| 20507 | 1026 | V1026Q | C | 8.1e-06 | 7.8e-06 | 1.1e-05 | 8.967E-06 | 1.767E-06 | 7.800E-06 | 1.100E-05 |
| 20508 | 1026 | V1026G | C | 8.9e-06 | 1e-05 | 6.7e-06 | 8.533E-06 | 1.680E-06 | 6.700E-06 | 1.000E-05 |
| 20509 | 1026 | V1026H | C | 7.7e-06 | 5.9e-06 | 3.5e-06 | 5.700E-06 | 2.107E-06 | 3.500E-06 | 7.700E-06 |
| 20510 | 1026 | V1026I | C | 5.1e-06 | 9.4e-06 | 6.9e-06 | 7.133E-06 | 2.159E-06 | 5.100E-06 | 9.400E-06 |
| 20511 | 1026 | V1026L | C | 6.6e-06 | 4.8e-06 | 1.4e-05 | 8.467E-06 | 4.876E-06 | 4.800E-06 | 1.400E-05 |
| 20512 | 1026 | V1026K | C | 8.6e-06 | 9.2e-06 | 7.2e-06 | 8.333E-06 | 1.026E-06 | 7.200E-06 | 9.200E-06 |
| 20513 | 1026 | V1026M | C | 9.4e-06 | 6.9e-06 | 9.7e-06 | 8.667E-06 | 1.537E-06 | 6.900E-06 | 9.700E-06 |
| 20514 | 1026 | V1026F | C | 5e-06 | 9.3e-06 | 5.8e-06 | 6.700E-06 | 2.287E-06 | 5.000E-06 | 9.300E-06 |
| 20515 | 1026 | V1026P | C | 6.2e-06 | 7e-06 | 5.2e-06 | 6.133E-06 | 9.018E-07 | 5.200E-06 | 7.000E-06 |
| 20516 | 1026 | V1026S | C | 6.9e-06 | 6.7e-06 | 1.2e-05 | 8.533E-06 | 3.004E-06 | 6.700E-06 | 1.200E-05 |
| 20517 | 1026 | V1026T | C | 4.8e-06 | 6.7e-06 | 6.8e-06 | 6.100E-06 | 1.127E-06 | 4.800E-06 | 6.800E-06 |
| 20518 | 1026 | V1026W | C | 7.1e-06 | 6.1e-06 | 6.9e-06 | 6.700E-06 | 5.292E-07 | 6.100E-06 | 7.100E-06 |
| 20519 | 1026 | V1026Y | C | 1.2e-05 | 7.3e-06 | 8.1e-06 | 9.133E-06 | 2.515E-06 | 7.300E-06 | 1.200E-05 |
| 20520 | 1026 | V1026V | C | 1.2e-05 | 6.3e-06 | 9.5e-06 | 9.267E-06 | 2.857E-06 | 6.300E-06 | 1.200E-05 |
| 20521 | 1027 | P1027A | C | 1e-05 | 1.4e-05 | 9.8e-06 | 1.127E-05 | 2.369E-06 | 9.800E-06 | 1.400E-05 |
| 20522 | 1027 | P1027R | C | 6.6e-06 | 7e-06 | 6.5e-06 | 6.700E-06 | 2.646E-07 | 6.500E-06 | 7.000E-06 |
| 20523 | 1027 | P1027N | C | 5.1e-06 | 9.2e-06 | 8.3e-06 | 7.533E-06 | 2.155E-06 | 5.100E-06 | 9.200E-06 |
| 20524 | 1027 | P1027D | C | 6.3e-06 | 7e-06 | 4.8e-06 | 6.033E-06 | 1.124E-06 | 4.800E-06 | 7.000E-06 |
| 20525 | 1027 | P1027C | C | 8.9e-06 | 8.3e-06 | 8e-06 | 8.400E-06 | 4.583E-07 | 8.000E-06 | 8.900E-06 |
| 20526 | 1027 | P1027E | C | 6.6e-06 | 8.1e-06 | 7.1e-06 | 7.267E-06 | 7.638E-07 | 6.600E-06 | 8.100E-06 |
| 20527 | 1027 | P1027Q | C | 7.5e-06 | 1.3e-05 | 8.3e-06 | 9.600E-06 | 2.972E-06 | 7.500E-06 | 1.300E-05 |
| 20528 | 1027 | P1027G | C | 1e-05 | 4.7e-06 | 8.5e-06 | 7.733E-06 | 2.732E-06 | 4.700E-06 | 1.000E-05 |
| 20529 | 1027 | P1027H | C | 6.6e-06 | 8.1e-06 | 5.8e-06 | 6.833E-06 | 1.168E-06 | 5.800E-06 | 8.100E-06 |
| 20530 | 1027 | P1027I | C | 6.7e-06 | 7e-06 | 8.6e-06 | 7.433E-06 | 1.021E-06 | 6.700E-06 | 8.600E-06 |
| 20531 | 1027 | P1027L | C | 1e-05 | 5.2e-06 | 8.2e-06 | 7.800E-06 | 2.425E-06 | 5.200E-06 | 1.000E-05 |
| 20532 | 1027 | P1027K | C | 6.5e-06 | 8.6e-06 | 1.8e-05 | 1.103E-05 | 6.124E-06 | 6.500E-06 | 1.800E-05 |
| 20533 | 1027 | P1027M | C | 6.4e-06 | 9.2e-06 | 1.2e-05 | 9.200E-06 | 2.800E-06 | 6.400E-06 | 1.200E-05 |
| 20534 | 1027 | P1027F | C | 9.3e-06 | 5.5e-06 | 5.8e-06 | 6.867E-06 | 2.113E-06 | 5.500E-06 | 9.300E-06 |
| 20535 | 1027 | P1027P | C | 1.3e-05 | 1.1e-05 | 6.6e-06 | 1.020E-05 | 3.274E-06 | 6.600E-06 | 1.300E-05 |
| 20536 | 1027 | P1027S | C | 8.8e-06 | 4.8e-06 | 6.2e-06 | 6.600E-06 | 2.030E-06 | 4.800E-06 | 8.800E-06 |
| 20537 | 1027 | P1027T | C | 9.5e-06 | 6e-06 | 1.2e-05 | 9.167E-06 | 3.014E-06 | 6.000E-06 | 1.200E-05 |
| 20538 | 1027 | P1027W | C | 6.9e-06 | 1.2e-05 | 6.3e-06 | 8.400E-06 | 3.132E-06 | 6.300E-06 | 1.200E-05 |
| 20539 | 1027 | P1027Y | C | 6.3e-06 | 6.7e-06 | 6.1e-06 | 6.367E-06 | 3.055E-07 | 6.100E-06 | 6.700E-06 |
| 20540 | 1027 | P1027V | C | 4.3e-06 | 8.4e-06 | 8.7e-06 | 7.133E-06 | 2.458E-06 | 4.300E-06 | 8.700E-06 |
| 20541 | 1028 | S1028A | C | 1.2e-05 | 6.5e-06 | 8.6e-06 | 9.033E-06 | 2.775E-06 | 6.500E-06 | 1.200E-05 |
| 20542 | 1028 | S1028R | C | 7.2e-06 | 8.9e-06 | 7e-06 | 7.700E-06 | 1.044E-06 | 7.000E-06 | 8.900E-06 |
| 20543 | 1028 | S1028N | C | 5.1e-06 | 9e-06 | 8.1e-06 | 7.400E-06 | 2.042E-06 | 5.100E-06 | 9.000E-06 |
| 20544 | 1028 | S1028D | C | 9.5e-06 | 7e-06 | 9.6e-06 | 8.700E-06 | 1.473E-06 | 7.000E-06 | 9.600E-06 |
| 20545 | 1028 | S1028C | C | 9.9e-06 | 9.7e-06 | 8.9e-06 | 9.500E-06 | 5.292E-07 | 8.900E-06 | 9.900E-06 |
| 20546 | 1028 | S1028E | C | 9.9e-06 | 6.8e-06 | 4.6e-06 | 7.100E-06 | 2.663E-06 | 4.600E-06 | 9.900E-06 |
| 20547 | 1028 | S1028Q | C | 9.1e-06 | 6.9e-06 | 4.2e-06 | 6.733E-06 | 2.454E-06 | 4.200E-06 | 9.100E-06 |
| 20548 | 1028 | S1028G | C | 7.1e-06 | 7.6e-06 | 6.2e-06 | 6.967E-06 | 7.095E-07 | 6.200E-06 | 7.600E-06 |
| 20549 | 1028 | S1028H | C | 1.4e-05 | 7.4e-06 | 9.3e-06 | 1.023E-05 | 3.398E-06 | 7.400E-06 | 1.400E-05 |
| 20550 | 1028 | S1028I | C | 8.3e-06 | 4.8e-06 | 1e-05 | 7.700E-06 | 2.651E-06 | 4.800E-06 | 1.000E-05 |
| 20551 | 1028 | S1028L | C | 1e-05 | 4.9e-06 | 1.4e-05 | 9.633E-06 | 4.561E-06 | 4.900E-06 | 1.400E-05 |
| 20552 | 1028 | S1028K | C | 1.3e-05 | 8.7e-06 | 7.8e-06 | 9.833E-06 | 2.779E-06 | 7.800E-06 | 1.300E-05 |
| 20553 | 1028 | S1028M | C | 9.3e-06 | 5.2e-06 | 8.7e-06 | 7.733E-06 | 2.214E-06 | 5.200E-06 | 9.300E-06 |
| 20554 | 1028 | S1028F | C | 1e-05 | 8.1e-06 | 7.2e-06 | 8.433E-06 | 1.429E-06 | 7.200E-06 | 1.000E-05 |
| 20555 | 1028 | S1028P | C | 4.7e-06 | 1.2e-05 | 5.2e-06 | 7.300E-06 | 4.078E-06 | 4.700E-06 | 1.200E-05 |
| 20556 | 1028 | S1028S | C | 7.2e-06 | 5.3e-06 | 1.4e-05 | 8.833E-06 | 4.574E-06 | 5.300E-06 | 1.400E-05 |
| 20557 | 1028 | S1028T | C | 9.5e-06 | 6.5e-06 | 1.1e-05 | 9.000E-06 | 2.291E-06 | 6.500E-06 | 1.100E-05 |
| 20558 | 1028 | S1028W | C | 5.8e-06 | 9.4e-06 | 1.3e-05 | 9.400E-06 | 3.600E-06 | 5.800E-06 | 1.300E-05 |
| 20559 | 1028 | S1028Y | C | 8.6e-06 | 1.4e-05 | 7.6e-06 | 1.007E-05 | 3.443E-06 | 7.600E-06 | 1.400E-05 |
| 20560 | 1028 | S1028V | C | 1.1e-05 | 1e-05 | 5.1e-06 | 8.700E-06 | 3.158E-06 | 5.100E-06 | 1.100E-05 |
| 20561 | 1029 | S1029A | C | 1.1e-05 | 1.2e-05 | 8e-06 | 1.033E-05 | 2.082E-06 | 8.000E-06 | 1.200E-05 |
| 20562 | 1029 | S1029R | C | 1.1e-05 | 8.7e-06 | 7.3e-06 | 9.000E-06 | 1.868E-06 | 7.300E-06 | 1.100E-05 |
| 20563 | 1029 | S1029N | C | 8.8e-06 | 1e-05 | 5.5e-06 | 8.100E-06 | 2.330E-06 | 5.500E-06 | 1.000E-05 |
| 20564 | 1029 | S1029D | C | 7e-06 | 6.2e-06 | 4.1e-06 | 5.767E-06 | 1.498E-06 | 4.100E-06 | 7.000E-06 |
| 20565 | 1029 | S1029C | C | 6.5e-06 | 1e-05 | 3.8e-06 | 6.767E-06 | 3.109E-06 | 3.800E-06 | 1.000E-05 |
| 20566 | 1029 | S1029E | C | 4.5e-06 | 6.4e-06 | 6.9e-06 | 5.933E-06 | 1.266E-06 | 4.500E-06 | 6.900E-06 |
| 20567 | 1029 | S1029Q | C | 1.3e-05 | 1.3e-05 | 9.2e-06 | 1.173E-05 | 2.194E-06 | 9.200E-06 | 1.300E-05 |
| 20568 | 1029 | S1029G | C | 9.8e-06 | 7.5e-06 | 1.2e-05 | 9.767E-06 | 2.250E-06 | 7.500E-06 | 1.200E-05 |
| 20569 | 1029 | S1029H | C | 7e-06 | 8.9e-06 | 8.4e-06 | 8.100E-06 | 9.849E-07 | 7.000E-06 | 8.900E-06 |

| | | | | | | | | | | |
|---|---|---|---|---|---|---|---|---|---|---|
| 20570 | 1029 | S1029I | C | 8.5e-06 | 9.3e-06 | 5.2e-06 | 7.667E-06 | 2.173E-06 | 5.200E-06 | 9.300E-06 |
| 20571 | 1029 | S1029L | C | 6.6e-06 | 1.1e-05 | 8.9e-06 | 8.833E-06 | 2.201E-06 | 6.600E-06 | 1.100E-05 |
| 20572 | 1029 | S1029K | C | 5.3e-06 | 6.4e-06 | 6.6e-06 | 6.100E-06 | 7.000E-07 | 5.300E-06 | 6.600E-06 |
| 20573 | 1029 | S1029M | C | 9.7e-06 | 7.4e-06 | 6.9e-06 | 8.000E-06 | 1.493E-06 | 6.900E-06 | 9.700E-06 |
| 20574 | 1029 | S1029F | C | 7.1e-06 | 1.3e-05 | 1.3e-05 | 1.103E-05 | 3.406E-06 | 7.100E-06 | 1.300E-05 |
| 20575 | 1029 | S1029P | C | 6.1e-06 | 1e-05 | 9.5e-06 | 8.533E-06 | 2.122E-06 | 6.100E-06 | 1.000E-05 |
| 20576 | 1029 | S1029S | C | 8.3e-06 | 9.5e-06 | 8.1e-06 | 8.633E-06 | 7.572E-07 | 8.100E-06 | 9.500E-06 |
| 20577 | 1029 | S1029T | C | 6.7e-06 | 1.2e-05 | 9.2e-06 | 9.300E-06 | 2.651E-06 | 6.700E-06 | 1.200E-05 |
| 20578 | 1029 | S1029W | C | 5.7e-06 | 7.9e-06 | 7.5e-06 | 7.033E-06 | 1.172E-06 | 5.700E-06 | 7.900E-06 |
| 20579 | 1029 | S1029Y | C | 6.9e-06 | 8e-06 | 9.9e-06 | 8.267E-06 | 1.518E-06 | 6.900E-06 | 9.900E-06 |
| 20580 | 1029 | S1029V | C | 8.7e-06 | 6.2e-06 | 7.2e-06 | 7.367E-06 | 1.258E-06 | 6.200E-06 | 8.700E-06 |
| 20581 | 1030 | S1030A | C | 1.2e-05 | 1.1e-05 | 9.3e-06 | 1.077E-05 | 1.365E-06 | 9.300E-06 | 1.200E-05 |
| 20582 | 1030 | S1030R | C | 9.8e-06 | 6.6e-06 | 6.7e-06 | 7.700E-06 | 1.819E-06 | 6.600E-06 | 9.800E-06 |
| 20583 | 1030 | S1030N | C | 7.6e-06 | 9.3e-06 | 5e-06 | 7.300E-06 | 2.166E-06 | 5.000E-06 | 9.300E-06 |
| 20584 | 1030 | S1030D | C | 5.5e-06 | 6.3e-06 | 4.9e-06 | 5.567E-06 | 7.024E-07 | 4.900E-06 | 6.300E-06 |
| 20585 | 1030 | S1030C | C | 1.5e-05 | 8.7e-06 | 1e-05 | 1.123E-05 | 3.326E-06 | 8.700E-06 | 1.500E-05 |
| 20586 | 1030 | S1030E | C | 7.1e-06 | 6.8e-06 | 6.3e-06 | 6.733E-06 | 4.041E-07 | 6.300E-06 | 7.100E-06 |
| 20587 | 1030 | S1030Q | C | 6.7e-06 | 9.5e-06 | 8.1e-06 | 8.100E-06 | 1.400E-06 | 6.700E-06 | 9.500E-06 |
| 20588 | 1030 | S1030G | C | 6.5e-06 | 7.2e-06 | 7.2e-06 | 6.967E-06 | 4.041E-07 | 6.500E-06 | 7.200E-06 |
| 20589 | 1030 | S1030H | C | 6.8e-06 | 1.1e-05 | 9.4e-06 | 9.067E-06 | 2.120E-06 | 6.800E-06 | 1.100E-05 |
| 20590 | 1030 | S1030I | C | 6.2e-06 | 1.3e-05 | 8.7e-06 | 9.300E-06 | 3.439E-06 | 6.200E-06 | 1.300E-05 |
| 20591 | 1030 | S1030L | C | 7.4e-06 | 5.9e-06 | 8.9e-06 | 7.400E-06 | 1.500E-06 | 5.900E-06 | 8.900E-06 |
| 20592 | 1030 | S1030K | C | 8.3e-06 | 6.5e-06 | 5.2e-06 | 6.667E-06 | 1.557E-06 | 5.200E-06 | 8.300E-06 |
| 20593 | 1030 | S1030M | C | 1e-05 | 1.1e-05 | 8.6e-06 | 9.867E-06 | 1.206E-06 | 8.600E-06 | 1.100E-05 |
| 20594 | 1030 | S1030F | C | 6.2e-06 | 7.2e-06 | 9.6e-06 | 7.667E-06 | 1.747E-06 | 6.200E-06 | 9.600E-06 |
| 20595 | 1030 | S1030P | C | 6.8e-06 | 6.5e-06 | 6.2e-06 | 6.500E-06 | 3.000E-07 | 6.200E-06 | 6.800E-06 |
| 20596 | 1030 | S1030S | C | 6.6e-06 | 1e-05 | 3.9e-06 | 6.833E-06 | 3.057E-06 | 3.900E-06 | 1.000E-05 |
| 20597 | 1030 | S1030T | C | 6.4e-06 | 7.2e-06 | 1.6e-05 | 9.867E-06 | 5.327E-06 | 6.400E-06 | 1.600E-05 |
| 20598 | 1030 | S1030W | C | 1.5e-05 | 1e-05 | 7.1e-06 | 1.070E-05 | 3.996E-06 | 7.100E-06 | 1.500E-05 |
| 20599 | 1030 | S1030Y | C | 8.4e-06 | 9.7e-06 | 1e-05 | 9.367E-06 | 8.505E-07 | 8.400E-06 | 1.000E-05 |
| 20600 | 1030 | S1030V | C | 1.3e-05 | 8.1e-06 | 9.5e-06 | 1.020E-05 | 2.524E-06 | 8.100E-06 | 1.300E-05 |
| 20601 | 1031 | L1031A | C | 9.5e-06 | 9.3e-06 | 4.7e-06 | 7.833E-06 | 2.715E-06 | 4.700E-06 | 9.500E-06 |
| 20602 | 1031 | L1031R | C | 1.2e-05 | 9.7e-06 | 5.9e-06 | 9.200E-06 | 3.081E-06 | 5.900E-06 | 1.200E-05 |
| 20603 | 1031 | L1031N | C | 8.8e-06 | 8.9e-06 | 1.3e-05 | 1.023E-05 | 2.397E-06 | 8.800E-06 | 1.300E-05 |
| 20604 | 1031 | L1031D | C | 5.8e-06 | 6.9e-06 | 8.2e-06 | 6.967E-06 | 1.201E-06 | 5.800E-06 | 8.200E-06 |
| 20605 | 1031 | L1031C | C | 8.8e-06 | 7.5e-06 | 6.1e-06 | 7.467E-06 | 1.350E-06 | 6.100E-06 | 8.800E-06 |
| 20606 | 1031 | L1031E | C | 8.5e-06 | 4.6e-06 | 6.2e-06 | 6.433E-06 | 1.960E-06 | 4.600E-06 | 8.500E-06 |
| 20607 | 1031 | L1031Q | C | 5.5e-06 | 7.4e-06 | 6.6e-06 | 6.500E-06 | 9.539E-07 | 5.500E-06 | 7.400E-06 |
| 20608 | 1031 | L1031G | C | 4.7e-06 | 1.1e-05 | 6.8e-06 | 7.500E-06 | 3.208E-06 | 4.700E-06 | 1.100E-05 |
| 20609 | 1031 | L1031H | C | 3.9e-06 | 7.9e-06 | 6.9e-06 | 6.233E-06 | 2.082E-06 | 3.900E-06 | 7.900E-06 |
| 20610 | 1031 | L1031I | C | 1.3e-05 | 9e-06 | 1e-05 | 1.067E-05 | 2.082E-06 | 9.000E-06 | 1.300E-05 |
| 20611 | 1031 | L1031L | C | 8.8e-06 | 6.8e-06 | 5.3e-06 | 6.967E-06 | 1.756E-06 | 5.300E-06 | 8.800E-06 |
| 20612 | 1031 | L1031K | C | 6.6e-06 | 3.3e-06 | 1.2e-05 | 7.300E-06 | 4.392E-06 | 3.300E-06 | 1.200E-05 |
| 20613 | 1031 | L1031M | C | 6.9e-06 | 5e-06 | 1.3e-05 | 8.300E-06 | 4.180E-06 | 5.000E-06 | 1.300E-05 |
| 20614 | 1031 | L1031F | C | 4.3e-06 | 9.6e-06 | 8.6e-06 | 7.500E-06 | 2.816E-06 | 4.300E-06 | 9.600E-06 |
| 20615 | 1031 | L1031P | C | 7.7e-06 | 5.5e-06 | 1.1e-05 | 8.067E-06 | 2.768E-06 | 5.500E-06 | 1.100E-05 |
| 20616 | 1031 | L1031S | C | 9.5e-06 | 7.8e-06 | 9.6e-06 | 8.967E-06 | 1.012E-06 | 7.800E-06 | 9.600E-06 |
| 20617 | 1031 | L1031T | C | 8.3e-06 | 6.2e-06 | 4.7e-06 | 6.400E-06 | 1.808E-06 | 4.700E-06 | 8.300E-06 |
| 20618 | 1031 | L1031W | C | 7e-06 | 6.2e-06 | 5.6e-06 | 6.267E-06 | 7.024E-07 | 5.600E-06 | 7.000E-06 |
| 20619 | 1031 | L1031Y | C | 6.6e-06 | 7.1e-06 | 8.1e-06 | 7.267E-06 | 7.638E-07 | 6.600E-06 | 8.100E-06 |
| 20620 | 1031 | L1031V | C | 1e-05 | 6.8e-06 | 7.4e-06 | 8.067E-06 | 1.701E-06 | 6.800E-06 | 1.000E-05 |
| 20621 | 1032 | G1032A | C | 3.4e-06 | 9.1e-06 | 1.4e-05 | 8.833E-06 | 5.305E-06 | 3.400E-06 | 1.400E-05 |
| 20622 | 1032 | G1032R | C | 1.1e-05 | 7.1e-06 | 4.8e-06 | 7.633E-06 | 3.134E-06 | 4.800E-06 | 1.100E-05 |
| 20623 | 1032 | G1032N | C | 4.8e-06 | 1.3e-05 | 8.7e-06 | 8.833E-06 | 4.102E-06 | 4.800E-06 | 1.300E-05 |
| 20624 | 1032 | G1032D | C | 9.3e-06 | 9e-06 | 1.8e-05 | 1.210E-05 | 5.112E-06 | 9.000E-06 | 1.800E-05 |
| 20625 | 1032 | G1032C | C | 5.3e-06 | 9.4e-06 | 7.2e-06 | 7.300E-06 | 2.052E-06 | 5.300E-06 | 9.400E-06 |
| 20626 | 1032 | G1032E | C | 7.7e-06 | 6.6e-06 | 6.5e-06 | 6.933E-06 | 6.658E-07 | 6.500E-06 | 7.700E-06 |
| 20627 | 1032 | G1032Q | C | 6.5e-06 | 1.3e-05 | 6.7e-06 | 8.733E-06 | 3.696E-06 | 6.500E-06 | 1.300E-05 |
| 20628 | 1032 | G1032G | C | 6.9e-06 | 8.1e-06 | 5.8e-06 | 6.933E-06 | 1.150E-06 | 5.800E-06 | 8.100E-06 |
| 20629 | 1032 | G1032H | C | 6.7e-06 | 5.5e-06 | 5.3e-06 | 5.833E-06 | 7.572E-07 | 5.300E-06 | 6.700E-06 |
| 20630 | 1032 | G1032I | C | 9.4e-06 | 7.2e-06 | 6.9e-06 | 7.833E-06 | 1.365E-06 | 6.900E-06 | 9.400E-06 |
| 20631 | 1032 | G1032L | C | 6.7e-06 | 5.4e-06 | 6.1e-06 | 6.067E-06 | 6.506E-07 | 5.400E-06 | 6.700E-06 |
| 20632 | 1032 | G1032K | C | 5e-06 | 7.5e-06 | 6.3e-06 | 6.267E-06 | 1.250E-06 | 5.000E-06 | 7.500E-06 |
| 20633 | 1032 | G1032M | C | 7.8e-06 | 6.5e-06 | 6.8e-06 | 7.033E-06 | 6.807E-07 | 6.500E-06 | 7.800E-06 |
| 20634 | 1032 | G1032F | C | 9.6e-06 | 1.7e-05 | 8.7e-06 | 1.177E-05 | 4.554E-06 | 8.700E-06 | 1.700E-05 |
| 20635 | 1032 | G1032P | C | 8.9e-06 | 5e-06 | 7.8e-06 | 7.233E-06 | 2.011E-06 | 5.000E-06 | 8.900E-06 |
| 20636 | 1032 | G1032S | C | 7.3e-06 | 9.9e-06 | 6.9e-06 | 8.033E-06 | 1.629E-06 | 6.900E-06 | 9.900E-06 |
| 20637 | 1032 | G1032T | C | 6.8e-06 | 7.6e-06 | 1.1e-05 | 8.467E-06 | 2.230E-06 | 6.800E-06 | 1.100E-05 |
| 20638 | 1032 | G1032W | C | 6.5e-06 | 4.3e-06 | 4.8e-06 | 5.200E-06 | 1.153E-06 | 4.300E-06 | 6.500E-06 |
| 20639 | 1032 | G1032Y | C | 8.3e-06 | 1.1e-05 | 1.1e-05 | 1.010E-05 | 1.559E-06 | 8.300E-06 | 1.100E-05 |
| 20640 | 1032 | G1032V | C | 6.4e-06 | 6.5e-06 | 8.2e-06 | 7.033E-06 | 1.012E-06 | 6.400E-06 | 8.200E-06 |
| 20641 | 1033 | T1033A | C | 9.4e-06 | 6.3e-06 | 1.1e-05 | 8.900E-06 | 2.390E-06 | 6.300E-06 | 1.100E-05 |
| 20642 | 1033 | T1033R | C | 5.7e-06 | 3.8e-06 | 5e-06 | 4.833E-06 | 9.609E-07 | 3.800E-06 | 5.700E-06 |
| 20643 | 1033 | T1033N | C | 6.4e-06 | 5.7e-06 | 5.9e-06 | 6.000E-06 | 3.606E-07 | 5.700E-06 | 6.400E-06 |
| 20644 | 1033 | T1033D | C | 1.2e-05 | 7.4e-06 | 1.1e-05 | 1.013E-05 | 2.419E-06 | 7.400E-06 | 1.200E-05 |
| 20645 | 1033 | T1033C | C | 8.2e-06 | 6.7e-06 | 7.9e-06 | 7.600E-06 | 7.937E-07 | 6.700E-06 | 8.200E-06 |
| 20646 | 1033 | T1033E | C | 8.4e-06 | 1e-05 | 5.7e-06 | 8.033E-06 | 2.173E-06 | 5.700E-06 | 1.000E-05 |
| 20647 | 1033 | T1033Q | C | 7.9e-06 | 8e-06 | 7.9e-06 | 7.933E-06 | 5.774E-08 | 7.900E-06 | 8.000E-06 |
| 20648 | 1033 | T1033G | C | 9.6e-06 | 1.5e-05 | 9.7e-06 | 1.143E-05 | 3.089E-06 | 9.600E-06 | 1.500E-05 |
| 20649 | 1033 | T1033H | C | 1.3e-05 | 7.4e-06 | 1.3e-05 | 1.113E-05 | 3.233E-06 | 7.400E-06 | 1.300E-05 |
| 20650 | 1033 | T1033I | C | 1.2e-05 | 7.6e-06 | 1.1e-05 | 1.020E-05 | 2.307E-06 | 7.600E-06 | 1.200E-05 |
| 20651 | 1033 | T1033L | C | 1.3e-05 | 1.3e-05 | 7.3e-06 | 1.110E-05 | 3.291E-06 | 7.300E-06 | 1.300E-05 |
| 20652 | 1033 | T1033K | C | 6.5e-06 | 8.2e-06 | 4.8e-06 | 6.500E-06 | 1.700E-06 | 4.800E-06 | 8.200E-06 |
| 20653 | 1033 | T1033M | C | 7e-06 | 1e-05 | 6.8e-06 | 7.933E-06 | 1.793E-06 | 6.800E-06 | 1.000E-05 |
| 20654 | 1033 | T1033F | C | 8e-06 | 6.9e-06 | 5.3e-06 | 6.733E-06 | 1.358E-06 | 5.300E-06 | 8.000E-06 |

| | | | | | | | | | | |
|---|---|---|---|---|---|---|---|---|---|---|
| 20655 | 1033 | T1033P | C | 7.7e-06 | 1.6e-05 | 7.1e-06 | 1.027E-05 | 4.974E-06 | 7.100E-06 | 1.600E-05 |
| 20656 | 1033 | T1033S | C | 6.7e-06 | 8.4e-06 | 6.3e-06 | 7.133E-06 | 1.115E-06 | 6.300E-06 | 8.400E-06 |
| 20657 | 1033 | T1033T | C | 7.2e-06 | 5.5e-06 | 5.6e-06 | 6.100E-06 | 9.539E-07 | 5.500E-06 | 7.200E-06 |
| 20658 | 1033 | T1033W | C | 9.3e-06 | 1e-05 | 5.3e-06 | 8.200E-06 | 2.536E-06 | 5.300E-06 | 1.000E-05 |
| 20659 | 1033 | T1033Y | C | 9.6e-06 | 5.4e-06 | 5.5e-06 | 6.833E-06 | 2.397E-06 | 5.400E-06 | 9.600E-06 |
| 20660 | 1033 | T1033V | C | 7.4e-06 | 6.6e-06 | 1.3e-05 | 9.000E-06 | 3.487E-06 | 6.600E-06 | 1.300E-05 |
| 20661 | 1034 | Q1034A | C | 1.1e-05 | 9.3e-06 | 8.3e-06 | 9.533E-06 | 1.365E-06 | 8.300E-06 | 1.100E-05 |
| 20662 | 1034 | Q1034R | C | 1.2e-05 | 9.4e-06 | 7.8e-06 | 9.733E-06 | 2.120E-06 | 7.800E-06 | 1.200E-05 |
| 20663 | 1034 | Q1034N | C | 1.2e-05 | 4.2e-06 | 7.1e-06 | 7.767E-06 | 3.943E-06 | 4.200E-06 | 1.200E-05 |
| 20664 | 1034 | Q1034D | C | 6.9e-06 | 7.7e-06 | 1.8e-05 | 1.087E-05 | 6.191E-06 | 6.900E-06 | 1.800E-05 |
| 20665 | 1034 | Q1034C | C | 6.6e-06 | 1e-05 | 8.1e-06 | 8.233E-06 | 1.704E-06 | 6.600E-06 | 1.000E-05 |
| 20666 | 1034 | Q1034E | C | 7e-06 | 1.4e-05 | 9e-06 | 1.000E-05 | 3.606E-06 | 7.000E-06 | 1.400E-05 |
| 20667 | 1034 | Q1034Q | C | 7.9e-06 | 5.3e-06 | 6.7e-06 | 6.633E-06 | 1.301E-06 | 5.300E-06 | 7.900E-06 |
| 20668 | 1034 | Q1034G | C | 1.1e-05 | 1.1e-05 | 1.3e-05 | 1.167E-05 | 1.155E-06 | 1.100E-05 | 1.300E-05 |
| 20669 | 1034 | Q1034H | C | 6.7e-06 | 6.7e-06 | 8.4e-06 | 7.267E-06 | 9.815E-07 | 6.700E-06 | 8.400E-06 |
| 20670 | 1034 | Q1034I | C | 1e-05 | 1.5e-05 | 9.5e-06 | 1.150E-05 | 3.041E-06 | 9.500E-06 | 1.500E-05 |
| 20671 | 1034 | Q1034L | C | 5.4e-06 | 8e-06 | 1.3e-05 | 8.800E-06 | 3.863E-06 | 5.400E-06 | 1.300E-05 |
| 20672 | 1034 | Q1034K | C | 9.3e-06 | 7.3e-06 | 5.3e-06 | 7.300E-06 | 2.000E-06 | 5.300E-06 | 9.300E-06 |
| 20673 | 1034 | Q1034M | C | 7.1e-06 | 1.2e-05 | 7.1e-06 | 8.733E-06 | 2.829E-06 | 7.100E-06 | 1.200E-05 |
| 20674 | 1034 | Q1034F | C | 9.7e-06 | 9.9e-06 | 8.6e-06 | 9.400E-06 | 7.000E-07 | 8.600E-06 | 9.900E-06 |
| 20675 | 1034 | Q1034P | C | 5.2e-06 | 6.1e-06 | 9.8e-06 | 7.033E-06 | 2.438E-06 | 5.200E-06 | 9.800E-06 |
| 20676 | 1034 | Q1034S | C | 1e-05 | 7e-06 | 1e-05 | 9.000E-06 | 1.732E-06 | 7.000E-06 | 1.000E-05 |
| 20677 | 1034 | Q1034T | C | 4.1e-06 | 9.7e-06 | 1e-05 | 7.933E-06 | 3.323E-06 | 4.100E-06 | 1.000E-05 |
| 20678 | 1034 | Q1034W | C | 8.8e-06 | 1.1e-05 | 6.9e-06 | 8.900E-06 | 2.052E-06 | 6.900E-06 | 1.100E-05 |
| 20679 | 1034 | Q1034Y | C | 1.6e-05 | 7.3e-06 | 6.1e-06 | 9.800E-06 | 5.403E-06 | 6.100E-06 | 1.600E-05 |
| 20680 | 1034 | Q1034V | C | 9.2e-06 | 4.9e-06 | 7.7e-06 | 7.267E-06 | 2.183E-06 | 4.900E-06 | 9.200E-06 |
| 20681 | 1035 | T1035A | C | 6.8e-06 | 1.6e-05 | 8.3e-06 | 1.037E-05 | 4.936E-06 | 6.800E-06 | 1.600E-05 |
| 20682 | 1035 | T1035R | C | 9.4e-06 | 1.2e-05 | 4.9e-06 | 8.767E-06 | 3.592E-06 | 4.900E-06 | 1.200E-05 |
| 20683 | 1035 | T1035N | C | 5.9e-06 | 7.6e-06 | 1.3e-05 | 8.833E-06 | 3.707E-06 | 5.900E-06 | 1.300E-05 |
| 20684 | 1035 | T1035D | C | 7.9e-06 | 9.7e-06 | 7.9e-06 | 8.500E-06 | 1.039E-06 | 7.900E-06 | 9.700E-06 |
| 20685 | 1035 | T1035C | C | 1e-05 | 9.8e-06 | 7.3e-06 | 9.033E-06 | 1.504E-06 | 7.300E-06 | 1.000E-05 |
| 20686 | 1035 | T1035E | C | 8.7e-06 | 1.4e-05 | 5.5e-06 | 9.400E-06 | 4.293E-06 | 5.500E-06 | 1.400E-05 |
| 20687 | 1035 | T1035Q | C | 6.5e-06 | 1.4e-05 | 9.9e-06 | 1.013E-05 | 3.755E-06 | 6.500E-06 | 1.400E-05 |
| 20688 | 1035 | T1035G | C | 4.9e-06 | 9.8e-06 | 5.7e-06 | 6.800E-06 | 2.629E-06 | 4.900E-06 | 9.800E-06 |
| 20689 | 1035 | T1035H | C | 9.9e-06 | 6.4e-06 | 7.1e-06 | 7.800E-06 | 1.852E-06 | 6.400E-06 | 9.900E-06 |
| 20690 | 1035 | T1035I | C | 1.1e-05 | 6.8e-06 | 8.9e-06 | 8.900E-06 | 2.100E-06 | 6.800E-06 | 1.100E-05 |
| 20691 | 1035 | T1035L | C | 5.5e-06 | 6.1e-06 | 7.4e-06 | 6.333E-06 | 9.713E-07 | 5.500E-06 | 7.400E-06 |
| 20692 | 1035 | T1035K | C | 9.6e-06 | 1e-05 | 4.8e-06 | 8.133E-06 | 2.894E-06 | 4.800E-06 | 1.000E-05 |
| 20693 | 1035 | T1035M | C | 5.3e-06 | 9.5e-06 | 7.5e-06 | 7.433E-06 | 2.101E-06 | 5.300E-06 | 9.500E-06 |
| 20694 | 1035 | T1035F | C | 5.6e-06 | 7.8e-06 | 5.2e-06 | 6.200E-06 | 1.400E-06 | 5.200E-06 | 7.800E-06 |
| 20695 | 1035 | T1035P | C | 9.9e-06 | 9.3e-06 | 8.3e-06 | 9.167E-06 | 8.083E-07 | 8.300E-06 | 9.900E-06 |
| 20696 | 1035 | T1035S | C | 5.6e-06 | 1.3e-05 | 6.4e-06 | 8.333E-06 | 4.061E-06 | 5.600E-06 | 1.300E-05 |
| 20697 | 1035 | T1035T | C | 1.6e-05 | 1.2e-05 | 5.9e-06 | 1.130E-05 | 5.086E-06 | 5.900E-06 | 1.600E-05 |
| 20698 | 1035 | T1035W | C | 8.9e-06 | 6.6e-06 | 7.3e-06 | 7.600E-06 | 1.179E-06 | 6.600E-06 | 8.900E-06 |
| 20699 | 1035 | T1035Y | C | 5e-06 | 6.4e-06 | 1e-05 | 7.133E-06 | 2.579E-06 | 5.000E-06 | 1.000E-05 |
| 20700 | 1035 | T1035V | C | 4.3e-06 | 6.2e-06 | 6.2e-06 | 5.567E-06 | 1.097E-06 | 4.300E-06 | 6.200E-06 |
| 20701 | 1036 | Y1036A | C | 6.8e-06 | 1.7e-05 | 6.7e-06 | 1.017E-05 | 5.918E-06 | 6.700E-06 | 1.700E-05 |
| 20702 | 1036 | Y1036R | C | 1.1e-05 | 5.7e-06 | 6.3e-06 | 7.667E-06 | 2.902E-06 | 5.700E-06 | 1.100E-05 |
| 20703 | 1036 | Y1036N | C | 8e-06 | 4.8e-06 | 9.6e-06 | 7.467E-06 | 2.444E-06 | 4.800E-06 | 9.600E-06 |
| 20704 | 1036 | Y1036D | C | 7.4e-06 | 9.4e-06 | 8.1e-06 | 8.300E-06 | 1.015E-06 | 7.400E-06 | 9.400E-06 |
| 20705 | 1036 | Y1036C | C | 9.5e-06 | 5.9e-06 | 7.1e-06 | 7.500E-06 | 1.833E-06 | 5.900E-06 | 9.500E-06 |
| 20706 | 1036 | Y1036E | C | 9.5e-06 | 9.8e-06 | 1e-05 | 9.767E-06 | 2.517E-07 | 9.500E-06 | 1.000E-05 |
| 20707 | 1036 | Y1036Q | C | 1.5e-05 | 9.8e-06 | 1e-05 | 1.160E-05 | 2.946E-06 | 9.800E-06 | 1.500E-05 |
| 20708 | 1036 | Y1036G | C | 1e-05 | 6.6e-06 | 7.5e-06 | 8.033E-06 | 1.762E-06 | 6.600E-06 | 1.000E-05 |
| 20709 | 1036 | Y1036H | C | 1.1e-05 | 6.9e-06 | 9.9e-06 | 9.267E-06 | 2.122E-06 | 6.900E-06 | 1.100E-05 |
| 20710 | 1036 | Y1036I | C | 6.3e-06 | 9.8e-06 | 8.8e-06 | 8.300E-06 | 1.803E-06 | 6.300E-06 | 9.800E-06 |
| 20711 | 1036 | Y1036L | C | 7.2e-06 | 6.5e-06 | 5.2e-06 | 6.300E-06 | 1.015E-06 | 5.200E-06 | 7.200E-06 |
| 20712 | 1036 | Y1036K | C | 7.2e-06 | 1.1e-05 | 4.5e-06 | 7.567E-06 | 3.265E-06 | 4.500E-06 | 1.100E-05 |
| 20713 | 1036 | Y1036M | C | 1.1e-05 | 5.9e-06 | 7.4e-06 | 8.100E-06 | 2.621E-06 | 5.900E-06 | 1.100E-05 |
| 20714 | 1036 | Y1036F | C | 6.5e-06 | 6.6e-06 | 6.8e-06 | 6.633E-06 | 1.528E-07 | 6.500E-06 | 6.800E-06 |
| 20715 | 1036 | Y1036P | C | 8e-06 | 6e-06 | 7.2e-06 | 7.067E-06 | 1.007E-06 | 6.000E-06 | 8.000E-06 |
| 20716 | 1036 | Y1036S | C | 7.3e-06 | 1e-05 | 4.4e-06 | 7.233E-06 | 2.801E-06 | 4.400E-06 | 1.000E-05 |
| 20717 | 1036 | Y1036T | C | 9.4e-06 | 1e-05 | 5.9e-06 | 8.433E-06 | 2.214E-06 | 5.900E-06 | 1.000E-05 |
| 20718 | 1036 | Y1036W | C | 8.2e-06 | 8.1e-06 | 9.4e-06 | 8.567E-06 | 7.234E-07 | 8.100E-06 | 9.400E-06 |
| 20719 | 1036 | Y1036Y | C | 1.1e-05 | 5.2e-06 | 1.1e-05 | 9.067E-06 | 3.349E-06 | 5.200E-06 | 1.100E-05 |
| 20720 | 1036 | Y1036V | C | 7.7e-06 | 1.3e-05 | 5.2e-06 | 8.633E-06 | 3.983E-06 | 5.200E-06 | 1.300E-05 |
| 20721 | 1037 | I1037A | C | 6.4e-06 | 7.5e-06 | 6.3e-06 | 6.733E-06 | 6.658E-07 | 6.300E-06 | 7.500E-06 |
| 20722 | 1037 | I1037R | C | 6e-06 | 8.8e-06 | 4.8e-06 | 6.533E-06 | 2.053E-06 | 4.800E-06 | 8.800E-06 |
| 20723 | 1037 | I1037N | C | 9.4e-06 | 5.3e-06 | 8.7e-06 | 7.800E-06 | 2.193E-06 | 5.300E-06 | 9.400E-06 |
| 20724 | 1037 | I1037D | C | 8.5e-06 | 8.4e-06 | 8.3e-06 | 8.400E-06 | 1.000E-07 | 8.300E-06 | 8.500E-06 |
| 20725 | 1037 | I1037C | C | 5.1e-06 | 8.2e-06 | 1.8e-05 | 1.043E-05 | 6.734E-06 | 5.100E-06 | 1.800E-05 |
| 20726 | 1037 | I1037E | C | 1.1e-05 | 5.7e-06 | 5.1e-06 | 7.267E-06 | 3.247E-06 | 5.100E-06 | 1.100E-05 |
| 20727 | 1037 | I1037Q | C | 5.1e-06 | 1.3e-05 | 8.8e-06 | 8.967E-06 | 3.953E-06 | 5.100E-06 | 1.300E-05 |
| 20728 | 1037 | I1037G | C | 6.3e-06 | 9.8e-06 | 8.8e-06 | 8.300E-06 | 1.803E-06 | 6.300E-06 | 9.800E-06 |
| 20729 | 1037 | I1037H | C | 1e-05 | 7e-06 | 6.3e-06 | 7.767E-06 | 1.966E-06 | 6.300E-06 | 1.000E-05 |
| 20730 | 1037 | I1037I | C | 9.7e-06 | 6.9e-06 | 9.5e-06 | 8.700E-06 | 1.562E-06 | 6.900E-06 | 9.700E-06 |
| 20731 | 1037 | I1037L | C | 5.7e-06 | 6.7e-06 | 1.3e-05 | 8.467E-06 | 3.958E-06 | 5.700E-06 | 1.300E-05 |
| 20732 | 1037 | I1037K | C | 1.2e-05 | 6.3e-06 | 5e-06 | 7.767E-06 | 3.723E-06 | 5.000E-06 | 1.200E-05 |
| 20733 | 1037 | I1037M | C | 7.5e-06 | 3.3e-06 | 8.2e-06 | 6.333E-06 | 2.650E-06 | 3.300E-06 | 8.200E-06 |
| 20734 | 1037 | I1037F | C | 1.6e-05 | 1e-05 | 1.1e-05 | 1.233E-05 | 3.215E-06 | 1.000E-05 | 1.600E-05 |
| 20735 | 1037 | I1037P | C | 1.2e-05 | 8.4e-06 | 4.4e-06 | 8.267E-06 | 3.802E-06 | 4.400E-06 | 1.200E-05 |
| 20736 | 1037 | I1037S | C | 8.3e-06 | 3.5e-06 | 9.2e-06 | 7.000E-06 | 3.064E-06 | 3.500E-06 | 9.200E-06 |
| 20737 | 1037 | I1037T | C | 6.9e-06 | 7.6e-06 | 7.1e-06 | 7.200E-06 | 3.606E-07 | 6.900E-06 | 7.600E-06 |
| 20738 | 1037 | I1037W | C | 8.7e-06 | 6.3e-06 | 5.8e-06 | 6.933E-06 | 1.550E-06 | 5.800E-06 | 8.700E-06 |
| 20739 | 1037 | I1037Y | C | 9e-06 | 9.5e-06 | 6.6e-06 | 8.367E-06 | 1.550E-06 | 6.600E-06 | 9.500E-06 |

| | | | | | | | | | | |
|---|---|---|---|---|---|---|---|---|---|---|
| 20740 | 1037 | I1037V | C | 7.3e-06 | 7.3e-06 | 1.3e-05 | 9.200E-06 | 3.291E-06 | 7.300E-06 | 1.300E-05 |
| 20741 | 1038 | C1038A | C | 4.4e-06 | 7e-06 | 8.9e-06 | 6.767E-06 | 2.259E-06 | 4.400E-06 | 8.900E-06 |
| 20742 | 1038 | C1038R | C | 3.5e-06 | 8.8e-06 | 1e-05 | 7.433E-06 | 3.459E-06 | 3.500E-06 | 1.000E-05 |
| 20743 | 1038 | C1038N | C | 4.7e-06 | 1.4e-05 | 7.1e-06 | 8.600E-06 | 4.828E-06 | 4.700E-06 | 1.400E-05 |
| 20744 | 1038 | C1038D | C | 6.4e-06 | 8.1e-06 | 4.4e-06 | 6.300E-06 | 1.852E-06 | 4.400E-06 | 8.100E-06 |
| 20745 | 1038 | C1038C | C | 6.2e-06 | 6.5e-06 | 9.2e-06 | 7.300E-06 | 1.652E-06 | 6.200E-06 | 9.200E-06 |
| 20746 | 1038 | C1038E | C | 7.2e-06 | 1.3e-05 | 4.7e-06 | 8.300E-06 | 4.258E-06 | 4.700E-06 | 1.300E-05 |
| 20747 | 1038 | C1038Q | C | 9.9e-06 | 8.9e-06 | 6.9e-06 | 8.567E-06 | 1.528E-06 | 6.900E-06 | 9.900E-06 |
| 20748 | 1038 | C1038G | C | 9.8e-06 | 7.2e-06 | 5.3e-06 | 7.433E-06 | 2.259E-06 | 5.300E-06 | 9.800E-06 |
| 20749 | 1038 | C1038H | C | 7.1e-06 | 9.7e-06 | 7.2e-06 | 8.000E-06 | 1.473E-06 | 7.100E-06 | 9.700E-06 |
| 20750 | 1038 | C1038I | C | 6.4e-06 | 1e-05 | 1.3e-05 | 9.800E-06 | 3.305E-06 | 6.400E-06 | 1.300E-05 |
| 20751 | 1038 | C1038L | C | 1e-05 | 5.3e-06 | 1.2e-05 | 9.100E-06 | 3.439E-06 | 5.300E-06 | 1.200E-05 |
| 20752 | 1038 | C1038K | C | 5.4e-06 | 7.3e-06 | 8.5e-06 | 7.067E-06 | 1.563E-06 | 5.400E-06 | 8.500E-06 |
| 20753 | 1038 | C1038M | C | 7.4e-06 | 1.1e-05 | 8.7e-06 | 9.033E-06 | 1.823E-06 | 7.400E-06 | 1.100E-05 |
| 20754 | 1038 | C1038F | C | 7.1e-06 | 7e-06 | 6.2e-06 | 6.767E-06 | 4.933E-07 | 6.200E-06 | 7.100E-06 |
| 20755 | 1038 | C1038P | C | 6.4e-06 | 1.2e-05 | 1.3e-05 | 1.047E-05 | 3.557E-06 | 6.400E-06 | 1.300E-05 |
| 20756 | 1038 | C1038S | C | 7.1e-06 | 1.3e-05 | 6.7e-06 | 8.933E-06 | 3.528E-06 | 6.700E-06 | 1.300E-05 |
| 20757 | 1038 | C1038T | C | 6.8e-06 | 9.6e-06 | 5.6e-06 | 7.333E-06 | 2.053E-06 | 5.600E-06 | 9.600E-06 |
| 20758 | 1038 | C1038W | C | 9.9e-06 | 8.4e-06 | 5.9e-06 | 8.067E-06 | 2.021E-06 | 5.900E-06 | 9.900E-06 |
| 20759 | 1038 | C1038Y | C | 6.1e-06 | 5.7e-06 | 1.1e-05 | 7.600E-06 | 2.951E-06 | 5.700E-06 | 1.100E-05 |
| 20760 | 1038 | C1038V | C | 6.9e-06 | 6.6e-06 | 8.3e-06 | 7.267E-06 | 9.074E-07 | 6.600E-06 | 8.300E-06 |
| 20761 | 1039 | N1039A | C | 8.2e-06 | 5.5e-06 | 5.1e-06 | 6.267E-06 | 1.686E-06 | 5.100E-06 | 8.200E-06 |
| 20762 | 1039 | N1039R | C | 7.2e-06 | 7.2e-06 | 6.8e-06 | 7.067E-06 | 2.309E-07 | 6.800E-06 | 7.200E-06 |
| 20763 | 1039 | N1039N | C | 5.2e-06 | 9.6e-06 | 6.8e-06 | 7.200E-06 | 2.227E-06 | 5.200E-06 | 9.600E-06 |
| 20764 | 1039 | N1039D | C | 7.2e-06 | 1.2e-05 | 9.1e-06 | 9.433E-06 | 2.417E-06 | 7.200E-06 | 1.200E-05 |
| 20765 | 1039 | N1039C | C | 8.7e-06 | 8.5e-06 | 9.2e-06 | 8.800E-06 | 3.606E-07 | 8.500E-06 | 9.200E-06 |
| 20766 | 1039 | N1039E | C | 6.4e-06 | 7.4e-06 | 7e-06 | 6.933E-06 | 5.033E-07 | 6.400E-06 | 7.400E-06 |
| 20767 | 1039 | N1039Q | C | 7.1e-06 | 1.2e-05 | 8.9e-06 | 9.333E-06 | 2.479E-06 | 7.100E-06 | 1.200E-05 |
| 20768 | 1039 | N1039G | C | 7.3e-06 | 5.4e-06 | 6.7e-06 | 6.467E-06 | 9.713E-07 | 5.400E-06 | 7.300E-06 |
| 20769 | 1039 | N1039H | C | 1e-05 | 5.7e-06 | 9.4e-06 | 8.367E-06 | 2.329E-06 | 5.700E-06 | 1.000E-05 |
| 20770 | 1039 | N1039I | C | 8.1e-06 | 8.6e-06 | 1.3e-05 | 9.900E-06 | 2.696E-06 | 8.100E-06 | 1.300E-05 |
| 20771 | 1039 | N1039L | C | 1.2e-05 | 7.5e-06 | 8.2e-06 | 9.233E-06 | 2.421E-06 | 7.500E-06 | 1.200E-05 |
| 20772 | 1039 | N1039K | C | 9e-06 | 3e-06 | 1e-05 | 7.333E-06 | 3.786E-06 | 3.000E-06 | 1.000E-05 |
| 20773 | 1039 | N1039M | C | 1.1e-05 | 8.9e-06 | 9.8e-06 | 9.900E-06 | 1.054E-06 | 8.900E-06 | 1.100E-05 |
| 20774 | 1039 | N1039F | C | 7e-06 | 1.2e-05 | 7.5e-06 | 8.833E-06 | 2.754E-06 | 7.000E-06 | 1.200E-05 |
| 20775 | 1039 | N1039P | C | 9.5e-06 | 8.3e-06 | 4.4e-06 | 7.400E-06 | 2.666E-06 | 4.400E-06 | 9.500E-06 |
| 20776 | 1039 | N1039S | C | 9e-06 | 4.6e-06 | 4.9e-06 | 6.167E-06 | 2.458E-06 | 4.600E-06 | 9.000E-06 |
| 20777 | 1039 | N1039T | C | 8.8e-06 | 7.3e-06 | 8.3e-06 | 8.133E-06 | 7.638E-07 | 7.300E-06 | 8.800E-06 |
| 20778 | 1039 | N1039W | C | 5.9e-06 | 6.1e-06 | 6.8e-06 | 6.267E-06 | 4.726E-07 | 5.900E-06 | 6.800E-06 |
| 20779 | 1039 | N1039Y | C | 9.4e-06 | 1.1e-05 | 5.8e-06 | 8.733E-06 | 2.663E-06 | 5.800E-06 | 1.100E-05 |
| 20780 | 1039 | N1039V | C | 8.3e-06 | 4.6e-06 | 7.8e-06 | 6.900E-06 | 2.007E-06 | 4.600E-06 | 8.300E-06 |
| 20781 | 1040 | V1040A | C | 8.4e-06 | 5.7e-06 | 1e-05 | 8.033E-06 | 2.173E-06 | 5.700E-06 | 1.000E-05 |
| 20782 | 1040 | V1040R | C | 5.4e-06 | 8.5e-06 | 7e-06 | 6.967E-06 | 1.550E-06 | 5.400E-06 | 8.500E-06 |
| 20783 | 1040 | V1040N | C | 5.7e-06 | 7.5e-06 | 6e-06 | 6.400E-06 | 9.644E-07 | 5.700E-06 | 7.500E-06 |
| 20784 | 1040 | V1040D | C | 9e-06 | 1.1e-05 | 4.3e-06 | 8.100E-06 | 3.439E-06 | 4.300E-06 | 1.100E-05 |
| 20785 | 1040 | V1040C | C | 3.4e-06 | 9.3e-06 | 6.4e-06 | 6.367E-06 | 2.950E-06 | 3.400E-06 | 9.300E-06 |
| 20786 | 1040 | V1040E | C | 1e-05 | 8.9e-06 | 6.5e-06 | 8.467E-06 | 1.790E-06 | 6.500E-06 | 1.000E-05 |
| 20787 | 1040 | V1040Q | C | 3.7e-06 | 3.5e-06 | 5e-06 | 4.067E-06 | 8.145E-07 | 3.500E-06 | 5.000E-06 |
| 20788 | 1040 | V1040G | C | 1.6e-05 | 6.6e-06 | 6.3e-06 | 9.633E-06 | 5.516E-06 | 6.300E-06 | 1.600E-05 |
| 20789 | 1040 | V1040H | C | 6.3e-06 | 7.9e-06 | 8.1e-06 | 7.433E-06 | 9.866E-07 | 6.300E-06 | 8.100E-06 |
| 20790 | 1040 | V1040I | C | 7.2e-06 | 1.1e-05 | 7.3e-06 | 8.500E-06 | 2.166E-06 | 7.200E-06 | 1.100E-05 |
| 20791 | 1040 | V1040L | C | 9.3e-06 | 7.7e-06 | 6.3e-06 | 7.767E-06 | 1.501E-06 | 6.300E-06 | 9.300E-06 |
| 20792 | 1040 | V1040K | C | 1.1e-05 | 1.1e-05 | 9.7e-06 | 1.057E-05 | 7.506E-07 | 9.700E-06 | 1.100E-05 |
| 20793 | 1040 | V1040M | C | 9.6e-06 | 9.4e-06 | 8.2e-06 | 9.067E-06 | 7.572E-07 | 8.200E-06 | 9.600E-06 |
| 20794 | 1040 | V1040F | C | 9.8e-06 | 1e-05 | 9.3e-06 | 9.700E-06 | 3.606E-07 | 9.300E-06 | 1.000E-05 |
| 20795 | 1040 | V1040P | C | 8.4e-06 | 7.4e-06 | 9e-06 | 8.267E-06 | 8.083E-07 | 7.400E-06 | 9.000E-06 |
| 20796 | 1040 | V1040S | C | 7.5e-06 | 8.3e-06 | 5.8e-06 | 7.200E-06 | 1.277E-06 | 5.800E-06 | 8.300E-06 |
| 20797 | 1040 | V1040T | C | 1e-05 | 6.4e-06 | 8.2e-06 | 8.200E-06 | 1.800E-06 | 6.400E-06 | 1.000E-05 |
| 20798 | 1040 | V1040W | C | 8.3e-06 | 6.6e-06 | 5.9e-06 | 6.933E-06 | 1.234E-06 | 5.900E-06 | 8.300E-06 |
| 20799 | 1040 | V1040Y | C | 7.1e-06 | 1.1e-05 | 9.2e-06 | 9.100E-06 | 1.952E-06 | 7.100E-06 | 1.100E-05 |
| 20800 | 1040 | V1040V | C | 9.7e-06 | 1.1e-05 | 6.9e-06 | 9.200E-06 | 2.095E-06 | 6.900E-06 | 1.100E-05 |
| 20801 | 1041 | N1041A | C | 3.6e-06 | 5.8e-06 | 1e-05 | 6.467E-06 | 3.252E-06 | 3.600E-06 | 1.000E-05 |
| 20802 | 1041 | N1041R | C | 1.7e-05 | 1e-05 | 1e-05 | 1.233E-05 | 4.041E-06 | 1.000E-05 | 1.700E-05 |
| 20803 | 1041 | N1041N | C | 7.1e-06 | 8.6e-06 | 9.8e-06 | 8.500E-06 | 1.353E-06 | 7.100E-06 | 9.800E-06 |
| 20804 | 1041 | N1041D | C | 9.2e-06 | 1.1e-05 | 8.2e-06 | 9.467E-06 | 1.419E-06 | 8.200E-06 | 1.100E-05 |
| 20805 | 1041 | N1041C | C | 7.2e-06 | 4.5e-06 | 5.5e-06 | 5.733E-06 | 1.365E-06 | 4.500E-06 | 7.200E-06 |
| 20806 | 1041 | N1041E | C | 6.9e-06 | 8.1e-06 | 9e-06 | 8.000E-06 | 1.054E-06 | 6.900E-06 | 9.000E-06 |
| 20807 | 1041 | N1041Q | C | 6.3e-06 | 5e-06 | 9.5e-06 | 6.933E-06 | 2.316E-06 | 5.000E-06 | 9.500E-06 |
| 20808 | 1041 | N1041G | C | 6.8e-06 | 6.5e-06 | 1.9e-05 | 1.077E-05 | 7.132E-06 | 6.500E-06 | 1.900E-05 |
| 20809 | 1041 | N1041H | C | 7.5e-06 | 8.5e-06 | 5.8e-06 | 7.267E-06 | 1.365E-06 | 5.800E-06 | 8.500E-06 |
| 20810 | 1041 | N1041I | C | 8e-06 | 1e-05 | 1e-05 | 9.333E-06 | 1.155E-06 | 8.000E-06 | 1.000E-05 |
| 20811 | 1041 | N1041L | C | 4.7e-06 | 7.6e-06 | 1e-05 | 7.433E-06 | 2.654E-06 | 4.700E-06 | 1.000E-05 |
| 20812 | 1041 | N1041K | C | 9.3e-06 | 6.8e-06 | 8.6e-06 | 8.233E-06 | 1.290E-06 | 6.800E-06 | 9.300E-06 |
| 20813 | 1041 | N1041M | C | 1.5e-05 | 5.7e-06 | 6.8e-06 | 9.167E-06 | 5.082E-06 | 5.700E-06 | 1.500E-05 |
| 20814 | 1041 | N1041F | C | 8e-06 | 8e-06 | 7.6e-06 | 7.867E-06 | 2.309E-07 | 7.600E-06 | 8.000E-06 |
| 20815 | 1041 | N1041P | C | 6.1e-06 | 8.3e-06 | 7.2e-06 | 7.200E-06 | 1.100E-06 | 6.100E-06 | 8.300E-06 |
| 20816 | 1041 | N1041S | C | 5.2e-06 | 7.3e-06 | 3.8e-06 | 5.433E-06 | 1.762E-06 | 3.800E-06 | 7.300E-06 |
| 20817 | 1041 | N1041T | C | 1e-05 | 6.8e-06 | 7.4e-06 | 8.067E-06 | 1.701E-06 | 6.800E-06 | 1.000E-05 |
| 20818 | 1041 | N1041W | C | 9.5e-06 | 8.3e-06 | 6.2e-06 | 8.000E-06 | 1.670E-06 | 6.200E-06 | 9.500E-06 |
| 20819 | 1041 | N1041Y | C | 8.2e-06 | 5.8e-06 | 3.3e-06 | 5.767E-06 | 2.450E-06 | 3.300E-06 | 8.200E-06 |
| 20820 | 1041 | N1041V | C | 9.5e-06 | 8.4e-06 | 4.8e-06 | 7.567E-06 | 2.458E-06 | 4.800E-06 | 9.500E-06 |
| 20821 | 1042 | H1042A | C | 6e-06 | 8.8e-06 | 8.1e-06 | 7.633E-06 | 1.457E-06 | 6.000E-06 | 8.800E-06 |
| 20822 | 1042 | H1042R | C | 1.3e-05 | 1.2e-05 | 9.5e-06 | 1.150E-05 | 1.803E-06 | 9.500E-06 | 1.300E-05 |
| 20823 | 1042 | H1042N | C | 9.2e-06 | 1.4e-05 | 1e-05 | 1.107E-05 | 2.572E-06 | 9.200E-06 | 1.400E-05 |
| 20824 | 1042 | H1042D | C | 4.9e-06 | 8.7e-06 | 4.2e-06 | 5.933E-06 | 2.421E-06 | 4.200E-06 | 8.700E-06 |

| | | | | | | | | | | |
|---|---|---|---|---|---|---|---|---|---|---|
| 20825 | 1042 | H1042C | C | 9.1e-06 | 6e-06 | 1.2e-05 | 9.033E-06 | 3.001E-06 | 6.000E-06 | 1.200E-05 |
| 20826 | 1042 | H1042E | C | 6.1e-06 | 9.1e-06 | 6.6e-06 | 7.267E-06 | 1.607E-06 | 6.100E-06 | 9.100E-06 |
| 20827 | 1042 | H1042Q | C | 8e-06 | 1.4e-05 | 6.8e-06 | 9.600E-06 | 3.857E-06 | 6.800E-06 | 1.400E-05 |
| 20828 | 1042 | H1042G | C | 6.7e-06 | 6.9e-06 | 8.3e-06 | 7.300E-06 | 8.718E-07 | 6.700E-06 | 8.300E-06 |
| 20829 | 1042 | H1042H | C | 1.4e-05 | 1e-05 | 7.8e-06 | 1.060E-05 | 3.143E-06 | 7.800E-06 | 1.400E-05 |
| 20830 | 1042 | H1042I | C | 7.7e-06 | 6.7e-06 | 7e-06 | 7.133E-06 | 5.132E-07 | 6.700E-06 | 7.700E-06 |
| 20831 | 1042 | H1042L | C | 6.6e-06 | 7.5e-06 | 7.5e-06 | 7.200E-06 | 5.196E-07 | 6.600E-06 | 7.500E-06 |
| 20832 | 1042 | H1042K | C | 6.4e-06 | 8.6e-06 | 9.2e-06 | 8.067E-06 | 1.474E-06 | 6.400E-06 | 9.200E-06 |
| 20833 | 1042 | H1042M | C | 6.4e-06 | 6.3e-06 | 8.6e-06 | 7.100E-06 | 1.300E-06 | 6.300E-06 | 8.600E-06 |
| 20834 | 1042 | H1042F | C | 1.1e-05 | 7.8e-06 | 5.2e-06 | 8.000E-06 | 2.905E-06 | 5.200E-06 | 1.100E-05 |
| 20835 | 1042 | H1042P | C | 6.8e-06 | 9.1e-06 | 4.5e-06 | 6.800E-06 | 2.300E-06 | 4.500E-06 | 9.100E-06 |
| 20836 | 1042 | H1042S | C | 6.5e-06 | 9.9e-06 | 5.5e-06 | 7.300E-06 | 2.307E-06 | 5.500E-06 | 9.900E-06 |
| 20837 | 1042 | H1042T | C | 7.4e-06 | 1e-05 | 1.3e-05 | 1.013E-05 | 2.802E-06 | 7.400E-06 | 1.300E-05 |
| 20838 | 1042 | H1042W | C | 6.1e-06 | 1.7e-05 | 7.7e-06 | 1.027E-05 | 5.886E-06 | 6.100E-06 | 1.700E-05 |
| 20839 | 1042 | H1042Y | C | 6.6e-06 | 5e-06 | 1e-05 | 7.200E-06 | 2.553E-06 | 5.000E-06 | 1.000E-05 |
| 20840 | 1042 | H1042V | C | 9.2e-06 | 1.4e-05 | 4.8e-06 | 9.333E-06 | 4.601E-06 | 4.800E-06 | 1.400E-05 |
| 20841 | 1043 | K1043A | C | 7e-06 | 5e-06 | 8.7e-06 | 6.900E-06 | 1.852E-06 | 5.000E-06 | 8.700E-06 |
| 20842 | 1043 | K1043R | C | 7.1e-06 | 8.8e-06 | 7.1e-06 | 7.667E-06 | 9.815E-07 | 7.100E-06 | 8.800E-06 |
| 20843 | 1043 | K1043N | C | 7.5e-06 | 6.2e-06 | 9.3e-06 | 7.667E-06 | 1.557E-06 | 6.200E-06 | 9.300E-06 |
| 20844 | 1043 | K1043D | C | 9.7e-06 | 6.2e-06 | 5.4e-06 | 7.100E-06 | 2.287E-06 | 5.400E-06 | 9.700E-06 |
| 20845 | 1043 | K1043C | C | 1.4e-05 | 6.9e-06 | 8e-06 | 9.633E-06 | 3.821E-06 | 6.900E-06 | 1.400E-05 |
| 20846 | 1043 | K1043E | C | 6e-06 | 7.7e-06 | 9.7e-06 | 7.800E-06 | 1.852E-06 | 6.000E-06 | 9.700E-06 |
| 20847 | 1043 | K1043Q | C | 9.7e-06 | 1.1e-05 | 6.2e-06 | 8.967E-06 | 2.483E-06 | 6.200E-06 | 1.100E-05 |
| 20848 | 1043 | K1043G | C | 4.1e-06 | 7.4e-06 | 7.9e-06 | 6.467E-06 | 2.065E-06 | 4.100E-06 | 7.900E-06 |
| 20849 | 1043 | K1043H | C | 6.5e-06 | 9.1e-06 | 9.2e-06 | 8.267E-06 | 1.531E-06 | 6.500E-06 | 9.200E-06 |
| 20850 | 1043 | K1043I | C | 5.6e-06 | 1.3e-05 | 6.8e-06 | 8.467E-06 | 3.972E-06 | 5.600E-06 | 1.300E-05 |
| 20851 | 1043 | K1043L | C | 5.2e-06 | 1.3e-05 | 4.6e-06 | 7.600E-06 | 4.686E-06 | 4.600E-06 | 1.300E-05 |
| 20852 | 1043 | K1043K | C | 4.4e-06 | 5.3e-06 | 6.7e-06 | 5.467E-06 | 1.159E-06 | 4.400E-06 | 6.700E-06 |
| 20853 | 1043 | K1043M | C | 8.3e-06 | 5.8e-06 | 4.5e-06 | 6.200E-06 | 1.931E-06 | 4.500E-06 | 8.300E-06 |
| 20854 | 1043 | K1043F | C | 4.3e-06 | 5.6e-06 | 7.7e-06 | 5.867E-06 | 1.716E-06 | 4.300E-06 | 7.700E-06 |
| 20855 | 1043 | K1043P | C | 4.7e-06 | 5.2e-06 | 6.4e-06 | 5.433E-06 | 8.737E-07 | 4.700E-06 | 6.400E-06 |
| 20856 | 1043 | K1043S | C | 7e-06 | 5.6e-06 | 7.7e-06 | 6.767E-06 | 1.069E-06 | 5.600E-06 | 7.700E-06 |
| 20857 | 1043 | K1043T | C | 5.9e-06 | 9.9e-06 | 4.3e-06 | 6.700E-06 | 2.884E-06 | 4.300E-06 | 9.900E-06 |
| 20858 | 1043 | K1043W | C | 9.1e-06 | 6.6e-06 | 9.3e-06 | 8.333E-06 | 1.504E-06 | 6.600E-06 | 9.300E-06 |
| 20859 | 1043 | K1043Y | C | 6.8e-06 | 9.5e-06 | 4.5e-06 | 6.933E-06 | 2.503E-06 | 4.500E-06 | 9.500E-06 |
| 20860 | 1043 | K1043V | C | 6.5e-06 | 6.1e-06 | 8.4e-06 | 7.000E-06 | 1.229E-06 | 6.100E-06 | 8.400E-06 |
| 20861 | 1044 | P1044A | C | 9e-06 | 1.3e-05 | 9.7e-06 | 1.057E-05 | 2.136E-06 | 9.000E-06 | 1.300E-05 |
| 20862 | 1044 | P1044R | C | 6e-06 | 9.5e-06 | 7.2e-06 | 7.567E-06 | 1.779E-06 | 6.000E-06 | 9.500E-06 |
| 20863 | 1044 | P1044N | C | 6.6e-06 | 6.3e-06 | 6e-06 | 6.300E-06 | 3.000E-07 | 6.000E-06 | 6.600E-06 |
| 20864 | 1044 | P1044D | C | 6.2e-06 | 1.1e-05 | 8.1e-06 | 8.433E-06 | 2.417E-06 | 6.200E-06 | 1.100E-05 |
| 20865 | 1044 | P1044C | C | 6.7e-06 | 1.3e-05 | 8.4e-06 | 9.367E-06 | 3.259E-06 | 6.700E-06 | 1.300E-05 |
| 20866 | 1044 | P1044E | C | 9.9e-06 | 6.6e-06 | 5e-06 | 7.167E-06 | 2.499E-06 | 5.000E-06 | 9.900E-06 |
| 20867 | 1044 | P1044Q | C | 1.2e-05 | 5.5e-06 | 8.2e-06 | 8.567E-06 | 3.265E-06 | 5.500E-06 | 1.200E-05 |
| 20868 | 1044 | P1044G | C | 1.3e-05 | 6.6e-06 | 5.3e-06 | 8.300E-06 | 4.122E-06 | 5.300E-06 | 1.300E-05 |
| 20869 | 1044 | P1044H | C | 6.7e-06 | 6.4e-06 | 4.7e-06 | 5.933E-06 | 1.079E-06 | 4.700E-06 | 6.700E-06 |
| 20870 | 1044 | P1044I | C | 6.3e-06 | 6.5e-06 | 8e-06 | 6.933E-06 | 9.292E-07 | 6.300E-06 | 8.000E-06 |
| 20871 | 1044 | P1044L | C | 8.5e-06 | 1.2e-05 | 1e-05 | 1.017E-05 | 1.756E-06 | 8.500E-06 | 1.200E-05 |
| 20872 | 1044 | P1044K | C | 6e-06 | 8.4e-06 | 6.5e-06 | 6.967E-06 | 1.266E-06 | 6.000E-06 | 8.400E-06 |
| 20873 | 1044 | P1044M | C | 9.9e-06 | 7e-06 | 1e-05 | 8.967E-06 | 1.704E-06 | 7.000E-06 | 1.000E-05 |
| 20874 | 1044 | P1044F | C | 6.6e-06 | 4.9e-06 | 8.2e-06 | 6.567E-06 | 1.650E-06 | 4.900E-06 | 8.200E-06 |
| 20875 | 1044 | P1044P | C | 6.9e-06 | 5.1e-06 | 7.7e-06 | 6.567E-06 | 1.332E-06 | 5.100E-06 | 7.700E-06 |
| 20876 | 1044 | P1044S | C | 7e-06 | 7.3e-06 | 8.8e-06 | 7.700E-06 | 9.644E-07 | 7.000E-06 | 8.800E-06 |
| 20877 | 1044 | P1044T | C | 7.5e-06 | 6.9e-06 | 7.5e-06 | 7.300E-06 | 3.464E-07 | 6.900E-06 | 7.500E-06 |
| 20878 | 1044 | P1044W | C | 7.2e-06 | 4e-06 | 6.6e-06 | 5.933E-06 | 1.701E-06 | 4.000E-06 | 7.200E-06 |
| 20879 | 1044 | P1044Y | C | 1.2e-05 | 5.9e-06 | 4.8e-06 | 7.567E-06 | 3.879E-06 | 4.800E-06 | 1.200E-05 |
| 20880 | 1044 | P1044V | C | 5.2e-06 | 6.3e-06 | 6.1e-06 | 5.867E-06 | 5.859E-07 | 5.200E-06 | 6.300E-06 |
| 20881 | 1045 | S1045A | C | 6.6e-06 | 9.8e-06 | 6.8e-06 | 7.733E-06 | 1.793E-06 | 6.600E-06 | 9.800E-06 |
| 20882 | 1045 | S1045R | C | 1e-05 | 6.2e-06 | 8.2e-06 | 8.133E-06 | 1.901E-06 | 6.200E-06 | 1.000E-05 |
| 20883 | 1045 | S1045N | C | 1.5e-05 | 1e-05 | 9.2e-06 | 1.140E-05 | 3.143E-06 | 9.200E-06 | 1.500E-05 |
| 20884 | 1045 | S1045D | C | 5.6e-06 | 7.9e-06 | 4.6e-06 | 6.033E-06 | 1.692E-06 | 4.600E-06 | 7.900E-06 |
| 20885 | 1045 | S1045C | C | 8.2e-06 | 1.3e-05 | 6.6e-06 | 9.267E-06 | 3.331E-06 | 6.600E-06 | 1.300E-05 |
| 20886 | 1045 | S1045E | C | 6.5e-06 | 6.2e-06 | 8.2e-06 | 6.967E-06 | 1.079E-06 | 6.200E-06 | 8.200E-06 |
| 20887 | 1045 | S1045Q | C | 5.4e-06 | 5.2e-06 | 5e-06 | 5.200E-06 | 2.000E-07 | 5.000E-06 | 5.400E-06 |
| 20888 | 1045 | S1045G | C | 8.8e-06 | 9.5e-06 | 6.4e-06 | 8.233E-06 | 1.626E-06 | 6.400E-06 | 9.500E-06 |
| 20889 | 1045 | S1045H | C | 7.5e-06 | 8.9e-06 | 7.9e-06 | 8.100E-06 | 7.211E-07 | 7.500E-06 | 8.900E-06 |
| 20890 | 1045 | S1045I | C | 1.1e-05 | 1.2e-05 | 7.9e-06 | 1.030E-05 | 2.138E-06 | 7.900E-06 | 1.200E-05 |
| 20891 | 1045 | S1045L | C | 1.1e-05 | 1.1e-05 | 7.5e-06 | 9.833E-06 | 2.021E-06 | 7.500E-06 | 1.100E-05 |
| 20892 | 1045 | S1045K | C | 7.1e-06 | 9.9e-06 | 1.1e-05 | 9.333E-06 | 2.011E-06 | 7.100E-06 | 1.100E-05 |
| 20893 | 1045 | S1045M | C | 7.6e-06 | 8.9e-06 | 4.6e-06 | 7.033E-06 | 2.205E-06 | 4.600E-06 | 8.900E-06 |
| 20894 | 1045 | S1045F | C | 1.4e-05 | 8.9e-06 | 7.8e-06 | 1.023E-05 | 3.308E-06 | 7.800E-06 | 1.400E-05 |
| 20895 | 1045 | S1045P | C | 9.5e-06 | 1.3e-05 | 7.3e-06 | 9.933E-06 | 2.875E-06 | 7.300E-06 | 1.300E-05 |
| 20896 | 1045 | S1045S | C | 6e-06 | 1.1e-05 | 4.7e-06 | 7.233E-06 | 3.326E-06 | 4.700E-06 | 1.100E-05 |
| 20897 | 1045 | S1045T | C | 1.4e-05 | 9.6e-06 | 3.6e-06 | 9.067E-06 | 5.220E-06 | 3.600E-06 | 1.400E-05 |
| 20898 | 1045 | S1045W | C | 8.9e-06 | 1.2e-05 | 5.1e-06 | 8.667E-06 | 3.456E-06 | 5.100E-06 | 1.200E-05 |
| 20899 | 1045 | S1045Y | C | 9.8e-06 | 1.2e-05 | 1.9e-05 | 1.360E-05 | 4.804E-06 | 9.800E-06 | 1.900E-05 |
| 20900 | 1045 | S1045V | C | 9.5e-06 | 9.4e-06 | 6.4e-06 | 8.433E-06 | 1.762E-06 | 6.400E-06 | 9.500E-06 |
| 20901 | 1046 | N1046A | C | 8.2e-06 | 6.6e-06 | 5.7e-06 | 6.833E-06 | 1.266E-06 | 5.700E-06 | 8.200E-06 |
| 20902 | 1046 | N1046R | C | 1.3e-05 | 6.2e-06 | 6.6e-06 | 8.600E-06 | 3.816E-06 | 6.200E-06 | 1.300E-05 |
| 20903 | 1046 | N1046N | C | 9.5e-06 | 6.3e-06 | 6.5e-06 | 7.433E-06 | 1.793E-06 | 6.300E-06 | 9.500E-06 |
| 20904 | 1046 | N1046D | C | 5.8e-06 | 7.8e-06 | 9.7e-06 | 7.767E-06 | 1.950E-06 | 5.800E-06 | 9.700E-06 |
| 20905 | 1046 | N1046C | C | 4.4e-06 | 6.7e-06 | 1.2e-05 | 7.700E-06 | 3.897E-06 | 4.400E-06 | 1.200E-05 |
| 20906 | 1046 | N1046E | C | 6.7e-06 | 9.5e-06 | 7.3e-06 | 7.833E-06 | 1.474E-06 | 6.700E-06 | 9.500E-06 |
| 20907 | 1046 | N1046Q | C | 6.4e-06 | 1e-05 | 7.4e-06 | 7.933E-06 | 1.858E-06 | 6.400E-06 | 1.000E-05 |
| 20908 | 1046 | N1046G | C | 1.3e-05 | 1e-05 | 7.1e-06 | 1.003E-05 | 2.950E-06 | 7.100E-06 | 1.300E-05 |
| 20909 | 1046 | N1046H | C | 1.1e-05 | 8.7e-06 | 9.8e-06 | 9.833E-06 | 1.150E-06 | 8.700E-06 | 1.100E-05 |

| | | | | | | | | | | |
|---|---|---|---|---|---|---|---|---|---|---|
| 20910 | 1046 | N1046I | C | 1e-05 | 6.9e-06 | 5e-06 | 7.300E-06 | 2.524E-06 | 5.000E-06 | 1.000E-05 |
| 20911 | 1046 | N1046L | C | 1.8e-05 | 1e-05 | 7e-06 | 1.167E-05 | 5.686E-06 | 7.000E-06 | 1.800E-05 |
| 20912 | 1046 | N1046K | C | 6.6e-06 | 1.5e-05 | 6.6e-06 | 9.400E-06 | 4.850E-06 | 6.600E-06 | 1.500E-05 |
| 20913 | 1046 | N1046M | C | 5.4e-06 | 6.1e-06 | 8.5e-06 | 6.667E-06 | 1.626E-06 | 5.400E-06 | 8.500E-06 |
| 20914 | 1046 | N1046F | C | 1.1e-05 | 1.4e-05 | 6.3e-06 | 1.043E-05 | 3.881E-06 | 6.300E-06 | 1.400E-05 |
| 20915 | 1046 | N1046P | C | 6.3e-06 | 8.8e-06 | 8.2e-06 | 7.767E-06 | 1.305E-06 | 6.300E-06 | 8.800E-06 |
| 20916 | 1046 | N1046S | C | 5.7e-06 | 6.1e-06 | 5.1e-06 | 5.633E-06 | 5.033E-07 | 5.100E-06 | 6.100E-06 |
| 20917 | 1046 | N1046T | C | 4.3e-06 | 1e-05 | 3.7e-06 | 6.000E-06 | 3.477E-06 | 3.700E-06 | 1.000E-05 |
| 20918 | 1046 | N1046W | C | 1e-05 | 4.5e-06 | 6.9e-06 | 7.133E-06 | 2.757E-06 | 4.500E-06 | 1.000E-05 |
| 20919 | 1046 | N1046Y | C | 9.2e-06 | 8.3e-06 | 5.1e-06 | 7.533E-06 | 2.155E-06 | 5.100E-06 | 9.200E-06 |
| 20920 | 1046 | N1046V | C | 1.3e-05 | 1.6e-05 | 9.7e-06 | 1.290E-05 | 3.151E-06 | 9.700E-06 | 1.600E-05 |
| 20921 | 1047 | T1047A | C | 6.7e-06 | 1e-05 | 1.1e-05 | 9.233E-06 | 2.250E-06 | 6.700E-06 | 1.100E-05 |
| 20922 | 1047 | T1047R | C | 7.6e-06 | 7.3e-06 | 5.1e-06 | 6.667E-06 | 1.365E-06 | 5.100E-06 | 7.600E-06 |
| 20923 | 1047 | T1047N | C | 1.1e-05 | 8.7e-06 | 7.6e-06 | 9.100E-06 | 1.735E-06 | 7.600E-06 | 1.100E-05 |
| 20924 | 1047 | T1047D | C | 9.9e-06 | 5.9e-06 | 4.6e-06 | 6.800E-06 | 2.762E-06 | 4.600E-06 | 9.900E-06 |
| 20925 | 1047 | T1047C | C | 9.2e-06 | 9.3e-06 | 7.4e-06 | 8.633E-06 | 1.069E-06 | 7.400E-06 | 9.300E-06 |
| 20926 | 1047 | T1047E | C | 9e-06 | 8.8e-06 | 8.8e-06 | 8.867E-06 | 1.155E-07 | 8.800E-06 | 9.000E-06 |
| 20927 | 1047 | T1047Q | C | 6.9e-06 | 9.7e-06 | 9.6e-06 | 8.733E-06 | 1.589E-06 | 6.900E-06 | 9.700E-06 |
| 20928 | 1047 | T1047G | C | 6.1e-06 | 5.3e-06 | 4.6e-06 | 5.333E-06 | 7.506E-07 | 4.600E-06 | 6.100E-06 |
| 20929 | 1047 | T1047H | C | 6.8e-06 | 4.8e-06 | 4.1e-06 | 5.233E-06 | 1.401E-06 | 4.100E-06 | 6.800E-06 |
| 20930 | 1047 | T1047I | C | 5.1e-06 | 1.1e-05 | 7.3e-06 | 7.800E-06 | 2.982E-06 | 5.100E-06 | 1.100E-05 |
| 20931 | 1047 | T1047L | C | 1.1e-05 | 8.4e-06 | 7.4e-06 | 8.933E-06 | 1.858E-06 | 7.400E-06 | 1.100E-05 |
| 20932 | 1047 | T1047K | C | 6.1e-06 | 8.8e-06 | 8.4e-06 | 7.767E-06 | 1.457E-06 | 6.100E-06 | 8.800E-06 |
| 20933 | 1047 | T1047M | C | 8.5e-06 | 1.1e-05 | 1.4e-05 | 1.117E-05 | 2.754E-06 | 8.500E-06 | 1.400E-05 |
| 20934 | 1047 | T1047F | C | 5.2e-06 | 6e-06 | 8e-06 | 6.400E-06 | 1.442E-06 | 5.200E-06 | 8.000E-06 |
| 20935 | 1047 | T1047P | C | 9.9e-06 | 8.9e-06 | 9.8e-06 | 9.533E-06 | 5.508E-07 | 8.900E-06 | 9.900E-06 |
| 20936 | 1047 | T1047S | C | 9.7e-06 | 8.7e-06 | 1e-05 | 9.467E-06 | 6.807E-07 | 8.700E-06 | 1.000E-05 |
| 20937 | 1047 | T1047T | C | 4.1e-06 | 9e-06 | 7.7e-06 | 6.933E-06 | 2.538E-06 | 4.100E-06 | 9.000E-06 |
| 20938 | 1047 | T1047W | C | 6.6e-06 | 6.7e-06 | 8.2e-06 | 7.167E-06 | 8.963E-07 | 6.600E-06 | 8.200E-06 |
| 20939 | 1047 | T1047Y | C | 8.2e-06 | 5.7e-06 | 7.3e-06 | 7.067E-06 | 1.266E-06 | 5.700E-06 | 8.200E-06 |
| 20940 | 1047 | T1047V | C | 3.9e-06 | 4.7e-06 | 3.3e-06 | 3.967E-06 | 7.024E-07 | 3.300E-06 | 4.700E-06 |
| 20941 | 1048 | K1048A | C | 9.3e-06 | 1e-05 | 9.3e-06 | 9.533E-06 | 4.041E-07 | 9.300E-06 | 1.000E-05 |
| 20942 | 1048 | K1048R | C | 7.7e-06 | 8.5e-06 | 7.5e-06 | 7.900E-06 | 5.292E-07 | 7.500E-06 | 8.500E-06 |
| 20943 | 1048 | K1048N | C | 6.2e-06 | 1.4e-05 | 1e-05 | 1.007E-05 | 3.900E-06 | 6.200E-06 | 1.400E-05 |
| 20944 | 1048 | K1048D | C | 6.5e-06 | 9.7e-06 | 6.1e-06 | 7.433E-06 | 1.973E-06 | 6.100E-06 | 9.700E-06 |
| 20945 | 1048 | K1048C | C | 7e-06 | 8.4e-06 | 8.2e-06 | 7.867E-06 | 7.572E-07 | 7.000E-06 | 8.400E-06 |
| 20946 | 1048 | K1048E | C | 1.4e-05 | 8.4e-06 | 8e-06 | 1.013E-05 | 3.355E-06 | 8.000E-06 | 1.400E-05 |
| 20947 | 1048 | K1048Q | C | 8.5e-06 | 8.1e-06 | 6.2e-06 | 7.600E-06 | 1.229E-06 | 6.200E-06 | 8.500E-06 |
| 20948 | 1048 | K1048G | C | 7e-06 | 8.2e-06 | 6.6e-06 | 7.267E-06 | 8.327E-07 | 6.600E-06 | 8.200E-06 |
| 20949 | 1048 | K1048H | C | 8.9e-06 | 7.4e-06 | 6.2e-06 | 7.500E-06 | 1.353E-06 | 6.200E-06 | 8.900E-06 |
| 20950 | 1048 | K1048I | C | 9.6e-06 | 6.2e-06 | 5.9e-06 | 7.233E-06 | 2.055E-06 | 5.900E-06 | 9.600E-06 |
| 20951 | 1048 | K1048L | C | 5.2e-06 | 9e-06 | 7.1e-06 | 7.100E-06 | 1.900E-06 | 5.200E-06 | 9.000E-06 |
| 20952 | 1048 | K1048K | C | 1.4e-05 | 8.2e-06 | 8.6e-06 | 1.027E-05 | 3.239E-06 | 8.200E-06 | 1.400E-05 |
| 20953 | 1048 | K1048M | C | 8.5e-06 | 5e-06 | 6.1e-06 | 6.533E-06 | 1.790E-06 | 5.000E-06 | 8.500E-06 |
| 20954 | 1048 | K1048F | C | 4.9e-06 | 7.3e-06 | 8.5e-06 | 6.900E-06 | 1.833E-06 | 4.900E-06 | 8.500E-06 |
| 20955 | 1048 | K1048P | C | 1e-05 | 8.7e-06 | 5.5e-06 | 8.067E-06 | 2.316E-06 | 5.500E-06 | 1.000E-05 |
| 20956 | 1048 | K1048S | C | 7.4e-06 | 8.9e-06 | 7.2e-06 | 7.833E-06 | 9.292E-07 | 7.200E-06 | 8.900E-06 |
| 20957 | 1048 | K1048T | C | 8.9e-06 | 9.3e-06 | 1.4e-05 | 1.073E-05 | 2.836E-06 | 8.900E-06 | 1.400E-05 |
| 20958 | 1048 | K1048W | C | 7.9e-06 | 6.5e-06 | 7.6e-06 | 7.333E-06 | 7.371E-07 | 6.500E-06 | 7.900E-06 |
| 20959 | 1048 | K1048Y | C | 7.7e-06 | 6.5e-06 | 8.3e-06 | 7.500E-06 | 9.165E-07 | 6.500E-06 | 8.300E-06 |
| 20960 | 1048 | K1048V | C | 6.3e-06 | 1e-05 | 1.3e-05 | 9.767E-06 | 3.356E-06 | 6.300E-06 | 1.300E-05 |
| 20961 | 1049 | V1049A | C | 8.7e-06 | 6.3e-06 | 1e-05 | 8.333E-06 | 1.877E-06 | 6.300E-06 | 1.000E-05 |
| 20962 | 1049 | V1049R | C | 1.4e-05 | 7.7e-06 | 8.7e-06 | 1.013E-05 | 3.386E-06 | 7.700E-06 | 1.400E-05 |
| 20963 | 1049 | V1049N | C | 5.4e-06 | 2.6e-06 | 6.5e-06 | 4.833E-06 | 2.011E-06 | 2.600E-06 | 6.500E-06 |
| 20964 | 1049 | V1049D | C | 5e-06 | 6e-06 | 2e-05 | 1.033E-05 | 8.386E-06 | 5.000E-06 | 2.000E-05 |
| 20965 | 1049 | V1049C | C | 1.2e-05 | 7.1e-06 | 5.3e-06 | 8.133E-06 | 3.467E-06 | 5.300E-06 | 1.200E-05 |
| 20966 | 1049 | V1049E | C | 7.7e-06 | 7.5e-06 | 1.2e-05 | 9.067E-06 | 2.542E-06 | 7.500E-06 | 1.200E-05 |
| 20967 | 1049 | V1049Q | C | 7.7e-06 | 7.7e-06 | 8.1e-06 | 7.833E-06 | 2.309E-07 | 7.700E-06 | 8.100E-06 |
| 20968 | 1049 | V1049G | C | 1.2e-05 | 1.4e-05 | 5.9e-06 | 1.063E-05 | 4.219E-06 | 5.900E-06 | 1.400E-05 |
| 20969 | 1049 | V1049H | C | 1.1e-05 | 4e-06 | 9.4e-06 | 8.133E-06 | 3.668E-06 | 4.000E-06 | 1.100E-05 |
| 20970 | 1049 | V1049I | C | 7.4e-06 | 8.2e-06 | 3.7e-06 | 6.433E-06 | 2.401E-06 | 3.700E-06 | 8.200E-06 |
| 20971 | 1049 | V1049L | C | 9.4e-06 | 1.5e-05 | 3.8e-06 | 9.400E-06 | 5.600E-06 | 3.800E-06 | 1.500E-05 |
| 20972 | 1049 | V1049K | C | 8.2e-06 | 4.8e-06 | 6.2e-06 | 6.400E-06 | 1.709E-06 | 4.800E-06 | 8.200E-06 |
| 20973 | 1049 | V1049M | C | 9e-06 | 4.4e-06 | 1.9e-05 | 1.080E-05 | 7.465E-06 | 4.400E-06 | 1.900E-05 |
| 20974 | 1049 | V1049F | C | 4.6e-06 | 5.6e-06 | 1e-05 | 6.733E-06 | 2.873E-06 | 4.600E-06 | 1.000E-05 |
| 20975 | 1049 | V1049P | C | 4.7e-06 | 9.5e-06 | 6.3e-06 | 6.833E-06 | 2.444E-06 | 4.700E-06 | 9.500E-06 |
| 20976 | 1049 | V1049S | C | 8.5e-06 | 7.1e-06 | 5.8e-06 | 7.133E-06 | 1.350E-06 | 5.800E-06 | 8.500E-06 |
| 20977 | 1049 | V1049T | C | 6.6e-06 | 5.9e-06 | 7.9e-06 | 6.800E-06 | 1.015E-06 | 5.900E-06 | 7.900E-06 |
| 20978 | 1049 | V1049W | C | 3.4e-06 | 1.2e-05 | 4e-06 | 6.467E-06 | 4.801E-06 | 3.400E-06 | 1.200E-05 |
| 20979 | 1049 | V1049Y | C | 6e-06 | 4.8e-06 | 6.2e-06 | 5.667E-06 | 7.572E-07 | 4.800E-06 | 6.200E-06 |
| 20980 | 1049 | V1049V | C | 9.1e-06 | 8.1e-06 | 5.7e-06 | 7.633E-06 | 1.747E-06 | 5.700E-06 | 9.100E-06 |
| 20981 | 1050 | D1050A | C | 9.7e-06 | 3.6e-06 | 5.6e-06 | 6.300E-06 | 3.110E-06 | 3.600E-06 | 9.700E-06 |
| 20982 | 1050 | D1050R | C | 8.3e-06 | 9.4e-06 | 7.2e-06 | 8.300E-06 | 1.100E-06 | 7.200E-06 | 9.400E-06 |
| 20983 | 1050 | D1050N | C | 5e-06 | 8.2e-06 | 9.9e-06 | 7.700E-06 | 2.488E-06 | 5.000E-06 | 9.900E-06 |
| 20984 | 1050 | D1050D | C | 9.2e-06 | 9.8e-06 | 8.2e-06 | 9.067E-06 | 8.083E-07 | 8.200E-06 | 9.800E-06 |
| 20985 | 1050 | D1050C | C | 1e-05 | 1e-05 | 7.3e-06 | 9.100E-06 | 1.559E-06 | 7.300E-06 | 1.000E-05 |
| 20986 | 1050 | D1050E | C | 5.4e-06 | 5.2e-06 | 7.2e-06 | 5.933E-06 | 1.102E-06 | 5.200E-06 | 7.200E-06 |
| 20987 | 1050 | D1050Q | C | 8.2e-06 | 6.3e-06 | 1.5e-05 | 9.833E-06 | 4.574E-06 | 6.300E-06 | 1.500E-05 |
| 20988 | 1050 | D1050G | C | 6.8e-06 | 9.8e-06 | 8.6e-06 | 8.400E-06 | 1.510E-06 | 6.800E-06 | 9.800E-06 |
| 20989 | 1050 | D1050H | C | 1e-05 | 6.2e-06 | 7.6e-06 | 7.933E-06 | 1.922E-06 | 6.200E-06 | 1.000E-05 |
| 20990 | 1050 | D1050I | C | 7.4e-06 | 5.3e-06 | 5.1e-06 | 5.933E-06 | 1.274E-06 | 5.100E-06 | 7.400E-06 |
| 20991 | 1050 | D1050L | C | 8.8e-06 | 7.7e-06 | 6.2e-06 | 7.567E-06 | 1.305E-06 | 6.200E-06 | 8.800E-06 |
| 20992 | 1050 | D1050K | C | 5.2e-06 | 5.2e-06 | 4.3e-06 | 4.900E-06 | 5.196E-07 | 4.300E-06 | 5.200E-06 |
| 20993 | 1050 | D1050M | C | 9.2e-06 | 6.6e-06 | 9.3e-06 | 8.367E-06 | 1.531E-06 | 6.600E-06 | 9.300E-06 |
| 20994 | 1050 | D1050F | C | 4.8e-06 | 1.1e-05 | 9.4e-06 | 8.400E-06 | 3.219E-06 | 4.800E-06 | 1.100E-05 |

| | | | | | | | | | | |
|---|---|---|---|---|---|---|---|---|---|---|
| 20995 | 1050 | D1050P | C | 1.4e-05 | 6.5e-06 | 1.4e-05 | 1.150E-05 | 4.330E-06 | 6.500E-06 | 1.400E-05 |
| 20996 | 1050 | D1050S | C | 7.6e-06 | 9.3e-06 | 8.2e-06 | 8.367E-06 | 8.622E-07 | 7.600E-06 | 9.300E-06 |
| 20997 | 1050 | D1050T | C | 8.9e-06 | 7.6e-06 | 9e-06 | 8.500E-06 | 7.810E-07 | 7.600E-06 | 9.000E-06 |
| 20998 | 1050 | D1050W | C | 6.3e-06 | 6.5e-06 | 8.9e-06 | 7.233E-06 | 1.447E-06 | 6.300E-06 | 8.900E-06 |
| 20999 | 1050 | D1050Y | C | 7.6e-06 | 9.6e-06 | 1.2e-05 | 9.733E-06 | 2.203E-06 | 7.600E-06 | 1.200E-05 |
| 21000 | 1050 | D1050V | C | 9e-06 | 4.8e-06 | 1e-05 | 7.933E-06 | 2.759E-06 | 4.800E-06 | 1.000E-05 |
| 21001 | 1051 | K1051A | C | 7e-06 | 7.6e-06 | 6.4e-06 | 7.000E-06 | 6.000E-07 | 6.400E-06 | 7.600E-06 |
| 21002 | 1051 | K1051R | C | 7.4e-06 | 1e-05 | 1.4e-05 | 1.047E-05 | 3.325E-06 | 7.400E-06 | 1.400E-05 |
| 21003 | 1051 | K1051N | C | 1.2e-05 | 7.6e-06 | 7.8e-06 | 9.133E-06 | 2.485E-06 | 7.600E-06 | 1.200E-05 |
| 21004 | 1051 | K1051D | C | 1.3e-05 | 5.9e-06 | 9.6e-06 | 9.500E-06 | 3.551E-06 | 5.900E-06 | 1.300E-05 |
| 21005 | 1051 | K1051C | C | 5.4e-06 | 1.2e-05 | 1.4e-05 | 1.047E-05 | 4.500E-06 | 5.400E-06 | 1.400E-05 |
| 21006 | 1051 | K1051E | C | 6.3e-06 | 9.6e-06 | 6.5e-06 | 7.467E-06 | 1.850E-06 | 6.300E-06 | 9.600E-06 |
| 21007 | 1051 | K1051Q | C | 1.1e-05 | 7.4e-06 | 6.1e-06 | 8.167E-06 | 2.538E-06 | 6.100E-06 | 1.100E-05 |
| 21008 | 1051 | K1051G | C | 9.4e-06 | 1e-05 | 5e-06 | 8.133E-06 | 2.730E-06 | 5.000E-06 | 1.000E-05 |
| 21009 | 1051 | K1051H | C | 8.9e-06 | 6.4e-06 | 8.8e-06 | 8.033E-06 | 1.415E-06 | 6.400E-06 | 8.900E-06 |
| 21010 | 1051 | K1051I | C | 9.3e-06 | 1e-05 | 1e-05 | 9.767E-06 | 4.041E-07 | 9.300E-06 | 1.000E-05 |
| 21011 | 1051 | K1051L | C | 1.1e-05 | 9.2e-06 | 9.6e-06 | 9.933E-06 | 9.452E-07 | 9.200E-06 | 1.100E-05 |
| 21012 | 1051 | K1051K | C | 7.3e-06 | 1.1e-05 | 7.8e-06 | 8.700E-06 | 2.007E-06 | 7.300E-06 | 1.100E-05 |
| 21013 | 1051 | K1051M | C | 4.7e-06 | 6.3e-06 | 6.1e-06 | 5.700E-06 | 8.718E-07 | 4.700E-06 | 6.300E-06 |
| 21014 | 1051 | K1051F | C | 6.1e-06 | 4e-06 | 4.3e-06 | 4.800E-06 | 1.136E-06 | 4.000E-06 | 6.100E-06 |
| 21015 | 1051 | K1051P | C | 6.7e-06 | 8.2e-06 | 6.5e-06 | 7.133E-06 | 9.292E-07 | 6.500E-06 | 8.200E-06 |
| 21016 | 1051 | K1051S | C | 9.1e-06 | 1e-05 | 7.2e-06 | 8.767E-06 | 1.429E-06 | 7.200E-06 | 1.000E-05 |
| 21017 | 1051 | K1051T | C | 1e-05 | 4.7e-06 | 8.7e-06 | 7.800E-06 | 2.762E-06 | 4.700E-06 | 1.000E-05 |
| 21018 | 1051 | K1051W | C | 5.4e-06 | 1.2e-05 | 9.6e-06 | 9.000E-06 | 3.341E-06 | 5.400E-06 | 1.200E-05 |
| 21019 | 1051 | K1051Y | C | 8.1e-06 | 8.2e-06 | 1.1e-05 | 9.100E-06 | 1.646E-06 | 8.100E-06 | 1.100E-05 |
| 21020 | 1051 | K1051V | C | 7e-06 | 6.3e-06 | 6.9e-06 | 6.733E-06 | 3.786E-07 | 6.300E-06 | 7.000E-06 |
| 21021 | 1052 | K1052A | C | 6.6e-06 | 1.7e-05 | 9.2e-06 | 1.093E-05 | 5.412E-06 | 6.600E-06 | 1.700E-05 |
| 21022 | 1052 | K1052R | C | 1.3e-05 | 6.5e-06 | 9.6e-06 | 9.700E-06 | 3.251E-06 | 6.500E-06 | 1.300E-05 |
| 21023 | 1052 | K1052N | C | 8e-06 | 1.2e-05 | 4.6e-06 | 8.200E-06 | 3.704E-06 | 4.600E-06 | 1.200E-05 |
| 21024 | 1052 | K1052D | C | 5.6e-06 | 7.9e-06 | 6.9e-06 | 6.800E-06 | 1.153E-06 | 5.600E-06 | 7.900E-06 |
| 21025 | 1052 | K1052C | C | 6.3e-06 | 8.3e-06 | 8.2e-06 | 7.600E-06 | 1.127E-06 | 6.300E-06 | 8.300E-06 |
| 21026 | 1052 | K1052E | C | 9.5e-06 | 6.8e-06 | 6.5e-06 | 7.600E-06 | 1.652E-06 | 6.500E-06 | 9.500E-06 |
| 21027 | 1052 | K1052Q | C | 1.3e-05 | 8.7e-06 | 4.4e-06 | 8.700E-06 | 4.300E-06 | 4.400E-06 | 1.300E-05 |
| 21028 | 1052 | K1052G | C | 1.1e-05 | 4.8e-06 | 9.3e-06 | 8.367E-06 | 3.204E-06 | 4.800E-06 | 1.100E-05 |
| 21029 | 1052 | K1052H | C | 6.9e-06 | 7.1e-06 | 9.3e-06 | 7.767E-06 | 1.332E-06 | 6.900E-06 | 9.300E-06 |
| 21030 | 1052 | K1052I | C | 6.7e-06 | 5.3e-06 | 9.5e-06 | 7.167E-06 | 2.139E-06 | 5.300E-06 | 9.500E-06 |
| 21031 | 1052 | K1052L | C | 1.2e-05 | 1e-05 | 6.6e-06 | 9.533E-06 | 2.730E-06 | 6.600E-06 | 1.200E-05 |
| 21032 | 1052 | K1052K | C | 4e-06 | 7.7e-06 | 1e-05 | 7.233E-06 | 3.027E-06 | 4.000E-06 | 1.000E-05 |
| 21033 | 1052 | K1052M | C | 6.2e-06 | 1e-05 | 1.5e-05 | 1.040E-05 | 4.414E-06 | 6.200E-06 | 1.500E-05 |
| 21034 | 1052 | K1052F | C | 5.7e-06 | 1e-05 | 9.6e-06 | 8.433E-06 | 2.376E-06 | 5.700E-06 | 1.000E-05 |
| 21035 | 1052 | K1052P | C | 9.4e-06 | 6.8e-06 | 9.6e-06 | 8.600E-06 | 1.562E-06 | 6.800E-06 | 9.600E-06 |
| 21036 | 1052 | K1052S | C | 1.1e-05 | 1.3e-05 | 6.6e-06 | 1.020E-05 | 3.274E-06 | 6.600E-06 | 1.300E-05 |
| 21037 | 1052 | K1052T | C | 9.9e-06 | 7.4e-06 | 1e-05 | 9.100E-06 | 1.473E-06 | 7.400E-06 | 1.000E-05 |
| 21038 | 1052 | K1052W | C | 5.7e-06 | 3.8e-06 | 9.7e-06 | 6.400E-06 | 3.012E-06 | 3.800E-06 | 9.700E-06 |
| 21039 | 1052 | K1052Y | C | 1.1e-05 | 5e-06 | 5.2e-06 | 7.067E-06 | 3.408E-06 | 5.000E-06 | 1.100E-05 |
| 21040 | 1052 | K1052V | C | 8.4e-06 | 5.6e-06 | 8.4e-06 | 7.467E-06 | 1.617E-06 | 5.600E-06 | 8.400E-06 |
| 21041 | 1053 | V1053A | C | 6.1e-06 | 7.7e-06 | 7.6e-06 | 7.133E-06 | 8.963E-07 | 6.100E-06 | 7.700E-06 |
| 21042 | 1053 | V1053R | C | 6.3e-06 | 8e-06 | 9.4e-06 | 7.900E-06 | 1.552E-06 | 6.300E-06 | 9.400E-06 |
| 21043 | 1053 | V1053N | C | 1.2e-05 | 5.6e-06 | 5.7e-06 | 7.767E-06 | 3.667E-06 | 5.600E-06 | 1.200E-05 |
| 21044 | 1053 | V1053D | C | 9e-06 | 6.1e-06 | 6.5e-06 | 7.200E-06 | 1.572E-06 | 6.100E-06 | 9.000E-06 |
| 21045 | 1053 | V1053C | C | 9.3e-06 | 4.8e-06 | 7.7e-06 | 7.267E-06 | 2.281E-06 | 4.800E-06 | 9.300E-06 |
| 21046 | 1053 | V1053E | C | 8.2e-06 | 4.3e-06 | 6.5e-06 | 6.333E-06 | 1.955E-06 | 4.300E-06 | 8.200E-06 |
| 21047 | 1053 | V1053Q | C | 4.3e-06 | 8.5e-06 | 8e-06 | 6.933E-06 | 2.294E-06 | 4.300E-06 | 8.500E-06 |
| 21048 | 1053 | V1053G | C | 8.4e-06 | 1e-05 | 9.6e-06 | 9.333E-06 | 8.327E-07 | 8.400E-06 | 1.000E-05 |
| 21049 | 1053 | V1053H | C | 1.2e-05 | 6.9e-06 | 6e-06 | 8.300E-06 | 3.236E-06 | 6.000E-06 | 1.200E-05 |
| 21050 | 1053 | V1053I | C | 6.6e-06 | 8.4e-06 | 3e-05 | 1.500E-05 | 1.302E-05 | 6.600E-06 | 3.000E-05 |
| 21051 | 1053 | V1053L | C | 9.2e-06 | 6.3e-06 | 9.8e-06 | 8.433E-06 | 1.872E-06 | 6.300E-06 | 9.800E-06 |
| 21052 | 1053 | V1053K | C | 5.1e-06 | 6.5e-06 | 9.9e-06 | 7.167E-06 | 2.468E-06 | 5.100E-06 | 9.900E-06 |
| 21053 | 1053 | V1053M | C | 6.7e-06 | 6.4e-06 | 4.8e-06 | 5.967E-06 | 1.021E-06 | 4.800E-06 | 6.700E-06 |
| 21054 | 1053 | V1053F | C | 6e-06 | 1.1e-05 | 5e-06 | 7.333E-06 | 3.215E-06 | 5.000E-06 | 1.100E-05 |
| 21055 | 1053 | V1053P | C | 2.2e-05 | 1.1e-05 | 5.3e-06 | 1.277E-05 | 8.489E-06 | 5.300E-06 | 2.200E-05 |
| 21056 | 1053 | V1053S | C | 8.3e-06 | 6.2e-06 | 7.5e-06 | 7.333E-06 | 1.060E-06 | 6.200E-06 | 8.300E-06 |
| 21057 | 1053 | V1053T | C | 1.2e-05 | 1e-05 | 6.4e-06 | 9.467E-06 | 2.838E-06 | 6.400E-06 | 1.200E-05 |
| 21058 | 1053 | V1053W | C | 9.7e-06 | 9.3e-06 | 4.6e-06 | 7.867E-06 | 2.836E-06 | 4.600E-06 | 9.700E-06 |
| 21059 | 1053 | V1053Y | C | 6.9e-06 | 5.3e-06 | 1e-05 | 7.400E-06 | 2.390E-06 | 5.300E-06 | 1.000E-05 |
| 21060 | 1053 | V1053V | C | 1.5e-05 | 1.9e-05 | 7.8e-06 | 1.393E-05 | 5.676E-06 | 7.800E-06 | 1.900E-05 |
| 21061 | 1054 | E1054A | C | 9.7e-06 | 1.1e-05 | 7e-06 | 9.233E-06 | 2.040E-06 | 7.000E-06 | 1.100E-05 |
| 21062 | 1054 | E1054R | C | 6.2e-06 | 9e-06 | 7.9e-06 | 7.700E-06 | 1.411E-06 | 6.200E-06 | 9.000E-06 |
| 21063 | 1054 | E1054N | C | 9.7e-06 | 1e-05 | 6.7e-06 | 8.800E-06 | 1.825E-06 | 6.700E-06 | 1.000E-05 |
| 21064 | 1054 | E1054D | C | 1.7e-05 | 4.5e-06 | 1.3e-05 | 1.150E-05 | 6.384E-06 | 4.500E-06 | 1.700E-05 |
| 21065 | 1054 | E1054C | C | 6.4e-06 | 8.2e-06 | 6.6e-06 | 7.067E-06 | 9.866E-07 | 6.400E-06 | 8.200E-06 |
| 21066 | 1054 | E1054E | C | 8.6e-06 | 9.9e-06 | 1.1e-05 | 9.833E-06 | 1.201E-06 | 8.600E-06 | 1.100E-05 |
| 21067 | 1054 | E1054Q | C | 9e-06 | 7.6e-06 | 1.5e-05 | 1.053E-05 | 3.931E-06 | 7.600E-06 | 1.500E-05 |
| 21068 | 1054 | E1054G | C | 4.9e-06 | 1.3e-05 | 1.1e-05 | 9.633E-06 | 4.219E-06 | 4.900E-06 | 1.300E-05 |
| 21069 | 1054 | E1054H | C | 1e-05 | 1.3e-05 | 8.2e-06 | 1.040E-05 | 2.425E-06 | 8.200E-06 | 1.300E-05 |
| 21070 | 1054 | E1054I | C | 7e-06 | 9e-06 | 7.1e-06 | 7.700E-06 | 1.127E-06 | 7.000E-06 | 9.000E-06 |
| 21071 | 1054 | E1054L | C | 7.1e-06 | 1.3e-05 | 8.2e-06 | 9.433E-06 | 3.137E-06 | 7.100E-06 | 1.300E-05 |
| 21072 | 1054 | E1054K | C | 9.2e-06 | 5.1e-06 | 7.6e-06 | 7.300E-06 | 2.066E-06 | 5.100E-06 | 9.200E-06 |
| 21073 | 1054 | E1054M | C | 6.7e-06 | 4.2e-06 | 8.2e-06 | 6.367E-06 | 2.021E-06 | 4.200E-06 | 8.200E-06 |
| 21074 | 1054 | E1054F | C | 9.6e-06 | 6.9e-06 | 9.5e-06 | 8.667E-06 | 1.531E-06 | 6.900E-06 | 9.600E-06 |
| 21075 | 1054 | E1054P | C | 6.3e-06 | 9.2e-06 | 9.2e-06 | 8.233E-06 | 1.674E-06 | 6.300E-06 | 9.200E-06 |
| 21076 | 1054 | E1054S | C | 1.2e-05 | 1e-05 | 7.4e-06 | 9.800E-06 | 2.307E-06 | 7.400E-06 | 1.200E-05 |
| 21077 | 1054 | E1054T | C | 1e-05 | 8.3e-06 | 8.5e-06 | 8.933E-06 | 9.292E-07 | 8.300E-06 | 1.000E-05 |
| 21078 | 1054 | E1054W | C | 8.3e-06 | 1.8e-05 | 6.2e-06 | 1.083E-05 | 6.295E-06 | 6.200E-06 | 1.800E-05 |
| 21079 | 1054 | E1054Y | C | 1e-05 | 3.1e-06 | 4.8e-06 | 5.967E-06 | 3.595E-06 | 3.100E-06 | 1.000E-05 |

| | | | | | | | | | | |
|---|---|---|---|---|---|---|---|---|---|---|
| 21080 | 1054 | E1054V | C | 5.7e-06 | 9.9e-06 | 6.5e-06 | 7.367E-06 | 2.230E-06 | 5.700E-06 | 9.900E-06 |
| 21081 | 1055 | P1055A | C | 6.5e-06 | 1e-05 | 1.4e-05 | 1.017E-05 | 3.753E-06 | 6.500E-06 | 1.400E-05 |
| 21082 | 1055 | P1055R | C | 1.1e-05 | 5.1e-06 | 4.6e-06 | 6.900E-06 | 3.559E-06 | 4.600E-06 | 1.100E-05 |
| 21083 | 1055 | P1055N | C | 6.5e-06 | 9.7e-06 | 3.3e-06 | 6.500E-06 | 3.200E-06 | 3.300E-06 | 9.700E-06 |
| 21084 | 1055 | P1055D | C | 7.3e-06 | 1.1e-05 | 8.9e-06 | 9.067E-06 | 1.856E-06 | 7.300E-06 | 1.100E-05 |
| 21085 | 1055 | P1055C | C | 6.3e-06 | 6.8e-06 | 1.2e-05 | 8.367E-06 | 3.156E-06 | 6.300E-06 | 1.200E-05 |
| 21086 | 1055 | P1055E | C | 9.3e-06 | 5.9e-06 | 8.8e-06 | 8.000E-06 | 1.836E-06 | 5.900E-06 | 9.300E-06 |
| 21087 | 1055 | P1055Q | C | 1.5e-05 | 9.9e-06 | 4.3e-06 | 9.733E-06 | 5.352E-06 | 4.300E-06 | 1.500E-05 |
| 21088 | 1055 | P1055G | C | 9.2e-06 | 8e-06 | 6.8e-06 | 8.000E-06 | 1.200E-06 | 6.800E-06 | 9.200E-06 |
| 21089 | 1055 | P1055H | C | 8.4e-06 | 1.2e-05 | 5.4e-06 | 8.600E-06 | 3.305E-06 | 5.400E-06 | 1.200E-05 |
| 21090 | 1055 | P1055I | C | 5.6e-06 | 9.5e-06 | 1.3e-05 | 9.367E-06 | 3.702E-06 | 5.600E-06 | 1.300E-05 |
| 21091 | 1055 | P1055L | C | 9e-06 | 5.3e-06 | 8.4e-06 | 7.567E-06 | 1.986E-06 | 5.300E-06 | 9.000E-06 |
| 21092 | 1055 | P1055K | C | 9.2e-06 | 8.5e-06 | 9.7e-06 | 9.133E-06 | 6.028E-07 | 8.500E-06 | 9.700E-06 |
| 21093 | 1055 | P1055M | C | 3.6e-06 | 1.2e-05 | 1.9e-05 | 1.153E-05 | 7.711E-06 | 3.600E-06 | 1.900E-05 |
| 21094 | 1055 | P1055F | C | 9e-06 | 1.3e-05 | 7.4e-06 | 9.800E-06 | 2.884E-06 | 7.400E-06 | 1.300E-05 |
| 21095 | 1055 | P1055P | C | 4.5e-06 | 5.7e-06 | 9.1e-06 | 6.433E-06 | 2.386E-06 | 4.500E-06 | 9.100E-06 |
| 21096 | 1055 | P1055S | C | 1.5e-05 | 4.7e-06 | 1e-05 | 9.900E-06 | 5.151E-06 | 4.700E-06 | 1.500E-05 |
| 21097 | 1055 | P1055T | C | 1.1e-05 | 5.8e-06 | 1.4e-05 | 1.027E-05 | 4.149E-06 | 5.800E-06 | 1.400E-05 |
| 21098 | 1055 | P1055W | C | 6.5e-06 | 4.6e-06 | 6.5e-06 | 5.867E-06 | 1.097E-06 | 4.600E-06 | 6.500E-06 |
| 21099 | 1055 | P1055Y | C | 8.1e-06 | 4.7e-06 | 1e-05 | 7.600E-06 | 2.685E-06 | 4.700E-06 | 1.000E-05 |
| 21100 | 1055 | P1055V | C | 6.7e-06 | 7.2e-06 | 7.5e-06 | 7.133E-06 | 4.041E-07 | 6.700E-06 | 7.500E-06 |
| 21101 | 1056 | K1056A | C | 9.4e-06 | 6.1e-06 | 5.1e-06 | 6.867E-06 | 2.250E-06 | 5.100E-06 | 9.400E-06 |
| 21102 | 1056 | K1056R | C | 7e-06 | 7.4e-06 | 5.8e-06 | 6.733E-06 | 8.327E-07 | 5.800E-06 | 7.400E-06 |
| 21103 | 1056 | K1056N | C | 4.8e-06 | 6.7e-06 | 8.3e-06 | 6.600E-06 | 1.752E-06 | 4.800E-06 | 8.300E-06 |
| 21104 | 1056 | K1056D | C | 3.6e-06 | 6e-06 | 7.1e-06 | 5.567E-06 | 1.790E-06 | 3.600E-06 | 7.100E-06 |
| 21105 | 1056 | K1056C | C | 6.6e-06 | 7.4e-06 | 9.9e-06 | 7.967E-06 | 1.721E-06 | 6.600E-06 | 9.900E-06 |
| 21106 | 1056 | K1056E | C | 9.5e-06 | 8.6e-06 | 3.8e-06 | 7.300E-06 | 3.064E-06 | 3.800E-06 | 9.500E-06 |
| 21107 | 1056 | K1056Q | C | 8.4e-06 | 8.4e-06 | 9.3e-06 | 8.700E-06 | 5.196E-07 | 8.400E-06 | 9.300E-06 |
| 21108 | 1056 | K1056G | C | 9.5e-06 | 6e-06 | 7.9e-06 | 7.800E-06 | 1.752E-06 | 6.000E-06 | 9.500E-06 |
| 21109 | 1056 | K1056H | C | 6.8e-06 | 8.2e-06 | 5.6e-06 | 6.867E-06 | 1.301E-06 | 5.600E-06 | 8.200E-06 |
| 21110 | 1056 | K1056I | C | 8e-06 | 7.8e-06 | 9.3e-06 | 8.367E-06 | 8.145E-07 | 7.800E-06 | 9.300E-06 |
| 21111 | 1056 | K1056L | C | 5.5e-06 | 8.8e-06 | 5.1e-06 | 6.467E-06 | 2.031E-06 | 5.100E-06 | 8.800E-06 |
| 21112 | 1056 | K1056K | C | 5.8e-06 | 6.2e-06 | 8.5e-06 | 6.833E-06 | 1.457E-06 | 5.800E-06 | 8.500E-06 |
| 21113 | 1056 | K1056M | C | 7.2e-06 | 6.7e-06 | 8.4e-06 | 7.433E-06 | 8.737E-07 | 6.700E-06 | 8.400E-06 |
| 21114 | 1056 | K1056F | C | 6e-06 | 7.7e-06 | 5.2e-06 | 6.300E-06 | 1.277E-06 | 5.200E-06 | 7.700E-06 |
| 21115 | 1056 | K1056P | C | 1.2e-05 | 5.3e-06 | 9.8e-06 | 9.033E-06 | 3.415E-06 | 5.300E-06 | 1.200E-05 |
| 21116 | 1056 | K1056S | C | 1.1e-05 | 9.7e-06 | 1.3e-05 | 1.123E-05 | 1.662E-06 | 9.700E-06 | 1.300E-05 |
| 21117 | 1056 | K1056T | C | 8.4e-06 | 4.9e-06 | 5.7e-06 | 6.333E-06 | 1.834E-06 | 4.900E-06 | 8.400E-06 |
| 21118 | 1056 | K1056W | C | 5.9e-06 | 9.8e-06 | 1.5e-05 | 1.023E-05 | 4.565E-06 | 5.900E-06 | 1.500E-05 |
| 21119 | 1056 | K1056Y | C | 9.1e-06 | 4.3e-06 | 8.5e-06 | 7.300E-06 | 2.615E-06 | 4.300E-06 | 9.100E-06 |
| 21120 | 1056 | K1056V | C | 6e-06 | 4.4e-06 | 5.6e-06 | 5.333E-06 | 8.327E-07 | 4.400E-06 | 6.000E-06 |
| 21121 | 1057 | S1057A | C | 1.3e-05 | 1e-05 | 9.8e-06 | 1.093E-05 | 1.793E-06 | 9.800E-06 | 1.300E-05 |
| 21122 | 1057 | S1057R | C | 1.1e-05 | 8.5e-06 | 7.7e-06 | 9.067E-06 | 1.721E-06 | 7.700E-06 | 1.100E-05 |
| 21123 | 1057 | S1057N | C | 9.7e-06 | 6.1e-06 | 6.8e-06 | 7.533E-06 | 1.909E-06 | 6.100E-06 | 9.700E-06 |
| 21124 | 1057 | S1057D | C | 7.3e-06 | 5.3e-06 | 1e-05 | 7.533E-06 | 2.359E-06 | 5.300E-06 | 1.000E-05 |
| 21125 | 1057 | S1057C | C | 9.5e-06 | 9.4e-06 | 3.8e-06 | 7.567E-06 | 3.262E-06 | 3.800E-06 | 9.500E-06 |
| 21126 | 1057 | S1057E | C | 1.7e-05 | 9.5e-06 | 8.3e-06 | 1.160E-05 | 4.715E-06 | 8.300E-06 | 1.700E-05 |
| 21127 | 1057 | S1057Q | C | 7.7e-06 | 1e-05 | 5.5e-06 | 7.733E-06 | 2.250E-06 | 5.500E-06 | 1.000E-05 |
| 21128 | 1057 | S1057G | C | 7.5e-06 | 4.8e-06 | 7.1e-06 | 6.467E-06 | 1.457E-06 | 4.800E-06 | 7.500E-06 |
| 21129 | 1057 | S1057H | C | 9.8e-06 | 9.1e-06 | 9.3e-06 | 9.400E-06 | 3.606E-07 | 9.100E-06 | 9.800E-06 |
| 21130 | 1057 | S1057I | C | 6.6e-06 | 8.9e-06 | 9e-06 | 8.167E-06 | 1.358E-06 | 6.600E-06 | 9.000E-06 |
| 21131 | 1057 | S1057L | C | 3.5e-06 | 7.7e-06 | 1.2e-05 | 7.733E-06 | 4.250E-06 | 3.500E-06 | 1.200E-05 |
| 21132 | 1057 | S1057K | C | 7e-06 | 1e-05 | 1e-05 | 9.000E-06 | 1.732E-06 | 7.000E-06 | 1.000E-05 |
| 21133 | 1057 | S1057M | C | 6.6e-06 | 8e-06 | 6.7e-06 | 7.100E-06 | 7.810E-07 | 6.600E-06 | 8.000E-06 |
| 21134 | 1057 | S1057F | C | 1.4e-05 | 5.7e-06 | 8.6e-06 | 9.433E-06 | 4.212E-06 | 5.700E-06 | 1.400E-05 |
| 21135 | 1057 | S1057P | C | 8.3e-06 | 8.3e-06 | 1.4e-05 | 1.020E-05 | 3.291E-06 | 8.300E-06 | 1.400E-05 |
| 21136 | 1057 | S1057S | C | 5.6e-06 | 1.2e-05 | 1e-05 | 9.200E-06 | 3.274E-06 | 5.600E-06 | 1.200E-05 |
| 21137 | 1057 | S1057T | C | 4.4e-06 | 5.4e-06 | 8.1e-06 | 5.967E-06 | 1.914E-06 | 4.400E-06 | 8.100E-06 |
| 21138 | 1057 | S1057W | C | 6.7e-06 | 6.6e-06 | 1.2e-05 | 8.433E-06 | 3.089E-06 | 6.600E-06 | 1.200E-05 |
| 21139 | 1057 | S1057Y | C | 5.4e-06 | 2e-05 | 9.7e-06 | 1.170E-05 | 7.503E-06 | 5.400E-06 | 2.000E-05 |
| 21140 | 1057 | S1057V | C | 4.9e-06 | 3.3e-06 | 7.7e-06 | 5.300E-06 | 2.227E-06 | 3.300E-06 | 7.700E-06 |
| 21141 | 1058 | C1058A | C | 1e-05 | 1e-05 | 7.1e-06 | 9.033E-06 | 1.674E-06 | 7.100E-06 | 1.000E-05 |
| 21142 | 1058 | C1058R | C | 7e-06 | 1.1e-05 | 1.2e-05 | 1.000E-05 | 2.646E-06 | 7.000E-06 | 1.200E-05 |
| 21143 | 1058 | C1058N | C | 9.2e-06 | 9.7e-06 | 6e-06 | 8.300E-06 | 2.007E-06 | 6.000E-06 | 9.700E-06 |
| 21144 | 1058 | C1058D | C | 6.1e-06 | 6e-06 | 1e-05 | 7.367E-06 | 2.281E-06 | 6.000E-06 | 1.000E-05 |
| 21145 | 1058 | C1058C | C | 4.6e-06 | 1.4e-05 | 6.2e-06 | 8.267E-06 | 5.029E-06 | 4.600E-06 | 1.400E-05 |
| 21146 | 1058 | C1058E | C | 9.8e-06 | 4.1e-06 | 9.6e-06 | 7.833E-06 | 3.235E-06 | 4.100E-06 | 9.800E-06 |
| 21147 | 1058 | C1058Q | C | 9.6e-06 | 1.9e-05 | 6.7e-06 | 1.177E-05 | 6.430E-06 | 6.700E-06 | 1.900E-05 |
| 21148 | 1058 | C1058G | C | 5e-06 | 6.2e-06 | 9.5e-06 | 6.900E-06 | 2.330E-06 | 5.000E-06 | 9.500E-06 |
| 21149 | 1058 | C1058H | C | 5.4e-06 | 7.6e-06 | 8.9e-06 | 7.300E-06 | 1.769E-06 | 5.400E-06 | 8.900E-06 |
| 21150 | 1058 | C1058I | C | 1.2e-05 | 9.1e-06 | 7.8e-06 | 9.633E-06 | 2.150E-06 | 7.800E-06 | 1.200E-05 |
| 21151 | 1058 | C1058L | C | 9.7e-06 | 6.9e-06 | 1.2e-05 | 9.533E-06 | 2.554E-06 | 6.900E-06 | 1.200E-05 |
| 21152 | 1058 | C1058K | C | 7.3e-06 | 6.3e-06 | 1.1e-05 | 8.200E-06 | 2.476E-06 | 6.300E-06 | 1.100E-05 |
| 21153 | 1058 | C1058M | C | 1.3e-05 | 1.9e-05 | 1.1e-05 | 1.433E-05 | 4.163E-06 | 1.100E-05 | 1.900E-05 |
| 21154 | 1058 | C1058F | C | 1e-05 | 4.8e-06 | 3.7e-06 | 6.167E-06 | 3.365E-06 | 3.700E-06 | 1.000E-05 |
| 21155 | 1058 | C1058P | C | 7.5e-06 | 2.2e-06 | 9.2e-06 | 6.300E-06 | 3.651E-06 | 2.200E-06 | 9.200E-06 |
| 21156 | 1058 | C1058S | C | 5.5e-06 | 6.7e-06 | 4.9e-06 | 5.700E-06 | 9.165E-07 | 4.900E-06 | 6.700E-06 |
| 21157 | 1058 | C1058T | C | 9.2e-06 | 7.6e-06 | 8.8e-06 | 8.533E-06 | 8.327E-07 | 7.600E-06 | 9.200E-06 |
| 21158 | 1058 | C1058W | C | 1e-05 | 8.7e-06 | 1.3e-05 | 1.057E-05 | 2.205E-06 | 8.700E-06 | 1.300E-05 |
| 21159 | 1058 | C1058Y | C | 6.8e-06 | 1e-05 | 6.3e-06 | 7.700E-06 | 2.007E-06 | 6.300E-06 | 1.000E-05 |
| 21160 | 1058 | C1058V | C | 8e-06 | 8.1e-06 | 1.1e-05 | 9.033E-06 | 1.704E-06 | 8.000E-06 | 1.100E-05 |
| 21161 | 1059 | D1059A | D | 4.5e-06 | 1.1e-05 | 6.7e-06 | 7.400E-06 | 3.306E-06 | 4.500E-06 | 1.100E-05 |
| 21162 | 1059 | D1059R | D | 1.2e-05 | 1.3e-05 | 7e-06 | 1.067E-05 | 3.215E-06 | 7.000E-06 | 1.300E-05 |
| 21163 | 1059 | D1059N | D | 4.2e-06 | 5.8e-06 | 4.8e-06 | 4.933E-06 | 8.083E-07 | 4.200E-06 | 5.800E-06 |
| 21164 | 1059 | D1059D | D | 1.2e-05 | 6.8e-06 | 1e-05 | 9.600E-06 | 2.623E-06 | 6.800E-06 | 1.200E-05 |

| | | | | | | | | | | |
|---|---|---|---|---|---|---|---|---|---|---|
| 21165 | 1059 | D1059C | D | 6.2e-06 | 5.1e-06 | 7.3e-06 | 6.200E-06 | 1.100E-06 | 5.100E-06 | 7.300E-06 |
| 21166 | 1059 | D1059E | D | 6.4e-06 | 6.3e-06 | 9.1e-06 | 7.267E-06 | 1.589E-06 | 6.300E-06 | 9.100E-06 |
| 21167 | 1059 | D1059Q | D | 1.1e-05 | 1.2e-05 | 8.6e-06 | 1.053E-05 | 1.747E-06 | 8.600E-06 | 1.200E-05 |
| 21168 | 1059 | D1059G | D | 6.5e-06 | 1.4e-05 | 6.4e-06 | 8.967E-06 | 4.359E-06 | 6.400E-06 | 1.400E-05 |
| 21169 | 1059 | D1059H | D | 9.3e-06 | 9.8e-06 | 9.7e-06 | 9.600E-06 | 2.646E-07 | 9.300E-06 | 9.800E-06 |
| 21170 | 1059 | D1059I | D | 1.3e-05 | 9.5e-06 | 6.2e-06 | 9.567E-06 | 3.400E-06 | 6.200E-06 | 1.300E-05 |
| 21171 | 1059 | D1059L | D | 7.6e-06 | 8.6e-06 | 7.1e-06 | 7.767E-06 | 7.638E-07 | 7.100E-06 | 8.600E-06 |
| 21172 | 1059 | D1059K | D | 6.1e-06 | 8e-06 | 9.4e-06 | 7.833E-06 | 1.656E-06 | 6.100E-06 | 9.400E-06 |
| 21173 | 1059 | D1059M | D | 5.5e-06 | 9.4e-06 | 8.6e-06 | 7.833E-06 | 2.060E-06 | 5.500E-06 | 9.400E-06 |
| 21174 | 1059 | D1059F | D | 5.9e-06 | 3.8e-06 | 1.5e-05 | 8.233E-06 | 5.953E-06 | 3.800E-06 | 1.500E-05 |
| 21175 | 1059 | D1059P | D | 5.1e-06 | 1.2e-05 | 7.3e-06 | 8.133E-06 | 3.525E-06 | 5.100E-06 | 1.200E-05 |
| 21176 | 1059 | D1059S | D | 8.6e-06 | 8.8e-06 | 6.7e-06 | 8.033E-06 | 1.159E-06 | 6.700E-06 | 8.800E-06 |
| 21177 | 1059 | D1059T | D | 1.2e-05 | 9e-06 | 9e-06 | 1.000E-05 | 1.732E-06 | 9.000E-06 | 1.200E-05 |
| 21178 | 1059 | D1059W | D | 9.8e-06 | 1.2e-05 | 6.9e-06 | 9.567E-06 | 2.558E-06 | 6.900E-06 | 1.200E-05 |
| 21179 | 1059 | D1059Y | D | 6.4e-06 | 1.2e-05 | 9.1e-06 | 9.167E-06 | 2.801E-06 | 6.400E-06 | 1.200E-05 |
| 21180 | 1059 | D1059V | D | 8.3e-06 | 6.4e-06 | 7.9e-06 | 7.533E-06 | 1.002E-06 | 6.400E-06 | 8.300E-06 |
| 21181 | 1060 | I1060A | D | 1.8e-05 | 7.5e-06 | 7.8e-06 | 1.110E-05 | 5.977E-06 | 7.500E-06 | 1.800E-05 |
| 21182 | 1060 | I1060R | D | 8.3e-06 | 9.7e-06 | 9.7e-06 | 9.233E-06 | 8.083E-07 | 8.300E-06 | 9.700E-06 |
| 21183 | 1060 | I1060N | D | 6.9e-06 | 1.2e-05 | 5.7e-06 | 8.200E-06 | 3.345E-06 | 5.700E-06 | 1.200E-05 |
| 21184 | 1060 | I1060D | D | 6.7e-06 | 4.3e-06 | 9.9e-06 | 6.967E-06 | 2.810E-06 | 4.300E-06 | 9.900E-06 |
| 21185 | 1060 | I1060C | D | 5.8e-06 | 1e-05 | 1.3e-05 | 9.600E-06 | 3.617E-06 | 5.800E-06 | 1.300E-05 |
| 21186 | 1060 | I1060E | D | 6.8e-06 | 9.9e-06 | 6.5e-06 | 7.733E-06 | 1.882E-06 | 6.500E-06 | 9.900E-06 |
| 21187 | 1060 | I1060Q | D | 5.5e-06 | 6.5e-06 | 4.6e-06 | 5.533E-06 | 9.504E-07 | 4.600E-06 | 6.500E-06 |
| 21188 | 1060 | I1060G | D | 9e-06 | 9.2e-06 | 6.7e-06 | 8.300E-06 | 1.389E-06 | 6.700E-06 | 9.200E-06 |
| 21189 | 1060 | I1060H | D | 1e-05 | 6.9e-06 | 6.4e-06 | 7.767E-06 | 1.950E-06 | 6.400E-06 | 1.000E-05 |
| 21190 | 1060 | I1060I | D | 1.2e-05 | 1.1e-05 | 9e-06 | 1.067E-05 | 1.528E-06 | 9.000E-06 | 1.200E-05 |
| 21191 | 1060 | I1060L | D | 1.1e-05 | 9.8e-06 | 1.2e-05 | 1.093E-05 | 1.102E-06 | 9.800E-06 | 1.200E-05 |
| 21192 | 1060 | I1060K | D | 8e-06 | 5.7e-06 | 1e-05 | 7.900E-06 | 2.152E-06 | 5.700E-06 | 1.000E-05 |
| 21193 | 1060 | I1060M | D | 4.7e-06 | 9e-06 | 4.2e-06 | 5.967E-06 | 2.639E-06 | 4.200E-06 | 9.000E-06 |
| 21194 | 1060 | I1060F | D | 9.4e-06 | 6.1e-06 | 8.1e-06 | 7.867E-06 | 1.662E-06 | 6.100E-06 | 9.400E-06 |
| 21195 | 1060 | I1060P | D | 8.8e-06 | 1.4e-05 | 5.1e-06 | 9.300E-06 | 4.471E-06 | 5.100E-06 | 1.400E-05 |
| 21196 | 1060 | I1060S | D | 6.8e-06 | 7.5e-06 | 7.8e-06 | 7.367E-06 | 5.132E-07 | 6.800E-06 | 7.800E-06 |
| 21197 | 1060 | I1060T | D | 1.1e-05 | 5.2e-06 | 9e-06 | 8.400E-06 | 2.946E-06 | 5.200E-06 | 1.100E-05 |
| 21198 | 1060 | I1060W | D | 5.1e-06 | 1.2e-05 | 7.9e-06 | 8.333E-06 | 3.470E-06 | 5.100E-06 | 1.200E-05 |
| 21199 | 1060 | I1060Y | D | 5.1e-06 | 6.5e-06 | 9.5e-06 | 7.033E-06 | 2.248E-06 | 5.100E-06 | 9.500E-06 |
| 21200 | 1060 | I1060V | D | 8.9e-06 | 7.7e-06 | 6.3e-06 | 7.633E-06 | 1.301E-06 | 6.300E-06 | 8.900E-06 |
| 21201 | 1061 | Q1061A | D | 1e-05 | 6.6e-06 | 8e-06 | 8.200E-06 | 1.709E-06 | 6.600E-06 | 1.000E-05 |
| 21202 | 1061 | Q1061R | D | 7e-06 | 5.2e-06 | 1.3e-05 | 8.400E-06 | 4.084E-06 | 5.200E-06 | 1.300E-05 |
| 21203 | 1061 | Q1061N | D | 9.7e-06 | 6.3e-06 | 8.8e-06 | 8.267E-06 | 1.762E-06 | 6.300E-06 | 9.700E-06 |
| 21204 | 1061 | Q1061D | D | 6.8e-06 | 1.1e-05 | 7.7e-06 | 8.500E-06 | 2.211E-06 | 6.800E-06 | 1.100E-05 |
| 21205 | 1061 | Q1061C | D | 9.1e-06 | 6.2e-06 | 8.7e-06 | 8.000E-06 | 1.572E-06 | 6.200E-06 | 9.100E-06 |
| 21206 | 1061 | Q1061E | D | 6.1e-06 | 9e-06 | 7.2e-06 | 7.433E-06 | 1.464E-06 | 6.100E-06 | 9.000E-06 |
| 21207 | 1061 | Q1061Q | D | 6.2e-06 | 4.6e-06 | 7.9e-06 | 6.233E-06 | 1.650E-06 | 4.600E-06 | 7.900E-06 |
| 21208 | 1061 | Q1061G | D | 7.8e-06 | 6.3e-06 | 5.6e-06 | 6.567E-06 | 1.124E-06 | 5.600E-06 | 7.800E-06 |
| 21209 | 1061 | Q1061H | D | 1e-05 | 6.1e-06 | 1.1e-05 | 9.033E-06 | 2.589E-06 | 6.100E-06 | 1.100E-05 |
| 21210 | 1061 | Q1061I | D | 7.4e-06 | 8.5e-06 | 1e-05 | 8.633E-06 | 1.305E-06 | 7.400E-06 | 1.000E-05 |
| 21211 | 1061 | Q1061L | D | 6.3e-06 | 6.4e-06 | 9.3e-06 | 7.333E-06 | 1.704E-06 | 6.300E-06 | 9.300E-06 |
| 21212 | 1061 | Q1061K | D | 6.6e-06 | 1e-05 | 1.2e-05 | 9.533E-06 | 2.730E-06 | 6.600E-06 | 1.200E-05 |
| 21213 | 1061 | Q1061M | D | 5.5e-06 | 9.5e-06 | 6.3e-06 | 7.100E-06 | 2.117E-06 | 5.500E-06 | 9.500E-06 |
| 21214 | 1061 | Q1061F | D | 6.4e-06 | 7.1e-06 | 9.3e-06 | 7.600E-06 | 1.513E-06 | 6.400E-06 | 9.300E-06 |
| 21215 | 1061 | Q1061P | D | 6.3e-06 | 7.3e-06 | 1.3e-05 | 8.867E-06 | 3.614E-06 | 6.300E-06 | 1.300E-05 |
| 21216 | 1061 | Q1061S | D | 7.3e-06 | 9.9e-06 | 9.7e-06 | 8.967E-06 | 1.447E-06 | 7.300E-06 | 9.900E-06 |
| 21217 | 1061 | Q1061T | D | 8.2e-06 | 5e-06 | 7.6e-06 | 6.933E-06 | 1.701E-06 | 5.000E-06 | 8.200E-06 |
| 21218 | 1061 | Q1061W | D | 6.7e-06 | 9.8e-06 | 5.7e-06 | 7.400E-06 | 2.138E-06 | 5.700E-06 | 9.800E-06 |
| 21219 | 1061 | Q1061Y | D | 7.5e-06 | 5.6e-06 | 6.7e-06 | 6.600E-06 | 9.539E-07 | 5.600E-06 | 7.500E-06 |
| 21220 | 1061 | Q1061V | D | 1.1e-05 | 9.9e-06 | 8.5e-06 | 9.800E-06 | 1.253E-06 | 8.500E-06 | 1.100E-05 |
| 21221 | 1062 | M1062A | D | 6.9e-06 | 9.2e-06 | 1.1e-05 | 9.033E-06 | 2.055E-06 | 6.900E-06 | 1.100E-05 |
| 21222 | 1062 | M1062R | D | 3.3e-06 | 8.1e-06 | 7.2e-06 | 6.200E-06 | 2.551E-06 | 3.300E-06 | 8.100E-06 |
| 21223 | 1062 | M1062N | D | 9e-06 | 1.1e-05 | 1.3e-05 | 1.100E-05 | 2.000E-06 | 9.000E-06 | 1.300E-05 |
| 21224 | 1062 | M1062D | D | 9.1e-06 | 9.8e-06 | 6.9e-06 | 8.600E-06 | 1.513E-06 | 6.900E-06 | 9.800E-06 |
| 21225 | 1062 | M1062C | D | 7.7e-06 | 6.1e-06 | 7.9e-06 | 7.233E-06 | 9.866E-07 | 6.100E-06 | 7.900E-06 |
| 21226 | 1062 | M1062E | D | 9.2e-06 | 1.2e-05 | 4e-06 | 8.400E-06 | 4.060E-06 | 4.000E-06 | 1.200E-05 |
| 21227 | 1062 | M1062Q | D | 8.9e-06 | 4.5e-06 | 1e-05 | 7.800E-06 | 2.910E-06 | 4.500E-06 | 1.000E-05 |
| 21228 | 1062 | M1062G | D | 5.7e-06 | 9.4e-06 | 7.3e-06 | 7.467E-06 | 1.856E-06 | 5.700E-06 | 9.400E-06 |
| 21229 | 1062 | M1062H | D | 5.3e-06 | 9.5e-06 | 8.1e-06 | 7.633E-06 | 2.139E-06 | 5.300E-06 | 9.500E-06 |
| 21230 | 1062 | M1062I | D | 1.2e-05 | 9.4e-06 | 7.5e-06 | 9.633E-06 | 2.259E-06 | 7.500E-06 | 1.200E-05 |
| 21231 | 1062 | M1062L | D | 7.3e-06 | 6.4e-06 | 4.8e-06 | 6.167E-06 | 1.266E-06 | 4.800E-06 | 7.300E-06 |
| 21232 | 1062 | M1062K | D | 1.1e-05 | 7.5e-06 | 9.4e-06 | 9.300E-06 | 1.752E-06 | 7.500E-06 | 1.100E-05 |
| 21233 | 1062 | M1062M | D | 6.3e-06 | 1e-05 | 7e-06 | 7.767E-06 | 1.966E-06 | 6.300E-06 | 1.000E-05 |
| 21234 | 1062 | M1062F | D | 4.7e-06 | 9.5e-06 | 9.9e-06 | 8.033E-06 | 2.894E-06 | 4.700E-06 | 9.900E-06 |
| 21235 | 1062 | M1062P | D | 9.9e-06 | 6.4e-06 | 7e-06 | 7.767E-06 | 1.872E-06 | 6.400E-06 | 9.900E-06 |
| 21236 | 1062 | M1062S | D | 6.5e-06 | 6.1e-06 | 9.5e-06 | 7.367E-06 | 1.858E-06 | 6.100E-06 | 9.500E-06 |
| 21237 | 1062 | M1062T | D | 4.8e-06 | 6.2e-06 | 8.4e-06 | 6.467E-06 | 1.815E-06 | 4.800E-06 | 8.400E-06 |
| 21238 | 1062 | M1062W | D | 7.8e-06 | 6.9e-06 | 9.8e-06 | 8.167E-06 | 1.484E-06 | 6.900E-06 | 9.800E-06 |
| 21239 | 1062 | M1062Y | D | 4.9e-06 | 8e-06 | 8.8e-06 | 7.233E-06 | 2.060E-06 | 4.900E-06 | 8.800E-06 |
| 21240 | 1062 | M1062V | D | 1.1e-05 | 6.5e-06 | 6.2e-06 | 7.900E-06 | 2.689E-06 | 6.200E-06 | 1.100E-05 |
| 21241 | 1063 | T1063A | D | 7.5e-06 | 7.1e-06 | 9.2e-06 | 7.933E-06 | 1.115E-06 | 7.100E-06 | 9.200E-06 |
| 21242 | 1063 | T1063R | D | 6.5e-06 | 1.4e-05 | 9.1e-06 | 9.867E-06 | 3.808E-06 | 6.500E-06 | 1.400E-05 |
| 21243 | 1063 | T1063N | D | 1.1e-05 | 7.7e-06 | 4.5e-06 | 7.733E-06 | 3.250E-06 | 4.500E-06 | 1.100E-05 |
| 21244 | 1063 | T1063D | D | 8.7e-06 | 1e-05 | 6.4e-06 | 8.367E-06 | 1.823E-06 | 6.400E-06 | 1.000E-05 |
| 21245 | 1063 | T1063C | D | 7.5e-06 | 9.6e-06 | 6.9e-06 | 8.000E-06 | 1.418E-06 | 6.900E-06 | 9.600E-06 |
| 21246 | 1063 | T1063E | D | 1.1e-05 | 6.2e-06 | 4.1e-06 | 7.100E-06 | 3.537E-06 | 4.100E-06 | 1.100E-05 |
| 21247 | 1063 | T1063Q | D | 8.7e-06 | 8.8e-06 | 9.9e-06 | 9.133E-06 | 6.658E-07 | 8.700E-06 | 9.900E-06 |
| 21248 | 1063 | T1063G | D | 9e-06 | 8e-06 | 1.4e-05 | 1.033E-05 | 3.215E-06 | 8.000E-06 | 1.400E-05 |
| 21249 | 1063 | T1063H | D | 4.4e-06 | 1.2e-05 | 5.6e-06 | 7.333E-06 | 4.086E-06 | 4.400E-06 | 1.200E-05 |

| | | | | | | | | | | |
|---|---|---|---|---|---|---|---|---|---|---|
| 21250 | 1063 | T1063I | D | 6.9e-06 | 4.8e-06 | 8.8e-06 | 6.833E-06 | 2.001E-06 | 4.800E-06 | 8.800E-06 |
| 21251 | 1063 | T1063L | D | 8.1e-06 | 4.7e-06 | 1.2e-05 | 8.267E-06 | 3.653E-06 | 4.700E-06 | 1.200E-05 |
| 21252 | 1063 | T1063K | D | 8.5e-06 | 4.8e-06 | 9.8e-06 | 7.700E-06 | 2.594E-06 | 4.800E-06 | 9.800E-06 |
| 21253 | 1063 | T1063M | D | 8.7e-06 | 1.2e-05 | 7.9e-06 | 9.533E-06 | 2.173E-06 | 7.900E-06 | 1.200E-05 |
| 21254 | 1063 | T1063F | D | 1e-05 | 1.2e-05 | 6e-06 | 9.333E-06 | 3.055E-06 | 6.000E-06 | 1.200E-05 |
| 21255 | 1063 | T1063P | D | 1.2e-05 | 5e-06 | 8.1e-06 | 8.367E-06 | 3.508E-06 | 5.000E-06 | 1.200E-05 |
| 21256 | 1063 | T1063S | D | 5.9e-06 | 5.5e-06 | 7.3e-06 | 6.233E-06 | 9.452E-07 | 5.500E-06 | 7.300E-06 |
| 21257 | 1063 | T1063T | D | 5.8e-06 | 9.6e-06 | 9.7e-06 | 8.367E-06 | 2.223E-06 | 5.800E-06 | 9.700E-06 |
| 21258 | 1063 | T1063W | D | 8.6e-06 | 9.6e-06 | 5.5e-06 | 7.900E-06 | 2.138E-06 | 5.500E-06 | 9.600E-06 |
| 21259 | 1063 | T1063Y | D | 9.3e-06 | 9.2e-06 | 5.7e-06 | 8.067E-06 | 2.050E-06 | 5.700E-06 | 9.300E-06 |
| 21260 | 1063 | T1063V | D | 1.1e-05 | 1e-05 | 7.9e-06 | 9.633E-06 | 1.582E-06 | 7.900E-06 | 1.100E-05 |
| 21261 | 1064 | Q1064A | D | 6.9e-06 | 6.7e-06 | 6.5e-06 | 6.700E-06 | 2.000E-07 | 6.500E-06 | 6.900E-06 |
| 21262 | 1064 | Q1064R | D | 6.5e-06 | 9.8e-06 | 6e-06 | 7.433E-06 | 2.065E-06 | 6.000E-06 | 9.800E-06 |
| 21263 | 1064 | Q1064N | D | 6.5e-06 | 5e-06 | 5.4e-06 | 5.633E-06 | 7.767E-07 | 5.000E-06 | 6.500E-06 |
| 21264 | 1064 | Q1064D | D | 6.5e-06 | 8.6e-06 | 5.7e-06 | 6.933E-06 | 1.498E-06 | 5.700E-06 | 8.600E-06 |
| 21265 | 1064 | Q1064C | D | 9.5e-06 | 7.1e-06 | 4.9e-06 | 7.167E-06 | 2.301E-06 | 4.900E-06 | 9.500E-06 |
| 21266 | 1064 | Q1064E | D | 4.5e-06 | 7.7e-06 | 1.1e-05 | 7.733E-06 | 3.250E-06 | 4.500E-06 | 1.100E-05 |
| 21267 | 1064 | Q1064Q | D | 1.2e-05 | 1e-05 | 6.9e-06 | 9.633E-06 | 2.570E-06 | 6.900E-06 | 1.200E-05 |
| 21268 | 1064 | Q1064G | D | 9.8e-06 | 6e-06 | 7.6e-06 | 7.800E-06 | 1.908E-06 | 6.000E-06 | 9.800E-06 |
| 21269 | 1064 | Q1064H | D | 7.5e-06 | 5.7e-06 | 6.4e-06 | 6.533E-06 | 9.074E-07 | 5.700E-06 | 7.500E-06 |
| 21270 | 1064 | Q1064I | D | 1.2e-05 | 9.3e-06 | 1.1e-05 | 1.077E-05 | 1.365E-06 | 9.300E-06 | 1.200E-05 |
| 21271 | 1064 | Q1064L | D | 1.4e-05 | 1e-05 | 6.1e-06 | 1.003E-05 | 3.950E-06 | 6.100E-06 | 1.400E-05 |
| 21272 | 1064 | Q1064K | D | 1.3e-05 | 8e-06 | 1e-05 | 1.033E-05 | 2.517E-06 | 8.000E-06 | 1.300E-05 |
| 21273 | 1064 | Q1064M | D | 5.1e-06 | 7.8e-06 | 5e-06 | 5.967E-06 | 1.589E-06 | 5.000E-06 | 7.800E-06 |
| 21274 | 1064 | Q1064F | D | 6.2e-06 | 9.7e-06 | 9.1e-06 | 8.333E-06 | 1.872E-06 | 6.200E-06 | 9.700E-06 |
| 21275 | 1064 | Q1064P | D | 9.4e-06 | 9e-06 | 4.8e-06 | 7.733E-06 | 2.548E-06 | 4.800E-06 | 9.400E-06 |
| 21276 | 1064 | Q1064S | D | 6.2e-06 | 5.2e-06 | 8.3e-06 | 6.567E-06 | 1.582E-06 | 5.200E-06 | 8.300E-06 |
| 21277 | 1064 | Q1064T | D | 1.6e-05 | 5.6e-06 | 7.3e-06 | 9.633E-06 | 5.579E-06 | 5.600E-06 | 1.600E-05 |
| 21278 | 1064 | Q1064W | D | 1.2e-05 | 8.8e-06 | 9.7e-06 | 1.017E-05 | 1.650E-06 | 8.800E-06 | 1.200E-05 |
| 21279 | 1064 | Q1064Y | D | 4.7e-06 | 7e-06 | 8.2e-06 | 6.633E-06 | 1.779E-06 | 4.700E-06 | 8.200E-06 |
| 21280 | 1064 | Q1064V | D | 7.4e-06 | 6.6e-06 | 1e-05 | 8.000E-06 | 1.778E-06 | 6.600E-06 | 1.000E-05 |
| 21281 | 1065 | S1065A | D | 1e-05 | 6.1e-06 | 4.5e-06 | 6.867E-06 | 2.829E-06 | 4.500E-06 | 1.000E-05 |
| 21282 | 1065 | S1065R | D | 1e-05 | 5.4e-06 | 8.5e-06 | 7.967E-06 | 2.346E-06 | 5.400E-06 | 1.000E-05 |
| 21283 | 1065 | S1065N | D | 7.8e-06 | 5.2e-06 | 8.7e-06 | 7.233E-06 | 1.818E-06 | 5.200E-06 | 8.700E-06 |
| 21284 | 1065 | S1065D | D | 4.8e-06 | 8.2e-06 | 1e-05 | 7.667E-06 | 2.641E-06 | 4.800E-06 | 1.000E-05 |
| 21285 | 1065 | S1065C | D | 6.1e-06 | 1.6e-05 | 6.8e-06 | 9.633E-06 | 5.525E-06 | 6.100E-06 | 1.600E-05 |
| 21286 | 1065 | S1065E | D | 1.2e-05 | 6.6e-06 | 4.6e-06 | 7.733E-06 | 3.828E-06 | 4.600E-06 | 1.200E-05 |
| 21287 | 1065 | S1065Q | D | 9.7e-06 | 1.1e-05 | 8.6e-06 | 9.767E-06 | 1.201E-06 | 8.600E-06 | 1.100E-05 |
| 21288 | 1065 | S1065G | D | 3.6e-06 | 4.6e-06 | 9.6e-06 | 5.933E-06 | 3.215E-06 | 3.600E-06 | 9.600E-06 |
| 21289 | 1065 | S1065H | D | 6.1e-06 | 7.9e-06 | 1e-05 | 8.000E-06 | 1.952E-06 | 6.100E-06 | 1.000E-05 |
| 21290 | 1065 | S1065I | D | 1e-05 | 6.1e-06 | 9e-06 | 8.367E-06 | 2.026E-06 | 6.100E-06 | 1.000E-05 |
| 21291 | 1065 | S1065L | D | 1.1e-05 | 1.4e-05 | 7.7e-06 | 1.090E-05 | 3.151E-06 | 7.700E-06 | 1.400E-05 |
| 21292 | 1065 | S1065K | D | 9.5e-06 | 8.9e-06 | 1.1e-05 | 9.800E-06 | 1.082E-06 | 8.900E-06 | 1.100E-05 |
| 21293 | 1065 | S1065M | D | 6.5e-06 | 1e-05 | 6.5e-06 | 7.667E-06 | 2.021E-06 | 6.500E-06 | 1.000E-05 |
| 21294 | 1065 | S1065F | D | 9.6e-06 | 8.3e-06 | 6.5e-06 | 8.133E-06 | 1.557E-06 | 6.500E-06 | 9.600E-06 |
| 21295 | 1065 | S1065P | D | 5e-06 | 1.1e-05 | 1.2e-05 | 9.333E-06 | 3.786E-06 | 5.000E-06 | 1.200E-05 |
| 21296 | 1065 | S1065S | D | 1.2e-05 | 8.6e-06 | 5.5e-06 | 8.700E-06 | 3.251E-06 | 5.500E-06 | 1.200E-05 |
| 21297 | 1065 | S1065T | D | 4.8e-06 | 4.7e-06 | 4.8e-06 | 4.767E-06 | 5.774E-08 | 4.700E-06 | 4.800E-06 |
| 21298 | 1065 | S1065W | D | 1.4e-05 | 6.2e-06 | 9.8e-06 | 1.000E-05 | 3.904E-06 | 6.200E-06 | 1.400E-05 |
| 21299 | 1065 | S1065Y | D | 1e-05 | 1.1e-05 | 6.3e-06 | 9.100E-06 | 2.476E-06 | 6.300E-06 | 1.100E-05 |
| 21300 | 1065 | S1065V | D | 1.1e-05 | 5.7e-06 | 6.2e-06 | 7.633E-06 | 2.926E-06 | 5.700E-06 | 1.100E-05 |
| 21301 | 1066 | P1066A | D | 6e-06 | 8e-06 | 9.4e-06 | 7.800E-06 | 1.709E-06 | 6.000E-06 | 9.400E-06 |
| 21302 | 1066 | P1066R | D | 1e-05 | 5.8e-06 | 1.3e-05 | 9.600E-06 | 3.617E-06 | 5.800E-06 | 1.300E-05 |
| 21303 | 1066 | P1066N | D | 1e-05 | 8.5e-06 | 7.4e-06 | 8.633E-06 | 1.305E-06 | 7.400E-06 | 1.000E-05 |
| 21304 | 1066 | P1066D | D | 1.2e-05 | 9.4e-06 | 5.5e-06 | 8.967E-06 | 3.272E-06 | 5.500E-06 | 1.200E-05 |
| 21305 | 1066 | P1066C | D | 5.2e-06 | 5.4e-06 | 5.3e-06 | 5.300E-06 | 1.000E-07 | 5.200E-06 | 5.400E-06 |
| 21306 | 1066 | P1066E | D | 9.2e-06 | 9.3e-06 | 1.3e-05 | 1.050E-05 | 2.166E-06 | 9.200E-06 | 1.300E-05 |
| 21307 | 1066 | P1066Q | D | 5.8e-06 | 9.2e-06 | 7.7e-06 | 7.567E-06 | 1.704E-06 | 5.800E-06 | 9.200E-06 |
| 21308 | 1066 | P1066G | D | 8.6e-06 | 1.2e-05 | 7.2e-06 | 9.267E-06 | 2.468E-06 | 7.200E-06 | 1.200E-05 |
| 21309 | 1066 | P1066H | D | 1.2e-05 | 5.6e-06 | 5.2e-06 | 7.600E-06 | 3.816E-06 | 5.200E-06 | 1.200E-05 |
| 21310 | 1066 | P1066I | D | 7.4e-06 | 6.6e-06 | 7.9e-06 | 7.300E-06 | 6.557E-07 | 6.600E-06 | 7.900E-06 |
| 21311 | 1066 | P1066L | D | 1.3e-05 | 8e-06 | 1.1e-05 | 1.067E-05 | 2.517E-06 | 8.000E-06 | 1.300E-05 |
| 21312 | 1066 | P1066K | D | 3.6e-06 | 5.5e-06 | 4.8e-06 | 4.633E-06 | 9.609E-07 | 3.600E-06 | 5.500E-06 |
| 21313 | 1066 | P1066M | D | 7.8e-06 | 9.8e-06 | 2.9e-05 | 1.553E-05 | 1.171E-05 | 7.800E-06 | 2.900E-05 |
| 21314 | 1066 | P1066F | D | 5e-06 | 5.4e-06 | 1.1e-05 | 7.133E-06 | 3.355E-06 | 5.000E-06 | 1.100E-05 |
| 21315 | 1066 | P1066P | D | 9.4e-06 | 7.4e-06 | 7.5e-06 | 8.100E-06 | 1.127E-06 | 7.400E-06 | 9.400E-06 |
| 21316 | 1066 | P1066S | D | 1e-05 | 6.7e-06 | 1.8e-05 | 1.157E-05 | 5.811E-06 | 6.700E-06 | 1.800E-05 |
| 21317 | 1066 | P1066T | D | 9.3e-06 | 3.6e-06 | 1.1e-05 | 7.967E-06 | 3.876E-06 | 3.600E-06 | 1.100E-05 |
| 21318 | 1066 | P1066W | D | 1.1e-05 | 9e-06 | 8.9e-06 | 9.633E-06 | 1.185E-06 | 8.900E-06 | 1.100E-05 |
| 21319 | 1066 | P1066Y | D | 1.3e-05 | 9.3e-06 | 1e-05 | 1.077E-05 | 1.966E-06 | 9.300E-06 | 1.300E-05 |
| 21320 | 1066 | P1066V | D | 8.6e-06 | 7.3e-06 | 1.3e-05 | 9.633E-06 | 2.987E-06 | 7.300E-06 | 1.300E-05 |
| 21321 | 1067 | S1067A | D | 7.2e-06 | 8.1e-06 | 1.3e-05 | 9.433E-06 | 3.121E-06 | 7.200E-06 | 1.300E-05 |
| 21322 | 1067 | S1067R | D | 1e-05 | 8e-06 | 7.1e-06 | 8.367E-06 | 1.484E-06 | 7.100E-06 | 1.000E-05 |
| 21323 | 1067 | S1067N | D | 6.7e-06 | 5.2e-06 | 6.1e-06 | 6.000E-06 | 7.550E-07 | 5.200E-06 | 6.700E-06 |
| 21324 | 1067 | S1067D | D | 1.5e-05 | 9.3e-06 | 6.8e-06 | 1.037E-05 | 4.203E-06 | 6.800E-06 | 1.500E-05 |
| 21325 | 1067 | S1067C | D | 8e-06 | 7.5e-06 | 7.2e-06 | 7.567E-06 | 4.041E-07 | 7.200E-06 | 8.000E-06 |
| 21326 | 1067 | S1067E | D | 9.9e-06 | 6.4e-06 | 5.2e-06 | 7.167E-06 | 2.442E-06 | 5.200E-06 | 9.900E-06 |
| 21327 | 1067 | S1067Q | D | 1e-05 | 5.4e-06 | 8.2e-06 | 7.867E-06 | 2.318E-06 | 5.400E-06 | 1.000E-05 |
| 21328 | 1067 | S1067G | D | 3.5e-06 | 6e-06 | 6.1e-06 | 5.200E-06 | 1.473E-06 | 3.500E-06 | 6.100E-06 |
| 21329 | 1067 | S1067H | D | 8.5e-06 | 7.9e-06 | 8.1e-06 | 8.167E-06 | 3.055E-07 | 7.900E-06 | 8.500E-06 |
| 21330 | 1067 | S1067I | D | 9.5e-06 | 5.9e-06 | 4.9e-06 | 6.767E-06 | 2.419E-06 | 4.900E-06 | 9.500E-06 |
| 21331 | 1067 | S1067L | D | 5.1e-06 | 5.2e-06 | 6.8e-06 | 5.700E-06 | 9.539E-07 | 5.100E-06 | 6.800E-06 |
| 21332 | 1067 | S1067K | D | 5.8e-06 | 7.7e-06 | 6.3e-06 | 6.600E-06 | 9.849E-07 | 5.800E-06 | 7.700E-06 |
| 21333 | 1067 | S1067M | D | 6.5e-06 | 7.3e-06 | 8.9e-06 | 7.567E-06 | 1.222E-06 | 6.500E-06 | 8.900E-06 |
| 21334 | 1067 | S1067F | D | 1e-05 | 9.1e-06 | 6.7e-06 | 8.600E-06 | 1.706E-06 | 6.700E-06 | 1.000E-05 |

| | | | | | | | | | | |
|---|---|---|---|---|---|---|---|---|---|---|
| 21335 | 1067 | S1067P | D | 4.3e-06 | 1.1e-05 | 1.3e-05 | 9.433E-06 | 4.557E-06 | 4.300E-06 | 1.300E-05 |
| 21336 | 1067 | S1067S | D | 7.7e-06 | 7.5e-06 | 1.5e-05 | 1.007E-05 | 4.274E-06 | 7.500E-06 | 1.500E-05 |
| 21337 | 1067 | S1067T | D | 8.8e-06 | 8.2e-06 | 7.1e-06 | 8.033E-06 | 8.622E-07 | 7.100E-06 | 8.800E-06 |
| 21338 | 1067 | S1067W | D | 9.7e-06 | 9.9e-06 | 1e-05 | 9.867E-06 | 1.528E-07 | 9.700E-06 | 1.000E-05 |
| 21339 | 1067 | S1067Y | D | 8.9e-06 | 5.6e-06 | 6.1e-06 | 6.867E-06 | 1.779E-06 | 5.600E-06 | 8.900E-06 |
| 21340 | 1067 | S1067V | D | 1.1e-05 | 6.8e-06 | 9.6e-06 | 9.133E-06 | 2.139E-06 | 6.800E-06 | 1.100E-05 |
| 21341 | 1068 | S1068A | D | 1.3e-05 | 4.3e-06 | 8.7e-06 | 8.667E-06 | 4.350E-06 | 4.300E-06 | 1.300E-05 |
| 21342 | 1068 | S1068R | D | 8.4e-06 | 9.9e-06 | 4.5e-06 | 7.600E-06 | 2.787E-06 | 4.500E-06 | 9.900E-06 |
| 21343 | 1068 | S1068N | D | 9.2e-06 | 7.7e-06 | 7.2e-06 | 8.033E-06 | 1.041E-06 | 7.200E-06 | 9.200E-06 |
| 21344 | 1068 | S1068D | D | 7.7e-06 | 1.2e-05 | 7.1e-06 | 8.933E-06 | 2.673E-06 | 7.100E-06 | 1.200E-05 |
| 21345 | 1068 | S1068C | D | 9.2e-06 | 7.8e-06 | 5.3e-06 | 7.433E-06 | 1.976E-06 | 5.300E-06 | 9.200E-06 |
| 21346 | 1068 | S1068E | D | 1.1e-05 | 9.9e-06 | 9.6e-06 | 1.017E-05 | 7.371E-07 | 9.600E-06 | 1.100E-05 |
| 21347 | 1068 | S1068Q | D | 6.7e-06 | 5.4e-06 | 9.5e-06 | 7.200E-06 | 2.095E-06 | 5.400E-06 | 9.500E-06 |
| 21348 | 1068 | S1068G | D | 1e-05 | 1.1e-05 | 7.7e-06 | 9.567E-06 | 1.692E-06 | 7.700E-06 | 1.100E-05 |
| 21349 | 1068 | S1068H | D | 1.3e-05 | 1.4e-05 | 5e-06 | 1.067E-05 | 4.933E-06 | 5.000E-06 | 1.400E-05 |
| 21350 | 1068 | S1068I | D | 1.2e-05 | 5.9e-06 | 8.7e-06 | 8.867E-06 | 3.053E-06 | 5.900E-06 | 1.200E-05 |
| 21351 | 1068 | S1068L | D | 7.3e-06 | 7.3e-06 | 7.4e-06 | 7.333E-06 | 5.774E-08 | 7.300E-06 | 7.400E-06 |
| 21352 | 1068 | S1068K | D | 1.3e-05 | 6.4e-06 | 1.5e-05 | 1.147E-05 | 4.500E-06 | 6.400E-06 | 1.500E-05 |
| 21353 | 1068 | S1068M | D | 8.4e-06 | 7.7e-06 | 6.5e-06 | 7.533E-06 | 9.609E-07 | 6.500E-06 | 8.400E-06 |
| 21354 | 1068 | S1068F | D | 8.9e-06 | 9.2e-06 | 7.9e-06 | 8.667E-06 | 6.807E-07 | 7.900E-06 | 9.200E-06 |
| 21355 | 1068 | S1068P | D | 1.3e-05 | 9e-06 | 1.6e-05 | 1.267E-05 | 3.512E-06 | 9.000E-06 | 1.600E-05 |
| 21356 | 1068 | S1068S | D | 9e-06 | 4.9e-06 | 6.9e-06 | 6.933E-06 | 2.050E-06 | 4.900E-06 | 9.000E-06 |
| 21357 | 1068 | S1068T | D | 1.5e-05 | 6.7e-06 | 9.8e-06 | 1.050E-05 | 4.194E-06 | 6.700E-06 | 1.500E-05 |
| 21358 | 1068 | S1068W | D | 1.3e-05 | 7.5e-06 | 1.7e-05 | 1.250E-05 | 4.770E-06 | 7.500E-06 | 1.700E-05 |
| 21359 | 1068 | S1068Y | D | 9.2e-06 | 1.2e-05 | 4.6e-06 | 8.600E-06 | 3.736E-06 | 4.600E-06 | 1.200E-05 |
| 21360 | 1068 | S1068V | D | 8.4e-06 | 6.3e-06 | 1e-05 | 8.233E-06 | 1.856E-06 | 6.300E-06 | 1.000E-05 |
| 21361 | 1069 | L1069A | D | 9e-06 | 5.9e-06 | 1.4e-05 | 9.633E-06 | 4.087E-06 | 5.900E-06 | 1.400E-05 |
| 21362 | 1069 | L1069R | D | 4.6e-06 | 5.7e-06 | 1.1e-05 | 7.100E-06 | 3.422E-06 | 4.600E-06 | 1.100E-05 |
| 21363 | 1069 | L1069N | D | 7.4e-06 | 5.6e-06 | 6.5e-06 | 6.500E-06 | 9.000E-07 | 5.600E-06 | 7.400E-06 |
| 21364 | 1069 | L1069D | D | 1.4e-05 | 4.8e-06 | 7.2e-06 | 8.667E-06 | 4.772E-06 | 4.800E-06 | 1.400E-05 |
| 21365 | 1069 | L1069C | D | 1.2e-05 | 8.6e-06 | 4.2e-06 | 8.267E-06 | 3.911E-06 | 4.200E-06 | 1.200E-05 |
| 21366 | 1069 | L1069E | D | 1.4e-05 | 6.4e-06 | 1.3e-05 | 1.113E-05 | 4.130E-06 | 6.400E-06 | 1.400E-05 |
| 21367 | 1069 | L1069Q | D | 9.3e-06 | 7.4e-06 | 6.5e-06 | 7.733E-06 | 1.429E-06 | 6.500E-06 | 9.300E-06 |
| 21368 | 1069 | L1069G | D | 3.4e-06 | 7.3e-06 | 1.1e-05 | 7.233E-06 | 3.800E-06 | 3.400E-06 | 1.100E-05 |
| 21369 | 1069 | L1069H | D | 5.3e-06 | 6.6e-06 | 3.3e-06 | 5.067E-06 | 1.662E-06 | 3.300E-06 | 6.600E-06 |
| 21370 | 1069 | L1069I | D | 8.9e-06 | 4.5e-06 | 5e-06 | 6.133E-06 | 2.409E-06 | 4.500E-06 | 8.900E-06 |
| 21371 | 1069 | L1069L | D | 7.7e-06 | 9.4e-06 | 7e-06 | 8.033E-06 | 1.234E-06 | 7.000E-06 | 9.400E-06 |
| 21372 | 1069 | L1069K | D | 5.5e-06 | 1.3e-05 | 8.9e-06 | 9.133E-06 | 3.755E-06 | 5.500E-06 | 1.300E-05 |
| 21373 | 1069 | L1069M | D | 7.6e-06 | 9.1e-06 | 8.5e-06 | 8.400E-06 | 7.550E-07 | 7.600E-06 | 9.100E-06 |
| 21374 | 1069 | L1069F | D | 6.9e-06 | 4.6e-06 | 1.3e-05 | 8.167E-06 | 4.341E-06 | 4.600E-06 | 1.300E-05 |
| 21375 | 1069 | L1069P | D | 8.2e-06 | 5.5e-06 | 6.1e-06 | 6.600E-06 | 1.418E-06 | 5.500E-06 | 8.200E-06 |
| 21376 | 1069 | L1069S | D | 6.8e-06 | 4.9e-06 | 9.7e-06 | 7.133E-06 | 2.417E-06 | 4.900E-06 | 9.700E-06 |
| 21377 | 1069 | L1069T | D | 5.5e-06 | 6.9e-06 | 5e-06 | 5.800E-06 | 9.849E-07 | 5.000E-06 | 6.900E-06 |
| 21378 | 1069 | L1069W | D | 1.1e-05 | 9e-06 | 8.1e-06 | 9.367E-06 | 1.484E-06 | 8.100E-06 | 1.100E-05 |
| 21379 | 1069 | L1069Y | D | 5.9e-06 | 6e-06 | 5.6e-06 | 5.833E-06 | 2.082E-07 | 5.600E-06 | 6.000E-06 |
| 21380 | 1069 | L1069V | D | 9e-06 | 6.7e-06 | 7.7e-06 | 7.800E-06 | 1.153E-06 | 6.700E-06 | 9.000E-06 |
| 21381 | 1070 | S1070A | D | 5.3e-06 | 6.7e-06 | 8e-06 | 6.667E-06 | 1.350E-06 | 5.300E-06 | 8.000E-06 |
| 21382 | 1070 | S1070R | D | 1.1e-05 | 9.4e-06 | 9.4e-06 | 9.933E-06 | 9.238E-07 | 9.400E-06 | 1.100E-05 |
| 21383 | 1070 | S1070N | D | 1e-05 | 7.8e-06 | 5.2e-06 | 7.667E-06 | 2.403E-06 | 5.200E-06 | 1.000E-05 |
| 21384 | 1070 | S1070D | D | 1.3e-05 | 1.1e-05 | 9.9e-06 | 1.130E-05 | 1.572E-06 | 9.900E-06 | 1.300E-05 |
| 21385 | 1070 | S1070C | D | 7.7e-06 | 5.5e-06 | 6.3e-06 | 6.500E-06 | 1.114E-06 | 5.500E-06 | 7.700E-06 |
| 21386 | 1070 | S1070E | D | 5.3e-06 | 1.1e-05 | 1.4e-05 | 1.010E-05 | 4.419E-06 | 5.300E-06 | 1.400E-05 |
| 21387 | 1070 | S1070Q | D | 7e-06 | 1.1e-05 | 6.5e-06 | 8.167E-06 | 2.466E-06 | 6.500E-06 | 1.100E-05 |
| 21388 | 1070 | S1070G | D | 1.1e-05 | 4.6e-06 | 9.9e-06 | 8.500E-06 | 3.422E-06 | 4.600E-06 | 1.100E-05 |
| 21389 | 1070 | S1070H | D | 7.8e-06 | 8.1e-06 | 9.1e-06 | 8.333E-06 | 6.807E-07 | 7.800E-06 | 9.100E-06 |
| 21390 | 1070 | S1070I | D | 1.1e-05 | 6.6e-06 | 8.1e-06 | 8.567E-06 | 2.237E-06 | 6.600E-06 | 1.100E-05 |
| 21391 | 1070 | S1070L | D | 1.1e-05 | 8.6e-06 | 1.2e-05 | 1.053E-05 | 1.747E-06 | 8.600E-06 | 1.200E-05 |
| 21392 | 1070 | S1070K | D | 1.1e-05 | 4.6e-06 | 1.4e-05 | 9.867E-06 | 4.801E-06 | 4.600E-06 | 1.400E-05 |
| 21393 | 1070 | S1070M | D | 1.1e-05 | 9.3e-06 | 8.1e-06 | 9.467E-06 | 1.457E-06 | 8.100E-06 | 1.100E-05 |
| 21394 | 1070 | S1070F | D | 8.8e-06 | 7.7e-06 | 8.7e-06 | 8.400E-06 | 6.083E-07 | 7.700E-06 | 8.800E-06 |
| 21395 | 1070 | S1070P | D | 8.6e-06 | 6.9e-06 | 9.6e-06 | 8.367E-06 | 1.365E-06 | 6.900E-06 | 9.600E-06 |
| 21396 | 1070 | S1070S | D | 8.7e-06 | 6.7e-06 | 8.3e-06 | 7.900E-06 | 1.058E-06 | 6.700E-06 | 8.700E-06 |
| 21397 | 1070 | S1070T | D | 1.1e-05 | 9.7e-06 | 1.2e-05 | 1.090E-05 | 1.153E-06 | 9.700E-06 | 1.200E-05 |
| 21398 | 1070 | S1070W | D | 9.9e-06 | 5e-06 | 6.8e-06 | 7.233E-06 | 2.479E-06 | 5.000E-06 | 9.900E-06 |
| 21399 | 1070 | S1070Y | D | 8.9e-06 | 1.1e-05 | 7.5e-06 | 9.133E-06 | 1.762E-06 | 7.500E-06 | 1.100E-05 |
| 21400 | 1070 | S1070V | D | 7.3e-06 | 1.4e-05 | 1.2e-05 | 1.110E-05 | 3.439E-06 | 7.300E-06 | 1.400E-05 |
| 21401 | 1071 | A1071A | D | 4.7e-06 | 5.5e-06 | 8.6e-06 | 6.267E-06 | 2.060E-06 | 4.700E-06 | 8.600E-06 |
| 21402 | 1071 | A1071R | D | 6.9e-06 | 1e-05 | 6.6e-06 | 7.833E-06 | 1.882E-06 | 6.600E-06 | 1.000E-05 |
| 21403 | 1071 | A1071N | D | 6.2e-06 | 1.1e-05 | 6.1e-06 | 7.767E-06 | 2.801E-06 | 6.100E-06 | 1.100E-05 |
| 21404 | 1071 | A1071D | D | 8.4e-06 | 6.6e-06 | 1.6e-05 | 1.033E-05 | 4.989E-06 | 6.600E-06 | 1.600E-05 |
| 21405 | 1071 | A1071C | D | 7.7e-06 | 6.2e-06 | 7.5e-06 | 7.133E-06 | 8.145E-07 | 6.200E-06 | 7.700E-06 |
| 21406 | 1071 | A1071E | D | 1.3e-05 | 9.1e-06 | 9.9e-06 | 1.067E-05 | 2.060E-06 | 9.100E-06 | 1.300E-05 |
| 21407 | 1071 | A1071Q | D | 8.6e-06 | 6.7e-06 | 6.4e-06 | 7.233E-06 | 1.193E-06 | 6.400E-06 | 8.600E-06 |
| 21408 | 1071 | A1071G | D | 1.1e-05 | 5.4e-06 | 5.9e-06 | 7.433E-06 | 3.099E-06 | 5.400E-06 | 1.100E-05 |
| 21409 | 1071 | A1071H | D | 6.1e-06 | 6.4e-06 | 4.4e-06 | 5.633E-06 | 1.079E-06 | 4.400E-06 | 6.400E-06 |
| 21410 | 1071 | A1071I | D | 8e-06 | 7.2e-06 | 6.2e-06 | 7.133E-06 | 9.018E-07 | 6.200E-06 | 8.000E-06 |
| 21411 | 1071 | A1071L | D | 9.7e-06 | 1e-05 | 7e-06 | 8.900E-06 | 1.652E-06 | 7.000E-06 | 1.000E-05 |
| 21412 | 1071 | A1071K | D | 6.5e-06 | 6.3e-06 | 9.5e-06 | 7.433E-06 | 1.793E-06 | 6.300E-06 | 9.500E-06 |
| 21413 | 1071 | A1071M | D | 5e-06 | 1.8e-05 | 1.4e-05 | 1.233E-05 | 6.658E-06 | 5.000E-06 | 1.800E-05 |
| 21414 | 1071 | A1071F | D | 5.8e-06 | 5.9e-06 | 6.5e-06 | 6.067E-06 | 3.786E-07 | 5.800E-06 | 6.500E-06 |
| 21415 | 1071 | A1071P | D | 5.9e-06 | 4.3e-06 | 1.1e-05 | 7.067E-06 | 3.499E-06 | 4.300E-06 | 1.100E-05 |
| 21416 | 1071 | A1071S | D | 1.5e-05 | 3.1e-06 | 8.5e-06 | 8.867E-06 | 5.958E-06 | 3.100E-06 | 1.500E-05 |
| 21417 | 1071 | A1071T | D | 1.3e-05 | 9.5e-06 | 7.9e-06 | 1.013E-05 | 2.608E-06 | 7.900E-06 | 1.300E-05 |
| 21418 | 1071 | A1071W | D | 8.2e-06 | 6.1e-06 | 4.8e-06 | 6.367E-06 | 1.716E-06 | 4.800E-06 | 8.200E-06 |
| 21419 | 1071 | A1071Y | D | 6.2e-06 | 6.3e-06 | 7.2e-06 | 6.567E-06 | 5.508E-07 | 6.200E-06 | 7.200E-06 |

| | | | | | | | | | | |
|---|---|---|---|---|---|---|---|---|---|---|
| 21420 | 1071 | A1071V | D | 8.6e-06 | 6e-06 | 7.5e-06 | 7.367E-06 | 1.305E-06 | 6.000E-06 | 8.600E-06 |
| 21421 | 1072 | S1072A | D | 6.8e-06 | 6.5e-06 | 1.1e-05 | 8.100E-06 | 2.516E-06 | 6.500E-06 | 1.100E-05 |
| 21422 | 1072 | S1072R | D | 9.3e-06 | 6e-06 | 6.1e-06 | 7.133E-06 | 1.877E-06 | 6.000E-06 | 9.300E-06 |
| 21423 | 1072 | S1072N | D | 1.1e-05 | 9.5e-06 | 5.7e-06 | 8.733E-06 | 2.732E-06 | 5.700E-06 | 1.100E-05 |
| 21424 | 1072 | S1072D | D | 1.1e-05 | 7.7e-06 | 9.5e-06 | 9.400E-06 | 1.652E-06 | 7.700E-06 | 1.100E-05 |
| 21425 | 1072 | S1072C | D | 9.7e-06 | 4.8e-06 | 5.2e-06 | 6.567E-06 | 2.721E-06 | 4.800E-06 | 9.700E-06 |
| 21426 | 1072 | S1072E | D | 8.7e-06 | 8e-06 | 1.3e-05 | 9.900E-06 | 2.707E-06 | 8.000E-06 | 1.300E-05 |
| 21427 | 1072 | S1072Q | D | 8.6e-06 | 3.1e-06 | 8.5e-06 | 6.733E-06 | 3.147E-06 | 3.100E-06 | 8.600E-06 |
| 21428 | 1072 | S1072G | D | 1.4e-05 | 9.7e-06 | 1.1e-05 | 1.157E-05 | 2.205E-06 | 9.700E-06 | 1.400E-05 |
| 21429 | 1072 | S1072H | D | 9.6e-06 | 9.3e-06 | 9.2e-06 | 9.367E-06 | 2.082E-07 | 9.200E-06 | 9.600E-06 |
| 21430 | 1072 | S1072I | D | 4.8e-06 | 9.4e-06 | 7.3e-06 | 7.167E-06 | 2.303E-06 | 4.800E-06 | 9.400E-06 |
| 21431 | 1072 | S1072L | D | 6.1e-06 | 6.9e-06 | 6.6e-06 | 6.533E-06 | 4.041E-07 | 6.100E-06 | 6.900E-06 |
| 21432 | 1072 | S1072K | D | 6.7e-06 | 6.2e-06 | 1.2e-05 | 8.300E-06 | 3.214E-06 | 6.200E-06 | 1.200E-05 |
| 21433 | 1072 | S1072M | D | 8.7e-06 | 6.6e-06 | 9.1e-06 | 8.133E-06 | 1.343E-06 | 6.600E-06 | 9.100E-06 |
| 21434 | 1072 | S1072F | D | 1.3e-05 | 1.4e-05 | 4.6e-06 | 1.053E-05 | 5.163E-06 | 4.600E-06 | 1.400E-05 |
| 21435 | 1072 | S1072P | D | 6e-06 | 1.4e-05 | 6.3e-06 | 8.767E-06 | 4.535E-06 | 6.000E-06 | 1.400E-05 |
| 21436 | 1072 | S1072S | D | 8.7e-06 | 8e-06 | 5.1e-06 | 7.267E-06 | 1.909E-06 | 5.100E-06 | 8.700E-06 |
| 21437 | 1072 | S1072T | D | 6.6e-06 | 9.1e-06 | 4.4e-06 | 6.700E-06 | 2.352E-06 | 4.400E-06 | 9.100E-06 |
| 21438 | 1072 | S1072W | D | 1.2e-05 | 1e-05 | 1.3e-05 | 1.167E-05 | 1.528E-06 | 1.000E-05 | 1.300E-05 |
| 21439 | 1072 | S1072Y | D | 5.8e-06 | 7.2e-06 | 1.3e-05 | 8.667E-06 | 3.818E-06 | 5.800E-06 | 1.300E-05 |
| 21440 | 1072 | S1072V | D | 7.7e-06 | 9.7e-06 | 8.8e-06 | 8.733E-06 | 1.002E-06 | 7.700E-06 | 9.700E-06 |
| 21441 | 1073 | V1073A | D | 9.7e-06 | 6.1e-06 | 7.8e-06 | 7.867E-06 | 1.801E-06 | 6.100E-06 | 9.700E-06 |
| 21442 | 1073 | V1073R | D | 7.4e-06 | 8.2e-06 | 5.1e-06 | 6.900E-06 | 1.609E-06 | 5.100E-06 | 8.200E-06 |
| 21443 | 1073 | V1073N | D | 6.2e-06 | 8e-06 | 4.7e-06 | 6.300E-06 | 1.652E-06 | 4.700E-06 | 8.000E-06 |
| 21444 | 1073 | V1073D | D | 7e-06 | 1.2e-05 | 8.2e-06 | 9.067E-06 | 2.610E-06 | 7.000E-06 | 1.200E-05 |
| 21445 | 1073 | V1073C | D | 7.6e-06 | 1.2e-05 | 8.5e-06 | 9.367E-06 | 2.325E-06 | 7.600E-06 | 1.200E-05 |
| 21446 | 1073 | V1073E | D | 8.1e-06 | 7.9e-06 | 8.2e-06 | 8.067E-06 | 1.528E-07 | 7.900E-06 | 8.200E-06 |
| 21447 | 1073 | V1073Q | D | 6.1e-06 | 8.4e-06 | 1.4e-05 | 9.500E-06 | 4.063E-06 | 6.100E-06 | 1.400E-05 |
| 21448 | 1073 | V1073G | D | 8e-06 | 6.6e-06 | 7.4e-06 | 7.333E-06 | 7.024E-07 | 6.600E-06 | 8.000E-06 |
| 21449 | 1073 | V1073H | D | 1e-05 | 7.2e-06 | 8.7e-06 | 8.633E-06 | 1.401E-06 | 7.200E-06 | 1.000E-05 |
| 21450 | 1073 | V1073I | D | 9.3e-06 | 7.9e-06 | 1.2e-05 | 9.733E-06 | 2.084E-06 | 7.900E-06 | 1.200E-05 |
| 21451 | 1073 | V1073L | D | 6.6e-06 | 9.3e-06 | 9.5e-06 | 8.467E-06 | 1.620E-06 | 6.600E-06 | 9.500E-06 |
| 21452 | 1073 | V1073K | D | 9.3e-06 | 6.2e-06 | 9.6e-06 | 8.367E-06 | 1.882E-06 | 6.200E-06 | 9.600E-06 |
| 21453 | 1073 | V1073M | D | 9.7e-06 | 7.1e-06 | 6.4e-06 | 7.733E-06 | 1.739E-06 | 6.400E-06 | 9.700E-06 |
| 21454 | 1073 | V1073F | D | 6e-06 | 1.1e-05 | 1.1e-05 | 9.333E-06 | 2.887E-06 | 6.000E-06 | 1.100E-05 |
| 21455 | 1073 | V1073P | D | 8.9e-06 | 5.9e-06 | 2e-05 | 1.160E-05 | 7.428E-06 | 5.900E-06 | 2.000E-05 |
| 21456 | 1073 | V1073S | D | 1.1e-05 | 1.1e-05 | 1e-05 | 1.067E-05 | 5.774E-07 | 1.000E-05 | 1.100E-05 |
| 21457 | 1073 | V1073T | D | 4.7e-06 | 7.1e-06 | 7.4e-06 | 6.400E-06 | 1.480E-06 | 4.700E-06 | 7.400E-06 |
| 21458 | 1073 | V1073W | D | 6e-06 | 1e-05 | 1e-05 | 8.667E-06 | 2.309E-06 | 6.000E-06 | 1.000E-05 |
| 21459 | 1073 | V1073Y | D | 9.1e-06 | 8.6e-06 | 5.6e-06 | 7.767E-06 | 1.893E-06 | 5.600E-06 | 9.100E-06 |
| 21460 | 1073 | V1073V | D | 9.4e-06 | 9.2e-06 | 1.2e-05 | 1.020E-05 | 1.562E-06 | 9.200E-06 | 1.200E-05 |
| 21461 | 1074 | G1074A | D | 1.5e-05 | 1.1e-05 | 4.5e-06 | 1.017E-05 | 5.299E-06 | 4.500E-06 | 1.500E-05 |
| 21462 | 1074 | G1074R | D | 5e-06 | 1.4e-05 | 8e-06 | 9.000E-06 | 4.583E-06 | 5.000E-06 | 1.400E-05 |
| 21463 | 1074 | G1074N | D | 8.4e-06 | 7.4e-06 | 7e-06 | 7.600E-06 | 7.211E-07 | 7.000E-06 | 8.400E-06 |
| 21464 | 1074 | G1074D | D | 7.7e-06 | 9.9e-06 | 8e-06 | 8.533E-06 | 1.193E-06 | 7.700E-06 | 9.900E-06 |
| 21465 | 1074 | G1074C | D | 1.2e-05 | 7.8e-06 | 6e-06 | 8.600E-06 | 3.079E-06 | 6.000E-06 | 1.200E-05 |
| 21466 | 1074 | G1074E | D | 9.5e-06 | 9.3e-06 | 8.7e-06 | 9.167E-06 | 4.163E-07 | 8.700E-06 | 9.500E-06 |
| 21467 | 1074 | G1074Q | D | 7.2e-06 | 4.6e-06 | 4.7e-06 | 5.500E-06 | 1.473E-06 | 4.600E-06 | 7.200E-06 |
| 21468 | 1074 | G1074G | D | 4.7e-06 | 8.9e-06 | 8.2e-06 | 7.267E-06 | 2.250E-06 | 4.700E-06 | 8.900E-06 |
| 21469 | 1074 | G1074H | D | 6.9e-06 | 1.1e-05 | 9e-06 | 8.967E-06 | 2.050E-06 | 6.900E-06 | 1.100E-05 |
| 21470 | 1074 | G1074I | D | 9.3e-06 | 8.6e-06 | 8e-06 | 8.633E-06 | 6.506E-07 | 8.000E-06 | 9.300E-06 |
| 21471 | 1074 | G1074L | D | 9.4e-06 | 8.5e-06 | 7.8e-06 | 8.567E-06 | 8.021E-07 | 7.800E-06 | 9.400E-06 |
| 21472 | 1074 | G1074K | D | 5.8e-06 | 8.7e-06 | 1.1e-05 | 8.500E-06 | 2.606E-06 | 5.800E-06 | 1.100E-05 |
| 21473 | 1074 | G1074M | D | 7.2e-06 | 6.9e-06 | 1.1e-05 | 8.367E-06 | 2.285E-06 | 6.900E-06 | 1.100E-05 |
| 21474 | 1074 | G1074F | D | 5.9e-06 | 4.6e-06 | 9.1e-06 | 6.533E-06 | 2.316E-06 | 4.600E-06 | 9.100E-06 |
| 21475 | 1074 | G1074P | D | 6.6e-06 | 6.4e-06 | 5.3e-06 | 6.100E-06 | 7.000E-07 | 5.300E-06 | 6.600E-06 |
| 21476 | 1074 | G1074S | D | 8.3e-06 | 5e-06 | 5.4e-06 | 6.233E-06 | 1.801E-06 | 5.000E-06 | 8.300E-06 |
| 21477 | 1074 | G1074T | D | 7.5e-06 | 6.1e-06 | 9.7e-06 | 7.767E-06 | 1.815E-06 | 6.100E-06 | 9.700E-06 |
| 21478 | 1074 | G1074W | D | 8.5e-06 | 7.4e-06 | 1.1e-05 | 8.967E-06 | 1.845E-06 | 7.400E-06 | 1.100E-05 |
| 21479 | 1074 | G1074Y | D | 6.9e-06 | 5.9e-06 | 1.2e-05 | 8.267E-06 | 3.272E-06 | 5.900E-06 | 1.200E-05 |
| 21480 | 1074 | G1074V | D | 9.3e-06 | 9.1e-06 | 6.7e-06 | 8.367E-06 | 1.447E-06 | 6.700E-06 | 9.300E-06 |
| 21481 | 1075 | D1075A | D | 9.9e-06 | 4.4e-06 | 5.6e-06 | 6.633E-06 | 2.892E-06 | 4.400E-06 | 9.900E-06 |
| 21482 | 1075 | D1075R | D | 6.7e-06 | 6.7e-06 | 6.1e-06 | 6.500E-06 | 3.464E-07 | 6.100E-06 | 6.700E-06 |
| 21483 | 1075 | D1075N | D | 1.1e-05 | 1.4e-05 | 8.1e-06 | 1.103E-05 | 2.950E-06 | 8.100E-06 | 1.400E-05 |
| 21484 | 1075 | D1075D | D | 7.4e-06 | 1.3e-05 | 8.5e-06 | 9.633E-06 | 2.967E-06 | 7.400E-06 | 1.300E-05 |
| 21485 | 1075 | D1075C | D | 7.8e-06 | 3.4e-06 | 8.1e-06 | 6.433E-06 | 2.631E-06 | 3.400E-06 | 8.100E-06 |
| 21486 | 1075 | D1075E | D | 9.1e-06 | 9.3e-06 | 6.7e-06 | 8.367E-06 | 1.447E-06 | 6.700E-06 | 9.300E-06 |
| 21487 | 1075 | D1075Q | D | 8.5e-06 | 5.5e-06 | 9.1e-06 | 7.700E-06 | 1.929E-06 | 5.500E-06 | 9.100E-06 |
| 21488 | 1075 | D1075G | D | 6.8e-06 | 1.3e-05 | 7.5e-06 | 9.100E-06 | 3.396E-06 | 6.800E-06 | 1.300E-05 |
| 21489 | 1075 | D1075H | D | 7.1e-06 | 5.7e-06 | 7e-06 | 6.600E-06 | 7.810E-07 | 5.700E-06 | 7.100E-06 |
| 21490 | 1075 | D1075I | D | 6.5e-06 | 8.5e-06 | 5.7e-06 | 6.900E-06 | 1.442E-06 | 5.700E-06 | 8.500E-06 |
| 21491 | 1075 | D1075L | D | 4.6e-06 | 1.4e-05 | 7.1e-06 | 8.567E-06 | 4.869E-06 | 4.600E-06 | 1.400E-05 |
| 21492 | 1075 | D1075K | D | 6.5e-06 | 6.2e-06 | 6.1e-06 | 6.267E-06 | 2.082E-07 | 6.100E-06 | 6.500E-06 |
| 21493 | 1075 | D1075M | D | 7.9e-06 | 5.4e-06 | 8.1e-06 | 7.133E-06 | 1.504E-06 | 5.400E-06 | 8.100E-06 |
| 21494 | 1075 | D1075F | D | 8.8e-06 | 8.7e-06 | 7e-06 | 8.167E-06 | 1.012E-06 | 7.000E-06 | 8.800E-06 |
| 21495 | 1075 | D1075P | D | 5.3e-06 | 1.3e-05 | 5.7e-06 | 8.000E-06 | 4.335E-06 | 5.300E-06 | 1.300E-05 |
| 21496 | 1075 | D1075S | D | 6.5e-06 | 9e-06 | 8.1e-06 | 7.867E-06 | 1.266E-06 | 6.500E-06 | 9.000E-06 |
| 21497 | 1075 | D1075T | D | 6.4e-06 | 1.3e-05 | 9.8e-06 | 9.733E-06 | 3.301E-06 | 6.400E-06 | 1.300E-05 |
| 21498 | 1075 | D1075W | D | 4.7e-06 | 5.3e-06 | 7.6e-06 | 5.867E-06 | 1.531E-06 | 4.700E-06 | 7.600E-06 |
| 21499 | 1075 | D1075Y | D | 6.4e-06 | 1.2e-05 | 7e-06 | 8.467E-06 | 3.075E-06 | 6.400E-06 | 1.200E-05 |
| 21500 | 1075 | D1075V | D | 9e-06 | 1.2e-05 | 8.9e-06 | 9.967E-06 | 1.762E-06 | 8.900E-06 | 1.200E-05 |
| 21501 | 1076 | R1076A | D | 5.8e-06 | 7e-06 | 1e-05 | 7.600E-06 | 2.163E-06 | 5.800E-06 | 1.000E-05 |
| 21502 | 1076 | R1076R | D | 5e-06 | 6.3e-06 | 6.8e-06 | 6.033E-06 | 9.292E-07 | 5.000E-06 | 6.800E-06 |
| 21503 | 1076 | R1076N | D | 8.8e-06 | 5.2e-06 | 8.8e-06 | 7.600E-06 | 2.078E-06 | 5.200E-06 | 8.800E-06 |
| 21504 | 1076 | R1076D | D | 6e-06 | 5.7e-06 | 1.1e-05 | 7.567E-06 | 2.977E-06 | 5.700E-06 | 1.100E-05 |

| | | | | | | | | | | |
|---|---|---|---|---|---|---|---|---|---|---|
| 21505 | 1076 | R1076C | D | 7.5e-06 | 1.1e-05 | 6.9e-06 | 8.467E-06 | 2.214E-06 | 6.900E-06 | 1.100E-05 |
| 21506 | 1076 | R1076E | D | 6.9e-06 | 1.2e-05 | 4.7e-06 | 7.867E-06 | 3.745E-06 | 4.700E-06 | 1.200E-05 |
| 21507 | 1076 | R1076Q | D | 5.3e-06 | 6.4e-06 | 7.8e-06 | 6.500E-06 | 1.253E-06 | 5.300E-06 | 7.800E-06 |
| 21508 | 1076 | R1076G | D | 6.3e-06 | 8.2e-06 | 1.3e-05 | 9.167E-06 | 3.453E-06 | 6.300E-06 | 1.300E-05 |
| 21509 | 1076 | R1076H | D | 3.7e-06 | 6.1e-06 | 7e-06 | 5.600E-06 | 1.706E-06 | 3.700E-06 | 7.000E-06 |
| 21510 | 1076 | R1076I | D | 3.6e-06 | 8.4e-06 | 1e-05 | 7.333E-06 | 3.331E-06 | 3.600E-06 | 1.000E-05 |
| 21511 | 1076 | R1076L | D | 6.4e-06 | 1.3e-05 | 7.2e-06 | 8.867E-06 | 3.602E-06 | 6.400E-06 | 1.300E-05 |
| 21512 | 1076 | R1076K | D | 8.8e-06 | 9.3e-06 | 6.2e-06 | 8.100E-06 | 1.664E-06 | 6.200E-06 | 9.300E-06 |
| 21513 | 1076 | R1076M | D | 5.8e-06 | 7.6e-06 | 1.3e-05 | 8.800E-06 | 3.747E-06 | 5.800E-06 | 1.300E-05 |
| 21514 | 1076 | R1076F | D | 8.7e-06 | 4.5e-06 | 6.7e-06 | 6.633E-06 | 2.101E-06 | 4.500E-06 | 8.700E-06 |
| 21515 | 1076 | R1076P | D | 6.3e-06 | 9.3e-06 | 8e-06 | 7.867E-06 | 1.504E-06 | 6.300E-06 | 9.300E-06 |
| 21516 | 1076 | R1076S | D | 9.3e-06 | 1.3e-05 | 2e-05 | 1.410E-05 | 5.434E-06 | 9.300E-06 | 2.000E-05 |
| 21517 | 1076 | R1076T | D | 6.4e-06 | 8e-06 | 5.5e-06 | 6.633E-06 | 1.266E-06 | 5.500E-06 | 8.000E-06 |
| 21518 | 1076 | R1076W | D | 9.3e-06 | 9.9e-06 | 6.2e-06 | 8.467E-06 | 1.986E-06 | 6.200E-06 | 9.900E-06 |
| 21519 | 1076 | R1076Y | D | 4.8e-06 | 6.5e-06 | 1.2e-05 | 7.767E-06 | 3.763E-06 | 4.800E-06 | 1.200E-05 |
| 21520 | 1076 | R1076V | D | 1e-05 | 1.1e-05 | 4.9e-06 | 8.633E-06 | 3.272E-06 | 4.900E-06 | 1.100E-05 |
| 21521 | 1077 | V1077A | D | 5.2e-06 | 8.2e-06 | 8.2e-06 | 7.200E-06 | 1.732E-06 | 5.200E-06 | 8.200E-06 |
| 21522 | 1077 | V1077R | D | 7.4e-06 | 4.7e-06 | 7.7e-06 | 6.600E-06 | 1.652E-06 | 4.700E-06 | 7.700E-06 |
| 21523 | 1077 | V1077N | D | 5.2e-06 | 9.4e-06 | 9.9e-06 | 8.167E-06 | 2.581E-06 | 5.200E-06 | 9.900E-06 |
| 21524 | 1077 | V1077D | D | 7e-06 | 6.8e-06 | 4.2e-06 | 6.000E-06 | 1.562E-06 | 4.200E-06 | 7.000E-06 |
| 21525 | 1077 | V1077C | D | 6.5e-06 | 6.5e-06 | 6.5e-06 | 6.500E-06 | 0.000E+00 | 6.500E-06 | 6.500E-06 |
| 21526 | 1077 | V1077E | D | 6.1e-06 | 8.6e-06 | 8.5e-06 | 7.733E-06 | 1.415E-06 | 6.100E-06 | 8.600E-06 |
| 21527 | 1077 | V1077Q | D | 5.9e-06 | 5.9e-06 | 7e-06 | 6.267E-06 | 6.351E-07 | 5.900E-06 | 7.000E-06 |
| 21528 | 1077 | V1077G | D | 5.1e-06 | 7.8e-06 | 9.3e-06 | 7.400E-06 | 2.128E-06 | 5.100E-06 | 9.300E-06 |
| 21529 | 1077 | V1077H | D | 8.2e-06 | 9.8e-06 | 4.7e-06 | 7.567E-06 | 2.608E-06 | 4.700E-06 | 9.800E-06 |
| 21530 | 1077 | V1077I | D | 5.1e-06 | 6.5e-06 | 6.7e-06 | 6.100E-06 | 8.718E-07 | 5.100E-06 | 6.700E-06 |
| 21531 | 1077 | V1077L | D | 1.2e-05 | 1.3e-05 | 6.7e-06 | 1.057E-05 | 3.386E-06 | 6.700E-06 | 1.300E-05 |
| 21532 | 1077 | V1077K | D | 7.3e-06 | 1e-05 | 1e-05 | 9.100E-06 | 1.559E-06 | 7.300E-06 | 1.000E-05 |
| 21533 | 1077 | V1077M | D | 7e-06 | 1.2e-05 | 7.9e-06 | 8.967E-06 | 2.665E-06 | 7.000E-06 | 1.200E-05 |
| 21534 | 1077 | V1077F | D | 1.2e-05 | 6.9e-06 | 9.5e-06 | 9.467E-06 | 2.550E-06 | 6.900E-06 | 1.200E-05 |
| 21535 | 1077 | V1077P | D | 1.4e-05 | 6.9e-06 | 6.3e-06 | 9.067E-06 | 4.283E-06 | 6.300E-06 | 1.400E-05 |
| 21536 | 1077 | V1077S | D | 9.9e-06 | 5.4e-06 | 8.8e-06 | 8.033E-06 | 2.346E-06 | 5.400E-06 | 9.900E-06 |
| 21537 | 1077 | V1077T | D | 8.3e-06 | 1.2e-05 | 7.5e-06 | 9.267E-06 | 2.401E-06 | 7.500E-06 | 1.200E-05 |
| 21538 | 1077 | V1077W | D | 4.5e-06 | 6.3e-06 | 6.7e-06 | 5.833E-06 | 1.172E-06 | 4.500E-06 | 6.700E-06 |
| 21539 | 1077 | V1077Y | D | 8.6e-06 | 1e-05 | 4.9e-06 | 7.833E-06 | 2.635E-06 | 4.900E-06 | 1.000E-05 |
| 21540 | 1077 | V1077V | D | 6.9e-06 | 1.5e-05 | 5.5e-06 | 9.133E-06 | 5.129E-06 | 5.500E-06 | 1.500E-05 |
| 21541 | 1078 | T1078A | D | 7.5e-06 | 1.2e-05 | 7.8e-06 | 9.100E-06 | 2.516E-06 | 7.500E-06 | 1.200E-05 |
| 21542 | 1078 | T1078R | D | 1.2e-05 | 8.3e-06 | 6.9e-06 | 9.067E-06 | 2.635E-06 | 6.900E-06 | 1.200E-05 |
| 21543 | 1078 | T1078N | D | 8.2e-06 | 9.2e-06 | 1e-05 | 9.133E-06 | 9.018E-07 | 8.200E-06 | 1.000E-05 |
| 21544 | 1078 | T1078D | D | 1.3e-05 | 7.6e-06 | 1.2e-05 | 1.087E-05 | 2.873E-06 | 7.600E-06 | 1.300E-05 |
| 21545 | 1078 | T1078C | D | 8.4e-06 | 5.2e-06 | 1.4e-05 | 9.200E-06 | 4.454E-06 | 5.200E-06 | 1.400E-05 |
| 21546 | 1078 | T1078E | D | 1.5e-05 | 7.1e-06 | 9.5e-06 | 1.053E-05 | 4.050E-06 | 7.100E-06 | 1.500E-05 |
| 21547 | 1078 | T1078Q | D | 6e-06 | 3.1e-06 | 9.3e-06 | 6.133E-06 | 3.102E-06 | 3.100E-06 | 9.300E-06 |
| 21548 | 1078 | T1078G | D | 1.5e-05 | 6.5e-06 | 9.9e-06 | 1.047E-05 | 4.278E-06 | 6.500E-06 | 1.500E-05 |
| 21549 | 1078 | T1078H | D | 9.4e-06 | 9.1e-06 | 6.5e-06 | 8.333E-06 | 1.595E-06 | 6.500E-06 | 9.400E-06 |
| 21550 | 1078 | T1078I | D | 6.3e-06 | 9.6e-06 | 4.6e-06 | 6.833E-06 | 2.542E-06 | 4.600E-06 | 9.600E-06 |
| 21551 | 1078 | T1078L | D | 8e-06 | 7.4e-06 | 5.6e-06 | 7.000E-06 | 1.249E-06 | 5.600E-06 | 8.000E-06 |
| 21552 | 1078 | T1078K | D | 5.3e-06 | 9.5e-06 | 4.5e-06 | 6.433E-06 | 2.686E-06 | 4.500E-06 | 9.500E-06 |
| 21553 | 1078 | T1078M | D | 9.3e-06 | 7.1e-06 | 5.4e-06 | 7.267E-06 | 1.955E-06 | 5.400E-06 | 9.300E-06 |
| 21554 | 1078 | T1078F | D | 6.2e-06 | 7.9e-06 | 9.7e-06 | 7.933E-06 | 1.750E-06 | 6.200E-06 | 9.700E-06 |
| 21555 | 1078 | T1078P | D | 1.2e-05 | 9.6e-06 | 1.3e-05 | 1.153E-05 | 1.747E-06 | 9.600E-06 | 1.300E-05 |
| 21556 | 1078 | T1078S | D | 8.5e-06 | 9.7e-06 | 5.7e-06 | 7.967E-06 | 2.053E-06 | 5.700E-06 | 9.700E-06 |
| 21557 | 1078 | T1078T | D | 6e-06 | 8.9e-06 | 9.5e-06 | 8.133E-06 | 1.872E-06 | 6.000E-06 | 9.500E-06 |
| 21558 | 1078 | T1078W | D | 5.5e-06 | 8.7e-06 | 4.8e-06 | 6.333E-06 | 2.079E-06 | 4.800E-06 | 8.700E-06 |
| 21559 | 1078 | T1078Y | D | 1.2e-05 | 4.7e-06 | 4.8e-06 | 7.167E-06 | 4.186E-06 | 4.700E-06 | 1.200E-05 |
| 21560 | 1078 | T1078V | D | 1.2e-05 | 4.1e-06 | 1.1e-05 | 9.033E-06 | 4.302E-06 | 4.100E-06 | 1.200E-05 |
| 21561 | 1079 | I1079A | D | 9.7e-06 | 4.6e-06 | 6e-06 | 6.767E-06 | 2.635E-06 | 4.600E-06 | 9.700E-06 |
| 21562 | 1079 | I1079R | D | 7.3e-06 | 1.3e-05 | 1e-05 | 1.010E-05 | 2.851E-06 | 7.300E-06 | 1.300E-05 |
| 21563 | 1079 | I1079N | D | 1e-05 | 7.9e-06 | 8.1e-06 | 8.667E-06 | 1.159E-06 | 7.900E-06 | 1.000E-05 |
| 21564 | 1079 | I1079D | D | 5.7e-06 | 3.4e-06 | 4.6e-06 | 4.567E-06 | 1.150E-06 | 3.400E-06 | 5.700E-06 |
| 21565 | 1079 | I1079C | D | 6.4e-06 | 8.6e-06 | 8.9e-06 | 7.967E-06 | 1.365E-06 | 6.400E-06 | 8.900E-06 |
| 21566 | 1079 | I1079E | D | 1.3e-05 | 8.5e-06 | 7e-06 | 9.500E-06 | 3.122E-06 | 7.000E-06 | 1.300E-05 |
| 21567 | 1079 | I1079Q | D | 8.5e-06 | 5.8e-06 | 6e-06 | 6.767E-06 | 1.504E-06 | 5.800E-06 | 8.500E-06 |
| 21568 | 1079 | I1079G | D | 1.1e-05 | 6.5e-06 | 7.1e-06 | 8.200E-06 | 2.443E-06 | 6.500E-06 | 1.100E-05 |
| 21569 | 1079 | I1079H | D | 1.2e-05 | 5.3e-06 | 1.1e-05 | 9.433E-06 | 3.614E-06 | 5.300E-06 | 1.200E-05 |
| 21570 | 1079 | I1079I | D | 5.5e-06 | 1.2e-05 | 9.2e-06 | 8.900E-06 | 3.260E-06 | 5.500E-06 | 1.200E-05 |
| 21571 | 1079 | I1079L | D | 8.5e-06 | 7.4e-06 | 1.1e-05 | 8.967E-06 | 1.845E-06 | 7.400E-06 | 1.100E-05 |
| 21572 | 1079 | I1079K | D | 1e-05 | 5.3e-06 | 1e-05 | 8.433E-06 | 2.714E-06 | 5.300E-06 | 1.000E-05 |
| 21573 | 1079 | I1079M | D | 5.6e-06 | 1e-05 | 7.4e-06 | 7.667E-06 | 2.212E-06 | 5.600E-06 | 1.000E-05 |
| 21574 | 1079 | I1079F | D | 1e-05 | 8.8e-06 | 3.1e-06 | 7.300E-06 | 3.686E-06 | 3.100E-06 | 1.000E-05 |
| 21575 | 1079 | I1079P | D | 1.1e-05 | 5.7e-06 | 7.1e-06 | 7.933E-06 | 2.747E-06 | 5.700E-06 | 1.100E-05 |
| 21576 | 1079 | I1079S | D | 7.4e-06 | 5.9e-06 | 6.3e-06 | 6.533E-06 | 7.767E-07 | 5.900E-06 | 7.400E-06 |
| 21577 | 1079 | I1079T | D | 6.7e-06 | 1.1e-05 | 9.9e-06 | 9.200E-06 | 2.234E-06 | 6.700E-06 | 1.100E-05 |
| 21578 | 1079 | I1079W | D | 5.4e-06 | 7.9e-06 | 8.5e-06 | 7.267E-06 | 1.644E-06 | 5.400E-06 | 8.500E-06 |
| 21579 | 1079 | I1079Y | D | 8.4e-06 | 7.3e-06 | 1.7e-05 | 1.090E-05 | 5.311E-06 | 7.300E-06 | 1.700E-05 |
| 21580 | 1079 | I1079V | D | 8.8e-06 | 7.2e-06 | 5.1e-06 | 7.033E-06 | 1.856E-06 | 5.100E-06 | 8.800E-06 |
| 21581 | 1080 | T1080A | D | 9e-06 | 7.9e-06 | 5.8e-06 | 7.567E-06 | 1.626E-06 | 5.800E-06 | 9.000E-06 |
| 21582 | 1080 | T1080R | D | 6.3e-06 | 1.5e-05 | 1.3e-05 | 1.143E-05 | 4.557E-06 | 6.300E-06 | 1.500E-05 |
| 21583 | 1080 | T1080N | D | 7.9e-06 | 1.1e-05 | 7.2e-06 | 8.700E-06 | 2.022E-06 | 7.200E-06 | 1.100E-05 |
| 21584 | 1080 | T1080D | D | 3.5e-06 | 6.8e-06 | 8.7e-06 | 6.333E-06 | 2.631E-06 | 3.500E-06 | 8.700E-06 |
| 21585 | 1080 | T1080C | D | 1.3e-05 | 1.4e-05 | 9.4e-06 | 1.213E-05 | 2.419E-06 | 9.400E-06 | 1.400E-05 |
| 21586 | 1080 | T1080E | D | 6.6e-06 | 6.9e-06 | 6.2e-06 | 6.567E-06 | 3.512E-07 | 6.200E-06 | 6.900E-06 |
| 21587 | 1080 | T1080Q | D | 1.3e-05 | 6.5e-06 | 1e-05 | 9.833E-06 | 3.253E-06 | 6.500E-06 | 1.300E-05 |
| 21588 | 1080 | T1080G | D | 1.3e-05 | 9.4e-06 | 9e-06 | 1.047E-05 | 2.203E-06 | 9.000E-06 | 1.300E-05 |
| 21589 | 1080 | T1080H | D | 1.1e-05 | 1.5e-05 | 5.7e-06 | 1.057E-05 | 4.665E-06 | 5.700E-06 | 1.500E-05 |

| | | | | | | | | | | |
|---|---|---|---|---|---|---|---|---|---|---|
| 21590 | 1080 | T1080I | D | 4.6e-06 | 7.2e-06 | 1.3e-05 | 8.267E-06 | 4.300E-06 | 4.600E-06 | 1.300E-05 |
| 21591 | 1080 | T1080L | D | 6.9e-06 | 7.9e-06 | 1e-05 | 8.267E-06 | 1.582E-06 | 6.900E-06 | 1.000E-05 |
| 21592 | 1080 | T1080K | D | 5.7e-06 | 9.4e-06 | 6.9e-06 | 7.333E-06 | 1.888E-06 | 5.700E-06 | 9.400E-06 |
| 21593 | 1080 | T1080M | D | 7.6e-06 | 5.3e-06 | 7.1e-06 | 6.667E-06 | 1.210E-06 | 5.300E-06 | 7.600E-06 |
| 21594 | 1080 | T1080F | D | 8.7e-06 | 8.2e-06 | 5.9e-06 | 7.600E-06 | 1.493E-06 | 5.900E-06 | 8.700E-06 |
| 21595 | 1080 | T1080P | D | 6.5e-06 | 7.4e-06 | 6.5e-06 | 6.800E-06 | 5.196E-07 | 6.500E-06 | 7.400E-06 |
| 21596 | 1080 | T1080S | D | 6.3e-06 | 6e-06 | 6.6e-06 | 6.300E-06 | 3.000E-07 | 6.000E-06 | 6.600E-06 |
| 21597 | 1080 | T1080T | D | 4e-06 | 7.6e-06 | 7e-06 | 6.200E-06 | 1.929E-06 | 4.000E-06 | 7.600E-06 |
| 21598 | 1080 | T1080W | D | 1e-05 | 4.9e-06 | 1.2e-05 | 8.967E-06 | 3.661E-06 | 4.900E-06 | 1.200E-05 |
| 21599 | 1080 | T1080Y | D | 7.3e-06 | 1.1e-05 | 6.7e-06 | 8.333E-06 | 2.329E-06 | 6.700E-06 | 1.100E-05 |
| 21600 | 1080 | T1080V | D | 6.1e-06 | 1e-05 | 9.5e-06 | 8.533E-06 | 2.122E-06 | 6.100E-06 | 1.000E-05 |
| 21601 | 1081 | C1081A | D | 5.4e-06 | 1.4e-05 | 9.5e-06 | 9.633E-06 | 4.302E-06 | 5.400E-06 | 1.400E-05 |
| 21602 | 1081 | C1081R | D | 9.5e-06 | 5.8e-06 | 6.3e-06 | 7.200E-06 | 2.007E-06 | 5.800E-06 | 9.500E-06 |
| 21603 | 1081 | C1081N | D | 7.3e-06 | 1.1e-05 | 1e-05 | 9.433E-06 | 1.914E-06 | 7.300E-06 | 1.100E-05 |
| 21604 | 1081 | C1081D | D | 1e-05 | 7.3e-06 | 5.9e-06 | 7.733E-06 | 2.084E-06 | 5.900E-06 | 1.000E-05 |
| 21605 | 1081 | C1081C | D | 8.3e-06 | 6.2e-06 | 6e-06 | 6.833E-06 | 1.274E-06 | 6.000E-06 | 8.300E-06 |
| 21606 | 1081 | C1081E | D | 1.1e-05 | 8.4e-06 | 4.1e-06 | 7.833E-06 | 3.485E-06 | 4.100E-06 | 1.100E-05 |
| 21607 | 1081 | C1081Q | D | 6.1e-06 | 5.7e-06 | 5.1e-06 | 5.633E-06 | 5.033E-07 | 5.100E-06 | 6.100E-06 |
| 21608 | 1081 | C1081G | D | 7.2e-06 | 7.9e-06 | 4.9e-06 | 6.667E-06 | 1.570E-06 | 4.900E-06 | 7.900E-06 |
| 21609 | 1081 | C1081H | D | 6.8e-06 | 1.1e-05 | 9.8e-06 | 9.200E-06 | 2.163E-06 | 6.800E-06 | 1.100E-05 |
| 21610 | 1081 | C1081I | D | 1e-05 | 5.6e-06 | 8.8e-06 | 8.133E-06 | 2.274E-06 | 5.600E-06 | 1.000E-05 |
| 21611 | 1081 | C1081L | D | 1.1e-05 | 9.5e-06 | 1.4e-05 | 1.150E-05 | 2.291E-06 | 9.500E-06 | 1.400E-05 |
| 21612 | 1081 | C1081K | D | 5.6e-06 | 9.5e-06 | 8.2e-06 | 7.767E-06 | 1.986E-06 | 5.600E-06 | 9.500E-06 |
| 21613 | 1081 | C1081M | D | 1.1e-05 | 5.5e-06 | 5e-06 | 7.167E-06 | 3.329E-06 | 5.000E-06 | 1.100E-05 |
| 21614 | 1081 | C1081F | D | 8.6e-06 | 4.7e-06 | 7.5e-06 | 6.933E-06 | 2.011E-06 | 4.700E-06 | 8.600E-06 |
| 21615 | 1081 | C1081P | D | 8.4e-06 | 9.7e-06 | 1.2e-05 | 1.003E-05 | 1.823E-06 | 8.400E-06 | 1.200E-05 |
| 21616 | 1081 | C1081S | D | 6.7e-06 | 8e-06 | 1.3e-05 | 9.233E-06 | 3.326E-06 | 6.700E-06 | 1.300E-05 |
| 21617 | 1081 | C1081T | D | 1e-05 | 6.5e-06 | 4.8e-06 | 7.100E-06 | 2.651E-06 | 4.800E-06 | 1.000E-05 |
| 21618 | 1081 | C1081W | D | 1.3e-05 | 5.9e-06 | 8.5e-06 | 9.133E-06 | 3.592E-06 | 5.900E-06 | 1.300E-05 |
| 21619 | 1081 | C1081Y | D | 1.2e-05 | 8.9e-06 | 7.7e-06 | 9.533E-06 | 2.219E-06 | 7.700E-06 | 1.200E-05 |
| 21620 | 1081 | C1081V | D | 5.2e-06 | 6.4e-06 | 6.9e-06 | 6.167E-06 | 8.737E-07 | 5.200E-06 | 6.900E-06 |
| 21621 | 1082 | R1082A | D | 7e-06 | 1.5e-05 | 7.7e-06 | 9.900E-06 | 4.431E-06 | 7.000E-06 | 1.500E-05 |
| 21622 | 1082 | R1082R | D | 6.1e-06 | 8.9e-06 | 9.5e-06 | 8.167E-06 | 1.815E-06 | 6.100E-06 | 9.500E-06 |
| 21623 | 1082 | R1082N | D | 1.4e-05 | 1.2e-05 | 7.6e-06 | 1.120E-05 | 3.274E-06 | 7.600E-06 | 1.400E-05 |
| 21624 | 1082 | R1082D | D | 8.2e-06 | 7.8e-06 | 9.4e-06 | 8.467E-06 | 8.327E-07 | 7.800E-06 | 9.400E-06 |
| 21625 | 1082 | R1082C | D | 6.9e-06 | 8.3e-06 | 5.1e-06 | 6.767E-06 | 1.604E-06 | 5.100E-06 | 8.300E-06 |
| 21626 | 1082 | R1082E | D | 1.2e-05 | 5.3e-06 | 5.3e-06 | 7.533E-06 | 3.868E-06 | 5.300E-06 | 1.200E-05 |
| 21627 | 1082 | R1082Q | D | 7.9e-06 | 8.3e-06 | 7.2e-06 | 7.800E-06 | 5.568E-07 | 7.200E-06 | 8.300E-06 |
| 21628 | 1082 | R1082G | D | 7.8e-06 | 9.2e-06 | 9.1e-06 | 8.700E-06 | 7.810E-07 | 7.800E-06 | 9.200E-06 |
| 21629 | 1082 | R1082H | D | 1.2e-05 | 7.9e-06 | 1.3e-05 | 1.097E-05 | 2.702E-06 | 7.900E-06 | 1.300E-05 |
| 21630 | 1082 | R1082I | D | 8.6e-06 | 1.4e-05 | 1.3e-05 | 1.187E-05 | 2.873E-06 | 8.600E-06 | 1.400E-05 |
| 21631 | 1082 | R1082L | D | 6.4e-06 | 8.3e-06 | 6e-06 | 6.900E-06 | 1.229E-06 | 6.000E-06 | 8.300E-06 |
| 21632 | 1082 | R1082K | D | 6.3e-06 | 7.2e-06 | 8.7e-06 | 7.400E-06 | 1.212E-06 | 6.300E-06 | 8.700E-06 |
| 21633 | 1082 | R1082M | D | 6.3e-06 | 7.8e-06 | 5.9e-06 | 6.667E-06 | 1.002E-06 | 5.900E-06 | 7.800E-06 |
| 21634 | 1082 | R1082F | D | 8.4e-06 | 1.1e-05 | 2e-05 | 1.313E-05 | 6.087E-06 | 8.400E-06 | 2.000E-05 |
| 21635 | 1082 | R1082P | D | 4.5e-06 | 5.8e-06 | 1.6e-05 | 8.767E-06 | 6.298E-06 | 4.500E-06 | 1.600E-05 |
| 21636 | 1082 | R1082S | D | 6.1e-06 | 5.8e-06 | 3.4e-06 | 5.100E-06 | 1.480E-06 | 3.400E-06 | 6.100E-06 |
| 21637 | 1082 | R1082T | D | 6.5e-06 | 8.4e-06 | 7.3e-06 | 7.400E-06 | 9.539E-07 | 6.500E-06 | 8.400E-06 |
| 21638 | 1082 | R1082W | D | 4.4e-06 | 7.6e-06 | 9.5e-06 | 7.167E-06 | 2.577E-06 | 4.400E-06 | 9.500E-06 |
| 21639 | 1082 | R1082Y | D | 9.5e-06 | 1.8e-05 | 4.9e-06 | 1.080E-05 | 6.646E-06 | 4.900E-06 | 1.800E-05 |
| 21640 | 1082 | R1082V | D | 5.2e-06 | 1e-05 | 1.4e-05 | 9.733E-06 | 4.406E-06 | 5.200E-06 | 1.400E-05 |
| 21641 | 1083 | A1083A | D | 8.8e-06 | 1.1e-05 | 1e-05 | 9.933E-06 | 1.102E-06 | 8.800E-06 | 1.100E-05 |
| 21642 | 1083 | A1083R | D | 8.7e-06 | 7e-06 | 8.2e-06 | 7.967E-06 | 8.737E-07 | 7.000E-06 | 8.700E-06 |
| 21643 | 1083 | A1083N | D | 8.2e-06 | 7.4e-06 | 1.1e-05 | 8.867E-06 | 1.890E-06 | 7.400E-06 | 1.100E-05 |
| 21644 | 1083 | A1083D | D | 5.9e-06 | 9.7e-06 | 6.5e-06 | 7.367E-06 | 2.043E-06 | 5.900E-06 | 9.700E-06 |
| 21645 | 1083 | A1083C | D | 9.5e-06 | 6.4e-06 | 7.4e-06 | 7.767E-06 | 1.582E-06 | 6.400E-06 | 9.500E-06 |
| 21646 | 1083 | A1083E | D | 4.7e-06 | 6.6e-06 | 9.8e-06 | 7.033E-06 | 2.577E-06 | 4.700E-06 | 9.800E-06 |
| 21647 | 1083 | A1083Q | D | 6.5e-06 | 9.9e-06 | 6e-06 | 7.467E-06 | 2.122E-06 | 6.000E-06 | 9.900E-06 |
| 21648 | 1083 | A1083G | D | 8.6e-06 | 6.4e-06 | 6.7e-06 | 7.233E-06 | 1.193E-06 | 6.400E-06 | 8.600E-06 |
| 21649 | 1083 | A1083H | D | 4.6e-06 | 7.4e-06 | 8.7e-06 | 6.900E-06 | 2.095E-06 | 4.600E-06 | 8.700E-06 |
| 21650 | 1083 | A1083I | D | 6.2e-06 | 6.5e-06 | 1e-05 | 7.567E-06 | 2.113E-06 | 6.200E-06 | 1.000E-05 |
| 21651 | 1083 | A1083L | D | 1.5e-05 | 9.2e-06 | 9.6e-06 | 1.127E-05 | 3.239E-06 | 9.200E-06 | 1.500E-05 |
| 21652 | 1083 | A1083K | D | 6.6e-06 | 1e-05 | 1.4e-05 | 1.020E-05 | 3.704E-06 | 6.600E-06 | 1.400E-05 |
| 21653 | 1083 | A1083M | D | 9.1e-06 | 9.8e-06 | 5.9e-06 | 8.267E-06 | 2.079E-06 | 5.900E-06 | 9.800E-06 |
| 21654 | 1083 | A1083F | D | 1.3e-05 | 1.8e-05 | 1.4e-05 | 1.500E-05 | 2.646E-06 | 1.300E-05 | 1.800E-05 |
| 21655 | 1083 | A1083P | D | 1.1e-05 | 6.3e-06 | 9.3e-06 | 8.867E-06 | 2.380E-06 | 6.300E-06 | 1.100E-05 |
| 21656 | 1083 | A1083S | D | 1.1e-05 | 8.1e-06 | 7.4e-06 | 8.833E-06 | 1.909E-06 | 7.400E-06 | 1.100E-05 |
| 21657 | 1083 | A1083T | D | 1.8e-05 | 5.6e-06 | 6e-06 | 9.867E-06 | 7.047E-06 | 5.600E-06 | 1.800E-05 |
| 21658 | 1083 | A1083W | D | 5e-06 | 8.8e-06 | 9.6e-06 | 7.800E-06 | 2.458E-06 | 5.000E-06 | 9.600E-06 |
| 21659 | 1083 | A1083Y | D | 4.5e-06 | 9.5e-06 | 5.5e-06 | 6.500E-06 | 2.646E-06 | 4.500E-06 | 9.500E-06 |
| 21660 | 1083 | A1083V | D | 9.8e-06 | 5.4e-06 | 1e-05 | 8.400E-06 | 2.600E-06 | 5.400E-06 | 1.000E-05 |
| 21661 | 1084 | S1084A | D | 1.4e-05 | 8e-06 | 1.1e-05 | 1.100E-05 | 3.000E-06 | 8.000E-06 | 1.400E-05 |
| 21662 | 1084 | S1084R | D | 1.1e-05 | 6.6e-06 | 4e-06 | 7.200E-06 | 3.538E-06 | 4.000E-06 | 1.100E-05 |
| 21663 | 1084 | S1084N | D | 9.6e-06 | 7.3e-06 | 6.4e-06 | 7.767E-06 | 1.650E-06 | 6.400E-06 | 9.600E-06 |
| 21664 | 1084 | S1084D | D | 6.5e-06 | 1.2e-05 | 9.6e-06 | 9.367E-06 | 2.757E-06 | 6.500E-06 | 1.200E-05 |
| 21665 | 1084 | S1084C | D | 1.1e-05 | 1e-05 | 5.6e-06 | 8.867E-06 | 2.873E-06 | 5.600E-06 | 1.100E-05 |
| 21666 | 1084 | S1084E | D | 1.4e-05 | 1.2e-05 | 1e-05 | 1.200E-05 | 2.000E-06 | 1.000E-05 | 1.400E-05 |
| 21667 | 1084 | S1084Q | D | 9.3e-06 | 4.8e-06 | 8.8e-06 | 7.633E-06 | 2.466E-06 | 4.800E-06 | 9.300E-06 |
| 21668 | 1084 | S1084G | D | 1e-05 | 5.9e-06 | 6.8e-06 | 7.567E-06 | 2.155E-06 | 5.900E-06 | 1.000E-05 |
| 21669 | 1084 | S1084H | D | 8.9e-06 | 8e-06 | 7e-06 | 7.967E-06 | 9.504E-07 | 7.000E-06 | 8.900E-06 |
| 21670 | 1084 | S1084I | D | 7.5e-06 | 9e-06 | 6.7e-06 | 7.733E-06 | 1.168E-06 | 6.700E-06 | 9.000E-06 |
| 21671 | 1084 | S1084L | D | 1.1e-05 | 9.7e-06 | 6.8e-06 | 9.167E-06 | 2.150E-06 | 6.800E-06 | 1.100E-05 |
| 21672 | 1084 | S1084K | D | 1.3e-05 | 4.8e-06 | 9.2e-06 | 9.000E-06 | 4.104E-06 | 4.800E-06 | 1.300E-05 |
| 21673 | 1084 | S1084M | D | 5.6e-06 | 6.1e-06 | 1.2e-05 | 7.900E-06 | 3.559E-06 | 5.600E-06 | 1.200E-05 |
| 21674 | 1084 | S1084F | D | 5.7e-06 | 1.8e-05 | 8.9e-06 | 1.087E-05 | 6.381E-06 | 5.700E-06 | 1.800E-05 |

| | | | | | | | | | | |
|---|---|---|---|---|---|---|---|---|---|---|
| 21675 | 1084 | S1084P | D | 9.6e-06 | 6.1e-06 | 8.5e-06 | 8.067E-06 | 1.790E-06 | 6.100E-06 | 9.600E-06 |
| 21676 | 1084 | S1084S | D | 1.5e-05 | 9.1e-06 | 9.6e-06 | 1.123E-05 | 3.272E-06 | 9.100E-06 | 1.500E-05 |
| 21677 | 1084 | S1084T | D | 1e-05 | 2.9e-06 | 7.7e-06 | 6.867E-06 | 3.623E-06 | 2.900E-06 | 1.000E-05 |
| 21678 | 1084 | S1084W | D | 6.8e-06 | 9.2e-06 | 4.7e-06 | 6.900E-06 | 2.252E-06 | 4.700E-06 | 9.200E-06 |
| 21679 | 1084 | S1084Y | D | 1.1e-05 | 4.5e-06 | 9.6e-06 | 8.367E-06 | 3.421E-06 | 4.500E-06 | 1.100E-05 |
| 21680 | 1084 | S1084V | D | 7.1e-06 | 8.5e-06 | 7e-06 | 7.533E-06 | 8.386E-07 | 7.000E-06 | 8.500E-06 |
| 21681 | 1085 | Q1085A | D | 5.4e-06 | 1.2e-05 | 1.3e-05 | 1.013E-05 | 4.130E-06 | 5.400E-06 | 1.300E-05 |
| 21682 | 1085 | Q1085R | D | 4.8e-06 | 1.3e-05 | 8.4e-06 | 8.733E-06 | 4.110E-06 | 4.800E-06 | 1.300E-05 |
| 21683 | 1085 | Q1085N | D | 9.9e-06 | 1.2e-05 | 6.8e-06 | 9.567E-06 | 2.616E-06 | 6.800E-06 | 1.200E-05 |
| 21684 | 1085 | Q1085D | D | 1.1e-05 | 9.9e-06 | 1.1e-05 | 1.063E-05 | 6.351E-07 | 9.900E-06 | 1.100E-05 |
| 21685 | 1085 | Q1085C | D | 8.2e-06 | 6.9e-06 | 1.1e-05 | 8.700E-06 | 2.095E-06 | 6.900E-06 | 1.100E-05 |
| 21686 | 1085 | Q1085E | D | 7.2e-06 | 1e-05 | 1.1e-05 | 9.400E-06 | 1.970E-06 | 7.200E-06 | 1.100E-05 |
| 21687 | 1085 | Q1085Q | D | 8e-06 | 4.6e-06 | 1.1e-05 | 7.867E-06 | 3.202E-06 | 4.600E-06 | 1.100E-05 |
| 21688 | 1085 | Q1085G | D | 7e-06 | 8.7e-06 | 5.6e-06 | 7.100E-06 | 1.552E-06 | 5.600E-06 | 8.700E-06 |
| 21689 | 1085 | Q1085H | D | 5.6e-06 | 8.3e-06 | 6e-06 | 6.633E-06 | 1.457E-06 | 5.600E-06 | 8.300E-06 |
| 21690 | 1085 | Q1085I | D | 1e-05 | 1.5e-05 | 6.6e-06 | 1.053E-05 | 4.225E-06 | 6.600E-06 | 1.500E-05 |
| 21691 | 1085 | Q1085L | D | 9.1e-06 | 9.9e-06 | 5.9e-06 | 8.300E-06 | 2.117E-06 | 5.900E-06 | 9.900E-06 |
| 21692 | 1085 | Q1085K | D | 4.7e-06 | 4.1e-06 | 1.4e-05 | 7.600E-06 | 5.551E-06 | 4.100E-06 | 1.400E-05 |
| 21693 | 1085 | Q1085M | D | 6.9e-06 | 5.7e-06 | 6.3e-06 | 6.300E-06 | 6.000E-07 | 5.700E-06 | 6.900E-06 |
| 21694 | 1085 | Q1085F | D | 9e-06 | 7e-06 | 7.1e-06 | 7.700E-06 | 1.127E-06 | 7.000E-06 | 9.000E-06 |
| 21695 | 1085 | Q1085P | D | 8.2e-06 | 1.6e-05 | 1e-05 | 1.140E-05 | 4.084E-06 | 8.200E-06 | 1.600E-05 |
| 21696 | 1085 | Q1085S | D | 8.4e-06 | 1.4e-05 | 6.3e-06 | 9.567E-06 | 3.980E-06 | 6.300E-06 | 1.400E-05 |
| 21697 | 1085 | Q1085T | D | 5e-06 | 9.2e-06 | 9.7e-06 | 7.967E-06 | 2.581E-06 | 5.000E-06 | 9.700E-06 |
| 21698 | 1085 | Q1085W | D | 6.4e-06 | 5.9e-06 | 1.8e-05 | 1.010E-05 | 6.846E-06 | 5.900E-06 | 1.800E-05 |
| 21699 | 1085 | Q1085Y | D | 6e-06 | 5.8e-06 | 1.1e-05 | 7.600E-06 | 2.946E-06 | 5.800E-06 | 1.100E-05 |
| 21700 | 1085 | Q1085V | D | 7.7e-06 | 8.9e-06 | 7.9e-06 | 8.167E-06 | 6.429E-07 | 7.700E-06 | 8.900E-06 |
| 21701 | 1086 | D1086A | D | 7e-06 | 5.9e-06 | 6.5e-06 | 6.467E-06 | 5.508E-07 | 5.900E-06 | 7.000E-06 |
| 21702 | 1086 | D1086R | D | 1.2e-05 | 6e-06 | 6e-06 | 8.000E-06 | 3.464E-06 | 6.000E-06 | 1.200E-05 |
| 21703 | 1086 | D1086N | D | 9.7e-06 | 9.7e-06 | 8.3e-06 | 9.233E-06 | 8.083E-07 | 8.300E-06 | 9.700E-06 |
| 21704 | 1086 | D1086D | D | 8.5e-06 | 1e-05 | 8.1e-06 | 8.867E-06 | 1.002E-06 | 8.100E-06 | 1.000E-05 |
| 21705 | 1086 | D1086C | D | 8.2e-06 | 5.4e-06 | 8.4e-06 | 7.333E-06 | 1.677E-06 | 5.400E-06 | 8.400E-06 |
| 21706 | 1086 | D1086E | D | 5e-06 | 5.9e-06 | 6.4e-06 | 5.767E-06 | 7.095E-07 | 5.000E-06 | 6.400E-06 |
| 21707 | 1086 | D1086Q | D | 1e-05 | 7.6e-06 | 6.5e-06 | 8.033E-06 | 1.790E-06 | 6.500E-06 | 1.000E-05 |
| 21708 | 1086 | D1086G | D | 9.1e-06 | 9.8e-06 | 4.8e-06 | 7.900E-06 | 2.707E-06 | 4.800E-06 | 9.800E-06 |
| 21709 | 1086 | D1086H | D | 6.4e-06 | 1.4e-05 | 7.3e-06 | 9.233E-06 | 4.153E-06 | 6.400E-06 | 1.400E-05 |
| 21710 | 1086 | D1086I | D | 1.4e-05 | 1e-05 | 6.5e-06 | 1.017E-05 | 3.753E-06 | 6.500E-06 | 1.400E-05 |
| 21711 | 1086 | D1086L | D | 7.5e-06 | 7.6e-06 | 8.3e-06 | 7.800E-06 | 4.359E-07 | 7.500E-06 | 8.300E-06 |
| 21712 | 1086 | D1086K | D | 6.7e-06 | 7.1e-06 | 3.9e-06 | 5.900E-06 | 1.744E-06 | 3.900E-06 | 7.100E-06 |
| 21713 | 1086 | D1086M | D | 7.7e-06 | 8e-06 | 1.2e-05 | 9.233E-06 | 2.401E-06 | 7.700E-06 | 1.200E-05 |
| 21714 | 1086 | D1086F | D | 9.4e-06 | 7.7e-06 | 8.9e-06 | 8.667E-06 | 8.737E-07 | 7.700E-06 | 9.400E-06 |
| 21715 | 1086 | D1086P | D | 6e-06 | 9.8e-06 | 8e-06 | 7.933E-06 | 1.901E-06 | 6.000E-06 | 9.800E-06 |
| 21716 | 1086 | D1086S | D | 5.7e-06 | 1.3e-05 | 8.7e-06 | 9.133E-06 | 3.669E-06 | 5.700E-06 | 1.300E-05 |
| 21717 | 1086 | D1086T | D | 5.4e-06 | 4.1e-06 | 8.8e-06 | 6.100E-06 | 2.427E-06 | 4.100E-06 | 8.800E-06 |
| 21718 | 1086 | D1086W | D | 7.9e-06 | 1.3e-05 | 6.6e-06 | 9.167E-06 | 3.383E-06 | 6.600E-06 | 1.300E-05 |
| 21719 | 1086 | D1086Y | D | 1e-05 | 4.8e-06 | 8.7e-06 | 7.833E-06 | 2.706E-06 | 4.800E-06 | 1.000E-05 |
| 21720 | 1086 | D1086V | D | 1e-05 | 8.6e-06 | 7.8e-06 | 8.800E-06 | 1.114E-06 | 7.800E-06 | 1.000E-05 |
| 21721 | 1087 | V1087A | D | 5.1e-06 | 1.5e-05 | 8.2e-06 | 9.433E-06 | 5.064E-06 | 5.100E-06 | 1.500E-05 |
| 21722 | 1087 | V1087R | D | 8.6e-06 | 1.1e-05 | 6.8e-06 | 8.800E-06 | 2.107E-06 | 6.800E-06 | 1.100E-05 |
| 21723 | 1087 | V1087N | D | 8.2e-06 | 9.4e-06 | 7.8e-06 | 8.467E-06 | 8.327E-07 | 7.800E-06 | 9.400E-06 |
| 21724 | 1087 | V1087D | D | 1e-05 | 5.2e-06 | 9.7e-06 | 8.300E-06 | 2.689E-06 | 5.200E-06 | 1.000E-05 |
| 21725 | 1087 | V1087C | D | 4.4e-06 | 5.1e-06 | 6.6e-06 | 5.367E-06 | 1.124E-06 | 4.400E-06 | 6.600E-06 |
| 21726 | 1087 | V1087E | D | 6.2e-06 | 4.9e-06 | 8.1e-06 | 6.400E-06 | 1.609E-06 | 4.900E-06 | 8.100E-06 |
| 21727 | 1087 | V1087Q | D | 6.3e-06 | 6.2e-06 | 4.4e-06 | 5.633E-06 | 1.069E-06 | 4.400E-06 | 6.300E-06 |
| 21728 | 1087 | V1087G | D | 1.1e-05 | 5e-06 | 6.7e-06 | 7.567E-06 | 3.092E-06 | 5.000E-06 | 1.100E-05 |
| 21729 | 1087 | V1087H | D | 8.2e-06 | 5.3e-06 | 4.6e-06 | 6.033E-06 | 1.909E-06 | 4.600E-06 | 8.200E-06 |
| 21730 | 1087 | V1087I | D | 5.4e-06 | 6.8e-06 | 7e-06 | 6.400E-06 | 8.718E-07 | 5.400E-06 | 7.000E-06 |
| 21731 | 1087 | V1087L | D | 6.5e-06 | 5.5e-06 | 1e-05 | 7.333E-06 | 2.363E-06 | 5.500E-06 | 1.000E-05 |
| 21732 | 1087 | V1087K | D | 6.1e-06 | 7.3e-06 | 1.1e-05 | 8.133E-06 | 2.554E-06 | 6.100E-06 | 1.100E-05 |
| 21733 | 1087 | V1087M | D | 9.4e-06 | 5.4e-06 | 9.5e-06 | 8.100E-06 | 2.339E-06 | 5.400E-06 | 9.500E-06 |
| 21734 | 1087 | V1087F | D | 8.8e-06 | 1.1e-05 | 5.6e-06 | 8.467E-06 | 2.715E-06 | 5.600E-06 | 1.100E-05 |
| 21735 | 1087 | V1087P | D | 2.1e-05 | 7.1e-06 | 6.1e-06 | 1.140E-05 | 8.329E-06 | 6.100E-06 | 2.100E-05 |
| 21736 | 1087 | V1087S | D | 7.6e-06 | 5.8e-06 | 3.9e-06 | 5.767E-06 | 1.850E-06 | 3.900E-06 | 7.600E-06 |
| 21737 | 1087 | V1087T | D | 5.4e-06 | 4.7e-06 | 6.9e-06 | 5.667E-06 | 1.124E-06 | 4.700E-06 | 6.900E-06 |
| 21738 | 1087 | V1087W | D | 4.2e-06 | 6.7e-06 | 6e-06 | 5.633E-06 | 1.290E-06 | 4.200E-06 | 6.700E-06 |
| 21739 | 1087 | V1087Y | D | 9.6e-06 | 9.8e-06 | 6.2e-06 | 8.533E-06 | 2.023E-06 | 6.200E-06 | 9.800E-06 |
| 21740 | 1087 | V1087V | D | 1.3e-05 | 9.9e-06 | 6.9e-06 | 9.933E-06 | 3.050E-06 | 6.900E-06 | 1.300E-05 |
| 21741 | 1088 | N1088A | D | 1.4e-05 | 9.1e-06 | 5.1e-06 | 9.400E-06 | 4.458E-06 | 5.100E-06 | 1.400E-05 |
| 21742 | 1088 | N1088R | D | 1.1e-05 | 8.2e-06 | 7.2e-06 | 8.800E-06 | 1.970E-06 | 7.200E-06 | 1.100E-05 |
| 21743 | 1088 | N1088N | D | 9.5e-06 | 6e-06 | 5.3e-06 | 6.933E-06 | 2.250E-06 | 5.300E-06 | 9.500E-06 |
| 21744 | 1088 | N1088D | D | 6.9e-06 | 5.9e-06 | 3.3e-06 | 5.367E-06 | 1.858E-06 | 3.300E-06 | 6.900E-06 |
| 21745 | 1088 | N1088C | D | 1.2e-05 | 8.6e-06 | 6.3e-06 | 8.967E-06 | 2.868E-06 | 6.300E-06 | 1.200E-05 |
| 21746 | 1088 | N1088E | D | 1.4e-05 | 5.6e-06 | 3.1e-06 | 7.567E-06 | 5.710E-06 | 3.100E-06 | 1.400E-05 |
| 21747 | 1088 | N1088Q | D | 1.4e-05 | 5.8e-06 | 8.6e-06 | 9.467E-06 | 4.168E-06 | 5.800E-06 | 1.400E-05 |
| 21748 | 1088 | N1088G | D | 1.4e-05 | 6.5e-06 | 2.1e-05 | 1.383E-05 | 7.251E-06 | 6.500E-06 | 2.100E-05 |
| 21749 | 1088 | N1088H | D | 8e-06 | 1e-05 | 5.5e-06 | 7.833E-06 | 2.255E-06 | 5.500E-06 | 1.000E-05 |
| 21750 | 1088 | N1088I | D | 5.4e-06 | 8e-06 | 1e-05 | 7.800E-06 | 2.307E-06 | 5.400E-06 | 1.000E-05 |
| 21751 | 1088 | N1088L | D | 4.2e-06 | 9.6e-06 | 6.8e-06 | 6.867E-06 | 2.701E-06 | 4.200E-06 | 9.600E-06 |
| 21752 | 1088 | N1088K | D | 5.1e-06 | 7.6e-06 | 9.5e-06 | 7.400E-06 | 2.207E-06 | 5.100E-06 | 9.500E-06 |
| 21753 | 1088 | N1088M | D | 5.1e-06 | 1.1e-05 | 9.9e-06 | 8.667E-06 | 3.137E-06 | 5.100E-06 | 1.100E-05 |
| 21754 | 1088 | N1088F | D | 1.1e-05 | 7.1e-06 | 9.4e-06 | 9.167E-06 | 1.960E-06 | 7.100E-06 | 1.100E-05 |
| 21755 | 1088 | N1088P | D | 1.7e-05 | 8.4e-06 | 8.7e-06 | 1.137E-05 | 4.881E-06 | 8.400E-06 | 1.700E-05 |
| 21756 | 1088 | N1088S | D | 1.4e-05 | 1.1e-05 | 6.4e-06 | 1.047E-05 | 3.828E-06 | 6.400E-06 | 1.400E-05 |
| 21757 | 1088 | N1088T | D | 7e-06 | 9.8e-06 | 1.4e-05 | 1.027E-05 | 3.523E-06 | 7.000E-06 | 1.400E-05 |
| 21758 | 1088 | N1088W | D | 5.9e-06 | 9.2e-06 | 7.2e-06 | 7.433E-06 | 1.662E-06 | 5.900E-06 | 9.200E-06 |
| 21759 | 1088 | N1088Y | D | 1.5e-05 | 9.2e-06 | 5.8e-06 | 1.000E-05 | 4.652E-06 | 5.800E-06 | 1.500E-05 |

| | | | | | | | | | | |
|---|---|---|---|---|---|---|---|---|---|---|
| 21760 | 1088 | N1088V | D | 9e-06 | 1.2e-05 | 5.6e-06 | 8.867E-06 | 3.202E-06 | 5.600E-06 | 1.200E-05 |
| 21761 | 1089 | T1089A | D | 6.3e-06 | 5.9e-06 | 6.8e-06 | 6.333E-06 | 4.509E-07 | 5.900E-06 | 6.800E-06 |
| 21762 | 1089 | T1089R | D | 8.6e-06 | 4.8e-06 | 6.7e-06 | 6.700E-06 | 1.900E-06 | 4.800E-06 | 8.600E-06 |
| 21763 | 1089 | T1089N | D | 5.3e-06 | 1.6e-05 | 9.3e-06 | 1.020E-05 | 5.406E-06 | 5.300E-06 | 1.600E-05 |
| 21764 | 1089 | T1089D | D | 7.3e-06 | 7.5e-06 | 7.7e-06 | 7.500E-06 | 2.000E-07 | 7.300E-06 | 7.700E-06 |
| 21765 | 1089 | T1089C | D | 6.2e-06 | 7.1e-06 | 6.1e-06 | 6.467E-06 | 5.508E-07 | 6.100E-06 | 7.100E-06 |
| 21766 | 1089 | T1089E | D | 5.5e-06 | 8.5e-06 | 9.7e-06 | 7.900E-06 | 2.163E-06 | 5.500E-06 | 9.700E-06 |
| 21767 | 1089 | T1089Q | D | 7.1e-06 | 9.9e-06 | 4.3e-06 | 7.100E-06 | 2.800E-06 | 4.300E-06 | 9.900E-06 |
| 21768 | 1089 | T1089G | D | 7.5e-06 | 7.3e-06 | 9.3e-06 | 8.033E-06 | 1.102E-06 | 7.300E-06 | 9.300E-06 |
| 21769 | 1089 | T1089H | D | 6e-06 | 7.5e-06 | 7.4e-06 | 6.967E-06 | 8.386E-07 | 6.000E-06 | 7.500E-06 |
| 21770 | 1089 | T1089I | D | 4.6e-06 | 6.3e-06 | 6.8e-06 | 5.900E-06 | 1.153E-06 | 4.600E-06 | 6.800E-06 |
| 21771 | 1089 | T1089L | D | 5.5e-06 | 6e-06 | 4.9e-06 | 5.467E-06 | 5.508E-07 | 4.900E-06 | 6.000E-06 |
| 21772 | 1089 | T1089K | D | 8e-06 | 7.1e-06 | 9.3e-06 | 8.133E-06 | 1.106E-06 | 7.100E-06 | 9.300E-06 |
| 21773 | 1089 | T1089M | D | 5.1e-06 | 3.2e-06 | 4.5e-06 | 4.267E-06 | 9.713E-07 | 3.200E-06 | 5.100E-06 |
| 21774 | 1089 | T1089F | D | 6.5e-06 | 4.8e-06 | 6.5e-06 | 5.933E-06 | 9.815E-07 | 4.800E-06 | 6.500E-06 |
| 21775 | 1089 | T1089P | D | 7.4e-06 | 1e-05 | 7e-06 | 8.133E-06 | 1.629E-06 | 7.000E-06 | 1.000E-05 |
| 21776 | 1089 | T1089S | D | 1.2e-05 | 7.9e-06 | 7.8e-06 | 9.233E-06 | 2.397E-06 | 7.800E-06 | 1.200E-05 |
| 21777 | 1089 | T1089T | D | 7.1e-06 | 8.2e-06 | 8.4e-06 | 7.900E-06 | 7.000E-07 | 7.100E-06 | 8.400E-06 |
| 21778 | 1089 | T1089W | D | 8.5e-06 | 6.6e-06 | 9.3e-06 | 8.133E-06 | 1.387E-06 | 6.600E-06 | 9.300E-06 |
| 21779 | 1089 | T1089Y | D | 7.1e-06 | 6.4e-06 | 1e-05 | 7.833E-06 | 1.909E-06 | 6.400E-06 | 1.000E-05 |
| 21780 | 1089 | T1089V | D | 1e-05 | 7e-06 | 4.6e-06 | 7.200E-06 | 2.706E-06 | 4.600E-06 | 1.000E-05 |
| 21781 | 1090 | A1090A | D | 1e-05 | 8.8e-06 | 7e-06 | 8.600E-06 | 1.510E-06 | 7.000E-06 | 1.000E-05 |
| 21782 | 1090 | A1090R | D | 2.8e-06 | 1.7e-06 | 2.2e-06 | 2.233E-06 | 5.508E-07 | 1.700E-06 | 2.800E-06 |
| 21783 | 1090 | A1090N | D | 2.9e-06 | 5.9e-06 | 2.8e-06 | 3.867E-06 | 1.762E-06 | 2.800E-06 | 5.900E-06 |
| 21784 | 1090 | A1090D | D | 4e-06 | 3.8e-06 | 8.9e-06 | 5.567E-06 | 2.888E-06 | 3.800E-06 | 8.900E-06 |
| 21785 | 1090 | A1090C | D | 8.3e-06 | 5.8e-06 | 6.8e-06 | 6.967E-06 | 1.258E-06 | 5.800E-06 | 8.300E-06 |
| 21786 | 1090 | A1090E | D | 5.1e-06 | 4.7e-06 | 5.9e-06 | 5.233E-06 | 6.110E-07 | 4.700E-06 | 5.900E-06 |
| 21787 | 1090 | A1090Q | D | 9.8e-06 | 1.3e-05 | 4.7e-06 | 9.167E-06 | 4.186E-06 | 4.700E-06 | 1.300E-05 |
| 21788 | 1090 | A1090G | D | 6e-06 | 7.9e-06 | 9.4e-06 | 7.767E-06 | 1.704E-06 | 6.000E-06 | 9.400E-06 |
| 21789 | 1090 | A1090H | D | 2e-06 | 3.8e-06 | 1.6e-06 | 2.467E-06 | 1.172E-06 | 1.600E-06 | 3.800E-06 |
| 21790 | 1090 | A1090I | D | 4.5e-06 | 6.4e-06 | 5.2e-06 | 5.367E-06 | 9.609E-07 | 4.500E-06 | 6.400E-06 |
| 21791 | 1090 | A1090L | D | 5.1e-06 | 7.7e-06 | 2.4e-06 | 5.067E-06 | 2.650E-06 | 2.400E-06 | 7.700E-06 |
| 21792 | 1090 | A1090K | D | 6e-06 | 3e-06 | 3e-06 | 4.000E-06 | 1.732E-06 | 3.000E-06 | 6.000E-06 |
| 21793 | 1090 | A1090M | D | 3.1e-06 | 2.9e-06 | 5.1e-06 | 3.700E-06 | 1.217E-06 | 2.900E-06 | 5.100E-06 |
| 21794 | 1090 | A1090F | D | 3.4e-06 | 3.5e-06 | 4.5e-06 | 3.800E-06 | 6.083E-07 | 3.400E-06 | 4.500E-06 |
| 21795 | 1090 | A1090P | D | 1.3e-05 | 9.1e-06 | 6.8e-06 | 9.633E-06 | 3.134E-06 | 6.800E-06 | 1.300E-05 |
| 21796 | 1090 | A1090S | D | 3.2e-06 | 3.2e-06 | 4.2e-06 | 3.533E-06 | 5.774E-07 | 3.200E-06 | 4.200E-06 |
| 21797 | 1090 | A1090T | D | 4.9e-06 | 1.2e-05 | 4.1e-06 | 7.000E-06 | 4.349E-06 | 4.100E-06 | 1.200E-05 |
| 21798 | 1090 | A1090W | D | 2.5e-06 | 2.9e-06 | 3e-06 | 2.800E-06 | 2.646E-07 | 2.500E-06 | 3.000E-06 |
| 21799 | 1090 | A1090Y | D | 4.7e-06 | 2.9e-06 | 4.3e-06 | 3.967E-06 | 9.452E-07 | 2.900E-06 | 4.700E-06 |
| 21800 | 1090 | A1090V | D | 6.9e-06 | 8.2e-06 | 3.1e-06 | 6.067E-06 | 2.650E-06 | 3.100E-06 | 8.200E-06 |
| 21801 | 1091 | V1091A | D | 7e-06 | 1.1e-05 | 4.6e-06 | 7.533E-06 | 3.233E-06 | 4.600E-06 | 1.100E-05 |
| 21802 | 1091 | V1091R | D | 6.4e-06 | 6.9e-06 | 8.2e-06 | 7.167E-06 | 9.292E-07 | 6.400E-06 | 8.200E-06 |
| 21803 | 1091 | V1091N | D | 1.3e-05 | 5.2e-06 | 9.5e-06 | 9.233E-06 | 3.907E-06 | 5.200E-06 | 1.300E-05 |
| 21804 | 1091 | V1091D | D | 6.8e-06 | 6.6e-06 | 8.8e-06 | 7.400E-06 | 1.217E-06 | 6.600E-06 | 8.800E-06 |
| 21805 | 1091 | V1091C | D | 7.8e-06 | 6.3e-06 | 4.4e-06 | 6.167E-06 | 1.704E-06 | 4.400E-06 | 7.800E-06 |
| 21806 | 1091 | V1091E | D | 4.2e-06 | 8.8e-06 | 1.1e-05 | 8.000E-06 | 3.470E-06 | 4.200E-06 | 1.100E-05 |
| 21807 | 1091 | V1091Q | D | 5.3e-06 | 8.7e-06 | 1.2e-05 | 8.667E-06 | 3.350E-06 | 5.300E-06 | 1.200E-05 |
| 21808 | 1091 | V1091G | D | 6.7e-06 | 7.4e-06 | 6.8e-06 | 6.967E-06 | 3.786E-07 | 6.700E-06 | 7.400E-06 |
| 21809 | 1091 | V1091H | D | 6.1e-06 | 6.7e-06 | 5.6e-06 | 6.133E-06 | 5.508E-07 | 5.600E-06 | 6.700E-06 |
| 21810 | 1091 | V1091I | D | 1.1e-05 | 5.7e-06 | 9e-06 | 8.567E-06 | 2.676E-06 | 5.700E-06 | 1.100E-05 |
| 21811 | 1091 | V1091L | D | 6.7e-06 | 9.3e-06 | 6.2e-06 | 7.400E-06 | 1.664E-06 | 6.200E-06 | 9.300E-06 |
| 21812 | 1091 | V1091K | D | 9.1e-06 | 4.9e-06 | 4.5e-06 | 6.167E-06 | 2.548E-06 | 4.500E-06 | 9.100E-06 |
| 21813 | 1091 | V1091M | D | 6.8e-06 | 7.1e-06 | 1.4e-05 | 9.300E-06 | 4.073E-06 | 6.800E-06 | 1.400E-05 |
| 21814 | 1091 | V1091F | D | 9.9e-06 | 8e-06 | 1.3e-05 | 1.030E-05 | 2.524E-06 | 8.000E-06 | 1.300E-05 |
| 21815 | 1091 | V1091P | D | 6.3e-06 | 7.2e-06 | 9.1e-06 | 7.533E-06 | 1.429E-06 | 6.300E-06 | 9.100E-06 |
| 21816 | 1091 | V1091S | D | 1e-05 | 5.9e-06 | 1.7e-05 | 1.097E-05 | 5.613E-06 | 5.900E-06 | 1.700E-05 |
| 21817 | 1091 | V1091T | D | 7.8e-06 | 5.2e-06 | 6.6e-06 | 6.533E-06 | 1.301E-06 | 5.200E-06 | 7.800E-06 |
| 21818 | 1091 | V1091W | D | 1e-05 | 7.7e-06 | 1.2e-05 | 9.900E-06 | 2.152E-06 | 7.700E-06 | 1.200E-05 |
| 21819 | 1091 | V1091Y | D | 1.3e-05 | 7e-06 | 1.3e-05 | 1.100E-05 | 3.464E-06 | 7.000E-06 | 1.300E-05 |
| 21820 | 1091 | V1091V | D | 7.6e-06 | 6.3e-06 | 8.5e-06 | 7.467E-06 | 1.106E-06 | 6.300E-06 | 8.500E-06 |
| 21821 | 1092 | A1092A | D | 7.3e-06 | 8.7e-06 | 5.3e-06 | 7.100E-06 | 1.709E-06 | 5.300E-06 | 8.700E-06 |
| 21822 | 1092 | A1092R | D | 9.2e-06 | 1.1e-05 | 9.8e-06 | 1.000E-05 | 9.165E-07 | 9.200E-06 | 1.100E-05 |
| 21823 | 1092 | A1092N | D | 8e-06 | 5.8e-06 | 5e-06 | 6.267E-06 | 1.553E-06 | 5.000E-06 | 8.000E-06 |
| 21824 | 1092 | A1092D | D | 6e-06 | 8.3e-06 | 6.9e-06 | 7.067E-06 | 1.159E-06 | 6.000E-06 | 8.300E-06 |
| 21825 | 1092 | A1092C | D | 6.5e-06 | 6.9e-06 | 1.3e-05 | 8.800E-06 | 3.643E-06 | 6.500E-06 | 1.300E-05 |
| 21826 | 1092 | A1092E | D | 7.7e-06 | 6e-06 | 7.1e-06 | 6.933E-06 | 8.622E-07 | 6.000E-06 | 7.700E-06 |
| 21827 | 1092 | A1092Q | D | 6.8e-06 | 4.9e-06 | 1.2e-05 | 7.900E-06 | 3.676E-06 | 4.900E-06 | 1.200E-05 |
| 21828 | 1092 | A1092G | D | 5.8e-06 | 4.9e-06 | 8.7e-06 | 6.467E-06 | 1.986E-06 | 4.900E-06 | 8.700E-06 |
| 21829 | 1092 | A1092H | D | 6.3e-06 | 4.8e-06 | 9.4e-06 | 6.833E-06 | 2.346E-06 | 4.800E-06 | 9.400E-06 |
| 21830 | 1092 | A1092I | D | 4.4e-06 | 5.1e-06 | 6.1e-06 | 5.200E-06 | 8.544E-07 | 4.400E-06 | 6.100E-06 |
| 21831 | 1092 | A1092L | D | 6e-06 | 5.4e-06 | 7.8e-06 | 6.400E-06 | 1.249E-06 | 5.400E-06 | 7.800E-06 |
| 21832 | 1092 | A1092K | D | 4.1e-06 | 1.1e-05 | 8.8e-06 | 7.967E-06 | 3.525E-06 | 4.100E-06 | 1.100E-05 |
| 21833 | 1092 | A1092M | D | 9.3e-06 | 3.3e-06 | 1.2e-05 | 8.200E-06 | 4.453E-06 | 3.300E-06 | 1.200E-05 |
| 21834 | 1092 | A1092F | D | 8.8e-06 | 7e-06 | 8.8e-06 | 8.200E-06 | 1.039E-06 | 7.000E-06 | 8.800E-06 |
| 21835 | 1092 | A1092P | D | 8.8e-06 | 8.8e-06 | 9.4e-06 | 9.000E-06 | 3.464E-07 | 8.800E-06 | 9.400E-06 |
| 21836 | 1092 | A1092S | D | 8.8e-06 | 8.5e-06 | 1.8e-05 | 1.177E-05 | 5.400E-06 | 8.500E-06 | 1.800E-05 |
| 21837 | 1092 | A1092T | D | 1e-05 | 5.5e-06 | 6.6e-06 | 7.367E-06 | 2.346E-06 | 5.500E-06 | 1.000E-05 |
| 21838 | 1092 | A1092W | D | 9e-06 | 5.5e-06 | 9e-06 | 7.833E-06 | 2.021E-06 | 5.500E-06 | 9.000E-06 |
| 21839 | 1092 | A1092Y | D | 9e-06 | 7.9e-06 | 8.3e-06 | 8.400E-06 | 5.568E-07 | 7.900E-06 | 9.000E-06 |
| 21840 | 1092 | A1092V | D | 5.8e-06 | 7.9e-06 | 5.9e-06 | 6.533E-06 | 1.185E-06 | 5.800E-06 | 7.900E-06 |
| 21841 | 1093 | W1093A | D | 6.9e-06 | 1.4e-05 | 1.4e-05 | 1.163E-05 | 4.099E-06 | 6.900E-06 | 1.400E-05 |
| 21842 | 1093 | W1093R | D | 9.7e-06 | 8.2e-06 | 1.2e-05 | 9.967E-06 | 1.914E-06 | 8.200E-06 | 1.200E-05 |
| 21843 | 1093 | W1093N | D | 8.8e-06 | 1e-05 | 5.4e-06 | 8.067E-06 | 2.386E-06 | 5.400E-06 | 1.000E-05 |
| 21844 | 1093 | W1093D | D | 7e-06 | 6.7e-06 | 8.8e-06 | 7.500E-06 | 1.136E-06 | 6.700E-06 | 8.800E-06 |

| | | | | | | | | | | |
|---|---|---|---|---|---|---|---|---|---|---|
| 21845 | 1093 | W1093C | D | 7.3e-06 | 6.6e-06 | 1e-05 | 7.967E-06 | 1.795E-06 | 6.600E-06 | 1.000E-05 |
| 21846 | 1093 | W1093E | D | 6.8e-06 | 1e-05 | 1.1e-05 | 9.267E-06 | 2.194E-06 | 6.800E-06 | 1.100E-05 |
| 21847 | 1093 | W1093Q | D | 8.5e-06 | 6.2e-06 | 4.4e-06 | 6.367E-06 | 2.055E-06 | 4.400E-06 | 8.500E-06 |
| 21848 | 1093 | W1093G | D | 6.9e-06 | 1.1e-05 | 5.6e-06 | 7.833E-06 | 2.818E-06 | 5.600E-06 | 1.100E-05 |
| 21849 | 1093 | W1093H | D | 6.6e-06 | 6.8e-06 | 9.4e-06 | 7.600E-06 | 1.562E-06 | 6.600E-06 | 9.400E-06 |
| 21850 | 1093 | W1093I | D | 8.5e-06 | 8e-06 | 1e-05 | 8.833E-06 | 1.041E-06 | 8.000E-06 | 1.000E-05 |
| 21851 | 1093 | W1093L | D | 9.3e-06 | 7.7e-06 | 4.1e-06 | 7.033E-06 | 2.663E-06 | 4.100E-06 | 9.300E-06 |
| 21852 | 1093 | W1093K | D | 1.4e-05 | 8.3e-06 | 7.1e-06 | 9.800E-06 | 3.686E-06 | 7.100E-06 | 1.400E-05 |
| 21853 | 1093 | W1093M | D | 7.4e-06 | 8.6e-06 | 8.5e-06 | 8.167E-06 | 6.658E-07 | 7.400E-06 | 8.600E-06 |
| 21854 | 1093 | W1093F | D | 8.9e-06 | 4.6e-06 | 9.6e-06 | 7.700E-06 | 2.707E-06 | 4.600E-06 | 9.600E-06 |
| 21855 | 1093 | W1093P | D | 9.1e-06 | 6.8e-06 | 6.5e-06 | 7.467E-06 | 1.422E-06 | 6.500E-06 | 9.100E-06 |
| 21856 | 1093 | W1093S | D | 9.5e-06 | 7e-06 | 1e-05 | 8.833E-06 | 1.607E-06 | 7.000E-06 | 1.000E-05 |
| 21857 | 1093 | W1093T | D | 1.1e-05 | 9.6e-06 | 7.7e-06 | 9.433E-06 | 1.656E-06 | 7.700E-06 | 1.100E-05 |
| 21858 | 1093 | W1093W | D | 4.7e-06 | 9.4e-06 | 6.9e-06 | 7.000E-06 | 2.352E-06 | 4.700E-06 | 9.400E-06 |
| 21859 | 1093 | W1093Y | D | 1e-05 | 8.3e-06 | 9.9e-06 | 9.400E-06 | 9.539E-07 | 8.300E-06 | 1.000E-05 |
| 21860 | 1093 | W1093V | D | 8.2e-06 | 8.3e-06 | 7.2e-06 | 7.900E-06 | 6.083E-07 | 7.200E-06 | 8.300E-06 |
| 21861 | 1094 | Y1094A | D | 8.8e-06 | 5.9e-06 | 1.2e-05 | 8.900E-06 | 3.051E-06 | 5.900E-06 | 1.200E-05 |
| 21862 | 1094 | Y1094R | D | 6.6e-06 | 8.8e-06 | 1.2e-05 | 9.133E-06 | 2.715E-06 | 6.600E-06 | 1.200E-05 |
| 21863 | 1094 | Y1094N | D | 8.5e-06 | 6.6e-06 | 4e-06 | 6.367E-06 | 2.259E-06 | 4.000E-06 | 8.500E-06 |
| 21864 | 1094 | Y1094D | D | 1.1e-05 | 1.2e-05 | 4.4e-06 | 9.133E-06 | 4.130E-06 | 4.400E-06 | 1.200E-05 |
| 21865 | 1094 | Y1094C | D | 1.1e-05 | 7.8e-06 | 6.7e-06 | 8.500E-06 | 2.234E-06 | 6.700E-06 | 1.100E-05 |
| 21866 | 1094 | Y1094E | D | 7.4e-06 | 5.2e-06 | 7.2e-06 | 6.600E-06 | 1.217E-06 | 5.200E-06 | 7.400E-06 |
| 21867 | 1094 | Y1094Q | D | 7.1e-06 | 7e-06 | 1.3e-05 | 9.033E-06 | 3.436E-06 | 7.000E-06 | 1.300E-05 |
| 21868 | 1094 | Y1094G | D | 4.4e-06 | 1.2e-05 | 5.7e-06 | 7.367E-06 | 4.065E-06 | 4.400E-06 | 1.200E-05 |
| 21869 | 1094 | Y1094H | D | 9.5e-06 | 5.1e-06 | 6.7e-06 | 7.100E-06 | 2.227E-06 | 5.100E-06 | 9.500E-06 |
| 21870 | 1094 | Y1094I | D | 1.3e-05 | 6.5e-06 | 8e-06 | 9.167E-06 | 3.403E-06 | 6.500E-06 | 1.300E-05 |
| 21871 | 1094 | Y1094L | D | 6.7e-06 | 6.8e-06 | 9e-06 | 7.500E-06 | 1.300E-06 | 6.700E-06 | 9.000E-06 |
| 21872 | 1094 | Y1094K | D | 1e-05 | 8.4e-06 | 6.3e-06 | 8.233E-06 | 1.856E-06 | 6.300E-06 | 1.000E-05 |
| 21873 | 1094 | Y1094M | D | 7.6e-06 | 1.2e-05 | 4.7e-06 | 8.100E-06 | 3.676E-06 | 4.700E-06 | 1.200E-05 |
| 21874 | 1094 | Y1094F | D | 8.7e-06 | 5.5e-06 | 8.8e-06 | 7.667E-06 | 1.877E-06 | 5.500E-06 | 8.800E-06 |
| 21875 | 1094 | Y1094P | D | 7.7e-06 | 7.2e-06 | 6.8e-06 | 7.233E-06 | 4.509E-07 | 6.800E-06 | 7.700E-06 |
| 21876 | 1094 | Y1094S | D | 7.2e-06 | 8.3e-06 | 2e-05 | 1.183E-05 | 7.094E-06 | 7.200E-06 | 2.000E-05 |
| 21877 | 1094 | Y1094T | D | 8.7e-06 | 4.7e-06 | 4.4e-06 | 5.933E-06 | 2.401E-06 | 4.400E-06 | 8.700E-06 |
| 21878 | 1094 | Y1094W | D | 9.7e-06 | 5.4e-06 | 1e-05 | 8.367E-06 | 2.574E-06 | 5.400E-06 | 1.000E-05 |
| 21879 | 1094 | Y1094Y | D | 8.3e-06 | 5.2e-06 | 8e-06 | 7.167E-06 | 1.710E-06 | 5.200E-06 | 8.300E-06 |
| 21880 | 1094 | Y1094V | D | 7.7e-06 | 5.9e-06 | 8.6e-06 | 7.400E-06 | 1.375E-06 | 5.900E-06 | 8.600E-06 |
| 21881 | 1095 | Q1095A | D | 9e-06 | 7.1e-06 | 5.5e-06 | 7.200E-06 | 1.752E-06 | 5.500E-06 | 9.000E-06 |
| 21882 | 1095 | Q1095R | D | 7.2e-06 | 1.2e-05 | 8e-06 | 9.067E-06 | 2.572E-06 | 7.200E-06 | 1.200E-05 |
| 21883 | 1095 | Q1095N | D | 7.2e-06 | 9.6e-06 | 5e-06 | 7.267E-06 | 2.301E-06 | 5.000E-06 | 9.600E-06 |
| 21884 | 1095 | Q1095D | D | 1.4e-05 | 5.1e-06 | 6.1e-06 | 8.400E-06 | 4.875E-06 | 5.100E-06 | 1.400E-05 |
| 21885 | 1095 | Q1095C | D | 9.2e-06 | 1.2e-05 | 6.7e-06 | 9.300E-06 | 2.651E-06 | 6.700E-06 | 1.200E-05 |
| 21886 | 1095 | Q1095E | D | 1.1e-05 | 1.1e-05 | 9.4e-06 | 1.047E-05 | 9.238E-07 | 9.400E-06 | 1.100E-05 |
| 21887 | 1095 | Q1095Q | D | 8.5e-06 | 7.4e-06 | 5.7e-06 | 7.200E-06 | 1.411E-06 | 5.700E-06 | 8.500E-06 |
| 21888 | 1095 | Q1095G | D | 6.7e-06 | 6.8e-06 | 8.9e-06 | 7.467E-06 | 1.242E-06 | 6.700E-06 | 8.900E-06 |
| 21889 | 1095 | Q1095H | D | 7.6e-06 | 1.5e-05 | 5.7e-06 | 9.433E-06 | 4.914E-06 | 5.700E-06 | 1.500E-05 |
| 21890 | 1095 | Q1095I | D | 9.2e-06 | 9.4e-06 | 4.8e-06 | 7.800E-06 | 2.600E-06 | 4.800E-06 | 9.400E-06 |
| 21891 | 1095 | Q1095L | D | 1e-05 | 1.1e-05 | 4.8e-06 | 8.600E-06 | 3.329E-06 | 4.800E-06 | 1.100E-05 |
| 21892 | 1095 | Q1095K | D | 1.2e-05 | 6.8e-06 | 7.6e-06 | 8.800E-06 | 2.800E-06 | 6.800E-06 | 1.200E-05 |
| 21893 | 1095 | Q1095M | D | 4.3e-06 | 7.4e-06 | 8.2e-06 | 6.633E-06 | 2.060E-06 | 4.300E-06 | 8.200E-06 |
| 21894 | 1095 | Q1095F | D | 1e-05 | 4.9e-06 | 1.1e-05 | 8.633E-06 | 3.272E-06 | 4.900E-06 | 1.100E-05 |
| 21895 | 1095 | Q1095P | D | 8e-06 | 1e-05 | 1.1e-05 | 9.667E-06 | 1.528E-06 | 8.000E-06 | 1.100E-05 |
| 21896 | 1095 | Q1095S | D | 7.1e-06 | 1e-05 | 8.2e-06 | 8.433E-06 | 1.464E-06 | 7.100E-06 | 1.000E-05 |
| 21897 | 1095 | Q1095T | D | 5.9e-06 | 6.4e-06 | 7e-06 | 6.433E-06 | 5.508E-07 | 5.900E-06 | 7.000E-06 |
| 21898 | 1095 | Q1095W | D | 4.9e-06 | 8e-06 | 8.4e-06 | 7.100E-06 | 1.916E-06 | 4.900E-06 | 8.400E-06 |
| 21899 | 1095 | Q1095Y | D | 1.4e-05 | 1.1e-05 | 5.2e-06 | 1.007E-05 | 4.474E-06 | 5.200E-06 | 1.400E-05 |
| 21900 | 1095 | Q1095V | D | 6.9e-06 | 9.9e-06 | 8.7e-06 | 8.500E-06 | 1.510E-06 | 6.900E-06 | 9.900E-06 |
| 21901 | 1096 | Q1096A | D | 4.9e-06 | 6.7e-06 | 1.3e-05 | 8.200E-06 | 4.253E-06 | 4.900E-06 | 1.300E-05 |
| 21902 | 1096 | Q1096R | D | 8.3e-06 | 9.7e-06 | 1.4e-05 | 1.067E-05 | 2.970E-06 | 8.300E-06 | 1.400E-05 |
| 21903 | 1096 | Q1096N | D | 1.2e-05 | 4.8e-06 | 5.4e-06 | 7.400E-06 | 3.995E-06 | 4.800E-06 | 1.200E-05 |
| 21904 | 1096 | Q1096D | D | 4.7e-06 | 5.9e-06 | 6.7e-06 | 5.767E-06 | 1.007E-06 | 4.700E-06 | 6.700E-06 |
| 21905 | 1096 | Q1096C | D | 6.3e-06 | 1.5e-05 | 1.4e-05 | 1.177E-05 | 4.761E-06 | 6.300E-06 | 1.500E-05 |
| 21906 | 1096 | Q1096E | D | 6.9e-06 | 9.6e-06 | 4.7e-06 | 7.067E-06 | 2.454E-06 | 4.700E-06 | 9.600E-06 |
| 21907 | 1096 | Q1096Q | D | 1.2e-05 | 9.4e-06 | 5.8e-06 | 9.067E-06 | 3.113E-06 | 5.800E-06 | 1.200E-05 |
| 21908 | 1096 | Q1096G | D | 5.1e-06 | 6.1e-06 | 3.9e-06 | 5.033E-06 | 1.102E-06 | 3.900E-06 | 6.100E-06 |
| 21909 | 1096 | Q1096H | D | 4.9e-06 | 7.6e-06 | 7.8e-06 | 6.767E-06 | 1.620E-06 | 4.900E-06 | 7.800E-06 |
| 21910 | 1096 | Q1096I | D | 1e-05 | 4.1e-06 | 1.2e-05 | 8.700E-06 | 4.107E-06 | 4.100E-06 | 1.200E-05 |
| 21911 | 1096 | Q1096L | D | 7.7e-06 | 1.3e-05 | 7.3e-06 | 9.333E-06 | 3.182E-06 | 7.300E-06 | 1.300E-05 |
| 21912 | 1096 | Q1096K | D | 1e-05 | 7.3e-06 | 6.7e-06 | 8.000E-06 | 1.758E-06 | 6.700E-06 | 1.000E-05 |
| 21913 | 1096 | Q1096M | D | 1.2e-05 | 7.8e-06 | 1.5e-05 | 1.160E-05 | 3.617E-06 | 7.800E-06 | 1.500E-05 |
| 21914 | 1096 | Q1096F | D | 1.3e-05 | 1e-05 | 6.3e-06 | 9.767E-06 | 3.356E-06 | 6.300E-06 | 1.300E-05 |
| 21915 | 1096 | Q1096P | D | 6.7e-06 | 5e-06 | 1e-05 | 7.233E-06 | 2.542E-06 | 5.000E-06 | 1.000E-05 |
| 21916 | 1096 | Q1096S | D | 7.3e-06 | 9.6e-06 | 6.7e-06 | 7.867E-06 | 1.531E-06 | 6.700E-06 | 9.600E-06 |
| 21917 | 1096 | Q1096T | D | 9.5e-06 | 4.5e-06 | 1.4e-05 | 9.333E-06 | 4.752E-06 | 4.500E-06 | 1.400E-05 |
| 21918 | 1096 | Q1096W | D | 6.3e-06 | 7.5e-06 | 4.2e-06 | 6.000E-06 | 1.670E-06 | 4.200E-06 | 7.500E-06 |
| 21919 | 1096 | Q1096Y | D | 4e-06 | 1e-05 | 3.7e-06 | 5.900E-06 | 3.554E-06 | 3.700E-06 | 1.000E-05 |
| 21920 | 1096 | Q1096V | D | 1.1e-05 | 7.7e-06 | 4.7e-06 | 7.800E-06 | 3.151E-06 | 4.700E-06 | 1.100E-05 |
| 21921 | 1097 | K1097A | D | 4.6e-06 | 7.6e-06 | 7.2e-06 | 6.467E-06 | 1.629E-06 | 4.600E-06 | 7.600E-06 |
| 21922 | 1097 | K1097R | D | 1.3e-05 | 1.2e-05 | 1.4e-05 | 1.300E-05 | 1.000E-06 | 1.200E-05 | 1.400E-05 |
| 21923 | 1097 | K1097N | D | 7.3e-06 | 1e-05 | 5.2e-06 | 7.500E-06 | 2.406E-06 | 5.200E-06 | 1.000E-05 |
| 21924 | 1097 | K1097D | D | 1.2e-05 | 8.2e-06 | 6e-06 | 8.733E-06 | 3.035E-06 | 6.000E-06 | 1.200E-05 |
| 21925 | 1097 | K1097C | D | 7.9e-06 | 9.6e-06 | 1.1e-05 | 9.500E-06 | 1.552E-06 | 7.900E-06 | 1.100E-05 |
| 21926 | 1097 | K1097E | D | 6.4e-06 | 1.1e-05 | 9.7e-06 | 9.033E-06 | 2.371E-06 | 6.400E-06 | 1.100E-05 |
| 21927 | 1097 | K1097Q | D | 9.7e-06 | 1.1e-05 | 8.8e-06 | 9.833E-06 | 1.106E-06 | 8.800E-06 | 1.100E-05 |
| 21928 | 1097 | K1097G | D | 6.6e-06 | 1.4e-05 | 8e-06 | 9.533E-06 | 3.931E-06 | 6.600E-06 | 1.400E-05 |
| 21929 | 1097 | K1097H | D | 9.5e-06 | 8.7e-06 | 1e-05 | 9.400E-06 | 6.557E-07 | 8.700E-06 | 1.000E-05 |

| | | | | | | | | | | |
|---|---|---|---|---|---|---|---|---|---|---|
| 21930 | 1097 | K1097I | D | 8e-06 | 5.2e-06 | 6.1e-06 | 6.433E-06 | 1.429E-06 | 5.200E-06 | 8.000E-06 |
| 21931 | 1097 | K1097L | D | 7.5e-06 | 6.9e-06 | 6.9e-06 | 7.100E-06 | 3.464E-07 | 6.900E-06 | 7.500E-06 |
| 21932 | 1097 | K1097K | D | 7.8e-06 | 7e-06 | 9.4e-06 | 8.067E-06 | 1.222E-06 | 7.000E-06 | 9.400E-06 |
| 21933 | 1097 | K1097M | D | 9.4e-06 | 1e-05 | 1.1e-05 | 1.013E-05 | 8.083E-07 | 9.400E-06 | 1.100E-05 |
| 21934 | 1097 | K1097F | D | 1.3e-05 | 1.1e-05 | 5.5e-06 | 9.833E-06 | 3.884E-06 | 5.500E-06 | 1.300E-05 |
| 21935 | 1097 | K1097P | D | 4.6e-06 | 9.1e-06 | 7e-06 | 6.900E-06 | 2.252E-06 | 4.600E-06 | 9.100E-06 |
| 21936 | 1097 | K1097S | D | 1.1e-05 | 6.7e-06 | 1e-05 | 9.233E-06 | 2.250E-06 | 6.700E-06 | 1.100E-05 |
| 21937 | 1097 | K1097T | D | 6.4e-06 | 1.4e-05 | 5e-06 | 8.467E-06 | 4.843E-06 | 5.000E-06 | 1.400E-05 |
| 21938 | 1097 | K1097W | D | 7.1e-06 | 7.2e-06 | 1.3e-05 | 9.100E-06 | 3.378E-06 | 7.100E-06 | 1.300E-05 |
| 21939 | 1097 | K1097Y | D | 8.7e-06 | 8.5e-06 | 4.6e-06 | 7.267E-06 | 2.312E-06 | 4.600E-06 | 8.700E-06 |
| 21940 | 1097 | K1097V | D | 6.2e-06 | 9.5e-06 | 4.7e-06 | 6.800E-06 | 2.456E-06 | 4.700E-06 | 9.500E-06 |
| 21941 | 1098 | P1098A | D | 5.7e-06 | 7e-06 | 8.8e-06 | 7.167E-06 | 1.557E-06 | 5.700E-06 | 8.800E-06 |
| 21942 | 1098 | P1098R | D | 6.7e-06 | 9.2e-06 | 7.1e-06 | 7.667E-06 | 1.343E-06 | 6.700E-06 | 9.200E-06 |
| 21943 | 1098 | P1098N | D | 7.6e-06 | 8.9e-06 | 9.8e-06 | 8.767E-06 | 1.106E-06 | 7.600E-06 | 9.800E-06 |
| 21944 | 1098 | P1098D | D | 1.5e-05 | 7.8e-06 | 9.2e-06 | 1.067E-05 | 3.818E-06 | 7.800E-06 | 1.500E-05 |
| 21945 | 1098 | P1098C | D | 9.1e-06 | 7e-06 | 6.8e-06 | 7.633E-06 | 1.274E-06 | 6.800E-06 | 9.100E-06 |
| 21946 | 1098 | P1098E | D | 6.4e-06 | 8.5e-06 | 9.9e-06 | 8.267E-06 | 1.762E-06 | 6.400E-06 | 9.900E-06 |
| 21947 | 1098 | P1098Q | D | 7.8e-06 | 6.5e-06 | 8.5e-06 | 7.600E-06 | 1.015E-06 | 6.500E-06 | 8.500E-06 |
| 21948 | 1098 | P1098G | D | 7.9e-06 | 1.5e-05 | 5.8e-06 | 9.567E-06 | 4.821E-06 | 5.800E-06 | 1.500E-05 |
| 21949 | 1098 | P1098H | D | 5.3e-06 | 4.1e-06 | 8.2e-06 | 5.867E-06 | 2.108E-06 | 4.100E-06 | 8.200E-06 |
| 21950 | 1098 | P1098I | D | 4.4e-06 | 9.8e-06 | 8.4e-06 | 7.533E-06 | 2.802E-06 | 4.400E-06 | 9.800E-06 |
| 21951 | 1098 | P1098L | D | 1.1e-05 | 8.8e-06 | 6.4e-06 | 8.733E-06 | 2.301E-06 | 6.400E-06 | 1.100E-05 |
| 21952 | 1098 | P1098K | D | 1e-05 | 5.8e-06 | 8.2e-06 | 8.000E-06 | 2.107E-06 | 5.800E-06 | 1.000E-05 |
| 21953 | 1098 | P1098M | D | 8.5e-06 | 1.2e-05 | 6.8e-06 | 9.100E-06 | 2.651E-06 | 6.800E-06 | 1.200E-05 |
| 21954 | 1098 | P1098F | D | 6.2e-06 | 6.5e-06 | 7.7e-06 | 6.800E-06 | 7.937E-07 | 6.200E-06 | 7.700E-06 |
| 21955 | 1098 | P1098P | D | 8.5e-06 | 6.7e-06 | 7.4e-06 | 7.533E-06 | 9.074E-07 | 6.700E-06 | 8.500E-06 |
| 21956 | 1098 | P1098S | D | 9.4e-06 | 9.6e-06 | 6.6e-06 | 8.533E-06 | 1.677E-06 | 6.600E-06 | 9.600E-06 |
| 21957 | 1098 | P1098T | D | 4.8e-06 | 8.5e-06 | 1.2e-05 | 8.433E-06 | 3.600E-06 | 4.800E-06 | 1.200E-05 |
| 21958 | 1098 | P1098W | D | 9.5e-06 | 6e-06 | 8.9e-06 | 8.133E-06 | 1.872E-06 | 6.000E-06 | 9.500E-06 |
| 21959 | 1098 | P1098Y | D | 6.4e-06 | 1.3e-05 | 6e-06 | 8.467E-06 | 3.931E-06 | 6.000E-06 | 1.300E-05 |
| 21960 | 1098 | P1098V | D | 9.9e-06 | 6.4e-06 | 1.4e-05 | 1.010E-05 | 3.804E-06 | 6.400E-06 | 1.400E-05 |
| 21961 | 1099 | G1099A | D | 1e-05 | 8e-06 | 6.1e-06 | 8.033E-06 | 1.950E-06 | 6.100E-06 | 1.000E-05 |
| 21962 | 1099 | G1099R | D | 9.5e-06 | 4.9e-06 | 1.4e-05 | 9.467E-06 | 4.550E-06 | 4.900E-06 | 1.400E-05 |
| 21963 | 1099 | G1099N | D | 4.4e-06 | 1.1e-05 | 8.3e-06 | 7.900E-06 | 3.318E-06 | 4.400E-06 | 1.100E-05 |
| 21964 | 1099 | G1099D | D | 6.4e-06 | 1e-05 | 6.7e-06 | 7.700E-06 | 1.997E-06 | 6.400E-06 | 1.000E-05 |
| 21965 | 1099 | G1099C | D | 1.4e-05 | 7.5e-06 | 5.6e-06 | 9.033E-06 | 4.405E-06 | 5.600E-06 | 1.400E-05 |
| 21966 | 1099 | G1099E | D | 1.2e-05 | 6.5e-06 | 1.1e-05 | 9.833E-06 | 2.930E-06 | 6.500E-06 | 1.200E-05 |
| 21967 | 1099 | G1099Q | D | 1.1e-05 | 6.1e-06 | 1.4e-05 | 1.037E-05 | 3.988E-06 | 6.100E-06 | 1.400E-05 |
| 21968 | 1099 | G1099G | D | 5.9e-06 | 1.4e-05 | 1e-05 | 9.967E-06 | 4.050E-06 | 5.900E-06 | 1.400E-05 |
| 21969 | 1099 | G1099H | D | 1.4e-05 | 1.4e-05 | 4.8e-06 | 1.093E-05 | 5.312E-06 | 4.800E-06 | 1.400E-05 |
| 21970 | 1099 | G1099I | D | 7e-06 | 1.4e-05 | 6.3e-06 | 9.100E-06 | 4.258E-06 | 6.300E-06 | 1.400E-05 |
| 21971 | 1099 | G1099L | D | 4.5e-06 | 1.3e-05 | 9.6e-06 | 9.033E-06 | 4.278E-06 | 4.500E-06 | 1.300E-05 |
| 21972 | 1099 | G1099K | D | 7.8e-06 | 6.2e-06 | 1e-05 | 8.000E-06 | 1.908E-06 | 6.200E-06 | 1.000E-05 |
| 21973 | 1099 | G1099M | D | 6.8e-06 | 5.6e-06 | 8e-06 | 6.800E-06 | 1.200E-06 | 5.600E-06 | 8.000E-06 |
| 21974 | 1099 | G1099F | D | 7.2e-06 | 5.7e-06 | 4.8e-06 | 5.900E-06 | 1.212E-06 | 4.800E-06 | 7.200E-06 |
| 21975 | 1099 | G1099P | D | 3.2e-06 | 7e-06 | 1.3e-05 | 7.733E-06 | 4.941E-06 | 3.200E-06 | 1.300E-05 |
| 21976 | 1099 | G1099S | D | 1e-05 | 8.9e-06 | 4.9e-06 | 7.933E-06 | 2.684E-06 | 4.900E-06 | 1.000E-05 |
| 21977 | 1099 | G1099T | D | 6.4e-06 | 9e-06 | 9.9e-06 | 8.433E-06 | 1.818E-06 | 6.400E-06 | 9.900E-06 |
| 21978 | 1099 | G1099W | D | 5.5e-06 | 4.7e-06 | 1e-05 | 6.733E-06 | 2.857E-06 | 4.700E-06 | 1.000E-05 |
| 21979 | 1099 | G1099Y | D | 7.4e-06 | 7.1e-06 | 6.6e-06 | 7.033E-06 | 4.041E-07 | 6.600E-06 | 7.400E-06 |
| 21980 | 1099 | G1099V | D | 1.8e-05 | 6.1e-06 | 4.7e-06 | 9.600E-06 | 7.308E-06 | 4.700E-06 | 1.800E-05 |
| 21981 | 1100 | K1100A | D | 1.5e-05 | 9.2e-06 | 8e-06 | 1.073E-05 | 3.743E-06 | 8.000E-06 | 1.500E-05 |
| 21982 | 1100 | K1100R | D | 8.9e-06 | 6.5e-06 | 1.3e-05 | 9.467E-06 | 3.287E-06 | 6.500E-06 | 1.300E-05 |
| 21983 | 1100 | K1100N | D | 7.1e-06 | 4.4e-06 | 6.7e-06 | 6.067E-06 | 1.457E-06 | 4.400E-06 | 7.100E-06 |
| 21984 | 1100 | K1100D | D | 1.1e-05 | 5e-06 | 5.2e-06 | 7.067E-06 | 3.408E-06 | 5.000E-06 | 1.100E-05 |
| 21985 | 1100 | K1100C | D | 7.5e-06 | 6.1e-06 | 4.8e-06 | 6.133E-06 | 1.350E-06 | 4.800E-06 | 7.500E-06 |
| 21986 | 1100 | K1100E | D | 6.5e-06 | 7.7e-06 | 1.1e-05 | 8.400E-06 | 2.330E-06 | 6.500E-06 | 1.100E-05 |
| 21987 | 1100 | K1100Q | D | 8.5e-06 | 1e-05 | 9e-06 | 9.167E-06 | 7.638E-07 | 8.500E-06 | 1.000E-05 |
| 21988 | 1100 | K1100G | D | 5.5e-06 | 6.8e-06 | 3.5e-06 | 5.267E-06 | 1.662E-06 | 3.500E-06 | 6.800E-06 |
| 21989 | 1100 | K1100H | D | 5.3e-06 | 6.2e-06 | 7.7e-06 | 6.400E-06 | 1.212E-06 | 5.300E-06 | 7.700E-06 |
| 21990 | 1100 | K1100I | D | 9.6e-06 | 4.7e-06 | 5.7e-06 | 6.667E-06 | 2.589E-06 | 4.700E-06 | 9.600E-06 |
| 21991 | 1100 | K1100L | D | 6.7e-06 | 5.2e-06 | 1.1e-05 | 7.633E-06 | 3.011E-06 | 5.200E-06 | 1.100E-05 |
| 21992 | 1100 | K1100K | D | 4.5e-06 | 6.9e-06 | 1e-05 | 7.133E-06 | 2.757E-06 | 4.500E-06 | 1.000E-05 |
| 21993 | 1100 | K1100M | D | 3.4e-06 | 6.1e-06 | 1.5e-05 | 8.167E-06 | 6.070E-06 | 3.400E-06 | 1.500E-05 |
| 21994 | 1100 | K1100F | D | 4.6e-06 | 1.1e-05 | 1.2e-05 | 9.200E-06 | 4.015E-06 | 4.600E-06 | 1.200E-05 |
| 21995 | 1100 | K1100P | D | 7e-06 | 5.3e-06 | 6.8e-06 | 6.367E-06 | 9.292E-07 | 5.300E-06 | 7.000E-06 |
| 21996 | 1100 | K1100S | D | 9.9e-06 | 5.6e-06 | 8.4e-06 | 7.967E-06 | 2.183E-06 | 5.600E-06 | 9.900E-06 |
| 21997 | 1100 | K1100T | D | 6.3e-06 | 7e-06 | 7.1e-06 | 6.800E-06 | 4.359E-07 | 6.300E-06 | 7.100E-06 |
| 21998 | 1100 | K1100W | D | 1.3e-05 | 9.1e-06 | 9.2e-06 | 1.043E-05 | 2.223E-06 | 9.100E-06 | 1.300E-05 |
| 21999 | 1100 | K1100Y | D | 9.6e-06 | 8e-06 | 1.4e-05 | 1.053E-05 | 3.107E-06 | 8.000E-06 | 1.400E-05 |
| 22000 | 1100 | K1100V | D | 8e-06 | 1e-05 | 9.7e-06 | 9.233E-06 | 1.079E-06 | 8.000E-06 | 1.000E-05 |
| 22001 | 1101 | A1101A | D | 5.2e-06 | 1.1e-05 | 1.1e-05 | 9.067E-06 | 3.349E-06 | 5.200E-06 | 1.100E-05 |
| 22002 | 1101 | A1101R | D | 8.4e-06 | 1.3e-05 | 7.5e-06 | 9.633E-06 | 2.950E-06 | 7.500E-06 | 1.300E-05 |
| 22003 | 1101 | A1101N | D | 6.3e-06 | 4.9e-06 | 9.6e-06 | 6.933E-06 | 2.413E-06 | 4.900E-06 | 9.600E-06 |
| 22004 | 1101 | A1101D | D | 4.7e-06 | 1.2e-05 | 6.6e-06 | 7.767E-06 | 3.787E-06 | 4.700E-06 | 1.200E-05 |
| 22005 | 1101 | A1101C | D | 9.1e-06 | 5.3e-06 | 1.3e-05 | 9.133E-06 | 3.850E-06 | 5.300E-06 | 1.300E-05 |
| 22006 | 1101 | A1101E | D | 5e-06 | 6.6e-06 | 9.1e-06 | 6.900E-06 | 2.066E-06 | 5.000E-06 | 9.100E-06 |
| 22007 | 1101 | A1101Q | D | 9.3e-06 | 7.4e-06 | 7.6e-06 | 8.100E-06 | 1.044E-06 | 7.400E-06 | 9.300E-06 |
| 22008 | 1101 | A1101G | D | 1.3e-05 | 9.8e-06 | 1.1e-05 | 1.127E-05 | 1.617E-06 | 9.800E-06 | 1.300E-05 |
| 22009 | 1101 | A1101H | D | 8.4e-06 | 4.3e-06 | 1.3e-05 | 8.567E-06 | 4.352E-06 | 4.300E-06 | 1.300E-05 |
| 22010 | 1101 | A1101I | D | 8.6e-06 | 6.1e-06 | 5.7e-06 | 6.800E-06 | 1.572E-06 | 5.700E-06 | 8.600E-06 |
| 22011 | 1101 | A1101L | D | 5.9e-06 | 7e-06 | 9.5e-06 | 7.467E-06 | 1.845E-06 | 5.900E-06 | 9.500E-06 |
| 22012 | 1101 | A1101K | D | 7e-06 | 6.4e-06 | 1.3e-05 | 8.800E-06 | 3.650E-06 | 6.400E-06 | 1.300E-05 |
| 22013 | 1101 | A1101M | D | 5.9e-06 | 6.2e-06 | 6.2e-06 | 6.100E-06 | 1.732E-07 | 5.900E-06 | 6.200E-06 |
| 22014 | 1101 | A1101F | D | 6.8e-06 | 1.1e-05 | 6.7e-06 | 8.167E-06 | 2.454E-06 | 6.700E-06 | 1.100E-05 |

| | | | | | | | | | | |
|---|---|---|---|---|---|---|---|---|---|---|
| 22015 | 1101 | A1101P | D | 8.4e-06 | 7.2e-06 | 1.7e-05 | 1.087E-05 | 5.345E-06 | 7.200E-06 | 1.700E-05 |
| 22016 | 1101 | A1101S | D | 7.1e-06 | 4.3e-06 | 8.5e-06 | 6.633E-06 | 2.139E-06 | 4.300E-06 | 8.500E-06 |
| 22017 | 1101 | A1101T | D | 9.3e-06 | 8.1e-06 | 8.2e-06 | 8.533E-06 | 6.658E-07 | 8.100E-06 | 9.300E-06 |
| 22018 | 1101 | A1101W | D | 7.6e-06 | 7.6e-06 | 7.6e-06 | 7.600E-06 | 1.037E-21 | 7.600E-06 | 7.600E-06 |
| 22019 | 1101 | A1101Y | D | 6.9e-06 | 1e-05 | 1.1e-05 | 9.300E-06 | 2.138E-06 | 6.900E-06 | 1.100E-05 |
| 22020 | 1101 | A1101V | D | 8.3e-06 | 1.4e-05 | 6.4e-06 | 9.567E-06 | 3.955E-06 | 6.400E-06 | 1.400E-05 |
| 22021 | 1102 | P1102A | D | 7.2e-06 | 7.1e-06 | 9.2e-06 | 7.833E-06 | 1.185E-06 | 7.100E-06 | 9.200E-06 |
| 22022 | 1102 | P1102R | D | 1.6e-05 | 9.2e-06 | 7.5e-06 | 1.090E-05 | 4.498E-06 | 7.500E-06 | 1.600E-05 |
| 22023 | 1102 | P1102N | D | 6.5e-06 | 7.5e-06 | 6.7e-06 | 6.900E-06 | 5.292E-07 | 6.500E-06 | 7.500E-06 |
| 22024 | 1102 | P1102D | D | 6.5e-06 | 6.1e-06 | 1e-05 | 7.533E-06 | 2.146E-06 | 6.100E-06 | 1.000E-05 |
| 22025 | 1102 | P1102C | D | 8.2e-06 | 1.5e-05 | 8.1e-06 | 1.043E-05 | 3.955E-06 | 8.100E-06 | 1.500E-05 |
| 22026 | 1102 | P1102E | D | 1e-05 | 6.2e-06 | 4.7e-06 | 6.967E-06 | 2.732E-06 | 4.700E-06 | 1.000E-05 |
| 22027 | 1102 | P1102Q | D | 1.2e-05 | 1.1e-05 | 9e-06 | 1.067E-05 | 1.528E-06 | 9.000E-06 | 1.200E-05 |
| 22028 | 1102 | P1102G | D | 5.9e-06 | 6.6e-06 | 1.1e-05 | 7.833E-06 | 2.765E-06 | 5.900E-06 | 1.100E-05 |
| 22029 | 1102 | P1102H | D | 7.3e-06 | 6.5e-06 | 1e-05 | 7.933E-06 | 1.834E-06 | 6.500E-06 | 1.000E-05 |
| 22030 | 1102 | P1102I | D | 6.5e-06 | 7.7e-06 | 9.1e-06 | 7.767E-06 | 1.301E-06 | 6.500E-06 | 9.100E-06 |
| 22031 | 1102 | P1102L | D | 5.6e-06 | 6.9e-06 | 7.7e-06 | 6.733E-06 | 1.060E-06 | 5.600E-06 | 7.700E-06 |
| 22032 | 1102 | P1102K | D | 9.1e-06 | 7.7e-06 | 7.1e-06 | 7.967E-06 | 1.026E-06 | 7.100E-06 | 9.100E-06 |
| 22033 | 1102 | P1102M | D | 9.4e-06 | 6.6e-06 | 5.4e-06 | 7.133E-06 | 2.053E-06 | 5.400E-06 | 9.400E-06 |
| 22034 | 1102 | P1102F | D | 7.5e-06 | 8.4e-06 | 1e-05 | 8.633E-06 | 1.266E-06 | 7.500E-06 | 1.000E-05 |
| 22035 | 1102 | P1102P | D | 7.5e-06 | 8.9e-06 | 5e-06 | 7.133E-06 | 1.976E-06 | 5.000E-06 | 8.900E-06 |
| 22036 | 1102 | P1102S | D | 6e-06 | 8.8e-06 | 8.3e-06 | 7.700E-06 | 1.493E-06 | 6.000E-06 | 8.800E-06 |
| 22037 | 1102 | P1102T | D | 1.4e-05 | 6.9e-06 | 4.5e-06 | 8.467E-06 | 4.940E-06 | 4.500E-06 | 1.400E-05 |
| 22038 | 1102 | P1102W | D | 1e-05 | 1.2e-05 | 1.2e-05 | 1.133E-05 | 1.155E-06 | 1.000E-05 | 1.200E-05 |
| 22039 | 1102 | P1102Y | D | 6e-06 | 1e-05 | 9.3e-06 | 8.433E-06 | 2.136E-06 | 6.000E-06 | 1.000E-05 |
| 22040 | 1102 | P1102V | D | 5.3e-06 | 7.7e-06 | 3.3e-06 | 5.433E-06 | 2.203E-06 | 3.300E-06 | 7.700E-06 |
| 22041 | 1103 | K1103A | D | 9.4e-06 | 5.6e-06 | 6.6e-06 | 7.200E-06 | 1.970E-06 | 5.600E-06 | 9.400E-06 |
| 22042 | 1103 | K1103R | D | 6.9e-06 | 5e-06 | 9.8e-06 | 7.233E-06 | 2.417E-06 | 5.000E-06 | 9.800E-06 |
| 22043 | 1103 | K1103N | D | 8.4e-06 | 7.8e-06 | 9.9e-06 | 8.700E-06 | 1.082E-06 | 7.800E-06 | 9.900E-06 |
| 22044 | 1103 | K1103D | D | 1e-05 | 4.5e-06 | 9.2e-06 | 7.900E-06 | 2.972E-06 | 4.500E-06 | 1.000E-05 |
| 22045 | 1103 | K1103C | D | 4.5e-06 | 1.3e-05 | 5.3e-06 | 7.600E-06 | 4.694E-06 | 4.500E-06 | 1.300E-05 |
| 22046 | 1103 | K1103E | D | 7.3e-06 | 7.1e-06 | 9.7e-06 | 8.033E-06 | 1.447E-06 | 7.100E-06 | 9.700E-06 |
| 22047 | 1103 | K1103Q | D | 4.5e-06 | 1.3e-05 | 8.8e-06 | 8.767E-06 | 4.250E-06 | 4.500E-06 | 1.300E-05 |
| 22048 | 1103 | K1103G | D | 8e-06 | 6.8e-06 | 4.1e-06 | 6.300E-06 | 1.997E-06 | 4.100E-06 | 8.000E-06 |
| 22049 | 1103 | K1103H | D | 5.9e-06 | 6.5e-06 | 8.6e-06 | 7.000E-06 | 1.418E-06 | 5.900E-06 | 8.600E-06 |
| 22050 | 1103 | K1103I | D | 1e-05 | 8e-06 | 6.5e-06 | 8.167E-06 | 1.756E-06 | 6.500E-06 | 1.000E-05 |
| 22051 | 1103 | K1103L | D | 5.2e-06 | 6.8e-06 | 1.3e-05 | 8.333E-06 | 4.120E-06 | 5.200E-06 | 1.300E-05 |
| 22052 | 1103 | K1103K | D | 9e-06 | 7.3e-06 | 9.4e-06 | 8.567E-06 | 1.115E-06 | 7.300E-06 | 9.400E-06 |
| 22053 | 1103 | K1103M | D | 1.3e-05 | 1.3e-05 | 8.2e-06 | 1.140E-05 | 2.771E-06 | 8.200E-06 | 1.300E-05 |
| 22054 | 1103 | K1103F | D | 8.5e-06 | 5.2e-06 | 1.4e-05 | 9.233E-06 | 4.446E-06 | 5.200E-06 | 1.400E-05 |
| 22055 | 1103 | K1103P | D | 7.1e-06 | 1.2e-05 | 1.1e-05 | 1.003E-05 | 2.589E-06 | 7.100E-06 | 1.200E-05 |
| 22056 | 1103 | K1103S | D | 9.8e-06 | 5.9e-06 | 1.1e-05 | 8.900E-06 | 2.666E-06 | 5.900E-06 | 1.100E-05 |
| 22057 | 1103 | K1103T | D | 8.7e-06 | 6.5e-06 | 9e-06 | 8.067E-06 | 1.365E-06 | 6.500E-06 | 9.000E-06 |
| 22058 | 1103 | K1103W | D | 1.1e-05 | 7.4e-06 | 9.9e-06 | 9.433E-06 | 1.845E-06 | 7.400E-06 | 1.100E-05 |
| 22059 | 1103 | K1103Y | D | 1.2e-05 | 7.9e-06 | 5.6e-06 | 8.500E-06 | 3.242E-06 | 5.600E-06 | 1.200E-05 |
| 22060 | 1103 | K1103V | D | 7.6e-06 | 6.7e-06 | 7.9e-06 | 7.400E-06 | 6.245E-07 | 6.700E-06 | 7.900E-06 |
| 22061 | 1104 | L1104A | D | 6.7e-06 | 4.4e-06 | 1.9e-05 | 1.003E-05 | 7.850E-06 | 4.400E-06 | 1.900E-05 |
| 22062 | 1104 | L1104R | D | 1.3e-05 | 1.8e-05 | 6.6e-06 | 1.253E-05 | 5.714E-06 | 6.600E-06 | 1.800E-05 |
| 22063 | 1104 | L1104N | D | 6.2e-06 | 5.2e-06 | 6.5e-06 | 5.967E-06 | 6.807E-07 | 5.200E-06 | 6.500E-06 |
| 22064 | 1104 | L1104D | D | 1.9e-05 | 5.7e-06 | 4.3e-06 | 9.667E-06 | 8.113E-06 | 4.300E-06 | 1.900E-05 |
| 22065 | 1104 | L1104C | D | 9.3e-06 | 1e-05 | 9.8e-06 | 9.700E-06 | 3.606E-07 | 9.300E-06 | 1.000E-05 |
| 22066 | 1104 | L1104E | D | 9.7e-06 | 1.1e-05 | 7.9e-06 | 9.533E-06 | 1.557E-06 | 7.900E-06 | 1.100E-05 |
| 22067 | 1104 | L1104Q | D | 7.7e-06 | 8.3e-06 | 7.3e-06 | 7.767E-06 | 5.033E-07 | 7.300E-06 | 8.300E-06 |
| 22068 | 1104 | L1104G | D | 1.5e-05 | 8.4e-06 | 6.1e-06 | 9.833E-06 | 4.620E-06 | 6.100E-06 | 1.500E-05 |
| 22069 | 1104 | L1104H | D | 7.4e-06 | 9.3e-06 | 7.7e-06 | 8.133E-06 | 1.021E-06 | 7.400E-06 | 9.300E-06 |
| 22070 | 1104 | L1104I | D | 8.3e-06 | 5.7e-06 | 9.4e-06 | 7.800E-06 | 1.900E-06 | 5.700E-06 | 9.400E-06 |
| 22071 | 1104 | L1104L | D | 8.8e-06 | 8.4e-06 | 4.9e-06 | 7.367E-06 | 2.146E-06 | 4.900E-06 | 8.800E-06 |
| 22072 | 1104 | L1104K | D | 7.5e-06 | 6.7e-06 | 1e-05 | 8.067E-06 | 1.721E-06 | 6.700E-06 | 1.000E-05 |
| 22073 | 1104 | L1104M | D | 4.4e-06 | 9.2e-06 | 5e-06 | 6.200E-06 | 2.615E-06 | 4.400E-06 | 9.200E-06 |
| 22074 | 1104 | L1104F | D | 8.7e-06 | 8e-06 | 6.6e-06 | 7.767E-06 | 1.069E-06 | 6.600E-06 | 8.700E-06 |
| 22075 | 1104 | L1104P | D | 8.1e-06 | 1.2e-05 | 7.8e-06 | 9.300E-06 | 2.343E-06 | 7.800E-06 | 1.200E-05 |
| 22076 | 1104 | L1104S | D | 8e-06 | 1.1e-05 | 8.2e-06 | 9.067E-06 | 1.677E-06 | 8.000E-06 | 1.100E-05 |
| 22077 | 1104 | L1104T | D | 7.9e-06 | 1.3e-05 | 3.8e-06 | 8.233E-06 | 4.609E-06 | 3.800E-06 | 1.300E-05 |
| 22078 | 1104 | L1104W | D | 9.2e-06 | 1.3e-05 | 6.7e-06 | 9.633E-06 | 3.172E-06 | 6.700E-06 | 1.300E-05 |
| 22079 | 1104 | L1104Y | D | 6.6e-06 | 9.9e-06 | 8.3e-06 | 8.267E-06 | 1.650E-06 | 6.600E-06 | 9.900E-06 |
| 22080 | 1104 | L1104V | D | 4.1e-06 | 8.6e-06 | 7.1e-06 | 6.600E-06 | 2.291E-06 | 4.100E-06 | 8.600E-06 |
| 22081 | 1105 | L1105A | D | 8e-06 | 1.2e-05 | 3.9e-06 | 7.967E-06 | 4.050E-06 | 3.900E-06 | 1.200E-05 |
| 22082 | 1105 | L1105R | D | 7.1e-06 | 1.2e-05 | 7e-06 | 8.700E-06 | 2.858E-06 | 7.000E-06 | 1.200E-05 |
| 22083 | 1105 | L1105N | D | 7.4e-06 | 5.4e-06 | 1.1e-05 | 7.933E-06 | 2.838E-06 | 5.400E-06 | 1.100E-05 |
| 22084 | 1105 | L1105D | D | 1.3e-05 | 7.2e-06 | 7.7e-06 | 9.300E-06 | 3.214E-06 | 7.200E-06 | 1.300E-05 |
| 22085 | 1105 | L1105C | D | 1.4e-05 | 9.4e-06 | 5.4e-06 | 9.600E-06 | 4.303E-06 | 5.400E-06 | 1.400E-05 |
| 22086 | 1105 | L1105E | D | 8.6e-06 | 5.6e-06 | 6.3e-06 | 6.833E-06 | 1.570E-06 | 5.600E-06 | 8.600E-06 |
| 22087 | 1105 | L1105Q | D | 5e-06 | 7.8e-06 | 5.1e-06 | 5.967E-06 | 1.589E-06 | 5.000E-06 | 7.800E-06 |
| 22088 | 1105 | L1105G | D | 7.2e-06 | 9.3e-06 | 7.2e-06 | 7.900E-06 | 1.212E-06 | 7.200E-06 | 9.300E-06 |
| 22089 | 1105 | L1105H | D | 5.9e-06 | 5.9e-06 | 7.2e-06 | 6.333E-06 | 7.506E-07 | 5.900E-06 | 7.200E-06 |
| 22090 | 1105 | L1105I | D | 1.3e-05 | 4.9e-06 | 4e-06 | 7.300E-06 | 4.957E-06 | 4.000E-06 | 1.300E-05 |
| 22091 | 1105 | L1105L | D | 1.7e-05 | 7.3e-06 | 7e-06 | 1.043E-05 | 5.689E-06 | 7.000E-06 | 1.700E-05 |
| 22092 | 1105 | L1105K | D | 6.2e-06 | 9.9e-06 | 7.4e-06 | 7.833E-06 | 1.888E-06 | 6.200E-06 | 9.900E-06 |
| 22093 | 1105 | L1105M | D | 1.4e-05 | 1.2e-05 | 7.4e-06 | 1.113E-05 | 3.384E-06 | 7.400E-06 | 1.400E-05 |
| 22094 | 1105 | L1105F | D | 7.1e-06 | 6.1e-06 | 6.8e-06 | 6.667E-06 | 5.132E-07 | 6.100E-06 | 7.100E-06 |
| 22095 | 1105 | L1105P | D | 1.1e-05 | 6.6e-06 | 5.9e-06 | 7.833E-06 | 2.765E-06 | 5.900E-06 | 1.100E-05 |
| 22096 | 1105 | L1105S | D | 7.4e-06 | 6.8e-06 | 9.1e-06 | 7.767E-06 | 1.193E-06 | 6.800E-06 | 9.100E-06 |
| 22097 | 1105 | L1105T | D | 7.8e-06 | 6e-06 | 1.1e-05 | 8.267E-06 | 2.532E-06 | 6.000E-06 | 1.100E-05 |
| 22098 | 1105 | L1105W | D | 6e-06 | 8.4e-06 | 8e-06 | 7.467E-06 | 1.286E-06 | 6.000E-06 | 8.400E-06 |
| 22099 | 1105 | L1105Y | D | 5.8e-06 | 8e-06 | 6.6e-06 | 6.800E-06 | 1.114E-06 | 5.800E-06 | 8.000E-06 |

| | | | | | | | | | | |
|---|---|---|---|---|---|---|---|---|---|---|
| 22100 | 1105 | L1105V | D | 8.3e-06 | 9.7e-06 | 1.1e-05 | 9.667E-06 | 1.350E-06 | 8.300E-06 | 1.100E-05 |
| 22101 | 1106 | I1106A | D | 9.3e-06 | 1.3e-05 | 6.6e-06 | 9.633E-06 | 3.213E-06 | 6.600E-06 | 1.300E-05 |
| 22102 | 1106 | I1106R | D | 1.3e-05 | 6.9e-06 | 1.2e-05 | 1.063E-05 | 3.272E-06 | 6.900E-06 | 1.300E-05 |
| 22103 | 1106 | I1106N | D | 6.8e-06 | 1.4e-05 | 6e-06 | 8.933E-06 | 4.406E-06 | 6.000E-06 | 1.400E-05 |
| 22104 | 1106 | I1106D | D | 7.8e-06 | 6.4e-06 | 6.3e-06 | 6.833E-06 | 8.386E-07 | 6.300E-06 | 7.800E-06 |
| 22105 | 1106 | I1106C | D | 5e-06 | 8e-06 | 6.4e-06 | 6.467E-06 | 1.501E-06 | 5.000E-06 | 8.000E-06 |
| 22106 | 1106 | I1106E | D | 5.3e-06 | 1.2e-05 | 1.1e-05 | 9.433E-06 | 3.614E-06 | 5.300E-06 | 1.200E-05 |
| 22107 | 1106 | I1106Q | D | 8.8e-06 | 7.3e-06 | 4.4e-06 | 6.833E-06 | 2.237E-06 | 4.400E-06 | 8.800E-06 |
| 22108 | 1106 | I1106G | D | 7.8e-06 | 7.2e-06 | 1.2e-05 | 9.000E-06 | 2.615E-06 | 7.200E-06 | 1.200E-05 |
| 22109 | 1106 | I1106H | D | 1.1e-05 | 8.4e-06 | 1.3e-05 | 1.080E-05 | 2.307E-06 | 8.400E-06 | 1.300E-05 |
| 22110 | 1106 | I1106I | D | 1.4e-05 | 7.7e-06 | 9.4e-06 | 1.037E-05 | 3.259E-06 | 7.700E-06 | 1.400E-05 |
| 22111 | 1106 | I1106L | D | 6.1e-06 | 1.2e-05 | 1.5e-05 | 1.103E-05 | 4.528E-06 | 6.100E-06 | 1.500E-05 |
| 22112 | 1106 | I1106K | D | 9.4e-06 | 5.1e-06 | 5.9e-06 | 6.800E-06 | 2.287E-06 | 5.100E-06 | 9.400E-06 |
| 22113 | 1106 | I1106M | D | 6.5e-06 | 7.7e-06 | 5.4e-06 | 6.533E-06 | 1.150E-06 | 5.400E-06 | 7.700E-06 |
| 22114 | 1106 | I1106F | D | 8e-06 | 7.4e-06 | 6.2e-06 | 7.200E-06 | 9.165E-07 | 6.200E-06 | 8.000E-06 |
| 22115 | 1106 | I1106P | D | 6.7e-06 | 8.5e-06 | 8.5e-06 | 7.900E-06 | 1.039E-06 | 6.700E-06 | 8.500E-06 |
| 22116 | 1106 | I1106S | D | 5.2e-06 | 7.6e-06 | 7.1e-06 | 6.633E-06 | 1.266E-06 | 5.200E-06 | 7.600E-06 |
| 22117 | 1106 | I1106T | D | 9.5e-06 | 1.2e-05 | 8.5e-06 | 1.000E-05 | 1.803E-06 | 8.500E-06 | 1.200E-05 |
| 22118 | 1106 | I1106W | D | 1.2e-05 | 5.6e-06 | 8.1e-06 | 8.567E-06 | 3.225E-06 | 5.600E-06 | 1.200E-05 |
| 22119 | 1106 | I1106Y | D | 5.9e-06 | 6.2e-06 | 9.3e-06 | 7.133E-06 | 1.882E-06 | 5.900E-06 | 9.300E-06 |
| 22120 | 1106 | I1106V | D | 9.7e-06 | 4.4e-06 | 9.7e-06 | 7.933E-06 | 3.060E-06 | 4.400E-06 | 9.700E-06 |
| 22121 | 1107 | Y1107A | D | 5.9e-06 | 9.5e-06 | 9.4e-06 | 8.267E-06 | 2.050E-06 | 5.900E-06 | 9.500E-06 |
| 22122 | 1107 | Y1107R | D | 7.8e-06 | 6.3e-06 | 9.5e-06 | 7.867E-06 | 1.601E-06 | 6.300E-06 | 9.500E-06 |
| 22123 | 1107 | Y1107N | D | 1.3e-05 | 9.2e-06 | 8.1e-06 | 1.010E-05 | 2.571E-06 | 8.100E-06 | 1.300E-05 |
| 22124 | 1107 | Y1107D | D | 6.2e-06 | 9e-06 | 1.4e-05 | 9.733E-06 | 3.951E-06 | 6.200E-06 | 1.400E-05 |
| 22125 | 1107 | Y1107C | D | 1.3e-05 | 5.6e-06 | 5e-06 | 7.867E-06 | 4.456E-06 | 5.000E-06 | 1.300E-05 |
| 22126 | 1107 | Y1107E | D | 1.3e-05 | 1.4e-05 | 7.8e-06 | 1.160E-05 | 3.329E-06 | 7.800E-06 | 1.400E-05 |
| 22127 | 1107 | Y1107Q | D | 6.1e-06 | 8.2e-06 | 7.9e-06 | 7.400E-06 | 1.136E-06 | 6.100E-06 | 8.200E-06 |
| 22128 | 1107 | Y1107G | D | 1.2e-05 | 6e-06 | 5.1e-06 | 7.700E-06 | 3.751E-06 | 5.100E-06 | 1.200E-05 |
| 22129 | 1107 | Y1107H | D | 6.7e-06 | 7.8e-06 | 9e-06 | 7.833E-06 | 1.150E-06 | 6.700E-06 | 9.000E-06 |
| 22130 | 1107 | Y1107I | D | 7e-06 | 1.1e-05 | 8e-06 | 8.667E-06 | 2.082E-06 | 7.000E-06 | 1.100E-05 |
| 22131 | 1107 | Y1107L | D | 3.1e-06 | 6.4e-06 | 5.6e-06 | 5.033E-06 | 1.721E-06 | 3.100E-06 | 6.400E-06 |
| 22132 | 1107 | Y1107K | D | 5.8e-06 | 4.6e-06 | 5.5e-06 | 5.300E-06 | 6.245E-07 | 4.600E-06 | 5.800E-06 |
| 22133 | 1107 | Y1107M | D | 8.9e-06 | 5.8e-06 | 8.5e-06 | 7.733E-06 | 1.686E-06 | 5.800E-06 | 8.900E-06 |
| 22134 | 1107 | Y1107F | D | 1.2e-05 | 1e-05 | 1.5e-05 | 1.233E-05 | 2.517E-06 | 1.000E-05 | 1.500E-05 |
| 22135 | 1107 | Y1107P | D | 1.4e-05 | 1e-05 | 1e-05 | 1.133E-05 | 2.309E-06 | 1.000E-05 | 1.400E-05 |
| 22136 | 1107 | Y1107S | D | 4.7e-06 | 4.4e-06 | 9.2e-06 | 6.100E-06 | 2.689E-06 | 4.400E-06 | 9.200E-06 |
| 22137 | 1107 | Y1107T | D | 8.5e-06 | 5e-06 | 5.7e-06 | 6.400E-06 | 1.852E-06 | 5.000E-06 | 8.500E-06 |
| 22138 | 1107 | Y1107W | D | 7.1e-06 | 8.3e-06 | 6e-06 | 7.133E-06 | 1.150E-06 | 6.000E-06 | 8.300E-06 |
| 22139 | 1107 | Y1107Y | D | 4.3e-06 | 6.8e-06 | 7e-06 | 6.033E-06 | 1.504E-06 | 4.300E-06 | 7.000E-06 |
| 22140 | 1107 | Y1107V | D | 1.2e-05 | 8e-06 | 7.9e-06 | 9.300E-06 | 2.339E-06 | 7.900E-06 | 1.200E-05 |
| 22141 | 1108 | S1108A | D | 7.2e-06 | 6.1e-06 | 4.5e-06 | 5.933E-06 | 1.358E-06 | 4.500E-06 | 7.200E-06 |
| 22142 | 1108 | S1108R | D | 8.1e-06 | 4.3e-06 | 4.7e-06 | 5.700E-06 | 2.088E-06 | 4.300E-06 | 8.100E-06 |
| 22143 | 1108 | S1108N | D | 4.8e-06 | 6.2e-06 | 4.4e-06 | 5.133E-06 | 9.452E-07 | 4.400E-06 | 6.200E-06 |
| 22144 | 1108 | S1108D | D | 1.2e-05 | 1e-05 | 5.9e-06 | 9.300E-06 | 3.110E-06 | 5.900E-06 | 1.200E-05 |
| 22145 | 1108 | S1108C | D | 4.4e-06 | 4.5e-06 | 4.2e-06 | 4.367E-06 | 1.528E-07 | 4.200E-06 | 4.500E-06 |
| 22146 | 1108 | S1108E | D | 1e-05 | 8.7e-06 | 4.7e-06 | 7.800E-06 | 2.762E-06 | 4.700E-06 | 1.000E-05 |
| 22147 | 1108 | S1108Q | D | 5.3e-06 | 8.8e-06 | 5.1e-06 | 6.400E-06 | 2.081E-06 | 5.100E-06 | 8.800E-06 |
| 22148 | 1108 | S1108G | D | 4.5e-06 | 5.3e-06 | 7.7e-06 | 5.833E-06 | 1.665E-06 | 4.500E-06 | 7.700E-06 |
| 22149 | 1108 | S1108H | D | 1e-05 | 4e-06 | 5.2e-06 | 6.400E-06 | 3.175E-06 | 4.000E-06 | 1.000E-05 |
| 22150 | 1108 | S1108I | D | 6.9e-06 | 4e-06 | 8.4e-06 | 6.433E-06 | 2.237E-06 | 4.000E-06 | 8.400E-06 |
| 22151 | 1108 | S1108L | D | 3.7e-06 | 5.7e-06 | 6.6e-06 | 5.333E-06 | 1.484E-06 | 3.700E-06 | 6.600E-06 |
| 22152 | 1108 | S1108K | D | 6.5e-06 | 3.6e-06 | 7e-06 | 5.700E-06 | 1.836E-06 | 3.600E-06 | 7.000E-06 |
| 22153 | 1108 | S1108M | D | 9.2e-06 | 4.9e-06 | 6e-06 | 6.700E-06 | 2.234E-06 | 4.900E-06 | 9.200E-06 |
| 22154 | 1108 | S1108F | D | 8.2e-06 | 4.4e-06 | 3.6e-06 | 5.400E-06 | 2.458E-06 | 3.600E-06 | 8.200E-06 |
| 22155 | 1108 | S1108P | D | 4.7e-06 | 5.2e-06 | 1.6e-05 | 8.633E-06 | 6.385E-06 | 4.700E-06 | 1.600E-05 |
| 22156 | 1108 | S1108S | D | 8.8e-06 | 9.8e-06 | 8.1e-06 | 8.900E-06 | 8.544E-07 | 8.100E-06 | 9.800E-06 |
| 22157 | 1108 | S1108T | D | 6.1e-06 | 7.1e-06 | 8.7e-06 | 7.300E-06 | 1.311E-06 | 6.100E-06 | 8.700E-06 |
| 22158 | 1108 | S1108W | D | 5.9e-06 | 5.3e-06 | 5.7e-06 | 5.633E-06 | 3.055E-07 | 5.300E-06 | 5.900E-06 |
| 22159 | 1108 | S1108Y | D | 6.7e-06 | 4.8e-06 | 6.8e-06 | 6.100E-06 | 1.127E-06 | 4.800E-06 | 6.800E-06 |
| 22160 | 1108 | S1108V | D | 7.2e-06 | 7.5e-06 | 6.9e-06 | 7.200E-06 | 3.000E-07 | 6.900E-06 | 7.500E-06 |
| 22161 | 1109 | A1109A | D | 7e-06 | 7.5e-06 | 8.5e-06 | 7.667E-06 | 7.638E-07 | 7.000E-06 | 8.500E-06 |
| 22162 | 1109 | A1109R | D | 9.5e-06 | 4.9e-06 | 5e-06 | 6.467E-06 | 2.627E-06 | 4.900E-06 | 9.500E-06 |
| 22163 | 1109 | A1109N | D | 9e-06 | 1.2e-05 | 7.9e-06 | 9.633E-06 | 2.122E-06 | 7.900E-06 | 1.200E-05 |
| 22164 | 1109 | A1109D | D | 9.3e-06 | 9e-06 | 6.9e-06 | 8.400E-06 | 1.308E-06 | 6.900E-06 | 9.300E-06 |
| 22165 | 1109 | A1109C | D | 9.1e-06 | 9.7e-06 | 5.2e-06 | 8.000E-06 | 2.443E-06 | 5.200E-06 | 9.700E-06 |
| 22166 | 1109 | A1109E | D | 6.7e-06 | 8.2e-06 | 5.2e-06 | 6.700E-06 | 1.500E-06 | 5.200E-06 | 8.200E-06 |
| 22167 | 1109 | A1109Q | D | 4e-06 | 4.5e-06 | 5.7e-06 | 4.733E-06 | 8.737E-07 | 4.000E-06 | 5.700E-06 |
| 22168 | 1109 | A1109G | D | 7.9e-06 | 1.2e-05 | 9.5e-06 | 9.800E-06 | 2.066E-06 | 7.900E-06 | 1.200E-05 |
| 22169 | 1109 | A1109H | D | 1.3e-05 | 4.6e-06 | 6.4e-06 | 8.000E-06 | 4.423E-06 | 4.600E-06 | 1.300E-05 |
| 22170 | 1109 | A1109I | D | 6.3e-06 | 1.3e-05 | 5.4e-06 | 8.233E-06 | 4.153E-06 | 5.400E-06 | 1.300E-05 |
| 22171 | 1109 | A1109L | D | 4.3e-06 | 8.9e-06 | 5.4e-06 | 6.200E-06 | 2.402E-06 | 4.300E-06 | 8.900E-06 |
| 22172 | 1109 | A1109K | D | 3.5e-06 | 6.6e-06 | 8.2e-06 | 6.100E-06 | 2.390E-06 | 3.500E-06 | 8.200E-06 |
| 22173 | 1109 | A1109M | D | 5.5e-06 | 1.2e-05 | 9.1e-06 | 8.867E-06 | 3.256E-06 | 5.500E-06 | 1.200E-05 |
| 22174 | 1109 | A1109F | D | 4.3e-06 | 6.8e-06 | 7e-06 | 6.033E-06 | 1.504E-06 | 4.300E-06 | 7.000E-06 |
| 22175 | 1109 | A1109P | D | 5.4e-06 | 3.4e-06 | 1e-05 | 6.267E-06 | 3.384E-06 | 3.400E-06 | 1.000E-05 |
| 22176 | 1109 | A1109S | D | 1.5e-05 | 6.3e-06 | 1e-05 | 1.043E-05 | 4.366E-06 | 6.300E-06 | 1.500E-05 |
| 22177 | 1109 | A1109T | D | 1.4e-05 | 6.5e-06 | 1.1e-05 | 1.050E-05 | 3.775E-06 | 6.500E-06 | 1.400E-05 |
| 22178 | 1109 | A1109W | D | 4.8e-06 | 8.8e-06 | 9.8e-06 | 7.800E-06 | 2.646E-06 | 4.800E-06 | 9.800E-06 |
| 22179 | 1109 | A1109Y | D | 6.2e-06 | 9.2e-06 | 6.4e-06 | 7.267E-06 | 1.677E-06 | 6.200E-06 | 9.200E-06 |
| 22180 | 1109 | A1109V | D | 7.3e-06 | 5.6e-06 | 7.9e-06 | 6.933E-06 | 1.193E-06 | 5.600E-06 | 7.900E-06 |
| 22181 | 1110 | S1110A | D | 1e-05 | 6.8e-06 | 7.5e-06 | 8.100E-06 | 1.682E-06 | 6.800E-06 | 1.000E-05 |
| 22182 | 1110 | S1110R | D | 9.4e-06 | 4.9e-06 | 9.9e-06 | 8.067E-06 | 2.754E-06 | 4.900E-06 | 9.900E-06 |
| 22183 | 1110 | S1110N | D | 6e-06 | 1.2e-05 | 1.1e-05 | 9.667E-06 | 3.215E-06 | 6.000E-06 | 1.200E-05 |
| 22184 | 1110 | S1110D | D | 1.1e-05 | 6.8e-06 | 8.1e-06 | 8.633E-06 | 2.150E-06 | 6.800E-06 | 1.100E-05 |

| | | | | | | | | | | |
|---|---|---|---|---|---|---|---|---|---|---|
| 22185 | 1110 | S1110C | D | 1.2e-05 | 6.4e-06 | 1.2e-05 | 1.013E-05 | 3.233E-06 | 6.400E-06 | 1.200E-05 |
| 22186 | 1110 | S1110E | D | 1.2e-05 | 5.7e-06 | 6.6e-06 | 8.100E-06 | 3.407E-06 | 5.700E-06 | 1.200E-05 |
| 22187 | 1110 | S1110Q | D | 8.6e-06 | 6.2e-06 | 9.6e-06 | 8.133E-06 | 1.747E-06 | 6.200E-06 | 9.600E-06 |
| 22188 | 1110 | S1110G | D | 6.6e-06 | 4.7e-06 | 7.6e-06 | 6.300E-06 | 1.473E-06 | 4.700E-06 | 7.600E-06 |
| 22189 | 1110 | S1110H | D | 1.3e-05 | 8.3e-06 | 5.9e-06 | 9.067E-06 | 3.612E-06 | 5.900E-06 | 1.300E-05 |
| 22190 | 1110 | S1110I | D | 1.1e-05 | 1.3e-05 | 8.9e-06 | 1.097E-05 | 2.050E-06 | 8.900E-06 | 1.300E-05 |
| 22191 | 1110 | S1110L | D | 1.2e-05 | 1.3e-05 | 5.5e-06 | 1.017E-05 | 4.072E-06 | 5.500E-06 | 1.300E-05 |
| 22192 | 1110 | S1110K | D | 4.6e-06 | 5.3e-06 | 1.4e-05 | 7.967E-06 | 5.237E-06 | 4.600E-06 | 1.400E-05 |
| 22193 | 1110 | S1110M | D | 9.1e-06 | 1.1e-05 | 4.8e-06 | 8.300E-06 | 3.176E-06 | 4.800E-06 | 1.100E-05 |
| 22194 | 1110 | S1110F | D | 6.3e-06 | 9.9e-06 | 8.4e-06 | 8.200E-06 | 1.808E-06 | 6.300E-06 | 9.900E-06 |
| 22195 | 1110 | S1110P | D | 6.4e-06 | 9.2e-06 | 1.1e-05 | 8.867E-06 | 2.318E-06 | 6.400E-06 | 1.100E-05 |
| 22196 | 1110 | S1110S | D | 7.5e-06 | 7.2e-06 | 9.4e-06 | 8.033E-06 | 1.193E-06 | 7.200E-06 | 9.400E-06 |
| 22197 | 1110 | S1110T | D | 6.2e-06 | 6.6e-06 | 1.1e-05 | 7.933E-06 | 2.663E-06 | 6.200E-06 | 1.100E-05 |
| 22198 | 1110 | S1110W | D | 9.7e-06 | 4.4e-06 | 9.1e-06 | 7.733E-06 | 2.902E-06 | 4.400E-06 | 9.700E-06 |
| 22199 | 1110 | S1110Y | D | 6.4e-06 | 8.3e-06 | 7e-06 | 7.233E-06 | 9.713E-07 | 6.400E-06 | 8.300E-06 |
| 22200 | 1110 | S1110V | D | 1e-05 | 5.4e-06 | 1.2e-05 | 9.133E-06 | 3.384E-06 | 5.400E-06 | 1.200E-05 |
| 22201 | 1111 | F1111A | D | 6.7e-06 | 9.6e-06 | 1.3e-05 | 9.767E-06 | 3.153E-06 | 6.700E-06 | 1.300E-05 |
| 22202 | 1111 | F1111R | D | 3.9e-06 | 4.4e-06 | 5.2e-06 | 4.500E-06 | 6.557E-07 | 3.900E-06 | 5.200E-06 |
| 22203 | 1111 | F1111N | D | 2e-06 | 2.1e-06 | 2.2e-06 | 2.100E-06 | 1.000E-07 | 2.000E-06 | 2.200E-06 |
| 22204 | 1111 | F1111D | D | 7.6e-06 | 3e-06 | 4.8e-06 | 5.133E-06 | 2.318E-06 | 3.000E-06 | 7.600E-06 |
| 22205 | 1111 | F1111C | D | 4.8e-06 | 6.3e-06 | 9e-06 | 6.700E-06 | 2.128E-06 | 4.800E-06 | 9.000E-06 |
| 22206 | 1111 | F1111E | D | 6.6e-06 | 4.3e-06 | 4.3e-06 | 5.067E-06 | 1.328E-06 | 4.300E-06 | 6.600E-06 |
| 22207 | 1111 | F1111Q | D | 3.1e-06 | 4.9e-06 | 7.6e-06 | 5.200E-06 | 2.265E-06 | 3.100E-06 | 7.600E-06 |
| 22208 | 1111 | F1111G | D | 4.7e-06 | 7.2e-06 | 8.9e-06 | 6.933E-06 | 2.113E-06 | 4.700E-06 | 8.900E-06 |
| 22209 | 1111 | F1111H | D | 6.5e-06 | 3.6e-06 | 4.2e-06 | 4.767E-06 | 1.531E-06 | 3.600E-06 | 6.500E-06 |
| 22210 | 1111 | F1111I | D | 8.1e-06 | 9.2e-06 | 6e-06 | 7.767E-06 | 1.626E-06 | 6.000E-06 | 9.200E-06 |
| 22211 | 1111 | F1111L | D | 7.3e-06 | 6.5e-06 | 8.6e-06 | 7.467E-06 | 1.060E-06 | 6.500E-06 | 8.600E-06 |
| 22212 | 1111 | F1111K | D | 4.4e-06 | 6.4e-06 | 5.3e-06 | 5.367E-06 | 1.002E-06 | 4.400E-06 | 6.400E-06 |
| 22213 | 1111 | F1111M | D | 1.1e-05 | 5e-06 | 5.3e-06 | 7.100E-06 | 3.381E-06 | 5.000E-06 | 1.100E-05 |
| 22214 | 1111 | F1111F | D | 1e-05 | 9.3e-06 | 8.8e-06 | 9.367E-06 | 6.028E-07 | 8.800E-06 | 1.000E-05 |
| 22215 | 1111 | F1111P | D | 8e-06 | 9.2e-06 | 1e-05 | 9.067E-06 | 1.007E-06 | 8.000E-06 | 1.000E-05 |
| 22216 | 1111 | F1111S | D | 8.4e-06 | 6e-06 | 8e-06 | 7.467E-06 | 1.286E-06 | 6.000E-06 | 8.400E-06 |
| 22217 | 1111 | F1111T | D | 5.2e-06 | 1.3e-05 | 5.9e-06 | 8.033E-06 | 4.315E-06 | 5.200E-06 | 1.300E-05 |
| 22218 | 1111 | F1111W | D | 5.8e-06 | 6.2e-06 | 1e-05 | 7.333E-06 | 2.318E-06 | 5.800E-06 | 1.000E-05 |
| 22219 | 1111 | F1111Y | D | 1.4e-05 | 5.8e-06 | 1e-05 | 9.933E-06 | 4.100E-06 | 5.800E-06 | 1.400E-05 |
| 22220 | 1111 | F1111V | D | 4.4e-06 | 9.3e-06 | 9.6e-06 | 7.767E-06 | 2.919E-06 | 4.400E-06 | 9.600E-06 |
| 22221 | 1112 | L1112A | D | 4.8e-06 | 1.8e-05 | 4.8e-06 | 9.200E-06 | 7.621E-06 | 4.800E-06 | 1.800E-05 |
| 22222 | 1112 | L1112R | D | 9.9e-06 | 7.6e-06 | 6.3e-06 | 7.933E-06 | 1.823E-06 | 6.300E-06 | 9.900E-06 |
| 22223 | 1112 | L1112N | D | 9.3e-06 | 1.3e-05 | 1.1e-05 | 1.110E-05 | 1.852E-06 | 9.300E-06 | 1.300E-05 |
| 22224 | 1112 | L1112D | D | 1.2e-05 | 4.4e-06 | 7.8e-06 | 8.067E-06 | 3.807E-06 | 4.400E-06 | 1.200E-05 |
| 22225 | 1112 | L1112C | D | 3.8e-06 | 6.8e-06 | 8.8e-06 | 6.467E-06 | 2.517E-06 | 3.800E-06 | 8.800E-06 |
| 22226 | 1112 | L1112E | D | 9.9e-06 | 8.4e-06 | 7.9e-06 | 8.733E-06 | 1.041E-06 | 7.900E-06 | 9.900E-06 |
| 22227 | 1112 | L1112Q | D | 9.3e-06 | 8.5e-06 | 9.8e-06 | 9.200E-06 | 6.557E-07 | 8.500E-06 | 9.800E-06 |
| 22228 | 1112 | L1112G | D | 9.7e-06 | 1.4e-05 | 8.1e-06 | 1.060E-05 | 3.051E-06 | 8.100E-06 | 1.400E-05 |
| 22229 | 1112 | L1112H | D | 5.9e-06 | 1.3e-05 | 6.1e-06 | 8.333E-06 | 4.043E-06 | 5.900E-06 | 1.300E-05 |
| 22230 | 1112 | L1112I | D | 1.1e-05 | 7.6e-06 | 6.7e-06 | 8.433E-06 | 2.268E-06 | 6.700E-06 | 1.100E-05 |
| 22231 | 1112 | L1112L | D | 9.2e-06 | 6.3e-06 | 9.3e-06 | 8.267E-06 | 1.704E-06 | 6.300E-06 | 9.300E-06 |
| 22232 | 1112 | L1112K | D | 9.2e-06 | 9.2e-06 | 9.2e-06 | 9.200E-06 | 0.000E+00 | 9.200E-06 | 9.200E-06 |
| 22233 | 1112 | L1112M | D | 5.7e-06 | 1.3e-05 | 9.4e-06 | 9.367E-06 | 3.650E-06 | 5.700E-06 | 1.300E-05 |
| 22234 | 1112 | L1112F | D | 1.1e-05 | 1.2e-05 | 1.1e-05 | 1.133E-05 | 5.774E-07 | 1.100E-05 | 1.200E-05 |
| 22235 | 1112 | L1112P | D | 6.5e-06 | 5.5e-06 | 9.1e-06 | 7.033E-06 | 1.858E-06 | 5.500E-06 | 9.100E-06 |
| 22236 | 1112 | L1112S | D | 5.8e-06 | 1.4e-05 | 6.5e-06 | 8.767E-06 | 4.546E-06 | 5.800E-06 | 1.400E-05 |
| 22237 | 1112 | L1112T | D | 8.5e-06 | 6.4e-06 | 1e-05 | 8.300E-06 | 1.808E-06 | 6.400E-06 | 1.000E-05 |
| 22238 | 1112 | L1112W | D | 5.7e-06 | 7.9e-06 | 7.3e-06 | 6.967E-06 | 1.137E-06 | 5.700E-06 | 7.900E-06 |
| 22239 | 1112 | L1112Y | D | 6.2e-06 | 5.9e-06 | 9.7e-06 | 7.267E-06 | 2.113E-06 | 5.900E-06 | 9.700E-06 |
| 22240 | 1112 | L1112V | D | 8.8e-06 | 9.1e-06 | 8.6e-06 | 8.833E-06 | 2.517E-07 | 8.600E-06 | 9.100E-06 |
| 22241 | 1113 | Y1113A | D | 7.8e-06 | 6.7e-06 | 1e-05 | 8.167E-06 | 1.680E-06 | 6.700E-06 | 1.000E-05 |
| 22242 | 1113 | Y1113R | D | 5.7e-06 | 6e-06 | 5.3e-06 | 5.667E-06 | 3.512E-07 | 5.300E-06 | 6.000E-06 |
| 22243 | 1113 | Y1113N | D | 8.7e-06 | 1.1e-05 | 8.6e-06 | 9.433E-06 | 1.358E-06 | 8.600E-06 | 1.100E-05 |
| 22244 | 1113 | Y1113D | D | 8.9e-06 | 5e-06 | 6.5e-06 | 6.800E-06 | 1.967E-06 | 5.000E-06 | 8.900E-06 |
| 22245 | 1113 | Y1113C | D | 6.8e-06 | 7.6e-06 | 6.7e-06 | 7.033E-06 | 4.933E-07 | 6.700E-06 | 7.600E-06 |
| 22246 | 1113 | Y1113E | D | 1e-05 | 5.6e-06 | 8.8e-06 | 8.133E-06 | 2.274E-06 | 5.600E-06 | 1.000E-05 |
| 22247 | 1113 | Y1113Q | D | 6.2e-06 | 1.3e-05 | 3.7e-06 | 7.633E-06 | 4.813E-06 | 3.700E-06 | 1.300E-05 |
| 22248 | 1113 | Y1113G | D | 8e-06 | 5.2e-06 | 9.2e-06 | 7.467E-06 | 2.053E-06 | 5.200E-06 | 9.200E-06 |
| 22249 | 1113 | Y1113H | D | 1.1e-05 | 6.4e-06 | 4.4e-06 | 7.267E-06 | 3.384E-06 | 4.400E-06 | 1.100E-05 |
| 22250 | 1113 | Y1113I | D | 9.6e-06 | 7.1e-06 | 8.9e-06 | 8.533E-06 | 1.290E-06 | 7.100E-06 | 9.600E-06 |
| 22251 | 1113 | Y1113L | D | 9.5e-06 | 7.5e-06 | 8.5e-06 | 8.500E-06 | 1.000E-06 | 7.500E-06 | 9.500E-06 |
| 22252 | 1113 | Y1113K | D | 3.3e-06 | 9.8e-06 | 1.2e-05 | 8.367E-06 | 4.524E-06 | 3.300E-06 | 1.200E-05 |
| 22253 | 1113 | Y1113M | D | 1e-05 | 1.4e-05 | 8.8e-06 | 1.093E-05 | 2.723E-06 | 8.800E-06 | 1.400E-05 |
| 22254 | 1113 | Y1113F | D | 5.9e-06 | 9.1e-06 | 9.9e-06 | 8.300E-06 | 2.117E-06 | 5.900E-06 | 9.900E-06 |
| 22255 | 1113 | Y1113P | D | 1.1e-05 | 6e-06 | 5.9e-06 | 7.633E-06 | 2.916E-06 | 5.900E-06 | 1.100E-05 |
| 22256 | 1113 | Y1113S | D | 6.6e-06 | 8.7e-06 | 9.9e-06 | 8.400E-06 | 1.670E-06 | 6.600E-06 | 9.900E-06 |
| 22257 | 1113 | Y1113T | D | 9.1e-06 | 6.9e-06 | 8.1e-06 | 8.033E-06 | 1.102E-06 | 6.900E-06 | 9.100E-06 |
| 22258 | 1113 | Y1113W | D | 6.7e-06 | 8.1e-06 | 8.7e-06 | 7.833E-06 | 1.026E-06 | 6.700E-06 | 8.700E-06 |
| 22259 | 1113 | Y1113Y | D | 8.4e-06 | 5.4e-06 | 7e-06 | 6.933E-06 | 1.501E-06 | 5.400E-06 | 8.400E-06 |
| 22260 | 1113 | Y1113V | D | 1.1e-05 | 9.4e-06 | 1.3e-05 | 1.113E-05 | 1.804E-06 | 9.400E-06 | 1.300E-05 |
| 22261 | 1114 | S1114A | D | 1.1e-05 | 8.3e-06 | 1.4e-05 | 1.110E-05 | 2.851E-06 | 8.300E-06 | 1.400E-05 |
| 22262 | 1114 | S1114R | D | 5.3e-06 | 2.2e-05 | 1.1e-05 | 1.277E-05 | 8.489E-06 | 5.300E-06 | 2.200E-05 |
| 22263 | 1114 | S1114N | D | 9.2e-06 | 7.7e-06 | 1.8e-05 | 1.163E-05 | 5.564E-06 | 7.700E-06 | 1.800E-05 |
| 22264 | 1114 | S1114D | D | 8.9e-06 | 1.1e-05 | 1.1e-05 | 1.030E-05 | 1.212E-06 | 8.900E-06 | 1.100E-05 |
| 22265 | 1114 | S1114C | D | 1.3e-05 | 1.7e-05 | 1.8e-05 | 1.600E-05 | 2.646E-06 | 1.300E-05 | 1.800E-05 |
| 22266 | 1114 | S1114E | D | 5.9e-06 | 1.2e-05 | 1e-05 | 9.300E-06 | 3.110E-06 | 5.900E-06 | 1.200E-05 |
| 22267 | 1114 | S1114Q | D | 8.9e-06 | 6.7e-06 | 2e-05 | 1.187E-05 | 7.129E-06 | 6.700E-06 | 2.000E-05 |
| 22268 | 1114 | S1114G | D | 1.1e-05 | 1.5e-05 | 1.1e-05 | 1.233E-05 | 2.309E-06 | 1.100E-05 | 1.500E-05 |
| 22269 | 1114 | S1114H | D | 8.7e-06 | 1.7e-05 | 7.5e-06 | 1.107E-05 | 5.173E-06 | 7.500E-06 | 1.700E-05 |

| | | | | | | | | | | |
|---|---|---|---|---|---|---|---|---|---|---|
| 22270 | 1114 | S1114I | D | 1.1e-05 | 1.7e-05 | 1.5e-05 | 1.433E-05 | 3.055E-06 | 1.100E-05 | 1.700E-05 |
| 22271 | 1114 | S1114L | D | 1.9e-05 | 8.3e-06 | 1.2e-05 | 1.310E-05 | 5.434E-06 | 8.300E-06 | 1.900E-05 |
| 22272 | 1114 | S1114K | D | 7.4e-06 | 6.3e-06 | 9.9e-06 | 7.867E-06 | 1.845E-06 | 6.300E-06 | 9.900E-06 |
| 22273 | 1114 | S1114M | D | 6.6e-06 | 6.3e-06 | 1.2e-05 | 8.300E-06 | 3.208E-06 | 6.300E-06 | 1.200E-05 |
| 22274 | 1114 | S1114F | D | 4.8e-06 | 6.5e-06 | 1.4e-05 | 8.433E-06 | 4.895E-06 | 4.800E-06 | 1.400E-05 |
| 22275 | 1114 | S1114P | D | 1.1e-05 | 1.9e-05 | 1.2e-05 | 1.400E-05 | 4.359E-06 | 1.100E-05 | 1.900E-05 |
| 22276 | 1114 | S1114S | D | 6.4e-06 | 9.4e-06 | 7.9e-06 | 7.900E-06 | 1.500E-06 | 6.400E-06 | 9.400E-06 |
| 22277 | 1114 | S1114T | D | 8.2e-06 | 7.2e-06 | 6.5e-06 | 7.300E-06 | 8.544E-07 | 6.500E-06 | 8.200E-06 |
| 22278 | 1114 | S1114W | D | 1e-05 | 6.3e-06 | 7.5e-06 | 7.933E-06 | 1.888E-06 | 6.300E-06 | 1.000E-05 |
| 22279 | 1114 | S1114Y | D | 1.3e-05 | 5.5e-06 | 1.3e-05 | 1.050E-05 | 4.330E-06 | 5.500E-06 | 1.300E-05 |
| 22280 | 1114 | S1114V | D | 7.1e-06 | 8.6e-06 | 6.7e-06 | 7.467E-06 | 1.002E-06 | 6.700E-06 | 8.600E-06 |
| 22281 | 1115 | G1115A | D | 1.1e-05 | 1.3e-05 | 9.6e-06 | 1.120E-05 | 1.709E-06 | 9.600E-06 | 1.300E-05 |
| 22282 | 1115 | G1115R | D | 6.2e-06 | 8.7e-06 | 6.8e-06 | 7.233E-06 | 1.305E-06 | 6.200E-06 | 8.700E-06 |
| 22283 | 1115 | G1115N | D | 7.5e-06 | 3.3e-06 | 7.6e-06 | 6.133E-06 | 2.454E-06 | 3.300E-06 | 7.600E-06 |
| 22284 | 1115 | G1115D | D | 7e-06 | 1.1e-05 | 1e-05 | 9.333E-06 | 2.082E-06 | 7.000E-06 | 1.100E-05 |
| 22285 | 1115 | G1115C | D | 7.1e-06 | 9e-06 | 1.2e-05 | 9.367E-06 | 2.470E-06 | 7.100E-06 | 1.200E-05 |
| 22286 | 1115 | G1115E | D | 6.8e-06 | 7.2e-06 | 1e-05 | 8.000E-06 | 1.744E-06 | 6.800E-06 | 1.000E-05 |
| 22287 | 1115 | G1115Q | D | 8.4e-06 | 6.7e-06 | 4.1e-06 | 6.400E-06 | 2.166E-06 | 4.100E-06 | 8.400E-06 |
| 22288 | 1115 | G1115G | D | 1.2e-05 | 9.6e-06 | 1.5e-05 | 1.220E-05 | 2.706E-06 | 9.600E-06 | 1.500E-05 |
| 22289 | 1115 | G1115H | D | 8.2e-06 | 7.4e-06 | 1e-05 | 8.533E-06 | 1.332E-06 | 7.400E-06 | 1.000E-05 |
| 22290 | 1115 | G1115I | D | 4.8e-06 | 9.3e-06 | 8e-06 | 7.367E-06 | 2.316E-06 | 4.800E-06 | 9.300E-06 |
| 22291 | 1115 | G1115L | D | 6.2e-06 | 1.4e-05 | 1.1e-05 | 1.040E-05 | 3.934E-06 | 6.200E-06 | 1.400E-05 |
| 22292 | 1115 | G1115K | D | 1.2e-05 | 5.7e-06 | 4.7e-06 | 7.467E-06 | 3.958E-06 | 4.700E-06 | 1.200E-05 |
| 22293 | 1115 | G1115M | D | 9.5e-06 | 8.6e-06 | 6.9e-06 | 8.333E-06 | 1.320E-06 | 6.900E-06 | 9.500E-06 |
| 22294 | 1115 | G1115F | D | 8.2e-06 | 4.6e-06 | 1.3e-05 | 8.600E-06 | 4.214E-06 | 4.600E-06 | 1.300E-05 |
| 22295 | 1115 | G1115P | D | 8.1e-06 | 9.7e-06 | 6.6e-06 | 8.133E-06 | 1.550E-06 | 6.600E-06 | 9.700E-06 |
| 22296 | 1115 | G1115S | D | 1e-05 | 1.3e-05 | 5.2e-06 | 9.400E-06 | 3.934E-06 | 5.200E-06 | 1.300E-05 |
| 22297 | 1115 | G1115T | D | 5.6e-06 | 6.4e-06 | 8.3e-06 | 6.767E-06 | 1.387E-06 | 5.600E-06 | 8.300E-06 |
| 22298 | 1115 | G1115W | D | 7.3e-06 | 9.6e-06 | 4.5e-06 | 7.133E-06 | 2.554E-06 | 4.500E-06 | 9.600E-06 |
| 22299 | 1115 | G1115Y | D | 8.9e-06 | 6.8e-06 | 1.1e-05 | 8.900E-06 | 2.100E-06 | 6.800E-06 | 1.100E-05 |
| 22300 | 1115 | G1115V | D | 6.8e-06 | 8.8e-06 | 5.8e-06 | 7.133E-06 | 1.528E-06 | 5.800E-06 | 8.800E-06 |
| 22301 | 1116 | V1116A | D | 1.3e-05 | 1.2e-05 | 7e-06 | 1.067E-05 | 3.215E-06 | 7.000E-06 | 1.300E-05 |
| 22302 | 1116 | V1116R | D | 8.1e-06 | 8e-06 | 1e-05 | 8.700E-06 | 1.127E-06 | 8.000E-06 | 1.000E-05 |
| 22303 | 1116 | V1116N | D | 1.1e-05 | 1.3e-05 | 4.6e-06 | 9.533E-06 | 4.388E-06 | 4.600E-06 | 1.300E-05 |
| 22304 | 1116 | V1116D | D | 7.1e-06 | 9.7e-06 | 6.3e-06 | 7.700E-06 | 1.778E-06 | 6.300E-06 | 9.700E-06 |
| 22305 | 1116 | V1116C | D | 6.3e-06 | 1.3e-05 | 8.3e-06 | 9.200E-06 | 3.439E-06 | 6.300E-06 | 1.300E-05 |
| 22306 | 1116 | V1116E | D | 5.6e-06 | 8.7e-06 | 1e-05 | 8.100E-06 | 2.261E-06 | 5.600E-06 | 1.000E-05 |
| 22307 | 1116 | V1116Q | D | 7e-06 | 9.3e-06 | 7.3e-06 | 7.867E-06 | 1.250E-06 | 7.000E-06 | 9.300E-06 |
| 22308 | 1116 | V1116G | D | 1.1e-05 | 1e-05 | 1.1e-05 | 1.067E-05 | 5.774E-07 | 1.000E-05 | 1.100E-05 |
| 22309 | 1116 | V1116H | D | 9.9e-06 | 6.5e-06 | 8.3e-06 | 8.233E-06 | 1.701E-06 | 6.500E-06 | 9.900E-06 |
| 22310 | 1116 | V1116I | D | 6.1e-06 | 9.6e-06 | 1.1e-05 | 8.900E-06 | 2.524E-06 | 6.100E-06 | 1.100E-05 |
| 22311 | 1116 | V1116L | D | 5.4e-06 | 9e-06 | 8.5e-06 | 7.633E-06 | 1.950E-06 | 5.400E-06 | 9.000E-06 |
| 22312 | 1116 | V1116K | D | 7.1e-06 | 1e-05 | 8.3e-06 | 8.467E-06 | 1.457E-06 | 7.100E-06 | 1.000E-05 |
| 22313 | 1116 | V1116M | D | 1e-05 | 6.7e-06 | 4.6e-06 | 7.100E-06 | 2.722E-06 | 4.600E-06 | 1.000E-05 |
| 22314 | 1116 | V1116F | D | 6.7e-06 | 1e-05 | 7.2e-06 | 7.967E-06 | 1.779E-06 | 6.700E-06 | 1.000E-05 |
| 22315 | 1116 | V1116P | D | 5.1e-06 | 6.2e-06 | 9e-06 | 6.767E-06 | 2.011E-06 | 5.100E-06 | 9.000E-06 |
| 22316 | 1116 | V1116S | D | 6.3e-06 | 6.6e-06 | 6.5e-06 | 6.467E-06 | 1.528E-07 | 6.300E-06 | 6.600E-06 |
| 22317 | 1116 | V1116T | D | 1.2e-05 | 1e-05 | 9e-06 | 1.033E-05 | 1.528E-06 | 9.000E-06 | 1.200E-05 |
| 22318 | 1116 | V1116W | D | 6.9e-06 | 6.2e-06 | 6.7e-06 | 6.600E-06 | 3.606E-07 | 6.200E-06 | 6.900E-06 |
| 22319 | 1116 | V1116Y | D | 6.8e-06 | 6.1e-06 | 4.9e-06 | 5.933E-06 | 9.609E-07 | 4.900E-06 | 6.800E-06 |
| 22320 | 1116 | V1116V | D | 1.5e-05 | 5.8e-06 | 3.6e-06 | 8.133E-06 | 6.048E-06 | 3.600E-06 | 1.500E-05 |
| 22321 | 1117 | P1117A | D | 6.7e-06 | 9.7e-06 | 8.7e-06 | 8.367E-06 | 1.528E-06 | 6.700E-06 | 9.700E-06 |
| 22322 | 1117 | P1117R | D | 3.2e-06 | 7.6e-06 | 1e-05 | 6.933E-06 | 3.449E-06 | 3.200E-06 | 1.000E-05 |
| 22323 | 1117 | P1117N | D | 6.9e-06 | 1.3e-05 | 8.4e-06 | 9.433E-06 | 3.179E-06 | 6.900E-06 | 1.300E-05 |
| 22324 | 1117 | P1117D | D | 6.1e-06 | 7.6e-06 | 6.2e-06 | 6.633E-06 | 8.386E-07 | 6.100E-06 | 7.600E-06 |
| 22325 | 1117 | P1117C | D | 7e-06 | 8.6e-06 | 6.8e-06 | 7.467E-06 | 9.866E-07 | 6.800E-06 | 8.600E-06 |
| 22326 | 1117 | P1117E | D | 4.8e-06 | 3.2e-06 | 5.9e-06 | 4.633E-06 | 1.358E-06 | 3.200E-06 | 5.900E-06 |
| 22327 | 1117 | P1117Q | D | 1.4e-05 | 8.6e-06 | 4.6e-06 | 9.067E-06 | 4.717E-06 | 4.600E-06 | 1.400E-05 |
| 22328 | 1117 | P1117G | D | 1.3e-05 | 8.2e-06 | 8.4e-06 | 9.867E-06 | 2.715E-06 | 8.200E-06 | 1.300E-05 |
| 22329 | 1117 | P1117H | D | 8.3e-06 | 5.2e-06 | 6.7e-06 | 6.733E-06 | 1.550E-06 | 5.200E-06 | 8.300E-06 |
| 22330 | 1117 | P1117I | D | 6.4e-06 | 4.7e-06 | 5e-06 | 5.367E-06 | 9.074E-07 | 4.700E-06 | 6.400E-06 |
| 22331 | 1117 | P1117L | D | 6.6e-06 | 8.3e-06 | 8.1e-06 | 7.667E-06 | 9.292E-07 | 6.600E-06 | 8.300E-06 |
| 22332 | 1117 | P1117K | D | 6.5e-06 | 6.3e-06 | 4.1e-06 | 5.633E-06 | 1.332E-06 | 4.100E-06 | 6.500E-06 |
| 22333 | 1117 | P1117M | D | 9.6e-06 | 4.6e-06 | 9.4e-06 | 7.867E-06 | 2.831E-06 | 4.600E-06 | 9.600E-06 |
| 22334 | 1117 | P1117F | D | 7.2e-06 | 7.6e-06 | 9.8e-06 | 8.200E-06 | 1.400E-06 | 7.200E-06 | 9.800E-06 |
| 22335 | 1117 | P1117P | D | 1.2e-05 | 7.4e-06 | 1.4e-05 | 1.113E-05 | 3.384E-06 | 7.400E-06 | 1.400E-05 |
| 22336 | 1117 | P1117S | D | 5.3e-06 | 1e-05 | 6.6e-06 | 7.300E-06 | 2.427E-06 | 5.300E-06 | 1.000E-05 |
| 22337 | 1117 | P1117T | D | 5.5e-06 | 1.4e-05 | 5.5e-06 | 8.333E-06 | 4.907E-06 | 5.500E-06 | 1.400E-05 |
| 22338 | 1117 | P1117W | D | 1.2e-05 | 9.7e-06 | 9.3e-06 | 1.033E-05 | 1.457E-06 | 9.300E-06 | 1.200E-05 |
| 22339 | 1117 | P1117Y | D | 5.4e-06 | 6.7e-06 | 1.1e-05 | 7.700E-06 | 2.931E-06 | 5.400E-06 | 1.100E-05 |
| 22340 | 1117 | P1117V | D | 1e-05 | 1.8e-05 | 5.2e-06 | 1.107E-05 | 6.466E-06 | 5.200E-06 | 1.800E-05 |
| 22341 | 1118 | S1118A | D | 5.6e-06 | 1.4e-05 | 9e-06 | 9.533E-06 | 4.225E-06 | 5.600E-06 | 1.400E-05 |
| 22342 | 1118 | S1118R | D | 9.8e-06 | 6.8e-06 | 9.7e-06 | 8.767E-06 | 1.704E-06 | 6.800E-06 | 9.800E-06 |
| 22343 | 1118 | S1118N | D | 1.1e-05 | 5.8e-06 | 8.6e-06 | 8.467E-06 | 2.603E-06 | 5.800E-06 | 1.100E-05 |
| 22344 | 1118 | S1118D | D | 6e-06 | 8.2e-06 | 8.2e-06 | 7.467E-06 | 1.270E-06 | 6.000E-06 | 8.200E-06 |
| 22345 | 1118 | S1118C | D | 6.5e-06 | 4.7e-06 | 5.8e-06 | 5.667E-06 | 9.074E-07 | 4.700E-06 | 6.500E-06 |
| 22346 | 1118 | S1118E | D | 9.3e-06 | 1.1e-05 | 1.1e-05 | 1.043E-05 | 9.815E-07 | 9.300E-06 | 1.100E-05 |
| 22347 | 1118 | S1118Q | D | 1e-05 | 9.8e-06 | 7.4e-06 | 9.067E-06 | 1.447E-06 | 7.400E-06 | 1.000E-05 |
| 22348 | 1118 | S1118G | D | 9.7e-06 | 4.7e-06 | 7.3e-06 | 7.233E-06 | 2.501E-06 | 4.700E-06 | 9.700E-06 |
| 22349 | 1118 | S1118H | D | 9.6e-06 | 9.4e-06 | 5.9e-06 | 8.300E-06 | 2.081E-06 | 5.900E-06 | 9.600E-06 |
| 22350 | 1118 | S1118I | D | 9e-06 | 9.4e-06 | 1.1e-05 | 9.800E-06 | 1.058E-06 | 9.000E-06 | 1.100E-05 |
| 22351 | 1118 | S1118L | D | 9.6e-06 | 3.8e-06 | 8e-06 | 7.133E-06 | 2.996E-06 | 3.800E-06 | 9.600E-06 |
| 22352 | 1118 | S1118K | D | 5.6e-06 | 8.1e-06 | 6.1e-06 | 6.600E-06 | 1.323E-06 | 5.600E-06 | 8.100E-06 |
| 22353 | 1118 | S1118M | D | 6.8e-06 | 6.8e-06 | 3.8e-06 | 5.800E-06 | 1.732E-06 | 3.800E-06 | 6.800E-06 |
| 22354 | 1118 | S1118F | D | 1.1e-05 | 1.3e-05 | 5.9e-06 | 9.967E-06 | 3.661E-06 | 5.900E-06 | 1.300E-05 |

| | | | | | | | | | | |
|---|---|---|---|---|---|---|---|---|---|---|
| 22355 | 1118 | S1118P | D | 9.5e-06 | 1.1e-05 | 6.7e-06 | 9.067E-06 | 2.183E-06 | 6.700E-06 | 1.100E-05 |
| 22356 | 1118 | S1118S | D | 6.6e-06 | 5.7e-06 | 7.9e-06 | 6.733E-06 | 1.106E-06 | 5.700E-06 | 7.900E-06 |
| 22357 | 1118 | S1118T | D | 6.4e-06 | 8.7e-06 | 6.7e-06 | 7.267E-06 | 1.250E-06 | 6.400E-06 | 8.700E-06 |
| 22358 | 1118 | S1118W | D | 1.3e-05 | 9e-06 | 4.9e-06 | 8.967E-06 | 4.050E-06 | 4.900E-06 | 1.300E-05 |
| 22359 | 1118 | S1118Y | D | 9e-06 | 8.6e-06 | 9.6e-06 | 9.067E-06 | 5.033E-07 | 8.600E-06 | 9.600E-06 |
| 22360 | 1118 | S1118V | D | 7.8e-06 | 1e-05 | 1.1e-05 | 9.600E-06 | 1.637E-06 | 7.800E-06 | 1.100E-05 |
| 22361 | 1119 | R1119A | D | 1.1e-05 | 5.4e-06 | 9.5e-06 | 8.633E-06 | 2.899E-06 | 5.400E-06 | 1.100E-05 |
| 22362 | 1119 | R1119R | D | 9.8e-06 | 6.8e-06 | 8.3e-06 | 8.300E-06 | 1.500E-06 | 6.800E-06 | 9.800E-06 |
| 22363 | 1119 | R1119N | D | 6.5e-06 | 6.4e-06 | 8.4e-06 | 7.100E-06 | 1.127E-06 | 6.400E-06 | 8.400E-06 |
| 22364 | 1119 | R1119D | D | 7.1e-06 | 8.5e-06 | 4.9e-06 | 6.833E-06 | 1.815E-06 | 4.900E-06 | 8.500E-06 |
| 22365 | 1119 | R1119C | D | 6.4e-06 | 5.5e-06 | 1e-05 | 7.300E-06 | 2.381E-06 | 5.500E-06 | 1.000E-05 |
| 22366 | 1119 | R1119E | D | 1.1e-05 | 1.3e-05 | 9.2e-06 | 1.107E-05 | 1.901E-06 | 9.200E-06 | 1.300E-05 |
| 22367 | 1119 | R1119Q | D | 1.3e-05 | 1.3e-05 | 1.3e-05 | 1.300E-05 | 0.000E+00 | 1.300E-05 | 1.300E-05 |
| 22368 | 1119 | R1119G | D | 1.3e-05 | 5.8e-06 | 1.2e-05 | 1.027E-05 | 3.900E-06 | 5.800E-06 | 1.300E-05 |
| 22369 | 1119 | R1119H | D | 7.8e-06 | 6.8e-06 | 8.6e-06 | 7.733E-06 | 9.018E-07 | 6.800E-06 | 8.600E-06 |
| 22370 | 1119 | R1119I | D | 1.1e-05 | 1.3e-05 | 7.9e-06 | 1.063E-05 | 2.570E-06 | 7.900E-06 | 1.300E-05 |
| 22371 | 1119 | R1119L | D | 7.3e-06 | 7.5e-06 | 1e-05 | 8.267E-06 | 1.504E-06 | 7.300E-06 | 1.000E-05 |
| 22372 | 1119 | R1119K | D | 6.6e-06 | 7.4e-06 | 1.1e-05 | 8.333E-06 | 2.344E-06 | 6.600E-06 | 1.100E-05 |
| 22373 | 1119 | R1119M | D | 7e-06 | 7.2e-06 | 9.1e-06 | 7.767E-06 | 1.159E-06 | 7.000E-06 | 9.100E-06 |
| 22374 | 1119 | R1119F | D | 6e-06 | 6.4e-06 | 8.2e-06 | 6.867E-06 | 1.172E-06 | 6.000E-06 | 8.200E-06 |
| 22375 | 1119 | R1119P | D | 6e-06 | 9.6e-06 | 9.1e-06 | 8.233E-06 | 1.950E-06 | 6.000E-06 | 9.600E-06 |
| 22376 | 1119 | R1119S | D | 1.3e-05 | 1.3e-05 | 9.8e-06 | 1.193E-05 | 1.848E-06 | 9.800E-06 | 1.300E-05 |
| 22377 | 1119 | R1119T | D | 1.3e-05 | 6.9e-06 | 1.1e-05 | 1.030E-05 | 3.110E-06 | 6.900E-06 | 1.300E-05 |
| 22378 | 1119 | R1119W | D | 6.8e-06 | 1.3e-05 | 5.5e-06 | 8.433E-06 | 4.008E-06 | 5.500E-06 | 1.300E-05 |
| 22379 | 1119 | R1119Y | D | 6.3e-06 | 6.8e-06 | 8.3e-06 | 7.133E-06 | 1.041E-06 | 6.300E-06 | 8.300E-06 |
| 22380 | 1119 | R1119V | D | 6.6e-06 | 1.2e-05 | 1.1e-05 | 9.867E-06 | 2.873E-06 | 6.600E-06 | 1.200E-05 |
| 22381 | 1120 | F1120A | D | 5.4e-06 | 1.1e-05 | 5.4e-06 | 7.267E-06 | 3.233E-06 | 5.400E-06 | 1.100E-05 |
| 22382 | 1120 | F1120R | D | 1.1e-05 | 7.9e-06 | 6.8e-06 | 8.567E-06 | 2.178E-06 | 6.800E-06 | 1.100E-05 |
| 22383 | 1120 | F1120N | D | 1.4e-05 | 1.9e-05 | 9.5e-06 | 1.417E-05 | 4.752E-06 | 9.500E-06 | 1.900E-05 |
| 22384 | 1120 | F1120D | D | 5.9e-06 | 9.4e-06 | 9.9e-06 | 8.400E-06 | 2.179E-06 | 5.900E-06 | 9.900E-06 |
| 22385 | 1120 | F1120C | D | 7.5e-06 | 1.2e-05 | 9.2e-06 | 9.567E-06 | 2.272E-06 | 7.500E-06 | 1.200E-05 |
| 22386 | 1120 | F1120E | D | 7e-06 | 6.1e-06 | 5.5e-06 | 6.200E-06 | 7.550E-07 | 5.500E-06 | 7.000E-06 |
| 22387 | 1120 | F1120Q | D | 7.7e-06 | 6.1e-06 | 8.4e-06 | 7.400E-06 | 1.179E-06 | 6.100E-06 | 8.400E-06 |
| 22388 | 1120 | F1120G | D | 9.4e-06 | 1e-05 | 8.1e-06 | 9.167E-06 | 9.713E-07 | 8.100E-06 | 1.000E-05 |
| 22389 | 1120 | F1120H | D | 6.3e-06 | 6.9e-06 | 6.9e-06 | 6.700E-06 | 3.464E-07 | 6.300E-06 | 6.900E-06 |
| 22390 | 1120 | F1120I | D | 5.7e-06 | 4.7e-06 | 6.1e-06 | 5.500E-06 | 7.211E-07 | 4.700E-06 | 6.100E-06 |
| 22391 | 1120 | F1120L | D | 9.7e-06 | 1.1e-05 | 8e-06 | 9.567E-06 | 1.504E-06 | 8.000E-06 | 1.100E-05 |
| 22392 | 1120 | F1120K | D | 4.8e-06 | 4e-06 | 1.2e-05 | 6.933E-06 | 4.406E-06 | 4.000E-06 | 1.200E-05 |
| 22393 | 1120 | F1120M | D | 5.3e-06 | 7.7e-06 | 8.3e-06 | 7.100E-06 | 1.587E-06 | 5.300E-06 | 8.300E-06 |
| 22394 | 1120 | F1120F | D | 9.1e-06 | 1.1e-05 | 4.4e-06 | 8.167E-06 | 3.398E-06 | 4.400E-06 | 1.100E-05 |
| 22395 | 1120 | F1120P | D | 5.2e-06 | 1.1e-05 | 8.1e-06 | 8.100E-06 | 2.900E-06 | 5.200E-06 | 1.100E-05 |
| 22396 | 1120 | F1120S | D | 8.5e-06 | 7.3e-06 | 8.2e-06 | 8.000E-06 | 6.245E-07 | 7.300E-06 | 8.500E-06 |
| 22397 | 1120 | F1120T | D | 8.7e-06 | 5.8e-06 | 6.9e-06 | 7.133E-06 | 1.464E-06 | 5.800E-06 | 8.700E-06 |
| 22398 | 1120 | F1120W | D | 9.7e-06 | 9.4e-06 | 6.4e-06 | 8.500E-06 | 1.825E-06 | 6.400E-06 | 9.700E-06 |
| 22399 | 1120 | F1120Y | D | 1e-05 | 6.4e-06 | 8.3e-06 | 8.233E-06 | 1.801E-06 | 6.400E-06 | 1.000E-05 |
| 22400 | 1120 | F1120V | D | 5.4e-06 | 5.5e-06 | 5e-06 | 5.300E-06 | 2.646E-07 | 5.000E-06 | 5.500E-06 |
| 22401 | 1121 | S1121A | D | 7.8e-06 | 6.4e-06 | 1.1e-05 | 8.400E-06 | 2.358E-06 | 6.400E-06 | 1.100E-05 |
| 22402 | 1121 | S1121R | D | 1.3e-05 | 9.8e-06 | 9.3e-06 | 1.070E-05 | 2.007E-06 | 9.300E-06 | 1.300E-05 |
| 22403 | 1121 | S1121N | D | 1e-05 | 8.4e-06 | 2.9e-06 | 7.100E-06 | 3.724E-06 | 2.900E-06 | 1.000E-05 |
| 22404 | 1121 | S1121D | D | 1e-05 | 4.6e-06 | 1.2e-05 | 8.867E-06 | 3.828E-06 | 4.600E-06 | 1.200E-05 |
| 22405 | 1121 | S1121C | D | 5.7e-06 | 7.2e-06 | 7.9e-06 | 6.933E-06 | 1.124E-06 | 5.700E-06 | 7.900E-06 |
| 22406 | 1121 | S1121E | D | 7.4e-06 | 5.9e-06 | 1.8e-05 | 1.043E-05 | 6.596E-06 | 5.900E-06 | 1.800E-05 |
| 22407 | 1121 | S1121Q | D | 5.4e-06 | 8.9e-06 | 7.4e-06 | 7.233E-06 | 1.756E-06 | 5.400E-06 | 8.900E-06 |
| 22408 | 1121 | S1121G | D | 7.4e-06 | 7.7e-06 | 8.1e-06 | 7.733E-06 | 3.512E-07 | 7.400E-06 | 8.100E-06 |
| 22409 | 1121 | S1121H | D | 7e-06 | 8.4e-06 | 7.3e-06 | 7.567E-06 | 7.371E-07 | 7.000E-06 | 8.400E-06 |
| 22410 | 1121 | S1121I | D | 9.6e-06 | 5e-06 | 8.5e-06 | 7.700E-06 | 2.402E-06 | 5.000E-06 | 9.600E-06 |
| 22411 | 1121 | S1121L | D | 7.3e-06 | 6.6e-06 | 7.5e-06 | 7.133E-06 | 4.726E-07 | 6.600E-06 | 7.500E-06 |
| 22412 | 1121 | S1121K | D | 5.9e-06 | 1.1e-05 | 9.6e-06 | 8.833E-06 | 2.635E-06 | 5.900E-06 | 1.100E-05 |
| 22413 | 1121 | S1121M | D | 8.9e-06 | 8.1e-06 | 1.2e-05 | 9.667E-06 | 2.060E-06 | 8.100E-06 | 1.200E-05 |
| 22414 | 1121 | S1121F | D | 7.2e-06 | 7.4e-06 | 1.3e-05 | 9.200E-06 | 3.292E-06 | 7.200E-06 | 1.300E-05 |
| 22415 | 1121 | S1121P | D | 1.3e-05 | 1.1e-05 | 5e-06 | 9.667E-06 | 4.163E-06 | 5.000E-06 | 1.300E-05 |
| 22416 | 1121 | S1121S | D | 9.7e-06 | 9.9e-06 | 8.8e-06 | 9.467E-06 | 5.859E-07 | 8.800E-06 | 9.900E-06 |
| 22417 | 1121 | S1121T | D | 8.5e-06 | 8e-06 | 1e-05 | 8.833E-06 | 1.041E-06 | 8.000E-06 | 1.000E-05 |
| 22418 | 1121 | S1121W | D | 8.4e-06 | 9.8e-06 | 8.2e-06 | 8.800E-06 | 8.718E-07 | 8.200E-06 | 9.800E-06 |
| 22419 | 1121 | S1121Y | D | 7.1e-06 | 4.6e-06 | 7e-06 | 6.233E-06 | 1.415E-06 | 4.600E-06 | 7.100E-06 |
| 22420 | 1121 | S1121V | D | 1.1e-05 | 1.2e-05 | 9.8e-06 | 1.093E-05 | 1.102E-06 | 9.800E-06 | 1.200E-05 |
| 22421 | 1122 | G1122A | D | 8.9e-06 | 6.6e-06 | 8.4e-06 | 7.967E-06 | 1.210E-06 | 6.600E-06 | 8.900E-06 |
| 22422 | 1122 | G1122R | D | 6.9e-06 | 8.2e-06 | 9.5e-06 | 8.200E-06 | 1.300E-06 | 6.900E-06 | 9.500E-06 |
| 22423 | 1122 | G1122N | D | 9.9e-06 | 7e-06 | 5.7e-06 | 7.533E-06 | 2.150E-06 | 5.700E-06 | 9.900E-06 |
| 22424 | 1122 | G1122D | D | 5.8e-06 | 7.1e-06 | 6.3e-06 | 6.400E-06 | 6.557E-07 | 5.800E-06 | 7.100E-06 |
| 22425 | 1122 | G1122C | D | 1.2e-05 | 9.5e-06 | 8.6e-06 | 1.003E-05 | 1.762E-06 | 8.600E-06 | 1.200E-05 |
| 22426 | 1122 | G1122E | D | 9.4e-06 | 1.6e-05 | 1.1e-05 | 1.213E-05 | 3.443E-06 | 9.400E-06 | 1.600E-05 |
| 22427 | 1122 | G1122Q | D | 1.2e-05 | 8.4e-06 | 8.9e-06 | 9.767E-06 | 1.950E-06 | 8.400E-06 | 1.200E-05 |
| 22428 | 1122 | G1122G | D | 8.2e-06 | 6.2e-06 | 7.1e-06 | 7.167E-06 | 1.002E-06 | 6.200E-06 | 8.200E-06 |
| 22429 | 1122 | G1122H | D | 4.4e-06 | 1e-05 | 5.6e-06 | 6.667E-06 | 2.948E-06 | 4.400E-06 | 1.000E-05 |
| 22430 | 1122 | G1122I | D | 4.9e-06 | 7.4e-06 | 1.4e-05 | 8.767E-06 | 4.701E-06 | 4.900E-06 | 1.400E-05 |
| 22431 | 1122 | G1122L | D | 9.8e-06 | 1.3e-05 | 9.3e-06 | 1.070E-05 | 2.007E-06 | 9.300E-06 | 1.300E-05 |
| 22432 | 1122 | G1122K | D | 5.3e-06 | 5.8e-06 | 5.2e-06 | 5.433E-06 | 3.215E-07 | 5.200E-06 | 5.800E-06 |
| 22433 | 1122 | G1122M | D | 9.3e-06 | 1.8e-05 | 5.8e-06 | 1.103E-05 | 6.282E-06 | 5.800E-06 | 1.800E-05 |
| 22434 | 1122 | G1122F | D | 1.4e-05 | 6.7e-06 | 1.1e-05 | 1.057E-05 | 3.669E-06 | 6.700E-06 | 1.400E-05 |
| 22435 | 1122 | G1122P | D | 9.6e-06 | 6.4e-06 | 6.9e-06 | 7.633E-06 | 1.721E-06 | 6.400E-06 | 9.600E-06 |
| 22436 | 1122 | G1122S | D | 6e-06 | 4.5e-06 | 9.1e-06 | 6.533E-06 | 2.346E-06 | 4.500E-06 | 9.100E-06 |
| 22437 | 1122 | G1122T | D | 1e-05 | 7.8e-06 | 1.1e-05 | 9.600E-06 | 1.637E-06 | 7.800E-06 | 1.100E-05 |
| 22438 | 1122 | G1122W | D | 7.3e-06 | 6.1e-06 | 9e-06 | 7.467E-06 | 1.457E-06 | 6.100E-06 | 9.000E-06 |
| 22439 | 1122 | G1122Y | D | 1.6e-05 | 8.6e-06 | 7.3e-06 | 1.063E-05 | 4.693E-06 | 7.300E-06 | 1.600E-05 |

| | | | | | | | | | | |
|---|---|---|---|---|---|---|---|---|---|---|
| 22440 | 1122 | G1122V | D | 6.4e-06 | 7.8e-06 | 1e-05 | 8.067E-06 | 1.815E-06 | 6.400E-06 | 1.000E-05 |
| 22441 | 1123 | S1123A | D | 8.9e-06 | 6.1e-06 | 6.7e-06 | 7.233E-06 | 1.474E-06 | 6.100E-06 | 8.900E-06 |
| 22442 | 1123 | S1123R | D | 8.6e-06 | 8.8e-06 | 8.3e-06 | 8.567E-06 | 2.517E-07 | 8.300E-06 | 8.800E-06 |
| 22443 | 1123 | S1123N | D | 9.5e-06 | 1e-05 | 6.2e-06 | 8.567E-06 | 2.065E-06 | 6.200E-06 | 1.000E-05 |
| 22444 | 1123 | S1123D | D | 7.8e-06 | 7.9e-06 | 6.3e-06 | 7.333E-06 | 8.963E-07 | 6.300E-06 | 7.900E-06 |
| 22445 | 1123 | S1123C | D | 1.1e-05 | 7.9e-06 | 8.5e-06 | 9.133E-06 | 1.644E-06 | 7.900E-06 | 1.100E-05 |
| 22446 | 1123 | S1123E | D | 5.3e-06 | 7e-06 | 1.2e-05 | 8.100E-06 | 3.483E-06 | 5.300E-06 | 1.200E-05 |
| 22447 | 1123 | S1123Q | D | 7.4e-06 | 5.4e-06 | 8.2e-06 | 7.000E-06 | 1.442E-06 | 5.400E-06 | 8.200E-06 |
| 22448 | 1123 | S1123G | D | 4.3e-06 | 1.4e-05 | 1.6e-05 | 1.143E-05 | 6.258E-06 | 4.300E-06 | 1.600E-05 |
| 22449 | 1123 | S1123H | D | 1.1e-05 | 8.7e-06 | 1.1e-05 | 1.023E-05 | 1.328E-06 | 8.700E-06 | 1.100E-05 |
| 22450 | 1123 | S1123I | D | 1.1e-05 | 5.5e-06 | 3.8e-06 | 6.767E-06 | 3.763E-06 | 3.800E-06 | 1.100E-05 |
| 22451 | 1123 | S1123L | D | 8.9e-06 | 9.2e-06 | 5.9e-06 | 8.000E-06 | 1.825E-06 | 5.900E-06 | 9.200E-06 |
| 22452 | 1123 | S1123K | D | 9.9e-06 | 9.2e-06 | 7.2e-06 | 8.767E-06 | 1.401E-06 | 7.200E-06 | 9.900E-06 |
| 22453 | 1123 | S1123M | D | 1.5e-05 | 6.3e-06 | 7.4e-06 | 9.567E-06 | 4.737E-06 | 6.300E-06 | 1.500E-05 |
| 22454 | 1123 | S1123F | D | 5e-06 | 7.3e-06 | 5.5e-06 | 5.933E-06 | 1.210E-06 | 5.000E-06 | 7.300E-06 |
| 22455 | 1123 | S1123P | D | 7.4e-06 | 8.4e-06 | 7.8e-06 | 7.867E-06 | 5.033E-07 | 7.400E-06 | 8.400E-06 |
| 22456 | 1123 | S1123S | D | 6.4e-06 | 1e-05 | 7.7e-06 | 8.033E-06 | 1.823E-06 | 6.400E-06 | 1.000E-05 |
| 22457 | 1123 | S1123T | D | 6.2e-06 | 4.8e-06 | 7.1e-06 | 6.033E-06 | 1.159E-06 | 4.800E-06 | 7.100E-06 |
| 22458 | 1123 | S1123W | D | 7.8e-06 | 7.2e-06 | 1e-05 | 8.333E-06 | 1.474E-06 | 7.200E-06 | 1.000E-05 |
| 22459 | 1123 | S1123Y | D | 5.8e-06 | 8.9e-06 | 6.8e-06 | 7.167E-06 | 1.582E-06 | 5.800E-06 | 8.900E-06 |
| 22460 | 1123 | S1123V | D | 8.3e-06 | 8.6e-06 | 6.9e-06 | 7.933E-06 | 9.074E-07 | 6.900E-06 | 8.600E-06 |
| 22461 | 1124 | R1124A | D | 8.4e-06 | 5e-06 | 6e-06 | 6.467E-06 | 1.747E-06 | 5.000E-06 | 8.400E-06 |
| 22462 | 1124 | R1124R | D | 7.8e-06 | 1e-05 | 8.4e-06 | 8.733E-06 | 1.137E-06 | 7.800E-06 | 1.000E-05 |
| 22463 | 1124 | R1124N | D | 1.4e-05 | 9.2e-06 | 3.6e-06 | 8.933E-06 | 5.205E-06 | 3.600E-06 | 1.400E-05 |
| 22464 | 1124 | R1124D | D | 4.8e-06 | 1.3e-05 | 7.5e-06 | 8.433E-06 | 4.179E-06 | 4.800E-06 | 1.300E-05 |
| 22465 | 1124 | R1124C | D | 8.1e-06 | 4.6e-06 | 7.8e-06 | 6.833E-06 | 1.940E-06 | 4.600E-06 | 8.100E-06 |
| 22466 | 1124 | R1124E | D | 1e-05 | 1e-05 | 6.2e-06 | 8.733E-06 | 2.194E-06 | 6.200E-06 | 1.000E-05 |
| 22467 | 1124 | R1124Q | D | 6.4e-06 | 1.1e-05 | 4e-06 | 7.133E-06 | 3.557E-06 | 4.000E-06 | 1.100E-05 |
| 22468 | 1124 | R1124G | D | 7.8e-06 | 1.6e-05 | 9.7e-06 | 1.117E-05 | 4.292E-06 | 7.800E-06 | 1.600E-05 |
| 22469 | 1124 | R1124H | D | 8.8e-06 | 6.7e-06 | 6.8e-06 | 7.433E-06 | 1.185E-06 | 6.700E-06 | 8.800E-06 |
| 22470 | 1124 | R1124I | D | 6.8e-06 | 1e-05 | 7.7e-06 | 8.167E-06 | 1.650E-06 | 6.800E-06 | 1.000E-05 |
| 22471 | 1124 | R1124L | D | 7.3e-06 | 7.6e-06 | 4.9e-06 | 6.600E-06 | 1.480E-06 | 4.900E-06 | 7.600E-06 |
| 22472 | 1124 | R1124K | D | 6.9e-06 | 8.3e-06 | 4.1e-06 | 6.433E-06 | 2.139E-06 | 4.100E-06 | 8.300E-06 |
| 22473 | 1124 | R1124M | D | 1.2e-05 | 8.9e-06 | 1.2e-05 | 1.097E-05 | 1.790E-06 | 8.900E-06 | 1.200E-05 |
| 22474 | 1124 | R1124F | D | 6.4e-06 | 5e-06 | 5.1e-06 | 5.500E-06 | 7.810E-07 | 5.000E-06 | 6.400E-06 |
| 22475 | 1124 | R1124P | D | 1.4e-05 | 8.8e-06 | 9.6e-06 | 1.080E-05 | 2.800E-06 | 8.800E-06 | 1.400E-05 |
| 22476 | 1124 | R1124S | D | 9.2e-06 | 6.3e-06 | 5.1e-06 | 6.867E-06 | 2.108E-06 | 5.100E-06 | 9.200E-06 |
| 22477 | 1124 | R1124T | D | 7.1e-06 | 5.5e-06 | 8.4e-06 | 7.000E-06 | 1.453E-06 | 5.500E-06 | 8.400E-06 |
| 22478 | 1124 | R1124W | D | 6e-06 | 1.2e-05 | 8.6e-06 | 8.867E-06 | 3.009E-06 | 6.000E-06 | 1.200E-05 |
| 22479 | 1124 | R1124Y | D | 6.9e-06 | 6.9e-06 | 1.2e-05 | 8.600E-06 | 2.944E-06 | 6.900E-06 | 1.200E-05 |
| 22480 | 1124 | R1124V | D | 5.6e-06 | 4.7e-06 | 9.9e-06 | 6.733E-06 | 2.779E-06 | 4.700E-06 | 9.900E-06 |
| 22481 | 1125 | S1125A | D | 8.7e-06 | 1.1e-05 | 1.1e-05 | 1.023E-05 | 1.328E-06 | 8.700E-06 | 1.100E-05 |
| 22482 | 1125 | S1125R | D | 7.2e-06 | 1e-05 | 9.3e-06 | 8.833E-06 | 1.457E-06 | 7.200E-06 | 1.000E-05 |
| 22483 | 1125 | S1125N | D | 9.7e-06 | 1e-05 | 7.7e-06 | 9.133E-06 | 1.250E-06 | 7.700E-06 | 1.000E-05 |
| 22484 | 1125 | S1125D | D | 6.3e-06 | 6.5e-06 | 1.3e-05 | 8.600E-06 | 3.812E-06 | 6.300E-06 | 1.300E-05 |
| 22485 | 1125 | S1125C | D | 1e-05 | 6.9e-06 | 9.5e-06 | 8.800E-06 | 1.664E-06 | 6.900E-06 | 1.000E-05 |
| 22486 | 1125 | S1125E | D | 9.6e-06 | 6.7e-06 | 1.1e-05 | 9.100E-06 | 2.193E-06 | 6.700E-06 | 1.100E-05 |
| 22487 | 1125 | S1125Q | D | 1e-05 | 1.3e-05 | 3.6e-06 | 8.867E-06 | 4.801E-06 | 3.600E-06 | 1.300E-05 |
| 22488 | 1125 | S1125G | D | 7.3e-06 | 7e-06 | 5e-06 | 6.433E-06 | 1.250E-06 | 5.000E-06 | 7.300E-06 |
| 22489 | 1125 | S1125H | D | 5.3e-06 | 6.4e-06 | 9.7e-06 | 7.133E-06 | 2.290E-06 | 5.300E-06 | 9.700E-06 |
| 22490 | 1125 | S1125I | D | 5.8e-06 | 8.8e-06 | 6e-06 | 6.867E-06 | 1.677E-06 | 5.800E-06 | 8.800E-06 |
| 22491 | 1125 | S1125L | D | 7.3e-06 | 1e-05 | 7.7e-06 | 8.333E-06 | 1.457E-06 | 7.300E-06 | 1.000E-05 |
| 22492 | 1125 | S1125K | D | 6e-06 | 3.8e-06 | 7.7e-06 | 5.833E-06 | 1.955E-06 | 3.800E-06 | 7.700E-06 |
| 22493 | 1125 | S1125M | D | 9.1e-06 | 8e-06 | 8.6e-06 | 8.567E-06 | 5.508E-07 | 8.000E-06 | 9.100E-06 |
| 22494 | 1125 | S1125F | D | 7.6e-06 | 7e-06 | 1.3e-05 | 9.200E-06 | 3.305E-06 | 7.000E-06 | 1.300E-05 |
| 22495 | 1125 | S1125P | D | 6.2e-06 | 1.3e-05 | 6.4e-06 | 8.533E-06 | 3.870E-06 | 6.200E-06 | 1.300E-05 |
| 22496 | 1125 | S1125S | D | 9.2e-06 | 4.8e-06 | 8.1e-06 | 7.367E-06 | 2.290E-06 | 4.800E-06 | 9.200E-06 |
| 22497 | 1125 | S1125T | D | 1.1e-05 | 6.4e-06 | 8.2e-06 | 8.533E-06 | 2.318E-06 | 6.400E-06 | 1.100E-05 |
| 22498 | 1125 | S1125W | D | 5e-06 | 6.6e-06 | 5.5e-06 | 5.700E-06 | 8.185E-07 | 5.000E-06 | 6.600E-06 |
| 22499 | 1125 | S1125Y | D | 9.2e-06 | 5.3e-06 | 9.6e-06 | 8.033E-06 | 2.376E-06 | 5.300E-06 | 9.600E-06 |
| 22500 | 1125 | S1125V | D | 5.7e-06 | 4.5e-06 | 5.9e-06 | 5.367E-06 | 7.572E-07 | 4.500E-06 | 5.900E-06 |
| 22501 | 1126 | G1126A | D | 8.9e-06 | 8.4e-06 | 9.6e-06 | 8.967E-06 | 6.028E-07 | 8.400E-06 | 9.600E-06 |
| 22502 | 1126 | G1126R | D | 4e-06 | 9.8e-06 | 8.9e-06 | 7.567E-06 | 3.121E-06 | 4.000E-06 | 9.800E-06 |
| 22503 | 1126 | G1126N | D | 6.9e-06 | 5.8e-06 | 1e-05 | 7.567E-06 | 2.178E-06 | 5.800E-06 | 1.000E-05 |
| 22504 | 1126 | G1126D | D | 4.5e-06 | 6.9e-06 | 4e-06 | 5.133E-06 | 1.550E-06 | 4.000E-06 | 6.900E-06 |
| 22505 | 1126 | G1126C | D | 1.2e-05 | 9e-06 | 9.5e-06 | 1.017E-05 | 1.607E-06 | 9.000E-06 | 1.200E-05 |
| 22506 | 1126 | G1126E | D | 6.3e-06 | 9.7e-06 | 7e-06 | 7.667E-06 | 1.795E-06 | 6.300E-06 | 9.700E-06 |
| 22507 | 1126 | G1126Q | D | 5.6e-06 | 1.1e-05 | 9.6e-06 | 8.733E-06 | 2.802E-06 | 5.600E-06 | 1.100E-05 |
| 22508 | 1126 | G1126G | D | 9.7e-06 | 1.4e-05 | 9.3e-06 | 1.100E-05 | 2.606E-06 | 9.300E-06 | 1.400E-05 |
| 22509 | 1126 | G1126H | D | 5.4e-06 | 8.2e-06 | 9.4e-06 | 7.667E-06 | 2.053E-06 | 5.400E-06 | 9.400E-06 |
| 22510 | 1126 | G1126I | D | 7.1e-06 | 9.7e-06 | 9.2e-06 | 8.667E-06 | 1.380E-06 | 7.100E-06 | 9.700E-06 |
| 22511 | 1126 | G1126L | D | 9e-06 | 6e-06 | 8.9e-06 | 7.967E-06 | 1.704E-06 | 6.000E-06 | 9.000E-06 |
| 22512 | 1126 | G1126K | D | 4.6e-06 | 9.8e-06 | 4.5e-06 | 6.300E-06 | 3.032E-06 | 4.500E-06 | 9.800E-06 |
| 22513 | 1126 | G1126M | D | 1e-05 | 5.2e-06 | 8.6e-06 | 7.933E-06 | 2.468E-06 | 5.200E-06 | 1.000E-05 |
| 22514 | 1126 | G1126F | D | 6.3e-06 | 4.6e-06 | 6.1e-06 | 5.667E-06 | 9.292E-07 | 4.600E-06 | 6.300E-06 |
| 22515 | 1126 | G1126P | D | 1.5e-05 | 9.6e-06 | 4.7e-06 | 9.767E-06 | 5.152E-06 | 4.700E-06 | 1.500E-05 |
| 22516 | 1126 | G1126S | D | 4.3e-06 | 8.8e-06 | 7.1e-06 | 6.733E-06 | 2.272E-06 | 4.300E-06 | 8.800E-06 |
| 22517 | 1126 | G1126T | D | 9.8e-06 | 1.1e-05 | 1.1e-05 | 1.060E-05 | 6.928E-07 | 9.800E-06 | 1.100E-05 |
| 22518 | 1126 | G1126W | D | 4.8e-06 | 8.6e-06 | 7.7e-06 | 7.033E-06 | 1.986E-06 | 4.800E-06 | 8.600E-06 |
| 22519 | 1126 | G1126Y | D | 8.3e-06 | 6.5e-06 | 9.2e-06 | 8.000E-06 | 1.375E-06 | 6.500E-06 | 9.200E-06 |
| 22520 | 1126 | G1126V | D | 7.1e-06 | 5.4e-06 | 6.3e-06 | 6.267E-06 | 8.505E-07 | 5.400E-06 | 7.100E-06 |
| 22521 | 1127 | T1127A | D | 4.7e-06 | 6.3e-06 | 5.9e-06 | 5.633E-06 | 8.327E-07 | 4.700E-06 | 6.300E-06 |
| 22522 | 1127 | T1127R | D | 6.5e-06 | 9.8e-06 | 1.1e-05 | 9.100E-06 | 2.330E-06 | 6.500E-06 | 1.100E-05 |
| 22523 | 1127 | T1127N | D | 1e-05 | 9.4e-06 | 5.6e-06 | 8.333E-06 | 2.386E-06 | 5.600E-06 | 1.000E-05 |
| 22524 | 1127 | T1127D | D | 5.4e-06 | 1.1e-05 | 9.1e-06 | 8.500E-06 | 2.848E-06 | 5.400E-06 | 1.100E-05 |

| | | | | | | | | | | |
|---|---|---|---|---|---|---|---|---|---|---|
| 22525 | 1127 | T1127C | D | 7.4e-06 | 7e-06 | 5.5e-06 | 6.633E-06 | 1.002E-06 | 5.500E-06 | 7.400E-06 |
| 22526 | 1127 | T1127E | D | 8.1e-06 | 7.6e-06 | 1e-05 | 8.567E-06 | 1.266E-06 | 7.600E-06 | 1.000E-05 |
| 22527 | 1127 | T1127Q | D | 8.5e-06 | 8e-06 | 8.9e-06 | 8.467E-06 | 4.509E-07 | 8.000E-06 | 8.900E-06 |
| 22528 | 1127 | T1127G | D | 6.7e-06 | 8.3e-06 | 1.4e-05 | 9.667E-06 | 3.837E-06 | 6.700E-06 | 1.400E-05 |
| 22529 | 1127 | T1127H | D | 7e-06 | 7.4e-06 | 8.1e-06 | 7.500E-06 | 5.568E-07 | 7.000E-06 | 8.100E-06 |
| 22530 | 1127 | T1127I | D | 7.7e-06 | 6.6e-06 | 4.8e-06 | 6.367E-06 | 1.464E-06 | 4.800E-06 | 7.700E-06 |
| 22531 | 1127 | T1127L | D | 8.5e-06 | 3.8e-06 | 9.2e-06 | 7.167E-06 | 2.937E-06 | 3.800E-06 | 9.200E-06 |
| 22532 | 1127 | T1127K | D | 4.5e-06 | 9.9e-06 | 5.8e-06 | 6.733E-06 | 2.818E-06 | 4.500E-06 | 9.900E-06 |
| 22533 | 1127 | T1127M | D | 1.2e-05 | 1.2e-05 | 9.8e-06 | 1.127E-05 | 1.270E-06 | 9.800E-06 | 1.200E-05 |
| 22534 | 1127 | T1127F | D | 1.5e-05 | 9.6e-06 | 9.3e-06 | 1.130E-05 | 3.208E-06 | 9.300E-06 | 1.500E-05 |
| 22535 | 1127 | T1127P | D | 1e-05 | 8e-06 | 1.2e-05 | 1.000E-05 | 2.000E-06 | 8.000E-06 | 1.200E-05 |
| 22536 | 1127 | T1127S | D | 9.8e-06 | 5.5e-06 | 5.3e-06 | 6.867E-06 | 2.542E-06 | 5.300E-06 | 9.800E-06 |
| 22537 | 1127 | T1127T | D | 7.6e-06 | 6.7e-06 | 1.4e-05 | 9.433E-06 | 3.980E-06 | 6.700E-06 | 1.400E-05 |
| 22538 | 1127 | T1127W | D | 6.7e-06 | 8.1e-06 | 4.8e-06 | 6.533E-06 | 1.656E-06 | 4.800E-06 | 8.100E-06 |
| 22539 | 1127 | T1127Y | D | 6.1e-06 | 9.1e-06 | 9.8e-06 | 8.333E-06 | 1.966E-06 | 6.100E-06 | 9.800E-06 |
| 22540 | 1127 | T1127V | D | 7.8e-06 | 1.2e-05 | 7.1e-06 | 8.967E-06 | 2.650E-06 | 7.100E-06 | 1.200E-05 |
| 22541 | 1128 | D1128A | D | 9.6e-06 | 9.5e-06 | 8.9e-06 | 9.333E-06 | 3.786E-07 | 8.900E-06 | 9.600E-06 |
| 22542 | 1128 | D1128R | D | 7e-06 | 8.7e-06 | 1.1e-05 | 8.900E-06 | 2.007E-06 | 7.000E-06 | 1.100E-05 |
| 22543 | 1128 | D1128N | D | 7.3e-06 | 9.4e-06 | 6.7e-06 | 7.800E-06 | 1.418E-06 | 6.700E-06 | 9.400E-06 |
| 22544 | 1128 | D1128D | D | 9.2e-06 | 4.7e-06 | 8.4e-06 | 7.433E-06 | 2.401E-06 | 4.700E-06 | 9.200E-06 |
| 22545 | 1128 | D1128C | D | 8.5e-06 | 7.3e-06 | 1.4e-05 | 9.933E-06 | 3.573E-06 | 7.300E-06 | 1.400E-05 |
| 22546 | 1128 | D1128E | D | 9e-06 | 6.8e-06 | 8.5e-06 | 8.100E-06 | 1.153E-06 | 6.800E-06 | 9.000E-06 |
| 22547 | 1128 | D1128Q | D | 5.6e-06 | 7.5e-06 | 5.2e-06 | 6.100E-06 | 1.229E-06 | 5.200E-06 | 7.500E-06 |
| 22548 | 1128 | D1128G | D | 1.1e-05 | 6.7e-06 | 1.3e-05 | 1.023E-05 | 3.219E-06 | 6.700E-06 | 1.300E-05 |
| 22549 | 1128 | D1128H | D | 7.6e-06 | 1.1e-05 | 8.8e-06 | 9.133E-06 | 1.724E-06 | 7.600E-06 | 1.100E-05 |
| 22550 | 1128 | D1128I | D | 6.3e-06 | 6.9e-06 | 7.6e-06 | 6.933E-06 | 6.506E-07 | 6.300E-06 | 7.600E-06 |
| 22551 | 1128 | D1128L | D | 1e-05 | 7.1e-06 | 8.5e-06 | 8.533E-06 | 1.450E-06 | 7.100E-06 | 1.000E-05 |
| 22552 | 1128 | D1128K | D | 9.4e-06 | 6.7e-06 | 9.8e-06 | 8.633E-06 | 1.686E-06 | 6.700E-06 | 9.800E-06 |
| 22553 | 1128 | D1128M | D | 1.5e-05 | 7.6e-06 | 8.7e-06 | 1.043E-05 | 3.993E-06 | 7.600E-06 | 1.500E-05 |
| 22554 | 1128 | D1128F | D | 4.5e-06 | 1.1e-05 | 6.5e-06 | 7.333E-06 | 3.329E-06 | 4.500E-06 | 1.100E-05 |
| 22555 | 1128 | D1128P | D | 7.2e-06 | 6.4e-06 | 6e-06 | 6.533E-06 | 6.110E-07 | 6.000E-06 | 7.200E-06 |
| 22556 | 1128 | D1128S | D | 1e-05 | 5.6e-06 | 8.1e-06 | 7.900E-06 | 2.207E-06 | 5.600E-06 | 1.000E-05 |
| 22557 | 1128 | D1128T | D | 7.2e-06 | 6.7e-06 | 6e-06 | 6.633E-06 | 6.028E-07 | 6.000E-06 | 7.200E-06 |
| 22558 | 1128 | D1128W | D | 7.4e-06 | 6.3e-06 | 7.9e-06 | 7.200E-06 | 8.185E-07 | 6.300E-06 | 7.900E-06 |
| 22559 | 1128 | D1128Y | D | 9.1e-06 | 6.5e-06 | 3.7e-06 | 6.433E-06 | 2.701E-06 | 3.700E-06 | 9.100E-06 |
| 22560 | 1128 | D1128V | D | 6.3e-06 | 6.3e-06 | 1e-05 | 7.533E-06 | 2.136E-06 | 6.300E-06 | 1.000E-05 |
| 22561 | 1129 | F1129A | D | 5.4e-06 | 8.9e-06 | 8.6e-06 | 7.633E-06 | 1.940E-06 | 5.400E-06 | 8.900E-06 |
| 22562 | 1129 | F1129R | D | 1e-05 | 6e-06 | 6.8e-06 | 7.600E-06 | 2.117E-06 | 6.000E-06 | 1.000E-05 |
| 22563 | 1129 | F1129N | D | 5.6e-06 | 7e-06 | 5.1e-06 | 5.900E-06 | 9.849E-07 | 5.100E-06 | 7.000E-06 |
| 22564 | 1129 | F1129D | D | 6.7e-06 | 9.8e-06 | 6.3e-06 | 7.600E-06 | 1.916E-06 | 6.300E-06 | 9.800E-06 |
| 22565 | 1129 | F1129C | D | 1.2e-05 | 8.3e-06 | 9.4e-06 | 9.900E-06 | 1.900E-06 | 8.300E-06 | 1.200E-05 |
| 22566 | 1129 | F1129E | D | 9.3e-06 | 9.3e-06 | 8.7e-06 | 9.100E-06 | 3.464E-07 | 8.700E-06 | 9.300E-06 |
| 22567 | 1129 | F1129Q | D | 5.2e-06 | 7.1e-06 | 6.9e-06 | 6.400E-06 | 1.044E-06 | 5.200E-06 | 7.100E-06 |
| 22568 | 1129 | F1129G | D | 7.3e-06 | 7.3e-06 | 8.7e-06 | 7.767E-06 | 8.083E-07 | 7.300E-06 | 8.700E-06 |
| 22569 | 1129 | F1129H | D | 6.3e-06 | 1.3e-05 | 9.5e-06 | 9.600E-06 | 3.351E-06 | 6.300E-06 | 1.300E-05 |
| 22570 | 1129 | F1129I | D | 1.4e-05 | 1.4e-05 | 8.4e-06 | 1.213E-05 | 3.233E-06 | 8.400E-06 | 1.400E-05 |
| 22571 | 1129 | F1129L | D | 5.3e-06 | 5.7e-06 | 7.2e-06 | 6.067E-06 | 1.002E-06 | 5.300E-06 | 7.200E-06 |
| 22572 | 1129 | F1129K | D | 6.6e-06 | 7.5e-06 | 6.5e-06 | 6.867E-06 | 5.508E-07 | 6.500E-06 | 7.500E-06 |
| 22573 | 1129 | F1129M | D | 4.6e-06 | 5.5e-06 | 6.6e-06 | 5.567E-06 | 1.002E-06 | 4.600E-06 | 6.600E-06 |
| 22574 | 1129 | F1129F | D | 5.8e-06 | 5.3e-06 | 8.4e-06 | 6.500E-06 | 1.664E-06 | 5.300E-06 | 8.400E-06 |
| 22575 | 1129 | F1129P | D | 1.1e-05 | 9e-06 | 1.1e-05 | 1.033E-05 | 1.155E-06 | 9.000E-06 | 1.100E-05 |
| 22576 | 1129 | F1129S | D | 9.8e-06 | 9.1e-06 | 6.6e-06 | 8.500E-06 | 1.682E-06 | 6.600E-06 | 9.800E-06 |
| 22577 | 1129 | F1129T | D | 6.2e-06 | 1e-05 | 1.1e-05 | 9.067E-06 | 2.532E-06 | 6.200E-06 | 1.100E-05 |
| 22578 | 1129 | F1129W | D | 3.9e-06 | 9.3e-06 | 9.3e-06 | 7.500E-06 | 3.118E-06 | 3.900E-06 | 9.300E-06 |
| 22579 | 1129 | F1129Y | D | 7.8e-06 | 6.8e-06 | 7.1e-06 | 7.233E-06 | 5.132E-07 | 6.800E-06 | 7.800E-06 |
| 22580 | 1129 | F1129V | D | 9.3e-06 | 9.3e-06 | 9e-06 | 9.200E-06 | 1.732E-07 | 9.000E-06 | 9.300E-06 |
| 22581 | 1130 | T1130A | D | 8e-06 | 4.8e-06 | 4.9e-06 | 5.900E-06 | 1.819E-06 | 4.800E-06 | 8.000E-06 |
| 22582 | 1130 | T1130R | D | 7.1e-06 | 9.2e-06 | 6.9e-06 | 7.733E-06 | 1.274E-06 | 6.900E-06 | 9.200E-06 |
| 22583 | 1130 | T1130N | D | 8.6e-06 | 7.7e-06 | 1.1e-05 | 9.100E-06 | 1.706E-06 | 7.700E-06 | 1.100E-05 |
| 22584 | 1130 | T1130D | D | 8.6e-06 | 6.7e-06 | 4.4e-06 | 6.567E-06 | 2.103E-06 | 4.400E-06 | 8.600E-06 |
| 22585 | 1130 | T1130C | D | 6.7e-06 | 1.3e-05 | 9.6e-06 | 9.767E-06 | 3.153E-06 | 6.700E-06 | 1.300E-05 |
| 22586 | 1130 | T1130E | D | 6.8e-06 | 8.5e-06 | 1.1e-05 | 8.767E-06 | 2.113E-06 | 6.800E-06 | 1.100E-05 |
| 22587 | 1130 | T1130Q | D | 1e-05 | 1.2e-05 | 7.4e-06 | 9.800E-06 | 2.307E-06 | 7.400E-06 | 1.200E-05 |
| 22588 | 1130 | T1130G | D | 6.2e-06 | 1.2e-05 | 1.2e-05 | 1.007E-05 | 3.349E-06 | 6.200E-06 | 1.200E-05 |
| 22589 | 1130 | T1130H | D | 1.3e-05 | 4.4e-06 | 1e-05 | 9.133E-06 | 4.365E-06 | 4.400E-06 | 1.300E-05 |
| 22590 | 1130 | T1130I | D | 9.2e-06 | 5.5e-06 | 8.4e-06 | 7.700E-06 | 1.947E-06 | 5.500E-06 | 9.200E-06 |
| 22591 | 1130 | T1130L | D | 9.3e-06 | 7.8e-06 | 5.7e-06 | 7.600E-06 | 1.808E-06 | 5.700E-06 | 9.300E-06 |
| 22592 | 1130 | T1130K | D | 1.1e-05 | 6.7e-06 | 8.2e-06 | 8.633E-06 | 2.183E-06 | 6.700E-06 | 1.100E-05 |
| 22593 | 1130 | T1130M | D | 6.5e-06 | 4.8e-06 | 6.9e-06 | 6.067E-06 | 1.115E-06 | 4.800E-06 | 6.900E-06 |
| 22594 | 1130 | T1130F | D | 1e-05 | 4.8e-06 | 9.7e-06 | 8.167E-06 | 2.919E-06 | 4.800E-06 | 1.000E-05 |
| 22595 | 1130 | T1130P | D | 1e-05 | 7e-06 | 6.2e-06 | 7.733E-06 | 2.003E-06 | 6.200E-06 | 1.000E-05 |
| 22596 | 1130 | T1130S | D | 7.8e-06 | 4.4e-06 | 1.3e-05 | 8.400E-06 | 4.331E-06 | 4.400E-06 | 1.300E-05 |
| 22597 | 1130 | T1130T | D | 5.5e-06 | 6.6e-06 | 1.1e-05 | 7.700E-06 | 2.910E-06 | 5.500E-06 | 1.100E-05 |
| 22598 | 1130 | T1130W | D | 1.4e-05 | 7.1e-06 | 3.9e-06 | 8.333E-06 | 5.162E-06 | 3.900E-06 | 1.400E-05 |
| 22599 | 1130 | T1130Y | D | 6.8e-06 | 9.7e-06 | 6.3e-06 | 7.600E-06 | 1.836E-06 | 6.300E-06 | 9.700E-06 |
| 22600 | 1130 | T1130V | D | 1.3e-05 | 6.1e-06 | 4.5e-06 | 7.867E-06 | 4.517E-06 | 4.500E-06 | 1.300E-05 |
| 22601 | 1131 | L1131A | D | 7.4e-06 | 7.6e-06 | 8.9e-06 | 7.967E-06 | 8.145E-07 | 7.400E-06 | 8.900E-06 |
| 22602 | 1131 | L1131R | D | 1e-05 | 1e-05 | 2.1e-06 | 7.367E-06 | 4.561E-06 | 2.100E-06 | 1.000E-05 |
| 22603 | 1131 | L1131N | D | 4.1e-06 | 1e-05 | 8.2e-06 | 7.433E-06 | 3.024E-06 | 4.100E-06 | 1.000E-05 |
| 22604 | 1131 | L1131D | D | 6.2e-06 | 5.7e-06 | 1.4e-05 | 8.633E-06 | 4.654E-06 | 5.700E-06 | 1.400E-05 |
| 22605 | 1131 | L1131C | D | 1e-05 | 8.3e-06 | 9.9e-06 | 9.400E-06 | 9.539E-07 | 8.300E-06 | 1.000E-05 |
| 22606 | 1131 | L1131E | D | 4.1e-06 | 1.1e-05 | 9.6e-06 | 8.233E-06 | 3.647E-06 | 4.100E-06 | 1.100E-05 |
| 22607 | 1131 | L1131Q | D | 9e-06 | 7.5e-06 | 1.3e-05 | 9.833E-06 | 2.843E-06 | 7.500E-06 | 1.300E-05 |
| 22608 | 1131 | L1131G | D | 9.6e-06 | 5.1e-06 | 6.7e-06 | 7.133E-06 | 2.281E-06 | 5.100E-06 | 9.600E-06 |
| 22609 | 1131 | L1131H | D | 1.1e-05 | 6.4e-06 | 6.3e-06 | 7.900E-06 | 2.685E-06 | 6.300E-06 | 1.100E-05 |

| | | | | | | | | | | |
|---|---|---|---|---|---|---|---|---|---|---|
| 22610 | 1131 | L1131I | D | 6.1e-06 | 1.4e-05 | 1e-05 | 1.003E-05 | 3.950E-06 | 6.100E-06 | 1.400E-05 |
| 22611 | 1131 | L1131L | D | 7.1e-06 | 6.4e-06 | 9.1e-06 | 7.533E-06 | 1.401E-06 | 6.400E-06 | 9.100E-06 |
| 22612 | 1131 | L1131K | D | 1.3e-05 | 6.4e-06 | 5.4e-06 | 8.267E-06 | 4.130E-06 | 5.400E-06 | 1.300E-05 |
| 22613 | 1131 | L1131M | D | 5e-06 | 6.9e-06 | 1.2e-05 | 7.967E-06 | 3.620E-06 | 5.000E-06 | 1.200E-05 |
| 22614 | 1131 | L1131F | D | 7.9e-06 | 6.7e-06 | 1.2e-05 | 8.867E-06 | 2.779E-06 | 6.700E-06 | 1.200E-05 |
| 22615 | 1131 | L1131P | D | 7.9e-06 | 6.8e-06 | 1.3e-05 | 9.233E-06 | 3.308E-06 | 6.800E-06 | 1.300E-05 |
| 22616 | 1131 | L1131S | D | 7e-06 | 7.1e-06 | 9.6e-06 | 7.900E-06 | 1.473E-06 | 7.000E-06 | 9.600E-06 |
| 22617 | 1131 | L1131T | D | 9.3e-06 | 9.1e-06 | 9.7e-06 | 9.367E-06 | 3.055E-07 | 9.100E-06 | 9.700E-06 |
| 22618 | 1131 | L1131W | D | 8e-06 | 1e-05 | 5e-06 | 7.667E-06 | 2.517E-06 | 5.000E-06 | 1.000E-05 |
| 22619 | 1131 | L1131Y | D | 7.4e-06 | 3.8e-06 | 5.6e-06 | 5.600E-06 | 1.800E-06 | 3.800E-06 | 7.400E-06 |
| 22620 | 1131 | L1131V | D | 6e-06 | 7.5e-06 | 6e-06 | 6.500E-06 | 8.660E-07 | 6.000E-06 | 7.500E-06 |
| 22621 | 1132 | T1132A | D | 7.9e-06 | 1.2e-05 | 4.3e-06 | 8.067E-06 | 3.853E-06 | 4.300E-06 | 1.200E-05 |
| 22622 | 1132 | T1132R | D | 9.8e-06 | 9.2e-06 | 1e-05 | 9.667E-06 | 4.163E-07 | 9.200E-06 | 1.000E-05 |
| 22623 | 1132 | T1132N | D | 9.1e-06 | 7.2e-06 | 1.2e-05 | 9.433E-06 | 2.417E-06 | 7.200E-06 | 1.200E-05 |
| 22624 | 1132 | T1132D | D | 7.2e-06 | 5.9e-06 | 8.5e-06 | 7.200E-06 | 1.300E-06 | 5.900E-06 | 8.500E-06 |
| 22625 | 1132 | T1132C | D | 1.2e-05 | 5.8e-06 | 9.2e-06 | 9.000E-06 | 3.105E-06 | 5.800E-06 | 1.200E-05 |
| 22626 | 1132 | T1132E | D | 9.5e-06 | 7.2e-06 | 6.1e-06 | 7.600E-06 | 1.735E-06 | 6.100E-06 | 9.500E-06 |
| 22627 | 1132 | T1132Q | D | 1e-05 | 4.2e-06 | 1.3e-05 | 9.067E-06 | 4.474E-06 | 4.200E-06 | 1.300E-05 |
| 22628 | 1132 | T1132G | D | 1.3e-05 | 9e-06 | 7.7e-06 | 9.900E-06 | 2.762E-06 | 7.700E-06 | 1.300E-05 |
| 22629 | 1132 | T1132H | D | 4.4e-06 | 9.6e-06 | 6e-06 | 6.667E-06 | 2.663E-06 | 4.400E-06 | 9.600E-06 |
| 22630 | 1132 | T1132I | D | 1e-05 | 9.7e-06 | 7.7e-06 | 9.133E-06 | 1.250E-06 | 7.700E-06 | 1.000E-05 |
| 22631 | 1132 | T1132L | D | 9e-06 | 8.5e-06 | 1.3e-05 | 1.017E-05 | 2.466E-06 | 8.500E-06 | 1.300E-05 |
| 22632 | 1132 | T1132K | D | 8.1e-06 | 1e-05 | 8.8e-06 | 8.967E-06 | 9.609E-07 | 8.100E-06 | 1.000E-05 |
| 22633 | 1132 | T1132M | D | 1.4e-05 | 6e-06 | 5.8e-06 | 8.600E-06 | 4.678E-06 | 5.800E-06 | 1.400E-05 |
| 22634 | 1132 | T1132F | D | 4.6e-06 | 5.6e-06 | 1e-05 | 6.733E-06 | 2.873E-06 | 4.600E-06 | 1.000E-05 |
| 22635 | 1132 | T1132P | D | 4.3e-06 | 7.3e-06 | 6.5e-06 | 6.033E-06 | 1.553E-06 | 4.300E-06 | 7.300E-06 |
| 22636 | 1132 | T1132S | D | 8.7e-06 | 7.7e-06 | 6.2e-06 | 7.533E-06 | 1.258E-06 | 6.200E-06 | 8.700E-06 |
| 22637 | 1132 | T1132T | D | 9.6e-06 | 9.5e-06 | 8.1e-06 | 9.067E-06 | 8.386E-07 | 8.100E-06 | 9.600E-06 |
| 22638 | 1132 | T1132W | D | 8.6e-06 | 9.6e-06 | 5.8e-06 | 8.000E-06 | 1.970E-06 | 5.800E-06 | 9.600E-06 |
| 22639 | 1132 | T1132Y | D | 1.2e-05 | 4.4e-06 | 7.4e-06 | 7.933E-06 | 3.828E-06 | 4.400E-06 | 1.200E-05 |
| 22640 | 1132 | T1132V | D | 9.7e-06 | 1e-05 | 9.2e-06 | 9.633E-06 | 4.041E-07 | 9.200E-06 | 1.000E-05 |
| 22641 | 1133 | I1133A | D | 7.1e-06 | 5.8e-06 | 6.5e-06 | 6.467E-06 | 6.506E-07 | 5.800E-06 | 7.100E-06 |
| 22642 | 1133 | I1133R | D | 3.8e-06 | 6.2e-06 | 6.5e-06 | 5.500E-06 | 1.480E-06 | 3.800E-06 | 6.500E-06 |
| 22643 | 1133 | I1133N | D | 9.9e-06 | 1.1e-05 | 4.7e-06 | 8.533E-06 | 3.365E-06 | 4.700E-06 | 1.100E-05 |
| 22644 | 1133 | I1133D | D | 6.5e-06 | 1.2e-05 | 1.6e-05 | 1.150E-05 | 4.770E-06 | 6.500E-06 | 1.600E-05 |
| 22645 | 1133 | I1133C | D | 1e-05 | 7.1e-06 | 9.7e-06 | 8.933E-06 | 1.595E-06 | 7.100E-06 | 1.000E-05 |
| 22646 | 1133 | I1133E | D | 5.2e-06 | 9.9e-06 | 7.8e-06 | 7.633E-06 | 2.354E-06 | 5.200E-06 | 9.900E-06 |
| 22647 | 1133 | I1133Q | D | 3.7e-06 | 7.2e-06 | 6.6e-06 | 5.833E-06 | 1.872E-06 | 3.700E-06 | 7.200E-06 |
| 22648 | 1133 | I1133G | D | 7.6e-06 | 1.2e-05 | 8.1e-06 | 9.233E-06 | 2.409E-06 | 7.600E-06 | 1.200E-05 |
| 22649 | 1133 | I1133H | D | 7.3e-06 | 6.9e-06 | 7.2e-06 | 7.133E-06 | 2.082E-07 | 6.900E-06 | 7.300E-06 |
| 22650 | 1133 | I1133I | D | 6.5e-06 | 9.6e-06 | 5.1e-06 | 7.067E-06 | 2.303E-06 | 5.100E-06 | 9.600E-06 |
| 22651 | 1133 | I1133L | D | 7.5e-06 | 5.9e-06 | 5.3e-06 | 6.233E-06 | 1.137E-06 | 5.300E-06 | 7.500E-06 |
| 22652 | 1133 | I1133K | D | 5.4e-06 | 5.9e-06 | 8.3e-06 | 6.533E-06 | 1.550E-06 | 5.400E-06 | 8.300E-06 |
| 22653 | 1133 | I1133M | D | 5.5e-06 | 7.7e-06 | 6.2e-06 | 6.467E-06 | 1.124E-06 | 5.500E-06 | 7.700E-06 |
| 22654 | 1133 | I1133F | D | 7.6e-06 | 4.3e-06 | 6.8e-06 | 6.233E-06 | 1.721E-06 | 4.300E-06 | 7.600E-06 |
| 22655 | 1133 | I1133P | D | 6.5e-06 | 4.4e-06 | 8.6e-06 | 6.500E-06 | 2.100E-06 | 4.400E-06 | 8.600E-06 |
| 22656 | 1133 | I1133S | D | 6.9e-06 | 4.7e-06 | 1.3e-05 | 8.200E-06 | 4.300E-06 | 4.700E-06 | 1.300E-05 |
| 22657 | 1133 | I1133T | D | 6.6e-06 | 9.6e-06 | 7.8e-06 | 8.000E-06 | 1.510E-06 | 6.600E-06 | 9.600E-06 |
| 22658 | 1133 | I1133W | D | 7.2e-06 | 4.5e-06 | 8.9e-06 | 6.867E-06 | 2.219E-06 | 4.500E-06 | 8.900E-06 |
| 22659 | 1133 | I1133Y | D | 8.8e-06 | 7.5e-06 | 6.9e-06 | 7.733E-06 | 9.713E-07 | 6.900E-06 | 8.800E-06 |
| 22660 | 1133 | I1133V | D | 1e-05 | 1.4e-05 | 6.1e-06 | 1.003E-05 | 3.950E-06 | 6.100E-06 | 1.400E-05 |
| 22661 | 1134 | S1134A | D | 8.5e-06 | 8.2e-06 | 1e-05 | 8.900E-06 | 9.644E-07 | 8.200E-06 | 1.000E-05 |
| 22662 | 1134 | S1134R | D | 5.5e-06 | 5.3e-06 | 8.9e-06 | 6.567E-06 | 2.023E-06 | 5.300E-06 | 8.900E-06 |
| 22663 | 1134 | S1134N | D | 7.5e-06 | 1.3e-05 | 4.3e-06 | 8.267E-06 | 4.400E-06 | 4.300E-06 | 1.300E-05 |
| 22664 | 1134 | S1134D | D | 7.1e-06 | 5.1e-06 | 1.3e-05 | 8.400E-06 | 4.107E-06 | 5.100E-06 | 1.300E-05 |
| 22665 | 1134 | S1134C | D | 8.6e-06 | 1.4e-05 | 1.2e-05 | 1.153E-05 | 2.730E-06 | 8.600E-06 | 1.400E-05 |
| 22666 | 1134 | S1134E | D | 1.3e-05 | 6.1e-06 | 1.1e-05 | 1.003E-05 | 3.550E-06 | 6.100E-06 | 1.300E-05 |
| 22667 | 1134 | S1134Q | D | 9.3e-06 | 7.4e-06 | 9.9e-06 | 8.867E-06 | 1.305E-06 | 7.400E-06 | 9.900E-06 |
| 22668 | 1134 | S1134G | D | 6e-06 | 9.6e-06 | 9.3e-06 | 8.300E-06 | 1.997E-06 | 6.000E-06 | 9.600E-06 |
| 22669 | 1134 | S1134H | D | 1e-05 | 1.2e-05 | 1.7e-05 | 1.300E-05 | 3.606E-06 | 1.000E-05 | 1.700E-05 |
| 22670 | 1134 | S1134I | D | 5.4e-06 | 1.1e-05 | 5e-06 | 7.133E-06 | 3.355E-06 | 5.000E-06 | 1.100E-05 |
| 22671 | 1134 | S1134L | D | 9.7e-06 | 5.5e-06 | 5.9e-06 | 7.033E-06 | 2.318E-06 | 5.500E-06 | 9.700E-06 |
| 22672 | 1134 | S1134K | D | 6.1e-06 | 1.9e-05 | 8e-06 | 1.103E-05 | 6.964E-06 | 6.100E-06 | 1.900E-05 |
| 22673 | 1134 | S1134M | D | 7.8e-06 | 7.5e-06 | 7.1e-06 | 7.467E-06 | 3.512E-07 | 7.100E-06 | 7.800E-06 |
| 22674 | 1134 | S1134F | D | 8.7e-06 | 8.4e-06 | 4.9e-06 | 7.333E-06 | 2.113E-06 | 4.900E-06 | 8.700E-06 |
| 22675 | 1134 | S1134P | D | 8.5e-06 | 6.6e-06 | 3.6e-06 | 6.233E-06 | 2.470E-06 | 3.600E-06 | 8.500E-06 |
| 22676 | 1134 | S1134S | D | 6.5e-06 | 1e-05 | 1.2e-05 | 9.500E-06 | 2.784E-06 | 6.500E-06 | 1.200E-05 |
| 22677 | 1134 | S1134T | D | 7.8e-06 | 3.4e-06 | 7.2e-06 | 6.133E-06 | 2.386E-06 | 3.400E-06 | 7.800E-06 |
| 22678 | 1134 | S1134W | D | 9.3e-06 | 7.6e-06 | 9.3e-06 | 8.733E-06 | 9.815E-07 | 7.600E-06 | 9.300E-06 |
| 22679 | 1134 | S1134Y | D | 6.3e-06 | 9.1e-06 | 5.1e-06 | 6.833E-06 | 2.053E-06 | 5.100E-06 | 9.100E-06 |
| 22680 | 1134 | S1134V | D | 9.8e-06 | 9.2e-06 | 7.3e-06 | 8.767E-06 | 1.305E-06 | 7.300E-06 | 9.800E-06 |
| 22681 | 1135 | S1135A | D | 4.8e-06 | 5.3e-06 | 8.2e-06 | 6.100E-06 | 1.836E-06 | 4.800E-06 | 8.200E-06 |
| 22682 | 1135 | S1135R | D | 8.9e-06 | 9.1e-06 | 8e-06 | 8.667E-06 | 5.859E-07 | 8.000E-06 | 9.100E-06 |
| 22683 | 1135 | S1135N | D | 5.3e-06 | 1.1e-05 | 1.3e-05 | 9.767E-06 | 3.995E-06 | 5.300E-06 | 1.300E-05 |
| 22684 | 1135 | S1135D | D | 1.1e-05 | 5e-06 | 1.5e-05 | 1.033E-05 | 5.033E-06 | 5.000E-06 | 1.500E-05 |
| 22685 | 1135 | S1135C | D | 6.4e-06 | 5.8e-06 | 1.4e-05 | 8.733E-06 | 4.571E-06 | 5.800E-06 | 1.400E-05 |
| 22686 | 1135 | S1135E | D | 5.5e-06 | 5.5e-06 | 5e-06 | 5.333E-06 | 2.887E-07 | 5.000E-06 | 5.500E-06 |
| 22687 | 1135 | S1135Q | D | 5.5e-06 | 5.9e-06 | 1e-05 | 7.133E-06 | 2.491E-06 | 5.500E-06 | 1.000E-05 |
| 22688 | 1135 | S1135G | D | 8.6e-06 | 1.4e-05 | 6.7e-06 | 9.767E-06 | 3.787E-06 | 6.700E-06 | 1.400E-05 |
| 22689 | 1135 | S1135H | D | 6.1e-06 | 7.9e-06 | 6.2e-06 | 6.733E-06 | 1.012E-06 | 6.100E-06 | 7.900E-06 |
| 22690 | 1135 | S1135I | D | 5.5e-06 | 1.2e-05 | 4.8e-06 | 7.433E-06 | 3.970E-06 | 4.800E-06 | 1.200E-05 |
| 22691 | 1135 | S1135L | D | 5.4e-06 | 6.7e-06 | 1.4e-05 | 8.700E-06 | 4.636E-06 | 5.400E-06 | 1.400E-05 |
| 22692 | 1135 | S1135K | D | 9.7e-06 | 9.5e-06 | 5.9e-06 | 8.367E-06 | 2.139E-06 | 5.900E-06 | 9.700E-06 |
| 22693 | 1135 | S1135M | D | 1e-05 | 6.6e-06 | 7.9e-06 | 8.167E-06 | 1.716E-06 | 6.600E-06 | 1.000E-05 |
| 22694 | 1135 | S1135F | D | 8.7e-06 | 9.4e-06 | 4.8e-06 | 7.633E-06 | 2.479E-06 | 4.800E-06 | 9.400E-06 |

| | | | | | | | | | | |
|---|---|---|---|---|---|---|---|---|---|---|
| 22695 | 1135 | S1135P | D | 7e-06 | 9.6e-06 | 9.2e-06 | 8.600E-06 | 1.400E-06 | 7.000E-06 | 9.600E-06 |
| 22696 | 1135 | S1135S | D | 7.7e-06 | 4.6e-06 | 4.9e-06 | 5.733E-06 | 1.710E-06 | 4.600E-06 | 7.700E-06 |
| 22697 | 1135 | S1135T | D | 1.1e-05 | 7.2e-06 | 9.4e-06 | 9.200E-06 | 1.908E-06 | 7.200E-06 | 1.100E-05 |
| 22698 | 1135 | S1135W | D | 7.3e-06 | 4.8e-06 | 4.2e-06 | 5.433E-06 | 1.644E-06 | 4.200E-06 | 7.300E-06 |
| 22699 | 1135 | S1135Y | D | 4.9e-06 | 7.8e-06 | 1.2e-05 | 8.233E-06 | 3.570E-06 | 4.900E-06 | 1.200E-05 |
| 22700 | 1135 | S1135V | D | 1e-05 | 4.9e-06 | 3.9e-06 | 6.267E-06 | 3.272E-06 | 3.900E-06 | 1.000E-05 |
| 22701 | 1136 | L1136A | D | 7.7e-06 | 1.1e-05 | 1e-05 | 9.567E-06 | 1.692E-06 | 7.700E-06 | 1.100E-05 |
| 22702 | 1136 | L1136R | D | 8.3e-06 | 7.5e-06 | 8.5e-06 | 8.100E-06 | 5.292E-07 | 7.500E-06 | 8.500E-06 |
| 22703 | 1136 | L1136N | D | 1.3e-05 | 6.5e-06 | 8.5e-06 | 9.333E-06 | 3.329E-06 | 6.500E-06 | 1.300E-05 |
| 22704 | 1136 | L1136D | D | 8.1e-06 | 5.3e-06 | 1.3e-05 | 8.800E-06 | 3.897E-06 | 5.300E-06 | 1.300E-05 |
| 22705 | 1136 | L1136C | D | 9.3e-06 | 4.9e-06 | 1.2e-05 | 8.733E-06 | 3.584E-06 | 4.900E-06 | 1.200E-05 |
| 22706 | 1136 | L1136E | D | 8.7e-06 | 7.1e-06 | 1e-05 | 8.600E-06 | 1.453E-06 | 7.100E-06 | 1.000E-05 |
| 22707 | 1136 | L1136Q | D | 4.8e-06 | 6.9e-06 | 6.6e-06 | 6.100E-06 | 1.136E-06 | 4.800E-06 | 6.900E-06 |
| 22708 | 1136 | L1136G | D | 6.1e-06 | 7.7e-06 | 6.5e-06 | 6.767E-06 | 8.327E-07 | 6.100E-06 | 7.700E-06 |
| 22709 | 1136 | L1136H | D | 9.2e-06 | 8.2e-06 | 6.4e-06 | 7.933E-06 | 1.419E-06 | 6.400E-06 | 9.200E-06 |
| 22710 | 1136 | L1136I | D | 1.3e-05 | 6.4e-06 | 1.3e-05 | 1.080E-05 | 3.811E-06 | 6.400E-06 | 1.300E-05 |
| 22711 | 1136 | L1136L | D | 4.4e-06 | 8.3e-06 | 5.1e-06 | 5.933E-06 | 2.079E-06 | 4.400E-06 | 8.300E-06 |
| 22712 | 1136 | L1136K | D | 1e-05 | 6.9e-06 | 1.1e-05 | 9.300E-06 | 2.138E-06 | 6.900E-06 | 1.100E-05 |
| 22713 | 1136 | L1136M | D | 7.2e-06 | 7.1e-06 | 5.9e-06 | 6.733E-06 | 7.234E-07 | 5.900E-06 | 7.200E-06 |
| 22714 | 1136 | L1136F | D | 5e-06 | 4.5e-06 | 6.6e-06 | 5.367E-06 | 1.097E-06 | 4.500E-06 | 6.600E-06 |
| 22715 | 1136 | L1136P | D | 1e-05 | 6.9e-06 | 9.7e-06 | 8.867E-06 | 1.710E-06 | 6.900E-06 | 1.000E-05 |
| 22716 | 1136 | L1136S | D | 1.1e-05 | 7.3e-06 | 4.5e-06 | 7.600E-06 | 3.260E-06 | 4.500E-06 | 1.100E-05 |
| 22717 | 1136 | L1136T | D | 9.6e-06 | 5.6e-06 | 8.2e-06 | 7.800E-06 | 2.030E-06 | 5.600E-06 | 9.600E-06 |
| 22718 | 1136 | L1136W | D | 8.6e-06 | 6.4e-06 | 7e-06 | 7.333E-06 | 1.137E-06 | 6.400E-06 | 8.600E-06 |
| 22719 | 1136 | L1136Y | D | 4.9e-06 | 6.6e-06 | 1e-05 | 7.167E-06 | 2.597E-06 | 4.900E-06 | 1.000E-05 |
| 22720 | 1136 | L1136V | D | 5.6e-06 | 7.2e-06 | 1.4e-05 | 8.933E-06 | 4.460E-06 | 5.600E-06 | 1.400E-05 |
| 22721 | 1137 | Q1137A | D | 5.1e-06 | 9e-06 | 9.4e-06 | 7.833E-06 | 2.376E-06 | 5.100E-06 | 9.400E-06 |
| 22722 | 1137 | Q1137R | D | 6.9e-06 | 1e-05 | 1e-05 | 8.967E-06 | 1.790E-06 | 6.900E-06 | 1.000E-05 |
| 22723 | 1137 | Q1137N | D | 9.3e-06 | 9.1e-06 | 6.3e-06 | 8.233E-06 | 1.677E-06 | 6.300E-06 | 9.300E-06 |
| 22724 | 1137 | Q1137D | D | 1e-05 | 1.7e-05 | 8.4e-06 | 1.180E-05 | 4.574E-06 | 8.400E-06 | 1.700E-05 |
| 22725 | 1137 | Q1137C | D | 8.2e-06 | 6.6e-06 | 9.5e-06 | 8.100E-06 | 1.453E-06 | 6.600E-06 | 9.500E-06 |
| 22726 | 1137 | Q1137E | D | 9.8e-06 | 1e-05 | 7.5e-06 | 9.100E-06 | 1.389E-06 | 7.500E-06 | 1.000E-05 |
| 22727 | 1137 | Q1137Q | D | 7.2e-06 | 8.6e-06 | 9.4e-06 | 8.400E-06 | 1.114E-06 | 7.200E-06 | 9.400E-06 |
| 22728 | 1137 | Q1137G | D | 5.8e-06 | 8.7e-06 | 7.2e-06 | 7.233E-06 | 1.450E-06 | 5.800E-06 | 8.700E-06 |
| 22729 | 1137 | Q1137H | D | 1.2e-05 | 7.6e-06 | 7e-06 | 8.867E-06 | 2.730E-06 | 7.000E-06 | 1.200E-05 |
| 22730 | 1137 | Q1137I | D | 1e-05 | 9.1e-06 | 1.4e-05 | 1.103E-05 | 2.608E-06 | 9.100E-06 | 1.400E-05 |
| 22731 | 1137 | Q1137L | D | 9.2e-06 | 6.3e-06 | 1e-05 | 8.500E-06 | 1.947E-06 | 6.300E-06 | 1.000E-05 |
| 22732 | 1137 | Q1137K | D | 8.7e-06 | 5.4e-06 | 6.4e-06 | 6.833E-06 | 1.692E-06 | 5.400E-06 | 8.700E-06 |
| 22733 | 1137 | Q1137M | D | 7.5e-06 | 7.4e-06 | 5e-06 | 6.633E-06 | 1.415E-06 | 5.000E-06 | 7.500E-06 |
| 22734 | 1137 | Q1137F | D | 6.4e-06 | 8.2e-06 | 9.3e-06 | 7.967E-06 | 1.464E-06 | 6.400E-06 | 9.300E-06 |
| 22735 | 1137 | Q1137P | D | 6.3e-06 | 4.2e-06 | 1.2e-05 | 7.500E-06 | 4.036E-06 | 4.200E-06 | 1.200E-05 |
| 22736 | 1137 | Q1137S | D | 5.2e-06 | 8.8e-06 | 9.3e-06 | 7.767E-06 | 2.237E-06 | 5.200E-06 | 9.300E-06 |
| 22737 | 1137 | Q1137T | D | 1.7e-05 | 8e-06 | 5.3e-06 | 1.010E-05 | 6.126E-06 | 5.300E-06 | 1.700E-05 |
| 22738 | 1137 | Q1137W | D | 3.9e-06 | 8.1e-06 | 1.4e-05 | 8.667E-06 | 5.074E-06 | 3.900E-06 | 1.400E-05 |
| 22739 | 1137 | Q1137Y | D | 6.7e-06 | 6.8e-06 | 5.2e-06 | 6.233E-06 | 8.963E-07 | 5.200E-06 | 6.800E-06 |
| 22740 | 1137 | Q1137V | D | 9.7e-06 | 1.3e-05 | 1.4e-05 | 1.223E-05 | 2.250E-06 | 9.700E-06 | 1.400E-05 |
| 22741 | 1138 | P1138A | D | 7.3e-06 | 6.6e-06 | 1.3e-05 | 8.967E-06 | 3.510E-06 | 6.600E-06 | 1.300E-05 |
| 22742 | 1138 | P1138R | D | 9.4e-06 | 1.3e-05 | 8.1e-06 | 1.017E-05 | 2.538E-06 | 8.100E-06 | 1.300E-05 |
| 22743 | 1138 | P1138N | D | 5.8e-06 | 5.8e-06 | 6.1e-06 | 5.900E-06 | 1.732E-07 | 5.800E-06 | 6.100E-06 |
| 22744 | 1138 | P1138D | D | 4.9e-06 | 9.2e-06 | 4.6e-06 | 6.233E-06 | 2.574E-06 | 4.600E-06 | 9.200E-06 |
| 22745 | 1138 | P1138C | D | 1.1e-05 | 5.5e-06 | 5.3e-06 | 7.267E-06 | 3.235E-06 | 5.300E-06 | 1.100E-05 |
| 22746 | 1138 | P1138E | D | 4.8e-06 | 1e-05 | 5.8e-06 | 6.867E-06 | 2.759E-06 | 4.800E-06 | 1.000E-05 |
| 22747 | 1138 | P1138Q | D | 6.8e-06 | 7.1e-06 | 4.5e-06 | 6.133E-06 | 1.422E-06 | 4.500E-06 | 7.100E-06 |
| 22748 | 1138 | P1138G | D | 5.4e-06 | 6.5e-06 | 1.2e-05 | 7.967E-06 | 3.536E-06 | 5.400E-06 | 1.200E-05 |
| 22749 | 1138 | P1138H | D | 7.7e-06 | 5.9e-06 | 5.2e-06 | 6.267E-06 | 1.290E-06 | 5.200E-06 | 7.700E-06 |
| 22750 | 1138 | P1138I | D | 6.2e-06 | 9.3e-06 | 9.6e-06 | 8.367E-06 | 1.882E-06 | 6.200E-06 | 9.600E-06 |
| 22751 | 1138 | P1138L | D | 1.1e-05 | 1e-05 | 9.2e-06 | 1.007E-05 | 9.018E-07 | 9.200E-06 | 1.100E-05 |
| 22752 | 1138 | P1138K | D | 6.9e-06 | 1e-05 | 9.7e-06 | 8.867E-06 | 1.710E-06 | 6.900E-06 | 1.000E-05 |
| 22753 | 1138 | P1138M | D | 9.3e-06 | 9.7e-06 | 7.1e-06 | 8.700E-06 | 1.400E-06 | 7.100E-06 | 9.700E-06 |
| 22754 | 1138 | P1138F | D | 6.9e-06 | 6.7e-06 | 6.7e-06 | 6.767E-06 | 1.155E-07 | 6.700E-06 | 6.900E-06 |
| 22755 | 1138 | P1138P | D | 5.9e-06 | 7.1e-06 | 9.9e-06 | 7.633E-06 | 2.053E-06 | 5.900E-06 | 9.900E-06 |
| 22756 | 1138 | P1138S | D | 1e-05 | 1.2e-05 | 5.6e-06 | 9.200E-06 | 3.274E-06 | 5.600E-06 | 1.200E-05 |
| 22757 | 1138 | P1138T | D | 6.1e-06 | 1.2e-05 | 8.3e-06 | 8.800E-06 | 2.982E-06 | 6.100E-06 | 1.200E-05 |
| 22758 | 1138 | P1138W | D | 9e-06 | 1.1e-05 | 7.4e-06 | 9.133E-06 | 1.804E-06 | 7.400E-06 | 1.100E-05 |
| 22759 | 1138 | P1138Y | D | 6.2e-06 | 1e-05 | 7.3e-06 | 7.833E-06 | 1.955E-06 | 6.200E-06 | 1.000E-05 |
| 22760 | 1138 | P1138V | D | 8.3e-06 | 1.4e-05 | 8.5e-06 | 1.027E-05 | 3.235E-06 | 8.300E-06 | 1.400E-05 |
| 22761 | 1139 | E1139A | D | 9.3e-06 | 7.1e-06 | 6e-06 | 7.467E-06 | 1.680E-06 | 6.000E-06 | 9.300E-06 |
| 22762 | 1139 | E1139R | D | 9.7e-06 | 9e-06 | 6.7e-06 | 8.467E-06 | 1.570E-06 | 6.700E-06 | 9.700E-06 |
| 22763 | 1139 | E1139N | D | 6.5e-06 | 5.4e-06 | 6.1e-06 | 6.000E-06 | 5.568E-07 | 5.400E-06 | 6.500E-06 |
| 22764 | 1139 | E1139D | D | 5.7e-06 | 6.7e-06 | 6.4e-06 | 6.267E-06 | 5.132E-07 | 5.700E-06 | 6.700E-06 |
| 22765 | 1139 | E1139C | D | 8.5e-06 | 7.2e-06 | 1.2e-05 | 9.233E-06 | 2.483E-06 | 7.200E-06 | 1.200E-05 |
| 22766 | 1139 | E1139E | D | 1.3e-05 | 6.6e-06 | 8e-06 | 9.200E-06 | 3.365E-06 | 6.600E-06 | 1.300E-05 |
| 22767 | 1139 | E1139Q | D | 6e-06 | 6.8e-06 | 6e-06 | 6.267E-06 | 4.619E-07 | 6.000E-06 | 6.800E-06 |
| 22768 | 1139 | E1139G | D | 1.3e-05 | 9e-06 | 7.7e-06 | 9.900E-06 | 2.762E-06 | 7.700E-06 | 1.300E-05 |
| 22769 | 1139 | E1139H | D | 1.1e-05 | 7.5e-06 | 5.4e-06 | 7.967E-06 | 2.829E-06 | 5.400E-06 | 1.100E-05 |
| 22770 | 1139 | E1139I | D | 8.4e-06 | 1e-05 | 7e-06 | 8.467E-06 | 1.501E-06 | 7.000E-06 | 1.000E-05 |
| 22771 | 1139 | E1139L | D | 1.5e-05 | 1.3e-05 | 8.3e-06 | 1.210E-05 | 3.439E-06 | 8.300E-06 | 1.500E-05 |
| 22772 | 1139 | E1139K | D | 9e-06 | 5.2e-06 | 9.6e-06 | 7.933E-06 | 2.386E-06 | 5.200E-06 | 9.600E-06 |
| 22773 | 1139 | E1139M | D | 1.4e-05 | 6.8e-06 | 8.8e-06 | 9.867E-06 | 3.717E-06 | 6.800E-06 | 1.400E-05 |
| 22774 | 1139 | E1139F | D | 8e-06 | 7.4e-06 | 9.7e-06 | 8.367E-06 | 1.193E-06 | 7.400E-06 | 9.700E-06 |
| 22775 | 1139 | E1139P | D | 1.3e-05 | 7.2e-06 | 1e-05 | 1.007E-05 | 2.901E-06 | 7.200E-06 | 1.300E-05 |
| 22776 | 1139 | E1139S | D | 7.1e-06 | 9.9e-06 | 9.2e-06 | 8.733E-06 | 1.457E-06 | 7.100E-06 | 9.900E-06 |
| 22777 | 1139 | E1139T | D | 6.4e-06 | 8.1e-06 | 7.3e-06 | 7.267E-06 | 8.505E-07 | 6.400E-06 | 8.100E-06 |
| 22778 | 1139 | E1139W | D | 7.5e-06 | 6.9e-06 | 9.7e-06 | 8.033E-06 | 1.474E-06 | 6.900E-06 | 9.700E-06 |
| 22779 | 1139 | E1139Y | D | 4.9e-06 | 8.3e-06 | 1.2e-05 | 8.400E-06 | 3.551E-06 | 4.900E-06 | 1.200E-05 |

| | | | | | | | | | | |
|---|---|---|---|---|---|---|---|---|---|---|
| 22780 | 1139 | E1139V | D | 6.6e-06 | 9.1e-06 | 9.5e-06 | 8.400E-06 | 1.572E-06 | 6.600E-06 | 9.500E-06 |
| 22781 | 1140 | D1140A | D | 1.2e-05 | 7.4e-06 | 7e-06 | 8.800E-06 | 2.778E-06 | 7.000E-06 | 1.200E-05 |
| 22782 | 1140 | D1140R | D | 9.9e-06 | 6.4e-06 | 4.4e-06 | 6.900E-06 | 2.784E-06 | 4.400E-06 | 9.900E-06 |
| 22783 | 1140 | D1140N | D | 7.4e-06 | 5e-06 | 6.7e-06 | 6.367E-06 | 1.234E-06 | 5.000E-06 | 7.400E-06 |
| 22784 | 1140 | D1140D | D | 8.2e-06 | 9.3e-06 | 6.4e-06 | 7.967E-06 | 1.464E-06 | 6.400E-06 | 9.300E-06 |
| 22785 | 1140 | D1140C | D | 9e-06 | 1.9e-05 | 7.4e-06 | 1.180E-05 | 6.286E-06 | 7.400E-06 | 1.900E-05 |
| 22786 | 1140 | D1140E | D | 8.1e-06 | 6e-06 | 6e-06 | 6.700E-06 | 1.212E-06 | 6.000E-06 | 8.100E-06 |
| 22787 | 1140 | D1140Q | D | 8.8e-06 | 1.3e-05 | 5.6e-06 | 9.133E-06 | 3.711E-06 | 5.600E-06 | 1.300E-05 |
| 22788 | 1140 | D1140G | D | 4.5e-06 | 8.2e-06 | 8e-06 | 6.900E-06 | 2.081E-06 | 4.500E-06 | 8.200E-06 |
| 22789 | 1140 | D1140H | D | 8e-06 | 3.9e-06 | 5.7e-06 | 5.867E-06 | 2.055E-06 | 3.900E-06 | 8.000E-06 |
| 22790 | 1140 | D1140I | D | 7.4e-06 | 1e-05 | 8.1e-06 | 8.500E-06 | 1.345E-06 | 7.400E-06 | 1.000E-05 |
| 22791 | 1140 | D1140L | D | 9.8e-06 | 7.5e-06 | 7.7e-06 | 8.333E-06 | 1.274E-06 | 7.500E-06 | 9.800E-06 |
| 22792 | 1140 | D1140K | D | 9.5e-06 | 1.3e-05 | 1.4e-05 | 1.217E-05 | 2.363E-06 | 9.500E-06 | 1.400E-05 |
| 22793 | 1140 | D1140M | D | 1.3e-05 | 7.4e-06 | 5.5e-06 | 8.633E-06 | 3.899E-06 | 5.500E-06 | 1.300E-05 |
| 22794 | 1140 | D1140F | D | 8.9e-06 | 1e-05 | 5.6e-06 | 8.167E-06 | 2.290E-06 | 5.600E-06 | 1.000E-05 |
| 22795 | 1140 | D1140P | D | 5.6e-06 | 1.2e-05 | 8e-06 | 8.533E-06 | 3.233E-06 | 5.600E-06 | 1.200E-05 |
| 22796 | 1140 | D1140S | D | 7e-06 | 5.9e-06 | 1.4e-05 | 8.967E-06 | 4.394E-06 | 5.900E-06 | 1.400E-05 |
| 22797 | 1140 | D1140T | D | 8.1e-06 | 1e-05 | 9.2e-06 | 9.100E-06 | 9.539E-07 | 8.100E-06 | 1.000E-05 |
| 22798 | 1140 | D1140W | D | 4.2e-06 | 5.9e-06 | 4.7e-06 | 4.933E-06 | 8.737E-07 | 4.200E-06 | 5.900E-06 |
| 22799 | 1140 | D1140Y | D | 1e-05 | 6.5e-06 | 5.3e-06 | 7.267E-06 | 2.442E-06 | 5.300E-06 | 1.000E-05 |
| 22800 | 1140 | D1140V | D | 7.9e-06 | 7.8e-06 | 1.3e-05 | 9.567E-06 | 2.974E-06 | 7.800E-06 | 1.300E-05 |
| 22801 | 1141 | F1141A | D | 9.5e-06 | 8.8e-06 | 8.8e-06 | 9.033E-06 | 4.041E-07 | 8.800E-06 | 9.500E-06 |
| 22802 | 1141 | F1141R | D | 1.1e-05 | 4.7e-06 | 6.8e-06 | 7.500E-06 | 3.208E-06 | 4.700E-06 | 1.100E-05 |
| 22803 | 1141 | F1141N | D | 7e-06 | 7.2e-06 | 6.9e-06 | 7.033E-06 | 1.528E-07 | 6.900E-06 | 7.200E-06 |
| 22804 | 1141 | F1141D | D | 6.3e-06 | 5.3e-06 | 1e-05 | 7.200E-06 | 2.476E-06 | 5.300E-06 | 1.000E-05 |
| 22805 | 1141 | F1141C | D | 8.9e-06 | 8.2e-06 | 7.6e-06 | 8.233E-06 | 6.506E-07 | 7.600E-06 | 8.900E-06 |
| 22806 | 1141 | F1141E | D | 6.2e-06 | 6.7e-06 | 6.8e-06 | 6.567E-06 | 3.215E-07 | 6.200E-06 | 6.800E-06 |
| 22807 | 1141 | F1141Q | D | 8.7e-06 | 1.3e-05 | 8e-06 | 9.900E-06 | 2.707E-06 | 8.000E-06 | 1.300E-05 |
| 22808 | 1141 | F1141G | D | 6.2e-06 | 8.2e-06 | 9.4e-06 | 7.933E-06 | 1.617E-06 | 6.200E-06 | 9.400E-06 |
| 22809 | 1141 | F1141H | D | 6.3e-06 | 8.6e-06 | 8.4e-06 | 7.767E-06 | 1.274E-06 | 6.300E-06 | 8.600E-06 |
| 22810 | 1141 | F1141I | D | 9.6e-06 | 8.2e-06 | 7.6e-06 | 8.467E-06 | 1.026E-06 | 7.600E-06 | 9.600E-06 |
| 22811 | 1141 | F1141L | D | 9.1e-06 | 7.8e-06 | 8.5e-06 | 8.467E-06 | 6.506E-07 | 7.800E-06 | 9.100E-06 |
| 22812 | 1141 | F1141K | D | 1.3e-05 | 1.1e-05 | 6.2e-06 | 1.007E-05 | 3.495E-06 | 6.200E-06 | 1.300E-05 |
| 22813 | 1141 | F1141M | D | 5.5e-06 | 7.4e-06 | 7.1e-06 | 6.667E-06 | 1.021E-06 | 5.500E-06 | 7.400E-06 |
| 22814 | 1141 | F1141F | D | 1.2e-05 | 1.1e-05 | 4.1e-06 | 9.033E-06 | 4.302E-06 | 4.100E-06 | 1.200E-05 |
| 22815 | 1141 | F1141P | D | 9.2e-06 | 8.6e-06 | 1.1e-05 | 9.600E-06 | 1.249E-06 | 8.600E-06 | 1.100E-05 |
| 22816 | 1141 | F1141S | D | 8.9e-06 | 1.2e-05 | 6.4e-06 | 9.100E-06 | 2.805E-06 | 6.400E-06 | 1.200E-05 |
| 22817 | 1141 | F1141T | D | 7.9e-06 | 4.4e-06 | 1e-05 | 7.433E-06 | 2.829E-06 | 4.400E-06 | 1.000E-05 |
| 22818 | 1141 | F1141W | D | 9.9e-06 | 5.4e-06 | 4.1e-06 | 6.467E-06 | 3.044E-06 | 4.100E-06 | 9.900E-06 |
| 22819 | 1141 | F1141Y | D | 7.6e-06 | 9.1e-06 | 8e-06 | 8.233E-06 | 7.767E-07 | 7.600E-06 | 9.100E-06 |
| 22820 | 1141 | F1141V | D | 9.1e-06 | 8e-06 | 7.8e-06 | 8.300E-06 | 7.000E-07 | 7.800E-06 | 9.100E-06 |
| 22821 | 1142 | A1142A | D | 9e-06 | 6.2e-06 | 6.3e-06 | 7.167E-06 | 1.589E-06 | 6.200E-06 | 9.000E-06 |
| 22822 | 1142 | A1142R | D | 5.3e-06 | 1.4e-05 | 6.6e-06 | 8.633E-06 | 4.693E-06 | 5.300E-06 | 1.400E-05 |
| 22823 | 1142 | A1142N | D | 6.3e-06 | 4.9e-06 | 6.4e-06 | 5.867E-06 | 8.386E-07 | 4.900E-06 | 6.400E-06 |
| 22824 | 1142 | A1142D | D | 6.1e-06 | 3.5e-06 | 6.6e-06 | 5.400E-06 | 1.664E-06 | 3.500E-06 | 6.600E-06 |
| 22825 | 1142 | A1142C | D | 8e-06 | 9.8e-06 | 1.1e-05 | 9.600E-06 | 1.510E-06 | 8.000E-06 | 1.100E-05 |
| 22826 | 1142 | A1142E | D | 6.8e-06 | 7.7e-06 | 1.1e-05 | 8.500E-06 | 2.211E-06 | 6.800E-06 | 1.100E-05 |
| 22827 | 1142 | A1142Q | D | 6.4e-06 | 9.3e-06 | 9.6e-06 | 8.433E-06 | 1.767E-06 | 6.400E-06 | 9.600E-06 |
| 22828 | 1142 | A1142G | D | 8.9e-06 | 7.5e-06 | 1.1e-05 | 9.133E-06 | 1.762E-06 | 7.500E-06 | 1.100E-05 |
| 22829 | 1142 | A1142H | D | 1.2e-05 | 7e-06 | 1e-05 | 9.667E-06 | 2.517E-06 | 7.000E-06 | 1.200E-05 |
| 22830 | 1142 | A1142I | D | 6.9e-06 | 7.7e-06 | 4.6e-06 | 6.400E-06 | 1.609E-06 | 4.600E-06 | 7.700E-06 |
| 22831 | 1142 | A1142L | D | 6.7e-06 | 5.8e-06 | 8.6e-06 | 7.033E-06 | 1.429E-06 | 5.800E-06 | 8.600E-06 |
| 22832 | 1142 | A1142K | D | 4.7e-06 | 5.4e-06 | 7.1e-06 | 5.733E-06 | 1.234E-06 | 4.700E-06 | 7.100E-06 |
| 22833 | 1142 | A1142M | D | 9.9e-06 | 7.4e-06 | 8.6e-06 | 8.633E-06 | 1.250E-06 | 7.400E-06 | 9.900E-06 |
| 22834 | 1142 | A1142F | D | 1e-05 | 8.3e-06 | 7.6e-06 | 8.633E-06 | 1.234E-06 | 7.600E-06 | 1.000E-05 |
| 22835 | 1142 | A1142P | D | 8.3e-06 | 7.3e-06 | 9.1e-06 | 8.233E-06 | 9.018E-07 | 7.300E-06 | 9.100E-06 |
| 22836 | 1142 | A1142S | D | 4.3e-06 | 7e-06 | 1.1e-05 | 7.433E-06 | 3.371E-06 | 4.300E-06 | 1.100E-05 |
| 22837 | 1142 | A1142T | D | 6.3e-06 | 8.6e-06 | 1.3e-05 | 9.300E-06 | 3.404E-06 | 6.300E-06 | 1.300E-05 |
| 22838 | 1142 | A1142W | D | 1e-05 | 1e-05 | 1.1e-05 | 1.033E-05 | 5.774E-07 | 1.000E-05 | 1.100E-05 |
| 22839 | 1142 | A1142Y | D | 8.9e-06 | 1.5e-05 | 1.1e-05 | 1.163E-05 | 3.099E-06 | 8.900E-06 | 1.500E-05 |
| 22840 | 1142 | A1142V | D | 7e-06 | 8.1e-06 | 1e-05 | 8.367E-06 | 1.518E-06 | 7.000E-06 | 1.000E-05 |
| 22841 | 1143 | T1143A | D | 6.9e-06 | 1.4e-05 | 5.6e-06 | 8.833E-06 | 4.521E-06 | 5.600E-06 | 1.400E-05 |
| 22842 | 1143 | T1143R | D | 8.3e-06 | 3.8e-06 | 9.6e-06 | 7.233E-06 | 3.044E-06 | 3.800E-06 | 9.600E-06 |
| 22843 | 1143 | T1143N | D | 6.7e-06 | 6.8e-06 | 9.9e-06 | 7.800E-06 | 1.819E-06 | 6.700E-06 | 9.900E-06 |
| 22844 | 1143 | T1143D | D | 1e-05 | 6.8e-06 | 9.1e-06 | 8.633E-06 | 1.650E-06 | 6.800E-06 | 1.000E-05 |
| 22845 | 1143 | T1143C | D | 6.4e-06 | 1.1e-05 | 6.4e-06 | 7.933E-06 | 2.656E-06 | 6.400E-06 | 1.100E-05 |
| 22846 | 1143 | T1143E | D | 9.1e-06 | 7.9e-06 | 9.1e-06 | 8.700E-06 | 6.928E-07 | 7.900E-06 | 9.100E-06 |
| 22847 | 1143 | T1143Q | D | 7.4e-06 | 4e-06 | 1.2e-05 | 7.800E-06 | 4.015E-06 | 4.000E-06 | 1.200E-05 |
| 22848 | 1143 | T1143G | D | 6.7e-06 | 1.5e-05 | 6.4e-06 | 9.367E-06 | 4.881E-06 | 6.400E-06 | 1.500E-05 |
| 22849 | 1143 | T1143H | D | 9.8e-06 | 8.6e-06 | 6.3e-06 | 8.233E-06 | 1.779E-06 | 6.300E-06 | 9.800E-06 |
| 22850 | 1143 | T1143I | D | 6.9e-06 | 1.3e-05 | 6.5e-06 | 8.800E-06 | 3.643E-06 | 6.500E-06 | 1.300E-05 |
| 22851 | 1143 | T1143L | D | 5.1e-06 | 8.5e-06 | 1.4e-05 | 9.200E-06 | 4.491E-06 | 5.100E-06 | 1.400E-05 |
| 22852 | 1143 | T1143K | D | 9.3e-06 | 5.6e-06 | 7.8e-06 | 7.567E-06 | 1.861E-06 | 5.600E-06 | 9.300E-06 |
| 22853 | 1143 | T1143M | D | 1.4e-05 | 9.9e-06 | 6.5e-06 | 1.013E-05 | 3.755E-06 | 6.500E-06 | 1.400E-05 |
| 22854 | 1143 | T1143F | D | 1.5e-05 | 8.9e-06 | 1.3e-05 | 1.230E-05 | 3.110E-06 | 8.900E-06 | 1.500E-05 |
| 22855 | 1143 | T1143P | D | 8.9e-06 | 7.9e-06 | 6.6e-06 | 7.800E-06 | 1.153E-06 | 6.600E-06 | 8.900E-06 |
| 22856 | 1143 | T1143S | D | 1.8e-05 | 8e-06 | 9.8e-06 | 1.193E-05 | 5.330E-06 | 8.000E-06 | 1.800E-05 |
| 22857 | 1143 | T1143T | D | 6.7e-06 | 1.3e-05 | 6.9e-06 | 8.867E-06 | 3.581E-06 | 6.700E-06 | 1.300E-05 |
| 22858 | 1143 | T1143W | D | 8.5e-06 | 1.2e-05 | 6.3e-06 | 8.933E-06 | 2.875E-06 | 6.300E-06 | 1.200E-05 |
| 22859 | 1143 | T1143Y | D | 7.7e-06 | 5e-06 | 1.3e-05 | 8.567E-06 | 4.070E-06 | 5.000E-06 | 1.300E-05 |
| 22860 | 1143 | T1143V | D | 5.4e-06 | 1e-05 | 6.6e-06 | 7.333E-06 | 2.386E-06 | 5.400E-06 | 1.000E-05 |
| 22861 | 1144 | Y1144A | D | 1.1e-05 | 1.2e-05 | 1.1e-05 | 1.133E-05 | 5.774E-07 | 1.100E-05 | 1.200E-05 |
| 22862 | 1144 | Y1144R | D | 6.3e-06 | 9.8e-06 | 7.4e-06 | 7.833E-06 | 1.790E-06 | 6.300E-06 | 9.800E-06 |
| 22863 | 1144 | Y1144N | D | 8.7e-06 | 6.9e-06 | 6.9e-06 | 7.500E-06 | 1.039E-06 | 6.900E-06 | 8.700E-06 |
| 22864 | 1144 | Y1144D | D | 9.6e-06 | 1e-05 | 1.1e-05 | 1.020E-05 | 7.211E-07 | 9.600E-06 | 1.100E-05 |

| | | | | | | | | | | |
|---|---|---|---|---|---|---|---|---|---|---|
| 22865 | 1144 | Y1144C | D | 9.4e-06 | 1.1e-05 | 3.5e-06 | 7.967E-06 | 3.950E-06 | 3.500E-06 | 1.100E-05 |
| 22866 | 1144 | Y1144E | D | 4.1e-06 | 1e-05 | 8.5e-06 | 7.533E-06 | 3.066E-06 | 4.100E-06 | 1.000E-05 |
| 22867 | 1144 | Y1144Q | D | 1.1e-05 | 1.3e-05 | 1.9e-05 | 1.433E-05 | 4.163E-06 | 1.100E-05 | 1.900E-05 |
| 22868 | 1144 | Y1144G | D | 5.2e-06 | 7.9e-06 | 9.7e-06 | 7.600E-06 | 2.265E-06 | 5.200E-06 | 9.700E-06 |
| 22869 | 1144 | Y1144H | D | 5.7e-06 | 1.1e-05 | 5.7e-06 | 7.467E-06 | 3.060E-06 | 5.700E-06 | 1.100E-05 |
| 22870 | 1144 | Y1144I | D | 8.1e-06 | 9.5e-06 | 6.9e-06 | 8.167E-06 | 1.301E-06 | 6.900E-06 | 9.500E-06 |
| 22871 | 1144 | Y1144L | D | 9.8e-06 | 8.4e-06 | 8e-06 | 8.733E-06 | 9.452E-07 | 8.000E-06 | 9.800E-06 |
| 22872 | 1144 | Y1144K | D | 4.5e-06 | 1.3e-05 | 6.9e-06 | 8.133E-06 | 4.382E-06 | 4.500E-06 | 1.300E-05 |
| 22873 | 1144 | Y1144M | D | 6.6e-06 | 1e-05 | 9.5e-06 | 8.700E-06 | 1.836E-06 | 6.600E-06 | 1.000E-05 |
| 22874 | 1144 | Y1144F | D | 7.6e-06 | 1.1e-05 | 1.4e-05 | 1.087E-05 | 3.202E-06 | 7.600E-06 | 1.400E-05 |
| 22875 | 1144 | Y1144P | D | 6.6e-06 | 1.4e-05 | 6.1e-06 | 8.900E-06 | 4.424E-06 | 6.100E-06 | 1.400E-05 |
| 22876 | 1144 | Y1144S | D | 1.2e-05 | 7.3e-06 | 9.6e-06 | 9.633E-06 | 2.350E-06 | 7.300E-06 | 1.200E-05 |
| 22877 | 1144 | Y1144T | D | 8e-06 | 6.6e-06 | 8.6e-06 | 7.733E-06 | 1.026E-06 | 6.600E-06 | 8.600E-06 |
| 22878 | 1144 | Y1144W | D | 4.6e-06 | 3.1e-06 | 9.4e-06 | 5.700E-06 | 3.291E-06 | 3.100E-06 | 9.400E-06 |
| 22879 | 1144 | Y1144Y | D | 4.7e-06 | 6.5e-06 | 4.9e-06 | 5.367E-06 | 9.866E-07 | 4.700E-06 | 6.500E-06 |
| 22880 | 1144 | Y1144V | D | 9.5e-06 | 1.5e-05 | 9e-06 | 1.117E-05 | 3.329E-06 | 9.000E-06 | 1.500E-05 |
| 22881 | 1145 | Y1145A | D | 7.3e-06 | 1.4e-05 | 9.4e-06 | 1.023E-05 | 3.427E-06 | 7.300E-06 | 1.400E-05 |
| 22882 | 1145 | Y1145R | D | 6.3e-06 | 6.6e-06 | 5.9e-06 | 6.267E-06 | 3.512E-07 | 5.900E-06 | 6.600E-06 |
| 22883 | 1145 | Y1145N | D | 1.1e-05 | 6.7e-06 | 7.4e-06 | 8.367E-06 | 2.307E-06 | 6.700E-06 | 1.100E-05 |
| 22884 | 1145 | Y1145D | D | 1.2e-05 | 6.9e-06 | 8.1e-06 | 9.000E-06 | 2.666E-06 | 6.900E-06 | 1.200E-05 |
| 22885 | 1145 | Y1145C | D | 8.1e-06 | 8.8e-06 | 5.1e-06 | 7.333E-06 | 1.966E-06 | 5.100E-06 | 8.800E-06 |
| 22886 | 1145 | Y1145E | D | 6.5e-06 | 8.3e-06 | 8.8e-06 | 7.867E-06 | 1.210E-06 | 6.500E-06 | 8.800E-06 |
| 22887 | 1145 | Y1145Q | D | 6.2e-06 | 9.8e-06 | 6.9e-06 | 7.633E-06 | 1.909E-06 | 6.200E-06 | 9.800E-06 |
| 22888 | 1145 | Y1145G | D | 1.1e-05 | 4.4e-06 | 8e-06 | 7.800E-06 | 3.305E-06 | 4.400E-06 | 1.100E-05 |
| 22889 | 1145 | Y1145H | D | 4.2e-06 | 9.9e-06 | 5.7e-06 | 6.600E-06 | 2.955E-06 | 4.200E-06 | 9.900E-06 |
| 22890 | 1145 | Y1145I | D | 1.2e-05 | 1e-05 | 9e-06 | 1.033E-05 | 1.528E-06 | 9.000E-06 | 1.200E-05 |
| 22891 | 1145 | Y1145L | D | 8.7e-06 | 7.2e-06 | 4e-06 | 6.633E-06 | 2.401E-06 | 4.000E-06 | 8.700E-06 |
| 22892 | 1145 | Y1145K | D | 4.6e-06 | 6.6e-06 | 7.5e-06 | 6.233E-06 | 1.484E-06 | 4.600E-06 | 7.500E-06 |
| 22893 | 1145 | Y1145M | D | 7.1e-06 | 7.6e-06 | 7.7e-06 | 7.467E-06 | 3.215E-07 | 7.100E-06 | 7.700E-06 |
| 22894 | 1145 | Y1145F | D | 8.5e-06 | 5e-06 | 5.6e-06 | 6.367E-06 | 1.872E-06 | 5.000E-06 | 8.500E-06 |
| 22895 | 1145 | Y1145P | D | 7.7e-06 | 7e-06 | 1.1e-05 | 8.567E-06 | 2.136E-06 | 7.000E-06 | 1.100E-05 |
| 22896 | 1145 | Y1145S | D | 6.8e-06 | 8.7e-06 | 8.2e-06 | 7.900E-06 | 9.849E-07 | 6.800E-06 | 8.700E-06 |
| 22897 | 1145 | Y1145T | D | 8.7e-06 | 9.1e-06 | 8.7e-06 | 8.833E-06 | 2.309E-07 | 8.700E-06 | 9.100E-06 |
| 22898 | 1145 | Y1145W | D | 5.1e-06 | 6.9e-06 | 6.4e-06 | 6.133E-06 | 9.292E-07 | 5.100E-06 | 6.900E-06 |
| 22899 | 1145 | Y1145Y | D | 4.9e-06 | 9e-06 | 5.1e-06 | 6.333E-06 | 2.312E-06 | 4.900E-06 | 9.000E-06 |
| 22900 | 1145 | Y1145V | D | 1.5e-05 | 1.2e-05 | 9e-06 | 1.200E-05 | 3.000E-06 | 9.000E-06 | 1.500E-05 |
| 22901 | 1146 | C1146A | D | 7.2e-06 | 5.3e-06 | 9.9e-06 | 7.467E-06 | 2.312E-06 | 5.300E-06 | 9.900E-06 |
| 22902 | 1146 | C1146R | D | 9e-06 | 8.9e-06 | 4.9e-06 | 7.600E-06 | 2.339E-06 | 4.900E-06 | 9.000E-06 |
| 22903 | 1146 | C1146N | D | 6.1e-06 | 9.7e-06 | 1.2e-05 | 9.267E-06 | 2.974E-06 | 6.100E-06 | 1.200E-05 |
| 22904 | 1146 | C1146D | D | 5.9e-06 | 9.6e-06 | 6.7e-06 | 7.400E-06 | 1.947E-06 | 5.900E-06 | 9.600E-06 |
| 22905 | 1146 | C1146C | D | 1.2e-05 | 8.5e-06 | 8.4e-06 | 9.633E-06 | 2.050E-06 | 8.400E-06 | 1.200E-05 |
| 22906 | 1146 | C1146E | D | 5.1e-06 | 1.3e-05 | 1e-05 | 9.367E-06 | 3.988E-06 | 5.100E-06 | 1.300E-05 |
| 22907 | 1146 | C1146Q | D | 6.9e-06 | 8.4e-06 | 5.8e-06 | 7.033E-06 | 1.305E-06 | 5.800E-06 | 8.400E-06 |
| 22908 | 1146 | C1146G | D | 8.8e-06 | 6.4e-06 | 1.5e-05 | 1.007E-05 | 4.438E-06 | 6.400E-06 | 1.500E-05 |
| 22909 | 1146 | C1146H | D | 1.3e-05 | 7.2e-06 | 6.3e-06 | 8.833E-06 | 3.636E-06 | 6.300E-06 | 1.300E-05 |
| 22910 | 1146 | C1146I | D | 7.5e-06 | 7.7e-06 | 1.1e-05 | 8.733E-06 | 1.966E-06 | 7.500E-06 | 1.100E-05 |
| 22911 | 1146 | C1146L | D | 7.4e-06 | 9e-06 | 1.2e-05 | 9.467E-06 | 2.335E-06 | 7.400E-06 | 1.200E-05 |
| 22912 | 1146 | C1146K | D | 4.7e-06 | 6.1e-06 | 9.3e-06 | 6.700E-06 | 2.358E-06 | 4.700E-06 | 9.300E-06 |
| 22913 | 1146 | C1146M | D | 6.5e-06 | 4.5e-06 | 8.4e-06 | 6.467E-06 | 1.950E-06 | 4.500E-06 | 8.400E-06 |
| 22914 | 1146 | C1146F | D | 7.9e-06 | 6.9e-06 | 1.4e-05 | 9.600E-06 | 3.843E-06 | 6.900E-06 | 1.400E-05 |
| 22915 | 1146 | C1146P | D | 4.4e-06 | 4.3e-06 | 8.1e-06 | 5.600E-06 | 2.166E-06 | 4.300E-06 | 8.100E-06 |
| 22916 | 1146 | C1146S | D | 7.9e-06 | 5.6e-06 | 5.9e-06 | 6.467E-06 | 1.250E-06 | 5.600E-06 | 7.900E-06 |
| 22917 | 1146 | C1146T | D | 9.6e-06 | 5.2e-06 | 8.9e-06 | 7.900E-06 | 2.364E-06 | 5.200E-06 | 9.600E-06 |
| 22918 | 1146 | C1146W | D | 1.2e-05 | 8e-06 | 6.6e-06 | 8.867E-06 | 2.802E-06 | 6.600E-06 | 1.200E-05 |
| 22919 | 1146 | C1146Y | D | 9.7e-06 | 7.7e-06 | 7.3e-06 | 8.233E-06 | 1.286E-06 | 7.300E-06 | 9.700E-06 |
| 22920 | 1146 | C1146V | D | 9.5e-06 | 1.3e-05 | 4.7e-06 | 9.067E-06 | 4.167E-06 | 4.700E-06 | 1.300E-05 |
| 22921 | 1147 | Q1147A | D | 7.6e-06 | 2.9e-06 | 9.2e-06 | 6.567E-06 | 3.275E-06 | 2.900E-06 | 9.200E-06 |
| 22922 | 1147 | Q1147R | D | 5.6e-06 | 1.3e-05 | 1.2e-05 | 1.020E-05 | 4.015E-06 | 5.600E-06 | 1.300E-05 |
| 22923 | 1147 | Q1147N | D | 5.5e-06 | 8.4e-06 | 6.2e-06 | 6.700E-06 | 1.513E-06 | 5.500E-06 | 8.400E-06 |
| 22924 | 1147 | Q1147D | D | 9.4e-06 | 2e-05 | 9.5e-06 | 1.297E-05 | 6.091E-06 | 9.400E-06 | 2.000E-05 |
| 22925 | 1147 | Q1147C | D | 1.3e-05 | 1e-05 | 6.5e-06 | 9.833E-06 | 3.253E-06 | 6.500E-06 | 1.300E-05 |
| 22926 | 1147 | Q1147E | D | 9.1e-06 | 4.5e-06 | 6.4e-06 | 6.667E-06 | 2.312E-06 | 4.500E-06 | 9.100E-06 |
| 22927 | 1147 | Q1147Q | D | 6.1e-06 | 5.4e-06 | 4.9e-06 | 5.467E-06 | 6.028E-07 | 4.900E-06 | 6.100E-06 |
| 22928 | 1147 | Q1147G | D | 1.1e-05 | 6.2e-06 | 1.1e-05 | 9.400E-06 | 2.771E-06 | 6.200E-06 | 1.100E-05 |
| 22929 | 1147 | Q1147H | D | 5.9e-06 | 8.2e-06 | 6.6e-06 | 6.900E-06 | 1.179E-06 | 5.900E-06 | 8.200E-06 |
| 22930 | 1147 | Q1147I | D | 5.5e-06 | 7.1e-06 | 6.3e-06 | 6.300E-06 | 8.000E-07 | 5.500E-06 | 7.100E-06 |
| 22931 | 1147 | Q1147L | D | 5.4e-06 | 6.2e-06 | 4.3e-06 | 5.300E-06 | 9.539E-07 | 4.300E-06 | 6.200E-06 |
| 22932 | 1147 | Q1147K | D | 6.6e-06 | 7.5e-06 | 8.9e-06 | 7.667E-06 | 1.159E-06 | 6.600E-06 | 8.900E-06 |
| 22933 | 1147 | Q1147M | D | 1.4e-05 | 5.4e-06 | 8.7e-06 | 9.367E-06 | 4.339E-06 | 5.400E-06 | 1.400E-05 |
| 22934 | 1147 | Q1147F | D | 1.1e-05 | 6.5e-06 | 6.4e-06 | 7.967E-06 | 2.627E-06 | 6.400E-06 | 1.100E-05 |
| 22935 | 1147 | Q1147P | D | 8.4e-06 | 9.1e-06 | 9.6e-06 | 9.033E-06 | 6.028E-07 | 8.400E-06 | 9.600E-06 |
| 22936 | 1147 | Q1147S | D | 9.4e-06 | 4.6e-06 | 9.3e-06 | 7.767E-06 | 2.743E-06 | 4.600E-06 | 9.400E-06 |
| 22937 | 1147 | Q1147T | D | 9.1e-06 | 5.3e-06 | 6e-06 | 6.800E-06 | 2.022E-06 | 5.300E-06 | 9.100E-06 |
| 22938 | 1147 | Q1147W | D | 9.6e-06 | 6.9e-06 | 3.6e-06 | 6.700E-06 | 3.005E-06 | 3.600E-06 | 9.600E-06 |
| 22939 | 1147 | Q1147Y | D | 6.5e-06 | 7.9e-06 | 4.4e-06 | 6.267E-06 | 1.762E-06 | 4.400E-06 | 7.900E-06 |
| 22940 | 1147 | Q1147V | D | 7.2e-06 | 4.2e-06 | 7e-06 | 6.133E-06 | 1.677E-06 | 4.200E-06 | 7.200E-06 |
| 22941 | 1148 | Q1148A | D | 6.6e-06 | 5.7e-06 | 7.2e-06 | 6.500E-06 | 7.550E-07 | 5.700E-06 | 7.200E-06 |
| 22942 | 1148 | Q1148R | D | 7.1e-06 | 7.8e-06 | 9.7e-06 | 8.200E-06 | 1.345E-06 | 7.100E-06 | 9.700E-06 |
| 22943 | 1148 | Q1148N | D | 6.6e-06 | 3.5e-06 | 9.2e-06 | 6.433E-06 | 2.854E-06 | 3.500E-06 | 9.200E-06 |
| 22944 | 1148 | Q1148D | D | 4.8e-06 | 7.9e-06 | 7.3e-06 | 6.667E-06 | 1.644E-06 | 4.800E-06 | 7.900E-06 |
| 22945 | 1148 | Q1148C | D | 5.3e-06 | 7.2e-06 | 9.9e-06 | 7.467E-06 | 2.312E-06 | 5.300E-06 | 9.900E-06 |
| 22946 | 1148 | Q1148E | D | 7.1e-06 | 5e-06 | 6.2e-06 | 6.100E-06 | 1.054E-06 | 5.000E-06 | 7.100E-06 |
| 22947 | 1148 | Q1148Q | D | 4e-06 | 6.8e-06 | 1.4e-05 | 8.267E-06 | 5.159E-06 | 4.000E-06 | 1.400E-05 |
| 22948 | 1148 | Q1148G | D | 8e-06 | 1.2e-05 | 5.9e-06 | 8.633E-06 | 3.099E-06 | 5.900E-06 | 1.200E-05 |
| 22949 | 1148 | Q1148H | D | 1e-05 | 1.3e-05 | 1.1e-05 | 1.133E-05 | 1.528E-06 | 1.000E-05 | 1.300E-05 |

| | | | | | | | | | | |
|---|---|---|---|---|---|---|---|---|---|---|
| 22950 | 1148 | Q1148I | D | 3.9e-06 | 6.1e-06 | 9.6e-06 | 6.533E-06 | 2.875E-06 | 3.900E-06 | 9.600E-06 |
| 22951 | 1148 | Q1148L | D | 1.4e-05 | 9.1e-06 | 6e-06 | 9.700E-06 | 4.034E-06 | 6.000E-06 | 1.400E-05 |
| 22952 | 1148 | Q1148K | D | 8.4e-06 | 1.3e-05 | 5.8e-06 | 9.067E-06 | 3.646E-06 | 5.800E-06 | 1.300E-05 |
| 22953 | 1148 | Q1148M | D | 1.3e-05 | 1.2e-05 | 9.3e-06 | 1.143E-05 | 1.914E-06 | 9.300E-06 | 1.300E-05 |
| 22954 | 1148 | Q1148F | D | 5.6e-06 | 9.5e-06 | 1.1e-05 | 8.700E-06 | 2.787E-06 | 5.600E-06 | 1.100E-05 |
| 22955 | 1148 | Q1148P | D | 5.2e-06 | 9.8e-06 | 6.1e-06 | 7.033E-06 | 2.438E-06 | 5.200E-06 | 9.800E-06 |
| 22956 | 1148 | Q1148S | D | 3.9e-06 | 5.5e-06 | 1.1e-05 | 6.800E-06 | 3.724E-06 | 3.900E-06 | 1.100E-05 |
| 22957 | 1148 | Q1148T | D | 1.4e-05 | 6.7e-06 | 9.9e-06 | 1.020E-05 | 3.659E-06 | 6.700E-06 | 1.400E-05 |
| 22958 | 1148 | Q1148W | D | 1.3e-05 | 9.6e-06 | 6.1e-06 | 9.567E-06 | 3.450E-06 | 6.100E-06 | 1.300E-05 |
| 22959 | 1148 | Q1148Y | D | 7.8e-06 | 8.8e-06 | 9.2e-06 | 8.600E-06 | 7.211E-07 | 7.800E-06 | 9.200E-06 |
| 22960 | 1148 | Q1148V | D | 8e-06 | 7.7e-06 | 6.9e-06 | 7.533E-06 | 5.686E-07 | 6.900E-06 | 8.000E-06 |
| 22961 | 1149 | H1149A | D | 1.4e-05 | 7.1e-06 | 3.2e-05 | 1.770E-05 | 1.286E-05 | 7.100E-06 | 3.200E-05 |
| 22962 | 1149 | H1149R | D | 5.5e-06 | 5.8e-06 | 5.9e-06 | 5.733E-06 | 2.082E-07 | 5.500E-06 | 5.900E-06 |
| 22963 | 1149 | H1149N | D | 4.2e-06 | 4.2e-06 | 5e-06 | 4.467E-06 | 4.619E-07 | 4.200E-06 | 5.000E-06 |
| 22964 | 1149 | H1149D | D | 8e-06 | 7.6e-06 | 7e-06 | 7.533E-06 | 5.033E-07 | 7.000E-06 | 8.000E-06 |
| 22965 | 1149 | H1149C | D | 9.5e-06 | 9.9e-06 | 9.1e-06 | 9.500E-06 | 4.000E-07 | 9.100E-06 | 9.900E-06 |
| 22966 | 1149 | H1149E | D | 9.8e-06 | 9e-06 | 5.9e-06 | 8.233E-06 | 2.060E-06 | 5.900E-06 | 9.800E-06 |
| 22967 | 1149 | H1149Q | D | 5.1e-06 | 7.2e-06 | 7.4e-06 | 6.567E-06 | 1.274E-06 | 5.100E-06 | 7.400E-06 |
| 22968 | 1149 | H1149G | D | 1.2e-05 | 1.3e-05 | 1.5e-05 | 1.333E-05 | 1.528E-06 | 1.200E-05 | 1.500E-05 |
| 22969 | 1149 | H1149H | D | 1.4e-05 | 6.6e-06 | 1.3e-05 | 1.120E-05 | 4.015E-06 | 6.600E-06 | 1.400E-05 |
| 22970 | 1149 | H1149I | D | 1.2e-05 | 1.4e-05 | 1.5e-05 | 1.367E-05 | 1.528E-06 | 1.200E-05 | 1.500E-05 |
| 22971 | 1149 | H1149L | D | 9.2e-06 | 1.9e-05 | 8.4e-06 | 1.220E-05 | 5.903E-06 | 8.400E-06 | 1.900E-05 |
| 22972 | 1149 | H1149K | D | 1.4e-05 | 9.8e-06 | 1.2e-05 | 1.193E-05 | 2.101E-06 | 9.800E-06 | 1.400E-05 |
| 22973 | 1149 | H1149M | D | 1.1e-05 | 6.5e-06 | 1.7e-05 | 1.150E-05 | 5.268E-06 | 6.500E-06 | 1.700E-05 |
| 22974 | 1149 | H1149F | D | 1e-05 | 7.9e-06 | 1.5e-05 | 1.097E-05 | 3.647E-06 | 7.900E-06 | 1.500E-05 |
| 22975 | 1149 | H1149P | D | 1.4e-05 | 2.2e-05 | 8.3e-06 | 1.477E-05 | 6.882E-06 | 8.300E-06 | 2.200E-05 |
| 22976 | 1149 | H1149S | D | 3.8e-06 | 6.8e-06 | 4.4e-06 | 5.000E-06 | 1.587E-06 | 3.800E-06 | 6.800E-06 |
| 22977 | 1149 | H1149T | D | 8.3e-06 | 5.4e-06 | 3.8e-06 | 5.833E-06 | 2.281E-06 | 3.800E-06 | 8.300E-06 |
| 22978 | 1149 | H1149W | D | 1.1e-05 | 9.9e-06 | 1.9e-05 | 1.330E-05 | 4.967E-06 | 9.900E-06 | 1.900E-05 |
| 22979 | 1149 | H1149Y | D | 6.7e-06 | 9.8e-06 | 1.3e-05 | 9.833E-06 | 3.150E-06 | 6.700E-06 | 1.300E-05 |
| 22980 | 1149 | H1149V | D | 7.2e-06 | 1.2e-05 | 1.9e-05 | 1.273E-05 | 5.934E-06 | 7.200E-06 | 1.900E-05 |
| 22981 | 1150 | Y1150A | D | 6.5e-06 | 9.8e-06 | 8.8e-06 | 8.367E-06 | 1.692E-06 | 6.500E-06 | 9.800E-06 |
| 22982 | 1150 | Y1150R | D | 1e-05 | 4.1e-06 | 1.1e-05 | 8.367E-06 | 3.729E-06 | 4.100E-06 | 1.100E-05 |
| 22983 | 1150 | Y1150N | D | 5.9e-06 | 5.8e-06 | 5.6e-06 | 5.767E-06 | 1.528E-07 | 5.600E-06 | 5.900E-06 |
| 22984 | 1150 | Y1150D | D | 4.9e-06 | 4.7e-06 | 9.7e-06 | 6.433E-06 | 2.831E-06 | 4.700E-06 | 9.700E-06 |
| 22985 | 1150 | Y1150C | D | 8.3e-06 | 1.4e-05 | 6.6e-06 | 9.633E-06 | 3.876E-06 | 6.600E-06 | 1.400E-05 |
| 22986 | 1150 | Y1150E | D | 7.5e-06 | 7.3e-06 | 4.9e-06 | 6.567E-06 | 1.447E-06 | 4.900E-06 | 7.500E-06 |
| 22987 | 1150 | Y1150Q | D | 4.7e-06 | 8e-06 | 5.4e-06 | 6.033E-06 | 1.739E-06 | 4.700E-06 | 8.000E-06 |
| 22988 | 1150 | Y1150G | D | 1.3e-05 | 8.7e-06 | 5.6e-06 | 9.100E-06 | 3.716E-06 | 5.600E-06 | 1.300E-05 |
| 22989 | 1150 | Y1150H | D | 7e-06 | 7.7e-06 | 5.8e-06 | 6.833E-06 | 9.609E-07 | 5.800E-06 | 7.700E-06 |
| 22990 | 1150 | Y1150I | D | 1.2e-05 | 7.6e-06 | 1.1e-05 | 1.020E-05 | 2.307E-06 | 7.600E-06 | 1.200E-05 |
| 22991 | 1150 | Y1150L | D | 6.5e-06 | 1.4e-05 | 1.2e-05 | 1.083E-05 | 3.884E-06 | 6.500E-06 | 1.400E-05 |
| 22992 | 1150 | Y1150K | D | 5.3e-06 | 6.5e-06 | 5.4e-06 | 5.733E-06 | 6.658E-07 | 5.300E-06 | 6.500E-06 |
| 22993 | 1150 | Y1150M | D | 6.5e-06 | 6.7e-06 | 9.9e-06 | 7.700E-06 | 1.908E-06 | 6.500E-06 | 9.900E-06 |
| 22994 | 1150 | Y1150F | D | 9.7e-06 | 9.4e-06 | 6e-06 | 8.367E-06 | 2.055E-06 | 6.000E-06 | 9.700E-06 |
| 22995 | 1150 | Y1150P | D | 9.8e-06 | 9.7e-06 | 1.2e-05 | 1.050E-05 | 1.300E-06 | 9.700E-06 | 1.200E-05 |
| 22996 | 1150 | Y1150S | D | 3.3e-06 | 4.3e-06 | 5e-06 | 4.200E-06 | 8.544E-07 | 3.300E-06 | 5.000E-06 |
| 22997 | 1150 | Y1150T | D | 3.8e-06 | 5.1e-06 | 7.4e-06 | 5.433E-06 | 1.823E-06 | 3.800E-06 | 7.400E-06 |
| 22998 | 1150 | Y1150W | D | 9.9e-06 | 5e-06 | 1e-05 | 8.300E-06 | 2.858E-06 | 5.000E-06 | 1.000E-05 |
| 22999 | 1150 | Y1150Y | D | 4.8e-06 | 9.4e-06 | 4.4e-06 | 6.200E-06 | 2.778E-06 | 4.400E-06 | 9.400E-06 |
| 23000 | 1150 | Y1150V | D | 8.8e-06 | 6.9e-06 | 6.8e-06 | 7.500E-06 | 1.127E-06 | 6.800E-06 | 8.800E-06 |
| 23001 | 1151 | T1151A | D | 6.7e-06 | 8.3e-06 | 1.4e-05 | 9.667E-06 | 3.837E-06 | 6.700E-06 | 1.400E-05 |
| 23002 | 1151 | T1151R | D | 6.1e-06 | 7.9e-06 | 4.2e-06 | 6.067E-06 | 1.850E-06 | 4.200E-06 | 7.900E-06 |
| 23003 | 1151 | T1151N | D | 7.3e-06 | 6.5e-06 | 6.4e-06 | 6.733E-06 | 4.933E-07 | 6.400E-06 | 7.300E-06 |
| 23004 | 1151 | T1151D | D | 8.5e-06 | 7e-06 | 5.6e-06 | 7.033E-06 | 1.450E-06 | 5.600E-06 | 8.500E-06 |
| 23005 | 1151 | T1151C | D | 5.4e-06 | 1.1e-05 | 5.3e-06 | 7.233E-06 | 3.262E-06 | 5.300E-06 | 1.100E-05 |
| 23006 | 1151 | T1151E | D | 9.1e-06 | 6.9e-06 | 8e-06 | 8.000E-06 | 1.100E-06 | 6.900E-06 | 9.100E-06 |
| 23007 | 1151 | T1151Q | D | 5e-06 | 4.3e-06 | 5.9e-06 | 5.067E-06 | 8.021E-07 | 4.300E-06 | 5.900E-06 |
| 23008 | 1151 | T1151G | D | 7.2e-06 | 1.4e-05 | 1.3e-05 | 1.140E-05 | 3.672E-06 | 7.200E-06 | 1.400E-05 |
| 23009 | 1151 | T1151H | D | 6.3e-06 | 5.7e-06 | 7.2e-06 | 6.400E-06 | 7.550E-07 | 5.700E-06 | 7.200E-06 |
| 23010 | 1151 | T1151I | D | 1.7e-05 | 1e-05 | 4.2e-06 | 1.040E-05 | 6.409E-06 | 4.200E-06 | 1.700E-05 |
| 23011 | 1151 | T1151L | D | 7.1e-06 | 8.6e-06 | 8.4e-06 | 8.033E-06 | 8.145E-07 | 7.100E-06 | 8.600E-06 |
| 23012 | 1151 | T1151K | D | 2e-05 | 7.5e-06 | 8.9e-06 | 1.213E-05 | 6.849E-06 | 7.500E-06 | 2.000E-05 |
| 23013 | 1151 | T1151M | D | 7e-06 | 6.2e-06 | 8.3e-06 | 7.167E-06 | 1.060E-06 | 6.200E-06 | 8.300E-06 |
| 23014 | 1151 | T1151F | D | 8.7e-06 | 6e-06 | 6.2e-06 | 6.967E-06 | 1.504E-06 | 6.000E-06 | 8.700E-06 |
| 23015 | 1151 | T1151P | D | 1.2e-05 | 8.8e-06 | 8.4e-06 | 9.733E-06 | 1.973E-06 | 8.400E-06 | 1.200E-05 |
| 23016 | 1151 | T1151S | D | 7.7e-06 | 7.5e-06 | 8.6e-06 | 7.933E-06 | 5.859E-07 | 7.500E-06 | 8.600E-06 |
| 23017 | 1151 | T1151T | D | 1.1e-05 | 9.2e-06 | 9.2e-06 | 9.800E-06 | 1.039E-06 | 9.200E-06 | 1.100E-05 |
| 23018 | 1151 | T1151W | D | 5.8e-06 | 1.1e-05 | 5.8e-06 | 7.533E-06 | 3.002E-06 | 5.800E-06 | 1.100E-05 |
| 23019 | 1151 | T1151Y | D | 9.3e-06 | 1.4e-05 | 8.4e-06 | 1.057E-05 | 3.007E-06 | 8.400E-06 | 1.400E-05 |
| 23020 | 1151 | T1151V | D | 8.7e-06 | 1.1e-05 | 5.3e-06 | 8.333E-06 | 2.868E-06 | 5.300E-06 | 1.100E-05 |
| 23021 | 1152 | T1152A | D | 1.3e-05 | 1.1e-05 | 1.1e-05 | 1.167E-05 | 1.155E-06 | 1.100E-05 | 1.300E-05 |
| 23022 | 1152 | T1152R | D | 9.2e-06 | 5.4e-06 | 8e-06 | 7.533E-06 | 1.943E-06 | 5.400E-06 | 9.200E-06 |
| 23023 | 1152 | T1152N | D | 7.7e-06 | 9.1e-06 | 8.2e-06 | 8.333E-06 | 7.095E-07 | 7.700E-06 | 9.100E-06 |
| 23024 | 1152 | T1152D | D | 9.8e-06 | 6.9e-06 | 8.9e-06 | 8.533E-06 | 1.484E-06 | 6.900E-06 | 9.800E-06 |
| 23025 | 1152 | T1152C | D | 1.3e-05 | 1.1e-05 | 8.7e-06 | 1.090E-05 | 2.152E-06 | 8.700E-06 | 1.300E-05 |
| 23026 | 1152 | T1152E | D | 5.1e-06 | 5.1e-06 | 8.9e-06 | 6.367E-06 | 2.194E-06 | 5.100E-06 | 8.900E-06 |
| 23027 | 1152 | T1152Q | D | 8.1e-06 | 3.6e-06 | 6.6e-06 | 6.100E-06 | 2.291E-06 | 3.600E-06 | 8.100E-06 |
| 23028 | 1152 | T1152G | D | 9.4e-06 | 1.1e-05 | 1.2e-05 | 1.080E-05 | 1.311E-06 | 9.400E-06 | 1.200E-05 |
| 23029 | 1152 | T1152H | D | 6.1e-06 | 5.8e-06 | 1e-05 | 7.300E-06 | 2.343E-06 | 5.800E-06 | 1.000E-05 |
| 23030 | 1152 | T1152I | D | 8.5e-06 | 6e-06 | 7e-06 | 7.167E-06 | 1.258E-06 | 6.000E-06 | 8.500E-06 |
| 23031 | 1152 | T1152L | D | 5.1e-06 | 1.2e-05 | 9e-06 | 8.700E-06 | 3.460E-06 | 5.100E-06 | 1.200E-05 |
| 23032 | 1152 | T1152K | D | 5.1e-06 | 6.5e-06 | 8.9e-06 | 6.833E-06 | 1.922E-06 | 5.100E-06 | 8.900E-06 |
| 23033 | 1152 | T1152M | D | 5.4e-06 | 5.5e-06 | 6.2e-06 | 5.700E-06 | 4.359E-07 | 5.400E-06 | 6.200E-06 |
| 23034 | 1152 | T1152F | D | 7e-06 | 6.2e-06 | 6e-06 | 6.400E-06 | 5.292E-07 | 6.000E-06 | 7.000E-06 |

| | | | | | | | | | | |
|---|---|---|---|---|---|---|---|---|---|---|
| 23035 | 1152 | T1152P | D | 8e-06 | 7.1e-06 | 1.3e-05 | 9.367E-06 | 3.179E-06 | 7.100E-06 | 1.300E-05 |
| 23036 | 1152 | T1152S | D | 6.2e-06 | 5.3e-06 | 6.7e-06 | 6.067E-06 | 7.095E-07 | 5.300E-06 | 6.700E-06 |
| 23037 | 1152 | T1152T | D | 4.7e-06 | 8.2e-06 | 8.9e-06 | 7.267E-06 | 2.250E-06 | 4.700E-06 | 8.900E-06 |
| 23038 | 1152 | T1152W | D | 8.2e-06 | 5.9e-06 | 1.1e-05 | 8.367E-06 | 2.554E-06 | 5.900E-06 | 1.100E-05 |
| 23039 | 1152 | T1152Y | D | 1.2e-05 | 5.4e-06 | 9e-06 | 8.800E-06 | 3.305E-06 | 5.400E-06 | 1.200E-05 |
| 23040 | 1152 | T1152V | D | 7.4e-06 | 6.8e-06 | 6.5e-06 | 6.900E-06 | 4.583E-07 | 6.500E-06 | 7.400E-06 |
| 23041 | 1153 | P1153A | D | 8.5e-06 | 1.3e-05 | 8.3e-06 | 9.933E-06 | 2.658E-06 | 8.300E-06 | 1.300E-05 |
| 23042 | 1153 | P1153R | D | 9.8e-06 | 6.3e-06 | 4.5e-06 | 6.867E-06 | 2.695E-06 | 4.500E-06 | 9.800E-06 |
| 23043 | 1153 | P1153N | D | 1.9e-05 | 9.5e-06 | 5.9e-06 | 1.147E-05 | 6.768E-06 | 5.900E-06 | 1.900E-05 |
| 23044 | 1153 | P1153D | D | 6.7e-06 | 7.2e-06 | 9e-06 | 7.633E-06 | 1.210E-06 | 6.700E-06 | 9.000E-06 |
| 23045 | 1153 | P1153C | D | 5.8e-06 | 4.8e-06 | 6.7e-06 | 5.767E-06 | 9.504E-07 | 4.800E-06 | 6.700E-06 |
| 23046 | 1153 | P1153E | D | 6.9e-06 | 1.4e-05 | 4.7e-06 | 8.533E-06 | 4.860E-06 | 4.700E-06 | 1.400E-05 |
| 23047 | 1153 | P1153Q | D | 6.5e-06 | 1.1e-05 | 1e-05 | 9.167E-06 | 2.363E-06 | 6.500E-06 | 1.100E-05 |
| 23048 | 1153 | P1153G | D | 5.4e-06 | 9e-06 | 1e-05 | 8.133E-06 | 2.419E-06 | 5.400E-06 | 1.000E-05 |
| 23049 | 1153 | P1153H | D | 6.2e-06 | 1.5e-05 | 7e-06 | 9.400E-06 | 4.866E-06 | 6.200E-06 | 1.500E-05 |
| 23050 | 1153 | P1153I | D | 6.8e-06 | 1e-05 | 6.6e-06 | 7.800E-06 | 1.908E-06 | 6.600E-06 | 1.000E-05 |
| 23051 | 1153 | P1153L | D | 6.3e-06 | 7.5e-06 | 6.6e-06 | 6.800E-06 | 6.245E-07 | 6.300E-06 | 7.500E-06 |
| 23052 | 1153 | P1153K | D | 1.4e-05 | 9.2e-06 | 8.4e-06 | 1.053E-05 | 3.029E-06 | 8.400E-06 | 1.400E-05 |
| 23053 | 1153 | P1153M | D | 1.1e-05 | 1e-05 | 1e-05 | 1.033E-05 | 5.774E-07 | 1.000E-05 | 1.100E-05 |
| 23054 | 1153 | P1153F | D | 3.9e-06 | 6.7e-06 | 7.1e-06 | 5.900E-06 | 1.744E-06 | 3.900E-06 | 7.100E-06 |
| 23055 | 1153 | P1153P | D | 1e-05 | 1.2e-05 | 5.8e-06 | 9.267E-06 | 3.164E-06 | 5.800E-06 | 1.200E-05 |
| 23056 | 1153 | P1153S | D | 7.1e-06 | 5.5e-06 | 6.1e-06 | 6.233E-06 | 8.083E-07 | 5.500E-06 | 7.100E-06 |
| 23057 | 1153 | P1153T | D | 6.1e-06 | 8.5e-06 | 5.8e-06 | 6.800E-06 | 1.480E-06 | 5.800E-06 | 8.500E-06 |
| 23058 | 1153 | P1153W | D | 8.9e-06 | 7.8e-06 | 1.2e-05 | 9.567E-06 | 2.178E-06 | 7.800E-06 | 1.200E-05 |
| 23059 | 1153 | P1153Y | D | 5.5e-06 | 1.2e-05 | 8.2e-06 | 8.567E-06 | 3.265E-06 | 5.500E-06 | 1.200E-05 |
| 23060 | 1153 | P1153V | D | 5e-06 | 1e-05 | 9.4e-06 | 8.133E-06 | 2.730E-06 | 5.000E-06 | 1.000E-05 |
| 23061 | 1154 | P1154A | D | 5.9e-06 | 6.6e-06 | 7.3e-06 | 6.600E-06 | 7.000E-07 | 5.900E-06 | 7.300E-06 |
| 23062 | 1154 | P1154R | D | 4e-06 | 6.9e-06 | 4.1e-06 | 5.000E-06 | 1.646E-06 | 4.000E-06 | 6.900E-06 |
| 23063 | 1154 | P1154N | D | 6.7e-06 | 9.8e-06 | 8.8e-06 | 8.433E-06 | 1.582E-06 | 6.700E-06 | 9.800E-06 |
| 23064 | 1154 | P1154D | D | 7.4e-06 | 7.7e-06 | 6.9e-06 | 7.333E-06 | 4.041E-07 | 6.900E-06 | 7.700E-06 |
| 23065 | 1154 | P1154C | D | 9e-06 | 5.9e-06 | 5.6e-06 | 6.833E-06 | 1.882E-06 | 5.600E-06 | 9.000E-06 |
| 23066 | 1154 | P1154E | D | 9.5e-06 | 6.9e-06 | 1.2e-05 | 9.467E-06 | 2.550E-06 | 6.900E-06 | 1.200E-05 |
| 23067 | 1154 | P1154Q | D | 1.3e-05 | 2.8e-06 | 7.6e-06 | 7.800E-06 | 5.103E-06 | 2.800E-06 | 1.300E-05 |
| 23068 | 1154 | P1154G | D | 7.7e-06 | 9.1e-06 | 9.1e-06 | 8.633E-06 | 8.083E-07 | 7.700E-06 | 9.100E-06 |
| 23069 | 1154 | P1154H | D | 5.5e-06 | 3.8e-06 | 1e-05 | 6.433E-06 | 3.204E-06 | 3.800E-06 | 1.000E-05 |
| 23070 | 1154 | P1154I | D | 7.8e-06 | 1.4e-05 | 1.4e-05 | 1.193E-05 | 3.580E-06 | 7.800E-06 | 1.400E-05 |
| 23071 | 1154 | P1154L | D | 7.9e-06 | 1.2e-05 | 7.6e-06 | 9.167E-06 | 2.458E-06 | 7.600E-06 | 1.200E-05 |
| 23072 | 1154 | P1154K | D | 7e-06 | 5.2e-06 | 4e-06 | 5.400E-06 | 1.510E-06 | 4.000E-06 | 7.000E-06 |
| 23073 | 1154 | P1154M | D | 7.3e-06 | 1.4e-05 | 8.2e-06 | 9.833E-06 | 3.636E-06 | 7.300E-06 | 1.400E-05 |
| 23074 | 1154 | P1154F | D | 9.9e-06 | 1.2e-05 | 6.6e-06 | 9.500E-06 | 2.722E-06 | 6.600E-06 | 1.200E-05 |
| 23075 | 1154 | P1154P | D | 6.1e-06 | 1.3e-05 | 1.3e-05 | 1.070E-05 | 3.984E-06 | 6.100E-06 | 1.300E-05 |
| 23076 | 1154 | P1154S | D | 1.1e-05 | 6.7e-06 | 1.2e-05 | 9.900E-06 | 2.816E-06 | 6.700E-06 | 1.200E-05 |
| 23077 | 1154 | P1154T | D | 9.6e-06 | 5.8e-06 | 5.8e-06 | 7.067E-06 | 2.194E-06 | 5.800E-06 | 9.600E-06 |
| 23078 | 1154 | P1154W | D | 9.6e-06 | 9.8e-06 | 1.3e-05 | 1.080E-05 | 1.908E-06 | 9.600E-06 | 1.300E-05 |
| 23079 | 1154 | P1154Y | D | 6.7e-06 | 5.5e-06 | 5.5e-06 | 5.900E-06 | 6.928E-07 | 5.500E-06 | 6.700E-06 |
| 23080 | 1154 | P1154V | D | 1.3e-05 | 6e-06 | 6e-06 | 8.333E-06 | 4.041E-06 | 6.000E-06 | 1.300E-05 |
| 23081 | 1155 | T1155A | D | 8.9e-06 | 7.7e-06 | 9.3e-06 | 8.633E-06 | 8.327E-07 | 7.700E-06 | 9.300E-06 |
| 23082 | 1155 | T1155R | D | 9.4e-06 | 7.3e-06 | 6.4e-06 | 7.700E-06 | 1.539E-06 | 6.400E-06 | 9.400E-06 |
| 23083 | 1155 | T1155N | D | 6.7e-06 | 9.1e-06 | 4.6e-06 | 6.800E-06 | 2.252E-06 | 4.600E-06 | 9.100E-06 |
| 23084 | 1155 | T1155D | D | 9.7e-06 | 5.6e-06 | 1.1e-05 | 8.767E-06 | 2.818E-06 | 5.600E-06 | 1.100E-05 |
| 23085 | 1155 | T1155C | D | 6.4e-06 | 5.8e-06 | 4.7e-06 | 5.633E-06 | 8.622E-07 | 4.700E-06 | 6.400E-06 |
| 23086 | 1155 | T1155E | D | 6.3e-06 | 1e-05 | 5.2e-06 | 7.167E-06 | 2.515E-06 | 5.200E-06 | 1.000E-05 |
| 23087 | 1155 | T1155Q | D | 1.6e-05 | 7.7e-06 | 1.2e-05 | 1.190E-05 | 4.151E-06 | 7.700E-06 | 1.600E-05 |
| 23088 | 1155 | T1155G | D | 1e-05 | 7.4e-06 | 1e-05 | 9.133E-06 | 1.501E-06 | 7.400E-06 | 1.000E-05 |
| 23089 | 1155 | T1155H | D | 8.1e-06 | 7.1e-06 | 8.4e-06 | 7.867E-06 | 6.807E-07 | 7.100E-06 | 8.400E-06 |
| 23090 | 1155 | T1155I | D | 6.4e-06 | 9.9e-06 | 1.2e-05 | 9.433E-06 | 2.829E-06 | 6.400E-06 | 1.200E-05 |
| 23091 | 1155 | T1155L | D | 7e-06 | 8.5e-06 | 6.1e-06 | 7.200E-06 | 1.212E-06 | 6.100E-06 | 8.500E-06 |
| 23092 | 1155 | T1155K | D | 5.5e-06 | 6.4e-06 | 7.7e-06 | 6.533E-06 | 1.106E-06 | 5.500E-06 | 7.700E-06 |
| 23093 | 1155 | T1155M | D | 5.8e-06 | 1.1e-05 | 1.2e-05 | 9.600E-06 | 3.329E-06 | 5.800E-06 | 1.200E-05 |
| 23094 | 1155 | T1155F | D | 8.3e-06 | 5.5e-06 | 1.3e-05 | 8.933E-06 | 3.790E-06 | 5.500E-06 | 1.300E-05 |
| 23095 | 1155 | T1155P | D | 7e-06 | 5.2e-06 | 9.5e-06 | 7.233E-06 | 2.159E-06 | 5.200E-06 | 9.500E-06 |
| 23096 | 1155 | T1155S | D | 8.7e-06 | 5.9e-06 | 6.7e-06 | 7.100E-06 | 1.442E-06 | 5.900E-06 | 8.700E-06 |
| 23097 | 1155 | T1155T | D | 1.3e-05 | 8.3e-06 | 4.9e-06 | 8.733E-06 | 4.067E-06 | 4.900E-06 | 1.300E-05 |
| 23098 | 1155 | T1155W | D | 1.2e-05 | 6.8e-06 | 5.4e-06 | 8.067E-06 | 3.478E-06 | 5.400E-06 | 1.200E-05 |
| 23099 | 1155 | T1155Y | D | 1.1e-05 | 6.7e-06 | 6.4e-06 | 8.033E-06 | 2.574E-06 | 6.400E-06 | 1.100E-05 |
| 23100 | 1155 | T1155V | D | 9.5e-06 | 1e-05 | 9.2e-06 | 9.567E-06 | 4.041E-07 | 9.200E-06 | 1.000E-05 |
| 23101 | 1156 | F1156A | D | 1.7e-05 | 7.1e-06 | 8e-06 | 1.070E-05 | 5.474E-06 | 7.100E-06 | 1.700E-05 |
| 23102 | 1156 | F1156R | D | 9.6e-06 | 7e-06 | 6.1e-06 | 7.567E-06 | 1.818E-06 | 6.100E-06 | 9.600E-06 |
| 23103 | 1156 | F1156N | D | 4.1e-06 | 6.8e-06 | 7.7e-06 | 6.200E-06 | 1.873E-06 | 4.100E-06 | 7.700E-06 |
| 23104 | 1156 | F1156D | D | 1.1e-05 | 1.1e-05 | 5.4e-06 | 9.133E-06 | 3.233E-06 | 5.400E-06 | 1.100E-05 |
| 23105 | 1156 | F1156C | D | 8.3e-06 | 1.2e-05 | 8.8e-06 | 9.700E-06 | 2.007E-06 | 8.300E-06 | 1.200E-05 |
| 23106 | 1156 | F1156E | D | 1.1e-05 | 7e-06 | 4.6e-06 | 7.533E-06 | 3.233E-06 | 4.600E-06 | 1.100E-05 |
| 23107 | 1156 | F1156Q | D | 9.1e-06 | 4.9e-06 | 1.1e-05 | 8.333E-06 | 3.121E-06 | 4.900E-06 | 1.100E-05 |
| 23108 | 1156 | F1156G | D | 6.3e-06 | 6.8e-06 | 5.3e-06 | 6.133E-06 | 7.638E-07 | 5.300E-06 | 6.800E-06 |
| 23109 | 1156 | F1156H | D | 1e-05 | 7e-06 | 4.6e-06 | 7.200E-06 | 2.706E-06 | 4.600E-06 | 1.000E-05 |
| 23110 | 1156 | F1156I | D | 1.3e-05 | 1.2e-05 | 7.8e-06 | 1.093E-05 | 2.759E-06 | 7.800E-06 | 1.300E-05 |
| 23111 | 1156 | F1156L | D | 4.7e-06 | 5.7e-06 | 7.8e-06 | 6.067E-06 | 1.582E-06 | 4.700E-06 | 7.800E-06 |
| 23112 | 1156 | F1156K | D | 6e-06 | 6.3e-06 | 3.1e-06 | 5.133E-06 | 1.767E-06 | 3.100E-06 | 6.300E-06 |
| 23113 | 1156 | F1156M | D | 1e-05 | 1.1e-05 | 8e-06 | 9.667E-06 | 1.528E-06 | 8.000E-06 | 1.100E-05 |
| 23114 | 1156 | F1156F | D | 8.9e-06 | 9.3e-06 | 8.2e-06 | 8.800E-06 | 5.568E-07 | 8.200E-06 | 9.300E-06 |
| 23115 | 1156 | F1156P | D | 9.4e-06 | 1.3e-05 | 1.1e-05 | 1.113E-05 | 1.804E-06 | 9.400E-06 | 1.300E-05 |
| 23116 | 1156 | F1156S | D | 6.6e-06 | 6.5e-06 | 1e-05 | 7.700E-06 | 1.992E-06 | 6.500E-06 | 1.000E-05 |
| 23117 | 1156 | F1156T | D | 8.7e-06 | 5.8e-06 | 4.6e-06 | 6.367E-06 | 2.108E-06 | 4.600E-06 | 8.700E-06 |
| 23118 | 1156 | F1156W | D | 8.3e-06 | 8.9e-06 | 7.5e-06 | 8.233E-06 | 7.024E-07 | 7.500E-06 | 8.900E-06 |
| 23119 | 1156 | F1156Y | D | 1.3e-05 | 4.8e-06 | 1e-05 | 9.267E-06 | 4.149E-06 | 4.800E-06 | 1.300E-05 |

| | | | | | | | | | | |
|---|---|---|---|---|---|---|---|---|---|---|
| 23120 | 1156 | F1156V | D | 8.1e-06 | 8e-06 | 7.9e-06 | 8.000E-06 | 1.000E-07 | 7.900E-06 | 8.100E-06 |
| 23121 | 1157 | G1157A | D | 1.1e-05 | 9.9e-06 | 6e-06 | 8.967E-06 | 2.627E-06 | 6.000E-06 | 1.100E-05 |
| 23122 | 1157 | G1157R | D | 1.7e-05 | 5.1e-06 | 7.5e-06 | 9.867E-06 | 6.293E-06 | 5.100E-06 | 1.700E-05 |
| 23123 | 1157 | G1157N | D | 1.2e-05 | 7.1e-06 | 7.8e-06 | 8.967E-06 | 2.650E-06 | 7.100E-06 | 1.200E-05 |
| 23124 | 1157 | G1157D | D | 1.5e-05 | 7.6e-06 | 4.8e-06 | 9.133E-06 | 5.270E-06 | 4.800E-06 | 1.500E-05 |
| 23125 | 1157 | G1157C | D | 5.4e-06 | 1.8e-05 | 7.7e-06 | 1.037E-05 | 6.710E-06 | 5.400E-06 | 1.800E-05 |
| 23126 | 1157 | G1157E | D | 7.3e-06 | 1e-05 | 5.1e-06 | 7.467E-06 | 2.454E-06 | 5.100E-06 | 1.000E-05 |
| 23127 | 1157 | G1157Q | D | 1e-05 | 1.2e-05 | 9.4e-06 | 1.047E-05 | 1.361E-06 | 9.400E-06 | 1.200E-05 |
| 23128 | 1157 | G1157G | D | 6.9e-06 | 8.2e-06 | 1.3e-05 | 9.367E-06 | 3.213E-06 | 6.900E-06 | 1.300E-05 |
| 23129 | 1157 | G1157H | D | 7.5e-06 | 5.6e-06 | 9.9e-06 | 7.667E-06 | 2.155E-06 | 5.600E-06 | 9.900E-06 |
| 23130 | 1157 | G1157I | D | 9e-06 | 6.9e-06 | 7.3e-06 | 7.733E-06 | 1.115E-06 | 6.900E-06 | 9.000E-06 |
| 23131 | 1157 | G1157L | D | 7.1e-06 | 9.4e-06 | 6.3e-06 | 7.600E-06 | 1.609E-06 | 6.300E-06 | 9.400E-06 |
| 23132 | 1157 | G1157K | D | 6.6e-06 | 8.9e-06 | 6.3e-06 | 7.267E-06 | 1.422E-06 | 6.300E-06 | 8.900E-06 |
| 23133 | 1157 | G1157M | D | 2.8e-06 | 1.3e-05 | 1.1e-05 | 8.933E-06 | 5.405E-06 | 2.800E-06 | 1.300E-05 |
| 23134 | 1157 | G1157F | D | 4.7e-06 | 6.9e-06 | 6.1e-06 | 5.900E-06 | 1.114E-06 | 4.700E-06 | 6.900E-06 |
| 23135 | 1157 | G1157P | D | 5.5e-06 | 4.8e-06 | 1.2e-05 | 7.433E-06 | 3.970E-06 | 4.800E-06 | 1.200E-05 |
| 23136 | 1157 | G1157S | D | 9.1e-06 | 6.3e-06 | 7e-06 | 7.467E-06 | 1.457E-06 | 6.300E-06 | 9.100E-06 |
| 23137 | 1157 | G1157T | D | 6.4e-06 | 9.4e-06 | 1.8e-05 | 1.127E-05 | 6.021E-06 | 6.400E-06 | 1.800E-05 |
| 23138 | 1157 | G1157W | D | 9.6e-06 | 1.3e-05 | 1.3e-05 | 1.187E-05 | 1.963E-06 | 9.600E-06 | 1.300E-05 |
| 23139 | 1157 | G1157Y | D | 6.4e-06 | 6.9e-06 | 1.2e-05 | 8.433E-06 | 3.099E-06 | 6.400E-06 | 1.200E-05 |
| 23140 | 1157 | G1157V | D | 6.8e-06 | 1.4e-05 | 1.2e-05 | 1.093E-05 | 3.717E-06 | 6.800E-06 | 1.400E-05 |
| 23141 | 1158 | Q1158A | D | 9.3e-06 | 1.4e-05 | 1e-05 | 1.110E-05 | 2.536E-06 | 9.300E-06 | 1.400E-05 |
| 23142 | 1158 | Q1158R | D | 6.7e-06 | 6.1e-06 | 5e-06 | 5.933E-06 | 8.622E-07 | 5.000E-06 | 6.700E-06 |
| 23143 | 1158 | Q1158N | D | 6.2e-06 | 6.9e-06 | 8e-06 | 7.033E-06 | 9.074E-07 | 6.200E-06 | 8.000E-06 |
| 23144 | 1158 | Q1158D | D | 7.9e-06 | 6.1e-06 | 1e-05 | 8.000E-06 | 1.952E-06 | 6.100E-06 | 1.000E-05 |
| 23145 | 1158 | Q1158C | D | 9.5e-06 | 9.3e-06 | 5.7e-06 | 8.167E-06 | 2.139E-06 | 5.700E-06 | 9.500E-06 |
| 23146 | 1158 | Q1158E | D | 8.5e-06 | 5.3e-06 | 1.6e-05 | 9.933E-06 | 5.492E-06 | 5.300E-06 | 1.600E-05 |
| 23147 | 1158 | Q1158Q | D | 6.1e-06 | 6.5e-06 | 6.1e-06 | 6.233E-06 | 2.309E-07 | 6.100E-06 | 6.500E-06 |
| 23148 | 1158 | Q1158G | D | 1.4e-05 | 9.2e-06 | 6.3e-06 | 9.833E-06 | 3.889E-06 | 6.300E-06 | 1.400E-05 |
| 23149 | 1158 | Q1158H | D | 5.2e-06 | 4.2e-06 | 9.5e-06 | 6.300E-06 | 2.816E-06 | 4.200E-06 | 9.500E-06 |
| 23150 | 1158 | Q1158I | D | 9.9e-06 | 5.4e-06 | 8.1e-06 | 7.800E-06 | 2.265E-06 | 5.400E-06 | 9.900E-06 |
| 23151 | 1158 | Q1158L | D | 4.7e-06 | 8.2e-06 | 8.3e-06 | 7.067E-06 | 2.050E-06 | 4.700E-06 | 8.300E-06 |
| 23152 | 1158 | Q1158K | D | 7.9e-06 | 8.8e-06 | 4.7e-06 | 7.133E-06 | 2.155E-06 | 4.700E-06 | 8.800E-06 |
| 23153 | 1158 | Q1158M | D | 7.8e-06 | 6.5e-06 | 1e-05 | 8.100E-06 | 1.769E-06 | 6.500E-06 | 1.000E-05 |
| 23154 | 1158 | Q1158F | D | 9.3e-06 | 7.5e-06 | 8.9e-06 | 8.567E-06 | 9.452E-07 | 7.500E-06 | 9.300E-06 |
| 23155 | 1158 | Q1158P | D | 1e-05 | 1.5e-05 | 4.7e-06 | 9.900E-06 | 5.151E-06 | 4.700E-06 | 1.500E-05 |
| 23156 | 1158 | Q1158S | D | 9e-06 | 6.4e-06 | 8.6e-06 | 8.000E-06 | 1.400E-06 | 6.400E-06 | 9.000E-06 |
| 23157 | 1158 | Q1158T | D | 5.7e-06 | 3.3e-06 | 5.7e-06 | 4.900E-06 | 1.386E-06 | 3.300E-06 | 5.700E-06 |
| 23158 | 1158 | Q1158W | D | 7.1e-06 | 6.6e-06 | 7.6e-06 | 7.100E-06 | 5.000E-07 | 6.600E-06 | 7.600E-06 |
| 23159 | 1158 | Q1158Y | D | 1e-05 | 4.4e-06 | 5.9e-06 | 6.767E-06 | 2.899E-06 | 4.400E-06 | 1.000E-05 |
| 23160 | 1158 | Q1158V | D | 8.5e-06 | 1e-05 | 1.2e-05 | 1.017E-05 | 1.756E-06 | 8.500E-06 | 1.200E-05 |
| 23161 | 1159 | G1159A | D | 8.4e-06 | 8.9e-06 | 1.1e-05 | 9.433E-06 | 1.380E-06 | 8.400E-06 | 1.100E-05 |
| 23162 | 1159 | G1159R | D | 9.5e-06 | 5.4e-06 | 6.8e-06 | 7.233E-06 | 2.084E-06 | 5.400E-06 | 9.500E-06 |
| 23163 | 1159 | G1159N | D | 5.4e-06 | 6.1e-06 | 5.5e-06 | 5.667E-06 | 3.786E-07 | 5.400E-06 | 6.100E-06 |
| 23164 | 1159 | G1159D | D | 8.9e-06 | 8.4e-06 | 4.4e-06 | 7.233E-06 | 2.466E-06 | 4.400E-06 | 8.900E-06 |
| 23165 | 1159 | G1159C | D | 9.2e-06 | 9.7e-06 | 9e-06 | 9.300E-06 | 3.606E-07 | 9.000E-06 | 9.700E-06 |
| 23166 | 1159 | G1159E | D | 6.7e-06 | 7.6e-06 | 1.2e-05 | 8.767E-06 | 2.836E-06 | 6.700E-06 | 1.200E-05 |
| 23167 | 1159 | G1159Q | D | 3.9e-06 | 1.2e-05 | 1.2e-05 | 9.300E-06 | 4.677E-06 | 3.900E-06 | 1.200E-05 |
| 23168 | 1159 | G1159G | D | 1.4e-05 | 1e-05 | 8.4e-06 | 1.080E-05 | 2.884E-06 | 8.400E-06 | 1.400E-05 |
| 23169 | 1159 | G1159H | D | 6.7e-06 | 5.4e-06 | 6.6e-06 | 6.233E-06 | 7.234E-07 | 5.400E-06 | 6.700E-06 |
| 23170 | 1159 | G1159I | D | 8.8e-06 | 9.2e-06 | 9.5e-06 | 9.167E-06 | 3.512E-07 | 8.800E-06 | 9.500E-06 |
| 23171 | 1159 | G1159L | D | 1.2e-05 | 8.8e-06 | 9.2e-06 | 1.000E-05 | 1.744E-06 | 8.800E-06 | 1.200E-05 |
| 23172 | 1159 | G1159K | D | 6e-06 | 6e-06 | 1.1e-05 | 7.667E-06 | 2.887E-06 | 6.000E-06 | 1.100E-05 |
| 23173 | 1159 | G1159M | D | 5.4e-06 | 6e-06 | 9.6e-06 | 7.000E-06 | 2.272E-06 | 5.400E-06 | 9.600E-06 |
| 23174 | 1159 | G1159F | D | 6.2e-06 | 5e-06 | 6.3e-06 | 5.833E-06 | 7.234E-07 | 5.000E-06 | 6.300E-06 |
| 23175 | 1159 | G1159P | D | 6.4e-06 | 6.1e-06 | 7.9e-06 | 6.800E-06 | 9.644E-07 | 6.100E-06 | 7.900E-06 |
| 23176 | 1159 | G1159S | D | 4.4e-06 | 6.8e-06 | 9.7e-06 | 6.967E-06 | 2.654E-06 | 4.400E-06 | 9.700E-06 |
| 23177 | 1159 | G1159T | D | 1e-05 | 9e-06 | 8.4e-06 | 9.133E-06 | 8.083E-07 | 8.400E-06 | 1.000E-05 |
| 23178 | 1159 | G1159W | D | 6.3e-06 | 8.9e-06 | 8.2e-06 | 7.800E-06 | 1.345E-06 | 6.300E-06 | 8.900E-06 |
| 23179 | 1159 | G1159Y | D | 1.4e-05 | 1.3e-05 | 5.4e-06 | 1.080E-05 | 4.703E-06 | 5.400E-06 | 1.400E-05 |
| 23180 | 1159 | G1159V | D | 6.9e-06 | 5.7e-06 | 9.4e-06 | 7.333E-06 | 1.888E-06 | 5.700E-06 | 9.400E-06 |
| 23181 | 1160 | T1160A | D | 6.4e-06 | 8.5e-06 | 9.2e-06 | 8.033E-06 | 1.457E-06 | 6.400E-06 | 9.200E-06 |
| 23182 | 1160 | T1160R | D | 1e-05 | 1.1e-05 | 1.2e-05 | 1.100E-05 | 1.000E-06 | 1.000E-05 | 1.200E-05 |
| 23183 | 1160 | T1160N | D | 6.8e-06 | 1.1e-05 | 8.8e-06 | 8.867E-06 | 2.101E-06 | 6.800E-06 | 1.100E-05 |
| 23184 | 1160 | T1160D | D | 8.7e-06 | 1.4e-05 | 6.8e-06 | 9.833E-06 | 3.731E-06 | 6.800E-06 | 1.400E-05 |
| 23185 | 1160 | T1160C | D | 7.7e-06 | 1.3e-05 | 6.3e-06 | 9.000E-06 | 3.534E-06 | 6.300E-06 | 1.300E-05 |
| 23186 | 1160 | T1160E | D | 6.6e-06 | 8e-06 | 6.5e-06 | 7.033E-06 | 8.386E-07 | 6.500E-06 | 8.000E-06 |
| 23187 | 1160 | T1160Q | D | 7.2e-06 | 6.3e-06 | 1.3e-05 | 8.833E-06 | 3.636E-06 | 6.300E-06 | 1.300E-05 |
| 23188 | 1160 | T1160G | D | 1.4e-05 | 1.4e-05 | 7e-06 | 1.167E-05 | 4.041E-06 | 7.000E-06 | 1.400E-05 |
| 23189 | 1160 | T1160H | D | 8.5e-06 | 3.2e-06 | 8e-06 | 6.567E-06 | 2.926E-06 | 3.200E-06 | 8.500E-06 |
| 23190 | 1160 | T1160I | D | 5.6e-06 | 6e-06 | 4.8e-06 | 5.467E-06 | 6.110E-07 | 4.800E-06 | 6.000E-06 |
| 23191 | 1160 | T1160L | D | 1.1e-05 | 1.1e-05 | 1e-05 | 1.067E-05 | 5.774E-07 | 1.000E-05 | 1.100E-05 |
| 23192 | 1160 | T1160K | D | 7.1e-06 | 1.3e-05 | 8.3e-06 | 9.467E-06 | 3.118E-06 | 7.100E-06 | 1.300E-05 |
| 23193 | 1160 | T1160M | D | 5.6e-06 | 1.8e-05 | 8.1e-06 | 1.057E-05 | 6.558E-06 | 5.600E-06 | 1.800E-05 |
| 23194 | 1160 | T1160F | D | 8.6e-06 | 4.9e-06 | 7.1e-06 | 6.867E-06 | 1.861E-06 | 4.900E-06 | 8.600E-06 |
| 23195 | 1160 | T1160P | D | 4.6e-06 | 9.4e-06 | 8.2e-06 | 7.400E-06 | 2.498E-06 | 4.600E-06 | 9.400E-06 |
| 23196 | 1160 | T1160S | D | 6.4e-06 | 8.4e-06 | 8.2e-06 | 7.667E-06 | 1.102E-06 | 6.400E-06 | 8.400E-06 |
| 23197 | 1160 | T1160T | D | 5.1e-06 | 7.2e-06 | 1e-05 | 7.433E-06 | 2.458E-06 | 5.100E-06 | 1.000E-05 |
| 23198 | 1160 | T1160W | D | 1.1e-05 | 9.5e-06 | 9.4e-06 | 9.967E-06 | 8.963E-07 | 9.400E-06 | 1.100E-05 |
| 23199 | 1160 | T1160Y | D | 6.4e-06 | 6.2e-06 | 8.6e-06 | 7.067E-06 | 1.332E-06 | 6.200E-06 | 8.600E-06 |
| 23200 | 1160 | T1160V | D | 5.3e-06 | 1.5e-05 | 5.4e-06 | 8.567E-06 | 5.572E-06 | 5.300E-06 | 1.500E-05 |
| 23201 | 1161 | K1161A | D | 4.9e-06 | 1.3e-05 | 8.5e-06 | 8.800E-06 | 4.058E-06 | 4.900E-06 | 1.300E-05 |
| 23202 | 1161 | K1161R | D | 8.6e-06 | 9.6e-06 | 6.4e-06 | 8.200E-06 | 1.637E-06 | 6.400E-06 | 9.600E-06 |
| 23203 | 1161 | K1161N | D | 6.7e-06 | 7.3e-06 | 9.4e-06 | 7.800E-06 | 1.418E-06 | 6.700E-06 | 9.400E-06 |
| 23204 | 1161 | K1161D | D | 6.5e-06 | 4.6e-06 | 5.1e-06 | 5.400E-06 | 9.849E-07 | 4.600E-06 | 6.500E-06 |

| | | | | | | | | | | |
|---|---|---|---|---|---|---|---|---|---|---|
| 23205 | 1161 | K1161C | D | 6.5e-06 | 5.1e-06 | 4.8e-06 | 5.467E-06 | 9.074E-07 | 4.800E-06 | 6.500E-06 |
| 23206 | 1161 | K1161E | D | 7.1e-06 | 6.6e-06 | 8.1e-06 | 7.267E-06 | 7.638E-07 | 6.600E-06 | 8.100E-06 |
| 23207 | 1161 | K1161Q | D | 6.4e-06 | 4.4e-06 | 1.2e-05 | 7.600E-06 | 3.940E-06 | 4.400E-06 | 1.200E-05 |
| 23208 | 1161 | K1161G | D | 1.2e-05 | 8.7e-06 | 7.1e-06 | 9.267E-06 | 2.499E-06 | 7.100E-06 | 1.200E-05 |
| 23209 | 1161 | K1161H | D | 4e-06 | 9.3e-06 | 1.2e-05 | 8.433E-06 | 4.070E-06 | 4.000E-06 | 1.200E-05 |
| 23210 | 1161 | K1161I | D | 8e-06 | 6.7e-06 | 1.3e-05 | 9.233E-06 | 3.326E-06 | 6.700E-06 | 1.300E-05 |
| 23211 | 1161 | K1161L | D | 6.6e-06 | 1e-05 | 1.1e-05 | 9.200E-06 | 2.307E-06 | 6.600E-06 | 1.100E-05 |
| 23212 | 1161 | K1161K | D | 1e-05 | 8.9e-06 | 4.7e-06 | 7.867E-06 | 2.797E-06 | 4.700E-06 | 1.000E-05 |
| 23213 | 1161 | K1161M | D | 1.3e-05 | 1e-05 | 8.6e-06 | 1.053E-05 | 2.248E-06 | 8.600E-06 | 1.300E-05 |
| 23214 | 1161 | K1161F | D | 6.4e-06 | 1.4e-05 | 5.2e-06 | 8.533E-06 | 4.772E-06 | 5.200E-06 | 1.400E-05 |
| 23215 | 1161 | K1161P | D | 7e-06 | 4.8e-06 | 1e-05 | 7.267E-06 | 2.610E-06 | 4.800E-06 | 1.000E-05 |
| 23216 | 1161 | K1161S | D | 9e-06 | 8.9e-06 | 7.2e-06 | 8.367E-06 | 1.012E-06 | 7.200E-06 | 9.000E-06 |
| 23217 | 1161 | K1161T | D | 7.6e-06 | 6.2e-06 | 1.1e-05 | 8.267E-06 | 2.468E-06 | 6.200E-06 | 1.100E-05 |
| 23218 | 1161 | K1161W | D | 5e-06 | 6.1e-06 | 7.5e-06 | 6.200E-06 | 1.253E-06 | 5.000E-06 | 7.500E-06 |
| 23219 | 1161 | K1161Y | D | 1.5e-05 | 1.1e-05 | 4.6e-06 | 1.020E-05 | 5.246E-06 | 4.600E-06 | 1.500E-05 |
| 23220 | 1161 | K1161V | D | 1.1e-05 | 4.4e-06 | 4.7e-06 | 6.700E-06 | 3.727E-06 | 4.400E-06 | 1.100E-05 |
| 23221 | 1162 | V1162A | D | 9.8e-06 | 6.5e-06 | 9.6e-06 | 8.633E-06 | 1.850E-06 | 6.500E-06 | 9.800E-06 |
| 23222 | 1162 | V1162R | D | 4.8e-06 | 8.9e-06 | 5.1e-06 | 6.267E-06 | 2.285E-06 | 4.800E-06 | 8.900E-06 |
| 23223 | 1162 | V1162N | D | 6.5e-06 | 1.2e-05 | 3.9e-06 | 7.467E-06 | 4.136E-06 | 3.900E-06 | 1.200E-05 |
| 23224 | 1162 | V1162D | D | 6.2e-06 | 9.5e-06 | 8.3e-06 | 8.000E-06 | 1.670E-06 | 6.200E-06 | 9.500E-06 |
| 23225 | 1162 | V1162C | D | 1e-05 | 9.9e-06 | 8.7e-06 | 9.533E-06 | 7.234E-07 | 8.700E-06 | 1.000E-05 |
| 23226 | 1162 | V1162E | D | 8.9e-06 | 9.7e-06 | 6.9e-06 | 8.500E-06 | 1.442E-06 | 6.900E-06 | 9.700E-06 |
| 23227 | 1162 | V1162Q | D | 7.2e-06 | 8.1e-06 | 4.1e-06 | 6.467E-06 | 2.098E-06 | 4.100E-06 | 8.100E-06 |
| 23228 | 1162 | V1162G | D | 8.2e-06 | 1.2e-05 | 4.8e-06 | 8.333E-06 | 3.602E-06 | 4.800E-06 | 1.200E-05 |
| 23229 | 1162 | V1162H | D | 5.9e-06 | 6.7e-06 | 6.2e-06 | 6.267E-06 | 4.041E-07 | 5.900E-06 | 6.700E-06 |
| 23230 | 1162 | V1162I | D | 1.2e-05 | 1.2e-05 | 8.2e-06 | 1.073E-05 | 2.194E-06 | 8.200E-06 | 1.200E-05 |
| 23231 | 1162 | V1162L | D | 8e-06 | 6.4e-06 | 1.1e-05 | 8.467E-06 | 2.335E-06 | 6.400E-06 | 1.100E-05 |
| 23232 | 1162 | V1162K | D | 3.7e-06 | 4.6e-06 | 6.8e-06 | 5.033E-06 | 1.595E-06 | 3.700E-06 | 6.800E-06 |
| 23233 | 1162 | V1162M | D | 1.4e-05 | 1e-05 | 1.1e-05 | 1.167E-05 | 2.082E-06 | 1.000E-05 | 1.400E-05 |
| 23234 | 1162 | V1162F | D | 1.4e-05 | 1e-05 | 6e-06 | 1.000E-05 | 4.000E-06 | 6.000E-06 | 1.400E-05 |
| 23235 | 1162 | V1162P | D | 6.4e-06 | 6.1e-06 | 1.1e-05 | 7.833E-06 | 2.747E-06 | 6.100E-06 | 1.100E-05 |
| 23236 | 1162 | V1162S | D | 8.4e-06 | 1.1e-05 | 6.9e-06 | 8.767E-06 | 2.074E-06 | 6.900E-06 | 1.100E-05 |
| 23237 | 1162 | V1162T | D | 5.9e-06 | 4.8e-06 | 1.7e-05 | 9.233E-06 | 6.749E-06 | 4.800E-06 | 1.700E-05 |
| 23238 | 1162 | V1162W | D | 4.8e-06 | 8.1e-06 | 8.8e-06 | 7.233E-06 | 2.136E-06 | 4.800E-06 | 8.800E-06 |
| 23239 | 1162 | V1162Y | D | 1.1e-05 | 6.6e-06 | 1.3e-05 | 1.020E-05 | 3.274E-06 | 6.600E-06 | 1.300E-05 |
| 23240 | 1162 | V1162V | D | 1e-05 | 5.4e-06 | 5.8e-06 | 7.067E-06 | 2.548E-06 | 5.400E-06 | 1.000E-05 |
| 23241 | 1163 | E1163A | D | 6.1e-06 | 3.3e-06 | 8.3e-06 | 5.900E-06 | 2.506E-06 | 3.300E-06 | 8.300E-06 |
| 23242 | 1163 | E1163R | D | 5.3e-06 | 7.9e-06 | 1e-05 | 7.733E-06 | 2.354E-06 | 5.300E-06 | 1.000E-05 |
| 23243 | 1163 | E1163N | D | 7.2e-06 | 7.2e-06 | 6.5e-06 | 6.967E-06 | 4.041E-07 | 6.500E-06 | 7.200E-06 |
| 23244 | 1163 | E1163D | D | 7.6e-06 | 9e-06 | 9.2e-06 | 8.600E-06 | 8.718E-07 | 7.600E-06 | 9.200E-06 |
| 23245 | 1163 | E1163C | D | 5.9e-06 | 6e-06 | 5.4e-06 | 5.767E-06 | 3.215E-07 | 5.400E-06 | 6.000E-06 |
| 23246 | 1163 | E1163E | D | 9.1e-06 | 9.5e-06 | 9e-06 | 9.200E-06 | 2.646E-07 | 9.000E-06 | 9.500E-06 |
| 23247 | 1163 | E1163Q | D | 9.3e-06 | 9.3e-06 | 6.7e-06 | 8.433E-06 | 1.501E-06 | 6.700E-06 | 9.300E-06 |
| 23248 | 1163 | E1163G | D | 6e-06 | 7.8e-06 | 6.3e-06 | 6.700E-06 | 9.644E-07 | 6.000E-06 | 7.800E-06 |
| 23249 | 1163 | E1163H | D | 9.6e-06 | 6.7e-06 | 5.8e-06 | 7.367E-06 | 1.986E-06 | 5.800E-06 | 9.600E-06 |
| 23250 | 1163 | E1163I | D | 8e-06 | 1.2e-05 | 6.7e-06 | 8.900E-06 | 2.762E-06 | 6.700E-06 | 1.200E-05 |
| 23251 | 1163 | E1163L | D | 1.3e-05 | 9.6e-06 | 1.1e-05 | 1.120E-05 | 1.709E-06 | 9.600E-06 | 1.300E-05 |
| 23252 | 1163 | E1163K | D | 1.2e-05 | 6.8e-06 | 7.5e-06 | 8.767E-06 | 2.822E-06 | 6.800E-06 | 1.200E-05 |
| 23253 | 1163 | E1163M | D | 7.5e-06 | 5.8e-06 | 8.9e-06 | 7.400E-06 | 1.552E-06 | 5.800E-06 | 8.900E-06 |
| 23254 | 1163 | E1163F | D | 7.6e-06 | 1.1e-05 | 7.1e-06 | 8.567E-06 | 2.122E-06 | 7.100E-06 | 1.100E-05 |
| 23255 | 1163 | E1163P | D | 9.3e-06 | 8.6e-06 | 4.8e-06 | 7.567E-06 | 2.421E-06 | 4.800E-06 | 9.300E-06 |
| 23256 | 1163 | E1163S | D | 6.4e-06 | 4.9e-06 | 4.8e-06 | 5.367E-06 | 8.963E-07 | 4.800E-06 | 6.400E-06 |
| 23257 | 1163 | E1163T | D | 7.1e-06 | 7.3e-06 | 1.8e-05 | 1.080E-05 | 6.236E-06 | 7.100E-06 | 1.800E-05 |
| 23258 | 1163 | E1163W | D | 4.9e-06 | 4.6e-06 | 6.5e-06 | 5.333E-06 | 1.021E-06 | 4.600E-06 | 6.500E-06 |
| 23259 | 1163 | E1163Y | D | 8.5e-06 | 1.1e-05 | 6.2e-06 | 8.567E-06 | 2.401E-06 | 6.200E-06 | 1.100E-05 |
| 23260 | 1163 | E1163V | D | 4.9e-06 | 5.5e-06 | 9.2e-06 | 6.533E-06 | 2.329E-06 | 4.900E-06 | 9.200E-06 |
| 23261 | 1164 | I1164A | D | 7.5e-06 | 8.3e-06 | 5.5e-06 | 7.100E-06 | 1.442E-06 | 5.500E-06 | 8.300E-06 |
| 23262 | 1164 | I1164R | D | 5.5e-06 | 5.1e-06 | 6.8e-06 | 5.800E-06 | 8.888E-07 | 5.100E-06 | 6.800E-06 |
| 23263 | 1164 | I1164N | D | 9.1e-06 | 8.7e-06 | 1.6e-05 | 1.127E-05 | 4.104E-06 | 8.700E-06 | 1.600E-05 |
| 23264 | 1164 | I1164D | D | 9.5e-06 | 9.5e-06 | 6.3e-06 | 8.433E-06 | 1.848E-06 | 6.300E-06 | 9.500E-06 |
| 23265 | 1164 | I1164C | D | 6.3e-06 | 8.3e-06 | 8.1e-06 | 7.567E-06 | 1.102E-06 | 6.300E-06 | 8.300E-06 |
| 23266 | 1164 | I1164E | D | 4.4e-06 | 6e-06 | 9.8e-06 | 6.733E-06 | 2.774E-06 | 4.400E-06 | 9.800E-06 |
| 23267 | 1164 | I1164Q | D | 7.5e-06 | 7.3e-06 | 5.5e-06 | 6.767E-06 | 1.102E-06 | 5.500E-06 | 7.500E-06 |
| 23268 | 1164 | I1164G | D | 1e-05 | 6.2e-06 | 1e-05 | 8.733E-06 | 2.194E-06 | 6.200E-06 | 1.000E-05 |
| 23269 | 1164 | I1164H | D | 6.5e-06 | 9.7e-06 | 7.3e-06 | 7.833E-06 | 1.665E-06 | 6.500E-06 | 9.700E-06 |
| 23270 | 1164 | I1164I | D | 1.1e-05 | 4.3e-06 | 7.5e-06 | 7.600E-06 | 3.351E-06 | 4.300E-06 | 1.100E-05 |
| 23271 | 1164 | I1164L | D | 9.4e-06 | 9.5e-06 | 9.2e-06 | 9.367E-06 | 1.528E-07 | 9.200E-06 | 9.500E-06 |
| 23272 | 1164 | I1164K | D | 8.9e-06 | 8.5e-06 | 1e-05 | 9.133E-06 | 7.767E-07 | 8.500E-06 | 1.000E-05 |
| 23273 | 1164 | I1164M | D | 6.6e-06 | 7.2e-06 | 8.1e-06 | 7.300E-06 | 7.550E-07 | 6.600E-06 | 8.100E-06 |
| 23274 | 1164 | I1164F | D | 1.5e-05 | 5.6e-06 | 1e-05 | 1.020E-05 | 4.703E-06 | 5.600E-06 | 1.500E-05 |
| 23275 | 1164 | I1164P | D | 1.2e-05 | 6.3e-06 | 9.7e-06 | 9.333E-06 | 2.868E-06 | 6.300E-06 | 1.200E-05 |
| 23276 | 1164 | I1164S | D | 5e-06 | 6.3e-06 | 1.2e-05 | 7.767E-06 | 3.723E-06 | 5.000E-06 | 1.200E-05 |
| 23277 | 1164 | I1164T | D | 9.8e-06 | 9.9e-06 | 6.1e-06 | 8.600E-06 | 2.166E-06 | 6.100E-06 | 9.900E-06 |
| 23278 | 1164 | I1164W | D | 6.8e-06 | 6.2e-06 | 5.4e-06 | 6.133E-06 | 7.024E-07 | 5.400E-06 | 6.800E-06 |
| 23279 | 1164 | I1164Y | D | 3.9e-06 | 6.4e-06 | 7.8e-06 | 6.033E-06 | 1.976E-06 | 3.900E-06 | 7.800E-06 |
| 23280 | 1164 | I1164V | D | 9.6e-06 | 5.9e-06 | 1e-05 | 8.500E-06 | 2.261E-06 | 5.900E-06 | 1.000E-05 |
| 23281 | 1165 | K1165A | D | 5.8e-06 | 1.5e-05 | 9.9e-06 | 1.023E-05 | 4.609E-06 | 5.800E-06 | 1.500E-05 |
| 23282 | 1165 | K1165R | D | 5.9e-06 | 1.1e-05 | 1e-05 | 8.967E-06 | 2.702E-06 | 5.900E-06 | 1.100E-05 |
| 23283 | 1165 | K1165N | D | 7.7e-06 | 7.2e-06 | 7.5e-06 | 7.467E-06 | 2.517E-07 | 7.200E-06 | 7.700E-06 |
| 23284 | 1165 | K1165D | D | 6.9e-06 | 7.7e-06 | 9.6e-06 | 8.067E-06 | 1.387E-06 | 6.900E-06 | 9.600E-06 |
| 23285 | 1165 | K1165C | D | 1.3e-05 | 6e-06 | 9.3e-06 | 9.433E-06 | 3.502E-06 | 6.000E-06 | 1.300E-05 |
| 23286 | 1165 | K1165E | D | 5e-06 | 5.8e-06 | 6.6e-06 | 5.800E-06 | 8.000E-07 | 5.000E-06 | 6.600E-06 |
| 23287 | 1165 | K1165Q | D | 7.5e-06 | 1.1e-05 | 5.6e-06 | 8.033E-06 | 2.739E-06 | 5.600E-06 | 1.100E-05 |
| 23288 | 1165 | K1165G | D | 7e-06 | 1e-05 | 9.6e-06 | 8.867E-06 | 1.629E-06 | 7.000E-06 | 1.000E-05 |
| 23289 | 1165 | K1165H | D | 6.8e-06 | 8.8e-06 | 1.3e-05 | 9.533E-06 | 3.164E-06 | 6.800E-06 | 1.300E-05 |

| | | | | | | | | | | |
|---|---|---|---|---|---|---|---|---|---|---|
| 23290 | 1165 | K1165I | D | 8e-06 | 4.2e-06 | 1e-05 | 7.400E-06 | 2.946E-06 | 4.200E-06 | 1.000E-05 |
| 23291 | 1165 | K1165L | D | 8.1e-06 | 8.4e-06 | 4.2e-06 | 6.900E-06 | 2.343E-06 | 4.200E-06 | 8.400E-06 |
| 23292 | 1165 | K1165K | D | 8.5e-06 | 7.5e-06 | 5.2e-06 | 7.067E-06 | 1.692E-06 | 5.200E-06 | 8.500E-06 |
| 23293 | 1165 | K1165M | D | 1.2e-05 | 9.8e-06 | 4.4e-06 | 8.733E-06 | 3.911E-06 | 4.400E-06 | 1.200E-05 |
| 23294 | 1165 | K1165F | D | 9.1e-06 | 1.1e-05 | 7e-06 | 9.033E-06 | 2.001E-06 | 7.000E-06 | 1.100E-05 |
| 23295 | 1165 | K1165P | D | 5.2e-06 | 5.6e-06 | 6.1e-06 | 5.633E-06 | 4.509E-07 | 5.200E-06 | 6.100E-06 |
| 23296 | 1165 | K1165S | D | 6.7e-06 | 7e-06 | 9.3e-06 | 7.667E-06 | 1.422E-06 | 6.700E-06 | 9.300E-06 |
| 23297 | 1165 | K1165T | D | 8e-06 | 9.7e-06 | 1.2e-05 | 9.900E-06 | 2.007E-06 | 8.000E-06 | 1.200E-05 |
| 23298 | 1165 | K1165W | D | 5.6e-06 | 6.7e-06 | 4.3e-06 | 5.533E-06 | 1.201E-06 | 4.300E-06 | 6.700E-06 |
| 23299 | 1165 | K1165Y | D | 1.1e-05 | 7e-06 | 7.8e-06 | 8.600E-06 | 2.117E-06 | 7.000E-06 | 1.100E-05 |
| 23300 | 1165 | K1165V | D | 4.7e-06 | 9.6e-06 | 9.2e-06 | 7.833E-06 | 2.721E-06 | 4.700E-06 | 9.600E-06 |
| 23301 | 1166 | R1166A | D | 8.2e-06 | 4.8e-06 | 9.1e-06 | 7.367E-06 | 2.268E-06 | 4.800E-06 | 9.100E-06 |
| 23302 | 1166 | R1166R | D | 9.3e-06 | 9.3e-06 | 8e-06 | 8.867E-06 | 7.506E-07 | 8.000E-06 | 9.300E-06 |
| 23303 | 1166 | R1166N | D | 6.8e-06 | 6.6e-06 | 6.6e-06 | 6.667E-06 | 1.155E-07 | 6.600E-06 | 6.800E-06 |
| 23304 | 1166 | R1166D | D | 8.5e-06 | 8.4e-06 | 5.5e-06 | 7.467E-06 | 1.704E-06 | 5.500E-06 | 8.500E-06 |
| 23305 | 1166 | R1166C | D | 7e-06 | 9.7e-06 | 5.5e-06 | 7.400E-06 | 2.128E-06 | 5.500E-06 | 9.700E-06 |
| 23306 | 1166 | R1166E | D | 6.1e-06 | 8e-06 | 1.1e-05 | 8.367E-06 | 2.470E-06 | 6.100E-06 | 1.100E-05 |
| 23307 | 1166 | R1166Q | D | 8.1e-06 | 1.1e-05 | 7.8e-06 | 8.967E-06 | 1.767E-06 | 7.800E-06 | 1.100E-05 |
| 23308 | 1166 | R1166G | D | 5.7e-06 | 8.9e-06 | 1.2e-05 | 8.867E-06 | 3.150E-06 | 5.700E-06 | 1.200E-05 |
| 23309 | 1166 | R1166H | D | 5e-06 | 9.2e-06 | 9.3e-06 | 7.833E-06 | 2.454E-06 | 5.000E-06 | 9.300E-06 |
| 23310 | 1166 | R1166I | D | 1.6e-05 | 6.8e-06 | 1e-05 | 1.093E-05 | 4.670E-06 | 6.800E-06 | 1.600E-05 |
| 23311 | 1166 | R1166L | D | 8.9e-06 | 2.1e-05 | 5.5e-06 | 1.180E-05 | 8.147E-06 | 5.500E-06 | 2.100E-05 |
| 23312 | 1166 | R1166K | D | 5.5e-06 | 6.9e-06 | 1.6e-05 | 9.467E-06 | 5.701E-06 | 5.500E-06 | 1.600E-05 |
| 23313 | 1166 | R1166M | D | 4.8e-06 | 9.1e-06 | 8.5e-06 | 7.467E-06 | 2.329E-06 | 4.800E-06 | 9.100E-06 |
| 23314 | 1166 | R1166F | D | 9.7e-06 | 5.5e-06 | 4.6e-06 | 6.600E-06 | 2.722E-06 | 4.600E-06 | 9.700E-06 |
| 23315 | 1166 | R1166P | D | 7.6e-06 | 1.3e-05 | 8e-06 | 9.533E-06 | 3.009E-06 | 7.600E-06 | 1.300E-05 |
| 23316 | 1166 | R1166S | D | 3.6e-06 | 7.4e-06 | 9.6e-06 | 6.867E-06 | 3.035E-06 | 3.600E-06 | 9.600E-06 |
| 23317 | 1166 | R1166T | D | 9.6e-06 | 1.3e-05 | 1e-05 | 1.087E-05 | 1.858E-06 | 9.600E-06 | 1.300E-05 |
| 23318 | 1166 | R1166W | D | 6.5e-06 | 1.3e-05 | 1.2e-05 | 1.050E-05 | 3.500E-06 | 6.500E-06 | 1.300E-05 |
| 23319 | 1166 | R1166Y | D | 4.9e-06 | 7e-06 | 6.4e-06 | 6.100E-06 | 1.082E-06 | 4.900E-06 | 7.000E-06 |
| 23320 | 1166 | R1166V | D | 7.4e-06 | 1e-05 | 5.9e-06 | 7.767E-06 | 2.074E-06 | 5.900E-06 | 1.000E-05 |
| 23321 | 1167 | T1167A | D | 1.2e-05 | 3.3e-06 | 6.7e-06 | 7.333E-06 | 4.384E-06 | 3.300E-06 | 1.200E-05 |
| 23322 | 1167 | T1167R | D | 7.1e-06 | 8.6e-06 | 9.4e-06 | 8.367E-06 | 1.168E-06 | 7.100E-06 | 9.400E-06 |
| 23323 | 1167 | T1167N | D | 8.6e-06 | 2.9e-06 | 6.8e-06 | 6.100E-06 | 2.914E-06 | 2.900E-06 | 8.600E-06 |
| 23324 | 1167 | T1167D | D | 1e-05 | 9.7e-06 | 6.7e-06 | 8.800E-06 | 1.825E-06 | 6.700E-06 | 1.000E-05 |
| 23325 | 1167 | T1167C | D | 1e-05 | 7.9e-06 | 7.2e-06 | 8.367E-06 | 1.457E-06 | 7.200E-06 | 1.000E-05 |
| 23326 | 1167 | T1167E | D | 6.9e-06 | 7.9e-06 | 9.3e-06 | 8.033E-06 | 1.206E-06 | 6.900E-06 | 9.300E-06 |
| 23327 | 1167 | T1167Q | D | 5.5e-06 | 9.4e-06 | 7.3e-06 | 7.400E-06 | 1.952E-06 | 5.500E-06 | 9.400E-06 |
| 23328 | 1167 | T1167G | D | 1.1e-05 | 7.6e-06 | 5.2e-06 | 7.933E-06 | 2.914E-06 | 5.200E-06 | 1.100E-05 |
| 23329 | 1167 | T1167H | D | 9e-06 | 1e-05 | 6.6e-06 | 8.533E-06 | 1.747E-06 | 6.600E-06 | 1.000E-05 |
| 23330 | 1167 | T1167I | D | 7e-06 | 9.2e-06 | 4.8e-06 | 7.000E-06 | 2.200E-06 | 4.800E-06 | 9.200E-06 |
| 23331 | 1167 | T1167L | D | 6.7e-06 | 6.6e-06 | 6.5e-06 | 6.600E-06 | 1.000E-07 | 6.500E-06 | 6.700E-06 |
| 23332 | 1167 | T1167K | D | 1.1e-05 | 7.8e-06 | 5.6e-06 | 8.133E-06 | 2.715E-06 | 5.600E-06 | 1.100E-05 |
| 23333 | 1167 | T1167M | D | 9.9e-06 | 7.8e-06 | 1e-05 | 9.233E-06 | 1.242E-06 | 7.800E-06 | 1.000E-05 |
| 23334 | 1167 | T1167F | D | 6.4e-06 | 1.4e-05 | 5.3e-06 | 8.567E-06 | 4.737E-06 | 5.300E-06 | 1.400E-05 |
| 23335 | 1167 | T1167P | D | 6.6e-06 | 9e-06 | 9.4e-06 | 8.333E-06 | 1.514E-06 | 6.600E-06 | 9.400E-06 |
| 23336 | 1167 | T1167S | D | 6.2e-06 | 1e-05 | 7.2e-06 | 7.800E-06 | 1.970E-06 | 6.200E-06 | 1.000E-05 |
| 23337 | 1167 | T1167T | D | 9.1e-06 | 9.6e-06 | 1.2e-05 | 1.023E-05 | 1.550E-06 | 9.100E-06 | 1.200E-05 |
| 23338 | 1167 | T1167W | D | 9.9e-06 | 7.5e-06 | 5.6e-06 | 7.667E-06 | 2.155E-06 | 5.600E-06 | 9.900E-06 |
| 23339 | 1167 | T1167Y | D | 8.1e-06 | 3.2e-06 | 8.5e-06 | 6.600E-06 | 2.951E-06 | 3.200E-06 | 8.500E-06 |
| 23340 | 1167 | T1167V | D | 8.7e-06 | 9.2e-06 | 7.5e-06 | 8.467E-06 | 8.737E-07 | 7.500E-06 | 9.200E-06 |
| 23341 | 1168 | V1168A | D | 1.1e-05 | 1.6e-05 | 1e-05 | 1.233E-05 | 3.215E-06 | 1.000E-05 | 1.600E-05 |
| 23342 | 1168 | V1168R | D | 6.9e-06 | 5.9e-06 | 9.4e-06 | 7.400E-06 | 1.803E-06 | 5.900E-06 | 9.400E-06 |
| 23343 | 1168 | V1168N | D | 9.6e-06 | 7.9e-06 | 9e-06 | 8.833E-06 | 8.622E-07 | 7.900E-06 | 9.600E-06 |
| 23344 | 1168 | V1168D | D | 1.1e-05 | 7.1e-06 | 4.6e-06 | 7.567E-06 | 3.225E-06 | 4.600E-06 | 1.100E-05 |
| 23345 | 1168 | V1168C | D | 5.9e-06 | 8e-06 | 6.9e-06 | 6.933E-06 | 1.050E-06 | 5.900E-06 | 8.000E-06 |
| 23346 | 1168 | V1168E | D | 1.3e-05 | 9.6e-06 | 1.2e-05 | 1.153E-05 | 1.747E-06 | 9.600E-06 | 1.300E-05 |
| 23347 | 1168 | V1168Q | D | 1.5e-05 | 5.4e-06 | 8.5e-06 | 9.633E-06 | 4.899E-06 | 5.400E-06 | 1.500E-05 |
| 23348 | 1168 | V1168G | D | 8.7e-06 | 6.7e-06 | 5.9e-06 | 7.100E-06 | 1.442E-06 | 5.900E-06 | 8.700E-06 |
| 23349 | 1168 | V1168H | D | 9.3e-06 | 4.4e-06 | 1.3e-05 | 8.900E-06 | 4.314E-06 | 4.400E-06 | 1.300E-05 |
| 23350 | 1168 | V1168I | D | 5.5e-06 | 3.7e-06 | 1e-05 | 6.400E-06 | 3.245E-06 | 3.700E-06 | 1.000E-05 |
| 23351 | 1168 | V1168L | D | 1.2e-05 | 1.2e-05 | 6.7e-06 | 1.023E-05 | 3.060E-06 | 6.700E-06 | 1.200E-05 |
| 23352 | 1168 | V1168K | D | 8e-06 | 5.2e-06 | 1.1e-05 | 8.067E-06 | 2.901E-06 | 5.200E-06 | 1.100E-05 |
| 23353 | 1168 | V1168M | D | 8.7e-06 | 9.4e-06 | 7.9e-06 | 8.667E-06 | 7.506E-07 | 7.900E-06 | 9.400E-06 |
| 23354 | 1168 | V1168F | D | 6.7e-06 | 5.9e-06 | 8.9e-06 | 7.167E-06 | 1.553E-06 | 5.900E-06 | 8.900E-06 |
| 23355 | 1168 | V1168P | D | 1.3e-05 | 8.8e-06 | 6.3e-06 | 9.367E-06 | 3.386E-06 | 6.300E-06 | 1.300E-05 |
| 23356 | 1168 | V1168S | D | 1.3e-05 | 8.2e-06 | 5.1e-06 | 8.767E-06 | 3.980E-06 | 5.100E-06 | 1.300E-05 |
| 23357 | 1168 | V1168T | D | 7.3e-06 | 9.6e-06 | 5.5e-06 | 7.467E-06 | 2.055E-06 | 5.500E-06 | 9.600E-06 |
| 23358 | 1168 | V1168W | D | 1.4e-05 | 1.6e-05 | 8.4e-06 | 1.280E-05 | 3.940E-06 | 8.400E-06 | 1.600E-05 |
| 23359 | 1168 | V1168Y | D | 1.2e-05 | 1.2e-05 | 6.9e-06 | 1.030E-05 | 2.944E-06 | 6.900E-06 | 1.200E-05 |
| 23360 | 1168 | V1168V | D | 7.5e-06 | 1.7e-05 | 7.2e-06 | 1.057E-05 | 5.573E-06 | 7.200E-06 | 1.700E-05 |
| 23361 | 1169 | A1169A | D | 9.5e-06 | 1.3e-05 | 1.3e-05 | 1.183E-05 | 2.021E-06 | 9.500E-06 | 1.300E-05 |
| 23362 | 1169 | A1169R | D | 6.1e-06 | 5.2e-06 | 8.9e-06 | 6.733E-06 | 1.930E-06 | 5.200E-06 | 8.900E-06 |
| 23363 | 1169 | A1169N | D | 3.1e-06 | 8.5e-06 | 4.3e-06 | 5.300E-06 | 2.835E-06 | 3.100E-06 | 8.500E-06 |
| 23364 | 1169 | A1169D | D | 1.1e-05 | 6.2e-06 | 1.3e-05 | 1.007E-05 | 3.495E-06 | 6.200E-06 | 1.300E-05 |
| 23365 | 1169 | A1169C | D | 8e-06 | 7.6e-06 | 1e-05 | 8.533E-06 | 1.286E-06 | 7.600E-06 | 1.000E-05 |
| 23366 | 1169 | A1169E | D | 9.1e-06 | 7e-06 | 9.8e-06 | 8.633E-06 | 1.457E-06 | 7.000E-06 | 9.800E-06 |
| 23367 | 1169 | A1169Q | D | 7.7e-06 | 6.1e-06 | 5.9e-06 | 6.567E-06 | 9.866E-07 | 5.900E-06 | 7.700E-06 |
| 23368 | 1169 | A1169G | D | 7e-06 | 8.5e-06 | 9.3e-06 | 8.267E-06 | 1.168E-06 | 7.000E-06 | 9.300E-06 |
| 23369 | 1169 | A1169H | D | 1.2e-05 | 6.5e-06 | 9.4e-06 | 9.300E-06 | 2.751E-06 | 6.500E-06 | 1.200E-05 |
| 23370 | 1169 | A1169I | D | 5.9e-06 | 5e-06 | 8.8e-06 | 6.567E-06 | 1.986E-06 | 5.000E-06 | 8.800E-06 |
| 23371 | 1169 | A1169L | D | 1.4e-05 | 4.6e-06 | 8.1e-06 | 8.900E-06 | 4.751E-06 | 4.600E-06 | 1.400E-05 |
| 23372 | 1169 | A1169K | D | 9e-06 | 5.8e-06 | 1.1e-05 | 8.600E-06 | 2.623E-06 | 5.800E-06 | 1.100E-05 |
| 23373 | 1169 | A1169M | D | 1e-05 | 9.8e-06 | 5.4e-06 | 8.400E-06 | 2.600E-06 | 5.400E-06 | 1.000E-05 |
| 23374 | 1169 | A1169F | D | 9.6e-06 | 8.6e-06 | 9.6e-06 | 9.267E-06 | 5.774E-07 | 8.600E-06 | 9.600E-06 |

| | | | | | | | | | | |
|---|---|---|---|---|---|---|---|---|---|---|
| 23375 | 1169 | A1169P | D | 5.5e-06 | 7e-06 | 7.9e-06 | 6.800E-06 | 1.212E-06 | 5.500E-06 | 7.900E-06 |
| 23376 | 1169 | A1169S | D | 8.5e-06 | 8.1e-06 | 3.3e-06 | 6.633E-06 | 2.894E-06 | 3.300E-06 | 8.500E-06 |
| 23377 | 1169 | A1169T | D | 7.4e-06 | 7.1e-06 | 6.8e-06 | 7.100E-06 | 3.000E-07 | 6.800E-06 | 7.400E-06 |
| 23378 | 1169 | A1169W | D | 9.5e-06 | 6.1e-06 | 5.8e-06 | 7.133E-06 | 2.055E-06 | 5.800E-06 | 9.500E-06 |
| 23379 | 1169 | A1169Y | D | 7.7e-06 | 4.2e-06 | 1.2e-05 | 7.967E-06 | 3.907E-06 | 4.200E-06 | 1.200E-05 |
| 23380 | 1169 | A1169V | D | 5.3e-06 | 6.9e-06 | 6.2e-06 | 6.133E-06 | 8.021E-07 | 5.300E-06 | 6.900E-06 |
| 23381 | 1170 | A1170A | D | 7.1e-06 | 6.4e-06 | 1.2e-05 | 8.500E-06 | 3.051E-06 | 6.400E-06 | 1.200E-05 |
| 23382 | 1170 | A1170R | D | 6.4e-06 | 6.8e-06 | 8.9e-06 | 7.367E-06 | 1.343E-06 | 6.400E-06 | 8.900E-06 |
| 23383 | 1170 | A1170N | D | 5.7e-06 | 8.6e-06 | 8.8e-06 | 7.700E-06 | 1.735E-06 | 5.700E-06 | 8.800E-06 |
| 23384 | 1170 | A1170D | D | 6.9e-06 | 7.5e-06 | 7.1e-06 | 7.167E-06 | 3.055E-07 | 6.900E-06 | 7.500E-06 |
| 23385 | 1170 | A1170C | D | 9.3e-06 | 8.8e-06 | 8.9e-06 | 9.000E-06 | 2.646E-07 | 8.800E-06 | 9.300E-06 |
| 23386 | 1170 | A1170E | D | 8.3e-06 | 6.9e-06 | 6.5e-06 | 7.233E-06 | 9.452E-07 | 6.500E-06 | 8.300E-06 |
| 23387 | 1170 | A1170Q | D | 4.7e-06 | 7.5e-06 | 9.4e-06 | 7.200E-06 | 2.364E-06 | 4.700E-06 | 9.400E-06 |
| 23388 | 1170 | A1170G | D | 9.9e-06 | 7.7e-06 | 4.6e-06 | 7.400E-06 | 2.663E-06 | 4.600E-06 | 9.900E-06 |
| 23389 | 1170 | A1170H | D | 6.5e-06 | 8.1e-06 | 9.8e-06 | 8.133E-06 | 1.650E-06 | 6.500E-06 | 9.800E-06 |
| 23390 | 1170 | A1170I | D | 7.5e-06 | 4.3e-06 | 6.4e-06 | 6.067E-06 | 1.626E-06 | 4.300E-06 | 7.500E-06 |
| 23391 | 1170 | A1170L | D | 8.8e-06 | 3.6e-06 | 6.5e-06 | 6.300E-06 | 2.606E-06 | 3.600E-06 | 8.800E-06 |
| 23392 | 1170 | A1170K | D | 8.2e-06 | 7.3e-06 | 7.1e-06 | 7.533E-06 | 5.859E-07 | 7.100E-06 | 8.200E-06 |
| 23393 | 1170 | A1170M | D | 1.7e-05 | 7.1e-06 | 5.3e-06 | 9.800E-06 | 6.300E-06 | 5.300E-06 | 1.700E-05 |
| 23394 | 1170 | A1170F | D | 8.6e-06 | 5.6e-06 | 1e-05 | 8.067E-06 | 2.248E-06 | 5.600E-06 | 1.000E-05 |
| 23395 | 1170 | A1170P | D | 7.1e-06 | 5.8e-06 | 1.3e-05 | 8.633E-06 | 3.837E-06 | 5.800E-06 | 1.300E-05 |
| 23396 | 1170 | A1170S | D | 6e-06 | 9.5e-06 | 1.9e-05 | 1.150E-05 | 6.727E-06 | 6.000E-06 | 1.900E-05 |
| 23397 | 1170 | A1170T | D | 1.3e-05 | 5.1e-06 | 7.9e-06 | 8.667E-06 | 4.005E-06 | 5.100E-06 | 1.300E-05 |
| 23398 | 1170 | A1170W | D | 5.2e-06 | 5.7e-06 | 7.5e-06 | 6.133E-06 | 1.210E-06 | 5.200E-06 | 7.500E-06 |
| 23399 | 1170 | A1170Y | D | 6.1e-06 | 7.8e-06 | 6.4e-06 | 6.767E-06 | 9.074E-07 | 6.100E-06 | 7.800E-06 |
| 23400 | 1170 | A1170V | D | 5.1e-06 | 4.7e-06 | 9e-06 | 6.267E-06 | 2.376E-06 | 4.700E-06 | 9.000E-06 |
| 23401 | 1171 | P1171A | D | 6.5e-06 | 1e-05 | 8.8e-06 | 8.433E-06 | 1.779E-06 | 6.500E-06 | 1.000E-05 |
| 23402 | 1171 | P1171R | D | 3.9e-06 | 6.1e-06 | 5.7e-06 | 5.233E-06 | 1.172E-06 | 3.900E-06 | 6.100E-06 |
| 23403 | 1171 | P1171N | D | 1.2e-05 | 4.3e-06 | 1.5e-05 | 1.043E-05 | 5.519E-06 | 4.300E-06 | 1.500E-05 |
| 23404 | 1171 | P1171D | D | 1.1e-05 | 7.2e-06 | 5.9e-06 | 8.033E-06 | 2.650E-06 | 5.900E-06 | 1.100E-05 |
| 23405 | 1171 | P1171C | D | 6e-06 | 4e-06 | 5.2e-06 | 5.067E-06 | 1.007E-06 | 4.000E-06 | 6.000E-06 |
| 23406 | 1171 | P1171E | D | 1e-05 | 7.7e-06 | 7.4e-06 | 8.367E-06 | 1.422E-06 | 7.400E-06 | 1.000E-05 |
| 23407 | 1171 | P1171Q | D | 9.7e-06 | 7.4e-06 | 4.7e-06 | 7.267E-06 | 2.503E-06 | 4.700E-06 | 9.700E-06 |
| 23408 | 1171 | P1171G | D | 8e-06 | 6.5e-06 | 1.4e-05 | 9.500E-06 | 3.969E-06 | 6.500E-06 | 1.400E-05 |
| 23409 | 1171 | P1171H | D | 7.9e-06 | 4.6e-06 | 1.2e-05 | 8.167E-06 | 3.707E-06 | 4.600E-06 | 1.200E-05 |
| 23410 | 1171 | P1171I | D | 1.1e-05 | 1.3e-05 | 6e-06 | 1.000E-05 | 3.606E-06 | 6.000E-06 | 1.300E-05 |
| 23411 | 1171 | P1171L | D | 5.7e-06 | 9e-06 | 1e-05 | 8.233E-06 | 2.250E-06 | 5.700E-06 | 1.000E-05 |
| 23412 | 1171 | P1171K | D | 7e-06 | 8.9e-06 | 5.3e-06 | 7.067E-06 | 1.801E-06 | 5.300E-06 | 8.900E-06 |
| 23413 | 1171 | P1171M | D | 1.3e-05 | 8.8e-06 | 9.2e-06 | 1.033E-05 | 2.318E-06 | 8.800E-06 | 1.300E-05 |
| 23414 | 1171 | P1171F | D | 5.6e-06 | 6.6e-06 | 6.7e-06 | 6.300E-06 | 6.083E-07 | 5.600E-06 | 6.700E-06 |
| 23415 | 1171 | P1171P | D | 6.2e-06 | 6.7e-06 | 6e-06 | 6.300E-06 | 3.606E-07 | 6.000E-06 | 6.700E-06 |
| 23416 | 1171 | P1171S | D | 6.2e-06 | 1e-05 | 1e-05 | 8.733E-06 | 2.194E-06 | 6.200E-06 | 1.000E-05 |
| 23417 | 1171 | P1171T | D | 7.2e-06 | 7.4e-06 | 4.6e-06 | 6.400E-06 | 1.562E-06 | 4.600E-06 | 7.400E-06 |
| 23418 | 1171 | P1171W | D | 7.8e-06 | 5.7e-06 | 1.2e-05 | 8.500E-06 | 3.208E-06 | 5.700E-06 | 1.200E-05 |
| 23419 | 1171 | P1171Y | D | 5.6e-06 | 8.3e-06 | 6e-06 | 6.633E-06 | 1.457E-06 | 5.600E-06 | 8.300E-06 |
| 23420 | 1171 | P1171V | D | 6.6e-06 | 6.6e-06 | 7.6e-06 | 6.933E-06 | 5.774E-07 | 6.600E-06 | 7.600E-06 |
| 23421 | 1172 | S1172A | D | 5.2e-06 | 9.8e-06 | 9.4e-06 | 8.133E-06 | 2.548E-06 | 5.200E-06 | 9.800E-06 |
| 23422 | 1172 | S1172R | D | 4.8e-06 | 8.7e-06 | 9.5e-06 | 7.667E-06 | 2.515E-06 | 4.800E-06 | 9.500E-06 |
| 23423 | 1172 | S1172N | D | 1.3e-05 | 6.7e-06 | 9.4e-06 | 9.700E-06 | 3.161E-06 | 6.700E-06 | 1.300E-05 |
| 23424 | 1172 | S1172D | D | 1.2e-05 | 9.4e-06 | 7.4e-06 | 9.600E-06 | 2.307E-06 | 7.400E-06 | 1.200E-05 |
| 23425 | 1172 | S1172C | D | 9.1e-06 | 9.6e-06 | 7.7e-06 | 8.800E-06 | 9.849E-07 | 7.700E-06 | 9.600E-06 |
| 23426 | 1172 | S1172E | D | 8.9e-06 | 6.1e-06 | 1.2e-05 | 9.000E-06 | 2.951E-06 | 6.100E-06 | 1.200E-05 |
| 23427 | 1172 | S1172Q | D | 5e-06 | 6.8e-06 | 5.6e-06 | 5.800E-06 | 9.165E-07 | 5.000E-06 | 6.800E-06 |
| 23428 | 1172 | S1172G | D | 4.1e-06 | 5.7e-06 | 7.8e-06 | 5.867E-06 | 1.856E-06 | 4.100E-06 | 7.800E-06 |
| 23429 | 1172 | S1172H | D | 5.8e-06 | 1e-05 | 8.5e-06 | 8.100E-06 | 2.128E-06 | 5.800E-06 | 1.000E-05 |
| 23430 | 1172 | S1172I | D | 7.8e-06 | 1e-05 | 8.2e-06 | 8.667E-06 | 1.172E-06 | 7.800E-06 | 1.000E-05 |
| 23431 | 1172 | S1172L | D | 6.6e-06 | 6.3e-06 | 3.3e-06 | 5.400E-06 | 1.825E-06 | 3.300E-06 | 6.600E-06 |
| 23432 | 1172 | S1172K | D | 7.7e-06 | 5.4e-06 | 9.7e-06 | 7.600E-06 | 2.152E-06 | 5.400E-06 | 9.700E-06 |
| 23433 | 1172 | S1172M | D | 6.3e-06 | 6e-06 | 5.5e-06 | 5.933E-06 | 4.041E-07 | 5.500E-06 | 6.300E-06 |
| 23434 | 1172 | S1172F | D | 6.2e-06 | 1.4e-05 | 7.2e-06 | 9.133E-06 | 4.244E-06 | 6.200E-06 | 1.400E-05 |
| 23435 | 1172 | S1172P | D | 7e-06 | 1e-05 | 8.6e-06 | 8.533E-06 | 1.501E-06 | 7.000E-06 | 1.000E-05 |
| 23436 | 1172 | S1172S | D | 5.7e-06 | 6.2e-06 | 1.1e-05 | 7.633E-06 | 2.926E-06 | 5.700E-06 | 1.100E-05 |
| 23437 | 1172 | S1172T | D | 9.2e-06 | 5.8e-06 | 7.3e-06 | 7.433E-06 | 1.704E-06 | 5.800E-06 | 9.200E-06 |
| 23438 | 1172 | S1172W | D | 1.1e-05 | 5.2e-06 | 1.3e-05 | 9.733E-06 | 4.051E-06 | 5.200E-06 | 1.300E-05 |
| 23439 | 1172 | S1172Y | D | 6.3e-06 | 5.8e-06 | 9.9e-06 | 7.333E-06 | 2.237E-06 | 5.800E-06 | 9.900E-06 |
| 23440 | 1172 | S1172V | D | 9e-06 | 8.9e-06 | 7.2e-06 | 8.367E-06 | 1.012E-06 | 7.200E-06 | 9.000E-06 |
| 23441 | 1173 | V1173A | D | 6e-06 | 1.3e-05 | 8.1e-06 | 9.033E-06 | 3.592E-06 | 6.000E-06 | 1.300E-05 |
| 23442 | 1173 | V1173R | D | 7.9e-06 | 8.8e-06 | 6.9e-06 | 7.867E-06 | 9.504E-07 | 6.900E-06 | 8.800E-06 |
| 23443 | 1173 | V1173N | D | 1.3e-05 | 8.2e-06 | 8.2e-06 | 9.800E-06 | 2.771E-06 | 8.200E-06 | 1.300E-05 |
| 23444 | 1173 | V1173D | D | 8.4e-06 | 7.4e-06 | 1.1e-05 | 8.933E-06 | 1.858E-06 | 7.400E-06 | 1.100E-05 |
| 23445 | 1173 | V1173C | D | 8.6e-06 | 7.4e-06 | 7e-06 | 7.667E-06 | 8.327E-07 | 7.000E-06 | 8.600E-06 |
| 23446 | 1173 | V1173E | D | 7.2e-06 | 6.7e-06 | 1.7e-05 | 1.030E-05 | 5.808E-06 | 6.700E-06 | 1.700E-05 |
| 23447 | 1173 | V1173Q | D | 1.3e-05 | 5.8e-06 | 9e-06 | 9.267E-06 | 3.607E-06 | 5.800E-06 | 1.300E-05 |
| 23448 | 1173 | V1173G | D | 1.1e-05 | 1.1e-05 | 6.2e-06 | 9.400E-06 | 2.771E-06 | 6.200E-06 | 1.100E-05 |
| 23449 | 1173 | V1173H | D | 5.1e-06 | 7.9e-06 | 8.1e-06 | 7.033E-06 | 1.677E-06 | 5.100E-06 | 8.100E-06 |
| 23450 | 1173 | V1173I | D | 6.2e-06 | 7.5e-06 | 1.2e-05 | 8.567E-06 | 3.044E-06 | 6.200E-06 | 1.200E-05 |
| 23451 | 1173 | V1173L | D | 6.1e-06 | 8.1e-06 | 5.7e-06 | 6.633E-06 | 1.286E-06 | 5.700E-06 | 8.100E-06 |
| 23452 | 1173 | V1173K | D | 6.6e-06 | 8.6e-06 | 8.1e-06 | 7.767E-06 | 1.041E-06 | 6.600E-06 | 8.600E-06 |
| 23453 | 1173 | V1173M | D | 5.7e-06 | 6.3e-06 | 9.2e-06 | 7.067E-06 | 1.872E-06 | 5.700E-06 | 9.200E-06 |
| 23454 | 1173 | V1173F | D | 4e-06 | 9.6e-06 | 7.7e-06 | 7.100E-06 | 2.848E-06 | 4.000E-06 | 9.600E-06 |
| 23455 | 1173 | V1173P | D | 1.1e-05 | 3.2e-06 | 1.3e-05 | 9.067E-06 | 5.178E-06 | 3.200E-06 | 1.300E-05 |
| 23456 | 1173 | V1173S | D | 1e-05 | 1.2e-05 | 5.7e-06 | 9.233E-06 | 3.219E-06 | 5.700E-06 | 1.200E-05 |
| 23457 | 1173 | V1173T | D | 6.8e-06 | 6.1e-06 | 7.3e-06 | 6.733E-06 | 6.028E-07 | 6.100E-06 | 7.300E-06 |
| 23458 | 1173 | V1173W | D | 9.7e-06 | 8.2e-06 | 7.3e-06 | 8.400E-06 | 1.212E-06 | 7.300E-06 | 9.700E-06 |
| 23459 | 1173 | V1173Y | D | 6e-06 | 6.3e-06 | 7.7e-06 | 6.667E-06 | 9.074E-07 | 6.000E-06 | 7.700E-06 |

| | | | | | | | | | | |
|---|---|---|---|---|---|---|---|---|---|---|
| 23460 | 1173 | V1173V | D | 1.3e-05 | 5.2e-06 | 5.5e-06 | 7.900E-06 | 4.419E-06 | 5.200E-06 | 1.300E-05 |
| 23461 | 1174 | F1174A | D | 8.8e-06 | 5e-06 | 6.9e-06 | 6.900E-06 | 1.900E-06 | 5.000E-06 | 8.800E-06 |
| 23462 | 1174 | F1174R | D | 6.7e-06 | 7.2e-06 | 6.7e-06 | 6.867E-06 | 2.887E-07 | 6.700E-06 | 7.200E-06 |
| 23463 | 1174 | F1174N | D | 8.4e-06 | 6.5e-06 | 9.9e-06 | 8.267E-06 | 1.704E-06 | 6.500E-06 | 9.900E-06 |
| 23464 | 1174 | F1174D | D | 8.4e-06 | 9.3e-06 | 9.7e-06 | 9.133E-06 | 6.658E-07 | 8.400E-06 | 9.700E-06 |
| 23465 | 1174 | F1174C | D | 9.3e-06 | 6.6e-06 | 1.6e-05 | 1.063E-05 | 4.840E-06 | 6.600E-06 | 1.600E-05 |
| 23466 | 1174 | F1174E | D | 9.3e-06 | 6.5e-06 | 9.5e-06 | 8.433E-06 | 1.677E-06 | 6.500E-06 | 9.500E-06 |
| 23467 | 1174 | F1174Q | D | 7e-06 | 6.1e-06 | 5.4e-06 | 6.167E-06 | 8.021E-07 | 5.400E-06 | 7.000E-06 |
| 23468 | 1174 | F1174G | D | 5.1e-06 | 9.2e-06 | 6.9e-06 | 7.067E-06 | 2.055E-06 | 5.100E-06 | 9.200E-06 |
| 23469 | 1174 | F1174H | D | 7.5e-06 | 7.8e-06 | 6.5e-06 | 7.267E-06 | 6.807E-07 | 6.500E-06 | 7.800E-06 |
| 23470 | 1174 | F1174I | D | 5.4e-06 | 5.4e-06 | 8.5e-06 | 6.433E-06 | 1.790E-06 | 5.400E-06 | 8.500E-06 |
| 23471 | 1174 | F1174L | D | 1e-05 | 5.9e-06 | 9.1e-06 | 8.333E-06 | 2.155E-06 | 5.900E-06 | 1.000E-05 |
| 23472 | 1174 | F1174K | D | 6.3e-06 | 7.2e-06 | 6.4e-06 | 6.633E-06 | 4.933E-07 | 6.300E-06 | 7.200E-06 |
| 23473 | 1174 | F1174M | D | 3.4e-06 | 7.9e-06 | 8.6e-06 | 6.633E-06 | 2.822E-06 | 3.400E-06 | 8.600E-06 |
| 23474 | 1174 | F1174F | D | 6.2e-06 | 1.2e-05 | 1.2e-05 | 1.007E-05 | 3.349E-06 | 6.200E-06 | 1.200E-05 |
| 23475 | 1174 | F1174P | D | 3.5e-06 | 1.2e-05 | 6.9e-06 | 7.467E-06 | 4.278E-06 | 3.500E-06 | 1.200E-05 |
| 23476 | 1174 | F1174S | D | 6.7e-06 | 1.1e-05 | 1.2e-05 | 9.900E-06 | 2.816E-06 | 6.700E-06 | 1.200E-05 |
| 23477 | 1174 | F1174T | D | 1.2e-05 | 7.3e-06 | 6.5e-06 | 8.600E-06 | 2.972E-06 | 6.500E-06 | 1.200E-05 |
| 23478 | 1174 | F1174W | D | 1.7e-05 | 9.5e-06 | 1.9e-05 | 1.517E-05 | 5.008E-06 | 9.500E-06 | 1.900E-05 |
| 23479 | 1174 | F1174Y | D | 1.3e-05 | 4.5e-06 | 5.4e-06 | 7.633E-06 | 4.669E-06 | 4.500E-06 | 1.300E-05 |
| 23480 | 1174 | F1174V | D | 7.9e-06 | 7.7e-06 | 5.6e-06 | 7.067E-06 | 1.274E-06 | 5.600E-06 | 7.900E-06 |
| 23481 | 1175 | I1175A | D | 6.9e-06 | 8.1e-06 | 1.1e-05 | 8.667E-06 | 2.108E-06 | 6.900E-06 | 1.100E-05 |
| 23482 | 1175 | I1175R | D | 4.3e-06 | 5.6e-06 | 7.2e-06 | 5.700E-06 | 1.453E-06 | 4.300E-06 | 7.200E-06 |
| 23483 | 1175 | I1175N | D | 9.5e-06 | 9e-06 | 6.2e-06 | 8.233E-06 | 1.779E-06 | 6.200E-06 | 9.500E-06 |
| 23484 | 1175 | I1175D | D | 6.4e-06 | 6.8e-06 | 6.9e-06 | 6.700E-06 | 2.646E-07 | 6.400E-06 | 6.900E-06 |
| 23485 | 1175 | I1175C | D | 7.1e-06 | 7.2e-06 | 1.1e-05 | 8.433E-06 | 2.223E-06 | 7.100E-06 | 1.100E-05 |
| 23486 | 1175 | I1175E | D | 7.9e-06 | 1.8e-05 | 7.6e-06 | 1.117E-05 | 5.920E-06 | 7.600E-06 | 1.800E-05 |
| 23487 | 1175 | I1175Q | D | 7.7e-06 | 8.3e-06 | 3.2e-06 | 6.400E-06 | 2.787E-06 | 3.200E-06 | 8.300E-06 |
| 23488 | 1175 | I1175G | D | 8.9e-06 | 1.1e-05 | 4.9e-06 | 8.267E-06 | 3.099E-06 | 4.900E-06 | 1.100E-05 |
| 23489 | 1175 | I1175H | D | 6.9e-06 | 7.2e-06 | 7.1e-06 | 7.067E-06 | 1.528E-07 | 6.900E-06 | 7.200E-06 |
| 23490 | 1175 | I1175I | D | 8.5e-06 | 9.5e-06 | 9.3e-06 | 9.100E-06 | 5.292E-07 | 8.500E-06 | 9.500E-06 |
| 23491 | 1175 | I1175L | D | 5.9e-06 | 9.1e-06 | 8.6e-06 | 7.867E-06 | 1.721E-06 | 5.900E-06 | 9.100E-06 |
| 23492 | 1175 | I1175K | D | 8.3e-06 | 7.3e-06 | 6.5e-06 | 7.367E-06 | 9.018E-07 | 6.500E-06 | 8.300E-06 |
| 23493 | 1175 | I1175M | D | 6.5e-06 | 1.2e-05 | 6.9e-06 | 8.467E-06 | 3.066E-06 | 6.500E-06 | 1.200E-05 |
| 23494 | 1175 | I1175F | D | 7.4e-06 | 4.6e-06 | 8.6e-06 | 6.867E-06 | 2.053E-06 | 4.600E-06 | 8.600E-06 |
| 23495 | 1175 | I1175P | D | 9.8e-06 | 2e-05 | 6.7e-06 | 1.217E-05 | 6.959E-06 | 6.700E-06 | 2.000E-05 |
| 23496 | 1175 | I1175S | D | 4.7e-06 | 8.4e-06 | 6.4e-06 | 6.500E-06 | 1.852E-06 | 4.700E-06 | 8.400E-06 |
| 23497 | 1175 | I1175T | D | 8.2e-06 | 1.3e-05 | 9.1e-06 | 1.010E-05 | 2.551E-06 | 8.200E-06 | 1.300E-05 |
| 23498 | 1175 | I1175W | D | 5e-06 | 6.8e-06 | 6.8e-06 | 6.200E-06 | 1.039E-06 | 5.000E-06 | 6.800E-06 |
| 23499 | 1175 | I1175Y | D | 7.6e-06 | 9.9e-06 | 6.3e-06 | 7.933E-06 | 1.823E-06 | 6.300E-06 | 9.900E-06 |
| 23500 | 1175 | I1175V | D | 5.8e-06 | 8.2e-06 | 7e-06 | 7.000E-06 | 1.200E-06 | 5.800E-06 | 8.200E-06 |
| 23501 | 1176 | F1176A | D | 1.2e-05 | 9.6e-06 | 9.6e-06 | 1.040E-05 | 1.386E-06 | 9.600E-06 | 1.200E-05 |
| 23502 | 1176 | F1176R | D | 1e-05 | 5.2e-06 | 1e-05 | 8.400E-06 | 2.771E-06 | 5.200E-06 | 1.000E-05 |
| 23503 | 1176 | F1176N | D | 6.3e-06 | 6.4e-06 | 4.8e-06 | 5.833E-06 | 8.963E-07 | 4.800E-06 | 6.400E-06 |
| 23504 | 1176 | F1176D | D | 5.3e-06 | 1.1e-05 | 1.2e-05 | 9.433E-06 | 3.614E-06 | 5.300E-06 | 1.200E-05 |
| 23505 | 1176 | F1176C | D | 1.3e-05 | 7.8e-06 | 6.5e-06 | 9.100E-06 | 3.439E-06 | 6.500E-06 | 1.300E-05 |
| 23506 | 1176 | F1176E | D | 9.6e-06 | 9.5e-06 | 9.5e-06 | 9.533E-06 | 5.774E-08 | 9.500E-06 | 9.600E-06 |
| 23507 | 1176 | F1176Q | D | 1.3e-05 | 4.1e-06 | 5.1e-06 | 7.400E-06 | 4.875E-06 | 4.100E-06 | 1.300E-05 |
| 23508 | 1176 | F1176G | D | 8.9e-06 | 7.4e-06 | 8.1e-06 | 8.133E-06 | 7.506E-07 | 7.400E-06 | 8.900E-06 |
| 23509 | 1176 | F1176H | D | 7.3e-06 | 7.5e-06 | 8.9e-06 | 7.900E-06 | 8.718E-07 | 7.300E-06 | 8.900E-06 |
| 23510 | 1176 | F1176I | D | 1.3e-05 | 5.6e-06 | 7.8e-06 | 8.800E-06 | 3.800E-06 | 5.600E-06 | 1.300E-05 |
| 23511 | 1176 | F1176L | D | 9.4e-06 | 6.1e-06 | 8.9e-06 | 8.133E-06 | 1.779E-06 | 6.100E-06 | 9.400E-06 |
| 23512 | 1176 | F1176K | D | 5.3e-06 | 9e-06 | 6.2e-06 | 6.833E-06 | 1.930E-06 | 5.300E-06 | 9.000E-06 |
| 23513 | 1176 | F1176M | D | 6.6e-06 | 5.8e-06 | 9.4e-06 | 7.267E-06 | 1.890E-06 | 5.800E-06 | 9.400E-06 |
| 23514 | 1176 | F1176F | D | 6.3e-06 | 1.1e-05 | 8.1e-06 | 8.467E-06 | 2.371E-06 | 6.300E-06 | 1.100E-05 |
| 23515 | 1176 | F1176P | D | 5.4e-06 | 8.7e-06 | 6.6e-06 | 6.900E-06 | 1.670E-06 | 5.400E-06 | 8.700E-06 |
| 23516 | 1176 | F1176S | D | 8.9e-06 | 6.4e-06 | 9.7e-06 | 8.333E-06 | 1.721E-06 | 6.400E-06 | 9.700E-06 |
| 23517 | 1176 | F1176T | D | 6.6e-06 | 8.1e-06 | 1.4e-05 | 9.567E-06 | 3.912E-06 | 6.600E-06 | 1.400E-05 |
| 23518 | 1176 | F1176W | D | 8.8e-06 | 1.2e-05 | 1.1e-05 | 1.060E-05 | 1.637E-06 | 8.800E-06 | 1.200E-05 |
| 23519 | 1176 | F1176Y | D | 9.7e-06 | 8.8e-06 | 9.4e-06 | 9.300E-06 | 4.583E-07 | 8.800E-06 | 9.700E-06 |
| 23520 | 1176 | F1176V | D | 1.3e-05 | 5.9e-06 | 5.2e-06 | 8.033E-06 | 4.315E-06 | 5.200E-06 | 1.300E-05 |
| 23521 | 1177 | P1177A | D | 1e-05 | 1.3e-05 | 8.2e-06 | 1.040E-05 | 2.425E-06 | 8.200E-06 | 1.300E-05 |
| 23522 | 1177 | P1177R | D | 9.5e-06 | 7.8e-06 | 7.8e-06 | 8.367E-06 | 9.815E-07 | 7.800E-06 | 9.500E-06 |
| 23523 | 1177 | P1177N | D | 8.9e-06 | 5.9e-06 | 6.3e-06 | 7.033E-06 | 1.629E-06 | 5.900E-06 | 8.900E-06 |
| 23524 | 1177 | P1177D | D | 7e-06 | 6.6e-06 | 8.3e-06 | 7.300E-06 | 8.888E-07 | 6.600E-06 | 8.300E-06 |
| 23525 | 1177 | P1177C | D | 1e-05 | 9.2e-06 | 7.2e-06 | 8.800E-06 | 1.442E-06 | 7.200E-06 | 1.000E-05 |
| 23526 | 1177 | P1177E | D | 1.3e-05 | 6.2e-06 | 5.1e-06 | 8.100E-06 | 4.279E-06 | 5.100E-06 | 1.300E-05 |
| 23527 | 1177 | P1177Q | D | 7.6e-06 | 7.3e-06 | 6.9e-06 | 7.267E-06 | 3.512E-07 | 6.900E-06 | 7.600E-06 |
| 23528 | 1177 | P1177G | D | 1.2e-05 | 3.9e-06 | 6.1e-06 | 7.333E-06 | 4.188E-06 | 3.900E-06 | 1.200E-05 |
| 23529 | 1177 | P1177H | D | 9.1e-06 | 9e-06 | 5.7e-06 | 7.933E-06 | 1.935E-06 | 5.700E-06 | 9.100E-06 |
| 23530 | 1177 | P1177I | D | 8.7e-06 | 1e-05 | 8.1e-06 | 8.933E-06 | 9.713E-07 | 8.100E-06 | 1.000E-05 |
| 23531 | 1177 | P1177L | D | 6.9e-06 | 1.2e-05 | 6e-06 | 8.300E-06 | 3.236E-06 | 6.000E-06 | 1.200E-05 |
| 23532 | 1177 | P1177K | D | 7.6e-06 | 6.9e-06 | 1.1e-05 | 8.500E-06 | 2.193E-06 | 6.900E-06 | 1.100E-05 |
| 23533 | 1177 | P1177M | D | 9.1e-06 | 4.9e-06 | 7.4e-06 | 7.133E-06 | 2.113E-06 | 4.900E-06 | 9.100E-06 |
| 23534 | 1177 | P1177F | D | 4.8e-06 | 7e-06 | 7.5e-06 | 6.433E-06 | 1.436E-06 | 4.800E-06 | 7.500E-06 |
| 23535 | 1177 | P1177P | D | 1.1e-05 | 1.3e-05 | 1.4e-05 | 1.267E-05 | 1.528E-06 | 1.100E-05 | 1.400E-05 |
| 23536 | 1177 | P1177S | D | 9.8e-06 | 7e-06 | 7.4e-06 | 8.067E-06 | 1.514E-06 | 7.000E-06 | 9.800E-06 |
| 23537 | 1177 | P1177T | D | 9.6e-06 | 9.9e-06 | 6.8e-06 | 8.767E-06 | 1.710E-06 | 6.800E-06 | 9.900E-06 |
| 23538 | 1177 | P1177W | D | 9.1e-06 | 8.8e-06 | 7.1e-06 | 8.333E-06 | 1.079E-06 | 7.100E-06 | 9.100E-06 |
| 23539 | 1177 | P1177Y | D | 1.2e-05 | 8.6e-06 | 4.4e-06 | 8.333E-06 | 3.807E-06 | 4.400E-06 | 1.200E-05 |
| 23540 | 1177 | P1177V | D | 8.2e-06 | 7.9e-06 | 7.1e-06 | 7.733E-06 | 5.686E-07 | 7.100E-06 | 8.200E-06 |
| 23541 | 1178 | P1178A | D | 1.2e-05 | 6.4e-06 | 1.3e-05 | 1.047E-05 | 3.557E-06 | 6.400E-06 | 1.300E-05 |
| 23542 | 1178 | P1178R | D | 8e-06 | 6.8e-06 | 4.9e-06 | 6.567E-06 | 1.563E-06 | 4.900E-06 | 8.000E-06 |
| 23543 | 1178 | P1178N | D | 7.2e-06 | 7.1e-06 | 6.5e-06 | 6.933E-06 | 3.786E-07 | 6.500E-06 | 7.200E-06 |
| 23544 | 1178 | P1178D | D | 6.8e-06 | 1.4e-05 | 6.9e-06 | 9.233E-06 | 4.128E-06 | 6.800E-06 | 1.400E-05 |

| | | | | | | | | | | |
|---|---|---|---|---|---|---|---|---|---|---|
| 23545 | 1178 | P1178C | D | 8.2e-06 | 8.6e-06 | 5.9e-06 | 7.567E-06 | 1.457E-06 | 5.900E-06 | 8.600E-06 |
| 23546 | 1178 | P1178E | D | 6.2e-06 | 7.9e-06 | 9.7e-06 | 7.933E-06 | 1.750E-06 | 6.200E-06 | 9.700E-06 |
| 23547 | 1178 | P1178Q | D | 9.1e-06 | 6.4e-06 | 6.3e-06 | 7.267E-06 | 1.589E-06 | 6.300E-06 | 9.100E-06 |
| 23548 | 1178 | P1178G | D | 5.3e-06 | 6.6e-06 | 6e-06 | 5.967E-06 | 6.506E-07 | 5.300E-06 | 6.600E-06 |
| 23549 | 1178 | P1178H | D | 9.4e-06 | 1.5e-05 | 9.4e-06 | 1.127E-05 | 3.233E-06 | 9.400E-06 | 1.500E-05 |
| 23550 | 1178 | P1178I | D | 5.3e-06 | 9.4e-06 | 4.6e-06 | 6.433E-06 | 2.593E-06 | 4.600E-06 | 9.400E-06 |
| 23551 | 1178 | P1178L | D | 6.3e-06 | 8.9e-06 | 1e-05 | 8.400E-06 | 1.900E-06 | 6.300E-06 | 1.000E-05 |
| 23552 | 1178 | P1178K | D | 4.9e-06 | 9e-06 | 5.2e-06 | 6.367E-06 | 2.285E-06 | 4.900E-06 | 9.000E-06 |
| 23553 | 1178 | P1178M | D | 6.6e-06 | 1e-05 | 5.6e-06 | 7.400E-06 | 2.307E-06 | 5.600E-06 | 1.000E-05 |
| 23554 | 1178 | P1178F | D | 6.4e-06 | 6.5e-06 | 5.6e-06 | 6.167E-06 | 4.933E-07 | 5.600E-06 | 6.500E-06 |
| 23555 | 1178 | P1178P | D | 9.3e-06 | 9.5e-06 | 8.7e-06 | 9.167E-06 | 4.163E-07 | 8.700E-06 | 9.500E-06 |
| 23556 | 1178 | P1178S | D | 8.7e-06 | 7.2e-06 | 4.5e-06 | 6.800E-06 | 2.128E-06 | 4.500E-06 | 8.700E-06 |
| 23557 | 1178 | P1178T | D | 5.1e-06 | 1.5e-05 | 8.1e-06 | 9.400E-06 | 5.076E-06 | 5.100E-06 | 1.500E-05 |
| 23558 | 1178 | P1178W | D | 1.1e-05 | 9e-06 | 9e-06 | 9.667E-06 | 1.155E-06 | 9.000E-06 | 1.100E-05 |
| 23559 | 1178 | P1178Y | D | 1.6e-05 | 7.6e-06 | 8e-06 | 1.053E-05 | 4.738E-06 | 7.600E-06 | 1.600E-05 |
| 23560 | 1178 | P1178V | D | 8.9e-06 | 7.1e-06 | 5.4e-06 | 7.133E-06 | 1.750E-06 | 5.400E-06 | 8.900E-06 |
| 23561 | 1179 | S1179A | D | 8.7e-06 | 7.1e-06 | 4.1e-06 | 6.633E-06 | 2.335E-06 | 4.100E-06 | 8.700E-06 |
| 23562 | 1179 | S1179R | D | 9.6e-06 | 9.9e-06 | 7.4e-06 | 8.967E-06 | 1.365E-06 | 7.400E-06 | 9.900E-06 |
| 23563 | 1179 | S1179N | D | 7.1e-06 | 4.9e-06 | 7.5e-06 | 6.500E-06 | 1.400E-06 | 4.900E-06 | 7.500E-06 |
| 23564 | 1179 | S1179D | D | 1.1e-05 | 6.2e-06 | 7.3e-06 | 8.167E-06 | 2.515E-06 | 6.200E-06 | 1.100E-05 |
| 23565 | 1179 | S1179C | D | 1.2e-05 | 8.4e-06 | 5.3e-06 | 8.567E-06 | 3.353E-06 | 5.300E-06 | 1.200E-05 |
| 23566 | 1179 | S1179E | D | 8e-06 | 7.8e-06 | 8.3e-06 | 8.033E-06 | 2.517E-07 | 7.800E-06 | 8.300E-06 |
| 23567 | 1179 | S1179Q | D | 7.8e-06 | 9.4e-06 | 1e-05 | 9.067E-06 | 1.137E-06 | 7.800E-06 | 1.000E-05 |
| 23568 | 1179 | S1179G | D | 8.4e-06 | 9.2e-06 | 9.6e-06 | 9.067E-06 | 6.110E-07 | 8.400E-06 | 9.600E-06 |
| 23569 | 1179 | S1179H | D | 9e-06 | 5.8e-06 | 6.1e-06 | 6.967E-06 | 1.767E-06 | 5.800E-06 | 9.000E-06 |
| 23570 | 1179 | S1179I | D | 9.3e-06 | 5.5e-06 | 5.4e-06 | 6.733E-06 | 2.223E-06 | 5.400E-06 | 9.300E-06 |
| 23571 | 1179 | S1179L | D | 9.3e-06 | 7.3e-06 | 9.9e-06 | 8.833E-06 | 1.361E-06 | 7.300E-06 | 9.900E-06 |
| 23572 | 1179 | S1179K | D | 8.2e-06 | 6.8e-06 | 5.1e-06 | 6.700E-06 | 1.552E-06 | 5.100E-06 | 8.200E-06 |
| 23573 | 1179 | S1179M | D | 6.5e-06 | 7.2e-06 | 8.8e-06 | 7.500E-06 | 1.179E-06 | 6.500E-06 | 8.800E-06 |
| 23574 | 1179 | S1179F | D | 1e-05 | 5.6e-06 | 6.8e-06 | 7.467E-06 | 2.274E-06 | 5.600E-06 | 1.000E-05 |
| 23575 | 1179 | S1179P | D | 1.2e-05 | 8.2e-06 | 6.7e-06 | 8.967E-06 | 2.732E-06 | 6.700E-06 | 1.200E-05 |
| 23576 | 1179 | S1179S | D | 4.5e-06 | 9.5e-06 | 1e-05 | 8.000E-06 | 3.041E-06 | 4.500E-06 | 1.000E-05 |
| 23577 | 1179 | S1179T | D | 1.3e-05 | 7.9e-06 | 8.2e-06 | 9.700E-06 | 2.862E-06 | 7.900E-06 | 1.300E-05 |
| 23578 | 1179 | S1179W | D | 9.6e-06 | 5.6e-06 | 8.7e-06 | 7.967E-06 | 2.098E-06 | 5.600E-06 | 9.600E-06 |
| 23579 | 1179 | S1179Y | D | 7.1e-06 | 8.5e-06 | 4.5e-06 | 6.700E-06 | 2.030E-06 | 4.500E-06 | 8.500E-06 |
| 23580 | 1179 | S1179V | D | 5.7e-06 | 6e-06 | 6.8e-06 | 6.167E-06 | 5.686E-07 | 5.700E-06 | 6.800E-06 |
| 23581 | 1180 | D1180A | D | 5.1e-06 | 5e-06 | 6.3e-06 | 5.467E-06 | 7.234E-07 | 5.000E-06 | 6.300E-06 |
| 23582 | 1180 | D1180R | D | 7.6e-06 | 8.7e-06 | 5.1e-06 | 7.133E-06 | 1.845E-06 | 5.100E-06 | 8.700E-06 |
| 23583 | 1180 | D1180N | D | 9.9e-06 | 1.7e-05 | 9e-06 | 1.197E-05 | 4.382E-06 | 9.000E-06 | 1.700E-05 |
| 23584 | 1180 | D1180D | D | 9.8e-06 | 9.2e-06 | 6.6e-06 | 8.533E-06 | 1.701E-06 | 6.600E-06 | 9.800E-06 |
| 23585 | 1180 | D1180C | D | 3.9e-06 | 5.8e-06 | 7.9e-06 | 5.867E-06 | 2.001E-06 | 3.900E-06 | 7.900E-06 |
| 23586 | 1180 | D1180E | D | 6.7e-06 | 8.9e-06 | 7.7e-06 | 7.767E-06 | 1.102E-06 | 6.700E-06 | 8.900E-06 |
| 23587 | 1180 | D1180Q | D | 5.1e-06 | 1.1e-05 | 7.3e-06 | 7.800E-06 | 2.982E-06 | 5.100E-06 | 1.100E-05 |
| 23588 | 1180 | D1180G | D | 9.7e-06 | 4.9e-06 | 8.2e-06 | 7.600E-06 | 2.456E-06 | 4.900E-06 | 9.700E-06 |
| 23589 | 1180 | D1180H | D | 6.5e-06 | 6.5e-06 | 4.3e-06 | 5.767E-06 | 1.270E-06 | 4.300E-06 | 6.500E-06 |
| 23590 | 1180 | D1180I | D | 7e-06 | 6.7e-06 | 6.5e-06 | 6.733E-06 | 2.517E-07 | 6.500E-06 | 7.000E-06 |
| 23591 | 1180 | D1180L | D | 8.3e-06 | 1e-05 | 1e-05 | 9.433E-06 | 9.815E-07 | 8.300E-06 | 1.000E-05 |
| 23592 | 1180 | D1180K | D | 5e-06 | 4.5e-06 | 6.4e-06 | 5.300E-06 | 9.849E-07 | 4.500E-06 | 6.400E-06 |
| 23593 | 1180 | D1180M | D | 9.6e-06 | 9.3e-06 | 1.1e-05 | 9.967E-06 | 9.074E-07 | 9.300E-06 | 1.100E-05 |
| 23594 | 1180 | D1180F | D | 7.6e-06 | 7.9e-06 | 9.5e-06 | 8.333E-06 | 1.021E-06 | 7.600E-06 | 9.500E-06 |
| 23595 | 1180 | D1180P | D | 5.1e-06 | 6.8e-06 | 9.8e-06 | 7.233E-06 | 2.380E-06 | 5.100E-06 | 9.800E-06 |
| 23596 | 1180 | D1180S | D | 6.9e-06 | 4.8e-06 | 1e-05 | 7.233E-06 | 2.616E-06 | 4.800E-06 | 1.000E-05 |
| 23597 | 1180 | D1180T | D | 8.5e-06 | 8.5e-06 | 9.1e-06 | 8.700E-06 | 3.464E-07 | 8.500E-06 | 9.100E-06 |
| 23598 | 1180 | D1180W | D | 5.8e-06 | 8.1e-06 | 6.3e-06 | 6.733E-06 | 1.210E-06 | 5.800E-06 | 8.100E-06 |
| 23599 | 1180 | D1180Y | D | 7.7e-06 | 5.4e-06 | 7.3e-06 | 6.800E-06 | 1.229E-06 | 5.400E-06 | 7.700E-06 |
| 23600 | 1180 | D1180V | D | 6.6e-06 | 9.1e-06 | 8.3e-06 | 8.000E-06 | 1.277E-06 | 6.600E-06 | 9.100E-06 |
| 23601 | 1181 | E1181A | D | 1.7e-05 | 8.2e-06 | 6.2e-06 | 1.047E-05 | 5.746E-06 | 6.200E-06 | 1.700E-05 |
| 23602 | 1181 | E1181R | D | 7.1e-06 | 7.3e-06 | 1.3e-05 | 9.133E-06 | 3.350E-06 | 7.100E-06 | 1.300E-05 |
| 23603 | 1181 | E1181N | D | 7.6e-06 | 1.3e-05 | 8.7e-06 | 9.767E-06 | 2.854E-06 | 7.600E-06 | 1.300E-05 |
| 23604 | 1181 | E1181D | D | 5.5e-06 | 5.6e-06 | 6.9e-06 | 6.000E-06 | 7.810E-07 | 5.500E-06 | 6.900E-06 |
| 23605 | 1181 | E1181C | D | 7.3e-06 | 6.8e-06 | 4.5e-06 | 6.200E-06 | 1.493E-06 | 4.500E-06 | 7.300E-06 |
| 23606 | 1181 | E1181E | D | 8.7e-06 | 6.4e-06 | 7.3e-06 | 7.467E-06 | 1.159E-06 | 6.400E-06 | 8.700E-06 |
| 23607 | 1181 | E1181Q | D | 8.2e-06 | 8.4e-06 | 7.6e-06 | 8.067E-06 | 4.163E-07 | 7.600E-06 | 8.400E-06 |
| 23608 | 1181 | E1181G | D | 4.7e-06 | 6.2e-06 | 1.1e-05 | 7.300E-06 | 3.291E-06 | 4.700E-06 | 1.100E-05 |
| 23609 | 1181 | E1181H | D | 6.7e-06 | 5.8e-06 | 7.7e-06 | 6.733E-06 | 9.504E-07 | 5.800E-06 | 7.700E-06 |
| 23610 | 1181 | E1181I | D | 6.9e-06 | 5.6e-06 | 6.9e-06 | 6.467E-06 | 7.506E-07 | 5.600E-06 | 6.900E-06 |
| 23611 | 1181 | E1181L | D | 5.2e-06 | 7.5e-06 | 5.8e-06 | 6.167E-06 | 1.193E-06 | 5.200E-06 | 7.500E-06 |
| 23612 | 1181 | E1181K | D | 4.9e-06 | 8e-06 | 9.8e-06 | 7.567E-06 | 2.479E-06 | 4.900E-06 | 9.800E-06 |
| 23613 | 1181 | E1181M | D | 7.8e-06 | 8.3e-06 | 6.7e-06 | 7.600E-06 | 8.185E-07 | 6.700E-06 | 8.300E-06 |
| 23614 | 1181 | E1181F | D | 1.1e-05 | 8e-06 | 1e-05 | 9.667E-06 | 1.528E-06 | 8.000E-06 | 1.100E-05 |
| 23615 | 1181 | E1181P | D | 5.5e-06 | 8.9e-06 | 6.6e-06 | 7.000E-06 | 1.735E-06 | 5.500E-06 | 8.900E-06 |
| 23616 | 1181 | E1181S | D | 7e-06 | 1.1e-05 | 6.4e-06 | 8.133E-06 | 2.501E-06 | 6.400E-06 | 1.100E-05 |
| 23617 | 1181 | E1181T | D | 1.1e-05 | 6.8e-06 | 5.8e-06 | 7.867E-06 | 2.759E-06 | 5.800E-06 | 1.100E-05 |
| 23618 | 1181 | E1181W | D | 9e-06 | 1.1e-05 | 6.2e-06 | 8.733E-06 | 2.411E-06 | 6.200E-06 | 1.100E-05 |
| 23619 | 1181 | E1181Y | D | 6.8e-06 | 7.6e-06 | 9e-06 | 7.800E-06 | 1.114E-06 | 6.800E-06 | 9.000E-06 |
| 23620 | 1181 | E1181V | D | 6.3e-06 | 5.1e-06 | 1.3e-05 | 8.133E-06 | 4.257E-06 | 5.100E-06 | 1.300E-05 |
| 23621 | 1182 | Q1182A | D | 8.2e-06 | 9.3e-06 | 8.9e-06 | 8.800E-06 | 5.568E-07 | 8.200E-06 | 9.300E-06 |
| 23622 | 1182 | Q1182R | D | 1.1e-05 | 1.3e-05 | 5.9e-06 | 9.967E-06 | 3.661E-06 | 5.900E-06 | 1.300E-05 |
| 23623 | 1182 | Q1182N | D | 6.7e-06 | 1.1e-05 | 6e-06 | 7.900E-06 | 2.707E-06 | 6.000E-06 | 1.100E-05 |
| 23624 | 1182 | Q1182D | D | 1.3e-05 | 5.6e-06 | 8.2e-06 | 8.933E-06 | 3.754E-06 | 5.600E-06 | 1.300E-05 |
| 23625 | 1182 | Q1182C | D | 9.1e-06 | 9.5e-06 | 1.4e-05 | 1.087E-05 | 2.721E-06 | 9.100E-06 | 1.400E-05 |
| 23626 | 1182 | Q1182E | D | 7.6e-06 | 2.7e-06 | 6.9e-06 | 5.733E-06 | 2.650E-06 | 2.700E-06 | 7.600E-06 |
| 23627 | 1182 | Q1182Q | D | 8.7e-06 | 5.2e-06 | 5.4e-06 | 6.433E-06 | 1.966E-06 | 5.200E-06 | 8.700E-06 |
| 23628 | 1182 | Q1182G | D | 5.3e-06 | 5.9e-06 | 6.8e-06 | 6.000E-06 | 7.550E-07 | 5.300E-06 | 6.800E-06 |
| 23629 | 1182 | Q1182H | D | 7.6e-06 | 7.3e-06 | 4.9e-06 | 6.600E-06 | 1.480E-06 | 4.900E-06 | 7.600E-06 |

| | | | | | | | | | | |
|---|---|---|---|---|---|---|---|---|---|---|
| 23630 | 1182 | Q1182I | D | 6.5e-06 | 1e-05 | 1.1e-05 | 9.167E-06 | 2.363E-06 | 6.500E-06 | 1.100E-05 |
| 23631 | 1182 | Q1182L | D | 5.5e-06 | 7.2e-06 | 7.8e-06 | 6.833E-06 | 1.193E-06 | 5.500E-06 | 7.800E-06 |
| 23632 | 1182 | Q1182K | D | 6e-06 | 9.6e-06 | 4.5e-06 | 6.700E-06 | 2.621E-06 | 4.500E-06 | 9.600E-06 |
| 23633 | 1182 | Q1182M | D | 1e-05 | 5.6e-06 | 7.7e-06 | 7.767E-06 | 2.201E-06 | 5.600E-06 | 1.000E-05 |
| 23634 | 1182 | Q1182F | D | 7.3e-06 | 5.9e-06 | 7.5e-06 | 6.900E-06 | 8.718E-07 | 5.900E-06 | 7.500E-06 |
| 23635 | 1182 | Q1182P | D | 1e-05 | 5.3e-06 | 8.4e-06 | 7.900E-06 | 2.390E-06 | 5.300E-06 | 1.000E-05 |
| 23636 | 1182 | Q1182S | D | 1.3e-05 | 1.4e-05 | 5.6e-06 | 1.087E-05 | 4.588E-06 | 5.600E-06 | 1.400E-05 |
| 23637 | 1182 | Q1182T | D | 9.6e-06 | 8.5e-06 | 7e-06 | 8.367E-06 | 1.305E-06 | 7.000E-06 | 9.600E-06 |
| 23638 | 1182 | Q1182W | D | 8.1e-06 | 1.5e-05 | 9.5e-06 | 1.087E-05 | 3.647E-06 | 8.100E-06 | 1.500E-05 |
| 23639 | 1182 | Q1182Y | D | 5.6e-06 | 8.8e-06 | 6.5e-06 | 6.967E-06 | 1.650E-06 | 5.600E-06 | 8.800E-06 |
| 23640 | 1182 | Q1182V | D | 1e-05 | 8.3e-06 | 4.7e-06 | 7.667E-06 | 2.706E-06 | 4.700E-06 | 1.000E-05 |
| 23641 | 1183 | L1183A | D | 6.4e-06 | 9e-06 | 3.6e-06 | 6.333E-06 | 2.701E-06 | 3.600E-06 | 9.000E-06 |
| 23642 | 1183 | L1183R | D | 1e-05 | 4.8e-06 | 4.8e-06 | 6.533E-06 | 3.002E-06 | 4.800E-06 | 1.000E-05 |
| 23643 | 1183 | L1183N | D | 6.8e-06 | 9.7e-06 | 9.1e-06 | 8.533E-06 | 1.531E-06 | 6.800E-06 | 9.700E-06 |
| 23644 | 1183 | L1183D | D | 9.9e-06 | 8.1e-06 | 1.2e-05 | 1.000E-05 | 1.952E-06 | 8.100E-06 | 1.200E-05 |
| 23645 | 1183 | L1183C | D | 5.1e-06 | 8.5e-06 | 7.3e-06 | 6.967E-06 | 1.724E-06 | 5.100E-06 | 8.500E-06 |
| 23646 | 1183 | L1183E | D | 4.5e-06 | 6.2e-06 | 1.3e-05 | 7.900E-06 | 4.498E-06 | 4.500E-06 | 1.300E-05 |
| 23647 | 1183 | L1183Q | D | 1.8e-05 | 8.6e-06 | 8.6e-06 | 1.173E-05 | 5.427E-06 | 8.600E-06 | 1.800E-05 |
| 23648 | 1183 | L1183G | D | 8.9e-06 | 1.6e-05 | 4.7e-06 | 9.867E-06 | 5.712E-06 | 4.700E-06 | 1.600E-05 |
| 23649 | 1183 | L1183H | D | 6.7e-06 | 8.8e-06 | 1.1e-05 | 8.833E-06 | 2.150E-06 | 6.700E-06 | 1.100E-05 |
| 23650 | 1183 | L1183I | D | 1.2e-05 | 9.4e-06 | 9.5e-06 | 1.030E-05 | 1.473E-06 | 9.400E-06 | 1.200E-05 |
| 23651 | 1183 | L1183L | D | 9.4e-06 | 1.4e-05 | 1.2e-05 | 1.180E-05 | 2.307E-06 | 9.400E-06 | 1.400E-05 |
| 23652 | 1183 | L1183K | D | 8.6e-06 | 1.2e-05 | 5.2e-06 | 8.600E-06 | 3.400E-06 | 5.200E-06 | 1.200E-05 |
| 23653 | 1183 | L1183M | D | 5.3e-06 | 7.7e-06 | 6.1e-06 | 6.367E-06 | 1.222E-06 | 5.300E-06 | 7.700E-06 |
| 23654 | 1183 | L1183F | D | 1e-05 | 7.2e-06 | 9.6e-06 | 8.933E-06 | 1.514E-06 | 7.200E-06 | 1.000E-05 |
| 23655 | 1183 | L1183P | D | 8.6e-06 | 1.2e-05 | 9.6e-06 | 1.007E-05 | 1.747E-06 | 8.600E-06 | 1.200E-05 |
| 23656 | 1183 | L1183S | D | 9.1e-06 | 1.1e-05 | 8.6e-06 | 9.567E-06 | 1.266E-06 | 8.600E-06 | 1.100E-05 |
| 23657 | 1183 | L1183T | D | 1e-05 | 5.9e-06 | 1.5e-05 | 1.030E-05 | 4.557E-06 | 5.900E-06 | 1.500E-05 |
| 23658 | 1183 | L1183W | D | 8.6e-06 | 7.9e-06 | 8.7e-06 | 8.400E-06 | 4.359E-07 | 7.900E-06 | 8.700E-06 |
| 23659 | 1183 | L1183Y | D | 1.4e-05 | 6.5e-06 | 8.8e-06 | 9.767E-06 | 3.842E-06 | 6.500E-06 | 1.400E-05 |
| 23660 | 1183 | L1183V | D | 4.5e-06 | 6.9e-06 | 6.9e-06 | 6.100E-06 | 1.386E-06 | 4.500E-06 | 6.900E-06 |
| 23661 | 1184 | K1184A | D | 7.9e-06 | 8.7e-06 | 1e-05 | 8.867E-06 | 1.060E-06 | 7.900E-06 | 1.000E-05 |
| 23662 | 1184 | K1184R | D | 7.6e-06 | 7.1e-06 | 9.5e-06 | 8.067E-06 | 1.266E-06 | 7.100E-06 | 9.500E-06 |
| 23663 | 1184 | K1184N | D | 7.2e-06 | 6.2e-06 | 5.9e-06 | 6.433E-06 | 6.807E-07 | 5.900E-06 | 7.200E-06 |
| 23664 | 1184 | K1184D | D | 7e-06 | 1.6e-05 | 6.9e-06 | 9.967E-06 | 5.225E-06 | 6.900E-06 | 1.600E-05 |
| 23665 | 1184 | K1184C | D | 4.5e-06 | 8.8e-06 | 4.4e-06 | 5.900E-06 | 2.512E-06 | 4.400E-06 | 8.800E-06 |
| 23666 | 1184 | K1184E | D | 9.2e-06 | 9.7e-06 | 8.4e-06 | 9.100E-06 | 6.557E-07 | 8.400E-06 | 9.700E-06 |
| 23667 | 1184 | K1184Q | D | 7.2e-06 | 5.4e-06 | 9.6e-06 | 7.400E-06 | 2.107E-06 | 5.400E-06 | 9.600E-06 |
| 23668 | 1184 | K1184G | D | 4.8e-06 | 1.2e-05 | 1e-05 | 8.933E-06 | 3.717E-06 | 4.800E-06 | 1.200E-05 |
| 23669 | 1184 | K1184H | D | 4.3e-06 | 7.7e-06 | 6.1e-06 | 6.033E-06 | 1.701E-06 | 4.300E-06 | 7.700E-06 |
| 23670 | 1184 | K1184I | D | 6.6e-06 | 1.9e-05 | 6.2e-06 | 1.060E-05 | 7.277E-06 | 6.200E-06 | 1.900E-05 |
| 23671 | 1184 | K1184L | D | 1.1e-05 | 1.1e-05 | 9.4e-06 | 1.047E-05 | 9.238E-07 | 9.400E-06 | 1.100E-05 |
| 23672 | 1184 | K1184K | D | 6.6e-06 | 6.7e-06 | 7.6e-06 | 6.967E-06 | 5.508E-07 | 6.600E-06 | 7.600E-06 |
| 23673 | 1184 | K1184M | D | 9.5e-06 | 3.7e-06 | 5.1e-06 | 6.100E-06 | 3.027E-06 | 3.700E-06 | 9.500E-06 |
| 23674 | 1184 | K1184F | D | 7e-06 | 6.4e-06 | 5e-06 | 6.133E-06 | 1.026E-06 | 5.000E-06 | 7.000E-06 |
| 23675 | 1184 | K1184P | D | 1.5e-05 | 6.8e-06 | 9.2e-06 | 1.033E-05 | 4.216E-06 | 6.800E-06 | 1.500E-05 |
| 23676 | 1184 | K1184S | D | 6e-06 | 6.6e-06 | 4.9e-06 | 5.833E-06 | 8.622E-07 | 4.900E-06 | 6.600E-06 |
| 23677 | 1184 | K1184T | D | 1.1e-05 | 7.9e-06 | 1.3e-05 | 1.063E-05 | 2.570E-06 | 7.900E-06 | 1.300E-05 |
| 23678 | 1184 | K1184W | D | 1.4e-05 | 7.6e-06 | 8.6e-06 | 1.007E-05 | 3.443E-06 | 7.600E-06 | 1.400E-05 |
| 23679 | 1184 | K1184Y | D | 5.5e-06 | 5.3e-06 | 1.3e-05 | 7.933E-06 | 4.389E-06 | 5.300E-06 | 1.300E-05 |
| 23680 | 1184 | K1184V | D | 4.6e-06 | 6.6e-06 | 6.2e-06 | 5.800E-06 | 1.058E-06 | 4.600E-06 | 6.600E-06 |
| 23681 | 1185 | S1185A | D | 1.4e-05 | 6.5e-06 | 6.9e-06 | 9.133E-06 | 4.219E-06 | 6.500E-06 | 1.400E-05 |
| 23682 | 1185 | S1185R | D | 9.6e-06 | 9.5e-06 | 7.7e-06 | 8.933E-06 | 1.069E-06 | 7.700E-06 | 9.600E-06 |
| 23683 | 1185 | S1185N | D | 1.5e-05 | 4.4e-06 | 1.7e-05 | 1.213E-05 | 6.772E-06 | 4.400E-06 | 1.700E-05 |
| 23684 | 1185 | S1185D | D | 1.1e-05 | 6.1e-06 | 1.1e-05 | 9.367E-06 | 2.829E-06 | 6.100E-06 | 1.100E-05 |
| 23685 | 1185 | S1185C | D | 4.9e-06 | 1.2e-05 | 6.6e-06 | 7.833E-06 | 3.707E-06 | 4.900E-06 | 1.200E-05 |
| 23686 | 1185 | S1185E | D | 6.9e-06 | 9.1e-06 | 1.1e-05 | 9.000E-06 | 2.052E-06 | 6.900E-06 | 1.100E-05 |
| 23687 | 1185 | S1185Q | D | 8e-06 | 6.8e-06 | 8e-06 | 7.600E-06 | 6.928E-07 | 6.800E-06 | 8.000E-06 |
| 23688 | 1185 | S1185G | D | 9.7e-06 | 6.2e-06 | 9.4e-06 | 8.433E-06 | 1.940E-06 | 6.200E-06 | 9.700E-06 |
| 23689 | 1185 | S1185H | D | 5e-06 | 6.6e-06 | 6.7e-06 | 6.100E-06 | 9.539E-07 | 5.000E-06 | 6.700E-06 |
| 23690 | 1185 | S1185I | D | 7.3e-06 | 5.2e-06 | 8.4e-06 | 6.967E-06 | 1.626E-06 | 5.200E-06 | 8.400E-06 |
| 23691 | 1185 | S1185L | D | 1.5e-05 | 5.6e-06 | 1.1e-05 | 1.053E-05 | 4.717E-06 | 5.600E-06 | 1.500E-05 |
| 23692 | 1185 | S1185K | D | 7.8e-06 | 1.1e-05 | 5.4e-06 | 8.067E-06 | 2.810E-06 | 5.400E-06 | 1.100E-05 |
| 23693 | 1185 | S1185M | D | 6.4e-06 | 1e-05 | 5.8e-06 | 7.400E-06 | 2.272E-06 | 5.800E-06 | 1.000E-05 |
| 23694 | 1185 | S1185F | D | 5.7e-06 | 7.9e-06 | 7.2e-06 | 6.933E-06 | 1.124E-06 | 5.700E-06 | 7.900E-06 |
| 23695 | 1185 | S1185P | D | 9.7e-06 | 1.2e-05 | 8.9e-06 | 1.020E-05 | 1.609E-06 | 8.900E-06 | 1.200E-05 |
| 23696 | 1185 | S1185S | D | 5.4e-06 | 9.6e-06 | 9e-06 | 8.000E-06 | 2.272E-06 | 5.400E-06 | 9.600E-06 |
| 23697 | 1185 | S1185T | D | 1.3e-05 | 7.6e-06 | 4.2e-06 | 8.267E-06 | 4.438E-06 | 4.200E-06 | 1.300E-05 |
| 23698 | 1185 | S1185W | D | 7.5e-06 | 8.9e-06 | 6.6e-06 | 7.667E-06 | 1.159E-06 | 6.600E-06 | 8.900E-06 |
| 23699 | 1185 | S1185Y | D | 9.1e-06 | 7.7e-06 | 8.7e-06 | 8.500E-06 | 7.211E-07 | 7.700E-06 | 9.100E-06 |
| 23700 | 1185 | S1185V | D | 7.7e-06 | 6.2e-06 | 1.6e-05 | 9.967E-06 | 5.279E-06 | 6.200E-06 | 1.600E-05 |
| 23701 | 1186 | G1186A | D | 8.8e-06 | 6.4e-06 | 1.5e-05 | 1.007E-05 | 4.438E-06 | 6.400E-06 | 1.500E-05 |
| 23702 | 1186 | G1186R | D | 5.2e-06 | 7.9e-06 | 1.3e-05 | 8.700E-06 | 3.961E-06 | 5.200E-06 | 1.300E-05 |
| 23703 | 1186 | G1186N | D | 1.3e-05 | 1.3e-05 | 8.8e-06 | 1.160E-05 | 2.425E-06 | 8.800E-06 | 1.300E-05 |
| 23704 | 1186 | G1186D | D | 5.8e-06 | 4.4e-06 | 9.7e-06 | 6.633E-06 | 2.747E-06 | 4.400E-06 | 9.700E-06 |
| 23705 | 1186 | G1186C | D | 6.1e-06 | 9.1e-06 | 6.9e-06 | 7.367E-06 | 1.553E-06 | 6.100E-06 | 9.100E-06 |
| 23706 | 1186 | G1186E | D | 6.8e-06 | 8.9e-06 | 8.6e-06 | 8.100E-06 | 1.136E-06 | 6.800E-06 | 8.900E-06 |
| 23707 | 1186 | G1186Q | D | 7.3e-06 | 5.3e-06 | 7.4e-06 | 6.667E-06 | 1.185E-06 | 5.300E-06 | 7.400E-06 |
| 23708 | 1186 | G1186G | D | 6.9e-06 | 5.5e-06 | 5.4e-06 | 5.933E-06 | 8.386E-07 | 5.400E-06 | 6.900E-06 |
| 23709 | 1186 | G1186H | D | 6.9e-06 | 9.9e-06 | 9e-06 | 8.600E-06 | 1.539E-06 | 6.900E-06 | 9.900E-06 |
| 23710 | 1186 | G1186I | D | 9.6e-06 | 5.6e-06 | 1.2e-05 | 9.067E-06 | 3.233E-06 | 5.600E-06 | 1.200E-05 |
| 23711 | 1186 | G1186L | D | 1.2e-05 | 7e-06 | 9.4e-06 | 9.467E-06 | 2.501E-06 | 7.000E-06 | 1.200E-05 |
| 23712 | 1186 | G1186K | D | 1.3e-05 | 7.3e-06 | 7.8e-06 | 9.367E-06 | 3.156E-06 | 7.300E-06 | 1.300E-05 |
| 23713 | 1186 | G1186M | D | 6.1e-06 | 9.3e-06 | 7.8e-06 | 7.733E-06 | 1.601E-06 | 6.100E-06 | 9.300E-06 |
| 23714 | 1186 | G1186F | D | 9.8e-06 | 9e-06 | 7.2e-06 | 8.667E-06 | 1.332E-06 | 7.200E-06 | 9.800E-06 |

| | | | | | | | | | | |
|---|---|---|---|---|---|---|---|---|---|---|
| 23715 | 1186 | G1186P | D | 9e-06 | 7.2e-06 | 6.8e-06 | 7.667E-06 | 1.172E-06 | 6.800E-06 | 9.000E-06 |
| 23716 | 1186 | G1186S | D | 8.1e-06 | 6.6e-06 | 9e-06 | 7.900E-06 | 1.212E-06 | 6.600E-06 | 9.000E-06 |
| 23717 | 1186 | G1186T | D | 1.7e-05 | 5.2e-06 | 1.3e-05 | 1.173E-05 | 6.001E-06 | 5.200E-06 | 1.700E-05 |
| 23718 | 1186 | G1186W | D | 5.4e-06 | 7e-06 | 5.8e-06 | 6.067E-06 | 8.327E-07 | 5.400E-06 | 7.000E-06 |
| 23719 | 1186 | G1186Y | D | 1.1e-05 | 6.7e-06 | 7.9e-06 | 8.533E-06 | 2.219E-06 | 6.700E-06 | 1.100E-05 |
| 23720 | 1186 | G1186V | D | 7.7e-06 | 3.5e-06 | 8.9e-06 | 6.700E-06 | 2.835E-06 | 3.500E-06 | 8.900E-06 |
| 23721 | 1187 | T1187A | D | 1.1e-05 | 8.5e-06 | 5.4e-06 | 8.300E-06 | 2.805E-06 | 5.400E-06 | 1.100E-05 |
| 23722 | 1187 | T1187R | D | 1e-05 | 1.3e-05 | 5.7e-06 | 9.567E-06 | 3.669E-06 | 5.700E-06 | 1.300E-05 |
| 23723 | 1187 | T1187N | D | 1.2e-05 | 9.7e-06 | 9.5e-06 | 1.040E-05 | 1.389E-06 | 9.500E-06 | 1.200E-05 |
| 23724 | 1187 | T1187D | D | 8.6e-06 | 6.6e-06 | 9.5e-06 | 8.233E-06 | 1.484E-06 | 6.600E-06 | 9.500E-06 |
| 23725 | 1187 | T1187C | D | 4.7e-06 | 6.2e-06 | 5.8e-06 | 5.567E-06 | 7.767E-07 | 4.700E-06 | 6.200E-06 |
| 23726 | 1187 | T1187E | D | 6.4e-06 | 6.6e-06 | 7.7e-06 | 6.900E-06 | 7.000E-07 | 6.400E-06 | 7.700E-06 |
| 23727 | 1187 | T1187Q | D | 7e-06 | 6.7e-06 | 6.4e-06 | 6.700E-06 | 3.000E-07 | 6.400E-06 | 7.000E-06 |
| 23728 | 1187 | T1187G | D | 9.7e-06 | 7.1e-06 | 7.6e-06 | 8.133E-06 | 1.380E-06 | 7.100E-06 | 9.700E-06 |
| 23729 | 1187 | T1187H | D | 9.8e-06 | 6.3e-06 | 1.3e-05 | 9.700E-06 | 3.351E-06 | 6.300E-06 | 1.300E-05 |
| 23730 | 1187 | T1187I | D | 4.5e-06 | 4.8e-06 | 7.2e-06 | 5.500E-06 | 1.480E-06 | 4.500E-06 | 7.200E-06 |
| 23731 | 1187 | T1187L | D | 1.1e-05 | 5.3e-06 | 1.1e-05 | 9.100E-06 | 3.291E-06 | 5.300E-06 | 1.100E-05 |
| 23732 | 1187 | T1187K | D | 9.1e-06 | 8.7e-06 | 1.2e-05 | 9.933E-06 | 1.801E-06 | 8.700E-06 | 1.200E-05 |
| 23733 | 1187 | T1187M | D | 8.3e-06 | 1.1e-05 | 8.5e-06 | 9.267E-06 | 1.504E-06 | 8.300E-06 | 1.100E-05 |
| 23734 | 1187 | T1187F | D | 7e-06 | 6.5e-06 | 1e-05 | 7.833E-06 | 1.893E-06 | 6.500E-06 | 1.000E-05 |
| 23735 | 1187 | T1187P | D | 9.3e-06 | 9.2e-06 | 1.4e-05 | 1.083E-05 | 2.743E-06 | 9.200E-06 | 1.400E-05 |
| 23736 | 1187 | T1187S | D | 1.4e-05 | 8e-06 | 5.6e-06 | 9.200E-06 | 4.327E-06 | 5.600E-06 | 1.400E-05 |
| 23737 | 1187 | T1187T | D | 9.9e-06 | 8.2e-06 | 1.4e-05 | 1.070E-05 | 2.982E-06 | 8.200E-06 | 1.400E-05 |
| 23738 | 1187 | T1187W | D | 1.2e-05 | 5.7e-06 | 9.3e-06 | 9.000E-06 | 3.161E-06 | 5.700E-06 | 1.200E-05 |
| 23739 | 1187 | T1187Y | D | 8.1e-06 | 1e-05 | 5.3e-06 | 7.800E-06 | 2.364E-06 | 5.300E-06 | 1.000E-05 |
| 23740 | 1187 | T1187V | D | 4.4e-06 | 6.5e-06 | 9.7e-06 | 6.867E-06 | 2.669E-06 | 4.400E-06 | 9.700E-06 |
| 23741 | 1188 | A1188A | D | 8.1e-06 | 8.2e-06 | 7e-06 | 7.767E-06 | 6.658E-07 | 7.000E-06 | 8.200E-06 |
| 23742 | 1188 | A1188R | D | 8.5e-06 | 7.3e-06 | 6.3e-06 | 7.367E-06 | 1.102E-06 | 6.300E-06 | 8.500E-06 |
| 23743 | 1188 | A1188N | D | 5.5e-06 | 6e-06 | 6.8e-06 | 6.100E-06 | 6.557E-07 | 5.500E-06 | 6.800E-06 |
| 23744 | 1188 | A1188D | D | 6.3e-06 | 1.5e-05 | 9.4e-06 | 1.023E-05 | 4.409E-06 | 6.300E-06 | 1.500E-05 |
| 23745 | 1188 | A1188C | D | 6.6e-06 | 6.9e-06 | 8.5e-06 | 7.333E-06 | 1.021E-06 | 6.600E-06 | 8.500E-06 |
| 23746 | 1188 | A1188E | D | 1.2e-05 | 6.2e-06 | 7.5e-06 | 8.567E-06 | 3.044E-06 | 6.200E-06 | 1.200E-05 |
| 23747 | 1188 | A1188Q | D | 9.4e-06 | 7.4e-06 | 8.8e-06 | 8.533E-06 | 1.026E-06 | 7.400E-06 | 9.400E-06 |
| 23748 | 1188 | A1188G | D | 1e-05 | 8.2e-06 | 1.4e-05 | 1.073E-05 | 2.969E-06 | 8.200E-06 | 1.400E-05 |
| 23749 | 1188 | A1188H | D | 9.6e-06 | 5.6e-06 | 9.7e-06 | 8.300E-06 | 2.339E-06 | 5.600E-06 | 9.700E-06 |
| 23750 | 1188 | A1188I | D | 8.2e-06 | 7.1e-06 | 1.2e-05 | 9.100E-06 | 2.571E-06 | 7.100E-06 | 1.200E-05 |
| 23751 | 1188 | A1188L | D | 4.9e-06 | 9.5e-06 | 7.4e-06 | 7.267E-06 | 2.303E-06 | 4.900E-06 | 9.500E-06 |
| 23752 | 1188 | A1188K | D | 5.2e-06 | 8e-06 | 8.4e-06 | 7.200E-06 | 1.744E-06 | 5.200E-06 | 8.400E-06 |
| 23753 | 1188 | A1188M | D | 1.1e-05 | 9.9e-06 | 7.8e-06 | 9.567E-06 | 1.626E-06 | 7.800E-06 | 1.100E-05 |
| 23754 | 1188 | A1188F | D | 9.8e-06 | 7.2e-06 | 5.5e-06 | 7.500E-06 | 2.166E-06 | 5.500E-06 | 9.800E-06 |
| 23755 | 1188 | A1188P | D | 5.5e-06 | 8e-06 | 6.7e-06 | 6.733E-06 | 1.250E-06 | 5.500E-06 | 8.000E-06 |
| 23756 | 1188 | A1188S | D | 1.1e-05 | 1.3e-05 | 1.2e-05 | 1.200E-05 | 1.000E-06 | 1.100E-05 | 1.300E-05 |
| 23757 | 1188 | A1188T | D | 8.9e-06 | 8.7e-06 | 7.1e-06 | 8.233E-06 | 9.866E-07 | 7.100E-06 | 8.900E-06 |
| 23758 | 1188 | A1188W | D | 7.3e-06 | 6.4e-06 | 7.6e-06 | 7.100E-06 | 6.245E-07 | 6.400E-06 | 7.600E-06 |
| 23759 | 1188 | A1188Y | D | 5.8e-06 | 5.3e-06 | 9.1e-06 | 6.733E-06 | 2.065E-06 | 5.300E-06 | 9.100E-06 |
| 23760 | 1188 | A1188V | D | 7e-06 | 7.3e-06 | 6.9e-06 | 7.067E-06 | 2.082E-07 | 6.900E-06 | 7.300E-06 |
| 23761 | 1189 | S1189A | D | 1e-05 | 3.3e-06 | 1.1e-05 | 8.100E-06 | 4.187E-06 | 3.300E-06 | 1.100E-05 |
| 23762 | 1189 | S1189R | D | 6.7e-06 | 9.3e-06 | 9.5e-06 | 8.500E-06 | 1.562E-06 | 6.700E-06 | 9.500E-06 |
| 23763 | 1189 | S1189N | D | 9.5e-06 | 5e-06 | 1.3e-05 | 9.167E-06 | 4.010E-06 | 5.000E-06 | 1.300E-05 |
| 23764 | 1189 | S1189D | D | 8.5e-06 | 9e-06 | 6.5e-06 | 8.000E-06 | 1.323E-06 | 6.500E-06 | 9.000E-06 |
| 23765 | 1189 | S1189C | D | 1.3e-05 | 1.2e-05 | 9e-06 | 1.133E-05 | 2.082E-06 | 9.000E-06 | 1.300E-05 |
| 23766 | 1189 | S1189E | D | 1.2e-05 | 6.7e-06 | 1.2e-05 | 1.023E-05 | 3.060E-06 | 6.700E-06 | 1.200E-05 |
| 23767 | 1189 | S1189Q | D | 7.4e-06 | 8.9e-06 | 7.4e-06 | 7.900E-06 | 8.660E-07 | 7.400E-06 | 8.900E-06 |
| 23768 | 1189 | S1189G | D | 8.1e-06 | 9.8e-06 | 7.7e-06 | 8.533E-06 | 1.115E-06 | 7.700E-06 | 9.800E-06 |
| 23769 | 1189 | S1189H | D | 6.2e-06 | 8.2e-06 | 1.6e-05 | 1.013E-05 | 5.178E-06 | 6.200E-06 | 1.600E-05 |
| 23770 | 1189 | S1189I | D | 1.3e-05 | 9.2e-06 | 8.9e-06 | 1.037E-05 | 2.285E-06 | 8.900E-06 | 1.300E-05 |
| 23771 | 1189 | S1189L | D | 3.8e-06 | 8.4e-06 | 8.2e-06 | 6.800E-06 | 2.600E-06 | 3.800E-06 | 8.400E-06 |
| 23772 | 1189 | S1189K | D | 9.1e-06 | 9e-06 | 6.7e-06 | 8.267E-06 | 1.358E-06 | 6.700E-06 | 9.100E-06 |
| 23773 | 1189 | S1189M | D | 4.5e-06 | 8.2e-06 | 6.2e-06 | 6.300E-06 | 1.852E-06 | 4.500E-06 | 8.200E-06 |
| 23774 | 1189 | S1189F | D | 1.3e-05 | 6.4e-06 | 4.9e-06 | 8.100E-06 | 4.309E-06 | 4.900E-06 | 1.300E-05 |
| 23775 | 1189 | S1189P | D | 9.1e-06 | 7.5e-06 | 6.2e-06 | 7.600E-06 | 1.453E-06 | 6.200E-06 | 9.100E-06 |
| 23776 | 1189 | S1189S | D | 8.2e-06 | 6.3e-06 | 7.9e-06 | 7.467E-06 | 1.021E-06 | 6.300E-06 | 8.200E-06 |
| 23777 | 1189 | S1189T | D | 2.8e-06 | 8.3e-06 | 6.6e-06 | 5.900E-06 | 2.816E-06 | 2.800E-06 | 8.300E-06 |
| 23778 | 1189 | S1189W | D | 6.9e-06 | 1.4e-05 | 1e-05 | 1.030E-05 | 3.559E-06 | 6.900E-06 | 1.400E-05 |
| 23779 | 1189 | S1189Y | D | 9.7e-06 | 9.9e-06 | 8.2e-06 | 9.267E-06 | 9.292E-07 | 8.200E-06 | 9.900E-06 |
| 23780 | 1189 | S1189V | D | 1.3e-05 | 8e-06 | 4.7e-06 | 8.567E-06 | 4.179E-06 | 4.700E-06 | 1.300E-05 |
| 23781 | 1190 | V1190A | D | 1.2e-05 | 5.7e-06 | 8.2e-06 | 8.633E-06 | 3.172E-06 | 5.700E-06 | 1.200E-05 |
| 23782 | 1190 | V1190R | D | 6.3e-06 | 7.3e-06 | 9.1e-06 | 7.567E-06 | 1.419E-06 | 6.300E-06 | 9.100E-06 |
| 23783 | 1190 | V1190N | D | 9.3e-06 | 6.1e-06 | 8.2e-06 | 7.867E-06 | 1.626E-06 | 6.100E-06 | 9.300E-06 |
| 23784 | 1190 | V1190D | D | 1.1e-05 | 8.1e-06 | 1.1e-05 | 1.003E-05 | 1.674E-06 | 8.100E-06 | 1.100E-05 |
| 23785 | 1190 | V1190C | D | 7.6e-06 | 1.2e-05 | 8.9e-06 | 9.500E-06 | 2.261E-06 | 7.600E-06 | 1.200E-05 |
| 23786 | 1190 | V1190E | D | 1.2e-05 | 6.2e-06 | 9.3e-06 | 9.167E-06 | 2.902E-06 | 6.200E-06 | 1.200E-05 |
| 23787 | 1190 | V1190Q | D | 6e-06 | 7.9e-06 | 7e-06 | 6.967E-06 | 9.504E-07 | 6.000E-06 | 7.900E-06 |
| 23788 | 1190 | V1190G | D | 9.5e-06 | 4.5e-06 | 3.7e-06 | 5.900E-06 | 3.143E-06 | 3.700E-06 | 9.500E-06 |
| 23789 | 1190 | V1190H | D | 7e-06 | 6.3e-06 | 1.4e-05 | 9.100E-06 | 4.258E-06 | 6.300E-06 | 1.400E-05 |
| 23790 | 1190 | V1190I | D | 8.5e-06 | 7.8e-06 | 1.5e-05 | 1.043E-05 | 3.970E-06 | 7.800E-06 | 1.500E-05 |
| 23791 | 1190 | V1190L | D | 7e-06 | 8.7e-06 | 9.2e-06 | 8.300E-06 | 1.153E-06 | 7.000E-06 | 9.200E-06 |
| 23792 | 1190 | V1190K | D | 7.1e-06 | 6.5e-06 | 9.2e-06 | 7.600E-06 | 1.418E-06 | 6.500E-06 | 9.200E-06 |
| 23793 | 1190 | V1190M | D | 1.5e-05 | 9.5e-06 | 6.4e-06 | 1.030E-05 | 4.355E-06 | 6.400E-06 | 1.500E-05 |
| 23794 | 1190 | V1190F | D | 1.3e-05 | 4.1e-06 | 8.2e-06 | 8.433E-06 | 4.455E-06 | 4.100E-06 | 1.300E-05 |
| 23795 | 1190 | V1190P | D | 1e-05 | 6.9e-06 | 7.9e-06 | 8.267E-06 | 1.582E-06 | 6.900E-06 | 1.000E-05 |
| 23796 | 1190 | V1190S | D | 9.1e-06 | 1.2e-05 | 1.4e-05 | 1.170E-05 | 2.464E-06 | 9.100E-06 | 1.400E-05 |
| 23797 | 1190 | V1190T | D | 1.2e-05 | 3.9e-06 | 1e-05 | 8.633E-06 | 4.219E-06 | 3.900E-06 | 1.200E-05 |
| 23798 | 1190 | V1190W | D | 8.6e-06 | 4.1e-06 | 8.9e-06 | 7.200E-06 | 2.689E-06 | 4.100E-06 | 8.900E-06 |
| 23799 | 1190 | V1190Y | D | 1.2e-05 | 8e-06 | 9.4e-06 | 9.800E-06 | 2.030E-06 | 8.000E-06 | 1.200E-05 |

| | | | | | | | | | | |
|---|---|---|---|---|---|---|---|---|---|---|
| 23800 | 1190 | V1190V | D | 7.3e-06 | 4.7e-06 | 7.2e-06 | 6.400E-06 | 1.473E-06 | 4.700E-06 | 7.300E-06 |
| 23801 | 1191 | V1191A | D | 9.8e-06 | 8.6e-06 | 8.4e-06 | 8.933E-06 | 7.572E-07 | 8.400E-06 | 9.800E-06 |
| 23802 | 1191 | V1191R | D | 6.4e-06 | 8.7e-06 | 6.9e-06 | 7.333E-06 | 1.210E-06 | 6.400E-06 | 8.700E-06 |
| 23803 | 1191 | V1191N | D | 6.7e-06 | 1.1e-05 | 4.7e-06 | 7.467E-06 | 3.219E-06 | 4.700E-06 | 1.100E-05 |
| 23804 | 1191 | V1191D | D | 9.6e-06 | 4.9e-06 | 1e-05 | 8.167E-06 | 2.836E-06 | 4.900E-06 | 1.000E-05 |
| 23805 | 1191 | V1191C | D | 8.8e-06 | 6.1e-06 | 7.7e-06 | 7.533E-06 | 1.358E-06 | 6.100E-06 | 8.800E-06 |
| 23806 | 1191 | V1191E | D | 6.1e-06 | 5.2e-06 | 6.1e-06 | 5.800E-06 | 5.196E-07 | 5.200E-06 | 6.100E-06 |
| 23807 | 1191 | V1191Q | D | 7.9e-06 | 4.7e-06 | 8.8e-06 | 7.133E-06 | 2.155E-06 | 4.700E-06 | 8.800E-06 |
| 23808 | 1191 | V1191G | D | 1e-05 | 9.9e-06 | 7e-06 | 8.967E-06 | 1.704E-06 | 7.000E-06 | 1.000E-05 |
| 23809 | 1191 | V1191H | D | 7.8e-06 | 3.2e-06 | 4.8e-06 | 5.267E-06 | 2.335E-06 | 3.200E-06 | 7.800E-06 |
| 23810 | 1191 | V1191I | D | 6e-06 | 1.3e-05 | 4.8e-06 | 7.933E-06 | 4.429E-06 | 4.800E-06 | 1.300E-05 |
| 23811 | 1191 | V1191L | D | 8.7e-06 | 1.1e-05 | 9.4e-06 | 9.700E-06 | 1.179E-06 | 8.700E-06 | 1.100E-05 |
| 23812 | 1191 | V1191K | D | 4.6e-06 | 9.2e-06 | 9.5e-06 | 7.767E-06 | 2.747E-06 | 4.600E-06 | 9.500E-06 |
| 23813 | 1191 | V1191M | D | 8.8e-06 | 9.1e-06 | 1.4e-05 | 1.063E-05 | 2.919E-06 | 8.800E-06 | 1.400E-05 |
| 23814 | 1191 | V1191F | D | 7.4e-06 | 6.6e-06 | 8.1e-06 | 7.367E-06 | 7.506E-07 | 6.600E-06 | 8.100E-06 |
| 23815 | 1191 | V1191P | D | 9.1e-06 | 6.8e-06 | 9.3e-06 | 8.400E-06 | 1.389E-06 | 6.800E-06 | 9.300E-06 |
| 23816 | 1191 | V1191S | D | 9.9e-06 | 9.6e-06 | 8.9e-06 | 9.467E-06 | 5.132E-07 | 8.900E-06 | 9.900E-06 |
| 23817 | 1191 | V1191T | D | 6.3e-06 | 9.4e-06 | 9.7e-06 | 8.467E-06 | 1.882E-06 | 6.300E-06 | 9.700E-06 |
| 23818 | 1191 | V1191W | D | 6.6e-06 | 6.6e-06 | 7.6e-06 | 6.933E-06 | 5.774E-07 | 6.600E-06 | 7.600E-06 |
| 23819 | 1191 | V1191Y | D | 3.2e-06 | 6.5e-06 | 6.9e-06 | 5.533E-06 | 2.031E-06 | 3.200E-06 | 6.900E-06 |
| 23820 | 1191 | V1191V | D | 6.4e-06 | 9.1e-06 | 1e-05 | 8.500E-06 | 1.873E-06 | 6.400E-06 | 1.000E-05 |
| 23821 | 1192 | C1192A | D | 6.3e-06 | 6.9e-06 | 1.1e-05 | 8.067E-06 | 2.558E-06 | 6.300E-06 | 1.100E-05 |
| 23822 | 1192 | C1192R | D | 6.7e-06 | 7.1e-06 | 9.7e-06 | 7.833E-06 | 1.629E-06 | 6.700E-06 | 9.700E-06 |
| 23823 | 1192 | C1192N | D | 4.4e-06 | 8.9e-06 | 1.3e-05 | 8.767E-06 | 4.302E-06 | 4.400E-06 | 1.300E-05 |
| 23824 | 1192 | C1192D | D | 7.8e-06 | 4.9e-06 | 1.4e-05 | 8.900E-06 | 4.649E-06 | 4.900E-06 | 1.400E-05 |
| 23825 | 1192 | C1192C | D | 9.1e-06 | 1e-05 | 9.4e-06 | 9.500E-06 | 4.583E-07 | 9.100E-06 | 1.000E-05 |
| 23826 | 1192 | C1192E | D | 1e-05 | 9.7e-06 | 6.3e-06 | 8.667E-06 | 2.055E-06 | 6.300E-06 | 1.000E-05 |
| 23827 | 1192 | C1192Q | D | 6.4e-06 | 1.3e-05 | 6.2e-06 | 8.533E-06 | 3.870E-06 | 6.200E-06 | 1.300E-05 |
| 23828 | 1192 | C1192G | D | 9.4e-06 | 1.5e-05 | 5.9e-06 | 1.010E-05 | 4.590E-06 | 5.900E-06 | 1.500E-05 |
| 23829 | 1192 | C1192H | D | 8.3e-06 | 9e-06 | 7.5e-06 | 8.267E-06 | 7.506E-07 | 7.500E-06 | 9.000E-06 |
| 23830 | 1192 | C1192I | D | 5.3e-06 | 6e-06 | 7.6e-06 | 6.300E-06 | 1.179E-06 | 5.300E-06 | 7.600E-06 |
| 23831 | 1192 | C1192L | D | 7.3e-06 | 1.5e-05 | 8.8e-06 | 1.037E-05 | 4.082E-06 | 7.300E-06 | 1.500E-05 |
| 23832 | 1192 | C1192K | D | 1e-05 | 4.8e-06 | 7.7e-06 | 7.500E-06 | 2.606E-06 | 4.800E-06 | 1.000E-05 |
| 23833 | 1192 | C1192M | D | 7.8e-06 | 8.8e-06 | 6.7e-06 | 7.767E-06 | 1.050E-06 | 6.700E-06 | 8.800E-06 |
| 23834 | 1192 | C1192F | D | 8.6e-06 | 9.5e-06 | 5.5e-06 | 7.867E-06 | 2.098E-06 | 5.500E-06 | 9.500E-06 |
| 23835 | 1192 | C1192P | D | 8.9e-06 | 8.9e-06 | 7.3e-06 | 8.367E-06 | 9.238E-07 | 7.300E-06 | 8.900E-06 |
| 23836 | 1192 | C1192S | D | 9.9e-06 | 5.7e-06 | 1.1e-05 | 8.867E-06 | 2.797E-06 | 5.700E-06 | 1.100E-05 |
| 23837 | 1192 | C1192T | D | 6.2e-06 | 7.2e-06 | 1.8e-05 | 1.047E-05 | 6.543E-06 | 6.200E-06 | 1.800E-05 |
| 23838 | 1192 | C1192W | D | 7.5e-06 | 1.2e-05 | 8.3e-06 | 9.267E-06 | 2.401E-06 | 7.500E-06 | 1.200E-05 |
| 23839 | 1192 | C1192Y | D | 1.1e-05 | 5.2e-06 | 6.7e-06 | 7.633E-06 | 3.011E-06 | 5.200E-06 | 1.100E-05 |
| 23840 | 1192 | C1192V | D | 1.2e-05 | 1.1e-05 | 1.2e-05 | 1.167E-05 | 5.774E-07 | 1.100E-05 | 1.200E-05 |
| 23841 | 1193 | L1193A | D | 7.7e-06 | 7.1e-06 | 4.2e-06 | 6.333E-06 | 1.872E-06 | 4.200E-06 | 7.700E-06 |
| 23842 | 1193 | L1193R | D | 7.7e-06 | 5.5e-06 | 1.5e-05 | 9.400E-06 | 4.973E-06 | 5.500E-06 | 1.500E-05 |
| 23843 | 1193 | L1193N | D | 9.7e-06 | 6.1e-06 | 1.2e-05 | 9.267E-06 | 2.974E-06 | 6.100E-06 | 1.200E-05 |
| 23844 | 1193 | L1193D | D | 8.9e-06 | 9.4e-06 | 6.8e-06 | 8.367E-06 | 1.380E-06 | 6.800E-06 | 9.400E-06 |
| 23845 | 1193 | L1193C | D | 4.8e-06 | 6.1e-06 | 1.1e-05 | 7.300E-06 | 3.270E-06 | 4.800E-06 | 1.100E-05 |
| 23846 | 1193 | L1193E | D | 8.8e-06 | 7.2e-06 | 1.3e-05 | 9.667E-06 | 2.996E-06 | 7.200E-06 | 1.300E-05 |
| 23847 | 1193 | L1193Q | D | 1.1e-05 | 9.2e-06 | 6.1e-06 | 8.767E-06 | 2.479E-06 | 6.100E-06 | 1.100E-05 |
| 23848 | 1193 | L1193G | D | 6.5e-06 | 1.3e-05 | 1e-05 | 9.833E-06 | 3.253E-06 | 6.500E-06 | 1.300E-05 |
| 23849 | 1193 | L1193H | D | 7.4e-06 | 6e-06 | 8.9e-06 | 7.433E-06 | 1.450E-06 | 6.000E-06 | 8.900E-06 |
| 23850 | 1193 | L1193I | D | 8e-06 | 5.9e-06 | 1.1e-05 | 8.300E-06 | 2.563E-06 | 5.900E-06 | 1.100E-05 |
| 23851 | 1193 | L1193L | D | 9.7e-06 | 5.3e-06 | 8.6e-06 | 7.867E-06 | 2.290E-06 | 5.300E-06 | 9.700E-06 |
| 23852 | 1193 | L1193K | D | 1e-05 | 5.5e-06 | 1.1e-05 | 8.833E-06 | 2.930E-06 | 5.500E-06 | 1.100E-05 |
| 23853 | 1193 | L1193M | D | 1.4e-05 | 5e-06 | 5.9e-06 | 8.300E-06 | 4.957E-06 | 5.000E-06 | 1.400E-05 |
| 23854 | 1193 | L1193F | D | 8.7e-06 | 1.4e-05 | 6.4e-06 | 9.700E-06 | 3.897E-06 | 6.400E-06 | 1.400E-05 |
| 23855 | 1193 | L1193P | D | 7.1e-06 | 9.5e-06 | 6.9e-06 | 7.833E-06 | 1.447E-06 | 6.900E-06 | 9.500E-06 |
| 23856 | 1193 | L1193S | D | 1e-05 | 8.1e-06 | 8.6e-06 | 8.900E-06 | 9.849E-07 | 8.100E-06 | 1.000E-05 |
| 23857 | 1193 | L1193T | D | 5.1e-06 | 9.9e-06 | 6.2e-06 | 7.067E-06 | 2.515E-06 | 5.100E-06 | 9.900E-06 |
| 23858 | 1193 | L1193W | D | 1.4e-05 | 9.9e-06 | 4.8e-06 | 9.567E-06 | 4.609E-06 | 4.800E-06 | 1.400E-05 |
| 23859 | 1193 | L1193Y | D | 1.8e-05 | 7.3e-06 | 5.8e-06 | 1.037E-05 | 6.653E-06 | 5.800E-06 | 1.800E-05 |
| 23860 | 1193 | L1193V | D | 8.1e-06 | 7.3e-06 | 5e-06 | 6.800E-06 | 1.609E-06 | 5.000E-06 | 8.100E-06 |
| 23861 | 1194 | L1194A | D | 7.7e-06 | 5.6e-06 | 7e-06 | 6.767E-06 | 1.069E-06 | 5.600E-06 | 7.700E-06 |
| 23862 | 1194 | L1194R | D | 7.5e-06 | 5.5e-06 | 9.6e-06 | 7.533E-06 | 2.050E-06 | 5.500E-06 | 9.600E-06 |
| 23863 | 1194 | L1194N | D | 6.9e-06 | 1.3e-05 | 1e-05 | 9.967E-06 | 3.050E-06 | 6.900E-06 | 1.300E-05 |
| 23864 | 1194 | L1194D | D | 1e-05 | 1.2e-05 | 1.7e-05 | 1.300E-05 | 3.606E-06 | 1.000E-05 | 1.700E-05 |
| 23865 | 1194 | L1194C | D | 8e-06 | 5.4e-06 | 9e-06 | 7.467E-06 | 1.858E-06 | 5.400E-06 | 9.000E-06 |
| 23866 | 1194 | L1194E | D | 9.1e-06 | 1.2e-05 | 6.1e-06 | 9.067E-06 | 2.950E-06 | 6.100E-06 | 1.200E-05 |
| 23867 | 1194 | L1194Q | D | 6e-06 | 8.9e-06 | 6.7e-06 | 7.200E-06 | 1.513E-06 | 6.000E-06 | 8.900E-06 |
| 23868 | 1194 | L1194G | D | 1.3e-05 | 5.5e-06 | 4.7e-06 | 7.733E-06 | 4.579E-06 | 4.700E-06 | 1.300E-05 |
| 23869 | 1194 | L1194H | D | 7e-06 | 1.1e-05 | 7.6e-06 | 8.533E-06 | 2.157E-06 | 7.000E-06 | 1.100E-05 |
| 23870 | 1194 | L1194I | D | 8.8e-06 | 1e-05 | 9.4e-06 | 9.400E-06 | 6.000E-07 | 8.800E-06 | 1.000E-05 |
| 23871 | 1194 | L1194L | D | 8.6e-06 | 1e-05 | 8.9e-06 | 9.167E-06 | 7.371E-07 | 8.600E-06 | 1.000E-05 |
| 23872 | 1194 | L1194K | D | 1e-05 | 4e-06 | 9.8e-06 | 7.933E-06 | 3.408E-06 | 4.000E-06 | 1.000E-05 |
| 23873 | 1194 | L1194M | D | 7.1e-06 | 7.5e-06 | 6.5e-06 | 7.033E-06 | 5.033E-07 | 6.500E-06 | 7.500E-06 |
| 23874 | 1194 | L1194F | D | 1e-05 | 4.8e-06 | 8.1e-06 | 7.633E-06 | 2.631E-06 | 4.800E-06 | 1.000E-05 |
| 23875 | 1194 | L1194P | D | 5.6e-06 | 6.8e-06 | 7.7e-06 | 6.700E-06 | 1.054E-06 | 5.600E-06 | 7.700E-06 |
| 23876 | 1194 | L1194S | D | 6.5e-06 | 8.8e-06 | 7.2e-06 | 7.500E-06 | 1.179E-06 | 6.500E-06 | 8.800E-06 |
| 23877 | 1194 | L1194T | D | 8.5e-06 | 6.4e-06 | 1.2e-05 | 8.967E-06 | 2.829E-06 | 6.400E-06 | 1.200E-05 |
| 23878 | 1194 | L1194W | D | 1e-05 | 1.3e-05 | 8.3e-06 | 1.043E-05 | 2.380E-06 | 8.300E-06 | 1.300E-05 |
| 23879 | 1194 | L1194Y | D | 8e-06 | 6.5e-06 | 6.2e-06 | 6.900E-06 | 9.644E-07 | 6.200E-06 | 8.000E-06 |
| 23880 | 1194 | L1194V | D | 8e-06 | 1.9e-05 | 8.9e-06 | 1.197E-05 | 6.108E-06 | 8.000E-06 | 1.900E-05 |
| 23881 | 1195 | N1195A | D | 6.1e-06 | 9.4e-06 | 6.8e-06 | 7.433E-06 | 1.739E-06 | 6.100E-06 | 9.400E-06 |
| 23882 | 1195 | N1195R | D | 1.2e-05 | 7.5e-06 | 8.5e-06 | 9.333E-06 | 2.363E-06 | 7.500E-06 | 1.200E-05 |
| 23883 | 1195 | N1195N | D | 5e-06 | 7.1e-06 | 9.6e-06 | 7.233E-06 | 2.303E-06 | 5.000E-06 | 9.600E-06 |
| 23884 | 1195 | N1195D | D | 1.1e-05 | 1.1e-05 | 7.8e-06 | 9.933E-06 | 1.848E-06 | 7.800E-06 | 1.100E-05 |

| | | | | | | | | | | |
|---|---|---|---|---|---|---|---|---|---|---|
| 23885 | 1195 | N1195C | D | 1.3e-05 | 1.4e-05 | 1.5e-05 | 1.400E-05 | 1.000E-06 | 1.300E-05 | 1.500E-05 |
| 23886 | 1195 | N1195E | D | 7.8e-06 | 1.1e-05 | 4.3e-06 | 7.700E-06 | 3.351E-06 | 4.300E-06 | 1.100E-05 |
| 23887 | 1195 | N1195Q | D | 6e-06 | 5.4e-06 | 6e-06 | 5.800E-06 | 3.464E-07 | 5.400E-06 | 6.000E-06 |
| 23888 | 1195 | N1195G | D | 7e-06 | 1e-05 | 5.9e-06 | 7.633E-06 | 2.122E-06 | 5.900E-06 | 1.000E-05 |
| 23889 | 1195 | N1195H | D | 6.1e-06 | 4.4e-06 | 9.5e-06 | 6.667E-06 | 2.597E-06 | 4.400E-06 | 9.500E-06 |
| 23890 | 1195 | N1195I | D | 1.4e-05 | 9.2e-06 | 8.6e-06 | 1.060E-05 | 2.960E-06 | 8.600E-06 | 1.400E-05 |
| 23891 | 1195 | N1195L | D | 7.8e-06 | 8.4e-06 | 1.3e-05 | 9.733E-06 | 2.845E-06 | 7.800E-06 | 1.300E-05 |
| 23892 | 1195 | N1195K | D | 1.3e-05 | 5.4e-06 | 7.1e-06 | 8.500E-06 | 3.989E-06 | 5.400E-06 | 1.300E-05 |
| 23893 | 1195 | N1195M | D | 6.6e-06 | 9.2e-06 | 9.8e-06 | 8.533E-06 | 1.701E-06 | 6.600E-06 | 9.800E-06 |
| 23894 | 1195 | N1195F | D | 1.3e-05 | 7e-06 | 1.8e-05 | 1.267E-05 | 5.508E-06 | 7.000E-06 | 1.800E-05 |
| 23895 | 1195 | N1195P | D | 1.3e-05 | 5.7e-06 | 5.7e-06 | 8.133E-06 | 4.215E-06 | 5.700E-06 | 1.300E-05 |
| 23896 | 1195 | N1195S | D | 6.4e-06 | 1.3e-05 | 1.1e-05 | 1.013E-05 | 3.384E-06 | 6.400E-06 | 1.300E-05 |
| 23897 | 1195 | N1195T | D | 5.9e-06 | 5.3e-06 | 8.4e-06 | 6.533E-06 | 1.644E-06 | 5.300E-06 | 8.400E-06 |
| 23898 | 1195 | N1195W | D | 6.6e-06 | 9.1e-06 | 6.6e-06 | 7.433E-06 | 1.443E-06 | 6.600E-06 | 9.100E-06 |
| 23899 | 1195 | N1195Y | D | 9e-06 | 5.7e-06 | 1.1e-05 | 8.567E-06 | 2.676E-06 | 5.700E-06 | 1.100E-05 |
| 23900 | 1195 | N1195V | D | 6.6e-06 | 1.4e-05 | 6.3e-06 | 8.967E-06 | 4.362E-06 | 6.300E-06 | 1.400E-05 |
| 23901 | 1196 | N1196A | D | 8.9e-06 | 9.2e-06 | 1.1e-05 | 9.700E-06 | 1.136E-06 | 8.900E-06 | 1.100E-05 |
| 23902 | 1196 | N1196R | D | 6e-06 | 8e-06 | 1.2e-05 | 8.667E-06 | 3.055E-06 | 6.000E-06 | 1.200E-05 |
| 23903 | 1196 | N1196N | D | 7.8e-06 | 1.1e-05 | 7.1e-06 | 8.633E-06 | 2.079E-06 | 7.100E-06 | 1.100E-05 |
| 23904 | 1196 | N1196D | D | 5e-06 | 6.4e-06 | 8.9e-06 | 6.767E-06 | 1.976E-06 | 5.000E-06 | 8.900E-06 |
| 23905 | 1196 | N1196C | D | 4.6e-06 | 5.7e-06 | 7.7e-06 | 6.000E-06 | 1.572E-06 | 4.600E-06 | 7.700E-06 |
| 23906 | 1196 | N1196E | D | 9.1e-06 | 8.8e-06 | 1.2e-05 | 9.967E-06 | 1.767E-06 | 8.800E-06 | 1.200E-05 |
| 23907 | 1196 | N1196Q | D | 6.5e-06 | 8e-06 | 6.7e-06 | 7.067E-06 | 8.145E-07 | 6.500E-06 | 8.000E-06 |
| 23908 | 1196 | N1196G | D | 4.5e-06 | 6.6e-06 | 7.8e-06 | 6.300E-06 | 1.670E-06 | 4.500E-06 | 7.800E-06 |
| 23909 | 1196 | N1196H | D | 9.6e-06 | 5.2e-06 | 5.9e-06 | 6.900E-06 | 2.364E-06 | 5.200E-06 | 9.600E-06 |
| 23910 | 1196 | N1196I | D | 1e-05 | 1.3e-05 | 1.4e-05 | 1.233E-05 | 2.082E-06 | 1.000E-05 | 1.400E-05 |
| 23911 | 1196 | N1196L | D | 8.9e-06 | 8.4e-06 | 9.1e-06 | 8.800E-06 | 3.606E-07 | 8.400E-06 | 9.100E-06 |
| 23912 | 1196 | N1196K | D | 1.2e-05 | 9.4e-06 | 5.9e-06 | 9.100E-06 | 3.061E-06 | 5.900E-06 | 1.200E-05 |
| 23913 | 1196 | N1196M | D | 9.9e-06 | 6.3e-06 | 7.2e-06 | 7.800E-06 | 1.873E-06 | 6.300E-06 | 9.900E-06 |
| 23914 | 1196 | N1196F | D | 1e-05 | 6.8e-06 | 7.3e-06 | 8.033E-06 | 1.721E-06 | 6.800E-06 | 1.000E-05 |
| 23915 | 1196 | N1196P | D | 4.9e-06 | 7.2e-06 | 8.9e-06 | 7.000E-06 | 2.007E-06 | 4.900E-06 | 8.900E-06 |
| 23916 | 1196 | N1196S | D | 6.9e-06 | 9.5e-06 | 5.6e-06 | 7.333E-06 | 1.986E-06 | 5.600E-06 | 9.500E-06 |
| 23917 | 1196 | N1196T | D | 9.5e-06 | 1.2e-05 | 7.5e-06 | 9.667E-06 | 2.255E-06 | 7.500E-06 | 1.200E-05 |
| 23918 | 1196 | N1196W | D | 1.1e-05 | 6.8e-06 | 6.5e-06 | 8.100E-06 | 2.516E-06 | 6.500E-06 | 1.100E-05 |
| 23919 | 1196 | N1196Y | D | 7.5e-06 | 8e-06 | 5.9e-06 | 7.133E-06 | 1.097E-06 | 5.900E-06 | 8.000E-06 |
| 23920 | 1196 | N1196V | D | 6.6e-06 | 7e-06 | 7e-06 | 6.867E-06 | 2.309E-07 | 6.600E-06 | 7.000E-06 |
| 23921 | 1197 | F1197A | D | 4.2e-06 | 6.1e-06 | 1e-05 | 6.767E-06 | 2.957E-06 | 4.200E-06 | 1.000E-05 |
| 23922 | 1197 | F1197R | D | 3e-06 | 1.2e-05 | 1e-05 | 8.333E-06 | 4.726E-06 | 3.000E-06 | 1.200E-05 |
| 23923 | 1197 | F1197N | D | 1.1e-05 | 6.2e-06 | 1.2e-05 | 9.733E-06 | 3.101E-06 | 6.200E-06 | 1.200E-05 |
| 23924 | 1197 | F1197D | D | 6.3e-06 | 9.6e-06 | 8.3e-06 | 8.067E-06 | 1.662E-06 | 6.300E-06 | 9.600E-06 |
| 23925 | 1197 | F1197C | D | 6.1e-06 | 8.3e-06 | 7.7e-06 | 7.367E-06 | 1.137E-06 | 6.100E-06 | 8.300E-06 |
| 23926 | 1197 | F1197E | D | 1.2e-05 | 9.2e-06 | 1.4e-05 | 1.173E-05 | 2.411E-06 | 9.200E-06 | 1.400E-05 |
| 23927 | 1197 | F1197Q | D | 6.2e-06 | 1.2e-05 | 8.6e-06 | 8.933E-06 | 2.914E-06 | 6.200E-06 | 1.200E-05 |
| 23928 | 1197 | F1197G | D | 8.7e-06 | 8.6e-06 | 7.9e-06 | 8.400E-06 | 4.359E-07 | 7.900E-06 | 8.700E-06 |
| 23929 | 1197 | F1197H | D | 1.2e-05 | 6.4e-06 | 8.8e-06 | 9.067E-06 | 2.810E-06 | 6.400E-06 | 1.200E-05 |
| 23930 | 1197 | F1197I | D | 5.1e-06 | 5.7e-06 | 1.1e-05 | 7.267E-06 | 3.247E-06 | 5.100E-06 | 1.100E-05 |
| 23931 | 1197 | F1197L | D | 6.8e-06 | 4.8e-06 | 3.8e-06 | 5.133E-06 | 1.528E-06 | 3.800E-06 | 6.800E-06 |
| 23932 | 1197 | F1197K | D | 6.6e-06 | 6.2e-06 | 7.7e-06 | 6.833E-06 | 7.767E-07 | 6.200E-06 | 7.700E-06 |
| 23933 | 1197 | F1197M | D | 9.1e-06 | 9.3e-06 | 5.2e-06 | 7.867E-06 | 2.312E-06 | 5.200E-06 | 9.300E-06 |
| 23934 | 1197 | F1197F | D | 9.3e-06 | 6.8e-06 | 8.9e-06 | 8.333E-06 | 1.343E-06 | 6.800E-06 | 9.300E-06 |
| 23935 | 1197 | F1197P | D | 7.5e-06 | 7e-06 | 3.9e-06 | 6.133E-06 | 1.950E-06 | 3.900E-06 | 7.500E-06 |
| 23936 | 1197 | F1197S | D | 4.7e-06 | 7.2e-06 | 6.8e-06 | 6.233E-06 | 1.343E-06 | 4.700E-06 | 7.200E-06 |
| 23937 | 1197 | F1197T | D | 9.5e-06 | 5.2e-06 | 6.8e-06 | 7.167E-06 | 2.173E-06 | 5.200E-06 | 9.500E-06 |
| 23938 | 1197 | F1197W | D | 4.6e-06 | 1e-05 | 9.9e-06 | 8.167E-06 | 3.089E-06 | 4.600E-06 | 1.000E-05 |
| 23939 | 1197 | F1197Y | D | 1.3e-05 | 1.5e-05 | 3.3e-06 | 1.043E-05 | 6.258E-06 | 3.300E-06 | 1.500E-05 |
| 23940 | 1197 | F1197V | D | 5.2e-06 | 1.1e-05 | 9.5e-06 | 8.567E-06 | 3.011E-06 | 5.200E-06 | 1.100E-05 |
| 23941 | 1198 | Y1198A | D | 7.7e-06 | 1.3e-05 | 6.9e-06 | 9.200E-06 | 3.315E-06 | 6.900E-06 | 1.300E-05 |
| 23942 | 1198 | Y1198R | D | 7.6e-06 | 8.1e-06 | 1.5e-05 | 1.023E-05 | 4.136E-06 | 7.600E-06 | 1.500E-05 |
| 23943 | 1198 | Y1198N | D | 4e-06 | 7.1e-06 | 6.5e-06 | 5.867E-06 | 1.644E-06 | 4.000E-06 | 7.100E-06 |
| 23944 | 1198 | Y1198D | D | 4.8e-06 | 1.1e-05 | 6.4e-06 | 7.400E-06 | 3.219E-06 | 4.800E-06 | 1.100E-05 |
| 23945 | 1198 | Y1198C | D | 1.1e-05 | 7.9e-06 | 6.5e-06 | 8.467E-06 | 2.303E-06 | 6.500E-06 | 1.100E-05 |
| 23946 | 1198 | Y1198E | D | 8.9e-06 | 8.4e-06 | 1.1e-05 | 9.433E-06 | 1.380E-06 | 8.400E-06 | 1.100E-05 |
| 23947 | 1198 | Y1198Q | D | 5.7e-06 | 9.6e-06 | 5.2e-06 | 6.833E-06 | 2.409E-06 | 5.200E-06 | 9.600E-06 |
| 23948 | 1198 | Y1198G | D | 4.7e-06 | 9e-06 | 9.2e-06 | 7.633E-06 | 2.542E-06 | 4.700E-06 | 9.200E-06 |
| 23949 | 1198 | Y1198H | D | 1.2e-05 | 6.6e-06 | 6.3e-06 | 8.300E-06 | 3.208E-06 | 6.300E-06 | 1.200E-05 |
| 23950 | 1198 | Y1198I | D | 4e-06 | 6.8e-06 | 1.2e-05 | 7.600E-06 | 4.060E-06 | 4.000E-06 | 1.200E-05 |
| 23951 | 1198 | Y1198L | D | 1.1e-05 | 1.1e-05 | 6.9e-06 | 9.633E-06 | 2.367E-06 | 6.900E-06 | 1.100E-05 |
| 23952 | 1198 | Y1198K | D | 8.9e-06 | 8.1e-06 | 7.6e-06 | 8.200E-06 | 6.557E-07 | 7.600E-06 | 8.900E-06 |
| 23953 | 1198 | Y1198M | D | 1e-05 | 6.8e-06 | 1.1e-05 | 9.267E-06 | 2.194E-06 | 6.800E-06 | 1.100E-05 |
| 23954 | 1198 | Y1198F | D | 9.3e-06 | 7.7e-06 | 7.5e-06 | 8.167E-06 | 9.866E-07 | 7.500E-06 | 9.300E-06 |
| 23955 | 1198 | Y1198P | D | 7.5e-06 | 7e-06 | 9.6e-06 | 8.033E-06 | 1.380E-06 | 7.000E-06 | 9.600E-06 |
| 23956 | 1198 | Y1198S | D | 1.4e-05 | 4.6e-06 | 7.7e-06 | 8.767E-06 | 4.790E-06 | 4.600E-06 | 1.400E-05 |
| 23957 | 1198 | Y1198T | D | 6.5e-06 | 8.4e-06 | 5.5e-06 | 6.800E-06 | 1.473E-06 | 5.500E-06 | 8.400E-06 |
| 23958 | 1198 | Y1198W | D | 4e-06 | 5e-06 | 8.7e-06 | 5.900E-06 | 2.476E-06 | 4.000E-06 | 8.700E-06 |
| 23959 | 1198 | Y1198Y | D | 1.3e-05 | 8.3e-06 | 9.9e-06 | 1.040E-05 | 2.390E-06 | 8.300E-06 | 1.300E-05 |
| 23960 | 1198 | Y1198V | D | 6.2e-06 | 7.4e-06 | 9.3e-06 | 7.633E-06 | 1.563E-06 | 6.200E-06 | 9.300E-06 |
| 23961 | 1199 | P1199A | D | 1.2e-05 | 4.6e-06 | 6.3e-06 | 7.633E-06 | 3.876E-06 | 4.600E-06 | 1.200E-05 |
| 23962 | 1199 | P1199R | D | 8e-06 | 7.6e-06 | 7.6e-06 | 7.733E-06 | 2.309E-07 | 7.600E-06 | 8.000E-06 |
| 23963 | 1199 | P1199N | D | 1.1e-05 | 7.3e-06 | 9.2e-06 | 9.167E-06 | 1.850E-06 | 7.300E-06 | 1.100E-05 |
| 23964 | 1199 | P1199D | D | 8e-06 | 9.2e-06 | 5.3e-06 | 7.500E-06 | 1.997E-06 | 5.300E-06 | 9.200E-06 |
| 23965 | 1199 | P1199C | D | 6.8e-06 | 4.3e-06 | 9.3e-06 | 6.800E-06 | 2.500E-06 | 4.300E-06 | 9.300E-06 |
| 23966 | 1199 | P1199E | D | 9.4e-06 | 7e-06 | 1.6e-05 | 1.080E-05 | 4.660E-06 | 7.000E-06 | 1.600E-05 |
| 23967 | 1199 | P1199Q | D | 8.1e-06 | 7.5e-06 | 9.8e-06 | 8.467E-06 | 1.193E-06 | 7.500E-06 | 9.800E-06 |
| 23968 | 1199 | P1199G | D | 6.7e-06 | 6.4e-06 | 1e-05 | 7.700E-06 | 1.997E-06 | 6.400E-06 | 1.000E-05 |
| 23969 | 1199 | P1199H | D | 6.5e-06 | 4.4e-06 | 6.9e-06 | 5.933E-06 | 1.343E-06 | 4.400E-06 | 6.900E-06 |

| | | | | | | | | | | |
|---|---|---|---|---|---|---|---|---|---|---|
| 23970 | 1199 | P1199I | D | 6.2e-06 | 8.2e-06 | 6.5e-06 | 6.967E-06 | 1.079E-06 | 6.200E-06 | 8.200E-06 |
| 23971 | 1199 | P1199L | D | 7.8e-06 | 8.4e-06 | 7.7e-06 | 7.967E-06 | 3.786E-07 | 7.700E-06 | 8.400E-06 |
| 23972 | 1199 | P1199K | D | 4.6e-06 | 6.9e-06 | 6e-06 | 5.833E-06 | 1.159E-06 | 4.600E-06 | 6.900E-06 |
| 23973 | 1199 | P1199M | D | 7.2e-06 | 1.1e-05 | 5.3e-06 | 7.833E-06 | 2.902E-06 | 5.300E-06 | 1.100E-05 |
| 23974 | 1199 | P1199F | D | 1.4e-05 | 4.7e-06 | 5.1e-06 | 7.933E-06 | 5.258E-06 | 4.700E-06 | 1.400E-05 |
| 23975 | 1199 | P1199P | D | 8.1e-06 | 1e-05 | 1.1e-05 | 9.700E-06 | 1.473E-06 | 8.100E-06 | 1.100E-05 |
| 23976 | 1199 | P1199S | D | 7.2e-06 | 9.9e-06 | 7.2e-06 | 8.100E-06 | 1.559E-06 | 7.200E-06 | 9.900E-06 |
| 23977 | 1199 | P1199T | D | 7.5e-06 | 4.8e-06 | 9.2e-06 | 7.167E-06 | 2.219E-06 | 4.800E-06 | 9.200E-06 |
| 23978 | 1199 | P1199W | D | 8.7e-06 | 1.5e-05 | 5.9e-06 | 9.867E-06 | 4.661E-06 | 5.900E-06 | 1.500E-05 |
| 23979 | 1199 | P1199Y | D | 4e-06 | 8.1e-06 | 4.2e-06 | 5.433E-06 | 2.312E-06 | 4.000E-06 | 8.100E-06 |
| 23980 | 1199 | P1199V | D | 1.2e-05 | 7e-06 | 4e-06 | 7.667E-06 | 4.041E-06 | 4.000E-06 | 1.200E-05 |
| 23981 | 1200 | R1200A | D | 3.8e-06 | 1.3e-05 | 6.7e-06 | 7.833E-06 | 4.704E-06 | 3.800E-06 | 1.300E-05 |
| 23982 | 1200 | R1200R | D | 9.7e-06 | 6.3e-06 | 8.2e-06 | 8.067E-06 | 1.704E-06 | 6.300E-06 | 9.700E-06 |
| 23983 | 1200 | R1200N | D | 6.9e-06 | 5.2e-06 | 6.5e-06 | 6.200E-06 | 8.888E-07 | 5.200E-06 | 6.900E-06 |
| 23984 | 1200 | R1200D | D | 1.3e-05 | 9.7e-06 | 1.2e-05 | 1.157E-05 | 1.692E-06 | 9.700E-06 | 1.300E-05 |
| 23985 | 1200 | R1200C | D | 1.3e-05 | 3.3e-06 | 6.9e-06 | 7.733E-06 | 4.903E-06 | 3.300E-06 | 1.300E-05 |
| 23986 | 1200 | R1200E | D | 6.9e-06 | 6e-06 | 1.2e-05 | 8.300E-06 | 3.236E-06 | 6.000E-06 | 1.200E-05 |
| 23987 | 1200 | R1200Q | D | 9.4e-06 | 7.5e-06 | 6.7e-06 | 7.867E-06 | 1.387E-06 | 6.700E-06 | 9.400E-06 |
| 23988 | 1200 | R1200G | D | 8.3e-06 | 5.1e-06 | 9.8e-06 | 7.733E-06 | 2.401E-06 | 5.100E-06 | 9.800E-06 |
| 23989 | 1200 | R1200H | D | 5.4e-06 | 6.9e-06 | 9.3e-06 | 7.200E-06 | 1.967E-06 | 5.400E-06 | 9.300E-06 |
| 23990 | 1200 | R1200I | D | 6e-06 | 7.4e-06 | 9.1e-06 | 7.500E-06 | 1.552E-06 | 6.000E-06 | 9.100E-06 |
| 23991 | 1200 | R1200L | D | 9.8e-06 | 8.7e-06 | 6.4e-06 | 8.300E-06 | 1.735E-06 | 6.400E-06 | 9.800E-06 |
| 23992 | 1200 | R1200K | D | 8.5e-06 | 6.9e-06 | 1.3e-05 | 9.467E-06 | 3.163E-06 | 6.900E-06 | 1.300E-05 |
| 23993 | 1200 | R1200M | D | 6.7e-06 | 8.7e-06 | 5e-06 | 6.800E-06 | 1.852E-06 | 5.000E-06 | 8.700E-06 |
| 23994 | 1200 | R1200F | D | 9.2e-06 | 8.3e-06 | 5.5e-06 | 7.667E-06 | 1.930E-06 | 5.500E-06 | 9.200E-06 |
| 23995 | 1200 | R1200P | D | 7.1e-06 | 7.6e-06 | 8.9e-06 | 7.867E-06 | 9.292E-07 | 7.100E-06 | 8.900E-06 |
| 23996 | 1200 | R1200S | D | 8.2e-06 | 8.1e-06 | 1.4e-05 | 1.010E-05 | 3.378E-06 | 8.100E-06 | 1.400E-05 |
| 23997 | 1200 | R1200T | D | 9.2e-06 | 9.4e-06 | 8.9e-06 | 9.167E-06 | 2.517E-07 | 8.900E-06 | 9.400E-06 |
| 23998 | 1200 | R1200W | D | 7.2e-06 | 6e-06 | 7.1e-06 | 6.767E-06 | 6.658E-07 | 6.000E-06 | 7.200E-06 |
| 23999 | 1200 | R1200Y | D | 1.1e-05 | 5.6e-06 | 1.4e-05 | 1.020E-05 | 4.257E-06 | 5.600E-06 | 1.400E-05 |
| 24000 | 1200 | R1200V | D | 9.1e-06 | 1e-05 | 1.1e-05 | 1.003E-05 | 9.504E-07 | 9.100E-06 | 1.100E-05 |
| 24001 | 1201 | E1201A | D | 8e-06 | 7.1e-06 | 1e-05 | 8.367E-06 | 1.484E-06 | 7.100E-06 | 1.000E-05 |
| 24002 | 1201 | E1201R | D | 7.6e-06 | 1.1e-05 | 7.9e-06 | 8.833E-06 | 1.882E-06 | 7.600E-06 | 1.100E-05 |
| 24003 | 1201 | E1201N | D | 5.8e-06 | 7.1e-06 | 7.5e-06 | 6.800E-06 | 8.888E-07 | 5.800E-06 | 7.500E-06 |
| 24004 | 1201 | E1201D | D | 1e-05 | 8.6e-06 | 6.2e-06 | 8.267E-06 | 1.922E-06 | 6.200E-06 | 1.000E-05 |
| 24005 | 1201 | E1201C | D | 9.5e-06 | 5.5e-06 | 8.5e-06 | 7.833E-06 | 2.082E-06 | 5.500E-06 | 9.500E-06 |
| 24006 | 1201 | E1201E | D | 1e-05 | 6.2e-06 | 9.4e-06 | 8.533E-06 | 2.043E-06 | 6.200E-06 | 1.000E-05 |
| 24007 | 1201 | E1201Q | D | 1e-05 | 1.4e-05 | 8.7e-06 | 1.090E-05 | 2.762E-06 | 8.700E-06 | 1.400E-05 |
| 24008 | 1201 | E1201G | D | 4.8e-06 | 6.8e-06 | 5.1e-06 | 5.567E-06 | 1.079E-06 | 4.800E-06 | 6.800E-06 |
| 24009 | 1201 | E1201H | D | 1.2e-05 | 1.1e-05 | 8.7e-06 | 1.057E-05 | 1.692E-06 | 8.700E-06 | 1.200E-05 |
| 24010 | 1201 | E1201I | D | 9.4e-06 | 9.6e-06 | 7e-06 | 8.667E-06 | 1.447E-06 | 7.000E-06 | 9.600E-06 |
| 24011 | 1201 | E1201L | D | 6.9e-06 | 9.3e-06 | 9.7e-06 | 8.633E-06 | 1.514E-06 | 6.900E-06 | 9.700E-06 |
| 24012 | 1201 | E1201K | D | 6.9e-06 | 6e-06 | 8e-06 | 6.967E-06 | 1.002E-06 | 6.000E-06 | 8.000E-06 |
| 24013 | 1201 | E1201M | D | 7.4e-06 | 6.1e-06 | 8.9e-06 | 7.467E-06 | 1.401E-06 | 6.100E-06 | 8.900E-06 |
| 24014 | 1201 | E1201F | D | 9.1e-06 | 4.8e-06 | 6.9e-06 | 6.933E-06 | 2.150E-06 | 4.800E-06 | 9.100E-06 |
| 24015 | 1201 | E1201P | D | 8.7e-06 | 4.9e-06 | 7.1e-06 | 6.900E-06 | 1.908E-06 | 4.900E-06 | 8.700E-06 |
| 24016 | 1201 | E1201S | D | 9.4e-06 | 1.1e-05 | 6.1e-06 | 8.833E-06 | 2.499E-06 | 6.100E-06 | 1.100E-05 |
| 24017 | 1201 | E1201T | D | 1.3e-05 | 1.3e-05 | 6e-06 | 1.067E-05 | 4.041E-06 | 6.000E-06 | 1.300E-05 |
| 24018 | 1201 | E1201W | D | 6.4e-06 | 1.4e-05 | 8.6e-06 | 9.667E-06 | 3.911E-06 | 6.400E-06 | 1.400E-05 |
| 24019 | 1201 | E1201Y | D | 8.7e-06 | 1.8e-05 | 8e-06 | 1.157E-05 | 5.582E-06 | 8.000E-06 | 1.800E-05 |
| 24020 | 1201 | E1201V | D | 7.6e-06 | 9e-06 | 9.3e-06 | 8.633E-06 | 9.074E-07 | 7.600E-06 | 9.300E-06 |
| 24021 | 1202 | A1202A | D | 9.7e-06 | 1e-05 | 3.4e-06 | 7.700E-06 | 3.727E-06 | 3.400E-06 | 1.000E-05 |
| 24022 | 1202 | A1202R | D | 5.5e-06 | 9.7e-06 | 5.2e-06 | 6.800E-06 | 2.516E-06 | 5.200E-06 | 9.700E-06 |
| 24023 | 1202 | A1202N | D | 6.7e-06 | 8.8e-06 | 7e-06 | 7.500E-06 | 1.136E-06 | 6.700E-06 | 8.800E-06 |
| 24024 | 1202 | A1202D | D | 1.1e-05 | 9.6e-06 | 9.4e-06 | 1.000E-05 | 8.718E-07 | 9.400E-06 | 1.100E-05 |
| 24025 | 1202 | A1202C | D | 7.4e-06 | 9e-06 | 1.2e-05 | 9.467E-06 | 2.335E-06 | 7.400E-06 | 1.200E-05 |
| 24026 | 1202 | A1202E | D | 4.8e-06 | 7.1e-06 | 9.7e-06 | 7.200E-06 | 2.452E-06 | 4.800E-06 | 9.700E-06 |
| 24027 | 1202 | A1202Q | D | 5.2e-06 | 6.7e-06 | 1.1e-05 | 7.633E-06 | 3.011E-06 | 5.200E-06 | 1.100E-05 |
| 24028 | 1202 | A1202G | D | 6.2e-06 | 1.4e-05 | 8.5e-06 | 9.567E-06 | 4.008E-06 | 6.200E-06 | 1.400E-05 |
| 24029 | 1202 | A1202H | D | 5.3e-06 | 5.9e-06 | 1.2e-05 | 7.733E-06 | 3.707E-06 | 5.300E-06 | 1.200E-05 |
| 24030 | 1202 | A1202I | D | 9.9e-06 | 6.7e-06 | 1e-05 | 8.867E-06 | 1.877E-06 | 6.700E-06 | 1.000E-05 |
| 24031 | 1202 | A1202L | D | 6.2e-06 | 8.9e-06 | 5e-06 | 6.700E-06 | 1.997E-06 | 5.000E-06 | 8.900E-06 |
| 24032 | 1202 | A1202K | D | 9.8e-06 | 1.3e-05 | 9.3e-06 | 1.070E-05 | 2.007E-06 | 9.300E-06 | 1.300E-05 |
| 24033 | 1202 | A1202M | D | 1.1e-05 | 5.7e-06 | 8.4e-06 | 8.367E-06 | 2.650E-06 | 5.700E-06 | 1.100E-05 |
| 24034 | 1202 | A1202F | D | 6.8e-06 | 9e-06 | 5e-06 | 6.933E-06 | 2.003E-06 | 5.000E-06 | 9.000E-06 |
| 24035 | 1202 | A1202P | D | 7.2e-06 | 5.9e-06 | 5e-06 | 6.033E-06 | 1.106E-06 | 5.000E-06 | 7.200E-06 |
| 24036 | 1202 | A1202S | D | 9.3e-06 | 4.3e-06 | 6.9e-06 | 6.833E-06 | 2.501E-06 | 4.300E-06 | 9.300E-06 |
| 24037 | 1202 | A1202T | D | 6.4e-06 | 1.1e-05 | 7.8e-06 | 8.400E-06 | 2.358E-06 | 6.400E-06 | 1.100E-05 |
| 24038 | 1202 | A1202W | D | 8.5e-06 | 6.6e-06 | 6.1e-06 | 7.067E-06 | 1.266E-06 | 6.100E-06 | 8.500E-06 |
| 24039 | 1202 | A1202Y | D | 1.2e-05 | 5.9e-06 | 7.2e-06 | 8.367E-06 | 3.213E-06 | 5.900E-06 | 1.200E-05 |
| 24040 | 1202 | A1202V | D | 9.6e-06 | 8e-06 | 8.7e-06 | 8.767E-06 | 8.021E-07 | 8.000E-06 | 9.600E-06 |
| 24041 | 1203 | K1203A | D | 6e-06 | 1.3e-05 | 8e-06 | 9.000E-06 | 3.606E-06 | 6.000E-06 | 1.300E-05 |
| 24042 | 1203 | K1203R | D | 9.7e-06 | 9.3e-06 | 8.8e-06 | 9.267E-06 | 4.509E-07 | 8.800E-06 | 9.700E-06 |
| 24043 | 1203 | K1203N | D | 9.9e-06 | 6e-06 | 7.9e-06 | 7.933E-06 | 1.950E-06 | 6.000E-06 | 9.900E-06 |
| 24044 | 1203 | K1203D | D | 8.5e-06 | 1.1e-05 | 6.1e-06 | 8.533E-06 | 2.450E-06 | 6.100E-06 | 1.100E-05 |
| 24045 | 1203 | K1203C | D | 9e-06 | 5.3e-06 | 5.3e-06 | 6.533E-06 | 2.136E-06 | 5.300E-06 | 9.000E-06 |
| 24046 | 1203 | K1203E | D | 9.7e-06 | 9.5e-06 | 1.1e-05 | 1.007E-05 | 8.145E-07 | 9.500E-06 | 1.100E-05 |
| 24047 | 1203 | K1203Q | D | 4e-06 | 6.4e-06 | 8.1e-06 | 6.167E-06 | 2.060E-06 | 4.000E-06 | 8.100E-06 |
| 24048 | 1203 | K1203G | D | 6.6e-06 | 8.1e-06 | 8.9e-06 | 7.867E-06 | 1.168E-06 | 6.600E-06 | 8.900E-06 |
| 24049 | 1203 | K1203H | D | 9.4e-06 | 7.6e-06 | 6.5e-06 | 7.833E-06 | 1.464E-06 | 6.500E-06 | 9.400E-06 |
| 24050 | 1203 | K1203I | D | 3.8e-06 | 9.1e-06 | 9.2e-06 | 7.367E-06 | 3.089E-06 | 3.800E-06 | 9.200E-06 |
| 24051 | 1203 | K1203L | D | 5.2e-06 | 1e-05 | 8.3e-06 | 7.833E-06 | 2.434E-06 | 5.200E-06 | 1.000E-05 |
| 24052 | 1203 | K1203K | D | 6.2e-06 | 9.6e-06 | 3.8e-06 | 6.533E-06 | 2.914E-06 | 3.800E-06 | 9.600E-06 |
| 24053 | 1203 | K1203M | D | 9.5e-06 | 1.2e-05 | 8e-06 | 9.833E-06 | 2.021E-06 | 8.000E-06 | 1.200E-05 |
| 24054 | 1203 | K1203F | D | 7.7e-06 | 1e-05 | 1.5e-05 | 1.090E-05 | 3.732E-06 | 7.700E-06 | 1.500E-05 |

| | | | | | | | | | | |
|---|---|---|---|---|---|---|---|---|---|---|
| 24055 | 1203 | K1203P | D | 6.9e-06 | 1e-05 | 1.1e-05 | 9.300E-06 | 2.138E-06 | 6.900E-06 | 1.100E-05 |
| 24056 | 1203 | K1203S | D | 6.7e-06 | 1.4e-05 | 1.2e-05 | 1.090E-05 | 3.772E-06 | 6.700E-06 | 1.400E-05 |
| 24057 | 1203 | K1203T | D | 9.8e-06 | 4.2e-06 | 8.2e-06 | 7.400E-06 | 2.884E-06 | 4.200E-06 | 9.800E-06 |
| 24058 | 1203 | K1203W | D | 4.8e-06 | 3.5e-06 | 4.9e-06 | 4.400E-06 | 7.810E-07 | 3.500E-06 | 4.900E-06 |
| 24059 | 1203 | K1203Y | D | 1.1e-05 | 5.8e-06 | 8.1e-06 | 8.300E-06 | 2.606E-06 | 5.800E-06 | 1.100E-05 |
| 24060 | 1203 | K1203V | D | 7e-06 | 1.2e-05 | 4.5e-06 | 7.833E-06 | 3.819E-06 | 4.500E-06 | 1.200E-05 |
| 24061 | 1204 | V1204A | D | 6.8e-06 | 6.4e-06 | 1.2e-05 | 8.400E-06 | 3.124E-06 | 6.400E-06 | 1.200E-05 |
| 24062 | 1204 | V1204R | D | 6.5e-06 | 1e-05 | 6.6e-06 | 7.700E-06 | 1.992E-06 | 6.500E-06 | 1.000E-05 |
| 24063 | 1204 | V1204N | D | 6.3e-06 | 1.4e-05 | 6e-06 | 8.767E-06 | 4.535E-06 | 6.000E-06 | 1.400E-05 |
| 24064 | 1204 | V1204D | D | 1.1e-05 | 9.9e-06 | 1e-05 | 1.030E-05 | 6.083E-07 | 9.900E-06 | 1.100E-05 |
| 24065 | 1204 | V1204C | D | 8.6e-06 | 5.2e-06 | 1e-05 | 7.933E-06 | 2.468E-06 | 5.200E-06 | 1.000E-05 |
| 24066 | 1204 | V1204E | D | 7.8e-06 | 5.1e-06 | 5.4e-06 | 6.100E-06 | 1.480E-06 | 5.100E-06 | 7.800E-06 |
| 24067 | 1204 | V1204Q | D | 9.1e-06 | 8.2e-06 | 6e-06 | 7.767E-06 | 1.595E-06 | 6.000E-06 | 9.100E-06 |
| 24068 | 1204 | V1204G | D | 7.3e-06 | 9.8e-06 | 9.5e-06 | 8.867E-06 | 1.365E-06 | 7.300E-06 | 9.800E-06 |
| 24069 | 1204 | V1204H | D | 6.3e-06 | 9.9e-06 | 8.3e-06 | 8.167E-06 | 1.804E-06 | 6.300E-06 | 9.900E-06 |
| 24070 | 1204 | V1204I | D | 1.1e-05 | 8.4e-06 | 6.4e-06 | 8.600E-06 | 2.307E-06 | 6.400E-06 | 1.100E-05 |
| 24071 | 1204 | V1204L | D | 4.8e-06 | 6.2e-06 | 9.1e-06 | 6.700E-06 | 2.193E-06 | 4.800E-06 | 9.100E-06 |
| 24072 | 1204 | V1204K | D | 5.2e-06 | 6.7e-06 | 7.7e-06 | 6.533E-06 | 1.258E-06 | 5.200E-06 | 7.700E-06 |
| 24073 | 1204 | V1204M | D | 7.6e-06 | 5.8e-06 | 1.1e-05 | 8.133E-06 | 2.641E-06 | 5.800E-06 | 1.100E-05 |
| 24074 | 1204 | V1204F | D | 8.1e-06 | 1.2e-05 | 8.8e-06 | 9.633E-06 | 2.079E-06 | 8.100E-06 | 1.200E-05 |
| 24075 | 1204 | V1204P | D | 1.6e-05 | 6.3e-06 | 7.2e-06 | 9.833E-06 | 5.359E-06 | 6.300E-06 | 1.600E-05 |
| 24076 | 1204 | V1204S | D | 5.1e-06 | 6.9e-06 | 6e-06 | 6.000E-06 | 9.000E-07 | 5.100E-06 | 6.900E-06 |
| 24077 | 1204 | V1204T | D | 4.5e-06 | 9.2e-06 | 7e-06 | 6.900E-06 | 2.352E-06 | 4.500E-06 | 9.200E-06 |
| 24078 | 1204 | V1204W | D | 7e-06 | 9.1e-06 | 1.2e-05 | 9.367E-06 | 2.511E-06 | 7.000E-06 | 1.200E-05 |
| 24079 | 1204 | V1204Y | D | 6.2e-06 | 8.7e-06 | 5.5e-06 | 6.800E-06 | 1.682E-06 | 5.500E-06 | 8.700E-06 |
| 24080 | 1204 | V1204V | D | 5.8e-06 | 1.4e-05 | 1.4e-05 | 1.127E-05 | 4.734E-06 | 5.800E-06 | 1.400E-05 |
| 24081 | 1205 | Q1205A | D | 7.3e-06 | 8.9e-06 | 4.3e-06 | 6.833E-06 | 2.335E-06 | 4.300E-06 | 8.900E-06 |
| 24082 | 1205 | Q1205R | D | 8.4e-06 | 5.5e-06 | 1.3e-05 | 8.967E-06 | 3.782E-06 | 5.500E-06 | 1.300E-05 |
| 24083 | 1205 | Q1205N | D | 6.4e-06 | 6.8e-06 | 8.4e-06 | 7.200E-06 | 1.058E-06 | 6.400E-06 | 8.400E-06 |
| 24084 | 1205 | Q1205D | D | 5.5e-06 | 5e-06 | 7.5e-06 | 6.000E-06 | 1.323E-06 | 5.000E-06 | 7.500E-06 |
| 24085 | 1205 | Q1205C | D | 8.2e-06 | 1e-05 | 9.5e-06 | 9.233E-06 | 9.292E-07 | 8.200E-06 | 1.000E-05 |
| 24086 | 1205 | Q1205E | D | 9.3e-06 | 9.3e-06 | 9.8e-06 | 9.467E-06 | 2.887E-07 | 9.300E-06 | 9.800E-06 |
| 24087 | 1205 | Q1205Q | D | 8.7e-06 | 9.6e-06 | 1.3e-05 | 1.043E-05 | 2.268E-06 | 8.700E-06 | 1.300E-05 |
| 24088 | 1205 | Q1205G | D | 1.2e-05 | 8.2e-06 | 6.9e-06 | 9.033E-06 | 2.650E-06 | 6.900E-06 | 1.200E-05 |
| 24089 | 1205 | Q1205H | D | 9.7e-06 | 8.4e-06 | 7.6e-06 | 8.567E-06 | 1.060E-06 | 7.600E-06 | 9.700E-06 |
| 24090 | 1205 | Q1205I | D | 1.3e-05 | 6.2e-06 | 4.8e-06 | 8.000E-06 | 4.386E-06 | 4.800E-06 | 1.300E-05 |
| 24091 | 1205 | Q1205L | D | 1e-05 | 9.2e-06 | 9e-06 | 9.400E-06 | 5.292E-07 | 9.000E-06 | 1.000E-05 |
| 24092 | 1205 | Q1205K | D | 1.2e-05 | 6.3e-06 | 7.4e-06 | 8.567E-06 | 3.024E-06 | 6.300E-06 | 1.200E-05 |
| 24093 | 1205 | Q1205M | D | 6.7e-06 | 7.8e-06 | 7.8e-06 | 7.433E-06 | 6.351E-07 | 6.700E-06 | 7.800E-06 |
| 24094 | 1205 | Q1205F | D | 8.5e-06 | 6.4e-06 | 6.2e-06 | 7.033E-06 | 1.274E-06 | 6.200E-06 | 8.500E-06 |
| 24095 | 1205 | Q1205P | D | 4.7e-06 | 1.3e-05 | 7e-06 | 8.233E-06 | 4.285E-06 | 4.700E-06 | 1.300E-05 |
| 24096 | 1205 | Q1205S | D | 9.8e-06 | 6.5e-06 | 7.4e-06 | 7.900E-06 | 1.706E-06 | 6.500E-06 | 9.800E-06 |
| 24097 | 1205 | Q1205T | D | 9e-06 | 5.3e-06 | 4.5e-06 | 6.267E-06 | 2.401E-06 | 4.500E-06 | 9.000E-06 |
| 24098 | 1205 | Q1205W | D | 5.8e-06 | 5.5e-06 | 8.2e-06 | 6.500E-06 | 1.480E-06 | 5.500E-06 | 8.200E-06 |
| 24099 | 1205 | Q1205Y | D | 9.6e-06 | 8.4e-06 | 6e-06 | 8.000E-06 | 1.833E-06 | 6.000E-06 | 9.600E-06 |
| 24100 | 1205 | Q1205V | D | 5.8e-06 | 7e-06 | 8.5e-06 | 7.100E-06 | 1.353E-06 | 5.800E-06 | 8.500E-06 |
| 24101 | 1206 | W1206A | D | 7.9e-06 | 8.8e-06 | 9.8e-06 | 8.833E-06 | 9.504E-07 | 7.900E-06 | 9.800E-06 |
| 24102 | 1206 | W1206R | D | 6.9e-06 | 6.6e-06 | 6.8e-06 | 6.767E-06 | 1.528E-07 | 6.600E-06 | 6.900E-06 |
| 24103 | 1206 | W1206N | D | 8.1e-06 | 5.1e-06 | 8e-06 | 7.067E-06 | 1.704E-06 | 5.100E-06 | 8.100E-06 |
| 24104 | 1206 | W1206D | D | 7.8e-06 | 9.5e-06 | 8.3e-06 | 8.533E-06 | 8.737E-07 | 7.800E-06 | 9.500E-06 |
| 24105 | 1206 | W1206C | D | 5.8e-06 | 1e-05 | 1e-05 | 8.600E-06 | 2.425E-06 | 5.800E-06 | 1.000E-05 |
| 24106 | 1206 | W1206E | D | 9.9e-06 | 8.3e-06 | 8.7e-06 | 8.967E-06 | 8.327E-07 | 8.300E-06 | 9.900E-06 |
| 24107 | 1206 | W1206Q | D | 6.1e-06 | 8.5e-06 | 6.7e-06 | 7.100E-06 | 1.249E-06 | 6.100E-06 | 8.500E-06 |
| 24108 | 1206 | W1206G | D | 1.3e-05 | 9e-06 | 6e-06 | 9.333E-06 | 3.512E-06 | 6.000E-06 | 1.300E-05 |
| 24109 | 1206 | W1206H | D | 4.5e-06 | 8.9e-06 | 8.2e-06 | 7.200E-06 | 2.364E-06 | 4.500E-06 | 8.900E-06 |
| 24110 | 1206 | W1206I | D | 5.9e-06 | 1.2e-05 | 6.6e-06 | 8.167E-06 | 3.338E-06 | 5.900E-06 | 1.200E-05 |
| 24111 | 1206 | W1206L | D | 8.2e-06 | 4.6e-06 | 9.4e-06 | 7.400E-06 | 2.498E-06 | 4.600E-06 | 9.400E-06 |
| 24112 | 1206 | W1206K | D | 6.8e-06 | 1.2e-05 | 8.6e-06 | 9.133E-06 | 2.641E-06 | 6.800E-06 | 1.200E-05 |
| 24113 | 1206 | W1206M | D | 1.2e-05 | 8.8e-06 | 9.6e-06 | 1.013E-05 | 1.665E-06 | 8.800E-06 | 1.200E-05 |
| 24114 | 1206 | W1206F | D | 1.3e-05 | 7.6e-06 | 7.8e-06 | 9.467E-06 | 3.062E-06 | 7.600E-06 | 1.300E-05 |
| 24115 | 1206 | W1206P | D | 5.4e-06 | 1e-05 | 1.2e-05 | 9.133E-06 | 3.384E-06 | 5.400E-06 | 1.200E-05 |
| 24116 | 1206 | W1206S | D | 7.5e-06 | 4.2e-06 | 1.1e-05 | 7.567E-06 | 3.400E-06 | 4.200E-06 | 1.100E-05 |
| 24117 | 1206 | W1206T | D | 9e-06 | 8.7e-06 | 5.3e-06 | 7.667E-06 | 2.055E-06 | 5.300E-06 | 9.000E-06 |
| 24118 | 1206 | W1206W | D | 6.6e-06 | 8.1e-06 | 1.1e-05 | 8.567E-06 | 2.237E-06 | 6.600E-06 | 1.100E-05 |
| 24119 | 1206 | W1206Y | D | 5.4e-06 | 5.8e-06 | 4.3e-06 | 5.167E-06 | 7.767E-07 | 4.300E-06 | 5.800E-06 |
| 24120 | 1206 | W1206V | D | 5.6e-06 | 8.6e-06 | 6.1e-06 | 6.767E-06 | 1.607E-06 | 5.600E-06 | 8.600E-06 |
| 24121 | 1207 | K1207A | D | 6.4e-06 | 7.3e-06 | 5.2e-06 | 6.300E-06 | 1.054E-06 | 5.200E-06 | 7.300E-06 |
| 24122 | 1207 | K1207R | D | 8.3e-06 | 3.6e-06 | 7.5e-06 | 6.467E-06 | 2.515E-06 | 3.600E-06 | 8.300E-06 |
| 24123 | 1207 | K1207N | D | 5.9e-06 | 1e-05 | 4.9e-06 | 6.933E-06 | 2.702E-06 | 4.900E-06 | 1.000E-05 |
| 24124 | 1207 | K1207D | D | 9.3e-06 | 1.4e-05 | 1e-05 | 1.110E-05 | 2.536E-06 | 9.300E-06 | 1.400E-05 |
| 24125 | 1207 | K1207C | D | 8.9e-06 | 9.8e-06 | 9.9e-06 | 9.533E-06 | 5.508E-07 | 8.900E-06 | 9.900E-06 |
| 24126 | 1207 | K1207E | D | 7.8e-06 | 1.3e-05 | 6.9e-06 | 9.233E-06 | 3.293E-06 | 6.900E-06 | 1.300E-05 |
| 24127 | 1207 | K1207Q | D | 9.3e-06 | 1.3e-05 | 6.6e-06 | 9.633E-06 | 3.213E-06 | 6.600E-06 | 1.300E-05 |
| 24128 | 1207 | K1207G | D | 9.5e-06 | 1.3e-05 | 5.3e-06 | 9.267E-06 | 3.855E-06 | 5.300E-06 | 1.300E-05 |
| 24129 | 1207 | K1207H | D | 5e-06 | 5.7e-06 | 7.1e-06 | 5.933E-06 | 1.069E-06 | 5.000E-06 | 7.100E-06 |
| 24130 | 1207 | K1207I | D | 1.2e-05 | 5.8e-06 | 5.3e-06 | 7.700E-06 | 3.732E-06 | 5.300E-06 | 1.200E-05 |
| 24131 | 1207 | K1207L | D | 2e-05 | 7.2e-06 | 1.2e-05 | 1.307E-05 | 6.466E-06 | 7.200E-06 | 2.000E-05 |
| 24132 | 1207 | K1207K | D | 4.5e-06 | 1.2e-05 | 6.2e-06 | 7.567E-06 | 3.932E-06 | 4.500E-06 | 1.200E-05 |
| 24133 | 1207 | K1207M | D | 1.6e-05 | 7.2e-06 | 4.8e-06 | 9.333E-06 | 5.897E-06 | 4.800E-06 | 1.600E-05 |
| 24134 | 1207 | K1207F | D | 5.8e-06 | 8.6e-06 | 5.7e-06 | 6.700E-06 | 1.646E-06 | 5.700E-06 | 8.600E-06 |
| 24135 | 1207 | K1207P | D | 6.5e-06 | 7.1e-06 | 7.4e-06 | 7.000E-06 | 4.583E-07 | 6.500E-06 | 7.400E-06 |
| 24136 | 1207 | K1207S | D | 6.6e-06 | 9.2e-06 | 8.3e-06 | 8.033E-06 | 1.320E-06 | 6.600E-06 | 9.200E-06 |
| 24137 | 1207 | K1207T | D | 6.2e-06 | 9.4e-06 | 6.5e-06 | 7.367E-06 | 1.767E-06 | 6.200E-06 | 9.400E-06 |
| 24138 | 1207 | K1207W | D | 9.9e-06 | 6.9e-06 | 6e-06 | 7.600E-06 | 2.042E-06 | 6.000E-06 | 9.900E-06 |
| 24139 | 1207 | K1207Y | D | 8.4e-06 | 7.3e-06 | 6.2e-06 | 7.300E-06 | 1.100E-06 | 6.200E-06 | 8.400E-06 |

| | | | | | | | | | | |
|---|---|---|---|---|---|---|---|---|---|---|
| 24140 | 1207 | K1207V | D | 8.6e-06 | 8.7e-06 | 6.3e-06 | 7.867E-06 | 1.358E-06 | 6.300E-06 | 8.700E-06 |
| 24141 | 1208 | V1208A | D | 4.2e-06 | 7.6e-06 | 9.9e-06 | 7.233E-06 | 2.868E-06 | 4.200E-06 | 9.900E-06 |
| 24142 | 1208 | V1208R | D | 7.9e-06 | 6.3e-06 | 6.3e-06 | 6.833E-06 | 9.238E-07 | 6.300E-06 | 7.900E-06 |
| 24143 | 1208 | V1208N | D | 7.1e-06 | 1.1e-05 | 9.2e-06 | 9.100E-06 | 1.952E-06 | 7.100E-06 | 1.100E-05 |
| 24144 | 1208 | V1208D | D | 1.4e-05 | 8.6e-06 | 8.9e-06 | 1.050E-05 | 3.035E-06 | 8.600E-06 | 1.400E-05 |
| 24145 | 1208 | V1208C | D | 8.8e-06 | 6e-06 | 5.2e-06 | 6.667E-06 | 1.890E-06 | 5.200E-06 | 8.800E-06 |
| 24146 | 1208 | V1208E | D | 1.3e-05 | 6.6e-06 | 5.7e-06 | 8.433E-06 | 3.980E-06 | 5.700E-06 | 1.300E-05 |
| 24147 | 1208 | V1208Q | D | 6.2e-06 | 7e-06 | 1.3e-05 | 8.733E-06 | 3.717E-06 | 6.200E-06 | 1.300E-05 |
| 24148 | 1208 | V1208G | D | 9.9e-06 | 6.8e-06 | 5.6e-06 | 7.433E-06 | 2.219E-06 | 5.600E-06 | 9.900E-06 |
| 24149 | 1208 | V1208H | D | 7.6e-06 | 8.3e-06 | 1.5e-05 | 1.030E-05 | 4.085E-06 | 7.600E-06 | 1.500E-05 |
| 24150 | 1208 | V1208I | D | 8.7e-06 | 9.3e-06 | 8.7e-06 | 8.900E-06 | 3.464E-07 | 8.700E-06 | 9.300E-06 |
| 24151 | 1208 | V1208L | D | 9.8e-06 | 6.9e-06 | 6.1e-06 | 7.600E-06 | 1.947E-06 | 6.100E-06 | 9.800E-06 |
| 24152 | 1208 | V1208K | D | 9.7e-06 | 1e-05 | 1.9e-05 | 1.290E-05 | 5.285E-06 | 9.700E-06 | 1.900E-05 |
| 24153 | 1208 | V1208M | D | 8.3e-06 | 8.2e-06 | 1.1e-05 | 9.167E-06 | 1.589E-06 | 8.200E-06 | 1.100E-05 |
| 24154 | 1208 | V1208F | D | 6.2e-06 | 5.6e-06 | 9.2e-06 | 7.000E-06 | 1.929E-06 | 5.600E-06 | 9.200E-06 |
| 24155 | 1208 | V1208P | D | 4.3e-06 | 6e-06 | 4.8e-06 | 5.033E-06 | 8.737E-07 | 4.300E-06 | 6.000E-06 |
| 24156 | 1208 | V1208S | D | 9.3e-06 | 7.8e-06 | 6.4e-06 | 7.833E-06 | 1.450E-06 | 6.400E-06 | 9.300E-06 |
| 24157 | 1208 | V1208T | D | 6.2e-06 | 7.8e-06 | 8.6e-06 | 7.533E-06 | 1.222E-06 | 6.200E-06 | 8.600E-06 |
| 24158 | 1208 | V1208W | D | 4.6e-06 | 7.2e-06 | 3.9e-06 | 5.233E-06 | 1.739E-06 | 3.900E-06 | 7.200E-06 |
| 24159 | 1208 | V1208Y | D | 8.7e-06 | 7.2e-06 | 8.4e-06 | 8.100E-06 | 7.937E-07 | 7.200E-06 | 8.700E-06 |
| 24160 | 1208 | V1208V | D | 6.8e-06 | 1e-05 | 7e-06 | 7.933E-06 | 1.793E-06 | 6.800E-06 | 1.000E-05 |
| 24161 | 1209 | D1209A | D | 7.6e-06 | 6.2e-06 | 1.1e-05 | 8.267E-06 | 2.468E-06 | 6.200E-06 | 1.100E-05 |
| 24162 | 1209 | D1209R | D | 6.6e-06 | 5.6e-06 | 5.8e-06 | 6.000E-06 | 5.292E-07 | 5.600E-06 | 6.600E-06 |
| 24163 | 1209 | D1209N | D | 7.2e-06 | 9.9e-06 | 8.4e-06 | 8.500E-06 | 1.353E-06 | 7.200E-06 | 9.900E-06 |
| 24164 | 1209 | D1209D | D | 9.4e-06 | 1.6e-05 | 4.6e-06 | 1.000E-05 | 5.724E-06 | 4.600E-06 | 1.600E-05 |
| 24165 | 1209 | D1209C | D | 6e-06 | 1.1e-05 | 8.9e-06 | 8.633E-06 | 2.511E-06 | 6.000E-06 | 1.100E-05 |
| 24166 | 1209 | D1209E | D | 1.3e-05 | 6.9e-06 | 9e-06 | 9.633E-06 | 3.099E-06 | 6.900E-06 | 1.300E-05 |
| 24167 | 1209 | D1209Q | D | 9.9e-06 | 9.1e-06 | 1.5e-05 | 1.133E-05 | 3.201E-06 | 9.100E-06 | 1.500E-05 |
| 24168 | 1209 | D1209G | D | 6.9e-06 | 8.1e-06 | 6.2e-06 | 7.067E-06 | 9.609E-07 | 6.200E-06 | 8.100E-06 |
| 24169 | 1209 | D1209H | D | 7.7e-06 | 5.6e-06 | 4.4e-06 | 5.900E-06 | 1.670E-06 | 4.400E-06 | 7.700E-06 |
| 24170 | 1209 | D1209I | D | 9.1e-06 | 5.5e-06 | 8.6e-06 | 7.733E-06 | 1.950E-06 | 5.500E-06 | 9.100E-06 |
| 24171 | 1209 | D1209L | D | 1.3e-05 | 9.5e-06 | 8.5e-06 | 1.033E-05 | 2.363E-06 | 8.500E-06 | 1.300E-05 |
| 24172 | 1209 | D1209K | D | 9.3e-06 | 8.1e-06 | 8.5e-06 | 8.633E-06 | 6.110E-07 | 8.100E-06 | 9.300E-06 |
| 24173 | 1209 | D1209M | D | 5.1e-06 | 9.1e-06 | 5.1e-06 | 6.433E-06 | 2.309E-06 | 5.100E-06 | 9.100E-06 |
| 24174 | 1209 | D1209F | D | 7.2e-06 | 7.6e-06 | 1.3e-05 | 9.267E-06 | 3.239E-06 | 7.200E-06 | 1.300E-05 |
| 24175 | 1209 | D1209P | D | 6.3e-06 | 6.4e-06 | 1.2e-05 | 8.233E-06 | 3.262E-06 | 6.300E-06 | 1.200E-05 |
| 24176 | 1209 | D1209S | D | 4.5e-06 | 7.3e-06 | 5.3e-06 | 5.700E-06 | 1.442E-06 | 4.500E-06 | 7.300E-06 |
| 24177 | 1209 | D1209T | D | 1.2e-05 | 6.1e-06 | 9.1e-06 | 9.067E-06 | 2.950E-06 | 6.100E-06 | 1.200E-05 |
| 24178 | 1209 | D1209W | D | 1e-05 | 1.2e-05 | 3.6e-06 | 8.533E-06 | 4.388E-06 | 3.600E-06 | 1.200E-05 |
| 24179 | 1209 | D1209Y | D | 6.5e-06 | 4.6e-06 | 9.4e-06 | 6.833E-06 | 2.417E-06 | 4.600E-06 | 9.400E-06 |
| 24180 | 1209 | D1209V | D | 7.9e-06 | 6.4e-06 | 6.2e-06 | 6.833E-06 | 9.292E-07 | 6.200E-06 | 7.900E-06 |
| 24181 | 1210 | N1210A | D | 6.9e-06 | 8.7e-06 | 1.2e-05 | 9.200E-06 | 2.587E-06 | 6.900E-06 | 1.200E-05 |
| 24182 | 1210 | N1210R | D | 4.9e-06 | 9.9e-06 | 1e-05 | 8.267E-06 | 2.916E-06 | 4.900E-06 | 1.000E-05 |
| 24183 | 1210 | N1210N | D | 5.5e-06 | 7.5e-06 | 8.9e-06 | 7.300E-06 | 1.709E-06 | 5.500E-06 | 8.900E-06 |
| 24184 | 1210 | N1210D | D | 1e-05 | 5.9e-06 | 6.7e-06 | 7.533E-06 | 2.173E-06 | 5.900E-06 | 1.000E-05 |
| 24185 | 1210 | N1210C | D | 9.7e-06 | 9.5e-06 | 7.6e-06 | 8.933E-06 | 1.159E-06 | 7.600E-06 | 9.700E-06 |
| 24186 | 1210 | N1210E | D | 7.4e-06 | 6.2e-06 | 1.2e-05 | 8.533E-06 | 3.062E-06 | 6.200E-06 | 1.200E-05 |
| 24187 | 1210 | N1210Q | D | 8.9e-06 | 5e-06 | 1.3e-05 | 8.967E-06 | 4.000E-06 | 5.000E-06 | 1.300E-05 |
| 24188 | 1210 | N1210G | D | 1.1e-05 | 5.1e-06 | 5.4e-06 | 7.167E-06 | 3.323E-06 | 5.100E-06 | 1.100E-05 |
| 24189 | 1210 | N1210H | D | 1.3e-05 | 1e-05 | 6.9e-06 | 9.967E-06 | 3.050E-06 | 6.900E-06 | 1.300E-05 |
| 24190 | 1210 | N1210I | D | 9.5e-06 | 9.7e-06 | 7.5e-06 | 8.900E-06 | 1.217E-06 | 7.500E-06 | 9.700E-06 |
| 24191 | 1210 | N1210L | D | 4.9e-06 | 6.7e-06 | 1.1e-05 | 7.533E-06 | 3.134E-06 | 4.900E-06 | 1.100E-05 |
| 24192 | 1210 | N1210K | D | 9.9e-06 | 1e-05 | 7.8e-06 | 9.233E-06 | 1.242E-06 | 7.800E-06 | 1.000E-05 |
| 24193 | 1210 | N1210M | D | 6.9e-06 | 1.3e-05 | 1.3e-05 | 1.097E-05 | 3.522E-06 | 6.900E-06 | 1.300E-05 |
| 24194 | 1210 | N1210F | D | 8.2e-06 | 6.9e-06 | 1e-05 | 8.367E-06 | 1.557E-06 | 6.900E-06 | 1.000E-05 |
| 24195 | 1210 | N1210P | D | 1.1e-05 | 4.8e-06 | 1e-05 | 8.600E-06 | 3.329E-06 | 4.800E-06 | 1.100E-05 |
| 24196 | 1210 | N1210S | D | 5.1e-06 | 9.6e-06 | 4.5e-06 | 6.400E-06 | 2.787E-06 | 4.500E-06 | 9.600E-06 |
| 24197 | 1210 | N1210T | D | 7e-06 | 1e-05 | 6.4e-06 | 7.800E-06 | 1.929E-06 | 6.400E-06 | 1.000E-05 |
| 24198 | 1210 | N1210W | D | 1.2e-05 | 1e-05 | 6.7e-06 | 9.567E-06 | 2.676E-06 | 6.700E-06 | 1.200E-05 |
| 24199 | 1210 | N1210Y | D | 7e-06 | 9.7e-06 | 8.9e-06 | 8.533E-06 | 1.387E-06 | 7.000E-06 | 9.700E-06 |
| 24200 | 1210 | N1210V | D | 7.4e-06 | 7.2e-06 | 6.5e-06 | 7.033E-06 | 4.726E-07 | 6.500E-06 | 7.400E-06 |
| 24201 | 1211 | A1211A | D | 7.5e-06 | 9.4e-06 | 7.4e-06 | 8.100E-06 | 1.127E-06 | 7.400E-06 | 9.400E-06 |
| 24202 | 1211 | A1211R | D | 6.5e-06 | 1.3e-05 | 7.2e-06 | 8.900E-06 | 3.568E-06 | 6.500E-06 | 1.300E-05 |
| 24203 | 1211 | A1211N | D | 6.6e-06 | 9.7e-06 | 5.8e-06 | 7.367E-06 | 2.060E-06 | 5.800E-06 | 9.700E-06 |
| 24204 | 1211 | A1211D | D | 1.2e-05 | 1e-05 | 1.4e-05 | 1.200E-05 | 2.000E-06 | 1.000E-05 | 1.400E-05 |
| 24205 | 1211 | A1211C | D | 6.7e-06 | 1.3e-05 | 6.2e-06 | 8.633E-06 | 3.790E-06 | 6.200E-06 | 1.300E-05 |
| 24206 | 1211 | A1211E | D | 1.3e-05 | 5.8e-06 | 1.1e-05 | 9.933E-06 | 3.717E-06 | 5.800E-06 | 1.300E-05 |
| 24207 | 1211 | A1211Q | D | 1.1e-05 | 9.1e-06 | 8.8e-06 | 9.633E-06 | 1.193E-06 | 8.800E-06 | 1.100E-05 |
| 24208 | 1211 | A1211G | D | 6.1e-06 | 8.8e-06 | 1e-05 | 8.300E-06 | 1.997E-06 | 6.100E-06 | 1.000E-05 |
| 24209 | 1211 | A1211H | D | 7e-06 | 1e-05 | 5.8e-06 | 7.600E-06 | 2.163E-06 | 5.800E-06 | 1.000E-05 |
| 24210 | 1211 | A1211I | D | 8.1e-06 | 5.6e-06 | 9.4e-06 | 7.700E-06 | 1.931E-06 | 5.600E-06 | 9.400E-06 |
| 24211 | 1211 | A1211L | D | 9.1e-06 | 8.3e-06 | 7.5e-06 | 8.300E-06 | 8.000E-07 | 7.500E-06 | 9.100E-06 |
| 24212 | 1211 | A1211K | D | 8.2e-06 | 1.2e-05 | 9.3e-06 | 9.833E-06 | 1.955E-06 | 8.200E-06 | 1.200E-05 |
| 24213 | 1211 | A1211M | D | 1.1e-05 | 8.4e-06 | 6.4e-06 | 8.600E-06 | 2.307E-06 | 6.400E-06 | 1.100E-05 |
| 24214 | 1211 | A1211F | D | 7.4e-06 | 6.5e-06 | 7.1e-06 | 7.000E-06 | 4.583E-07 | 6.500E-06 | 7.400E-06 |
| 24215 | 1211 | A1211P | D | 1e-05 | 9.4e-06 | 9.9e-06 | 9.767E-06 | 3.215E-07 | 9.400E-06 | 1.000E-05 |
| 24216 | 1211 | A1211S | D | 7.1e-06 | 1.2e-05 | 1.2e-05 | 1.037E-05 | 2.829E-06 | 7.100E-06 | 1.200E-05 |
| 24217 | 1211 | A1211T | D | 4.3e-06 | 6.6e-06 | 9.2e-06 | 6.700E-06 | 2.452E-06 | 4.300E-06 | 9.200E-06 |
| 24218 | 1211 | A1211W | D | 6.6e-06 | 5.4e-06 | 6.5e-06 | 6.167E-06 | 6.658E-07 | 5.400E-06 | 6.600E-06 |
| 24219 | 1211 | A1211Y | D | 8.9e-06 | 4.7e-06 | 4.4e-06 | 6.000E-06 | 2.516E-06 | 4.400E-06 | 8.900E-06 |
| 24220 | 1211 | A1211V | D | 9e-06 | 6.8e-06 | 6.2e-06 | 7.333E-06 | 1.474E-06 | 6.200E-06 | 9.000E-06 |
| 24221 | 1212 | L1212A | D | 1.4e-05 | 8e-06 | 5.5e-06 | 9.167E-06 | 4.368E-06 | 5.500E-06 | 1.400E-05 |
| 24222 | 1212 | L1212R | D | 8.5e-06 | 6.4e-06 | 5.1e-06 | 6.667E-06 | 1.716E-06 | 5.100E-06 | 8.500E-06 |
| 24223 | 1212 | L1212N | D | 5.1e-06 | 7.5e-06 | 9.6e-06 | 7.400E-06 | 2.252E-06 | 5.100E-06 | 9.600E-06 |
| 24224 | 1212 | L1212D | D | 7.1e-06 | 5.6e-06 | 1.5e-05 | 9.233E-06 | 5.050E-06 | 5.600E-06 | 1.500E-05 |

| | | | | | | | | | | |
|---|---|---|---|---|---|---|---|---|---|---|
| 24225 | 1212 | L1212C | D | 6.6e-06 | 6.3e-06 | 9.2e-06 | 7.367E-06 | 1.595E-06 | 6.300E-06 | 9.200E-06 |
| 24226 | 1212 | L1212E | D | 6.4e-06 | 1.4e-05 | 6.6e-06 | 9.000E-06 | 4.331E-06 | 6.400E-06 | 1.400E-05 |
| 24227 | 1212 | L1212Q | D | 4.7e-06 | 6.5e-06 | 8.2e-06 | 6.467E-06 | 1.750E-06 | 4.700E-06 | 8.200E-06 |
| 24228 | 1212 | L1212G | D | 6.3e-06 | 1e-05 | 1.1e-05 | 9.100E-06 | 2.476E-06 | 6.300E-06 | 1.100E-05 |
| 24229 | 1212 | L1212H | D | 1e-05 | 9.8e-06 | 1.1e-05 | 1.027E-05 | 6.429E-07 | 9.800E-06 | 1.100E-05 |
| 24230 | 1212 | L1212I | D | 1.2e-05 | 8.5e-06 | 6.4e-06 | 8.967E-06 | 2.829E-06 | 6.400E-06 | 1.200E-05 |
| 24231 | 1212 | L1212L | D | 6.7e-06 | 9.4e-06 | 8.4e-06 | 8.167E-06 | 1.365E-06 | 6.700E-06 | 9.400E-06 |
| 24232 | 1212 | L1212K | D | 9.8e-06 | 9.7e-06 | 7e-06 | 8.833E-06 | 1.589E-06 | 7.000E-06 | 9.800E-06 |
| 24233 | 1212 | L1212M | D | 6.9e-06 | 8.6e-06 | 6.3e-06 | 7.267E-06 | 1.193E-06 | 6.300E-06 | 8.600E-06 |
| 24234 | 1212 | L1212F | D | 8.6e-06 | 8.6e-06 | 9.2e-06 | 8.800E-06 | 3.464E-07 | 8.600E-06 | 9.200E-06 |
| 24235 | 1212 | L1212P | D | 9.9e-06 | 5.9e-06 | 7.8e-06 | 7.867E-06 | 2.001E-06 | 5.900E-06 | 9.900E-06 |
| 24236 | 1212 | L1212S | D | 5.3e-06 | 7.5e-06 | 5.6e-06 | 6.133E-06 | 1.193E-06 | 5.300E-06 | 7.500E-06 |
| 24237 | 1212 | L1212T | D | 5.8e-06 | 8.5e-06 | 7.2e-06 | 7.167E-06 | 1.350E-06 | 5.800E-06 | 8.500E-06 |
| 24238 | 1212 | L1212W | D | 4.7e-06 | 9.8e-06 | 9.7e-06 | 8.067E-06 | 2.916E-06 | 4.700E-06 | 9.800E-06 |
| 24239 | 1212 | L1212Y | D | 4.4e-06 | 7.5e-06 | 6.9e-06 | 6.267E-06 | 1.644E-06 | 4.400E-06 | 7.500E-06 |
| 24240 | 1212 | L1212V | D | 7.7e-06 | 1.3e-05 | 9.8e-06 | 1.017E-05 | 2.669E-06 | 7.700E-06 | 1.300E-05 |
| 24241 | 1213 | Q1213A | D | 9.6e-06 | 9e-06 | 1e-05 | 9.533E-06 | 5.033E-07 | 9.000E-06 | 1.000E-05 |
| 24242 | 1213 | Q1213R | D | 6.5e-06 | 8.9e-06 | 5.4e-06 | 6.933E-06 | 1.790E-06 | 5.400E-06 | 8.900E-06 |
| 24243 | 1213 | Q1213N | D | 6.8e-06 | 1e-05 | 7.3e-06 | 8.033E-06 | 1.721E-06 | 6.800E-06 | 1.000E-05 |
| 24244 | 1213 | Q1213D | D | 5e-06 | 9.4e-06 | 6.4e-06 | 6.933E-06 | 2.248E-06 | 5.000E-06 | 9.400E-06 |
| 24245 | 1213 | Q1213C | D | 4.6e-06 | 7.5e-06 | 9.1e-06 | 7.067E-06 | 2.281E-06 | 4.600E-06 | 9.100E-06 |
| 24246 | 1213 | Q1213E | D | 1.4e-05 | 5.7e-06 | 9.9e-06 | 9.867E-06 | 4.150E-06 | 5.700E-06 | 1.400E-05 |
| 24247 | 1213 | Q1213Q | D | 1e-05 | 1.4e-05 | 6.8e-06 | 1.027E-05 | 3.607E-06 | 6.800E-06 | 1.400E-05 |
| 24248 | 1213 | Q1213G | D | 8.5e-06 | 8e-06 | 8.1e-06 | 8.200E-06 | 2.646E-07 | 8.000E-06 | 8.500E-06 |
| 24249 | 1213 | Q1213H | D | 7.5e-06 | 7.8e-06 | 1e-05 | 8.433E-06 | 1.365E-06 | 7.500E-06 | 1.000E-05 |
| 24250 | 1213 | Q1213I | D | 9.4e-06 | 8.4e-06 | 5.3e-06 | 7.700E-06 | 2.138E-06 | 5.300E-06 | 9.400E-06 |
| 24251 | 1213 | Q1213L | D | 1.3e-05 | 7.4e-06 | 8.9e-06 | 9.767E-06 | 2.899E-06 | 7.400E-06 | 1.300E-05 |
| 24252 | 1213 | Q1213K | D | 5.3e-06 | 5.8e-06 | 1e-05 | 7.033E-06 | 2.581E-06 | 5.300E-06 | 1.000E-05 |
| 24253 | 1213 | Q1213M | D | 5.1e-06 | 1.1e-05 | 9.7e-06 | 8.600E-06 | 3.100E-06 | 5.100E-06 | 1.100E-05 |
| 24254 | 1213 | Q1213F | D | 1.5e-05 | 8.9e-06 | 6.2e-06 | 1.003E-05 | 4.508E-06 | 6.200E-06 | 1.500E-05 |
| 24255 | 1213 | Q1213P | D | 8.9e-06 | 1e-05 | 9.6e-06 | 9.500E-06 | 5.568E-07 | 8.900E-06 | 1.000E-05 |
| 24256 | 1213 | Q1213S | D | 5.2e-06 | 9.8e-06 | 7.5e-06 | 7.500E-06 | 2.300E-06 | 5.200E-06 | 9.800E-06 |
| 24257 | 1213 | Q1213T | D | 6.1e-06 | 1e-05 | 5.4e-06 | 7.167E-06 | 2.479E-06 | 5.400E-06 | 1.000E-05 |
| 24258 | 1213 | Q1213W | D | 1.4e-05 | 8.6e-06 | 5.6e-06 | 9.400E-06 | 4.257E-06 | 5.600E-06 | 1.400E-05 |
| 24259 | 1213 | Q1213Y | D | 7.7e-06 | 1.1e-05 | 5.8e-06 | 8.167E-06 | 2.631E-06 | 5.800E-06 | 1.100E-05 |
| 24260 | 1213 | Q1213V | D | 9.7e-06 | 1.3e-05 | 1.2e-05 | 1.157E-05 | 1.692E-06 | 9.700E-06 | 1.300E-05 |
| 24261 | 1214 | S1214A | D | 6.7e-06 | 1.4e-05 | 1e-05 | 1.023E-05 | 3.656E-06 | 6.700E-06 | 1.400E-05 |
| 24262 | 1214 | S1214R | D | 7.7e-06 | 7.2e-06 | 1e-05 | 8.300E-06 | 1.493E-06 | 7.200E-06 | 1.000E-05 |
| 24263 | 1214 | S1214N | D | 7.9e-06 | 1.5e-05 | 9.8e-06 | 1.090E-05 | 3.676E-06 | 7.900E-06 | 1.500E-05 |
| 24264 | 1214 | S1214D | D | 1.1e-05 | 7.3e-06 | 1.2e-05 | 1.010E-05 | 2.476E-06 | 7.300E-06 | 1.200E-05 |
| 24265 | 1214 | S1214C | D | 5.6e-06 | 6.4e-06 | 8.5e-06 | 6.833E-06 | 1.498E-06 | 5.600E-06 | 8.500E-06 |
| 24266 | 1214 | S1214E | D | 1.1e-05 | 1e-05 | 9.5e-06 | 1.017E-05 | 7.638E-07 | 9.500E-06 | 1.100E-05 |
| 24267 | 1214 | S1214Q | D | 7.9e-06 | 6.7e-06 | 8.6e-06 | 7.733E-06 | 9.609E-07 | 6.700E-06 | 8.600E-06 |
| 24268 | 1214 | S1214G | D | 9.4e-06 | 5e-06 | 3.1e-06 | 5.833E-06 | 3.232E-06 | 3.100E-06 | 9.400E-06 |
| 24269 | 1214 | S1214H | D | 8.2e-06 | 1.2e-05 | 9.1e-06 | 9.767E-06 | 1.986E-06 | 8.200E-06 | 1.200E-05 |
| 24270 | 1214 | S1214I | D | 8.2e-06 | 1e-05 | 5.9e-06 | 8.033E-06 | 2.055E-06 | 5.900E-06 | 1.000E-05 |
| 24271 | 1214 | S1214L | D | 8.2e-06 | 1e-05 | 1e-05 | 9.400E-06 | 1.039E-06 | 8.200E-06 | 1.000E-05 |
| 24272 | 1214 | S1214K | D | 6.1e-06 | 1.4e-05 | 6e-06 | 8.700E-06 | 4.590E-06 | 6.000E-06 | 1.400E-05 |
| 24273 | 1214 | S1214M | D | 6.3e-06 | 5.7e-06 | 1.5e-05 | 9.000E-06 | 5.205E-06 | 5.700E-06 | 1.500E-05 |
| 24274 | 1214 | S1214F | D | 1.3e-05 | 5.6e-06 | 1.1e-05 | 9.867E-06 | 3.828E-06 | 5.600E-06 | 1.300E-05 |
| 24275 | 1214 | S1214P | D | 6.4e-06 | 5.5e-06 | 8.9e-06 | 6.933E-06 | 1.762E-06 | 5.500E-06 | 8.900E-06 |
| 24276 | 1214 | S1214S | D | 8.5e-06 | 9e-06 | 6.3e-06 | 7.933E-06 | 1.436E-06 | 6.300E-06 | 9.000E-06 |
| 24277 | 1214 | S1214T | D | 9.7e-06 | 1.1e-05 | 8.5e-06 | 9.733E-06 | 1.250E-06 | 8.500E-06 | 1.100E-05 |
| 24278 | 1214 | S1214W | D | 1.5e-05 | 1.1e-05 | 1.1e-05 | 1.233E-05 | 2.309E-06 | 1.100E-05 | 1.500E-05 |
| 24279 | 1214 | S1214Y | D | 1.1e-05 | 5.2e-06 | 1.2e-05 | 9.400E-06 | 3.672E-06 | 5.200E-06 | 1.200E-05 |
| 24280 | 1214 | S1214V | D | 8.2e-06 | 8.9e-06 | 5.5e-06 | 7.533E-06 | 1.795E-06 | 5.500E-06 | 8.900E-06 |
| 24281 | 1215 | G1215A | D | 5.6e-06 | 3.9e-06 | 1e-05 | 6.500E-06 | 3.148E-06 | 3.900E-06 | 1.000E-05 |
| 24282 | 1215 | G1215R | D | 6.8e-06 | 8.3e-06 | 6.1e-06 | 7.067E-06 | 1.124E-06 | 6.100E-06 | 8.300E-06 |
| 24283 | 1215 | G1215N | D | 5.2e-06 | 1.2e-05 | 6.6e-06 | 7.933E-06 | 3.591E-06 | 5.200E-06 | 1.200E-05 |
| 24284 | 1215 | G1215D | D | 1e-05 | 6.7e-06 | 4.5e-06 | 7.067E-06 | 2.768E-06 | 4.500E-06 | 1.000E-05 |
| 24285 | 1215 | G1215C | D | 5.8e-06 | 1.2e-05 | 7.9e-06 | 8.567E-06 | 3.153E-06 | 5.800E-06 | 1.200E-05 |
| 24286 | 1215 | G1215E | D | 1e-05 | 3.4e-06 | 6.2e-06 | 6.533E-06 | 3.313E-06 | 3.400E-06 | 1.000E-05 |
| 24287 | 1215 | G1215Q | D | 6.3e-06 | 1e-05 | 1e-05 | 8.767E-06 | 2.136E-06 | 6.300E-06 | 1.000E-05 |
| 24288 | 1215 | G1215G | D | 4.9e-06 | 7.1e-06 | 9.2e-06 | 7.067E-06 | 2.150E-06 | 4.900E-06 | 9.200E-06 |
| 24289 | 1215 | G1215H | D | 1.1e-05 | 1.2e-05 | 1e-05 | 1.100E-05 | 1.000E-06 | 1.000E-05 | 1.200E-05 |
| 24290 | 1215 | G1215I | D | 1.3e-05 | 7.9e-06 | 1.1e-05 | 1.063E-05 | 2.570E-06 | 7.900E-06 | 1.300E-05 |
| 24291 | 1215 | G1215L | D | 6.8e-06 | 6e-06 | 1.1e-05 | 7.933E-06 | 2.686E-06 | 6.000E-06 | 1.100E-05 |
| 24292 | 1215 | G1215K | D | 1.3e-05 | 5.7e-06 | 7.1e-06 | 8.600E-06 | 3.874E-06 | 5.700E-06 | 1.300E-05 |
| 24293 | 1215 | G1215M | D | 6.6e-06 | 9.8e-06 | 5.3e-06 | 7.233E-06 | 2.316E-06 | 5.300E-06 | 9.800E-06 |
| 24294 | 1215 | G1215F | D | 5.5e-06 | 6.9e-06 | 6.7e-06 | 6.367E-06 | 7.572E-07 | 5.500E-06 | 6.900E-06 |
| 24295 | 1215 | G1215P | D | 6.7e-06 | 9.3e-06 | 7e-06 | 7.667E-06 | 1.422E-06 | 6.700E-06 | 9.300E-06 |
| 24296 | 1215 | G1215S | D | 4.9e-06 | 7.4e-06 | 6.1e-06 | 6.133E-06 | 1.250E-06 | 4.900E-06 | 7.400E-06 |
| 24297 | 1215 | G1215T | D | 1.1e-05 | 7.4e-06 | 5.9e-06 | 8.100E-06 | 2.621E-06 | 5.900E-06 | 1.100E-05 |
| 24298 | 1215 | G1215W | D | 8.7e-06 | 7.3e-06 | 4.6e-06 | 6.867E-06 | 2.084E-06 | 4.600E-06 | 8.700E-06 |
| 24299 | 1215 | G1215Y | D | 6.4e-06 | 9.1e-06 | 6.5e-06 | 7.333E-06 | 1.531E-06 | 6.400E-06 | 9.100E-06 |
| 24300 | 1215 | G1215V | D | 1.2e-05 | 4.4e-06 | 6.9e-06 | 7.767E-06 | 3.873E-06 | 4.400E-06 | 1.200E-05 |
| 24301 | 1216 | N1216A | D | 1.2e-05 | 9.2e-06 | 9e-06 | 1.007E-05 | 1.677E-06 | 9.000E-06 | 1.200E-05 |
| 24302 | 1216 | N1216R | D | 3.1e-06 | 6.7e-06 | 6.4e-06 | 5.400E-06 | 1.997E-06 | 3.100E-06 | 6.700E-06 |
| 24303 | 1216 | N1216N | D | 8.6e-06 | 6.6e-06 | 6e-06 | 7.067E-06 | 1.361E-06 | 6.000E-06 | 8.600E-06 |
| 24304 | 1216 | N1216D | D | 1.1e-05 | 1e-05 | 2.6e-06 | 7.867E-06 | 4.588E-06 | 2.600E-06 | 1.100E-05 |
| 24305 | 1216 | N1216C | D | 9.8e-06 | 9.3e-06 | 1.3e-05 | 1.070E-05 | 2.007E-06 | 9.300E-06 | 1.300E-05 |
| 24306 | 1216 | N1216E | D | 9.2e-06 | 6.3e-06 | 6.1e-06 | 7.200E-06 | 1.735E-06 | 6.100E-06 | 9.200E-06 |
| 24307 | 1216 | N1216Q | D | 9.8e-06 | 1.1e-05 | 4.7e-06 | 8.500E-06 | 3.345E-06 | 4.700E-06 | 1.100E-05 |
| 24308 | 1216 | N1216G | D | 4.4e-06 | 8.6e-06 | 9.4e-06 | 7.467E-06 | 2.686E-06 | 4.400E-06 | 9.400E-06 |
| 24309 | 1216 | N1216H | D | 6.9e-06 | 5.9e-06 | 1.2e-05 | 8.267E-06 | 3.272E-06 | 5.900E-06 | 1.200E-05 |

| | | | | | | | | | | |
|---|---|---|---|---|---|---|---|---|---|---|
| 24310 | 1216 | N1216I | D | 6.8e-06 | 9e-06 | 8.4e-06 | 8.067E-06 | 1.137E-06 | 6.800E-06 | 9.000E-06 |
| 24311 | 1216 | N1216L | D | 6.1e-06 | 6.2e-06 | 1.3e-05 | 8.433E-06 | 3.955E-06 | 6.100E-06 | 1.300E-05 |
| 24312 | 1216 | N1216K | D | 9.1e-06 | 9.7e-06 | 9.2e-06 | 9.333E-06 | 3.215E-07 | 9.100E-06 | 9.700E-06 |
| 24313 | 1216 | N1216M | D | 7.1e-06 | 6.9e-06 | 8.6e-06 | 7.533E-06 | 9.292E-07 | 6.900E-06 | 8.600E-06 |
| 24314 | 1216 | N1216F | D | 9.9e-06 | 5.6e-06 | 4.8e-06 | 6.767E-06 | 2.743E-06 | 4.800E-06 | 9.900E-06 |
| 24315 | 1216 | N1216P | D | 6.2e-06 | 1.1e-05 | 6.6e-06 | 7.933E-06 | 2.663E-06 | 6.200E-06 | 1.100E-05 |
| 24316 | 1216 | N1216S | D | 9.3e-06 | 5.7e-06 | 1.3e-05 | 9.333E-06 | 3.650E-06 | 5.700E-06 | 1.300E-05 |
| 24317 | 1216 | N1216T | D | 5.9e-06 | 5.5e-06 | 9.7e-06 | 7.033E-06 | 2.318E-06 | 5.500E-06 | 9.700E-06 |
| 24318 | 1216 | N1216W | D | 1.1e-05 | 5.8e-06 | 7e-06 | 7.933E-06 | 2.723E-06 | 5.800E-06 | 1.100E-05 |
| 24319 | 1216 | N1216Y | D | 1.1e-05 | 6.1e-06 | 9.8e-06 | 8.967E-06 | 2.554E-06 | 6.100E-06 | 1.100E-05 |
| 24320 | 1216 | N1216V | D | 9.6e-06 | 6.9e-06 | 1.2e-05 | 9.500E-06 | 2.551E-06 | 6.900E-06 | 1.200E-05 |
| 24321 | 1217 | S1217A | D | 6.6e-06 | 1.5e-05 | 9.1e-06 | 1.023E-05 | 4.313E-06 | 6.600E-06 | 1.500E-05 |
| 24322 | 1217 | S1217R | D | 7.1e-06 | 6.7e-06 | 6.8e-06 | 6.867E-06 | 2.082E-07 | 6.700E-06 | 7.100E-06 |
| 24323 | 1217 | S1217N | D | 1e-05 | 6e-06 | 6e-06 | 7.333E-06 | 2.309E-06 | 6.000E-06 | 1.000E-05 |
| 24324 | 1217 | S1217D | D | 1.2e-05 | 6.6e-06 | 4.5e-06 | 7.700E-06 | 3.869E-06 | 4.500E-06 | 1.200E-05 |
| 24325 | 1217 | S1217C | D | 1.5e-05 | 4.8e-06 | 8.3e-06 | 9.367E-06 | 5.183E-06 | 4.800E-06 | 1.500E-05 |
| 24326 | 1217 | S1217E | D | 9.8e-06 | 8.2e-06 | 6.2e-06 | 8.067E-06 | 1.804E-06 | 6.200E-06 | 9.800E-06 |
| 24327 | 1217 | S1217Q | D | 6e-06 | 6.6e-06 | 1.9e-05 | 1.053E-05 | 7.338E-06 | 6.000E-06 | 1.900E-05 |
| 24328 | 1217 | S1217G | D | 9.1e-06 | 1e-05 | 5.1e-06 | 8.067E-06 | 2.608E-06 | 5.100E-06 | 1.000E-05 |
| 24329 | 1217 | S1217H | D | 1e-05 | 6.3e-06 | 8.4e-06 | 8.233E-06 | 1.856E-06 | 6.300E-06 | 1.000E-05 |
| 24330 | 1217 | S1217I | D | 7.2e-06 | 4.6e-06 | 7e-06 | 6.267E-06 | 1.447E-06 | 4.600E-06 | 7.200E-06 |
| 24331 | 1217 | S1217L | D | 6.1e-06 | 7.3e-06 | 1.8e-05 | 1.047E-05 | 6.552E-06 | 6.100E-06 | 1.800E-05 |
| 24332 | 1217 | S1217K | D | 1.1e-05 | 8.3e-06 | 6.8e-06 | 8.700E-06 | 2.128E-06 | 6.800E-06 | 1.100E-05 |
| 24333 | 1217 | S1217M | D | 6.3e-06 | 4.9e-06 | 1.4e-05 | 8.400E-06 | 4.900E-06 | 4.900E-06 | 1.400E-05 |
| 24334 | 1217 | S1217F | D | 6e-06 | 1.3e-05 | 1.4e-05 | 1.100E-05 | 4.359E-06 | 6.000E-06 | 1.400E-05 |
| 24335 | 1217 | S1217P | D | 5.6e-06 | 1.1e-05 | 4.1e-06 | 6.900E-06 | 3.629E-06 | 4.100E-06 | 1.100E-05 |
| 24336 | 1217 | S1217S | D | 1.1e-05 | 5.4e-06 | 1.3e-05 | 9.800E-06 | 3.940E-06 | 5.400E-06 | 1.300E-05 |
| 24337 | 1217 | S1217T | D | 7.8e-06 | 5.6e-06 | 1e-05 | 7.800E-06 | 2.200E-06 | 5.600E-06 | 1.000E-05 |
| 24338 | 1217 | S1217W | D | 1.4e-05 | 6.7e-06 | 1.3e-05 | 1.123E-05 | 3.958E-06 | 6.700E-06 | 1.400E-05 |
| 24339 | 1217 | S1217Y | D | 9.7e-06 | 6.6e-06 | 1e-05 | 8.767E-06 | 1.882E-06 | 6.600E-06 | 1.000E-05 |
| 24340 | 1217 | S1217V | D | 9.5e-06 | 6.4e-06 | 9.1e-06 | 8.333E-06 | 1.686E-06 | 6.400E-06 | 9.500E-06 |
| 24341 | 1218 | Q1218A | D | 1.1e-05 | 9.3e-06 | 8.3e-06 | 9.533E-06 | 1.365E-06 | 8.300E-06 | 1.100E-05 |
| 24342 | 1218 | Q1218R | D | 9.4e-06 | 6.3e-06 | 6.8e-06 | 7.500E-06 | 1.664E-06 | 6.300E-06 | 9.400E-06 |
| 24343 | 1218 | Q1218N | D | 6.9e-06 | 9.1e-06 | 5.3e-06 | 7.100E-06 | 1.908E-06 | 5.300E-06 | 9.100E-06 |
| 24344 | 1218 | Q1218D | D | 8.4e-06 | 9.5e-06 | 9.7e-06 | 9.200E-06 | 7.000E-07 | 8.400E-06 | 9.700E-06 |
| 24345 | 1218 | Q1218C | D | 1.2e-05 | 3.6e-06 | 1.1e-05 | 8.867E-06 | 4.588E-06 | 3.600E-06 | 1.200E-05 |
| 24346 | 1218 | Q1218E | D | 6e-06 | 1e-05 | 9.6e-06 | 8.533E-06 | 2.203E-06 | 6.000E-06 | 1.000E-05 |
| 24347 | 1218 | Q1218Q | D | 8.2e-06 | 7.7e-06 | 8.5e-06 | 8.133E-06 | 4.041E-07 | 7.700E-06 | 8.500E-06 |
| 24348 | 1218 | Q1218G | D | 6.3e-06 | 7.3e-06 | 1e-05 | 7.867E-06 | 1.914E-06 | 6.300E-06 | 1.000E-05 |
| 24349 | 1218 | Q1218H | D | 5.8e-06 | 6.7e-06 | 7e-06 | 6.500E-06 | 6.245E-07 | 5.800E-06 | 7.000E-06 |
| 24350 | 1218 | Q1218I | D | 8.9e-06 | 7e-06 | 1.2e-05 | 9.300E-06 | 2.524E-06 | 7.000E-06 | 1.200E-05 |
| 24351 | 1218 | Q1218L | D | 7.4e-06 | 8.1e-06 | 7.1e-06 | 7.533E-06 | 5.132E-07 | 7.100E-06 | 8.100E-06 |
| 24352 | 1218 | Q1218K | D | 3.3e-06 | 6.9e-06 | 8.5e-06 | 6.233E-06 | 2.663E-06 | 3.300E-06 | 8.500E-06 |
| 24353 | 1218 | Q1218M | D | 7.5e-06 | 8.2e-06 | 6.8e-06 | 7.500E-06 | 7.000E-07 | 6.800E-06 | 8.200E-06 |
| 24354 | 1218 | Q1218F | D | 9e-06 | 9.4e-06 | 1.3e-05 | 1.047E-05 | 2.203E-06 | 9.000E-06 | 1.300E-05 |
| 24355 | 1218 | Q1218P | D | 6.2e-06 | 8.9e-06 | 1.1e-05 | 8.700E-06 | 2.406E-06 | 6.200E-06 | 1.100E-05 |
| 24356 | 1218 | Q1218S | D | 5e-06 | 5.8e-06 | 1.3e-05 | 7.933E-06 | 4.406E-06 | 5.000E-06 | 1.300E-05 |
| 24357 | 1218 | Q1218T | D | 9.7e-06 | 6.1e-06 | 5.6e-06 | 7.133E-06 | 2.237E-06 | 5.600E-06 | 9.700E-06 |
| 24358 | 1218 | Q1218W | D | 8.8e-06 | 9e-06 | 9.2e-06 | 9.000E-06 | 2.000E-07 | 8.800E-06 | 9.200E-06 |
| 24359 | 1218 | Q1218Y | D | 6e-06 | 8.1e-06 | 1.4e-05 | 9.367E-06 | 4.148E-06 | 6.000E-06 | 1.400E-05 |
| 24360 | 1218 | Q1218V | D | 5.3e-06 | 5e-06 | 7.2e-06 | 5.833E-06 | 1.193E-06 | 5.000E-06 | 7.200E-06 |
| 24361 | 1219 | E1219A | D | 9.8e-06 | 8.4e-06 | 4.5e-06 | 7.567E-06 | 2.747E-06 | 4.500E-06 | 9.800E-06 |
| 24362 | 1219 | E1219R | D | 7.4e-06 | 6.9e-06 | 6e-06 | 6.767E-06 | 7.095E-07 | 6.000E-06 | 7.400E-06 |
| 24363 | 1219 | E1219N | D | 6.1e-06 | 6.9e-06 | 7.7e-06 | 6.900E-06 | 8.000E-07 | 6.100E-06 | 7.700E-06 |
| 24364 | 1219 | E1219D | D | 6.2e-06 | 9.3e-06 | 1.3e-05 | 9.500E-06 | 3.404E-06 | 6.200E-06 | 1.300E-05 |
| 24365 | 1219 | E1219C | D | 1.5e-05 | 8.4e-06 | 9.9e-06 | 1.110E-05 | 3.460E-06 | 8.400E-06 | 1.500E-05 |
| 24366 | 1219 | E1219E | D | 1e-05 | 5.5e-06 | 1.2e-05 | 9.167E-06 | 3.329E-06 | 5.500E-06 | 1.200E-05 |
| 24367 | 1219 | E1219Q | D | 7.8e-06 | 4.1e-06 | 7.8e-06 | 6.567E-06 | 2.136E-06 | 4.100E-06 | 7.800E-06 |
| 24368 | 1219 | E1219G | D | 1.4e-05 | 1.2e-05 | 9.6e-06 | 1.187E-05 | 2.203E-06 | 9.600E-06 | 1.400E-05 |
| 24369 | 1219 | E1219H | D | 7e-06 | 7.5e-06 | 8.1e-06 | 7.533E-06 | 5.508E-07 | 7.000E-06 | 8.100E-06 |
| 24370 | 1219 | E1219I | D | 8.1e-06 | 8.3e-06 | 6.6e-06 | 7.667E-06 | 9.292E-07 | 6.600E-06 | 8.300E-06 |
| 24371 | 1219 | E1219L | D | 5.3e-06 | 1.3e-05 | 5.6e-06 | 7.967E-06 | 4.362E-06 | 5.300E-06 | 1.300E-05 |
| 24372 | 1219 | E1219K | D | 6.2e-06 | 8.1e-06 | 4e-06 | 6.100E-06 | 2.052E-06 | 4.000E-06 | 8.100E-06 |
| 24373 | 1219 | E1219M | D | 1.7e-05 | 9.2e-06 | 6.3e-06 | 1.083E-05 | 5.534E-06 | 6.300E-06 | 1.700E-05 |
| 24374 | 1219 | E1219F | D | 7e-06 | 1.1e-05 | 7.4e-06 | 8.467E-06 | 2.203E-06 | 7.000E-06 | 1.100E-05 |
| 24375 | 1219 | E1219P | D | 9.3e-06 | 9e-06 | 7.8e-06 | 8.700E-06 | 7.937E-07 | 7.800E-06 | 9.300E-06 |
| 24376 | 1219 | E1219S | D | 9.3e-06 | 6.4e-06 | 7.5e-06 | 7.733E-06 | 1.464E-06 | 6.400E-06 | 9.300E-06 |
| 24377 | 1219 | E1219T | D | 6.9e-06 | 6.1e-06 | 1.1e-05 | 8.000E-06 | 2.629E-06 | 6.100E-06 | 1.100E-05 |
| 24378 | 1219 | E1219W | D | 9.6e-06 | 1.6e-05 | 7e-06 | 1.087E-05 | 4.632E-06 | 7.000E-06 | 1.600E-05 |
| 24379 | 1219 | E1219Y | D | 5.1e-06 | 1.2e-05 | 6.5e-06 | 7.867E-06 | 3.647E-06 | 5.100E-06 | 1.200E-05 |
| 24380 | 1219 | E1219V | D | 5.1e-06 | 5e-06 | 3.5e-06 | 4.533E-06 | 8.963E-07 | 3.500E-06 | 5.100E-06 |
| 24381 | 1220 | S1220A | D | 7e-06 | 1.1e-05 | 6.6e-06 | 8.200E-06 | 2.433E-06 | 6.600E-06 | 1.100E-05 |
| 24382 | 1220 | S1220R | D | 1.3e-05 | 9.7e-06 | 7.2e-06 | 9.967E-06 | 2.909E-06 | 7.200E-06 | 1.300E-05 |
| 24383 | 1220 | S1220N | D | 1.2e-05 | 5e-06 | 6.2e-06 | 7.733E-06 | 3.743E-06 | 5.000E-06 | 1.200E-05 |
| 24384 | 1220 | S1220D | D | 6.6e-06 | 6.3e-06 | 1.2e-05 | 8.300E-06 | 3.208E-06 | 6.300E-06 | 1.200E-05 |
| 24385 | 1220 | S1220C | D | 1.2e-05 | 9e-06 | 5.1e-06 | 8.700E-06 | 3.460E-06 | 5.100E-06 | 1.200E-05 |
| 24386 | 1220 | S1220E | D | 9.5e-06 | 1.2e-05 | 3.3e-06 | 8.267E-06 | 4.479E-06 | 3.300E-06 | 1.200E-05 |
| 24387 | 1220 | S1220Q | D | 6.7e-06 | 4.9e-06 | 9.3e-06 | 6.967E-06 | 2.212E-06 | 4.900E-06 | 9.300E-06 |
| 24388 | 1220 | S1220G | D | 7.4e-06 | 4.4e-06 | 7e-06 | 6.267E-06 | 1.629E-06 | 4.400E-06 | 7.400E-06 |
| 24389 | 1220 | S1220H | D | 6.5e-06 | 8.7e-06 | 8.7e-06 | 7.967E-06 | 1.270E-06 | 6.500E-06 | 8.700E-06 |
| 24390 | 1220 | S1220I | D | 6.6e-06 | 7e-06 | 7.7e-06 | 7.100E-06 | 5.568E-07 | 6.600E-06 | 7.700E-06 |
| 24391 | 1220 | S1220L | D | 1.1e-05 | 6.4e-06 | 1.1e-05 | 9.467E-06 | 2.656E-06 | 6.400E-06 | 1.100E-05 |
| 24392 | 1220 | S1220K | D | 6.7e-06 | 6.6e-06 | 1.1e-05 | 8.100E-06 | 2.512E-06 | 6.600E-06 | 1.100E-05 |
| 24393 | 1220 | S1220M | D | 9.7e-06 | 6.8e-06 | 4.9e-06 | 7.133E-06 | 2.417E-06 | 4.900E-06 | 9.700E-06 |
| 24394 | 1220 | S1220F | D | 6.1e-06 | 7.8e-06 | 1e-05 | 7.967E-06 | 1.955E-06 | 6.100E-06 | 1.000E-05 |

| | | | | | | | | | | |
|---|---|---|---|---|---|---|---|---|---|---|
| 24395 | 1220 | S1220P | D | 1.6e-05 | 9.1e-06 | 1.1e-05 | 1.203E-05 | 3.564E-06 | 9.100E-06 | 1.600E-05 |
| 24396 | 1220 | S1220S | D | 5.9e-06 | 7.2e-06 | 7.7e-06 | 6.933E-06 | 9.292E-07 | 5.900E-06 | 7.700E-06 |
| 24397 | 1220 | S1220T | D | 5.2e-06 | 8.8e-06 | 8.6e-06 | 7.533E-06 | 2.023E-06 | 5.200E-06 | 8.800E-06 |
| 24398 | 1220 | S1220W | D | 9.2e-06 | 1.2e-05 | 1.1e-05 | 1.073E-05 | 1.419E-06 | 9.200E-06 | 1.200E-05 |
| 24399 | 1220 | S1220Y | D | 8.5e-06 | 8.1e-06 | 5.8e-06 | 7.467E-06 | 1.457E-06 | 5.800E-06 | 8.500E-06 |
| 24400 | 1220 | S1220V | D | 5e-06 | 8.7e-06 | 9.4e-06 | 7.700E-06 | 2.364E-06 | 5.000E-06 | 9.400E-06 |
| 24401 | 1221 | V1221A | D | 4.4e-06 | 1e-05 | 9.4e-06 | 7.933E-06 | 3.075E-06 | 4.400E-06 | 1.000E-05 |
| 24402 | 1221 | V1221R | D | 9.9e-06 | 8.3e-06 | 8e-06 | 8.733E-06 | 1.021E-06 | 8.000E-06 | 9.900E-06 |
| 24403 | 1221 | V1221N | D | 8.3e-06 | 1.3e-05 | 5.8e-06 | 9.033E-06 | 3.656E-06 | 5.800E-06 | 1.300E-05 |
| 24404 | 1221 | V1221D | D | 6.7e-06 | 5.7e-06 | 5.9e-06 | 6.100E-06 | 5.292E-07 | 5.700E-06 | 6.700E-06 |
| 24405 | 1221 | V1221C | D | 8.5e-06 | 5.1e-06 | 9.4e-06 | 7.667E-06 | 2.268E-06 | 5.100E-06 | 9.400E-06 |
| 24406 | 1221 | V1221E | D | 9.9e-06 | 6.1e-06 | 7.7e-06 | 7.900E-06 | 1.908E-06 | 6.100E-06 | 9.900E-06 |
| 24407 | 1221 | V1221Q | D | 9e-06 | 7.9e-06 | 1e-05 | 8.967E-06 | 1.050E-06 | 7.900E-06 | 1.000E-05 |
| 24408 | 1221 | V1221G | D | 5.2e-06 | 7.4e-06 | 7.9e-06 | 6.833E-06 | 1.436E-06 | 5.200E-06 | 7.900E-06 |
| 24409 | 1221 | V1221H | D | 8.6e-06 | 1.6e-05 | 1.2e-05 | 1.220E-05 | 3.704E-06 | 8.600E-06 | 1.600E-05 |
| 24410 | 1221 | V1221I | D | 5.8e-06 | 5.7e-06 | 7.6e-06 | 6.367E-06 | 1.069E-06 | 5.700E-06 | 7.600E-06 |
| 24411 | 1221 | V1221L | D | 8.2e-06 | 5.8e-06 | 5e-06 | 6.333E-06 | 1.665E-06 | 5.000E-06 | 8.200E-06 |
| 24412 | 1221 | V1221K | D | 7.9e-06 | 1.3e-05 | 9.2e-06 | 1.003E-05 | 2.650E-06 | 7.900E-06 | 1.300E-05 |
| 24413 | 1221 | V1221M | D | 9e-06 | 8.8e-06 | 1.2e-05 | 9.933E-06 | 1.793E-06 | 8.800E-06 | 1.200E-05 |
| 24414 | 1221 | V1221F | D | 9.7e-06 | 6.2e-06 | 1.1e-05 | 8.967E-06 | 2.483E-06 | 6.200E-06 | 1.100E-05 |
| 24415 | 1221 | V1221P | D | 4.5e-06 | 6.5e-06 | 6.3e-06 | 5.767E-06 | 1.102E-06 | 4.500E-06 | 6.500E-06 |
| 24416 | 1221 | V1221S | D | 4.3e-06 | 9.3e-06 | 6.4e-06 | 6.667E-06 | 2.511E-06 | 4.300E-06 | 9.300E-06 |
| 24417 | 1221 | V1221T | D | 8e-06 | 4.6e-06 | 5.3e-06 | 5.967E-06 | 1.795E-06 | 4.600E-06 | 8.000E-06 |
| 24418 | 1221 | V1221W | D | 8.3e-06 | 3.1e-06 | 9.3e-06 | 6.900E-06 | 3.329E-06 | 3.100E-06 | 9.300E-06 |
| 24419 | 1221 | V1221Y | D | 5.5e-06 | 5.5e-06 | 8e-06 | 6.333E-06 | 1.443E-06 | 5.500E-06 | 8.000E-06 |
| 24420 | 1221 | V1221V | D | 7.6e-06 | 7.4e-06 | 6e-06 | 7.000E-06 | 8.718E-07 | 6.000E-06 | 7.600E-06 |
| 24421 | 1222 | T1222A | D | 6.1e-06 | 7.7e-06 | 1e-05 | 7.933E-06 | 1.960E-06 | 6.100E-06 | 1.000E-05 |
| 24422 | 1222 | T1222R | D | 8e-06 | 1e-05 | 8.3e-06 | 8.767E-06 | 1.079E-06 | 8.000E-06 | 1.000E-05 |
| 24423 | 1222 | T1222N | D | 1e-05 | 7.3e-06 | 6.5e-06 | 7.933E-06 | 1.834E-06 | 6.500E-06 | 1.000E-05 |
| 24424 | 1222 | T1222D | D | 7.2e-06 | 1.6e-05 | 7.2e-06 | 1.013E-05 | 5.081E-06 | 7.200E-06 | 1.600E-05 |
| 24425 | 1222 | T1222C | D | 8.2e-06 | 8.5e-06 | 4.1e-06 | 6.933E-06 | 2.458E-06 | 4.100E-06 | 8.500E-06 |
| 24426 | 1222 | T1222E | D | 8.9e-06 | 9.9e-06 | 8.6e-06 | 9.133E-06 | 6.807E-07 | 8.600E-06 | 9.900E-06 |
| 24427 | 1222 | T1222Q | D | 7.5e-06 | 1.3e-05 | 1.2e-05 | 1.083E-05 | 2.930E-06 | 7.500E-06 | 1.300E-05 |
| 24428 | 1222 | T1222G | D | 7.8e-06 | 1.1e-05 | 8e-06 | 8.933E-06 | 1.793E-06 | 7.800E-06 | 1.100E-05 |
| 24429 | 1222 | T1222H | D | 1.1e-05 | 8e-06 | 6.7e-06 | 8.567E-06 | 2.205E-06 | 6.700E-06 | 1.100E-05 |
| 24430 | 1222 | T1222I | D | 7.5e-06 | 7.2e-06 | 1.4e-05 | 9.567E-06 | 3.842E-06 | 7.200E-06 | 1.400E-05 |
| 24431 | 1222 | T1222L | D | 7.7e-06 | 1.3e-05 | 4.8e-06 | 8.500E-06 | 4.158E-06 | 4.800E-06 | 1.300E-05 |
| 24432 | 1222 | T1222K | D | 1e-05 | 7.9e-06 | 5e-06 | 7.633E-06 | 2.511E-06 | 5.000E-06 | 1.000E-05 |
| 24433 | 1222 | T1222M | D | 1.1e-05 | 7.9e-06 | 5.3e-06 | 8.067E-06 | 2.854E-06 | 5.300E-06 | 1.100E-05 |
| 24434 | 1222 | T1222F | D | 9.5e-06 | 9.3e-06 | 8.7e-06 | 9.167E-06 | 4.163E-07 | 8.700E-06 | 9.500E-06 |
| 24435 | 1222 | T1222P | D | 7.8e-06 | 1.2e-05 | 7.4e-06 | 9.067E-06 | 2.548E-06 | 7.400E-06 | 1.200E-05 |
| 24436 | 1222 | T1222S | D | 9.5e-06 | 9.9e-06 | 8.5e-06 | 9.300E-06 | 7.211E-07 | 8.500E-06 | 9.900E-06 |
| 24437 | 1222 | T1222T | D | 8.6e-06 | 6.2e-06 | 1.1e-05 | 8.600E-06 | 2.400E-06 | 6.200E-06 | 1.100E-05 |
| 24438 | 1222 | T1222W | D | 6.9e-06 | 1.4e-05 | 5e-06 | 8.633E-06 | 4.744E-06 | 5.000E-06 | 1.400E-05 |
| 24439 | 1222 | T1222Y | D | 6.4e-06 | 1.3e-05 | 8e-06 | 9.133E-06 | 3.443E-06 | 6.400E-06 | 1.300E-05 |
| 24440 | 1222 | T1222V | D | 5.7e-06 | 1.2e-05 | 7e-06 | 8.233E-06 | 3.326E-06 | 5.700E-06 | 1.200E-05 |
| 24441 | 1223 | E1223A | D | 4.6e-06 | 5.8e-06 | 8.8e-06 | 6.400E-06 | 2.163E-06 | 4.600E-06 | 8.800E-06 |
| 24442 | 1223 | E1223R | D | 8e-06 | 8.2e-06 | 9.6e-06 | 8.600E-06 | 8.718E-07 | 8.000E-06 | 9.600E-06 |
| 24443 | 1223 | E1223N | D | 8.9e-06 | 6.8e-06 | 6.1e-06 | 7.267E-06 | 1.457E-06 | 6.100E-06 | 8.900E-06 |
| 24444 | 1223 | E1223D | D | 9.3e-06 | 1.3e-05 | 9e-06 | 1.043E-05 | 2.228E-06 | 9.000E-06 | 1.300E-05 |
| 24445 | 1223 | E1223C | D | 1.2e-05 | 7.3e-06 | 8.2e-06 | 9.167E-06 | 2.495E-06 | 7.300E-06 | 1.200E-05 |
| 24446 | 1223 | E1223E | D | 6.5e-06 | 7.9e-06 | 6.7e-06 | 7.033E-06 | 7.572E-07 | 6.500E-06 | 7.900E-06 |
| 24447 | 1223 | E1223Q | D | 5.1e-06 | 6.8e-06 | 1.1e-05 | 7.633E-06 | 3.037E-06 | 5.100E-06 | 1.100E-05 |
| 24448 | 1223 | E1223G | D | 9.3e-06 | 7.7e-06 | 6.7e-06 | 7.900E-06 | 1.311E-06 | 6.700E-06 | 9.300E-06 |
| 24449 | 1223 | E1223H | D | 6.3e-06 | 6.4e-06 | 6.3e-06 | 6.333E-06 | 5.774E-08 | 6.300E-06 | 6.400E-06 |
| 24450 | 1223 | E1223I | D | 7.9e-06 | 1e-05 | 6.5e-06 | 8.133E-06 | 1.762E-06 | 6.500E-06 | 1.000E-05 |
| 24451 | 1223 | E1223L | D | 5.1e-06 | 1.3e-05 | 8.3e-06 | 8.800E-06 | 3.974E-06 | 5.100E-06 | 1.300E-05 |
| 24452 | 1223 | E1223K | D | 6.6e-06 | 8.7e-06 | 5.1e-06 | 6.800E-06 | 1.808E-06 | 5.100E-06 | 8.700E-06 |
| 24453 | 1223 | E1223M | D | 1e-05 | 9.8e-06 | 1e-05 | 9.933E-06 | 1.155E-07 | 9.800E-06 | 1.000E-05 |
| 24454 | 1223 | E1223F | D | 5.6e-06 | 5e-06 | 9.7e-06 | 6.767E-06 | 2.558E-06 | 5.000E-06 | 9.700E-06 |
| 24455 | 1223 | E1223P | D | 7.7e-06 | 8.3e-06 | 5.9e-06 | 7.300E-06 | 1.249E-06 | 5.900E-06 | 8.300E-06 |
| 24456 | 1223 | E1223S | D | 7.1e-06 | 6e-06 | 5e-06 | 6.033E-06 | 1.050E-06 | 5.000E-06 | 7.100E-06 |
| 24457 | 1223 | E1223T | D | 5.9e-06 | 5e-06 | 3.5e-06 | 4.800E-06 | 1.212E-06 | 3.500E-06 | 5.900E-06 |
| 24458 | 1223 | E1223W | D | 9.6e-06 | 8.4e-06 | 5.7e-06 | 7.900E-06 | 1.997E-06 | 5.700E-06 | 9.600E-06 |
| 24459 | 1223 | E1223Y | D | 6.5e-06 | 7.2e-06 | 6.3e-06 | 6.667E-06 | 4.726E-07 | 6.300E-06 | 7.200E-06 |
| 24460 | 1223 | E1223V | D | 8.4e-06 | 8.7e-06 | 7.9e-06 | 8.333E-06 | 4.041E-07 | 7.900E-06 | 8.700E-06 |
| 24461 | 1224 | Q1224A | D | 7.6e-06 | 5.1e-06 | 8.9e-06 | 7.200E-06 | 1.931E-06 | 5.100E-06 | 8.900E-06 |
| 24462 | 1224 | Q1224R | D | 5.8e-06 | 9.4e-06 | 9.6e-06 | 8.267E-06 | 2.139E-06 | 5.800E-06 | 9.600E-06 |
| 24463 | 1224 | Q1224N | D | 1e-05 | 5.9e-06 | 7.1e-06 | 7.667E-06 | 2.108E-06 | 5.900E-06 | 1.000E-05 |
| 24464 | 1224 | Q1224D | D | 7.9e-06 | 1.3e-05 | 7.8e-06 | 9.567E-06 | 2.974E-06 | 7.800E-06 | 1.300E-05 |
| 24465 | 1224 | Q1224C | D | 1.2e-05 | 6.3e-06 | 6.5e-06 | 8.267E-06 | 3.235E-06 | 6.300E-06 | 1.200E-05 |
| 24466 | 1224 | Q1224E | D | 1.3e-05 | 1.4e-05 | 8.9e-06 | 1.197E-05 | 2.702E-06 | 8.900E-06 | 1.400E-05 |
| 24467 | 1224 | Q1224Q | D | 1e-05 | 1.1e-05 | 9.3e-06 | 1.010E-05 | 8.544E-07 | 9.300E-06 | 1.100E-05 |
| 24468 | 1224 | Q1224G | D | 6.4e-06 | 5.9e-06 | 9.9e-06 | 7.400E-06 | 2.179E-06 | 5.900E-06 | 9.900E-06 |
| 24469 | 1224 | Q1224H | D | 1.7e-05 | 1.7e-05 | 5.2e-06 | 1.307E-05 | 6.813E-06 | 5.200E-06 | 1.700E-05 |
| 24470 | 1224 | Q1224I | D | 9.3e-06 | 7.5e-06 | 1e-05 | 8.933E-06 | 1.290E-06 | 7.500E-06 | 1.000E-05 |
| 24471 | 1224 | Q1224L | D | 1.4e-05 | 6.6e-06 | 7.3e-06 | 9.300E-06 | 4.085E-06 | 6.600E-06 | 1.400E-05 |
| 24472 | 1224 | Q1224K | D | 1.1e-05 | 7.2e-06 | 6.2e-06 | 8.133E-06 | 2.532E-06 | 6.200E-06 | 1.100E-05 |
| 24473 | 1224 | Q1224M | D | 9.6e-06 | 6.9e-06 | 5.7e-06 | 7.400E-06 | 1.997E-06 | 5.700E-06 | 9.600E-06 |
| 24474 | 1224 | Q1224F | D | 8e-06 | 8.6e-06 | 9.9e-06 | 8.833E-06 | 9.713E-07 | 8.000E-06 | 9.900E-06 |
| 24475 | 1224 | Q1224P | D | 1.4e-05 | 1e-05 | 5.9e-06 | 9.967E-06 | 4.050E-06 | 5.900E-06 | 1.400E-05 |
| 24476 | 1224 | Q1224S | D | 8.9e-06 | 7.7e-06 | 8.6e-06 | 8.400E-06 | 6.245E-07 | 7.700E-06 | 8.900E-06 |
| 24477 | 1224 | Q1224T | D | 7.9e-06 | 9.6e-06 | 7.6e-06 | 8.367E-06 | 1.079E-06 | 7.600E-06 | 9.600E-06 |
| 24478 | 1224 | Q1224W | D | 9.5e-06 | 5.8e-06 | 4.8e-06 | 6.700E-06 | 2.476E-06 | 4.800E-06 | 9.500E-06 |
| 24479 | 1224 | Q1224Y | D | 6.5e-06 | 8.7e-06 | 6.6e-06 | 7.267E-06 | 1.242E-06 | 6.500E-06 | 8.700E-06 |

| | | | | | | | | | | |
|---|---|---|---|---|---|---|---|---|---|---|
| 24480 | 1224 | Q1224V | D | 1.9e-05 | 1e-05 | 6e-06 | 1.167E-05 | 6.658E-06 | 6.000E-06 | 1.900E-05 |
| 24481 | 1225 | D1225A | D | 6.5e-06 | 8.5e-06 | 4.2e-06 | 6.400E-06 | 2.152E-06 | 4.200E-06 | 8.500E-06 |
| 24482 | 1225 | D1225R | D | 5.7e-06 | 6.4e-06 | 7.4e-06 | 6.500E-06 | 8.544E-07 | 5.700E-06 | 7.400E-06 |
| 24483 | 1225 | D1225N | D | 1.3e-05 | 9.1e-06 | 5.3e-06 | 9.133E-06 | 3.850E-06 | 5.300E-06 | 1.300E-05 |
| 24484 | 1225 | D1225D | D | 1.8e-05 | 7e-06 | 8e-06 | 1.100E-05 | 6.083E-06 | 7.000E-06 | 1.800E-05 |
| 24485 | 1225 | D1225C | D | 8.4e-06 | 9.4e-06 | 1.2e-05 | 9.933E-06 | 1.858E-06 | 8.400E-06 | 1.200E-05 |
| 24486 | 1225 | D1225E | D | 9.4e-06 | 5.5e-06 | 5.9e-06 | 6.933E-06 | 2.146E-06 | 5.500E-06 | 9.400E-06 |
| 24487 | 1225 | D1225Q | D | 7.4e-06 | 6.2e-06 | 7.4e-06 | 7.000E-06 | 6.928E-07 | 6.200E-06 | 7.400E-06 |
| 24488 | 1225 | D1225G | D | 8.2e-06 | 7.5e-06 | 3.4e-06 | 6.367E-06 | 2.593E-06 | 3.400E-06 | 8.200E-06 |
| 24489 | 1225 | D1225H | D | 9.2e-06 | 6.2e-06 | 8.3e-06 | 7.900E-06 | 1.539E-06 | 6.200E-06 | 9.200E-06 |
| 24490 | 1225 | D1225I | D | 7.4e-06 | 5.1e-06 | 9.3e-06 | 7.267E-06 | 2.103E-06 | 5.100E-06 | 9.300E-06 |
| 24491 | 1225 | D1225L | D | 6e-06 | 5.5e-06 | 7.9e-06 | 6.467E-06 | 1.266E-06 | 5.500E-06 | 7.900E-06 |
| 24492 | 1225 | D1225K | D | 1.3e-05 | 5.9e-06 | 6.8e-06 | 8.567E-06 | 3.866E-06 | 5.900E-06 | 1.300E-05 |
| 24493 | 1225 | D1225M | D | 1.3e-05 | 8.4e-06 | 1e-05 | 1.047E-05 | 2.335E-06 | 8.400E-06 | 1.300E-05 |
| 24494 | 1225 | D1225F | D | 8.3e-06 | 5.6e-06 | 6.8e-06 | 6.900E-06 | 1.353E-06 | 5.600E-06 | 8.300E-06 |
| 24495 | 1225 | D1225P | D | 6.1e-06 | 6.6e-06 | 7.3e-06 | 6.667E-06 | 6.028E-07 | 6.100E-06 | 7.300E-06 |
| 24496 | 1225 | D1225S | D | 6.9e-06 | 8.9e-06 | 7.7e-06 | 7.833E-06 | 1.007E-06 | 6.900E-06 | 8.900E-06 |
| 24497 | 1225 | D1225T | D | 8.6e-06 | 1e-05 | 1.1e-05 | 9.867E-06 | 1.206E-06 | 8.600E-06 | 1.100E-05 |
| 24498 | 1225 | D1225W | D | 9.9e-06 | 5.4e-06 | 7e-06 | 7.433E-06 | 2.281E-06 | 5.400E-06 | 9.900E-06 |
| 24499 | 1225 | D1225Y | D | 8.9e-06 | 6.2e-06 | 1.3e-05 | 9.367E-06 | 3.424E-06 | 6.200E-06 | 1.300E-05 |
| 24500 | 1225 | D1225V | D | 1.2e-05 | 6.2e-06 | 6.8e-06 | 8.333E-06 | 3.190E-06 | 6.200E-06 | 1.200E-05 |
| 24501 | 1226 | S1226A | D | 7.8e-06 | 1.3e-05 | 6.5e-06 | 9.100E-06 | 3.439E-06 | 6.500E-06 | 1.300E-05 |
| 24502 | 1226 | S1226R | D | 9.2e-06 | 6.7e-06 | 9e-06 | 8.300E-06 | 1.389E-06 | 6.700E-06 | 9.200E-06 |
| 24503 | 1226 | S1226N | D | 5.8e-06 | 7.1e-06 | 1e-05 | 7.633E-06 | 2.150E-06 | 5.800E-06 | 1.000E-05 |
| 24504 | 1226 | S1226D | D | 6.3e-06 | 1e-05 | 1.2e-05 | 9.433E-06 | 2.892E-06 | 6.300E-06 | 1.200E-05 |
| 24505 | 1226 | S1226C | D | 6.5e-06 | 7e-06 | 6.3e-06 | 6.600E-06 | 3.606E-07 | 6.300E-06 | 7.000E-06 |
| 24506 | 1226 | S1226E | D | 1e-05 | 7.4e-06 | 1.2e-05 | 9.800E-06 | 2.307E-06 | 7.400E-06 | 1.200E-05 |
| 24507 | 1226 | S1226Q | D | 7.6e-06 | 1.6e-05 | 4.3e-06 | 9.300E-06 | 6.032E-06 | 4.300E-06 | 1.600E-05 |
| 24508 | 1226 | S1226G | D | 1e-05 | 7.8e-06 | 8e-06 | 8.600E-06 | 1.217E-06 | 7.800E-06 | 1.000E-05 |
| 24509 | 1226 | S1226H | D | 1e-05 | 5.8e-06 | 1.5e-05 | 1.027E-05 | 4.606E-06 | 5.800E-06 | 1.500E-05 |
| 24510 | 1226 | S1226I | D | 4.8e-06 | 9.6e-06 | 7.5e-06 | 7.300E-06 | 2.406E-06 | 4.800E-06 | 9.600E-06 |
| 24511 | 1226 | S1226L | D | 9e-06 | 9.8e-06 | 1.2e-05 | 1.027E-05 | 1.553E-06 | 9.000E-06 | 1.200E-05 |
| 24512 | 1226 | S1226K | D | 6.1e-06 | 1.3e-05 | 6.5e-06 | 8.533E-06 | 3.873E-06 | 6.100E-06 | 1.300E-05 |
| 24513 | 1226 | S1226M | D | 1.4e-05 | 8.8e-06 | 5.9e-06 | 9.567E-06 | 4.104E-06 | 5.900E-06 | 1.400E-05 |
| 24514 | 1226 | S1226F | D | 7e-06 | 8.7e-06 | 9.5e-06 | 8.400E-06 | 1.277E-06 | 7.000E-06 | 9.500E-06 |
| 24515 | 1226 | S1226P | D | 8.6e-06 | 1.2e-05 | 1e-05 | 1.020E-05 | 1.709E-06 | 8.600E-06 | 1.200E-05 |
| 24516 | 1226 | S1226S | D | 4.6e-06 | 1.3e-05 | 4.5e-06 | 7.367E-06 | 4.879E-06 | 4.500E-06 | 1.300E-05 |
| 24517 | 1226 | S1226T | D | 1.7e-05 | 6.2e-06 | 7e-06 | 1.007E-05 | 6.018E-06 | 6.200E-06 | 1.700E-05 |
| 24518 | 1226 | S1226W | D | 7.1e-06 | 4.6e-06 | 1.3e-05 | 8.233E-06 | 4.313E-06 | 4.600E-06 | 1.300E-05 |
| 24519 | 1226 | S1226Y | D | 9.9e-06 | 1.4e-05 | 7.5e-06 | 1.047E-05 | 3.287E-06 | 7.500E-06 | 1.400E-05 |
| 24520 | 1226 | S1226V | D | 6e-06 | 8.6e-06 | 1.1e-05 | 8.533E-06 | 2.501E-06 | 6.000E-06 | 1.100E-05 |
| 24521 | 1227 | K1227A | D | 6.1e-06 | 1.1e-05 | 8.8e-06 | 8.633E-06 | 2.454E-06 | 6.100E-06 | 1.100E-05 |
| 24522 | 1227 | K1227R | D | 8e-06 | 5.8e-06 | 1.2e-05 | 8.600E-06 | 3.143E-06 | 5.800E-06 | 1.200E-05 |
| 24523 | 1227 | K1227N | D | 8.6e-06 | 3.8e-06 | 8.8e-06 | 7.067E-06 | 2.831E-06 | 3.800E-06 | 8.800E-06 |
| 24524 | 1227 | K1227D | D | 1.7e-05 | 8.1e-06 | 1.1e-05 | 1.203E-05 | 4.539E-06 | 8.100E-06 | 1.700E-05 |
| 24525 | 1227 | K1227C | D | 8.3e-06 | 8.6e-06 | 6.8e-06 | 7.900E-06 | 9.644E-07 | 6.800E-06 | 8.600E-06 |
| 24526 | 1227 | K1227E | D | 7e-06 | 1.2e-05 | 8.7e-06 | 9.233E-06 | 2.542E-06 | 7.000E-06 | 1.200E-05 |
| 24527 | 1227 | K1227Q | D | 9e-06 | 7.4e-06 | 4.9e-06 | 7.100E-06 | 2.066E-06 | 4.900E-06 | 9.000E-06 |
| 24528 | 1227 | K1227G | D | 3.4e-06 | 7.2e-06 | 8.1e-06 | 6.233E-06 | 2.495E-06 | 3.400E-06 | 8.100E-06 |
| 24529 | 1227 | K1227H | D | 1.3e-05 | 9.2e-06 | 1.5e-05 | 1.240E-05 | 2.946E-06 | 9.200E-06 | 1.500E-05 |
| 24530 | 1227 | K1227I | D | 7.1e-06 | 7.5e-06 | 7.6e-06 | 7.400E-06 | 2.646E-07 | 7.100E-06 | 7.600E-06 |
| 24531 | 1227 | K1227L | D | 9.7e-06 | 5.9e-06 | 1e-05 | 8.533E-06 | 2.285E-06 | 5.900E-06 | 1.000E-05 |
| 24532 | 1227 | K1227K | D | 8.4e-06 | 5.9e-06 | 7.9e-06 | 7.400E-06 | 1.323E-06 | 5.900E-06 | 8.400E-06 |
| 24533 | 1227 | K1227M | D | 5.4e-06 | 1.1e-05 | 8.8e-06 | 8.400E-06 | 2.821E-06 | 5.400E-06 | 1.100E-05 |
| 24534 | 1227 | K1227F | D | 6.3e-06 | 8.1e-06 | 7.2e-06 | 7.200E-06 | 9.000E-07 | 6.300E-06 | 8.100E-06 |
| 24535 | 1227 | K1227P | D | 1e-05 | 7.5e-06 | 8e-06 | 8.500E-06 | 1.323E-06 | 7.500E-06 | 1.000E-05 |
| 24536 | 1227 | K1227S | D | 8.8e-06 | 1.4e-05 | 6.3e-06 | 9.700E-06 | 3.928E-06 | 6.300E-06 | 1.400E-05 |
| 24537 | 1227 | K1227T | D | 6.5e-06 | 7.2e-06 | 8.7e-06 | 7.467E-06 | 1.124E-06 | 6.500E-06 | 8.700E-06 |
| 24538 | 1227 | K1227W | D | 8.5e-06 | 9.8e-06 | 9.1e-06 | 9.133E-06 | 6.506E-07 | 8.500E-06 | 9.800E-06 |
| 24539 | 1227 | K1227Y | D | 8.3e-06 | 8e-06 | 1.3e-05 | 9.767E-06 | 2.804E-06 | 8.000E-06 | 1.300E-05 |
| 24540 | 1227 | K1227V | D | 8.9e-06 | 9.5e-06 | 8.2e-06 | 8.867E-06 | 6.506E-07 | 8.200E-06 | 9.500E-06 |
| 24541 | 1228 | D1228A | D | 1e-05 | 9.3e-06 | 7.7e-06 | 9.000E-06 | 1.179E-06 | 7.700E-06 | 1.000E-05 |
| 24542 | 1228 | D1228R | D | 6.6e-06 | 7e-06 | 9.1e-06 | 7.567E-06 | 1.343E-06 | 6.600E-06 | 9.100E-06 |
| 24543 | 1228 | D1228N | D | 1e-05 | 7.5e-06 | 9.7e-06 | 9.067E-06 | 1.365E-06 | 7.500E-06 | 1.000E-05 |
| 24544 | 1228 | D1228D | D | 7.9e-06 | 7.7e-06 | 8.4e-06 | 8.000E-06 | 3.606E-07 | 7.700E-06 | 8.400E-06 |
| 24545 | 1228 | D1228C | D | 5.7e-06 | 9.1e-06 | 7.4e-06 | 7.400E-06 | 1.700E-06 | 5.700E-06 | 9.100E-06 |
| 24546 | 1228 | D1228E | D | 5.2e-06 | 1.2e-05 | 6.3e-06 | 7.833E-06 | 3.650E-06 | 5.200E-06 | 1.200E-05 |
| 24547 | 1228 | D1228Q | D | 8.4e-06 | 8.9e-06 | 8.7e-06 | 8.667E-06 | 2.517E-07 | 8.400E-06 | 8.900E-06 |
| 24548 | 1228 | D1228G | D | 5.5e-06 | 9.8e-06 | 8.2e-06 | 7.833E-06 | 2.173E-06 | 5.500E-06 | 9.800E-06 |
| 24549 | 1228 | D1228H | D | 7.1e-06 | 6.4e-06 | 6.7e-06 | 6.733E-06 | 3.512E-07 | 6.400E-06 | 7.100E-06 |
| 24550 | 1228 | D1228I | D | 6.2e-06 | 9.2e-06 | 1e-05 | 8.467E-06 | 2.003E-06 | 6.200E-06 | 1.000E-05 |
| 24551 | 1228 | D1228L | D | 6.4e-06 | 8.6e-06 | 5.5e-06 | 6.833E-06 | 1.595E-06 | 5.500E-06 | 8.600E-06 |
| 24552 | 1228 | D1228K | D | 8.9e-06 | 7.5e-06 | 6.8e-06 | 7.733E-06 | 1.069E-06 | 6.800E-06 | 8.900E-06 |
| 24553 | 1228 | D1228M | D | 1.2e-05 | 5.1e-06 | 9.3e-06 | 8.800E-06 | 3.477E-06 | 5.100E-06 | 1.200E-05 |
| 24554 | 1228 | D1228F | D | 7.4e-06 | 7.8e-06 | 5.8e-06 | 7.000E-06 | 1.058E-06 | 5.800E-06 | 7.800E-06 |
| 24555 | 1228 | D1228P | D | 9.8e-06 | 1.1e-05 | 9.5e-06 | 1.010E-05 | 7.937E-07 | 9.500E-06 | 1.100E-05 |
| 24556 | 1228 | D1228S | D | 3.2e-06 | 5.4e-06 | 6.2e-06 | 4.933E-06 | 1.553E-06 | 3.200E-06 | 6.200E-06 |
| 24557 | 1228 | D1228T | D | 6.3e-06 | 9.8e-06 | 6.4e-06 | 7.500E-06 | 1.992E-06 | 6.300E-06 | 9.800E-06 |
| 24558 | 1228 | D1228W | D | 9e-06 | 1.4e-05 | 4.5e-06 | 9.167E-06 | 4.752E-06 | 4.500E-06 | 1.400E-05 |
| 24559 | 1228 | D1228Y | D | 7.2e-06 | 1.3e-05 | 1e-05 | 1.007E-05 | 2.901E-06 | 7.200E-06 | 1.300E-05 |
| 24560 | 1228 | D1228V | D | 1.1e-05 | 1e-05 | 6e-06 | 9.000E-06 | 2.646E-06 | 6.000E-06 | 1.100E-05 |
| 24561 | 1229 | S1229A | D | 8.9e-06 | 7.6e-06 | 8.1e-06 | 8.200E-06 | 6.557E-07 | 7.600E-06 | 8.900E-06 |
| 24562 | 1229 | S1229R | D | 7.3e-06 | 6.3e-06 | 6.4e-06 | 6.667E-06 | 5.508E-07 | 6.300E-06 | 7.300E-06 |
| 24563 | 1229 | S1229N | D | 7.9e-06 | 1e-05 | 6.2e-06 | 8.033E-06 | 1.904E-06 | 6.200E-06 | 1.000E-05 |
| 24564 | 1229 | S1229D | D | 5.9e-06 | 7e-06 | 1.2e-05 | 8.300E-06 | 3.251E-06 | 5.900E-06 | 1.200E-05 |

| | | | | | | | | | | |
|---|---|---|---|---|---|---|---|---|---|---|
| 24565 | 1229 | S1229C | D | 1.2e-05 | 9.4e-06 | 8.5e-06 | 9.967E-06 | 1.818E-06 | 8.500E-06 | 1.200E-05 |
| 24566 | 1229 | S1229E | D | 8.2e-06 | 4.1e-06 | 9.8e-06 | 7.367E-06 | 2.940E-06 | 4.100E-06 | 9.800E-06 |
| 24567 | 1229 | S1229Q | D | 8.6e-06 | 1.2e-05 | 6.7e-06 | 9.100E-06 | 2.685E-06 | 6.700E-06 | 1.200E-05 |
| 24568 | 1229 | S1229G | D | 8.2e-06 | 8.8e-06 | 6.5e-06 | 7.833E-06 | 1.193E-06 | 6.500E-06 | 8.800E-06 |
| 24569 | 1229 | S1229H | D | 1.5e-05 | 5.2e-06 | 1.4e-05 | 1.140E-05 | 5.393E-06 | 5.200E-06 | 1.500E-05 |
| 24570 | 1229 | S1229I | D | 5e-06 | 6.7e-06 | 4.5e-06 | 5.400E-06 | 1.153E-06 | 4.500E-06 | 6.700E-06 |
| 24571 | 1229 | S1229L | D | 6e-06 | 6.3e-06 | 1.2e-05 | 8.100E-06 | 3.381E-06 | 6.000E-06 | 1.200E-05 |
| 24572 | 1229 | S1229K | D | 1e-05 | 6.9e-06 | 8.2e-06 | 8.367E-06 | 1.557E-06 | 6.900E-06 | 1.000E-05 |
| 24573 | 1229 | S1229M | D | 9.8e-06 | 1.2e-05 | 1.5e-05 | 1.227E-05 | 2.610E-06 | 9.800E-06 | 1.500E-05 |
| 24574 | 1229 | S1229F | D | 7.8e-06 | 1.2e-05 | 7.4e-06 | 9.067E-06 | 2.548E-06 | 7.400E-06 | 1.200E-05 |
| 24575 | 1229 | S1229P | D | 5e-06 | 7.3e-06 | 7.9e-06 | 6.733E-06 | 1.531E-06 | 5.000E-06 | 7.900E-06 |
| 24576 | 1229 | S1229S | D | 1.3e-05 | 9.4e-06 | 1.2e-05 | 1.147E-05 | 1.858E-06 | 9.400E-06 | 1.300E-05 |
| 24577 | 1229 | S1229T | D | 8.8e-06 | 5.6e-06 | 8.2e-06 | 7.533E-06 | 1.701E-06 | 5.600E-06 | 8.800E-06 |
| 24578 | 1229 | S1229W | D | 5.3e-06 | 9.6e-06 | 6.7e-06 | 7.200E-06 | 2.193E-06 | 5.300E-06 | 9.600E-06 |
| 24579 | 1229 | S1229Y | D | 8.5e-06 | 4.6e-06 | 5.9e-06 | 6.333E-06 | 1.986E-06 | 4.600E-06 | 8.500E-06 |
| 24580 | 1229 | S1229V | D | 1.1e-05 | 1.2e-05 | 9.2e-06 | 1.073E-05 | 1.419E-06 | 9.200E-06 | 1.200E-05 |
| 24581 | 1230 | T1230A | D | 1.5e-05 | 7.8e-06 | 7.4e-06 | 1.007E-05 | 4.277E-06 | 7.400E-06 | 1.500E-05 |
| 24582 | 1230 | T1230R | D | 7.4e-06 | 8.5e-06 | 8.1e-06 | 8.000E-06 | 5.568E-07 | 7.400E-06 | 8.500E-06 |
| 24583 | 1230 | T1230N | D | 5e-06 | 5.6e-06 | 8.6e-06 | 6.400E-06 | 1.929E-06 | 5.000E-06 | 8.600E-06 |
| 24584 | 1230 | T1230D | D | 6.4e-06 | 6.2e-06 | 8.2e-06 | 6.933E-06 | 1.102E-06 | 6.200E-06 | 8.200E-06 |
| 24585 | 1230 | T1230C | D | 9.8e-06 | 8.7e-06 | 9.9e-06 | 9.467E-06 | 6.658E-07 | 8.700E-06 | 9.900E-06 |
| 24586 | 1230 | T1230E | D | 6.3e-06 | 1e-05 | 7.5e-06 | 7.933E-06 | 1.888E-06 | 6.300E-06 | 1.000E-05 |
| 24587 | 1230 | T1230Q | D | 4.8e-06 | 9.5e-06 | 7.7e-06 | 7.333E-06 | 2.371E-06 | 4.800E-06 | 9.500E-06 |
| 24588 | 1230 | T1230G | D | 5.8e-06 | 5.1e-06 | 1.1e-05 | 7.300E-06 | 3.223E-06 | 5.100E-06 | 1.100E-05 |
| 24589 | 1230 | T1230H | D | 8.6e-06 | 8.1e-06 | 9.7e-06 | 8.800E-06 | 8.185E-07 | 8.100E-06 | 9.700E-06 |
| 24590 | 1230 | T1230I | D | 9.3e-06 | 9.6e-06 | 6.3e-06 | 8.400E-06 | 1.825E-06 | 6.300E-06 | 9.600E-06 |
| 24591 | 1230 | T1230L | D | 1.7e-05 | 6.9e-06 | 9.9e-06 | 1.127E-05 | 5.187E-06 | 6.900E-06 | 1.700E-05 |
| 24592 | 1230 | T1230K | D | 1.1e-05 | 7.1e-06 | 1.2e-05 | 1.003E-05 | 2.589E-06 | 7.100E-06 | 1.200E-05 |
| 24593 | 1230 | T1230M | D | 9.2e-06 | 6.5e-06 | 7.8e-06 | 7.833E-06 | 1.350E-06 | 6.500E-06 | 9.200E-06 |
| 24594 | 1230 | T1230F | D | 5e-06 | 7.8e-06 | 7.4e-06 | 6.733E-06 | 1.514E-06 | 5.000E-06 | 7.800E-06 |
| 24595 | 1230 | T1230P | D | 9.9e-06 | 9.2e-06 | 5.9e-06 | 8.333E-06 | 2.136E-06 | 5.900E-06 | 9.900E-06 |
| 24596 | 1230 | T1230S | D | 1.1e-05 | 7.3e-06 | 6.8e-06 | 8.367E-06 | 2.294E-06 | 6.800E-06 | 1.100E-05 |
| 24597 | 1230 | T1230T | D | 8.3e-06 | 7.1e-06 | 5.9e-06 | 7.100E-06 | 1.200E-06 | 5.900E-06 | 8.300E-06 |
| 24598 | 1230 | T1230W | D | 6.1e-06 | 8e-06 | 1.2e-05 | 8.700E-06 | 3.012E-06 | 6.100E-06 | 1.200E-05 |
| 24599 | 1230 | T1230Y | D | 2.1e-05 | 5.2e-06 | 5.1e-06 | 1.043E-05 | 9.151E-06 | 5.100E-06 | 2.100E-05 |
| 24600 | 1230 | T1230V | D | 1.1e-05 | 6.9e-06 | 9.6e-06 | 9.167E-06 | 2.084E-06 | 6.900E-06 | 1.100E-05 |
| 24601 | 1231 | Y1231A | D | 1.6e-05 | 1.3e-05 | 1.1e-05 | 1.333E-05 | 2.517E-06 | 1.100E-05 | 1.600E-05 |
| 24602 | 1231 | Y1231R | D | 4.9e-06 | 9.5e-06 | 9.8e-06 | 8.067E-06 | 2.747E-06 | 4.900E-06 | 9.800E-06 |
| 24603 | 1231 | Y1231N | D | 9e-06 | 9.5e-06 | 1.4e-05 | 1.083E-05 | 2.754E-06 | 9.000E-06 | 1.400E-05 |
| 24604 | 1231 | Y1231D | D | 8.4e-06 | 4.7e-06 | 5.8e-06 | 6.300E-06 | 1.900E-06 | 4.700E-06 | 8.400E-06 |
| 24605 | 1231 | Y1231C | D | 8.9e-06 | 8.1e-06 | 1e-05 | 9.000E-06 | 9.539E-07 | 8.100E-06 | 1.000E-05 |
| 24606 | 1231 | Y1231E | D | 9.9e-06 | 4.5e-06 | 6e-06 | 6.800E-06 | 2.787E-06 | 4.500E-06 | 9.900E-06 |
| 24607 | 1231 | Y1231Q | D | 8.1e-06 | 1.1e-05 | 8.3e-06 | 9.133E-06 | 1.620E-06 | 8.100E-06 | 1.100E-05 |
| 24608 | 1231 | Y1231G | D | 1e-05 | 6.6e-06 | 5.3e-06 | 7.300E-06 | 2.427E-06 | 5.300E-06 | 1.000E-05 |
| 24609 | 1231 | Y1231H | D | 1.3e-05 | 9.6e-06 | 5.9e-06 | 9.500E-06 | 3.551E-06 | 5.900E-06 | 1.300E-05 |
| 24610 | 1231 | Y1231I | D | 6.4e-06 | 8.7e-06 | 6.3e-06 | 7.133E-06 | 1.358E-06 | 6.300E-06 | 8.700E-06 |
| 24611 | 1231 | Y1231L | D | 1.2e-05 | 8.8e-06 | 8.7e-06 | 9.833E-06 | 1.877E-06 | 8.700E-06 | 1.200E-05 |
| 24612 | 1231 | Y1231K | D | 7.6e-06 | 1e-05 | 4.8e-06 | 7.467E-06 | 2.603E-06 | 4.800E-06 | 1.000E-05 |
| 24613 | 1231 | Y1231M | D | 7.4e-06 | 9.5e-06 | 1.1e-05 | 9.300E-06 | 1.808E-06 | 7.400E-06 | 1.100E-05 |
| 24614 | 1231 | Y1231F | D | 1.3e-05 | 5.5e-06 | 6.8e-06 | 8.433E-06 | 4.008E-06 | 5.500E-06 | 1.300E-05 |
| 24615 | 1231 | Y1231P | D | 4.8e-06 | 6.9e-06 | 1.4e-05 | 8.567E-06 | 4.821E-06 | 4.800E-06 | 1.400E-05 |
| 24616 | 1231 | Y1231S | D | 5.2e-06 | 4.9e-06 | 8e-06 | 6.033E-06 | 1.710E-06 | 4.900E-06 | 8.000E-06 |
| 24617 | 1231 | Y1231T | D | 4e-06 | 8.2e-06 | 6.3e-06 | 6.167E-06 | 2.103E-06 | 4.000E-06 | 8.200E-06 |
| 24618 | 1231 | Y1231W | D | 8.3e-06 | 4.6e-06 | 7.9e-06 | 6.933E-06 | 2.031E-06 | 4.600E-06 | 8.300E-06 |
| 24619 | 1231 | Y1231Y | D | 7.5e-06 | 6.5e-06 | 6.2e-06 | 6.733E-06 | 6.807E-07 | 6.200E-06 | 7.500E-06 |
| 24620 | 1231 | Y1231V | D | 1.5e-05 | 9.9e-06 | 7.9e-06 | 1.093E-05 | 3.661E-06 | 7.900E-06 | 1.500E-05 |
| 24621 | 1232 | S1232A | D | 9.3e-06 | 9.3e-06 | 3.6e-06 | 7.400E-06 | 3.291E-06 | 3.600E-06 | 9.300E-06 |
| 24622 | 1232 | S1232R | D | 6.3e-06 | 6.9e-06 | 8.2e-06 | 7.133E-06 | 9.713E-07 | 6.300E-06 | 8.200E-06 |
| 24623 | 1232 | S1232N | D | 6e-06 | 5.4e-06 | 6.4e-06 | 5.933E-06 | 5.033E-07 | 5.400E-06 | 6.400E-06 |
| 24624 | 1232 | S1232D | D | 1e-05 | 9.3e-06 | 1e-05 | 9.767E-06 | 4.041E-07 | 9.300E-06 | 1.000E-05 |
| 24625 | 1232 | S1232C | D | 9e-06 | 8.4e-06 | 3.9e-06 | 7.100E-06 | 2.787E-06 | 3.900E-06 | 9.000E-06 |
| 24626 | 1232 | S1232E | D | 6.9e-06 | 6.1e-06 | 5e-06 | 6.000E-06 | 9.539E-07 | 5.000E-06 | 6.900E-06 |
| 24627 | 1232 | S1232Q | D | 8.7e-06 | 4.2e-06 | 9.6e-06 | 7.500E-06 | 2.893E-06 | 4.200E-06 | 9.600E-06 |
| 24628 | 1232 | S1232G | D | 9.3e-06 | 7e-06 | 8e-06 | 8.100E-06 | 1.153E-06 | 7.000E-06 | 9.300E-06 |
| 24629 | 1232 | S1232H | D | 1.3e-05 | 9.7e-06 | 1.3e-05 | 1.190E-05 | 1.905E-06 | 9.700E-06 | 1.300E-05 |
| 24630 | 1232 | S1232I | D | 6.4e-06 | 7.3e-06 | 1.2e-05 | 8.567E-06 | 3.007E-06 | 6.400E-06 | 1.200E-05 |
| 24631 | 1232 | S1232L | D | 1.5e-05 | 7.4e-06 | 1e-05 | 1.080E-05 | 3.863E-06 | 7.400E-06 | 1.500E-05 |
| 24632 | 1232 | S1232K | D | 8.1e-06 | 8.3e-06 | 1e-05 | 8.800E-06 | 1.044E-06 | 8.100E-06 | 1.000E-05 |
| 24633 | 1232 | S1232M | D | 9.1e-06 | 9e-06 | 7.5e-06 | 8.533E-06 | 8.963E-07 | 7.500E-06 | 9.100E-06 |
| 24634 | 1232 | S1232F | D | 6.4e-06 | 5.6e-06 | 8.2e-06 | 6.733E-06 | 1.332E-06 | 5.600E-06 | 8.200E-06 |
| 24635 | 1232 | S1232P | D | 7.5e-06 | 4e-06 | 1.1e-05 | 7.500E-06 | 3.500E-06 | 4.000E-06 | 1.100E-05 |
| 24636 | 1232 | S1232S | D | 1.1e-05 | 7.4e-06 | 9e-06 | 9.133E-06 | 1.804E-06 | 7.400E-06 | 1.100E-05 |
| 24637 | 1232 | S1232T | D | 7.9e-06 | 4.5e-06 | 9.1e-06 | 7.167E-06 | 2.386E-06 | 4.500E-06 | 9.100E-06 |
| 24638 | 1232 | S1232W | D | 7.6e-06 | 7.6e-06 | 8.1e-06 | 7.767E-06 | 2.887E-07 | 7.600E-06 | 8.100E-06 |
| 24639 | 1232 | S1232Y | D | 6.9e-06 | 7.6e-06 | 3.5e-06 | 6.000E-06 | 2.193E-06 | 3.500E-06 | 7.600E-06 |
| 24640 | 1232 | S1232V | D | 1.2e-05 | 1.2e-05 | 7.5e-06 | 1.050E-05 | 2.598E-06 | 7.500E-06 | 1.200E-05 |
| 24641 | 1233 | L1233A | D | 6.1e-06 | 8.9e-06 | 8.3e-06 | 7.767E-06 | 1.474E-06 | 6.100E-06 | 8.900E-06 |
| 24642 | 1233 | L1233R | D | 6.2e-06 | 1e-05 | 1.3e-05 | 9.733E-06 | 3.408E-06 | 6.200E-06 | 1.300E-05 |
| 24643 | 1233 | L1233N | D | 4.4e-06 | 8.3e-06 | 6e-06 | 6.233E-06 | 1.960E-06 | 4.400E-06 | 8.300E-06 |
| 24644 | 1233 | L1233D | D | 4.4e-06 | 5.3e-06 | 8.1e-06 | 5.933E-06 | 1.930E-06 | 4.400E-06 | 8.100E-06 |
| 24645 | 1233 | L1233C | D | 5.5e-06 | 6.7e-06 | 4.1e-06 | 5.433E-06 | 1.301E-06 | 4.100E-06 | 6.700E-06 |
| 24646 | 1233 | L1233E | D | 1.5e-05 | 1e-05 | 1.3e-05 | 1.267E-05 | 2.517E-06 | 1.000E-05 | 1.500E-05 |
| 24647 | 1233 | L1233Q | D | 7.7e-06 | 7.1e-06 | 6.8e-06 | 7.200E-06 | 4.583E-07 | 6.800E-06 | 7.700E-06 |
| 24648 | 1233 | L1233G | D | 6.8e-06 | 7.3e-06 | 9.1e-06 | 7.733E-06 | 1.210E-06 | 6.800E-06 | 9.100E-06 |
| 24649 | 1233 | L1233H | D | 8.2e-06 | 5.7e-06 | 5.3e-06 | 6.400E-06 | 1.572E-06 | 5.300E-06 | 8.200E-06 |

| | | | | | | | | | | |
|---|---|---|---|---|---|---|---|---|---|---|
| 24650 | 1233 | L1233I | D | 5.7e-06 | 1.3e-05 | 1.3e-05 | 1.057E-05 | 4.215E-06 | 5.700E-06 | 1.300E-05 |
| 24651 | 1233 | L1233L | D | 6.6e-06 | 9.8e-06 | 6.9e-06 | 7.767E-06 | 1.767E-06 | 6.600E-06 | 9.800E-06 |
| 24652 | 1233 | L1233K | D | 1.1e-05 | 8.9e-06 | 8.7e-06 | 9.533E-06 | 1.274E-06 | 8.700E-06 | 1.100E-05 |
| 24653 | 1233 | L1233M | D | 9.9e-06 | 1e-05 | 8.5e-06 | 9.467E-06 | 8.386E-07 | 8.500E-06 | 1.000E-05 |
| 24654 | 1233 | L1233F | D | 1e-05 | 6.9e-06 | 6.2e-06 | 7.700E-06 | 2.022E-06 | 6.200E-06 | 1.000E-05 |
| 24655 | 1233 | L1233P | D | 9.3e-06 | 7.7e-06 | 4.9e-06 | 7.300E-06 | 2.227E-06 | 4.900E-06 | 9.300E-06 |
| 24656 | 1233 | L1233S | D | 7.3e-06 | 7.4e-06 | 1.1e-05 | 8.567E-06 | 2.108E-06 | 7.300E-06 | 1.100E-05 |
| 24657 | 1233 | L1233T | D | 8.3e-06 | 7.5e-06 | 7.5e-06 | 7.767E-06 | 4.619E-07 | 7.500E-06 | 8.300E-06 |
| 24658 | 1233 | L1233W | D | 7e-06 | 7.3e-06 | 8.8e-06 | 7.700E-06 | 9.644E-07 | 7.000E-06 | 8.800E-06 |
| 24659 | 1233 | L1233Y | D | 6.7e-06 | 1.4e-05 | 8.4e-06 | 9.700E-06 | 3.820E-06 | 6.700E-06 | 1.400E-05 |
| 24660 | 1233 | L1233V | D | 9e-06 | 1e-05 | 1.7e-05 | 1.200E-05 | 4.359E-06 | 9.000E-06 | 1.700E-05 |
| 24661 | 1234 | S1234A | D | 1.4e-05 | 1.1e-05 | 5.3e-06 | 1.010E-05 | 4.419E-06 | 5.300E-06 | 1.400E-05 |
| 24662 | 1234 | S1234R | D | 9.7e-06 | 1e-05 | 7.5e-06 | 9.067E-06 | 1.365E-06 | 7.500E-06 | 1.000E-05 |
| 24663 | 1234 | S1234N | D | 1.4e-05 | 1.1e-05 | 8.3e-06 | 1.110E-05 | 2.851E-06 | 8.300E-06 | 1.400E-05 |
| 24664 | 1234 | S1234D | D | 9.5e-06 | 7e-06 | 1.2e-05 | 9.500E-06 | 2.500E-06 | 7.000E-06 | 1.200E-05 |
| 24665 | 1234 | S1234C | D | 6.4e-06 | 1e-05 | 5.3e-06 | 7.233E-06 | 2.458E-06 | 5.300E-06 | 1.000E-05 |
| 24666 | 1234 | S1234E | D | 1.1e-05 | 6.6e-06 | 1.1e-05 | 9.533E-06 | 2.540E-06 | 6.600E-06 | 1.100E-05 |
| 24667 | 1234 | S1234Q | D | 8.2e-06 | 9.9e-06 | 6.8e-06 | 8.300E-06 | 1.552E-06 | 6.800E-06 | 9.900E-06 |
| 24668 | 1234 | S1234G | D | 5.9e-06 | 1.4e-05 | 8.4e-06 | 9.433E-06 | 4.148E-06 | 5.900E-06 | 1.400E-05 |
| 24669 | 1234 | S1234H | D | 5.5e-06 | 1.1e-05 | 1.5e-05 | 1.050E-05 | 4.770E-06 | 5.500E-06 | 1.500E-05 |
| 24670 | 1234 | S1234I | D | 4.1e-06 | 7.6e-06 | 9.4e-06 | 7.033E-06 | 2.695E-06 | 4.100E-06 | 9.400E-06 |
| 24671 | 1234 | S1234L | D | 6.7e-06 | 9.6e-06 | 7.5e-06 | 7.933E-06 | 1.498E-06 | 6.700E-06 | 9.600E-06 |
| 24672 | 1234 | S1234K | D | 7.4e-06 | 1e-05 | 4.7e-06 | 7.367E-06 | 2.650E-06 | 4.700E-06 | 1.000E-05 |
| 24673 | 1234 | S1234M | D | 4.3e-06 | 4.8e-06 | 8.5e-06 | 5.867E-06 | 2.294E-06 | 4.300E-06 | 8.500E-06 |
| 24674 | 1234 | S1234F | D | 1.1e-05 | 4.8e-06 | 6.7e-06 | 7.500E-06 | 3.176E-06 | 4.800E-06 | 1.100E-05 |
| 24675 | 1234 | S1234P | D | 7.5e-06 | 9.5e-06 | 1.1e-05 | 9.333E-06 | 1.756E-06 | 7.500E-06 | 1.100E-05 |
| 24676 | 1234 | S1234S | D | 6.7e-06 | 9e-06 | 5.2e-06 | 6.967E-06 | 1.914E-06 | 5.200E-06 | 9.000E-06 |
| 24677 | 1234 | S1234T | D | 6.7e-06 | 7.9e-06 | 6.9e-06 | 7.167E-06 | 6.429E-07 | 6.700E-06 | 7.900E-06 |
| 24678 | 1234 | S1234W | D | 6.8e-06 | 6.6e-06 | 1.4e-05 | 9.133E-06 | 4.216E-06 | 6.600E-06 | 1.400E-05 |
| 24679 | 1234 | S1234Y | D | 4.6e-06 | 9.7e-06 | 7.7e-06 | 7.333E-06 | 2.570E-06 | 4.600E-06 | 9.700E-06 |
| 24680 | 1234 | S1234V | D | 7.6e-06 | 1.3e-05 | 1.4e-05 | 1.153E-05 | 3.443E-06 | 7.600E-06 | 1.400E-05 |
| 24681 | 1235 | S1235A | D | 1.2e-05 | 1.1e-05 | 5.2e-06 | 9.400E-06 | 3.672E-06 | 5.200E-06 | 1.200E-05 |
| 24682 | 1235 | S1235R | D | 1.1e-05 | 6.9e-06 | 4.7e-06 | 7.533E-06 | 3.197E-06 | 4.700E-06 | 1.100E-05 |
| 24683 | 1235 | S1235N | D | 2.1e-05 | 9.8e-06 | 6.7e-06 | 1.250E-05 | 7.523E-06 | 6.700E-06 | 2.100E-05 |
| 24684 | 1235 | S1235D | D | 8.2e-06 | 9.6e-06 | 5.1e-06 | 7.633E-06 | 2.303E-06 | 5.100E-06 | 9.600E-06 |
| 24685 | 1235 | S1235C | D | 1.2e-05 | 4.6e-06 | 8.3e-06 | 8.300E-06 | 3.700E-06 | 4.600E-06 | 1.200E-05 |
| 24686 | 1235 | S1235E | D | 9e-06 | 4.2e-06 | 1.4e-05 | 9.067E-06 | 4.900E-06 | 4.200E-06 | 1.400E-05 |
| 24687 | 1235 | S1235Q | D | 8.8e-06 | 4.7e-06 | 9.8e-06 | 7.767E-06 | 2.702E-06 | 4.700E-06 | 9.800E-06 |
| 24688 | 1235 | S1235G | D | 5.5e-06 | 9.6e-06 | 1e-05 | 8.367E-06 | 2.491E-06 | 5.500E-06 | 1.000E-05 |
| 24689 | 1235 | S1235H | D | 4.9e-06 | 9.1e-06 | 1.8e-05 | 1.067E-05 | 6.689E-06 | 4.900E-06 | 1.800E-05 |
| 24690 | 1235 | S1235I | D | 1.1e-05 | 6.9e-06 | 1.4e-05 | 1.063E-05 | 3.564E-06 | 6.900E-06 | 1.400E-05 |
| 24691 | 1235 | S1235L | D | 9.2e-06 | 5.5e-06 | 1.2e-05 | 8.900E-06 | 3.260E-06 | 5.500E-06 | 1.200E-05 |
| 24692 | 1235 | S1235K | D | 6.8e-06 | 1.2e-05 | 8.7e-06 | 9.167E-06 | 2.631E-06 | 6.800E-06 | 1.200E-05 |
| 24693 | 1235 | S1235M | D | 1.4e-05 | 7.5e-06 | 9.4e-06 | 1.030E-05 | 3.342E-06 | 7.500E-06 | 1.400E-05 |
| 24694 | 1235 | S1235F | D | 5.1e-06 | 8.6e-06 | 4.8e-06 | 6.167E-06 | 2.113E-06 | 4.800E-06 | 8.600E-06 |
| 24695 | 1235 | S1235P | D | 5.9e-06 | 6.5e-06 | 6.8e-06 | 6.400E-06 | 4.583E-07 | 5.900E-06 | 6.800E-06 |
| 24696 | 1235 | S1235S | D | 9.6e-06 | 6.2e-06 | 1.4e-05 | 9.933E-06 | 3.911E-06 | 6.200E-06 | 1.400E-05 |
| 24697 | 1235 | S1235T | D | 1e-05 | 1.9e-05 | 1.2e-05 | 1.367E-05 | 4.726E-06 | 1.000E-05 | 1.900E-05 |
| 24698 | 1235 | S1235W | D | 4e-06 | 6.9e-06 | 9e-06 | 6.633E-06 | 2.511E-06 | 4.000E-06 | 9.000E-06 |
| 24699 | 1235 | S1235Y | D | 1e-05 | 7.5e-06 | 6.9e-06 | 8.133E-06 | 1.644E-06 | 6.900E-06 | 1.000E-05 |
| 24700 | 1235 | S1235V | D | 5.6e-06 | 1.3e-05 | 6.4e-06 | 8.333E-06 | 4.061E-06 | 5.600E-06 | 1.300E-05 |
| 24701 | 1236 | T1236A | D | 9.1e-06 | 5.8e-06 | 6.4e-06 | 7.100E-06 | 1.758E-06 | 5.800E-06 | 9.100E-06 |
| 24702 | 1236 | T1236R | D | 9.8e-06 | 7.2e-06 | 8.6e-06 | 8.533E-06 | 1.301E-06 | 7.200E-06 | 9.800E-06 |
| 24703 | 1236 | T1236N | D | 5.3e-06 | 8e-06 | 1.2e-05 | 8.433E-06 | 3.371E-06 | 5.300E-06 | 1.200E-05 |
| 24704 | 1236 | T1236D | D | 9.8e-06 | 5.7e-06 | 1.3e-05 | 9.500E-06 | 3.659E-06 | 5.700E-06 | 1.300E-05 |
| 24705 | 1236 | T1236C | D | 9.8e-06 | 6.8e-06 | 7.9e-06 | 8.167E-06 | 1.518E-06 | 6.800E-06 | 9.800E-06 |
| 24706 | 1236 | T1236E | D | 9.9e-06 | 8.4e-06 | 1.2e-05 | 1.010E-05 | 1.808E-06 | 8.400E-06 | 1.200E-05 |
| 24707 | 1236 | T1236Q | D | 6.8e-06 | 1e-05 | 5.8e-06 | 7.533E-06 | 2.194E-06 | 5.800E-06 | 1.000E-05 |
| 24708 | 1236 | T1236G | D | 1.3e-05 | 8.6e-06 | 6.7e-06 | 9.433E-06 | 3.232E-06 | 6.700E-06 | 1.300E-05 |
| 24709 | 1236 | T1236H | D | 9.4e-06 | 6.4e-06 | 4.3e-06 | 6.700E-06 | 2.563E-06 | 4.300E-06 | 9.400E-06 |
| 24710 | 1236 | T1236I | D | 5.8e-06 | 7.3e-06 | 8.3e-06 | 7.133E-06 | 1.258E-06 | 5.800E-06 | 8.300E-06 |
| 24711 | 1236 | T1236L | D | 8.8e-06 | 6.2e-06 | 9.8e-06 | 8.267E-06 | 1.858E-06 | 6.200E-06 | 9.800E-06 |
| 24712 | 1236 | T1236K | D | 7.2e-06 | 5.3e-06 | 5.6e-06 | 6.033E-06 | 1.021E-06 | 5.300E-06 | 7.200E-06 |
| 24713 | 1236 | T1236M | D | 1e-05 | 9.6e-06 | 4.7e-06 | 8.100E-06 | 2.951E-06 | 4.700E-06 | 1.000E-05 |
| 24714 | 1236 | T1236F | D | 6.5e-06 | 1e-05 | 1e-05 | 8.833E-06 | 2.021E-06 | 6.500E-06 | 1.000E-05 |
| 24715 | 1236 | T1236P | D | 9.4e-06 | 9.6e-06 | 6.7e-06 | 8.567E-06 | 1.620E-06 | 6.700E-06 | 9.600E-06 |
| 24716 | 1236 | T1236S | D | 8.8e-06 | 9.7e-06 | 7.6e-06 | 8.700E-06 | 1.054E-06 | 7.600E-06 | 9.700E-06 |
| 24717 | 1236 | T1236T | D | 1e-05 | 6.3e-06 | 1.4e-05 | 1.010E-05 | 3.851E-06 | 6.300E-06 | 1.400E-05 |
| 24718 | 1236 | T1236W | D | 9.6e-06 | 5.6e-06 | 6.7e-06 | 7.300E-06 | 2.066E-06 | 5.600E-06 | 9.600E-06 |
| 24719 | 1236 | T1236Y | D | 4.9e-06 | 8.7e-06 | 4.4e-06 | 6.000E-06 | 2.352E-06 | 4.400E-06 | 8.700E-06 |
| 24720 | 1236 | T1236V | D | 1e-05 | 7.1e-06 | 8.9e-06 | 8.667E-06 | 1.464E-06 | 7.100E-06 | 1.000E-05 |
| 24721 | 1237 | L1237A | D | 1e-05 | 8.6e-06 | 8.9e-06 | 9.167E-06 | 7.371E-07 | 8.600E-06 | 1.000E-05 |
| 24722 | 1237 | L1237R | D | 7.2e-06 | 4.6e-06 | 9.7e-06 | 7.167E-06 | 2.550E-06 | 4.600E-06 | 9.700E-06 |
| 24723 | 1237 | L1237N | D | 8.7e-06 | 9.2e-06 | 6.6e-06 | 8.167E-06 | 1.380E-06 | 6.600E-06 | 9.200E-06 |
| 24724 | 1237 | L1237D | D | 5.4e-06 | 5.9e-06 | 1.4e-05 | 8.433E-06 | 4.827E-06 | 5.400E-06 | 1.400E-05 |
| 24725 | 1237 | L1237C | D | 6.8e-06 | 8.8e-06 | 9.1e-06 | 8.233E-06 | 1.250E-06 | 6.800E-06 | 9.100E-06 |
| 24726 | 1237 | L1237E | D | 4.4e-06 | 1.2e-05 | 8.4e-06 | 8.267E-06 | 3.802E-06 | 4.400E-06 | 1.200E-05 |
| 24727 | 1237 | L1237Q | D | 1.1e-05 | 1e-05 | 7.5e-06 | 9.500E-06 | 1.803E-06 | 7.500E-06 | 1.100E-05 |
| 24728 | 1237 | L1237G | D | 9.4e-06 | 9.3e-06 | 7.1e-06 | 8.600E-06 | 1.300E-06 | 7.100E-06 | 9.400E-06 |
| 24729 | 1237 | L1237H | D | 4.3e-06 | 1.2e-05 | 1.1e-05 | 9.100E-06 | 4.187E-06 | 4.300E-06 | 1.200E-05 |
| 24730 | 1237 | L1237I | D | 8.1e-06 | 9.5e-06 | 7e-06 | 8.200E-06 | 1.253E-06 | 7.000E-06 | 9.500E-06 |
| 24731 | 1237 | L1237L | D | 7.1e-06 | 1.4e-05 | 8.5e-06 | 9.867E-06 | 3.647E-06 | 7.100E-06 | 1.400E-05 |
| 24732 | 1237 | L1237K | D | 8.5e-06 | 1.2e-05 | 6.4e-06 | 8.967E-06 | 2.829E-06 | 6.400E-06 | 1.200E-05 |
| 24733 | 1237 | L1237M | D | 1.1e-05 | 1.1e-05 | 6.7e-06 | 9.567E-06 | 2.483E-06 | 6.700E-06 | 1.100E-05 |
| 24734 | 1237 | L1237F | D | 7.5e-06 | 4.2e-06 | 6e-06 | 5.900E-06 | 1.652E-06 | 4.200E-06 | 7.500E-06 |

| | | | | | | | | | | |
|---|---|---|---|---|---|---|---|---|---|---|
| 24735 | 1237 | L1237P | D | 9.6e-06 | 6.2e-06 | 7e-06 | 7.600E-06 | 1.778E-06 | 6.200E-06 | 9.600E-06 |
| 24736 | 1237 | L1237S | D | 6.2e-06 | 6.5e-06 | 4.3e-06 | 5.667E-06 | 1.193E-06 | 4.300E-06 | 6.500E-06 |
| 24737 | 1237 | L1237T | D | 1.2e-05 | 9.9e-06 | 8.5e-06 | 1.013E-05 | 1.762E-06 | 8.500E-06 | 1.200E-05 |
| 24738 | 1237 | L1237W | D | 6.4e-06 | 5.9e-06 | 6.5e-06 | 6.267E-06 | 3.215E-07 | 5.900E-06 | 6.500E-06 |
| 24739 | 1237 | L1237Y | D | 1e-05 | 1.2e-05 | 6.6e-06 | 9.533E-06 | 2.730E-06 | 6.600E-06 | 1.200E-05 |
| 24740 | 1237 | L1237V | D | 9.4e-06 | 8.2e-06 | 1.4e-05 | 1.053E-05 | 3.062E-06 | 8.200E-06 | 1.400E-05 |
| 24741 | 1238 | T1238A | D | 1.1e-05 | 1.6e-05 | 4.8e-06 | 1.060E-05 | 5.611E-06 | 4.800E-06 | 1.600E-05 |
| 24742 | 1238 | T1238R | D | 5.1e-06 | 1.2e-05 | 9.6e-06 | 8.900E-06 | 3.503E-06 | 5.100E-06 | 1.200E-05 |
| 24743 | 1238 | T1238N | D | 7.2e-06 | 9.1e-06 | 1.1e-05 | 9.100E-06 | 1.900E-06 | 7.200E-06 | 1.100E-05 |
| 24744 | 1238 | T1238D | D | 8.5e-06 | 7.6e-06 | 1e-05 | 8.700E-06 | 1.212E-06 | 7.600E-06 | 1.000E-05 |
| 24745 | 1238 | T1238C | D | 1.5e-05 | 9.5e-06 | 8.6e-06 | 1.103E-05 | 3.465E-06 | 8.600E-06 | 1.500E-05 |
| 24746 | 1238 | T1238E | D | 7.3e-06 | 8.2e-06 | 1.1e-05 | 8.833E-06 | 1.930E-06 | 7.300E-06 | 1.100E-05 |
| 24747 | 1238 | T1238Q | D | 7.1e-06 | 1.4e-05 | 1.4e-05 | 1.170E-05 | 3.984E-06 | 7.100E-06 | 1.400E-05 |
| 24748 | 1238 | T1238G | D | 7.6e-06 | 4.9e-06 | 6.2e-06 | 6.233E-06 | 1.350E-06 | 4.900E-06 | 7.600E-06 |
| 24749 | 1238 | T1238H | D | 4.3e-06 | 6.1e-06 | 7.5e-06 | 5.967E-06 | 1.604E-06 | 4.300E-06 | 7.500E-06 |
| 24750 | 1238 | T1238I | D | 7.5e-06 | 5.9e-06 | 9.8e-06 | 7.733E-06 | 1.960E-06 | 5.900E-06 | 9.800E-06 |
| 24751 | 1238 | T1238L | D | 1.1e-05 | 7.3e-06 | 8.4e-06 | 8.900E-06 | 1.900E-06 | 7.300E-06 | 1.100E-05 |
| 24752 | 1238 | T1238K | D | 6.1e-06 | 5.8e-06 | 4.9e-06 | 5.600E-06 | 6.245E-07 | 4.900E-06 | 6.100E-06 |
| 24753 | 1238 | T1238M | D | 9.5e-06 | 7.1e-06 | 1.1e-05 | 9.200E-06 | 1.967E-06 | 7.100E-06 | 1.100E-05 |
| 24754 | 1238 | T1238F | D | 7.1e-06 | 1.3e-05 | 7.6e-06 | 9.233E-06 | 3.272E-06 | 7.100E-06 | 1.300E-05 |
| 24755 | 1238 | T1238P | D | 5e-06 | 1.2e-05 | 1e-05 | 9.000E-06 | 3.606E-06 | 5.000E-06 | 1.200E-05 |
| 24756 | 1238 | T1238S | D | 8.8e-06 | 7.6e-06 | 5.7e-06 | 7.367E-06 | 1.563E-06 | 5.700E-06 | 8.800E-06 |
| 24757 | 1238 | T1238T | D | 9.1e-06 | 8.4e-06 | 6.4e-06 | 7.967E-06 | 1.401E-06 | 6.400E-06 | 9.100E-06 |
| 24758 | 1238 | T1238W | D | 6.3e-06 | 5.7e-06 | 6.1e-06 | 6.033E-06 | 3.055E-07 | 5.700E-06 | 6.300E-06 |
| 24759 | 1238 | T1238Y | D | 7e-06 | 9.2e-06 | 7.5e-06 | 7.900E-06 | 1.153E-06 | 7.000E-06 | 9.200E-06 |
| 24760 | 1238 | T1238V | D | 5.3e-06 | 6.5e-06 | 5.1e-06 | 5.633E-06 | 7.572E-07 | 5.100E-06 | 6.500E-06 |
| 24761 | 1239 | L1239A | D | 6.6e-06 | 7e-06 | 8.5e-06 | 7.367E-06 | 1.002E-06 | 6.600E-06 | 8.500E-06 |
| 24762 | 1239 | L1239R | D | 1.1e-05 | 8.6e-06 | 1.1e-05 | 1.020E-05 | 1.386E-06 | 8.600E-06 | 1.100E-05 |
| 24763 | 1239 | L1239N | D | 5.9e-06 | 7.6e-06 | 1.3e-05 | 8.833E-06 | 3.707E-06 | 5.900E-06 | 1.300E-05 |
| 24764 | 1239 | L1239D | D | 9.3e-06 | 4.5e-06 | 1e-05 | 7.933E-06 | 2.994E-06 | 4.500E-06 | 1.000E-05 |
| 24765 | 1239 | L1239C | D | 1.5e-05 | 6.9e-06 | 9.3e-06 | 1.040E-05 | 4.161E-06 | 6.900E-06 | 1.500E-05 |
| 24766 | 1239 | L1239E | D | 6e-06 | 9.1e-06 | 6.1e-06 | 7.067E-06 | 1.762E-06 | 6.000E-06 | 9.100E-06 |
| 24767 | 1239 | L1239Q | D | 6.3e-06 | 5.6e-06 | 9e-06 | 6.967E-06 | 1.795E-06 | 5.600E-06 | 9.000E-06 |
| 24768 | 1239 | L1239G | D | 1.2e-05 | 1.3e-05 | 1.4e-05 | 1.300E-05 | 1.000E-06 | 1.200E-05 | 1.400E-05 |
| 24769 | 1239 | L1239H | D | 5.7e-06 | 8.7e-06 | 8.9e-06 | 7.767E-06 | 1.793E-06 | 5.700E-06 | 8.900E-06 |
| 24770 | 1239 | L1239I | D | 6.1e-06 | 9e-06 | 8.1e-06 | 7.733E-06 | 1.484E-06 | 6.100E-06 | 9.000E-06 |
| 24771 | 1239 | L1239L | D | 1e-05 | 4.3e-06 | 1e-05 | 8.100E-06 | 3.291E-06 | 4.300E-06 | 1.000E-05 |
| 24772 | 1239 | L1239K | D | 7.4e-06 | 5.8e-06 | 7.2e-06 | 6.800E-06 | 8.718E-07 | 5.800E-06 | 7.400E-06 |
| 24773 | 1239 | L1239M | D | 9.3e-06 | 6.6e-06 | 4.5e-06 | 6.800E-06 | 2.406E-06 | 4.500E-06 | 9.300E-06 |
| 24774 | 1239 | L1239F | D | 6.6e-06 | 1e-05 | 6.2e-06 | 7.600E-06 | 2.088E-06 | 6.200E-06 | 1.000E-05 |
| 24775 | 1239 | L1239P | D | 5.8e-06 | 9.8e-06 | 1.2e-05 | 9.200E-06 | 3.143E-06 | 5.800E-06 | 1.200E-05 |
| 24776 | 1239 | L1239S | D | 7e-06 | 6e-06 | 1e-05 | 7.667E-06 | 2.082E-06 | 6.000E-06 | 1.000E-05 |
| 24777 | 1239 | L1239T | D | 8.1e-06 | 6.8e-06 | 9.7e-06 | 8.200E-06 | 1.453E-06 | 6.800E-06 | 9.700E-06 |
| 24778 | 1239 | L1239W | D | 6.5e-06 | 5.9e-06 | 1.2e-05 | 8.133E-06 | 3.362E-06 | 5.900E-06 | 1.200E-05 |
| 24779 | 1239 | L1239Y | D | 3.5e-06 | 6.8e-06 | 8.6e-06 | 6.300E-06 | 2.587E-06 | 3.500E-06 | 8.600E-06 |
| 24780 | 1239 | L1239V | D | 9.2e-06 | 7e-06 | 7.1e-06 | 7.767E-06 | 1.242E-06 | 7.000E-06 | 9.200E-06 |
| 24781 | 1240 | S1240A | D | 7.1e-06 | 4e-06 | 6.2e-06 | 5.767E-06 | 1.595E-06 | 4.000E-06 | 7.100E-06 |
| 24782 | 1240 | S1240R | D | 7.2e-06 | 9.6e-06 | 6.8e-06 | 7.867E-06 | 1.514E-06 | 6.800E-06 | 9.600E-06 |
| 24783 | 1240 | S1240N | D | 5.3e-06 | 4.7e-06 | 5.9e-06 | 5.300E-06 | 6.000E-07 | 4.700E-06 | 5.900E-06 |
| 24784 | 1240 | S1240D | D | 1e-05 | 8.7e-06 | 1.1e-05 | 9.900E-06 | 1.153E-06 | 8.700E-06 | 1.100E-05 |
| 24785 | 1240 | S1240C | D | 1.5e-05 | 8.4e-06 | 9.4e-06 | 1.093E-05 | 3.557E-06 | 8.400E-06 | 1.500E-05 |
| 24786 | 1240 | S1240E | D | 1e-05 | 1e-05 | 7.2e-06 | 9.067E-06 | 1.617E-06 | 7.200E-06 | 1.000E-05 |
| 24787 | 1240 | S1240Q | D | 8.4e-06 | 6.7e-06 | 1e-05 | 8.367E-06 | 1.650E-06 | 6.700E-06 | 1.000E-05 |
| 24788 | 1240 | S1240G | D | 7.4e-06 | 9.3e-06 | 8.8e-06 | 8.500E-06 | 9.849E-07 | 7.400E-06 | 9.300E-06 |
| 24789 | 1240 | S1240H | D | 9.7e-06 | 7.4e-06 | 7.8e-06 | 8.300E-06 | 1.229E-06 | 7.400E-06 | 9.700E-06 |
| 24790 | 1240 | S1240I | D | 8.2e-06 | 9.5e-06 | 1.3e-05 | 1.023E-05 | 2.483E-06 | 8.200E-06 | 1.300E-05 |
| 24791 | 1240 | S1240L | D | 8.7e-06 | 1e-05 | 1.4e-05 | 1.090E-05 | 2.762E-06 | 8.700E-06 | 1.400E-05 |
| 24792 | 1240 | S1240K | D | 9.5e-06 | 7.7e-06 | 4.4e-06 | 7.200E-06 | 2.587E-06 | 4.400E-06 | 9.500E-06 |
| 24793 | 1240 | S1240M | D | 9.5e-06 | 1e-05 | 6.7e-06 | 8.733E-06 | 1.779E-06 | 6.700E-06 | 1.000E-05 |
| 24794 | 1240 | S1240F | D | 9.7e-06 | 7.8e-06 | 8.5e-06 | 8.667E-06 | 9.609E-07 | 7.800E-06 | 9.700E-06 |
| 24795 | 1240 | S1240P | D | 7.3e-06 | 6.3e-06 | 6.3e-06 | 6.633E-06 | 5.774E-07 | 6.300E-06 | 7.300E-06 |
| 24796 | 1240 | S1240S | D | 4.5e-06 | 1e-05 | 1.1e-05 | 8.500E-06 | 3.500E-06 | 4.500E-06 | 1.100E-05 |
| 24797 | 1240 | S1240T | D | 4.7e-06 | 5.1e-06 | 9.7e-06 | 6.500E-06 | 2.778E-06 | 4.700E-06 | 9.700E-06 |
| 24798 | 1240 | S1240W | D | 5e-06 | 7.6e-06 | 9.1e-06 | 7.233E-06 | 2.074E-06 | 5.000E-06 | 9.100E-06 |
| 24799 | 1240 | S1240Y | D | 6.8e-06 | 6.1e-06 | 7.2e-06 | 6.700E-06 | 5.568E-07 | 6.100E-06 | 7.200E-06 |
| 24800 | 1240 | S1240V | D | 5.8e-06 | 1.3e-05 | 1.3e-05 | 1.060E-05 | 4.157E-06 | 5.800E-06 | 1.300E-05 |
| 24801 | 1241 | K1241A | D | 1.2e-05 | 6.5e-06 | 1.1e-05 | 9.833E-06 | 2.930E-06 | 6.500E-06 | 1.200E-05 |
| 24802 | 1241 | K1241R | D | 5.2e-06 | 5.8e-06 | 7.3e-06 | 6.100E-06 | 1.082E-06 | 5.200E-06 | 7.300E-06 |
| 24803 | 1241 | K1241N | D | 1.3e-05 | 6.9e-06 | 6.5e-06 | 8.800E-06 | 3.643E-06 | 6.500E-06 | 1.300E-05 |
| 24804 | 1241 | K1241D | D | 1.2e-05 | 4.6e-06 | 8.9e-06 | 8.500E-06 | 3.716E-06 | 4.600E-06 | 1.200E-05 |
| 24805 | 1241 | K1241C | D | 7.9e-06 | 7e-06 | 6.4e-06 | 7.100E-06 | 7.550E-07 | 6.400E-06 | 7.900E-06 |
| 24806 | 1241 | K1241E | D | 7.9e-06 | 1e-05 | 1.1e-05 | 9.633E-06 | 1.582E-06 | 7.900E-06 | 1.100E-05 |
| 24807 | 1241 | K1241Q | D | 8e-06 | 6.2e-06 | 1.1e-05 | 8.400E-06 | 2.425E-06 | 6.200E-06 | 1.100E-05 |
| 24808 | 1241 | K1241G | D | 8.2e-06 | 8.4e-06 | 7.6e-06 | 8.067E-06 | 4.163E-07 | 7.600E-06 | 8.400E-06 |
| 24809 | 1241 | K1241H | D | 7.7e-06 | 1.3e-05 | 8.4e-06 | 9.700E-06 | 2.879E-06 | 7.700E-06 | 1.300E-05 |
| 24810 | 1241 | K1241I | D | 1.3e-05 | 4.8e-06 | 9.1e-06 | 8.967E-06 | 4.102E-06 | 4.800E-06 | 1.300E-05 |
| 24811 | 1241 | K1241L | D | 1e-05 | 1e-05 | 1.1e-05 | 1.033E-05 | 5.774E-07 | 1.000E-05 | 1.100E-05 |
| 24812 | 1241 | K1241K | D | 1.2e-05 | 8e-06 | 4.7e-06 | 8.233E-06 | 3.656E-06 | 4.700E-06 | 1.200E-05 |
| 24813 | 1241 | K1241M | D | 7.6e-06 | 5.3e-06 | 6.5e-06 | 6.467E-06 | 1.150E-06 | 5.300E-06 | 7.600E-06 |
| 24814 | 1241 | K1241F | D | 7.6e-06 | 7.1e-06 | 5.9e-06 | 6.867E-06 | 8.737E-07 | 5.900E-06 | 7.600E-06 |
| 24815 | 1241 | K1241P | D | 5.1e-06 | 7.3e-06 | 8.5e-06 | 6.967E-06 | 1.724E-06 | 5.100E-06 | 8.500E-06 |
| 24816 | 1241 | K1241S | D | 8e-06 | 7.7e-06 | 9.3e-06 | 8.333E-06 | 8.505E-07 | 7.700E-06 | 9.300E-06 |
| 24817 | 1241 | K1241T | D | 1.6e-05 | 9.5e-06 | 4.9e-06 | 1.013E-05 | 5.577E-06 | 4.900E-06 | 1.600E-05 |
| 24818 | 1241 | K1241W | D | 8.8e-06 | 5.7e-06 | 8e-06 | 7.500E-06 | 1.609E-06 | 5.700E-06 | 8.800E-06 |
| 24819 | 1241 | K1241Y | D | 6.1e-06 | 7e-06 | 8.1e-06 | 7.067E-06 | 1.002E-06 | 6.100E-06 | 8.100E-06 |

| | | | | | | | | | | |
|---|---|---|---|---|---|---|---|---|---|---|
| 24820 | 1241 | K1241V | D | 7.7e-06 | 8.7e-06 | 7.6e-06 | 8.000E-06 | 6.083E-07 | 7.600E-06 | 8.700E-06 |
| 24821 | 1242 | A1242A | D | 6e-06 | 1e-05 | 5.6e-06 | 7.200E-06 | 2.433E-06 | 5.600E-06 | 1.000E-05 |
| 24822 | 1242 | A1242R | D | 7.4e-06 | 6.6e-06 | 9.9e-06 | 7.967E-06 | 1.721E-06 | 6.600E-06 | 9.900E-06 |
| 24823 | 1242 | A1242N | D | 8.1e-06 | 8.7e-06 | 5.9e-06 | 7.567E-06 | 1.474E-06 | 5.900E-06 | 8.700E-06 |
| 24824 | 1242 | A1242D | D | 8.7e-06 | 6.6e-06 | 5.5e-06 | 6.933E-06 | 1.626E-06 | 5.500E-06 | 8.700E-06 |
| 24825 | 1242 | A1242C | D | 4.1e-06 | 7.1e-06 | 1.3e-05 | 8.067E-06 | 4.528E-06 | 4.100E-06 | 1.300E-05 |
| 24826 | 1242 | A1242E | D | 1.2e-05 | 1.3e-05 | 7.6e-06 | 1.087E-05 | 2.873E-06 | 7.600E-06 | 1.300E-05 |
| 24827 | 1242 | A1242Q | D | 9.1e-06 | 5e-06 | 7.4e-06 | 7.167E-06 | 2.060E-06 | 5.000E-06 | 9.100E-06 |
| 24828 | 1242 | A1242G | D | 5.3e-06 | 1e-05 | 6.7e-06 | 7.333E-06 | 2.413E-06 | 5.300E-06 | 1.000E-05 |
| 24829 | 1242 | A1242H | D | 8.6e-06 | 4.9e-06 | 9.3e-06 | 7.600E-06 | 2.364E-06 | 4.900E-06 | 9.300E-06 |
| 24830 | 1242 | A1242I | D | 1.1e-05 | 6.2e-06 | 1.3e-05 | 1.007E-05 | 3.495E-06 | 6.200E-06 | 1.300E-05 |
| 24831 | 1242 | A1242L | D | 9.2e-06 | 5.2e-06 | 7.4e-06 | 7.267E-06 | 2.003E-06 | 5.200E-06 | 9.200E-06 |
| 24832 | 1242 | A1242K | D | 7.3e-06 | 1.4e-05 | 6.4e-06 | 9.233E-06 | 4.153E-06 | 6.400E-06 | 1.400E-05 |
| 24833 | 1242 | A1242M | D | 5.6e-06 | 7.1e-06 | 8.1e-06 | 6.933E-06 | 1.258E-06 | 5.600E-06 | 8.100E-06 |
| 24834 | 1242 | A1242F | D | 6.1e-06 | 9.1e-06 | 4.1e-06 | 6.433E-06 | 2.517E-06 | 4.100E-06 | 9.100E-06 |
| 24835 | 1242 | A1242P | D | 7e-06 | 1e-05 | 1.3e-05 | 1.000E-05 | 3.000E-06 | 7.000E-06 | 1.300E-05 |
| 24836 | 1242 | A1242S | D | 1.9e-05 | 9.7e-06 | 7.1e-06 | 1.193E-05 | 6.256E-06 | 7.100E-06 | 1.900E-05 |
| 24837 | 1242 | A1242T | D | 3.2e-06 | 8.7e-06 | 9.1e-06 | 7.000E-06 | 3.297E-06 | 3.200E-06 | 9.100E-06 |
| 24838 | 1242 | A1242W | D | 9.8e-06 | 9.1e-06 | 6.8e-06 | 8.567E-06 | 1.570E-06 | 6.800E-06 | 9.800E-06 |
| 24839 | 1242 | A1242Y | D | 6.8e-06 | 1.2e-05 | 9.9e-06 | 9.567E-06 | 2.616E-06 | 6.800E-06 | 1.200E-05 |
| 24840 | 1242 | A1242V | D | 9.9e-06 | 7.4e-06 | 9.7e-06 | 9.000E-06 | 1.389E-06 | 7.400E-06 | 9.900E-06 |
| 24841 | 1243 | D1243A | D | 1e-05 | 6.4e-06 | 6.6e-06 | 7.667E-06 | 2.023E-06 | 6.400E-06 | 1.000E-05 |
| 24842 | 1243 | D1243R | D | 8.4e-06 | 7.3e-06 | 5e-06 | 6.900E-06 | 1.735E-06 | 5.000E-06 | 8.400E-06 |
| 24843 | 1243 | D1243N | D | 6e-06 | 1.2e-05 | 1.2e-05 | 1.000E-05 | 3.464E-06 | 6.000E-06 | 1.200E-05 |
| 24844 | 1243 | D1243D | D | 4e-06 | 9e-06 | 6.6e-06 | 6.533E-06 | 2.501E-06 | 4.000E-06 | 9.000E-06 |
| 24845 | 1243 | D1243C | D | 5.7e-06 | 8.2e-06 | 1e-05 | 7.967E-06 | 2.159E-06 | 5.700E-06 | 1.000E-05 |
| 24846 | 1243 | D1243E | D | 5.3e-06 | 5.3e-06 | 1.2e-05 | 7.533E-06 | 3.868E-06 | 5.300E-06 | 1.200E-05 |
| 24847 | 1243 | D1243Q | D | 6.7e-06 | 1.1e-05 | 5.5e-06 | 7.733E-06 | 2.892E-06 | 5.500E-06 | 1.100E-05 |
| 24848 | 1243 | D1243G | D | 1e-05 | 6.7e-06 | 9e-06 | 8.567E-06 | 1.692E-06 | 6.700E-06 | 1.000E-05 |
| 24849 | 1243 | D1243H | D | 6.2e-06 | 9.2e-06 | 5e-06 | 6.800E-06 | 2.163E-06 | 5.000E-06 | 9.200E-06 |
| 24850 | 1243 | D1243I | D | 9.9e-06 | 7.2e-06 | 6.3e-06 | 7.800E-06 | 1.873E-06 | 6.300E-06 | 9.900E-06 |
| 24851 | 1243 | D1243L | D | 6.5e-06 | 8.7e-06 | 9.3e-06 | 8.167E-06 | 1.474E-06 | 6.500E-06 | 9.300E-06 |
| 24852 | 1243 | D1243K | D | 7.8e-06 | 8.7e-06 | 6.5e-06 | 7.667E-06 | 1.106E-06 | 6.500E-06 | 8.700E-06 |
| 24853 | 1243 | D1243M | D | 9.7e-06 | 9e-06 | 1e-05 | 9.567E-06 | 5.132E-07 | 9.000E-06 | 1.000E-05 |
| 24854 | 1243 | D1243F | D | 7.3e-06 | 9.2e-06 | 1.7e-05 | 1.117E-05 | 5.140E-06 | 7.300E-06 | 1.700E-05 |
| 24855 | 1243 | D1243P | D | 6.1e-06 | 7.9e-06 | 1.3e-05 | 9.000E-06 | 3.579E-06 | 6.100E-06 | 1.300E-05 |
| 24856 | 1243 | D1243S | D | 1.3e-05 | 4.9e-06 | 8.8e-06 | 8.900E-06 | 4.051E-06 | 4.900E-06 | 1.300E-05 |
| 24857 | 1243 | D1243T | D | 5.3e-06 | 7.3e-06 | 1.3e-05 | 8.533E-06 | 3.995E-06 | 5.300E-06 | 1.300E-05 |
| 24858 | 1243 | D1243W | D | 8.5e-06 | 6.5e-06 | 1.1e-05 | 8.667E-06 | 2.255E-06 | 6.500E-06 | 1.100E-05 |
| 24859 | 1243 | D1243Y | D | 1.4e-05 | 6.9e-06 | 7.3e-06 | 9.400E-06 | 3.989E-06 | 6.900E-06 | 1.400E-05 |
| 24860 | 1243 | D1243V | D | 5e-06 | 6.8e-06 | 7.5e-06 | 6.433E-06 | 1.290E-06 | 5.000E-06 | 7.500E-06 |
| 24861 | 1244 | Y1244A | D | 9.4e-06 | 6.1e-06 | 1.3e-05 | 9.500E-06 | 3.451E-06 | 6.100E-06 | 1.300E-05 |
| 24862 | 1244 | Y1244R | D | 8e-06 | 6.6e-06 | 8.9e-06 | 7.833E-06 | 1.159E-06 | 6.600E-06 | 8.900E-06 |
| 24863 | 1244 | Y1244N | D | 9.6e-06 | 8.5e-06 | 9.3e-06 | 9.133E-06 | 5.686E-07 | 8.500E-06 | 9.600E-06 |
| 24864 | 1244 | Y1244D | D | 4.7e-06 | 7.8e-06 | 9.5e-06 | 7.333E-06 | 2.434E-06 | 4.700E-06 | 9.500E-06 |
| 24865 | 1244 | Y1244C | D | 1.1e-05 | 4.6e-06 | 1e-05 | 8.533E-06 | 3.443E-06 | 4.600E-06 | 1.100E-05 |
| 24866 | 1244 | Y1244E | D | 7.8e-06 | 1.6e-05 | 1e-05 | 1.127E-05 | 4.244E-06 | 7.800E-06 | 1.600E-05 |
| 24867 | 1244 | Y1244Q | D | 8.4e-06 | 7.1e-06 | 7.5e-06 | 7.667E-06 | 6.658E-07 | 7.100E-06 | 8.400E-06 |
| 24868 | 1244 | Y1244G | D | 9.6e-06 | 6e-06 | 1.3e-05 | 9.533E-06 | 3.500E-06 | 6.000E-06 | 1.300E-05 |
| 24869 | 1244 | Y1244H | D | 1e-05 | 6.4e-06 | 7.3e-06 | 7.900E-06 | 1.873E-06 | 6.400E-06 | 1.000E-05 |
| 24870 | 1244 | Y1244I | D | 4.8e-06 | 1.4e-05 | 6.4e-06 | 8.400E-06 | 4.915E-06 | 4.800E-06 | 1.400E-05 |
| 24871 | 1244 | Y1244L | D | 5.3e-06 | 6.7e-06 | 6.8e-06 | 6.267E-06 | 8.386E-07 | 5.300E-06 | 6.800E-06 |
| 24872 | 1244 | Y1244K | D | 1.5e-05 | 9.1e-06 | 6.4e-06 | 1.017E-05 | 4.398E-06 | 6.400E-06 | 1.500E-05 |
| 24873 | 1244 | Y1244M | D | 7e-06 | 6.5e-06 | 1e-05 | 7.833E-06 | 1.893E-06 | 6.500E-06 | 1.000E-05 |
| 24874 | 1244 | Y1244F | D | 7.3e-06 | 6.5e-06 | 8.1e-06 | 7.300E-06 | 8.000E-07 | 6.500E-06 | 8.100E-06 |
| 24875 | 1244 | Y1244P | D | 4.6e-06 | 7.3e-06 | 8.8e-06 | 6.900E-06 | 2.128E-06 | 4.600E-06 | 8.800E-06 |
| 24876 | 1244 | Y1244S | D | 6.5e-06 | 1.3e-05 | 6.1e-06 | 8.533E-06 | 3.873E-06 | 6.100E-06 | 1.300E-05 |
| 24877 | 1244 | Y1244T | D | 1.2e-05 | 9.9e-06 | 8e-06 | 9.967E-06 | 2.001E-06 | 8.000E-06 | 1.200E-05 |
| 24878 | 1244 | Y1244W | D | 5e-06 | 6.5e-06 | 8.3e-06 | 6.600E-06 | 1.652E-06 | 5.000E-06 | 8.300E-06 |
| 24879 | 1244 | Y1244Y | D | 5.6e-06 | 8.3e-06 | 6.1e-06 | 6.667E-06 | 1.436E-06 | 5.600E-06 | 8.300E-06 |
| 24880 | 1244 | Y1244V | D | 6.8e-06 | 6.3e-06 | 6.9e-06 | 6.667E-06 | 3.215E-07 | 6.300E-06 | 6.900E-06 |
| 24881 | 1245 | E1245A | D | 9.2e-06 | 6.9e-06 | 6.4e-06 | 7.500E-06 | 1.493E-06 | 6.400E-06 | 9.200E-06 |
| 24882 | 1245 | E1245R | D | 4.9e-06 | 2.4e-06 | 1.1e-05 | 6.100E-06 | 4.424E-06 | 2.400E-06 | 1.100E-05 |
| 24883 | 1245 | E1245N | D | 6.1e-06 | 3.5e-06 | 1.1e-05 | 6.867E-06 | 3.808E-06 | 3.500E-06 | 1.100E-05 |
| 24884 | 1245 | E1245D | D | 8.4e-06 | 6.1e-06 | 1e-05 | 8.167E-06 | 1.960E-06 | 6.100E-06 | 1.000E-05 |
| 24885 | 1245 | E1245C | D | 6.6e-06 | 5.4e-06 | 7.2e-06 | 6.400E-06 | 9.165E-07 | 5.400E-06 | 7.200E-06 |
| 24886 | 1245 | E1245E | D | 5.1e-06 | 9.9e-06 | 9.4e-06 | 8.133E-06 | 2.639E-06 | 5.100E-06 | 9.900E-06 |
| 24887 | 1245 | E1245Q | D | 8.5e-06 | 6.5e-06 | 6.4e-06 | 7.133E-06 | 1.185E-06 | 6.400E-06 | 8.500E-06 |
| 24888 | 1245 | E1245G | D | 1e-05 | 9.1e-06 | 7.1e-06 | 8.733E-06 | 1.484E-06 | 7.100E-06 | 1.000E-05 |
| 24889 | 1245 | E1245H | D | 7.5e-06 | 5.4e-06 | 9.5e-06 | 7.467E-06 | 2.050E-06 | 5.400E-06 | 9.500E-06 |
| 24890 | 1245 | E1245I | D | 5.3e-06 | 5.4e-06 | 8.1e-06 | 6.267E-06 | 1.589E-06 | 5.300E-06 | 8.100E-06 |
| 24891 | 1245 | E1245L | D | 1e-05 | 5.7e-06 | 5.8e-06 | 7.167E-06 | 2.454E-06 | 5.700E-06 | 1.000E-05 |
| 24892 | 1245 | E1245K | D | 1.1e-05 | 6e-06 | 1e-05 | 9.000E-06 | 2.646E-06 | 6.000E-06 | 1.100E-05 |
| 24893 | 1245 | E1245M | D | 1e-05 | 1.4e-05 | 9.3e-06 | 1.110E-05 | 2.536E-06 | 9.300E-06 | 1.400E-05 |
| 24894 | 1245 | E1245F | D | 9.7e-06 | 6.6e-06 | 9.3e-06 | 8.533E-06 | 1.686E-06 | 6.600E-06 | 9.700E-06 |
| 24895 | 1245 | E1245P | D | 4.8e-06 | 7.2e-06 | 8.4e-06 | 6.800E-06 | 1.833E-06 | 4.800E-06 | 8.400E-06 |
| 24896 | 1245 | E1245S | D | 6.2e-06 | 1e-05 | 6.5e-06 | 7.567E-06 | 2.113E-06 | 6.200E-06 | 1.000E-05 |
| 24897 | 1245 | E1245T | D | 7.1e-06 | 9.4e-06 | 1.1e-05 | 9.167E-06 | 1.960E-06 | 7.100E-06 | 1.100E-05 |
| 24898 | 1245 | E1245W | D | 6.8e-06 | 7.4e-06 | 1.4e-05 | 9.400E-06 | 3.995E-06 | 6.800E-06 | 1.400E-05 |
| 24899 | 1245 | E1245Y | D | 6.8e-06 | 9.7e-06 | 7.3e-06 | 7.933E-06 | 1.550E-06 | 6.800E-06 | 9.700E-06 |
| 24900 | 1245 | E1245V | D | 5.5e-06 | 7.8e-06 | 9.8e-06 | 7.700E-06 | 2.152E-06 | 5.500E-06 | 9.800E-06 |
| 24901 | 1246 | K1246A | D | 5.7e-06 | 7e-06 | 6.7e-06 | 6.467E-06 | 6.807E-07 | 5.700E-06 | 7.000E-06 |
| 24902 | 1246 | K1246R | D | 6.7e-06 | 1e-05 | 1.3e-05 | 9.900E-06 | 3.151E-06 | 6.700E-06 | 1.300E-05 |
| 24903 | 1246 | K1246N | D | 8.3e-06 | 8.1e-06 | 4.6e-06 | 7.000E-06 | 2.081E-06 | 4.600E-06 | 8.300E-06 |
| 24904 | 1246 | K1246D | D | 7.5e-06 | 9.2e-06 | 1.5e-05 | 1.057E-05 | 3.932E-06 | 7.500E-06 | 1.500E-05 |

| | | | | | | | | | | |
|---|---|---|---|---|---|---|---|---|---|---|
| 24905 | 1246 | K1246C | D | 7.3e-06 | 4.9e-06 | 1.3e-05 | 8.400E-06 | 4.161E-06 | 4.900E-06 | 1.300E-05 |
| 24906 | 1246 | K1246E | D | 7.6e-06 | 6.9e-06 | 6.7e-06 | 7.067E-06 | 4.726E-07 | 6.700E-06 | 7.600E-06 |
| 24907 | 1246 | K1246Q | D | 1.5e-05 | 8e-06 | 7e-06 | 1.000E-05 | 4.359E-06 | 7.000E-06 | 1.500E-05 |
| 24908 | 1246 | K1246G | D | 7e-06 | 6.7e-06 | 1e-05 | 7.900E-06 | 1.825E-06 | 6.700E-06 | 1.000E-05 |
| 24909 | 1246 | K1246H | D | 4.9e-06 | 4.3e-06 | 7e-06 | 5.400E-06 | 1.418E-06 | 4.300E-06 | 7.000E-06 |
| 24910 | 1246 | K1246I | D | 4.8e-06 | 1.1e-05 | 9e-06 | 8.267E-06 | 3.164E-06 | 4.800E-06 | 1.100E-05 |
| 24911 | 1246 | K1246L | D | 7.6e-06 | 1.1e-05 | 5.3e-06 | 7.967E-06 | 2.868E-06 | 5.300E-06 | 1.100E-05 |
| 24912 | 1246 | K1246K | D | 8.6e-06 | 1e-05 | 9.8e-06 | 9.467E-06 | 7.572E-07 | 8.600E-06 | 1.000E-05 |
| 24913 | 1246 | K1246M | D | 6.2e-06 | 6.1e-06 | 6.4e-06 | 6.233E-06 | 1.528E-07 | 6.100E-06 | 6.400E-06 |
| 24914 | 1246 | K1246F | D | 1.2e-05 | 1e-05 | 1.3e-05 | 1.167E-05 | 1.528E-06 | 1.000E-05 | 1.300E-05 |
| 24915 | 1246 | K1246P | D | 9.7e-06 | 3e-06 | 6e-06 | 6.233E-06 | 3.356E-06 | 3.000E-06 | 9.700E-06 |
| 24916 | 1246 | K1246S | D | 4.2e-06 | 9.5e-06 | 1e-05 | 7.900E-06 | 3.214E-06 | 4.200E-06 | 1.000E-05 |
| 24917 | 1246 | K1246T | D | 6.5e-06 | 7.9e-06 | 6.4e-06 | 6.933E-06 | 8.386E-07 | 6.400E-06 | 7.900E-06 |
| 24918 | 1246 | K1246W | D | 8.9e-06 | 1e-05 | 1.1e-05 | 9.967E-06 | 1.050E-06 | 8.900E-06 | 1.100E-05 |
| 24919 | 1246 | K1246Y | D | 5e-06 | 5.7e-06 | 7.2e-06 | 5.967E-06 | 1.124E-06 | 5.000E-06 | 7.200E-06 |
| 24920 | 1246 | K1246V | D | 8e-06 | 1.1e-05 | 4.6e-06 | 7.867E-06 | 3.202E-06 | 4.600E-06 | 1.100E-05 |
| 24921 | 1247 | H1247A | D | 1.3e-05 | 4.8e-06 | 5.7e-06 | 7.833E-06 | 4.497E-06 | 4.800E-06 | 1.300E-05 |
| 24922 | 1247 | H1247R | D | 7e-06 | 1.1e-05 | 3.7e-06 | 7.233E-06 | 3.656E-06 | 3.700E-06 | 1.100E-05 |
| 24923 | 1247 | H1247N | D | 9.3e-06 | 5.7e-06 | 7.2e-06 | 7.400E-06 | 1.808E-06 | 5.700E-06 | 9.300E-06 |
| 24924 | 1247 | H1247D | D | 9.2e-06 | 8.6e-06 | 1e-05 | 9.267E-06 | 7.024E-07 | 8.600E-06 | 1.000E-05 |
| 24925 | 1247 | H1247C | D | 4.6e-06 | 6.5e-06 | 6.2e-06 | 5.767E-06 | 1.021E-06 | 4.600E-06 | 6.500E-06 |
| 24926 | 1247 | H1247E | D | 1.2e-05 | 1e-05 | 5.6e-06 | 9.200E-06 | 3.274E-06 | 5.600E-06 | 1.200E-05 |
| 24927 | 1247 | H1247Q | D | 7.3e-06 | 1.2e-05 | 5.4e-06 | 8.233E-06 | 3.398E-06 | 5.400E-06 | 1.200E-05 |
| 24928 | 1247 | H1247G | D | 1.1e-05 | 1.3e-05 | 5.8e-06 | 9.933E-06 | 3.717E-06 | 5.800E-06 | 1.300E-05 |
| 24929 | 1247 | H1247H | D | 5.2e-06 | 4.8e-06 | 8.8e-06 | 6.267E-06 | 2.203E-06 | 4.800E-06 | 8.800E-06 |
| 24930 | 1247 | H1247I | D | 8.5e-06 | 7.4e-06 | 5.2e-06 | 7.033E-06 | 1.680E-06 | 5.200E-06 | 8.500E-06 |
| 24931 | 1247 | H1247L | D | 7.3e-06 | 6.8e-06 | 7.1e-06 | 7.067E-06 | 2.517E-07 | 6.800E-06 | 7.300E-06 |
| 24932 | 1247 | H1247K | D | 6.1e-06 | 3.5e-06 | 6.9e-06 | 5.500E-06 | 1.778E-06 | 3.500E-06 | 6.900E-06 |
| 24933 | 1247 | H1247M | D | 1.3e-05 | 9.7e-06 | 1.1e-05 | 1.123E-05 | 1.662E-06 | 9.700E-06 | 1.300E-05 |
| 24934 | 1247 | H1247F | D | 5.9e-06 | 5e-06 | 9.3e-06 | 6.733E-06 | 2.268E-06 | 5.000E-06 | 9.300E-06 |
| 24935 | 1247 | H1247P | D | 4.7e-06 | 6.9e-06 | 1.1e-05 | 7.533E-06 | 3.197E-06 | 4.700E-06 | 1.100E-05 |
| 24936 | 1247 | H1247S | D | 7.2e-06 | 1.1e-05 | 5.7e-06 | 7.967E-06 | 2.732E-06 | 5.700E-06 | 1.100E-05 |
| 24937 | 1247 | H1247T | D | 9.3e-06 | 6e-06 | 6.7e-06 | 7.333E-06 | 1.739E-06 | 6.000E-06 | 9.300E-06 |
| 24938 | 1247 | H1247W | D | 7.1e-06 | 1.4e-05 | 6.8e-06 | 9.300E-06 | 4.073E-06 | 6.800E-06 | 1.400E-05 |
| 24939 | 1247 | H1247Y | D | 5.5e-06 | 4.5e-06 | 6.1e-06 | 5.367E-06 | 8.083E-07 | 4.500E-06 | 6.100E-06 |
| 24940 | 1247 | H1247V | D | 9.3e-06 | 6.8e-06 | 5.8e-06 | 7.300E-06 | 1.803E-06 | 5.800E-06 | 9.300E-06 |
| 24941 | 1248 | K1248A | D | 5.7e-06 | 7.9e-06 | 6.8e-06 | 6.800E-06 | 1.100E-06 | 5.700E-06 | 7.900E-06 |
| 24942 | 1248 | K1248R | D | 9.6e-06 | 1.3e-05 | 6.2e-06 | 9.600E-06 | 3.400E-06 | 6.200E-06 | 1.300E-05 |
| 24943 | 1248 | K1248N | D | 1e-05 | 5.3e-06 | 8.9e-06 | 8.067E-06 | 2.458E-06 | 5.300E-06 | 1.000E-05 |
| 24944 | 1248 | K1248D | D | 5.4e-06 | 6.6e-06 | 6.2e-06 | 6.067E-06 | 6.110E-07 | 5.400E-06 | 6.600E-06 |
| 24945 | 1248 | K1248C | D | 8.6e-06 | 9.7e-06 | 1.7e-05 | 1.177E-05 | 4.565E-06 | 8.600E-06 | 1.700E-05 |
| 24946 | 1248 | K1248E | D | 7.3e-06 | 7e-06 | 9.2e-06 | 7.833E-06 | 1.193E-06 | 7.000E-06 | 9.200E-06 |
| 24947 | 1248 | K1248Q | D | 8.5e-06 | 6.9e-06 | 6.9e-06 | 7.433E-06 | 9.238E-07 | 6.900E-06 | 8.500E-06 |
| 24948 | 1248 | K1248G | D | 1.2e-05 | 1e-05 | 8e-06 | 1.000E-05 | 2.000E-06 | 8.000E-06 | 1.200E-05 |
| 24949 | 1248 | K1248H | D | 6.6e-06 | 5.5e-06 | 2.8e-06 | 4.967E-06 | 1.955E-06 | 2.800E-06 | 6.600E-06 |
| 24950 | 1248 | K1248I | D | 9.3e-06 | 8.3e-06 | 8e-06 | 8.533E-06 | 6.807E-07 | 8.000E-06 | 9.300E-06 |
| 24951 | 1248 | K1248L | D | 6.1e-06 | 1.3e-05 | 8.3e-06 | 9.133E-06 | 3.525E-06 | 6.100E-06 | 1.300E-05 |
| 24952 | 1248 | K1248K | D | 7.8e-06 | 5.3e-06 | 4e-06 | 5.700E-06 | 1.931E-06 | 4.000E-06 | 7.800E-06 |
| 24953 | 1248 | K1248M | D | 9.6e-06 | 6.7e-06 | 1.5e-05 | 1.043E-05 | 4.212E-06 | 6.700E-06 | 1.500E-05 |
| 24954 | 1248 | K1248F | D | 5.8e-06 | 8.9e-06 | 7.5e-06 | 7.400E-06 | 1.552E-06 | 5.800E-06 | 8.900E-06 |
| 24955 | 1248 | K1248P | D | 7.9e-06 | 6.3e-06 | 6.9e-06 | 7.033E-06 | 8.083E-07 | 6.300E-06 | 7.900E-06 |
| 24956 | 1248 | K1248S | D | 4e-06 | 7.2e-06 | 6.3e-06 | 5.833E-06 | 1.650E-06 | 4.000E-06 | 7.200E-06 |
| 24957 | 1248 | K1248T | D | 1.1e-05 | 7.4e-06 | 8.8e-06 | 9.067E-06 | 1.815E-06 | 7.400E-06 | 1.100E-05 |
| 24958 | 1248 | K1248W | D | 5.3e-06 | 1.1e-05 | 1.6e-05 | 1.077E-05 | 5.354E-06 | 5.300E-06 | 1.600E-05 |
| 24959 | 1248 | K1248Y | D | 9e-06 | 9e-06 | 7.3e-06 | 8.433E-06 | 9.815E-07 | 7.300E-06 | 9.000E-06 |
| 24960 | 1248 | K1248V | D | 1.4e-05 | 6.7e-06 | 6.7e-06 | 9.133E-06 | 4.215E-06 | 6.700E-06 | 1.400E-05 |
| 24961 | 1249 | V1249A | D | 7.6e-06 | 1.1e-05 | 7.8e-06 | 8.800E-06 | 1.908E-06 | 7.600E-06 | 1.100E-05 |
| 24962 | 1249 | V1249R | D | 8.8e-06 | 9.4e-06 | 1.1e-05 | 9.733E-06 | 1.137E-06 | 8.800E-06 | 1.100E-05 |
| 24963 | 1249 | V1249N | D | 6.6e-06 | 7.1e-06 | 8.7e-06 | 7.467E-06 | 1.097E-06 | 6.600E-06 | 8.700E-06 |
| 24964 | 1249 | V1249D | D | 8.8e-06 | 6.7e-06 | 7e-06 | 7.500E-06 | 1.136E-06 | 6.700E-06 | 8.800E-06 |
| 24965 | 1249 | V1249C | D | 9.1e-06 | 6.2e-06 | 7e-06 | 7.433E-06 | 1.498E-06 | 6.200E-06 | 9.100E-06 |
| 24966 | 1249 | V1249E | D | 1.2e-05 | 7.9e-06 | 6.6e-06 | 8.833E-06 | 2.818E-06 | 6.600E-06 | 1.200E-05 |
| 24967 | 1249 | V1249Q | D | 9.5e-06 | 8.9e-06 | 8.2e-06 | 8.867E-06 | 6.506E-07 | 8.200E-06 | 9.500E-06 |
| 24968 | 1249 | V1249G | D | 7.6e-06 | 5.3e-06 | 6.4e-06 | 6.433E-06 | 1.150E-06 | 5.300E-06 | 7.600E-06 |
| 24969 | 1249 | V1249H | D | 6e-06 | 6.9e-06 | 6.6e-06 | 6.500E-06 | 4.583E-07 | 6.000E-06 | 6.900E-06 |
| 24970 | 1249 | V1249I | D | 6.7e-06 | 7.4e-06 | 3.1e-06 | 5.733E-06 | 2.307E-06 | 3.100E-06 | 7.400E-06 |
| 24971 | 1249 | V1249L | D | 9.8e-06 | 1.2e-05 | 5.2e-06 | 9.000E-06 | 3.470E-06 | 5.200E-06 | 1.200E-05 |
| 24972 | 1249 | V1249K | D | 7.4e-06 | 6.8e-06 | 5.8e-06 | 6.667E-06 | 8.083E-07 | 5.800E-06 | 7.400E-06 |
| 24973 | 1249 | V1249M | D | 1.4e-05 | 6.6e-06 | 1e-05 | 1.020E-05 | 3.704E-06 | 6.600E-06 | 1.400E-05 |
| 24974 | 1249 | V1249F | D | 8.4e-06 | 1.2e-05 | 7.6e-06 | 9.333E-06 | 2.344E-06 | 7.600E-06 | 1.200E-05 |
| 24975 | 1249 | V1249P | D | 1.2e-05 | 1.3e-05 | 8.4e-06 | 1.113E-05 | 2.419E-06 | 8.400E-06 | 1.300E-05 |
| 24976 | 1249 | V1249S | D | 1.1e-05 | 5.8e-06 | 8.1e-06 | 8.300E-06 | 2.606E-06 | 5.800E-06 | 1.100E-05 |
| 24977 | 1249 | V1249T | D | 1.3e-05 | 4.7e-06 | 5.2e-06 | 7.633E-06 | 4.654E-06 | 4.700E-06 | 1.300E-05 |
| 24978 | 1249 | V1249W | D | 7.2e-06 | 1.4e-05 | 5.1e-06 | 8.767E-06 | 4.652E-06 | 5.100E-06 | 1.400E-05 |
| 24979 | 1249 | V1249Y | D | 5.7e-06 | 9.8e-06 | 4.1e-06 | 6.533E-06 | 2.940E-06 | 4.100E-06 | 9.800E-06 |
| 24980 | 1249 | V1249V | D | 7.7e-06 | 9.5e-06 | 6.5e-06 | 7.900E-06 | 1.510E-06 | 6.500E-06 | 9.500E-06 |
| 24981 | 1250 | Y1250A | D | 9.3e-06 | 3.9e-06 | 1.1e-05 | 8.067E-06 | 3.707E-06 | 3.900E-06 | 1.100E-05 |
| 24982 | 1250 | Y1250R | D | 9.8e-06 | 7.7e-06 | 9e-06 | 8.833E-06 | 1.060E-06 | 7.700E-06 | 9.800E-06 |
| 24983 | 1250 | Y1250N | D | 6.8e-06 | 5.6e-06 | 6.7e-06 | 6.367E-06 | 6.658E-07 | 5.600E-06 | 6.800E-06 |
| 24984 | 1250 | Y1250D | D | 7.2e-06 | 6.6e-06 | 1.3e-05 | 8.933E-06 | 3.535E-06 | 6.600E-06 | 1.300E-05 |
| 24985 | 1250 | Y1250C | D | 1e-05 | 5.5e-06 | 8.9e-06 | 8.133E-06 | 2.346E-06 | 5.500E-06 | 1.000E-05 |
| 24986 | 1250 | Y1250E | D | 5.1e-06 | 7.9e-06 | 1.4e-05 | 9.000E-06 | 4.551E-06 | 5.100E-06 | 1.400E-05 |
| 24987 | 1250 | Y1250Q | D | 8.4e-06 | 4.8e-06 | 8.1e-06 | 7.100E-06 | 1.997E-06 | 4.800E-06 | 8.400E-06 |
| 24988 | 1250 | Y1250G | D | 7.9e-06 | 9.1e-06 | 6.4e-06 | 7.800E-06 | 1.353E-06 | 6.400E-06 | 9.100E-06 |
| 24989 | 1250 | Y1250H | D | 1e-05 | 7.8e-06 | 6.5e-06 | 8.100E-06 | 1.769E-06 | 6.500E-06 | 1.000E-05 |

| | | | | | | | | | | |
|---|---|---|---|---|---|---|---|---|---|---|
| 24990 | 1250 | Y1250I | D | 5.6e-06 | 4.4e-06 | 1.5e-05 | 8.333E-06 | 5.805E-06 | 4.400E-06 | 1.500E-05 |
| 24991 | 1250 | Y1250L | D | 6.2e-06 | 7.4e-06 | 3.7e-06 | 5.767E-06 | 1.888E-06 | 3.700E-06 | 7.400E-06 |
| 24992 | 1250 | Y1250K | D | 7.2e-06 | 1e-05 | 6.5e-06 | 7.900E-06 | 1.852E-06 | 6.500E-06 | 1.000E-05 |
| 24993 | 1250 | Y1250M | D | 7.6e-06 | 1.2e-05 | 9.5e-06 | 9.700E-06 | 2.207E-06 | 7.600E-06 | 1.200E-05 |
| 24994 | 1250 | Y1250F | D | 1.1e-05 | 9.9e-06 | 1.1e-05 | 1.063E-05 | 6.351E-07 | 9.900E-06 | 1.100E-05 |
| 24995 | 1250 | Y1250P | D | 4.5e-06 | 6.6e-06 | 9.3e-06 | 6.800E-06 | 2.406E-06 | 4.500E-06 | 9.300E-06 |
| 24996 | 1250 | Y1250S | D | 7.5e-06 | 7.8e-06 | 6.9e-06 | 7.400E-06 | 4.583E-07 | 6.900E-06 | 7.800E-06 |
| 24997 | 1250 | Y1250T | D | 1.2e-05 | 7.8e-06 | 5.1e-06 | 8.300E-06 | 3.477E-06 | 5.100E-06 | 1.200E-05 |
| 24998 | 1250 | Y1250W | D | 8.8e-06 | 5.3e-06 | 5.7e-06 | 6.600E-06 | 1.916E-06 | 5.300E-06 | 8.800E-06 |
| 24999 | 1250 | Y1250Y | D | 9.3e-06 | 9.5e-06 | 6.4e-06 | 8.400E-06 | 1.735E-06 | 6.400E-06 | 9.500E-06 |
| 25000 | 1250 | Y1250V | D | 1.3e-05 | 4.8e-06 | 9.4e-06 | 9.067E-06 | 4.110E-06 | 4.800E-06 | 1.300E-05 |
| 25001 | 1251 | A1251A | D | 8.3e-06 | 8.6e-06 | 1e-05 | 8.967E-06 | 9.074E-07 | 8.300E-06 | 1.000E-05 |
| 25002 | 1251 | A1251R | D | 6.6e-06 | 1.3e-05 | 1.2e-05 | 1.053E-05 | 3.443E-06 | 6.600E-06 | 1.300E-05 |
| 25003 | 1251 | A1251N | D | 1.1e-05 | 1.2e-05 | 6.4e-06 | 9.800E-06 | 2.987E-06 | 6.400E-06 | 1.200E-05 |
| 25004 | 1251 | A1251D | D | 4.9e-06 | 6.9e-06 | 1.1e-05 | 7.600E-06 | 3.110E-06 | 4.900E-06 | 1.100E-05 |
| 25005 | 1251 | A1251C | D | 8.8e-06 | 9e-06 | 5.1e-06 | 7.633E-06 | 2.196E-06 | 5.100E-06 | 9.000E-06 |
| 25006 | 1251 | A1251E | D | 1e-05 | 5.9e-06 | 4.8e-06 | 6.900E-06 | 2.740E-06 | 4.800E-06 | 1.000E-05 |
| 25007 | 1251 | A1251Q | D | 6.8e-06 | 5.7e-06 | 7.8e-06 | 6.767E-06 | 1.050E-06 | 5.700E-06 | 7.800E-06 |
| 25008 | 1251 | A1251G | D | 1.1e-05 | 1.4e-05 | 5e-06 | 1.000E-05 | 4.583E-06 | 5.000E-06 | 1.400E-05 |
| 25009 | 1251 | A1251H | D | 6.6e-06 | 7.9e-06 | 1.2e-05 | 8.833E-06 | 2.818E-06 | 6.600E-06 | 1.200E-05 |
| 25010 | 1251 | A1251I | D | 9.2e-06 | 1.5e-05 | 8.7e-06 | 1.097E-05 | 3.502E-06 | 8.700E-06 | 1.500E-05 |
| 25011 | 1251 | A1251L | D | 9.3e-06 | 7.4e-06 | 7.3e-06 | 8.000E-06 | 1.127E-06 | 7.300E-06 | 9.300E-06 |
| 25012 | 1251 | A1251K | D | 1.1e-05 | 3.7e-06 | 7.6e-06 | 7.433E-06 | 3.653E-06 | 3.700E-06 | 1.100E-05 |
| 25013 | 1251 | A1251M | D | 6.7e-06 | 9.7e-06 | 6.3e-06 | 7.567E-06 | 1.858E-06 | 6.300E-06 | 9.700E-06 |
| 25014 | 1251 | A1251F | D | 1.2e-05 | 1.2e-05 | 1.3e-05 | 1.233E-05 | 5.774E-07 | 1.200E-05 | 1.300E-05 |
| 25015 | 1251 | A1251P | D | 5.7e-06 | 9.7e-06 | 6.1e-06 | 7.167E-06 | 2.203E-06 | 5.700E-06 | 9.700E-06 |
| 25016 | 1251 | A1251S | D | 8.9e-06 | 8.8e-06 | 4.9e-06 | 7.533E-06 | 2.281E-06 | 4.900E-06 | 8.900E-06 |
| 25017 | 1251 | A1251T | D | 7e-06 | 8.7e-06 | 6.8e-06 | 7.500E-06 | 1.044E-06 | 6.800E-06 | 8.700E-06 |
| 25018 | 1251 | A1251W | D | 7.7e-06 | 7.1e-06 | 1.2e-05 | 8.933E-06 | 2.673E-06 | 7.100E-06 | 1.200E-05 |
| 25019 | 1251 | A1251Y | D | 1.5e-05 | 9e-06 | 6.3e-06 | 1.010E-05 | 4.453E-06 | 6.300E-06 | 1.500E-05 |
| 25020 | 1251 | A1251V | D | 3.7e-06 | 5.4e-06 | 1.1e-05 | 6.700E-06 | 3.820E-06 | 3.700E-06 | 1.100E-05 |
| 25021 | 1252 | C1252A | D | 5.4e-06 | 5.2e-06 | 8.2e-06 | 6.267E-06 | 1.677E-06 | 5.200E-06 | 8.200E-06 |
| 25022 | 1252 | C1252R | D | 5.7e-06 | 7.9e-06 | 8.3e-06 | 7.300E-06 | 1.400E-06 | 5.700E-06 | 8.300E-06 |
| 25023 | 1252 | C1252N | D | 7.7e-06 | 4.8e-06 | 7.7e-06 | 6.733E-06 | 1.674E-06 | 4.800E-06 | 7.700E-06 |
| 25024 | 1252 | C1252D | D | 6e-06 | 9.1e-06 | 1e-05 | 8.367E-06 | 2.098E-06 | 6.000E-06 | 1.000E-05 |
| 25025 | 1252 | C1252C | D | 1e-05 | 8.3e-06 | 5.6e-06 | 7.967E-06 | 2.219E-06 | 5.600E-06 | 1.000E-05 |
| 25026 | 1252 | C1252E | D | 1.1e-05 | 1.3e-05 | 8.3e-06 | 1.077E-05 | 2.359E-06 | 8.300E-06 | 1.300E-05 |
| 25027 | 1252 | C1252Q | D | 7.2e-06 | 7.2e-06 | 4.9e-06 | 6.433E-06 | 1.328E-06 | 4.900E-06 | 7.200E-06 |
| 25028 | 1252 | C1252G | D | 7.8e-06 | 5.2e-06 | 7.5e-06 | 6.833E-06 | 1.422E-06 | 5.200E-06 | 7.800E-06 |
| 25029 | 1252 | C1252H | D | 6.4e-06 | 6.7e-06 | 1.2e-05 | 8.367E-06 | 3.150E-06 | 6.400E-06 | 1.200E-05 |
| 25030 | 1252 | C1252I | D | 1.4e-05 | 9.8e-06 | 1.3e-05 | 1.227E-05 | 2.194E-06 | 9.800E-06 | 1.400E-05 |
| 25031 | 1252 | C1252L | D | 8.1e-06 | 7.3e-06 | 3.1e-06 | 6.167E-06 | 2.686E-06 | 3.100E-06 | 8.100E-06 |
| 25032 | 1252 | C1252K | D | 8.5e-06 | 7.4e-06 | 1e-05 | 8.633E-06 | 1.305E-06 | 7.400E-06 | 1.000E-05 |
| 25033 | 1252 | C1252M | D | 9.1e-06 | 4.4e-06 | 6.4e-06 | 6.633E-06 | 2.359E-06 | 4.400E-06 | 9.100E-06 |
| 25034 | 1252 | C1252F | D | 4e-06 | 5.7e-06 | 1.3e-05 | 7.567E-06 | 4.782E-06 | 4.000E-06 | 1.300E-05 |
| 25035 | 1252 | C1252P | D | 6.4e-06 | 7.7e-06 | 7.9e-06 | 7.333E-06 | 8.145E-07 | 6.400E-06 | 7.900E-06 |
| 25036 | 1252 | C1252S | D | 1.4e-05 | 8.7e-06 | 5.6e-06 | 9.433E-06 | 4.248E-06 | 5.600E-06 | 1.400E-05 |
| 25037 | 1252 | C1252T | D | 1.3e-05 | 1.4e-05 | 9.6e-06 | 1.220E-05 | 2.307E-06 | 9.600E-06 | 1.400E-05 |
| 25038 | 1252 | C1252W | D | 4.9e-06 | 1.1e-05 | 7.2e-06 | 7.700E-06 | 3.081E-06 | 4.900E-06 | 1.100E-05 |
| 25039 | 1252 | C1252Y | D | 8.4e-06 | 1e-05 | 6.9e-06 | 8.433E-06 | 1.550E-06 | 6.900E-06 | 1.000E-05 |
| 25040 | 1252 | C1252V | D | 9.5e-06 | 1.3e-05 | 7e-06 | 9.833E-06 | 3.014E-06 | 7.000E-06 | 1.300E-05 |
| 25041 | 1253 | E1253A | D | 6.8e-06 | 6.6e-06 | 5.5e-06 | 6.300E-06 | 7.000E-07 | 5.500E-06 | 6.800E-06 |
| 25042 | 1253 | E1253R | D | 1.3e-05 | 5e-06 | 9.7e-06 | 9.233E-06 | 4.020E-06 | 5.000E-06 | 1.300E-05 |
| 25043 | 1253 | E1253N | D | 8.6e-06 | 5.5e-06 | 6.5e-06 | 6.867E-06 | 1.582E-06 | 5.500E-06 | 8.600E-06 |
| 25044 | 1253 | E1253D | D | 9.7e-06 | 5.7e-06 | 6.9e-06 | 7.433E-06 | 2.053E-06 | 5.700E-06 | 9.700E-06 |
| 25045 | 1253 | E1253C | D | 1.1e-05 | 9.6e-06 | 7.8e-06 | 9.467E-06 | 1.604E-06 | 7.800E-06 | 1.100E-05 |
| 25046 | 1253 | E1253E | D | 6.8e-06 | 8.9e-06 | 5.1e-06 | 6.933E-06 | 1.904E-06 | 5.100E-06 | 8.900E-06 |
| 25047 | 1253 | E1253Q | D | 9.1e-06 | 1.3e-05 | 7.1e-06 | 9.733E-06 | 3.001E-06 | 7.100E-06 | 1.300E-05 |
| 25048 | 1253 | E1253G | D | 1.3e-05 | 4.2e-06 | 8.9e-06 | 8.700E-06 | 4.403E-06 | 4.200E-06 | 1.300E-05 |
| 25049 | 1253 | E1253H | D | 9.9e-06 | 7e-06 | 9.8e-06 | 8.900E-06 | 1.646E-06 | 7.000E-06 | 9.900E-06 |
| 25050 | 1253 | E1253I | D | 9.9e-06 | 1.1e-05 | 5e-06 | 8.633E-06 | 3.194E-06 | 5.000E-06 | 1.100E-05 |
| 25051 | 1253 | E1253L | D | 9.1e-06 | 7.5e-06 | 7.3e-06 | 7.967E-06 | 9.866E-07 | 7.300E-06 | 9.100E-06 |
| 25052 | 1253 | E1253K | D | 1.1e-05 | 1.1e-05 | 1.1e-05 | 1.100E-05 | 2.075E-21 | 1.100E-05 | 1.100E-05 |
| 25053 | 1253 | E1253M | D | 4.6e-06 | 6.5e-06 | 7.5e-06 | 6.200E-06 | 1.473E-06 | 4.600E-06 | 7.500E-06 |
| 25054 | 1253 | E1253F | D | 4.7e-06 | 1.1e-05 | 9.6e-06 | 8.433E-06 | 3.308E-06 | 4.700E-06 | 1.100E-05 |
| 25055 | 1253 | E1253P | D | 7.3e-06 | 9.3e-06 | 8.9e-06 | 8.500E-06 | 1.058E-06 | 7.300E-06 | 9.300E-06 |
| 25056 | 1253 | E1253S | D | 8.2e-06 | 4.7e-06 | 1.1e-05 | 7.967E-06 | 3.156E-06 | 4.700E-06 | 1.100E-05 |
| 25057 | 1253 | E1253T | D | 6.9e-06 | 7.4e-06 | 6.2e-06 | 6.833E-06 | 6.028E-07 | 6.200E-06 | 7.400E-06 |
| 25058 | 1253 | E1253W | D | 1.3e-05 | 5.8e-06 | 8.4e-06 | 9.067E-06 | 3.646E-06 | 5.800E-06 | 1.300E-05 |
| 25059 | 1253 | E1253Y | D | 1.7e-05 | 6e-06 | 6.3e-06 | 9.767E-06 | 6.266E-06 | 6.000E-06 | 1.700E-05 |
| 25060 | 1253 | E1253V | D | 6.6e-06 | 7.4e-06 | 8.3e-06 | 7.433E-06 | 8.505E-07 | 6.600E-06 | 8.300E-06 |
| 25061 | 1254 | V1254A | D | 6.7e-06 | 1.1e-05 | 5.9e-06 | 7.867E-06 | 2.743E-06 | 5.900E-06 | 1.100E-05 |
| 25062 | 1254 | V1254R | D | 9.8e-06 | 4.9e-06 | 5.7e-06 | 6.800E-06 | 2.629E-06 | 4.900E-06 | 9.800E-06 |
| 25063 | 1254 | V1254N | D | 9.1e-06 | 5.2e-06 | 6.2e-06 | 6.833E-06 | 2.026E-06 | 5.200E-06 | 9.100E-06 |
| 25064 | 1254 | V1254D | D | 7.5e-06 | 8.6e-06 | 7.3e-06 | 7.800E-06 | 7.000E-07 | 7.300E-06 | 8.600E-06 |
| 25065 | 1254 | V1254C | D | 1.1e-05 | 6.1e-06 | 9.3e-06 | 8.800E-06 | 2.488E-06 | 6.100E-06 | 1.100E-05 |
| 25066 | 1254 | V1254E | D | 5.7e-06 | 7.3e-06 | 7e-06 | 6.667E-06 | 8.505E-07 | 5.700E-06 | 7.300E-06 |
| 25067 | 1254 | V1254Q | D | 7.1e-06 | 8.7e-06 | 4.5e-06 | 6.767E-06 | 2.120E-06 | 4.500E-06 | 8.700E-06 |
| 25068 | 1254 | V1254G | D | 9.2e-06 | 1.2e-05 | 9.6e-06 | 1.027E-05 | 1.514E-06 | 9.200E-06 | 1.200E-05 |
| 25069 | 1254 | V1254H | D | 9.9e-06 | 5.1e-06 | 8.8e-06 | 7.933E-06 | 2.515E-06 | 5.100E-06 | 9.900E-06 |
| 25070 | 1254 | V1254I | D | 7.2e-06 | 6.1e-06 | 1.1e-05 | 8.100E-06 | 2.571E-06 | 6.100E-06 | 1.100E-05 |
| 25071 | 1254 | V1254L | D | 6.2e-06 | 8.5e-06 | 5.8e-06 | 6.833E-06 | 1.457E-06 | 5.800E-06 | 8.500E-06 |
| 25072 | 1254 | V1254K | D | 5.1e-06 | 5.9e-06 | 1.3e-05 | 8.000E-06 | 4.349E-06 | 5.100E-06 | 1.300E-05 |
| 25073 | 1254 | V1254M | D | 6.6e-06 | 8.2e-06 | 9.9e-06 | 8.233E-06 | 1.650E-06 | 6.600E-06 | 9.900E-06 |
| 25074 | 1254 | V1254F | D | 5.4e-06 | 7e-06 | 1e-05 | 7.467E-06 | 2.335E-06 | 5.400E-06 | 1.000E-05 |

| | | | | | | | | | | |
|---|---|---|---|---|---|---|---|---|---|---|
| 25075 | 1254 | V1254P | D | 7.1e-06 | 8.2e-06 | 1.2e-05 | 9.100E-06 | 2.571E-06 | 7.100E-06 | 1.200E-05 |
| 25076 | 1254 | V1254S | D | 5.8e-06 | 5.4e-06 | 7.6e-06 | 6.267E-06 | 1.172E-06 | 5.400E-06 | 7.600E-06 |
| 25077 | 1254 | V1254T | D | 7e-06 | 4.7e-06 | 5.2e-06 | 5.633E-06 | 1.210E-06 | 4.700E-06 | 7.000E-06 |
| 25078 | 1254 | V1254W | D | 6e-06 | 1.3e-05 | 1.4e-05 | 1.100E-05 | 4.359E-06 | 6.000E-06 | 1.400E-05 |
| 25079 | 1254 | V1254Y | D | 4.6e-06 | 7.2e-06 | 1e-05 | 7.267E-06 | 2.701E-06 | 4.600E-06 | 1.000E-05 |
| 25080 | 1254 | V1254V | D | 3.1e-06 | 8.3e-06 | 1e-05 | 7.133E-06 | 3.595E-06 | 3.100E-06 | 1.000E-05 |
| 25081 | 1255 | T1255A | D | 8.9e-06 | 5.1e-06 | 1.1e-05 | 8.333E-06 | 2.991E-06 | 5.100E-06 | 1.100E-05 |
| 25082 | 1255 | T1255R | D | 6.1e-06 | 9e-06 | 8.6e-06 | 7.900E-06 | 1.572E-06 | 6.100E-06 | 9.000E-06 |
| 25083 | 1255 | T1255N | D | 7.3e-06 | 9.4e-06 | 1.6e-05 | 1.090E-05 | 4.540E-06 | 7.300E-06 | 1.600E-05 |
| 25084 | 1255 | T1255D | D | 5.6e-06 | 9.4e-06 | 9.5e-06 | 8.167E-06 | 2.223E-06 | 5.600E-06 | 9.500E-06 |
| 25085 | 1255 | T1255C | D | 9.5e-06 | 1.5e-05 | 1e-05 | 1.150E-05 | 3.041E-06 | 9.500E-06 | 1.500E-05 |
| 25086 | 1255 | T1255E | D | 4.4e-06 | 9.7e-06 | 1.5e-05 | 9.700E-06 | 5.300E-06 | 4.400E-06 | 1.500E-05 |
| 25087 | 1255 | T1255Q | D | 8.4e-06 | 9.4e-06 | 4.4e-06 | 7.400E-06 | 2.646E-06 | 4.400E-06 | 9.400E-06 |
| 25088 | 1255 | T1255G | D | 8e-06 | 1.2e-05 | 7e-06 | 9.000E-06 | 2.646E-06 | 7.000E-06 | 1.200E-05 |
| 25089 | 1255 | T1255H | D | 1.3e-05 | 1e-05 | 5.1e-06 | 9.367E-06 | 3.988E-06 | 5.100E-06 | 1.300E-05 |
| 25090 | 1255 | T1255I | D | 1.1e-05 | 5.2e-06 | 7.2e-06 | 7.800E-06 | 2.946E-06 | 5.200E-06 | 1.100E-05 |
| 25091 | 1255 | T1255L | D | 1e-05 | 8.7e-06 | 1.6e-05 | 1.157E-05 | 3.894E-06 | 8.700E-06 | 1.600E-05 |
| 25092 | 1255 | T1255K | D | 8.9e-06 | 4.2e-06 | 9.1e-06 | 7.400E-06 | 2.773E-06 | 4.200E-06 | 9.100E-06 |
| 25093 | 1255 | T1255M | D | 6e-06 | 9.8e-06 | 4.5e-06 | 6.767E-06 | 2.732E-06 | 4.500E-06 | 9.800E-06 |
| 25094 | 1255 | T1255F | D | 9.7e-06 | 6.2e-06 | 8.2e-06 | 8.033E-06 | 1.756E-06 | 6.200E-06 | 9.700E-06 |
| 25095 | 1255 | T1255P | D | 6.4e-06 | 4.9e-06 | 7.9e-06 | 6.400E-06 | 1.500E-06 | 4.900E-06 | 7.900E-06 |
| 25096 | 1255 | T1255S | D | 8.6e-06 | 6.5e-06 | 7.7e-06 | 7.600E-06 | 1.054E-06 | 6.500E-06 | 8.600E-06 |
| 25097 | 1255 | T1255T | D | 7.7e-06 | 7.9e-06 | 9e-06 | 8.200E-06 | 7.000E-07 | 7.700E-06 | 9.000E-06 |
| 25098 | 1255 | T1255W | D | 8.2e-06 | 1.1e-05 | 8.4e-06 | 9.200E-06 | 1.562E-06 | 8.200E-06 | 1.100E-05 |
| 25099 | 1255 | T1255Y | D | 6.8e-06 | 5.5e-06 | 6.1e-06 | 6.133E-06 | 6.506E-07 | 5.500E-06 | 6.800E-06 |
| 25100 | 1255 | T1255V | D | 6.7e-06 | 8.4e-06 | 1.2e-05 | 9.033E-06 | 2.706E-06 | 6.700E-06 | 1.200E-05 |
| 25101 | 1256 | H1256A | D | 1.2e-05 | 9.9e-06 | 6.8e-06 | 9.567E-06 | 2.616E-06 | 6.800E-06 | 1.200E-05 |
| 25102 | 1256 | H1256R | D | 9.6e-06 | 9.1e-06 | 5.1e-06 | 7.933E-06 | 2.466E-06 | 5.100E-06 | 9.600E-06 |
| 25103 | 1256 | H1256N | D | 8e-06 | 9.1e-06 | 6.6e-06 | 7.900E-06 | 1.253E-06 | 6.600E-06 | 9.100E-06 |
| 25104 | 1256 | H1256D | D | 7.9e-06 | 9.4e-06 | 6.7e-06 | 8.000E-06 | 1.353E-06 | 6.700E-06 | 9.400E-06 |
| 25105 | 1256 | H1256C | D | 7.4e-06 | 1e-05 | 1e-05 | 9.133E-06 | 1.501E-06 | 7.400E-06 | 1.000E-05 |
| 25106 | 1256 | H1256E | D | 5.8e-06 | 9.3e-06 | 1.1e-05 | 8.700E-06 | 2.651E-06 | 5.800E-06 | 1.100E-05 |
| 25107 | 1256 | H1256Q | D | 5e-06 | 1.3e-05 | 9.4e-06 | 9.133E-06 | 4.007E-06 | 5.000E-06 | 1.300E-05 |
| 25108 | 1256 | H1256G | D | 7.7e-06 | 1.5e-05 | 5.9e-06 | 9.533E-06 | 4.819E-06 | 5.900E-06 | 1.500E-05 |
| 25109 | 1256 | H1256H | D | 5.4e-06 | 6.6e-06 | 1.2e-05 | 8.000E-06 | 3.516E-06 | 5.400E-06 | 1.200E-05 |
| 25110 | 1256 | H1256I | D | 7.7e-06 | 5.9e-06 | 5.7e-06 | 6.433E-06 | 1.102E-06 | 5.700E-06 | 7.700E-06 |
| 25111 | 1256 | H1256L | D | 9.3e-06 | 7.5e-06 | 5.2e-06 | 7.333E-06 | 2.055E-06 | 5.200E-06 | 9.300E-06 |
| 25112 | 1256 | H1256K | D | 9e-06 | 6.8e-06 | 5.3e-06 | 7.033E-06 | 1.861E-06 | 5.300E-06 | 9.000E-06 |
| 25113 | 1256 | H1256M | D | 5.9e-06 | 7.2e-06 | 1.3e-05 | 8.700E-06 | 3.780E-06 | 5.900E-06 | 1.300E-05 |
| 25114 | 1256 | H1256F | D | 9.8e-06 | 7.1e-06 | 6.2e-06 | 7.700E-06 | 1.873E-06 | 6.200E-06 | 9.800E-06 |
| 25115 | 1256 | H1256P | D | 1.4e-05 | 1.1e-05 | 1.7e-05 | 1.400E-05 | 3.000E-06 | 1.100E-05 | 1.700E-05 |
| 25116 | 1256 | H1256S | D | 9.5e-06 | 6.2e-06 | 7.4e-06 | 7.700E-06 | 1.670E-06 | 6.200E-06 | 9.500E-06 |
| 25117 | 1256 | H1256T | D | 9.4e-06 | 4.9e-06 | 5.2e-06 | 6.500E-06 | 2.516E-06 | 4.900E-06 | 9.400E-06 |
| 25118 | 1256 | H1256W | D | 6.9e-06 | 7.2e-06 | 6e-06 | 6.700E-06 | 6.245E-07 | 6.000E-06 | 7.200E-06 |
| 25119 | 1256 | H1256Y | D | 2e-05 | 7.3e-06 | 8.8e-06 | 1.203E-05 | 6.940E-06 | 7.300E-06 | 2.000E-05 |
| 25120 | 1256 | H1256V | D | 5.3e-06 | 9.1e-06 | 5.6e-06 | 6.667E-06 | 2.113E-06 | 5.300E-06 | 9.100E-06 |
| 25121 | 1257 | Q1257A | D | 1e-05 | 1.3e-05 | 8.6e-06 | 1.053E-05 | 2.248E-06 | 8.600E-06 | 1.300E-05 |
| 25122 | 1257 | Q1257R | D | 4.6e-06 | 1.1e-05 | 5.7e-06 | 7.100E-06 | 3.422E-06 | 4.600E-06 | 1.100E-05 |
| 25123 | 1257 | Q1257N | D | 6.2e-06 | 9.8e-06 | 8.5e-06 | 8.167E-06 | 1.823E-06 | 6.200E-06 | 9.800E-06 |
| 25124 | 1257 | Q1257D | D | 5.7e-06 | 7.8e-06 | 8.9e-06 | 7.467E-06 | 1.626E-06 | 5.700E-06 | 8.900E-06 |
| 25125 | 1257 | Q1257C | D | 6.2e-06 | 5.5e-06 | 8.5e-06 | 6.733E-06 | 1.570E-06 | 5.500E-06 | 8.500E-06 |
| 25126 | 1257 | Q1257E | D | 5.4e-06 | 4.4e-06 | 7.6e-06 | 5.800E-06 | 1.637E-06 | 4.400E-06 | 7.600E-06 |
| 25127 | 1257 | Q1257Q | D | 1.2e-05 | 4.9e-06 | 5.5e-06 | 7.467E-06 | 3.937E-06 | 4.900E-06 | 1.200E-05 |
| 25128 | 1257 | Q1257G | D | 1.1e-05 | 1.2e-05 | 5.7e-06 | 9.567E-06 | 3.386E-06 | 5.700E-06 | 1.200E-05 |
| 25129 | 1257 | Q1257H | D | 8.8e-06 | 9.6e-06 | 4.9e-06 | 7.767E-06 | 2.515E-06 | 4.900E-06 | 9.600E-06 |
| 25130 | 1257 | Q1257I | D | 7.1e-06 | 9.2e-06 | 8.1e-06 | 8.133E-06 | 1.050E-06 | 7.100E-06 | 9.200E-06 |
| 25131 | 1257 | Q1257L | D | 6.6e-06 | 4.7e-06 | 5.2e-06 | 5.500E-06 | 9.849E-07 | 4.700E-06 | 6.600E-06 |
| 25132 | 1257 | Q1257K | D | 1e-05 | 1e-05 | 5.9e-06 | 8.633E-06 | 2.367E-06 | 5.900E-06 | 1.000E-05 |
| 25133 | 1257 | Q1257M | D | 9.3e-06 | 1.3e-05 | 6.7e-06 | 9.667E-06 | 3.166E-06 | 6.700E-06 | 1.300E-05 |
| 25134 | 1257 | Q1257F | D | 1.1e-05 | 9.6e-06 | 7.8e-06 | 9.467E-06 | 1.604E-06 | 7.800E-06 | 1.100E-05 |
| 25135 | 1257 | Q1257P | D | 1.1e-05 | 1e-05 | 6.3e-06 | 9.100E-06 | 2.476E-06 | 6.300E-06 | 1.100E-05 |
| 25136 | 1257 | Q1257S | D | 8.8e-06 | 4.5e-06 | 7.2e-06 | 6.833E-06 | 2.173E-06 | 4.500E-06 | 8.800E-06 |
| 25137 | 1257 | Q1257T | D | 5.1e-06 | 7e-06 | 7.8e-06 | 6.633E-06 | 1.387E-06 | 5.100E-06 | 7.800E-06 |
| 25138 | 1257 | Q1257W | D | 8.1e-06 | 6.5e-06 | 5.1e-06 | 6.567E-06 | 1.501E-06 | 5.100E-06 | 8.100E-06 |
| 25139 | 1257 | Q1257Y | D | 6.8e-06 | 5.7e-06 | 5.7e-06 | 6.067E-06 | 6.351E-07 | 5.700E-06 | 6.800E-06 |
| 25140 | 1257 | Q1257V | D | 1.4e-05 | 9.4e-06 | 9.1e-06 | 1.083E-05 | 2.747E-06 | 9.100E-06 | 1.400E-05 |
| 25141 | 1258 | G1258A | D | 8.5e-06 | 6.9e-06 | 6.1e-06 | 7.167E-06 | 1.222E-06 | 6.100E-06 | 8.500E-06 |
| 25142 | 1258 | G1258R | D | 1.2e-05 | 5e-06 | 7.4e-06 | 8.133E-06 | 3.557E-06 | 5.000E-06 | 1.200E-05 |
| 25143 | 1258 | G1258N | D | 6.3e-06 | 8.9e-06 | 9.2e-06 | 8.133E-06 | 1.595E-06 | 6.300E-06 | 9.200E-06 |
| 25144 | 1258 | G1258D | D | 7.8e-06 | 9.6e-06 | 9.2e-06 | 8.867E-06 | 9.452E-07 | 7.800E-06 | 9.600E-06 |
| 25145 | 1258 | G1258C | D | 1.7e-05 | 1.2e-05 | 1.4e-05 | 1.433E-05 | 2.517E-06 | 1.200E-05 | 1.700E-05 |
| 25146 | 1258 | G1258E | D | 1.5e-05 | 5e-06 | 1.6e-05 | 1.200E-05 | 6.083E-06 | 5.000E-06 | 1.600E-05 |
| 25147 | 1258 | G1258Q | D | 9.4e-06 | 9.1e-06 | 5.6e-06 | 8.033E-06 | 2.113E-06 | 5.600E-06 | 9.400E-06 |
| 25148 | 1258 | G1258G | D | 1.1e-05 | 9.3e-06 | 1.3e-05 | 1.110E-05 | 1.852E-06 | 9.300E-06 | 1.300E-05 |
| 25149 | 1258 | G1258H | D | 9.9e-06 | 6.7e-06 | 9.5e-06 | 8.700E-06 | 1.744E-06 | 6.700E-06 | 9.900E-06 |
| 25150 | 1258 | G1258I | D | 5.2e-06 | 7.7e-06 | 6.6e-06 | 6.500E-06 | 1.253E-06 | 5.200E-06 | 7.700E-06 |
| 25151 | 1258 | G1258L | D | 8.4e-06 | 8.8e-06 | 5.1e-06 | 7.433E-06 | 2.031E-06 | 5.100E-06 | 8.800E-06 |
| 25152 | 1258 | G1258K | D | 6.7e-06 | 5.4e-06 | 9.3e-06 | 7.133E-06 | 1.986E-06 | 5.400E-06 | 9.300E-06 |
| 25153 | 1258 | G1258M | D | 1.2e-05 | 9.7e-06 | 7.1e-06 | 9.600E-06 | 2.452E-06 | 7.100E-06 | 1.200E-05 |
| 25154 | 1258 | G1258F | D | 9e-06 | 4.2e-06 | 7.8e-06 | 7.000E-06 | 2.498E-06 | 4.200E-06 | 9.000E-06 |
| 25155 | 1258 | G1258P | D | 1.3e-05 | 5.4e-06 | 7.4e-06 | 8.600E-06 | 3.940E-06 | 5.400E-06 | 1.300E-05 |
| 25156 | 1258 | G1258S | D | 7.6e-06 | 1e-05 | 3.3e-06 | 6.967E-06 | 3.395E-06 | 3.300E-06 | 1.000E-05 |
| 25157 | 1258 | G1258T | D | 7e-06 | 9.6e-06 | 1.1e-05 | 9.200E-06 | 2.030E-06 | 7.000E-06 | 1.100E-05 |
| 25158 | 1258 | G1258W | D | 6.9e-06 | 8.7e-06 | 5.9e-06 | 7.167E-06 | 1.419E-06 | 5.900E-06 | 8.700E-06 |
| 25159 | 1258 | G1258Y | D | 6.1e-06 | 9.2e-06 | 5.9e-06 | 7.067E-06 | 1.850E-06 | 5.900E-06 | 9.200E-06 |

| | | | | | | | | | | |
|---|---|---|---|---|---|---|---|---|---|---|
| 25160 | 1258 | G1258V | D | 1e-05 | 9.1e-06 | 4.7e-06 | 7.933E-06 | 2.836E-06 | 4.700E-06 | 1.000E-05 |
| 25161 | 1259 | L1259A | D | 7.3e-06 | 7.3e-06 | 9.6e-06 | 8.067E-06 | 1.328E-06 | 7.300E-06 | 9.600E-06 |
| 25162 | 1259 | L1259R | D | 8.6e-06 | 9.4e-06 | 1.1e-05 | 9.667E-06 | 1.222E-06 | 8.600E-06 | 1.100E-05 |
| 25163 | 1259 | L1259N | D | 6.6e-06 | 6.3e-06 | 7.2e-06 | 6.700E-06 | 4.583E-07 | 6.300E-06 | 7.200E-06 |
| 25164 | 1259 | L1259D | D | 7.3e-06 | 6.6e-06 | 6.3e-06 | 6.733E-06 | 5.132E-07 | 6.300E-06 | 7.300E-06 |
| 25165 | 1259 | L1259C | D | 9.2e-06 | 6.9e-06 | 9.7e-06 | 8.600E-06 | 1.493E-06 | 6.900E-06 | 9.700E-06 |
| 25166 | 1259 | L1259E | D | 9.9e-06 | 9.6e-06 | 4.8e-06 | 8.100E-06 | 2.862E-06 | 4.800E-06 | 9.900E-06 |
| 25167 | 1259 | L1259Q | D | 4.1e-06 | 6.6e-06 | 8.4e-06 | 6.367E-06 | 2.159E-06 | 4.100E-06 | 8.400E-06 |
| 25168 | 1259 | L1259G | D | 1.3e-05 | 9.4e-06 | 5.2e-06 | 9.200E-06 | 3.904E-06 | 5.200E-06 | 1.300E-05 |
| 25169 | 1259 | L1259H | D | 6.2e-06 | 9.8e-06 | 6.4e-06 | 7.467E-06 | 2.023E-06 | 6.200E-06 | 9.800E-06 |
| 25170 | 1259 | L1259I | D | 9.6e-06 | 1.1e-05 | 7.1e-06 | 9.233E-06 | 1.976E-06 | 7.100E-06 | 1.100E-05 |
| 25171 | 1259 | L1259L | D | 8e-06 | 1e-05 | 9.7e-06 | 9.233E-06 | 1.079E-06 | 8.000E-06 | 1.000E-05 |
| 25172 | 1259 | L1259K | D | 6.2e-06 | 9.9e-06 | 6.5e-06 | 7.533E-06 | 2.055E-06 | 6.200E-06 | 9.900E-06 |
| 25173 | 1259 | L1259M | D | 9.4e-06 | 6.3e-06 | 1.1e-05 | 8.900E-06 | 2.390E-06 | 6.300E-06 | 1.100E-05 |
| 25174 | 1259 | L1259F | D | 6.7e-06 | 6.7e-06 | 1.4e-05 | 9.133E-06 | 4.215E-06 | 6.700E-06 | 1.400E-05 |
| 25175 | 1259 | L1259P | D | 6.7e-06 | 9.2e-06 | 5.6e-06 | 7.167E-06 | 1.845E-06 | 5.600E-06 | 9.200E-06 |
| 25176 | 1259 | L1259S | D | 7.5e-06 | 4.8e-06 | 5.5e-06 | 5.933E-06 | 1.401E-06 | 4.800E-06 | 7.500E-06 |
| 25177 | 1259 | L1259T | D | 5.5e-06 | 9.9e-06 | 1e-05 | 8.467E-06 | 2.570E-06 | 5.500E-06 | 1.000E-05 |
| 25178 | 1259 | L1259W | D | 5.9e-06 | 9.3e-06 | 7.3e-06 | 7.500E-06 | 1.709E-06 | 5.900E-06 | 9.300E-06 |
| 25179 | 1259 | L1259Y | D | 8.4e-06 | 6.6e-06 | 1.3e-05 | 9.333E-06 | 3.301E-06 | 6.600E-06 | 1.300E-05 |
| 25180 | 1259 | L1259V | D | 1.4e-05 | 1.1e-05 | 6.6e-06 | 1.053E-05 | 3.722E-06 | 6.600E-06 | 1.400E-05 |
| 25181 | 1260 | S1260A | D | 8.1e-06 | 7e-06 | 8.9e-06 | 8.000E-06 | 9.539E-07 | 7.000E-06 | 8.900E-06 |
| 25182 | 1260 | S1260R | D | 1e-05 | 7.6e-06 | 4.4e-06 | 7.333E-06 | 2.810E-06 | 4.400E-06 | 1.000E-05 |
| 25183 | 1260 | S1260N | D | 6.4e-06 | 5.3e-06 | 9.2e-06 | 6.967E-06 | 2.011E-06 | 5.300E-06 | 9.200E-06 |
| 25184 | 1260 | S1260D | D | 5.2e-06 | 4e-06 | 4.7e-06 | 4.633E-06 | 6.028E-07 | 4.000E-06 | 5.200E-06 |
| 25185 | 1260 | S1260C | D | 6.4e-06 | 7.7e-06 | 7e-06 | 7.033E-06 | 6.506E-07 | 6.400E-06 | 7.700E-06 |
| 25186 | 1260 | S1260E | D | 9.8e-06 | 9.7e-06 | 6.2e-06 | 8.567E-06 | 2.050E-06 | 6.200E-06 | 9.800E-06 |
| 25187 | 1260 | S1260Q | D | 6.9e-06 | 7.8e-06 | 4.8e-06 | 6.500E-06 | 1.539E-06 | 4.800E-06 | 7.800E-06 |
| 25188 | 1260 | S1260G | D | 9.3e-06 | 6.7e-06 | 6.9e-06 | 7.633E-06 | 1.447E-06 | 6.700E-06 | 9.300E-06 |
| 25189 | 1260 | S1260H | D | 6.6e-06 | 9.4e-06 | 9.7e-06 | 8.567E-06 | 1.710E-06 | 6.600E-06 | 9.700E-06 |
| 25190 | 1260 | S1260I | D | 8.3e-06 | 1.3e-05 | 6.7e-06 | 9.333E-06 | 3.275E-06 | 6.700E-06 | 1.300E-05 |
| 25191 | 1260 | S1260L | D | 1.3e-05 | 8e-06 | 7e-06 | 9.333E-06 | 3.215E-06 | 7.000E-06 | 1.300E-05 |
| 25192 | 1260 | S1260K | D | 8.3e-06 | 5.7e-06 | 6.9e-06 | 6.967E-06 | 1.301E-06 | 5.700E-06 | 8.300E-06 |
| 25193 | 1260 | S1260M | D | 9e-06 | 5.3e-06 | 3.4e-06 | 5.900E-06 | 2.848E-06 | 3.400E-06 | 9.000E-06 |
| 25194 | 1260 | S1260F | D | 9.5e-06 | 5.1e-06 | 7.2e-06 | 7.267E-06 | 2.201E-06 | 5.100E-06 | 9.500E-06 |
| 25195 | 1260 | S1260P | D | 1.3e-05 | 5.7e-06 | 9.6e-06 | 9.433E-06 | 3.653E-06 | 5.700E-06 | 1.300E-05 |
| 25196 | 1260 | S1260S | D | 1.5e-05 | 6.4e-06 | 1.1e-05 | 1.080E-05 | 4.303E-06 | 6.400E-06 | 1.500E-05 |
| 25197 | 1260 | S1260T | D | 8.3e-06 | 6.2e-06 | 6.9e-06 | 7.133E-06 | 1.069E-06 | 6.200E-06 | 8.300E-06 |
| 25198 | 1260 | S1260W | D | 7.4e-06 | 1.3e-05 | 9.7e-06 | 1.003E-05 | 2.815E-06 | 7.400E-06 | 1.300E-05 |
| 25199 | 1260 | S1260Y | D | 7.9e-06 | 4.8e-06 | 7.8e-06 | 6.833E-06 | 1.762E-06 | 4.800E-06 | 7.900E-06 |
| 25200 | 1260 | S1260V | D | 3.8e-06 | 8.7e-06 | 1.2e-05 | 8.167E-06 | 4.126E-06 | 3.800E-06 | 1.200E-05 |
| 25201 | 1261 | S1261A | D | 1.1e-05 | 7.7e-06 | 1e-05 | 9.567E-06 | 1.692E-06 | 7.700E-06 | 1.100E-05 |
| 25202 | 1261 | S1261R | D | 1.4e-05 | 7.3e-06 | 7.3e-06 | 9.533E-06 | 3.868E-06 | 7.300E-06 | 1.400E-05 |
| 25203 | 1261 | S1261N | D | 4.6e-06 | 6.3e-06 | 4.4e-06 | 5.100E-06 | 1.044E-06 | 4.400E-06 | 6.300E-06 |
| 25204 | 1261 | S1261D | D | 6.7e-06 | 6e-06 | 1e-05 | 7.567E-06 | 2.136E-06 | 6.000E-06 | 1.000E-05 |
| 25205 | 1261 | S1261C | D | 1e-05 | 8.5e-06 | 7.8e-06 | 8.767E-06 | 1.124E-06 | 7.800E-06 | 1.000E-05 |
| 25206 | 1261 | S1261E | D | 7.4e-06 | 6.3e-06 | 6.4e-06 | 6.700E-06 | 6.083E-07 | 6.300E-06 | 7.400E-06 |
| 25207 | 1261 | S1261Q | D | 7.7e-06 | 8.6e-06 | 7.7e-06 | 8.000E-06 | 5.196E-07 | 7.700E-06 | 8.600E-06 |
| 25208 | 1261 | S1261G | D | 1.1e-05 | 6.9e-06 | 7.8e-06 | 8.567E-06 | 2.155E-06 | 6.900E-06 | 1.100E-05 |
| 25209 | 1261 | S1261H | D | 6.8e-06 | 1.4e-05 | 7.4e-06 | 9.400E-06 | 3.995E-06 | 6.800E-06 | 1.400E-05 |
| 25210 | 1261 | S1261I | D | 7.4e-06 | 4.7e-06 | 9e-06 | 7.033E-06 | 2.173E-06 | 4.700E-06 | 9.000E-06 |
| 25211 | 1261 | S1261L | D | 6.9e-06 | 9.1e-06 | 1e-05 | 8.667E-06 | 1.595E-06 | 6.900E-06 | 1.000E-05 |
| 25212 | 1261 | S1261K | D | 7.1e-06 | 1e-05 | 6.9e-06 | 8.000E-06 | 1.735E-06 | 6.900E-06 | 1.000E-05 |
| 25213 | 1261 | S1261M | D | 7.7e-06 | 7e-06 | 8e-06 | 7.567E-06 | 5.132E-07 | 7.000E-06 | 8.000E-06 |
| 25214 | 1261 | S1261F | D | 8.5e-06 | 1.5e-05 | 9.7e-06 | 1.107E-05 | 3.459E-06 | 8.500E-06 | 1.500E-05 |
| 25215 | 1261 | S1261P | D | 1e-05 | 9.5e-06 | 7.2e-06 | 8.900E-06 | 1.493E-06 | 7.200E-06 | 1.000E-05 |
| 25216 | 1261 | S1261S | D | 4.8e-06 | 9.7e-06 | 9.3e-06 | 7.933E-06 | 2.721E-06 | 4.800E-06 | 9.700E-06 |
| 25217 | 1261 | S1261T | D | 8.7e-06 | 7.7e-06 | 1.1e-05 | 9.133E-06 | 1.692E-06 | 7.700E-06 | 1.100E-05 |
| 25218 | 1261 | S1261W | D | 9.3e-06 | 9.2e-06 | 4.6e-06 | 7.700E-06 | 2.685E-06 | 4.600E-06 | 9.300E-06 |
| 25219 | 1261 | S1261Y | D | 4.5e-06 | 9e-06 | 8.5e-06 | 7.333E-06 | 2.466E-06 | 4.500E-06 | 9.000E-06 |
| 25220 | 1261 | S1261V | D | 8.7e-06 | 7.8e-06 | 1.2e-05 | 9.500E-06 | 2.211E-06 | 7.800E-06 | 1.200E-05 |
| 25221 | 1262 | P1262A | D | 5e-06 | 9.2e-06 | 7.9e-06 | 7.367E-06 | 2.150E-06 | 5.000E-06 | 9.200E-06 |
| 25222 | 1262 | P1262R | D | 9.2e-06 | 4.6e-06 | 4.9e-06 | 6.233E-06 | 2.574E-06 | 4.600E-06 | 9.200E-06 |
| 25223 | 1262 | P1262N | D | 1.5e-05 | 6.4e-06 | 1.3e-05 | 1.147E-05 | 4.500E-06 | 6.400E-06 | 1.500E-05 |
| 25224 | 1262 | P1262D | D | 9.5e-06 | 6.5e-06 | 1.8e-05 | 1.133E-05 | 5.965E-06 | 6.500E-06 | 1.800E-05 |
| 25225 | 1262 | P1262C | D | 7.6e-06 | 7.5e-06 | 6.5e-06 | 7.200E-06 | 6.083E-07 | 6.500E-06 | 7.600E-06 |
| 25226 | 1262 | P1262E | D | 6.5e-06 | 1.1e-05 | 9.6e-06 | 9.033E-06 | 2.303E-06 | 6.500E-06 | 1.100E-05 |
| 25227 | 1262 | P1262Q | D | 6.3e-06 | 6.8e-06 | 5.1e-06 | 6.067E-06 | 8.737E-07 | 5.100E-06 | 6.800E-06 |
| 25228 | 1262 | P1262G | D | 9.9e-06 | 1.1e-05 | 1.6e-05 | 1.230E-05 | 3.251E-06 | 9.900E-06 | 1.600E-05 |
| 25229 | 1262 | P1262H | D | 1.4e-05 | 6.3e-06 | 5.6e-06 | 8.633E-06 | 4.661E-06 | 5.600E-06 | 1.400E-05 |
| 25230 | 1262 | P1262I | D | 1.3e-05 | 6.7e-06 | 7.5e-06 | 9.067E-06 | 3.430E-06 | 6.700E-06 | 1.300E-05 |
| 25231 | 1262 | P1262L | D | 7.7e-06 | 4.6e-06 | 8.8e-06 | 7.033E-06 | 2.178E-06 | 4.600E-06 | 8.800E-06 |
| 25232 | 1262 | P1262K | D | 6e-06 | 9.8e-06 | 5.1e-06 | 6.967E-06 | 2.495E-06 | 5.100E-06 | 9.800E-06 |
| 25233 | 1262 | P1262M | D | 8.6e-06 | 9e-06 | 8e-06 | 8.533E-06 | 5.033E-07 | 8.000E-06 | 9.000E-06 |
| 25234 | 1262 | P1262F | D | 6.8e-06 | 9.1e-06 | 6.3e-06 | 7.400E-06 | 1.493E-06 | 6.300E-06 | 9.100E-06 |
| 25235 | 1262 | P1262P | D | 5.1e-06 | 6.9e-06 | 8.2e-06 | 6.733E-06 | 1.557E-06 | 5.100E-06 | 8.200E-06 |
| 25236 | 1262 | P1262S | D | 7.3e-06 | 6.4e-06 | 6.3e-06 | 6.667E-06 | 5.508E-07 | 6.300E-06 | 7.300E-06 |
| 25237 | 1262 | P1262T | D | 8.7e-06 | 9.8e-06 | 6.4e-06 | 8.300E-06 | 1.735E-06 | 6.400E-06 | 9.800E-06 |
| 25238 | 1262 | P1262W | D | 9.1e-06 | 8.9e-06 | 9.2e-06 | 9.067E-06 | 1.528E-07 | 8.900E-06 | 9.200E-06 |
| 25239 | 1262 | P1262Y | D | 3.5e-06 | 1.2e-05 | 8.9e-06 | 8.133E-06 | 4.302E-06 | 3.500E-06 | 1.200E-05 |
| 25240 | 1262 | P1262V | D | 1.1e-05 | 5.4e-06 | 6.4e-06 | 7.600E-06 | 2.987E-06 | 5.400E-06 | 1.100E-05 |
| 25241 | 1263 | V1263A | D | 5.7e-06 | 5.5e-06 | 7.4e-06 | 6.200E-06 | 1.044E-06 | 5.500E-06 | 7.400E-06 |
| 25242 | 1263 | V1263R | D | 1e-05 | 1.1e-05 | 4.4e-06 | 8.467E-06 | 3.557E-06 | 4.400E-06 | 1.100E-05 |
| 25243 | 1263 | V1263N | D | 9.6e-06 | 1.3e-05 | 7.2e-06 | 9.933E-06 | 2.914E-06 | 7.200E-06 | 1.300E-05 |
| 25244 | 1263 | V1263D | D | 6.4e-06 | 5.5e-06 | 6.6e-06 | 6.167E-06 | 5.859E-07 | 5.500E-06 | 6.600E-06 |

| | | | | | | | | | | |
|---|---|---|---|---|---|---|---|---|---|---|
| 25245 | 1263 | V1263C | D | 9.6e-06 | 8.8e-06 | 5.8e-06 | 8.067E-06 | 2.003E-06 | 5.800E-06 | 9.600E-06 |
| 25246 | 1263 | V1263E | D | 9.7e-06 | 6.6e-06 | 9.6e-06 | 8.633E-06 | 1.762E-06 | 6.600E-06 | 9.700E-06 |
| 25247 | 1263 | V1263Q | D | 6.3e-06 | 5.6e-06 | 9.6e-06 | 7.167E-06 | 2.136E-06 | 5.600E-06 | 9.600E-06 |
| 25248 | 1263 | V1263G | D | 6.3e-06 | 9.2e-06 | 7e-06 | 7.500E-06 | 1.513E-06 | 6.300E-06 | 9.200E-06 |
| 25249 | 1263 | V1263H | D | 8.2e-06 | 1.3e-05 | 1e-05 | 1.040E-05 | 2.425E-06 | 8.200E-06 | 1.300E-05 |
| 25250 | 1263 | V1263I | D | 7.7e-06 | 8.9e-06 | 9.8e-06 | 8.800E-06 | 1.054E-06 | 7.700E-06 | 9.800E-06 |
| 25251 | 1263 | V1263L | D | 4.6e-06 | 5.2e-06 | 7.6e-06 | 5.800E-06 | 1.587E-06 | 4.600E-06 | 7.600E-06 |
| 25252 | 1263 | V1263K | D | 8.3e-06 | 1.3e-05 | 8.7e-06 | 1.000E-05 | 2.606E-06 | 8.300E-06 | 1.300E-05 |
| 25253 | 1263 | V1263M | D | 8.8e-06 | 8.6e-06 | 8.1e-06 | 8.500E-06 | 3.606E-07 | 8.100E-06 | 8.800E-06 |
| 25254 | 1263 | V1263F | D | 6.6e-06 | 5.3e-06 | 1.3e-05 | 8.300E-06 | 4.122E-06 | 5.300E-06 | 1.300E-05 |
| 25255 | 1263 | V1263P | D | 6.2e-06 | 6.2e-06 | 7.8e-06 | 6.733E-06 | 9.238E-07 | 6.200E-06 | 7.800E-06 |
| 25256 | 1263 | V1263S | D | 1.5e-05 | 9e-06 | 7.8e-06 | 1.060E-05 | 3.857E-06 | 7.800E-06 | 1.500E-05 |
| 25257 | 1263 | V1263T | D | 7.9e-06 | 6.9e-06 | 6.5e-06 | 7.100E-06 | 7.211E-07 | 6.500E-06 | 7.900E-06 |
| 25258 | 1263 | V1263W | D | 8.4e-06 | 8.3e-06 | 6.6e-06 | 7.767E-06 | 1.012E-06 | 6.600E-06 | 8.400E-06 |
| 25259 | 1263 | V1263Y | D | 1.2e-05 | 1.2e-05 | 6.5e-06 | 1.017E-05 | 3.175E-06 | 6.500E-06 | 1.200E-05 |
| 25260 | 1263 | V1263V | D | 5.5e-06 | 9.1e-06 | 7.6e-06 | 7.400E-06 | 1.808E-06 | 5.500E-06 | 9.100E-06 |
| 25261 | 1264 | T1264A | D | 1.3e-05 | 1.2e-05 | 7e-06 | 1.067E-05 | 3.215E-06 | 7.000E-06 | 1.300E-05 |
| 25262 | 1264 | T1264R | D | 8.2e-06 | 1.3e-05 | 1.1e-05 | 1.073E-05 | 2.411E-06 | 8.200E-06 | 1.300E-05 |
| 25263 | 1264 | T1264N | D | 7.6e-06 | 6.1e-06 | 5e-06 | 6.233E-06 | 1.305E-06 | 5.000E-06 | 7.600E-06 |
| 25264 | 1264 | T1264D | D | 7.5e-06 | 7.4e-06 | 1.1e-05 | 8.633E-06 | 2.050E-06 | 7.400E-06 | 1.100E-05 |
| 25265 | 1264 | T1264C | D | 4.4e-06 | 1.1e-05 | 8.2e-06 | 7.867E-06 | 3.313E-06 | 4.400E-06 | 1.100E-05 |
| 25266 | 1264 | T1264E | D | 9.1e-06 | 6.2e-06 | 1.2e-05 | 9.100E-06 | 2.900E-06 | 6.200E-06 | 1.200E-05 |
| 25267 | 1264 | T1264Q | D | 5.2e-06 | 9.4e-06 | 6.9e-06 | 7.167E-06 | 2.113E-06 | 5.200E-06 | 9.400E-06 |
| 25268 | 1264 | T1264G | D | 7.2e-06 | 1.1e-05 | 7.9e-06 | 8.700E-06 | 2.022E-06 | 7.200E-06 | 1.100E-05 |
| 25269 | 1264 | T1264H | D | 6.4e-06 | 5.5e-06 | 9.6e-06 | 7.167E-06 | 2.155E-06 | 5.500E-06 | 9.600E-06 |
| 25270 | 1264 | T1264I | D | 7.8e-06 | 7.2e-06 | 8e-06 | 7.667E-06 | 4.163E-07 | 7.200E-06 | 8.000E-06 |
| 25271 | 1264 | T1264L | D | 5.9e-06 | 1.1e-05 | 6.7e-06 | 7.867E-06 | 2.743E-06 | 5.900E-06 | 1.100E-05 |
| 25272 | 1264 | T1264K | D | 9.8e-06 | 9.7e-06 | 6.4e-06 | 8.633E-06 | 1.935E-06 | 6.400E-06 | 9.800E-06 |
| 25273 | 1264 | T1264M | D | 7.2e-06 | 7.8e-06 | 1.4e-05 | 9.667E-06 | 3.765E-06 | 7.200E-06 | 1.400E-05 |
| 25274 | 1264 | T1264F | D | 6.6e-06 | 7.2e-06 | 9.9e-06 | 7.900E-06 | 1.758E-06 | 6.600E-06 | 9.900E-06 |
| 25275 | 1264 | T1264P | D | 1e-05 | 1e-05 | 1e-05 | 1.000E-05 | 0.000E+00 | 1.000E-05 | 1.000E-05 |
| 25276 | 1264 | T1264S | D | 7e-06 | 6.9e-06 | 8.8e-06 | 7.567E-06 | 1.069E-06 | 6.900E-06 | 8.800E-06 |
| 25277 | 1264 | T1264T | D | 9.5e-06 | 7.8e-06 | 6.8e-06 | 8.033E-06 | 1.365E-06 | 6.800E-06 | 9.500E-06 |
| 25278 | 1264 | T1264W | D | 7.8e-06 | 7e-06 | 9.7e-06 | 8.167E-06 | 1.387E-06 | 7.000E-06 | 9.700E-06 |
| 25279 | 1264 | T1264Y | D | 7e-06 | 1e-05 | 1.1e-05 | 9.333E-06 | 2.082E-06 | 7.000E-06 | 1.100E-05 |
| 25280 | 1264 | T1264V | D | 4.7e-06 | 5e-06 | 6.7e-06 | 5.467E-06 | 1.079E-06 | 4.700E-06 | 6.700E-06 |
| 25281 | 1265 | K1265A | D | 5.5e-06 | 6.1e-06 | 6.7e-06 | 6.100E-06 | 6.000E-07 | 5.500E-06 | 6.700E-06 |
| 25282 | 1265 | K1265R | D | 9.5e-06 | 7e-06 | 1.1e-05 | 9.167E-06 | 2.021E-06 | 7.000E-06 | 1.100E-05 |
| 25283 | 1265 | K1265N | D | 1.1e-05 | 5.4e-06 | 1.2e-05 | 9.467E-06 | 3.557E-06 | 5.400E-06 | 1.200E-05 |
| 25284 | 1265 | K1265D | D | 7.9e-06 | 8.5e-06 | 9.7e-06 | 8.700E-06 | 9.165E-07 | 7.900E-06 | 9.700E-06 |
| 25285 | 1265 | K1265C | D | 9.6e-06 | 5.5e-06 | 8.9e-06 | 8.000E-06 | 2.193E-06 | 5.500E-06 | 9.600E-06 |
| 25286 | 1265 | K1265E | D | 5.4e-06 | 7.3e-06 | 9.8e-06 | 7.500E-06 | 2.207E-06 | 5.400E-06 | 9.800E-06 |
| 25287 | 1265 | K1265Q | D | 1.1e-05 | 7.9e-06 | 6.9e-06 | 8.600E-06 | 2.138E-06 | 6.900E-06 | 1.100E-05 |
| 25288 | 1265 | K1265G | D | 1.2e-05 | 6.1e-06 | 9.8e-06 | 9.300E-06 | 2.982E-06 | 6.100E-06 | 1.200E-05 |
| 25289 | 1265 | K1265H | D | 8.5e-06 | 6.4e-06 | 1.1e-05 | 8.633E-06 | 2.303E-06 | 6.400E-06 | 1.100E-05 |
| 25290 | 1265 | K1265I | D | 1.2e-05 | 6e-06 | 8.2e-06 | 8.733E-06 | 3.035E-06 | 6.000E-06 | 1.200E-05 |
| 25291 | 1265 | K1265L | D | 5e-06 | 1.1e-05 | 1.2e-05 | 9.333E-06 | 3.786E-06 | 5.000E-06 | 1.200E-05 |
| 25292 | 1265 | K1265K | D | 1.3e-05 | 8.1e-06 | 3.4e-06 | 8.167E-06 | 4.800E-06 | 3.400E-06 | 1.300E-05 |
| 25293 | 1265 | K1265M | D | 4.9e-06 | 9.9e-06 | 1.4e-05 | 9.600E-06 | 4.557E-06 | 4.900E-06 | 1.400E-05 |
| 25294 | 1265 | K1265F | D | 1e-05 | 8.3e-06 | 1.3e-05 | 1.043E-05 | 2.380E-06 | 8.300E-06 | 1.300E-05 |
| 25295 | 1265 | K1265P | D | 7.3e-06 | 8.4e-06 | 5.1e-06 | 6.933E-06 | 1.680E-06 | 5.100E-06 | 8.400E-06 |
| 25296 | 1265 | K1265S | D | 5.8e-06 | 5.5e-06 | 1.4e-05 | 8.433E-06 | 4.823E-06 | 5.500E-06 | 1.400E-05 |
| 25297 | 1265 | K1265T | D | 7.1e-06 | 8.9e-06 | 7.9e-06 | 7.967E-06 | 9.018E-07 | 7.100E-06 | 8.900E-06 |
| 25298 | 1265 | K1265W | D | 1.1e-05 | 6.5e-06 | 7.8e-06 | 8.433E-06 | 2.316E-06 | 6.500E-06 | 1.100E-05 |
| 25299 | 1265 | K1265Y | D | 6e-06 | 9.3e-06 | 1.4e-05 | 9.767E-06 | 4.020E-06 | 6.000E-06 | 1.400E-05 |
| 25300 | 1265 | K1265V | D | 9.8e-06 | 1.1e-05 | 7.2e-06 | 9.333E-06 | 1.943E-06 | 7.200E-06 | 1.100E-05 |
| 25301 | 1266 | S1266A | D | 7e-06 | 9.1e-06 | 6.6e-06 | 7.567E-06 | 1.343E-06 | 6.600E-06 | 9.100E-06 |
| 25302 | 1266 | S1266R | D | 6.7e-06 | 9e-06 | 9e-06 | 8.233E-06 | 1.328E-06 | 6.700E-06 | 9.000E-06 |
| 25303 | 1266 | S1266N | D | 8.1e-06 | 7e-06 | 6e-06 | 7.033E-06 | 1.050E-06 | 6.000E-06 | 8.100E-06 |
| 25304 | 1266 | S1266D | D | 7.2e-06 | 7.8e-06 | 7.1e-06 | 7.367E-06 | 3.786E-07 | 7.100E-06 | 7.800E-06 |
| 25305 | 1266 | S1266C | D | 1.1e-05 | 4.8e-06 | 7.7e-06 | 7.833E-06 | 3.102E-06 | 4.800E-06 | 1.100E-05 |
| 25306 | 1266 | S1266E | D | 7.9e-06 | 6.2e-06 | 4.7e-06 | 6.267E-06 | 1.601E-06 | 4.700E-06 | 7.900E-06 |
| 25307 | 1266 | S1266Q | D | 6.7e-06 | 7e-06 | 5.9e-06 | 6.533E-06 | 5.686E-07 | 5.900E-06 | 7.000E-06 |
| 25308 | 1266 | S1266G | D | 9.6e-06 | 1e-05 | 6.5e-06 | 8.700E-06 | 1.916E-06 | 6.500E-06 | 1.000E-05 |
| 25309 | 1266 | S1266H | D | 9.3e-06 | 7.5e-06 | 6.6e-06 | 7.800E-06 | 1.375E-06 | 6.600E-06 | 9.300E-06 |
| 25310 | 1266 | S1266I | D | 9.1e-06 | 4.5e-06 | 1.2e-05 | 8.533E-06 | 3.782E-06 | 4.500E-06 | 1.200E-05 |
| 25311 | 1266 | S1266L | D | 6.6e-06 | 8.6e-06 | 9.6e-06 | 8.267E-06 | 1.528E-06 | 6.600E-06 | 9.600E-06 |
| 25312 | 1266 | S1266K | D | 6.9e-06 | 1e-05 | 6.7e-06 | 7.867E-06 | 1.850E-06 | 6.700E-06 | 1.000E-05 |
| 25313 | 1266 | S1266M | D | 1.2e-05 | 1.1e-05 | 1.2e-05 | 1.167E-05 | 5.774E-07 | 1.100E-05 | 1.200E-05 |
| 25314 | 1266 | S1266F | D | 5.8e-06 | 9.8e-06 | 5.2e-06 | 6.933E-06 | 2.501E-06 | 5.200E-06 | 9.800E-06 |
| 25315 | 1266 | S1266P | D | 7.2e-06 | 5.8e-06 | 7.2e-06 | 6.733E-06 | 8.083E-07 | 5.800E-06 | 7.200E-06 |
| 25316 | 1266 | S1266S | D | 6e-06 | 5.9e-06 | 1e-05 | 7.300E-06 | 2.339E-06 | 5.900E-06 | 1.000E-05 |
| 25317 | 1266 | S1266T | D | 6.6e-06 | 9.3e-06 | 4.4e-06 | 6.767E-06 | 2.454E-06 | 4.400E-06 | 9.300E-06 |
| 25318 | 1266 | S1266W | D | 5e-06 | 9.9e-06 | 6.8e-06 | 7.233E-06 | 2.479E-06 | 5.000E-06 | 9.900E-06 |
| 25319 | 1266 | S1266Y | D | 6.6e-06 | 3.9e-06 | 8.1e-06 | 6.200E-06 | 2.128E-06 | 3.900E-06 | 8.100E-06 |
| 25320 | 1266 | S1266V | D | 3.4e-06 | 1e-05 | 7.8e-06 | 7.067E-06 | 3.361E-06 | 3.400E-06 | 1.000E-05 |
| 25321 | 1267 | F1267A | D | 4.1e-06 | 4.8e-06 | 5.8e-06 | 4.900E-06 | 8.544E-07 | 4.100E-06 | 5.800E-06 |
| 25322 | 1267 | F1267R | D | 1.1e-05 | 9.2e-06 | 5.6e-06 | 8.600E-06 | 2.750E-06 | 5.600E-06 | 1.100E-05 |
| 25323 | 1267 | F1267N | D | 8.3e-06 | 5.7e-06 | 7.8e-06 | 7.267E-06 | 1.380E-06 | 5.700E-06 | 8.300E-06 |
| 25324 | 1267 | F1267D | D | 8.4e-06 | 8e-06 | 9.2e-06 | 8.533E-06 | 6.110E-07 | 8.000E-06 | 9.200E-06 |
| 25325 | 1267 | F1267C | D | 5.4e-06 | 8.8e-06 | 5.9e-06 | 6.700E-06 | 1.836E-06 | 5.400E-06 | 8.800E-06 |
| 25326 | 1267 | F1267E | D | 8e-06 | 1.9e-05 | 9.4e-06 | 1.213E-05 | 5.988E-06 | 8.000E-06 | 1.900E-05 |
| 25327 | 1267 | F1267Q | D | 6.3e-06 | 1.2e-05 | 1.4e-05 | 1.077E-05 | 3.995E-06 | 6.300E-06 | 1.400E-05 |
| 25328 | 1267 | F1267G | D | 8.2e-06 | 1.3e-05 | 4.9e-06 | 8.700E-06 | 4.073E-06 | 4.900E-06 | 1.300E-05 |
| 25329 | 1267 | F1267H | D | 5.3e-06 | 5.5e-06 | 4.8e-06 | 5.200E-06 | 3.606E-07 | 4.800E-06 | 5.500E-06 |

| | | | | | | | | | | |
|---|---|---|---|---|---|---|---|---|---|---|
| 25330 | 1267 | F1267I | D | 8e-06 | 6.4e-06 | 6.8e-06 | 7.067E-06 | 8.327E-07 | 6.400E-06 | 8.000E-06 |
| 25331 | 1267 | F1267L | D | 9.3e-06 | 1.3e-05 | 9.5e-06 | 1.060E-05 | 2.081E-06 | 9.300E-06 | 1.300E-05 |
| 25332 | 1267 | F1267K | D | 4.2e-06 | 6.5e-06 | 5.4e-06 | 5.367E-06 | 1.150E-06 | 4.200E-06 | 6.500E-06 |
| 25333 | 1267 | F1267M | D | 4.4e-06 | 8.3e-06 | 6.8e-06 | 6.500E-06 | 1.967E-06 | 4.400E-06 | 8.300E-06 |
| 25334 | 1267 | F1267F | D | 7.9e-06 | 1.2e-05 | 1.2e-05 | 1.063E-05 | 2.367E-06 | 7.900E-06 | 1.200E-05 |
| 25335 | 1267 | F1267P | D | 4.5e-06 | 5.5e-06 | 9.8e-06 | 6.600E-06 | 2.816E-06 | 4.500E-06 | 9.800E-06 |
| 25336 | 1267 | F1267S | D | 6.3e-06 | 8.4e-06 | 1.5e-05 | 9.900E-06 | 4.540E-06 | 6.300E-06 | 1.500E-05 |
| 25337 | 1267 | F1267T | D | 1.7e-05 | 1.1e-05 | 1e-05 | 1.267E-05 | 3.786E-06 | 1.000E-05 | 1.700E-05 |
| 25338 | 1267 | F1267W | D | 8.5e-06 | 1.2e-05 | 8e-06 | 9.500E-06 | 2.179E-06 | 8.000E-06 | 1.200E-05 |
| 25339 | 1267 | F1267Y | D | 8e-06 | 6.7e-06 | 1.2e-05 | 8.900E-06 | 2.762E-06 | 6.700E-06 | 1.200E-05 |
| 25340 | 1267 | F1267V | D | 7.7e-06 | 6.2e-06 | 1.1e-05 | 8.300E-06 | 2.456E-06 | 6.200E-06 | 1.100E-05 |
| 25341 | 1268 | N1268A | D | 1.2e-05 | 4.2e-06 | 7.3e-06 | 7.833E-06 | 3.927E-06 | 4.200E-06 | 1.200E-05 |
| 25342 | 1268 | N1268R | D | 5.9e-06 | 1.3e-05 | 1.5e-05 | 1.130E-05 | 4.782E-06 | 5.900E-06 | 1.500E-05 |
| 25343 | 1268 | N1268N | D | 6.9e-06 | 7.1e-06 | 4.4e-06 | 6.133E-06 | 1.504E-06 | 4.400E-06 | 7.100E-06 |
| 25344 | 1268 | N1268D | D | 7.9e-06 | 6.7e-06 | 6.8e-06 | 7.133E-06 | 6.658E-07 | 6.700E-06 | 7.900E-06 |
| 25345 | 1268 | N1268C | D | 1.4e-05 | 1.1e-05 | 5.9e-06 | 1.030E-05 | 4.095E-06 | 5.900E-06 | 1.400E-05 |
| 25346 | 1268 | N1268E | D | 1.5e-05 | 9.8e-06 | 6.3e-06 | 1.037E-05 | 4.378E-06 | 6.300E-06 | 1.500E-05 |
| 25347 | 1268 | N1268Q | D | 1.2e-05 | 3.4e-06 | 7e-06 | 7.467E-06 | 4.319E-06 | 3.400E-06 | 1.200E-05 |
| 25348 | 1268 | N1268G | D | 6.4e-06 | 5.7e-06 | 6.7e-06 | 6.267E-06 | 5.132E-07 | 5.700E-06 | 6.700E-06 |
| 25349 | 1268 | N1268H | D | 6.8e-06 | 5.4e-06 | 9.7e-06 | 7.300E-06 | 2.193E-06 | 5.400E-06 | 9.700E-06 |
| 25350 | 1268 | N1268I | D | 5e-06 | 7e-06 | 7.7e-06 | 6.567E-06 | 1.401E-06 | 5.000E-06 | 7.700E-06 |
| 25351 | 1268 | N1268L | D | 9.6e-06 | 1e-05 | 5.8e-06 | 8.467E-06 | 2.318E-06 | 5.800E-06 | 1.000E-05 |
| 25352 | 1268 | N1268K | D | 9.8e-06 | 8.1e-06 | 8.3e-06 | 8.733E-06 | 9.292E-07 | 8.100E-06 | 9.800E-06 |
| 25353 | 1268 | N1268M | D | 1e-05 | 1.1e-05 | 7.5e-06 | 9.500E-06 | 1.803E-06 | 7.500E-06 | 1.100E-05 |
| 25354 | 1268 | N1268F | D | 9.1e-06 | 9.7e-06 | 7.5e-06 | 8.767E-06 | 1.137E-06 | 7.500E-06 | 9.700E-06 |
| 25355 | 1268 | N1268P | D | 5.1e-06 | 1.1e-05 | 8.1e-06 | 8.067E-06 | 2.950E-06 | 5.100E-06 | 1.100E-05 |
| 25356 | 1268 | N1268S | D | 6.9e-06 | 8.1e-06 | 5.8e-06 | 6.933E-06 | 1.150E-06 | 5.800E-06 | 8.100E-06 |
| 25357 | 1268 | N1268T | D | 8.2e-06 | 9e-06 | 8.8e-06 | 8.667E-06 | 4.163E-07 | 8.200E-06 | 9.000E-06 |
| 25358 | 1268 | N1268W | D | 8.2e-06 | 8.5e-06 | 1e-05 | 8.900E-06 | 9.644E-07 | 8.200E-06 | 1.000E-05 |
| 25359 | 1268 | N1268Y | D | 8e-06 | 1.7e-05 | 6.2e-06 | 1.040E-05 | 5.786E-06 | 6.200E-06 | 1.700E-05 |
| 25360 | 1268 | N1268V | D | 9.1e-06 | 1.2e-05 | 8.5e-06 | 9.867E-06 | 1.872E-06 | 8.500E-06 | 1.200E-05 |
| 25361 | 1269 | R1269A | D | 9.2e-06 | 1.1e-05 | 1e-05 | 1.007E-05 | 9.018E-07 | 9.200E-06 | 1.100E-05 |
| 25362 | 1269 | R1269R | D | 8.4e-06 | 8e-06 | 8.1e-06 | 8.167E-06 | 2.082E-07 | 8.000E-06 | 8.400E-06 |
| 25363 | 1269 | R1269N | D | 9.2e-06 | 3.2e-06 | 8.9e-06 | 7.100E-06 | 3.381E-06 | 3.200E-06 | 9.200E-06 |
| 25364 | 1269 | R1269D | D | 5.4e-06 | 8e-06 | 6.3e-06 | 6.567E-06 | 1.320E-06 | 5.400E-06 | 8.000E-06 |
| 25365 | 1269 | R1269C | D | 4.9e-06 | 1.3e-05 | 8.9e-06 | 8.933E-06 | 4.050E-06 | 4.900E-06 | 1.300E-05 |
| 25366 | 1269 | R1269E | D | 8.4e-06 | 8.9e-06 | 1e-05 | 9.100E-06 | 8.185E-07 | 8.400E-06 | 1.000E-05 |
| 25367 | 1269 | R1269Q | D | 5.3e-06 | 9.8e-06 | 6.8e-06 | 7.300E-06 | 2.291E-06 | 5.300E-06 | 9.800E-06 |
| 25368 | 1269 | R1269G | D | 1e-05 | 8.5e-06 | 8e-06 | 8.833E-06 | 1.041E-06 | 8.000E-06 | 1.000E-05 |
| 25369 | 1269 | R1269H | D | 7e-06 | 8.2e-06 | 8.5e-06 | 7.900E-06 | 7.937E-07 | 7.000E-06 | 8.500E-06 |
| 25370 | 1269 | R1269I | D | 7.9e-06 | 5.4e-06 | 8.3e-06 | 7.200E-06 | 1.572E-06 | 5.400E-06 | 8.300E-06 |
| 25371 | 1269 | R1269L | D | 9.6e-06 | 6.5e-06 | 1.3e-05 | 9.700E-06 | 3.251E-06 | 6.500E-06 | 1.300E-05 |
| 25372 | 1269 | R1269K | D | 3.3e-06 | 7.6e-06 | 9e-06 | 6.633E-06 | 2.970E-06 | 3.300E-06 | 9.000E-06 |
| 25373 | 1269 | R1269M | D | 7.7e-06 | 7.8e-06 | 4.9e-06 | 6.800E-06 | 1.646E-06 | 4.900E-06 | 7.800E-06 |
| 25374 | 1269 | R1269F | D | 5.4e-06 | 8.3e-06 | 1e-05 | 7.900E-06 | 2.326E-06 | 5.400E-06 | 1.000E-05 |
| 25375 | 1269 | R1269P | D | 8.3e-06 | 1.1e-05 | 7.3e-06 | 8.867E-06 | 1.914E-06 | 7.300E-06 | 1.100E-05 |
| 25376 | 1269 | R1269S | D | 9.6e-06 | 1.2e-05 | 5.4e-06 | 9.000E-06 | 3.341E-06 | 5.400E-06 | 1.200E-05 |
| 25377 | 1269 | R1269T | D | 1.3e-05 | 1.6e-05 | 1.3e-05 | 1.400E-05 | 1.732E-06 | 1.300E-05 | 1.600E-05 |
| 25378 | 1269 | R1269W | D | 5.9e-06 | 6.4e-06 | 1.1e-05 | 7.767E-06 | 2.811E-06 | 5.900E-06 | 1.100E-05 |
| 25379 | 1269 | R1269Y | D | 8.8e-06 | 6.7e-06 | 1e-05 | 8.500E-06 | 1.670E-06 | 6.700E-06 | 1.000E-05 |
| 25380 | 1269 | R1269V | D | 6.6e-06 | 5.7e-06 | 7.5e-06 | 6.600E-06 | 9.000E-07 | 5.700E-06 | 7.500E-06 |
| 25381 | 1270 | G1270A | D | 7.7e-06 | 6.3e-06 | 9.4e-06 | 7.800E-06 | 1.552E-06 | 6.300E-06 | 9.400E-06 |
| 25382 | 1270 | G1270R | D | 6.8e-06 | 3.5e-06 | 9.3e-06 | 6.533E-06 | 2.909E-06 | 3.500E-06 | 9.300E-06 |
| 25383 | 1270 | G1270N | D | 5e-06 | 1.1e-05 | 8.7e-06 | 8.233E-06 | 3.027E-06 | 5.000E-06 | 1.100E-05 |
| 25384 | 1270 | G1270D | D | 9e-06 | 9.8e-06 | 7.4e-06 | 8.733E-06 | 1.222E-06 | 7.400E-06 | 9.800E-06 |
| 25385 | 1270 | G1270C | D | 9.4e-06 | 1.1e-05 | 7e-06 | 9.133E-06 | 2.013E-06 | 7.000E-06 | 1.100E-05 |
| 25386 | 1270 | G1270E | D | 1.3e-05 | 4.5e-06 | 4.6e-06 | 7.367E-06 | 4.879E-06 | 4.500E-06 | 1.300E-05 |
| 25387 | 1270 | G1270Q | D | 8.4e-06 | 9.3e-06 | 7.5e-06 | 8.400E-06 | 9.000E-07 | 7.500E-06 | 9.300E-06 |
| 25388 | 1270 | G1270G | D | 9.2e-06 | 8.3e-06 | 7.5e-06 | 8.333E-06 | 8.505E-07 | 7.500E-06 | 9.200E-06 |
| 25389 | 1270 | G1270H | D | 9.4e-06 | 3.9e-06 | 8.7e-06 | 7.333E-06 | 2.994E-06 | 3.900E-06 | 9.400E-06 |
| 25390 | 1270 | G1270I | D | 6.6e-06 | 9.9e-06 | 1.1e-05 | 9.167E-06 | 2.290E-06 | 6.600E-06 | 1.100E-05 |
| 25391 | 1270 | G1270L | D | 6.5e-06 | 1.2e-05 | 6.3e-06 | 8.267E-06 | 3.235E-06 | 6.300E-06 | 1.200E-05 |
| 25392 | 1270 | G1270K | D | 8.3e-06 | 4.6e-06 | 6.3e-06 | 6.400E-06 | 1.852E-06 | 4.600E-06 | 8.300E-06 |
| 25393 | 1270 | G1270M | D | 5.7e-06 | 7e-06 | 5.4e-06 | 6.033E-06 | 8.505E-07 | 5.400E-06 | 7.000E-06 |
| 25394 | 1270 | G1270F | D | 8.7e-06 | 7.6e-06 | 9.8e-06 | 8.700E-06 | 1.100E-06 | 7.600E-06 | 9.800E-06 |
| 25395 | 1270 | G1270P | D | 5.4e-06 | 7.7e-06 | 5.1e-06 | 6.067E-06 | 1.422E-06 | 5.100E-06 | 7.700E-06 |
| 25396 | 1270 | G1270S | D | 8.4e-06 | 7e-06 | 7.5e-06 | 7.633E-06 | 7.095E-07 | 7.000E-06 | 8.400E-06 |
| 25397 | 1270 | G1270T | D | 4.9e-06 | 6.9e-06 | 1.3e-05 | 8.267E-06 | 4.219E-06 | 4.900E-06 | 1.300E-05 |
| 25398 | 1270 | G1270W | D | 9.1e-06 | 7.7e-06 | 9e-06 | 8.600E-06 | 7.810E-07 | 7.700E-06 | 9.100E-06 |
| 25399 | 1270 | G1270Y | D | 4.8e-06 | 1.4e-05 | 6.1e-06 | 8.300E-06 | 4.979E-06 | 4.800E-06 | 1.400E-05 |
| 25400 | 1270 | G1270V | D | 1.5e-05 | 7.7e-06 | 9.9e-06 | 1.087E-05 | 3.745E-06 | 7.700E-06 | 1.500E-05 |
| 25401 | 1271 | E1271A | D | 8.7e-06 | 1.2e-05 | 6.8e-06 | 9.167E-06 | 2.631E-06 | 6.800E-06 | 1.200E-05 |
| 25402 | 1271 | E1271R | D | 5.8e-06 | 1.1e-05 | 8.2e-06 | 8.333E-06 | 2.603E-06 | 5.800E-06 | 1.100E-05 |
| 25403 | 1271 | E1271N | D | 7.8e-06 | 7.6e-06 | 6.4e-06 | 7.267E-06 | 7.572E-07 | 6.400E-06 | 7.800E-06 |
| 25404 | 1271 | E1271D | D | 6.5e-06 | 6.4e-06 | 8.4e-06 | 7.100E-06 | 1.127E-06 | 6.400E-06 | 8.400E-06 |
| 25405 | 1271 | E1271C | D | 7.4e-06 | 8.2e-06 | 7.7e-06 | 7.767E-06 | 4.041E-07 | 7.400E-06 | 8.200E-06 |
| 25406 | 1271 | E1271E | D | 4.5e-06 | 1.1e-05 | 1.1e-05 | 8.833E-06 | 3.753E-06 | 4.500E-06 | 1.100E-05 |
| 25407 | 1271 | E1271Q | D | 5e-06 | 1.4e-05 | 1e-05 | 9.667E-06 | 4.509E-06 | 5.000E-06 | 1.400E-05 |
| 25408 | 1271 | E1271G | D | 4.7e-06 | 6.4e-06 | 1.2e-05 | 7.700E-06 | 3.820E-06 | 4.700E-06 | 1.200E-05 |
| 25409 | 1271 | E1271H | D | 6.2e-06 | 6.5e-06 | 7.2e-06 | 6.633E-06 | 5.132E-07 | 6.200E-06 | 7.200E-06 |
| 25410 | 1271 | E1271I | D | 7.3e-06 | 9.6e-06 | 8.2e-06 | 8.367E-06 | 1.159E-06 | 7.300E-06 | 9.600E-06 |
| 25411 | 1271 | E1271L | D | 4.8e-06 | 8.4e-06 | 6.8e-06 | 6.667E-06 | 1.804E-06 | 4.800E-06 | 8.400E-06 |
| 25412 | 1271 | E1271K | D | 5.4e-06 | 1e-05 | 6.6e-06 | 7.333E-06 | 2.386E-06 | 5.400E-06 | 1.000E-05 |
| 25413 | 1271 | E1271M | D | 4.6e-06 | 5.9e-06 | 4.4e-06 | 4.967E-06 | 8.145E-07 | 4.400E-06 | 5.900E-06 |
| 25414 | 1271 | E1271F | D | 9.3e-06 | 1.1e-05 | 8.2e-06 | 9.500E-06 | 1.411E-06 | 8.200E-06 | 1.100E-05 |

| | | | | | | | | | | |
|---|---|---|---|---|---|---|---|---|---|---|
| 25415 | 1271 | E1271P | D | 6e-06 | 5.9e-06 | 9.1e-06 | 7.000E-06 | 1.819E-06 | 5.900E-06 | 9.100E-06 |
| 25416 | 1271 | E1271S | D | 1.1e-05 | 6.5e-06 | 6.4e-06 | 7.967E-06 | 2.627E-06 | 6.400E-06 | 1.100E-05 |
| 25417 | 1271 | E1271T | D | 4.7e-06 | 7.7e-06 | 7.3e-06 | 6.567E-06 | 1.629E-06 | 4.700E-06 | 7.700E-06 |
| 25418 | 1271 | E1271W | D | 7.3e-06 | 9.9e-06 | 5.1e-06 | 7.433E-06 | 2.403E-06 | 5.100E-06 | 9.900E-06 |
| 25419 | 1271 | E1271Y | D | 8.2e-06 | 5.4e-06 | 8.6e-06 | 7.400E-06 | 1.744E-06 | 5.400E-06 | 8.600E-06 |
| 25420 | 1271 | E1271V | D | 9.2e-06 | 4.1e-06 | 8.9e-06 | 7.400E-06 | 2.862E-06 | 4.100E-06 | 9.200E-06 |
| 25421 | 1272 | C1272A | D | 4.6e-06 | 1e-05 | 1.1e-05 | 8.533E-06 | 3.443E-06 | 4.600E-06 | 1.100E-05 |
| 25422 | 1272 | C1272R | D | 1.3e-05 | 8.3e-06 | 5.2e-06 | 8.833E-06 | 3.927E-06 | 5.200E-06 | 1.300E-05 |
| 25423 | 1272 | C1272N | D | 9.7e-06 | 7.4e-06 | 4.7e-06 | 7.267E-06 | 2.503E-06 | 4.700E-06 | 9.700E-06 |
| 25424 | 1272 | C1272D | D | 3.8e-06 | 8e-06 | 1.4e-05 | 8.600E-06 | 5.126E-06 | 3.800E-06 | 1.400E-05 |
| 25425 | 1272 | C1272C | D | 5.2e-06 | 8.7e-06 | 6.2e-06 | 6.700E-06 | 1.803E-06 | 5.200E-06 | 8.700E-06 |
| 25426 | 1272 | C1272E | D | 3.4e-06 | 5.6e-06 | 1.2e-05 | 7.000E-06 | 4.468E-06 | 3.400E-06 | 1.200E-05 |
| 25427 | 1272 | C1272Q | D | 3.8e-06 | 6.8e-06 | 7.1e-06 | 5.900E-06 | 1.825E-06 | 3.800E-06 | 7.100E-06 |
| 25428 | 1272 | C1272G | D | 5e-06 | 9.7e-06 | 1e-05 | 8.233E-06 | 2.804E-06 | 5.000E-06 | 1.000E-05 |
| 25429 | 1272 | C1272H | D | 9.5e-06 | 7.1e-06 | 4.2e-06 | 6.933E-06 | 2.654E-06 | 4.200E-06 | 9.500E-06 |
| 25430 | 1272 | C1272I | D | 4.4e-06 | 7.6e-06 | 5.9e-06 | 5.967E-06 | 1.601E-06 | 4.400E-06 | 7.600E-06 |
| 25431 | 1272 | C1272L | D | 7.1e-06 | 6.6e-06 | 7.6e-06 | 7.100E-06 | 5.000E-07 | 6.600E-06 | 7.600E-06 |
| 25432 | 1272 | C1272K | D | 8.5e-06 | 6.9e-06 | 8e-06 | 7.800E-06 | 8.185E-07 | 6.900E-06 | 8.500E-06 |
| 25433 | 1272 | C1272M | D | 5e-06 | 9.1e-06 | 1.1e-05 | 8.367E-06 | 3.066E-06 | 5.000E-06 | 1.100E-05 |
| 25434 | 1272 | C1272F | D | 8.1e-06 | 1.5e-05 | 7e-06 | 1.003E-05 | 4.336E-06 | 7.000E-06 | 1.500E-05 |
| 25435 | 1272 | C1272P | D | 1.2e-05 | 5e-06 | 6.8e-06 | 7.933E-06 | 3.635E-06 | 5.000E-06 | 1.200E-05 |
| 25436 | 1272 | C1272S | D | 9.9e-06 | 8.9e-06 | 1.2e-05 | 1.027E-05 | 1.582E-06 | 8.900E-06 | 1.200E-05 |
| 25437 | 1272 | C1272T | D | 5.8e-06 | 5.5e-06 | 7.9e-06 | 6.400E-06 | 1.308E-06 | 5.500E-06 | 7.900E-06 |
| 25438 | 1272 | C1272W | D | 1.8e-05 | 6.5e-06 | 9.3e-06 | 1.127E-05 | 5.997E-06 | 6.500E-06 | 1.800E-05 |
| 25439 | 1272 | C1272Y | D | 7.2e-06 | 6.9e-06 | 5.6e-06 | 6.567E-06 | 8.505E-07 | 5.600E-06 | 7.200E-06 |
| 25440 | 1272 | C1272V | D | 9.7e-06 | 9.5e-06 | 1.1e-05 | 1.007E-05 | 8.145E-07 | 9.500E-06 | 1.100E-05 |
| 25441 | 1273 | E1273A | E | 5.2e-06 | 6.3e-06 | 7.9e-06 | 6.467E-06 | 1.358E-06 | 5.200E-06 | 7.900E-06 |
| 25442 | 1273 | E1273R | E | 9.4e-06 | 4.8e-06 | 1.1e-05 | 8.400E-06 | 3.219E-06 | 4.800E-06 | 1.100E-05 |
| 25443 | 1273 | E1273N | E | 8.5e-06 | 5.4e-06 | 1.3e-05 | 8.967E-06 | 3.821E-06 | 5.400E-06 | 1.300E-05 |
| 25444 | 1273 | E1273D | E | 5.5e-06 | 9.1e-06 | 4.8e-06 | 6.467E-06 | 2.307E-06 | 4.800E-06 | 9.100E-06 |
| 25445 | 1273 | E1273C | E | 8.5e-06 | 7.3e-06 | 5.5e-06 | 7.100E-06 | 1.510E-06 | 5.500E-06 | 8.500E-06 |
| 25446 | 1273 | E1273E | E | 4.9e-06 | 6.7e-06 | 9.9e-06 | 7.167E-06 | 2.532E-06 | 4.900E-06 | 9.900E-06 |
| 25447 | 1273 | E1273Q | E | 7.3e-06 | 6.7e-06 | 1e-05 | 8.000E-06 | 1.758E-06 | 6.700E-06 | 1.000E-05 |
| 25448 | 1273 | E1273G | E | 7e-06 | 7.3e-06 | 7.9e-06 | 7.400E-06 | 4.583E-07 | 7.000E-06 | 7.900E-06 |
| 25449 | 1273 | E1273H | E | 7.1e-06 | 6.7e-06 | 5.5e-06 | 6.433E-06 | 8.327E-07 | 5.500E-06 | 7.100E-06 |
| 25450 | 1273 | E1273I | E | 1.4e-05 | 9.4e-06 | 7.6e-06 | 1.033E-05 | 3.301E-06 | 7.600E-06 | 1.400E-05 |
| 25451 | 1273 | E1273L | E | 1.2e-05 | 7.3e-06 | 1.3e-05 | 1.077E-05 | 3.044E-06 | 7.300E-06 | 1.300E-05 |
| 25452 | 1273 | E1273K | E | 8.1e-06 | 7.1e-06 | 8.7e-06 | 7.967E-06 | 8.083E-07 | 7.100E-06 | 8.700E-06 |
| 25453 | 1273 | E1273M | E | 6.9e-06 | 5.4e-06 | 5.8e-06 | 6.033E-06 | 7.767E-07 | 5.400E-06 | 6.900E-06 |
| 25454 | 1273 | E1273F | E | 1e-05 | 6.6e-06 | 4.6e-06 | 7.067E-06 | 2.730E-06 | 4.600E-06 | 1.000E-05 |
| 25455 | 1273 | E1273P | E | 6.5e-06 | 1.2e-05 | 1.2e-05 | 1.017E-05 | 3.175E-06 | 6.500E-06 | 1.200E-05 |
| 25456 | 1273 | E1273S | E | 7.2e-06 | 9e-06 | 8.5e-06 | 8.233E-06 | 9.292E-07 | 7.200E-06 | 9.000E-06 |
| 25457 | 1273 | E1273T | E | 7e-06 | 8.4e-06 | 6.6e-06 | 7.333E-06 | 9.452E-07 | 6.600E-06 | 8.400E-06 |
| 25458 | 1273 | E1273W | E | 6.7e-06 | 1e-05 | 1.1e-05 | 9.233E-06 | 2.250E-06 | 6.700E-06 | 1.100E-05 |
| 25459 | 1273 | E1273Y | E | 5.4e-06 | 7.6e-06 | 3.2e-06 | 5.400E-06 | 2.200E-06 | 3.200E-06 | 7.600E-06 |
| 25460 | 1273 | E1273V | E | 1e-05 | 1.4e-05 | 9.2e-06 | 1.107E-05 | 2.572E-06 | 9.200E-06 | 1.400E-05 |
| 25461 | 1274 | V1274A | E | 9.8e-06 | 4.7e-06 | 8e-06 | 7.500E-06 | 2.587E-06 | 4.700E-06 | 9.800E-06 |
| 25462 | 1274 | V1274R | E | 9.7e-06 | 1.1e-05 | 1e-05 | 1.023E-05 | 6.807E-07 | 9.700E-06 | 1.100E-05 |
| 25463 | 1274 | V1274N | E | 1.1e-05 | 8.1e-06 | 9.4e-06 | 9.500E-06 | 1.453E-06 | 8.100E-06 | 1.100E-05 |
| 25464 | 1274 | V1274D | E | 7.4e-06 | 7e-06 | 8.5e-06 | 7.633E-06 | 7.767E-07 | 7.000E-06 | 8.500E-06 |
| 25465 | 1274 | V1274C | E | 7.1e-06 | 9.5e-06 | 6.3e-06 | 7.633E-06 | 1.665E-06 | 6.300E-06 | 9.500E-06 |
| 25466 | 1274 | V1274E | E | 6.2e-06 | 6.5e-06 | 6.5e-06 | 6.400E-06 | 1.732E-07 | 6.200E-06 | 6.500E-06 |
| 25467 | 1274 | V1274Q | E | 8.9e-06 | 5.3e-06 | 7.1e-06 | 7.100E-06 | 1.800E-06 | 5.300E-06 | 8.900E-06 |
| 25468 | 1274 | V1274G | E | 9.5e-06 | 1.4e-05 | 1e-05 | 1.117E-05 | 2.466E-06 | 9.500E-06 | 1.400E-05 |
| 25469 | 1274 | V1274H | E | 9.5e-06 | 8.8e-06 | 8.2e-06 | 8.833E-06 | 6.506E-07 | 8.200E-06 | 9.500E-06 |
| 25470 | 1274 | V1274I | E | 1e-05 | 1.3e-05 | 5.1e-06 | 9.367E-06 | 3.988E-06 | 5.100E-06 | 1.300E-05 |
| 25471 | 1274 | V1274L | E | 9.9e-06 | 7.5e-06 | 1.3e-05 | 1.013E-05 | 2.757E-06 | 7.500E-06 | 1.300E-05 |
| 25472 | 1274 | V1274K | E | 7.9e-06 | 4.9e-06 | 6.9e-06 | 6.567E-06 | 1.528E-06 | 4.900E-06 | 7.900E-06 |
| 25473 | 1274 | V1274M | E | 1.1e-05 | 6.3e-06 | 1.1e-05 | 9.433E-06 | 2.714E-06 | 6.300E-06 | 1.100E-05 |
| 25474 | 1274 | V1274F | E | 4.9e-06 | 9e-06 | 9.5e-06 | 7.800E-06 | 2.524E-06 | 4.900E-06 | 9.500E-06 |
| 25475 | 1274 | V1274P | E | 9.9e-06 | 1.4e-05 | 1e-05 | 1.130E-05 | 2.339E-06 | 9.900E-06 | 1.400E-05 |
| 25476 | 1274 | V1274S | E | 9.8e-06 | 4.8e-06 | 5.1e-06 | 6.567E-06 | 2.804E-06 | 4.800E-06 | 9.800E-06 |
| 25477 | 1274 | V1274T | E | 8.7e-06 | 9.2e-06 | 6.9e-06 | 8.267E-06 | 1.210E-06 | 6.900E-06 | 9.200E-06 |
| 25478 | 1274 | V1274W | E | 1.5e-05 | 8.4e-06 | 6e-06 | 9.800E-06 | 4.660E-06 | 6.000E-06 | 1.500E-05 |
| 25479 | 1274 | V1274Y | E | 8e-06 | 6e-06 | 5.2e-06 | 6.400E-06 | 1.442E-06 | 5.200E-06 | 8.000E-06 |
| 25480 | 1274 | V1274V | E | 2.4e-05 | 6.7e-06 | 7e-06 | 1.257E-05 | 9.903E-06 | 6.700E-06 | 2.400E-05 |
| 25481 | 1275 | Q1275A | E | 8.4e-06 | 4.7e-06 | 8.1e-06 | 7.067E-06 | 2.055E-06 | 4.700E-06 | 8.400E-06 |
| 25482 | 1275 | Q1275R | E | 7.9e-06 | 9.9e-06 | 8.4e-06 | 8.733E-06 | 1.041E-06 | 7.900E-06 | 9.900E-06 |
| 25483 | 1275 | Q1275N | E | 4.7e-06 | 1.4e-05 | 5.6e-06 | 8.100E-06 | 5.129E-06 | 4.700E-06 | 1.400E-05 |
| 25484 | 1275 | Q1275D | E | 1.8e-05 | 1.1e-05 | 6.6e-06 | 1.187E-05 | 5.749E-06 | 6.600E-06 | 1.800E-05 |
| 25485 | 1275 | Q1275C | E | 7.2e-06 | 7.7e-06 | 6.2e-06 | 7.033E-06 | 7.638E-07 | 6.200E-06 | 7.700E-06 |
| 25486 | 1275 | Q1275E | E | 1.4e-05 | 1e-05 | 1.3e-05 | 1.233E-05 | 2.082E-06 | 1.000E-05 | 1.400E-05 |
| 25487 | 1275 | Q1275Q | E | 8.6e-06 | 6.2e-06 | 5.8e-06 | 6.867E-06 | 1.514E-06 | 5.800E-06 | 8.600E-06 |
| 25488 | 1275 | Q1275G | E | 6.2e-06 | 1.4e-05 | 7.9e-06 | 9.367E-06 | 4.102E-06 | 6.200E-06 | 1.400E-05 |
| 25489 | 1275 | Q1275H | E | 9.9e-06 | 4.9e-06 | 9.2e-06 | 8.000E-06 | 2.707E-06 | 4.900E-06 | 9.900E-06 |
| 25490 | 1275 | Q1275I | E | 8.7e-06 | 8.4e-06 | 9.6e-06 | 8.900E-06 | 6.245E-07 | 8.400E-06 | 9.600E-06 |
| 25491 | 1275 | Q1275L | E | 9.4e-06 | 1.4e-05 | 9.3e-06 | 1.090E-05 | 2.685E-06 | 9.300E-06 | 1.400E-05 |
| 25492 | 1275 | Q1275K | E | 1.4e-05 | 3.5e-06 | 9.2e-06 | 8.900E-06 | 5.256E-06 | 3.500E-06 | 1.400E-05 |
| 25493 | 1275 | Q1275M | E | 9.8e-06 | 8.9e-06 | 5.9e-06 | 8.200E-06 | 2.042E-06 | 5.900E-06 | 9.800E-06 |
| 25494 | 1275 | Q1275F | E | 1.6e-05 | 7.6e-06 | 6.1e-06 | 9.900E-06 | 5.336E-06 | 6.100E-06 | 1.600E-05 |
| 25495 | 1275 | Q1275P | E | 4.7e-06 | 9.5e-06 | 6.3e-06 | 6.833E-06 | 2.444E-06 | 4.700E-06 | 9.500E-06 |
| 25496 | 1275 | Q1275S | E | 9.8e-06 | 4.7e-06 | 8.1e-06 | 7.533E-06 | 2.597E-06 | 4.700E-06 | 9.800E-06 |
| 25497 | 1275 | Q1275T | E | 9.5e-06 | 6.8e-06 | 9.1e-06 | 8.467E-06 | 1.457E-06 | 6.800E-06 | 9.500E-06 |
| 25498 | 1275 | Q1275W | E | 1e-05 | 7.1e-06 | 1.1e-05 | 9.367E-06 | 2.026E-06 | 7.100E-06 | 1.100E-05 |
| 25499 | 1275 | Q1275Y | E | 1.2e-05 | 8.5e-06 | 6.8e-06 | 9.100E-06 | 2.651E-06 | 6.800E-06 | 1.200E-05 |

| | | | | | | | | | | |
|---|---|---|---|---|---|---|---|---|---|---|
| 25500 | 1275 | Q1275V | E | 8.2e-06 | 1.2e-05 | 7.7e-06 | 9.300E-06 | 2.352E-06 | 7.700E-06 | 1.200E-05 |
| 25501 | 1276 | L1276A | E | 4.3e-06 | 9e-06 | 1e-05 | 7.767E-06 | 3.044E-06 | 4.300E-06 | 1.000E-05 |
| 25502 | 1276 | L1276R | E | 7.4e-06 | 5.4e-06 | 7.1e-06 | 6.633E-06 | 1.079E-06 | 5.400E-06 | 7.400E-06 |
| 25503 | 1276 | L1276N | E | 8.6e-06 | 5.8e-06 | 8.9e-06 | 7.767E-06 | 1.710E-06 | 5.800E-06 | 8.900E-06 |
| 25504 | 1276 | L1276D | E | 9.8e-06 | 8.1e-06 | 9.4e-06 | 9.100E-06 | 8.888E-07 | 8.100E-06 | 9.800E-06 |
| 25505 | 1276 | L1276C | E | 6.2e-06 | 1e-05 | 1.3e-05 | 9.733E-06 | 3.408E-06 | 6.200E-06 | 1.300E-05 |
| 25506 | 1276 | L1276E | E | 7.7e-06 | 1.4e-05 | 1e-05 | 1.057E-05 | 3.188E-06 | 7.700E-06 | 1.400E-05 |
| 25507 | 1276 | L1276Q | E | 8.8e-06 | 8.8e-06 | 1e-05 | 9.200E-06 | 6.928E-07 | 8.800E-06 | 1.000E-05 |
| 25508 | 1276 | L1276G | E | 8.8e-06 | 4.2e-06 | 1.2e-05 | 8.333E-06 | 3.921E-06 | 4.200E-06 | 1.200E-05 |
| 25509 | 1276 | L1276H | E | 9.8e-06 | 9.7e-06 | 6.9e-06 | 8.800E-06 | 1.646E-06 | 6.900E-06 | 9.800E-06 |
| 25510 | 1276 | L1276I | E | 7.2e-06 | 7e-06 | 8.2e-06 | 7.467E-06 | 6.429E-07 | 7.000E-06 | 8.200E-06 |
| 25511 | 1276 | L1276L | E | 6.6e-06 | 7.2e-06 | 1e-05 | 7.933E-06 | 1.815E-06 | 6.600E-06 | 1.000E-05 |
| 25512 | 1276 | L1276K | E | 9.3e-06 | 1.4e-05 | 5.9e-06 | 9.733E-06 | 4.067E-06 | 5.900E-06 | 1.400E-05 |
| 25513 | 1276 | L1276M | E | 9e-06 | 8e-06 | 9.4e-06 | 8.800E-06 | 7.211E-07 | 8.000E-06 | 9.400E-06 |
| 25514 | 1276 | L1276F | E | 8.1e-06 | 5.5e-06 | 9.3e-06 | 7.633E-06 | 1.943E-06 | 5.500E-06 | 9.300E-06 |
| 25515 | 1276 | L1276P | E | 5.4e-06 | 6.2e-06 | 5.6e-06 | 5.733E-06 | 4.163E-07 | 5.400E-06 | 6.200E-06 |
| 25516 | 1276 | L1276S | E | 9.7e-06 | 5.6e-06 | 8.3e-06 | 7.867E-06 | 2.084E-06 | 5.600E-06 | 9.700E-06 |
| 25517 | 1276 | L1276T | E | 7.3e-06 | 4.6e-06 | 9.9e-06 | 7.267E-06 | 2.650E-06 | 4.600E-06 | 9.900E-06 |
| 25518 | 1276 | L1276W | E | 1.1e-05 | 8.9e-06 | 5.1e-06 | 8.333E-06 | 2.991E-06 | 5.100E-06 | 1.100E-05 |
| 25519 | 1276 | L1276Y | E | 5.7e-06 | 8.4e-06 | 8.9e-06 | 7.667E-06 | 1.721E-06 | 5.700E-06 | 8.900E-06 |
| 25520 | 1276 | L1276V | E | 9.3e-06 | 9.1e-06 | 1.3e-05 | 1.047E-05 | 2.196E-06 | 9.100E-06 | 1.300E-05 |
| 25521 | 1277 | V1277A | E | 9.3e-06 | 9.5e-06 | 6.6e-06 | 8.467E-06 | 1.620E-06 | 6.600E-06 | 9.500E-06 |
| 25522 | 1277 | V1277R | E | 1.2e-05 | 8.8e-06 | 5.4e-06 | 8.733E-06 | 3.301E-06 | 5.400E-06 | 1.200E-05 |
| 25523 | 1277 | V1277N | E | 8.7e-06 | 9.1e-06 | 7e-06 | 8.267E-06 | 1.115E-06 | 7.000E-06 | 9.100E-06 |
| 25524 | 1277 | V1277D | E | 9.2e-06 | 7.7e-06 | 8.9e-06 | 8.600E-06 | 7.937E-07 | 7.700E-06 | 9.200E-06 |
| 25525 | 1277 | V1277C | E | 1.4e-05 | 9.3e-06 | 8.5e-06 | 1.060E-05 | 2.972E-06 | 8.500E-06 | 1.400E-05 |
| 25526 | 1277 | V1277E | E | 8.4e-06 | 5.7e-06 | 1.1e-05 | 8.367E-06 | 2.650E-06 | 5.700E-06 | 1.100E-05 |
| 25527 | 1277 | V1277Q | E | 6.5e-06 | 6.3e-06 | 6.9e-06 | 6.567E-06 | 3.055E-07 | 6.300E-06 | 6.900E-06 |
| 25528 | 1277 | V1277G | E | 5.5e-06 | 5.9e-06 | 4.7e-06 | 5.367E-06 | 6.110E-07 | 4.700E-06 | 5.900E-06 |
| 25529 | 1277 | V1277H | E | 1.2e-05 | 7e-06 | 8.9e-06 | 9.300E-06 | 2.524E-06 | 7.000E-06 | 1.200E-05 |
| 25530 | 1277 | V1277I | E | 9e-06 | 8.7e-06 | 1e-05 | 9.233E-06 | 6.807E-07 | 8.700E-06 | 1.000E-05 |
| 25531 | 1277 | V1277L | E | 1e-05 | 7.6e-06 | 1.1e-05 | 9.533E-06 | 1.747E-06 | 7.600E-06 | 1.100E-05 |
| 25532 | 1277 | V1277K | E | 1.4e-05 | 6.9e-06 | 9.7e-06 | 1.020E-05 | 3.576E-06 | 6.900E-06 | 1.400E-05 |
| 25533 | 1277 | V1277M | E | 1e-05 | 9.5e-06 | 6.2e-06 | 8.567E-06 | 2.065E-06 | 6.200E-06 | 1.000E-05 |
| 25534 | 1277 | V1277F | E | 6.6e-06 | 6.3e-06 | 9.3e-06 | 7.400E-06 | 1.652E-06 | 6.300E-06 | 9.300E-06 |
| 25535 | 1277 | V1277P | E | 1.4e-05 | 6.3e-06 | 8.8e-06 | 9.700E-06 | 3.928E-06 | 6.300E-06 | 1.400E-05 |
| 25536 | 1277 | V1277S | E | 5.9e-06 | 6.4e-06 | 5.2e-06 | 5.833E-06 | 6.028E-07 | 5.200E-06 | 6.400E-06 |
| 25537 | 1277 | V1277T | E | 5.6e-06 | 6e-06 | 7e-06 | 6.200E-06 | 7.211E-07 | 5.600E-06 | 7.000E-06 |
| 25538 | 1277 | V1277W | E | 9.4e-06 | 1e-05 | 7.4e-06 | 8.933E-06 | 1.361E-06 | 7.400E-06 | 1.000E-05 |
| 25539 | 1277 | V1277Y | E | 8.3e-06 | 9.8e-06 | 7.9e-06 | 8.667E-06 | 1.002E-06 | 7.900E-06 | 9.800E-06 |
| 25540 | 1277 | V1277V | E | 6.2e-06 | 7.7e-06 | 9.1e-06 | 7.667E-06 | 1.450E-06 | 6.200E-06 | 9.100E-06 |
| 25541 | 1278 | E1278A | E | 5.1e-06 | 1.2e-05 | 9e-06 | 8.700E-06 | 3.460E-06 | 5.100E-06 | 1.200E-05 |
| 25542 | 1278 | E1278R | E | 9.4e-06 | 1.3e-05 | 6.9e-06 | 9.767E-06 | 3.066E-06 | 6.900E-06 | 1.300E-05 |
| 25543 | 1278 | E1278N | E | 1e-05 | 1.2e-05 | 8.3e-06 | 1.010E-05 | 1.852E-06 | 8.300E-06 | 1.200E-05 |
| 25544 | 1278 | E1278D | E | 1e-05 | 4.3e-06 | 1e-05 | 8.100E-06 | 3.291E-06 | 4.300E-06 | 1.000E-05 |
| 25545 | 1278 | E1278C | E | 7.6e-06 | 9.8e-06 | 7.9e-06 | 8.433E-06 | 1.193E-06 | 7.600E-06 | 9.800E-06 |
| 25546 | 1278 | E1278E | E | 3.8e-06 | 5.8e-06 | 6.9e-06 | 5.500E-06 | 1.572E-06 | 3.800E-06 | 6.900E-06 |
| 25547 | 1278 | E1278Q | E | 5.7e-06 | 9e-06 | 6.5e-06 | 7.067E-06 | 1.721E-06 | 5.700E-06 | 9.000E-06 |
| 25548 | 1278 | E1278G | E | 1.2e-05 | 7.3e-06 | 6.6e-06 | 8.633E-06 | 2.937E-06 | 6.600E-06 | 1.200E-05 |
| 25549 | 1278 | E1278H | E | 2.1e-05 | 6.3e-06 | 5.2e-06 | 1.083E-05 | 8.822E-06 | 5.200E-06 | 2.100E-05 |
| 25550 | 1278 | E1278I | E | 4.8e-06 | 7.9e-06 | 7.2e-06 | 6.633E-06 | 1.626E-06 | 4.800E-06 | 7.900E-06 |
| 25551 | 1278 | E1278L | E | 5.8e-06 | 9.9e-06 | 7.9e-06 | 7.867E-06 | 2.050E-06 | 5.800E-06 | 9.900E-06 |
| 25552 | 1278 | E1278K | E | 9.7e-06 | 9e-06 | 1.2e-05 | 1.023E-05 | 1.570E-06 | 9.000E-06 | 1.200E-05 |
| 25553 | 1278 | E1278M | E | 8.8e-06 | 3.9e-06 | 7.8e-06 | 6.833E-06 | 2.589E-06 | 3.900E-06 | 8.800E-06 |
| 25554 | 1278 | E1278F | E | 9.8e-06 | 7.6e-06 | 9e-06 | 8.800E-06 | 1.114E-06 | 7.600E-06 | 9.800E-06 |
| 25555 | 1278 | E1278P | E | 7.1e-06 | 3.2e-06 | 7.5e-06 | 5.933E-06 | 2.376E-06 | 3.200E-06 | 7.500E-06 |
| 25556 | 1278 | E1278S | E | 3.8e-06 | 8.3e-06 | 9.2e-06 | 7.100E-06 | 2.893E-06 | 3.800E-06 | 9.200E-06 |
| 25557 | 1278 | E1278T | E | 1.1e-05 | 2.1e-05 | 7.3e-06 | 1.310E-05 | 7.087E-06 | 7.300E-06 | 2.100E-05 |
| 25558 | 1278 | E1278W | E | 4.9e-06 | 1e-05 | 8.9e-06 | 7.933E-06 | 2.684E-06 | 4.900E-06 | 1.000E-05 |
| 25559 | 1278 | E1278Y | E | 1.1e-05 | 1.1e-05 | 8.4e-06 | 1.013E-05 | 1.501E-06 | 8.400E-06 | 1.100E-05 |
| 25560 | 1278 | E1278V | E | 6.8e-06 | 5.2e-06 | 8e-06 | 6.667E-06 | 1.405E-06 | 5.200E-06 | 8.000E-06 |
| 25561 | 1279 | S1279A | E | 8.6e-06 | 1e-05 | 9.7e-06 | 9.433E-06 | 7.371E-07 | 8.600E-06 | 1.000E-05 |
| 25562 | 1279 | S1279R | E | 1.3e-05 | 8.1e-06 | 6.7e-06 | 9.267E-06 | 3.308E-06 | 6.700E-06 | 1.300E-05 |
| 25563 | 1279 | S1279N | E | 1.7e-05 | 9.4e-06 | 6.6e-06 | 1.100E-05 | 5.381E-06 | 6.600E-06 | 1.700E-05 |
| 25564 | 1279 | S1279D | E | 9e-06 | 5.8e-06 | 1e-05 | 8.267E-06 | 2.194E-06 | 5.800E-06 | 1.000E-05 |
| 25565 | 1279 | S1279C | E | 6.2e-06 | 5.3e-06 | 1.3e-05 | 8.167E-06 | 4.210E-06 | 5.300E-06 | 1.300E-05 |
| 25566 | 1279 | S1279E | E | 6.3e-06 | 8.6e-06 | 6.7e-06 | 7.200E-06 | 1.229E-06 | 6.300E-06 | 8.600E-06 |
| 25567 | 1279 | S1279Q | E | 6.9e-06 | 7.3e-06 | 1.1e-05 | 8.400E-06 | 2.261E-06 | 6.900E-06 | 1.100E-05 |
| 25568 | 1279 | S1279G | E | 8.7e-06 | 9.7e-06 | 6.3e-06 | 8.233E-06 | 1.747E-06 | 6.300E-06 | 9.700E-06 |
| 25569 | 1279 | S1279H | E | 5.7e-06 | 7.1e-06 | 1.2e-05 | 8.267E-06 | 3.308E-06 | 5.700E-06 | 1.200E-05 |
| 25570 | 1279 | S1279I | E | 7.1e-06 | 6.4e-06 | 9.5e-06 | 7.667E-06 | 1.626E-06 | 6.400E-06 | 9.500E-06 |
| 25571 | 1279 | S1279L | E | 7.2e-06 | 8e-06 | 4.7e-06 | 6.633E-06 | 1.721E-06 | 4.700E-06 | 8.000E-06 |
| 25572 | 1279 | S1279K | E | 5.8e-06 | 7.9e-06 | 6.1e-06 | 6.600E-06 | 1.136E-06 | 5.800E-06 | 7.900E-06 |
| 25573 | 1279 | S1279M | E | 7.7e-06 | 1.1e-05 | 7.5e-06 | 8.733E-06 | 1.966E-06 | 7.500E-06 | 1.100E-05 |
| 25574 | 1279 | S1279F | E | 6.3e-06 | 5e-06 | 6.3e-06 | 5.867E-06 | 7.506E-07 | 5.000E-06 | 6.300E-06 |
| 25575 | 1279 | S1279P | E | 1.1e-05 | 5.8e-06 | 1.1e-05 | 9.267E-06 | 3.002E-06 | 5.800E-06 | 1.100E-05 |
| 25576 | 1279 | S1279S | E | 1.1e-05 | 7.1e-06 | 6.9e-06 | 8.333E-06 | 2.312E-06 | 6.900E-06 | 1.100E-05 |
| 25577 | 1279 | S1279T | E | 8.1e-06 | 1.2e-05 | 9.6e-06 | 9.900E-06 | 1.967E-06 | 8.100E-06 | 1.200E-05 |
| 25578 | 1279 | S1279W | E | 1.2e-05 | 1e-05 | 6.3e-06 | 9.433E-06 | 2.892E-06 | 6.300E-06 | 1.200E-05 |
| 25579 | 1279 | S1279Y | E | 1.2e-05 | 8.1e-06 | 7e-06 | 9.033E-06 | 2.627E-06 | 7.000E-06 | 1.200E-05 |
| 25580 | 1279 | S1279V | E | 1.1e-05 | 1e-05 | 5.9e-06 | 8.967E-06 | 2.702E-06 | 5.900E-06 | 1.100E-05 |
| 25581 | 1280 | G1280A | E | 5.4e-06 | 4.3e-06 | 7.3e-06 | 5.667E-06 | 1.518E-06 | 4.300E-06 | 7.300E-06 |
| 25582 | 1280 | G1280R | E | 4.6e-06 | 1.3e-05 | 7.1e-06 | 8.233E-06 | 4.313E-06 | 4.600E-06 | 1.300E-05 |
| 25583 | 1280 | G1280N | E | 4.4e-06 | 8.3e-06 | 7.1e-06 | 6.600E-06 | 1.997E-06 | 4.400E-06 | 8.300E-06 |
| 25584 | 1280 | G1280D | E | 6.5e-06 | 5.2e-06 | 1.7e-05 | 9.567E-06 | 6.470E-06 | 5.200E-06 | 1.700E-05 |

| | | | | | | | | | | |
|---|---|---|---|---|---|---|---|---|---|---|
| 25585 | 1280 | G1280C | E | 9.2e-06 | 6.7e-06 | 9.7e-06 | 8.533E-06 | 1.607E-06 | 6.700E-06 | 9.700E-06 |
| 25586 | 1280 | G1280E | E | 1.2e-05 | 5.5e-06 | 9.3e-06 | 8.933E-06 | 3.265E-06 | 5.500E-06 | 1.200E-05 |
| 25587 | 1280 | G1280Q | E | 7.6e-06 | 8.2e-06 | 9.4e-06 | 8.400E-06 | 9.165E-07 | 7.600E-06 | 9.400E-06 |
| 25588 | 1280 | G1280G | E | 9.6e-06 | 1e-05 | 8.7e-06 | 9.433E-06 | 6.658E-07 | 8.700E-06 | 1.000E-05 |
| 25589 | 1280 | G1280H | E | 1e-05 | 6.6e-06 | 8e-06 | 8.200E-06 | 1.709E-06 | 6.600E-06 | 1.000E-05 |
| 25590 | 1280 | G1280I | E | 7.3e-06 | 5.3e-06 | 8.8e-06 | 7.133E-06 | 1.756E-06 | 5.300E-06 | 8.800E-06 |
| 25591 | 1280 | G1280L | E | 8e-06 | 8.4e-06 | 9.6e-06 | 8.667E-06 | 8.327E-07 | 8.000E-06 | 9.600E-06 |
| 25592 | 1280 | G1280K | E | 1.1e-05 | 6e-06 | 8.4e-06 | 8.467E-06 | 2.501E-06 | 6.000E-06 | 1.100E-05 |
| 25593 | 1280 | G1280M | E | 1e-05 | 9.9e-06 | 5.8e-06 | 8.567E-06 | 2.397E-06 | 5.800E-06 | 1.000E-05 |
| 25594 | 1280 | G1280F | E | 7.1e-06 | 7.5e-06 | 5.8e-06 | 6.800E-06 | 8.888E-07 | 5.800E-06 | 7.500E-06 |
| 25595 | 1280 | G1280P | E | 8.4e-06 | 5.5e-06 | 3.8e-06 | 5.900E-06 | 2.326E-06 | 3.800E-06 | 8.400E-06 |
| 25596 | 1280 | G1280S | E | 4.6e-06 | 8.6e-06 | 6.7e-06 | 6.633E-06 | 2.001E-06 | 4.600E-06 | 8.600E-06 |
| 25597 | 1280 | G1280T | E | 4.6e-06 | 5.5e-06 | 6.1e-06 | 5.400E-06 | 7.550E-07 | 4.600E-06 | 6.100E-06 |
| 25598 | 1280 | G1280W | E | 1.3e-05 | 5.9e-06 | 8.2e-06 | 9.033E-06 | 3.623E-06 | 5.900E-06 | 1.300E-05 |
| 25599 | 1280 | G1280Y | E | 7e-06 | 1e-05 | 9.4e-06 | 8.800E-06 | 1.587E-06 | 7.000E-06 | 1.000E-05 |
| 25600 | 1280 | G1280V | E | 8.5e-06 | 5.1e-06 | 8.2e-06 | 7.267E-06 | 1.882E-06 | 5.100E-06 | 8.500E-06 |
| 25601 | 1281 | G1281A | E | 1.2e-05 | 6.6e-06 | 8.6e-06 | 9.067E-06 | 2.730E-06 | 6.600E-06 | 1.200E-05 |
| 25602 | 1281 | G1281R | E | 1.3e-05 | 7e-06 | 6.5e-06 | 8.833E-06 | 3.617E-06 | 6.500E-06 | 1.300E-05 |
| 25603 | 1281 | G1281N | E | 1e-05 | 7.4e-06 | 1.1e-05 | 9.467E-06 | 1.858E-06 | 7.400E-06 | 1.100E-05 |
| 25604 | 1281 | G1281D | E | 7.7e-06 | 9.3e-06 | 1.1e-05 | 9.333E-06 | 1.650E-06 | 7.700E-06 | 1.100E-05 |
| 25605 | 1281 | G1281C | E | 6.3e-06 | 5.2e-06 | 6.2e-06 | 5.900E-06 | 6.083E-07 | 5.200E-06 | 6.300E-06 |
| 25606 | 1281 | G1281E | E | 6.3e-06 | 4.5e-06 | 1.3e-05 | 7.933E-06 | 4.479E-06 | 4.500E-06 | 1.300E-05 |
| 25607 | 1281 | G1281Q | E | 1.3e-05 | 5.3e-06 | 1.6e-05 | 1.143E-05 | 5.519E-06 | 5.300E-06 | 1.600E-05 |
| 25608 | 1281 | G1281G | E | 8.8e-06 | 7.7e-06 | 4.6e-06 | 7.033E-06 | 2.178E-06 | 4.600E-06 | 8.800E-06 |
| 25609 | 1281 | G1281H | E | 7e-06 | 6.9e-06 | 1e-05 | 7.967E-06 | 1.762E-06 | 6.900E-06 | 1.000E-05 |
| 25610 | 1281 | G1281I | E | 9.1e-06 | 5.4e-06 | 5.7e-06 | 6.733E-06 | 2.055E-06 | 5.400E-06 | 9.100E-06 |
| 25611 | 1281 | G1281L | E | 7.2e-06 | 8.2e-06 | 7.7e-06 | 7.700E-06 | 5.000E-07 | 7.200E-06 | 8.200E-06 |
| 25612 | 1281 | G1281K | E | 9.9e-06 | 9.8e-06 | 8.6e-06 | 9.433E-06 | 7.234E-07 | 8.600E-06 | 9.900E-06 |
| 25613 | 1281 | G1281M | E | 6.5e-06 | 1.1e-05 | 9.1e-06 | 8.867E-06 | 2.259E-06 | 6.500E-06 | 1.100E-05 |
| 25614 | 1281 | G1281F | E | 1e-05 | 1.3e-05 | 6.4e-06 | 9.800E-06 | 3.305E-06 | 6.400E-06 | 1.300E-05 |
| 25615 | 1281 | G1281P | E | 6.1e-06 | 8.9e-06 | 8.5e-06 | 7.833E-06 | 1.514E-06 | 6.100E-06 | 8.900E-06 |
| 25616 | 1281 | G1281S | E | 3.8e-06 | 4.4e-06 | 4.9e-06 | 4.367E-06 | 5.508E-07 | 3.800E-06 | 4.900E-06 |
| 25617 | 1281 | G1281T | E | 9e-06 | 9.4e-06 | 9.6e-06 | 9.333E-06 | 3.055E-07 | 9.000E-06 | 9.600E-06 |
| 25618 | 1281 | G1281W | E | 8.6e-06 | 7e-06 | 8.6e-06 | 8.067E-06 | 9.238E-07 | 7.000E-06 | 8.600E-06 |
| 25619 | 1281 | G1281Y | E | 1.4e-05 | 7e-06 | 8.3e-06 | 9.767E-06 | 3.723E-06 | 7.000E-06 | 1.400E-05 |
| 25620 | 1281 | G1281V | E | 4.7e-06 | 4.9e-06 | 8e-06 | 5.867E-06 | 1.850E-06 | 4.700E-06 | 8.000E-06 |
| 25621 | 1282 | G1282A | E | 6.2e-06 | 9.7e-06 | 1e-05 | 8.633E-06 | 2.113E-06 | 6.200E-06 | 1.000E-05 |
| 25622 | 1282 | G1282R | E | 9.1e-06 | 6.8e-06 | 4e-06 | 6.633E-06 | 2.554E-06 | 4.000E-06 | 9.100E-06 |
| 25623 | 1282 | G1282N | E | 7.8e-06 | 1e-05 | 7.8e-06 | 8.533E-06 | 1.270E-06 | 7.800E-06 | 1.000E-05 |
| 25624 | 1282 | G1282D | E | 1e-05 | 8.8e-06 | 6.2e-06 | 8.333E-06 | 1.943E-06 | 6.200E-06 | 1.000E-05 |
| 25625 | 1282 | G1282C | E | 8.2e-06 | 6.6e-06 | 6.6e-06 | 7.133E-06 | 9.238E-07 | 6.600E-06 | 8.200E-06 |
| 25626 | 1282 | G1282E | E | 6.9e-06 | 5.7e-06 | 9e-06 | 7.200E-06 | 1.670E-06 | 5.700E-06 | 9.000E-06 |
| 25627 | 1282 | G1282Q | E | 6.6e-06 | 1.1e-05 | 8.7e-06 | 8.767E-06 | 2.201E-06 | 6.600E-06 | 1.100E-05 |
| 25628 | 1282 | G1282G | E | 1.2e-05 | 1e-05 | 5.4e-06 | 9.133E-06 | 3.384E-06 | 5.400E-06 | 1.200E-05 |
| 25629 | 1282 | G1282H | E | 8.5e-06 | 8.1e-06 | 1.4e-05 | 1.020E-05 | 3.297E-06 | 8.100E-06 | 1.400E-05 |
| 25630 | 1282 | G1282I | E | 6.9e-06 | 7.2e-06 | 9.5e-06 | 7.867E-06 | 1.422E-06 | 6.900E-06 | 9.500E-06 |
| 25631 | 1282 | G1282L | E | 1.3e-05 | 9.2e-06 | 9.3e-06 | 1.050E-05 | 2.166E-06 | 9.200E-06 | 1.300E-05 |
| 25632 | 1282 | G1282K | E | 5.1e-06 | 9.7e-06 | 7.1e-06 | 7.300E-06 | 2.307E-06 | 5.100E-06 | 9.700E-06 |
| 25633 | 1282 | G1282M | E | 1.1e-05 | 1.2e-05 | 8.6e-06 | 1.053E-05 | 1.747E-06 | 8.600E-06 | 1.200E-05 |
| 25634 | 1282 | G1282F | E | 8.9e-06 | 9.3e-06 | 6.1e-06 | 8.100E-06 | 1.744E-06 | 6.100E-06 | 9.300E-06 |
| 25635 | 1282 | G1282P | E | 7.5e-06 | 6e-06 | 9.6e-06 | 7.700E-06 | 1.808E-06 | 6.000E-06 | 9.600E-06 |
| 25636 | 1282 | G1282S | E | 1.1e-05 | 1.3e-05 | 6e-06 | 1.000E-05 | 3.606E-06 | 6.000E-06 | 1.300E-05 |
| 25637 | 1282 | G1282T | E | 5.6e-06 | 7.7e-06 | 1.3e-05 | 8.767E-06 | 3.814E-06 | 5.600E-06 | 1.300E-05 |
| 25638 | 1282 | G1282W | E | 6.3e-06 | 1.1e-05 | 8.2e-06 | 8.500E-06 | 2.364E-06 | 6.300E-06 | 1.100E-05 |
| 25639 | 1282 | G1282Y | E | 7.5e-06 | 1.3e-05 | 9.8e-06 | 1.010E-05 | 2.762E-06 | 7.500E-06 | 1.300E-05 |
| 25640 | 1282 | G1282V | E | 5.9e-06 | 4.6e-06 | 8.2e-06 | 6.233E-06 | 1.823E-06 | 4.600E-06 | 8.200E-06 |
| 25641 | 1283 | L1283A | E | 8.1e-06 | 6.8e-06 | 4.5e-06 | 6.467E-06 | 1.823E-06 | 4.500E-06 | 8.100E-06 |
| 25642 | 1283 | L1283R | E | 8.5e-06 | 7.4e-06 | 5.7e-06 | 7.200E-06 | 1.411E-06 | 5.700E-06 | 8.500E-06 |
| 25643 | 1283 | L1283N | E | 8.4e-06 | 4.7e-06 | 9.5e-06 | 7.533E-06 | 2.515E-06 | 4.700E-06 | 9.500E-06 |
| 25644 | 1283 | L1283D | E | 7.5e-06 | 8.8e-06 | 7.6e-06 | 7.967E-06 | 7.234E-07 | 7.500E-06 | 8.800E-06 |
| 25645 | 1283 | L1283C | E | 2.9e-06 | 6.8e-06 | 1.2e-05 | 7.233E-06 | 4.565E-06 | 2.900E-06 | 1.200E-05 |
| 25646 | 1283 | L1283E | E | 7.5e-06 | 6.5e-06 | 4.9e-06 | 6.300E-06 | 1.311E-06 | 4.900E-06 | 7.500E-06 |
| 25647 | 1283 | L1283Q | E | 9.4e-06 | 8e-06 | 7.4e-06 | 8.267E-06 | 1.026E-06 | 7.400E-06 | 9.400E-06 |
| 25648 | 1283 | L1283G | E | 6.8e-06 | 6.6e-06 | 1.3e-05 | 8.800E-06 | 3.639E-06 | 6.600E-06 | 1.300E-05 |
| 25649 | 1283 | L1283H | E | 8.2e-06 | 1.4e-05 | 1.3e-05 | 1.173E-05 | 3.101E-06 | 8.200E-06 | 1.400E-05 |
| 25650 | 1283 | L1283I | E | 1.2e-05 | 4.9e-06 | 8.4e-06 | 8.433E-06 | 3.550E-06 | 4.900E-06 | 1.200E-05 |
| 25651 | 1283 | L1283L | E | 3.7e-06 | 9.2e-06 | 8.6e-06 | 7.167E-06 | 3.017E-06 | 3.700E-06 | 9.200E-06 |
| 25652 | 1283 | L1283K | E | 1.1e-05 | 9.6e-06 | 8.1e-06 | 9.567E-06 | 1.450E-06 | 8.100E-06 | 1.100E-05 |
| 25653 | 1283 | L1283M | E | 7.2e-06 | 8.9e-06 | 8.8e-06 | 8.300E-06 | 9.539E-07 | 7.200E-06 | 8.900E-06 |
| 25654 | 1283 | L1283F | E | 7.7e-06 | 6.5e-06 | 8.9e-06 | 7.700E-06 | 1.200E-06 | 6.500E-06 | 8.900E-06 |
| 25655 | 1283 | L1283P | E | 4.9e-06 | 4.8e-06 | 6.3e-06 | 5.333E-06 | 8.386E-07 | 4.800E-06 | 6.300E-06 |
| 25656 | 1283 | L1283S | E | 4.3e-06 | 1e-05 | 7.8e-06 | 7.367E-06 | 2.875E-06 | 4.300E-06 | 1.000E-05 |
| 25657 | 1283 | L1283T | E | 6.4e-06 | 6.6e-06 | 6.8e-06 | 6.600E-06 | 2.000E-07 | 6.400E-06 | 6.800E-06 |
| 25658 | 1283 | L1283W | E | 1.7e-05 | 1.2e-05 | 1.4e-05 | 1.433E-05 | 2.517E-06 | 1.200E-05 | 1.700E-05 |
| 25659 | 1283 | L1283Y | E | 9.1e-06 | 6.3e-06 | 4.7e-06 | 6.700E-06 | 2.227E-06 | 4.700E-06 | 9.100E-06 |
| 25660 | 1283 | L1283V | E | 6.9e-06 | 7.4e-06 | 6.8e-06 | 7.033E-06 | 3.215E-07 | 6.800E-06 | 7.400E-06 |
| 25661 | 1284 | V1284A | E | 7.5e-06 | 1.1e-05 | 6.8e-06 | 8.433E-06 | 2.250E-06 | 6.800E-06 | 1.100E-05 |
| 25662 | 1284 | V1284R | E | 8.3e-06 | 7.5e-06 | 7.3e-06 | 7.700E-06 | 5.292E-07 | 7.300E-06 | 8.300E-06 |
| 25663 | 1284 | V1284N | E | 1.1e-05 | 8.8e-06 | 4.4e-06 | 8.067E-06 | 3.361E-06 | 4.400E-06 | 1.100E-05 |
| 25664 | 1284 | V1284D | E | 6.2e-06 | 9.7e-06 | 7.3e-06 | 7.733E-06 | 1.790E-06 | 6.200E-06 | 9.700E-06 |
| 25665 | 1284 | V1284C | E | 9.6e-06 | 8.5e-06 | 8.7e-06 | 8.933E-06 | 5.859E-07 | 8.500E-06 | 9.600E-06 |
| 25666 | 1284 | V1284E | E | 1.1e-05 | 5.1e-06 | 4.9e-06 | 7.000E-06 | 3.466E-06 | 4.900E-06 | 1.100E-05 |
| 25667 | 1284 | V1284Q | E | 1.5e-05 | 9.1e-06 | 8.3e-06 | 1.080E-05 | 3.659E-06 | 8.300E-06 | 1.500E-05 |
| 25668 | 1284 | V1284G | E | 9.7e-06 | 6.9e-06 | 7.1e-06 | 7.900E-06 | 1.562E-06 | 6.900E-06 | 9.700E-06 |
| 25669 | 1284 | V1284H | E | 4.7e-06 | 8e-06 | 5.5e-06 | 6.067E-06 | 1.721E-06 | 4.700E-06 | 8.000E-06 |

| | | | | | | | | | | |
|---|---|---|---|---|---|---|---|---|---|---|
| 25670 | 1284 | V1284I | E | 1e-05 | 9.4e-06 | 7.6e-06 | 9.000E-06 | 1.249E-06 | 7.600E-06 | 1.000E-05 |
| 25671 | 1284 | V1284L | E | 4.8e-06 | 7.3e-06 | 6.2e-06 | 6.100E-06 | 1.253E-06 | 4.800E-06 | 7.300E-06 |
| 25672 | 1284 | V1284K | E | 6.1e-06 | 9.8e-06 | 4.1e-06 | 6.667E-06 | 2.892E-06 | 4.100E-06 | 9.800E-06 |
| 25673 | 1284 | V1284M | E | 5.6e-06 | 5.4e-06 | 1.4e-05 | 8.333E-06 | 4.908E-06 | 5.400E-06 | 1.400E-05 |
| 25674 | 1284 | V1284F | E | 4.4e-06 | 1.3e-05 | 6.5e-06 | 7.967E-06 | 4.484E-06 | 4.400E-06 | 1.300E-05 |
| 25675 | 1284 | V1284P | E | 9.4e-06 | 6.5e-06 | 1e-05 | 8.633E-06 | 1.872E-06 | 6.500E-06 | 1.000E-05 |
| 25676 | 1284 | V1284S | E | 3.2e-06 | 9.4e-06 | 9.2e-06 | 7.267E-06 | 3.523E-06 | 3.200E-06 | 9.400E-06 |
| 25677 | 1284 | V1284T | E | 9.5e-06 | 6.7e-06 | 1e-05 | 8.733E-06 | 1.779E-06 | 6.700E-06 | 1.000E-05 |
| 25678 | 1284 | V1284W | E | 6.6e-06 | 1.4e-05 | 9.6e-06 | 1.007E-05 | 3.722E-06 | 6.600E-06 | 1.400E-05 |
| 25679 | 1284 | V1284Y | E | 6.8e-06 | 7.7e-06 | 9.9e-06 | 8.133E-06 | 1.595E-06 | 6.800E-06 | 9.900E-06 |
| 25680 | 1284 | V1284V | E | 8e-06 | 5.5e-06 | 1.1e-05 | 8.167E-06 | 2.754E-06 | 5.500E-06 | 1.100E-05 |
| 25681 | 1285 | Q1285A | E | 6.4e-06 | 4.9e-06 | 9.1e-06 | 6.800E-06 | 2.128E-06 | 4.900E-06 | 9.100E-06 |
| 25682 | 1285 | Q1285R | E | 8e-06 | 9.5e-06 | 6.6e-06 | 8.033E-06 | 1.450E-06 | 6.600E-06 | 9.500E-06 |
| 25683 | 1285 | Q1285N | E | 6.7e-06 | 3.9e-06 | 1e-05 | 6.867E-06 | 3.053E-06 | 3.900E-06 | 1.000E-05 |
| 25684 | 1285 | Q1285D | E | 7.9e-06 | 1e-05 | 6.3e-06 | 8.067E-06 | 1.856E-06 | 6.300E-06 | 1.000E-05 |
| 25685 | 1285 | Q1285C | E | 7.7e-06 | 8.5e-06 | 5.8e-06 | 7.333E-06 | 1.387E-06 | 5.800E-06 | 8.500E-06 |
| 25686 | 1285 | Q1285E | E | 1e-05 | 4.4e-06 | 6.8e-06 | 7.067E-06 | 2.810E-06 | 4.400E-06 | 1.000E-05 |
| 25687 | 1285 | Q1285Q | E | 7.2e-06 | 6.7e-06 | 6.9e-06 | 6.933E-06 | 2.517E-07 | 6.700E-06 | 7.200E-06 |
| 25688 | 1285 | Q1285G | E | 5.6e-06 | 9.6e-06 | 6.6e-06 | 7.267E-06 | 2.082E-06 | 5.600E-06 | 9.600E-06 |
| 25689 | 1285 | Q1285H | E | 1.9e-05 | 8.2e-06 | 7.6e-06 | 1.160E-05 | 6.416E-06 | 7.600E-06 | 1.900E-05 |
| 25690 | 1285 | Q1285I | E | 6.7e-06 | 6.3e-06 | 8.4e-06 | 7.133E-06 | 1.115E-06 | 6.300E-06 | 8.400E-06 |
| 25691 | 1285 | Q1285L | E | 5e-06 | 7.8e-06 | 1.2e-05 | 8.267E-06 | 3.523E-06 | 5.000E-06 | 1.200E-05 |
| 25692 | 1285 | Q1285K | E | 5.7e-06 | 6.7e-06 | 9.1e-06 | 7.167E-06 | 1.747E-06 | 5.700E-06 | 9.100E-06 |
| 25693 | 1285 | Q1285M | E | 4.3e-06 | 7.1e-06 | 1.5e-05 | 8.800E-06 | 5.549E-06 | 4.300E-06 | 1.500E-05 |
| 25694 | 1285 | Q1285F | E | 4.8e-06 | 5.3e-06 | 6.2e-06 | 5.433E-06 | 7.095E-07 | 4.800E-06 | 6.200E-06 |
| 25695 | 1285 | Q1285P | E | 1.3e-05 | 1.3e-05 | 5.3e-06 | 1.043E-05 | 4.446E-06 | 5.300E-06 | 1.300E-05 |
| 25696 | 1285 | Q1285S | E | 5.7e-06 | 5.3e-06 | 7.7e-06 | 6.233E-06 | 1.286E-06 | 5.300E-06 | 7.700E-06 |
| 25697 | 1285 | Q1285T | E | 8.4e-06 | 7.8e-06 | 7.3e-06 | 7.833E-06 | 5.508E-07 | 7.300E-06 | 8.400E-06 |
| 25698 | 1285 | Q1285W | E | 9.9e-06 | 1.2e-05 | 9.2e-06 | 1.037E-05 | 1.457E-06 | 9.200E-06 | 1.200E-05 |
| 25699 | 1285 | Q1285Y | E | 9e-06 | 8.4e-06 | 8.9e-06 | 8.767E-06 | 3.215E-07 | 8.400E-06 | 9.000E-06 |
| 25700 | 1285 | Q1285V | E | 5e-06 | 1.1e-05 | 1.5e-05 | 1.033E-05 | 5.033E-06 | 5.000E-06 | 1.500E-05 |
| 25701 | 1286 | P1286A | E | 3.9e-06 | 1.3e-05 | 1e-05 | 8.967E-06 | 4.637E-06 | 3.900E-06 | 1.300E-05 |
| 25702 | 1286 | P1286R | E | 6.2e-06 | 4.7e-06 | 5.9e-06 | 5.600E-06 | 7.937E-07 | 4.700E-06 | 6.200E-06 |
| 25703 | 1286 | P1286N | E | 1.1e-05 | 9.1e-06 | 5.5e-06 | 8.533E-06 | 2.793E-06 | 5.500E-06 | 1.100E-05 |
| 25704 | 1286 | P1286D | E | 7.4e-06 | 1.2e-05 | 6.8e-06 | 8.733E-06 | 2.845E-06 | 6.800E-06 | 1.200E-05 |
| 25705 | 1286 | P1286C | E | 9.1e-06 | 3.9e-06 | 8.9e-06 | 7.300E-06 | 2.946E-06 | 3.900E-06 | 9.100E-06 |
| 25706 | 1286 | P1286E | E | 7.7e-06 | 6.9e-06 | 6.5e-06 | 7.033E-06 | 6.110E-07 | 6.500E-06 | 7.700E-06 |
| 25707 | 1286 | P1286Q | E | 6.3e-06 | 1.6e-05 | 6e-06 | 9.433E-06 | 5.689E-06 | 6.000E-06 | 1.600E-05 |
| 25708 | 1286 | P1286G | E | 1.1e-05 | 5.8e-06 | 4e-06 | 6.933E-06 | 3.635E-06 | 4.000E-06 | 1.100E-05 |
| 25709 | 1286 | P1286H | E | 5.9e-06 | 7.9e-06 | 7.9e-06 | 7.233E-06 | 1.155E-06 | 5.900E-06 | 7.900E-06 |
| 25710 | 1286 | P1286I | E | 4.6e-06 | 1.1e-05 | 2e-05 | 1.187E-05 | 7.736E-06 | 4.600E-06 | 2.000E-05 |
| 25711 | 1286 | P1286L | E | 6.9e-06 | 1.7e-05 | 6.7e-06 | 1.020E-05 | 5.890E-06 | 6.700E-06 | 1.700E-05 |
| 25712 | 1286 | P1286K | E | 9.7e-06 | 5.5e-06 | 1e-05 | 8.400E-06 | 2.516E-06 | 5.500E-06 | 1.000E-05 |
| 25713 | 1286 | P1286M | E | 1.2e-05 | 9.2e-06 | 6.4e-06 | 9.200E-06 | 2.800E-06 | 6.400E-06 | 1.200E-05 |
| 25714 | 1286 | P1286F | E | 6.9e-06 | 9.2e-06 | 5e-06 | 7.033E-06 | 2.103E-06 | 5.000E-06 | 9.200E-06 |
| 25715 | 1286 | P1286P | E | 7.9e-06 | 7.7e-06 | 4.2e-06 | 6.600E-06 | 2.081E-06 | 4.200E-06 | 7.900E-06 |
| 25716 | 1286 | P1286S | E | 1.2e-05 | 7.1e-06 | 9.6e-06 | 9.567E-06 | 2.450E-06 | 7.100E-06 | 1.200E-05 |
| 25717 | 1286 | P1286T | E | 8.6e-06 | 6.1e-06 | 5.8e-06 | 6.833E-06 | 1.537E-06 | 5.800E-06 | 8.600E-06 |
| 25718 | 1286 | P1286W | E | 5.4e-06 | 8.7e-06 | 7.8e-06 | 7.300E-06 | 1.706E-06 | 5.400E-06 | 8.700E-06 |
| 25719 | 1286 | P1286Y | E | 7.8e-06 | 1.2e-05 | 6.9e-06 | 8.900E-06 | 2.722E-06 | 6.900E-06 | 1.200E-05 |
| 25720 | 1286 | P1286V | E | 4.7e-06 | 6.7e-06 | 9e-06 | 6.800E-06 | 2.152E-06 | 4.700E-06 | 9.000E-06 |
| 25721 | 1287 | G1287A | E | 1e-05 | 1.1e-05 | 7e-06 | 9.333E-06 | 2.082E-06 | 7.000E-06 | 1.100E-05 |
| 25722 | 1287 | G1287R | E | 6.8e-06 | 7.2e-06 | 7.8e-06 | 7.267E-06 | 5.033E-07 | 6.800E-06 | 7.800E-06 |
| 25723 | 1287 | G1287N | E | 1e-05 | 9.6e-06 | 5.7e-06 | 8.433E-06 | 2.376E-06 | 5.700E-06 | 1.000E-05 |
| 25724 | 1287 | G1287D | E | 4.6e-06 | 8.7e-06 | 1.5e-05 | 9.433E-06 | 5.239E-06 | 4.600E-06 | 1.500E-05 |
| 25725 | 1287 | G1287C | E | 1.1e-05 | 5.8e-06 | 1.3e-05 | 9.933E-06 | 3.717E-06 | 5.800E-06 | 1.300E-05 |
| 25726 | 1287 | G1287E | E | 1.1e-05 | 5.6e-06 | 5.6e-06 | 7.400E-06 | 3.118E-06 | 5.600E-06 | 1.100E-05 |
| 25727 | 1287 | G1287Q | E | 7.4e-06 | 1.1e-05 | 6e-06 | 8.133E-06 | 2.579E-06 | 6.000E-06 | 1.100E-05 |
| 25728 | 1287 | G1287G | E | 6.8e-06 | 5.8e-06 | 9.3e-06 | 7.300E-06 | 1.803E-06 | 5.800E-06 | 9.300E-06 |
| 25729 | 1287 | G1287H | E | 9.1e-06 | 8.1e-06 | 7e-06 | 8.067E-06 | 1.050E-06 | 7.000E-06 | 9.100E-06 |
| 25730 | 1287 | G1287I | E | 9.3e-06 | 5e-06 | 7.7e-06 | 7.333E-06 | 2.173E-06 | 5.000E-06 | 9.300E-06 |
| 25731 | 1287 | G1287L | E | 1e-05 | 8.7e-06 | 3.8e-06 | 7.500E-06 | 3.270E-06 | 3.800E-06 | 1.000E-05 |
| 25732 | 1287 | G1287K | E | 7.7e-06 | 1.1e-05 | 7.5e-06 | 8.733E-06 | 1.966E-06 | 7.500E-06 | 1.100E-05 |
| 25733 | 1287 | G1287M | E | 7.8e-06 | 7.2e-06 | 5.4e-06 | 6.800E-06 | 1.249E-06 | 5.400E-06 | 7.800E-06 |
| 25734 | 1287 | G1287F | E | 4.6e-06 | 8.2e-06 | 6.8e-06 | 6.533E-06 | 1.815E-06 | 4.600E-06 | 8.200E-06 |
| 25735 | 1287 | G1287P | E | 5.2e-06 | 7.3e-06 | 8.5e-06 | 7.000E-06 | 1.670E-06 | 5.200E-06 | 8.500E-06 |
| 25736 | 1287 | G1287S | E | 4.9e-06 | 9.8e-06 | 4.8e-06 | 6.500E-06 | 2.858E-06 | 4.800E-06 | 9.800E-06 |
| 25737 | 1287 | G1287T | E | 5e-06 | 6.9e-06 | 9.7e-06 | 7.200E-06 | 2.364E-06 | 5.000E-06 | 9.700E-06 |
| 25738 | 1287 | G1287W | E | 6.1e-06 | 1.1e-05 | 7.2e-06 | 8.100E-06 | 2.571E-06 | 6.100E-06 | 1.100E-05 |
| 25739 | 1287 | G1287Y | E | 6.2e-06 | 1.4e-05 | 1.3e-05 | 1.107E-05 | 4.244E-06 | 6.200E-06 | 1.400E-05 |
| 25740 | 1287 | G1287V | E | 7e-06 | 6.8e-06 | 6.9e-06 | 6.900E-06 | 1.000E-07 | 6.800E-06 | 7.000E-06 |
| 25741 | 1288 | G1288A | E | 1.1e-05 | 7.5e-06 | 7.3e-06 | 8.600E-06 | 2.081E-06 | 7.300E-06 | 1.100E-05 |
| 25742 | 1288 | G1288R | E | 8.5e-06 | 6.6e-06 | 8e-06 | 7.700E-06 | 9.849E-07 | 6.600E-06 | 8.500E-06 |
| 25743 | 1288 | G1288N | E | 8.5e-06 | 4.9e-06 | 8.6e-06 | 7.333E-06 | 2.108E-06 | 4.900E-06 | 8.600E-06 |
| 25744 | 1288 | G1288D | E | 7.2e-06 | 9.4e-06 | 4.9e-06 | 7.167E-06 | 2.250E-06 | 4.900E-06 | 9.400E-06 |
| 25745 | 1288 | G1288C | E | 4.7e-06 | 1.1e-05 | 9e-06 | 8.233E-06 | 3.219E-06 | 4.700E-06 | 1.100E-05 |
| 25746 | 1288 | G1288E | E | 7e-06 | 5.9e-06 | 7.6e-06 | 6.833E-06 | 8.622E-07 | 5.900E-06 | 7.600E-06 |
| 25747 | 1288 | G1288Q | E | 1.2e-05 | 4.6e-06 | 7.3e-06 | 7.967E-06 | 3.745E-06 | 4.600E-06 | 1.200E-05 |
| 25748 | 1288 | G1288G | E | 1e-05 | 5.7e-06 | 7.7e-06 | 7.800E-06 | 2.152E-06 | 5.700E-06 | 1.000E-05 |
| 25749 | 1288 | G1288H | E | 7.5e-06 | 4.7e-06 | 6.2e-06 | 6.133E-06 | 1.401E-06 | 4.700E-06 | 7.500E-06 |
| 25750 | 1288 | G1288I | E | 8e-06 | 1.1e-05 | 9.9e-06 | 9.633E-06 | 1.518E-06 | 8.000E-06 | 1.100E-05 |
| 25751 | 1288 | G1288L | E | 5.8e-06 | 8.1e-06 | 8.6e-06 | 7.500E-06 | 1.493E-06 | 5.800E-06 | 8.600E-06 |
| 25752 | 1288 | G1288K | E | 8.5e-06 | 5.9e-06 | 6.9e-06 | 7.100E-06 | 1.311E-06 | 5.900E-06 | 8.500E-06 |
| 25753 | 1288 | G1288M | E | 5.5e-06 | 1.5e-05 | 9.3e-06 | 9.933E-06 | 4.782E-06 | 5.500E-06 | 1.500E-05 |
| 25754 | 1288 | G1288F | E | 4.5e-06 | 4.3e-06 | 7.1e-06 | 5.300E-06 | 1.562E-06 | 4.300E-06 | 7.100E-06 |

| | | | | | | | | | | |
|---|---|---|---|---|---|---|---|---|---|---|
| 25755 | 1288 | G1288P | E | 1.1e-05 | 1.4e-05 | 9.8e-06 | 1.160E-05 | 2.163E-06 | 9.800E-06 | 1.400E-05 |
| 25756 | 1288 | G1288S | E | 9.2e-06 | 8.7e-06 | 1e-05 | 9.300E-06 | 6.557E-07 | 8.700E-06 | 1.000E-05 |
| 25757 | 1288 | G1288T | E | 6.8e-06 | 6.9e-06 | 1.3e-05 | 8.900E-06 | 3.551E-06 | 6.800E-06 | 1.300E-05 |
| 25758 | 1288 | G1288W | E | 9.2e-06 | 6.6e-06 | 4.9e-06 | 6.900E-06 | 2.166E-06 | 4.900E-06 | 9.200E-06 |
| 25759 | 1288 | G1288Y | E | 3.6e-06 | 9e-06 | 1.1e-05 | 7.867E-06 | 3.828E-06 | 3.600E-06 | 1.100E-05 |
| 25760 | 1288 | G1288V | E | 8e-06 | 9.4e-06 | 4.1e-06 | 7.167E-06 | 2.747E-06 | 4.100E-06 | 9.400E-06 |
| 25761 | 1289 | S1289A | E | 6.6e-06 | 8.9e-06 | 1.5e-05 | 1.017E-05 | 4.341E-06 | 6.600E-06 | 1.500E-05 |
| 25762 | 1289 | S1289R | E | 9.1e-06 | 1.2e-05 | 6e-06 | 9.033E-06 | 3.001E-06 | 6.000E-06 | 1.200E-05 |
| 25763 | 1289 | S1289N | E | 6.7e-06 | 8.6e-06 | 6.2e-06 | 7.167E-06 | 1.266E-06 | 6.200E-06 | 8.600E-06 |
| 25764 | 1289 | S1289D | E | 7.5e-06 | 1.4e-05 | 1.1e-05 | 1.083E-05 | 3.253E-06 | 7.500E-06 | 1.400E-05 |
| 25765 | 1289 | S1289C | E | 6.3e-06 | 1e-05 | 1.2e-05 | 9.433E-06 | 2.892E-06 | 6.300E-06 | 1.200E-05 |
| 25766 | 1289 | S1289E | E | 7e-06 | 8.1e-06 | 4.6e-06 | 6.567E-06 | 1.790E-06 | 4.600E-06 | 8.100E-06 |
| 25767 | 1289 | S1289Q | E | 6.4e-06 | 1.3e-05 | 3.3e-06 | 7.567E-06 | 4.954E-06 | 3.300E-06 | 1.300E-05 |
| 25768 | 1289 | S1289G | E | 1.4e-05 | 9.3e-06 | 9.2e-06 | 1.083E-05 | 2.743E-06 | 9.200E-06 | 1.400E-05 |
| 25769 | 1289 | S1289H | E | 4.5e-06 | 1.5e-05 | 4.8e-06 | 8.100E-06 | 5.977E-06 | 4.500E-06 | 1.500E-05 |
| 25770 | 1289 | S1289I | E | 4.9e-06 | 4.9e-06 | 1.1e-05 | 6.933E-06 | 3.522E-06 | 4.900E-06 | 1.100E-05 |
| 25771 | 1289 | S1289L | E | 7.7e-06 | 5.5e-06 | 8.2e-06 | 7.133E-06 | 1.436E-06 | 5.500E-06 | 8.200E-06 |
| 25772 | 1289 | S1289K | E | 1.4e-05 | 8.6e-06 | 5.6e-06 | 9.400E-06 | 4.257E-06 | 5.600E-06 | 1.400E-05 |
| 25773 | 1289 | S1289M | E | 7.5e-06 | 8.5e-06 | 1.2e-05 | 9.333E-06 | 2.363E-06 | 7.500E-06 | 1.200E-05 |
| 25774 | 1289 | S1289F | E | 1.4e-05 | 7.6e-06 | 8.5e-06 | 1.003E-05 | 3.465E-06 | 7.600E-06 | 1.400E-05 |
| 25775 | 1289 | S1289P | E | 7.1e-06 | 5.7e-06 | 6.9e-06 | 6.567E-06 | 7.572E-07 | 5.700E-06 | 7.100E-06 |
| 25776 | 1289 | S1289S | E | 1.1e-05 | 7.9e-06 | 1.7e-05 | 1.197E-05 | 4.626E-06 | 7.900E-06 | 1.700E-05 |
| 25777 | 1289 | S1289T | E | 1.1e-05 | 1.2e-05 | 6.2e-06 | 9.733E-06 | 3.101E-06 | 6.200E-06 | 1.200E-05 |
| 25778 | 1289 | S1289W | E | 9.6e-06 | 9.5e-06 | 9.4e-06 | 9.500E-06 | 1.000E-07 | 9.400E-06 | 9.600E-06 |
| 25779 | 1289 | S1289Y | E | 9.7e-06 | 7.7e-06 | 6.5e-06 | 7.967E-06 | 1.617E-06 | 6.500E-06 | 9.700E-06 |
| 25780 | 1289 | S1289V | E | 4.7e-06 | 8.3e-06 | 8.7e-06 | 7.233E-06 | 2.203E-06 | 4.700E-06 | 8.700E-06 |
| 25781 | 1290 | L1290A | E | 7.5e-06 | 5e-06 | 7.8e-06 | 6.767E-06 | 1.537E-06 | 5.000E-06 | 7.800E-06 |
| 25782 | 1290 | L1290R | E | 1.1e-05 | 6.1e-06 | 6.3e-06 | 7.800E-06 | 2.773E-06 | 6.100E-06 | 1.100E-05 |
| 25783 | 1290 | L1290N | E | 1.1e-05 | 7.8e-06 | 5e-06 | 7.933E-06 | 3.002E-06 | 5.000E-06 | 1.100E-05 |
| 25784 | 1290 | L1290D | E | 8.8e-06 | 6.8e-06 | 5.9e-06 | 7.167E-06 | 1.484E-06 | 5.900E-06 | 8.800E-06 |
| 25785 | 1290 | L1290C | E | 6.4e-06 | 6.5e-06 | 1e-05 | 7.633E-06 | 2.050E-06 | 6.400E-06 | 1.000E-05 |
| 25786 | 1290 | L1290E | E | 5e-06 | 9.3e-06 | 1.3e-05 | 9.100E-06 | 4.004E-06 | 5.000E-06 | 1.300E-05 |
| 25787 | 1290 | L1290Q | E | 4.9e-06 | 5.2e-06 | 1.4e-05 | 8.033E-06 | 5.169E-06 | 4.900E-06 | 1.400E-05 |
| 25788 | 1290 | L1290G | E | 2.2e-05 | 8.3e-06 | 5.7e-06 | 1.200E-05 | 8.757E-06 | 5.700E-06 | 2.200E-05 |
| 25789 | 1290 | L1290H | E | 6.1e-06 | 6.6e-06 | 3.4e-06 | 5.367E-06 | 1.721E-06 | 3.400E-06 | 6.600E-06 |
| 25790 | 1290 | L1290I | E | 6.6e-06 | 6.1e-06 | 8.2e-06 | 6.967E-06 | 1.097E-06 | 6.100E-06 | 8.200E-06 |
| 25791 | 1290 | L1290L | E | 6.3e-06 | 1e-05 | 1.1e-05 | 9.100E-06 | 2.476E-06 | 6.300E-06 | 1.100E-05 |
| 25792 | 1290 | L1290K | E | 8.1e-06 | 8.8e-06 | 5.4e-06 | 7.433E-06 | 1.795E-06 | 5.400E-06 | 8.800E-06 |
| 25793 | 1290 | L1290M | E | 5.9e-06 | 8e-06 | 8.2e-06 | 7.367E-06 | 1.274E-06 | 5.900E-06 | 8.200E-06 |
| 25794 | 1290 | L1290F | E | 7.7e-06 | 9e-06 | 9.2e-06 | 8.633E-06 | 8.145E-07 | 7.700E-06 | 9.200E-06 |
| 25795 | 1290 | L1290P | E | 6.3e-06 | 5.4e-06 | 7.5e-06 | 6.400E-06 | 1.054E-06 | 5.400E-06 | 7.500E-06 |
| 25796 | 1290 | L1290S | E | 6e-06 | 1.1e-05 | 6.5e-06 | 7.833E-06 | 2.754E-06 | 6.000E-06 | 1.100E-05 |
| 25797 | 1290 | L1290T | E | 7.7e-06 | 6.8e-06 | 4.1e-06 | 6.200E-06 | 1.873E-06 | 4.100E-06 | 7.700E-06 |
| 25798 | 1290 | L1290W | E | 1e-05 | 7.2e-06 | 1.2e-05 | 9.733E-06 | 2.411E-06 | 7.200E-06 | 1.200E-05 |
| 25799 | 1290 | L1290Y | E | 1.8e-05 | 1.7e-05 | 1.4e-05 | 1.633E-05 | 2.082E-06 | 1.400E-05 | 1.800E-05 |
| 25800 | 1290 | L1290V | E | 6.4e-06 | 7.4e-06 | 1e-05 | 7.933E-06 | 1.858E-06 | 6.400E-06 | 1.000E-05 |
| 25801 | 1291 | R1291A | E | 1.2e-05 | 4.2e-06 | 4.9e-06 | 7.033E-06 | 4.315E-06 | 4.200E-06 | 1.200E-05 |
| 25802 | 1291 | R1291R | E | 4.9e-06 | 1e-05 | 9.5e-06 | 8.133E-06 | 2.811E-06 | 4.900E-06 | 1.000E-05 |
| 25803 | 1291 | R1291N | E | 5.9e-06 | 5.4e-06 | 8.6e-06 | 6.633E-06 | 1.721E-06 | 5.400E-06 | 8.600E-06 |
| 25804 | 1291 | R1291D | E | 1.2e-05 | 4.6e-06 | 4.8e-06 | 7.133E-06 | 4.216E-06 | 4.600E-06 | 1.200E-05 |
| 25805 | 1291 | R1291C | E | 2.9e-06 | 6.4e-06 | 8.9e-06 | 6.067E-06 | 3.014E-06 | 2.900E-06 | 8.900E-06 |
| 25806 | 1291 | R1291E | E | 7.8e-06 | 9.3e-06 | 7.7e-06 | 8.267E-06 | 8.963E-07 | 7.700E-06 | 9.300E-06 |
| 25807 | 1291 | R1291Q | E | 1.2e-05 | 7.3e-06 | 3.5e-06 | 7.600E-06 | 4.258E-06 | 3.500E-06 | 1.200E-05 |
| 25808 | 1291 | R1291G | E | 1e-05 | 5.2e-06 | 8e-06 | 7.733E-06 | 2.411E-06 | 5.200E-06 | 1.000E-05 |
| 25809 | 1291 | R1291H | E | 1.3e-05 | 7.3e-06 | 8.9e-06 | 9.733E-06 | 2.940E-06 | 7.300E-06 | 1.300E-05 |
| 25810 | 1291 | R1291I | E | 1.4e-05 | 9.3e-06 | 8e-06 | 1.043E-05 | 3.156E-06 | 8.000E-06 | 1.400E-05 |
| 25811 | 1291 | R1291L | E | 7.8e-06 | 9.2e-06 | 7.2e-06 | 8.067E-06 | 1.026E-06 | 7.200E-06 | 9.200E-06 |
| 25812 | 1291 | R1291K | E | 9e-06 | 6.6e-06 | 1e-05 | 8.533E-06 | 1.747E-06 | 6.600E-06 | 1.000E-05 |
| 25813 | 1291 | R1291M | E | 6.1e-06 | 7.7e-06 | 7.4e-06 | 7.067E-06 | 8.505E-07 | 6.100E-06 | 7.700E-06 |
| 25814 | 1291 | R1291F | E | 7.7e-06 | 4.9e-06 | 5.8e-06 | 6.133E-06 | 1.429E-06 | 4.900E-06 | 7.700E-06 |
| 25815 | 1291 | R1291P | E | 1e-05 | 8.1e-06 | 5.4e-06 | 7.833E-06 | 2.312E-06 | 5.400E-06 | 1.000E-05 |
| 25816 | 1291 | R1291S | E | 4.7e-06 | 4.7e-06 | 1.3e-05 | 7.467E-06 | 4.792E-06 | 4.700E-06 | 1.300E-05 |
| 25817 | 1291 | R1291T | E | 9.4e-06 | 9.6e-06 | 1.1e-05 | 1.000E-05 | 8.718E-07 | 9.400E-06 | 1.100E-05 |
| 25818 | 1291 | R1291W | E | 5.4e-06 | 6.5e-06 | 8.7e-06 | 6.867E-06 | 1.680E-06 | 5.400E-06 | 8.700E-06 |
| 25819 | 1291 | R1291Y | E | 4.4e-06 | 5.9e-06 | 7e-06 | 5.767E-06 | 1.305E-06 | 4.400E-06 | 7.000E-06 |
| 25820 | 1291 | R1291V | E | 6.6e-06 | 1e-05 | 7.3e-06 | 7.967E-06 | 1.795E-06 | 6.600E-06 | 1.000E-05 |
| 25821 | 1292 | L1292A | E | 4.9e-06 | 9.3e-06 | 4.6e-06 | 6.267E-06 | 2.631E-06 | 4.600E-06 | 9.300E-06 |
| 25822 | 1292 | L1292R | E | 6.9e-06 | 9.5e-06 | 8.7e-06 | 8.367E-06 | 1.332E-06 | 6.900E-06 | 9.500E-06 |
| 25823 | 1292 | L1292N | E | 1.2e-05 | 5.2e-06 | 5.1e-06 | 7.433E-06 | 3.955E-06 | 5.100E-06 | 1.200E-05 |
| 25824 | 1292 | L1292D | E | 6.9e-06 | 5.1e-06 | 7.4e-06 | 6.467E-06 | 1.210E-06 | 5.100E-06 | 7.400E-06 |
| 25825 | 1292 | L1292C | E | 8.6e-06 | 5.7e-06 | 5.5e-06 | 6.600E-06 | 1.735E-06 | 5.500E-06 | 8.600E-06 |
| 25826 | 1292 | L1292E | E | 9.2e-06 | 1.2e-05 | 6.1e-06 | 9.100E-06 | 2.951E-06 | 6.100E-06 | 1.200E-05 |
| 25827 | 1292 | L1292Q | E | 1e-05 | 6e-06 | 8.1e-06 | 8.033E-06 | 2.001E-06 | 6.000E-06 | 1.000E-05 |
| 25828 | 1292 | L1292G | E | 6e-06 | 5.6e-06 | 9.2e-06 | 6.933E-06 | 1.973E-06 | 5.600E-06 | 9.200E-06 |
| 25829 | 1292 | L1292H | E | 6.1e-06 | 4.9e-06 | 7e-06 | 6.000E-06 | 1.054E-06 | 4.900E-06 | 7.000E-06 |
| 25830 | 1292 | L1292I | E | 6.7e-06 | 7.7e-06 | 1.3e-05 | 9.133E-06 | 3.386E-06 | 6.700E-06 | 1.300E-05 |
| 25831 | 1292 | L1292L | E | 8.6e-06 | 3.9e-06 | 9.3e-06 | 7.267E-06 | 2.937E-06 | 3.900E-06 | 9.300E-06 |
| 25832 | 1292 | L1292K | E | 4.7e-06 | 3.9e-06 | 9e-06 | 5.867E-06 | 2.743E-06 | 3.900E-06 | 9.000E-06 |
| 25833 | 1292 | L1292M | E | 6.6e-06 | 1.2e-05 | 8.4e-06 | 9.000E-06 | 2.750E-06 | 6.600E-06 | 1.200E-05 |
| 25834 | 1292 | L1292F | E | 7.9e-06 | 8.6e-06 | 8.1e-06 | 8.200E-06 | 3.606E-07 | 7.900E-06 | 8.600E-06 |
| 25835 | 1292 | L1292P | E | 6.6e-06 | 7e-06 | 9.7e-06 | 7.767E-06 | 1.686E-06 | 6.600E-06 | 9.700E-06 |
| 25836 | 1292 | L1292S | E | 6e-06 | 8.6e-06 | 5.2e-06 | 6.600E-06 | 1.778E-06 | 5.200E-06 | 8.600E-06 |
| 25837 | 1292 | L1292T | E | 1e-05 | 7.9e-06 | 9.1e-06 | 9.000E-06 | 1.054E-06 | 7.900E-06 | 1.000E-05 |
| 25838 | 1292 | L1292W | E | 7.3e-06 | 1e-05 | 6.5e-06 | 7.933E-06 | 1.834E-06 | 6.500E-06 | 1.000E-05 |
| 25839 | 1292 | L1292Y | E | 6.7e-06 | 7.2e-06 | 5.9e-06 | 6.600E-06 | 6.557E-07 | 5.900E-06 | 7.200E-06 |

| | | | | | | | | | | |
|---|---|---|---|---|---|---|---|---|---|---|
| 25840 | 1292 | L1292V | E | 9.2e-06 | 1.3e-05 | 1.3e-05 | 1.173E-05 | 2.194E-06 | 9.200E-06 | 1.300E-05 |
| 25841 | 1293 | S1293A | E | 9.6e-06 | 1.2e-05 | 7.3e-06 | 9.633E-06 | 2.350E-06 | 7.300E-06 | 1.200E-05 |
| 25842 | 1293 | S1293R | E | 5.2e-06 | 8.2e-06 | 8.4e-06 | 7.267E-06 | 1.793E-06 | 5.200E-06 | 8.400E-06 |
| 25843 | 1293 | S1293N | E | 6e-06 | 9.3e-06 | 8.6e-06 | 7.967E-06 | 1.739E-06 | 6.000E-06 | 9.300E-06 |
| 25844 | 1293 | S1293D | E | 8.9e-06 | 9.1e-06 | 3.2e-06 | 7.067E-06 | 3.350E-06 | 3.200E-06 | 9.100E-06 |
| 25845 | 1293 | S1293C | E | 7.9e-06 | 5.3e-06 | 5.7e-06 | 6.300E-06 | 1.400E-06 | 5.300E-06 | 7.900E-06 |
| 25846 | 1293 | S1293E | E | 7.8e-06 | 7.3e-06 | 3.9e-06 | 6.333E-06 | 2.122E-06 | 3.900E-06 | 7.800E-06 |
| 25847 | 1293 | S1293Q | E | 9.1e-06 | 2.7e-06 | 7.4e-06 | 6.400E-06 | 3.315E-06 | 2.700E-06 | 9.100E-06 |
| 25848 | 1293 | S1293G | E | 1.5e-05 | 1.2e-05 | 4.9e-06 | 1.063E-05 | 5.187E-06 | 4.900E-06 | 1.500E-05 |
| 25849 | 1293 | S1293H | E | 8.1e-06 | 1.2e-05 | 8.5e-06 | 9.533E-06 | 2.146E-06 | 8.100E-06 | 1.200E-05 |
| 25850 | 1293 | S1293I | E | 9e-06 | 6.8e-06 | 7.2e-06 | 7.667E-06 | 1.172E-06 | 6.800E-06 | 9.000E-06 |
| 25851 | 1293 | S1293L | E | 5.2e-06 | 4.8e-06 | 7.6e-06 | 5.867E-06 | 1.514E-06 | 4.800E-06 | 7.600E-06 |
| 25852 | 1293 | S1293K | E | 8.5e-06 | 5.3e-06 | 7e-06 | 6.933E-06 | 1.601E-06 | 5.300E-06 | 8.500E-06 |
| 25853 | 1293 | S1293M | E | 5.3e-06 | 4.6e-06 | 1.1e-05 | 6.967E-06 | 3.510E-06 | 4.600E-06 | 1.100E-05 |
| 25854 | 1293 | S1293F | E | 8.4e-06 | 1.2e-05 | 9.1e-06 | 9.833E-06 | 1.909E-06 | 8.400E-06 | 1.200E-05 |
| 25855 | 1293 | S1293P | E | 1.5e-05 | 9.8e-06 | 6.2e-06 | 1.033E-05 | 4.424E-06 | 6.200E-06 | 1.500E-05 |
| 25856 | 1293 | S1293S | E | 1e-05 | 4.9e-06 | 6.1e-06 | 7.000E-06 | 2.666E-06 | 4.900E-06 | 1.000E-05 |
| 25857 | 1293 | S1293T | E | 6.1e-06 | 1.2e-05 | 3.7e-06 | 7.267E-06 | 4.271E-06 | 3.700E-06 | 1.200E-05 |
| 25858 | 1293 | S1293W | E | 1.1e-05 | 5.1e-06 | 7.8e-06 | 7.967E-06 | 2.954E-06 | 5.100E-06 | 1.100E-05 |
| 25859 | 1293 | S1293Y | E | 1.4e-05 | 8.8e-06 | 1.1e-05 | 1.127E-05 | 2.610E-06 | 8.800E-06 | 1.400E-05 |
| 25860 | 1293 | S1293V | E | 8.6e-06 | 9e-06 | 8.9e-06 | 8.833E-06 | 2.082E-07 | 8.600E-06 | 9.000E-06 |
| 25861 | 1294 | C1294A | E | 6e-06 | 1.4e-05 | 6.8e-06 | 8.933E-06 | 4.406E-06 | 6.000E-06 | 1.400E-05 |
| 25862 | 1294 | C1294R | E | 7.1e-06 | 1.2e-05 | 8.5e-06 | 9.200E-06 | 2.524E-06 | 7.100E-06 | 1.200E-05 |
| 25863 | 1294 | C1294N | E | 6.6e-06 | 6.8e-06 | 4.9e-06 | 6.100E-06 | 1.044E-06 | 4.900E-06 | 6.800E-06 |
| 25864 | 1294 | C1294D | E | 9.3e-06 | 1.3e-05 | 9.4e-06 | 1.057E-05 | 2.108E-06 | 9.300E-06 | 1.300E-05 |
| 25865 | 1294 | C1294C | E | 7e-06 | 6.5e-06 | 4.9e-06 | 6.133E-06 | 1.097E-06 | 4.900E-06 | 7.000E-06 |
| 25866 | 1294 | C1294E | E | 6e-06 | 5.1e-06 | 6.4e-06 | 5.833E-06 | 6.658E-07 | 5.100E-06 | 6.400E-06 |
| 25867 | 1294 | C1294Q | E | 4.9e-06 | 1.1e-05 | 7.2e-06 | 7.700E-06 | 3.081E-06 | 4.900E-06 | 1.100E-05 |
| 25868 | 1294 | C1294G | E | 1.4e-05 | 5.3e-06 | 9.2e-06 | 9.500E-06 | 4.358E-06 | 5.300E-06 | 1.400E-05 |
| 25869 | 1294 | C1294H | E | 4.6e-06 | 1e-05 | 7.6e-06 | 7.400E-06 | 2.706E-06 | 4.600E-06 | 1.000E-05 |
| 25870 | 1294 | C1294I | E | 1.2e-05 | 1.2e-05 | 5.1e-06 | 9.700E-06 | 3.984E-06 | 5.100E-06 | 1.200E-05 |
| 25871 | 1294 | C1294L | E | 4.7e-06 | 8.3e-06 | 6.5e-06 | 6.500E-06 | 1.800E-06 | 4.700E-06 | 8.300E-06 |
| 25872 | 1294 | C1294K | E | 1e-05 | 6.6e-06 | 4.6e-06 | 7.067E-06 | 2.730E-06 | 4.600E-06 | 1.000E-05 |
| 25873 | 1294 | C1294M | E | 6.9e-06 | 8.5e-06 | 1e-05 | 8.467E-06 | 1.550E-06 | 6.900E-06 | 1.000E-05 |
| 25874 | 1294 | C1294F | E | 8.7e-06 | 8.4e-06 | 6e-06 | 7.700E-06 | 1.480E-06 | 6.000E-06 | 8.700E-06 |
| 25875 | 1294 | C1294P | E | 4.8e-06 | 8.5e-06 | 3.4e-06 | 5.567E-06 | 2.635E-06 | 3.400E-06 | 8.500E-06 |
| 25876 | 1294 | C1294S | E | 6.9e-06 | 1e-05 | 6.8e-06 | 7.900E-06 | 1.819E-06 | 6.800E-06 | 1.000E-05 |
| 25877 | 1294 | C1294T | E | 8.5e-06 | 7.1e-06 | 9e-06 | 8.200E-06 | 9.849E-07 | 7.100E-06 | 9.000E-06 |
| 25878 | 1294 | C1294W | E | 6.2e-06 | 5e-06 | 1.1e-05 | 7.400E-06 | 3.175E-06 | 5.000E-06 | 1.100E-05 |
| 25879 | 1294 | C1294Y | E | 1.2e-05 | 8.8e-06 | 6.4e-06 | 9.067E-06 | 2.810E-06 | 6.400E-06 | 1.200E-05 |
| 25880 | 1294 | C1294V | E | 9.8e-06 | 1.8e-05 | 1.3e-05 | 1.360E-05 | 4.133E-06 | 9.800E-06 | 1.800E-05 |
| 25881 | 1295 | A1295A | E | 1.3e-05 | 9.9e-06 | 6.4e-06 | 9.767E-06 | 3.302E-06 | 6.400E-06 | 1.300E-05 |
| 25882 | 1295 | A1295R | E | 4.6e-06 | 4.4e-06 | 9.9e-06 | 6.300E-06 | 3.119E-06 | 4.400E-06 | 9.900E-06 |
| 25883 | 1295 | A1295N | E | 6.2e-06 | 1.4e-05 | 5.8e-06 | 8.667E-06 | 4.623E-06 | 5.800E-06 | 1.400E-05 |
| 25884 | 1295 | A1295D | E | 1.2e-05 | 7e-06 | 5e-06 | 8.000E-06 | 3.606E-06 | 5.000E-06 | 1.200E-05 |
| 25885 | 1295 | A1295C | E | 9e-06 | 6.1e-06 | 6.3e-06 | 7.133E-06 | 1.620E-06 | 6.100E-06 | 9.000E-06 |
| 25886 | 1295 | A1295E | E | 1.2e-05 | 9.3e-06 | 9e-06 | 1.010E-05 | 1.652E-06 | 9.000E-06 | 1.200E-05 |
| 25887 | 1295 | A1295Q | E | 6.3e-06 | 8.4e-06 | 9.4e-06 | 8.033E-06 | 1.582E-06 | 6.300E-06 | 9.400E-06 |
| 25888 | 1295 | A1295G | E | 1.1e-05 | 7.4e-06 | 7.9e-06 | 8.767E-06 | 1.950E-06 | 7.400E-06 | 1.100E-05 |
| 25889 | 1295 | A1295H | E | 1.6e-05 | 6.1e-06 | 1.4e-05 | 1.203E-05 | 5.235E-06 | 6.100E-06 | 1.600E-05 |
| 25890 | 1295 | A1295I | E | 9.4e-06 | 6.1e-06 | 9.1e-06 | 8.200E-06 | 1.825E-06 | 6.100E-06 | 9.400E-06 |
| 25891 | 1295 | A1295L | E | 8e-06 | 5e-06 | 1.1e-05 | 8.000E-06 | 3.000E-06 | 5.000E-06 | 1.100E-05 |
| 25892 | 1295 | A1295K | E | 6.6e-06 | 6.2e-06 | 1.1e-05 | 7.933E-06 | 2.663E-06 | 6.200E-06 | 1.100E-05 |
| 25893 | 1295 | A1295M | E | 1.1e-05 | 8.7e-06 | 5e-06 | 8.233E-06 | 3.027E-06 | 5.000E-06 | 1.100E-05 |
| 25894 | 1295 | A1295F | E | 6.1e-06 | 1e-05 | 8.2e-06 | 8.100E-06 | 1.952E-06 | 6.100E-06 | 1.000E-05 |
| 25895 | 1295 | A1295P | E | 1.3e-05 | 1.4e-05 | 5.2e-06 | 1.073E-05 | 4.818E-06 | 5.200E-06 | 1.400E-05 |
| 25896 | 1295 | A1295S | E | 7.1e-06 | 8e-06 | 7.2e-06 | 7.433E-06 | 4.933E-07 | 7.100E-06 | 8.000E-06 |
| 25897 | 1295 | A1295T | E | 1.3e-05 | 6.4e-06 | 1.3e-05 | 1.080E-05 | 3.811E-06 | 6.400E-06 | 1.300E-05 |
| 25898 | 1295 | A1295W | E | 9e-06 | 7.1e-06 | 9.5e-06 | 8.533E-06 | 1.266E-06 | 7.100E-06 | 9.500E-06 |
| 25899 | 1295 | A1295Y | E | 1e-05 | 5.5e-06 | 9.7e-06 | 8.400E-06 | 2.516E-06 | 5.500E-06 | 1.000E-05 |
| 25900 | 1295 | A1295V | E | 7.6e-06 | 8.7e-06 | 1.1e-05 | 9.100E-06 | 1.735E-06 | 7.600E-06 | 1.100E-05 |
| 25901 | 1296 | A1296A | E | 7.4e-06 | 9.5e-06 | 6.5e-06 | 7.800E-06 | 1.539E-06 | 6.500E-06 | 9.500E-06 |
| 25902 | 1296 | A1296R | E | 5.5e-06 | 1.1e-05 | 3.4e-06 | 6.633E-06 | 3.925E-06 | 3.400E-06 | 1.100E-05 |
| 25903 | 1296 | A1296N | E | 4.5e-06 | 4e-06 | 6.4e-06 | 4.967E-06 | 1.266E-06 | 4.000E-06 | 6.400E-06 |
| 25904 | 1296 | A1296D | E | 1e-05 | 1.2e-05 | 1.1e-05 | 1.100E-05 | 1.000E-06 | 1.000E-05 | 1.200E-05 |
| 25905 | 1296 | A1296C | E | 9.1e-06 | 7.6e-06 | 5.4e-06 | 7.367E-06 | 1.861E-06 | 5.400E-06 | 9.100E-06 |
| 25906 | 1296 | A1296E | E | 4.9e-06 | 1.4e-05 | 5.2e-06 | 8.033E-06 | 5.169E-06 | 4.900E-06 | 1.400E-05 |
| 25907 | 1296 | A1296Q | E | 6.3e-06 | 8e-06 | 1.1e-05 | 8.433E-06 | 2.380E-06 | 6.300E-06 | 1.100E-05 |
| 25908 | 1296 | A1296G | E | 7.2e-06 | 5.4e-06 | 1.2e-05 | 8.200E-06 | 3.412E-06 | 5.400E-06 | 1.200E-05 |
| 25909 | 1296 | A1296H | E | 4.6e-06 | 6.3e-06 | 6.4e-06 | 5.767E-06 | 1.012E-06 | 4.600E-06 | 6.400E-06 |
| 25910 | 1296 | A1296I | E | 1.1e-05 | 1e-05 | 1e-05 | 1.033E-05 | 5.774E-07 | 1.000E-05 | 1.100E-05 |
| 25911 | 1296 | A1296L | E | 9.8e-06 | 1.3e-05 | 9.5e-06 | 1.077E-05 | 1.940E-06 | 9.500E-06 | 1.300E-05 |
| 25912 | 1296 | A1296K | E | 4.2e-06 | 9.5e-06 | 9e-06 | 7.567E-06 | 2.926E-06 | 4.200E-06 | 9.500E-06 |
| 25913 | 1296 | A1296M | E | 7.6e-06 | 1.3e-05 | 6.3e-06 | 8.967E-06 | 3.553E-06 | 6.300E-06 | 1.300E-05 |
| 25914 | 1296 | A1296F | E | 5e-06 | 1.2e-05 | 4.5e-06 | 7.167E-06 | 4.193E-06 | 4.500E-06 | 1.200E-05 |
| 25915 | 1296 | A1296P | E | 7e-06 | 4.4e-06 | 5.9e-06 | 5.767E-06 | 1.305E-06 | 4.400E-06 | 7.000E-06 |
| 25916 | 1296 | A1296S | E | 5.5e-06 | 4.6e-06 | 6.1e-06 | 5.400E-06 | 7.550E-07 | 4.600E-06 | 6.100E-06 |
| 25917 | 1296 | A1296T | E | 8.3e-06 | 8.5e-06 | 6.3e-06 | 7.700E-06 | 1.217E-06 | 6.300E-06 | 8.500E-06 |
| 25918 | 1296 | A1296W | E | 7.2e-06 | 8.6e-06 | 1e-05 | 8.600E-06 | 1.400E-06 | 7.200E-06 | 1.000E-05 |
| 25919 | 1296 | A1296Y | E | 1.9e-05 | 4.9e-06 | 8.7e-06 | 1.087E-05 | 7.295E-06 | 4.900E-06 | 1.900E-05 |
| 25920 | 1296 | A1296V | E | 8.9e-06 | 7.1e-06 | 5.7e-06 | 7.233E-06 | 1.604E-06 | 5.700E-06 | 8.900E-06 |
| 25921 | 1297 | S1297A | E | 5.8e-06 | 1.1e-05 | 5e-06 | 7.267E-06 | 3.258E-06 | 5.000E-06 | 1.100E-05 |
| 25922 | 1297 | S1297R | E | 7.7e-06 | 7.1e-06 | 6.5e-06 | 7.100E-06 | 6.000E-07 | 6.500E-06 | 7.700E-06 |
| 25923 | 1297 | S1297N | E | 8.8e-06 | 9.1e-06 | 9.1e-06 | 9.000E-06 | 1.732E-07 | 8.800E-06 | 9.100E-06 |
| 25924 | 1297 | S1297D | E | 6.5e-06 | 4e-06 | 1.1e-05 | 7.167E-06 | 3.547E-06 | 4.000E-06 | 1.100E-05 |

| | | | | | | | | | | |
|---|---|---|---|---|---|---|---|---|---|---|
| 25925 | 1297 | S1297C | E | 1e-05 | 6.2e-06 | 4.4e-06 | 6.867E-06 | 2.859E-06 | 4.400E-06 | 1.000E-05 |
| 25926 | 1297 | S1297E | E | 5.7e-06 | 1.4e-05 | 9.3e-06 | 9.667E-06 | 4.162E-06 | 5.700E-06 | 1.400E-05 |
| 25927 | 1297 | S1297Q | E | 7.2e-06 | 6.1e-06 | 7.2e-06 | 6.833E-06 | 6.351E-07 | 6.100E-06 | 7.200E-06 |
| 25928 | 1297 | S1297G | E | 7.1e-06 | 4.7e-06 | 3.7e-06 | 5.167E-06 | 1.747E-06 | 3.700E-06 | 7.100E-06 |
| 25929 | 1297 | S1297H | E | 6.9e-06 | 1e-05 | 5.1e-06 | 7.333E-06 | 2.479E-06 | 5.100E-06 | 1.000E-05 |
| 25930 | 1297 | S1297I | E | 6e-06 | 5.9e-06 | 7e-06 | 6.300E-06 | 6.083E-07 | 5.900E-06 | 7.000E-06 |
| 25931 | 1297 | S1297L | E | 5.6e-06 | 9e-06 | 1.4e-05 | 9.533E-06 | 4.225E-06 | 5.600E-06 | 1.400E-05 |
| 25932 | 1297 | S1297K | E | 1e-05 | 1.2e-05 | 4.3e-06 | 8.767E-06 | 3.995E-06 | 4.300E-06 | 1.200E-05 |
| 25933 | 1297 | S1297M | E | 8.3e-06 | 4.8e-06 | 9.4e-06 | 7.500E-06 | 2.402E-06 | 4.800E-06 | 9.400E-06 |
| 25934 | 1297 | S1297F | E | 1.2e-05 | 1e-05 | 1e-05 | 1.067E-05 | 1.155E-06 | 1.000E-05 | 1.200E-05 |
| 25935 | 1297 | S1297P | E | 8.8e-06 | 1.1e-05 | 8.1e-06 | 9.300E-06 | 1.513E-06 | 8.100E-06 | 1.100E-05 |
| 25936 | 1297 | S1297S | E | 7.6e-06 | 9.5e-06 | 1e-05 | 9.033E-06 | 1.266E-06 | 7.600E-06 | 1.000E-05 |
| 25937 | 1297 | S1297T | E | 8.2e-06 | 9.3e-06 | 9.3e-06 | 8.933E-06 | 6.351E-07 | 8.200E-06 | 9.300E-06 |
| 25938 | 1297 | S1297W | E | 8.2e-06 | 8.7e-06 | 7.5e-06 | 8.133E-06 | 6.028E-07 | 7.500E-06 | 8.700E-06 |
| 25939 | 1297 | S1297Y | E | 9.5e-06 | 7.4e-06 | 9e-06 | 8.633E-06 | 1.097E-06 | 7.400E-06 | 9.500E-06 |
| 25940 | 1297 | S1297V | E | 1.1e-05 | 7.1e-06 | 7.3e-06 | 8.467E-06 | 2.196E-06 | 7.100E-06 | 1.100E-05 |
| 25941 | 1298 | G1298A | E | 5.5e-06 | 8.5e-06 | 3.9e-06 | 5.967E-06 | 2.335E-06 | 3.900E-06 | 8.500E-06 |
| 25942 | 1298 | G1298R | E | 7.2e-06 | 4.3e-06 | 6.6e-06 | 6.033E-06 | 1.531E-06 | 4.300E-06 | 7.200E-06 |
| 25943 | 1298 | G1298N | E | 9.7e-06 | 6.7e-06 | 8.6e-06 | 8.333E-06 | 1.518E-06 | 6.700E-06 | 9.700E-06 |
| 25944 | 1298 | G1298D | E | 5.9e-06 | 5.3e-06 | 7.6e-06 | 6.267E-06 | 1.193E-06 | 5.300E-06 | 7.600E-06 |
| 25945 | 1298 | G1298C | E | 4e-06 | 4.3e-06 | 9.4e-06 | 5.900E-06 | 3.035E-06 | 4.000E-06 | 9.400E-06 |
| 25946 | 1298 | G1298E | E | 5e-06 | 5.1e-06 | 7.8e-06 | 5.967E-06 | 1.589E-06 | 5.000E-06 | 7.800E-06 |
| 25947 | 1298 | G1298Q | E | 7.9e-06 | 7.5e-06 | 1.7e-05 | 1.080E-05 | 5.373E-06 | 7.500E-06 | 1.700E-05 |
| 25948 | 1298 | G1298G | E | 1e-05 | 6e-06 | 6.4e-06 | 7.467E-06 | 2.203E-06 | 6.000E-06 | 1.000E-05 |
| 25949 | 1298 | G1298H | E | 7.4e-06 | 1.1e-05 | 6.3e-06 | 8.233E-06 | 2.458E-06 | 6.300E-06 | 1.100E-05 |
| 25950 | 1298 | G1298I | E | 6.4e-06 | 6.3e-06 | 6e-06 | 6.233E-06 | 2.082E-07 | 6.000E-06 | 6.400E-06 |
| 25951 | 1298 | G1298L | E | 6.8e-06 | 4.7e-06 | 8.6e-06 | 6.700E-06 | 1.952E-06 | 4.700E-06 | 8.600E-06 |
| 25952 | 1298 | G1298K | E | 6.4e-06 | 8.4e-06 | 9.6e-06 | 8.133E-06 | 1.617E-06 | 6.400E-06 | 9.600E-06 |
| 25953 | 1298 | G1298M | E | 1e-05 | 7.7e-06 | 9.6e-06 | 9.100E-06 | 1.229E-06 | 7.700E-06 | 1.000E-05 |
| 25954 | 1298 | G1298F | E | 6e-06 | 6.5e-06 | 6.9e-06 | 6.467E-06 | 4.509E-07 | 6.000E-06 | 6.900E-06 |
| 25955 | 1298 | G1298P | E | 1e-05 | 8.6e-06 | 8.3e-06 | 8.967E-06 | 9.074E-07 | 8.300E-06 | 1.000E-05 |
| 25956 | 1298 | G1298S | E | 8.4e-06 | 9e-06 | 9.1e-06 | 8.833E-06 | 3.786E-07 | 8.400E-06 | 9.100E-06 |
| 25957 | 1298 | G1298T | E | 3.7e-06 | 1.4e-05 | 1.4e-05 | 1.057E-05 | 5.947E-06 | 3.700E-06 | 1.400E-05 |
| 25958 | 1298 | G1298W | E | 1.3e-05 | 1.8e-05 | 5.7e-06 | 1.223E-05 | 6.186E-06 | 5.700E-06 | 1.800E-05 |
| 25959 | 1298 | G1298Y | E | 1e-05 | 6.2e-06 | 5.2e-06 | 7.133E-06 | 2.532E-06 | 5.200E-06 | 1.000E-05 |
| 25960 | 1298 | G1298V | E | 4.5e-06 | 4.7e-06 | 6.3e-06 | 5.167E-06 | 9.866E-07 | 4.500E-06 | 6.300E-06 |
| 25961 | 1299 | F1299A | E | 4.9e-06 | 4.8e-06 | 5.9e-06 | 5.200E-06 | 6.083E-07 | 4.800E-06 | 5.900E-06 |
| 25962 | 1299 | F1299R | E | 1.3e-05 | 9.2e-06 | 7.9e-06 | 1.003E-05 | 2.650E-06 | 7.900E-06 | 1.300E-05 |
| 25963 | 1299 | F1299N | E | 9.1e-06 | 9.1e-06 | 7.3e-06 | 8.500E-06 | 1.039E-06 | 7.300E-06 | 9.100E-06 |
| 25964 | 1299 | F1299D | E | 7.2e-06 | 6.8e-06 | 1.2e-05 | 8.667E-06 | 2.894E-06 | 6.800E-06 | 1.200E-05 |
| 25965 | 1299 | F1299C | E | 6.8e-06 | 9.9e-06 | 7.5e-06 | 8.067E-06 | 1.626E-06 | 6.800E-06 | 9.900E-06 |
| 25966 | 1299 | F1299E | E | 7.2e-06 | 7.7e-06 | 9.1e-06 | 8.000E-06 | 9.849E-07 | 7.200E-06 | 9.100E-06 |
| 25967 | 1299 | F1299Q | E | 6.1e-06 | 8.8e-06 | 1.2e-05 | 8.967E-06 | 2.954E-06 | 6.100E-06 | 1.200E-05 |
| 25968 | 1299 | F1299G | E | 6.7e-06 | 6e-06 | 6.9e-06 | 6.533E-06 | 4.726E-07 | 6.000E-06 | 6.900E-06 |
| 25969 | 1299 | F1299H | E | 1e-05 | 9.2e-06 | 9.4e-06 | 9.533E-06 | 4.163E-07 | 9.200E-06 | 1.000E-05 |
| 25970 | 1299 | F1299I | E | 6.3e-06 | 7.1e-06 | 8.2e-06 | 7.200E-06 | 9.539E-07 | 6.300E-06 | 8.200E-06 |
| 25971 | 1299 | F1299L | E | 5.8e-06 | 9.4e-06 | 5.6e-06 | 6.933E-06 | 2.139E-06 | 5.600E-06 | 9.400E-06 |
| 25972 | 1299 | F1299K | E | 4.4e-06 | 8.8e-06 | 5.6e-06 | 6.267E-06 | 2.274E-06 | 4.400E-06 | 8.800E-06 |
| 25973 | 1299 | F1299M | E | 9e-06 | 6.8e-06 | 9.6e-06 | 8.467E-06 | 1.474E-06 | 6.800E-06 | 9.600E-06 |
| 25974 | 1299 | F1299F | E | 1.4e-05 | 4.9e-06 | 4.5e-06 | 7.800E-06 | 5.373E-06 | 4.500E-06 | 1.400E-05 |
| 25975 | 1299 | F1299P | E | 1.1e-05 | 7.1e-06 | 7.5e-06 | 8.533E-06 | 2.146E-06 | 7.100E-06 | 1.100E-05 |
| 25976 | 1299 | F1299S | E | 1.6e-05 | 1.1e-05 | 4.6e-06 | 1.053E-05 | 5.714E-06 | 4.600E-06 | 1.600E-05 |
| 25977 | 1299 | F1299T | E | 9.4e-06 | 5.6e-06 | 9.6e-06 | 8.200E-06 | 2.254E-06 | 5.600E-06 | 9.600E-06 |
| 25978 | 1299 | F1299W | E | 1.1e-05 | 6.9e-06 | 9.4e-06 | 9.100E-06 | 2.066E-06 | 6.900E-06 | 1.100E-05 |
| 25979 | 1299 | F1299Y | E | 9.4e-06 | 8.9e-06 | 1.2e-05 | 1.010E-05 | 1.664E-06 | 8.900E-06 | 1.200E-05 |
| 25980 | 1299 | F1299V | E | 6.9e-06 | 5.2e-06 | 1e-05 | 7.367E-06 | 2.434E-06 | 5.200E-06 | 1.000E-05 |
| 25981 | 1300 | N1300A | E | 8.9e-06 | 8.4e-06 | 6.1e-06 | 7.800E-06 | 1.493E-06 | 6.100E-06 | 8.900E-06 |
| 25982 | 1300 | N1300R | E | 6.5e-06 | 4.2e-06 | 1e-05 | 6.900E-06 | 2.921E-06 | 4.200E-06 | 1.000E-05 |
| 25983 | 1300 | N1300N | E | 9.5e-06 | 1e-05 | 4.4e-06 | 7.967E-06 | 3.099E-06 | 4.400E-06 | 1.000E-05 |
| 25984 | 1300 | N1300D | E | 6.4e-06 | 8.8e-06 | 6.6e-06 | 7.267E-06 | 1.332E-06 | 6.400E-06 | 8.800E-06 |
| 25985 | 1300 | N1300C | E | 6.4e-06 | 7.6e-06 | 7.8e-06 | 7.267E-06 | 7.572E-07 | 6.400E-06 | 7.800E-06 |
| 25986 | 1300 | N1300E | E | 1.2e-05 | 8.7e-06 | 6.3e-06 | 9.000E-06 | 2.862E-06 | 6.300E-06 | 1.200E-05 |
| 25987 | 1300 | N1300Q | E | 1.3e-05 | 1e-05 | 7.5e-06 | 1.017E-05 | 2.754E-06 | 7.500E-06 | 1.300E-05 |
| 25988 | 1300 | N1300G | E | 8.2e-06 | 9.2e-06 | 7.8e-06 | 8.400E-06 | 7.211E-07 | 7.800E-06 | 9.200E-06 |
| 25989 | 1300 | N1300H | E | 9.7e-06 | 6.7e-06 | 8.5e-06 | 8.300E-06 | 1.510E-06 | 6.700E-06 | 9.700E-06 |
| 25990 | 1300 | N1300I | E | 1.1e-05 | 8e-06 | 4.7e-06 | 7.900E-06 | 3.151E-06 | 4.700E-06 | 1.100E-05 |
| 25991 | 1300 | N1300L | E | 7.8e-06 | 1.1e-05 | 4.7e-06 | 7.833E-06 | 3.150E-06 | 4.700E-06 | 1.100E-05 |
| 25992 | 1300 | N1300K | E | 1e-05 | 7.3e-06 | 9.1e-06 | 8.800E-06 | 1.375E-06 | 7.300E-06 | 1.000E-05 |
| 25993 | 1300 | N1300M | E | 4.7e-06 | 1.4e-05 | 9.6e-06 | 9.433E-06 | 4.652E-06 | 4.700E-06 | 1.400E-05 |
| 25994 | 1300 | N1300F | E | 7.9e-06 | 7.2e-06 | 1.4e-05 | 9.700E-06 | 3.740E-06 | 7.200E-06 | 1.400E-05 |
| 25995 | 1300 | N1300P | E | 7e-06 | 7.2e-06 | 1e-05 | 8.067E-06 | 1.677E-06 | 7.000E-06 | 1.000E-05 |
| 25996 | 1300 | N1300S | E | 1e-05 | 4.8e-06 | 8.3e-06 | 7.700E-06 | 2.651E-06 | 4.800E-06 | 1.000E-05 |
| 25997 | 1300 | N1300T | E | 6.9e-06 | 1.1e-05 | 8.3e-06 | 8.733E-06 | 2.084E-06 | 6.900E-06 | 1.100E-05 |
| 25998 | 1300 | N1300W | E | 1e-05 | 7.6e-06 | 1.1e-05 | 9.533E-06 | 1.747E-06 | 7.600E-06 | 1.100E-05 |
| 25999 | 1300 | N1300Y | E | 7.4e-06 | 9.7e-06 | 1.1e-05 | 9.367E-06 | 1.823E-06 | 7.400E-06 | 1.100E-05 |
| 26000 | 1300 | N1300V | E | 6.4e-06 | 1.5e-05 | 6.7e-06 | 9.367E-06 | 4.881E-06 | 6.400E-06 | 1.500E-05 |
| 26001 | 1301 | I1301A | E | 1.3e-05 | 1.1e-05 | 1.4e-05 | 1.267E-05 | 1.528E-06 | 1.100E-05 | 1.400E-05 |
| 26002 | 1301 | I1301R | E | 1e-05 | 7.6e-06 | 6.2e-06 | 7.933E-06 | 1.922E-06 | 6.200E-06 | 1.000E-05 |
| 26003 | 1301 | I1301N | E | 1.1e-05 | 6.3e-06 | 1.5e-05 | 1.077E-05 | 4.355E-06 | 6.300E-06 | 1.500E-05 |
| 26004 | 1301 | I1301D | E | 5.2e-06 | 1.1e-05 | 8.1e-06 | 8.100E-06 | 2.900E-06 | 5.200E-06 | 1.100E-05 |
| 26005 | 1301 | I1301C | E | 7e-06 | 4.4e-06 | 8.4e-06 | 6.600E-06 | 2.030E-06 | 4.400E-06 | 8.400E-06 |
| 26006 | 1301 | I1301E | E | 1.1e-05 | 8e-06 | 7.1e-06 | 8.700E-06 | 2.042E-06 | 7.100E-06 | 1.100E-05 |
| 26007 | 1301 | I1301Q | E | 1.2e-05 | 7e-06 | 1e-05 | 9.667E-06 | 2.517E-06 | 7.000E-06 | 1.200E-05 |
| 26008 | 1301 | I1301G | E | 5.1e-06 | 8.9e-06 | 9.8e-06 | 7.933E-06 | 2.495E-06 | 5.100E-06 | 9.800E-06 |
| 26009 | 1301 | I1301H | E | 6.7e-06 | 5.5e-06 | 9e-06 | 7.067E-06 | 1.779E-06 | 5.500E-06 | 9.000E-06 |

| | | | | | | | | | | |
|---|---|---|---|---|---|---|---|---|---|---|
| 26010 | 1301 | I1301I | E | 7.3e-06 | 1.2e-05 | 4.8e-06 | 8.033E-06 | 3.656E-06 | 4.800E-06 | 1.200E-05 |
| 26011 | 1301 | I1301L | E | 1e-05 | 3.2e-06 | 1e-05 | 7.733E-06 | 3.926E-06 | 3.200E-06 | 1.000E-05 |
| 26012 | 1301 | I1301K | E | 1.2e-05 | 4.5e-06 | 6.5e-06 | 7.667E-06 | 3.884E-06 | 4.500E-06 | 1.200E-05 |
| 26013 | 1301 | I1301M | E | 1.1e-05 | 1e-05 | 7.8e-06 | 9.600E-06 | 1.637E-06 | 7.800E-06 | 1.100E-05 |
| 26014 | 1301 | I1301F | E | 5.9e-06 | 1.5e-05 | 8.3e-06 | 9.733E-06 | 4.716E-06 | 5.900E-06 | 1.500E-05 |
| 26015 | 1301 | I1301P | E | 5.1e-06 | 8.5e-06 | 4e-06 | 5.867E-06 | 2.346E-06 | 4.000E-06 | 8.500E-06 |
| 26016 | 1301 | I1301S | E | 6.7e-06 | 7.1e-06 | 5.7e-06 | 6.500E-06 | 7.211E-07 | 5.700E-06 | 7.100E-06 |
| 26017 | 1301 | I1301T | E | 6.9e-06 | 1.4e-05 | 1.4e-05 | 1.163E-05 | 4.099E-06 | 6.900E-06 | 1.400E-05 |
| 26018 | 1301 | I1301W | E | 1.1e-05 | 1.1e-05 | 8.3e-06 | 1.010E-05 | 1.559E-06 | 8.300E-06 | 1.100E-05 |
| 26019 | 1301 | I1301Y | E | 6e-06 | 9.9e-06 | 1.4e-05 | 9.967E-06 | 4.000E-06 | 6.000E-06 | 1.400E-05 |
| 26020 | 1301 | I1301V | E | 8.3e-06 | 1e-05 | 9.7e-06 | 9.333E-06 | 9.074E-07 | 8.300E-06 | 1.000E-05 |
| 26021 | 1302 | K1302A | E | 1.1e-05 | 6.3e-06 | 9e-06 | 8.767E-06 | 2.359E-06 | 6.300E-06 | 1.100E-05 |
| 26022 | 1302 | K1302R | E | 1.4e-05 | 7e-06 | 9.6e-06 | 1.020E-05 | 3.538E-06 | 7.000E-06 | 1.400E-05 |
| 26023 | 1302 | K1302N | E | 7.5e-06 | 3e-06 | 7.1e-06 | 5.867E-06 | 2.491E-06 | 3.000E-06 | 7.500E-06 |
| 26024 | 1302 | K1302D | E | 7e-06 | 1.5e-05 | 1e-05 | 1.067E-05 | 4.041E-06 | 7.000E-06 | 1.500E-05 |
| 26025 | 1302 | K1302C | E | 1.3e-05 | 8.4e-06 | 4.5e-06 | 8.633E-06 | 4.255E-06 | 4.500E-06 | 1.300E-05 |
| 26026 | 1302 | K1302E | E | 7.1e-06 | 8.7e-06 | 7.5e-06 | 7.767E-06 | 8.327E-07 | 7.100E-06 | 8.700E-06 |
| 26027 | 1302 | K1302Q | E | 8.1e-06 | 4.6e-06 | 7.5e-06 | 6.733E-06 | 1.872E-06 | 4.600E-06 | 8.100E-06 |
| 26028 | 1302 | K1302G | E | 1e-05 | 4.5e-06 | 9.7e-06 | 8.067E-06 | 3.092E-06 | 4.500E-06 | 1.000E-05 |
| 26029 | 1302 | K1302H | E | 7.1e-06 | 6.7e-06 | 9.8e-06 | 7.867E-06 | 1.686E-06 | 6.700E-06 | 9.800E-06 |
| 26030 | 1302 | K1302I | E | 3.6e-06 | 6.6e-06 | 4.1e-06 | 4.767E-06 | 1.607E-06 | 3.600E-06 | 6.600E-06 |
| 26031 | 1302 | K1302L | E | 1e-05 | 1.3e-05 | 4.6e-06 | 9.200E-06 | 4.257E-06 | 4.600E-06 | 1.300E-05 |
| 26032 | 1302 | K1302K | E | 1.5e-05 | 6.3e-06 | 5.6e-06 | 8.967E-06 | 5.237E-06 | 5.600E-06 | 1.500E-05 |
| 26033 | 1302 | K1302M | E | 8.2e-06 | 4e-06 | 8.5e-06 | 6.900E-06 | 2.516E-06 | 4.000E-06 | 8.500E-06 |
| 26034 | 1302 | K1302F | E | 6.6e-06 | 7.3e-06 | 8.2e-06 | 7.367E-06 | 8.021E-07 | 6.600E-06 | 8.200E-06 |
| 26035 | 1302 | K1302P | E | 7.4e-06 | 1.7e-05 | 7.6e-06 | 1.067E-05 | 5.486E-06 | 7.400E-06 | 1.700E-05 |
| 26036 | 1302 | K1302S | E | 9.1e-06 | 6.5e-06 | 6.3e-06 | 7.300E-06 | 1.562E-06 | 6.300E-06 | 9.100E-06 |
| 26037 | 1302 | K1302T | E | 7.2e-06 | 7.1e-06 | 6.8e-06 | 7.033E-06 | 2.082E-07 | 6.800E-06 | 7.200E-06 |
| 26038 | 1302 | K1302W | E | 4.3e-06 | 9.2e-06 | 6.4e-06 | 6.633E-06 | 2.458E-06 | 4.300E-06 | 9.200E-06 |
| 26039 | 1302 | K1302Y | E | 1e-05 | 4.9e-06 | 4.9e-06 | 6.600E-06 | 2.944E-06 | 4.900E-06 | 1.000E-05 |
| 26040 | 1302 | K1302V | E | 6.3e-06 | 6.2e-06 | 9.4e-06 | 7.300E-06 | 1.819E-06 | 6.200E-06 | 9.400E-06 |
| 26041 | 1303 | D1303A | E | 8.2e-06 | 1.3e-05 | 4.3e-06 | 8.500E-06 | 4.358E-06 | 4.300E-06 | 1.300E-05 |
| 26042 | 1303 | D1303R | E | 4.3e-06 | 9.6e-06 | 8.8e-06 | 7.567E-06 | 2.857E-06 | 4.300E-06 | 9.600E-06 |
| 26043 | 1303 | D1303N | E | 8.8e-06 | 9.7e-06 | 7.9e-06 | 8.800E-06 | 9.000E-07 | 7.900E-06 | 9.700E-06 |
| 26044 | 1303 | D1303D | E | 8.8e-06 | 6.8e-06 | 3.4e-06 | 6.333E-06 | 2.730E-06 | 3.400E-06 | 8.800E-06 |
| 26045 | 1303 | D1303C | E | 6e-06 | 4.3e-06 | 2e-05 | 1.010E-05 | 8.616E-06 | 4.300E-06 | 2.000E-05 |
| 26046 | 1303 | D1303E | E | 5.2e-06 | 5.8e-06 | 1.1e-05 | 7.333E-06 | 3.190E-06 | 5.200E-06 | 1.100E-05 |
| 26047 | 1303 | D1303Q | E | 9.1e-06 | 6.2e-06 | 4.8e-06 | 6.700E-06 | 2.193E-06 | 4.800E-06 | 9.100E-06 |
| 26048 | 1303 | D1303G | E | 1.1e-05 | 7.7e-06 | 9.1e-06 | 9.267E-06 | 1.656E-06 | 7.700E-06 | 1.100E-05 |
| 26049 | 1303 | D1303H | E | 7.5e-06 | 1.3e-05 | 9.1e-06 | 9.867E-06 | 2.829E-06 | 7.500E-06 | 1.300E-05 |
| 26050 | 1303 | D1303I | E | 5.4e-06 | 4.4e-06 | 1e-05 | 6.600E-06 | 2.987E-06 | 4.400E-06 | 1.000E-05 |
| 26051 | 1303 | D1303L | E | 1e-05 | 5.8e-06 | 4.6e-06 | 6.800E-06 | 2.835E-06 | 4.600E-06 | 1.000E-05 |
| 26052 | 1303 | D1303K | E | 9.7e-06 | 1.2e-05 | 5.2e-06 | 8.967E-06 | 3.459E-06 | 5.200E-06 | 1.200E-05 |
| 26053 | 1303 | D1303M | E | 9.4e-06 | 1e-05 | 4.1e-06 | 7.833E-06 | 3.247E-06 | 4.100E-06 | 1.000E-05 |
| 26054 | 1303 | D1303F | E | 1.2e-05 | 8.5e-06 | 6.4e-06 | 8.967E-06 | 2.829E-06 | 6.400E-06 | 1.200E-05 |
| 26055 | 1303 | D1303P | E | 8.4e-06 | 5.2e-06 | 9.8e-06 | 7.800E-06 | 2.358E-06 | 5.200E-06 | 9.800E-06 |
| 26056 | 1303 | D1303S | E | 5.8e-06 | 6.4e-06 | 6.4e-06 | 6.200E-06 | 3.464E-07 | 5.800E-06 | 6.400E-06 |
| 26057 | 1303 | D1303T | E | 1.2e-05 | 6.1e-06 | 6e-06 | 8.033E-06 | 3.436E-06 | 6.000E-06 | 1.200E-05 |
| 26058 | 1303 | D1303W | E | 6.8e-06 | 7.5e-06 | 7.7e-06 | 7.333E-06 | 4.726E-07 | 6.800E-06 | 7.700E-06 |
| 26059 | 1303 | D1303Y | E | 6.3e-06 | 7.2e-06 | 9.6e-06 | 7.700E-06 | 1.706E-06 | 6.300E-06 | 9.600E-06 |
| 26060 | 1303 | D1303V | E | 1e-05 | 2.3e-06 | 8e-06 | 6.767E-06 | 3.995E-06 | 2.300E-06 | 1.000E-05 |
| 26061 | 1304 | T1304A | E | 1.3e-05 | 4.8e-06 | 5.1e-06 | 7.633E-06 | 4.650E-06 | 4.800E-06 | 1.300E-05 |
| 26062 | 1304 | T1304R | E | 1e-05 | 6.7e-06 | 3.6e-06 | 6.767E-06 | 3.201E-06 | 3.600E-06 | 1.000E-05 |
| 26063 | 1304 | T1304N | E | 9.6e-06 | 3.6e-06 | 8.6e-06 | 7.267E-06 | 3.215E-06 | 3.600E-06 | 9.600E-06 |
| 26064 | 1304 | T1304D | E | 5.7e-06 | 1.5e-05 | 6e-06 | 8.900E-06 | 5.285E-06 | 5.700E-06 | 1.500E-05 |
| 26065 | 1304 | T1304C | E | 6.3e-06 | 1e-05 | 1.2e-05 | 9.433E-06 | 2.892E-06 | 6.300E-06 | 1.200E-05 |
| 26066 | 1304 | T1304E | E | 1e-05 | 8.1e-06 | 5.3e-06 | 7.800E-06 | 2.364E-06 | 5.300E-06 | 1.000E-05 |
| 26067 | 1304 | T1304Q | E | 4.9e-06 | 5.5e-06 | 1.2e-05 | 7.467E-06 | 3.937E-06 | 4.900E-06 | 1.200E-05 |
| 26068 | 1304 | T1304G | E | 1.2e-05 | 6.9e-06 | 5.9e-06 | 8.267E-06 | 3.272E-06 | 5.900E-06 | 1.200E-05 |
| 26069 | 1304 | T1304H | E | 8.3e-06 | 8.1e-06 | 7.3e-06 | 7.900E-06 | 5.292E-07 | 7.300E-06 | 8.300E-06 |
| 26070 | 1304 | T1304I | E | 8.4e-06 | 7.1e-06 | 5.6e-06 | 7.033E-06 | 1.401E-06 | 5.600E-06 | 8.400E-06 |
| 26071 | 1304 | T1304L | E | 6.9e-06 | 8.9e-06 | 9.5e-06 | 8.433E-06 | 1.361E-06 | 6.900E-06 | 9.500E-06 |
| 26072 | 1304 | T1304K | E | 1e-05 | 4.6e-06 | 5.3e-06 | 6.633E-06 | 2.937E-06 | 4.600E-06 | 1.000E-05 |
| 26073 | 1304 | T1304M | E | 5.6e-06 | 6.6e-06 | 1.4e-05 | 8.733E-06 | 4.588E-06 | 5.600E-06 | 1.400E-05 |
| 26074 | 1304 | T1304F | E | 5.6e-06 | 7.4e-06 | 7.4e-06 | 6.800E-06 | 1.039E-06 | 5.600E-06 | 7.400E-06 |
| 26075 | 1304 | T1304P | E | 1.3e-05 | 3.8e-06 | 5.8e-06 | 7.533E-06 | 4.839E-06 | 3.800E-06 | 1.300E-05 |
| 26076 | 1304 | T1304S | E | 1.2e-05 | 1.3e-05 | 7.6e-06 | 1.087E-05 | 2.873E-06 | 7.600E-06 | 1.300E-05 |
| 26077 | 1304 | T1304T | E | 9.3e-06 | 7.7e-06 | 7.9e-06 | 8.300E-06 | 8.718E-07 | 7.700E-06 | 9.300E-06 |
| 26078 | 1304 | T1304W | E | 5.4e-06 | 1.4e-05 | 9.6e-06 | 9.667E-06 | 4.300E-06 | 5.400E-06 | 1.400E-05 |
| 26079 | 1304 | T1304Y | E | 8.3e-06 | 7.9e-06 | 7e-06 | 7.733E-06 | 6.658E-07 | 7.000E-06 | 8.300E-06 |
| 26080 | 1304 | T1304V | E | 5.7e-06 | 6.4e-06 | 3.3e-06 | 5.133E-06 | 1.626E-06 | 3.300E-06 | 6.400E-06 |
| 26081 | 1305 | Y1305A | E | 8.7e-06 | 1.1e-05 | 8e-06 | 9.233E-06 | 1.570E-06 | 8.000E-06 | 1.100E-05 |
| 26082 | 1305 | Y1305R | E | 7.1e-06 | 6.9e-06 | 3.4e-06 | 5.800E-06 | 2.081E-06 | 3.400E-06 | 7.100E-06 |
| 26083 | 1305 | Y1305N | E | 1.3e-05 | 9.8e-06 | 7.3e-06 | 1.003E-05 | 2.857E-06 | 7.300E-06 | 1.300E-05 |
| 26084 | 1305 | Y1305D | E | 4.8e-06 | 4.7e-06 | 4.8e-06 | 4.767E-06 | 5.774E-08 | 4.700E-06 | 4.800E-06 |
| 26085 | 1305 | Y1305C | E | 1e-05 | 3.3e-06 | 6e-06 | 6.433E-06 | 3.371E-06 | 3.300E-06 | 1.000E-05 |
| 26086 | 1305 | Y1305E | E | 6.8e-06 | 9.8e-06 | 5.6e-06 | 7.400E-06 | 2.163E-06 | 5.600E-06 | 9.800E-06 |
| 26087 | 1305 | Y1305Q | E | 7.8e-06 | 4.6e-06 | 1.2e-05 | 8.133E-06 | 3.711E-06 | 4.600E-06 | 1.200E-05 |
| 26088 | 1305 | Y1305G | E | 9.5e-06 | 7.4e-06 | 4.8e-06 | 7.233E-06 | 2.354E-06 | 4.800E-06 | 9.500E-06 |
| 26089 | 1305 | Y1305H | E | 9.6e-06 | 4.7e-06 | 1.2e-05 | 8.767E-06 | 3.721E-06 | 4.700E-06 | 1.200E-05 |
| 26090 | 1305 | Y1305I | E | 8.2e-06 | 9.7e-06 | 1.4e-05 | 1.063E-05 | 3.011E-06 | 8.200E-06 | 1.400E-05 |
| 26091 | 1305 | Y1305L | E | 6.4e-06 | 5e-06 | 8.3e-06 | 6.567E-06 | 1.656E-06 | 5.000E-06 | 8.300E-06 |
| 26092 | 1305 | Y1305K | E | 7.8e-06 | 5.4e-06 | 5.7e-06 | 6.300E-06 | 1.308E-06 | 5.400E-06 | 7.800E-06 |
| 26093 | 1305 | Y1305M | E | 7.8e-06 | 7.3e-06 | 9.3e-06 | 8.133E-06 | 1.041E-06 | 7.300E-06 | 9.300E-06 |
| 26094 | 1305 | Y1305F | E | 1.1e-05 | 8.4e-06 | 4.7e-06 | 8.033E-06 | 3.166E-06 | 4.700E-06 | 1.100E-05 |

| | | | | | | | | | | |
|---|---|---|---|---|---|---|---|---|---|---|
| 26095 | 1305 | Y1305P | E | 9.4e-06 | 6.9e-06 | 1.3e-05 | 9.767E-06 | 3.066E-06 | 6.900E-06 | 1.300E-05 |
| 26096 | 1305 | Y1305S | E | 8.3e-06 | 7.9e-06 | 6.6e-06 | 7.600E-06 | 8.888E-07 | 6.600E-06 | 8.300E-06 |
| 26097 | 1305 | Y1305T | E | 6.7e-06 | 8.8e-06 | 9.7e-06 | 8.400E-06 | 1.539E-06 | 6.700E-06 | 9.700E-06 |
| 26098 | 1305 | Y1305W | E | 8.3e-06 | 9.9e-06 | 1.6e-05 | 1.140E-05 | 4.063E-06 | 8.300E-06 | 1.600E-05 |
| 26099 | 1305 | Y1305Y | E | 5.2e-06 | 1.4e-05 | 9.2e-06 | 9.467E-06 | 4.406E-06 | 5.200E-06 | 1.400E-05 |
| 26100 | 1305 | Y1305V | E | 7.7e-06 | 4.3e-06 | 7.1e-06 | 6.367E-06 | 1.815E-06 | 4.300E-06 | 7.700E-06 |
| 26101 | 1306 | I1306A | E | 5.8e-06 | 5.9e-06 | 6e-06 | 5.900E-06 | 1.000E-07 | 5.800E-06 | 6.000E-06 |
| 26102 | 1306 | I1306R | E | 1e-05 | 4.8e-06 | 5.1e-06 | 6.633E-06 | 2.919E-06 | 4.800E-06 | 1.000E-05 |
| 26103 | 1306 | I1306N | E | 1.2e-05 | 7.8e-06 | 1.6e-05 | 1.193E-05 | 4.100E-06 | 7.800E-06 | 1.600E-05 |
| 26104 | 1306 | I1306D | E | 8e-06 | 5.6e-06 | 8.6e-06 | 7.400E-06 | 1.587E-06 | 5.600E-06 | 8.600E-06 |
| 26105 | 1306 | I1306C | E | 9.1e-06 | 9.2e-06 | 9.9e-06 | 9.400E-06 | 4.359E-07 | 9.100E-06 | 9.900E-06 |
| 26106 | 1306 | I1306E | E | 6.6e-06 | 1.2e-05 | 1.1e-05 | 9.867E-06 | 2.873E-06 | 6.600E-06 | 1.200E-05 |
| 26107 | 1306 | I1306Q | E | 7.8e-06 | 7.1e-06 | 4.8e-06 | 6.567E-06 | 1.570E-06 | 4.800E-06 | 7.800E-06 |
| 26108 | 1306 | I1306G | E | 1.3e-05 | 7.7e-06 | 4.7e-06 | 8.467E-06 | 4.203E-06 | 4.700E-06 | 1.300E-05 |
| 26109 | 1306 | I1306H | E | 7.1e-06 | 9.2e-06 | 9.9e-06 | 8.733E-06 | 1.457E-06 | 7.100E-06 | 9.900E-06 |
| 26110 | 1306 | I1306I | E | 7.6e-06 | 9.2e-06 | 1.5e-05 | 1.060E-05 | 3.894E-06 | 7.600E-06 | 1.500E-05 |
| 26111 | 1306 | I1306L | E | 5.8e-06 | 8.2e-06 | 1.3e-05 | 9.000E-06 | 3.666E-06 | 5.800E-06 | 1.300E-05 |
| 26112 | 1306 | I1306K | E | 6.7e-06 | 6.4e-06 | 1e-05 | 7.700E-06 | 1.997E-06 | 6.400E-06 | 1.000E-05 |
| 26113 | 1306 | I1306M | E | 6.9e-06 | 9.9e-06 | 1e-05 | 8.933E-06 | 1.762E-06 | 6.900E-06 | 1.000E-05 |
| 26114 | 1306 | I1306F | E | 5.8e-06 | 5.4e-06 | 1.2e-05 | 7.733E-06 | 3.700E-06 | 5.400E-06 | 1.200E-05 |
| 26115 | 1306 | I1306P | E | 1.1e-05 | 1.1e-05 | 7.5e-06 | 9.833E-06 | 2.021E-06 | 7.500E-06 | 1.100E-05 |
| 26116 | 1306 | I1306S | E | 6.2e-06 | 5.9e-06 | 9.9e-06 | 7.333E-06 | 2.228E-06 | 5.900E-06 | 9.900E-06 |
| 26117 | 1306 | I1306T | E | 5.6e-06 | 5.3e-06 | 4.7e-06 | 5.200E-06 | 4.583E-07 | 4.700E-06 | 5.600E-06 |
| 26118 | 1306 | I1306W | E | 6.8e-06 | 6.6e-06 | 1.1e-05 | 8.133E-06 | 2.485E-06 | 6.600E-06 | 1.100E-05 |
| 26119 | 1306 | I1306Y | E | 8.3e-06 | 8.5e-06 | 6.4e-06 | 7.733E-06 | 1.159E-06 | 6.400E-06 | 8.500E-06 |
| 26120 | 1306 | I1306V | E | 6.7e-06 | 1.4e-05 | 6.4e-06 | 9.033E-06 | 4.304E-06 | 6.400E-06 | 1.400E-05 |
| 26121 | 1307 | H1307A | E | 8.2e-06 | 1.1e-05 | 7.9e-06 | 9.033E-06 | 1.710E-06 | 7.900E-06 | 1.100E-05 |
| 26122 | 1307 | H1307R | E | 9.5e-06 | 1.3e-05 | 6.6e-06 | 9.700E-06 | 3.205E-06 | 6.600E-06 | 1.300E-05 |
| 26123 | 1307 | H1307N | E | 6.7e-06 | 7.4e-06 | 6.7e-06 | 6.933E-06 | 4.041E-07 | 6.700E-06 | 7.400E-06 |
| 26124 | 1307 | H1307D | E | 6.8e-06 | 6.1e-06 | 7.4e-06 | 6.767E-06 | 6.506E-07 | 6.100E-06 | 7.400E-06 |
| 26125 | 1307 | H1307C | E | 6.3e-06 | 8e-06 | 8.9e-06 | 7.733E-06 | 1.320E-06 | 6.300E-06 | 8.900E-06 |
| 26126 | 1307 | H1307E | E | 6.8e-06 | 1e-05 | 1.2e-05 | 9.600E-06 | 2.623E-06 | 6.800E-06 | 1.200E-05 |
| 26127 | 1307 | H1307Q | E | 8.8e-06 | 1.3e-05 | 5.1e-06 | 8.967E-06 | 3.953E-06 | 5.100E-06 | 1.300E-05 |
| 26128 | 1307 | H1307G | E | 1e-05 | 5.7e-06 | 1e-05 | 8.567E-06 | 2.483E-06 | 5.700E-06 | 1.000E-05 |
| 26129 | 1307 | H1307H | E | 1.1e-05 | 1.1e-05 | 1.4e-05 | 1.200E-05 | 1.732E-06 | 1.100E-05 | 1.400E-05 |
| 26130 | 1307 | H1307I | E | 1.2e-05 | 7.2e-06 | 9.1e-06 | 9.433E-06 | 2.417E-06 | 7.200E-06 | 1.200E-05 |
| 26131 | 1307 | H1307L | E | 9.2e-06 | 5.7e-06 | 1.8e-05 | 1.097E-05 | 6.337E-06 | 5.700E-06 | 1.800E-05 |
| 26132 | 1307 | H1307K | E | 6.6e-06 | 8.1e-06 | 1.1e-05 | 8.567E-06 | 2.237E-06 | 6.600E-06 | 1.100E-05 |
| 26133 | 1307 | H1307M | E | 1.4e-05 | 5.4e-06 | 7.9e-06 | 9.100E-06 | 4.424E-06 | 5.400E-06 | 1.400E-05 |
| 26134 | 1307 | H1307F | E | 6.1e-06 | 7e-06 | 1e-05 | 7.700E-06 | 2.042E-06 | 6.100E-06 | 1.000E-05 |
| 26135 | 1307 | H1307P | E | 5.8e-06 | 5.9e-06 | 6.5e-06 | 6.067E-06 | 3.786E-07 | 5.800E-06 | 6.500E-06 |
| 26136 | 1307 | H1307S | E | 9.5e-06 | 7.5e-06 | 6.4e-06 | 7.800E-06 | 1.572E-06 | 6.400E-06 | 9.500E-06 |
| 26137 | 1307 | H1307T | E | 9.3e-06 | 3.7e-06 | 6e-06 | 6.333E-06 | 2.815E-06 | 3.700E-06 | 9.300E-06 |
| 26138 | 1307 | H1307W | E | 3e-06 | 3.1e-06 | 7.5e-06 | 4.533E-06 | 2.570E-06 | 3.000E-06 | 7.500E-06 |
| 26139 | 1307 | H1307Y | E | 5.6e-06 | 6.9e-06 | 1.3e-05 | 8.500E-06 | 3.951E-06 | 5.600E-06 | 1.300E-05 |
| 26140 | 1307 | H1307V | E | 1.2e-05 | 6.7e-06 | 9.7e-06 | 9.467E-06 | 2.658E-06 | 6.700E-06 | 1.200E-05 |
| 26141 | 1308 | W1308A | E | 1.4e-05 | 5.5e-06 | 7.5e-06 | 9.000E-06 | 4.444E-06 | 5.500E-06 | 1.400E-05 |
| 26142 | 1308 | W1308R | E | 9e-06 | 1.2e-05 | 1.9e-05 | 1.333E-05 | 5.132E-06 | 9.000E-06 | 1.900E-05 |
| 26143 | 1308 | W1308N | E | 4.2e-06 | 8.9e-06 | 9.8e-06 | 7.633E-06 | 3.007E-06 | 4.200E-06 | 9.800E-06 |
| 26144 | 1308 | W1308D | E | 9.1e-06 | 7.4e-06 | 7.3e-06 | 7.933E-06 | 1.012E-06 | 7.300E-06 | 9.100E-06 |
| 26145 | 1308 | W1308C | E | 5.6e-06 | 7.7e-06 | 8.9e-06 | 7.400E-06 | 1.670E-06 | 5.600E-06 | 8.900E-06 |
| 26146 | 1308 | W1308E | E | 5.3e-06 | 8.3e-06 | 4.9e-06 | 6.167E-06 | 1.858E-06 | 4.900E-06 | 8.300E-06 |
| 26147 | 1308 | W1308Q | E | 5.4e-06 | 1e-05 | 1.3e-05 | 9.467E-06 | 3.828E-06 | 5.400E-06 | 1.300E-05 |
| 26148 | 1308 | W1308G | E | 9.7e-06 | 1.3e-05 | 1.2e-05 | 1.157E-05 | 1.692E-06 | 9.700E-06 | 1.300E-05 |
| 26149 | 1308 | W1308H | E | 8.2e-06 | 1.3e-05 | 9.8e-06 | 1.033E-05 | 2.444E-06 | 8.200E-06 | 1.300E-05 |
| 26150 | 1308 | W1308I | E | 4.9e-06 | 5.7e-06 | 8.8e-06 | 6.467E-06 | 2.060E-06 | 4.900E-06 | 8.800E-06 |
| 26151 | 1308 | W1308L | E | 7.8e-06 | 8.2e-06 | 9.1e-06 | 8.367E-06 | 6.658E-07 | 7.800E-06 | 9.100E-06 |
| 26152 | 1308 | W1308K | E | 7.4e-06 | 1.1e-05 | 6.3e-06 | 8.233E-06 | 2.458E-06 | 6.300E-06 | 1.100E-05 |
| 26153 | 1308 | W1308M | E | 6e-06 | 7.5e-06 | 9.3e-06 | 7.600E-06 | 1.652E-06 | 6.000E-06 | 9.300E-06 |
| 26154 | 1308 | W1308F | E | 4.1e-06 | 6.1e-06 | 9.2e-06 | 6.467E-06 | 2.570E-06 | 4.100E-06 | 9.200E-06 |
| 26155 | 1308 | W1308P | E | 1.2e-05 | 9.2e-06 | 8.1e-06 | 9.767E-06 | 2.011E-06 | 8.100E-06 | 1.200E-05 |
| 26156 | 1308 | W1308S | E | 6.8e-06 | 9.1e-06 | 1e-05 | 8.633E-06 | 1.650E-06 | 6.800E-06 | 1.000E-05 |
| 26157 | 1308 | W1308T | E | 6.4e-06 | 5.3e-06 | 4.6e-06 | 5.433E-06 | 9.074E-07 | 4.600E-06 | 6.400E-06 |
| 26158 | 1308 | W1308W | E | 9.1e-06 | 9e-06 | 6.5e-06 | 8.200E-06 | 1.473E-06 | 6.500E-06 | 9.100E-06 |
| 26159 | 1308 | W1308Y | E | 7.1e-06 | 6.2e-06 | 6.9e-06 | 6.733E-06 | 4.726E-07 | 6.200E-06 | 7.100E-06 |
| 26160 | 1308 | W1308V | E | 5.1e-06 | 1.2e-05 | 6.8e-06 | 7.967E-06 | 3.595E-06 | 5.100E-06 | 1.200E-05 |
| 26161 | 1309 | V1309A | E | 7.2e-06 | 5.9e-06 | 7.5e-06 | 6.867E-06 | 8.505E-07 | 5.900E-06 | 7.500E-06 |
| 26162 | 1309 | V1309R | E | 7.7e-06 | 4.4e-06 | 1e-05 | 7.367E-06 | 2.815E-06 | 4.400E-06 | 1.000E-05 |
| 26163 | 1309 | V1309N | E | 1.1e-05 | 6.3e-06 | 7.4e-06 | 8.233E-06 | 2.458E-06 | 6.300E-06 | 1.100E-05 |
| 26164 | 1309 | V1309D | E | 8.9e-06 | 7.3e-06 | 1e-05 | 8.733E-06 | 1.358E-06 | 7.300E-06 | 1.000E-05 |
| 26165 | 1309 | V1309C | E | 5.4e-06 | 8.6e-06 | 9.1e-06 | 7.700E-06 | 2.007E-06 | 5.400E-06 | 9.100E-06 |
| 26166 | 1309 | V1309E | E | 1.2e-05 | 6.1e-06 | 7.6e-06 | 8.567E-06 | 3.066E-06 | 6.100E-06 | 1.200E-05 |
| 26167 | 1309 | V1309Q | E | 6.9e-06 | 9.6e-06 | 8.4e-06 | 8.300E-06 | 1.353E-06 | 6.900E-06 | 9.600E-06 |
| 26168 | 1309 | V1309G | E | 8.5e-06 | 1e-05 | 1e-05 | 9.500E-06 | 8.660E-07 | 8.500E-06 | 1.000E-05 |
| 26169 | 1309 | V1309H | E | 9e-06 | 6.7e-06 | 1.2e-05 | 9.233E-06 | 2.658E-06 | 6.700E-06 | 1.200E-05 |
| 26170 | 1309 | V1309I | E | 4.1e-06 | 1.1e-05 | 5.6e-06 | 6.900E-06 | 3.629E-06 | 4.100E-06 | 1.100E-05 |
| 26171 | 1309 | V1309L | E | 1e-05 | 5.5e-06 | 6.3e-06 | 7.267E-06 | 2.401E-06 | 5.500E-06 | 1.000E-05 |
| 26172 | 1309 | V1309K | E | 4.6e-06 | 6.4e-06 | 6.6e-06 | 5.867E-06 | 1.102E-06 | 4.600E-06 | 6.600E-06 |
| 26173 | 1309 | V1309M | E | 6.9e-06 | 1e-05 | 1.5e-05 | 1.063E-05 | 4.087E-06 | 6.900E-06 | 1.500E-05 |
| 26174 | 1309 | V1309F | E | 4.3e-06 | 7.8e-06 | 6e-06 | 6.033E-06 | 1.750E-06 | 4.300E-06 | 7.800E-06 |
| 26175 | 1309 | V1309P | E | 8.4e-06 | 9.9e-06 | 1e-05 | 9.433E-06 | 8.963E-07 | 8.400E-06 | 1.000E-05 |
| 26176 | 1309 | V1309S | E | 7e-06 | 9.6e-06 | 6.4e-06 | 7.667E-06 | 1.701E-06 | 6.400E-06 | 9.600E-06 |
| 26177 | 1309 | V1309T | E | 8.4e-06 | 9.7e-06 | 7.2e-06 | 8.433E-06 | 1.250E-06 | 7.200E-06 | 9.700E-06 |
| 26178 | 1309 | V1309W | E | 4.1e-06 | 1.1e-05 | 1.8e-05 | 1.103E-05 | 6.950E-06 | 4.100E-06 | 1.800E-05 |
| 26179 | 1309 | V1309Y | E | 6.2e-06 | 3.7e-06 | 1.4e-05 | 7.967E-06 | 5.372E-06 | 3.700E-06 | 1.400E-05 |

| | | | | | | | | | | |
|---|---|---|---|---|---|---|---|---|---|---|
| 26180 | 1309 | V1309V | E | 7.1e-06 | 6.6e-06 | 8.4e-06 | 7.367E-06 | 9.292E-07 | 6.600E-06 | 8.400E-06 |
| 26181 | 1310 | R1310A | E | 5.4e-06 | 1.5e-05 | 8.4e-06 | 9.600E-06 | 4.911E-06 | 5.400E-06 | 1.500E-05 |
| 26182 | 1310 | R1310R | E | 6.7e-06 | 4.3e-06 | 1.1e-05 | 7.333E-06 | 3.395E-06 | 4.300E-06 | 1.100E-05 |
| 26183 | 1310 | R1310N | E | 6.4e-06 | 1.1e-05 | 1.1e-05 | 9.467E-06 | 2.656E-06 | 6.400E-06 | 1.100E-05 |
| 26184 | 1310 | R1310D | E | 6.5e-06 | 1e-05 | 1.1e-05 | 9.167E-06 | 2.363E-06 | 6.500E-06 | 1.100E-05 |
| 26185 | 1310 | R1310C | E | 7.2e-06 | 4.3e-06 | 1.1e-05 | 7.500E-06 | 3.360E-06 | 4.300E-06 | 1.100E-05 |
| 26186 | 1310 | R1310E | E | 6.9e-06 | 8.3e-06 | 9.3e-06 | 8.167E-06 | 1.206E-06 | 6.900E-06 | 9.300E-06 |
| 26187 | 1310 | R1310Q | E | 6.8e-06 | 8.9e-06 | 1.1e-05 | 8.900E-06 | 2.100E-06 | 6.800E-06 | 1.100E-05 |
| 26188 | 1310 | R1310G | E | 5.3e-06 | 6.2e-06 | 7.4e-06 | 6.300E-06 | 1.054E-06 | 5.300E-06 | 7.400E-06 |
| 26189 | 1310 | R1310H | E | 6.6e-06 | 8.7e-06 | 4.7e-06 | 6.667E-06 | 2.001E-06 | 4.700E-06 | 8.700E-06 |
| 26190 | 1310 | R1310I | E | 4.5e-06 | 7.4e-06 | 5.9e-06 | 5.933E-06 | 1.450E-06 | 4.500E-06 | 7.400E-06 |
| 26191 | 1310 | R1310L | E | 1.1e-05 | 9.7e-06 | 7.8e-06 | 9.500E-06 | 1.609E-06 | 7.800E-06 | 1.100E-05 |
| 26192 | 1310 | R1310K | E | 8.4e-06 | 1.3e-05 | 8.4e-06 | 9.933E-06 | 2.656E-06 | 8.400E-06 | 1.300E-05 |
| 26193 | 1310 | R1310M | E | 5.3e-06 | 7e-06 | 6.7e-06 | 6.333E-06 | 9.074E-07 | 5.300E-06 | 7.000E-06 |
| 26194 | 1310 | R1310F | E | 7.1e-06 | 9.4e-06 | 5.1e-06 | 7.200E-06 | 2.152E-06 | 5.100E-06 | 9.400E-06 |
| 26195 | 1310 | R1310P | E | 9.4e-06 | 9.8e-06 | 9.1e-06 | 9.433E-06 | 3.512E-07 | 9.100E-06 | 9.800E-06 |
| 26196 | 1310 | R1310S | E | 7e-06 | 9.2e-06 | 7.3e-06 | 7.833E-06 | 1.193E-06 | 7.000E-06 | 9.200E-06 |
| 26197 | 1310 | R1310T | E | 6.7e-06 | 1.3e-05 | 1.3e-05 | 1.090E-05 | 3.637E-06 | 6.700E-06 | 1.300E-05 |
| 26198 | 1310 | R1310W | E | 9.1e-06 | 6.7e-06 | 9e-06 | 8.267E-06 | 1.358E-06 | 6.700E-06 | 9.100E-06 |
| 26199 | 1310 | R1310Y | E | 7.9e-06 | 8.6e-06 | 4.7e-06 | 7.067E-06 | 2.079E-06 | 4.700E-06 | 8.600E-06 |
| 26200 | 1310 | R1310V | E | 6.4e-06 | 1.3e-05 | 6.3e-06 | 8.567E-06 | 3.840E-06 | 6.300E-06 | 1.300E-05 |
| 26201 | 1311 | Q1311A | E | 6.1e-06 | 7.3e-06 | 8.2e-06 | 7.200E-06 | 1.054E-06 | 6.100E-06 | 8.200E-06 |
| 26202 | 1311 | Q1311R | E | 9e-06 | 8.9e-06 | 6.4e-06 | 8.100E-06 | 1.473E-06 | 6.400E-06 | 9.000E-06 |
| 26203 | 1311 | Q1311N | E | 6.2e-06 | 8.9e-06 | 1.3e-05 | 9.367E-06 | 3.424E-06 | 6.200E-06 | 1.300E-05 |
| 26204 | 1311 | Q1311D | E | 6.7e-06 | 1.3e-05 | 8.1e-06 | 9.267E-06 | 3.308E-06 | 6.700E-06 | 1.300E-05 |
| 26205 | 1311 | Q1311C | E | 9.8e-06 | 1.4e-05 | 8.4e-06 | 1.073E-05 | 2.914E-06 | 8.400E-06 | 1.400E-05 |
| 26206 | 1311 | Q1311E | E | 1.4e-05 | 7.1e-06 | 8.5e-06 | 9.867E-06 | 3.647E-06 | 7.100E-06 | 1.400E-05 |
| 26207 | 1311 | Q1311Q | E | 5.4e-06 | 9.8e-06 | 5.2e-06 | 6.800E-06 | 2.600E-06 | 5.200E-06 | 9.800E-06 |
| 26208 | 1311 | Q1311G | E | 7e-06 | 8.6e-06 | 4.9e-06 | 6.833E-06 | 1.856E-06 | 4.900E-06 | 8.600E-06 |
| 26209 | 1311 | Q1311H | E | 8.2e-06 | 6e-06 | 6.3e-06 | 6.833E-06 | 1.193E-06 | 6.000E-06 | 8.200E-06 |
| 26210 | 1311 | Q1311I | E | 8e-06 | 6.7e-06 | 1e-05 | 8.233E-06 | 1.662E-06 | 6.700E-06 | 1.000E-05 |
| 26211 | 1311 | Q1311L | E | 8.4e-06 | 5.8e-06 | 5.9e-06 | 6.700E-06 | 1.473E-06 | 5.800E-06 | 8.400E-06 |
| 26212 | 1311 | Q1311K | E | 9e-06 | 6.7e-06 | 8.5e-06 | 8.067E-06 | 1.210E-06 | 6.700E-06 | 9.000E-06 |
| 26213 | 1311 | Q1311M | E | 8.5e-06 | 5.7e-06 | 1.6e-05 | 1.007E-05 | 5.326E-06 | 5.700E-06 | 1.600E-05 |
| 26214 | 1311 | Q1311F | E | 8.4e-06 | 8.9e-06 | 9.1e-06 | 8.800E-06 | 3.606E-07 | 8.400E-06 | 9.100E-06 |
| 26215 | 1311 | Q1311P | E | 9.5e-06 | 1.2e-05 | 7.6e-06 | 9.700E-06 | 2.207E-06 | 7.600E-06 | 1.200E-05 |
| 26216 | 1311 | Q1311S | E | 7.2e-06 | 9.4e-06 | 1.1e-05 | 9.200E-06 | 1.908E-06 | 7.200E-06 | 1.100E-05 |
| 26217 | 1311 | Q1311T | E | 9.1e-06 | 1.1e-05 | 1.1e-05 | 1.037E-05 | 1.097E-06 | 9.100E-06 | 1.100E-05 |
| 26218 | 1311 | Q1311W | E | 9.1e-06 | 1.3e-05 | 7.6e-06 | 9.900E-06 | 2.787E-06 | 7.600E-06 | 1.300E-05 |
| 26219 | 1311 | Q1311Y | E | 5e-06 | 9.2e-06 | 1.1e-05 | 8.400E-06 | 3.079E-06 | 5.000E-06 | 1.100E-05 |
| 26220 | 1311 | Q1311V | E | 9.3e-06 | 8.9e-06 | 5.4e-06 | 7.867E-06 | 2.146E-06 | 5.400E-06 | 9.300E-06 |
| 26221 | 1312 | A1312A | E | 8.5e-06 | 6.7e-06 | 6.5e-06 | 7.233E-06 | 1.102E-06 | 6.500E-06 | 8.500E-06 |
| 26222 | 1312 | A1312R | E | 8e-06 | 6.7e-06 | 1.2e-05 | 8.900E-06 | 2.762E-06 | 6.700E-06 | 1.200E-05 |
| 26223 | 1312 | A1312N | E | 6.3e-06 | 8.8e-06 | 6.9e-06 | 7.333E-06 | 1.305E-06 | 6.300E-06 | 8.800E-06 |
| 26224 | 1312 | A1312D | E | 5.7e-06 | 1.1e-05 | 1.2e-05 | 9.567E-06 | 3.386E-06 | 5.700E-06 | 1.200E-05 |
| 26225 | 1312 | A1312C | E | 4.7e-06 | 5.8e-06 | 9.1e-06 | 6.533E-06 | 2.290E-06 | 4.700E-06 | 9.100E-06 |
| 26226 | 1312 | A1312E | E | 7.2e-06 | 6.8e-06 | 7e-06 | 7.000E-06 | 2.000E-07 | 6.800E-06 | 7.200E-06 |
| 26227 | 1312 | A1312Q | E | 9.2e-06 | 9.3e-06 | 9.7e-06 | 9.400E-06 | 2.646E-07 | 9.200E-06 | 9.700E-06 |
| 26228 | 1312 | A1312G | E | 5.1e-06 | 9.3e-06 | 7.6e-06 | 7.333E-06 | 2.113E-06 | 5.100E-06 | 9.300E-06 |
| 26229 | 1312 | A1312H | E | 6.3e-06 | 8.6e-06 | 1.2e-05 | 8.967E-06 | 2.868E-06 | 6.300E-06 | 1.200E-05 |
| 26230 | 1312 | A1312I | E | 9.9e-06 | 9.5e-06 | 9.4e-06 | 9.600E-06 | 2.646E-07 | 9.400E-06 | 9.900E-06 |
| 26231 | 1312 | A1312L | E | 8.3e-06 | 4.7e-06 | 7.4e-06 | 6.800E-06 | 1.873E-06 | 4.700E-06 | 8.300E-06 |
| 26232 | 1312 | A1312K | E | 8.4e-06 | 6.6e-06 | 9e-06 | 8.000E-06 | 1.249E-06 | 6.600E-06 | 9.000E-06 |
| 26233 | 1312 | A1312M | E | 7.7e-06 | 9.6e-06 | 7.9e-06 | 8.400E-06 | 1.044E-06 | 7.700E-06 | 9.600E-06 |
| 26234 | 1312 | A1312F | E | 7.5e-06 | 8.1e-06 | 5e-06 | 6.867E-06 | 1.644E-06 | 5.000E-06 | 8.100E-06 |
| 26235 | 1312 | A1312P | E | 8.6e-06 | 8.5e-06 | 6.6e-06 | 7.900E-06 | 1.127E-06 | 6.600E-06 | 8.600E-06 |
| 26236 | 1312 | A1312S | E | 6.9e-06 | 6.4e-06 | 1e-05 | 7.767E-06 | 1.950E-06 | 6.400E-06 | 1.000E-05 |
| 26237 | 1312 | A1312T | E | 8e-06 | 8.9e-06 | 6.8e-06 | 7.900E-06 | 1.054E-06 | 6.800E-06 | 8.900E-06 |
| 26238 | 1312 | A1312W | E | 8.8e-06 | 7.2e-06 | 9.5e-06 | 8.500E-06 | 1.179E-06 | 7.200E-06 | 9.500E-06 |
| 26239 | 1312 | A1312Y | E | 4.8e-06 | 8.7e-06 | 7.6e-06 | 7.033E-06 | 2.011E-06 | 4.800E-06 | 8.700E-06 |
| 26240 | 1312 | A1312V | E | 9.9e-06 | 4.8e-06 | 9e-06 | 7.900E-06 | 2.722E-06 | 4.800E-06 | 9.900E-06 |
| 26241 | 1313 | P1313A | E | 5.2e-06 | 9.5e-06 | 1.3e-05 | 9.233E-06 | 3.907E-06 | 5.200E-06 | 1.300E-05 |
| 26242 | 1313 | P1313R | E | 9.2e-06 | 7.4e-06 | 1.5e-05 | 1.053E-05 | 3.972E-06 | 7.400E-06 | 1.500E-05 |
| 26243 | 1313 | P1313N | E | 9.4e-06 | 7.7e-06 | 1.1e-05 | 9.367E-06 | 1.650E-06 | 7.700E-06 | 1.100E-05 |
| 26244 | 1313 | P1313D | E | 7e-06 | 1e-05 | 7.5e-06 | 8.167E-06 | 1.607E-06 | 7.000E-06 | 1.000E-05 |
| 26245 | 1313 | P1313C | E | 1e-05 | 1.3e-05 | 5.5e-06 | 9.500E-06 | 3.775E-06 | 5.500E-06 | 1.300E-05 |
| 26246 | 1313 | P1313E | E | 1.2e-05 | 8.8e-06 | 5.9e-06 | 8.900E-06 | 3.051E-06 | 5.900E-06 | 1.200E-05 |
| 26247 | 1313 | P1313Q | E | 8.2e-06 | 9.1e-06 | 7.7e-06 | 8.333E-06 | 7.095E-07 | 7.700E-06 | 9.100E-06 |
| 26248 | 1313 | P1313G | E | 9.5e-06 | 9.1e-06 | 9.6e-06 | 9.400E-06 | 2.646E-07 | 9.100E-06 | 9.600E-06 |
| 26249 | 1313 | P1313H | E | 1e-05 | 8.6e-06 | 1e-05 | 9.533E-06 | 8.083E-07 | 8.600E-06 | 1.000E-05 |
| 26250 | 1313 | P1313I | E | 6.9e-06 | 9.4e-06 | 1.3e-05 | 9.767E-06 | 3.066E-06 | 6.900E-06 | 1.300E-05 |
| 26251 | 1313 | P1313L | E | 5e-06 | 8.6e-06 | 7.1e-06 | 6.900E-06 | 1.808E-06 | 5.000E-06 | 8.600E-06 |
| 26252 | 1313 | P1313K | E | 4.9e-06 | 6e-06 | 1.9e-05 | 9.967E-06 | 7.842E-06 | 4.900E-06 | 1.900E-05 |
| 26253 | 1313 | P1313M | E | 1.3e-05 | 1e-05 | 5.6e-06 | 9.533E-06 | 3.722E-06 | 5.600E-06 | 1.300E-05 |
| 26254 | 1313 | P1313F | E | 6.3e-06 | 1.1e-05 | 5.7e-06 | 7.667E-06 | 2.902E-06 | 5.700E-06 | 1.100E-05 |
| 26255 | 1313 | P1313P | E | 8.8e-06 | 6.1e-06 | 8.2e-06 | 7.700E-06 | 1.418E-06 | 6.100E-06 | 8.800E-06 |
| 26256 | 1313 | P1313S | E | 9.1e-06 | 7.9e-06 | 7.4e-06 | 8.133E-06 | 8.737E-07 | 7.400E-06 | 9.100E-06 |
| 26257 | 1313 | P1313T | E | 6.6e-06 | 6.7e-06 | 5e-06 | 6.100E-06 | 9.539E-07 | 5.000E-06 | 6.700E-06 |
| 26258 | 1313 | P1313W | E | 9.7e-06 | 7.7e-06 | 8.4e-06 | 8.600E-06 | 1.015E-06 | 7.700E-06 | 9.700E-06 |
| 26259 | 1313 | P1313Y | E | 5e-06 | 6e-06 | 7.8e-06 | 6.267E-06 | 1.419E-06 | 5.000E-06 | 7.800E-06 |
| 26260 | 1313 | P1313V | E | 7.9e-06 | 9.1e-06 | 1.4e-05 | 1.033E-05 | 3.232E-06 | 7.900E-06 | 1.400E-05 |
| 26261 | 1314 | G1314A | E | 8.8e-06 | 5.3e-06 | 1.1e-05 | 8.367E-06 | 2.875E-06 | 5.300E-06 | 1.100E-05 |
| 26262 | 1314 | G1314R | E | 5.5e-06 | 6e-06 | 8.5e-06 | 6.667E-06 | 1.607E-06 | 5.500E-06 | 8.500E-06 |
| 26263 | 1314 | G1314N | E | 6.1e-06 | 8.1e-06 | 6e-06 | 6.733E-06 | 1.185E-06 | 6.000E-06 | 8.100E-06 |
| 26264 | 1314 | G1314D | E | 1.3e-05 | 6.6e-06 | 9.3e-06 | 9.633E-06 | 3.213E-06 | 6.600E-06 | 1.300E-05 |

| | | | | | | | | | | |
|---|---|---|---|---|---|---|---|---|---|---|
| 26265 | 1314 | G1314C | E | 6.3e-06 | 9.5e-06 | 8.8e-06 | 8.200E-06 | 1.682E-06 | 6.300E-06 | 9.500E-06 |
| 26266 | 1314 | G1314E | E | 5.4e-06 | 1.1e-05 | 1.2e-05 | 9.467E-06 | 3.557E-06 | 5.400E-06 | 1.200E-05 |
| 26267 | 1314 | G1314Q | E | 1.6e-05 | 9e-06 | 4.6e-06 | 9.867E-06 | 5.749E-06 | 4.600E-06 | 1.600E-05 |
| 26268 | 1314 | G1314G | E | 7.8e-06 | 1.3e-05 | 7.4e-06 | 9.400E-06 | 3.124E-06 | 7.400E-06 | 1.300E-05 |
| 26269 | 1314 | G1314H | E | 6.2e-06 | 8.8e-06 | 4.3e-06 | 6.433E-06 | 2.259E-06 | 4.300E-06 | 8.800E-06 |
| 26270 | 1314 | G1314I | E | 1.3e-05 | 6.3e-06 | 7.8e-06 | 9.033E-06 | 3.516E-06 | 6.300E-06 | 1.300E-05 |
| 26271 | 1314 | G1314L | E | 8.9e-06 | 8.4e-06 | 9.2e-06 | 8.833E-06 | 4.041E-07 | 8.400E-06 | 9.200E-06 |
| 26272 | 1314 | G1314K | E | 8.1e-06 | 6.7e-06 | 9.4e-06 | 8.067E-06 | 1.350E-06 | 6.700E-06 | 9.400E-06 |
| 26273 | 1314 | G1314M | E | 9.3e-06 | 1e-05 | 1.3e-05 | 1.077E-05 | 1.966E-06 | 9.300E-06 | 1.300E-05 |
| 26274 | 1314 | G1314F | E | 9.8e-06 | 7.5e-06 | 6.1e-06 | 7.800E-06 | 1.868E-06 | 6.100E-06 | 9.800E-06 |
| 26275 | 1314 | G1314P | E | 8.2e-06 | 8.1e-06 | 1.1e-05 | 9.100E-06 | 1.646E-06 | 8.100E-06 | 1.100E-05 |
| 26276 | 1314 | G1314S | E | 9.1e-06 | 6.8e-06 | 8.1e-06 | 8.000E-06 | 1.153E-06 | 6.800E-06 | 9.100E-06 |
| 26277 | 1314 | G1314T | E | 5.4e-06 | 1e-05 | 6.8e-06 | 7.400E-06 | 2.358E-06 | 5.400E-06 | 1.000E-05 |
| 26278 | 1314 | G1314W | E | 9.6e-06 | 3.8e-06 | 8e-06 | 7.133E-06 | 2.996E-06 | 3.800E-06 | 9.600E-06 |
| 26279 | 1314 | G1314Y | E | 6.2e-06 | 7.3e-06 | 1.2e-05 | 8.500E-06 | 3.081E-06 | 6.200E-06 | 1.200E-05 |
| 26280 | 1314 | G1314V | E | 9.4e-06 | 1.1e-05 | 9.4e-06 | 9.933E-06 | 9.238E-07 | 9.400E-06 | 1.100E-05 |
| 26281 | 1315 | K1315A | E | 3.9e-06 | 6.2e-06 | 6.9e-06 | 5.667E-06 | 1.570E-06 | 3.900E-06 | 6.900E-06 |
| 26282 | 1315 | K1315R | E | 1.9e-05 | 9e-06 | 5.3e-06 | 1.110E-05 | 7.087E-06 | 5.300E-06 | 1.900E-05 |
| 26283 | 1315 | K1315N | E | 9.6e-06 | 1.1e-05 | 9.7e-06 | 1.010E-05 | 7.810E-07 | 9.600E-06 | 1.100E-05 |
| 26284 | 1315 | K1315D | E | 1.4e-05 | 7.2e-06 | 7e-06 | 9.400E-06 | 3.985E-06 | 7.000E-06 | 1.400E-05 |
| 26285 | 1315 | K1315C | E | 5.7e-06 | 7.3e-06 | 1.7e-05 | 1.000E-05 | 6.115E-06 | 5.700E-06 | 1.700E-05 |
| 26286 | 1315 | K1315E | E | 9.5e-06 | 1e-05 | 1.1e-05 | 1.017E-05 | 7.638E-07 | 9.500E-06 | 1.100E-05 |
| 26287 | 1315 | K1315Q | E | 1.4e-05 | 1.1e-05 | 9.2e-06 | 1.140E-05 | 2.425E-06 | 9.200E-06 | 1.400E-05 |
| 26288 | 1315 | K1315G | E | 9.3e-06 | 8.4e-06 | 1.2e-05 | 9.900E-06 | 1.873E-06 | 8.400E-06 | 1.200E-05 |
| 26289 | 1315 | K1315H | E | 1.2e-05 | 7.5e-06 | 9.2e-06 | 9.567E-06 | 2.272E-06 | 7.500E-06 | 1.200E-05 |
| 26290 | 1315 | K1315I | E | 8.5e-06 | 7.8e-06 | 1e-05 | 8.767E-06 | 1.124E-06 | 7.800E-06 | 1.000E-05 |
| 26291 | 1315 | K1315L | E | 1.6e-05 | 6.7e-06 | 7.8e-06 | 1.017E-05 | 5.082E-06 | 6.700E-06 | 1.600E-05 |
| 26292 | 1315 | K1315K | E | 4.5e-06 | 4.8e-06 | 5.7e-06 | 5.000E-06 | 6.245E-07 | 4.500E-06 | 5.700E-06 |
| 26293 | 1315 | K1315M | E | 2e-05 | 4.5e-06 | 1.1e-05 | 1.183E-05 | 7.784E-06 | 4.500E-06 | 2.000E-05 |
| 26294 | 1315 | K1315F | E | 1.2e-05 | 6.6e-06 | 1e-05 | 9.533E-06 | 2.730E-06 | 6.600E-06 | 1.200E-05 |
| 26295 | 1315 | K1315P | E | 1.2e-05 | 6.4e-06 | 1.4e-05 | 1.080E-05 | 3.940E-06 | 6.400E-06 | 1.400E-05 |
| 26296 | 1315 | K1315S | E | 7.7e-06 | 7e-06 | 9.2e-06 | 7.967E-06 | 1.124E-06 | 7.000E-06 | 9.200E-06 |
| 26297 | 1315 | K1315T | E | 1.1e-05 | 5.7e-06 | 4.8e-06 | 7.167E-06 | 3.350E-06 | 4.800E-06 | 1.100E-05 |
| 26298 | 1315 | K1315W | E | 7.7e-06 | 6.8e-06 | 4.9e-06 | 6.467E-06 | 1.429E-06 | 4.900E-06 | 7.700E-06 |
| 26299 | 1315 | K1315Y | E | 7.6e-06 | 6.5e-06 | 7.6e-06 | 7.233E-06 | 6.351E-07 | 6.500E-06 | 7.600E-06 |
| 26300 | 1315 | K1315V | E | 3.2e-06 | 7.4e-06 | 4.6e-06 | 5.067E-06 | 2.139E-06 | 3.200E-06 | 7.400E-06 |
| 26301 | 1316 | G1316A | E | 4.5e-06 | 6.8e-06 | 5e-06 | 5.433E-06 | 1.210E-06 | 4.500E-06 | 6.800E-06 |
| 26302 | 1316 | G1316R | E | 8.9e-06 | 5.7e-06 | 6.7e-06 | 7.100E-06 | 1.637E-06 | 5.700E-06 | 8.900E-06 |
| 26303 | 1316 | G1316N | E | 6.3e-06 | 5.6e-06 | 9e-06 | 6.967E-06 | 1.795E-06 | 5.600E-06 | 9.000E-06 |
| 26304 | 1316 | G1316D | E | 1.3e-05 | 9.6e-06 | 8.7e-06 | 1.043E-05 | 2.268E-06 | 8.700E-06 | 1.300E-05 |
| 26305 | 1316 | G1316C | E | 9.6e-06 | 5.5e-06 | 5.4e-06 | 6.833E-06 | 2.397E-06 | 5.400E-06 | 9.600E-06 |
| 26306 | 1316 | G1316E | E | 9e-06 | 5e-06 | 6.6e-06 | 6.867E-06 | 2.013E-06 | 5.000E-06 | 9.000E-06 |
| 26307 | 1316 | G1316Q | E | 8.8e-06 | 5.6e-06 | 7.6e-06 | 7.333E-06 | 1.617E-06 | 5.600E-06 | 8.800E-06 |
| 26308 | 1316 | G1316G | E | 5.6e-06 | 9.6e-06 | 1.1e-05 | 8.733E-06 | 2.802E-06 | 5.600E-06 | 1.100E-05 |
| 26309 | 1316 | G1316H | E | 1.3e-05 | 4.5e-06 | 8.9e-06 | 8.800E-06 | 4.251E-06 | 4.500E-06 | 1.300E-05 |
| 26310 | 1316 | G1316I | E | 9.5e-06 | 7.3e-06 | 1e-05 | 8.933E-06 | 1.436E-06 | 7.300E-06 | 1.000E-05 |
| 26311 | 1316 | G1316L | E | 4.7e-06 | 7.8e-06 | 6.1e-06 | 6.200E-06 | 1.552E-06 | 4.700E-06 | 7.800E-06 |
| 26312 | 1316 | G1316K | E | 7.6e-06 | 1.2e-05 | 6.2e-06 | 8.600E-06 | 3.027E-06 | 6.200E-06 | 1.200E-05 |
| 26313 | 1316 | G1316M | E | 4.9e-06 | 1.2e-05 | 5e-06 | 7.300E-06 | 4.071E-06 | 4.900E-06 | 1.200E-05 |
| 26314 | 1316 | G1316F | E | 1.1e-05 | 4.8e-06 | 6.5e-06 | 7.433E-06 | 3.204E-06 | 4.800E-06 | 1.100E-05 |
| 26315 | 1316 | G1316P | E | 9e-06 | 1e-05 | 8.6e-06 | 9.200E-06 | 7.211E-07 | 8.600E-06 | 1.000E-05 |
| 26316 | 1316 | G1316S | E | 9.7e-06 | 9.3e-06 | 9.4e-06 | 9.467E-06 | 2.082E-07 | 9.300E-06 | 9.700E-06 |
| 26317 | 1316 | G1316T | E | 9.9e-06 | 6.1e-06 | 9.8e-06 | 8.600E-06 | 2.166E-06 | 6.100E-06 | 9.900E-06 |
| 26318 | 1316 | G1316W | E | 9.9e-06 | 1e-05 | 1.2e-05 | 1.063E-05 | 1.185E-06 | 9.900E-06 | 1.200E-05 |
| 26319 | 1316 | G1316Y | E | 8e-06 | 4.9e-06 | 5.7e-06 | 6.200E-06 | 1.609E-06 | 4.900E-06 | 8.000E-06 |
| 26320 | 1316 | G1316V | E | 7.8e-06 | 1.3e-05 | 6.2e-06 | 9.000E-06 | 3.555E-06 | 6.200E-06 | 1.300E-05 |
| 26321 | 1317 | L1317A | E | 9.5e-06 | 6e-06 | 1.1e-05 | 8.833E-06 | 2.566E-06 | 6.000E-06 | 1.100E-05 |
| 26322 | 1317 | L1317R | E | 1.1e-05 | 7.5e-06 | 3.7e-06 | 7.400E-06 | 3.651E-06 | 3.700E-06 | 1.100E-05 |
| 26323 | 1317 | L1317N | E | 1.2e-05 | 6.6e-06 | 5.8e-06 | 8.133E-06 | 3.372E-06 | 5.800E-06 | 1.200E-05 |
| 26324 | 1317 | L1317D | E | 5.8e-06 | 7.5e-06 | 6e-06 | 6.433E-06 | 9.292E-07 | 5.800E-06 | 7.500E-06 |
| 26325 | 1317 | L1317C | E | 1.2e-05 | 5.8e-06 | 5.9e-06 | 7.900E-06 | 3.551E-06 | 5.800E-06 | 1.200E-05 |
| 26326 | 1317 | L1317E | E | 1.2e-05 | 5.7e-06 | 4.6e-06 | 7.433E-06 | 3.993E-06 | 4.600E-06 | 1.200E-05 |
| 26327 | 1317 | L1317Q | E | 1.4e-05 | 7.6e-06 | 5.4e-06 | 9.000E-06 | 4.468E-06 | 5.400E-06 | 1.400E-05 |
| 26328 | 1317 | L1317G | E | 6.9e-06 | 6.2e-06 | 4e-06 | 5.700E-06 | 1.513E-06 | 4.000E-06 | 6.900E-06 |
| 26329 | 1317 | L1317H | E | 6.1e-06 | 9.3e-06 | 5e-06 | 6.800E-06 | 2.234E-06 | 5.000E-06 | 9.300E-06 |
| 26330 | 1317 | L1317I | E | 9.8e-06 | 1.5e-05 | 9e-06 | 1.127E-05 | 3.258E-06 | 9.000E-06 | 1.500E-05 |
| 26331 | 1317 | L1317L | E | 1.2e-05 | 8.4e-06 | 1.1e-05 | 1.047E-05 | 1.858E-06 | 8.400E-06 | 1.200E-05 |
| 26332 | 1317 | L1317K | E | 7.4e-06 | 1.5e-05 | 7.7e-06 | 1.003E-05 | 4.304E-06 | 7.400E-06 | 1.500E-05 |
| 26333 | 1317 | L1317M | E | 8.1e-06 | 5.2e-06 | 4.3e-06 | 5.867E-06 | 1.986E-06 | 4.300E-06 | 8.100E-06 |
| 26334 | 1317 | L1317F | E | 9.3e-06 | 7.7e-06 | 1.1e-05 | 9.333E-06 | 1.650E-06 | 7.700E-06 | 1.100E-05 |
| 26335 | 1317 | L1317P | E | 1.4e-05 | 6.7e-06 | 8.9e-06 | 9.867E-06 | 3.745E-06 | 6.700E-06 | 1.400E-05 |
| 26336 | 1317 | L1317S | E | 8.9e-06 | 8.3e-06 | 8.9e-06 | 8.700E-06 | 3.464E-07 | 8.300E-06 | 8.900E-06 |
| 26337 | 1317 | L1317T | E | 6.3e-06 | 1.3e-05 | 1.1e-05 | 1.010E-05 | 3.439E-06 | 6.300E-06 | 1.300E-05 |
| 26338 | 1317 | L1317W | E | 9.2e-06 | 6.6e-06 | 8.1e-06 | 7.967E-06 | 1.305E-06 | 6.600E-06 | 9.200E-06 |
| 26339 | 1317 | L1317Y | E | 1.3e-05 | 6e-06 | 8.1e-06 | 9.033E-06 | 3.592E-06 | 6.000E-06 | 1.300E-05 |
| 26340 | 1317 | L1317V | E | 1.2e-05 | 7.7e-06 | 8.4e-06 | 9.367E-06 | 2.307E-06 | 7.700E-06 | 1.200E-05 |
| 26341 | 1318 | E1318A | E | 7.2e-06 | 4.4e-06 | 6.2e-06 | 5.933E-06 | 1.419E-06 | 4.400E-06 | 7.200E-06 |
| 26342 | 1318 | E1318R | E | 6e-06 | 4.5e-06 | 8.1e-06 | 6.200E-06 | 1.808E-06 | 4.500E-06 | 8.100E-06 |
| 26343 | 1318 | E1318N | E | 1.1e-05 | 9.2e-06 | 4.4e-06 | 8.200E-06 | 3.412E-06 | 4.400E-06 | 1.100E-05 |
| 26344 | 1318 | E1318D | E | 7.5e-06 | 9.5e-06 | 1.1e-05 | 9.333E-06 | 1.756E-06 | 7.500E-06 | 1.100E-05 |
| 26345 | 1318 | E1318C | E | 8.5e-06 | 8.3e-06 | 1.1e-05 | 9.267E-06 | 1.504E-06 | 8.300E-06 | 1.100E-05 |
| 26346 | 1318 | E1318E | E | 7.6e-06 | 9.7e-06 | 5.2e-06 | 7.500E-06 | 2.252E-06 | 5.200E-06 | 9.700E-06 |
| 26347 | 1318 | E1318Q | E | 8.1e-06 | 7.5e-06 | 8e-06 | 7.867E-06 | 3.215E-07 | 7.500E-06 | 8.100E-06 |
| 26348 | 1318 | E1318G | E | 8.2e-06 | 9.2e-06 | 1e-05 | 9.133E-06 | 9.018E-07 | 8.200E-06 | 1.000E-05 |
| 26349 | 1318 | E1318H | E | 6.2e-06 | 1.2e-05 | 5.5e-06 | 7.900E-06 | 3.568E-06 | 5.500E-06 | 1.200E-05 |

| | | | | | | | | | | |
|---|---|---|---|---|---|---|---|---|---|---|
| 26350 | 1318 | E1318I | E | 5.2e-06 | 1.7e-05 | 1e-05 | 1.073E-05 | 5.934E-06 | 5.200E-06 | 1.700E-05 |
| 26351 | 1318 | E1318L | E | 1e-05 | 5e-06 | 6.5e-06 | 7.167E-06 | 2.566E-06 | 5.000E-06 | 1.000E-05 |
| 26352 | 1318 | E1318K | E | 1.1e-05 | 5.4e-06 | 7.6e-06 | 8.000E-06 | 2.821E-06 | 5.400E-06 | 1.100E-05 |
| 26353 | 1318 | E1318M | E | 1.2e-05 | 5.1e-06 | 6.8e-06 | 7.967E-06 | 3.595E-06 | 5.100E-06 | 1.200E-05 |
| 26354 | 1318 | E1318F | E | 5.5e-06 | 6.8e-06 | 9.3e-06 | 7.200E-06 | 1.931E-06 | 5.500E-06 | 9.300E-06 |
| 26355 | 1318 | E1318P | E | 4.7e-06 | 5.1e-06 | 6.5e-06 | 5.433E-06 | 9.452E-07 | 4.700E-06 | 6.500E-06 |
| 26356 | 1318 | E1318S | E | 1.1e-05 | 9.1e-06 | 5.8e-06 | 8.633E-06 | 2.631E-06 | 5.800E-06 | 1.100E-05 |
| 26357 | 1318 | E1318T | E | 6.1e-06 | 7.5e-06 | 6e-06 | 6.533E-06 | 8.386E-07 | 6.000E-06 | 7.500E-06 |
| 26358 | 1318 | E1318W | E | 8.2e-06 | 9.6e-06 | 7.5e-06 | 8.433E-06 | 1.069E-06 | 7.500E-06 | 9.600E-06 |
| 26359 | 1318 | E1318Y | E | 6.9e-06 | 5.6e-06 | 8.3e-06 | 6.933E-06 | 1.350E-06 | 5.600E-06 | 8.300E-06 |
| 26360 | 1318 | E1318V | E | 6.8e-06 | 4.8e-06 | 8.8e-06 | 6.800E-06 | 2.000E-06 | 4.800E-06 | 8.800E-06 |
| 26361 | 1319 | W1319A | E | 1.2e-05 | 8.7e-06 | 6.7e-06 | 9.133E-06 | 2.676E-06 | 6.700E-06 | 1.200E-05 |
| 26362 | 1319 | W1319R | E | 9.1e-06 | 8.5e-06 | 1.1e-05 | 9.533E-06 | 1.305E-06 | 8.500E-06 | 1.100E-05 |
| 26363 | 1319 | W1319N | E | 7.5e-06 | 1e-05 | 7.6e-06 | 8.367E-06 | 1.415E-06 | 7.500E-06 | 1.000E-05 |
| 26364 | 1319 | W1319D | E | 1.2e-05 | 7.2e-06 | 9.2e-06 | 9.467E-06 | 2.411E-06 | 7.200E-06 | 1.200E-05 |
| 26365 | 1319 | W1319C | E | 4.8e-06 | 3.8e-06 | 3.9e-06 | 4.167E-06 | 5.508E-07 | 3.800E-06 | 4.800E-06 |
| 26366 | 1319 | W1319E | E | 1.2e-05 | 6.9e-06 | 5.7e-06 | 8.200E-06 | 3.345E-06 | 5.700E-06 | 1.200E-05 |
| 26367 | 1319 | W1319Q | E | 5.5e-06 | 4.4e-06 | 1.1e-05 | 6.967E-06 | 3.536E-06 | 4.400E-06 | 1.100E-05 |
| 26368 | 1319 | W1319G | E | 4.6e-06 | 1.3e-05 | 6.8e-06 | 8.133E-06 | 4.356E-06 | 4.600E-06 | 1.300E-05 |
| 26369 | 1319 | W1319H | E | 8.8e-06 | 4.4e-06 | 7.8e-06 | 7.000E-06 | 2.307E-06 | 4.400E-06 | 8.800E-06 |
| 26370 | 1319 | W1319I | E | 1e-05 | 1.3e-05 | 1.4e-05 | 1.233E-05 | 2.082E-06 | 1.000E-05 | 1.400E-05 |
| 26371 | 1319 | W1319L | E | 9.3e-06 | 4.6e-06 | 8.3e-06 | 7.400E-06 | 2.476E-06 | 4.600E-06 | 9.300E-06 |
| 26372 | 1319 | W1319K | E | 6.5e-06 | 1.2e-05 | 1.1e-05 | 9.833E-06 | 2.930E-06 | 6.500E-06 | 1.200E-05 |
| 26373 | 1319 | W1319M | E | 5.8e-06 | 1.3e-05 | 1.4e-05 | 1.093E-05 | 4.474E-06 | 5.800E-06 | 1.400E-05 |
| 26374 | 1319 | W1319F | E | 7.3e-06 | 1.1e-05 | 9.7e-06 | 9.333E-06 | 1.877E-06 | 7.300E-06 | 1.100E-05 |
| 26375 | 1319 | W1319P | E | 4.6e-06 | 1e-05 | 8.9e-06 | 7.833E-06 | 2.854E-06 | 4.600E-06 | 1.000E-05 |
| 26376 | 1319 | W1319S | E | 8.8e-06 | 1e-05 | 9.4e-06 | 9.400E-06 | 6.000E-07 | 8.800E-06 | 1.000E-05 |
| 26377 | 1319 | W1319T | E | 7e-06 | 6.6e-06 | 9.5e-06 | 7.700E-06 | 1.572E-06 | 6.600E-06 | 9.500E-06 |
| 26378 | 1319 | W1319W | E | 5e-06 | 8e-06 | 7.3e-06 | 6.767E-06 | 1.570E-06 | 5.000E-06 | 8.000E-06 |
| 26379 | 1319 | W1319Y | E | 1.4e-05 | 6.1e-06 | 7.3e-06 | 9.133E-06 | 4.257E-06 | 6.100E-06 | 1.400E-05 |
| 26380 | 1319 | W1319V | E | 1e-05 | 1.4e-05 | 1.2e-05 | 1.200E-05 | 2.000E-06 | 1.000E-05 | 1.400E-05 |
| 26381 | 1320 | V1320A | E | 7.1e-06 | 5.1e-06 | 7e-06 | 6.400E-06 | 1.127E-06 | 5.100E-06 | 7.100E-06 |
| 26382 | 1320 | V1320R | E | 1.4e-05 | 9.3e-06 | 6.4e-06 | 9.900E-06 | 3.835E-06 | 6.400E-06 | 1.400E-05 |
| 26383 | 1320 | V1320N | E | 8.7e-06 | 1e-05 | 6.2e-06 | 8.300E-06 | 1.931E-06 | 6.200E-06 | 1.000E-05 |
| 26384 | 1320 | V1320D | E | 1.4e-05 | 7.2e-06 | 8.6e-06 | 9.933E-06 | 3.591E-06 | 7.200E-06 | 1.400E-05 |
| 26385 | 1320 | V1320C | E | 7.1e-06 | 6.2e-06 | 4.7e-06 | 6.000E-06 | 1.212E-06 | 4.700E-06 | 7.100E-06 |
| 26386 | 1320 | V1320E | E | 7.1e-06 | 6.3e-06 | 7.4e-06 | 6.933E-06 | 5.686E-07 | 6.300E-06 | 7.400E-06 |
| 26387 | 1320 | V1320Q | E | 9.5e-06 | 9.9e-06 | 1.7e-05 | 1.213E-05 | 4.219E-06 | 9.500E-06 | 1.700E-05 |
| 26388 | 1320 | V1320G | E | 4.9e-06 | 1.3e-05 | 8.4e-06 | 8.767E-06 | 4.062E-06 | 4.900E-06 | 1.300E-05 |
| 26389 | 1320 | V1320H | E | 1.1e-05 | 6e-06 | 8.7e-06 | 8.567E-06 | 2.503E-06 | 6.000E-06 | 1.100E-05 |
| 26390 | 1320 | V1320I | E | 1.3e-05 | 1.3e-05 | 5.7e-06 | 1.057E-05 | 4.215E-06 | 5.700E-06 | 1.300E-05 |
| 26391 | 1320 | V1320L | E | 1.3e-05 | 1.2e-05 | 6.4e-06 | 1.047E-05 | 3.557E-06 | 6.400E-06 | 1.300E-05 |
| 26392 | 1320 | V1320K | E | 1.2e-05 | 8.3e-06 | 6.3e-06 | 8.867E-06 | 2.892E-06 | 6.300E-06 | 1.200E-05 |
| 26393 | 1320 | V1320M | E | 8.3e-06 | 5.4e-06 | 8.4e-06 | 7.367E-06 | 1.704E-06 | 5.400E-06 | 8.400E-06 |
| 26394 | 1320 | V1320F | E | 1.2e-05 | 7.2e-06 | 5.6e-06 | 8.267E-06 | 3.331E-06 | 5.600E-06 | 1.200E-05 |
| 26395 | 1320 | V1320P | E | 9.4e-06 | 5e-06 | 5.5e-06 | 6.633E-06 | 2.409E-06 | 5.000E-06 | 9.400E-06 |
| 26396 | 1320 | V1320S | E | 1.1e-05 | 6.6e-06 | 9.1e-06 | 8.900E-06 | 2.207E-06 | 6.600E-06 | 1.100E-05 |
| 26397 | 1320 | V1320T | E | 1.2e-05 | 8.6e-06 | 1.2e-05 | 1.087E-05 | 1.963E-06 | 8.600E-06 | 1.200E-05 |
| 26398 | 1320 | V1320W | E | 6.5e-06 | 1.2e-05 | 8.5e-06 | 9.000E-06 | 2.784E-06 | 6.500E-06 | 1.200E-05 |
| 26399 | 1320 | V1320Y | E | 8.1e-06 | 5.6e-06 | 9.8e-06 | 7.833E-06 | 2.113E-06 | 5.600E-06 | 9.800E-06 |
| 26400 | 1320 | V1320V | E | 1.1e-05 | 8.8e-06 | 5.4e-06 | 8.400E-06 | 2.821E-06 | 5.400E-06 | 1.100E-05 |
| 26401 | 1321 | A1321A | E | 4.6e-06 | 4.5e-06 | 9.6e-06 | 6.233E-06 | 2.916E-06 | 4.500E-06 | 9.600E-06 |
| 26402 | 1321 | A1321R | E | 8.4e-06 | 1.3e-05 | 5.9e-06 | 9.100E-06 | 3.601E-06 | 5.900E-06 | 1.300E-05 |
| 26403 | 1321 | A1321N | E | 5.4e-06 | 5.9e-06 | 8.9e-06 | 6.733E-06 | 1.893E-06 | 5.400E-06 | 8.900E-06 |
| 26404 | 1321 | A1321D | E | 8.2e-06 | 8.6e-06 | 9.1e-06 | 8.633E-06 | 4.509E-07 | 8.200E-06 | 9.100E-06 |
| 26405 | 1321 | A1321C | E | 9.2e-06 | 4.1e-06 | 1.4e-05 | 9.100E-06 | 4.951E-06 | 4.100E-06 | 1.400E-05 |
| 26406 | 1321 | A1321E | E | 7.2e-06 | 9.2e-06 | 4.8e-06 | 7.067E-06 | 2.203E-06 | 4.800E-06 | 9.200E-06 |
| 26407 | 1321 | A1321Q | E | 7.7e-06 | 4.9e-06 | 8.3e-06 | 6.967E-06 | 1.815E-06 | 4.900E-06 | 8.300E-06 |
| 26408 | 1321 | A1321G | E | 6.7e-06 | 1.3e-05 | 5.9e-06 | 8.533E-06 | 3.889E-06 | 5.900E-06 | 1.300E-05 |
| 26409 | 1321 | A1321H | E | 9.9e-06 | 5.2e-06 | 1.3e-05 | 9.367E-06 | 3.927E-06 | 5.200E-06 | 1.300E-05 |
| 26410 | 1321 | A1321I | E | 6.3e-06 | 8.1e-06 | 9.3e-06 | 7.900E-06 | 1.510E-06 | 6.300E-06 | 9.300E-06 |
| 26411 | 1321 | A1321L | E | 4.9e-06 | 9e-06 | 5.5e-06 | 6.467E-06 | 2.214E-06 | 4.900E-06 | 9.000E-06 |
| 26412 | 1321 | A1321K | E | 6.6e-06 | 7.4e-06 | 1e-05 | 8.000E-06 | 1.778E-06 | 6.600E-06 | 1.000E-05 |
| 26413 | 1321 | A1321M | E | 8.8e-06 | 1.1e-05 | 6.7e-06 | 8.833E-06 | 2.150E-06 | 6.700E-06 | 1.100E-05 |
| 26414 | 1321 | A1321F | E | 1.1e-05 | 1.2e-05 | 1.1e-05 | 1.133E-05 | 5.774E-07 | 1.100E-05 | 1.200E-05 |
| 26415 | 1321 | A1321P | E | 9.1e-06 | 7.5e-06 | 6.1e-06 | 7.567E-06 | 1.501E-06 | 6.100E-06 | 9.100E-06 |
| 26416 | 1321 | A1321S | E | 9.5e-06 | 7.7e-06 | 7.9e-06 | 8.367E-06 | 9.866E-07 | 7.700E-06 | 9.500E-06 |
| 26417 | 1321 | A1321T | E | 5.6e-06 | 3.6e-06 | 1.9e-05 | 9.400E-06 | 8.374E-06 | 3.600E-06 | 1.900E-05 |
| 26418 | 1321 | A1321W | E | 4.8e-06 | 3.4e-06 | 8.5e-06 | 5.567E-06 | 2.635E-06 | 3.400E-06 | 8.500E-06 |
| 26419 | 1321 | A1321Y | E | 8.6e-06 | 5.3e-06 | 5.2e-06 | 6.367E-06 | 1.935E-06 | 5.200E-06 | 8.600E-06 |
| 26420 | 1321 | A1321V | E | 5.2e-06 | 9.7e-06 | 1.3e-05 | 9.300E-06 | 3.915E-06 | 5.200E-06 | 1.300E-05 |
| 26421 | 1322 | R1322A | E | 7.3e-06 | 1.8e-05 | 1.4e-05 | 1.310E-05 | 5.406E-06 | 7.300E-06 | 1.800E-05 |
| 26422 | 1322 | R1322R | E | 4.8e-06 | 5.6e-06 | 6e-06 | 5.467E-06 | 6.110E-07 | 4.800E-06 | 6.000E-06 |
| 26423 | 1322 | R1322N | E | 6.9e-06 | 5.2e-06 | 9.1e-06 | 7.067E-06 | 1.955E-06 | 5.200E-06 | 9.100E-06 |
| 26424 | 1322 | R1322D | E | 4.1e-06 | 7e-06 | 6.1e-06 | 5.733E-06 | 1.484E-06 | 4.100E-06 | 7.000E-06 |
| 26425 | 1322 | R1322C | E | 1.2e-05 | 1.1e-05 | 7.2e-06 | 1.007E-05 | 2.532E-06 | 7.200E-06 | 1.200E-05 |
| 26426 | 1322 | R1322E | E | 7.8e-06 | 8.8e-06 | 7.1e-06 | 7.900E-06 | 8.544E-07 | 7.100E-06 | 8.800E-06 |
| 26427 | 1322 | R1322Q | E | 4e-06 | 1.4e-05 | 5.9e-06 | 7.967E-06 | 5.311E-06 | 4.000E-06 | 1.400E-05 |
| 26428 | 1322 | R1322G | E | 7.9e-06 | 4.4e-06 | 5.4e-06 | 5.900E-06 | 1.803E-06 | 4.400E-06 | 7.900E-06 |
| 26429 | 1322 | R1322H | E | 6.7e-06 | 3.2e-06 | 6.6e-06 | 5.500E-06 | 1.992E-06 | 3.200E-06 | 6.700E-06 |
| 26430 | 1322 | R1322I | E | 7.2e-06 | 5.1e-06 | 1.2e-05 | 8.100E-06 | 3.537E-06 | 5.100E-06 | 1.200E-05 |
| 26431 | 1322 | R1322L | E | 9.5e-06 | 5.6e-06 | 4.8e-06 | 6.633E-06 | 2.515E-06 | 4.800E-06 | 9.500E-06 |
| 26432 | 1322 | R1322K | E | 9e-06 | 1.3e-05 | 9.1e-06 | 1.037E-05 | 2.281E-06 | 9.000E-06 | 1.300E-05 |
| 26433 | 1322 | R1322M | E | 9.7e-06 | 7.8e-06 | 9.9e-06 | 9.133E-06 | 1.159E-06 | 7.800E-06 | 9.900E-06 |
| 26434 | 1322 | R1322F | E | 1.1e-05 | 6.6e-06 | 9.2e-06 | 8.933E-06 | 2.212E-06 | 6.600E-06 | 1.100E-05 |

| | | | | | | | | | | |
|---|---|---|---|---|---|---|---|---|---|---|
| 26435 | 1322 | R1322P | E | 8.6e-06 | 1.2e-05 | 6.1e-06 | 8.900E-06 | 2.961E-06 | 6.100E-06 | 1.200E-05 |
| 26436 | 1322 | R1322S | E | 5.4e-06 | 8.4e-06 | 1.3e-05 | 8.933E-06 | 3.828E-06 | 5.400E-06 | 1.300E-05 |
| 26437 | 1322 | R1322T | E | 8.6e-06 | 4.4e-06 | 6.6e-06 | 6.533E-06 | 2.101E-06 | 4.400E-06 | 8.600E-06 |
| 26438 | 1322 | R1322W | E | 6.5e-06 | 4.7e-06 | 7.1e-06 | 6.100E-06 | 1.249E-06 | 4.700E-06 | 7.100E-06 |
| 26439 | 1322 | R1322Y | E | 8.3e-06 | 9.1e-06 | 9.2e-06 | 8.867E-06 | 4.933E-07 | 8.300E-06 | 9.200E-06 |
| 26440 | 1322 | R1322V | E | 6.2e-06 | 8.9e-06 | 1.2e-05 | 9.033E-06 | 2.902E-06 | 6.200E-06 | 1.200E-05 |
| 26441 | 1323 | I1323A | E | 7.3e-06 | 5.7e-06 | 8.7e-06 | 7.233E-06 | 1.501E-06 | 5.700E-06 | 8.700E-06 |
| 26442 | 1323 | I1323R | E | 1.2e-05 | 4.9e-06 | 7e-06 | 7.967E-06 | 3.647E-06 | 4.900E-06 | 1.200E-05 |
| 26443 | 1323 | I1323N | E | 6.7e-06 | 9.1e-06 | 1.1e-05 | 8.933E-06 | 2.155E-06 | 6.700E-06 | 1.100E-05 |
| 26444 | 1323 | I1323D | E | 4.6e-06 | 1.1e-05 | 1.6e-05 | 1.053E-05 | 5.714E-06 | 4.600E-06 | 1.600E-05 |
| 26445 | 1323 | I1323C | E | 5.4e-06 | 8.5e-06 | 5.1e-06 | 6.333E-06 | 1.882E-06 | 5.100E-06 | 8.500E-06 |
| 26446 | 1323 | I1323E | E | 9.4e-06 | 1.4e-05 | 1.3e-05 | 1.213E-05 | 2.419E-06 | 9.400E-06 | 1.400E-05 |
| 26447 | 1323 | I1323Q | E | 6.3e-06 | 7.3e-06 | 4.3e-06 | 5.967E-06 | 1.528E-06 | 4.300E-06 | 7.300E-06 |
| 26448 | 1323 | I1323G | E | 1.4e-05 | 7.4e-06 | 7.1e-06 | 9.500E-06 | 3.900E-06 | 7.100E-06 | 1.400E-05 |
| 26449 | 1323 | I1323H | E | 9.7e-06 | 4.6e-06 | 4.8e-06 | 6.367E-06 | 2.888E-06 | 4.600E-06 | 9.700E-06 |
| 26450 | 1323 | I1323I | E | 9.4e-06 | 9.7e-06 | 5.9e-06 | 8.333E-06 | 2.113E-06 | 5.900E-06 | 9.700E-06 |
| 26451 | 1323 | I1323L | E | 9.1e-06 | 1e-05 | 5.3e-06 | 8.133E-06 | 2.495E-06 | 5.300E-06 | 1.000E-05 |
| 26452 | 1323 | I1323K | E | 7.5e-06 | 1.1e-05 | 1.5e-05 | 1.117E-05 | 3.753E-06 | 7.500E-06 | 1.500E-05 |
| 26453 | 1323 | I1323M | E | 5.5e-06 | 4.3e-06 | 6.9e-06 | 5.567E-06 | 1.301E-06 | 4.300E-06 | 6.900E-06 |
| 26454 | 1323 | I1323F | E | 4.7e-06 | 6.6e-06 | 1.1e-05 | 7.433E-06 | 3.232E-06 | 4.700E-06 | 1.100E-05 |
| 26455 | 1323 | I1323P | E | 1e-05 | 4.9e-06 | 9.1e-06 | 8.000E-06 | 2.722E-06 | 4.900E-06 | 1.000E-05 |
| 26456 | 1323 | I1323S | E | 5.1e-06 | 9.3e-06 | 6.3e-06 | 6.900E-06 | 2.163E-06 | 5.100E-06 | 9.300E-06 |
| 26457 | 1323 | I1323T | E | 7.1e-06 | 1e-05 | 1.3e-05 | 1.003E-05 | 2.950E-06 | 7.100E-06 | 1.300E-05 |
| 26458 | 1323 | I1323W | E | 9.8e-06 | 8.6e-06 | 6.1e-06 | 8.167E-06 | 1.888E-06 | 6.100E-06 | 9.800E-06 |
| 26459 | 1323 | I1323Y | E | 6.9e-06 | 8.6e-06 | 6.2e-06 | 7.233E-06 | 1.234E-06 | 6.200E-06 | 8.600E-06 |
| 26460 | 1323 | I1323V | E | 1.3e-05 | 1.3e-05 | 5e-06 | 1.033E-05 | 4.619E-06 | 5.000E-06 | 1.300E-05 |
| 26461 | 1324 | Y1324A | E | 6e-06 | 9.7e-06 | 6.7e-06 | 7.467E-06 | 1.966E-06 | 6.000E-06 | 9.700E-06 |
| 26462 | 1324 | Y1324R | E | 5.5e-06 | 8.7e-06 | 6.4e-06 | 6.867E-06 | 1.650E-06 | 5.500E-06 | 8.700E-06 |
| 26463 | 1324 | Y1324N | E | 6.1e-06 | 1.3e-05 | 1e-05 | 9.700E-06 | 3.460E-06 | 6.100E-06 | 1.300E-05 |
| 26464 | 1324 | Y1324D | E | 9.2e-06 | 9.3e-06 | 1.4e-05 | 1.083E-05 | 2.743E-06 | 9.200E-06 | 1.400E-05 |
| 26465 | 1324 | Y1324C | E | 1.3e-05 | 1e-05 | 6.8e-06 | 9.933E-06 | 3.101E-06 | 6.800E-06 | 1.300E-05 |
| 26466 | 1324 | Y1324E | E | 4.1e-06 | 7e-06 | 9.5e-06 | 6.867E-06 | 2.702E-06 | 4.100E-06 | 9.500E-06 |
| 26467 | 1324 | Y1324Q | E | 6.3e-06 | 1.2e-05 | 1.1e-05 | 9.767E-06 | 3.044E-06 | 6.300E-06 | 1.200E-05 |
| 26468 | 1324 | Y1324G | E | 6.6e-06 | 6.6e-06 | 6.9e-06 | 6.700E-06 | 1.732E-07 | 6.600E-06 | 6.900E-06 |
| 26469 | 1324 | Y1324H | E | 9.6e-06 | 1.3e-05 | 6.6e-06 | 9.733E-06 | 3.202E-06 | 6.600E-06 | 1.300E-05 |
| 26470 | 1324 | Y1324I | E | 8.5e-06 | 1.2e-05 | 1.1e-05 | 1.050E-05 | 1.803E-06 | 8.500E-06 | 1.200E-05 |
| 26471 | 1324 | Y1324L | E | 8.7e-06 | 4.7e-06 | 5.1e-06 | 6.167E-06 | 2.203E-06 | 4.700E-06 | 8.700E-06 |
| 26472 | 1324 | Y1324K | E | 7.4e-06 | 9.9e-06 | 5e-06 | 7.433E-06 | 2.450E-06 | 5.000E-06 | 9.900E-06 |
| 26473 | 1324 | Y1324M | E | 1.2e-05 | 7.3e-06 | 5.9e-06 | 8.400E-06 | 3.195E-06 | 5.900E-06 | 1.200E-05 |
| 26474 | 1324 | Y1324F | E | 1.5e-05 | 1.1e-05 | 1.3e-05 | 1.300E-05 | 2.000E-06 | 1.100E-05 | 1.500E-05 |
| 26475 | 1324 | Y1324P | E | 6.3e-06 | 5.1e-06 | 6e-06 | 5.800E-06 | 6.245E-07 | 5.100E-06 | 6.300E-06 |
| 26476 | 1324 | Y1324S | E | 5.1e-06 | 7e-06 | 6.6e-06 | 6.233E-06 | 1.002E-06 | 5.100E-06 | 7.000E-06 |
| 26477 | 1324 | Y1324T | E | 1e-05 | 6.8e-06 | 9.4e-06 | 8.733E-06 | 1.701E-06 | 6.800E-06 | 1.000E-05 |
| 26478 | 1324 | Y1324W | E | 1.1e-05 | 4e-06 | 9.3e-06 | 8.100E-06 | 3.651E-06 | 4.000E-06 | 1.100E-05 |
| 26479 | 1324 | Y1324Y | E | 5.2e-06 | 8.8e-06 | 8.6e-06 | 7.533E-06 | 2.023E-06 | 5.200E-06 | 8.800E-06 |
| 26480 | 1324 | Y1324V | E | 4.2e-06 | 6.2e-06 | 9.6e-06 | 6.667E-06 | 2.730E-06 | 4.200E-06 | 9.600E-06 |
| 26481 | 1325 | P1325A | E | 8.9e-06 | 1e-05 | 1e-05 | 9.633E-06 | 6.351E-07 | 8.900E-06 | 1.000E-05 |
| 26482 | 1325 | P1325R | E | 7.1e-06 | 8e-06 | 9.5e-06 | 8.200E-06 | 1.212E-06 | 7.100E-06 | 9.500E-06 |
| 26483 | 1325 | P1325N | E | 8.2e-06 | 4.4e-06 | 1.2e-05 | 8.200E-06 | 3.800E-06 | 4.400E-06 | 1.200E-05 |
| 26484 | 1325 | P1325D | E | 6.5e-06 | 5.9e-06 | 6.2e-06 | 6.200E-06 | 3.000E-07 | 5.900E-06 | 6.500E-06 |
| 26485 | 1325 | P1325C | E | 9e-06 | 6.4e-06 | 6.2e-06 | 7.200E-06 | 1.562E-06 | 6.200E-06 | 9.000E-06 |
| 26486 | 1325 | P1325E | E | 5.3e-06 | 9.8e-06 | 6.9e-06 | 7.333E-06 | 2.281E-06 | 5.300E-06 | 9.800E-06 |
| 26487 | 1325 | P1325Q | E | 8.3e-06 | 9.3e-06 | 7.9e-06 | 8.500E-06 | 7.211E-07 | 7.900E-06 | 9.300E-06 |
| 26488 | 1325 | P1325G | E | 6e-06 | 1.8e-05 | 7.3e-06 | 1.043E-05 | 6.585E-06 | 6.000E-06 | 1.800E-05 |
| 26489 | 1325 | P1325H | E | 8.6e-06 | 5.4e-06 | 8.6e-06 | 7.533E-06 | 1.848E-06 | 5.400E-06 | 8.600E-06 |
| 26490 | 1325 | P1325I | E | 8.1e-06 | 7.2e-06 | 9.9e-06 | 8.400E-06 | 1.375E-06 | 7.200E-06 | 9.900E-06 |
| 26491 | 1325 | P1325L | E | 1.2e-05 | 8.9e-06 | 6.2e-06 | 9.033E-06 | 2.902E-06 | 6.200E-06 | 1.200E-05 |
| 26492 | 1325 | P1325K | E | 5.8e-06 | 6.1e-06 | 7e-06 | 6.300E-06 | 6.245E-07 | 5.800E-06 | 7.000E-06 |
| 26493 | 1325 | P1325M | E | 1e-05 | 5.4e-06 | 6.4e-06 | 7.267E-06 | 2.419E-06 | 5.400E-06 | 1.000E-05 |
| 26494 | 1325 | P1325F | E | 4e-06 | 8.2e-06 | 9e-06 | 7.067E-06 | 2.686E-06 | 4.000E-06 | 9.000E-06 |
| 26495 | 1325 | P1325P | E | 1.1e-05 | 8.6e-06 | 7.9e-06 | 9.167E-06 | 1.626E-06 | 7.900E-06 | 1.100E-05 |
| 26496 | 1325 | P1325S | E | 1e-05 | 5e-06 | 5.7e-06 | 6.900E-06 | 2.707E-06 | 5.000E-06 | 1.000E-05 |
| 26497 | 1325 | P1325T | E | 6.4e-06 | 7.6e-06 | 8.3e-06 | 7.433E-06 | 9.609E-07 | 6.400E-06 | 8.300E-06 |
| 26498 | 1325 | P1325W | E | 7.4e-06 | 7e-06 | 7e-06 | 7.133E-06 | 2.309E-07 | 7.000E-06 | 7.400E-06 |
| 26499 | 1325 | P1325Y | E | 1.2e-05 | 7.2e-06 | 1.3e-05 | 1.073E-05 | 3.101E-06 | 7.200E-06 | 1.300E-05 |
| 26500 | 1325 | P1325V | E | 5.6e-06 | 9.4e-06 | 5.7e-06 | 6.900E-06 | 2.166E-06 | 5.600E-06 | 9.400E-06 |
| 26501 | 1326 | T1326A | E | 6.6e-06 | 1.3e-05 | 8e-06 | 9.200E-06 | 3.365E-06 | 6.600E-06 | 1.300E-05 |
| 26502 | 1326 | T1326R | E | 4.9e-06 | 6.2e-06 | 4.6e-06 | 5.233E-06 | 8.505E-07 | 4.600E-06 | 6.200E-06 |
| 26503 | 1326 | T1326N | E | 9.2e-06 | 6.1e-06 | 6.1e-06 | 7.133E-06 | 1.790E-06 | 6.100E-06 | 9.200E-06 |
| 26504 | 1326 | T1326D | E | 9.8e-06 | 7.3e-06 | 1e-05 | 9.033E-06 | 1.504E-06 | 7.300E-06 | 1.000E-05 |
| 26505 | 1326 | T1326C | E | 6.9e-06 | 1.2e-05 | 6.9e-06 | 8.600E-06 | 2.944E-06 | 6.900E-06 | 1.200E-05 |
| 26506 | 1326 | T1326E | E | 7.3e-06 | 1.3e-05 | 6.5e-06 | 8.933E-06 | 3.544E-06 | 6.500E-06 | 1.300E-05 |
| 26507 | 1326 | T1326Q | E | 8.2e-06 | 9.6e-06 | 1e-05 | 9.267E-06 | 9.452E-07 | 8.200E-06 | 1.000E-05 |
| 26508 | 1326 | T1326G | E | 1e-05 | 8.7e-06 | 4.1e-06 | 7.600E-06 | 3.100E-06 | 4.100E-06 | 1.000E-05 |
| 26509 | 1326 | T1326H | E | 1.4e-05 | 1.1e-05 | 4.9e-06 | 9.967E-06 | 4.637E-06 | 4.900E-06 | 1.400E-05 |
| 26510 | 1326 | T1326I | E | 6.1e-06 | 6.3e-06 | 7.2e-06 | 6.533E-06 | 5.859E-07 | 6.100E-06 | 7.200E-06 |
| 26511 | 1326 | T1326L | E | 4.8e-06 | 6.8e-06 | 8.7e-06 | 6.767E-06 | 1.950E-06 | 4.800E-06 | 8.700E-06 |
| 26512 | 1326 | T1326K | E | 6.8e-06 | 7.5e-06 | 9.2e-06 | 7.833E-06 | 1.234E-06 | 6.800E-06 | 9.200E-06 |
| 26513 | 1326 | T1326M | E | 6.9e-06 | 1.1e-05 | 1.4e-05 | 1.063E-05 | 3.564E-06 | 6.900E-06 | 1.400E-05 |
| 26514 | 1326 | T1326F | E | 7.6e-06 | 1.3e-05 | 9.4e-06 | 1.000E-05 | 2.750E-06 | 7.600E-06 | 1.300E-05 |
| 26515 | 1326 | T1326P | E | 9.7e-06 | 1.1e-05 | 7e-06 | 9.233E-06 | 2.040E-06 | 7.000E-06 | 1.100E-05 |
| 26516 | 1326 | T1326S | E | 6.3e-06 | 8.2e-06 | 5.2e-06 | 6.567E-06 | 1.518E-06 | 5.200E-06 | 8.200E-06 |
| 26517 | 1326 | T1326T | E | 5.6e-06 | 7.6e-06 | 1.4e-05 | 9.067E-06 | 4.388E-06 | 5.600E-06 | 1.400E-05 |
| 26518 | 1326 | T1326W | E | 6.9e-06 | 7.5e-06 | 7.6e-06 | 7.333E-06 | 3.786E-07 | 6.900E-06 | 7.600E-06 |
| 26519 | 1326 | T1326Y | E | 8.1e-06 | 7.7e-06 | 9.4e-06 | 8.400E-06 | 8.888E-07 | 7.700E-06 | 9.400E-06 |

| | | | | | | | | | | |
|---|---|---|---|---|---|---|---|---|---|---|
| 26520 | 1326 | T1326V | E | 7.6e-06 | 4.7e-06 | 1.1e-05 | 7.767E-06 | 3.153E-06 | 4.700E-06 | 1.100E-05 |
| 26521 | 1327 | N1327A | E | 4.6e-06 | 4.1e-06 | 8.1e-06 | 5.600E-06 | 2.179E-06 | 4.100E-06 | 8.100E-06 |
| 26522 | 1327 | N1327R | E | 6.9e-06 | 9.6e-06 | 6.8e-06 | 7.767E-06 | 1.589E-06 | 6.800E-06 | 9.600E-06 |
| 26523 | 1327 | N1327N | E | 6e-06 | 1.4e-05 | 9.4e-06 | 9.800E-06 | 4.015E-06 | 6.000E-06 | 1.400E-05 |
| 26524 | 1327 | N1327D | E | 1.1e-05 | 8e-06 | 1.4e-05 | 1.100E-05 | 3.000E-06 | 8.000E-06 | 1.400E-05 |
| 26525 | 1327 | N1327C | E | 5.2e-06 | 6.7e-06 | 4.7e-06 | 5.533E-06 | 1.041E-06 | 4.700E-06 | 6.700E-06 |
| 26526 | 1327 | N1327E | E | 9.8e-06 | 8.8e-06 | 9.7e-06 | 9.433E-06 | 5.508E-07 | 8.800E-06 | 9.800E-06 |
| 26527 | 1327 | N1327Q | E | 8e-06 | 9e-06 | 7.5e-06 | 8.167E-06 | 7.638E-07 | 7.500E-06 | 9.000E-06 |
| 26528 | 1327 | N1327G | E | 8.9e-06 | 7.7e-06 | 1.2e-05 | 9.533E-06 | 2.219E-06 | 7.700E-06 | 1.200E-05 |
| 26529 | 1327 | N1327H | E | 1.1e-05 | 7.8e-06 | 1.2e-05 | 1.027E-05 | 2.194E-06 | 7.800E-06 | 1.200E-05 |
| 26530 | 1327 | N1327I | E | 6.9e-06 | 9.4e-06 | 9.2e-06 | 8.500E-06 | 1.389E-06 | 6.900E-06 | 9.400E-06 |
| 26531 | 1327 | N1327L | E | 4.5e-06 | 8.9e-06 | 1.1e-05 | 8.133E-06 | 3.317E-06 | 4.500E-06 | 1.100E-05 |
| 26532 | 1327 | N1327K | E | 7.7e-06 | 5.5e-06 | 1e-05 | 7.733E-06 | 2.250E-06 | 5.500E-06 | 1.000E-05 |
| 26533 | 1327 | N1327M | E | 7.2e-06 | 1.3e-05 | 9.6e-06 | 9.933E-06 | 2.914E-06 | 7.200E-06 | 1.300E-05 |
| 26534 | 1327 | N1327F | E | 1.2e-05 | 7.8e-06 | 1e-05 | 9.933E-06 | 2.101E-06 | 7.800E-06 | 1.200E-05 |
| 26535 | 1327 | N1327P | E | 6.7e-06 | 7e-06 | 6.6e-06 | 6.767E-06 | 2.082E-07 | 6.600E-06 | 7.000E-06 |
| 26536 | 1327 | N1327S | E | 1.1e-05 | 9.9e-06 | 4.9e-06 | 8.600E-06 | 3.251E-06 | 4.900E-06 | 1.100E-05 |
| 26537 | 1327 | N1327T | E | 9.4e-06 | 5.1e-06 | 8.2e-06 | 7.567E-06 | 2.219E-06 | 5.100E-06 | 9.400E-06 |
| 26538 | 1327 | N1327W | E | 1.1e-05 | 7e-06 | 5.4e-06 | 7.800E-06 | 2.884E-06 | 5.400E-06 | 1.100E-05 |
| 26539 | 1327 | N1327Y | E | 9e-06 | 1.2e-05 | 1.1e-05 | 1.067E-05 | 1.528E-06 | 9.000E-06 | 1.200E-05 |
| 26540 | 1327 | N1327V | E | 7.6e-06 | 8.3e-06 | 8.2e-06 | 8.033E-06 | 3.786E-07 | 7.600E-06 | 8.300E-06 |
| 26541 | 1328 | G1328A | E | 7.2e-06 | 1.2e-05 | 6.9e-06 | 8.700E-06 | 2.862E-06 | 6.900E-06 | 1.200E-05 |
| 26542 | 1328 | G1328R | E | 7.5e-06 | 1e-05 | 1.5e-05 | 1.083E-05 | 3.819E-06 | 7.500E-06 | 1.500E-05 |
| 26543 | 1328 | G1328N | E | 5.3e-06 | 1e-05 | 6.9e-06 | 7.400E-06 | 2.390E-06 | 5.300E-06 | 1.000E-05 |
| 26544 | 1328 | G1328D | E | 9.3e-06 | 7.8e-06 | 9.4e-06 | 8.833E-06 | 8.963E-07 | 7.800E-06 | 9.400E-06 |
| 26545 | 1328 | G1328C | E | 1.3e-05 | 8.8e-06 | 7.2e-06 | 9.667E-06 | 2.996E-06 | 7.200E-06 | 1.300E-05 |
| 26546 | 1328 | G1328E | E | 4.2e-06 | 5.1e-06 | 4.6e-06 | 4.633E-06 | 4.509E-07 | 4.200E-06 | 5.100E-06 |
| 26547 | 1328 | G1328Q | E | 6.2e-06 | 1.1e-05 | 6.9e-06 | 8.033E-06 | 2.593E-06 | 6.200E-06 | 1.100E-05 |
| 26548 | 1328 | G1328G | E | 6.3e-06 | 9.4e-06 | 7.3e-06 | 7.667E-06 | 1.582E-06 | 6.300E-06 | 9.400E-06 |
| 26549 | 1328 | G1328H | E | 7.2e-06 | 6e-06 | 5.6e-06 | 6.267E-06 | 8.327E-07 | 5.600E-06 | 7.200E-06 |
| 26550 | 1328 | G1328I | E | 5.3e-06 | 1e-05 | 4.8e-06 | 6.700E-06 | 2.869E-06 | 4.800E-06 | 1.000E-05 |
| 26551 | 1328 | G1328L | E | 7.6e-06 | 4.6e-06 | 1.4e-05 | 8.733E-06 | 4.801E-06 | 4.600E-06 | 1.400E-05 |
| 26552 | 1328 | G1328K | E | 1.1e-05 | 9.9e-06 | 8.3e-06 | 9.733E-06 | 1.358E-06 | 8.300E-06 | 1.100E-05 |
| 26553 | 1328 | G1328M | E | 1.1e-05 | 6.2e-06 | 1e-05 | 9.067E-06 | 2.532E-06 | 6.200E-06 | 1.100E-05 |
| 26554 | 1328 | G1328F | E | 7.8e-06 | 5.4e-06 | 1.2e-05 | 8.400E-06 | 3.341E-06 | 5.400E-06 | 1.200E-05 |
| 26555 | 1328 | G1328P | E | 9.3e-06 | 1e-05 | 9.8e-06 | 9.700E-06 | 3.606E-07 | 9.300E-06 | 1.000E-05 |
| 26556 | 1328 | G1328S | E | 6.9e-06 | 8.5e-06 | 8e-06 | 7.800E-06 | 8.185E-07 | 6.900E-06 | 8.500E-06 |
| 26557 | 1328 | G1328T | E | 6.8e-06 | 6.4e-06 | 5.1e-06 | 6.100E-06 | 8.888E-07 | 5.100E-06 | 6.800E-06 |
| 26558 | 1328 | G1328W | E | 8.1e-06 | 9.2e-06 | 9e-06 | 8.767E-06 | 5.859E-07 | 8.100E-06 | 9.200E-06 |
| 26559 | 1328 | G1328Y | E | 4.7e-06 | 8e-06 | 9.6e-06 | 7.433E-06 | 2.499E-06 | 4.700E-06 | 9.600E-06 |
| 26560 | 1328 | G1328V | E | 1.3e-05 | 1.2e-05 | 6.9e-06 | 1.063E-05 | 3.272E-06 | 6.900E-06 | 1.300E-05 |
| 26561 | 1329 | Y1329A | E | 9.9e-06 | 1.3e-05 | 1.1e-05 | 1.130E-05 | 1.572E-06 | 9.900E-06 | 1.300E-05 |
| 26562 | 1329 | Y1329R | E | 6.4e-06 | 6.3e-06 | 6.2e-06 | 6.300E-06 | 1.000E-07 | 6.200E-06 | 6.400E-06 |
| 26563 | 1329 | Y1329N | E | 8.5e-06 | 8.8e-06 | 4.4e-06 | 7.233E-06 | 2.458E-06 | 4.400E-06 | 8.800E-06 |
| 26564 | 1329 | Y1329D | E | 8.8e-06 | 9.2e-06 | 1.4e-05 | 1.067E-05 | 2.894E-06 | 8.800E-06 | 1.400E-05 |
| 26565 | 1329 | Y1329C | E | 5.6e-06 | 7.7e-06 | 1.3e-05 | 8.767E-06 | 3.814E-06 | 5.600E-06 | 1.300E-05 |
| 26566 | 1329 | Y1329E | E | 7.7e-06 | 6.2e-06 | 6.8e-06 | 6.900E-06 | 7.550E-07 | 6.200E-06 | 7.700E-06 |
| 26567 | 1329 | Y1329Q | E | 6.4e-06 | 5.6e-06 | 1.2e-05 | 8.000E-06 | 3.487E-06 | 5.600E-06 | 1.200E-05 |
| 26568 | 1329 | Y1329G | E | 7.2e-06 | 9.9e-06 | 6.6e-06 | 7.900E-06 | 1.758E-06 | 6.600E-06 | 9.900E-06 |
| 26569 | 1329 | Y1329H | E | 9.8e-06 | 7.9e-06 | 8.6e-06 | 8.767E-06 | 9.609E-07 | 7.900E-06 | 9.800E-06 |
| 26570 | 1329 | Y1329I | E | 6.8e-06 | 4.5e-06 | 5.3e-06 | 5.533E-06 | 1.168E-06 | 4.500E-06 | 6.800E-06 |
| 26571 | 1329 | Y1329L | E | 8.7e-06 | 8.6e-06 | 8.5e-06 | 8.600E-06 | 1.000E-07 | 8.500E-06 | 8.700E-06 |
| 26572 | 1329 | Y1329K | E | 8.4e-06 | 7.8e-06 | 4.8e-06 | 7.000E-06 | 1.929E-06 | 4.800E-06 | 8.400E-06 |
| 26573 | 1329 | Y1329M | E | 7.3e-06 | 6.5e-06 | 7.4e-06 | 7.067E-06 | 4.933E-07 | 6.500E-06 | 7.400E-06 |
| 26574 | 1329 | Y1329F | E | 1.2e-05 | 5e-06 | 8.9e-06 | 8.633E-06 | 3.508E-06 | 5.000E-06 | 1.200E-05 |
| 26575 | 1329 | Y1329P | E | 1.9e-05 | 6.4e-06 | 3.4e-06 | 9.600E-06 | 8.278E-06 | 3.400E-06 | 1.900E-05 |
| 26576 | 1329 | Y1329S | E | 5e-06 | 1.1e-05 | 7.5e-06 | 7.833E-06 | 3.014E-06 | 5.000E-06 | 1.100E-05 |
| 26577 | 1329 | Y1329T | E | 9.9e-06 | 8.3e-06 | 9.8e-06 | 9.333E-06 | 8.963E-07 | 8.300E-06 | 9.900E-06 |
| 26578 | 1329 | Y1329W | E | 7.1e-06 | 8.1e-06 | 1.3e-05 | 9.400E-06 | 3.158E-06 | 7.100E-06 | 1.300E-05 |
| 26579 | 1329 | Y1329Y | E | 9.2e-06 | 9e-06 | 8.6e-06 | 8.933E-06 | 3.055E-07 | 8.600E-06 | 9.200E-06 |
| 26580 | 1329 | Y1329V | E | 9.5e-06 | 7.2e-06 | 1.1e-05 | 9.233E-06 | 1.914E-06 | 7.200E-06 | 1.100E-05 |
| 26581 | 1330 | T1330A | E | 6.6e-06 | 1.4e-05 | 1e-05 | 1.020E-05 | 3.704E-06 | 6.600E-06 | 1.400E-05 |
| 26582 | 1330 | T1330R | E | 6.2e-06 | 7.8e-06 | 8.1e-06 | 7.367E-06 | 1.021E-06 | 6.200E-06 | 8.100E-06 |
| 26583 | 1330 | T1330N | E | 7.8e-06 | 8.8e-06 | 1.3e-05 | 9.867E-06 | 2.759E-06 | 7.800E-06 | 1.300E-05 |
| 26584 | 1330 | T1330D | E | 7e-06 | 8.9e-06 | 7e-06 | 7.633E-06 | 1.097E-06 | 7.000E-06 | 8.900E-06 |
| 26585 | 1330 | T1330C | E | 4.9e-06 | 6.8e-06 | 1e-05 | 7.233E-06 | 2.577E-06 | 4.900E-06 | 1.000E-05 |
| 26586 | 1330 | T1330E | E | 1.2e-05 | 5.4e-06 | 1.1e-05 | 9.467E-06 | 3.557E-06 | 5.400E-06 | 1.200E-05 |
| 26587 | 1330 | T1330Q | E | 8e-06 | 7.4e-06 | 1.1e-05 | 8.800E-06 | 1.929E-06 | 7.400E-06 | 1.100E-05 |
| 26588 | 1330 | T1330G | E | 6.9e-06 | 1.4e-05 | 1e-05 | 1.030E-05 | 3.559E-06 | 6.900E-06 | 1.400E-05 |
| 26589 | 1330 | T1330H | E | 7.9e-06 | 8e-06 | 9e-06 | 8.300E-06 | 6.083E-07 | 7.900E-06 | 9.000E-06 |
| 26590 | 1330 | T1330I | E | 1.3e-05 | 9.2e-06 | 8.2e-06 | 1.013E-05 | 2.532E-06 | 8.200E-06 | 1.300E-05 |
| 26591 | 1330 | T1330L | E | 1.1e-05 | 5.7e-06 | 6.7e-06 | 7.800E-06 | 2.816E-06 | 5.700E-06 | 1.100E-05 |
| 26592 | 1330 | T1330K | E | 6.7e-06 | 9.2e-06 | 8.3e-06 | 8.067E-06 | 1.266E-06 | 6.700E-06 | 9.200E-06 |
| 26593 | 1330 | T1330M | E | 5.8e-06 | 5.7e-06 | 1.3e-05 | 8.167E-06 | 4.186E-06 | 5.700E-06 | 1.300E-05 |
| 26594 | 1330 | T1330F | E | 8.1e-06 | 4.2e-06 | 8.6e-06 | 6.967E-06 | 2.409E-06 | 4.200E-06 | 8.600E-06 |
| 26595 | 1330 | T1330P | E | 7.5e-06 | 1e-05 | 8.8e-06 | 8.767E-06 | 1.250E-06 | 7.500E-06 | 1.000E-05 |
| 26596 | 1330 | T1330S | E | 1.2e-05 | 1.3e-05 | 6.5e-06 | 1.050E-05 | 3.500E-06 | 6.500E-06 | 1.300E-05 |
| 26597 | 1330 | T1330T | E | 1.1e-05 | 6.2e-06 | 1e-05 | 9.067E-06 | 2.532E-06 | 6.200E-06 | 1.100E-05 |
| 26598 | 1330 | T1330W | E | 1.3e-05 | 3.7e-06 | 9.1e-06 | 8.600E-06 | 4.670E-06 | 3.700E-06 | 1.300E-05 |
| 26599 | 1330 | T1330Y | E | 9.3e-06 | 6.8e-06 | 1e-05 | 8.700E-06 | 1.682E-06 | 6.800E-06 | 1.000E-05 |
| 26600 | 1330 | T1330V | E | 1.1e-05 | 8.2e-06 | 7.8e-06 | 9.000E-06 | 1.744E-06 | 7.800E-06 | 1.100E-05 |
| 26601 | 1331 | R1331A | E | 6.3e-06 | 6.8e-06 | 6.1e-06 | 6.400E-06 | 3.606E-07 | 6.100E-06 | 6.800E-06 |
| 26602 | 1331 | R1331R | E | 5.8e-06 | 8.9e-06 | 9.2e-06 | 7.967E-06 | 1.882E-06 | 5.800E-06 | 9.200E-06 |
| 26603 | 1331 | R1331N | E | 1.2e-05 | 5.2e-06 | 6e-06 | 7.733E-06 | 3.717E-06 | 5.200E-06 | 1.200E-05 |
| 26604 | 1331 | R1331D | E | 1.3e-05 | 7e-06 | 8.1e-06 | 9.367E-06 | 3.194E-06 | 7.000E-06 | 1.300E-05 |

| | | | | | | | | | | |
|---|---|---|---|---|---|---|---|---|---|---|
| 26605 | 1331 | R1331C | E | 4.9e-06 | 9.4e-06 | 8.9e-06 | 7.733E-06 | 2.466E-06 | 4.900E-06 | 9.400E-06 |
| 26606 | 1331 | R1331E | E | 1.2e-05 | 8.6e-06 | 6.9e-06 | 9.167E-06 | 2.597E-06 | 6.900E-06 | 1.200E-05 |
| 26607 | 1331 | R1331Q | E | 7e-06 | 6.1e-06 | 4.3e-06 | 5.800E-06 | 1.375E-06 | 4.300E-06 | 7.000E-06 |
| 26608 | 1331 | R1331G | E | 6.5e-06 | 1.1e-05 | 8.3e-06 | 8.600E-06 | 2.265E-06 | 6.500E-06 | 1.100E-05 |
| 26609 | 1331 | R1331H | E | 1.3e-05 | 5.4e-06 | 1e-05 | 9.467E-06 | 3.828E-06 | 5.400E-06 | 1.300E-05 |
| 26610 | 1331 | R1331I | E | 1.1e-05 | 1.6e-05 | 5e-06 | 1.067E-05 | 5.508E-06 | 5.000E-06 | 1.600E-05 |
| 26611 | 1331 | R1331L | E | 1.5e-05 | 5e-06 | 6.4e-06 | 8.800E-06 | 5.415E-06 | 5.000E-06 | 1.500E-05 |
| 26612 | 1331 | R1331K | E | 9.1e-06 | 9e-06 | 8.8e-06 | 8.967E-06 | 1.528E-07 | 8.800E-06 | 9.100E-06 |
| 26613 | 1331 | R1331M | E | 1e-05 | 8.1e-06 | 9.6e-06 | 9.233E-06 | 1.002E-06 | 8.100E-06 | 1.000E-05 |
| 26614 | 1331 | R1331F | E | 1.3e-05 | 8.9e-06 | 6.8e-06 | 9.567E-06 | 3.153E-06 | 6.800E-06 | 1.300E-05 |
| 26615 | 1331 | R1331P | E | 8.5e-06 | 6.8e-06 | 1.5e-05 | 1.010E-05 | 4.328E-06 | 6.800E-06 | 1.500E-05 |
| 26616 | 1331 | R1331S | E | 1e-05 | 7e-06 | 7.6e-06 | 8.200E-06 | 1.587E-06 | 7.000E-06 | 1.000E-05 |
| 26617 | 1331 | R1331T | E | 4.5e-06 | 9.8e-06 | 1.3e-05 | 9.100E-06 | 4.293E-06 | 4.500E-06 | 1.300E-05 |
| 26618 | 1331 | R1331W | E | 1.3e-05 | 1e-05 | 1.4e-05 | 1.233E-05 | 2.082E-06 | 1.000E-05 | 1.400E-05 |
| 26619 | 1331 | R1331Y | E | 6.6e-06 | 1.3e-05 | 9.4e-06 | 9.667E-06 | 3.208E-06 | 6.600E-06 | 1.300E-05 |
| 26620 | 1331 | R1331V | E | 5.4e-06 | 5.4e-06 | 1.2e-05 | 7.600E-06 | 3.811E-06 | 5.400E-06 | 1.200E-05 |
| 26621 | 1332 | Y1332A | E | 6e-06 | 5.2e-06 | 1.4e-05 | 8.400E-06 | 4.866E-06 | 5.200E-06 | 1.400E-05 |
| 26622 | 1332 | Y1332R | E | 1.3e-05 | 3.8e-06 | 1e-05 | 8.933E-06 | 4.692E-06 | 3.800E-06 | 1.300E-05 |
| 26623 | 1332 | Y1332N | E | 7.9e-06 | 4.6e-06 | 9e-06 | 7.167E-06 | 2.290E-06 | 4.600E-06 | 9.000E-06 |
| 26624 | 1332 | Y1332D | E | 4.2e-06 | 9.2e-06 | 7e-06 | 6.800E-06 | 2.506E-06 | 4.200E-06 | 9.200E-06 |
| 26625 | 1332 | Y1332C | E | 6.5e-06 | 9.5e-06 | 1.1e-05 | 9.000E-06 | 2.291E-06 | 6.500E-06 | 1.100E-05 |
| 26626 | 1332 | Y1332E | E | 5.8e-06 | 1.4e-05 | 9.9e-06 | 9.900E-06 | 4.100E-06 | 5.800E-06 | 1.400E-05 |
| 26627 | 1332 | Y1332Q | E | 4.2e-06 | 6.2e-06 | 1.1e-05 | 7.133E-06 | 3.495E-06 | 4.200E-06 | 1.100E-05 |
| 26628 | 1332 | Y1332G | E | 1.1e-05 | 8.5e-06 | 5.6e-06 | 8.367E-06 | 2.702E-06 | 5.600E-06 | 1.100E-05 |
| 26629 | 1332 | Y1332H | E | 5.7e-06 | 8.1e-06 | 5.3e-06 | 6.367E-06 | 1.514E-06 | 5.300E-06 | 8.100E-06 |
| 26630 | 1332 | Y1332I | E | 7e-06 | 1e-05 | 9.9e-06 | 8.967E-06 | 1.704E-06 | 7.000E-06 | 1.000E-05 |
| 26631 | 1332 | Y1332L | E | 3.3e-06 | 7e-06 | 7.2e-06 | 5.833E-06 | 2.196E-06 | 3.300E-06 | 7.200E-06 |
| 26632 | 1332 | Y1332K | E | 1.3e-05 | 7.8e-06 | 6e-06 | 8.933E-06 | 3.635E-06 | 6.000E-06 | 1.300E-05 |
| 26633 | 1332 | Y1332M | E | 8.4e-06 | 1.2e-05 | 8.1e-06 | 9.500E-06 | 2.170E-06 | 8.100E-06 | 1.200E-05 |
| 26634 | 1332 | Y1332F | E | 7.5e-06 | 1.2e-05 | 5.7e-06 | 8.400E-06 | 3.245E-06 | 5.700E-06 | 1.200E-05 |
| 26635 | 1332 | Y1332P | E | 1.1e-05 | 1.4e-05 | 1.3e-05 | 1.267E-05 | 1.528E-06 | 1.100E-05 | 1.400E-05 |
| 26636 | 1332 | Y1332S | E | 1.1e-05 | 1.3e-05 | 4.7e-06 | 9.567E-06 | 4.332E-06 | 4.700E-06 | 1.300E-05 |
| 26637 | 1332 | Y1332T | E | 1.1e-05 | 6.3e-06 | 8e-06 | 8.433E-06 | 2.380E-06 | 6.300E-06 | 1.100E-05 |
| 26638 | 1332 | Y1332W | E | 9.9e-06 | 2.1e-05 | 6.5e-06 | 1.247E-05 | 7.583E-06 | 6.500E-06 | 2.100E-05 |
| 26639 | 1332 | Y1332Y | E | 6.1e-06 | 1.2e-05 | 1.8e-05 | 1.203E-05 | 5.950E-06 | 6.100E-06 | 1.800E-05 |
| 26640 | 1332 | Y1332V | E | 6e-06 | 6.4e-06 | 8.1e-06 | 6.833E-06 | 1.115E-06 | 6.000E-06 | 8.100E-06 |
| 26641 | 1333 | A1333A | E | 8.9e-06 | 4.4e-06 | 9.3e-06 | 7.533E-06 | 2.721E-06 | 4.400E-06 | 9.300E-06 |
| 26642 | 1333 | A1333R | E | 6.9e-06 | 6.7e-06 | 7.2e-06 | 6.933E-06 | 2.517E-07 | 6.700E-06 | 7.200E-06 |
| 26643 | 1333 | A1333N | E | 8.8e-06 | 7.2e-06 | 1.3e-05 | 9.667E-06 | 2.996E-06 | 7.200E-06 | 1.300E-05 |
| 26644 | 1333 | A1333D | E | 8.5e-06 | 4.5e-06 | 7e-06 | 6.667E-06 | 2.021E-06 | 4.500E-06 | 8.500E-06 |
| 26645 | 1333 | A1333C | E | 5.8e-06 | 5.4e-06 | 5.4e-06 | 5.533E-06 | 2.309E-07 | 5.400E-06 | 5.800E-06 |
| 26646 | 1333 | A1333E | E | 1.2e-05 | 9.7e-06 | 7.9e-06 | 9.867E-06 | 2.055E-06 | 7.900E-06 | 1.200E-05 |
| 26647 | 1333 | A1333Q | E | 7.3e-06 | 6.2e-06 | 6.8e-06 | 6.767E-06 | 5.508E-07 | 6.200E-06 | 7.300E-06 |
| 26648 | 1333 | A1333G | E | 1.3e-05 | 7.6e-06 | 9.9e-06 | 1.017E-05 | 2.710E-06 | 7.600E-06 | 1.300E-05 |
| 26649 | 1333 | A1333H | E | 6.4e-06 | 6.3e-06 | 6.4e-06 | 6.367E-06 | 5.774E-08 | 6.300E-06 | 6.400E-06 |
| 26650 | 1333 | A1333I | E | 9.7e-06 | 4.6e-06 | 1.2e-05 | 8.767E-06 | 3.787E-06 | 4.600E-06 | 1.200E-05 |
| 26651 | 1333 | A1333L | E | 6.3e-06 | 9.4e-06 | 5.1e-06 | 6.933E-06 | 2.219E-06 | 5.100E-06 | 9.400E-06 |
| 26652 | 1333 | A1333K | E | 1.3e-05 | 7.7e-06 | 1.1e-05 | 1.057E-05 | 2.676E-06 | 7.700E-06 | 1.300E-05 |
| 26653 | 1333 | A1333M | E | 7.4e-06 | 6.1e-06 | 9.9e-06 | 7.800E-06 | 1.931E-06 | 6.100E-06 | 9.900E-06 |
| 26654 | 1333 | A1333F | E | 9.6e-06 | 8.6e-06 | 1e-05 | 9.400E-06 | 7.211E-07 | 8.600E-06 | 1.000E-05 |
| 26655 | 1333 | A1333P | E | 6.3e-06 | 6.6e-06 | 9.4e-06 | 7.433E-06 | 1.710E-06 | 6.300E-06 | 9.400E-06 |
| 26656 | 1333 | A1333S | E | 4.6e-06 | 6.4e-06 | 8.4e-06 | 6.467E-06 | 1.901E-06 | 4.600E-06 | 8.400E-06 |
| 26657 | 1333 | A1333T | E | 4.3e-06 | 9.4e-06 | 6.1e-06 | 6.600E-06 | 2.587E-06 | 4.300E-06 | 9.400E-06 |
| 26658 | 1333 | A1333W | E | 9.1e-06 | 8.5e-06 | 6.1e-06 | 7.900E-06 | 1.587E-06 | 6.100E-06 | 9.100E-06 |
| 26659 | 1333 | A1333Y | E | 5.2e-06 | 5e-06 | 9.2e-06 | 6.467E-06 | 2.369E-06 | 5.000E-06 | 9.200E-06 |
| 26660 | 1333 | A1333V | E | 1.2e-05 | 4.4e-06 | 1.1e-05 | 9.133E-06 | 4.130E-06 | 4.400E-06 | 1.200E-05 |
| 26661 | 1334 | D1334A | E | 5.6e-06 | 1.2e-05 | 1.1e-05 | 9.533E-06 | 3.443E-06 | 5.600E-06 | 1.200E-05 |
| 26662 | 1334 | D1334R | E | 6.4e-06 | 5e-06 | 5.8e-06 | 5.733E-06 | 7.024E-07 | 5.000E-06 | 6.400E-06 |
| 26663 | 1334 | D1334N | E | 8.2e-06 | 7.8e-06 | 5.9e-06 | 7.300E-06 | 1.229E-06 | 5.900E-06 | 8.200E-06 |
| 26664 | 1334 | D1334D | E | 8.1e-06 | 9.2e-06 | 8.6e-06 | 8.633E-06 | 5.508E-07 | 8.100E-06 | 9.200E-06 |
| 26665 | 1334 | D1334C | E | 4.4e-06 | 1.2e-05 | 1e-05 | 8.800E-06 | 3.940E-06 | 4.400E-06 | 1.200E-05 |
| 26666 | 1334 | D1334E | E | 1.1e-05 | 5.1e-06 | 6.5e-06 | 7.533E-06 | 3.083E-06 | 5.100E-06 | 1.100E-05 |
| 26667 | 1334 | D1334Q | E | 3e-06 | 6.1e-06 | 7.1e-06 | 5.400E-06 | 2.138E-06 | 3.000E-06 | 7.100E-06 |
| 26668 | 1334 | D1334G | E | 4.8e-06 | 9.2e-06 | 6.4e-06 | 6.800E-06 | 2.227E-06 | 4.800E-06 | 9.200E-06 |
| 26669 | 1334 | D1334H | E | 9.9e-06 | 9.3e-06 | 6.2e-06 | 8.467E-06 | 1.986E-06 | 6.200E-06 | 9.900E-06 |
| 26670 | 1334 | D1334I | E | 1.5e-05 | 4.9e-06 | 6.8e-06 | 8.900E-06 | 5.367E-06 | 4.900E-06 | 1.500E-05 |
| 26671 | 1334 | D1334L | E | 4.1e-06 | 7.2e-06 | 8.1e-06 | 6.467E-06 | 2.098E-06 | 4.100E-06 | 8.100E-06 |
| 26672 | 1334 | D1334K | E | 6.9e-06 | 8.3e-06 | 9.8e-06 | 8.333E-06 | 1.450E-06 | 6.900E-06 | 9.800E-06 |
| 26673 | 1334 | D1334M | E | 1.1e-05 | 9.1e-06 | 1.3e-05 | 1.103E-05 | 1.950E-06 | 9.100E-06 | 1.300E-05 |
| 26674 | 1334 | D1334F | E | 8.3e-06 | 8.7e-06 | 9e-06 | 8.667E-06 | 3.512E-07 | 8.300E-06 | 9.000E-06 |
| 26675 | 1334 | D1334P | E | 7.8e-06 | 5.6e-06 | 5.6e-06 | 6.333E-06 | 1.270E-06 | 5.600E-06 | 7.800E-06 |
| 26676 | 1334 | D1334S | E | 3.2e-06 | 6.3e-06 | 4.6e-06 | 4.700E-06 | 1.552E-06 | 3.200E-06 | 6.300E-06 |
| 26677 | 1334 | D1334T | E | 1.3e-05 | 1.2e-05 | 7.5e-06 | 1.083E-05 | 2.930E-06 | 7.500E-06 | 1.300E-05 |
| 26678 | 1334 | D1334W | E | 9.8e-06 | 6.8e-06 | 5.1e-06 | 7.233E-06 | 2.380E-06 | 5.100E-06 | 9.800E-06 |
| 26679 | 1334 | D1334Y | E | 4.6e-06 | 9.1e-06 | 5.5e-06 | 6.400E-06 | 2.381E-06 | 4.600E-06 | 9.100E-06 |
| 26680 | 1334 | D1334V | E | 1.2e-05 | 7.3e-06 | 8.7e-06 | 9.333E-06 | 2.413E-06 | 7.300E-06 | 1.200E-05 |
| 26681 | 1335 | S1335A | E | 8.8e-06 | 1e-05 | 8.2e-06 | 9.000E-06 | 9.165E-07 | 8.200E-06 | 1.000E-05 |
| 26682 | 1335 | S1335R | E | 6.5e-06 | 6.6e-06 | 7.2e-06 | 6.767E-06 | 3.786E-07 | 6.500E-06 | 7.200E-06 |
| 26683 | 1335 | S1335N | E | 4.1e-06 | 4.7e-06 | 8.8e-06 | 5.867E-06 | 2.558E-06 | 4.100E-06 | 8.800E-06 |
| 26684 | 1335 | S1335D | E | 9.3e-06 | 1.4e-05 | 5.6e-06 | 9.633E-06 | 4.210E-06 | 5.600E-06 | 1.400E-05 |
| 26685 | 1335 | S1335C | E | 9.3e-06 | 6.9e-06 | 9.5e-06 | 8.567E-06 | 1.447E-06 | 6.900E-06 | 9.500E-06 |
| 26686 | 1335 | S1335E | E | 5.1e-06 | 6.7e-06 | 7.2e-06 | 6.333E-06 | 1.097E-06 | 5.100E-06 | 7.200E-06 |
| 26687 | 1335 | S1335Q | E | 1.2e-05 | 4.4e-06 | 1.5e-05 | 1.047E-05 | 5.464E-06 | 4.400E-06 | 1.500E-05 |
| 26688 | 1335 | S1335G | E | 9.9e-06 | 8.9e-06 | 7.1e-06 | 8.633E-06 | 1.419E-06 | 7.100E-06 | 9.900E-06 |
| 26689 | 1335 | S1335H | E | 7.5e-06 | 8.5e-06 | 7.9e-06 | 7.967E-06 | 5.033E-07 | 7.500E-06 | 8.500E-06 |

| | | | | | | | | | | |
|---|---|---|---|---|---|---|---|---|---|---|
| 26690 | 1335 | S1335I | E | 7.7e-06 | 4.6e-06 | 7.4e-06 | 6.567E-06 | 1.710E-06 | 4.600E-06 | 7.700E-06 |
| 26691 | 1335 | S1335L | E | 1.1e-05 | 8.5e-06 | 5e-06 | 8.167E-06 | 3.014E-06 | 5.000E-06 | 1.100E-05 |
| 26692 | 1335 | S1335K | E | 8.4e-06 | 9.8e-06 | 1.3e-05 | 1.040E-05 | 2.358E-06 | 8.400E-06 | 1.300E-05 |
| 26693 | 1335 | S1335M | E | 6.9e-06 | 8.7e-06 | 1.3e-05 | 9.533E-06 | 3.134E-06 | 6.900E-06 | 1.300E-05 |
| 26694 | 1335 | S1335F | E | 8.3e-06 | 8.8e-06 | 6.4e-06 | 7.833E-06 | 1.266E-06 | 6.400E-06 | 8.800E-06 |
| 26695 | 1335 | S1335P | E | 6.5e-06 | 5.3e-06 | 6.7e-06 | 6.167E-06 | 7.572E-07 | 5.300E-06 | 6.700E-06 |
| 26696 | 1335 | S1335S | E | 7e-06 | 6.4e-06 | 6e-06 | 6.467E-06 | 5.033E-07 | 6.000E-06 | 7.000E-06 |
| 26697 | 1335 | S1335T | E | 8.6e-06 | 6.8e-06 | 5.2e-06 | 6.867E-06 | 1.701E-06 | 5.200E-06 | 8.600E-06 |
| 26698 | 1335 | S1335W | E | 8.3e-06 | 8.7e-06 | 8.8e-06 | 8.600E-06 | 2.646E-07 | 8.300E-06 | 8.800E-06 |
| 26699 | 1335 | S1335Y | E | 8.1e-06 | 1.4e-05 | 8.5e-06 | 1.020E-05 | 3.297E-06 | 8.100E-06 | 1.400E-05 |
| 26700 | 1335 | S1335V | E | 5.4e-06 | 6.6e-06 | 3.8e-06 | 5.267E-06 | 1.405E-06 | 3.800E-06 | 6.600E-06 |
| 26701 | 1336 | V1336A | E | 6e-06 | 7.5e-06 | 5.6e-06 | 6.367E-06 | 1.002E-06 | 5.600E-06 | 7.500E-06 |
| 26702 | 1336 | V1336R | E | 7.7e-06 | 1.1e-05 | 9.8e-06 | 9.500E-06 | 1.670E-06 | 7.700E-06 | 1.100E-05 |
| 26703 | 1336 | V1336N | E | 5.6e-06 | 1.2e-05 | 7e-06 | 8.200E-06 | 3.365E-06 | 5.600E-06 | 1.200E-05 |
| 26704 | 1336 | V1336D | E | 8.9e-06 | 9.7e-06 | 5.6e-06 | 8.067E-06 | 2.173E-06 | 5.600E-06 | 9.700E-06 |
| 26705 | 1336 | V1336C | E | 8e-06 | 4.4e-06 | 7e-06 | 6.467E-06 | 1.858E-06 | 4.400E-06 | 8.000E-06 |
| 26706 | 1336 | V1336E | E | 7.2e-06 | 8.3e-06 | 6.2e-06 | 7.233E-06 | 1.050E-06 | 6.200E-06 | 8.300E-06 |
| 26707 | 1336 | V1336Q | E | 6.6e-06 | 5.8e-06 | 7.5e-06 | 6.633E-06 | 8.505E-07 | 5.800E-06 | 7.500E-06 |
| 26708 | 1336 | V1336G | E | 7.5e-06 | 4.9e-06 | 6.8e-06 | 6.400E-06 | 1.345E-06 | 4.900E-06 | 7.500E-06 |
| 26709 | 1336 | V1336H | E | 6.3e-06 | 5.6e-06 | 5.9e-06 | 5.933E-06 | 3.512E-07 | 5.600E-06 | 6.300E-06 |
| 26710 | 1336 | V1336I | E | 8.5e-06 | 7.9e-06 | 1.1e-05 | 9.133E-06 | 1.644E-06 | 7.900E-06 | 1.100E-05 |
| 26711 | 1336 | V1336L | E | 6.7e-06 | 7.4e-06 | 7.7e-06 | 7.267E-06 | 5.132E-07 | 6.700E-06 | 7.700E-06 |
| 26712 | 1336 | V1336K | E | 8.5e-06 | 5.7e-06 | 1.2e-05 | 8.733E-06 | 3.156E-06 | 5.700E-06 | 1.200E-05 |
| 26713 | 1336 | V1336M | E | 9.3e-06 | 6.1e-06 | 8.8e-06 | 8.067E-06 | 1.721E-06 | 6.100E-06 | 9.300E-06 |
| 26714 | 1336 | V1336F | E | 6.8e-06 | 7.9e-06 | 4.8e-06 | 6.500E-06 | 1.572E-06 | 4.800E-06 | 7.900E-06 |
| 26715 | 1336 | V1336P | E | 9.1e-06 | 6.8e-06 | 8.5e-06 | 8.133E-06 | 1.193E-06 | 6.800E-06 | 9.100E-06 |
| 26716 | 1336 | V1336S | E | 4.5e-06 | 6.7e-06 | 1e-05 | 7.067E-06 | 2.768E-06 | 4.500E-06 | 1.000E-05 |
| 26717 | 1336 | V1336T | E | 4.7e-06 | 5.3e-06 | 7.7e-06 | 5.900E-06 | 1.587E-06 | 4.700E-06 | 7.700E-06 |
| 26718 | 1336 | V1336W | E | 1.3e-05 | 6.3e-06 | 9.7e-06 | 9.667E-06 | 3.350E-06 | 6.300E-06 | 1.300E-05 |
| 26719 | 1336 | V1336Y | E | 6.6e-06 | 6.9e-06 | 6.1e-06 | 6.533E-06 | 4.041E-07 | 6.100E-06 | 6.900E-06 |
| 26720 | 1336 | V1336V | E | 9.7e-06 | 6.8e-06 | 1.2e-05 | 9.500E-06 | 2.606E-06 | 6.800E-06 | 1.200E-05 |
| 26721 | 1337 | K1337A | E | 7.1e-06 | 8.5e-06 | 5.8e-06 | 7.133E-06 | 1.350E-06 | 5.800E-06 | 8.500E-06 |
| 26722 | 1337 | K1337R | E | 1.1e-05 | 9.3e-06 | 7.8e-06 | 9.367E-06 | 1.601E-06 | 7.800E-06 | 1.100E-05 |
| 26723 | 1337 | K1337N | E | 5.2e-06 | 6.9e-06 | 8.4e-06 | 6.833E-06 | 1.601E-06 | 5.200E-06 | 8.400E-06 |
| 26724 | 1337 | K1337D | E | 6.3e-06 | 4.5e-06 | 8e-06 | 6.267E-06 | 1.750E-06 | 4.500E-06 | 8.000E-06 |
| 26725 | 1337 | K1337C | E | 1.2e-05 | 1.1e-05 | 7.4e-06 | 1.013E-05 | 2.419E-06 | 7.400E-06 | 1.200E-05 |
| 26726 | 1337 | K1337E | E | 1e-05 | 5.5e-06 | 6.7e-06 | 7.400E-06 | 2.330E-06 | 5.500E-06 | 1.000E-05 |
| 26727 | 1337 | K1337Q | E | 5.8e-06 | 8.7e-06 | 8e-06 | 7.500E-06 | 1.513E-06 | 5.800E-06 | 8.700E-06 |
| 26728 | 1337 | K1337G | E | 3.7e-06 | 9.3e-06 | 1.4e-05 | 9.000E-06 | 5.157E-06 | 3.700E-06 | 1.400E-05 |
| 26729 | 1337 | K1337H | E | 4.9e-06 | 1e-05 | 6.4e-06 | 7.100E-06 | 2.621E-06 | 4.900E-06 | 1.000E-05 |
| 26730 | 1337 | K1337I | E | 7.9e-06 | 7.3e-06 | 1e-05 | 8.400E-06 | 1.418E-06 | 7.300E-06 | 1.000E-05 |
| 26731 | 1337 | K1337L | E | 7.1e-06 | 3.9e-06 | 1.3e-05 | 8.000E-06 | 4.616E-06 | 3.900E-06 | 1.300E-05 |
| 26732 | 1337 | K1337K | E | 8.7e-06 | 8.3e-06 | 8.4e-06 | 8.467E-06 | 2.082E-07 | 8.300E-06 | 8.700E-06 |
| 26733 | 1337 | K1337M | E | 6.1e-06 | 6.5e-06 | 7.5e-06 | 6.700E-06 | 7.211E-07 | 6.100E-06 | 7.500E-06 |
| 26734 | 1337 | K1337F | E | 5e-06 | 9.3e-06 | 8.4e-06 | 7.567E-06 | 2.268E-06 | 5.000E-06 | 9.300E-06 |
| 26735 | 1337 | K1337P | E | 1.3e-05 | 9.2e-06 | 9.4e-06 | 1.053E-05 | 2.139E-06 | 9.200E-06 | 1.300E-05 |
| 26736 | 1337 | K1337S | E | 6.3e-06 | 7.8e-06 | 8.6e-06 | 7.567E-06 | 1.168E-06 | 6.300E-06 | 8.600E-06 |
| 26737 | 1337 | K1337T | E | 1.1e-05 | 4.6e-06 | 7.5e-06 | 7.700E-06 | 3.205E-06 | 4.600E-06 | 1.100E-05 |
| 26738 | 1337 | K1337W | E | 9.8e-06 | 9.3e-06 | 1.4e-05 | 1.103E-05 | 2.581E-06 | 9.300E-06 | 1.400E-05 |
| 26739 | 1337 | K1337Y | E | 6.5e-06 | 3.1e-06 | 8.1e-06 | 5.900E-06 | 2.553E-06 | 3.100E-06 | 8.100E-06 |
| 26740 | 1337 | K1337V | E | 3.9e-06 | 6.4e-06 | 6.2e-06 | 5.500E-06 | 1.389E-06 | 3.900E-06 | 6.400E-06 |
| 26741 | 1338 | G1338A | E | 7.6e-06 | 7.2e-06 | 7.7e-06 | 7.500E-06 | 2.646E-07 | 7.200E-06 | 7.700E-06 |
| 26742 | 1338 | G1338R | E | 6.8e-06 | 6.6e-06 | 9.8e-06 | 7.733E-06 | 1.793E-06 | 6.600E-06 | 9.800E-06 |
| 26743 | 1338 | G1338N | E | 5e-06 | 4.6e-06 | 8.4e-06 | 6.000E-06 | 2.088E-06 | 4.600E-06 | 8.400E-06 |
| 26744 | 1338 | G1338D | E | 4.9e-06 | 6.6e-06 | 5.2e-06 | 5.567E-06 | 9.074E-07 | 4.900E-06 | 6.600E-06 |
| 26745 | 1338 | G1338C | E | 4.2e-06 | 9.1e-06 | 8.4e-06 | 7.233E-06 | 2.650E-06 | 4.200E-06 | 9.100E-06 |
| 26746 | 1338 | G1338E | E | 6.4e-06 | 1.7e-05 | 4.4e-06 | 9.267E-06 | 6.772E-06 | 4.400E-06 | 1.700E-05 |
| 26747 | 1338 | G1338Q | E | 1.3e-05 | 4.6e-06 | 6.7e-06 | 8.100E-06 | 4.371E-06 | 4.600E-06 | 1.300E-05 |
| 26748 | 1338 | G1338G | E | 6.8e-06 | 7.3e-06 | 5.2e-06 | 6.433E-06 | 1.097E-06 | 5.200E-06 | 7.300E-06 |
| 26749 | 1338 | G1338H | E | 8.3e-06 | 4.9e-06 | 6.4e-06 | 6.533E-06 | 1.704E-06 | 4.900E-06 | 8.300E-06 |
| 26750 | 1338 | G1338I | E | 6.1e-06 | 4.9e-06 | 9.9e-06 | 6.967E-06 | 2.610E-06 | 4.900E-06 | 9.900E-06 |
| 26751 | 1338 | G1338L | E | 9.7e-06 | 1.1e-05 | 8.6e-06 | 9.767E-06 | 1.201E-06 | 8.600E-06 | 1.100E-05 |
| 26752 | 1338 | G1338K | E | 6.4e-06 | 6.4e-06 | 8.3e-06 | 7.033E-06 | 1.097E-06 | 6.400E-06 | 8.300E-06 |
| 26753 | 1338 | G1338M | E | 4.3e-06 | 4.7e-06 | 1.5e-05 | 8.000E-06 | 6.065E-06 | 4.300E-06 | 1.500E-05 |
| 26754 | 1338 | G1338F | E | 9.8e-06 | 6.7e-06 | 7.3e-06 | 7.933E-06 | 1.644E-06 | 6.700E-06 | 9.800E-06 |
| 26755 | 1338 | G1338P | E | 7.7e-06 | 4.1e-06 | 5.4e-06 | 5.733E-06 | 1.823E-06 | 4.100E-06 | 7.700E-06 |
| 26756 | 1338 | G1338S | E | 5.8e-06 | 4.6e-06 | 9.4e-06 | 6.600E-06 | 2.498E-06 | 4.600E-06 | 9.400E-06 |
| 26757 | 1338 | G1338T | E | 7.5e-06 | 4.7e-06 | 6.6e-06 | 6.267E-06 | 1.429E-06 | 4.700E-06 | 7.500E-06 |
| 26758 | 1338 | G1338W | E | 7.6e-06 | 9.3e-06 | 9.6e-06 | 8.833E-06 | 1.079E-06 | 7.600E-06 | 9.600E-06 |
| 26759 | 1338 | G1338Y | E | 6.3e-06 | 6.5e-06 | 8.2e-06 | 7.000E-06 | 1.044E-06 | 6.300E-06 | 8.200E-06 |
| 26760 | 1338 | G1338V | E | 9.3e-06 | 8.2e-06 | 8.1e-06 | 8.533E-06 | 6.658E-07 | 8.100E-06 | 9.300E-06 |
| 26761 | 1339 | R1339A | E | 6.4e-06 | 8.2e-06 | 8.1e-06 | 7.567E-06 | 1.012E-06 | 6.400E-06 | 8.200E-06 |
| 26762 | 1339 | R1339R | E | 6.3e-06 | 6.5e-06 | 6.2e-06 | 6.333E-06 | 1.528E-07 | 6.200E-06 | 6.500E-06 |
| 26763 | 1339 | R1339N | E | 7.7e-06 | 6.4e-06 | 8.4e-06 | 7.500E-06 | 1.015E-06 | 6.400E-06 | 8.400E-06 |
| 26764 | 1339 | R1339D | E | 4.5e-06 | 1.1e-05 | 4.7e-06 | 6.733E-06 | 3.696E-06 | 4.500E-06 | 1.100E-05 |
| 26765 | 1339 | R1339C | E | 7.1e-06 | 5.1e-06 | 1.3e-05 | 8.400E-06 | 4.107E-06 | 5.100E-06 | 1.300E-05 |
| 26766 | 1339 | R1339E | E | 5.1e-06 | 4.6e-06 | 9.4e-06 | 6.367E-06 | 2.639E-06 | 4.600E-06 | 9.400E-06 |
| 26767 | 1339 | R1339Q | E | 1e-05 | 5.9e-06 | 7.7e-06 | 7.867E-06 | 2.055E-06 | 5.900E-06 | 1.000E-05 |
| 26768 | 1339 | R1339G | E | 4.6e-06 | 8.9e-06 | 1.3e-05 | 8.833E-06 | 4.200E-06 | 4.600E-06 | 1.300E-05 |
| 26769 | 1339 | R1339H | E | 5.8e-06 | 6.4e-06 | 5.6e-06 | 5.933E-06 | 4.163E-07 | 5.600E-06 | 6.400E-06 |
| 26770 | 1339 | R1339I | E | 9.1e-06 | 8.1e-06 | 1.3e-05 | 1.007E-05 | 2.589E-06 | 8.100E-06 | 1.300E-05 |
| 26771 | 1339 | R1339L | E | 7.6e-06 | 5e-06 | 9.8e-06 | 7.467E-06 | 2.403E-06 | 5.000E-06 | 9.800E-06 |
| 26772 | 1339 | R1339K | E | 9e-06 | 8.1e-06 | 5.6e-06 | 7.567E-06 | 1.762E-06 | 5.600E-06 | 9.000E-06 |
| 26773 | 1339 | R1339M | E | 1.2e-05 | 1e-05 | 5.3e-06 | 9.100E-06 | 3.439E-06 | 5.300E-06 | 1.200E-05 |
| 26774 | 1339 | R1339F | E | 4.9e-06 | 1e-05 | 9.5e-06 | 8.133E-06 | 2.811E-06 | 4.900E-06 | 1.000E-05 |

| | | | | | | | | | | |
|---|---|---|---|---|---|---|---|---|---|---|
| 26775 | 1339 | R1339P | E | 7.1e-06 | 1.1e-05 | 6.8e-06 | 8.300E-06 | 2.343E-06 | 6.800E-06 | 1.100E-05 |
| 26776 | 1339 | R1339S | E | 1.4e-05 | 8.2e-06 | 7e-06 | 9.733E-06 | 3.743E-06 | 7.000E-06 | 1.400E-05 |
| 26777 | 1339 | R1339T | E | 7.3e-06 | 5.1e-06 | 7.1e-06 | 6.500E-06 | 1.217E-06 | 5.100E-06 | 7.300E-06 |
| 26778 | 1339 | R1339W | E | 8e-06 | 6e-06 | 3.6e-06 | 5.867E-06 | 2.203E-06 | 3.600E-06 | 8.000E-06 |
| 26779 | 1339 | R1339Y | E | 8e-06 | 6.6e-06 | 9.6e-06 | 8.067E-06 | 1.501E-06 | 6.600E-06 | 9.600E-06 |
| 26780 | 1339 | R1339V | E | 6e-06 | 7.5e-06 | 5.8e-06 | 6.433E-06 | 9.292E-07 | 5.800E-06 | 7.500E-06 |
| 26781 | 1340 | F1340A | E | 6.3e-06 | 9.3e-06 | 9.8e-06 | 8.467E-06 | 1.893E-06 | 6.300E-06 | 9.800E-06 |
| 26782 | 1340 | F1340R | E | 7.5e-06 | 1e-05 | 3.1e-06 | 6.867E-06 | 3.493E-06 | 3.100E-06 | 1.000E-05 |
| 26783 | 1340 | F1340N | E | 7.2e-06 | 4.5e-06 | 7.9e-06 | 6.533E-06 | 1.795E-06 | 4.500E-06 | 7.900E-06 |
| 26784 | 1340 | F1340D | E | 9.3e-06 | 8.2e-06 | 1.3e-05 | 1.017E-05 | 2.515E-06 | 8.200E-06 | 1.300E-05 |
| 26785 | 1340 | F1340C | E | 6e-06 | 9.1e-06 | 5.3e-06 | 6.800E-06 | 2.022E-06 | 5.300E-06 | 9.100E-06 |
| 26786 | 1340 | F1340E | E | 5.5e-06 | 9.4e-06 | 9e-06 | 7.967E-06 | 2.146E-06 | 5.500E-06 | 9.400E-06 |
| 26787 | 1340 | F1340Q | E | 8.5e-06 | 5.8e-06 | 1.1e-05 | 8.433E-06 | 2.601E-06 | 5.800E-06 | 1.100E-05 |
| 26788 | 1340 | F1340G | E | 3.4e-06 | 1.4e-05 | 1.1e-05 | 9.467E-06 | 5.464E-06 | 3.400E-06 | 1.400E-05 |
| 26789 | 1340 | F1340H | E | 9.6e-06 | 9.7e-06 | 9.4e-06 | 9.567E-06 | 1.528E-07 | 9.400E-06 | 9.700E-06 |
| 26790 | 1340 | F1340I | E | 6.1e-06 | 1.1e-05 | 1.2e-05 | 9.700E-06 | 3.158E-06 | 6.100E-06 | 1.200E-05 |
| 26791 | 1340 | F1340L | E | 1e-05 | 5.1e-06 | 1.2e-05 | 9.033E-06 | 3.550E-06 | 5.100E-06 | 1.200E-05 |
| 26792 | 1340 | F1340K | E | 9.7e-06 | 9.3e-06 | 1.1e-05 | 1.000E-05 | 8.888E-07 | 9.300E-06 | 1.100E-05 |
| 26793 | 1340 | F1340M | E | 7.7e-06 | 1.1e-05 | 5.6e-06 | 8.100E-06 | 2.722E-06 | 5.600E-06 | 1.100E-05 |
| 26794 | 1340 | F1340F | E | 9.4e-06 | 1.6e-05 | 9.3e-06 | 1.157E-05 | 3.840E-06 | 9.300E-06 | 1.600E-05 |
| 26795 | 1340 | F1340P | E | 5.9e-06 | 8.9e-06 | 7.8e-06 | 7.533E-06 | 1.518E-06 | 5.900E-06 | 8.900E-06 |
| 26796 | 1340 | F1340S | E | 6.1e-06 | 7.1e-06 | 7.3e-06 | 6.833E-06 | 6.429E-07 | 6.100E-06 | 7.300E-06 |
| 26797 | 1340 | F1340T | E | 1.1e-05 | 7.1e-06 | 1.1e-05 | 9.700E-06 | 2.252E-06 | 7.100E-06 | 1.100E-05 |
| 26798 | 1340 | F1340W | E | 5.7e-06 | 7.4e-06 | 6.5e-06 | 6.533E-06 | 8.505E-07 | 5.700E-06 | 7.400E-06 |
| 26799 | 1340 | F1340Y | E | 8.5e-06 | 9.8e-06 | 1.1e-05 | 9.767E-06 | 1.250E-06 | 8.500E-06 | 1.100E-05 |
| 26800 | 1340 | F1340V | E | 6.1e-06 | 1e-05 | 1.3e-05 | 9.700E-06 | 3.460E-06 | 6.100E-06 | 1.300E-05 |
| 26801 | 1341 | T1341A | E | 9.1e-06 | 5.5e-06 | 9.1e-06 | 7.900E-06 | 2.078E-06 | 5.500E-06 | 9.100E-06 |
| 26802 | 1341 | T1341R | E | 9.6e-06 | 8.5e-06 | 8.2e-06 | 8.767E-06 | 7.371E-07 | 8.200E-06 | 9.600E-06 |
| 26803 | 1341 | T1341N | E | 8.1e-06 | 5.9e-06 | 7e-06 | 7.000E-06 | 1.100E-06 | 5.900E-06 | 8.100E-06 |
| 26804 | 1341 | T1341D | E | 6.6e-06 | 1.6e-05 | 5e-06 | 9.200E-06 | 5.943E-06 | 5.000E-06 | 1.600E-05 |
| 26805 | 1341 | T1341C | E | 1e-05 | 1.5e-05 | 9.9e-06 | 1.163E-05 | 2.916E-06 | 9.900E-06 | 1.500E-05 |
| 26806 | 1341 | T1341E | E | 1.1e-05 | 7.1e-06 | 1.3e-05 | 1.037E-05 | 3.001E-06 | 7.100E-06 | 1.300E-05 |
| 26807 | 1341 | T1341Q | E | 9.3e-06 | 1.1e-05 | 7.5e-06 | 9.267E-06 | 1.750E-06 | 7.500E-06 | 1.100E-05 |
| 26808 | 1341 | T1341G | E | 9.7e-06 | 3.6e-06 | 5.9e-06 | 6.400E-06 | 3.081E-06 | 3.600E-06 | 9.700E-06 |
| 26809 | 1341 | T1341H | E | 1.2e-05 | 8.5e-06 | 1.3e-05 | 1.117E-05 | 2.363E-06 | 8.500E-06 | 1.300E-05 |
| 26810 | 1341 | T1341I | E | 8.5e-06 | 8.6e-06 | 6.4e-06 | 7.833E-06 | 1.242E-06 | 6.400E-06 | 8.600E-06 |
| 26811 | 1341 | T1341L | E | 1.3e-05 | 7.1e-06 | 1e-05 | 1.003E-05 | 2.950E-06 | 7.100E-06 | 1.300E-05 |
| 26812 | 1341 | T1341K | E | 1.4e-05 | 7e-06 | 8.5e-06 | 9.833E-06 | 3.686E-06 | 7.000E-06 | 1.400E-05 |
| 26813 | 1341 | T1341M | E | 6e-06 | 1.4e-05 | 6.8e-06 | 8.933E-06 | 4.406E-06 | 6.000E-06 | 1.400E-05 |
| 26814 | 1341 | T1341F | E | 9.3e-06 | 9e-06 | 6.1e-06 | 8.133E-06 | 1.767E-06 | 6.100E-06 | 9.300E-06 |
| 26815 | 1341 | T1341P | E | 1.1e-05 | 4.7e-06 | 5.6e-06 | 7.100E-06 | 3.407E-06 | 4.700E-06 | 1.100E-05 |
| 26816 | 1341 | T1341S | E | 5.1e-06 | 9.2e-06 | 8.2e-06 | 7.500E-06 | 2.138E-06 | 5.100E-06 | 9.200E-06 |
| 26817 | 1341 | T1341T | E | 6.2e-06 | 4.8e-06 | 4.1e-06 | 5.033E-06 | 1.069E-06 | 4.100E-06 | 6.200E-06 |
| 26818 | 1341 | T1341W | E | 5.2e-06 | 5.4e-06 | 5.1e-06 | 5.233E-06 | 1.528E-07 | 5.100E-06 | 5.400E-06 |
| 26819 | 1341 | T1341Y | E | 1.3e-05 | 8.6e-06 | 1.5e-05 | 1.220E-05 | 3.274E-06 | 8.600E-06 | 1.500E-05 |
| 26820 | 1341 | T1341V | E | 8.4e-06 | 1.2e-05 | 8.8e-06 | 9.733E-06 | 1.973E-06 | 8.400E-06 | 1.200E-05 |
| 26821 | 1342 | I1342A | E | 5.6e-06 | 8e-06 | 7.8e-06 | 7.133E-06 | 1.332E-06 | 5.600E-06 | 8.000E-06 |
| 26822 | 1342 | I1342R | E | 7.7e-06 | 8.1e-06 | 5.8e-06 | 7.200E-06 | 1.229E-06 | 5.800E-06 | 8.100E-06 |
| 26823 | 1342 | I1342N | E | 6.7e-06 | 6.4e-06 | 5.9e-06 | 6.333E-06 | 4.041E-07 | 5.900E-06 | 6.700E-06 |
| 26824 | 1342 | I1342D | E | 4.2e-06 | 6.5e-06 | 9.7e-06 | 6.800E-06 | 2.762E-06 | 4.200E-06 | 9.700E-06 |
| 26825 | 1342 | I1342C | E | 4.4e-06 | 8.8e-06 | 1.1e-05 | 8.067E-06 | 3.361E-06 | 4.400E-06 | 1.100E-05 |
| 26826 | 1342 | I1342E | E | 3.6e-06 | 9.4e-06 | 1.2e-05 | 8.333E-06 | 4.300E-06 | 3.600E-06 | 1.200E-05 |
| 26827 | 1342 | I1342Q | E | 1e-05 | 1.3e-05 | 6e-06 | 9.667E-06 | 3.512E-06 | 6.000E-06 | 1.300E-05 |
| 26828 | 1342 | I1342G | E | 1.4e-05 | 1.3e-05 | 4.2e-06 | 1.040E-05 | 5.393E-06 | 4.200E-06 | 1.400E-05 |
| 26829 | 1342 | I1342H | E | 7.8e-06 | 6.4e-06 | 6.1e-06 | 6.767E-06 | 9.074E-07 | 6.100E-06 | 7.800E-06 |
| 26830 | 1342 | I1342I | E | 1e-05 | 6.4e-06 | 5.7e-06 | 7.367E-06 | 2.307E-06 | 5.700E-06 | 1.000E-05 |
| 26831 | 1342 | I1342L | E | 6.8e-06 | 9e-06 | 6.7e-06 | 7.500E-06 | 1.300E-06 | 6.700E-06 | 9.000E-06 |
| 26832 | 1342 | I1342K | E | 8.4e-06 | 6.5e-06 | 9.5e-06 | 8.133E-06 | 1.518E-06 | 6.500E-06 | 9.500E-06 |
| 26833 | 1342 | I1342M | E | 7.3e-06 | 1.1e-05 | 5.4e-06 | 7.900E-06 | 2.848E-06 | 5.400E-06 | 1.100E-05 |
| 26834 | 1342 | I1342F | E | 6.3e-06 | 1.4e-05 | 5.5e-06 | 8.600E-06 | 4.694E-06 | 5.500E-06 | 1.400E-05 |
| 26835 | 1342 | I1342P | E | 6.6e-06 | 1e-05 | 3.9e-06 | 6.833E-06 | 3.057E-06 | 3.900E-06 | 1.000E-05 |
| 26836 | 1342 | I1342S | E | 5.6e-06 | 6.6e-06 | 6.3e-06 | 6.167E-06 | 5.132E-07 | 5.600E-06 | 6.600E-06 |
| 26837 | 1342 | I1342T | E | 3.8e-06 | 1e-05 | 4.2e-06 | 6.000E-06 | 3.470E-06 | 3.800E-06 | 1.000E-05 |
| 26838 | 1342 | I1342W | E | 6.4e-06 | 7.6e-06 | 9.9e-06 | 7.967E-06 | 1.779E-06 | 6.400E-06 | 9.900E-06 |
| 26839 | 1342 | I1342Y | E | 8.6e-06 | 1.2e-05 | 5.3e-06 | 8.633E-06 | 3.350E-06 | 5.300E-06 | 1.200E-05 |
| 26840 | 1342 | I1342V | E | 7.1e-06 | 5.8e-06 | 1.1e-05 | 7.967E-06 | 2.706E-06 | 5.800E-06 | 1.100E-05 |
| 26841 | 1343 | S1343A | E | 9.5e-06 | 9.9e-06 | 8.8e-06 | 9.400E-06 | 5.568E-07 | 8.800E-06 | 9.900E-06 |
| 26842 | 1343 | S1343R | E | 1.1e-05 | 9.7e-06 | 1.3e-05 | 1.123E-05 | 1.662E-06 | 9.700E-06 | 1.300E-05 |
| 26843 | 1343 | S1343N | E | 9.7e-06 | 1e-05 | 5.9e-06 | 8.533E-06 | 2.285E-06 | 5.900E-06 | 1.000E-05 |
| 26844 | 1343 | S1343D | E | 1e-05 | 7.1e-06 | 1.5e-05 | 1.070E-05 | 3.996E-06 | 7.100E-06 | 1.500E-05 |
| 26845 | 1343 | S1343C | E | 7.4e-06 | 8e-06 | 8.8e-06 | 8.067E-06 | 7.024E-07 | 7.400E-06 | 8.800E-06 |
| 26846 | 1343 | S1343E | E | 9.4e-06 | 1e-05 | 1.3e-05 | 1.080E-05 | 1.929E-06 | 9.400E-06 | 1.300E-05 |
| 26847 | 1343 | S1343Q | E | 8.5e-06 | 1.2e-05 | 6.5e-06 | 9.000E-06 | 2.784E-06 | 6.500E-06 | 1.200E-05 |
| 26848 | 1343 | S1343G | E | 9.5e-06 | 8.8e-06 | 4.9e-06 | 7.733E-06 | 2.479E-06 | 4.900E-06 | 9.500E-06 |
| 26849 | 1343 | S1343H | E | 7.9e-06 | 7.1e-06 | 6.9e-06 | 7.300E-06 | 5.292E-07 | 6.900E-06 | 7.900E-06 |
| 26850 | 1343 | S1343I | E | 6.4e-06 | 7.1e-06 | 1.3e-05 | 8.833E-06 | 3.625E-06 | 6.400E-06 | 1.300E-05 |
| 26851 | 1343 | S1343L | E | 1.5e-05 | 1.4e-05 | 9.1e-06 | 1.270E-05 | 3.158E-06 | 9.100E-06 | 1.500E-05 |
| 26852 | 1343 | S1343K | E | 7.1e-06 | 1.3e-05 | 1e-05 | 1.003E-05 | 2.950E-06 | 7.100E-06 | 1.300E-05 |
| 26853 | 1343 | S1343M | E | 5.9e-06 | 9.6e-06 | 8.2e-06 | 7.900E-06 | 1.868E-06 | 5.900E-06 | 9.600E-06 |
| 26854 | 1343 | S1343F | E | 7.7e-06 | 6.3e-06 | 6.2e-06 | 6.733E-06 | 8.386E-07 | 6.200E-06 | 7.700E-06 |
| 26855 | 1343 | S1343P | E | 5.5e-06 | 1.2e-05 | 7.1e-06 | 8.200E-06 | 3.387E-06 | 5.500E-06 | 1.200E-05 |
| 26856 | 1343 | S1343S | E | 8.4e-06 | 6.8e-06 | 2.7e-06 | 5.967E-06 | 2.940E-06 | 2.700E-06 | 8.400E-06 |
| 26857 | 1343 | S1343T | E | 1e-05 | 7.9e-06 | 7.5e-06 | 8.467E-06 | 1.343E-06 | 7.500E-06 | 1.000E-05 |
| 26858 | 1343 | S1343W | E | 8.2e-06 | 7.7e-06 | 8.4e-06 | 8.100E-06 | 3.606E-07 | 7.700E-06 | 8.400E-06 |
| 26859 | 1343 | S1343Y | E | 6e-06 | 6.3e-06 | 4.8e-06 | 5.700E-06 | 7.937E-07 | 4.800E-06 | 6.300E-06 |

| | | | | | | | | | | |
|---|---|---|---|---|---|---|---|---|---|---|
| 26860 | 1343 | S1343V | E | 1e-05 | 6.1e-06 | 7.2e-06 | 7.767E-06 | 2.011E-06 | 6.100E-06 | 1.000E-05 |
| 26861 | 1344 | A1344A | E | 9.3e-06 | 6.1e-06 | 5.8e-06 | 7.067E-06 | 1.940E-06 | 5.800E-06 | 9.300E-06 |
| 26862 | 1344 | A1344R | E | 5.5e-06 | 1.3e-05 | 9.4e-06 | 9.300E-06 | 3.751E-06 | 5.500E-06 | 1.300E-05 |
| 26863 | 1344 | A1344N | E | 9.1e-06 | 7.4e-06 | 5.7e-06 | 7.400E-06 | 1.700E-06 | 5.700E-06 | 9.100E-06 |
| 26864 | 1344 | A1344D | E | 6.9e-06 | 8.4e-06 | 6.6e-06 | 7.300E-06 | 9.644E-07 | 6.600E-06 | 8.400E-06 |
| 26865 | 1344 | A1344C | E | 7.4e-06 | 4.6e-06 | 9.3e-06 | 7.100E-06 | 2.364E-06 | 4.600E-06 | 9.300E-06 |
| 26866 | 1344 | A1344E | E | 4.7e-06 | 6.8e-06 | 4.8e-06 | 5.433E-06 | 1.185E-06 | 4.700E-06 | 6.800E-06 |
| 26867 | 1344 | A1344Q | E | 7.3e-06 | 9.3e-06 | 5.3e-06 | 7.300E-06 | 2.000E-06 | 5.300E-06 | 9.300E-06 |
| 26868 | 1344 | A1344G | E | 6.8e-06 | 7.8e-06 | 7.1e-06 | 7.233E-06 | 5.132E-07 | 6.800E-06 | 7.800E-06 |
| 26869 | 1344 | A1344H | E | 7.2e-06 | 9.1e-06 | 1e-05 | 8.767E-06 | 1.429E-06 | 7.200E-06 | 1.000E-05 |
| 26870 | 1344 | A1344I | E | 9.9e-06 | 8.9e-06 | 1.2e-05 | 1.027E-05 | 1.582E-06 | 8.900E-06 | 1.200E-05 |
| 26871 | 1344 | A1344L | E | 8.7e-06 | 1.4e-05 | 9.3e-06 | 1.067E-05 | 2.902E-06 | 8.700E-06 | 1.400E-05 |
| 26872 | 1344 | A1344K | E | 6.6e-06 | 9.5e-06 | 6.1e-06 | 7.400E-06 | 1.836E-06 | 6.100E-06 | 9.500E-06 |
| 26873 | 1344 | A1344M | E | 6.1e-06 | 5.6e-06 | 7.4e-06 | 6.367E-06 | 9.292E-07 | 5.600E-06 | 7.400E-06 |
| 26874 | 1344 | A1344F | E | 5.6e-06 | 7e-06 | 7.9e-06 | 6.833E-06 | 1.159E-06 | 5.600E-06 | 7.900E-06 |
| 26875 | 1344 | A1344P | E | 5.4e-06 | 1e-05 | 9.5e-06 | 8.300E-06 | 2.524E-06 | 5.400E-06 | 1.000E-05 |
| 26876 | 1344 | A1344S | E | 1.3e-05 | 5.9e-06 | 1e-05 | 9.633E-06 | 3.564E-06 | 5.900E-06 | 1.300E-05 |
| 26877 | 1344 | A1344T | E | 4.5e-06 | 7.4e-06 | 8e-06 | 6.633E-06 | 1.872E-06 | 4.500E-06 | 8.000E-06 |
| 26878 | 1344 | A1344W | E | 7e-06 | 9.5e-06 | 7.4e-06 | 7.967E-06 | 1.343E-06 | 7.000E-06 | 9.500E-06 |
| 26879 | 1344 | A1344Y | E | 9.5e-06 | 8.8e-06 | 8.8e-06 | 9.033E-06 | 4.041E-07 | 8.800E-06 | 9.500E-06 |
| 26880 | 1344 | A1344V | E | 9.2e-06 | 9.5e-06 | 9e-06 | 9.233E-06 | 2.517E-07 | 9.000E-06 | 9.500E-06 |
| 26881 | 1345 | D1345A | E | 4.7e-06 | 9.9e-06 | 5.8e-06 | 6.800E-06 | 2.740E-06 | 4.700E-06 | 9.900E-06 |
| 26882 | 1345 | D1345R | E | 8.5e-06 | 6.6e-06 | 1.1e-05 | 8.700E-06 | 2.207E-06 | 6.600E-06 | 1.100E-05 |
| 26883 | 1345 | D1345N | E | 1.1e-05 | 6.1e-06 | 7.9e-06 | 8.333E-06 | 2.479E-06 | 6.100E-06 | 1.100E-05 |
| 26884 | 1345 | D1345D | E | 5.6e-06 | 1.1e-05 | 6.4e-06 | 7.667E-06 | 2.914E-06 | 5.600E-06 | 1.100E-05 |
| 26885 | 1345 | D1345C | E | 6.9e-06 | 5.6e-06 | 8e-06 | 6.833E-06 | 1.201E-06 | 5.600E-06 | 8.000E-06 |
| 26886 | 1345 | D1345E | E | 9.7e-06 | 4.4e-06 | 5e-06 | 6.367E-06 | 2.902E-06 | 4.400E-06 | 9.700E-06 |
| 26887 | 1345 | D1345Q | E | 8.4e-06 | 7.1e-06 | 6.6e-06 | 7.367E-06 | 9.292E-07 | 6.600E-06 | 8.400E-06 |
| 26888 | 1345 | D1345G | E | 1.1e-05 | 8.7e-06 | 9.4e-06 | 9.700E-06 | 1.179E-06 | 8.700E-06 | 1.100E-05 |
| 26889 | 1345 | D1345H | E | 1.1e-05 | 5.6e-06 | 7.3e-06 | 7.967E-06 | 2.761E-06 | 5.600E-06 | 1.100E-05 |
| 26890 | 1345 | D1345I | E | 1e-05 | 9.1e-06 | 8.2e-06 | 9.100E-06 | 9.000E-07 | 8.200E-06 | 1.000E-05 |
| 26891 | 1345 | D1345L | E | 8e-06 | 4.9e-06 | 9.2e-06 | 7.367E-06 | 2.219E-06 | 4.900E-06 | 9.200E-06 |
| 26892 | 1345 | D1345K | E | 6.9e-06 | 7.6e-06 | 7.5e-06 | 7.333E-06 | 3.786E-07 | 6.900E-06 | 7.600E-06 |
| 26893 | 1345 | D1345M | E | 9.2e-06 | 8.9e-06 | 8.9e-06 | 9.000E-06 | 1.732E-07 | 8.900E-06 | 9.200E-06 |
| 26894 | 1345 | D1345F | E | 5e-06 | 4.8e-06 | 1.4e-05 | 7.933E-06 | 5.255E-06 | 4.800E-06 | 1.400E-05 |
| 26895 | 1345 | D1345P | E | 7.7e-06 | 9.3e-06 | 5.5e-06 | 7.500E-06 | 1.908E-06 | 5.500E-06 | 9.300E-06 |
| 26896 | 1345 | D1345S | E | 9e-06 | 1.4e-05 | 6.3e-06 | 9.767E-06 | 3.907E-06 | 6.300E-06 | 1.400E-05 |
| 26897 | 1345 | D1345T | E | 1.8e-05 | 5.3e-06 | 6.7e-06 | 1.000E-05 | 6.963E-06 | 5.300E-06 | 1.800E-05 |
| 26898 | 1345 | D1345W | E | 1.7e-05 | 6.2e-06 | 6.4e-06 | 9.867E-06 | 6.178E-06 | 6.200E-06 | 1.700E-05 |
| 26899 | 1345 | D1345Y | E | 6.2e-06 | 8.8e-06 | 6.3e-06 | 7.100E-06 | 1.473E-06 | 6.200E-06 | 8.800E-06 |
| 26900 | 1345 | D1345V | E | 5.8e-06 | 7.8e-06 | 6.4e-06 | 6.667E-06 | 1.026E-06 | 5.800E-06 | 7.800E-06 |
| 26901 | 1346 | T1346A | E | 1.3e-05 | 8.4e-06 | 9.4e-06 | 1.027E-05 | 2.419E-06 | 8.400E-06 | 1.300E-05 |
| 26902 | 1346 | T1346R | E | 6.4e-06 | 9.9e-06 | 1.2e-05 | 9.433E-06 | 2.829E-06 | 6.400E-06 | 1.200E-05 |
| 26903 | 1346 | T1346N | E | 4.6e-06 | 9.2e-06 | 9.5e-06 | 7.767E-06 | 2.747E-06 | 4.600E-06 | 9.500E-06 |
| 26904 | 1346 | T1346D | E | 7.7e-06 | 1.1e-05 | 9.1e-06 | 9.267E-06 | 1.656E-06 | 7.700E-06 | 1.100E-05 |
| 26905 | 1346 | T1346C | E | 5e-06 | 9.3e-06 | 2.3e-06 | 5.533E-06 | 3.530E-06 | 2.300E-06 | 9.300E-06 |
| 26906 | 1346 | T1346E | E | 8.8e-06 | 7.1e-06 | 7.5e-06 | 7.800E-06 | 8.888E-07 | 7.100E-06 | 8.800E-06 |
| 26907 | 1346 | T1346Q | E | 1.3e-05 | 4.5e-06 | 5e-06 | 7.500E-06 | 4.770E-06 | 4.500E-06 | 1.300E-05 |
| 26908 | 1346 | T1346G | E | 5.1e-06 | 9.3e-06 | 8e-06 | 7.467E-06 | 2.150E-06 | 5.100E-06 | 9.300E-06 |
| 26909 | 1346 | T1346H | E | 1.2e-05 | 9.3e-06 | 4.8e-06 | 8.700E-06 | 3.637E-06 | 4.800E-06 | 1.200E-05 |
| 26910 | 1346 | T1346I | E | 1.7e-05 | 4.9e-06 | 5.7e-06 | 9.200E-06 | 6.767E-06 | 4.900E-06 | 1.700E-05 |
| 26911 | 1346 | T1346L | E | 6.9e-06 | 7.2e-06 | 1.1e-05 | 8.367E-06 | 2.285E-06 | 6.900E-06 | 1.100E-05 |
| 26912 | 1346 | T1346K | E | 9.2e-06 | 5.5e-06 | 1.3e-05 | 9.233E-06 | 3.750E-06 | 5.500E-06 | 1.300E-05 |
| 26913 | 1346 | T1346M | E | 1.1e-05 | 9.8e-06 | 8.4e-06 | 9.733E-06 | 1.301E-06 | 8.400E-06 | 1.100E-05 |
| 26914 | 1346 | T1346F | E | 6.6e-06 | 7.5e-06 | 7.4e-06 | 7.167E-06 | 4.933E-07 | 6.600E-06 | 7.500E-06 |
| 26915 | 1346 | T1346P | E | 5e-06 | 5.5e-06 | 4e-06 | 4.833E-06 | 7.638E-07 | 4.000E-06 | 5.500E-06 |
| 26916 | 1346 | T1346S | E | 9e-06 | 6.6e-06 | 7e-06 | 7.533E-06 | 1.286E-06 | 6.600E-06 | 9.000E-06 |
| 26917 | 1346 | T1346T | E | 6.1e-06 | 6.7e-06 | 5.7e-06 | 6.167E-06 | 5.033E-07 | 5.700E-06 | 6.700E-06 |
| 26918 | 1346 | T1346W | E | 8.4e-06 | 9.2e-06 | 6.8e-06 | 8.133E-06 | 1.222E-06 | 6.800E-06 | 9.200E-06 |
| 26919 | 1346 | T1346Y | E | 1e-05 | 9e-06 | 1e-05 | 9.667E-06 | 5.774E-07 | 9.000E-06 | 1.000E-05 |
| 26920 | 1346 | T1346V | E | 8.3e-06 | 1.2e-05 | 1.1e-05 | 1.043E-05 | 1.914E-06 | 8.300E-06 | 1.200E-05 |
| 26921 | 1347 | S1347A | E | 6.1e-06 | 5.9e-06 | 9.6e-06 | 7.200E-06 | 2.081E-06 | 5.900E-06 | 9.600E-06 |
| 26922 | 1347 | S1347R | E | 4.9e-06 | 9.7e-06 | 6.8e-06 | 7.133E-06 | 2.417E-06 | 4.900E-06 | 9.700E-06 |
| 26923 | 1347 | S1347N | E | 7.5e-06 | 5.4e-06 | 9.1e-06 | 7.333E-06 | 1.856E-06 | 5.400E-06 | 9.100E-06 |
| 26924 | 1347 | S1347D | E | 9.3e-06 | 6.8e-06 | 5.4e-06 | 7.167E-06 | 1.976E-06 | 5.400E-06 | 9.300E-06 |
| 26925 | 1347 | S1347C | E | 1e-05 | 1e-05 | 8.1e-06 | 9.367E-06 | 1.097E-06 | 8.100E-06 | 1.000E-05 |
| 26926 | 1347 | S1347E | E | 7.7e-06 | 6e-06 | 5.6e-06 | 6.433E-06 | 1.115E-06 | 5.600E-06 | 7.700E-06 |
| 26927 | 1347 | S1347Q | E | 1e-05 | 6.2e-06 | 4.6e-06 | 6.933E-06 | 2.774E-06 | 4.600E-06 | 1.000E-05 |
| 26928 | 1347 | S1347G | E | 7.1e-06 | 1e-05 | 5.5e-06 | 7.533E-06 | 2.281E-06 | 5.500E-06 | 1.000E-05 |
| 26929 | 1347 | S1347H | E | 6.8e-06 | 6.6e-06 | 6.3e-06 | 6.567E-06 | 2.517E-07 | 6.300E-06 | 6.800E-06 |
| 26930 | 1347 | S1347I | E | 8.4e-06 | 9.6e-06 | 4.7e-06 | 7.567E-06 | 2.554E-06 | 4.700E-06 | 9.600E-06 |
| 26931 | 1347 | S1347L | E | 6.2e-06 | 6.2e-06 | 8.9e-06 | 7.100E-06 | 1.559E-06 | 6.200E-06 | 8.900E-06 |
| 26932 | 1347 | S1347K | E | 1.2e-05 | 6.8e-06 | 1.1e-05 | 9.933E-06 | 2.759E-06 | 6.800E-06 | 1.200E-05 |
| 26933 | 1347 | S1347M | E | 6.6e-06 | 8.3e-06 | 6.1e-06 | 7.000E-06 | 1.153E-06 | 6.100E-06 | 8.300E-06 |
| 26934 | 1347 | S1347F | E | 6.4e-06 | 7.9e-06 | 7.2e-06 | 7.167E-06 | 7.506E-07 | 6.400E-06 | 7.900E-06 |
| 26935 | 1347 | S1347P | E | 1e-05 | 9.6e-06 | 7.6e-06 | 9.067E-06 | 1.286E-06 | 7.600E-06 | 1.000E-05 |
| 26936 | 1347 | S1347S | E | 9.8e-06 | 1e-05 | 7.7e-06 | 9.167E-06 | 1.274E-06 | 7.700E-06 | 1.000E-05 |
| 26937 | 1347 | S1347T | E | 5.2e-06 | 6.6e-06 | 1.3e-05 | 8.267E-06 | 4.159E-06 | 5.200E-06 | 1.300E-05 |
| 26938 | 1347 | S1347W | E | 1.7e-05 | 8.9e-06 | 8.4e-06 | 1.143E-05 | 4.827E-06 | 8.400E-06 | 1.700E-05 |
| 26939 | 1347 | S1347Y | E | 1.4e-05 | 3.5e-06 | 9.7e-06 | 9.067E-06 | 5.279E-06 | 3.500E-06 | 1.400E-05 |
| 26940 | 1347 | S1347V | E | 5.5e-06 | 6.9e-06 | 6.9e-06 | 6.433E-06 | 8.083E-07 | 5.500E-06 | 6.900E-06 |
| 26941 | 1348 | K1348A | E | 7.3e-06 | 1e-05 | 9.3e-06 | 8.867E-06 | 1.401E-06 | 7.300E-06 | 1.000E-05 |
| 26942 | 1348 | K1348R | E | 5.8e-06 | 1e-05 | 9.1e-06 | 8.300E-06 | 2.211E-06 | 5.800E-06 | 1.000E-05 |
| 26943 | 1348 | K1348N | E | 6.8e-06 | 6.9e-06 | 7.4e-06 | 7.033E-06 | 3.215E-07 | 6.800E-06 | 7.400E-06 |
| 26944 | 1348 | K1348D | E | 6.9e-06 | 1.2e-05 | 7.5e-06 | 8.800E-06 | 2.787E-06 | 6.900E-06 | 1.200E-05 |

| | | | | | | | | | | |
|---|---|---|---|---|---|---|---|---|---|---|
| 26945 | 1348 | K1348C | E | 8e-06 | 7.7e-06 | 6.8e-06 | 7.500E-06 | 6.245E-07 | 6.800E-06 | 8.000E-06 |
| 26946 | 1348 | K1348E | E | 4.1e-06 | 8.5e-06 | 1.2e-05 | 8.200E-06 | 3.959E-06 | 4.100E-06 | 1.200E-05 |
| 26947 | 1348 | K1348Q | E | 8.8e-06 | 1.4e-05 | 1e-05 | 1.093E-05 | 2.723E-06 | 8.800E-06 | 1.400E-05 |
| 26948 | 1348 | K1348G | E | 1.1e-05 | 8e-06 | 8.6e-06 | 9.200E-06 | 1.587E-06 | 8.000E-06 | 1.100E-05 |
| 26949 | 1348 | K1348H | E | 6.7e-06 | 7.9e-06 | 9.7e-06 | 8.100E-06 | 1.510E-06 | 6.700E-06 | 9.700E-06 |
| 26950 | 1348 | K1348I | E | 5.1e-06 | 6.6e-06 | 8.3e-06 | 6.667E-06 | 1.601E-06 | 5.100E-06 | 8.300E-06 |
| 26951 | 1348 | K1348L | E | 7.8e-06 | 5.5e-06 | 9.3e-06 | 7.533E-06 | 1.914E-06 | 5.500E-06 | 9.300E-06 |
| 26952 | 1348 | K1348K | E | 1e-05 | 9.5e-06 | 6.5e-06 | 8.667E-06 | 1.893E-06 | 6.500E-06 | 1.000E-05 |
| 26953 | 1348 | K1348M | E | 1.3e-05 | 7.3e-06 | 2.7e-06 | 7.667E-06 | 5.160E-06 | 2.700E-06 | 1.300E-05 |
| 26954 | 1348 | K1348F | E | 8.1e-06 | 8.7e-06 | 5.9e-06 | 7.567E-06 | 1.474E-06 | 5.900E-06 | 8.700E-06 |
| 26955 | 1348 | K1348P | E | 1.3e-05 | 8.6e-06 | 6.5e-06 | 9.367E-06 | 3.317E-06 | 6.500E-06 | 1.300E-05 |
| 26956 | 1348 | K1348S | E | 8.9e-06 | 6.7e-06 | 6.3e-06 | 7.300E-06 | 1.400E-06 | 6.300E-06 | 8.900E-06 |
| 26957 | 1348 | K1348T | E | 8.1e-06 | 9.5e-06 | 1e-05 | 9.200E-06 | 9.849E-07 | 8.100E-06 | 1.000E-05 |
| 26958 | 1348 | K1348W | E | 8.7e-06 | 1.2e-05 | 1.2e-05 | 1.090E-05 | 1.905E-06 | 8.700E-06 | 1.200E-05 |
| 26959 | 1348 | K1348Y | E | 9.1e-06 | 8.4e-06 | 3.8e-06 | 7.100E-06 | 2.879E-06 | 3.800E-06 | 9.100E-06 |
| 26960 | 1348 | K1348V | E | 1.2e-05 | 1.4e-05 | 9.9e-06 | 1.197E-05 | 2.050E-06 | 9.900E-06 | 1.400E-05 |
| 26961 | 1349 | N1349A | E | 6.8e-06 | 4.6e-06 | 5.1e-06 | 5.500E-06 | 1.153E-06 | 4.600E-06 | 6.800E-06 |
| 26962 | 1349 | N1349R | E | 1.1e-05 | 1.2e-05 | 6.6e-06 | 9.867E-06 | 2.873E-06 | 6.600E-06 | 1.200E-05 |
| 26963 | 1349 | N1349N | E | 6.5e-06 | 9e-06 | 1e-05 | 8.500E-06 | 1.803E-06 | 6.500E-06 | 1.000E-05 |
| 26964 | 1349 | N1349D | E | 6.3e-06 | 8.3e-06 | 5.7e-06 | 6.767E-06 | 1.361E-06 | 5.700E-06 | 8.300E-06 |
| 26965 | 1349 | N1349C | E | 5.4e-06 | 5.1e-06 | 9.2e-06 | 6.567E-06 | 2.285E-06 | 5.100E-06 | 9.200E-06 |
| 26966 | 1349 | N1349E | E | 6.4e-06 | 6.8e-06 | 1e-05 | 7.733E-06 | 1.973E-06 | 6.400E-06 | 1.000E-05 |
| 26967 | 1349 | N1349Q | E | 1.2e-05 | 5.5e-06 | 1.4e-05 | 1.050E-05 | 4.444E-06 | 5.500E-06 | 1.400E-05 |
| 26968 | 1349 | N1349G | E | 7.9e-06 | 1.2e-05 | 5.4e-06 | 8.433E-06 | 3.332E-06 | 5.400E-06 | 1.200E-05 |
| 26969 | 1349 | N1349H | E | 7.1e-06 | 1.1e-05 | 6e-06 | 8.033E-06 | 2.627E-06 | 6.000E-06 | 1.100E-05 |
| 26970 | 1349 | N1349I | E | 4.8e-06 | 4.9e-06 | 1.1e-05 | 6.900E-06 | 3.551E-06 | 4.800E-06 | 1.100E-05 |
| 26971 | 1349 | N1349L | E | 7.3e-06 | 7.6e-06 | 7.2e-06 | 7.367E-06 | 2.082E-07 | 7.200E-06 | 7.600E-06 |
| 26972 | 1349 | N1349K | E | 8.4e-06 | 6.5e-06 | 1.1e-05 | 8.633E-06 | 2.259E-06 | 6.500E-06 | 1.100E-05 |
| 26973 | 1349 | N1349M | E | 6.4e-06 | 1.2e-05 | 8.9e-06 | 9.100E-06 | 2.805E-06 | 6.400E-06 | 1.200E-05 |
| 26974 | 1349 | N1349F | E | 1.5e-05 | 6.2e-06 | 4.9e-06 | 8.700E-06 | 5.495E-06 | 4.900E-06 | 1.500E-05 |
| 26975 | 1349 | N1349P | E | 8.6e-06 | 8.6e-06 | 8.5e-06 | 8.567E-06 | 5.774E-08 | 8.500E-06 | 8.600E-06 |
| 26976 | 1349 | N1349S | E | 7.4e-06 | 9e-06 | 9.8e-06 | 8.733E-06 | 1.222E-06 | 7.400E-06 | 9.800E-06 |
| 26977 | 1349 | N1349T | E | 7.7e-06 | 1e-05 | 8.2e-06 | 8.633E-06 | 1.210E-06 | 7.700E-06 | 1.000E-05 |
| 26978 | 1349 | N1349W | E | 8.2e-06 | 6.5e-06 | 8.5e-06 | 7.733E-06 | 1.079E-06 | 6.500E-06 | 8.500E-06 |
| 26979 | 1349 | N1349Y | E | 5.8e-06 | 1.4e-05 | 8.6e-06 | 9.467E-06 | 4.168E-06 | 5.800E-06 | 1.400E-05 |
| 26980 | 1349 | N1349V | E | 1e-05 | 8e-06 | 7.1e-06 | 8.367E-06 | 1.484E-06 | 7.100E-06 | 1.000E-05 |
| 26981 | 1350 | T1350A | E | 6e-06 | 5.6e-06 | 5.8e-06 | 5.800E-06 | 2.000E-07 | 5.600E-06 | 6.000E-06 |
| 26982 | 1350 | T1350R | E | 5.9e-06 | 4.8e-06 | 7.1e-06 | 5.933E-06 | 1.150E-06 | 4.800E-06 | 7.100E-06 |
| 26983 | 1350 | T1350N | E | 1.1e-05 | 6.7e-06 | 6.2e-06 | 7.967E-06 | 2.639E-06 | 6.200E-06 | 1.100E-05 |
| 26984 | 1350 | T1350D | E | 5.8e-06 | 6.1e-06 | 6.6e-06 | 6.167E-06 | 4.041E-07 | 5.800E-06 | 6.600E-06 |
| 26985 | 1350 | T1350C | E | 8.7e-06 | 4.9e-06 | 1.3e-05 | 8.867E-06 | 4.053E-06 | 4.900E-06 | 1.300E-05 |
| 26986 | 1350 | T1350E | E | 9.4e-06 | 6.9e-06 | 6.4e-06 | 7.567E-06 | 1.607E-06 | 6.400E-06 | 9.400E-06 |
| 26987 | 1350 | T1350Q | E | 7.7e-06 | 7e-06 | 8.7e-06 | 7.800E-06 | 8.544E-07 | 7.000E-06 | 8.700E-06 |
| 26988 | 1350 | T1350G | E | 6.6e-06 | 4.3e-06 | 5.5e-06 | 5.467E-06 | 1.150E-06 | 4.300E-06 | 6.600E-06 |
| 26989 | 1350 | T1350H | E | 7e-06 | 9.6e-06 | 4.8e-06 | 7.133E-06 | 2.403E-06 | 4.800E-06 | 9.600E-06 |
| 26990 | 1350 | T1350I | E | 4.4e-06 | 6.8e-06 | 1.3e-05 | 8.067E-06 | 4.438E-06 | 4.400E-06 | 1.300E-05 |
| 26991 | 1350 | T1350L | E | 4.9e-06 | 8.1e-06 | 1.5e-05 | 9.333E-06 | 5.162E-06 | 4.900E-06 | 1.500E-05 |
| 26992 | 1350 | T1350K | E | 8.1e-06 | 4.6e-06 | 9.5e-06 | 7.400E-06 | 2.524E-06 | 4.600E-06 | 9.500E-06 |
| 26993 | 1350 | T1350M | E | 8.5e-06 | 9.8e-06 | 7.7e-06 | 8.667E-06 | 1.060E-06 | 7.700E-06 | 9.800E-06 |
| 26994 | 1350 | T1350F | E | 5.5e-06 | 1e-05 | 1e-05 | 8.500E-06 | 2.598E-06 | 5.500E-06 | 1.000E-05 |
| 26995 | 1350 | T1350P | E | 9.5e-06 | 1.3e-05 | 7.9e-06 | 1.013E-05 | 2.608E-06 | 7.900E-06 | 1.300E-05 |
| 26996 | 1350 | T1350S | E | 1.3e-05 | 6.6e-06 | 5.7e-06 | 8.433E-06 | 3.980E-06 | 5.700E-06 | 1.300E-05 |
| 26997 | 1350 | T1350T | E | 1.3e-05 | 3.1e-06 | 5.1e-06 | 7.067E-06 | 5.235E-06 | 3.100E-06 | 1.300E-05 |
| 26998 | 1350 | T1350W | E | 6.7e-06 | 8.6e-06 | 1e-05 | 8.433E-06 | 1.656E-06 | 6.700E-06 | 1.000E-05 |
| 26999 | 1350 | T1350Y | E | 6.6e-06 | 1.1e-05 | 1.5e-05 | 1.087E-05 | 4.202E-06 | 6.600E-06 | 1.500E-05 |
| 27000 | 1350 | T1350V | E | 5.5e-06 | 8.7e-06 | 1.3e-05 | 9.067E-06 | 3.763E-06 | 5.500E-06 | 1.300E-05 |
| 27001 | 1351 | A1351A | E | 6.5e-06 | 8.6e-06 | 1.8e-05 | 1.103E-05 | 6.124E-06 | 6.500E-06 | 1.800E-05 |
| 27002 | 1351 | A1351R | E | 7.4e-06 | 1e-05 | 6.6e-06 | 8.000E-06 | 1.778E-06 | 6.600E-06 | 1.000E-05 |
| 27003 | 1351 | A1351N | E | 9.3e-06 | 9.4e-06 | 5.8e-06 | 8.167E-06 | 2.050E-06 | 5.800E-06 | 9.400E-06 |
| 27004 | 1351 | A1351D | E | 1e-05 | 1e-05 | 9.2e-06 | 9.733E-06 | 4.619E-07 | 9.200E-06 | 1.000E-05 |
| 27005 | 1351 | A1351C | E | 7.3e-06 | 6.2e-06 | 8.9e-06 | 7.467E-06 | 1.358E-06 | 6.200E-06 | 8.900E-06 |
| 27006 | 1351 | A1351E | E | 8.1e-06 | 6.1e-06 | 1.4e-05 | 9.400E-06 | 4.107E-06 | 6.100E-06 | 1.400E-05 |
| 27007 | 1351 | A1351Q | E | 1.1e-05 | 6.4e-06 | 6.3e-06 | 7.900E-06 | 2.685E-06 | 6.300E-06 | 1.100E-05 |
| 27008 | 1351 | A1351G | E | 1.2e-05 | 7.9e-06 | 1.1e-05 | 1.030E-05 | 2.138E-06 | 7.900E-06 | 1.200E-05 |
| 27009 | 1351 | A1351H | E | 1e-05 | 5.9e-06 | 7.9e-06 | 7.933E-06 | 2.050E-06 | 5.900E-06 | 1.000E-05 |
| 27010 | 1351 | A1351I | E | 7.5e-06 | 1.3e-05 | 9.4e-06 | 9.967E-06 | 2.793E-06 | 7.500E-06 | 1.300E-05 |
| 27011 | 1351 | A1351L | E | 1.2e-05 | 3.3e-06 | 1e-05 | 8.433E-06 | 4.557E-06 | 3.300E-06 | 1.200E-05 |
| 27012 | 1351 | A1351K | E | 6.1e-06 | 4.6e-06 | 1e-05 | 6.900E-06 | 2.787E-06 | 4.600E-06 | 1.000E-05 |
| 27013 | 1351 | A1351M | E | 1.4e-05 | 6.8e-06 | 6.5e-06 | 9.100E-06 | 4.246E-06 | 6.500E-06 | 1.400E-05 |
| 27014 | 1351 | A1351F | E | 8e-06 | 1.4e-05 | 8.5e-06 | 1.017E-05 | 3.329E-06 | 8.000E-06 | 1.400E-05 |
| 27015 | 1351 | A1351P | E | 7e-06 | 7.2e-06 | 8.7e-06 | 7.633E-06 | 9.292E-07 | 7.000E-06 | 8.700E-06 |
| 27016 | 1351 | A1351S | E | 4.7e-06 | 7.4e-06 | 1.2e-05 | 8.033E-06 | 3.691E-06 | 4.700E-06 | 1.200E-05 |
| 27017 | 1351 | A1351T | E | 8.5e-06 | 8.6e-06 | 1.3e-05 | 1.003E-05 | 2.570E-06 | 8.500E-06 | 1.300E-05 |
| 27018 | 1351 | A1351W | E | 8.2e-06 | 6.7e-06 | 4.6e-06 | 6.500E-06 | 1.808E-06 | 4.600E-06 | 8.200E-06 |
| 27019 | 1351 | A1351Y | E | 6.4e-06 | 4.5e-06 | 8.6e-06 | 6.500E-06 | 2.052E-06 | 4.500E-06 | 8.600E-06 |
| 27020 | 1351 | A1351V | E | 4.4e-06 | 1.1e-05 | 5.2e-06 | 6.867E-06 | 3.602E-06 | 4.400E-06 | 1.100E-05 |
| 27021 | 1352 | Y1352A | E | 8.6e-06 | 6.2e-06 | 6.3e-06 | 7.033E-06 | 1.358E-06 | 6.200E-06 | 8.600E-06 |
| 27022 | 1352 | Y1352R | E | 9.1e-06 | 7.3e-06 | 6.7e-06 | 7.700E-06 | 1.249E-06 | 6.700E-06 | 9.100E-06 |
| 27023 | 1352 | Y1352N | E | 7.8e-06 | 6.4e-06 | 7.2e-06 | 7.133E-06 | 7.024E-07 | 6.400E-06 | 7.800E-06 |
| 27024 | 1352 | Y1352D | E | 6.1e-06 | 6.3e-06 | 5.2e-06 | 5.867E-06 | 5.859E-07 | 5.200E-06 | 6.300E-06 |
| 27025 | 1352 | Y1352C | E | 1.5e-05 | 9.1e-06 | 1.1e-05 | 1.170E-05 | 3.012E-06 | 9.100E-06 | 1.500E-05 |
| 27026 | 1352 | Y1352E | E | 6.6e-06 | 5.1e-06 | 8.2e-06 | 6.633E-06 | 1.550E-06 | 5.100E-06 | 8.200E-06 |
| 27027 | 1352 | Y1352Q | E | 1.3e-05 | 1.1e-05 | 6.7e-06 | 1.023E-05 | 3.219E-06 | 6.700E-06 | 1.300E-05 |
| 27028 | 1352 | Y1352G | E | 7e-06 | 7.2e-06 | 1.2e-05 | 8.733E-06 | 2.831E-06 | 7.000E-06 | 1.200E-05 |
| 27029 | 1352 | Y1352H | E | 8.8e-06 | 5.9e-06 | 7.9e-06 | 7.533E-06 | 1.484E-06 | 5.900E-06 | 8.800E-06 |

| | | | | | | | | | | |
|---|---|---|---|---|---|---|---|---|---|---|
| 27030 | 1352 | Y1352I | E | 9e-06 | 9.8e-06 | 6.8e-06 | 8.533E-06 | 1.553E-06 | 6.800E-06 | 9.800E-06 |
| 27031 | 1352 | Y1352L | E | 8.7e-06 | 8.1e-06 | 4.5e-06 | 7.100E-06 | 2.272E-06 | 4.500E-06 | 8.700E-06 |
| 27032 | 1352 | Y1352K | E | 5.1e-06 | 9.9e-06 | 1e-05 | 8.333E-06 | 2.801E-06 | 5.100E-06 | 1.000E-05 |
| 27033 | 1352 | Y1352M | E | 5.8e-06 | 4.8e-06 | 7.1e-06 | 5.900E-06 | 1.153E-06 | 4.800E-06 | 7.100E-06 |
| 27034 | 1352 | Y1352F | E | 6.1e-06 | 5.1e-06 | 7.1e-06 | 6.100E-06 | 1.000E-06 | 5.100E-06 | 7.100E-06 |
| 27035 | 1352 | Y1352P | E | 8.6e-06 | 6.3e-06 | 4.6e-06 | 6.500E-06 | 2.007E-06 | 4.600E-06 | 8.600E-06 |
| 27036 | 1352 | Y1352S | E | 6.1e-06 | 9e-06 | 8.8e-06 | 7.967E-06 | 1.620E-06 | 6.100E-06 | 9.000E-06 |
| 27037 | 1352 | Y1352T | E | 9e-06 | 7e-06 | 6.3e-06 | 7.433E-06 | 1.401E-06 | 6.300E-06 | 9.000E-06 |
| 27038 | 1352 | Y1352W | E | 8.5e-06 | 8.8e-06 | 6.5e-06 | 7.933E-06 | 1.250E-06 | 6.500E-06 | 8.800E-06 |
| 27039 | 1352 | Y1352Y | E | 9.9e-06 | 4.4e-06 | 1.3e-05 | 9.100E-06 | 4.355E-06 | 4.400E-06 | 1.300E-05 |
| 27040 | 1352 | Y1352V | E | 8.7e-06 | 8.1e-06 | 6.7e-06 | 7.833E-06 | 1.026E-06 | 6.700E-06 | 8.700E-06 |
| 27041 | 1353 | L1353A | E | 1.1e-05 | 5.3e-06 | 4.8e-06 | 7.033E-06 | 3.444E-06 | 4.800E-06 | 1.100E-05 |
| 27042 | 1353 | L1353R | E | 7.6e-06 | 7.3e-06 | 6.2e-06 | 7.033E-06 | 7.371E-07 | 6.200E-06 | 7.600E-06 |
| 27043 | 1353 | L1353N | E | 1.4e-05 | 5.5e-06 | 9.2e-06 | 9.567E-06 | 4.262E-06 | 5.500E-06 | 1.400E-05 |
| 27044 | 1353 | L1353D | E | 6.8e-06 | 4.8e-06 | 1.2e-05 | 7.867E-06 | 3.717E-06 | 4.800E-06 | 1.200E-05 |
| 27045 | 1353 | L1353C | E | 6.3e-06 | 4.5e-06 | 3.1e-06 | 4.633E-06 | 1.604E-06 | 3.100E-06 | 6.300E-06 |
| 27046 | 1353 | L1353E | E | 8e-06 | 5.8e-06 | 1.1e-05 | 8.267E-06 | 2.610E-06 | 5.800E-06 | 1.100E-05 |
| 27047 | 1353 | L1353Q | E | 8.4e-06 | 6.2e-06 | 5.7e-06 | 6.767E-06 | 1.436E-06 | 5.700E-06 | 8.400E-06 |
| 27048 | 1353 | L1353G | E | 5.3e-06 | 6.5e-06 | 8.4e-06 | 6.733E-06 | 1.563E-06 | 5.300E-06 | 8.400E-06 |
| 27049 | 1353 | L1353H | E | 8.6e-06 | 9.4e-06 | 1e-05 | 9.333E-06 | 7.024E-07 | 8.600E-06 | 1.000E-05 |
| 27050 | 1353 | L1353I | E | 1e-05 | 6.6e-06 | 7.7e-06 | 8.100E-06 | 1.735E-06 | 6.600E-06 | 1.000E-05 |
| 27051 | 1353 | L1353L | E | 1.5e-05 | 4.6e-06 | 6.6e-06 | 8.733E-06 | 5.518E-06 | 4.600E-06 | 1.500E-05 |
| 27052 | 1353 | L1353K | E | 7.5e-06 | 5.1e-06 | 5.2e-06 | 5.933E-06 | 1.358E-06 | 5.100E-06 | 7.500E-06 |
| 27053 | 1353 | L1353M | E | 1e-05 | 5.1e-06 | 6.9e-06 | 7.333E-06 | 2.479E-06 | 5.100E-06 | 1.000E-05 |
| 27054 | 1353 | L1353F | E | 7.3e-06 | 4.7e-06 | 1.1e-05 | 7.667E-06 | 3.166E-06 | 4.700E-06 | 1.100E-05 |
| 27055 | 1353 | L1353P | E | 6.7e-06 | 7.6e-06 | 6e-06 | 6.767E-06 | 8.021E-07 | 6.000E-06 | 7.600E-06 |
| 27056 | 1353 | L1353S | E | 8.1e-06 | 8.1e-06 | 1.1e-05 | 9.067E-06 | 1.674E-06 | 8.100E-06 | 1.100E-05 |
| 27057 | 1353 | L1353T | E | 8.8e-06 | 5.6e-06 | 6.3e-06 | 6.900E-06 | 1.682E-06 | 5.600E-06 | 8.800E-06 |
| 27058 | 1353 | L1353W | E | 5.7e-06 | 9.3e-06 | 9.2e-06 | 8.067E-06 | 2.050E-06 | 5.700E-06 | 9.300E-06 |
| 27059 | 1353 | L1353Y | E | 1e-05 | 5.9e-06 | 8.8e-06 | 8.233E-06 | 2.108E-06 | 5.900E-06 | 1.000E-05 |
| 27060 | 1353 | L1353V | E | 7.2e-06 | 9.8e-06 | 9.6e-06 | 8.867E-06 | 1.447E-06 | 7.200E-06 | 9.800E-06 |
| 27061 | 1354 | Q1354A | E | 1.3e-05 | 7.7e-06 | 4.7e-06 | 8.467E-06 | 4.203E-06 | 4.700E-06 | 1.300E-05 |
| 27062 | 1354 | Q1354R | E | 3.7e-06 | 1.2e-05 | 1.1e-05 | 8.900E-06 | 4.531E-06 | 3.700E-06 | 1.200E-05 |
| 27063 | 1354 | Q1354N | E | 9.2e-06 | 7.8e-06 | 7e-06 | 8.000E-06 | 1.114E-06 | 7.000E-06 | 9.200E-06 |
| 27064 | 1354 | Q1354D | E | 9.3e-06 | 6.4e-06 | 5.5e-06 | 7.067E-06 | 1.986E-06 | 5.500E-06 | 9.300E-06 |
| 27065 | 1354 | Q1354C | E | 1.1e-05 | 7.7e-06 | 5.2e-06 | 7.967E-06 | 2.909E-06 | 5.200E-06 | 1.100E-05 |
| 27066 | 1354 | Q1354E | E | 1.3e-05 | 9.5e-06 | 4.6e-06 | 9.033E-06 | 4.219E-06 | 4.600E-06 | 1.300E-05 |
| 27067 | 1354 | Q1354Q | E | 6.5e-06 | 6.2e-06 | 9.4e-06 | 7.367E-06 | 1.767E-06 | 6.200E-06 | 9.400E-06 |
| 27068 | 1354 | Q1354G | E | 6.5e-06 | 6.5e-06 | 9.1e-06 | 7.367E-06 | 1.501E-06 | 6.500E-06 | 9.100E-06 |
| 27069 | 1354 | Q1354H | E | 7.4e-06 | 7.2e-06 | 5e-06 | 6.533E-06 | 1.332E-06 | 5.000E-06 | 7.400E-06 |
| 27070 | 1354 | Q1354I | E | 7.5e-06 | 1e-05 | 6.7e-06 | 8.067E-06 | 1.721E-06 | 6.700E-06 | 1.000E-05 |
| 27071 | 1354 | Q1354L | E | 1.1e-05 | 4.8e-06 | 7.8e-06 | 7.867E-06 | 3.101E-06 | 4.800E-06 | 1.100E-05 |
| 27072 | 1354 | Q1354K | E | 9e-06 | 5.5e-06 | 7.7e-06 | 7.400E-06 | 1.769E-06 | 5.500E-06 | 9.000E-06 |
| 27073 | 1354 | Q1354M | E | 5e-06 | 5.5e-06 | 5.1e-06 | 5.200E-06 | 2.646E-07 | 5.000E-06 | 5.500E-06 |
| 27074 | 1354 | Q1354F | E | 8.7e-06 | 9.5e-06 | 1.2e-05 | 1.007E-05 | 1.721E-06 | 8.700E-06 | 1.200E-05 |
| 27075 | 1354 | Q1354P | E | 9.8e-06 | 1.2e-05 | 6.7e-06 | 9.500E-06 | 2.663E-06 | 6.700E-06 | 1.200E-05 |
| 27076 | 1354 | Q1354S | E | 6.9e-06 | 6.4e-06 | 7.8e-06 | 7.033E-06 | 7.095E-07 | 6.400E-06 | 7.800E-06 |
| 27077 | 1354 | Q1354T | E | 6.9e-06 | 6.1e-06 | 5.3e-06 | 6.100E-06 | 8.000E-07 | 5.300E-06 | 6.900E-06 |
| 27078 | 1354 | Q1354W | E | 9.2e-06 | 5.2e-06 | 7.5e-06 | 7.300E-06 | 2.007E-06 | 5.200E-06 | 9.200E-06 |
| 27079 | 1354 | Q1354Y | E | 4.8e-06 | 9.4e-06 | 7.6e-06 | 7.267E-06 | 2.318E-06 | 4.800E-06 | 9.400E-06 |
| 27080 | 1354 | Q1354V | E | 4.4e-06 | 1.1e-05 | 9.1e-06 | 8.167E-06 | 3.398E-06 | 4.400E-06 | 1.100E-05 |
| 27081 | 1355 | M1355A | E | 7.6e-06 | 8.9e-06 | 8.1e-06 | 8.200E-06 | 6.557E-07 | 7.600E-06 | 8.900E-06 |
| 27082 | 1355 | M1355R | E | 6.6e-06 | 1.2e-05 | 7.1e-06 | 8.567E-06 | 2.984E-06 | 6.600E-06 | 1.200E-05 |
| 27083 | 1355 | M1355N | E | 1e-05 | 1.4e-05 | 5.2e-06 | 9.733E-06 | 4.406E-06 | 5.200E-06 | 1.400E-05 |
| 27084 | 1355 | M1355D | E | 6.6e-06 | 1.4e-05 | 7e-06 | 9.200E-06 | 4.162E-06 | 6.600E-06 | 1.400E-05 |
| 27085 | 1355 | M1355C | E | 1.4e-05 | 5.3e-06 | 6.4e-06 | 8.567E-06 | 4.737E-06 | 5.300E-06 | 1.400E-05 |
| 27086 | 1355 | M1355E | E | 7.8e-06 | 1.5e-05 | 6.5e-06 | 9.767E-06 | 4.579E-06 | 6.500E-06 | 1.500E-05 |
| 27087 | 1355 | M1355Q | E | 7.3e-06 | 4.4e-06 | 1.2e-05 | 7.900E-06 | 3.835E-06 | 4.400E-06 | 1.200E-05 |
| 27088 | 1355 | M1355G | E | 1.5e-05 | 9e-06 | 8.5e-06 | 1.083E-05 | 3.617E-06 | 8.500E-06 | 1.500E-05 |
| 27089 | 1355 | M1355H | E | 5e-06 | 1e-05 | 4.9e-06 | 6.633E-06 | 2.916E-06 | 4.900E-06 | 1.000E-05 |
| 27090 | 1355 | M1355I | E | 7.4e-06 | 9e-06 | 5.8e-06 | 7.400E-06 | 1.600E-06 | 5.800E-06 | 9.000E-06 |
| 27091 | 1355 | M1355L | E | 1.2e-05 | 5.1e-06 | 5.9e-06 | 7.667E-06 | 3.774E-06 | 5.100E-06 | 1.200E-05 |
| 27092 | 1355 | M1355K | E | 7.3e-06 | 1.2e-05 | 1.2e-05 | 1.043E-05 | 2.714E-06 | 7.300E-06 | 1.200E-05 |
| 27093 | 1355 | M1355M | E | 8.4e-06 | 7.6e-06 | 6.9e-06 | 7.633E-06 | 7.506E-07 | 6.900E-06 | 8.400E-06 |
| 27094 | 1355 | M1355F | E | 6e-06 | 4.4e-06 | 1e-05 | 6.800E-06 | 2.884E-06 | 4.400E-06 | 1.000E-05 |
| 27095 | 1355 | M1355P | E | 8.3e-06 | 1.3e-05 | 6.9e-06 | 9.400E-06 | 3.195E-06 | 6.900E-06 | 1.300E-05 |
| 27096 | 1355 | M1355S | E | 9.2e-06 | 9.4e-06 | 8e-06 | 8.867E-06 | 7.572E-07 | 8.000E-06 | 9.400E-06 |
| 27097 | 1355 | M1355T | E | 8e-06 | 8.4e-06 | 7.6e-06 | 8.000E-06 | 4.000E-07 | 7.600E-06 | 8.400E-06 |
| 27098 | 1355 | M1355W | E | 1.2e-05 | 5.8e-06 | 9.3e-06 | 9.033E-06 | 3.109E-06 | 5.800E-06 | 1.200E-05 |
| 27099 | 1355 | M1355Y | E | 4.8e-06 | 6.3e-06 | 1e-05 | 7.033E-06 | 2.676E-06 | 4.800E-06 | 1.000E-05 |
| 27100 | 1355 | M1355V | E | 5.4e-06 | 1.1e-05 | 1e-05 | 8.800E-06 | 2.987E-06 | 5.400E-06 | 1.100E-05 |
| 27101 | 1356 | N1356A | E | 9.8e-06 | 1.9e-05 | 1.4e-05 | 1.427E-05 | 4.606E-06 | 9.800E-06 | 1.900E-05 |
| 27102 | 1356 | N1356R | E | 7.6e-06 | 6.6e-06 | 7.2e-06 | 7.133E-06 | 5.033E-07 | 6.600E-06 | 7.600E-06 |
| 27103 | 1356 | N1356N | E | 6.9e-06 | 9.2e-06 | 7.1e-06 | 7.733E-06 | 1.274E-06 | 6.900E-06 | 9.200E-06 |
| 27104 | 1356 | N1356D | E | 8.6e-06 | 1.4e-05 | 1e-05 | 1.087E-05 | 2.802E-06 | 8.600E-06 | 1.400E-05 |
| 27105 | 1356 | N1356C | E | 8.5e-06 | 5.2e-06 | 7e-06 | 6.900E-06 | 1.652E-06 | 5.200E-06 | 8.500E-06 |
| 27106 | 1356 | N1356E | E | 8e-06 | 9.9e-06 | 6.2e-06 | 8.033E-06 | 1.850E-06 | 6.200E-06 | 9.900E-06 |
| 27107 | 1356 | N1356Q | E | 1.5e-05 | 6.4e-06 | 4.4e-06 | 8.600E-06 | 5.632E-06 | 4.400E-06 | 1.500E-05 |
| 27108 | 1356 | N1356G | E | 7e-06 | 8.2e-06 | 7e-06 | 7.400E-06 | 6.928E-07 | 7.000E-06 | 8.200E-06 |
| 27109 | 1356 | N1356H | E | 6.6e-06 | 9.5e-06 | 6.7e-06 | 7.600E-06 | 1.646E-06 | 6.600E-06 | 9.500E-06 |
| 27110 | 1356 | N1356I | E | 6.2e-06 | 1.1e-05 | 1.4e-05 | 1.040E-05 | 3.934E-06 | 6.200E-06 | 1.400E-05 |
| 27111 | 1356 | N1356L | E | 6.1e-06 | 9.9e-06 | 7.3e-06 | 7.767E-06 | 1.943E-06 | 6.100E-06 | 9.900E-06 |
| 27112 | 1356 | N1356K | E | 9.7e-06 | 9.9e-06 | 7.1e-06 | 8.900E-06 | 1.562E-06 | 7.100E-06 | 9.900E-06 |
| 27113 | 1356 | N1356M | E | 7.8e-06 | 7.5e-06 | 8.8e-06 | 8.033E-06 | 6.807E-07 | 7.500E-06 | 8.800E-06 |
| 27114 | 1356 | N1356F | E | 7.9e-06 | 1.2e-05 | 1.3e-05 | 1.097E-05 | 2.702E-06 | 7.900E-06 | 1.300E-05 |

| | | | | | | | | | | |
|---|---|---|---|---|---|---|---|---|---|---|
| 27115 | 1356 | N1356P | E | 1.2e-05 | 7.2e-06 | 8.4e-06 | 9.200E-06 | 2.498E-06 | 7.200E-06 | 1.200E-05 |
| 27116 | 1356 | N1356S | E | 6.2e-06 | 9.9e-06 | 8.6e-06 | 8.233E-06 | 1.877E-06 | 6.200E-06 | 9.900E-06 |
| 27117 | 1356 | N1356T | E | 1e-05 | 8.4e-06 | 1.1e-05 | 9.800E-06 | 1.311E-06 | 8.400E-06 | 1.100E-05 |
| 27118 | 1356 | N1356W | E | 6.2e-06 | 7.4e-06 | 8.3e-06 | 7.300E-06 | 1.054E-06 | 6.200E-06 | 8.300E-06 |
| 27119 | 1356 | N1356Y | E | 8.7e-06 | 8.9e-06 | 7.5e-06 | 8.367E-06 | 7.572E-07 | 7.500E-06 | 8.900E-06 |
| 27120 | 1356 | N1356V | E | 6e-06 | 5.4e-06 | 5e-06 | 5.467E-06 | 5.033E-07 | 5.000E-06 | 6.000E-06 |
| 27121 | 1357 | S1357A | E | 7.3e-06 | 6.2e-06 | 7.5e-06 | 7.000E-06 | 7.000E-07 | 6.200E-06 | 7.500E-06 |
| 27122 | 1357 | S1357R | E | 1.3e-05 | 4.5e-06 | 8.2e-06 | 8.567E-06 | 4.262E-06 | 4.500E-06 | 1.300E-05 |
| 27123 | 1357 | S1357N | E | 1.2e-05 | 7.7e-06 | 5.9e-06 | 8.533E-06 | 3.134E-06 | 5.900E-06 | 1.200E-05 |
| 27124 | 1357 | S1357D | E | 9.5e-06 | 9.2e-06 | 1.1e-05 | 9.900E-06 | 9.644E-07 | 9.200E-06 | 1.100E-05 |
| 27125 | 1357 | S1357C | E | 7.3e-06 | 1.3e-05 | 7e-06 | 9.100E-06 | 3.381E-06 | 7.000E-06 | 1.300E-05 |
| 27126 | 1357 | S1357E | E | 6.4e-06 | 8.5e-06 | 1e-05 | 8.300E-06 | 1.808E-06 | 6.400E-06 | 1.000E-05 |
| 27127 | 1357 | S1357Q | E | 6.8e-06 | 4.7e-06 | 6.6e-06 | 6.033E-06 | 1.159E-06 | 4.700E-06 | 6.800E-06 |
| 27128 | 1357 | S1357G | E | 8.2e-06 | 6.8e-06 | 1.1e-05 | 8.667E-06 | 2.139E-06 | 6.800E-06 | 1.100E-05 |
| 27129 | 1357 | S1357H | E | 1.1e-05 | 5.9e-06 | 7.1e-06 | 8.000E-06 | 2.666E-06 | 5.900E-06 | 1.100E-05 |
| 27130 | 1357 | S1357I | E | 9.4e-06 | 6.2e-06 | 7.4e-06 | 7.667E-06 | 1.617E-06 | 6.200E-06 | 9.400E-06 |
| 27131 | 1357 | S1357L | E | 6.1e-06 | 9.4e-06 | 9e-06 | 8.167E-06 | 1.801E-06 | 6.100E-06 | 9.400E-06 |
| 27132 | 1357 | S1357K | E | 4.7e-06 | 9.4e-06 | 1.1e-05 | 8.367E-06 | 3.275E-06 | 4.700E-06 | 1.100E-05 |
| 27133 | 1357 | S1357M | E | 6.6e-06 | 5e-06 | 9.5e-06 | 7.033E-06 | 2.281E-06 | 5.000E-06 | 9.500E-06 |
| 27134 | 1357 | S1357F | E | 1.1e-05 | 1.5e-05 | 1e-05 | 1.200E-05 | 2.646E-06 | 1.000E-05 | 1.500E-05 |
| 27135 | 1357 | S1357P | E | 6.8e-06 | 9.3e-06 | 7.1e-06 | 7.733E-06 | 1.365E-06 | 6.800E-06 | 9.300E-06 |
| 27136 | 1357 | S1357S | E | 7.4e-06 | 9.8e-06 | 1.1e-05 | 9.400E-06 | 1.833E-06 | 7.400E-06 | 1.100E-05 |
| 27137 | 1357 | S1357T | E | 9.4e-06 | 9.7e-06 | 4.7e-06 | 7.933E-06 | 2.804E-06 | 4.700E-06 | 9.700E-06 |
| 27138 | 1357 | S1357W | E | 8.6e-06 | 1.2e-05 | 1.3e-05 | 1.120E-05 | 2.307E-06 | 8.600E-06 | 1.300E-05 |
| 27139 | 1357 | S1357Y | E | 1.1e-05 | 4.8e-06 | 8.2e-06 | 8.000E-06 | 3.105E-06 | 4.800E-06 | 1.100E-05 |
| 27140 | 1357 | S1357V | E | 1.3e-05 | 6.3e-06 | 8.5e-06 | 9.267E-06 | 3.415E-06 | 6.300E-06 | 1.300E-05 |
| 27141 | 1358 | L1358A | E | 6.7e-06 | 6.7e-06 | 1e-05 | 7.800E-06 | 1.905E-06 | 6.700E-06 | 1.000E-05 |
| 27142 | 1358 | L1358R | E | 9.9e-06 | 1e-05 | 8.6e-06 | 9.500E-06 | 7.810E-07 | 8.600E-06 | 1.000E-05 |
| 27143 | 1358 | L1358N | E | 9.6e-06 | 5.9e-06 | 5.6e-06 | 7.033E-06 | 2.228E-06 | 5.600E-06 | 9.600E-06 |
| 27144 | 1358 | L1358D | E | 6.9e-06 | 5.8e-06 | 7e-06 | 6.567E-06 | 6.658E-07 | 5.800E-06 | 7.000E-06 |
| 27145 | 1358 | L1358C | E | 6.1e-06 | 9.3e-06 | 8.2e-06 | 7.867E-06 | 1.626E-06 | 6.100E-06 | 9.300E-06 |
| 27146 | 1358 | L1358E | E | 5.7e-06 | 4.9e-06 | 1.3e-05 | 7.867E-06 | 4.464E-06 | 4.900E-06 | 1.300E-05 |
| 27147 | 1358 | L1358Q | E | 7.3e-06 | 1.3e-05 | 6.4e-06 | 8.900E-06 | 3.579E-06 | 6.400E-06 | 1.300E-05 |
| 27148 | 1358 | L1358G | E | 4.6e-06 | 1.3e-05 | 6.6e-06 | 8.067E-06 | 4.388E-06 | 4.600E-06 | 1.300E-05 |
| 27149 | 1358 | L1358H | E | 1.2e-05 | 5.6e-06 | 6e-06 | 7.867E-06 | 3.585E-06 | 5.600E-06 | 1.200E-05 |
| 27150 | 1358 | L1358I | E | 1.2e-05 | 8.5e-06 | 5.1e-06 | 8.533E-06 | 3.450E-06 | 5.100E-06 | 1.200E-05 |
| 27151 | 1358 | L1358L | E | 6.6e-06 | 1.2e-05 | 9.6e-06 | 9.400E-06 | 2.706E-06 | 6.600E-06 | 1.200E-05 |
| 27152 | 1358 | L1358K | E | 6.5e-06 | 8.3e-06 | 8e-06 | 7.600E-06 | 9.644E-07 | 6.500E-06 | 8.300E-06 |
| 27153 | 1358 | L1358M | E | 4.9e-06 | 8.8e-06 | 5.5e-06 | 6.400E-06 | 2.100E-06 | 4.900E-06 | 8.800E-06 |
| 27154 | 1358 | L1358F | E | 6.1e-06 | 8.2e-06 | 6.2e-06 | 6.833E-06 | 1.185E-06 | 6.100E-06 | 8.200E-06 |
| 27155 | 1358 | L1358P | E | 9.2e-06 | 9.6e-06 | 1.5e-05 | 1.127E-05 | 3.239E-06 | 9.200E-06 | 1.500E-05 |
| 27156 | 1358 | L1358S | E | 8.3e-06 | 7.2e-06 | 6.3e-06 | 7.267E-06 | 1.002E-06 | 6.300E-06 | 8.300E-06 |
| 27157 | 1358 | L1358T | E | 1.3e-05 | 8.7e-06 | 1.4e-05 | 1.190E-05 | 2.816E-06 | 8.700E-06 | 1.400E-05 |
| 27158 | 1358 | L1358W | E | 1e-05 | 5.7e-06 | 5.7e-06 | 7.133E-06 | 2.483E-06 | 5.700E-06 | 1.000E-05 |
| 27159 | 1358 | L1358Y | E | 5.9e-06 | 8e-06 | 7.3e-06 | 7.067E-06 | 1.069E-06 | 5.900E-06 | 8.000E-06 |
| 27160 | 1358 | L1358V | E | 6.5e-06 | 4.6e-06 | 7e-06 | 6.033E-06 | 1.266E-06 | 4.600E-06 | 7.000E-06 |
| 27161 | 1359 | R1359A | E | 6.7e-06 | 8.5e-06 | 1.6e-05 | 1.040E-05 | 4.933E-06 | 6.700E-06 | 1.600E-05 |
| 27162 | 1359 | R1359R | E | 6.6e-06 | 7.1e-06 | 8.6e-06 | 7.433E-06 | 1.041E-06 | 6.600E-06 | 8.600E-06 |
| 27163 | 1359 | R1359N | E | 5e-06 | 7.2e-06 | 1e-05 | 7.400E-06 | 2.506E-06 | 5.000E-06 | 1.000E-05 |
| 27164 | 1359 | R1359D | E | 8.4e-06 | 8.3e-06 | 9.1e-06 | 8.600E-06 | 4.359E-07 | 8.300E-06 | 9.100E-06 |
| 27165 | 1359 | R1359C | E | 9.1e-06 | 8.9e-06 | 6.9e-06 | 8.300E-06 | 1.217E-06 | 6.900E-06 | 9.100E-06 |
| 27166 | 1359 | R1359E | E | 1.2e-05 | 1.4e-05 | 7.2e-06 | 1.107E-05 | 3.495E-06 | 7.200E-06 | 1.400E-05 |
| 27167 | 1359 | R1359Q | E | 1.4e-05 | 1.5e-05 | 7.8e-06 | 1.227E-05 | 3.900E-06 | 7.800E-06 | 1.500E-05 |
| 27168 | 1359 | R1359G | E | 1e-05 | 7.6e-06 | 4.9e-06 | 7.500E-06 | 2.551E-06 | 4.900E-06 | 1.000E-05 |
| 27169 | 1359 | R1359H | E | 8.5e-06 | 8.7e-06 | 1.3e-05 | 1.007E-05 | 2.542E-06 | 8.500E-06 | 1.300E-05 |
| 27170 | 1359 | R1359I | E | 7.1e-06 | 9.4e-06 | 4.9e-06 | 7.133E-06 | 2.250E-06 | 4.900E-06 | 9.400E-06 |
| 27171 | 1359 | R1359L | E | 8.8e-06 | 6.4e-06 | 9.8e-06 | 8.333E-06 | 1.747E-06 | 6.400E-06 | 9.800E-06 |
| 27172 | 1359 | R1359K | E | 1.4e-05 | 9.2e-06 | 9.3e-06 | 1.083E-05 | 2.743E-06 | 9.200E-06 | 1.400E-05 |
| 27173 | 1359 | R1359M | E | 1e-05 | 7.1e-06 | 1.3e-05 | 1.003E-05 | 2.950E-06 | 7.100E-06 | 1.300E-05 |
| 27174 | 1359 | R1359F | E | 5e-06 | 9.5e-06 | 1e-05 | 8.167E-06 | 2.754E-06 | 5.000E-06 | 1.000E-05 |
| 27175 | 1359 | R1359P | E | 5.9e-06 | 1.2e-05 | 5.6e-06 | 7.833E-06 | 3.612E-06 | 5.600E-06 | 1.200E-05 |
| 27176 | 1359 | R1359S | E | 1.5e-05 | 9e-06 | 4.8e-06 | 9.600E-06 | 5.126E-06 | 4.800E-06 | 1.500E-05 |
| 27177 | 1359 | R1359T | E | 6.6e-06 | 6.2e-06 | 8.7e-06 | 7.167E-06 | 1.343E-06 | 6.200E-06 | 8.700E-06 |
| 27178 | 1359 | R1359W | E | 6.5e-06 | 4.9e-06 | 9.3e-06 | 6.900E-06 | 2.227E-06 | 4.900E-06 | 9.300E-06 |
| 27179 | 1359 | R1359Y | E | 6.6e-06 | 9e-06 | 7.9e-06 | 7.833E-06 | 1.201E-06 | 6.600E-06 | 9.000E-06 |
| 27180 | 1359 | R1359V | E | 2.8e-06 | 5.8e-06 | 9.9e-06 | 6.167E-06 | 3.564E-06 | 2.800E-06 | 9.900E-06 |
| 27181 | 1360 | A1360A | E | 1.2e-05 | 5.4e-06 | 5.4e-06 | 7.600E-06 | 3.811E-06 | 5.400E-06 | 1.200E-05 |
| 27182 | 1360 | A1360R | E | 6.3e-06 | 9.6e-06 | 8.6e-06 | 8.167E-06 | 1.692E-06 | 6.300E-06 | 9.600E-06 |
| 27183 | 1360 | A1360N | E | 5.6e-06 | 6.6e-06 | 8.5e-06 | 6.900E-06 | 1.473E-06 | 5.600E-06 | 8.500E-06 |
| 27184 | 1360 | A1360D | E | 6.2e-06 | 7.9e-06 | 7.2e-06 | 7.100E-06 | 8.544E-07 | 6.200E-06 | 7.900E-06 |
| 27185 | 1360 | A1360C | E | 6e-06 | 7.9e-06 | 1.3e-05 | 8.967E-06 | 3.620E-06 | 6.000E-06 | 1.300E-05 |
| 27186 | 1360 | A1360E | E | 8.6e-06 | 6.5e-06 | 1.1e-05 | 8.700E-06 | 2.252E-06 | 6.500E-06 | 1.100E-05 |
| 27187 | 1360 | A1360Q | E | 9.6e-06 | 9e-06 | 4.2e-06 | 7.600E-06 | 2.960E-06 | 4.200E-06 | 9.600E-06 |
| 27188 | 1360 | A1360G | E | 3.8e-06 | 7.3e-06 | 7.4e-06 | 6.167E-06 | 2.050E-06 | 3.800E-06 | 7.400E-06 |
| 27189 | 1360 | A1360H | E | 8.2e-06 | 5.4e-06 | 1.1e-05 | 8.200E-06 | 2.800E-06 | 5.400E-06 | 1.100E-05 |
| 27190 | 1360 | A1360I | E | 7.7e-06 | 4.9e-06 | 8.4e-06 | 7.000E-06 | 1.852E-06 | 4.900E-06 | 8.400E-06 |
| 27191 | 1360 | A1360L | E | 6.7e-06 | 5.6e-06 | 3.8e-06 | 5.367E-06 | 1.464E-06 | 3.800E-06 | 6.700E-06 |
| 27192 | 1360 | A1360K | E | 8.9e-06 | 9.3e-06 | 8.8e-06 | 9.000E-06 | 2.646E-07 | 8.800E-06 | 9.300E-06 |
| 27193 | 1360 | A1360M | E | 6.8e-06 | 1.1e-05 | 5.2e-06 | 7.667E-06 | 2.996E-06 | 5.200E-06 | 1.100E-05 |
| 27194 | 1360 | A1360F | E | 8.1e-06 | 7.1e-06 | 1.2e-05 | 9.067E-06 | 2.589E-06 | 7.100E-06 | 1.200E-05 |
| 27195 | 1360 | A1360P | E | 8.6e-06 | 4.9e-06 | 6.7e-06 | 6.733E-06 | 1.850E-06 | 4.900E-06 | 8.600E-06 |
| 27196 | 1360 | A1360S | E | 2e-05 | 6.4e-06 | 1.2e-05 | 1.280E-05 | 6.835E-06 | 6.400E-06 | 2.000E-05 |
| 27197 | 1360 | A1360T | E | 9e-06 | 8.7e-06 | 8.2e-06 | 8.633E-06 | 4.041E-07 | 8.200E-06 | 9.000E-06 |
| 27198 | 1360 | A1360W | E | 9.9e-06 | 7e-06 | 6.5e-06 | 7.800E-06 | 1.836E-06 | 6.500E-06 | 9.900E-06 |
| 27199 | 1360 | A1360Y | E | 6.9e-06 | 5.9e-06 | 6.7e-06 | 6.500E-06 | 5.292E-07 | 5.900E-06 | 6.900E-06 |

| | | | | | | | | | | |
|---|---|---|---|---|---|---|---|---|---|---|
| 27200 | 1360 | A1360V | E | 7.6e-06 | 1.1e-05 | 8.8e-06 | 9.133E-06 | 1.724E-06 | 7.600E-06 | 1.100E-05 |
| 27201 | 1361 | E1361A | E | 8.8e-06 | 1.3e-05 | 5.4e-06 | 9.067E-06 | 3.807E-06 | 5.400E-06 | 1.300E-05 |
| 27202 | 1361 | E1361R | E | 7.3e-06 | 6.3e-06 | 8.2e-06 | 7.267E-06 | 9.504E-07 | 6.300E-06 | 8.200E-06 |
| 27203 | 1361 | E1361N | E | 6.2e-06 | 9e-06 | 4.8e-06 | 6.667E-06 | 2.139E-06 | 4.800E-06 | 9.000E-06 |
| 27204 | 1361 | E1361D | E | 5.6e-06 | 1.2e-05 | 1.1e-05 | 9.533E-06 | 3.443E-06 | 5.600E-06 | 1.200E-05 |
| 27205 | 1361 | E1361C | E | 5.8e-06 | 4.9e-06 | 1.2e-05 | 7.567E-06 | 3.866E-06 | 4.900E-06 | 1.200E-05 |
| 27206 | 1361 | E1361E | E | 5.4e-06 | 8.8e-06 | 5.9e-06 | 6.700E-06 | 1.836E-06 | 5.400E-06 | 8.800E-06 |
| 27207 | 1361 | E1361Q | E | 5.6e-06 | 9.2e-06 | 1.1e-05 | 8.600E-06 | 2.750E-06 | 5.600E-06 | 1.100E-05 |
| 27208 | 1361 | E1361G | E | 1.2e-05 | 5.4e-06 | 6.3e-06 | 7.900E-06 | 3.579E-06 | 5.400E-06 | 1.200E-05 |
| 27209 | 1361 | E1361H | E | 9.5e-06 | 7.8e-06 | 8.6e-06 | 8.633E-06 | 8.505E-07 | 7.800E-06 | 9.500E-06 |
| 27210 | 1361 | E1361I | E | 6.4e-06 | 1.2e-05 | 6.9e-06 | 8.433E-06 | 3.099E-06 | 6.400E-06 | 1.200E-05 |
| 27211 | 1361 | E1361L | E | 8.1e-06 | 8.9e-06 | 9.4e-06 | 8.800E-06 | 6.557E-07 | 8.100E-06 | 9.400E-06 |
| 27212 | 1361 | E1361K | E | 8.2e-06 | 1.3e-05 | 5.5e-06 | 8.900E-06 | 3.799E-06 | 5.500E-06 | 1.300E-05 |
| 27213 | 1361 | E1361M | E | 1.3e-05 | 1.4e-05 | 4.3e-06 | 1.043E-05 | 5.335E-06 | 4.300E-06 | 1.400E-05 |
| 27214 | 1361 | E1361F | E | 5.8e-06 | 9.9e-06 | 6.7e-06 | 7.467E-06 | 2.155E-06 | 5.800E-06 | 9.900E-06 |
| 27215 | 1361 | E1361P | E | 5.6e-06 | 1.4e-05 | 5.9e-06 | 8.500E-06 | 4.766E-06 | 5.600E-06 | 1.400E-05 |
| 27216 | 1361 | E1361S | E | 5.3e-06 | 6.1e-06 | 4.6e-06 | 5.333E-06 | 7.506E-07 | 4.600E-06 | 6.100E-06 |
| 27217 | 1361 | E1361T | E | 7.5e-06 | 9e-06 | 4.2e-06 | 6.900E-06 | 2.456E-06 | 4.200E-06 | 9.000E-06 |
| 27218 | 1361 | E1361W | E | 7.1e-06 | 9.5e-06 | 9.7e-06 | 8.767E-06 | 1.447E-06 | 7.100E-06 | 9.700E-06 |
| 27219 | 1361 | E1361Y | E | 5e-06 | 9.2e-06 | 9.7e-06 | 7.967E-06 | 2.581E-06 | 5.000E-06 | 9.700E-06 |
| 27220 | 1361 | E1361V | E | 1.2e-05 | 4.8e-06 | 6.5e-06 | 7.767E-06 | 3.763E-06 | 4.800E-06 | 1.200E-05 |
| 27221 | 1362 | D1362A | E | 9.1e-06 | 9.3e-06 | 9.1e-06 | 9.167E-06 | 1.155E-07 | 9.100E-06 | 9.300E-06 |
| 27222 | 1362 | D1362R | E | 1.4e-05 | 9.2e-06 | 6.3e-06 | 9.833E-06 | 3.889E-06 | 6.300E-06 | 1.400E-05 |
| 27223 | 1362 | D1362N | E | 4.4e-06 | 8.4e-06 | 6.3e-06 | 6.367E-06 | 2.001E-06 | 4.400E-06 | 8.400E-06 |
| 27224 | 1362 | D1362D | E | 6.6e-06 | 6.9e-06 | 5.2e-06 | 6.233E-06 | 9.074E-07 | 5.200E-06 | 6.900E-06 |
| 27225 | 1362 | D1362C | E | 1.1e-05 | 8.8e-06 | 6.5e-06 | 8.767E-06 | 2.250E-06 | 6.500E-06 | 1.100E-05 |
| 27226 | 1362 | D1362E | E | 9.7e-06 | 1.4e-05 | 7.3e-06 | 1.033E-05 | 3.395E-06 | 7.300E-06 | 1.400E-05 |
| 27227 | 1362 | D1362Q | E | 6.9e-06 | 4.7e-06 | 6e-06 | 5.867E-06 | 1.106E-06 | 4.700E-06 | 6.900E-06 |
| 27228 | 1362 | D1362G | E | 9.8e-06 | 9.6e-06 | 9.2e-06 | 9.533E-06 | 3.055E-07 | 9.200E-06 | 9.800E-06 |
| 27229 | 1362 | D1362H | E | 7.3e-06 | 8.8e-06 | 4.4e-06 | 6.833E-06 | 2.237E-06 | 4.400E-06 | 8.800E-06 |
| 27230 | 1362 | D1362I | E | 9.4e-06 | 7.6e-06 | 4.3e-06 | 7.100E-06 | 2.587E-06 | 4.300E-06 | 9.400E-06 |
| 27231 | 1362 | D1362L | E | 7.9e-06 | 1.2e-05 | 6.2e-06 | 8.700E-06 | 2.982E-06 | 6.200E-06 | 1.200E-05 |
| 27232 | 1362 | D1362K | E | 9.2e-06 | 1.3e-05 | 4.4e-06 | 8.867E-06 | 4.310E-06 | 4.400E-06 | 1.300E-05 |
| 27233 | 1362 | D1362M | E | 9.4e-06 | 1.4e-05 | 9.4e-06 | 1.093E-05 | 2.656E-06 | 9.400E-06 | 1.400E-05 |
| 27234 | 1362 | D1362F | E | 8.5e-06 | 1.4e-05 | 1.4e-05 | 1.217E-05 | 3.175E-06 | 8.500E-06 | 1.400E-05 |
| 27235 | 1362 | D1362P | E | 5.7e-06 | 1e-05 | 8.3e-06 | 8.000E-06 | 2.166E-06 | 5.700E-06 | 1.000E-05 |
| 27236 | 1362 | D1362S | E | 6.4e-06 | 7.1e-06 | 6.6e-06 | 6.700E-06 | 3.606E-07 | 6.400E-06 | 7.100E-06 |
| 27237 | 1362 | D1362T | E | 6.2e-06 | 7.3e-06 | 4.7e-06 | 6.067E-06 | 1.305E-06 | 4.700E-06 | 7.300E-06 |
| 27238 | 1362 | D1362W | E | 3.9e-06 | 5.7e-06 | 7.4e-06 | 5.667E-06 | 1.750E-06 | 3.900E-06 | 7.400E-06 |
| 27239 | 1362 | D1362Y | E | 3.8e-06 | 9.9e-06 | 7.7e-06 | 7.133E-06 | 3.089E-06 | 3.800E-06 | 9.900E-06 |
| 27240 | 1362 | D1362V | E | 5.1e-06 | 9.3e-06 | 9.1e-06 | 7.833E-06 | 2.369E-06 | 5.100E-06 | 9.300E-06 |
| 27241 | 1363 | T1363A | E | 4.9e-06 | 1.1e-05 | 7.8e-06 | 7.900E-06 | 3.051E-06 | 4.900E-06 | 1.100E-05 |
| 27242 | 1363 | T1363R | E | 8.5e-06 | 9.8e-06 | 5.1e-06 | 7.800E-06 | 2.427E-06 | 5.100E-06 | 9.800E-06 |
| 27243 | 1363 | T1363N | E | 1.1e-05 | 6.4e-06 | 7.5e-06 | 8.300E-06 | 2.402E-06 | 6.400E-06 | 1.100E-05 |
| 27244 | 1363 | T1363D | E | 5.7e-06 | 6.5e-06 | 1.1e-05 | 7.733E-06 | 2.857E-06 | 5.700E-06 | 1.100E-05 |
| 27245 | 1363 | T1363C | E | 5.6e-06 | 6.4e-06 | 7.5e-06 | 6.500E-06 | 9.539E-07 | 5.600E-06 | 7.500E-06 |
| 27246 | 1363 | T1363E | E | 1.7e-05 | 7.9e-06 | 8.9e-06 | 1.127E-05 | 4.990E-06 | 7.900E-06 | 1.700E-05 |
| 27247 | 1363 | T1363Q | E | 8.9e-06 | 1.3e-05 | 5.4e-06 | 9.100E-06 | 3.804E-06 | 5.400E-06 | 1.300E-05 |
| 27248 | 1363 | T1363G | E | 5e-06 | 6.7e-06 | 5.7e-06 | 5.800E-06 | 8.544E-07 | 5.000E-06 | 6.700E-06 |
| 27249 | 1363 | T1363H | E | 6.8e-06 | 7.6e-06 | 7.9e-06 | 7.433E-06 | 5.686E-07 | 6.800E-06 | 7.900E-06 |
| 27250 | 1363 | T1363I | E | 4.5e-06 | 4.8e-06 | 5.9e-06 | 5.067E-06 | 7.371E-07 | 4.500E-06 | 5.900E-06 |
| 27251 | 1363 | T1363L | E | 1.1e-05 | 7.9e-06 | 8e-06 | 8.967E-06 | 1.762E-06 | 7.900E-06 | 1.100E-05 |
| 27252 | 1363 | T1363K | E | 7.2e-06 | 5.6e-06 | 8.5e-06 | 7.100E-06 | 1.453E-06 | 5.600E-06 | 8.500E-06 |
| 27253 | 1363 | T1363M | E | 4.7e-06 | 7e-06 | 6.9e-06 | 6.200E-06 | 1.300E-06 | 4.700E-06 | 7.000E-06 |
| 27254 | 1363 | T1363F | E | 9.3e-06 | 1.3e-05 | 7.5e-06 | 9.933E-06 | 2.804E-06 | 7.500E-06 | 1.300E-05 |
| 27255 | 1363 | T1363P | E | 8.5e-06 | 6.5e-06 | 1.3e-05 | 9.333E-06 | 3.329E-06 | 6.500E-06 | 1.300E-05 |
| 27256 | 1363 | T1363S | E | 9.3e-06 | 1.1e-05 | 9.6e-06 | 9.967E-06 | 9.074E-07 | 9.300E-06 | 1.100E-05 |
| 27257 | 1363 | T1363T | E | 4.3e-06 | 6.7e-06 | 1e-05 | 7.000E-06 | 2.862E-06 | 4.300E-06 | 1.000E-05 |
| 27258 | 1363 | T1363W | E | 1.2e-05 | 5.5e-06 | 7.8e-06 | 8.433E-06 | 3.296E-06 | 5.500E-06 | 1.200E-05 |
| 27259 | 1363 | T1363Y | E | 8.3e-06 | 8.1e-06 | 8.1e-06 | 8.167E-06 | 1.155E-07 | 8.100E-06 | 8.300E-06 |
| 27260 | 1363 | T1363V | E | 1.4e-05 | 7.7e-06 | 7.6e-06 | 9.767E-06 | 3.667E-06 | 7.600E-06 | 1.400E-05 |
| 27261 | 1364 | A1364A | E | 9.5e-06 | 6.5e-06 | 9.8e-06 | 8.600E-06 | 1.825E-06 | 6.500E-06 | 9.800E-06 |
| 27262 | 1364 | A1364R | E | 1e-05 | 6.8e-06 | 6.9e-06 | 7.900E-06 | 1.819E-06 | 6.800E-06 | 1.000E-05 |
| 27263 | 1364 | A1364N | E | 5.9e-06 | 7.2e-06 | 5.6e-06 | 6.233E-06 | 8.505E-07 | 5.600E-06 | 7.200E-06 |
| 27264 | 1364 | A1364D | E | 6.9e-06 | 5.7e-06 | 4.4e-06 | 5.667E-06 | 1.250E-06 | 4.400E-06 | 6.900E-06 |
| 27265 | 1364 | A1364C | E | 6.6e-06 | 7.1e-06 | 7.4e-06 | 7.033E-06 | 4.041E-07 | 6.600E-06 | 7.400E-06 |
| 27266 | 1364 | A1364E | E | 5.6e-06 | 9.3e-06 | 6.9e-06 | 7.267E-06 | 1.877E-06 | 5.600E-06 | 9.300E-06 |
| 27267 | 1364 | A1364Q | E | 6.9e-06 | 6.9e-06 | 6.4e-06 | 6.733E-06 | 2.887E-07 | 6.400E-06 | 6.900E-06 |
| 27268 | 1364 | A1364G | E | 6.1e-06 | 6.1e-06 | 4.6e-06 | 5.600E-06 | 8.660E-07 | 4.600E-06 | 6.100E-06 |
| 27269 | 1364 | A1364H | E | 9.2e-06 | 4.7e-06 | 5.7e-06 | 6.533E-06 | 2.363E-06 | 4.700E-06 | 9.200E-06 |
| 27270 | 1364 | A1364I | E | 4.9e-06 | 2e-05 | 7.8e-06 | 1.090E-05 | 8.013E-06 | 4.900E-06 | 2.000E-05 |
| 27271 | 1364 | A1364L | E | 1e-05 | 4.7e-06 | 7.3e-06 | 7.333E-06 | 2.650E-06 | 4.700E-06 | 1.000E-05 |
| 27272 | 1364 | A1364K | E | 5.6e-06 | 6.5e-06 | 8.9e-06 | 7.000E-06 | 1.706E-06 | 5.600E-06 | 8.900E-06 |
| 27273 | 1364 | A1364M | E | 8.4e-06 | 5.9e-06 | 5.5e-06 | 6.600E-06 | 1.572E-06 | 5.500E-06 | 8.400E-06 |
| 27274 | 1364 | A1364F | E | 1.2e-05 | 4.9e-06 | 8e-06 | 8.300E-06 | 3.559E-06 | 4.900E-06 | 1.200E-05 |
| 27275 | 1364 | A1364P | E | 1e-05 | 6.3e-06 | 7.2e-06 | 7.833E-06 | 1.930E-06 | 6.300E-06 | 1.000E-05 |
| 27276 | 1364 | A1364S | E | 6.2e-06 | 8.1e-06 | 9.1e-06 | 7.800E-06 | 1.473E-06 | 6.200E-06 | 9.100E-06 |
| 27277 | 1364 | A1364T | E | 9.4e-06 | 8.9e-06 | 8.3e-06 | 8.867E-06 | 5.508E-07 | 8.300E-06 | 9.400E-06 |
| 27278 | 1364 | A1364W | E | 6.3e-06 | 7.3e-06 | 1.6e-05 | 9.867E-06 | 5.335E-06 | 6.300E-06 | 1.600E-05 |
| 27279 | 1364 | A1364Y | E | 8.5e-06 | 6e-06 | 6.9e-06 | 7.133E-06 | 1.266E-06 | 6.000E-06 | 8.500E-06 |
| 27280 | 1364 | A1364V | E | 9.3e-06 | 1.3e-05 | 5.3e-06 | 9.200E-06 | 3.851E-06 | 5.300E-06 | 1.300E-05 |
| 27281 | 1365 | V1365A | E | 7.3e-06 | 6.7e-06 | 8.3e-06 | 7.433E-06 | 8.083E-07 | 6.700E-06 | 8.300E-06 |
| 27282 | 1365 | V1365R | E | 9.1e-06 | 5.3e-06 | 1.3e-05 | 9.133E-06 | 3.850E-06 | 5.300E-06 | 1.300E-05 |
| 27283 | 1365 | V1365N | E | 7.5e-06 | 4.5e-06 | 5.7e-06 | 5.900E-06 | 1.510E-06 | 4.500E-06 | 7.500E-06 |
| 27284 | 1365 | V1365D | E | 7e-06 | 7.3e-06 | 6.2e-06 | 6.833E-06 | 5.686E-07 | 6.200E-06 | 7.300E-06 |

| | | | | | | | | | | |
|---|---|---|---|---|---|---|---|---|---|---|
| 27285 | 1365 | V1365C | E | 7.2e-06 | 1.3e-05 | 3.5e-06 | 7.900E-06 | 4.789E-06 | 3.500E-06 | 1.300E-05 |
| 27286 | 1365 | V1365E | E | 4.6e-06 | 5.4e-06 | 7e-06 | 5.667E-06 | 1.222E-06 | 4.600E-06 | 7.000E-06 |
| 27287 | 1365 | V1365Q | E | 6.2e-06 | 5.2e-06 | 9e-06 | 6.800E-06 | 1.970E-06 | 5.200E-06 | 9.000E-06 |
| 27288 | 1365 | V1365G | E | 1e-05 | 1.4e-05 | 8.3e-06 | 1.077E-05 | 2.926E-06 | 8.300E-06 | 1.400E-05 |
| 27289 | 1365 | V1365H | E | 9.2e-06 | 1.3e-05 | 7.6e-06 | 9.933E-06 | 2.774E-06 | 7.600E-06 | 1.300E-05 |
| 27290 | 1365 | V1365I | E | 3.9e-06 | 8.8e-06 | 7e-06 | 6.567E-06 | 2.479E-06 | 3.900E-06 | 8.800E-06 |
| 27291 | 1365 | V1365L | E | 1e-05 | 9.1e-06 | 1.3e-05 | 1.070E-05 | 2.042E-06 | 9.100E-06 | 1.300E-05 |
| 27292 | 1365 | V1365K | E | 9.3e-06 | 1.3e-05 | 8e-06 | 1.010E-05 | 2.594E-06 | 8.000E-06 | 1.300E-05 |
| 27293 | 1365 | V1365M | E | 6.8e-06 | 7.7e-06 | 1.1e-05 | 8.500E-06 | 2.211E-06 | 6.800E-06 | 1.100E-05 |
| 27294 | 1365 | V1365F | E | 4.5e-06 | 1.1e-05 | 1.4e-05 | 9.833E-06 | 4.856E-06 | 4.500E-06 | 1.400E-05 |
| 27295 | 1365 | V1365P | E | 1.3e-05 | 1.2e-05 | 8.5e-06 | 1.117E-05 | 2.363E-06 | 8.500E-06 | 1.300E-05 |
| 27296 | 1365 | V1365S | E | 6.6e-06 | 9.1e-06 | 8.5e-06 | 8.067E-06 | 1.305E-06 | 6.600E-06 | 9.100E-06 |
| 27297 | 1365 | V1365T | E | 9.2e-06 | 6.1e-06 | 7.4e-06 | 7.567E-06 | 1.557E-06 | 6.100E-06 | 9.200E-06 |
| 27298 | 1365 | V1365W | E | 7e-06 | 6.2e-06 | 9.8e-06 | 7.667E-06 | 1.890E-06 | 6.200E-06 | 9.800E-06 |
| 27299 | 1365 | V1365Y | E | 4.6e-06 | 6.6e-06 | 6.8e-06 | 6.000E-06 | 1.217E-06 | 4.600E-06 | 6.800E-06 |
| 27300 | 1365 | V1365V | E | 1.1e-05 | 9.2e-06 | 1.1e-05 | 1.040E-05 | 1.039E-06 | 9.200E-06 | 1.100E-05 |
| 27301 | 1366 | Y1366A | E | 7.6e-06 | 9.1e-06 | 9.3e-06 | 8.667E-06 | 9.292E-07 | 7.600E-06 | 9.300E-06 |
| 27302 | 1366 | Y1366R | E | 7.1e-06 | 1e-05 | 6.6e-06 | 7.900E-06 | 1.836E-06 | 6.600E-06 | 1.000E-05 |
| 27303 | 1366 | Y1366N | E | 9.9e-06 | 9.8e-06 | 3.5e-06 | 7.733E-06 | 3.667E-06 | 3.500E-06 | 9.900E-06 |
| 27304 | 1366 | Y1366D | E | 5.9e-06 | 1.1e-05 | 6.3e-06 | 7.733E-06 | 2.836E-06 | 5.900E-06 | 1.100E-05 |
| 27305 | 1366 | Y1366C | E | 1e-05 | 8.4e-06 | 1e-05 | 9.467E-06 | 9.238E-07 | 8.400E-06 | 1.000E-05 |
| 27306 | 1366 | Y1366E | E | 9.2e-06 | 5.6e-06 | 1.1e-05 | 8.600E-06 | 2.750E-06 | 5.600E-06 | 1.100E-05 |
| 27307 | 1366 | Y1366Q | E | 7e-06 | 1e-05 | 7.4e-06 | 8.133E-06 | 1.629E-06 | 7.000E-06 | 1.000E-05 |
| 27308 | 1366 | Y1366G | E | 1.1e-05 | 8.5e-06 | 9.2e-06 | 9.567E-06 | 1.290E-06 | 8.500E-06 | 1.100E-05 |
| 27309 | 1366 | Y1366H | E | 7.5e-06 | 8.6e-06 | 7.7e-06 | 7.933E-06 | 5.859E-07 | 7.500E-06 | 8.600E-06 |
| 27310 | 1366 | Y1366I | E | 8.6e-06 | 1e-05 | 4.6e-06 | 7.733E-06 | 2.802E-06 | 4.600E-06 | 1.000E-05 |
| 27311 | 1366 | Y1366L | E | 4.8e-06 | 1.5e-05 | 5e-06 | 8.267E-06 | 5.832E-06 | 4.800E-06 | 1.500E-05 |
| 27312 | 1366 | Y1366K | E | 8.2e-06 | 5.6e-06 | 7.5e-06 | 7.100E-06 | 1.345E-06 | 5.600E-06 | 8.200E-06 |
| 27313 | 1366 | Y1366M | E | 5.3e-06 | 9.3e-06 | 4.5e-06 | 6.367E-06 | 2.572E-06 | 4.500E-06 | 9.300E-06 |
| 27314 | 1366 | Y1366F | E | 1.1e-05 | 1.6e-05 | 8.2e-06 | 1.173E-05 | 3.951E-06 | 8.200E-06 | 1.600E-05 |
| 27315 | 1366 | Y1366P | E | 1e-05 | 1.3e-05 | 1e-05 | 1.100E-05 | 1.732E-06 | 1.000E-05 | 1.300E-05 |
| 27316 | 1366 | Y1366S | E | 7.1e-06 | 6.2e-06 | 9.2e-06 | 7.500E-06 | 1.539E-06 | 6.200E-06 | 9.200E-06 |
| 27317 | 1366 | Y1366T | E | 7.5e-06 | 9.7e-06 | 7.1e-06 | 8.100E-06 | 1.400E-06 | 7.100E-06 | 9.700E-06 |
| 27318 | 1366 | Y1366W | E | 9.6e-06 | 1.1e-05 | 8.4e-06 | 9.667E-06 | 1.301E-06 | 8.400E-06 | 1.100E-05 |
| 27319 | 1366 | Y1366Y | E | 9.3e-06 | 9.5e-06 | 9.9e-06 | 9.567E-06 | 3.055E-07 | 9.300E-06 | 9.900E-06 |
| 27320 | 1366 | Y1366V | E | 1e-05 | 1.6e-05 | 8.1e-06 | 1.137E-05 | 4.124E-06 | 8.100E-06 | 1.600E-05 |
| 27321 | 1367 | Y1367A | E | 4.8e-06 | 6.7e-06 | 9e-06 | 6.833E-06 | 2.103E-06 | 4.800E-06 | 9.000E-06 |
| 27322 | 1367 | Y1367R | E | 9.1e-06 | 9.3e-06 | 7.4e-06 | 8.600E-06 | 1.044E-06 | 7.400E-06 | 9.300E-06 |
| 27323 | 1367 | Y1367N | E | 8.7e-06 | 4.6e-06 | 9.3e-06 | 7.533E-06 | 2.558E-06 | 4.600E-06 | 9.300E-06 |
| 27324 | 1367 | Y1367D | E | 6.1e-06 | 5.8e-06 | 9e-06 | 6.967E-06 | 1.767E-06 | 5.800E-06 | 9.000E-06 |
| 27325 | 1367 | Y1367C | E | 4.6e-06 | 7.6e-06 | 6.5e-06 | 6.233E-06 | 1.518E-06 | 4.600E-06 | 7.600E-06 |
| 27326 | 1367 | Y1367E | E | 1.3e-05 | 1.3e-05 | 1.1e-05 | 1.233E-05 | 1.155E-06 | 1.100E-05 | 1.300E-05 |
| 27327 | 1367 | Y1367Q | E | 1.1e-05 | 5.2e-06 | 1.2e-05 | 9.400E-06 | 3.672E-06 | 5.200E-06 | 1.200E-05 |
| 27328 | 1367 | Y1367G | E | 9.5e-06 | 4.9e-06 | 9.1e-06 | 7.833E-06 | 2.548E-06 | 4.900E-06 | 9.500E-06 |
| 27329 | 1367 | Y1367H | E | 7.8e-06 | 7.2e-06 | 1.2e-05 | 9.000E-06 | 2.615E-06 | 7.200E-06 | 1.200E-05 |
| 27330 | 1367 | Y1367I | E | 9.6e-06 | 1e-05 | 7.5e-06 | 9.033E-06 | 1.343E-06 | 7.500E-06 | 1.000E-05 |
| 27331 | 1367 | Y1367L | E | 6.3e-06 | 8.2e-06 | 1.3e-05 | 9.167E-06 | 3.453E-06 | 6.300E-06 | 1.300E-05 |
| 27332 | 1367 | Y1367K | E | 8.9e-06 | 1.3e-05 | 5.8e-06 | 9.233E-06 | 3.612E-06 | 5.800E-06 | 1.300E-05 |
| 27333 | 1367 | Y1367M | E | 1.3e-05 | 4.5e-06 | 8.2e-06 | 8.567E-06 | 4.262E-06 | 4.500E-06 | 1.300E-05 |
| 27334 | 1367 | Y1367F | E | 6.3e-06 | 6.8e-06 | 4.8e-06 | 5.967E-06 | 1.041E-06 | 4.800E-06 | 6.800E-06 |
| 27335 | 1367 | Y1367P | E | 6.9e-06 | 6.6e-06 | 4.8e-06 | 6.100E-06 | 1.136E-06 | 4.800E-06 | 6.900E-06 |
| 27336 | 1367 | Y1367S | E | 6.6e-06 | 6.6e-06 | 8.5e-06 | 7.233E-06 | 1.097E-06 | 6.600E-06 | 8.500E-06 |
| 27337 | 1367 | Y1367T | E | 1e-05 | 6.1e-06 | 7.7e-06 | 7.933E-06 | 1.960E-06 | 6.100E-06 | 1.000E-05 |
| 27338 | 1367 | Y1367W | E | 6.1e-06 | 5.9e-06 | 9.4e-06 | 7.133E-06 | 1.966E-06 | 5.900E-06 | 9.400E-06 |
| 27339 | 1367 | Y1367Y | E | 1.3e-05 | 5e-06 | 7.9e-06 | 8.633E-06 | 4.050E-06 | 5.000E-06 | 1.300E-05 |
| 27340 | 1367 | Y1367V | E | 9.6e-06 | 1e-05 | 6e-06 | 8.533E-06 | 2.203E-06 | 6.000E-06 | 1.000E-05 |
| 27341 | 1368 | C1368A | E | 5.5e-06 | 6.3e-06 | 7e-06 | 6.267E-06 | 7.506E-07 | 5.500E-06 | 7.000E-06 |
| 27342 | 1368 | C1368R | E | 6.1e-06 | 7.1e-06 | 4.8e-06 | 6.000E-06 | 1.153E-06 | 4.800E-06 | 7.100E-06 |
| 27343 | 1368 | C1368N | E | 6.4e-06 | 6.3e-06 | 8.9e-06 | 7.200E-06 | 1.473E-06 | 6.300E-06 | 8.900E-06 |
| 27344 | 1368 | C1368D | E | 6e-06 | 9.2e-06 | 5.3e-06 | 6.833E-06 | 2.079E-06 | 5.300E-06 | 9.200E-06 |
| 27345 | 1368 | C1368C | E | 5.1e-06 | 5.2e-06 | 4.9e-06 | 5.067E-06 | 1.528E-07 | 4.900E-06 | 5.200E-06 |
| 27346 | 1368 | C1368E | E | 6.5e-06 | 7e-06 | 7.1e-06 | 6.867E-06 | 3.215E-07 | 6.500E-06 | 7.100E-06 |
| 27347 | 1368 | C1368Q | E | 8.3e-06 | 6.5e-06 | 6.5e-06 | 7.100E-06 | 1.039E-06 | 6.500E-06 | 8.300E-06 |
| 27348 | 1368 | C1368G | E | 8.8e-06 | 6.5e-06 | 3.7e-06 | 6.333E-06 | 2.554E-06 | 3.700E-06 | 8.800E-06 |
| 27349 | 1368 | C1368H | E | 7e-06 | 6.9e-06 | 6.5e-06 | 6.800E-06 | 2.646E-07 | 6.500E-06 | 7.000E-06 |
| 27350 | 1368 | C1368I | E | 5.5e-06 | 6.3e-06 | 9.5e-06 | 7.100E-06 | 2.117E-06 | 5.500E-06 | 9.500E-06 |
| 27351 | 1368 | C1368L | E | 5.1e-06 | 1.3e-05 | 6.7e-06 | 8.267E-06 | 4.177E-06 | 5.100E-06 | 1.300E-05 |
| 27352 | 1368 | C1368K | E | 6.8e-06 | 8.5e-06 | 8.4e-06 | 7.900E-06 | 9.539E-07 | 6.800E-06 | 8.500E-06 |
| 27353 | 1368 | C1368M | E | 1.2e-05 | 8.9e-06 | 5.5e-06 | 8.800E-06 | 3.251E-06 | 5.500E-06 | 1.200E-05 |
| 27354 | 1368 | C1368F | E | 6.4e-06 | 9.8e-06 | 6.6e-06 | 7.600E-06 | 1.908E-06 | 6.400E-06 | 9.800E-06 |
| 27355 | 1368 | C1368P | E | 8.1e-06 | 6e-06 | 1.1e-05 | 8.367E-06 | 2.511E-06 | 6.000E-06 | 1.100E-05 |
| 27356 | 1368 | C1368S | E | 8.1e-06 | 8.1e-06 | 8e-06 | 8.067E-06 | 5.774E-08 | 8.000E-06 | 8.100E-06 |
| 27357 | 1368 | C1368T | E | 1.4e-05 | 9.1e-06 | 9.9e-06 | 1.100E-05 | 2.629E-06 | 9.100E-06 | 1.400E-05 |
| 27358 | 1368 | C1368W | E | 9.8e-06 | 1.4e-05 | 9.3e-06 | 1.103E-05 | 2.581E-06 | 9.300E-06 | 1.400E-05 |
| 27359 | 1368 | C1368Y | E | 7.9e-06 | 1.5e-05 | 7.9e-06 | 1.027E-05 | 4.099E-06 | 7.900E-06 | 1.500E-05 |
| 27360 | 1368 | C1368V | E | 4.6e-06 | 6.3e-06 | 6.7e-06 | 5.867E-06 | 1.115E-06 | 4.600E-06 | 6.700E-06 |
| 27361 | 1369 | S1369A | E | 5.6e-06 | 6.2e-06 | 8.3e-06 | 6.700E-06 | 1.418E-06 | 5.600E-06 | 8.300E-06 |
| 27362 | 1369 | S1369R | E | 7.6e-06 | 9.7e-06 | 9.2e-06 | 8.833E-06 | 1.097E-06 | 7.600E-06 | 9.700E-06 |
| 27363 | 1369 | S1369N | E | 1e-05 | 9.4e-06 | 9.7e-06 | 9.700E-06 | 3.000E-07 | 9.400E-06 | 1.000E-05 |
| 27364 | 1369 | S1369D | E | 6.1e-06 | 4.2e-06 | 8.4e-06 | 6.233E-06 | 2.103E-06 | 4.200E-06 | 8.400E-06 |
| 27365 | 1369 | S1369C | E | 8.9e-06 | 1.1e-05 | 9.8e-06 | 9.900E-06 | 1.054E-06 | 8.900E-06 | 1.100E-05 |
| 27366 | 1369 | S1369E | E | 1e-05 | 7.8e-06 | 1.1e-05 | 9.600E-06 | 1.637E-06 | 7.800E-06 | 1.100E-05 |
| 27367 | 1369 | S1369Q | E | 6e-06 | 1e-05 | 1.5e-05 | 1.033E-05 | 4.509E-06 | 6.000E-06 | 1.500E-05 |
| 27368 | 1369 | S1369G | E | 1.3e-05 | 1.1e-05 | 7.5e-06 | 1.050E-05 | 2.784E-06 | 7.500E-06 | 1.300E-05 |
| 27369 | 1369 | S1369H | E | 5.1e-06 | 1.5e-05 | 6.8e-06 | 8.967E-06 | 5.294E-06 | 5.100E-06 | 1.500E-05 |

| | | | | | | | | | | |
|---|---|---|---|---|---|---|---|---|---|---|
| 27370 | 1369 | S1369I | E | 9.6e-06 | 1.1e-05 | 7.7e-06 | 9.433E-06 | 1.656E-06 | 7.700E-06 | 1.100E-05 |
| 27371 | 1369 | S1369L | E | 9.3e-06 | 9.1e-06 | 4.7e-06 | 7.700E-06 | 2.600E-06 | 4.700E-06 | 9.300E-06 |
| 27372 | 1369 | S1369K | E | 4.9e-06 | 4e-06 | 8.1e-06 | 5.667E-06 | 2.155E-06 | 4.000E-06 | 8.100E-06 |
| 27373 | 1369 | S1369M | E | 8.4e-06 | 9.8e-06 | 6.3e-06 | 8.167E-06 | 1.762E-06 | 6.300E-06 | 9.800E-06 |
| 27374 | 1369 | S1369F | E | 6.6e-06 | 9.3e-06 | 7.6e-06 | 7.833E-06 | 1.365E-06 | 6.600E-06 | 9.300E-06 |
| 27375 | 1369 | S1369P | E | 6.8e-06 | 9.6e-06 | 1.3e-05 | 9.800E-06 | 3.105E-06 | 6.800E-06 | 1.300E-05 |
| 27376 | 1369 | S1369S | E | 5.4e-06 | 6.9e-06 | 4.3e-06 | 5.533E-06 | 1.305E-06 | 4.300E-06 | 6.900E-06 |
| 27377 | 1369 | S1369T | E | 9.7e-06 | 1.2e-05 | 5.5e-06 | 9.067E-06 | 3.296E-06 | 5.500E-06 | 1.200E-05 |
| 27378 | 1369 | S1369W | E | 1.4e-05 | 8e-06 | 6.8e-06 | 9.600E-06 | 3.857E-06 | 6.800E-06 | 1.400E-05 |
| 27379 | 1369 | S1369Y | E | 9.7e-06 | 7.5e-06 | 6.8e-06 | 8.000E-06 | 1.513E-06 | 6.800E-06 | 9.700E-06 |
| 27380 | 1369 | S1369V | E | 1.3e-05 | 1e-05 | 1e-05 | 1.100E-05 | 1.732E-06 | 1.000E-05 | 1.300E-05 |
| 27381 | 1370 | R1370A | E | 5.2e-06 | 5.9e-06 | 5.5e-06 | 5.533E-06 | 3.512E-07 | 5.200E-06 | 5.900E-06 |
| 27382 | 1370 | R1370R | E | 1.4e-05 | 1.2e-05 | 1e-05 | 1.200E-05 | 2.000E-06 | 1.000E-05 | 1.400E-05 |
| 27383 | 1370 | R1370N | E | 1.3e-05 | 6.4e-06 | 8.3e-06 | 9.233E-06 | 3.398E-06 | 6.400E-06 | 1.300E-05 |
| 27384 | 1370 | R1370D | E | 8.1e-06 | 6.6e-06 | 5.6e-06 | 6.767E-06 | 1.258E-06 | 5.600E-06 | 8.100E-06 |
| 27385 | 1370 | R1370C | E | 6.1e-06 | 8.1e-06 | 1.3e-05 | 9.067E-06 | 3.550E-06 | 6.100E-06 | 1.300E-05 |
| 27386 | 1370 | R1370E | E | 5.7e-06 | 5e-06 | 9.1e-06 | 6.600E-06 | 2.193E-06 | 5.000E-06 | 9.100E-06 |
| 27387 | 1370 | R1370Q | E | 6.9e-06 | 5.7e-06 | 7.7e-06 | 6.767E-06 | 1.007E-06 | 5.700E-06 | 7.700E-06 |
| 27388 | 1370 | R1370G | E | 7e-06 | 5.3e-06 | 1.4e-05 | 8.767E-06 | 4.611E-06 | 5.300E-06 | 1.400E-05 |
| 27389 | 1370 | R1370H | E | 9.7e-06 | 7.6e-06 | 6.2e-06 | 7.833E-06 | 1.762E-06 | 6.200E-06 | 9.700E-06 |
| 27390 | 1370 | R1370I | E | 6.3e-06 | 7.2e-06 | 1e-05 | 7.833E-06 | 1.930E-06 | 6.300E-06 | 1.000E-05 |
| 27391 | 1370 | R1370L | E | 8.4e-06 | 6.6e-06 | 1.4e-05 | 9.667E-06 | 3.859E-06 | 6.600E-06 | 1.400E-05 |
| 27392 | 1370 | R1370K | E | 9.3e-06 | 1.3e-05 | 8.7e-06 | 1.033E-05 | 2.329E-06 | 8.700E-06 | 1.300E-05 |
| 27393 | 1370 | R1370M | E | 5.2e-06 | 6.5e-06 | 6.1e-06 | 5.933E-06 | 6.658E-07 | 5.200E-06 | 6.500E-06 |
| 27394 | 1370 | R1370F | E | 5.2e-06 | 8e-06 | 1.2e-05 | 8.400E-06 | 3.418E-06 | 5.200E-06 | 1.200E-05 |
| 27395 | 1370 | R1370P | E | 8.7e-06 | 7.6e-06 | 1.3e-05 | 9.767E-06 | 2.854E-06 | 7.600E-06 | 1.300E-05 |
| 27396 | 1370 | R1370S | E | 9.2e-06 | 6.6e-06 | 9.1e-06 | 8.300E-06 | 1.473E-06 | 6.600E-06 | 9.200E-06 |
| 27397 | 1370 | R1370T | E | 4.4e-06 | 4.6e-06 | 1.4e-05 | 7.667E-06 | 5.486E-06 | 4.400E-06 | 1.400E-05 |
| 27398 | 1370 | R1370W | E | 4.8e-06 | 8.6e-06 | 1.3e-05 | 8.800E-06 | 4.104E-06 | 4.800E-06 | 1.300E-05 |
| 27399 | 1370 | R1370Y | E | 9.6e-06 | 6.5e-06 | 5.3e-06 | 7.133E-06 | 2.219E-06 | 5.300E-06 | 9.600E-06 |
| 27400 | 1370 | R1370V | E | 8.8e-06 | 7.5e-06 | 7.5e-06 | 7.933E-06 | 7.506E-07 | 7.500E-06 | 8.800E-06 |
| 27401 | 1371 | W1371A | E | 7.6e-06 | 1.2e-05 | 8.6e-06 | 9.400E-06 | 2.307E-06 | 7.600E-06 | 1.200E-05 |
| 27402 | 1371 | W1371R | E | 6.7e-06 | 6.6e-06 | 1.2e-05 | 8.433E-06 | 3.089E-06 | 6.600E-06 | 1.200E-05 |
| 27403 | 1371 | W1371N | E | 6.5e-06 | 4.8e-06 | 5.9e-06 | 5.733E-06 | 8.622E-07 | 4.800E-06 | 6.500E-06 |
| 27404 | 1371 | W1371D | E | 6.5e-06 | 7.5e-06 | 9.9e-06 | 7.967E-06 | 1.747E-06 | 6.500E-06 | 9.900E-06 |
| 27405 | 1371 | W1371C | E | 8.9e-06 | 7.6e-06 | 4.7e-06 | 7.067E-06 | 2.150E-06 | 4.700E-06 | 8.900E-06 |
| 27406 | 1371 | W1371E | E | 8.8e-06 | 3.3e-06 | 5.4e-06 | 5.833E-06 | 2.775E-06 | 3.300E-06 | 8.800E-06 |
| 27407 | 1371 | W1371Q | E | 1.2e-05 | 6.9e-06 | 9.4e-06 | 9.433E-06 | 2.550E-06 | 6.900E-06 | 1.200E-05 |
| 27408 | 1371 | W1371G | E | 6.6e-06 | 5.9e-06 | 9.1e-06 | 7.200E-06 | 1.682E-06 | 5.900E-06 | 9.100E-06 |
| 27409 | 1371 | W1371H | E | 7.3e-06 | 4.1e-06 | 1.1e-05 | 7.467E-06 | 3.453E-06 | 4.100E-06 | 1.100E-05 |
| 27410 | 1371 | W1371I | E | 7.2e-06 | 7.1e-06 | 1.2e-05 | 8.767E-06 | 2.801E-06 | 7.100E-06 | 1.200E-05 |
| 27411 | 1371 | W1371L | E | 6.1e-06 | 1.2e-05 | 1e-05 | 9.367E-06 | 3.001E-06 | 6.100E-06 | 1.200E-05 |
| 27412 | 1371 | W1371K | E | 1.5e-05 | 1e-05 | 6.6e-06 | 1.053E-05 | 4.225E-06 | 6.600E-06 | 1.500E-05 |
| 27413 | 1371 | W1371M | E | 7.1e-06 | 1.2e-05 | 9.2e-06 | 9.433E-06 | 2.458E-06 | 7.100E-06 | 1.200E-05 |
| 27414 | 1371 | W1371F | E | 8.8e-06 | 9.2e-06 | 6.5e-06 | 8.167E-06 | 1.457E-06 | 6.500E-06 | 9.200E-06 |
| 27415 | 1371 | W1371P | E | 9.7e-06 | 7.4e-06 | 6.4e-06 | 7.833E-06 | 1.692E-06 | 6.400E-06 | 9.700E-06 |
| 27416 | 1371 | W1371S | E | 9.7e-06 | 7.3e-06 | 6.8e-06 | 7.933E-06 | 1.550E-06 | 6.800E-06 | 9.700E-06 |
| 27417 | 1371 | W1371T | E | 7.3e-06 | 1.2e-05 | 6.4e-06 | 8.567E-06 | 3.007E-06 | 6.400E-06 | 1.200E-05 |
| 27418 | 1371 | W1371W | E | 5.2e-06 | 5e-06 | 9.6e-06 | 6.600E-06 | 2.600E-06 | 5.000E-06 | 9.600E-06 |
| 27419 | 1371 | W1371Y | E | 4.8e-06 | 5e-06 | 8e-06 | 5.933E-06 | 1.793E-06 | 4.800E-06 | 8.000E-06 |
| 27420 | 1371 | W1371V | E | 6.1e-06 | 6.7e-06 | 9.6e-06 | 7.467E-06 | 1.872E-06 | 6.100E-06 | 9.600E-06 |
| 27421 | 1372 | G1372A | E | 6.1e-06 | 8.8e-06 | 4.8e-06 | 6.567E-06 | 2.040E-06 | 4.800E-06 | 8.800E-06 |
| 27422 | 1372 | G1372R | E | 1e-05 | 1.2e-05 | 5.9e-06 | 9.300E-06 | 3.110E-06 | 5.900E-06 | 1.200E-05 |
| 27423 | 1372 | G1372N | E | 8.9e-06 | 8.3e-06 | 6.8e-06 | 8.000E-06 | 1.082E-06 | 6.800E-06 | 8.900E-06 |
| 27424 | 1372 | G1372D | E | 5.3e-06 | 6.2e-06 | 1.2e-05 | 7.833E-06 | 3.636E-06 | 5.300E-06 | 1.200E-05 |
| 27425 | 1372 | G1372C | E | 6.6e-06 | 1.2e-05 | 1e-05 | 9.533E-06 | 2.730E-06 | 6.600E-06 | 1.200E-05 |
| 27426 | 1372 | G1372E | E | 1.2e-05 | 9.3e-06 | 6.4e-06 | 9.233E-06 | 2.801E-06 | 6.400E-06 | 1.200E-05 |
| 27427 | 1372 | G1372Q | E | 8.9e-06 | 7.8e-06 | 9.1e-06 | 8.600E-06 | 7.000E-07 | 7.800E-06 | 9.100E-06 |
| 27428 | 1372 | G1372G | E | 5.8e-06 | 4e-06 | 6.7e-06 | 5.500E-06 | 1.375E-06 | 4.000E-06 | 6.700E-06 |
| 27429 | 1372 | G1372H | E | 6.7e-06 | 7e-06 | 8.4e-06 | 7.367E-06 | 9.074E-07 | 6.700E-06 | 8.400E-06 |
| 27430 | 1372 | G1372I | E | 6.5e-06 | 9.5e-06 | 4.5e-06 | 6.833E-06 | 2.517E-06 | 4.500E-06 | 9.500E-06 |
| 27431 | 1372 | G1372L | E | 7.1e-06 | 7.2e-06 | 4.7e-06 | 6.333E-06 | 1.415E-06 | 4.700E-06 | 7.200E-06 |
| 27432 | 1372 | G1372K | E | 9.5e-06 | 1.1e-05 | 6.8e-06 | 9.100E-06 | 2.128E-06 | 6.800E-06 | 1.100E-05 |
| 27433 | 1372 | G1372M | E | 9.7e-06 | 6.9e-06 | 1.3e-05 | 9.867E-06 | 3.053E-06 | 6.900E-06 | 1.300E-05 |
| 27434 | 1372 | G1372F | E | 8.8e-06 | 8.2e-06 | 1.1e-05 | 9.333E-06 | 1.474E-06 | 8.200E-06 | 1.100E-05 |
| 27435 | 1372 | G1372P | E | 7.7e-06 | 7.6e-06 | 8.2e-06 | 7.833E-06 | 3.215E-07 | 7.600E-06 | 8.200E-06 |
| 27436 | 1372 | G1372S | E | 8.3e-06 | 6.8e-06 | 6.5e-06 | 7.200E-06 | 9.644E-07 | 6.500E-06 | 8.300E-06 |
| 27437 | 1372 | G1372T | E | 1.1e-05 | 4.8e-06 | 8.3e-06 | 8.033E-06 | 3.109E-06 | 4.800E-06 | 1.100E-05 |
| 27438 | 1372 | G1372W | E | 8.7e-06 | 1.8e-05 | 6.3e-06 | 1.100E-05 | 6.180E-06 | 6.300E-06 | 1.800E-05 |
| 27439 | 1372 | G1372Y | E | 6.2e-06 | 5.7e-06 | 6e-06 | 5.967E-06 | 2.517E-07 | 5.700E-06 | 6.200E-06 |
| 27440 | 1372 | G1372V | E | 5.6e-06 | 9e-06 | 9e-06 | 7.867E-06 | 1.963E-06 | 5.600E-06 | 9.000E-06 |
| 27441 | 1373 | G1373A | E | 1.3e-05 | 6.8e-06 | 1e-05 | 9.933E-06 | 3.101E-06 | 6.800E-06 | 1.300E-05 |
| 27442 | 1373 | G1373R | E | 5.7e-06 | 6.1e-06 | 5.7e-06 | 5.833E-06 | 2.309E-07 | 5.700E-06 | 6.100E-06 |
| 27443 | 1373 | G1373N | E | 9.2e-06 | 5.1e-06 | 5.6e-06 | 6.633E-06 | 2.237E-06 | 5.100E-06 | 9.200E-06 |
| 27444 | 1373 | G1373D | E | 8.9e-06 | 6.3e-06 | 1.3e-05 | 9.400E-06 | 3.378E-06 | 6.300E-06 | 1.300E-05 |
| 27445 | 1373 | G1373C | E | 9.7e-06 | 6.8e-06 | 7.3e-06 | 7.933E-06 | 1.550E-06 | 6.800E-06 | 9.700E-06 |
| 27446 | 1373 | G1373E | E | 1.6e-05 | 5.9e-06 | 5.5e-06 | 9.133E-06 | 5.950E-06 | 5.500E-06 | 1.600E-05 |
| 27447 | 1373 | G1373Q | E | 9.9e-06 | 4.9e-06 | 4.3e-06 | 6.367E-06 | 3.075E-06 | 4.300E-06 | 9.900E-06 |
| 27448 | 1373 | G1373G | E | 6.8e-06 | 9.5e-06 | 7e-06 | 7.767E-06 | 1.504E-06 | 6.800E-06 | 9.500E-06 |
| 27449 | 1373 | G1373H | E | 1.1e-05 | 9.6e-06 | 8e-06 | 9.533E-06 | 1.501E-06 | 8.000E-06 | 1.100E-05 |
| 27450 | 1373 | G1373I | E | 9.3e-06 | 9.8e-06 | 9.7e-06 | 9.600E-06 | 2.646E-07 | 9.300E-06 | 9.800E-06 |
| 27451 | 1373 | G1373L | E | 1.2e-05 | 5.2e-06 | 1.9e-05 | 1.207E-05 | 6.900E-06 | 5.200E-06 | 1.900E-05 |
| 27452 | 1373 | G1373K | E | 1e-05 | 8.8e-06 | 1.1e-05 | 9.933E-06 | 1.102E-06 | 8.800E-06 | 1.100E-05 |
| 27453 | 1373 | G1373M | E | 6.7e-06 | 1.9e-05 | 7.6e-06 | 1.110E-05 | 6.856E-06 | 6.700E-06 | 1.900E-05 |
| 27454 | 1373 | G1373F | E | 7.6e-06 | 1.2e-05 | 5.2e-06 | 8.267E-06 | 3.449E-06 | 5.200E-06 | 1.200E-05 |

| | | | | | | | | | | |
|---|---|---|---|---|---|---|---|---|---|---|
| 27455 | 1373 | G1373P | E | 8.9e-06 | 6.5e-06 | 1.3e-05 | 9.467E-06 | 3.287E-06 | 6.500E-06 | 1.300E-05 |
| 27456 | 1373 | G1373S | E | 1.2e-05 | 7.9e-06 | 7.2e-06 | 9.033E-06 | 2.593E-06 | 7.200E-06 | 1.200E-05 |
| 27457 | 1373 | G1373T | E | 1.4e-05 | 5.6e-06 | 4e-06 | 7.867E-06 | 5.372E-06 | 4.000E-06 | 1.400E-05 |
| 27458 | 1373 | G1373W | E | 4.3e-06 | 1.4e-05 | 7.8e-06 | 8.700E-06 | 4.912E-06 | 4.300E-06 | 1.400E-05 |
| 27459 | 1373 | G1373Y | E | 6.8e-06 | 9.6e-06 | 6.4e-06 | 7.600E-06 | 1.744E-06 | 6.400E-06 | 9.600E-06 |
| 27460 | 1373 | G1373V | E | 5.5e-06 | 7.6e-06 | 7e-06 | 6.700E-06 | 1.082E-06 | 5.500E-06 | 7.600E-06 |
| 27461 | 1374 | D1374A | E | 1.1e-05 | 9.5e-06 | 1.5e-05 | 1.183E-05 | 2.843E-06 | 9.500E-06 | 1.500E-05 |
| 27462 | 1374 | D1374R | E | 8.1e-06 | 5.3e-06 | 5.9e-06 | 6.433E-06 | 1.474E-06 | 5.300E-06 | 8.100E-06 |
| 27463 | 1374 | D1374N | E | 1.3e-05 | 1.1e-05 | 9e-06 | 1.100E-05 | 2.000E-06 | 9.000E-06 | 1.300E-05 |
| 27464 | 1374 | D1374D | E | 8.7e-06 | 6e-06 | 6.9e-06 | 7.200E-06 | 1.375E-06 | 6.000E-06 | 8.700E-06 |
| 27465 | 1374 | D1374C | E | 1.1e-05 | 3.8e-06 | 6.5e-06 | 7.100E-06 | 3.637E-06 | 3.800E-06 | 1.100E-05 |
| 27466 | 1374 | D1374E | E | 6.6e-06 | 1.3e-05 | 8.8e-06 | 9.467E-06 | 3.252E-06 | 6.600E-06 | 1.300E-05 |
| 27467 | 1374 | D1374Q | E | 7.2e-06 | 1.2e-05 | 9.8e-06 | 9.667E-06 | 2.403E-06 | 7.200E-06 | 1.200E-05 |
| 27468 | 1374 | D1374G | E | 5.8e-06 | 6.2e-06 | 9.8e-06 | 7.267E-06 | 2.203E-06 | 5.800E-06 | 9.800E-06 |
| 27469 | 1374 | D1374H | E | 7.2e-06 | 4.6e-06 | 1.6e-05 | 9.267E-06 | 5.974E-06 | 4.600E-06 | 1.600E-05 |
| 27470 | 1374 | D1374I | E | 7e-06 | 8e-06 | 8.7e-06 | 7.900E-06 | 8.544E-07 | 7.000E-06 | 8.700E-06 |
| 27471 | 1374 | D1374L | E | 9.2e-06 | 4.8e-06 | 1.5e-05 | 9.667E-06 | 5.116E-06 | 4.800E-06 | 1.500E-05 |
| 27472 | 1374 | D1374K | E | 6.2e-06 | 9.4e-06 | 9.4e-06 | 8.333E-06 | 1.848E-06 | 6.200E-06 | 9.400E-06 |
| 27473 | 1374 | D1374M | E | 5.2e-06 | 9.4e-06 | 1.2e-05 | 8.867E-06 | 3.431E-06 | 5.200E-06 | 1.200E-05 |
| 27474 | 1374 | D1374F | E | 4.2e-06 | 7.1e-06 | 1e-05 | 7.100E-06 | 2.900E-06 | 4.200E-06 | 1.000E-05 |
| 27475 | 1374 | D1374P | E | 1.7e-05 | 1.1e-05 | 6.5e-06 | 1.150E-05 | 5.268E-06 | 6.500E-06 | 1.700E-05 |
| 27476 | 1374 | D1374S | E | 1e-05 | 6.9e-06 | 7.2e-06 | 8.033E-06 | 1.710E-06 | 6.900E-06 | 1.000E-05 |
| 27477 | 1374 | D1374T | E | 1.2e-05 | 6.7e-06 | 9.9e-06 | 9.533E-06 | 2.669E-06 | 6.700E-06 | 1.200E-05 |
| 27478 | 1374 | D1374W | E | 4.6e-06 | 4.5e-06 | 9.4e-06 | 6.167E-06 | 2.801E-06 | 4.500E-06 | 9.400E-06 |
| 27479 | 1374 | D1374Y | E | 9.8e-06 | 7.4e-06 | 9.9e-06 | 9.033E-06 | 1.415E-06 | 7.400E-06 | 9.900E-06 |
| 27480 | 1374 | D1374V | E | 5e-06 | 1.3e-05 | 6.9e-06 | 8.300E-06 | 4.180E-06 | 5.000E-06 | 1.300E-05 |
| 27481 | 1375 | G1375A | E | 6e-06 | 7.5e-06 | 6.4e-06 | 6.633E-06 | 7.767E-07 | 6.000E-06 | 7.500E-06 |
| 27482 | 1375 | G1375R | E | 8.4e-06 | 8.4e-06 | 1.3e-05 | 9.933E-06 | 2.656E-06 | 8.400E-06 | 1.300E-05 |
| 27483 | 1375 | G1375N | E | 7.4e-06 | 6.2e-06 | 6.8e-06 | 6.800E-06 | 6.000E-07 | 6.200E-06 | 7.400E-06 |
| 27484 | 1375 | G1375D | E | 8.3e-06 | 7e-06 | 9.2e-06 | 8.167E-06 | 1.106E-06 | 7.000E-06 | 9.200E-06 |
| 27485 | 1375 | G1375C | E | 4.7e-06 | 9e-06 | 5.5e-06 | 6.400E-06 | 2.287E-06 | 4.700E-06 | 9.000E-06 |
| 27486 | 1375 | G1375E | E | 6.7e-06 | 1.2e-05 | 7.3e-06 | 8.667E-06 | 2.902E-06 | 6.700E-06 | 1.200E-05 |
| 27487 | 1375 | G1375Q | E | 1.1e-05 | 5.8e-06 | 1.2e-05 | 9.600E-06 | 3.329E-06 | 5.800E-06 | 1.200E-05 |
| 27488 | 1375 | G1375G | E | 5.4e-06 | 1.3e-05 | 1e-05 | 9.467E-06 | 3.828E-06 | 5.400E-06 | 1.300E-05 |
| 27489 | 1375 | G1375H | E | 6.6e-06 | 5.9e-06 | 9.3e-06 | 7.267E-06 | 1.795E-06 | 5.900E-06 | 9.300E-06 |
| 27490 | 1375 | G1375I | E | 5.5e-06 | 6.4e-06 | 9.6e-06 | 7.167E-06 | 2.155E-06 | 5.500E-06 | 9.600E-06 |
| 27491 | 1375 | G1375L | E | 5.7e-06 | 1.4e-05 | 5.8e-06 | 8.500E-06 | 4.763E-06 | 5.700E-06 | 1.400E-05 |
| 27492 | 1375 | G1375K | E | 8.1e-06 | 9.7e-06 | 6.6e-06 | 8.133E-06 | 1.550E-06 | 6.600E-06 | 9.700E-06 |
| 27493 | 1375 | G1375M | E | 7e-06 | 1.3e-05 | 8.3e-06 | 9.433E-06 | 3.156E-06 | 7.000E-06 | 1.300E-05 |
| 27494 | 1375 | G1375F | E | 6.6e-06 | 5e-06 | 1.3e-05 | 8.200E-06 | 4.233E-06 | 5.000E-06 | 1.300E-05 |
| 27495 | 1375 | G1375P | E | 8.4e-06 | 6.5e-06 | 9.6e-06 | 8.167E-06 | 1.563E-06 | 6.500E-06 | 9.600E-06 |
| 27496 | 1375 | G1375S | E | 5.2e-06 | 6e-06 | 1.2e-05 | 7.733E-06 | 3.717E-06 | 5.200E-06 | 1.200E-05 |
| 27497 | 1375 | G1375T | E | 8e-06 | 8.1e-06 | 9.2e-06 | 8.433E-06 | 6.658E-07 | 8.000E-06 | 9.200E-06 |
| 27498 | 1375 | G1375W | E | 8.9e-06 | 8.1e-06 | 5.3e-06 | 7.433E-06 | 1.890E-06 | 5.300E-06 | 8.900E-06 |
| 27499 | 1375 | G1375Y | E | 6.3e-06 | 9.6e-06 | 5.1e-06 | 7.000E-06 | 2.330E-06 | 5.100E-06 | 9.600E-06 |
| 27500 | 1375 | G1375V | E | 6.3e-06 | 7e-06 | 5.5e-06 | 6.267E-06 | 7.506E-07 | 5.500E-06 | 7.000E-06 |
| 27501 | 1376 | F1376A | E | 6.6e-06 | 7.1e-06 | 1.2e-05 | 8.567E-06 | 2.984E-06 | 6.600E-06 | 1.200E-05 |
| 27502 | 1376 | F1376R | E | 6.3e-06 | 1.7e-05 | 6.5e-06 | 9.933E-06 | 6.121E-06 | 6.300E-06 | 1.700E-05 |
| 27503 | 1376 | F1376N | E | 1e-05 | 1.3e-05 | 1.5e-05 | 1.267E-05 | 2.517E-06 | 1.000E-05 | 1.500E-05 |
| 27504 | 1376 | F1376D | E | 1e-05 | 1.1e-05 | 7.8e-06 | 9.600E-06 | 1.637E-06 | 7.800E-06 | 1.100E-05 |
| 27505 | 1376 | F1376C | E | 6.7e-06 | 9.5e-06 | 1e-05 | 8.733E-06 | 1.779E-06 | 6.700E-06 | 1.000E-05 |
| 27506 | 1376 | F1376E | E | 9.6e-06 | 1e-05 | 6.2e-06 | 8.600E-06 | 2.088E-06 | 6.200E-06 | 1.000E-05 |
| 27507 | 1376 | F1376Q | E | 7.8e-06 | 9.8e-06 | 3.4e-06 | 7.000E-06 | 3.274E-06 | 3.400E-06 | 9.800E-06 |
| 27508 | 1376 | F1376G | E | 8.7e-06 | 5.1e-06 | 8.9e-06 | 7.567E-06 | 2.139E-06 | 5.100E-06 | 8.900E-06 |
| 27509 | 1376 | F1376H | E | 9.1e-06 | 7.6e-06 | 7.2e-06 | 7.967E-06 | 1.002E-06 | 7.200E-06 | 9.100E-06 |
| 27510 | 1376 | F1376I | E | 7.4e-06 | 1.3e-05 | 8.5e-06 | 9.633E-06 | 2.967E-06 | 7.400E-06 | 1.300E-05 |
| 27511 | 1376 | F1376L | E | 6.5e-06 | 1.6e-05 | 5.1e-06 | 9.200E-06 | 5.930E-06 | 5.100E-06 | 1.600E-05 |
| 27512 | 1376 | F1376K | E | 6.6e-06 | 9.8e-06 | 4.9e-06 | 7.100E-06 | 2.488E-06 | 4.900E-06 | 9.800E-06 |
| 27513 | 1376 | F1376M | E | 1.6e-05 | 9e-06 | 6.7e-06 | 1.057E-05 | 4.844E-06 | 6.700E-06 | 1.600E-05 |
| 27514 | 1376 | F1376F | E | 8.7e-06 | 4.6e-06 | 6.7e-06 | 6.667E-06 | 2.050E-06 | 4.600E-06 | 8.700E-06 |
| 27515 | 1376 | F1376P | E | 1.7e-05 | 1.2e-05 | 1.1e-05 | 1.333E-05 | 3.215E-06 | 1.100E-05 | 1.700E-05 |
| 27516 | 1376 | F1376S | E | 6.9e-06 | 8.9e-06 | 6.7e-06 | 7.500E-06 | 1.217E-06 | 6.700E-06 | 8.900E-06 |
| 27517 | 1376 | F1376T | E | 6.6e-06 | 7.8e-06 | 1.1e-05 | 8.467E-06 | 2.274E-06 | 6.600E-06 | 1.100E-05 |
| 27518 | 1376 | F1376W | E | 6.1e-06 | 1.1e-05 | 9e-06 | 8.700E-06 | 2.464E-06 | 6.100E-06 | 1.100E-05 |
| 27519 | 1376 | F1376Y | E | 6.5e-06 | 1e-05 | 8.3e-06 | 8.267E-06 | 1.750E-06 | 6.500E-06 | 1.000E-05 |
| 27520 | 1376 | F1376V | E | 1.1e-05 | 4.7e-06 | 6.5e-06 | 7.400E-06 | 3.245E-06 | 4.700E-06 | 1.100E-05 |
| 27521 | 1377 | Y1377A | E | 5.5e-06 | 8.7e-06 | 1.2e-05 | 8.733E-06 | 3.250E-06 | 5.500E-06 | 1.200E-05 |
| 27522 | 1377 | Y1377R | E | 6.9e-06 | 4.6e-06 | 1.3e-05 | 8.167E-06 | 4.341E-06 | 4.600E-06 | 1.300E-05 |
| 27523 | 1377 | Y1377N | E | 8.7e-06 | 9.3e-06 | 9.9e-06 | 9.300E-06 | 6.000E-07 | 8.700E-06 | 9.900E-06 |
| 27524 | 1377 | Y1377D | E | 7.8e-06 | 6.6e-06 | 5.1e-06 | 6.500E-06 | 1.353E-06 | 5.100E-06 | 7.800E-06 |
| 27525 | 1377 | Y1377C | E | 7.9e-06 | 5e-06 | 8.5e-06 | 7.133E-06 | 1.872E-06 | 5.000E-06 | 8.500E-06 |
| 27526 | 1377 | Y1377E | E | 1.2e-05 | 5.4e-06 | 9.4e-06 | 8.933E-06 | 3.325E-06 | 5.400E-06 | 1.200E-05 |
| 27527 | 1377 | Y1377Q | E | 6.5e-06 | 7e-06 | 7.4e-06 | 6.967E-06 | 4.509E-07 | 6.500E-06 | 7.400E-06 |
| 27528 | 1377 | Y1377G | E | 6.4e-06 | 6.6e-06 | 7.9e-06 | 6.967E-06 | 8.145E-07 | 6.400E-06 | 7.900E-06 |
| 27529 | 1377 | Y1377H | E | 6.4e-06 | 7.3e-06 | 8e-06 | 7.233E-06 | 8.021E-07 | 6.400E-06 | 8.000E-06 |
| 27530 | 1377 | Y1377I | E | 1.3e-05 | 5.5e-06 | 6.1e-06 | 8.200E-06 | 4.168E-06 | 5.500E-06 | 1.300E-05 |
| 27531 | 1377 | Y1377L | E | 1.6e-05 | 1e-05 | 6.2e-06 | 1.073E-05 | 4.941E-06 | 6.200E-06 | 1.600E-05 |
| 27532 | 1377 | Y1377K | E | 8.7e-06 | 7e-06 | 6.4e-06 | 7.367E-06 | 1.193E-06 | 6.400E-06 | 8.700E-06 |
| 27533 | 1377 | Y1377M | E | 6.8e-06 | 5.4e-06 | 9.6e-06 | 7.267E-06 | 2.139E-06 | 5.400E-06 | 9.600E-06 |
| 27534 | 1377 | Y1377F | E | 9.2e-06 | 8.5e-06 | 6.7e-06 | 8.133E-06 | 1.290E-06 | 6.700E-06 | 9.200E-06 |
| 27535 | 1377 | Y1377P | E | 7.1e-06 | 1.5e-05 | 8.8e-06 | 1.030E-05 | 4.158E-06 | 7.100E-06 | 1.500E-05 |
| 27536 | 1377 | Y1377S | E | 1e-05 | 6.8e-06 | 8e-06 | 8.267E-06 | 1.617E-06 | 6.800E-06 | 1.000E-05 |
| 27537 | 1377 | Y1377T | E | 5.2e-06 | 8.8e-06 | 7.6e-06 | 7.200E-06 | 1.833E-06 | 5.200E-06 | 8.800E-06 |
| 27538 | 1377 | Y1377W | E | 8.2e-06 | 8.6e-06 | 6.5e-06 | 7.767E-06 | 1.115E-06 | 6.500E-06 | 8.600E-06 |
| 27539 | 1377 | Y1377Y | E | 7.4e-06 | 9.5e-06 | 6.3e-06 | 7.733E-06 | 1.626E-06 | 6.300E-06 | 9.500E-06 |

| | | | | | | | | | | |
|---|---|---|---|---|---|---|---|---|---|---|
| 27540 | 1377 | Y1377V | E | 1.6e-05 | 1.4e-05 | 9.8e-06 | 1.327E-05 | 3.164E-06 | 9.800E-06 | 1.600E-05 |
| 27541 | 1378 | A1378A | E | 5.5e-06 | 1.3e-05 | 5.8e-06 | 8.100E-06 | 4.246E-06 | 5.500E-06 | 1.300E-05 |
| 27542 | 1378 | A1378R | E | 8.1e-06 | 5.9e-06 | 7.1e-06 | 7.033E-06 | 1.102E-06 | 5.900E-06 | 8.100E-06 |
| 27543 | 1378 | A1378N | E | 6.3e-06 | 7.6e-06 | 7e-06 | 6.967E-06 | 6.506E-07 | 6.300E-06 | 7.600E-06 |
| 27544 | 1378 | A1378D | E | 4.7e-06 | 9.5e-06 | 6e-06 | 6.733E-06 | 2.483E-06 | 4.700E-06 | 9.500E-06 |
| 27545 | 1378 | A1378C | E | 5.4e-06 | 5e-06 | 8.2e-06 | 6.200E-06 | 1.744E-06 | 5.000E-06 | 8.200E-06 |
| 27546 | 1378 | A1378E | E | 8.2e-06 | 5.3e-06 | 6.1e-06 | 6.533E-06 | 1.498E-06 | 5.300E-06 | 8.200E-06 |
| 27547 | 1378 | A1378Q | E | 1.3e-05 | 9.6e-06 | 7.1e-06 | 9.900E-06 | 2.961E-06 | 7.100E-06 | 1.300E-05 |
| 27548 | 1378 | A1378G | E | 5.9e-06 | 7.6e-06 | 6.3e-06 | 6.600E-06 | 8.888E-07 | 5.900E-06 | 7.600E-06 |
| 27549 | 1378 | A1378H | E | 7.9e-06 | 9.1e-06 | 4.4e-06 | 7.133E-06 | 2.442E-06 | 4.400E-06 | 9.100E-06 |
| 27550 | 1378 | A1378I | E | 6.8e-06 | 8e-06 | 9.1e-06 | 7.967E-06 | 1.150E-06 | 6.800E-06 | 9.100E-06 |
| 27551 | 1378 | A1378L | E | 1.1e-05 | 6.6e-06 | 7.9e-06 | 8.500E-06 | 2.261E-06 | 6.600E-06 | 1.100E-05 |
| 27552 | 1378 | A1378K | E | 7.5e-06 | 6.9e-06 | 8.8e-06 | 7.733E-06 | 9.713E-07 | 6.900E-06 | 8.800E-06 |
| 27553 | 1378 | A1378M | E | 6.2e-06 | 8.3e-06 | 4.7e-06 | 6.400E-06 | 1.808E-06 | 4.700E-06 | 8.300E-06 |
| 27554 | 1378 | A1378F | E | 4.9e-06 | 8.7e-06 | 1e-05 | 7.867E-06 | 2.650E-06 | 4.900E-06 | 1.000E-05 |
| 27555 | 1378 | A1378P | E | 9.8e-06 | 6.7e-06 | 7.7e-06 | 8.067E-06 | 1.582E-06 | 6.700E-06 | 9.800E-06 |
| 27556 | 1378 | A1378S | E | 8e-06 | 9.4e-06 | 7.7e-06 | 8.367E-06 | 9.074E-07 | 7.700E-06 | 9.400E-06 |
| 27557 | 1378 | A1378T | E | 5e-06 | 1.2e-05 | 1.3e-05 | 1.000E-05 | 4.359E-06 | 5.000E-06 | 1.300E-05 |
| 27558 | 1378 | A1378W | E | 4.5e-06 | 1.2e-05 | 1.1e-05 | 9.167E-06 | 4.072E-06 | 4.500E-06 | 1.200E-05 |
| 27559 | 1378 | A1378Y | E | 9.4e-06 | 8.4e-06 | 8.1e-06 | 8.633E-06 | 6.807E-07 | 8.100E-06 | 9.400E-06 |
| 27560 | 1378 | A1378V | E | 5.9e-06 | 8.2e-06 | 1.3e-05 | 9.033E-06 | 3.623E-06 | 5.900E-06 | 1.300E-05 |
| 27561 | 1379 | M1379A | E | 7.6e-06 | 6.4e-06 | 9.2e-06 | 7.733E-06 | 1.405E-06 | 6.400E-06 | 9.200E-06 |
| 27562 | 1379 | M1379R | E | 9.8e-06 | 1.3e-05 | 1.2e-05 | 1.160E-05 | 1.637E-06 | 9.800E-06 | 1.300E-05 |
| 27563 | 1379 | M1379N | E | 1.4e-05 | 1e-05 | 1e-05 | 1.133E-05 | 2.309E-06 | 1.000E-05 | 1.400E-05 |
| 27564 | 1379 | M1379D | E | 7.5e-06 | 6.5e-06 | 5.5e-06 | 6.500E-06 | 1.000E-06 | 5.500E-06 | 7.500E-06 |
| 27565 | 1379 | M1379C | E | 8.3e-06 | 8.4e-06 | 9.1e-06 | 8.600E-06 | 4.359E-07 | 8.300E-06 | 9.100E-06 |
| 27566 | 1379 | M1379E | E | 2e-05 | 1e-05 | 5.9e-06 | 1.197E-05 | 7.253E-06 | 5.900E-06 | 2.000E-05 |
| 27567 | 1379 | M1379Q | E | 6.3e-06 | 9.8e-06 | 1.4e-05 | 1.003E-05 | 3.855E-06 | 6.300E-06 | 1.400E-05 |
| 27568 | 1379 | M1379G | E | 8.9e-06 | 1.2e-05 | 7.2e-06 | 9.367E-06 | 2.434E-06 | 7.200E-06 | 1.200E-05 |
| 27569 | 1379 | M1379H | E | 7.3e-06 | 6.3e-06 | 7.5e-06 | 7.033E-06 | 6.429E-07 | 6.300E-06 | 7.500E-06 |
| 27570 | 1379 | M1379I | E | 6.7e-06 | 1.2e-05 | 1.2e-05 | 1.023E-05 | 3.060E-06 | 6.700E-06 | 1.200E-05 |
| 27571 | 1379 | M1379L | E | 1.3e-05 | 9.1e-06 | 6.6e-06 | 9.567E-06 | 3.225E-06 | 6.600E-06 | 1.300E-05 |
| 27572 | 1379 | M1379K | E | 9.1e-06 | 1.4e-05 | 6.7e-06 | 9.933E-06 | 3.721E-06 | 6.700E-06 | 1.400E-05 |
| 27573 | 1379 | M1379M | E | 9.5e-06 | 8.2e-06 | 8.5e-06 | 8.733E-06 | 6.807E-07 | 8.200E-06 | 9.500E-06 |
| 27574 | 1379 | M1379F | E | 1.3e-05 | 1.3e-05 | 9.5e-06 | 1.183E-05 | 2.021E-06 | 9.500E-06 | 1.300E-05 |
| 27575 | 1379 | M1379P | E | 8.2e-06 | 5e-06 | 9.5e-06 | 7.567E-06 | 2.316E-06 | 5.000E-06 | 9.500E-06 |
| 27576 | 1379 | M1379S | E | 9.8e-06 | 6.5e-06 | 8.5e-06 | 8.267E-06 | 1.662E-06 | 6.500E-06 | 9.800E-06 |
| 27577 | 1379 | M1379T | E | 6.7e-06 | 4.3e-06 | 8.8e-06 | 6.600E-06 | 2.252E-06 | 4.300E-06 | 8.800E-06 |
| 27578 | 1379 | M1379W | E | 8.6e-06 | 9.8e-06 | 6e-06 | 8.133E-06 | 1.943E-06 | 6.000E-06 | 9.800E-06 |
| 27579 | 1379 | M1379Y | E | 8.3e-06 | 1.4e-05 | 9.6e-06 | 1.063E-05 | 2.987E-06 | 8.300E-06 | 1.400E-05 |
| 27580 | 1379 | M1379V | E | 4.3e-06 | 1.5e-05 | 1.5e-05 | 1.143E-05 | 6.178E-06 | 4.300E-06 | 1.500E-05 |
| 27581 | 1380 | D1380A | E | 8.9e-06 | 8.2e-06 | 1.1e-05 | 9.367E-06 | 1.457E-06 | 8.200E-06 | 1.100E-05 |
| 27582 | 1380 | D1380R | E | 9.3e-06 | 6.3e-06 | 6e-06 | 7.200E-06 | 1.825E-06 | 6.000E-06 | 9.300E-06 |
| 27583 | 1380 | D1380N | E | 1.5e-05 | 7.2e-06 | 7.6e-06 | 9.933E-06 | 4.392E-06 | 7.200E-06 | 1.500E-05 |
| 27584 | 1380 | D1380D | E | 9.7e-06 | 8.3e-06 | 9.8e-06 | 9.267E-06 | 8.386E-07 | 8.300E-06 | 9.800E-06 |
| 27585 | 1380 | D1380C | E | 8.7e-06 | 6.7e-06 | 9.6e-06 | 8.333E-06 | 1.484E-06 | 6.700E-06 | 9.600E-06 |
| 27586 | 1380 | D1380E | E | 8.4e-06 | 9.7e-06 | 5.3e-06 | 7.800E-06 | 2.261E-06 | 5.300E-06 | 9.700E-06 |
| 27587 | 1380 | D1380Q | E | 7e-06 | 6.5e-06 | 1e-05 | 7.833E-06 | 1.893E-06 | 6.500E-06 | 1.000E-05 |
| 27588 | 1380 | D1380G | E | 1.3e-05 | 5.9e-06 | 7e-06 | 8.633E-06 | 3.821E-06 | 5.900E-06 | 1.300E-05 |
| 27589 | 1380 | D1380H | E | 6.5e-06 | 5.1e-06 | 1.3e-05 | 8.200E-06 | 4.215E-06 | 5.100E-06 | 1.300E-05 |
| 27590 | 1380 | D1380I | E | 1.3e-05 | 9.2e-06 | 6.8e-06 | 9.667E-06 | 3.126E-06 | 6.800E-06 | 1.300E-05 |
| 27591 | 1380 | D1380L | E | 7.6e-06 | 7.3e-06 | 1.3e-05 | 9.300E-06 | 3.208E-06 | 7.300E-06 | 1.300E-05 |
| 27592 | 1380 | D1380K | E | 9.2e-06 | 5.6e-06 | 6.6e-06 | 7.133E-06 | 1.858E-06 | 5.600E-06 | 9.200E-06 |
| 27593 | 1380 | D1380M | E | 4.8e-06 | 6.9e-06 | 7.1e-06 | 6.267E-06 | 1.274E-06 | 4.800E-06 | 7.100E-06 |
| 27594 | 1380 | D1380F | E | 6.1e-06 | 8e-06 | 1.3e-05 | 9.033E-06 | 3.564E-06 | 6.100E-06 | 1.300E-05 |
| 27595 | 1380 | D1380P | E | 9.3e-06 | 7.9e-06 | 9.3e-06 | 8.833E-06 | 8.083E-07 | 7.900E-06 | 9.300E-06 |
| 27596 | 1380 | D1380S | E | 6.9e-06 | 5.2e-06 | 8.1e-06 | 6.733E-06 | 1.457E-06 | 5.200E-06 | 8.100E-06 |
| 27597 | 1380 | D1380T | E | 6.2e-06 | 7.2e-06 | 7.6e-06 | 7.000E-06 | 7.211E-07 | 6.200E-06 | 7.600E-06 |
| 27598 | 1380 | D1380W | E | 7.1e-06 | 8.6e-06 | 6.5e-06 | 7.400E-06 | 1.082E-06 | 6.500E-06 | 8.600E-06 |
| 27599 | 1380 | D1380Y | E | 4.1e-06 | 5.4e-06 | 4.9e-06 | 4.800E-06 | 6.557E-07 | 4.100E-06 | 5.400E-06 |
| 27600 | 1380 | D1380V | E | 6.2e-06 | 6.2e-06 | 6.6e-06 | 6.333E-06 | 2.309E-07 | 6.200E-06 | 6.600E-06 |
| 27601 | 1381 | Y1381A | E | 4.9e-06 | 4.4e-06 | 1e-05 | 6.433E-06 | 3.099E-06 | 4.400E-06 | 1.000E-05 |
| 27602 | 1381 | Y1381R | E | 7.1e-06 | 7e-06 | 9.7e-06 | 7.933E-06 | 1.531E-06 | 7.000E-06 | 9.700E-06 |
| 27603 | 1381 | Y1381N | E | 4.8e-06 | 5.9e-06 | 7.8e-06 | 6.167E-06 | 1.518E-06 | 4.800E-06 | 7.800E-06 |
| 27604 | 1381 | Y1381D | E | 8.7e-06 | 7.4e-06 | 1.1e-05 | 9.033E-06 | 1.823E-06 | 7.400E-06 | 1.100E-05 |
| 27605 | 1381 | Y1381C | E | 4.9e-06 | 1e-05 | 7.9e-06 | 7.600E-06 | 2.563E-06 | 4.900E-06 | 1.000E-05 |
| 27606 | 1381 | Y1381E | E | 6.3e-06 | 1e-05 | 6.7e-06 | 7.667E-06 | 2.031E-06 | 6.300E-06 | 1.000E-05 |
| 27607 | 1381 | Y1381Q | E | 6.7e-06 | 6.8e-06 | 6.8e-06 | 6.767E-06 | 5.774E-08 | 6.700E-06 | 6.800E-06 |
| 27608 | 1381 | Y1381G | E | 7.5e-06 | 8.3e-06 | 1e-05 | 8.600E-06 | 1.277E-06 | 7.500E-06 | 1.000E-05 |
| 27609 | 1381 | Y1381H | E | 1.1e-05 | 1.1e-05 | 9.3e-06 | 1.043E-05 | 9.815E-07 | 9.300E-06 | 1.100E-05 |
| 27610 | 1381 | Y1381I | E | 1.8e-05 | 7.5e-06 | 6e-06 | 1.050E-05 | 6.538E-06 | 6.000E-06 | 1.800E-05 |
| 27611 | 1381 | Y1381L | E | 1.1e-05 | 4.5e-06 | 8.5e-06 | 8.000E-06 | 3.279E-06 | 4.500E-06 | 1.100E-05 |
| 27612 | 1381 | Y1381K | E | 7e-06 | 9.2e-06 | 6.9e-06 | 7.700E-06 | 1.300E-06 | 6.900E-06 | 9.200E-06 |
| 27613 | 1381 | Y1381M | E | 7.2e-06 | 1.4e-05 | 1.3e-05 | 1.140E-05 | 3.672E-06 | 7.200E-06 | 1.400E-05 |
| 27614 | 1381 | Y1381F | E | 4.4e-06 | 9.1e-06 | 5.4e-06 | 6.300E-06 | 2.476E-06 | 4.400E-06 | 9.100E-06 |
| 27615 | 1381 | Y1381P | E | 8.3e-06 | 6e-06 | 4.3e-06 | 6.200E-06 | 2.007E-06 | 4.300E-06 | 8.300E-06 |
| 27616 | 1381 | Y1381S | E | 9.2e-06 | 1.3e-05 | 5.5e-06 | 9.233E-06 | 3.750E-06 | 5.500E-06 | 1.300E-05 |
| 27617 | 1381 | Y1381T | E | 8.8e-06 | 7.9e-06 | 9.5e-06 | 8.733E-06 | 8.021E-07 | 7.900E-06 | 9.500E-06 |
| 27618 | 1381 | Y1381W | E | 6.2e-06 | 7.6e-06 | 6.9e-06 | 6.900E-06 | 7.000E-07 | 6.200E-06 | 7.600E-06 |
| 27619 | 1381 | Y1381Y | E | 4.5e-06 | 5.4e-06 | 5.2e-06 | 5.033E-06 | 4.726E-07 | 4.500E-06 | 5.400E-06 |
| 27620 | 1381 | Y1381V | E | 1.5e-05 | 7.6e-06 | 9.7e-06 | 1.077E-05 | 3.814E-06 | 7.600E-06 | 1.500E-05 |
| 27621 | 1382 | W1382A | E | 6.3e-06 | 8.4e-06 | 6.3e-06 | 7.000E-06 | 1.212E-06 | 6.300E-06 | 8.400E-06 |
| 27622 | 1382 | W1382R | E | 1.4e-05 | 8.8e-06 | 7.4e-06 | 1.007E-05 | 3.478E-06 | 7.400E-06 | 1.400E-05 |
| 27623 | 1382 | W1382N | E | 1.1e-05 | 7.9e-06 | 7.1e-06 | 8.667E-06 | 2.060E-06 | 7.100E-06 | 1.100E-05 |
| 27624 | 1382 | W1382D | E | 8.6e-06 | 1.1e-05 | 8.2e-06 | 9.267E-06 | 1.514E-06 | 8.200E-06 | 1.100E-05 |

| | | | | | | | | | | |
|---|---|---|---|---|---|---|---|---|---|---|
| 27625 | 1382 | W1382C | E | 5.3e-06 | 1e-05 | 1.2e-05 | 9.100E-06 | 3.439E-06 | 5.300E-06 | 1.200E-05 |
| 27626 | 1382 | W1382E | E | 1e-05 | 5.9e-06 | 1e-05 | 8.633E-06 | 2.367E-06 | 5.900E-06 | 1.000E-05 |
| 27627 | 1382 | W1382Q | E | 7.1e-06 | 7.4e-06 | 5e-06 | 6.500E-06 | 1.308E-06 | 5.000E-06 | 7.400E-06 |
| 27628 | 1382 | W1382G | E | 1.9e-05 | 9.4e-06 | 8.7e-06 | 1.237E-05 | 5.755E-06 | 8.700E-06 | 1.900E-05 |
| 27629 | 1382 | W1382H | E | 9.7e-06 | 8.3e-06 | 9.7e-06 | 9.233E-06 | 8.083E-07 | 8.300E-06 | 9.700E-06 |
| 27630 | 1382 | W1382I | E | 6e-06 | 3.5e-06 | 7.4e-06 | 5.633E-06 | 1.976E-06 | 3.500E-06 | 7.400E-06 |
| 27631 | 1382 | W1382L | E | 1.3e-05 | 6.3e-06 | 1.3e-05 | 1.077E-05 | 3.868E-06 | 6.300E-06 | 1.300E-05 |
| 27632 | 1382 | W1382K | E | 1e-05 | 6.7e-06 | 1e-05 | 8.900E-06 | 1.905E-06 | 6.700E-06 | 1.000E-05 |
| 27633 | 1382 | W1382M | E | 6.9e-06 | 8.3e-06 | 6.5e-06 | 7.233E-06 | 9.452E-07 | 6.500E-06 | 8.300E-06 |
| 27634 | 1382 | W1382F | E | 8.6e-06 | 1.2e-05 | 5.8e-06 | 8.800E-06 | 3.105E-06 | 5.800E-06 | 1.200E-05 |
| 27635 | 1382 | W1382P | E | 7.7e-06 | 9.1e-06 | 1.1e-05 | 9.267E-06 | 1.656E-06 | 7.700E-06 | 1.100E-05 |
| 27636 | 1382 | W1382S | E | 6.7e-06 | 1.3e-05 | 6.2e-06 | 8.633E-06 | 3.790E-06 | 6.200E-06 | 1.300E-05 |
| 27637 | 1382 | W1382T | E | 1.3e-05 | 7.7e-06 | 9.5e-06 | 1.007E-05 | 2.695E-06 | 7.700E-06 | 1.300E-05 |
| 27638 | 1382 | W1382W | E | 1.2e-05 | 6.6e-06 | 7.6e-06 | 8.733E-06 | 2.873E-06 | 6.600E-06 | 1.200E-05 |
| 27639 | 1382 | W1382Y | E | 4.5e-06 | 1.2e-05 | 6.6e-06 | 7.700E-06 | 3.869E-06 | 4.500E-06 | 1.200E-05 |
| 27640 | 1382 | W1382V | E | 8e-06 | 9e-06 | 1e-05 | 9.000E-06 | 1.000E-06 | 8.000E-06 | 1.000E-05 |
| 27641 | 1383 | G1383A | E | 5.6e-06 | 4.4e-06 | 6.4e-06 | 5.467E-06 | 1.007E-06 | 4.400E-06 | 6.400E-06 |
| 27642 | 1383 | G1383R | E | 7.6e-06 | 5.5e-06 | 5.9e-06 | 6.333E-06 | 1.115E-06 | 5.500E-06 | 7.600E-06 |
| 27643 | 1383 | G1383N | E | 5.7e-06 | 7.8e-06 | 9.1e-06 | 7.533E-06 | 1.716E-06 | 5.700E-06 | 9.100E-06 |
| 27644 | 1383 | G1383D | E | 1e-05 | 8.4e-06 | 9.6e-06 | 9.333E-06 | 8.327E-07 | 8.400E-06 | 1.000E-05 |
| 27645 | 1383 | G1383C | E | 6.9e-06 | 1.2e-05 | 9.3e-06 | 9.400E-06 | 2.551E-06 | 6.900E-06 | 1.200E-05 |
| 27646 | 1383 | G1383E | E | 7e-06 | 1e-05 | 6.1e-06 | 7.700E-06 | 2.042E-06 | 6.100E-06 | 1.000E-05 |
| 27647 | 1383 | G1383Q | E | 5.9e-06 | 1.2e-05 | 1e-05 | 9.300E-06 | 3.110E-06 | 5.900E-06 | 1.200E-05 |
| 27648 | 1383 | G1383G | E | 8.3e-06 | 4.1e-06 | 8.7e-06 | 7.033E-06 | 2.548E-06 | 4.100E-06 | 8.700E-06 |
| 27649 | 1383 | G1383H | E | 7.2e-06 | 1.3e-05 | 8.5e-06 | 9.567E-06 | 3.044E-06 | 7.200E-06 | 1.300E-05 |
| 27650 | 1383 | G1383I | E | 4.3e-06 | 1.1e-05 | 9e-06 | 8.100E-06 | 3.439E-06 | 4.300E-06 | 1.100E-05 |
| 27651 | 1383 | G1383L | E | 1.3e-05 | 3.1e-06 | 4.8e-06 | 6.967E-06 | 5.294E-06 | 3.100E-06 | 1.300E-05 |
| 27652 | 1383 | G1383K | E | 5.7e-06 | 6.3e-06 | 6e-06 | 6.000E-06 | 3.000E-07 | 5.700E-06 | 6.300E-06 |
| 27653 | 1383 | G1383M | E | 7.2e-06 | 8.3e-06 | 7.2e-06 | 7.567E-06 | 6.351E-07 | 7.200E-06 | 8.300E-06 |
| 27654 | 1383 | G1383F | E | 6.7e-06 | 6.9e-06 | 7.6e-06 | 7.067E-06 | 4.726E-07 | 6.700E-06 | 7.600E-06 |
| 27655 | 1383 | G1383P | E | 1e-05 | 6.7e-06 | 9.5e-06 | 8.733E-06 | 1.779E-06 | 6.700E-06 | 1.000E-05 |
| 27656 | 1383 | G1383S | E | 6.1e-06 | 6.6e-06 | 6.5e-06 | 6.400E-06 | 2.646E-07 | 6.100E-06 | 6.600E-06 |
| 27657 | 1383 | G1383T | E | 9.3e-06 | 1.2e-05 | 5.8e-06 | 9.033E-06 | 3.109E-06 | 5.800E-06 | 1.200E-05 |
| 27658 | 1383 | G1383W | E | 4.8e-06 | 9.6e-06 | 6.3e-06 | 6.900E-06 | 2.456E-06 | 4.800E-06 | 9.600E-06 |
| 27659 | 1383 | G1383Y | E | 1e-05 | 6.1e-06 | 7.2e-06 | 7.767E-06 | 2.011E-06 | 6.100E-06 | 1.000E-05 |
| 27660 | 1383 | G1383V | E | 6.9e-06 | 8.3e-06 | 1e-05 | 8.400E-06 | 1.552E-06 | 6.900E-06 | 1.000E-05 |
| 27661 | 1384 | Q1384A | E | 6.3e-06 | 1.1e-05 | 6.3e-06 | 7.867E-06 | 2.714E-06 | 6.300E-06 | 1.100E-05 |
| 27662 | 1384 | Q1384R | E | 1.1e-05 | 7.4e-06 | 8.5e-06 | 8.967E-06 | 1.845E-06 | 7.400E-06 | 1.100E-05 |
| 27663 | 1384 | Q1384N | E | 5.4e-06 | 6e-06 | 1.4e-05 | 8.467E-06 | 4.801E-06 | 5.400E-06 | 1.400E-05 |
| 27664 | 1384 | Q1384D | E | 7.8e-06 | 6.8e-06 | 6.5e-06 | 7.033E-06 | 6.807E-07 | 6.500E-06 | 7.800E-06 |
| 27665 | 1384 | Q1384C | E | 8.4e-06 | 1.1e-05 | 6.9e-06 | 8.767E-06 | 2.074E-06 | 6.900E-06 | 1.100E-05 |
| 27666 | 1384 | Q1384E | E | 7.9e-06 | 6.6e-06 | 9.6e-06 | 8.033E-06 | 1.504E-06 | 6.600E-06 | 9.600E-06 |
| 27667 | 1384 | Q1384Q | E | 9.5e-06 | 7e-06 | 1.1e-05 | 9.167E-06 | 2.021E-06 | 7.000E-06 | 1.100E-05 |
| 27668 | 1384 | Q1384G | E | 6.3e-06 | 8.2e-06 | 8.2e-06 | 7.567E-06 | 1.097E-06 | 6.300E-06 | 8.200E-06 |
| 27669 | 1384 | Q1384H | E | 6.5e-06 | 1.3e-05 | 1.3e-05 | 1.083E-05 | 3.753E-06 | 6.500E-06 | 1.300E-05 |
| 27670 | 1384 | Q1384I | E | 9.6e-06 | 9e-06 | 7.4e-06 | 8.667E-06 | 1.137E-06 | 7.400E-06 | 9.600E-06 |
| 27671 | 1384 | Q1384L | E | 1.2e-05 | 7.5e-06 | 6.5e-06 | 8.667E-06 | 2.930E-06 | 6.500E-06 | 1.200E-05 |
| 27672 | 1384 | Q1384K | E | 1.3e-05 | 8.7e-06 | 8e-06 | 9.900E-06 | 2.707E-06 | 8.000E-06 | 1.300E-05 |
| 27673 | 1384 | Q1384M | E | 8e-06 | 3.5e-06 | 8.7e-06 | 6.733E-06 | 2.822E-06 | 3.500E-06 | 8.700E-06 |
| 27674 | 1384 | Q1384F | E | 1e-05 | 5e-06 | 8.5e-06 | 7.833E-06 | 2.566E-06 | 5.000E-06 | 1.000E-05 |
| 27675 | 1384 | Q1384P | E | 6e-06 | 6.9e-06 | 7.8e-06 | 6.900E-06 | 9.000E-07 | 6.000E-06 | 7.800E-06 |
| 27676 | 1384 | Q1384S | E | 8.7e-06 | 8.8e-06 | 1.3e-05 | 1.017E-05 | 2.454E-06 | 8.700E-06 | 1.300E-05 |
| 27677 | 1384 | Q1384T | E | 6.8e-06 | 4.6e-06 | 7e-06 | 6.133E-06 | 1.332E-06 | 4.600E-06 | 7.000E-06 |
| 27678 | 1384 | Q1384W | E | 9.6e-06 | 1e-05 | 5e-06 | 8.200E-06 | 2.778E-06 | 5.000E-06 | 1.000E-05 |
| 27679 | 1384 | Q1384Y | E | 8e-06 | 6.7e-06 | 4.5e-06 | 6.400E-06 | 1.769E-06 | 4.500E-06 | 8.000E-06 |
| 27680 | 1384 | Q1384V | E | 8.1e-06 | 8.7e-06 | 5.8e-06 | 7.533E-06 | 1.531E-06 | 5.800E-06 | 8.700E-06 |
| 27681 | 1385 | G1385A | E | 6.2e-06 | 6.6e-06 | 1.3e-05 | 8.600E-06 | 3.816E-06 | 6.200E-06 | 1.300E-05 |
| 27682 | 1385 | G1385R | E | 5.9e-06 | 9.8e-06 | 7e-06 | 7.567E-06 | 2.011E-06 | 5.900E-06 | 9.800E-06 |
| 27683 | 1385 | G1385N | E | 9.8e-06 | 7.5e-06 | 6.5e-06 | 7.933E-06 | 1.692E-06 | 6.500E-06 | 9.800E-06 |
| 27684 | 1385 | G1385D | E | 4.8e-06 | 5.2e-06 | 6.8e-06 | 5.600E-06 | 1.058E-06 | 4.800E-06 | 6.800E-06 |
| 27685 | 1385 | G1385C | E | 8.5e-06 | 6.4e-06 | 8.4e-06 | 7.767E-06 | 1.185E-06 | 6.400E-06 | 8.500E-06 |
| 27686 | 1385 | G1385E | E | 9.6e-06 | 9.7e-06 | 7.3e-06 | 8.867E-06 | 1.358E-06 | 7.300E-06 | 9.700E-06 |
| 27687 | 1385 | G1385Q | E | 5.1e-06 | 4.9e-06 | 8.8e-06 | 6.267E-06 | 2.196E-06 | 4.900E-06 | 8.800E-06 |
| 27688 | 1385 | G1385G | E | 6.5e-06 | 1.3e-05 | 1.1e-05 | 1.017E-05 | 3.329E-06 | 6.500E-06 | 1.300E-05 |
| 27689 | 1385 | G1385H | E | 8e-06 | 6.6e-06 | 4.8e-06 | 6.467E-06 | 1.604E-06 | 4.800E-06 | 8.000E-06 |
| 27690 | 1385 | G1385I | E | 9.2e-06 | 1.2e-05 | 9.3e-06 | 1.017E-05 | 1.589E-06 | 9.200E-06 | 1.200E-05 |
| 27691 | 1385 | G1385L | E | 1.3e-05 | 1.2e-05 | 5.7e-06 | 1.023E-05 | 3.958E-06 | 5.700E-06 | 1.300E-05 |
| 27692 | 1385 | G1385K | E | 6.3e-06 | 6.5e-06 | 1.2e-05 | 8.267E-06 | 3.235E-06 | 6.300E-06 | 1.200E-05 |
| 27693 | 1385 | G1385M | E | 8.5e-06 | 1.2e-05 | 7.9e-06 | 9.467E-06 | 2.214E-06 | 7.900E-06 | 1.200E-05 |
| 27694 | 1385 | G1385F | E | 4.7e-06 | 5.5e-06 | 9.5e-06 | 6.567E-06 | 2.572E-06 | 4.700E-06 | 9.500E-06 |
| 27695 | 1385 | G1385P | E | 1.2e-05 | 7.8e-06 | 4.4e-06 | 8.067E-06 | 3.807E-06 | 4.400E-06 | 1.200E-05 |
| 27696 | 1385 | G1385S | E | 7.3e-06 | 1e-05 | 4e-06 | 7.100E-06 | 3.005E-06 | 4.000E-06 | 1.000E-05 |
| 27697 | 1385 | G1385T | E | 9.7e-06 | 1e-05 | 1.3e-05 | 1.090E-05 | 1.825E-06 | 9.700E-06 | 1.300E-05 |
| 27698 | 1385 | G1385W | E | 1.1e-05 | 8e-06 | 4.4e-06 | 7.800E-06 | 3.305E-06 | 4.400E-06 | 1.100E-05 |
| 27699 | 1385 | G1385Y | E | 7.8e-06 | 6.1e-06 | 1.4e-05 | 9.300E-06 | 4.158E-06 | 6.100E-06 | 1.400E-05 |
| 27700 | 1385 | G1385V | E | 1.3e-05 | 3.8e-06 | 7.5e-06 | 8.100E-06 | 4.629E-06 | 3.800E-06 | 1.300E-05 |
| 27701 | 1386 | T1386A | E | 1.1e-05 | 2.9e-06 | 6.6e-06 | 6.833E-06 | 4.055E-06 | 2.900E-06 | 1.100E-05 |
| 27702 | 1386 | T1386R | E | 1.2e-05 | 9.4e-06 | 6.2e-06 | 9.200E-06 | 2.905E-06 | 6.200E-06 | 1.200E-05 |
| 27703 | 1386 | T1386N | E | 8.8e-06 | 9.9e-06 | 8.5e-06 | 9.067E-06 | 7.371E-07 | 8.500E-06 | 9.900E-06 |
| 27704 | 1386 | T1386D | E | 8.4e-06 | 6.8e-06 | 6.4e-06 | 7.200E-06 | 1.058E-06 | 6.400E-06 | 8.400E-06 |
| 27705 | 1386 | T1386C | E | 5.1e-06 | 8.6e-06 | 7.8e-06 | 7.167E-06 | 1.834E-06 | 5.100E-06 | 8.600E-06 |
| 27706 | 1386 | T1386E | E | 1.1e-05 | 4.5e-06 | 7.7e-06 | 7.733E-06 | 3.250E-06 | 4.500E-06 | 1.100E-05 |
| 27707 | 1386 | T1386Q | E | 6.4e-06 | 1.2e-05 | 7.7e-06 | 8.700E-06 | 2.931E-06 | 6.400E-06 | 1.200E-05 |
| 27708 | 1386 | T1386G | E | 1e-05 | 9.9e-06 | 7.7e-06 | 9.200E-06 | 1.300E-06 | 7.700E-06 | 1.000E-05 |
| 27709 | 1386 | T1386H | E | 1.3e-05 | 8.9e-06 | 7.5e-06 | 9.800E-06 | 2.858E-06 | 7.500E-06 | 1.300E-05 |

| | | | | | | | | | | |
|---|---|---|---|---|---|---|---|---|---|---|
| 27710 | 1386 | T1386I | E | 7.3e-06 | 6.3e-06 | 7.3e-06 | 6.967E-06 | 5.774E-07 | 6.300E-06 | 7.300E-06 |
| 27711 | 1386 | T1386L | E | 9.4e-06 | 1.4e-05 | 1.2e-05 | 1.180E-05 | 2.307E-06 | 9.400E-06 | 1.400E-05 |
| 27712 | 1386 | T1386K | E | 6.5e-06 | 1.2e-05 | 8.1e-06 | 8.867E-06 | 2.829E-06 | 6.500E-06 | 1.200E-05 |
| 27713 | 1386 | T1386M | E | 9.8e-06 | 9e-06 | 8.3e-06 | 9.033E-06 | 7.506E-07 | 8.300E-06 | 9.800E-06 |
| 27714 | 1386 | T1386F | E | 8.7e-06 | 8.3e-06 | 5e-06 | 7.333E-06 | 2.031E-06 | 5.000E-06 | 8.700E-06 |
| 27715 | 1386 | T1386P | E | 4.9e-06 | 6.8e-06 | 9.6e-06 | 7.100E-06 | 2.364E-06 | 4.900E-06 | 9.600E-06 |
| 27716 | 1386 | T1386S | E | 5.7e-06 | 4.7e-06 | 5.6e-06 | 5.333E-06 | 5.508E-07 | 4.700E-06 | 5.700E-06 |
| 27717 | 1386 | T1386T | E | 1.1e-05 | 5.2e-06 | 6.7e-06 | 7.633E-06 | 3.011E-06 | 5.200E-06 | 1.100E-05 |
| 27718 | 1386 | T1386W | E | 9.5e-06 | 1e-05 | 6.8e-06 | 8.767E-06 | 1.721E-06 | 6.800E-06 | 1.000E-05 |
| 27719 | 1386 | T1386Y | E | 5.5e-06 | 7.2e-06 | 5.9e-06 | 6.200E-06 | 8.888E-07 | 5.500E-06 | 7.200E-06 |
| 27720 | 1386 | T1386V | E | 1e-05 | 9.1e-06 | 6.4e-06 | 8.500E-06 | 1.873E-06 | 6.400E-06 | 1.000E-05 |
| 27721 | 1387 | L1387A | E | 1.1e-05 | 1.3e-05 | 1.1e-05 | 1.167E-05 | 1.155E-06 | 1.100E-05 | 1.300E-05 |
| 27722 | 1387 | L1387R | E | 6.4e-06 | 6.6e-06 | 8.6e-06 | 7.200E-06 | 1.217E-06 | 6.400E-06 | 8.600E-06 |
| 27723 | 1387 | L1387N | E | 5e-06 | 1.8e-05 | 7.5e-06 | 1.017E-05 | 6.898E-06 | 5.000E-06 | 1.800E-05 |
| 27724 | 1387 | L1387D | E | 1.3e-05 | 6.1e-06 | 1.3e-05 | 1.070E-05 | 3.984E-06 | 6.100E-06 | 1.300E-05 |
| 27725 | 1387 | L1387C | E | 6.4e-06 | 8.4e-06 | 6.1e-06 | 6.967E-06 | 1.250E-06 | 6.100E-06 | 8.400E-06 |
| 27726 | 1387 | L1387E | E | 9e-06 | 8.8e-06 | 9.4e-06 | 9.067E-06 | 3.055E-07 | 8.800E-06 | 9.400E-06 |
| 27727 | 1387 | L1387Q | E | 7.3e-06 | 7.7e-06 | 5.5e-06 | 6.833E-06 | 1.172E-06 | 5.500E-06 | 7.700E-06 |
| 27728 | 1387 | L1387G | E | 1e-05 | 7.7e-06 | 9.2e-06 | 8.967E-06 | 1.168E-06 | 7.700E-06 | 1.000E-05 |
| 27729 | 1387 | L1387H | E | 5.3e-06 | 6.7e-06 | 5.9e-06 | 5.967E-06 | 7.024E-07 | 5.300E-06 | 6.700E-06 |
| 27730 | 1387 | L1387I | E | 9.8e-06 | 8.4e-06 | 5e-06 | 7.733E-06 | 2.468E-06 | 5.000E-06 | 9.800E-06 |
| 27731 | 1387 | L1387L | E | 4.9e-06 | 5.3e-06 | 7.6e-06 | 5.933E-06 | 1.457E-06 | 4.900E-06 | 7.600E-06 |
| 27732 | 1387 | L1387K | E | 9.2e-06 | 9.7e-06 | 6.8e-06 | 8.567E-06 | 1.550E-06 | 6.800E-06 | 9.700E-06 |
| 27733 | 1387 | L1387M | E | 5.8e-06 | 1.1e-05 | 1.4e-05 | 1.027E-05 | 4.149E-06 | 5.800E-06 | 1.400E-05 |
| 27734 | 1387 | L1387F | E | 6.3e-06 | 1.4e-05 | 8.1e-06 | 9.467E-06 | 4.028E-06 | 6.300E-06 | 1.400E-05 |
| 27735 | 1387 | L1387P | E | 5.5e-06 | 8.7e-06 | 6.8e-06 | 7.000E-06 | 1.609E-06 | 5.500E-06 | 8.700E-06 |
| 27736 | 1387 | L1387S | E | 4.7e-06 | 7e-06 | 9e-06 | 6.900E-06 | 2.152E-06 | 4.700E-06 | 9.000E-06 |
| 27737 | 1387 | L1387T | E | 6.5e-06 | 5.2e-06 | 7.2e-06 | 6.300E-06 | 1.015E-06 | 5.200E-06 | 7.200E-06 |
| 27738 | 1387 | L1387W | E | 4.8e-06 | 9.2e-06 | 5.8e-06 | 6.600E-06 | 2.307E-06 | 4.800E-06 | 9.200E-06 |
| 27739 | 1387 | L1387Y | E | 5.4e-06 | 9.6e-06 | 8.5e-06 | 7.833E-06 | 2.178E-06 | 5.400E-06 | 9.600E-06 |
| 27740 | 1387 | L1387V | E | 1.3e-05 | 1.2e-05 | 8.1e-06 | 1.103E-05 | 2.589E-06 | 8.100E-06 | 1.300E-05 |
| 27741 | 1388 | V1388A | E | 7.7e-06 | 7.5e-06 | 9.3e-06 | 8.167E-06 | 9.866E-07 | 7.500E-06 | 9.300E-06 |
| 27742 | 1388 | V1388R | E | 7e-06 | 9.1e-06 | 6.7e-06 | 7.600E-06 | 1.308E-06 | 6.700E-06 | 9.100E-06 |
| 27743 | 1388 | V1388N | E | 9.9e-06 | 6.7e-06 | 8.6e-06 | 8.400E-06 | 1.609E-06 | 6.700E-06 | 9.900E-06 |
| 27744 | 1388 | V1388D | E | 8.2e-06 | 5.9e-06 | 4.1e-06 | 6.067E-06 | 2.055E-06 | 4.100E-06 | 8.200E-06 |
| 27745 | 1388 | V1388C | E | 6.5e-06 | 6.4e-06 | 6.3e-06 | 6.400E-06 | 1.000E-07 | 6.300E-06 | 6.500E-06 |
| 27746 | 1388 | V1388E | E | 1.3e-05 | 7.4e-06 | 4.9e-06 | 8.433E-06 | 4.148E-06 | 4.900E-06 | 1.300E-05 |
| 27747 | 1388 | V1388Q | E | 8.2e-06 | 1e-05 | 7.2e-06 | 8.467E-06 | 1.419E-06 | 7.200E-06 | 1.000E-05 |
| 27748 | 1388 | V1388G | E | 6.7e-06 | 7.6e-06 | 1.3e-05 | 9.100E-06 | 3.407E-06 | 6.700E-06 | 1.300E-05 |
| 27749 | 1388 | V1388H | E | 6.2e-06 | 4.4e-06 | 1.3e-05 | 7.867E-06 | 4.536E-06 | 4.400E-06 | 1.300E-05 |
| 27750 | 1388 | V1388I | E | 7.2e-06 | 1.3e-05 | 5.8e-06 | 8.667E-06 | 3.818E-06 | 5.800E-06 | 1.300E-05 |
| 27751 | 1388 | V1388L | E | 1.2e-05 | 1.7e-05 | 7.7e-06 | 1.223E-05 | 4.654E-06 | 7.700E-06 | 1.700E-05 |
| 27752 | 1388 | V1388K | E | 1.1e-05 | 6.3e-06 | 8.8e-06 | 8.700E-06 | 2.352E-06 | 6.300E-06 | 1.100E-05 |
| 27753 | 1388 | V1388M | E | 5.7e-06 | 7.1e-06 | 6.7e-06 | 6.500E-06 | 7.211E-07 | 5.700E-06 | 7.100E-06 |
| 27754 | 1388 | V1388F | E | 8.6e-06 | 1e-05 | 8.3e-06 | 8.967E-06 | 9.074E-07 | 8.300E-06 | 1.000E-05 |
| 27755 | 1388 | V1388P | E | 5.7e-06 | 7.2e-06 | 9.3e-06 | 7.400E-06 | 1.808E-06 | 5.700E-06 | 9.300E-06 |
| 27756 | 1388 | V1388S | E | 6.1e-06 | 8.3e-06 | 5.5e-06 | 6.633E-06 | 1.474E-06 | 5.500E-06 | 8.300E-06 |
| 27757 | 1388 | V1388T | E | 1.3e-05 | 6e-06 | 6.6e-06 | 8.533E-06 | 3.880E-06 | 6.000E-06 | 1.300E-05 |
| 27758 | 1388 | V1388W | E | 1.1e-05 | 9.1e-06 | 6.5e-06 | 8.867E-06 | 2.259E-06 | 6.500E-06 | 1.100E-05 |
| 27759 | 1388 | V1388Y | E | 7.6e-06 | 5.7e-06 | 8.2e-06 | 7.167E-06 | 1.305E-06 | 5.700E-06 | 8.200E-06 |
| 27760 | 1388 | V1388V | E | 1.2e-05 | 7.7e-06 | 8.4e-06 | 9.367E-06 | 2.307E-06 | 7.700E-06 | 1.200E-05 |
| 27761 | 1389 | T1389A | E | 5.2e-06 | 1.2e-05 | 7.7e-06 | 8.300E-06 | 3.439E-06 | 5.200E-06 | 1.200E-05 |
| 27762 | 1389 | T1389R | E | 1.2e-05 | 5.9e-06 | 8.8e-06 | 8.900E-06 | 3.051E-06 | 5.900E-06 | 1.200E-05 |
| 27763 | 1389 | T1389N | E | 9.8e-06 | 5.3e-06 | 5e-06 | 6.700E-06 | 2.689E-06 | 5.000E-06 | 9.800E-06 |
| 27764 | 1389 | T1389D | E | 1.5e-05 | 7.8e-06 | 5.2e-06 | 9.333E-06 | 5.077E-06 | 5.200E-06 | 1.500E-05 |
| 27765 | 1389 | T1389C | E | 6.9e-06 | 8e-06 | 9e-06 | 7.967E-06 | 1.050E-06 | 6.900E-06 | 9.000E-06 |
| 27766 | 1389 | T1389E | E | 1.3e-05 | 9.6e-06 | 8.7e-06 | 1.043E-05 | 2.268E-06 | 8.700E-06 | 1.300E-05 |
| 27767 | 1389 | T1389Q | E | 1.2e-05 | 6.8e-06 | 5.3e-06 | 8.033E-06 | 3.516E-06 | 5.300E-06 | 1.200E-05 |
| 27768 | 1389 | T1389G | E | 5.3e-06 | 9e-06 | 8.3e-06 | 7.533E-06 | 1.966E-06 | 5.300E-06 | 9.000E-06 |
| 27769 | 1389 | T1389H | E | 1e-05 | 6.6e-06 | 4.3e-06 | 6.967E-06 | 2.868E-06 | 4.300E-06 | 1.000E-05 |
| 27770 | 1389 | T1389I | E | 7.1e-06 | 1e-05 | 9.4e-06 | 8.833E-06 | 1.531E-06 | 7.100E-06 | 1.000E-05 |
| 27771 | 1389 | T1389L | E | 1e-05 | 7.6e-06 | 7.6e-06 | 8.400E-06 | 1.386E-06 | 7.600E-06 | 1.000E-05 |
| 27772 | 1389 | T1389K | E | 7.6e-06 | 6e-06 | 5.4e-06 | 6.333E-06 | 1.137E-06 | 5.400E-06 | 7.600E-06 |
| 27773 | 1389 | T1389M | E | 1.3e-05 | 1.3e-05 | 1.3e-05 | 1.300E-05 | 0.000E+00 | 1.300E-05 | 1.300E-05 |
| 27774 | 1389 | T1389F | E | 7.6e-06 | 7.2e-06 | 1.1e-05 | 8.600E-06 | 2.088E-06 | 7.200E-06 | 1.100E-05 |
| 27775 | 1389 | T1389P | E | 1e-05 | 6.3e-06 | 7.4e-06 | 7.900E-06 | 1.900E-06 | 6.300E-06 | 1.000E-05 |
| 27776 | 1389 | T1389S | E | 6.4e-06 | 5.8e-06 | 3.3e-06 | 5.167E-06 | 1.644E-06 | 3.300E-06 | 6.400E-06 |
| 27777 | 1389 | T1389T | E | 8.6e-06 | 6.7e-06 | 6.5e-06 | 7.267E-06 | 1.159E-06 | 6.500E-06 | 8.600E-06 |
| 27778 | 1389 | T1389W | E | 6.9e-06 | 8.8e-06 | 6.4e-06 | 7.367E-06 | 1.266E-06 | 6.400E-06 | 8.800E-06 |
| 27779 | 1389 | T1389Y | E | 5.8e-06 | 1.2e-05 | 9.2e-06 | 9.000E-06 | 3.105E-06 | 5.800E-06 | 1.200E-05 |
| 27780 | 1389 | T1389V | E | 1e-05 | 8.1e-06 | 1.3e-05 | 1.037E-05 | 2.470E-06 | 8.100E-06 | 1.300E-05 |
| 27781 | 1390 | V1390A | E | 8.6e-06 | 1.5e-05 | 7.6e-06 | 1.040E-05 | 4.015E-06 | 7.600E-06 | 1.500E-05 |
| 27782 | 1390 | V1390R | E | 6e-06 | 9.6e-06 | 6.3e-06 | 7.300E-06 | 1.997E-06 | 6.000E-06 | 9.600E-06 |
| 27783 | 1390 | V1390N | E | 3.5e-06 | 1.1e-05 | 6.3e-06 | 6.933E-06 | 3.790E-06 | 3.500E-06 | 1.100E-05 |
| 27784 | 1390 | V1390D | E | 9e-06 | 4.9e-06 | 1.1e-05 | 8.300E-06 | 3.110E-06 | 4.900E-06 | 1.100E-05 |
| 27785 | 1390 | V1390C | E | 8.2e-06 | 1.3e-05 | 1.2e-05 | 1.107E-05 | 2.532E-06 | 8.200E-06 | 1.300E-05 |
| 27786 | 1390 | V1390E | E | 6.3e-06 | 6.5e-06 | 8.1e-06 | 6.967E-06 | 9.866E-07 | 6.300E-06 | 8.100E-06 |
| 27787 | 1390 | V1390Q | E | 8.1e-06 | 1.5e-05 | 9.8e-06 | 1.097E-05 | 3.595E-06 | 8.100E-06 | 1.500E-05 |
| 27788 | 1390 | V1390G | E | 1.2e-05 | 4.8e-06 | 4.5e-06 | 7.100E-06 | 4.246E-06 | 4.500E-06 | 1.200E-05 |
| 27789 | 1390 | V1390H | E | 7.5e-06 | 5e-06 | 9.7e-06 | 7.400E-06 | 2.352E-06 | 5.000E-06 | 9.700E-06 |
| 27790 | 1390 | V1390I | E | 6e-06 | 6.3e-06 | 8.5e-06 | 6.933E-06 | 1.365E-06 | 6.000E-06 | 8.500E-06 |
| 27791 | 1390 | V1390L | E | 8.2e-06 | 8.8e-06 | 1.2e-05 | 9.667E-06 | 2.043E-06 | 8.200E-06 | 1.200E-05 |
| 27792 | 1390 | V1390K | E | 1.3e-05 | 1.2e-05 | 5.5e-06 | 1.017E-05 | 4.072E-06 | 5.500E-06 | 1.300E-05 |
| 27793 | 1390 | V1390M | E | 9.2e-06 | 1.1e-05 | 4.6e-06 | 8.267E-06 | 3.301E-06 | 4.600E-06 | 1.100E-05 |
| 27794 | 1390 | V1390F | E | 8.9e-06 | 6.6e-06 | 9.3e-06 | 8.267E-06 | 1.457E-06 | 6.600E-06 | 9.300E-06 |

| | | | | | | | | | | |
|---|---|---|---|---|---|---|---|---|---|---|
| 27795 | 1390 | V1390P | E | 8.9e-06 | 5e-06 | 8.9e-06 | 7.600E-06 | 2.252E-06 | 5.000E-06 | 8.900E-06 |
| 27796 | 1390 | V1390S | E | 4.3e-06 | 7.3e-06 | 9.8e-06 | 7.133E-06 | 2.754E-06 | 4.300E-06 | 9.800E-06 |
| 27797 | 1390 | V1390T | E | 6e-06 | 6.3e-06 | 4.5e-06 | 5.600E-06 | 9.644E-07 | 4.500E-06 | 6.300E-06 |
| 27798 | 1390 | V1390W | E | 6.4e-06 | 7.6e-06 | 1.1e-05 | 8.333E-06 | 2.386E-06 | 6.400E-06 | 1.100E-05 |
| 27799 | 1390 | V1390Y | E | 9.3e-06 | 7e-06 | 1e-05 | 8.767E-06 | 1.570E-06 | 7.000E-06 | 1.000E-05 |
| 27800 | 1390 | V1390V | E | 3.4e-06 | 5.8e-06 | 8.2e-06 | 5.800E-06 | 2.400E-06 | 3.400E-06 | 8.200E-06 |
| 27801 | 1391 | S1391A | E | 1.5e-05 | 6.9e-06 | 3.3e-06 | 8.400E-06 | 5.992E-06 | 3.300E-06 | 1.500E-05 |
| 27802 | 1391 | S1391R | E | 6e-06 | 8.5e-06 | 1.3e-05 | 9.167E-06 | 3.547E-06 | 6.000E-06 | 1.300E-05 |
| 27803 | 1391 | S1391N | E | 6.4e-06 | 4.4e-06 | 8.7e-06 | 6.500E-06 | 2.152E-06 | 4.400E-06 | 8.700E-06 |
| 27804 | 1391 | S1391D | E | 9.9e-06 | 1.4e-05 | 9.8e-06 | 1.123E-05 | 2.397E-06 | 9.800E-06 | 1.400E-05 |
| 27805 | 1391 | S1391C | E | 7.7e-06 | 7e-06 | 6.6e-06 | 7.100E-06 | 5.568E-07 | 6.600E-06 | 7.700E-06 |
| 27806 | 1391 | S1391E | E | 1.1e-05 | 4e-06 | 8.8e-06 | 7.933E-06 | 3.580E-06 | 4.000E-06 | 1.100E-05 |
| 27807 | 1391 | S1391Q | E | 8.5e-06 | 1.3e-05 | 5.6e-06 | 9.033E-06 | 3.729E-06 | 5.600E-06 | 1.300E-05 |
| 27808 | 1391 | S1391G | E | 7.4e-06 | 1.4e-05 | 7e-06 | 9.467E-06 | 3.931E-06 | 7.000E-06 | 1.400E-05 |
| 27809 | 1391 | S1391H | E | 1.3e-05 | 8.1e-06 | 7.8e-06 | 9.633E-06 | 2.919E-06 | 7.800E-06 | 1.300E-05 |
| 27810 | 1391 | S1391I | E | 7.3e-06 | 9.6e-06 | 7.9e-06 | 8.267E-06 | 1.193E-06 | 7.300E-06 | 9.600E-06 |
| 27811 | 1391 | S1391L | E | 7e-06 | 6e-06 | 5.9e-06 | 6.300E-06 | 6.083E-07 | 5.900E-06 | 7.000E-06 |
| 27812 | 1391 | S1391K | E | 9e-06 | 5.2e-06 | 7.5e-06 | 7.233E-06 | 1.914E-06 | 5.200E-06 | 9.000E-06 |
| 27813 | 1391 | S1391M | E | 5.4e-06 | 7.2e-06 | 1e-05 | 7.533E-06 | 2.318E-06 | 5.400E-06 | 1.000E-05 |
| 27814 | 1391 | S1391F | E | 9.7e-06 | 6.6e-06 | 1.4e-05 | 1.010E-05 | 3.716E-06 | 6.600E-06 | 1.400E-05 |
| 27815 | 1391 | S1391P | E | 5.3e-06 | 6.1e-06 | 5.1e-06 | 5.500E-06 | 5.292E-07 | 5.100E-06 | 6.100E-06 |
| 27816 | 1391 | S1391S | E | 6.7e-06 | 8.4e-06 | 9.7e-06 | 8.267E-06 | 1.504E-06 | 6.700E-06 | 9.700E-06 |
| 27817 | 1391 | S1391T | E | 8.2e-06 | 5.7e-06 | 9.8e-06 | 7.900E-06 | 2.066E-06 | 5.700E-06 | 9.800E-06 |
| 27818 | 1391 | S1391W | E | 5.5e-06 | 4.8e-06 | 1.2e-05 | 7.433E-06 | 3.970E-06 | 4.800E-06 | 1.200E-05 |
| 27819 | 1391 | S1391Y | E | 8.7e-06 | 4.4e-06 | 1.5e-05 | 9.367E-06 | 5.331E-06 | 4.400E-06 | 1.500E-05 |
| 27820 | 1391 | S1391V | E | 8e-06 | 1e-05 | 8.3e-06 | 8.767E-06 | 1.079E-06 | 8.000E-06 | 1.000E-05 |
| 27821 | 1392 | S1392A | E | 6.8e-06 | 4.8e-06 | 4.3e-06 | 5.300E-06 | 1.323E-06 | 4.300E-06 | 6.800E-06 |
| 27822 | 1392 | S1392R | E | 8.9e-06 | 9e-06 | 5.8e-06 | 7.900E-06 | 1.819E-06 | 5.800E-06 | 9.000E-06 |
| 27823 | 1392 | S1392N | E | 1.2e-05 | 6.3e-06 | 9.8e-06 | 9.367E-06 | 2.875E-06 | 6.300E-06 | 1.200E-05 |
| 27824 | 1392 | S1392D | E | 1e-05 | 1.2e-05 | 7.5e-06 | 9.833E-06 | 2.255E-06 | 7.500E-06 | 1.200E-05 |
| 27825 | 1392 | S1392C | E | 5.5e-06 | 3.9e-06 | 7.6e-06 | 5.667E-06 | 1.856E-06 | 3.900E-06 | 7.600E-06 |
| 27826 | 1392 | S1392E | E | 5.7e-06 | 8.6e-06 | 7.1e-06 | 7.133E-06 | 1.450E-06 | 5.700E-06 | 8.600E-06 |
| 27827 | 1392 | S1392Q | E | 8.6e-06 | 7.5e-06 | 5.5e-06 | 7.200E-06 | 1.572E-06 | 5.500E-06 | 8.600E-06 |
| 27828 | 1392 | S1392G | E | 5.2e-06 | 6.6e-06 | 9e-06 | 6.933E-06 | 1.922E-06 | 5.200E-06 | 9.000E-06 |
| 27829 | 1392 | S1392H | E | 7.6e-06 | 4.9e-06 | 5.9e-06 | 6.133E-06 | 1.365E-06 | 4.900E-06 | 7.600E-06 |
| 27830 | 1392 | S1392I | E | 7.3e-06 | 9.2e-06 | 8.3e-06 | 8.267E-06 | 9.504E-07 | 7.300E-06 | 9.200E-06 |
| 27831 | 1392 | S1392L | E | 9.6e-06 | 9.6e-06 | 9.2e-06 | 9.467E-06 | 2.309E-07 | 9.200E-06 | 9.600E-06 |
| 27832 | 1392 | S1392K | E | 1e-05 | 8.3e-06 | 9.9e-06 | 9.400E-06 | 9.539E-07 | 8.300E-06 | 1.000E-05 |
| 27833 | 1392 | S1392M | E | 9.3e-06 | 4.9e-06 | 9.2e-06 | 7.800E-06 | 2.512E-06 | 4.900E-06 | 9.300E-06 |
| 27834 | 1392 | S1392F | E | 1.4e-05 | 6.9e-06 | 1.3e-05 | 1.130E-05 | 3.843E-06 | 6.900E-06 | 1.400E-05 |
| 27835 | 1392 | S1392P | E | 7.3e-06 | 8.3e-06 | 8.7e-06 | 8.100E-06 | 7.211E-07 | 7.300E-06 | 8.700E-06 |
| 27836 | 1392 | S1392S | E | 7.7e-06 | 8.3e-06 | 5.9e-06 | 7.300E-06 | 1.249E-06 | 5.900E-06 | 8.300E-06 |
| 27837 | 1392 | S1392T | E | 5.1e-06 | 6.6e-06 | 3.2e-06 | 4.967E-06 | 1.704E-06 | 3.200E-06 | 6.600E-06 |
| 27838 | 1392 | S1392W | E | 9.1e-06 | 9.5e-06 | 5.2e-06 | 7.933E-06 | 2.376E-06 | 5.200E-06 | 9.500E-06 |
| 27839 | 1392 | S1392Y | E | 6.5e-06 | 1.1e-05 | 9.8e-06 | 9.100E-06 | 2.330E-06 | 6.500E-06 | 1.100E-05 |
| 27840 | 1392 | S1392V | E | 9.1e-06 | 9e-06 | 7e-06 | 8.367E-06 | 1.185E-06 | 7.000E-06 | 9.100E-06 |
| 27841 | 1393 | A1393A | E | 1.3e-05 | 4.4e-06 | 6.2e-06 | 7.867E-06 | 4.536E-06 | 4.400E-06 | 1.300E-05 |
| 27842 | 1393 | A1393R | E | 5.6e-06 | 9.6e-06 | 8.7e-06 | 7.967E-06 | 2.098E-06 | 5.600E-06 | 9.600E-06 |
| 27843 | 1393 | A1393N | E | 6.5e-06 | 7.3e-06 | 6.7e-06 | 6.833E-06 | 4.163E-07 | 6.500E-06 | 7.300E-06 |
| 27844 | 1393 | A1393D | E | 7.5e-06 | 1.1e-05 | 9.5e-06 | 9.333E-06 | 1.756E-06 | 7.500E-06 | 1.100E-05 |
| 27845 | 1393 | A1393C | E | 1.3e-05 | 5.8e-06 | 6.5e-06 | 8.433E-06 | 3.970E-06 | 5.800E-06 | 1.300E-05 |
| 27846 | 1393 | A1393E | E | 7.1e-06 | 1.8e-05 | 5.1e-06 | 1.007E-05 | 6.943E-06 | 5.100E-06 | 1.800E-05 |
| 27847 | 1393 | A1393Q | E | 1.1e-05 | 6.1e-06 | 1.4e-05 | 1.037E-05 | 3.988E-06 | 6.100E-06 | 1.400E-05 |
| 27848 | 1393 | A1393G | E | 1e-05 | 1e-05 | 9.8e-06 | 9.933E-06 | 1.155E-07 | 9.800E-06 | 1.000E-05 |
| 27849 | 1393 | A1393H | E | 5.6e-06 | 1.9e-05 | 6.6e-06 | 1.040E-05 | 7.465E-06 | 5.600E-06 | 1.900E-05 |
| 27850 | 1393 | A1393I | E | 9.6e-06 | 6.8e-06 | 6.1e-06 | 7.500E-06 | 1.852E-06 | 6.100E-06 | 9.600E-06 |
| 27851 | 1393 | A1393L | E | 4.7e-06 | 6.4e-06 | 4.3e-06 | 5.133E-06 | 1.115E-06 | 4.300E-06 | 6.400E-06 |
| 27852 | 1393 | A1393K | E | 5e-06 | 1e-05 | 9.8e-06 | 8.267E-06 | 2.831E-06 | 5.000E-06 | 1.000E-05 |
| 27853 | 1393 | A1393M | E | 5.5e-06 | 7.6e-06 | 5e-06 | 6.033E-06 | 1.380E-06 | 5.000E-06 | 7.600E-06 |
| 27854 | 1393 | A1393F | E | 8.4e-06 | 7.4e-06 | 8.4e-06 | 8.067E-06 | 5.774E-07 | 7.400E-06 | 8.400E-06 |
| 27855 | 1393 | A1393P | E | 3.8e-06 | 3.8e-06 | 6.3e-06 | 4.633E-06 | 1.443E-06 | 3.800E-06 | 6.300E-06 |
| 27856 | 1393 | A1393S | E | 1e-05 | 5.6e-06 | 1.1e-05 | 8.867E-06 | 2.873E-06 | 5.600E-06 | 1.100E-05 |
| 27857 | 1393 | A1393T | E | 6.6e-06 | 1.2e-05 | 6.7e-06 | 8.433E-06 | 3.089E-06 | 6.600E-06 | 1.200E-05 |
| 27858 | 1393 | A1393W | E | 9.3e-06 | 8.3e-06 | 7.2e-06 | 8.267E-06 | 1.050E-06 | 7.200E-06 | 9.300E-06 |
| 27859 | 1393 | A1393Y | E | 9.5e-06 | 6.5e-06 | 9.4e-06 | 8.467E-06 | 1.704E-06 | 6.500E-06 | 9.500E-06 |
| 27860 | 1393 | A1393V | E | 8.6e-06 | 8.5e-06 | 9.3e-06 | 8.800E-06 | 4.359E-07 | 8.500E-06 | 9.300E-06 |
| 27861 | 1394 | S1394A | E | 7.4e-06 | 9.8e-06 | 9.6e-06 | 8.933E-06 | 1.332E-06 | 7.400E-06 | 9.800E-06 |
| 27862 | 1394 | S1394R | E | 6e-06 | 5.2e-06 | 7.6e-06 | 6.267E-06 | 1.222E-06 | 5.200E-06 | 7.600E-06 |
| 27863 | 1394 | S1394N | E | 6.9e-06 | 6.7e-06 | 1.3e-05 | 8.867E-06 | 3.581E-06 | 6.700E-06 | 1.300E-05 |
| 27864 | 1394 | S1394D | E | 6.4e-06 | 5.9e-06 | 6.8e-06 | 6.367E-06 | 4.509E-07 | 5.900E-06 | 6.800E-06 |
| 27865 | 1394 | S1394C | E | 6.1e-06 | 9.9e-06 | 5.7e-06 | 7.233E-06 | 2.318E-06 | 5.700E-06 | 9.900E-06 |
| 27866 | 1394 | S1394E | E | 4.6e-06 | 9e-06 | 1.1e-05 | 8.200E-06 | 3.274E-06 | 4.600E-06 | 1.100E-05 |
| 27867 | 1394 | S1394Q | E | 7.4e-06 | 1.1e-05 | 6.4e-06 | 8.267E-06 | 2.419E-06 | 6.400E-06 | 1.100E-05 |
| 27868 | 1394 | S1394G | E | 9.9e-06 | 5.2e-06 | 6.5e-06 | 7.200E-06 | 2.427E-06 | 5.200E-06 | 9.900E-06 |
| 27869 | 1394 | S1394H | E | 9.4e-06 | 9.9e-06 | 9.4e-06 | 9.567E-06 | 2.887E-07 | 9.400E-06 | 9.900E-06 |
| 27870 | 1394 | S1394I | E | 8.6e-06 | 6.8e-06 | 3.8e-06 | 6.400E-06 | 2.425E-06 | 3.800E-06 | 8.600E-06 |
| 27871 | 1394 | S1394L | E | 6e-06 | 7.5e-06 | 9.7e-06 | 7.733E-06 | 1.861E-06 | 6.000E-06 | 9.700E-06 |
| 27872 | 1394 | S1394K | E | 1.3e-05 | 7.5e-06 | 6.8e-06 | 9.100E-06 | 3.396E-06 | 6.800E-06 | 1.300E-05 |
| 27873 | 1394 | S1394M | E | 5.7e-06 | 7.8e-06 | 4.1e-06 | 5.867E-06 | 1.856E-06 | 4.100E-06 | 7.800E-06 |
| 27874 | 1394 | S1394F | E | 7.7e-06 | 7.4e-06 | 7.7e-06 | 7.600E-06 | 1.732E-07 | 7.400E-06 | 7.700E-06 |
| 27875 | 1394 | S1394P | E | 5.9e-06 | 1e-05 | 8.8e-06 | 8.233E-06 | 2.108E-06 | 5.900E-06 | 1.000E-05 |
| 27876 | 1394 | S1394S | E | 6.4e-06 | 1.3e-05 | 1.4e-05 | 1.113E-05 | 4.130E-06 | 6.400E-06 | 1.400E-05 |
| 27877 | 1394 | S1394T | E | 1.3e-05 | 5.3e-06 | 4.5e-06 | 7.600E-06 | 4.694E-06 | 4.500E-06 | 1.300E-05 |
| 27878 | 1394 | S1394W | E | 7.5e-06 | 6.1e-06 | 1e-05 | 7.867E-06 | 1.976E-06 | 6.100E-06 | 1.000E-05 |
| 27879 | 1394 | S1394Y | E | 6.2e-06 | 9.2e-06 | 8.4e-06 | 7.933E-06 | 1.553E-06 | 6.200E-06 | 9.200E-06 |

| | | | | | | | | | | |
|---|---|---|---|---|---|---|---|---|---|---|
| 27880 | 1394 | S1394V | E | 7.8e-06 | 6.4e-06 | 1e-05 | 8.067E-06 | 1.815E-06 | 6.400E-06 | 1.000E-05 |
| 27881 | 1395 | T1395A | E | 1.7e-05 | 8.5e-06 | 6.2e-06 | 1.057E-05 | 5.689E-06 | 6.200E-06 | 1.700E-05 |
| 27882 | 1395 | T1395R | E | 8.4e-06 | 9.6e-06 | 1e-05 | 9.333E-06 | 8.327E-07 | 8.400E-06 | 1.000E-05 |
| 27883 | 1395 | T1395N | E | 9.6e-06 | 5.8e-06 | 8.2e-06 | 7.867E-06 | 1.922E-06 | 5.800E-06 | 9.600E-06 |
| 27884 | 1395 | T1395D | E | 8.3e-06 | 1.4e-05 | 1.2e-05 | 1.143E-05 | 2.892E-06 | 8.300E-06 | 1.400E-05 |
| 27885 | 1395 | T1395C | E | 4.8e-06 | 8e-06 | 6.2e-06 | 6.333E-06 | 1.604E-06 | 4.800E-06 | 8.000E-06 |
| 27886 | 1395 | T1395E | E | 8.7e-06 | 7.4e-06 | 6.5e-06 | 7.533E-06 | 1.106E-06 | 6.500E-06 | 8.700E-06 |
| 27887 | 1395 | T1395Q | E | 5.6e-06 | 9.1e-06 | 5.5e-06 | 6.733E-06 | 2.050E-06 | 5.500E-06 | 9.100E-06 |
| 27888 | 1395 | T1395G | E | 4.2e-06 | 4.4e-06 | 5.8e-06 | 4.800E-06 | 8.718E-07 | 4.200E-06 | 5.800E-06 |
| 27889 | 1395 | T1395H | E | 8.8e-06 | 7.7e-06 | 1.3e-05 | 9.833E-06 | 2.797E-06 | 7.700E-06 | 1.300E-05 |
| 27890 | 1395 | T1395I | E | 6.4e-06 | 7.3e-06 | 6.9e-06 | 6.867E-06 | 4.509E-07 | 6.400E-06 | 7.300E-06 |
| 27891 | 1395 | T1395L | E | 9.2e-06 | 9.9e-06 | 6.9e-06 | 8.667E-06 | 1.570E-06 | 6.900E-06 | 9.900E-06 |
| 27892 | 1395 | T1395K | E | 4.6e-06 | 6.2e-06 | 9.7e-06 | 6.833E-06 | 2.608E-06 | 4.600E-06 | 9.700E-06 |
| 27893 | 1395 | T1395M | E | 6.4e-06 | 5.9e-06 | 1.5e-05 | 9.100E-06 | 5.116E-06 | 5.900E-06 | 1.500E-05 |
| 27894 | 1395 | T1395F | E | 6.9e-06 | 5.5e-06 | 4.7e-06 | 5.700E-06 | 1.114E-06 | 4.700E-06 | 6.900E-06 |
| 27895 | 1395 | T1395P | E | 9.4e-06 | 8.4e-06 | 1.5e-05 | 1.093E-05 | 3.557E-06 | 8.400E-06 | 1.500E-05 |
| 27896 | 1395 | T1395S | E | 8.9e-06 | 6.2e-06 | 5.6e-06 | 6.900E-06 | 1.758E-06 | 5.600E-06 | 8.900E-06 |
| 27897 | 1395 | T1395T | E | 9.3e-06 | 1.4e-05 | 2.4e-06 | 8.567E-06 | 5.835E-06 | 2.400E-06 | 1.400E-05 |
| 27898 | 1395 | T1395W | E | 6.1e-06 | 7.8e-06 | 9.1e-06 | 7.667E-06 | 1.504E-06 | 6.100E-06 | 9.100E-06 |
| 27899 | 1395 | T1395Y | E | 9e-06 | 1e-05 | 8.7e-06 | 9.233E-06 | 6.807E-07 | 8.700E-06 | 1.000E-05 |
| 27900 | 1395 | T1395V | E | 6.4e-06 | 7.1e-06 | 8.8e-06 | 7.433E-06 | 1.234E-06 | 6.400E-06 | 8.800E-06 |
| 27901 | 1396 | K1396A | E | 5.6e-06 | 7.5e-06 | 5.2e-06 | 6.100E-06 | 1.229E-06 | 5.200E-06 | 7.500E-06 |
| 27902 | 1396 | K1396R | E | 6e-06 | 9.3e-06 | 9.1e-06 | 8.133E-06 | 1.850E-06 | 6.000E-06 | 9.300E-06 |
| 27903 | 1396 | K1396N | E | 9.8e-06 | 9.2e-06 | 5.9e-06 | 8.300E-06 | 2.100E-06 | 5.900E-06 | 9.800E-06 |
| 27904 | 1396 | K1396D | E | 8.9e-06 | 6.2e-06 | 2.1e-05 | 1.203E-05 | 7.882E-06 | 6.200E-06 | 2.100E-05 |
| 27905 | 1396 | K1396C | E | 1.3e-05 | 5e-06 | 6.9e-06 | 8.300E-06 | 4.180E-06 | 5.000E-06 | 1.300E-05 |
| 27906 | 1396 | K1396E | E | 1.8e-05 | 6.1e-06 | 6.5e-06 | 1.020E-05 | 6.758E-06 | 6.100E-06 | 1.800E-05 |
| 27907 | 1396 | K1396Q | E | 7.7e-06 | 9e-06 | 9.5e-06 | 8.733E-06 | 9.292E-07 | 7.700E-06 | 9.500E-06 |
| 27908 | 1396 | K1396G | E | 4.3e-06 | 9e-06 | 1.5e-05 | 9.433E-06 | 5.363E-06 | 4.300E-06 | 1.500E-05 |
| 27909 | 1396 | K1396H | E | 7.4e-06 | 7.8e-06 | 5e-06 | 6.733E-06 | 1.514E-06 | 5.000E-06 | 7.800E-06 |
| 27910 | 1396 | K1396I | E | 1.4e-05 | 3.4e-06 | 7.4e-06 | 8.267E-06 | 5.353E-06 | 3.400E-06 | 1.400E-05 |
| 27911 | 1396 | K1396L | E | 8.6e-06 | 1.6e-05 | 6.2e-06 | 1.027E-05 | 5.108E-06 | 6.200E-06 | 1.600E-05 |
| 27912 | 1396 | K1396K | E | 7.2e-06 | 5.8e-06 | 8.6e-06 | 7.200E-06 | 1.400E-06 | 5.800E-06 | 8.600E-06 |
| 27913 | 1396 | K1396M | E | 9.5e-06 | 5.4e-06 | 8.9e-06 | 7.933E-06 | 2.214E-06 | 5.400E-06 | 9.500E-06 |
| 27914 | 1396 | K1396F | E | 7.2e-06 | 6.1e-06 | 1.1e-05 | 8.100E-06 | 2.571E-06 | 6.100E-06 | 1.100E-05 |
| 27915 | 1396 | K1396P | E | 6.3e-06 | 8.3e-06 | 8e-06 | 7.533E-06 | 1.079E-06 | 6.300E-06 | 8.300E-06 |
| 27916 | 1396 | K1396S | E | 8.7e-06 | 9.5e-06 | 1e-05 | 9.400E-06 | 6.557E-07 | 8.700E-06 | 1.000E-05 |
| 27917 | 1396 | K1396T | E | 5.6e-06 | 1.3e-05 | 9.4e-06 | 9.333E-06 | 3.700E-06 | 5.600E-06 | 1.300E-05 |
| 27918 | 1396 | K1396W | E | 6.8e-06 | 4.5e-06 | 8.5e-06 | 6.600E-06 | 2.007E-06 | 4.500E-06 | 8.500E-06 |
| 27919 | 1396 | K1396Y | E | 5.8e-06 | 9.4e-06 | 9.4e-06 | 8.200E-06 | 2.078E-06 | 5.800E-06 | 9.400E-06 |
| 27920 | 1396 | K1396V | E | 4.6e-06 | 8.1e-06 | 6.4e-06 | 6.367E-06 | 1.750E-06 | 4.600E-06 | 8.100E-06 |
| 27921 | 1397 | G1397A | E | 7.8e-06 | 8.1e-06 | 8.7e-06 | 8.200E-06 | 4.583E-07 | 7.800E-06 | 8.700E-06 |
| 27922 | 1397 | G1397R | E | 8e-06 | 1.5e-05 | 9.4e-06 | 1.080E-05 | 3.704E-06 | 8.000E-06 | 1.500E-05 |
| 27923 | 1397 | G1397N | E | 1e-05 | 9.4e-06 | 6.5e-06 | 8.633E-06 | 1.872E-06 | 6.500E-06 | 1.000E-05 |
| 27924 | 1397 | G1397D | E | 8.9e-06 | 9.7e-06 | 5.1e-06 | 7.900E-06 | 2.458E-06 | 5.100E-06 | 9.700E-06 |
| 27925 | 1397 | G1397C | E | 9.5e-06 | 6.5e-06 | 8.6e-06 | 8.200E-06 | 1.539E-06 | 6.500E-06 | 9.500E-06 |
| 27926 | 1397 | G1397E | E | 7e-06 | 4e-06 | 1.6e-05 | 9.000E-06 | 6.245E-06 | 4.000E-06 | 1.600E-05 |
| 27927 | 1397 | G1397Q | E | 9.4e-06 | 6e-06 | 9.5e-06 | 8.300E-06 | 1.992E-06 | 6.000E-06 | 9.500E-06 |
| 27928 | 1397 | G1397G | E | 8.3e-06 | 7.9e-06 | 9.7e-06 | 8.633E-06 | 9.452E-07 | 7.900E-06 | 9.700E-06 |
| 27929 | 1397 | G1397H | E | 6.1e-06 | 7.5e-06 | 1e-05 | 7.867E-06 | 1.976E-06 | 6.100E-06 | 1.000E-05 |
| 27930 | 1397 | G1397I | E | 8.2e-06 | 6.8e-06 | 1.2e-05 | 9.000E-06 | 2.691E-06 | 6.800E-06 | 1.200E-05 |
| 27931 | 1397 | G1397L | E | 1.5e-05 | 7.8e-06 | 1.4e-05 | 1.227E-05 | 3.900E-06 | 7.800E-06 | 1.500E-05 |
| 27932 | 1397 | G1397K | E | 6.6e-06 | 6.3e-06 | 9.4e-06 | 7.433E-06 | 1.710E-06 | 6.300E-06 | 9.400E-06 |
| 27933 | 1397 | G1397M | E | 5.7e-06 | 7.1e-06 | 6.2e-06 | 6.333E-06 | 7.095E-07 | 5.700E-06 | 7.100E-06 |
| 27934 | 1397 | G1397F | E | 8.4e-06 | 9.3e-06 | 7.6e-06 | 8.433E-06 | 8.505E-07 | 7.600E-06 | 9.300E-06 |
| 27935 | 1397 | G1397P | E | 3.7e-06 | 6.2e-06 | 6.2e-06 | 5.367E-06 | 1.443E-06 | 3.700E-06 | 6.200E-06 |
| 27936 | 1397 | G1397S | E | 6.6e-06 | 4.8e-06 | 6.2e-06 | 5.867E-06 | 9.452E-07 | 4.800E-06 | 6.600E-06 |
| 27937 | 1397 | G1397T | E | 5.6e-06 | 6.2e-06 | 1.1e-05 | 7.600E-06 | 2.960E-06 | 5.600E-06 | 1.100E-05 |
| 27938 | 1397 | G1397W | E | 9.8e-06 | 1e-05 | 6.4e-06 | 8.733E-06 | 2.023E-06 | 6.400E-06 | 1.000E-05 |
| 27939 | 1397 | G1397Y | E | 9.5e-06 | 9.7e-06 | 1.3e-05 | 1.073E-05 | 1.966E-06 | 9.500E-06 | 1.300E-05 |
| 27940 | 1397 | G1397V | E | 6.2e-06 | 8.9e-06 | 1.5e-05 | 1.003E-05 | 4.508E-06 | 6.200E-06 | 1.500E-05 |
| 27941 | 1398 | P1398A | E | 7e-06 | 8.7e-06 | 6.8e-06 | 7.500E-06 | 1.044E-06 | 6.800E-06 | 8.700E-06 |
| 27942 | 1398 | P1398R | E | 4.1e-06 | 6.3e-06 | 1.3e-05 | 7.800E-06 | 4.636E-06 | 4.100E-06 | 1.300E-05 |
| 27943 | 1398 | P1398N | E | 9e-06 | 7.4e-06 | 1.4e-05 | 1.013E-05 | 3.443E-06 | 7.400E-06 | 1.400E-05 |
| 27944 | 1398 | P1398D | E | 1.1e-05 | 8.9e-06 | 9.9e-06 | 9.933E-06 | 1.050E-06 | 8.900E-06 | 1.100E-05 |
| 27945 | 1398 | P1398C | E | 1.3e-05 | 5.4e-06 | 1.1e-05 | 9.800E-06 | 3.940E-06 | 5.400E-06 | 1.300E-05 |
| 27946 | 1398 | P1398E | E | 5.9e-06 | 5.3e-06 | 7e-06 | 6.067E-06 | 8.622E-07 | 5.300E-06 | 7.000E-06 |
| 27947 | 1398 | P1398Q | E | 4.1e-06 | 5.8e-06 | 1.1e-05 | 6.967E-06 | 3.595E-06 | 4.100E-06 | 1.100E-05 |
| 27948 | 1398 | P1398G | E | 6.9e-06 | 8.2e-06 | 1.2e-05 | 9.033E-06 | 2.650E-06 | 6.900E-06 | 1.200E-05 |
| 27949 | 1398 | P1398H | E | 7.4e-06 | 6.6e-06 | 7.1e-06 | 7.033E-06 | 4.041E-07 | 6.600E-06 | 7.400E-06 |
| 27950 | 1398 | P1398I | E | 7.5e-06 | 1e-05 | 6.1e-06 | 7.867E-06 | 1.976E-06 | 6.100E-06 | 1.000E-05 |
| 27951 | 1398 | P1398L | E | 8.2e-06 | 6.5e-06 | 1.3e-05 | 9.233E-06 | 3.371E-06 | 6.500E-06 | 1.300E-05 |
| 27952 | 1398 | P1398K | E | 7e-06 | 1.3e-05 | 1.1e-05 | 1.033E-05 | 3.055E-06 | 7.000E-06 | 1.300E-05 |
| 27953 | 1398 | P1398M | E | 9.8e-06 | 1e-05 | 8.7e-06 | 9.500E-06 | 7.000E-07 | 8.700E-06 | 1.000E-05 |
| 27954 | 1398 | P1398F | E | 8.9e-06 | 7.1e-06 | 6.2e-06 | 7.400E-06 | 1.375E-06 | 6.200E-06 | 8.900E-06 |
| 27955 | 1398 | P1398P | E | 8.2e-06 | 6.6e-06 | 9.4e-06 | 8.067E-06 | 1.405E-06 | 6.600E-06 | 9.400E-06 |
| 27956 | 1398 | P1398S | E | 8.8e-06 | 7.5e-06 | 5.8e-06 | 7.367E-06 | 1.504E-06 | 5.800E-06 | 8.800E-06 |
| 27957 | 1398 | P1398T | E | 6.6e-06 | 7.9e-06 | 9.6e-06 | 8.033E-06 | 1.504E-06 | 6.600E-06 | 9.600E-06 |
| 27958 | 1398 | P1398W | E | 9.4e-06 | 5.2e-06 | 8.7e-06 | 7.767E-06 | 2.250E-06 | 5.200E-06 | 9.400E-06 |
| 27959 | 1398 | P1398Y | E | 7.2e-06 | 5.6e-06 | 6.3e-06 | 6.367E-06 | 8.021E-07 | 5.600E-06 | 7.200E-06 |
| 27960 | 1398 | P1398V | E | 7.5e-06 | 8.7e-06 | 7.1e-06 | 7.767E-06 | 8.327E-07 | 7.100E-06 | 8.700E-06 |
| 27961 | 1399 | S1399A | E | 1e-05 | 4.3e-06 | 1e-05 | 8.100E-06 | 3.291E-06 | 4.300E-06 | 1.000E-05 |
| 27962 | 1399 | S1399R | E | 9.8e-06 | 9.8e-06 | 6.6e-06 | 8.733E-06 | 1.848E-06 | 6.600E-06 | 9.800E-06 |
| 27963 | 1399 | S1399N | E | 1e-05 | 6.4e-06 | 1.3e-05 | 9.800E-06 | 3.305E-06 | 6.400E-06 | 1.300E-05 |
| 27964 | 1399 | S1399D | E | 6.1e-06 | 6.4e-06 | 5.1e-06 | 5.867E-06 | 6.807E-07 | 5.100E-06 | 6.400E-06 |

| | | | | | | | | | | |
|---|---|---|---|---|---|---|---|---|---|---|
| 27965 | 1399 | S1399C | E | 7.3e-06 | 8.7e-06 | 1e-05 | 8.667E-06 | 1.350E-06 | 7.300E-06 | 1.000E-05 |
| 27966 | 1399 | S1399E | E | 5.9e-06 | 6.8e-06 | 5.6e-06 | 6.100E-06 | 6.245E-07 | 5.600E-06 | 6.800E-06 |
| 27967 | 1399 | S1399Q | E | 6.3e-06 | 8.8e-06 | 1e-05 | 8.367E-06 | 1.888E-06 | 6.300E-06 | 1.000E-05 |
| 27968 | 1399 | S1399G | E | 1.1e-05 | 6.3e-06 | 8.9e-06 | 8.733E-06 | 2.354E-06 | 6.300E-06 | 1.100E-05 |
| 27969 | 1399 | S1399H | E | 6.8e-06 | 5.1e-06 | 9.8e-06 | 7.233E-06 | 2.380E-06 | 5.100E-06 | 9.800E-06 |
| 27970 | 1399 | S1399I | E | 6.2e-06 | 1.1e-05 | 6.6e-06 | 7.933E-06 | 2.663E-06 | 6.200E-06 | 1.100E-05 |
| 27971 | 1399 | S1399L | E | 1.3e-05 | 4.3e-06 | 1.1e-05 | 9.433E-06 | 4.557E-06 | 4.300E-06 | 1.300E-05 |
| 27972 | 1399 | S1399K | E | 5.7e-06 | 6.3e-06 | 9.9e-06 | 7.300E-06 | 2.272E-06 | 5.700E-06 | 9.900E-06 |
| 27973 | 1399 | S1399M | E | 8.9e-06 | 1.3e-05 | 8.3e-06 | 1.007E-05 | 2.558E-06 | 8.300E-06 | 1.300E-05 |
| 27974 | 1399 | S1399F | E | 7.3e-06 | 7.3e-06 | 9.9e-06 | 8.167E-06 | 1.501E-06 | 7.300E-06 | 9.900E-06 |
| 27975 | 1399 | S1399P | E | 8.1e-06 | 5e-06 | 5.4e-06 | 6.167E-06 | 1.686E-06 | 5.000E-06 | 8.100E-06 |
| 27976 | 1399 | S1399S | E | 1.8e-05 | 6.7e-06 | 4.3e-06 | 9.667E-06 | 7.316E-06 | 4.300E-06 | 1.800E-05 |
| 27977 | 1399 | S1399T | E | 9.4e-06 | 8.7e-06 | 1.4e-05 | 1.070E-05 | 2.879E-06 | 8.700E-06 | 1.400E-05 |
| 27978 | 1399 | S1399W | E | 6.9e-06 | 7.5e-06 | 8.5e-06 | 7.633E-06 | 8.083E-07 | 6.900E-06 | 8.500E-06 |
| 27979 | 1399 | S1399Y | E | 1.6e-05 | 1e-05 | 9.3e-06 | 1.177E-05 | 3.683E-06 | 9.300E-06 | 1.600E-05 |
| 27980 | 1399 | S1399V | E | 6.9e-06 | 6.9e-06 | 1.3e-05 | 8.933E-06 | 3.522E-06 | 6.900E-06 | 1.300E-05 |
| 27981 | 1400 | V1400A | E | 1.2e-05 | 5.7e-06 | 5.5e-06 | 7.733E-06 | 3.696E-06 | 5.500E-06 | 1.200E-05 |
| 27982 | 1400 | V1400R | E | 8.2e-06 | 4e-06 | 1.3e-05 | 8.400E-06 | 4.503E-06 | 4.000E-06 | 1.300E-05 |
| 27983 | 1400 | V1400N | E | 6.1e-06 | 8.4e-06 | 7e-06 | 7.167E-06 | 1.159E-06 | 6.100E-06 | 8.400E-06 |
| 27984 | 1400 | V1400D | E | 7.2e-06 | 1e-05 | 4.6e-06 | 7.267E-06 | 2.701E-06 | 4.600E-06 | 1.000E-05 |
| 27985 | 1400 | V1400C | E | 7.9e-06 | 4.8e-06 | 1.3e-05 | 8.567E-06 | 4.140E-06 | 4.800E-06 | 1.300E-05 |
| 27986 | 1400 | V1400E | E | 1e-05 | 7.5e-06 | 9e-06 | 8.833E-06 | 1.258E-06 | 7.500E-06 | 1.000E-05 |
| 27987 | 1400 | V1400Q | E | 5.3e-06 | 5.8e-06 | 6.6e-06 | 5.900E-06 | 6.557E-07 | 5.300E-06 | 6.600E-06 |
| 27988 | 1400 | V1400G | E | 7.1e-06 | 8.8e-06 | 1e-05 | 8.633E-06 | 1.457E-06 | 7.100E-06 | 1.000E-05 |
| 27989 | 1400 | V1400H | E | 1.4e-05 | 8.6e-06 | 6.9e-06 | 9.833E-06 | 3.707E-06 | 6.900E-06 | 1.400E-05 |
| 27990 | 1400 | V1400I | E | 1.4e-05 | 6.2e-06 | 7.9e-06 | 9.367E-06 | 4.102E-06 | 6.200E-06 | 1.400E-05 |
| 27991 | 1400 | V1400L | E | 1.1e-05 | 8.2e-06 | 1.2e-05 | 1.040E-05 | 1.970E-06 | 8.200E-06 | 1.200E-05 |
| 27992 | 1400 | V1400K | E | 8.5e-06 | 7.2e-06 | 8e-06 | 7.900E-06 | 6.557E-07 | 7.200E-06 | 8.500E-06 |
| 27993 | 1400 | V1400M | E | 3.7e-06 | 1.3e-05 | 6.3e-06 | 7.667E-06 | 4.798E-06 | 3.700E-06 | 1.300E-05 |
| 27994 | 1400 | V1400F | E | 6.9e-06 | 6.7e-06 | 7.6e-06 | 7.067E-06 | 4.726E-07 | 6.700E-06 | 7.600E-06 |
| 27995 | 1400 | V1400P | E | 7.3e-06 | 6.2e-06 | 8.6e-06 | 7.367E-06 | 1.201E-06 | 6.200E-06 | 8.600E-06 |
| 27996 | 1400 | V1400S | E | 6.9e-06 | 8.2e-06 | 8.6e-06 | 7.900E-06 | 8.888E-07 | 6.900E-06 | 8.600E-06 |
| 27997 | 1400 | V1400T | E | 6.5e-06 | 6.4e-06 | 4.7e-06 | 5.867E-06 | 1.012E-06 | 4.700E-06 | 6.500E-06 |
| 27998 | 1400 | V1400W | E | 6.4e-06 | 4.2e-06 | 6.8e-06 | 5.800E-06 | 1.400E-06 | 4.200E-06 | 6.800E-06 |
| 27999 | 1400 | V1400Y | E | 5.7e-06 | 6.5e-06 | 9.3e-06 | 7.167E-06 | 1.890E-06 | 5.700E-06 | 9.300E-06 |
| 28000 | 1400 | V1400V | E | 6.1e-06 | 6.7e-06 | 5.2e-06 | 6.000E-06 | 7.550E-07 | 5.200E-06 | 6.700E-06 |
| 28001 | 1401 | F1401A | E | 7.4e-06 | 5.6e-06 | 8.4e-06 | 7.133E-06 | 1.419E-06 | 5.600E-06 | 8.400E-06 |
| 28002 | 1401 | F1401R | E | 6.5e-06 | 1.1e-05 | 9.5e-06 | 9.000E-06 | 2.291E-06 | 6.500E-06 | 1.100E-05 |
| 28003 | 1401 | F1401N | E | 1.3e-05 | 8.3e-06 | 1.4e-05 | 1.177E-05 | 3.044E-06 | 8.300E-06 | 1.400E-05 |
| 28004 | 1401 | F1401D | E | 1.3e-05 | 9.6e-06 | 1.2e-05 | 1.153E-05 | 1.747E-06 | 9.600E-06 | 1.300E-05 |
| 28005 | 1401 | F1401C | E | 4.5e-06 | 6.6e-06 | 7.3e-06 | 6.133E-06 | 1.457E-06 | 4.500E-06 | 7.300E-06 |
| 28006 | 1401 | F1401E | E | 7.4e-06 | 1e-05 | 4.8e-06 | 7.400E-06 | 2.600E-06 | 4.800E-06 | 1.000E-05 |
| 28007 | 1401 | F1401Q | E | 1.4e-05 | 9.3e-06 | 6.2e-06 | 9.833E-06 | 3.927E-06 | 6.200E-06 | 1.400E-05 |
| 28008 | 1401 | F1401G | E | 7.7e-06 | 6.2e-06 | 1e-05 | 7.967E-06 | 1.914E-06 | 6.200E-06 | 1.000E-05 |
| 28009 | 1401 | F1401H | E | 4.5e-06 | 8.9e-06 | 1.3e-05 | 8.800E-06 | 4.251E-06 | 4.500E-06 | 1.300E-05 |
| 28010 | 1401 | F1401I | E | 1.3e-05 | 6.8e-06 | 8.8e-06 | 9.533E-06 | 3.164E-06 | 6.800E-06 | 1.300E-05 |
| 28011 | 1401 | F1401L | E | 4.5e-06 | 4.9e-06 | 4.5e-06 | 4.633E-06 | 2.309E-07 | 4.500E-06 | 4.900E-06 |
| 28012 | 1401 | F1401K | E | 9.7e-06 | 8.5e-06 | 9.5e-06 | 9.233E-06 | 6.429E-07 | 8.500E-06 | 9.700E-06 |
| 28013 | 1401 | F1401M | E | 5e-06 | 7e-06 | 5.3e-06 | 5.767E-06 | 1.079E-06 | 5.000E-06 | 7.000E-06 |
| 28014 | 1401 | F1401F | E | 1.9e-05 | 6.7e-06 | 8.6e-06 | 1.143E-05 | 6.621E-06 | 6.700E-06 | 1.900E-05 |
| 28015 | 1401 | F1401P | E | 6.5e-06 | 9.6e-06 | 1.1e-05 | 9.033E-06 | 2.303E-06 | 6.500E-06 | 1.100E-05 |
| 28016 | 1401 | F1401S | E | 9.3e-06 | 5.4e-06 | 9.6e-06 | 8.100E-06 | 2.343E-06 | 5.400E-06 | 9.600E-06 |
| 28017 | 1401 | F1401T | E | 7e-06 | 5.4e-06 | 6e-06 | 6.133E-06 | 8.083E-07 | 5.400E-06 | 7.000E-06 |
| 28018 | 1401 | F1401W | E | 6e-06 | 5.6e-06 | 5.6e-06 | 5.733E-06 | 2.309E-07 | 5.600E-06 | 6.000E-06 |
| 28019 | 1401 | F1401Y | E | 5e-06 | 9.4e-06 | 8e-06 | 7.467E-06 | 2.248E-06 | 5.000E-06 | 9.400E-06 |
| 28020 | 1401 | F1401V | E | 1e-05 | 9.7e-06 | 9.7e-06 | 9.800E-06 | 1.732E-07 | 9.700E-06 | 1.000E-05 |
| 28021 | 1402 | P1402A | E | 5.8e-06 | 6e-06 | 1e-05 | 7.267E-06 | 2.369E-06 | 5.800E-06 | 1.000E-05 |
| 28022 | 1402 | P1402R | E | 6.7e-06 | 1.1e-05 | 9.4e-06 | 9.033E-06 | 2.173E-06 | 6.700E-06 | 1.100E-05 |
| 28023 | 1402 | P1402N | E | 8e-06 | 6.5e-06 | 1.1e-05 | 8.500E-06 | 2.291E-06 | 6.500E-06 | 1.100E-05 |
| 28024 | 1402 | P1402D | E | 6.3e-06 | 7.2e-06 | 8.8e-06 | 7.433E-06 | 1.266E-06 | 6.300E-06 | 8.800E-06 |
| 28025 | 1402 | P1402C | E | 7.1e-06 | 7.5e-06 | 7.9e-06 | 7.500E-06 | 4.000E-07 | 7.100E-06 | 7.900E-06 |
| 28026 | 1402 | P1402E | E | 6.3e-06 | 9.7e-06 | 8e-06 | 8.000E-06 | 1.700E-06 | 6.300E-06 | 9.700E-06 |
| 28027 | 1402 | P1402Q | E | 1.3e-05 | 7.6e-06 | 6e-06 | 8.867E-06 | 3.668E-06 | 6.000E-06 | 1.300E-05 |
| 28028 | 1402 | P1402G | E | 7.4e-06 | 7.3e-06 | 9.1e-06 | 7.933E-06 | 1.012E-06 | 7.300E-06 | 9.100E-06 |
| 28029 | 1402 | P1402H | E | 5.4e-06 | 1.5e-05 | 8e-06 | 9.467E-06 | 4.965E-06 | 5.400E-06 | 1.500E-05 |
| 28030 | 1402 | P1402I | E | 1.1e-05 | 1.1e-05 | 9e-06 | 1.033E-05 | 1.155E-06 | 9.000E-06 | 1.100E-05 |
| 28031 | 1402 | P1402L | E | 6.9e-06 | 5.7e-06 | 5.5e-06 | 6.033E-06 | 7.572E-07 | 5.500E-06 | 6.900E-06 |
| 28032 | 1402 | P1402K | E | 1.2e-05 | 4.5e-06 | 6.7e-06 | 7.733E-06 | 3.855E-06 | 4.500E-06 | 1.200E-05 |
| 28033 | 1402 | P1402M | E | 1.2e-05 | 8.8e-06 | 5.6e-06 | 8.800E-06 | 3.200E-06 | 5.600E-06 | 1.200E-05 |
| 28034 | 1402 | P1402F | E | 5.9e-06 | 7e-06 | 4.8e-06 | 5.900E-06 | 1.100E-06 | 4.800E-06 | 7.000E-06 |
| 28035 | 1402 | P1402P | E | 9e-06 | 8.9e-06 | 1.2e-05 | 9.967E-06 | 1.762E-06 | 8.900E-06 | 1.200E-05 |
| 28036 | 1402 | P1402S | E | 9.8e-06 | 5.3e-06 | 6.4e-06 | 7.167E-06 | 2.346E-06 | 5.300E-06 | 9.800E-06 |
| 28037 | 1402 | P1402T | E | 6.6e-06 | 1.8e-05 | 9.3e-06 | 1.130E-05 | 5.957E-06 | 6.600E-06 | 1.800E-05 |
| 28038 | 1402 | P1402W | E | 1.2e-05 | 6.1e-06 | 1.1e-05 | 9.700E-06 | 3.158E-06 | 6.100E-06 | 1.200E-05 |
| 28039 | 1402 | P1402Y | E | 1.1e-05 | 7.8e-06 | 6.2e-06 | 8.333E-06 | 2.444E-06 | 6.200E-06 | 1.100E-05 |
| 28040 | 1402 | P1402V | E | 5e-06 | 8.9e-06 | 8e-06 | 7.300E-06 | 2.042E-06 | 5.000E-06 | 8.900E-06 |
| 28041 | 1403 | L1403A | E | 1.3e-05 | 1.1e-05 | 9.5e-06 | 1.117E-05 | 1.756E-06 | 9.500E-06 | 1.300E-05 |
| 28042 | 1403 | L1403R | E | 6.5e-06 | 6.4e-06 | 9.6e-06 | 7.500E-06 | 1.819E-06 | 6.400E-06 | 9.600E-06 |
| 28043 | 1403 | L1403N | E | 1.2e-05 | 6e-06 | 9.8e-06 | 9.267E-06 | 3.035E-06 | 6.000E-06 | 1.200E-05 |
| 28044 | 1403 | L1403D | E | 4.3e-06 | 6.8e-06 | 8.7e-06 | 6.600E-06 | 2.207E-06 | 4.300E-06 | 8.700E-06 |
| 28045 | 1403 | L1403C | E | 1.5e-05 | 4.7e-06 | 1.3e-05 | 1.090E-05 | 5.462E-06 | 4.700E-06 | 1.500E-05 |
| 28046 | 1403 | L1403E | E | 8.2e-06 | 6e-06 | 5.4e-06 | 6.533E-06 | 1.474E-06 | 5.400E-06 | 8.200E-06 |
| 28047 | 1403 | L1403Q | E | 6.4e-06 | 6.7e-06 | 4.7e-06 | 5.933E-06 | 1.079E-06 | 4.700E-06 | 6.700E-06 |
| 28048 | 1403 | L1403G | E | 8.4e-06 | 9.3e-06 | 6.8e-06 | 8.167E-06 | 1.266E-06 | 6.800E-06 | 9.300E-06 |
| 28049 | 1403 | L1403H | E | 1e-05 | 1.2e-05 | 8.3e-06 | 1.010E-05 | 1.852E-06 | 8.300E-06 | 1.200E-05 |

| | | | | | | | | | | |
|---|---|---|---|---|---|---|---|---|---|---|
| 28050 | 1403 | L1403I | E | 5.3e-06 | 5.1e-06 | 1.3e-05 | 7.800E-06 | 4.504E-06 | 5.100E-06 | 1.300E-05 |
| 28051 | 1403 | L1403L | E | 1.8e-05 | 8.2e-06 | 6.7e-06 | 1.097E-05 | 6.137E-06 | 6.700E-06 | 1.800E-05 |
| 28052 | 1403 | L1403K | E | 9.3e-06 | 1e-05 | 1.6e-05 | 1.177E-05 | 3.683E-06 | 9.300E-06 | 1.600E-05 |
| 28053 | 1403 | L1403M | E | 9.2e-06 | 7.5e-06 | 8.8e-06 | 8.500E-06 | 8.888E-07 | 7.500E-06 | 9.200E-06 |
| 28054 | 1403 | L1403F | E | 9.2e-06 | 6e-06 | 6.2e-06 | 7.133E-06 | 1.793E-06 | 6.000E-06 | 9.200E-06 |
| 28055 | 1403 | L1403P | E | 7.4e-06 | 1.4e-05 | 6.7e-06 | 9.367E-06 | 4.028E-06 | 6.700E-06 | 1.400E-05 |
| 28056 | 1403 | L1403S | E | 6.4e-06 | 8.2e-06 | 6.5e-06 | 7.033E-06 | 1.012E-06 | 6.400E-06 | 8.200E-06 |
| 28057 | 1403 | L1403T | E | 1e-05 | 7.6e-06 | 8e-06 | 8.533E-06 | 1.286E-06 | 7.600E-06 | 1.000E-05 |
| 28058 | 1403 | L1403W | E | 6.7e-06 | 1.4e-05 | 9.1e-06 | 9.933E-06 | 3.721E-06 | 6.700E-06 | 1.400E-05 |
| 28059 | 1403 | L1403Y | E | 6.4e-06 | 6.7e-06 | 2e-05 | 1.103E-05 | 7.767E-06 | 6.400E-06 | 2.000E-05 |
| 28060 | 1403 | L1403V | E | 9.3e-06 | 7.4e-06 | 6.3e-06 | 7.667E-06 | 1.518E-06 | 6.300E-06 | 9.300E-06 |
| 28061 | 1404 | A1404A | E | 5.4e-06 | 5.2e-06 | 5.6e-06 | 5.400E-06 | 2.000E-07 | 5.200E-06 | 5.600E-06 |
| 28062 | 1404 | A1404R | E | 6.5e-06 | 6.9e-06 | 1.3e-05 | 8.800E-06 | 3.643E-06 | 6.500E-06 | 1.300E-05 |
| 28063 | 1404 | A1404N | E | 4.8e-06 | 9.1e-06 | 7.8e-06 | 7.233E-06 | 2.205E-06 | 4.800E-06 | 9.100E-06 |
| 28064 | 1404 | A1404D | E | 1.1e-05 | 9.7e-06 | 1e-05 | 1.023E-05 | 6.807E-07 | 9.700E-06 | 1.100E-05 |
| 28065 | 1404 | A1404C | E | 6.6e-06 | 1.3e-05 | 7e-06 | 8.867E-06 | 3.585E-06 | 6.600E-06 | 1.300E-05 |
| 28066 | 1404 | A1404E | E | 8.9e-06 | 4.6e-06 | 6.3e-06 | 6.600E-06 | 2.166E-06 | 4.600E-06 | 8.900E-06 |
| 28067 | 1404 | A1404Q | E | 6.1e-06 | 6.2e-06 | 8.9e-06 | 7.067E-06 | 1.589E-06 | 6.100E-06 | 8.900E-06 |
| 28068 | 1404 | A1404G | E | 5.5e-06 | 8.6e-06 | 4.6e-06 | 6.233E-06 | 2.098E-06 | 4.600E-06 | 8.600E-06 |
| 28069 | 1404 | A1404H | E | 9.5e-06 | 6.1e-06 | 1.8e-05 | 1.120E-05 | 6.129E-06 | 6.100E-06 | 1.800E-05 |
| 28070 | 1404 | A1404I | E | 5.2e-06 | 5e-06 | 9.2e-06 | 6.467E-06 | 2.369E-06 | 5.000E-06 | 9.200E-06 |
| 28071 | 1404 | A1404L | E | 9.2e-06 | 4.9e-06 | 1e-05 | 8.033E-06 | 2.743E-06 | 4.900E-06 | 1.000E-05 |
| 28072 | 1404 | A1404K | E | 1.1e-05 | 7.4e-06 | 7.7e-06 | 8.700E-06 | 1.997E-06 | 7.400E-06 | 1.100E-05 |
| 28073 | 1404 | A1404M | E | 1.2e-05 | 9e-06 | 7.8e-06 | 9.600E-06 | 2.163E-06 | 7.800E-06 | 1.200E-05 |
| 28074 | 1404 | A1404F | E | 5.4e-06 | 7.8e-06 | 7.2e-06 | 6.800E-06 | 1.249E-06 | 5.400E-06 | 7.800E-06 |
| 28075 | 1404 | A1404P | E | 9.1e-06 | 1.2e-05 | 8.7e-06 | 9.933E-06 | 1.801E-06 | 8.700E-06 | 1.200E-05 |
| 28076 | 1404 | A1404S | E | 9.1e-06 | 1e-05 | 6.5e-06 | 8.533E-06 | 1.818E-06 | 6.500E-06 | 1.000E-05 |
| 28077 | 1404 | A1404T | E | 8.6e-06 | 6.5e-06 | 9.1e-06 | 8.067E-06 | 1.380E-06 | 6.500E-06 | 9.100E-06 |
| 28078 | 1404 | A1404W | E | 8e-06 | 8.7e-06 | 6.1e-06 | 7.600E-06 | 1.345E-06 | 6.100E-06 | 8.700E-06 |
| 28079 | 1404 | A1404Y | E | 4.2e-06 | 6.9e-06 | 9.7e-06 | 6.933E-06 | 2.750E-06 | 4.200E-06 | 9.700E-06 |
| 28080 | 1404 | A1404V | E | 9.2e-06 | 1e-05 | 4.2e-06 | 7.800E-06 | 3.143E-06 | 4.200E-06 | 1.000E-05 |
| 28081 | 1405 | P1405A | E | 8.3e-06 | 9.3e-06 | 1.1e-05 | 9.533E-06 | 1.365E-06 | 8.300E-06 | 1.100E-05 |
| 28082 | 1405 | P1405R | E | 6.2e-06 | 1.4e-05 | 4.8e-06 | 8.333E-06 | 4.957E-06 | 4.800E-06 | 1.400E-05 |
| 28083 | 1405 | P1405N | E | 8.2e-06 | 8.5e-06 | 9.8e-06 | 8.833E-06 | 8.505E-07 | 8.200E-06 | 9.800E-06 |
| 28084 | 1405 | P1405D | E | 1e-05 | 5.6e-06 | 9.6e-06 | 8.400E-06 | 2.433E-06 | 5.600E-06 | 1.000E-05 |
| 28085 | 1405 | P1405C | E | 6.3e-06 | 1.4e-05 | 8.4e-06 | 9.567E-06 | 3.980E-06 | 6.300E-06 | 1.400E-05 |
| 28086 | 1405 | P1405E | E | 1.4e-05 | 7.9e-06 | 4.6e-06 | 8.833E-06 | 4.769E-06 | 4.600E-06 | 1.400E-05 |
| 28087 | 1405 | P1405Q | E | 4.8e-06 | 8.4e-06 | 7.7e-06 | 6.967E-06 | 1.909E-06 | 4.800E-06 | 8.400E-06 |
| 28088 | 1405 | P1405G | E | 1.4e-05 | 8.7e-06 | 6.8e-06 | 9.833E-06 | 3.731E-06 | 6.800E-06 | 1.400E-05 |
| 28089 | 1405 | P1405H | E | 9.8e-06 | 4.6e-06 | 5.1e-06 | 6.500E-06 | 2.869E-06 | 4.600E-06 | 9.800E-06 |
| 28090 | 1405 | P1405I | E | 6.9e-06 | 8.3e-06 | 5.8e-06 | 7.000E-06 | 1.253E-06 | 5.800E-06 | 8.300E-06 |
| 28091 | 1405 | P1405L | E | 8.1e-06 | 9.6e-06 | 9.7e-06 | 9.133E-06 | 8.963E-07 | 8.100E-06 | 9.700E-06 |
| 28092 | 1405 | P1405K | E | 7e-06 | 5.9e-06 | 4.4e-06 | 5.767E-06 | 1.305E-06 | 4.400E-06 | 7.000E-06 |
| 28093 | 1405 | P1405M | E | 1e-05 | 8.3e-06 | 8.8e-06 | 9.033E-06 | 8.737E-07 | 8.300E-06 | 1.000E-05 |
| 28094 | 1405 | P1405F | E | 6.2e-06 | 1.1e-05 | 7.2e-06 | 8.133E-06 | 2.532E-06 | 6.200E-06 | 1.100E-05 |
| 28095 | 1405 | P1405P | E | 7.8e-06 | 3.8e-06 | 5.4e-06 | 5.667E-06 | 2.013E-06 | 3.800E-06 | 7.800E-06 |
| 28096 | 1405 | P1405S | E | 6.6e-06 | 5.2e-06 | 6.7e-06 | 6.167E-06 | 8.386E-07 | 5.200E-06 | 6.700E-06 |
| 28097 | 1405 | P1405T | E | 6.6e-06 | 1.3e-05 | 7.7e-06 | 9.100E-06 | 3.422E-06 | 6.600E-06 | 1.300E-05 |
| 28098 | 1405 | P1405W | E | 7e-06 | 9.6e-06 | 7.2e-06 | 7.933E-06 | 1.447E-06 | 7.000E-06 | 9.600E-06 |
| 28099 | 1405 | P1405Y | E | 1.1e-05 | 5.4e-06 | 4.9e-06 | 7.100E-06 | 3.387E-06 | 4.900E-06 | 1.100E-05 |
| 28100 | 1405 | P1405V | E | 6.3e-06 | 1.5e-05 | 4.7e-06 | 8.667E-06 | 5.543E-06 | 4.700E-06 | 1.500E-05 |
| 28101 | 1406 | S1406A | E | 6.9e-06 | 1e-05 | 7.5e-06 | 8.133E-06 | 1.644E-06 | 6.900E-06 | 1.000E-05 |
| 28102 | 1406 | S1406R | E | 6.6e-06 | 8e-06 | 4.2e-06 | 6.267E-06 | 1.922E-06 | 4.200E-06 | 8.000E-06 |
| 28103 | 1406 | S1406N | E | 4.2e-06 | 9.4e-06 | 7.7e-06 | 7.100E-06 | 2.651E-06 | 4.200E-06 | 9.400E-06 |
| 28104 | 1406 | S1406D | E | 8.6e-06 | 1e-05 | 7.9e-06 | 8.833E-06 | 1.069E-06 | 7.900E-06 | 1.000E-05 |
| 28105 | 1406 | S1406C | E | 1e-05 | 7.9e-06 | 7.7e-06 | 8.533E-06 | 1.274E-06 | 7.700E-06 | 1.000E-05 |
| 28106 | 1406 | S1406E | E | 8.1e-06 | 1e-05 | 8.1e-06 | 8.733E-06 | 1.097E-06 | 8.100E-06 | 1.000E-05 |
| 28107 | 1406 | S1406Q | E | 1.3e-05 | 9.5e-06 | 7.5e-06 | 1.000E-05 | 2.784E-06 | 7.500E-06 | 1.300E-05 |
| 28108 | 1406 | S1406G | E | 1.4e-05 | 9.6e-06 | 8.3e-06 | 1.063E-05 | 2.987E-06 | 8.300E-06 | 1.400E-05 |
| 28109 | 1406 | S1406H | E | 8.7e-06 | 6.9e-06 | 8.2e-06 | 7.933E-06 | 9.292E-07 | 6.900E-06 | 8.700E-06 |
| 28110 | 1406 | S1406I | E | 9e-06 | 1.1e-05 | 9.4e-06 | 9.800E-06 | 1.058E-06 | 9.000E-06 | 1.100E-05 |
| 28111 | 1406 | S1406L | E | 8.1e-06 | 7.7e-06 | 8.9e-06 | 8.233E-06 | 6.110E-07 | 7.700E-06 | 8.900E-06 |
| 28112 | 1406 | S1406K | E | 2e-05 | 8.1e-06 | 9.4e-06 | 1.250E-05 | 6.528E-06 | 8.100E-06 | 2.000E-05 |
| 28113 | 1406 | S1406M | E | 8.6e-06 | 4e-06 | 5.2e-06 | 5.933E-06 | 2.386E-06 | 4.000E-06 | 8.600E-06 |
| 28114 | 1406 | S1406F | E | 5.8e-06 | 7e-06 | 8.1e-06 | 6.967E-06 | 1.150E-06 | 5.800E-06 | 8.100E-06 |
| 28115 | 1406 | S1406P | E | 5.3e-06 | 2e-05 | 4.7e-06 | 1.000E-05 | 8.665E-06 | 4.700E-06 | 2.000E-05 |
| 28116 | 1406 | S1406S | E | 4.8e-06 | 8.5e-06 | 4.1e-06 | 5.800E-06 | 2.364E-06 | 4.100E-06 | 8.500E-06 |
| 28117 | 1406 | S1406T | E | 6.9e-06 | 3.7e-06 | 1e-05 | 6.867E-06 | 3.150E-06 | 3.700E-06 | 1.000E-05 |
| 28118 | 1406 | S1406W | E | 6.8e-06 | 6.1e-06 | 8.4e-06 | 7.100E-06 | 1.179E-06 | 6.100E-06 | 8.400E-06 |
| 28119 | 1406 | S1406Y | E | 8.4e-06 | 9.3e-06 | 5.8e-06 | 7.833E-06 | 1.818E-06 | 5.800E-06 | 9.300E-06 |
| 28120 | 1406 | S1406V | E | 8.1e-06 | 9.1e-06 | 7e-06 | 8.067E-06 | 1.050E-06 | 7.000E-06 | 9.100E-06 |
| 28121 | 1407 | S1407A | E | 5.2e-06 | 5e-06 | 7.2e-06 | 5.800E-06 | 1.217E-06 | 5.000E-06 | 7.200E-06 |
| 28122 | 1407 | S1407R | E | 9.1e-06 | 5.8e-06 | 8.8e-06 | 7.900E-06 | 1.825E-06 | 5.800E-06 | 9.100E-06 |
| 28123 | 1407 | S1407N | E | 9.2e-06 | 6.1e-06 | 6.9e-06 | 7.400E-06 | 1.609E-06 | 6.100E-06 | 9.200E-06 |
| 28124 | 1407 | S1407D | E | 4.7e-06 | 1.5e-05 | 7.2e-06 | 8.967E-06 | 5.372E-06 | 4.700E-06 | 1.500E-05 |
| 28125 | 1407 | S1407C | E | 5e-06 | 5.3e-06 | 8.2e-06 | 6.167E-06 | 1.767E-06 | 5.000E-06 | 8.200E-06 |
| 28126 | 1407 | S1407E | E | 1.1e-05 | 6.7e-06 | 7.1e-06 | 8.267E-06 | 2.376E-06 | 6.700E-06 | 1.100E-05 |
| 28127 | 1407 | S1407Q | E | 6.4e-06 | 7.3e-06 | 7.2e-06 | 6.967E-06 | 4.933E-07 | 6.400E-06 | 7.300E-06 |
| 28128 | 1407 | S1407G | E | 6.7e-06 | 8.5e-06 | 9.6e-06 | 8.267E-06 | 1.464E-06 | 6.700E-06 | 9.600E-06 |
| 28129 | 1407 | S1407H | E | 7.3e-06 | 9e-06 | 4.5e-06 | 6.933E-06 | 2.272E-06 | 4.500E-06 | 9.000E-06 |
| 28130 | 1407 | S1407I | E | 7.9e-06 | 1e-05 | 6.3e-06 | 8.067E-06 | 1.856E-06 | 6.300E-06 | 1.000E-05 |
| 28131 | 1407 | S1407L | E | 8.9e-06 | 6.2e-06 | 1.2e-05 | 9.033E-06 | 2.902E-06 | 6.200E-06 | 1.200E-05 |
| 28132 | 1407 | S1407K | E | 1e-05 | 9.8e-06 | 6.4e-06 | 8.733E-06 | 2.023E-06 | 6.400E-06 | 1.000E-05 |
| 28133 | 1407 | S1407M | E | 8.5e-06 | 1.1e-05 | 8.1e-06 | 9.200E-06 | 1.572E-06 | 8.100E-06 | 1.100E-05 |
| 28134 | 1407 | S1407F | E | 1.1e-05 | 6.1e-06 | 9.2e-06 | 8.767E-06 | 2.479E-06 | 6.100E-06 | 1.100E-05 |

| | | | | | | | | | | |
|---|---|---|---|---|---|---|---|---|---|---|
| 28135 | 1407 | S1407P | E | 9.2e-06 | 5.9e-06 | 6.3e-06 | 7.133E-06 | 1.801E-06 | 5.900E-06 | 9.200E-06 |
| 28136 | 1407 | S1407S | E | 1.9e-05 | 9.8e-06 | 9.6e-06 | 1.280E-05 | 5.370E-06 | 9.600E-06 | 1.900E-05 |
| 28137 | 1407 | S1407T | E | 1e-05 | 9.5e-06 | 1.4e-05 | 1.117E-05 | 2.466E-06 | 9.500E-06 | 1.400E-05 |
| 28138 | 1407 | S1407W | E | 5.7e-06 | 5.9e-06 | 6.4e-06 | 6.000E-06 | 3.606E-07 | 5.700E-06 | 6.400E-06 |
| 28139 | 1407 | S1407Y | E | 7.4e-06 | 6.3e-06 | 8.6e-06 | 7.433E-06 | 1.150E-06 | 6.300E-06 | 8.600E-06 |
| 28140 | 1407 | S1407V | E | 6e-06 | 9.3e-06 | 6.4e-06 | 7.233E-06 | 1.801E-06 | 6.000E-06 | 9.300E-06 |
| 28141 | 1408 | K1408A | E | 6.4e-06 | 9e-06 | 9.5e-06 | 8.300E-06 | 1.664E-06 | 6.400E-06 | 9.500E-06 |
| 28142 | 1408 | K1408R | E | 7.8e-06 | 8.8e-06 | 7.6e-06 | 8.067E-06 | 6.429E-07 | 7.600E-06 | 8.800E-06 |
| 28143 | 1408 | K1408N | E | 5.1e-06 | 7.9e-06 | 9.9e-06 | 7.633E-06 | 2.411E-06 | 5.100E-06 | 9.900E-06 |
| 28144 | 1408 | K1408D | E | 7.6e-06 | 5.6e-06 | 6.6e-06 | 6.600E-06 | 1.000E-06 | 5.600E-06 | 7.600E-06 |
| 28145 | 1408 | K1408C | E | 1.2e-05 | 6.4e-06 | 5.6e-06 | 8.000E-06 | 3.487E-06 | 5.600E-06 | 1.200E-05 |
| 28146 | 1408 | K1408E | E | 1.2e-05 | 1e-05 | 5e-06 | 9.000E-06 | 3.606E-06 | 5.000E-06 | 1.200E-05 |
| 28147 | 1408 | K1408Q | E | 8.2e-06 | 9.6e-06 | 1.8e-05 | 1.193E-05 | 5.300E-06 | 8.200E-06 | 1.800E-05 |
| 28148 | 1408 | K1408G | E | 3.5e-06 | 4.9e-06 | 1e-05 | 6.133E-06 | 3.421E-06 | 3.500E-06 | 1.000E-05 |
| 28149 | 1408 | K1408H | E | 9.8e-06 | 8.1e-06 | 9.8e-06 | 9.233E-06 | 9.815E-07 | 8.100E-06 | 9.800E-06 |
| 28150 | 1408 | K1408I | E | 1.1e-05 | 5e-06 | 1e-05 | 8.667E-06 | 3.215E-06 | 5.000E-06 | 1.100E-05 |
| 28151 | 1408 | K1408L | E | 8.7e-06 | 9.8e-06 | 8e-06 | 8.833E-06 | 9.074E-07 | 8.000E-06 | 9.800E-06 |
| 28152 | 1408 | K1408K | E | 5.6e-06 | 5.2e-06 | 9.7e-06 | 6.833E-06 | 2.491E-06 | 5.200E-06 | 9.700E-06 |
| 28153 | 1408 | K1408M | E | 6.1e-06 | 6.9e-06 | 6.4e-06 | 6.467E-06 | 4.041E-07 | 6.100E-06 | 6.900E-06 |
| 28154 | 1408 | K1408F | E | 7.2e-06 | 9.9e-06 | 4.8e-06 | 7.300E-06 | 2.551E-06 | 4.800E-06 | 9.900E-06 |
| 28155 | 1408 | K1408P | E | 9.2e-06 | 6.4e-06 | 1.3e-05 | 9.533E-06 | 3.313E-06 | 6.400E-06 | 1.300E-05 |
| 28156 | 1408 | K1408S | E | 6.7e-06 | 4.3e-06 | 1.1e-05 | 7.333E-06 | 3.395E-06 | 4.300E-06 | 1.100E-05 |
| 28157 | 1408 | K1408T | E | 8e-06 | 9.7e-06 | 8.9e-06 | 8.867E-06 | 8.505E-07 | 8.000E-06 | 9.700E-06 |
| 28158 | 1408 | K1408W | E | 6e-06 | 4.5e-06 | 8.4e-06 | 6.300E-06 | 1.967E-06 | 4.500E-06 | 8.400E-06 |
| 28159 | 1408 | K1408Y | E | 6.1e-06 | 5e-06 | 1.3e-05 | 8.033E-06 | 4.336E-06 | 5.000E-06 | 1.300E-05 |
| 28160 | 1408 | K1408V | E | 4.5e-06 | 9.3e-06 | 1.4e-05 | 9.267E-06 | 4.750E-06 | 4.500E-06 | 1.400E-05 |
| 28161 | 1409 | S1409A | E | 1.7e-05 | 6.4e-06 | 5.8e-06 | 9.733E-06 | 6.300E-06 | 5.800E-06 | 1.700E-05 |
| 28162 | 1409 | S1409R | E | 1.3e-05 | 8.1e-06 | 7.7e-06 | 9.600E-06 | 2.951E-06 | 7.700E-06 | 1.300E-05 |
| 28163 | 1409 | S1409N | E | 6.7e-06 | 7e-06 | 7e-06 | 6.900E-06 | 1.732E-07 | 6.700E-06 | 7.000E-06 |
| 28164 | 1409 | S1409D | E | 6.3e-06 | 6.3e-06 | 7.8e-06 | 6.800E-06 | 8.660E-07 | 6.300E-06 | 7.800E-06 |
| 28165 | 1409 | S1409C | E | 1.2e-05 | 6.7e-06 | 6e-06 | 8.233E-06 | 3.281E-06 | 6.000E-06 | 1.200E-05 |
| 28166 | 1409 | S1409E | E | 1.4e-05 | 1.1e-05 | 9.5e-06 | 1.150E-05 | 2.291E-06 | 9.500E-06 | 1.400E-05 |
| 28167 | 1409 | S1409Q | E | 3.9e-06 | 5.5e-06 | 5e-06 | 4.800E-06 | 8.185E-07 | 3.900E-06 | 5.500E-06 |
| 28168 | 1409 | S1409G | E | 6.6e-06 | 6.7e-06 | 6.3e-06 | 6.533E-06 | 2.082E-07 | 6.300E-06 | 6.700E-06 |
| 28169 | 1409 | S1409H | E | 1.2e-05 | 2e-05 | 7.2e-06 | 1.307E-05 | 6.466E-06 | 7.200E-06 | 2.000E-05 |
| 28170 | 1409 | S1409I | E | 7.7e-06 | 1e-05 | 6.6e-06 | 8.100E-06 | 1.735E-06 | 6.600E-06 | 1.000E-05 |
| 28171 | 1409 | S1409L | E | 6.7e-06 | 7.7e-06 | 6.3e-06 | 6.900E-06 | 7.211E-07 | 6.300E-06 | 7.700E-06 |
| 28172 | 1409 | S1409K | E | 8.1e-06 | 8e-06 | 8.4e-06 | 8.167E-06 | 2.082E-07 | 8.000E-06 | 8.400E-06 |
| 28173 | 1409 | S1409M | E | 9.6e-06 | 7.1e-06 | 7.1e-06 | 7.933E-06 | 1.443E-06 | 7.100E-06 | 9.600E-06 |
| 28174 | 1409 | S1409F | E | 4.4e-06 | 6.5e-06 | 5.3e-06 | 5.400E-06 | 1.054E-06 | 4.400E-06 | 6.500E-06 |
| 28175 | 1409 | S1409P | E | 1.1e-05 | 8.5e-06 | 9.8e-06 | 9.767E-06 | 1.250E-06 | 8.500E-06 | 1.100E-05 |
| 28176 | 1409 | S1409S | E | 6.6e-06 | 6.4e-06 | 9.7e-06 | 7.567E-06 | 1.850E-06 | 6.400E-06 | 9.700E-06 |
| 28177 | 1409 | S1409T | E | 7.8e-06 | 6e-06 | 8e-06 | 7.267E-06 | 1.102E-06 | 6.000E-06 | 8.000E-06 |
| 28178 | 1409 | S1409W | E | 1.1e-05 | 7.6e-06 | 8.1e-06 | 8.900E-06 | 1.836E-06 | 7.600E-06 | 1.100E-05 |
| 28179 | 1409 | S1409Y | E | 7.7e-06 | 7.1e-06 | 7e-06 | 7.267E-06 | 3.786E-07 | 7.000E-06 | 7.700E-06 |
| 28180 | 1409 | S1409V | E | 4.5e-06 | 7.4e-06 | 4.8e-06 | 5.567E-06 | 1.595E-06 | 4.500E-06 | 7.400E-06 |
| 28181 | 1410 | T1410A | E | 6.5e-06 | 5.4e-06 | 1.6e-05 | 9.300E-06 | 5.828E-06 | 5.400E-06 | 1.600E-05 |
| 28182 | 1410 | T1410R | E | 6.7e-06 | 1.4e-05 | 1.3e-05 | 1.123E-05 | 3.958E-06 | 6.700E-06 | 1.400E-05 |
| 28183 | 1410 | T1410N | E | 9.3e-06 | 7.1e-06 | 1.2e-05 | 9.467E-06 | 2.454E-06 | 7.100E-06 | 1.200E-05 |
| 28184 | 1410 | T1410D | E | 1.1e-05 | 8e-06 | 1.4e-05 | 1.100E-05 | 3.000E-06 | 8.000E-06 | 1.400E-05 |
| 28185 | 1410 | T1410C | E | 1.7e-05 | 1.3e-05 | 7.4e-06 | 1.247E-05 | 4.822E-06 | 7.400E-06 | 1.700E-05 |
| 28186 | 1410 | T1410E | E | 4.8e-06 | 1.2e-05 | 1.7e-05 | 1.127E-05 | 6.133E-06 | 4.800E-06 | 1.700E-05 |
| 28187 | 1410 | T1410Q | E | 9.8e-06 | 8.9e-06 | 1e-05 | 9.567E-06 | 5.859E-07 | 8.900E-06 | 1.000E-05 |
| 28188 | 1410 | T1410G | E | 8e-06 | 4.5e-06 | 6.3e-06 | 6.267E-06 | 1.750E-06 | 4.500E-06 | 8.000E-06 |
| 28189 | 1410 | T1410H | E | 9.9e-06 | 7.2e-06 | 6.8e-06 | 7.967E-06 | 1.686E-06 | 6.800E-06 | 9.900E-06 |
| 28190 | 1410 | T1410I | E | 2.1e-05 | 7.4e-06 | 1e-05 | 1.280E-05 | 7.219E-06 | 7.400E-06 | 2.100E-05 |
| 28191 | 1410 | T1410L | E | 6.7e-06 | 9.7e-06 | 4.8e-06 | 7.067E-06 | 2.470E-06 | 4.800E-06 | 9.700E-06 |
| 28192 | 1410 | T1410K | E | 5e-06 | 1.4e-05 | 1.3e-05 | 1.067E-05 | 4.933E-06 | 5.000E-06 | 1.400E-05 |
| 28193 | 1410 | T1410M | E | 1.3e-05 | 4.9e-06 | 6.5e-06 | 8.133E-06 | 4.290E-06 | 4.900E-06 | 1.300E-05 |
| 28194 | 1410 | T1410F | E | 1.3e-05 | 5.8e-06 | 8.2e-06 | 9.000E-06 | 3.666E-06 | 5.800E-06 | 1.300E-05 |
| 28195 | 1410 | T1410P | E | 1e-05 | 6.7e-06 | 5.8e-06 | 7.500E-06 | 2.211E-06 | 5.800E-06 | 1.000E-05 |
| 28196 | 1410 | T1410S | E | 1.3e-05 | 7.4e-06 | 1.1e-05 | 1.047E-05 | 2.838E-06 | 7.400E-06 | 1.300E-05 |
| 28197 | 1410 | T1410T | E | 1.2e-05 | 9.3e-06 | 6e-06 | 9.100E-06 | 3.005E-06 | 6.000E-06 | 1.200E-05 |
| 28198 | 1410 | T1410W | E | 6.7e-06 | 1.1e-05 | 6.7e-06 | 8.133E-06 | 2.483E-06 | 6.700E-06 | 1.100E-05 |
| 28199 | 1410 | T1410Y | E | 4.4e-06 | 8.3e-06 | 1.5e-05 | 9.233E-06 | 5.361E-06 | 4.400E-06 | 1.500E-05 |
| 28200 | 1410 | T1410V | E | 6.7e-06 | 8.1e-06 | 8.8e-06 | 7.867E-06 | 1.069E-06 | 6.700E-06 | 8.800E-06 |
| 28201 | 1411 | S1411A | E | 8.4e-06 | 1e-05 | 6.8e-06 | 8.400E-06 | 1.600E-06 | 6.800E-06 | 1.000E-05 |
| 28202 | 1411 | S1411R | E | 7e-06 | 5.7e-06 | 8.9e-06 | 7.200E-06 | 1.609E-06 | 5.700E-06 | 8.900E-06 |
| 28203 | 1411 | S1411N | E | 6.2e-06 | 1.4e-05 | 7.3e-06 | 9.167E-06 | 4.222E-06 | 6.200E-06 | 1.400E-05 |
| 28204 | 1411 | S1411D | E | 8.7e-06 | 4.2e-06 | 7.2e-06 | 6.700E-06 | 2.291E-06 | 4.200E-06 | 8.700E-06 |
| 28205 | 1411 | S1411C | E | 5.2e-06 | 5.9e-06 | 7.3e-06 | 6.133E-06 | 1.069E-06 | 5.200E-06 | 7.300E-06 |
| 28206 | 1411 | S1411E | E | 1.1e-05 | 6.8e-06 | 9.2e-06 | 9.000E-06 | 2.107E-06 | 6.800E-06 | 1.100E-05 |
| 28207 | 1411 | S1411Q | E | 1.1e-05 | 6.2e-06 | 5.5e-06 | 7.567E-06 | 2.994E-06 | 5.500E-06 | 1.100E-05 |
| 28208 | 1411 | S1411G | E | 5.6e-06 | 6.2e-06 | 1e-05 | 7.267E-06 | 2.386E-06 | 5.600E-06 | 1.000E-05 |
| 28209 | 1411 | S1411H | E | 9.3e-06 | 8.8e-06 | 8.2e-06 | 8.767E-06 | 5.508E-07 | 8.200E-06 | 9.300E-06 |
| 28210 | 1411 | S1411I | E | 7.3e-06 | 1.1e-05 | 4.6e-06 | 7.633E-06 | 3.213E-06 | 4.600E-06 | 1.100E-05 |
| 28211 | 1411 | S1411L | E | 8.1e-06 | 1e-05 | 8.5e-06 | 8.867E-06 | 1.002E-06 | 8.100E-06 | 1.000E-05 |
| 28212 | 1411 | S1411K | E | 8.8e-06 | 1.5e-05 | 5.7e-06 | 9.833E-06 | 4.735E-06 | 5.700E-06 | 1.500E-05 |
| 28213 | 1411 | S1411M | E | 1.4e-05 | 5.4e-06 | 6e-06 | 8.467E-06 | 4.801E-06 | 5.400E-06 | 1.400E-05 |
| 28214 | 1411 | S1411F | E | 9.6e-06 | 8.2e-06 | 1.7e-05 | 1.160E-05 | 4.729E-06 | 8.200E-06 | 1.700E-05 |
| 28215 | 1411 | S1411P | E | 8.6e-06 | 3.9e-06 | 9.8e-06 | 7.433E-06 | 3.118E-06 | 3.900E-06 | 9.800E-06 |
| 28216 | 1411 | S1411S | E | 5.8e-06 | 1.1e-05 | 1e-05 | 8.933E-06 | 2.759E-06 | 5.800E-06 | 1.100E-05 |
| 28217 | 1411 | S1411T | E | 6.8e-06 | 9.6e-06 | 5.6e-06 | 7.333E-06 | 2.053E-06 | 5.600E-06 | 9.600E-06 |
| 28218 | 1411 | S1411W | E | 6.6e-06 | 6.1e-06 | 8.9e-06 | 7.200E-06 | 1.493E-06 | 6.100E-06 | 8.900E-06 |
| 28219 | 1411 | S1411Y | E | 4.9e-06 | 1.1e-05 | 7.5e-06 | 7.800E-06 | 3.061E-06 | 4.900E-06 | 1.100E-05 |

| | | | | | | | | | | |
|---|---|---|---|---|---|---|---|---|---|---|
| 28220 | 1411 | S1411V | E | 5.8e-06 | 8.2e-06 | 7.6e-06 | 7.200E-06 | 1.249E-06 | 5.800E-06 | 8.200E-06 |
| 28221 | 1412 | G1412A | E | 6.7e-06 | 9.6e-06 | 6e-06 | 7.433E-06 | 1.909E-06 | 6.000E-06 | 9.600E-06 |
| 28222 | 1412 | G1412R | E | 6.7e-06 | 9.5e-06 | 6.7e-06 | 7.633E-06 | 1.617E-06 | 6.700E-06 | 9.500E-06 |
| 28223 | 1412 | G1412N | E | 4.1e-06 | 1.4e-05 | 5.9e-06 | 8.000E-06 | 5.274E-06 | 4.100E-06 | 1.400E-05 |
| 28224 | 1412 | G1412D | E | 5.3e-06 | 1e-05 | 6.6e-06 | 7.300E-06 | 2.427E-06 | 5.300E-06 | 1.000E-05 |
| 28225 | 1412 | G1412C | E | 8.6e-06 | 9.9e-06 | 7.1e-06 | 8.533E-06 | 1.401E-06 | 7.100E-06 | 9.900E-06 |
| 28226 | 1412 | G1412E | E | 1e-05 | 5e-06 | 6.8e-06 | 7.267E-06 | 2.532E-06 | 5.000E-06 | 1.000E-05 |
| 28227 | 1412 | G1412Q | E | 1.1e-05 | 6.5e-06 | 6.2e-06 | 7.900E-06 | 2.689E-06 | 6.200E-06 | 1.100E-05 |
| 28228 | 1412 | G1412G | E | 9.2e-06 | 4.8e-06 | 1.2e-05 | 8.667E-06 | 3.630E-06 | 4.800E-06 | 1.200E-05 |
| 28229 | 1412 | G1412H | E | 4.5e-06 | 6.5e-06 | 8.8e-06 | 6.600E-06 | 2.152E-06 | 4.500E-06 | 8.800E-06 |
| 28230 | 1412 | G1412I | E | 3.2e-06 | 6.5e-06 | 6.3e-06 | 5.333E-06 | 1.850E-06 | 3.200E-06 | 6.500E-06 |
| 28231 | 1412 | G1412L | E | 7.1e-06 | 8e-06 | 8e-06 | 7.700E-06 | 5.196E-07 | 7.100E-06 | 8.000E-06 |
| 28232 | 1412 | G1412K | E | 6.2e-06 | 9.5e-06 | 5.4e-06 | 7.033E-06 | 2.173E-06 | 5.400E-06 | 9.500E-06 |
| 28233 | 1412 | G1412M | E | 9.8e-06 | 7.6e-06 | 4.3e-06 | 7.233E-06 | 2.768E-06 | 4.300E-06 | 9.800E-06 |
| 28234 | 1412 | G1412F | E | 5.4e-06 | 7.8e-06 | 8e-06 | 7.067E-06 | 1.447E-06 | 5.400E-06 | 8.000E-06 |
| 28235 | 1412 | G1412P | E | 5.5e-06 | 6.2e-06 | 4.8e-06 | 5.500E-06 | 7.000E-07 | 4.800E-06 | 6.200E-06 |
| 28236 | 1412 | G1412S | E | 7.3e-06 | 5.8e-06 | 1.2e-05 | 8.367E-06 | 3.235E-06 | 5.800E-06 | 1.200E-05 |
| 28237 | 1412 | G1412T | E | 5.5e-06 | 8.4e-06 | 4e-06 | 5.967E-06 | 2.237E-06 | 4.000E-06 | 8.400E-06 |
| 28238 | 1412 | G1412W | E | 6.4e-06 | 1.3e-05 | 6.5e-06 | 8.633E-06 | 3.782E-06 | 6.400E-06 | 1.300E-05 |
| 28239 | 1412 | G1412Y | E | 3.2e-06 | 1.2e-05 | 7.4e-06 | 7.533E-06 | 4.402E-06 | 3.200E-06 | 1.200E-05 |
| 28240 | 1412 | G1412V | E | 4.4e-06 | 8.1e-06 | 8.6e-06 | 7.033E-06 | 2.294E-06 | 4.400E-06 | 8.600E-06 |
| 28241 | 1413 | G1413A | E | 6.2e-06 | 1.4e-05 | 8.1e-06 | 9.433E-06 | 4.067E-06 | 6.200E-06 | 1.400E-05 |
| 28242 | 1413 | G1413R | E | 9.5e-06 | 7e-06 | 8.9e-06 | 8.467E-06 | 1.305E-06 | 7.000E-06 | 9.500E-06 |
| 28243 | 1413 | G1413N | E | 7.1e-06 | 6.2e-06 | 9.8e-06 | 7.700E-06 | 1.873E-06 | 6.200E-06 | 9.800E-06 |
| 28244 | 1413 | G1413D | E | 9.8e-06 | 1.2e-05 | 8.1e-06 | 9.967E-06 | 1.955E-06 | 8.100E-06 | 1.200E-05 |
| 28245 | 1413 | G1413C | E | 9.2e-06 | 7.6e-06 | 1.1e-05 | 9.267E-06 | 1.701E-06 | 7.600E-06 | 1.100E-05 |
| 28246 | 1413 | G1413E | E | 1.5e-05 | 9.5e-06 | 6.3e-06 | 1.027E-05 | 4.400E-06 | 6.300E-06 | 1.500E-05 |
| 28247 | 1413 | G1413Q | E | 7.7e-06 | 4.4e-06 | 6.8e-06 | 6.300E-06 | 1.706E-06 | 4.400E-06 | 7.700E-06 |
| 28248 | 1413 | G1413G | E | 4.5e-06 | 7.9e-06 | 9.6e-06 | 7.333E-06 | 2.597E-06 | 4.500E-06 | 9.600E-06 |
| 28249 | 1413 | G1413H | E | 6e-06 | 6.9e-06 | 6.6e-06 | 6.500E-06 | 4.583E-07 | 6.000E-06 | 6.900E-06 |
| 28250 | 1413 | G1413I | E | 7.7e-06 | 8.1e-06 | 7e-06 | 7.600E-06 | 5.568E-07 | 7.000E-06 | 8.100E-06 |
| 28251 | 1413 | G1413L | E | 1.2e-05 | 4.8e-06 | 6.9e-06 | 7.900E-06 | 3.703E-06 | 4.800E-06 | 1.200E-05 |
| 28252 | 1413 | G1413K | E | 7.5e-06 | 4.7e-06 | 6.2e-06 | 6.133E-06 | 1.401E-06 | 4.700E-06 | 7.500E-06 |
| 28253 | 1413 | G1413M | E | 7.2e-06 | 5.9e-06 | 1.4e-05 | 9.033E-06 | 4.350E-06 | 5.900E-06 | 1.400E-05 |
| 28254 | 1413 | G1413F | E | 4.8e-06 | 9.1e-06 | 1.1e-05 | 8.300E-06 | 3.176E-06 | 4.800E-06 | 1.100E-05 |
| 28255 | 1413 | G1413P | E | 1.9e-05 | 6.5e-06 | 9.1e-06 | 1.153E-05 | 6.596E-06 | 6.500E-06 | 1.900E-05 |
| 28256 | 1413 | G1413S | E | 5.1e-06 | 6.5e-06 | 7.6e-06 | 6.400E-06 | 1.253E-06 | 5.100E-06 | 7.600E-06 |
| 28257 | 1413 | G1413T | E | 1.1e-05 | 1.2e-05 | 4.4e-06 | 9.133E-06 | 4.130E-06 | 4.400E-06 | 1.200E-05 |
| 28258 | 1413 | G1413W | E | 5.5e-06 | 9.9e-06 | 5.9e-06 | 7.100E-06 | 2.433E-06 | 5.500E-06 | 9.900E-06 |
| 28259 | 1413 | G1413Y | E | 8.2e-06 | 1e-05 | 6.3e-06 | 8.167E-06 | 1.850E-06 | 6.300E-06 | 1.000E-05 |
| 28260 | 1413 | G1413V | E | 4.8e-06 | 6.7e-06 | 6.6e-06 | 6.033E-06 | 1.069E-06 | 4.800E-06 | 6.700E-06 |
| 28261 | 1414 | T1414A | E | 7.1e-06 | 6e-06 | 9.1e-06 | 7.400E-06 | 1.572E-06 | 6.000E-06 | 9.100E-06 |
| 28262 | 1414 | T1414R | E | 6.5e-06 | 8e-06 | 6.1e-06 | 6.867E-06 | 1.002E-06 | 6.100E-06 | 8.000E-06 |
| 28263 | 1414 | T1414N | E | 7.5e-06 | 6.8e-06 | 6.1e-06 | 6.800E-06 | 7.000E-07 | 6.100E-06 | 7.500E-06 |
| 28264 | 1414 | T1414D | E | 1e-05 | 7.6e-06 | 9.9e-06 | 9.167E-06 | 1.358E-06 | 7.600E-06 | 1.000E-05 |
| 28265 | 1414 | T1414C | E | 6.9e-06 | 5.8e-06 | 9e-06 | 7.233E-06 | 1.626E-06 | 5.800E-06 | 9.000E-06 |
| 28266 | 1414 | T1414E | E | 1.1e-05 | 1.1e-05 | 5.9e-06 | 9.300E-06 | 2.944E-06 | 5.900E-06 | 1.100E-05 |
| 28267 | 1414 | T1414Q | E | 9.3e-06 | 7.2e-06 | 1.4e-05 | 1.017E-05 | 3.482E-06 | 7.200E-06 | 1.400E-05 |
| 28268 | 1414 | T1414G | E | 7.9e-06 | 6.4e-06 | 5.9e-06 | 6.733E-06 | 1.041E-06 | 5.900E-06 | 7.900E-06 |
| 28269 | 1414 | T1414H | E | 9.5e-06 | 6.7e-06 | 1.4e-05 | 1.007E-05 | 3.683E-06 | 6.700E-06 | 1.400E-05 |
| 28270 | 1414 | T1414I | E | 1.1e-05 | 1.3e-05 | 7.3e-06 | 1.043E-05 | 2.892E-06 | 7.300E-06 | 1.300E-05 |
| 28271 | 1414 | T1414L | E | 6.8e-06 | 8.4e-06 | 1.1e-05 | 8.733E-06 | 2.120E-06 | 6.800E-06 | 1.100E-05 |
| 28272 | 1414 | T1414K | E | 1.4e-05 | 8.3e-06 | 9.8e-06 | 1.070E-05 | 2.955E-06 | 8.300E-06 | 1.400E-05 |
| 28273 | 1414 | T1414M | E | 5.7e-06 | 8.2e-06 | 8.8e-06 | 7.567E-06 | 1.644E-06 | 5.700E-06 | 8.800E-06 |
| 28274 | 1414 | T1414F | E | 6.6e-06 | 8.3e-06 | 9.4e-06 | 8.100E-06 | 1.411E-06 | 6.600E-06 | 9.400E-06 |
| 28275 | 1414 | T1414P | E | 8.5e-06 | 1.1e-05 | 7.1e-06 | 8.867E-06 | 1.976E-06 | 7.100E-06 | 1.100E-05 |
| 28276 | 1414 | T1414S | E | 6.2e-06 | 4.9e-06 | 9.5e-06 | 6.867E-06 | 2.371E-06 | 4.900E-06 | 9.500E-06 |
| 28277 | 1414 | T1414T | E | 6.3e-06 | 1.2e-05 | 5.4e-06 | 7.900E-06 | 3.579E-06 | 5.400E-06 | 1.200E-05 |
| 28278 | 1414 | T1414W | E | 8.2e-06 | 4.7e-06 | 6.2e-06 | 6.367E-06 | 1.756E-06 | 4.700E-06 | 8.200E-06 |
| 28279 | 1414 | T1414Y | E | 8.4e-06 | 9.7e-06 | 8.9e-06 | 9.000E-06 | 6.557E-07 | 8.400E-06 | 9.700E-06 |
| 28280 | 1414 | T1414V | E | 8.9e-06 | 5.9e-06 | 1.4e-05 | 9.600E-06 | 4.095E-06 | 5.900E-06 | 1.400E-05 |
| 28281 | 1415 | A1415A | E | 1.2e-05 | 6.5e-06 | 1.9e-05 | 1.250E-05 | 6.265E-06 | 6.500E-06 | 1.900E-05 |
| 28282 | 1415 | A1415R | E | 6.5e-06 | 8.1e-06 | 6.3e-06 | 6.967E-06 | 9.866E-07 | 6.300E-06 | 8.100E-06 |
| 28283 | 1415 | A1415N | E | 7e-06 | 7.7e-06 | 1e-05 | 8.233E-06 | 1.570E-06 | 7.000E-06 | 1.000E-05 |
| 28284 | 1415 | A1415D | E | 5.8e-06 | 6.4e-06 | 6.5e-06 | 6.233E-06 | 3.786E-07 | 5.800E-06 | 6.500E-06 |
| 28285 | 1415 | A1415C | E | 7.5e-06 | 9.9e-06 | 5.2e-06 | 7.533E-06 | 2.350E-06 | 5.200E-06 | 9.900E-06 |
| 28286 | 1415 | A1415E | E | 6.4e-06 | 6.3e-06 | 7.9e-06 | 6.867E-06 | 8.963E-07 | 6.300E-06 | 7.900E-06 |
| 28287 | 1415 | A1415Q | E | 6.2e-06 | 5.2e-06 | 8.2e-06 | 6.533E-06 | 1.528E-06 | 5.200E-06 | 8.200E-06 |
| 28288 | 1415 | A1415G | E | 8.2e-06 | 8.9e-06 | 5.1e-06 | 7.400E-06 | 2.022E-06 | 5.100E-06 | 8.900E-06 |
| 28289 | 1415 | A1415H | E | 8.6e-06 | 6.1e-06 | 6.8e-06 | 7.167E-06 | 1.290E-06 | 6.100E-06 | 8.600E-06 |
| 28290 | 1415 | A1415I | E | 6.2e-06 | 1e-05 | 8.1e-06 | 8.100E-06 | 1.900E-06 | 6.200E-06 | 1.000E-05 |
| 28291 | 1415 | A1415L | E | 5.9e-06 | 9.5e-06 | 4.9e-06 | 6.767E-06 | 2.419E-06 | 4.900E-06 | 9.500E-06 |
| 28292 | 1415 | A1415K | E | 8.2e-06 | 1e-05 | 6.4e-06 | 8.200E-06 | 1.800E-06 | 6.400E-06 | 1.000E-05 |
| 28293 | 1415 | A1415M | E | 1.2e-05 | 9.4e-06 | 1.4e-05 | 1.180E-05 | 2.307E-06 | 9.400E-06 | 1.400E-05 |
| 28294 | 1415 | A1415F | E | 9.9e-06 | 8.1e-06 | 6.8e-06 | 8.267E-06 | 1.557E-06 | 6.800E-06 | 9.900E-06 |
| 28295 | 1415 | A1415P | E | 5e-06 | 5.7e-06 | 6.5e-06 | 5.733E-06 | 7.506E-07 | 5.000E-06 | 6.500E-06 |
| 28296 | 1415 | A1415S | E | 6.6e-06 | 9.7e-06 | 5.5e-06 | 7.267E-06 | 2.178E-06 | 5.500E-06 | 9.700E-06 |
| 28297 | 1415 | A1415T | E | 7.5e-06 | 1e-05 | 7.9e-06 | 8.467E-06 | 1.343E-06 | 7.500E-06 | 1.000E-05 |
| 28298 | 1415 | A1415W | E | 9.3e-06 | 6.3e-06 | 4.8e-06 | 6.800E-06 | 2.291E-06 | 4.800E-06 | 9.300E-06 |
| 28299 | 1415 | A1415Y | E | 1.1e-05 | 6.1e-06 | 1e-05 | 9.033E-06 | 2.589E-06 | 6.100E-06 | 1.100E-05 |
| 28300 | 1415 | A1415V | E | 1.2e-05 | 6.5e-06 | 4.8e-06 | 7.767E-06 | 3.763E-06 | 4.800E-06 | 1.200E-05 |
| 28301 | 1416 | A1416A | E | 8.4e-06 | 8.3e-06 | 1.2e-05 | 9.567E-06 | 2.108E-06 | 8.300E-06 | 1.200E-05 |
| 28302 | 1416 | A1416R | E | 9.4e-06 | 9.3e-06 | 6.6e-06 | 8.433E-06 | 1.589E-06 | 6.600E-06 | 9.400E-06 |
| 28303 | 1416 | A1416N | E | 8.9e-06 | 7.5e-06 | 1.1e-05 | 9.133E-06 | 1.762E-06 | 7.500E-06 | 1.100E-05 |
| 28304 | 1416 | A1416D | E | 6e-06 | 8.3e-06 | 7.7e-06 | 7.333E-06 | 1.193E-06 | 6.000E-06 | 8.300E-06 |

| | | | | | | | | | | |
|---|---|---|---|---|---|---|---|---|---|---|
| 28305 | 1416 | A1416C | E | 8.5e-06 | 1.1e-05 | 9.3e-06 | 9.600E-06 | 1.277E-06 | 8.500E-06 | 1.100E-05 |
| 28306 | 1416 | A1416E | E | 1.2e-05 | 8.7e-06 | 6.7e-06 | 9.133E-06 | 2.676E-06 | 6.700E-06 | 1.200E-05 |
| 28307 | 1416 | A1416Q | E | 7.4e-06 | 7e-06 | 7e-06 | 7.133E-06 | 2.309E-07 | 7.000E-06 | 7.400E-06 |
| 28308 | 1416 | A1416G | E | 8.7e-06 | 9.4e-06 | 9.6e-06 | 9.233E-06 | 4.726E-07 | 8.700E-06 | 9.600E-06 |
| 28309 | 1416 | A1416H | E | 9e-06 | 4.6e-06 | 7.9e-06 | 7.167E-06 | 2.290E-06 | 4.600E-06 | 9.000E-06 |
| 28310 | 1416 | A1416I | E | 3.4e-06 | 6.5e-06 | 8.4e-06 | 6.100E-06 | 2.524E-06 | 3.400E-06 | 8.400E-06 |
| 28311 | 1416 | A1416L | E | 9.8e-06 | 6.9e-06 | 4.7e-06 | 7.133E-06 | 2.558E-06 | 4.700E-06 | 9.800E-06 |
| 28312 | 1416 | A1416K | E | 4.5e-06 | 6.1e-06 | 1.1e-05 | 7.200E-06 | 3.387E-06 | 4.500E-06 | 1.100E-05 |
| 28313 | 1416 | A1416M | E | 1.2e-05 | 7e-06 | 1e-05 | 9.667E-06 | 2.517E-06 | 7.000E-06 | 1.200E-05 |
| 28314 | 1416 | A1416F | E | 1e-05 | 8.3e-06 | 4.9e-06 | 7.733E-06 | 2.597E-06 | 4.900E-06 | 1.000E-05 |
| 28315 | 1416 | A1416P | E | 4.9e-06 | 6.2e-06 | 9e-06 | 6.700E-06 | 2.095E-06 | 4.900E-06 | 9.000E-06 |
| 28316 | 1416 | A1416S | E | 7.1e-06 | 1.3e-05 | 1.4e-05 | 1.137E-05 | 3.729E-06 | 7.100E-06 | 1.400E-05 |
| 28317 | 1416 | A1416T | E | 7.8e-06 | 6e-06 | 1.2e-05 | 8.600E-06 | 3.079E-06 | 6.000E-06 | 1.200E-05 |
| 28318 | 1416 | A1416W | E | 1e-05 | 6.8e-06 | 6.7e-06 | 7.833E-06 | 1.877E-06 | 6.700E-06 | 1.000E-05 |
| 28319 | 1416 | A1416Y | E | 7.1e-06 | 7.5e-06 | 4.8e-06 | 6.467E-06 | 1.457E-06 | 4.800E-06 | 7.500E-06 |
| 28320 | 1416 | A1416V | E | 1.3e-05 | 7e-06 | 1.4e-05 | 1.133E-05 | 3.786E-06 | 7.000E-06 | 1.400E-05 |
| 28321 | 1417 | L1417A | E | 1.4e-05 | 4.7e-06 | 8.1e-06 | 8.933E-06 | 4.706E-06 | 4.700E-06 | 1.400E-05 |
| 28322 | 1417 | L1417R | E | 6.9e-06 | 1.2e-05 | 6.6e-06 | 8.500E-06 | 3.035E-06 | 6.600E-06 | 1.200E-05 |
| 28323 | 1417 | L1417N | E | 7e-06 | 6.9e-06 | 7e-06 | 6.967E-06 | 5.774E-08 | 6.900E-06 | 7.000E-06 |
| 28324 | 1417 | L1417D | E | 5e-06 | 1.2e-05 | 6.7e-06 | 7.900E-06 | 3.651E-06 | 5.000E-06 | 1.200E-05 |
| 28325 | 1417 | L1417C | E | 6.2e-06 | 1e-05 | 8.8e-06 | 8.333E-06 | 1.943E-06 | 6.200E-06 | 1.000E-05 |
| 28326 | 1417 | L1417E | E | 8.4e-06 | 1.1e-05 | 5.4e-06 | 8.267E-06 | 2.802E-06 | 5.400E-06 | 1.100E-05 |
| 28327 | 1417 | L1417Q | E | 1.1e-05 | 1.4e-05 | 2.8e-06 | 9.267E-06 | 5.798E-06 | 2.800E-06 | 1.400E-05 |
| 28328 | 1417 | L1417G | E | 5.7e-06 | 1e-05 | 8.9e-06 | 8.200E-06 | 2.234E-06 | 5.700E-06 | 1.000E-05 |
| 28329 | 1417 | L1417H | E | 7.4e-06 | 6.7e-06 | 3.6e-06 | 5.900E-06 | 2.022E-06 | 3.600E-06 | 7.400E-06 |
| 28330 | 1417 | L1417I | E | 1.3e-05 | 4.5e-06 | 1.6e-05 | 1.117E-05 | 5.965E-06 | 4.500E-06 | 1.600E-05 |
| 28331 | 1417 | L1417L | E | 1.1e-05 | 4.6e-06 | 1.3e-05 | 9.533E-06 | 4.388E-06 | 4.600E-06 | 1.300E-05 |
| 28332 | 1417 | L1417K | E | 1.2e-05 | 5.2e-06 | 6.8e-06 | 8.000E-06 | 3.555E-06 | 5.200E-06 | 1.200E-05 |
| 28333 | 1417 | L1417M | E | 5e-06 | 1e-05 | 5.2e-06 | 6.733E-06 | 2.831E-06 | 5.000E-06 | 1.000E-05 |
| 28334 | 1417 | L1417F | E | 9.4e-06 | 5.6e-06 | 9.5e-06 | 8.167E-06 | 2.223E-06 | 5.600E-06 | 9.500E-06 |
| 28335 | 1417 | L1417P | E | 5.6e-06 | 6.2e-06 | 5.3e-06 | 5.700E-06 | 4.583E-07 | 5.300E-06 | 6.200E-06 |
| 28336 | 1417 | L1417S | E | 8e-06 | 5.6e-06 | 5.7e-06 | 6.433E-06 | 1.358E-06 | 5.600E-06 | 8.000E-06 |
| 28337 | 1417 | L1417T | E | 6.5e-06 | 9e-06 | 7.1e-06 | 7.533E-06 | 1.305E-06 | 6.500E-06 | 9.000E-06 |
| 28338 | 1417 | L1417W | E | 8.9e-06 | 7.4e-06 | 7e-06 | 7.767E-06 | 1.002E-06 | 7.000E-06 | 8.900E-06 |
| 28339 | 1417 | L1417Y | E | 5.9e-06 | 6e-06 | 8.8e-06 | 6.900E-06 | 1.646E-06 | 5.900E-06 | 8.800E-06 |
| 28340 | 1417 | L1417V | E | 1.1e-05 | 6.4e-06 | 1.3e-05 | 1.013E-05 | 3.384E-06 | 6.400E-06 | 1.300E-05 |
| 28341 | 1418 | G1418A | E | 1.5e-05 | 6.5e-06 | 1.2e-05 | 1.117E-05 | 4.311E-06 | 6.500E-06 | 1.500E-05 |
| 28342 | 1418 | G1418R | E | 6.1e-06 | 6.2e-06 | 8e-06 | 6.767E-06 | 1.069E-06 | 6.100E-06 | 8.000E-06 |
| 28343 | 1418 | G1418N | E | 9.5e-06 | 7.5e-06 | 5.4e-06 | 7.467E-06 | 2.050E-06 | 5.400E-06 | 9.500E-06 |
| 28344 | 1418 | G1418D | E | 7.8e-06 | 9e-06 | 9.2e-06 | 8.667E-06 | 7.572E-07 | 7.800E-06 | 9.200E-06 |
| 28345 | 1418 | G1418C | E | 6.3e-06 | 7.7e-06 | 8.5e-06 | 7.500E-06 | 1.114E-06 | 6.300E-06 | 8.500E-06 |
| 28346 | 1418 | G1418E | E | 9.5e-06 | 7.8e-06 | 1.2e-05 | 9.767E-06 | 2.113E-06 | 7.800E-06 | 1.200E-05 |
| 28347 | 1418 | G1418Q | E | 9.1e-06 | 8e-06 | 1.2e-05 | 9.700E-06 | 2.066E-06 | 8.000E-06 | 1.200E-05 |
| 28348 | 1418 | G1418G | E | 7.9e-06 | 6.6e-06 | 7.8e-06 | 7.433E-06 | 7.234E-07 | 6.600E-06 | 7.900E-06 |
| 28349 | 1418 | G1418H | E | 8.2e-06 | 7.1e-06 | 5.8e-06 | 7.033E-06 | 1.201E-06 | 5.800E-06 | 8.200E-06 |
| 28350 | 1418 | G1418I | E | 7.4e-06 | 6.4e-06 | 1e-05 | 7.933E-06 | 1.858E-06 | 6.400E-06 | 1.000E-05 |
| 28351 | 1418 | G1418L | E | 5.1e-06 | 1e-05 | 9.1e-06 | 8.067E-06 | 2.608E-06 | 5.100E-06 | 1.000E-05 |
| 28352 | 1418 | G1418K | E | 9.6e-06 | 5.8e-06 | 6e-06 | 7.133E-06 | 2.139E-06 | 5.800E-06 | 9.600E-06 |
| 28353 | 1418 | G1418M | E | 4.7e-06 | 6.7e-06 | 6.3e-06 | 5.900E-06 | 1.058E-06 | 4.700E-06 | 6.700E-06 |
| 28354 | 1418 | G1418F | E | 1.4e-05 | 8.5e-06 | 1.5e-05 | 1.250E-05 | 3.500E-06 | 8.500E-06 | 1.500E-05 |
| 28355 | 1418 | G1418P | E | 6.3e-06 | 1.3e-05 | 4.6e-06 | 7.967E-06 | 4.441E-06 | 4.600E-06 | 1.300E-05 |
| 28356 | 1418 | G1418S | E | 1.4e-05 | 6e-06 | 8.2e-06 | 9.400E-06 | 4.133E-06 | 6.000E-06 | 1.400E-05 |
| 28357 | 1418 | G1418T | E | 1.3e-05 | 6.5e-06 | 5e-06 | 8.167E-06 | 4.252E-06 | 5.000E-06 | 1.300E-05 |
| 28358 | 1418 | G1418W | E | 6.9e-06 | 9.2e-06 | 1.5e-05 | 1.037E-05 | 4.174E-06 | 6.900E-06 | 1.500E-05 |
| 28359 | 1418 | G1418Y | E | 9.8e-06 | 1.4e-05 | 5.8e-06 | 9.867E-06 | 4.100E-06 | 5.800E-06 | 1.400E-05 |
| 28360 | 1418 | G1418V | E | 9.8e-06 | 6.8e-06 | 1.2e-05 | 9.533E-06 | 2.610E-06 | 6.800E-06 | 1.200E-05 |
| 28361 | 1419 | C1419A | E | 7.8e-06 | 9.8e-06 | 8.2e-06 | 8.600E-06 | 1.058E-06 | 7.800E-06 | 9.800E-06 |
| 28362 | 1419 | C1419R | E | 7.4e-06 | 9.7e-06 | 1.4e-05 | 1.037E-05 | 3.350E-06 | 7.400E-06 | 1.400E-05 |
| 28363 | 1419 | C1419N | E | 9.3e-06 | 5.1e-06 | 1.1e-05 | 8.467E-06 | 3.037E-06 | 5.100E-06 | 1.100E-05 |
| 28364 | 1419 | C1419D | E | 8.9e-06 | 8.3e-06 | 6.5e-06 | 7.900E-06 | 1.249E-06 | 6.500E-06 | 8.900E-06 |
| 28365 | 1419 | C1419C | E | 9.3e-06 | 5.1e-06 | 6.6e-06 | 7.000E-06 | 2.128E-06 | 5.100E-06 | 9.300E-06 |
| 28366 | 1419 | C1419E | E | 9.6e-06 | 1.4e-05 | 1.4e-05 | 1.253E-05 | 2.540E-06 | 9.600E-06 | 1.400E-05 |
| 28367 | 1419 | C1419Q | E | 8e-06 | 6e-06 | 6.8e-06 | 6.933E-06 | 1.007E-06 | 6.000E-06 | 8.000E-06 |
| 28368 | 1419 | C1419G | E | 1e-05 | 6.1e-06 | 1.4e-05 | 1.003E-05 | 3.950E-06 | 6.100E-06 | 1.400E-05 |
| 28369 | 1419 | C1419H | E | 9e-06 | 9.5e-06 | 5.4e-06 | 7.967E-06 | 2.237E-06 | 5.400E-06 | 9.500E-06 |
| 28370 | 1419 | C1419I | E | 7.3e-06 | 1e-05 | 7e-06 | 8.100E-06 | 1.652E-06 | 7.000E-06 | 1.000E-05 |
| 28371 | 1419 | C1419L | E | 8.5e-06 | 4.5e-06 | 8.4e-06 | 7.133E-06 | 2.281E-06 | 4.500E-06 | 8.500E-06 |
| 28372 | 1419 | C1419K | E | 6.6e-06 | 4.2e-06 | 7.4e-06 | 6.067E-06 | 1.665E-06 | 4.200E-06 | 7.400E-06 |
| 28373 | 1419 | C1419M | E | 1.7e-05 | 5.9e-06 | 6.1e-06 | 9.667E-06 | 6.352E-06 | 5.900E-06 | 1.700E-05 |
| 28374 | 1419 | C1419F | E | 1e-05 | 6.9e-06 | 8.1e-06 | 8.333E-06 | 1.563E-06 | 6.900E-06 | 1.000E-05 |
| 28375 | 1419 | C1419P | E | 1e-05 | 9.6e-06 | 9.9e-06 | 9.833E-06 | 2.082E-07 | 9.600E-06 | 1.000E-05 |
| 28376 | 1419 | C1419S | E | 5.7e-06 | 7.2e-06 | 8.5e-06 | 7.133E-06 | 1.401E-06 | 5.700E-06 | 8.500E-06 |
| 28377 | 1419 | C1419T | E | 9.9e-06 | 6e-06 | 6.5e-06 | 7.467E-06 | 2.122E-06 | 6.000E-06 | 9.900E-06 |
| 28378 | 1419 | C1419W | E | 1.4e-05 | 6.9e-06 | 4.8e-06 | 8.567E-06 | 4.821E-06 | 4.800E-06 | 1.400E-05 |
| 28379 | 1419 | C1419Y | E | 4.3e-06 | 1.2e-05 | 5.4e-06 | 7.233E-06 | 4.165E-06 | 4.300E-06 | 1.200E-05 |
| 28380 | 1419 | C1419V | E | 6e-06 | 5.3e-06 | 7.2e-06 | 6.167E-06 | 9.609E-07 | 5.300E-06 | 7.200E-06 |
| 28381 | 1420 | L1420A | E | 6e-06 | 5.6e-06 | 8.9e-06 | 6.833E-06 | 1.801E-06 | 5.600E-06 | 8.900E-06 |
| 28382 | 1420 | L1420R | E | 8.6e-06 | 7.8e-06 | 6.8e-06 | 7.733E-06 | 9.018E-07 | 6.800E-06 | 8.600E-06 |
| 28383 | 1420 | L1420N | E | 5.9e-06 | 9.3e-06 | 9.9e-06 | 8.367E-06 | 2.157E-06 | 5.900E-06 | 9.900E-06 |
| 28384 | 1420 | L1420D | E | 9.2e-06 | 6.1e-06 | 9.1e-06 | 8.133E-06 | 1.762E-06 | 6.100E-06 | 9.200E-06 |
| 28385 | 1420 | L1420C | E | 4.1e-06 | 5e-06 | 7e-06 | 5.367E-06 | 1.484E-06 | 4.100E-06 | 7.000E-06 |
| 28386 | 1420 | L1420E | E | 6.6e-06 | 4.3e-06 | 3.3e-06 | 4.733E-06 | 1.692E-06 | 3.300E-06 | 6.600E-06 |
| 28387 | 1420 | L1420Q | E | 6.7e-06 | 1.1e-05 | 4.1e-06 | 7.267E-06 | 3.485E-06 | 4.100E-06 | 1.100E-05 |
| 28388 | 1420 | L1420G | E | 7.6e-06 | 6.2e-06 | 8.8e-06 | 7.533E-06 | 1.301E-06 | 6.200E-06 | 8.800E-06 |
| 28389 | 1420 | L1420H | E | 6e-06 | 1.1e-05 | 9.4e-06 | 8.800E-06 | 2.553E-06 | 6.000E-06 | 1.100E-05 |

| | | | | | | | | | | |
|---|---|---|---|---|---|---|---|---|---|---|
| 28390 | 1420 | L1420I | E | 5.7e-06 | 6.8e-06 | 1e-05 | 7.500E-06 | 2.234E-06 | 5.700E-06 | 1.000E-05 |
| 28391 | 1420 | L1420L | E | 1.2e-05 | 5.8e-06 | 5.4e-06 | 7.733E-06 | 3.700E-06 | 5.400E-06 | 1.200E-05 |
| 28392 | 1420 | L1420K | E | 6.4e-06 | 1.4e-05 | 6.4e-06 | 8.933E-06 | 4.388E-06 | 6.400E-06 | 1.400E-05 |
| 28393 | 1420 | L1420M | E | 1.2e-05 | 9.4e-06 | 5.9e-06 | 9.100E-06 | 3.061E-06 | 5.900E-06 | 1.200E-05 |
| 28394 | 1420 | L1420F | E | 8.7e-06 | 6.8e-06 | 6.5e-06 | 7.333E-06 | 1.193E-06 | 6.500E-06 | 8.700E-06 |
| 28395 | 1420 | L1420P | E | 7.4e-06 | 6.4e-06 | 1.2e-05 | 8.600E-06 | 2.987E-06 | 6.400E-06 | 1.200E-05 |
| 28396 | 1420 | L1420S | E | 5.4e-06 | 1.2e-05 | 7.5e-06 | 8.300E-06 | 3.372E-06 | 5.400E-06 | 1.200E-05 |
| 28397 | 1420 | L1420T | E | 9.5e-06 | 6.8e-06 | 6.3e-06 | 7.533E-06 | 1.721E-06 | 6.300E-06 | 9.500E-06 |
| 28398 | 1420 | L1420W | E | 9.5e-06 | 7.8e-06 | 6.2e-06 | 7.833E-06 | 1.650E-06 | 6.200E-06 | 9.500E-06 |
| 28399 | 1420 | L1420Y | E | 1.1e-05 | 1.2e-05 | 8.1e-06 | 1.037E-05 | 2.026E-06 | 8.100E-06 | 1.200E-05 |
| 28400 | 1420 | L1420V | E | 9.5e-06 | 6.2e-06 | 6.5e-06 | 7.400E-06 | 1.825E-06 | 6.200E-06 | 9.500E-06 |
| 28401 | 1421 | V1421A | E | 1.1e-05 | 1e-05 | 5.6e-06 | 8.867E-06 | 2.873E-06 | 5.600E-06 | 1.100E-05 |
| 28402 | 1421 | V1421R | E | 1e-05 | 6.5e-06 | 7.3e-06 | 7.933E-06 | 1.834E-06 | 6.500E-06 | 1.000E-05 |
| 28403 | 1421 | V1421N | E | 7.2e-06 | 7.6e-06 | 1e-05 | 8.267E-06 | 1.514E-06 | 7.200E-06 | 1.000E-05 |
| 28404 | 1421 | V1421D | E | 1e-05 | 7.8e-06 | 7.4e-06 | 8.400E-06 | 1.400E-06 | 7.400E-06 | 1.000E-05 |
| 28405 | 1421 | V1421C | E | 9.2e-06 | 3.8e-06 | 4.7e-06 | 5.900E-06 | 2.893E-06 | 3.800E-06 | 9.200E-06 |
| 28406 | 1421 | V1421E | E | 9e-06 | 9.9e-06 | 1e-05 | 9.633E-06 | 5.508E-07 | 9.000E-06 | 1.000E-05 |
| 28407 | 1421 | V1421Q | E | 9.6e-06 | 7.8e-06 | 9.1e-06 | 8.833E-06 | 9.292E-07 | 7.800E-06 | 9.600E-06 |
| 28408 | 1421 | V1421G | E | 8.3e-06 | 4e-06 | 1.1e-05 | 7.767E-06 | 3.530E-06 | 4.000E-06 | 1.100E-05 |
| 28409 | 1421 | V1421H | E | 7.6e-06 | 4.4e-06 | 7.6e-06 | 6.533E-06 | 1.848E-06 | 4.400E-06 | 7.600E-06 |
| 28410 | 1421 | V1421I | E | 5.3e-06 | 6.8e-06 | 3.9e-06 | 5.333E-06 | 1.450E-06 | 3.900E-06 | 6.800E-06 |
| 28411 | 1421 | V1421L | E | 7.6e-06 | 8.2e-06 | 7.2e-06 | 7.667E-06 | 5.033E-07 | 7.200E-06 | 8.200E-06 |
| 28412 | 1421 | V1421K | E | 9.4e-06 | 6.5e-06 | 8.3e-06 | 8.067E-06 | 1.464E-06 | 6.500E-06 | 9.400E-06 |
| 28413 | 1421 | V1421M | E | 6.2e-06 | 5.2e-06 | 1.5e-05 | 8.800E-06 | 5.393E-06 | 5.200E-06 | 1.500E-05 |
| 28414 | 1421 | V1421F | E | 1.2e-05 | 1.1e-05 | 5.3e-06 | 9.433E-06 | 3.614E-06 | 5.300E-06 | 1.200E-05 |
| 28415 | 1421 | V1421P | E | 9.7e-06 | 5.8e-06 | 4.3e-06 | 6.600E-06 | 2.787E-06 | 4.300E-06 | 9.700E-06 |
| 28416 | 1421 | V1421S | E | 6.8e-06 | 8e-06 | 9.4e-06 | 8.067E-06 | 1.301E-06 | 6.800E-06 | 9.400E-06 |
| 28417 | 1421 | V1421T | E | 3.6e-06 | 4.6e-06 | 7.1e-06 | 5.100E-06 | 1.803E-06 | 3.600E-06 | 7.100E-06 |
| 28418 | 1421 | V1421W | E | 1.3e-05 | 6.5e-06 | 4.6e-06 | 8.033E-06 | 4.405E-06 | 4.600E-06 | 1.300E-05 |
| 28419 | 1421 | V1421Y | E | 8.6e-06 | 1.4e-05 | 1.1e-05 | 1.120E-05 | 2.706E-06 | 8.600E-06 | 1.400E-05 |
| 28420 | 1421 | V1421V | E | 1.4e-05 | 5.2e-06 | 1.2e-05 | 1.040E-05 | 4.613E-06 | 5.200E-06 | 1.400E-05 |
| 28421 | 1422 | K1422A | E | 8.2e-06 | 4.8e-06 | 7e-06 | 6.667E-06 | 1.724E-06 | 4.800E-06 | 8.200E-06 |
| 28422 | 1422 | K1422R | E | 9.6e-06 | 8.1e-06 | 1.2e-05 | 9.900E-06 | 1.967E-06 | 8.100E-06 | 1.200E-05 |
| 28423 | 1422 | K1422N | E | 8.6e-06 | 7.7e-06 | 8.4e-06 | 8.233E-06 | 4.726E-07 | 7.700E-06 | 8.600E-06 |
| 28424 | 1422 | K1422D | E | 6.4e-06 | 7.3e-06 | 9.2e-06 | 7.633E-06 | 1.429E-06 | 6.400E-06 | 9.200E-06 |
| 28425 | 1422 | K1422C | E | 7.1e-06 | 1.3e-05 | 1.4e-05 | 1.137E-05 | 3.729E-06 | 7.100E-06 | 1.400E-05 |
| 28426 | 1422 | K1422E | E | 9e-06 | 6.6e-06 | 6.8e-06 | 7.467E-06 | 1.332E-06 | 6.600E-06 | 9.000E-06 |
| 28427 | 1422 | K1422Q | E | 9.7e-06 | 8.7e-06 | 7.6e-06 | 8.667E-06 | 1.050E-06 | 7.600E-06 | 9.700E-06 |
| 28428 | 1422 | K1422G | E | 7.9e-06 | 9e-06 | 9.7e-06 | 8.867E-06 | 9.074E-07 | 7.900E-06 | 9.700E-06 |
| 28429 | 1422 | K1422H | E | 4.4e-06 | 5.9e-06 | 1e-05 | 6.767E-06 | 2.899E-06 | 4.400E-06 | 1.000E-05 |
| 28430 | 1422 | K1422I | E | 1e-05 | 8.9e-06 | 9.5e-06 | 9.467E-06 | 5.508E-07 | 8.900E-06 | 1.000E-05 |
| 28431 | 1422 | K1422L | E | 6.2e-06 | 8.7e-06 | 7.8e-06 | 7.567E-06 | 1.266E-06 | 6.200E-06 | 8.700E-06 |
| 28432 | 1422 | K1422K | E | 7.5e-06 | 7.2e-06 | 1.6e-05 | 1.023E-05 | 4.996E-06 | 7.200E-06 | 1.600E-05 |
| 28433 | 1422 | K1422M | E | 1.1e-05 | 7.4e-06 | 1.4e-05 | 1.080E-05 | 3.305E-06 | 7.400E-06 | 1.400E-05 |
| 28434 | 1422 | K1422F | E | 5.1e-06 | 9e-06 | 7.6e-06 | 7.233E-06 | 1.976E-06 | 5.100E-06 | 9.000E-06 |
| 28435 | 1422 | K1422P | E | 3.8e-06 | 7.6e-06 | 6.6e-06 | 6.000E-06 | 1.970E-06 | 3.800E-06 | 7.600E-06 |
| 28436 | 1422 | K1422S | E | 9.2e-06 | 4.8e-06 | 4.6e-06 | 6.200E-06 | 2.600E-06 | 4.600E-06 | 9.200E-06 |
| 28437 | 1422 | K1422T | E | 1.1e-05 | 8.7e-06 | 3.5e-06 | 7.733E-06 | 3.842E-06 | 3.500E-06 | 1.100E-05 |
| 28438 | 1422 | K1422W | E | 9.4e-06 | 5.2e-06 | 6.2e-06 | 6.933E-06 | 2.194E-06 | 5.200E-06 | 9.400E-06 |
| 28439 | 1422 | K1422Y | E | 1e-05 | 8.8e-06 | 3.9e-06 | 7.567E-06 | 3.232E-06 | 3.900E-06 | 1.000E-05 |
| 28440 | 1422 | K1422V | E | 8.2e-06 | 7e-06 | 8.3e-06 | 7.833E-06 | 7.234E-07 | 7.000E-06 | 8.300E-06 |
| 28441 | 1423 | D1423A | E | 5.9e-06 | 6.8e-06 | 6.6e-06 | 6.433E-06 | 4.726E-07 | 5.900E-06 | 6.800E-06 |
| 28442 | 1423 | D1423R | E | 6.6e-06 | 4.8e-06 | 9e-06 | 6.800E-06 | 2.107E-06 | 4.800E-06 | 9.000E-06 |
| 28443 | 1423 | D1423N | E | 1.1e-05 | 1.3e-05 | 5.1e-06 | 9.700E-06 | 4.107E-06 | 5.100E-06 | 1.300E-05 |
| 28444 | 1423 | D1423D | E | 4.5e-06 | 8.8e-06 | 9.2e-06 | 7.500E-06 | 2.606E-06 | 4.500E-06 | 9.200E-06 |
| 28445 | 1423 | D1423C | E | 1.2e-05 | 7.8e-06 | 6.4e-06 | 8.733E-06 | 2.914E-06 | 6.400E-06 | 1.200E-05 |
| 28446 | 1423 | D1423E | E | 4.1e-06 | 5.9e-06 | 8.4e-06 | 6.133E-06 | 2.159E-06 | 4.100E-06 | 8.400E-06 |
| 28447 | 1423 | D1423Q | E | 1.2e-05 | 1.1e-05 | 8.9e-06 | 1.063E-05 | 1.582E-06 | 8.900E-06 | 1.200E-05 |
| 28448 | 1423 | D1423G | E | 5.4e-06 | 8.9e-06 | 8.2e-06 | 7.500E-06 | 1.852E-06 | 5.400E-06 | 8.900E-06 |
| 28449 | 1423 | D1423H | E | 1e-05 | 9.3e-06 | 6.6e-06 | 8.633E-06 | 1.795E-06 | 6.600E-06 | 1.000E-05 |
| 28450 | 1423 | D1423I | E | 6.5e-06 | 7.1e-06 | 9e-06 | 7.533E-06 | 1.305E-06 | 6.500E-06 | 9.000E-06 |
| 28451 | 1423 | D1423L | E | 5.1e-06 | 7.6e-06 | 5.9e-06 | 6.200E-06 | 1.277E-06 | 5.100E-06 | 7.600E-06 |
| 28452 | 1423 | D1423K | E | 8.3e-06 | 9.5e-06 | 1.1e-05 | 9.600E-06 | 1.353E-06 | 8.300E-06 | 1.100E-05 |
| 28453 | 1423 | D1423M | E | 9.7e-06 | 6.5e-06 | 9.7e-06 | 8.633E-06 | 1.848E-06 | 6.500E-06 | 9.700E-06 |
| 28454 | 1423 | D1423F | E | 9e-06 | 6.7e-06 | 7e-06 | 7.567E-06 | 1.250E-06 | 6.700E-06 | 9.000E-06 |
| 28455 | 1423 | D1423P | E | 8.8e-06 | 7.8e-06 | 1.2e-05 | 9.533E-06 | 2.194E-06 | 7.800E-06 | 1.200E-05 |
| 28456 | 1423 | D1423S | E | 8.4e-06 | 6.8e-06 | 8.6e-06 | 7.933E-06 | 9.866E-07 | 6.800E-06 | 8.600E-06 |
| 28457 | 1423 | D1423T | E | 6.4e-06 | 8.9e-06 | 3.9e-06 | 6.400E-06 | 2.500E-06 | 3.900E-06 | 8.900E-06 |
| 28458 | 1423 | D1423W | E | 9.1e-06 | 5.5e-06 | 4.3e-06 | 6.300E-06 | 2.498E-06 | 4.300E-06 | 9.100E-06 |
| 28459 | 1423 | D1423Y | E | 9.4e-06 | 5.1e-06 | 1.4e-05 | 9.500E-06 | 4.451E-06 | 5.100E-06 | 1.400E-05 |
| 28460 | 1423 | D1423V | E | 5.3e-06 | 8.9e-06 | 5.2e-06 | 6.467E-06 | 2.108E-06 | 5.200E-06 | 8.900E-06 |
| 28461 | 1424 | Y1424A | E | 5.5e-06 | 6.9e-06 | 7.2e-06 | 6.533E-06 | 9.074E-07 | 5.500E-06 | 7.200E-06 |
| 28462 | 1424 | Y1424R | E | 1.3e-05 | 9.6e-06 | 7.7e-06 | 1.010E-05 | 2.685E-06 | 7.700E-06 | 1.300E-05 |
| 28463 | 1424 | Y1424N | E | 1.3e-05 | 1.1e-05 | 1.6e-05 | 1.333E-05 | 2.517E-06 | 1.100E-05 | 1.600E-05 |
| 28464 | 1424 | Y1424D | E | 7.2e-06 | 6.6e-06 | 5.7e-06 | 6.500E-06 | 7.550E-07 | 5.700E-06 | 7.200E-06 |
| 28465 | 1424 | Y1424C | E | 5.2e-06 | 7.2e-06 | 1.3e-05 | 8.467E-06 | 4.051E-06 | 5.200E-06 | 1.300E-05 |
| 28466 | 1424 | Y1424E | E | 9.3e-06 | 8.5e-06 | 5.1e-06 | 7.633E-06 | 2.230E-06 | 5.100E-06 | 9.300E-06 |
| 28467 | 1424 | Y1424Q | E | 6.2e-06 | 7.4e-06 | 7.5e-06 | 7.033E-06 | 7.234E-07 | 6.200E-06 | 7.500E-06 |
| 28468 | 1424 | Y1424G | E | 6.6e-06 | 1e-05 | 6.4e-06 | 7.667E-06 | 2.023E-06 | 6.400E-06 | 1.000E-05 |
| 28469 | 1424 | Y1424H | E | 9.1e-06 | 8.4e-06 | 9.5e-06 | 9.000E-06 | 5.568E-07 | 8.400E-06 | 9.500E-06 |
| 28470 | 1424 | Y1424I | E | 8.3e-06 | 1.1e-05 | 6.4e-06 | 8.567E-06 | 2.312E-06 | 6.400E-06 | 1.100E-05 |
| 28471 | 1424 | Y1424L | E | 1.1e-05 | 1.1e-05 | 6.3e-06 | 9.433E-06 | 2.714E-06 | 6.300E-06 | 1.100E-05 |
| 28472 | 1424 | Y1424K | E | 5.1e-06 | 7.8e-06 | 9.5e-06 | 7.467E-06 | 2.219E-06 | 5.100E-06 | 9.500E-06 |
| 28473 | 1424 | Y1424M | E | 4.5e-06 | 5e-06 | 6.5e-06 | 5.333E-06 | 1.041E-06 | 4.500E-06 | 6.500E-06 |
| 28474 | 1424 | Y1424F | E | 9.1e-06 | 7.1e-06 | 5.4e-06 | 7.200E-06 | 1.852E-06 | 5.400E-06 | 9.100E-06 |

| | | | | | | | | | | |
|---|---|---|---|---|---|---|---|---|---|---|
| 28475 | 1424 | Y1424P | E | 1.3e-05 | 6.3e-06 | 6.3e-06 | 8.533E-06 | 3.868E-06 | 6.300E-06 | 1.300E-05 |
| 28476 | 1424 | Y1424S | E | 1.3e-05 | 1e-05 | 7e-06 | 1.000E-05 | 3.000E-06 | 7.000E-06 | 1.300E-05 |
| 28477 | 1424 | Y1424T | E | 1.3e-05 | 6.9e-06 | 7.3e-06 | 9.067E-06 | 3.412E-06 | 6.900E-06 | 1.300E-05 |
| 28478 | 1424 | Y1424W | E | 4.5e-06 | 9.2e-06 | 8.8e-06 | 7.500E-06 | 2.606E-06 | 4.500E-06 | 9.200E-06 |
| 28479 | 1424 | Y1424Y | E | 9e-06 | 6.6e-06 | 1.3e-05 | 9.533E-06 | 3.233E-06 | 6.600E-06 | 1.300E-05 |
| 28480 | 1424 | Y1424V | E | 8.6e-06 | 8e-06 | 6.8e-06 | 7.800E-06 | 9.165E-07 | 6.800E-06 | 8.600E-06 |
| 28481 | 1425 | F1425A | E | 9.1e-06 | 5.9e-06 | 6.5e-06 | 7.167E-06 | 1.701E-06 | 5.900E-06 | 9.100E-06 |
| 28482 | 1425 | F1425R | E | 6.3e-06 | 1.2e-05 | 9.5e-06 | 9.267E-06 | 2.857E-06 | 6.300E-06 | 1.200E-05 |
| 28483 | 1425 | F1425N | E | 1.3e-05 | 1.1e-05 | 8.4e-06 | 1.080E-05 | 2.307E-06 | 8.400E-06 | 1.300E-05 |
| 28484 | 1425 | F1425D | E | 1.3e-05 | 1.2e-05 | 6.9e-06 | 1.063E-05 | 3.272E-06 | 6.900E-06 | 1.300E-05 |
| 28485 | 1425 | F1425C | E | 1.2e-05 | 9.2e-06 | 8.6e-06 | 9.933E-06 | 1.815E-06 | 8.600E-06 | 1.200E-05 |
| 28486 | 1425 | F1425E | E | 7.2e-06 | 6.5e-06 | 9.9e-06 | 7.867E-06 | 1.795E-06 | 6.500E-06 | 9.900E-06 |
| 28487 | 1425 | F1425Q | E | 6.9e-06 | 7.9e-06 | 5.2e-06 | 6.667E-06 | 1.365E-06 | 5.200E-06 | 7.900E-06 |
| 28488 | 1425 | F1425G | E | 1.2e-05 | 6.7e-06 | 5.4e-06 | 8.033E-06 | 3.496E-06 | 5.400E-06 | 1.200E-05 |
| 28489 | 1425 | F1425H | E | 1e-05 | 7.9e-06 | 9.2e-06 | 9.033E-06 | 1.060E-06 | 7.900E-06 | 1.000E-05 |
| 28490 | 1425 | F1425I | E | 1.4e-05 | 7e-06 | 9.6e-06 | 1.020E-05 | 3.538E-06 | 7.000E-06 | 1.400E-05 |
| 28491 | 1425 | F1425L | E | 9.1e-06 | 5.2e-06 | 7e-06 | 7.100E-06 | 1.952E-06 | 5.200E-06 | 9.100E-06 |
| 28492 | 1425 | F1425K | E | 1.2e-05 | 6.7e-06 | 9.2e-06 | 9.300E-06 | 2.651E-06 | 6.700E-06 | 1.200E-05 |
| 28493 | 1425 | F1425M | E | 1.2e-05 | 4.1e-06 | 4.6e-06 | 6.900E-06 | 4.424E-06 | 4.100E-06 | 1.200E-05 |
| 28494 | 1425 | F1425F | E | 1.2e-05 | 6.8e-06 | 9.3e-06 | 9.367E-06 | 2.601E-06 | 6.800E-06 | 1.200E-05 |
| 28495 | 1425 | F1425P | E | 1.4e-05 | 9.4e-06 | 7.3e-06 | 1.023E-05 | 3.427E-06 | 7.300E-06 | 1.400E-05 |
| 28496 | 1425 | F1425S | E | 5.8e-06 | 6e-06 | 1e-05 | 7.267E-06 | 2.369E-06 | 5.800E-06 | 1.000E-05 |
| 28497 | 1425 | F1425T | E | 4.4e-06 | 9.4e-06 | 6.5e-06 | 6.767E-06 | 2.511E-06 | 4.400E-06 | 9.400E-06 |
| 28498 | 1425 | F1425W | E | 3.9e-06 | 6e-06 | 7.2e-06 | 5.700E-06 | 1.670E-06 | 3.900E-06 | 7.200E-06 |
| 28499 | 1425 | F1425Y | E | 8.5e-06 | 6.7e-06 | 5e-06 | 6.733E-06 | 1.750E-06 | 5.000E-06 | 8.500E-06 |
| 28500 | 1425 | F1425V | E | 1.2e-05 | 8.4e-06 | 6.7e-06 | 9.033E-06 | 2.706E-06 | 6.700E-06 | 1.200E-05 |
| 28501 | 1426 | P1426A | E | 6.9e-06 | 1.4e-05 | 1e-05 | 1.030E-05 | 3.559E-06 | 6.900E-06 | 1.400E-05 |
| 28502 | 1426 | P1426R | E | 6.7e-06 | 4.1e-06 | 9.3e-06 | 6.700E-06 | 2.600E-06 | 4.100E-06 | 9.300E-06 |
| 28503 | 1426 | P1426N | E | 4.4e-06 | 8.3e-06 | 9.4e-06 | 7.367E-06 | 2.627E-06 | 4.400E-06 | 9.400E-06 |
| 28504 | 1426 | P1426D | E | 6.6e-06 | 9.2e-06 | 1.2e-05 | 9.267E-06 | 2.701E-06 | 6.600E-06 | 1.200E-05 |
| 28505 | 1426 | P1426C | E | 8e-06 | 8.8e-06 | 1.2e-05 | 9.600E-06 | 2.117E-06 | 8.000E-06 | 1.200E-05 |
| 28506 | 1426 | P1426E | E | 7.3e-06 | 1e-05 | 7.6e-06 | 8.300E-06 | 1.480E-06 | 7.300E-06 | 1.000E-05 |
| 28507 | 1426 | P1426Q | E | 5.2e-06 | 1.6e-05 | 4.9e-06 | 8.700E-06 | 6.324E-06 | 4.900E-06 | 1.600E-05 |
| 28508 | 1426 | P1426G | E | 5.8e-06 | 9.2e-06 | 9.2e-06 | 8.067E-06 | 1.963E-06 | 5.800E-06 | 9.200E-06 |
| 28509 | 1426 | P1426H | E | 5.2e-06 | 7.7e-06 | 4.9e-06 | 5.933E-06 | 1.537E-06 | 4.900E-06 | 7.700E-06 |
| 28510 | 1426 | P1426I | E | 9.3e-06 | 7.6e-06 | 5.4e-06 | 7.433E-06 | 1.955E-06 | 5.400E-06 | 9.300E-06 |
| 28511 | 1426 | P1426L | E | 1.1e-05 | 9.5e-06 | 6.4e-06 | 8.967E-06 | 2.346E-06 | 6.400E-06 | 1.100E-05 |
| 28512 | 1426 | P1426K | E | 9.1e-06 | 1e-05 | 7.1e-06 | 8.733E-06 | 1.484E-06 | 7.100E-06 | 1.000E-05 |
| 28513 | 1426 | P1426M | E | 6.9e-06 | 6.7e-06 | 9.9e-06 | 7.833E-06 | 1.793E-06 | 6.700E-06 | 9.900E-06 |
| 28514 | 1426 | P1426F | E | 8.9e-06 | 9.7e-06 | 8.2e-06 | 8.933E-06 | 7.506E-07 | 8.200E-06 | 9.700E-06 |
| 28515 | 1426 | P1426P | E | 7e-06 | 6.4e-06 | 4.7e-06 | 6.033E-06 | 1.193E-06 | 4.700E-06 | 7.000E-06 |
| 28516 | 1426 | P1426S | E | 6.5e-06 | 7.7e-06 | 8.8e-06 | 7.667E-06 | 1.150E-06 | 6.500E-06 | 8.800E-06 |
| 28517 | 1426 | P1426T | E | 8.7e-06 | 7.2e-06 | 9.8e-06 | 8.567E-06 | 1.305E-06 | 7.200E-06 | 9.800E-06 |
| 28518 | 1426 | P1426W | E | 8.6e-06 | 8.9e-06 | 1.2e-05 | 9.833E-06 | 1.882E-06 | 8.600E-06 | 1.200E-05 |
| 28519 | 1426 | P1426Y | E | 6.6e-06 | 1.1e-05 | 8.2e-06 | 8.600E-06 | 2.227E-06 | 6.600E-06 | 1.100E-05 |
| 28520 | 1426 | P1426V | E | 6.8e-06 | 9.1e-06 | 1.1e-05 | 8.967E-06 | 2.103E-06 | 6.800E-06 | 1.100E-05 |
| 28521 | 1427 | E1427A | E | 7.5e-06 | 6.6e-06 | 1.4e-05 | 9.367E-06 | 4.038E-06 | 6.600E-06 | 1.400E-05 |
| 28522 | 1427 | E1427R | E | 5.3e-06 | 6.5e-06 | 8.3e-06 | 6.700E-06 | 1.510E-06 | 5.300E-06 | 8.300E-06 |
| 28523 | 1427 | E1427N | E | 4.9e-06 | 1.3e-05 | 5.5e-06 | 7.800E-06 | 4.513E-06 | 4.900E-06 | 1.300E-05 |
| 28524 | 1427 | E1427D | E | 1.4e-05 | 6.7e-06 | 1e-05 | 1.023E-05 | 3.656E-06 | 6.700E-06 | 1.400E-05 |
| 28525 | 1427 | E1427C | E | 9.6e-06 | 6.2e-06 | 1.3e-05 | 9.600E-06 | 3.400E-06 | 6.200E-06 | 1.300E-05 |
| 28526 | 1427 | E1427E | E | 8.2e-06 | 6.3e-06 | 6.4e-06 | 6.967E-06 | 1.069E-06 | 6.300E-06 | 8.200E-06 |
| 28527 | 1427 | E1427Q | E | 9.9e-06 | 1.2e-05 | 6.3e-06 | 9.400E-06 | 2.883E-06 | 6.300E-06 | 1.200E-05 |
| 28528 | 1427 | E1427G | E | 4.7e-06 | 6.4e-06 | 9.6e-06 | 6.900E-06 | 2.488E-06 | 4.700E-06 | 9.600E-06 |
| 28529 | 1427 | E1427H | E | 8.9e-06 | 1.1e-05 | 1.1e-05 | 1.030E-05 | 1.212E-06 | 8.900E-06 | 1.100E-05 |
| 28530 | 1427 | E1427I | E | 7.8e-06 | 5e-06 | 9.6e-06 | 7.467E-06 | 2.318E-06 | 5.000E-06 | 9.600E-06 |
| 28531 | 1427 | E1427L | E | 5.1e-06 | 6.8e-06 | 8.4e-06 | 6.767E-06 | 1.650E-06 | 5.100E-06 | 8.400E-06 |
| 28532 | 1427 | E1427K | E | 7.1e-06 | 6.6e-06 | 5.7e-06 | 6.467E-06 | 7.095E-07 | 5.700E-06 | 7.100E-06 |
| 28533 | 1427 | E1427M | E | 1.8e-05 | 6.6e-06 | 1.3e-05 | 1.253E-05 | 5.714E-06 | 6.600E-06 | 1.800E-05 |
| 28534 | 1427 | E1427F | E | 5.3e-06 | 9.1e-06 | 1.7e-05 | 1.047E-05 | 5.969E-06 | 5.300E-06 | 1.700E-05 |
| 28535 | 1427 | E1427P | E | 1.2e-05 | 9.2e-06 | 1.2e-05 | 1.107E-05 | 1.617E-06 | 9.200E-06 | 1.200E-05 |
| 28536 | 1427 | E1427S | E | 8.4e-06 | 8.6e-06 | 7.5e-06 | 8.167E-06 | 5.859E-07 | 7.500E-06 | 8.600E-06 |
| 28537 | 1427 | E1427T | E | 4.1e-06 | 4.4e-06 | 5.5e-06 | 4.667E-06 | 7.371E-07 | 4.100E-06 | 5.500E-06 |
| 28538 | 1427 | E1427W | E | 5.3e-06 | 5e-06 | 1.3e-05 | 7.767E-06 | 4.535E-06 | 5.000E-06 | 1.300E-05 |
| 28539 | 1427 | E1427Y | E | 8e-06 | 5e-06 | 8.7e-06 | 7.233E-06 | 1.966E-06 | 5.000E-06 | 8.700E-06 |
| 28540 | 1427 | E1427V | E | 5.6e-06 | 6.9e-06 | 9.1e-06 | 7.200E-06 | 1.769E-06 | 5.600E-06 | 9.100E-06 |
| 28541 | 1428 | P1428A | E | 6.7e-06 | 5.4e-06 | 4.4e-06 | 5.500E-06 | 1.153E-06 | 4.400E-06 | 6.700E-06 |
| 28542 | 1428 | P1428R | E | 1.2e-05 | 8.4e-06 | 6.7e-06 | 9.033E-06 | 2.706E-06 | 6.700E-06 | 1.200E-05 |
| 28543 | 1428 | P1428N | E | 9.3e-06 | 1.2e-05 | 7.9e-06 | 9.733E-06 | 2.084E-06 | 7.900E-06 | 1.200E-05 |
| 28544 | 1428 | P1428D | E | 9.6e-06 | 6.2e-06 | 9e-06 | 8.267E-06 | 1.815E-06 | 6.200E-06 | 9.600E-06 |
| 28545 | 1428 | P1428C | E | 7.9e-06 | 9.1e-06 | 1.1e-05 | 9.333E-06 | 1.563E-06 | 7.900E-06 | 1.100E-05 |
| 28546 | 1428 | P1428E | E | 7.6e-06 | 7.2e-06 | 9.3e-06 | 8.033E-06 | 1.115E-06 | 7.200E-06 | 9.300E-06 |
| 28547 | 1428 | P1428Q | E | 1.3e-05 | 4.3e-06 | 6.3e-06 | 7.867E-06 | 4.557E-06 | 4.300E-06 | 1.300E-05 |
| 28548 | 1428 | P1428G | E | 5.5e-06 | 9.1e-06 | 4.8e-06 | 6.467E-06 | 2.307E-06 | 4.800E-06 | 9.100E-06 |
| 28549 | 1428 | P1428H | E | 9.8e-06 | 6.5e-06 | 8.6e-06 | 8.300E-06 | 1.670E-06 | 6.500E-06 | 9.800E-06 |
| 28550 | 1428 | P1428I | E | 5.9e-06 | 7.8e-06 | 8.3e-06 | 7.333E-06 | 1.266E-06 | 5.900E-06 | 8.300E-06 |
| 28551 | 1428 | P1428L | E | 9.7e-06 | 5.3e-06 | 4.5e-06 | 6.500E-06 | 2.800E-06 | 4.500E-06 | 9.700E-06 |
| 28552 | 1428 | P1428K | E | 5.4e-06 | 1e-05 | 9.9e-06 | 8.433E-06 | 2.627E-06 | 5.400E-06 | 1.000E-05 |
| 28553 | 1428 | P1428M | E | 5.9e-06 | 1e-05 | 9.2e-06 | 8.367E-06 | 2.173E-06 | 5.900E-06 | 1.000E-05 |
| 28554 | 1428 | P1428F | E | 9.6e-06 | 1.3e-05 | 5.4e-06 | 9.333E-06 | 3.807E-06 | 5.400E-06 | 1.300E-05 |
| 28555 | 1428 | P1428P | E | 5.7e-06 | 9.2e-06 | 1.5e-05 | 9.967E-06 | 4.697E-06 | 5.700E-06 | 1.500E-05 |
| 28556 | 1428 | P1428S | E | 6.4e-06 | 6.2e-06 | 1.1e-05 | 7.867E-06 | 2.715E-06 | 6.200E-06 | 1.100E-05 |
| 28557 | 1428 | P1428T | E | 8.8e-06 | 7.7e-06 | 9.5e-06 | 8.667E-06 | 9.074E-07 | 7.700E-06 | 9.500E-06 |
| 28558 | 1428 | P1428W | E | 7.1e-06 | 9.3e-06 | 8.3e-06 | 8.233E-06 | 1.102E-06 | 7.100E-06 | 9.300E-06 |
| 28559 | 1428 | P1428Y | E | 7.7e-06 | 1e-05 | 5.3e-06 | 7.667E-06 | 2.350E-06 | 5.300E-06 | 1.000E-05 |

| | | | | | | | | | | |
|---|---|---|---|---|---|---|---|---|---|---|
| 28560 | 1428 | P1428V | E | 6.1e-06 | 7.6e-06 | 9.6e-06 | 7.767E-06 | 1.756E-06 | 6.100E-06 | 9.600E-06 |
| 28561 | 1429 | V1429A | E | 9.8e-06 | 1e-05 | 8.2e-06 | 9.333E-06 | 9.866E-07 | 8.200E-06 | 1.000E-05 |
| 28562 | 1429 | V1429R | E | 8.1e-06 | 7.8e-06 | 9.3e-06 | 8.400E-06 | 7.937E-07 | 7.800E-06 | 9.300E-06 |
| 28563 | 1429 | V1429N | E | 9.2e-06 | 7e-06 | 8.1e-06 | 8.100E-06 | 1.100E-06 | 7.000E-06 | 9.200E-06 |
| 28564 | 1429 | V1429D | E | 6.4e-06 | 1.2e-05 | 3.9e-06 | 7.433E-06 | 4.148E-06 | 3.900E-06 | 1.200E-05 |
| 28565 | 1429 | V1429C | E | 6.3e-06 | 3.1e-06 | 3.9e-06 | 4.433E-06 | 1.665E-06 | 3.100E-06 | 6.300E-06 |
| 28566 | 1429 | V1429E | E | 5.7e-06 | 1e-05 | 1.1e-05 | 8.900E-06 | 2.816E-06 | 5.700E-06 | 1.100E-05 |
| 28567 | 1429 | V1429Q | E | 7.4e-06 | 9.5e-06 | 4.6e-06 | 7.167E-06 | 2.458E-06 | 4.600E-06 | 9.500E-06 |
| 28568 | 1429 | V1429G | E | 8.5e-06 | 7.3e-06 | 5.7e-06 | 7.167E-06 | 1.405E-06 | 5.700E-06 | 8.500E-06 |
| 28569 | 1429 | V1429H | E | 6.1e-06 | 9e-06 | 8.7e-06 | 7.933E-06 | 1.595E-06 | 6.100E-06 | 9.000E-06 |
| 28570 | 1429 | V1429I | E | 9.6e-06 | 6.6e-06 | 1.3e-05 | 9.733E-06 | 3.202E-06 | 6.600E-06 | 1.300E-05 |
| 28571 | 1429 | V1429L | E | 6e-06 | 4.6e-06 | 9.4e-06 | 6.667E-06 | 2.468E-06 | 4.600E-06 | 9.400E-06 |
| 28572 | 1429 | V1429K | E | 8e-06 | 6.8e-06 | 1.3e-05 | 9.267E-06 | 3.288E-06 | 6.800E-06 | 1.300E-05 |
| 28573 | 1429 | V1429M | E | 7.3e-06 | 6.4e-06 | 5.4e-06 | 6.367E-06 | 9.504E-07 | 5.400E-06 | 7.300E-06 |
| 28574 | 1429 | V1429F | E | 8e-06 | 6.4e-06 | 1.2e-05 | 8.800E-06 | 2.884E-06 | 6.400E-06 | 1.200E-05 |
| 28575 | 1429 | V1429P | E | 5e-06 | 9.8e-06 | 8.7e-06 | 7.833E-06 | 2.515E-06 | 5.000E-06 | 9.800E-06 |
| 28576 | 1429 | V1429S | E | 7.4e-06 | 1.1e-05 | 8.3e-06 | 8.900E-06 | 1.873E-06 | 7.400E-06 | 1.100E-05 |
| 28577 | 1429 | V1429T | E | 7.8e-06 | 7.3e-06 | 7.2e-06 | 7.433E-06 | 3.215E-07 | 7.200E-06 | 7.800E-06 |
| 28578 | 1429 | V1429W | E | 1.4e-05 | 5.4e-06 | 8.8e-06 | 9.400E-06 | 4.331E-06 | 5.400E-06 | 1.400E-05 |
| 28579 | 1429 | V1429Y | E | 7.8e-06 | 1e-05 | 1.2e-05 | 9.933E-06 | 2.101E-06 | 7.800E-06 | 1.200E-05 |
| 28580 | 1429 | V1429V | E | 6.3e-06 | 1e-05 | 6.5e-06 | 7.600E-06 | 2.081E-06 | 6.300E-06 | 1.000E-05 |
| 28581 | 1430 | T1430A | E | 8.8e-06 | 4.7e-06 | 7.4e-06 | 6.967E-06 | 2.084E-06 | 4.700E-06 | 8.800E-06 |
| 28582 | 1430 | T1430R | E | 6.1e-06 | 5.9e-06 | 1.4e-05 | 8.667E-06 | 4.620E-06 | 5.900E-06 | 1.400E-05 |
| 28583 | 1430 | T1430N | E | 6.6e-06 | 6.6e-06 | 9e-06 | 7.400E-06 | 1.386E-06 | 6.600E-06 | 9.000E-06 |
| 28584 | 1430 | T1430D | E | 9.4e-06 | 7e-06 | 1.5e-05 | 1.047E-05 | 4.105E-06 | 7.000E-06 | 1.500E-05 |
| 28585 | 1430 | T1430C | E | 6.2e-06 | 5.7e-06 | 5.8e-06 | 5.900E-06 | 2.646E-07 | 5.700E-06 | 6.200E-06 |
| 28586 | 1430 | T1430E | E | 9e-06 | 6.7e-06 | 8.2e-06 | 7.967E-06 | 1.168E-06 | 6.700E-06 | 9.000E-06 |
| 28587 | 1430 | T1430Q | E | 6e-06 | 9e-06 | 7.8e-06 | 7.600E-06 | 1.510E-06 | 6.000E-06 | 9.000E-06 |
| 28588 | 1430 | T1430G | E | 4e-06 | 6.8e-06 | 7.4e-06 | 6.067E-06 | 1.815E-06 | 4.000E-06 | 7.400E-06 |
| 28589 | 1430 | T1430H | E | 6.9e-06 | 1e-05 | 4.7e-06 | 7.200E-06 | 2.663E-06 | 4.700E-06 | 1.000E-05 |
| 28590 | 1430 | T1430I | E | 9.9e-06 | 1e-05 | 4.3e-06 | 8.067E-06 | 3.262E-06 | 4.300E-06 | 1.000E-05 |
| 28591 | 1430 | T1430L | E | 1e-05 | 6.7e-06 | 7e-06 | 7.900E-06 | 1.825E-06 | 6.700E-06 | 1.000E-05 |
| 28592 | 1430 | T1430K | E | 6.3e-06 | 7.8e-06 | 1.1e-05 | 8.367E-06 | 2.401E-06 | 6.300E-06 | 1.100E-05 |
| 28593 | 1430 | T1430M | E | 8.8e-06 | 5.8e-06 | 7.1e-06 | 7.233E-06 | 1.504E-06 | 5.800E-06 | 8.800E-06 |
| 28594 | 1430 | T1430F | E | 7.4e-06 | 1e-05 | 7.3e-06 | 8.233E-06 | 1.531E-06 | 7.300E-06 | 1.000E-05 |
| 28595 | 1430 | T1430P | E | 9.4e-06 | 9.4e-06 | 1e-05 | 9.600E-06 | 3.464E-07 | 9.400E-06 | 1.000E-05 |
| 28596 | 1430 | T1430S | E | 7.8e-06 | 5e-06 | 8.2e-06 | 7.000E-06 | 1.744E-06 | 5.000E-06 | 8.200E-06 |
| 28597 | 1430 | T1430T | E | 8.4e-06 | 7.4e-06 | 1e-05 | 8.600E-06 | 1.311E-06 | 7.400E-06 | 1.000E-05 |
| 28598 | 1430 | T1430W | E | 9.3e-06 | 8.4e-06 | 6.7e-06 | 8.133E-06 | 1.320E-06 | 6.700E-06 | 9.300E-06 |
| 28599 | 1430 | T1430Y | E | 6.2e-06 | 6.1e-06 | 6.6e-06 | 6.300E-06 | 2.646E-07 | 6.100E-06 | 6.600E-06 |
| 28600 | 1430 | T1430V | E | 8.5e-06 | 9.3e-06 | 1.2e-05 | 9.933E-06 | 1.834E-06 | 8.500E-06 | 1.200E-05 |
| 28601 | 1431 | V1431A | E | 5.6e-06 | 8.4e-06 | 6.8e-06 | 6.933E-06 | 1.405E-06 | 5.600E-06 | 8.400E-06 |
| 28602 | 1431 | V1431R | E | 7.2e-06 | 5.7e-06 | 8.8e-06 | 7.233E-06 | 1.550E-06 | 5.700E-06 | 8.800E-06 |
| 28603 | 1431 | V1431N | E | 9.4e-06 | 3.1e-06 | 5.7e-06 | 6.067E-06 | 3.166E-06 | 3.100E-06 | 9.400E-06 |
| 28604 | 1431 | V1431D | E | 6.3e-06 | 9.1e-06 | 7.8e-06 | 7.733E-06 | 1.401E-06 | 6.300E-06 | 9.100E-06 |
| 28605 | 1431 | V1431C | E | 1.2e-05 | 6.4e-06 | 5.4e-06 | 7.933E-06 | 3.557E-06 | 5.400E-06 | 1.200E-05 |
| 28606 | 1431 | V1431E | E | 6.4e-06 | 6.4e-06 | 6.5e-06 | 6.433E-06 | 5.774E-08 | 6.400E-06 | 6.500E-06 |
| 28607 | 1431 | V1431Q | E | 8.5e-06 | 6.7e-06 | 9.6e-06 | 8.267E-06 | 1.464E-06 | 6.700E-06 | 9.600E-06 |
| 28608 | 1431 | V1431G | E | 4.7e-06 | 4.9e-06 | 7.8e-06 | 5.800E-06 | 1.735E-06 | 4.700E-06 | 7.800E-06 |
| 28609 | 1431 | V1431H | E | 6.8e-06 | 3.3e-06 | 7e-06 | 5.700E-06 | 2.081E-06 | 3.300E-06 | 7.000E-06 |
| 28610 | 1431 | V1431I | E | 8.2e-06 | 1e-05 | 8.9e-06 | 9.033E-06 | 9.074E-07 | 8.200E-06 | 1.000E-05 |
| 28611 | 1431 | V1431L | E | 8.1e-06 | 1.4e-05 | 4.7e-06 | 8.933E-06 | 4.706E-06 | 4.700E-06 | 1.400E-05 |
| 28612 | 1431 | V1431K | E | 8.9e-06 | 1.4e-05 | 8.3e-06 | 1.040E-05 | 3.132E-06 | 8.300E-06 | 1.400E-05 |
| 28613 | 1431 | V1431M | E | 9.1e-06 | 8.9e-06 | 6.5e-06 | 8.167E-06 | 1.447E-06 | 6.500E-06 | 9.100E-06 |
| 28614 | 1431 | V1431F | E | 7.7e-06 | 6.9e-06 | 4.8e-06 | 6.467E-06 | 1.498E-06 | 4.800E-06 | 7.700E-06 |
| 28615 | 1431 | V1431P | E | 1.1e-05 | 4.5e-06 | 1.1e-05 | 8.833E-06 | 3.753E-06 | 4.500E-06 | 1.100E-05 |
| 28616 | 1431 | V1431S | E | 1e-05 | 7.5e-06 | 7.9e-06 | 8.467E-06 | 1.343E-06 | 7.500E-06 | 1.000E-05 |
| 28617 | 1431 | V1431T | E | 8.5e-06 | 6.1e-06 | 1.2e-05 | 8.867E-06 | 2.967E-06 | 6.100E-06 | 1.200E-05 |
| 28618 | 1431 | V1431W | E | 6.6e-06 | 5.9e-06 | 6.5e-06 | 6.333E-06 | 3.786E-07 | 5.900E-06 | 6.600E-06 |
| 28619 | 1431 | V1431Y | E | 8.9e-06 | 5.7e-06 | 8.5e-06 | 7.700E-06 | 1.744E-06 | 5.700E-06 | 8.900E-06 |
| 28620 | 1431 | V1431V | E | 7e-06 | 9.3e-06 | 8.3e-06 | 8.200E-06 | 1.153E-06 | 7.000E-06 | 9.300E-06 |
| 28621 | 1432 | S1432A | E | 9.3e-06 | 9.5e-06 | 6.2e-06 | 8.333E-06 | 1.850E-06 | 6.200E-06 | 9.500E-06 |
| 28622 | 1432 | S1432R | E | 3.5e-06 | 7.8e-06 | 9.3e-06 | 6.867E-06 | 3.011E-06 | 3.500E-06 | 9.300E-06 |
| 28623 | 1432 | S1432N | E | 1.3e-05 | 5.2e-06 | 6.1e-06 | 8.100E-06 | 4.267E-06 | 5.200E-06 | 1.300E-05 |
| 28624 | 1432 | S1432D | E | 9.3e-06 | 4.8e-06 | 5.3e-06 | 6.467E-06 | 2.466E-06 | 4.800E-06 | 9.300E-06 |
| 28625 | 1432 | S1432C | E | 1.2e-05 | 8.3e-06 | 1.6e-05 | 1.210E-05 | 3.851E-06 | 8.300E-06 | 1.600E-05 |
| 28626 | 1432 | S1432E | E | 6.2e-06 | 1.1e-05 | 4.6e-06 | 7.267E-06 | 3.331E-06 | 4.600E-06 | 1.100E-05 |
| 28627 | 1432 | S1432Q | E | 5.3e-06 | 6e-06 | 5.5e-06 | 5.600E-06 | 3.606E-07 | 5.300E-06 | 6.000E-06 |
| 28628 | 1432 | S1432G | E | 7.9e-06 | 8e-06 | 8.5e-06 | 8.133E-06 | 3.215E-07 | 7.900E-06 | 8.500E-06 |
| 28629 | 1432 | S1432H | E | 7.4e-06 | 6.4e-06 | 5.1e-06 | 6.300E-06 | 1.153E-06 | 5.100E-06 | 7.400E-06 |
| 28630 | 1432 | S1432I | E | 8.5e-06 | 1.4e-05 | 1.5e-05 | 1.250E-05 | 3.500E-06 | 8.500E-06 | 1.500E-05 |
| 28631 | 1432 | S1432L | E | 7.6e-06 | 6e-06 | 6.4e-06 | 6.667E-06 | 8.327E-07 | 6.000E-06 | 7.600E-06 |
| 28632 | 1432 | S1432K | E | 8.3e-06 | 1.2e-05 | 8e-06 | 9.433E-06 | 2.228E-06 | 8.000E-06 | 1.200E-05 |
| 28633 | 1432 | S1432M | E | 1e-05 | 9.4e-06 | 6.9e-06 | 8.767E-06 | 1.644E-06 | 6.900E-06 | 1.000E-05 |
| 28634 | 1432 | S1432F | E | 7.8e-06 | 7.6e-06 | 1.4e-05 | 9.800E-06 | 3.639E-06 | 7.600E-06 | 1.400E-05 |
| 28635 | 1432 | S1432P | E | 4.9e-06 | 8e-06 | 6.9e-06 | 6.600E-06 | 1.572E-06 | 4.900E-06 | 8.000E-06 |
| 28636 | 1432 | S1432S | E | 7.6e-06 | 6e-06 | 8e-06 | 7.200E-06 | 1.058E-06 | 6.000E-06 | 8.000E-06 |
| 28637 | 1432 | S1432T | E | 9.9e-06 | 6e-06 | 6.8e-06 | 7.567E-06 | 2.060E-06 | 6.000E-06 | 9.900E-06 |
| 28638 | 1432 | S1432W | E | 4.8e-06 | 5.1e-06 | 6.6e-06 | 5.500E-06 | 9.644E-07 | 4.800E-06 | 6.600E-06 |
| 28639 | 1432 | S1432Y | E | 7.1e-06 | 7.9e-06 | 1e-05 | 8.333E-06 | 1.498E-06 | 7.100E-06 | 1.000E-05 |
| 28640 | 1432 | S1432V | E | 5.8e-06 | 6e-06 | 8.1e-06 | 6.633E-06 | 1.274E-06 | 5.800E-06 | 8.100E-06 |
| 28641 | 1433 | W1433A | E | 9e-06 | 6.2e-06 | 9.2e-06 | 8.133E-06 | 1.677E-06 | 6.200E-06 | 9.200E-06 |
| 28642 | 1433 | W1433R | E | 5.9e-06 | 1.1e-05 | 6.8e-06 | 7.900E-06 | 2.722E-06 | 5.900E-06 | 1.100E-05 |
| 28643 | 1433 | W1433N | E | 7.8e-06 | 8.2e-06 | 8.6e-06 | 8.200E-06 | 4.000E-07 | 7.800E-06 | 8.600E-06 |
| 28644 | 1433 | W1433D | E | 1.4e-05 | 6.7e-06 | 7.2e-06 | 9.300E-06 | 4.078E-06 | 6.700E-06 | 1.400E-05 |

| | | | | | | | | | | |
|---|---|---|---|---|---|---|---|---|---|---|
| 28645 | 1433 | W1433C | E | 4.3e-06 | 6.7e-06 | 1.1e-05 | 7.333E-06 | 3.395E-06 | 4.300E-06 | 1.100E-05 |
| 28646 | 1433 | W1433E | E | 1.1e-05 | 7.4e-06 | 8.3e-06 | 8.900E-06 | 1.873E-06 | 7.400E-06 | 1.100E-05 |
| 28647 | 1433 | W1433Q | E | 1.1e-05 | 1.1e-05 | 6.2e-06 | 9.400E-06 | 2.771E-06 | 6.200E-06 | 1.100E-05 |
| 28648 | 1433 | W1433G | E | 6.2e-06 | 9.8e-06 | 8.5e-06 | 8.167E-06 | 1.823E-06 | 6.200E-06 | 9.800E-06 |
| 28649 | 1433 | W1433H | E | 8e-06 | 8.9e-06 | 7.2e-06 | 8.033E-06 | 8.505E-07 | 7.200E-06 | 8.900E-06 |
| 28650 | 1433 | W1433I | E | 1e-05 | 7.1e-06 | 1e-05 | 9.033E-06 | 1.674E-06 | 7.100E-06 | 1.000E-05 |
| 28651 | 1433 | W1433L | E | 8.1e-06 | 7.9e-06 | 1.4e-05 | 1.000E-05 | 3.466E-06 | 7.900E-06 | 1.400E-05 |
| 28652 | 1433 | W1433K | E | 1e-05 | 6.8e-06 | 1e-05 | 8.933E-06 | 1.848E-06 | 6.800E-06 | 1.000E-05 |
| 28653 | 1433 | W1433M | E | 1.1e-05 | 8.3e-06 | 9.9e-06 | 9.733E-06 | 1.358E-06 | 8.300E-06 | 1.100E-05 |
| 28654 | 1433 | W1433F | E | 8.4e-06 | 7.5e-06 | 6.3e-06 | 7.400E-06 | 1.054E-06 | 6.300E-06 | 8.400E-06 |
| 28655 | 1433 | W1433P | E | 6.2e-06 | 9.5e-06 | 8.7e-06 | 8.133E-06 | 1.721E-06 | 6.200E-06 | 9.500E-06 |
| 28656 | 1433 | W1433S | E | 5.9e-06 | 1.4e-05 | 5.9e-06 | 8.600E-06 | 4.677E-06 | 5.900E-06 | 1.400E-05 |
| 28657 | 1433 | W1433T | E | 1.1e-05 | 5.1e-06 | 1.3e-05 | 9.700E-06 | 4.107E-06 | 5.100E-06 | 1.300E-05 |
| 28658 | 1433 | W1433W | E | 6.3e-06 | 8.4e-06 | 5.7e-06 | 6.800E-06 | 1.418E-06 | 5.700E-06 | 8.400E-06 |
| 28659 | 1433 | W1433Y | E | 7.2e-06 | 9.8e-06 | 7.8e-06 | 8.267E-06 | 1.361E-06 | 7.200E-06 | 9.800E-06 |
| 28660 | 1433 | W1433V | E | 7.6e-06 | 5.8e-06 | 9.6e-06 | 7.667E-06 | 1.901E-06 | 5.800E-06 | 9.600E-06 |
| 28661 | 1434 | N1434A | E | 1e-05 | 1.2e-05 | 1e-05 | 1.067E-05 | 1.155E-06 | 1.000E-05 | 1.200E-05 |
| 28662 | 1434 | N1434R | E | 8.3e-06 | 4.4e-06 | 4.3e-06 | 5.667E-06 | 2.281E-06 | 4.300E-06 | 8.300E-06 |
| 28663 | 1434 | N1434N | E | 9.8e-06 | 4.5e-06 | 7.1e-06 | 7.133E-06 | 2.650E-06 | 4.500E-06 | 9.800E-06 |
| 28664 | 1434 | N1434D | E | 1e-05 | 8.9e-06 | 7.3e-06 | 8.733E-06 | 1.358E-06 | 7.300E-06 | 1.000E-05 |
| 28665 | 1434 | N1434C | E | 7.2e-06 | 4.8e-06 | 9.5e-06 | 7.167E-06 | 2.350E-06 | 4.800E-06 | 9.500E-06 |
| 28666 | 1434 | N1434E | E | 7.1e-06 | 9.6e-06 | 6.5e-06 | 7.733E-06 | 1.644E-06 | 6.500E-06 | 9.600E-06 |
| 28667 | 1434 | N1434Q | E | 1.3e-05 | 1e-05 | 9.3e-06 | 1.077E-05 | 1.966E-06 | 9.300E-06 | 1.300E-05 |
| 28668 | 1434 | N1434G | E | 1e-05 | 6.7e-06 | 6.3e-06 | 7.667E-06 | 2.031E-06 | 6.300E-06 | 1.000E-05 |
| 28669 | 1434 | N1434H | E | 6e-06 | 9.3e-06 | 9.6e-06 | 8.300E-06 | 1.997E-06 | 6.000E-06 | 9.600E-06 |
| 28670 | 1434 | N1434I | E | 1.1e-05 | 8.9e-06 | 1.1e-05 | 1.030E-05 | 1.212E-06 | 8.900E-06 | 1.100E-05 |
| 28671 | 1434 | N1434L | E | 5.8e-06 | 8.8e-06 | 6.3e-06 | 6.967E-06 | 1.607E-06 | 5.800E-06 | 8.800E-06 |
| 28672 | 1434 | N1434K | E | 1e-05 | 8.8e-06 | 5.1e-06 | 7.967E-06 | 2.554E-06 | 5.100E-06 | 1.000E-05 |
| 28673 | 1434 | N1434M | E | 5.8e-06 | 6.9e-06 | 1.1e-05 | 7.900E-06 | 2.740E-06 | 5.800E-06 | 1.100E-05 |
| 28674 | 1434 | N1434F | E | 9.4e-06 | 5.9e-06 | 3.7e-06 | 6.333E-06 | 2.875E-06 | 3.700E-06 | 9.400E-06 |
| 28675 | 1434 | N1434P | E | 9.9e-06 | 7.8e-06 | 7.4e-06 | 8.367E-06 | 1.343E-06 | 7.400E-06 | 9.900E-06 |
| 28676 | 1434 | N1434S | E | 7.6e-06 | 5e-06 | 8.7e-06 | 7.100E-06 | 1.900E-06 | 5.000E-06 | 8.700E-06 |
| 28677 | 1434 | N1434T | E | 7.1e-06 | 6.2e-06 | 1e-05 | 7.767E-06 | 1.986E-06 | 6.200E-06 | 1.000E-05 |
| 28678 | 1434 | N1434W | E | 6e-06 | 1.4e-05 | 8.2e-06 | 9.400E-06 | 4.133E-06 | 6.000E-06 | 1.400E-05 |
| 28679 | 1434 | N1434Y | E | 6.7e-06 | 1e-05 | 5.9e-06 | 7.533E-06 | 2.173E-06 | 5.900E-06 | 1.000E-05 |
| 28680 | 1434 | N1434V | E | 9.7e-06 | 6.9e-06 | 5.4e-06 | 7.333E-06 | 2.183E-06 | 5.400E-06 | 9.700E-06 |
| 28681 | 1435 | S1435A | E | 8.4e-06 | 4.6e-06 | 9.9e-06 | 7.633E-06 | 2.732E-06 | 4.600E-06 | 9.900E-06 |
| 28682 | 1435 | S1435R | E | 7.3e-06 | 6.8e-06 | 6.7e-06 | 6.933E-06 | 3.215E-07 | 6.700E-06 | 7.300E-06 |
| 28683 | 1435 | S1435N | E | 1.8e-05 | 9.8e-06 | 4.2e-06 | 1.067E-05 | 6.941E-06 | 4.200E-06 | 1.800E-05 |
| 28684 | 1435 | S1435D | E | 9.3e-06 | 6.5e-06 | 8e-06 | 7.933E-06 | 1.401E-06 | 6.500E-06 | 9.300E-06 |
| 28685 | 1435 | S1435C | E | 1.5e-05 | 1e-05 | 9.9e-06 | 1.163E-05 | 2.916E-06 | 9.900E-06 | 1.500E-05 |
| 28686 | 1435 | S1435E | E | 7.7e-06 | 7.4e-06 | 6.5e-06 | 7.200E-06 | 6.245E-07 | 6.500E-06 | 7.700E-06 |
| 28687 | 1435 | S1435Q | E | 4.5e-06 | 5.7e-06 | 1.2e-05 | 7.400E-06 | 4.029E-06 | 4.500E-06 | 1.200E-05 |
| 28688 | 1435 | S1435G | E | 6.5e-06 | 9.8e-06 | 1.2e-05 | 9.433E-06 | 2.768E-06 | 6.500E-06 | 1.200E-05 |
| 28689 | 1435 | S1435H | E | 9.9e-06 | 5.7e-06 | 9.1e-06 | 8.233E-06 | 2.230E-06 | 5.700E-06 | 9.900E-06 |
| 28690 | 1435 | S1435I | E | 7.3e-06 | 4.7e-06 | 1.4e-05 | 8.667E-06 | 4.798E-06 | 4.700E-06 | 1.400E-05 |
| 28691 | 1435 | S1435L | E | 6.3e-06 | 1.3e-05 | 9.4e-06 | 9.567E-06 | 3.353E-06 | 6.300E-06 | 1.300E-05 |
| 28692 | 1435 | S1435K | E | 5.8e-06 | 9.7e-06 | 4.6e-06 | 6.700E-06 | 2.666E-06 | 4.600E-06 | 9.700E-06 |
| 28693 | 1435 | S1435M | E | 1.1e-05 | 7.9e-06 | 1.1e-05 | 9.967E-06 | 1.790E-06 | 7.900E-06 | 1.100E-05 |
| 28694 | 1435 | S1435F | E | 9.4e-06 | 5.5e-06 | 6.9e-06 | 7.267E-06 | 1.976E-06 | 5.500E-06 | 9.400E-06 |
| 28695 | 1435 | S1435P | E | 9.5e-06 | 1.2e-05 | 5.7e-06 | 9.067E-06 | 3.172E-06 | 5.700E-06 | 1.200E-05 |
| 28696 | 1435 | S1435S | E | 6.4e-06 | 9.8e-06 | 7e-06 | 7.733E-06 | 1.815E-06 | 6.400E-06 | 9.800E-06 |
| 28697 | 1435 | S1435T | E | 1.1e-05 | 4.5e-06 | 7.9e-06 | 7.800E-06 | 3.251E-06 | 4.500E-06 | 1.100E-05 |
| 28698 | 1435 | S1435W | E | 7.5e-06 | 1.3e-05 | 6e-06 | 8.833E-06 | 3.686E-06 | 6.000E-06 | 1.300E-05 |
| 28699 | 1435 | S1435Y | E | 6.8e-06 | 7.3e-06 | 1.2e-05 | 8.700E-06 | 2.869E-06 | 6.800E-06 | 1.200E-05 |
| 28700 | 1435 | S1435V | E | 6.5e-06 | 9.6e-06 | 8.5e-06 | 8.200E-06 | 1.572E-06 | 6.500E-06 | 9.600E-06 |
| 28701 | 1436 | G1436A | E | 7.2e-06 | 9.8e-06 | 6.6e-06 | 7.867E-06 | 1.701E-06 | 6.600E-06 | 9.800E-06 |
| 28702 | 1436 | G1436R | E | 2.6e-06 | 6.6e-06 | 1.1e-05 | 6.733E-06 | 4.202E-06 | 2.600E-06 | 1.100E-05 |
| 28703 | 1436 | G1436N | E | 5.2e-06 | 1.4e-05 | 8.4e-06 | 9.200E-06 | 4.454E-06 | 5.200E-06 | 1.400E-05 |
| 28704 | 1436 | G1436D | E | 7.1e-06 | 1.2e-05 | 8.3e-06 | 9.133E-06 | 2.554E-06 | 7.100E-06 | 1.200E-05 |
| 28705 | 1436 | G1436C | E | 9.6e-06 | 9.6e-06 | 8.3e-06 | 9.167E-06 | 7.506E-07 | 8.300E-06 | 9.600E-06 |
| 28706 | 1436 | G1436E | E | 6.6e-06 | 8.1e-06 | 1.4e-05 | 9.567E-06 | 3.912E-06 | 6.600E-06 | 1.400E-05 |
| 28707 | 1436 | G1436Q | E | 9.6e-06 | 9e-06 | 9.4e-06 | 9.333E-06 | 3.055E-07 | 9.000E-06 | 9.600E-06 |
| 28708 | 1436 | G1436G | E | 1.1e-05 | 9.1e-06 | 7.5e-06 | 9.200E-06 | 1.752E-06 | 7.500E-06 | 1.100E-05 |
| 28709 | 1436 | G1436H | E | 1e-05 | 9.4e-06 | 9.2e-06 | 9.533E-06 | 4.163E-07 | 9.200E-06 | 1.000E-05 |
| 28710 | 1436 | G1436I | E | 5.1e-06 | 6.6e-06 | 9.1e-06 | 6.933E-06 | 2.021E-06 | 5.100E-06 | 9.100E-06 |
| 28711 | 1436 | G1436L | E | 1.1e-05 | 9.3e-06 | 6e-06 | 8.767E-06 | 2.542E-06 | 6.000E-06 | 1.100E-05 |
| 28712 | 1436 | G1436K | E | 2.7e-06 | 4.4e-06 | 7.5e-06 | 4.867E-06 | 2.434E-06 | 2.700E-06 | 7.500E-06 |
| 28713 | 1436 | G1436M | E | 1e-05 | 1.1e-05 | 1.2e-05 | 1.100E-05 | 1.000E-06 | 1.000E-05 | 1.200E-05 |
| 28714 | 1436 | G1436F | E | 1.1e-05 | 6.3e-06 | 1.1e-05 | 9.433E-06 | 2.714E-06 | 6.300E-06 | 1.100E-05 |
| 28715 | 1436 | G1436P | E | 1.2e-05 | 9.8e-06 | 6.8e-06 | 9.533E-06 | 2.610E-06 | 6.800E-06 | 1.200E-05 |
| 28716 | 1436 | G1436S | E | 9.3e-06 | 7.5e-06 | 9.1e-06 | 8.633E-06 | 9.866E-07 | 7.500E-06 | 9.300E-06 |
| 28717 | 1436 | G1436T | E | 9.5e-06 | 4.9e-06 | 9.7e-06 | 8.033E-06 | 2.715E-06 | 4.900E-06 | 9.700E-06 |
| 28718 | 1436 | G1436W | E | 8.6e-06 | 1.2e-05 | 7.3e-06 | 9.300E-06 | 2.427E-06 | 7.300E-06 | 1.200E-05 |
| 28719 | 1436 | G1436Y | E | 5.5e-06 | 4.4e-06 | 6.3e-06 | 5.400E-06 | 9.539E-07 | 4.400E-06 | 6.300E-06 |
| 28720 | 1436 | G1436V | E | 5.1e-06 | 1.3e-05 | 8e-06 | 8.700E-06 | 3.996E-06 | 5.100E-06 | 1.300E-05 |
| 28721 | 1437 | A1437A | E | 1.4e-05 | 1.1e-05 | 6.3e-06 | 1.043E-05 | 3.881E-06 | 6.300E-06 | 1.400E-05 |
| 28722 | 1437 | A1437R | E | 7.3e-06 | 1.1e-05 | 5.8e-06 | 8.033E-06 | 2.676E-06 | 5.800E-06 | 1.100E-05 |
| 28723 | 1437 | A1437N | E | 5.7e-06 | 1.3e-05 | 6.8e-06 | 8.500E-06 | 3.936E-06 | 5.700E-06 | 1.300E-05 |
| 28724 | 1437 | A1437D | E | 6.2e-06 | 1.3e-05 | 8.5e-06 | 9.233E-06 | 3.459E-06 | 6.200E-06 | 1.300E-05 |
| 28725 | 1437 | A1437C | E | 5e-06 | 7.3e-06 | 5.5e-06 | 5.933E-06 | 1.210E-06 | 5.000E-06 | 7.300E-06 |
| 28726 | 1437 | A1437E | E | 6e-06 | 6.8e-06 | 8.7e-06 | 7.167E-06 | 1.387E-06 | 6.000E-06 | 8.700E-06 |
| 28727 | 1437 | A1437Q | E | 9.8e-06 | 4.6e-06 | 8.9e-06 | 7.767E-06 | 2.779E-06 | 4.600E-06 | 9.800E-06 |
| 28728 | 1437 | A1437G | E | 6.3e-06 | 8.9e-06 | 7.4e-06 | 7.533E-06 | 1.305E-06 | 6.300E-06 | 8.900E-06 |
| 28729 | 1437 | A1437H | E | 6.6e-06 | 6.3e-06 | 4.8e-06 | 5.900E-06 | 9.644E-07 | 4.800E-06 | 6.600E-06 |

| | | | | | | | | | | |
|---|---|---|---|---|---|---|---|---|---|---|
| 28730 | 1437 | A1437I | E | 1.1e-05 | 4.9e-06 | 7.3e-06 | 7.733E-06 | 3.073E-06 | 4.900E-06 | 1.100E-05 |
| 28731 | 1437 | A1437L | E | 7.8e-06 | 7.2e-06 | 6.4e-06 | 7.133E-06 | 7.024E-07 | 6.400E-06 | 7.800E-06 |
| 28732 | 1437 | A1437K | E | 6.7e-06 | 9.4e-06 | 1.9e-05 | 1.170E-05 | 6.465E-06 | 6.700E-06 | 1.900E-05 |
| 28733 | 1437 | A1437M | E | 6.2e-06 | 5.7e-06 | 8.4e-06 | 6.767E-06 | 1.436E-06 | 5.700E-06 | 8.400E-06 |
| 28734 | 1437 | A1437F | E | 3.2e-06 | 8.8e-06 | 1.3e-05 | 8.333E-06 | 4.917E-06 | 3.200E-06 | 1.300E-05 |
| 28735 | 1437 | A1437P | E | 1.3e-05 | 1.4e-05 | 4.8e-06 | 1.060E-05 | 5.048E-06 | 4.800E-06 | 1.400E-05 |
| 28736 | 1437 | A1437S | E | 4.1e-06 | 9.8e-06 | 7.6e-06 | 7.167E-06 | 2.875E-06 | 4.100E-06 | 9.800E-06 |
| 28737 | 1437 | A1437T | E | 8.1e-06 | 6.6e-06 | 9.6e-06 | 8.100E-06 | 1.500E-06 | 6.600E-06 | 9.600E-06 |
| 28738 | 1437 | A1437W | E | 5.7e-06 | 1e-05 | 5.8e-06 | 7.167E-06 | 2.454E-06 | 5.700E-06 | 1.000E-05 |
| 28739 | 1437 | A1437Y | E | 5.6e-06 | 9.4e-06 | 1e-05 | 8.333E-06 | 2.386E-06 | 5.600E-06 | 1.000E-05 |
| 28740 | 1437 | A1437V | E | 1e-05 | 5.7e-06 | 6.1e-06 | 7.267E-06 | 2.376E-06 | 5.700E-06 | 1.000E-05 |
| 28741 | 1438 | L1438A | E | 4.9e-06 | 1e-05 | 7.1e-06 | 7.333E-06 | 2.558E-06 | 4.900E-06 | 1.000E-05 |
| 28742 | 1438 | L1438R | E | 5.8e-06 | 6.1e-06 | 6.4e-06 | 6.100E-06 | 3.000E-07 | 5.800E-06 | 6.400E-06 |
| 28743 | 1438 | L1438N | E | 7.3e-06 | 7.8e-06 | 7.3e-06 | 7.467E-06 | 2.887E-07 | 7.300E-06 | 7.800E-06 |
| 28744 | 1438 | L1438D | E | 8.1e-06 | 7.3e-06 | 1e-05 | 8.467E-06 | 1.387E-06 | 7.300E-06 | 1.000E-05 |
| 28745 | 1438 | L1438C | E | 8.5e-06 | 1.7e-05 | 1.1e-05 | 1.217E-05 | 4.368E-06 | 8.500E-06 | 1.700E-05 |
| 28746 | 1438 | L1438E | E | 1.2e-05 | 6e-06 | 1e-05 | 9.333E-06 | 3.055E-06 | 6.000E-06 | 1.200E-05 |
| 28747 | 1438 | L1438Q | E | 5.4e-06 | 9.1e-06 | 9.8e-06 | 8.100E-06 | 2.364E-06 | 5.400E-06 | 9.800E-06 |
| 28748 | 1438 | L1438G | E | 8e-06 | 5.5e-06 | 7.8e-06 | 7.100E-06 | 1.389E-06 | 5.500E-06 | 8.000E-06 |
| 28749 | 1438 | L1438H | E | 7.8e-06 | 4.7e-06 | 8.7e-06 | 7.067E-06 | 2.098E-06 | 4.700E-06 | 8.700E-06 |
| 28750 | 1438 | L1438I | E | 8.5e-06 | 8.1e-06 | 4.5e-06 | 7.033E-06 | 2.203E-06 | 4.500E-06 | 8.500E-06 |
| 28751 | 1438 | L1438L | E | 5e-06 | 7.6e-06 | 7.4e-06 | 6.667E-06 | 1.447E-06 | 5.000E-06 | 7.600E-06 |
| 28752 | 1438 | L1438K | E | 7.8e-06 | 1.2e-05 | 9.1e-06 | 9.633E-06 | 2.150E-06 | 7.800E-06 | 1.200E-05 |
| 28753 | 1438 | L1438M | E | 5.5e-06 | 4.8e-06 | 7.1e-06 | 5.800E-06 | 1.179E-06 | 4.800E-06 | 7.100E-06 |
| 28754 | 1438 | L1438F | E | 7e-06 | 7.8e-06 | 1.3e-05 | 9.267E-06 | 3.258E-06 | 7.000E-06 | 1.300E-05 |
| 28755 | 1438 | L1438P | E | 6.3e-06 | 5.6e-06 | 6.3e-06 | 6.067E-06 | 4.041E-07 | 5.600E-06 | 6.300E-06 |
| 28756 | 1438 | L1438S | E | 9.4e-06 | 2.1e-05 | 6.4e-06 | 1.227E-05 | 7.711E-06 | 6.400E-06 | 2.100E-05 |
| 28757 | 1438 | L1438T | E | 7.2e-06 | 6e-06 | 8.5e-06 | 7.233E-06 | 1.250E-06 | 6.000E-06 | 8.500E-06 |
| 28758 | 1438 | L1438W | E | 6.7e-06 | 7.6e-06 | 5.1e-06 | 6.467E-06 | 1.266E-06 | 5.100E-06 | 7.600E-06 |
| 28759 | 1438 | L1438Y | E | 7.6e-06 | 5.4e-06 | 9.6e-06 | 7.533E-06 | 2.101E-06 | 5.400E-06 | 9.600E-06 |
| 28760 | 1438 | L1438V | E | 7.2e-06 | 7.1e-06 | 9.5e-06 | 7.933E-06 | 1.358E-06 | 7.100E-06 | 9.500E-06 |
| 28761 | 1439 | T1439A | E | 1.3e-05 | 5.9e-06 | 7.4e-06 | 8.767E-06 | 3.742E-06 | 5.900E-06 | 1.300E-05 |
| 28762 | 1439 | T1439R | E | 6.3e-06 | 4.8e-06 | 5.9e-06 | 5.667E-06 | 7.767E-07 | 4.800E-06 | 6.300E-06 |
| 28763 | 1439 | T1439N | E | 7.4e-06 | 9.7e-06 | 6.9e-06 | 8.000E-06 | 1.493E-06 | 6.900E-06 | 9.700E-06 |
| 28764 | 1439 | T1439D | E | 5.4e-06 | 1.1e-05 | 7.1e-06 | 7.833E-06 | 2.871E-06 | 5.400E-06 | 1.100E-05 |
| 28765 | 1439 | T1439C | E | 7.9e-06 | 9.6e-06 | 9.1e-06 | 8.867E-06 | 8.737E-07 | 7.900E-06 | 9.600E-06 |
| 28766 | 1439 | T1439E | E | 9.4e-06 | 6.8e-06 | 9.2e-06 | 8.467E-06 | 1.447E-06 | 6.800E-06 | 9.400E-06 |
| 28767 | 1439 | T1439Q | E | 8.7e-06 | 6.1e-06 | 7.1e-06 | 7.300E-06 | 1.311E-06 | 6.100E-06 | 8.700E-06 |
| 28768 | 1439 | T1439G | E | 1e-05 | 9.1e-06 | 1e-05 | 9.700E-06 | 5.196E-07 | 9.100E-06 | 1.000E-05 |
| 28769 | 1439 | T1439H | E | 7.5e-06 | 4e-06 | 6.6e-06 | 6.033E-06 | 1.818E-06 | 4.000E-06 | 7.500E-06 |
| 28770 | 1439 | T1439I | E | 6.1e-06 | 4.8e-06 | 4.6e-06 | 5.167E-06 | 8.145E-07 | 4.600E-06 | 6.100E-06 |
| 28771 | 1439 | T1439L | E | 9.8e-06 | 5.3e-06 | 1e-05 | 8.367E-06 | 2.658E-06 | 5.300E-06 | 1.000E-05 |
| 28772 | 1439 | T1439K | E | 1.1e-05 | 7.5e-06 | 9.5e-06 | 9.333E-06 | 1.756E-06 | 7.500E-06 | 1.100E-05 |
| 28773 | 1439 | T1439M | E | 8.7e-06 | 6.2e-06 | 6.5e-06 | 7.133E-06 | 1.365E-06 | 6.200E-06 | 8.700E-06 |
| 28774 | 1439 | T1439F | E | 6.6e-06 | 1.3e-05 | 6.7e-06 | 8.767E-06 | 3.667E-06 | 6.600E-06 | 1.300E-05 |
| 28775 | 1439 | T1439P | E | 6.8e-06 | 6.6e-06 | 8.8e-06 | 7.400E-06 | 1.217E-06 | 6.600E-06 | 8.800E-06 |
| 28776 | 1439 | T1439S | E | 8.3e-06 | 1.1e-05 | 1.1e-05 | 1.010E-05 | 1.559E-06 | 8.300E-06 | 1.100E-05 |
| 28777 | 1439 | T1439T | E | 9.6e-06 | 1.4e-05 | 1.5e-05 | 1.287E-05 | 2.873E-06 | 9.600E-06 | 1.500E-05 |
| 28778 | 1439 | T1439W | E | 9.3e-06 | 7.9e-06 | 4.8e-06 | 7.333E-06 | 2.303E-06 | 4.800E-06 | 9.300E-06 |
| 28779 | 1439 | T1439Y | E | 7.5e-06 | 8.1e-06 | 1.1e-05 | 8.867E-06 | 1.872E-06 | 7.500E-06 | 1.100E-05 |
| 28780 | 1439 | T1439V | E | 9.2e-06 | 7.7e-06 | 9e-06 | 8.633E-06 | 8.145E-07 | 7.700E-06 | 9.200E-06 |
| 28781 | 1440 | S1440A | E | 8.2e-06 | 1.3e-05 | 1.2e-05 | 1.107E-05 | 2.532E-06 | 8.200E-06 | 1.300E-05 |
| 28782 | 1440 | S1440R | E | 6.2e-06 | 1e-05 | 9e-06 | 8.400E-06 | 1.970E-06 | 6.200E-06 | 1.000E-05 |
| 28783 | 1440 | S1440N | E | 9.2e-06 | 5e-06 | 6.8e-06 | 7.000E-06 | 2.107E-06 | 5.000E-06 | 9.200E-06 |
| 28784 | 1440 | S1440D | E | 9.4e-06 | 1.1e-05 | 5.1e-06 | 8.500E-06 | 3.051E-06 | 5.100E-06 | 1.100E-05 |
| 28785 | 1440 | S1440C | E | 5.7e-06 | 1.1e-05 | 9.3e-06 | 8.667E-06 | 2.706E-06 | 5.700E-06 | 1.100E-05 |
| 28786 | 1440 | S1440E | E | 6.8e-06 | 6e-06 | 6.1e-06 | 6.300E-06 | 4.359E-07 | 6.000E-06 | 6.800E-06 |
| 28787 | 1440 | S1440Q | E | 4.3e-06 | 9.7e-06 | 7.4e-06 | 7.133E-06 | 2.710E-06 | 4.300E-06 | 9.700E-06 |
| 28788 | 1440 | S1440G | E | 7.2e-06 | 1.1e-05 | 1.1e-05 | 9.733E-06 | 2.194E-06 | 7.200E-06 | 1.100E-05 |
| 28789 | 1440 | S1440H | E | 7.1e-06 | 6.1e-06 | 9.3e-06 | 7.500E-06 | 1.637E-06 | 6.100E-06 | 9.300E-06 |
| 28790 | 1440 | S1440I | E | 4e-06 | 1e-05 | 2e-05 | 1.133E-05 | 8.083E-06 | 4.000E-06 | 2.000E-05 |
| 28791 | 1440 | S1440L | E | 7.7e-06 | 9.6e-06 | 1.1e-05 | 9.433E-06 | 1.656E-06 | 7.700E-06 | 1.100E-05 |
| 28792 | 1440 | S1440K | E | 8.9e-06 | 6.2e-06 | 9.9e-06 | 8.333E-06 | 1.914E-06 | 6.200E-06 | 9.900E-06 |
| 28793 | 1440 | S1440M | E | 7.9e-06 | 7e-06 | 9.3e-06 | 8.067E-06 | 1.159E-06 | 7.000E-06 | 9.300E-06 |
| 28794 | 1440 | S1440F | E | 7.4e-06 | 7e-06 | 5e-06 | 6.467E-06 | 1.286E-06 | 5.000E-06 | 7.400E-06 |
| 28795 | 1440 | S1440P | E | 1.8e-05 | 9.9e-06 | 5.6e-06 | 1.117E-05 | 6.296E-06 | 5.600E-06 | 1.800E-05 |
| 28796 | 1440 | S1440S | E | 7.8e-06 | 1.1e-05 | 4e-06 | 7.600E-06 | 3.504E-06 | 4.000E-06 | 1.100E-05 |
| 28797 | 1440 | S1440T | E | 5.8e-06 | 6.3e-06 | 7.4e-06 | 6.500E-06 | 8.185E-07 | 5.800E-06 | 7.400E-06 |
| 28798 | 1440 | S1440W | E | 1e-05 | 7.9e-06 | 4.1e-06 | 7.333E-06 | 2.991E-06 | 4.100E-06 | 1.000E-05 |
| 28799 | 1440 | S1440Y | E | 5.2e-06 | 9.6e-06 | 8.3e-06 | 7.700E-06 | 2.261E-06 | 5.200E-06 | 9.600E-06 |
| 28800 | 1440 | S1440V | E | 9.3e-06 | 1.5e-05 | 8e-06 | 1.077E-05 | 3.723E-06 | 8.000E-06 | 1.500E-05 |
| 28801 | 1441 | G1441A | E | 5.2e-06 | 9.1e-06 | 9.1e-06 | 7.800E-06 | 2.252E-06 | 5.200E-06 | 9.100E-06 |
| 28802 | 1441 | G1441R | E | 1e-05 | 1.3e-05 | 9.8e-06 | 1.093E-05 | 1.793E-06 | 9.800E-06 | 1.300E-05 |
| 28803 | 1441 | G1441N | E | 1.3e-05 | 1.2e-05 | 6.1e-06 | 1.037E-05 | 3.729E-06 | 6.100E-06 | 1.300E-05 |
| 28804 | 1441 | G1441D | E | 8.8e-06 | 8.9e-06 | 9.3e-06 | 9.000E-06 | 2.646E-07 | 8.800E-06 | 9.300E-06 |
| 28805 | 1441 | G1441C | E | 8.7e-06 | 9.3e-06 | 8.3e-06 | 8.767E-06 | 5.033E-07 | 8.300E-06 | 9.300E-06 |
| 28806 | 1441 | G1441E | E | 5.2e-06 | 3.6e-06 | 6.6e-06 | 5.133E-06 | 1.501E-06 | 3.600E-06 | 6.600E-06 |
| 28807 | 1441 | G1441Q | E | 9.1e-06 | 1.1e-05 | 7.3e-06 | 9.133E-06 | 1.850E-06 | 7.300E-06 | 1.100E-05 |
| 28808 | 1441 | G1441G | E | 9.6e-06 | 7.8e-06 | 7e-06 | 8.133E-06 | 1.332E-06 | 7.000E-06 | 9.600E-06 |
| 28809 | 1441 | G1441H | E | 8e-06 | 7.9e-06 | 7e-06 | 7.633E-06 | 5.508E-07 | 7.000E-06 | 8.000E-06 |
| 28810 | 1441 | G1441I | E | 7e-06 | 6.9e-06 | 6.1e-06 | 6.667E-06 | 4.933E-07 | 6.100E-06 | 7.000E-06 |
| 28811 | 1441 | G1441L | E | 5.7e-06 | 1.3e-05 | 1.3e-05 | 1.057E-05 | 4.215E-06 | 5.700E-06 | 1.300E-05 |
| 28812 | 1441 | G1441K | E | 6.7e-06 | 8.8e-06 | 6.4e-06 | 7.300E-06 | 1.308E-06 | 6.400E-06 | 8.800E-06 |
| 28813 | 1441 | G1441M | E | 9.1e-06 | 5.1e-06 | 1.2e-05 | 8.733E-06 | 3.465E-06 | 5.100E-06 | 1.200E-05 |
| 28814 | 1441 | G1441F | E | 1.3e-05 | 2.3e-05 | 7.5e-06 | 1.450E-05 | 7.858E-06 | 7.500E-06 | 2.300E-05 |

| | | | | | | | | | | |
|---|---|---|---|---|---|---|---|---|---|---|
| 28815 | 1441 | G1441P | E | 1.3e-05 | 9.3e-06 | 6.4e-06 | 9.567E-06 | 3.308E-06 | 6.400E-06 | 1.300E-05 |
| 28816 | 1441 | G1441S | E | 8.8e-06 | 7.1e-06 | 4.8e-06 | 6.900E-06 | 2.007E-06 | 4.800E-06 | 8.800E-06 |
| 28817 | 1441 | G1441T | E | 6.9e-06 | 5.8e-06 | 1.2e-05 | 8.233E-06 | 3.308E-06 | 5.800E-06 | 1.200E-05 |
| 28818 | 1441 | G1441W | E | 6.2e-06 | 4.3e-06 | 4.9e-06 | 5.133E-06 | 9.713E-07 | 4.300E-06 | 6.200E-06 |
| 28819 | 1441 | G1441Y | E | 8.6e-06 | 6.2e-06 | 1e-05 | 8.267E-06 | 1.922E-06 | 6.200E-06 | 1.000E-05 |
| 28820 | 1441 | G1441V | E | 1e-05 | 9.6e-06 | 8.5e-06 | 9.367E-06 | 7.767E-07 | 8.500E-06 | 1.000E-05 |
| 28821 | 1442 | V1442A | E | 5.4e-06 | 7.3e-06 | 7.2e-06 | 6.633E-06 | 1.069E-06 | 5.400E-06 | 7.300E-06 |
| 28822 | 1442 | V1442R | E | 9e-06 | 8.2e-06 | 1.1e-05 | 9.400E-06 | 1.442E-06 | 8.200E-06 | 1.100E-05 |
| 28823 | 1442 | V1442N | E | 7.9e-06 | 5.5e-06 | 1.2e-05 | 8.467E-06 | 3.287E-06 | 5.500E-06 | 1.200E-05 |
| 28824 | 1442 | V1442D | E | 6.3e-06 | 1.3e-05 | 1.1e-05 | 1.010E-05 | 3.439E-06 | 6.300E-06 | 1.300E-05 |
| 28825 | 1442 | V1442C | E | 6.3e-06 | 5.4e-06 | 5.3e-06 | 5.667E-06 | 5.508E-07 | 5.300E-06 | 6.300E-06 |
| 28826 | 1442 | V1442E | E | 1e-05 | 9.3e-06 | 1.2e-05 | 1.043E-05 | 1.401E-06 | 9.300E-06 | 1.200E-05 |
| 28827 | 1442 | V1442Q | E | 6.8e-06 | 7e-06 | 1.2e-05 | 8.600E-06 | 2.946E-06 | 6.800E-06 | 1.200E-05 |
| 28828 | 1442 | V1442G | E | 9.7e-06 | 6.4e-06 | 1.2e-05 | 9.367E-06 | 2.815E-06 | 6.400E-06 | 1.200E-05 |
| 28829 | 1442 | V1442H | E | 4.4e-06 | 7.4e-06 | 4.3e-06 | 5.367E-06 | 1.762E-06 | 4.300E-06 | 7.400E-06 |
| 28830 | 1442 | V1442I | E | 8.1e-06 | 8.9e-06 | 8.5e-06 | 8.500E-06 | 4.000E-07 | 8.100E-06 | 8.900E-06 |
| 28831 | 1442 | V1442L | E | 1.1e-05 | 5.1e-06 | 9.4e-06 | 8.500E-06 | 3.051E-06 | 5.100E-06 | 1.100E-05 |
| 28832 | 1442 | V1442K | E | 6.6e-06 | 9.8e-06 | 5.8e-06 | 7.400E-06 | 2.117E-06 | 5.800E-06 | 9.800E-06 |
| 28833 | 1442 | V1442M | E | 5.7e-06 | 1e-05 | 6.4e-06 | 7.367E-06 | 2.307E-06 | 5.700E-06 | 1.000E-05 |
| 28834 | 1442 | V1442F | E | 4.3e-06 | 3.5e-06 | 1.5e-05 | 7.600E-06 | 6.421E-06 | 3.500E-06 | 1.500E-05 |
| 28835 | 1442 | V1442P | E | 1e-05 | 7.6e-06 | 1e-05 | 9.200E-06 | 1.386E-06 | 7.600E-06 | 1.000E-05 |
| 28836 | 1442 | V1442S | E | 4.3e-06 | 8.8e-06 | 1.2e-05 | 8.367E-06 | 3.868E-06 | 4.300E-06 | 1.200E-05 |
| 28837 | 1442 | V1442T | E | 7e-06 | 9.4e-06 | 7.8e-06 | 8.067E-06 | 1.222E-06 | 7.000E-06 | 9.400E-06 |
| 28838 | 1442 | V1442W | E | 1.1e-05 | 4.9e-06 | 4.5e-06 | 6.800E-06 | 3.643E-06 | 4.500E-06 | 1.100E-05 |
| 28839 | 1442 | V1442Y | E | 6.8e-06 | 8.3e-06 | 1.2e-05 | 9.033E-06 | 2.676E-06 | 6.800E-06 | 1.200E-05 |
| 28840 | 1442 | V1442V | E | 6.1e-06 | 9.2e-06 | 7.6e-06 | 7.633E-06 | 1.550E-06 | 6.100E-06 | 9.200E-06 |
| 28841 | 1443 | H1443A | E | 1e-05 | 1.2e-05 | 1.1e-05 | 1.100E-05 | 1.000E-06 | 1.000E-05 | 1.200E-05 |
| 28842 | 1443 | H1443R | E | 1.3e-05 | 4.8e-06 | 7.5e-06 | 8.433E-06 | 4.179E-06 | 4.800E-06 | 1.300E-05 |
| 28843 | 1443 | H1443N | E | 9.3e-06 | 5.4e-06 | 4.7e-06 | 6.467E-06 | 2.479E-06 | 4.700E-06 | 9.300E-06 |
| 28844 | 1443 | H1443D | E | 1.1e-05 | 6.2e-06 | 6e-06 | 7.733E-06 | 2.831E-06 | 6.000E-06 | 1.100E-05 |
| 28845 | 1443 | H1443C | E | 9e-06 | 5.5e-06 | 1.2e-05 | 8.833E-06 | 3.253E-06 | 5.500E-06 | 1.200E-05 |
| 28846 | 1443 | H1443E | E | 1e-05 | 7.7e-06 | 8.1e-06 | 8.600E-06 | 1.229E-06 | 7.700E-06 | 1.000E-05 |
| 28847 | 1443 | H1443Q | E | 9.6e-06 | 1.1e-05 | 7.8e-06 | 9.467E-06 | 1.604E-06 | 7.800E-06 | 1.100E-05 |
| 28848 | 1443 | H1443G | E | 8.3e-06 | 9.5e-06 | 9.8e-06 | 9.200E-06 | 7.937E-07 | 8.300E-06 | 9.800E-06 |
| 28849 | 1443 | H1443H | E | 5.5e-06 | 1e-05 | 5.5e-06 | 7.000E-06 | 2.598E-06 | 5.500E-06 | 1.000E-05 |
| 28850 | 1443 | H1443I | E | 8.5e-06 | 6.4e-06 | 9.5e-06 | 8.133E-06 | 1.582E-06 | 6.400E-06 | 9.500E-06 |
| 28851 | 1443 | H1443L | E | 7.6e-06 | 6.8e-06 | 4.4e-06 | 6.267E-06 | 1.665E-06 | 4.400E-06 | 7.600E-06 |
| 28852 | 1443 | H1443K | E | 9.7e-06 | 6.9e-06 | 9.2e-06 | 8.600E-06 | 1.493E-06 | 6.900E-06 | 9.700E-06 |
| 28853 | 1443 | H1443M | E | 9.6e-06 | 7e-06 | 8.6e-06 | 8.400E-06 | 1.311E-06 | 7.000E-06 | 9.600E-06 |
| 28854 | 1443 | H1443F | E | 9.9e-06 | 9e-06 | 7.5e-06 | 8.800E-06 | 1.212E-06 | 7.500E-06 | 9.900E-06 |
| 28855 | 1443 | H1443P | E | 8.5e-06 | 6.1e-06 | 1.4e-05 | 9.533E-06 | 4.050E-06 | 6.100E-06 | 1.400E-05 |
| 28856 | 1443 | H1443S | E | 4.8e-06 | 4.6e-06 | 7.7e-06 | 5.700E-06 | 1.735E-06 | 4.600E-06 | 7.700E-06 |
| 28857 | 1443 | H1443T | E | 6.1e-06 | 2.7e-06 | 8.9e-06 | 5.900E-06 | 3.105E-06 | 2.700E-06 | 8.900E-06 |
| 28858 | 1443 | H1443W | E | 9.9e-06 | 4.6e-06 | 1.2e-05 | 8.833E-06 | 3.814E-06 | 4.600E-06 | 1.200E-05 |
| 28859 | 1443 | H1443Y | E | 6.8e-06 | 6.9e-06 | 5.7e-06 | 6.467E-06 | 6.658E-07 | 5.700E-06 | 6.900E-06 |
| 28860 | 1443 | H1443V | E | 6.9e-06 | 4e-06 | 1e-05 | 6.967E-06 | 3.001E-06 | 4.000E-06 | 1.000E-05 |
| 28861 | 1444 | T1444A | E | 9.9e-06 | 1.4e-05 | 8.9e-06 | 1.093E-05 | 2.702E-06 | 8.900E-06 | 1.400E-05 |
| 28862 | 1444 | T1444R | E | 1e-05 | 9.8e-06 | 7.3e-06 | 9.033E-06 | 1.504E-06 | 7.300E-06 | 1.000E-05 |
| 28863 | 1444 | T1444N | E | 1.3e-05 | 8.8e-06 | 9.1e-06 | 1.030E-05 | 2.343E-06 | 8.800E-06 | 1.300E-05 |
| 28864 | 1444 | T1444D | E | 1.1e-05 | 8.7e-06 | 1e-05 | 9.900E-06 | 1.153E-06 | 8.700E-06 | 1.100E-05 |
| 28865 | 1444 | T1444C | E | 7.8e-06 | 3.7e-06 | 1e-05 | 7.167E-06 | 3.197E-06 | 3.700E-06 | 1.000E-05 |
| 28866 | 1444 | T1444E | E | 6.3e-06 | 7.7e-06 | 7.3e-06 | 7.100E-06 | 7.211E-07 | 6.300E-06 | 7.700E-06 |
| 28867 | 1444 | T1444Q | E | 7e-06 | 6.5e-06 | 5.6e-06 | 6.367E-06 | 7.095E-07 | 5.600E-06 | 7.000E-06 |
| 28868 | 1444 | T1444G | E | 7.6e-06 | 1.3e-05 | 9.7e-06 | 1.010E-05 | 2.722E-06 | 7.600E-06 | 1.300E-05 |
| 28869 | 1444 | T1444H | E | 6.3e-06 | 8.8e-06 | 6.4e-06 | 7.167E-06 | 1.415E-06 | 6.300E-06 | 8.800E-06 |
| 28870 | 1444 | T1444I | E | 4.6e-06 | 1e-05 | 7.2e-06 | 7.267E-06 | 2.701E-06 | 4.600E-06 | 1.000E-05 |
| 28871 | 1444 | T1444L | E | 5e-06 | 1.1e-05 | 1.3e-05 | 9.667E-06 | 4.163E-06 | 5.000E-06 | 1.300E-05 |
| 28872 | 1444 | T1444K | E | 3.3e-06 | 4.7e-06 | 1.2e-05 | 6.667E-06 | 4.672E-06 | 3.300E-06 | 1.200E-05 |
| 28873 | 1444 | T1444M | E | 1e-05 | 7.1e-06 | 9.3e-06 | 8.800E-06 | 1.513E-06 | 7.100E-06 | 1.000E-05 |
| 28874 | 1444 | T1444F | E | 9.6e-06 | 6.3e-06 | 5e-06 | 6.967E-06 | 2.371E-06 | 5.000E-06 | 9.600E-06 |
| 28875 | 1444 | T1444P | E | 1.3e-05 | 7e-06 | 9.6e-06 | 9.867E-06 | 3.009E-06 | 7.000E-06 | 1.300E-05 |
| 28876 | 1444 | T1444S | E | 7.2e-06 | 7.3e-06 | 1.8e-05 | 1.083E-05 | 6.207E-06 | 7.200E-06 | 1.800E-05 |
| 28877 | 1444 | T1444T | E | 5.6e-06 | 1e-05 | 6.7e-06 | 7.433E-06 | 2.290E-06 | 5.600E-06 | 1.000E-05 |
| 28878 | 1444 | T1444W | E | 7.8e-06 | 5.2e-06 | 8.5e-06 | 7.167E-06 | 1.739E-06 | 5.200E-06 | 8.500E-06 |
| 28879 | 1444 | T1444Y | E | 1e-05 | 1e-05 | 1.1e-05 | 1.033E-05 | 5.774E-07 | 1.000E-05 | 1.100E-05 |
| 28880 | 1444 | T1444V | E | 8.9e-06 | 6.8e-06 | 1.3e-05 | 9.567E-06 | 3.153E-06 | 6.800E-06 | 1.300E-05 |
| 28881 | 1445 | F1445A | E | 1.4e-05 | 1.2e-05 | 1.1e-05 | 1.233E-05 | 1.528E-06 | 1.100E-05 | 1.400E-05 |
| 28882 | 1445 | F1445R | E | 6.8e-06 | 4.5e-06 | 6.5e-06 | 5.933E-06 | 1.250E-06 | 4.500E-06 | 6.800E-06 |
| 28883 | 1445 | F1445N | E | 8.3e-06 | 1.1e-05 | 8.3e-06 | 9.200E-06 | 1.559E-06 | 8.300E-06 | 1.100E-05 |
| 28884 | 1445 | F1445D | E | 6.5e-06 | 6.7e-06 | 1e-05 | 7.733E-06 | 1.966E-06 | 6.500E-06 | 1.000E-05 |
| 28885 | 1445 | F1445C | E | 6.1e-06 | 4.8e-06 | 5.2e-06 | 5.367E-06 | 6.658E-07 | 4.800E-06 | 6.100E-06 |
| 28886 | 1445 | F1445E | E | 6e-06 | 6.9e-06 | 7.4e-06 | 6.767E-06 | 7.095E-07 | 6.000E-06 | 7.400E-06 |
| 28887 | 1445 | F1445Q | E | 5.5e-06 | 8.8e-06 | 9.5e-06 | 7.933E-06 | 2.136E-06 | 5.500E-06 | 9.500E-06 |
| 28888 | 1445 | F1445G | E | 6.1e-06 | 6.5e-06 | 9.3e-06 | 7.300E-06 | 1.744E-06 | 6.100E-06 | 9.300E-06 |
| 28889 | 1445 | F1445H | E | 6.3e-06 | 1.4e-05 | 7.3e-06 | 9.200E-06 | 4.187E-06 | 6.300E-06 | 1.400E-05 |
| 28890 | 1445 | F1445I | E | 6.5e-06 | 8.6e-06 | 7.3e-06 | 7.467E-06 | 1.060E-06 | 6.500E-06 | 8.600E-06 |
| 28891 | 1445 | F1445L | E | 7.5e-06 | 1.3e-05 | 8.5e-06 | 9.667E-06 | 2.930E-06 | 7.500E-06 | 1.300E-05 |
| 28892 | 1445 | F1445K | E | 4.5e-06 | 8.8e-06 | 6.5e-06 | 6.600E-06 | 2.152E-06 | 4.500E-06 | 8.800E-06 |
| 28893 | 1445 | F1445M | E | 4.7e-06 | 7.6e-06 | 7.8e-06 | 6.700E-06 | 1.735E-06 | 4.700E-06 | 7.800E-06 |
| 28894 | 1445 | F1445F | E | 6.9e-06 | 6.2e-06 | 9.7e-06 | 7.600E-06 | 1.852E-06 | 6.200E-06 | 9.700E-06 |
| 28895 | 1445 | F1445P | E | 4.8e-06 | 7.9e-06 | 9.2e-06 | 7.300E-06 | 2.261E-06 | 4.800E-06 | 9.200E-06 |
| 28896 | 1445 | F1445S | E | 7.8e-06 | 7e-06 | 8.7e-06 | 7.833E-06 | 8.505E-07 | 7.000E-06 | 8.700E-06 |
| 28897 | 1445 | F1445T | E | 8.5e-06 | 9.9e-06 | 7e-06 | 8.467E-06 | 1.450E-06 | 7.000E-06 | 9.900E-06 |
| 28898 | 1445 | F1445W | E | 6.4e-06 | 6.7e-06 | 9.2e-06 | 7.433E-06 | 1.537E-06 | 6.400E-06 | 9.200E-06 |
| 28899 | 1445 | F1445Y | E | 6.5e-06 | 1.7e-05 | 5.7e-06 | 9.733E-06 | 6.306E-06 | 5.700E-06 | 1.700E-05 |

| | | | | | | | | | | |
|---|---|---|---|---|---|---|---|---|---|---|
| 28900 | 1445 | F1445V | E | 9.6e-06 | 1.2e-05 | 7.8e-06 | 9.800E-06 | 2.107E-06 | 7.800E-06 | 1.200E-05 |
| 28901 | 1446 | P1446A | E | 1e-05 | 5.8e-06 | 7.2e-06 | 7.667E-06 | 2.139E-06 | 5.800E-06 | 1.000E-05 |
| 28902 | 1446 | P1446R | E | 7e-06 | 4.6e-06 | 8.8e-06 | 6.800E-06 | 2.107E-06 | 4.600E-06 | 8.800E-06 |
| 28903 | 1446 | P1446N | E | 1.2e-05 | 8e-06 | 1e-05 | 1.000E-05 | 2.000E-06 | 8.000E-06 | 1.200E-05 |
| 28904 | 1446 | P1446D | E | 6.8e-06 | 4.8e-06 | 9.9e-06 | 7.167E-06 | 2.570E-06 | 4.800E-06 | 9.900E-06 |
| 28905 | 1446 | P1446C | E | 9.4e-06 | 8.7e-06 | 4.6e-06 | 7.567E-06 | 2.593E-06 | 4.600E-06 | 9.400E-06 |
| 28906 | 1446 | P1446E | E | 1.2e-05 | 1.2e-05 | 6.5e-06 | 1.017E-05 | 3.175E-06 | 6.500E-06 | 1.200E-05 |
| 28907 | 1446 | P1446Q | E | 8.9e-06 | 6.8e-06 | 6.9e-06 | 7.533E-06 | 1.185E-06 | 6.800E-06 | 8.900E-06 |
| 28908 | 1446 | P1446G | E | 8.5e-06 | 9.3e-06 | 1.2e-05 | 9.933E-06 | 1.834E-06 | 8.500E-06 | 1.200E-05 |
| 28909 | 1446 | P1446H | E | 4.4e-06 | 9.1e-06 | 9.2e-06 | 7.567E-06 | 2.743E-06 | 4.400E-06 | 9.200E-06 |
| 28910 | 1446 | P1446I | E | 1.2e-05 | 9e-06 | 5.9e-06 | 8.967E-06 | 3.050E-06 | 5.900E-06 | 1.200E-05 |
| 28911 | 1446 | P1446L | E | 7e-06 | 7.9e-06 | 1.1e-05 | 8.633E-06 | 2.098E-06 | 7.000E-06 | 1.100E-05 |
| 28912 | 1446 | P1446K | E | 9e-06 | 9e-06 | 8.1e-06 | 8.700E-06 | 5.196E-07 | 8.100E-06 | 9.000E-06 |
| 28913 | 1446 | P1446M | E | 6.4e-06 | 9.5e-06 | 9.3e-06 | 8.400E-06 | 1.735E-06 | 6.400E-06 | 9.500E-06 |
| 28914 | 1446 | P1446F | E | 1e-05 | 9.3e-06 | 5.6e-06 | 8.300E-06 | 2.364E-06 | 5.600E-06 | 1.000E-05 |
| 28915 | 1446 | P1446P | E | 4.8e-06 | 4.8e-06 | 9.7e-06 | 6.433E-06 | 2.829E-06 | 4.800E-06 | 9.700E-06 |
| 28916 | 1446 | P1446S | E | 7e-06 | 9.7e-06 | 5.3e-06 | 7.333E-06 | 2.219E-06 | 5.300E-06 | 9.700E-06 |
| 28917 | 1446 | P1446T | E | 4.1e-06 | 1.1e-05 | 4.5e-06 | 6.533E-06 | 3.873E-06 | 4.100E-06 | 1.100E-05 |
| 28918 | 1446 | P1446W | E | 6.1e-06 | 9e-06 | 7.8e-06 | 7.633E-06 | 1.457E-06 | 6.100E-06 | 9.000E-06 |
| 28919 | 1446 | P1446Y | E | 6.8e-06 | 7.7e-06 | 4.5e-06 | 6.333E-06 | 1.650E-06 | 4.500E-06 | 7.700E-06 |
| 28920 | 1446 | P1446V | E | 8.2e-06 | 5.6e-06 | 4.5e-06 | 6.100E-06 | 1.900E-06 | 4.500E-06 | 8.200E-06 |
| 28921 | 1447 | A1447A | E | 8.4e-06 | 7.2e-06 | 6.3e-06 | 7.300E-06 | 1.054E-06 | 6.300E-06 | 8.400E-06 |
| 28922 | 1447 | A1447R | E | 7.9e-06 | 7.8e-06 | 6.7e-06 | 7.467E-06 | 6.658E-07 | 6.700E-06 | 7.900E-06 |
| 28923 | 1447 | A1447N | E | 9.3e-06 | 6e-06 | 9.9e-06 | 8.400E-06 | 2.100E-06 | 6.000E-06 | 9.900E-06 |
| 28924 | 1447 | A1447D | E | 5.8e-06 | 9e-06 | 9.9e-06 | 8.233E-06 | 2.155E-06 | 5.800E-06 | 9.900E-06 |
| 28925 | 1447 | A1447C | E | 9.6e-06 | 6.3e-06 | 1e-05 | 8.633E-06 | 2.031E-06 | 6.300E-06 | 1.000E-05 |
| 28926 | 1447 | A1447E | E | 7.6e-06 | 7.1e-06 | 4.8e-06 | 6.500E-06 | 1.493E-06 | 4.800E-06 | 7.600E-06 |
| 28927 | 1447 | A1447Q | E | 1.2e-05 | 1.2e-05 | 6.8e-06 | 1.027E-05 | 3.002E-06 | 6.800E-06 | 1.200E-05 |
| 28928 | 1447 | A1447G | E | 1e-05 | 1.4e-05 | 1.2e-05 | 1.200E-05 | 2.000E-06 | 1.000E-05 | 1.400E-05 |
| 28929 | 1447 | A1447H | E | 8e-06 | 7.3e-06 | 6.5e-06 | 7.267E-06 | 7.506E-07 | 6.500E-06 | 8.000E-06 |
| 28930 | 1447 | A1447I | E | 1.2e-05 | 5.5e-06 | 9.6e-06 | 9.033E-06 | 3.287E-06 | 5.500E-06 | 1.200E-05 |
| 28931 | 1447 | A1447L | E | 1e-05 | 8.8e-06 | 8e-06 | 8.933E-06 | 1.007E-06 | 8.000E-06 | 1.000E-05 |
| 28932 | 1447 | A1447K | E | 6.6e-06 | 6.1e-06 | 6.2e-06 | 6.300E-06 | 2.646E-07 | 6.100E-06 | 6.600E-06 |
| 28933 | 1447 | A1447M | E | 5.7e-06 | 1e-05 | 1.1e-05 | 8.900E-06 | 2.816E-06 | 5.700E-06 | 1.100E-05 |
| 28934 | 1447 | A1447F | E | 6.6e-06 | 8.2e-06 | 7.4e-06 | 7.400E-06 | 8.000E-07 | 6.600E-06 | 8.200E-06 |
| 28935 | 1447 | A1447P | E | 5.8e-06 | 7.1e-06 | 9.5e-06 | 7.467E-06 | 1.877E-06 | 5.800E-06 | 9.500E-06 |
| 28936 | 1447 | A1447S | E | 6.3e-06 | 1.4e-05 | 9.4e-06 | 9.900E-06 | 3.874E-06 | 6.300E-06 | 1.400E-05 |
| 28937 | 1447 | A1447T | E | 1.1e-05 | 1.3e-05 | 8.8e-06 | 1.093E-05 | 2.101E-06 | 8.800E-06 | 1.300E-05 |
| 28938 | 1447 | A1447W | E | 7.3e-06 | 1e-05 | 7e-06 | 8.100E-06 | 1.652E-06 | 7.000E-06 | 1.000E-05 |
| 28939 | 1447 | A1447Y | E | 1.7e-05 | 7.2e-06 | 5.2e-06 | 9.800E-06 | 6.315E-06 | 5.200E-06 | 1.700E-05 |
| 28940 | 1447 | A1447V | E | 7.3e-06 | 5.7e-06 | 8.3e-06 | 7.100E-06 | 1.311E-06 | 5.700E-06 | 8.300E-06 |
| 28941 | 1448 | V1448A | E | 6e-06 | 8.9e-06 | 9.1e-06 | 8.000E-06 | 1.735E-06 | 6.000E-06 | 9.100E-06 |
| 28942 | 1448 | V1448R | E | 8.7e-06 | 6e-06 | 5e-06 | 6.567E-06 | 1.914E-06 | 5.000E-06 | 8.700E-06 |
| 28943 | 1448 | V1448N | E | 8.5e-06 | 9.8e-06 | 7.9e-06 | 8.733E-06 | 9.713E-07 | 7.900E-06 | 9.800E-06 |
| 28944 | 1448 | V1448D | E | 7.5e-06 | 1.3e-05 | 6.6e-06 | 9.033E-06 | 3.465E-06 | 6.600E-06 | 1.300E-05 |
| 28945 | 1448 | V1448C | E | 7.9e-06 | 9.1e-06 | 9.5e-06 | 8.833E-06 | 8.327E-07 | 7.900E-06 | 9.500E-06 |
| 28946 | 1448 | V1448E | E | 1.3e-05 | 7.1e-06 | 1e-05 | 1.003E-05 | 2.950E-06 | 7.100E-06 | 1.300E-05 |
| 28947 | 1448 | V1448Q | E | 4.8e-06 | 6e-06 | 1.3e-05 | 7.933E-06 | 4.429E-06 | 4.800E-06 | 1.300E-05 |
| 28948 | 1448 | V1448G | E | 8.7e-06 | 1.2e-05 | 5.8e-06 | 8.833E-06 | 3.102E-06 | 5.800E-06 | 1.200E-05 |
| 28949 | 1448 | V1448H | E | 1.3e-05 | 6.7e-06 | 8.5e-06 | 9.400E-06 | 3.245E-06 | 6.700E-06 | 1.300E-05 |
| 28950 | 1448 | V1448I | E | 5.5e-06 | 5.1e-06 | 4.6e-06 | 5.067E-06 | 4.509E-07 | 4.600E-06 | 5.500E-06 |
| 28951 | 1448 | V1448L | E | 6.4e-06 | 5.5e-06 | 8.2e-06 | 6.700E-06 | 1.375E-06 | 5.500E-06 | 8.200E-06 |
| 28952 | 1448 | V1448K | E | 6.1e-06 | 8.2e-06 | 5.8e-06 | 6.700E-06 | 1.308E-06 | 5.800E-06 | 8.200E-06 |
| 28953 | 1448 | V1448M | E | 1e-05 | 9.1e-06 | 1.1e-05 | 1.003E-05 | 9.504E-07 | 9.100E-06 | 1.100E-05 |
| 28954 | 1448 | V1448F | E | 8.1e-06 | 1e-05 | 6.3e-06 | 8.133E-06 | 1.850E-06 | 6.300E-06 | 1.000E-05 |
| 28955 | 1448 | V1448P | E | 1.1e-05 | 9.9e-06 | 5.9e-06 | 8.933E-06 | 2.684E-06 | 5.900E-06 | 1.100E-05 |
| 28956 | 1448 | V1448S | E | 7.1e-06 | 6.6e-06 | 4.7e-06 | 6.133E-06 | 1.266E-06 | 4.700E-06 | 7.100E-06 |
| 28957 | 1448 | V1448T | E | 8.6e-06 | 7.6e-06 | 1.3e-05 | 9.733E-06 | 2.873E-06 | 7.600E-06 | 1.300E-05 |
| 28958 | 1448 | V1448W | E | 1.1e-05 | 5.9e-06 | 6.3e-06 | 7.733E-06 | 2.836E-06 | 5.900E-06 | 1.100E-05 |
| 28959 | 1448 | V1448Y | E | 9e-06 | 5.2e-06 | 8.2e-06 | 7.467E-06 | 2.003E-06 | 5.200E-06 | 9.000E-06 |
| 28960 | 1448 | V1448V | E | 6.7e-06 | 9.7e-06 | 9.6e-06 | 8.667E-06 | 1.704E-06 | 6.700E-06 | 9.700E-06 |
| 28961 | 1449 | L1449A | E | 6.2e-06 | 4.4e-06 | 1e-05 | 6.867E-06 | 2.859E-06 | 4.400E-06 | 1.000E-05 |
| 28962 | 1449 | L1449R | E | 6.9e-06 | 9.3e-06 | 7e-06 | 7.733E-06 | 1.358E-06 | 6.900E-06 | 9.300E-06 |
| 28963 | 1449 | L1449N | E | 7.7e-06 | 8.8e-06 | 7.1e-06 | 7.867E-06 | 8.622E-07 | 7.100E-06 | 8.800E-06 |
| 28964 | 1449 | L1449D | E | 7.2e-06 | 4.5e-06 | 5.3e-06 | 5.667E-06 | 1.387E-06 | 4.500E-06 | 7.200E-06 |
| 28965 | 1449 | L1449C | E | 1.2e-05 | 8.2e-06 | 6.4e-06 | 8.867E-06 | 2.859E-06 | 6.400E-06 | 1.200E-05 |
| 28966 | 1449 | L1449E | E | 6.6e-06 | 8e-06 | 5.1e-06 | 6.567E-06 | 1.450E-06 | 5.100E-06 | 8.000E-06 |
| 28967 | 1449 | L1449Q | E | 9.3e-06 | 4.5e-06 | 4.3e-06 | 6.033E-06 | 2.831E-06 | 4.300E-06 | 9.300E-06 |
| 28968 | 1449 | L1449G | E | 8.7e-06 | 9.2e-06 | 6.7e-06 | 8.200E-06 | 1.323E-06 | 6.700E-06 | 9.200E-06 |
| 28969 | 1449 | L1449H | E | 6.4e-06 | 5.3e-06 | 8.9e-06 | 6.867E-06 | 1.845E-06 | 5.300E-06 | 8.900E-06 |
| 28970 | 1449 | L1449I | E | 1.1e-05 | 6.7e-06 | 8.6e-06 | 8.767E-06 | 2.155E-06 | 6.700E-06 | 1.100E-05 |
| 28971 | 1449 | L1449L | E | 7e-06 | 1.1e-05 | 1.3e-05 | 1.033E-05 | 3.055E-06 | 7.000E-06 | 1.300E-05 |
| 28972 | 1449 | L1449K | E | 6.9e-06 | 7.3e-06 | 9.8e-06 | 8.000E-06 | 1.572E-06 | 6.900E-06 | 9.800E-06 |
| 28973 | 1449 | L1449M | E | 9.5e-06 | 6.3e-06 | 5.7e-06 | 7.167E-06 | 2.043E-06 | 5.700E-06 | 9.500E-06 |
| 28974 | 1449 | L1449F | E | 5.9e-06 | 6.6e-06 | 8.1e-06 | 6.867E-06 | 1.124E-06 | 5.900E-06 | 8.100E-06 |
| 28975 | 1449 | L1449P | E | 9.6e-06 | 1.3e-05 | 1.2e-05 | 1.153E-05 | 1.747E-06 | 9.600E-06 | 1.300E-05 |
| 28976 | 1449 | L1449S | E | 6.8e-06 | 1.4e-05 | 8.8e-06 | 9.867E-06 | 3.717E-06 | 6.800E-06 | 1.400E-05 |
| 28977 | 1449 | L1449T | E | 7.8e-06 | 4.5e-06 | 8e-06 | 6.767E-06 | 1.966E-06 | 4.500E-06 | 8.000E-06 |
| 28978 | 1449 | L1449W | E | 1.3e-05 | 9.4e-06 | 5.3e-06 | 9.233E-06 | 3.853E-06 | 5.300E-06 | 1.300E-05 |
| 28979 | 1449 | L1449Y | E | 8.9e-06 | 8e-06 | 4.4e-06 | 7.100E-06 | 2.381E-06 | 4.400E-06 | 8.900E-06 |
| 28980 | 1449 | L1449V | E | 6.4e-06 | 9.9e-06 | 5.9e-06 | 7.400E-06 | 2.179E-06 | 5.900E-06 | 9.900E-06 |
| 28981 | 1450 | Q1450A | E | 1.3e-05 | 8e-06 | 9.6e-06 | 1.020E-05 | 2.553E-06 | 8.000E-06 | 1.300E-05 |
| 28982 | 1450 | Q1450R | E | 6.5e-06 | 1.1e-05 | 6.6e-06 | 8.033E-06 | 2.570E-06 | 6.500E-06 | 1.100E-05 |
| 28983 | 1450 | Q1450N | E | 8.3e-06 | 6e-06 | 8.6e-06 | 7.633E-06 | 1.422E-06 | 6.000E-06 | 8.600E-06 |
| 28984 | 1450 | Q1450D | E | 1.4e-05 | 9.8e-06 | 5e-06 | 9.600E-06 | 4.503E-06 | 5.000E-06 | 1.400E-05 |

| | | | | | | | | | | |
|---|---|---|---|---|---|---|---|---|---|---|
| 28985 | 1450 | Q1450C | E | 4.5e-06 | 9.6e-06 | 8.2e-06 | 7.433E-06 | 2.635E-06 | 4.500E-06 | 9.600E-06 |
| 28986 | 1450 | Q1450E | E | 5.2e-06 | 6.7e-06 | 6.1e-06 | 6.000E-06 | 7.550E-07 | 5.200E-06 | 6.700E-06 |
| 28987 | 1450 | Q1450Q | E | 6.8e-06 | 9.3e-06 | 1.2e-05 | 9.367E-06 | 2.601E-06 | 6.800E-06 | 1.200E-05 |
| 28988 | 1450 | Q1450G | E | 1e-05 | 9.9e-06 | 4.3e-06 | 8.067E-06 | 3.262E-06 | 4.300E-06 | 1.000E-05 |
| 28989 | 1450 | Q1450H | E | 8.9e-06 | 8.8e-06 | 8.5e-06 | 8.733E-06 | 2.082E-07 | 8.500E-06 | 8.900E-06 |
| 28990 | 1450 | Q1450I | E | 1e-05 | 7.8e-06 | 8e-06 | 8.600E-06 | 1.217E-06 | 7.800E-06 | 1.000E-05 |
| 28991 | 1450 | Q1450L | E | 7.9e-06 | 6.7e-06 | 5.1e-06 | 6.567E-06 | 1.405E-06 | 5.100E-06 | 7.900E-06 |
| 28992 | 1450 | Q1450K | E | 6.5e-06 | 5.5e-06 | 1.6e-05 | 9.333E-06 | 5.795E-06 | 5.500E-06 | 1.600E-05 |
| 28993 | 1450 | Q1450M | E | 7e-06 | 8.4e-06 | 4.9e-06 | 6.767E-06 | 1.762E-06 | 4.900E-06 | 8.400E-06 |
| 28994 | 1450 | Q1450F | E | 1.5e-05 | 1.1e-05 | 1.3e-05 | 1.300E-05 | 2.000E-06 | 1.100E-05 | 1.500E-05 |
| 28995 | 1450 | Q1450P | E | 1.3e-05 | 7.4e-06 | 6.6e-06 | 9.000E-06 | 3.487E-06 | 6.600E-06 | 1.300E-05 |
| 28996 | 1450 | Q1450S | E | 6.5e-06 | 1e-05 | 7.1e-06 | 7.867E-06 | 1.872E-06 | 6.500E-06 | 1.000E-05 |
| 28997 | 1450 | Q1450T | E | 7.1e-06 | 6.6e-06 | 5e-06 | 6.233E-06 | 1.097E-06 | 5.000E-06 | 7.100E-06 |
| 28998 | 1450 | Q1450W | E | 6.7e-06 | 6.9e-06 | 7.7e-06 | 7.100E-06 | 5.292E-07 | 6.700E-06 | 7.700E-06 |
| 28999 | 1450 | Q1450Y | E | 8.8e-06 | 9.2e-06 | 5e-06 | 7.667E-06 | 2.318E-06 | 5.000E-06 | 9.200E-06 |
| 29000 | 1450 | Q1450V | E | 7.2e-06 | 9.6e-06 | 5.7e-06 | 7.500E-06 | 1.967E-06 | 5.700E-06 | 9.600E-06 |
| 29001 | 1451 | S1451A | E | 5e-06 | 1.5e-05 | 6.5e-06 | 8.833E-06 | 5.393E-06 | 5.000E-06 | 1.500E-05 |
| 29002 | 1451 | S1451R | E | 6.2e-06 | 9.3e-06 | 6.8e-06 | 7.433E-06 | 1.644E-06 | 6.200E-06 | 9.300E-06 |
| 29003 | 1451 | S1451N | E | 7e-06 | 9.8e-06 | 6e-06 | 7.600E-06 | 1.970E-06 | 6.000E-06 | 9.800E-06 |
| 29004 | 1451 | S1451D | E | 6.6e-06 | 8.2e-06 | 7.2e-06 | 7.333E-06 | 8.083E-07 | 6.600E-06 | 8.200E-06 |
| 29005 | 1451 | S1451C | E | 7.2e-06 | 1e-05 | 1.3e-05 | 1.007E-05 | 2.901E-06 | 7.200E-06 | 1.300E-05 |
| 29006 | 1451 | S1451E | E | 1.9e-05 | 6.7e-06 | 9.3e-06 | 1.167E-05 | 6.483E-06 | 6.700E-06 | 1.900E-05 |
| 29007 | 1451 | S1451Q | E | 9.3e-06 | 8.4e-06 | 2e-05 | 1.257E-05 | 6.453E-06 | 8.400E-06 | 2.000E-05 |
| 29008 | 1451 | S1451G | E | 4.5e-06 | 8.4e-06 | 5.7e-06 | 6.200E-06 | 1.997E-06 | 4.500E-06 | 8.400E-06 |
| 29009 | 1451 | S1451H | E | 4.1e-06 | 6.5e-06 | 6.5e-06 | 5.700E-06 | 1.386E-06 | 4.100E-06 | 6.500E-06 |
| 29010 | 1451 | S1451I | E | 1.3e-05 | 7.1e-06 | 1.4e-05 | 1.137E-05 | 3.729E-06 | 7.100E-06 | 1.400E-05 |
| 29011 | 1451 | S1451L | E | 8.9e-06 | 1.1e-05 | 7.4e-06 | 9.100E-06 | 1.808E-06 | 7.400E-06 | 1.100E-05 |
| 29012 | 1451 | S1451K | E | 5.7e-06 | 8.4e-06 | 8.9e-06 | 7.667E-06 | 1.721E-06 | 5.700E-06 | 8.900E-06 |
| 29013 | 1451 | S1451M | E | 5.5e-06 | 8e-06 | 1e-05 | 7.833E-06 | 2.255E-06 | 5.500E-06 | 1.000E-05 |
| 29014 | 1451 | S1451F | E | 6.5e-06 | 1e-05 | 4.7e-06 | 7.067E-06 | 2.695E-06 | 4.700E-06 | 1.000E-05 |
| 29015 | 1451 | S1451P | E | 6.9e-06 | 1e-05 | 7.8e-06 | 8.233E-06 | 1.595E-06 | 6.900E-06 | 1.000E-05 |
| 29016 | 1451 | S1451S | E | 1.1e-05 | 1.3e-05 | 7.3e-06 | 1.043E-05 | 2.892E-06 | 7.300E-06 | 1.300E-05 |
| 29017 | 1451 | S1451T | E | 4e-06 | 5.7e-06 | 1.3e-05 | 7.567E-06 | 4.782E-06 | 4.000E-06 | 1.300E-05 |
| 29018 | 1451 | S1451W | E | 1.1e-05 | 6e-06 | 1.1e-05 | 9.333E-06 | 2.887E-06 | 6.000E-06 | 1.100E-05 |
| 29019 | 1451 | S1451Y | E | 1.2e-05 | 1e-05 | 8.9e-06 | 1.030E-05 | 1.572E-06 | 8.900E-06 | 1.200E-05 |
| 29020 | 1451 | S1451V | E | 1.1e-05 | 5.5e-06 | 1.4e-05 | 1.017E-05 | 4.311E-06 | 5.500E-06 | 1.400E-05 |
| 29021 | 1452 | S1452A | E | 1.3e-05 | 6.3e-06 | 8.7e-06 | 9.333E-06 | 3.395E-06 | 6.300E-06 | 1.300E-05 |
| 29022 | 1452 | S1452R | E | 5.8e-06 | 1.1e-05 | 6.4e-06 | 7.733E-06 | 2.845E-06 | 5.800E-06 | 1.100E-05 |
| 29023 | 1452 | S1452N | E | 6.6e-06 | 6.4e-06 | 6.1e-06 | 6.367E-06 | 2.517E-07 | 6.100E-06 | 6.600E-06 |
| 29024 | 1452 | S1452D | E | 5.8e-06 | 6.9e-06 | 1e-05 | 7.567E-06 | 2.178E-06 | 5.800E-06 | 1.000E-05 |
| 29025 | 1452 | S1452C | E | 7.4e-06 | 1.1e-05 | 5.4e-06 | 7.933E-06 | 2.838E-06 | 5.400E-06 | 1.100E-05 |
| 29026 | 1452 | S1452E | E | 6.4e-06 | 1e-05 | 8.3e-06 | 8.233E-06 | 1.801E-06 | 6.400E-06 | 1.000E-05 |
| 29027 | 1452 | S1452Q | E | 9.7e-06 | 1.8e-05 | 7.3e-06 | 1.167E-05 | 5.615E-06 | 7.300E-06 | 1.800E-05 |
| 29028 | 1452 | S1452G | E | 9.2e-06 | 1.1e-05 | 9.1e-06 | 9.767E-06 | 1.069E-06 | 9.100E-06 | 1.100E-05 |
| 29029 | 1452 | S1452H | E | 8.8e-06 | 7.1e-06 | 8.4e-06 | 8.100E-06 | 8.888E-07 | 7.100E-06 | 8.800E-06 |
| 29030 | 1452 | S1452I | E | 1.1e-05 | 1e-05 | 9.6e-06 | 1.020E-05 | 7.211E-07 | 9.600E-06 | 1.100E-05 |
| 29031 | 1452 | S1452L | E | 6.3e-06 | 5.3e-06 | 4.8e-06 | 5.467E-06 | 7.638E-07 | 4.800E-06 | 6.300E-06 |
| 29032 | 1452 | S1452K | E | 7e-06 | 9.3e-06 | 9.5e-06 | 8.600E-06 | 1.389E-06 | 7.000E-06 | 9.500E-06 |
| 29033 | 1452 | S1452M | E | 1.3e-05 | 6.6e-06 | 6.1e-06 | 8.567E-06 | 3.848E-06 | 6.100E-06 | 1.300E-05 |
| 29034 | 1452 | S1452F | E | 1e-05 | 8.9e-06 | 6.4e-06 | 8.433E-06 | 1.845E-06 | 6.400E-06 | 1.000E-05 |
| 29035 | 1452 | S1452P | E | 5.4e-06 | 1.4e-05 | 5.8e-06 | 8.400E-06 | 4.854E-06 | 5.400E-06 | 1.400E-05 |
| 29036 | 1452 | S1452S | E | 6.6e-06 | 1.5e-05 | 5.6e-06 | 9.067E-06 | 5.163E-06 | 5.600E-06 | 1.500E-05 |
| 29037 | 1452 | S1452T | E | 6.2e-06 | 5.1e-06 | 8.2e-06 | 6.500E-06 | 1.572E-06 | 5.100E-06 | 8.200E-06 |
| 29038 | 1452 | S1452W | E | 7.2e-06 | 1.4e-05 | 8.7e-06 | 9.967E-06 | 3.573E-06 | 7.200E-06 | 1.400E-05 |
| 29039 | 1452 | S1452Y | E | 8.9e-06 | 8.5e-06 | 5.8e-06 | 7.733E-06 | 1.686E-06 | 5.800E-06 | 8.900E-06 |
| 29040 | 1452 | S1452V | E | 9.4e-06 | 6.5e-06 | 1e-05 | 8.633E-06 | 1.872E-06 | 6.500E-06 | 1.000E-05 |
| 29041 | 1453 | G1453A | E | 9.1e-06 | 6.7e-06 | 9.1e-06 | 8.300E-06 | 1.386E-06 | 6.700E-06 | 9.100E-06 |
| 29042 | 1453 | G1453R | E | 7.7e-06 | 8.1e-06 | 1.8e-05 | 1.127E-05 | 5.835E-06 | 7.700E-06 | 1.800E-05 |
| 29043 | 1453 | G1453N | E | 5.4e-06 | 7.4e-06 | 9.5e-06 | 7.433E-06 | 2.050E-06 | 5.400E-06 | 9.500E-06 |
| 29044 | 1453 | G1453D | E | 7.7e-06 | 4.6e-06 | 1.2e-05 | 8.100E-06 | 3.716E-06 | 4.600E-06 | 1.200E-05 |
| 29045 | 1453 | G1453C | E | 1.2e-05 | 8.2e-06 | 8.8e-06 | 9.667E-06 | 2.043E-06 | 8.200E-06 | 1.200E-05 |
| 29046 | 1453 | G1453E | E | 7.2e-06 | 8.3e-06 | 5.5e-06 | 7.000E-06 | 1.411E-06 | 5.500E-06 | 8.300E-06 |
| 29047 | 1453 | G1453Q | E | 7.8e-06 | 3.5e-06 | 1.4e-05 | 8.433E-06 | 5.279E-06 | 3.500E-06 | 1.400E-05 |
| 29048 | 1453 | G1453G | E | 4.4e-06 | 9e-06 | 5.3e-06 | 6.233E-06 | 2.438E-06 | 4.400E-06 | 9.000E-06 |
| 29049 | 1453 | G1453H | E | 6.4e-06 | 8.6e-06 | 4.1e-06 | 6.367E-06 | 2.250E-06 | 4.100E-06 | 8.600E-06 |
| 29050 | 1453 | G1453I | E | 8.4e-06 | 9.6e-06 | 1.9e-05 | 1.233E-05 | 5.805E-06 | 8.400E-06 | 1.900E-05 |
| 29051 | 1453 | G1453L | E | 4.9e-06 | 6.1e-06 | 7.3e-06 | 6.100E-06 | 1.200E-06 | 4.900E-06 | 7.300E-06 |
| 29052 | 1453 | G1453K | E | 7.4e-06 | 5.6e-06 | 1.1e-05 | 8.000E-06 | 2.750E-06 | 5.600E-06 | 1.100E-05 |
| 29053 | 1453 | G1453M | E | 9.2e-06 | 1.2e-05 | 6.3e-06 | 9.167E-06 | 2.850E-06 | 6.300E-06 | 1.200E-05 |
| 29054 | 1453 | G1453F | E | 1.4e-05 | 9e-06 | 9.2e-06 | 1.073E-05 | 2.831E-06 | 9.000E-06 | 1.400E-05 |
| 29055 | 1453 | G1453P | E | 9.4e-06 | 8.6e-06 | 4.9e-06 | 7.633E-06 | 2.401E-06 | 4.900E-06 | 9.400E-06 |
| 29056 | 1453 | G1453S | E | 4.3e-06 | 9.5e-06 | 1.1e-05 | 8.267E-06 | 3.516E-06 | 4.300E-06 | 1.100E-05 |
| 29057 | 1453 | G1453T | E | 5.5e-06 | 1.2e-05 | 6.4e-06 | 7.967E-06 | 3.522E-06 | 5.500E-06 | 1.200E-05 |
| 29058 | 1453 | G1453W | E | 8.4e-06 | 9.4e-06 | 6e-06 | 7.933E-06 | 1.747E-06 | 6.000E-06 | 9.400E-06 |
| 29059 | 1453 | G1453Y | E | 7.2e-06 | 8.1e-06 | 6.8e-06 | 7.367E-06 | 6.658E-07 | 6.800E-06 | 8.100E-06 |
| 29060 | 1453 | G1453V | E | 9.1e-06 | 6.5e-06 | 1.3e-05 | 9.533E-06 | 3.272E-06 | 6.500E-06 | 1.300E-05 |
| 29061 | 1454 | L1454A | E | 5.5e-06 | 6.2e-06 | 7.1e-06 | 6.267E-06 | 8.021E-07 | 5.500E-06 | 7.100E-06 |
| 29062 | 1454 | L1454R | E | 6.8e-06 | 1.2e-05 | 9.8e-06 | 9.533E-06 | 2.610E-06 | 6.800E-06 | 1.200E-05 |
| 29063 | 1454 | L1454N | E | 4.9e-06 | 6.3e-06 | 8.9e-06 | 6.700E-06 | 2.030E-06 | 4.900E-06 | 8.900E-06 |
| 29064 | 1454 | L1454D | E | 1.1e-05 | 1.3e-05 | 6.5e-06 | 1.017E-05 | 3.329E-06 | 6.500E-06 | 1.300E-05 |
| 29065 | 1454 | L1454C | E | 1.1e-05 | 9.8e-06 | 1.2e-05 | 1.093E-05 | 1.102E-06 | 9.800E-06 | 1.200E-05 |
| 29066 | 1454 | L1454E | E | 1.2e-05 | 6.1e-06 | 8.9e-06 | 9.000E-06 | 2.951E-06 | 6.100E-06 | 1.200E-05 |
| 29067 | 1454 | L1454Q | E | 8.8e-06 | 4.4e-06 | 8.5e-06 | 7.233E-06 | 2.458E-06 | 4.400E-06 | 8.800E-06 |
| 29068 | 1454 | L1454G | E | 7.2e-06 | 6.7e-06 | 1.6e-05 | 9.967E-06 | 5.231E-06 | 6.700E-06 | 1.600E-05 |
| 29069 | 1454 | L1454H | E | 7.6e-06 | 5.1e-06 | 5.2e-06 | 5.967E-06 | 1.415E-06 | 5.100E-06 | 7.600E-06 |

| | | | | | | | | | | |
|---|---|---|---|---|---|---|---|---|---|---|
| 29070 | 1454 | L1454I | E | 5.3e-06 | 5.2e-06 | 7.6e-06 | 6.033E-06 | 1.358E-06 | 5.200E-06 | 7.600E-06 |
| 29071 | 1454 | L1454L | E | 9.1e-06 | 9.3e-06 | 1.1e-05 | 9.800E-06 | 1.044E-06 | 9.100E-06 | 1.100E-05 |
| 29072 | 1454 | L1454K | E | 5.9e-06 | 7.7e-06 | 7.2e-06 | 6.933E-06 | 9.292E-07 | 5.900E-06 | 7.700E-06 |
| 29073 | 1454 | L1454M | E | 4.4e-06 | 4.9e-06 | 9.4e-06 | 6.233E-06 | 2.754E-06 | 4.400E-06 | 9.400E-06 |
| 29074 | 1454 | L1454F | E | 5.2e-06 | 1e-05 | 9e-06 | 8.067E-06 | 2.532E-06 | 5.200E-06 | 1.000E-05 |
| 29075 | 1454 | L1454P | E | 1.1e-05 | 9.6e-06 | 6e-06 | 8.867E-06 | 2.579E-06 | 6.000E-06 | 1.100E-05 |
| 29076 | 1454 | L1454S | E | 7.6e-06 | 8.8e-06 | 5e-06 | 7.133E-06 | 1.943E-06 | 5.000E-06 | 8.800E-06 |
| 29077 | 1454 | L1454T | E | 5.7e-06 | 8.7e-06 | 6.8e-06 | 7.067E-06 | 1.518E-06 | 5.700E-06 | 8.700E-06 |
| 29078 | 1454 | L1454W | E | 1e-05 | 5.6e-06 | 5.8e-06 | 7.133E-06 | 2.485E-06 | 5.600E-06 | 1.000E-05 |
| 29079 | 1454 | L1454Y | E | 9.4e-06 | 8.1e-06 | 6.4e-06 | 7.967E-06 | 1.504E-06 | 6.400E-06 | 9.400E-06 |
| 29080 | 1454 | L1454V | E | 5.2e-06 | 6.4e-06 | 7.1e-06 | 6.233E-06 | 9.609E-07 | 5.200E-06 | 7.100E-06 |
| 29081 | 1455 | Y1455A | E | 5e-06 | 7.5e-06 | 8.9e-06 | 7.133E-06 | 1.976E-06 | 5.000E-06 | 8.900E-06 |
| 29082 | 1455 | Y1455R | E | 6.1e-06 | 6.2e-06 | 1.3e-05 | 8.433E-06 | 3.955E-06 | 6.100E-06 | 1.300E-05 |
| 29083 | 1455 | Y1455N | E | 9.9e-06 | 9.7e-06 | 8.2e-06 | 9.267E-06 | 9.292E-07 | 8.200E-06 | 9.900E-06 |
| 29084 | 1455 | Y1455D | E | 1.2e-05 | 1.1e-05 | 6.3e-06 | 9.767E-06 | 3.044E-06 | 6.300E-06 | 1.200E-05 |
| 29085 | 1455 | Y1455C | E | 1.1e-05 | 6.3e-06 | 7.3e-06 | 8.200E-06 | 2.476E-06 | 6.300E-06 | 1.100E-05 |
| 29086 | 1455 | Y1455E | E | 1.4e-05 | 6e-06 | 9.2e-06 | 9.733E-06 | 4.027E-06 | 6.000E-06 | 1.400E-05 |
| 29087 | 1455 | Y1455Q | E | 8.8e-06 | 8.3e-06 | 4.6e-06 | 7.233E-06 | 2.294E-06 | 4.600E-06 | 8.800E-06 |
| 29088 | 1455 | Y1455G | E | 3.2e-06 | 1.3e-05 | 7.1e-06 | 7.767E-06 | 4.934E-06 | 3.200E-06 | 1.300E-05 |
| 29089 | 1455 | Y1455H | E | 1.2e-05 | 5.1e-06 | 5.2e-06 | 7.433E-06 | 3.955E-06 | 5.100E-06 | 1.200E-05 |
| 29090 | 1455 | Y1455I | E | 1.3e-05 | 9e-06 | 8.1e-06 | 1.003E-05 | 2.608E-06 | 8.100E-06 | 1.300E-05 |
| 29091 | 1455 | Y1455L | E | 8.6e-06 | 6.7e-06 | 9.7e-06 | 8.333E-06 | 1.518E-06 | 6.700E-06 | 9.700E-06 |
| 29092 | 1455 | Y1455K | E | 7.6e-06 | 7.5e-06 | 1e-05 | 8.367E-06 | 1.415E-06 | 7.500E-06 | 1.000E-05 |
| 29093 | 1455 | Y1455M | E | 7.9e-06 | 9e-06 | 9.8e-06 | 8.900E-06 | 9.539E-07 | 7.900E-06 | 9.800E-06 |
| 29094 | 1455 | Y1455F | E | 5.7e-06 | 8.2e-06 | 5.7e-06 | 6.533E-06 | 1.443E-06 | 5.700E-06 | 8.200E-06 |
| 29095 | 1455 | Y1455P | E | 9.4e-06 | 3.4e-06 | 8.9e-06 | 7.233E-06 | 3.329E-06 | 3.400E-06 | 9.400E-06 |
| 29096 | 1455 | Y1455S | E | 8e-06 | 8.5e-06 | 7.8e-06 | 8.100E-06 | 3.606E-07 | 7.800E-06 | 8.500E-06 |
| 29097 | 1455 | Y1455T | E | 9.1e-06 | 7.9e-06 | 6e-06 | 7.667E-06 | 1.563E-06 | 6.000E-06 | 9.100E-06 |
| 29098 | 1455 | Y1455W | E | 4.4e-06 | 1.1e-05 | 4.6e-06 | 6.667E-06 | 3.754E-06 | 4.400E-06 | 1.100E-05 |
| 29099 | 1455 | Y1455Y | E | 7.5e-06 | 3.9e-06 | 9.7e-06 | 7.033E-06 | 2.928E-06 | 3.900E-06 | 9.700E-06 |
| 29100 | 1455 | Y1455V | E | 1.2e-05 | 1e-05 | 7.7e-06 | 9.900E-06 | 2.152E-06 | 7.700E-06 | 1.200E-05 |
| 29101 | 1456 | S1456A | E | 1.2e-05 | 1.3e-05 | 4.7e-06 | 9.900E-06 | 4.531E-06 | 4.700E-06 | 1.300E-05 |
| 29102 | 1456 | S1456R | E | 1.4e-05 | 5.9e-06 | 8.9e-06 | 9.600E-06 | 4.095E-06 | 5.900E-06 | 1.400E-05 |
| 29103 | 1456 | S1456N | E | 8.3e-06 | 8.7e-06 | 8.8e-06 | 8.600E-06 | 2.646E-07 | 8.300E-06 | 8.800E-06 |
| 29104 | 1456 | S1456D | E | 9.8e-06 | 5.5e-06 | 1.2e-05 | 9.100E-06 | 3.306E-06 | 5.500E-06 | 1.200E-05 |
| 29105 | 1456 | S1456C | E | 5.7e-06 | 9.8e-06 | 7.2e-06 | 7.567E-06 | 2.074E-06 | 5.700E-06 | 9.800E-06 |
| 29106 | 1456 | S1456E | E | 8.4e-06 | 6.6e-06 | 1.4e-05 | 9.667E-06 | 3.859E-06 | 6.600E-06 | 1.400E-05 |
| 29107 | 1456 | S1456Q | E | 4.9e-06 | 9.7e-06 | 1.4e-05 | 9.533E-06 | 4.552E-06 | 4.900E-06 | 1.400E-05 |
| 29108 | 1456 | S1456G | E | 7.9e-06 | 5.3e-06 | 1.3e-05 | 8.733E-06 | 3.917E-06 | 5.300E-06 | 1.300E-05 |
| 29109 | 1456 | S1456H | E | 6.9e-06 | 7.5e-06 | 4.3e-06 | 6.233E-06 | 1.701E-06 | 4.300E-06 | 7.500E-06 |
| 29110 | 1456 | S1456I | E | 8.9e-06 | 9.5e-06 | 6.6e-06 | 8.333E-06 | 1.531E-06 | 6.600E-06 | 9.500E-06 |
| 29111 | 1456 | S1456L | E | 6.1e-06 | 7.1e-06 | 8.2e-06 | 7.133E-06 | 1.050E-06 | 6.100E-06 | 8.200E-06 |
| 29112 | 1456 | S1456K | E | 7.3e-06 | 8.9e-06 | 5.6e-06 | 7.267E-06 | 1.650E-06 | 5.600E-06 | 8.900E-06 |
| 29113 | 1456 | S1456M | E | 1e-05 | 9e-06 | 9.3e-06 | 9.433E-06 | 5.132E-07 | 9.000E-06 | 1.000E-05 |
| 29114 | 1456 | S1456F | E | 1e-05 | 3.9e-06 | 6.9e-06 | 6.933E-06 | 3.050E-06 | 3.900E-06 | 1.000E-05 |
| 29115 | 1456 | S1456P | E | 1.1e-05 | 1.5e-05 | 9.9e-06 | 1.197E-05 | 2.684E-06 | 9.900E-06 | 1.500E-05 |
| 29116 | 1456 | S1456S | E | 7e-06 | 1e-05 | 7.8e-06 | 8.267E-06 | 1.553E-06 | 7.000E-06 | 1.000E-05 |
| 29117 | 1456 | S1456T | E | 6.3e-06 | 6.6e-06 | 8.6e-06 | 7.167E-06 | 1.250E-06 | 6.300E-06 | 8.600E-06 |
| 29118 | 1456 | S1456W | E | 1e-05 | 7.3e-06 | 6.9e-06 | 8.067E-06 | 1.686E-06 | 6.900E-06 | 1.000E-05 |
| 29119 | 1456 | S1456Y | E | 5.9e-06 | 6.5e-06 | 1.1e-05 | 7.800E-06 | 2.787E-06 | 5.900E-06 | 1.100E-05 |
| 29120 | 1456 | S1456V | E | 7.3e-06 | 9.5e-06 | 8.7e-06 | 8.500E-06 | 1.114E-06 | 7.300E-06 | 9.500E-06 |
| 29121 | 1457 | L1457A | E | 9.8e-06 | 4.8e-06 | 6.7e-06 | 7.100E-06 | 2.524E-06 | 4.800E-06 | 9.800E-06 |
| 29122 | 1457 | L1457R | E | 1.3e-05 | 8.7e-06 | 7.2e-06 | 9.633E-06 | 3.011E-06 | 7.200E-06 | 1.300E-05 |
| 29123 | 1457 | L1457N | E | 5.8e-06 | 1e-05 | 8.7e-06 | 8.167E-06 | 2.150E-06 | 5.800E-06 | 1.000E-05 |
| 29124 | 1457 | L1457D | E | 6.3e-06 | 1.1e-05 | 1e-05 | 9.100E-06 | 2.476E-06 | 6.300E-06 | 1.100E-05 |
| 29125 | 1457 | L1457C | E | 6.6e-06 | 9.5e-06 | 4.5e-06 | 6.867E-06 | 2.511E-06 | 4.500E-06 | 9.500E-06 |
| 29126 | 1457 | L1457E | E | 3.1e-06 | 8.7e-06 | 8.1e-06 | 6.633E-06 | 3.075E-06 | 3.100E-06 | 8.700E-06 |
| 29127 | 1457 | L1457Q | E | 5.3e-06 | 9.3e-06 | 5.1e-06 | 6.567E-06 | 2.369E-06 | 5.100E-06 | 9.300E-06 |
| 29128 | 1457 | L1457G | E | 6.5e-06 | 6.2e-06 | 7e-06 | 6.567E-06 | 4.041E-07 | 6.200E-06 | 7.000E-06 |
| 29129 | 1457 | L1457H | E | 5.8e-06 | 8e-06 | 6.5e-06 | 6.767E-06 | 1.124E-06 | 5.800E-06 | 8.000E-06 |
| 29130 | 1457 | L1457I | E | 1.2e-05 | 9.4e-06 | 7.7e-06 | 9.700E-06 | 2.166E-06 | 7.700E-06 | 1.200E-05 |
| 29131 | 1457 | L1457L | E | 1.1e-05 | 8e-06 | 7.5e-06 | 8.833E-06 | 1.893E-06 | 7.500E-06 | 1.100E-05 |
| 29132 | 1457 | L1457K | E | 9.4e-06 | 8.7e-06 | 6.5e-06 | 8.200E-06 | 1.513E-06 | 6.500E-06 | 9.400E-06 |
| 29133 | 1457 | L1457M | E | 1.4e-05 | 5e-06 | 4.6e-06 | 7.867E-06 | 5.315E-06 | 4.600E-06 | 1.400E-05 |
| 29134 | 1457 | L1457F | E | 6.9e-06 | 7.8e-06 | 7.7e-06 | 7.467E-06 | 4.933E-07 | 6.900E-06 | 7.800E-06 |
| 29135 | 1457 | L1457P | E | 6.4e-06 | 9.1e-06 | 7.8e-06 | 7.767E-06 | 1.350E-06 | 6.400E-06 | 9.100E-06 |
| 29136 | 1457 | L1457S | E | 7.4e-06 | 1.4e-05 | 7.9e-06 | 9.767E-06 | 3.675E-06 | 7.400E-06 | 1.400E-05 |
| 29137 | 1457 | L1457T | E | 8.9e-06 | 8.1e-06 | 6.6e-06 | 7.867E-06 | 1.168E-06 | 6.600E-06 | 8.900E-06 |
| 29138 | 1457 | L1457W | E | 8.8e-06 | 6.3e-06 | 7.1e-06 | 7.400E-06 | 1.277E-06 | 6.300E-06 | 8.800E-06 |
| 29139 | 1457 | L1457Y | E | 4.9e-06 | 8.4e-06 | 6.7e-06 | 6.667E-06 | 1.750E-06 | 4.900E-06 | 8.400E-06 |
| 29140 | 1457 | L1457V | E | 1.1e-05 | 1.3e-05 | 1.1e-05 | 1.167E-05 | 1.155E-06 | 1.100E-05 | 1.300E-05 |
| 29141 | 1458 | S1458A | E | 8.8e-06 | 5.7e-06 | 7.5e-06 | 7.333E-06 | 1.557E-06 | 5.700E-06 | 8.800E-06 |
| 29142 | 1458 | S1458R | E | 1.1e-05 | 1.6e-05 | 8.9e-06 | 1.197E-05 | 3.647E-06 | 8.900E-06 | 1.600E-05 |
| 29143 | 1458 | S1458N | E | 9.3e-06 | 5.2e-06 | 9.5e-06 | 8.000E-06 | 2.427E-06 | 5.200E-06 | 9.500E-06 |
| 29144 | 1458 | S1458D | E | 7.3e-06 | 8e-06 | 9.4e-06 | 8.233E-06 | 1.069E-06 | 7.300E-06 | 9.400E-06 |
| 29145 | 1458 | S1458C | E | 8.1e-06 | 7.2e-06 | 7.1e-06 | 7.467E-06 | 5.508E-07 | 7.100E-06 | 8.100E-06 |
| 29146 | 1458 | S1458E | E | 9.1e-06 | 6.3e-06 | 1.1e-05 | 8.800E-06 | 2.364E-06 | 6.300E-06 | 1.100E-05 |
| 29147 | 1458 | S1458Q | E | 6.9e-06 | 1.4e-05 | 4.7e-06 | 8.533E-06 | 4.860E-06 | 4.700E-06 | 1.400E-05 |
| 29148 | 1458 | S1458G | E | 8.8e-06 | 6.7e-06 | 6.2e-06 | 7.233E-06 | 1.380E-06 | 6.200E-06 | 8.800E-06 |
| 29149 | 1458 | S1458H | E | 5.9e-06 | 6.3e-06 | 7.4e-06 | 6.533E-06 | 7.767E-07 | 5.900E-06 | 7.400E-06 |
| 29150 | 1458 | S1458I | E | 7.6e-06 | 6.2e-06 | 9.2e-06 | 7.667E-06 | 1.501E-06 | 6.200E-06 | 9.200E-06 |
| 29151 | 1458 | S1458L | E | 1e-05 | 1.4e-05 | 1e-05 | 1.133E-05 | 2.309E-06 | 1.000E-05 | 1.400E-05 |
| 29152 | 1458 | S1458K | E | 8.8e-06 | 6.9e-06 | 1e-05 | 8.567E-06 | 1.563E-06 | 6.900E-06 | 1.000E-05 |
| 29153 | 1458 | S1458M | E | 7e-06 | 7.2e-06 | 6.2e-06 | 6.800E-06 | 5.292E-07 | 6.200E-06 | 7.200E-06 |
| 29154 | 1458 | S1458F | E | 6.8e-06 | 1.5e-05 | 6.1e-06 | 9.300E-06 | 4.949E-06 | 6.100E-06 | 1.500E-05 |

| | | | | | | | | | | |
|---|---|---|---|---|---|---|---|---|---|---|
| 29155 | 1458 | S1458P | E | 1.4e-05 | 8.6e-06 | 5.8e-06 | 9.467E-06 | 4.168E-06 | 5.800E-06 | 1.400E-05 |
| 29156 | 1458 | S1458S | E | 6.2e-06 | 1.3e-05 | 1e-05 | 9.733E-06 | 3.408E-06 | 6.200E-06 | 1.300E-05 |
| 29157 | 1458 | S1458T | E | 9.6e-06 | 9.3e-06 | 7e-06 | 8.633E-06 | 1.422E-06 | 7.000E-06 | 9.600E-06 |
| 29158 | 1458 | S1458W | E | 8.2e-06 | 8.7e-06 | 7.7e-06 | 8.200E-06 | 5.000E-07 | 7.700E-06 | 8.700E-06 |
| 29159 | 1458 | S1458Y | E | 7.9e-06 | 1.5e-05 | 6.8e-06 | 9.900E-06 | 4.451E-06 | 6.800E-06 | 1.500E-05 |
| 29160 | 1458 | S1458V | E | 4.5e-06 | 9.1e-06 | 9.7e-06 | 7.767E-06 | 2.845E-06 | 4.500E-06 | 9.700E-06 |
| 29161 | 1459 | S1459A | E | 1.2e-05 | 1e-05 | 6.5e-06 | 9.500E-06 | 2.784E-06 | 6.500E-06 | 1.200E-05 |
| 29162 | 1459 | S1459R | E | 9.6e-06 | 5.9e-06 | 4.6e-06 | 6.700E-06 | 2.594E-06 | 4.600E-06 | 9.600E-06 |
| 29163 | 1459 | S1459N | E | 9.3e-06 | 7.2e-06 | 7.9e-06 | 8.133E-06 | 1.069E-06 | 7.200E-06 | 9.300E-06 |
| 29164 | 1459 | S1459D | E | 6.8e-06 | 4.9e-06 | 9.8e-06 | 7.167E-06 | 2.470E-06 | 4.900E-06 | 9.800E-06 |
| 29165 | 1459 | S1459C | E | 1.1e-05 | 1.4e-05 | 6.9e-06 | 1.063E-05 | 3.564E-06 | 6.900E-06 | 1.400E-05 |
| 29166 | 1459 | S1459E | E | 8.6e-06 | 7e-06 | 7.1e-06 | 7.567E-06 | 8.963E-07 | 7.000E-06 | 8.600E-06 |
| 29167 | 1459 | S1459Q | E | 6.1e-06 | 1.2e-05 | 1e-05 | 9.367E-06 | 3.001E-06 | 6.100E-06 | 1.200E-05 |
| 29168 | 1459 | S1459G | E | 5.9e-06 | 6.5e-06 | 8.6e-06 | 7.000E-06 | 1.418E-06 | 5.900E-06 | 8.600E-06 |
| 29169 | 1459 | S1459H | E | 1.3e-05 | 9.2e-06 | 1.3e-05 | 1.173E-05 | 2.194E-06 | 9.200E-06 | 1.300E-05 |
| 29170 | 1459 | S1459I | E | 7.9e-06 | 6.3e-06 | 1.2e-05 | 8.733E-06 | 2.940E-06 | 6.300E-06 | 1.200E-05 |
| 29171 | 1459 | S1459L | E | 5.4e-06 | 7.8e-06 | 7.2e-06 | 6.800E-06 | 1.249E-06 | 5.400E-06 | 7.800E-06 |
| 29172 | 1459 | S1459K | E | 6.5e-06 | 6.8e-06 | 9.1e-06 | 7.467E-06 | 1.422E-06 | 6.500E-06 | 9.100E-06 |
| 29173 | 1459 | S1459M | E | 7.9e-06 | 5.2e-06 | 5.6e-06 | 6.233E-06 | 1.457E-06 | 5.200E-06 | 7.900E-06 |
| 29174 | 1459 | S1459F | E | 4.5e-06 | 5.2e-06 | 9.7e-06 | 6.467E-06 | 2.822E-06 | 4.500E-06 | 9.700E-06 |
| 29175 | 1459 | S1459P | E | 5.5e-06 | 1.3e-05 | 7.8e-06 | 8.767E-06 | 3.842E-06 | 5.500E-06 | 1.300E-05 |
| 29176 | 1459 | S1459S | E | 1.4e-05 | 9.8e-06 | 5e-06 | 9.600E-06 | 4.503E-06 | 5.000E-06 | 1.400E-05 |
| 29177 | 1459 | S1459T | E | 1.4e-05 | 9.8e-06 | 4.7e-06 | 9.500E-06 | 4.657E-06 | 4.700E-06 | 1.400E-05 |
| 29178 | 1459 | S1459W | E | 4.6e-06 | 8e-06 | 8.1e-06 | 6.900E-06 | 1.992E-06 | 4.600E-06 | 8.100E-06 |
| 29179 | 1459 | S1459Y | E | 9.3e-06 | 6.5e-06 | 4.8e-06 | 6.867E-06 | 2.272E-06 | 4.800E-06 | 9.300E-06 |
| 29180 | 1459 | S1459V | E | 7.6e-06 | 8.9e-06 | 6.8e-06 | 7.767E-06 | 1.060E-06 | 6.800E-06 | 8.900E-06 |
| 29181 | 1460 | V1460A | E | 9.8e-06 | 7.4e-06 | 1e-05 | 9.067E-06 | 1.447E-06 | 7.400E-06 | 1.000E-05 |
| 29182 | 1460 | V1460R | E | 7.9e-06 | 9.7e-06 | 5.9e-06 | 7.833E-06 | 1.901E-06 | 5.900E-06 | 9.700E-06 |
| 29183 | 1460 | V1460N | E | 7.8e-06 | 8.8e-06 | 1e-05 | 8.867E-06 | 1.102E-06 | 7.800E-06 | 1.000E-05 |
| 29184 | 1460 | V1460D | E | 1.9e-05 | 1.2e-05 | 9.8e-06 | 1.360E-05 | 4.804E-06 | 9.800E-06 | 1.900E-05 |
| 29185 | 1460 | V1460C | E | 9.2e-06 | 9.5e-06 | 3e-06 | 7.233E-06 | 3.669E-06 | 3.000E-06 | 9.500E-06 |
| 29186 | 1460 | V1460E | E | 1.2e-05 | 6.4e-06 | 6.4e-06 | 8.267E-06 | 3.233E-06 | 6.400E-06 | 1.200E-05 |
| 29187 | 1460 | V1460Q | E | 3.3e-06 | 5.2e-06 | 4e-06 | 4.167E-06 | 9.609E-07 | 3.300E-06 | 5.200E-06 |
| 29188 | 1460 | V1460G | E | 8.1e-06 | 6.4e-06 | 7.4e-06 | 7.300E-06 | 8.544E-07 | 6.400E-06 | 8.100E-06 |
| 29189 | 1460 | V1460H | E | 1.2e-05 | 9.2e-06 | 7.9e-06 | 9.700E-06 | 2.095E-06 | 7.900E-06 | 1.200E-05 |
| 29190 | 1460 | V1460I | E | 6.4e-06 | 5.9e-06 | 4e-06 | 5.433E-06 | 1.266E-06 | 4.000E-06 | 6.400E-06 |
| 29191 | 1460 | V1460L | E | 6.7e-06 | 6.8e-06 | 7.1e-06 | 6.867E-06 | 2.082E-07 | 6.700E-06 | 7.100E-06 |
| 29192 | 1460 | V1460K | E | 6.3e-06 | 7e-06 | 5.4e-06 | 6.233E-06 | 8.021E-07 | 5.400E-06 | 7.000E-06 |
| 29193 | 1460 | V1460M | E | 8.2e-06 | 5.3e-06 | 8.2e-06 | 7.233E-06 | 1.674E-06 | 5.300E-06 | 8.200E-06 |
| 29194 | 1460 | V1460F | E | 8.7e-06 | 3.1e-06 | 1.1e-05 | 7.600E-06 | 4.063E-06 | 3.100E-06 | 1.100E-05 |
| 29195 | 1460 | V1460P | E | 7e-06 | 7.7e-06 | 1e-05 | 8.233E-06 | 1.570E-06 | 7.000E-06 | 1.000E-05 |
| 29196 | 1460 | V1460S | E | 8.8e-06 | 8.3e-06 | 9.2e-06 | 8.767E-06 | 4.509E-07 | 8.300E-06 | 9.200E-06 |
| 29197 | 1460 | V1460T | E | 6.1e-06 | 9.5e-06 | 5.8e-06 | 7.133E-06 | 2.055E-06 | 5.800E-06 | 9.500E-06 |
| 29198 | 1460 | V1460W | E | 6.1e-06 | 9.9e-06 | 1.1e-05 | 9.000E-06 | 2.571E-06 | 6.100E-06 | 1.100E-05 |
| 29199 | 1460 | V1460Y | E | 6.1e-06 | 7e-06 | 7.6e-06 | 6.900E-06 | 7.550E-07 | 6.100E-06 | 7.600E-06 |
| 29200 | 1460 | V1460V | E | 5.8e-06 | 6e-06 | 9.3e-06 | 7.033E-06 | 1.966E-06 | 5.800E-06 | 9.300E-06 |
| 29201 | 1461 | V1461A | E | 8e-06 | 6.9e-06 | 5.2e-06 | 6.700E-06 | 1.411E-06 | 5.200E-06 | 8.000E-06 |
| 29202 | 1461 | V1461R | E | 6.8e-06 | 1.3e-05 | 1e-05 | 9.933E-06 | 3.101E-06 | 6.800E-06 | 1.300E-05 |
| 29203 | 1461 | V1461N | E | 1.2e-05 | 1.6e-05 | 1.1e-05 | 1.300E-05 | 2.646E-06 | 1.100E-05 | 1.600E-05 |
| 29204 | 1461 | V1461D | E | 6.6e-06 | 7e-06 | 9.6e-06 | 7.733E-06 | 1.629E-06 | 6.600E-06 | 9.600E-06 |
| 29205 | 1461 | V1461C | E | 7.5e-06 | 1.3e-05 | 8.8e-06 | 9.767E-06 | 2.875E-06 | 7.500E-06 | 1.300E-05 |
| 29206 | 1461 | V1461E | E | 7.4e-06 | 7.5e-06 | 8.4e-06 | 7.767E-06 | 5.508E-07 | 7.400E-06 | 8.400E-06 |
| 29207 | 1461 | V1461Q | E | 4.7e-06 | 1.8e-05 | 6.5e-06 | 9.733E-06 | 7.215E-06 | 4.700E-06 | 1.800E-05 |
| 29208 | 1461 | V1461G | E | 1.3e-05 | 5.4e-06 | 6e-06 | 8.133E-06 | 4.225E-06 | 5.400E-06 | 1.300E-05 |
| 29209 | 1461 | V1461H | E | 5.2e-06 | 5.8e-06 | 9.5e-06 | 6.833E-06 | 2.329E-06 | 5.200E-06 | 9.500E-06 |
| 29210 | 1461 | V1461I | E | 8.1e-06 | 5.2e-06 | 9.5e-06 | 7.600E-06 | 2.193E-06 | 5.200E-06 | 9.500E-06 |
| 29211 | 1461 | V1461L | E | 7.2e-06 | 6.4e-06 | 4.6e-06 | 6.067E-06 | 1.332E-06 | 4.600E-06 | 7.200E-06 |
| 29212 | 1461 | V1461K | E | 9.1e-06 | 6.8e-06 | 4.5e-06 | 6.800E-06 | 2.300E-06 | 4.500E-06 | 9.100E-06 |
| 29213 | 1461 | V1461M | E | 8.5e-06 | 7e-06 | 1.4e-05 | 9.833E-06 | 3.686E-06 | 7.000E-06 | 1.400E-05 |
| 29214 | 1461 | V1461F | E | 8.8e-06 | 8.1e-06 | 5.8e-06 | 7.567E-06 | 1.570E-06 | 5.800E-06 | 8.800E-06 |
| 29215 | 1461 | V1461P | E | 9.3e-06 | 8.7e-06 | 5.8e-06 | 7.933E-06 | 1.872E-06 | 5.800E-06 | 9.300E-06 |
| 29216 | 1461 | V1461S | E | 9.1e-06 | 1e-05 | 9.9e-06 | 9.667E-06 | 4.933E-07 | 9.100E-06 | 1.000E-05 |
| 29217 | 1461 | V1461T | E | 7.3e-06 | 6.9e-06 | 6.2e-06 | 6.800E-06 | 5.568E-07 | 6.200E-06 | 7.300E-06 |
| 29218 | 1461 | V1461W | E | 6.5e-06 | 1.2e-05 | 6.6e-06 | 8.367E-06 | 3.147E-06 | 6.500E-06 | 1.200E-05 |
| 29219 | 1461 | V1461Y | E | 6.8e-06 | 6.8e-06 | 4.7e-06 | 6.100E-06 | 1.212E-06 | 4.700E-06 | 6.800E-06 |
| 29220 | 1461 | V1461V | E | 8e-06 | 6.6e-06 | 9.8e-06 | 8.133E-06 | 1.604E-06 | 6.600E-06 | 9.800E-06 |
| 29221 | 1462 | T1462A | E | 1.7e-05 | 1.1e-05 | 8.3e-06 | 1.210E-05 | 4.453E-06 | 8.300E-06 | 1.700E-05 |
| 29222 | 1462 | T1462R | E | 1.1e-05 | 7.8e-06 | 6.9e-06 | 8.567E-06 | 2.155E-06 | 6.900E-06 | 1.100E-05 |
| 29223 | 1462 | T1462N | E | 7.5e-06 | 5.9e-06 | 8.4e-06 | 7.267E-06 | 1.266E-06 | 5.900E-06 | 8.400E-06 |
| 29224 | 1462 | T1462D | E | 1.6e-05 | 5.5e-06 | 6.2e-06 | 9.233E-06 | 5.871E-06 | 5.500E-06 | 1.600E-05 |
| 29225 | 1462 | T1462C | E | 6.7e-06 | 8.9e-06 | 9.4e-06 | 8.333E-06 | 1.436E-06 | 6.700E-06 | 9.400E-06 |
| 29226 | 1462 | T1462E | E | 5.5e-06 | 6.8e-06 | 7.7e-06 | 6.667E-06 | 1.106E-06 | 5.500E-06 | 7.700E-06 |
| 29227 | 1462 | T1462Q | E | 6.5e-06 | 9.2e-06 | 1e-05 | 8.567E-06 | 1.834E-06 | 6.500E-06 | 1.000E-05 |
| 29228 | 1462 | T1462G | E | 7e-06 | 1.5e-05 | 1.1e-05 | 1.100E-05 | 4.000E-06 | 7.000E-06 | 1.500E-05 |
| 29229 | 1462 | T1462H | E | 5e-06 | 4.6e-06 | 9.9e-06 | 6.500E-06 | 2.951E-06 | 4.600E-06 | 9.900E-06 |
| 29230 | 1462 | T1462I | E | 1.3e-05 | 8.9e-06 | 9.6e-06 | 1.050E-05 | 2.193E-06 | 8.900E-06 | 1.300E-05 |
| 29231 | 1462 | T1462L | E | 8.4e-06 | 1.5e-05 | 1.3e-05 | 1.213E-05 | 3.384E-06 | 8.400E-06 | 1.500E-05 |
| 29232 | 1462 | T1462K | E | 7.3e-06 | 8.5e-06 | 5e-06 | 6.933E-06 | 1.779E-06 | 5.000E-06 | 8.500E-06 |
| 29233 | 1462 | T1462M | E | 7.2e-06 | 1.3e-05 | 7.3e-06 | 9.167E-06 | 3.320E-06 | 7.200E-06 | 1.300E-05 |
| 29234 | 1462 | T1462F | E | 1.2e-05 | 1.3e-05 | 9e-06 | 1.133E-05 | 2.082E-06 | 9.000E-06 | 1.300E-05 |
| 29235 | 1462 | T1462P | E | 1e-05 | 1e-05 | 7.7e-06 | 9.233E-06 | 1.328E-06 | 7.700E-06 | 1.000E-05 |
| 29236 | 1462 | T1462S | E | 5.4e-06 | 1.3e-05 | 7e-06 | 8.467E-06 | 4.007E-06 | 5.400E-06 | 1.300E-05 |
| 29237 | 1462 | T1462T | E | 4.1e-06 | 8.9e-06 | 6.6e-06 | 6.533E-06 | 2.401E-06 | 4.100E-06 | 8.900E-06 |
| 29238 | 1462 | T1462W | E | 1.2e-05 | 5.6e-06 | 1.3e-05 | 1.020E-05 | 4.015E-06 | 5.600E-06 | 1.300E-05 |
| 29239 | 1462 | T1462Y | E | 9.1e-06 | 9.7e-06 | 7.2e-06 | 8.667E-06 | 1.305E-06 | 7.200E-06 | 9.700E-06 |

| | | | | | | | | | | |
|---|---|---|---|---|---|---|---|---|---|---|
| 29240 | 1462 | T1462V | E | 7.2e-06 | 1.1e-05 | 1.3e-05 | 1.040E-05 | 2.946E-06 | 7.200E-06 | 1.300E-05 |
| 29241 | 1463 | V1463A | E | 7.3e-06 | 1.2e-05 | 7.7e-06 | 9.000E-06 | 2.606E-06 | 7.300E-06 | 1.200E-05 |
| 29242 | 1463 | V1463R | E | 9.1e-06 | 7.9e-06 | 8.5e-06 | 8.500E-06 | 6.000E-07 | 7.900E-06 | 9.100E-06 |
| 29243 | 1463 | V1463N | E | 5.4e-06 | 1e-05 | 1.7e-05 | 1.080E-05 | 5.841E-06 | 5.400E-06 | 1.700E-05 |
| 29244 | 1463 | V1463D | E | 1.5e-05 | 9.5e-06 | 8.6e-06 | 1.103E-05 | 3.465E-06 | 8.600E-06 | 1.500E-05 |
| 29245 | 1463 | V1463C | E | 8.4e-06 | 5.2e-06 | 9.2e-06 | 7.600E-06 | 2.117E-06 | 5.200E-06 | 9.200E-06 |
| 29246 | 1463 | V1463E | E | 1.2e-05 | 6.3e-06 | 8.6e-06 | 8.967E-06 | 2.868E-06 | 6.300E-06 | 1.200E-05 |
| 29247 | 1463 | V1463Q | E | 1.3e-05 | 1.2e-05 | 6.3e-06 | 1.043E-05 | 3.614E-06 | 6.300E-06 | 1.300E-05 |
| 29248 | 1463 | V1463G | E | 1e-05 | 6.3e-06 | 9.5e-06 | 8.600E-06 | 2.007E-06 | 6.300E-06 | 1.000E-05 |
| 29249 | 1463 | V1463H | E | 7.7e-06 | 8.2e-06 | 9.1e-06 | 8.333E-06 | 7.095E-07 | 7.700E-06 | 9.100E-06 |
| 29250 | 1463 | V1463I | E | 7.4e-06 | 7e-06 | 4.1e-06 | 6.167E-06 | 1.801E-06 | 4.100E-06 | 7.400E-06 |
| 29251 | 1463 | V1463L | E | 6.2e-06 | 9.5e-06 | 7.1e-06 | 7.600E-06 | 1.706E-06 | 6.200E-06 | 9.500E-06 |
| 29252 | 1463 | V1463K | E | 6.3e-06 | 7.2e-06 | 1.3e-05 | 8.833E-06 | 3.636E-06 | 6.300E-06 | 1.300E-05 |
| 29253 | 1463 | V1463M | E | 9.1e-06 | 9.8e-06 | 6.4e-06 | 8.433E-06 | 1.795E-06 | 6.400E-06 | 9.800E-06 |
| 29254 | 1463 | V1463F | E | 5.2e-06 | 7.4e-06 | 1.1e-05 | 7.867E-06 | 2.928E-06 | 5.200E-06 | 1.100E-05 |
| 29255 | 1463 | V1463P | E | 9.4e-06 | 9.8e-06 | 9e-06 | 9.400E-06 | 4.000E-07 | 9.000E-06 | 9.800E-06 |
| 29256 | 1463 | V1463S | E | 6.3e-06 | 1.3e-05 | 8.5e-06 | 9.267E-06 | 3.415E-06 | 6.300E-06 | 1.300E-05 |
| 29257 | 1463 | V1463T | E | 7.1e-06 | 6.1e-06 | 7.6e-06 | 6.933E-06 | 7.638E-07 | 6.100E-06 | 7.600E-06 |
| 29258 | 1463 | V1463W | E | 9.3e-06 | 7.3e-06 | 6.6e-06 | 7.733E-06 | 1.401E-06 | 6.600E-06 | 9.300E-06 |
| 29259 | 1463 | V1463Y | E | 8.1e-06 | 9.2e-06 | 6.5e-06 | 7.933E-06 | 1.358E-06 | 6.500E-06 | 9.200E-06 |
| 29260 | 1463 | V1463V | E | 7.1e-06 | 1e-05 | 9.5e-06 | 8.867E-06 | 1.550E-06 | 7.100E-06 | 1.000E-05 |
| 29261 | 1464 | P1464A | E | 1e-05 | 5.5e-06 | 6.5e-06 | 7.333E-06 | 2.363E-06 | 5.500E-06 | 1.000E-05 |
| 29262 | 1464 | P1464R | E | 1.3e-05 | 1.3e-05 | 1e-05 | 1.200E-05 | 1.732E-06 | 1.000E-05 | 1.300E-05 |
| 29263 | 1464 | P1464N | E | 1.5e-05 | 9.1e-06 | 4.9e-06 | 9.667E-06 | 5.074E-06 | 4.900E-06 | 1.500E-05 |
| 29264 | 1464 | P1464D | E | 7.9e-06 | 6.8e-06 | 1e-05 | 8.233E-06 | 1.626E-06 | 6.800E-06 | 1.000E-05 |
| 29265 | 1464 | P1464C | E | 4.6e-06 | 9.4e-06 | 6.2e-06 | 6.733E-06 | 2.444E-06 | 4.600E-06 | 9.400E-06 |
| 29266 | 1464 | P1464E | E | 6.2e-06 | 1.3e-05 | 6.4e-06 | 8.533E-06 | 3.870E-06 | 6.200E-06 | 1.300E-05 |
| 29267 | 1464 | P1464Q | E | 1.2e-05 | 8.7e-06 | 1.1e-05 | 1.057E-05 | 1.692E-06 | 8.700E-06 | 1.200E-05 |
| 29268 | 1464 | P1464G | E | 7e-06 | 7.7e-06 | 1.4e-05 | 9.567E-06 | 3.855E-06 | 7.000E-06 | 1.400E-05 |
| 29269 | 1464 | P1464H | E | 9.1e-06 | 6.6e-06 | 1e-05 | 8.567E-06 | 1.762E-06 | 6.600E-06 | 1.000E-05 |
| 29270 | 1464 | P1464I | E | 7.5e-06 | 4.6e-06 | 4.8e-06 | 5.633E-06 | 1.620E-06 | 4.600E-06 | 7.500E-06 |
| 29271 | 1464 | P1464L | E | 7e-06 | 6.9e-06 | 9.1e-06 | 7.667E-06 | 1.242E-06 | 6.900E-06 | 9.100E-06 |
| 29272 | 1464 | P1464K | E | 7.3e-06 | 1.2e-05 | 8.9e-06 | 9.400E-06 | 2.390E-06 | 7.300E-06 | 1.200E-05 |
| 29273 | 1464 | P1464M | E | 5e-06 | 8.4e-06 | 6.8e-06 | 6.733E-06 | 1.701E-06 | 5.000E-06 | 8.400E-06 |
| 29274 | 1464 | P1464F | E | 3.9e-06 | 5.8e-06 | 1.2e-05 | 7.233E-06 | 4.236E-06 | 3.900E-06 | 1.200E-05 |
| 29275 | 1464 | P1464P | E | 1.4e-05 | 1.1e-05 | 1.3e-05 | 1.267E-05 | 1.528E-06 | 1.100E-05 | 1.400E-05 |
| 29276 | 1464 | P1464S | E | 4.7e-06 | 1.3e-05 | 7e-06 | 8.233E-06 | 4.285E-06 | 4.700E-06 | 1.300E-05 |
| 29277 | 1464 | P1464T | E | 5.9e-06 | 7.5e-06 | 5.5e-06 | 6.300E-06 | 1.058E-06 | 5.500E-06 | 7.500E-06 |
| 29278 | 1464 | P1464W | E | 5.3e-06 | 5.4e-06 | 9.9e-06 | 6.867E-06 | 2.627E-06 | 5.300E-06 | 9.900E-06 |
| 29279 | 1464 | P1464Y | E | 6.1e-06 | 9.1e-06 | 8.8e-06 | 8.000E-06 | 1.652E-06 | 6.100E-06 | 9.100E-06 |
| 29280 | 1464 | P1464V | E | 6.2e-06 | 9.3e-06 | 7e-06 | 7.500E-06 | 1.609E-06 | 6.200E-06 | 9.300E-06 |
| 29281 | 1465 | S1465A | E | 1e-05 | 7.6e-06 | 3.9e-06 | 7.167E-06 | 3.073E-06 | 3.900E-06 | 1.000E-05 |
| 29282 | 1465 | S1465R | E | 7e-06 | 1.2e-05 | 8.2e-06 | 9.067E-06 | 2.610E-06 | 7.000E-06 | 1.200E-05 |
| 29283 | 1465 | S1465N | E | 9.8e-06 | 8.6e-06 | 1.3e-05 | 1.047E-05 | 2.274E-06 | 8.600E-06 | 1.300E-05 |
| 29284 | 1465 | S1465D | E | 5.8e-06 | 7.1e-06 | 1.1e-05 | 7.967E-06 | 2.706E-06 | 5.800E-06 | 1.100E-05 |
| 29285 | 1465 | S1465C | E | 7.7e-06 | 5.6e-06 | 6.9e-06 | 6.733E-06 | 1.060E-06 | 5.600E-06 | 7.700E-06 |
| 29286 | 1465 | S1465E | E | 1.5e-05 | 6.6e-06 | 8.5e-06 | 1.003E-05 | 4.405E-06 | 6.600E-06 | 1.500E-05 |
| 29287 | 1465 | S1465Q | E | 9.8e-06 | 8.9e-06 | 6.3e-06 | 8.333E-06 | 1.818E-06 | 6.300E-06 | 9.800E-06 |
| 29288 | 1465 | S1465G | E | 8.3e-06 | 7.2e-06 | 1.3e-05 | 9.500E-06 | 3.081E-06 | 7.200E-06 | 1.300E-05 |
| 29289 | 1465 | S1465H | E | 5.5e-06 | 1.1e-05 | 1.3e-05 | 9.833E-06 | 3.884E-06 | 5.500E-06 | 1.300E-05 |
| 29290 | 1465 | S1465I | E | 1e-05 | 1.3e-05 | 8.4e-06 | 1.047E-05 | 2.335E-06 | 8.400E-06 | 1.300E-05 |
| 29291 | 1465 | S1465L | E | 8.7e-06 | 7.7e-06 | 9.2e-06 | 8.533E-06 | 7.638E-07 | 7.700E-06 | 9.200E-06 |
| 29292 | 1465 | S1465K | E | 5.9e-06 | 7.7e-06 | 9.7e-06 | 7.767E-06 | 1.901E-06 | 5.900E-06 | 9.700E-06 |
| 29293 | 1465 | S1465M | E | 6.9e-06 | 4.8e-06 | 6.7e-06 | 6.133E-06 | 1.159E-06 | 4.800E-06 | 6.900E-06 |
| 29294 | 1465 | S1465F | E | 6.7e-06 | 5.9e-06 | 5.9e-06 | 6.167E-06 | 4.619E-07 | 5.900E-06 | 6.700E-06 |
| 29295 | 1465 | S1465P | E | 6.6e-06 | 9.8e-06 | 6.6e-06 | 7.667E-06 | 1.848E-06 | 6.600E-06 | 9.800E-06 |
| 29296 | 1465 | S1465S | E | 9.3e-06 | 8.1e-06 | 6.6e-06 | 8.000E-06 | 1.353E-06 | 6.600E-06 | 9.300E-06 |
| 29297 | 1465 | S1465T | E | 9e-06 | 1.4e-05 | 8.8e-06 | 1.060E-05 | 2.946E-06 | 8.800E-06 | 1.400E-05 |
| 29298 | 1465 | S1465W | E | 8.8e-06 | 5.3e-06 | 1.3e-05 | 9.033E-06 | 3.855E-06 | 5.300E-06 | 1.300E-05 |
| 29299 | 1465 | S1465Y | E | 5e-06 | 4.7e-06 | 1.8e-05 | 9.233E-06 | 7.594E-06 | 4.700E-06 | 1.800E-05 |
| 29300 | 1465 | S1465V | E | 4.5e-06 | 6.5e-06 | 7.1e-06 | 6.033E-06 | 1.361E-06 | 4.500E-06 | 7.100E-06 |
| 29301 | 1466 | S1466A | E | 1.1e-05 | 1e-05 | 5e-06 | 8.667E-06 | 3.215E-06 | 5.000E-06 | 1.100E-05 |
| 29302 | 1466 | S1466R | E | 5.3e-06 | 4.8e-06 | 7.4e-06 | 5.833E-06 | 1.380E-06 | 4.800E-06 | 7.400E-06 |
| 29303 | 1466 | S1466N | E | 9.1e-06 | 7.2e-06 | 8.8e-06 | 8.367E-06 | 1.021E-06 | 7.200E-06 | 9.100E-06 |
| 29304 | 1466 | S1466D | E | 6.3e-06 | 5.1e-06 | 1e-05 | 7.133E-06 | 2.554E-06 | 5.100E-06 | 1.000E-05 |
| 29305 | 1466 | S1466C | E | 7.7e-06 | 6.4e-06 | 1.1e-05 | 8.367E-06 | 2.371E-06 | 6.400E-06 | 1.100E-05 |
| 29306 | 1466 | S1466E | E | 1e-05 | 8.8e-06 | 1.3e-05 | 1.060E-05 | 2.163E-06 | 8.800E-06 | 1.300E-05 |
| 29307 | 1466 | S1466Q | E | 1.3e-05 | 7.3e-06 | 3.6e-06 | 7.967E-06 | 4.735E-06 | 3.600E-06 | 1.300E-05 |
| 29308 | 1466 | S1466G | E | 6.3e-06 | 5.4e-06 | 6e-06 | 5.900E-06 | 4.583E-07 | 5.400E-06 | 6.300E-06 |
| 29309 | 1466 | S1466H | E | 1.9e-05 | 8.2e-06 | 5.3e-06 | 1.083E-05 | 7.220E-06 | 5.300E-06 | 1.900E-05 |
| 29310 | 1466 | S1466I | E | 5.8e-06 | 5.1e-06 | 5.9e-06 | 5.600E-06 | 4.359E-07 | 5.100E-06 | 5.900E-06 |
| 29311 | 1466 | S1466L | E | 4.8e-06 | 7.7e-06 | 5.1e-06 | 5.867E-06 | 1.595E-06 | 4.800E-06 | 7.700E-06 |
| 29312 | 1466 | S1466K | E | 1.1e-05 | 1.4e-05 | 4.2e-06 | 9.733E-06 | 5.021E-06 | 4.200E-06 | 1.400E-05 |
| 29313 | 1466 | S1466M | E | 8.4e-06 | 1e-05 | 1.1e-05 | 9.800E-06 | 1.311E-06 | 8.400E-06 | 1.100E-05 |
| 29314 | 1466 | S1466F | E | 5.5e-06 | 9.6e-06 | 1.4e-05 | 9.700E-06 | 4.251E-06 | 5.500E-06 | 1.400E-05 |
| 29315 | 1466 | S1466P | E | 1.1e-05 | 7.3e-06 | 7.7e-06 | 8.667E-06 | 2.031E-06 | 7.300E-06 | 1.100E-05 |
| 29316 | 1466 | S1466S | E | 6e-06 | 7e-06 | 4.2e-06 | 5.733E-06 | 1.419E-06 | 4.200E-06 | 7.000E-06 |
| 29317 | 1466 | S1466T | E | 4.8e-06 | 5e-06 | 5.6e-06 | 5.133E-06 | 4.163E-07 | 4.800E-06 | 5.600E-06 |
| 29318 | 1466 | S1466W | E | 7.8e-06 | 6.4e-06 | 7.4e-06 | 7.200E-06 | 7.211E-07 | 6.400E-06 | 7.800E-06 |
| 29319 | 1466 | S1466Y | E | 8.1e-06 | 1e-05 | 6.4e-06 | 8.167E-06 | 1.801E-06 | 6.400E-06 | 1.000E-05 |
| 29320 | 1466 | S1466V | E | 4.3e-06 | 9.4e-06 | 1e-05 | 7.900E-06 | 3.132E-06 | 4.300E-06 | 1.000E-05 |
| 29321 | 1467 | S1467A | E | 1.1e-05 | 6e-06 | 4.5e-06 | 7.167E-06 | 3.403E-06 | 4.500E-06 | 1.100E-05 |
| 29322 | 1467 | S1467R | E | 8.8e-06 | 9.6e-06 | 9.4e-06 | 9.267E-06 | 4.163E-07 | 8.800E-06 | 9.600E-06 |
| 29323 | 1467 | S1467N | E | 9.6e-06 | 8.8e-06 | 1.1e-05 | 9.800E-06 | 1.114E-06 | 8.800E-06 | 1.100E-05 |
| 29324 | 1467 | S1467D | E | 9e-06 | 6.4e-06 | 9.1e-06 | 8.167E-06 | 1.531E-06 | 6.400E-06 | 9.100E-06 |

| | | | | | | | | | | |
|---|---|---|---|---|---|---|---|---|---|---|
| 29325 | 1467 | S1467C | E | 6.7e-06 | 9.7e-06 | 7.5e-06 | 7.967E-06 | 1.553E-06 | 6.700E-06 | 9.700E-06 |
| 29326 | 1467 | S1467E | E | 5.8e-06 | 6.5e-06 | 6e-06 | 6.100E-06 | 3.606E-07 | 5.800E-06 | 6.500E-06 |
| 29327 | 1467 | S1467Q | E | 9.1e-06 | 1.1e-05 | 7e-06 | 9.033E-06 | 2.001E-06 | 7.000E-06 | 1.100E-05 |
| 29328 | 1467 | S1467G | E | 6.6e-06 | 4.9e-06 | 9.9e-06 | 7.133E-06 | 2.542E-06 | 4.900E-06 | 9.900E-06 |
| 29329 | 1467 | S1467H | E | 5.7e-06 | 9.2e-06 | 6.5e-06 | 7.133E-06 | 1.834E-06 | 5.700E-06 | 9.200E-06 |
| 29330 | 1467 | S1467I | E | 1e-05 | 6.4e-06 | 5.2e-06 | 7.200E-06 | 2.498E-06 | 5.200E-06 | 1.000E-05 |
| 29331 | 1467 | S1467L | E | 5.7e-06 | 7.8e-06 | 6.7e-06 | 6.733E-06 | 1.050E-06 | 5.700E-06 | 7.800E-06 |
| 29332 | 1467 | S1467K | E | 8.4e-06 | 4.5e-06 | 1.1e-05 | 7.967E-06 | 3.272E-06 | 4.500E-06 | 1.100E-05 |
| 29333 | 1467 | S1467M | E | 6.4e-06 | 5.9e-06 | 4.3e-06 | 5.533E-06 | 1.097E-06 | 4.300E-06 | 6.400E-06 |
| 29334 | 1467 | S1467F | E | 8.3e-06 | 9.2e-06 | 6.9e-06 | 8.133E-06 | 1.159E-06 | 6.900E-06 | 9.200E-06 |
| 29335 | 1467 | S1467P | E | 9.6e-06 | 1e-05 | 5.8e-06 | 8.467E-06 | 2.318E-06 | 5.800E-06 | 1.000E-05 |
| 29336 | 1467 | S1467S | E | 9.6e-06 | 9.8e-06 | 9.2e-06 | 9.533E-06 | 3.055E-07 | 9.200E-06 | 9.800E-06 |
| 29337 | 1467 | S1467T | E | 7.5e-06 | 5.8e-06 | 9.5e-06 | 7.600E-06 | 1.852E-06 | 5.800E-06 | 9.500E-06 |
| 29338 | 1467 | S1467W | E | 8.3e-06 | 2.1e-05 | 9.9e-06 | 1.307E-05 | 6.917E-06 | 8.300E-06 | 2.100E-05 |
| 29339 | 1467 | S1467Y | E | 1e-05 | 7.5e-06 | 4.3e-06 | 7.267E-06 | 2.857E-06 | 4.300E-06 | 1.000E-05 |
| 29340 | 1467 | S1467V | E | 8.6e-06 | 7.2e-06 | 1e-05 | 8.600E-06 | 1.400E-06 | 7.200E-06 | 1.000E-05 |
| 29341 | 1468 | L1468A | E | 7.6e-06 | 7.5e-06 | 9.1e-06 | 8.067E-06 | 8.963E-07 | 7.500E-06 | 9.100E-06 |
| 29342 | 1468 | L1468R | E | 8.7e-06 | 6.7e-06 | 6.1e-06 | 7.167E-06 | 1.361E-06 | 6.100E-06 | 8.700E-06 |
| 29343 | 1468 | L1468N | E | 1.3e-05 | 1.2e-05 | 7.4e-06 | 1.080E-05 | 2.987E-06 | 7.400E-06 | 1.300E-05 |
| 29344 | 1468 | L1468D | E | 7.8e-06 | 1.1e-05 | 9e-06 | 9.267E-06 | 1.617E-06 | 7.800E-06 | 1.100E-05 |
| 29345 | 1468 | L1468C | E | 9e-06 | 9.3e-06 | 1e-05 | 9.433E-06 | 5.132E-07 | 9.000E-06 | 1.000E-05 |
| 29346 | 1468 | L1468E | E | 1e-05 | 7.1e-06 | 1e-05 | 9.033E-06 | 1.674E-06 | 7.100E-06 | 1.000E-05 |
| 29347 | 1468 | L1468Q | E | 5.8e-06 | 8.9e-06 | 6.4e-06 | 7.033E-06 | 1.644E-06 | 5.800E-06 | 8.900E-06 |
| 29348 | 1468 | L1468G | E | 6.2e-06 | 9.7e-06 | 1.3e-05 | 9.633E-06 | 3.400E-06 | 6.200E-06 | 1.300E-05 |
| 29349 | 1468 | L1468H | E | 9.9e-06 | 6.3e-06 | 1.2e-05 | 9.400E-06 | 2.883E-06 | 6.300E-06 | 1.200E-05 |
| 29350 | 1468 | L1468I | E | 1e-05 | 9.9e-06 | 1e-05 | 9.967E-06 | 5.774E-08 | 9.900E-06 | 1.000E-05 |
| 29351 | 1468 | L1468L | E | 1.4e-05 | 6.5e-06 | 8.5e-06 | 9.667E-06 | 3.884E-06 | 6.500E-06 | 1.400E-05 |
| 29352 | 1468 | L1468K | E | 1.5e-05 | 1.2e-05 | 1.3e-05 | 1.333E-05 | 1.528E-06 | 1.200E-05 | 1.500E-05 |
| 29353 | 1468 | L1468M | E | 8e-06 | 5.1e-06 | 1.3e-05 | 8.700E-06 | 3.996E-06 | 5.100E-06 | 1.300E-05 |
| 29354 | 1468 | L1468F | E | 1.1e-05 | 7.7e-06 | 4.3e-06 | 7.667E-06 | 3.350E-06 | 4.300E-06 | 1.100E-05 |
| 29355 | 1468 | L1468P | E | 5e-06 | 9.6e-06 | 6.9e-06 | 7.167E-06 | 2.312E-06 | 5.000E-06 | 9.600E-06 |
| 29356 | 1468 | L1468S | E | 7.2e-06 | 1e-05 | 9.7e-06 | 8.967E-06 | 1.537E-06 | 7.200E-06 | 1.000E-05 |
| 29357 | 1468 | L1468T | E | 7.1e-06 | 7.1e-06 | 7.8e-06 | 7.333E-06 | 4.041E-07 | 7.100E-06 | 7.800E-06 |
| 29358 | 1468 | L1468W | E | 8.6e-06 | 1.2e-05 | 6.6e-06 | 9.067E-06 | 2.730E-06 | 6.600E-06 | 1.200E-05 |
| 29359 | 1468 | L1468Y | E | 8.4e-06 | 7.9e-06 | 4.2e-06 | 6.833E-06 | 2.294E-06 | 4.200E-06 | 8.400E-06 |
| 29360 | 1468 | L1468V | E | 6.1e-06 | 1.1e-05 | 6.8e-06 | 7.967E-06 | 2.650E-06 | 6.100E-06 | 1.100E-05 |
| 29361 | 1469 | G1469A | E | 8.3e-06 | 9.2e-06 | 4.7e-06 | 7.400E-06 | 2.381E-06 | 4.700E-06 | 9.200E-06 |
| 29362 | 1469 | G1469R | E | 9.4e-06 | 8.8e-06 | 1.8e-05 | 1.207E-05 | 5.147E-06 | 8.800E-06 | 1.800E-05 |
| 29363 | 1469 | G1469N | E | 7.3e-06 | 7.8e-06 | 8e-06 | 7.700E-06 | 3.606E-07 | 7.300E-06 | 8.000E-06 |
| 29364 | 1469 | G1469D | E | 3.4e-06 | 1.1e-05 | 1.3e-05 | 9.133E-06 | 5.065E-06 | 3.400E-06 | 1.300E-05 |
| 29365 | 1469 | G1469C | E | 6.2e-06 | 6.9e-06 | 7.2e-06 | 6.767E-06 | 5.132E-07 | 6.200E-06 | 7.200E-06 |
| 29366 | 1469 | G1469E | E | 1.2e-05 | 9.2e-06 | 1.1e-05 | 1.073E-05 | 1.419E-06 | 9.200E-06 | 1.200E-05 |
| 29367 | 1469 | G1469Q | E | 7.3e-06 | 1.3e-05 | 4.6e-06 | 8.300E-06 | 4.288E-06 | 4.600E-06 | 1.300E-05 |
| 29368 | 1469 | G1469G | E | 1.3e-05 | 1.1e-05 | 1.2e-05 | 1.200E-05 | 1.000E-06 | 1.100E-05 | 1.300E-05 |
| 29369 | 1469 | G1469H | E | 7.1e-06 | 1e-05 | 8.2e-06 | 8.433E-06 | 1.464E-06 | 7.100E-06 | 1.000E-05 |
| 29370 | 1469 | G1469I | E | 6.5e-06 | 7.9e-06 | 7.7e-06 | 7.367E-06 | 7.572E-07 | 6.500E-06 | 7.900E-06 |
| 29371 | 1469 | G1469L | E | 9e-06 | 1.3e-05 | 5.9e-06 | 9.300E-06 | 3.559E-06 | 5.900E-06 | 1.300E-05 |
| 29372 | 1469 | G1469K | E | 6.3e-06 | 4.7e-06 | 8.4e-06 | 6.467E-06 | 1.856E-06 | 4.700E-06 | 8.400E-06 |
| 29373 | 1469 | G1469M | E | 9.6e-06 | 5.4e-06 | 8.9e-06 | 7.967E-06 | 2.250E-06 | 5.400E-06 | 9.600E-06 |
| 29374 | 1469 | G1469F | E | 9.6e-06 | 9.6e-06 | 7.7e-06 | 8.967E-06 | 1.097E-06 | 7.700E-06 | 9.600E-06 |
| 29375 | 1469 | G1469P | E | 8.9e-06 | 7.3e-06 | 7.2e-06 | 7.800E-06 | 9.539E-07 | 7.200E-06 | 8.900E-06 |
| 29376 | 1469 | G1469S | E | 7.8e-06 | 1.4e-05 | 8.6e-06 | 1.013E-05 | 3.372E-06 | 7.800E-06 | 1.400E-05 |
| 29377 | 1469 | G1469T | E | 9.8e-06 | 1.1e-05 | 1e-05 | 1.027E-05 | 6.429E-07 | 9.800E-06 | 1.100E-05 |
| 29378 | 1469 | G1469W | E | 9.4e-06 | 9.4e-06 | 3.8e-06 | 7.533E-06 | 3.233E-06 | 3.800E-06 | 9.400E-06 |
| 29379 | 1469 | G1469Y | E | 4.5e-06 | 9.7e-06 | 8.8e-06 | 7.667E-06 | 2.779E-06 | 4.500E-06 | 9.700E-06 |
| 29380 | 1469 | G1469V | E | 1e-05 | 6.9e-06 | 7.7e-06 | 8.200E-06 | 1.609E-06 | 6.900E-06 | 1.000E-05 |
| 29381 | 1470 | T1470A | E | 5.8e-06 | 1.2e-05 | 3.4e-06 | 7.067E-06 | 4.438E-06 | 3.400E-06 | 1.200E-05 |
| 29382 | 1470 | T1470R | E | 6.7e-06 | 7.8e-06 | 7.8e-06 | 7.433E-06 | 6.351E-07 | 6.700E-06 | 7.800E-06 |
| 29383 | 1470 | T1470N | E | 5.2e-06 | 5.8e-06 | 8.9e-06 | 6.633E-06 | 1.986E-06 | 5.200E-06 | 8.900E-06 |
| 29384 | 1470 | T1470D | E | 5.7e-06 | 7.9e-06 | 8.7e-06 | 7.433E-06 | 1.553E-06 | 5.700E-06 | 8.700E-06 |
| 29385 | 1470 | T1470C | E | 6.6e-06 | 4.7e-06 | 9.9e-06 | 7.067E-06 | 2.631E-06 | 4.700E-06 | 9.900E-06 |
| 29386 | 1470 | T1470E | E | 6.7e-06 | 7.9e-06 | 5.5e-06 | 6.700E-06 | 1.200E-06 | 5.500E-06 | 7.900E-06 |
| 29387 | 1470 | T1470Q | E | 1.1e-05 | 6.2e-06 | 1e-05 | 9.067E-06 | 2.532E-06 | 6.200E-06 | 1.100E-05 |
| 29388 | 1470 | T1470G | E | 9.2e-06 | 8e-06 | 7.9e-06 | 8.367E-06 | 7.234E-07 | 7.900E-06 | 9.200E-06 |
| 29389 | 1470 | T1470H | E | 6.8e-06 | 7e-06 | 4.6e-06 | 6.133E-06 | 1.332E-06 | 4.600E-06 | 7.000E-06 |
| 29390 | 1470 | T1470I | E | 6e-06 | 9.1e-06 | 6.2e-06 | 7.100E-06 | 1.735E-06 | 6.000E-06 | 9.100E-06 |
| 29391 | 1470 | T1470L | E | 6.3e-06 | 1.3e-05 | 1.4e-05 | 1.110E-05 | 4.187E-06 | 6.300E-06 | 1.400E-05 |
| 29392 | 1470 | T1470K | E | 7.6e-06 | 6.5e-06 | 1.1e-05 | 8.367E-06 | 2.346E-06 | 6.500E-06 | 1.100E-05 |
| 29393 | 1470 | T1470M | E | 1.2e-05 | 5.2e-06 | 8.3e-06 | 8.500E-06 | 3.404E-06 | 5.200E-06 | 1.200E-05 |
| 29394 | 1470 | T1470F | E | 1e-05 | 5.1e-06 | 7.9e-06 | 7.667E-06 | 2.458E-06 | 5.100E-06 | 1.000E-05 |
| 29395 | 1470 | T1470P | E | 6.7e-06 | 6.7e-06 | 1.1e-05 | 8.133E-06 | 2.483E-06 | 6.700E-06 | 1.100E-05 |
| 29396 | 1470 | T1470S | E | 7.2e-06 | 9e-06 | 9.4e-06 | 8.533E-06 | 1.172E-06 | 7.200E-06 | 9.400E-06 |
| 29397 | 1470 | T1470T | E | 9.6e-06 | 1.3e-05 | 9.1e-06 | 1.057E-05 | 2.122E-06 | 9.100E-06 | 1.300E-05 |
| 29398 | 1470 | T1470W | E | 5.8e-06 | 4.5e-06 | 8.1e-06 | 6.133E-06 | 1.823E-06 | 4.500E-06 | 8.100E-06 |
| 29399 | 1470 | T1470Y | E | 9.6e-06 | 8.5e-06 | 6.6e-06 | 8.233E-06 | 1.518E-06 | 6.600E-06 | 9.600E-06 |
| 29400 | 1470 | T1470V | E | 1.2e-05 | 6.8e-06 | 6.8e-06 | 8.533E-06 | 3.002E-06 | 6.800E-06 | 1.200E-05 |
| 29401 | 1471 | Q1471A | E | 1.4e-05 | 5.3e-06 | 9.9e-06 | 9.733E-06 | 4.352E-06 | 5.300E-06 | 1.400E-05 |
| 29402 | 1471 | Q1471R | E | 6e-06 | 7.6e-06 | 6.6e-06 | 6.733E-06 | 8.083E-07 | 6.000E-06 | 7.600E-06 |
| 29403 | 1471 | Q1471N | E | 5.3e-06 | 4.1e-06 | 1.1e-05 | 6.800E-06 | 3.686E-06 | 4.100E-06 | 1.100E-05 |
| 29404 | 1471 | Q1471D | E | 8.4e-06 | 7.1e-06 | 9.8e-06 | 8.433E-06 | 1.350E-06 | 7.100E-06 | 9.800E-06 |
| 29405 | 1471 | Q1471C | E | 6.3e-06 | 9.5e-06 | 5.9e-06 | 7.233E-06 | 1.973E-06 | 5.900E-06 | 9.500E-06 |
| 29406 | 1471 | Q1471E | E | 8.1e-06 | 9.7e-06 | 6.1e-06 | 7.967E-06 | 1.804E-06 | 6.100E-06 | 9.700E-06 |
| 29407 | 1471 | Q1471Q | E | 6.5e-06 | 8.4e-06 | 9.3e-06 | 8.067E-06 | 1.429E-06 | 6.500E-06 | 9.300E-06 |
| 29408 | 1471 | Q1471G | E | 5.6e-06 | 8.2e-06 | 9.4e-06 | 7.733E-06 | 1.943E-06 | 5.600E-06 | 9.400E-06 |
| 29409 | 1471 | Q1471H | E | 9.7e-06 | 5.9e-06 | 6.7e-06 | 7.433E-06 | 2.003E-06 | 5.900E-06 | 9.700E-06 |

| | | | | | | | | | | |
|---|---|---|---|---|---|---|---|---|---|---|
| 29410 | 1471 | Q1471I | E | 8.4e-06 | 9.7e-06 | 9.5e-06 | 9.200E-06 | 7.000E-07 | 8.400E-06 | 9.700E-06 |
| 29411 | 1471 | Q1471L | E | 9.8e-06 | 1.1e-05 | 8.6e-06 | 9.800E-06 | 1.200E-06 | 8.600E-06 | 1.100E-05 |
| 29412 | 1471 | Q1471K | E | 8.5e-06 | 6.4e-06 | 1.1e-05 | 8.633E-06 | 2.303E-06 | 6.400E-06 | 1.100E-05 |
| 29413 | 1471 | Q1471M | E | 1.5e-05 | 9.6e-06 | 5.9e-06 | 1.017E-05 | 4.576E-06 | 5.900E-06 | 1.500E-05 |
| 29414 | 1471 | Q1471F | E | 6.2e-06 | 6.3e-06 | 5e-06 | 5.833E-06 | 7.234E-07 | 5.000E-06 | 6.300E-06 |
| 29415 | 1471 | Q1471P | E | 7e-06 | 8.4e-06 | 4.7e-06 | 6.700E-06 | 1.868E-06 | 4.700E-06 | 8.400E-06 |
| 29416 | 1471 | Q1471S | E | 8.5e-06 | 1.9e-05 | 6.9e-06 | 1.147E-05 | 6.573E-06 | 6.900E-06 | 1.900E-05 |
| 29417 | 1471 | Q1471T | E | 4.2e-06 | 8.5e-06 | 6.9e-06 | 6.533E-06 | 2.173E-06 | 4.200E-06 | 8.500E-06 |
| 29418 | 1471 | Q1471W | E | 5.2e-06 | 9.7e-06 | 1.6e-05 | 1.030E-05 | 5.425E-06 | 5.200E-06 | 1.600E-05 |
| 29419 | 1471 | Q1471Y | E | 6.8e-06 | 4.5e-06 | 9.3e-06 | 6.867E-06 | 2.401E-06 | 4.500E-06 | 9.300E-06 |
| 29420 | 1471 | Q1471V | E | 5.7e-06 | 7.1e-06 | 8.3e-06 | 7.033E-06 | 1.301E-06 | 5.700E-06 | 8.300E-06 |
| 29421 | 1472 | T1472A | E | 5e-06 | 4.6e-06 | 9.9e-06 | 6.500E-06 | 2.951E-06 | 4.600E-06 | 9.900E-06 |
| 29422 | 1472 | T1472R | E | 9.8e-06 | 9.6e-06 | 4.8e-06 | 8.067E-06 | 2.831E-06 | 4.800E-06 | 9.800E-06 |
| 29423 | 1472 | T1472N | E | 4.6e-06 | 9e-06 | 7.6e-06 | 7.067E-06 | 2.248E-06 | 4.600E-06 | 9.000E-06 |
| 29424 | 1472 | T1472D | E | 7.8e-06 | 9.2e-06 | 8.7e-06 | 8.567E-06 | 7.095E-07 | 7.800E-06 | 9.200E-06 |
| 29425 | 1472 | T1472C | E | 6.2e-06 | 6.6e-06 | 9e-06 | 7.267E-06 | 1.514E-06 | 6.200E-06 | 9.000E-06 |
| 29426 | 1472 | T1472E | E | 1e-05 | 8.9e-06 | 7.1e-06 | 8.667E-06 | 1.464E-06 | 7.100E-06 | 1.000E-05 |
| 29427 | 1472 | T1472Q | E | 9.4e-06 | 7.3e-06 | 8.1e-06 | 8.267E-06 | 1.060E-06 | 7.300E-06 | 9.400E-06 |
| 29428 | 1472 | T1472G | E | 6.3e-06 | 8.6e-06 | 9.6e-06 | 8.167E-06 | 1.692E-06 | 6.300E-06 | 9.600E-06 |
| 29429 | 1472 | T1472H | E | 4.1e-06 | 6.8e-06 | 6.7e-06 | 5.867E-06 | 1.531E-06 | 4.100E-06 | 6.800E-06 |
| 29430 | 1472 | T1472I | E | 1.4e-05 | 6.8e-06 | 8.8e-06 | 9.867E-06 | 3.717E-06 | 6.800E-06 | 1.400E-05 |
| 29431 | 1472 | T1472L | E | 5.9e-06 | 6.2e-06 | 9.5e-06 | 7.200E-06 | 1.997E-06 | 5.900E-06 | 9.500E-06 |
| 29432 | 1472 | T1472K | E | 6.4e-06 | 1e-05 | 1e-05 | 8.800E-06 | 2.078E-06 | 6.400E-06 | 1.000E-05 |
| 29433 | 1472 | T1472M | E | 6.5e-06 | 1.2e-05 | 1e-05 | 9.500E-06 | 2.784E-06 | 6.500E-06 | 1.200E-05 |
| 29434 | 1472 | T1472F | E | 1.3e-05 | 5.3e-06 | 7.2e-06 | 8.500E-06 | 4.011E-06 | 5.300E-06 | 1.300E-05 |
| 29435 | 1472 | T1472P | E | 5.8e-06 | 8.7e-06 | 6.6e-06 | 7.033E-06 | 1.498E-06 | 5.800E-06 | 8.700E-06 |
| 29436 | 1472 | T1472S | E | 1e-05 | 8.8e-06 | 8e-06 | 8.933E-06 | 1.007E-06 | 8.000E-06 | 1.000E-05 |
| 29437 | 1472 | T1472T | E | 1.4e-05 | 5.8e-06 | 1.3e-05 | 1.093E-05 | 4.474E-06 | 5.800E-06 | 1.400E-05 |
| 29438 | 1472 | T1472W | E | 1.1e-05 | 1.3e-05 | 8.5e-06 | 1.083E-05 | 2.255E-06 | 8.500E-06 | 1.300E-05 |
| 29439 | 1472 | T1472Y | E | 8.4e-06 | 1e-05 | 1.4e-05 | 1.080E-05 | 2.884E-06 | 8.400E-06 | 1.400E-05 |
| 29440 | 1472 | T1472V | E | 1.3e-05 | 1.2e-05 | 7.6e-06 | 1.087E-05 | 2.873E-06 | 7.600E-06 | 1.300E-05 |
| 29441 | 1473 | Y1473A | E | 1.5e-05 | 6.4e-06 | 8.1e-06 | 9.833E-06 | 4.554E-06 | 6.400E-06 | 1.500E-05 |
| 29442 | 1473 | Y1473R | E | 9.3e-06 | 7.6e-06 | 7e-06 | 7.967E-06 | 1.193E-06 | 7.000E-06 | 9.300E-06 |
| 29443 | 1473 | Y1473N | E | 6.4e-06 | 6.8e-06 | 6.9e-06 | 6.700E-06 | 2.646E-07 | 6.400E-06 | 6.900E-06 |
| 29444 | 1473 | Y1473D | E | 6.3e-06 | 5.6e-06 | 7.9e-06 | 6.600E-06 | 1.179E-06 | 5.600E-06 | 7.900E-06 |
| 29445 | 1473 | Y1473C | E | 7.5e-06 | 1.1e-05 | 4.4e-06 | 7.633E-06 | 3.302E-06 | 4.400E-06 | 1.100E-05 |
| 29446 | 1473 | Y1473E | E | 6.5e-06 | 7.3e-06 | 8.8e-06 | 7.533E-06 | 1.168E-06 | 6.500E-06 | 8.800E-06 |
| 29447 | 1473 | Y1473Q | E | 6.5e-06 | 5.6e-06 | 7e-06 | 6.367E-06 | 7.095E-07 | 5.600E-06 | 7.000E-06 |
| 29448 | 1473 | Y1473G | E | 1.3e-05 | 8e-06 | 6.8e-06 | 9.267E-06 | 3.288E-06 | 6.800E-06 | 1.300E-05 |
| 29449 | 1473 | Y1473H | E | 1.4e-05 | 1.1e-05 | 9.8e-06 | 1.160E-05 | 2.163E-06 | 9.800E-06 | 1.400E-05 |
| 29450 | 1473 | Y1473I | E | 4.3e-06 | 4.5e-06 | 5.4e-06 | 4.733E-06 | 5.859E-07 | 4.300E-06 | 5.400E-06 |
| 29451 | 1473 | Y1473L | E | 6.1e-06 | 1.1e-05 | 1.4e-05 | 1.037E-05 | 3.988E-06 | 6.100E-06 | 1.400E-05 |
| 29452 | 1473 | Y1473K | E | 9.5e-06 | 1.4e-05 | 1.1e-05 | 1.150E-05 | 2.291E-06 | 9.500E-06 | 1.400E-05 |
| 29453 | 1473 | Y1473M | E | 1.4e-05 | 7.1e-06 | 7.9e-06 | 9.667E-06 | 3.774E-06 | 7.100E-06 | 1.400E-05 |
| 29454 | 1473 | Y1473F | E | 6.7e-06 | 8.5e-06 | 5.1e-06 | 6.767E-06 | 1.701E-06 | 5.100E-06 | 8.500E-06 |
| 29455 | 1473 | Y1473P | E | 8.9e-06 | 9.3e-06 | 7.7e-06 | 8.633E-06 | 8.327E-07 | 7.700E-06 | 9.300E-06 |
| 29456 | 1473 | Y1473S | E | 9.7e-06 | 8.1e-06 | 7.2e-06 | 8.333E-06 | 1.266E-06 | 7.200E-06 | 9.700E-06 |
| 29457 | 1473 | Y1473T | E | 6.8e-06 | 7.2e-06 | 6.2e-06 | 6.733E-06 | 5.033E-07 | 6.200E-06 | 7.200E-06 |
| 29458 | 1473 | Y1473W | E | 8e-06 | 1.1e-05 | 8.9e-06 | 9.300E-06 | 1.539E-06 | 8.000E-06 | 1.100E-05 |
| 29459 | 1473 | Y1473Y | E | 9.6e-06 | 9.4e-06 | 1e-05 | 9.667E-06 | 3.055E-07 | 9.400E-06 | 1.000E-05 |
| 29460 | 1473 | Y1473V | E | 4.5e-06 | 6.9e-06 | 7.2e-06 | 6.200E-06 | 1.480E-06 | 4.500E-06 | 7.200E-06 |
| 29461 | 1474 | I1474A | E | 8.2e-06 | 9.1e-06 | 5.1e-06 | 7.467E-06 | 2.098E-06 | 5.100E-06 | 9.100E-06 |
| 29462 | 1474 | I1474R | E | 6.5e-06 | 5.6e-06 | 1.3e-05 | 8.367E-06 | 4.038E-06 | 5.600E-06 | 1.300E-05 |
| 29463 | 1474 | I1474N | E | 9.1e-06 | 1.1e-05 | 5e-06 | 8.367E-06 | 3.066E-06 | 5.000E-06 | 1.100E-05 |
| 29464 | 1474 | I1474D | E | 6e-06 | 8.1e-06 | 1e-05 | 8.033E-06 | 2.001E-06 | 6.000E-06 | 1.000E-05 |
| 29465 | 1474 | I1474C | E | 8.2e-06 | 9e-06 | 5.8e-06 | 7.667E-06 | 1.665E-06 | 5.800E-06 | 9.000E-06 |
| 29466 | 1474 | I1474E | E | 7e-06 | 1e-05 | 4.1e-06 | 7.033E-06 | 2.950E-06 | 4.100E-06 | 1.000E-05 |
| 29467 | 1474 | I1474Q | E | 6.6e-06 | 1e-05 | 7.4e-06 | 8.000E-06 | 1.778E-06 | 6.600E-06 | 1.000E-05 |
| 29468 | 1474 | I1474G | E | 6.7e-06 | 7.3e-06 | 6.5e-06 | 6.833E-06 | 4.163E-07 | 6.500E-06 | 7.300E-06 |
| 29469 | 1474 | I1474H | E | 4.2e-06 | 8.4e-06 | 1.4e-05 | 8.867E-06 | 4.917E-06 | 4.200E-06 | 1.400E-05 |
| 29470 | 1474 | I1474I | E | 1.1e-05 | 8.8e-06 | 7.5e-06 | 9.100E-06 | 1.769E-06 | 7.500E-06 | 1.100E-05 |
| 29471 | 1474 | I1474L | E | 9.3e-06 | 1.4e-05 | 5e-06 | 9.433E-06 | 4.501E-06 | 5.000E-06 | 1.400E-05 |
| 29472 | 1474 | I1474K | E | 8.2e-06 | 7.5e-06 | 6.6e-06 | 7.433E-06 | 8.021E-07 | 6.600E-06 | 8.200E-06 |
| 29473 | 1474 | I1474M | E | 7.9e-06 | 6.7e-06 | 7.4e-06 | 7.333E-06 | 6.028E-07 | 6.700E-06 | 7.900E-06 |
| 29474 | 1474 | I1474F | E | 4.6e-06 | 1e-05 | 8.3e-06 | 7.633E-06 | 2.761E-06 | 4.600E-06 | 1.000E-05 |
| 29475 | 1474 | I1474P | E | 1.5e-05 | 5.7e-06 | 1.1e-05 | 1.057E-05 | 4.665E-06 | 5.700E-06 | 1.500E-05 |
| 29476 | 1474 | I1474S | E | 1.3e-05 | 7.1e-06 | 6.5e-06 | 8.867E-06 | 3.592E-06 | 6.500E-06 | 1.300E-05 |
| 29477 | 1474 | I1474T | E | 4.4e-06 | 7.4e-06 | 6.5e-06 | 6.100E-06 | 1.539E-06 | 4.400E-06 | 7.400E-06 |
| 29478 | 1474 | I1474W | E | 4.9e-06 | 5.8e-06 | 8.5e-06 | 6.400E-06 | 1.873E-06 | 4.900E-06 | 8.500E-06 |
| 29479 | 1474 | I1474Y | E | 9.3e-06 | 5.7e-06 | 1.5e-05 | 1.000E-05 | 4.689E-06 | 5.700E-06 | 1.500E-05 |
| 29480 | 1474 | I1474V | E | 1.3e-05 | 6.9e-06 | 4.4e-06 | 8.100E-06 | 4.424E-06 | 4.400E-06 | 1.300E-05 |
| 29481 | 1475 | C1475A | E | 5.5e-06 | 1.1e-05 | 1.3e-05 | 9.833E-06 | 3.884E-06 | 5.500E-06 | 1.300E-05 |
| 29482 | 1475 | C1475R | E | 7.6e-06 | 9.2e-06 | 6.7e-06 | 7.833E-06 | 1.266E-06 | 6.700E-06 | 9.200E-06 |
| 29483 | 1475 | C1475N | E | 3.8e-06 | 5.2e-06 | 6.1e-06 | 5.033E-06 | 1.159E-06 | 3.800E-06 | 6.100E-06 |
| 29484 | 1475 | C1475D | E | 7.2e-06 | 3.7e-06 | 1.1e-05 | 7.300E-06 | 3.651E-06 | 3.700E-06 | 1.100E-05 |
| 29485 | 1475 | C1475C | E | 7e-06 | 7e-06 | 9.7e-06 | 7.900E-06 | 1.559E-06 | 7.000E-06 | 9.700E-06 |
| 29486 | 1475 | C1475E | E | 1.1e-05 | 1e-05 | 8.9e-06 | 9.967E-06 | 1.050E-06 | 8.900E-06 | 1.100E-05 |
| 29487 | 1475 | C1475Q | E | 1.2e-05 | 5.5e-06 | 6.4e-06 | 7.967E-06 | 3.522E-06 | 5.500E-06 | 1.200E-05 |
| 29488 | 1475 | C1475G | E | 7.9e-06 | 6.8e-06 | 5.5e-06 | 6.733E-06 | 1.201E-06 | 5.500E-06 | 7.900E-06 |
| 29489 | 1475 | C1475H | E | 5.4e-06 | 6.4e-06 | 4.8e-06 | 5.533E-06 | 8.083E-07 | 4.800E-06 | 6.400E-06 |
| 29490 | 1475 | C1475I | E | 9.1e-06 | 6.7e-06 | 1.1e-05 | 8.933E-06 | 2.155E-06 | 6.700E-06 | 1.100E-05 |
| 29491 | 1475 | C1475L | E | 6.3e-06 | 5.9e-06 | 4.5e-06 | 5.567E-06 | 9.452E-07 | 4.500E-06 | 6.300E-06 |
| 29492 | 1475 | C1475K | E | 7.5e-06 | 9e-06 | 7e-06 | 7.833E-06 | 1.041E-06 | 7.000E-06 | 9.000E-06 |
| 29493 | 1475 | C1475M | E | 9.9e-06 | 9.2e-06 | 4.9e-06 | 8.000E-06 | 2.707E-06 | 4.900E-06 | 9.900E-06 |
| 29494 | 1475 | C1475F | E | 4.6e-06 | 9.5e-06 | 1.3e-05 | 9.033E-06 | 4.219E-06 | 4.600E-06 | 1.300E-05 |

| | | | | | | | | | | |
|---|---|---|---|---|---|---|---|---|---|---|
| 29495 | 1475 | C1475P | E | 9.3e-06 | 1.3e-05 | 9.5e-06 | 1.060E-05 | 2.081E-06 | 9.300E-06 | 1.300E-05 |
| 29496 | 1475 | C1475S | E | 9.4e-06 | 9.7e-06 | 8.3e-06 | 9.133E-06 | 7.371E-07 | 8.300E-06 | 9.700E-06 |
| 29497 | 1475 | C1475T | E | 1e-05 | 9.3e-06 | 5e-06 | 8.100E-06 | 2.707E-06 | 5.000E-06 | 1.000E-05 |
| 29498 | 1475 | C1475W | E | 6.5e-06 | 9.9e-06 | 1.1e-05 | 9.133E-06 | 2.346E-06 | 6.500E-06 | 1.100E-05 |
| 29499 | 1475 | C1475Y | E | 9.2e-06 | 9.8e-06 | 1.3e-05 | 1.067E-05 | 2.043E-06 | 9.200E-06 | 1.300E-05 |
| 29500 | 1475 | C1475V | E | 9.7e-06 | 8.2e-06 | 7.3e-06 | 8.400E-06 | 1.212E-06 | 7.300E-06 | 9.700E-06 |
| 29501 | 1476 | N1476A | E | 5.3e-06 | 7.4e-06 | 8.1e-06 | 6.933E-06 | 1.457E-06 | 5.300E-06 | 8.100E-06 |
| 29502 | 1476 | N1476R | E | 1.1e-05 | 5.4e-06 | 1.4e-05 | 1.013E-05 | 4.365E-06 | 5.400E-06 | 1.400E-05 |
| 29503 | 1476 | N1476N | E | 6.5e-06 | 8.6e-06 | 8.8e-06 | 7.967E-06 | 1.274E-06 | 6.500E-06 | 8.800E-06 |
| 29504 | 1476 | N1476D | E | 5.5e-06 | 1.3e-05 | 6.3e-06 | 8.267E-06 | 4.119E-06 | 5.500E-06 | 1.300E-05 |
| 29505 | 1476 | N1476C | E | 9e-06 | 7.8e-06 | 7.3e-06 | 8.033E-06 | 8.737E-07 | 7.300E-06 | 9.000E-06 |
| 29506 | 1476 | N1476E | E | 8.8e-06 | 9.1e-06 | 8.9e-06 | 8.933E-06 | 1.528E-07 | 8.800E-06 | 9.100E-06 |
| 29507 | 1476 | N1476Q | E | 8.8e-06 | 9.4e-06 | 4.8e-06 | 7.667E-06 | 2.501E-06 | 4.800E-06 | 9.400E-06 |
| 29508 | 1476 | N1476G | E | 6.4e-06 | 8.1e-06 | 6.1e-06 | 6.867E-06 | 1.079E-06 | 6.100E-06 | 8.100E-06 |
| 29509 | 1476 | N1476H | E | 9.3e-06 | 6.8e-06 | 6.9e-06 | 7.667E-06 | 1.415E-06 | 6.800E-06 | 9.300E-06 |
| 29510 | 1476 | N1476I | E | 9.4e-06 | 7.8e-06 | 6.1e-06 | 7.767E-06 | 1.650E-06 | 6.100E-06 | 9.400E-06 |
| 29511 | 1476 | N1476L | E | 7.2e-06 | 5.9e-06 | 1.3e-05 | 8.700E-06 | 3.780E-06 | 5.900E-06 | 1.300E-05 |
| 29512 | 1476 | N1476K | E | 9.6e-06 | 6.6e-06 | 1.1e-05 | 9.067E-06 | 2.248E-06 | 6.600E-06 | 1.100E-05 |
| 29513 | 1476 | N1476M | E | 1.5e-05 | 5.1e-06 | 1.5e-05 | 1.170E-05 | 5.716E-06 | 5.100E-06 | 1.500E-05 |
| 29514 | 1476 | N1476F | E | 8.8e-06 | 5.9e-06 | 7.4e-06 | 7.367E-06 | 1.450E-06 | 5.900E-06 | 8.800E-06 |
| 29515 | 1476 | N1476P | E | 7.5e-06 | 7.5e-06 | 1.5e-05 | 1.000E-05 | 4.330E-06 | 7.500E-06 | 1.500E-05 |
| 29516 | 1476 | N1476S | E | 9.1e-06 | 6.4e-06 | 7.1e-06 | 7.533E-06 | 1.401E-06 | 6.400E-06 | 9.100E-06 |
| 29517 | 1476 | N1476T | E | 1.2e-05 | 1.3e-05 | 5.5e-06 | 1.017E-05 | 4.072E-06 | 5.500E-06 | 1.300E-05 |
| 29518 | 1476 | N1476W | E | 8e-06 | 7.3e-06 | 6.4e-06 | 7.233E-06 | 8.021E-07 | 6.400E-06 | 8.000E-06 |
| 29519 | 1476 | N1476Y | E | 9.1e-06 | 9.4e-06 | 4.8e-06 | 7.767E-06 | 2.574E-06 | 4.800E-06 | 9.400E-06 |
| 29520 | 1476 | N1476V | E | 1.1e-05 | 1.4e-05 | 5.2e-06 | 1.007E-05 | 4.474E-06 | 5.200E-06 | 1.400E-05 |
| 29521 | 1477 | V1477A | E | 5.9e-06 | 1e-05 | 1.6e-05 | 1.063E-05 | 5.080E-06 | 5.900E-06 | 1.600E-05 |
| 29522 | 1477 | V1477R | E | 6.3e-06 | 6.8e-06 | 7.2e-06 | 6.767E-06 | 4.509E-07 | 6.300E-06 | 7.200E-06 |
| 29523 | 1477 | V1477N | E | 6.8e-06 | 4.7e-06 | 1.2e-05 | 7.833E-06 | 3.758E-06 | 4.700E-06 | 1.200E-05 |
| 29524 | 1477 | V1477D | E | 1.4e-05 | 1.2e-05 | 4.8e-06 | 1.027E-05 | 4.839E-06 | 4.800E-06 | 1.400E-05 |
| 29525 | 1477 | V1477C | E | 8.7e-06 | 6.9e-06 | 1.1e-05 | 8.867E-06 | 2.055E-06 | 6.900E-06 | 1.100E-05 |
| 29526 | 1477 | V1477E | E | 5.7e-06 | 9.1e-06 | 9.8e-06 | 8.200E-06 | 2.193E-06 | 5.700E-06 | 9.800E-06 |
| 29527 | 1477 | V1477Q | E | 8.6e-06 | 6.4e-06 | 8.5e-06 | 7.833E-06 | 1.242E-06 | 6.400E-06 | 8.600E-06 |
| 29528 | 1477 | V1477G | E | 6.6e-06 | 7.5e-06 | 9.9e-06 | 8.000E-06 | 1.706E-06 | 6.600E-06 | 9.900E-06 |
| 29529 | 1477 | V1477H | E | 7.2e-06 | 8.4e-06 | 5.8e-06 | 7.133E-06 | 1.301E-06 | 5.800E-06 | 8.400E-06 |
| 29530 | 1477 | V1477I | E | 1.1e-05 | 7.7e-06 | 8.3e-06 | 9.000E-06 | 1.758E-06 | 7.700E-06 | 1.100E-05 |
| 29531 | 1477 | V1477L | E | 1.2e-05 | 8.1e-06 | 9.3e-06 | 9.800E-06 | 1.997E-06 | 8.100E-06 | 1.200E-05 |
| 29532 | 1477 | V1477K | E | 9.4e-06 | 7e-06 | 8e-06 | 8.133E-06 | 1.206E-06 | 7.000E-06 | 9.400E-06 |
| 29533 | 1477 | V1477M | E | 8.6e-06 | 8.4e-06 | 9.3e-06 | 8.767E-06 | 4.726E-07 | 8.400E-06 | 9.300E-06 |
| 29534 | 1477 | V1477F | E | 6.7e-06 | 7.4e-06 | 9.7e-06 | 7.933E-06 | 1.570E-06 | 6.700E-06 | 9.700E-06 |
| 29535 | 1477 | V1477P | E | 5.4e-06 | 8.8e-06 | 1e-05 | 8.067E-06 | 2.386E-06 | 5.400E-06 | 1.000E-05 |
| 29536 | 1477 | V1477S | E | 8.9e-06 | 1.4e-05 | 6.6e-06 | 9.833E-06 | 3.787E-06 | 6.600E-06 | 1.400E-05 |
| 29537 | 1477 | V1477T | E | 5.1e-06 | 1.7e-05 | 1e-05 | 1.070E-05 | 5.981E-06 | 5.100E-06 | 1.700E-05 |
| 29538 | 1477 | V1477W | E | 5.8e-06 | 7.7e-06 | 7.8e-06 | 7.100E-06 | 1.127E-06 | 5.800E-06 | 7.800E-06 |
| 29539 | 1477 | V1477Y | E | 7.7e-06 | 6.8e-06 | 8.5e-06 | 7.667E-06 | 8.505E-07 | 6.800E-06 | 8.500E-06 |
| 29540 | 1477 | V1477V | E | 8.6e-06 | 8.6e-06 | 1e-05 | 9.067E-06 | 8.083E-07 | 8.600E-06 | 1.000E-05 |
| 29541 | 1478 | N1478A | E | 5.3e-06 | 7.4e-06 | 1.4e-05 | 8.900E-06 | 4.540E-06 | 5.300E-06 | 1.400E-05 |
| 29542 | 1478 | N1478R | E | 6.3e-06 | 9.7e-06 | 9e-06 | 8.333E-06 | 1.795E-06 | 6.300E-06 | 9.700E-06 |
| 29543 | 1478 | N1478N | E | 1.1e-05 | 9.7e-06 | 1e-05 | 1.023E-05 | 6.807E-07 | 9.700E-06 | 1.100E-05 |
| 29544 | 1478 | N1478D | E | 6.7e-06 | 8.8e-06 | 7.7e-06 | 7.733E-06 | 1.050E-06 | 6.700E-06 | 8.800E-06 |
| 29545 | 1478 | N1478C | E | 8.7e-06 | 8e-06 | 7e-06 | 7.900E-06 | 8.544E-07 | 7.000E-06 | 8.700E-06 |
| 29546 | 1478 | N1478E | E | 6.8e-06 | 6.9e-06 | 8.7e-06 | 7.467E-06 | 1.069E-06 | 6.800E-06 | 8.700E-06 |
| 29547 | 1478 | N1478Q | E | 1.1e-05 | 7.9e-06 | 6.3e-06 | 8.400E-06 | 2.390E-06 | 6.300E-06 | 1.100E-05 |
| 29548 | 1478 | N1478G | E | 6.6e-06 | 9e-06 | 8.1e-06 | 7.900E-06 | 1.212E-06 | 6.600E-06 | 9.000E-06 |
| 29549 | 1478 | N1478H | E | 1.1e-05 | 9e-06 | 6.1e-06 | 8.700E-06 | 2.464E-06 | 6.100E-06 | 1.100E-05 |
| 29550 | 1478 | N1478I | E | 8.7e-06 | 9.6e-06 | 9.2e-06 | 9.167E-06 | 4.509E-07 | 8.700E-06 | 9.600E-06 |
| 29551 | 1478 | N1478L | E | 1.4e-05 | 8.2e-06 | 1.1e-05 | 1.107E-05 | 2.901E-06 | 8.200E-06 | 1.400E-05 |
| 29552 | 1478 | N1478K | E | 7e-06 | 8.4e-06 | 1.5e-05 | 1.013E-05 | 4.272E-06 | 7.000E-06 | 1.500E-05 |
| 29553 | 1478 | N1478M | E | 5.5e-06 | 9.5e-06 | 9.5e-06 | 8.167E-06 | 2.309E-06 | 5.500E-06 | 9.500E-06 |
| 29554 | 1478 | N1478F | E | 9.4e-06 | 8e-06 | 7e-06 | 8.133E-06 | 1.206E-06 | 7.000E-06 | 9.400E-06 |
| 29555 | 1478 | N1478P | E | 5.5e-06 | 8.3e-06 | 6e-06 | 6.600E-06 | 1.493E-06 | 5.500E-06 | 8.300E-06 |
| 29556 | 1478 | N1478S | E | 9.8e-06 | 1.1e-05 | 8.2e-06 | 9.667E-06 | 1.405E-06 | 8.200E-06 | 1.100E-05 |
| 29557 | 1478 | N1478T | E | 9.2e-06 | 4.7e-06 | 8.8e-06 | 7.567E-06 | 2.491E-06 | 4.700E-06 | 9.200E-06 |
| 29558 | 1478 | N1478W | E | 1.1e-05 | 9.8e-06 | 5e-06 | 8.600E-06 | 3.175E-06 | 5.000E-06 | 1.100E-05 |
| 29559 | 1478 | N1478Y | E | 1.3e-05 | 6.4e-06 | 7.9e-06 | 9.100E-06 | 3.460E-06 | 6.400E-06 | 1.300E-05 |
| 29560 | 1478 | N1478V | E | 1e-05 | 9.1e-06 | 5.2e-06 | 8.100E-06 | 2.551E-06 | 5.200E-06 | 1.000E-05 |
| 29561 | 1479 | H1479A | E | 1.2e-05 | 6.5e-06 | 6.1e-06 | 8.200E-06 | 3.297E-06 | 6.100E-06 | 1.200E-05 |
| 29562 | 1479 | H1479R | E | 7.9e-06 | 8.5e-06 | 1.8e-05 | 1.147E-05 | 5.666E-06 | 7.900E-06 | 1.800E-05 |
| 29563 | 1479 | H1479N | E | 6.5e-06 | 9.9e-06 | 7.3e-06 | 7.900E-06 | 1.778E-06 | 6.500E-06 | 9.900E-06 |
| 29564 | 1479 | H1479D | E | 4.9e-06 | 7.3e-06 | 9.4e-06 | 7.200E-06 | 2.252E-06 | 4.900E-06 | 9.400E-06 |
| 29565 | 1479 | H1479C | E | 7.4e-06 | 4.9e-06 | 5.6e-06 | 5.967E-06 | 1.290E-06 | 4.900E-06 | 7.400E-06 |
| 29566 | 1479 | H1479E | E | 6.8e-06 | 6.7e-06 | 8.6e-06 | 7.367E-06 | 1.069E-06 | 6.700E-06 | 8.600E-06 |
| 29567 | 1479 | H1479Q | E | 4.6e-06 | 4.3e-06 | 5.5e-06 | 4.800E-06 | 6.245E-07 | 4.300E-06 | 5.500E-06 |
| 29568 | 1479 | H1479G | E | 7.4e-06 | 4.5e-06 | 6.2e-06 | 6.033E-06 | 1.457E-06 | 4.500E-06 | 7.400E-06 |
| 29569 | 1479 | H1479H | E | 6.9e-06 | 9.7e-06 | 5.5e-06 | 7.367E-06 | 2.139E-06 | 5.500E-06 | 9.700E-06 |
| 29570 | 1479 | H1479I | E | 2.9e-06 | 7.1e-06 | 5.3e-06 | 5.100E-06 | 2.107E-06 | 2.900E-06 | 7.100E-06 |
| 29571 | 1479 | H1479L | E | 1.1e-05 | 4.6e-06 | 6.5e-06 | 7.367E-06 | 3.287E-06 | 4.600E-06 | 1.100E-05 |
| 29572 | 1479 | H1479K | E | 5.1e-06 | 7.7e-06 | 8.6e-06 | 7.133E-06 | 1.818E-06 | 5.100E-06 | 8.600E-06 |
| 29573 | 1479 | H1479M | E | 5.9e-06 | 1.3e-05 | 1.6e-05 | 1.163E-05 | 5.187E-06 | 5.900E-06 | 1.600E-05 |
| 29574 | 1479 | H1479F | E | 1e-05 | 7.9e-06 | 4.9e-06 | 7.600E-06 | 2.563E-06 | 4.900E-06 | 1.000E-05 |
| 29575 | 1479 | H1479P | E | 6.2e-06 | 8.9e-06 | 5.2e-06 | 6.767E-06 | 1.914E-06 | 5.200E-06 | 8.900E-06 |
| 29576 | 1479 | H1479S | E | 6.7e-06 | 8.8e-06 | 6.7e-06 | 7.400E-06 | 1.212E-06 | 6.700E-06 | 8.800E-06 |
| 29577 | 1479 | H1479T | E | 8.6e-06 | 6.8e-06 | 6e-06 | 7.133E-06 | 1.332E-06 | 6.000E-06 | 8.600E-06 |
| 29578 | 1479 | H1479W | E | 4.8e-06 | 4.5e-06 | 7e-06 | 5.433E-06 | 1.365E-06 | 4.500E-06 | 7.000E-06 |
| 29579 | 1479 | H1479Y | E | 8e-06 | 4e-06 | 8.7e-06 | 6.900E-06 | 2.536E-06 | 4.000E-06 | 8.700E-06 |

| | | | | | | | | | | |
|---|---|---|---|---|---|---|---|---|---|---|
| 29580 | 1479 | H1479V | E | 6.6e-06 | 8e-06 | 1.3e-05 | 9.200E-06 | 3.365E-06 | 6.600E-06 | 1.300E-05 |
| 29581 | 1480 | K1480A | E | 6.6e-06 | 9.7e-06 | 6.2e-06 | 7.500E-06 | 1.916E-06 | 6.200E-06 | 9.700E-06 |
| 29582 | 1480 | K1480R | E | 1.2e-05 | 9.3e-06 | 6.6e-06 | 9.300E-06 | 2.700E-06 | 6.600E-06 | 1.200E-05 |
| 29583 | 1480 | K1480N | E | 8.6e-06 | 7.8e-06 | 6.9e-06 | 7.767E-06 | 8.505E-07 | 6.900E-06 | 8.600E-06 |
| 29584 | 1480 | K1480D | E | 7.2e-06 | 6.4e-06 | 8.5e-06 | 7.367E-06 | 1.060E-06 | 6.400E-06 | 8.500E-06 |
| 29585 | 1480 | K1480C | E | 1.4e-05 | 1.2e-05 | 1e-05 | 1.200E-05 | 2.000E-06 | 1.000E-05 | 1.400E-05 |
| 29586 | 1480 | K1480E | E | 6.2e-06 | 1.3e-05 | 7.5e-06 | 8.900E-06 | 3.610E-06 | 6.200E-06 | 1.300E-05 |
| 29587 | 1480 | K1480Q | E | 9.3e-06 | 8.9e-06 | 1.4e-05 | 1.073E-05 | 2.836E-06 | 8.900E-06 | 1.400E-05 |
| 29588 | 1480 | K1480G | E | 6.5e-06 | 9.9e-06 | 8.5e-06 | 8.300E-06 | 1.709E-06 | 6.500E-06 | 9.900E-06 |
| 29589 | 1480 | K1480H | E | 7.3e-06 | 8.7e-06 | 1e-05 | 8.667E-06 | 1.350E-06 | 7.300E-06 | 1.000E-05 |
| 29590 | 1480 | K1480I | E | 7.2e-06 | 6.5e-06 | 6.7e-06 | 6.800E-06 | 3.606E-07 | 6.500E-06 | 7.200E-06 |
| 29591 | 1480 | K1480L | E | 6.5e-06 | 6.4e-06 | 7.4e-06 | 6.767E-06 | 5.508E-07 | 6.400E-06 | 7.400E-06 |
| 29592 | 1480 | K1480K | E | 8.6e-06 | 8.3e-06 | 7.1e-06 | 8.000E-06 | 7.937E-07 | 7.100E-06 | 8.600E-06 |
| 29593 | 1480 | K1480M | E | 1.3e-05 | 5.8e-06 | 1.2e-05 | 1.027E-05 | 3.900E-06 | 5.800E-06 | 1.300E-05 |
| 29594 | 1480 | K1480F | E | 9.7e-06 | 6.7e-06 | 9.9e-06 | 8.767E-06 | 1.793E-06 | 6.700E-06 | 9.900E-06 |
| 29595 | 1480 | K1480P | E | 1.2e-05 | 1e-05 | 5.9e-06 | 9.300E-06 | 3.110E-06 | 5.900E-06 | 1.200E-05 |
| 29596 | 1480 | K1480S | E | 7.4e-06 | 7.8e-06 | 3.8e-06 | 6.333E-06 | 2.203E-06 | 3.800E-06 | 7.800E-06 |
| 29597 | 1480 | K1480T | E | 8.1e-06 | 8.9e-06 | 6.7e-06 | 7.900E-06 | 1.114E-06 | 6.700E-06 | 8.900E-06 |
| 29598 | 1480 | K1480W | E | 1.4e-05 | 9.6e-06 | 1.1e-05 | 1.153E-05 | 2.248E-06 | 9.600E-06 | 1.400E-05 |
| 29599 | 1480 | K1480Y | E | 4.7e-06 | 1.3e-05 | 6.3e-06 | 8.000E-06 | 4.403E-06 | 4.700E-06 | 1.300E-05 |
| 29600 | 1480 | K1480V | E | 5.9e-06 | 1.1e-05 | 6.2e-06 | 7.700E-06 | 2.862E-06 | 5.900E-06 | 1.100E-05 |
| 29601 | 1481 | P1481A | E | 1.2e-05 | 9.8e-06 | 6.6e-06 | 9.467E-06 | 2.715E-06 | 6.600E-06 | 1.200E-05 |
| 29602 | 1481 | P1481R | E | 1.3e-05 | 9e-06 | 7.5e-06 | 9.833E-06 | 2.843E-06 | 7.500E-06 | 1.300E-05 |
| 29603 | 1481 | P1481N | E | 7.6e-06 | 9.2e-06 | 6.9e-06 | 7.900E-06 | 1.179E-06 | 6.900E-06 | 9.200E-06 |
| 29604 | 1481 | P1481D | E | 8.5e-06 | 1.2e-05 | 5.7e-06 | 8.733E-06 | 3.156E-06 | 5.700E-06 | 1.200E-05 |
| 29605 | 1481 | P1481C | E | 9.5e-06 | 9.2e-06 | 1.3e-05 | 1.057E-05 | 2.113E-06 | 9.200E-06 | 1.300E-05 |
| 29606 | 1481 | P1481E | E | 1.2e-05 | 5.7e-06 | 6.8e-06 | 8.167E-06 | 3.365E-06 | 5.700E-06 | 1.200E-05 |
| 29607 | 1481 | P1481Q | E | 5.6e-06 | 1.1e-05 | 1.3e-05 | 9.867E-06 | 3.828E-06 | 5.600E-06 | 1.300E-05 |
| 29608 | 1481 | P1481G | E | 7.6e-06 | 8.4e-06 | 8.1e-06 | 8.033E-06 | 4.041E-07 | 7.600E-06 | 8.400E-06 |
| 29609 | 1481 | P1481H | E | 7.8e-06 | 7e-06 | 5e-06 | 6.600E-06 | 1.442E-06 | 5.000E-06 | 7.800E-06 |
| 29610 | 1481 | P1481I | E | 9.2e-06 | 1.1e-05 | 6.6e-06 | 8.933E-06 | 2.212E-06 | 6.600E-06 | 1.100E-05 |
| 29611 | 1481 | P1481L | E | 6.8e-06 | 8.7e-06 | 6.1e-06 | 7.200E-06 | 1.345E-06 | 6.100E-06 | 8.700E-06 |
| 29612 | 1481 | P1481K | E | 6.6e-06 | 6.5e-06 | 8.2e-06 | 7.100E-06 | 9.539E-07 | 6.500E-06 | 8.200E-06 |
| 29613 | 1481 | P1481M | E | 6.5e-06 | 6.1e-06 | 6.4e-06 | 6.333E-06 | 2.082E-07 | 6.100E-06 | 6.500E-06 |
| 29614 | 1481 | P1481F | E | 9.7e-06 | 8.8e-06 | 7.5e-06 | 8.667E-06 | 1.106E-06 | 7.500E-06 | 9.700E-06 |
| 29615 | 1481 | P1481P | E | 7.2e-06 | 7.6e-06 | 1.4e-05 | 9.600E-06 | 3.816E-06 | 7.200E-06 | 1.400E-05 |
| 29616 | 1481 | P1481S | E | 7.8e-06 | 7.2e-06 | 6.8e-06 | 7.267E-06 | 5.033E-07 | 6.800E-06 | 7.800E-06 |
| 29617 | 1481 | P1481T | E | 7.4e-06 | 9.2e-06 | 5.8e-06 | 7.467E-06 | 1.701E-06 | 5.800E-06 | 9.200E-06 |
| 29618 | 1481 | P1481W | E | 7.9e-06 | 6.4e-06 | 9.8e-06 | 8.033E-06 | 1.704E-06 | 6.400E-06 | 9.800E-06 |
| 29619 | 1481 | P1481Y | E | 6e-06 | 8.3e-06 | 8.2e-06 | 7.500E-06 | 1.300E-06 | 6.000E-06 | 8.300E-06 |
| 29620 | 1481 | P1481V | E | 9.6e-06 | 6.2e-06 | 1.5e-05 | 1.027E-05 | 4.438E-06 | 6.200E-06 | 1.500E-05 |
| 29621 | 1482 | S1482A | E | 4.3e-06 | 1.5e-05 | 7.2e-06 | 8.833E-06 | 5.534E-06 | 4.300E-06 | 1.500E-05 |
| 29622 | 1482 | S1482R | E | 1.1e-05 | 6.9e-06 | 6.9e-06 | 8.267E-06 | 2.367E-06 | 6.900E-06 | 1.100E-05 |
| 29623 | 1482 | S1482N | E | 1.3e-05 | 7.3e-06 | 1.4e-05 | 1.143E-05 | 3.614E-06 | 7.300E-06 | 1.400E-05 |
| 29624 | 1482 | S1482D | E | 7.9e-06 | 6.7e-06 | 7.9e-06 | 7.500E-06 | 6.928E-07 | 6.700E-06 | 7.900E-06 |
| 29625 | 1482 | S1482C | E | 8.9e-06 | 9.2e-06 | 4.2e-06 | 7.433E-06 | 2.804E-06 | 4.200E-06 | 9.200E-06 |
| 29626 | 1482 | S1482E | E | 7e-06 | 9.5e-06 | 1.9e-05 | 1.183E-05 | 6.331E-06 | 7.000E-06 | 1.900E-05 |
| 29627 | 1482 | S1482Q | E | 6.6e-06 | 1.1e-05 | 5.5e-06 | 7.700E-06 | 2.910E-06 | 5.500E-06 | 1.100E-05 |
| 29628 | 1482 | S1482G | E | 9.5e-06 | 7.5e-06 | 7.3e-06 | 8.100E-06 | 1.217E-06 | 7.300E-06 | 9.500E-06 |
| 29629 | 1482 | S1482H | E | 1.6e-05 | 1.1e-05 | 1e-05 | 1.233E-05 | 3.215E-06 | 1.000E-05 | 1.600E-05 |
| 29630 | 1482 | S1482I | E | 6.7e-06 | 5.5e-06 | 8.9e-06 | 7.033E-06 | 1.724E-06 | 5.500E-06 | 8.900E-06 |
| 29631 | 1482 | S1482L | E | 7.5e-06 | 4.9e-06 | 6.5e-06 | 6.300E-06 | 1.311E-06 | 4.900E-06 | 7.500E-06 |
| 29632 | 1482 | S1482K | E | 8.6e-06 | 1.4e-05 | 8.2e-06 | 1.027E-05 | 3.239E-06 | 8.200E-06 | 1.400E-05 |
| 29633 | 1482 | S1482M | E | 1.8e-05 | 4.8e-06 | 4.7e-06 | 9.167E-06 | 7.650E-06 | 4.700E-06 | 1.800E-05 |
| 29634 | 1482 | S1482F | E | 1.2e-05 | 9.6e-06 | 1.4e-05 | 1.187E-05 | 2.203E-06 | 9.600E-06 | 1.400E-05 |
| 29635 | 1482 | S1482P | E | 1.3e-05 | 9.5e-06 | 1.1e-05 | 1.117E-05 | 1.756E-06 | 9.500E-06 | 1.300E-05 |
| 29636 | 1482 | S1482S | E | 6.9e-06 | 6.5e-06 | 8e-06 | 7.133E-06 | 7.767E-07 | 6.500E-06 | 8.000E-06 |
| 29637 | 1482 | S1482T | E | 1e-05 | 9.3e-06 | 4.8e-06 | 8.033E-06 | 2.822E-06 | 4.800E-06 | 1.000E-05 |
| 29638 | 1482 | S1482W | E | 5.1e-06 | 4.7e-06 | 5.2e-06 | 5.000E-06 | 2.646E-07 | 4.700E-06 | 5.200E-06 |
| 29639 | 1482 | S1482Y | E | 1e-05 | 8.3e-06 | 8e-06 | 8.767E-06 | 1.079E-06 | 8.000E-06 | 1.000E-05 |
| 29640 | 1482 | S1482V | E | 5.6e-06 | 7.5e-06 | 1e-05 | 7.700E-06 | 2.207E-06 | 5.600E-06 | 1.000E-05 |
| 29641 | 1483 | N1483A | E | 8.7e-06 | 1.3e-05 | 6.5e-06 | 9.400E-06 | 3.306E-06 | 6.500E-06 | 1.300E-05 |
| 29642 | 1483 | N1483R | E | 1.3e-05 | 5.1e-06 | 4.8e-06 | 7.633E-06 | 4.650E-06 | 4.800E-06 | 1.300E-05 |
| 29643 | 1483 | N1483N | E | 1.1e-05 | 4.6e-06 | 5.8e-06 | 7.133E-06 | 3.402E-06 | 4.600E-06 | 1.100E-05 |
| 29644 | 1483 | N1483D | E | 1.5e-05 | 6.5e-06 | 2.3e-06 | 7.933E-06 | 6.470E-06 | 2.300E-06 | 1.500E-05 |
| 29645 | 1483 | N1483C | E | 1.4e-05 | 8.8e-06 | 7.9e-06 | 1.023E-05 | 3.293E-06 | 7.900E-06 | 1.400E-05 |
| 29646 | 1483 | N1483E | E | 1e-05 | 6.5e-06 | 8.8e-06 | 8.433E-06 | 1.779E-06 | 6.500E-06 | 1.000E-05 |
| 29647 | 1483 | N1483Q | E | 6.4e-06 | 1.4e-05 | 1.3e-05 | 1.113E-05 | 4.130E-06 | 6.400E-06 | 1.400E-05 |
| 29648 | 1483 | N1483G | E | 8e-06 | 9.9e-06 | 1e-05 | 9.300E-06 | 1.127E-06 | 8.000E-06 | 1.000E-05 |
| 29649 | 1483 | N1483H | E | 5.6e-06 | 8e-06 | 1e-05 | 7.867E-06 | 2.203E-06 | 5.600E-06 | 1.000E-05 |
| 29650 | 1483 | N1483I | E | 5.2e-06 | 5e-06 | 6.4e-06 | 5.533E-06 | 7.572E-07 | 5.000E-06 | 6.400E-06 |
| 29651 | 1483 | N1483L | E | 6.7e-06 | 6.4e-06 | 9.2e-06 | 7.433E-06 | 1.537E-06 | 6.400E-06 | 9.200E-06 |
| 29652 | 1483 | N1483K | E | 9.4e-06 | 1e-05 | 8.8e-06 | 9.400E-06 | 6.000E-07 | 8.800E-06 | 1.000E-05 |
| 29653 | 1483 | N1483M | E | 7e-06 | 8.1e-06 | 1.4e-05 | 9.700E-06 | 3.764E-06 | 7.000E-06 | 1.400E-05 |
| 29654 | 1483 | N1483F | E | 4.8e-06 | 5.9e-06 | 9.4e-06 | 6.700E-06 | 2.402E-06 | 4.800E-06 | 9.400E-06 |
| 29655 | 1483 | N1483P | E | 1.3e-05 | 6.6e-06 | 6.2e-06 | 8.600E-06 | 3.816E-06 | 6.200E-06 | 1.300E-05 |
| 29656 | 1483 | N1483S | E | 9.6e-06 | 8.1e-06 | 1.3e-05 | 1.023E-05 | 2.511E-06 | 8.100E-06 | 1.300E-05 |
| 29657 | 1483 | N1483T | E | 7.1e-06 | 6.9e-06 | 6.5e-06 | 6.833E-06 | 3.055E-07 | 6.500E-06 | 7.100E-06 |
| 29658 | 1483 | N1483W | E | 8.9e-06 | 9.9e-06 | 1.2e-05 | 1.027E-05 | 1.582E-06 | 8.900E-06 | 1.200E-05 |
| 29659 | 1483 | N1483Y | E | 1.3e-05 | 1.2e-05 | 1.2e-05 | 1.233E-05 | 5.774E-07 | 1.200E-05 | 1.300E-05 |
| 29660 | 1483 | N1483V | E | 6.5e-06 | 1e-05 | 7.2e-06 | 7.900E-06 | 1.852E-06 | 6.500E-06 | 1.000E-05 |
| 29661 | 1484 | T1484A | E | 5.3e-06 | 7.2e-06 | 6.7e-06 | 6.400E-06 | 9.849E-07 | 5.300E-06 | 7.200E-06 |
| 29662 | 1484 | T1484R | E | 9.6e-06 | 7.6e-06 | 6e-06 | 7.733E-06 | 1.804E-06 | 6.000E-06 | 9.600E-06 |
| 29663 | 1484 | T1484N | E | 1.2e-05 | 4.3e-06 | 7.8e-06 | 8.033E-06 | 3.855E-06 | 4.300E-06 | 1.200E-05 |
| 29664 | 1484 | T1484D | E | 7.5e-06 | 5.1e-06 | 5.3e-06 | 5.967E-06 | 1.332E-06 | 5.100E-06 | 7.500E-06 |

| | | | | | | | | | | |
|---|---|---|---|---|---|---|---|---|---|---|
| 29665 | 1484 | T1484C | E | 6.9e-06 | 1.3e-05 | 7.5e-06 | 9.133E-06 | 3.362E-06 | 6.900E-06 | 1.300E-05 |
| 29666 | 1484 | T1484E | E | 6e-06 | 5.2e-06 | 8.8e-06 | 6.667E-06 | 1.890E-06 | 5.200E-06 | 8.800E-06 |
| 29667 | 1484 | T1484Q | E | 7.9e-06 | 1.2e-05 | 7.4e-06 | 9.100E-06 | 2.524E-06 | 7.400E-06 | 1.200E-05 |
| 29668 | 1484 | T1484G | E | 1.2e-05 | 1.1e-05 | 6.9e-06 | 9.967E-06 | 2.702E-06 | 6.900E-06 | 1.200E-05 |
| 29669 | 1484 | T1484H | E | 1e-05 | 1.1e-05 | 1.1e-05 | 1.067E-05 | 5.774E-07 | 1.000E-05 | 1.100E-05 |
| 29670 | 1484 | T1484I | E | 1.1e-05 | 1e-05 | 9.5e-06 | 1.017E-05 | 7.638E-07 | 9.500E-06 | 1.100E-05 |
| 29671 | 1484 | T1484L | E | 4.7e-06 | 9.6e-06 | 7.9e-06 | 7.400E-06 | 2.488E-06 | 4.700E-06 | 9.600E-06 |
| 29672 | 1484 | T1484K | E | 7.7e-06 | 1.8e-05 | 9.9e-06 | 1.187E-05 | 5.424E-06 | 7.700E-06 | 1.800E-05 |
| 29673 | 1484 | T1484M | E | 1.1e-05 | 7.2e-06 | 4.9e-06 | 7.700E-06 | 3.081E-06 | 4.900E-06 | 1.100E-05 |
| 29674 | 1484 | T1484F | E | 9.5e-06 | 1.1e-05 | 9.8e-06 | 1.010E-05 | 7.937E-07 | 9.500E-06 | 1.100E-05 |
| 29675 | 1484 | T1484P | E | 7e-06 | 8.6e-06 | 5.6e-06 | 7.067E-06 | 1.501E-06 | 5.600E-06 | 8.600E-06 |
| 29676 | 1484 | T1484S | E | 1.2e-05 | 6.9e-06 | 1.2e-05 | 1.030E-05 | 2.944E-06 | 6.900E-06 | 1.200E-05 |
| 29677 | 1484 | T1484T | E | 7.9e-06 | 1.1e-05 | 9e-06 | 9.300E-06 | 1.572E-06 | 7.900E-06 | 1.100E-05 |
| 29678 | 1484 | T1484W | E | 3.3e-06 | 6.4e-06 | 8.9e-06 | 6.200E-06 | 2.805E-06 | 3.300E-06 | 8.900E-06 |
| 29679 | 1484 | T1484Y | E | 1.3e-05 | 8.9e-06 | 1e-05 | 1.063E-05 | 2.122E-06 | 8.900E-06 | 1.300E-05 |
| 29680 | 1484 | T1484V | E | 6.6e-06 | 1.1e-05 | 5.5e-06 | 7.700E-06 | 2.910E-06 | 5.500E-06 | 1.100E-05 |
| 29681 | 1485 | K1485A | E | 8.1e-06 | 5e-06 | 9.3e-06 | 7.467E-06 | 2.219E-06 | 5.000E-06 | 9.300E-06 |
| 29682 | 1485 | K1485R | E | 7e-06 | 1e-05 | 6.6e-06 | 7.867E-06 | 1.858E-06 | 6.600E-06 | 1.000E-05 |
| 29683 | 1485 | K1485N | E | 5.6e-06 | 1.3e-05 | 1e-05 | 9.533E-06 | 3.722E-06 | 5.600E-06 | 1.300E-05 |
| 29684 | 1485 | K1485D | E | 9.1e-06 | 7.6e-06 | 6.7e-06 | 7.800E-06 | 1.212E-06 | 6.700E-06 | 9.100E-06 |
| 29685 | 1485 | K1485C | E | 7.3e-06 | 9.2e-06 | 4.5e-06 | 7.000E-06 | 2.364E-06 | 4.500E-06 | 9.200E-06 |
| 29686 | 1485 | K1485E | E | 1.9e-05 | 6.7e-06 | 8.7e-06 | 1.147E-05 | 6.600E-06 | 6.700E-06 | 1.900E-05 |
| 29687 | 1485 | K1485Q | E | 7.8e-06 | 1.1e-05 | 4.5e-06 | 7.767E-06 | 3.250E-06 | 4.500E-06 | 1.100E-05 |
| 29688 | 1485 | K1485G | E | 1.1e-05 | 1.7e-05 | 7.6e-06 | 1.187E-05 | 4.760E-06 | 7.600E-06 | 1.700E-05 |
| 29689 | 1485 | K1485H | E | 9e-06 | 9.7e-06 | 9.5e-06 | 9.400E-06 | 3.606E-07 | 9.000E-06 | 9.700E-06 |
| 29690 | 1485 | K1485I | E | 1.4e-05 | 5e-06 | 5.2e-06 | 8.067E-06 | 5.139E-06 | 5.000E-06 | 1.400E-05 |
| 29691 | 1485 | K1485L | E | 9.9e-06 | 7.5e-06 | 1.2e-05 | 9.800E-06 | 2.252E-06 | 7.500E-06 | 1.200E-05 |
| 29692 | 1485 | K1485K | E | 7e-06 | 4.7e-06 | 6.4e-06 | 6.033E-06 | 1.193E-06 | 4.700E-06 | 7.000E-06 |
| 29693 | 1485 | K1485M | E | 7.1e-06 | 9e-06 | 9.4e-06 | 8.500E-06 | 1.229E-06 | 7.100E-06 | 9.400E-06 |
| 29694 | 1485 | K1485F | E | 4.7e-06 | 5.9e-06 | 7.6e-06 | 6.067E-06 | 1.457E-06 | 4.700E-06 | 7.600E-06 |
| 29695 | 1485 | K1485P | E | 1.2e-05 | 4.8e-06 | 7.8e-06 | 8.200E-06 | 3.617E-06 | 4.800E-06 | 1.200E-05 |
| 29696 | 1485 | K1485S | E | 7.4e-06 | 6.1e-06 | 9.2e-06 | 7.567E-06 | 1.557E-06 | 6.100E-06 | 9.200E-06 |
| 29697 | 1485 | K1485T | E | 1e-05 | 8.7e-06 | 6.5e-06 | 8.400E-06 | 1.769E-06 | 6.500E-06 | 1.000E-05 |
| 29698 | 1485 | K1485W | E | 4.5e-06 | 7.1e-06 | 7.7e-06 | 6.433E-06 | 1.701E-06 | 4.500E-06 | 7.700E-06 |
| 29699 | 1485 | K1485Y | E | 4.4e-06 | 9.5e-06 | 8.6e-06 | 7.500E-06 | 2.722E-06 | 4.400E-06 | 9.500E-06 |
| 29700 | 1485 | K1485V | E | 6e-06 | 6.6e-06 | 1e-05 | 7.533E-06 | 2.157E-06 | 6.000E-06 | 1.000E-05 |
| 29701 | 1486 | V1486A | E | 4.5e-06 | 5.9e-06 | 6.5e-06 | 5.633E-06 | 1.026E-06 | 4.500E-06 | 6.500E-06 |
| 29702 | 1486 | V1486R | E | 8.7e-06 | 1.2e-05 | 7.5e-06 | 9.400E-06 | 2.330E-06 | 7.500E-06 | 1.200E-05 |
| 29703 | 1486 | V1486N | E | 4.6e-06 | 1.3e-05 | 9e-06 | 8.867E-06 | 4.202E-06 | 4.600E-06 | 1.300E-05 |
| 29704 | 1486 | V1486D | E | 6.5e-06 | 1.4e-05 | 5.1e-06 | 8.533E-06 | 4.786E-06 | 5.100E-06 | 1.400E-05 |
| 29705 | 1486 | V1486C | E | 8.9e-06 | 1.4e-05 | 9.4e-06 | 1.077E-05 | 2.811E-06 | 8.900E-06 | 1.400E-05 |
| 29706 | 1486 | V1486E | E | 7.9e-06 | 1.1e-05 | 9.2e-06 | 9.367E-06 | 1.557E-06 | 7.900E-06 | 1.100E-05 |
| 29707 | 1486 | V1486Q | E | 8.1e-06 | 9.9e-06 | 5.4e-06 | 7.800E-06 | 2.265E-06 | 5.400E-06 | 9.900E-06 |
| 29708 | 1486 | V1486G | E | 8.7e-06 | 1.3e-05 | 9.4e-06 | 1.037E-05 | 2.307E-06 | 8.700E-06 | 1.300E-05 |
| 29709 | 1486 | V1486H | E | 7.6e-06 | 7.8e-06 | 9.9e-06 | 8.433E-06 | 1.274E-06 | 7.600E-06 | 9.900E-06 |
| 29710 | 1486 | V1486I | E | 6.2e-06 | 6e-06 | 8.3e-06 | 6.833E-06 | 1.274E-06 | 6.000E-06 | 8.300E-06 |
| 29711 | 1486 | V1486L | E | 1e-05 | 8.9e-06 | 7.3e-06 | 8.733E-06 | 1.358E-06 | 7.300E-06 | 1.000E-05 |
| 29712 | 1486 | V1486K | E | 6.5e-06 | 1.2e-05 | 8.7e-06 | 9.067E-06 | 2.768E-06 | 6.500E-06 | 1.200E-05 |
| 29713 | 1486 | V1486M | E | 6.9e-06 | 1.1e-05 | 8.8e-06 | 8.900E-06 | 2.052E-06 | 6.900E-06 | 1.100E-05 |
| 29714 | 1486 | V1486F | E | 1.5e-05 | 1.3e-05 | 6.7e-06 | 1.157E-05 | 4.332E-06 | 6.700E-06 | 1.500E-05 |
| 29715 | 1486 | V1486P | E | 9.5e-06 | 1.3e-05 | 8.2e-06 | 1.023E-05 | 2.483E-06 | 8.200E-06 | 1.300E-05 |
| 29716 | 1486 | V1486S | E | 6.3e-06 | 7.6e-06 | 6.6e-06 | 6.833E-06 | 6.807E-07 | 6.300E-06 | 7.600E-06 |
| 29717 | 1486 | V1486T | E | 1.1e-05 | 9.4e-06 | 9.3e-06 | 9.900E-06 | 9.539E-07 | 9.300E-06 | 1.100E-05 |
| 29718 | 1486 | V1486W | E | 6.2e-06 | 6.1e-06 | 1.2e-05 | 8.100E-06 | 3.378E-06 | 6.100E-06 | 1.200E-05 |
| 29719 | 1486 | V1486Y | E | 1.2e-05 | 1.3e-05 | 4.3e-06 | 9.767E-06 | 4.761E-06 | 4.300E-06 | 1.300E-05 |
| 29720 | 1486 | V1486V | E | 1.2e-05 | 5.8e-06 | 1.4e-05 | 1.060E-05 | 4.276E-06 | 5.800E-06 | 1.400E-05 |
| 29721 | 1487 | D1487A | E | 6.3e-06 | 7e-06 | 6.3e-06 | 6.533E-06 | 4.041E-07 | 6.300E-06 | 7.000E-06 |
| 29722 | 1487 | D1487R | E | 6.4e-06 | 4.5e-06 | 7.2e-06 | 6.033E-06 | 1.387E-06 | 4.500E-06 | 7.200E-06 |
| 29723 | 1487 | D1487N | E | 1.3e-05 | 1.3e-05 | 1.4e-05 | 1.333E-05 | 5.774E-07 | 1.300E-05 | 1.400E-05 |
| 29724 | 1487 | D1487D | E | 1.2e-05 | 9.4e-06 | 7.2e-06 | 9.533E-06 | 2.403E-06 | 7.200E-06 | 1.200E-05 |
| 29725 | 1487 | D1487C | E | 6.7e-06 | 6.5e-06 | 6.1e-06 | 6.433E-06 | 3.055E-07 | 6.100E-06 | 6.700E-06 |
| 29726 | 1487 | D1487E | E | 3.5e-06 | 6e-06 | 1.2e-05 | 7.167E-06 | 4.368E-06 | 3.500E-06 | 1.200E-05 |
| 29727 | 1487 | D1487Q | E | 9.7e-06 | 7.1e-06 | 6.3e-06 | 7.700E-06 | 1.778E-06 | 6.300E-06 | 9.700E-06 |
| 29728 | 1487 | D1487G | E | 1.4e-05 | 6.3e-06 | 5e-06 | 8.433E-06 | 4.864E-06 | 5.000E-06 | 1.400E-05 |
| 29729 | 1487 | D1487H | E | 1e-05 | 9.2e-06 | 6.3e-06 | 8.500E-06 | 1.947E-06 | 6.300E-06 | 1.000E-05 |
| 29730 | 1487 | D1487I | E | 9.1e-06 | 7.5e-06 | 1e-05 | 8.867E-06 | 1.266E-06 | 7.500E-06 | 1.000E-05 |
| 29731 | 1487 | D1487L | E | 8.8e-06 | 6.7e-06 | 1.1e-05 | 8.833E-06 | 2.150E-06 | 6.700E-06 | 1.100E-05 |
| 29732 | 1487 | D1487K | E | 8.8e-06 | 1.1e-05 | 8.2e-06 | 9.333E-06 | 1.474E-06 | 8.200E-06 | 1.100E-05 |
| 29733 | 1487 | D1487M | E | 1.3e-05 | 9.8e-06 | 3.8e-06 | 8.867E-06 | 4.670E-06 | 3.800E-06 | 1.300E-05 |
| 29734 | 1487 | D1487F | E | 8.9e-06 | 7.7e-06 | 7.6e-06 | 8.067E-06 | 7.234E-07 | 7.600E-06 | 8.900E-06 |
| 29735 | 1487 | D1487P | E | 6.7e-06 | 6.6e-06 | 5.9e-06 | 6.400E-06 | 4.359E-07 | 5.900E-06 | 6.700E-06 |
| 29736 | 1487 | D1487S | E | 8.6e-06 | 1.5e-05 | 9.3e-06 | 1.097E-05 | 3.510E-06 | 8.600E-06 | 1.500E-05 |
| 29737 | 1487 | D1487T | E | 5.7e-06 | 7.8e-06 | 6.4e-06 | 6.633E-06 | 1.069E-06 | 5.700E-06 | 7.800E-06 |
| 29738 | 1487 | D1487W | E | 6.3e-06 | 4.3e-06 | 1.8e-05 | 9.533E-06 | 7.400E-06 | 4.300E-06 | 1.800E-05 |
| 29739 | 1487 | D1487Y | E | 9.4e-06 | 1.2e-05 | 8.5e-06 | 9.967E-06 | 1.818E-06 | 8.500E-06 | 1.200E-05 |
| 29740 | 1487 | D1487V | E | 1e-05 | 4.4e-06 | 7.7e-06 | 7.367E-06 | 2.815E-06 | 4.400E-06 | 1.000E-05 |
| 29741 | 1488 | K1488A | E | 9.3e-06 | 1.3e-05 | 7.8e-06 | 1.003E-05 | 2.676E-06 | 7.800E-06 | 1.300E-05 |
| 29742 | 1488 | K1488R | E | 1.4e-05 | 8.9e-06 | 6.5e-06 | 9.800E-06 | 3.830E-06 | 6.500E-06 | 1.400E-05 |
| 29743 | 1488 | K1488N | E | 7.7e-06 | 1e-05 | 4.6e-06 | 7.433E-06 | 2.710E-06 | 4.600E-06 | 1.000E-05 |
| 29744 | 1488 | K1488D | E | 6.4e-06 | 1.1e-05 | 9.7e-06 | 9.033E-06 | 2.371E-06 | 6.400E-06 | 1.100E-05 |
| 29745 | 1488 | K1488C | E | 5.2e-06 | 9.7e-06 | 8.5e-06 | 7.800E-06 | 2.330E-06 | 5.200E-06 | 9.700E-06 |
| 29746 | 1488 | K1488E | E | 1.1e-05 | 7.4e-06 | 1.3e-05 | 1.047E-05 | 2.838E-06 | 7.400E-06 | 1.300E-05 |
| 29747 | 1488 | K1488Q | E | 1.1e-05 | 7e-06 | 6.4e-06 | 8.133E-06 | 2.501E-06 | 6.400E-06 | 1.100E-05 |
| 29748 | 1488 | K1488G | E | 7.8e-06 | 8.2e-06 | 1.4e-05 | 1.000E-05 | 3.470E-06 | 7.800E-06 | 1.400E-05 |
| 29749 | 1488 | K1488H | E | 8.3e-06 | 4.4e-06 | 7.6e-06 | 6.767E-06 | 2.079E-06 | 4.400E-06 | 8.300E-06 |

| | | | | | | | | | | |
|---|---|---|---|---|---|---|---|---|---|---|
| 29750 | 1488 | K1488I | E | 1.1e-05 | 1.2e-05 | 7.3e-06 | 1.010E-05 | 2.476E-06 | 7.300E-06 | 1.200E-05 |
| 29751 | 1488 | K1488L | E | 5e-06 | 1.2e-05 | 1.7e-05 | 1.133E-05 | 6.028E-06 | 5.000E-06 | 1.700E-05 |
| 29752 | 1488 | K1488K | E | 6.8e-06 | 5.6e-06 | 7e-06 | 6.467E-06 | 7.572E-07 | 5.600E-06 | 7.000E-06 |
| 29753 | 1488 | K1488M | E | 4.4e-06 | 6.2e-06 | 1.2e-05 | 7.533E-06 | 3.972E-06 | 4.400E-06 | 1.200E-05 |
| 29754 | 1488 | K1488F | E | 8e-06 | 1.1e-05 | 1e-05 | 9.667E-06 | 1.528E-06 | 8.000E-06 | 1.100E-05 |
| 29755 | 1488 | K1488P | E | 8.1e-06 | 7.4e-06 | 6.5e-06 | 7.333E-06 | 8.021E-07 | 6.500E-06 | 8.100E-06 |
| 29756 | 1488 | K1488S | E | 6.5e-06 | 6.4e-06 | 9.5e-06 | 7.467E-06 | 1.762E-06 | 6.400E-06 | 9.500E-06 |
| 29757 | 1488 | K1488T | E | 7.2e-06 | 5.3e-06 | 8.5e-06 | 7.000E-06 | 1.609E-06 | 5.300E-06 | 8.500E-06 |
| 29758 | 1488 | K1488W | E | 5.3e-06 | 6.5e-06 | 9.6e-06 | 7.133E-06 | 2.219E-06 | 5.300E-06 | 9.600E-06 |
| 29759 | 1488 | K1488Y | E | 7.4e-06 | 7.3e-06 | 7.8e-06 | 7.500E-06 | 2.646E-07 | 7.300E-06 | 7.800E-06 |
| 29760 | 1488 | K1488V | E | 1.3e-05 | 6.6e-06 | 9.3e-06 | 9.633E-06 | 3.213E-06 | 6.600E-06 | 1.300E-05 |
| 29761 | 1489 | K1489A | E | 8.6e-06 | 1.1e-05 | 7.2e-06 | 8.933E-06 | 1.922E-06 | 7.200E-06 | 1.100E-05 |
| 29762 | 1489 | K1489R | E | 7.4e-06 | 8.8e-06 | 5e-06 | 7.067E-06 | 1.922E-06 | 5.000E-06 | 8.800E-06 |
| 29763 | 1489 | K1489N | E | 6.9e-06 | 4.2e-06 | 5.8e-06 | 5.633E-06 | 1.358E-06 | 4.200E-06 | 6.900E-06 |
| 29764 | 1489 | K1489D | E | 1.1e-05 | 1.2e-05 | 8.3e-06 | 1.043E-05 | 1.914E-06 | 8.300E-06 | 1.200E-05 |
| 29765 | 1489 | K1489C | E | 7.9e-06 | 6.7e-06 | 9.8e-06 | 8.133E-06 | 1.563E-06 | 6.700E-06 | 9.800E-06 |
| 29766 | 1489 | K1489E | E | 4.1e-06 | 4.5e-06 | 6.6e-06 | 5.067E-06 | 1.343E-06 | 4.100E-06 | 6.600E-06 |
| 29767 | 1489 | K1489Q | E | 9.8e-06 | 1.2e-05 | 8.5e-06 | 1.010E-05 | 1.769E-06 | 8.500E-06 | 1.200E-05 |
| 29768 | 1489 | K1489G | E | 5.8e-06 | 9.8e-06 | 9.7e-06 | 8.433E-06 | 2.281E-06 | 5.800E-06 | 9.800E-06 |
| 29769 | 1489 | K1489H | E | 8.1e-06 | 8.5e-06 | 8e-06 | 8.200E-06 | 2.646E-07 | 8.000E-06 | 8.500E-06 |
| 29770 | 1489 | K1489I | E | 6.6e-06 | 9.7e-06 | 7.2e-06 | 7.833E-06 | 1.644E-06 | 6.600E-06 | 9.700E-06 |
| 29771 | 1489 | K1489L | E | 6.5e-06 | 1e-05 | 8.8e-06 | 8.433E-06 | 1.779E-06 | 6.500E-06 | 1.000E-05 |
| 29772 | 1489 | K1489K | E | 6.5e-06 | 1.2e-05 | 1.1e-05 | 9.833E-06 | 2.930E-06 | 6.500E-06 | 1.200E-05 |
| 29773 | 1489 | K1489M | E | 1e-05 | 4.7e-06 | 7.1e-06 | 7.267E-06 | 2.654E-06 | 4.700E-06 | 1.000E-05 |
| 29774 | 1489 | K1489F | E | 7.1e-06 | 8e-06 | 7.5e-06 | 7.533E-06 | 4.509E-07 | 7.100E-06 | 8.000E-06 |
| 29775 | 1489 | K1489P | E | 1.4e-05 | 8.2e-06 | 6.4e-06 | 9.533E-06 | 3.972E-06 | 6.400E-06 | 1.400E-05 |
| 29776 | 1489 | K1489S | E | 7.5e-06 | 8.5e-06 | 7.8e-06 | 7.933E-06 | 5.132E-07 | 7.500E-06 | 8.500E-06 |
| 29777 | 1489 | K1489T | E | 5.9e-06 | 8.7e-06 | 7.8e-06 | 7.467E-06 | 1.429E-06 | 5.900E-06 | 8.700E-06 |
| 29778 | 1489 | K1489W | E | 6.1e-06 | 6.6e-06 | 8.2e-06 | 6.967E-06 | 1.097E-06 | 6.100E-06 | 8.200E-06 |
| 29779 | 1489 | K1489Y | E | 1e-05 | 6.4e-06 | 1e-05 | 8.800E-06 | 2.078E-06 | 6.400E-06 | 1.000E-05 |
| 29780 | 1489 | K1489V | E | 6.5e-06 | 6.4e-06 | 6.6e-06 | 6.500E-06 | 1.000E-07 | 6.400E-06 | 6.600E-06 |
| 29781 | 1490 | V1490A | E | 6.8e-06 | 9.6e-06 | 8.6e-06 | 8.333E-06 | 1.419E-06 | 6.800E-06 | 9.600E-06 |
| 29782 | 1490 | V1490R | E | 7.8e-06 | 4.6e-06 | 9.4e-06 | 7.267E-06 | 2.444E-06 | 4.600E-06 | 9.400E-06 |
| 29783 | 1490 | V1490N | E | 7.5e-06 | 3.9e-06 | 1.2e-05 | 7.800E-06 | 4.058E-06 | 3.900E-06 | 1.200E-05 |
| 29784 | 1490 | V1490D | E | 8e-06 | 9.3e-06 | 9.7e-06 | 9.000E-06 | 8.888E-07 | 8.000E-06 | 9.700E-06 |
| 29785 | 1490 | V1490C | E | 1.2e-05 | 5.7e-06 | 1e-05 | 9.233E-06 | 3.219E-06 | 5.700E-06 | 1.200E-05 |
| 29786 | 1490 | V1490E | E | 1.2e-05 | 9.3e-06 | 4.6e-06 | 8.633E-06 | 3.745E-06 | 4.600E-06 | 1.200E-05 |
| 29787 | 1490 | V1490Q | E | 9.1e-06 | 5.4e-06 | 5.3e-06 | 6.600E-06 | 2.166E-06 | 5.300E-06 | 9.100E-06 |
| 29788 | 1490 | V1490G | E | 6.5e-06 | 7.4e-06 | 9.7e-06 | 7.867E-06 | 1.650E-06 | 6.500E-06 | 9.700E-06 |
| 29789 | 1490 | V1490H | E | 1.3e-05 | 6.3e-06 | 5.8e-06 | 8.367E-06 | 4.020E-06 | 5.800E-06 | 1.300E-05 |
| 29790 | 1490 | V1490I | E | 9.3e-06 | 9.5e-06 | 1.3e-05 | 1.060E-05 | 2.081E-06 | 9.300E-06 | 1.300E-05 |
| 29791 | 1490 | V1490L | E | 8.1e-06 | 6.2e-06 | 9.7e-06 | 8.000E-06 | 1.752E-06 | 6.200E-06 | 9.700E-06 |
| 29792 | 1490 | V1490K | E | 6.3e-06 | 7.8e-06 | 1.3e-05 | 9.033E-06 | 3.516E-06 | 6.300E-06 | 1.300E-05 |
| 29793 | 1490 | V1490M | E | 7.4e-06 | 5.2e-06 | 6.8e-06 | 6.467E-06 | 1.137E-06 | 5.200E-06 | 7.400E-06 |
| 29794 | 1490 | V1490F | E | 4.7e-06 | 6.9e-06 | 9.3e-06 | 6.967E-06 | 2.301E-06 | 4.700E-06 | 9.300E-06 |
| 29795 | 1490 | V1490P | E | 9.4e-06 | 5.9e-06 | 1.1e-05 | 8.767E-06 | 2.608E-06 | 5.900E-06 | 1.100E-05 |
| 29796 | 1490 | V1490S | E | 4.7e-06 | 5.9e-06 | 8.2e-06 | 6.267E-06 | 1.779E-06 | 4.700E-06 | 8.200E-06 |
| 29797 | 1490 | V1490T | E | 1.2e-05 | 1e-05 | 9.3e-06 | 1.043E-05 | 1.401E-06 | 9.300E-06 | 1.200E-05 |
| 29798 | 1490 | V1490W | E | 4.6e-06 | 9.5e-06 | 8.2e-06 | 7.433E-06 | 2.538E-06 | 4.600E-06 | 9.500E-06 |
| 29799 | 1490 | V1490Y | E | 7.6e-06 | 9.7e-06 | 6.6e-06 | 7.967E-06 | 1.582E-06 | 6.600E-06 | 9.700E-06 |
| 29800 | 1490 | V1490V | E | 9.1e-06 | 1.3e-05 | 7.2e-06 | 9.767E-06 | 2.957E-06 | 7.200E-06 | 1.300E-05 |
| 29801 | 1491 | E1491A | E | 1.3e-05 | 1.4e-05 | 1.2e-05 | 1.300E-05 | 1.000E-06 | 1.200E-05 | 1.400E-05 |
| 29802 | 1491 | E1491R | E | 7e-06 | 6.9e-06 | 6.2e-06 | 6.700E-06 | 4.359E-07 | 6.200E-06 | 7.000E-06 |
| 29803 | 1491 | E1491N | E | 8.6e-06 | 5.9e-06 | 9e-06 | 7.833E-06 | 1.686E-06 | 5.900E-06 | 9.000E-06 |
| 29804 | 1491 | E1491D | E | 1.4e-05 | 5.8e-06 | 6.3e-06 | 8.700E-06 | 4.597E-06 | 5.800E-06 | 1.400E-05 |
| 29805 | 1491 | E1491C | E | 5.4e-06 | 6e-06 | 6.5e-06 | 5.967E-06 | 5.508E-07 | 5.400E-06 | 6.500E-06 |
| 29806 | 1491 | E1491E | E | 5.8e-06 | 7.3e-06 | 6.1e-06 | 6.400E-06 | 7.937E-07 | 5.800E-06 | 7.300E-06 |
| 29807 | 1491 | E1491Q | E | 4.8e-06 | 6.2e-06 | 5.1e-06 | 5.367E-06 | 7.371E-07 | 4.800E-06 | 6.200E-06 |
| 29808 | 1491 | E1491G | E | 6.9e-06 | 7.1e-06 | 1e-05 | 8.000E-06 | 1.735E-06 | 6.900E-06 | 1.000E-05 |
| 29809 | 1491 | E1491H | E | 8.1e-06 | 6.3e-06 | 6.5e-06 | 6.967E-06 | 9.866E-07 | 6.300E-06 | 8.100E-06 |
| 29810 | 1491 | E1491I | E | 9e-06 | 5.6e-06 | 8.9e-06 | 7.833E-06 | 1.935E-06 | 5.600E-06 | 9.000E-06 |
| 29811 | 1491 | E1491L | E | 8.6e-06 | 9.5e-06 | 8.7e-06 | 8.933E-06 | 4.933E-07 | 8.600E-06 | 9.500E-06 |
| 29812 | 1491 | E1491K | E | 7e-06 | 6.7e-06 | 9.3e-06 | 7.667E-06 | 1.422E-06 | 6.700E-06 | 9.300E-06 |
| 29813 | 1491 | E1491M | E | 9.4e-06 | 4.7e-06 | 1e-05 | 8.033E-06 | 2.902E-06 | 4.700E-06 | 1.000E-05 |
| 29814 | 1491 | E1491F | E | 5.5e-06 | 7.8e-06 | 1.8e-05 | 1.043E-05 | 6.653E-06 | 5.500E-06 | 1.800E-05 |
| 29815 | 1491 | E1491P | E | 6.2e-06 | 7.7e-06 | 7.7e-06 | 7.200E-06 | 8.660E-07 | 6.200E-06 | 7.700E-06 |
| 29816 | 1491 | E1491S | E | 8.7e-06 | 5e-06 | 7.1e-06 | 6.933E-06 | 1.856E-06 | 5.000E-06 | 8.700E-06 |
| 29817 | 1491 | E1491T | E | 8.6e-06 | 9.7e-06 | 8.8e-06 | 9.033E-06 | 5.859E-07 | 8.600E-06 | 9.700E-06 |
| 29818 | 1491 | E1491W | E | 6.7e-06 | 9.1e-06 | 8.2e-06 | 8.000E-06 | 1.212E-06 | 6.700E-06 | 9.100E-06 |
| 29819 | 1491 | E1491Y | E | 8.3e-06 | 1e-05 | 1.2e-05 | 1.010E-05 | 1.852E-06 | 8.300E-06 | 1.200E-05 |
| 29820 | 1491 | E1491V | E | 8.2e-06 | 6.2e-06 | 9.1e-06 | 7.833E-06 | 1.484E-06 | 6.200E-06 | 9.100E-06 |
| 29821 | 1492 | P1492A | E | 9.5e-06 | 1e-05 | 9.8e-06 | 9.767E-06 | 2.517E-07 | 9.500E-06 | 1.000E-05 |
| 29822 | 1492 | P1492R | E | 8.6e-06 | 5.8e-06 | 8.7e-06 | 7.700E-06 | 1.646E-06 | 5.800E-06 | 8.700E-06 |
| 29823 | 1492 | P1492N | E | 7.3e-06 | 3.4e-06 | 8.4e-06 | 6.367E-06 | 2.627E-06 | 3.400E-06 | 8.400E-06 |
| 29824 | 1492 | P1492D | E | 9.2e-06 | 7e-06 | 5.3e-06 | 7.167E-06 | 1.955E-06 | 5.300E-06 | 9.200E-06 |
| 29825 | 1492 | P1492C | E | 1.5e-05 | 8.9e-06 | 8.1e-06 | 1.067E-05 | 3.774E-06 | 8.100E-06 | 1.500E-05 |
| 29826 | 1492 | P1492E | E | 1.3e-05 | 4.2e-06 | 7.4e-06 | 8.200E-06 | 4.454E-06 | 4.200E-06 | 1.300E-05 |
| 29827 | 1492 | P1492Q | E | 5.7e-06 | 5.1e-06 | 4.6e-06 | 5.133E-06 | 5.508E-07 | 4.600E-06 | 5.700E-06 |
| 29828 | 1492 | P1492G | E | 1.3e-05 | 5e-06 | 1.1e-05 | 9.667E-06 | 4.163E-06 | 5.000E-06 | 1.300E-05 |
| 29829 | 1492 | P1492H | E | 6.9e-06 | 9.6e-06 | 7.1e-06 | 7.867E-06 | 1.504E-06 | 6.900E-06 | 9.600E-06 |
| 29830 | 1492 | P1492I | E | 1.1e-05 | 5.4e-06 | 7.3e-06 | 7.900E-06 | 2.848E-06 | 5.400E-06 | 1.100E-05 |
| 29831 | 1492 | P1492L | E | 5.6e-06 | 7e-06 | 6.5e-06 | 6.367E-06 | 7.095E-07 | 5.600E-06 | 7.000E-06 |
| 29832 | 1492 | P1492K | E | 7.7e-06 | 9.3e-06 | 4.6e-06 | 7.200E-06 | 2.390E-06 | 4.600E-06 | 9.300E-06 |
| 29833 | 1492 | P1492M | E | 8.6e-06 | 9e-06 | 8.5e-06 | 8.700E-06 | 2.646E-07 | 8.500E-06 | 9.000E-06 |
| 29834 | 1492 | P1492F | E | 4.9e-06 | 5.1e-06 | 1e-05 | 6.667E-06 | 2.888E-06 | 4.900E-06 | 1.000E-05 |

| | | | | | | | | | | |
|---|---|---|---|---|---|---|---|---|---|---|
| 29835 | 1492 | P1492P | E | 9.3e-06 | 9e-06 | 7.5e-06 | 8.600E-06 | 9.644E-07 | 7.500E-06 | 9.300E-06 |
| 29836 | 1492 | P1492S | E | 4.5e-06 | 7.6e-06 | 5e-06 | 5.700E-06 | 1.664E-06 | 4.500E-06 | 7.600E-06 |
| 29837 | 1492 | P1492T | E | 3.3e-06 | 9.6e-06 | 8.4e-06 | 7.100E-06 | 3.345E-06 | 3.300E-06 | 9.600E-06 |
| 29838 | 1492 | P1492W | E | 7e-06 | 6e-06 | 6.2e-06 | 6.400E-06 | 5.292E-07 | 6.000E-06 | 7.000E-06 |
| 29839 | 1492 | P1492Y | E | 8.3e-06 | 8.3e-06 | 9.9e-06 | 8.833E-06 | 9.238E-07 | 8.300E-06 | 9.900E-06 |
| 29840 | 1492 | P1492V | E | 1.1e-05 | 1.2e-05 | 1.1e-05 | 1.133E-05 | 5.774E-07 | 1.100E-05 | 1.200E-05 |

Table 5: A scalable Her2-Trastuzumab-Pertuzumab binding affinity landscape. In this table, **ResID** represents the amino acid residue number of PDB entry 6OGE, while **Muta** represents the site-specific missense mutation introduced into the 1492 amino acid residues in PDB entry 6OGE, **Min**, **Max**, **Mean** and **Std** represent the minimum, the maximum, the average and the standard deviation of the $K_d$ values calculated using Prodigy, respectively.

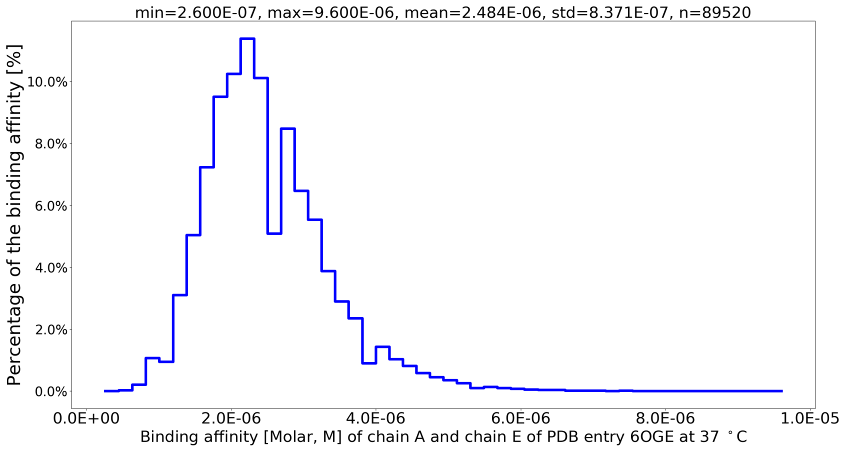


Figure 4: A histogram of the distribution pattern of the Her2-Trastuzumab-Pertuzumab binding affinities between chain A and chain E (listed in Table 1) of PDB entry 6OGE with one site-specific missense mutation.

| No. | ResID | Muta | ChainID | $K_{d1}$ (M) | $K_{d2}$ (M) | $K_{d3}$ (M) | Mean (M) | Std (M) | Min (M) | Max (M) |
|---|---|---|---|---|---|---|---|---|---|---|
| 1 | 1 | T1A | A | 3.1e-06 | 3.1e-06 | 3.1e-06 | 3.100E-06 | 5.187E-22 | 3.100E-06 | 3.100E-06 |
| 2 | 1 | T1R | A | 1.8e-06 | 1.8e-06 | 1.8e-06 | 1.800E-06 | 0.000E+00 | 1.800E-06 | 1.800E-06 |
| 3 | 1 | T1N | A | 2.2e-06 | 2.2e-06 | 2.2e-06 | 2.200E-06 | 0.000E+00 | 2.200E-06 | 2.200E-06 |
| 4 | 1 | T1D | A | 2.9e-06 | 2.9e-06 | 2.9e-06 | 2.900E-06 | 5.187E-22 | 2.900E-06 | 2.900E-06 |
| 5 | 1 | T1C | A | 2.3e-06 | 2.3e-06 | 2.3e-06 | 2.300E-06 | 0.000E+00 | 2.300E-06 | 2.300E-06 |
| 6 | 1 | T1E | A | 1.8e-06 | 1.8e-06 | 1.8e-06 | 1.800E-06 | 0.000E+00 | 1.800E-06 | 1.800E-06 |
| 7 | 1 | T1Q | A | 2.7e-06 | 2.7e-06 | 2.7e-06 | 2.700E-06 | 0.000E+00 | 2.700E-06 | 2.700E-06 |
| 8 | 1 | T1G | A | 2.1e-06 | 2.1e-06 | 2.1e-06 | 2.100E-06 | 0.000E+00 | 2.100E-06 | 2.100E-06 |
| 9 | 1 | T1H | A | 2.4e-06 | 2.4e-06 | 2.4e-06 | 2.400E-06 | 0.000E+00 | 2.400E-06 | 2.400E-06 |
| 10 | 1 | T1I | A | 5.2e-06 | 5.2e-06 | 5.2e-06 | 5.200E-06 | 0.000E+00 | 5.200E-06 | 5.200E-06 |
| 11 | 1 | T1L | A | 2.6e-06 | 2.6e-06 | 2.6e-06 | 2.600E-06 | 0.000E+00 | 2.600E-06 | 2.600E-06 |
| 12 | 1 | T1K | A | 1.4e-06 | 1.4e-06 | 1.4e-06 | 1.400E-06 | 0.000E+00 | 1.400E-06 | 1.400E-06 |
| 13 | 1 | T1M | A | 2e-06 | 2e-06 | 2e-06 | 2.000E-06 | 0.000E+00 | 2.000E-06 | 2.000E-06 |
| 14 | 1 | T1F | A | 5.2e-06 | 5.2e-06 | 5.2e-06 | 5.200E-06 | 0.000E+00 | 5.200E-06 | 5.200E-06 |
| 15 | 1 | T1P | A | 3e-06 | 3e-06 | 3e-06 | 3.000E-06 | 0.000E+00 | 3.000E-06 | 3.000E-06 |
| 16 | 1 | T1S | A | 3.4e-06 | 3.4e-06 | 3.4e-06 | 3.400E-06 | 0.000E+00 | 3.400E-06 | 3.400E-06 |
| 17 | 1 | T1T | A | 3.1e-06 | 3.1e-06 | 3.1e-06 | 3.100E-06 | 5.187E-22 | 3.100E-06 | 3.100E-06 |
| 18 | 1 | T1W | A | 1.7e-06 | 1.7e-06 | 1.7e-06 | 1.700E-06 | 0.000E+00 | 1.700E-06 | 1.700E-06 |
| 19 | 1 | T1Y | A | 2.7e-06 | 2.7e-06 | 2.7e-06 | 2.700E-06 | 0.000E+00 | 2.700E-06 | 2.700E-06 |
| 20 | 1 | T1V | A | 3.1e-06 | 3.1e-06 | 3.1e-06 | 3.100E-06 | 5.187E-22 | 3.100E-06 | 3.100E-06 |
| 21 | 2 | Q2A | A | 3.1e-06 | 3.1e-06 | 3.1e-06 | 3.100E-06 | 5.187E-22 | 3.100E-06 | 3.100E-06 |
| 22 | 2 | Q2R | A | 2.6e-06 | 2.6e-06 | 2.6e-06 | 2.600E-06 | 0.000E+00 | 2.600E-06 | 2.600E-06 |
| 23 | 2 | Q2N | A | 3.5e-06 | 3.5e-06 | 3.5e-06 | 3.500E-06 | 0.000E+00 | 3.500E-06 | 3.500E-06 |
| 24 | 2 | Q2D | A | 4.1e-06 | 4.1e-06 | 4.1e-06 | 4.100E-06 | 0.000E+00 | 4.100E-06 | 4.100E-06 |
| 25 | 2 | Q2C | A | 1.7e-06 | 3.7e-06 | 3.7e-06 | 3.033E-06 | 1.155E-06 | 1.700E-06 | 3.700E-06 |
| 26 | 2 | Q2E | A | 1.8e-06 | 2.7e-06 | 2.7e-06 | 2.400E-06 | 5.196E-07 | 1.800E-06 | 2.700E-06 |
| 27 | 2 | Q2Q | A | 3e-06 | 1.3e-06 | 2.6e-06 | 2.300E-06 | 8.888E-07 | 1.300E-06 | 3.000E-06 |
| 28 | 2 | Q2G | A | 2.4e-06 | 2.6e-06 | 2.7e-06 | 2.567E-06 | 1.528E-07 | 2.400E-06 | 2.700E-06 |
| 29 | 2 | Q2H | A | 2.1e-06 | 3.9e-06 | 3.9e-06 | 3.300E-06 | 1.039E-06 | 2.100E-06 | 3.900E-06 |
| 30 | 2 | Q2I | A | 1.2e-06 | 2.1e-06 | 1.2e-06 | 1.500E-06 | 5.196E-07 | 1.200E-06 | 2.100E-06 |
| 31 | 2 | Q2L | A | 3.5e-06 | 4.2e-06 | 3.5e-06 | 3.733E-06 | 4.041E-07 | 3.500E-06 | 4.200E-06 |
| 32 | 2 | Q2K | A | 1.8e-06 | 1.8e-06 | 2.9e-06 | 2.167E-06 | 6.351E-07 | 1.800E-06 | 2.900E-06 |
| 33 | 2 | Q2M | A | 3.7e-06 | 3.7e-06 | 2.3e-06 | 3.233E-06 | 8.083E-07 | 2.300E-06 | 3.700E-06 |
| 34 | 2 | Q2F | A | 2.2e-06 | 2.1e-06 | 2.2e-06 | 2.167E-06 | 5.774E-08 | 2.100E-06 | 2.200E-06 |
| 35 | 2 | Q2P | A | 3.9e-06 | 3e-06 | 3.9e-06 | 3.600E-06 | 5.196E-07 | 3.000E-06 | 3.900E-06 |
| 36 | 2 | Q2S | A | 2.7e-06 | 2.7e-06 | 2.7e-06 | 2.700E-06 | 0.000E+00 | 2.700E-06 | 2.700E-06 |
| 37 | 2 | Q2T | A | 3.8e-06 | 2e-06 | 1.4e-06 | 2.400E-06 | 1.249E-06 | 1.400E-06 | 3.800E-06 |
| 38 | 2 | Q2W | A | 2.1e-06 | 1.7e-06 | 1.9e-06 | 1.900E-06 | 2.000E-07 | 1.700E-06 | 2.100E-06 |
| 39 | 2 | Q2Y | A | 2.7e-06 | 1.4e-06 | 5.3e-06 | 3.133E-06 | 1.986E-06 | 1.400E-06 | 5.300E-06 |
| 40 | 2 | Q2V | A | 1.6e-06 | 1.2e-06 | 1.2e-06 | 1.333E-06 | 2.309E-07 | 1.200E-06 | 1.600E-06 |
| 41 | 3 | V3A | A | 3e-06 | 3e-06 | 2.2e-06 | 2.733E-06 | 4.619E-07 | 2.200E-06 | 3.000E-06 |
| 42 | 3 | V3R | A | 2.6e-06 | 4.6e-06 | 4.6e-06 | 3.933E-06 | 1.155E-06 | 2.600E-06 | 4.600E-06 |
| 43 | 3 | V3N | A | 3.3e-06 | 3.3e-06 | 3.3e-06 | 3.300E-06 | 0.000E+00 | 3.300E-06 | 3.300E-06 |
| 44 | 3 | V3D | A | 2.2e-06 | 2.2e-06 | 2.2e-06 | 2.200E-06 | 0.000E+00 | 2.200E-06 | 2.200E-06 |
| 45 | 3 | V3C | A | 2.5e-06 | 2.5e-06 | 2.3e-06 | 2.433E-06 | 1.155E-07 | 2.300E-06 | 2.500E-06 |
| 46 | 3 | V3E | A | 2e-06 | 2e-06 | 2.3e-06 | 2.100E-06 | 1.732E-07 | 2.000E-06 | 2.300E-06 |
| 47 | 3 | V3Q | A | 2.9e-06 | 2.9e-06 | 3.8e-06 | 3.200E-06 | 5.196E-07 | 2.900E-06 | 3.800E-06 |
| 48 | 3 | V3G | A | 1.5e-06 | 1.5e-06 | 1.5e-06 | 1.500E-06 | 0.000E+00 | 1.500E-06 | 1.500E-06 |
| 49 | 3 | V3H | A | 3.1e-06 | 1.9e-06 | 2e-06 | 2.333E-06 | 6.658E-07 | 1.900E-06 | 3.100E-06 |
| 50 | 3 | V3I | A | 1.3e-06 | 3e-06 | 1.3e-06 | 1.867E-06 | 9.815E-07 | 1.300E-06 | 3.000E-06 |
| 51 | 3 | V3L | A | 1.9e-06 | 2.1e-06 | 2.1e-06 | 2.033E-06 | 1.155E-07 | 1.900E-06 | 2.100E-06 |
| 52 | 3 | V3K | A | 2.7e-06 | 2.6e-06 | 2.2e-06 | 2.500E-06 | 2.646E-07 | 2.200E-06 | 2.700E-06 |
| 53 | 3 | V3M | A | 2.7e-06 | 1.6e-06 | 2.3e-06 | 2.200E-06 | 5.568E-07 | 1.600E-06 | 2.700E-06 |
| 54 | 3 | V3F | A | 2.6e-06 | 1.1e-06 | 3.3e-06 | 2.333E-06 | 1.124E-06 | 1.100E-06 | 3.300E-06 |
| 55 | 3 | V3P | A | 2.4e-06 | 1.9e-06 | 1.9e-06 | 2.067E-06 | 2.887E-07 | 1.900E-06 | 2.400E-06 |
| 56 | 3 | V3S | A | 2.3e-06 | 2.3e-06 | 2.2e-06 | 2.267E-06 | 5.774E-08 | 2.200E-06 | 2.300E-06 |
| 57 | 3 | V3T | A | 2.1e-06 | 1.9e-06 | 2e-06 | 2.000E-06 | 1.000E-07 | 1.900E-06 | 2.100E-06 |
| 58 | 3 | V3W | A | 3e-06 | 3e-06 | 2.6e-06 | 2.867E-06 | 2.309E-07 | 2.600E-06 | 3.000E-06 |
| 59 | 3 | V3Y | A | 1.6e-06 | 2.2e-06 | 3.1e-06 | 2.300E-06 | 7.550E-07 | 1.600E-06 | 3.100E-06 |
| 60 | 3 | V3V | A | 2.1e-06 | 1.9e-06 | 1.6e-06 | 1.867E-06 | 2.517E-07 | 1.600E-06 | 2.100E-06 |
| 61 | 4 | C4A | A | 1.6e-06 | 3.5e-06 | 1.8e-06 | 2.300E-06 | 1.044E-06 | 1.600E-06 | 3.500E-06 |
| 62 | 4 | C4R | A | 2.7e-06 | 2.4e-06 | 1.9e-06 | 2.333E-06 | 4.041E-07 | 1.900E-06 | 2.700E-06 |
| 63 | 4 | C4N | A | 2.7e-06 | 3.4e-06 | 2.4e-06 | 2.833E-06 | 5.132E-07 | 2.400E-06 | 3.400E-06 |
| 64 | 4 | C4D | A | 2.8e-06 | 2.4e-06 | 2.5e-06 | 2.567E-06 | 2.082E-07 | 2.400E-06 | 2.800E-06 |
| 65 | 4 | C4C | A | 1.8e-06 | 1.8e-06 | 1.8e-06 | 1.800E-06 | 0.000E+00 | 1.800E-06 | 1.800E-06 |
| 66 | 4 | C4E | A | 2.1e-06 | 2.4e-06 | 2.2e-06 | 2.233E-06 | 1.528E-07 | 2.100E-06 | 2.400E-06 |
| 67 | 4 | C4Q | A | 1.3e-06 | 2.2e-06 | 1.4e-06 | 1.633E-06 | 4.933E-07 | 1.300E-06 | 2.200E-06 |
| 68 | 4 | C4G | A | 4.1e-06 | 2.5e-06 | 2.3e-06 | 2.967E-06 | 9.866E-07 | 2.300E-06 | 4.100E-06 |
| 69 | 4 | C4H | A | 2.4e-06 | 2.3e-06 | 1.5e-06 | 2.067E-06 | 4.933E-07 | 1.500E-06 | 2.400E-06 |
| 70 | 4 | C4I | A | 2.3e-06 | 2.3e-06 | 3.5e-06 | 2.700E-06 | 6.928E-07 | 2.300E-06 | 3.500E-06 |
| 71 | 4 | C4L | A | 2.1e-06 | 2.2e-06 | 2.2e-06 | 2.167E-06 | 5.774E-08 | 2.100E-06 | 2.200E-06 |
| 72 | 4 | C4K | A | 2.1e-06 | 2.1e-06 | 3.6e-06 | 2.600E-06 | 8.660E-07 | 2.100E-06 | 3.600E-06 |
| 73 | 4 | C4M | A | 2.2e-06 | 1.9e-06 | 2e-06 | 2.033E-06 | 1.528E-07 | 1.900E-06 | 2.200E-06 |
| 74 | 4 | C4F | A | 2.1e-06 | 2.8e-06 | 3.5e-06 | 2.800E-06 | 7.000E-07 | 2.100E-06 | 3.500E-06 |
| 75 | 4 | C4P | A | 2.7e-06 | 2e-06 | 2.9e-06 | 2.533E-06 | 4.726E-07 | 2.000E-06 | 2.900E-06 |
| 76 | 4 | C4S | A | 4.5e-06 | 1.7e-06 | 1.9e-06 | 2.700E-06 | 1.562E-06 | 1.700E-06 | 4.500E-06 |
| 77 | 4 | C4T | A | 1.6e-06 | 2.8e-06 | 3.1e-06 | 2.500E-06 | 7.937E-07 | 1.600E-06 | 3.100E-06 |
| 78 | 4 | C4W | A | 2.2e-06 | 2.8e-06 | 1.9e-06 | 2.300E-06 | 4.583E-07 | 1.900E-06 | 2.800E-06 |
| 79 | 4 | C4Y | A | 2.1e-06 | 2.3e-06 | 1.8e-06 | 2.067E-06 | 2.517E-07 | 1.800E-06 | 2.300E-06 |
| 80 | 4 | C4V | A | 4.2e-06 | 1.9e-06 | 2.6e-06 | 2.900E-06 | 1.179E-06 | 1.900E-06 | 4.200E-06 |
| 81 | 5 | T5A | A | 1.9e-06 | 2.6e-06 | 2.8e-06 | 2.433E-06 | 4.726E-07 | 1.900E-06 | 2.800E-06 |
| 82 | 5 | T5R | A | 2.2e-06 | 2.9e-06 | 3.1e-06 | 2.733E-06 | 4.726E-07 | 2.200E-06 | 3.100E-06 |
| 83 | 5 | T5N | A | 2.2e-06 | 4.6e-06 | 2.1e-06 | 2.967E-06 | 1.415E-06 | 2.100E-06 | 4.600E-06 |
| 84 | 5 | T5D | A | 3.3e-06 | 2e-06 | 2e-06 | 2.433E-06 | 7.506E-07 | 2.000E-06 | 3.300E-06 |

| | | | | | | | | | | |
|---|---|---|---|---|---|---|---|---|---|---|
| 85 | 5 | T5C | A | 2.1e-06 | 2.6e-06 | 1.8e-06 | 2.167E-06 | 4.041E-07 | 1.800E-06 | 2.600E-06 |
| 86 | 5 | T5E | A | 4.1e-06 | 3.6e-06 | 2.8e-06 | 3.500E-06 | 6.557E-07 | 2.800E-06 | 4.100E-06 |
| 87 | 5 | T5Q | A | 2.2e-06 | 1.6e-06 | 2.2e-06 | 2.000E-06 | 3.464E-07 | 1.600E-06 | 2.200E-06 |
| 88 | 5 | T5G | A | 2.9e-06 | 2.2e-06 | 3e-06 | 2.700E-06 | 4.359E-07 | 2.200E-06 | 3.000E-06 |
| 89 | 5 | T5H | A | 3.5e-06 | 3e-06 | 7.9e-06 | 4.800E-06 | 2.696E-06 | 3.000E-06 | 7.900E-06 |
| 90 | 5 | T5I | A | 2.8e-06 | 3.7e-06 | 2.2e-06 | 2.900E-06 | 7.550E-07 | 2.200E-06 | 3.700E-06 |
| 91 | 5 | T5L | A | 2.9e-06 | 2.9e-06 | 2.1e-06 | 2.633E-06 | 4.619E-07 | 2.100E-06 | 2.900E-06 |
| 92 | 5 | T5K | A | 1.5e-06 | 3.1e-06 | 2e-06 | 2.200E-06 | 8.185E-07 | 1.500E-06 | 3.100E-06 |
| 93 | 5 | T5M | A | 2.1e-06 | 1.9e-06 | 2.3e-06 | 2.100E-06 | 2.000E-07 | 1.900E-06 | 2.300E-06 |
| 94 | 5 | T5F | A | 9.1e-07 | 1.7e-06 | 1.7e-06 | 1.437E-06 | 4.561E-07 | 9.100E-07 | 1.700E-06 |
| 95 | 5 | T5P | A | 3.4e-06 | 2.8e-06 | 3.1e-06 | 3.100E-06 | 3.000E-07 | 2.800E-06 | 3.400E-06 |
| 96 | 5 | T5S | A | 1.6e-06 | 1.8e-06 | 2.7e-06 | 2.033E-06 | 5.859E-07 | 1.600E-06 | 2.700E-06 |
| 97 | 5 | T5T | A | 1.4e-06 | 2.5e-06 | 2.4e-06 | 2.100E-06 | 6.083E-07 | 1.400E-06 | 2.500E-06 |
| 98 | 5 | T5W | A | 2.1e-06 | 4e-06 | 1.6e-06 | 2.567E-06 | 1.266E-06 | 1.600E-06 | 4.000E-06 |
| 99 | 5 | T5Y | A | 1.8e-06 | 2e-06 | 3.7e-06 | 2.500E-06 | 1.044E-06 | 1.800E-06 | 3.700E-06 |
| 100 | 5 | T5V | A | 2.9e-06 | 2.4e-06 | 2.3e-06 | 2.533E-06 | 3.215E-07 | 2.300E-06 | 2.900E-06 |
| 101 | 6 | G6A | A | 1.7e-06 | 1.7e-06 | 2.1e-06 | 1.833E-06 | 2.309E-07 | 1.700E-06 | 2.100E-06 |
| 102 | 6 | G6R | A | 3.9e-06 | 2.2e-06 | 2.5e-06 | 2.867E-06 | 9.074E-07 | 2.200E-06 | 3.900E-06 |
| 103 | 6 | G6N | A | 2.6e-06 | 1.8e-06 | 4.9e-06 | 3.100E-06 | 1.609E-06 | 1.800E-06 | 4.900E-06 |
| 104 | 6 | G6D | A | 2.3e-06 | 1.5e-06 | 2e-06 | 1.933E-06 | 4.041E-07 | 1.500E-06 | 2.300E-06 |
| 105 | 6 | G6C | A | 1.6e-06 | 1.3e-06 | 3.2e-06 | 2.033E-06 | 1.021E-06 | 1.300E-06 | 3.200E-06 |
| 106 | 6 | G6E | A | 1.4e-06 | 3.7e-06 | 2.2e-06 | 2.433E-06 | 1.168E-06 | 1.400E-06 | 3.700E-06 |
| 107 | 6 | G6Q | A | 2.3e-06 | 5e-06 | 2.8e-06 | 3.367E-06 | 1.436E-06 | 2.300E-06 | 5.000E-06 |
| 108 | 6 | G6G | A | 2.3e-06 | 1.4e-06 | 1.3e-06 | 1.667E-06 | 5.508E-07 | 1.300E-06 | 2.300E-06 |
| 109 | 6 | G6H | A | 3e-06 | 1.3e-06 | 2.5e-06 | 2.267E-06 | 8.737E-07 | 1.300E-06 | 3.000E-06 |
| 110 | 6 | G6I | A | 2.3e-06 | 1.4e-06 | 1.8e-06 | 1.833E-06 | 4.509E-07 | 1.400E-06 | 2.300E-06 |
| 111 | 6 | G6L | A | 3.2e-06 | 3.3e-06 | 3e-06 | 3.167E-06 | 1.528E-07 | 3.000E-06 | 3.300E-06 |
| 112 | 6 | G6K | A | 1.8e-06 | 2.3e-06 | 2.8e-06 | 2.300E-06 | 5.000E-07 | 1.800E-06 | 2.800E-06 |
| 113 | 6 | G6M | A | 2.9e-06 | 2.3e-06 | 2.8e-06 | 2.667E-06 | 3.215E-07 | 2.300E-06 | 2.900E-06 |
| 114 | 6 | G6F | A | 4.5e-06 | 1.9e-06 | 3.1e-06 | 3.167E-06 | 1.301E-06 | 1.900E-06 | 4.500E-06 |
| 115 | 6 | G6P | A | 2e-06 | 2.8e-06 | 1.6e-06 | 2.133E-06 | 6.110E-07 | 1.600E-06 | 2.800E-06 |
| 116 | 6 | G6S | A | 3.5e-06 | 2.5e-06 | 2e-06 | 2.667E-06 | 7.638E-07 | 2.000E-06 | 3.500E-06 |
| 117 | 6 | G6T | A | 3.5e-06 | 3e-06 | 4.5e-06 | 3.667E-06 | 7.638E-07 | 3.000E-06 | 4.500E-06 |
| 118 | 6 | G6W | A | 3.4e-06 | 2.5e-06 | 4.2e-06 | 3.367E-06 | 8.505E-07 | 2.500E-06 | 4.200E-06 |
| 119 | 6 | G6Y | A | 2.8e-06 | 3.5e-06 | 3.6e-06 | 3.300E-06 | 4.359E-07 | 2.800E-06 | 3.600E-06 |
| 120 | 6 | G6V | A | 2.3e-06 | 3e-06 | 2e-06 | 2.433E-06 | 5.132E-07 | 2.000E-06 | 3.000E-06 |
| 121 | 7 | T7A | A | 3.2e-06 | 2.5e-06 | 3.2e-06 | 2.967E-06 | 4.041E-07 | 2.500E-06 | 3.200E-06 |
| 122 | 7 | T7R | A | 1.5e-06 | 3.1e-06 | 2.6e-06 | 2.400E-06 | 8.185E-07 | 1.500E-06 | 3.100E-06 |
| 123 | 7 | T7N | A | 1.7e-06 | 1e-06 | 1.7e-06 | 1.467E-06 | 4.041E-07 | 1.000E-06 | 1.700E-06 |
| 124 | 7 | T7D | A | 5.5e-06 | 2.1e-06 | 2.1e-06 | 3.233E-06 | 1.963E-06 | 2.100E-06 | 5.500E-06 |
| 125 | 7 | T7C | A | 2.8e-06 | 4.3e-06 | 2.5e-06 | 3.200E-06 | 9.644E-07 | 2.500E-06 | 4.300E-06 |
| 126 | 7 | T7E | A | 1.8e-06 | 3.2e-06 | 2.1e-06 | 2.367E-06 | 7.371E-07 | 1.800E-06 | 3.200E-06 |
| 127 | 7 | T7Q | A | 2.7e-06 | 1.8e-06 | 3.8e-06 | 2.767E-06 | 1.002E-06 | 1.800E-06 | 3.800E-06 |
| 128 | 7 | T7G | A | 1.3e-06 | 2.2e-06 | 1.9e-06 | 1.800E-06 | 4.583E-07 | 1.300E-06 | 2.200E-06 |
| 129 | 7 | T7H | A | 2.7e-06 | 3.8e-06 | 3.8e-06 | 3.433E-06 | 6.351E-07 | 2.700E-06 | 3.800E-06 |
| 130 | 7 | T7I | A | 3.2e-06 | 2.1e-06 | 3.6e-06 | 2.967E-06 | 7.767E-07 | 2.100E-06 | 3.600E-06 |
| 131 | 7 | T7L | A | 4.8e-06 | 2.1e-06 | 2.3e-06 | 3.067E-06 | 1.504E-06 | 2.100E-06 | 4.800E-06 |
| 132 | 7 | T7K | A | 2.5e-06 | 1.4e-06 | 2.8e-06 | 2.233E-06 | 7.371E-07 | 1.400E-06 | 2.800E-06 |
| 133 | 7 | T7M | A | 2.9e-06 | 2.6e-06 | 2e-06 | 2.500E-06 | 4.583E-07 | 2.000E-06 | 2.900E-06 |
| 134 | 7 | T7F | A | 1.2e-06 | 1.7e-06 | 2.8e-06 | 1.900E-06 | 8.185E-07 | 1.200E-06 | 2.800E-06 |
| 135 | 7 | T7P | A | 2.5e-06 | 1.6e-06 | 1.6e-06 | 1.900E-06 | 5.196E-07 | 1.600E-06 | 2.500E-06 |
| 136 | 7 | T7S | A | 1.9e-06 | 2.7e-06 | 2.7e-06 | 2.433E-06 | 4.619E-07 | 1.900E-06 | 2.700E-06 |
| 137 | 7 | T7T | A | 2.2e-06 | 2.8e-06 | 3.1e-06 | 2.700E-06 | 4.583E-07 | 2.200E-06 | 3.100E-06 |
| 138 | 7 | T7W | A | 1.9e-06 | 2e-06 | 2.3e-06 | 2.067E-06 | 2.082E-07 | 1.900E-06 | 2.300E-06 |
| 139 | 7 | T7Y | A | 3.4e-06 | 2e-06 | 3e-06 | 2.800E-06 | 7.211E-07 | 2.000E-06 | 3.400E-06 |
| 140 | 7 | T7V | A | 1.8e-06 | 2.8e-06 | 1.5e-06 | 2.033E-06 | 6.807E-07 | 1.500E-06 | 2.800E-06 |
| 141 | 8 | D8A | A | 1.4e-06 | 2.5e-06 | 1.8e-06 | 1.900E-06 | 5.568E-07 | 1.400E-06 | 2.500E-06 |
| 142 | 8 | D8R | A | 2.5e-06 | 3.4e-06 | 3.4e-06 | 3.100E-06 | 5.196E-07 | 2.500E-06 | 3.400E-06 |
| 143 | 8 | D8N | A | 3.2e-06 | 2.9e-06 | 1.8e-06 | 2.633E-06 | 7.371E-07 | 1.800E-06 | 3.200E-06 |
| 144 | 8 | D8D | A | 1.8e-06 | 1.7e-06 | 2.2e-06 | 1.900E-06 | 2.646E-07 | 1.700E-06 | 2.200E-06 |
| 145 | 8 | D8C | A | 2.4e-06 | 3.3e-06 | 1.9e-06 | 2.533E-06 | 7.095E-07 | 1.900E-06 | 3.300E-06 |
| 146 | 8 | D8E | A | 2.2e-06 | 2.1e-06 | 2.5e-06 | 2.267E-06 | 2.082E-07 | 2.100E-06 | 2.500E-06 |
| 147 | 8 | D8Q | A | 1.6e-06 | 2.4e-06 | 2e-06 | 2.000E-06 | 4.000E-07 | 1.600E-06 | 2.400E-06 |
| 148 | 8 | D8G | A | 2.6e-06 | 1.9e-06 | 4.3e-06 | 2.933E-06 | 1.234E-06 | 1.900E-06 | 4.300E-06 |
| 149 | 8 | D8H | A | 1.5e-06 | 2.1e-06 | 1.2e-06 | 1.600E-06 | 4.583E-07 | 1.200E-06 | 2.100E-06 |
| 150 | 8 | D8I | A | 2e-06 | 3.8e-06 | 2.4e-06 | 2.733E-06 | 9.452E-07 | 2.000E-06 | 3.800E-06 |
| 151 | 8 | D8L | A | 3.4e-06 | 1.9e-06 | 2.2e-06 | 2.500E-06 | 7.937E-07 | 1.900E-06 | 3.400E-06 |
| 152 | 8 | D8K | A | 1.2e-06 | 2.7e-06 | 2.7e-06 | 2.200E-06 | 8.660E-07 | 1.200E-06 | 2.700E-06 |
| 153 | 8 | D8M | A | 2.6e-06 | 2.6e-06 | 1.1e-06 | 2.100E-06 | 8.660E-07 | 1.100E-06 | 2.600E-06 |
| 154 | 8 | D8F | A | 2.6e-06 | 1.5e-06 | 3.6e-06 | 2.567E-06 | 1.050E-06 | 1.500E-06 | 3.600E-06 |
| 155 | 8 | D8P | A | 2.4e-06 | 2.2e-06 | 3.2e-06 | 2.600E-06 | 5.292E-07 | 2.200E-06 | 3.200E-06 |
| 156 | 8 | D8S | A | 2.5e-06 | 2.3e-06 | 2.6e-06 | 2.467E-06 | 1.528E-07 | 2.300E-06 | 2.600E-06 |
| 157 | 8 | D8T | A | 3.1e-06 | 2.6e-06 | 1.9e-06 | 2.533E-06 | 6.028E-07 | 1.900E-06 | 3.100E-06 |
| 158 | 8 | D8W | A | 1.9e-06 | 3.2e-06 | 2.3e-06 | 2.467E-06 | 6.658E-07 | 1.900E-06 | 3.200E-06 |
| 159 | 8 | D8Y | A | 1.9e-06 | 2e-06 | 2.5e-06 | 2.133E-06 | 3.215E-07 | 1.900E-06 | 2.500E-06 |
| 160 | 8 | D8V | A | 3.5e-06 | 1.5e-06 | 4.2e-06 | 3.067E-06 | 1.401E-06 | 1.500E-06 | 4.200E-06 |
| 161 | 9 | M9A | A | 2.3e-06 | 1.2e-06 | 1.7e-06 | 1.733E-06 | 5.508E-07 | 1.200E-06 | 2.300E-06 |
| 162 | 9 | M9R | A | 2.9e-06 | 2.1e-06 | 2.7e-06 | 2.567E-06 | 4.163E-07 | 2.100E-06 | 2.900E-06 |
| 163 | 9 | M9N | A | 1.7e-06 | 1.9e-06 | 2.5e-06 | 2.033E-06 | 4.163E-07 | 1.700E-06 | 2.500E-06 |
| 164 | 9 | M9D | A | 2e-06 | 1.9e-06 | 2.3e-06 | 2.067E-06 | 2.082E-07 | 1.900E-06 | 2.300E-06 |
| 165 | 9 | M9C | A | 2.5e-06 | 2.7e-06 | 1.8e-06 | 2.333E-06 | 4.726E-07 | 1.800E-06 | 2.700E-06 |
| 166 | 9 | M9E | A | 3.4e-06 | 2.2e-06 | 2.3e-06 | 2.633E-06 | 6.658E-07 | 2.200E-06 | 3.400E-06 |
| 167 | 9 | M9Q | A | 2.5e-06 | 2.4e-06 | 9.7e-07 | 1.957E-06 | 8.559E-07 | 9.700E-07 | 2.500E-06 |
| 168 | 9 | M9G | A | 1.9e-06 | 9.2e-07 | 1.4e-06 | 1.407E-06 | 4.900E-07 | 9.200E-07 | 1.900E-06 |
| 169 | 9 | M9H | A | 1.7e-06 | 1.7e-06 | 2.2e-06 | 1.867E-06 | 2.887E-07 | 1.700E-06 | 2.200E-06 |

| | | | | | | | | | | |
|---|---|---|---|---|---|---|---|---|---|---|
| 170 | 9 | M9I | A | 2e-06 | 2.5e-06 | 1.6e-06 | 2.033E-06 | 4.509E-07 | 1.600E-06 | 2.500E-06 |
| 171 | 9 | M9L | A | 2.2e-06 | 4e-06 | 3.5e-06 | 3.233E-06 | 9.292E-07 | 2.200E-06 | 4.000E-06 |
| 172 | 9 | M9K | A | 1.4e-06 | 1.6e-06 | 3e-06 | 2.000E-06 | 8.718E-07 | 1.400E-06 | 3.000E-06 |
| 173 | 9 | M9M | A | 3.9e-06 | 3.3e-06 | 1.6e-06 | 2.933E-06 | 1.193E-06 | 1.600E-06 | 3.900E-06 |
| 174 | 9 | M9F | A | 2e-06 | 3.1e-06 | 2.4e-06 | 2.500E-06 | 5.568E-07 | 2.000E-06 | 3.100E-06 |
| 175 | 9 | M9P | A | 1.9e-06 | 2e-06 | 2.9e-06 | 2.267E-06 | 5.508E-07 | 1.900E-06 | 2.900E-06 |
| 176 | 9 | M9S | A | 2.5e-06 | 2.3e-06 | 9e-07 | 1.900E-06 | 8.718E-07 | 9.000E-07 | 2.500E-06 |
| 177 | 9 | M9T | A | 4.4e-06 | 2.8e-06 | 2.7e-06 | 3.300E-06 | 9.539E-07 | 2.700E-06 | 4.400E-06 |
| 178 | 9 | M9W | A | 2.1e-06 | 2.3e-06 | 2.2e-06 | 2.200E-06 | 1.000E-07 | 2.100E-06 | 2.300E-06 |
| 179 | 9 | M9Y | A | 3.5e-06 | 1.9e-06 | 1.7e-06 | 2.367E-06 | 9.866E-07 | 1.700E-06 | 3.500E-06 |
| 180 | 9 | M9V | A | 2.2e-06 | 1.6e-06 | 3.8e-06 | 2.533E-06 | 1.137E-06 | 1.600E-06 | 3.800E-06 |
| 181 | 10 | K10A | A | 2.3e-06 | 1.5e-06 | 2.3e-06 | 2.033E-06 | 4.619E-07 | 1.500E-06 | 2.300E-06 |
| 182 | 10 | K10R | A | 3e-06 | 1.8e-06 | 2.1e-06 | 2.300E-06 | 6.245E-07 | 1.800E-06 | 3.000E-06 |
| 183 | 10 | K10N | A | 2.1e-06 | 2.3e-06 | 2.7e-06 | 2.367E-06 | 3.055E-07 | 2.100E-06 | 2.700E-06 |
| 184 | 10 | K10D | A | 1.3e-06 | 3.8e-06 | 1.5e-06 | 2.200E-06 | 1.389E-06 | 1.300E-06 | 3.800E-06 |
| 185 | 10 | K10C | A | 2.1e-06 | 8.6e-07 | 1.8e-06 | 1.587E-06 | 6.469E-07 | 8.600E-07 | 2.100E-06 |
| 186 | 10 | K10E | A | 2.3e-06 | 2.5e-06 | 1.9e-06 | 2.233E-06 | 3.055E-07 | 1.900E-06 | 2.500E-06 |
| 187 | 10 | K10Q | A | 2.2e-06 | 2.1e-06 | 2.8e-06 | 2.367E-06 | 3.786E-07 | 2.100E-06 | 2.800E-06 |
| 188 | 10 | K10G | A | 3.4e-06 | 1.6e-06 | 2.8e-06 | 2.600E-06 | 9.165E-07 | 1.600E-06 | 3.400E-06 |
| 189 | 10 | K10H | A | 1.9e-06 | 2.2e-06 | 3.5e-06 | 2.533E-06 | 8.505E-07 | 1.900E-06 | 3.500E-06 |
| 190 | 10 | K10I | A | 3.8e-06 | 2.8e-06 | 1.5e-06 | 2.700E-06 | 1.153E-06 | 1.500E-06 | 3.800E-06 |
| 191 | 10 | K10L | A | 1.5e-06 | 3e-06 | 1.9e-06 | 2.133E-06 | 7.767E-07 | 1.500E-06 | 3.000E-06 |
| 192 | 10 | K10K | A | 2e-06 | 4e-06 | 2e-06 | 2.667E-06 | 1.155E-06 | 2.000E-06 | 4.000E-06 |
| 193 | 10 | K10M | A | 4e-06 | 1.2e-06 | 1.5e-06 | 2.233E-06 | 1.537E-06 | 1.200E-06 | 4.000E-06 |
| 194 | 10 | K10F | A | 2e-06 | 4.6e-06 | 1.8e-06 | 2.800E-06 | 1.562E-06 | 1.800E-06 | 4.600E-06 |
| 195 | 10 | K10P | A | 2.5e-06 | 2.2e-06 | 2.1e-06 | 2.267E-06 | 2.082E-07 | 2.100E-06 | 2.500E-06 |
| 196 | 10 | K10S | A | 1.3e-06 | 3.4e-06 | 1.5e-06 | 2.067E-06 | 1.159E-06 | 1.300E-06 | 3.400E-06 |
| 197 | 10 | K10T | A | 2.2e-06 | 2.1e-06 | 2.7e-06 | 2.333E-06 | 3.215E-07 | 2.100E-06 | 2.700E-06 |
| 198 | 10 | K10W | A | 2.6e-06 | 2.3e-06 | 3e-06 | 2.633E-06 | 3.512E-07 | 2.300E-06 | 3.000E-06 |
| 199 | 10 | K10Y | A | 1.7e-06 | 1.6e-06 | 2e-06 | 1.767E-06 | 2.082E-07 | 1.600E-06 | 2.000E-06 |
| 200 | 10 | K10V | A | 3e-06 | 2.2e-06 | 4.2e-06 | 3.133E-06 | 1.007E-06 | 2.200E-06 | 4.200E-06 |
| 201 | 11 | L11A | A | 2.7e-06 | 2.7e-06 | 1.7e-06 | 2.367E-06 | 5.774E-07 | 1.700E-06 | 2.700E-06 |
| 202 | 11 | L11R | A | 1.1e-06 | 2.1e-06 | 2.7e-06 | 1.967E-06 | 8.083E-07 | 1.100E-06 | 2.700E-06 |
| 203 | 11 | L11N | A | 2.5e-06 | 4.1e-06 | 1.8e-06 | 2.800E-06 | 1.179E-06 | 1.800E-06 | 4.100E-06 |
| 204 | 11 | L11D | A | 2.6e-06 | 2.3e-06 | 2.1e-06 | 2.333E-06 | 2.517E-07 | 2.100E-06 | 2.600E-06 |
| 205 | 11 | L11C | A | 3.1e-06 | 4e-06 | 1.7e-06 | 2.933E-06 | 1.159E-06 | 1.700E-06 | 4.000E-06 |
| 206 | 11 | L11E | A | 1.2e-06 | 1.6e-06 | 2.9e-06 | 1.900E-06 | 8.888E-07 | 1.200E-06 | 2.900E-06 |
| 207 | 11 | L11Q | A | 2.1e-06 | 3e-06 | 3.2e-06 | 2.767E-06 | 5.859E-07 | 2.100E-06 | 3.200E-06 |
| 208 | 11 | L11G | A | 2.4e-06 | 3e-06 | 2.8e-06 | 2.733E-06 | 3.055E-07 | 2.400E-06 | 3.000E-06 |
| 209 | 11 | L11H | A | 2.1e-06 | 3.1e-06 | 4e-06 | 3.067E-06 | 9.504E-07 | 2.100E-06 | 4.000E-06 |
| 210 | 11 | L11I | A | 2.2e-06 | 2.4e-06 | 1.5e-06 | 2.033E-06 | 4.726E-07 | 1.500E-06 | 2.400E-06 |
| 211 | 11 | L11L | A | 2.3e-06 | 1.9e-06 | 2.8e-06 | 2.333E-06 | 4.509E-07 | 1.900E-06 | 2.800E-06 |
| 212 | 11 | L11K | A | 2.9e-06 | 2.4e-06 | 2.4e-06 | 2.567E-06 | 2.887E-07 | 2.400E-06 | 2.900E-06 |
| 213 | 11 | L11M | A | 3.1e-06 | 2.4e-06 | 2.7e-06 | 2.733E-06 | 3.512E-07 | 2.400E-06 | 3.100E-06 |
| 214 | 11 | L11F | A | 3.3e-06 | 2.4e-06 | 2e-06 | 2.567E-06 | 6.658E-07 | 2.000E-06 | 3.300E-06 |
| 215 | 11 | L11P | A | 3.1e-06 | 2.6e-06 | 1.9e-06 | 2.533E-06 | 6.028E-07 | 1.900E-06 | 3.100E-06 |
| 216 | 11 | L11S | A | 3.2e-06 | 2.8e-06 | 3.8e-06 | 3.267E-06 | 5.033E-07 | 2.800E-06 | 3.800E-06 |
| 217 | 11 | L11T | A | 2e-06 | 4.4e-06 | 2e-06 | 2.800E-06 | 1.386E-06 | 2.000E-06 | 4.400E-06 |
| 218 | 11 | L11W | A | 1.9e-06 | 1.2e-06 | 3e-06 | 2.033E-06 | 9.074E-07 | 1.200E-06 | 3.000E-06 |
| 219 | 11 | L11Y | A | 1.6e-06 | 1.8e-06 | 3.1e-06 | 2.167E-06 | 8.145E-07 | 1.600E-06 | 3.100E-06 |
| 220 | 11 | L11V | A | 1.8e-06 | 2.7e-06 | 2.8e-06 | 2.433E-06 | 5.508E-07 | 1.800E-06 | 2.800E-06 |
| 221 | 12 | R12A | A | 1.1e-06 | 1.3e-06 | 2.3e-06 | 1.567E-06 | 6.429E-07 | 1.100E-06 | 2.300E-06 |
| 222 | 12 | R12R | A | 1.6e-06 | 2e-06 | 2.2e-06 | 1.933E-06 | 3.055E-07 | 1.600E-06 | 2.200E-06 |
| 223 | 12 | R12N | A | 2.2e-06 | 1.8e-06 | 3e-06 | 2.333E-06 | 6.110E-07 | 1.800E-06 | 3.000E-06 |
| 224 | 12 | R12D | A | 2.7e-06 | 2.9e-06 | 2.3e-06 | 2.633E-06 | 3.055E-07 | 2.300E-06 | 2.900E-06 |
| 225 | 12 | R12C | A | 1.7e-06 | 3.3e-06 | 2.4e-06 | 2.467E-06 | 8.021E-07 | 1.700E-06 | 3.300E-06 |
| 226 | 12 | R12E | A | 2.2e-06 | 2.2e-06 | 3.4e-06 | 2.600E-06 | 6.928E-07 | 2.200E-06 | 3.400E-06 |
| 227 | 12 | R12Q | A | 2.2e-06 | 3.5e-06 | 2.6e-06 | 2.767E-06 | 6.658E-07 | 2.200E-06 | 3.500E-06 |
| 228 | 12 | R12G | A | 1.5e-06 | 2.1e-06 | 2.5e-06 | 2.033E-06 | 5.033E-07 | 1.500E-06 | 2.500E-06 |
| 229 | 12 | R12H | A | 2.9e-06 | 2.7e-06 | 2.8e-06 | 2.800E-06 | 1.000E-07 | 2.700E-06 | 2.900E-06 |
| 230 | 12 | R12I | A | 1.8e-06 | 2.8e-06 | 2.5e-06 | 2.367E-06 | 5.132E-07 | 1.800E-06 | 2.800E-06 |
| 231 | 12 | R12L | A | 2.7e-06 | 1.4e-06 | 2.2e-06 | 2.100E-06 | 6.557E-07 | 1.400E-06 | 2.700E-06 |
| 232 | 12 | R12K | A | 3.9e-06 | 1.8e-06 | 2.8e-06 | 2.833E-06 | 1.050E-06 | 1.800E-06 | 3.900E-06 |
| 233 | 12 | R12M | A | 1.8e-06 | 3.7e-06 | 2e-06 | 2.500E-06 | 1.044E-06 | 1.800E-06 | 3.700E-06 |
| 234 | 12 | R12F | A | 2.7e-06 | 3.9e-06 | 2.1e-06 | 2.900E-06 | 9.165E-07 | 2.100E-06 | 3.900E-06 |
| 235 | 12 | R12P | A | 2.4e-06 | 1.9e-06 | 2.2e-06 | 2.167E-06 | 2.517E-07 | 1.900E-06 | 2.400E-06 |
| 236 | 12 | R12S | A | 2.4e-06 | 3e-06 | 1.9e-06 | 2.433E-06 | 5.508E-07 | 1.900E-06 | 3.000E-06 |
| 237 | 12 | R12T | A | 2.2e-06 | 1.5e-06 | 3.3e-06 | 2.333E-06 | 9.074E-07 | 1.500E-06 | 3.300E-06 |
| 238 | 12 | R12W | A | 2.2e-06 | 2.2e-06 | 1.8e-06 | 2.067E-06 | 2.309E-07 | 1.800E-06 | 2.200E-06 |
| 239 | 12 | R12Y | A | 1.6e-06 | 3.5e-06 | 2.5e-06 | 2.533E-06 | 9.504E-07 | 1.600E-06 | 3.500E-06 |
| 240 | 12 | R12V | A | 1.5e-06 | 2.4e-06 | 3e-06 | 2.300E-06 | 7.550E-07 | 1.500E-06 | 3.000E-06 |
| 241 | 13 | L13A | A | 2.8e-06 | 1.2e-06 | 2.6e-06 | 2.200E-06 | 8.718E-07 | 1.200E-06 | 2.800E-06 |
| 242 | 13 | L13R | A | 2e-06 | 2.4e-06 | 1.7e-06 | 2.033E-06 | 3.512E-07 | 1.700E-06 | 2.400E-06 |
| 243 | 13 | L13N | A | 2.8e-06 | 2e-06 | 2.1e-06 | 2.300E-06 | 4.359E-07 | 2.000E-06 | 2.800E-06 |
| 244 | 13 | L13D | A | 2.5e-06 | 2.8e-06 | 2.2e-06 | 2.500E-06 | 3.000E-07 | 2.200E-06 | 2.800E-06 |
| 245 | 13 | L13C | A | 3.8e-06 | 1.4e-06 | 2.2e-06 | 2.467E-06 | 1.222E-06 | 1.400E-06 | 3.800E-06 |
| 246 | 13 | L13E | A | 2.4e-06 | 2.2e-06 | 3.4e-06 | 2.667E-06 | 6.429E-07 | 2.200E-06 | 3.400E-06 |
| 247 | 13 | L13Q | A | 3.1e-06 | 2.4e-06 | 2.4e-06 | 2.633E-06 | 4.041E-07 | 2.400E-06 | 3.100E-06 |
| 248 | 13 | L13G | A | 1.4e-06 | 2.1e-06 | 2e-06 | 1.833E-06 | 3.786E-07 | 1.400E-06 | 2.100E-06 |
| 249 | 13 | L13H | A | 2.9e-06 | 1.8e-06 | 1.2e-06 | 1.967E-06 | 8.622E-07 | 1.200E-06 | 2.900E-06 |
| 250 | 13 | L13I | A | 3.1e-06 | 2.4e-06 | 4.6e-06 | 3.367E-06 | 1.124E-06 | 2.400E-06 | 4.600E-06 |
| 251 | 13 | L13L | A | 3.9e-06 | 2.4e-06 | 2.7e-06 | 3.000E-06 | 7.937E-07 | 2.400E-06 | 3.900E-06 |
| 252 | 13 | L13K | A | 5.6e-06 | 1.9e-06 | 3.2e-06 | 3.567E-06 | 1.877E-06 | 1.900E-06 | 5.600E-06 |
| 253 | 13 | L13M | A | 2.7e-06 | 4.8e-06 | 2e-06 | 3.167E-06 | 1.457E-06 | 2.000E-06 | 4.800E-06 |
| 254 | 13 | L13F | A | 3.6e-06 | 1.9e-06 | 1.9e-06 | 2.467E-06 | 9.815E-07 | 1.900E-06 | 3.600E-06 |

| | | | | | | | | | | |
|---|---|---|---|---|---|---|---|---|---|---|
| 255 | 13 | L13P | A | 1.3e-06 | 2.8e-06 | 3.4e-06 | 2.500E-06 | 1.082E-06 | 1.300E-06 | 3.400E-06 |
| 256 | 13 | L13S | A | 2e-06 | 2.3e-06 | 2.4e-06 | 2.233E-06 | 2.082E-07 | 2.000E-06 | 2.400E-06 |
| 257 | 13 | L13T | A | 2.4e-06 | 1.1e-06 | 2.1e-06 | 1.867E-06 | 6.807E-07 | 1.100E-06 | 2.400E-06 |
| 258 | 13 | L13W | A | 2.8e-06 | 4.1e-06 | 2.5e-06 | 3.133E-06 | 8.505E-07 | 2.500E-06 | 4.100E-06 |
| 259 | 13 | L13Y | A | 4.4e-06 | 3e-06 | 2.6e-06 | 3.333E-06 | 9.452E-07 | 2.600E-06 | 4.400E-06 |
| 260 | 13 | L13V | A | 1.8e-06 | 5.6e-06 | 2.4e-06 | 3.267E-06 | 2.043E-06 | 1.800E-06 | 5.600E-06 |
| 261 | 14 | P14A | A | 2.3e-06 | 2e-06 | 1.8e-06 | 2.033E-06 | 2.517E-07 | 1.800E-06 | 2.300E-06 |
| 262 | 14 | P14R | A | 1.7e-06 | 2.6e-06 | 3e-06 | 2.433E-06 | 6.658E-07 | 1.700E-06 | 3.000E-06 |
| 263 | 14 | P14N | A | 3e-06 | 1.8e-06 | 2.5e-06 | 2.433E-06 | 6.028E-07 | 1.800E-06 | 3.000E-06 |
| 264 | 14 | P14D | A | 2e-06 | 2.4e-06 | 2.8e-06 | 2.400E-06 | 4.000E-07 | 2.000E-06 | 2.800E-06 |
| 265 | 14 | P14C | A | 8.4e-07 | 1.3e-06 | 2.5e-06 | 1.547E-06 | 8.570E-07 | 8.400E-07 | 2.500E-06 |
| 266 | 14 | P14E | A | 3.1e-06 | 2.5e-06 | 2.6e-06 | 2.733E-06 | 3.215E-07 | 2.500E-06 | 3.100E-06 |
| 267 | 14 | P14Q | A | 1.3e-06 | 2.1e-06 | 1.9e-06 | 1.767E-06 | 4.163E-07 | 1.300E-06 | 2.100E-06 |
| 268 | 14 | P14G | A | 2.1e-06 | 4e-06 | 2e-06 | 2.700E-06 | 1.127E-06 | 2.000E-06 | 4.000E-06 |
| 269 | 14 | P14H | A | 2.6e-06 | 1.4e-06 | 2e-06 | 2.000E-06 | 6.000E-07 | 1.400E-06 | 2.600E-06 |
| 270 | 14 | P14I | A | 2.3e-06 | 3.3e-06 | 5.3e-06 | 3.633E-06 | 1.528E-06 | 2.300E-06 | 5.300E-06 |
| 271 | 14 | P14L | A | 2.1e-06 | 1.8e-06 | 2.6e-06 | 2.167E-06 | 4.041E-07 | 1.800E-06 | 2.600E-06 |
| 272 | 14 | P14K | A | 2.6e-06 | 2.7e-06 | 3.2e-06 | 2.833E-06 | 3.215E-07 | 2.600E-06 | 3.200E-06 |
| 273 | 14 | P14M | A | 3e-06 | 2.8e-06 | 3.3e-06 | 3.033E-06 | 2.517E-07 | 2.800E-06 | 3.300E-06 |
| 274 | 14 | P14F | A | 1.9e-06 | 3.1e-06 | 1.9e-06 | 2.300E-06 | 6.928E-07 | 1.900E-06 | 3.100E-06 |
| 275 | 14 | P14P | A | 2.6e-06 | 1.6e-06 | 2.1e-06 | 2.100E-06 | 5.000E-07 | 1.600E-06 | 2.600E-06 |
| 276 | 14 | P14S | A | 1.5e-06 | 3.8e-06 | 1.9e-06 | 2.400E-06 | 1.229E-06 | 1.500E-06 | 3.800E-06 |
| 277 | 14 | P14T | A | 1.6e-06 | 2.7e-06 | 2.1e-06 | 2.133E-06 | 5.508E-07 | 1.600E-06 | 2.700E-06 |
| 278 | 14 | P14W | A | 2.2e-06 | 2.6e-06 | 9.2e-07 | 1.907E-06 | 8.776E-07 | 9.200E-07 | 2.600E-06 |
| 279 | 14 | P14Y | A | 2.2e-06 | 9.3e-07 | 2.1e-06 | 1.743E-06 | 7.061E-07 | 9.300E-07 | 2.200E-06 |
| 280 | 14 | P14V | A | 2.4e-06 | 1.9e-06 | 3.5e-06 | 2.600E-06 | 8.185E-07 | 1.900E-06 | 3.500E-06 |
| 281 | 15 | A15A | A | 3.1e-06 | 4.3e-06 | 2.1e-06 | 3.167E-06 | 1.102E-06 | 2.100E-06 | 4.300E-06 |
| 282 | 15 | A15R | A | 1.8e-06 | 2.5e-06 | 3.1e-06 | 2.467E-06 | 6.506E-07 | 1.800E-06 | 3.100E-06 |
| 283 | 15 | A15N | A | 2.8e-06 | 2.6e-06 | 1.8e-06 | 2.400E-06 | 5.292E-07 | 1.800E-06 | 2.800E-06 |
| 284 | 15 | A15D | A | 1.8e-06 | 1.7e-06 | 2.9e-06 | 2.133E-06 | 6.658E-07 | 1.700E-06 | 2.900E-06 |
| 285 | 15 | A15C | A | 1.9e-06 | 2.3e-06 | 2.3e-06 | 2.167E-06 | 2.309E-07 | 1.900E-06 | 2.300E-06 |
| 286 | 15 | A15E | A | 2.9e-06 | 1.7e-06 | 2.4e-06 | 2.333E-06 | 6.028E-07 | 1.700E-06 | 2.900E-06 |
| 287 | 15 | A15Q | A | 3.2e-06 | 2.6e-06 | 3e-06 | 2.933E-06 | 3.055E-07 | 2.600E-06 | 3.200E-06 |
| 288 | 15 | A15G | A | 1.3e-06 | 2.7e-06 | 3e-06 | 2.333E-06 | 9.074E-07 | 1.300E-06 | 3.000E-06 |
| 289 | 15 | A15H | A | 2.3e-06 | 2.1e-06 | 2.8e-06 | 2.400E-06 | 3.606E-07 | 2.100E-06 | 2.800E-06 |
| 290 | 15 | A15I | A | 1.2e-06 | 3.3e-06 | 4.4e-06 | 2.967E-06 | 1.626E-06 | 1.200E-06 | 4.400E-06 |
| 291 | 15 | A15L | A | 2.3e-06 | 2.5e-06 | 1.6e-06 | 2.133E-06 | 4.726E-07 | 1.600E-06 | 2.500E-06 |
| 292 | 15 | A15K | A | 2.4e-06 | 2.2e-06 | 2.8e-06 | 2.467E-06 | 3.055E-07 | 2.200E-06 | 2.800E-06 |
| 293 | 15 | A15M | A | 2.8e-06 | 2.8e-06 | 2.4e-06 | 2.667E-06 | 2.309E-07 | 2.400E-06 | 2.800E-06 |
| 294 | 15 | A15F | A | 3.2e-06 | 3.2e-06 | 1.6e-06 | 2.667E-06 | 9.238E-07 | 1.600E-06 | 3.200E-06 |
| 295 | 15 | A15P | A | 2.9e-06 | 2.3e-06 | 1.5e-06 | 2.233E-06 | 7.024E-07 | 1.500E-06 | 2.900E-06 |
| 296 | 15 | A15S | A | 1.6e-06 | 2.6e-06 | 2.1e-06 | 2.100E-06 | 5.000E-07 | 1.600E-06 | 2.600E-06 |
| 297 | 15 | A15T | A | 2.2e-06 | 1.9e-06 | 1.7e-06 | 1.933E-06 | 2.517E-07 | 1.700E-06 | 2.200E-06 |
| 298 | 15 | A15W | A | 1.8e-06 | 1.8e-06 | 3.2e-06 | 2.267E-06 | 8.083E-07 | 1.800E-06 | 3.200E-06 |
| 299 | 15 | A15Y | A | 2e-06 | 1.4e-06 | 2.2e-06 | 1.867E-06 | 4.163E-07 | 1.400E-06 | 2.200E-06 |
| 300 | 15 | A15V | A | 2.4e-06 | 3.4e-06 | 1.5e-06 | 2.433E-06 | 9.504E-07 | 1.500E-06 | 3.400E-06 |
| 301 | 16 | S16A | A | 1.7e-06 | 2.1e-06 | 3.2e-06 | 2.333E-06 | 7.767E-07 | 1.700E-06 | 3.200E-06 |
| 302 | 16 | S16R | A | 2.2e-06 | 9.5e-07 | 3.8e-06 | 2.317E-06 | 1.429E-06 | 9.500E-07 | 3.800E-06 |
| 303 | 16 | S16N | A | 1.8e-06 | 1.7e-06 | 2.8e-06 | 2.100E-06 | 6.083E-07 | 1.700E-06 | 2.800E-06 |
| 304 | 16 | S16D | A | 2.2e-06 | 2.1e-06 | 2.4e-06 | 2.233E-06 | 1.528E-07 | 2.100E-06 | 2.400E-06 |
| 305 | 16 | S16C | A | 1.2e-06 | 1.7e-06 | 2.7e-06 | 1.867E-06 | 7.638E-07 | 1.200E-06 | 2.700E-06 |
| 306 | 16 | S16E | A | 2.2e-06 | 3.9e-06 | 1.5e-06 | 2.533E-06 | 1.234E-06 | 1.500E-06 | 3.900E-06 |
| 307 | 16 | S16Q | A | 2.3e-06 | 2.1e-06 | 2e-06 | 2.133E-06 | 1.528E-07 | 2.000E-06 | 2.300E-06 |
| 308 | 16 | S16G | A | 3e-06 | 2.7e-06 | 1.2e-06 | 2.300E-06 | 9.644E-07 | 1.200E-06 | 3.000E-06 |
| 309 | 16 | S16H | A | 4.6e-06 | 2.4e-06 | 2.7e-06 | 3.233E-06 | 1.193E-06 | 2.400E-06 | 4.600E-06 |
| 310 | 16 | S16I | A | 4.3e-06 | 2.9e-06 | 1.8e-06 | 3.000E-06 | 1.253E-06 | 1.800E-06 | 4.300E-06 |
| 311 | 16 | S16L | A | 2.7e-06 | 2.1e-06 | 2.7e-06 | 2.500E-06 | 3.464E-07 | 2.100E-06 | 2.700E-06 |
| 312 | 16 | S16K | A | 2.7e-06 | 3.7e-06 | 2.1e-06 | 2.833E-06 | 8.083E-07 | 2.100E-06 | 3.700E-06 |
| 313 | 16 | S16M | A | 2.9e-06 | 2.3e-06 | 2.8e-06 | 2.667E-06 | 3.215E-07 | 2.300E-06 | 2.900E-06 |
| 314 | 16 | S16F | A | 2.5e-06 | 2e-06 | 1.1e-06 | 1.867E-06 | 7.095E-07 | 1.100E-06 | 2.500E-06 |
| 315 | 16 | S16P | A | 1.9e-06 | 2.6e-06 | 1.4e-06 | 1.967E-06 | 6.028E-07 | 1.400E-06 | 2.600E-06 |
| 316 | 16 | S16S | A | 3.3e-06 | 1.3e-06 | 2.4e-06 | 2.333E-06 | 1.002E-06 | 1.300E-06 | 3.300E-06 |
| 317 | 16 | S16T | A | 1.9e-06 | 2.7e-06 | 2.5e-06 | 2.367E-06 | 4.163E-07 | 1.900E-06 | 2.700E-06 |
| 318 | 16 | S16W | A | 2.2e-06 | 1.9e-06 | 1.8e-06 | 1.967E-06 | 2.082E-07 | 1.800E-06 | 2.200E-06 |
| 319 | 16 | S16Y | A | 2.7e-06 | 1.7e-06 | 2.2e-06 | 2.200E-06 | 5.000E-07 | 1.700E-06 | 2.700E-06 |
| 320 | 16 | S16V | A | 2.1e-06 | 4.1e-06 | 1.2e-06 | 2.467E-06 | 1.484E-06 | 1.200E-06 | 4.100E-06 |
| 321 | 17 | P17A | A | 2e-06 | 3.2e-06 | 4.9e-06 | 3.367E-06 | 1.457E-06 | 2.000E-06 | 4.900E-06 |
| 322 | 17 | P17R | A | 1.9e-06 | 1.5e-06 | 3.1e-06 | 2.167E-06 | 8.327E-07 | 1.500E-06 | 3.100E-06 |
| 323 | 17 | P17N | A | 2.5e-06 | 1.8e-06 | 1.8e-06 | 2.033E-06 | 4.041E-07 | 1.800E-06 | 2.500E-06 |
| 324 | 17 | P17D | A | 2.3e-06 | 2.6e-06 | 2.6e-06 | 2.500E-06 | 1.732E-07 | 2.300E-06 | 2.600E-06 |
| 325 | 17 | P17C | A | 3e-06 | 2.6e-06 | 4.2e-06 | 3.267E-06 | 8.327E-07 | 2.600E-06 | 4.200E-06 |
| 326 | 17 | P17E | A | 2.1e-06 | 2.6e-06 | 2.8e-06 | 2.500E-06 | 3.606E-07 | 2.100E-06 | 2.800E-06 |
| 327 | 17 | P17Q | A | 2.5e-06 | 2.3e-06 | 2.3e-06 | 2.367E-06 | 1.155E-07 | 2.300E-06 | 2.500E-06 |
| 328 | 17 | P17G | A | 2.9e-06 | 1.5e-06 | 2.2e-06 | 2.200E-06 | 7.000E-07 | 1.500E-06 | 2.900E-06 |
| 329 | 17 | P17H | A | 2.6e-06 | 2.3e-06 | 1.5e-06 | 2.133E-06 | 5.686E-07 | 1.500E-06 | 2.600E-06 |
| 330 | 17 | P17I | A | 1.9e-06 | 1.7e-06 | 1.5e-06 | 1.700E-06 | 2.000E-07 | 1.500E-06 | 1.900E-06 |
| 331 | 17 | P17L | A | 2.2e-06 | 2.2e-06 | 2.1e-06 | 2.167E-06 | 5.774E-08 | 2.100E-06 | 2.200E-06 |
| 332 | 17 | P17K | A | 2.3e-06 | 1.7e-06 | 1.7e-06 | 1.900E-06 | 3.464E-07 | 1.700E-06 | 2.300E-06 |
| 333 | 17 | P17M | A | 2.1e-06 | 3.9e-06 | 2.3e-06 | 2.767E-06 | 9.866E-07 | 2.100E-06 | 3.900E-06 |
| 334 | 17 | P17F | A | 2.9e-06 | 2.2e-06 | 2.3e-06 | 2.467E-06 | 3.786E-07 | 2.200E-06 | 2.900E-06 |
| 335 | 17 | P17P | A | 3e-06 | 2.4e-06 | 2e-06 | 2.467E-06 | 5.033E-07 | 2.000E-06 | 3.000E-06 |
| 336 | 17 | P17S | A | 2.3e-06 | 2.2e-06 | 2.4e-06 | 2.300E-06 | 1.000E-07 | 2.200E-06 | 2.400E-06 |
| 337 | 17 | P17T | A | 2e-06 | 2.2e-06 | 2.5e-06 | 2.233E-06 | 2.517E-07 | 2.000E-06 | 2.500E-06 |
| 338 | 17 | P17W | A | 2.7e-06 | 1.5e-06 | 1.4e-06 | 1.867E-06 | 7.234E-07 | 1.400E-06 | 2.700E-06 |
| 339 | 17 | P17Y | A | 2.4e-06 | 1.8e-06 | 2.5e-06 | 2.233E-06 | 3.786E-07 | 1.800E-06 | 2.500E-06 |

| | | | | | | | | | | |
|---|---|---|---|---|---|---|---|---|---|---|
| 340 | 17 | P17V | A | 3.6e-06 | 2.9e-06 | 1.9e-06 | 2.800E-06 | 8.544E-07 | 1.900E-06 | 3.600E-06 |
| 341 | 18 | E18A | A | 2.4e-06 | 3.1e-06 | 2.5e-06 | 2.667E-06 | 3.786E-07 | 2.400E-06 | 3.100E-06 |
| 342 | 18 | E18R | A | 1.7e-06 | 1.9e-06 | 2.9e-06 | 2.167E-06 | 6.429E-07 | 1.700E-06 | 2.900E-06 |
| 343 | 18 | E18N | A | 2e-06 | 2.9e-06 | 1.3e-06 | 2.067E-06 | 8.021E-07 | 1.300E-06 | 2.900E-06 |
| 344 | 18 | E18D | A | 3e-06 | 1.7e-06 | 2.7e-06 | 2.467E-06 | 6.807E-07 | 1.700E-06 | 3.000E-06 |
| 345 | 18 | E18C | A | 2.5e-06 | 2.8e-06 | 3.3e-06 | 2.867E-06 | 4.041E-07 | 2.500E-06 | 3.300E-06 |
| 346 | 18 | E18E | A | 3.5e-06 | 2.2e-06 | 2.9e-06 | 2.867E-06 | 6.506E-07 | 2.200E-06 | 3.500E-06 |
| 347 | 18 | E18Q | A | 1.8e-06 | 1.5e-06 | 3e-06 | 2.100E-06 | 7.937E-07 | 1.500E-06 | 3.000E-06 |
| 348 | 18 | E18G | A | 2.6e-06 | 2.9e-06 | 2.1e-06 | 2.533E-06 | 4.041E-07 | 2.100E-06 | 2.900E-06 |
| 349 | 18 | E18H | A | 2.9e-06 | 2.3e-06 | 2e-06 | 2.400E-06 | 4.583E-07 | 2.000E-06 | 2.900E-06 |
| 350 | 18 | E18I | A | 2.3e-06 | 1.9e-06 | 2.5e-06 | 2.233E-06 | 3.055E-07 | 1.900E-06 | 2.500E-06 |
| 351 | 18 | E18L | A | 3e-06 | 2.2e-06 | 2.6e-06 | 2.600E-06 | 4.000E-07 | 2.200E-06 | 3.000E-06 |
| 352 | 18 | E18K | A | 1.6e-06 | 3e-06 | 1.8e-06 | 2.133E-06 | 7.572E-07 | 1.600E-06 | 3.000E-06 |
| 353 | 18 | E18M | A | 2e-06 | 1.9e-06 | 3.3e-06 | 2.400E-06 | 7.810E-07 | 1.900E-06 | 3.300E-06 |
| 354 | 18 | E18F | A | 3e-06 | 3.2e-06 | 3e-06 | 3.067E-06 | 1.155E-07 | 3.000E-06 | 3.200E-06 |
| 355 | 18 | E18P | A | 1.5e-06 | 2.5e-06 | 3.3e-06 | 2.433E-06 | 9.018E-07 | 1.500E-06 | 3.300E-06 |
| 356 | 18 | E18S | A | 2.6e-06 | 1.3e-06 | 3.3e-06 | 2.400E-06 | 1.015E-06 | 1.300E-06 | 3.300E-06 |
| 357 | 18 | E18T | A | 2.1e-06 | 1.5e-06 | 3.5e-06 | 2.367E-06 | 1.026E-06 | 1.500E-06 | 3.500E-06 |
| 358 | 18 | E18W | A | 1.9e-06 | 2.8e-06 | 8.8e-07 | 1.860E-06 | 9.606E-07 | 8.800E-07 | 2.800E-06 |
| 359 | 18 | E18Y | A | 1.3e-06 | 3.6e-06 | 3e-06 | 2.633E-06 | 1.193E-06 | 1.300E-06 | 3.600E-06 |
| 360 | 18 | E18V | A | 2.4e-06 | 5.1e-06 | 2.6e-06 | 3.367E-06 | 1.504E-06 | 2.400E-06 | 5.100E-06 |
| 361 | 19 | T19A | A | 2.3e-06 | 2.9e-06 | 4.1e-06 | 3.100E-06 | 9.165E-07 | 2.300E-06 | 4.100E-06 |
| 362 | 19 | T19R | A | 2.5e-06 | 2.3e-06 | 2.9e-06 | 2.567E-06 | 3.055E-07 | 2.300E-06 | 2.900E-06 |
| 363 | 19 | T19N | A | 2.6e-06 | 3.4e-06 | 2.6e-06 | 2.867E-06 | 4.619E-07 | 2.600E-06 | 3.400E-06 |
| 364 | 19 | T19D | A | 2.2e-06 | 3e-06 | 2.1e-06 | 2.433E-06 | 4.933E-07 | 2.100E-06 | 3.000E-06 |
| 365 | 19 | T19C | A | 2.8e-06 | 3.4e-06 | 2.6e-06 | 2.933E-06 | 4.163E-07 | 2.600E-06 | 3.400E-06 |
| 366 | 19 | T19E | A | 1.8e-06 | 3.7e-06 | 1.8e-06 | 2.433E-06 | 1.097E-06 | 1.800E-06 | 3.700E-06 |
| 367 | 19 | T19Q | A | 2.2e-06 | 1.4e-06 | 3.7e-06 | 2.433E-06 | 1.168E-06 | 1.400E-06 | 3.700E-06 |
| 368 | 19 | T19G | A | 2.6e-06 | 2.6e-06 | 2.8e-06 | 2.667E-06 | 1.155E-07 | 2.600E-06 | 2.800E-06 |
| 369 | 19 | T19H | A | 1.8e-06 | 1.9e-06 | 2.2e-06 | 1.967E-06 | 2.082E-07 | 1.800E-06 | 2.200E-06 |
| 370 | 19 | T19I | A | 2.3e-06 | 2.7e-06 | 2.3e-06 | 2.433E-06 | 2.309E-07 | 2.300E-06 | 2.700E-06 |
| 371 | 19 | T19L | A | 3e-06 | 4.1e-06 | 2.7e-06 | 3.267E-06 | 7.371E-07 | 2.700E-06 | 4.100E-06 |
| 372 | 19 | T19K | A | 2.3e-06 | 1.9e-06 | 2.1e-06 | 2.100E-06 | 2.000E-07 | 1.900E-06 | 2.300E-06 |
| 373 | 19 | T19M | A | 2.1e-06 | 3.6e-06 | 2.5e-06 | 2.733E-06 | 7.767E-07 | 2.100E-06 | 3.600E-06 |
| 374 | 19 | T19F | A | 2.9e-06 | 2.2e-06 | 3.5e-06 | 2.867E-06 | 6.506E-07 | 2.200E-06 | 3.500E-06 |
| 375 | 19 | T19P | A | 2.2e-06 | 2.5e-06 | 4.2e-06 | 2.967E-06 | 1.079E-06 | 2.200E-06 | 4.200E-06 |
| 376 | 19 | T19S | A | 1.4e-06 | 3.1e-06 | 2e-06 | 2.167E-06 | 8.622E-07 | 1.400E-06 | 3.100E-06 |
| 377 | 19 | T19T | A | 2.2e-06 | 1.3e-06 | 3.5e-06 | 2.333E-06 | 1.106E-06 | 1.300E-06 | 3.500E-06 |
| 378 | 19 | T19W | A | 1.6e-06 | 4.2e-06 | 2.1e-06 | 2.633E-06 | 1.380E-06 | 1.600E-06 | 4.200E-06 |
| 379 | 19 | T19Y | A | 3.7e-06 | 2.1e-06 | 3e-06 | 2.933E-06 | 8.021E-07 | 2.100E-06 | 3.700E-06 |
| 380 | 19 | T19V | A | 2.3e-06 | 2.3e-06 | 2.6e-06 | 2.400E-06 | 1.732E-07 | 2.300E-06 | 2.600E-06 |
| 381 | 20 | H20A | A | 1.9e-06 | 2e-06 | 4.1e-06 | 2.667E-06 | 1.242E-06 | 1.900E-06 | 4.100E-06 |
| 382 | 20 | H20R | A | 2.8e-06 | 2e-06 | 2e-06 | 2.267E-06 | 4.619E-07 | 2.000E-06 | 2.800E-06 |
| 383 | 20 | H20N | A | 2.5e-06 | 3.9e-06 | 3e-06 | 3.133E-06 | 7.095E-07 | 2.500E-06 | 3.900E-06 |
| 384 | 20 | H20D | A | 4.1e-06 | 2.1e-06 | 2.7e-06 | 2.967E-06 | 1.026E-06 | 2.100E-06 | 4.100E-06 |
| 385 | 20 | H20C | A | 2.2e-06 | 5.6e-06 | 2.9e-06 | 3.567E-06 | 1.795E-06 | 2.200E-06 | 5.600E-06 |
| 386 | 20 | H20E | A | 2.4e-06 | 2e-06 | 3.8e-06 | 2.733E-06 | 9.452E-07 | 2.000E-06 | 3.800E-06 |
| 387 | 20 | H20Q | A | 2.5e-06 | 3.3e-06 | 2.1e-06 | 2.633E-06 | 6.110E-07 | 2.100E-06 | 3.300E-06 |
| 388 | 20 | H20G | A | 3.2e-06 | 2.8e-06 | 5.9e-06 | 3.967E-06 | 1.686E-06 | 2.800E-06 | 5.900E-06 |
| 389 | 20 | H20H | A | 3e-06 | 2.2e-06 | 3.7e-06 | 2.967E-06 | 7.506E-07 | 2.200E-06 | 3.700E-06 |
| 390 | 20 | H20I | A | 2.5e-06 | 1.3e-06 | 2.2e-06 | 2.000E-06 | 6.245E-07 | 1.300E-06 | 2.500E-06 |
| 391 | 20 | H20L | A | 2.3e-06 | 3.9e-06 | 3.5e-06 | 3.233E-06 | 8.327E-07 | 2.300E-06 | 3.900E-06 |
| 392 | 20 | H20K | A | 4e-06 | 2.6e-06 | 2.3e-06 | 2.967E-06 | 9.074E-07 | 2.300E-06 | 4.000E-06 |
| 393 | 20 | H20M | A | 1.5e-06 | 2.9e-06 | 2.1e-06 | 2.167E-06 | 7.024E-07 | 1.500E-06 | 2.900E-06 |
| 394 | 20 | H20F | A | 4.7e-06 | 1.4e-06 | 2.9e-06 | 3.000E-06 | 1.652E-06 | 1.400E-06 | 4.700E-06 |
| 395 | 20 | H20P | A | 3.3e-06 | 2.5e-06 | 2.4e-06 | 2.733E-06 | 4.933E-07 | 2.400E-06 | 3.300E-06 |
| 396 | 20 | H20S | A | 2.3e-06 | 1.8e-06 | 3e-06 | 2.367E-06 | 6.028E-07 | 1.800E-06 | 3.000E-06 |
| 397 | 20 | H20T | A | 2.2e-06 | 1.8e-06 | 2e-06 | 2.000E-06 | 2.000E-07 | 1.800E-06 | 2.200E-06 |
| 398 | 20 | H20W | A | 2.7e-06 | 4e-06 | 2.4e-06 | 3.033E-06 | 8.505E-07 | 2.400E-06 | 4.000E-06 |
| 399 | 20 | H20Y | A | 1.8e-06 | 2.3e-06 | 2e-06 | 2.033E-06 | 2.517E-07 | 1.800E-06 | 2.300E-06 |
| 400 | 20 | H20V | A | 3.5e-06 | 2.7e-06 | 3.2e-06 | 3.133E-06 | 4.041E-07 | 2.700E-06 | 3.500E-06 |
| 401 | 21 | L21A | A | 2.4e-06 | 2.9e-06 | 1.5e-06 | 2.267E-06 | 7.095E-07 | 1.500E-06 | 2.900E-06 |
| 402 | 21 | L21R | A | 2.2e-06 | 3.1e-06 | 2.3e-06 | 2.533E-06 | 4.933E-07 | 2.200E-06 | 3.100E-06 |
| 403 | 21 | L21N | A | 1.3e-06 | 1.9e-06 | 2.2e-06 | 1.800E-06 | 4.583E-07 | 1.300E-06 | 2.200E-06 |
| 404 | 21 | L21D | A | 2.2e-06 | 2.6e-06 | 2.5e-06 | 2.433E-06 | 2.082E-07 | 2.200E-06 | 2.600E-06 |
| 405 | 21 | L21C | A | 2.2e-06 | 2.6e-06 | 3.1e-06 | 2.633E-06 | 4.509E-07 | 2.200E-06 | 3.100E-06 |
| 406 | 21 | L21E | A | 2.6e-06 | 1.4e-06 | 2e-06 | 2.000E-06 | 6.000E-07 | 1.400E-06 | 2.600E-06 |
| 407 | 21 | L21Q | A | 2e-06 | 2.5e-06 | 2.1e-06 | 2.200E-06 | 2.646E-07 | 2.000E-06 | 2.500E-06 |
| 408 | 21 | L21G | A | 2.1e-06 | 3.1e-06 | 1.4e-06 | 2.200E-06 | 8.544E-07 | 1.400E-06 | 3.100E-06 |
| 409 | 21 | L21H | A | 3.5e-06 | 2.3e-06 | 1.6e-06 | 2.467E-06 | 9.609E-07 | 1.600E-06 | 3.500E-06 |
| 410 | 21 | L21I | A | 3.2e-06 | 2.1e-06 | 2.7e-06 | 2.667E-06 | 5.508E-07 | 2.100E-06 | 3.200E-06 |
| 411 | 21 | L21L | A | 2.7e-06 | 4e-06 | 2.2e-06 | 2.967E-06 | 9.292E-07 | 2.200E-06 | 4.000E-06 |
| 412 | 21 | L21K | A | 3e-06 | 2.3e-06 | 2.6e-06 | 2.633E-06 | 3.512E-07 | 2.300E-06 | 3.000E-06 |
| 413 | 21 | L21M | A | 1.9e-06 | 2.7e-06 | 2.2e-06 | 2.267E-06 | 4.041E-07 | 1.900E-06 | 2.700E-06 |
| 414 | 21 | L21F | A | 2.4e-06 | 2.5e-06 | 2.8e-06 | 2.567E-06 | 2.082E-07 | 2.400E-06 | 2.800E-06 |
| 415 | 21 | L21P | A | 2.2e-06 | 2.7e-06 | 2.5e-06 | 2.467E-06 | 2.517E-07 | 2.200E-06 | 2.700E-06 |
| 416 | 21 | L21S | A | 1.5e-06 | 2.7e-06 | 1.9e-06 | 2.033E-06 | 6.110E-07 | 1.500E-06 | 2.700E-06 |
| 417 | 21 | L21T | A | 3.1e-06 | 1.1e-06 | 2.2e-06 | 2.133E-06 | 1.002E-06 | 1.100E-06 | 3.100E-06 |
| 418 | 21 | L21W | A | 2e-06 | 2.6e-06 | 2.4e-06 | 2.333E-06 | 3.055E-07 | 2.000E-06 | 2.600E-06 |
| 419 | 21 | L21Y | A | 3e-06 | 3.1e-06 | 2.7e-06 | 2.933E-06 | 2.082E-07 | 2.700E-06 | 3.100E-06 |
| 420 | 21 | L21V | A | 4.1e-06 | 1.1e-06 | 3.8e-06 | 3.000E-06 | 1.652E-06 | 1.100E-06 | 4.100E-06 |
| 421 | 22 | D22A | A | 2.4e-06 | 2.4e-06 | 5.1e-06 | 3.300E-06 | 1.559E-06 | 2.400E-06 | 5.100E-06 |
| 422 | 22 | D22R | A | 1.7e-06 | 3.4e-06 | 2.9e-06 | 2.667E-06 | 8.737E-07 | 1.700E-06 | 3.400E-06 |
| 423 | 22 | D22N | A | 1.9e-06 | 2.1e-06 | 3.3e-06 | 2.433E-06 | 7.572E-07 | 1.900E-06 | 3.300E-06 |
| 424 | 22 | D22D | A | 1.8e-06 | 1.9e-06 | 2.3e-06 | 2.000E-06 | 2.646E-07 | 1.800E-06 | 2.300E-06 |

| | | | | | | | | | | |
|---|---|---|---|---|---|---|---|---|---|---|
| 425 | 22 | D22C | A | 1.9e-06 | 3.4e-06 | 2.4e-06 | 2.567E-06 | 7.638E-07 | 1.900E-06 | 3.400E-06 |
| 426 | 22 | D22E | A | 1.8e-06 | 1.8e-06 | 3.6e-06 | 2.400E-06 | 1.039E-06 | 1.800E-06 | 3.600E-06 |
| 427 | 22 | D22Q | A | 2e-06 | 1.4e-06 | 2.6e-06 | 2.000E-06 | 6.000E-07 | 1.400E-06 | 2.600E-06 |
| 428 | 22 | D22G | A | 2.7e-06 | 2.9e-06 | 4.5e-06 | 3.367E-06 | 9.866E-07 | 2.700E-06 | 4.500E-06 |
| 429 | 22 | D22H | A | 2.1e-06 | 5.9e-06 | 2.2e-06 | 3.400E-06 | 2.166E-06 | 2.100E-06 | 5.900E-06 |
| 430 | 22 | D22I | A | 1.7e-06 | 3.6e-06 | 2.3e-06 | 2.533E-06 | 9.713E-07 | 1.700E-06 | 3.600E-06 |
| 431 | 22 | D22L | A | 1.9e-06 | 1.5e-06 | 2.2e-06 | 1.867E-06 | 3.512E-07 | 1.500E-06 | 2.200E-06 |
| 432 | 22 | D22K | A | 2e-06 | 2.1e-06 | 1.8e-06 | 1.967E-06 | 1.528E-07 | 1.800E-06 | 2.100E-06 |
| 433 | 22 | D22M | A | 3.4e-06 | 2e-06 | 3.8e-06 | 3.067E-06 | 9.452E-07 | 2.000E-06 | 3.800E-06 |
| 434 | 22 | D22F | A | 4e-06 | 1.3e-06 | 1.3e-06 | 2.200E-06 | 1.559E-06 | 1.300E-06 | 4.000E-06 |
| 435 | 22 | D22P | A | 2.1e-06 | 2.7e-06 | 2.3e-06 | 2.367E-06 | 3.055E-07 | 2.100E-06 | 2.700E-06 |
| 436 | 22 | D22S | A | 1.3e-06 | 3.7e-06 | 1.9e-06 | 2.300E-06 | 1.249E-06 | 1.300E-06 | 3.700E-06 |
| 437 | 22 | D22T | A | 2.6e-06 | 3e-06 | 2e-06 | 2.533E-06 | 5.033E-07 | 2.000E-06 | 3.000E-06 |
| 438 | 22 | D22W | A | 1.5e-06 | 3.5e-06 | 2.8e-06 | 2.600E-06 | 1.015E-06 | 1.500E-06 | 3.500E-06 |
| 439 | 22 | D22Y | A | 2.5e-06 | 3.5e-06 | 3.1e-06 | 3.033E-06 | 5.033E-07 | 2.500E-06 | 3.500E-06 |
| 440 | 22 | D22V | A | 2.8e-06 | 2.7e-06 | 2.1e-06 | 2.533E-06 | 3.786E-07 | 2.100E-06 | 2.800E-06 |
| 441 | 23 | M23A | A | 2.5e-06 | 1.7e-06 | 2.5e-06 | 2.233E-06 | 4.619E-07 | 1.700E-06 | 2.500E-06 |
| 442 | 23 | M23R | A | 4.5e-06 | 2e-06 | 1.4e-06 | 2.633E-06 | 1.644E-06 | 1.400E-06 | 4.500E-06 |
| 443 | 23 | M23N | A | 2.2e-06 | 2e-06 | 1.5e-06 | 1.900E-06 | 3.606E-07 | 1.500E-06 | 2.200E-06 |
| 444 | 23 | M23D | A | 2.4e-06 | 2e-06 | 2.2e-06 | 2.200E-06 | 2.000E-07 | 2.000E-06 | 2.400E-06 |
| 445 | 23 | M23C | A | 2.6e-06 | 4.7e-06 | 1.8e-06 | 3.033E-06 | 1.498E-06 | 1.800E-06 | 4.700E-06 |
| 446 | 23 | M23E | A | 3.9e-06 | 3.1e-06 | 2.5e-06 | 3.167E-06 | 7.024E-07 | 2.500E-06 | 3.900E-06 |
| 447 | 23 | M23Q | A | 4.9e-06 | 4.2e-06 | 2.1e-06 | 3.733E-06 | 1.457E-06 | 2.100E-06 | 4.900E-06 |
| 448 | 23 | M23G | A | 4.9e-06 | 3.8e-06 | 3.4e-06 | 4.033E-06 | 7.767E-07 | 3.400E-06 | 4.900E-06 |
| 449 | 23 | M23H | A | 3.4e-06 | 4.1e-06 | 1.4e-06 | 2.967E-06 | 1.401E-06 | 1.400E-06 | 4.100E-06 |
| 450 | 23 | M23I | A | 2.4e-06 | 3.3e-06 | 2.3e-06 | 2.667E-06 | 5.508E-07 | 2.300E-06 | 3.300E-06 |
| 451 | 23 | M23L | A | 3.5e-06 | 3.1e-06 | 2.2e-06 | 2.933E-06 | 6.658E-07 | 2.200E-06 | 3.500E-06 |
| 452 | 23 | M23K | A | 2.9e-06 | 2e-06 | 2.6e-06 | 2.500E-06 | 4.583E-07 | 2.000E-06 | 2.900E-06 |
| 453 | 23 | M23M | A | 2.8e-06 | 1.9e-06 | 2.8e-06 | 2.500E-06 | 5.196E-07 | 1.900E-06 | 2.800E-06 |
| 454 | 23 | M23F | A | 4.5e-06 | 2e-06 | 2.7e-06 | 3.067E-06 | 1.290E-06 | 2.000E-06 | 4.500E-06 |
| 455 | 23 | M23P | A | 2.1e-06 | 3.2e-06 | 2.1e-06 | 2.467E-06 | 6.351E-07 | 2.100E-06 | 3.200E-06 |
| 456 | 23 | M23S | A | 4.7e-06 | 4.1e-06 | 3.8e-06 | 4.200E-06 | 4.583E-07 | 3.800E-06 | 4.700E-06 |
| 457 | 23 | M23T | A | 1.7e-06 | 2.2e-06 | 3.1e-06 | 2.333E-06 | 7.095E-07 | 1.700E-06 | 3.100E-06 |
| 458 | 23 | M23W | A | 3.4e-06 | 2.8e-06 | 2.6e-06 | 2.933E-06 | 4.163E-07 | 2.600E-06 | 3.400E-06 |
| 459 | 23 | M23Y | A | 6.2e-07 | 1.6e-06 | 2.9e-06 | 1.707E-06 | 1.144E-06 | 6.200E-07 | 2.900E-06 |
| 460 | 23 | M23V | A | 2.3e-06 | 1.4e-06 | 2.7e-06 | 2.133E-06 | 6.658E-07 | 1.400E-06 | 2.700E-06 |
| 461 | 24 | L24A | A | 3.9e-06 | 1.5e-06 | 2.3e-06 | 2.567E-06 | 1.222E-06 | 1.500E-06 | 3.900E-06 |
| 462 | 24 | L24R | A | 2.6e-06 | 2.5e-06 | 1e-06 | 2.033E-06 | 8.963E-07 | 1.000E-06 | 2.600E-06 |
| 463 | 24 | L24N | A | 2e-06 | 2e-06 | 3.7e-06 | 2.567E-06 | 9.815E-07 | 2.000E-06 | 3.700E-06 |
| 464 | 24 | L24D | A | 2.1e-06 | 1.7e-06 | 3.3e-06 | 2.367E-06 | 8.327E-07 | 1.700E-06 | 3.300E-06 |
| 465 | 24 | L24C | A | 1.9e-06 | 5e-06 | 2e-06 | 2.967E-06 | 1.762E-06 | 1.900E-06 | 5.000E-06 |
| 466 | 24 | L24E | A | 3.6e-06 | 1.8e-06 | 2.8e-06 | 2.733E-06 | 9.018E-07 | 1.800E-06 | 3.600E-06 |
| 467 | 24 | L24Q | A | 2.7e-06 | 2e-06 | 2.9e-06 | 2.533E-06 | 4.726E-07 | 2.000E-06 | 2.900E-06 |
| 468 | 24 | L24G | A | 1.9e-06 | 2.5e-06 | 2e-06 | 2.133E-06 | 3.215E-07 | 1.900E-06 | 2.500E-06 |
| 469 | 24 | L24H | A | 4e-06 | 2.9e-06 | 2.8e-06 | 3.233E-06 | 6.658E-07 | 2.800E-06 | 4.000E-06 |
| 470 | 24 | L24I | A | 1.8e-06 | 1.6e-06 | 3.2e-06 | 2.200E-06 | 8.718E-07 | 1.600E-06 | 3.200E-06 |
| 471 | 24 | L24L | A | 9.5e-07 | 1.4e-06 | 2.2e-06 | 1.517E-06 | 6.331E-07 | 9.500E-07 | 2.200E-06 |
| 472 | 24 | L24K | A | 2.7e-06 | 1.8e-06 | 1.3e-06 | 1.933E-06 | 7.095E-07 | 1.300E-06 | 2.700E-06 |
| 473 | 24 | L24M | A | 1.4e-06 | 3.2e-06 | 2e-06 | 2.200E-06 | 9.165E-07 | 1.400E-06 | 3.200E-06 |
| 474 | 24 | L24F | A | 2e-06 | 2e-06 | 1.3e-06 | 1.767E-06 | 4.041E-07 | 1.300E-06 | 2.000E-06 |
| 475 | 24 | L24P | A | 2.5e-06 | 2.8e-06 | 3.5e-06 | 2.933E-06 | 5.132E-07 | 2.500E-06 | 3.500E-06 |
| 476 | 24 | L24S | A | 1.2e-06 | 2.6e-06 | 1.9e-06 | 1.900E-06 | 7.000E-07 | 1.200E-06 | 2.600E-06 |
| 477 | 24 | L24T | A | 3.5e-06 | 3.4e-06 | 3.6e-06 | 3.500E-06 | 1.000E-07 | 3.400E-06 | 3.600E-06 |
| 478 | 24 | L24W | A | 4.3e-06 | 2.6e-06 | 2.9e-06 | 3.267E-06 | 9.074E-07 | 2.600E-06 | 4.300E-06 |
| 479 | 24 | L24Y | A | 2.3e-06 | 3.3e-06 | 3.2e-06 | 2.933E-06 | 5.508E-07 | 2.300E-06 | 3.300E-06 |
| 480 | 24 | L24V | A | 3.1e-06 | 2.2e-06 | 5e-06 | 3.433E-06 | 1.429E-06 | 2.200E-06 | 5.000E-06 |
| 481 | 25 | R25A | A | 1.9e-06 | 1.3e-06 | 2.6e-06 | 1.933E-06 | 6.506E-07 | 1.300E-06 | 2.600E-06 |
| 482 | 25 | R25R | A | 3.7e-06 | 2e-06 | 1.9e-06 | 2.533E-06 | 1.012E-06 | 1.900E-06 | 3.700E-06 |
| 483 | 25 | R25N | A | 2.4e-06 | 2.1e-06 | 1.7e-06 | 2.067E-06 | 3.512E-07 | 1.700E-06 | 2.400E-06 |
| 484 | 25 | R25D | A | 2e-06 | 2.5e-06 | 1.9e-06 | 2.133E-06 | 3.215E-07 | 1.900E-06 | 2.500E-06 |
| 485 | 25 | R25C | A | 2.1e-06 | 2.9e-06 | 1.6e-06 | 2.200E-06 | 6.557E-07 | 1.600E-06 | 2.900E-06 |
| 486 | 25 | R25E | A | 3.4e-06 | 1.3e-06 | 2.4e-06 | 2.367E-06 | 1.050E-06 | 1.300E-06 | 3.400E-06 |
| 487 | 25 | R25Q | A | 1.2e-06 | 1.4e-06 | 2.8e-06 | 1.800E-06 | 8.718E-07 | 1.200E-06 | 2.800E-06 |
| 488 | 25 | R25G | A | 2e-06 | 2e-06 | 2.4e-06 | 2.133E-06 | 2.309E-07 | 2.000E-06 | 2.400E-06 |
| 489 | 25 | R25H | A | 2.5e-06 | 2.9e-06 | 2.2e-06 | 2.533E-06 | 3.512E-07 | 2.200E-06 | 2.900E-06 |
| 490 | 25 | R25I | A | 1.9e-06 | 2e-06 | 2.4e-06 | 2.100E-06 | 2.646E-07 | 1.900E-06 | 2.400E-06 |
| 491 | 25 | R25L | A | 2.4e-06 | 2.1e-06 | 2.1e-06 | 2.200E-06 | 1.732E-07 | 2.100E-06 | 2.400E-06 |
| 492 | 25 | R25K | A | 1.4e-06 | 2.1e-06 | 3.4e-06 | 2.300E-06 | 1.015E-06 | 1.400E-06 | 3.400E-06 |
| 493 | 25 | R25M | A | 1.9e-06 | 4.8e-06 | 2.3e-06 | 3.000E-06 | 1.572E-06 | 1.900E-06 | 4.800E-06 |
| 494 | 25 | R25F | A | 1.8e-06 | 4e-06 | 2.6e-06 | 2.800E-06 | 1.114E-06 | 1.800E-06 | 4.000E-06 |
| 495 | 25 | R25P | A | 2.7e-06 | 2.8e-06 | 3.6e-06 | 3.033E-06 | 4.933E-07 | 2.700E-06 | 3.600E-06 |
| 496 | 25 | R25S | A | 1.3e-06 | 1.4e-06 | 2.3e-06 | 1.667E-06 | 5.508E-07 | 1.300E-06 | 2.300E-06 |
| 497 | 25 | R25T | A | 2.6e-06 | 3.1e-06 | 2.6e-06 | 2.767E-06 | 2.887E-07 | 2.600E-06 | 3.100E-06 |
| 498 | 25 | R25W | A | 2.9e-06 | 2.7e-06 | 3e-06 | 2.867E-06 | 1.528E-07 | 2.700E-06 | 3.000E-06 |
| 499 | 25 | R25Y | A | 2.1e-06 | 2.2e-06 | 1.8e-06 | 2.033E-06 | 2.082E-07 | 1.800E-06 | 2.200E-06 |
| 500 | 25 | R25V | A | 4e-06 | 2.9e-06 | 2.9e-06 | 3.267E-06 | 6.351E-07 | 2.900E-06 | 4.000E-06 |
| 501 | 26 | H26A | A | 6e-06 | 2.6e-06 | 2.4e-06 | 3.667E-06 | 2.023E-06 | 2.400E-06 | 6.000E-06 |
| 502 | 26 | H26R | A | 2.6e-06 | 2.3e-06 | 2e-06 | 2.300E-06 | 3.000E-07 | 2.000E-06 | 2.600E-06 |
| 503 | 26 | H26N | A | 1.7e-06 | 4.3e-06 | 1.4e-06 | 2.467E-06 | 1.595E-06 | 1.400E-06 | 4.300E-06 |
| 504 | 26 | H26D | A | 2.5e-06 | 1.6e-06 | 4e-06 | 2.700E-06 | 1.212E-06 | 1.600E-06 | 4.000E-06 |
| 505 | 26 | H26C | A | 4.7e-06 | 2e-06 | 4e-06 | 3.567E-06 | 1.401E-06 | 2.000E-06 | 4.700E-06 |
| 506 | 26 | H26E | A | 2.4e-06 | 2.8e-06 | 1.9e-06 | 2.367E-06 | 4.509E-07 | 1.900E-06 | 2.800E-06 |
| 507 | 26 | H26Q | A | 2.3e-06 | 1.5e-06 | 1.7e-06 | 1.833E-06 | 4.163E-07 | 1.500E-06 | 2.300E-06 |
| 508 | 26 | H26G | A | 3.4e-06 | 3.2e-06 | 3.9e-06 | 3.500E-06 | 3.606E-07 | 3.200E-06 | 3.900E-06 |
| 509 | 26 | H26H | A | 1.7e-06 | 3.7e-06 | 2.2e-06 | 2.533E-06 | 1.041E-06 | 1.700E-06 | 3.700E-06 |

| | | | | | | | | | | |
|---|---|---|---|---|---|---|---|---|---|---|
| 510 | 26 | H26I | A | 2.3e-06 | 1.2e-06 | 2.8e-06 | 2.100E-06 | 8.185E-07 | 1.200E-06 | 2.800E-06 |
| 511 | 26 | H26L | A | 1.5e-06 | 2.3e-06 | 1.7e-06 | 1.833E-06 | 4.163E-07 | 1.500E-06 | 2.300E-06 |
| 512 | 26 | H26K | A | 2.3e-06 | 1.7e-06 | 2.3e-06 | 2.100E-06 | 3.464E-07 | 1.700E-06 | 2.300E-06 |
| 513 | 26 | H26M | A | 1.6e-06 | 2.7e-06 | 1.5e-06 | 1.933E-06 | 6.658E-07 | 1.500E-06 | 2.700E-06 |
| 514 | 26 | H26F | A | 3.3e-06 | 3.1e-06 | 1.9e-06 | 2.767E-06 | 7.572E-07 | 1.900E-06 | 3.300E-06 |
| 515 | 26 | H26P | A | 3.6e-06 | 2.3e-06 | 1.9e-06 | 2.600E-06 | 8.888E-07 | 1.900E-06 | 3.600E-06 |
| 516 | 26 | H26S | A | 1.8e-06 | 3.8e-06 | 2.5e-06 | 2.700E-06 | 1.015E-06 | 1.800E-06 | 3.800E-06 |
| 517 | 26 | H26T | A | 1.8e-06 | 2.1e-06 | 1.4e-06 | 1.767E-06 | 3.512E-07 | 1.400E-06 | 2.100E-06 |
| 518 | 26 | H26W | A | 4.2e-06 | 2.5e-06 | 2.6e-06 | 3.100E-06 | 9.539E-07 | 2.500E-06 | 4.200E-06 |
| 519 | 26 | H26Y | A | 2.5e-06 | 3e-06 | 2.8e-06 | 2.767E-06 | 2.517E-07 | 2.500E-06 | 3.000E-06 |
| 520 | 26 | H26V | A | 2.7e-06 | 2.2e-06 | 1.8e-06 | 2.233E-06 | 4.509E-07 | 1.800E-06 | 2.700E-06 |
| 521 | 27 | L27A | A | 1.9e-06 | 1.4e-06 | 3.8e-06 | 2.367E-06 | 1.266E-06 | 1.400E-06 | 3.800E-06 |
| 522 | 27 | L27R | A | 2.8e-06 | 3.6e-06 | 1.9e-06 | 2.767E-06 | 8.505E-07 | 1.900E-06 | 3.600E-06 |
| 523 | 27 | L27N | A | 3.1e-06 | 1.5e-06 | 4.2e-06 | 2.933E-06 | 1.358E-06 | 1.500E-06 | 4.200E-06 |
| 524 | 27 | L27D | A | 1.7e-06 | 3.2e-06 | 2.4e-06 | 2.433E-06 | 7.506E-07 | 1.700E-06 | 3.200E-06 |
| 525 | 27 | L27C | A | 2e-06 | 2.7e-06 | 1.6e-06 | 2.100E-06 | 5.568E-07 | 1.600E-06 | 2.700E-06 |
| 526 | 27 | L27E | A | 5.6e-06 | 1.9e-06 | 3.7e-06 | 3.733E-06 | 1.850E-06 | 1.900E-06 | 5.600E-06 |
| 527 | 27 | L27Q | A | 3.2e-06 | 2.3e-06 | 1.4e-06 | 2.300E-06 | 9.000E-07 | 1.400E-06 | 3.200E-06 |
| 528 | 27 | L27G | A | 1.5e-06 | 1.8e-06 | 3.2e-06 | 2.167E-06 | 9.074E-07 | 1.500E-06 | 3.200E-06 |
| 529 | 27 | L27H | A | 2.1e-06 | 2.5e-06 | 1.8e-06 | 2.133E-06 | 3.512E-07 | 1.800E-06 | 2.500E-06 |
| 530 | 27 | L27I | A | 3.4e-06 | 2.3e-06 | 1.9e-06 | 2.533E-06 | 7.767E-07 | 1.900E-06 | 3.400E-06 |
| 531 | 27 | L27L | A | 3.5e-06 | 2.8e-06 | 2.5e-06 | 2.933E-06 | 5.132E-07 | 2.500E-06 | 3.500E-06 |
| 532 | 27 | L27K | A | 1.3e-06 | 2e-06 | 1.7e-06 | 1.667E-06 | 3.512E-07 | 1.300E-06 | 2.000E-06 |
| 533 | 27 | L27M | A | 1.9e-06 | 2e-06 | 2.2e-06 | 2.033E-06 | 1.528E-07 | 1.900E-06 | 2.200E-06 |
| 534 | 27 | L27F | A | 1.9e-06 | 1.3e-06 | 4.2e-06 | 2.467E-06 | 1.531E-06 | 1.300E-06 | 4.200E-06 |
| 535 | 27 | L27P | A | 4.9e-06 | 1.8e-06 | 1.7e-06 | 2.800E-06 | 1.819E-06 | 1.700E-06 | 4.900E-06 |
| 536 | 27 | L27S | A | 3.1e-06 | 3e-06 | 3.2e-06 | 3.100E-06 | 1.000E-07 | 3.000E-06 | 3.200E-06 |
| 537 | 27 | L27T | A | 3.5e-06 | 5.6e-06 | 2.6e-06 | 3.900E-06 | 1.539E-06 | 2.600E-06 | 5.600E-06 |
| 538 | 27 | L27W | A | 3.2e-06 | 2.5e-06 | 2.9e-06 | 2.867E-06 | 3.512E-07 | 2.500E-06 | 3.200E-06 |
| 539 | 27 | L27Y | A | 1.9e-06 | 2e-06 | 3.5e-06 | 2.467E-06 | 8.963E-07 | 1.900E-06 | 3.500E-06 |
| 540 | 27 | L27V | A | 2e-06 | 2.4e-06 | 1.8e-06 | 2.067E-06 | 3.055E-07 | 1.800E-06 | 2.400E-06 |
| 541 | 28 | Y28A | A | 3.2e-06 | 3.6e-06 | 3.5e-06 | 3.433E-06 | 2.082E-07 | 3.200E-06 | 3.600E-06 |
| 542 | 28 | Y28R | A | 3.2e-06 | 2.3e-06 | 2.9e-06 | 2.800E-06 | 4.583E-07 | 2.300E-06 | 3.200E-06 |
| 543 | 28 | Y28N | A | 2.3e-06 | 2.4e-06 | 2.3e-06 | 2.333E-06 | 5.774E-08 | 2.300E-06 | 2.400E-06 |
| 544 | 28 | Y28D | A | 2.9e-06 | 2.4e-06 | 3.7e-06 | 3.000E-06 | 6.557E-07 | 2.400E-06 | 3.700E-06 |
| 545 | 28 | Y28C | A | 3.5e-06 | 2.7e-06 | 1.4e-06 | 2.533E-06 | 1.060E-06 | 1.400E-06 | 3.500E-06 |
| 546 | 28 | Y28E | A | 2.3e-06 | 5.1e-06 | 2.4e-06 | 3.267E-06 | 1.589E-06 | 2.300E-06 | 5.100E-06 |
| 547 | 28 | Y28Q | A | 2.9e-06 | 2.5e-06 | 2.4e-06 | 2.600E-06 | 2.646E-07 | 2.400E-06 | 2.900E-06 |
| 548 | 28 | Y28G | A | 3e-06 | 1.3e-06 | 2.3e-06 | 2.200E-06 | 8.544E-07 | 1.300E-06 | 3.000E-06 |
| 549 | 28 | Y28H | A | 1.6e-06 | 2.4e-06 | 2.4e-06 | 2.133E-06 | 4.619E-07 | 1.600E-06 | 2.400E-06 |
| 550 | 28 | Y28I | A | 3.1e-06 | 3.5e-06 | 3.6e-06 | 3.400E-06 | 2.646E-07 | 3.100E-06 | 3.600E-06 |
| 551 | 28 | Y28L | A | 2.4e-06 | 2.5e-06 | 1.6e-06 | 2.167E-06 | 4.933E-07 | 1.600E-06 | 2.500E-06 |
| 552 | 28 | Y28K | A | 3.7e-06 | 1.3e-06 | 2.5e-06 | 2.500E-06 | 1.200E-06 | 1.300E-06 | 3.700E-06 |
| 553 | 28 | Y28M | A | 2.9e-06 | 2.3e-06 | 3e-06 | 2.733E-06 | 3.786E-07 | 2.300E-06 | 3.000E-06 |
| 554 | 28 | Y28F | A | 1.9e-06 | 2.8e-06 | 2.1e-06 | 2.267E-06 | 4.726E-07 | 1.900E-06 | 2.800E-06 |
| 555 | 28 | Y28P | A | 4e-06 | 2.6e-06 | 3.1e-06 | 3.233E-06 | 7.095E-07 | 2.600E-06 | 4.000E-06 |
| 556 | 28 | Y28S | A | 1.7e-06 | 2.7e-06 | 1.6e-06 | 2.000E-06 | 6.083E-07 | 1.600E-06 | 2.700E-06 |
| 557 | 28 | Y28T | A | 1.5e-06 | 2e-06 | 2e-06 | 1.833E-06 | 2.887E-07 | 1.500E-06 | 2.000E-06 |
| 558 | 28 | Y28W | A | 1.5e-06 | 2.1e-06 | 2.5e-06 | 2.033E-06 | 5.033E-07 | 1.500E-06 | 2.500E-06 |
| 559 | 28 | Y28Y | A | 1.9e-06 | 3e-06 | 1.2e-06 | 2.033E-06 | 9.074E-07 | 1.200E-06 | 3.000E-06 |
| 560 | 28 | Y28V | A | 1.9e-06 | 2.8e-06 | 3.3e-06 | 2.667E-06 | 7.095E-07 | 1.900E-06 | 3.300E-06 |
| 561 | 29 | Q29A | A | 2.5e-06 | 1e-06 | 4.6e-06 | 2.700E-06 | 1.808E-06 | 1.000E-06 | 4.600E-06 |
| 562 | 29 | Q29R | A | 3.4e-06 | 2.2e-06 | 2.2e-06 | 2.600E-06 | 6.928E-07 | 2.200E-06 | 3.400E-06 |
| 563 | 29 | Q29N | A | 4.6e-06 | 2.7e-06 | 2.3e-06 | 3.200E-06 | 1.229E-06 | 2.300E-06 | 4.600E-06 |
| 564 | 29 | Q29D | A | 2.1e-06 | 3.6e-06 | 2.2e-06 | 2.633E-06 | 8.386E-07 | 2.100E-06 | 3.600E-06 |
| 565 | 29 | Q29C | A | 3.6e-06 | 2.1e-06 | 1.1e-06 | 2.267E-06 | 1.258E-06 | 1.100E-06 | 3.600E-06 |
| 566 | 29 | Q29E | A | 2.7e-06 | 3.1e-06 | 1.8e-06 | 2.533E-06 | 6.658E-07 | 1.800E-06 | 3.100E-06 |
| 567 | 29 | Q29Q | A | 1.8e-06 | 3.2e-06 | 1.7e-06 | 2.233E-06 | 8.386E-07 | 1.700E-06 | 3.200E-06 |
| 568 | 29 | Q29G | A | 1.8e-06 | 3.3e-06 | 1.6e-06 | 2.233E-06 | 9.292E-07 | 1.600E-06 | 3.300E-06 |
| 569 | 29 | Q29H | A | 3e-06 | 2.3e-06 | 2.9e-06 | 2.733E-06 | 3.786E-07 | 2.300E-06 | 3.000E-06 |
| 570 | 29 | Q29I | A | 3.1e-06 | 2.3e-06 | 1.9e-06 | 2.433E-06 | 6.110E-07 | 1.900E-06 | 3.100E-06 |
| 571 | 29 | Q29L | A | 3.3e-06 | 3.1e-06 | 2.6e-06 | 3.000E-06 | 3.606E-07 | 2.600E-06 | 3.300E-06 |
| 572 | 29 | Q29K | A | 2.1e-06 | 1.8e-06 | 1.9e-06 | 1.933E-06 | 1.528E-07 | 1.800E-06 | 2.100E-06 |
| 573 | 29 | Q29M | A | 5.4e-06 | 1.3e-06 | 2.7e-06 | 3.133E-06 | 2.084E-06 | 1.300E-06 | 5.400E-06 |
| 574 | 29 | Q29F | A | 2.4e-06 | 2.2e-06 | 3.3e-06 | 2.633E-06 | 5.859E-07 | 2.200E-06 | 3.300E-06 |
| 575 | 29 | Q29P | A | 3.3e-06 | 3.8e-06 | 2.3e-06 | 3.133E-06 | 7.638E-07 | 2.300E-06 | 3.800E-06 |
| 576 | 29 | Q29S | A | 9.4e-07 | 3.3e-06 | 1.7e-06 | 1.980E-06 | 1.205E-06 | 9.400E-07 | 3.300E-06 |
| 577 | 29 | Q29T | A | 3.6e-06 | 2.1e-06 | 2.2e-06 | 2.633E-06 | 8.386E-07 | 2.100E-06 | 3.600E-06 |
| 578 | 29 | Q29W | A | 2.8e-06 | 4.1e-06 | 2.8e-06 | 3.233E-06 | 7.506E-07 | 2.800E-06 | 4.100E-06 |
| 579 | 29 | Q29Y | A | 1.9e-06 | 2.2e-06 | 2e-06 | 2.033E-06 | 1.528E-07 | 1.900E-06 | 2.200E-06 |
| 580 | 29 | Q29V | A | 3.5e-06 | 1.5e-06 | 3.2e-06 | 2.733E-06 | 1.079E-06 | 1.500E-06 | 3.500E-06 |
| 581 | 30 | G30A | A | 3.6e-06 | 1.6e-06 | 3e-06 | 2.733E-06 | 1.026E-06 | 1.600E-06 | 3.600E-06 |
| 582 | 30 | G30R | A | 2.5e-06 | 2.5e-06 | 1.7e-06 | 2.233E-06 | 4.619E-07 | 1.700E-06 | 2.500E-06 |
| 583 | 30 | G30N | A | 3.4e-06 | 2e-06 | 2.7e-06 | 2.700E-06 | 7.000E-07 | 2.000E-06 | 3.400E-06 |
| 584 | 30 | G30D | A | 1.9e-06 | 2.2e-06 | 5.4e-06 | 3.167E-06 | 1.940E-06 | 1.900E-06 | 5.400E-06 |
| 585 | 30 | G30C | A | 1.8e-06 | 3e-06 | 1.7e-06 | 2.167E-06 | 7.234E-07 | 1.700E-06 | 3.000E-06 |
| 586 | 30 | G30E | A | 1.5e-06 | 3.2e-06 | 2.6e-06 | 2.433E-06 | 8.622E-07 | 1.500E-06 | 3.200E-06 |
| 587 | 30 | G30Q | A | 1.6e-06 | 2.5e-06 | 1.6e-06 | 1.900E-06 | 5.196E-07 | 1.600E-06 | 2.500E-06 |
| 588 | 30 | G30G | A | 3.1e-06 | 1.8e-06 | 3.3e-06 | 2.733E-06 | 8.145E-07 | 1.800E-06 | 3.300E-06 |
| 589 | 30 | G30H | A | 1.2e-06 | 2.5e-06 | 2.6e-06 | 2.100E-06 | 7.810E-07 | 1.200E-06 | 2.600E-06 |
| 590 | 30 | G30I | A | 2.2e-06 | 1.5e-06 | 3.7e-06 | 2.467E-06 | 1.124E-06 | 1.500E-06 | 3.700E-06 |
| 591 | 30 | G30L | A | 1.9e-06 | 4.1e-06 | 2.3e-06 | 2.767E-06 | 1.172E-06 | 1.900E-06 | 4.100E-06 |
| 592 | 30 | G30K | A | 3e-06 | 2.2e-06 | 2.9e-06 | 2.700E-06 | 4.359E-07 | 2.200E-06 | 3.000E-06 |
| 593 | 30 | G30M | A | 2.5e-06 | 1.7e-06 | 2.6e-06 | 2.267E-06 | 4.933E-07 | 1.700E-06 | 2.600E-06 |
| 594 | 30 | G30F | A | 3.2e-06 | 2.1e-06 | 3.2e-06 | 2.833E-06 | 6.351E-07 | 2.100E-06 | 3.200E-06 |

| | | | | | | | | | | |
|---|---|---|---|---|---|---|---|---|---|---|
| 595 | 30 | G30P | A | 2.8e-06 | 1.6e-06 | 2.6e-06 | 2.333E-06 | 6.429E-07 | 1.600E-06 | 2.800E-06 |
| 596 | 30 | G30S | A | 1.4e-06 | 3.6e-06 | 2.9e-06 | 2.633E-06 | 1.124E-06 | 1.400E-06 | 3.600E-06 |
| 597 | 30 | G30T | A | 2.2e-06 | 2.2e-06 | 3.5e-06 | 2.633E-06 | 7.506E-07 | 2.200E-06 | 3.500E-06 |
| 598 | 30 | G30W | A | 2.7e-06 | 2.4e-06 | 2.2e-06 | 2.433E-06 | 2.517E-07 | 2.200E-06 | 2.700E-06 |
| 599 | 30 | G30Y | A | 2.5e-06 | 2e-06 | 1.2e-06 | 1.900E-06 | 6.557E-07 | 1.200E-06 | 2.500E-06 |
| 600 | 30 | G30V | A | 2.4e-06 | 2.4e-06 | 2.1e-06 | 2.300E-06 | 1.732E-07 | 2.100E-06 | 2.400E-06 |
| 601 | 31 | C31A | A | 2.6e-06 | 2.5e-06 | 2.2e-06 | 2.433E-06 | 2.082E-07 | 2.200E-06 | 2.600E-06 |
| 602 | 31 | C31R | A | 3.6e-06 | 1.2e-06 | 3.1e-06 | 2.633E-06 | 1.266E-06 | 1.200E-06 | 3.600E-06 |
| 603 | 31 | C31N | A | 2.9e-06 | 2.6e-06 | 1.8e-06 | 2.433E-06 | 5.686E-07 | 1.800E-06 | 2.900E-06 |
| 604 | 31 | C31D | A | 2.1e-06 | 1.9e-06 | 3.5e-06 | 2.500E-06 | 8.718E-07 | 1.900E-06 | 3.500E-06 |
| 605 | 31 | C31C | A | 3e-06 | 2.6e-06 | 2.1e-06 | 2.567E-06 | 4.509E-07 | 2.100E-06 | 3.000E-06 |
| 606 | 31 | C31E | A | 1.9e-06 | 2.6e-06 | 2.7e-06 | 2.400E-06 | 4.359E-07 | 1.900E-06 | 2.700E-06 |
| 607 | 31 | C31Q | A | 1.1e-06 | 2.7e-06 | 2.7e-06 | 2.167E-06 | 9.238E-07 | 1.100E-06 | 2.700E-06 |
| 608 | 31 | C31G | A | 2e-06 | 2.1e-06 | 2.3e-06 | 2.133E-06 | 1.528E-07 | 2.000E-06 | 2.300E-06 |
| 609 | 31 | C31H | A | 3.1e-06 | 2.1e-06 | 4.5e-06 | 3.233E-06 | 1.206E-06 | 2.100E-06 | 4.500E-06 |
| 610 | 31 | C31I | A | 3.8e-06 | 2.1e-06 | 2.3e-06 | 2.733E-06 | 9.292E-07 | 2.100E-06 | 3.800E-06 |
| 611 | 31 | C31L | A | 2.8e-06 | 3.1e-06 | 2.5e-06 | 2.800E-06 | 3.000E-07 | 2.500E-06 | 3.100E-06 |
| 612 | 31 | C31K | A | 2.5e-06 | 2.7e-06 | 1.8e-06 | 2.333E-06 | 4.726E-07 | 1.800E-06 | 2.700E-06 |
| 613 | 31 | C31M | A | 1.9e-06 | 3.7e-06 | 3e-06 | 2.867E-06 | 9.074E-07 | 1.900E-06 | 3.700E-06 |
| 614 | 31 | C31F | A | 4.1e-06 | 1.4e-06 | 1.9e-06 | 2.467E-06 | 1.436E-06 | 1.400E-06 | 4.100E-06 |
| 615 | 31 | C31P | A | 2.1e-06 | 2.7e-06 | 3.6e-06 | 2.800E-06 | 7.550E-07 | 2.100E-06 | 3.600E-06 |
| 616 | 31 | C31S | A | 1.9e-06 | 2e-06 | 1.9e-06 | 1.933E-06 | 5.774E-08 | 1.900E-06 | 2.000E-06 |
| 617 | 31 | C31T | A | 3.4e-06 | 1.7e-06 | 1.8e-06 | 2.300E-06 | 9.539E-07 | 1.700E-06 | 3.400E-06 |
| 618 | 31 | C31W | A | 2.8e-06 | 2.8e-06 | 2.8e-06 | 2.800E-06 | 0.000E+00 | 2.800E-06 | 2.800E-06 |
| 619 | 31 | C31Y | A | 2.7e-06 | 1.6e-06 | 2.5e-06 | 2.267E-06 | 5.859E-07 | 1.600E-06 | 2.700E-06 |
| 620 | 31 | C31V | A | 2e-06 | 2.1e-06 | 2.5e-06 | 2.200E-06 | 2.646E-07 | 2.000E-06 | 2.500E-06 |
| 621 | 32 | Q32A | A | 1.6e-06 | 4.5e-06 | 2.4e-06 | 2.833E-06 | 1.498E-06 | 1.600E-06 | 4.500E-06 |
| 622 | 32 | Q32R | A | 2.5e-06 | 1e-06 | 2.2e-06 | 1.900E-06 | 7.937E-07 | 1.000E-06 | 2.500E-06 |
| 623 | 32 | Q32N | A | 1.8e-06 | 1.5e-06 | 2.7e-06 | 2.000E-06 | 6.245E-07 | 1.500E-06 | 2.700E-06 |
| 624 | 32 | Q32D | A | 5.6e-06 | 2.6e-06 | 4e-06 | 4.067E-06 | 1.501E-06 | 2.600E-06 | 5.600E-06 |
| 625 | 32 | Q32C | A | 1.3e-06 | 2.9e-06 | 1.1e-06 | 1.767E-06 | 9.866E-07 | 1.100E-06 | 2.900E-06 |
| 626 | 32 | Q32E | A | 3.6e-06 | 4.5e-06 | 2.5e-06 | 3.533E-06 | 1.002E-06 | 2.500E-06 | 4.500E-06 |
| 627 | 32 | Q32Q | A | 1.9e-06 | 2.3e-06 | 3.5e-06 | 2.567E-06 | 8.327E-07 | 1.900E-06 | 3.500E-06 |
| 628 | 32 | Q32G | A | 3e-06 | 1.5e-06 | 3.3e-06 | 2.600E-06 | 9.644E-07 | 1.500E-06 | 3.300E-06 |
| 629 | 32 | Q32H | A | 3.7e-06 | 2e-06 | 3e-06 | 2.900E-06 | 8.544E-07 | 2.000E-06 | 3.700E-06 |
| 630 | 32 | Q32I | A | 2.8e-06 | 1.8e-06 | 2.4e-06 | 2.333E-06 | 5.033E-07 | 1.800E-06 | 2.800E-06 |
| 631 | 32 | Q32L | A | 2.5e-06 | 2.3e-06 | 3.2e-06 | 2.667E-06 | 4.726E-07 | 2.300E-06 | 3.200E-06 |
| 632 | 32 | Q32K | A | 1.9e-06 | 2.1e-06 | 3.6e-06 | 2.533E-06 | 9.292E-07 | 1.900E-06 | 3.600E-06 |
| 633 | 32 | Q32M | A | 2.5e-06 | 1.8e-06 | 1.4e-06 | 1.900E-06 | 5.568E-07 | 1.400E-06 | 2.500E-06 |
| 634 | 32 | Q32F | A | 3e-06 | 2.2e-06 | 2.9e-06 | 2.700E-06 | 4.359E-07 | 2.200E-06 | 3.000E-06 |
| 635 | 32 | Q32P | A | 1.7e-06 | 3.4e-06 | 2.1e-06 | 2.400E-06 | 8.888E-07 | 1.700E-06 | 3.400E-06 |
| 636 | 32 | Q32S | A | 2.3e-06 | 2.2e-06 | 1.7e-06 | 2.067E-06 | 3.215E-07 | 1.700E-06 | 2.300E-06 |
| 637 | 32 | Q32T | A | 2.2e-06 | 2.2e-06 | 2.2e-06 | 2.200E-06 | 0.000E+00 | 2.200E-06 | 2.200E-06 |
| 638 | 32 | Q32W | A | 3.6e-06 | 3.4e-06 | 9.3e-07 | 2.643E-06 | 1.487E-06 | 9.300E-07 | 3.600E-06 |
| 639 | 32 | Q32Y | A | 1.8e-06 | 2.5e-06 | 2.6e-06 | 2.300E-06 | 4.359E-07 | 1.800E-06 | 2.600E-06 |
| 640 | 32 | Q32V | A | 3.3e-06 | 1.9e-06 | 2e-06 | 2.400E-06 | 7.810E-07 | 1.900E-06 | 3.300E-06 |
| 641 | 33 | V33A | A | 1.5e-06 | 3.7e-06 | 2.6e-06 | 2.600E-06 | 1.100E-06 | 1.500E-06 | 3.700E-06 |
| 642 | 33 | V33R | A | 2.7e-06 | 9.8e-07 | 2.6e-06 | 2.093E-06 | 9.655E-07 | 9.800E-07 | 2.700E-06 |
| 643 | 33 | V33N | A | 1.8e-06 | 2.5e-06 | 2.2e-06 | 2.167E-06 | 3.512E-07 | 1.800E-06 | 2.500E-06 |
| 644 | 33 | V33D | A | 2.1e-06 | 2.1e-06 | 1.6e-06 | 1.933E-06 | 2.887E-07 | 1.600E-06 | 2.100E-06 |
| 645 | 33 | V33C | A | 3.4e-06 | 1.7e-06 | 1e-06 | 2.033E-06 | 1.234E-06 | 1.000E-06 | 3.400E-06 |
| 646 | 33 | V33E | A | 2.2e-06 | 3.1e-06 | 1.4e-06 | 2.233E-06 | 8.505E-07 | 1.400E-06 | 3.100E-06 |
| 647 | 33 | V33Q | A | 3.1e-06 | 3.3e-06 | 2.4e-06 | 2.933E-06 | 4.726E-07 | 2.400E-06 | 3.300E-06 |
| 648 | 33 | V33G | A | 2.1e-06 | 1.2e-06 | 4.1e-06 | 2.467E-06 | 1.484E-06 | 1.200E-06 | 4.100E-06 |
| 649 | 33 | V33H | A | 3.5e-06 | 2.2e-06 | 2.2e-06 | 2.633E-06 | 7.506E-07 | 2.200E-06 | 3.500E-06 |
| 650 | 33 | V33I | A | 2.3e-06 | 2.3e-06 | 2.3e-06 | 2.300E-06 | 0.000E+00 | 2.300E-06 | 2.300E-06 |
| 651 | 33 | V33L | A | 1.3e-06 | 2.2e-06 | 2.6e-06 | 2.033E-06 | 6.658E-07 | 1.300E-06 | 2.600E-06 |
| 652 | 33 | V33K | A | 2.4e-06 | 2.8e-06 | 2e-06 | 2.400E-06 | 4.000E-07 | 2.000E-06 | 2.800E-06 |
| 653 | 33 | V33M | A | 1.7e-06 | 1.9e-06 | 1e-06 | 1.533E-06 | 4.726E-07 | 1.000E-06 | 1.900E-06 |
| 654 | 33 | V33F | A | 2.4e-06 | 3.1e-06 | 2.7e-06 | 2.733E-06 | 3.512E-07 | 2.400E-06 | 3.100E-06 |
| 655 | 33 | V33P | A | 1.9e-06 | 1.7e-06 | 4.6e-06 | 2.733E-06 | 1.620E-06 | 1.700E-06 | 4.600E-06 |
| 656 | 33 | V33S | A | 2.6e-06 | 1.6e-06 | 2.1e-06 | 2.100E-06 | 5.000E-07 | 1.600E-06 | 2.600E-06 |
| 657 | 33 | V33T | A | 2.2e-06 | 1.7e-06 | 2.1e-06 | 2.000E-06 | 2.646E-07 | 1.700E-06 | 2.200E-06 |
| 658 | 33 | V33W | A | 1.8e-06 | 2.1e-06 | 3.2e-06 | 2.367E-06 | 7.371E-07 | 1.800E-06 | 3.200E-06 |
| 659 | 33 | V33Y | A | 1.5e-06 | 2.2e-06 | 1.3e-06 | 1.667E-06 | 4.726E-07 | 1.300E-06 | 2.200E-06 |
| 660 | 33 | V33V | A | 2.7e-06 | 2.5e-06 | 2.6e-06 | 2.600E-06 | 1.000E-07 | 2.500E-06 | 2.700E-06 |
| 661 | 34 | V34A | A | 3e-06 | 3.9e-06 | 4.2e-06 | 3.700E-06 | 6.245E-07 | 3.000E-06 | 4.200E-06 |
| 662 | 34 | V34R | A | 4.2e-06 | 1.8e-06 | 4.9e-06 | 3.633E-06 | 1.626E-06 | 1.800E-06 | 4.900E-06 |
| 663 | 34 | V34N | A | 3.5e-06 | 1.9e-06 | 2.5e-06 | 2.633E-06 | 8.083E-07 | 1.900E-06 | 3.500E-06 |
| 664 | 34 | V34D | A | 3.1e-06 | 1.8e-06 | 2.4e-06 | 2.433E-06 | 6.506E-07 | 1.800E-06 | 3.100E-06 |
| 665 | 34 | V34C | A | 2.6e-06 | 3.9e-06 | 1.6e-06 | 2.700E-06 | 1.153E-06 | 1.600E-06 | 3.900E-06 |
| 666 | 34 | V34E | A | 1.8e-06 | 3.8e-06 | 4.2e-06 | 3.267E-06 | 1.286E-06 | 1.800E-06 | 4.200E-06 |
| 667 | 34 | V34Q | A | 2e-06 | 2.7e-06 | 2.2e-06 | 2.300E-06 | 3.606E-07 | 2.000E-06 | 2.700E-06 |
| 668 | 34 | V34G | A | 2.8e-06 | 1.6e-06 | 2.7e-06 | 2.367E-06 | 6.658E-07 | 1.600E-06 | 2.800E-06 |
| 669 | 34 | V34H | A | 2.3e-06 | 2.8e-06 | 2.6e-06 | 2.567E-06 | 2.517E-07 | 2.300E-06 | 2.800E-06 |
| 670 | 34 | V34I | A | 3.4e-06 | 2.9e-06 | 1.8e-06 | 2.700E-06 | 8.185E-07 | 1.800E-06 | 3.400E-06 |
| 671 | 34 | V34L | A | 1.4e-06 | 2.1e-06 | 2.7e-06 | 2.067E-06 | 6.506E-07 | 1.400E-06 | 2.700E-06 |
| 672 | 34 | V34K | A | 3e-06 | 2.4e-06 | 4e-06 | 3.133E-06 | 8.083E-07 | 2.400E-06 | 4.000E-06 |
| 673 | 34 | V34M | A | 2.2e-06 | 1.7e-06 | 1.9e-06 | 1.933E-06 | 2.517E-07 | 1.700E-06 | 2.200E-06 |
| 674 | 34 | V34F | A | 3.3e-06 | 2.1e-06 | 3.6e-06 | 3.000E-06 | 7.937E-07 | 2.100E-06 | 3.600E-06 |
| 675 | 34 | V34P | A | 2e-06 | 1.8e-06 | 1.4e-06 | 1.733E-06 | 3.055E-07 | 1.400E-06 | 2.000E-06 |
| 676 | 34 | V34S | A | 1.4e-06 | 2.8e-06 | 2.4e-06 | 2.200E-06 | 7.211E-07 | 1.400E-06 | 2.800E-06 |
| 677 | 34 | V34T | A | 2.7e-06 | 2.9e-06 | 1.4e-06 | 2.333E-06 | 8.145E-07 | 1.400E-06 | 2.900E-06 |
| 678 | 34 | V34W | A | 2.5e-06 | 1.2e-06 | 1.7e-06 | 1.800E-06 | 6.557E-07 | 1.200E-06 | 2.500E-06 |
| 679 | 34 | V34Y | A | 1.4e-06 | 3.5e-06 | 3.6e-06 | 2.833E-06 | 1.242E-06 | 1.400E-06 | 3.600E-06 |

| | | | | | | | | | | |
|---|---|---|---|---|---|---|---|---|---|---|
| 680 | 34 | V34V | A | 2.5e-06 | 1.6e-06 | 3.5e-06 | 2.533E-06 | 9.504E-07 | 1.600E-06 | 3.500E-06 |
| 681 | 35 | Q35A | A | 2.9e-06 | 2.7e-06 | 2.5e-06 | 2.700E-06 | 2.000E-07 | 2.500E-06 | 2.900E-06 |
| 682 | 35 | Q35R | A | 2.6e-06 | 3.4e-06 | 1.8e-06 | 2.600E-06 | 8.000E-07 | 1.800E-06 | 3.400E-06 |
| 683 | 35 | Q35N | A | 2.3e-06 | 2.7e-06 | 3.1e-06 | 2.700E-06 | 4.000E-07 | 2.300E-06 | 3.100E-06 |
| 684 | 35 | Q35D | A | 3.1e-06 | 3e-06 | 4.3e-06 | 3.467E-06 | 7.234E-07 | 3.000E-06 | 4.300E-06 |
| 685 | 35 | Q35C | A | 5e-06 | 2.6e-06 | 4.6e-06 | 4.067E-06 | 1.286E-06 | 2.600E-06 | 5.000E-06 |
| 686 | 35 | Q35E | A | 1.8e-06 | 3e-06 | 1.3e-06 | 2.033E-06 | 8.737E-07 | 1.300E-06 | 3.000E-06 |
| 687 | 35 | Q35Q | A | 2.5e-06 | 2.9e-06 | 1.3e-06 | 2.233E-06 | 8.327E-07 | 1.300E-06 | 2.900E-06 |
| 688 | 35 | Q35G | A | 2.6e-06 | 2.8e-06 | 2.2e-06 | 2.533E-06 | 3.055E-07 | 2.200E-06 | 2.800E-06 |
| 689 | 35 | Q35H | A | 2.8e-06 | 3.4e-06 | 1.7e-06 | 2.633E-06 | 8.622E-07 | 1.700E-06 | 3.400E-06 |
| 690 | 35 | Q35I | A | 3.3e-06 | 2.1e-06 | 1.9e-06 | 2.433E-06 | 7.572E-07 | 1.900E-06 | 3.300E-06 |
| 691 | 35 | Q35L | A | 2.8e-06 | 1.6e-06 | 2.1e-06 | 2.167E-06 | 6.028E-07 | 1.600E-06 | 2.800E-06 |
| 692 | 35 | Q35K | A | 2.1e-06 | 2e-06 | 2.6e-06 | 2.233E-06 | 3.215E-07 | 2.000E-06 | 2.600E-06 |
| 693 | 35 | Q35M | A | 1.8e-06 | 1.4e-06 | 1.4e-06 | 1.533E-06 | 2.309E-07 | 1.400E-06 | 1.800E-06 |
| 694 | 35 | Q35F | A | 3.9e-06 | 2.9e-06 | 2.4e-06 | 3.067E-06 | 7.638E-07 | 2.400E-06 | 3.900E-06 |
| 695 | 35 | Q35P | A | 4.3e-06 | 1.8e-06 | 1.9e-06 | 2.667E-06 | 1.415E-06 | 1.800E-06 | 4.300E-06 |
| 696 | 35 | Q35S | A | 2.8e-06 | 3.7e-06 | 2.2e-06 | 2.900E-06 | 7.550E-07 | 2.200E-06 | 3.700E-06 |
| 697 | 35 | Q35T | A | 1.9e-06 | 2.6e-06 | 3e-06 | 2.500E-06 | 5.568E-07 | 1.900E-06 | 3.000E-06 |
| 698 | 35 | Q35W | A | 2e-06 | 3.2e-06 | 3.6e-06 | 2.933E-06 | 8.327E-07 | 2.000E-06 | 3.600E-06 |
| 699 | 35 | Q35Y | A | 2.6e-06 | 4.1e-06 | 2.7e-06 | 3.133E-06 | 8.386E-07 | 2.600E-06 | 4.100E-06 |
| 700 | 35 | Q35V | A | 2.9e-06 | 2.1e-06 | 1.1e-06 | 2.033E-06 | 9.018E-07 | 1.100E-06 | 2.900E-06 |
| 701 | 36 | G36A | A | 2.7e-06 | 4.2e-06 | 3.9e-06 | 3.600E-06 | 7.937E-07 | 2.700E-06 | 4.200E-06 |
| 702 | 36 | G36R | A | 1.3e-06 | 1.4e-06 | 2.9e-06 | 1.867E-06 | 8.963E-07 | 1.300E-06 | 2.900E-06 |
| 703 | 36 | G36N | A | 1.4e-06 | 3.3e-06 | 2.2e-06 | 2.300E-06 | 9.539E-07 | 1.400E-06 | 3.300E-06 |
| 704 | 36 | G36D | A | 2.3e-06 | 2.6e-06 | 2.6e-06 | 2.500E-06 | 1.732E-07 | 2.300E-06 | 2.600E-06 |
| 705 | 36 | G36C | A | 2.5e-06 | 1.8e-06 | 2.3e-06 | 2.200E-06 | 3.606E-07 | 1.800E-06 | 2.500E-06 |
| 706 | 36 | G36E | A | 2.6e-06 | 3.5e-06 | 2.9e-06 | 3.000E-06 | 4.583E-07 | 2.600E-06 | 3.500E-06 |
| 707 | 36 | G36Q | A | 2.1e-06 | 4.3e-06 | 2.1e-06 | 2.833E-06 | 1.270E-06 | 2.100E-06 | 4.300E-06 |
| 708 | 36 | G36G | A | 2.4e-06 | 1.9e-06 | 2.2e-06 | 2.167E-06 | 2.517E-07 | 1.900E-06 | 2.400E-06 |
| 709 | 36 | G36H | A | 2.7e-06 | 1.7e-06 | 2.3e-06 | 2.233E-06 | 5.033E-07 | 1.700E-06 | 2.700E-06 |
| 710 | 36 | G36I | A | 3.6e-06 | 2.6e-06 | 2.1e-06 | 2.767E-06 | 7.638E-07 | 2.100E-06 | 3.600E-06 |
| 711 | 36 | G36L | A | 2.6e-06 | 2e-06 | 2.3e-06 | 2.300E-06 | 3.000E-07 | 2.000E-06 | 2.600E-06 |
| 712 | 36 | G36K | A | 3.1e-06 | 2.7e-06 | 3.2e-06 | 3.000E-06 | 2.646E-07 | 2.700E-06 | 3.200E-06 |
| 713 | 36 | G36M | A | 3.7e-06 | 2.1e-06 | 1.7e-06 | 2.500E-06 | 1.058E-06 | 1.700E-06 | 3.700E-06 |
| 714 | 36 | G36F | A | 3.1e-06 | 2.2e-06 | 3.1e-06 | 2.800E-06 | 5.196E-07 | 2.200E-06 | 3.100E-06 |
| 715 | 36 | G36P | A | 2.3e-06 | 1.4e-06 | 3.1e-06 | 2.267E-06 | 8.505E-07 | 1.400E-06 | 3.100E-06 |
| 716 | 36 | G36S | A | 1.8e-06 | 3e-06 | 1.8e-06 | 2.200E-06 | 6.928E-07 | 1.800E-06 | 3.000E-06 |
| 717 | 36 | G36T | A | 2.1e-06 | 1.6e-06 | 2.8e-06 | 2.167E-06 | 6.028E-07 | 1.600E-06 | 2.800E-06 |
| 718 | 36 | G36W | A | 1.6e-06 | 1.8e-06 | 1.4e-06 | 1.600E-06 | 2.000E-07 | 1.400E-06 | 1.800E-06 |
| 719 | 36 | G36Y | A | 3.2e-06 | 3.2e-06 | 2.1e-06 | 2.833E-06 | 6.351E-07 | 2.100E-06 | 3.200E-06 |
| 720 | 36 | G36V | A | 2.7e-06 | 1.7e-06 | 2.6e-06 | 2.333E-06 | 5.508E-07 | 1.700E-06 | 2.700E-06 |
| 721 | 37 | N37A | A | 2.6e-06 | 2.5e-06 | 1.9e-06 | 2.333E-06 | 3.786E-07 | 1.900E-06 | 2.600E-06 |
| 722 | 37 | N37R | A | 3.7e-06 | 1.1e-06 | 3.1e-06 | 2.633E-06 | 1.361E-06 | 1.100E-06 | 3.700E-06 |
| 723 | 37 | N37N | A | 2.2e-06 | 1.8e-06 | 2.9e-06 | 2.300E-06 | 5.568E-07 | 1.800E-06 | 2.900E-06 |
| 724 | 37 | N37D | A | 2.6e-06 | 2.5e-06 | 1.9e-06 | 2.333E-06 | 3.786E-07 | 1.900E-06 | 2.600E-06 |
| 725 | 37 | N37C | A | 2.6e-06 | 3.3e-06 | 3.7e-06 | 3.200E-06 | 5.568E-07 | 2.600E-06 | 3.700E-06 |
| 726 | 37 | N37E | A | 3.5e-06 | 1.5e-06 | 3.7e-06 | 2.900E-06 | 1.217E-06 | 1.500E-06 | 3.700E-06 |
| 727 | 37 | N37Q | A | 2.2e-06 | 2.6e-06 | 1.6e-06 | 2.133E-06 | 5.033E-07 | 1.600E-06 | 2.600E-06 |
| 728 | 37 | N37G | A | 1.5e-06 | 2.5e-06 | 2.6e-06 | 2.200E-06 | 6.083E-07 | 1.500E-06 | 2.600E-06 |
| 729 | 37 | N37H | A | 3.5e-06 | 1.9e-06 | 2.4e-06 | 2.600E-06 | 8.185E-07 | 1.900E-06 | 3.500E-06 |
| 730 | 37 | N37I | A | 1.3e-06 | 2.3e-06 | 3.2e-06 | 2.267E-06 | 9.504E-07 | 1.300E-06 | 3.200E-06 |
| 731 | 37 | N37L | A | 1.4e-06 | 1.8e-06 | 2.3e-06 | 1.833E-06 | 4.509E-07 | 1.400E-06 | 2.300E-06 |
| 732 | 37 | N37K | A | 1.3e-06 | 1.5e-06 | 2.6e-06 | 1.800E-06 | 7.000E-07 | 1.300E-06 | 2.600E-06 |
| 733 | 37 | N37M | A | 3.1e-06 | 2.6e-06 | 2.6e-06 | 2.767E-06 | 2.887E-07 | 2.600E-06 | 3.100E-06 |
| 734 | 37 | N37F | A | 2.5e-06 | 1.6e-06 | 2.2e-06 | 2.100E-06 | 4.583E-07 | 1.600E-06 | 2.500E-06 |
| 735 | 37 | N37P | A | 2.3e-06 | 4e-06 | 1.8e-06 | 2.700E-06 | 1.153E-06 | 1.800E-06 | 4.000E-06 |
| 736 | 37 | N37S | A | 2.9e-06 | 2.5e-06 | 2.8e-06 | 2.733E-06 | 2.082E-07 | 2.500E-06 | 2.900E-06 |
| 737 | 37 | N37T | A | 2.9e-06 | 2.2e-06 | 2.7e-06 | 2.600E-06 | 3.606E-07 | 2.200E-06 | 2.900E-06 |
| 738 | 37 | N37W | A | 1.9e-06 | 2.5e-06 | 1.5e-06 | 1.967E-06 | 5.033E-07 | 1.500E-06 | 2.500E-06 |
| 739 | 37 | N37Y | A | 1.2e-06 | 2.5e-06 | 2.7e-06 | 2.133E-06 | 8.145E-07 | 1.200E-06 | 2.700E-06 |
| 740 | 37 | N37V | A | 3.8e-06 | 1.2e-06 | 5.2e-06 | 3.400E-06 | 2.030E-06 | 1.200E-06 | 5.200E-06 |
| 741 | 38 | L38A | A | 1.5e-06 | 1.9e-06 | 3.3e-06 | 2.233E-06 | 9.452E-07 | 1.500E-06 | 3.300E-06 |
| 742 | 38 | L38R | A | 1.8e-06 | 3.3e-06 | 1.9e-06 | 2.333E-06 | 8.386E-07 | 1.800E-06 | 3.300E-06 |
| 743 | 38 | L38N | A | 1.6e-06 | 3.1e-06 | 1.8e-06 | 2.167E-06 | 8.145E-07 | 1.600E-06 | 3.100E-06 |
| 744 | 38 | L38D | A | 1.4e-06 | 1.6e-06 | 4.3e-06 | 2.433E-06 | 1.620E-06 | 1.400E-06 | 4.300E-06 |
| 745 | 38 | L38C | A | 4.1e-06 | 4.8e-06 | 2.2e-06 | 3.700E-06 | 1.345E-06 | 2.200E-06 | 4.800E-06 |
| 746 | 38 | L38E | A | 1.9e-06 | 2.5e-06 | 3.9e-06 | 2.767E-06 | 1.026E-06 | 1.900E-06 | 3.900E-06 |
| 747 | 38 | L38Q | A | 3e-06 | 3.7e-06 | 3.4e-06 | 3.367E-06 | 3.512E-07 | 3.000E-06 | 3.700E-06 |
| 748 | 38 | L38G | A | 3.9e-06 | 2.8e-06 | 1.6e-06 | 2.767E-06 | 1.150E-06 | 1.600E-06 | 3.900E-06 |
| 749 | 38 | L38H | A | 1.5e-06 | 1.4e-06 | 2e-06 | 1.633E-06 | 3.215E-07 | 1.400E-06 | 2.000E-06 |
| 750 | 38 | L38I | A | 2.4e-06 | 1.5e-06 | 2.5e-06 | 2.133E-06 | 5.508E-07 | 1.500E-06 | 2.500E-06 |
| 751 | 38 | L38L | A | 2.8e-06 | 3.2e-06 | 1.4e-06 | 2.467E-06 | 9.452E-07 | 1.400E-06 | 3.200E-06 |
| 752 | 38 | L38K | A | 3.2e-06 | 1.9e-06 | 1e-06 | 2.033E-06 | 1.106E-06 | 1.000E-06 | 3.200E-06 |
| 753 | 38 | L38M | A | 2.1e-06 | 1.2e-06 | 4.8e-06 | 2.700E-06 | 1.873E-06 | 1.200E-06 | 4.800E-06 |
| 754 | 38 | L38F | A | 3.2e-06 | 2.1e-06 | 1.6e-06 | 2.300E-06 | 8.185E-07 | 1.600E-06 | 3.200E-06 |
| 755 | 38 | L38P | A | 1.8e-06 | 1.8e-06 | 2.5e-06 | 2.033E-06 | 4.041E-07 | 1.800E-06 | 2.500E-06 |
| 756 | 38 | L38S | A | 3.1e-06 | 4.7e-06 | 1.5e-06 | 3.100E-06 | 1.600E-06 | 1.500E-06 | 4.700E-06 |
| 757 | 38 | L38T | A | 2.2e-06 | 1.8e-06 | 2.6e-06 | 2.200E-06 | 4.000E-07 | 1.800E-06 | 2.600E-06 |
| 758 | 38 | L38W | A | 4.3e-06 | 2.8e-06 | 3e-06 | 3.367E-06 | 8.145E-07 | 2.800E-06 | 4.300E-06 |
| 759 | 38 | L38Y | A | 2.7e-06 | 2.5e-06 | 3.3e-06 | 2.833E-06 | 4.163E-07 | 2.500E-06 | 3.300E-06 |
| 760 | 38 | L38V | A | 2.4e-06 | 4.1e-06 | 2.6e-06 | 3.033E-06 | 9.292E-07 | 2.400E-06 | 4.100E-06 |
| 761 | 39 | E39A | A | 2.5e-06 | 2.5e-06 | 2e-06 | 2.333E-06 | 2.887E-07 | 2.000E-06 | 2.500E-06 |
| 762 | 39 | E39R | A | 3e-06 | 1.8e-06 | 2.4e-06 | 2.400E-06 | 6.000E-07 | 1.800E-06 | 3.000E-06 |
| 763 | 39 | E39N | A | 2.4e-06 | 2.6e-06 | 3.4e-06 | 2.800E-06 | 5.292E-07 | 2.400E-06 | 3.400E-06 |
| 764 | 39 | E39D | A | 4.1e-06 | 3.3e-06 | 1.5e-06 | 2.967E-06 | 1.332E-06 | 1.500E-06 | 4.100E-06 |

| | | | | | | | | | | |
|---|---|---|---|---|---|---|---|---|---|---|
| 765 | 39 | E39C | A | 2.2e-06 | 2.4e-06 | 1.5e-06 | 2.033E-06 | 4.726E-07 | 1.500E-06 | 2.400E-06 |
| 766 | 39 | E39E | A | 2.1e-06 | 2.8e-06 | 2.1e-06 | 2.333E-06 | 4.041E-07 | 2.100E-06 | 2.800E-06 |
| 767 | 39 | E39Q | A | 3.7e-06 | 2.5e-06 | 3.1e-06 | 3.100E-06 | 6.000E-07 | 2.500E-06 | 3.700E-06 |
| 768 | 39 | E39G | A | 3.4e-06 | 9.8e-07 | 4.1e-06 | 2.827E-06 | 1.637E-06 | 9.800E-07 | 4.100E-06 |
| 769 | 39 | E39H | A | 2.9e-06 | 2.5e-06 | 2.5e-06 | 2.633E-06 | 2.309E-07 | 2.500E-06 | 2.900E-06 |
| 770 | 39 | E39I | A | 5.1e-06 | 2.6e-06 | 2.5e-06 | 3.400E-06 | 1.473E-06 | 2.500E-06 | 5.100E-06 |
| 771 | 39 | E39L | A | 1.3e-06 | 2.2e-06 | 2.3e-06 | 1.933E-06 | 5.508E-07 | 1.300E-06 | 2.300E-06 |
| 772 | 39 | E39K | A | 2.3e-06 | 2.5e-06 | 2.3e-06 | 2.367E-06 | 1.155E-07 | 2.300E-06 | 2.500E-06 |
| 773 | 39 | E39M | A | 1.9e-06 | 2e-06 | 2e-06 | 1.967E-06 | 5.774E-08 | 1.900E-06 | 2.000E-06 |
| 774 | 39 | E39F | A | 1.5e-06 | 3.5e-06 | 3.2e-06 | 2.733E-06 | 1.079E-06 | 1.500E-06 | 3.500E-06 |
| 775 | 39 | E39P | A | 2.3e-06 | 2e-06 | 2e-06 | 2.100E-06 | 1.732E-07 | 2.000E-06 | 2.300E-06 |
| 776 | 39 | E39S | A | 2.2e-06 | 2.5e-06 | 1.2e-06 | 1.967E-06 | 6.807E-07 | 1.200E-06 | 2.500E-06 |
| 777 | 39 | E39T | A | 2.2e-06 | 1.3e-06 | 1.5e-06 | 1.667E-06 | 4.726E-07 | 1.300E-06 | 2.200E-06 |
| 778 | 39 | E39W | A | 1.7e-06 | 2.5e-06 | 2.5e-06 | 2.233E-06 | 4.619E-07 | 1.700E-06 | 2.500E-06 |
| 779 | 39 | E39Y | A | 1.6e-06 | 2.4e-06 | 1.4e-06 | 1.800E-06 | 5.292E-07 | 1.400E-06 | 2.400E-06 |
| 780 | 39 | E39V | A | 3.4e-06 | 1e-06 | 2.2e-06 | 2.200E-06 | 1.200E-06 | 1.000E-06 | 3.400E-06 |
| 781 | 40 | L40A | A | 2.4e-06 | 2.8e-06 | 3.1e-06 | 2.767E-06 | 3.512E-07 | 2.400E-06 | 3.100E-06 |
| 782 | 40 | L40R | A | 2e-06 | 2.3e-06 | 4.5e-06 | 2.933E-06 | 1.365E-06 | 2.000E-06 | 4.500E-06 |
| 783 | 40 | L40N | A | 1.5e-06 | 1.6e-06 | 2.3e-06 | 1.800E-06 | 4.359E-07 | 1.500E-06 | 2.300E-06 |
| 784 | 40 | L40D | A | 3.4e-06 | 3.1e-06 | 2.3e-06 | 2.933E-06 | 5.686E-07 | 2.300E-06 | 3.400E-06 |
| 785 | 40 | L40C | A | 2.6e-06 | 1.1e-06 | 2.5e-06 | 2.067E-06 | 8.386E-07 | 1.100E-06 | 2.600E-06 |
| 786 | 40 | L40E | A | 1.4e-06 | 2.2e-06 | 2.5e-06 | 2.033E-06 | 5.686E-07 | 1.400E-06 | 2.500E-06 |
| 787 | 40 | L40Q | A | 3.9e-06 | 3e-06 | 4.3e-06 | 3.733E-06 | 6.658E-07 | 3.000E-06 | 4.300E-06 |
| 788 | 40 | L40G | A | 3.3e-06 | 1.7e-06 | 1.6e-06 | 2.200E-06 | 9.539E-07 | 1.600E-06 | 3.300E-06 |
| 789 | 40 | L40H | A | 4.1e-06 | 1.8e-06 | 2.9e-06 | 2.933E-06 | 1.150E-06 | 1.800E-06 | 4.100E-06 |
| 790 | 40 | L40I | A | 2.5e-06 | 2e-06 | 2.4e-06 | 2.300E-06 | 2.646E-07 | 2.000E-06 | 2.500E-06 |
| 791 | 40 | L40L | A | 2.9e-06 | 2.6e-06 | 1.9e-06 | 2.467E-06 | 5.132E-07 | 1.900E-06 | 2.900E-06 |
| 792 | 40 | L40K | A | 2.1e-06 | 2.2e-06 | 1.8e-06 | 2.033E-06 | 2.082E-07 | 1.800E-06 | 2.200E-06 |
| 793 | 40 | L40M | A | 1.8e-06 | 2.3e-06 | 2.2e-06 | 2.100E-06 | 2.646E-07 | 1.800E-06 | 2.300E-06 |
| 794 | 40 | L40F | A | 2.8e-06 | 2.3e-06 | 2e-06 | 2.367E-06 | 4.041E-07 | 2.000E-06 | 2.800E-06 |
| 795 | 40 | L40P | A | 1.7e-06 | 3.1e-06 | 2.5e-06 | 2.433E-06 | 7.024E-07 | 1.700E-06 | 3.100E-06 |
| 796 | 40 | L40S | A | 1.6e-06 | 1.4e-06 | 2.3e-06 | 1.767E-06 | 4.726E-07 | 1.400E-06 | 2.300E-06 |
| 797 | 40 | L40T | A | 2e-06 | 2.2e-06 | 3.3e-06 | 2.500E-06 | 7.000E-07 | 2.000E-06 | 3.300E-06 |
| 798 | 40 | L40W | A | 1.9e-06 | 2e-06 | 3.1e-06 | 2.333E-06 | 6.658E-07 | 1.900E-06 | 3.100E-06 |
| 799 | 40 | L40Y | A | 1.6e-06 | 2.3e-06 | 2.7e-06 | 2.200E-06 | 5.568E-07 | 1.600E-06 | 2.700E-06 |
| 800 | 40 | L40V | A | 2.6e-06 | 2.1e-06 | 1.9e-06 | 2.200E-06 | 3.606E-07 | 1.900E-06 | 2.600E-06 |
| 801 | 41 | T41A | A | 3.2e-06 | 2.6e-06 | 1.7e-06 | 2.500E-06 | 7.550E-07 | 1.700E-06 | 3.200E-06 |
| 802 | 41 | T41R | A | 2.1e-06 | 1.9e-06 | 2.4e-06 | 2.133E-06 | 2.517E-07 | 1.900E-06 | 2.400E-06 |
| 803 | 41 | T41N | A | 3.3e-06 | 1.2e-06 | 3.7e-06 | 2.733E-06 | 1.343E-06 | 1.200E-06 | 3.700E-06 |
| 804 | 41 | T41D | A | 2.8e-06 | 1.9e-06 | 1.7e-06 | 2.133E-06 | 5.859E-07 | 1.700E-06 | 2.800E-06 |
| 805 | 41 | T41C | A | 3.3e-06 | 2.5e-06 | 1.7e-06 | 2.500E-06 | 8.000E-07 | 1.700E-06 | 3.300E-06 |
| 806 | 41 | T41E | A | 4e-06 | 8.5e-07 | 2.9e-06 | 2.583E-06 | 1.599E-06 | 8.500E-07 | 4.000E-06 |
| 807 | 41 | T41Q | A | 2.3e-06 | 3.9e-06 | 1.3e-06 | 2.500E-06 | 1.311E-06 | 1.300E-06 | 3.900E-06 |
| 808 | 41 | T41G | A | 2e-06 | 2.3e-06 | 1.9e-06 | 2.067E-06 | 2.082E-07 | 1.900E-06 | 2.300E-06 |
| 809 | 41 | T41H | A | 2.8e-06 | 4.3e-06 | 4.8e-06 | 3.967E-06 | 1.041E-06 | 2.800E-06 | 4.800E-06 |
| 810 | 41 | T41I | A | 1.6e-06 | 2.6e-06 | 2.3e-06 | 2.167E-06 | 5.132E-07 | 1.600E-06 | 2.600E-06 |
| 811 | 41 | T41L | A | 2.2e-06 | 3.9e-06 | 2.9e-06 | 3.000E-06 | 8.544E-07 | 2.200E-06 | 3.900E-06 |
| 812 | 41 | T41K | A | 3.3e-06 | 1.9e-06 | 9e-06 | 4.733E-06 | 3.761E-06 | 1.900E-06 | 9.000E-06 |
| 813 | 41 | T41M | A | 2.1e-06 | 1.5e-06 | 2.2e-06 | 1.933E-06 | 3.786E-07 | 1.500E-06 | 2.200E-06 |
| 814 | 41 | T41F | A | 2.1e-06 | 2.3e-06 | 1.3e-06 | 1.900E-06 | 5.292E-07 | 1.300E-06 | 2.300E-06 |
| 815 | 41 | T41P | A | 1.7e-06 | 2.2e-06 | 2.3e-06 | 2.067E-06 | 3.215E-07 | 1.700E-06 | 2.300E-06 |
| 816 | 41 | T41S | A | 2.1e-06 | 4.7e-06 | 1.4e-06 | 2.733E-06 | 1.739E-06 | 1.400E-06 | 4.700E-06 |
| 817 | 41 | T41T | A | 1.3e-06 | 1.1e-06 | 1.8e-06 | 1.400E-06 | 3.606E-07 | 1.100E-06 | 1.800E-06 |
| 818 | 41 | T41W | A | 2.5e-06 | 3.4e-06 | 1.6e-06 | 2.500E-06 | 9.000E-07 | 1.600E-06 | 3.400E-06 |
| 819 | 41 | T41Y | A | 3.3e-06 | 2.5e-06 | 2.5e-06 | 2.767E-06 | 4.619E-07 | 2.500E-06 | 3.300E-06 |
| 820 | 41 | T41V | A | 1.5e-06 | 2.2e-06 | 2.6e-06 | 2.100E-06 | 5.568E-07 | 1.500E-06 | 2.600E-06 |
| 821 | 42 | Y42A | A | 2.5e-06 | 4.6e-06 | 3.4e-06 | 3.500E-06 | 1.054E-06 | 2.500E-06 | 4.600E-06 |
| 822 | 42 | Y42R | A | 3.2e-06 | 1.8e-06 | 2.5e-06 | 2.500E-06 | 7.000E-07 | 1.800E-06 | 3.200E-06 |
| 823 | 42 | Y42N | A | 2.1e-06 | 2e-06 | 2.1e-06 | 2.067E-06 | 5.774E-08 | 2.000E-06 | 2.100E-06 |
| 824 | 42 | Y42D | A | 2e-06 | 2.2e-06 | 2.4e-06 | 2.200E-06 | 2.000E-07 | 2.000E-06 | 2.400E-06 |
| 825 | 42 | Y42C | A | 3.4e-06 | 1.8e-06 | 2.8e-06 | 2.667E-06 | 8.083E-07 | 1.800E-06 | 3.400E-06 |
| 826 | 42 | Y42E | A | 1.3e-06 | 2.3e-06 | 2.3e-06 | 1.967E-06 | 5.774E-07 | 1.300E-06 | 2.300E-06 |
| 827 | 42 | Y42Q | A | 3e-06 | 3.3e-06 | 2.7e-06 | 3.000E-06 | 3.000E-07 | 2.700E-06 | 3.300E-06 |
| 828 | 42 | Y42G | A | 2.8e-06 | 2.6e-06 | 5.1e-06 | 3.500E-06 | 1.389E-06 | 2.600E-06 | 5.100E-06 |
| 829 | 42 | Y42H | A | 4.1e-06 | 3.5e-06 | 1.4e-06 | 3.000E-06 | 1.418E-06 | 1.400E-06 | 4.100E-06 |
| 830 | 42 | Y42I | A | 2.6e-06 | 1e-06 | 2.2e-06 | 1.933E-06 | 8.327E-07 | 1.000E-06 | 2.600E-06 |
| 831 | 42 | Y42L | A | 1.8e-06 | 3.1e-06 | 3.1e-06 | 2.667E-06 | 7.506E-07 | 1.800E-06 | 3.100E-06 |
| 832 | 42 | Y42K | A | 1.8e-06 | 1.4e-06 | 2.2e-06 | 1.800E-06 | 4.000E-07 | 1.400E-06 | 2.200E-06 |
| 833 | 42 | Y42M | A | 2.2e-06 | 1.4e-06 | 2.4e-06 | 2.000E-06 | 5.292E-07 | 1.400E-06 | 2.400E-06 |
| 834 | 42 | Y42F | A | 2.5e-06 | 2.1e-06 | 8.5e-07 | 1.817E-06 | 8.607E-07 | 8.500E-07 | 2.500E-06 |
| 835 | 42 | Y42P | A | 2.3e-06 | 1.5e-06 | 1.7e-06 | 1.833E-06 | 4.163E-07 | 1.500E-06 | 2.300E-06 |
| 836 | 42 | Y42S | A | 2.4e-06 | 2.2e-06 | 1.5e-06 | 2.033E-06 | 4.726E-07 | 1.500E-06 | 2.400E-06 |
| 837 | 42 | Y42T | A | 2.4e-06 | 1.6e-06 | 2.5e-06 | 2.167E-06 | 4.933E-07 | 1.600E-06 | 2.500E-06 |
| 838 | 42 | Y42W | A | 2.7e-06 | 2e-06 | 3.6e-06 | 2.767E-06 | 8.021E-07 | 2.000E-06 | 3.600E-06 |
| 839 | 42 | Y42Y | A | 1.8e-06 | 2.2e-06 | 2e-06 | 2.000E-06 | 2.000E-07 | 1.800E-06 | 2.200E-06 |
| 840 | 42 | Y42V | A | 2e-06 | 2e-06 | 3.2e-06 | 2.400E-06 | 6.928E-07 | 2.000E-06 | 3.200E-06 |
| 841 | 43 | L43A | A | 1.5e-06 | 2.7e-06 | 4.6e-06 | 2.933E-06 | 1.563E-06 | 1.500E-06 | 4.600E-06 |
| 842 | 43 | L43R | A | 2.6e-06 | 2.2e-06 | 1.6e-06 | 2.133E-06 | 5.033E-07 | 1.600E-06 | 2.600E-06 |
| 843 | 43 | L43N | A | 4.8e-06 | 3e-06 | 1.7e-06 | 3.167E-06 | 1.557E-06 | 1.700E-06 | 4.800E-06 |
| 844 | 43 | L43D | A | 2.2e-06 | 3.4e-06 | 2.3e-06 | 2.633E-06 | 6.658E-07 | 2.200E-06 | 3.400E-06 |
| 845 | 43 | L43C | A | 2.9e-06 | 2.9e-06 | 6e-06 | 3.933E-06 | 1.790E-06 | 2.900E-06 | 6.000E-06 |
| 846 | 43 | L43E | A | 2.4e-06 | 3.2e-06 | 3e-06 | 2.867E-06 | 4.163E-07 | 2.400E-06 | 3.200E-06 |
| 847 | 43 | L43Q | A | 2.3e-06 | 2.9e-06 | 1.4e-06 | 2.200E-06 | 7.550E-07 | 1.400E-06 | 2.900E-06 |
| 848 | 43 | L43G | A | 3.2e-06 | 1.6e-06 | 3.9e-06 | 2.900E-06 | 1.179E-06 | 1.600E-06 | 3.900E-06 |
| 849 | 43 | L43H | A | 2.2e-06 | 2.7e-06 | 3.7e-06 | 2.867E-06 | 7.638E-07 | 2.200E-06 | 3.700E-06 |

| | | | | | | | | | | |
|---|---|---|---|---|---|---|---|---|---|---|
| 850 | 43 | L43I | A | 3.2e-06 | 2.6e-06 | 2.6e-06 | 2.800E-06 | 3.464E-07 | 2.600E-06 | 3.200E-06 |
| 851 | 43 | L43L | A | 2e-06 | 4.7e-06 | 3.2e-06 | 3.300E-06 | 1.353E-06 | 2.000E-06 | 4.700E-06 |
| 852 | 43 | L43K | A | 2.2e-06 | 1.6e-06 | 3.1e-06 | 2.300E-06 | 7.550E-07 | 1.600E-06 | 3.100E-06 |
| 853 | 43 | L43M | A | 2.3e-06 | 2.7e-06 | 3.3e-06 | 2.767E-06 | 5.033E-07 | 2.300E-06 | 3.300E-06 |
| 854 | 43 | L43F | A | 1.3e-06 | 2.3e-06 | 3.7e-06 | 2.433E-06 | 1.206E-06 | 1.300E-06 | 3.700E-06 |
| 855 | 43 | L43P | A | 1.8e-06 | 2.5e-06 | 1.5e-06 | 1.933E-06 | 5.132E-07 | 1.500E-06 | 2.500E-06 |
| 856 | 43 | L43S | A | 2.8e-06 | 1.8e-06 | 1.8e-06 | 2.133E-06 | 5.774E-07 | 1.800E-06 | 2.800E-06 |
| 857 | 43 | L43T | A | 4.4e-06 | 2e-06 | 2.9e-06 | 3.100E-06 | 1.212E-06 | 2.000E-06 | 4.400E-06 |
| 858 | 43 | L43W | A | 2.1e-06 | 2.2e-06 | 2.3e-06 | 2.200E-06 | 1.000E-07 | 2.100E-06 | 2.300E-06 |
| 859 | 43 | L43Y | A | 2.4e-06 | 1.8e-06 | 6e-06 | 3.400E-06 | 2.272E-06 | 1.800E-06 | 6.000E-06 |
| 860 | 43 | L43V | A | 1.9e-06 | 3.3e-06 | 5.9e-06 | 3.700E-06 | 2.030E-06 | 1.900E-06 | 5.900E-06 |
| 861 | 44 | P44A | A | 3.4e-06 | 1.8e-06 | 2.4e-06 | 2.533E-06 | 8.083E-07 | 1.800E-06 | 3.400E-06 |
| 862 | 44 | P44R | A | 1.5e-06 | 2.5e-06 | 2.5e-06 | 2.167E-06 | 5.774E-07 | 1.500E-06 | 2.500E-06 |
| 863 | 44 | P44N | A | 2.3e-06 | 1.6e-06 | 1.9e-06 | 1.933E-06 | 3.512E-07 | 1.600E-06 | 2.300E-06 |
| 864 | 44 | P44D | A | 1.9e-06 | 2.5e-06 | 3.3e-06 | 2.567E-06 | 7.024E-07 | 1.900E-06 | 3.300E-06 |
| 865 | 44 | P44C | A | 2.6e-06 | 2.6e-06 | 2.2e-06 | 2.467E-06 | 2.309E-07 | 2.200E-06 | 2.600E-06 |
| 866 | 44 | P44E | A | 1.8e-06 | 2e-06 | 1.8e-06 | 1.867E-06 | 1.155E-07 | 1.800E-06 | 2.000E-06 |
| 867 | 44 | P44Q | A | 2.8e-06 | 4.9e-06 | 1.7e-06 | 3.133E-06 | 1.626E-06 | 1.700E-06 | 4.900E-06 |
| 868 | 44 | P44G | A | 1.9e-06 | 1.8e-06 | 2.2e-06 | 1.967E-06 | 2.082E-07 | 1.800E-06 | 2.200E-06 |
| 869 | 44 | P44H | A | 2.5e-06 | 3.5e-06 | 2.5e-06 | 2.833E-06 | 5.774E-07 | 2.500E-06 | 3.500E-06 |
| 870 | 44 | P44I | A | 1.6e-06 | 2.4e-06 | 3.1e-06 | 2.367E-06 | 7.506E-07 | 1.600E-06 | 3.100E-06 |
| 871 | 44 | P44L | A | 2.7e-06 | 2.3e-06 | 2.8e-06 | 2.600E-06 | 2.646E-07 | 2.300E-06 | 2.800E-06 |
| 872 | 44 | P44K | A | 2.3e-06 | 2.9e-06 | 2.3e-06 | 2.500E-06 | 3.464E-07 | 2.300E-06 | 2.900E-06 |
| 873 | 44 | P44M | A | 2.5e-06 | 2.5e-06 | 3.3e-06 | 2.767E-06 | 4.619E-07 | 2.500E-06 | 3.300E-06 |
| 874 | 44 | P44F | A | 2.1e-06 | 1.8e-06 | 5.3e-06 | 3.067E-06 | 1.940E-06 | 1.800E-06 | 5.300E-06 |
| 875 | 44 | P44P | A | 1.3e-06 | 1.8e-06 | 1.9e-06 | 1.667E-06 | 3.215E-07 | 1.300E-06 | 1.900E-06 |
| 876 | 44 | P44S | A | 2.5e-06 | 4e-06 | 1.7e-06 | 2.733E-06 | 1.168E-06 | 1.700E-06 | 4.000E-06 |
| 877 | 44 | P44T | A | 1.9e-06 | 1.7e-06 | 2.3e-06 | 1.967E-06 | 3.055E-07 | 1.700E-06 | 2.300E-06 |
| 878 | 44 | P44W | A | 1.7e-06 | 1.9e-06 | 1.9e-06 | 1.833E-06 | 1.155E-07 | 1.700E-06 | 1.900E-06 |
| 879 | 44 | P44Y | A | 5.2e-06 | 5.3e-06 | 3.3e-06 | 4.600E-06 | 1.127E-06 | 3.300E-06 | 5.300E-06 |
| 880 | 44 | P44V | A | 3.9e-06 | 1.6e-06 | 1.8e-06 | 2.433E-06 | 1.274E-06 | 1.600E-06 | 3.900E-06 |
| 881 | 45 | T45A | A | 2.8e-06 | 2e-06 | 2.3e-06 | 2.367E-06 | 4.041E-07 | 2.000E-06 | 2.800E-06 |
| 882 | 45 | T45R | A | 2.6e-06 | 1.7e-06 | 2.9e-06 | 2.400E-06 | 6.245E-07 | 1.700E-06 | 2.900E-06 |
| 883 | 45 | T45N | A | 2.3e-06 | 2.1e-06 | 3.2e-06 | 2.533E-06 | 5.859E-07 | 2.100E-06 | 3.200E-06 |
| 884 | 45 | T45D | A | 2.2e-06 | 3e-06 | 2.6e-06 | 2.600E-06 | 4.000E-07 | 2.200E-06 | 3.000E-06 |
| 885 | 45 | T45C | A | 2.2e-06 | 3.2e-06 | 1.6e-06 | 2.333E-06 | 8.083E-07 | 1.600E-06 | 3.200E-06 |
| 886 | 45 | T45E | A | 2.4e-06 | 1.6e-06 | 2.1e-06 | 2.033E-06 | 4.041E-07 | 1.600E-06 | 2.400E-06 |
| 887 | 45 | T45Q | A | 1.8e-06 | 1.6e-06 | 2.7e-06 | 2.033E-06 | 5.859E-07 | 1.600E-06 | 2.700E-06 |
| 888 | 45 | T45G | A | 2.7e-06 | 4.6e-06 | 2.4e-06 | 3.233E-06 | 1.193E-06 | 2.400E-06 | 4.600E-06 |
| 889 | 45 | T45H | A | 1.9e-06 | 3.5e-06 | 2.1e-06 | 2.500E-06 | 8.718E-07 | 1.900E-06 | 3.500E-06 |
| 890 | 45 | T45I | A | 4.2e-06 | 1.8e-06 | 2.1e-06 | 2.700E-06 | 1.308E-06 | 1.800E-06 | 4.200E-06 |
| 891 | 45 | T45L | A | 1.8e-06 | 1.6e-06 | 1.8e-06 | 1.733E-06 | 1.155E-07 | 1.600E-06 | 1.800E-06 |
| 892 | 45 | T45K | A | 3.3e-06 | 2.1e-06 | 2.4e-06 | 2.600E-06 | 6.245E-07 | 2.100E-06 | 3.300E-06 |
| 893 | 45 | T45M | A | 5.1e-06 | 2.9e-06 | 1.9e-06 | 3.300E-06 | 1.637E-06 | 1.900E-06 | 5.100E-06 |
| 894 | 45 | T45F | A | 1.8e-06 | 2.9e-06 | 2.1e-06 | 2.267E-06 | 5.686E-07 | 1.800E-06 | 2.900E-06 |
| 895 | 45 | T45P | A | 2.2e-06 | 2.2e-06 | 2e-06 | 2.133E-06 | 1.155E-07 | 2.000E-06 | 2.200E-06 |
| 896 | 45 | T45S | A | 2.3e-06 | 2.4e-06 | 2.5e-06 | 2.400E-06 | 1.000E-07 | 2.300E-06 | 2.500E-06 |
| 897 | 45 | T45T | A | 1.8e-06 | 2.7e-06 | 1.9e-06 | 2.133E-06 | 4.933E-07 | 1.800E-06 | 2.700E-06 |
| 898 | 45 | T45W | A | 1.5e-06 | 2.5e-06 | 2.1e-06 | 2.033E-06 | 5.033E-07 | 1.500E-06 | 2.500E-06 |
| 899 | 45 | T45Y | A | 1.8e-06 | 1.5e-06 | 2e-06 | 1.767E-06 | 2.517E-07 | 1.500E-06 | 2.000E-06 |
| 900 | 45 | T45V | A | 9.2e-07 | 3.7e-06 | 2.8e-06 | 2.473E-06 | 1.418E-06 | 9.200E-07 | 3.700E-06 |
| 901 | 46 | N46A | A | 1.6e-06 | 4.1e-06 | 2.3e-06 | 2.667E-06 | 1.290E-06 | 1.600E-06 | 4.100E-06 |
| 902 | 46 | N46R | A | 3.9e-06 | 3.2e-06 | 3.7e-06 | 3.600E-06 | 3.606E-07 | 3.200E-06 | 3.900E-06 |
| 903 | 46 | N46N | A | 2.5e-06 | 2.8e-06 | 3.4e-06 | 2.900E-06 | 4.583E-07 | 2.500E-06 | 3.400E-06 |
| 904 | 46 | N46D | A | 3e-06 | 1.9e-06 | 1.4e-06 | 2.100E-06 | 8.185E-07 | 1.400E-06 | 3.000E-06 |
| 905 | 46 | N46C | A | 5e-06 | 3.4e-06 | 2.7e-06 | 3.700E-06 | 1.179E-06 | 2.700E-06 | 5.000E-06 |
| 906 | 46 | N46E | A | 1.5e-06 | 2.6e-06 | 2.6e-06 | 2.233E-06 | 6.351E-07 | 1.500E-06 | 2.600E-06 |
| 907 | 46 | N46Q | A | 3.5e-06 | 3.6e-06 | 1.7e-06 | 2.933E-06 | 1.069E-06 | 1.700E-06 | 3.600E-06 |
| 908 | 46 | N46G | A | 2.7e-06 | 3.2e-06 | 1.4e-06 | 2.433E-06 | 9.292E-07 | 1.400E-06 | 3.200E-06 |
| 909 | 46 | N46H | A | 1.5e-06 | 1.9e-06 | 2.7e-06 | 2.033E-06 | 6.110E-07 | 1.500E-06 | 2.700E-06 |
| 910 | 46 | N46I | A | 3e-06 | 3.2e-06 | 2.3e-06 | 2.833E-06 | 4.726E-07 | 2.300E-06 | 3.200E-06 |
| 911 | 46 | N46L | A | 2.2e-06 | 3.6e-06 | 9.2e-07 | 2.240E-06 | 1.340E-06 | 9.200E-07 | 3.600E-06 |
| 912 | 46 | N46K | A | 3.6e-06 | 3.3e-06 | 2.9e-06 | 3.267E-06 | 3.512E-07 | 2.900E-06 | 3.600E-06 |
| 913 | 46 | N46M | A | 3.5e-06 | 2.2e-06 | 2.6e-06 | 2.767E-06 | 6.658E-07 | 2.200E-06 | 3.500E-06 |
| 914 | 46 | N46F | A | 2.9e-06 | 3.2e-06 | 3.6e-06 | 3.233E-06 | 3.512E-07 | 2.900E-06 | 3.600E-06 |
| 915 | 46 | N46P | A | 1.8e-06 | 2.8e-06 | 2.6e-06 | 2.400E-06 | 5.292E-07 | 1.800E-06 | 2.800E-06 |
| 916 | 46 | N46S | A | 1.7e-06 | 2.1e-06 | 1.5e-06 | 1.767E-06 | 3.055E-07 | 1.500E-06 | 2.100E-06 |
| 917 | 46 | N46T | A | 2e-06 | 2.2e-06 | 2.1e-06 | 2.100E-06 | 1.000E-07 | 2.000E-06 | 2.200E-06 |
| 918 | 46 | N46W | A | 4.2e-06 | 3.1e-06 | 1.5e-06 | 2.933E-06 | 1.358E-06 | 1.500E-06 | 4.200E-06 |
| 919 | 46 | N46Y | A | 2.6e-06 | 1.5e-06 | 1.7e-06 | 1.933E-06 | 5.859E-07 | 1.500E-06 | 2.600E-06 |
| 920 | 46 | N46V | A | 1.9e-06 | 2.5e-06 | 2.5e-06 | 2.300E-06 | 3.464E-07 | 1.900E-06 | 2.500E-06 |
| 921 | 47 | A47A | A | 2.3e-06 | 2.2e-06 | 2.2e-06 | 2.233E-06 | 5.774E-08 | 2.200E-06 | 2.300E-06 |
| 922 | 47 | A47R | A | 2.2e-06 | 1.6e-06 | 2e-06 | 1.933E-06 | 3.055E-07 | 1.600E-06 | 2.200E-06 |
| 923 | 47 | A47N | A | 2.7e-06 | 2.9e-06 | 2.5e-06 | 2.700E-06 | 2.000E-07 | 2.500E-06 | 2.900E-06 |
| 924 | 47 | A47D | A | 2.1e-06 | 3.9e-06 | 2.4e-06 | 2.800E-06 | 9.644E-07 | 2.100E-06 | 3.900E-06 |
| 925 | 47 | A47C | A | 2.6e-06 | 1.4e-06 | 2.4e-06 | 2.133E-06 | 6.429E-07 | 1.400E-06 | 2.600E-06 |
| 926 | 47 | A47E | A | 1.7e-06 | 2.1e-06 | 2.4e-06 | 2.067E-06 | 3.512E-07 | 1.700E-06 | 2.400E-06 |
| 927 | 47 | A47Q | A | 1.7e-06 | 4.4e-06 | 2.3e-06 | 2.800E-06 | 1.418E-06 | 1.700E-06 | 4.400E-06 |
| 928 | 47 | A47G | A | 2.2e-06 | 2.4e-06 | 2.5e-06 | 2.367E-06 | 1.528E-07 | 2.200E-06 | 2.500E-06 |
| 929 | 47 | A47H | A | 3.7e-06 | 3.6e-06 | 2.7e-06 | 3.333E-06 | 5.508E-07 | 2.700E-06 | 3.700E-06 |
| 930 | 47 | A47I | A | 2.6e-06 | 2.2e-06 | 3e-06 | 2.600E-06 | 4.000E-07 | 2.200E-06 | 3.000E-06 |
| 931 | 47 | A47L | A | 4.6e-06 | 3e-06 | 2.1e-06 | 3.233E-06 | 1.266E-06 | 2.100E-06 | 4.600E-06 |
| 932 | 47 | A47K | A | 1.8e-06 | 3e-06 | 2.4e-06 | 2.400E-06 | 6.000E-07 | 1.800E-06 | 3.000E-06 |
| 933 | 47 | A47M | A | 2.3e-06 | 1.8e-06 | 2.4e-06 | 2.167E-06 | 3.215E-07 | 1.800E-06 | 2.400E-06 |
| 934 | 47 | A47F | A | 1.6e-06 | 3.1e-06 | 1.1e-06 | 1.933E-06 | 1.041E-06 | 1.100E-06 | 3.100E-06 |

| | | | | | | | | | | |
|---|---|---|---|---|---|---|---|---|---|---|
| 935 | 47 | A47P | A | 3.2e-06 | 1.7e-06 | 3.5e-06 | 2.800E-06 | 9.644E-07 | 1.700E-06 | 3.500E-06 |
| 936 | 47 | A47S | A | 2.1e-06 | 3.6e-06 | 2.7e-06 | 2.800E-06 | 7.550E-07 | 2.100E-06 | 3.600E-06 |
| 937 | 47 | A47T | A | 3.7e-06 | 2.4e-06 | 4.6e-06 | 3.567E-06 | 1.106E-06 | 2.400E-06 | 4.600E-06 |
| 938 | 47 | A47W | A | 3.2e-06 | 2.3e-06 | 1.6e-06 | 2.367E-06 | 8.021E-07 | 1.600E-06 | 3.200E-06 |
| 939 | 47 | A47Y | A | 2.9e-06 | 1.1e-06 | 2e-06 | 2.000E-06 | 9.000E-07 | 1.100E-06 | 2.900E-06 |
| 940 | 47 | A47V | A | 1.9e-06 | 3.4e-06 | 2.5e-06 | 2.600E-06 | 7.550E-07 | 1.900E-06 | 3.400E-06 |
| 941 | 48 | S48A | A | 2.5e-06 | 1.8e-06 | 3e-06 | 2.433E-06 | 6.028E-07 | 1.800E-06 | 3.000E-06 |
| 942 | 48 | S48R | A | 1.5e-06 | 3.6e-06 | 3.9e-06 | 3.000E-06 | 1.308E-06 | 1.500E-06 | 3.900E-06 |
| 943 | 48 | S48N | A | 3e-06 | 1.7e-06 | 2.6e-06 | 2.433E-06 | 6.658E-07 | 1.700E-06 | 3.000E-06 |
| 944 | 48 | S48D | A | 7e-06 | 2.6e-06 | 3.1e-06 | 4.233E-06 | 2.409E-06 | 2.600E-06 | 7.000E-06 |
| 945 | 48 | S48C | A | 2.3e-06 | 1.9e-06 | 1.3e-06 | 1.833E-06 | 5.033E-07 | 1.300E-06 | 2.300E-06 |
| 946 | 48 | S48E | A | 3.2e-06 | 4.7e-06 | 3.2e-06 | 3.700E-06 | 8.660E-07 | 3.200E-06 | 4.700E-06 |
| 947 | 48 | S48Q | A | 3.4e-06 | 3e-06 | 2.6e-06 | 3.000E-06 | 4.000E-07 | 2.600E-06 | 3.400E-06 |
| 948 | 48 | S48G | A | 2.1e-06 | 1.5e-06 | 2.2e-06 | 1.933E-06 | 3.786E-07 | 1.500E-06 | 2.200E-06 |
| 949 | 48 | S48H | A | 3e-06 | 1.7e-06 | 1.9e-06 | 2.200E-06 | 7.000E-07 | 1.700E-06 | 3.000E-06 |
| 950 | 48 | S48I | A | 1.5e-06 | 2e-06 | 1.2e-06 | 1.567E-06 | 4.041E-07 | 1.200E-06 | 2.000E-06 |
| 951 | 48 | S48L | A | 2.9e-06 | 3.2e-06 | 2.7e-06 | 2.933E-06 | 2.517E-07 | 2.700E-06 | 3.200E-06 |
| 952 | 48 | S48K | A | 1.6e-06 | 1.9e-06 | 1.6e-06 | 1.700E-06 | 1.732E-07 | 1.600E-06 | 1.900E-06 |
| 953 | 48 | S48M | A | 2.7e-06 | 4.2e-06 | 2.7e-06 | 3.200E-06 | 8.660E-07 | 2.700E-06 | 4.200E-06 |
| 954 | 48 | S48F | A | 2.7e-06 | 2.8e-06 | 2.2e-06 | 2.567E-06 | 3.215E-07 | 2.200E-06 | 2.800E-06 |
| 955 | 48 | S48P | A | 2.9e-06 | 2.9e-06 | 2.3e-06 | 2.700E-06 | 3.464E-07 | 2.300E-06 | 2.900E-06 |
| 956 | 48 | S48S | A | 1.9e-06 | 2.8e-06 | 2.3e-06 | 2.333E-06 | 4.509E-07 | 1.900E-06 | 2.800E-06 |
| 957 | 48 | S48T | A | 3.9e-06 | 2.1e-06 | 2e-06 | 2.667E-06 | 1.069E-06 | 2.000E-06 | 3.900E-06 |
| 958 | 48 | S48W | A | 2.6e-06 | 2.1e-06 | 2.9e-06 | 2.533E-06 | 4.041E-07 | 2.100E-06 | 2.900E-06 |
| 959 | 48 | S48Y | A | 3.7e-06 | 3e-06 | 2.2e-06 | 2.967E-06 | 7.506E-07 | 2.200E-06 | 3.700E-06 |
| 960 | 48 | S48V | A | 1.6e-06 | 2.2e-06 | 9.6e-07 | 1.587E-06 | 6.201E-07 | 9.600E-07 | 2.200E-06 |
| 961 | 49 | L49A | A | 2.6e-06 | 2e-06 | 3.1e-06 | 2.567E-06 | 5.508E-07 | 2.000E-06 | 3.100E-06 |
| 962 | 49 | L49R | A | 1.8e-06 | 1.5e-06 | 2.5e-06 | 1.933E-06 | 5.132E-07 | 1.500E-06 | 2.500E-06 |
| 963 | 49 | L49N | A | 2.4e-06 | 2.8e-06 | 2.2e-06 | 2.467E-06 | 3.055E-07 | 2.200E-06 | 2.800E-06 |
| 964 | 49 | L49D | A | 2.2e-06 | 2.6e-06 | 2.6e-06 | 2.467E-06 | 2.309E-07 | 2.200E-06 | 2.600E-06 |
| 965 | 49 | L49C | A | 3.2e-06 | 2.5e-06 | 2.5e-06 | 2.733E-06 | 4.041E-07 | 2.500E-06 | 3.200E-06 |
| 966 | 49 | L49E | A | 1.5e-06 | 1.6e-06 | 2.6e-06 | 1.900E-06 | 6.083E-07 | 1.500E-06 | 2.600E-06 |
| 967 | 49 | L49Q | A | 2.1e-06 | 3e-06 | 2.1e-06 | 2.400E-06 | 5.196E-07 | 2.100E-06 | 3.000E-06 |
| 968 | 49 | L49G | A | 1.8e-06 | 1.9e-06 | 2.7e-06 | 2.133E-06 | 4.933E-07 | 1.800E-06 | 2.700E-06 |
| 969 | 49 | L49H | A | 2.4e-06 | 2.2e-06 | 1.9e-06 | 2.167E-06 | 2.517E-07 | 1.900E-06 | 2.400E-06 |
| 970 | 49 | L49I | A | 2.4e-06 | 2e-06 | 1.8e-06 | 2.067E-06 | 3.055E-07 | 1.800E-06 | 2.400E-06 |
| 971 | 49 | L49L | A | 3e-06 | 2.4e-06 | 4.1e-06 | 3.167E-06 | 8.622E-07 | 2.400E-06 | 4.100E-06 |
| 972 | 49 | L49K | A | 1.9e-06 | 5.3e-06 | 2.8e-06 | 3.333E-06 | 1.762E-06 | 1.900E-06 | 5.300E-06 |
| 973 | 49 | L49M | A | 4.2e-06 | 1.7e-06 | 1.9e-06 | 2.600E-06 | 1.389E-06 | 1.700E-06 | 4.200E-06 |
| 974 | 49 | L49F | A | 1.8e-06 | 2.2e-06 | 3.5e-06 | 2.500E-06 | 8.888E-07 | 1.800E-06 | 3.500E-06 |
| 975 | 49 | L49P | A | 2.1e-06 | 3.2e-06 | 2.9e-06 | 2.733E-06 | 5.686E-07 | 2.100E-06 | 3.200E-06 |
| 976 | 49 | L49S | A | 2.7e-06 | 3.7e-06 | 2e-06 | 2.800E-06 | 8.544E-07 | 2.000E-06 | 3.700E-06 |
| 977 | 49 | L49T | A | 1.5e-06 | 1.8e-06 | 1.4e-06 | 1.567E-06 | 2.082E-07 | 1.400E-06 | 1.800E-06 |
| 978 | 49 | L49W | A | 2.1e-06 | 1.6e-06 | 2.3e-06 | 2.000E-06 | 3.606E-07 | 1.600E-06 | 2.300E-06 |
| 979 | 49 | L49Y | A | 1.6e-06 | 2.7e-06 | 2.6e-06 | 2.300E-06 | 6.083E-07 | 1.600E-06 | 2.700E-06 |
| 980 | 49 | L49V | A | 5.1e-06 | 4.2e-06 | 2.1e-06 | 3.800E-06 | 1.539E-06 | 2.100E-06 | 5.100E-06 |
| 981 | 50 | S50A | A | 2.2e-06 | 2.6e-06 | 3.6e-06 | 2.800E-06 | 7.211E-07 | 2.200E-06 | 3.600E-06 |
| 982 | 50 | S50R | A | 1.2e-06 | 2.1e-06 | 3.6e-06 | 2.300E-06 | 1.212E-06 | 1.200E-06 | 3.600E-06 |
| 983 | 50 | S50N | A | 2.8e-06 | 1.4e-06 | 2.7e-06 | 2.300E-06 | 7.810E-07 | 1.400E-06 | 2.800E-06 |
| 984 | 50 | S50D | A | 2.7e-06 | 2.1e-06 | 1.8e-06 | 2.200E-06 | 4.583E-07 | 1.800E-06 | 2.700E-06 |
| 985 | 50 | S50C | A | 1.6e-06 | 2.4e-06 | 1.9e-06 | 1.967E-06 | 4.041E-07 | 1.600E-06 | 2.400E-06 |
| 986 | 50 | S50E | A | 2.4e-06 | 1.8e-06 | 3.4e-06 | 2.533E-06 | 8.083E-07 | 1.800E-06 | 3.400E-06 |
| 987 | 50 | S50Q | A | 1.5e-06 | 4e-06 | 3.2e-06 | 2.900E-06 | 1.277E-06 | 1.500E-06 | 4.000E-06 |
| 988 | 50 | S50G | A | 2.6e-06 | 2.7e-06 | 2.5e-06 | 2.600E-06 | 1.000E-07 | 2.500E-06 | 2.700E-06 |
| 989 | 50 | S50H | A | 2.6e-06 | 1.6e-06 | 3.3e-06 | 2.500E-06 | 8.544E-07 | 1.600E-06 | 3.300E-06 |
| 990 | 50 | S50I | A | 4.2e-06 | 1.8e-06 | 2.4e-06 | 2.800E-06 | 1.249E-06 | 1.800E-06 | 4.200E-06 |
| 991 | 50 | S50L | A | 2.7e-06 | 2.8e-06 | 2e-06 | 2.500E-06 | 4.359E-07 | 2.000E-06 | 2.800E-06 |
| 992 | 50 | S50K | A | 1.8e-06 | 2.2e-06 | 3.1e-06 | 2.367E-06 | 6.658E-07 | 1.800E-06 | 3.100E-06 |
| 993 | 50 | S50M | A | 3.2e-06 | 1.9e-06 | 3e-06 | 2.700E-06 | 7.000E-07 | 1.900E-06 | 3.200E-06 |
| 994 | 50 | S50F | A | 1.6e-06 | 4.4e-06 | 2.7e-06 | 2.900E-06 | 1.411E-06 | 1.600E-06 | 4.400E-06 |
| 995 | 50 | S50P | A | 2.3e-06 | 2.8e-06 | 3.3e-06 | 2.800E-06 | 5.000E-07 | 2.300E-06 | 3.300E-06 |
| 996 | 50 | S50S | A | 2.4e-06 | 1.4e-06 | 1.8e-06 | 1.867E-06 | 5.033E-07 | 1.400E-06 | 2.400E-06 |
| 997 | 50 | S50T | A | 3.9e-06 | 2.6e-06 | 1.8e-06 | 2.767E-06 | 1.060E-06 | 1.800E-06 | 3.900E-06 |
| 998 | 50 | S50W | A | 1.6e-06 | 2.2e-06 | 2.9e-06 | 2.233E-06 | 6.506E-07 | 1.600E-06 | 2.900E-06 |
| 999 | 50 | S50Y | A | 2.1e-06 | 2.2e-06 | 4e-06 | 2.767E-06 | 1.069E-06 | 2.100E-06 | 4.000E-06 |
| 1000 | 50 | S50V | A | 1.9e-06 | 2.2e-06 | 1.9e-06 | 2.000E-06 | 1.732E-07 | 1.900E-06 | 2.200E-06 |
| 1001 | 51 | F51A | A | 1.4e-06 | 2.1e-06 | 2.6e-06 | 2.033E-06 | 6.028E-07 | 1.400E-06 | 2.600E-06 |
| 1002 | 51 | F51R | A | 3.8e-06 | 1e-06 | 2.6e-06 | 2.467E-06 | 1.405E-06 | 1.000E-06 | 3.800E-06 |
| 1003 | 51 | F51N | A | 3.8e-06 | 2.2e-06 | 3.2e-06 | 3.067E-06 | 8.083E-07 | 2.200E-06 | 3.800E-06 |
| 1004 | 51 | F51D | A | 2.3e-06 | 1.9e-06 | 4.5e-06 | 2.900E-06 | 1.400E-06 | 1.900E-06 | 4.500E-06 |
| 1005 | 51 | F51C | A | 2.4e-06 | 3.1e-06 | 2.5e-06 | 2.667E-06 | 3.786E-07 | 2.400E-06 | 3.100E-06 |
| 1006 | 51 | F51E | A | 2.6e-06 | 1.3e-06 | 2.6e-06 | 2.167E-06 | 7.506E-07 | 1.300E-06 | 2.600E-06 |
| 1007 | 51 | F51Q | A | 1.7e-06 | 1.9e-06 | 2.4e-06 | 2.000E-06 | 3.606E-07 | 1.700E-06 | 2.400E-06 |
| 1008 | 51 | F51G | A | 2.6e-06 | 1.9e-06 | 2.5e-06 | 2.333E-06 | 3.786E-07 | 1.900E-06 | 2.600E-06 |
| 1009 | 51 | F51H | A | 1.6e-06 | 2.5e-06 | 2.5e-06 | 2.200E-06 | 5.196E-07 | 1.600E-06 | 2.500E-06 |
| 1010 | 51 | F51I | A | 2.3e-06 | 3.2e-06 | 2.3e-06 | 2.600E-06 | 5.196E-07 | 2.300E-06 | 3.200E-06 |
| 1011 | 51 | F51L | A | 2.4e-06 | 2.5e-06 | 2.3e-06 | 2.400E-06 | 1.000E-07 | 2.300E-06 | 2.500E-06 |
| 1012 | 51 | F51K | A | 4.9e-06 | 4e-06 | 2.9e-06 | 3.933E-06 | 1.002E-06 | 2.900E-06 | 4.900E-06 |
| 1013 | 51 | F51M | A | 2.1e-06 | 1.2e-06 | 4.1e-06 | 2.467E-06 | 1.484E-06 | 1.200E-06 | 4.100E-06 |
| 1014 | 51 | F51F | A | 1.2e-06 | 1.5e-06 | 2.1e-06 | 1.600E-06 | 4.583E-07 | 1.200E-06 | 2.100E-06 |
| 1015 | 51 | F51P | A | 4e-06 | 3.1e-06 | 2.2e-06 | 3.100E-06 | 9.000E-07 | 2.200E-06 | 4.000E-06 |
| 1016 | 51 | F51S | A | 1.4e-06 | 2.5e-06 | 2.5e-06 | 2.133E-06 | 6.351E-07 | 1.400E-06 | 2.500E-06 |
| 1017 | 51 | F51T | A | 1.6e-06 | 1.5e-06 | 3.4e-06 | 2.167E-06 | 1.069E-06 | 1.500E-06 | 3.400E-06 |
| 1018 | 51 | F51W | A | 2.5e-06 | 2.5e-06 | 3.2e-06 | 2.733E-06 | 4.041E-07 | 2.500E-06 | 3.200E-06 |
| 1019 | 51 | F51Y | A | 3.1e-06 | 2.1e-06 | 3.6e-06 | 2.933E-06 | 7.638E-07 | 2.100E-06 | 3.600E-06 |

| | | | | | | | | | | |
|---|---|---|---|---|---|---|---|---|---|---|
| 1020 | 51 | F51V | A | 2.3e-06 | 1.2e-06 | 1.6e-06 | 1.700E-06 | 5.568E-07 | 1.200E-06 | 2.300E-06 |
| 1021 | 52 | L52A | A | 3e-06 | 2.2e-06 | 2.3e-06 | 2.500E-06 | 4.359E-07 | 2.200E-06 | 3.000E-06 |
| 1022 | 52 | L52R | A | 3.2e-06 | 2.3e-06 | 4.4e-06 | 3.300E-06 | 1.054E-06 | 2.300E-06 | 4.400E-06 |
| 1023 | 52 | L52N | A | 2.7e-06 | 2.4e-06 | 1.1e-06 | 2.067E-06 | 8.505E-07 | 1.100E-06 | 2.700E-06 |
| 1024 | 52 | L52D | A | 3.5e-06 | 2.6e-06 | 3.2e-06 | 3.100E-06 | 4.583E-07 | 2.600E-06 | 3.500E-06 |
| 1025 | 52 | L52C | A | 2.9e-06 | 1.6e-06 | 1.8e-06 | 2.100E-06 | 7.000E-07 | 1.600E-06 | 2.900E-06 |
| 1026 | 52 | L52E | A | 1.5e-06 | 2.4e-06 | 2.6e-06 | 2.167E-06 | 5.859E-07 | 1.500E-06 | 2.600E-06 |
| 1027 | 52 | L52Q | A | 2e-06 | 2.7e-06 | 3.4e-06 | 2.700E-06 | 7.000E-07 | 2.000E-06 | 3.400E-06 |
| 1028 | 52 | L52G | A | 2.2e-06 | 2.2e-06 | 5.4e-06 | 3.267E-06 | 1.848E-06 | 2.200E-06 | 5.400E-06 |
| 1029 | 52 | L52H | A | 3.9e-06 | 1.6e-06 | 2.7e-06 | 2.733E-06 | 1.150E-06 | 1.600E-06 | 3.900E-06 |
| 1030 | 52 | L52I | A | 2.2e-06 | 1.4e-06 | 2.5e-06 | 2.033E-06 | 5.686E-07 | 1.400E-06 | 2.500E-06 |
| 1031 | 52 | L52L | A | 4.2e-06 | 1.7e-06 | 3.8e-06 | 3.233E-06 | 1.343E-06 | 1.700E-06 | 4.200E-06 |
| 1032 | 52 | L52K | A | 1.5e-06 | 1.3e-06 | 1.6e-06 | 1.467E-06 | 1.528E-07 | 1.300E-06 | 1.600E-06 |
| 1033 | 52 | L52M | A | 2.1e-06 | 2.2e-06 | 3.8e-06 | 2.700E-06 | 9.539E-07 | 2.100E-06 | 3.800E-06 |
| 1034 | 52 | L52F | A | 2.8e-06 | 3.9e-06 | 2.5e-06 | 3.067E-06 | 7.371E-07 | 2.500E-06 | 3.900E-06 |
| 1035 | 52 | L52P | A | 1.6e-06 | 2.6e-06 | 2.5e-06 | 2.233E-06 | 5.508E-07 | 1.600E-06 | 2.600E-06 |
| 1036 | 52 | L52S | A | 2.6e-06 | 1.4e-06 | 1.8e-06 | 1.933E-06 | 6.110E-07 | 1.400E-06 | 2.600E-06 |
| 1037 | 52 | L52T | A | 2.7e-06 | 1.1e-06 | 1.7e-06 | 1.833E-06 | 8.083E-07 | 1.100E-06 | 2.700E-06 |
| 1038 | 52 | L52W | A | 2.5e-06 | 4.6e-06 | 2.8e-06 | 3.300E-06 | 1.136E-06 | 2.500E-06 | 4.600E-06 |
| 1039 | 52 | L52Y | A | 3.6e-06 | 2.5e-06 | 2.9e-06 | 3.000E-06 | 5.568E-07 | 2.500E-06 | 3.600E-06 |
| 1040 | 52 | L52V | A | 1.5e-06 | 2.6e-06 | 1.5e-06 | 1.867E-06 | 6.351E-07 | 1.500E-06 | 2.600E-06 |
| 1041 | 53 | Q53A | A | 4.1e-06 | 3.7e-06 | 3.9e-06 | 3.900E-06 | 2.000E-07 | 3.700E-06 | 4.100E-06 |
| 1042 | 53 | Q53R | A | 2.2e-06 | 2.4e-06 | 3.5e-06 | 2.700E-06 | 7.000E-07 | 2.200E-06 | 3.500E-06 |
| 1043 | 53 | Q53N | A | 2.7e-06 | 1.5e-06 | 2.6e-06 | 2.267E-06 | 6.658E-07 | 1.500E-06 | 2.700E-06 |
| 1044 | 53 | Q53D | A | 1.7e-06 | 1.4e-06 | 2.1e-06 | 1.733E-06 | 3.512E-07 | 1.400E-06 | 2.100E-06 |
| 1045 | 53 | Q53C | A | 1.7e-06 | 1.3e-06 | 3.4e-06 | 2.133E-06 | 1.115E-06 | 1.300E-06 | 3.400E-06 |
| 1046 | 53 | Q53E | A | 2.1e-06 | 2.8e-06 | 3.3e-06 | 2.733E-06 | 6.028E-07 | 2.100E-06 | 3.300E-06 |
| 1047 | 53 | Q53Q | A | 2.9e-06 | 3.3e-06 | 1.7e-06 | 2.633E-06 | 8.327E-07 | 1.700E-06 | 3.300E-06 |
| 1048 | 53 | Q53G | A | 1.2e-06 | 2e-06 | 2e-06 | 1.733E-06 | 4.619E-07 | 1.200E-06 | 2.000E-06 |
| 1049 | 53 | Q53H | A | 2e-06 | 1.8e-06 | 2.3e-06 | 2.033E-06 | 2.517E-07 | 1.800E-06 | 2.300E-06 |
| 1050 | 53 | Q53I | A | 1.3e-06 | 2.3e-06 | 2.8e-06 | 2.133E-06 | 7.638E-07 | 1.300E-06 | 2.800E-06 |
| 1051 | 53 | Q53L | A | 1.4e-06 | 1.7e-06 | 2e-06 | 1.700E-06 | 3.000E-07 | 1.400E-06 | 2.000E-06 |
| 1052 | 53 | Q53K | A | 1.7e-06 | 1.5e-06 | 2.4e-06 | 1.867E-06 | 4.726E-07 | 1.500E-06 | 2.400E-06 |
| 1053 | 53 | Q53M | A | 2.6e-06 | 1.9e-06 | 2.9e-06 | 2.467E-06 | 5.132E-07 | 1.900E-06 | 2.900E-06 |
| 1054 | 53 | Q53F | A | 2.7e-06 | 5.4e-06 | 1.8e-06 | 3.300E-06 | 1.873E-06 | 1.800E-06 | 5.400E-06 |
| 1055 | 53 | Q53P | A | 1.6e-06 | 1.7e-06 | 3.2e-06 | 2.167E-06 | 8.963E-07 | 1.600E-06 | 3.200E-06 |
| 1056 | 53 | Q53S | A | 3.3e-06 | 4.6e-06 | 2e-06 | 3.300E-06 | 1.300E-06 | 2.000E-06 | 4.600E-06 |
| 1057 | 53 | Q53T | A | 2.4e-06 | 1.8e-06 | 1.1e-06 | 1.767E-06 | 6.506E-07 | 1.100E-06 | 2.400E-06 |
| 1058 | 53 | Q53W | A | 2.9e-06 | 1.8e-06 | 3.6e-06 | 2.767E-06 | 9.074E-07 | 1.800E-06 | 3.600E-06 |
| 1059 | 53 | Q53Y | A | 2.1e-06 | 3.3e-06 | 2.3e-06 | 2.567E-06 | 6.429E-07 | 2.100E-06 | 3.300E-06 |
| 1060 | 53 | Q53V | A | 1.8e-06 | 2.8e-06 | 3.3e-06 | 2.633E-06 | 7.638E-07 | 1.800E-06 | 3.300E-06 |
| 1061 | 54 | D54A | A | 2.2e-06 | 2.1e-06 | 1.8e-06 | 2.033E-06 | 2.082E-07 | 1.800E-06 | 2.200E-06 |
| 1062 | 54 | D54R | A | 2.3e-06 | 2.9e-06 | 3e-06 | 2.733E-06 | 3.786E-07 | 2.300E-06 | 3.000E-06 |
| 1063 | 54 | D54N | A | 5.1e-06 | 1.9e-06 | 1.7e-06 | 2.900E-06 | 1.908E-06 | 1.700E-06 | 5.100E-06 |
| 1064 | 54 | D54D | A | 2.8e-06 | 2.4e-06 | 2.2e-06 | 2.467E-06 | 3.055E-07 | 2.200E-06 | 2.800E-06 |
| 1065 | 54 | D54C | A | 2.1e-06 | 2.6e-06 | 2.4e-06 | 2.367E-06 | 2.517E-07 | 2.100E-06 | 2.600E-06 |
| 1066 | 54 | D54E | A | 3e-06 | 3.2e-06 | 5.6e-06 | 3.933E-06 | 1.447E-06 | 3.000E-06 | 5.600E-06 |
| 1067 | 54 | D54Q | A | 2.3e-06 | 1.7e-06 | 2.5e-06 | 2.167E-06 | 4.163E-07 | 1.700E-06 | 2.500E-06 |
| 1068 | 54 | D54G | A | 3.1e-06 | 2.2e-06 | 2.5e-06 | 2.600E-06 | 4.583E-07 | 2.200E-06 | 3.100E-06 |
| 1069 | 54 | D54H | A | 2.2e-06 | 2e-06 | 4.5e-06 | 2.900E-06 | 1.389E-06 | 2.000E-06 | 4.500E-06 |
| 1070 | 54 | D54I | A | 1.8e-06 | 2.3e-06 | 2.2e-06 | 2.100E-06 | 2.646E-07 | 1.800E-06 | 2.300E-06 |
| 1071 | 54 | D54L | A | 4.5e-06 | 2.4e-06 | 2.3e-06 | 3.067E-06 | 1.242E-06 | 2.300E-06 | 4.500E-06 |
| 1072 | 54 | D54K | A | 1.7e-06 | 4.1e-06 | 1.9e-06 | 2.567E-06 | 1.332E-06 | 1.700E-06 | 4.100E-06 |
| 1073 | 54 | D54M | A | 1.8e-06 | 4.1e-06 | 2.8e-06 | 2.900E-06 | 1.153E-06 | 1.800E-06 | 4.100E-06 |
| 1074 | 54 | D54F | A | 1.9e-06 | 2.6e-06 | 9.5e-07 | 1.817E-06 | 8.282E-07 | 9.500E-07 | 2.600E-06 |
| 1075 | 54 | D54P | A | 1.4e-06 | 3.4e-06 | 2.4e-06 | 2.400E-06 | 1.000E-06 | 1.400E-06 | 3.400E-06 |
| 1076 | 54 | D54S | A | 1.8e-06 | 2.1e-06 | 2.2e-06 | 2.033E-06 | 2.082E-07 | 1.800E-06 | 2.200E-06 |
| 1077 | 54 | D54T | A | 3e-06 | 2e-06 | 3.3e-06 | 2.767E-06 | 6.807E-07 | 2.000E-06 | 3.300E-06 |
| 1078 | 54 | D54W | A | 2.3e-06 | 2.3e-06 | 2.2e-06 | 2.267E-06 | 5.774E-08 | 2.200E-06 | 2.300E-06 |
| 1079 | 54 | D54Y | A | 1.5e-06 | 2.2e-06 | 2.1e-06 | 1.933E-06 | 3.786E-07 | 1.500E-06 | 2.200E-06 |
| 1080 | 54 | D54V | A | 3.1e-06 | 3.4e-06 | 2.8e-06 | 3.100E-06 | 3.000E-07 | 2.800E-06 | 3.400E-06 |
| 1081 | 55 | I55A | A | 1.7e-06 | 1.9e-06 | 1.2e-06 | 1.600E-06 | 3.606E-07 | 1.200E-06 | 1.900E-06 |
| 1082 | 55 | I55R | A | 1.9e-06 | 2.8e-06 | 5.6e-06 | 3.433E-06 | 1.930E-06 | 1.900E-06 | 5.600E-06 |
| 1083 | 55 | I55N | A | 2.7e-06 | 5.8e-06 | 2.6e-06 | 3.700E-06 | 1.819E-06 | 2.600E-06 | 5.800E-06 |
| 1084 | 55 | I55D | A | 1.3e-06 | 3.1e-06 | 2.8e-06 | 2.400E-06 | 9.644E-07 | 1.300E-06 | 3.100E-06 |
| 1085 | 55 | I55C | A | 2.2e-06 | 2.1e-06 | 2.8e-06 | 2.367E-06 | 3.786E-07 | 2.100E-06 | 2.800E-06 |
| 1086 | 55 | I55E | A | 1.5e-06 | 1.9e-06 | 5.5e-06 | 2.967E-06 | 2.203E-06 | 1.500E-06 | 5.500E-06 |
| 1087 | 55 | I55Q | A | 1.8e-06 | 1.4e-06 | 3.1e-06 | 2.100E-06 | 8.888E-07 | 1.400E-06 | 3.100E-06 |
| 1088 | 55 | I55G | A | 2.6e-06 | 2.2e-06 | 2.1e-06 | 2.300E-06 | 2.646E-07 | 2.100E-06 | 2.600E-06 |
| 1089 | 55 | I55H | A | 3.5e-06 | 2.1e-06 | 3.5e-06 | 3.033E-06 | 8.083E-07 | 2.100E-06 | 3.500E-06 |
| 1090 | 55 | I55I | A | 6.8e-07 | 3.8e-06 | 1.9e-06 | 2.127E-06 | 1.572E-06 | 6.800E-07 | 3.800E-06 |
| 1091 | 55 | I55L | A | 2.6e-06 | 1.7e-06 | 2.2e-06 | 2.167E-06 | 4.509E-07 | 1.700E-06 | 2.600E-06 |
| 1092 | 55 | I55K | A | 2.2e-06 | 2.8e-06 | 2.3e-06 | 2.433E-06 | 3.215E-07 | 2.200E-06 | 2.800E-06 |
| 1093 | 55 | I55M | A | 1.2e-06 | 3e-06 | 1.9e-06 | 2.033E-06 | 9.074E-07 | 1.200E-06 | 3.000E-06 |
| 1094 | 55 | I55F | A | 1.5e-06 | 3.4e-06 | 2.3e-06 | 2.400E-06 | 9.539E-07 | 1.500E-06 | 3.400E-06 |
| 1095 | 55 | I55P | A | 4.1e-06 | 2.3e-06 | 3.5e-06 | 3.300E-06 | 9.165E-07 | 2.300E-06 | 4.100E-06 |
| 1096 | 55 | I55S | A | 2.1e-06 | 2.6e-06 | 1.9e-06 | 2.200E-06 | 3.606E-07 | 1.900E-06 | 2.600E-06 |
| 1097 | 55 | I55T | A | 1.4e-06 | 2.9e-06 | 2.2e-06 | 2.167E-06 | 7.506E-07 | 1.400E-06 | 2.900E-06 |
| 1098 | 55 | I55W | A | 2.5e-06 | 1.9e-06 | 2.2e-06 | 2.200E-06 | 3.000E-07 | 1.900E-06 | 2.500E-06 |
| 1099 | 55 | I55Y | A | 3e-06 | 1.2e-06 | 2.4e-06 | 2.200E-06 | 9.165E-07 | 1.200E-06 | 3.000E-06 |
| 1100 | 55 | I55V | A | 2.4e-06 | 1.5e-06 | 1.9e-06 | 1.933E-06 | 4.509E-07 | 1.500E-06 | 2.400E-06 |
| 1101 | 56 | Q56A | A | 3.4e-06 | 2.5e-06 | 2e-06 | 2.633E-06 | 7.095E-07 | 2.000E-06 | 3.400E-06 |
| 1102 | 56 | Q56R | A | 3e-06 | 2.3e-06 | 3.8e-06 | 3.033E-06 | 7.506E-07 | 2.300E-06 | 3.800E-06 |
| 1103 | 56 | Q56N | A | 2.1e-06 | 2.9e-06 | 2.3e-06 | 2.433E-06 | 4.163E-07 | 2.100E-06 | 2.900E-06 |
| 1104 | 56 | Q56D | A | 2e-06 | 4.5e-06 | 2.3e-06 | 2.933E-06 | 1.365E-06 | 2.000E-06 | 4.500E-06 |

| | | | | | | | | | | |
|---|---|---|---|---|---|---|---|---|---|---|
| 1105 | 56 | Q56C | A | 2.7e-06 | 1.8e-06 | 2.1e-06 | 2.200E-06 | 4.583E-07 | 1.800E-06 | 2.700E-06 |
| 1106 | 56 | Q56E | A | 2.3e-06 | 2.7e-06 | 1.7e-06 | 2.233E-06 | 5.033E-07 | 1.700E-06 | 2.700E-06 |
| 1107 | 56 | Q56Q | A | 1.6e-06 | 2e-06 | 3e-06 | 2.200E-06 | 7.211E-07 | 1.600E-06 | 3.000E-06 |
| 1108 | 56 | Q56G | A | 2.7e-06 | 2.7e-06 | 2.8e-06 | 2.733E-06 | 5.774E-08 | 2.700E-06 | 2.800E-06 |
| 1109 | 56 | Q56H | A | 1.7e-06 | 2.8e-06 | 3.3e-06 | 2.600E-06 | 8.185E-07 | 1.700E-06 | 3.300E-06 |
| 1110 | 56 | Q56I | A | 2.1e-06 | 1.7e-06 | 2.3e-06 | 2.033E-06 | 3.055E-07 | 1.700E-06 | 2.300E-06 |
| 1111 | 56 | Q56L | A | 2.3e-06 | 1.9e-06 | 3.1e-06 | 2.433E-06 | 6.110E-07 | 1.900E-06 | 3.100E-06 |
| 1112 | 56 | Q56K | A | 2.7e-06 | 2.6e-06 | 1.2e-06 | 2.167E-06 | 8.386E-07 | 1.200E-06 | 2.700E-06 |
| 1113 | 56 | Q56M | A | 2.5e-06 | 3.1e-06 | 1.3e-06 | 2.300E-06 | 9.165E-07 | 1.300E-06 | 3.100E-06 |
| 1114 | 56 | Q56F | A | 3.9e-06 | 3.8e-06 | 2e-06 | 3.233E-06 | 1.069E-06 | 2.000E-06 | 3.900E-06 |
| 1115 | 56 | Q56P | A | 2.6e-06 | 2.3e-06 | 2e-06 | 2.300E-06 | 3.000E-07 | 2.000E-06 | 2.600E-06 |
| 1116 | 56 | Q56S | A | 2.1e-06 | 4.9e-06 | 2.7e-06 | 3.233E-06 | 1.474E-06 | 2.100E-06 | 4.900E-06 |
| 1117 | 56 | Q56T | A | 1.6e-06 | 1.2e-06 | 1.1e-06 | 1.300E-06 | 2.646E-07 | 1.100E-06 | 1.600E-06 |
| 1118 | 56 | Q56W | A | 9.9e-07 | 1.3e-06 | 4.6e-06 | 2.297E-06 | 2.001E-06 | 9.900E-07 | 4.600E-06 |
| 1119 | 56 | Q56Y | A | 2.7e-06 | 3e-06 | 1.7e-06 | 2.467E-06 | 6.807E-07 | 1.700E-06 | 3.000E-06 |
| 1120 | 56 | Q56V | A | 1.6e-06 | 2.4e-06 | 1e-06 | 1.667E-06 | 7.024E-07 | 1.000E-06 | 2.400E-06 |
| 1121 | 57 | E57A | A | 3.2e-06 | 2.4e-06 | 1.9e-06 | 2.500E-06 | 6.557E-07 | 1.900E-06 | 3.200E-06 |
| 1122 | 57 | E57R | A | 2.3e-06 | 2.7e-06 | 3.3e-06 | 2.767E-06 | 5.033E-07 | 2.300E-06 | 3.300E-06 |
| 1123 | 57 | E57N | A | 2.1e-06 | 3.1e-06 | 2.9e-06 | 2.700E-06 | 5.292E-07 | 2.100E-06 | 3.100E-06 |
| 1124 | 57 | E57D | A | 1.2e-06 | 2e-06 | 1.8e-06 | 1.667E-06 | 4.163E-07 | 1.200E-06 | 2.000E-06 |
| 1125 | 57 | E57C | A | 1.8e-06 | 2.3e-06 | 1.5e-06 | 1.867E-06 | 4.041E-07 | 1.500E-06 | 2.300E-06 |
| 1126 | 57 | E57E | A | 1.8e-06 | 2.7e-06 | 1.4e-06 | 1.967E-06 | 6.658E-07 | 1.400E-06 | 2.700E-06 |
| 1127 | 57 | E57Q | A | 2.3e-06 | 1.9e-06 | 1.6e-06 | 1.933E-06 | 3.512E-07 | 1.600E-06 | 2.300E-06 |
| 1128 | 57 | E57G | A | 1.4e-06 | 3.3e-06 | 1.5e-06 | 2.067E-06 | 1.069E-06 | 1.400E-06 | 3.300E-06 |
| 1129 | 57 | E57H | A | 1.5e-06 | 2.8e-06 | 1.5e-06 | 1.933E-06 | 7.506E-07 | 1.500E-06 | 2.800E-06 |
| 1130 | 57 | E57I | A | 1.9e-06 | 2.7e-06 | 3.4e-06 | 2.667E-06 | 7.506E-07 | 1.900E-06 | 3.400E-06 |
| 1131 | 57 | E57L | A | 2.6e-06 | 2.8e-06 | 1.5e-06 | 2.300E-06 | 7.000E-07 | 1.500E-06 | 2.800E-06 |
| 1132 | 57 | E57K | A | 2.5e-06 | 4.2e-06 | 4.7e-06 | 3.800E-06 | 1.153E-06 | 2.500E-06 | 4.700E-06 |
| 1133 | 57 | E57M | A | 2.9e-06 | 1.5e-06 | 3.3e-06 | 2.567E-06 | 9.452E-07 | 1.500E-06 | 3.300E-06 |
| 1134 | 57 | E57F | A | 3.6e-06 | 3e-06 | 2.2e-06 | 2.933E-06 | 7.024E-07 | 2.200E-06 | 3.600E-06 |
| 1135 | 57 | E57P | A | 6.2e-07 | 3.8e-06 | 2.2e-06 | 2.207E-06 | 1.590E-06 | 6.200E-07 | 3.800E-06 |
| 1136 | 57 | E57S | A | 3.8e-06 | 6.5e-06 | 3.3e-06 | 4.533E-06 | 1.721E-06 | 3.300E-06 | 6.500E-06 |
| 1137 | 57 | E57T | A | 1.8e-06 | 2e-06 | 1.7e-06 | 1.833E-06 | 1.528E-07 | 1.700E-06 | 2.000E-06 |
| 1138 | 57 | E57W | A | 2.5e-06 | 3.2e-06 | 2e-06 | 2.567E-06 | 6.028E-07 | 2.000E-06 | 3.200E-06 |
| 1139 | 57 | E57Y | A | 2.3e-06 | 2.4e-06 | 3.2e-06 | 2.633E-06 | 4.933E-07 | 2.300E-06 | 3.200E-06 |
| 1140 | 57 | E57V | A | 2.5e-06 | 2.1e-06 | 3.1e-06 | 2.567E-06 | 5.033E-07 | 2.100E-06 | 3.100E-06 |
| 1141 | 58 | V58A | A | 1.7e-06 | 2.1e-06 | 3.6e-06 | 2.467E-06 | 1.002E-06 | 1.700E-06 | 3.600E-06 |
| 1142 | 58 | V58R | A | 3e-06 | 2.8e-06 | 2.3e-06 | 2.700E-06 | 3.606E-07 | 2.300E-06 | 3.000E-06 |
| 1143 | 58 | V58N | A | 3e-06 | 1.5e-06 | 2.3e-06 | 2.267E-06 | 7.506E-07 | 1.500E-06 | 3.000E-06 |
| 1144 | 58 | V58D | A | 3.2e-06 | 3.5e-06 | 2.5e-06 | 3.067E-06 | 5.132E-07 | 2.500E-06 | 3.500E-06 |
| 1145 | 58 | V58C | A | 2.6e-06 | 2.8e-06 | 2.4e-06 | 2.600E-06 | 2.000E-07 | 2.400E-06 | 2.800E-06 |
| 1146 | 58 | V58E | A | 2.1e-06 | 1.9e-06 | 3.7e-06 | 2.567E-06 | 9.866E-07 | 1.900E-06 | 3.700E-06 |
| 1147 | 58 | V58Q | A | 1.4e-06 | 1.8e-06 | 6.5e-06 | 3.233E-06 | 2.836E-06 | 1.400E-06 | 6.500E-06 |
| 1148 | 58 | V58G | A | 2e-06 | 2e-06 | 2.3e-06 | 2.100E-06 | 1.732E-07 | 2.000E-06 | 2.300E-06 |
| 1149 | 58 | V58H | A | 1.9e-06 | 2.2e-06 | 2.8e-06 | 2.300E-06 | 4.583E-07 | 1.900E-06 | 2.800E-06 |
| 1150 | 58 | V58I | A | 2.5e-06 | 2.9e-06 | 2.2e-06 | 2.533E-06 | 3.512E-07 | 2.200E-06 | 2.900E-06 |
| 1151 | 58 | V58L | A | 3.7e-06 | 1.2e-06 | 2.2e-06 | 2.367E-06 | 1.258E-06 | 1.200E-06 | 3.700E-06 |
| 1152 | 58 | V58K | A | 2.1e-06 | 1.5e-06 | 2.9e-06 | 2.167E-06 | 7.024E-07 | 1.500E-06 | 2.900E-06 |
| 1153 | 58 | V58M | A | 2.7e-06 | 3.1e-06 | 2.1e-06 | 2.633E-06 | 5.033E-07 | 2.100E-06 | 3.100E-06 |
| 1154 | 58 | V58F | A | 3.3e-06 | 3.1e-06 | 1.8e-06 | 2.733E-06 | 8.145E-07 | 1.800E-06 | 3.300E-06 |
| 1155 | 58 | V58P | A | 2e-06 | 2.3e-06 | 2.7e-06 | 2.333E-06 | 3.512E-07 | 2.000E-06 | 2.700E-06 |
| 1156 | 58 | V58S | A | 2.5e-06 | 2.1e-06 | 2.8e-06 | 2.467E-06 | 3.512E-07 | 2.100E-06 | 2.800E-06 |
| 1157 | 58 | V58T | A | 3.1e-06 | 3.5e-06 | 2.1e-06 | 2.900E-06 | 7.211E-07 | 2.100E-06 | 3.500E-06 |
| 1158 | 58 | V58W | A | 2.7e-06 | 1.4e-06 | 2.1e-06 | 2.067E-06 | 6.506E-07 | 1.400E-06 | 2.700E-06 |
| 1159 | 58 | V58Y | A | 8.3e-07 | 1.6e-06 | 1.6e-06 | 1.343E-06 | 4.446E-07 | 8.300E-07 | 1.600E-06 |
| 1160 | 58 | V58V | A | 2.2e-06 | 2.6e-06 | 2.6e-06 | 2.467E-06 | 2.309E-07 | 2.200E-06 | 2.600E-06 |
| 1161 | 59 | Q59A | A | 1.8e-06 | 3.3e-06 | 1.9e-06 | 2.333E-06 | 8.386E-07 | 1.800E-06 | 3.300E-06 |
| 1162 | 59 | Q59R | A | 4.4e-06 | 1.8e-06 | 1.4e-06 | 2.533E-06 | 1.629E-06 | 1.400E-06 | 4.400E-06 |
| 1163 | 59 | Q59N | A | 2.9e-06 | 1.9e-06 | 3.4e-06 | 2.733E-06 | 7.638E-07 | 1.900E-06 | 3.400E-06 |
| 1164 | 59 | Q59D | A | 2.1e-06 | 1.4e-06 | 1.6e-06 | 1.700E-06 | 3.606E-07 | 1.400E-06 | 2.100E-06 |
| 1165 | 59 | Q59C | A | 1.4e-06 | 2.2e-06 | 1.6e-06 | 1.733E-06 | 4.163E-07 | 1.400E-06 | 2.200E-06 |
| 1166 | 59 | Q59E | A | 4.1e-06 | 2.5e-06 | 4.2e-06 | 3.600E-06 | 9.539E-07 | 2.500E-06 | 4.200E-06 |
| 1167 | 59 | Q59Q | A | 2.4e-06 | 3.1e-06 | 2.7e-06 | 2.733E-06 | 3.512E-07 | 2.400E-06 | 3.100E-06 |
| 1168 | 59 | Q59G | A | 2.5e-06 | 1.6e-06 | 1.8e-06 | 1.967E-06 | 4.726E-07 | 1.600E-06 | 2.500E-06 |
| 1169 | 59 | Q59H | A | 1.7e-06 | 3.2e-06 | 3e-06 | 2.633E-06 | 8.145E-07 | 1.700E-06 | 3.200E-06 |
| 1170 | 59 | Q59I | A | 2.4e-06 | 1.4e-06 | 1.2e-06 | 1.667E-06 | 6.429E-07 | 1.200E-06 | 2.400E-06 |
| 1171 | 59 | Q59L | A | 2.4e-06 | 2.7e-06 | 2.4e-06 | 2.500E-06 | 1.732E-07 | 2.400E-06 | 2.700E-06 |
| 1172 | 59 | Q59K | A | 2.6e-06 | 2.3e-06 | 2e-06 | 2.300E-06 | 3.000E-07 | 2.000E-06 | 2.600E-06 |
| 1173 | 59 | Q59M | A | 2.7e-06 | 2.1e-06 | 4e-06 | 2.933E-06 | 9.713E-07 | 2.100E-06 | 4.000E-06 |
| 1174 | 59 | Q59F | A | 1.4e-06 | 2.4e-06 | 3.3e-06 | 2.367E-06 | 9.504E-07 | 1.400E-06 | 3.300E-06 |
| 1175 | 59 | Q59P | A | 1.7e-06 | 4e-06 | 1.4e-06 | 2.367E-06 | 1.422E-06 | 1.400E-06 | 4.000E-06 |
| 1176 | 59 | Q59S | A | 3.5e-06 | 2.2e-06 | 2.1e-06 | 2.600E-06 | 7.810E-07 | 2.100E-06 | 3.500E-06 |
| 1177 | 59 | Q59T | A | 2.4e-06 | 2.6e-06 | 1.2e-06 | 2.067E-06 | 7.572E-07 | 1.200E-06 | 2.600E-06 |
| 1178 | 59 | Q59W | A | 2.9e-06 | 1.5e-06 | 2.6e-06 | 2.333E-06 | 7.371E-07 | 1.500E-06 | 2.900E-06 |
| 1179 | 59 | Q59Y | A | 3.7e-06 | 2.8e-06 | 3.5e-06 | 3.333E-06 | 4.726E-07 | 2.800E-06 | 3.700E-06 |
| 1180 | 59 | Q59V | A | 1.6e-06 | 3.6e-06 | 3.9e-06 | 3.033E-06 | 1.250E-06 | 1.600E-06 | 3.900E-06 |
| 1181 | 60 | G60A | A | 2.2e-06 | 1.8e-06 | 3.4e-06 | 2.467E-06 | 8.327E-07 | 1.800E-06 | 3.400E-06 |
| 1182 | 60 | G60R | A | 3.4e-06 | 1.7e-06 | 1.1e-06 | 2.067E-06 | 1.193E-06 | 1.100E-06 | 3.400E-06 |
| 1183 | 60 | G60N | A | 1.7e-06 | 4.2e-06 | 3.9e-06 | 3.267E-06 | 1.365E-06 | 1.700E-06 | 4.200E-06 |
| 1184 | 60 | G60D | A | 3.1e-06 | 2e-06 | 2.7e-06 | 2.600E-06 | 5.568E-07 | 2.000E-06 | 3.100E-06 |
| 1185 | 60 | G60C | A | 1.7e-06 | 3.9e-06 | 2.3e-06 | 2.633E-06 | 1.137E-06 | 1.700E-06 | 3.900E-06 |
| 1186 | 60 | G60E | A | 1.5e-06 | 2.7e-06 | 2.4e-06 | 2.200E-06 | 6.245E-07 | 1.500E-06 | 2.700E-06 |
| 1187 | 60 | G60Q | A | 2.2e-06 | 2.7e-06 | 2e-06 | 2.300E-06 | 3.606E-07 | 2.000E-06 | 2.700E-06 |
| 1188 | 60 | G60G | A | 1.2e-06 | 1.7e-06 | 3.9e-06 | 2.267E-06 | 1.436E-06 | 1.200E-06 | 3.900E-06 |
| 1189 | 60 | G60H | A | 2.7e-06 | 2.3e-06 | 2.9e-06 | 2.633E-06 | 3.055E-07 | 2.300E-06 | 2.900E-06 |

| | | | | | | | | | | |
|---|---|---|---|---|---|---|---|---|---|---|
| 1190 | 60 | G60I | A | 1.8e-06 | 3.1e-06 | 3.3e-06 | 2.733E-06 | 8.145E-07 | 1.800E-06 | 3.300E-06 |
| 1191 | 60 | G60L | A | 2e-06 | 2.2e-06 | 2.1e-06 | 2.100E-06 | 1.000E-07 | 2.000E-06 | 2.200E-06 |
| 1192 | 60 | G60K | A | 3e-06 | 4.2e-06 | 2.1e-06 | 3.100E-06 | 1.054E-06 | 2.100E-06 | 4.200E-06 |
| 1193 | 60 | G60M | A | 2.4e-06 | 2.7e-06 | 4.4e-06 | 3.167E-06 | 1.079E-06 | 2.400E-06 | 4.400E-06 |
| 1194 | 60 | G60F | A | 2.7e-06 | 2e-06 | 1.3e-06 | 2.000E-06 | 7.000E-07 | 1.300E-06 | 2.700E-06 |
| 1195 | 60 | G60P | A | 3.3e-06 | 3.3e-06 | 2e-06 | 2.867E-06 | 7.506E-07 | 2.000E-06 | 3.300E-06 |
| 1196 | 60 | G60S | A | 1.6e-06 | 2.1e-06 | 1.7e-06 | 1.800E-06 | 2.646E-07 | 1.600E-06 | 2.100E-06 |
| 1197 | 60 | G60T | A | 2.6e-06 | 2e-06 | 9.6e-07 | 1.853E-06 | 8.298E-07 | 9.600E-07 | 2.600E-06 |
| 1198 | 60 | G60W | A | 1.3e-06 | 2.8e-06 | 2.9e-06 | 2.333E-06 | 8.963E-07 | 1.300E-06 | 2.900E-06 |
| 1199 | 60 | G60Y | A | 2.7e-06 | 2e-06 | 3.7e-06 | 2.800E-06 | 8.544E-07 | 2.000E-06 | 3.700E-06 |
| 1200 | 60 | G60V | A | 2.8e-06 | 2.3e-06 | 3.1e-06 | 2.733E-06 | 4.041E-07 | 2.300E-06 | 3.100E-06 |
| 1201 | 61 | Y61A | A | 2.6e-06 | 3.1e-06 | 1.5e-06 | 2.400E-06 | 8.185E-07 | 1.500E-06 | 3.100E-06 |
| 1202 | 61 | Y61R | A | 3e-06 | 3.1e-06 | 2.4e-06 | 2.833E-06 | 3.786E-07 | 2.400E-06 | 3.100E-06 |
| 1203 | 61 | Y61N | A | 1.7e-06 | 2.2e-06 | 1.8e-06 | 1.900E-06 | 2.646E-07 | 1.700E-06 | 2.200E-06 |
| 1204 | 61 | Y61D | A | 2.2e-06 | 2.6e-06 | 3e-06 | 2.600E-06 | 4.000E-07 | 2.200E-06 | 3.000E-06 |
| 1205 | 61 | Y61C | A | 4.8e-06 | 1.4e-06 | 3.3e-06 | 3.167E-06 | 1.704E-06 | 1.400E-06 | 4.800E-06 |
| 1206 | 61 | Y61E | A | 1.9e-06 | 3.7e-06 | 2.4e-06 | 2.667E-06 | 9.292E-07 | 1.900E-06 | 3.700E-06 |
| 1207 | 61 | Y61Q | A | 1.9e-06 | 2.4e-06 | 2.3e-06 | 2.200E-06 | 2.646E-07 | 1.900E-06 | 2.400E-06 |
| 1208 | 61 | Y61G | A | 2.1e-06 | 1.9e-06 | 2.1e-06 | 2.033E-06 | 1.155E-07 | 1.900E-06 | 2.100E-06 |
| 1209 | 61 | Y61H | A | 2.4e-06 | 2.8e-06 | 3.2e-06 | 2.800E-06 | 4.000E-07 | 2.400E-06 | 3.200E-06 |
| 1210 | 61 | Y61I | A | 1.8e-06 | 3.7e-06 | 2.4e-06 | 2.633E-06 | 9.713E-07 | 1.800E-06 | 3.700E-06 |
| 1211 | 61 | Y61L | A | 3.5e-06 | 1.7e-06 | 2.6e-06 | 2.600E-06 | 9.000E-07 | 1.700E-06 | 3.500E-06 |
| 1212 | 61 | Y61K | A | 1.9e-06 | 2.7e-06 | 3.6e-06 | 2.733E-06 | 8.505E-07 | 1.900E-06 | 3.600E-06 |
| 1213 | 61 | Y61M | A | 1.1e-06 | 3.3e-06 | 1.4e-06 | 1.933E-06 | 1.193E-06 | 1.100E-06 | 3.300E-06 |
| 1214 | 61 | Y61F | A | 2.4e-06 | 2.5e-06 | 2.1e-06 | 2.333E-06 | 2.082E-07 | 2.100E-06 | 2.500E-06 |
| 1215 | 61 | Y61P | A | 2e-06 | 2.5e-06 | 1.7e-06 | 2.067E-06 | 4.041E-07 | 1.700E-06 | 2.500E-06 |
| 1216 | 61 | Y61S | A | 1.3e-06 | 2.5e-06 | 2.3e-06 | 2.033E-06 | 6.429E-07 | 1.300E-06 | 2.500E-06 |
| 1217 | 61 | Y61T | A | 2.4e-06 | 2.1e-06 | 2.5e-06 | 2.333E-06 | 2.082E-07 | 2.100E-06 | 2.500E-06 |
| 1218 | 61 | Y61W | A | 3.4e-06 | 2.1e-06 | 1.9e-06 | 2.467E-06 | 8.145E-07 | 1.900E-06 | 3.400E-06 |
| 1219 | 61 | Y61Y | A | 1.2e-06 | 2.6e-06 | 4.5e-06 | 2.767E-06 | 1.656E-06 | 1.200E-06 | 4.500E-06 |
| 1220 | 61 | Y61V | A | 2e-06 | 1.1e-06 | 1.8e-06 | 1.633E-06 | 4.726E-07 | 1.100E-06 | 2.000E-06 |
| 1221 | 62 | V62A | A | 1.9e-06 | 2.1e-06 | 3.5e-06 | 2.500E-06 | 8.718E-07 | 1.900E-06 | 3.500E-06 |
| 1222 | 62 | V62R | A | 1.5e-06 | 2.7e-06 | 1.2e-06 | 1.800E-06 | 7.937E-07 | 1.200E-06 | 2.700E-06 |
| 1223 | 62 | V62N | A | 2.7e-06 | 1.9e-06 | 1.7e-06 | 2.100E-06 | 5.292E-07 | 1.700E-06 | 2.700E-06 |
| 1224 | 62 | V62D | A | 3.9e-06 | 1.7e-06 | 1.9e-06 | 2.500E-06 | 1.217E-06 | 1.700E-06 | 3.900E-06 |
| 1225 | 62 | V62C | A | 1.7e-06 | 2.5e-06 | 2.6e-06 | 2.267E-06 | 4.933E-07 | 1.700E-06 | 2.600E-06 |
| 1226 | 62 | V62E | A | 5.3e-06 | 2.7e-06 | 1.3e-06 | 3.100E-06 | 2.030E-06 | 1.300E-06 | 5.300E-06 |
| 1227 | 62 | V62Q | A | 3.2e-06 | 1.9e-06 | 2.3e-06 | 2.467E-06 | 6.658E-07 | 1.900E-06 | 3.200E-06 |
| 1228 | 62 | V62G | A | 3.1e-06 | 3.1e-06 | 2.8e-06 | 3.000E-06 | 1.732E-07 | 2.800E-06 | 3.100E-06 |
| 1229 | 62 | V62H | A | 1.3e-06 | 2.5e-06 | 2.7e-06 | 2.167E-06 | 7.572E-07 | 1.300E-06 | 2.700E-06 |
| 1230 | 62 | V62I | A | 2.1e-06 | 2e-06 | 2.1e-06 | 2.067E-06 | 5.774E-08 | 2.000E-06 | 2.100E-06 |
| 1231 | 62 | V62L | A | 4.9e-06 | 3.3e-06 | 2.4e-06 | 3.533E-06 | 1.266E-06 | 2.400E-06 | 4.900E-06 |
| 1232 | 62 | V62K | A | 1.6e-06 | 1.5e-06 | 2e-06 | 1.700E-06 | 2.646E-07 | 1.500E-06 | 2.000E-06 |
| 1233 | 62 | V62M | A | 2.4e-06 | 2e-06 | 2.9e-06 | 2.433E-06 | 4.509E-07 | 2.000E-06 | 2.900E-06 |
| 1234 | 62 | V62F | A | 1.5e-06 | 2.3e-06 | 2.8e-06 | 2.200E-06 | 6.557E-07 | 1.500E-06 | 2.800E-06 |
| 1235 | 62 | V62P | A | 2.9e-06 | 2.6e-06 | 2e-06 | 2.500E-06 | 4.583E-07 | 2.000E-06 | 2.900E-06 |
| 1236 | 62 | V62S | A | 2.5e-06 | 2.3e-06 | 2.5e-06 | 2.433E-06 | 1.155E-07 | 2.300E-06 | 2.500E-06 |
| 1237 | 62 | V62T | A | 1.5e-06 | 2.4e-06 | 2e-06 | 1.967E-06 | 4.509E-07 | 1.500E-06 | 2.400E-06 |
| 1238 | 62 | V62W | A | 1.3e-06 | 2.2e-06 | 2e-06 | 1.833E-06 | 4.726E-07 | 1.300E-06 | 2.200E-06 |
| 1239 | 62 | V62Y | A | 2.5e-06 | 2.6e-06 | 1.7e-06 | 2.267E-06 | 4.933E-07 | 1.700E-06 | 2.600E-06 |
| 1240 | 62 | V62V | A | 1.5e-06 | 2.1e-06 | 2.2e-06 | 1.933E-06 | 3.786E-07 | 1.500E-06 | 2.200E-06 |
| 1241 | 63 | L63A | A | 4.8e-06 | 2.6e-06 | 3.1e-06 | 3.500E-06 | 1.153E-06 | 2.600E-06 | 4.800E-06 |
| 1242 | 63 | L63R | A | 2.1e-06 | 1.9e-06 | 2.4e-06 | 2.133E-06 | 2.517E-07 | 1.900E-06 | 2.400E-06 |
| 1243 | 63 | L63N | A | 1.6e-06 | 3.1e-06 | 2e-06 | 2.233E-06 | 7.767E-07 | 1.600E-06 | 3.100E-06 |
| 1244 | 63 | L63D | A | 3.4e-06 | 9.2e-07 | 3.7e-06 | 2.673E-06 | 1.526E-06 | 9.200E-07 | 3.700E-06 |
| 1245 | 63 | L63C | A | 2.8e-06 | 1.8e-06 | 1.9e-06 | 2.167E-06 | 5.508E-07 | 1.800E-06 | 2.800E-06 |
| 1246 | 63 | L63E | A | 2.5e-06 | 2.5e-06 | 2.1e-06 | 2.367E-06 | 2.309E-07 | 2.100E-06 | 2.500E-06 |
| 1247 | 63 | L63Q | A | 3.8e-06 | 2.1e-06 | 1.8e-06 | 2.567E-06 | 1.079E-06 | 1.800E-06 | 3.800E-06 |
| 1248 | 63 | L63G | A | 3.3e-06 | 1.6e-06 | 2.6e-06 | 2.500E-06 | 8.544E-07 | 1.600E-06 | 3.300E-06 |
| 1249 | 63 | L63H | A | 2.1e-06 | 2.1e-06 | 2.6e-06 | 2.267E-06 | 2.887E-07 | 2.100E-06 | 2.600E-06 |
| 1250 | 63 | L63I | A | 2.6e-06 | 2.4e-06 | 1.9e-06 | 2.300E-06 | 3.606E-07 | 1.900E-06 | 2.600E-06 |
| 1251 | 63 | L63L | A | 2.9e-06 | 1.6e-06 | 2.9e-06 | 2.467E-06 | 7.506E-07 | 1.600E-06 | 2.900E-06 |
| 1252 | 63 | L63K | A | 2.2e-06 | 1.9e-06 | 1.7e-06 | 1.933E-06 | 2.517E-07 | 1.700E-06 | 2.200E-06 |
| 1253 | 63 | L63M | A | 3.3e-06 | 2.7e-06 | 2e-06 | 2.667E-06 | 6.506E-07 | 2.000E-06 | 3.300E-06 |
| 1254 | 63 | L63F | A | 1.5e-06 | 2.7e-06 | 2.1e-06 | 2.100E-06 | 6.000E-07 | 1.500E-06 | 2.700E-06 |
| 1255 | 63 | L63P | A | 2.6e-06 | 2.9e-06 | 3e-06 | 2.833E-06 | 2.082E-07 | 2.600E-06 | 3.000E-06 |
| 1256 | 63 | L63S | A | 2.6e-06 | 2.3e-06 | 4.2e-06 | 3.033E-06 | 1.021E-06 | 2.300E-06 | 4.200E-06 |
| 1257 | 63 | L63T | A | 3e-06 | 1.8e-06 | 2.2e-06 | 2.333E-06 | 6.110E-07 | 1.800E-06 | 3.000E-06 |
| 1258 | 63 | L63W | A | 2.8e-06 | 3.2e-06 | 2.4e-06 | 2.800E-06 | 4.000E-07 | 2.400E-06 | 3.200E-06 |
| 1259 | 63 | L63Y | A | 2.1e-06 | 2.9e-06 | 2e-06 | 2.333E-06 | 4.933E-07 | 2.000E-06 | 2.900E-06 |
| 1260 | 63 | L63V | A | 3.5e-06 | 2e-06 | 1.8e-06 | 2.433E-06 | 9.292E-07 | 1.800E-06 | 3.500E-06 |
| 1261 | 64 | I64A | A | 1.8e-06 | 2.7e-06 | 2.9e-06 | 2.467E-06 | 5.859E-07 | 1.800E-06 | 2.900E-06 |
| 1262 | 64 | I64R | A | 1.8e-06 | 2.2e-06 | 1.5e-06 | 1.833E-06 | 3.512E-07 | 1.500E-06 | 2.200E-06 |
| 1263 | 64 | I64N | A | 3.2e-06 | 2.2e-06 | 1.8e-06 | 2.400E-06 | 7.211E-07 | 1.800E-06 | 3.200E-06 |
| 1264 | 64 | I64D | A | 2e-06 | 1.2e-06 | 1.9e-06 | 1.700E-06 | 4.359E-07 | 1.200E-06 | 2.000E-06 |
| 1265 | 64 | I64C | A | 2.2e-06 | 2.4e-06 | 3e-06 | 2.533E-06 | 4.163E-07 | 2.200E-06 | 3.000E-06 |
| 1266 | 64 | I64E | A | 2e-06 | 4.8e-06 | 2.3e-06 | 3.033E-06 | 1.537E-06 | 2.000E-06 | 4.800E-06 |
| 1267 | 64 | I64Q | A | 1.9e-06 | 2.6e-06 | 1.7e-06 | 2.067E-06 | 4.726E-07 | 1.700E-06 | 2.600E-06 |
| 1268 | 64 | I64G | A | 2.6e-06 | 1.9e-06 | 4e-06 | 2.833E-06 | 1.069E-06 | 1.900E-06 | 4.000E-06 |
| 1269 | 64 | I64H | A | 2.7e-06 | 1.6e-06 | 2.5e-06 | 2.267E-06 | 5.859E-07 | 1.600E-06 | 2.700E-06 |
| 1270 | 64 | I64I | A | 3.3e-06 | 1.9e-06 | 3.3e-06 | 2.833E-06 | 8.083E-07 | 1.900E-06 | 3.300E-06 |
| 1271 | 64 | I64L | A | 2.4e-06 | 2.1e-06 | 2.3e-06 | 2.267E-06 | 1.528E-07 | 2.100E-06 | 2.400E-06 |
| 1272 | 64 | I64K | A | 1.5e-06 | 2.9e-06 | 2.3e-06 | 2.233E-06 | 7.024E-07 | 1.500E-06 | 2.900E-06 |
| 1273 | 64 | I64M | A | 2.9e-06 | 3.7e-06 | 2.7e-06 | 3.100E-06 | 5.292E-07 | 2.700E-06 | 3.700E-06 |
| 1274 | 64 | I64F | A | 1.8e-06 | 2.5e-06 | 2.4e-06 | 2.233E-06 | 3.786E-07 | 1.800E-06 | 2.500E-06 |

| | | | | | | | | | | |
|---|---|---|---|---|---|---|---|---|---|---|
| 1275 | 64 | I64P | A | 1.7e-06 | 2.1e-06 | 4.5e-06 | 2.767E-06 | 1.514E-06 | 1.700E-06 | 4.500E-06 |
| 1276 | 64 | I64S | A | 2.5e-06 | 4.7e-06 | 1.2e-06 | 2.800E-06 | 1.769E-06 | 1.200E-06 | 4.700E-06 |
| 1277 | 64 | I64T | A | 3.1e-06 | 2.1e-06 | 3.6e-06 | 2.933E-06 | 7.638E-07 | 2.100E-06 | 3.600E-06 |
| 1278 | 64 | I64W | A | 2.4e-06 | 1.9e-06 | 1.5e-06 | 1.933E-06 | 4.509E-07 | 1.500E-06 | 2.400E-06 |
| 1279 | 64 | I64Y | A | 1.5e-06 | 1.8e-06 | 1.2e-06 | 1.500E-06 | 3.000E-07 | 1.200E-06 | 1.800E-06 |
| 1280 | 64 | I64V | A | 1.7e-06 | 2e-06 | 2.8e-06 | 2.167E-06 | 5.686E-07 | 1.700E-06 | 2.800E-06 |
| 1281 | 65 | A65A | A | 2.6e-06 | 2.1e-06 | 2.6e-06 | 2.433E-06 | 2.887E-07 | 2.100E-06 | 2.600E-06 |
| 1282 | 65 | A65R | A | 2.7e-06 | 3.3e-06 | 2.1e-06 | 2.700E-06 | 6.000E-07 | 2.100E-06 | 3.300E-06 |
| 1283 | 65 | A65N | A | 1.4e-06 | 3e-06 | 2.8e-06 | 2.400E-06 | 8.718E-07 | 1.400E-06 | 3.000E-06 |
| 1284 | 65 | A65D | A | 1.5e-06 | 2.1e-06 | 3.5e-06 | 2.367E-06 | 1.026E-06 | 1.500E-06 | 3.500E-06 |
| 1285 | 65 | A65C | A | 2.2e-06 | 2.1e-06 | 2.5e-06 | 2.267E-06 | 2.082E-07 | 2.100E-06 | 2.500E-06 |
| 1286 | 65 | A65E | A | 3.1e-06 | 2.6e-06 | 2.2e-06 | 2.633E-06 | 4.509E-07 | 2.200E-06 | 3.100E-06 |
| 1287 | 65 | A65Q | A | 3.9e-06 | 2.1e-06 | 2.4e-06 | 2.800E-06 | 9.644E-07 | 2.100E-06 | 3.900E-06 |
| 1288 | 65 | A65G | A | 2.2e-06 | 1.8e-06 | 2.6e-06 | 2.200E-06 | 4.000E-07 | 1.800E-06 | 2.600E-06 |
| 1289 | 65 | A65H | A | 2.2e-06 | 1.6e-06 | 3.9e-06 | 2.567E-06 | 1.193E-06 | 1.600E-06 | 3.900E-06 |
| 1290 | 65 | A65I | A | 1.9e-06 | 1.6e-06 | 1e-06 | 1.500E-06 | 4.583E-07 | 1.000E-06 | 1.900E-06 |
| 1291 | 65 | A65L | A | 2e-06 | 2.8e-06 | 2.3e-06 | 2.367E-06 | 4.041E-07 | 2.000E-06 | 2.800E-06 |
| 1292 | 65 | A65K | A | 3.3e-06 | 2.6e-06 | 2.2e-06 | 2.700E-06 | 5.568E-07 | 2.200E-06 | 3.300E-06 |
| 1293 | 65 | A65M | A | 1e-06 | 2.1e-06 | 3.3e-06 | 2.133E-06 | 1.150E-06 | 1.000E-06 | 3.300E-06 |
| 1294 | 65 | A65F | A | 1.8e-06 | 1.9e-06 | 2.2e-06 | 1.967E-06 | 2.082E-07 | 1.800E-06 | 2.200E-06 |
| 1295 | 65 | A65P | A | 2.2e-06 | 1.8e-06 | 3e-06 | 2.333E-06 | 6.110E-07 | 1.800E-06 | 3.000E-06 |
| 1296 | 65 | A65S | A | 1.7e-06 | 2.8e-06 | 2.2e-06 | 2.233E-06 | 5.508E-07 | 1.700E-06 | 2.800E-06 |
| 1297 | 65 | A65T | A | 2.1e-06 | 1.1e-06 | 1.5e-06 | 1.567E-06 | 5.033E-07 | 1.100E-06 | 2.100E-06 |
| 1298 | 65 | A65W | A | 3.9e-06 | 1e-06 | 1.6e-06 | 2.167E-06 | 1.531E-06 | 1.000E-06 | 3.900E-06 |
| 1299 | 65 | A65Y | A | 2.4e-06 | 2.5e-06 | 1.8e-06 | 2.233E-06 | 3.786E-07 | 1.800E-06 | 2.500E-06 |
| 1300 | 65 | A65V | A | 2.9e-06 | 2e-06 | 3.4e-06 | 2.767E-06 | 7.095E-07 | 2.000E-06 | 3.400E-06 |
| 1301 | 66 | H66A | A | 2.7e-06 | 2.7e-06 | 1.9e-06 | 2.433E-06 | 4.619E-07 | 1.900E-06 | 2.700E-06 |
| 1302 | 66 | H66R | A | 2.4e-06 | 2.8e-06 | 1.4e-06 | 2.200E-06 | 7.211E-07 | 1.400E-06 | 2.800E-06 |
| 1303 | 66 | H66N | A | 2e-06 | 3.6e-06 | 1.4e-06 | 2.333E-06 | 1.137E-06 | 1.400E-06 | 3.600E-06 |
| 1304 | 66 | H66D | A | 2e-06 | 1.8e-06 | 2.3e-06 | 2.033E-06 | 2.517E-07 | 1.800E-06 | 2.300E-06 |
| 1305 | 66 | H66C | A | 2e-06 | 3.3e-06 | 2.2e-06 | 2.500E-06 | 7.000E-07 | 2.000E-06 | 3.300E-06 |
| 1306 | 66 | H66E | A | 1.7e-06 | 5.1e-06 | 2.5e-06 | 3.100E-06 | 1.778E-06 | 1.700E-06 | 5.100E-06 |
| 1307 | 66 | H66Q | A | 3.1e-06 | 3.5e-06 | 1.9e-06 | 2.833E-06 | 8.327E-07 | 1.900E-06 | 3.500E-06 |
| 1308 | 66 | H66G | A | 2.7e-06 | 2.2e-06 | 2e-06 | 2.300E-06 | 3.606E-07 | 2.000E-06 | 2.700E-06 |
| 1309 | 66 | H66H | A | 2.3e-06 | 4.2e-06 | 2.5e-06 | 3.000E-06 | 1.044E-06 | 2.300E-06 | 4.200E-06 |
| 1310 | 66 | H66I | A | 2.9e-06 | 2.6e-06 | 2.6e-06 | 2.700E-06 | 1.732E-07 | 2.600E-06 | 2.900E-06 |
| 1311 | 66 | H66L | A | 3e-06 | 1.1e-06 | 2.5e-06 | 2.200E-06 | 9.849E-07 | 1.100E-06 | 3.000E-06 |
| 1312 | 66 | H66K | A | 3.3e-06 | 3.1e-06 | 2.3e-06 | 2.900E-06 | 5.292E-07 | 2.300E-06 | 3.300E-06 |
| 1313 | 66 | H66M | A | 1.2e-06 | 1.6e-06 | 3.8e-06 | 2.200E-06 | 1.400E-06 | 1.200E-06 | 3.800E-06 |
| 1314 | 66 | H66F | A | 2e-06 | 2.5e-06 | 1.8e-06 | 2.100E-06 | 3.606E-07 | 1.800E-06 | 2.500E-06 |
| 1315 | 66 | H66P | A | 3.1e-06 | 2.1e-06 | 2.6e-06 | 2.600E-06 | 5.000E-07 | 2.100E-06 | 3.100E-06 |
| 1316 | 66 | H66S | A | 2.3e-06 | 1.6e-06 | 2.7e-06 | 2.200E-06 | 5.568E-07 | 1.600E-06 | 2.700E-06 |
| 1317 | 66 | H66T | A | 3.5e-06 | 4.7e-06 | 2.5e-06 | 3.567E-06 | 1.102E-06 | 2.500E-06 | 4.700E-06 |
| 1318 | 66 | H66W | A | 2.1e-06 | 2.3e-06 | 2.8e-06 | 2.400E-06 | 3.606E-07 | 2.100E-06 | 2.800E-06 |
| 1319 | 66 | H66Y | A | 4.5e-06 | 3.7e-06 | 2.9e-06 | 3.700E-06 | 8.000E-07 | 2.900E-06 | 4.500E-06 |
| 1320 | 66 | H66V | A | 2.6e-06 | 2.2e-06 | 1.7e-06 | 2.167E-06 | 4.509E-07 | 1.700E-06 | 2.600E-06 |
| 1321 | 67 | N67A | A | 3.7e-06 | 2.4e-06 | 4.1e-06 | 3.400E-06 | 8.888E-07 | 2.400E-06 | 4.100E-06 |
| 1322 | 67 | N67R | A | 4e-06 | 1.6e-06 | 1.8e-06 | 2.467E-06 | 1.332E-06 | 1.600E-06 | 4.000E-06 |
| 1323 | 67 | N67N | A | 3.6e-06 | 2.2e-06 | 3.2e-06 | 3.000E-06 | 7.211E-07 | 2.200E-06 | 3.600E-06 |
| 1324 | 67 | N67D | A | 1.6e-06 | 2.9e-06 | 4e-06 | 2.833E-06 | 1.201E-06 | 1.600E-06 | 4.000E-06 |
| 1325 | 67 | N67C | A | 1.6e-06 | 1.9e-06 | 3.2e-06 | 2.233E-06 | 8.505E-07 | 1.600E-06 | 3.200E-06 |
| 1326 | 67 | N67E | A | 1.5e-06 | 2.4e-06 | 3.9e-06 | 2.600E-06 | 1.212E-06 | 1.500E-06 | 3.900E-06 |
| 1327 | 67 | N67Q | A | 2.7e-06 | 1.4e-06 | 3e-06 | 2.367E-06 | 8.505E-07 | 1.400E-06 | 3.000E-06 |
| 1328 | 67 | N67G | A | 2e-06 | 2.2e-06 | 1.7e-06 | 1.967E-06 | 2.517E-07 | 1.700E-06 | 2.200E-06 |
| 1329 | 67 | N67H | A | 2.5e-06 | 3e-06 | 2.7e-06 | 2.733E-06 | 2.517E-07 | 2.500E-06 | 3.000E-06 |
| 1330 | 67 | N67I | A | 1.4e-06 | 2.2e-06 | 1.2e-06 | 1.600E-06 | 5.292E-07 | 1.200E-06 | 2.200E-06 |
| 1331 | 67 | N67L | A | 2.2e-06 | 2.6e-06 | 1.9e-06 | 2.233E-06 | 3.512E-07 | 1.900E-06 | 2.600E-06 |
| 1332 | 67 | N67K | A | 3.8e-06 | 3.5e-06 | 3.7e-06 | 3.667E-06 | 1.528E-07 | 3.500E-06 | 3.800E-06 |
| 1333 | 67 | N67M | A | 2.9e-06 | 2e-06 | 2.9e-06 | 2.600E-06 | 5.196E-07 | 2.000E-06 | 2.900E-06 |
| 1334 | 67 | N67F | A | 1.9e-06 | 1.9e-06 | 2.5e-06 | 2.100E-06 | 3.464E-07 | 1.900E-06 | 2.500E-06 |
| 1335 | 67 | N67P | A | 1.6e-06 | 4.4e-06 | 3.4e-06 | 3.133E-06 | 1.419E-06 | 1.600E-06 | 4.400E-06 |
| 1336 | 67 | N67S | A | 2e-06 | 2.9e-06 | 3.3e-06 | 2.733E-06 | 6.658E-07 | 2.000E-06 | 3.300E-06 |
| 1337 | 67 | N67T | A | 2.8e-06 | 2.4e-06 | 4.5e-06 | 3.233E-06 | 1.115E-06 | 2.400E-06 | 4.500E-06 |
| 1338 | 67 | N67W | A | 1.3e-06 | 2.5e-06 | 1.9e-06 | 1.900E-06 | 6.000E-07 | 1.300E-06 | 2.500E-06 |
| 1339 | 67 | N67Y | A | 3.3e-06 | 2.1e-06 | 2.5e-06 | 2.633E-06 | 6.110E-07 | 2.100E-06 | 3.300E-06 |
| 1340 | 67 | N67V | A | 2.7e-06 | 2.5e-06 | 2.6e-06 | 2.600E-06 | 1.000E-07 | 2.500E-06 | 2.700E-06 |
| 1341 | 68 | Q68A | A | 1.4e-06 | 2e-06 | 2.1e-06 | 1.833E-06 | 3.786E-07 | 1.400E-06 | 2.100E-06 |
| 1342 | 68 | Q68R | A | 2.9e-06 | 2.7e-06 | 3.1e-06 | 2.900E-06 | 2.000E-07 | 2.700E-06 | 3.100E-06 |
| 1343 | 68 | Q68N | A | 2.4e-06 | 3.7e-06 | 2.6e-06 | 2.900E-06 | 7.000E-07 | 2.400E-06 | 3.700E-06 |
| 1344 | 68 | Q68D | A | 1.6e-06 | 4.4e-06 | 2.9e-06 | 2.967E-06 | 1.401E-06 | 1.600E-06 | 4.400E-06 |
| 1345 | 68 | Q68C | A | 2.1e-06 | 2e-06 | 2.6e-06 | 2.233E-06 | 3.215E-07 | 2.000E-06 | 2.600E-06 |
| 1346 | 68 | Q68E | A | 3.1e-06 | 2e-06 | 2.4e-06 | 2.500E-06 | 5.568E-07 | 2.000E-06 | 3.100E-06 |
| 1347 | 68 | Q68Q | A | 2.9e-06 | 3.4e-06 | 2.7e-06 | 3.000E-06 | 3.606E-07 | 2.700E-06 | 3.400E-06 |
| 1348 | 68 | Q68G | A | 2.8e-06 | 2e-06 | 2.2e-06 | 2.333E-06 | 4.163E-07 | 2.000E-06 | 2.800E-06 |
| 1349 | 68 | Q68H | A | 2.4e-06 | 1.4e-06 | 2.3e-06 | 2.033E-06 | 5.508E-07 | 1.400E-06 | 2.400E-06 |
| 1350 | 68 | Q68I | A | 1.6e-06 | 3.5e-06 | 2e-06 | 2.367E-06 | 1.002E-06 | 1.600E-06 | 3.500E-06 |
| 1351 | 68 | Q68L | A | 1.8e-06 | 3e-06 | 2.9e-06 | 2.567E-06 | 6.658E-07 | 1.800E-06 | 3.000E-06 |
| 1352 | 68 | Q68K | A | 3.4e-06 | 2.7e-06 | 3.9e-06 | 3.333E-06 | 6.028E-07 | 2.700E-06 | 3.900E-06 |
| 1353 | 68 | Q68M | A | 1.2e-06 | 4.1e-06 | 2e-06 | 2.433E-06 | 1.498E-06 | 1.200E-06 | 4.100E-06 |
| 1354 | 68 | Q68F | A | 2.1e-06 | 2.4e-06 | 4.6e-06 | 3.033E-06 | 1.365E-06 | 2.100E-06 | 4.600E-06 |
| 1355 | 68 | Q68P | A | 1.7e-06 | 2.4e-06 | 1.2e-06 | 1.767E-06 | 6.028E-07 | 1.200E-06 | 2.400E-06 |
| 1356 | 68 | Q68S | A | 2.1e-06 | 2.9e-06 | 3.4e-06 | 2.800E-06 | 6.557E-07 | 2.100E-06 | 3.400E-06 |
| 1357 | 68 | Q68T | A | 2.2e-06 | 1.9e-06 | 2.1e-06 | 2.067E-06 | 1.528E-07 | 1.900E-06 | 2.200E-06 |
| 1358 | 68 | Q68W | A | 2.1e-06 | 2.3e-06 | 2e-06 | 2.133E-06 | 1.528E-07 | 2.000E-06 | 2.300E-06 |
| 1359 | 68 | Q68Y | A | 1.8e-06 | 2.1e-06 | 3.6e-06 | 2.500E-06 | 9.644E-07 | 1.800E-06 | 3.600E-06 |

| | | | | | | | | | | |
|---|---|---|---|---|---|---|---|---|---|---|
| 1360 | 68 | Q68V | A | 1.5e-06 | 2.9e-06 | 2.1e-06 | 2.167E-06 | 7.024E-07 | 1.500E-06 | 2.900E-06 |
| 1361 | 69 | V69A | A | 2.2e-06 | 3.1e-06 | 2.7e-06 | 2.667E-06 | 4.509E-07 | 2.200E-06 | 3.100E-06 |
| 1362 | 69 | V69R | A | 2e-06 | 2.7e-06 | 2.6e-06 | 2.433E-06 | 3.786E-07 | 2.000E-06 | 2.700E-06 |
| 1363 | 69 | V69N | A | 2.5e-06 | 1.5e-06 | 2.9e-06 | 2.300E-06 | 7.211E-07 | 1.500E-06 | 2.900E-06 |
| 1364 | 69 | V69D | A | 1.5e-06 | 2.5e-06 | 1.8e-06 | 1.933E-06 | 5.132E-07 | 1.500E-06 | 2.500E-06 |
| 1365 | 69 | V69C | A | 2.1e-06 | 2.5e-06 | 2.5e-06 | 2.367E-06 | 2.309E-07 | 2.100E-06 | 2.500E-06 |
| 1366 | 69 | V69E | A | 1.3e-06 | 2.8e-06 | 2.6e-06 | 2.233E-06 | 8.145E-07 | 1.300E-06 | 2.800E-06 |
| 1367 | 69 | V69Q | A | 1.5e-06 | 1.3e-06 | 6.2e-06 | 3.000E-06 | 2.773E-06 | 1.300E-06 | 6.200E-06 |
| 1368 | 69 | V69G | A | 1.8e-06 | 2.8e-06 | 2.3e-06 | 2.300E-06 | 5.000E-07 | 1.800E-06 | 2.800E-06 |
| 1369 | 69 | V69H | A | 3e-06 | 4.1e-06 | 2e-06 | 3.033E-06 | 1.050E-06 | 2.000E-06 | 4.100E-06 |
| 1370 | 69 | V69I | A | 2.4e-06 | 3.5e-06 | 2.2e-06 | 2.700E-06 | 7.000E-07 | 2.200E-06 | 3.500E-06 |
| 1371 | 69 | V69L | A | 1.8e-06 | 2.2e-06 | 2.7e-06 | 2.233E-06 | 4.509E-07 | 1.800E-06 | 2.700E-06 |
| 1372 | 69 | V69K | A | 2.7e-06 | 2.8e-06 | 2.3e-06 | 2.600E-06 | 2.646E-07 | 2.300E-06 | 2.800E-06 |
| 1373 | 69 | V69M | A | 2.9e-06 | 2e-06 | 3.7e-06 | 2.867E-06 | 8.505E-07 | 2.000E-06 | 3.700E-06 |
| 1374 | 69 | V69F | A | 1.6e-06 | 1.6e-06 | 2.5e-06 | 1.900E-06 | 5.196E-07 | 1.600E-06 | 2.500E-06 |
| 1375 | 69 | V69P | A | 1.3e-06 | 2.5e-06 | 1.1e-06 | 1.633E-06 | 7.572E-07 | 1.100E-06 | 2.500E-06 |
| 1376 | 69 | V69S | A | 2.6e-06 | 2.9e-06 | 2.3e-06 | 2.600E-06 | 3.000E-07 | 2.300E-06 | 2.900E-06 |
| 1377 | 69 | V69T | A | 2.4e-06 | 2.5e-06 | 4e-06 | 2.967E-06 | 8.963E-07 | 2.400E-06 | 4.000E-06 |
| 1378 | 69 | V69W | A | 3.7e-06 | 3.1e-06 | 2.3e-06 | 3.033E-06 | 7.024E-07 | 2.300E-06 | 3.700E-06 |
| 1379 | 69 | V69Y | A | 2.1e-06 | 3.4e-06 | 1.8e-06 | 2.433E-06 | 8.505E-07 | 1.800E-06 | 3.400E-06 |
| 1380 | 69 | V69V | A | 4.3e-06 | 2.3e-06 | 2.7e-06 | 3.100E-06 | 1.058E-06 | 2.300E-06 | 4.300E-06 |
| 1381 | 70 | R70A | A | 3.4e-06 | 2.9e-06 | 3e-06 | 3.100E-06 | 2.646E-07 | 2.900E-06 | 3.400E-06 |
| 1382 | 70 | R70R | A | 3.2e-06 | 2.3e-06 | 2.4e-06 | 2.633E-06 | 4.933E-07 | 2.300E-06 | 3.200E-06 |
| 1383 | 70 | R70N | A | 2.9e-06 | 2.9e-06 | 2e-06 | 2.600E-06 | 5.196E-07 | 2.000E-06 | 2.900E-06 |
| 1384 | 70 | R70D | A | 3.6e-06 | 1.2e-06 | 2.2e-06 | 2.333E-06 | 1.206E-06 | 1.200E-06 | 3.600E-06 |
| 1385 | 70 | R70C | A | 2.7e-06 | 3.7e-06 | 1.4e-06 | 2.600E-06 | 1.153E-06 | 1.400E-06 | 3.700E-06 |
| 1386 | 70 | R70E | A | 2.3e-06 | 3.2e-06 | 4e-06 | 3.167E-06 | 8.505E-07 | 2.300E-06 | 4.000E-06 |
| 1387 | 70 | R70Q | A | 1.8e-06 | 4.7e-06 | 4.1e-06 | 3.533E-06 | 1.531E-06 | 1.800E-06 | 4.700E-06 |
| 1388 | 70 | R70G | A | 3.3e-06 | 3.6e-06 | 2e-06 | 2.967E-06 | 8.505E-07 | 2.000E-06 | 3.600E-06 |
| 1389 | 70 | R70H | A | 2e-06 | 1.7e-06 | 2.4e-06 | 2.033E-06 | 3.512E-07 | 1.700E-06 | 2.400E-06 |
| 1390 | 70 | R70I | A | 2.6e-06 | 2.1e-06 | 3.8e-06 | 2.833E-06 | 8.737E-07 | 2.100E-06 | 3.800E-06 |
| 1391 | 70 | R70L | A | 2.3e-06 | 2.5e-06 | 9.1e-07 | 1.903E-06 | 8.660E-07 | 9.100E-07 | 2.500E-06 |
| 1392 | 70 | R70K | A | 3.9e-06 | 2e-06 | 2e-06 | 2.633E-06 | 1.097E-06 | 2.000E-06 | 3.900E-06 |
| 1393 | 70 | R70M | A | 2.3e-06 | 3.7e-06 | 2.3e-06 | 2.767E-06 | 8.083E-07 | 2.300E-06 | 3.700E-06 |
| 1394 | 70 | R70F | A | 1.6e-06 | 3e-06 | 2.9e-06 | 2.500E-06 | 7.810E-07 | 1.600E-06 | 3.000E-06 |
| 1395 | 70 | R70P | A | 2.1e-06 | 2.2e-06 | 4.9e-06 | 3.067E-06 | 1.589E-06 | 2.100E-06 | 4.900E-06 |
| 1396 | 70 | R70S | A | 3.8e-06 | 2.4e-06 | 1.9e-06 | 2.700E-06 | 9.849E-07 | 1.900E-06 | 3.800E-06 |
| 1397 | 70 | R70T | A | 3.7e-06 | 2.6e-06 | 2.9e-06 | 3.067E-06 | 5.686E-07 | 2.600E-06 | 3.700E-06 |
| 1398 | 70 | R70W | A | 2.9e-06 | 2.4e-06 | 2.5e-06 | 2.600E-06 | 2.646E-07 | 2.400E-06 | 2.900E-06 |
| 1399 | 70 | R70Y | A | 3e-06 | 1.4e-06 | 2.3e-06 | 2.233E-06 | 8.021E-07 | 1.400E-06 | 3.000E-06 |
| 1400 | 70 | R70V | A | 2.2e-06 | 1.8e-06 | 3.1e-06 | 2.367E-06 | 6.658E-07 | 1.800E-06 | 3.100E-06 |
| 1401 | 71 | Q71A | A | 2.7e-06 | 3.1e-06 | 2.4e-06 | 2.733E-06 | 3.512E-07 | 2.400E-06 | 3.100E-06 |
| 1402 | 71 | Q71R | A | 3.2e-06 | 2.2e-06 | 2.6e-06 | 2.667E-06 | 5.033E-07 | 2.200E-06 | 3.200E-06 |
| 1403 | 71 | Q71N | A | 3.1e-06 | 1.1e-06 | 2.7e-06 | 2.300E-06 | 1.058E-06 | 1.100E-06 | 3.100E-06 |
| 1404 | 71 | Q71D | A | 2.7e-06 | 2.3e-06 | 1.9e-06 | 2.300E-06 | 4.000E-07 | 1.900E-06 | 2.700E-06 |
| 1405 | 71 | Q71C | A | 1.8e-06 | 1.9e-06 | 2.5e-06 | 2.067E-06 | 3.786E-07 | 1.800E-06 | 2.500E-06 |
| 1406 | 71 | Q71E | A | 1.6e-06 | 1.6e-06 | 1.6e-06 | 1.600E-06 | 0.000E+00 | 1.600E-06 | 1.600E-06 |
| 1407 | 71 | Q71Q | A | 1.3e-06 | 2.9e-06 | 2.6e-06 | 2.267E-06 | 8.505E-07 | 1.300E-06 | 2.900E-06 |
| 1408 | 71 | Q71G | A | 1.9e-06 | 2.6e-06 | 1.9e-06 | 2.133E-06 | 4.041E-07 | 1.900E-06 | 2.600E-06 |
| 1409 | 71 | Q71H | A | 3.2e-06 | 4e-06 | 1.3e-06 | 2.833E-06 | 1.387E-06 | 1.300E-06 | 4.000E-06 |
| 1410 | 71 | Q71I | A | 2.5e-06 | 1.9e-06 | 2.1e-06 | 2.167E-06 | 3.055E-07 | 1.900E-06 | 2.500E-06 |
| 1411 | 71 | Q71L | A | 3.1e-06 | 2.6e-06 | 2.4e-06 | 2.700E-06 | 3.606E-07 | 2.400E-06 | 3.100E-06 |
| 1412 | 71 | Q71K | A | 2.1e-06 | 2.2e-06 | 2.9e-06 | 2.400E-06 | 4.359E-07 | 2.100E-06 | 2.900E-06 |
| 1413 | 71 | Q71M | A | 1.8e-06 | 2.8e-06 | 1.7e-06 | 2.100E-06 | 6.083E-07 | 1.700E-06 | 2.800E-06 |
| 1414 | 71 | Q71F | A | 1.5e-06 | 2.9e-06 | 3e-06 | 2.467E-06 | 8.386E-07 | 1.500E-06 | 3.000E-06 |
| 1415 | 71 | Q71P | A | 2.9e-06 | 2.2e-06 | 2.4e-06 | 2.500E-06 | 3.606E-07 | 2.200E-06 | 2.900E-06 |
| 1416 | 71 | Q71S | A | 2.2e-06 | 1.9e-06 | 1.1e-06 | 1.733E-06 | 5.686E-07 | 1.100E-06 | 2.200E-06 |
| 1417 | 71 | Q71T | A | 2.7e-06 | 3.2e-06 | 1.9e-06 | 2.600E-06 | 6.557E-07 | 1.900E-06 | 3.200E-06 |
| 1418 | 71 | Q71W | A | 2.8e-06 | 1.9e-06 | 2.9e-06 | 2.533E-06 | 5.508E-07 | 1.900E-06 | 2.900E-06 |
| 1419 | 71 | Q71Y | A | 1.5e-06 | 3.2e-06 | 2.2e-06 | 2.300E-06 | 8.544E-07 | 1.500E-06 | 3.200E-06 |
| 1420 | 71 | Q71V | A | 2.2e-06 | 1.6e-06 | 2.8e-06 | 2.200E-06 | 6.000E-07 | 1.600E-06 | 2.800E-06 |
| 1421 | 72 | V72A | A | 1.9e-06 | 1.8e-06 | 1.4e-06 | 1.700E-06 | 2.646E-07 | 1.400E-06 | 1.900E-06 |
| 1422 | 72 | V72R | A | 2.6e-06 | 1.6e-06 | 1.1e-06 | 1.767E-06 | 7.638E-07 | 1.100E-06 | 2.600E-06 |
| 1423 | 72 | V72N | A | 2.6e-06 | 2.9e-06 | 1.9e-06 | 2.467E-06 | 5.132E-07 | 1.900E-06 | 2.900E-06 |
| 1424 | 72 | V72D | A | 2.3e-06 | 1.9e-06 | 1.8e-06 | 2.000E-06 | 2.646E-07 | 1.800E-06 | 2.300E-06 |
| 1425 | 72 | V72C | A | 3e-06 | 1.7e-06 | 3.7e-06 | 2.800E-06 | 1.015E-06 | 1.700E-06 | 3.700E-06 |
| 1426 | 72 | V72E | A | 2.5e-06 | 2.4e-06 | 3.9e-06 | 2.933E-06 | 8.386E-07 | 2.400E-06 | 3.900E-06 |
| 1427 | 72 | V72Q | A | 3e-06 | 3.2e-06 | 2.3e-06 | 2.833E-06 | 4.726E-07 | 2.300E-06 | 3.200E-06 |
| 1428 | 72 | V72G | A | 6.1e-06 | 2.1e-06 | 4e-06 | 4.067E-06 | 2.001E-06 | 2.100E-06 | 6.100E-06 |
| 1429 | 72 | V72H | A | 3.8e-06 | 2.3e-06 | 3.5e-06 | 3.200E-06 | 7.937E-07 | 2.300E-06 | 3.800E-06 |
| 1430 | 72 | V72I | A | 2.5e-06 | 1.5e-06 | 2.2e-06 | 2.067E-06 | 5.132E-07 | 1.500E-06 | 2.500E-06 |
| 1431 | 72 | V72L | A | 2.7e-06 | 1.9e-06 | 2.2e-06 | 2.267E-06 | 4.041E-07 | 1.900E-06 | 2.700E-06 |
| 1432 | 72 | V72K | A | 1.4e-06 | 2.9e-06 | 2.5e-06 | 2.267E-06 | 7.767E-07 | 1.400E-06 | 2.900E-06 |
| 1433 | 72 | V72M | A | 2.6e-06 | 5e-06 | 1.1e-06 | 2.900E-06 | 1.967E-06 | 1.100E-06 | 5.000E-06 |
| 1434 | 72 | V72F | A | 1.2e-06 | 2e-06 | 2.2e-06 | 1.800E-06 | 5.292E-07 | 1.200E-06 | 2.200E-06 |
| 1435 | 72 | V72P | A | 1.7e-06 | 3.3e-06 | 2e-06 | 2.333E-06 | 8.505E-07 | 1.700E-06 | 3.300E-06 |
| 1436 | 72 | V72S | A | 3.2e-06 | 2.4e-06 | 2.1e-06 | 2.567E-06 | 5.686E-07 | 2.100E-06 | 3.200E-06 |
| 1437 | 72 | V72T | A | 2e-06 | 2.3e-06 | 1.6e-06 | 1.967E-06 | 3.512E-07 | 1.600E-06 | 2.300E-06 |
| 1438 | 72 | V72W | A | 2.8e-06 | 2.4e-06 | 2e-06 | 2.400E-06 | 4.000E-07 | 2.000E-06 | 2.800E-06 |
| 1439 | 72 | V72Y | A | 2.4e-06 | 2.3e-06 | 2e-06 | 2.233E-06 | 2.082E-07 | 2.000E-06 | 2.400E-06 |
| 1440 | 72 | V72V | A | 3.1e-06 | 2.4e-06 | 3.2e-06 | 2.900E-06 | 4.359E-07 | 2.400E-06 | 3.200E-06 |
| 1441 | 73 | P73A | A | 1.5e-06 | 2e-06 | 2.9e-06 | 2.133E-06 | 7.095E-07 | 1.500E-06 | 2.900E-06 |
| 1442 | 73 | P73R | A | 2.6e-06 | 2.9e-06 | 1.3e-06 | 2.267E-06 | 8.505E-07 | 1.300E-06 | 2.900E-06 |
| 1443 | 73 | P73N | A | 1.9e-06 | 1.8e-06 | 1.7e-06 | 1.800E-06 | 1.000E-07 | 1.700E-06 | 1.900E-06 |
| 1444 | 73 | P73D | A | 2e-06 | 1.6e-06 | 2.7e-06 | 2.100E-06 | 5.568E-07 | 1.600E-06 | 2.700E-06 |

| | | | | | | | | | | |
|---|---|---|---|---|---|---|---|---|---|---|
| 1445 | 73 | P73C | A | 2.1e-06 | 1.7e-06 | 1.7e-06 | 1.833E-06 | 2.309E-07 | 1.700E-06 | 2.100E-06 |
| 1446 | 73 | P73E | A | 3.4e-06 | 2.1e-06 | 1.7e-06 | 2.400E-06 | 8.888E-07 | 1.700E-06 | 3.400E-06 |
| 1447 | 73 | P73Q | A | 2e-06 | 3.8e-06 | 2.8e-06 | 2.867E-06 | 9.018E-07 | 2.000E-06 | 3.800E-06 |
| 1448 | 73 | P73G | A | 1.9e-06 | 3.4e-06 | 2.8e-06 | 2.700E-06 | 7.550E-07 | 1.900E-06 | 3.400E-06 |
| 1449 | 73 | P73H | A | 4.6e-06 | 2.7e-06 | 2.6e-06 | 3.300E-06 | 1.127E-06 | 2.600E-06 | 4.600E-06 |
| 1450 | 73 | P73I | A | 2.8e-06 | 3e-06 | 1.2e-06 | 2.333E-06 | 9.866E-07 | 1.200E-06 | 3.000E-06 |
| 1451 | 73 | P73L | A | 1.8e-06 | 4.8e-06 | 1.9e-06 | 2.833E-06 | 1.704E-06 | 1.800E-06 | 4.800E-06 |
| 1452 | 73 | P73K | A | 3.1e-06 | 2.4e-06 | 4.5e-06 | 3.333E-06 | 1.069E-06 | 2.400E-06 | 4.500E-06 |
| 1453 | 73 | P73M | A | 2e-06 | 2.6e-06 | 3.5e-06 | 2.700E-06 | 7.550E-07 | 2.000E-06 | 3.500E-06 |
| 1454 | 73 | P73F | A | 2.8e-06 | 1.8e-06 | 1.7e-06 | 2.100E-06 | 6.083E-07 | 1.700E-06 | 2.800E-06 |
| 1455 | 73 | P73P | A | 1.9e-06 | 2.2e-06 | 1.5e-06 | 1.867E-06 | 3.512E-07 | 1.500E-06 | 2.200E-06 |
| 1456 | 73 | P73S | A | 1.7e-06 | 3e-06 | 3.3e-06 | 2.667E-06 | 8.505E-07 | 1.700E-06 | 3.300E-06 |
| 1457 | 73 | P73T | A | 2.5e-06 | 3e-06 | 4.5e-06 | 3.333E-06 | 1.041E-06 | 2.500E-06 | 4.500E-06 |
| 1458 | 73 | P73W | A | 2e-06 | 2.6e-06 | 1.6e-06 | 2.067E-06 | 5.033E-07 | 1.600E-06 | 2.600E-06 |
| 1459 | 73 | P73Y | A | 2.7e-06 | 2.2e-06 | 3e-06 | 2.633E-06 | 4.041E-07 | 2.200E-06 | 3.000E-06 |
| 1460 | 73 | P73V | A | 3.3e-06 | 2.4e-06 | 2.3e-06 | 2.667E-06 | 5.508E-07 | 2.300E-06 | 3.300E-06 |
| 1461 | 74 | L74A | A | 1.6e-06 | 2.1e-06 | 4.7e-06 | 2.800E-06 | 1.664E-06 | 1.600E-06 | 4.700E-06 |
| 1462 | 74 | L74R | A | 1.8e-06 | 1.3e-06 | 3.1e-06 | 2.067E-06 | 9.292E-07 | 1.300E-06 | 3.100E-06 |
| 1463 | 74 | L74N | A | 2.4e-06 | 2.4e-06 | 1.1e-06 | 1.967E-06 | 7.506E-07 | 1.100E-06 | 2.400E-06 |
| 1464 | 74 | L74D | A | 3.7e-06 | 2.4e-06 | 2.1e-06 | 2.733E-06 | 8.505E-07 | 2.100E-06 | 3.700E-06 |
| 1465 | 74 | L74C | A | 1.4e-06 | 2.1e-06 | 2.3e-06 | 1.933E-06 | 4.726E-07 | 1.400E-06 | 2.300E-06 |
| 1466 | 74 | L74E | A | 3.1e-06 | 2.7e-06 | 2.2e-06 | 2.667E-06 | 4.509E-07 | 2.200E-06 | 3.100E-06 |
| 1467 | 74 | L74Q | A | 1.7e-06 | 1.8e-06 | 1.4e-06 | 1.633E-06 | 2.082E-07 | 1.400E-06 | 1.800E-06 |
| 1468 | 74 | L74G | A | 1.9e-06 | 2.7e-06 | 2.1e-06 | 2.233E-06 | 4.163E-07 | 1.900E-06 | 2.700E-06 |
| 1469 | 74 | L74H | A | 2.7e-06 | 2.7e-06 | 2.2e-06 | 2.533E-06 | 2.887E-07 | 2.200E-06 | 2.700E-06 |
| 1470 | 74 | L74I | A | 1.9e-06 | 2.3e-06 | 3e-06 | 2.400E-06 | 5.568E-07 | 1.900E-06 | 3.000E-06 |
| 1471 | 74 | L74L | A | 1.9e-06 | 2.4e-06 | 2.4e-06 | 2.233E-06 | 2.887E-07 | 1.900E-06 | 2.400E-06 |
| 1472 | 74 | L74K | A | 2.6e-06 | 1.9e-06 | 3e-06 | 2.500E-06 | 5.568E-07 | 1.900E-06 | 3.000E-06 |
| 1473 | 74 | L74M | A | 2.6e-06 | 3.2e-06 | 4.2e-06 | 3.333E-06 | 8.083E-07 | 2.600E-06 | 4.200E-06 |
| 1474 | 74 | L74F | A | 2.8e-06 | 2e-06 | 3.1e-06 | 2.633E-06 | 5.686E-07 | 2.000E-06 | 3.100E-06 |
| 1475 | 74 | L74P | A | 2.4e-06 | 1.4e-06 | 2.3e-06 | 2.033E-06 | 5.508E-07 | 1.400E-06 | 2.400E-06 |
| 1476 | 74 | L74S | A | 2.6e-06 | 1.7e-06 | 2.5e-06 | 2.267E-06 | 4.933E-07 | 1.700E-06 | 2.600E-06 |
| 1477 | 74 | L74T | A | 2.8e-06 | 3.3e-06 | 2.7e-06 | 2.933E-06 | 3.215E-07 | 2.700E-06 | 3.300E-06 |
| 1478 | 74 | L74W | A | 2.4e-06 | 2.7e-06 | 1.6e-06 | 2.233E-06 | 5.686E-07 | 1.600E-06 | 2.700E-06 |
| 1479 | 74 | L74Y | A | 2.6e-06 | 2.5e-06 | 2.6e-06 | 2.567E-06 | 5.774E-08 | 2.500E-06 | 2.600E-06 |
| 1480 | 74 | L74V | A | 3.1e-06 | 2e-06 | 2.9e-06 | 2.667E-06 | 5.859E-07 | 2.000E-06 | 3.100E-06 |
| 1481 | 75 | Q75A | A | 3.3e-06 | 2.2e-06 | 2.3e-06 | 2.600E-06 | 6.083E-07 | 2.200E-06 | 3.300E-06 |
| 1482 | 75 | Q75R | A | 2.2e-06 | 1.6e-06 | 2.3e-06 | 2.033E-06 | 3.786E-07 | 1.600E-06 | 2.300E-06 |
| 1483 | 75 | Q75N | A | 2.1e-06 | 2.1e-06 | 2.2e-06 | 2.133E-06 | 5.774E-08 | 2.100E-06 | 2.200E-06 |
| 1484 | 75 | Q75D | A | 1.8e-06 | 2.2e-06 | 1.1e-06 | 1.700E-06 | 5.568E-07 | 1.100E-06 | 2.200E-06 |
| 1485 | 75 | Q75C | A | 8.2e-07 | 1.8e-06 | 2.8e-06 | 1.807E-06 | 9.900E-07 | 8.200E-07 | 2.800E-06 |
| 1486 | 75 | Q75E | A | 2.3e-06 | 3.5e-06 | 2.7e-06 | 2.833E-06 | 6.110E-07 | 2.300E-06 | 3.500E-06 |
| 1487 | 75 | Q75Q | A | 2.5e-06 | 2.1e-06 | 2.1e-06 | 2.233E-06 | 2.309E-07 | 2.100E-06 | 2.500E-06 |
| 1488 | 75 | Q75G | A | 3.1e-06 | 2.4e-06 | 3.2e-06 | 2.900E-06 | 4.359E-07 | 2.400E-06 | 3.200E-06 |
| 1489 | 75 | Q75H | A | 2.3e-06 | 2.6e-06 | 3.6e-06 | 2.833E-06 | 6.807E-07 | 2.300E-06 | 3.600E-06 |
| 1490 | 75 | Q75I | A | 2.7e-06 | 2.8e-06 | 2.6e-06 | 2.700E-06 | 1.000E-07 | 2.600E-06 | 2.800E-06 |
| 1491 | 75 | Q75L | A | 4.9e-06 | 2.4e-06 | 2.8e-06 | 3.367E-06 | 1.343E-06 | 2.400E-06 | 4.900E-06 |
| 1492 | 75 | Q75K | A | 5.1e-06 | 2.9e-06 | 2.2e-06 | 3.400E-06 | 1.513E-06 | 2.200E-06 | 5.100E-06 |
| 1493 | 75 | Q75M | A | 2.4e-06 | 2.7e-06 | 2.4e-06 | 2.500E-06 | 1.732E-07 | 2.400E-06 | 2.700E-06 |
| 1494 | 75 | Q75F | A | 4.5e-06 | 2.1e-06 | 2.6e-06 | 3.067E-06 | 1.266E-06 | 2.100E-06 | 4.500E-06 |
| 1495 | 75 | Q75P | A | 1.8e-06 | 3.3e-06 | 2.8e-06 | 2.633E-06 | 7.638E-07 | 1.800E-06 | 3.300E-06 |
| 1496 | 75 | Q75S | A | 1.3e-06 | 1.9e-06 | 1.7e-06 | 1.633E-06 | 3.055E-07 | 1.300E-06 | 1.900E-06 |
| 1497 | 75 | Q75T | A | 2.4e-06 | 2.4e-06 | 2e-06 | 2.267E-06 | 2.309E-07 | 2.000E-06 | 2.400E-06 |
| 1498 | 75 | Q75W | A | 1.3e-06 | 2.2e-06 | 2.7e-06 | 2.067E-06 | 7.095E-07 | 1.300E-06 | 2.700E-06 |
| 1499 | 75 | Q75Y | A | 2.9e-06 | 3.9e-06 | 1.8e-06 | 2.867E-06 | 1.050E-06 | 1.800E-06 | 3.900E-06 |
| 1500 | 75 | Q75V | A | 1.9e-06 | 2.4e-06 | 2.4e-06 | 2.233E-06 | 2.887E-07 | 1.900E-06 | 2.400E-06 |
| 1501 | 76 | R76A | A | 1.3e-06 | 2.3e-06 | 4.3e-06 | 2.633E-06 | 1.528E-06 | 1.300E-06 | 4.300E-06 |
| 1502 | 76 | R76R | A | 2.6e-06 | 2.3e-06 | 2.2e-06 | 2.367E-06 | 2.082E-07 | 2.200E-06 | 2.600E-06 |
| 1503 | 76 | R76N | A | 3.2e-06 | 1.3e-06 | 1.6e-06 | 2.033E-06 | 1.021E-06 | 1.300E-06 | 3.200E-06 |
| 1504 | 76 | R76D | A | 4.8e-06 | 2.3e-06 | 2.3e-06 | 3.133E-06 | 1.443E-06 | 2.300E-06 | 4.800E-06 |
| 1505 | 76 | R76C | A | 1.2e-06 | 2.4e-06 | 1.3e-06 | 1.633E-06 | 6.658E-07 | 1.200E-06 | 2.400E-06 |
| 1506 | 76 | R76E | A | 3.9e-06 | 2.2e-06 | 1.4e-06 | 2.500E-06 | 1.277E-06 | 1.400E-06 | 3.900E-06 |
| 1507 | 76 | R76Q | A | 2.8e-06 | 1.6e-06 | 1.7e-06 | 2.033E-06 | 6.658E-07 | 1.600E-06 | 2.800E-06 |
| 1508 | 76 | R76G | A | 1.9e-06 | 2e-06 | 2.7e-06 | 2.200E-06 | 4.359E-07 | 1.900E-06 | 2.700E-06 |
| 1509 | 76 | R76H | A | 3.1e-06 | 2.3e-06 | 2e-06 | 2.467E-06 | 5.686E-07 | 2.000E-06 | 3.100E-06 |
| 1510 | 76 | R76I | A | 2.2e-06 | 3.5e-06 | 1.9e-06 | 2.533E-06 | 8.505E-07 | 1.900E-06 | 3.500E-06 |
| 1511 | 76 | R76L | A | 2.3e-06 | 2.6e-06 | 9.5e-07 | 1.950E-06 | 8.789E-07 | 9.500E-07 | 2.600E-06 |
| 1512 | 76 | R76K | A | 1.4e-06 | 2.4e-06 | 2.4e-06 | 2.067E-06 | 5.774E-07 | 1.400E-06 | 2.400E-06 |
| 1513 | 76 | R76M | A | 2.1e-06 | 2.6e-06 | 2.2e-06 | 2.300E-06 | 2.646E-07 | 2.100E-06 | 2.600E-06 |
| 1514 | 76 | R76F | A | 3.6e-06 | 2e-06 | 2.2e-06 | 2.600E-06 | 8.718E-07 | 2.000E-06 | 3.600E-06 |
| 1515 | 76 | R76P | A | 2.2e-06 | 2e-06 | 1.9e-06 | 2.033E-06 | 1.528E-07 | 1.900E-06 | 2.200E-06 |
| 1516 | 76 | R76S | A | 2.4e-06 | 2.7e-06 | 2.9e-06 | 2.667E-06 | 2.517E-07 | 2.400E-06 | 2.900E-06 |
| 1517 | 76 | R76T | A | 1.8e-06 | 1e-06 | 3.9e-06 | 2.233E-06 | 1.498E-06 | 1.000E-06 | 3.900E-06 |
| 1518 | 76 | R76W | A | 1.4e-06 | 1.8e-06 | 2.8e-06 | 2.000E-06 | 7.211E-07 | 1.400E-06 | 2.800E-06 |
| 1519 | 76 | R76Y | A | 3.1e-06 | 2.6e-06 | 1.8e-06 | 2.500E-06 | 6.557E-07 | 1.800E-06 | 3.100E-06 |
| 1520 | 76 | R76V | A | 4e-06 | 3.1e-06 | 2.3e-06 | 3.133E-06 | 8.505E-07 | 2.300E-06 | 4.000E-06 |
| 1521 | 77 | L77A | A | 3.3e-06 | 1.7e-06 | 1.6e-06 | 2.200E-06 | 9.539E-07 | 1.600E-06 | 3.300E-06 |
| 1522 | 77 | L77R | A | 2.7e-06 | 1.8e-06 | 2.3e-06 | 2.267E-06 | 4.509E-07 | 1.800E-06 | 2.700E-06 |
| 1523 | 77 | L77N | A | 2.3e-06 | 2.1e-06 | 1.8e-06 | 2.067E-06 | 2.517E-07 | 1.800E-06 | 2.300E-06 |
| 1524 | 77 | L77D | A | 1.5e-06 | 3.1e-06 | 3.6e-06 | 2.733E-06 | 1.097E-06 | 1.500E-06 | 3.600E-06 |
| 1525 | 77 | L77C | A | 1.7e-06 | 1.6e-06 | 2e-06 | 1.767E-06 | 2.082E-07 | 1.600E-06 | 2.000E-06 |
| 1526 | 77 | L77E | A | 4.1e-06 | 2.6e-06 | 2.5e-06 | 3.067E-06 | 8.963E-07 | 2.500E-06 | 4.100E-06 |
| 1527 | 77 | L77Q | A | 3e-06 | 3.1e-06 | 2.4e-06 | 2.833E-06 | 3.786E-07 | 2.400E-06 | 3.100E-06 |
| 1528 | 77 | L77G | A | 2.6e-06 | 3.3e-06 | 2.2e-06 | 2.700E-06 | 5.568E-07 | 2.200E-06 | 3.300E-06 |
| 1529 | 77 | L77H | A | 1e-06 | 2.3e-06 | 1.6e-06 | 1.633E-06 | 6.506E-07 | 1.000E-06 | 2.300E-06 |

| | | | | | | | | | | |
|---|---|---|---|---|---|---|---|---|---|---|
| 1530 | 77 | L77I | A | 4.1e-06 | 4e-06 | 2.7e-06 | 3.600E-06 | 7.810E-07 | 2.700E-06 | 4.100E-06 |
| 1531 | 77 | L77L | A | 3.7e-06 | 1.8e-06 | 2.7e-06 | 2.733E-06 | 9.504E-07 | 1.800E-06 | 3.700E-06 |
| 1532 | 77 | L77K | A | 3.6e-06 | 2.1e-06 | 1.9e-06 | 2.533E-06 | 9.292E-07 | 1.900E-06 | 3.600E-06 |
| 1533 | 77 | L77M | A | 2.5e-06 | 3e-06 | 2.4e-06 | 2.633E-06 | 3.215E-07 | 2.400E-06 | 3.000E-06 |
| 1534 | 77 | L77F | A | 2.3e-06 | 3.8e-06 | 2.4e-06 | 2.833E-06 | 8.386E-07 | 2.300E-06 | 3.800E-06 |
| 1535 | 77 | L77P | A | 2.8e-06 | 3e-06 | 2.9e-06 | 2.900E-06 | 1.000E-07 | 2.800E-06 | 3.000E-06 |
| 1536 | 77 | L77S | A | 1.9e-06 | 2.3e-06 | 4.5e-06 | 2.900E-06 | 1.400E-06 | 1.900E-06 | 4.500E-06 |
| 1537 | 77 | L77T | A | 2.8e-06 | 2.4e-06 | 1.8e-06 | 2.333E-06 | 5.033E-07 | 1.800E-06 | 2.800E-06 |
| 1538 | 77 | L77W | A | 1.5e-06 | 2.1e-06 | 2.2e-06 | 1.933E-06 | 3.786E-07 | 1.500E-06 | 2.200E-06 |
| 1539 | 77 | L77Y | A | 2.5e-06 | 1.3e-06 | 2e-06 | 1.933E-06 | 6.028E-07 | 1.300E-06 | 2.500E-06 |
| 1540 | 77 | L77V | A | 1.7e-06 | 2.6e-06 | 3.7e-06 | 2.667E-06 | 1.002E-06 | 1.700E-06 | 3.700E-06 |
| 1541 | 78 | R78A | A | 1.9e-06 | 1.5e-06 | 3.1e-06 | 2.167E-06 | 8.327E-07 | 1.500E-06 | 3.100E-06 |
| 1542 | 78 | R78R | A | 2.5e-06 | 4.1e-06 | 1.5e-06 | 2.700E-06 | 1.311E-06 | 1.500E-06 | 4.100E-06 |
| 1543 | 78 | R78N | A | 3e-06 | 2.3e-06 | 1.7e-06 | 2.333E-06 | 6.506E-07 | 1.700E-06 | 3.000E-06 |
| 1544 | 78 | R78D | A | 2.1e-06 | 1.7e-06 | 1.7e-06 | 1.833E-06 | 2.309E-07 | 1.700E-06 | 2.100E-06 |
| 1545 | 78 | R78C | A | 1.8e-06 | 2.2e-06 | 2.9e-06 | 2.300E-06 | 5.568E-07 | 1.800E-06 | 2.900E-06 |
| 1546 | 78 | R78E | A | 2.8e-06 | 1.1e-06 | 1.7e-06 | 1.867E-06 | 8.622E-07 | 1.100E-06 | 2.800E-06 |
| 1547 | 78 | R78Q | A | 5.2e-06 | 2.6e-06 | 2.2e-06 | 3.333E-06 | 1.629E-06 | 2.200E-06 | 5.200E-06 |
| 1548 | 78 | R78G | A | 2.1e-06 | 1.5e-06 | 3.3e-06 | 2.300E-06 | 9.165E-07 | 1.500E-06 | 3.300E-06 |
| 1549 | 78 | R78H | A | 3.3e-06 | 2.8e-06 | 1.8e-06 | 2.633E-06 | 7.638E-07 | 1.800E-06 | 3.300E-06 |
| 1550 | 78 | R78I | A | 1.8e-06 | 1.4e-06 | 2.6e-06 | 1.933E-06 | 6.110E-07 | 1.400E-06 | 2.600E-06 |
| 1551 | 78 | R78L | A | 2.9e-06 | 2.6e-06 | 2.9e-06 | 2.800E-06 | 1.732E-07 | 2.600E-06 | 2.900E-06 |
| 1552 | 78 | R78K | A | 3.5e-06 | 2.1e-06 | 1.7e-06 | 2.433E-06 | 9.452E-07 | 1.700E-06 | 3.500E-06 |
| 1553 | 78 | R78M | A | 2.7e-06 | 2.7e-06 | 2.7e-06 | 2.700E-06 | 0.000E+00 | 2.700E-06 | 2.700E-06 |
| 1554 | 78 | R78F | A | 3.1e-06 | 2.1e-06 | 3.1e-06 | 2.767E-06 | 5.774E-07 | 2.100E-06 | 3.100E-06 |
| 1555 | 78 | R78P | A | 2e-06 | 2.6e-06 | 2.8e-06 | 2.467E-06 | 4.163E-07 | 2.000E-06 | 2.800E-06 |
| 1556 | 78 | R78S | A | 2.1e-06 | 2.2e-06 | 3.6e-06 | 2.633E-06 | 8.386E-07 | 2.100E-06 | 3.600E-06 |
| 1557 | 78 | R78T | A | 2.7e-06 | 1.9e-06 | 2.5e-06 | 2.367E-06 | 4.163E-07 | 1.900E-06 | 2.700E-06 |
| 1558 | 78 | R78W | A | 4e-06 | 2.5e-06 | 1.4e-06 | 2.633E-06 | 1.305E-06 | 1.400E-06 | 4.000E-06 |
| 1559 | 78 | R78Y | A | 2e-06 | 2.2e-06 | 1.2e-06 | 1.800E-06 | 5.292E-07 | 1.200E-06 | 2.200E-06 |
| 1560 | 78 | R78V | A | 2.4e-06 | 1.9e-06 | 2.3e-06 | 2.200E-06 | 2.646E-07 | 1.900E-06 | 2.400E-06 |
| 1561 | 79 | I79A | A | 2e-06 | 2.3e-06 | 9.9e-07 | 1.763E-06 | 6.863E-07 | 9.900E-07 | 2.300E-06 |
| 1562 | 79 | I79R | A | 2.1e-06 | 1.5e-06 | 2.5e-06 | 2.033E-06 | 5.033E-07 | 1.500E-06 | 2.500E-06 |
| 1563 | 79 | I79N | A | 1.4e-06 | 3.4e-06 | 2.2e-06 | 2.333E-06 | 1.007E-06 | 1.400E-06 | 3.400E-06 |
| 1564 | 79 | I79D | A | 2.3e-06 | 1.5e-06 | 3.5e-06 | 2.433E-06 | 1.007E-06 | 1.500E-06 | 3.500E-06 |
| 1565 | 79 | I79C | A | 2.5e-06 | 2.1e-06 | 2.3e-06 | 2.300E-06 | 2.000E-07 | 2.100E-06 | 2.500E-06 |
| 1566 | 79 | I79E | A | 1.5e-06 | 1.7e-06 | 1.9e-06 | 1.700E-06 | 2.000E-07 | 1.500E-06 | 1.900E-06 |
| 1567 | 79 | I79Q | A | 2.9e-06 | 2.5e-06 | 1.7e-06 | 2.367E-06 | 6.110E-07 | 1.700E-06 | 2.900E-06 |
| 1568 | 79 | I79G | A | 2e-06 | 1.8e-06 | 3.6e-06 | 2.467E-06 | 9.866E-07 | 1.800E-06 | 3.600E-06 |
| 1569 | 79 | I79H | A | 1.6e-06 | 1.6e-06 | 1.7e-06 | 1.633E-06 | 5.774E-08 | 1.600E-06 | 1.700E-06 |
| 1570 | 79 | I79I | A | 1.9e-06 | 2.7e-06 | 2.2e-06 | 2.267E-06 | 4.041E-07 | 1.900E-06 | 2.700E-06 |
| 1571 | 79 | I79L | A | 4e-06 | 3.7e-06 | 1.8e-06 | 3.167E-06 | 1.193E-06 | 1.800E-06 | 4.000E-06 |
| 1572 | 79 | I79K | A | 2.1e-06 | 2.4e-06 | 2.1e-06 | 2.200E-06 | 1.732E-07 | 2.100E-06 | 2.400E-06 |
| 1573 | 79 | I79M | A | 3.7e-06 | 4e-06 | 1.9e-06 | 3.200E-06 | 1.136E-06 | 1.900E-06 | 4.000E-06 |
| 1574 | 79 | I79F | A | 3.2e-06 | 1.3e-06 | 1.9e-06 | 2.133E-06 | 9.713E-07 | 1.300E-06 | 3.200E-06 |
| 1575 | 79 | I79P | A | 3e-06 | 3e-06 | 2e-06 | 2.667E-06 | 5.774E-07 | 2.000E-06 | 3.000E-06 |
| 1576 | 79 | I79S | A | 2.7e-06 | 2.5e-06 | 2.7e-06 | 2.633E-06 | 1.155E-07 | 2.500E-06 | 2.700E-06 |
| 1577 | 79 | I79T | A | 2.8e-06 | 1.9e-06 | 1.1e-06 | 1.933E-06 | 8.505E-07 | 1.100E-06 | 2.800E-06 |
| 1578 | 79 | I79W | A | 3.6e-06 | 1.4e-06 | 4.4e-06 | 3.133E-06 | 1.553E-06 | 1.400E-06 | 4.400E-06 |
| 1579 | 79 | I79Y | A | 2.4e-06 | 1.6e-06 | 2.6e-06 | 2.200E-06 | 5.292E-07 | 1.600E-06 | 2.600E-06 |
| 1580 | 79 | I79V | A | 2.3e-06 | 1.7e-06 | 2.2e-06 | 2.067E-06 | 3.215E-07 | 1.700E-06 | 2.300E-06 |
| 1581 | 80 | V80A | A | 2.2e-06 | 2.3e-06 | 1.9e-06 | 2.133E-06 | 2.082E-07 | 1.900E-06 | 2.300E-06 |
| 1582 | 80 | V80R | A | 2.5e-06 | 2.5e-06 | 3.6e-06 | 2.867E-06 | 6.351E-07 | 2.500E-06 | 3.600E-06 |
| 1583 | 80 | V80N | A | 4.6e-06 | 3.5e-06 | 2.4e-06 | 3.500E-06 | 1.100E-06 | 2.400E-06 | 4.600E-06 |
| 1584 | 80 | V80D | A | 2.8e-06 | 2.5e-06 | 4e-06 | 3.100E-06 | 7.937E-07 | 2.500E-06 | 4.000E-06 |
| 1585 | 80 | V80C | A | 1.5e-06 | 3.4e-06 | 3.8e-06 | 2.900E-06 | 1.229E-06 | 1.500E-06 | 3.800E-06 |
| 1586 | 80 | V80E | A | 3.7e-06 | 2.3e-06 | 1.5e-06 | 2.500E-06 | 1.114E-06 | 1.500E-06 | 3.700E-06 |
| 1587 | 80 | V80Q | A | 2.3e-06 | 2.6e-06 | 3.1e-06 | 2.667E-06 | 4.041E-07 | 2.300E-06 | 3.100E-06 |
| 1588 | 80 | V80G | A | 2.2e-06 | 2.6e-06 | 2.5e-06 | 2.433E-06 | 2.082E-07 | 2.200E-06 | 2.600E-06 |
| 1589 | 80 | V80H | A | 1.8e-06 | 3.2e-06 | 1.9e-06 | 2.300E-06 | 7.810E-07 | 1.800E-06 | 3.200E-06 |
| 1590 | 80 | V80I | A | 1.9e-06 | 1.7e-06 | 1.9e-06 | 1.833E-06 | 1.155E-07 | 1.700E-06 | 1.900E-06 |
| 1591 | 80 | V80L | A | 3.9e-06 | 3.2e-06 | 2.7e-06 | 3.267E-06 | 6.028E-07 | 2.700E-06 | 3.900E-06 |
| 1592 | 80 | V80K | A | 4.3e-06 | 2.7e-06 | 1.4e-06 | 2.800E-06 | 1.453E-06 | 1.400E-06 | 4.300E-06 |
| 1593 | 80 | V80M | A | 3.6e-06 | 2.2e-06 | 1.6e-06 | 2.467E-06 | 1.026E-06 | 1.600E-06 | 3.600E-06 |
| 1594 | 80 | V80F | A | 5.1e-06 | 2.3e-06 | 2.2e-06 | 3.200E-06 | 1.646E-06 | 2.200E-06 | 5.100E-06 |
| 1595 | 80 | V80P | A | 3.3e-06 | 2.7e-06 | 2e-06 | 2.667E-06 | 6.506E-07 | 2.000E-06 | 3.300E-06 |
| 1596 | 80 | V80S | A | 9.9e-07 | 3e-06 | 3.2e-06 | 2.397E-06 | 1.222E-06 | 9.900E-07 | 3.200E-06 |
| 1597 | 80 | V80T | A | 3.2e-06 | 4.5e-06 | 1.7e-06 | 3.133E-06 | 1.401E-06 | 1.700E-06 | 4.500E-06 |
| 1598 | 80 | V80W | A | 1.4e-06 | 2.5e-06 | 2.5e-06 | 2.133E-06 | 6.351E-07 | 1.400E-06 | 2.500E-06 |
| 1599 | 80 | V80Y | A | 1e-06 | 3.7e-06 | 1.7e-06 | 2.133E-06 | 1.401E-06 | 1.000E-06 | 3.700E-06 |
| 1600 | 80 | V80V | A | 2.5e-06 | 2.3e-06 | 2.6e-06 | 2.467E-06 | 1.528E-07 | 2.300E-06 | 2.600E-06 |
| 1601 | 81 | R81A | A | 1.9e-06 | 2.6e-06 | 2e-06 | 2.167E-06 | 3.786E-07 | 1.900E-06 | 2.600E-06 |
| 1602 | 81 | R81R | A | 1.7e-06 | 1e-06 | 2.2e-06 | 1.633E-06 | 6.028E-07 | 1.000E-06 | 2.200E-06 |
| 1603 | 81 | R81N | A | 1.8e-06 | 2.6e-06 | 3e-06 | 2.467E-06 | 6.110E-07 | 1.800E-06 | 3.000E-06 |
| 1604 | 81 | R81D | A | 5.8e-06 | 2.8e-06 | 4.1e-06 | 4.233E-06 | 1.504E-06 | 2.800E-06 | 5.800E-06 |
| 1605 | 81 | R81C | A | 2.7e-06 | 3.4e-06 | 1.9e-06 | 2.667E-06 | 7.506E-07 | 1.900E-06 | 3.400E-06 |
| 1606 | 81 | R81E | A | 1.8e-06 | 3.3e-06 | 1.9e-06 | 2.333E-06 | 8.386E-07 | 1.800E-06 | 3.300E-06 |
| 1607 | 81 | R81Q | A | 1.5e-06 | 1.7e-06 | 1.9e-06 | 1.700E-06 | 2.000E-07 | 1.500E-06 | 1.900E-06 |
| 1608 | 81 | R81G | A | 1.8e-06 | 1.4e-06 | 4.2e-06 | 2.467E-06 | 1.514E-06 | 1.400E-06 | 4.200E-06 |
| 1609 | 81 | R81H | A | 2.1e-06 | 2.5e-06 | 1.7e-06 | 2.100E-06 | 4.000E-07 | 1.700E-06 | 2.500E-06 |
| 1610 | 81 | R81I | A | 2.2e-06 | 1.4e-06 | 3.4e-06 | 2.333E-06 | 1.007E-06 | 1.400E-06 | 3.400E-06 |
| 1611 | 81 | R81L | A | 2.7e-06 | 2.3e-06 | 1.4e-06 | 2.133E-06 | 6.658E-07 | 1.400E-06 | 2.700E-06 |
| 1612 | 81 | R81K | A | 2.5e-06 | 1.7e-06 | 2e-06 | 2.067E-06 | 4.041E-07 | 1.700E-06 | 2.500E-06 |
| 1613 | 81 | R81M | A | 2.8e-06 | 1.7e-06 | 2.5e-06 | 2.333E-06 | 5.686E-07 | 1.700E-06 | 2.800E-06 |
| 1614 | 81 | R81F | A | 3.2e-06 | 1.2e-06 | 2.5e-06 | 2.300E-06 | 1.015E-06 | 1.200E-06 | 3.200E-06 |

| | | | | | | | | | | |
|---|---|---|---|---|---|---|---|---|---|---|
| 1615 | 81 | R81P | A | 2.7e-06 | 3.2e-06 | 2.4e-06 | 2.767E-06 | 4.041E-07 | 2.400E-06 | 3.200E-06 |
| 1616 | 81 | R81S | A | 1.7e-06 | 2.1e-06 | 3.1e-06 | 2.300E-06 | 7.211E-07 | 1.700E-06 | 3.100E-06 |
| 1617 | 81 | R81T | A | 2.6e-06 | 2.7e-06 | 3.2e-06 | 2.833E-06 | 3.215E-07 | 2.600E-06 | 3.200E-06 |
| 1618 | 81 | R81W | A | 3.1e-06 | 2.5e-06 | 3.2e-06 | 2.933E-06 | 3.786E-07 | 2.500E-06 | 3.200E-06 |
| 1619 | 81 | R81Y | A | 2e-06 | 2e-06 | 2.2e-06 | 2.067E-06 | 1.155E-07 | 2.000E-06 | 2.200E-06 |
| 1620 | 81 | R81V | A | 1.8e-06 | 2.8e-06 | 3.1e-06 | 2.567E-06 | 6.807E-07 | 1.800E-06 | 3.100E-06 |
| 1621 | 82 | G82A | A | 2.9e-06 | 2.4e-06 | 2.5e-06 | 2.600E-06 | 2.646E-07 | 2.400E-06 | 2.900E-06 |
| 1622 | 82 | G82R | A | 2.1e-06 | 1.5e-06 | 2.7e-06 | 2.100E-06 | 6.000E-07 | 1.500E-06 | 2.700E-06 |
| 1623 | 82 | G82N | A | 2.6e-06 | 2.9e-06 | 2.9e-06 | 2.800E-06 | 1.732E-07 | 2.600E-06 | 2.900E-06 |
| 1624 | 82 | G82D | A | 2.1e-06 | 2.3e-06 | 5.1e-06 | 3.167E-06 | 1.677E-06 | 2.100E-06 | 5.100E-06 |
| 1625 | 82 | G82C | A | 2.7e-06 | 3.5e-06 | 2.6e-06 | 2.933E-06 | 4.933E-07 | 2.600E-06 | 3.500E-06 |
| 1626 | 82 | G82E | A | 2e-06 | 3.5e-06 | 5e-06 | 3.500E-06 | 1.500E-06 | 2.000E-06 | 5.000E-06 |
| 1627 | 82 | G82Q | A | 3.1e-06 | 3.3e-06 | 1.8e-06 | 2.733E-06 | 8.145E-07 | 1.800E-06 | 3.300E-06 |
| 1628 | 82 | G82G | A | 3.7e-06 | 3.6e-06 | 2.1e-06 | 3.133E-06 | 8.963E-07 | 2.100E-06 | 3.700E-06 |
| 1629 | 82 | G82H | A | 2.5e-06 | 1.8e-06 | 2.3e-06 | 2.200E-06 | 3.606E-07 | 1.800E-06 | 2.500E-06 |
| 1630 | 82 | G82I | A | 2.6e-06 | 4e-06 | 2.9e-06 | 3.167E-06 | 7.371E-07 | 2.600E-06 | 4.000E-06 |
| 1631 | 82 | G82L | A | 2e-06 | 2.2e-06 | 2.3e-06 | 2.167E-06 | 1.528E-07 | 2.000E-06 | 2.300E-06 |
| 1632 | 82 | G82K | A | 2.3e-06 | 3.5e-06 | 1.5e-06 | 2.433E-06 | 1.007E-06 | 1.500E-06 | 3.500E-06 |
| 1633 | 82 | G82M | A | 2.4e-06 | 2.1e-06 | 2.3e-06 | 2.267E-06 | 1.528E-07 | 2.100E-06 | 2.400E-06 |
| 1634 | 82 | G82F | A | 2.8e-06 | 5.3e-06 | 1.4e-06 | 3.167E-06 | 1.976E-06 | 1.400E-06 | 5.300E-06 |
| 1635 | 82 | G82P | A | 4.1e-06 | 2.2e-06 | 2.4e-06 | 2.900E-06 | 1.044E-06 | 2.200E-06 | 4.100E-06 |
| 1636 | 82 | G82S | A | 2.3e-06 | 1.4e-06 | 9.5e-07 | 1.550E-06 | 6.874E-07 | 9.500E-07 | 2.300E-06 |
| 1637 | 82 | G82T | A | 2e-06 | 2.8e-06 | 2.3e-06 | 2.367E-06 | 4.041E-07 | 2.000E-06 | 2.800E-06 |
| 1638 | 82 | G82W | A | 2.7e-06 | 2.3e-06 | 2e-06 | 2.333E-06 | 3.512E-07 | 2.000E-06 | 2.700E-06 |
| 1639 | 82 | G82Y | A | 2.9e-06 | 2.4e-06 | 2.1e-06 | 2.467E-06 | 4.041E-07 | 2.100E-06 | 2.900E-06 |
| 1640 | 82 | G82V | A | 2.3e-06 | 2.4e-06 | 3.9e-06 | 2.867E-06 | 8.963E-07 | 2.300E-06 | 3.900E-06 |
| 1641 | 83 | T83A | A | 3.8e-06 | 2.3e-06 | 2e-06 | 2.700E-06 | 9.644E-07 | 2.000E-06 | 3.800E-06 |
| 1642 | 83 | T83R | A | 2.6e-06 | 2.4e-06 | 2.4e-06 | 2.467E-06 | 1.155E-07 | 2.400E-06 | 2.600E-06 |
| 1643 | 83 | T83N | A | 2.4e-06 | 3.4e-06 | 3.2e-06 | 3.000E-06 | 5.292E-07 | 2.400E-06 | 3.400E-06 |
| 1644 | 83 | T83D | A | 3.1e-06 | 3.1e-06 | 2.3e-06 | 2.833E-06 | 4.619E-07 | 2.300E-06 | 3.100E-06 |
| 1645 | 83 | T83C | A | 2.5e-06 | 2.2e-06 | 2.7e-06 | 2.467E-06 | 2.517E-07 | 2.200E-06 | 2.700E-06 |
| 1646 | 83 | T83E | A | 2.8e-06 | 2.6e-06 | 2.7e-06 | 2.700E-06 | 1.000E-07 | 2.600E-06 | 2.800E-06 |
| 1647 | 83 | T83Q | A | 6.8e-07 | 2e-06 | 1.3e-06 | 1.327E-06 | 6.604E-07 | 6.800E-07 | 2.000E-06 |
| 1648 | 83 | T83G | A | 1.8e-06 | 2.5e-06 | 2.8e-06 | 2.367E-06 | 5.132E-07 | 1.800E-06 | 2.800E-06 |
| 1649 | 83 | T83H | A | 3e-06 | 4.4e-06 | 2.1e-06 | 3.167E-06 | 1.159E-06 | 2.100E-06 | 4.400E-06 |
| 1650 | 83 | T83I | A | 2.7e-06 | 2.4e-06 | 2.1e-06 | 2.400E-06 | 3.000E-07 | 2.100E-06 | 2.700E-06 |
| 1651 | 83 | T83L | A | 2.7e-06 | 1.7e-06 | 3.4e-06 | 2.600E-06 | 8.544E-07 | 1.700E-06 | 3.400E-06 |
| 1652 | 83 | T83K | A | 2e-06 | 3.2e-06 | 2.6e-06 | 2.600E-06 | 6.000E-07 | 2.000E-06 | 3.200E-06 |
| 1653 | 83 | T83M | A | 3.2e-06 | 1.8e-06 | 2e-06 | 2.333E-06 | 7.572E-07 | 1.800E-06 | 3.200E-06 |
| 1654 | 83 | T83F | A | 2e-06 | 1.3e-06 | 3.2e-06 | 2.167E-06 | 9.609E-07 | 1.300E-06 | 3.200E-06 |
| 1655 | 83 | T83P | A | 2e-06 | 2.8e-06 | 4.2e-06 | 3.000E-06 | 1.114E-06 | 2.000E-06 | 4.200E-06 |
| 1656 | 83 | T83S | A | 2.5e-06 | 3.1e-06 | 1.8e-06 | 2.467E-06 | 6.506E-07 | 1.800E-06 | 3.100E-06 |
| 1657 | 83 | T83T | A | 1.4e-06 | 2.5e-06 | 2.4e-06 | 2.100E-06 | 6.083E-07 | 1.400E-06 | 2.500E-06 |
| 1658 | 83 | T83W | A | 2.3e-06 | 2e-06 | 1.7e-06 | 2.000E-06 | 3.000E-07 | 1.700E-06 | 2.300E-06 |
| 1659 | 83 | T83Y | A | 3.6e-06 | 2.7e-06 | 2.3e-06 | 2.867E-06 | 6.658E-07 | 2.300E-06 | 3.600E-06 |
| 1660 | 83 | T83V | A | 2.8e-06 | 2.1e-06 | 3e-06 | 2.633E-06 | 4.726E-07 | 2.100E-06 | 3.000E-06 |
| 1661 | 84 | Q84A | A | 2.8e-06 | 3.2e-06 | 2.2e-06 | 2.733E-06 | 5.033E-07 | 2.200E-06 | 3.200E-06 |
| 1662 | 84 | Q84R | A | 2.2e-06 | 2.9e-06 | 2e-06 | 2.367E-06 | 4.726E-07 | 2.000E-06 | 2.900E-06 |
| 1663 | 84 | Q84N | A | 2.7e-06 | 1.9e-06 | 2e-06 | 2.200E-06 | 4.359E-07 | 1.900E-06 | 2.700E-06 |
| 1664 | 84 | Q84D | A | 2e-06 | 2.5e-06 | 1.9e-06 | 2.133E-06 | 3.215E-07 | 1.900E-06 | 2.500E-06 |
| 1665 | 84 | Q84C | A | 2.7e-06 | 3.9e-06 | 5e-06 | 3.867E-06 | 1.150E-06 | 2.700E-06 | 5.000E-06 |
| 1666 | 84 | Q84E | A | 3.1e-06 | 2.7e-06 | 3e-06 | 2.933E-06 | 2.082E-07 | 2.700E-06 | 3.100E-06 |
| 1667 | 84 | Q84Q | A | 1.7e-06 | 2.2e-06 | 9.4e-07 | 1.613E-06 | 6.345E-07 | 9.400E-07 | 2.200E-06 |
| 1668 | 84 | Q84G | A | 3.1e-06 | 3.3e-06 | 3.8e-06 | 3.400E-06 | 3.606E-07 | 3.100E-06 | 3.800E-06 |
| 1669 | 84 | Q84H | A | 2.5e-06 | 2.8e-06 | 3.5e-06 | 2.933E-06 | 5.132E-07 | 2.500E-06 | 3.500E-06 |
| 1670 | 84 | Q84I | A | 2.1e-06 | 2.8e-06 | 9.9e-07 | 1.963E-06 | 9.127E-07 | 9.900E-07 | 2.800E-06 |
| 1671 | 84 | Q84L | A | 2.3e-06 | 1.7e-06 | 2e-06 | 2.000E-06 | 3.000E-07 | 1.700E-06 | 2.300E-06 |
| 1672 | 84 | Q84K | A | 3e-06 | 1.6e-06 | 2.1e-06 | 2.233E-06 | 7.095E-07 | 1.600E-06 | 3.000E-06 |
| 1673 | 84 | Q84M | A | 3.7e-06 | 1.9e-06 | 2.1e-06 | 2.567E-06 | 9.866E-07 | 1.900E-06 | 3.700E-06 |
| 1674 | 84 | Q84F | A | 2.6e-06 | 2.4e-06 | 2.8e-06 | 2.600E-06 | 2.000E-07 | 2.400E-06 | 2.800E-06 |
| 1675 | 84 | Q84P | A | 2.8e-06 | 2.6e-06 | 1.9e-06 | 2.433E-06 | 4.726E-07 | 1.900E-06 | 2.800E-06 |
| 1676 | 84 | Q84S | A | 3.1e-06 | 2.1e-06 | 1.7e-06 | 2.300E-06 | 7.211E-07 | 1.700E-06 | 3.100E-06 |
| 1677 | 84 | Q84T | A | 2.3e-06 | 2.7e-06 | 2.4e-06 | 2.467E-06 | 2.082E-07 | 2.300E-06 | 2.700E-06 |
| 1678 | 84 | Q84W | A | 1.6e-06 | 2.5e-06 | 1.2e-06 | 1.767E-06 | 6.658E-07 | 1.200E-06 | 2.500E-06 |
| 1679 | 84 | Q84Y | A | 2.5e-06 | 2.2e-06 | 2.4e-06 | 2.367E-06 | 1.528E-07 | 2.200E-06 | 2.500E-06 |
| 1680 | 84 | Q84V | A | 2.6e-06 | 1.5e-06 | 3.6e-06 | 2.567E-06 | 1.050E-06 | 1.500E-06 | 3.600E-06 |
| 1681 | 85 | L85A | A | 2e-06 | 2.3e-06 | 1.7e-06 | 2.000E-06 | 3.000E-07 | 1.700E-06 | 2.300E-06 |
| 1682 | 85 | L85R | A | 2.7e-06 | 2.2e-06 | 2.3e-06 | 2.400E-06 | 2.646E-07 | 2.200E-06 | 2.700E-06 |
| 1683 | 85 | L85N | A | 2.6e-06 | 2.2e-06 | 4.6e-06 | 3.133E-06 | 1.286E-06 | 2.200E-06 | 4.600E-06 |
| 1684 | 85 | L85D | A | 3.5e-06 | 4.2e-06 | 4.7e-06 | 4.133E-06 | 6.028E-07 | 3.500E-06 | 4.700E-06 |
| 1685 | 85 | L85C | A | 2.1e-06 | 1.7e-06 | 2.9e-06 | 2.233E-06 | 6.110E-07 | 1.700E-06 | 2.900E-06 |
| 1686 | 85 | L85E | A | 3.5e-06 | 1.9e-06 | 2.8e-06 | 2.733E-06 | 8.021E-07 | 1.900E-06 | 3.500E-06 |
| 1687 | 85 | L85Q | A | 3e-06 | 2.3e-06 | 2.7e-06 | 2.667E-06 | 3.512E-07 | 2.300E-06 | 3.000E-06 |
| 1688 | 85 | L85G | A | 2.3e-06 | 2.7e-06 | 1.5e-06 | 2.167E-06 | 6.110E-07 | 1.500E-06 | 2.700E-06 |
| 1689 | 85 | L85H | A | 2e-06 | 4.1e-06 | 2.8e-06 | 2.967E-06 | 1.060E-06 | 2.000E-06 | 4.100E-06 |
| 1690 | 85 | L85I | A | 2e-06 | 2e-06 | 2.2e-06 | 2.067E-06 | 1.155E-07 | 2.000E-06 | 2.200E-06 |
| 1691 | 85 | L85L | A | 1.7e-06 | 1.9e-06 | 2.2e-06 | 1.933E-06 | 2.517E-07 | 1.700E-06 | 2.200E-06 |
| 1692 | 85 | L85K | A | 1.2e-06 | 1.1e-06 | 2.3e-06 | 1.533E-06 | 6.658E-07 | 1.100E-06 | 2.300E-06 |
| 1693 | 85 | L85M | A | 2.6e-06 | 1.9e-06 | 2.4e-06 | 2.300E-06 | 3.606E-07 | 1.900E-06 | 2.600E-06 |
| 1694 | 85 | L85F | A | 5.1e-06 | 1.5e-06 | 9.8e-07 | 2.527E-06 | 2.244E-06 | 9.800E-07 | 5.100E-06 |
| 1695 | 85 | L85P | A | 2.4e-06 | 3.3e-06 | 3.3e-06 | 3.000E-06 | 5.196E-07 | 2.400E-06 | 3.300E-06 |
| 1696 | 85 | L85S | A | 1.8e-06 | 1.8e-06 | 2.8e-06 | 2.133E-06 | 5.774E-07 | 1.800E-06 | 2.800E-06 |
| 1697 | 85 | L85T | A | 1.7e-06 | 2e-06 | 3.7e-06 | 2.467E-06 | 1.079E-06 | 1.700E-06 | 3.700E-06 |
| 1698 | 85 | L85W | A | 1.6e-06 | 1.8e-06 | 2.2e-06 | 1.867E-06 | 3.055E-07 | 1.600E-06 | 2.200E-06 |
| 1699 | 85 | L85Y | A | 2e-06 | 2.4e-06 | 2e-06 | 2.133E-06 | 2.309E-07 | 2.000E-06 | 2.400E-06 |

| | | | | | | | | | | |
|---|---|---|---|---|---|---|---|---|---|---|
| 1700 | 85 | L85V | A | 2.8e-06 | 3.1e-06 | 2.9e-06 | 2.933E-06 | 1.528E-07 | 2.800E-06 | 3.100E-06 |
| 1701 | 86 | F86A | A | 2.6e-06 | 3.5e-06 | 2.5e-06 | 2.867E-06 | 5.508E-07 | 2.500E-06 | 3.500E-06 |
| 1702 | 86 | F86R | A | 1.3e-06 | 1.6e-06 | 2.2e-06 | 1.700E-06 | 4.583E-07 | 1.300E-06 | 2.200E-06 |
| 1703 | 86 | F86N | A | 1.4e-06 | 1.4e-06 | 2e-06 | 1.600E-06 | 3.464E-07 | 1.400E-06 | 2.000E-06 |
| 1704 | 86 | F86D | A | 1.8e-06 | 2.4e-06 | 2.2e-06 | 2.133E-06 | 3.055E-07 | 1.800E-06 | 2.400E-06 |
| 1705 | 86 | F86C | A | 2.6e-06 | 2.1e-06 | 2.9e-06 | 2.533E-06 | 4.041E-07 | 2.100E-06 | 2.900E-06 |
| 1706 | 86 | F86E | A | 2.5e-06 | 2.4e-06 | 1.7e-06 | 2.200E-06 | 4.359E-07 | 1.700E-06 | 2.500E-06 |
| 1707 | 86 | F86Q | A | 2.3e-06 | 2.8e-06 | 3.4e-06 | 2.833E-06 | 5.508E-07 | 2.300E-06 | 3.400E-06 |
| 1708 | 86 | F86G | A | 3.5e-06 | 5.4e-06 | 1.3e-06 | 3.400E-06 | 2.052E-06 | 1.300E-06 | 5.400E-06 |
| 1709 | 86 | F86H | A | 2.7e-06 | 2e-06 | 2.3e-06 | 2.333E-06 | 3.512E-07 | 2.000E-06 | 2.700E-06 |
| 1710 | 86 | F86I | A | 2.5e-06 | 2.9e-06 | 2.9e-06 | 2.767E-06 | 2.309E-07 | 2.500E-06 | 2.900E-06 |
| 1711 | 86 | F86L | A | 3.5e-06 | 1.9e-06 | 1.8e-06 | 2.400E-06 | 9.539E-07 | 1.800E-06 | 3.500E-06 |
| 1712 | 86 | F86K | A | 2.5e-06 | 2e-06 | 2.4e-06 | 2.300E-06 | 2.646E-07 | 2.000E-06 | 2.500E-06 |
| 1713 | 86 | F86M | A | 1e-06 | 1.3e-06 | 2e-06 | 1.433E-06 | 5.132E-07 | 1.000E-06 | 2.000E-06 |
| 1714 | 86 | F86F | A | 2e-06 | 1.5e-06 | 2.2e-06 | 1.900E-06 | 3.606E-07 | 1.500E-06 | 2.200E-06 |
| 1715 | 86 | F86P | A | 2.2e-06 | 4.4e-06 | 1.7e-06 | 2.767E-06 | 1.436E-06 | 1.700E-06 | 4.400E-06 |
| 1716 | 86 | F86S | A | 2.1e-06 | 2e-06 | 2.3e-06 | 2.133E-06 | 1.528E-07 | 2.000E-06 | 2.300E-06 |
| 1717 | 86 | F86T | A | 3.4e-06 | 2e-06 | 1.5e-06 | 2.300E-06 | 9.849E-07 | 1.500E-06 | 3.400E-06 |
| 1718 | 86 | F86W | A | 1.7e-06 | 2.6e-06 | 2.1e-06 | 2.133E-06 | 4.509E-07 | 1.700E-06 | 2.600E-06 |
| 1719 | 86 | F86Y | A | 2.3e-06 | 1.7e-06 | 2.8e-06 | 2.267E-06 | 5.508E-07 | 1.700E-06 | 2.800E-06 |
| 1720 | 86 | F86V | A | 2.7e-06 | 1e-06 | 2.5e-06 | 2.067E-06 | 9.292E-07 | 1.000E-06 | 2.700E-06 |
| 1721 | 87 | E87A | A | 4e-06 | 1.4e-06 | 3.1e-06 | 2.833E-06 | 1.320E-06 | 1.400E-06 | 4.000E-06 |
| 1722 | 87 | E87R | A | 4.4e-06 | 3.5e-06 | 2.1e-06 | 3.333E-06 | 1.159E-06 | 2.100E-06 | 4.400E-06 |
| 1723 | 87 | E87N | A | 1.6e-06 | 4.1e-06 | 2.3e-06 | 2.667E-06 | 1.290E-06 | 1.600E-06 | 4.100E-06 |
| 1724 | 87 | E87D | A | 2.7e-06 | 3.2e-06 | 4.6e-06 | 3.500E-06 | 9.849E-07 | 2.700E-06 | 4.600E-06 |
| 1725 | 87 | E87C | A | 3.5e-06 | 3e-06 | 3.8e-06 | 3.433E-06 | 4.041E-07 | 3.000E-06 | 3.800E-06 |
| 1726 | 87 | E87E | A | 3.5e-06 | 2.4e-06 | 1.6e-06 | 2.500E-06 | 9.539E-07 | 1.600E-06 | 3.500E-06 |
| 1727 | 87 | E87Q | A | 1.3e-06 | 1.5e-06 | 1.9e-06 | 1.567E-06 | 3.055E-07 | 1.300E-06 | 1.900E-06 |
| 1728 | 87 | E87G | A | 2.4e-06 | 2.6e-06 | 3.3e-06 | 2.767E-06 | 4.726E-07 | 2.400E-06 | 3.300E-06 |
| 1729 | 87 | E87H | A | 2.8e-06 | 2.2e-06 | 2.9e-06 | 2.633E-06 | 3.786E-07 | 2.200E-06 | 2.900E-06 |
| 1730 | 87 | E87I | A | 2.5e-06 | 1.5e-06 | 2e-06 | 2.000E-06 | 5.000E-07 | 1.500E-06 | 2.500E-06 |
| 1731 | 87 | E87L | A | 2.1e-06 | 3.6e-06 | 2.2e-06 | 2.633E-06 | 8.386E-07 | 2.100E-06 | 3.600E-06 |
| 1732 | 87 | E87K | A | 3.8e-06 | 2.7e-06 | 1.7e-06 | 2.733E-06 | 1.050E-06 | 1.700E-06 | 3.800E-06 |
| 1733 | 87 | E87M | A | 1.2e-06 | 2.5e-06 | 3.2e-06 | 2.300E-06 | 1.015E-06 | 1.200E-06 | 3.200E-06 |
| 1734 | 87 | E87F | A | 2.8e-06 | 2.3e-06 | 2.5e-06 | 2.533E-06 | 2.517E-07 | 2.300E-06 | 2.800E-06 |
| 1735 | 87 | E87P | A | 3e-06 | 1.7e-06 | 4e-06 | 2.900E-06 | 1.153E-06 | 1.700E-06 | 4.000E-06 |
| 1736 | 87 | E87S | A | 2.7e-06 | 3.2e-06 | 1.7e-06 | 2.533E-06 | 7.638E-07 | 1.700E-06 | 3.200E-06 |
| 1737 | 87 | E87T | A | 2.4e-06 | 1.9e-06 | 2.7e-06 | 2.333E-06 | 4.041E-07 | 1.900E-06 | 2.700E-06 |
| 1738 | 87 | E87W | A | 2.5e-06 | 3e-06 | 2.1e-06 | 2.533E-06 | 4.509E-07 | 2.100E-06 | 3.000E-06 |
| 1739 | 87 | E87Y | A | 2.2e-06 | 2.5e-06 | 2.8e-06 | 2.500E-06 | 3.000E-07 | 2.200E-06 | 2.800E-06 |
| 1740 | 87 | E87V | A | 2.2e-06 | 1.4e-06 | 1.9e-06 | 1.833E-06 | 4.041E-07 | 1.400E-06 | 2.200E-06 |
| 1741 | 88 | D88A | A | 1.4e-06 | 2.2e-06 | 3.5e-06 | 2.367E-06 | 1.060E-06 | 1.400E-06 | 3.500E-06 |
| 1742 | 88 | D88R | A | 2.4e-06 | 2.4e-06 | 2.3e-06 | 2.367E-06 | 5.774E-08 | 2.300E-06 | 2.400E-06 |
| 1743 | 88 | D88N | A | 2e-06 | 1.2e-06 | 2.2e-06 | 1.800E-06 | 5.292E-07 | 1.200E-06 | 2.200E-06 |
| 1744 | 88 | D88D | A | 2.6e-06 | 1.4e-06 | 1.9e-06 | 1.967E-06 | 6.028E-07 | 1.400E-06 | 2.600E-06 |
| 1745 | 88 | D88C | A | 2.1e-06 | 2.4e-06 | 2.5e-06 | 2.333E-06 | 2.082E-07 | 2.100E-06 | 2.500E-06 |
| 1746 | 88 | D88E | A | 4.2e-06 | 5.3e-06 | 3.3e-06 | 4.267E-06 | 1.002E-06 | 3.300E-06 | 5.300E-06 |
| 1747 | 88 | D88Q | A | 4.5e-06 | 2.9e-06 | 3e-06 | 3.467E-06 | 8.963E-07 | 2.900E-06 | 4.500E-06 |
| 1748 | 88 | D88G | A | 3.9e-06 | 3.6e-06 | 3.2e-06 | 3.567E-06 | 3.512E-07 | 3.200E-06 | 3.900E-06 |
| 1749 | 88 | D88H | A | 2.3e-06 | 2e-06 | 3.3e-06 | 2.533E-06 | 6.807E-07 | 2.000E-06 | 3.300E-06 |
| 1750 | 88 | D88I | A | 2.6e-06 | 3e-06 | 2.8e-06 | 2.800E-06 | 2.000E-07 | 2.600E-06 | 3.000E-06 |
| 1751 | 88 | D88L | A | 2.5e-06 | 2.1e-06 | 1.6e-06 | 2.067E-06 | 4.509E-07 | 1.600E-06 | 2.500E-06 |
| 1752 | 88 | D88K | A | 1.9e-06 | 3e-06 | 1e-06 | 1.967E-06 | 1.002E-06 | 1.000E-06 | 3.000E-06 |
| 1753 | 88 | D88M | A | 5.1e-06 | 2.4e-06 | 2.8e-06 | 3.433E-06 | 1.457E-06 | 2.400E-06 | 5.100E-06 |
| 1754 | 88 | D88F | A | 2.5e-06 | 2.1e-06 | 2.9e-06 | 2.500E-06 | 4.000E-07 | 2.100E-06 | 2.900E-06 |
| 1755 | 88 | D88P | A | 2.8e-06 | 3.2e-06 | 2.9e-06 | 2.967E-06 | 2.082E-07 | 2.800E-06 | 3.200E-06 |
| 1756 | 88 | D88S | A | 3.7e-06 | 1.7e-06 | 2.2e-06 | 2.533E-06 | 1.041E-06 | 1.700E-06 | 3.700E-06 |
| 1757 | 88 | D88T | A | 2.2e-06 | 2.2e-06 | 2.8e-06 | 2.400E-06 | 3.464E-07 | 2.200E-06 | 2.800E-06 |
| 1758 | 88 | D88W | A | 1.4e-06 | 2.5e-06 | 3e-06 | 2.300E-06 | 8.185E-07 | 1.400E-06 | 3.000E-06 |
| 1759 | 88 | D88Y | A | 3.4e-06 | 2.6e-06 | 1.3e-06 | 2.433E-06 | 1.060E-06 | 1.300E-06 | 3.400E-06 |
| 1760 | 88 | D88V | A | 2.2e-06 | 1.7e-06 | 1.5e-06 | 1.800E-06 | 3.606E-07 | 1.500E-06 | 2.200E-06 |
| 1761 | 89 | N89A | A | 2.3e-06 | 3.1e-06 | 1.6e-06 | 2.333E-06 | 7.506E-07 | 1.600E-06 | 3.100E-06 |
| 1762 | 89 | N89R | A | 2.5e-06 | 2.2e-06 | 1.5e-06 | 2.067E-06 | 5.132E-07 | 1.500E-06 | 2.500E-06 |
| 1763 | 89 | N89N | A | 1.9e-06 | 2.4e-06 | 1.7e-06 | 2.000E-06 | 3.606E-07 | 1.700E-06 | 2.400E-06 |
| 1764 | 89 | N89D | A | 2.5e-06 | 2.5e-06 | 2.2e-06 | 2.400E-06 | 1.732E-07 | 2.200E-06 | 2.500E-06 |
| 1765 | 89 | N89C | A | 2.5e-06 | 1.6e-06 | 1.6e-06 | 1.900E-06 | 5.196E-07 | 1.600E-06 | 2.500E-06 |
| 1766 | 89 | N89E | A | 2.5e-06 | 5.2e-06 | 2.3e-06 | 3.333E-06 | 1.620E-06 | 2.300E-06 | 5.200E-06 |
| 1767 | 89 | N89Q | A | 2.8e-06 | 2.7e-06 | 2.6e-06 | 2.700E-06 | 1.000E-07 | 2.600E-06 | 2.800E-06 |
| 1768 | 89 | N89G | A | 1.7e-06 | 3.1e-06 | 2.9e-06 | 2.567E-06 | 7.572E-07 | 1.700E-06 | 3.100E-06 |
| 1769 | 89 | N89H | A | 2.8e-06 | 4e-06 | 2.4e-06 | 3.067E-06 | 8.327E-07 | 2.400E-06 | 4.000E-06 |
| 1770 | 89 | N89I | A | 2.1e-06 | 1.9e-06 | 4.5e-06 | 2.833E-06 | 1.447E-06 | 1.900E-06 | 4.500E-06 |
| 1771 | 89 | N89L | A | 3.4e-06 | 2.2e-06 | 2.7e-06 | 2.767E-06 | 6.028E-07 | 2.200E-06 | 3.400E-06 |
| 1772 | 89 | N89K | A | 2.3e-06 | 3.3e-06 | 2.9e-06 | 2.833E-06 | 5.033E-07 | 2.300E-06 | 3.300E-06 |
| 1773 | 89 | N89M | A | 5.5e-06 | 2.2e-06 | 2.1e-06 | 3.267E-06 | 1.935E-06 | 2.100E-06 | 5.500E-06 |
| 1774 | 89 | N89F | A | 1.6e-06 | 1.9e-06 | 2e-06 | 1.833E-06 | 2.082E-07 | 1.600E-06 | 2.000E-06 |
| 1775 | 89 | N89P | A | 1.4e-06 | 5.5e-06 | 1.8e-06 | 2.900E-06 | 2.261E-06 | 1.400E-06 | 5.500E-06 |
| 1776 | 89 | N89S | A | 1.3e-06 | 2.5e-06 | 2.5e-06 | 2.100E-06 | 6.928E-07 | 1.300E-06 | 2.500E-06 |
| 1777 | 89 | N89T | A | 2.1e-06 | 3.2e-06 | 4e-06 | 3.100E-06 | 9.539E-07 | 2.100E-06 | 4.000E-06 |
| 1778 | 89 | N89W | A | 1.3e-06 | 2.1e-06 | 3.5e-06 | 2.300E-06 | 1.114E-06 | 1.300E-06 | 3.500E-06 |
| 1779 | 89 | N89Y | A | 3.1e-06 | 3.5e-06 | 2.1e-06 | 2.900E-06 | 7.211E-07 | 2.100E-06 | 3.500E-06 |
| 1780 | 89 | N89V | A | 3.4e-06 | 2.2e-06 | 1.8e-06 | 2.467E-06 | 8.327E-07 | 1.800E-06 | 3.400E-06 |
| 1781 | 90 | Y90A | A | 1.4e-06 | 2.4e-06 | 2.4e-06 | 2.067E-06 | 5.774E-07 | 1.400E-06 | 2.400E-06 |
| 1782 | 90 | Y90R | A | 1.5e-06 | 2.9e-06 | 1.1e-06 | 1.833E-06 | 9.452E-07 | 1.100E-06 | 2.900E-06 |
| 1783 | 90 | Y90N | A | 1.7e-06 | 2e-06 | 1.4e-06 | 1.700E-06 | 3.000E-07 | 1.400E-06 | 2.000E-06 |
| 1784 | 90 | Y90D | A | 2.7e-06 | 2.4e-06 | 2.8e-06 | 2.633E-06 | 2.082E-07 | 2.400E-06 | 2.800E-06 |

| | | | | | | | | | | |
|---|---|---|---|---|---|---|---|---|---|---|
| 1785 | 90 | Y90C | A | 1.4e-06 | 2.1e-06 | 2.5e-06 | 2.000E-06 | 5.568E-07 | 1.400E-06 | 2.500E-06 |
| 1786 | 90 | Y90E | A | 1.7e-06 | 2e-06 | 1.7e-06 | 1.800E-06 | 1.732E-07 | 1.700E-06 | 2.000E-06 |
| 1787 | 90 | Y90Q | A | 3.2e-06 | 3e-06 | 2e-06 | 2.733E-06 | 6.429E-07 | 2.000E-06 | 3.200E-06 |
| 1788 | 90 | Y90G | A | 2.8e-06 | 3.8e-06 | 3.2e-06 | 3.267E-06 | 5.033E-07 | 2.800E-06 | 3.800E-06 |
| 1789 | 90 | Y90H | A | 3.1e-06 | 2.9e-06 | 1.3e-06 | 2.433E-06 | 9.866E-07 | 1.300E-06 | 3.100E-06 |
| 1790 | 90 | Y90I | A | 1.2e-06 | 2.2e-06 | 1.9e-06 | 1.767E-06 | 5.132E-07 | 1.200E-06 | 2.200E-06 |
| 1791 | 90 | Y90L | A | 1.5e-06 | 3.2e-06 | 3.8e-06 | 2.833E-06 | 1.193E-06 | 1.500E-06 | 3.800E-06 |
| 1792 | 90 | Y90K | A | 2.2e-06 | 2.7e-06 | 4.8e-06 | 3.233E-06 | 1.380E-06 | 2.200E-06 | 4.800E-06 |
| 1793 | 90 | Y90M | A | 2.1e-06 | 3.4e-06 | 3.1e-06 | 2.867E-06 | 6.807E-07 | 2.100E-06 | 3.400E-06 |
| 1794 | 90 | Y90F | A | 2.9e-06 | 1.7e-06 | 2.9e-06 | 2.500E-06 | 6.928E-07 | 1.700E-06 | 2.900E-06 |
| 1795 | 90 | Y90P | A | 2.7e-06 | 1.6e-06 | 1.6e-06 | 1.967E-06 | 6.351E-07 | 1.600E-06 | 2.700E-06 |
| 1796 | 90 | Y90S | A | 2.1e-06 | 1.8e-06 | 1.4e-06 | 1.767E-06 | 3.512E-07 | 1.400E-06 | 2.100E-06 |
| 1797 | 90 | Y90T | A | 2.8e-06 | 1.8e-06 | 2.6e-06 | 2.400E-06 | 5.292E-07 | 1.800E-06 | 2.800E-06 |
| 1798 | 90 | Y90W | A | 1.3e-06 | 2.5e-06 | 3.1e-06 | 2.300E-06 | 9.165E-07 | 1.300E-06 | 3.100E-06 |
| 1799 | 90 | Y90Y | A | 2.2e-06 | 1.9e-06 | 3.4e-06 | 2.500E-06 | 7.937E-07 | 1.900E-06 | 3.400E-06 |
| 1800 | 90 | Y90V | A | 1.7e-06 | 2.8e-06 | 1.5e-06 | 2.000E-06 | 7.000E-07 | 1.500E-06 | 2.800E-06 |
| 1801 | 91 | A91A | A | 2.7e-06 | 4.4e-06 | 9e-07 | 2.667E-06 | 1.750E-06 | 9.000E-07 | 4.400E-06 |
| 1802 | 91 | A91R | A | 3.8e-06 | 1.1e-06 | 3.2e-06 | 2.700E-06 | 1.418E-06 | 1.100E-06 | 3.800E-06 |
| 1803 | 91 | A91N | A | 4e-06 | 3.9e-06 | 2.5e-06 | 3.467E-06 | 8.386E-07 | 2.500E-06 | 4.000E-06 |
| 1804 | 91 | A91D | A | 3.8e-06 | 1.5e-06 | 1.5e-06 | 2.267E-06 | 1.328E-06 | 1.500E-06 | 3.800E-06 |
| 1805 | 91 | A91C | A | 1.2e-06 | 1.6e-06 | 2.2e-06 | 1.667E-06 | 5.033E-07 | 1.200E-06 | 2.200E-06 |
| 1806 | 91 | A91E | A | 2.3e-06 | 2.1e-06 | 1.6e-06 | 2.000E-06 | 3.606E-07 | 1.600E-06 | 2.300E-06 |
| 1807 | 91 | A91Q | A | 1.9e-06 | 1.3e-06 | 2.3e-06 | 1.833E-06 | 5.033E-07 | 1.300E-06 | 2.300E-06 |
| 1808 | 91 | A91G | A | 2.6e-06 | 2.5e-06 | 3.8e-06 | 2.967E-06 | 7.234E-07 | 2.500E-06 | 3.800E-06 |
| 1809 | 91 | A91H | A | 3.1e-06 | 2.1e-06 | 3e-06 | 2.733E-06 | 5.508E-07 | 2.100E-06 | 3.100E-06 |
| 1810 | 91 | A91I | A | 2.4e-06 | 1.5e-06 | 2.5e-06 | 2.133E-06 | 5.508E-07 | 1.500E-06 | 2.500E-06 |
| 1811 | 91 | A91L | A | 3.3e-06 | 1.5e-06 | 3.1e-06 | 2.633E-06 | 9.866E-07 | 1.500E-06 | 3.300E-06 |
| 1812 | 91 | A91K | A | 3e-06 | 2.7e-06 | 2.4e-06 | 2.700E-06 | 3.000E-07 | 2.400E-06 | 3.000E-06 |
| 1813 | 91 | A91M | A | 1.4e-06 | 3.4e-06 | 1.8e-06 | 2.200E-06 | 1.058E-06 | 1.400E-06 | 3.400E-06 |
| 1814 | 91 | A91F | A | 2.7e-06 | 2.2e-06 | 3e-06 | 2.633E-06 | 4.041E-07 | 2.200E-06 | 3.000E-06 |
| 1815 | 91 | A91P | A | 1.7e-06 | 3.2e-06 | 1.4e-06 | 2.100E-06 | 9.644E-07 | 1.400E-06 | 3.200E-06 |
| 1816 | 91 | A91S | A | 1.7e-06 | 2.2e-06 | 3.4e-06 | 2.433E-06 | 8.737E-07 | 1.700E-06 | 3.400E-06 |
| 1817 | 91 | A91T | A | 1.7e-06 | 1.8e-06 | 4.5e-06 | 2.667E-06 | 1.589E-06 | 1.700E-06 | 4.500E-06 |
| 1818 | 91 | A91W | A | 3.9e-06 | 1.8e-06 | 1.9e-06 | 2.533E-06 | 1.185E-06 | 1.800E-06 | 3.900E-06 |
| 1819 | 91 | A91Y | A | 2.5e-06 | 2.9e-06 | 3e-06 | 2.800E-06 | 2.646E-07 | 2.500E-06 | 3.000E-06 |
| 1820 | 91 | A91V | A | 2.9e-06 | 3.3e-06 | 2.4e-06 | 2.867E-06 | 4.509E-07 | 2.400E-06 | 3.300E-06 |
| 1821 | 92 | L92A | A | 2.9e-06 | 3.8e-06 | 1.8e-06 | 2.833E-06 | 1.002E-06 | 1.800E-06 | 3.800E-06 |
| 1822 | 92 | L92R | A | 2.2e-06 | 1.9e-06 | 3.3e-06 | 2.467E-06 | 7.371E-07 | 1.900E-06 | 3.300E-06 |
| 1823 | 92 | L92N | A | 2.8e-06 | 1.7e-06 | 2.2e-06 | 2.233E-06 | 5.508E-07 | 1.700E-06 | 2.800E-06 |
| 1824 | 92 | L92D | A | 2.5e-06 | 1.8e-06 | 1.4e-06 | 1.900E-06 | 5.568E-07 | 1.400E-06 | 2.500E-06 |
| 1825 | 92 | L92C | A | 2.7e-06 | 2.7e-06 | 3.2e-06 | 2.867E-06 | 2.887E-07 | 2.700E-06 | 3.200E-06 |
| 1826 | 92 | L92E | A | 2.1e-06 | 2.8e-06 | 3.3e-06 | 2.733E-06 | 6.028E-07 | 2.100E-06 | 3.300E-06 |
| 1827 | 92 | L92Q | A | 2.5e-06 | 2.1e-06 | 2.4e-06 | 2.333E-06 | 2.082E-07 | 2.100E-06 | 2.500E-06 |
| 1828 | 92 | L92G | A | 2.3e-06 | 3.1e-06 | 3e-06 | 2.800E-06 | 4.359E-07 | 2.300E-06 | 3.100E-06 |
| 1829 | 92 | L92H | A | 2.5e-06 | 1.9e-06 | 2.5e-06 | 2.300E-06 | 3.464E-07 | 1.900E-06 | 2.500E-06 |
| 1830 | 92 | L92I | A | 3.9e-06 | 2.1e-06 | 2.8e-06 | 2.933E-06 | 9.074E-07 | 2.100E-06 | 3.900E-06 |
| 1831 | 92 | L92L | A | 1.8e-06 | 2.4e-06 | 2.3e-06 | 2.167E-06 | 3.215E-07 | 1.800E-06 | 2.400E-06 |
| 1832 | 92 | L92K | A | 3.1e-06 | 1.8e-06 | 4.4e-06 | 3.100E-06 | 1.300E-06 | 1.800E-06 | 4.400E-06 |
| 1833 | 92 | L92M | A | 2.6e-06 | 3e-06 | 2.1e-06 | 2.567E-06 | 4.509E-07 | 2.100E-06 | 3.000E-06 |
| 1834 | 92 | L92F | A | 1.6e-06 | 1.2e-06 | 2.7e-06 | 1.833E-06 | 7.767E-07 | 1.200E-06 | 2.700E-06 |
| 1835 | 92 | L92P | A | 2.6e-06 | 2e-06 | 2e-06 | 2.200E-06 | 3.464E-07 | 2.000E-06 | 2.600E-06 |
| 1836 | 92 | L92S | A | 2.4e-06 | 1.9e-06 | 2.4e-06 | 2.233E-06 | 2.887E-07 | 1.900E-06 | 2.400E-06 |
| 1837 | 92 | L92T | A | 1.9e-06 | 2.4e-06 | 3.1e-06 | 2.467E-06 | 6.028E-07 | 1.900E-06 | 3.100E-06 |
| 1838 | 92 | L92W | A | 1.5e-06 | 3.9e-06 | 2.5e-06 | 2.633E-06 | 1.206E-06 | 1.500E-06 | 3.900E-06 |
| 1839 | 92 | L92Y | A | 2.9e-06 | 1.9e-06 | 2.9e-06 | 2.567E-06 | 5.774E-07 | 1.900E-06 | 2.900E-06 |
| 1840 | 92 | L92V | A | 3.7e-06 | 2.7e-06 | 2.4e-06 | 2.933E-06 | 6.807E-07 | 2.400E-06 | 3.700E-06 |
| 1841 | 93 | A93A | A | 3.8e-06 | 2.5e-06 | 2.9e-06 | 3.067E-06 | 6.658E-07 | 2.500E-06 | 3.800E-06 |
| 1842 | 93 | A93R | A | 1.1e-06 | 3.9e-06 | 1.8e-06 | 2.267E-06 | 1.457E-06 | 1.100E-06 | 3.900E-06 |
| 1843 | 93 | A93N | A | 2.1e-06 | 2.1e-06 | 2.2e-06 | 2.133E-06 | 5.774E-08 | 2.100E-06 | 2.200E-06 |
| 1844 | 93 | A93D | A | 2.1e-06 | 3.8e-06 | 2.4e-06 | 2.767E-06 | 9.074E-07 | 2.100E-06 | 3.800E-06 |
| 1845 | 93 | A93C | A | 2.4e-06 | 2.9e-06 | 2.6e-06 | 2.633E-06 | 2.517E-07 | 2.400E-06 | 2.900E-06 |
| 1846 | 93 | A93E | A | 1.5e-06 | 2.6e-06 | 4.6e-06 | 2.900E-06 | 1.572E-06 | 1.500E-06 | 4.600E-06 |
| 1847 | 93 | A93Q | A | 2.1e-06 | 1.9e-06 | 2.3e-06 | 2.100E-06 | 2.000E-07 | 1.900E-06 | 2.300E-06 |
| 1848 | 93 | A93G | A | 2.8e-06 | 1.4e-06 | 1.6e-06 | 1.933E-06 | 7.572E-07 | 1.400E-06 | 2.800E-06 |
| 1849 | 93 | A93H | A | 2.4e-06 | 3.2e-06 | 3.5e-06 | 3.033E-06 | 5.686E-07 | 2.400E-06 | 3.500E-06 |
| 1850 | 93 | A93I | A | 1.4e-06 | 3.1e-06 | 1.3e-06 | 1.933E-06 | 1.012E-06 | 1.300E-06 | 3.100E-06 |
| 1851 | 93 | A93L | A | 1.8e-06 | 2.2e-06 | 1.7e-06 | 1.900E-06 | 2.646E-07 | 1.700E-06 | 2.200E-06 |
| 1852 | 93 | A93K | A | 2.4e-06 | 2.6e-06 | 3e-06 | 2.667E-06 | 3.055E-07 | 2.400E-06 | 3.000E-06 |
| 1853 | 93 | A93M | A | 2.8e-06 | 2.9e-06 | 3.6e-06 | 3.100E-06 | 4.359E-07 | 2.800E-06 | 3.600E-06 |
| 1854 | 93 | A93F | A | 1.8e-06 | 2.6e-06 | 2.1e-06 | 2.167E-06 | 4.041E-07 | 1.800E-06 | 2.600E-06 |
| 1855 | 93 | A93P | A | 3.7e-06 | 4.5e-06 | 2.8e-06 | 3.667E-06 | 8.505E-07 | 2.800E-06 | 4.500E-06 |
| 1856 | 93 | A93S | A | 1.7e-06 | 1.9e-06 | 2.8e-06 | 2.133E-06 | 5.859E-07 | 1.700E-06 | 2.800E-06 |
| 1857 | 93 | A93T | A | 1.9e-06 | 2.8e-06 | 1.7e-06 | 2.133E-06 | 5.859E-07 | 1.700E-06 | 2.800E-06 |
| 1858 | 93 | A93W | A | 3.3e-06 | 3e-06 | 3.7e-06 | 3.333E-06 | 3.512E-07 | 3.000E-06 | 3.700E-06 |
| 1859 | 93 | A93Y | A | 3.1e-06 | 2.9e-06 | 2.9e-06 | 2.967E-06 | 1.155E-07 | 2.900E-06 | 3.100E-06 |
| 1860 | 93 | A93V | A | 2.4e-06 | 1.7e-06 | 2.2e-06 | 2.100E-06 | 3.606E-07 | 1.700E-06 | 2.400E-06 |
| 1861 | 94 | V94A | A | 3e-06 | 2.7e-06 | 3.9e-06 | 3.200E-06 | 6.245E-07 | 2.700E-06 | 3.900E-06 |
| 1862 | 94 | V94R | A | 1.8e-06 | 1.5e-06 | 2.3e-06 | 1.867E-06 | 4.041E-07 | 1.500E-06 | 2.300E-06 |
| 1863 | 94 | V94N | A | 4e-06 | 1.7e-06 | 1.4e-06 | 2.367E-06 | 1.422E-06 | 1.400E-06 | 4.000E-06 |
| 1864 | 94 | V94D | A | 2.8e-06 | 3.3e-06 | 3.5e-06 | 3.200E-06 | 3.606E-07 | 2.800E-06 | 3.500E-06 |
| 1865 | 94 | V94C | A | 3.5e-06 | 4.4e-06 | 2.3e-06 | 3.400E-06 | 1.054E-06 | 2.300E-06 | 4.400E-06 |
| 1866 | 94 | V94E | A | 2.4e-06 | 2.6e-06 | 2.9e-06 | 2.633E-06 | 2.517E-07 | 2.400E-06 | 2.900E-06 |
| 1867 | 94 | V94Q | A | 1.8e-06 | 1.6e-06 | 3.9e-06 | 2.433E-06 | 1.274E-06 | 1.600E-06 | 3.900E-06 |
| 1868 | 94 | V94G | A | 3.5e-06 | 3e-06 | 1.7e-06 | 2.733E-06 | 9.292E-07 | 1.700E-06 | 3.500E-06 |
| 1869 | 94 | V94H | A | 2.6e-06 | 1.7e-06 | 2.2e-06 | 2.167E-06 | 4.509E-07 | 1.700E-06 | 2.600E-06 |

| | | | | | | | | | | |
|---|---|---|---|---|---|---|---|---|---|---|
| 1870 | 94 | V94I | A | 1.4e-06 | 1.6e-06 | 1.6e-06 | 1.533E-06 | 1.155E-07 | 1.400E-06 | 1.600E-06 |
| 1871 | 94 | V94L | A | 1.7e-06 | 1e-06 | 2.4e-06 | 1.700E-06 | 7.000E-07 | 1.000E-06 | 2.400E-06 |
| 1872 | 94 | V94K | A | 2.1e-06 | 2.2e-06 | 1.2e-06 | 1.833E-06 | 5.508E-07 | 1.200E-06 | 2.200E-06 |
| 1873 | 94 | V94M | A | 2.6e-06 | 3.3e-06 | 1.2e-06 | 2.367E-06 | 1.069E-06 | 1.200E-06 | 3.300E-06 |
| 1874 | 94 | V94F | A | 2e-06 | 2.8e-06 | 2.1e-06 | 2.300E-06 | 4.359E-07 | 2.000E-06 | 2.800E-06 |
| 1875 | 94 | V94P | A | 1.7e-06 | 3.8e-06 | 2.3e-06 | 2.600E-06 | 1.082E-06 | 1.700E-06 | 3.800E-06 |
| 1876 | 94 | V94S | A | 2.7e-06 | 2.2e-06 | 2.9e-06 | 2.600E-06 | 3.606E-07 | 2.200E-06 | 2.900E-06 |
| 1877 | 94 | V94T | A | 3.2e-06 | 4.6e-06 | 2.5e-06 | 3.433E-06 | 1.069E-06 | 2.500E-06 | 4.600E-06 |
| 1878 | 94 | V94W | A | 1.6e-06 | 1.6e-06 | 2.1e-06 | 1.767E-06 | 2.887E-07 | 1.600E-06 | 2.100E-06 |
| 1879 | 94 | V94Y | A | 1.8e-06 | 2.8e-06 | 2.4e-06 | 2.333E-06 | 5.033E-07 | 1.800E-06 | 2.800E-06 |
| 1880 | 94 | V94V | A | 2e-06 | 2.1e-06 | 2.8e-06 | 2.300E-06 | 4.359E-07 | 2.000E-06 | 2.800E-06 |
| 1881 | 95 | L95A | A | 3.1e-06 | 1.9e-06 | 2.7e-06 | 2.567E-06 | 6.110E-07 | 1.900E-06 | 3.100E-06 |
| 1882 | 95 | L95R | A | 2.2e-06 | 3.3e-06 | 2.3e-06 | 2.600E-06 | 6.083E-07 | 2.200E-06 | 3.300E-06 |
| 1883 | 95 | L95N | A | 2.1e-06 | 2e-06 | 3.8e-06 | 2.633E-06 | 1.012E-06 | 2.000E-06 | 3.800E-06 |
| 1884 | 95 | L95D | A | 3e-06 | 2.9e-06 | 1.4e-06 | 2.433E-06 | 8.963E-07 | 1.400E-06 | 3.000E-06 |
| 1885 | 95 | L95C | A | 2.8e-06 | 1.8e-06 | 4.2e-06 | 2.933E-06 | 1.206E-06 | 1.800E-06 | 4.200E-06 |
| 1886 | 95 | L95E | A | 2.3e-06 | 2.6e-06 | 3.3e-06 | 2.733E-06 | 5.132E-07 | 2.300E-06 | 3.300E-06 |
| 1887 | 95 | L95Q | A | 1.9e-06 | 3.3e-06 | 2.7e-06 | 2.633E-06 | 7.024E-07 | 1.900E-06 | 3.300E-06 |
| 1888 | 95 | L95G | A | 2.9e-06 | 2.1e-06 | 2.3e-06 | 2.433E-06 | 4.163E-07 | 2.100E-06 | 2.900E-06 |
| 1889 | 95 | L95H | A | 2.3e-06 | 2.3e-06 | 2e-06 | 2.200E-06 | 1.732E-07 | 2.000E-06 | 2.300E-06 |
| 1890 | 95 | L95I | A | 3.2e-06 | 1.9e-06 | 2.2e-06 | 2.433E-06 | 6.807E-07 | 1.900E-06 | 3.200E-06 |
| 1891 | 95 | L95L | A | 2.9e-06 | 1.8e-06 | 2.4e-06 | 2.367E-06 | 5.508E-07 | 1.800E-06 | 2.900E-06 |
| 1892 | 95 | L95K | A | 2e-06 | 2.4e-06 | 1.4e-06 | 1.933E-06 | 5.033E-07 | 1.400E-06 | 2.400E-06 |
| 1893 | 95 | L95M | A | 2.4e-06 | 2.1e-06 | 1.7e-06 | 2.067E-06 | 3.512E-07 | 1.700E-06 | 2.400E-06 |
| 1894 | 95 | L95F | A | 2.3e-06 | 2.7e-06 | 2.4e-06 | 2.467E-06 | 2.082E-07 | 2.300E-06 | 2.700E-06 |
| 1895 | 95 | L95P | A | 2e-06 | 4.4e-06 | 2.6e-06 | 3.000E-06 | 1.249E-06 | 2.000E-06 | 4.400E-06 |
| 1896 | 95 | L95S | A | 3.3e-06 | 2.6e-06 | 3.9e-06 | 3.267E-06 | 6.506E-07 | 2.600E-06 | 3.900E-06 |
| 1897 | 95 | L95T | A | 1.5e-06 | 1.9e-06 | 2.8e-06 | 2.067E-06 | 6.658E-07 | 1.500E-06 | 2.800E-06 |
| 1898 | 95 | L95W | A | 1.5e-06 | 1.2e-06 | 1.6e-06 | 1.433E-06 | 2.082E-07 | 1.200E-06 | 1.600E-06 |
| 1899 | 95 | L95Y | A | 2.3e-06 | 1.2e-06 | 1.8e-06 | 1.767E-06 | 5.508E-07 | 1.200E-06 | 2.300E-06 |
| 1900 | 95 | L95V | A | 1.8e-06 | 4.5e-06 | 2.2e-06 | 2.833E-06 | 1.457E-06 | 1.800E-06 | 4.500E-06 |
| 1901 | 96 | D96A | A | 2.2e-06 | 2.2e-06 | 2.6e-06 | 2.333E-06 | 2.309E-07 | 2.200E-06 | 2.600E-06 |
| 1902 | 96 | D96R | A | 2.3e-06 | 2.3e-06 | 2.2e-06 | 2.267E-06 | 5.774E-08 | 2.200E-06 | 2.300E-06 |
| 1903 | 96 | D96N | A | 2.3e-06 | 1.9e-06 | 2.5e-06 | 2.233E-06 | 3.055E-07 | 1.900E-06 | 2.500E-06 |
| 1904 | 96 | D96D | A | 2.2e-06 | 2.7e-06 | 2.3e-06 | 2.400E-06 | 2.646E-07 | 2.200E-06 | 2.700E-06 |
| 1905 | 96 | D96C | A | 2.8e-06 | 1.8e-06 | 2e-06 | 2.200E-06 | 5.292E-07 | 1.800E-06 | 2.800E-06 |
| 1906 | 96 | D96E | A | 1.9e-06 | 2.2e-06 | 2.4e-06 | 2.167E-06 | 2.517E-07 | 1.900E-06 | 2.400E-06 |
| 1907 | 96 | D96Q | A | 1.8e-06 | 1.4e-06 | 2.5e-06 | 1.900E-06 | 5.568E-07 | 1.400E-06 | 2.500E-06 |
| 1908 | 96 | D96G | A | 3e-06 | 2.1e-06 | 1.8e-06 | 2.300E-06 | 6.245E-07 | 1.800E-06 | 3.000E-06 |
| 1909 | 96 | D96H | A | 1.6e-06 | 1.9e-06 | 1.9e-06 | 1.800E-06 | 1.732E-07 | 1.600E-06 | 1.900E-06 |
| 1910 | 96 | D96I | A | 3.3e-06 | 2.7e-06 | 3.2e-06 | 3.067E-06 | 3.215E-07 | 2.700E-06 | 3.300E-06 |
| 1911 | 96 | D96L | A | 3.7e-06 | 2.7e-06 | 2.6e-06 | 3.000E-06 | 6.083E-07 | 2.600E-06 | 3.700E-06 |
| 1912 | 96 | D96K | A | 1.8e-06 | 3.8e-06 | 2.3e-06 | 2.633E-06 | 1.041E-06 | 1.800E-06 | 3.800E-06 |
| 1913 | 96 | D96M | A | 2.6e-06 | 1.8e-06 | 2e-06 | 2.133E-06 | 4.163E-07 | 1.800E-06 | 2.600E-06 |
| 1914 | 96 | D96F | A | 2.2e-06 | 1.9e-06 | 1.7e-06 | 1.933E-06 | 2.517E-07 | 1.700E-06 | 2.200E-06 |
| 1915 | 96 | D96P | A | 1.8e-06 | 4.5e-06 | 2.1e-06 | 2.800E-06 | 1.480E-06 | 1.800E-06 | 4.500E-06 |
| 1916 | 96 | D96S | A | 1.9e-06 | 1.5e-06 | 1.7e-06 | 1.700E-06 | 2.000E-07 | 1.500E-06 | 1.900E-06 |
| 1917 | 96 | D96T | A | 2.1e-06 | 3.4e-06 | 2.6e-06 | 2.700E-06 | 6.557E-07 | 2.100E-06 | 3.400E-06 |
| 1918 | 96 | D96W | A | 2.3e-06 | 2.2e-06 | 2.4e-06 | 2.300E-06 | 1.000E-07 | 2.200E-06 | 2.400E-06 |
| 1919 | 96 | D96Y | A | 1.7e-06 | 2e-06 | 4e-06 | 2.567E-06 | 1.250E-06 | 1.700E-06 | 4.000E-06 |
| 1920 | 96 | D96V | A | 2.3e-06 | 4.4e-06 | 2.1e-06 | 2.933E-06 | 1.274E-06 | 2.100E-06 | 4.400E-06 |
| 1921 | 97 | N97A | A | 1.9e-06 | 2e-06 | 2.9e-06 | 2.267E-06 | 5.508E-07 | 1.900E-06 | 2.900E-06 |
| 1922 | 97 | N97R | A | 1.4e-06 | 3.4e-06 | 4e-06 | 2.933E-06 | 1.361E-06 | 1.400E-06 | 4.000E-06 |
| 1923 | 97 | N97N | A | 3.4e-06 | 2.6e-06 | 1e-06 | 2.333E-06 | 1.222E-06 | 1.000E-06 | 3.400E-06 |
| 1924 | 97 | N97D | A | 1.8e-06 | 2.6e-06 | 1.6e-06 | 2.000E-06 | 5.292E-07 | 1.600E-06 | 2.600E-06 |
| 1925 | 97 | N97C | A | 2.8e-06 | 3.1e-06 | 3e-06 | 2.967E-06 | 1.528E-07 | 2.800E-06 | 3.100E-06 |
| 1926 | 97 | N97E | A | 1.8e-06 | 2.6e-06 | 3.2e-06 | 2.533E-06 | 7.024E-07 | 1.800E-06 | 3.200E-06 |
| 1927 | 97 | N97Q | A | 2.1e-06 | 1.9e-06 | 2.7e-06 | 2.233E-06 | 4.163E-07 | 1.900E-06 | 2.700E-06 |
| 1928 | 97 | N97G | A | 2.6e-06 | 3.4e-06 | 3.1e-06 | 3.033E-06 | 4.041E-07 | 2.600E-06 | 3.400E-06 |
| 1929 | 97 | N97H | A | 2e-06 | 1.9e-06 | 2.3e-06 | 2.067E-06 | 2.082E-07 | 1.900E-06 | 2.300E-06 |
| 1930 | 97 | N97I | A | 2.9e-06 | 2.4e-06 | 2.1e-06 | 2.467E-06 | 4.041E-07 | 2.100E-06 | 2.900E-06 |
| 1931 | 97 | N97L | A | 2.2e-06 | 3.7e-06 | 1e-06 | 2.300E-06 | 1.353E-06 | 1.000E-06 | 3.700E-06 |
| 1932 | 97 | N97K | A | 4.5e-06 | 3e-06 | 3.2e-06 | 3.567E-06 | 8.145E-07 | 3.000E-06 | 4.500E-06 |
| 1933 | 97 | N97M | A | 2.5e-06 | 2.4e-06 | 3.4e-06 | 2.767E-06 | 5.508E-07 | 2.400E-06 | 3.400E-06 |
| 1934 | 97 | N97F | A | 3e-06 | 2.4e-06 | 2.9e-06 | 2.767E-06 | 3.215E-07 | 2.400E-06 | 3.000E-06 |
| 1935 | 97 | N97P | A | 3.5e-06 | 1.7e-06 | 3.2e-06 | 2.800E-06 | 9.644E-07 | 1.700E-06 | 3.500E-06 |
| 1936 | 97 | N97S | A | 2.6e-06 | 2.9e-06 | 2.8e-06 | 2.767E-06 | 1.528E-07 | 2.600E-06 | 2.900E-06 |
| 1937 | 97 | N97T | A | 2.5e-06 | 3e-06 | 1e-06 | 2.167E-06 | 1.041E-06 | 1.000E-06 | 3.000E-06 |
| 1938 | 97 | N97W | A | 1.6e-06 | 2.2e-06 | 2.4e-06 | 2.067E-06 | 4.163E-07 | 1.600E-06 | 2.400E-06 |
| 1939 | 97 | N97Y | A | 2.3e-06 | 1.5e-06 | 2.6e-06 | 2.133E-06 | 5.686E-07 | 1.500E-06 | 2.600E-06 |
| 1940 | 97 | N97V | A | 2.4e-06 | 2.6e-06 | 1.6e-06 | 2.200E-06 | 5.292E-07 | 1.600E-06 | 2.600E-06 |
| 1941 | 98 | G98A | A | 3e-06 | 1.6e-06 | 2.5e-06 | 2.367E-06 | 7.095E-07 | 1.600E-06 | 3.000E-06 |
| 1942 | 98 | G98R | A | 1.9e-06 | 2.3e-06 | 2.3e-06 | 2.167E-06 | 2.309E-07 | 1.900E-06 | 2.300E-06 |
| 1943 | 98 | G98N | A | 1.3e-06 | 2.2e-06 | 2.1e-06 | 1.867E-06 | 4.933E-07 | 1.300E-06 | 2.200E-06 |
| 1944 | 98 | G98D | A | 3.4e-06 | 2.9e-06 | 2.5e-06 | 2.933E-06 | 4.509E-07 | 2.500E-06 | 3.400E-06 |
| 1945 | 98 | G98C | A | 1.6e-06 | 2.9e-06 | 2.4e-06 | 2.300E-06 | 6.557E-07 | 1.600E-06 | 2.900E-06 |
| 1946 | 98 | G98E | A | 2.3e-06 | 3.8e-06 | 3e-06 | 3.033E-06 | 7.506E-07 | 2.300E-06 | 3.800E-06 |
| 1947 | 98 | G98Q | A | 2.6e-06 | 1.6e-06 | 2.4e-06 | 2.200E-06 | 5.292E-07 | 1.600E-06 | 2.600E-06 |
| 1948 | 98 | G98G | A | 2.8e-06 | 3.1e-06 | 2.6e-06 | 2.833E-06 | 2.517E-07 | 2.600E-06 | 3.100E-06 |
| 1949 | 98 | G98H | A | 2.1e-06 | 2.2e-06 | 1.9e-06 | 2.067E-06 | 1.528E-07 | 1.900E-06 | 2.200E-06 |
| 1950 | 98 | G98I | A | 3.3e-06 | 3.5e-06 | 1.8e-06 | 2.867E-06 | 9.292E-07 | 1.800E-06 | 3.500E-06 |
| 1951 | 98 | G98L | A | 2.7e-06 | 1.9e-06 | 2.2e-06 | 2.267E-06 | 4.041E-07 | 1.900E-06 | 2.700E-06 |
| 1952 | 98 | G98K | A | 2.4e-06 | 1.7e-06 | 2.7e-06 | 2.267E-06 | 5.132E-07 | 1.700E-06 | 2.700E-06 |
| 1953 | 98 | G98M | A | 2.4e-06 | 1.3e-06 | 1.1e-06 | 1.600E-06 | 7.000E-07 | 1.100E-06 | 2.400E-06 |
| 1954 | 98 | G98F | A | 2.3e-06 | 1.7e-06 | 2e-06 | 2.000E-06 | 3.000E-07 | 1.700E-06 | 2.300E-06 |

| | | | | | | | | | | |
|---|---|---|---|---|---|---|---|---|---|---|
| 1955 | 98 | G98P | A | 2.5e-06 | 1.9e-06 | 4.8e-06 | 3.067E-06 | 1.531E-06 | 1.900E-06 | 4.800E-06 |
| 1956 | 98 | G98S | A | 3.1e-06 | 2.5e-06 | 3.3e-06 | 2.967E-06 | 4.163E-07 | 2.500E-06 | 3.300E-06 |
| 1957 | 98 | G98T | A | 1.4e-06 | 2.3e-06 | 1.1e-06 | 1.600E-06 | 6.245E-07 | 1.100E-06 | 2.300E-06 |
| 1958 | 98 | G98W | A | 2.7e-06 | 4.6e-06 | 2e-06 | 3.100E-06 | 1.345E-06 | 2.000E-06 | 4.600E-06 |
| 1959 | 98 | G98Y | A | 1.5e-06 | 1.8e-06 | 2e-06 | 1.767E-06 | 2.517E-07 | 1.500E-06 | 2.000E-06 |
| 1960 | 98 | G98V | A | 1.2e-06 | 1.8e-06 | 2.9e-06 | 1.967E-06 | 8.622E-07 | 1.200E-06 | 2.900E-06 |
| 1961 | 99 | D99A | A | 2.1e-06 | 4.2e-06 | 2.4e-06 | 2.900E-06 | 1.136E-06 | 2.100E-06 | 4.200E-06 |
| 1962 | 99 | D99R | A | 2.6e-06 | 1.7e-06 | 1.9e-06 | 2.067E-06 | 4.726E-07 | 1.700E-06 | 2.600E-06 |
| 1963 | 99 | D99N | A | 3.6e-06 | 2e-06 | 1.6e-06 | 2.400E-06 | 1.058E-06 | 1.600E-06 | 3.600E-06 |
| 1964 | 99 | D99D | A | 2.3e-06 | 1.7e-06 | 1.6e-06 | 1.867E-06 | 3.786E-07 | 1.600E-06 | 2.300E-06 |
| 1965 | 99 | D99C | A | 2e-06 | 2.4e-06 | 1.1e-06 | 1.833E-06 | 6.658E-07 | 1.100E-06 | 2.400E-06 |
| 1966 | 99 | D99E | A | 2e-06 | 2e-06 | 1.5e-06 | 1.833E-06 | 2.887E-07 | 1.500E-06 | 2.000E-06 |
| 1967 | 99 | D99Q | A | 1.7e-06 | 2.1e-06 | 4.1e-06 | 2.633E-06 | 1.286E-06 | 1.700E-06 | 4.100E-06 |
| 1968 | 99 | D99G | A | 5.4e-06 | 3.3e-06 | 1.9e-06 | 3.533E-06 | 1.762E-06 | 1.900E-06 | 5.400E-06 |
| 1969 | 99 | D99H | A | 2.8e-06 | 2.3e-06 | 2.4e-06 | 2.500E-06 | 2.646E-07 | 2.300E-06 | 2.800E-06 |
| 1970 | 99 | D99I | A | 3.2e-06 | 2.3e-06 | 2.5e-06 | 2.667E-06 | 4.726E-07 | 2.300E-06 | 3.200E-06 |
| 1971 | 99 | D99L | A | 1.1e-06 | 2.8e-06 | 3.1e-06 | 2.333E-06 | 1.079E-06 | 1.100E-06 | 3.100E-06 |
| 1972 | 99 | D99K | A | 2.4e-06 | 2.9e-06 | 2.1e-06 | 2.467E-06 | 4.041E-07 | 2.100E-06 | 2.900E-06 |
| 1973 | 99 | D99M | A | 4.1e-06 | 2e-06 | 3.7e-06 | 3.267E-06 | 1.115E-06 | 2.000E-06 | 4.100E-06 |
| 1974 | 99 | D99F | A | 4.2e-06 | 3e-06 | 3.6e-06 | 3.600E-06 | 6.000E-07 | 3.000E-06 | 4.200E-06 |
| 1975 | 99 | D99P | A | 2e-06 | 1.7e-06 | 1.7e-06 | 1.800E-06 | 1.732E-07 | 1.700E-06 | 2.000E-06 |
| 1976 | 99 | D99S | A | 1.8e-06 | 2.2e-06 | 2e-06 | 2.000E-06 | 2.000E-07 | 1.800E-06 | 2.200E-06 |
| 1977 | 99 | D99T | A | 1.2e-06 | 1.5e-06 | 1.2e-06 | 1.300E-06 | 1.732E-07 | 1.200E-06 | 1.500E-06 |
| 1978 | 99 | D99W | A | 1.8e-06 | 1.4e-06 | 1.9e-06 | 1.700E-06 | 2.646E-07 | 1.400E-06 | 1.900E-06 |
| 1979 | 99 | D99Y | A | 1.7e-06 | 2.1e-06 | 2.6e-06 | 2.133E-06 | 4.509E-07 | 1.700E-06 | 2.600E-06 |
| 1980 | 99 | D99V | A | 3.7e-06 | 2.5e-06 | 2.9e-06 | 3.033E-06 | 6.110E-07 | 2.500E-06 | 3.700E-06 |
| 1981 | 100 | P100A | A | 2.8e-06 | 2.3e-06 | 1.8e-06 | 2.300E-06 | 5.000E-07 | 1.800E-06 | 2.800E-06 |
| 1982 | 100 | P100R | A | 2.3e-06 | 1.5e-06 | 2.6e-06 | 2.133E-06 | 5.686E-07 | 1.500E-06 | 2.600E-06 |
| 1983 | 100 | P100N | A | 1.9e-06 | 2.3e-06 | 1.8e-06 | 2.000E-06 | 2.646E-07 | 1.800E-06 | 2.300E-06 |
| 1984 | 100 | P100D | A | 2.3e-06 | 1.8e-06 | 2.5e-06 | 2.200E-06 | 3.606E-07 | 1.800E-06 | 2.500E-06 |
| 1985 | 100 | P100C | A | 4.3e-06 | 1.8e-06 | 1.8e-06 | 2.633E-06 | 1.443E-06 | 1.800E-06 | 4.300E-06 |
| 1986 | 100 | P100E | A | 2e-06 | 1.9e-06 | 1.5e-06 | 1.800E-06 | 2.646E-07 | 1.500E-06 | 2.000E-06 |
| 1987 | 100 | P100Q | A | 3.2e-06 | 2.1e-06 | 4.5e-06 | 3.267E-06 | 1.201E-06 | 2.100E-06 | 4.500E-06 |
| 1988 | 100 | P100G | A | 4.5e-06 | 2.9e-06 | 1.6e-06 | 3.000E-06 | 1.453E-06 | 1.600E-06 | 4.500E-06 |
| 1989 | 100 | P100H | A | 1.9e-06 | 2.6e-06 | 4.2e-06 | 2.900E-06 | 1.179E-06 | 1.900E-06 | 4.200E-06 |
| 1990 | 100 | P100I | A | 2.8e-06 | 2.2e-06 | 1.5e-06 | 2.167E-06 | 6.506E-07 | 1.500E-06 | 2.800E-06 |
| 1991 | 100 | P100L | A | 2e-06 | 1.6e-06 | 4e-06 | 2.533E-06 | 1.286E-06 | 1.600E-06 | 4.000E-06 |
| 1992 | 100 | P100K | A | 1.7e-06 | 2.6e-06 | 3.2e-06 | 2.500E-06 | 7.550E-07 | 1.700E-06 | 3.200E-06 |
| 1993 | 100 | P100M | A | 2.2e-06 | 2.2e-06 | 1.9e-06 | 2.100E-06 | 1.732E-07 | 1.900E-06 | 2.200E-06 |
| 1994 | 100 | P100F | A | 1.8e-06 | 2.6e-06 | 1.7e-06 | 2.033E-06 | 4.933E-07 | 1.700E-06 | 2.600E-06 |
| 1995 | 100 | P100P | A | 3.1e-06 | 4e-06 | 1.9e-06 | 3.000E-06 | 1.054E-06 | 1.900E-06 | 4.000E-06 |
| 1996 | 100 | P100S | A | 2.3e-06 | 3.8e-06 | 2.5e-06 | 2.867E-06 | 8.145E-07 | 2.300E-06 | 3.800E-06 |
| 1997 | 100 | P100T | A | 2.5e-06 | 2.9e-06 | 4.6e-06 | 3.333E-06 | 1.115E-06 | 2.500E-06 | 4.600E-06 |
| 1998 | 100 | P100W | A | 2.3e-06 | 3.2e-06 | 1.9e-06 | 2.467E-06 | 6.658E-07 | 1.900E-06 | 3.200E-06 |
| 1999 | 100 | P100Y | A | 1.4e-06 | 1.6e-06 | 2.5e-06 | 1.833E-06 | 5.859E-07 | 1.400E-06 | 2.500E-06 |
| 2000 | 100 | P100V | A | 2.9e-06 | 2.2e-06 | 3e-06 | 2.700E-06 | 4.359E-07 | 2.200E-06 | 3.000E-06 |
| 2001 | 101 | L101A | A | 2.8e-06 | 2.4e-06 | 2.9e-06 | 2.700E-06 | 2.646E-07 | 2.400E-06 | 2.900E-06 |
| 2002 | 101 | L101R | A | 2.9e-06 | 2.9e-06 | 1.7e-06 | 2.500E-06 | 6.928E-07 | 1.700E-06 | 2.900E-06 |
| 2003 | 101 | L101N | A | 2.7e-06 | 2.2e-06 | 3.9e-06 | 2.933E-06 | 8.737E-07 | 2.200E-06 | 3.900E-06 |
| 2004 | 101 | L101D | A | 2.3e-06 | 2.1e-06 | 1.2e-06 | 1.867E-06 | 5.859E-07 | 1.200E-06 | 2.300E-06 |
| 2005 | 101 | L101C | A | 2.2e-06 | 3.5e-06 | 2.4e-06 | 2.700E-06 | 7.000E-07 | 2.200E-06 | 3.500E-06 |
| 2006 | 101 | L101E | A | 2.4e-06 | 1.9e-06 | 2.1e-06 | 2.133E-06 | 2.517E-07 | 1.900E-06 | 2.400E-06 |
| 2007 | 101 | L101Q | A | 2.5e-06 | 2.8e-06 | 1.9e-06 | 2.400E-06 | 4.583E-07 | 1.900E-06 | 2.800E-06 |
| 2008 | 101 | L101G | A | 4e-06 | 2.2e-06 | 2.6e-06 | 2.933E-06 | 9.452E-07 | 2.200E-06 | 4.000E-06 |
| 2009 | 101 | L101H | A | 3e-06 | 2.1e-06 | 2.7e-06 | 2.600E-06 | 4.583E-07 | 2.100E-06 | 3.000E-06 |
| 2010 | 101 | L101I | A | 1.6e-06 | 3.3e-06 | 2.2e-06 | 2.367E-06 | 8.622E-07 | 1.600E-06 | 3.300E-06 |
| 2011 | 101 | L101L | A | 1.4e-06 | 2.1e-06 | 2.3e-06 | 1.933E-06 | 4.726E-07 | 1.400E-06 | 2.300E-06 |
| 2012 | 101 | L101K | A | 5.1e-06 | 1.5e-06 | 2.2e-06 | 2.933E-06 | 1.909E-06 | 1.500E-06 | 5.100E-06 |
| 2013 | 101 | L101M | A | 3e-06 | 2.3e-06 | 2.3e-06 | 2.533E-06 | 4.041E-07 | 2.300E-06 | 3.000E-06 |
| 2014 | 101 | L101F | A | 1.4e-06 | 2.9e-06 | 1.3e-06 | 1.867E-06 | 8.963E-07 | 1.300E-06 | 2.900E-06 |
| 2015 | 101 | L101P | A | 2.4e-06 | 3e-06 | 2.9e-06 | 2.767E-06 | 3.215E-07 | 2.400E-06 | 3.000E-06 |
| 2016 | 101 | L101S | A | 2.7e-06 | 1.2e-06 | 3.7e-06 | 2.533E-06 | 1.258E-06 | 1.200E-06 | 3.700E-06 |
| 2017 | 101 | L101T | A | 2.9e-06 | 1.9e-06 | 2.8e-06 | 2.533E-06 | 5.508E-07 | 1.900E-06 | 2.900E-06 |
| 2018 | 101 | L101W | A | 2.6e-06 | 1.8e-06 | 1.7e-06 | 2.033E-06 | 4.933E-07 | 1.700E-06 | 2.600E-06 |
| 2019 | 101 | L101Y | A | 1.9e-06 | 3.3e-06 | 2.3e-06 | 2.500E-06 | 7.211E-07 | 1.900E-06 | 3.300E-06 |
| 2020 | 101 | L101V | A | 1.6e-06 | 3.1e-06 | 3e-06 | 2.567E-06 | 8.386E-07 | 1.600E-06 | 3.100E-06 |
| 2021 | 102 | N102A | A | 3.9e-06 | 2e-06 | 3.3e-06 | 3.067E-06 | 9.713E-07 | 2.000E-06 | 3.900E-06 |
| 2022 | 102 | N102R | A | 2.5e-06 | 2.2e-06 | 3.8e-06 | 2.833E-06 | 8.505E-07 | 2.200E-06 | 3.800E-06 |
| 2023 | 102 | N102N | A | 2.3e-06 | 8.4e-07 | 2.7e-06 | 1.947E-06 | 9.790E-07 | 8.400E-07 | 2.700E-06 |
| 2024 | 102 | N102D | A | 2.7e-06 | 3.4e-06 | 2.5e-06 | 2.867E-06 | 4.726E-07 | 2.500E-06 | 3.400E-06 |
| 2025 | 102 | N102C | A | 3.3e-06 | 1.3e-06 | 2.7e-06 | 2.433E-06 | 1.026E-06 | 1.300E-06 | 3.300E-06 |
| 2026 | 102 | N102E | A | 3.1e-06 | 1.4e-06 | 3.2e-06 | 2.567E-06 | 1.012E-06 | 1.400E-06 | 3.200E-06 |
| 2027 | 102 | N102Q | A | 2.6e-06 | 2.2e-06 | 5.4e-06 | 3.400E-06 | 1.744E-06 | 2.200E-06 | 5.400E-06 |
| 2028 | 102 | N102G | A | 3.6e-06 | 2.7e-06 | 2.6e-06 | 2.967E-06 | 5.508E-07 | 2.600E-06 | 3.600E-06 |
| 2029 | 102 | N102H | A | 2.6e-06 | 1.9e-06 | 2.5e-06 | 2.333E-06 | 3.786E-07 | 1.900E-06 | 2.600E-06 |
| 2030 | 102 | N102I | A | 2.7e-06 | 1.9e-06 | 3e-06 | 2.533E-06 | 5.686E-07 | 1.900E-06 | 3.000E-06 |
| 2031 | 102 | N102L | A | 3.7e-06 | 2.2e-06 | 1.5e-06 | 2.467E-06 | 1.124E-06 | 1.500E-06 | 3.700E-06 |
| 2032 | 102 | N102K | A | 1e-06 | 2.9e-06 | 2.1e-06 | 2.000E-06 | 9.539E-07 | 1.000E-06 | 2.900E-06 |
| 2033 | 102 | N102M | A | 2.6e-06 | 2.3e-06 | 2.5e-06 | 2.467E-06 | 1.528E-07 | 2.300E-06 | 2.600E-06 |
| 2034 | 102 | N102F | A | 2e-06 | 3.5e-06 | 2.7e-06 | 2.733E-06 | 7.506E-07 | 2.000E-06 | 3.500E-06 |
| 2035 | 102 | N102P | A | 2.2e-06 | 2.3e-06 | 2e-06 | 2.167E-06 | 1.528E-07 | 2.000E-06 | 2.300E-06 |
| 2036 | 102 | N102S | A | 1.9e-06 | 2.9e-06 | 1e-06 | 1.933E-06 | 9.504E-07 | 1.000E-06 | 2.900E-06 |
| 2037 | 102 | N102T | A | 1.9e-06 | 5.2e-06 | 1.6e-06 | 2.900E-06 | 1.997E-06 | 1.600E-06 | 5.200E-06 |
| 2038 | 102 | N102W | A | 2.6e-06 | 4.4e-06 | 2.5e-06 | 3.167E-06 | 1.069E-06 | 2.500E-06 | 4.400E-06 |
| 2039 | 102 | N102Y | A | 3.7e-06 | 1.9e-06 | 2.2e-06 | 2.600E-06 | 9.644E-07 | 1.900E-06 | 3.700E-06 |

| | | | | | | | | | | |
|---|---|---|---|---|---|---|---|---|---|---|
| 2040 | 102 | N102V | A | 2.1e-06 | 1.6e-06 | 5.2e-06 | 2.967E-06 | 1.950E-06 | 1.600E-06 | 5.200E-06 |
| 2041 | 103 | N103A | A | 2.2e-06 | 1.8e-06 | 2.1e-06 | 2.033E-06 | 2.082E-07 | 1.800E-06 | 2.200E-06 |
| 2042 | 103 | N103R | A | 2.8e-06 | 3.8e-06 | 2.1e-06 | 2.900E-06 | 8.544E-07 | 2.100E-06 | 3.800E-06 |
| 2043 | 103 | N103N | A | 1.9e-06 | 1.5e-06 | 2.5e-06 | 1.967E-06 | 5.033E-07 | 1.500E-06 | 2.500E-06 |
| 2044 | 103 | N103D | A | 1.9e-06 | 2.6e-06 | 2.5e-06 | 2.333E-06 | 3.786E-07 | 1.900E-06 | 2.600E-06 |
| 2045 | 103 | N103C | A | 2.5e-06 | 2.4e-06 | 2.4e-06 | 2.433E-06 | 5.774E-08 | 2.400E-06 | 2.500E-06 |
| 2046 | 103 | N103E | A | 2e-06 | 3.5e-06 | 2.3e-06 | 2.600E-06 | 7.937E-07 | 2.000E-06 | 3.500E-06 |
| 2047 | 103 | N103Q | A | 3.5e-06 | 1.6e-06 | 2.9e-06 | 2.667E-06 | 9.713E-07 | 1.600E-06 | 3.500E-06 |
| 2048 | 103 | N103G | A | 2.9e-06 | 1.7e-06 | 2.2e-06 | 2.267E-06 | 6.028E-07 | 1.700E-06 | 2.900E-06 |
| 2049 | 103 | N103H | A | 2.3e-06 | 1.9e-06 | 1.9e-06 | 2.033E-06 | 2.309E-07 | 1.900E-06 | 2.300E-06 |
| 2050 | 103 | N103I | A | 2.1e-06 | 4e-06 | 2e-06 | 2.700E-06 | 1.127E-06 | 2.000E-06 | 4.000E-06 |
| 2051 | 103 | N103L | A | 2.1e-06 | 2.8e-06 | 2.5e-06 | 2.467E-06 | 3.512E-07 | 2.100E-06 | 2.800E-06 |
| 2052 | 103 | N103K | A | 1.8e-06 | 2.7e-06 | 1.4e-06 | 1.967E-06 | 6.658E-07 | 1.400E-06 | 2.700E-06 |
| 2053 | 103 | N103M | A | 2.3e-06 | 3.7e-06 | 3.3e-06 | 3.100E-06 | 7.211E-07 | 2.300E-06 | 3.700E-06 |
| 2054 | 103 | N103F | A | 3.8e-06 | 2.2e-06 | 1.8e-06 | 2.600E-06 | 1.058E-06 | 1.800E-06 | 3.800E-06 |
| 2055 | 103 | N103P | A | 2.5e-06 | 2.2e-06 | 1.9e-06 | 2.200E-06 | 3.000E-07 | 1.900E-06 | 2.500E-06 |
| 2056 | 103 | N103S | A | 2.4e-06 | 2.8e-06 | 2.9e-06 | 2.700E-06 | 2.646E-07 | 2.400E-06 | 2.900E-06 |
| 2057 | 103 | N103T | A | 2.6e-06 | 1.3e-06 | 2.1e-06 | 2.000E-06 | 6.557E-07 | 1.300E-06 | 2.600E-06 |
| 2058 | 103 | N103W | A | 2e-06 | 2.5e-06 | 2.5e-06 | 2.333E-06 | 2.887E-07 | 2.000E-06 | 2.500E-06 |
| 2059 | 103 | N103Y | A | 2.2e-06 | 2.7e-06 | 3.9e-06 | 2.933E-06 | 8.737E-07 | 2.200E-06 | 3.900E-06 |
| 2060 | 103 | N103V | A | 3.4e-06 | 3.1e-06 | 2.2e-06 | 2.900E-06 | 6.245E-07 | 2.200E-06 | 3.400E-06 |
| 2061 | 104 | T104A | A | 3.1e-06 | 1.9e-06 | 3e-06 | 2.667E-06 | 6.658E-07 | 1.900E-06 | 3.100E-06 |
| 2062 | 104 | T104R | A | 1.8e-06 | 3.1e-06 | 4e-06 | 2.967E-06 | 1.106E-06 | 1.800E-06 | 4.000E-06 |
| 2063 | 104 | T104N | A | 8.3e-07 | 2.3e-06 | 1.1e-06 | 1.410E-06 | 7.825E-07 | 8.300E-07 | 2.300E-06 |
| 2064 | 104 | T104D | A | 3.5e-06 | 1.6e-06 | 2.3e-06 | 2.467E-06 | 9.609E-07 | 1.600E-06 | 3.500E-06 |
| 2065 | 104 | T104C | A | 9.5e-07 | 2.5e-06 | 1.9e-06 | 1.783E-06 | 7.816E-07 | 9.500E-07 | 2.500E-06 |
| 2066 | 104 | T104E | A | 2.1e-06 | 1.5e-06 | 1.8e-06 | 1.800E-06 | 3.000E-07 | 1.500E-06 | 2.100E-06 |
| 2067 | 104 | T104Q | A | 2.1e-06 | 2.3e-06 | 2.2e-06 | 2.200E-06 | 1.000E-07 | 2.100E-06 | 2.300E-06 |
| 2068 | 104 | T104G | A | 4.3e-06 | 2.3e-06 | 4.3e-06 | 3.633E-06 | 1.155E-06 | 2.300E-06 | 4.300E-06 |
| 2069 | 104 | T104H | A | 2.4e-06 | 3.4e-06 | 3.1e-06 | 2.967E-06 | 5.132E-07 | 2.400E-06 | 3.400E-06 |
| 2070 | 104 | T104I | A | 1.3e-06 | 2.4e-06 | 3.1e-06 | 2.267E-06 | 9.074E-07 | 1.300E-06 | 3.100E-06 |
| 2071 | 104 | T104L | A | 2.3e-06 | 2.5e-06 | 2.8e-06 | 2.533E-06 | 2.517E-07 | 2.300E-06 | 2.800E-06 |
| 2072 | 104 | T104K | A | 1.3e-06 | 3.1e-06 | 2.8e-06 | 2.400E-06 | 9.644E-07 | 1.300E-06 | 3.100E-06 |
| 2073 | 104 | T104M | A | 1.5e-06 | 1.8e-06 | 1.2e-06 | 1.500E-06 | 3.000E-07 | 1.200E-06 | 1.800E-06 |
| 2074 | 104 | T104F | A | 1.2e-06 | 2.9e-06 | 3.3e-06 | 2.467E-06 | 1.115E-06 | 1.200E-06 | 3.300E-06 |
| 2075 | 104 | T104P | A | 1.4e-06 | 2.5e-06 | 2.5e-06 | 2.133E-06 | 6.351E-07 | 1.400E-06 | 2.500E-06 |
| 2076 | 104 | T104S | A | 2.7e-06 | 1.8e-06 | 2.7e-06 | 2.400E-06 | 5.196E-07 | 1.800E-06 | 2.700E-06 |
| 2077 | 104 | T104T | A | 1.2e-06 | 3.2e-06 | 2.2e-06 | 2.200E-06 | 1.000E-06 | 1.200E-06 | 3.200E-06 |
| 2078 | 104 | T104W | A | 3.5e-06 | 1.4e-06 | 2.8e-06 | 2.567E-06 | 1.069E-06 | 1.400E-06 | 3.500E-06 |
| 2079 | 104 | T104Y | A | 3e-06 | 3.1e-06 | 1.9e-06 | 2.667E-06 | 6.658E-07 | 1.900E-06 | 3.100E-06 |
| 2080 | 104 | T104V | A | 2.6e-06 | 3.5e-06 | 2.8e-06 | 2.967E-06 | 4.726E-07 | 2.600E-06 | 3.500E-06 |
| 2081 | 105 | T105A | A | 1.1e-06 | 2.1e-06 | 1.2e-06 | 1.467E-06 | 5.508E-07 | 1.100E-06 | 2.100E-06 |
| 2082 | 105 | T105R | A | 2.6e-06 | 2.4e-06 | 2.5e-06 | 2.500E-06 | 1.000E-07 | 2.400E-06 | 2.600E-06 |
| 2083 | 105 | T105N | A | 2.4e-06 | 2.2e-06 | 2.7e-06 | 2.433E-06 | 2.517E-07 | 2.200E-06 | 2.700E-06 |
| 2084 | 105 | T105D | A | 2.9e-06 | 2.6e-06 | 2.1e-06 | 2.533E-06 | 4.041E-07 | 2.100E-06 | 2.900E-06 |
| 2085 | 105 | T105C | A | 2.2e-06 | 3.4e-06 | 2.3e-06 | 2.633E-06 | 6.658E-07 | 2.200E-06 | 3.400E-06 |
| 2086 | 105 | T105E | A | 3.3e-06 | 5.5e-06 | 2.5e-06 | 3.767E-06 | 1.553E-06 | 2.500E-06 | 5.500E-06 |
| 2087 | 105 | T105Q | A | 3.1e-06 | 2e-06 | 4.1e-06 | 3.067E-06 | 1.050E-06 | 2.000E-06 | 4.100E-06 |
| 2088 | 105 | T105G | A | 2.6e-06 | 2.2e-06 | 3.6e-06 | 2.800E-06 | 7.211E-07 | 2.200E-06 | 3.600E-06 |
| 2089 | 105 | T105H | A | 2.1e-06 | 3.9e-06 | 2.6e-06 | 2.867E-06 | 9.292E-07 | 2.100E-06 | 3.900E-06 |
| 2090 | 105 | T105I | A | 9.5e-07 | 1.4e-06 | 1.6e-06 | 1.317E-06 | 3.329E-07 | 9.500E-07 | 1.600E-06 |
| 2091 | 105 | T105L | A | 4.5e-06 | 3.2e-06 | 1.3e-06 | 3.000E-06 | 1.609E-06 | 1.300E-06 | 4.500E-06 |
| 2092 | 105 | T105K | A | 3.3e-06 | 2.4e-06 | 3e-06 | 2.900E-06 | 4.583E-07 | 2.400E-06 | 3.300E-06 |
| 2093 | 105 | T105M | A | 1.8e-06 | 3.9e-06 | 2.4e-06 | 2.700E-06 | 1.082E-06 | 1.800E-06 | 3.900E-06 |
| 2094 | 105 | T105F | A | 1.6e-06 | 3.1e-06 | 4.1e-06 | 2.933E-06 | 1.258E-06 | 1.600E-06 | 4.100E-06 |
| 2095 | 105 | T105P | A | 2.3e-06 | 1.9e-06 | 3.4e-06 | 2.533E-06 | 7.767E-07 | 1.900E-06 | 3.400E-06 |
| 2096 | 105 | T105S | A | 3.9e-06 | 3.3e-06 | 2.1e-06 | 3.100E-06 | 9.165E-07 | 2.100E-06 | 3.900E-06 |
| 2097 | 105 | T105T | A | 1.9e-06 | 2.5e-06 | 2.2e-06 | 2.200E-06 | 3.000E-07 | 1.900E-06 | 2.500E-06 |
| 2098 | 105 | T105W | A | 2.1e-06 | 2.3e-06 | 3.3e-06 | 2.567E-06 | 6.429E-07 | 2.100E-06 | 3.300E-06 |
| 2099 | 105 | T105Y | A | 2.2e-06 | 5.6e-06 | 1.5e-06 | 3.100E-06 | 2.193E-06 | 1.500E-06 | 5.600E-06 |
| 2100 | 105 | T105V | A | 3.6e-06 | 2.3e-06 | 2.5e-06 | 2.800E-06 | 7.000E-07 | 2.300E-06 | 3.600E-06 |
| 2101 | 106 | P106A | A | 1.8e-06 | 3.3e-06 | 1.1e-06 | 2.067E-06 | 1.124E-06 | 1.100E-06 | 3.300E-06 |
| 2102 | 106 | P106R | A | 3.4e-06 | 2.1e-06 | 2.5e-06 | 2.667E-06 | 6.658E-07 | 2.100E-06 | 3.400E-06 |
| 2103 | 106 | P106N | A | 2.7e-06 | 2.7e-06 | 1.5e-06 | 2.300E-06 | 6.928E-07 | 1.500E-06 | 2.700E-06 |
| 2104 | 106 | P106D | A | 2.8e-06 | 1.7e-06 | 2.6e-06 | 2.367E-06 | 5.859E-07 | 1.700E-06 | 2.800E-06 |
| 2105 | 106 | P106C | A | 2.3e-06 | 2.2e-06 | 1.9e-06 | 2.133E-06 | 2.082E-07 | 1.900E-06 | 2.300E-06 |
| 2106 | 106 | P106E | A | 4.6e-06 | 1.8e-06 | 2.2e-06 | 2.867E-06 | 1.514E-06 | 1.800E-06 | 4.600E-06 |
| 2107 | 106 | P106Q | A | 2.8e-06 | 2e-06 | 3.1e-06 | 2.633E-06 | 5.686E-07 | 2.000E-06 | 3.100E-06 |
| 2108 | 106 | P106G | A | 3e-06 | 2.8e-06 | 2.2e-06 | 2.667E-06 | 4.163E-07 | 2.200E-06 | 3.000E-06 |
| 2109 | 106 | P106H | A | 1.5e-06 | 2.3e-06 | 2.8e-06 | 2.200E-06 | 6.557E-07 | 1.500E-06 | 2.800E-06 |
| 2110 | 106 | P106I | A | 2.6e-06 | 2.5e-06 | 3.5e-06 | 2.867E-06 | 5.508E-07 | 2.500E-06 | 3.500E-06 |
| 2111 | 106 | P106L | A | 3e-06 | 1.9e-06 | 1.8e-06 | 2.233E-06 | 6.658E-07 | 1.800E-06 | 3.000E-06 |
| 2112 | 106 | P106K | A | 2.5e-06 | 3.3e-06 | 2.3e-06 | 2.700E-06 | 5.292E-07 | 2.300E-06 | 3.300E-06 |
| 2113 | 106 | P106M | A | 2.7e-06 | 1.7e-06 | 2.1e-06 | 2.167E-06 | 5.033E-07 | 1.700E-06 | 2.700E-06 |
| 2114 | 106 | P106F | A | 4.8e-06 | 2e-06 | 3e-06 | 3.267E-06 | 1.419E-06 | 2.000E-06 | 4.800E-06 |
| 2115 | 106 | P106P | A | 2.1e-06 | 5.3e-06 | 3e-06 | 3.467E-06 | 1.650E-06 | 2.100E-06 | 5.300E-06 |
| 2116 | 106 | P106S | A | 3.6e-06 | 1.8e-06 | 2.4e-06 | 2.600E-06 | 9.165E-07 | 1.800E-06 | 3.600E-06 |
| 2117 | 106 | P106T | A | 2e-06 | 3.4e-06 | 2.8e-06 | 2.733E-06 | 7.024E-07 | 2.000E-06 | 3.400E-06 |
| 2118 | 106 | P106W | A | 4.4e-06 | 2.9e-06 | 3.1e-06 | 3.467E-06 | 8.145E-07 | 2.900E-06 | 4.400E-06 |
| 2119 | 106 | P106Y | A | 2.7e-06 | 4e-06 | 2e-06 | 2.900E-06 | 1.015E-06 | 2.000E-06 | 4.000E-06 |
| 2120 | 106 | P106V | A | 3.2e-06 | 2.6e-06 | 2.8e-06 | 2.867E-06 | 3.055E-07 | 2.600E-06 | 3.200E-06 |
| 2121 | 107 | V107A | A | 3.8e-06 | 4.6e-06 | 2.4e-06 | 3.600E-06 | 1.114E-06 | 2.400E-06 | 4.600E-06 |
| 2122 | 107 | V107R | A | 1.1e-06 | 3.2e-06 | 2.8e-06 | 2.367E-06 | 1.115E-06 | 1.100E-06 | 3.200E-06 |
| 2123 | 107 | V107N | A | 2.3e-06 | 2.6e-06 | 1.8e-06 | 2.233E-06 | 4.041E-07 | 1.800E-06 | 2.600E-06 |
| 2124 | 107 | V107D | A | 4.2e-06 | 2.7e-06 | 2.3e-06 | 3.067E-06 | 1.002E-06 | 2.300E-06 | 4.200E-06 |

| | | | | | | | | | | |
|---|---|---|---|---|---|---|---|---|---|---|
| 2125 | 107 | V107C | A | 3.9e-06 | 1.8e-06 | 1.5e-06 | 2.400E-06 | 1.308E-06 | 1.500E-06 | 3.900E-06 |
| 2126 | 107 | V107E | A | 5.3e-06 | 4.3e-06 | 1.3e-06 | 3.633E-06 | 2.082E-06 | 1.300E-06 | 5.300E-06 |
| 2127 | 107 | V107Q | A | 1.6e-06 | 4.2e-06 | 1.8e-06 | 2.533E-06 | 1.447E-06 | 1.600E-06 | 4.200E-06 |
| 2128 | 107 | V107G | A | 2.7e-06 | 1.3e-06 | 4.6e-06 | 2.867E-06 | 1.656E-06 | 1.300E-06 | 4.600E-06 |
| 2129 | 107 | V107H | A | 2.3e-06 | 1.3e-06 | 4e-06 | 2.533E-06 | 1.365E-06 | 1.300E-06 | 4.000E-06 |
| 2130 | 107 | V107I | A | 3.9e-06 | 2.1e-06 | 2.8e-06 | 2.933E-06 | 9.074E-07 | 2.100E-06 | 3.900E-06 |
| 2131 | 107 | V107L | A | 2.3e-06 | 3e-06 | 1.6e-06 | 2.300E-06 | 7.000E-07 | 1.600E-06 | 3.000E-06 |
| 2132 | 107 | V107K | A | 3.1e-06 | 1.8e-06 | 2.5e-06 | 2.467E-06 | 6.506E-07 | 1.800E-06 | 3.100E-06 |
| 2133 | 107 | V107M | A | 2.1e-06 | 5e-06 | 3.6e-06 | 3.567E-06 | 1.450E-06 | 2.100E-06 | 5.000E-06 |
| 2134 | 107 | V107F | A | 3.4e-06 | 1.6e-06 | 3.5e-06 | 2.833E-06 | 1.069E-06 | 1.600E-06 | 3.500E-06 |
| 2135 | 107 | V107P | A | 2.3e-06 | 6e-06 | 2.8e-06 | 3.700E-06 | 2.007E-06 | 2.300E-06 | 6.000E-06 |
| 2136 | 107 | V107S | A | 2.1e-06 | 2.6e-06 | 1.8e-06 | 2.167E-06 | 4.041E-07 | 1.800E-06 | 2.600E-06 |
| 2137 | 107 | V107T | A | 2.9e-06 | 1.3e-06 | 1.7e-06 | 1.967E-06 | 8.327E-07 | 1.300E-06 | 2.900E-06 |
| 2138 | 107 | V107W | A | 3.3e-06 | 2.2e-06 | 2.8e-06 | 2.767E-06 | 5.508E-07 | 2.200E-06 | 3.300E-06 |
| 2139 | 107 | V107Y | A | 2.1e-06 | 1.4e-06 | 1.8e-06 | 1.767E-06 | 3.512E-07 | 1.400E-06 | 2.100E-06 |
| 2140 | 107 | V107V | A | 2.6e-06 | 2.6e-06 | 2.4e-06 | 2.533E-06 | 1.155E-07 | 2.400E-06 | 2.600E-06 |
| 2141 | 108 | T108A | A | 2.6e-06 | 3.8e-06 | 3.6e-06 | 3.333E-06 | 6.429E-07 | 2.600E-06 | 3.800E-06 |
| 2142 | 108 | T108R | A | 3.9e-06 | 3.3e-06 | 1.2e-06 | 2.800E-06 | 1.418E-06 | 1.200E-06 | 3.900E-06 |
| 2143 | 108 | T108N | A | 9.2e-07 | 2.3e-06 | 1.8e-06 | 1.673E-06 | 6.987E-07 | 9.200E-07 | 2.300E-06 |
| 2144 | 108 | T108D | A | 1.8e-06 | 2.1e-06 | 2.1e-06 | 2.000E-06 | 1.732E-07 | 1.800E-06 | 2.100E-06 |
| 2145 | 108 | T108C | A | 1.6e-06 | 3.5e-06 | 2e-06 | 2.367E-06 | 1.002E-06 | 1.600E-06 | 3.500E-06 |
| 2146 | 108 | T108E | A | 2.1e-06 | 3.4e-06 | 3.1e-06 | 2.867E-06 | 6.807E-07 | 2.100E-06 | 3.400E-06 |
| 2147 | 108 | T108Q | A | 2.3e-06 | 2.3e-06 | 2.2e-06 | 2.267E-06 | 5.774E-08 | 2.200E-06 | 2.300E-06 |
| 2148 | 108 | T108G | A | 2.8e-06 | 1.4e-06 | 3.9e-06 | 2.700E-06 | 1.253E-06 | 1.400E-06 | 3.900E-06 |
| 2149 | 108 | T108H | A | 3.8e-06 | 2.7e-06 | 3.1e-06 | 3.200E-06 | 5.568E-07 | 2.700E-06 | 3.800E-06 |
| 2150 | 108 | T108I | A | 3.2e-06 | 1.8e-06 | 1.6e-06 | 2.200E-06 | 8.718E-07 | 1.600E-06 | 3.200E-06 |
| 2151 | 108 | T108L | A | 2e-06 | 1.8e-06 | 2.7e-06 | 2.167E-06 | 4.726E-07 | 1.800E-06 | 2.700E-06 |
| 2152 | 108 | T108K | A | 2.4e-06 | 2.8e-06 | 2.4e-06 | 2.533E-06 | 2.309E-07 | 2.400E-06 | 2.800E-06 |
| 2153 | 108 | T108M | A | 3e-06 | 2.7e-06 | 2.8e-06 | 2.833E-06 | 1.528E-07 | 2.700E-06 | 3.000E-06 |
| 2154 | 108 | T108F | A | 1.8e-06 | 2.5e-06 | 2.6e-06 | 2.300E-06 | 4.359E-07 | 1.800E-06 | 2.600E-06 |
| 2155 | 108 | T108P | A | 1.9e-06 | 2.1e-06 | 2.4e-06 | 2.133E-06 | 2.517E-07 | 1.900E-06 | 2.400E-06 |
| 2156 | 108 | T108S | A | 1.7e-06 | 4.1e-06 | 1.1e-06 | 2.300E-06 | 1.587E-06 | 1.100E-06 | 4.100E-06 |
| 2157 | 108 | T108T | A | 1.7e-06 | 2.9e-06 | 2.7e-06 | 2.433E-06 | 6.429E-07 | 1.700E-06 | 2.900E-06 |
| 2158 | 108 | T108W | A | 2.7e-06 | 2.3e-06 | 1.8e-06 | 2.267E-06 | 4.509E-07 | 1.800E-06 | 2.700E-06 |
| 2159 | 108 | T108Y | A | 2.8e-06 | 3.4e-06 | 3e-06 | 3.067E-06 | 3.055E-07 | 2.800E-06 | 3.400E-06 |
| 2160 | 108 | T108V | A | 2e-06 | 2.2e-06 | 2.8e-06 | 2.333E-06 | 4.163E-07 | 2.000E-06 | 2.800E-06 |
| 2161 | 109 | G109A | A | 2.2e-06 | 3.4e-06 | 3.6e-06 | 3.067E-06 | 7.572E-07 | 2.200E-06 | 3.600E-06 |
| 2162 | 109 | G109R | A | 9.3e-07 | 3.9e-06 | 2e-06 | 2.277E-06 | 1.504E-06 | 9.300E-07 | 3.900E-06 |
| 2163 | 109 | G109N | A | 2.3e-06 | 2.3e-06 | 3.1e-06 | 2.567E-06 | 4.619E-07 | 2.300E-06 | 3.100E-06 |
| 2164 | 109 | G109D | A | 3.6e-06 | 2.7e-06 | 2.2e-06 | 2.833E-06 | 7.095E-07 | 2.200E-06 | 3.600E-06 |
| 2165 | 109 | G109C | A | 2.6e-06 | 2e-06 | 2.5e-06 | 2.367E-06 | 3.215E-07 | 2.000E-06 | 2.600E-06 |
| 2166 | 109 | G109E | A | 2.4e-06 | 3e-06 | 2.5e-06 | 2.633E-06 | 3.215E-07 | 2.400E-06 | 3.000E-06 |
| 2167 | 109 | G109Q | A | 2.4e-06 | 2.1e-06 | 1.8e-06 | 2.100E-06 | 3.000E-07 | 1.800E-06 | 2.400E-06 |
| 2168 | 109 | G109G | A | 3e-06 | 1.1e-06 | 1.5e-06 | 1.867E-06 | 1.002E-06 | 1.100E-06 | 3.000E-06 |
| 2169 | 109 | G109H | A | 2.4e-06 | 1.4e-06 | 2.2e-06 | 2.000E-06 | 5.292E-07 | 1.400E-06 | 2.400E-06 |
| 2170 | 109 | G109I | A | 3.1e-06 | 2.1e-06 | 2.2e-06 | 2.467E-06 | 5.508E-07 | 2.100E-06 | 3.100E-06 |
| 2171 | 109 | G109L | A | 2.8e-06 | 1.8e-06 | 2.7e-06 | 2.433E-06 | 5.508E-07 | 1.800E-06 | 2.800E-06 |
| 2172 | 109 | G109K | A | 3.1e-06 | 1.9e-06 | 1.5e-06 | 2.167E-06 | 8.327E-07 | 1.500E-06 | 3.100E-06 |
| 2173 | 109 | G109M | A | 2.4e-06 | 2.6e-06 | 1.9e-06 | 2.300E-06 | 3.606E-07 | 1.900E-06 | 2.600E-06 |
| 2174 | 109 | G109F | A | 2.7e-06 | 2.8e-06 | 1.9e-06 | 2.467E-06 | 4.933E-07 | 1.900E-06 | 2.800E-06 |
| 2175 | 109 | G109P | A | 2.5e-06 | 2.5e-06 | 2.4e-06 | 2.467E-06 | 5.774E-08 | 2.400E-06 | 2.500E-06 |
| 2176 | 109 | G109S | A | 1.9e-06 | 2.6e-06 | 1.9e-06 | 2.133E-06 | 4.041E-07 | 1.900E-06 | 2.600E-06 |
| 2177 | 109 | G109T | A | 2.3e-06 | 3.5e-06 | 3.8e-06 | 3.200E-06 | 7.937E-07 | 2.300E-06 | 3.800E-06 |
| 2178 | 109 | G109W | A | 3.8e-06 | 1.9e-06 | 4.6e-06 | 3.433E-06 | 1.387E-06 | 1.900E-06 | 4.600E-06 |
| 2179 | 109 | G109Y | A | 1.6e-06 | 2e-06 | 2.2e-06 | 1.933E-06 | 3.055E-07 | 1.600E-06 | 2.200E-06 |
| 2180 | 109 | G109V | A | 2.3e-06 | 2.6e-06 | 2.8e-06 | 2.567E-06 | 2.517E-07 | 2.300E-06 | 2.800E-06 |
| 2181 | 110 | A110A | A | 3e-06 | 2e-06 | 2.6e-06 | 2.533E-06 | 5.033E-07 | 2.000E-06 | 3.000E-06 |
| 2182 | 110 | A110R | A | 2.5e-06 | 2.7e-06 | 1.9e-06 | 2.367E-06 | 4.163E-07 | 1.900E-06 | 2.700E-06 |
| 2183 | 110 | A110N | A | 3.4e-06 | 3.6e-06 | 2.2e-06 | 3.067E-06 | 7.572E-07 | 2.200E-06 | 3.600E-06 |
| 2184 | 110 | A110D | A | 4.1e-06 | 1.7e-06 | 2.7e-06 | 2.833E-06 | 1.206E-06 | 1.700E-06 | 4.100E-06 |
| 2185 | 110 | A110C | A | 3.7e-06 | 1.5e-06 | 3.1e-06 | 2.767E-06 | 1.137E-06 | 1.500E-06 | 3.700E-06 |
| 2186 | 110 | A110E | A | 2.5e-06 | 2.4e-06 | 1.7e-06 | 2.200E-06 | 4.359E-07 | 1.700E-06 | 2.500E-06 |
| 2187 | 110 | A110Q | A | 1.9e-06 | 2.3e-06 | 2.1e-06 | 2.100E-06 | 2.000E-07 | 1.900E-06 | 2.300E-06 |
| 2188 | 110 | A110G | A | 2.8e-06 | 2.9e-06 | 2.2e-06 | 2.633E-06 | 3.786E-07 | 2.200E-06 | 2.900E-06 |
| 2189 | 110 | A110H | A | 2.3e-06 | 1.8e-06 | 3.1e-06 | 2.400E-06 | 6.557E-07 | 1.800E-06 | 3.100E-06 |
| 2190 | 110 | A110I | A | 2e-06 | 4.3e-06 | 2.1e-06 | 2.800E-06 | 1.300E-06 | 2.000E-06 | 4.300E-06 |
| 2191 | 110 | A110L | A | 2.5e-06 | 3.3e-06 | 3e-06 | 2.933E-06 | 4.041E-07 | 2.500E-06 | 3.300E-06 |
| 2192 | 110 | A110K | A | 1.2e-06 | 1.3e-06 | 2.1e-06 | 1.533E-06 | 4.933E-07 | 1.200E-06 | 2.100E-06 |
| 2193 | 110 | A110M | A | 1.5e-06 | 2.4e-06 | 1.5e-06 | 1.800E-06 | 5.196E-07 | 1.500E-06 | 2.400E-06 |
| 2194 | 110 | A110F | A | 2.3e-06 | 3e-06 | 1.4e-06 | 2.233E-06 | 8.021E-07 | 1.400E-06 | 3.000E-06 |
| 2195 | 110 | A110P | A | 2.2e-06 | 3.4e-06 | 2.8e-06 | 2.800E-06 | 6.000E-07 | 2.200E-06 | 3.400E-06 |
| 2196 | 110 | A110S | A | 3.2e-06 | 2.3e-06 | 2.3e-06 | 2.600E-06 | 5.196E-07 | 2.300E-06 | 3.200E-06 |
| 2197 | 110 | A110T | A | 2.1e-06 | 3.3e-06 | 2.2e-06 | 2.533E-06 | 6.658E-07 | 2.100E-06 | 3.300E-06 |
| 2198 | 110 | A110W | A | 2.8e-06 | 1.6e-06 | 1.3e-06 | 1.900E-06 | 7.937E-07 | 1.300E-06 | 2.800E-06 |
| 2199 | 110 | A110Y | A | 3e-06 | 3.6e-06 | 1.9e-06 | 2.833E-06 | 8.622E-07 | 1.900E-06 | 3.600E-06 |
| 2200 | 110 | A110V | A | 2.9e-06 | 1.7e-06 | 1.8e-06 | 2.133E-06 | 6.658E-07 | 1.700E-06 | 2.900E-06 |
| 2201 | 111 | S111A | A | 4.1e-06 | 2.8e-06 | 1.9e-06 | 2.933E-06 | 1.106E-06 | 1.900E-06 | 4.100E-06 |
| 2202 | 111 | S111R | A | 2.1e-06 | 2.7e-06 | 2.2e-06 | 2.333E-06 | 3.215E-07 | 2.100E-06 | 2.700E-06 |
| 2203 | 111 | S111N | A | 3.1e-06 | 1.6e-06 | 3.1e-06 | 2.600E-06 | 8.660E-07 | 1.600E-06 | 3.100E-06 |
| 2204 | 111 | S111D | A | 2.5e-06 | 2.8e-06 | 1.5e-06 | 2.267E-06 | 6.807E-07 | 1.500E-06 | 2.800E-06 |
| 2205 | 111 | S111C | A | 2.6e-06 | 2.3e-06 | 2.5e-06 | 2.467E-06 | 1.528E-07 | 2.300E-06 | 2.600E-06 |
| 2206 | 111 | S111E | A | 1.9e-06 | 1.2e-06 | 2.2e-06 | 1.767E-06 | 5.132E-07 | 1.200E-06 | 2.200E-06 |
| 2207 | 111 | S111Q | A | 4.8e-06 | 2.3e-06 | 1.8e-06 | 2.967E-06 | 1.607E-06 | 1.800E-06 | 4.800E-06 |
| 2208 | 111 | S111G | A | 2.5e-06 | 1.6e-06 | 2e-06 | 2.033E-06 | 4.509E-07 | 1.600E-06 | 2.500E-06 |
| 2209 | 111 | S111H | A | 2e-06 | 2.7e-06 | 9.9e-07 | 1.897E-06 | 8.597E-07 | 9.900E-07 | 2.700E-06 |

| | | | | | | | | | | |
|---|---|---|---|---|---|---|---|---|---|---|
| 2210 | 111 | S111I | A | 3.5e-06 | 2.5e-06 | 2.1e-06 | 2.700E-06 | 7.211E-07 | 2.100E-06 | 3.500E-06 |
| 2211 | 111 | S111L | A | 3.2e-06 | 1.3e-06 | 1.5e-06 | 2.000E-06 | 1.044E-06 | 1.300E-06 | 3.200E-06 |
| 2212 | 111 | S111K | A | 2e-06 | 1.7e-06 | 1.4e-06 | 1.700E-06 | 3.000E-07 | 1.400E-06 | 2.000E-06 |
| 2213 | 111 | S111M | A | 3.2e-06 | 8.9e-07 | 1.8e-06 | 1.963E-06 | 1.164E-06 | 8.900E-07 | 3.200E-06 |
| 2214 | 111 | S111F | A | 2.5e-06 | 3.1e-06 | 1.7e-06 | 2.433E-06 | 7.024E-07 | 1.700E-06 | 3.100E-06 |
| 2215 | 111 | S111P | A | 2.6e-06 | 2.1e-06 | 2.9e-06 | 2.533E-06 | 4.041E-07 | 2.100E-06 | 2.900E-06 |
| 2216 | 111 | S111S | A | 1.4e-06 | 2.3e-06 | 2.1e-06 | 1.933E-06 | 4.726E-07 | 1.400E-06 | 2.300E-06 |
| 2217 | 111 | S111T | A | 1.9e-06 | 2.3e-06 | 2.6e-06 | 2.267E-06 | 3.512E-07 | 1.900E-06 | 2.600E-06 |
| 2218 | 111 | S111W | A | 2.8e-06 | 1.7e-06 | 2.4e-06 | 2.300E-06 | 5.568E-07 | 1.700E-06 | 2.800E-06 |
| 2219 | 111 | S111Y | A | 1.9e-06 | 3.6e-06 | 1.4e-06 | 2.300E-06 | 1.153E-06 | 1.400E-06 | 3.600E-06 |
| 2220 | 111 | S111V | A | 2.7e-06 | 2.2e-06 | 3e-06 | 2.633E-06 | 4.041E-07 | 2.200E-06 | 3.000E-06 |
| 2221 | 112 | P112A | A | 3.1e-06 | 8.4e-07 | 1.5e-06 | 1.813E-06 | 1.162E-06 | 8.400E-07 | 3.100E-06 |
| 2222 | 112 | P112R | A | 1.7e-06 | 1.9e-06 | 2.2e-06 | 1.933E-06 | 2.517E-07 | 1.700E-06 | 2.200E-06 |
| 2223 | 112 | P112N | A | 1.7e-06 | 2.3e-06 | 2.3e-06 | 2.100E-06 | 3.464E-07 | 1.700E-06 | 2.300E-06 |
| 2224 | 112 | P112D | A | 2.1e-06 | 1.9e-06 | 2.1e-06 | 2.033E-06 | 1.155E-07 | 1.900E-06 | 2.100E-06 |
| 2225 | 112 | P112C | A | 4e-06 | 2.4e-06 | 2.2e-06 | 2.867E-06 | 9.866E-07 | 2.200E-06 | 4.000E-06 |
| 2226 | 112 | P112E | A | 5.5e-06 | 2.6e-06 | 1.4e-06 | 3.167E-06 | 2.108E-06 | 1.400E-06 | 5.500E-06 |
| 2227 | 112 | P112Q | A | 3.4e-06 | 1.8e-06 | 2.9e-06 | 2.700E-06 | 8.185E-07 | 1.800E-06 | 3.400E-06 |
| 2228 | 112 | P112G | A | 2.5e-06 | 2.1e-06 | 1.4e-06 | 2.000E-06 | 5.568E-07 | 1.400E-06 | 2.500E-06 |
| 2229 | 112 | P112H | A | 1.8e-06 | 4.1e-06 | 2.6e-06 | 2.833E-06 | 1.168E-06 | 1.800E-06 | 4.100E-06 |
| 2230 | 112 | P112I | A | 1.8e-06 | 1.9e-06 | 2.1e-06 | 1.933E-06 | 1.528E-07 | 1.800E-06 | 2.100E-06 |
| 2231 | 112 | P112L | A | 1.9e-06 | 1.9e-06 | 2e-06 | 1.933E-06 | 5.774E-08 | 1.900E-06 | 2.000E-06 |
| 2232 | 112 | P112K | A | 3.5e-06 | 2.5e-06 | 2.7e-06 | 2.900E-06 | 5.292E-07 | 2.500E-06 | 3.500E-06 |
| 2233 | 112 | P112M | A | 2e-06 | 2.4e-06 | 1.9e-06 | 2.100E-06 | 2.646E-07 | 1.900E-06 | 2.400E-06 |
| 2234 | 112 | P112F | A | 2.7e-06 | 3.1e-06 | 4.7e-06 | 3.500E-06 | 1.058E-06 | 2.700E-06 | 4.700E-06 |
| 2235 | 112 | P112P | A | 1.7e-06 | 9.9e-07 | 3.2e-06 | 1.963E-06 | 1.128E-06 | 9.900E-07 | 3.200E-06 |
| 2236 | 112 | P112S | A | 3.2e-06 | 2.1e-06 | 2e-06 | 2.433E-06 | 6.658E-07 | 2.000E-06 | 3.200E-06 |
| 2237 | 112 | P112T | A | 3.3e-06 | 2.6e-06 | 3.3e-06 | 3.067E-06 | 4.041E-07 | 2.600E-06 | 3.300E-06 |
| 2238 | 112 | P112W | A | 2.9e-06 | 3e-06 | 2e-06 | 2.633E-06 | 5.508E-07 | 2.000E-06 | 3.000E-06 |
| 2239 | 112 | P112Y | A | 2.4e-06 | 2.3e-06 | 4.5e-06 | 3.067E-06 | 1.242E-06 | 2.300E-06 | 4.500E-06 |
| 2240 | 112 | P112V | A | 7.7e-07 | 2.5e-06 | 2.5e-06 | 1.923E-06 | 9.988E-07 | 7.700E-07 | 2.500E-06 |
| 2241 | 113 | G113A | A | 2.3e-06 | 2.8e-06 | 2.4e-06 | 2.500E-06 | 2.646E-07 | 2.300E-06 | 2.800E-06 |
| 2242 | 113 | G113R | A | 2.1e-06 | 2.9e-06 | 2.4e-06 | 2.467E-06 | 4.041E-07 | 2.100E-06 | 2.900E-06 |
| 2243 | 113 | G113N | A | 1.7e-06 | 3e-06 | 2.2e-06 | 2.300E-06 | 6.557E-07 | 1.700E-06 | 3.000E-06 |
| 2244 | 113 | G113D | A | 4.9e-06 | 4.9e-06 | 1.5e-06 | 3.767E-06 | 1.963E-06 | 1.500E-06 | 4.900E-06 |
| 2245 | 113 | G113C | A | 2.9e-06 | 3.9e-06 | 3.1e-06 | 3.300E-06 | 5.292E-07 | 2.900E-06 | 3.900E-06 |
| 2246 | 113 | G113E | A | 1.4e-06 | 8.7e-07 | 2.2e-06 | 1.490E-06 | 6.696E-07 | 8.700E-07 | 2.200E-06 |
| 2247 | 113 | G113Q | A | 5e-06 | 3e-06 | 2.4e-06 | 3.467E-06 | 1.361E-06 | 2.400E-06 | 5.000E-06 |
| 2248 | 113 | G113G | A | 2.1e-06 | 2.3e-06 | 1.2e-06 | 1.867E-06 | 5.859E-07 | 1.200E-06 | 2.300E-06 |
| 2249 | 113 | G113H | A | 1.6e-06 | 1.7e-06 | 4e-06 | 2.433E-06 | 1.358E-06 | 1.600E-06 | 4.000E-06 |
| 2250 | 113 | G113I | A | 2.6e-06 | 2.6e-06 | 3.5e-06 | 2.900E-06 | 5.196E-07 | 2.600E-06 | 3.500E-06 |
| 2251 | 113 | G113L | A | 2e-06 | 1e-06 | 2.6e-06 | 1.867E-06 | 8.083E-07 | 1.000E-06 | 2.600E-06 |
| 2252 | 113 | G113K | A | 1.3e-06 | 1.4e-06 | 2.2e-06 | 1.633E-06 | 4.933E-07 | 1.300E-06 | 2.200E-06 |
| 2253 | 113 | G113M | A | 2.4e-06 | 2.9e-06 | 2.5e-06 | 2.600E-06 | 2.646E-07 | 2.400E-06 | 2.900E-06 |
| 2254 | 113 | G113F | A | 2.7e-06 | 3.5e-06 | 1.2e-06 | 2.467E-06 | 1.168E-06 | 1.200E-06 | 3.500E-06 |
| 2255 | 113 | G113P | A | 1.9e-06 | 1.5e-06 | 2.4e-06 | 1.933E-06 | 4.509E-07 | 1.500E-06 | 2.400E-06 |
| 2256 | 113 | G113S | A | 2.9e-06 | 3.1e-06 | 2.8e-06 | 2.933E-06 | 1.528E-07 | 2.800E-06 | 3.100E-06 |
| 2257 | 113 | G113T | A | 3.9e-06 | 2.2e-06 | 2.9e-06 | 3.000E-06 | 8.544E-07 | 2.200E-06 | 3.900E-06 |
| 2258 | 113 | G113W | A | 2.7e-06 | 2.3e-06 | 2.1e-06 | 2.367E-06 | 3.055E-07 | 2.100E-06 | 2.700E-06 |
| 2259 | 113 | G113Y | A | 2.2e-06 | 2.6e-06 | 2.1e-06 | 2.300E-06 | 2.646E-07 | 2.100E-06 | 2.600E-06 |
| 2260 | 113 | G113V | A | 3e-06 | 2.8e-06 | 1.8e-06 | 2.533E-06 | 6.429E-07 | 1.800E-06 | 3.000E-06 |
| 2261 | 114 | G114A | A | 1.7e-06 | 3e-06 | 1.4e-06 | 2.033E-06 | 8.505E-07 | 1.400E-06 | 3.000E-06 |
| 2262 | 114 | G114R | A | 3.1e-06 | 2.7e-06 | 3.8e-06 | 3.200E-06 | 5.568E-07 | 2.700E-06 | 3.800E-06 |
| 2263 | 114 | G114N | A | 2.3e-06 | 4.2e-06 | 4.5e-06 | 3.667E-06 | 1.193E-06 | 2.300E-06 | 4.500E-06 |
| 2264 | 114 | G114D | A | 2.1e-06 | 2.6e-06 | 1.7e-06 | 2.133E-06 | 4.509E-07 | 1.700E-06 | 2.600E-06 |
| 2265 | 114 | G114C | A | 1.8e-06 | 2e-06 | 1.9e-06 | 1.900E-06 | 1.000E-07 | 1.800E-06 | 2.000E-06 |
| 2266 | 114 | G114E | A | 2.7e-06 | 1.6e-06 | 3.2e-06 | 2.500E-06 | 8.185E-07 | 1.600E-06 | 3.200E-06 |
| 2267 | 114 | G114Q | A | 2.1e-06 | 1.9e-06 | 2.6e-06 | 2.200E-06 | 3.606E-07 | 1.900E-06 | 2.600E-06 |
| 2268 | 114 | G114G | A | 3.4e-06 | 3e-06 | 1.6e-06 | 2.667E-06 | 9.452E-07 | 1.600E-06 | 3.400E-06 |
| 2269 | 114 | G114H | A | 2.9e-06 | 2.4e-06 | 2.8e-06 | 2.700E-06 | 2.646E-07 | 2.400E-06 | 2.900E-06 |
| 2270 | 114 | G114I | A | 2.1e-06 | 2.7e-06 | 1.5e-06 | 2.100E-06 | 6.000E-07 | 1.500E-06 | 2.700E-06 |
| 2271 | 114 | G114L | A | 2.1e-06 | 1.5e-06 | 2.6e-06 | 2.067E-06 | 5.508E-07 | 1.500E-06 | 2.600E-06 |
| 2272 | 114 | G114K | A | 2.6e-06 | 1.7e-06 | 1.9e-06 | 2.067E-06 | 4.726E-07 | 1.700E-06 | 2.600E-06 |
| 2273 | 114 | G114M | A | 4.3e-06 | 2.1e-06 | 2.8e-06 | 3.067E-06 | 1.124E-06 | 2.100E-06 | 4.300E-06 |
| 2274 | 114 | G114F | A | 3.5e-06 | 2.2e-06 | 1.7e-06 | 2.467E-06 | 9.292E-07 | 1.700E-06 | 3.500E-06 |
| 2275 | 114 | G114P | A | 2.2e-06 | 1.9e-06 | 1.7e-06 | 1.933E-06 | 2.517E-07 | 1.700E-06 | 2.200E-06 |
| 2276 | 114 | G114S | A | 3.8e-06 | 1.3e-06 | 1.3e-06 | 2.133E-06 | 1.443E-06 | 1.300E-06 | 3.800E-06 |
| 2277 | 114 | G114T | A | 2.8e-06 | 2e-06 | 2.8e-06 | 2.533E-06 | 4.619E-07 | 2.000E-06 | 2.800E-06 |
| 2278 | 114 | G114W | A | 1.9e-06 | 2.3e-06 | 2.6e-06 | 2.267E-06 | 3.512E-07 | 1.900E-06 | 2.600E-06 |
| 2279 | 114 | G114Y | A | 2.3e-06 | 2.2e-06 | 2e-06 | 2.167E-06 | 1.528E-07 | 2.000E-06 | 2.300E-06 |
| 2280 | 114 | G114V | A | 1.2e-06 | 3.4e-06 | 1.6e-06 | 2.067E-06 | 1.172E-06 | 1.200E-06 | 3.400E-06 |
| 2281 | 115 | L115A | A | 2.1e-06 | 2.8e-06 | 3.4e-06 | 2.767E-06 | 6.506E-07 | 2.100E-06 | 3.400E-06 |
| 2282 | 115 | L115R | A | 3.4e-06 | 1.9e-06 | 2.4e-06 | 2.567E-06 | 7.638E-07 | 1.900E-06 | 3.400E-06 |
| 2283 | 115 | L115N | A | 3.6e-06 | 1.6e-06 | 3e-06 | 2.733E-06 | 1.026E-06 | 1.600E-06 | 3.600E-06 |
| 2284 | 115 | L115D | A | 2.4e-06 | 1.6e-06 | 2.9e-06 | 2.300E-06 | 6.557E-07 | 1.600E-06 | 2.900E-06 |
| 2285 | 115 | L115C | A | 3.4e-06 | 3.5e-06 | 3.1e-06 | 3.333E-06 | 2.082E-07 | 3.100E-06 | 3.500E-06 |
| 2286 | 115 | L115E | A | 2.2e-06 | 2.6e-06 | 1.4e-06 | 2.067E-06 | 6.110E-07 | 1.400E-06 | 2.600E-06 |
| 2287 | 115 | L115Q | A | 2.8e-06 | 2.8e-06 | 2e-06 | 2.533E-06 | 4.619E-07 | 2.000E-06 | 2.800E-06 |
| 2288 | 115 | L115G | A | 3e-06 | 2.8e-06 | 1.4e-06 | 2.400E-06 | 8.718E-07 | 1.400E-06 | 3.000E-06 |
| 2289 | 115 | L115H | A | 2.9e-06 | 3.1e-06 | 2.7e-06 | 2.900E-06 | 2.000E-07 | 2.700E-06 | 3.100E-06 |
| 2290 | 115 | L115I | A | 1.7e-06 | 1.2e-06 | 2.8e-06 | 1.900E-06 | 8.185E-07 | 1.200E-06 | 2.800E-06 |
| 2291 | 115 | L115L | A | 3.2e-06 | 1.4e-06 | 1.7e-06 | 2.100E-06 | 9.644E-07 | 1.400E-06 | 3.200E-06 |
| 2292 | 115 | L115K | A | 1.1e-06 | 2.1e-06 | 2.3e-06 | 1.833E-06 | 6.429E-07 | 1.100E-06 | 2.300E-06 |
| 2293 | 115 | L115M | A | 2.3e-06 | 2.6e-06 | 2.7e-06 | 2.533E-06 | 2.082E-07 | 2.300E-06 | 2.700E-06 |
| 2294 | 115 | L115F | A | 2.3e-06 | 3.1e-06 | 2.9e-06 | 2.767E-06 | 4.163E-07 | 2.300E-06 | 3.100E-06 |

| | | | | | | | | | | |
|---|---|---|---|---|---|---|---|---|---|---|
| 2295 | 115 | L115P | A | 2.2e-06 | 2.2e-06 | 3.8e-06 | 2.733E-06 | 9.238E-07 | 2.200E-06 | 3.800E-06 |
| 2296 | 115 | L115S | A | 2.6e-06 | 2.1e-06 | 1.6e-06 | 2.100E-06 | 5.000E-07 | 1.600E-06 | 2.600E-06 |
| 2297 | 115 | L115T | A | 2.3e-06 | 3.1e-06 | 1.9e-06 | 2.433E-06 | 6.110E-07 | 1.900E-06 | 3.100E-06 |
| 2298 | 115 | L115W | A | 1.6e-06 | 2.5e-06 | 1.6e-06 | 1.900E-06 | 5.196E-07 | 1.600E-06 | 2.500E-06 |
| 2299 | 115 | L115Y | A | 2e-06 | 1.6e-06 | 2.6e-06 | 2.067E-06 | 5.033E-07 | 1.600E-06 | 2.600E-06 |
| 2300 | 115 | L115V | A | 2.1e-06 | 1.7e-06 | 3.3e-06 | 2.367E-06 | 8.327E-07 | 1.700E-06 | 3.300E-06 |
| 2301 | 116 | R116A | A | 1.5e-06 | 2.1e-06 | 1.7e-06 | 1.767E-06 | 3.055E-07 | 1.500E-06 | 2.100E-06 |
| 2302 | 116 | R116R | A | 3.5e-06 | 1.4e-06 | 1.6e-06 | 2.167E-06 | 1.159E-06 | 1.400E-06 | 3.500E-06 |
| 2303 | 116 | R116N | A | 4.1e-06 | 1.9e-06 | 2e-06 | 2.667E-06 | 1.242E-06 | 1.900E-06 | 4.100E-06 |
| 2304 | 116 | R116D | A | 3.3e-06 | 1.8e-06 | 2.2e-06 | 2.433E-06 | 7.767E-07 | 1.800E-06 | 3.300E-06 |
| 2305 | 116 | R116C | A | 2.3e-06 | 2.1e-06 | 1.8e-06 | 2.067E-06 | 2.517E-07 | 1.800E-06 | 2.300E-06 |
| 2306 | 116 | R116E | A | 2.4e-06 | 2e-06 | 3.1e-06 | 2.500E-06 | 5.568E-07 | 2.000E-06 | 3.100E-06 |
| 2307 | 116 | R116Q | A | 2e-06 | 2.8e-06 | 3.4e-06 | 2.733E-06 | 7.024E-07 | 2.000E-06 | 3.400E-06 |
| 2308 | 116 | R116G | A | 2.6e-06 | 1.9e-06 | 2.7e-06 | 2.400E-06 | 4.359E-07 | 1.900E-06 | 2.700E-06 |
| 2309 | 116 | R116H | A | 1.1e-06 | 2.6e-06 | 2.1e-06 | 1.933E-06 | 7.638E-07 | 1.100E-06 | 2.600E-06 |
| 2310 | 116 | R116I | A | 1.4e-06 | 1.9e-06 | 2.4e-06 | 1.900E-06 | 5.000E-07 | 1.400E-06 | 2.400E-06 |
| 2311 | 116 | R116L | A | 2.1e-06 | 2.6e-06 | 1.6e-06 | 2.100E-06 | 5.000E-07 | 1.600E-06 | 2.600E-06 |
| 2312 | 116 | R116K | A | 3e-06 | 1.9e-06 | 2.3e-06 | 2.400E-06 | 5.568E-07 | 1.900E-06 | 3.000E-06 |
| 2313 | 116 | R116M | A | 1.1e-06 | 1.8e-06 | 2.5e-06 | 1.800E-06 | 7.000E-07 | 1.100E-06 | 2.500E-06 |
| 2314 | 116 | R116F | A | 4.2e-06 | 1.8e-06 | 2.5e-06 | 2.833E-06 | 1.234E-06 | 1.800E-06 | 4.200E-06 |
| 2315 | 116 | R116P | A | 2.2e-06 | 2.3e-06 | 2.1e-06 | 2.200E-06 | 1.000E-07 | 2.100E-06 | 2.300E-06 |
| 2316 | 116 | R116S | A | 3.5e-06 | 2.2e-06 | 2.3e-06 | 2.667E-06 | 7.234E-07 | 2.200E-06 | 3.500E-06 |
| 2317 | 116 | R116T | A | 2.4e-06 | 2.8e-06 | 4e-06 | 3.067E-06 | 8.327E-07 | 2.400E-06 | 4.000E-06 |
| 2318 | 116 | R116W | A | 2.7e-06 | 2.8e-06 | 3.8e-06 | 3.100E-06 | 6.083E-07 | 2.700E-06 | 3.800E-06 |
| 2319 | 116 | R116Y | A | 2.8e-06 | 2.9e-06 | 2.6e-06 | 2.767E-06 | 1.528E-07 | 2.600E-06 | 2.900E-06 |
| 2320 | 116 | R116V | A | 2.5e-06 | 3.8e-06 | 4e-06 | 3.433E-06 | 8.145E-07 | 2.500E-06 | 4.000E-06 |
| 2321 | 117 | E117A | A | 1.9e-06 | 2.3e-06 | 2.8e-06 | 2.333E-06 | 4.509E-07 | 1.900E-06 | 2.800E-06 |
| 2322 | 117 | E117R | A | 3.1e-06 | 2.4e-06 | 2.6e-06 | 2.700E-06 | 3.606E-07 | 2.400E-06 | 3.100E-06 |
| 2323 | 117 | E117N | A | 2.4e-06 | 3e-06 | 3e-06 | 2.800E-06 | 3.464E-07 | 2.400E-06 | 3.000E-06 |
| 2324 | 117 | E117D | A | 1.6e-06 | 1.9e-06 | 3e-06 | 2.167E-06 | 7.371E-07 | 1.600E-06 | 3.000E-06 |
| 2325 | 117 | E117C | A | 3.8e-06 | 6e-07 | 2e-06 | 2.133E-06 | 1.604E-06 | 6.000E-07 | 3.800E-06 |
| 2326 | 117 | E117E | A | 2.4e-06 | 1.8e-06 | 1.3e-06 | 1.833E-06 | 5.508E-07 | 1.300E-06 | 2.400E-06 |
| 2327 | 117 | E117Q | A | 2.5e-06 | 2.7e-06 | 2.2e-06 | 2.467E-06 | 2.517E-07 | 2.200E-06 | 2.700E-06 |
| 2328 | 117 | E117G | A | 2.2e-06 | 3.2e-06 | 3.9e-06 | 3.100E-06 | 8.544E-07 | 2.200E-06 | 3.900E-06 |
| 2329 | 117 | E117H | A | 3.5e-06 | 2.5e-06 | 1.9e-06 | 2.633E-06 | 8.083E-07 | 1.900E-06 | 3.500E-06 |
| 2330 | 117 | E117I | A | 1.9e-06 | 3.2e-06 | 1.5e-06 | 2.200E-06 | 8.888E-07 | 1.500E-06 | 3.200E-06 |
| 2331 | 117 | E117L | A | 2.2e-06 | 1.7e-06 | 1.9e-06 | 1.933E-06 | 2.517E-07 | 1.700E-06 | 2.200E-06 |
| 2332 | 117 | E117K | A | 1.4e-06 | 3.3e-06 | 2.3e-06 | 2.333E-06 | 9.504E-07 | 1.400E-06 | 3.300E-06 |
| 2333 | 117 | E117M | A | 1.9e-06 | 1.6e-06 | 3.3e-06 | 2.267E-06 | 9.074E-07 | 1.600E-06 | 3.300E-06 |
| 2334 | 117 | E117F | A | 4.3e-06 | 5.6e-06 | 2.6e-06 | 4.167E-06 | 1.504E-06 | 2.600E-06 | 5.600E-06 |
| 2335 | 117 | E117P | A | 2.1e-06 | 1.2e-06 | 2.7e-06 | 2.000E-06 | 7.550E-07 | 1.200E-06 | 2.700E-06 |
| 2336 | 117 | E117S | A | 2.5e-06 | 3.4e-06 | 2.8e-06 | 2.900E-06 | 4.583E-07 | 2.500E-06 | 3.400E-06 |
| 2337 | 117 | E117T | A | 1.2e-06 | 2.9e-06 | 2e-06 | 2.033E-06 | 8.505E-07 | 1.200E-06 | 2.900E-06 |
| 2338 | 117 | E117W | A | 3.1e-06 | 2.1e-06 | 2.4e-06 | 2.533E-06 | 5.132E-07 | 2.100E-06 | 3.100E-06 |
| 2339 | 117 | E117Y | A | 1.5e-06 | 2.7e-06 | 1.3e-06 | 1.833E-06 | 7.572E-07 | 1.300E-06 | 2.700E-06 |
| 2340 | 117 | E117V | A | 1.8e-06 | 4.4e-06 | 1.4e-06 | 2.533E-06 | 1.629E-06 | 1.400E-06 | 4.400E-06 |
| 2341 | 118 | L118A | A | 3.4e-06 | 3.3e-06 | 3.3e-06 | 3.333E-06 | 5.774E-08 | 3.300E-06 | 3.400E-06 |
| 2342 | 118 | L118R | A | 2.6e-06 | 2.4e-06 | 2e-06 | 2.333E-06 | 3.055E-07 | 2.000E-06 | 2.600E-06 |
| 2343 | 118 | L118N | A | 2.1e-06 | 3e-06 | 4.4e-06 | 3.167E-06 | 1.159E-06 | 2.100E-06 | 4.400E-06 |
| 2344 | 118 | L118D | A | 2.2e-06 | 3.2e-06 | 2.4e-06 | 2.600E-06 | 5.292E-07 | 2.200E-06 | 3.200E-06 |
| 2345 | 118 | L118C | A | 2.4e-06 | 2.6e-06 | 4.9e-06 | 3.300E-06 | 1.389E-06 | 2.400E-06 | 4.900E-06 |
| 2346 | 118 | L118E | A | 4.2e-06 | 2.3e-06 | 1.9e-06 | 2.800E-06 | 1.229E-06 | 1.900E-06 | 4.200E-06 |
| 2347 | 118 | L118Q | A | 2.4e-06 | 1.4e-06 | 2.6e-06 | 2.133E-06 | 6.429E-07 | 1.400E-06 | 2.600E-06 |
| 2348 | 118 | L118G | A | 5.2e-06 | 1.6e-06 | 2.3e-06 | 3.033E-06 | 1.909E-06 | 1.600E-06 | 5.200E-06 |
| 2349 | 118 | L118H | A | 2.8e-06 | 2.6e-06 | 1.2e-06 | 2.200E-06 | 8.718E-07 | 1.200E-06 | 2.800E-06 |
| 2350 | 118 | L118I | A | 3.1e-06 | 2.6e-06 | 2.7e-06 | 2.800E-06 | 2.646E-07 | 2.600E-06 | 3.100E-06 |
| 2351 | 118 | L118L | A | 4e-06 | 4.6e-06 | 1.9e-06 | 3.500E-06 | 1.418E-06 | 1.900E-06 | 4.600E-06 |
| 2352 | 118 | L118K | A | 3e-06 | 2.6e-06 | 2.8e-06 | 2.800E-06 | 2.000E-07 | 2.600E-06 | 3.000E-06 |
| 2353 | 118 | L118M | A | 1.9e-06 | 1.3e-06 | 2.7e-06 | 1.967E-06 | 7.024E-07 | 1.300E-06 | 2.700E-06 |
| 2354 | 118 | L118F | A | 2.5e-06 | 2.1e-06 | 2.9e-06 | 2.500E-06 | 4.000E-07 | 2.100E-06 | 2.900E-06 |
| 2355 | 118 | L118P | A | 3e-06 | 2.1e-06 | 2e-06 | 2.367E-06 | 5.508E-07 | 2.000E-06 | 3.000E-06 |
| 2356 | 118 | L118S | A | 2.8e-06 | 2.1e-06 | 2.5e-06 | 2.467E-06 | 3.512E-07 | 2.100E-06 | 2.800E-06 |
| 2357 | 118 | L118T | A | 1.9e-06 | 2.4e-06 | 2.3e-06 | 2.200E-06 | 2.646E-07 | 1.900E-06 | 2.400E-06 |
| 2358 | 118 | L118W | A | 3e-06 | 2.8e-06 | 1.5e-06 | 2.433E-06 | 8.145E-07 | 1.500E-06 | 3.000E-06 |
| 2359 | 118 | L118Y | A | 1.8e-06 | 3.3e-06 | 1.5e-06 | 2.200E-06 | 9.644E-07 | 1.500E-06 | 3.300E-06 |
| 2360 | 118 | L118V | A | 4.5e-06 | 2.2e-06 | 2.8e-06 | 3.167E-06 | 1.193E-06 | 2.200E-06 | 4.500E-06 |
| 2361 | 119 | Q119A | A | 2.2e-06 | 4e-06 | 1.8e-06 | 2.667E-06 | 1.172E-06 | 1.800E-06 | 4.000E-06 |
| 2362 | 119 | Q119R | A | 3.9e-06 | 6e-06 | 3.2e-06 | 4.367E-06 | 1.457E-06 | 3.200E-06 | 6.000E-06 |
| 2363 | 119 | Q119N | A | 1.6e-06 | 3.5e-06 | 3.6e-06 | 2.900E-06 | 1.127E-06 | 1.600E-06 | 3.600E-06 |
| 2364 | 119 | Q119D | A | 2.3e-06 | 4.8e-06 | 2.2e-06 | 3.100E-06 | 1.473E-06 | 2.200E-06 | 4.800E-06 |
| 2365 | 119 | Q119C | A | 2.6e-06 | 3e-06 | 2.4e-06 | 2.667E-06 | 3.055E-07 | 2.400E-06 | 3.000E-06 |
| 2366 | 119 | Q119E | A | 3.4e-06 | 1.8e-06 | 1.9e-06 | 2.367E-06 | 8.963E-07 | 1.800E-06 | 3.400E-06 |
| 2367 | 119 | Q119Q | A | 1.6e-06 | 2.1e-06 | 2.1e-06 | 1.933E-06 | 2.887E-07 | 1.600E-06 | 2.100E-06 |
| 2368 | 119 | Q119G | A | 2.8e-06 | 2.4e-06 | 1.8e-06 | 2.333E-06 | 5.033E-07 | 1.800E-06 | 2.800E-06 |
| 2369 | 119 | Q119H | A | 1.9e-06 | 2.4e-06 | 2.5e-06 | 2.267E-06 | 3.215E-07 | 1.900E-06 | 2.500E-06 |
| 2370 | 119 | Q119I | A | 2.9e-06 | 8.5e-07 | 5.2e-06 | 2.983E-06 | 2.176E-06 | 8.500E-07 | 5.200E-06 |
| 2371 | 119 | Q119L | A | 1.9e-06 | 1.7e-06 | 2.4e-06 | 2.000E-06 | 3.606E-07 | 1.700E-06 | 2.400E-06 |
| 2372 | 119 | Q119K | A | 2.8e-06 | 2.3e-06 | 1.4e-06 | 2.167E-06 | 7.095E-07 | 1.400E-06 | 2.800E-06 |
| 2373 | 119 | Q119M | A | 1.7e-06 | 2e-06 | 2.5e-06 | 2.067E-06 | 4.041E-07 | 1.700E-06 | 2.500E-06 |
| 2374 | 119 | Q119F | A | 2.4e-06 | 2.9e-06 | 2.9e-06 | 2.733E-06 | 2.887E-07 | 2.400E-06 | 2.900E-06 |
| 2375 | 119 | Q119P | A | 3.2e-06 | 1.4e-06 | 3.9e-06 | 2.833E-06 | 1.290E-06 | 1.400E-06 | 3.900E-06 |
| 2376 | 119 | Q119S | A | 2.1e-06 | 3.1e-06 | 1.9e-06 | 2.367E-06 | 6.429E-07 | 1.900E-06 | 3.100E-06 |
| 2377 | 119 | Q119T | A | 3.5e-06 | 1e-06 | 2.4e-06 | 2.300E-06 | 1.253E-06 | 1.000E-06 | 3.500E-06 |
| 2378 | 119 | Q119W | A | 4.1e-06 | 2.2e-06 | 1.8e-06 | 2.700E-06 | 1.229E-06 | 1.800E-06 | 4.100E-06 |
| 2379 | 119 | Q119Y | A | 2e-06 | 2.7e-06 | 1.4e-06 | 2.033E-06 | 6.506E-07 | 1.400E-06 | 2.700E-06 |

| | | | | | | | | | | |
|---|---|---|---|---|---|---|---|---|---|---|
| 2380 | 119 | Q119V | A | 2.8e-06 | 2e-06 | 2.4e-06 | 2.400E-06 | 4.000E-07 | 2.000E-06 | 2.800E-06 |
| 2381 | 120 | L120A | A | 1.7e-06 | 1.9e-06 | 1.6e-06 | 1.733E-06 | 1.528E-07 | 1.600E-06 | 1.900E-06 |
| 2382 | 120 | L120R | A | 3.8e-06 | 2.5e-06 | 2.8e-06 | 3.033E-06 | 6.807E-07 | 2.500E-06 | 3.800E-06 |
| 2383 | 120 | L120N | A | 2.2e-06 | 2.1e-06 | 2.2e-06 | 2.167E-06 | 5.774E-08 | 2.100E-06 | 2.200E-06 |
| 2384 | 120 | L120D | A | 2.9e-06 | 1.7e-06 | 1.9e-06 | 2.167E-06 | 6.429E-07 | 1.700E-06 | 2.900E-06 |
| 2385 | 120 | L120C | A | 3.1e-06 | 2.1e-06 | 1.8e-06 | 2.333E-06 | 6.807E-07 | 1.800E-06 | 3.100E-06 |
| 2386 | 120 | L120E | A | 1.9e-06 | 1.7e-06 | 2.6e-06 | 2.067E-06 | 4.726E-07 | 1.700E-06 | 2.600E-06 |
| 2387 | 120 | L120Q | A | 1.3e-06 | 1.6e-06 | 1.7e-06 | 1.533E-06 | 2.082E-07 | 1.300E-06 | 1.700E-06 |
| 2388 | 120 | L120G | A | 2e-06 | 2e-06 | 1.5e-06 | 1.833E-06 | 2.887E-07 | 1.500E-06 | 2.000E-06 |
| 2389 | 120 | L120H | A | 1.3e-06 | 3.1e-06 | 2.5e-06 | 2.300E-06 | 9.165E-07 | 1.300E-06 | 3.100E-06 |
| 2390 | 120 | L120I | A | 1e-06 | 3.2e-06 | 1.5e-06 | 1.900E-06 | 1.153E-06 | 1.000E-06 | 3.200E-06 |
| 2391 | 120 | L120L | A | 1.7e-06 | 2.9e-06 | 3.8e-06 | 2.800E-06 | 1.054E-06 | 1.700E-06 | 3.800E-06 |
| 2392 | 120 | L120K | A | 1e-06 | 2.8e-06 | 1.8e-06 | 1.867E-06 | 9.018E-07 | 1.000E-06 | 2.800E-06 |
| 2393 | 120 | L120M | A | 2.1e-06 | 1.8e-06 | 4.2e-06 | 2.700E-06 | 1.308E-06 | 1.800E-06 | 4.200E-06 |
| 2394 | 120 | L120F | A | 2.4e-06 | 2.7e-06 | 3.4e-06 | 2.833E-06 | 5.132E-07 | 2.400E-06 | 3.400E-06 |
| 2395 | 120 | L120P | A | 3.6e-06 | 3.2e-06 | 2.6e-06 | 3.133E-06 | 5.033E-07 | 2.600E-06 | 3.600E-06 |
| 2396 | 120 | L120S | A | 1.8e-06 | 2.6e-06 | 1.2e-06 | 1.867E-06 | 7.024E-07 | 1.200E-06 | 2.600E-06 |
| 2397 | 120 | L120T | A | 2.6e-06 | 2.2e-06 | 2.3e-06 | 2.367E-06 | 2.082E-07 | 2.200E-06 | 2.600E-06 |
| 2398 | 120 | L120W | A | 2.5e-06 | 2.1e-06 | 2.9e-06 | 2.500E-06 | 4.000E-07 | 2.100E-06 | 2.900E-06 |
| 2399 | 120 | L120Y | A | 3.3e-06 | 1.5e-06 | 1.7e-06 | 2.167E-06 | 9.866E-07 | 1.500E-06 | 3.300E-06 |
| 2400 | 120 | L120V | A | 2.8e-06 | 3.2e-06 | 1.1e-06 | 2.367E-06 | 1.115E-06 | 1.100E-06 | 3.200E-06 |
| 2401 | 121 | R121A | A | 1.7e-06 | 9e-07 | 2.3e-06 | 1.633E-06 | 7.024E-07 | 9.000E-07 | 2.300E-06 |
| 2402 | 121 | R121R | A | 2.3e-06 | 4e-06 | 2.1e-06 | 2.800E-06 | 1.044E-06 | 2.100E-06 | 4.000E-06 |
| 2403 | 121 | R121N | A | 1.8e-06 | 2.6e-06 | 2.2e-06 | 2.200E-06 | 4.000E-07 | 1.800E-06 | 2.600E-06 |
| 2404 | 121 | R121D | A | 2.2e-06 | 2.9e-06 | 1.9e-06 | 2.333E-06 | 5.132E-07 | 1.900E-06 | 2.900E-06 |
| 2405 | 121 | R121C | A | 1.6e-06 | 1.6e-06 | 1.8e-06 | 1.667E-06 | 1.155E-07 | 1.600E-06 | 1.800E-06 |
| 2406 | 121 | R121E | A | 1.6e-06 | 1.1e-06 | 4e-06 | 2.233E-06 | 1.550E-06 | 1.100E-06 | 4.000E-06 |
| 2407 | 121 | R121Q | A | 2e-06 | 3.4e-06 | 2.3e-06 | 2.567E-06 | 7.371E-07 | 2.000E-06 | 3.400E-06 |
| 2408 | 121 | R121G | A | 1.7e-06 | 1.7e-06 | 2.4e-06 | 1.933E-06 | 4.041E-07 | 1.700E-06 | 2.400E-06 |
| 2409 | 121 | R121H | A | 3e-06 | 2.7e-06 | 2.5e-06 | 2.733E-06 | 2.517E-07 | 2.500E-06 | 3.000E-06 |
| 2410 | 121 | R121I | A | 1.8e-06 | 2.4e-06 | 2.1e-06 | 2.100E-06 | 3.000E-07 | 1.800E-06 | 2.400E-06 |
| 2411 | 121 | R121L | A | 2.5e-06 | 2.4e-06 | 1.9e-06 | 2.267E-06 | 3.215E-07 | 1.900E-06 | 2.500E-06 |
| 2412 | 121 | R121K | A | 2.2e-06 | 3e-06 | 2.3e-06 | 2.500E-06 | 4.359E-07 | 2.200E-06 | 3.000E-06 |
| 2413 | 121 | R121M | A | 1.9e-06 | 2.6e-06 | 3.2e-06 | 2.567E-06 | 6.506E-07 | 1.900E-06 | 3.200E-06 |
| 2414 | 121 | R121F | A | 2e-06 | 3e-06 | 2.3e-06 | 2.433E-06 | 5.132E-07 | 2.000E-06 | 3.000E-06 |
| 2415 | 121 | R121P | A | 2.6e-06 | 2.6e-06 | 1.8e-06 | 2.333E-06 | 4.619E-07 | 1.800E-06 | 2.600E-06 |
| 2416 | 121 | R121S | A | 3.1e-06 | 2.6e-06 | 3.3e-06 | 3.000E-06 | 3.606E-07 | 2.600E-06 | 3.300E-06 |
| 2417 | 121 | R121T | A | 2.9e-06 | 1.8e-06 | 2.4e-06 | 2.367E-06 | 5.508E-07 | 1.800E-06 | 2.900E-06 |
| 2418 | 121 | R121W | A | 2.8e-06 | 2.7e-06 | 1.7e-06 | 2.400E-06 | 6.083E-07 | 1.700E-06 | 2.800E-06 |
| 2419 | 121 | R121Y | A | 4e-06 | 2.5e-06 | 2.4e-06 | 2.967E-06 | 8.963E-07 | 2.400E-06 | 4.000E-06 |
| 2420 | 121 | R121V | A | 2.7e-06 | 2.2e-06 | 3.3e-06 | 2.733E-06 | 5.508E-07 | 2.200E-06 | 3.300E-06 |
| 2421 | 122 | S122A | A | 1.3e-06 | 2.4e-06 | 2.1e-06 | 1.933E-06 | 5.686E-07 | 1.300E-06 | 2.400E-06 |
| 2422 | 122 | S122R | A | 2e-06 | 2.1e-06 | 3.4e-06 | 2.500E-06 | 7.810E-07 | 2.000E-06 | 3.400E-06 |
| 2423 | 122 | S122N | A | 2.4e-06 | 1.4e-06 | 1.1e-06 | 1.633E-06 | 6.807E-07 | 1.100E-06 | 2.400E-06 |
| 2424 | 122 | S122D | A | 2.8e-06 | 3e-06 | 2.7e-06 | 2.833E-06 | 1.528E-07 | 2.700E-06 | 3.000E-06 |
| 2425 | 122 | S122C | A | 1.8e-06 | 1e-06 | 3.6e-06 | 2.133E-06 | 1.332E-06 | 1.000E-06 | 3.600E-06 |
| 2426 | 122 | S122E | A | 1.4e-06 | 1.9e-06 | 1.7e-06 | 1.667E-06 | 2.517E-07 | 1.400E-06 | 1.900E-06 |
| 2427 | 122 | S122Q | A | 2.3e-06 | 1.8e-06 | 1.5e-06 | 1.867E-06 | 4.041E-07 | 1.500E-06 | 2.300E-06 |
| 2428 | 122 | S122G | A | 2.8e-06 | 2.6e-06 | 2.1e-06 | 2.500E-06 | 3.606E-07 | 2.100E-06 | 2.800E-06 |
| 2429 | 122 | S122H | A | 3.8e-06 | 1.6e-06 | 2.9e-06 | 2.767E-06 | 1.106E-06 | 1.600E-06 | 3.800E-06 |
| 2430 | 122 | S122I | A | 2.6e-06 | 2.3e-06 | 2.6e-06 | 2.500E-06 | 1.732E-07 | 2.300E-06 | 2.600E-06 |
| 2431 | 122 | S122L | A | 1.7e-06 | 2.3e-06 | 5.1e-06 | 3.033E-06 | 1.815E-06 | 1.700E-06 | 5.100E-06 |
| 2432 | 122 | S122K | A | 3.3e-06 | 3.5e-06 | 2.5e-06 | 3.100E-06 | 5.292E-07 | 2.500E-06 | 3.500E-06 |
| 2433 | 122 | S122M | A | 2.6e-06 | 3.6e-06 | 3.2e-06 | 3.133E-06 | 5.033E-07 | 2.600E-06 | 3.600E-06 |
| 2434 | 122 | S122F | A | 2.9e-06 | 3.2e-06 | 3.2e-06 | 3.100E-06 | 1.732E-07 | 2.900E-06 | 3.200E-06 |
| 2435 | 122 | S122P | A | 2.7e-06 | 2.3e-06 | 2e-06 | 2.333E-06 | 3.512E-07 | 2.000E-06 | 2.700E-06 |
| 2436 | 122 | S122S | A | 2.1e-06 | 1.5e-06 | 1.7e-06 | 1.767E-06 | 3.055E-07 | 1.500E-06 | 2.100E-06 |
| 2437 | 122 | S122T | A | 1.8e-06 | 1.9e-06 | 3.4e-06 | 2.367E-06 | 8.963E-07 | 1.800E-06 | 3.400E-06 |
| 2438 | 122 | S122W | A | 1.9e-06 | 2.9e-06 | 2.3e-06 | 2.367E-06 | 5.033E-07 | 1.900E-06 | 2.900E-06 |
| 2439 | 122 | S122Y | A | 3e-06 | 3.5e-06 | 3.1e-06 | 3.200E-06 | 2.646E-07 | 3.000E-06 | 3.500E-06 |
| 2440 | 122 | S122V | A | 2.4e-06 | 2.5e-06 | 2e-06 | 2.300E-06 | 2.646E-07 | 2.000E-06 | 2.500E-06 |
| 2441 | 123 | L123A | A | 5.2e-06 | 2.9e-06 | 2e-06 | 3.367E-06 | 1.650E-06 | 2.000E-06 | 5.200E-06 |
| 2442 | 123 | L123R | A | 2.7e-06 | 2.4e-06 | 2.7e-06 | 2.600E-06 | 1.732E-07 | 2.400E-06 | 2.700E-06 |
| 2443 | 123 | L123N | A | 1.1e-06 | 2.2e-06 | 2e-06 | 1.767E-06 | 5.859E-07 | 1.100E-06 | 2.200E-06 |
| 2444 | 123 | L123D | A | 3.3e-06 | 2.4e-06 | 2.6e-06 | 2.767E-06 | 4.726E-07 | 2.400E-06 | 3.300E-06 |
| 2445 | 123 | L123C | A | 2.1e-06 | 3e-06 | 2.7e-06 | 2.600E-06 | 4.583E-07 | 2.100E-06 | 3.000E-06 |
| 2446 | 123 | L123E | A | 1.5e-06 | 1.6e-06 | 1.4e-06 | 1.500E-06 | 1.000E-07 | 1.400E-06 | 1.600E-06 |
| 2447 | 123 | L123Q | A | 2.7e-06 | 3.6e-06 | 1.7e-06 | 2.667E-06 | 9.504E-07 | 1.700E-06 | 3.600E-06 |
| 2448 | 123 | L123G | A | 1.9e-06 | 2.2e-06 | 2.7e-06 | 2.267E-06 | 4.041E-07 | 1.900E-06 | 2.700E-06 |
| 2449 | 123 | L123H | A | 1.6e-06 | 3.3e-06 | 2.5e-06 | 2.467E-06 | 8.505E-07 | 1.600E-06 | 3.300E-06 |
| 2450 | 123 | L123I | A | 3.3e-06 | 2.2e-06 | 3e-06 | 2.833E-06 | 5.686E-07 | 2.200E-06 | 3.300E-06 |
| 2451 | 123 | L123L | A | 2.2e-06 | 2e-06 | 2.2e-06 | 2.133E-06 | 1.155E-07 | 2.000E-06 | 2.200E-06 |
| 2452 | 123 | L123K | A | 3.4e-06 | 2.2e-06 | 1.9e-06 | 2.500E-06 | 7.937E-07 | 1.900E-06 | 3.400E-06 |
| 2453 | 123 | L123M | A | 2.4e-06 | 3.2e-06 | 1.7e-06 | 2.433E-06 | 7.506E-07 | 1.700E-06 | 3.200E-06 |
| 2454 | 123 | L123F | A | 1.9e-06 | 2.2e-06 | 2.2e-06 | 2.100E-06 | 1.732E-07 | 1.900E-06 | 2.200E-06 |
| 2455 | 123 | L123P | A | 3.1e-06 | 5e-06 | 1.2e-06 | 3.100E-06 | 1.900E-06 | 1.200E-06 | 5.000E-06 |
| 2456 | 123 | L123S | A | 3.1e-06 | 4.3e-06 | 1.2e-06 | 2.867E-06 | 1.563E-06 | 1.200E-06 | 4.300E-06 |
| 2457 | 123 | L123T | A | 2.1e-06 | 3.7e-06 | 1.5e-06 | 2.433E-06 | 1.137E-06 | 1.500E-06 | 3.700E-06 |
| 2458 | 123 | L123W | A | 2e-06 | 2.6e-06 | 3.8e-06 | 2.800E-06 | 9.165E-07 | 2.000E-06 | 3.800E-06 |
| 2459 | 123 | L123Y | A | 5e-06 | 2.1e-06 | 2.1e-06 | 3.067E-06 | 1.674E-06 | 2.100E-06 | 5.000E-06 |
| 2460 | 123 | L123V | A | 1.9e-06 | 2.6e-06 | 2.2e-06 | 2.233E-06 | 3.512E-07 | 1.900E-06 | 2.600E-06 |
| 2461 | 124 | T124A | A | 2.3e-06 | 4.3e-06 | 3e-06 | 3.200E-06 | 1.015E-06 | 2.300E-06 | 4.300E-06 |
| 2462 | 124 | T124R | A | 2.4e-06 | 2.2e-06 | 2.6e-06 | 2.400E-06 | 2.000E-07 | 2.200E-06 | 2.600E-06 |
| 2463 | 124 | T124N | A | 1.8e-06 | 3.8e-06 | 3.3e-06 | 2.967E-06 | 1.041E-06 | 1.800E-06 | 3.800E-06 |
| 2464 | 124 | T124D | A | 2.4e-06 | 2.4e-06 | 3.5e-06 | 2.767E-06 | 6.351E-07 | 2.400E-06 | 3.500E-06 |

| | | | | | | | | | | |
|---|---|---|---|---|---|---|---|---|---|---|
| 2465 | 124 | T124C | A | 2.4e-06 | 3.1e-06 | 3.4e-06 | 2.967E-06 | 5.132E-07 | 2.400E-06 | 3.400E-06 |
| 2466 | 124 | T124E | A | 2.8e-06 | 1.8e-06 | 1.7e-06 | 2.100E-06 | 6.083E-07 | 1.700E-06 | 2.800E-06 |
| 2467 | 124 | T124Q | A | 2.9e-06 | 2e-06 | 2.8e-06 | 2.567E-06 | 4.933E-07 | 2.000E-06 | 2.900E-06 |
| 2468 | 124 | T124G | A | 3.2e-06 | 5.3e-06 | 1.8e-06 | 3.433E-06 | 1.762E-06 | 1.800E-06 | 5.300E-06 |
| 2469 | 124 | T124H | A | 3.8e-06 | 2.7e-06 | 1.8e-06 | 2.767E-06 | 1.002E-06 | 1.800E-06 | 3.800E-06 |
| 2470 | 124 | T124I | A | 1.6e-06 | 2.1e-06 | 2.9e-06 | 2.200E-06 | 6.557E-07 | 1.600E-06 | 2.900E-06 |
| 2471 | 124 | T124L | A | 2.1e-06 | 3.5e-06 | 1.9e-06 | 2.500E-06 | 8.718E-07 | 1.900E-06 | 3.500E-06 |
| 2472 | 124 | T124K | A | 1.5e-06 | 1.8e-06 | 1.5e-06 | 1.600E-06 | 1.732E-07 | 1.500E-06 | 1.800E-06 |
| 2473 | 124 | T124M | A | 4e-06 | 2.2e-06 | 2.9e-06 | 3.033E-06 | 9.074E-07 | 2.200E-06 | 4.000E-06 |
| 2474 | 124 | T124F | A | 2.6e-06 | 1.8e-06 | 2.2e-06 | 2.200E-06 | 4.000E-07 | 1.800E-06 | 2.600E-06 |
| 2475 | 124 | T124P | A | 1.6e-06 | 2.7e-06 | 2.8e-06 | 2.367E-06 | 6.658E-07 | 1.600E-06 | 2.800E-06 |
| 2476 | 124 | T124S | A | 1.8e-06 | 2.5e-06 | 3.2e-06 | 2.500E-06 | 7.000E-07 | 1.800E-06 | 3.200E-06 |
| 2477 | 124 | T124T | A | 2.2e-06 | 2e-06 | 2.4e-06 | 2.200E-06 | 2.000E-07 | 2.000E-06 | 2.400E-06 |
| 2478 | 124 | T124W | A | 2.5e-06 | 2.5e-06 | 2.6e-06 | 2.533E-06 | 5.774E-08 | 2.500E-06 | 2.600E-06 |
| 2479 | 124 | T124Y | A | 2.2e-06 | 2.2e-06 | 2.4e-06 | 2.267E-06 | 1.155E-07 | 2.200E-06 | 2.400E-06 |
| 2480 | 124 | T124V | A | 4.9e-06 | 2.2e-06 | 3.5e-06 | 3.533E-06 | 1.350E-06 | 2.200E-06 | 4.900E-06 |
| 2481 | 125 | E125A | A | 4.1e-06 | 2.1e-06 | 2.2e-06 | 2.800E-06 | 1.127E-06 | 2.100E-06 | 4.100E-06 |
| 2482 | 125 | E125R | A | 2.5e-06 | 1.1e-06 | 1.8e-06 | 1.800E-06 | 7.000E-07 | 1.100E-06 | 2.500E-06 |
| 2483 | 125 | E125N | A | 3.1e-06 | 3e-06 | 3.1e-06 | 3.067E-06 | 5.774E-08 | 3.000E-06 | 3.100E-06 |
| 2484 | 125 | E125D | A | 1.5e-06 | 1.7e-06 | 3e-06 | 2.067E-06 | 8.145E-07 | 1.500E-06 | 3.000E-06 |
| 2485 | 125 | E125C | A | 2.1e-06 | 3e-06 | 2.4e-06 | 2.500E-06 | 4.583E-07 | 2.100E-06 | 3.000E-06 |
| 2486 | 125 | E125E | A | 3.1e-06 | 2.3e-06 | 2e-06 | 2.467E-06 | 5.686E-07 | 2.000E-06 | 3.100E-06 |
| 2487 | 125 | E125Q | A | 2.4e-06 | 2.5e-06 | 2.5e-06 | 2.467E-06 | 5.774E-08 | 2.400E-06 | 2.500E-06 |
| 2488 | 125 | E125G | A | 2.7e-06 | 2.6e-06 | 2.9e-06 | 2.733E-06 | 1.528E-07 | 2.600E-06 | 2.900E-06 |
| 2489 | 125 | E125H | A | 3.1e-06 | 3.4e-06 | 2.2e-06 | 2.900E-06 | 6.245E-07 | 2.200E-06 | 3.400E-06 |
| 2490 | 125 | E125I | A | 2.6e-06 | 3.4e-06 | 2.4e-06 | 2.800E-06 | 5.292E-07 | 2.400E-06 | 3.400E-06 |
| 2491 | 125 | E125L | A | 2.8e-06 | 3.4e-06 | 2.1e-06 | 2.767E-06 | 6.506E-07 | 2.100E-06 | 3.400E-06 |
| 2492 | 125 | E125K | A | 2.6e-06 | 3.2e-06 | 2.5e-06 | 2.767E-06 | 3.786E-07 | 2.500E-06 | 3.200E-06 |
| 2493 | 125 | E125M | A | 1.5e-06 | 2.3e-06 | 2.2e-06 | 2.000E-06 | 4.359E-07 | 1.500E-06 | 2.300E-06 |
| 2494 | 125 | E125F | A | 1.7e-06 | 2e-06 | 2.3e-06 | 2.000E-06 | 3.000E-07 | 1.700E-06 | 2.300E-06 |
| 2495 | 125 | E125P | A | 2.3e-06 | 3.6e-06 | 4.1e-06 | 3.333E-06 | 9.292E-07 | 2.300E-06 | 4.100E-06 |
| 2496 | 125 | E125S | A | 1.6e-06 | 3.2e-06 | 2.5e-06 | 2.433E-06 | 8.021E-07 | 1.600E-06 | 3.200E-06 |
| 2497 | 125 | E125T | A | 3.2e-06 | 1.8e-06 | 2.3e-06 | 2.433E-06 | 7.095E-07 | 1.800E-06 | 3.200E-06 |
| 2498 | 125 | E125W | A | 2.8e-06 | 1.5e-06 | 1.8e-06 | 2.033E-06 | 6.807E-07 | 1.500E-06 | 2.800E-06 |
| 2499 | 125 | E125Y | A | 2.7e-06 | 1.1e-06 | 3.2e-06 | 2.333E-06 | 1.097E-06 | 1.100E-06 | 3.200E-06 |
| 2500 | 125 | E125V | A | 1.3e-06 | 1.5e-06 | 2.8e-06 | 1.867E-06 | 8.145E-07 | 1.300E-06 | 2.800E-06 |
| 2501 | 126 | I126A | A | 3.6e-06 | 1.9e-06 | 1.7e-06 | 2.400E-06 | 1.044E-06 | 1.700E-06 | 3.600E-06 |
| 2502 | 126 | I126R | A | 1.9e-06 | 2.7e-06 | 2.3e-06 | 2.300E-06 | 4.000E-07 | 1.900E-06 | 2.700E-06 |
| 2503 | 126 | I126N | A | 2.3e-06 | 2.4e-06 | 2.3e-06 | 2.333E-06 | 5.774E-08 | 2.300E-06 | 2.400E-06 |
| 2504 | 126 | I126D | A | 2.1e-06 | 3.6e-06 | 1.6e-06 | 2.433E-06 | 1.041E-06 | 1.600E-06 | 3.600E-06 |
| 2505 | 126 | I126C | A | 1.7e-06 | 3.3e-06 | 2e-06 | 2.333E-06 | 8.505E-07 | 1.700E-06 | 3.300E-06 |
| 2506 | 126 | I126E | A | 4.7e-06 | 2.7e-06 | 1.5e-06 | 2.967E-06 | 1.617E-06 | 1.500E-06 | 4.700E-06 |
| 2507 | 126 | I126Q | A | 2.1e-06 | 1.8e-06 | 3.2e-06 | 2.367E-06 | 7.371E-07 | 1.800E-06 | 3.200E-06 |
| 2508 | 126 | I126G | A | 2e-06 | 3.3e-06 | 2.3e-06 | 2.533E-06 | 6.807E-07 | 2.000E-06 | 3.300E-06 |
| 2509 | 126 | I126H | A | 1.4e-06 | 1.1e-06 | 3.8e-06 | 2.100E-06 | 1.480E-06 | 1.100E-06 | 3.800E-06 |
| 2510 | 126 | I126I | A | 1.6e-06 | 2.6e-06 | 1.9e-06 | 2.033E-06 | 5.132E-07 | 1.600E-06 | 2.600E-06 |
| 2511 | 126 | I126L | A | 2.8e-06 | 1.2e-06 | 2.9e-06 | 2.300E-06 | 9.539E-07 | 1.200E-06 | 2.900E-06 |
| 2512 | 126 | I126K | A | 2.2e-06 | 2.8e-06 | 2.8e-06 | 2.600E-06 | 3.464E-07 | 2.200E-06 | 2.800E-06 |
| 2513 | 126 | I126M | A | 2e-06 | 3.3e-06 | 3e-06 | 2.767E-06 | 6.807E-07 | 2.000E-06 | 3.300E-06 |
| 2514 | 126 | I126F | A | 2.7e-06 | 2.2e-06 | 2.4e-06 | 2.433E-06 | 2.517E-07 | 2.200E-06 | 2.700E-06 |
| 2515 | 126 | I126P | A | 3.4e-06 | 2.1e-06 | 1.3e-06 | 2.267E-06 | 1.060E-06 | 1.300E-06 | 3.400E-06 |
| 2516 | 126 | I126S | A | 1.8e-06 | 2.6e-06 | 3.9e-06 | 2.767E-06 | 1.060E-06 | 1.800E-06 | 3.900E-06 |
| 2517 | 126 | I126T | A | 2.2e-06 | 2.6e-06 | 1.9e-06 | 2.233E-06 | 3.512E-07 | 1.900E-06 | 2.600E-06 |
| 2518 | 126 | I126W | A | 2.2e-06 | 2.5e-06 | 2.5e-06 | 2.400E-06 | 1.732E-07 | 2.200E-06 | 2.500E-06 |
| 2519 | 126 | I126Y | A | 2.1e-06 | 3.4e-06 | 3.3e-06 | 2.933E-06 | 7.234E-07 | 2.100E-06 | 3.400E-06 |
| 2520 | 126 | I126V | A | 4e-06 | 1.3e-06 | 1.4e-06 | 2.233E-06 | 1.531E-06 | 1.300E-06 | 4.000E-06 |
| 2521 | 127 | L127A | A | 2e-06 | 1.6e-06 | 1.2e-06 | 1.600E-06 | 4.000E-07 | 1.200E-06 | 2.000E-06 |
| 2522 | 127 | L127R | A | 3.7e-06 | 1.7e-06 | 2.6e-06 | 2.667E-06 | 1.002E-06 | 1.700E-06 | 3.700E-06 |
| 2523 | 127 | L127N | A | 3.9e-06 | 2.7e-06 | 1.7e-06 | 2.767E-06 | 1.102E-06 | 1.700E-06 | 3.900E-06 |
| 2524 | 127 | L127D | A | 2.2e-06 | 2.4e-06 | 3.5e-06 | 2.700E-06 | 7.000E-07 | 2.200E-06 | 3.500E-06 |
| 2525 | 127 | L127C | A | 2.8e-06 | 2.6e-06 | 4.8e-06 | 3.400E-06 | 1.217E-06 | 2.600E-06 | 4.800E-06 |
| 2526 | 127 | L127E | A | 2.9e-06 | 2.5e-06 | 2.3e-06 | 2.567E-06 | 3.055E-07 | 2.300E-06 | 2.900E-06 |
| 2527 | 127 | L127Q | A | 2.9e-06 | 1.4e-06 | 2.2e-06 | 2.167E-06 | 7.506E-07 | 1.400E-06 | 2.900E-06 |
| 2528 | 127 | L127G | A | 2.6e-06 | 2.3e-06 | 4e-06 | 2.967E-06 | 9.074E-07 | 2.300E-06 | 4.000E-06 |
| 2529 | 127 | L127H | A | 2.5e-06 | 1.7e-06 | 1.8e-06 | 2.000E-06 | 4.359E-07 | 1.700E-06 | 2.500E-06 |
| 2530 | 127 | L127I | A | 4.4e-06 | 3.2e-06 | 2.4e-06 | 3.333E-06 | 1.007E-06 | 2.400E-06 | 4.400E-06 |
| 2531 | 127 | L127L | A | 1.9e-06 | 2.4e-06 | 2.3e-06 | 2.200E-06 | 2.646E-07 | 1.900E-06 | 2.400E-06 |
| 2532 | 127 | L127K | A | 1.8e-06 | 2.7e-06 | 1.7e-06 | 2.067E-06 | 5.508E-07 | 1.700E-06 | 2.700E-06 |
| 2533 | 127 | L127M | A | 2.7e-06 | 4.6e-06 | 1.8e-06 | 3.033E-06 | 1.429E-06 | 1.800E-06 | 4.600E-06 |
| 2534 | 127 | L127F | A | 2.9e-06 | 1.4e-06 | 2.5e-06 | 2.267E-06 | 7.767E-07 | 1.400E-06 | 2.900E-06 |
| 2535 | 127 | L127P | A | 4.1e-06 | 1.7e-06 | 4.4e-06 | 3.400E-06 | 1.480E-06 | 1.700E-06 | 4.400E-06 |
| 2536 | 127 | L127S | A | 1.7e-06 | 2.4e-06 | 1.5e-06 | 1.867E-06 | 4.726E-07 | 1.500E-06 | 2.400E-06 |
| 2537 | 127 | L127T | A | 4.7e-06 | 4.5e-06 | 2.1e-06 | 3.767E-06 | 1.447E-06 | 2.100E-06 | 4.700E-06 |
| 2538 | 127 | L127W | A | 3.1e-06 | 3.4e-06 | 1.9e-06 | 2.800E-06 | 7.937E-07 | 1.900E-06 | 3.400E-06 |
| 2539 | 127 | L127Y | A | 2.1e-06 | 2.1e-06 | 3.2e-06 | 2.467E-06 | 6.351E-07 | 2.100E-06 | 3.200E-06 |
| 2540 | 127 | L127V | A | 2.9e-06 | 1.3e-06 | 3.1e-06 | 2.433E-06 | 9.866E-07 | 1.300E-06 | 3.100E-06 |
| 2541 | 128 | K128A | A | 2.4e-06 | 1.5e-06 | 4.2e-06 | 2.700E-06 | 1.375E-06 | 1.500E-06 | 4.200E-06 |
| 2542 | 128 | K128R | A | 2.3e-06 | 2.9e-06 | 3.7e-06 | 2.967E-06 | 7.024E-07 | 2.300E-06 | 3.700E-06 |
| 2543 | 128 | K128N | A | 2.4e-06 | 2.4e-06 | 2.2e-06 | 2.333E-06 | 1.155E-07 | 2.200E-06 | 2.400E-06 |
| 2544 | 128 | K128D | A | 1.7e-06 | 2.3e-06 | 2.1e-06 | 2.033E-06 | 3.055E-07 | 1.700E-06 | 2.300E-06 |
| 2545 | 128 | K128C | A | 2.1e-06 | 2.2e-06 | 3.4e-06 | 2.567E-06 | 7.234E-07 | 2.100E-06 | 3.400E-06 |
| 2546 | 128 | K128E | A | 2.6e-06 | 2.3e-06 | 2.5e-06 | 2.467E-06 | 1.528E-07 | 2.300E-06 | 2.600E-06 |
| 2547 | 128 | K128Q | A | 2e-06 | 2.4e-06 | 1.6e-06 | 2.000E-06 | 4.000E-07 | 1.600E-06 | 2.400E-06 |
| 2548 | 128 | K128G | A | 1.9e-06 | 3.1e-06 | 2.1e-06 | 2.367E-06 | 6.429E-07 | 1.900E-06 | 3.100E-06 |
| 2549 | 128 | K128H | A | 2.2e-06 | 1.6e-06 | 3.9e-06 | 2.567E-06 | 1.193E-06 | 1.600E-06 | 3.900E-06 |

| | | | | | | | | | | |
|---|---|---|---|---|---|---|---|---|---|---|
| 2550 | 128 | K128I | A | 2e-06 | 3.5e-06 | 2e-06 | 2.500E-06 | 8.660E-07 | 2.000E-06 | 3.500E-06 |
| 2551 | 128 | K128L | A | 1.3e-06 | 3.2e-06 | 2.5e-06 | 2.333E-06 | 9.609E-07 | 1.300E-06 | 3.200E-06 |
| 2552 | 128 | K128K | A | 2e-06 | 2.5e-06 | 2.3e-06 | 2.267E-06 | 2.517E-07 | 2.000E-06 | 2.500E-06 |
| 2553 | 128 | K128M | A | 2.4e-06 | 2.9e-06 | 2.1e-06 | 2.467E-06 | 4.041E-07 | 2.100E-06 | 2.900E-06 |
| 2554 | 128 | K128F | A | 2.4e-06 | 1.9e-06 | 3e-06 | 2.433E-06 | 5.508E-07 | 1.900E-06 | 3.000E-06 |
| 2555 | 128 | K128P | A | 2.5e-06 | 3.5e-06 | 2.3e-06 | 2.767E-06 | 6.429E-07 | 2.300E-06 | 3.500E-06 |
| 2556 | 128 | K128S | A | 2.3e-06 | 1.9e-06 | 5e-06 | 3.067E-06 | 1.686E-06 | 1.900E-06 | 5.000E-06 |
| 2557 | 128 | K128T | A | 1.9e-06 | 3.5e-06 | 4e-06 | 3.133E-06 | 1.097E-06 | 1.900E-06 | 4.000E-06 |
| 2558 | 128 | K128W | A | 2.1e-06 | 2.6e-06 | 2.5e-06 | 2.400E-06 | 2.646E-07 | 2.100E-06 | 2.600E-06 |
| 2559 | 128 | K128Y | A | 2.2e-06 | 2.4e-06 | 1.8e-06 | 2.133E-06 | 3.055E-07 | 1.800E-06 | 2.400E-06 |
| 2560 | 128 | K128V | A | 1.6e-06 | 2.9e-06 | 3.3e-06 | 2.600E-06 | 8.888E-07 | 1.600E-06 | 3.300E-06 |
| 2561 | 129 | G129A | A | 1.7e-06 | 2e-06 | 2.4e-06 | 2.033E-06 | 3.512E-07 | 1.700E-06 | 2.400E-06 |
| 2562 | 129 | G129R | A | 1.7e-06 | 2e-06 | 2.7e-06 | 2.133E-06 | 5.132E-07 | 1.700E-06 | 2.700E-06 |
| 2563 | 129 | G129N | A | 4.1e-06 | 2.1e-06 | 3.1e-06 | 3.100E-06 | 1.000E-06 | 2.100E-06 | 4.100E-06 |
| 2564 | 129 | G129D | A | 2.4e-06 | 2.2e-06 | 2.3e-06 | 2.300E-06 | 1.000E-07 | 2.200E-06 | 2.400E-06 |
| 2565 | 129 | G129C | A | 2.8e-06 | 4.7e-06 | 2.3e-06 | 3.267E-06 | 1.266E-06 | 2.300E-06 | 4.700E-06 |
| 2566 | 129 | G129E | A | 3.4e-06 | 1.3e-06 | 3e-06 | 2.567E-06 | 1.115E-06 | 1.300E-06 | 3.400E-06 |
| 2567 | 129 | G129Q | A | 2e-06 | 1.7e-06 | 2.5e-06 | 2.067E-06 | 4.041E-07 | 1.700E-06 | 2.500E-06 |
| 2568 | 129 | G129G | A | 2.1e-06 | 4.1e-06 | 3e-06 | 3.067E-06 | 1.002E-06 | 2.100E-06 | 4.100E-06 |
| 2569 | 129 | G129H | A | 5.2e-06 | 9.4e-07 | 1.4e-06 | 2.513E-06 | 2.338E-06 | 9.400E-07 | 5.200E-06 |
| 2570 | 129 | G129I | A | 4.3e-06 | 2.2e-06 | 1.6e-06 | 2.700E-06 | 1.418E-06 | 1.600E-06 | 4.300E-06 |
| 2571 | 129 | G129L | A | 3.2e-06 | 2.8e-06 | 2.7e-06 | 2.900E-06 | 2.646E-07 | 2.700E-06 | 3.200E-06 |
| 2572 | 129 | G129K | A | 1.3e-06 | 3.3e-06 | 2.5e-06 | 2.367E-06 | 1.007E-06 | 1.300E-06 | 3.300E-06 |
| 2573 | 129 | G129M | A | 2.9e-06 | 2.1e-06 | 2.3e-06 | 2.433E-06 | 4.163E-07 | 2.100E-06 | 2.900E-06 |
| 2574 | 129 | G129F | A | 1.7e-06 | 4.5e-06 | 1.9e-06 | 2.700E-06 | 1.562E-06 | 1.700E-06 | 4.500E-06 |
| 2575 | 129 | G129P | A | 3.2e-06 | 2.7e-06 | 1.4e-06 | 2.433E-06 | 9.292E-07 | 1.400E-06 | 3.200E-06 |
| 2576 | 129 | G129S | A | 2.8e-06 | 1.9e-06 | 2.6e-06 | 2.433E-06 | 4.726E-07 | 1.900E-06 | 2.800E-06 |
| 2577 | 129 | G129T | A | 2.8e-06 | 2.5e-06 | 1.8e-06 | 2.367E-06 | 5.132E-07 | 1.800E-06 | 2.800E-06 |
| 2578 | 129 | G129W | A | 2.7e-06 | 2.9e-06 | 3.4e-06 | 3.000E-06 | 3.606E-07 | 2.700E-06 | 3.400E-06 |
| 2579 | 129 | G129Y | A | 2.9e-06 | 1.1e-06 | 2.6e-06 | 2.200E-06 | 9.644E-07 | 1.100E-06 | 2.900E-06 |
| 2580 | 129 | G129V | A | 2.7e-06 | 2.1e-06 | 2e-06 | 2.267E-06 | 3.786E-07 | 2.000E-06 | 2.700E-06 |
| 2581 | 130 | G130A | A | 2.5e-06 | 2.7e-06 | 2.2e-06 | 2.467E-06 | 2.517E-07 | 2.200E-06 | 2.700E-06 |
| 2582 | 130 | G130R | A | 2e-06 | 2.1e-06 | 2.4e-06 | 2.167E-06 | 2.082E-07 | 2.000E-06 | 2.400E-06 |
| 2583 | 130 | G130N | A | 1.5e-06 | 2.7e-06 | 2.1e-06 | 2.100E-06 | 6.000E-07 | 1.500E-06 | 2.700E-06 |
| 2584 | 130 | G130D | A | 6.3e-06 | 2.2e-06 | 1.9e-06 | 3.467E-06 | 2.458E-06 | 1.900E-06 | 6.300E-06 |
| 2585 | 130 | G130C | A | 3.2e-06 | 2.6e-06 | 2.6e-06 | 2.800E-06 | 3.464E-07 | 2.600E-06 | 3.200E-06 |
| 2586 | 130 | G130E | A | 2.5e-06 | 2.1e-06 | 2.6e-06 | 2.400E-06 | 2.646E-07 | 2.100E-06 | 2.600E-06 |
| 2587 | 130 | G130Q | A | 3.2e-06 | 2.9e-06 | 2.6e-06 | 2.900E-06 | 3.000E-07 | 2.600E-06 | 3.200E-06 |
| 2588 | 130 | G130G | A | 4e-06 | 3e-06 | 2.8e-06 | 3.267E-06 | 6.429E-07 | 2.800E-06 | 4.000E-06 |
| 2589 | 130 | G130H | A | 1.7e-06 | 2e-06 | 2.4e-06 | 2.033E-06 | 3.512E-07 | 1.700E-06 | 2.400E-06 |
| 2590 | 130 | G130I | A | 3e-06 | 3.9e-06 | 3.8e-06 | 3.567E-06 | 4.933E-07 | 3.000E-06 | 3.900E-06 |
| 2591 | 130 | G130L | A | 2.4e-06 | 2.7e-06 | 2.5e-06 | 2.533E-06 | 1.528E-07 | 2.400E-06 | 2.700E-06 |
| 2592 | 130 | G130K | A | 2.2e-06 | 3.5e-06 | 1.2e-06 | 2.300E-06 | 1.153E-06 | 1.200E-06 | 3.500E-06 |
| 2593 | 130 | G130M | A | 2.1e-06 | 2.1e-06 | 1.6e-06 | 1.933E-06 | 2.887E-07 | 1.600E-06 | 2.100E-06 |
| 2594 | 130 | G130F | A | 1.3e-06 | 2.8e-06 | 2.3e-06 | 2.133E-06 | 7.638E-07 | 1.300E-06 | 2.800E-06 |
| 2595 | 130 | G130P | A | 2.1e-06 | 2.7e-06 | 2.4e-06 | 2.400E-06 | 3.000E-07 | 2.100E-06 | 2.700E-06 |
| 2596 | 130 | G130S | A | 1.6e-06 | 2.9e-06 | 3.4e-06 | 2.633E-06 | 9.292E-07 | 1.600E-06 | 3.400E-06 |
| 2597 | 130 | G130T | A | 3.6e-06 | 3.2e-06 | 4e-06 | 3.600E-06 | 4.000E-07 | 3.200E-06 | 4.000E-06 |
| 2598 | 130 | G130W | A | 1.9e-06 | 1.6e-06 | 2.8e-06 | 2.100E-06 | 6.245E-07 | 1.600E-06 | 2.800E-06 |
| 2599 | 130 | G130Y | A | 2.2e-06 | 1.8e-06 | 2.6e-06 | 2.200E-06 | 4.000E-07 | 1.800E-06 | 2.600E-06 |
| 2600 | 130 | G130V | A | 1.4e-06 | 1.1e-06 | 3.1e-06 | 1.867E-06 | 1.079E-06 | 1.100E-06 | 3.100E-06 |
| 2601 | 131 | V131A | A | 2.9e-06 | 2e-06 | 3.5e-06 | 2.800E-06 | 7.550E-07 | 2.000E-06 | 3.500E-06 |
| 2602 | 131 | V131R | A | 1.4e-06 | 1.9e-06 | 1.8e-06 | 1.700E-06 | 2.646E-07 | 1.400E-06 | 1.900E-06 |
| 2603 | 131 | V131N | A | 1.6e-06 | 2.2e-06 | 4.3e-06 | 2.700E-06 | 1.418E-06 | 1.600E-06 | 4.300E-06 |
| 2604 | 131 | V131D | A | 3.1e-06 | 1.6e-06 | 2e-06 | 2.233E-06 | 7.767E-07 | 1.600E-06 | 3.100E-06 |
| 2605 | 131 | V131C | A | 1.2e-06 | 4.6e-06 | 2.8e-06 | 2.867E-06 | 1.701E-06 | 1.200E-06 | 4.600E-06 |
| 2606 | 131 | V131E | A | 2.2e-06 | 2.9e-06 | 2.5e-06 | 2.533E-06 | 3.512E-07 | 2.200E-06 | 2.900E-06 |
| 2607 | 131 | V131Q | A | 2.3e-06 | 2.2e-06 | 2.5e-06 | 2.333E-06 | 1.528E-07 | 2.200E-06 | 2.500E-06 |
| 2608 | 131 | V131G | A | 1.6e-06 | 2.2e-06 | 3e-06 | 2.267E-06 | 7.024E-07 | 1.600E-06 | 3.000E-06 |
| 2609 | 131 | V131H | A | 1.6e-06 | 1.9e-06 | 1.1e-06 | 1.533E-06 | 4.041E-07 | 1.100E-06 | 1.900E-06 |
| 2610 | 131 | V131I | A | 1.3e-06 | 3.4e-06 | 2.5e-06 | 2.400E-06 | 1.054E-06 | 1.300E-06 | 3.400E-06 |
| 2611 | 131 | V131L | A | 3.4e-06 | 3.2e-06 | 1.9e-06 | 2.833E-06 | 8.145E-07 | 1.900E-06 | 3.400E-06 |
| 2612 | 131 | V131K | A | 2.1e-06 | 3.6e-06 | 2.2e-06 | 2.633E-06 | 8.386E-07 | 2.100E-06 | 3.600E-06 |
| 2613 | 131 | V131M | A | 1.9e-06 | 3e-06 | 3.4e-06 | 2.767E-06 | 7.767E-07 | 1.900E-06 | 3.400E-06 |
| 2614 | 131 | V131F | A | 2.6e-06 | 2.5e-06 | 3.1e-06 | 2.733E-06 | 3.215E-07 | 2.500E-06 | 3.100E-06 |
| 2615 | 131 | V131P | A | 1.6e-06 | 2.8e-06 | 1.6e-06 | 2.000E-06 | 6.928E-07 | 1.600E-06 | 2.800E-06 |
| 2616 | 131 | V131S | A | 1.6e-06 | 2.2e-06 | 2.8e-06 | 2.200E-06 | 6.000E-07 | 1.600E-06 | 2.800E-06 |
| 2617 | 131 | V131T | A | 2.9e-06 | 2.6e-06 | 2.5e-06 | 2.667E-06 | 2.082E-07 | 2.500E-06 | 2.900E-06 |
| 2618 | 131 | V131W | A | 1.9e-06 | 2.1e-06 | 1.7e-06 | 1.900E-06 | 2.000E-07 | 1.700E-06 | 2.100E-06 |
| 2619 | 131 | V131Y | A | 1.9e-06 | 2.2e-06 | 1.5e-06 | 1.867E-06 | 3.512E-07 | 1.500E-06 | 2.200E-06 |
| 2620 | 131 | V131V | A | 2.8e-06 | 2.9e-06 | 2.2e-06 | 2.633E-06 | 3.786E-07 | 2.200E-06 | 2.900E-06 |
| 2621 | 132 | L132A | A | 1.9e-06 | 1.2e-06 | 4e-06 | 2.367E-06 | 1.457E-06 | 1.200E-06 | 4.000E-06 |
| 2622 | 132 | L132R | A | 2.4e-06 | 2.3e-06 | 2.3e-06 | 2.333E-06 | 5.774E-08 | 2.300E-06 | 2.400E-06 |
| 2623 | 132 | L132N | A | 4.9e-06 | 2.2e-06 | 1.6e-06 | 2.900E-06 | 1.758E-06 | 1.600E-06 | 4.900E-06 |
| 2624 | 132 | L132D | A | 2.2e-06 | 2e-06 | 3.1e-06 | 2.433E-06 | 5.859E-07 | 2.000E-06 | 3.100E-06 |
| 2625 | 132 | L132C | A | 3.1e-06 | 3.2e-06 | 2e-06 | 2.767E-06 | 6.658E-07 | 2.000E-06 | 3.200E-06 |
| 2626 | 132 | L132E | A | 2.8e-06 | 3e-06 | 1.8e-06 | 2.533E-06 | 6.429E-07 | 1.800E-06 | 3.000E-06 |
| 2627 | 132 | L132Q | A | 1.2e-06 | 2.7e-06 | 2.9e-06 | 2.267E-06 | 9.292E-07 | 1.200E-06 | 2.900E-06 |
| 2628 | 132 | L132G | A | 2.3e-06 | 2.9e-06 | 4.6e-06 | 3.267E-06 | 1.193E-06 | 2.300E-06 | 4.600E-06 |
| 2629 | 132 | L132H | A | 2.1e-06 | 2.5e-06 | 2.8e-06 | 2.467E-06 | 3.512E-07 | 2.100E-06 | 2.800E-06 |
| 2630 | 132 | L132I | A | 3.6e-06 | 2.9e-06 | 2.5e-06 | 3.000E-06 | 5.568E-07 | 2.500E-06 | 3.600E-06 |
| 2631 | 132 | L132L | A | 1.5e-06 | 1.8e-06 | 1.6e-06 | 1.633E-06 | 1.528E-07 | 1.500E-06 | 1.800E-06 |
| 2632 | 132 | L132K | A | 3.3e-06 | 1.4e-06 | 2.6e-06 | 2.433E-06 | 9.609E-07 | 1.400E-06 | 3.300E-06 |
| 2633 | 132 | L132M | A | 2.5e-06 | 3.8e-06 | 2.2e-06 | 2.833E-06 | 8.505E-07 | 2.200E-06 | 3.800E-06 |
| 2634 | 132 | L132F | A | 3.3e-06 | 2e-06 | 2.8e-06 | 2.700E-06 | 6.557E-07 | 2.000E-06 | 3.300E-06 |

| | | | | | | | | | | |
|---|---|---|---|---|---|---|---|---|---|---|
| 2635 | 132 | L132P | A | 3.5e-06 | 2.7e-06 | 3.3e-06 | 3.167E-06 | 4.163E-07 | 2.700E-06 | 3.500E-06 |
| 2636 | 132 | L132S | A | 2e-06 | 2.9e-06 | 2e-06 | 2.300E-06 | 5.196E-07 | 2.000E-06 | 2.900E-06 |
| 2637 | 132 | L132T | A | 5.6e-06 | 4e-06 | 1.1e-06 | 3.567E-06 | 2.281E-06 | 1.100E-06 | 5.600E-06 |
| 2638 | 132 | L132W | A | 1.8e-06 | 2.9e-06 | 3.1e-06 | 2.600E-06 | 7.000E-07 | 1.800E-06 | 3.100E-06 |
| 2639 | 132 | L132Y | A | 2.6e-06 | 2.6e-06 | 3.3e-06 | 2.833E-06 | 4.041E-07 | 2.600E-06 | 3.300E-06 |
| 2640 | 132 | L132V | A | 3.9e-06 | 2.5e-06 | 2.2e-06 | 2.867E-06 | 9.074E-07 | 2.200E-06 | 3.900E-06 |
| 2641 | 133 | I133A | A | 3.5e-06 | 3.4e-06 | 2e-06 | 2.967E-06 | 8.386E-07 | 2.000E-06 | 3.500E-06 |
| 2642 | 133 | I133R | A | 2.9e-06 | 3.3e-06 | 1.7e-06 | 2.633E-06 | 8.327E-07 | 1.700E-06 | 3.300E-06 |
| 2643 | 133 | I133N | A | 2.4e-06 | 1.9e-06 | 1.1e-06 | 1.800E-06 | 6.557E-07 | 1.100E-06 | 2.400E-06 |
| 2644 | 133 | I133D | A | 2.2e-06 | 1.4e-06 | 4.2e-06 | 2.600E-06 | 1.442E-06 | 1.400E-06 | 4.200E-06 |
| 2645 | 133 | I133C | A | 2e-06 | 6.6e-07 | 1.8e-06 | 1.487E-06 | 7.229E-07 | 6.600E-07 | 2.000E-06 |
| 2646 | 133 | I133E | A | 3.7e-06 | 3.6e-06 | 2.4e-06 | 3.233E-06 | 7.234E-07 | 2.400E-06 | 3.700E-06 |
| 2647 | 133 | I133Q | A | 4.1e-06 | 2.5e-06 | 3.2e-06 | 3.267E-06 | 8.021E-07 | 2.500E-06 | 4.100E-06 |
| 2648 | 133 | I133G | A | 1.1e-06 | 2.5e-06 | 2.2e-06 | 1.933E-06 | 7.371E-07 | 1.100E-06 | 2.500E-06 |
| 2649 | 133 | I133H | A | 1.7e-06 | 1.7e-06 | 2.1e-06 | 1.833E-06 | 2.309E-07 | 1.700E-06 | 2.100E-06 |
| 2650 | 133 | I133I | A | 2.6e-06 | 1.4e-06 | 1.9e-06 | 1.967E-06 | 6.028E-07 | 1.400E-06 | 2.600E-06 |
| 2651 | 133 | I133L | A | 9.3e-07 | 2e-06 | 2.4e-06 | 1.777E-06 | 7.600E-07 | 9.300E-07 | 2.400E-06 |
| 2652 | 133 | I133K | A | 3.5e-06 | 1.6e-06 | 2.7e-06 | 2.600E-06 | 9.539E-07 | 1.600E-06 | 3.500E-06 |
| 2653 | 133 | I133M | A | 1.7e-06 | 2.3e-06 | 2.9e-06 | 2.300E-06 | 6.000E-07 | 1.700E-06 | 2.900E-06 |
| 2654 | 133 | I133F | A | 3e-06 | 1.9e-06 | 2.3e-06 | 2.400E-06 | 5.568E-07 | 1.900E-06 | 3.000E-06 |
| 2655 | 133 | I133P | A | 3.7e-06 | 1.9e-06 | 1.9e-06 | 2.500E-06 | 1.039E-06 | 1.900E-06 | 3.700E-06 |
| 2656 | 133 | I133S | A | 3.2e-06 | 2.9e-06 | 1.7e-06 | 2.600E-06 | 7.937E-07 | 1.700E-06 | 3.200E-06 |
| 2657 | 133 | I133T | A | 2.1e-06 | 2.1e-06 | 1.6e-06 | 1.933E-06 | 2.887E-07 | 1.600E-06 | 2.100E-06 |
| 2658 | 133 | I133W | A | 2.6e-06 | 1.7e-06 | 3.6e-06 | 2.633E-06 | 9.504E-07 | 1.700E-06 | 3.600E-06 |
| 2659 | 133 | I133Y | A | 2.2e-06 | 2.1e-06 | 9.9e-07 | 1.763E-06 | 6.716E-07 | 9.900E-07 | 2.200E-06 |
| 2660 | 133 | I133V | A | 1.2e-06 | 4.5e-06 | 1.9e-06 | 2.533E-06 | 1.739E-06 | 1.200E-06 | 4.500E-06 |
| 2661 | 134 | Q134A | A | 2.4e-06 | 3.4e-06 | 2.3e-06 | 2.700E-06 | 6.083E-07 | 2.300E-06 | 3.400E-06 |
| 2662 | 134 | Q134R | A | 1.6e-06 | 3.6e-06 | 2.4e-06 | 2.533E-06 | 1.007E-06 | 1.600E-06 | 3.600E-06 |
| 2663 | 134 | Q134N | A | 2e-06 | 1.9e-06 | 2.2e-06 | 2.033E-06 | 1.528E-07 | 1.900E-06 | 2.200E-06 |
| 2664 | 134 | Q134D | A | 2.5e-06 | 2.9e-06 | 2.2e-06 | 2.533E-06 | 3.512E-07 | 2.200E-06 | 2.900E-06 |
| 2665 | 134 | Q134C | A | 1.9e-06 | 2.5e-06 | 1.6e-06 | 2.000E-06 | 4.583E-07 | 1.600E-06 | 2.500E-06 |
| 2666 | 134 | Q134E | A | 2.7e-06 | 2e-06 | 1.2e-06 | 1.967E-06 | 7.506E-07 | 1.200E-06 | 2.700E-06 |
| 2667 | 134 | Q134Q | A | 2.1e-06 | 2.5e-06 | 2.2e-06 | 2.267E-06 | 2.082E-07 | 2.100E-06 | 2.500E-06 |
| 2668 | 134 | Q134G | A | 3.5e-06 | 3.7e-06 | 2.6e-06 | 3.267E-06 | 5.859E-07 | 2.600E-06 | 3.700E-06 |
| 2669 | 134 | Q134H | A | 1.5e-06 | 2.8e-06 | 2.5e-06 | 2.267E-06 | 6.807E-07 | 1.500E-06 | 2.800E-06 |
| 2670 | 134 | Q134I | A | 2e-06 | 3.3e-06 | 2.3e-06 | 2.533E-06 | 6.807E-07 | 2.000E-06 | 3.300E-06 |
| 2671 | 134 | Q134L | A | 2.6e-06 | 2.3e-06 | 4.2e-06 | 3.033E-06 | 1.021E-06 | 2.300E-06 | 4.200E-06 |
| 2672 | 134 | Q134K | A | 2.4e-06 | 2.4e-06 | 2.8e-06 | 2.533E-06 | 2.309E-07 | 2.400E-06 | 2.800E-06 |
| 2673 | 134 | Q134M | A | 3.4e-06 | 1.4e-06 | 2.5e-06 | 2.433E-06 | 1.002E-06 | 1.400E-06 | 3.400E-06 |
| 2674 | 134 | Q134F | A | 2e-06 | 3.2e-06 | 3.8e-06 | 3.000E-06 | 9.165E-07 | 2.000E-06 | 3.800E-06 |
| 2675 | 134 | Q134P | A | 2.3e-06 | 2e-06 | 2e-06 | 2.100E-06 | 1.732E-07 | 2.000E-06 | 2.300E-06 |
| 2676 | 134 | Q134S | A | 2.7e-06 | 2.5e-06 | 2.8e-06 | 2.667E-06 | 1.528E-07 | 2.500E-06 | 2.800E-06 |
| 2677 | 134 | Q134T | A | 3.4e-06 | 2e-06 | 1.9e-06 | 2.433E-06 | 8.386E-07 | 1.900E-06 | 3.400E-06 |
| 2678 | 134 | Q134W | A | 2e-06 | 2.2e-06 | 3.2e-06 | 2.467E-06 | 6.429E-07 | 2.000E-06 | 3.200E-06 |
| 2679 | 134 | Q134Y | A | 2.4e-06 | 2.4e-06 | 3.1e-06 | 2.633E-06 | 4.041E-07 | 2.400E-06 | 3.100E-06 |
| 2680 | 134 | Q134V | A | 3.1e-06 | 1.4e-06 | 2.7e-06 | 2.400E-06 | 8.888E-07 | 1.400E-06 | 3.100E-06 |
| 2681 | 135 | R135A | A | 1.6e-06 | 2.6e-06 | 3.9e-06 | 2.700E-06 | 1.153E-06 | 1.600E-06 | 3.900E-06 |
| 2682 | 135 | R135R | A | 3.6e-06 | 2e-06 | 3.7e-06 | 3.100E-06 | 9.539E-07 | 2.000E-06 | 3.700E-06 |
| 2683 | 135 | R135N | A | 2.8e-06 | 1.5e-06 | 1.4e-06 | 1.900E-06 | 7.810E-07 | 1.400E-06 | 2.800E-06 |
| 2684 | 135 | R135D | A | 2e-06 | 2.1e-06 | 2.9e-06 | 2.333E-06 | 4.933E-07 | 2.000E-06 | 2.900E-06 |
| 2685 | 135 | R135C | A | 1.9e-06 | 9.1e-07 | 1.2e-06 | 1.337E-06 | 5.090E-07 | 9.100E-07 | 1.900E-06 |
| 2686 | 135 | R135E | A | 2.3e-06 | 1.7e-06 | 2.4e-06 | 2.133E-06 | 3.786E-07 | 1.700E-06 | 2.400E-06 |
| 2687 | 135 | R135Q | A | 2.4e-06 | 2.8e-06 | 4.5e-06 | 3.233E-06 | 1.115E-06 | 2.400E-06 | 4.500E-06 |
| 2688 | 135 | R135G | A | 1.3e-06 | 2.3e-06 | 1.4e-06 | 1.667E-06 | 5.508E-07 | 1.300E-06 | 2.300E-06 |
| 2689 | 135 | R135H | A | 2.3e-06 | 3.1e-06 | 2e-06 | 2.467E-06 | 5.686E-07 | 2.000E-06 | 3.100E-06 |
| 2690 | 135 | R135I | A | 2.3e-06 | 2.9e-06 | 2.5e-06 | 2.567E-06 | 3.055E-07 | 2.300E-06 | 2.900E-06 |
| 2691 | 135 | R135L | A | 2.4e-06 | 2.3e-06 | 3e-06 | 2.567E-06 | 3.786E-07 | 2.300E-06 | 3.000E-06 |
| 2692 | 135 | R135K | A | 2.9e-06 | 1.5e-06 | 1.6e-06 | 2.000E-06 | 7.810E-07 | 1.500E-06 | 2.900E-06 |
| 2693 | 135 | R135M | A | 1.7e-06 | 1.1e-06 | 2.3e-06 | 1.700E-06 | 6.000E-07 | 1.100E-06 | 2.300E-06 |
| 2694 | 135 | R135F | A | 2.7e-06 | 3.2e-06 | 2.9e-06 | 2.933E-06 | 2.517E-07 | 2.700E-06 | 3.200E-06 |
| 2695 | 135 | R135P | A | 1.9e-06 | 2.3e-06 | 2.6e-06 | 2.267E-06 | 3.512E-07 | 1.900E-06 | 2.600E-06 |
| 2696 | 135 | R135S | A | 1.9e-06 | 3.5e-06 | 3.1e-06 | 2.833E-06 | 8.327E-07 | 1.900E-06 | 3.500E-06 |
| 2697 | 135 | R135T | A | 2.8e-06 | 2.4e-06 | 3.3e-06 | 2.833E-06 | 4.509E-07 | 2.400E-06 | 3.300E-06 |
| 2698 | 135 | R135W | A | 2.8e-06 | 3.3e-06 | 2.4e-06 | 2.833E-06 | 4.509E-07 | 2.400E-06 | 3.300E-06 |
| 2699 | 135 | R135Y | A | 3.2e-06 | 2.3e-06 | 2.3e-06 | 2.600E-06 | 5.196E-07 | 2.300E-06 | 3.200E-06 |
| 2700 | 135 | R135V | A | 3.4e-06 | 2.1e-06 | 2.5e-06 | 2.667E-06 | 6.658E-07 | 2.100E-06 | 3.400E-06 |
| 2701 | 136 | N136A | A | 1.7e-06 | 1.9e-06 | 2.3e-06 | 1.967E-06 | 3.055E-07 | 1.700E-06 | 2.300E-06 |
| 2702 | 136 | N136R | A | 3e-06 | 2.1e-06 | 1.9e-06 | 2.333E-06 | 5.859E-07 | 1.900E-06 | 3.000E-06 |
| 2703 | 136 | N136N | A | 1.3e-06 | 2.3e-06 | 1.4e-06 | 1.667E-06 | 5.508E-07 | 1.300E-06 | 2.300E-06 |
| 2704 | 136 | N136D | A | 1.8e-06 | 2.8e-06 | 4.5e-06 | 3.033E-06 | 1.365E-06 | 1.800E-06 | 4.500E-06 |
| 2705 | 136 | N136C | A | 3.3e-06 | 3.2e-06 | 2.5e-06 | 3.000E-06 | 4.359E-07 | 2.500E-06 | 3.300E-06 |
| 2706 | 136 | N136E | A | 1.8e-06 | 2.4e-06 | 3e-06 | 2.400E-06 | 6.000E-07 | 1.800E-06 | 3.000E-06 |
| 2707 | 136 | N136Q | A | 1.8e-06 | 2.1e-06 | 2.7e-06 | 2.200E-06 | 4.583E-07 | 1.800E-06 | 2.700E-06 |
| 2708 | 136 | N136G | A | 1.8e-06 | 2e-06 | 3.4e-06 | 2.400E-06 | 8.718E-07 | 1.800E-06 | 3.400E-06 |
| 2709 | 136 | N136H | A | 1.9e-06 | 2.4e-06 | 4.1e-06 | 2.800E-06 | 1.153E-06 | 1.900E-06 | 4.100E-06 |
| 2710 | 136 | N136I | A | 3e-06 | 5.8e-06 | 2.1e-06 | 3.633E-06 | 1.930E-06 | 2.100E-06 | 5.800E-06 |
| 2711 | 136 | N136L | A | 1.8e-06 | 2.2e-06 | 2.1e-06 | 2.033E-06 | 2.082E-07 | 1.800E-06 | 2.200E-06 |
| 2712 | 136 | N136K | A | 2.7e-06 | 2.7e-06 | 2.3e-06 | 2.567E-06 | 2.309E-07 | 2.300E-06 | 2.700E-06 |
| 2713 | 136 | N136M | A | 2.4e-06 | 3.5e-06 | 2.7e-06 | 2.867E-06 | 5.686E-07 | 2.400E-06 | 3.500E-06 |
| 2714 | 136 | N136F | A | 2.1e-06 | 2.7e-06 | 2.7e-06 | 2.500E-06 | 3.464E-07 | 2.100E-06 | 2.700E-06 |
| 2715 | 136 | N136P | A | 2.1e-06 | 1.3e-06 | 1.4e-06 | 1.600E-06 | 4.359E-07 | 1.300E-06 | 2.100E-06 |
| 2716 | 136 | N136S | A | 3.3e-06 | 2.3e-06 | 1.3e-06 | 2.300E-06 | 1.000E-06 | 1.300E-06 | 3.300E-06 |
| 2717 | 136 | N136T | A | 2.3e-06 | 1.8e-06 | 1.2e-06 | 1.767E-06 | 5.508E-07 | 1.200E-06 | 2.300E-06 |
| 2718 | 136 | N136W | A | 2e-06 | 3e-06 | 3.5e-06 | 2.833E-06 | 7.638E-07 | 2.000E-06 | 3.500E-06 |
| 2719 | 136 | N136Y | A | 3.1e-06 | 2.5e-06 | 2.3e-06 | 2.633E-06 | 4.163E-07 | 2.300E-06 | 3.100E-06 |

| | | | | | | | | | | |
|---|---|---|---|---|---|---|---|---|---|---|
| 2720 | 136 | N136V | A | 2.1e-06 | 1.8e-06 | 2.5e-06 | 2.133E-06 | 3.512E-07 | 1.800E-06 | 2.500E-06 |
| 2721 | 137 | P137A | A | 2.4e-06 | 4e-06 | 2.1e-06 | 2.833E-06 | 1.021E-06 | 2.100E-06 | 4.000E-06 |
| 2722 | 137 | P137R | A | 2.8e-06 | 1.6e-06 | 1.1e-06 | 1.833E-06 | 8.737E-07 | 1.100E-06 | 2.800E-06 |
| 2723 | 137 | P137N | A | 1.3e-06 | 2.7e-06 | 2.1e-06 | 2.033E-06 | 7.024E-07 | 1.300E-06 | 2.700E-06 |
| 2724 | 137 | P137D | A | 2.5e-06 | 1.8e-06 | 3.1e-06 | 2.467E-06 | 6.506E-07 | 1.800E-06 | 3.100E-06 |
| 2725 | 137 | P137C | A | 2.3e-06 | 4.1e-06 | 2.2e-06 | 2.867E-06 | 1.069E-06 | 2.200E-06 | 4.100E-06 |
| 2726 | 137 | P137E | A | 2.4e-06 | 2.4e-06 | 1.6e-06 | 2.133E-06 | 4.619E-07 | 1.600E-06 | 2.400E-06 |
| 2727 | 137 | P137Q | A | 1.7e-06 | 3.8e-06 | 2.9e-06 | 2.800E-06 | 1.054E-06 | 1.700E-06 | 3.800E-06 |
| 2728 | 137 | P137G | A | 1.9e-06 | 2.2e-06 | 3.1e-06 | 2.400E-06 | 6.245E-07 | 1.900E-06 | 3.100E-06 |
| 2729 | 137 | P137H | A | 1.8e-06 | 2e-06 | 2.3e-06 | 2.033E-06 | 2.517E-07 | 1.800E-06 | 2.300E-06 |
| 2730 | 137 | P137I | A | 2e-06 | 2.3e-06 | 2e-06 | 2.100E-06 | 1.732E-07 | 2.000E-06 | 2.300E-06 |
| 2731 | 137 | P137L | A | 3.6e-06 | 2.5e-06 | 1.9e-06 | 2.667E-06 | 8.622E-07 | 1.900E-06 | 3.600E-06 |
| 2732 | 137 | P137K | A | 2.3e-06 | 2.2e-06 | 1.5e-06 | 2.000E-06 | 4.359E-07 | 1.500E-06 | 2.300E-06 |
| 2733 | 137 | P137M | A | 3.9e-06 | 2.9e-06 | 2.4e-06 | 3.067E-06 | 7.638E-07 | 2.400E-06 | 3.900E-06 |
| 2734 | 137 | P137F | A | 3.4e-06 | 2.8e-06 | 2.4e-06 | 2.867E-06 | 5.033E-07 | 2.400E-06 | 3.400E-06 |
| 2735 | 137 | P137P | A | 2.6e-06 | 3.1e-06 | 2.7e-06 | 2.800E-06 | 2.646E-07 | 2.600E-06 | 3.100E-06 |
| 2736 | 137 | P137S | A | 2.5e-06 | 1.5e-06 | 2.3e-06 | 2.100E-06 | 5.292E-07 | 1.500E-06 | 2.500E-06 |
| 2737 | 137 | P137T | A | 2.9e-06 | 2.1e-06 | 1.5e-06 | 2.167E-06 | 7.024E-07 | 1.500E-06 | 2.900E-06 |
| 2738 | 137 | P137W | A | 2.1e-06 | 2.4e-06 | 1.7e-06 | 2.067E-06 | 3.512E-07 | 1.700E-06 | 2.400E-06 |
| 2739 | 137 | P137Y | A | 2.7e-06 | 2.2e-06 | 5.7e-06 | 3.533E-06 | 1.893E-06 | 2.200E-06 | 5.700E-06 |
| 2740 | 137 | P137V | A | 2.1e-06 | 3e-06 | 2.2e-06 | 2.433E-06 | 4.933E-07 | 2.100E-06 | 3.000E-06 |
| 2741 | 138 | Q138A | A | 1.9e-06 | 2.1e-06 | 3.1e-06 | 2.367E-06 | 6.429E-07 | 1.900E-06 | 3.100E-06 |
| 2742 | 138 | Q138R | A | 1.2e-06 | 1.3e-06 | 1.6e-06 | 1.367E-06 | 2.082E-07 | 1.200E-06 | 1.600E-06 |
| 2743 | 138 | Q138N | A | 2.8e-06 | 2.5e-06 | 2.4e-06 | 2.567E-06 | 2.082E-07 | 2.400E-06 | 2.800E-06 |
| 2744 | 138 | Q138D | A | 2.5e-06 | 4.9e-06 | 2.1e-06 | 3.167E-06 | 1.514E-06 | 2.100E-06 | 4.900E-06 |
| 2745 | 138 | Q138C | A | 2.1e-06 | 3.1e-06 | 2.5e-06 | 2.567E-06 | 5.033E-07 | 2.100E-06 | 3.100E-06 |
| 2746 | 138 | Q138E | A | 2.7e-06 | 2.5e-06 | 3.7e-06 | 2.967E-06 | 6.429E-07 | 2.500E-06 | 3.700E-06 |
| 2747 | 138 | Q138Q | A | 5e-06 | 2.8e-06 | 3.8e-06 | 3.867E-06 | 1.102E-06 | 2.800E-06 | 5.000E-06 |
| 2748 | 138 | Q138G | A | 2.5e-06 | 2.2e-06 | 2e-06 | 2.233E-06 | 2.517E-07 | 2.000E-06 | 2.500E-06 |
| 2749 | 138 | Q138H | A | 2.7e-06 | 2e-06 | 2.4e-06 | 2.367E-06 | 3.512E-07 | 2.000E-06 | 2.700E-06 |
| 2750 | 138 | Q138I | A | 1.8e-06 | 3.5e-06 | 5.4e-06 | 3.567E-06 | 1.801E-06 | 1.800E-06 | 5.400E-06 |
| 2751 | 138 | Q138L | A | 2.3e-06 | 1.7e-06 | 3.2e-06 | 2.400E-06 | 7.550E-07 | 1.700E-06 | 3.200E-06 |
| 2752 | 138 | Q138K | A | 2.2e-06 | 2.4e-06 | 2.4e-06 | 2.333E-06 | 1.155E-07 | 2.200E-06 | 2.400E-06 |
| 2753 | 138 | Q138M | A | 3.3e-06 | 2.5e-06 | 2e-06 | 2.600E-06 | 6.557E-07 | 2.000E-06 | 3.300E-06 |
| 2754 | 138 | Q138F | A | 1.8e-06 | 1.9e-06 | 2.9e-06 | 2.200E-06 | 6.083E-07 | 1.800E-06 | 2.900E-06 |
| 2755 | 138 | Q138P | A | 3.1e-06 | 2.3e-06 | 2.4e-06 | 2.600E-06 | 4.359E-07 | 2.300E-06 | 3.100E-06 |
| 2756 | 138 | Q138S | A | 6e-06 | 2.8e-06 | 2.8e-06 | 3.867E-06 | 1.848E-06 | 2.800E-06 | 6.000E-06 |
| 2757 | 138 | Q138T | A | 2.3e-06 | 1.6e-06 | 3.2e-06 | 2.367E-06 | 8.021E-07 | 1.600E-06 | 3.200E-06 |
| 2758 | 138 | Q138W | A | 3e-06 | 2.2e-06 | 1.7e-06 | 2.300E-06 | 6.557E-07 | 1.700E-06 | 3.000E-06 |
| 2759 | 138 | Q138Y | A | 3e-06 | 3.8e-06 | 2.8e-06 | 3.200E-06 | 5.292E-07 | 2.800E-06 | 3.800E-06 |
| 2760 | 138 | Q138V | A | 1.4e-06 | 3.1e-06 | 3.4e-06 | 2.633E-06 | 1.079E-06 | 1.400E-06 | 3.400E-06 |
| 2761 | 139 | L139A | A | 3e-06 | 2.1e-06 | 2.2e-06 | 2.433E-06 | 4.933E-07 | 2.100E-06 | 3.000E-06 |
| 2762 | 139 | L139R | A | 1.5e-06 | 2.6e-06 | 2e-06 | 2.033E-06 | 5.508E-07 | 1.500E-06 | 2.600E-06 |
| 2763 | 139 | L139N | A | 1.1e-06 | 2.7e-06 | 1.4e-06 | 1.733E-06 | 8.505E-07 | 1.100E-06 | 2.700E-06 |
| 2764 | 139 | L139D | A | 3.8e-06 | 3e-06 | 3.1e-06 | 3.300E-06 | 4.359E-07 | 3.000E-06 | 3.800E-06 |
| 2765 | 139 | L139C | A | 3.5e-06 | 2.9e-06 | 1.8e-06 | 2.733E-06 | 8.622E-07 | 1.800E-06 | 3.500E-06 |
| 2766 | 139 | L139E | A | 2e-06 | 2.8e-06 | 2.7e-06 | 2.500E-06 | 4.359E-07 | 2.000E-06 | 2.800E-06 |
| 2767 | 139 | L139Q | A | 2.4e-06 | 1.8e-06 | 2e-06 | 2.067E-06 | 3.055E-07 | 1.800E-06 | 2.400E-06 |
| 2768 | 139 | L139G | A | 1.3e-06 | 1.9e-06 | 2.8e-06 | 2.000E-06 | 7.550E-07 | 1.300E-06 | 2.800E-06 |
| 2769 | 139 | L139H | A | 1.6e-06 | 2e-06 | 3.2e-06 | 2.267E-06 | 8.327E-07 | 1.600E-06 | 3.200E-06 |
| 2770 | 139 | L139I | A | 4.3e-06 | 2.6e-06 | 2.5e-06 | 3.133E-06 | 1.012E-06 | 2.500E-06 | 4.300E-06 |
| 2771 | 139 | L139L | A | 4.1e-06 | 2e-06 | 3.3e-06 | 3.133E-06 | 1.060E-06 | 2.000E-06 | 4.100E-06 |
| 2772 | 139 | L139K | A | 3.3e-06 | 1.9e-06 | 2e-06 | 2.400E-06 | 7.810E-07 | 1.900E-06 | 3.300E-06 |
| 2773 | 139 | L139M | A | 1.9e-06 | 2.7e-06 | 1.4e-06 | 2.000E-06 | 6.557E-07 | 1.400E-06 | 2.700E-06 |
| 2774 | 139 | L139F | A | 1.6e-06 | 2.3e-06 | 4.5e-06 | 2.800E-06 | 1.513E-06 | 1.600E-06 | 4.500E-06 |
| 2775 | 139 | L139P | A | 1.6e-06 | 3.5e-06 | 2.6e-06 | 2.567E-06 | 9.504E-07 | 1.600E-06 | 3.500E-06 |
| 2776 | 139 | L139S | A | 2.3e-06 | 2.6e-06 | 2.1e-06 | 2.333E-06 | 2.517E-07 | 2.100E-06 | 2.600E-06 |
| 2777 | 139 | L139T | A | 1.7e-06 | 2.5e-06 | 2.3e-06 | 2.167E-06 | 4.163E-07 | 1.700E-06 | 2.500E-06 |
| 2778 | 139 | L139W | A | 4e-06 | 2.4e-06 | 1.5e-06 | 2.633E-06 | 1.266E-06 | 1.500E-06 | 4.000E-06 |
| 2779 | 139 | L139Y | A | 2.1e-06 | 1.9e-06 | 1.3e-06 | 1.767E-06 | 4.163E-07 | 1.300E-06 | 2.100E-06 |
| 2780 | 139 | L139V | A | 1.6e-06 | 2.9e-06 | 1.8e-06 | 2.100E-06 | 7.000E-07 | 1.600E-06 | 2.900E-06 |
| 2781 | 140 | C140A | A | 2.3e-06 | 3.5e-06 | 1.3e-06 | 2.367E-06 | 1.102E-06 | 1.300E-06 | 3.500E-06 |
| 2782 | 140 | C140R | A | 2e-06 | 2.8e-06 | 2.8e-06 | 2.533E-06 | 4.619E-07 | 2.000E-06 | 2.800E-06 |
| 2783 | 140 | C140N | A | 3.2e-06 | 2.3e-06 | 2.4e-06 | 2.633E-06 | 4.933E-07 | 2.300E-06 | 3.200E-06 |
| 2784 | 140 | C140D | A | 3e-06 | 2.2e-06 | 4e-06 | 3.067E-06 | 9.018E-07 | 2.200E-06 | 4.000E-06 |
| 2785 | 140 | C140C | A | 2.8e-06 | 1.3e-06 | 3.5e-06 | 2.533E-06 | 1.124E-06 | 1.300E-06 | 3.500E-06 |
| 2786 | 140 | C140E | A | 2.3e-06 | 1.8e-06 | 2.8e-06 | 2.300E-06 | 5.000E-07 | 1.800E-06 | 2.800E-06 |
| 2787 | 140 | C140Q | A | 1.9e-06 | 1.8e-06 | 2.9e-06 | 2.200E-06 | 6.083E-07 | 1.800E-06 | 2.900E-06 |
| 2788 | 140 | C140G | A | 2.7e-06 | 2.5e-06 | 4.2e-06 | 3.133E-06 | 9.292E-07 | 2.500E-06 | 4.200E-06 |
| 2789 | 140 | C140H | A | 1.2e-06 | 2.5e-06 | 1.6e-06 | 1.767E-06 | 6.658E-07 | 1.200E-06 | 2.500E-06 |
| 2790 | 140 | C140I | A | 3e-06 | 4e-06 | 2.7e-06 | 3.233E-06 | 6.807E-07 | 2.700E-06 | 4.000E-06 |
| 2791 | 140 | C140L | A | 3.5e-06 | 2.8e-06 | 2.4e-06 | 2.900E-06 | 5.568E-07 | 2.400E-06 | 3.500E-06 |
| 2792 | 140 | C140K | A | 2.4e-06 | 1.1e-06 | 1.7e-06 | 1.733E-06 | 6.506E-07 | 1.100E-06 | 2.400E-06 |
| 2793 | 140 | C140M | A | 3e-06 | 1.6e-06 | 3e-06 | 2.533E-06 | 8.083E-07 | 1.600E-06 | 3.000E-06 |
| 2794 | 140 | C140F | A | 2.7e-06 | 3.7e-06 | 2.3e-06 | 2.900E-06 | 7.211E-07 | 2.300E-06 | 3.700E-06 |
| 2795 | 140 | C140P | A | 3.9e-06 | 1.6e-06 | 2.3e-06 | 2.600E-06 | 1.179E-06 | 1.600E-06 | 3.900E-06 |
| 2796 | 140 | C140S | A | 1.6e-06 | 1.8e-06 | 3.2e-06 | 2.200E-06 | 8.718E-07 | 1.600E-06 | 3.200E-06 |
| 2797 | 140 | C140T | A | 2.5e-06 | 2.2e-06 | 4.1e-06 | 2.933E-06 | 1.021E-06 | 2.200E-06 | 4.100E-06 |
| 2798 | 140 | C140W | A | 1.4e-06 | 2.9e-06 | 2.7e-06 | 2.333E-06 | 8.145E-07 | 1.400E-06 | 2.900E-06 |
| 2799 | 140 | C140Y | A | 3.4e-06 | 2.8e-06 | 2.4e-06 | 2.867E-06 | 5.033E-07 | 2.400E-06 | 3.400E-06 |
| 2800 | 140 | C140V | A | 2.8e-06 | 2e-06 | 1.7e-06 | 2.167E-06 | 5.686E-07 | 1.700E-06 | 2.800E-06 |
| 2801 | 141 | Y141A | A | 1.8e-06 | 1.9e-06 | 4.5e-06 | 2.733E-06 | 1.531E-06 | 1.800E-06 | 4.500E-06 |
| 2802 | 141 | Y141R | A | 1.6e-06 | 1.5e-06 | 2.7e-06 | 1.933E-06 | 6.658E-07 | 1.500E-06 | 2.700E-06 |
| 2803 | 141 | Y141N | A | 3.2e-06 | 4.1e-06 | 3.2e-06 | 3.500E-06 | 5.196E-07 | 3.200E-06 | 4.100E-06 |
| 2804 | 141 | Y141D | A | 2.6e-06 | 1.1e-06 | 1.9e-06 | 1.867E-06 | 7.506E-07 | 1.100E-06 | 2.600E-06 |

| | | | | | | | | | | |
|---|---|---|---|---|---|---|---|---|---|---|
| 2805 | 141 | Y141C | A | 1.6e-06 | 3.1e-06 | 1.6e-06 | 2.100E-06 | 8.660E-07 | 1.600E-06 | 3.100E-06 |
| 2806 | 141 | Y141E | A | 2.7e-06 | 1.9e-06 | 2.3e-06 | 2.300E-06 | 4.000E-07 | 1.900E-06 | 2.700E-06 |
| 2807 | 141 | Y141Q | A | 1.2e-06 | 2e-06 | 2.9e-06 | 2.033E-06 | 8.505E-07 | 1.200E-06 | 2.900E-06 |
| 2808 | 141 | Y141G | A | 2.3e-06 | 3.6e-06 | 2.4e-06 | 2.767E-06 | 7.234E-07 | 2.300E-06 | 3.600E-06 |
| 2809 | 141 | Y141H | A | 3.3e-06 | 3.2e-06 | 1.8e-06 | 2.767E-06 | 8.386E-07 | 1.800E-06 | 3.300E-06 |
| 2810 | 141 | Y141I | A | 3.8e-06 | 2.4e-06 | 3.9e-06 | 3.367E-06 | 8.386E-07 | 2.400E-06 | 3.900E-06 |
| 2811 | 141 | Y141L | A | 4.3e-06 | 2.4e-06 | 3.5e-06 | 3.400E-06 | 9.539E-07 | 2.400E-06 | 4.300E-06 |
| 2812 | 141 | Y141K | A | 1.2e-06 | 2.2e-06 | 2.4e-06 | 1.933E-06 | 6.429E-07 | 1.200E-06 | 2.400E-06 |
| 2813 | 141 | Y141M | A | 2.8e-06 | 2.3e-06 | 3.4e-06 | 2.833E-06 | 5.508E-07 | 2.300E-06 | 3.400E-06 |
| 2814 | 141 | Y141F | A | 3.3e-06 | 2e-06 | 2.9e-06 | 2.733E-06 | 6.658E-07 | 2.000E-06 | 3.300E-06 |
| 2815 | 141 | Y141P | A | 5.2e-06 | 2.2e-06 | 2e-06 | 3.133E-06 | 1.793E-06 | 2.000E-06 | 5.200E-06 |
| 2816 | 141 | Y141S | A | 2e-06 | 4e-06 | 2.1e-06 | 2.700E-06 | 1.127E-06 | 2.000E-06 | 4.000E-06 |
| 2817 | 141 | Y141T | A | 2.2e-06 | 2.8e-06 | 1.5e-06 | 2.167E-06 | 6.506E-07 | 1.500E-06 | 2.800E-06 |
| 2818 | 141 | Y141W | A | 3.1e-06 | 2e-06 | 3.1e-06 | 2.733E-06 | 6.351E-07 | 2.000E-06 | 3.100E-06 |
| 2819 | 141 | Y141Y | A | 3.3e-06 | 3.2e-06 | 4e-06 | 3.500E-06 | 4.359E-07 | 3.200E-06 | 4.000E-06 |
| 2820 | 141 | Y141V | A | 2.1e-06 | 4e-06 | 1.2e-06 | 2.433E-06 | 1.429E-06 | 1.200E-06 | 4.000E-06 |
| 2821 | 142 | Q142A | A | 2.2e-06 | 4.8e-06 | 2.8e-06 | 3.267E-06 | 1.361E-06 | 2.200E-06 | 4.800E-06 |
| 2822 | 142 | Q142R | A | 1.9e-06 | 2.3e-06 | 2.2e-06 | 2.133E-06 | 2.082E-07 | 1.900E-06 | 2.300E-06 |
| 2823 | 142 | Q142N | A | 2e-06 | 2.9e-06 | 1.5e-06 | 2.133E-06 | 7.095E-07 | 1.500E-06 | 2.900E-06 |
| 2824 | 142 | Q142D | A | 3.7e-06 | 2e-06 | 2.2e-06 | 2.633E-06 | 9.292E-07 | 2.000E-06 | 3.700E-06 |
| 2825 | 142 | Q142C | A | 3.9e-06 | 2.3e-06 | 3.3e-06 | 3.167E-06 | 8.083E-07 | 2.300E-06 | 3.900E-06 |
| 2826 | 142 | Q142E | A | 2.2e-06 | 2.3e-06 | 1.4e-06 | 1.967E-06 | 4.933E-07 | 1.400E-06 | 2.300E-06 |
| 2827 | 142 | Q142Q | A | 4.3e-06 | 2.4e-06 | 2e-06 | 2.900E-06 | 1.229E-06 | 2.000E-06 | 4.300E-06 |
| 2828 | 142 | Q142G | A | 2.1e-06 | 1.9e-06 | 2e-06 | 2.000E-06 | 1.000E-07 | 1.900E-06 | 2.100E-06 |
| 2829 | 142 | Q142H | A | 2.2e-06 | 2.1e-06 | 2.3e-06 | 2.200E-06 | 1.000E-07 | 2.100E-06 | 2.300E-06 |
| 2830 | 142 | Q142I | A | 2.9e-06 | 2e-06 | 1.6e-06 | 2.167E-06 | 6.658E-07 | 1.600E-06 | 2.900E-06 |
| 2831 | 142 | Q142L | A | 2.3e-06 | 2.1e-06 | 1.9e-06 | 2.100E-06 | 2.000E-07 | 1.900E-06 | 2.300E-06 |
| 2832 | 142 | Q142K | A | 2.4e-06 | 1.6e-06 | 2.2e-06 | 2.067E-06 | 4.163E-07 | 1.600E-06 | 2.400E-06 |
| 2833 | 142 | Q142M | A | 3.3e-06 | 2.7e-06 | 1.9e-06 | 2.633E-06 | 7.024E-07 | 1.900E-06 | 3.300E-06 |
| 2834 | 142 | Q142F | A | 1.8e-06 | 1.9e-06 | 2.1e-06 | 1.933E-06 | 1.528E-07 | 1.800E-06 | 2.100E-06 |
| 2835 | 142 | Q142P | A | 2.6e-06 | 2.9e-06 | 3.8e-06 | 3.100E-06 | 6.245E-07 | 2.600E-06 | 3.800E-06 |
| 2836 | 142 | Q142S | A | 1.6e-06 | 1.4e-06 | 3.9e-06 | 2.300E-06 | 1.389E-06 | 1.400E-06 | 3.900E-06 |
| 2837 | 142 | Q142T | A | 1.9e-06 | 2.4e-06 | 2.1e-06 | 2.133E-06 | 2.517E-07 | 1.900E-06 | 2.400E-06 |
| 2838 | 142 | Q142W | A | 3.3e-06 | 2.6e-06 | 3.3e-06 | 3.067E-06 | 4.041E-07 | 2.600E-06 | 3.300E-06 |
| 2839 | 142 | Q142Y | A | 2.5e-06 | 2.4e-06 | 4.1e-06 | 3.000E-06 | 9.539E-07 | 2.400E-06 | 4.100E-06 |
| 2840 | 142 | Q142V | A | 2.2e-06 | 3.8e-06 | 2.4e-06 | 2.800E-06 | 8.718E-07 | 2.200E-06 | 3.800E-06 |
| 2841 | 143 | D143A | A | 2.2e-06 | 2.5e-06 | 2.4e-06 | 2.367E-06 | 1.528E-07 | 2.200E-06 | 2.500E-06 |
| 2842 | 143 | D143R | A | 1.6e-06 | 3.7e-06 | 2.2e-06 | 2.500E-06 | 1.082E-06 | 1.600E-06 | 3.700E-06 |
| 2843 | 143 | D143N | A | 3.2e-06 | 2.3e-06 | 1.6e-06 | 2.367E-06 | 8.021E-07 | 1.600E-06 | 3.200E-06 |
| 2844 | 143 | D143D | A | 2.7e-06 | 1.7e-06 | 1.9e-06 | 2.100E-06 | 5.292E-07 | 1.700E-06 | 2.700E-06 |
| 2845 | 143 | D143C | A | 1.2e-06 | 2.6e-06 | 3.2e-06 | 2.333E-06 | 1.026E-06 | 1.200E-06 | 3.200E-06 |
| 2846 | 143 | D143E | A | 1.9e-06 | 2.4e-06 | 2.8e-06 | 2.367E-06 | 4.509E-07 | 1.900E-06 | 2.800E-06 |
| 2847 | 143 | D143Q | A | 2.1e-06 | 2e-06 | 2.8e-06 | 2.300E-06 | 4.359E-07 | 2.000E-06 | 2.800E-06 |
| 2848 | 143 | D143G | A | 2.8e-06 | 2.2e-06 | 3e-06 | 2.667E-06 | 4.163E-07 | 2.200E-06 | 3.000E-06 |
| 2849 | 143 | D143H | A | 3.3e-06 | 2.2e-06 | 2.4e-06 | 2.633E-06 | 5.859E-07 | 2.200E-06 | 3.300E-06 |
| 2850 | 143 | D143I | A | 2.5e-06 | 2.4e-06 | 2.7e-06 | 2.533E-06 | 1.528E-07 | 2.400E-06 | 2.700E-06 |
| 2851 | 143 | D143L | A | 4.2e-06 | 4.1e-06 | 2.9e-06 | 3.733E-06 | 7.234E-07 | 2.900E-06 | 4.200E-06 |
| 2852 | 143 | D143K | A | 2.2e-06 | 1.6e-06 | 3.6e-06 | 2.467E-06 | 1.026E-06 | 1.600E-06 | 3.600E-06 |
| 2853 | 143 | D143M | A | 2.5e-06 | 2.5e-06 | 2.6e-06 | 2.533E-06 | 5.774E-08 | 2.500E-06 | 2.600E-06 |
| 2854 | 143 | D143F | A | 1.7e-06 | 1.5e-06 | 1.7e-06 | 1.633E-06 | 1.155E-07 | 1.500E-06 | 1.700E-06 |
| 2855 | 143 | D143P | A | 1.3e-06 | 2.2e-06 | 1.9e-06 | 1.800E-06 | 4.583E-07 | 1.300E-06 | 2.200E-06 |
| 2856 | 143 | D143S | A | 2e-06 | 2.4e-06 | 1.6e-06 | 2.000E-06 | 4.000E-07 | 1.600E-06 | 2.400E-06 |
| 2857 | 143 | D143T | A | 2.2e-06 | 2.1e-06 | 2.7e-06 | 2.333E-06 | 3.215E-07 | 2.100E-06 | 2.700E-06 |
| 2858 | 143 | D143W | A | 4e-06 | 3.2e-06 | 2.2e-06 | 3.133E-06 | 9.018E-07 | 2.200E-06 | 4.000E-06 |
| 2859 | 143 | D143Y | A | 2e-06 | 3.3e-06 | 2.6e-06 | 2.633E-06 | 6.506E-07 | 2.000E-06 | 3.300E-06 |
| 2860 | 143 | D143V | A | 2e-06 | 2.7e-06 | 1.6e-06 | 2.100E-06 | 5.568E-07 | 1.600E-06 | 2.700E-06 |
| 2861 | 144 | T144A | A | 2e-06 | 1.8e-06 | 2.5e-06 | 2.100E-06 | 3.606E-07 | 1.800E-06 | 2.500E-06 |
| 2862 | 144 | T144R | A | 2e-06 | 3.4e-06 | 3e-06 | 2.800E-06 | 7.211E-07 | 2.000E-06 | 3.400E-06 |
| 2863 | 144 | T144N | A | 1.9e-06 | 1.9e-06 | 1.5e-06 | 1.767E-06 | 2.309E-07 | 1.500E-06 | 1.900E-06 |
| 2864 | 144 | T144D | A | 3.1e-06 | 3.5e-06 | 3.2e-06 | 3.267E-06 | 2.082E-07 | 3.100E-06 | 3.500E-06 |
| 2865 | 144 | T144C | A | 1.5e-06 | 2.8e-06 | 3.4e-06 | 2.567E-06 | 9.713E-07 | 1.500E-06 | 3.400E-06 |
| 2866 | 144 | T144E | A | 3.4e-06 | 1.7e-06 | 2.6e-06 | 2.567E-06 | 8.505E-07 | 1.700E-06 | 3.400E-06 |
| 2867 | 144 | T144Q | A | 2e-06 | 1.6e-06 | 1.3e-06 | 1.633E-06 | 3.512E-07 | 1.300E-06 | 2.000E-06 |
| 2868 | 144 | T144G | A | 4.2e-06 | 3e-06 | 4e-06 | 3.733E-06 | 6.429E-07 | 3.000E-06 | 4.200E-06 |
| 2869 | 144 | T144H | A | 4.6e-06 | 2.8e-06 | 1.9e-06 | 3.100E-06 | 1.375E-06 | 1.900E-06 | 4.600E-06 |
| 2870 | 144 | T144I | A | 2.4e-06 | 2.5e-06 | 2.6e-06 | 2.500E-06 | 1.000E-07 | 2.400E-06 | 2.600E-06 |
| 2871 | 144 | T144L | A | 1.9e-06 | 1.9e-06 | 1.9e-06 | 1.900E-06 | 2.593E-22 | 1.900E-06 | 1.900E-06 |
| 2872 | 144 | T144K | A | 2.1e-06 | 2.2e-06 | 3.5e-06 | 2.600E-06 | 7.810E-07 | 2.100E-06 | 3.500E-06 |
| 2873 | 144 | T144M | A | 5.3e-06 | 2e-06 | 1.9e-06 | 3.067E-06 | 1.935E-06 | 1.900E-06 | 5.300E-06 |
| 2874 | 144 | T144F | A | 2.6e-06 | 1.8e-06 | 1.7e-06 | 2.033E-06 | 4.933E-07 | 1.700E-06 | 2.600E-06 |
| 2875 | 144 | T144P | A | 2.8e-06 | 3.3e-06 | 1.6e-06 | 2.567E-06 | 8.737E-07 | 1.600E-06 | 3.300E-06 |
| 2876 | 144 | T144S | A | 2.9e-06 | 2.7e-06 | 2.8e-06 | 2.800E-06 | 1.000E-07 | 2.700E-06 | 2.900E-06 |
| 2877 | 144 | T144T | A | 1.1e-06 | 1.6e-06 | 2.1e-06 | 1.600E-06 | 5.000E-07 | 1.100E-06 | 2.100E-06 |
| 2878 | 144 | T144W | A | 2.9e-06 | 4e-06 | 2.1e-06 | 3.000E-06 | 9.539E-07 | 2.100E-06 | 4.000E-06 |
| 2879 | 144 | T144Y | A | 2.5e-06 | 3.6e-06 | 2e-06 | 2.700E-06 | 8.185E-07 | 2.000E-06 | 3.600E-06 |
| 2880 | 144 | T144V | A | 2.4e-06 | 1.7e-06 | 3.1e-06 | 2.400E-06 | 7.000E-07 | 1.700E-06 | 3.100E-06 |
| 2881 | 145 | I145A | A | 2.4e-06 | 1.4e-06 | 3.1e-06 | 2.300E-06 | 8.544E-07 | 1.400E-06 | 3.100E-06 |
| 2882 | 145 | I145R | A | 1.4e-06 | 2.9e-06 | 1.9e-06 | 2.067E-06 | 7.638E-07 | 1.400E-06 | 2.900E-06 |
| 2883 | 145 | I145N | A | 1.7e-06 | 4.6e-06 | 1.7e-06 | 2.667E-06 | 1.674E-06 | 1.700E-06 | 4.600E-06 |
| 2884 | 145 | I145D | A | 1.4e-06 | 2.8e-06 | 1.7e-06 | 1.967E-06 | 7.371E-07 | 1.400E-06 | 2.800E-06 |
| 2885 | 145 | I145C | A | 2.1e-06 | 1.6e-06 | 3.5e-06 | 2.400E-06 | 9.849E-07 | 1.600E-06 | 3.500E-06 |
| 2886 | 145 | I145E | A | 2.7e-06 | 3.9e-06 | 2.2e-06 | 2.933E-06 | 8.737E-07 | 2.200E-06 | 3.900E-06 |
| 2887 | 145 | I145Q | A | 1.8e-06 | 2.4e-06 | 2.3e-06 | 2.167E-06 | 3.215E-07 | 1.800E-06 | 2.400E-06 |
| 2888 | 145 | I145G | A | 2.6e-06 | 1.2e-06 | 3.4e-06 | 2.400E-06 | 1.114E-06 | 1.200E-06 | 3.400E-06 |
| 2889 | 145 | I145H | A | 1.8e-06 | 2.6e-06 | 3.7e-06 | 2.700E-06 | 9.539E-07 | 1.800E-06 | 3.700E-06 |

| | | | | | | | | | | |
|---|---|---|---|---|---|---|---|---|---|---|
| 2890 | 145 | I145I | A | 2.1e-06 | 2.1e-06 | 2.5e-06 | 2.233E-06 | 2.309E-07 | 2.100E-06 | 2.500E-06 |
| 2891 | 145 | I145L | A | 3.5e-06 | 1.3e-06 | 2.2e-06 | 2.333E-06 | 1.106E-06 | 1.300E-06 | 3.500E-06 |
| 2892 | 145 | I145K | A | 3.2e-06 | 3.1e-06 | 2.4e-06 | 2.900E-06 | 4.359E-07 | 2.400E-06 | 3.200E-06 |
| 2893 | 145 | I145M | A | 2.1e-06 | 1.3e-06 | 2.8e-06 | 2.067E-06 | 7.506E-07 | 1.300E-06 | 2.800E-06 |
| 2894 | 145 | I145F | A | 2.8e-06 | 4.5e-06 | 1.8e-06 | 3.033E-06 | 1.365E-06 | 1.800E-06 | 4.500E-06 |
| 2895 | 145 | I145P | A | 2.6e-06 | 2.7e-06 | 1.9e-06 | 2.400E-06 | 4.359E-07 | 1.900E-06 | 2.700E-06 |
| 2896 | 145 | I145S | A | 2.6e-06 | 1.5e-06 | 3.1e-06 | 2.400E-06 | 8.185E-07 | 1.500E-06 | 3.100E-06 |
| 2897 | 145 | I145T | A | 2.7e-06 | 2.4e-06 | 1.9e-06 | 2.333E-06 | 4.041E-07 | 1.900E-06 | 2.700E-06 |
| 2898 | 145 | I145W | A | 1.7e-06 | 2.8e-06 | 2e-06 | 2.167E-06 | 5.686E-07 | 1.700E-06 | 2.800E-06 |
| 2899 | 145 | I145Y | A | 1.8e-06 | 2e-06 | 2.9e-06 | 2.233E-06 | 5.859E-07 | 1.800E-06 | 2.900E-06 |
| 2900 | 145 | I145V | A | 2.7e-06 | 1.8e-06 | 2.4e-06 | 2.300E-06 | 4.583E-07 | 1.800E-06 | 2.700E-06 |
| 2901 | 146 | L146A | A | 2.5e-06 | 4.1e-06 | 2.2e-06 | 2.933E-06 | 1.021E-06 | 2.200E-06 | 4.100E-06 |
| 2902 | 146 | L146R | A | 5.3e-06 | 2.2e-06 | 2.3e-06 | 3.267E-06 | 1.762E-06 | 2.200E-06 | 5.300E-06 |
| 2903 | 146 | L146N | A | 1.6e-06 | 2e-06 | 2.4e-06 | 2.000E-06 | 4.000E-07 | 1.600E-06 | 2.400E-06 |
| 2904 | 146 | L146D | A | 2.6e-06 | 2.6e-06 | 3.1e-06 | 2.767E-06 | 2.887E-07 | 2.600E-06 | 3.100E-06 |
| 2905 | 146 | L146C | A | 1.8e-06 | 2.9e-06 | 1.5e-06 | 2.067E-06 | 7.371E-07 | 1.500E-06 | 2.900E-06 |
| 2906 | 146 | L146E | A | 1.5e-06 | 3e-06 | 2.1e-06 | 2.200E-06 | 7.550E-07 | 1.500E-06 | 3.000E-06 |
| 2907 | 146 | L146Q | A | 8e-07 | 1.5e-06 | 1.5e-06 | 1.267E-06 | 4.041E-07 | 8.000E-07 | 1.500E-06 |
| 2908 | 146 | L146G | A | 2.4e-06 | 2.9e-06 | 3.1e-06 | 2.800E-06 | 3.606E-07 | 2.400E-06 | 3.100E-06 |
| 2909 | 146 | L146H | A | 2.2e-06 | 1.4e-06 | 2e-06 | 1.867E-06 | 4.163E-07 | 1.400E-06 | 2.200E-06 |
| 2910 | 146 | L146I | A | 1.7e-06 | 3e-06 | 3.5e-06 | 2.733E-06 | 9.292E-07 | 1.700E-06 | 3.500E-06 |
| 2911 | 146 | L146L | A | 3.1e-06 | 1.5e-06 | 2.8e-06 | 2.467E-06 | 8.505E-07 | 1.500E-06 | 3.100E-06 |
| 2912 | 146 | L146K | A | 3.6e-06 | 3e-06 | 1.7e-06 | 2.767E-06 | 9.713E-07 | 1.700E-06 | 3.600E-06 |
| 2913 | 146 | L146M | A | 3.1e-06 | 1.7e-06 | 2.4e-06 | 2.400E-06 | 7.000E-07 | 1.700E-06 | 3.100E-06 |
| 2914 | 146 | L146F | A | 2.2e-06 | 3.9e-06 | 1.8e-06 | 2.633E-06 | 1.115E-06 | 1.800E-06 | 3.900E-06 |
| 2915 | 146 | L146P | A | 3.1e-06 | 3.2e-06 | 2.7e-06 | 3.000E-06 | 2.646E-07 | 2.700E-06 | 3.200E-06 |
| 2916 | 146 | L146S | A | 3.7e-06 | 2.6e-06 | 1.3e-06 | 2.533E-06 | 1.201E-06 | 1.300E-06 | 3.700E-06 |
| 2917 | 146 | L146T | A | 3.9e-06 | 3e-06 | 1.8e-06 | 2.900E-06 | 1.054E-06 | 1.800E-06 | 3.900E-06 |
| 2918 | 146 | L146W | A | 2.2e-06 | 1.2e-06 | 3.6e-06 | 2.333E-06 | 1.206E-06 | 1.200E-06 | 3.600E-06 |
| 2919 | 146 | L146Y | A | 1.1e-06 | 2.7e-06 | 3.3e-06 | 2.367E-06 | 1.137E-06 | 1.100E-06 | 3.300E-06 |
| 2920 | 146 | L146V | A | 2.8e-06 | 2.5e-06 | 2.7e-06 | 2.667E-06 | 1.528E-07 | 2.500E-06 | 2.800E-06 |
| 2921 | 147 | W147A | A | 2.4e-06 | 2.3e-06 | 3.2e-06 | 2.633E-06 | 4.933E-07 | 2.300E-06 | 3.200E-06 |
| 2922 | 147 | W147R | A | 1.8e-06 | 2.2e-06 | 2.9e-06 | 2.300E-06 | 5.568E-07 | 1.800E-06 | 2.900E-06 |
| 2923 | 147 | W147N | A | 2.5e-06 | 2.1e-06 | 3.1e-06 | 2.567E-06 | 5.033E-07 | 2.100E-06 | 3.100E-06 |
| 2924 | 147 | W147D | A | 1.6e-06 | 1.8e-06 | 2.8e-06 | 2.067E-06 | 6.429E-07 | 1.600E-06 | 2.800E-06 |
| 2925 | 147 | W147C | A | 1.8e-06 | 2.3e-06 | 2.5e-06 | 2.200E-06 | 3.606E-07 | 1.800E-06 | 2.500E-06 |
| 2926 | 147 | W147E | A | 1.9e-06 | 2.4e-06 | 1.2e-06 | 1.833E-06 | 6.028E-07 | 1.200E-06 | 2.400E-06 |
| 2927 | 147 | W147Q | A | 2.2e-06 | 1.2e-06 | 3.6e-06 | 2.333E-06 | 1.206E-06 | 1.200E-06 | 3.600E-06 |
| 2928 | 147 | W147G | A | 2.6e-06 | 3.2e-06 | 2.1e-06 | 2.633E-06 | 5.508E-07 | 2.100E-06 | 3.200E-06 |
| 2929 | 147 | W147H | A | 3.1e-06 | 2e-06 | 2.7e-06 | 2.600E-06 | 5.568E-07 | 2.000E-06 | 3.100E-06 |
| 2930 | 147 | W147I | A | 2.4e-06 | 3.2e-06 | 2.9e-06 | 2.833E-06 | 4.041E-07 | 2.400E-06 | 3.200E-06 |
| 2931 | 147 | W147L | A | 2.4e-06 | 2.6e-06 | 3.3e-06 | 2.767E-06 | 4.726E-07 | 2.400E-06 | 3.300E-06 |
| 2932 | 147 | W147K | A | 4.1e-06 | 2.2e-06 | 2.1e-06 | 2.800E-06 | 1.127E-06 | 2.100E-06 | 4.100E-06 |
| 2933 | 147 | W147M | A | 4e-06 | 2.4e-06 | 2.3e-06 | 2.900E-06 | 9.539E-07 | 2.300E-06 | 4.000E-06 |
| 2934 | 147 | W147F | A | 1.6e-06 | 2.1e-06 | 2.1e-06 | 1.933E-06 | 2.887E-07 | 1.600E-06 | 2.100E-06 |
| 2935 | 147 | W147P | A | 1.3e-06 | 2.2e-06 | 3e-06 | 2.167E-06 | 8.505E-07 | 1.300E-06 | 3.000E-06 |
| 2936 | 147 | W147S | A | 2.7e-06 | 2.1e-06 | 3.7e-06 | 2.833E-06 | 8.083E-07 | 2.100E-06 | 3.700E-06 |
| 2937 | 147 | W147T | A | 1.5e-06 | 8e-07 | 1.9e-06 | 1.400E-06 | 5.568E-07 | 8.000E-07 | 1.900E-06 |
| 2938 | 147 | W147W | A | 3.4e-06 | 1.1e-06 | 4e-06 | 2.833E-06 | 1.531E-06 | 1.100E-06 | 4.000E-06 |
| 2939 | 147 | W147Y | A | 1.5e-06 | 2.4e-06 | 2.4e-06 | 2.100E-06 | 5.196E-07 | 1.500E-06 | 2.400E-06 |
| 2940 | 147 | W147V | A | 2.1e-06 | 2.3e-06 | 1.6e-06 | 2.000E-06 | 3.606E-07 | 1.600E-06 | 2.300E-06 |
| 2941 | 148 | K148A | A | 4.4e-06 | 2.7e-06 | 2e-06 | 3.033E-06 | 1.234E-06 | 2.000E-06 | 4.400E-06 |
| 2942 | 148 | K148R | A | 2e-06 | 2.6e-06 | 2e-06 | 2.200E-06 | 3.464E-07 | 2.000E-06 | 2.600E-06 |
| 2943 | 148 | K148N | A | 1.3e-06 | 3.3e-06 | 2.5e-06 | 2.367E-06 | 1.007E-06 | 1.300E-06 | 3.300E-06 |
| 2944 | 148 | K148D | A | 2.9e-06 | 1.5e-06 | 2.6e-06 | 2.333E-06 | 7.371E-07 | 1.500E-06 | 2.900E-06 |
| 2945 | 148 | K148C | A | 3e-06 | 2.2e-06 | 2.6e-06 | 2.600E-06 | 4.000E-07 | 2.200E-06 | 3.000E-06 |
| 2946 | 148 | K148E | A | 2.2e-06 | 2.5e-06 | 4.8e-06 | 3.167E-06 | 1.422E-06 | 2.200E-06 | 4.800E-06 |
| 2947 | 148 | K148Q | A | 1.9e-06 | 2.3e-06 | 2.2e-06 | 2.133E-06 | 2.082E-07 | 1.900E-06 | 2.300E-06 |
| 2948 | 148 | K148G | A | 2.1e-06 | 2.2e-06 | 1.7e-06 | 2.000E-06 | 2.646E-07 | 1.700E-06 | 2.200E-06 |
| 2949 | 148 | K148H | A | 2.4e-06 | 2.2e-06 | 1.7e-06 | 2.100E-06 | 3.606E-07 | 1.700E-06 | 2.400E-06 |
| 2950 | 148 | K148I | A | 3e-06 | 1.5e-06 | 4.9e-06 | 3.133E-06 | 1.704E-06 | 1.500E-06 | 4.900E-06 |
| 2951 | 148 | K148L | A | 2e-06 | 1.8e-06 | 1.8e-06 | 1.867E-06 | 1.155E-07 | 1.800E-06 | 2.000E-06 |
| 2952 | 148 | K148K | A | 2.6e-06 | 1.6e-06 | 2.9e-06 | 2.367E-06 | 6.807E-07 | 1.600E-06 | 2.900E-06 |
| 2953 | 148 | K148M | A | 2.3e-06 | 1.4e-06 | 3e-06 | 2.233E-06 | 8.021E-07 | 1.400E-06 | 3.000E-06 |
| 2954 | 148 | K148F | A | 3.1e-06 | 2.8e-06 | 2.3e-06 | 2.733E-06 | 4.041E-07 | 2.300E-06 | 3.100E-06 |
| 2955 | 148 | K148P | A | 2.6e-06 | 2.3e-06 | 4.5e-06 | 3.133E-06 | 1.193E-06 | 2.300E-06 | 4.500E-06 |
| 2956 | 148 | K148S | A | 2.7e-06 | 2.9e-06 | 3e-06 | 2.867E-06 | 1.528E-07 | 2.700E-06 | 3.000E-06 |
| 2957 | 148 | K148T | A | 2.2e-06 | 2e-06 | 3.2e-06 | 2.467E-06 | 6.429E-07 | 2.000E-06 | 3.200E-06 |
| 2958 | 148 | K148W | A | 1.5e-06 | 2.8e-06 | 1.6e-06 | 1.967E-06 | 7.234E-07 | 1.500E-06 | 2.800E-06 |
| 2959 | 148 | K148Y | A | 2.3e-06 | 2.7e-06 | 2.2e-06 | 2.400E-06 | 2.646E-07 | 2.200E-06 | 2.700E-06 |
| 2960 | 148 | K148V | A | 1.9e-06 | 2.2e-06 | 2.8e-06 | 2.300E-06 | 4.583E-07 | 1.900E-06 | 2.800E-06 |
| 2961 | 149 | D149A | A | 1.6e-06 | 1.3e-06 | 3.2e-06 | 2.033E-06 | 1.021E-06 | 1.300E-06 | 3.200E-06 |
| 2962 | 149 | D149R | A | 2.9e-06 | 1.5e-06 | 2e-06 | 2.133E-06 | 7.095E-07 | 1.500E-06 | 2.900E-06 |
| 2963 | 149 | D149N | A | 2.4e-06 | 1.5e-06 | 1.3e-06 | 1.733E-06 | 5.859E-07 | 1.300E-06 | 2.400E-06 |
| 2964 | 149 | D149D | A | 3.4e-06 | 3.6e-06 | 3.1e-06 | 3.367E-06 | 2.517E-07 | 3.100E-06 | 3.600E-06 |
| 2965 | 149 | D149C | A | 2.2e-06 | 4.4e-06 | 1.6e-06 | 2.733E-06 | 1.474E-06 | 1.600E-06 | 4.400E-06 |
| 2966 | 149 | D149E | A | 2.1e-06 | 3.1e-06 | 2e-06 | 2.400E-06 | 6.083E-07 | 2.000E-06 | 3.100E-06 |
| 2967 | 149 | D149Q | A | 1.4e-06 | 3.5e-06 | 1.9e-06 | 2.267E-06 | 1.097E-06 | 1.400E-06 | 3.500E-06 |
| 2968 | 149 | D149G | A | 2.4e-06 | 2.8e-06 | 4.2e-06 | 3.133E-06 | 9.452E-07 | 2.400E-06 | 4.200E-06 |
| 2969 | 149 | D149H | A | 2.3e-06 | 2.5e-06 | 4.1e-06 | 2.967E-06 | 9.866E-07 | 2.300E-06 | 4.100E-06 |
| 2970 | 149 | D149I | A | 4.1e-06 | 2.8e-06 | 2.5e-06 | 3.133E-06 | 8.505E-07 | 2.500E-06 | 4.100E-06 |
| 2971 | 149 | D149L | A | 1.6e-06 | 3.1e-06 | 2.6e-06 | 2.433E-06 | 7.638E-07 | 1.600E-06 | 3.100E-06 |
| 2972 | 149 | D149K | A | 1.8e-06 | 4.2e-06 | 2.7e-06 | 2.900E-06 | 1.212E-06 | 1.800E-06 | 4.200E-06 |
| 2973 | 149 | D149M | A | 7.2e-07 | 2.4e-06 | 1.7e-06 | 1.607E-06 | 8.439E-07 | 7.200E-07 | 2.400E-06 |
| 2974 | 149 | D149F | A | 1.3e-06 | 1.8e-06 | 2.6e-06 | 1.900E-06 | 6.557E-07 | 1.300E-06 | 2.600E-06 |

| | | | | | | | | | | |
|---|---|---|---|---|---|---|---|---|---|---|
| 2975 | 149 | D149P | A | 2.9e-06 | 2.3e-06 | 2.1e-06 | 2.433E-06 | 4.163E-07 | 2.100E-06 | 2.900E-06 |
| 2976 | 149 | D149S | A | 2.5e-06 | 2.1e-06 | 3.4e-06 | 2.667E-06 | 6.658E-07 | 2.100E-06 | 3.400E-06 |
| 2977 | 149 | D149T | A | 5.1e-06 | 2.8e-06 | 2.9e-06 | 3.600E-06 | 1.300E-06 | 2.800E-06 | 5.100E-06 |
| 2978 | 149 | D149W | A | 2e-06 | 2.8e-06 | 1.9e-06 | 2.233E-06 | 4.933E-07 | 1.900E-06 | 2.800E-06 |
| 2979 | 149 | D149Y | A | 2.3e-06 | 1.5e-06 | 2.7e-06 | 2.167E-06 | 6.110E-07 | 1.500E-06 | 2.700E-06 |
| 2980 | 149 | D149V | A | 1.2e-06 | 2.4e-06 | 1.4e-06 | 1.667E-06 | 6.429E-07 | 1.200E-06 | 2.400E-06 |
| 2981 | 150 | I150A | A | 5.2e-06 | 1.9e-06 | 1.8e-06 | 2.967E-06 | 1.935E-06 | 1.800E-06 | 5.200E-06 |
| 2982 | 150 | I150R | A | 3.4e-06 | 3.8e-06 | 2.9e-06 | 3.367E-06 | 4.509E-07 | 2.900E-06 | 3.800E-06 |
| 2983 | 150 | I150N | A | 2.6e-06 | 8.4e-07 | 2.1e-06 | 1.847E-06 | 9.069E-07 | 8.400E-07 | 2.600E-06 |
| 2984 | 150 | I150D | A | 1.7e-06 | 2.2e-06 | 1.9e-06 | 1.933E-06 | 2.517E-07 | 1.700E-06 | 2.200E-06 |
| 2985 | 150 | I150C | A | 2.5e-06 | 1.9e-06 | 1.9e-06 | 2.100E-06 | 3.464E-07 | 1.900E-06 | 2.500E-06 |
| 2986 | 150 | I150E | A | 2.4e-06 | 1.8e-06 | 1.3e-06 | 1.833E-06 | 5.508E-07 | 1.300E-06 | 2.400E-06 |
| 2987 | 150 | I150Q | A | 2.3e-06 | 3.3e-06 | 2.6e-06 | 2.733E-06 | 5.132E-07 | 2.300E-06 | 3.300E-06 |
| 2988 | 150 | I150G | A | 3.1e-06 | 2e-06 | 1.4e-06 | 2.167E-06 | 8.622E-07 | 1.400E-06 | 3.100E-06 |
| 2989 | 150 | I150H | A | 2e-06 | 2.4e-06 | 2.5e-06 | 2.300E-06 | 2.646E-07 | 2.000E-06 | 2.500E-06 |
| 2990 | 150 | I150I | A | 2.5e-06 | 2.2e-06 | 2.7e-06 | 2.467E-06 | 2.517E-07 | 2.200E-06 | 2.700E-06 |
| 2991 | 150 | I150L | A | 3.2e-06 | 3.4e-06 | 3.9e-06 | 3.500E-06 | 3.606E-07 | 3.200E-06 | 3.900E-06 |
| 2992 | 150 | I150K | A | 1.9e-06 | 1.3e-06 | 2.2e-06 | 1.800E-06 | 4.583E-07 | 1.300E-06 | 2.200E-06 |
| 2993 | 150 | I150M | A | 1.1e-06 | 3.1e-06 | 4.4e-06 | 2.867E-06 | 1.662E-06 | 1.100E-06 | 4.400E-06 |
| 2994 | 150 | I150F | A | 1.3e-06 | 2.5e-06 | 1.6e-06 | 1.800E-06 | 6.245E-07 | 1.300E-06 | 2.500E-06 |
| 2995 | 150 | I150P | A | 2.2e-06 | 2.3e-06 | 2.2e-06 | 2.233E-06 | 5.774E-08 | 2.200E-06 | 2.300E-06 |
| 2996 | 150 | I150S | A | 3.1e-06 | 2.5e-06 | 2.6e-06 | 2.733E-06 | 3.215E-07 | 2.500E-06 | 3.100E-06 |
| 2997 | 150 | I150T | A | 2.3e-06 | 1.9e-06 | 1.7e-06 | 1.967E-06 | 3.055E-07 | 1.700E-06 | 2.300E-06 |
| 2998 | 150 | I150W | A | 2.7e-06 | 1.8e-06 | 2.8e-06 | 2.433E-06 | 5.508E-07 | 1.800E-06 | 2.800E-06 |
| 2999 | 150 | I150Y | A | 3.6e-06 | 3.1e-06 | 2.5e-06 | 3.067E-06 | 5.508E-07 | 2.500E-06 | 3.600E-06 |
| 3000 | 150 | I150V | A | 2.6e-06 | 2.6e-06 | 2.1e-06 | 2.433E-06 | 2.887E-07 | 2.100E-06 | 2.600E-06 |
| 3001 | 151 | F151A | A | 1.9e-06 | 2.5e-06 | 1.1e-06 | 1.833E-06 | 7.024E-07 | 1.100E-06 | 2.500E-06 |
| 3002 | 151 | F151R | A | 2.3e-06 | 3e-06 | 9.3e-07 | 2.077E-06 | 1.053E-06 | 9.300E-07 | 3.000E-06 |
| 3003 | 151 | F151N | A | 5.3e-06 | 1.9e-06 | 3.8e-06 | 3.667E-06 | 1.704E-06 | 1.900E-06 | 5.300E-06 |
| 3004 | 151 | F151D | A | 2e-06 | 2.4e-06 | 1.9e-06 | 2.100E-06 | 2.646E-07 | 1.900E-06 | 2.400E-06 |
| 3005 | 151 | F151C | A | 2e-06 | 1.8e-06 | 1.7e-06 | 1.833E-06 | 1.528E-07 | 1.700E-06 | 2.000E-06 |
| 3006 | 151 | F151E | A | 3.1e-06 | 1.7e-06 | 1.7e-06 | 2.167E-06 | 8.083E-07 | 1.700E-06 | 3.100E-06 |
| 3007 | 151 | F151Q | A | 3.6e-06 | 3.7e-06 | 3e-06 | 3.433E-06 | 3.786E-07 | 3.000E-06 | 3.700E-06 |
| 3008 | 151 | F151G | A | 1.8e-06 | 4.4e-06 | 2.3e-06 | 2.833E-06 | 1.380E-06 | 1.800E-06 | 4.400E-06 |
| 3009 | 151 | F151H | A | 2.7e-06 | 1.9e-06 | 2.4e-06 | 2.333E-06 | 4.041E-07 | 1.900E-06 | 2.700E-06 |
| 3010 | 151 | F151I | A | 4.1e-06 | 3.2e-06 | 2e-06 | 3.100E-06 | 1.054E-06 | 2.000E-06 | 4.100E-06 |
| 3011 | 151 | F151L | A | 3.4e-06 | 1.7e-06 | 1.6e-06 | 2.233E-06 | 1.012E-06 | 1.600E-06 | 3.400E-06 |
| 3012 | 151 | F151K | A | 2.9e-06 | 2.9e-06 | 2.3e-06 | 2.700E-06 | 3.464E-07 | 2.300E-06 | 2.900E-06 |
| 3013 | 151 | F151M | A | 2.9e-06 | 2.1e-06 | 2.7e-06 | 2.567E-06 | 4.163E-07 | 2.100E-06 | 2.900E-06 |
| 3014 | 151 | F151F | A | 2.5e-06 | 3e-06 | 1.7e-06 | 2.400E-06 | 6.557E-07 | 1.700E-06 | 3.000E-06 |
| 3015 | 151 | F151P | A | 2.9e-06 | 3e-06 | 1.2e-06 | 2.367E-06 | 1.012E-06 | 1.200E-06 | 3.000E-06 |
| 3016 | 151 | F151S | A | 1.9e-06 | 3.8e-06 | 1.9e-06 | 2.533E-06 | 1.097E-06 | 1.900E-06 | 3.800E-06 |
| 3017 | 151 | F151T | A | 3.8e-06 | 1.3e-06 | 1.6e-06 | 2.233E-06 | 1.365E-06 | 1.300E-06 | 3.800E-06 |
| 3018 | 151 | F151W | A | 2e-06 | 2.2e-06 | 4.3e-06 | 2.833E-06 | 1.274E-06 | 2.000E-06 | 4.300E-06 |
| 3019 | 151 | F151Y | A | 2.7e-06 | 1.3e-06 | 2.1e-06 | 2.033E-06 | 7.024E-07 | 1.300E-06 | 2.700E-06 |
| 3020 | 151 | F151V | A | 5.8e-06 | 3.8e-06 | 3.8e-06 | 4.467E-06 | 1.155E-06 | 3.800E-06 | 5.800E-06 |
| 3021 | 152 | H152A | A | 2.1e-06 | 1.6e-06 | 3.3e-06 | 2.333E-06 | 8.737E-07 | 1.600E-06 | 3.300E-06 |
| 3022 | 152 | H152R | A | 2.2e-06 | 3.5e-06 | 3.6e-06 | 3.100E-06 | 7.810E-07 | 2.200E-06 | 3.600E-06 |
| 3023 | 152 | H152N | A | 2.7e-06 | 1.7e-06 | 1.9e-06 | 2.100E-06 | 5.292E-07 | 1.700E-06 | 2.700E-06 |
| 3024 | 152 | H152D | A | 1.1e-06 | 1.6e-06 | 1.6e-06 | 1.433E-06 | 2.887E-07 | 1.100E-06 | 1.600E-06 |
| 3025 | 152 | H152C | A | 2.2e-06 | 1.6e-06 | 2.1e-06 | 1.967E-06 | 3.215E-07 | 1.600E-06 | 2.200E-06 |
| 3026 | 152 | H152E | A | 2.1e-06 | 2.3e-06 | 4e-06 | 2.800E-06 | 1.044E-06 | 2.100E-06 | 4.000E-06 |
| 3027 | 152 | H152Q | A | 2.1e-06 | 3.5e-06 | 6.9e-07 | 2.097E-06 | 1.405E-06 | 6.900E-07 | 3.500E-06 |
| 3028 | 152 | H152G | A | 1.6e-06 | 1.9e-06 | 1.9e-06 | 1.800E-06 | 1.732E-07 | 1.600E-06 | 1.900E-06 |
| 3029 | 152 | H152H | A | 2.1e-06 | 3.1e-06 | 1.3e-06 | 2.167E-06 | 9.018E-07 | 1.300E-06 | 3.100E-06 |
| 3030 | 152 | H152I | A | 1.3e-06 | 3.2e-06 | 2e-06 | 2.167E-06 | 9.609E-07 | 1.300E-06 | 3.200E-06 |
| 3031 | 152 | H152L | A | 1.7e-06 | 1.7e-06 | 1.9e-06 | 1.767E-06 | 1.155E-07 | 1.700E-06 | 1.900E-06 |
| 3032 | 152 | H152K | A | 1.5e-06 | 2e-06 | 2.1e-06 | 1.867E-06 | 3.215E-07 | 1.500E-06 | 2.100E-06 |
| 3033 | 152 | H152M | A | 1.7e-06 | 4.4e-06 | 3.2e-06 | 3.100E-06 | 1.353E-06 | 1.700E-06 | 4.400E-06 |
| 3034 | 152 | H152F | A | 1.9e-06 | 4e-06 | 2.1e-06 | 2.667E-06 | 1.159E-06 | 1.900E-06 | 4.000E-06 |
| 3035 | 152 | H152P | A | 4.3e-06 | 1.5e-06 | 1.6e-06 | 2.467E-06 | 1.589E-06 | 1.500E-06 | 4.300E-06 |
| 3036 | 152 | H152S | A | 2.1e-06 | 3.4e-06 | 2.4e-06 | 2.633E-06 | 6.807E-07 | 2.100E-06 | 3.400E-06 |
| 3037 | 152 | H152T | A | 2.4e-06 | 3.3e-06 | 1.3e-06 | 2.333E-06 | 1.002E-06 | 1.300E-06 | 3.300E-06 |
| 3038 | 152 | H152W | A | 3.1e-06 | 2.6e-06 | 1.8e-06 | 2.500E-06 | 6.557E-07 | 1.800E-06 | 3.100E-06 |
| 3039 | 152 | H152Y | A | 2.3e-06 | 1.9e-06 | 2e-06 | 2.067E-06 | 2.082E-07 | 1.900E-06 | 2.300E-06 |
| 3040 | 152 | H152V | A | 3e-06 | 3e-06 | 1.4e-06 | 2.467E-06 | 9.238E-07 | 1.400E-06 | 3.000E-06 |
| 3041 | 153 | K153A | A | 3.3e-06 | 2.6e-06 | 1.9e-06 | 2.600E-06 | 7.000E-07 | 1.900E-06 | 3.300E-06 |
| 3042 | 153 | K153R | A | 2.4e-06 | 2.4e-06 | 2.2e-06 | 2.333E-06 | 1.155E-07 | 2.200E-06 | 2.400E-06 |
| 3043 | 153 | K153N | A | 1.9e-06 | 4e-06 | 2.4e-06 | 2.767E-06 | 1.097E-06 | 1.900E-06 | 4.000E-06 |
| 3044 | 153 | K153D | A | 3.1e-06 | 1.1e-06 | 1.4e-06 | 1.867E-06 | 1.079E-06 | 1.100E-06 | 3.100E-06 |
| 3045 | 153 | K153C | A | 2.4e-06 | 2.3e-06 | 2.3e-06 | 2.333E-06 | 5.774E-08 | 2.300E-06 | 2.400E-06 |
| 3046 | 153 | K153E | A | 3.5e-06 | 2.5e-06 | 3.1e-06 | 3.033E-06 | 5.033E-07 | 2.500E-06 | 3.500E-06 |
| 3047 | 153 | K153Q | A | 2.4e-06 | 2.7e-06 | 2.2e-06 | 2.433E-06 | 2.517E-07 | 2.200E-06 | 2.700E-06 |
| 3048 | 153 | K153G | A | 2e-06 | 2.1e-06 | 3.4e-06 | 2.500E-06 | 7.810E-07 | 2.000E-06 | 3.400E-06 |
| 3049 | 153 | K153H | A | 1.9e-06 | 1.6e-06 | 1.9e-06 | 1.800E-06 | 1.732E-07 | 1.600E-06 | 1.900E-06 |
| 3050 | 153 | K153I | A | 1.5e-06 | 3.4e-06 | 2.8e-06 | 2.567E-06 | 9.713E-07 | 1.500E-06 | 3.400E-06 |
| 3051 | 153 | K153L | A | 1.7e-06 | 2.5e-06 | 2.6e-06 | 2.267E-06 | 4.933E-07 | 1.700E-06 | 2.600E-06 |
| 3052 | 153 | K153K | A | 1.8e-06 | 2.4e-06 | 2.3e-06 | 2.167E-06 | 3.215E-07 | 1.800E-06 | 2.400E-06 |
| 3053 | 153 | K153M | A | 2.8e-06 | 2.4e-06 | 2.7e-06 | 2.633E-06 | 2.082E-07 | 2.400E-06 | 2.800E-06 |
| 3054 | 153 | K153F | A | 1e-06 | 2.4e-06 | 2.6e-06 | 2.000E-06 | 8.718E-07 | 1.000E-06 | 2.600E-06 |
| 3055 | 153 | K153P | A | 1.5e-06 | 1.7e-06 | 4.6e-06 | 2.600E-06 | 1.735E-06 | 1.500E-06 | 4.600E-06 |
| 3056 | 153 | K153S | A | 2.2e-06 | 3.6e-06 | 2.7e-06 | 2.833E-06 | 7.095E-07 | 2.200E-06 | 3.600E-06 |
| 3057 | 153 | K153T | A | 1.7e-06 | 2.3e-06 | 3.8e-06 | 2.600E-06 | 1.082E-06 | 1.700E-06 | 3.800E-06 |
| 3058 | 153 | K153W | A | 3.3e-06 | 1.3e-06 | 1.1e-06 | 1.900E-06 | 1.217E-06 | 1.100E-06 | 3.300E-06 |
| 3059 | 153 | K153Y | A | 2.7e-06 | 2e-06 | 3.2e-06 | 2.633E-06 | 6.028E-07 | 2.000E-06 | 3.200E-06 |

| | | | | | | | | | | |
|---|---|---|---|---|---|---|---|---|---|---|
| 3060 | 153 | K153V | A | 1.5e-06 | 2.2e-06 | 2.7e-06 | 2.133E-06 | 6.028E-07 | 1.500E-06 | 2.700E-06 |
| 3061 | 154 | N154A | A | 1.7e-06 | 2e-06 | 2.9e-06 | 2.200E-06 | 6.245E-07 | 1.700E-06 | 2.900E-06 |
| 3062 | 154 | N154R | A | 2.9e-06 | 2.2e-06 | 2.5e-06 | 2.533E-06 | 3.512E-07 | 2.200E-06 | 2.900E-06 |
| 3063 | 154 | N154N | A | 4.2e-06 | 2.8e-06 | 4e-06 | 3.667E-06 | 7.572E-07 | 2.800E-06 | 4.200E-06 |
| 3064 | 154 | N154D | A | 3.1e-06 | 2.2e-06 | 3.5e-06 | 2.933E-06 | 6.658E-07 | 2.200E-06 | 3.500E-06 |
| 3065 | 154 | N154C | A | 2.6e-06 | 2.2e-06 | 2.7e-06 | 2.500E-06 | 2.646E-07 | 2.200E-06 | 2.700E-06 |
| 3066 | 154 | N154E | A | 1.7e-06 | 2.2e-06 | 2.6e-06 | 2.167E-06 | 4.509E-07 | 1.700E-06 | 2.600E-06 |
| 3067 | 154 | N154Q | A | 1.7e-06 | 2.6e-06 | 1.5e-06 | 1.933E-06 | 5.859E-07 | 1.500E-06 | 2.600E-06 |
| 3068 | 154 | N154G | A | 1.3e-06 | 3.7e-06 | 3.7e-06 | 2.900E-06 | 1.386E-06 | 1.300E-06 | 3.700E-06 |
| 3069 | 154 | N154H | A | 4.3e-06 | 2.8e-06 | 1.8e-06 | 2.967E-06 | 1.258E-06 | 1.800E-06 | 4.300E-06 |
| 3070 | 154 | N154I | A | 1.6e-06 | 1.6e-06 | 1.3e-06 | 1.500E-06 | 1.732E-07 | 1.300E-06 | 1.600E-06 |
| 3071 | 154 | N154L | A | 1.9e-06 | 1.9e-06 | 2.1e-06 | 1.967E-06 | 1.155E-07 | 1.900E-06 | 2.100E-06 |
| 3072 | 154 | N154K | A | 1.5e-06 | 1.5e-06 | 3.7e-06 | 2.233E-06 | 1.270E-06 | 1.500E-06 | 3.700E-06 |
| 3073 | 154 | N154M | A | 3.3e-06 | 1.9e-06 | 1.3e-06 | 2.167E-06 | 1.026E-06 | 1.300E-06 | 3.300E-06 |
| 3074 | 154 | N154F | A | 1.8e-06 | 1.9e-06 | 1.7e-06 | 1.800E-06 | 1.000E-07 | 1.700E-06 | 1.900E-06 |
| 3075 | 154 | N154P | A | 2.3e-06 | 2.8e-06 | 2.7e-06 | 2.600E-06 | 2.646E-07 | 2.300E-06 | 2.800E-06 |
| 3076 | 154 | N154S | A | 2.6e-06 | 1.5e-06 | 3e-06 | 2.367E-06 | 7.767E-07 | 1.500E-06 | 3.000E-06 |
| 3077 | 154 | N154T | A | 2.4e-06 | 1.5e-06 | 1.8e-06 | 1.900E-06 | 4.583E-07 | 1.500E-06 | 2.400E-06 |
| 3078 | 154 | N154W | A | 4.6e-06 | 1.8e-06 | 3.3e-06 | 3.233E-06 | 1.401E-06 | 1.800E-06 | 4.600E-06 |
| 3079 | 154 | N154Y | A | 2.8e-06 | 2e-06 | 2.5e-06 | 2.433E-06 | 4.041E-07 | 2.000E-06 | 2.800E-06 |
| 3080 | 154 | N154V | A | 2.3e-06 | 2.8e-06 | 2.9e-06 | 2.667E-06 | 3.215E-07 | 2.300E-06 | 2.900E-06 |
| 3081 | 155 | N155A | A | 3.4e-06 | 2.7e-06 | 1.4e-06 | 2.500E-06 | 1.015E-06 | 1.400E-06 | 3.400E-06 |
| 3082 | 155 | N155R | A | 1.2e-06 | 1.5e-06 | 2.2e-06 | 1.633E-06 | 5.132E-07 | 1.200E-06 | 2.200E-06 |
| 3083 | 155 | N155N | A | 1.9e-06 | 1.6e-06 | 2.2e-06 | 1.900E-06 | 3.000E-07 | 1.600E-06 | 2.200E-06 |
| 3084 | 155 | N155D | A | 1.5e-06 | 2.7e-06 | 1.9e-06 | 2.033E-06 | 6.110E-07 | 1.500E-06 | 2.700E-06 |
| 3085 | 155 | N155C | A | 2.5e-06 | 1.8e-06 | 1.3e-06 | 1.867E-06 | 6.028E-07 | 1.300E-06 | 2.500E-06 |
| 3086 | 155 | N155E | A | 4.6e-06 | 2.2e-06 | 2.6e-06 | 3.133E-06 | 1.286E-06 | 2.200E-06 | 4.600E-06 |
| 3087 | 155 | N155Q | A | 2.5e-06 | 2.6e-06 | 1.3e-06 | 2.133E-06 | 7.234E-07 | 1.300E-06 | 2.600E-06 |
| 3088 | 155 | N155G | A | 2.2e-06 | 2.2e-06 | 1.4e-06 | 1.933E-06 | 4.619E-07 | 1.400E-06 | 2.200E-06 |
| 3089 | 155 | N155H | A | 1.8e-06 | 3e-06 | 2.7e-06 | 2.500E-06 | 6.245E-07 | 1.800E-06 | 3.000E-06 |
| 3090 | 155 | N155I | A | 2.2e-06 | 3.9e-06 | 2e-06 | 2.700E-06 | 1.044E-06 | 2.000E-06 | 3.900E-06 |
| 3091 | 155 | N155L | A | 2.5e-06 | 2.2e-06 | 3.6e-06 | 2.767E-06 | 7.371E-07 | 2.200E-06 | 3.600E-06 |
| 3092 | 155 | N155K | A | 1.9e-06 | 1.9e-06 | 3e-06 | 2.267E-06 | 6.351E-07 | 1.900E-06 | 3.000E-06 |
| 3093 | 155 | N155M | A | 2.7e-06 | 1.3e-06 | 2e-06 | 2.000E-06 | 7.000E-07 | 1.300E-06 | 2.700E-06 |
| 3094 | 155 | N155F | A | 2.5e-06 | 4e-06 | 3.3e-06 | 3.267E-06 | 7.506E-07 | 2.500E-06 | 4.000E-06 |
| 3095 | 155 | N155P | A | 2.3e-06 | 2.4e-06 | 2.6e-06 | 2.433E-06 | 1.528E-07 | 2.300E-06 | 2.600E-06 |
| 3096 | 155 | N155S | A | 2.6e-06 | 2.4e-06 | 2.6e-06 | 2.533E-06 | 1.155E-07 | 2.400E-06 | 2.600E-06 |
| 3097 | 155 | N155T | A | 2.3e-06 | 2.2e-06 | 2.3e-06 | 2.267E-06 | 5.774E-08 | 2.200E-06 | 2.300E-06 |
| 3098 | 155 | N155W | A | 1.4e-06 | 1.9e-06 | 2.2e-06 | 1.833E-06 | 4.041E-07 | 1.400E-06 | 2.200E-06 |
| 3099 | 155 | N155Y | A | 2e-06 | 1.9e-06 | 1.7e-06 | 1.867E-06 | 1.528E-07 | 1.700E-06 | 2.000E-06 |
| 3100 | 155 | N155V | A | 1.4e-06 | 1.1e-06 | 3.9e-06 | 2.133E-06 | 1.537E-06 | 1.100E-06 | 3.900E-06 |
| 3101 | 156 | Q156A | A | 3.8e-06 | 2e-06 | 1.4e-06 | 2.400E-06 | 1.249E-06 | 1.400E-06 | 3.800E-06 |
| 3102 | 156 | Q156R | A | 2e-06 | 2.3e-06 | 2.6e-06 | 2.300E-06 | 3.000E-07 | 2.000E-06 | 2.600E-06 |
| 3103 | 156 | Q156N | A | 1.9e-06 | 2.3e-06 | 2.1e-06 | 2.100E-06 | 2.000E-07 | 1.900E-06 | 2.300E-06 |
| 3104 | 156 | Q156D | A | 2.7e-06 | 5.6e-06 | 1.9e-06 | 3.400E-06 | 1.947E-06 | 1.900E-06 | 5.600E-06 |
| 3105 | 156 | Q156C | A | 1.9e-06 | 2e-06 | 1.1e-06 | 1.667E-06 | 4.933E-07 | 1.100E-06 | 2.000E-06 |
| 3106 | 156 | Q156E | A | 1.9e-06 | 2.5e-06 | 2.3e-06 | 2.233E-06 | 3.055E-07 | 1.900E-06 | 2.500E-06 |
| 3107 | 156 | Q156Q | A | 2.1e-06 | 2.5e-06 | 1.6e-06 | 2.067E-06 | 4.509E-07 | 1.600E-06 | 2.500E-06 |
| 3108 | 156 | Q156G | A | 2.7e-06 | 3.3e-06 | 2.4e-06 | 2.800E-06 | 4.583E-07 | 2.400E-06 | 3.300E-06 |
| 3109 | 156 | Q156H | A | 3.2e-06 | 4.9e-06 | 2.6e-06 | 3.567E-06 | 1.193E-06 | 2.600E-06 | 4.900E-06 |
| 3110 | 156 | Q156I | A | 2.1e-06 | 2.2e-06 | 2.9e-06 | 2.400E-06 | 4.359E-07 | 2.100E-06 | 2.900E-06 |
| 3111 | 156 | Q156L | A | 1.9e-06 | 1.3e-06 | 1e-06 | 1.400E-06 | 4.583E-07 | 1.000E-06 | 1.900E-06 |
| 3112 | 156 | Q156K | A | 2.8e-06 | 3e-06 | 1.8e-06 | 2.533E-06 | 6.429E-07 | 1.800E-06 | 3.000E-06 |
| 3113 | 156 | Q156M | A | 2.4e-06 | 2.4e-06 | 2.3e-06 | 2.367E-06 | 5.774E-08 | 2.300E-06 | 2.400E-06 |
| 3114 | 156 | Q156F | A | 2.2e-06 | 1.9e-06 | 2e-06 | 2.033E-06 | 1.528E-07 | 1.900E-06 | 2.200E-06 |
| 3115 | 156 | Q156P | A | 3.1e-06 | 1.3e-06 | 1.6e-06 | 2.000E-06 | 9.644E-07 | 1.300E-06 | 3.100E-06 |
| 3116 | 156 | Q156S | A | 2.4e-06 | 2e-06 | 1.8e-06 | 2.067E-06 | 3.055E-07 | 1.800E-06 | 2.400E-06 |
| 3117 | 156 | Q156T | A | 3.6e-06 | 4e-06 | 2.4e-06 | 3.333E-06 | 8.327E-07 | 2.400E-06 | 4.000E-06 |
| 3118 | 156 | Q156W | A | 3.7e-06 | 1.9e-06 | 3.4e-06 | 3.000E-06 | 9.644E-07 | 1.900E-06 | 3.700E-06 |
| 3119 | 156 | Q156Y | A | 2e-06 | 2.5e-06 | 1.4e-06 | 1.967E-06 | 5.508E-07 | 1.400E-06 | 2.500E-06 |
| 3120 | 156 | Q156V | A | 8.4e-07 | 3e-06 | 2.6e-06 | 2.147E-06 | 1.149E-06 | 8.400E-07 | 3.000E-06 |
| 3121 | 157 | L157A | A | 3.1e-06 | 2.1e-06 | 2.2e-06 | 2.467E-06 | 5.508E-07 | 2.100E-06 | 3.100E-06 |
| 3122 | 157 | L157R | A | 2.5e-06 | 1.7e-06 | 2.1e-06 | 2.100E-06 | 4.000E-07 | 1.700E-06 | 2.500E-06 |
| 3123 | 157 | L157N | A | 2.1e-06 | 2.3e-06 | 1.7e-06 | 2.033E-06 | 3.055E-07 | 1.700E-06 | 2.300E-06 |
| 3124 | 157 | L157D | A | 2.4e-06 | 2.4e-06 | 3.2e-06 | 2.667E-06 | 4.619E-07 | 2.400E-06 | 3.200E-06 |
| 3125 | 157 | L157C | A | 1.4e-06 | 8.2e-07 | 1.2e-06 | 1.140E-06 | 2.946E-07 | 8.200E-07 | 1.400E-06 |
| 3126 | 157 | L157E | A | 2.9e-06 | 5.1e-06 | 2.6e-06 | 3.533E-06 | 1.365E-06 | 2.600E-06 | 5.100E-06 |
| 3127 | 157 | L157Q | A | 1.8e-06 | 2.3e-06 | 2.6e-06 | 2.233E-06 | 4.041E-07 | 1.800E-06 | 2.600E-06 |
| 3128 | 157 | L157G | A | 1.7e-06 | 3.5e-06 | 2.4e-06 | 2.533E-06 | 9.074E-07 | 1.700E-06 | 3.500E-06 |
| 3129 | 157 | L157H | A | 2.2e-06 | 2.6e-06 | 4.2e-06 | 3.000E-06 | 1.058E-06 | 2.200E-06 | 4.200E-06 |
| 3130 | 157 | L157I | A | 3.9e-06 | 3.2e-06 | 2.9e-06 | 3.333E-06 | 5.132E-07 | 2.900E-06 | 3.900E-06 |
| 3131 | 157 | L157L | A | 4.2e-06 | 1.6e-06 | 2.3e-06 | 2.700E-06 | 1.345E-06 | 1.600E-06 | 4.200E-06 |
| 3132 | 157 | L157K | A | 6.2e-06 | 1.5e-06 | 2.2e-06 | 3.300E-06 | 2.536E-06 | 1.500E-06 | 6.200E-06 |
| 3133 | 157 | L157M | A | 2.6e-06 | 3.1e-06 | 2.6e-06 | 2.767E-06 | 2.887E-07 | 2.600E-06 | 3.100E-06 |
| 3134 | 157 | L157F | A | 2.5e-06 | 3.2e-06 | 2.4e-06 | 2.700E-06 | 4.359E-07 | 2.400E-06 | 3.200E-06 |
| 3135 | 157 | L157P | A | 3e-06 | 2.5e-06 | 2.7e-06 | 2.733E-06 | 2.517E-07 | 2.500E-06 | 3.000E-06 |
| 3136 | 157 | L157S | A | 2e-06 | 2.5e-06 | 3.1e-06 | 2.533E-06 | 5.508E-07 | 2.000E-06 | 3.100E-06 |
| 3137 | 157 | L157T | A | 3.3e-06 | 2.3e-06 | 3.4e-06 | 3.000E-06 | 6.083E-07 | 2.300E-06 | 3.400E-06 |
| 3138 | 157 | L157W | A | 3.6e-06 | 1.3e-06 | 2e-06 | 2.300E-06 | 1.179E-06 | 1.300E-06 | 3.600E-06 |
| 3139 | 157 | L157Y | A | 2e-06 | 2.6e-06 | 2.5e-06 | 2.367E-06 | 3.215E-07 | 2.000E-06 | 2.600E-06 |
| 3140 | 157 | L157V | A | 1.4e-06 | 2.8e-06 | 1.3e-06 | 1.833E-06 | 8.386E-07 | 1.300E-06 | 2.800E-06 |
| 3141 | 158 | A158A | A | 3.5e-06 | 1.9e-06 | 2.3e-06 | 2.567E-06 | 8.327E-07 | 1.900E-06 | 3.500E-06 |
| 3142 | 158 | A158R | A | 1.9e-06 | 2e-06 | 1.9e-06 | 1.933E-06 | 5.774E-08 | 1.900E-06 | 2.000E-06 |
| 3143 | 158 | A158N | A | 4.2e-06 | 2e-06 | 1.1e-06 | 2.433E-06 | 1.595E-06 | 1.100E-06 | 4.200E-06 |
| 3144 | 158 | A158D | A | 3.5e-06 | 2e-06 | 2.9e-06 | 2.800E-06 | 7.550E-07 | 2.000E-06 | 3.500E-06 |

| | | | | | | | | | | |
|---|---|---|---|---|---|---|---|---|---|---|
| 3145 | 158 | A158C | A | 2.2e-06 | 3.1e-06 | 1.6e-06 | 2.300E-06 | 7.550E-07 | 1.600E-06 | 3.100E-06 |
| 3146 | 158 | A158E | A | 3.1e-06 | 1.8e-06 | 3.8e-06 | 2.900E-06 | 1.015E-06 | 1.800E-06 | 3.800E-06 |
| 3147 | 158 | A158Q | A | 1.3e-06 | 2.7e-06 | 1.6e-06 | 1.867E-06 | 7.371E-07 | 1.300E-06 | 2.700E-06 |
| 3148 | 158 | A158G | A | 2.2e-06 | 1.4e-06 | 2.3e-06 | 1.967E-06 | 4.933E-07 | 1.400E-06 | 2.300E-06 |
| 3149 | 158 | A158H | A | 1.4e-06 | 2.1e-06 | 2.2e-06 | 1.900E-06 | 4.359E-07 | 1.400E-06 | 2.200E-06 |
| 3150 | 158 | A158I | A | 2.1e-06 | 1.9e-06 | 2e-06 | 2.000E-06 | 1.000E-07 | 1.900E-06 | 2.100E-06 |
| 3151 | 158 | A158L | A | 2.7e-06 | 5.1e-06 | 2.1e-06 | 3.300E-06 | 1.587E-06 | 2.100E-06 | 5.100E-06 |
| 3152 | 158 | A158K | A | 1.4e-06 | 3e-06 | 3.1e-06 | 2.500E-06 | 9.539E-07 | 1.400E-06 | 3.100E-06 |
| 3153 | 158 | A158M | A | 7.3e-07 | 3.4e-06 | 2.7e-06 | 2.277E-06 | 1.384E-06 | 7.300E-07 | 3.400E-06 |
| 3154 | 158 | A158F | A | 2.2e-06 | 2.1e-06 | 2.4e-06 | 2.233E-06 | 1.528E-07 | 2.100E-06 | 2.400E-06 |
| 3155 | 158 | A158P | A | 1.7e-06 | 3.1e-06 | 2e-06 | 2.267E-06 | 7.371E-07 | 1.700E-06 | 3.100E-06 |
| 3156 | 158 | A158S | A | 3.8e-06 | 2.9e-06 | 1.4e-06 | 2.700E-06 | 1.212E-06 | 1.400E-06 | 3.800E-06 |
| 3157 | 158 | A158T | A | 3.6e-06 | 3.4e-06 | 2.7e-06 | 3.233E-06 | 4.726E-07 | 2.700E-06 | 3.600E-06 |
| 3158 | 158 | A158W | A | 2.2e-06 | 1.1e-06 | 1.8e-06 | 1.700E-06 | 5.568E-07 | 1.100E-06 | 2.200E-06 |
| 3159 | 158 | A158Y | A | 2.1e-06 | 2.4e-06 | 3.6e-06 | 2.700E-06 | 7.937E-07 | 2.100E-06 | 3.600E-06 |
| 3160 | 158 | A158V | A | 2.1e-06 | 1.9e-06 | 2.6e-06 | 2.200E-06 | 3.606E-07 | 1.900E-06 | 2.600E-06 |
| 3161 | 159 | L159A | A | 1.8e-06 | 3.2e-06 | 1.2e-06 | 2.067E-06 | 1.026E-06 | 1.200E-06 | 3.200E-06 |
| 3162 | 159 | L159R | A | 2.2e-06 | 2.4e-06 | 1.6e-06 | 2.067E-06 | 4.163E-07 | 1.600E-06 | 2.400E-06 |
| 3163 | 159 | L159N | A | 3.3e-06 | 2.5e-06 | 3.8e-06 | 3.200E-06 | 6.557E-07 | 2.500E-06 | 3.800E-06 |
| 3164 | 159 | L159D | A | 2.3e-06 | 2.2e-06 | 2.3e-06 | 2.267E-06 | 5.774E-08 | 2.200E-06 | 2.300E-06 |
| 3165 | 159 | L159C | A | 3.1e-06 | 2.1e-06 | 2.3e-06 | 2.500E-06 | 5.292E-07 | 2.100E-06 | 3.100E-06 |
| 3166 | 159 | L159E | A | 1.7e-06 | 3.3e-06 | 2.5e-06 | 2.500E-06 | 8.000E-07 | 1.700E-06 | 3.300E-06 |
| 3167 | 159 | L159Q | A | 2.2e-06 | 2.2e-06 | 1.4e-06 | 1.933E-06 | 4.619E-07 | 1.400E-06 | 2.200E-06 |
| 3168 | 159 | L159G | A | 1.4e-06 | 2.6e-06 | 3.3e-06 | 2.433E-06 | 9.609E-07 | 1.400E-06 | 3.300E-06 |
| 3169 | 159 | L159H | A | 1.8e-06 | 4.1e-06 | 2.9e-06 | 2.933E-06 | 1.150E-06 | 1.800E-06 | 4.100E-06 |
| 3170 | 159 | L159I | A | 2.7e-06 | 1.8e-06 | 2.9e-06 | 2.467E-06 | 5.859E-07 | 1.800E-06 | 2.900E-06 |
| 3171 | 159 | L159L | A | 1.7e-06 | 2.4e-06 | 1.9e-06 | 2.000E-06 | 3.606E-07 | 1.700E-06 | 2.400E-06 |
| 3172 | 159 | L159K | A | 2.3e-06 | 2.6e-06 | 2.5e-06 | 2.467E-06 | 1.528E-07 | 2.300E-06 | 2.600E-06 |
| 3173 | 159 | L159M | A | 2.2e-06 | 2.6e-06 | 2.8e-06 | 2.533E-06 | 3.055E-07 | 2.200E-06 | 2.800E-06 |
| 3174 | 159 | L159F | A | 3.3e-06 | 2.7e-06 | 2.8e-06 | 2.933E-06 | 3.215E-07 | 2.700E-06 | 3.300E-06 |
| 3175 | 159 | L159P | A | 1.9e-06 | 3.1e-06 | 1.5e-06 | 2.167E-06 | 8.327E-07 | 1.500E-06 | 3.100E-06 |
| 3176 | 159 | L159S | A | 3e-06 | 2e-06 | 2.7e-06 | 2.567E-06 | 5.132E-07 | 2.000E-06 | 3.000E-06 |
| 3177 | 159 | L159T | A | 1.7e-06 | 3e-06 | 1.9e-06 | 2.200E-06 | 7.000E-07 | 1.700E-06 | 3.000E-06 |
| 3178 | 159 | L159W | A | 2.4e-06 | 1.2e-06 | 2.1e-06 | 1.900E-06 | 6.245E-07 | 1.200E-06 | 2.400E-06 |
| 3179 | 159 | L159Y | A | 8.1e-07 | 2.3e-06 | 2.3e-06 | 1.803E-06 | 8.603E-07 | 8.100E-07 | 2.300E-06 |
| 3180 | 159 | L159V | A | 2.4e-06 | 1.6e-06 | 4.6e-06 | 2.867E-06 | 1.553E-06 | 1.600E-06 | 4.600E-06 |
| 3181 | 160 | T160A | A | 2.4e-06 | 2.9e-06 | 3.5e-06 | 2.933E-06 | 5.508E-07 | 2.400E-06 | 3.500E-06 |
| 3182 | 160 | T160R | A | 2.4e-06 | 2.1e-06 | 3.2e-06 | 2.567E-06 | 5.686E-07 | 2.100E-06 | 3.200E-06 |
| 3183 | 160 | T160N | A | 2.8e-06 | 4e-06 | 2.4e-06 | 3.067E-06 | 8.327E-07 | 2.400E-06 | 4.000E-06 |
| 3184 | 160 | T160D | A | 2.6e-06 | 1.6e-06 | 6.5e-06 | 3.567E-06 | 2.589E-06 | 1.600E-06 | 6.500E-06 |
| 3185 | 160 | T160C | A | 1.8e-06 | 2.6e-06 | 1.1e-06 | 1.833E-06 | 7.506E-07 | 1.100E-06 | 2.600E-06 |
| 3186 | 160 | T160E | A | 2.5e-06 | 2.2e-06 | 1.9e-06 | 2.200E-06 | 3.000E-07 | 1.900E-06 | 2.500E-06 |
| 3187 | 160 | T160Q | A | 1.8e-06 | 2.1e-06 | 2.6e-06 | 2.167E-06 | 4.041E-07 | 1.800E-06 | 2.600E-06 |
| 3188 | 160 | T160G | A | 4e-06 | 1.8e-06 | 2.6e-06 | 2.800E-06 | 1.114E-06 | 1.800E-06 | 4.000E-06 |
| 3189 | 160 | T160H | A | 1.6e-06 | 2e-06 | 1.9e-06 | 1.833E-06 | 2.082E-07 | 1.600E-06 | 2.000E-06 |
| 3190 | 160 | T160I | A | 2.9e-06 | 2.7e-06 | 3.6e-06 | 3.067E-06 | 4.726E-07 | 2.700E-06 | 3.600E-06 |
| 3191 | 160 | T160L | A | 2.5e-06 | 1.7e-06 | 2.3e-06 | 2.167E-06 | 4.163E-07 | 1.700E-06 | 2.500E-06 |
| 3192 | 160 | T160K | A | 1e-06 | 3.4e-06 | 3.6e-06 | 2.667E-06 | 1.447E-06 | 1.000E-06 | 3.600E-06 |
| 3193 | 160 | T160M | A | 2.2e-06 | 2.8e-06 | 2.4e-06 | 2.467E-06 | 3.055E-07 | 2.200E-06 | 2.800E-06 |
| 3194 | 160 | T160F | A | 1.9e-06 | 2.1e-06 | 3.3e-06 | 2.433E-06 | 7.572E-07 | 1.900E-06 | 3.300E-06 |
| 3195 | 160 | T160P | A | 2.4e-06 | 2.7e-06 | 2.7e-06 | 2.600E-06 | 1.732E-07 | 2.400E-06 | 2.700E-06 |
| 3196 | 160 | T160S | A | 3.4e-06 | 1.5e-06 | 2.7e-06 | 2.533E-06 | 9.609E-07 | 1.500E-06 | 3.400E-06 |
| 3197 | 160 | T160T | A | 2.4e-06 | 2.3e-06 | 3.2e-06 | 2.633E-06 | 4.933E-07 | 2.300E-06 | 3.200E-06 |
| 3198 | 160 | T160W | A | 2.7e-06 | 2.4e-06 | 1.4e-06 | 2.167E-06 | 6.807E-07 | 1.400E-06 | 2.700E-06 |
| 3199 | 160 | T160Y | A | 3.4e-06 | 1.9e-06 | 1.7e-06 | 2.333E-06 | 9.292E-07 | 1.700E-06 | 3.400E-06 |
| 3200 | 160 | T160V | A | 2.2e-06 | 2.4e-06 | 3.2e-06 | 2.600E-06 | 5.292E-07 | 2.200E-06 | 3.200E-06 |
| 3201 | 161 | L161A | A | 3.2e-06 | 3.5e-06 | 1.4e-06 | 2.700E-06 | 1.136E-06 | 1.400E-06 | 3.500E-06 |
| 3202 | 161 | L161R | A | 1.2e-06 | 3.1e-06 | 2e-06 | 2.100E-06 | 9.539E-07 | 1.200E-06 | 3.100E-06 |
| 3203 | 161 | L161N | A | 3.2e-06 | 2.4e-06 | 1.6e-06 | 2.400E-06 | 8.000E-07 | 1.600E-06 | 3.200E-06 |
| 3204 | 161 | L161D | A | 1.9e-06 | 2.4e-06 | 1.9e-06 | 2.067E-06 | 2.887E-07 | 1.900E-06 | 2.400E-06 |
| 3205 | 161 | L161C | A | 2.2e-06 | 2.2e-06 | 1.1e-06 | 1.833E-06 | 6.351E-07 | 1.100E-06 | 2.200E-06 |
| 3206 | 161 | L161E | A | 2.6e-06 | 1.3e-06 | 2.4e-06 | 2.100E-06 | 7.000E-07 | 1.300E-06 | 2.600E-06 |
| 3207 | 161 | L161Q | A | 1.8e-06 | 4.5e-06 | 2.7e-06 | 3.000E-06 | 1.375E-06 | 1.800E-06 | 4.500E-06 |
| 3208 | 161 | L161G | A | 1.9e-06 | 3.8e-06 | 5.3e-06 | 3.667E-06 | 1.704E-06 | 1.900E-06 | 5.300E-06 |
| 3209 | 161 | L161H | A | 1.4e-06 | 4.8e-06 | 2.6e-06 | 2.933E-06 | 1.724E-06 | 1.400E-06 | 4.800E-06 |
| 3210 | 161 | L161I | A | 1.1e-06 | 3.1e-06 | 3.5e-06 | 2.567E-06 | 1.286E-06 | 1.100E-06 | 3.500E-06 |
| 3211 | 161 | L161L | A | 2.7e-06 | 1.1e-06 | 2.2e-06 | 2.000E-06 | 8.185E-07 | 1.100E-06 | 2.700E-06 |
| 3212 | 161 | L161K | A | 3.5e-06 | 1.9e-06 | 2e-06 | 2.467E-06 | 8.963E-07 | 1.900E-06 | 3.500E-06 |
| 3213 | 161 | L161M | A | 1.7e-06 | 3.5e-06 | 2.6e-06 | 2.600E-06 | 9.000E-07 | 1.700E-06 | 3.500E-06 |
| 3214 | 161 | L161F | A | 2.3e-06 | 2.9e-06 | 2.7e-06 | 2.633E-06 | 3.055E-07 | 2.300E-06 | 2.900E-06 |
| 3215 | 161 | L161P | A | 2.2e-06 | 1.7e-06 | 5.6e-06 | 3.167E-06 | 2.122E-06 | 1.700E-06 | 5.600E-06 |
| 3216 | 161 | L161S | A | 2.2e-06 | 3e-06 | 3e-06 | 2.733E-06 | 4.619E-07 | 2.200E-06 | 3.000E-06 |
| 3217 | 161 | L161T | A | 2.3e-06 | 1.8e-06 | 1.8e-06 | 1.967E-06 | 2.887E-07 | 1.800E-06 | 2.300E-06 |
| 3218 | 161 | L161W | A | 2.6e-06 | 2.6e-06 | 2.1e-06 | 2.433E-06 | 2.887E-07 | 2.100E-06 | 2.600E-06 |
| 3219 | 161 | L161Y | A | 2.1e-06 | 2.4e-06 | 2.9e-06 | 2.467E-06 | 4.041E-07 | 2.100E-06 | 2.900E-06 |
| 3220 | 161 | L161V | A | 1.3e-06 | 2.6e-06 | 2.7e-06 | 2.200E-06 | 7.810E-07 | 1.300E-06 | 2.700E-06 |
| 3221 | 162 | I162A | A | 1.9e-06 | 1.9e-06 | 1.6e-06 | 1.800E-06 | 1.732E-07 | 1.600E-06 | 1.900E-06 |
| 3222 | 162 | I162R | A | 2e-06 | 2e-06 | 2.8e-06 | 2.267E-06 | 4.619E-07 | 2.000E-06 | 2.800E-06 |
| 3223 | 162 | I162N | A | 2.3e-06 | 1.8e-06 | 3.2e-06 | 2.433E-06 | 7.095E-07 | 1.800E-06 | 3.200E-06 |
| 3224 | 162 | I162D | A | 1.5e-06 | 1.3e-06 | 4.4e-06 | 2.400E-06 | 1.735E-06 | 1.300E-06 | 4.400E-06 |
| 3225 | 162 | I162C | A | 3.2e-06 | 1.9e-06 | 2.1e-06 | 2.400E-06 | 7.000E-07 | 1.900E-06 | 3.200E-06 |
| 3226 | 162 | I162E | A | 3.6e-06 | 2.3e-06 | 3.9e-06 | 3.267E-06 | 8.505E-07 | 2.300E-06 | 3.900E-06 |
| 3227 | 162 | I162Q | A | 2.7e-06 | 1.9e-06 | 2.7e-06 | 2.433E-06 | 4.619E-07 | 1.900E-06 | 2.700E-06 |
| 3228 | 162 | I162G | A | 2.2e-06 | 2.5e-06 | 1.2e-06 | 1.967E-06 | 6.807E-07 | 1.200E-06 | 2.500E-06 |
| 3229 | 162 | I162H | A | 2.6e-06 | 1.9e-06 | 2.3e-06 | 2.267E-06 | 3.512E-07 | 1.900E-06 | 2.600E-06 |

| | | | | | | | | | | |
|---|---|---|---|---|---|---|---|---|---|---|
| 3230 | 162 | I162I | A | 1.8e-06 | 2.4e-06 | 2.3e-06 | 2.167E-06 | 3.215E-07 | 1.800E-06 | 2.400E-06 |
| 3231 | 162 | I162L | A | 3.6e-06 | 3.3e-06 | 3e-06 | 3.300E-06 | 3.000E-07 | 3.000E-06 | 3.600E-06 |
| 3232 | 162 | I162K | A | 1.9e-06 | 1.9e-06 | 3.3e-06 | 2.367E-06 | 8.083E-07 | 1.900E-06 | 3.300E-06 |
| 3233 | 162 | I162M | A | 2.1e-06 | 3e-06 | 1.8e-06 | 2.300E-06 | 6.245E-07 | 1.800E-06 | 3.000E-06 |
| 3234 | 162 | I162F | A | 2.9e-06 | 2.5e-06 | 2.6e-06 | 2.667E-06 | 2.082E-07 | 2.500E-06 | 2.900E-06 |
| 3235 | 162 | I162P | A | 2.8e-06 | 1.4e-06 | 1.9e-06 | 2.033E-06 | 7.095E-07 | 1.400E-06 | 2.800E-06 |
| 3236 | 162 | I162S | A | 2.5e-06 | 1.9e-06 | 3.2e-06 | 2.533E-06 | 6.506E-07 | 1.900E-06 | 3.200E-06 |
| 3237 | 162 | I162T | A | 1.4e-06 | 2.4e-06 | 2.9e-06 | 2.233E-06 | 7.638E-07 | 1.400E-06 | 2.900E-06 |
| 3238 | 162 | I162W | A | 1.9e-06 | 1.2e-06 | 2.4e-06 | 1.833E-06 | 6.028E-07 | 1.200E-06 | 2.400E-06 |
| 3239 | 162 | I162Y | A | 1.4e-06 | 1.6e-06 | 2.1e-06 | 1.700E-06 | 3.606E-07 | 1.400E-06 | 2.100E-06 |
| 3240 | 162 | I162V | A | 2.2e-06 | 3.5e-06 | 4.9e-06 | 3.533E-06 | 1.350E-06 | 2.200E-06 | 4.900E-06 |
| 3241 | 163 | D163A | A | 3.1e-06 | 2.6e-06 | 2.4e-06 | 2.700E-06 | 3.606E-07 | 2.400E-06 | 3.100E-06 |
| 3242 | 163 | D163R | A | 2.5e-06 | 2.4e-06 | 2.5e-06 | 2.467E-06 | 5.774E-08 | 2.400E-06 | 2.500E-06 |
| 3243 | 163 | D163N | A | 2.8e-06 | 1.6e-06 | 2.5e-06 | 2.300E-06 | 6.245E-07 | 1.600E-06 | 2.800E-06 |
| 3244 | 163 | D163D | A | 1.4e-06 | 1.6e-06 | 2.6e-06 | 1.867E-06 | 6.429E-07 | 1.400E-06 | 2.600E-06 |
| 3245 | 163 | D163C | A | 2.7e-06 | 1.2e-06 | 3e-06 | 2.300E-06 | 9.644E-07 | 1.200E-06 | 3.000E-06 |
| 3246 | 163 | D163E | A | 2.5e-06 | 2.1e-06 | 3.8e-06 | 2.800E-06 | 8.888E-07 | 2.100E-06 | 3.800E-06 |
| 3247 | 163 | D163Q | A | 3e-06 | 3.2e-06 | 1.8e-06 | 2.667E-06 | 7.572E-07 | 1.800E-06 | 3.200E-06 |
| 3248 | 163 | D163G | A | 1.4e-06 | 3e-06 | 2.2e-06 | 2.200E-06 | 8.000E-07 | 1.400E-06 | 3.000E-06 |
| 3249 | 163 | D163H | A | 1.5e-06 | 3.6e-06 | 4.2e-06 | 3.100E-06 | 1.418E-06 | 1.500E-06 | 4.200E-06 |
| 3250 | 163 | D163I | A | 3.3e-06 | 2.6e-06 | 2.7e-06 | 2.867E-06 | 3.786E-07 | 2.600E-06 | 3.300E-06 |
| 3251 | 163 | D163L | A | 1.8e-06 | 1.3e-06 | 2.5e-06 | 1.867E-06 | 6.028E-07 | 1.300E-06 | 2.500E-06 |
| 3252 | 163 | D163K | A | 4.5e-06 | 3.2e-06 | 2.4e-06 | 3.367E-06 | 1.060E-06 | 2.400E-06 | 4.500E-06 |
| 3253 | 163 | D163M | A | 1.6e-06 | 1.7e-06 | 1.9e-06 | 1.733E-06 | 1.528E-07 | 1.600E-06 | 1.900E-06 |
| 3254 | 163 | D163F | A | 2.3e-06 | 2.6e-06 | 1.6e-06 | 2.167E-06 | 5.132E-07 | 1.600E-06 | 2.600E-06 |
| 3255 | 163 | D163P | A | 2.3e-06 | 2.2e-06 | 2.4e-06 | 2.300E-06 | 1.000E-07 | 2.200E-06 | 2.400E-06 |
| 3256 | 163 | D163S | A | 1.3e-06 | 1.3e-06 | 1.8e-06 | 1.467E-06 | 2.887E-07 | 1.300E-06 | 1.800E-06 |
| 3257 | 163 | D163T | A | 2.1e-06 | 1.5e-06 | 2.8e-06 | 2.133E-06 | 6.506E-07 | 1.500E-06 | 2.800E-06 |
| 3258 | 163 | D163W | A | 3.5e-06 | 1.8e-06 | 2.7e-06 | 2.667E-06 | 8.505E-07 | 1.800E-06 | 3.500E-06 |
| 3259 | 163 | D163Y | A | 1.5e-06 | 1.6e-06 | 2.6e-06 | 1.900E-06 | 6.083E-07 | 1.500E-06 | 2.600E-06 |
| 3260 | 163 | D163V | A | 2.4e-06 | 3e-06 | 2.3e-06 | 2.567E-06 | 3.786E-07 | 2.300E-06 | 3.000E-06 |
| 3261 | 164 | T164A | A | 2.5e-06 | 4.1e-06 | 3.6e-06 | 3.400E-06 | 8.185E-07 | 2.500E-06 | 4.100E-06 |
| 3262 | 164 | T164R | A | 4e-06 | 1.1e-06 | 1.9e-06 | 2.333E-06 | 1.498E-06 | 1.100E-06 | 4.000E-06 |
| 3263 | 164 | T164N | A | 2e-06 | 2.5e-06 | 2.1e-06 | 2.200E-06 | 2.646E-07 | 2.000E-06 | 2.500E-06 |
| 3264 | 164 | T164D | A | 1.2e-06 | 2.9e-06 | 2.6e-06 | 2.233E-06 | 9.074E-07 | 1.200E-06 | 2.900E-06 |
| 3265 | 164 | T164C | A | 4.4e-06 | 2.6e-06 | 2.4e-06 | 3.133E-06 | 1.102E-06 | 2.400E-06 | 4.400E-06 |
| 3266 | 164 | T164E | A | 1.3e-06 | 2.1e-06 | 2.5e-06 | 1.967E-06 | 6.110E-07 | 1.300E-06 | 2.500E-06 |
| 3267 | 164 | T164Q | A | 3e-06 | 1.7e-06 | 2.2e-06 | 2.300E-06 | 6.557E-07 | 1.700E-06 | 3.000E-06 |
| 3268 | 164 | T164G | A | 4.7e-06 | 2.8e-06 | 2.7e-06 | 3.400E-06 | 1.127E-06 | 2.700E-06 | 4.700E-06 |
| 3269 | 164 | T164H | A | 2.1e-06 | 1.8e-06 | 1.3e-06 | 1.733E-06 | 4.041E-07 | 1.300E-06 | 2.100E-06 |
| 3270 | 164 | T164I | A | 3.2e-06 | 2.7e-06 | 2e-06 | 2.633E-06 | 6.028E-07 | 2.000E-06 | 3.200E-06 |
| 3271 | 164 | T164L | A | 2.8e-06 | 3.9e-06 | 2.1e-06 | 2.933E-06 | 9.074E-07 | 2.100E-06 | 3.900E-06 |
| 3272 | 164 | T164K | A | 1.9e-06 | 2.8e-06 | 2.8e-06 | 2.500E-06 | 5.196E-07 | 1.900E-06 | 2.800E-06 |
| 3273 | 164 | T164M | A | 4.1e-06 | 5e-06 | 2.6e-06 | 3.900E-06 | 1.212E-06 | 2.600E-06 | 5.000E-06 |
| 3274 | 164 | T164F | A | 1.5e-06 | 2.8e-06 | 1.7e-06 | 2.000E-06 | 7.000E-07 | 1.500E-06 | 2.800E-06 |
| 3275 | 164 | T164P | A | 3.4e-06 | 1.4e-06 | 3.2e-06 | 2.667E-06 | 1.102E-06 | 1.400E-06 | 3.400E-06 |
| 3276 | 164 | T164S | A | 2.3e-06 | 2.5e-06 | 1.5e-06 | 2.100E-06 | 5.292E-07 | 1.500E-06 | 2.500E-06 |
| 3277 | 164 | T164T | A | 3.2e-06 | 1.5e-06 | 1.8e-06 | 2.167E-06 | 9.074E-07 | 1.500E-06 | 3.200E-06 |
| 3278 | 164 | T164W | A | 2.5e-06 | 2.3e-06 | 1.6e-06 | 2.133E-06 | 4.726E-07 | 1.600E-06 | 2.500E-06 |
| 3279 | 164 | T164Y | A | 2.5e-06 | 3.5e-06 | 3.5e-06 | 3.167E-06 | 5.774E-07 | 2.500E-06 | 3.500E-06 |
| 3280 | 164 | T164V | A | 2.5e-06 | 2.1e-06 | 2.3e-06 | 2.300E-06 | 2.000E-07 | 2.100E-06 | 2.500E-06 |
| 3281 | 165 | N165A | A | 3.3e-06 | 2.9e-06 | 1.6e-06 | 2.600E-06 | 8.888E-07 | 1.600E-06 | 3.300E-06 |
| 3282 | 165 | N165R | A | 3.1e-06 | 4.3e-06 | 1.6e-06 | 3.000E-06 | 1.353E-06 | 1.600E-06 | 4.300E-06 |
| 3283 | 165 | N165N | A | 1.3e-06 | 1.8e-06 | 2.7e-06 | 1.933E-06 | 7.095E-07 | 1.300E-06 | 2.700E-06 |
| 3284 | 165 | N165D | A | 3.5e-06 | 1.8e-06 | 2.5e-06 | 2.600E-06 | 8.544E-07 | 1.800E-06 | 3.500E-06 |
| 3285 | 165 | N165C | A | 2.5e-06 | 2.5e-06 | 2.3e-06 | 2.433E-06 | 1.155E-07 | 2.300E-06 | 2.500E-06 |
| 3286 | 165 | N165E | A | 4.5e-06 | 1.9e-06 | 1.4e-06 | 2.600E-06 | 1.664E-06 | 1.400E-06 | 4.500E-06 |
| 3287 | 165 | N165Q | A | 2.1e-06 | 2.5e-06 | 1.8e-06 | 2.133E-06 | 3.512E-07 | 1.800E-06 | 2.500E-06 |
| 3288 | 165 | N165G | A | 2.9e-06 | 2.7e-06 | 1.5e-06 | 2.367E-06 | 7.572E-07 | 1.500E-06 | 2.900E-06 |
| 3289 | 165 | N165H | A | 4.2e-06 | 2.4e-06 | 2.2e-06 | 2.933E-06 | 1.102E-06 | 2.200E-06 | 4.200E-06 |
| 3290 | 165 | N165I | A | 2.5e-06 | 9.2e-07 | 2.9e-06 | 2.107E-06 | 1.047E-06 | 9.200E-07 | 2.900E-06 |
| 3291 | 165 | N165L | A | 2.5e-06 | 2.1e-06 | 4e-06 | 2.867E-06 | 1.002E-06 | 2.100E-06 | 4.000E-06 |
| 3292 | 165 | N165K | A | 9.3e-07 | 2.2e-06 | 9e-07 | 1.343E-06 | 7.420E-07 | 9.000E-07 | 2.200E-06 |
| 3293 | 165 | N165M | A | 1.7e-06 | 2e-06 | 1.6e-06 | 1.767E-06 | 2.082E-07 | 1.600E-06 | 2.000E-06 |
| 3294 | 165 | N165F | A | 2.7e-06 | 2.5e-06 | 1.4e-06 | 2.200E-06 | 7.000E-07 | 1.400E-06 | 2.700E-06 |
| 3295 | 165 | N165P | A | 3.3e-06 | 2e-06 | 2.2e-06 | 2.500E-06 | 7.000E-07 | 2.000E-06 | 3.300E-06 |
| 3296 | 165 | N165S | A | 2.1e-06 | 2.4e-06 | 3e-06 | 2.500E-06 | 4.583E-07 | 2.100E-06 | 3.000E-06 |
| 3297 | 165 | N165T | A | 2.8e-06 | 5.8e-06 | 1.9e-06 | 3.500E-06 | 2.042E-06 | 1.900E-06 | 5.800E-06 |
| 3298 | 165 | N165W | A | 1.6e-06 | 3.5e-06 | 4.2e-06 | 3.100E-06 | 1.345E-06 | 1.600E-06 | 4.200E-06 |
| 3299 | 165 | N165Y | A | 3.7e-06 | 1.9e-06 | 1.2e-06 | 2.267E-06 | 1.290E-06 | 1.200E-06 | 3.700E-06 |
| 3300 | 165 | N165V | A | 2.2e-06 | 1.8e-06 | 2.3e-06 | 2.100E-06 | 2.646E-07 | 1.800E-06 | 2.300E-06 |
| 3301 | 166 | R166A | A | 1.8e-06 | 3.8e-06 | 4.8e-06 | 3.467E-06 | 1.528E-06 | 1.800E-06 | 4.800E-06 |
| 3302 | 166 | R166R | A | 1.6e-06 | 2.1e-06 | 1.8e-06 | 1.833E-06 | 2.517E-07 | 1.600E-06 | 2.100E-06 |
| 3303 | 166 | R166N | A | 2.6e-06 | 2.4e-06 | 1.8e-06 | 2.267E-06 | 4.163E-07 | 1.800E-06 | 2.600E-06 |
| 3304 | 166 | R166D | A | 2e-06 | 2.2e-06 | 6e-06 | 3.400E-06 | 2.254E-06 | 2.000E-06 | 6.000E-06 |
| 3305 | 166 | R166C | A | 2.7e-06 | 2.3e-06 | 2.5e-06 | 2.500E-06 | 2.000E-07 | 2.300E-06 | 2.700E-06 |
| 3306 | 166 | R166E | A | 2.7e-06 | 1.7e-06 | 2.4e-06 | 2.267E-06 | 5.132E-07 | 1.700E-06 | 2.700E-06 |
| 3307 | 166 | R166Q | A | 3e-06 | 1.2e-06 | 3.4e-06 | 2.533E-06 | 1.172E-06 | 1.200E-06 | 3.400E-06 |
| 3308 | 166 | R166G | A | 2.8e-06 | 3.5e-06 | 2.6e-06 | 2.967E-06 | 4.726E-07 | 2.600E-06 | 3.500E-06 |
| 3309 | 166 | R166H | A | 2e-06 | 1.6e-06 | 2.6e-06 | 2.067E-06 | 5.033E-07 | 1.600E-06 | 2.600E-06 |
| 3310 | 166 | R166I | A | 4.6e-06 | 2.3e-06 | 4.2e-06 | 3.700E-06 | 1.229E-06 | 2.300E-06 | 4.600E-06 |
| 3311 | 166 | R166L | A | 2.7e-06 | 2.7e-06 | 3e-06 | 2.800E-06 | 1.732E-07 | 2.700E-06 | 3.000E-06 |
| 3312 | 166 | R166K | A | 2.7e-06 | 2e-06 | 2.1e-06 | 2.267E-06 | 3.786E-07 | 2.000E-06 | 2.700E-06 |
| 3313 | 166 | R166M | A | 3.1e-06 | 2.7e-06 | 2.8e-06 | 2.867E-06 | 2.082E-07 | 2.700E-06 | 3.100E-06 |
| 3314 | 166 | R166F | A | 1.2e-06 | 1.2e-06 | 2.6e-06 | 1.667E-06 | 8.083E-07 | 1.200E-06 | 2.600E-06 |

| | | | | | | | | | | |
|---|---|---|---|---|---|---|---|---|---|---|
| 3315 | 166 | R166P | A | 1.6e-06 | 2.3e-06 | 3.1e-06 | 2.333E-06 | 7.506E-07 | 1.600E-06 | 3.100E-06 |
| 3316 | 166 | R166S | A | 4.3e-06 | 1.8e-06 | 3e-06 | 3.033E-06 | 1.250E-06 | 1.800E-06 | 4.300E-06 |
| 3317 | 166 | R166T | A | 2.8e-06 | 2.3e-06 | 1.9e-06 | 2.333E-06 | 4.509E-07 | 1.900E-06 | 2.800E-06 |
| 3318 | 166 | R166W | A | 3.3e-06 | 2.2e-06 | 3.3e-06 | 2.933E-06 | 6.351E-07 | 2.200E-06 | 3.300E-06 |
| 3319 | 166 | R166Y | A | 2.6e-06 | 2.3e-06 | 1.8e-06 | 2.233E-06 | 4.041E-07 | 1.800E-06 | 2.600E-06 |
| 3320 | 166 | R166V | A | 1.8e-06 | 2.5e-06 | 1.8e-06 | 2.033E-06 | 4.041E-07 | 1.800E-06 | 2.500E-06 |
| 3321 | 167 | S167A | A | 1.8e-06 | 2.5e-06 | 3e-06 | 2.433E-06 | 6.028E-07 | 1.800E-06 | 3.000E-06 |
| 3322 | 167 | S167R | A | 2e-06 | 2.9e-06 | 1.5e-06 | 2.133E-06 | 7.095E-07 | 1.500E-06 | 2.900E-06 |
| 3323 | 167 | S167N | A | 2.9e-06 | 1.4e-06 | 2e-06 | 2.100E-06 | 7.550E-07 | 1.400E-06 | 2.900E-06 |
| 3324 | 167 | S167D | A | 1.9e-06 | 4e-06 | 2.1e-06 | 2.667E-06 | 1.159E-06 | 1.900E-06 | 4.000E-06 |
| 3325 | 167 | S167C | A | 2.9e-06 | 4.4e-06 | 1.1e-06 | 2.800E-06 | 1.652E-06 | 1.100E-06 | 4.400E-06 |
| 3326 | 167 | S167E | A | 2.3e-06 | 2e-06 | 2.1e-06 | 2.133E-06 | 1.528E-07 | 2.000E-06 | 2.300E-06 |
| 3327 | 167 | S167Q | A | 2e-06 | 2.6e-06 | 2.2e-06 | 2.267E-06 | 3.055E-07 | 2.000E-06 | 2.600E-06 |
| 3328 | 167 | S167G | A | 2.6e-06 | 2.3e-06 | 3.7e-06 | 2.867E-06 | 7.371E-07 | 2.300E-06 | 3.700E-06 |
| 3329 | 167 | S167H | A | 2.3e-06 | 3.4e-06 | 1.6e-06 | 2.433E-06 | 9.074E-07 | 1.600E-06 | 3.400E-06 |
| 3330 | 167 | S167I | A | 2.7e-06 | 2.7e-06 | 2.2e-06 | 2.533E-06 | 2.887E-07 | 2.200E-06 | 2.700E-06 |
| 3331 | 167 | S167L | A | 2.7e-06 | 2.8e-06 | 2.2e-06 | 2.567E-06 | 3.215E-07 | 2.200E-06 | 2.800E-06 |
| 3332 | 167 | S167K | A | 3.6e-06 | 3.1e-06 | 2.6e-06 | 3.100E-06 | 5.000E-07 | 2.600E-06 | 3.600E-06 |
| 3333 | 167 | S167M | A | 2.7e-06 | 2.1e-06 | 5.2e-06 | 3.333E-06 | 1.644E-06 | 2.100E-06 | 5.200E-06 |
| 3334 | 167 | S167F | A | 2.1e-06 | 2.7e-06 | 1.8e-06 | 2.200E-06 | 4.583E-07 | 1.800E-06 | 2.700E-06 |
| 3335 | 167 | S167P | A | 3.3e-06 | 2.9e-06 | 2.2e-06 | 2.800E-06 | 5.568E-07 | 2.200E-06 | 3.300E-06 |
| 3336 | 167 | S167S | A | 4e-06 | 3.6e-06 | 4.5e-06 | 4.033E-06 | 4.509E-07 | 3.600E-06 | 4.500E-06 |
| 3337 | 167 | S167T | A | 1.1e-06 | 2.7e-06 | 2e-06 | 1.933E-06 | 8.021E-07 | 1.100E-06 | 2.700E-06 |
| 3338 | 167 | S167W | A | 4.6e-06 | 3.4e-06 | 2.5e-06 | 3.500E-06 | 1.054E-06 | 2.500E-06 | 4.600E-06 |
| 3339 | 167 | S167Y | A | 2.2e-06 | 1.3e-06 | 9.7e-07 | 1.490E-06 | 6.366E-07 | 9.700E-07 | 2.200E-06 |
| 3340 | 167 | S167V | A | 2.8e-06 | 2.6e-06 | 1.3e-06 | 2.233E-06 | 8.145E-07 | 1.300E-06 | 2.800E-06 |
| 3341 | 168 | R168A | A | 2.7e-06 | 3.7e-06 | 2.4e-06 | 2.933E-06 | 6.807E-07 | 2.400E-06 | 3.700E-06 |
| 3342 | 168 | R168R | A | 3.2e-06 | 6.4e-07 | 5.4e-06 | 3.080E-06 | 2.382E-06 | 6.400E-07 | 5.400E-06 |
| 3343 | 168 | R168N | A | 9e-07 | 1.7e-06 | 2.5e-06 | 1.700E-06 | 8.000E-07 | 9.000E-07 | 2.500E-06 |
| 3344 | 168 | R168D | A | 2.6e-06 | 2.2e-06 | 2.6e-06 | 2.467E-06 | 2.309E-07 | 2.200E-06 | 2.600E-06 |
| 3345 | 168 | R168C | A | 2.9e-06 | 2.9e-06 | 2.2e-06 | 2.667E-06 | 4.041E-07 | 2.200E-06 | 2.900E-06 |
| 3346 | 168 | R168E | A | 2.2e-06 | 3.1e-06 | 1.2e-06 | 2.167E-06 | 9.504E-07 | 1.200E-06 | 3.100E-06 |
| 3347 | 168 | R168Q | A | 2.6e-06 | 2.8e-06 | 2.6e-06 | 2.667E-06 | 1.155E-07 | 2.600E-06 | 2.800E-06 |
| 3348 | 168 | R168G | A | 3.6e-06 | 1.3e-06 | 2.4e-06 | 2.433E-06 | 1.150E-06 | 1.300E-06 | 3.600E-06 |
| 3349 | 168 | R168H | A | 2.5e-06 | 1.8e-06 | 1.1e-06 | 1.800E-06 | 7.000E-07 | 1.100E-06 | 2.500E-06 |
| 3350 | 168 | R168I | A | 2.1e-06 | 1.9e-06 | 1.4e-06 | 1.800E-06 | 3.606E-07 | 1.400E-06 | 2.100E-06 |
| 3351 | 168 | R168L | A | 2.8e-06 | 2.6e-06 | 2.4e-06 | 2.600E-06 | 2.000E-07 | 2.400E-06 | 2.800E-06 |
| 3352 | 168 | R168K | A | 2.8e-06 | 3.5e-06 | 8.1e-07 | 2.370E-06 | 1.396E-06 | 8.100E-07 | 3.500E-06 |
| 3353 | 168 | R168M | A | 2.3e-06 | 2.1e-06 | 3.6e-06 | 2.667E-06 | 8.145E-07 | 2.100E-06 | 3.600E-06 |
| 3354 | 168 | R168F | A | 2e-06 | 2.5e-06 | 3.2e-06 | 2.567E-06 | 6.028E-07 | 2.000E-06 | 3.200E-06 |
| 3355 | 168 | R168P | A | 1.5e-06 | 3.1e-06 | 1.1e-06 | 1.900E-06 | 1.058E-06 | 1.100E-06 | 3.100E-06 |
| 3356 | 168 | R168S | A | 2.6e-06 | 2.1e-06 | 1.6e-06 | 2.100E-06 | 5.000E-07 | 1.600E-06 | 2.600E-06 |
| 3357 | 168 | R168T | A | 1.3e-06 | 2.5e-06 | 2.8e-06 | 2.200E-06 | 7.937E-07 | 1.300E-06 | 2.800E-06 |
| 3358 | 168 | R168W | A | 3.1e-06 | 2.4e-06 | 2.2e-06 | 2.567E-06 | 4.726E-07 | 2.200E-06 | 3.100E-06 |
| 3359 | 168 | R168Y | A | 1.5e-06 | 1.9e-06 | 2.9e-06 | 2.100E-06 | 7.211E-07 | 1.500E-06 | 2.900E-06 |
| 3360 | 168 | R168V | A | 2.6e-06 | 2.1e-06 | 2.6e-06 | 2.433E-06 | 2.887E-07 | 2.100E-06 | 2.600E-06 |
| 3361 | 169 | A169A | A | 1.8e-06 | 1.6e-06 | 3e-06 | 2.133E-06 | 7.572E-07 | 1.600E-06 | 3.000E-06 |
| 3362 | 169 | A169R | A | 1.5e-06 | 1.9e-06 | 3.3e-06 | 2.233E-06 | 9.452E-07 | 1.500E-06 | 3.300E-06 |
| 3363 | 169 | A169N | A | 2.6e-06 | 3.9e-06 | 1.6e-06 | 2.700E-06 | 1.153E-06 | 1.600E-06 | 3.900E-06 |
| 3364 | 169 | A169D | A | 1.4e-06 | 2.4e-06 | 1.6e-06 | 1.800E-06 | 5.292E-07 | 1.400E-06 | 2.400E-06 |
| 3365 | 169 | A169C | A | 2.1e-06 | 2e-06 | 2.4e-06 | 2.167E-06 | 2.082E-07 | 2.000E-06 | 2.400E-06 |
| 3366 | 169 | A169E | A | 1.1e-06 | 3.3e-06 | 1.8e-06 | 2.067E-06 | 1.124E-06 | 1.100E-06 | 3.300E-06 |
| 3367 | 169 | A169Q | A | 1.8e-06 | 2.7e-06 | 2.8e-06 | 2.433E-06 | 5.508E-07 | 1.800E-06 | 2.800E-06 |
| 3368 | 169 | A169G | A | 1.5e-06 | 2.4e-06 | 2.2e-06 | 2.033E-06 | 4.726E-07 | 1.500E-06 | 2.400E-06 |
| 3369 | 169 | A169H | A | 1.2e-06 | 1.1e-06 | 4.7e-06 | 2.333E-06 | 2.050E-06 | 1.100E-06 | 4.700E-06 |
| 3370 | 169 | A169I | A | 1.7e-06 | 2.2e-06 | 2.3e-06 | 2.067E-06 | 3.215E-07 | 1.700E-06 | 2.300E-06 |
| 3371 | 169 | A169L | A | 9.2e-07 | 2.6e-06 | 2.3e-06 | 1.940E-06 | 8.960E-07 | 9.200E-07 | 2.600E-06 |
| 3372 | 169 | A169K | A | 1.7e-06 | 2.1e-06 | 3e-06 | 2.267E-06 | 6.658E-07 | 1.700E-06 | 3.000E-06 |
| 3373 | 169 | A169M | A | 2.1e-06 | 1.7e-06 | 2.6e-06 | 2.133E-06 | 4.509E-07 | 1.700E-06 | 2.600E-06 |
| 3374 | 169 | A169F | A | 1.8e-06 | 1.7e-06 | 1.1e-06 | 1.533E-06 | 3.786E-07 | 1.100E-06 | 1.800E-06 |
| 3375 | 169 | A169P | A | 2e-06 | 2.1e-06 | 1.6e-06 | 1.900E-06 | 2.646E-07 | 1.600E-06 | 2.100E-06 |
| 3376 | 169 | A169S | A | 1.6e-06 | 3.7e-06 | 2.9e-06 | 2.733E-06 | 1.060E-06 | 1.600E-06 | 3.700E-06 |
| 3377 | 169 | A169T | A | 2.2e-06 | 2.5e-06 | 2.2e-06 | 2.300E-06 | 1.732E-07 | 2.200E-06 | 2.500E-06 |
| 3378 | 169 | A169W | A | 1.5e-06 | 3.2e-06 | 1.7e-06 | 2.133E-06 | 9.292E-07 | 1.500E-06 | 3.200E-06 |
| 3379 | 169 | A169Y | A | 2.4e-06 | 1.5e-06 | 3.2e-06 | 2.367E-06 | 8.505E-07 | 1.500E-06 | 3.200E-06 |
| 3380 | 169 | A169V | A | 1.8e-06 | 2.2e-06 | 2.3e-06 | 2.100E-06 | 2.646E-07 | 1.800E-06 | 2.300E-06 |
| 3381 | 170 | C170A | A | 3.3e-06 | 2.5e-06 | 2.5e-06 | 2.767E-06 | 4.619E-07 | 2.500E-06 | 3.300E-06 |
| 3382 | 170 | C170R | A | 1.8e-06 | 1.3e-06 | 1.6e-06 | 1.567E-06 | 2.517E-07 | 1.300E-06 | 1.800E-06 |
| 3383 | 170 | C170N | A | 2.7e-06 | 2.5e-06 | 2.5e-06 | 2.567E-06 | 1.155E-07 | 2.500E-06 | 2.700E-06 |
| 3384 | 170 | C170D | A | 1.8e-06 | 3.1e-06 | 2.8e-06 | 2.567E-06 | 6.807E-07 | 1.800E-06 | 3.100E-06 |
| 3385 | 170 | C170C | A | 3.3e-06 | 2.7e-06 | 3.2e-06 | 3.067E-06 | 3.215E-07 | 2.700E-06 | 3.300E-06 |
| 3386 | 170 | C170E | A | 2.9e-06 | 1.6e-06 | 1.6e-06 | 2.033E-06 | 7.506E-07 | 1.600E-06 | 2.900E-06 |
| 3387 | 170 | C170Q | A | 2e-06 | 2.7e-06 | 2.9e-06 | 2.533E-06 | 4.726E-07 | 2.000E-06 | 2.900E-06 |
| 3388 | 170 | C170G | A | 1.6e-06 | 1.9e-06 | 1.5e-06 | 1.667E-06 | 2.082E-07 | 1.500E-06 | 1.900E-06 |
| 3389 | 170 | C170H | A | 2.1e-06 | 3.5e-06 | 3.2e-06 | 2.933E-06 | 7.371E-07 | 2.100E-06 | 3.500E-06 |
| 3390 | 170 | C170I | A | 2.5e-06 | 2.7e-06 | 1.3e-06 | 2.167E-06 | 7.572E-07 | 1.300E-06 | 2.700E-06 |
| 3391 | 170 | C170L | A | 1.3e-06 | 1.5e-06 | 2.5e-06 | 1.767E-06 | 6.429E-07 | 1.300E-06 | 2.500E-06 |
| 3392 | 170 | C170K | A | 2.3e-06 | 3.4e-06 | 4.2e-06 | 3.300E-06 | 9.539E-07 | 2.300E-06 | 4.200E-06 |
| 3393 | 170 | C170M | A | 7.8e-07 | 2.8e-06 | 3.3e-06 | 2.293E-06 | 1.334E-06 | 7.800E-07 | 3.300E-06 |
| 3394 | 170 | C170F | A | 2.2e-06 | 3e-06 | 1.4e-06 | 2.200E-06 | 8.000E-07 | 1.400E-06 | 3.000E-06 |
| 3395 | 170 | C170P | A | 1.7e-06 | 2.4e-06 | 3e-06 | 2.367E-06 | 6.506E-07 | 1.700E-06 | 3.000E-06 |
| 3396 | 170 | C170S | A | 2.8e-06 | 2.5e-06 | 1.9e-06 | 2.400E-06 | 4.583E-07 | 1.900E-06 | 2.800E-06 |
| 3397 | 170 | C170T | A | 3.9e-06 | 1.8e-06 | 3e-06 | 2.900E-06 | 1.054E-06 | 1.800E-06 | 3.900E-06 |
| 3398 | 170 | C170W | A | 1.1e-06 | 2.2e-06 | 2.5e-06 | 1.933E-06 | 7.371E-07 | 1.100E-06 | 2.500E-06 |
| 3399 | 170 | C170Y | A | 2.1e-06 | 2.8e-06 | 2.1e-06 | 2.333E-06 | 4.041E-07 | 2.100E-06 | 2.800E-06 |

| | | | | | | | | | | |
|---|---|---|---|---|---|---|---|---|---|---|
| 3400 | 170 | C170V | A | 2.8e-06 | 3e-06 | 3.6e-06 | 3.133E-06 | 4.163E-07 | 2.800E-06 | 3.600E-06 |
| 3401 | 171 | H171A | A | 1.2e-06 | 2.4e-06 | 1.7e-06 | 1.767E-06 | 6.028E-07 | 1.200E-06 | 2.400E-06 |
| 3402 | 171 | H171R | A | 4e-06 | 3.3e-06 | 2e-06 | 3.100E-06 | 1.015E-06 | 2.000E-06 | 4.000E-06 |
| 3403 | 171 | H171N | A | 3.4e-06 | 1.6e-06 | 4.6e-06 | 3.200E-06 | 1.510E-06 | 1.600E-06 | 4.600E-06 |
| 3404 | 171 | H171D | A | 2.6e-06 | 3.8e-06 | 1.9e-06 | 2.767E-06 | 9.609E-07 | 1.900E-06 | 3.800E-06 |
| 3405 | 171 | H171C | A | 1.6e-06 | 3.8e-06 | 1.6e-06 | 2.333E-06 | 1.270E-06 | 1.600E-06 | 3.800E-06 |
| 3406 | 171 | H171E | A | 2.8e-06 | 2.3e-06 | 3.4e-06 | 2.833E-06 | 5.508E-07 | 2.300E-06 | 3.400E-06 |
| 3407 | 171 | H171Q | A | 2.2e-06 | 1.4e-06 | 1.9e-06 | 1.833E-06 | 4.041E-07 | 1.400E-06 | 2.200E-06 |
| 3408 | 171 | H171G | A | 1.9e-06 | 4.6e-06 | 1.9e-06 | 2.800E-06 | 1.559E-06 | 1.900E-06 | 4.600E-06 |
| 3409 | 171 | H171H | A | 4.4e-06 | 2.2e-06 | 2.5e-06 | 3.033E-06 | 1.193E-06 | 2.200E-06 | 4.400E-06 |
| 3410 | 171 | H171I | A | 2.3e-06 | 2.7e-06 | 2.7e-06 | 2.567E-06 | 2.309E-07 | 2.300E-06 | 2.700E-06 |
| 3411 | 171 | H171L | A | 1.8e-06 | 7.9e-07 | 2.6e-06 | 1.730E-06 | 9.070E-07 | 7.900E-07 | 2.600E-06 |
| 3412 | 171 | H171K | A | 3.7e-06 | 2.2e-06 | 2.4e-06 | 2.767E-06 | 8.145E-07 | 2.200E-06 | 3.700E-06 |
| 3413 | 171 | H171M | A | 1.5e-06 | 2.3e-06 | 2.6e-06 | 2.133E-06 | 5.686E-07 | 1.500E-06 | 2.600E-06 |
| 3414 | 171 | H171F | A | 4.9e-06 | 2.2e-06 | 1.9e-06 | 3.000E-06 | 1.652E-06 | 1.900E-06 | 4.900E-06 |
| 3415 | 171 | H171P | A | 3e-06 | 4e-06 | 1.8e-06 | 2.933E-06 | 1.102E-06 | 1.800E-06 | 4.000E-06 |
| 3416 | 171 | H171S | A | 1.4e-06 | 2.1e-06 | 3.5e-06 | 2.333E-06 | 1.069E-06 | 1.400E-06 | 3.500E-06 |
| 3417 | 171 | H171T | A | 1.9e-06 | 2.9e-06 | 3.1e-06 | 2.633E-06 | 6.429E-07 | 1.900E-06 | 3.100E-06 |
| 3418 | 171 | H171W | A | 3.5e-06 | 2.5e-06 | 1.6e-06 | 2.533E-06 | 9.504E-07 | 1.600E-06 | 3.500E-06 |
| 3419 | 171 | H171Y | A | 2.6e-06 | 1.8e-06 | 2.4e-06 | 2.267E-06 | 4.163E-07 | 1.800E-06 | 2.600E-06 |
| 3420 | 171 | H171V | A | 1.6e-06 | 1.4e-06 | 2.3e-06 | 1.767E-06 | 4.726E-07 | 1.400E-06 | 2.300E-06 |
| 3421 | 172 | P172A | A | 1.6e-06 | 2.8e-06 | 2.5e-06 | 2.300E-06 | 6.245E-07 | 1.600E-06 | 2.800E-06 |
| 3422 | 172 | P172R | A | 3.5e-06 | 2.9e-06 | 2.2e-06 | 2.867E-06 | 6.506E-07 | 2.200E-06 | 3.500E-06 |
| 3423 | 172 | P172N | A | 5.9e-06 | 1.6e-06 | 3.9e-06 | 3.800E-06 | 2.152E-06 | 1.600E-06 | 5.900E-06 |
| 3424 | 172 | P172D | A | 1.1e-06 | 2.9e-06 | 4e-06 | 2.667E-06 | 1.464E-06 | 1.100E-06 | 4.000E-06 |
| 3425 | 172 | P172C | A | 1.2e-06 | 1.4e-06 | 1.6e-06 | 1.400E-06 | 2.000E-07 | 1.200E-06 | 1.600E-06 |
| 3426 | 172 | P172E | A | 2.9e-06 | 3.3e-06 | 4.1e-06 | 3.433E-06 | 6.110E-07 | 2.900E-06 | 4.100E-06 |
| 3427 | 172 | P172Q | A | 2.2e-06 | 2.4e-06 | 2.9e-06 | 2.500E-06 | 3.606E-07 | 2.200E-06 | 2.900E-06 |
| 3428 | 172 | P172G | A | 2.5e-06 | 2.7e-06 | 2.3e-06 | 2.500E-06 | 2.000E-07 | 2.300E-06 | 2.700E-06 |
| 3429 | 172 | P172H | A | 3.4e-06 | 1.9e-06 | 2.1e-06 | 2.467E-06 | 8.145E-07 | 1.900E-06 | 3.400E-06 |
| 3430 | 172 | P172I | A | 2.2e-06 | 3.1e-06 | 2.3e-06 | 2.533E-06 | 4.933E-07 | 2.200E-06 | 3.100E-06 |
| 3431 | 172 | P172L | A | 2e-06 | 1.7e-06 | 4.1e-06 | 2.600E-06 | 1.308E-06 | 1.700E-06 | 4.100E-06 |
| 3432 | 172 | P172K | A | 3.2e-06 | 2e-06 | 2.9e-06 | 2.700E-06 | 6.245E-07 | 2.000E-06 | 3.200E-06 |
| 3433 | 172 | P172M | A | 2e-06 | 1.9e-06 | 1.5e-06 | 1.800E-06 | 2.646E-07 | 1.500E-06 | 2.000E-06 |
| 3434 | 172 | P172F | A | 3.2e-06 | 1.6e-06 | 3e-06 | 2.600E-06 | 8.718E-07 | 1.600E-06 | 3.200E-06 |
| 3435 | 172 | P172P | A | 4.7e-06 | 2.7e-06 | 2.8e-06 | 3.400E-06 | 1.127E-06 | 2.700E-06 | 4.700E-06 |
| 3436 | 172 | P172S | A | 3.1e-06 | 2.8e-06 | 2.6e-06 | 2.833E-06 | 2.517E-07 | 2.600E-06 | 3.100E-06 |
| 3437 | 172 | P172T | A | 1.2e-06 | 1.8e-06 | 1.9e-06 | 1.633E-06 | 3.786E-07 | 1.200E-06 | 1.900E-06 |
| 3438 | 172 | P172W | A | 2.1e-06 | 3.3e-06 | 2.4e-06 | 2.600E-06 | 6.245E-07 | 2.100E-06 | 3.300E-06 |
| 3439 | 172 | P172Y | A | 1.4e-06 | 1.7e-06 | 2.5e-06 | 1.867E-06 | 5.686E-07 | 1.400E-06 | 2.500E-06 |
| 3440 | 172 | P172V | A | 1.9e-06 | 2e-06 | 2.1e-06 | 2.000E-06 | 1.000E-07 | 1.900E-06 | 2.100E-06 |
| 3441 | 173 | C173A | A | 2.9e-06 | 1.6e-06 | 3.3e-06 | 2.600E-06 | 8.888E-07 | 1.600E-06 | 3.300E-06 |
| 3442 | 173 | C173R | A | 1.5e-06 | 2.7e-06 | 1.5e-06 | 1.900E-06 | 6.928E-07 | 1.500E-06 | 2.700E-06 |
| 3443 | 173 | C173N | A | 3e-06 | 2.5e-06 | 3.3e-06 | 2.933E-06 | 4.041E-07 | 2.500E-06 | 3.300E-06 |
| 3444 | 173 | C173D | A | 1.9e-06 | 4.2e-06 | 2.1e-06 | 2.733E-06 | 1.274E-06 | 1.900E-06 | 4.200E-06 |
| 3445 | 173 | C173C | A | 2.9e-06 | 3e-06 | 2.2e-06 | 2.700E-06 | 4.359E-07 | 2.200E-06 | 3.000E-06 |
| 3446 | 173 | C173E | A | 2.9e-06 | 2.7e-06 | 2.7e-06 | 2.767E-06 | 1.155E-07 | 2.700E-06 | 2.900E-06 |
| 3447 | 173 | C173Q | A | 2.4e-06 | 3.8e-06 | 2.2e-06 | 2.800E-06 | 8.718E-07 | 2.200E-06 | 3.800E-06 |
| 3448 | 173 | C173G | A | 3.2e-06 | 2.4e-06 | 2.4e-06 | 2.667E-06 | 4.619E-07 | 2.400E-06 | 3.200E-06 |
| 3449 | 173 | C173H | A | 7e-06 | 1.8e-06 | 2.2e-06 | 3.667E-06 | 2.894E-06 | 1.800E-06 | 7.000E-06 |
| 3450 | 173 | C173I | A | 2.1e-06 | 2.7e-06 | 3.3e-06 | 2.700E-06 | 6.000E-07 | 2.100E-06 | 3.300E-06 |
| 3451 | 173 | C173L | A | 1.9e-06 | 1.3e-06 | 3.3e-06 | 2.167E-06 | 1.026E-06 | 1.300E-06 | 3.300E-06 |
| 3452 | 173 | C173K | A | 2.3e-06 | 1.6e-06 | 2.2e-06 | 2.033E-06 | 3.786E-07 | 1.600E-06 | 2.300E-06 |
| 3453 | 173 | C173M | A | 3.9e-06 | 4e-06 | 1.9e-06 | 3.267E-06 | 1.185E-06 | 1.900E-06 | 4.000E-06 |
| 3454 | 173 | C173F | A | 2.7e-06 | 2.1e-06 | 3.6e-06 | 2.800E-06 | 7.550E-07 | 2.100E-06 | 3.600E-06 |
| 3455 | 173 | C173P | A | 3.9e-06 | 1.7e-06 | 2.7e-06 | 2.767E-06 | 1.102E-06 | 1.700E-06 | 3.900E-06 |
| 3456 | 173 | C173S | A | 7.7e-07 | 1.4e-06 | 2e-06 | 1.390E-06 | 6.151E-07 | 7.700E-07 | 2.000E-06 |
| 3457 | 173 | C173T | A | 2.6e-06 | 2.2e-06 | 2.2e-06 | 2.333E-06 | 2.309E-07 | 2.200E-06 | 2.600E-06 |
| 3458 | 173 | C173W | A | 2.9e-06 | 3.8e-06 | 4.4e-06 | 3.700E-06 | 7.550E-07 | 2.900E-06 | 4.400E-06 |
| 3459 | 173 | C173Y | A | 3e-06 | 3.2e-06 | 4e-06 | 3.400E-06 | 5.292E-07 | 3.000E-06 | 4.000E-06 |
| 3460 | 173 | C173V | A | 1.8e-06 | 3.7e-06 | 2.5e-06 | 2.667E-06 | 9.609E-07 | 1.800E-06 | 3.700E-06 |
| 3461 | 174 | S174A | A | 3.1e-06 | 2.3e-06 | 2.6e-06 | 2.667E-06 | 4.041E-07 | 2.300E-06 | 3.100E-06 |
| 3462 | 174 | S174R | A | 2.7e-06 | 2e-06 | 1.5e-06 | 2.067E-06 | 6.028E-07 | 1.500E-06 | 2.700E-06 |
| 3463 | 174 | S174N | A | 2.8e-06 | 2.6e-06 | 3.1e-06 | 2.833E-06 | 2.517E-07 | 2.600E-06 | 3.100E-06 |
| 3464 | 174 | S174D | A | 2.4e-06 | 2.8e-06 | 2.2e-06 | 2.467E-06 | 3.055E-07 | 2.200E-06 | 2.800E-06 |
| 3465 | 174 | S174C | A | 2.3e-06 | 2.2e-06 | 2e-06 | 2.167E-06 | 1.528E-07 | 2.000E-06 | 2.300E-06 |
| 3466 | 174 | S174E | A | 1.7e-06 | 1.8e-06 | 2.2e-06 | 1.900E-06 | 2.646E-07 | 1.700E-06 | 2.200E-06 |
| 3467 | 174 | S174Q | A | 2.1e-06 | 3.6e-06 | 1.7e-06 | 2.467E-06 | 1.002E-06 | 1.700E-06 | 3.600E-06 |
| 3468 | 174 | S174G | A | 1.8e-06 | 8.4e-07 | 4.3e-06 | 2.313E-06 | 1.786E-06 | 8.400E-07 | 4.300E-06 |
| 3469 | 174 | S174H | A | 2.3e-06 | 1.8e-06 | 1.9e-06 | 2.000E-06 | 2.646E-07 | 1.800E-06 | 2.300E-06 |
| 3470 | 174 | S174I | A | 3.3e-06 | 2.1e-06 | 2.8e-06 | 2.733E-06 | 6.028E-07 | 2.100E-06 | 3.300E-06 |
| 3471 | 174 | S174L | A | 3e-06 | 3.1e-06 | 2.5e-06 | 2.867E-06 | 3.215E-07 | 2.500E-06 | 3.100E-06 |
| 3472 | 174 | S174K | A | 1.9e-06 | 1.9e-06 | 2.5e-06 | 2.100E-06 | 3.464E-07 | 1.900E-06 | 2.500E-06 |
| 3473 | 174 | S174M | A | 4.1e-06 | 1.9e-06 | 1.2e-06 | 2.400E-06 | 1.513E-06 | 1.200E-06 | 4.100E-06 |
| 3474 | 174 | S174F | A | 3.4e-06 | 3.8e-06 | 2.1e-06 | 3.100E-06 | 8.888E-07 | 2.100E-06 | 3.800E-06 |
| 3475 | 174 | S174P | A | 1.7e-06 | 3.1e-06 | 2.5e-06 | 2.433E-06 | 7.024E-07 | 1.700E-06 | 3.100E-06 |
| 3476 | 174 | S174S | A | 1.9e-06 | 4.3e-06 | 3e-06 | 3.067E-06 | 1.201E-06 | 1.900E-06 | 4.300E-06 |
| 3477 | 174 | S174T | A | 2.5e-06 | 1.6e-06 | 4e-06 | 2.700E-06 | 1.212E-06 | 1.600E-06 | 4.000E-06 |
| 3478 | 174 | S174W | A | 3.5e-06 | 2.1e-06 | 2.4e-06 | 2.667E-06 | 7.371E-07 | 2.100E-06 | 3.500E-06 |
| 3479 | 174 | S174Y | A | 1.5e-06 | 1.7e-06 | 2.6e-06 | 1.933E-06 | 5.859E-07 | 1.500E-06 | 2.600E-06 |
| 3480 | 174 | S174V | A | 2.3e-06 | 2.5e-06 | 3e-06 | 2.600E-06 | 3.606E-07 | 2.300E-06 | 3.000E-06 |
| 3481 | 175 | P175A | A | 3e-06 | 3.9e-06 | 1.7e-06 | 2.867E-06 | 1.106E-06 | 1.700E-06 | 3.900E-06 |
| 3482 | 175 | P175R | A | 1.9e-06 | 1.6e-06 | 2.3e-06 | 1.933E-06 | 3.512E-07 | 1.600E-06 | 2.300E-06 |
| 3483 | 175 | P175N | A | 2.1e-06 | 1.9e-06 | 1.2e-06 | 1.733E-06 | 4.726E-07 | 1.200E-06 | 2.100E-06 |
| 3484 | 175 | P175D | A | 2.3e-06 | 1.5e-06 | 3.4e-06 | 2.400E-06 | 9.539E-07 | 1.500E-06 | 3.400E-06 |

| | | | | | | | | | | |
|---|---|---|---|---|---|---|---|---|---|---|
| 3485 | 175 | P175C | A | 2.3e-06 | 3.3e-06 | 1.6e-06 | 2.400E-06 | 8.544E-07 | 1.600E-06 | 3.300E-06 |
| 3486 | 175 | P175E | A | 2.1e-06 | 4e-06 | 2.7e-06 | 2.933E-06 | 9.713E-07 | 2.100E-06 | 4.000E-06 |
| 3487 | 175 | P175Q | A | 2.1e-06 | 3.2e-06 | 3e-06 | 2.767E-06 | 5.859E-07 | 2.100E-06 | 3.200E-06 |
| 3488 | 175 | P175G | A | 1.8e-06 | 1.7e-06 | 1.7e-06 | 1.733E-06 | 5.774E-08 | 1.700E-06 | 1.800E-06 |
| 3489 | 175 | P175H | A | 3.1e-06 | 2e-06 | 2.1e-06 | 2.400E-06 | 6.083E-07 | 2.000E-06 | 3.100E-06 |
| 3490 | 175 | P175I | A | 4.8e-06 | 5.1e-06 | 2.1e-06 | 4.000E-06 | 1.652E-06 | 2.100E-06 | 5.100E-06 |
| 3491 | 175 | P175L | A | 1.3e-06 | 3e-06 | 3.4e-06 | 2.567E-06 | 1.115E-06 | 1.300E-06 | 3.400E-06 |
| 3492 | 175 | P175K | A | 7.6e-07 | 2e-06 | 2.4e-06 | 1.720E-06 | 8.551E-07 | 7.600E-07 | 2.400E-06 |
| 3493 | 175 | P175M | A | 4.1e-06 | 1.9e-06 | 2.6e-06 | 2.867E-06 | 1.124E-06 | 1.900E-06 | 4.100E-06 |
| 3494 | 175 | P175F | A | 3.3e-06 | 3.1e-06 | 1.7e-06 | 2.700E-06 | 8.718E-07 | 1.700E-06 | 3.300E-06 |
| 3495 | 175 | P175P | A | 1.8e-06 | 3.1e-06 | 2.4e-06 | 2.433E-06 | 6.506E-07 | 1.800E-06 | 3.100E-06 |
| 3496 | 175 | P175S | A | 1.2e-06 | 2e-06 | 3.1e-06 | 2.100E-06 | 9.539E-07 | 1.200E-06 | 3.100E-06 |
| 3497 | 175 | P175T | A | 2.8e-06 | 2.6e-06 | 1.3e-06 | 2.233E-06 | 8.145E-07 | 1.300E-06 | 2.800E-06 |
| 3498 | 175 | P175W | A | 2e-06 | 2.3e-06 | 2.4e-06 | 2.233E-06 | 2.082E-07 | 2.000E-06 | 2.400E-06 |
| 3499 | 175 | P175Y | A | 2.4e-06 | 1.2e-06 | 2.6e-06 | 2.067E-06 | 7.572E-07 | 1.200E-06 | 2.600E-06 |
| 3500 | 175 | P175V | A | 2.8e-06 | 2.3e-06 | 1.4e-06 | 2.167E-06 | 7.095E-07 | 1.400E-06 | 2.800E-06 |
| 3501 | 176 | M176A | A | 1.5e-06 | 3.1e-06 | 4.3e-06 | 2.967E-06 | 1.405E-06 | 1.500E-06 | 4.300E-06 |
| 3502 | 176 | M176R | A | 2.5e-06 | 2.5e-06 | 3e-06 | 2.667E-06 | 2.887E-07 | 2.500E-06 | 3.000E-06 |
| 3503 | 176 | M176N | A | 1.8e-06 | 2.1e-06 | 1.7e-06 | 1.867E-06 | 2.082E-07 | 1.700E-06 | 2.100E-06 |
| 3504 | 176 | M176D | A | 2.3e-06 | 2.2e-06 | 2e-06 | 2.167E-06 | 1.528E-07 | 2.000E-06 | 2.300E-06 |
| 3505 | 176 | M176C | A | 1.7e-06 | 3.2e-06 | 2.2e-06 | 2.367E-06 | 7.638E-07 | 1.700E-06 | 3.200E-06 |
| 3506 | 176 | M176E | A | 1.8e-06 | 1.2e-06 | 2.7e-06 | 1.900E-06 | 7.550E-07 | 1.200E-06 | 2.700E-06 |
| 3507 | 176 | M176Q | A | 3.4e-06 | 3.2e-06 | 2.8e-06 | 3.133E-06 | 3.055E-07 | 2.800E-06 | 3.400E-06 |
| 3508 | 176 | M176G | A | 1.4e-06 | 3e-06 | 1.6e-06 | 2.000E-06 | 8.718E-07 | 1.400E-06 | 3.000E-06 |
| 3509 | 176 | M176H | A | 4.1e-06 | 2.5e-06 | 2.6e-06 | 3.067E-06 | 8.963E-07 | 2.500E-06 | 4.100E-06 |
| 3510 | 176 | M176I | A | 1.5e-06 | 2.7e-06 | 2.8e-06 | 2.333E-06 | 7.234E-07 | 1.500E-06 | 2.800E-06 |
| 3511 | 176 | M176L | A | 2.4e-06 | 1.7e-06 | 1e-06 | 1.700E-06 | 7.000E-07 | 1.000E-06 | 2.400E-06 |
| 3512 | 176 | M176K | A | 1.9e-06 | 3e-06 | 1.9e-06 | 2.267E-06 | 6.351E-07 | 1.900E-06 | 3.000E-06 |
| 3513 | 176 | M176M | A | 1.1e-06 | 2.2e-06 | 1.8e-06 | 1.700E-06 | 5.568E-07 | 1.100E-06 | 2.200E-06 |
| 3514 | 176 | M176F | A | 2e-06 | 2.6e-06 | 1.5e-06 | 2.033E-06 | 5.508E-07 | 1.500E-06 | 2.600E-06 |
| 3515 | 176 | M176P | A | 3.1e-06 | 3.8e-06 | 2.7e-06 | 3.200E-06 | 5.568E-07 | 2.700E-06 | 3.800E-06 |
| 3516 | 176 | M176S | A | 2.3e-06 | 2.5e-06 | 2.5e-06 | 2.433E-06 | 1.155E-07 | 2.300E-06 | 2.500E-06 |
| 3517 | 176 | M176T | A | 3.3e-06 | 2.1e-06 | 2.5e-06 | 2.633E-06 | 6.110E-07 | 2.100E-06 | 3.300E-06 |
| 3518 | 176 | M176W | A | 2.2e-06 | 2.5e-06 | 1.9e-06 | 2.200E-06 | 3.000E-07 | 1.900E-06 | 2.500E-06 |
| 3519 | 176 | M176Y | A | 2.2e-06 | 1.7e-06 | 2.3e-06 | 2.067E-06 | 3.215E-07 | 1.700E-06 | 2.300E-06 |
| 3520 | 176 | M176V | A | 2.9e-06 | 2.4e-06 | 2.3e-06 | 2.533E-06 | 3.215E-07 | 2.300E-06 | 2.900E-06 |
| 3521 | 177 | C177A | A | 2.9e-06 | 4.4e-06 | 5.6e-06 | 4.300E-06 | 1.353E-06 | 2.900E-06 | 5.600E-06 |
| 3522 | 177 | C177R | A | 4.1e-06 | 3e-06 | 2.9e-06 | 3.333E-06 | 6.658E-07 | 2.900E-06 | 4.100E-06 |
| 3523 | 177 | C177N | A | 2.1e-06 | 3.9e-06 | 2.7e-06 | 2.900E-06 | 9.165E-07 | 2.100E-06 | 3.900E-06 |
| 3524 | 177 | C177D | A | 4.8e-06 | 1.8e-06 | 2.2e-06 | 2.933E-06 | 1.629E-06 | 1.800E-06 | 4.800E-06 |
| 3525 | 177 | C177C | A | 1.9e-06 | 1.1e-06 | 3.2e-06 | 2.067E-06 | 1.060E-06 | 1.100E-06 | 3.200E-06 |
| 3526 | 177 | C177E | A | 3.2e-06 | 3.4e-06 | 2.6e-06 | 3.067E-06 | 4.163E-07 | 2.600E-06 | 3.400E-06 |
| 3527 | 177 | C177Q | A | 3.7e-06 | 3.4e-06 | 3.2e-06 | 3.433E-06 | 2.517E-07 | 3.200E-06 | 3.700E-06 |
| 3528 | 177 | C177G | A | 4.8e-06 | 3.8e-06 | 2.5e-06 | 3.700E-06 | 1.153E-06 | 2.500E-06 | 4.800E-06 |
| 3529 | 177 | C177H | A | 7.1e-07 | 2.2e-06 | 2.4e-06 | 1.770E-06 | 9.234E-07 | 7.100E-07 | 2.400E-06 |
| 3530 | 177 | C177I | A | 2e-06 | 2.7e-06 | 2.5e-06 | 2.400E-06 | 3.606E-07 | 2.000E-06 | 2.700E-06 |
| 3531 | 177 | C177L | A | 2.7e-06 | 2.9e-06 | 2e-06 | 2.533E-06 | 4.726E-07 | 2.000E-06 | 2.900E-06 |
| 3532 | 177 | C177K | A | 3.6e-06 | 2.7e-06 | 2e-06 | 2.767E-06 | 8.021E-07 | 2.000E-06 | 3.600E-06 |
| 3533 | 177 | C177M | A | 4.1e-06 | 1.6e-06 | 2.4e-06 | 2.700E-06 | 1.277E-06 | 1.600E-06 | 4.100E-06 |
| 3534 | 177 | C177F | A | 4.5e-06 | 3.3e-06 | 2.8e-06 | 3.533E-06 | 8.737E-07 | 2.800E-06 | 4.500E-06 |
| 3535 | 177 | C177P | A | 2.3e-06 | 2.8e-06 | 1.9e-06 | 2.333E-06 | 4.509E-07 | 1.900E-06 | 2.800E-06 |
| 3536 | 177 | C177S | A | 9.4e-07 | 3.9e-06 | 1.8e-06 | 2.213E-06 | 1.523E-06 | 9.400E-07 | 3.900E-06 |
| 3537 | 177 | C177T | A | 3.2e-06 | 2.2e-06 | 2.4e-06 | 2.600E-06 | 5.292E-07 | 2.200E-06 | 3.200E-06 |
| 3538 | 177 | C177W | A | 4.5e-06 | 1.9e-06 | 1e-06 | 2.467E-06 | 1.818E-06 | 1.000E-06 | 4.500E-06 |
| 3539 | 177 | C177Y | A | 2.2e-06 | 1.3e-06 | 1.6e-06 | 1.700E-06 | 4.583E-07 | 1.300E-06 | 2.200E-06 |
| 3540 | 177 | C177V | A | 5.2e-06 | 1.9e-06 | 3e-06 | 3.367E-06 | 1.680E-06 | 1.900E-06 | 5.200E-06 |
| 3541 | 178 | K178A | A | 2e-06 | 1.9e-06 | 2.5e-06 | 2.133E-06 | 3.215E-07 | 1.900E-06 | 2.500E-06 |
| 3542 | 178 | K178R | A | 1.7e-06 | 1.4e-06 | 1.9e-06 | 1.667E-06 | 2.517E-07 | 1.400E-06 | 1.900E-06 |
| 3543 | 178 | K178N | A | 2.3e-06 | 8.5e-07 | 2.4e-06 | 1.850E-06 | 8.675E-07 | 8.500E-07 | 2.400E-06 |
| 3544 | 178 | K178D | A | 2e-06 | 2.4e-06 | 2e-06 | 2.133E-06 | 2.309E-07 | 2.000E-06 | 2.400E-06 |
| 3545 | 178 | K178C | A | 1.8e-06 | 2.5e-06 | 2.3e-06 | 2.200E-06 | 3.606E-07 | 1.800E-06 | 2.500E-06 |
| 3546 | 178 | K178E | A | 2.1e-06 | 1.5e-06 | 2.3e-06 | 1.967E-06 | 4.163E-07 | 1.500E-06 | 2.300E-06 |
| 3547 | 178 | K178Q | A | 1.9e-06 | 1.7e-06 | 2.6e-06 | 2.067E-06 | 4.726E-07 | 1.700E-06 | 2.600E-06 |
| 3548 | 178 | K178G | A | 2.9e-06 | 5.2e-06 | 1.6e-06 | 3.233E-06 | 1.823E-06 | 1.600E-06 | 5.200E-06 |
| 3549 | 178 | K178H | A | 3.5e-06 | 2.3e-06 | 4.8e-06 | 3.533E-06 | 1.250E-06 | 2.300E-06 | 4.800E-06 |
| 3550 | 178 | K178I | A | 3.2e-06 | 2e-06 | 2.2e-06 | 2.467E-06 | 6.429E-07 | 2.000E-06 | 3.200E-06 |
| 3551 | 178 | K178L | A | 5.4e-06 | 4e-06 | 3.1e-06 | 4.167E-06 | 1.159E-06 | 3.100E-06 | 5.400E-06 |
| 3552 | 178 | K178K | A | 2.6e-06 | 2.9e-06 | 2.6e-06 | 2.700E-06 | 1.732E-07 | 2.600E-06 | 2.900E-06 |
| 3553 | 178 | K178M | A | 2.4e-06 | 3e-06 | 1.8e-06 | 2.400E-06 | 6.000E-07 | 1.800E-06 | 3.000E-06 |
| 3554 | 178 | K178F | A | 8.6e-07 | 2e-06 | 1.9e-06 | 1.587E-06 | 6.313E-07 | 8.600E-07 | 2.000E-06 |
| 3555 | 178 | K178P | A | 2.9e-06 | 2.7e-06 | 2.3e-06 | 2.633E-06 | 3.055E-07 | 2.300E-06 | 2.900E-06 |
| 3556 | 178 | K178S | A | 3.7e-06 | 4.3e-06 | 2.7e-06 | 3.567E-06 | 8.083E-07 | 2.700E-06 | 4.300E-06 |
| 3557 | 178 | K178T | A | 3.4e-06 | 2.3e-06 | 3.2e-06 | 2.967E-06 | 5.859E-07 | 2.300E-06 | 3.400E-06 |
| 3558 | 178 | K178W | A | 2.1e-06 | 2e-06 | 3.5e-06 | 2.533E-06 | 8.386E-07 | 2.000E-06 | 3.500E-06 |
| 3559 | 178 | K178Y | A | 2.3e-06 | 2.3e-06 | 1.7e-06 | 2.100E-06 | 3.464E-07 | 1.700E-06 | 2.300E-06 |
| 3560 | 178 | K178V | A | 3e-06 | 2e-06 | 2.5e-06 | 2.500E-06 | 5.000E-07 | 2.000E-06 | 3.000E-06 |
| 3561 | 179 | G179A | A | 1.6e-06 | 2.3e-06 | 2e-06 | 1.967E-06 | 3.512E-07 | 1.600E-06 | 2.300E-06 |
| 3562 | 179 | G179R | A | 1.5e-06 | 1.9e-06 | 4.5e-06 | 2.633E-06 | 1.629E-06 | 1.500E-06 | 4.500E-06 |
| 3563 | 179 | G179N | A | 1.8e-06 | 2.3e-06 | 2.4e-06 | 2.167E-06 | 3.215E-07 | 1.800E-06 | 2.400E-06 |
| 3564 | 179 | G179D | A | 2.3e-06 | 2.2e-06 | 1.7e-06 | 2.067E-06 | 3.215E-07 | 1.700E-06 | 2.300E-06 |
| 3565 | 179 | G179C | A | 1.6e-06 | 1.7e-06 | 1.7e-06 | 1.667E-06 | 5.774E-08 | 1.600E-06 | 1.700E-06 |
| 3566 | 179 | G179E | A | 3.9e-06 | 5.2e-06 | 2.7e-06 | 3.933E-06 | 1.250E-06 | 2.700E-06 | 5.200E-06 |
| 3567 | 179 | G179Q | A | 3.7e-06 | 2.4e-06 | 2.2e-06 | 2.767E-06 | 8.145E-07 | 2.200E-06 | 3.700E-06 |
| 3568 | 179 | G179G | A | 1.7e-06 | 2.1e-06 | 1.2e-06 | 1.667E-06 | 4.509E-07 | 1.200E-06 | 2.100E-06 |
| 3569 | 179 | G179H | A | 2.6e-06 | 2.8e-06 | 2.4e-06 | 2.600E-06 | 2.000E-07 | 2.400E-06 | 2.800E-06 |

| | | | | | | | | | | |
|---|---|---|---|---|---|---|---|---|---|---|
| 3570 | 179 | G179I | A | 2.5e-06 | 1.6e-06 | 3e-06 | 2.367E-06 | 7.095E-07 | 1.600E-06 | 3.000E-06 |
| 3571 | 179 | G179L | A | 3.5e-06 | 3e-06 | 2.2e-06 | 2.900E-06 | 6.557E-07 | 2.200E-06 | 3.500E-06 |
| 3572 | 179 | G179K | A | 2.1e-06 | 1.9e-06 | 4.8e-06 | 2.933E-06 | 1.620E-06 | 1.900E-06 | 4.800E-06 |
| 3573 | 179 | G179M | A | 3.1e-06 | 2.4e-06 | 1.8e-06 | 2.433E-06 | 6.506E-07 | 1.800E-06 | 3.100E-06 |
| 3574 | 179 | G179F | A | 1.4e-06 | 3e-06 | 2.5e-06 | 2.300E-06 | 8.185E-07 | 1.400E-06 | 3.000E-06 |
| 3575 | 179 | G179P | A | 2.3e-06 | 3.5e-06 | 2.5e-06 | 2.767E-06 | 6.429E-07 | 2.300E-06 | 3.500E-06 |
| 3576 | 179 | G179S | A | 2.7e-06 | 2e-06 | 3.8e-06 | 2.833E-06 | 9.074E-07 | 2.000E-06 | 3.800E-06 |
| 3577 | 179 | G179T | A | 2e-06 | 2.9e-06 | 2.8e-06 | 2.567E-06 | 4.933E-07 | 2.000E-06 | 2.900E-06 |
| 3578 | 179 | G179W | A | 1e-06 | 2.1e-06 | 1.3e-06 | 1.467E-06 | 5.686E-07 | 1.000E-06 | 2.100E-06 |
| 3579 | 179 | G179Y | A | 1.3e-06 | 2e-06 | 1.8e-06 | 1.700E-06 | 3.606E-07 | 1.300E-06 | 2.000E-06 |
| 3580 | 179 | G179V | A | 1.6e-06 | 2.1e-06 | 3.5e-06 | 2.400E-06 | 9.849E-07 | 1.600E-06 | 3.500E-06 |
| 3581 | 180 | S180A | A | 1.8e-06 | 3.1e-06 | 2.7e-06 | 2.533E-06 | 6.658E-07 | 1.800E-06 | 3.100E-06 |
| 3582 | 180 | S180R | A | 2.1e-06 | 2.9e-06 | 3.8e-06 | 2.933E-06 | 8.505E-07 | 2.100E-06 | 3.800E-06 |
| 3583 | 180 | S180N | A | 2.6e-06 | 3.1e-06 | 2.3e-06 | 2.667E-06 | 4.041E-07 | 2.300E-06 | 3.100E-06 |
| 3584 | 180 | S180D | A | 4.9e-06 | 1.7e-06 | 3.3e-06 | 3.300E-06 | 1.600E-06 | 1.700E-06 | 4.900E-06 |
| 3585 | 180 | S180C | A | 3.7e-06 | 2.1e-06 | 2.4e-06 | 2.733E-06 | 8.505E-07 | 2.100E-06 | 3.700E-06 |
| 3586 | 180 | S180E | A | 2.1e-06 | 2.4e-06 | 4.2e-06 | 2.900E-06 | 1.136E-06 | 2.100E-06 | 4.200E-06 |
| 3587 | 180 | S180Q | A | 2.3e-06 | 2.2e-06 | 2.6e-06 | 2.367E-06 | 2.082E-07 | 2.200E-06 | 2.600E-06 |
| 3588 | 180 | S180G | A | 3.6e-06 | 2.1e-06 | 1.6e-06 | 2.433E-06 | 1.041E-06 | 1.600E-06 | 3.600E-06 |
| 3589 | 180 | S180H | A | 2.1e-06 | 2.1e-06 | 2.1e-06 | 2.100E-06 | 0.000E+00 | 2.100E-06 | 2.100E-06 |
| 3590 | 180 | S180I | A | 2.7e-06 | 9.9e-07 | 2.6e-06 | 2.097E-06 | 9.597E-07 | 9.900E-07 | 2.700E-06 |
| 3591 | 180 | S180L | A | 2.6e-06 | 2.4e-06 | 1.8e-06 | 2.267E-06 | 4.163E-07 | 1.800E-06 | 2.600E-06 |
| 3592 | 180 | S180K | A | 2.5e-06 | 2e-06 | 1.6e-06 | 2.033E-06 | 4.509E-07 | 1.600E-06 | 2.500E-06 |
| 3593 | 180 | S180M | A | 1.8e-06 | 2.4e-06 | 2.8e-06 | 2.333E-06 | 5.033E-07 | 1.800E-06 | 2.800E-06 |
| 3594 | 180 | S180F | A | 1.7e-06 | 4e-06 | 3.1e-06 | 2.933E-06 | 1.159E-06 | 1.700E-06 | 4.000E-06 |
| 3595 | 180 | S180P | A | 3.7e-06 | 2e-06 | 2.1e-06 | 2.600E-06 | 9.539E-07 | 2.000E-06 | 3.700E-06 |
| 3596 | 180 | S180S | A | 2.7e-06 | 3.1e-06 | 1.5e-06 | 2.433E-06 | 8.327E-07 | 1.500E-06 | 3.100E-06 |
| 3597 | 180 | S180T | A | 3.6e-06 | 2.4e-06 | 2.1e-06 | 2.700E-06 | 7.937E-07 | 2.100E-06 | 3.600E-06 |
| 3598 | 180 | S180W | A | 3.6e-06 | 3.2e-06 | 1.5e-06 | 2.767E-06 | 1.115E-06 | 1.500E-06 | 3.600E-06 |
| 3599 | 180 | S180Y | A | 2.9e-06 | 3.2e-06 | 1.9e-06 | 2.667E-06 | 6.807E-07 | 1.900E-06 | 3.200E-06 |
| 3600 | 180 | S180V | A | 2.6e-06 | 2.4e-06 | 1.6e-06 | 2.200E-06 | 5.292E-07 | 1.600E-06 | 2.600E-06 |
| 3601 | 181 | R181A | A | 3.9e-06 | 1.7e-06 | 3.1e-06 | 2.900E-06 | 1.114E-06 | 1.700E-06 | 3.900E-06 |
| 3602 | 181 | R181R | A | 2e-06 | 3.5e-06 | 3.4e-06 | 2.967E-06 | 8.386E-07 | 2.000E-06 | 3.500E-06 |
| 3603 | 181 | R181N | A | 2.1e-06 | 2.3e-06 | 1.9e-06 | 2.100E-06 | 2.000E-07 | 1.900E-06 | 2.300E-06 |
| 3604 | 181 | R181D | A | 3.8e-06 | 1e-06 | 2.9e-06 | 2.567E-06 | 1.429E-06 | 1.000E-06 | 3.800E-06 |
| 3605 | 181 | R181C | A | 3.4e-06 | 1.6e-06 | 3.2e-06 | 2.733E-06 | 9.866E-07 | 1.600E-06 | 3.400E-06 |
| 3606 | 181 | R181E | A | 3.9e-06 | 2.6e-06 | 3.6e-06 | 3.367E-06 | 6.807E-07 | 2.600E-06 | 3.900E-06 |
| 3607 | 181 | R181Q | A | 2.5e-06 | 1.7e-06 | 2.2e-06 | 2.133E-06 | 4.041E-07 | 1.700E-06 | 2.500E-06 |
| 3608 | 181 | R181G | A | 2.7e-06 | 4.5e-06 | 1.1e-06 | 2.767E-06 | 1.701E-06 | 1.100E-06 | 4.500E-06 |
| 3609 | 181 | R181H | A | 3.9e-06 | 3.1e-06 | 2.2e-06 | 3.067E-06 | 8.505E-07 | 2.200E-06 | 3.900E-06 |
| 3610 | 181 | R181I | A | 2.3e-06 | 2.3e-06 | 3.6e-06 | 2.733E-06 | 7.506E-07 | 2.300E-06 | 3.600E-06 |
| 3611 | 181 | R181L | A | 1.3e-06 | 2.3e-06 | 3.9e-06 | 2.500E-06 | 1.311E-06 | 1.300E-06 | 3.900E-06 |
| 3612 | 181 | R181K | A | 3.4e-06 | 2.9e-06 | 2.3e-06 | 2.867E-06 | 5.508E-07 | 2.300E-06 | 3.400E-06 |
| 3613 | 181 | R181M | A | 2.2e-06 | 4.3e-06 | 2.5e-06 | 3.000E-06 | 1.136E-06 | 2.200E-06 | 4.300E-06 |
| 3614 | 181 | R181F | A | 4.8e-06 | 2.1e-06 | 1.6e-06 | 2.833E-06 | 1.721E-06 | 1.600E-06 | 4.800E-06 |
| 3615 | 181 | R181P | A | 1.7e-06 | 1.4e-06 | 1.8e-06 | 1.633E-06 | 2.082E-07 | 1.400E-06 | 1.800E-06 |
| 3616 | 181 | R181S | A | 2.1e-06 | 1.9e-06 | 2.1e-06 | 2.033E-06 | 1.155E-07 | 1.900E-06 | 2.100E-06 |
| 3617 | 181 | R181T | A | 2.1e-06 | 3e-06 | 2.6e-06 | 2.567E-06 | 4.509E-07 | 2.100E-06 | 3.000E-06 |
| 3618 | 181 | R181W | A | 1.8e-06 | 2.2e-06 | 2.5e-06 | 2.167E-06 | 3.512E-07 | 1.800E-06 | 2.500E-06 |
| 3619 | 181 | R181Y | A | 2e-06 | 3.2e-06 | 2.7e-06 | 2.633E-06 | 6.028E-07 | 2.000E-06 | 3.200E-06 |
| 3620 | 181 | R181V | A | 2.7e-06 | 3.4e-06 | 3.2e-06 | 3.100E-06 | 3.606E-07 | 2.700E-06 | 3.400E-06 |
| 3621 | 182 | C182A | A | 1.7e-06 | 3.4e-06 | 1.6e-06 | 2.233E-06 | 1.012E-06 | 1.600E-06 | 3.400E-06 |
| 3622 | 182 | C182R | A | 3.8e-06 | 2.8e-06 | 1.7e-06 | 2.767E-06 | 1.050E-06 | 1.700E-06 | 3.800E-06 |
| 3623 | 182 | C182N | A | 1.5e-06 | 3.1e-06 | 2.2e-06 | 2.267E-06 | 8.021E-07 | 1.500E-06 | 3.100E-06 |
| 3624 | 182 | C182D | A | 2.7e-06 | 3.4e-06 | 2.1e-06 | 2.733E-06 | 6.506E-07 | 2.100E-06 | 3.400E-06 |
| 3625 | 182 | C182C | A | 2.9e-06 | 3.8e-06 | 2e-06 | 2.900E-06 | 9.000E-07 | 2.000E-06 | 3.800E-06 |
| 3626 | 182 | C182E | A | 1.9e-06 | 4.5e-06 | 2.7e-06 | 3.033E-06 | 1.332E-06 | 1.900E-06 | 4.500E-06 |
| 3627 | 182 | C182Q | A | 1.3e-06 | 4.4e-06 | 1.7e-06 | 2.467E-06 | 1.686E-06 | 1.300E-06 | 4.400E-06 |
| 3628 | 182 | C182G | A | 1.4e-06 | 2.6e-06 | 2.2e-06 | 2.067E-06 | 6.110E-07 | 1.400E-06 | 2.600E-06 |
| 3629 | 182 | C182H | A | 4.2e-06 | 1.4e-06 | 2.3e-06 | 2.633E-06 | 1.429E-06 | 1.400E-06 | 4.200E-06 |
| 3630 | 182 | C182I | A | 4.8e-06 | 3.5e-06 | 2.5e-06 | 3.600E-06 | 1.153E-06 | 2.500E-06 | 4.800E-06 |
| 3631 | 182 | C182L | A | 1.6e-06 | 2.6e-06 | 2.7e-06 | 2.300E-06 | 6.083E-07 | 1.600E-06 | 2.700E-06 |
| 3632 | 182 | C182K | A | 1.9e-06 | 3e-06 | 1.6e-06 | 2.167E-06 | 7.371E-07 | 1.600E-06 | 3.000E-06 |
| 3633 | 182 | C182M | A | 1.9e-06 | 3.7e-06 | 2.7e-06 | 2.767E-06 | 9.018E-07 | 1.900E-06 | 3.700E-06 |
| 3634 | 182 | C182F | A | 2.3e-06 | 2.7e-06 | 2.7e-06 | 2.567E-06 | 2.309E-07 | 2.300E-06 | 2.700E-06 |
| 3635 | 182 | C182P | A | 1.5e-06 | 3.5e-06 | 2.1e-06 | 2.367E-06 | 1.026E-06 | 1.500E-06 | 3.500E-06 |
| 3636 | 182 | C182S | A | 1.6e-06 | 2.4e-06 | 2.7e-06 | 2.233E-06 | 5.686E-07 | 1.600E-06 | 2.700E-06 |
| 3637 | 182 | C182T | A | 1.7e-06 | 2e-06 | 2.3e-06 | 2.000E-06 | 3.000E-07 | 1.700E-06 | 2.300E-06 |
| 3638 | 182 | C182W | A | 2.5e-06 | 3e-06 | 2.5e-06 | 2.667E-06 | 2.887E-07 | 2.500E-06 | 3.000E-06 |
| 3639 | 182 | C182Y | A | 2.3e-06 | 2.3e-06 | 1.9e-06 | 2.167E-06 | 2.309E-07 | 1.900E-06 | 2.300E-06 |
| 3640 | 182 | C182V | A | 5.4e-06 | 2.3e-06 | 2.3e-06 | 3.333E-06 | 1.790E-06 | 2.300E-06 | 5.400E-06 |
| 3641 | 183 | W183A | A | 3.3e-06 | 3.3e-06 | 1.8e-06 | 2.800E-06 | 8.660E-07 | 1.800E-06 | 3.300E-06 |
| 3642 | 183 | W183R | A | 1.8e-06 | 2.2e-06 | 2.4e-06 | 2.133E-06 | 3.055E-07 | 1.800E-06 | 2.400E-06 |
| 3643 | 183 | W183N | A | 2.2e-06 | 3.1e-06 | 2.6e-06 | 2.633E-06 | 4.509E-07 | 2.200E-06 | 3.100E-06 |
| 3644 | 183 | W183D | A | 2.7e-06 | 2.6e-06 | 2.3e-06 | 2.533E-06 | 2.082E-07 | 2.300E-06 | 2.700E-06 |
| 3645 | 183 | W183C | A | 1.9e-06 | 2.4e-06 | 2.8e-06 | 2.367E-06 | 4.509E-07 | 1.900E-06 | 2.800E-06 |
| 3646 | 183 | W183E | A | 4.3e-06 | 2.3e-06 | 1.7e-06 | 2.767E-06 | 1.361E-06 | 1.700E-06 | 4.300E-06 |
| 3647 | 183 | W183Q | A | 1.8e-06 | 1.8e-06 | 1.5e-06 | 1.700E-06 | 1.732E-07 | 1.500E-06 | 1.800E-06 |
| 3648 | 183 | W183G | A | 2.9e-06 | 2.9e-06 | 4.8e-06 | 3.533E-06 | 1.097E-06 | 2.900E-06 | 4.800E-06 |
| 3649 | 183 | W183H | A | 4e-06 | 2.2e-06 | 1.8e-06 | 2.667E-06 | 1.172E-06 | 1.800E-06 | 4.000E-06 |
| 3650 | 183 | W183I | A | 2.2e-06 | 4.3e-06 | 1.4e-06 | 2.633E-06 | 1.498E-06 | 1.400E-06 | 4.300E-06 |
| 3651 | 183 | W183L | A | 2e-06 | 1.7e-06 | 1.9e-06 | 1.867E-06 | 1.528E-07 | 1.700E-06 | 2.000E-06 |
| 3652 | 183 | W183K | A | 1.8e-06 | 3e-06 | 3.1e-06 | 2.633E-06 | 7.234E-07 | 1.800E-06 | 3.100E-06 |
| 3653 | 183 | W183M | A | 2.5e-06 | 6.9e-06 | 2.7e-06 | 4.033E-06 | 2.485E-06 | 2.500E-06 | 6.900E-06 |
| 3654 | 183 | W183F | A | 2.8e-06 | 2.4e-06 | 3e-06 | 2.733E-06 | 3.055E-07 | 2.400E-06 | 3.000E-06 |

| | | | | | | | | | | |
|---|---|---|---|---|---|---|---|---|---|---|
| 3655 | 183 | W183P | A | 2.3e-06 | 2.2e-06 | 1.2e-06 | 1.900E-06 | 6.083E-07 | 1.200E-06 | 2.300E-06 |
| 3656 | 183 | W183S | A | 2.1e-06 | 2.2e-06 | 3.4e-06 | 2.567E-06 | 7.234E-07 | 2.100E-06 | 3.400E-06 |
| 3657 | 183 | W183T | A | 2.8e-06 | 3.1e-06 | 1.9e-06 | 2.600E-06 | 6.245E-07 | 1.900E-06 | 3.100E-06 |
| 3658 | 183 | W183W | A | 1.7e-06 | 5.4e-06 | 3.3e-06 | 3.467E-06 | 1.856E-06 | 1.700E-06 | 5.400E-06 |
| 3659 | 183 | W183Y | A | 2.2e-06 | 2.3e-06 | 4.3e-06 | 2.933E-06 | 1.185E-06 | 2.200E-06 | 4.300E-06 |
| 3660 | 183 | W183V | A | 3.3e-06 | 2.7e-06 | 2.3e-06 | 2.767E-06 | 5.033E-07 | 2.300E-06 | 3.300E-06 |
| 3661 | 184 | G184A | A | 2.4e-06 | 2.2e-06 | 1.5e-06 | 2.033E-06 | 4.726E-07 | 1.500E-06 | 2.400E-06 |
| 3662 | 184 | G184R | A | 2.6e-06 | 2.5e-06 | 2.1e-06 | 2.400E-06 | 2.646E-07 | 2.100E-06 | 2.600E-06 |
| 3663 | 184 | G184N | A | 3.2e-06 | 5e-06 | 2.8e-06 | 3.667E-06 | 1.172E-06 | 2.800E-06 | 5.000E-06 |
| 3664 | 184 | G184D | A | 2.1e-06 | 2.1e-06 | 3.5e-06 | 2.567E-06 | 8.083E-07 | 2.100E-06 | 3.500E-06 |
| 3665 | 184 | G184C | A | 4.8e-06 | 2.9e-06 | 1.9e-06 | 3.200E-06 | 1.473E-06 | 1.900E-06 | 4.800E-06 |
| 3666 | 184 | G184E | A | 1.9e-06 | 2.2e-06 | 3.7e-06 | 2.600E-06 | 9.644E-07 | 1.900E-06 | 3.700E-06 |
| 3667 | 184 | G184Q | A | 2.1e-06 | 2.5e-06 | 2.5e-06 | 2.367E-06 | 2.309E-07 | 2.100E-06 | 2.500E-06 |
| 3668 | 184 | G184G | A | 9.9e-07 | 3.4e-06 | 2.8e-06 | 2.397E-06 | 1.255E-06 | 9.900E-07 | 3.400E-06 |
| 3669 | 184 | G184H | A | 1.8e-06 | 3.4e-06 | 1.8e-06 | 2.333E-06 | 9.238E-07 | 1.800E-06 | 3.400E-06 |
| 3670 | 184 | G184I | A | 2e-06 | 1.9e-06 | 4.8e-06 | 2.900E-06 | 1.646E-06 | 1.900E-06 | 4.800E-06 |
| 3671 | 184 | G184L | A | 2.5e-06 | 2.4e-06 | 3.2e-06 | 2.700E-06 | 4.359E-07 | 2.400E-06 | 3.200E-06 |
| 3672 | 184 | G184K | A | 3.1e-06 | 3.4e-06 | 4.6e-06 | 3.700E-06 | 7.937E-07 | 3.100E-06 | 4.600E-06 |
| 3673 | 184 | G184M | A | 4e-06 | 3.6e-06 | 1.9e-06 | 3.167E-06 | 1.115E-06 | 1.900E-06 | 4.000E-06 |
| 3674 | 184 | G184F | A | 2.3e-06 | 3e-06 | 2e-06 | 2.433E-06 | 5.132E-07 | 2.000E-06 | 3.000E-06 |
| 3675 | 184 | G184P | A | 2.5e-06 | 2.7e-06 | 2.3e-06 | 2.500E-06 | 2.000E-07 | 2.300E-06 | 2.700E-06 |
| 3676 | 184 | G184S | A | 1.8e-06 | 2.2e-06 | 2e-06 | 2.000E-06 | 2.000E-07 | 1.800E-06 | 2.200E-06 |
| 3677 | 184 | G184T | A | 2.8e-06 | 1.8e-06 | 2.3e-06 | 2.300E-06 | 5.000E-07 | 1.800E-06 | 2.800E-06 |
| 3678 | 184 | G184W | A | 3.4e-06 | 2.3e-06 | 3e-06 | 2.900E-06 | 5.568E-07 | 2.300E-06 | 3.400E-06 |
| 3679 | 184 | G184Y | A | 2.6e-06 | 3.1e-06 | 2.5e-06 | 2.733E-06 | 3.215E-07 | 2.500E-06 | 3.100E-06 |
| 3680 | 184 | G184V | A | 4.2e-06 | 2.2e-06 | 2.2e-06 | 2.867E-06 | 1.155E-06 | 2.200E-06 | 4.200E-06 |
| 3681 | 185 | E185A | A | 2.4e-06 | 2.8e-06 | 3.5e-06 | 2.900E-06 | 5.568E-07 | 2.400E-06 | 3.500E-06 |
| 3682 | 185 | E185R | A | 1.5e-06 | 4.9e-06 | 2.5e-06 | 2.967E-06 | 1.747E-06 | 1.500E-06 | 4.900E-06 |
| 3683 | 185 | E185N | A | 9.9e-07 | 3e-06 | 2.2e-06 | 2.063E-06 | 1.012E-06 | 9.900E-07 | 3.000E-06 |
| 3684 | 185 | E185D | A | 1.9e-06 | 2.4e-06 | 2.6e-06 | 2.300E-06 | 3.606E-07 | 1.900E-06 | 2.600E-06 |
| 3685 | 185 | E185C | A | 1.6e-06 | 2.5e-06 | 2.5e-06 | 2.200E-06 | 5.196E-07 | 1.600E-06 | 2.500E-06 |
| 3686 | 185 | E185E | A | 2.6e-06 | 1.9e-06 | 3.1e-06 | 2.533E-06 | 6.028E-07 | 1.900E-06 | 3.100E-06 |
| 3687 | 185 | E185Q | A | 2.9e-06 | 2.9e-06 | 2.2e-06 | 2.667E-06 | 4.041E-07 | 2.200E-06 | 2.900E-06 |
| 3688 | 185 | E185G | A | 2.3e-06 | 2.1e-06 | 4e-06 | 2.800E-06 | 1.044E-06 | 2.100E-06 | 4.000E-06 |
| 3689 | 185 | E185H | A | 1.7e-06 | 3.3e-06 | 2.7e-06 | 2.567E-06 | 8.083E-07 | 1.700E-06 | 3.300E-06 |
| 3690 | 185 | E185I | A | 2.9e-06 | 1.5e-06 | 2.4e-06 | 2.267E-06 | 7.095E-07 | 1.500E-06 | 2.900E-06 |
| 3691 | 185 | E185L | A | 3.2e-06 | 2.5e-06 | 2.2e-06 | 2.633E-06 | 5.132E-07 | 2.200E-06 | 3.200E-06 |
| 3692 | 185 | E185K | A | 2.4e-06 | 1.6e-06 | 2.1e-06 | 2.033E-06 | 4.041E-07 | 1.600E-06 | 2.400E-06 |
| 3693 | 185 | E185M | A | 5.1e-06 | 2.1e-06 | 2.6e-06 | 3.267E-06 | 1.607E-06 | 2.100E-06 | 5.100E-06 |
| 3694 | 185 | E185F | A | 1.9e-06 | 2.3e-06 | 1.7e-06 | 1.967E-06 | 3.055E-07 | 1.700E-06 | 2.300E-06 |
| 3695 | 185 | E185P | A | 2.7e-06 | 1.8e-06 | 3.2e-06 | 2.567E-06 | 7.095E-07 | 1.800E-06 | 3.200E-06 |
| 3696 | 185 | E185S | A | 2.6e-06 | 2e-06 | 1.8e-06 | 2.133E-06 | 4.163E-07 | 1.800E-06 | 2.600E-06 |
| 3697 | 185 | E185T | A | 2e-06 | 1.7e-06 | 1.7e-06 | 1.800E-06 | 1.732E-07 | 1.700E-06 | 2.000E-06 |
| 3698 | 185 | E185W | A | 2e-06 | 2.3e-06 | 1.3e-06 | 1.867E-06 | 5.132E-07 | 1.300E-06 | 2.300E-06 |
| 3699 | 185 | E185Y | A | 3.8e-06 | 3.6e-06 | 1.6e-06 | 3.000E-06 | 1.217E-06 | 1.600E-06 | 3.800E-06 |
| 3700 | 185 | E185V | A | 3.2e-06 | 2.7e-06 | 2.7e-06 | 2.867E-06 | 2.887E-07 | 2.700E-06 | 3.200E-06 |
| 3701 | 186 | S186A | A | 3.1e-06 | 1.6e-06 | 1.4e-06 | 2.033E-06 | 9.292E-07 | 1.400E-06 | 3.100E-06 |
| 3702 | 186 | S186R | A | 1.5e-06 | 3.1e-06 | 2.2e-06 | 2.267E-06 | 8.021E-07 | 1.500E-06 | 3.100E-06 |
| 3703 | 186 | S186N | A | 3e-06 | 4.5e-06 | 3.8e-06 | 3.767E-06 | 7.506E-07 | 3.000E-06 | 4.500E-06 |
| 3704 | 186 | S186D | A | 2.7e-06 | 2e-06 | 2.3e-06 | 2.333E-06 | 3.512E-07 | 2.000E-06 | 2.700E-06 |
| 3705 | 186 | S186C | A | 2.2e-06 | 1.8e-06 | 2.6e-06 | 2.200E-06 | 4.000E-07 | 1.800E-06 | 2.600E-06 |
| 3706 | 186 | S186E | A | 1.7e-06 | 2.7e-06 | 2.7e-06 | 2.367E-06 | 5.774E-07 | 1.700E-06 | 2.700E-06 |
| 3707 | 186 | S186Q | A | 1.7e-06 | 3e-06 | 2.6e-06 | 2.433E-06 | 6.658E-07 | 1.700E-06 | 3.000E-06 |
| 3708 | 186 | S186G | A | 4.9e-06 | 2.3e-06 | 3.6e-06 | 3.600E-06 | 1.300E-06 | 2.300E-06 | 4.900E-06 |
| 3709 | 186 | S186H | A | 2.4e-06 | 2.6e-06 | 2.5e-06 | 2.500E-06 | 1.000E-07 | 2.400E-06 | 2.600E-06 |
| 3710 | 186 | S186I | A | 1.9e-06 | 2.6e-06 | 2.9e-06 | 2.467E-06 | 5.132E-07 | 1.900E-06 | 2.900E-06 |
| 3711 | 186 | S186L | A | 2.4e-06 | 2.9e-06 | 2.4e-06 | 2.567E-06 | 2.887E-07 | 2.400E-06 | 2.900E-06 |
| 3712 | 186 | S186K | A | 2.7e-06 | 1.9e-06 | 3.1e-06 | 2.567E-06 | 6.110E-07 | 1.900E-06 | 3.100E-06 |
| 3713 | 186 | S186M | A | 2.3e-06 | 3e-06 | 3.8e-06 | 3.033E-06 | 7.506E-07 | 2.300E-06 | 3.800E-06 |
| 3714 | 186 | S186F | A | 3.5e-06 | 2.7e-06 | 2.3e-06 | 2.833E-06 | 6.110E-07 | 2.300E-06 | 3.500E-06 |
| 3715 | 186 | S186P | A | 1.1e-06 | 3.3e-06 | 2.2e-06 | 2.200E-06 | 1.100E-06 | 1.100E-06 | 3.300E-06 |
| 3716 | 186 | S186S | A | 2.3e-06 | 2.2e-06 | 1.9e-06 | 2.133E-06 | 2.082E-07 | 1.900E-06 | 2.300E-06 |
| 3717 | 186 | S186T | A | 2.1e-06 | 1.7e-06 | 1.6e-06 | 1.800E-06 | 2.646E-07 | 1.600E-06 | 2.100E-06 |
| 3718 | 186 | S186W | A | 3.4e-06 | 1.5e-06 | 1.5e-06 | 2.133E-06 | 1.097E-06 | 1.500E-06 | 3.400E-06 |
| 3719 | 186 | S186Y | A | 1.9e-06 | 2.4e-06 | 2.8e-06 | 2.367E-06 | 4.509E-07 | 1.900E-06 | 2.800E-06 |
| 3720 | 186 | S186V | A | 3.4e-06 | 1.9e-06 | 1.8e-06 | 2.367E-06 | 8.963E-07 | 1.800E-06 | 3.400E-06 |
| 3721 | 187 | S187A | A | 2e-06 | 2.4e-06 | 2.3e-06 | 2.233E-06 | 2.082E-07 | 2.000E-06 | 2.400E-06 |
| 3722 | 187 | S187R | A | 3.8e-06 | 2.9e-06 | 1.4e-06 | 2.700E-06 | 1.212E-06 | 1.400E-06 | 3.800E-06 |
| 3723 | 187 | S187N | A | 2.1e-06 | 2.1e-06 | 3.9e-06 | 2.700E-06 | 1.039E-06 | 2.100E-06 | 3.900E-06 |
| 3724 | 187 | S187D | A | 3.1e-06 | 1.7e-06 | 1.5e-06 | 2.100E-06 | 8.718E-07 | 1.500E-06 | 3.100E-06 |
| 3725 | 187 | S187C | A | 1.8e-06 | 4.4e-06 | 2.9e-06 | 3.033E-06 | 1.305E-06 | 1.800E-06 | 4.400E-06 |
| 3726 | 187 | S187E | A | 1.5e-06 | 1.6e-06 | 2.4e-06 | 1.833E-06 | 4.933E-07 | 1.500E-06 | 2.400E-06 |
| 3727 | 187 | S187Q | A | 2.2e-06 | 2.5e-06 | 2.2e-06 | 2.300E-06 | 1.732E-07 | 2.200E-06 | 2.500E-06 |
| 3728 | 187 | S187G | A | 3.5e-06 | 2.4e-06 | 3.2e-06 | 3.033E-06 | 5.686E-07 | 2.400E-06 | 3.500E-06 |
| 3729 | 187 | S187H | A | 2.8e-06 | 2.3e-06 | 2.2e-06 | 2.433E-06 | 3.215E-07 | 2.200E-06 | 2.800E-06 |
| 3730 | 187 | S187I | A | 3e-06 | 1.9e-06 | 5.4e-06 | 3.433E-06 | 1.790E-06 | 1.900E-06 | 5.400E-06 |
| 3731 | 187 | S187L | A | 2.2e-06 | 2.6e-06 | 2e-06 | 2.267E-06 | 3.055E-07 | 2.000E-06 | 2.600E-06 |
| 3732 | 187 | S187K | A | 1.5e-06 | 3.6e-06 | 2.8e-06 | 2.633E-06 | 1.060E-06 | 1.500E-06 | 3.600E-06 |
| 3733 | 187 | S187M | A | 1.9e-06 | 2.9e-06 | 3.3e-06 | 2.700E-06 | 7.211E-07 | 1.900E-06 | 3.300E-06 |
| 3734 | 187 | S187F | A | 2.6e-06 | 3.3e-06 | 2.8e-06 | 2.900E-06 | 3.606E-07 | 2.600E-06 | 3.300E-06 |
| 3735 | 187 | S187P | A | 2.7e-06 | 3e-06 | 2.2e-06 | 2.633E-06 | 4.041E-07 | 2.200E-06 | 3.000E-06 |
| 3736 | 187 | S187S | A | 2.1e-06 | 2.5e-06 | 2e-06 | 2.200E-06 | 2.646E-07 | 2.000E-06 | 2.500E-06 |
| 3737 | 187 | S187T | A | 2.3e-06 | 1.5e-06 | 3.2e-06 | 2.333E-06 | 8.505E-07 | 1.500E-06 | 3.200E-06 |
| 3738 | 187 | S187W | A | 2.1e-06 | 2.7e-06 | 1.4e-06 | 2.067E-06 | 6.506E-07 | 1.400E-06 | 2.700E-06 |
| 3739 | 187 | S187Y | A | 3e-06 | 3.1e-06 | 9.9e-07 | 2.363E-06 | 1.190E-06 | 9.900E-07 | 3.100E-06 |

| | | | | | | | | | | |
|---|---|---|---|---|---|---|---|---|---|---|
| 3740 | 187 | S187V | A | 1.4e-06 | 1.4e-06 | 1.5e-06 | 1.433E-06 | 5.774E-08 | 1.400E-06 | 1.500E-06 |
| 3741 | 188 | E188A | A | 4e-06 | 1.9e-06 | 3e-06 | 2.967E-06 | 1.050E-06 | 1.900E-06 | 4.000E-06 |
| 3742 | 188 | E188R | A | 2.4e-06 | 1.4e-06 | 1.4e-06 | 1.733E-06 | 5.774E-07 | 1.400E-06 | 2.400E-06 |
| 3743 | 188 | E188N | A | 3e-06 | 9.6e-07 | 2.6e-06 | 2.187E-06 | 1.081E-06 | 9.600E-07 | 3.000E-06 |
| 3744 | 188 | E188D | A | 2.5e-06 | 2.7e-06 | 2.8e-06 | 2.667E-06 | 1.528E-07 | 2.500E-06 | 2.800E-06 |
| 3745 | 188 | E188C | A | 2.8e-06 | 2.4e-06 | 1.8e-06 | 2.333E-06 | 5.033E-07 | 1.800E-06 | 2.800E-06 |
| 3746 | 188 | E188E | A | 2.6e-06 | 1.7e-06 | 1.9e-06 | 2.067E-06 | 4.726E-07 | 1.700E-06 | 2.600E-06 |
| 3747 | 188 | E188Q | A | 1.7e-06 | 3.2e-06 | 2.1e-06 | 2.333E-06 | 7.767E-07 | 1.700E-06 | 3.200E-06 |
| 3748 | 188 | E188G | A | 1.3e-06 | 4e-06 | 1.9e-06 | 2.400E-06 | 1.418E-06 | 1.300E-06 | 4.000E-06 |
| 3749 | 188 | E188H | A | 2.3e-06 | 1.4e-06 | 2.2e-06 | 1.967E-06 | 4.933E-07 | 1.400E-06 | 2.300E-06 |
| 3750 | 188 | E188I | A | 3.4e-06 | 2.1e-06 | 2e-06 | 2.500E-06 | 7.810E-07 | 2.000E-06 | 3.400E-06 |
| 3751 | 188 | E188L | A | 2.8e-06 | 3.2e-06 | 4.3e-06 | 3.433E-06 | 7.767E-07 | 2.800E-06 | 4.300E-06 |
| 3752 | 188 | E188K | A | 4.3e-06 | 2.1e-06 | 1.8e-06 | 2.733E-06 | 1.365E-06 | 1.800E-06 | 4.300E-06 |
| 3753 | 188 | E188M | A | 2e-06 | 3.4e-06 | 3.1e-06 | 2.833E-06 | 7.371E-07 | 2.000E-06 | 3.400E-06 |
| 3754 | 188 | E188F | A | 1.8e-06 | 3.7e-06 | 1.9e-06 | 2.467E-06 | 1.069E-06 | 1.800E-06 | 3.700E-06 |
| 3755 | 188 | E188P | A | 3.9e-06 | 2.4e-06 | 2.7e-06 | 3.000E-06 | 7.937E-07 | 2.400E-06 | 3.900E-06 |
| 3756 | 188 | E188S | A | 1.5e-06 | 3.4e-06 | 1.4e-06 | 2.100E-06 | 1.127E-06 | 1.400E-06 | 3.400E-06 |
| 3757 | 188 | E188T | A | 2e-06 | 8.6e-07 | 1.8e-06 | 1.553E-06 | 6.087E-07 | 8.600E-07 | 2.000E-06 |
| 3758 | 188 | E188W | A | 2.5e-06 | 5.1e-06 | 2.1e-06 | 3.233E-06 | 1.629E-06 | 2.100E-06 | 5.100E-06 |
| 3759 | 188 | E188Y | A | 1.6e-06 | 2.3e-06 | 2.7e-06 | 2.200E-06 | 5.568E-07 | 1.600E-06 | 2.700E-06 |
| 3760 | 188 | E188V | A | 1.3e-06 | 1.1e-06 | 3.2e-06 | 1.867E-06 | 1.159E-06 | 1.100E-06 | 3.200E-06 |
| 3761 | 189 | D189A | A | 1.1e-06 | 2.3e-06 | 2.1e-06 | 1.833E-06 | 6.429E-07 | 1.100E-06 | 2.300E-06 |
| 3762 | 189 | D189R | A | 1.9e-06 | 1.9e-06 | 2e-06 | 1.933E-06 | 5.774E-08 | 1.900E-06 | 2.000E-06 |
| 3763 | 189 | D189N | A | 2.8e-06 | 2.2e-06 | 2.6e-06 | 2.533E-06 | 3.055E-07 | 2.200E-06 | 2.800E-06 |
| 3764 | 189 | D189D | A | 4.7e-06 | 2.4e-06 | 2.3e-06 | 3.133E-06 | 1.358E-06 | 2.300E-06 | 4.700E-06 |
| 3765 | 189 | D189C | A | 2.3e-06 | 2.4e-06 | 1.6e-06 | 2.100E-06 | 4.359E-07 | 1.600E-06 | 2.400E-06 |
| 3766 | 189 | D189E | A | 2.2e-06 | 2e-06 | 8.7e-06 | 4.300E-06 | 3.812E-06 | 2.000E-06 | 8.700E-06 |
| 3767 | 189 | D189Q | A | 2.1e-06 | 1.6e-06 | 3.7e-06 | 2.467E-06 | 1.097E-06 | 1.600E-06 | 3.700E-06 |
| 3768 | 189 | D189G | A | 4.9e-06 | 3.6e-06 | 3.8e-06 | 4.100E-06 | 7.000E-07 | 3.600E-06 | 4.900E-06 |
| 3769 | 189 | D189H | A | 2.4e-06 | 5.1e-06 | 1.2e-06 | 2.900E-06 | 1.997E-06 | 1.200E-06 | 5.100E-06 |
| 3770 | 189 | D189I | A | 3.5e-06 | 3.8e-06 | 2e-06 | 3.100E-06 | 9.644E-07 | 2.000E-06 | 3.800E-06 |
| 3771 | 189 | D189L | A | 3.3e-06 | 3.2e-06 | 2.8e-06 | 3.100E-06 | 2.646E-07 | 2.800E-06 | 3.300E-06 |
| 3772 | 189 | D189K | A | 3.7e-06 | 2e-06 | 2.7e-06 | 2.800E-06 | 8.544E-07 | 2.000E-06 | 3.700E-06 |
| 3773 | 189 | D189M | A | 2e-06 | 2.4e-06 | 3.1e-06 | 2.500E-06 | 5.568E-07 | 2.000E-06 | 3.100E-06 |
| 3774 | 189 | D189F | A | 2.4e-06 | 2.8e-06 | 2.3e-06 | 2.500E-06 | 2.646E-07 | 2.300E-06 | 2.800E-06 |
| 3775 | 189 | D189P | A | 2.8e-06 | 2.5e-06 | 2.9e-06 | 2.733E-06 | 2.082E-07 | 2.500E-06 | 2.900E-06 |
| 3776 | 189 | D189S | A | 2.7e-06 | 1.9e-06 | 2.6e-06 | 2.400E-06 | 4.359E-07 | 1.900E-06 | 2.700E-06 |
| 3777 | 189 | D189T | A | 1.6e-06 | 1.5e-06 | 1.3e-06 | 1.467E-06 | 1.528E-07 | 1.300E-06 | 1.600E-06 |
| 3778 | 189 | D189W | A | 2.6e-06 | 2.7e-06 | 1.7e-06 | 2.333E-06 | 5.508E-07 | 1.700E-06 | 2.700E-06 |
| 3779 | 189 | D189Y | A | 1.6e-06 | 3e-06 | 2.6e-06 | 2.400E-06 | 7.211E-07 | 1.600E-06 | 3.000E-06 |
| 3780 | 189 | D189V | A | 1.7e-06 | 3.3e-06 | 3.4e-06 | 2.800E-06 | 9.539E-07 | 1.700E-06 | 3.400E-06 |
| 3781 | 190 | C190A | A | 3.5e-06 | 3.3e-06 | 4.2e-06 | 3.667E-06 | 4.726E-07 | 3.300E-06 | 4.200E-06 |
| 3782 | 190 | C190R | A | 3.3e-06 | 1.7e-06 | 2.6e-06 | 2.533E-06 | 8.021E-07 | 1.700E-06 | 3.300E-06 |
| 3783 | 190 | C190N | A | 2.3e-06 | 2.7e-06 | 3.2e-06 | 2.733E-06 | 4.509E-07 | 2.300E-06 | 3.200E-06 |
| 3784 | 190 | C190D | A | 2.1e-06 | 2e-06 | 2.8e-06 | 2.300E-06 | 4.359E-07 | 2.000E-06 | 2.800E-06 |
| 3785 | 190 | C190C | A | 2.5e-06 | 2.4e-06 | 3.7e-06 | 2.867E-06 | 7.234E-07 | 2.400E-06 | 3.700E-06 |
| 3786 | 190 | C190E | A | 2.4e-06 | 2.6e-06 | 3.2e-06 | 2.733E-06 | 4.163E-07 | 2.400E-06 | 3.200E-06 |
| 3787 | 190 | C190Q | A | 1.2e-06 | 2.5e-06 | 3.8e-06 | 2.500E-06 | 1.300E-06 | 1.200E-06 | 3.800E-06 |
| 3788 | 190 | C190G | A | 3.4e-06 | 2.4e-06 | 2e-06 | 2.600E-06 | 7.211E-07 | 2.000E-06 | 3.400E-06 |
| 3789 | 190 | C190H | A | 1.8e-06 | 2.3e-06 | 1.7e-06 | 1.933E-06 | 3.215E-07 | 1.700E-06 | 2.300E-06 |
| 3790 | 190 | C190I | A | 3.2e-06 | 2.6e-06 | 3.8e-06 | 3.200E-06 | 6.000E-07 | 2.600E-06 | 3.800E-06 |
| 3791 | 190 | C190L | A | 1.6e-06 | 2.4e-06 | 4.7e-06 | 2.900E-06 | 1.609E-06 | 1.600E-06 | 4.700E-06 |
| 3792 | 190 | C190K | A | 1.7e-06 | 1.6e-06 | 2.5e-06 | 1.933E-06 | 4.933E-07 | 1.600E-06 | 2.500E-06 |
| 3793 | 190 | C190M | A | 2.7e-06 | 2.5e-06 | 2.5e-06 | 2.567E-06 | 1.155E-07 | 2.500E-06 | 2.700E-06 |
| 3794 | 190 | C190F | A | 3.3e-06 | 4.3e-06 | 2.8e-06 | 3.467E-06 | 7.638E-07 | 2.800E-06 | 4.300E-06 |
| 3795 | 190 | C190P | A | 2.1e-06 | 2.4e-06 | 2.8e-06 | 2.433E-06 | 3.512E-07 | 2.100E-06 | 2.800E-06 |
| 3796 | 190 | C190S | A | 3.6e-06 | 1.5e-06 | 2e-06 | 2.367E-06 | 1.097E-06 | 1.500E-06 | 3.600E-06 |
| 3797 | 190 | C190T | A | 3.1e-06 | 1.8e-06 | 1.7e-06 | 2.200E-06 | 7.810E-07 | 1.700E-06 | 3.100E-06 |
| 3798 | 190 | C190W | A | 3e-06 | 1.8e-06 | 1.2e-06 | 2.000E-06 | 9.165E-07 | 1.200E-06 | 3.000E-06 |
| 3799 | 190 | C190Y | A | 2.4e-06 | 2.5e-06 | 2.2e-06 | 2.367E-06 | 1.528E-07 | 2.200E-06 | 2.500E-06 |
| 3800 | 190 | C190V | A | 2.4e-06 | 2.2e-06 | 3.1e-06 | 2.567E-06 | 4.726E-07 | 2.200E-06 | 3.100E-06 |
| 3801 | 191 | Q191A | A | 2.4e-06 | 3.2e-06 | 2.2e-06 | 2.600E-06 | 5.292E-07 | 2.200E-06 | 3.200E-06 |
| 3802 | 191 | Q191R | A | 2.2e-06 | 4.9e-06 | 1.6e-06 | 2.900E-06 | 1.758E-06 | 1.600E-06 | 4.900E-06 |
| 3803 | 191 | Q191N | A | 1.5e-06 | 1.1e-06 | 2.7e-06 | 1.767E-06 | 8.327E-07 | 1.100E-06 | 2.700E-06 |
| 3804 | 191 | Q191D | A | 3.6e-06 | 1.5e-06 | 8.2e-07 | 1.973E-06 | 1.449E-06 | 8.200E-07 | 3.600E-06 |
| 3805 | 191 | Q191C | A | 4.2e-06 | 2.2e-06 | 4e-06 | 3.467E-06 | 1.102E-06 | 2.200E-06 | 4.200E-06 |
| 3806 | 191 | Q191E | A | 2.9e-06 | 4.4e-06 | 2.4e-06 | 3.233E-06 | 1.041E-06 | 2.400E-06 | 4.400E-06 |
| 3807 | 191 | Q191Q | A | 2e-06 | 2.1e-06 | 3.1e-06 | 2.400E-06 | 6.083E-07 | 2.000E-06 | 3.100E-06 |
| 3808 | 191 | Q191G | A | 2.4e-06 | 2.3e-06 | 2.3e-06 | 2.333E-06 | 5.774E-08 | 2.300E-06 | 2.400E-06 |
| 3809 | 191 | Q191H | A | 3.7e-06 | 3.9e-06 | 1.4e-06 | 3.000E-06 | 1.389E-06 | 1.400E-06 | 3.900E-06 |
| 3810 | 191 | Q191I | A | 4.3e-06 | 2.3e-06 | 2.1e-06 | 2.900E-06 | 1.217E-06 | 2.100E-06 | 4.300E-06 |
| 3811 | 191 | Q191L | A | 2.4e-06 | 4e-06 | 2.5e-06 | 2.967E-06 | 8.963E-07 | 2.400E-06 | 4.000E-06 |
| 3812 | 191 | Q191K | A | 2.5e-06 | 2.1e-06 | 4.3e-06 | 2.967E-06 | 1.172E-06 | 2.100E-06 | 4.300E-06 |
| 3813 | 191 | Q191M | A | 2.1e-06 | 3.3e-06 | 3.8e-06 | 3.067E-06 | 8.737E-07 | 2.100E-06 | 3.800E-06 |
| 3814 | 191 | Q191F | A | 2.5e-06 | 4.3e-06 | 2.4e-06 | 3.067E-06 | 1.069E-06 | 2.400E-06 | 4.300E-06 |
| 3815 | 191 | Q191P | A | 2.4e-06 | 3.1e-06 | 1.7e-06 | 2.400E-06 | 7.000E-07 | 1.700E-06 | 3.100E-06 |
| 3816 | 191 | Q191S | A | 1.9e-06 | 1.7e-06 | 3.7e-06 | 2.433E-06 | 1.102E-06 | 1.700E-06 | 3.700E-06 |
| 3817 | 191 | Q191T | A | 2.8e-06 | 2.2e-06 | 1.4e-06 | 2.133E-06 | 7.024E-07 | 1.400E-06 | 2.800E-06 |
| 3818 | 191 | Q191W | A | 2.7e-06 | 1.9e-06 | 2.9e-06 | 2.500E-06 | 5.292E-07 | 1.900E-06 | 2.900E-06 |
| 3819 | 191 | Q191Y | A | 3.5e-06 | 1.8e-06 | 2.4e-06 | 2.567E-06 | 8.622E-07 | 1.800E-06 | 3.500E-06 |
| 3820 | 191 | Q191V | A | 2.7e-06 | 3.1e-06 | 2.1e-06 | 2.633E-06 | 5.033E-07 | 2.100E-06 | 3.100E-06 |
| 3821 | 192 | S192A | A | 4.1e-06 | 2.3e-06 | 2.4e-06 | 2.933E-06 | 1.012E-06 | 2.300E-06 | 4.100E-06 |
| 3822 | 192 | S192R | A | 2.5e-06 | 2.6e-06 | 2.8e-06 | 2.633E-06 | 1.528E-07 | 2.500E-06 | 2.800E-06 |
| 3823 | 192 | S192N | A | 2.3e-06 | 2.5e-06 | 2.3e-06 | 2.367E-06 | 1.155E-07 | 2.300E-06 | 2.500E-06 |
| 3824 | 192 | S192D | A | 2.8e-06 | 2.5e-06 | 9.5e-07 | 2.083E-06 | 9.929E-07 | 9.500E-07 | 2.800E-06 |

| | | | | | | | | | | |
|---|---|---|---|---|---|---|---|---|---|---|
| 3825 | 192 | S192C | A | 2.9e-06 | 3.4e-06 | 2.3e-06 | 2.867E-06 | 5.508E-07 | 2.300E-06 | 3.400E-06 |
| 3826 | 192 | S192E | A | 1.4e-06 | 3.2e-06 | 3e-06 | 2.533E-06 | 9.866E-07 | 1.400E-06 | 3.200E-06 |
| 3827 | 192 | S192Q | A | 2.2e-06 | 2.1e-06 | 1.8e-06 | 2.033E-06 | 2.082E-07 | 1.800E-06 | 2.200E-06 |
| 3828 | 192 | S192G | A | 1.1e-06 | 1.6e-06 | 2.5e-06 | 1.733E-06 | 7.095E-07 | 1.100E-06 | 2.500E-06 |
| 3829 | 192 | S192H | A | 2.7e-06 | 2.2e-06 | 2.4e-06 | 2.433E-06 | 2.517E-07 | 2.200E-06 | 2.700E-06 |
| 3830 | 192 | S192I | A | 1.9e-06 | 3.7e-06 | 2.6e-06 | 2.733E-06 | 9.074E-07 | 1.900E-06 | 3.700E-06 |
| 3831 | 192 | S192L | A | 2.7e-06 | 2.2e-06 | 2.2e-06 | 2.367E-06 | 2.887E-07 | 2.200E-06 | 2.700E-06 |
| 3832 | 192 | S192K | A | 1e-06 | 2.1e-06 | 3.9e-06 | 2.333E-06 | 1.464E-06 | 1.000E-06 | 3.900E-06 |
| 3833 | 192 | S192M | A | 1e-06 | 2.9e-06 | 2.4e-06 | 2.100E-06 | 9.849E-07 | 1.000E-06 | 2.900E-06 |
| 3834 | 192 | S192F | A | 2.9e-06 | 2.8e-06 | 2.8e-06 | 2.833E-06 | 5.774E-08 | 2.800E-06 | 2.900E-06 |
| 3835 | 192 | S192P | A | 2.7e-06 | 2.2e-06 | 1.4e-06 | 2.100E-06 | 6.557E-07 | 1.400E-06 | 2.700E-06 |
| 3836 | 192 | S192S | A | 2.4e-06 | 2.6e-06 | 1.8e-06 | 2.267E-06 | 4.163E-07 | 1.800E-06 | 2.600E-06 |
| 3837 | 192 | S192T | A | 1.5e-06 | 2.6e-06 | 1.9e-06 | 2.000E-06 | 5.568E-07 | 1.500E-06 | 2.600E-06 |
| 3838 | 192 | S192W | A | 3.2e-06 | 2.1e-06 | 2.1e-06 | 2.467E-06 | 6.351E-07 | 2.100E-06 | 3.200E-06 |
| 3839 | 192 | S192Y | A | 1.3e-06 | 2.2e-06 | 3.3e-06 | 2.267E-06 | 1.002E-06 | 1.300E-06 | 3.300E-06 |
| 3840 | 192 | S192V | A | 3e-06 | 2.2e-06 | 3.7e-06 | 2.967E-06 | 7.506E-07 | 2.200E-06 | 3.700E-06 |
| 3841 | 193 | L193A | A | 2.6e-06 | 3.1e-06 | 2.3e-06 | 2.667E-06 | 4.041E-07 | 2.300E-06 | 3.100E-06 |
| 3842 | 193 | L193R | A | 2e-06 | 2e-06 | 2.6e-06 | 2.200E-06 | 3.464E-07 | 2.000E-06 | 2.600E-06 |
| 3843 | 193 | L193N | A | 2e-06 | 2.4e-06 | 1.1e-06 | 1.833E-06 | 6.658E-07 | 1.100E-06 | 2.400E-06 |
| 3844 | 193 | L193D | A | 2.2e-06 | 1.7e-06 | 2.5e-06 | 2.133E-06 | 4.041E-07 | 1.700E-06 | 2.500E-06 |
| 3845 | 193 | L193C | A | 2.5e-06 | 2.6e-06 | 3.5e-06 | 2.867E-06 | 5.508E-07 | 2.500E-06 | 3.500E-06 |
| 3846 | 193 | L193E | A | 2.2e-06 | 2.5e-06 | 2.2e-06 | 2.300E-06 | 1.732E-07 | 2.200E-06 | 2.500E-06 |
| 3847 | 193 | L193Q | A | 2.2e-06 | 1.4e-06 | 2.8e-06 | 2.133E-06 | 7.024E-07 | 1.400E-06 | 2.800E-06 |
| 3848 | 193 | L193G | A | 3.3e-06 | 2.2e-06 | 2.3e-06 | 2.600E-06 | 6.083E-07 | 2.200E-06 | 3.300E-06 |
| 3849 | 193 | L193H | A | 2.3e-06 | 3.2e-06 | 2.2e-06 | 2.567E-06 | 5.508E-07 | 2.200E-06 | 3.200E-06 |
| 3850 | 193 | L193I | A | 2e-06 | 1.6e-06 | 1.9e-06 | 1.833E-06 | 2.082E-07 | 1.600E-06 | 2.000E-06 |
| 3851 | 193 | L193L | A | 2.3e-06 | 2.2e-06 | 2.4e-06 | 2.300E-06 | 1.000E-07 | 2.200E-06 | 2.400E-06 |
| 3852 | 193 | L193K | A | 1.4e-06 | 1.9e-06 | 1.8e-06 | 1.700E-06 | 2.646E-07 | 1.400E-06 | 1.900E-06 |
| 3853 | 193 | L193M | A | 1.4e-06 | 1.7e-06 | 2.2e-06 | 1.767E-06 | 4.041E-07 | 1.400E-06 | 2.200E-06 |
| 3854 | 193 | L193F | A | 2.3e-06 | 1.5e-06 | 2.9e-06 | 2.233E-06 | 7.024E-07 | 1.500E-06 | 2.900E-06 |
| 3855 | 193 | L193P | A | 2.2e-06 | 1.3e-06 | 2.7e-06 | 2.067E-06 | 7.095E-07 | 1.300E-06 | 2.700E-06 |
| 3856 | 193 | L193S | A | 3.4e-06 | 2.2e-06 | 2.6e-06 | 2.733E-06 | 6.110E-07 | 2.200E-06 | 3.400E-06 |
| 3857 | 193 | L193T | A | 2.8e-06 | 2.3e-06 | 4.7e-06 | 3.267E-06 | 1.266E-06 | 2.300E-06 | 4.700E-06 |
| 3858 | 193 | L193W | A | 2.8e-06 | 1.9e-06 | 2.6e-06 | 2.433E-06 | 4.726E-07 | 1.900E-06 | 2.800E-06 |
| 3859 | 193 | L193Y | A | 1.7e-06 | 1.1e-06 | 2.2e-06 | 1.667E-06 | 5.508E-07 | 1.100E-06 | 2.200E-06 |
| 3860 | 193 | L193V | A | 2.3e-06 | 3e-06 | 2.8e-06 | 2.700E-06 | 3.606E-07 | 2.300E-06 | 3.000E-06 |
| 3861 | 194 | T194A | A | 1.5e-06 | 2.5e-06 | 1.6e-06 | 1.867E-06 | 5.508E-07 | 1.500E-06 | 2.500E-06 |
| 3862 | 194 | T194R | A | 2.8e-06 | 4e-06 | 1.3e-06 | 2.700E-06 | 1.353E-06 | 1.300E-06 | 4.000E-06 |
| 3863 | 194 | T194N | A | 4.1e-06 | 1.7e-06 | 2.5e-06 | 2.767E-06 | 1.222E-06 | 1.700E-06 | 4.100E-06 |
| 3864 | 194 | T194D | A | 2.7e-06 | 1.8e-06 | 4.1e-06 | 2.867E-06 | 1.159E-06 | 1.800E-06 | 4.100E-06 |
| 3865 | 194 | T194C | A | 4e-06 | 2e-06 | 1.3e-06 | 2.433E-06 | 1.401E-06 | 1.300E-06 | 4.000E-06 |
| 3866 | 194 | T194E | A | 2.9e-06 | 7.7e-07 | 3.7e-06 | 2.457E-06 | 1.514E-06 | 7.700E-07 | 3.700E-06 |
| 3867 | 194 | T194Q | A | 2.5e-06 | 2.3e-06 | 2.6e-06 | 2.467E-06 | 1.528E-07 | 2.300E-06 | 2.600E-06 |
| 3868 | 194 | T194G | A | 1.9e-06 | 2.8e-06 | 2.3e-06 | 2.333E-06 | 4.509E-07 | 1.900E-06 | 2.800E-06 |
| 3869 | 194 | T194H | A | 1.4e-06 | 1.8e-06 | 2.4e-06 | 1.867E-06 | 5.033E-07 | 1.400E-06 | 2.400E-06 |
| 3870 | 194 | T194I | A | 2.6e-06 | 1.5e-06 | 2.2e-06 | 2.100E-06 | 5.568E-07 | 1.500E-06 | 2.600E-06 |
| 3871 | 194 | T194L | A | 1.9e-06 | 1.2e-06 | 2.3e-06 | 1.800E-06 | 5.568E-07 | 1.200E-06 | 2.300E-06 |
| 3872 | 194 | T194K | A | 2.3e-06 | 3.3e-06 | 2e-06 | 2.533E-06 | 6.807E-07 | 2.000E-06 | 3.300E-06 |
| 3873 | 194 | T194M | A | 2.2e-06 | 3e-06 | 3.5e-06 | 2.900E-06 | 6.557E-07 | 2.200E-06 | 3.500E-06 |
| 3874 | 194 | T194F | A | 1.8e-06 | 1.7e-06 | 3.1e-06 | 2.200E-06 | 7.810E-07 | 1.700E-06 | 3.100E-06 |
| 3875 | 194 | T194P | A | 2.8e-06 | 1.5e-06 | 1.6e-06 | 1.967E-06 | 7.234E-07 | 1.500E-06 | 2.800E-06 |
| 3876 | 194 | T194S | A | 2.5e-06 | 2.2e-06 | 1.3e-06 | 2.000E-06 | 6.245E-07 | 1.300E-06 | 2.500E-06 |
| 3877 | 194 | T194T | A | 2.6e-06 | 2e-06 | 2.4e-06 | 2.333E-06 | 3.055E-07 | 2.000E-06 | 2.600E-06 |
| 3878 | 194 | T194W | A | 2.4e-06 | 1.5e-06 | 2.7e-06 | 2.200E-06 | 6.245E-07 | 1.500E-06 | 2.700E-06 |
| 3879 | 194 | T194Y | A | 2.6e-06 | 1.6e-06 | 1.4e-06 | 1.867E-06 | 6.429E-07 | 1.400E-06 | 2.600E-06 |
| 3880 | 194 | T194V | A | 1.7e-06 | 1.7e-06 | 1.9e-06 | 1.767E-06 | 1.155E-07 | 1.700E-06 | 1.900E-06 |
| 3881 | 195 | R195A | A | 2.5e-06 | 1.6e-06 | 2e-06 | 2.033E-06 | 4.509E-07 | 1.600E-06 | 2.500E-06 |
| 3882 | 195 | R195R | A | 1.9e-06 | 1.9e-06 | 2.7e-06 | 2.167E-06 | 4.619E-07 | 1.900E-06 | 2.700E-06 |
| 3883 | 195 | R195N | A | 1.9e-06 | 3e-06 | 1.1e-06 | 2.000E-06 | 9.539E-07 | 1.100E-06 | 3.000E-06 |
| 3884 | 195 | R195D | A | 2.6e-06 | 1.5e-06 | 3.3e-06 | 2.467E-06 | 9.074E-07 | 1.500E-06 | 3.300E-06 |
| 3885 | 195 | R195C | A | 1.3e-06 | 1.9e-06 | 3.8e-06 | 2.333E-06 | 1.305E-06 | 1.300E-06 | 3.800E-06 |
| 3886 | 195 | R195E | A | 4e-06 | 3.8e-06 | 2.2e-06 | 3.333E-06 | 9.866E-07 | 2.200E-06 | 4.000E-06 |
| 3887 | 195 | R195Q | A | 2e-06 | 2.3e-06 | 1.3e-06 | 1.867E-06 | 5.132E-07 | 1.300E-06 | 2.300E-06 |
| 3888 | 195 | R195G | A | 3.9e-06 | 1.9e-06 | 3.6e-06 | 3.133E-06 | 1.079E-06 | 1.900E-06 | 3.900E-06 |
| 3889 | 195 | R195H | A | 3e-06 | 2e-06 | 1.4e-06 | 2.133E-06 | 8.083E-07 | 1.400E-06 | 3.000E-06 |
| 3890 | 195 | R195I | A | 2.4e-06 | 1.4e-06 | 3.4e-06 | 2.400E-06 | 1.000E-06 | 1.400E-06 | 3.400E-06 |
| 3891 | 195 | R195L | A | 2.6e-06 | 1.7e-06 | 4.3e-06 | 2.867E-06 | 1.320E-06 | 1.700E-06 | 4.300E-06 |
| 3892 | 195 | R195K | A | 1.8e-06 | 3.3e-06 | 2.5e-06 | 2.533E-06 | 7.506E-07 | 1.800E-06 | 3.300E-06 |
| 3893 | 195 | R195M | A | 3.1e-06 | 1.7e-06 | 2.1e-06 | 2.300E-06 | 7.211E-07 | 1.700E-06 | 3.100E-06 |
| 3894 | 195 | R195F | A | 1.8e-06 | 1.7e-06 | 2.7e-06 | 2.067E-06 | 5.508E-07 | 1.700E-06 | 2.700E-06 |
| 3895 | 195 | R195P | A | 9.6e-07 | 3e-06 | 2.6e-06 | 2.187E-06 | 1.081E-06 | 9.600E-07 | 3.000E-06 |
| 3896 | 195 | R195S | A | 2.8e-06 | 2.8e-06 | 2e-06 | 2.533E-06 | 4.619E-07 | 2.000E-06 | 2.800E-06 |
| 3897 | 195 | R195T | A | 9.5e-07 | 1.9e-06 | 2.4e-06 | 1.750E-06 | 7.365E-07 | 9.500E-07 | 2.400E-06 |
| 3898 | 195 | R195W | A | 1.7e-06 | 2.2e-06 | 3.8e-06 | 2.567E-06 | 1.097E-06 | 1.700E-06 | 3.800E-06 |
| 3899 | 195 | R195Y | A | 2.9e-06 | 1.6e-06 | 4.4e-06 | 2.967E-06 | 1.401E-06 | 1.600E-06 | 4.400E-06 |
| 3900 | 195 | R195V | A | 1.8e-06 | 2.6e-06 | 1.9e-06 | 2.100E-06 | 4.359E-07 | 1.800E-06 | 2.600E-06 |
| 3901 | 196 | T196A | A | 2.2e-06 | 2.6e-06 | 2.9e-06 | 2.567E-06 | 3.512E-07 | 2.200E-06 | 2.900E-06 |
| 3902 | 196 | T196R | A | 1.9e-06 | 2.5e-06 | 2.6e-06 | 2.333E-06 | 3.786E-07 | 1.900E-06 | 2.600E-06 |
| 3903 | 196 | T196N | A | 2.6e-06 | 2.3e-06 | 4.7e-06 | 3.200E-06 | 1.308E-06 | 2.300E-06 | 4.700E-06 |
| 3904 | 196 | T196D | A | 2.1e-06 | 2.2e-06 | 2.5e-06 | 2.267E-06 | 2.082E-07 | 2.100E-06 | 2.500E-06 |
| 3905 | 196 | T196C | A | 2.4e-06 | 2.9e-06 | 2.2e-06 | 2.500E-06 | 3.606E-07 | 2.200E-06 | 2.900E-06 |
| 3906 | 196 | T196E | A | 2.8e-06 | 2.3e-06 | 2.3e-06 | 2.467E-06 | 2.887E-07 | 2.300E-06 | 2.800E-06 |
| 3907 | 196 | T196Q | A | 2e-06 | 4.3e-06 | 2.6e-06 | 2.967E-06 | 1.193E-06 | 2.000E-06 | 4.300E-06 |
| 3908 | 196 | T196G | A | 2.4e-06 | 2.9e-06 | 3.4e-06 | 2.900E-06 | 5.000E-07 | 2.400E-06 | 3.400E-06 |
| 3909 | 196 | T196H | A | 1.8e-06 | 1.5e-06 | 1.3e-06 | 1.533E-06 | 2.517E-07 | 1.300E-06 | 1.800E-06 |

| | | | | | | | | | | |
|---|---|---|---|---|---|---|---|---|---|---|
| 3910 | 196 | T196I | A | 3.7e-06 | 2.2e-06 | 3.2e-06 | 3.033E-06 | 7.638E-07 | 2.200E-06 | 3.700E-06 |
| 3911 | 196 | T196L | A | 1.9e-06 | 1.8e-06 | 2.5e-06 | 2.067E-06 | 3.786E-07 | 1.800E-06 | 2.500E-06 |
| 3912 | 196 | T196K | A | 9.9e-07 | 2.5e-06 | 2.7e-06 | 2.063E-06 | 9.349E-07 | 9.900E-07 | 2.700E-06 |
| 3913 | 196 | T196M | A | 1.8e-06 | 1.9e-06 | 2.3e-06 | 2.000E-06 | 2.646E-07 | 1.800E-06 | 2.300E-06 |
| 3914 | 196 | T196F | A | 5.2e-06 | 2e-06 | 3.8e-06 | 3.667E-06 | 1.604E-06 | 2.000E-06 | 5.200E-06 |
| 3915 | 196 | T196P | A | 1.9e-06 | 2.3e-06 | 2.6e-06 | 2.267E-06 | 3.512E-07 | 1.900E-06 | 2.600E-06 |
| 3916 | 196 | T196S | A | 3e-06 | 2.1e-06 | 3.6e-06 | 2.900E-06 | 7.550E-07 | 2.100E-06 | 3.600E-06 |
| 3917 | 196 | T196T | A | 2.4e-06 | 2.6e-06 | 1.7e-06 | 2.233E-06 | 4.726E-07 | 1.700E-06 | 2.600E-06 |
| 3918 | 196 | T196W | A | 2e-06 | 3.9e-06 | 2.3e-06 | 2.733E-06 | 1.021E-06 | 2.000E-06 | 3.900E-06 |
| 3919 | 196 | T196Y | A | 2.3e-06 | 2.7e-06 | 2.8e-06 | 2.600E-06 | 2.646E-07 | 2.300E-06 | 2.800E-06 |
| 3920 | 196 | T196V | A | 1.6e-06 | 2.2e-06 | 1.8e-06 | 1.867E-06 | 3.055E-07 | 1.600E-06 | 2.200E-06 |
| 3921 | 197 | V197A | A | 2.1e-06 | 3.5e-06 | 1.8e-06 | 2.467E-06 | 9.074E-07 | 1.800E-06 | 3.500E-06 |
| 3922 | 197 | V197R | A | 1.4e-06 | 6.2e-07 | 2e-06 | 1.340E-06 | 6.920E-07 | 6.200E-07 | 2.000E-06 |
| 3923 | 197 | V197N | A | 3e-06 | 2.7e-06 | 3.3e-06 | 3.000E-06 | 3.000E-07 | 2.700E-06 | 3.300E-06 |
| 3924 | 197 | V197D | A | 1.6e-06 | 3.2e-06 | 3.3e-06 | 2.700E-06 | 9.539E-07 | 1.600E-06 | 3.300E-06 |
| 3925 | 197 | V197C | A | 1.3e-06 | 2.8e-06 | 1.7e-06 | 1.933E-06 | 7.767E-07 | 1.300E-06 | 2.800E-06 |
| 3926 | 197 | V197E | A | 2.7e-06 | 2.5e-06 | 1.7e-06 | 2.300E-06 | 5.292E-07 | 1.700E-06 | 2.700E-06 |
| 3927 | 197 | V197Q | A | 2.3e-06 | 1.5e-06 | 1.9e-06 | 1.900E-06 | 4.000E-07 | 1.500E-06 | 2.300E-06 |
| 3928 | 197 | V197G | A | 1.6e-06 | 2.9e-06 | 3.1e-06 | 2.533E-06 | 8.145E-07 | 1.600E-06 | 3.100E-06 |
| 3929 | 197 | V197H | A | 3.2e-06 | 4.1e-06 | 2.2e-06 | 3.167E-06 | 9.504E-07 | 2.200E-06 | 4.100E-06 |
| 3930 | 197 | V197I | A | 2.2e-06 | 1.9e-06 | 1.5e-06 | 1.867E-06 | 3.512E-07 | 1.500E-06 | 2.200E-06 |
| 3931 | 197 | V197L | A | 2.3e-06 | 2.3e-06 | 2.6e-06 | 2.400E-06 | 1.732E-07 | 2.300E-06 | 2.600E-06 |
| 3932 | 197 | V197K | A | 1.5e-06 | 2.8e-06 | 2.6e-06 | 2.300E-06 | 7.000E-07 | 1.500E-06 | 2.800E-06 |
| 3933 | 197 | V197M | A | 3.7e-06 | 1.4e-06 | 2e-06 | 2.367E-06 | 1.193E-06 | 1.400E-06 | 3.700E-06 |
| 3934 | 197 | V197F | A | 3.3e-06 | 2.1e-06 | 4.6e-06 | 3.333E-06 | 1.250E-06 | 2.100E-06 | 4.600E-06 |
| 3935 | 197 | V197P | A | 2.7e-06 | 2.6e-06 | 1.6e-06 | 2.300E-06 | 6.083E-07 | 1.600E-06 | 2.700E-06 |
| 3936 | 197 | V197S | A | 1.5e-06 | 1.2e-06 | 1.2e-06 | 1.300E-06 | 1.732E-07 | 1.200E-06 | 1.500E-06 |
| 3937 | 197 | V197T | A | 1.9e-06 | 2.1e-06 | 2.3e-06 | 2.100E-06 | 2.000E-07 | 1.900E-06 | 2.300E-06 |
| 3938 | 197 | V197W | A | 4.5e-06 | 2.4e-06 | 1.4e-06 | 2.767E-06 | 1.582E-06 | 1.400E-06 | 4.500E-06 |
| 3939 | 197 | V197Y | A | 3e-06 | 3.1e-06 | 3.2e-06 | 3.100E-06 | 1.000E-07 | 3.000E-06 | 3.200E-06 |
| 3940 | 197 | V197V | A | 3.2e-06 | 3.8e-06 | 2.1e-06 | 3.033E-06 | 8.622E-07 | 2.100E-06 | 3.800E-06 |
| 3941 | 198 | C198A | A | 1.5e-06 | 5.1e-06 | 1.3e-06 | 2.633E-06 | 2.139E-06 | 1.300E-06 | 5.100E-06 |
| 3942 | 198 | C198R | A | 2.3e-06 | 6.6e-06 | 1.3e-06 | 3.400E-06 | 2.816E-06 | 1.300E-06 | 6.600E-06 |
| 3943 | 198 | C198N | A | 3e-06 | 1.8e-06 | 2.5e-06 | 2.433E-06 | 6.028E-07 | 1.800E-06 | 3.000E-06 |
| 3944 | 198 | C198D | A | 3.5e-06 | 2e-06 | 1.3e-06 | 2.267E-06 | 1.124E-06 | 1.300E-06 | 3.500E-06 |
| 3945 | 198 | C198C | A | 2.5e-06 | 1.5e-06 | 2.3e-06 | 2.100E-06 | 5.292E-07 | 1.500E-06 | 2.500E-06 |
| 3946 | 198 | C198E | A | 1.9e-06 | 1.7e-06 | 2.9e-06 | 2.167E-06 | 6.429E-07 | 1.700E-06 | 2.900E-06 |
| 3947 | 198 | C198Q | A | 1.6e-06 | 1.9e-06 | 2e-06 | 1.833E-06 | 2.082E-07 | 1.600E-06 | 2.000E-06 |
| 3948 | 198 | C198G | A | 2.8e-06 | 3.7e-06 | 2.4e-06 | 2.967E-06 | 6.658E-07 | 2.400E-06 | 3.700E-06 |
| 3949 | 198 | C198H | A | 3.1e-06 | 3.9e-06 | 2.8e-06 | 3.267E-06 | 5.686E-07 | 2.800E-06 | 3.900E-06 |
| 3950 | 198 | C198I | A | 1.6e-06 | 2e-06 | 2.5e-06 | 2.033E-06 | 4.509E-07 | 1.600E-06 | 2.500E-06 |
| 3951 | 198 | C198L | A | 2.3e-06 | 2.6e-06 | 3.2e-06 | 2.700E-06 | 4.583E-07 | 2.300E-06 | 3.200E-06 |
| 3952 | 198 | C198K | A | 2.2e-06 | 2.2e-06 | 4.1e-06 | 2.833E-06 | 1.097E-06 | 2.200E-06 | 4.100E-06 |
| 3953 | 198 | C198M | A | 1.9e-06 | 2.2e-06 | 1.7e-06 | 1.933E-06 | 2.517E-07 | 1.700E-06 | 2.200E-06 |
| 3954 | 198 | C198F | A | 1.8e-06 | 2.6e-06 | 2.6e-06 | 2.333E-06 | 4.619E-07 | 1.800E-06 | 2.600E-06 |
| 3955 | 198 | C198P | A | 1.6e-06 | 3.1e-06 | 2.2e-06 | 2.300E-06 | 7.550E-07 | 1.600E-06 | 3.100E-06 |
| 3956 | 198 | C198S | A | 2.6e-06 | 1.9e-06 | 3.2e-06 | 2.567E-06 | 6.506E-07 | 1.900E-06 | 3.200E-06 |
| 3957 | 198 | C198T | A | 1.9e-06 | 3.9e-06 | 1.6e-06 | 2.467E-06 | 1.250E-06 | 1.600E-06 | 3.900E-06 |
| 3958 | 198 | C198W | A | 1.2e-06 | 1.7e-06 | 2.2e-06 | 1.700E-06 | 5.000E-07 | 1.200E-06 | 2.200E-06 |
| 3959 | 198 | C198Y | A | 2.2e-06 | 1.7e-06 | 2.8e-06 | 2.233E-06 | 5.508E-07 | 1.700E-06 | 2.800E-06 |
| 3960 | 198 | C198V | A | 2.7e-06 | 2.3e-06 | 1.7e-06 | 2.233E-06 | 5.033E-07 | 1.700E-06 | 2.700E-06 |
| 3961 | 199 | A199A | A | 2e-06 | 3.7e-06 | 2.1e-06 | 2.600E-06 | 9.539E-07 | 2.000E-06 | 3.700E-06 |
| 3962 | 199 | A199R | A | 2.4e-06 | 1.9e-06 | 1.9e-06 | 2.067E-06 | 2.887E-07 | 1.900E-06 | 2.400E-06 |
| 3963 | 199 | A199N | A | 2.4e-06 | 1.4e-06 | 2.6e-06 | 2.133E-06 | 6.429E-07 | 1.400E-06 | 2.600E-06 |
| 3964 | 199 | A199D | A | 2.6e-06 | 3.1e-06 | 6.4e-07 | 2.113E-06 | 1.300E-06 | 6.400E-07 | 3.100E-06 |
| 3965 | 199 | A199C | A | 2.7e-06 | 2.9e-06 | 3.9e-06 | 3.167E-06 | 6.429E-07 | 2.700E-06 | 3.900E-06 |
| 3966 | 199 | A199E | A | 2.3e-06 | 2e-06 | 3.6e-06 | 2.633E-06 | 8.505E-07 | 2.000E-06 | 3.600E-06 |
| 3967 | 199 | A199Q | A | 2.7e-06 | 8.3e-07 | 2.5e-06 | 2.010E-06 | 1.027E-06 | 8.300E-07 | 2.700E-06 |
| 3968 | 199 | A199G | A | 2.8e-06 | 1.7e-06 | 2.9e-06 | 2.467E-06 | 6.658E-07 | 1.700E-06 | 2.900E-06 |
| 3969 | 199 | A199H | A | 1.7e-06 | 3.8e-06 | 2.6e-06 | 2.700E-06 | 1.054E-06 | 1.700E-06 | 3.800E-06 |
| 3970 | 199 | A199I | A | 4.2e-06 | 2.7e-06 | 1.7e-06 | 2.867E-06 | 1.258E-06 | 1.700E-06 | 4.200E-06 |
| 3971 | 199 | A199L | A | 3.7e-06 | 3.3e-06 | 1.6e-06 | 2.867E-06 | 1.115E-06 | 1.600E-06 | 3.700E-06 |
| 3972 | 199 | A199K | A | 3.3e-06 | 3.2e-06 | 2.8e-06 | 3.100E-06 | 2.646E-07 | 2.800E-06 | 3.300E-06 |
| 3973 | 199 | A199M | A | 2.3e-06 | 1.7e-06 | 2.1e-06 | 2.033E-06 | 3.055E-07 | 1.700E-06 | 2.300E-06 |
| 3974 | 199 | A199F | A | 1.9e-06 | 2.1e-06 | 2.3e-06 | 2.100E-06 | 2.000E-07 | 1.900E-06 | 2.300E-06 |
| 3975 | 199 | A199P | A | 2.5e-06 | 3.2e-06 | 3.6e-06 | 3.100E-06 | 5.568E-07 | 2.500E-06 | 3.600E-06 |
| 3976 | 199 | A199S | A | 2.6e-06 | 3.7e-06 | 2.5e-06 | 2.933E-06 | 6.658E-07 | 2.500E-06 | 3.700E-06 |
| 3977 | 199 | A199T | A | 2e-06 | 1.7e-06 | 4.1e-06 | 2.600E-06 | 1.308E-06 | 1.700E-06 | 4.100E-06 |
| 3978 | 199 | A199W | A | 2e-06 | 1.9e-06 | 2.9e-06 | 2.267E-06 | 5.508E-07 | 1.900E-06 | 2.900E-06 |
| 3979 | 199 | A199Y | A | 2.2e-06 | 3.2e-06 | 2.2e-06 | 2.533E-06 | 5.774E-07 | 2.200E-06 | 3.200E-06 |
| 3980 | 199 | A199V | A | 3e-06 | 2.1e-06 | 3e-06 | 2.700E-06 | 5.196E-07 | 2.100E-06 | 3.000E-06 |
| 3981 | 200 | G200A | A | 2.1e-06 | 2.1e-06 | 2.5e-06 | 2.233E-06 | 2.309E-07 | 2.100E-06 | 2.500E-06 |
| 3982 | 200 | G200R | A | 1.7e-06 | 3.3e-06 | 1.7e-06 | 2.233E-06 | 9.238E-07 | 1.700E-06 | 3.300E-06 |
| 3983 | 200 | G200N | A | 2.2e-06 | 1.8e-06 | 2.2e-06 | 2.067E-06 | 2.309E-07 | 1.800E-06 | 2.200E-06 |
| 3984 | 200 | G200D | A | 3.1e-06 | 1.5e-06 | 4.6e-06 | 3.067E-06 | 1.550E-06 | 1.500E-06 | 4.600E-06 |
| 3985 | 200 | G200C | A | 1.4e-06 | 1.5e-06 | 1.6e-06 | 1.500E-06 | 1.000E-07 | 1.400E-06 | 1.600E-06 |
| 3986 | 200 | G200E | A | 2.4e-06 | 1.7e-06 | 2.7e-06 | 2.267E-06 | 5.132E-07 | 1.700E-06 | 2.700E-06 |
| 3987 | 200 | G200Q | A | 2.1e-06 | 1.2e-06 | 1.7e-06 | 1.667E-06 | 4.509E-07 | 1.200E-06 | 2.100E-06 |
| 3988 | 200 | G200G | A | 1.1e-06 | 3e-06 | 3e-06 | 2.367E-06 | 1.097E-06 | 1.100E-06 | 3.000E-06 |
| 3989 | 200 | G200H | A | 3.2e-06 | 2.5e-06 | 1.9e-06 | 2.533E-06 | 6.506E-07 | 1.900E-06 | 3.200E-06 |
| 3990 | 200 | G200I | A | 1.5e-06 | 2.6e-06 | 1.9e-06 | 2.000E-06 | 5.568E-07 | 1.500E-06 | 2.600E-06 |
| 3991 | 200 | G200L | A | 2.2e-06 | 2.9e-06 | 3.8e-06 | 2.967E-06 | 8.021E-07 | 2.200E-06 | 3.800E-06 |
| 3992 | 200 | G200K | A | 2.3e-06 | 2.1e-06 | 1.7e-06 | 2.033E-06 | 3.055E-07 | 1.700E-06 | 2.300E-06 |
| 3993 | 200 | G200M | A | 2.3e-06 | 4.8e-06 | 2.5e-06 | 3.200E-06 | 1.389E-06 | 2.300E-06 | 4.800E-06 |
| 3994 | 200 | G200F | A | 2.2e-06 | 3.3e-06 | 2e-06 | 2.500E-06 | 7.000E-07 | 2.000E-06 | 3.300E-06 |

| | | | | | | | | | | |
|---|---|---|---|---|---|---|---|---|---|---|
| 3995 | 200 | G200P | A | 3.3e-06 | 2.8e-06 | 3.8e-06 | 3.300E-06 | 5.000E-07 | 2.800E-06 | 3.800E-06 |
| 3996 | 200 | G200S | A | 3.8e-06 | 2.9e-06 | 1.2e-06 | 2.633E-06 | 1.320E-06 | 1.200E-06 | 3.800E-06 |
| 3997 | 200 | G200T | A | 3.1e-06 | 2.3e-06 | 1.8e-06 | 2.400E-06 | 6.557E-07 | 1.800E-06 | 3.100E-06 |
| 3998 | 200 | G200W | A | 1.8e-06 | 3.1e-06 | 2e-06 | 2.300E-06 | 7.000E-07 | 1.800E-06 | 3.100E-06 |
| 3999 | 200 | G200Y | A | 1.8e-06 | 2.8e-06 | 1.7e-06 | 2.100E-06 | 6.083E-07 | 1.700E-06 | 2.800E-06 |
| 4000 | 200 | G200V | A | 2e-06 | 1.8e-06 | 1.3e-06 | 1.700E-06 | 3.606E-07 | 1.300E-06 | 2.000E-06 |
| 4001 | 201 | G201A | A | 1.8e-06 | 3.6e-06 | 2.4e-06 | 2.600E-06 | 9.165E-07 | 1.800E-06 | 3.600E-06 |
| 4002 | 201 | G201R | A | 2.8e-06 | 1.6e-06 | 3.5e-06 | 2.633E-06 | 9.609E-07 | 1.600E-06 | 3.500E-06 |
| 4003 | 201 | G201N | A | 1.9e-06 | 3.3e-06 | 2.2e-06 | 2.467E-06 | 7.371E-07 | 1.900E-06 | 3.300E-06 |
| 4004 | 201 | G201D | A | 1.6e-06 | 1.4e-06 | 2.6e-06 | 1.867E-06 | 6.429E-07 | 1.400E-06 | 2.600E-06 |
| 4005 | 201 | G201C | A | 2.5e-06 | 9.7e-07 | 1e-06 | 1.490E-06 | 8.748E-07 | 9.700E-07 | 2.500E-06 |
| 4006 | 201 | G201E | A | 2e-06 | 2.9e-06 | 2.7e-06 | 2.533E-06 | 4.726E-07 | 2.000E-06 | 2.900E-06 |
| 4007 | 201 | G201Q | A | 5e-06 | 2.3e-06 | 8.9e-07 | 2.730E-06 | 2.088E-06 | 8.900E-07 | 5.000E-06 |
| 4008 | 201 | G201G | A | 2.2e-06 | 3e-06 | 1.6e-06 | 2.267E-06 | 7.024E-07 | 1.600E-06 | 3.000E-06 |
| 4009 | 201 | G201H | A | 3.4e-06 | 2.4e-06 | 9.9e-07 | 2.263E-06 | 1.211E-06 | 9.900E-07 | 3.400E-06 |
| 4010 | 201 | G201I | A | 3.4e-06 | 3.2e-06 | 2.1e-06 | 2.900E-06 | 7.000E-07 | 2.100E-06 | 3.400E-06 |
| 4011 | 201 | G201L | A | 1.6e-06 | 3e-06 | 3.1e-06 | 2.567E-06 | 8.386E-07 | 1.600E-06 | 3.100E-06 |
| 4012 | 201 | G201K | A | 3.1e-06 | 3e-06 | 3.5e-06 | 3.200E-06 | 2.646E-07 | 3.000E-06 | 3.500E-06 |
| 4013 | 201 | G201M | A | 2.4e-06 | 4.1e-06 | 3.1e-06 | 3.200E-06 | 8.544E-07 | 2.400E-06 | 4.100E-06 |
| 4014 | 201 | G201F | A | 2.7e-06 | 3.2e-06 | 2.4e-06 | 2.767E-06 | 4.041E-07 | 2.400E-06 | 3.200E-06 |
| 4015 | 201 | G201P | A | 1.7e-06 | 2.4e-06 | 1.9e-06 | 2.000E-06 | 3.606E-07 | 1.700E-06 | 2.400E-06 |
| 4016 | 201 | G201S | A | 2.3e-06 | 2.1e-06 | 2e-06 | 2.133E-06 | 1.528E-07 | 2.000E-06 | 2.300E-06 |
| 4017 | 201 | G201T | A | 2.5e-06 | 2.4e-06 | 1.9e-06 | 2.267E-06 | 3.215E-07 | 1.900E-06 | 2.500E-06 |
| 4018 | 201 | G201W | A | 3.2e-06 | 2.6e-06 | 2e-06 | 2.600E-06 | 6.000E-07 | 2.000E-06 | 3.200E-06 |
| 4019 | 201 | G201Y | A | 1.7e-06 | 2e-06 | 1.8e-06 | 1.833E-06 | 1.528E-07 | 1.700E-06 | 2.000E-06 |
| 4020 | 201 | G201V | A | 2.6e-06 | 2.7e-06 | 2.4e-06 | 2.567E-06 | 1.528E-07 | 2.400E-06 | 2.700E-06 |
| 4021 | 202 | C202A | A | 2.3e-06 | 3.8e-06 | 2.3e-06 | 2.800E-06 | 8.660E-07 | 2.300E-06 | 3.800E-06 |
| 4022 | 202 | C202R | A | 3.1e-06 | 2.6e-06 | 3.8e-06 | 3.167E-06 | 6.028E-07 | 2.600E-06 | 3.800E-06 |
| 4023 | 202 | C202N | A | 2.6e-06 | 2.6e-06 | 1.6e-06 | 2.267E-06 | 5.774E-07 | 1.600E-06 | 2.600E-06 |
| 4024 | 202 | C202D | A | 3.5e-06 | 2.1e-06 | 2.3e-06 | 2.633E-06 | 7.572E-07 | 2.100E-06 | 3.500E-06 |
| 4025 | 202 | C202C | A | 4e-06 | 3.3e-06 | 5.3e-06 | 4.200E-06 | 1.015E-06 | 3.300E-06 | 5.300E-06 |
| 4026 | 202 | C202E | A | 2.5e-06 | 2.4e-06 | 1.9e-06 | 2.267E-06 | 3.215E-07 | 1.900E-06 | 2.500E-06 |
| 4027 | 202 | C202Q | A | 1.2e-06 | 3.4e-06 | 1.7e-06 | 2.100E-06 | 1.153E-06 | 1.200E-06 | 3.400E-06 |
| 4028 | 202 | C202G | A | 2.2e-06 | 2.6e-06 | 2.5e-06 | 2.433E-06 | 2.082E-07 | 2.200E-06 | 2.600E-06 |
| 4029 | 202 | C202H | A | 3e-06 | 5.1e-06 | 2.5e-06 | 3.533E-06 | 1.380E-06 | 2.500E-06 | 5.100E-06 |
| 4030 | 202 | C202I | A | 3.3e-06 | 2.4e-06 | 2.7e-06 | 2.800E-06 | 4.583E-07 | 2.400E-06 | 3.300E-06 |
| 4031 | 202 | C202L | A | 3.5e-06 | 1.7e-06 | 2.7e-06 | 2.633E-06 | 9.018E-07 | 1.700E-06 | 3.500E-06 |
| 4032 | 202 | C202K | A | 1.8e-06 | 2.2e-06 | 1.6e-06 | 1.867E-06 | 3.055E-07 | 1.600E-06 | 2.200E-06 |
| 4033 | 202 | C202M | A | 2e-06 | 9.7e-07 | 2e-06 | 1.657E-06 | 5.947E-07 | 9.700E-07 | 2.000E-06 |
| 4034 | 202 | C202F | A | 3.1e-06 | 2.4e-06 | 1.9e-06 | 2.467E-06 | 6.028E-07 | 1.900E-06 | 3.100E-06 |
| 4035 | 202 | C202P | A | 3.7e-06 | 1.8e-06 | 1.9e-06 | 2.467E-06 | 1.069E-06 | 1.800E-06 | 3.700E-06 |
| 4036 | 202 | C202S | A | 2.4e-06 | 3.1e-06 | 1.7e-06 | 2.400E-06 | 7.000E-07 | 1.700E-06 | 3.100E-06 |
| 4037 | 202 | C202T | A | 1.7e-06 | 3.2e-06 | 1.7e-06 | 2.200E-06 | 8.660E-07 | 1.700E-06 | 3.200E-06 |
| 4038 | 202 | C202W | A | 3.5e-06 | 2.4e-06 | 1.9e-06 | 2.600E-06 | 8.185E-07 | 1.900E-06 | 3.500E-06 |
| 4039 | 202 | C202Y | A | 2.1e-06 | 1e-06 | 2.4e-06 | 1.833E-06 | 7.371E-07 | 1.000E-06 | 2.400E-06 |
| 4040 | 202 | C202V | A | 1.7e-06 | 2.2e-06 | 2.7e-06 | 2.200E-06 | 5.000E-07 | 1.700E-06 | 2.700E-06 |
| 4041 | 203 | A203A | A | 1.4e-06 | 1.7e-06 | 1.8e-06 | 1.633E-06 | 2.082E-07 | 1.400E-06 | 1.800E-06 |
| 4042 | 203 | A203R | A | 1.7e-06 | 1.5e-06 | 1.9e-06 | 1.700E-06 | 2.000E-07 | 1.500E-06 | 1.900E-06 |
| 4043 | 203 | A203N | A | 1.6e-06 | 1.6e-06 | 2.6e-06 | 1.933E-06 | 5.774E-07 | 1.600E-06 | 2.600E-06 |
| 4044 | 203 | A203D | A | 2.3e-06 | 3.7e-06 | 1.7e-06 | 2.567E-06 | 1.026E-06 | 1.700E-06 | 3.700E-06 |
| 4045 | 203 | A203C | A | 2.1e-06 | 2.8e-06 | 1.1e-06 | 2.000E-06 | 8.544E-07 | 1.100E-06 | 2.800E-06 |
| 4046 | 203 | A203E | A | 1.8e-06 | 2.4e-06 | 2.5e-06 | 2.233E-06 | 3.786E-07 | 1.800E-06 | 2.500E-06 |
| 4047 | 203 | A203Q | A | 9.1e-07 | 1.8e-06 | 1.7e-06 | 1.470E-06 | 4.875E-07 | 9.100E-07 | 1.800E-06 |
| 4048 | 203 | A203G | A | 6.2e-06 | 1.9e-06 | 2.2e-06 | 3.433E-06 | 2.401E-06 | 1.900E-06 | 6.200E-06 |
| 4049 | 203 | A203H | A | 4.2e-06 | 1.4e-06 | 3.5e-06 | 3.033E-06 | 1.457E-06 | 1.400E-06 | 4.200E-06 |
| 4050 | 203 | A203I | A | 2.9e-06 | 2.6e-06 | 2e-06 | 2.500E-06 | 4.583E-07 | 2.000E-06 | 2.900E-06 |
| 4051 | 203 | A203L | A | 6e-06 | 2.1e-06 | 2.9e-06 | 3.667E-06 | 2.060E-06 | 2.100E-06 | 6.000E-06 |
| 4052 | 203 | A203K | A | 1.6e-06 | 2.4e-06 | 2.8e-06 | 2.267E-06 | 6.110E-07 | 1.600E-06 | 2.800E-06 |
| 4053 | 203 | A203M | A | 2.6e-06 | 3.3e-06 | 2.6e-06 | 2.833E-06 | 4.041E-07 | 2.600E-06 | 3.300E-06 |
| 4054 | 203 | A203F | A | 2.3e-06 | 1.4e-06 | 3.8e-06 | 2.500E-06 | 1.212E-06 | 1.400E-06 | 3.800E-06 |
| 4055 | 203 | A203P | A | 2.5e-06 | 2.9e-06 | 2.9e-06 | 2.767E-06 | 2.309E-07 | 2.500E-06 | 2.900E-06 |
| 4056 | 203 | A203S | A | 3.5e-06 | 3.4e-06 | 2e-06 | 2.967E-06 | 8.386E-07 | 2.000E-06 | 3.500E-06 |
| 4057 | 203 | A203T | A | 2.9e-06 | 3.4e-06 | 2.4e-06 | 2.900E-06 | 5.000E-07 | 2.400E-06 | 3.400E-06 |
| 4058 | 203 | A203W | A | 6.3e-06 | 2.4e-06 | 1.6e-06 | 3.433E-06 | 2.515E-06 | 1.600E-06 | 6.300E-06 |
| 4059 | 203 | A203Y | A | 2.6e-06 | 2.2e-06 | 1.8e-06 | 2.200E-06 | 4.000E-07 | 1.800E-06 | 2.600E-06 |
| 4060 | 203 | A203V | A | 3.8e-06 | 3.8e-06 | 1.8e-06 | 3.133E-06 | 1.155E-06 | 1.800E-06 | 3.800E-06 |
| 4061 | 204 | R204A | A | 2.6e-06 | 2.2e-06 | 1.8e-06 | 2.200E-06 | 4.000E-07 | 1.800E-06 | 2.600E-06 |
| 4062 | 204 | R204R | A | 7.8e-07 | 1.1e-06 | 2e-06 | 1.293E-06 | 6.326E-07 | 7.800E-07 | 2.000E-06 |
| 4063 | 204 | R204N | A | 4.5e-06 | 2.3e-06 | 1.6e-06 | 2.800E-06 | 1.513E-06 | 1.600E-06 | 4.500E-06 |
| 4064 | 204 | R204D | A | 2.4e-06 | 1.5e-06 | 2.9e-06 | 2.267E-06 | 7.095E-07 | 1.500E-06 | 2.900E-06 |
| 4065 | 204 | R204C | A | 1.8e-06 | 1.6e-06 | 4.8e-06 | 2.733E-06 | 1.793E-06 | 1.600E-06 | 4.800E-06 |
| 4066 | 204 | R204E | A | 5.6e-06 | 2.8e-06 | 5.3e-06 | 4.567E-06 | 1.537E-06 | 2.800E-06 | 5.600E-06 |
| 4067 | 204 | R204Q | A | 2.3e-06 | 7.9e-07 | 3.1e-06 | 2.063E-06 | 1.173E-06 | 7.900E-07 | 3.100E-06 |
| 4068 | 204 | R204G | A | 2.6e-06 | 2.2e-06 | 3.1e-06 | 2.633E-06 | 4.509E-07 | 2.200E-06 | 3.100E-06 |
| 4069 | 204 | R204H | A | 1.6e-06 | 1.4e-06 | 3.4e-06 | 2.133E-06 | 1.102E-06 | 1.400E-06 | 3.400E-06 |
| 4070 | 204 | R204I | A | 2.3e-06 | 2.4e-06 | 2.9e-06 | 2.533E-06 | 3.215E-07 | 2.300E-06 | 2.900E-06 |
| 4071 | 204 | R204L | A | 3e-06 | 1.9e-06 | 2.1e-06 | 2.333E-06 | 5.859E-07 | 1.900E-06 | 3.000E-06 |
| 4072 | 204 | R204K | A | 1.8e-06 | 2.7e-06 | 2.3e-06 | 2.267E-06 | 4.509E-07 | 1.800E-06 | 2.700E-06 |
| 4073 | 204 | R204M | A | 2.1e-06 | 3e-06 | 2.7e-06 | 2.600E-06 | 4.583E-07 | 2.100E-06 | 3.000E-06 |
| 4074 | 204 | R204F | A | 3.4e-06 | 2.5e-06 | 3e-06 | 2.967E-06 | 4.509E-07 | 2.500E-06 | 3.400E-06 |
| 4075 | 204 | R204P | A | 3.1e-06 | 2.1e-06 | 3.2e-06 | 2.800E-06 | 6.083E-07 | 2.100E-06 | 3.200E-06 |
| 4076 | 204 | R204S | A | 2.2e-06 | 5.1e-06 | 1.6e-06 | 2.967E-06 | 1.872E-06 | 1.600E-06 | 5.100E-06 |
| 4077 | 204 | R204T | A | 3.5e-06 | 3.1e-06 | 2.1e-06 | 2.900E-06 | 7.211E-07 | 2.100E-06 | 3.500E-06 |
| 4078 | 204 | R204W | A | 3.2e-06 | 1.8e-06 | 3.7e-06 | 2.900E-06 | 9.849E-07 | 1.800E-06 | 3.700E-06 |
| 4079 | 204 | R204Y | A | 1.7e-06 | 1.6e-06 | 2.2e-06 | 1.833E-06 | 3.215E-07 | 1.600E-06 | 2.200E-06 |

| | | | | | | | | | | |
|---|---|---|---|---|---|---|---|---|---|---|
| 4080 | 204 | R204V | A | 3.4e-06 | 2.6e-06 | 4.9e-06 | 3.633E-06 | 1.168E-06 | 2.600E-06 | 4.900E-06 |
| 4081 | 205 | C205A | A | 1.6e-06 | 2.9e-06 | 1.6e-06 | 2.033E-06 | 7.506E-07 | 1.600E-06 | 2.900E-06 |
| 4082 | 205 | C205R | A | 2.9e-06 | 3.3e-06 | 2.8e-06 | 3.000E-06 | 2.646E-07 | 2.800E-06 | 3.300E-06 |
| 4083 | 205 | C205N | A | 3.1e-06 | 1.6e-06 | 4.1e-06 | 2.933E-06 | 1.258E-06 | 1.600E-06 | 4.100E-06 |
| 4084 | 205 | C205D | A | 1.8e-06 | 2.1e-06 | 1.3e-06 | 1.733E-06 | 4.041E-07 | 1.300E-06 | 2.100E-06 |
| 4085 | 205 | C205C | A | 2.8e-06 | 3.2e-06 | 2.7e-06 | 2.900E-06 | 2.646E-07 | 2.700E-06 | 3.200E-06 |
| 4086 | 205 | C205E | A | 3.3e-06 | 4.1e-06 | 3.2e-06 | 3.533E-06 | 4.933E-07 | 3.200E-06 | 4.100E-06 |
| 4087 | 205 | C205Q | A | 3.1e-06 | 1.7e-06 | 2.7e-06 | 2.500E-06 | 7.211E-07 | 1.700E-06 | 3.100E-06 |
| 4088 | 205 | C205G | A | 2.4e-06 | 2.7e-06 | 1.8e-06 | 2.300E-06 | 4.583E-07 | 1.800E-06 | 2.700E-06 |
| 4089 | 205 | C205H | A | 2.7e-06 | 1.9e-06 | 1.6e-06 | 2.067E-06 | 5.686E-07 | 1.600E-06 | 2.700E-06 |
| 4090 | 205 | C205I | A | 2.8e-06 | 3.8e-06 | 1.6e-06 | 2.733E-06 | 1.102E-06 | 1.600E-06 | 3.800E-06 |
| 4091 | 205 | C205L | A | 2.3e-06 | 2.9e-06 | 1.4e-06 | 2.200E-06 | 7.550E-07 | 1.400E-06 | 2.900E-06 |
| 4092 | 205 | C205K | A | 2.3e-06 | 1.3e-06 | 2.7e-06 | 2.100E-06 | 7.211E-07 | 1.300E-06 | 2.700E-06 |
| 4093 | 205 | C205M | A | 4e-06 | 2.1e-06 | 2.2e-06 | 2.767E-06 | 1.069E-06 | 2.100E-06 | 4.000E-06 |
| 4094 | 205 | C205F | A | 2.3e-06 | 2.9e-06 | 2.5e-06 | 2.567E-06 | 3.055E-07 | 2.300E-06 | 2.900E-06 |
| 4095 | 205 | C205P | A | 2.3e-06 | 2.1e-06 | 2.2e-06 | 2.200E-06 | 1.000E-07 | 2.100E-06 | 2.300E-06 |
| 4096 | 205 | C205S | A | 3.7e-06 | 1.1e-06 | 2.3e-06 | 2.367E-06 | 1.301E-06 | 1.100E-06 | 3.700E-06 |
| 4097 | 205 | C205T | A | 2.2e-06 | 2e-06 | 2.3e-06 | 2.167E-06 | 1.528E-07 | 2.000E-06 | 2.300E-06 |
| 4098 | 205 | C205W | A | 2.8e-06 | 4.4e-06 | 5.3e-06 | 4.167E-06 | 1.266E-06 | 2.800E-06 | 5.300E-06 |
| 4099 | 205 | C205Y | A | 3.9e-06 | 2.2e-06 | 3.7e-06 | 3.267E-06 | 9.292E-07 | 2.200E-06 | 3.900E-06 |
| 4100 | 205 | C205V | A | 3.3e-06 | 2e-06 | 1.5e-06 | 2.267E-06 | 9.292E-07 | 1.500E-06 | 3.300E-06 |
| 4101 | 206 | K206A | A | 1.9e-06 | 2.1e-06 | 1.7e-06 | 1.900E-06 | 2.000E-07 | 1.700E-06 | 2.100E-06 |
| 4102 | 206 | K206R | A | 2.9e-06 | 1.8e-06 | 1.9e-06 | 2.200E-06 | 6.083E-07 | 1.800E-06 | 2.900E-06 |
| 4103 | 206 | K206N | A | 2.3e-06 | 1.7e-06 | 4.5e-06 | 2.833E-06 | 1.474E-06 | 1.700E-06 | 4.500E-06 |
| 4104 | 206 | K206D | A | 1.9e-06 | 2.3e-06 | 1.6e-06 | 1.933E-06 | 3.512E-07 | 1.600E-06 | 2.300E-06 |
| 4105 | 206 | K206C | A | 1.5e-06 | 3.1e-06 | 1.7e-06 | 2.100E-06 | 8.718E-07 | 1.500E-06 | 3.100E-06 |
| 4106 | 206 | K206E | A | 2.3e-06 | 3.6e-06 | 1.7e-06 | 2.533E-06 | 9.713E-07 | 1.700E-06 | 3.600E-06 |
| 4107 | 206 | K206Q | A | 1.6e-06 | 4e-06 | 1.8e-06 | 2.467E-06 | 1.332E-06 | 1.600E-06 | 4.000E-06 |
| 4108 | 206 | K206G | A | 3.5e-06 | 2.2e-06 | 2.3e-06 | 2.667E-06 | 7.234E-07 | 2.200E-06 | 3.500E-06 |
| 4109 | 206 | K206H | A | 4.1e-06 | 2.4e-06 | 1.5e-06 | 2.667E-06 | 1.320E-06 | 1.500E-06 | 4.100E-06 |
| 4110 | 206 | K206I | A | 2.6e-06 | 2.1e-06 | 2.6e-06 | 2.433E-06 | 2.887E-07 | 2.100E-06 | 2.600E-06 |
| 4111 | 206 | K206L | A | 3e-06 | 3.4e-06 | 2.7e-06 | 3.033E-06 | 3.512E-07 | 2.700E-06 | 3.400E-06 |
| 4112 | 206 | K206K | A | 2e-06 | 3.5e-06 | 2.6e-06 | 2.700E-06 | 7.550E-07 | 2.000E-06 | 3.500E-06 |
| 4113 | 206 | K206M | A | 1.9e-06 | 2.6e-06 | 1.4e-06 | 1.967E-06 | 6.028E-07 | 1.400E-06 | 2.600E-06 |
| 4114 | 206 | K206F | A | 2.5e-06 | 2.9e-06 | 1.4e-06 | 2.267E-06 | 7.767E-07 | 1.400E-06 | 2.900E-06 |
| 4115 | 206 | K206P | A | 2.9e-06 | 2.4e-06 | 1.7e-06 | 2.333E-06 | 6.028E-07 | 1.700E-06 | 2.900E-06 |
| 4116 | 206 | K206S | A | 1.8e-06 | 2.8e-06 | 2.1e-06 | 2.233E-06 | 5.132E-07 | 1.800E-06 | 2.800E-06 |
| 4117 | 206 | K206T | A | 4.8e-06 | 2e-06 | 3.9e-06 | 3.567E-06 | 1.429E-06 | 2.000E-06 | 4.800E-06 |
| 4118 | 206 | K206W | A | 2.3e-06 | 4.7e-06 | 1.6e-06 | 2.867E-06 | 1.626E-06 | 1.600E-06 | 4.700E-06 |
| 4119 | 206 | K206Y | A | 1.6e-06 | 2.6e-06 | 2.2e-06 | 2.133E-06 | 5.033E-07 | 1.600E-06 | 2.600E-06 |
| 4120 | 206 | K206V | A | 1.3e-06 | 1.8e-06 | 1.9e-06 | 1.667E-06 | 3.215E-07 | 1.300E-06 | 1.900E-06 |
| 4121 | 207 | G207A | A | 4.1e-06 | 3.7e-06 | 1.5e-06 | 3.100E-06 | 1.400E-06 | 1.500E-06 | 4.100E-06 |
| 4122 | 207 | G207R | A | 1.7e-06 | 3e-06 | 1.9e-06 | 2.200E-06 | 7.000E-07 | 1.700E-06 | 3.000E-06 |
| 4123 | 207 | G207N | A | 2.1e-06 | 4.6e-06 | 2.7e-06 | 3.133E-06 | 1.305E-06 | 2.100E-06 | 4.600E-06 |
| 4124 | 207 | G207D | A | 3.1e-06 | 3.2e-06 | 2.9e-06 | 3.067E-06 | 1.528E-07 | 2.900E-06 | 3.200E-06 |
| 4125 | 207 | G207C | A | 2.2e-06 | 2.8e-06 | 1.6e-06 | 2.200E-06 | 6.000E-07 | 1.600E-06 | 2.800E-06 |
| 4126 | 207 | G207E | A | 2.6e-06 | 2.4e-06 | 2.3e-06 | 2.433E-06 | 1.528E-07 | 2.300E-06 | 2.600E-06 |
| 4127 | 207 | G207Q | A | 1.9e-06 | 3e-06 | 2.3e-06 | 2.400E-06 | 5.568E-07 | 1.900E-06 | 3.000E-06 |
| 4128 | 207 | G207G | A | 2.5e-06 | 2.5e-06 | 3.6e-06 | 2.867E-06 | 6.351E-07 | 2.500E-06 | 3.600E-06 |
| 4129 | 207 | G207H | A | 2.9e-06 | 3.3e-06 | 2.8e-06 | 3.000E-06 | 2.646E-07 | 2.800E-06 | 3.300E-06 |
| 4130 | 207 | G207I | A | 2e-06 | 3.2e-06 | 1.8e-06 | 2.333E-06 | 7.572E-07 | 1.800E-06 | 3.200E-06 |
| 4131 | 207 | G207L | A | 3.3e-06 | 2.4e-06 | 2.7e-06 | 2.800E-06 | 4.583E-07 | 2.400E-06 | 3.300E-06 |
| 4132 | 207 | G207K | A | 2.8e-06 | 2.8e-06 | 2.4e-06 | 2.667E-06 | 2.309E-07 | 2.400E-06 | 2.800E-06 |
| 4133 | 207 | G207M | A | 2.1e-06 | 1.5e-06 | 1.7e-06 | 1.767E-06 | 3.055E-07 | 1.500E-06 | 2.100E-06 |
| 4134 | 207 | G207F | A | 2.6e-06 | 3.3e-06 | 2e-06 | 2.633E-06 | 6.506E-07 | 2.000E-06 | 3.300E-06 |
| 4135 | 207 | G207P | A | 2.4e-06 | 2.2e-06 | 2.4e-06 | 2.333E-06 | 1.155E-07 | 2.200E-06 | 2.400E-06 |
| 4136 | 207 | G207S | A | 3.2e-06 | 1.7e-06 | 2.5e-06 | 2.467E-06 | 7.506E-07 | 1.700E-06 | 3.200E-06 |
| 4137 | 207 | G207T | A | 2e-06 | 2.4e-06 | 3.1e-06 | 2.500E-06 | 5.568E-07 | 2.000E-06 | 3.100E-06 |
| 4138 | 207 | G207W | A | 2.7e-06 | 1.4e-06 | 3.4e-06 | 2.500E-06 | 1.015E-06 | 1.400E-06 | 3.400E-06 |
| 4139 | 207 | G207Y | A | 3.6e-06 | 2.8e-06 | 2.4e-06 | 2.933E-06 | 6.110E-07 | 2.400E-06 | 3.600E-06 |
| 4140 | 207 | G207V | A | 2.9e-06 | 2.6e-06 | 1.3e-06 | 2.267E-06 | 8.505E-07 | 1.300E-06 | 2.900E-06 |
| 4141 | 208 | P208A | A | 2.8e-06 | 2e-06 | 3e-06 | 2.600E-06 | 5.292E-07 | 2.000E-06 | 3.000E-06 |
| 4142 | 208 | P208R | A | 2e-06 | 2.5e-06 | 1.6e-06 | 2.033E-06 | 4.509E-07 | 1.600E-06 | 2.500E-06 |
| 4143 | 208 | P208N | A | 2.6e-06 | 1.7e-06 | 3.5e-06 | 2.600E-06 | 9.000E-07 | 1.700E-06 | 3.500E-06 |
| 4144 | 208 | P208D | A | 3.5e-06 | 1.6e-06 | 2.5e-06 | 2.533E-06 | 9.504E-07 | 1.600E-06 | 3.500E-06 |
| 4145 | 208 | P208C | A | 3.8e-06 | 3.1e-06 | 3.1e-06 | 3.333E-06 | 4.041E-07 | 3.100E-06 | 3.800E-06 |
| 4146 | 208 | P208E | A | 2.7e-06 | 3.6e-06 | 1.4e-06 | 2.567E-06 | 1.106E-06 | 1.400E-06 | 3.600E-06 |
| 4147 | 208 | P208Q | A | 5.4e-06 | 3e-06 | 5.1e-06 | 4.500E-06 | 1.308E-06 | 3.000E-06 | 5.400E-06 |
| 4148 | 208 | P208G | A | 2e-06 | 2.7e-06 | 3.9e-06 | 2.867E-06 | 9.609E-07 | 2.000E-06 | 3.900E-06 |
| 4149 | 208 | P208H | A | 3.1e-06 | 2.2e-06 | 2.5e-06 | 2.600E-06 | 4.583E-07 | 2.200E-06 | 3.100E-06 |
| 4150 | 208 | P208I | A | 2.1e-06 | 2.6e-06 | 2e-06 | 2.233E-06 | 3.215E-07 | 2.000E-06 | 2.600E-06 |
| 4151 | 208 | P208L | A | 1.5e-06 | 1.8e-06 | 1.1e-06 | 1.467E-06 | 3.512E-07 | 1.100E-06 | 1.800E-06 |
| 4152 | 208 | P208K | A | 2.1e-06 | 3.1e-06 | 2.3e-06 | 2.500E-06 | 5.292E-07 | 2.100E-06 | 3.100E-06 |
| 4153 | 208 | P208M | A | 1.9e-06 | 1.8e-06 | 2.8e-06 | 2.167E-06 | 5.508E-07 | 1.800E-06 | 2.800E-06 |
| 4154 | 208 | P208F | A | 3.2e-06 | 2e-06 | 3.6e-06 | 2.933E-06 | 8.327E-07 | 2.000E-06 | 3.600E-06 |
| 4155 | 208 | P208P | A | 4.5e-06 | 3e-06 | 2.7e-06 | 3.400E-06 | 9.644E-07 | 2.700E-06 | 4.500E-06 |
| 4156 | 208 | P208S | A | 2.3e-06 | 2.8e-06 | 2.4e-06 | 2.500E-06 | 2.646E-07 | 2.300E-06 | 2.800E-06 |
| 4157 | 208 | P208T | A | 4.2e-06 | 2.8e-06 | 2.5e-06 | 3.167E-06 | 9.074E-07 | 2.500E-06 | 4.200E-06 |
| 4158 | 208 | P208W | A | 3.4e-06 | 2.9e-06 | 7.8e-07 | 2.360E-06 | 1.391E-06 | 7.800E-07 | 3.400E-06 |
| 4159 | 208 | P208Y | A | 2.6e-06 | 3e-06 | 3.6e-06 | 3.067E-06 | 5.033E-07 | 2.600E-06 | 3.600E-06 |
| 4160 | 208 | P208V | A | 1.6e-06 | 1.5e-06 | 1.7e-06 | 1.600E-06 | 1.000E-07 | 1.500E-06 | 1.700E-06 |
| 4161 | 209 | L209A | A | 1.3e-06 | 1.9e-06 | 2.1e-06 | 1.767E-06 | 4.163E-07 | 1.300E-06 | 2.100E-06 |
| 4162 | 209 | L209R | A | 2e-06 | 2.2e-06 | 2.3e-06 | 2.167E-06 | 1.528E-07 | 2.000E-06 | 2.300E-06 |
| 4163 | 209 | L209N | A | 4.4e-06 | 3.3e-06 | 3.2e-06 | 3.633E-06 | 6.658E-07 | 3.200E-06 | 4.400E-06 |
| 4164 | 209 | L209D | A | 2e-06 | 2.4e-06 | 2e-06 | 2.133E-06 | 2.309E-07 | 2.000E-06 | 2.400E-06 |

| | | | | | | | | | | |
|---|---|---|---|---|---|---|---|---|---|---|
| 4165 | 209 | L209C | A | 2.4e-06 | 2.4e-06 | 1.5e-06 | 2.100E-06 | 5.196E-07 | 1.500E-06 | 2.400E-06 |
| 4166 | 209 | L209E | A | 2e-06 | 1.4e-06 | 1.3e-06 | 1.567E-06 | 3.786E-07 | 1.300E-06 | 2.000E-06 |
| 4167 | 209 | L209Q | A | 3.6e-06 | 1.8e-06 | 1.3e-06 | 2.233E-06 | 1.210E-06 | 1.300E-06 | 3.600E-06 |
| 4168 | 209 | L209G | A | 1.7e-06 | 2.8e-06 | 1.7e-06 | 2.067E-06 | 6.351E-07 | 1.700E-06 | 2.800E-06 |
| 4169 | 209 | L209H | A | 4.8e-06 | 3.5e-06 | 2.6e-06 | 3.633E-06 | 1.106E-06 | 2.600E-06 | 4.800E-06 |
| 4170 | 209 | L209I | A | 5.9e-06 | 1.4e-06 | 2.3e-06 | 3.200E-06 | 2.381E-06 | 1.400E-06 | 5.900E-06 |
| 4171 | 209 | L209L | A | 1.3e-06 | 1.5e-06 | 2.2e-06 | 1.667E-06 | 4.726E-07 | 1.300E-06 | 2.200E-06 |
| 4172 | 209 | L209K | A | 2e-06 | 1.1e-06 | 3.3e-06 | 2.133E-06 | 1.106E-06 | 1.100E-06 | 3.300E-06 |
| 4173 | 209 | L209M | A | 2.5e-06 | 2.9e-06 | 1.6e-06 | 2.333E-06 | 6.658E-07 | 1.600E-06 | 2.900E-06 |
| 4174 | 209 | L209F | A | 1.8e-06 | 2.3e-06 | 3.2e-06 | 2.433E-06 | 7.095E-07 | 1.800E-06 | 3.200E-06 |
| 4175 | 209 | L209P | A | 2.6e-06 | 2.6e-06 | 2.8e-06 | 2.667E-06 | 1.155E-07 | 2.600E-06 | 2.800E-06 |
| 4176 | 209 | L209S | A | 1.4e-06 | 2.4e-06 | 1.2e-06 | 1.667E-06 | 6.429E-07 | 1.200E-06 | 2.400E-06 |
| 4177 | 209 | L209T | A | 2.4e-06 | 2.2e-06 | 3.3e-06 | 2.633E-06 | 5.859E-07 | 2.200E-06 | 3.300E-06 |
| 4178 | 209 | L209W | A | 2.1e-06 | 1.9e-06 | 3e-06 | 2.333E-06 | 5.859E-07 | 1.900E-06 | 3.000E-06 |
| 4179 | 209 | L209Y | A | 2.7e-06 | 1.6e-06 | 1.9e-06 | 2.067E-06 | 5.686E-07 | 1.600E-06 | 2.700E-06 |
| 4180 | 209 | L209V | A | 3.6e-06 | 2.1e-06 | 2.1e-06 | 2.600E-06 | 8.660E-07 | 2.100E-06 | 3.600E-06 |
| 4181 | 210 | P210A | A | 2.7e-06 | 1.5e-06 | 2e-06 | 2.067E-06 | 6.028E-07 | 1.500E-06 | 2.700E-06 |
| 4182 | 210 | P210R | A | 1.3e-06 | 1.2e-06 | 2.4e-06 | 1.633E-06 | 6.658E-07 | 1.200E-06 | 2.400E-06 |
| 4183 | 210 | P210N | A | 1.4e-06 | 1.7e-06 | 3.3e-06 | 2.133E-06 | 1.021E-06 | 1.400E-06 | 3.300E-06 |
| 4184 | 210 | P210D | A | 1.8e-06 | 2.3e-06 | 1.8e-06 | 1.967E-06 | 2.887E-07 | 1.800E-06 | 2.300E-06 |
| 4185 | 210 | P210C | A | 2.9e-06 | 1.5e-06 | 2.1e-06 | 2.167E-06 | 7.024E-07 | 1.500E-06 | 2.900E-06 |
| 4186 | 210 | P210E | A | 1.4e-06 | 3.7e-06 | 2.7e-06 | 2.600E-06 | 1.153E-06 | 1.400E-06 | 3.700E-06 |
| 4187 | 210 | P210Q | A | 2.7e-06 | 1.7e-06 | 2.2e-06 | 2.200E-06 | 5.000E-07 | 1.700E-06 | 2.700E-06 |
| 4188 | 210 | P210G | A | 2.1e-06 | 2.6e-06 | 2.6e-06 | 2.433E-06 | 2.887E-07 | 2.100E-06 | 2.600E-06 |
| 4189 | 210 | P210H | A | 1.5e-06 | 2e-06 | 1.4e-06 | 1.633E-06 | 3.215E-07 | 1.400E-06 | 2.000E-06 |
| 4190 | 210 | P210I | A | 2.5e-06 | 2.9e-06 | 3.4e-06 | 2.933E-06 | 4.509E-07 | 2.500E-06 | 3.400E-06 |
| 4191 | 210 | P210L | A | 2.1e-06 | 2.4e-06 | 2.7e-06 | 2.400E-06 | 3.000E-07 | 2.100E-06 | 2.700E-06 |
| 4192 | 210 | P210K | A | 3.8e-06 | 2e-06 | 1.7e-06 | 2.500E-06 | 1.136E-06 | 1.700E-06 | 3.800E-06 |
| 4193 | 210 | P210M | A | 3.4e-06 | 1.9e-06 | 3.6e-06 | 2.967E-06 | 9.292E-07 | 1.900E-06 | 3.600E-06 |
| 4194 | 210 | P210F | A | 2.6e-06 | 2.6e-06 | 1.2e-06 | 2.133E-06 | 8.083E-07 | 1.200E-06 | 2.600E-06 |
| 4195 | 210 | P210P | A | 1.9e-06 | 2.5e-06 | 2.4e-06 | 2.267E-06 | 3.215E-07 | 1.900E-06 | 2.500E-06 |
| 4196 | 210 | P210S | A | 2.1e-06 | 2.5e-06 | 1.8e-06 | 2.133E-06 | 3.512E-07 | 1.800E-06 | 2.500E-06 |
| 4197 | 210 | P210T | A | 1.3e-06 | 1.3e-06 | 4e-06 | 2.200E-06 | 1.559E-06 | 1.300E-06 | 4.000E-06 |
| 4198 | 210 | P210W | A | 5.8e-06 | 2e-06 | 2.7e-06 | 3.500E-06 | 2.022E-06 | 2.000E-06 | 5.800E-06 |
| 4199 | 210 | P210Y | A | 2.4e-06 | 2.3e-06 | 2.1e-06 | 2.267E-06 | 1.528E-07 | 2.100E-06 | 2.400E-06 |
| 4200 | 210 | P210V | A | 2.7e-06 | 2e-06 | 2.9e-06 | 2.533E-06 | 4.726E-07 | 2.000E-06 | 2.900E-06 |
| 4201 | 211 | T211A | A | 2.3e-06 | 2.7e-06 | 2.1e-06 | 2.367E-06 | 3.055E-07 | 2.100E-06 | 2.700E-06 |
| 4202 | 211 | T211R | A | 3.3e-06 | 1.6e-06 | 2.7e-06 | 2.533E-06 | 8.622E-07 | 1.600E-06 | 3.300E-06 |
| 4203 | 211 | T211N | A | 3.5e-06 | 2.1e-06 | 1.3e-06 | 2.300E-06 | 1.114E-06 | 1.300E-06 | 3.500E-06 |
| 4204 | 211 | T211D | A | 1.9e-06 | 3.4e-06 | 3.2e-06 | 2.833E-06 | 8.145E-07 | 1.900E-06 | 3.400E-06 |
| 4205 | 211 | T211C | A | 1.6e-06 | 3.5e-06 | 2.8e-06 | 2.633E-06 | 9.609E-07 | 1.600E-06 | 3.500E-06 |
| 4206 | 211 | T211E | A | 1.9e-06 | 1e-06 | 2.3e-06 | 1.733E-06 | 6.658E-07 | 1.000E-06 | 2.300E-06 |
| 4207 | 211 | T211Q | A | 2.4e-06 | 2.4e-06 | 2.1e-06 | 2.300E-06 | 1.732E-07 | 2.100E-06 | 2.400E-06 |
| 4208 | 211 | T211G | A | 2.8e-06 | 6.8e-06 | 1.7e-06 | 3.767E-06 | 2.684E-06 | 1.700E-06 | 6.800E-06 |
| 4209 | 211 | T211H | A | 3.5e-06 | 2.3e-06 | 1.6e-06 | 2.467E-06 | 9.609E-07 | 1.600E-06 | 3.500E-06 |
| 4210 | 211 | T211I | A | 3.5e-06 | 3e-06 | 2.6e-06 | 3.033E-06 | 4.509E-07 | 2.600E-06 | 3.500E-06 |
| 4211 | 211 | T211L | A | 2.5e-06 | 1.8e-06 | 2.7e-06 | 2.333E-06 | 4.726E-07 | 1.800E-06 | 2.700E-06 |
| 4212 | 211 | T211K | A | 1.1e-06 | 1.2e-06 | 1.9e-06 | 1.400E-06 | 4.359E-07 | 1.100E-06 | 1.900E-06 |
| 4213 | 211 | T211M | A | 5.6e-06 | 5.2e-06 | 2.6e-06 | 4.467E-06 | 1.629E-06 | 2.600E-06 | 5.600E-06 |
| 4214 | 211 | T211F | A | 3e-06 | 2.8e-06 | 3.7e-06 | 3.167E-06 | 4.726E-07 | 2.800E-06 | 3.700E-06 |
| 4215 | 211 | T211P | A | 1.6e-06 | 2.2e-06 | 3.1e-06 | 2.300E-06 | 7.550E-07 | 1.600E-06 | 3.100E-06 |
| 4216 | 211 | T211S | A | 2.2e-06 | 2.5e-06 | 2.9e-06 | 2.533E-06 | 3.512E-07 | 2.200E-06 | 2.900E-06 |
| 4217 | 211 | T211T | A | 2.8e-06 | 3e-06 | 3.6e-06 | 3.133E-06 | 4.163E-07 | 2.800E-06 | 3.600E-06 |
| 4218 | 211 | T211W | A | 2.1e-06 | 2e-06 | 2.9e-06 | 2.333E-06 | 4.933E-07 | 2.000E-06 | 2.900E-06 |
| 4219 | 211 | T211Y | A | 3e-06 | 3.5e-06 | 2.4e-06 | 2.967E-06 | 5.508E-07 | 2.400E-06 | 3.500E-06 |
| 4220 | 211 | T211V | A | 1.9e-06 | 2.9e-06 | 2e-06 | 2.267E-06 | 5.508E-07 | 1.900E-06 | 2.900E-06 |
| 4221 | 212 | D212A | A | 2e-06 | 2.2e-06 | 2.5e-06 | 2.233E-06 | 2.517E-07 | 2.000E-06 | 2.500E-06 |
| 4222 | 212 | D212R | A | 1.3e-06 | 4e-06 | 3.2e-06 | 2.833E-06 | 1.387E-06 | 1.300E-06 | 4.000E-06 |
| 4223 | 212 | D212N | A | 2.5e-06 | 3.7e-06 | 2.2e-06 | 2.800E-06 | 7.937E-07 | 2.200E-06 | 3.700E-06 |
| 4224 | 212 | D212D | A | 3.6e-06 | 3e-06 | 3.1e-06 | 3.233E-06 | 3.215E-07 | 3.000E-06 | 3.600E-06 |
| 4225 | 212 | D212C | A | 1.7e-06 | 1.8e-06 | 2.1e-06 | 1.867E-06 | 2.082E-07 | 1.700E-06 | 2.100E-06 |
| 4226 | 212 | D212E | A | 1.3e-06 | 2.2e-06 | 1.8e-06 | 1.767E-06 | 4.509E-07 | 1.300E-06 | 2.200E-06 |
| 4227 | 212 | D212Q | A | 2.6e-06 | 2e-06 | 2.5e-06 | 2.367E-06 | 3.215E-07 | 2.000E-06 | 2.600E-06 |
| 4228 | 212 | D212G | A | 2.1e-06 | 2.1e-06 | 1.6e-06 | 1.933E-06 | 2.887E-07 | 1.600E-06 | 2.100E-06 |
| 4229 | 212 | D212H | A | 2.6e-06 | 1.6e-06 | 2.4e-06 | 2.200E-06 | 5.292E-07 | 1.600E-06 | 2.600E-06 |
| 4230 | 212 | D212I | A | 2.5e-06 | 2.6e-06 | 1.4e-06 | 2.167E-06 | 6.658E-07 | 1.400E-06 | 2.600E-06 |
| 4231 | 212 | D212L | A | 2e-06 | 1.7e-06 | 2.9e-06 | 2.200E-06 | 6.245E-07 | 1.700E-06 | 2.900E-06 |
| 4232 | 212 | D212K | A | 2.9e-06 | 2.7e-06 | 1.6e-06 | 2.400E-06 | 7.000E-07 | 1.600E-06 | 2.900E-06 |
| 4233 | 212 | D212M | A | 2.9e-06 | 1e-06 | 1.2e-06 | 1.700E-06 | 1.044E-06 | 1.000E-06 | 2.900E-06 |
| 4234 | 212 | D212F | A | 2.2e-06 | 5.6e-06 | 2.8e-06 | 3.533E-06 | 1.815E-06 | 2.200E-06 | 5.600E-06 |
| 4235 | 212 | D212P | A | 2.6e-06 | 2.8e-06 | 2.5e-06 | 2.633E-06 | 1.528E-07 | 2.500E-06 | 2.800E-06 |
| 4236 | 212 | D212S | A | 1.5e-06 | 1.6e-06 | 2.3e-06 | 1.800E-06 | 4.359E-07 | 1.500E-06 | 2.300E-06 |
| 4237 | 212 | D212T | A | 2.9e-06 | 1.4e-06 | 2.3e-06 | 2.200E-06 | 7.550E-07 | 1.400E-06 | 2.900E-06 |
| 4238 | 212 | D212W | A | 3.1e-06 | 3e-06 | 3.3e-06 | 3.133E-06 | 1.528E-07 | 3.000E-06 | 3.300E-06 |
| 4239 | 212 | D212Y | A | 3.1e-06 | 3e-06 | 1.9e-06 | 2.667E-06 | 6.658E-07 | 1.900E-06 | 3.100E-06 |
| 4240 | 212 | D212V | A | 2.9e-06 | 4.1e-06 | 2.4e-06 | 3.133E-06 | 8.737E-07 | 2.400E-06 | 4.100E-06 |
| 4241 | 213 | C213A | A | 2.6e-06 | 3e-06 | 1.3e-06 | 2.300E-06 | 8.888E-07 | 1.300E-06 | 3.000E-06 |
| 4242 | 213 | C213R | A | 1.9e-06 | 3.1e-06 | 3e-06 | 2.667E-06 | 6.658E-07 | 1.900E-06 | 3.100E-06 |
| 4243 | 213 | C213N | A | 1.9e-06 | 2.5e-06 | 4.2e-06 | 2.867E-06 | 1.193E-06 | 1.900E-06 | 4.200E-06 |
| 4244 | 213 | C213D | A | 2.9e-06 | 2.3e-06 | 2.3e-06 | 2.500E-06 | 3.464E-07 | 2.300E-06 | 2.900E-06 |
| 4245 | 213 | C213C | A | 2.5e-06 | 3.6e-06 | 3.5e-06 | 3.200E-06 | 6.083E-07 | 2.500E-06 | 3.600E-06 |
| 4246 | 213 | C213E | A | 2.8e-06 | 2.4e-06 | 3e-06 | 2.733E-06 | 3.055E-07 | 2.400E-06 | 3.000E-06 |
| 4247 | 213 | C213Q | A | 2.1e-06 | 3.9e-06 | 1.4e-06 | 2.467E-06 | 1.290E-06 | 1.400E-06 | 3.900E-06 |
| 4248 | 213 | C213G | A | 3.8e-06 | 9.7e-07 | 2.5e-06 | 2.423E-06 | 1.417E-06 | 9.700E-07 | 3.800E-06 |
| 4249 | 213 | C213H | A | 1.9e-06 | 3.4e-06 | 2.2e-06 | 2.500E-06 | 7.937E-07 | 1.900E-06 | 3.400E-06 |

| | | | | | | | | | | |
|---|---|---|---|---|---|---|---|---|---|---|
| 4250 | 213 | C213I | A | 2.3e-06 | 2.2e-06 | 2.7e-06 | 2.400E-06 | 2.646E-07 | 2.200E-06 | 2.700E-06 |
| 4251 | 213 | C213L | A | 2.8e-06 | 2.3e-06 | 2.3e-06 | 2.467E-06 | 2.887E-07 | 2.300E-06 | 2.800E-06 |
| 4252 | 213 | C213K | A | 2.6e-06 | 3.4e-06 | 3.2e-06 | 3.067E-06 | 4.163E-07 | 2.600E-06 | 3.400E-06 |
| 4253 | 213 | C213M | A | 1.3e-06 | 2.9e-06 | 2e-06 | 2.067E-06 | 8.021E-07 | 1.300E-06 | 2.900E-06 |
| 4254 | 213 | C213F | A | 2.8e-06 | 1.8e-06 | 2.3e-06 | 2.300E-06 | 5.000E-07 | 1.800E-06 | 2.800E-06 |
| 4255 | 213 | C213P | A | 1.3e-06 | 2.6e-06 | 1.3e-06 | 1.733E-06 | 7.506E-07 | 1.300E-06 | 2.600E-06 |
| 4256 | 213 | C213S | A | 3.7e-06 | 3.1e-06 | 2.7e-06 | 3.167E-06 | 5.033E-07 | 2.700E-06 | 3.700E-06 |
| 4257 | 213 | C213T | A | 2.9e-06 | 2.5e-06 | 1.3e-06 | 2.233E-06 | 8.327E-07 | 1.300E-06 | 2.900E-06 |
| 4258 | 213 | C213W | A | 2.7e-06 | 1.1e-06 | 1.3e-06 | 1.700E-06 | 8.718E-07 | 1.100E-06 | 2.700E-06 |
| 4259 | 213 | C213Y | A | 1.7e-06 | 1.4e-06 | 1.5e-06 | 1.533E-06 | 1.528E-07 | 1.400E-06 | 1.700E-06 |
| 4260 | 213 | C213V | A | 2.5e-06 | 2.7e-06 | 1.4e-06 | 2.200E-06 | 7.000E-07 | 1.400E-06 | 2.700E-06 |
| 4261 | 214 | C214A | A | 2.6e-06 | 2.3e-06 | 1.8e-06 | 2.233E-06 | 4.041E-07 | 1.800E-06 | 2.600E-06 |
| 4262 | 214 | C214R | A | 2.4e-06 | 2.8e-06 | 3e-06 | 2.733E-06 | 3.055E-07 | 2.400E-06 | 3.000E-06 |
| 4263 | 214 | C214N | A | 2.9e-06 | 2.1e-06 | 3.3e-06 | 2.767E-06 | 6.110E-07 | 2.100E-06 | 3.300E-06 |
| 4264 | 214 | C214D | A | 2.3e-06 | 3.9e-06 | 3.8e-06 | 3.333E-06 | 8.963E-07 | 2.300E-06 | 3.900E-06 |
| 4265 | 214 | C214C | A | 2.1e-06 | 2.2e-06 | 2.6e-06 | 2.300E-06 | 2.646E-07 | 2.100E-06 | 2.600E-06 |
| 4266 | 214 | C214E | A | 1.8e-06 | 2.7e-06 | 4.8e-06 | 3.100E-06 | 1.539E-06 | 1.800E-06 | 4.800E-06 |
| 4267 | 214 | C214Q | A | 2.6e-06 | 2.3e-06 | 2.4e-06 | 2.433E-06 | 1.528E-07 | 2.300E-06 | 2.600E-06 |
| 4268 | 214 | C214G | A | 2.4e-06 | 3.4e-06 | 2.1e-06 | 2.633E-06 | 6.807E-07 | 2.100E-06 | 3.400E-06 |
| 4269 | 214 | C214H | A | 1.8e-06 | 2.8e-06 | 2.6e-06 | 2.400E-06 | 5.292E-07 | 1.800E-06 | 2.800E-06 |
| 4270 | 214 | C214I | A | 1.9e-06 | 2.7e-06 | 5e-06 | 3.200E-06 | 1.609E-06 | 1.900E-06 | 5.000E-06 |
| 4271 | 214 | C214L | A | 3.1e-06 | 3.7e-06 | 3.4e-06 | 3.400E-06 | 3.000E-07 | 3.100E-06 | 3.700E-06 |
| 4272 | 214 | C214K | A | 1.9e-06 | 2.4e-06 | 2.6e-06 | 2.300E-06 | 3.606E-07 | 1.900E-06 | 2.600E-06 |
| 4273 | 214 | C214M | A | 1.6e-06 | 4.4e-06 | 2.1e-06 | 2.700E-06 | 1.493E-06 | 1.600E-06 | 4.400E-06 |
| 4274 | 214 | C214F | A | 1.5e-06 | 2.1e-06 | 4.6e-06 | 2.733E-06 | 1.644E-06 | 1.500E-06 | 4.600E-06 |
| 4275 | 214 | C214P | A | 2.2e-06 | 2.8e-06 | 1.3e-06 | 2.100E-06 | 7.550E-07 | 1.300E-06 | 2.800E-06 |
| 4276 | 214 | C214S | A | 2.6e-06 | 2.2e-06 | 3.1e-06 | 2.633E-06 | 4.509E-07 | 2.200E-06 | 3.100E-06 |
| 4277 | 214 | C214T | A | 1.9e-06 | 2.5e-06 | 2.8e-06 | 2.400E-06 | 4.583E-07 | 1.900E-06 | 2.800E-06 |
| 4278 | 214 | C214W | A | 1.3e-06 | 1.8e-06 | 1.3e-06 | 1.467E-06 | 2.887E-07 | 1.300E-06 | 1.800E-06 |
| 4279 | 214 | C214Y | A | 3.2e-06 | 2.4e-06 | 3.4e-06 | 3.000E-06 | 5.292E-07 | 2.400E-06 | 3.400E-06 |
| 4280 | 214 | C214V | A | 2.1e-06 | 3.1e-06 | 1.6e-06 | 2.267E-06 | 7.638E-07 | 1.600E-06 | 3.100E-06 |
| 4281 | 215 | H215A | A | 1.5e-06 | 2.3e-06 | 2.7e-06 | 2.167E-06 | 6.110E-07 | 1.500E-06 | 2.700E-06 |
| 4282 | 215 | H215R | A | 1.8e-06 | 2.2e-06 | 2.3e-06 | 2.100E-06 | 2.646E-07 | 1.800E-06 | 2.300E-06 |
| 4283 | 215 | H215N | A | 2.8e-06 | 1.8e-06 | 3.2e-06 | 2.600E-06 | 7.211E-07 | 1.800E-06 | 3.200E-06 |
| 4284 | 215 | H215D | A | 1.3e-06 | 2.6e-06 | 2e-06 | 1.967E-06 | 6.506E-07 | 1.300E-06 | 2.600E-06 |
| 4285 | 215 | H215C | A | 2.4e-06 | 5e-06 | 2.3e-06 | 3.233E-06 | 1.531E-06 | 2.300E-06 | 5.000E-06 |
| 4286 | 215 | H215E | A | 4.6e-06 | 2.9e-06 | 3.9e-06 | 3.800E-06 | 8.544E-07 | 2.900E-06 | 4.600E-06 |
| 4287 | 215 | H215Q | A | 2.5e-06 | 2.3e-06 | 2.1e-06 | 2.300E-06 | 2.000E-07 | 2.100E-06 | 2.500E-06 |
| 4288 | 215 | H215G | A | 3.5e-06 | 3e-06 | 2.5e-06 | 3.000E-06 | 5.000E-07 | 2.500E-06 | 3.500E-06 |
| 4289 | 215 | H215H | A | 2.8e-06 | 3.9e-06 | 2.7e-06 | 3.133E-06 | 6.658E-07 | 2.700E-06 | 3.900E-06 |
| 4290 | 215 | H215I | A | 1.7e-06 | 2.8e-06 | 2.3e-06 | 2.267E-06 | 5.508E-07 | 1.700E-06 | 2.800E-06 |
| 4291 | 215 | H215L | A | 1.1e-06 | 2.5e-06 | 2.5e-06 | 2.033E-06 | 8.083E-07 | 1.100E-06 | 2.500E-06 |
| 4292 | 215 | H215K | A | 2.6e-06 | 2.3e-06 | 2.7e-06 | 2.533E-06 | 2.082E-07 | 2.300E-06 | 2.700E-06 |
| 4293 | 215 | H215M | A | 2.7e-06 | 2.3e-06 | 8.6e-07 | 1.953E-06 | 9.677E-07 | 8.600E-07 | 2.700E-06 |
| 4294 | 215 | H215F | A | 3.9e-06 | 2.4e-06 | 1.7e-06 | 2.667E-06 | 1.124E-06 | 1.700E-06 | 3.900E-06 |
| 4295 | 215 | H215P | A | 2.3e-06 | 2.2e-06 | 5.6e-06 | 3.367E-06 | 1.935E-06 | 2.200E-06 | 5.600E-06 |
| 4296 | 215 | H215S | A | 2.7e-06 | 1.7e-06 | 2.7e-06 | 2.367E-06 | 5.774E-07 | 1.700E-06 | 2.700E-06 |
| 4297 | 215 | H215T | A | 2.4e-06 | 2.4e-06 | 3.7e-06 | 2.833E-06 | 7.506E-07 | 2.400E-06 | 3.700E-06 |
| 4298 | 215 | H215W | A | 1.7e-06 | 9.8e-07 | 1.2e-06 | 1.293E-06 | 3.690E-07 | 9.800E-07 | 1.700E-06 |
| 4299 | 215 | H215Y | A | 3.2e-06 | 1.7e-06 | 4.2e-06 | 3.033E-06 | 1.258E-06 | 1.700E-06 | 4.200E-06 |
| 4300 | 215 | H215V | A | 2.2e-06 | 2.3e-06 | 3.2e-06 | 2.567E-06 | 5.508E-07 | 2.200E-06 | 3.200E-06 |
| 4301 | 216 | E216A | A | 2.5e-06 | 2.3e-06 | 1.7e-06 | 2.167E-06 | 4.163E-07 | 1.700E-06 | 2.500E-06 |
| 4302 | 216 | E216R | A | 2.9e-06 | 1.3e-06 | 2.6e-06 | 2.267E-06 | 8.505E-07 | 1.300E-06 | 2.900E-06 |
| 4303 | 216 | E216N | A | 1.1e-06 | 1.1e-06 | 3.1e-06 | 1.767E-06 | 1.155E-06 | 1.100E-06 | 3.100E-06 |
| 4304 | 216 | E216D | A | 1.9e-06 | 2e-06 | 3.3e-06 | 2.400E-06 | 7.810E-07 | 1.900E-06 | 3.300E-06 |
| 4305 | 216 | E216C | A | 1.9e-06 | 5.1e-06 | 2.6e-06 | 3.200E-06 | 1.682E-06 | 1.900E-06 | 5.100E-06 |
| 4306 | 216 | E216E | A | 2.1e-06 | 3.5e-06 | 1.8e-06 | 2.467E-06 | 9.074E-07 | 1.800E-06 | 3.500E-06 |
| 4307 | 216 | E216Q | A | 4e-06 | 3.2e-06 | 4e-06 | 3.733E-06 | 4.619E-07 | 3.200E-06 | 4.000E-06 |
| 4308 | 216 | E216G | A | 1.6e-06 | 1.9e-06 | 2.9e-06 | 2.133E-06 | 6.807E-07 | 1.600E-06 | 2.900E-06 |
| 4309 | 216 | E216H | A | 2.1e-06 | 4e-06 | 2.2e-06 | 2.767E-06 | 1.069E-06 | 2.100E-06 | 4.000E-06 |
| 4310 | 216 | E216I | A | 5.4e-06 | 4.7e-06 | 2.1e-06 | 4.067E-06 | 1.739E-06 | 2.100E-06 | 5.400E-06 |
| 4311 | 216 | E216L | A | 1.7e-06 | 1.5e-06 | 4.1e-06 | 2.433E-06 | 1.447E-06 | 1.500E-06 | 4.100E-06 |
| 4312 | 216 | E216K | A | 2.6e-06 | 3e-06 | 3.3e-06 | 2.967E-06 | 3.512E-07 | 2.600E-06 | 3.300E-06 |
| 4313 | 216 | E216M | A | 2.6e-06 | 3.6e-06 | 2e-06 | 2.733E-06 | 8.083E-07 | 2.000E-06 | 3.600E-06 |
| 4314 | 216 | E216F | A | 3.3e-06 | 1.6e-06 | 3.2e-06 | 2.700E-06 | 9.539E-07 | 1.600E-06 | 3.300E-06 |
| 4315 | 216 | E216P | A | 2.1e-06 | 2.2e-06 | 9.9e-07 | 1.763E-06 | 6.716E-07 | 9.900E-07 | 2.200E-06 |
| 4316 | 216 | E216S | A | 3.8e-06 | 1.7e-06 | 3.7e-06 | 3.067E-06 | 1.185E-06 | 1.700E-06 | 3.800E-06 |
| 4317 | 216 | E216T | A | 1.8e-06 | 2.4e-06 | 5.8e-06 | 3.333E-06 | 2.157E-06 | 1.800E-06 | 5.800E-06 |
| 4318 | 216 | E216W | A | 2.8e-06 | 1.9e-06 | 2.2e-06 | 2.300E-06 | 4.583E-07 | 1.900E-06 | 2.800E-06 |
| 4319 | 216 | E216Y | A | 2e-06 | 2.1e-06 | 4.4e-06 | 2.833E-06 | 1.358E-06 | 2.000E-06 | 4.400E-06 |
| 4320 | 216 | E216V | A | 2.9e-06 | 3e-06 | 2e-06 | 2.633E-06 | 5.508E-07 | 2.000E-06 | 3.000E-06 |
| 4321 | 217 | Q217A | A | 1.2e-06 | 2.2e-06 | 2.9e-06 | 2.100E-06 | 8.544E-07 | 1.200E-06 | 2.900E-06 |
| 4322 | 217 | Q217R | A | 2.1e-06 | 3.4e-06 | 3.5e-06 | 3.000E-06 | 7.810E-07 | 2.100E-06 | 3.500E-06 |
| 4323 | 217 | Q217N | A | 2.2e-06 | 2.7e-06 | 2.2e-06 | 2.367E-06 | 2.887E-07 | 2.200E-06 | 2.700E-06 |
| 4324 | 217 | Q217D | A | 1.7e-06 | 3e-06 | 1.1e-06 | 1.933E-06 | 9.713E-07 | 1.100E-06 | 3.000E-06 |
| 4325 | 217 | Q217C | A | 2.4e-06 | 1.1e-06 | 1.1e-06 | 1.533E-06 | 7.506E-07 | 1.100E-06 | 2.400E-06 |
| 4326 | 217 | Q217E | A | 2.5e-06 | 2e-06 | 2.5e-06 | 2.333E-06 | 2.887E-07 | 2.000E-06 | 2.500E-06 |
| 4327 | 217 | Q217Q | A | 2.6e-06 | 2.3e-06 | 3.4e-06 | 2.767E-06 | 5.686E-07 | 2.300E-06 | 3.400E-06 |
| 4328 | 217 | Q217G | A | 4.3e-06 | 2e-06 | 1.8e-06 | 2.700E-06 | 1.389E-06 | 1.800E-06 | 4.300E-06 |
| 4329 | 217 | Q217H | A | 1.2e-06 | 2.2e-06 | 3.9e-06 | 2.433E-06 | 1.365E-06 | 1.200E-06 | 3.900E-06 |
| 4330 | 217 | Q217I | A | 3.3e-06 | 2.6e-06 | 1.6e-06 | 2.500E-06 | 8.544E-07 | 1.600E-06 | 3.300E-06 |
| 4331 | 217 | Q217L | A | 2.1e-06 | 2.9e-06 | 2.1e-06 | 2.367E-06 | 4.619E-07 | 2.100E-06 | 2.900E-06 |
| 4332 | 217 | Q217K | A | 1.2e-06 | 1.3e-06 | 3.4e-06 | 1.967E-06 | 1.242E-06 | 1.200E-06 | 3.400E-06 |
| 4333 | 217 | Q217M | A | 2.5e-06 | 4e-06 | 2.3e-06 | 2.933E-06 | 9.292E-07 | 2.300E-06 | 4.000E-06 |
| 4334 | 217 | Q217F | A | 2.7e-06 | 3.2e-06 | 2.5e-06 | 2.800E-06 | 3.606E-07 | 2.500E-06 | 3.200E-06 |

| | | | | | | | | | | |
|---|---|---|---|---|---|---|---|---|---|---|
| 4335 | 217 | Q217P | A | 2.3e-06 | 1.7e-06 | 3e-06 | 2.333E-06 | 6.506E-07 | 1.700E-06 | 3.000E-06 |
| 4336 | 217 | Q217S | A | 1.9e-06 | 1.3e-06 | 1.1e-06 | 1.433E-06 | 4.163E-07 | 1.100E-06 | 1.900E-06 |
| 4337 | 217 | Q217T | A | 5.5e-06 | 2.1e-06 | 3.7e-06 | 3.767E-06 | 1.701E-06 | 2.100E-06 | 5.500E-06 |
| 4338 | 217 | Q217W | A | 2.2e-06 | 1.4e-06 | 2.6e-06 | 2.067E-06 | 6.110E-07 | 1.400E-06 | 2.600E-06 |
| 4339 | 217 | Q217Y | A | 2.8e-06 | 2e-06 | 2e-06 | 2.267E-06 | 4.619E-07 | 2.000E-06 | 2.800E-06 |
| 4340 | 217 | Q217V | A | 3.5e-06 | 1.5e-06 | 1.5e-06 | 2.167E-06 | 1.155E-06 | 1.500E-06 | 3.500E-06 |
| 4341 | 218 | C218A | A | 2.3e-06 | 2.5e-06 | 1.7e-06 | 2.167E-06 | 4.163E-07 | 1.700E-06 | 2.500E-06 |
| 4342 | 218 | C218R | A | 3.1e-06 | 1.2e-06 | 1.8e-06 | 2.033E-06 | 9.713E-07 | 1.200E-06 | 3.100E-06 |
| 4343 | 218 | C218N | A | 3.4e-06 | 2.6e-06 | 1.3e-06 | 2.433E-06 | 1.060E-06 | 1.300E-06 | 3.400E-06 |
| 4344 | 218 | C218D | A | 1.3e-06 | 2.2e-06 | 9.3e-07 | 1.477E-06 | 6.532E-07 | 9.300E-07 | 2.200E-06 |
| 4345 | 218 | C218C | A | 4.2e-06 | 2.2e-06 | 3.3e-06 | 3.233E-06 | 1.002E-06 | 2.200E-06 | 4.200E-06 |
| 4346 | 218 | C218E | A | 3.9e-06 | 1.3e-06 | 2.5e-06 | 2.567E-06 | 1.301E-06 | 1.300E-06 | 3.900E-06 |
| 4347 | 218 | C218Q | A | 1.7e-06 | 1.2e-06 | 3e-06 | 1.967E-06 | 9.292E-07 | 1.200E-06 | 3.000E-06 |
| 4348 | 218 | C218G | A | 1.9e-06 | 3.6e-06 | 2.2e-06 | 2.567E-06 | 9.074E-07 | 1.900E-06 | 3.600E-06 |
| 4349 | 218 | C218H | A | 2.3e-06 | 2.2e-06 | 1.8e-06 | 2.100E-06 | 2.646E-07 | 1.800E-06 | 2.300E-06 |
| 4350 | 218 | C218I | A | 3.3e-06 | 2.7e-06 | 1.6e-06 | 2.533E-06 | 8.622E-07 | 1.600E-06 | 3.300E-06 |
| 4351 | 218 | C218L | A | 1.3e-06 | 3.6e-06 | 1.7e-06 | 2.200E-06 | 1.229E-06 | 1.300E-06 | 3.600E-06 |
| 4352 | 218 | C218K | A | 3.1e-06 | 1.6e-06 | 1.7e-06 | 2.133E-06 | 8.386E-07 | 1.600E-06 | 3.100E-06 |
| 4353 | 218 | C218M | A | 2.4e-06 | 3.7e-06 | 1.8e-06 | 2.633E-06 | 9.713E-07 | 1.800E-06 | 3.700E-06 |
| 4354 | 218 | C218F | A | 1.7e-06 | 3.9e-06 | 2.2e-06 | 2.600E-06 | 1.153E-06 | 1.700E-06 | 3.900E-06 |
| 4355 | 218 | C218P | A | 2.5e-06 | 2.9e-06 | 1.9e-06 | 2.433E-06 | 5.033E-07 | 1.900E-06 | 2.900E-06 |
| 4356 | 218 | C218S | A | 3.1e-06 | 2.6e-06 | 2.1e-06 | 2.600E-06 | 5.000E-07 | 2.100E-06 | 3.100E-06 |
| 4357 | 218 | C218T | A | 1.2e-06 | 2.2e-06 | 2.9e-06 | 2.100E-06 | 8.544E-07 | 1.200E-06 | 2.900E-06 |
| 4358 | 218 | C218W | A | 4.1e-06 | 3.1e-06 | 3.4e-06 | 3.533E-06 | 5.132E-07 | 3.100E-06 | 4.100E-06 |
| 4359 | 218 | C218Y | A | 2.2e-06 | 2.3e-06 | 2.3e-06 | 2.267E-06 | 5.774E-08 | 2.200E-06 | 2.300E-06 |
| 4360 | 218 | C218V | A | 1.9e-06 | 2e-06 | 2.5e-06 | 2.133E-06 | 3.215E-07 | 1.900E-06 | 2.500E-06 |
| 4361 | 219 | A219A | A | 2.9e-06 | 1.8e-06 | 2.5e-06 | 2.400E-06 | 5.568E-07 | 1.800E-06 | 2.900E-06 |
| 4362 | 219 | A219R | A | 2.4e-06 | 1.7e-06 | 2.9e-06 | 2.333E-06 | 6.028E-07 | 1.700E-06 | 2.900E-06 |
| 4363 | 219 | A219N | A | 3.4e-06 | 3e-06 | 2.6e-06 | 3.000E-06 | 4.000E-07 | 2.600E-06 | 3.400E-06 |
| 4364 | 219 | A219D | A | 3.6e-06 | 3.8e-06 | 2.4e-06 | 3.267E-06 | 7.572E-07 | 2.400E-06 | 3.800E-06 |
| 4365 | 219 | A219C | A | 4.1e-06 | 2.6e-06 | 1.8e-06 | 2.833E-06 | 1.168E-06 | 1.800E-06 | 4.100E-06 |
| 4366 | 219 | A219E | A | 2.2e-06 | 1.4e-06 | 1.9e-06 | 1.833E-06 | 4.041E-07 | 1.400E-06 | 2.200E-06 |
| 4367 | 219 | A219Q | A | 1.5e-06 | 3.2e-06 | 2.9e-06 | 2.533E-06 | 9.074E-07 | 1.500E-06 | 3.200E-06 |
| 4368 | 219 | A219G | A | 2.2e-06 | 1.4e-06 | 4.6e-06 | 2.733E-06 | 1.665E-06 | 1.400E-06 | 4.600E-06 |
| 4369 | 219 | A219H | A | 2.1e-06 | 2.7e-06 | 2.4e-06 | 2.400E-06 | 3.000E-07 | 2.100E-06 | 2.700E-06 |
| 4370 | 219 | A219I | A | 2.4e-06 | 2.9e-06 | 4e-06 | 3.100E-06 | 8.185E-07 | 2.400E-06 | 4.000E-06 |
| 4371 | 219 | A219L | A | 1.4e-06 | 3.2e-06 | 2.8e-06 | 2.467E-06 | 9.452E-07 | 1.400E-06 | 3.200E-06 |
| 4372 | 219 | A219K | A | 2e-06 | 1.7e-06 | 4.5e-06 | 2.733E-06 | 1.537E-06 | 1.700E-06 | 4.500E-06 |
| 4373 | 219 | A219M | A | 2.3e-06 | 2.8e-06 | 2e-06 | 2.367E-06 | 4.041E-07 | 2.000E-06 | 2.800E-06 |
| 4374 | 219 | A219F | A | 2.4e-06 | 2.1e-06 | 2e-06 | 2.167E-06 | 2.082E-07 | 2.000E-06 | 2.400E-06 |
| 4375 | 219 | A219P | A | 1.9e-06 | 2.2e-06 | 2.7e-06 | 2.267E-06 | 4.041E-07 | 1.900E-06 | 2.700E-06 |
| 4376 | 219 | A219S | A | 2.5e-06 | 1.2e-06 | 2.1e-06 | 1.933E-06 | 6.658E-07 | 1.200E-06 | 2.500E-06 |
| 4377 | 219 | A219T | A | 4e-06 | 2.9e-06 | 3.1e-06 | 3.333E-06 | 5.859E-07 | 2.900E-06 | 4.000E-06 |
| 4378 | 219 | A219W | A | 3e-06 | 1.7e-06 | 1.8e-06 | 2.167E-06 | 7.234E-07 | 1.700E-06 | 3.000E-06 |
| 4379 | 219 | A219Y | A | 4.9e-06 | 3.6e-06 | 2.9e-06 | 3.800E-06 | 1.015E-06 | 2.900E-06 | 4.900E-06 |
| 4380 | 219 | A219V | A | 2.1e-06 | 2.6e-06 | 2.2e-06 | 2.300E-06 | 2.646E-07 | 2.100E-06 | 2.600E-06 |
| 4381 | 220 | A220A | A | 3.1e-06 | 2.4e-06 | 3.1e-06 | 2.867E-06 | 4.041E-07 | 2.400E-06 | 3.100E-06 |
| 4382 | 220 | A220R | A | 1.5e-06 | 2.9e-06 | 2.3e-06 | 2.233E-06 | 7.024E-07 | 1.500E-06 | 2.900E-06 |
| 4383 | 220 | A220N | A | 4.4e-06 | 2.4e-06 | 1.9e-06 | 2.900E-06 | 1.323E-06 | 1.900E-06 | 4.400E-06 |
| 4384 | 220 | A220D | A | 1.5e-06 | 1.9e-06 | 2.8e-06 | 2.067E-06 | 6.658E-07 | 1.500E-06 | 2.800E-06 |
| 4385 | 220 | A220C | A | 2.7e-06 | 1.3e-06 | 3.3e-06 | 2.433E-06 | 1.026E-06 | 1.300E-06 | 3.300E-06 |
| 4386 | 220 | A220E | A | 2.4e-06 | 1.9e-06 | 1.3e-06 | 1.867E-06 | 5.508E-07 | 1.300E-06 | 2.400E-06 |
| 4387 | 220 | A220Q | A | 2.2e-06 | 1.6e-06 | 2.6e-06 | 2.133E-06 | 5.033E-07 | 1.600E-06 | 2.600E-06 |
| 4388 | 220 | A220G | A | 2.4e-06 | 1e-06 | 2e-06 | 1.800E-06 | 7.211E-07 | 1.000E-06 | 2.400E-06 |
| 4389 | 220 | A220H | A | 2.7e-06 | 1.9e-06 | 1.8e-06 | 2.133E-06 | 4.933E-07 | 1.800E-06 | 2.700E-06 |
| 4390 | 220 | A220I | A | 2.6e-06 | 4.7e-06 | 2e-06 | 3.100E-06 | 1.418E-06 | 2.000E-06 | 4.700E-06 |
| 4391 | 220 | A220L | A | 1.4e-06 | 2.3e-06 | 1.9e-06 | 1.867E-06 | 4.509E-07 | 1.400E-06 | 2.300E-06 |
| 4392 | 220 | A220K | A | 2.2e-06 | 2.2e-06 | 2e-06 | 2.133E-06 | 1.155E-07 | 2.000E-06 | 2.200E-06 |
| 4393 | 220 | A220M | A | 1.6e-06 | 3.2e-06 | 3.7e-06 | 2.833E-06 | 1.097E-06 | 1.600E-06 | 3.700E-06 |
| 4394 | 220 | A220F | A | 1.7e-06 | 1.8e-06 | 2.8e-06 | 2.100E-06 | 6.083E-07 | 1.700E-06 | 2.800E-06 |
| 4395 | 220 | A220P | A | 1.9e-06 | 2.1e-06 | 3.3e-06 | 2.433E-06 | 7.572E-07 | 1.900E-06 | 3.300E-06 |
| 4396 | 220 | A220S | A | 2.6e-06 | 2.7e-06 | 5.4e-06 | 3.567E-06 | 1.589E-06 | 2.600E-06 | 5.400E-06 |
| 4397 | 220 | A220T | A | 3e-06 | 3.7e-06 | 2.2e-06 | 2.967E-06 | 7.506E-07 | 2.200E-06 | 3.700E-06 |
| 4398 | 220 | A220W | A | 2e-06 | 1.4e-06 | 3.4e-06 | 2.267E-06 | 1.026E-06 | 1.400E-06 | 3.400E-06 |
| 4399 | 220 | A220Y | A | 3.8e-06 | 3.1e-06 | 3.5e-06 | 3.467E-06 | 3.512E-07 | 3.100E-06 | 3.800E-06 |
| 4400 | 220 | A220V | A | 2.4e-06 | 2.8e-06 | 3.1e-06 | 2.767E-06 | 3.512E-07 | 2.400E-06 | 3.100E-06 |
| 4401 | 221 | G221A | A | 2.5e-06 | 1.2e-06 | 3.2e-06 | 2.300E-06 | 1.015E-06 | 1.200E-06 | 3.200E-06 |
| 4402 | 221 | G221R | A | 3.7e-06 | 2.5e-06 | 2e-06 | 2.733E-06 | 8.737E-07 | 2.000E-06 | 3.700E-06 |
| 4403 | 221 | G221N | A | 2.8e-06 | 2.6e-06 | 2.6e-06 | 2.667E-06 | 1.155E-07 | 2.600E-06 | 2.800E-06 |
| 4404 | 221 | G221D | A | 4.7e-06 | 2.4e-06 | 2.6e-06 | 3.233E-06 | 1.274E-06 | 2.400E-06 | 4.700E-06 |
| 4405 | 221 | G221C | A | 1.6e-06 | 3.5e-06 | 2.3e-06 | 2.467E-06 | 9.609E-07 | 1.600E-06 | 3.500E-06 |
| 4406 | 221 | G221E | A | 1.5e-06 | 3.2e-06 | 2e-06 | 2.233E-06 | 8.737E-07 | 1.500E-06 | 3.200E-06 |
| 4407 | 221 | G221Q | A | 4e-06 | 2.7e-06 | 3.6e-06 | 3.433E-06 | 6.658E-07 | 2.700E-06 | 4.000E-06 |
| 4408 | 221 | G221G | A | 2.1e-06 | 2.5e-06 | 2.2e-06 | 2.267E-06 | 2.082E-07 | 2.100E-06 | 2.500E-06 |
| 4409 | 221 | G221H | A | 3.3e-06 | 2.2e-06 | 2.7e-06 | 2.733E-06 | 5.508E-07 | 2.200E-06 | 3.300E-06 |
| 4410 | 221 | G221I | A | 2.8e-06 | 2.8e-06 | 4e-06 | 3.200E-06 | 6.928E-07 | 2.800E-06 | 4.000E-06 |
| 4411 | 221 | G221L | A | 3.2e-06 | 3.4e-06 | 3.1e-06 | 3.233E-06 | 1.528E-07 | 3.100E-06 | 3.400E-06 |
| 4412 | 221 | G221K | A | 3e-06 | 1.4e-06 | 2.6e-06 | 2.333E-06 | 8.327E-07 | 1.400E-06 | 3.000E-06 |
| 4413 | 221 | G221M | A | 2.6e-06 | 1.5e-06 | 2.3e-06 | 2.133E-06 | 5.686E-07 | 1.500E-06 | 2.600E-06 |
| 4414 | 221 | G221F | A | 2.7e-06 | 2.3e-06 | 1.7e-06 | 2.233E-06 | 5.033E-07 | 1.700E-06 | 2.700E-06 |
| 4415 | 221 | G221P | A | 2.6e-06 | 2.9e-06 | 1.6e-06 | 2.367E-06 | 6.807E-07 | 1.600E-06 | 2.900E-06 |
| 4416 | 221 | G221S | A | 3e-06 | 2.2e-06 | 2.5e-06 | 2.567E-06 | 4.041E-07 | 2.200E-06 | 3.000E-06 |
| 4417 | 221 | G221T | A | 2.7e-06 | 1.4e-06 | 2e-06 | 2.033E-06 | 6.506E-07 | 1.400E-06 | 2.700E-06 |
| 4418 | 221 | G221W | A | 2e-06 | 5e-06 | 1.6e-06 | 2.867E-06 | 1.858E-06 | 1.600E-06 | 5.000E-06 |
| 4419 | 221 | G221Y | A | 2.1e-06 | 2.6e-06 | 1.6e-06 | 2.100E-06 | 5.000E-07 | 1.600E-06 | 2.600E-06 |

| | | | | | | | | | | |
|---|---|---|---|---|---|---|---|---|---|---|
| 4420 | 221 | G221V | A | 4.6e-06 | 2.9e-06 | 2.4e-06 | 3.300E-06 | 1.153E-06 | 2.400E-06 | 4.600E-06 |
| 4421 | 222 | C222A | A | 1.9e-06 | 2.9e-06 | 2.4e-06 | 2.400E-06 | 5.000E-07 | 1.900E-06 | 2.900E-06 |
| 4422 | 222 | C222R | A | 2.5e-06 | 2.7e-06 | 3.1e-06 | 2.767E-06 | 3.055E-07 | 2.500E-06 | 3.100E-06 |
| 4423 | 222 | C222N | A | 5.9e-06 | 1.2e-06 | 1.8e-06 | 2.967E-06 | 2.558E-06 | 1.200E-06 | 5.900E-06 |
| 4424 | 222 | C222D | A | 1.3e-06 | 2.6e-06 | 2.9e-06 | 2.267E-06 | 8.505E-07 | 1.300E-06 | 2.900E-06 |
| 4425 | 222 | C222C | A | 2.1e-06 | 9.1e-07 | 1.7e-06 | 1.570E-06 | 6.056E-07 | 9.100E-07 | 2.100E-06 |
| 4426 | 222 | C222E | A | 3.5e-06 | 1.5e-06 | 1.7e-06 | 2.233E-06 | 1.102E-06 | 1.500E-06 | 3.500E-06 |
| 4427 | 222 | C222Q | A | 2.2e-06 | 3.3e-06 | 4.9e-06 | 3.467E-06 | 1.358E-06 | 2.200E-06 | 4.900E-06 |
| 4428 | 222 | C222G | A | 3.5e-06 | 2.8e-06 | 2e-06 | 2.767E-06 | 7.506E-07 | 2.000E-06 | 3.500E-06 |
| 4429 | 222 | C222H | A | 2.7e-06 | 2.7e-06 | 2.5e-06 | 2.633E-06 | 1.155E-07 | 2.500E-06 | 2.700E-06 |
| 4430 | 222 | C222I | A | 5.1e-06 | 1.9e-06 | 2.4e-06 | 3.133E-06 | 1.721E-06 | 1.900E-06 | 5.100E-06 |
| 4431 | 222 | C222L | A | 2.7e-06 | 3e-06 | 2.1e-06 | 2.600E-06 | 4.583E-07 | 2.100E-06 | 3.000E-06 |
| 4432 | 222 | C222K | A | 1.2e-06 | 4e-06 | 2.6e-06 | 2.600E-06 | 1.400E-06 | 1.200E-06 | 4.000E-06 |
| 4433 | 222 | C222M | A | 5.2e-06 | 2.2e-06 | 1.6e-06 | 3.000E-06 | 1.929E-06 | 1.600E-06 | 5.200E-06 |
| 4434 | 222 | C222F | A | 1.9e-06 | 2.9e-06 | 3e-06 | 2.600E-06 | 6.083E-07 | 1.900E-06 | 3.000E-06 |
| 4435 | 222 | C222P | A | 2.5e-06 | 1.2e-06 | 1.8e-06 | 1.833E-06 | 6.506E-07 | 1.200E-06 | 2.500E-06 |
| 4436 | 222 | C222S | A | 1.4e-06 | 2.6e-06 | 2.3e-06 | 2.100E-06 | 6.245E-07 | 1.400E-06 | 2.600E-06 |
| 4437 | 222 | C222T | A | 2.2e-06 | 2.9e-06 | 2e-06 | 2.367E-06 | 4.726E-07 | 2.000E-06 | 2.900E-06 |
| 4438 | 222 | C222W | A | 2.2e-06 | 2.1e-06 | 2.6e-06 | 2.300E-06 | 2.646E-07 | 2.100E-06 | 2.600E-06 |
| 4439 | 222 | C222Y | A | 1.4e-06 | 3.2e-06 | 1.1e-06 | 1.900E-06 | 1.136E-06 | 1.100E-06 | 3.200E-06 |
| 4440 | 222 | C222V | A | 1.7e-06 | 4.1e-06 | 1.3e-06 | 2.367E-06 | 1.514E-06 | 1.300E-06 | 4.100E-06 |
| 4441 | 223 | T223A | A | 1.4e-06 | 3.1e-06 | 2.4e-06 | 2.300E-06 | 8.544E-07 | 1.400E-06 | 3.100E-06 |
| 4442 | 223 | T223R | A | 2.8e-06 | 2.1e-06 | 2.5e-06 | 2.467E-06 | 3.512E-07 | 2.100E-06 | 2.800E-06 |
| 4443 | 223 | T223N | A | 3.3e-06 | 3.4e-06 | 2e-06 | 2.900E-06 | 7.810E-07 | 2.000E-06 | 3.400E-06 |
| 4444 | 223 | T223D | A | 4.8e-06 | 2.4e-06 | 2.4e-06 | 3.200E-06 | 1.386E-06 | 2.400E-06 | 4.800E-06 |
| 4445 | 223 | T223C | A | 2.2e-06 | 3.5e-06 | 1.9e-06 | 2.533E-06 | 8.505E-07 | 1.900E-06 | 3.500E-06 |
| 4446 | 223 | T223E | A | 2.2e-06 | 1.3e-06 | 2.3e-06 | 1.933E-06 | 5.508E-07 | 1.300E-06 | 2.300E-06 |
| 4447 | 223 | T223Q | A | 2.1e-06 | 1.7e-06 | 1.7e-06 | 1.833E-06 | 2.309E-07 | 1.700E-06 | 2.100E-06 |
| 4448 | 223 | T223G | A | 2.4e-06 | 2.1e-06 | 1.8e-06 | 2.100E-06 | 3.000E-07 | 1.800E-06 | 2.400E-06 |
| 4449 | 223 | T223H | A | 2.6e-06 | 1.7e-06 | 2.6e-06 | 2.300E-06 | 5.196E-07 | 1.700E-06 | 2.600E-06 |
| 4450 | 223 | T223I | A | 2.2e-06 | 3e-06 | 2.7e-06 | 2.633E-06 | 4.041E-07 | 2.200E-06 | 3.000E-06 |
| 4451 | 223 | T223L | A | 1.9e-06 | 1.9e-06 | 2.4e-06 | 2.067E-06 | 2.887E-07 | 1.900E-06 | 2.400E-06 |
| 4452 | 223 | T223K | A | 2e-06 | 1.7e-06 | 1.7e-06 | 1.800E-06 | 1.732E-07 | 1.700E-06 | 2.000E-06 |
| 4453 | 223 | T223M | A | 2.2e-06 | 1.6e-06 | 1.7e-06 | 1.833E-06 | 3.215E-07 | 1.600E-06 | 2.200E-06 |
| 4454 | 223 | T223F | A | 2.9e-06 | 4.6e-06 | 1.9e-06 | 3.133E-06 | 1.365E-06 | 1.900E-06 | 4.600E-06 |
| 4455 | 223 | T223P | A | 1.8e-06 | 3.8e-06 | 1.6e-06 | 2.400E-06 | 1.217E-06 | 1.600E-06 | 3.800E-06 |
| 4456 | 223 | T223S | A | 1.2e-06 | 1.8e-06 | 2e-06 | 1.667E-06 | 4.163E-07 | 1.200E-06 | 2.000E-06 |
| 4457 | 223 | T223T | A | 2e-06 | 1.7e-06 | 1.9e-06 | 1.867E-06 | 1.528E-07 | 1.700E-06 | 2.000E-06 |
| 4458 | 223 | T223W | A | 3.4e-06 | 2.6e-06 | 2.1e-06 | 2.700E-06 | 6.557E-07 | 2.100E-06 | 3.400E-06 |
| 4459 | 223 | T223Y | A | 2.1e-06 | 2.2e-06 | 2.5e-06 | 2.267E-06 | 2.082E-07 | 2.100E-06 | 2.500E-06 |
| 4460 | 223 | T223V | A | 2.6e-06 | 3.1e-06 | 2.7e-06 | 2.800E-06 | 2.646E-07 | 2.600E-06 | 3.100E-06 |
| 4461 | 224 | G224A | A | 2e-06 | 2.4e-06 | 2.8e-06 | 2.400E-06 | 4.000E-07 | 2.000E-06 | 2.800E-06 |
| 4462 | 224 | G224R | A | 2.6e-06 | 4.2e-06 | 1.4e-06 | 2.733E-06 | 1.405E-06 | 1.400E-06 | 4.200E-06 |
| 4463 | 224 | G224N | A | 3.3e-06 | 1.8e-06 | 2.7e-06 | 2.600E-06 | 7.550E-07 | 1.800E-06 | 3.300E-06 |
| 4464 | 224 | G224D | A | 1.3e-06 | 2.6e-06 | 2.6e-06 | 2.167E-06 | 7.506E-07 | 1.300E-06 | 2.600E-06 |
| 4465 | 224 | G224C | A | 1.2e-06 | 1.1e-06 | 2.4e-06 | 1.567E-06 | 7.234E-07 | 1.100E-06 | 2.400E-06 |
| 4466 | 224 | G224E | A | 2.9e-06 | 2.4e-06 | 2.1e-06 | 2.467E-06 | 4.041E-07 | 2.100E-06 | 2.900E-06 |
| 4467 | 224 | G224Q | A | 2.7e-06 | 1.4e-06 | 4.3e-06 | 2.800E-06 | 1.453E-06 | 1.400E-06 | 4.300E-06 |
| 4468 | 224 | G224G | A | 3e-06 | 1.6e-06 | 2.8e-06 | 2.467E-06 | 7.572E-07 | 1.600E-06 | 3.000E-06 |
| 4469 | 224 | G224H | A | 1.5e-06 | 1.7e-06 | 1.6e-06 | 1.600E-06 | 1.000E-07 | 1.500E-06 | 1.700E-06 |
| 4470 | 224 | G224I | A | 3e-06 | 2.5e-06 | 1.8e-06 | 2.433E-06 | 6.028E-07 | 1.800E-06 | 3.000E-06 |
| 4471 | 224 | G224L | A | 2.8e-06 | 2.7e-06 | 4.8e-06 | 3.433E-06 | 1.185E-06 | 2.700E-06 | 4.800E-06 |
| 4472 | 224 | G224K | A | 3.8e-06 | 3.5e-06 | 2.1e-06 | 3.133E-06 | 9.074E-07 | 2.100E-06 | 3.800E-06 |
| 4473 | 224 | G224M | A | 2.4e-06 | 2.6e-06 | 2.1e-06 | 2.367E-06 | 2.517E-07 | 2.100E-06 | 2.600E-06 |
| 4474 | 224 | G224F | A | 2.4e-06 | 2.2e-06 | 2.4e-06 | 2.333E-06 | 1.155E-07 | 2.200E-06 | 2.400E-06 |
| 4475 | 224 | G224P | A | 3.8e-06 | 2.5e-06 | 2.7e-06 | 3.000E-06 | 7.000E-07 | 2.500E-06 | 3.800E-06 |
| 4476 | 224 | G224S | A | 2.6e-06 | 1.6e-06 | 4e-06 | 2.733E-06 | 1.206E-06 | 1.600E-06 | 4.000E-06 |
| 4477 | 224 | G224T | A | 2e-06 | 2.7e-06 | 3.8e-06 | 2.833E-06 | 9.074E-07 | 2.000E-06 | 3.800E-06 |
| 4478 | 224 | G224W | A | 1.8e-06 | 5.5e-06 | 3.8e-06 | 3.700E-06 | 1.852E-06 | 1.800E-06 | 5.500E-06 |
| 4479 | 224 | G224Y | A | 2.8e-06 | 1.8e-06 | 9.3e-07 | 1.843E-06 | 9.358E-07 | 9.300E-07 | 2.800E-06 |
| 4480 | 224 | G224V | A | 2.4e-06 | 1.8e-06 | 3.7e-06 | 2.633E-06 | 9.713E-07 | 1.800E-06 | 3.700E-06 |
| 4481 | 225 | P225A | A | 2.4e-06 | 2.7e-06 | 2.6e-06 | 2.567E-06 | 1.528E-07 | 2.400E-06 | 2.700E-06 |
| 4482 | 225 | P225R | A | 2.4e-06 | 2.7e-06 | 2.1e-06 | 2.400E-06 | 3.000E-07 | 2.100E-06 | 2.700E-06 |
| 4483 | 225 | P225N | A | 2.4e-06 | 2.5e-06 | 2.1e-06 | 2.333E-06 | 2.082E-07 | 2.100E-06 | 2.500E-06 |
| 4484 | 225 | P225D | A | 3e-06 | 3.1e-06 | 2.6e-06 | 2.900E-06 | 2.646E-07 | 2.600E-06 | 3.100E-06 |
| 4485 | 225 | P225C | A | 3.6e-06 | 4e-06 | 2.5e-06 | 3.367E-06 | 7.767E-07 | 2.500E-06 | 4.000E-06 |
| 4486 | 225 | P225E | A | 2e-06 | 2.2e-06 | 3.2e-06 | 2.467E-06 | 6.429E-07 | 2.000E-06 | 3.200E-06 |
| 4487 | 225 | P225Q | A | 2.6e-06 | 8.7e-07 | 1.6e-06 | 1.690E-06 | 8.685E-07 | 8.700E-07 | 2.600E-06 |
| 4488 | 225 | P225G | A | 1.9e-06 | 3e-06 | 2.9e-06 | 2.600E-06 | 6.083E-07 | 1.900E-06 | 3.000E-06 |
| 4489 | 225 | P225H | A | 1.9e-06 | 3.1e-06 | 2.8e-06 | 2.600E-06 | 6.245E-07 | 1.900E-06 | 3.100E-06 |
| 4490 | 225 | P225I | A | 2.8e-06 | 1.9e-06 | 1.9e-06 | 2.200E-06 | 5.196E-07 | 1.900E-06 | 2.800E-06 |
| 4491 | 225 | P225L | A | 2.6e-06 | 2.1e-06 | 2.1e-06 | 2.267E-06 | 2.887E-07 | 2.100E-06 | 2.600E-06 |
| 4492 | 225 | P225K | A | 2.6e-06 | 2.5e-06 | 2.2e-06 | 2.433E-06 | 2.082E-07 | 2.200E-06 | 2.600E-06 |
| 4493 | 225 | P225M | A | 1.4e-06 | 1.3e-06 | 3.1e-06 | 1.933E-06 | 1.012E-06 | 1.300E-06 | 3.100E-06 |
| 4494 | 225 | P225F | A | 3.3e-06 | 2.1e-06 | 1.3e-06 | 2.233E-06 | 1.007E-06 | 1.300E-06 | 3.300E-06 |
| 4495 | 225 | P225P | A | 2.3e-06 | 2.6e-06 | 2.6e-06 | 2.500E-06 | 1.732E-07 | 2.300E-06 | 2.600E-06 |
| 4496 | 225 | P225S | A | 2.1e-06 | 3.1e-06 | 4.1e-06 | 3.100E-06 | 1.000E-06 | 2.100E-06 | 4.100E-06 |
| 4497 | 225 | P225T | A | 2.2e-06 | 2e-06 | 2.8e-06 | 2.333E-06 | 4.163E-07 | 2.000E-06 | 2.800E-06 |
| 4498 | 225 | P225W | A | 1.5e-06 | 2.5e-06 | 1.6e-06 | 1.867E-06 | 5.508E-07 | 1.500E-06 | 2.500E-06 |
| 4499 | 225 | P225Y | A | 2.6e-06 | 4.5e-06 | 2e-06 | 3.033E-06 | 1.305E-06 | 2.000E-06 | 4.500E-06 |
| 4500 | 225 | P225V | A | 2.5e-06 | 1.6e-06 | 2.1e-06 | 2.067E-06 | 4.509E-07 | 1.600E-06 | 2.500E-06 |
| 4501 | 226 | K226A | A | 4.5e-06 | 3e-06 | 2.3e-06 | 3.267E-06 | 1.124E-06 | 2.300E-06 | 4.500E-06 |
| 4502 | 226 | K226R | A | 2.4e-06 | 2.1e-06 | 3.9e-06 | 2.800E-06 | 9.644E-07 | 2.100E-06 | 3.900E-06 |
| 4503 | 226 | K226N | A | 1.1e-06 | 1.2e-06 | 1.9e-06 | 1.400E-06 | 4.359E-07 | 1.100E-06 | 1.900E-06 |
| 4504 | 226 | K226D | A | 3.2e-06 | 1.9e-06 | 4.1e-06 | 3.067E-06 | 1.106E-06 | 1.900E-06 | 4.100E-06 |

| | | | | | | | | | | |
|---|---|---|---|---|---|---|---|---|---|---|
| 4505 | 226 | K226C | A | 2.1e-06 | 1.9e-06 | 2.1e-06 | 2.033E-06 | 1.155E-07 | 1.900E-06 | 2.100E-06 |
| 4506 | 226 | K226E | A | 2.8e-06 | 2.2e-06 | 3.2e-06 | 2.733E-06 | 5.033E-07 | 2.200E-06 | 3.200E-06 |
| 4507 | 226 | K226Q | A | 1.8e-06 | 3.1e-06 | 2e-06 | 2.300E-06 | 7.000E-07 | 1.800E-06 | 3.100E-06 |
| 4508 | 226 | K226G | A | 4.3e-06 | 2.3e-06 | 2.5e-06 | 3.033E-06 | 1.102E-06 | 2.300E-06 | 4.300E-06 |
| 4509 | 226 | K226H | A | 8.3e-07 | 2.3e-06 | 3.4e-06 | 2.177E-06 | 1.289E-06 | 8.300E-07 | 3.400E-06 |
| 4510 | 226 | K226I | A | 2.8e-06 | 1.4e-06 | 2.9e-06 | 2.367E-06 | 8.386E-07 | 1.400E-06 | 2.900E-06 |
| 4511 | 226 | K226L | A | 2.3e-06 | 3.1e-06 | 2.5e-06 | 2.633E-06 | 4.163E-07 | 2.300E-06 | 3.100E-06 |
| 4512 | 226 | K226K | A | 2.4e-06 | 2.9e-06 | 1.6e-06 | 2.300E-06 | 6.557E-07 | 1.600E-06 | 2.900E-06 |
| 4513 | 226 | K226M | A | 1.7e-06 | 2.5e-06 | 2.3e-06 | 2.167E-06 | 4.163E-07 | 1.700E-06 | 2.500E-06 |
| 4514 | 226 | K226F | A | 2.1e-06 | 3.4e-06 | 1.9e-06 | 2.467E-06 | 8.145E-07 | 1.900E-06 | 3.400E-06 |
| 4515 | 226 | K226P | A | 2.2e-06 | 9.9e-07 | 3.2e-06 | 2.130E-06 | 1.107E-06 | 9.900E-07 | 3.200E-06 |
| 4516 | 226 | K226S | A | 3.4e-06 | 2.8e-06 | 1.6e-06 | 2.600E-06 | 9.165E-07 | 1.600E-06 | 3.400E-06 |
| 4517 | 226 | K226T | A | 3.3e-06 | 2.5e-06 | 2.7e-06 | 2.833E-06 | 4.163E-07 | 2.500E-06 | 3.300E-06 |
| 4518 | 226 | K226W | A | 2.7e-06 | 3.3e-06 | 3e-06 | 3.000E-06 | 3.000E-07 | 2.700E-06 | 3.300E-06 |
| 4519 | 226 | K226Y | A | 3.1e-06 | 1.9e-06 | 2.7e-06 | 2.567E-06 | 6.110E-07 | 1.900E-06 | 3.100E-06 |
| 4520 | 226 | K226V | A | 2e-06 | 1.8e-06 | 1.4e-06 | 1.733E-06 | 3.055E-07 | 1.400E-06 | 2.000E-06 |
| 4521 | 227 | H227A | A | 3e-06 | 2.6e-06 | 3.7e-06 | 3.100E-06 | 5.568E-07 | 2.600E-06 | 3.700E-06 |
| 4522 | 227 | H227R | A | 2.9e-06 | 1.4e-06 | 4e-06 | 2.767E-06 | 1.305E-06 | 1.400E-06 | 4.000E-06 |
| 4523 | 227 | H227N | A | 1.9e-06 | 2.4e-06 | 3e-06 | 2.433E-06 | 5.508E-07 | 1.900E-06 | 3.000E-06 |
| 4524 | 227 | H227D | A | 2.1e-06 | 1.9e-06 | 1.9e-06 | 1.967E-06 | 1.155E-07 | 1.900E-06 | 2.100E-06 |
| 4525 | 227 | H227C | A | 3.8e-06 | 1.3e-06 | 5.2e-06 | 3.433E-06 | 1.976E-06 | 1.300E-06 | 5.200E-06 |
| 4526 | 227 | H227E | A | 3.1e-06 | 1.9e-06 | 3.4e-06 | 2.800E-06 | 7.937E-07 | 1.900E-06 | 3.400E-06 |
| 4527 | 227 | H227Q | A | 2.3e-06 | 3.6e-06 | 1.2e-06 | 2.367E-06 | 1.201E-06 | 1.200E-06 | 3.600E-06 |
| 4528 | 227 | H227G | A | 2.2e-06 | 2.4e-06 | 2.8e-06 | 2.467E-06 | 3.055E-07 | 2.200E-06 | 2.800E-06 |
| 4529 | 227 | H227H | A | 2.9e-06 | 2.5e-06 | 1.7e-06 | 2.367E-06 | 6.110E-07 | 1.700E-06 | 2.900E-06 |
| 4530 | 227 | H227I | A | 2.7e-06 | 2.2e-06 | 2.3e-06 | 2.400E-06 | 2.646E-07 | 2.200E-06 | 2.700E-06 |
| 4531 | 227 | H227L | A | 1.5e-06 | 1.7e-06 | 2.7e-06 | 1.967E-06 | 6.429E-07 | 1.500E-06 | 2.700E-06 |
| 4532 | 227 | H227K | A | 1.5e-06 | 3.1e-06 | 2.9e-06 | 2.500E-06 | 8.718E-07 | 1.500E-06 | 3.100E-06 |
| 4533 | 227 | H227M | A | 1.6e-06 | 2.7e-06 | 2.5e-06 | 2.267E-06 | 5.859E-07 | 1.600E-06 | 2.700E-06 |
| 4534 | 227 | H227F | A | 1.8e-06 | 2.4e-06 | 1.9e-06 | 2.033E-06 | 3.215E-07 | 1.800E-06 | 2.400E-06 |
| 4535 | 227 | H227P | A | 5.2e-06 | 2.9e-06 | 2.6e-06 | 3.567E-06 | 1.422E-06 | 2.600E-06 | 5.200E-06 |
| 4536 | 227 | H227S | A | 1.9e-06 | 3e-06 | 2.5e-06 | 2.467E-06 | 5.508E-07 | 1.900E-06 | 3.000E-06 |
| 4537 | 227 | H227T | A | 1.5e-06 | 2.2e-06 | 2.9e-06 | 2.200E-06 | 7.000E-07 | 1.500E-06 | 2.900E-06 |
| 4538 | 227 | H227W | A | 3e-06 | 2.5e-06 | 2.8e-06 | 2.767E-06 | 2.517E-07 | 2.500E-06 | 3.000E-06 |
| 4539 | 227 | H227Y | A | 2.1e-06 | 4.7e-06 | 3.7e-06 | 3.500E-06 | 1.311E-06 | 2.100E-06 | 4.700E-06 |
| 4540 | 227 | H227V | A | 2.6e-06 | 2.7e-06 | 1.7e-06 | 2.333E-06 | 5.508E-07 | 1.700E-06 | 2.700E-06 |
| 4541 | 228 | S228A | A | 2.2e-06 | 1.9e-06 | 2.1e-06 | 2.067E-06 | 1.528E-07 | 1.900E-06 | 2.200E-06 |
| 4542 | 228 | S228R | A | 2e-06 | 3.1e-06 | 2.3e-06 | 2.467E-06 | 5.686E-07 | 2.000E-06 | 3.100E-06 |
| 4543 | 228 | S228N | A | 2.7e-06 | 2.8e-06 | 2.5e-06 | 2.667E-06 | 1.528E-07 | 2.500E-06 | 2.800E-06 |
| 4544 | 228 | S228D | A | 3.1e-06 | 1.9e-06 | 2.5e-06 | 2.500E-06 | 6.000E-07 | 1.900E-06 | 3.100E-06 |
| 4545 | 228 | S228C | A | 5.6e-06 | 2.5e-06 | 1.8e-06 | 3.300E-06 | 2.022E-06 | 1.800E-06 | 5.600E-06 |
| 4546 | 228 | S228E | A | 1.5e-06 | 1.7e-06 | 2.6e-06 | 1.933E-06 | 5.859E-07 | 1.500E-06 | 2.600E-06 |
| 4547 | 228 | S228Q | A | 2.6e-06 | 2.3e-06 | 1.1e-06 | 2.000E-06 | 7.937E-07 | 1.100E-06 | 2.600E-06 |
| 4548 | 228 | S228G | A | 4.2e-06 | 3.3e-06 | 2e-06 | 3.167E-06 | 1.106E-06 | 2.000E-06 | 4.200E-06 |
| 4549 | 228 | S228H | A | 1.2e-06 | 2.4e-06 | 2.6e-06 | 2.067E-06 | 7.572E-07 | 1.200E-06 | 2.600E-06 |
| 4550 | 228 | S228I | A | 3e-06 | 2.1e-06 | 2.2e-06 | 2.433E-06 | 4.933E-07 | 2.100E-06 | 3.000E-06 |
| 4551 | 228 | S228L | A | 3.1e-06 | 2.7e-06 | 1.8e-06 | 2.533E-06 | 6.658E-07 | 1.800E-06 | 3.100E-06 |
| 4552 | 228 | S228K | A | 1.9e-06 | 2.8e-06 | 2.6e-06 | 2.433E-06 | 4.726E-07 | 1.900E-06 | 2.800E-06 |
| 4553 | 228 | S228M | A | 1.7e-06 | 2.6e-06 | 1.6e-06 | 1.967E-06 | 5.508E-07 | 1.600E-06 | 2.600E-06 |
| 4554 | 228 | S228F | A | 1.6e-06 | 8.7e-07 | 2.7e-06 | 1.723E-06 | 9.212E-07 | 8.700E-07 | 2.700E-06 |
| 4555 | 228 | S228P | A | 3.4e-06 | 2.3e-06 | 1.6e-06 | 2.433E-06 | 9.074E-07 | 1.600E-06 | 3.400E-06 |
| 4556 | 228 | S228S | A | 3.5e-06 | 1.6e-06 | 1.8e-06 | 2.300E-06 | 1.044E-06 | 1.600E-06 | 3.500E-06 |
| 4557 | 228 | S228T | A | 2.1e-06 | 2.4e-06 | 2.9e-06 | 2.467E-06 | 4.041E-07 | 2.100E-06 | 2.900E-06 |
| 4558 | 228 | S228W | A | 2.4e-06 | 2.1e-06 | 3.7e-06 | 2.733E-06 | 8.505E-07 | 2.100E-06 | 3.700E-06 |
| 4559 | 228 | S228Y | A | 2.8e-06 | 2.2e-06 | 2.7e-06 | 2.567E-06 | 3.215E-07 | 2.200E-06 | 2.800E-06 |
| 4560 | 228 | S228V | A | 2.3e-06 | 2.4e-06 | 3.3e-06 | 2.667E-06 | 5.508E-07 | 2.300E-06 | 3.300E-06 |
| 4561 | 229 | D229A | A | 2.5e-06 | 2.2e-06 | 4.1e-06 | 2.933E-06 | 1.021E-06 | 2.200E-06 | 4.100E-06 |
| 4562 | 229 | D229R | A | 4.1e-06 | 1.5e-06 | 1e-06 | 2.200E-06 | 1.664E-06 | 1.000E-06 | 4.100E-06 |
| 4563 | 229 | D229N | A | 2.4e-06 | 3e-06 | 2e-06 | 2.467E-06 | 5.033E-07 | 2.000E-06 | 3.000E-06 |
| 4564 | 229 | D229D | A | 2.7e-06 | 2.3e-06 | 2.6e-06 | 2.533E-06 | 2.082E-07 | 2.300E-06 | 2.700E-06 |
| 4565 | 229 | D229C | A | 6e-06 | 1.6e-06 | 3e-06 | 3.533E-06 | 2.248E-06 | 1.600E-06 | 6.000E-06 |
| 4566 | 229 | D229E | A | 2.8e-06 | 3e-06 | 2.8e-06 | 2.867E-06 | 1.155E-07 | 2.800E-06 | 3.000E-06 |
| 4567 | 229 | D229Q | A | 1.4e-06 | 1.6e-06 | 1.9e-06 | 1.633E-06 | 2.517E-07 | 1.400E-06 | 1.900E-06 |
| 4568 | 229 | D229G | A | 2.4e-06 | 2.5e-06 | 2.6e-06 | 2.500E-06 | 1.000E-07 | 2.400E-06 | 2.600E-06 |
| 4569 | 229 | D229H | A | 1.2e-06 | 1.5e-06 | 2.4e-06 | 1.700E-06 | 6.245E-07 | 1.200E-06 | 2.400E-06 |
| 4570 | 229 | D229I | A | 2.1e-06 | 2.3e-06 | 3.2e-06 | 2.533E-06 | 5.859E-07 | 2.100E-06 | 3.200E-06 |
| 4571 | 229 | D229L | A | 5.2e-06 | 2.4e-06 | 1.8e-06 | 3.133E-06 | 1.815E-06 | 1.800E-06 | 5.200E-06 |
| 4572 | 229 | D229K | A | 3.1e-06 | 1.9e-06 | 1.5e-06 | 2.167E-06 | 8.327E-07 | 1.500E-06 | 3.100E-06 |
| 4573 | 229 | D229M | A | 1.2e-06 | 2.2e-06 | 1.4e-06 | 1.600E-06 | 5.292E-07 | 1.200E-06 | 2.200E-06 |
| 4574 | 229 | D229F | A | 2.5e-06 | 1.8e-06 | 2e-06 | 2.100E-06 | 3.606E-07 | 1.800E-06 | 2.500E-06 |
| 4575 | 229 | D229P | A | 3.2e-06 | 2.6e-06 | 2.8e-06 | 2.867E-06 | 3.055E-07 | 2.600E-06 | 3.200E-06 |
| 4576 | 229 | D229S | A | 2.4e-06 | 2.7e-06 | 2.7e-06 | 2.600E-06 | 1.732E-07 | 2.400E-06 | 2.700E-06 |
| 4577 | 229 | D229T | A | 3.1e-06 | 3.2e-06 | 3.2e-06 | 3.167E-06 | 5.774E-08 | 3.100E-06 | 3.200E-06 |
| 4578 | 229 | D229W | A | 3.3e-06 | 2.3e-06 | 2e-06 | 2.533E-06 | 6.807E-07 | 2.000E-06 | 3.300E-06 |
| 4579 | 229 | D229Y | A | 1.7e-06 | 2.1e-06 | 2.9e-06 | 2.233E-06 | 6.110E-07 | 1.700E-06 | 2.900E-06 |
| 4580 | 229 | D229V | A | 3.7e-06 | 3.2e-06 | 1.5e-06 | 2.800E-06 | 1.153E-06 | 1.500E-06 | 3.700E-06 |
| 4581 | 230 | C230A | A | 9.9e-07 | 1.2e-06 | 1.7e-06 | 1.297E-06 | 3.647E-07 | 9.900E-07 | 1.700E-06 |
| 4582 | 230 | C230R | A | 3.8e-06 | 2.4e-06 | 4e-06 | 3.400E-06 | 8.718E-07 | 2.400E-06 | 4.000E-06 |
| 4583 | 230 | C230N | A | 2.1e-06 | 1.9e-06 | 2.7e-06 | 2.233E-06 | 4.163E-07 | 1.900E-06 | 2.700E-06 |
| 4584 | 230 | C230D | A | 2.4e-06 | 3e-06 | 2.7e-06 | 2.700E-06 | 3.000E-07 | 2.400E-06 | 3.000E-06 |
| 4585 | 230 | C230C | A | 2e-06 | 3.6e-06 | 2.4e-06 | 2.667E-06 | 8.327E-07 | 2.000E-06 | 3.600E-06 |
| 4586 | 230 | C230E | A | 3.4e-06 | 2.8e-06 | 3.1e-06 | 3.100E-06 | 3.000E-07 | 2.800E-06 | 3.400E-06 |
| 4587 | 230 | C230Q | A | 2.4e-06 | 3.1e-06 | 2.1e-06 | 2.533E-06 | 5.132E-07 | 2.100E-06 | 3.100E-06 |
| 4588 | 230 | C230G | A | 3e-06 | 3.3e-06 | 1.9e-06 | 2.733E-06 | 7.371E-07 | 1.900E-06 | 3.300E-06 |
| 4589 | 230 | C230H | A | 1.7e-06 | 2.3e-06 | 1.7e-06 | 1.900E-06 | 3.464E-07 | 1.700E-06 | 2.300E-06 |

| | | | | | | | | | | |
|---|---|---|---|---|---|---|---|---|---|---|
| 4590 | 230 | C230I | A | 2.8e-06 | 1.5e-06 | 1.5e-06 | 1.933E-06 | 7.506E-07 | 1.500E-06 | 2.800E-06 |
| 4591 | 230 | C230L | A | 2.9e-06 | 2.7e-06 | 4.1e-06 | 3.233E-06 | 7.572E-07 | 2.700E-06 | 4.100E-06 |
| 4592 | 230 | C230K | A | 3.1e-06 | 3.1e-06 | 3.8e-06 | 3.333E-06 | 4.041E-07 | 3.100E-06 | 3.800E-06 |
| 4593 | 230 | C230M | A | 1.7e-06 | 2.5e-06 | 2.3e-06 | 2.167E-06 | 4.163E-07 | 1.700E-06 | 2.500E-06 |
| 4594 | 230 | C230F | A | 3.2e-06 | 2.5e-06 | 2.8e-06 | 2.833E-06 | 3.512E-07 | 2.500E-06 | 3.200E-06 |
| 4595 | 230 | C230P | A | 3.3e-06 | 2.6e-06 | 2.9e-06 | 2.933E-06 | 3.512E-07 | 2.600E-06 | 3.300E-06 |
| 4596 | 230 | C230S | A | 4.2e-06 | 2.9e-06 | 3.1e-06 | 3.400E-06 | 7.000E-07 | 2.900E-06 | 4.200E-06 |
| 4597 | 230 | C230T | A | 2e-06 | 1.3e-06 | 2.1e-06 | 1.800E-06 | 4.359E-07 | 1.300E-06 | 2.100E-06 |
| 4598 | 230 | C230W | A | 2.5e-06 | 2.7e-06 | 2.1e-06 | 2.433E-06 | 3.055E-07 | 2.100E-06 | 2.700E-06 |
| 4599 | 230 | C230Y | A | 2.1e-06 | 2.3e-06 | 2.1e-06 | 2.167E-06 | 1.155E-07 | 2.100E-06 | 2.300E-06 |
| 4600 | 230 | C230V | A | 4.1e-06 | 2.2e-06 | 1.2e-06 | 2.500E-06 | 1.473E-06 | 1.200E-06 | 4.100E-06 |
| 4601 | 231 | L231A | A | 3.4e-06 | 1.6e-06 | 2.6e-06 | 2.533E-06 | 9.018E-07 | 1.600E-06 | 3.400E-06 |
| 4602 | 231 | L231R | A | 2.3e-06 | 2.1e-06 | 2.5e-06 | 2.300E-06 | 2.000E-07 | 2.100E-06 | 2.500E-06 |
| 4603 | 231 | L231N | A | 2e-06 | 2.2e-06 | 2.8e-06 | 2.333E-06 | 4.163E-07 | 2.000E-06 | 2.800E-06 |
| 4604 | 231 | L231D | A | 2e-06 | 3.1e-06 | 2.2e-06 | 2.433E-06 | 5.859E-07 | 2.000E-06 | 3.100E-06 |
| 4605 | 231 | L231C | A | 2.8e-06 | 1.6e-06 | 3e-06 | 2.467E-06 | 7.572E-07 | 1.600E-06 | 3.000E-06 |
| 4606 | 231 | L231E | A | 2.3e-06 | 1.7e-06 | 1.6e-06 | 1.867E-06 | 3.786E-07 | 1.600E-06 | 2.300E-06 |
| 4607 | 231 | L231Q | A | 2.3e-06 | 5e-06 | 1.9e-06 | 3.067E-06 | 1.686E-06 | 1.900E-06 | 5.000E-06 |
| 4608 | 231 | L231G | A | 1.5e-06 | 2.7e-06 | 1.8e-06 | 2.000E-06 | 6.245E-07 | 1.500E-06 | 2.700E-06 |
| 4609 | 231 | L231H | A | 2.2e-06 | 3.3e-06 | 2.5e-06 | 2.667E-06 | 5.686E-07 | 2.200E-06 | 3.300E-06 |
| 4610 | 231 | L231I | A | 3e-06 | 2.3e-06 | 2e-06 | 2.433E-06 | 5.132E-07 | 2.000E-06 | 3.000E-06 |
| 4611 | 231 | L231L | A | 3.6e-06 | 2.4e-06 | 3.3e-06 | 3.100E-06 | 6.245E-07 | 2.400E-06 | 3.600E-06 |
| 4612 | 231 | L231K | A | 3.3e-06 | 1.3e-06 | 2.2e-06 | 2.267E-06 | 1.002E-06 | 1.300E-06 | 3.300E-06 |
| 4613 | 231 | L231M | A | 2e-06 | 3.2e-06 | 2.6e-06 | 2.600E-06 | 6.000E-07 | 2.000E-06 | 3.200E-06 |
| 4614 | 231 | L231F | A | 2.1e-06 | 2.9e-06 | 2.1e-06 | 2.367E-06 | 4.619E-07 | 2.100E-06 | 2.900E-06 |
| 4615 | 231 | L231P | A | 2.1e-06 | 2.2e-06 | 1.8e-06 | 2.033E-06 | 2.082E-07 | 1.800E-06 | 2.200E-06 |
| 4616 | 231 | L231S | A | 2.8e-06 | 2.5e-06 | 1.9e-06 | 2.400E-06 | 4.583E-07 | 1.900E-06 | 2.800E-06 |
| 4617 | 231 | L231T | A | 2.8e-06 | 1.6e-06 | 3.2e-06 | 2.533E-06 | 8.327E-07 | 1.600E-06 | 3.200E-06 |
| 4618 | 231 | L231W | A | 1.9e-06 | 2.2e-06 | 2.6e-06 | 2.233E-06 | 3.512E-07 | 1.900E-06 | 2.600E-06 |
| 4619 | 231 | L231Y | A | 1.8e-06 | 3.1e-06 | 2e-06 | 2.300E-06 | 7.000E-07 | 1.800E-06 | 3.100E-06 |
| 4620 | 231 | L231V | A | 3.1e-06 | 3.2e-06 | 2e-06 | 2.767E-06 | 6.658E-07 | 2.000E-06 | 3.200E-06 |
| 4621 | 232 | A232A | A | 2.8e-06 | 2.6e-06 | 2.2e-06 | 2.533E-06 | 3.055E-07 | 2.200E-06 | 2.800E-06 |
| 4622 | 232 | A232R | A | 3.2e-06 | 3e-06 | 2.8e-06 | 3.000E-06 | 2.000E-07 | 2.800E-06 | 3.200E-06 |
| 4623 | 232 | A232N | A | 1.8e-06 | 2.5e-06 | 2.3e-06 | 2.200E-06 | 3.606E-07 | 1.800E-06 | 2.500E-06 |
| 4624 | 232 | A232D | A | 3.3e-06 | 2e-06 | 2.2e-06 | 2.500E-06 | 7.000E-07 | 2.000E-06 | 3.300E-06 |
| 4625 | 232 | A232C | A | 3.3e-06 | 1.8e-06 | 1.3e-06 | 2.133E-06 | 1.041E-06 | 1.300E-06 | 3.300E-06 |
| 4626 | 232 | A232E | A | 3.1e-06 | 2.8e-06 | 2e-06 | 2.633E-06 | 5.686E-07 | 2.000E-06 | 3.100E-06 |
| 4627 | 232 | A232Q | A | 3.4e-06 | 2.9e-06 | 2.3e-06 | 2.867E-06 | 5.508E-07 | 2.300E-06 | 3.400E-06 |
| 4628 | 232 | A232G | A | 2.8e-06 | 3e-06 | 1.7e-06 | 2.500E-06 | 7.000E-07 | 1.700E-06 | 3.000E-06 |
| 4629 | 232 | A232H | A | 2.7e-06 | 2.1e-06 | 3.5e-06 | 2.767E-06 | 7.024E-07 | 2.100E-06 | 3.500E-06 |
| 4630 | 232 | A232I | A | 3.1e-06 | 3.6e-06 | 2e-06 | 2.900E-06 | 8.185E-07 | 2.000E-06 | 3.600E-06 |
| 4631 | 232 | A232L | A | 2.1e-06 | 3.9e-06 | 2e-06 | 2.667E-06 | 1.069E-06 | 2.000E-06 | 3.900E-06 |
| 4632 | 232 | A232K | A | 1.8e-06 | 3.4e-06 | 3.4e-06 | 2.867E-06 | 9.238E-07 | 1.800E-06 | 3.400E-06 |
| 4633 | 232 | A232M | A | 2.8e-06 | 2.5e-06 | 2.3e-06 | 2.533E-06 | 2.517E-07 | 2.300E-06 | 2.800E-06 |
| 4634 | 232 | A232F | A | 1.9e-06 | 1.6e-06 | 3.3e-06 | 2.267E-06 | 9.074E-07 | 1.600E-06 | 3.300E-06 |
| 4635 | 232 | A232P | A | 3.8e-06 | 2e-06 | 3.2e-06 | 3.000E-06 | 9.165E-07 | 2.000E-06 | 3.800E-06 |
| 4636 | 232 | A232S | A | 3.9e-06 | 2.7e-06 | 2.1e-06 | 2.900E-06 | 9.165E-07 | 2.100E-06 | 3.900E-06 |
| 4637 | 232 | A232T | A | 2.3e-06 | 2.5e-06 | 2.6e-06 | 2.467E-06 | 1.528E-07 | 2.300E-06 | 2.600E-06 |
| 4638 | 232 | A232W | A | 4.2e-06 | 2.1e-06 | 3.1e-06 | 3.133E-06 | 1.050E-06 | 2.100E-06 | 4.200E-06 |
| 4639 | 232 | A232Y | A | 3e-06 | 3.5e-06 | 2.1e-06 | 2.867E-06 | 7.095E-07 | 2.100E-06 | 3.500E-06 |
| 4640 | 232 | A232V | A | 1.7e-06 | 1.9e-06 | 2.5e-06 | 2.033E-06 | 4.163E-07 | 1.700E-06 | 2.500E-06 |
| 4641 | 233 | C233A | A | 2.5e-06 | 1.2e-06 | 2.5e-06 | 2.067E-06 | 7.506E-07 | 1.200E-06 | 2.500E-06 |
| 4642 | 233 | C233R | A | 1.8e-06 | 2.3e-06 | 2.8e-06 | 2.300E-06 | 5.000E-07 | 1.800E-06 | 2.800E-06 |
| 4643 | 233 | C233N | A | 3e-06 | 2.1e-06 | 2.5e-06 | 2.533E-06 | 4.509E-07 | 2.100E-06 | 3.000E-06 |
| 4644 | 233 | C233D | A | 2.4e-06 | 1.8e-06 | 2.1e-06 | 2.100E-06 | 3.000E-07 | 1.800E-06 | 2.400E-06 |
| 4645 | 233 | C233C | A | 2.5e-06 | 3e-06 | 2.9e-06 | 2.800E-06 | 2.646E-07 | 2.500E-06 | 3.000E-06 |
| 4646 | 233 | C233E | A | 4.5e-06 | 2.4e-06 | 2.6e-06 | 3.167E-06 | 1.159E-06 | 2.400E-06 | 4.500E-06 |
| 4647 | 233 | C233Q | A | 4.2e-06 | 2.2e-06 | 1.9e-06 | 2.767E-06 | 1.250E-06 | 1.900E-06 | 4.200E-06 |
| 4648 | 233 | C233G | A | 1.8e-06 | 1.9e-06 | 5.9e-06 | 3.200E-06 | 2.339E-06 | 1.800E-06 | 5.900E-06 |
| 4649 | 233 | C233H | A | 1.9e-06 | 3.1e-06 | 2.6e-06 | 2.533E-06 | 6.028E-07 | 1.900E-06 | 3.100E-06 |
| 4650 | 233 | C233I | A | 2.8e-06 | 2.6e-06 | 3.5e-06 | 2.967E-06 | 4.726E-07 | 2.600E-06 | 3.500E-06 |
| 4651 | 233 | C233L | A | 1.9e-06 | 2.7e-06 | 3e-06 | 2.533E-06 | 5.686E-07 | 1.900E-06 | 3.000E-06 |
| 4652 | 233 | C233K | A | 4.4e-06 | 2.1e-06 | 4.2e-06 | 3.567E-06 | 1.274E-06 | 2.100E-06 | 4.400E-06 |
| 4653 | 233 | C233M | A | 3.1e-06 | 2.4e-06 | 2.4e-06 | 2.633E-06 | 4.041E-07 | 2.400E-06 | 3.100E-06 |
| 4654 | 233 | C233F | A | 2.2e-06 | 3.4e-06 | 4.5e-06 | 3.367E-06 | 1.150E-06 | 2.200E-06 | 4.500E-06 |
| 4655 | 233 | C233P | A | 3.6e-06 | 3.1e-06 | 2.4e-06 | 3.033E-06 | 6.028E-07 | 2.400E-06 | 3.600E-06 |
| 4656 | 233 | C233S | A | 2e-06 | 3.1e-06 | 2.3e-06 | 2.467E-06 | 5.686E-07 | 2.000E-06 | 3.100E-06 |
| 4657 | 233 | C233T | A | 3.9e-06 | 2.3e-06 | 3.7e-06 | 3.300E-06 | 8.718E-07 | 2.300E-06 | 3.900E-06 |
| 4658 | 233 | C233W | A | 2.7e-06 | 2.7e-06 | 3.3e-06 | 2.900E-06 | 3.464E-07 | 2.700E-06 | 3.300E-06 |
| 4659 | 233 | C233Y | A | 4e-06 | 3e-06 | 2.2e-06 | 3.067E-06 | 9.018E-07 | 2.200E-06 | 4.000E-06 |
| 4660 | 233 | C233V | A | 2.8e-06 | 1.6e-06 | 2.9e-06 | 2.433E-06 | 7.234E-07 | 1.600E-06 | 2.900E-06 |
| 4661 | 234 | L234A | A | 2.3e-06 | 1.5e-06 | 3e-06 | 2.267E-06 | 7.506E-07 | 1.500E-06 | 3.000E-06 |
| 4662 | 234 | L234R | A | 3.1e-06 | 1.6e-06 | 2.2e-06 | 2.300E-06 | 7.550E-07 | 1.600E-06 | 3.100E-06 |
| 4663 | 234 | L234N | A | 2.3e-06 | 2.5e-06 | 3e-06 | 2.600E-06 | 3.606E-07 | 2.300E-06 | 3.000E-06 |
| 4664 | 234 | L234D | A | 6.3e-06 | 3.3e-06 | 1.8e-06 | 3.800E-06 | 2.291E-06 | 1.800E-06 | 6.300E-06 |
| 4665 | 234 | L234C | A | 2.1e-06 | 2.9e-06 | 2.6e-06 | 2.533E-06 | 4.041E-07 | 2.100E-06 | 2.900E-06 |
| 4666 | 234 | L234E | A | 2.5e-06 | 2.1e-06 | 3.1e-06 | 2.567E-06 | 5.033E-07 | 2.100E-06 | 3.100E-06 |
| 4667 | 234 | L234Q | A | 2.3e-06 | 2.4e-06 | 2.3e-06 | 2.333E-06 | 5.774E-08 | 2.300E-06 | 2.400E-06 |
| 4668 | 234 | L234G | A | 2.5e-06 | 3.8e-06 | 1.2e-06 | 2.500E-06 | 1.300E-06 | 1.200E-06 | 3.800E-06 |
| 4669 | 234 | L234H | A | 1.4e-06 | 2.4e-06 | 2.4e-06 | 2.067E-06 | 5.774E-07 | 1.400E-06 | 2.400E-06 |
| 4670 | 234 | L234I | A | 2.4e-06 | 3.4e-06 | 2.8e-06 | 2.867E-06 | 5.033E-07 | 2.400E-06 | 3.400E-06 |
| 4671 | 234 | L234L | A | 4e-06 | 2.9e-06 | 2.7e-06 | 3.200E-06 | 7.000E-07 | 2.700E-06 | 4.000E-06 |
| 4672 | 234 | L234K | A | 3e-06 | 2.1e-06 | 2.6e-06 | 2.567E-06 | 4.509E-07 | 2.100E-06 | 3.000E-06 |
| 4673 | 234 | L234M | A | 2.5e-06 | 2e-06 | 2.8e-06 | 2.433E-06 | 4.041E-07 | 2.000E-06 | 2.800E-06 |
| 4674 | 234 | L234F | A | 2.7e-06 | 4.1e-06 | 1.5e-06 | 2.767E-06 | 1.301E-06 | 1.500E-06 | 4.100E-06 |

| | | | | | | | | | | |
|---|---|---|---|---|---|---|---|---|---|---|
| 4675 | 234 | L234P | A | 3.2e-06 | 2.8e-06 | 1.8e-06 | 2.600E-06 | 7.211E-07 | 1.800E-06 | 3.200E-06 |
| 4676 | 234 | L234S | A | 3.9e-06 | 3e-06 | 2.3e-06 | 3.067E-06 | 8.021E-07 | 2.300E-06 | 3.900E-06 |
| 4677 | 234 | L234T | A | 2.3e-06 | 2.6e-06 | 3e-06 | 2.633E-06 | 3.512E-07 | 2.300E-06 | 3.000E-06 |
| 4678 | 234 | L234W | A | 2.4e-06 | 3.3e-06 | 2.9e-06 | 2.867E-06 | 4.509E-07 | 2.400E-06 | 3.300E-06 |
| 4679 | 234 | L234Y | A | 2.3e-06 | 1.9e-06 | 2.1e-06 | 2.100E-06 | 2.000E-07 | 1.900E-06 | 2.300E-06 |
| 4680 | 234 | L234V | A | 2.7e-06 | 1.8e-06 | 2.6e-06 | 2.367E-06 | 4.933E-07 | 1.800E-06 | 2.700E-06 |
| 4681 | 235 | H235A | A | 3.1e-06 | 2.2e-06 | 2.5e-06 | 2.600E-06 | 4.583E-07 | 2.200E-06 | 3.100E-06 |
| 4682 | 235 | H235R | A | 2.5e-06 | 1.5e-06 | 1.5e-06 | 1.833E-06 | 5.774E-07 | 1.500E-06 | 2.500E-06 |
| 4683 | 235 | H235N | A | 2.1e-06 | 2.4e-06 | 2.4e-06 | 2.300E-06 | 1.732E-07 | 2.100E-06 | 2.400E-06 |
| 4684 | 235 | H235D | A | 2.6e-06 | 1.9e-06 | 2.3e-06 | 2.267E-06 | 3.512E-07 | 1.900E-06 | 2.600E-06 |
| 4685 | 235 | H235C | A | 3.3e-06 | 3.5e-06 | 1.8e-06 | 2.867E-06 | 9.292E-07 | 1.800E-06 | 3.500E-06 |
| 4686 | 235 | H235E | A | 2.7e-06 | 2.6e-06 | 1.8e-06 | 2.367E-06 | 4.933E-07 | 1.800E-06 | 2.700E-06 |
| 4687 | 235 | H235Q | A | 1.4e-06 | 1.9e-06 | 1.8e-06 | 1.700E-06 | 2.646E-07 | 1.400E-06 | 1.900E-06 |
| 4688 | 235 | H235G | A | 2.4e-06 | 2.3e-06 | 2.4e-06 | 2.367E-06 | 5.774E-08 | 2.300E-06 | 2.400E-06 |
| 4689 | 235 | H235H | A | 3.8e-06 | 2.2e-06 | 2.4e-06 | 2.800E-06 | 8.718E-07 | 2.200E-06 | 3.800E-06 |
| 4690 | 235 | H235I | A | 2.7e-06 | 1.4e-06 | 2.5e-06 | 2.200E-06 | 7.000E-07 | 1.400E-06 | 2.700E-06 |
| 4691 | 235 | H235L | A | 1.9e-06 | 1.9e-06 | 2.6e-06 | 2.133E-06 | 4.041E-07 | 1.900E-06 | 2.600E-06 |
| 4692 | 235 | H235K | A | 2.3e-06 | 2.2e-06 | 1.5e-06 | 2.000E-06 | 4.359E-07 | 1.500E-06 | 2.300E-06 |
| 4693 | 235 | H235M | A | 4.2e-06 | 1.8e-06 | 2.1e-06 | 2.700E-06 | 1.308E-06 | 1.800E-06 | 4.200E-06 |
| 4694 | 235 | H235F | A | 3.7e-06 | 3e-06 | 2.2e-06 | 2.967E-06 | 7.506E-07 | 2.200E-06 | 3.700E-06 |
| 4695 | 235 | H235P | A | 1.7e-06 | 1.9e-06 | 1.4e-06 | 1.667E-06 | 2.517E-07 | 1.400E-06 | 1.900E-06 |
| 4696 | 235 | H235S | A | 1.8e-06 | 1.2e-06 | 2.6e-06 | 1.867E-06 | 7.024E-07 | 1.200E-06 | 2.600E-06 |
| 4697 | 235 | H235T | A | 2.2e-06 | 1.4e-06 | 2.9e-06 | 2.167E-06 | 7.506E-07 | 1.400E-06 | 2.900E-06 |
| 4698 | 235 | H235W | A | 3.1e-06 | 3.9e-06 | 4e-06 | 3.667E-06 | 4.933E-07 | 3.100E-06 | 4.000E-06 |
| 4699 | 235 | H235Y | A | 1.9e-06 | 3.6e-06 | 2.4e-06 | 2.633E-06 | 8.737E-07 | 1.900E-06 | 3.600E-06 |
| 4700 | 235 | H235V | A | 3.1e-06 | 1.6e-06 | 2.1e-06 | 2.267E-06 | 7.638E-07 | 1.600E-06 | 3.100E-06 |
| 4701 | 236 | F236A | A | 5.6e-06 | 2.7e-06 | 3.6e-06 | 3.967E-06 | 1.484E-06 | 2.700E-06 | 5.600E-06 |
| 4702 | 236 | F236R | A | 2.6e-06 | 2.4e-06 | 2.6e-06 | 2.533E-06 | 1.155E-07 | 2.400E-06 | 2.600E-06 |
| 4703 | 236 | F236N | A | 3.3e-06 | 3e-06 | 3.2e-06 | 3.167E-06 | 1.528E-07 | 3.000E-06 | 3.300E-06 |
| 4704 | 236 | F236D | A | 2.5e-06 | 1.8e-06 | 2.6e-06 | 2.300E-06 | 4.359E-07 | 1.800E-06 | 2.600E-06 |
| 4705 | 236 | F236C | A | 3.5e-06 | 2.1e-06 | 1.9e-06 | 2.500E-06 | 8.718E-07 | 1.900E-06 | 3.500E-06 |
| 4706 | 236 | F236E | A | 1.6e-06 | 1.9e-06 | 1.8e-06 | 1.767E-06 | 1.528E-07 | 1.600E-06 | 1.900E-06 |
| 4707 | 236 | F236Q | A | 2.2e-06 | 2.2e-06 | 1.4e-06 | 1.933E-06 | 4.619E-07 | 1.400E-06 | 2.200E-06 |
| 4708 | 236 | F236G | A | 2.8e-06 | 2.4e-06 | 3.2e-06 | 2.800E-06 | 4.000E-07 | 2.400E-06 | 3.200E-06 |
| 4709 | 236 | F236H | A | 2.5e-06 | 3.6e-06 | 3.5e-06 | 3.200E-06 | 6.083E-07 | 2.500E-06 | 3.600E-06 |
| 4710 | 236 | F236I | A | 3.5e-06 | 2.7e-06 | 1.6e-06 | 2.600E-06 | 9.539E-07 | 1.600E-06 | 3.500E-06 |
| 4711 | 236 | F236L | A | 2.6e-06 | 1.9e-06 | 2.2e-06 | 2.233E-06 | 3.512E-07 | 1.900E-06 | 2.600E-06 |
| 4712 | 236 | F236K | A | 2.5e-06 | 3.2e-06 | 2.5e-06 | 2.733E-06 | 4.041E-07 | 2.500E-06 | 3.200E-06 |
| 4713 | 236 | F236M | A | 4e-06 | 3.3e-06 | 2.6e-06 | 3.300E-06 | 7.000E-07 | 2.600E-06 | 4.000E-06 |
| 4714 | 236 | F236F | A | 2.4e-06 | 1.9e-06 | 1.9e-06 | 2.067E-06 | 2.887E-07 | 1.900E-06 | 2.400E-06 |
| 4715 | 236 | F236P | A | 1.8e-06 | 1.5e-06 | 2.7e-06 | 2.000E-06 | 6.245E-07 | 1.500E-06 | 2.700E-06 |
| 4716 | 236 | F236S | A | 2.7e-06 | 2e-06 | 2.1e-06 | 2.267E-06 | 3.786E-07 | 2.000E-06 | 2.700E-06 |
| 4717 | 236 | F236T | A | 2.2e-06 | 2.1e-06 | 1.9e-06 | 2.067E-06 | 1.528E-07 | 1.900E-06 | 2.200E-06 |
| 4718 | 236 | F236W | A | 3.5e-06 | 4.2e-06 | 2.1e-06 | 3.267E-06 | 1.069E-06 | 2.100E-06 | 4.200E-06 |
| 4719 | 236 | F236Y | A | 2.5e-06 | 1.8e-06 | 2.2e-06 | 2.167E-06 | 3.512E-07 | 1.800E-06 | 2.500E-06 |
| 4720 | 236 | F236V | A | 1.7e-06 | 2.3e-06 | 2.1e-06 | 2.033E-06 | 3.055E-07 | 1.700E-06 | 2.300E-06 |
| 4721 | 237 | N237A | A | 2.1e-06 | 2.7e-06 | 1.6e-06 | 2.133E-06 | 5.508E-07 | 1.600E-06 | 2.700E-06 |
| 4722 | 237 | N237R | A | 2.7e-06 | 1.4e-06 | 2e-06 | 2.033E-06 | 6.506E-07 | 1.400E-06 | 2.700E-06 |
| 4723 | 237 | N237N | A | 2.6e-06 | 2.9e-06 | 2.2e-06 | 2.567E-06 | 3.512E-07 | 2.200E-06 | 2.900E-06 |
| 4724 | 237 | N237D | A | 2.4e-06 | 1.7e-06 | 2.1e-06 | 2.067E-06 | 3.512E-07 | 1.700E-06 | 2.400E-06 |
| 4725 | 237 | N237C | A | 3.9e-06 | 3.3e-06 | 1.6e-06 | 2.933E-06 | 1.193E-06 | 1.600E-06 | 3.900E-06 |
| 4726 | 237 | N237E | A | 1.9e-06 | 2.8e-06 | 1.5e-06 | 2.067E-06 | 6.658E-07 | 1.500E-06 | 2.800E-06 |
| 4727 | 237 | N237Q | A | 2.4e-06 | 1.4e-06 | 3.1e-06 | 2.300E-06 | 8.544E-07 | 1.400E-06 | 3.100E-06 |
| 4728 | 237 | N237G | A | 3.4e-06 | 3.5e-06 | 2.4e-06 | 3.100E-06 | 6.083E-07 | 2.400E-06 | 3.500E-06 |
| 4729 | 237 | N237H | A | 2.7e-06 | 1.8e-06 | 1.5e-06 | 2.000E-06 | 6.245E-07 | 1.500E-06 | 2.700E-06 |
| 4730 | 237 | N237I | A | 2.7e-06 | 2.3e-06 | 3.1e-06 | 2.700E-06 | 4.000E-07 | 2.300E-06 | 3.100E-06 |
| 4731 | 237 | N237L | A | 1.8e-06 | 4.3e-06 | 1.6e-06 | 2.567E-06 | 1.504E-06 | 1.600E-06 | 4.300E-06 |
| 4732 | 237 | N237K | A | 2.7e-06 | 2.8e-06 | 1.7e-06 | 2.400E-06 | 6.083E-07 | 1.700E-06 | 2.800E-06 |
| 4733 | 237 | N237M | A | 2.1e-06 | 1.9e-06 | 3.2e-06 | 2.400E-06 | 7.000E-07 | 1.900E-06 | 3.200E-06 |
| 4734 | 237 | N237F | A | 9.4e-07 | 2.3e-06 | 2.1e-06 | 1.780E-06 | 7.343E-07 | 9.400E-07 | 2.300E-06 |
| 4735 | 237 | N237P | A | 2.2e-06 | 1.1e-06 | 2.6e-06 | 1.967E-06 | 7.767E-07 | 1.100E-06 | 2.600E-06 |
| 4736 | 237 | N237S | A | 2e-06 | 2.1e-06 | 3.9e-06 | 2.667E-06 | 1.069E-06 | 2.000E-06 | 3.900E-06 |
| 4737 | 237 | N237T | A | 2.6e-06 | 1.9e-06 | 3.4e-06 | 2.633E-06 | 7.506E-07 | 1.900E-06 | 3.400E-06 |
| 4738 | 237 | N237W | A | 3.5e-06 | 2.4e-06 | 3.3e-06 | 3.067E-06 | 5.859E-07 | 2.400E-06 | 3.500E-06 |
| 4739 | 237 | N237Y | A | 4.2e-06 | 1.9e-06 | 3.6e-06 | 3.233E-06 | 1.193E-06 | 1.900E-06 | 4.200E-06 |
| 4740 | 237 | N237V | A | 2.5e-06 | 4.1e-06 | 2.8e-06 | 3.133E-06 | 8.505E-07 | 2.500E-06 | 4.100E-06 |
| 4741 | 238 | H238A | A | 3.1e-06 | 1.5e-06 | 3.1e-06 | 2.567E-06 | 9.238E-07 | 1.500E-06 | 3.100E-06 |
| 4742 | 238 | H238R | A | 2.6e-06 | 3.1e-06 | 2e-06 | 2.567E-06 | 5.508E-07 | 2.000E-06 | 3.100E-06 |
| 4743 | 238 | H238N | A | 2.1e-06 | 3.5e-06 | 3e-06 | 2.867E-06 | 7.095E-07 | 2.100E-06 | 3.500E-06 |
| 4744 | 238 | H238D | A | 3.2e-06 | 2.4e-06 | 2.6e-06 | 2.733E-06 | 4.163E-07 | 2.400E-06 | 3.200E-06 |
| 4745 | 238 | H238C | A | 3.2e-06 | 2.1e-06 | 2.1e-06 | 2.467E-06 | 6.351E-07 | 2.100E-06 | 3.200E-06 |
| 4746 | 238 | H238E | A | 1.9e-06 | 2.5e-06 | 1.6e-06 | 2.000E-06 | 4.583E-07 | 1.600E-06 | 2.500E-06 |
| 4747 | 238 | H238Q | A | 3.2e-06 | 1.6e-06 | 2.4e-06 | 2.400E-06 | 8.000E-07 | 1.600E-06 | 3.200E-06 |
| 4748 | 238 | H238G | A | 2.2e-06 | 2.8e-06 | 3e-06 | 2.667E-06 | 4.163E-07 | 2.200E-06 | 3.000E-06 |
| 4749 | 238 | H238H | A | 3e-06 | 2.1e-06 | 3.1e-06 | 2.733E-06 | 5.508E-07 | 2.100E-06 | 3.100E-06 |
| 4750 | 238 | H238I | A | 2.3e-06 | 2e-06 | 3e-06 | 2.433E-06 | 5.132E-07 | 2.000E-06 | 3.000E-06 |
| 4751 | 238 | H238L | A | 2.8e-06 | 2.9e-06 | 3.8e-06 | 3.167E-06 | 5.508E-07 | 2.800E-06 | 3.800E-06 |
| 4752 | 238 | H238K | A | 1.6e-06 | 1.2e-06 | 3.6e-06 | 2.133E-06 | 1.286E-06 | 1.200E-06 | 3.600E-06 |
| 4753 | 238 | H238M | A | 2.5e-06 | 1.3e-06 | 3.1e-06 | 2.300E-06 | 9.165E-07 | 1.300E-06 | 3.100E-06 |
| 4754 | 238 | H238F | A | 2.2e-06 | 3e-06 | 2.6e-06 | 2.600E-06 | 4.000E-07 | 2.200E-06 | 3.000E-06 |
| 4755 | 238 | H238P | A | 3.3e-06 | 2.7e-06 | 2.8e-06 | 2.933E-06 | 3.215E-07 | 2.700E-06 | 3.300E-06 |
| 4756 | 238 | H238S | A | 2.2e-06 | 2.6e-06 | 2.3e-06 | 2.367E-06 | 2.082E-07 | 2.200E-06 | 2.600E-06 |
| 4757 | 238 | H238T | A | 1.4e-06 | 2e-06 | 2.3e-06 | 1.900E-06 | 4.583E-07 | 1.400E-06 | 2.300E-06 |
| 4758 | 238 | H238W | A | 1.3e-06 | 1.5e-06 | 2.5e-06 | 1.767E-06 | 6.429E-07 | 1.300E-06 | 2.500E-06 |
| 4759 | 238 | H238Y | A | 1.9e-06 | 4.6e-06 | 2.4e-06 | 2.967E-06 | 1.436E-06 | 1.900E-06 | 4.600E-06 |

| | | | | | | | | | | |
|---|---|---|---|---|---|---|---|---|---|---|
| 4760 | 238 | H238V | A | 4.6e-06 | 2.4e-06 | 2.5e-06 | 3.167E-06 | 1.242E-06 | 2.400E-06 | 4.600E-06 |
| 4761 | 239 | S239A | A | 1.5e-06 | 2.4e-06 | 3.6e-06 | 2.500E-06 | 1.054E-06 | 1.500E-06 | 3.600E-06 |
| 4762 | 239 | S239R | A | 4.1e-06 | 1.9e-06 | 3.6e-06 | 3.200E-06 | 1.153E-06 | 1.900E-06 | 4.100E-06 |
| 4763 | 239 | S239N | A | 1.8e-06 | 3.8e-06 | 1.8e-06 | 2.467E-06 | 1.155E-06 | 1.800E-06 | 3.800E-06 |
| 4764 | 239 | S239D | A | 2.8e-06 | 2.9e-06 | 2.5e-06 | 2.733E-06 | 2.082E-07 | 2.500E-06 | 2.900E-06 |
| 4765 | 239 | S239C | A | 2.9e-06 | 2.7e-06 | 3e-06 | 2.867E-06 | 1.528E-07 | 2.700E-06 | 3.000E-06 |
| 4766 | 239 | S239E | A | 2.6e-06 | 3.6e-06 | 2.7e-06 | 2.967E-06 | 5.508E-07 | 2.600E-06 | 3.600E-06 |
| 4767 | 239 | S239Q | A | 3.9e-06 | 2.5e-06 | 2.5e-06 | 2.967E-06 | 8.083E-07 | 2.500E-06 | 3.900E-06 |
| 4768 | 239 | S239G | A | 3e-06 | 2.7e-06 | 2.2e-06 | 2.633E-06 | 4.041E-07 | 2.200E-06 | 3.000E-06 |
| 4769 | 239 | S239H | A | 2.3e-06 | 4.1e-06 | 2.3e-06 | 2.900E-06 | 1.039E-06 | 2.300E-06 | 4.100E-06 |
| 4770 | 239 | S239I | A | 2e-06 | 2.7e-06 | 2.4e-06 | 2.367E-06 | 3.512E-07 | 2.000E-06 | 2.700E-06 |
| 4771 | 239 | S239L | A | 2.1e-06 | 2e-06 | 1.9e-06 | 2.000E-06 | 1.000E-07 | 1.900E-06 | 2.100E-06 |
| 4772 | 239 | S239K | A | 3e-06 | 3.7e-06 | 2.1e-06 | 2.933E-06 | 8.021E-07 | 2.100E-06 | 3.700E-06 |
| 4773 | 239 | S239M | A | 2.1e-06 | 2.4e-06 | 3.9e-06 | 2.800E-06 | 9.644E-07 | 2.100E-06 | 3.900E-06 |
| 4774 | 239 | S239F | A | 2.8e-06 | 3.4e-06 | 1.9e-06 | 2.700E-06 | 7.550E-07 | 1.900E-06 | 3.400E-06 |
| 4775 | 239 | S239P | A | 2.7e-06 | 4.6e-06 | 2.7e-06 | 3.333E-06 | 1.097E-06 | 2.700E-06 | 4.600E-06 |
| 4776 | 239 | S239S | A | 3.4e-06 | 2.5e-06 | 3.3e-06 | 3.067E-06 | 4.933E-07 | 2.500E-06 | 3.400E-06 |
| 4777 | 239 | S239T | A | 2.2e-06 | 2.7e-06 | 2.2e-06 | 2.367E-06 | 2.887E-07 | 2.200E-06 | 2.700E-06 |
| 4778 | 239 | S239W | A | 1.5e-06 | 2.7e-06 | 4e-06 | 2.733E-06 | 1.250E-06 | 1.500E-06 | 4.000E-06 |
| 4779 | 239 | S239Y | A | 2e-06 | 2.2e-06 | 1.9e-06 | 2.033E-06 | 1.528E-07 | 1.900E-06 | 2.200E-06 |
| 4780 | 239 | S239V | A | 4.6e-06 | 4.5e-06 | 2.7e-06 | 3.933E-06 | 1.069E-06 | 2.700E-06 | 4.600E-06 |
| 4781 | 240 | G240A | A | 1.6e-06 | 2.1e-06 | 4.4e-06 | 2.700E-06 | 1.493E-06 | 1.600E-06 | 4.400E-06 |
| 4782 | 240 | G240R | A | 1.5e-06 | 1.6e-06 | 1.6e-06 | 1.567E-06 | 5.774E-08 | 1.500E-06 | 1.600E-06 |
| 4783 | 240 | G240N | A | 1.6e-06 | 2.3e-06 | 1.9e-06 | 1.933E-06 | 3.512E-07 | 1.600E-06 | 2.300E-06 |
| 4784 | 240 | G240D | A | 1.4e-06 | 1.7e-06 | 2.8e-06 | 1.967E-06 | 7.371E-07 | 1.400E-06 | 2.800E-06 |
| 4785 | 240 | G240C | A | 2.1e-06 | 2.2e-06 | 2.5e-06 | 2.267E-06 | 2.082E-07 | 2.100E-06 | 2.500E-06 |
| 4786 | 240 | G240E | A | 2.6e-06 | 1.5e-06 | 1.9e-06 | 2.000E-06 | 5.568E-07 | 1.500E-06 | 2.600E-06 |
| 4787 | 240 | G240Q | A | 2.8e-06 | 2.1e-06 | 2.1e-06 | 2.333E-06 | 4.041E-07 | 2.100E-06 | 2.800E-06 |
| 4788 | 240 | G240G | A | 3.5e-06 | 1.4e-06 | 3.4e-06 | 2.767E-06 | 1.185E-06 | 1.400E-06 | 3.500E-06 |
| 4789 | 240 | G240H | A | 3.6e-06 | 1.8e-06 | 4.3e-06 | 3.233E-06 | 1.290E-06 | 1.800E-06 | 4.300E-06 |
| 4790 | 240 | G240I | A | 2.4e-06 | 1.5e-06 | 1.4e-06 | 1.767E-06 | 5.508E-07 | 1.400E-06 | 2.400E-06 |
| 4791 | 240 | G240L | A | 2.4e-06 | 2.2e-06 | 2.2e-06 | 2.267E-06 | 1.155E-07 | 2.200E-06 | 2.400E-06 |
| 4792 | 240 | G240K | A | 1.9e-06 | 9.7e-07 | 2.6e-06 | 1.823E-06 | 8.177E-07 | 9.700E-07 | 2.600E-06 |
| 4793 | 240 | G240M | A | 2.8e-06 | 2.6e-06 | 3.1e-06 | 2.833E-06 | 2.517E-07 | 2.600E-06 | 3.100E-06 |
| 4794 | 240 | G240F | A | 1.1e-06 | 1.9e-06 | 4.4e-06 | 2.467E-06 | 1.721E-06 | 1.100E-06 | 4.400E-06 |
| 4795 | 240 | G240P | A | 3.3e-06 | 1.2e-06 | 1.7e-06 | 2.067E-06 | 1.097E-06 | 1.200E-06 | 3.300E-06 |
| 4796 | 240 | G240S | A | 2.9e-06 | 2.2e-06 | 1.3e-06 | 2.133E-06 | 8.021E-07 | 1.300E-06 | 2.900E-06 |
| 4797 | 240 | G240T | A | 2.9e-06 | 2.6e-06 | 2e-06 | 2.500E-06 | 4.583E-07 | 2.000E-06 | 2.900E-06 |
| 4798 | 240 | G240W | A | 2e-06 | 2.6e-06 | 1.2e-06 | 1.933E-06 | 7.024E-07 | 1.200E-06 | 2.600E-06 |
| 4799 | 240 | G240Y | A | 1.1e-06 | 1.3e-06 | 2.9e-06 | 1.767E-06 | 9.866E-07 | 1.100E-06 | 2.900E-06 |
| 4800 | 240 | G240V | A | 2e-06 | 2e-06 | 3.8e-06 | 2.600E-06 | 1.039E-06 | 2.000E-06 | 3.800E-06 |
| 4801 | 241 | I241A | A | 2.3e-06 | 2e-06 | 4e-06 | 2.767E-06 | 1.079E-06 | 2.000E-06 | 4.000E-06 |
| 4802 | 241 | I241R | A | 1.7e-06 | 2e-06 | 3e-06 | 2.233E-06 | 6.807E-07 | 1.700E-06 | 3.000E-06 |
| 4803 | 241 | I241N | A | 1.9e-06 | 2.7e-06 | 2.8e-06 | 2.467E-06 | 4.933E-07 | 1.900E-06 | 2.800E-06 |
| 4804 | 241 | I241D | A | 2.2e-06 | 3e-06 | 2.3e-06 | 2.500E-06 | 4.359E-07 | 2.200E-06 | 3.000E-06 |
| 4805 | 241 | I241C | A | 1.8e-06 | 2.1e-06 | 4.1e-06 | 2.667E-06 | 1.250E-06 | 1.800E-06 | 4.100E-06 |
| 4806 | 241 | I241E | A | 1.2e-06 | 8.7e-07 | 3.1e-06 | 1.723E-06 | 1.204E-06 | 8.700E-07 | 3.100E-06 |
| 4807 | 241 | I241Q | A | 3.6e-06 | 1.5e-06 | 2.6e-06 | 2.567E-06 | 1.050E-06 | 1.500E-06 | 3.600E-06 |
| 4808 | 241 | I241G | A | 3.1e-06 | 6.5e-06 | 4.4e-06 | 4.667E-06 | 1.716E-06 | 3.100E-06 | 6.500E-06 |
| 4809 | 241 | I241H | A | 2.1e-06 | 1.9e-06 | 1.9e-06 | 1.967E-06 | 1.155E-07 | 1.900E-06 | 2.100E-06 |
| 4810 | 241 | I241I | A | 1.7e-06 | 2.2e-06 | 3.3e-06 | 2.400E-06 | 8.185E-07 | 1.700E-06 | 3.300E-06 |
| 4811 | 241 | I241L | A | 1.8e-06 | 1.6e-06 | 1.6e-06 | 1.667E-06 | 1.155E-07 | 1.600E-06 | 1.800E-06 |
| 4812 | 241 | I241K | A | 2.8e-06 | 2.7e-06 | 2.6e-06 | 2.700E-06 | 1.000E-07 | 2.600E-06 | 2.800E-06 |
| 4813 | 241 | I241M | A | 1.7e-06 | 3e-06 | 2.3e-06 | 2.333E-06 | 6.506E-07 | 1.700E-06 | 3.000E-06 |
| 4814 | 241 | I241F | A | 2.1e-06 | 2.8e-06 | 3.1e-06 | 2.667E-06 | 5.132E-07 | 2.100E-06 | 3.100E-06 |
| 4815 | 241 | I241P | A | 1.4e-06 | 2.3e-06 | 2.2e-06 | 1.967E-06 | 4.933E-07 | 1.400E-06 | 2.300E-06 |
| 4816 | 241 | I241S | A | 1.7e-06 | 3.1e-06 | 1.7e-06 | 2.167E-06 | 8.083E-07 | 1.700E-06 | 3.100E-06 |
| 4817 | 241 | I241T | A | 2.1e-06 | 4.3e-06 | 2e-06 | 2.800E-06 | 1.300E-06 | 2.000E-06 | 4.300E-06 |
| 4818 | 241 | I241W | A | 2.3e-06 | 1.5e-06 | 2e-06 | 1.933E-06 | 4.041E-07 | 1.500E-06 | 2.300E-06 |
| 4819 | 241 | I241Y | A | 2.9e-06 | 2.3e-06 | 2.2e-06 | 2.467E-06 | 3.786E-07 | 2.200E-06 | 2.900E-06 |
| 4820 | 241 | I241V | A | 2.3e-06 | 1.6e-06 | 3.6e-06 | 2.500E-06 | 1.015E-06 | 1.600E-06 | 3.600E-06 |
| 4821 | 242 | C242A | A | 2.3e-06 | 1.6e-06 | 1.1e-06 | 1.667E-06 | 6.028E-07 | 1.100E-06 | 2.300E-06 |
| 4822 | 242 | C242R | A | 1.9e-06 | 1.6e-06 | 1.4e-06 | 1.633E-06 | 2.517E-07 | 1.400E-06 | 1.900E-06 |
| 4823 | 242 | C242N | A | 1.9e-06 | 2.7e-06 | 1.5e-06 | 2.033E-06 | 6.110E-07 | 1.500E-06 | 2.700E-06 |
| 4824 | 242 | C242D | A | 3.9e-06 | 3.2e-06 | 2e-06 | 3.033E-06 | 9.609E-07 | 2.000E-06 | 3.900E-06 |
| 4825 | 242 | C242C | A | 8.8e-07 | 2.8e-06 | 2.4e-06 | 2.027E-06 | 1.013E-06 | 8.800E-07 | 2.800E-06 |
| 4826 | 242 | C242E | A | 2.8e-06 | 2.6e-06 | 2.6e-06 | 2.667E-06 | 1.155E-07 | 2.600E-06 | 2.800E-06 |
| 4827 | 242 | C242Q | A | 2.7e-06 | 1.3e-06 | 2.4e-06 | 2.133E-06 | 7.371E-07 | 1.300E-06 | 2.700E-06 |
| 4828 | 242 | C242G | A | 2.2e-06 | 2.9e-06 | 2.1e-06 | 2.400E-06 | 4.359E-07 | 2.100E-06 | 2.900E-06 |
| 4829 | 242 | C242H | A | 1.1e-06 | 3.1e-06 | 2.2e-06 | 2.133E-06 | 1.002E-06 | 1.100E-06 | 3.100E-06 |
| 4830 | 242 | C242I | A | 3.3e-06 | 2.3e-06 | 1.4e-06 | 2.333E-06 | 9.504E-07 | 1.400E-06 | 3.300E-06 |
| 4831 | 242 | C242L | A | 2.5e-06 | 2.1e-06 | 2.8e-06 | 2.467E-06 | 3.512E-07 | 2.100E-06 | 2.800E-06 |
| 4832 | 242 | C242K | A | 2.5e-06 | 1.9e-06 | 2.2e-06 | 2.200E-06 | 3.000E-07 | 1.900E-06 | 2.500E-06 |
| 4833 | 242 | C242M | A | 2.9e-06 | 1.8e-06 | 2.6e-06 | 2.433E-06 | 5.686E-07 | 1.800E-06 | 2.900E-06 |
| 4834 | 242 | C242F | A | 3.2e-06 | 1.1e-06 | 1.2e-06 | 1.833E-06 | 1.185E-06 | 1.100E-06 | 3.200E-06 |
| 4835 | 242 | C242P | A | 1.8e-06 | 1.8e-06 | 1.7e-06 | 1.767E-06 | 5.774E-08 | 1.700E-06 | 1.800E-06 |
| 4836 | 242 | C242S | A | 5.8e-06 | 1.9e-06 | 3.7e-06 | 3.800E-06 | 1.952E-06 | 1.900E-06 | 5.800E-06 |
| 4837 | 242 | C242T | A | 2.2e-06 | 2.1e-06 | 2.6e-06 | 2.300E-06 | 2.646E-07 | 2.100E-06 | 2.600E-06 |
| 4838 | 242 | C242W | A | 3e-06 | 2.5e-06 | 2e-06 | 2.500E-06 | 5.000E-07 | 2.000E-06 | 3.000E-06 |
| 4839 | 242 | C242Y | A | 2.8e-06 | 2.3e-06 | 2.9e-06 | 2.667E-06 | 3.215E-07 | 2.300E-06 | 2.900E-06 |
| 4840 | 242 | C242V | A | 1.5e-06 | 1.9e-06 | 2.8e-06 | 2.067E-06 | 6.658E-07 | 1.500E-06 | 2.800E-06 |
| 4841 | 243 | E243A | A | 2.5e-06 | 4.7e-06 | 2.6e-06 | 3.267E-06 | 1.242E-06 | 2.500E-06 | 4.700E-06 |
| 4842 | 243 | E243R | A | 2.5e-06 | 3.7e-06 | 2e-06 | 2.733E-06 | 8.737E-07 | 2.000E-06 | 3.700E-06 |
| 4843 | 243 | E243N | A | 9.6e-07 | 2.1e-06 | 3.9e-06 | 2.320E-06 | 1.482E-06 | 9.600E-07 | 3.900E-06 |
| 4844 | 243 | E243D | A | 2.7e-06 | 2.1e-06 | 2.3e-06 | 2.367E-06 | 3.055E-07 | 2.100E-06 | 2.700E-06 |

| | | | | | | | | | | |
|---|---|---|---|---|---|---|---|---|---|---|
| 4845 | 243 | E243C | A | 3.8e-06 | 2.3e-06 | 2.3e-06 | 2.800E-06 | 8.660E-07 | 2.300E-06 | 3.800E-06 |
| 4846 | 243 | E243E | A | 2.1e-06 | 2.9e-06 | 1.7e-06 | 2.233E-06 | 6.110E-07 | 1.700E-06 | 2.900E-06 |
| 4847 | 243 | E243Q | A | 1.2e-06 | 2.2e-06 | 3.3e-06 | 2.233E-06 | 1.050E-06 | 1.200E-06 | 3.300E-06 |
| 4848 | 243 | E243G | A | 3.1e-06 | 3.8e-06 | 1.2e-06 | 2.700E-06 | 1.345E-06 | 1.200E-06 | 3.800E-06 |
| 4849 | 243 | E243H | A | 1.3e-06 | 1.5e-06 | 3.2e-06 | 2.000E-06 | 1.044E-06 | 1.300E-06 | 3.200E-06 |
| 4850 | 243 | E243I | A | 2e-06 | 2.1e-06 | 1.6e-06 | 1.900E-06 | 2.646E-07 | 1.600E-06 | 2.100E-06 |
| 4851 | 243 | E243L | A | 2.8e-06 | 2.3e-06 | 4e-06 | 3.033E-06 | 8.737E-07 | 2.300E-06 | 4.000E-06 |
| 4852 | 243 | E243K | A | 2.4e-06 | 3e-06 | 2.3e-06 | 2.567E-06 | 3.786E-07 | 2.300E-06 | 3.000E-06 |
| 4853 | 243 | E243M | A | 1.7e-06 | 2.2e-06 | 2.7e-06 | 2.200E-06 | 5.000E-07 | 1.700E-06 | 2.700E-06 |
| 4854 | 243 | E243F | A | 2.9e-06 | 2.2e-06 | 2.2e-06 | 2.433E-06 | 4.041E-07 | 2.200E-06 | 2.900E-06 |
| 4855 | 243 | E243P | A | 3.7e-06 | 1.8e-06 | 2.5e-06 | 2.667E-06 | 9.609E-07 | 1.800E-06 | 3.700E-06 |
| 4856 | 243 | E243S | A | 2.3e-06 | 2.1e-06 | 1.8e-06 | 2.067E-06 | 2.517E-07 | 1.800E-06 | 2.300E-06 |
| 4857 | 243 | E243T | A | 1.9e-06 | 2e-06 | 3.7e-06 | 2.533E-06 | 1.012E-06 | 1.900E-06 | 3.700E-06 |
| 4858 | 243 | E243W | A | 2.8e-06 | 2.1e-06 | 2.5e-06 | 2.467E-06 | 3.512E-07 | 2.100E-06 | 2.800E-06 |
| 4859 | 243 | E243Y | A | 2.9e-06 | 2.8e-06 | 1.9e-06 | 2.533E-06 | 5.508E-07 | 1.900E-06 | 2.900E-06 |
| 4860 | 243 | E243V | A | 2.8e-06 | 4.2e-06 | 2.5e-06 | 3.167E-06 | 9.074E-07 | 2.500E-06 | 4.200E-06 |
| 4861 | 244 | L244A | A | 2.3e-06 | 2.9e-06 | 2.6e-06 | 2.600E-06 | 3.000E-07 | 2.300E-06 | 2.900E-06 |
| 4862 | 244 | L244R | A | 2.5e-06 | 1.8e-06 | 2.9e-06 | 2.400E-06 | 5.568E-07 | 1.800E-06 | 2.900E-06 |
| 4863 | 244 | L244N | A | 2.8e-06 | 2.2e-06 | 2.5e-06 | 2.500E-06 | 3.000E-07 | 2.200E-06 | 2.800E-06 |
| 4864 | 244 | L244D | A | 3.6e-06 | 1.5e-06 | 2.8e-06 | 2.633E-06 | 1.060E-06 | 1.500E-06 | 3.600E-06 |
| 4865 | 244 | L244C | A | 1.7e-06 | 2.9e-06 | 2.4e-06 | 2.333E-06 | 6.028E-07 | 1.700E-06 | 2.900E-06 |
| 4866 | 244 | L244E | A | 1.8e-06 | 2.1e-06 | 1.5e-06 | 1.800E-06 | 3.000E-07 | 1.500E-06 | 2.100E-06 |
| 4867 | 244 | L244Q | A | 3.8e-06 | 3.4e-06 | 2.5e-06 | 3.233E-06 | 6.658E-07 | 2.500E-06 | 3.800E-06 |
| 4868 | 244 | L244G | A | 2.7e-06 | 6.1e-07 | 2.2e-06 | 1.837E-06 | 1.091E-06 | 6.100E-07 | 2.700E-06 |
| 4869 | 244 | L244H | A | 2.6e-06 | 2.8e-06 | 2.5e-06 | 2.633E-06 | 1.528E-07 | 2.500E-06 | 2.800E-06 |
| 4870 | 244 | L244I | A | 3.2e-06 | 2.1e-06 | 2.3e-06 | 2.533E-06 | 5.859E-07 | 2.100E-06 | 3.200E-06 |
| 4871 | 244 | L244L | A | 3e-06 | 3.4e-06 | 3.3e-06 | 3.233E-06 | 2.082E-07 | 3.000E-06 | 3.400E-06 |
| 4872 | 244 | L244K | A | 1.7e-06 | 1.9e-06 | 2.5e-06 | 2.033E-06 | 4.163E-07 | 1.700E-06 | 2.500E-06 |
| 4873 | 244 | L244M | A | 2.8e-06 | 1.9e-06 | 2.2e-06 | 2.300E-06 | 4.583E-07 | 1.900E-06 | 2.800E-06 |
| 4874 | 244 | L244F | A | 5e-06 | 2.6e-06 | 1.8e-06 | 3.133E-06 | 1.665E-06 | 1.800E-06 | 5.000E-06 |
| 4875 | 244 | L244P | A | 3.9e-06 | 3.4e-06 | 3.3e-06 | 3.533E-06 | 3.215E-07 | 3.300E-06 | 3.900E-06 |
| 4876 | 244 | L244S | A | 2.2e-06 | 1.6e-06 | 1.6e-06 | 1.800E-06 | 3.464E-07 | 1.600E-06 | 2.200E-06 |
| 4877 | 244 | L244T | A | 2.5e-06 | 1.9e-06 | 1.7e-06 | 2.033E-06 | 4.163E-07 | 1.700E-06 | 2.500E-06 |
| 4878 | 244 | L244W | A | 1.9e-06 | 2.3e-06 | 2.1e-06 | 2.100E-06 | 2.000E-07 | 1.900E-06 | 2.300E-06 |
| 4879 | 244 | L244Y | A | 1.1e-06 | 3.4e-06 | 2.5e-06 | 2.333E-06 | 1.159E-06 | 1.100E-06 | 3.400E-06 |
| 4880 | 244 | L244V | A | 4e-06 | 2.4e-06 | 3.7e-06 | 3.367E-06 | 8.505E-07 | 2.400E-06 | 4.000E-06 |
| 4881 | 245 | H245A | A | 2.5e-06 | 2e-06 | 2.3e-06 | 2.267E-06 | 2.517E-07 | 2.000E-06 | 2.500E-06 |
| 4882 | 245 | H245R | A | 2.3e-06 | 2.3e-06 | 2.5e-06 | 2.367E-06 | 1.155E-07 | 2.300E-06 | 2.500E-06 |
| 4883 | 245 | H245N | A | 3.3e-06 | 1.3e-06 | 2.4e-06 | 2.333E-06 | 1.002E-06 | 1.300E-06 | 3.300E-06 |
| 4884 | 245 | H245D | A | 2.2e-06 | 2.1e-06 | 2.3e-06 | 2.200E-06 | 1.000E-07 | 2.100E-06 | 2.300E-06 |
| 4885 | 245 | H245C | A | 2.6e-06 | 2.1e-06 | 2.6e-06 | 2.433E-06 | 2.887E-07 | 2.100E-06 | 2.600E-06 |
| 4886 | 245 | H245E | A | 1.8e-06 | 2.1e-06 | 9.6e-07 | 1.620E-06 | 5.909E-07 | 9.600E-07 | 2.100E-06 |
| 4887 | 245 | H245Q | A | 2.6e-06 | 2.4e-06 | 1.7e-06 | 2.233E-06 | 4.726E-07 | 1.700E-06 | 2.600E-06 |
| 4888 | 245 | H245G | A | 2.7e-06 | 2.4e-06 | 1.9e-06 | 2.333E-06 | 4.041E-07 | 1.900E-06 | 2.700E-06 |
| 4889 | 245 | H245H | A | 1.5e-06 | 1.8e-06 | 1.9e-06 | 1.733E-06 | 2.082E-07 | 1.500E-06 | 1.900E-06 |
| 4890 | 245 | H245I | A | 1e-06 | 1.8e-06 | 3.2e-06 | 2.000E-06 | 1.114E-06 | 1.000E-06 | 3.200E-06 |
| 4891 | 245 | H245L | A | 4e-06 | 1.7e-06 | 2.7e-06 | 2.800E-06 | 1.153E-06 | 1.700E-06 | 4.000E-06 |
| 4892 | 245 | H245K | A | 2.3e-06 | 2.7e-06 | 2.6e-06 | 2.533E-06 | 2.082E-07 | 2.300E-06 | 2.700E-06 |
| 4893 | 245 | H245M | A | 2.2e-06 | 2.9e-06 | 3.6e-06 | 2.900E-06 | 7.000E-07 | 2.200E-06 | 3.600E-06 |
| 4894 | 245 | H245F | A | 2.2e-06 | 9.7e-07 | 2.6e-06 | 1.923E-06 | 8.495E-07 | 9.700E-07 | 2.600E-06 |
| 4895 | 245 | H245P | A | 2.5e-06 | 2.9e-06 | 2.4e-06 | 2.600E-06 | 2.646E-07 | 2.400E-06 | 2.900E-06 |
| 4896 | 245 | H245S | A | 1.8e-06 | 3e-06 | 2.2e-06 | 2.333E-06 | 6.110E-07 | 1.800E-06 | 3.000E-06 |
| 4897 | 245 | H245T | A | 1.8e-06 | 2.7e-06 | 3.1e-06 | 2.533E-06 | 6.658E-07 | 1.800E-06 | 3.100E-06 |
| 4898 | 245 | H245W | A | 3.1e-06 | 1.7e-06 | 4.1e-06 | 2.967E-06 | 1.206E-06 | 1.700E-06 | 4.100E-06 |
| 4899 | 245 | H245Y | A | 2.9e-06 | 2.8e-06 | 2.3e-06 | 2.667E-06 | 3.215E-07 | 2.300E-06 | 2.900E-06 |
| 4900 | 245 | H245V | A | 3.5e-06 | 2.4e-06 | 2.3e-06 | 2.733E-06 | 6.658E-07 | 2.300E-06 | 3.500E-06 |
| 4901 | 246 | C246A | A | 3.5e-06 | 2.4e-06 | 3.7e-06 | 3.200E-06 | 7.000E-07 | 2.400E-06 | 3.700E-06 |
| 4902 | 246 | C246R | A | 2.6e-06 | 2.5e-06 | 1.5e-06 | 2.200E-06 | 6.083E-07 | 1.500E-06 | 2.600E-06 |
| 4903 | 246 | C246N | A | 1.3e-06 | 1.8e-06 | 3e-06 | 2.033E-06 | 8.737E-07 | 1.300E-06 | 3.000E-06 |
| 4904 | 246 | C246D | A | 2.7e-06 | 3.6e-06 | 1.7e-06 | 2.667E-06 | 9.504E-07 | 1.700E-06 | 3.600E-06 |
| 4905 | 246 | C246C | A | 1.6e-06 | 3.7e-06 | 2e-06 | 2.433E-06 | 1.115E-06 | 1.600E-06 | 3.700E-06 |
| 4906 | 246 | C246E | A | 2.6e-06 | 2.3e-06 | 1.8e-06 | 2.233E-06 | 4.041E-07 | 1.800E-06 | 2.600E-06 |
| 4907 | 246 | C246Q | A | 2.2e-06 | 2.9e-06 | 6.4e-06 | 3.833E-06 | 2.250E-06 | 2.200E-06 | 6.400E-06 |
| 4908 | 246 | C246G | A | 1.8e-06 | 2.9e-06 | 3.1e-06 | 2.600E-06 | 7.000E-07 | 1.800E-06 | 3.100E-06 |
| 4909 | 246 | C246H | A | 1.7e-06 | 2e-06 | 2.6e-06 | 2.100E-06 | 4.583E-07 | 1.700E-06 | 2.600E-06 |
| 4910 | 246 | C246I | A | 3.7e-06 | 2.7e-06 | 2.2e-06 | 2.867E-06 | 7.638E-07 | 2.200E-06 | 3.700E-06 |
| 4911 | 246 | C246L | A | 1.6e-06 | 2e-06 | 3.1e-06 | 2.233E-06 | 7.767E-07 | 1.600E-06 | 3.100E-06 |
| 4912 | 246 | C246K | A | 2e-06 | 1.7e-06 | 3.5e-06 | 2.400E-06 | 9.644E-07 | 1.700E-06 | 3.500E-06 |
| 4913 | 246 | C246M | A | 4.5e-06 | 2.5e-06 | 3.1e-06 | 3.367E-06 | 1.026E-06 | 2.500E-06 | 4.500E-06 |
| 4914 | 246 | C246F | A | 1.8e-06 | 4.3e-06 | 2.5e-06 | 2.867E-06 | 1.290E-06 | 1.800E-06 | 4.300E-06 |
| 4915 | 246 | C246P | A | 1.8e-06 | 1.9e-06 | 2.3e-06 | 2.000E-06 | 2.646E-07 | 1.800E-06 | 2.300E-06 |
| 4916 | 246 | C246S | A | 2.4e-06 | 3.2e-06 | 2.6e-06 | 2.733E-06 | 4.163E-07 | 2.400E-06 | 3.200E-06 |
| 4917 | 246 | C246T | A | 2.2e-06 | 4.6e-06 | 2.3e-06 | 3.033E-06 | 1.358E-06 | 2.200E-06 | 4.600E-06 |
| 4918 | 246 | C246W | A | 3.2e-06 | 3.8e-06 | 2.6e-06 | 3.200E-06 | 6.000E-07 | 2.600E-06 | 3.800E-06 |
| 4919 | 246 | C246Y | A | 1.9e-06 | 2e-06 | 2.3e-06 | 2.067E-06 | 2.082E-07 | 1.900E-06 | 2.300E-06 |
| 4920 | 246 | C246V | A | 1.2e-06 | 1.7e-06 | 3.3e-06 | 2.067E-06 | 1.097E-06 | 1.200E-06 | 3.300E-06 |
| 4921 | 247 | P247A | A | 1.4e-06 | 3.3e-06 | 1.3e-06 | 2.000E-06 | 1.127E-06 | 1.300E-06 | 3.300E-06 |
| 4922 | 247 | P247R | A | 1.9e-06 | 3.3e-06 | 2.5e-06 | 2.567E-06 | 7.024E-07 | 1.900E-06 | 3.300E-06 |
| 4923 | 247 | P247N | A | 3.9e-06 | 3.1e-06 | 1.1e-06 | 2.700E-06 | 1.442E-06 | 1.100E-06 | 3.900E-06 |
| 4924 | 247 | P247D | A | 2.5e-06 | 3.3e-06 | 2.8e-06 | 2.867E-06 | 4.041E-07 | 2.500E-06 | 3.300E-06 |
| 4925 | 247 | P247C | A | 1.6e-06 | 4.6e-06 | 2.6e-06 | 2.933E-06 | 1.528E-06 | 1.600E-06 | 4.600E-06 |
| 4926 | 247 | P247E | A | 2.4e-06 | 1.5e-06 | 2.4e-06 | 2.100E-06 | 5.196E-07 | 1.500E-06 | 2.400E-06 |
| 4927 | 247 | P247Q | A | 2.9e-06 | 2.1e-06 | 1.6e-06 | 2.200E-06 | 6.557E-07 | 1.600E-06 | 2.900E-06 |
| 4928 | 247 | P247G | A | 1.8e-06 | 1.3e-06 | 2.3e-06 | 1.800E-06 | 5.000E-07 | 1.300E-06 | 2.300E-06 |
| 4929 | 247 | P247H | A | 2.9e-06 | 1.9e-06 | 2.5e-06 | 2.433E-06 | 5.033E-07 | 1.900E-06 | 2.900E-06 |

| | | | | | | | | | | |
|---|---|---|---|---|---|---|---|---|---|---|
| 4930 | 247 | P247I | A | 3.3e-06 | 3.4e-06 | 1.7e-06 | 2.800E-06 | 9.539E-07 | 1.700E-06 | 3.400E-06 |
| 4931 | 247 | P247L | A | 3.8e-06 | 2.1e-06 | 5.5e-06 | 3.800E-06 | 1.700E-06 | 2.100E-06 | 5.500E-06 |
| 4932 | 247 | P247K | A | 2.6e-06 | 3e-06 | 2.1e-06 | 2.567E-06 | 4.509E-07 | 2.100E-06 | 3.000E-06 |
| 4933 | 247 | P247M | A | 2.6e-06 | 3.4e-06 | 1.9e-06 | 2.633E-06 | 7.506E-07 | 1.900E-06 | 3.400E-06 |
| 4934 | 247 | P247F | A | 2.1e-06 | 2e-06 | 1.9e-06 | 2.000E-06 | 1.000E-07 | 1.900E-06 | 2.100E-06 |
| 4935 | 247 | P247P | A | 2.7e-06 | 2.6e-06 | 9.2e-07 | 2.073E-06 | 1.000E-06 | 9.200E-07 | 2.700E-06 |
| 4936 | 247 | P247S | A | 2.2e-06 | 1.7e-06 | 3.4e-06 | 2.433E-06 | 8.737E-07 | 1.700E-06 | 3.400E-06 |
| 4937 | 247 | P247T | A | 2.5e-06 | 2.3e-06 | 1.4e-06 | 2.067E-06 | 5.859E-07 | 1.400E-06 | 2.500E-06 |
| 4938 | 247 | P247W | A | 2.1e-06 | 2.6e-06 | 3.2e-06 | 2.633E-06 | 5.508E-07 | 2.100E-06 | 3.200E-06 |
| 4939 | 247 | P247Y | A | 2.2e-06 | 2.5e-06 | 1.7e-06 | 2.133E-06 | 4.041E-07 | 1.700E-06 | 2.500E-06 |
| 4940 | 247 | P247V | A | 2.5e-06 | 4e-06 | 2.4e-06 | 2.967E-06 | 8.963E-07 | 2.400E-06 | 4.000E-06 |
| 4941 | 248 | A248A | A | 3.4e-06 | 4e-06 | 1.9e-06 | 3.100E-06 | 1.082E-06 | 1.900E-06 | 4.000E-06 |
| 4942 | 248 | A248R | A | 2.2e-06 | 2.3e-06 | 2.8e-06 | 2.433E-06 | 3.215E-07 | 2.200E-06 | 2.800E-06 |
| 4943 | 248 | A248N | A | 2.1e-06 | 1.9e-06 | 1.9e-06 | 1.967E-06 | 1.155E-07 | 1.900E-06 | 2.100E-06 |
| 4944 | 248 | A248D | A | 1.6e-06 | 4.5e-06 | 2.7e-06 | 2.933E-06 | 1.464E-06 | 1.600E-06 | 4.500E-06 |
| 4945 | 248 | A248C | A | 1.8e-06 | 2e-06 | 4e-06 | 2.600E-06 | 1.217E-06 | 1.800E-06 | 4.000E-06 |
| 4946 | 248 | A248E | A | 2e-06 | 2.8e-06 | 1.9e-06 | 2.233E-06 | 4.933E-07 | 1.900E-06 | 2.800E-06 |
| 4947 | 248 | A248Q | A | 3.2e-06 | 1e-06 | 1.6e-06 | 1.933E-06 | 1.137E-06 | 1.000E-06 | 3.200E-06 |
| 4948 | 248 | A248G | A | 3e-06 | 2.8e-06 | 2.7e-06 | 2.833E-06 | 1.528E-07 | 2.700E-06 | 3.000E-06 |
| 4949 | 248 | A248H | A | 2.2e-06 | 1.8e-06 | 3.6e-06 | 2.533E-06 | 9.452E-07 | 1.800E-06 | 3.600E-06 |
| 4950 | 248 | A248I | A | 5.4e-06 | 2.2e-06 | 2.2e-06 | 3.267E-06 | 1.848E-06 | 2.200E-06 | 5.400E-06 |
| 4951 | 248 | A248L | A | 1.5e-06 | 1.8e-06 | 2.4e-06 | 1.900E-06 | 4.583E-07 | 1.500E-06 | 2.400E-06 |
| 4952 | 248 | A248K | A | 2.8e-06 | 3.6e-06 | 2.8e-06 | 3.067E-06 | 4.619E-07 | 2.800E-06 | 3.600E-06 |
| 4953 | 248 | A248M | A | 3.8e-06 | 3.9e-06 | 2.4e-06 | 3.367E-06 | 8.386E-07 | 2.400E-06 | 3.900E-06 |
| 4954 | 248 | A248F | A | 3.8e-06 | 1.6e-06 | 2.4e-06 | 2.600E-06 | 1.114E-06 | 1.600E-06 | 3.800E-06 |
| 4955 | 248 | A248P | A | 2.7e-06 | 2.8e-06 | 1.5e-06 | 2.333E-06 | 7.234E-07 | 1.500E-06 | 2.800E-06 |
| 4956 | 248 | A248S | A | 1.5e-06 | 1.8e-06 | 1.3e-06 | 1.533E-06 | 2.517E-07 | 1.300E-06 | 1.800E-06 |
| 4957 | 248 | A248T | A | 3.6e-06 | 3.2e-06 | 3.7e-06 | 3.500E-06 | 2.646E-07 | 3.200E-06 | 3.700E-06 |
| 4958 | 248 | A248W | A | 2.5e-06 | 2.6e-06 | 3.6e-06 | 2.900E-06 | 6.083E-07 | 2.500E-06 | 3.600E-06 |
| 4959 | 248 | A248Y | A | 4.3e-06 | 1.8e-06 | 1.9e-06 | 2.667E-06 | 1.415E-06 | 1.800E-06 | 4.300E-06 |
| 4960 | 248 | A248V | A | 2.6e-06 | 2.6e-06 | 3.6e-06 | 2.933E-06 | 5.774E-07 | 2.600E-06 | 3.600E-06 |
| 4961 | 249 | L249A | A | 1.2e-06 | 4.5e-06 | 3.3e-06 | 3.000E-06 | 1.670E-06 | 1.200E-06 | 4.500E-06 |
| 4962 | 249 | L249R | A | 1.2e-06 | 1.6e-06 | 3.4e-06 | 2.067E-06 | 1.172E-06 | 1.200E-06 | 3.400E-06 |
| 4963 | 249 | L249N | A | 1.4e-06 | 2.8e-06 | 3.1e-06 | 2.433E-06 | 9.074E-07 | 1.400E-06 | 3.100E-06 |
| 4964 | 249 | L249D | A | 3.6e-06 | 1.8e-06 | 2.5e-06 | 2.633E-06 | 9.074E-07 | 1.800E-06 | 3.600E-06 |
| 4965 | 249 | L249C | A | 1.9e-06 | 1.6e-06 | 4.8e-06 | 2.767E-06 | 1.767E-06 | 1.600E-06 | 4.800E-06 |
| 4966 | 249 | L249E | A | 3.8e-06 | 4.3e-06 | 3.5e-06 | 3.867E-06 | 4.041E-07 | 3.500E-06 | 4.300E-06 |
| 4967 | 249 | L249Q | A | 1.9e-06 | 2.7e-06 | 1.8e-06 | 2.133E-06 | 4.933E-07 | 1.800E-06 | 2.700E-06 |
| 4968 | 249 | L249G | A | 3e-06 | 1.7e-06 | 4.4e-06 | 3.033E-06 | 1.350E-06 | 1.700E-06 | 4.400E-06 |
| 4969 | 249 | L249H | A | 2e-06 | 3.2e-06 | 2e-06 | 2.400E-06 | 6.928E-07 | 2.000E-06 | 3.200E-06 |
| 4970 | 249 | L249I | A | 2.1e-06 | 3.9e-06 | 3.1e-06 | 3.033E-06 | 9.018E-07 | 2.100E-06 | 3.900E-06 |
| 4971 | 249 | L249L | A | 3.7e-06 | 2.6e-06 | 1.3e-06 | 2.533E-06 | 1.201E-06 | 1.300E-06 | 3.700E-06 |
| 4972 | 249 | L249K | A | 2.1e-06 | 1.9e-06 | 3.4e-06 | 2.467E-06 | 8.145E-07 | 1.900E-06 | 3.400E-06 |
| 4973 | 249 | L249M | A | 3.1e-06 | 3.1e-06 | 2.3e-06 | 2.833E-06 | 4.619E-07 | 2.300E-06 | 3.100E-06 |
| 4974 | 249 | L249F | A | 2.5e-06 | 1.7e-06 | 3.3e-06 | 2.500E-06 | 8.000E-07 | 1.700E-06 | 3.300E-06 |
| 4975 | 249 | L249P | A | 2.2e-06 | 2e-06 | 1.9e-06 | 2.033E-06 | 1.528E-07 | 1.900E-06 | 2.200E-06 |
| 4976 | 249 | L249S | A | 2e-06 | 1.4e-06 | 2.8e-06 | 2.067E-06 | 7.024E-07 | 1.400E-06 | 2.800E-06 |
| 4977 | 249 | L249T | A | 2.5e-06 | 1.3e-06 | 2.1e-06 | 1.967E-06 | 6.110E-07 | 1.300E-06 | 2.500E-06 |
| 4978 | 249 | L249W | A | 1.2e-06 | 1.9e-06 | 2.1e-06 | 1.733E-06 | 4.726E-07 | 1.200E-06 | 2.100E-06 |
| 4979 | 249 | L249Y | A | 1.7e-06 | 2.2e-06 | 1.8e-06 | 1.900E-06 | 2.646E-07 | 1.700E-06 | 2.200E-06 |
| 4980 | 249 | L249V | A | 2.6e-06 | 3.4e-06 | 1.4e-06 | 2.467E-06 | 1.007E-06 | 1.400E-06 | 3.400E-06 |
| 4981 | 250 | V250A | A | 2.1e-06 | 1.1e-06 | 2.4e-06 | 1.867E-06 | 6.807E-07 | 1.100E-06 | 2.400E-06 |
| 4982 | 250 | V250R | A | 3.3e-06 | 1.4e-06 | 2.5e-06 | 2.400E-06 | 9.539E-07 | 1.400E-06 | 3.300E-06 |
| 4983 | 250 | V250N | A | 3.1e-06 | 1.9e-06 | 2.2e-06 | 2.400E-06 | 6.245E-07 | 1.900E-06 | 3.100E-06 |
| 4984 | 250 | V250D | A | 2.9e-06 | 1.9e-06 | 4.2e-06 | 3.000E-06 | 1.153E-06 | 1.900E-06 | 4.200E-06 |
| 4985 | 250 | V250C | A | 1.5e-06 | 3.3e-06 | 2e-06 | 2.267E-06 | 9.292E-07 | 1.500E-06 | 3.300E-06 |
| 4986 | 250 | V250E | A | 3.4e-06 | 4.6e-06 | 1.4e-06 | 3.133E-06 | 1.617E-06 | 1.400E-06 | 4.600E-06 |
| 4987 | 250 | V250Q | A | 2.3e-06 | 3.5e-06 | 1.2e-06 | 2.333E-06 | 1.150E-06 | 1.200E-06 | 3.500E-06 |
| 4988 | 250 | V250G | A | 1.7e-06 | 2.4e-06 | 3.1e-06 | 2.400E-06 | 7.000E-07 | 1.700E-06 | 3.100E-06 |
| 4989 | 250 | V250H | A | 2.4e-06 | 1.4e-06 | 2.1e-06 | 1.967E-06 | 5.132E-07 | 1.400E-06 | 2.400E-06 |
| 4990 | 250 | V250I | A | 2.3e-06 | 1.9e-06 | 4.8e-06 | 3.000E-06 | 1.572E-06 | 1.900E-06 | 4.800E-06 |
| 4991 | 250 | V250L | A | 4.2e-06 | 2.6e-06 | 2.3e-06 | 3.033E-06 | 1.021E-06 | 2.300E-06 | 4.200E-06 |
| 4992 | 250 | V250K | A | 4.2e-06 | 2.7e-06 | 2.3e-06 | 3.067E-06 | 1.002E-06 | 2.300E-06 | 4.200E-06 |
| 4993 | 250 | V250M | A | 1.7e-06 | 4.3e-06 | 2.6e-06 | 2.867E-06 | 1.320E-06 | 1.700E-06 | 4.300E-06 |
| 4994 | 250 | V250F | A | 2.5e-06 | 2.8e-06 | 1.7e-06 | 2.333E-06 | 5.686E-07 | 1.700E-06 | 2.800E-06 |
| 4995 | 250 | V250P | A | 3.1e-06 | 2.6e-06 | 2.4e-06 | 2.700E-06 | 3.606E-07 | 2.400E-06 | 3.100E-06 |
| 4996 | 250 | V250S | A | 1.4e-06 | 1.8e-06 | 2.3e-06 | 1.833E-06 | 4.509E-07 | 1.400E-06 | 2.300E-06 |
| 4997 | 250 | V250T | A | 2.5e-06 | 2.7e-06 | 2.5e-06 | 2.567E-06 | 1.155E-07 | 2.500E-06 | 2.700E-06 |
| 4998 | 250 | V250W | A | 1.2e-06 | 1.9e-06 | 3.2e-06 | 2.100E-06 | 1.015E-06 | 1.200E-06 | 3.200E-06 |
| 4999 | 250 | V250Y | A | 1.2e-06 | 3.6e-06 | 2.4e-06 | 2.400E-06 | 1.200E-06 | 1.200E-06 | 3.600E-06 |
| 5000 | 250 | V250V | A | 2.7e-06 | 4.8e-06 | 3.3e-06 | 3.600E-06 | 1.082E-06 | 2.700E-06 | 4.800E-06 |
| 5001 | 251 | T251A | A | 2.9e-06 | 2.4e-06 | 1.3e-06 | 2.200E-06 | 8.185E-07 | 1.300E-06 | 2.900E-06 |
| 5002 | 251 | T251R | A | 3.2e-06 | 3.2e-06 | 2.1e-06 | 2.833E-06 | 6.351E-07 | 2.100E-06 | 3.200E-06 |
| 5003 | 251 | T251N | A | 2.3e-06 | 2.1e-06 | 3.2e-06 | 2.533E-06 | 5.859E-07 | 2.100E-06 | 3.200E-06 |
| 5004 | 251 | T251D | A | 3.1e-06 | 2.2e-06 | 4.8e-06 | 3.367E-06 | 1.320E-06 | 2.200E-06 | 4.800E-06 |
| 5005 | 251 | T251C | A | 3.8e-06 | 2.5e-06 | 2.8e-06 | 3.033E-06 | 6.807E-07 | 2.500E-06 | 3.800E-06 |
| 5006 | 251 | T251E | A | 1.2e-06 | 2.9e-06 | 2.1e-06 | 2.067E-06 | 8.505E-07 | 1.200E-06 | 2.900E-06 |
| 5007 | 251 | T251Q | A | 1.7e-06 | 3.6e-06 | 1.6e-06 | 2.300E-06 | 1.127E-06 | 1.600E-06 | 3.600E-06 |
| 5008 | 251 | T251G | A | 2.2e-06 | 3.8e-06 | 1e-06 | 2.333E-06 | 1.405E-06 | 1.000E-06 | 3.800E-06 |
| 5009 | 251 | T251H | A | 2.3e-06 | 2.6e-06 | 2.9e-06 | 2.600E-06 | 3.000E-07 | 2.300E-06 | 2.900E-06 |
| 5010 | 251 | T251I | A | 2.2e-06 | 1.1e-06 | 4.4e-06 | 2.567E-06 | 1.680E-06 | 1.100E-06 | 4.400E-06 |
| 5011 | 251 | T251L | A | 2.5e-06 | 2.3e-06 | 3.1e-06 | 2.633E-06 | 4.163E-07 | 2.300E-06 | 3.100E-06 |
| 5012 | 251 | T251K | A | 2.5e-06 | 2.4e-06 | 4.1e-06 | 3.000E-06 | 9.539E-07 | 2.400E-06 | 4.100E-06 |
| 5013 | 251 | T251M | A | 3.1e-06 | 1.4e-06 | 3.1e-06 | 2.533E-06 | 9.815E-07 | 1.400E-06 | 3.100E-06 |
| 5014 | 251 | T251F | A | 2.5e-06 | 3.8e-06 | 3.5e-06 | 3.267E-06 | 6.807E-07 | 2.500E-06 | 3.800E-06 |

| | | | | | | | | | | |
|---|---|---|---|---|---|---|---|---|---|---|
| 5015 | 251 | T251P | A | 3e-06 | 3.1e-06 | 2.4e-06 | 2.833E-06 | 3.786E-07 | 2.400E-06 | 3.100E-06 |
| 5016 | 251 | T251S | A | 2.1e-06 | 2.6e-06 | 2.1e-06 | 2.267E-06 | 2.887E-07 | 2.100E-06 | 2.600E-06 |
| 5017 | 251 | T251T | A | 2.3e-06 | 3.7e-06 | 2.3e-06 | 2.767E-06 | 8.083E-07 | 2.300E-06 | 3.700E-06 |
| 5018 | 251 | T251W | A | 4.7e-06 | 3.6e-06 | 2.7e-06 | 3.667E-06 | 1.002E-06 | 2.700E-06 | 4.700E-06 |
| 5019 | 251 | T251Y | A | 2.2e-06 | 3.3e-06 | 2.3e-06 | 2.600E-06 | 6.083E-07 | 2.200E-06 | 3.300E-06 |
| 5020 | 251 | T251V | A | 2.3e-06 | 2.3e-06 | 3.2e-06 | 2.600E-06 | 5.196E-07 | 2.300E-06 | 3.200E-06 |
| 5021 | 252 | Y252A | A | 1.8e-06 | 1.4e-06 | 2.1e-06 | 1.767E-06 | 3.512E-07 | 1.400E-06 | 2.100E-06 |
| 5022 | 252 | Y252R | A | 1.7e-06 | 1.8e-06 | 2.2e-06 | 1.900E-06 | 2.646E-07 | 1.700E-06 | 2.200E-06 |
| 5023 | 252 | Y252N | A | 1.8e-06 | 3e-06 | 2.1e-06 | 2.300E-06 | 6.245E-07 | 1.800E-06 | 3.000E-06 |
| 5024 | 252 | Y252D | A | 1.9e-06 | 2.9e-06 | 1.1e-06 | 1.967E-06 | 9.018E-07 | 1.100E-06 | 2.900E-06 |
| 5025 | 252 | Y252C | A | 2.2e-06 | 3.1e-06 | 1.3e-06 | 2.200E-06 | 9.000E-07 | 1.300E-06 | 3.100E-06 |
| 5026 | 252 | Y252E | A | 2.6e-06 | 2.5e-06 | 3.1e-06 | 2.733E-06 | 3.215E-07 | 2.500E-06 | 3.100E-06 |
| 5027 | 252 | Y252Q | A | 2e-06 | 1.7e-06 | 2.9e-06 | 2.200E-06 | 6.245E-07 | 1.700E-06 | 2.900E-06 |
| 5028 | 252 | Y252G | A | 3e-06 | 1.9e-06 | 2.3e-06 | 2.400E-06 | 5.568E-07 | 1.900E-06 | 3.000E-06 |
| 5029 | 252 | Y252H | A | 2.3e-06 | 2e-06 | 2e-06 | 2.100E-06 | 1.732E-07 | 2.000E-06 | 2.300E-06 |
| 5030 | 252 | Y252I | A | 2.9e-06 | 2.4e-06 | 2.4e-06 | 2.567E-06 | 2.887E-07 | 2.400E-06 | 2.900E-06 |
| 5031 | 252 | Y252L | A | 2.8e-06 | 2.7e-06 | 2.4e-06 | 2.633E-06 | 2.082E-07 | 2.400E-06 | 2.800E-06 |
| 5032 | 252 | Y252K | A | 2e-06 | 2e-06 | 2.5e-06 | 2.167E-06 | 2.887E-07 | 2.000E-06 | 2.500E-06 |
| 5033 | 252 | Y252M | A | 1.6e-06 | 1.8e-06 | 1.3e-06 | 1.567E-06 | 2.517E-07 | 1.300E-06 | 1.800E-06 |
| 5034 | 252 | Y252F | A | 2.7e-06 | 3.1e-06 | 2.8e-06 | 2.867E-06 | 2.082E-07 | 2.700E-06 | 3.100E-06 |
| 5035 | 252 | Y252P | A | 3.4e-06 | 1.5e-06 | 3.2e-06 | 2.700E-06 | 1.044E-06 | 1.500E-06 | 3.400E-06 |
| 5036 | 252 | Y252S | A | 1.7e-06 | 4.2e-06 | 3.1e-06 | 3.000E-06 | 1.253E-06 | 1.700E-06 | 4.200E-06 |
| 5037 | 252 | Y252T | A | 2.1e-06 | 3.2e-06 | 1.6e-06 | 2.300E-06 | 8.185E-07 | 1.600E-06 | 3.200E-06 |
| 5038 | 252 | Y252W | A | 2.5e-06 | 4.9e-06 | 2.4e-06 | 3.267E-06 | 1.415E-06 | 2.400E-06 | 4.900E-06 |
| 5039 | 252 | Y252Y | A | 1.5e-06 | 3.7e-06 | 1.5e-06 | 2.233E-06 | 1.270E-06 | 1.500E-06 | 3.700E-06 |
| 5040 | 252 | Y252V | A | 1.9e-06 | 2.5e-06 | 2.1e-06 | 2.167E-06 | 3.055E-07 | 1.900E-06 | 2.500E-06 |
| 5041 | 253 | N253A | A | 3.4e-06 | 2.3e-06 | 3.6e-06 | 3.100E-06 | 7.000E-07 | 2.300E-06 | 3.600E-06 |
| 5042 | 253 | N253R | A | 1.9e-06 | 3.1e-06 | 2e-06 | 2.333E-06 | 6.658E-07 | 1.900E-06 | 3.100E-06 |
| 5043 | 253 | N253N | A | 2.5e-06 | 2.8e-06 | 3.2e-06 | 2.833E-06 | 3.512E-07 | 2.500E-06 | 3.200E-06 |
| 5044 | 253 | N253D | A | 2.9e-06 | 2.6e-06 | 2.6e-06 | 2.700E-06 | 1.732E-07 | 2.600E-06 | 2.900E-06 |
| 5045 | 253 | N253C | A | 1.2e-06 | 2.4e-06 | 3.1e-06 | 2.233E-06 | 9.609E-07 | 1.200E-06 | 3.100E-06 |
| 5046 | 253 | N253E | A | 2.6e-06 | 2.1e-06 | 3.2e-06 | 2.633E-06 | 5.508E-07 | 2.100E-06 | 3.200E-06 |
| 5047 | 253 | N253Q | A | 4.1e-06 | 2.6e-06 | 4.4e-06 | 3.700E-06 | 9.644E-07 | 2.600E-06 | 4.400E-06 |
| 5048 | 253 | N253G | A | 3.1e-06 | 2.6e-06 | 1.9e-06 | 2.533E-06 | 6.028E-07 | 1.900E-06 | 3.100E-06 |
| 5049 | 253 | N253H | A | 2e-06 | 6e-06 | 4e-06 | 4.000E-06 | 2.000E-06 | 2.000E-06 | 6.000E-06 |
| 5050 | 253 | N253I | A | 1.7e-06 | 2.9e-06 | 3.3e-06 | 2.633E-06 | 8.327E-07 | 1.700E-06 | 3.300E-06 |
| 5051 | 253 | N253L | A | 3.2e-06 | 2.8e-06 | 3.2e-06 | 3.067E-06 | 2.309E-07 | 2.800E-06 | 3.200E-06 |
| 5052 | 253 | N253K | A | 2.3e-06 | 4.1e-06 | 9.4e-07 | 2.447E-06 | 1.585E-06 | 9.400E-07 | 4.100E-06 |
| 5053 | 253 | N253M | A | 2.6e-06 | 2.7e-06 | 3.4e-06 | 2.900E-06 | 4.359E-07 | 2.600E-06 | 3.400E-06 |
| 5054 | 253 | N253F | A | 2.7e-06 | 3.8e-06 | 2.2e-06 | 2.900E-06 | 8.185E-07 | 2.200E-06 | 3.800E-06 |
| 5055 | 253 | N253P | A | 2.8e-06 | 2.7e-06 | 2.4e-06 | 2.633E-06 | 2.082E-07 | 2.400E-06 | 2.800E-06 |
| 5056 | 253 | N253S | A | 2.6e-06 | 3.3e-06 | 4.5e-06 | 3.467E-06 | 9.609E-07 | 2.600E-06 | 4.500E-06 |
| 5057 | 253 | N253T | A | 3.6e-06 | 1.6e-06 | 3.3e-06 | 2.833E-06 | 1.079E-06 | 1.600E-06 | 3.600E-06 |
| 5058 | 253 | N253W | A | 1.3e-06 | 1.4e-06 | 2.2e-06 | 1.633E-06 | 4.933E-07 | 1.300E-06 | 2.200E-06 |
| 5059 | 253 | N253Y | A | 2.7e-06 | 1.7e-06 | 1.4e-06 | 1.933E-06 | 6.807E-07 | 1.400E-06 | 2.700E-06 |
| 5060 | 253 | N253V | A | 2.8e-06 | 3e-06 | 2.4e-06 | 2.733E-06 | 3.055E-07 | 2.400E-06 | 3.000E-06 |
| 5061 | 254 | T254A | A | 2.7e-06 | 5.6e-06 | 3.6e-06 | 3.967E-06 | 1.484E-06 | 2.700E-06 | 5.600E-06 |
| 5062 | 254 | T254R | A | 2.7e-06 | 1.2e-06 | 4.4e-06 | 2.767E-06 | 1.601E-06 | 1.200E-06 | 4.400E-06 |
| 5063 | 254 | T254N | A | 3.3e-06 | 2.6e-06 | 2.6e-06 | 2.833E-06 | 4.041E-07 | 2.600E-06 | 3.300E-06 |
| 5064 | 254 | T254D | A | 2.1e-06 | 2.7e-06 | 3.2e-06 | 2.667E-06 | 5.508E-07 | 2.100E-06 | 3.200E-06 |
| 5065 | 254 | T254C | A | 2.6e-06 | 3.2e-06 | 2e-06 | 2.600E-06 | 6.000E-07 | 2.000E-06 | 3.200E-06 |
| 5066 | 254 | T254E | A | 3e-06 | 3.6e-06 | 2.5e-06 | 3.033E-06 | 5.508E-07 | 2.500E-06 | 3.600E-06 |
| 5067 | 254 | T254Q | A | 2.4e-06 | 2.1e-06 | 1.9e-06 | 2.133E-06 | 2.517E-07 | 1.900E-06 | 2.400E-06 |
| 5068 | 254 | T254G | A | 2.4e-06 | 1.9e-06 | 2.4e-06 | 2.233E-06 | 2.887E-07 | 1.900E-06 | 2.400E-06 |
| 5069 | 254 | T254H | A | 2.8e-06 | 1.7e-06 | 1.7e-06 | 2.067E-06 | 6.351E-07 | 1.700E-06 | 2.800E-06 |
| 5070 | 254 | T254I | A | 2.2e-06 | 3.9e-06 | 2.9e-06 | 3.000E-06 | 8.544E-07 | 2.200E-06 | 3.900E-06 |
| 5071 | 254 | T254L | A | 1.9e-06 | 2.3e-06 | 4.1e-06 | 2.767E-06 | 1.172E-06 | 1.900E-06 | 4.100E-06 |
| 5072 | 254 | T254K | A | 1.8e-06 | 4e-06 | 2.9e-06 | 2.900E-06 | 1.100E-06 | 1.800E-06 | 4.000E-06 |
| 5073 | 254 | T254M | A | 2.2e-06 | 2.6e-06 | 2.1e-06 | 2.300E-06 | 2.646E-07 | 2.100E-06 | 2.600E-06 |
| 5074 | 254 | T254F | A | 3.8e-06 | 2.4e-06 | 3.2e-06 | 3.133E-06 | 7.024E-07 | 2.400E-06 | 3.800E-06 |
| 5075 | 254 | T254P | A | 2.3e-06 | 5.4e-06 | 3.1e-06 | 3.600E-06 | 1.609E-06 | 2.300E-06 | 5.400E-06 |
| 5076 | 254 | T254S | A | 2.3e-06 | 2.5e-06 | 3.4e-06 | 2.733E-06 | 5.859E-07 | 2.300E-06 | 3.400E-06 |
| 5077 | 254 | T254T | A | 1.8e-06 | 4.2e-06 | 1.3e-06 | 2.433E-06 | 1.550E-06 | 1.300E-06 | 4.200E-06 |
| 5078 | 254 | T254W | A | 2.3e-06 | 4.9e-06 | 2.6e-06 | 3.267E-06 | 1.422E-06 | 2.300E-06 | 4.900E-06 |
| 5079 | 254 | T254Y | A | 2.6e-06 | 1.7e-06 | 2.9e-06 | 2.400E-06 | 6.245E-07 | 1.700E-06 | 2.900E-06 |
| 5080 | 254 | T254V | A | 2.5e-06 | 2.2e-06 | 1.5e-06 | 2.067E-06 | 5.132E-07 | 1.500E-06 | 2.500E-06 |
| 5081 | 255 | D255A | A | 3e-06 | 3.4e-06 | 2.1e-06 | 2.833E-06 | 6.658E-07 | 2.100E-06 | 3.400E-06 |
| 5082 | 255 | D255R | A | 2.5e-06 | 2.7e-06 | 2.6e-06 | 2.600E-06 | 1.000E-07 | 2.500E-06 | 2.700E-06 |
| 5083 | 255 | D255N | A | 1.7e-06 | 2.9e-06 | 2.2e-06 | 2.267E-06 | 6.028E-07 | 1.700E-06 | 2.900E-06 |
| 5084 | 255 | D255D | A | 2.6e-06 | 2.5e-06 | 4.3e-06 | 3.133E-06 | 1.012E-06 | 2.500E-06 | 4.300E-06 |
| 5085 | 255 | D255C | A | 3e-06 | 3.1e-06 | 2.7e-06 | 2.933E-06 | 2.082E-07 | 2.700E-06 | 3.100E-06 |
| 5086 | 255 | D255E | A | 2.2e-06 | 1.8e-06 | 3.1e-06 | 2.367E-06 | 6.658E-07 | 1.800E-06 | 3.100E-06 |
| 5087 | 255 | D255Q | A | 2.2e-06 | 2.6e-06 | 2.7e-06 | 2.500E-06 | 2.646E-07 | 2.200E-06 | 2.700E-06 |
| 5088 | 255 | D255G | A | 2.5e-06 | 2.4e-06 | 3e-06 | 2.633E-06 | 3.215E-07 | 2.400E-06 | 3.000E-06 |
| 5089 | 255 | D255H | A | 2.2e-06 | 4.9e-06 | 2.2e-06 | 3.100E-06 | 1.559E-06 | 2.200E-06 | 4.900E-06 |
| 5090 | 255 | D255I | A | 2.7e-06 | 3.8e-06 | 3.3e-06 | 3.267E-06 | 5.508E-07 | 2.700E-06 | 3.800E-06 |
| 5091 | 255 | D255L | A | 1.9e-06 | 2.3e-06 | 1.5e-06 | 1.900E-06 | 4.000E-07 | 1.500E-06 | 2.300E-06 |
| 5092 | 255 | D255K | A | 2.4e-06 | 2e-06 | 4e-06 | 2.800E-06 | 1.058E-06 | 2.000E-06 | 4.000E-06 |
| 5093 | 255 | D255M | A | 2e-06 | 2.1e-06 | 2.1e-06 | 2.067E-06 | 5.774E-08 | 2.000E-06 | 2.100E-06 |
| 5094 | 255 | D255F | A | 2.2e-06 | 2.2e-06 | 1.5e-06 | 1.967E-06 | 4.041E-07 | 1.500E-06 | 2.200E-06 |
| 5095 | 255 | D255P | A | 1.7e-06 | 3e-06 | 1.8e-06 | 2.167E-06 | 7.234E-07 | 1.700E-06 | 3.000E-06 |
| 5096 | 255 | D255S | A | 2e-06 | 1.7e-06 | 2.3e-06 | 2.000E-06 | 3.000E-07 | 1.700E-06 | 2.300E-06 |
| 5097 | 255 | D255T | A | 1.6e-06 | 2.8e-06 | 2.9e-06 | 2.433E-06 | 7.234E-07 | 1.600E-06 | 2.900E-06 |
| 5098 | 255 | D255W | A | 2.5e-06 | 2.1e-06 | 2.7e-06 | 2.433E-06 | 3.055E-07 | 2.100E-06 | 2.700E-06 |
| 5099 | 255 | D255Y | A | 5.3e-06 | 2e-06 | 1.8e-06 | 3.033E-06 | 1.966E-06 | 1.800E-06 | 5.300E-06 |

| | | | | | | | | | | |
|---|---|---|---|---|---|---|---|---|---|---|
| 5100 | 255 | D255V | A | 2.7e-06 | 2.6e-06 | 2.2e-06 | 2.500E-06 | 2.646E-07 | 2.200E-06 | 2.700E-06 |
| 5101 | 256 | T256A | A | 1.9e-06 | 2.8e-06 | 1.7e-06 | 2.133E-06 | 5.859E-07 | 1.700E-06 | 2.800E-06 |
| 5102 | 256 | T256R | A | 1.9e-06 | 2.2e-06 | 2.5e-06 | 2.200E-06 | 3.000E-07 | 1.900E-06 | 2.500E-06 |
| 5103 | 256 | T256N | A | 2.7e-06 | 2.3e-06 | 1.7e-06 | 2.233E-06 | 5.033E-07 | 1.700E-06 | 2.700E-06 |
| 5104 | 256 | T256D | A | 1.5e-06 | 2.5e-06 | 2.7e-06 | 2.233E-06 | 6.429E-07 | 1.500E-06 | 2.700E-06 |
| 5105 | 256 | T256C | A | 1.8e-06 | 1.4e-06 | 2.3e-06 | 1.833E-06 | 4.509E-07 | 1.400E-06 | 2.300E-06 |
| 5106 | 256 | T256E | A | 1.8e-06 | 2.6e-06 | 3.2e-06 | 2.533E-06 | 7.024E-07 | 1.800E-06 | 3.200E-06 |
| 5107 | 256 | T256Q | A | 3.3e-06 | 1.5e-06 | 2.3e-06 | 2.367E-06 | 9.018E-07 | 1.500E-06 | 3.300E-06 |
| 5108 | 256 | T256G | A | 2.1e-06 | 1.5e-06 | 2e-06 | 1.867E-06 | 3.215E-07 | 1.500E-06 | 2.100E-06 |
| 5109 | 256 | T256H | A | 2.5e-06 | 3.7e-06 | 2.4e-06 | 2.867E-06 | 7.234E-07 | 2.400E-06 | 3.700E-06 |
| 5110 | 256 | T256I | A | 2e-06 | 3.5e-06 | 1.2e-06 | 2.233E-06 | 1.168E-06 | 1.200E-06 | 3.500E-06 |
| 5111 | 256 | T256L | A | 2.5e-06 | 1.3e-06 | 3.2e-06 | 2.333E-06 | 9.609E-07 | 1.300E-06 | 3.200E-06 |
| 5112 | 256 | T256K | A | 3e-06 | 1.6e-06 | 2.1e-06 | 2.233E-06 | 7.095E-07 | 1.600E-06 | 3.000E-06 |
| 5113 | 256 | T256M | A | 2.4e-06 | 1.6e-06 | 2.3e-06 | 2.100E-06 | 4.359E-07 | 1.600E-06 | 2.400E-06 |
| 5114 | 256 | T256F | A | 4.4e-06 | 2.4e-06 | 2e-06 | 2.933E-06 | 1.286E-06 | 2.000E-06 | 4.400E-06 |
| 5115 | 256 | T256P | A | 1.8e-06 | 2.5e-06 | 2e-06 | 2.100E-06 | 3.606E-07 | 1.800E-06 | 2.500E-06 |
| 5116 | 256 | T256S | A | 2.1e-06 | 1.8e-06 | 3.4e-06 | 2.433E-06 | 8.505E-07 | 1.800E-06 | 3.400E-06 |
| 5117 | 256 | T256T | A | 1.8e-06 | 1.9e-06 | 1.7e-06 | 1.800E-06 | 1.000E-07 | 1.700E-06 | 1.900E-06 |
| 5118 | 256 | T256W | A | 2.1e-06 | 2.1e-06 | 3.1e-06 | 2.433E-06 | 5.774E-07 | 2.100E-06 | 3.100E-06 |
| 5119 | 256 | T256Y | A | 1.3e-06 | 2.9e-06 | 3.2e-06 | 2.467E-06 | 1.021E-06 | 1.300E-06 | 3.200E-06 |
| 5120 | 256 | T256V | A | 1.9e-06 | 1.7e-06 | 4.5e-06 | 2.700E-06 | 1.562E-06 | 1.700E-06 | 4.500E-06 |
| 5121 | 257 | F257A | A | 2.1e-06 | 1.9e-06 | 2.5e-06 | 2.167E-06 | 3.055E-07 | 1.900E-06 | 2.500E-06 |
| 5122 | 257 | F257R | A | 1.9e-06 | 4.3e-06 | 2.3e-06 | 2.833E-06 | 1.286E-06 | 1.900E-06 | 4.300E-06 |
| 5123 | 257 | F257N | A | 2e-06 | 1.9e-06 | 2.7e-06 | 2.200E-06 | 4.359E-07 | 1.900E-06 | 2.700E-06 |
| 5124 | 257 | F257D | A | 3.6e-06 | 2.5e-06 | 3.2e-06 | 3.100E-06 | 5.568E-07 | 2.500E-06 | 3.600E-06 |
| 5125 | 257 | F257C | A | 2.4e-06 | 3.4e-06 | 1.8e-06 | 2.533E-06 | 8.083E-07 | 1.800E-06 | 3.400E-06 |
| 5126 | 257 | F257E | A | 2e-06 | 3.2e-06 | 1.9e-06 | 2.367E-06 | 7.234E-07 | 1.900E-06 | 3.200E-06 |
| 5127 | 257 | F257Q | A | 6.4e-07 | 3.4e-06 | 1.5e-06 | 1.847E-06 | 1.412E-06 | 6.400E-07 | 3.400E-06 |
| 5128 | 257 | F257G | A | 2.3e-06 | 1.8e-06 | 1.7e-06 | 1.933E-06 | 3.215E-07 | 1.700E-06 | 2.300E-06 |
| 5129 | 257 | F257H | A | 2.8e-06 | 3.3e-06 | 2.3e-06 | 2.800E-06 | 5.000E-07 | 2.300E-06 | 3.300E-06 |
| 5130 | 257 | F257I | A | 3.7e-06 | 2.5e-06 | 1.7e-06 | 2.633E-06 | 1.007E-06 | 1.700E-06 | 3.700E-06 |
| 5131 | 257 | F257L | A | 2.8e-06 | 2.6e-06 | 3.8e-06 | 3.067E-06 | 6.429E-07 | 2.600E-06 | 3.800E-06 |
| 5132 | 257 | F257K | A | 3.8e-06 | 1.4e-06 | 2.3e-06 | 2.500E-06 | 1.212E-06 | 1.400E-06 | 3.800E-06 |
| 5133 | 257 | F257M | A | 3.2e-06 | 1.8e-06 | 2e-06 | 2.333E-06 | 7.572E-07 | 1.800E-06 | 3.200E-06 |
| 5134 | 257 | F257F | A | 1.9e-06 | 2.7e-06 | 3.3e-06 | 2.633E-06 | 7.024E-07 | 1.900E-06 | 3.300E-06 |
| 5135 | 257 | F257P | A | 1.4e-06 | 2.4e-06 | 2.2e-06 | 2.000E-06 | 5.292E-07 | 1.400E-06 | 2.400E-06 |
| 5136 | 257 | F257S | A | 4.4e-06 | 2.7e-06 | 9.7e-07 | 2.690E-06 | 1.715E-06 | 9.700E-07 | 4.400E-06 |
| 5137 | 257 | F257T | A | 1.6e-06 | 1.8e-06 | 2e-06 | 1.800E-06 | 2.000E-07 | 1.600E-06 | 2.000E-06 |
| 5138 | 257 | F257W | A | 2.2e-06 | 1.6e-06 | 1.5e-06 | 1.767E-06 | 3.786E-07 | 1.500E-06 | 2.200E-06 |
| 5139 | 257 | F257Y | A | 2.4e-06 | 1.6e-06 | 3.1e-06 | 2.367E-06 | 7.506E-07 | 1.600E-06 | 3.100E-06 |
| 5140 | 257 | F257V | A | 1.6e-06 | 3.9e-06 | 2.2e-06 | 2.567E-06 | 1.193E-06 | 1.600E-06 | 3.900E-06 |
| 5141 | 258 | E258A | A | 7.8e-06 | 2.8e-06 | 2e-06 | 4.200E-06 | 3.143E-06 | 2.000E-06 | 7.800E-06 |
| 5142 | 258 | E258R | A | 1.6e-06 | 2.2e-06 | 3.1e-06 | 2.300E-06 | 7.550E-07 | 1.600E-06 | 3.100E-06 |
| 5143 | 258 | E258N | A | 2.4e-06 | 1.4e-06 | 2e-06 | 1.933E-06 | 5.033E-07 | 1.400E-06 | 2.400E-06 |
| 5144 | 258 | E258D | A | 2.3e-06 | 2.7e-06 | 3.1e-06 | 2.700E-06 | 4.000E-07 | 2.300E-06 | 3.100E-06 |
| 5145 | 258 | E258C | A | 2.1e-06 | 2.5e-06 | 2.5e-06 | 2.367E-06 | 2.309E-07 | 2.100E-06 | 2.500E-06 |
| 5146 | 258 | E258E | A | 1.6e-06 | 1.6e-06 | 1.4e-06 | 1.533E-06 | 1.155E-07 | 1.400E-06 | 1.600E-06 |
| 5147 | 258 | E258Q | A | 3.3e-06 | 6.7e-07 | 3.8e-06 | 2.590E-06 | 1.681E-06 | 6.700E-07 | 3.800E-06 |
| 5148 | 258 | E258G | A | 2.6e-06 | 3e-06 | 3.5e-06 | 3.033E-06 | 4.509E-07 | 2.600E-06 | 3.500E-06 |
| 5149 | 258 | E258H | A | 2.6e-06 | 3.8e-06 | 1.1e-06 | 2.500E-06 | 1.353E-06 | 1.100E-06 | 3.800E-06 |
| 5150 | 258 | E258I | A | 1.5e-06 | 3.2e-06 | 2.7e-06 | 2.467E-06 | 8.737E-07 | 1.500E-06 | 3.200E-06 |
| 5151 | 258 | E258L | A | 2.4e-06 | 1.8e-06 | 2.9e-06 | 2.367E-06 | 5.508E-07 | 1.800E-06 | 2.900E-06 |
| 5152 | 258 | E258K | A | 4.9e-06 | 2.3e-06 | 2.1e-06 | 3.100E-06 | 1.562E-06 | 2.100E-06 | 4.900E-06 |
| 5153 | 258 | E258M | A | 2.8e-06 | 2.4e-06 | 2.6e-06 | 2.600E-06 | 2.000E-07 | 2.400E-06 | 2.800E-06 |
| 5154 | 258 | E258F | A | 1.8e-06 | 3.3e-06 | 3.4e-06 | 2.833E-06 | 8.963E-07 | 1.800E-06 | 3.400E-06 |
| 5155 | 258 | E258P | A | 2.1e-06 | 2.6e-06 | 1.8e-06 | 2.167E-06 | 4.041E-07 | 1.800E-06 | 2.600E-06 |
| 5156 | 258 | E258S | A | 1.8e-06 | 1.6e-06 | 2.5e-06 | 1.967E-06 | 4.726E-07 | 1.600E-06 | 2.500E-06 |
| 5157 | 258 | E258T | A | 3.9e-06 | 3.1e-06 | 1.9e-06 | 2.967E-06 | 1.007E-06 | 1.900E-06 | 3.900E-06 |
| 5158 | 258 | E258W | A | 2.8e-06 | 1.3e-06 | 1.6e-06 | 1.900E-06 | 7.937E-07 | 1.300E-06 | 2.800E-06 |
| 5159 | 258 | E258Y | A | 3e-06 | 2.5e-06 | 2.9e-06 | 2.800E-06 | 2.646E-07 | 2.500E-06 | 3.000E-06 |
| 5160 | 258 | E258V | A | 2.2e-06 | 2.6e-06 | 2.3e-06 | 2.367E-06 | 2.082E-07 | 2.200E-06 | 2.600E-06 |
| 5161 | 259 | S259A | A | 1.7e-06 | 2.5e-06 | 1.9e-06 | 2.033E-06 | 4.163E-07 | 1.700E-06 | 2.500E-06 |
| 5162 | 259 | S259R | A | 4.2e-06 | 3.3e-06 | 2.5e-06 | 3.333E-06 | 8.505E-07 | 2.500E-06 | 4.200E-06 |
| 5163 | 259 | S259N | A | 2.5e-06 | 3.8e-06 | 1.7e-06 | 2.667E-06 | 1.060E-06 | 1.700E-06 | 3.800E-06 |
| 5164 | 259 | S259D | A | 3.5e-06 | 4.4e-06 | 2.9e-06 | 3.600E-06 | 7.550E-07 | 2.900E-06 | 4.400E-06 |
| 5165 | 259 | S259C | A | 3.5e-06 | 2.1e-06 | 2.9e-06 | 2.833E-06 | 7.024E-07 | 2.100E-06 | 3.500E-06 |
| 5166 | 259 | S259E | A | 3.1e-06 | 2e-06 | 2.1e-06 | 2.400E-06 | 6.083E-07 | 2.000E-06 | 3.100E-06 |
| 5167 | 259 | S259Q | A | 2.6e-06 | 3.7e-06 | 1.8e-06 | 2.700E-06 | 9.539E-07 | 1.800E-06 | 3.700E-06 |
| 5168 | 259 | S259G | A | 3.4e-06 | 2e-06 | 3.1e-06 | 2.833E-06 | 7.371E-07 | 2.000E-06 | 3.400E-06 |
| 5169 | 259 | S259H | A | 1.5e-06 | 1.8e-06 | 2e-06 | 1.767E-06 | 2.517E-07 | 1.500E-06 | 2.000E-06 |
| 5170 | 259 | S259I | A | 2.6e-06 | 8.4e-07 | 2.6e-06 | 2.013E-06 | 1.016E-06 | 8.400E-07 | 2.600E-06 |
| 5171 | 259 | S259L | A | 2.1e-06 | 3.2e-06 | 3e-06 | 2.767E-06 | 5.859E-07 | 2.100E-06 | 3.200E-06 |
| 5172 | 259 | S259K | A | 2.6e-06 | 1.1e-06 | 1.6e-06 | 1.767E-06 | 7.638E-07 | 1.100E-06 | 2.600E-06 |
| 5173 | 259 | S259M | A | 2.1e-06 | 1.9e-06 | 2.6e-06 | 2.200E-06 | 3.606E-07 | 1.900E-06 | 2.600E-06 |
| 5174 | 259 | S259F | A | 2.2e-06 | 2.3e-06 | 1.8e-06 | 2.100E-06 | 2.646E-07 | 1.800E-06 | 2.300E-06 |
| 5175 | 259 | S259P | A | 3.2e-06 | 2.1e-06 | 2.7e-06 | 2.667E-06 | 5.508E-07 | 2.100E-06 | 3.200E-06 |
| 5176 | 259 | S259S | A | 2.6e-06 | 2.6e-06 | 2.3e-06 | 2.500E-06 | 1.732E-07 | 2.300E-06 | 2.600E-06 |
| 5177 | 259 | S259T | A | 1.6e-06 | 3.7e-06 | 2.1e-06 | 2.467E-06 | 1.097E-06 | 1.600E-06 | 3.700E-06 |
| 5178 | 259 | S259W | A | 1.4e-06 | 2.2e-06 | 1.8e-06 | 1.800E-06 | 4.000E-07 | 1.400E-06 | 2.200E-06 |
| 5179 | 259 | S259Y | A | 2.9e-06 | 1.9e-06 | 2.1e-06 | 2.300E-06 | 5.292E-07 | 1.900E-06 | 2.900E-06 |
| 5180 | 259 | S259V | A | 3.5e-06 | 2.2e-06 | 3.3e-06 | 3.000E-06 | 7.000E-07 | 2.200E-06 | 3.500E-06 |
| 5181 | 260 | M260A | A | 2e-06 | 2.4e-06 | 2.7e-06 | 2.367E-06 | 3.512E-07 | 2.000E-06 | 2.700E-06 |
| 5182 | 260 | M260R | A | 2.6e-06 | 2e-06 | 1.2e-06 | 1.933E-06 | 7.024E-07 | 1.200E-06 | 2.600E-06 |
| 5183 | 260 | M260N | A | 2.5e-06 | 3e-06 | 2e-06 | 2.500E-06 | 5.000E-07 | 2.000E-06 | 3.000E-06 |
| 5184 | 260 | M260D | A | 1.9e-06 | 2.6e-06 | 1.5e-06 | 2.000E-06 | 5.568E-07 | 1.500E-06 | 2.600E-06 |

| | | | | | | | | | | |
|---|---|---|---|---|---|---|---|---|---|---|
| 5185 | 260 | M260C | A | 1.8e-06 | 1.4e-06 | 2.3e-06 | 1.833E-06 | 4.509E-07 | 1.400E-06 | 2.300E-06 |
| 5186 | 260 | M260E | A | 1.8e-06 | 3.4e-06 | 2.2e-06 | 2.467E-06 | 8.327E-07 | 1.800E-06 | 3.400E-06 |
| 5187 | 260 | M260Q | A | 1.9e-06 | 2e-06 | 3e-06 | 2.300E-06 | 6.083E-07 | 1.900E-06 | 3.000E-06 |
| 5188 | 260 | M260G | A | 1.4e-06 | 2.4e-06 | 2.1e-06 | 1.967E-06 | 5.132E-07 | 1.400E-06 | 2.400E-06 |
| 5189 | 260 | M260H | A | 3.3e-06 | 4.8e-06 | 2.8e-06 | 3.633E-06 | 1.041E-06 | 2.800E-06 | 4.800E-06 |
| 5190 | 260 | M260I | A | 1.6e-06 | 1.9e-06 | 2.1e-06 | 1.867E-06 | 2.517E-07 | 1.600E-06 | 2.100E-06 |
| 5191 | 260 | M260L | A | 3.4e-06 | 1.6e-06 | 2.5e-06 | 2.500E-06 | 9.000E-07 | 1.600E-06 | 3.400E-06 |
| 5192 | 260 | M260K | A | 2e-06 | 2.7e-06 | 2.1e-06 | 2.267E-06 | 3.786E-07 | 2.000E-06 | 2.700E-06 |
| 5193 | 260 | M260M | A | 3.2e-06 | 5e-06 | 1.3e-06 | 3.167E-06 | 1.850E-06 | 1.300E-06 | 5.000E-06 |
| 5194 | 260 | M260F | A | 8.9e-07 | 2.1e-06 | 2.6e-06 | 1.863E-06 | 8.792E-07 | 8.900E-07 | 2.600E-06 |
| 5195 | 260 | M260P | A | 2.5e-06 | 2.3e-06 | 2.1e-06 | 2.300E-06 | 2.000E-07 | 2.100E-06 | 2.500E-06 |
| 5196 | 260 | M260S | A | 2.2e-06 | 2.7e-06 | 3e-06 | 2.633E-06 | 4.041E-07 | 2.200E-06 | 3.000E-06 |
| 5197 | 260 | M260T | A | 2.8e-06 | 2.4e-06 | 2.6e-06 | 2.600E-06 | 2.000E-07 | 2.400E-06 | 2.800E-06 |
| 5198 | 260 | M260W | A | 1e-06 | 3.7e-06 | 2e-06 | 2.233E-06 | 1.365E-06 | 1.000E-06 | 3.700E-06 |
| 5199 | 260 | M260Y | A | 5.5e-06 | 2.7e-06 | 2.2e-06 | 3.467E-06 | 1.779E-06 | 2.200E-06 | 5.500E-06 |
| 5200 | 260 | M260V | A | 1.4e-06 | 2.8e-06 | 4e-06 | 2.733E-06 | 1.301E-06 | 1.400E-06 | 4.000E-06 |
| 5201 | 261 | P261A | A | 3.3e-06 | 2.4e-06 | 2.9e-06 | 2.867E-06 | 4.509E-07 | 2.400E-06 | 3.300E-06 |
| 5202 | 261 | P261R | A | 1.1e-06 | 2.4e-06 | 1.8e-06 | 1.767E-06 | 6.506E-07 | 1.100E-06 | 2.400E-06 |
| 5203 | 261 | P261N | A | 1.8e-06 | 2e-06 | 9.3e-07 | 1.577E-06 | 5.689E-07 | 9.300E-07 | 2.000E-06 |
| 5204 | 261 | P261D | A | 2.7e-06 | 2.3e-06 | 1.9e-06 | 2.300E-06 | 4.000E-07 | 1.900E-06 | 2.700E-06 |
| 5205 | 261 | P261C | A | 1.7e-06 | 2.7e-06 | 3.3e-06 | 2.567E-06 | 8.083E-07 | 1.700E-06 | 3.300E-06 |
| 5206 | 261 | P261E | A | 3.2e-06 | 2.5e-06 | 3.7e-06 | 3.133E-06 | 6.028E-07 | 2.500E-06 | 3.700E-06 |
| 5207 | 261 | P261Q | A | 2.8e-06 | 2.6e-06 | 3.8e-06 | 3.067E-06 | 6.429E-07 | 2.600E-06 | 3.800E-06 |
| 5208 | 261 | P261G | A | 2.2e-06 | 1.9e-06 | 3.2e-06 | 2.433E-06 | 6.807E-07 | 1.900E-06 | 3.200E-06 |
| 5209 | 261 | P261H | A | 2.7e-06 | 2.6e-06 | 9.7e-07 | 2.090E-06 | 9.712E-07 | 9.700E-07 | 2.700E-06 |
| 5210 | 261 | P261I | A | 1.9e-06 | 2.7e-06 | 3.2e-06 | 2.600E-06 | 6.557E-07 | 1.900E-06 | 3.200E-06 |
| 5211 | 261 | P261L | A | 2.4e-06 | 2.5e-06 | 2.6e-06 | 2.500E-06 | 1.000E-07 | 2.400E-06 | 2.600E-06 |
| 5212 | 261 | P261K | A | 3.6e-06 | 2.4e-06 | 2.5e-06 | 2.833E-06 | 6.658E-07 | 2.400E-06 | 3.600E-06 |
| 5213 | 261 | P261M | A | 2.9e-06 | 2.7e-06 | 1.6e-06 | 2.400E-06 | 7.000E-07 | 1.600E-06 | 2.900E-06 |
| 5214 | 261 | P261F | A | 1.8e-06 | 2.1e-06 | 2.5e-06 | 2.133E-06 | 3.512E-07 | 1.800E-06 | 2.500E-06 |
| 5215 | 261 | P261P | A | 3.6e-06 | 2.6e-06 | 2e-06 | 2.733E-06 | 8.083E-07 | 2.000E-06 | 3.600E-06 |
| 5216 | 261 | P261S | A | 1.9e-06 | 2.5e-06 | 2.3e-06 | 2.233E-06 | 3.055E-07 | 1.900E-06 | 2.500E-06 |
| 5217 | 261 | P261T | A | 1.5e-06 | 3.2e-06 | 3.3e-06 | 2.667E-06 | 1.012E-06 | 1.500E-06 | 3.300E-06 |
| 5218 | 261 | P261W | A | 1.3e-06 | 2.7e-06 | 3.1e-06 | 2.367E-06 | 9.452E-07 | 1.300E-06 | 3.100E-06 |
| 5219 | 261 | P261Y | A | 2.1e-06 | 1.8e-06 | 2.6e-06 | 2.167E-06 | 4.041E-07 | 1.800E-06 | 2.600E-06 |
| 5220 | 261 | P261V | A | 3e-06 | 2.6e-06 | 2.1e-06 | 2.567E-06 | 4.509E-07 | 2.100E-06 | 3.000E-06 |
| 5221 | 262 | N262A | A | 2.2e-06 | 2.8e-06 | 1e-06 | 2.000E-06 | 9.165E-07 | 1.000E-06 | 2.800E-06 |
| 5222 | 262 | N262R | A | 2.5e-06 | 2.1e-06 | 1.5e-06 | 2.033E-06 | 5.033E-07 | 1.500E-06 | 2.500E-06 |
| 5223 | 262 | N262N | A | 2.4e-06 | 3e-06 | 2.7e-06 | 2.700E-06 | 3.000E-07 | 2.400E-06 | 3.000E-06 |
| 5224 | 262 | N262D | A | 2.3e-06 | 2.8e-06 | 3.3e-06 | 2.800E-06 | 5.000E-07 | 2.300E-06 | 3.300E-06 |
| 5225 | 262 | N262C | A | 3.2e-06 | 3.2e-06 | 1.7e-06 | 2.700E-06 | 8.660E-07 | 1.700E-06 | 3.200E-06 |
| 5226 | 262 | N262E | A | 2e-06 | 1.9e-06 | 1.7e-06 | 1.867E-06 | 1.528E-07 | 1.700E-06 | 2.000E-06 |
| 5227 | 262 | N262Q | A | 1.7e-06 | 2.3e-06 | 2e-06 | 2.000E-06 | 3.000E-07 | 1.700E-06 | 2.300E-06 |
| 5228 | 262 | N262G | A | 8.4e-07 | 2.1e-06 | 1.9e-06 | 1.613E-06 | 6.772E-07 | 8.400E-07 | 2.100E-06 |
| 5229 | 262 | N262H | A | 2.6e-06 | 2.3e-06 | 1.8e-06 | 2.233E-06 | 4.041E-07 | 1.800E-06 | 2.600E-06 |
| 5230 | 262 | N262I | A | 2.7e-06 | 2.3e-06 | 2.3e-06 | 2.433E-06 | 2.309E-07 | 2.300E-06 | 2.700E-06 |
| 5231 | 262 | N262L | A | 2.5e-06 | 2.5e-06 | 2.2e-06 | 2.400E-06 | 1.732E-07 | 2.200E-06 | 2.500E-06 |
| 5232 | 262 | N262K | A | 2.6e-06 | 2.6e-06 | 1.8e-06 | 2.333E-06 | 4.619E-07 | 1.800E-06 | 2.600E-06 |
| 5233 | 262 | N262M | A | 1.9e-06 | 2.8e-06 | 2.2e-06 | 2.300E-06 | 4.583E-07 | 1.900E-06 | 2.800E-06 |
| 5234 | 262 | N262F | A | 2.5e-06 | 2.9e-06 | 2.7e-06 | 2.700E-06 | 2.000E-07 | 2.500E-06 | 2.900E-06 |
| 5235 | 262 | N262P | A | 3.1e-06 | 2.2e-06 | 1.8e-06 | 2.367E-06 | 6.658E-07 | 1.800E-06 | 3.100E-06 |
| 5236 | 262 | N262S | A | 2.5e-06 | 2.9e-06 | 3.4e-06 | 2.933E-06 | 4.509E-07 | 2.500E-06 | 3.400E-06 |
| 5237 | 262 | N262T | A | 3.3e-06 | 5.8e-06 | 1.3e-06 | 3.467E-06 | 2.255E-06 | 1.300E-06 | 5.800E-06 |
| 5238 | 262 | N262W | A | 2.6e-06 | 2.7e-06 | 2.9e-06 | 2.733E-06 | 1.528E-07 | 2.600E-06 | 2.900E-06 |
| 5239 | 262 | N262Y | A | 1.9e-06 | 1.5e-06 | 3.4e-06 | 2.267E-06 | 1.002E-06 | 1.500E-06 | 3.400E-06 |
| 5240 | 262 | N262V | A | 2.6e-06 | 3.1e-06 | 4.8e-06 | 3.500E-06 | 1.153E-06 | 2.600E-06 | 4.800E-06 |
| 5241 | 263 | P263A | A | 2.9e-06 | 2.4e-06 | 1.9e-06 | 2.400E-06 | 5.000E-07 | 1.900E-06 | 2.900E-06 |
| 5242 | 263 | P263R | A | 2e-06 | 2.6e-06 | 1.1e-06 | 1.900E-06 | 7.550E-07 | 1.100E-06 | 2.600E-06 |
| 5243 | 263 | P263N | A | 1.9e-06 | 2e-06 | 2e-06 | 1.967E-06 | 5.774E-08 | 1.900E-06 | 2.000E-06 |
| 5244 | 263 | P263D | A | 2.7e-06 | 1.2e-06 | 1.9e-06 | 1.933E-06 | 7.506E-07 | 1.200E-06 | 2.700E-06 |
| 5245 | 263 | P263C | A | 2.4e-06 | 1.4e-06 | 2.6e-06 | 2.133E-06 | 6.429E-07 | 1.400E-06 | 2.600E-06 |
| 5246 | 263 | P263E | A | 2.5e-06 | 1.9e-06 | 2.2e-06 | 2.200E-06 | 3.000E-07 | 1.900E-06 | 2.500E-06 |
| 5247 | 263 | P263Q | A | 3e-06 | 1.9e-06 | 1.7e-06 | 2.200E-06 | 7.000E-07 | 1.700E-06 | 3.000E-06 |
| 5248 | 263 | P263G | A | 1.9e-06 | 3e-06 | 1.8e-06 | 2.233E-06 | 6.658E-07 | 1.800E-06 | 3.000E-06 |
| 5249 | 263 | P263H | A | 4.4e-06 | 2.7e-06 | 1.5e-06 | 2.867E-06 | 1.457E-06 | 1.500E-06 | 4.400E-06 |
| 5250 | 263 | P263I | A | 1.5e-06 | 4.4e-06 | 2.2e-06 | 2.700E-06 | 1.513E-06 | 1.500E-06 | 4.400E-06 |
| 5251 | 263 | P263L | A | 3e-06 | 2.3e-06 | 5.2e-06 | 3.500E-06 | 1.513E-06 | 2.300E-06 | 5.200E-06 |
| 5252 | 263 | P263K | A | 2.9e-06 | 1.5e-06 | 4.8e-06 | 3.067E-06 | 1.656E-06 | 1.500E-06 | 4.800E-06 |
| 5253 | 263 | P263M | A | 3.3e-06 | 2.3e-06 | 4.1e-06 | 3.233E-06 | 9.018E-07 | 2.300E-06 | 4.100E-06 |
| 5254 | 263 | P263F | A | 3.2e-06 | 1.6e-06 | 2.2e-06 | 2.333E-06 | 8.083E-07 | 1.600E-06 | 3.200E-06 |
| 5255 | 263 | P263P | A | 1e-06 | 1.9e-06 | 2.3e-06 | 1.733E-06 | 6.658E-07 | 1.000E-06 | 2.300E-06 |
| 5256 | 263 | P263S | A | 4.7e-06 | 2e-06 | 2.6e-06 | 3.100E-06 | 1.418E-06 | 2.000E-06 | 4.700E-06 |
| 5257 | 263 | P263T | A | 1.1e-06 | 2.4e-06 | 3.4e-06 | 2.300E-06 | 1.153E-06 | 1.100E-06 | 3.400E-06 |
| 5258 | 263 | P263W | A | 1.2e-06 | 2.2e-06 | 2.5e-06 | 1.967E-06 | 6.807E-07 | 1.200E-06 | 2.500E-06 |
| 5259 | 263 | P263Y | A | 4e-06 | 2.7e-06 | 2.7e-06 | 3.133E-06 | 7.506E-07 | 2.700E-06 | 4.000E-06 |
| 5260 | 263 | P263V | A | 3.3e-06 | 4.7e-06 | 1.9e-06 | 3.300E-06 | 1.400E-06 | 1.900E-06 | 4.700E-06 |
| 5261 | 264 | E264A | A | 2.5e-06 | 1.7e-06 | 3.1e-06 | 2.433E-06 | 7.024E-07 | 1.700E-06 | 3.100E-06 |
| 5262 | 264 | E264R | A | 2.8e-06 | 2.3e-06 | 2.6e-06 | 2.567E-06 | 2.517E-07 | 2.300E-06 | 2.800E-06 |
| 5263 | 264 | E264N | A | 3.1e-06 | 4e-06 | 1.9e-06 | 3.000E-06 | 1.054E-06 | 1.900E-06 | 4.000E-06 |
| 5264 | 264 | E264D | A | 2.5e-06 | 2.2e-06 | 3e-06 | 2.567E-06 | 4.041E-07 | 2.200E-06 | 3.000E-06 |
| 5265 | 264 | E264C | A | 2.8e-06 | 1.4e-06 | 2.8e-06 | 2.333E-06 | 8.083E-07 | 1.400E-06 | 2.800E-06 |
| 5266 | 264 | E264E | A | 3.5e-06 | 1.8e-06 | 2.3e-06 | 2.533E-06 | 8.737E-07 | 1.800E-06 | 3.500E-06 |
| 5267 | 264 | E264Q | A | 2.3e-06 | 1.6e-06 | 2.7e-06 | 2.200E-06 | 5.568E-07 | 1.600E-06 | 2.700E-06 |
| 5268 | 264 | E264G | A | 2.4e-06 | 2.4e-06 | 2.8e-06 | 2.533E-06 | 2.309E-07 | 2.400E-06 | 2.800E-06 |
| 5269 | 264 | E264H | A | 2.8e-06 | 4.2e-06 | 2.5e-06 | 3.167E-06 | 9.074E-07 | 2.500E-06 | 4.200E-06 |

| | | | | | | | | | | |
|---|---|---|---|---|---|---|---|---|---|---|
| 5270 | 264 | E264I | A | 2.6e-06 | 2.2e-06 | 1.5e-06 | 2.100E-06 | 5.568E-07 | 1.500E-06 | 2.600E-06 |
| 5271 | 264 | E264L | A | 2.5e-06 | 1.9e-06 | 2.5e-06 | 2.300E-06 | 3.464E-07 | 1.900E-06 | 2.500E-06 |
| 5272 | 264 | E264K | A | 2.7e-06 | 2.4e-06 | 3.2e-06 | 2.767E-06 | 4.041E-07 | 2.400E-06 | 3.200E-06 |
| 5273 | 264 | E264M | A | 2.7e-06 | 1.7e-06 | 2.5e-06 | 2.300E-06 | 5.292E-07 | 1.700E-06 | 2.700E-06 |
| 5274 | 264 | E264F | A | 1.9e-06 | 1.4e-06 | 2.5e-06 | 1.933E-06 | 5.508E-07 | 1.400E-06 | 2.500E-06 |
| 5275 | 264 | E264P | A | 2.4e-06 | 3.8e-06 | 4.3e-06 | 3.500E-06 | 9.849E-07 | 2.400E-06 | 4.300E-06 |
| 5276 | 264 | E264S | A | 2.4e-06 | 2.9e-06 | 2e-06 | 2.433E-06 | 4.509E-07 | 2.000E-06 | 2.900E-06 |
| 5277 | 264 | E264T | A | 2.6e-06 | 2.8e-06 | 3.6e-06 | 3.000E-06 | 5.292E-07 | 2.600E-06 | 3.600E-06 |
| 5278 | 264 | E264W | A | 1.3e-06 | 1.4e-06 | 3.4e-06 | 2.033E-06 | 1.185E-06 | 1.300E-06 | 3.400E-06 |
| 5279 | 264 | E264Y | A | 2.8e-06 | 1.6e-06 | 2.5e-06 | 2.300E-06 | 6.245E-07 | 1.600E-06 | 2.800E-06 |
| 5280 | 264 | E264V | A | 3.6e-06 | 3.9e-06 | 1.4e-06 | 2.967E-06 | 1.365E-06 | 1.400E-06 | 3.900E-06 |
| 5281 | 265 | G265A | A | 3.7e-06 | 2.3e-06 | 2.4e-06 | 2.800E-06 | 7.810E-07 | 2.300E-06 | 3.700E-06 |
| 5282 | 265 | G265R | A | 3.6e-06 | 1.7e-06 | 2.7e-06 | 2.667E-06 | 9.504E-07 | 1.700E-06 | 3.600E-06 |
| 5283 | 265 | G265N | A | 1.8e-06 | 1.8e-06 | 3.9e-06 | 2.500E-06 | 1.212E-06 | 1.800E-06 | 3.900E-06 |
| 5284 | 265 | G265D | A | 2.8e-06 | 1.5e-06 | 2e-06 | 2.100E-06 | 6.557E-07 | 1.500E-06 | 2.800E-06 |
| 5285 | 265 | G265C | A | 3.4e-06 | 4.9e-06 | 2.2e-06 | 3.500E-06 | 1.353E-06 | 2.200E-06 | 4.900E-06 |
| 5286 | 265 | G265E | A | 3.9e-06 | 2.2e-06 | 1.5e-06 | 2.533E-06 | 1.234E-06 | 1.500E-06 | 3.900E-06 |
| 5287 | 265 | G265Q | A | 9e-07 | 1.1e-06 | 2.9e-06 | 1.633E-06 | 1.102E-06 | 9.000E-07 | 2.900E-06 |
| 5288 | 265 | G265G | A | 2.3e-06 | 2e-06 | 4.1e-06 | 2.800E-06 | 1.136E-06 | 2.000E-06 | 4.100E-06 |
| 5289 | 265 | G265H | A | 2.4e-06 | 2.7e-06 | 2.2e-06 | 2.433E-06 | 2.517E-07 | 2.200E-06 | 2.700E-06 |
| 5290 | 265 | G265I | A | 2.4e-06 | 2.6e-06 | 3.7e-06 | 2.900E-06 | 7.000E-07 | 2.400E-06 | 3.700E-06 |
| 5291 | 265 | G265L | A | 1.6e-06 | 3.4e-06 | 4.6e-06 | 3.200E-06 | 1.510E-06 | 1.600E-06 | 4.600E-06 |
| 5292 | 265 | G265K | A | 2e-06 | 2.9e-06 | 1.8e-06 | 2.233E-06 | 5.859E-07 | 1.800E-06 | 2.900E-06 |
| 5293 | 265 | G265M | A | 3.4e-06 | 2.2e-06 | 2.1e-06 | 2.567E-06 | 7.234E-07 | 2.100E-06 | 3.400E-06 |
| 5294 | 265 | G265F | A | 2.9e-06 | 1.5e-06 | 1.7e-06 | 2.033E-06 | 7.572E-07 | 1.500E-06 | 2.900E-06 |
| 5295 | 265 | G265P | A | 1.6e-06 | 2.7e-06 | 2.2e-06 | 2.167E-06 | 5.508E-07 | 1.600E-06 | 2.700E-06 |
| 5296 | 265 | G265S | A | 4e-06 | 3.2e-06 | 2.4e-06 | 3.200E-06 | 8.000E-07 | 2.400E-06 | 4.000E-06 |
| 5297 | 265 | G265T | A | 2.3e-06 | 2.8e-06 | 3.8e-06 | 2.967E-06 | 7.638E-07 | 2.300E-06 | 3.800E-06 |
| 5298 | 265 | G265W | A | 2.1e-06 | 2.6e-06 | 2.1e-06 | 2.267E-06 | 2.887E-07 | 2.100E-06 | 2.600E-06 |
| 5299 | 265 | G265Y | A | 3.5e-06 | 2.3e-06 | 2.5e-06 | 2.767E-06 | 6.429E-07 | 2.300E-06 | 3.500E-06 |
| 5300 | 265 | G265V | A | 2.6e-06 | 3.4e-06 | 1.8e-06 | 2.600E-06 | 8.000E-07 | 1.800E-06 | 3.400E-06 |
| 5301 | 266 | R266A | A | 2.4e-06 | 3e-06 | 1.5e-06 | 2.300E-06 | 7.550E-07 | 1.500E-06 | 3.000E-06 |
| 5302 | 266 | R266R | A | 4.3e-06 | 3e-06 | 2.6e-06 | 3.300E-06 | 8.888E-07 | 2.600E-06 | 4.300E-06 |
| 5303 | 266 | R266N | A | 3.1e-06 | 3.9e-06 | 4e-06 | 3.667E-06 | 4.933E-07 | 3.100E-06 | 4.000E-06 |
| 5304 | 266 | R266D | A | 2.8e-06 | 4.5e-06 | 2.5e-06 | 3.267E-06 | 1.079E-06 | 2.500E-06 | 4.500E-06 |
| 5305 | 266 | R266C | A | 2e-06 | 1.7e-06 | 2.1e-06 | 1.933E-06 | 2.082E-07 | 1.700E-06 | 2.100E-06 |
| 5306 | 266 | R266E | A | 2.7e-06 | 2.4e-06 | 3.5e-06 | 2.867E-06 | 5.686E-07 | 2.400E-06 | 3.500E-06 |
| 5307 | 266 | R266Q | A | 1.8e-06 | 2.1e-06 | 2.1e-06 | 2.000E-06 | 1.732E-07 | 1.800E-06 | 2.100E-06 |
| 5308 | 266 | R266G | A | 2.7e-06 | 2e-06 | 2.2e-06 | 2.300E-06 | 3.606E-07 | 2.000E-06 | 2.700E-06 |
| 5309 | 266 | R266H | A | 4.9e-06 | 2.2e-06 | 2.2e-06 | 3.100E-06 | 1.559E-06 | 2.200E-06 | 4.900E-06 |
| 5310 | 266 | R266I | A | 2.4e-06 | 1.3e-06 | 1.4e-06 | 1.700E-06 | 6.083E-07 | 1.300E-06 | 2.400E-06 |
| 5311 | 266 | R266L | A | 2.6e-06 | 1.6e-06 | 3.7e-06 | 2.633E-06 | 1.050E-06 | 1.600E-06 | 3.700E-06 |
| 5312 | 266 | R266K | A | 5e-06 | 1.6e-06 | 3.1e-06 | 3.233E-06 | 1.704E-06 | 1.600E-06 | 5.000E-06 |
| 5313 | 266 | R266M | A | 3.5e-06 | 1.5e-06 | 1.1e-06 | 2.033E-06 | 1.286E-06 | 1.100E-06 | 3.500E-06 |
| 5314 | 266 | R266F | A | 2.2e-06 | 3.2e-06 | 1.5e-06 | 2.300E-06 | 8.544E-07 | 1.500E-06 | 3.200E-06 |
| 5315 | 266 | R266P | A | 1.2e-06 | 2.1e-06 | 2.5e-06 | 1.933E-06 | 6.658E-07 | 1.200E-06 | 2.500E-06 |
| 5316 | 266 | R266S | A | 2e-06 | 1.9e-06 | 1.8e-06 | 1.900E-06 | 1.000E-07 | 1.800E-06 | 2.000E-06 |
| 5317 | 266 | R266T | A | 1.7e-06 | 2.5e-06 | 2.5e-06 | 2.233E-06 | 4.619E-07 | 1.700E-06 | 2.500E-06 |
| 5318 | 266 | R266W | A | 2e-06 | 4e-06 | 1.9e-06 | 2.633E-06 | 1.185E-06 | 1.900E-06 | 4.000E-06 |
| 5319 | 266 | R266Y | A | 1.6e-06 | 2.2e-06 | 2.4e-06 | 2.067E-06 | 4.163E-07 | 1.600E-06 | 2.400E-06 |
| 5320 | 266 | R266V | A | 2.3e-06 | 2.2e-06 | 3.3e-06 | 2.600E-06 | 6.083E-07 | 2.200E-06 | 3.300E-06 |
| 5321 | 267 | Y267A | A | 3.2e-06 | 1.4e-06 | 3.2e-06 | 2.600E-06 | 1.039E-06 | 1.400E-06 | 3.200E-06 |
| 5322 | 267 | Y267R | A | 1.5e-06 | 3.3e-06 | 1.4e-06 | 2.067E-06 | 1.069E-06 | 1.400E-06 | 3.300E-06 |
| 5323 | 267 | Y267N | A | 1.1e-06 | 2.9e-06 | 3.5e-06 | 2.500E-06 | 1.249E-06 | 1.100E-06 | 3.500E-06 |
| 5324 | 267 | Y267D | A | 2.1e-06 | 1.9e-06 | 4.2e-06 | 2.733E-06 | 1.274E-06 | 1.900E-06 | 4.200E-06 |
| 5325 | 267 | Y267C | A | 1e-06 | 2.9e-06 | 2.3e-06 | 2.067E-06 | 9.713E-07 | 1.000E-06 | 2.900E-06 |
| 5326 | 267 | Y267E | A | 2.2e-06 | 2.6e-06 | 3.3e-06 | 2.700E-06 | 5.568E-07 | 2.200E-06 | 3.300E-06 |
| 5327 | 267 | Y267Q | A | 2.4e-06 | 3.6e-06 | 2.5e-06 | 2.833E-06 | 6.658E-07 | 2.400E-06 | 3.600E-06 |
| 5328 | 267 | Y267G | A | 1.9e-06 | 1.2e-06 | 1.4e-06 | 1.500E-06 | 3.606E-07 | 1.200E-06 | 1.900E-06 |
| 5329 | 267 | Y267H | A | 2.1e-06 | 2.6e-06 | 2.3e-06 | 2.333E-06 | 2.517E-07 | 2.100E-06 | 2.600E-06 |
| 5330 | 267 | Y267I | A | 2.1e-06 | 1.2e-06 | 3.9e-06 | 2.400E-06 | 1.375E-06 | 1.200E-06 | 3.900E-06 |
| 5331 | 267 | Y267L | A | 1.9e-06 | 3.4e-06 | 1.9e-06 | 2.400E-06 | 8.660E-07 | 1.900E-06 | 3.400E-06 |
| 5332 | 267 | Y267K | A | 2.8e-06 | 3.3e-06 | 1.6e-06 | 2.567E-06 | 8.737E-07 | 1.600E-06 | 3.300E-06 |
| 5333 | 267 | Y267M | A | 2e-06 | 2.5e-06 | 4.5e-06 | 3.000E-06 | 1.323E-06 | 2.000E-06 | 4.500E-06 |
| 5334 | 267 | Y267F | A | 1.8e-06 | 2.6e-06 | 3.3e-06 | 2.567E-06 | 7.506E-07 | 1.800E-06 | 3.300E-06 |
| 5335 | 267 | Y267P | A | 3.7e-06 | 3.2e-06 | 2.7e-06 | 3.200E-06 | 5.000E-07 | 2.700E-06 | 3.700E-06 |
| 5336 | 267 | Y267S | A | 3.3e-06 | 1.5e-06 | 2.8e-06 | 2.533E-06 | 9.292E-07 | 1.500E-06 | 3.300E-06 |
| 5337 | 267 | Y267T | A | 1.5e-06 | 1.3e-06 | 2.4e-06 | 1.733E-06 | 5.859E-07 | 1.300E-06 | 2.400E-06 |
| 5338 | 267 | Y267W | A | 2.5e-06 | 2e-06 | 2.5e-06 | 2.333E-06 | 2.887E-07 | 2.000E-06 | 2.500E-06 |
| 5339 | 267 | Y267Y | A | 1.7e-06 | 6.8e-07 | 8e-07 | 1.060E-06 | 5.575E-07 | 6.800E-07 | 1.700E-06 |
| 5340 | 267 | Y267V | A | 2.8e-06 | 4.3e-06 | 3.1e-06 | 3.400E-06 | 7.937E-07 | 2.800E-06 | 4.300E-06 |
| 5341 | 268 | T268A | A | 3.5e-06 | 2.2e-06 | 1.7e-06 | 2.467E-06 | 9.292E-07 | 1.700E-06 | 3.500E-06 |
| 5342 | 268 | T268R | A | 1.5e-06 | 3.8e-06 | 1.3e-06 | 2.200E-06 | 1.389E-06 | 1.300E-06 | 3.800E-06 |
| 5343 | 268 | T268N | A | 2.3e-06 | 2.8e-06 | 2.5e-06 | 2.533E-06 | 2.517E-07 | 2.300E-06 | 2.800E-06 |
| 5344 | 268 | T268D | A | 2e-06 | 2.7e-06 | 2.6e-06 | 2.433E-06 | 3.786E-07 | 2.000E-06 | 2.700E-06 |
| 5345 | 268 | T268C | A | 2.8e-06 | 3.4e-06 | 2.6e-06 | 2.933E-06 | 4.163E-07 | 2.600E-06 | 3.400E-06 |
| 5346 | 268 | T268E | A | 4.9e-06 | 3e-06 | 2.5e-06 | 3.467E-06 | 1.266E-06 | 2.500E-06 | 4.900E-06 |
| 5347 | 268 | T268Q | A | 3e-06 | 3.7e-06 | 2.4e-06 | 3.033E-06 | 6.506E-07 | 2.400E-06 | 3.700E-06 |
| 5348 | 268 | T268G | A | 2.4e-06 | 1.5e-06 | 2.7e-06 | 2.200E-06 | 6.245E-07 | 1.500E-06 | 2.700E-06 |
| 5349 | 268 | T268H | A | 3.4e-06 | 1.4e-06 | 2.3e-06 | 2.367E-06 | 1.002E-06 | 1.400E-06 | 3.400E-06 |
| 5350 | 268 | T268I | A | 2e-06 | 2.7e-06 | 2e-06 | 2.233E-06 | 4.041E-07 | 2.000E-06 | 2.700E-06 |
| 5351 | 268 | T268L | A | 1.8e-06 | 2.3e-06 | 1.9e-06 | 2.000E-06 | 2.646E-07 | 1.800E-06 | 2.300E-06 |
| 5352 | 268 | T268K | A | 2.2e-06 | 3.7e-06 | 2.3e-06 | 2.733E-06 | 8.386E-07 | 2.200E-06 | 3.700E-06 |
| 5353 | 268 | T268M | A | 1.9e-06 | 2e-06 | 1.7e-06 | 1.867E-06 | 1.528E-07 | 1.700E-06 | 2.000E-06 |
| 5354 | 268 | T268F | A | 1.9e-06 | 1.7e-06 | 2.2e-06 | 1.933E-06 | 2.517E-07 | 1.700E-06 | 2.200E-06 |

| | | | | | | | | | | |
|---|---|---|---|---|---|---|---|---|---|---|
| 5355 | 268 | T268P | A | 2e-06 | 2e-06 | 1.9e-06 | 1.967E-06 | 5.774E-08 | 1.900E-06 | 2.000E-06 |
| 5356 | 268 | T268S | A | 2.1e-06 | 3.1e-06 | 2e-06 | 2.400E-06 | 6.083E-07 | 2.000E-06 | 3.100E-06 |
| 5357 | 268 | T268T | A | 2.5e-06 | 1.9e-06 | 4.3e-06 | 2.900E-06 | 1.249E-06 | 1.900E-06 | 4.300E-06 |
| 5358 | 268 | T268W | A | 1.9e-06 | 1.6e-06 | 1.7e-06 | 1.733E-06 | 1.528E-07 | 1.600E-06 | 1.900E-06 |
| 5359 | 268 | T268Y | A | 2.4e-06 | 2.8e-06 | 1.4e-06 | 2.200E-06 | 7.211E-07 | 1.400E-06 | 2.800E-06 |
| 5360 | 268 | T268V | A | 3.2e-06 | 2e-06 | 4.4e-06 | 3.200E-06 | 1.200E-06 | 2.000E-06 | 4.400E-06 |
| 5361 | 269 | F269A | A | 2.3e-06 | 1.8e-06 | 1.3e-06 | 1.800E-06 | 5.000E-07 | 1.300E-06 | 2.300E-06 |
| 5362 | 269 | F269R | A | 1.7e-06 | 2.3e-06 | 3.1e-06 | 2.367E-06 | 7.024E-07 | 1.700E-06 | 3.100E-06 |
| 5363 | 269 | F269N | A | 3e-06 | 2.1e-06 | 2.8e-06 | 2.633E-06 | 4.726E-07 | 2.100E-06 | 3.000E-06 |
| 5364 | 269 | F269D | A | 2.3e-06 | 1.2e-06 | 1.8e-06 | 1.767E-06 | 5.508E-07 | 1.200E-06 | 2.300E-06 |
| 5365 | 269 | F269C | A | 2.4e-06 | 2.5e-06 | 2e-06 | 2.300E-06 | 2.646E-07 | 2.000E-06 | 2.500E-06 |
| 5366 | 269 | F269E | A | 2.2e-06 | 3.2e-06 | 3.3e-06 | 2.900E-06 | 6.083E-07 | 2.200E-06 | 3.300E-06 |
| 5367 | 269 | F269Q | A | 3.4e-06 | 1.4e-06 | 1.8e-06 | 2.200E-06 | 1.058E-06 | 1.400E-06 | 3.400E-06 |
| 5368 | 269 | F269G | A | 3.2e-06 | 4.2e-06 | 2.1e-06 | 3.167E-06 | 1.050E-06 | 2.100E-06 | 4.200E-06 |
| 5369 | 269 | F269H | A | 1.9e-06 | 3.5e-06 | 1.4e-06 | 2.267E-06 | 1.097E-06 | 1.400E-06 | 3.500E-06 |
| 5370 | 269 | F269I | A | 3.2e-06 | 2.4e-06 | 3.1e-06 | 2.900E-06 | 4.359E-07 | 2.400E-06 | 3.200E-06 |
| 5371 | 269 | F269L | A | 1.7e-06 | 2.4e-06 | 1.8e-06 | 1.967E-06 | 3.786E-07 | 1.700E-06 | 2.400E-06 |
| 5372 | 269 | F269K | A | 1.3e-06 | 2e-06 | 2.3e-06 | 1.867E-06 | 5.132E-07 | 1.300E-06 | 2.300E-06 |
| 5373 | 269 | F269M | A | 2e-06 | 4.6e-06 | 2.2e-06 | 2.933E-06 | 1.447E-06 | 2.000E-06 | 4.600E-06 |
| 5374 | 269 | F269F | A | 1.4e-06 | 2.6e-06 | 2.2e-06 | 2.067E-06 | 6.110E-07 | 1.400E-06 | 2.600E-06 |
| 5375 | 269 | F269P | A | 3.1e-06 | 1.7e-06 | 2.3e-06 | 2.367E-06 | 7.024E-07 | 1.700E-06 | 3.100E-06 |
| 5376 | 269 | F269S | A | 1.2e-06 | 1.5e-06 | 2.8e-06 | 1.833E-06 | 8.505E-07 | 1.200E-06 | 2.800E-06 |
| 5377 | 269 | F269T | A | 2.6e-06 | 2.7e-06 | 1.9e-06 | 2.400E-06 | 4.359E-07 | 1.900E-06 | 2.700E-06 |
| 5378 | 269 | F269W | A | 1.3e-06 | 2e-06 | 2.6e-06 | 1.967E-06 | 6.506E-07 | 1.300E-06 | 2.600E-06 |
| 5379 | 269 | F269Y | A | 2.1e-06 | 2.8e-06 | 1.7e-06 | 2.200E-06 | 5.568E-07 | 1.700E-06 | 2.800E-06 |
| 5380 | 269 | F269V | A | 3.2e-06 | 2.1e-06 | 5.1e-06 | 3.467E-06 | 1.518E-06 | 2.100E-06 | 5.100E-06 |
| 5381 | 270 | G270A | A | 1.2e-06 | 3.8e-06 | 2.9e-06 | 2.633E-06 | 1.320E-06 | 1.200E-06 | 3.800E-06 |
| 5382 | 270 | G270R | A | 1.8e-06 | 3.4e-06 | 1.8e-06 | 2.333E-06 | 9.238E-07 | 1.800E-06 | 3.400E-06 |
| 5383 | 270 | G270N | A | 2.2e-06 | 3.3e-06 | 1.9e-06 | 2.467E-06 | 7.371E-07 | 1.900E-06 | 3.300E-06 |
| 5384 | 270 | G270D | A | 2.5e-06 | 2.2e-06 | 4.2e-06 | 2.967E-06 | 1.079E-06 | 2.200E-06 | 4.200E-06 |
| 5385 | 270 | G270C | A | 2.4e-06 | 2.2e-06 | 1.9e-06 | 2.167E-06 | 2.517E-07 | 1.900E-06 | 2.400E-06 |
| 5386 | 270 | G270E | A | 2.2e-06 | 1.4e-06 | 4.7e-06 | 2.767E-06 | 1.721E-06 | 1.400E-06 | 4.700E-06 |
| 5387 | 270 | G270Q | A | 2.4e-06 | 2.2e-06 | 3.7e-06 | 2.767E-06 | 8.145E-07 | 2.200E-06 | 3.700E-06 |
| 5388 | 270 | G270G | A | 4e-06 | 2.6e-06 | 1.8e-06 | 2.800E-06 | 1.114E-06 | 1.800E-06 | 4.000E-06 |
| 5389 | 270 | G270H | A | 1.7e-06 | 1.5e-06 | 2.2e-06 | 1.800E-06 | 3.606E-07 | 1.500E-06 | 2.200E-06 |
| 5390 | 270 | G270I | A | 2.5e-06 | 2.9e-06 | 1.3e-06 | 2.233E-06 | 8.327E-07 | 1.300E-06 | 2.900E-06 |
| 5391 | 270 | G270L | A | 1.6e-06 | 2e-06 | 2.9e-06 | 2.167E-06 | 6.658E-07 | 1.600E-06 | 2.900E-06 |
| 5392 | 270 | G270K | A | 1.1e-06 | 3.3e-06 | 1e-06 | 1.800E-06 | 1.300E-06 | 1.000E-06 | 3.300E-06 |
| 5393 | 270 | G270M | A | 2.4e-06 | 1.9e-06 | 2.1e-06 | 2.133E-06 | 2.517E-07 | 1.900E-06 | 2.400E-06 |
| 5394 | 270 | G270F | A | 2.6e-06 | 3.3e-06 | 2.5e-06 | 2.800E-06 | 4.359E-07 | 2.500E-06 | 3.300E-06 |
| 5395 | 270 | G270P | A | 1.6e-06 | 2.6e-06 | 1.6e-06 | 1.933E-06 | 5.774E-07 | 1.600E-06 | 2.600E-06 |
| 5396 | 270 | G270S | A | 4.3e-07 | 2.5e-06 | 2.1e-06 | 1.677E-06 | 1.098E-06 | 4.300E-07 | 2.500E-06 |
| 5397 | 270 | G270T | A | 1.7e-06 | 2.9e-06 | 3.9e-06 | 2.833E-06 | 1.102E-06 | 1.700E-06 | 3.900E-06 |
| 5398 | 270 | G270W | A | 3.8e-06 | 2e-06 | 2e-06 | 2.600E-06 | 1.039E-06 | 2.000E-06 | 3.800E-06 |
| 5399 | 270 | G270Y | A | 3.3e-06 | 1.8e-06 | 2.2e-06 | 2.433E-06 | 7.767E-07 | 1.800E-06 | 3.300E-06 |
| 5400 | 270 | G270V | A | 2.3e-06 | 9.1e-07 | 3.8e-06 | 2.337E-06 | 1.445E-06 | 9.100E-07 | 3.800E-06 |
| 5401 | 271 | A271A | A | 1.5e-06 | 1.6e-06 | 2.1e-06 | 1.733E-06 | 3.215E-07 | 1.500E-06 | 2.100E-06 |
| 5402 | 271 | A271R | A | 8.6e-06 | 2.5e-06 | 2.8e-06 | 4.633E-06 | 3.439E-06 | 2.500E-06 | 8.600E-06 |
| 5403 | 271 | A271N | A | 1.8e-06 | 2.3e-06 | 2.3e-06 | 2.133E-06 | 2.887E-07 | 1.800E-06 | 2.300E-06 |
| 5404 | 271 | A271D | A | 2.7e-06 | 1.8e-06 | 1.3e-06 | 1.933E-06 | 7.095E-07 | 1.300E-06 | 2.700E-06 |
| 5405 | 271 | A271C | A | 1.9e-06 | 2.4e-06 | 2.6e-06 | 2.300E-06 | 3.606E-07 | 1.900E-06 | 2.600E-06 |
| 5406 | 271 | A271E | A | 1.8e-06 | 1.9e-06 | 2.6e-06 | 2.100E-06 | 4.359E-07 | 1.800E-06 | 2.600E-06 |
| 5407 | 271 | A271Q | A | 3.2e-06 | 1.6e-06 | 2.9e-06 | 2.567E-06 | 8.505E-07 | 1.600E-06 | 3.200E-06 |
| 5408 | 271 | A271G | A | 2.3e-06 | 2.9e-06 | 2.6e-06 | 2.600E-06 | 3.000E-07 | 2.300E-06 | 2.900E-06 |
| 5409 | 271 | A271H | A | 2.7e-06 | 3.4e-06 | 2.8e-06 | 2.967E-06 | 3.786E-07 | 2.700E-06 | 3.400E-06 |
| 5410 | 271 | A271I | A | 3.2e-06 | 2e-06 | 2.8e-06 | 2.667E-06 | 6.110E-07 | 2.000E-06 | 3.200E-06 |
| 5411 | 271 | A271L | A | 2.4e-06 | 2.3e-06 | 4.3e-06 | 3.000E-06 | 1.127E-06 | 2.300E-06 | 4.300E-06 |
| 5412 | 271 | A271K | A | 2.7e-06 | 1.9e-06 | 2.3e-06 | 2.300E-06 | 4.000E-07 | 1.900E-06 | 2.700E-06 |
| 5413 | 271 | A271M | A | 1.9e-06 | 2.2e-06 | 1.8e-06 | 1.967E-06 | 2.082E-07 | 1.800E-06 | 2.200E-06 |
| 5414 | 271 | A271F | A | 3.5e-06 | 3.1e-06 | 3.4e-06 | 3.333E-06 | 2.082E-07 | 3.100E-06 | 3.500E-06 |
| 5415 | 271 | A271P | A | 2.2e-06 | 2.5e-06 | 2.1e-06 | 2.267E-06 | 2.082E-07 | 2.100E-06 | 2.500E-06 |
| 5416 | 271 | A271S | A | 2.5e-06 | 4.3e-06 | 2.9e-06 | 3.233E-06 | 9.452E-07 | 2.500E-06 | 4.300E-06 |
| 5417 | 271 | A271T | A | 2.7e-06 | 2.7e-06 | 3.4e-06 | 2.933E-06 | 4.041E-07 | 2.700E-06 | 3.400E-06 |
| 5418 | 271 | A271W | A | 2.4e-06 | 2e-06 | 2.2e-06 | 2.200E-06 | 2.000E-07 | 2.000E-06 | 2.400E-06 |
| 5419 | 271 | A271Y | A | 4e-06 | 2.5e-06 | 1.7e-06 | 2.733E-06 | 1.168E-06 | 1.700E-06 | 4.000E-06 |
| 5420 | 271 | A271V | A | 1.7e-06 | 1.9e-06 | 2.9e-06 | 2.167E-06 | 6.429E-07 | 1.700E-06 | 2.900E-06 |
| 5421 | 272 | S272A | A | 4e-06 | 2.8e-06 | 2.5e-06 | 3.100E-06 | 7.937E-07 | 2.500E-06 | 4.000E-06 |
| 5422 | 272 | S272R | A | 2.4e-06 | 2.3e-06 | 3.2e-06 | 2.633E-06 | 4.933E-07 | 2.300E-06 | 3.200E-06 |
| 5423 | 272 | S272N | A | 2.1e-06 | 3.2e-06 | 4.4e-06 | 3.233E-06 | 1.150E-06 | 2.100E-06 | 4.400E-06 |
| 5424 | 272 | S272D | A | 4.4e-06 | 2.2e-06 | 3.2e-06 | 3.267E-06 | 1.102E-06 | 2.200E-06 | 4.400E-06 |
| 5425 | 272 | S272C | A | 3.3e-06 | 2.3e-06 | 2.6e-06 | 2.733E-06 | 5.132E-07 | 2.300E-06 | 3.300E-06 |
| 5426 | 272 | S272E | A | 2.1e-06 | 2.9e-06 | 3.3e-06 | 2.767E-06 | 6.110E-07 | 2.100E-06 | 3.300E-06 |
| 5427 | 272 | S272Q | A | 3.2e-06 | 2.1e-06 | 2.8e-06 | 2.700E-06 | 5.568E-07 | 2.100E-06 | 3.200E-06 |
| 5428 | 272 | S272G | A | 2.7e-06 | 2.1e-06 | 1.9e-06 | 2.233E-06 | 4.163E-07 | 1.900E-06 | 2.700E-06 |
| 5429 | 272 | S272H | A | 3.3e-06 | 2.8e-06 | 3.9e-06 | 3.333E-06 | 5.508E-07 | 2.800E-06 | 3.900E-06 |
| 5430 | 272 | S272I | A | 3.5e-06 | 2.8e-06 | 2.7e-06 | 3.000E-06 | 4.359E-07 | 2.700E-06 | 3.500E-06 |
| 5431 | 272 | S272L | A | 2.1e-06 | 2.7e-06 | 1.7e-06 | 2.167E-06 | 5.033E-07 | 1.700E-06 | 2.700E-06 |
| 5432 | 272 | S272K | A | 3.4e-06 | 3.6e-06 | 3.5e-06 | 3.500E-06 | 1.000E-07 | 3.400E-06 | 3.600E-06 |
| 5433 | 272 | S272M | A | 2e-06 | 2.9e-06 | 2.1e-06 | 2.333E-06 | 4.933E-07 | 2.000E-06 | 2.900E-06 |
| 5434 | 272 | S272F | A | 3.6e-06 | 2.1e-06 | 2.3e-06 | 2.667E-06 | 8.145E-07 | 2.100E-06 | 3.600E-06 |
| 5435 | 272 | S272P | A | 1.3e-06 | 2.4e-06 | 2e-06 | 1.900E-06 | 5.568E-07 | 1.300E-06 | 2.400E-06 |
| 5436 | 272 | S272S | A | 1.2e-06 | 2.7e-06 | 1.1e-06 | 1.667E-06 | 8.963E-07 | 1.100E-06 | 2.700E-06 |
| 5437 | 272 | S272T | A | 2.1e-06 | 4e-06 | 1.9e-06 | 2.667E-06 | 1.159E-06 | 1.900E-06 | 4.000E-06 |
| 5438 | 272 | S272W | A | 2.3e-06 | 1.9e-06 | 5.2e-06 | 3.133E-06 | 1.801E-06 | 1.900E-06 | 5.200E-06 |
| 5439 | 272 | S272Y | A | 2.5e-06 | 1.4e-06 | 2.8e-06 | 2.233E-06 | 7.371E-07 | 1.400E-06 | 2.800E-06 |

| | | | | | | | | | | |
|---|---|---|---|---|---|---|---|---|---|---|
| 5440 | 272 | S272V | A | 5e-06 | 2.2e-06 | 3.4e-06 | 3.533E-06 | 1.405E-06 | 2.200E-06 | 5.000E-06 |
| 5441 | 273 | C273A | A | 1.6e-06 | 2.9e-06 | 2e-06 | 2.167E-06 | 6.658E-07 | 1.600E-06 | 2.900E-06 |
| 5442 | 273 | C273R | A | 1.6e-06 | 2.1e-06 | 1.8e-06 | 1.833E-06 | 2.517E-07 | 1.600E-06 | 2.100E-06 |
| 5443 | 273 | C273N | A | 2.1e-06 | 2.5e-06 | 2.3e-06 | 2.300E-06 | 2.000E-07 | 2.100E-06 | 2.500E-06 |
| 5444 | 273 | C273D | A | 2.6e-06 | 2e-06 | 1.7e-06 | 2.100E-06 | 4.583E-07 | 1.700E-06 | 2.600E-06 |
| 5445 | 273 | C273C | A | 1.8e-06 | 2.6e-06 | 2.4e-06 | 2.267E-06 | 4.163E-07 | 1.800E-06 | 2.600E-06 |
| 5446 | 273 | C273E | A | 2.6e-06 | 2e-06 | 2.7e-06 | 2.433E-06 | 3.786E-07 | 2.000E-06 | 2.700E-06 |
| 5447 | 273 | C273Q | A | 2.3e-06 | 1.6e-06 | 1.1e-06 | 1.667E-06 | 6.028E-07 | 1.100E-06 | 2.300E-06 |
| 5448 | 273 | C273G | A | 1.7e-06 | 1.8e-06 | 2.6e-06 | 2.033E-06 | 4.933E-07 | 1.700E-06 | 2.600E-06 |
| 5449 | 273 | C273H | A | 1.4e-06 | 2.3e-06 | 1.9e-06 | 1.867E-06 | 4.509E-07 | 1.400E-06 | 2.300E-06 |
| 5450 | 273 | C273I | A | 1.8e-06 | 2.8e-06 | 4.8e-06 | 3.133E-06 | 1.528E-06 | 1.800E-06 | 4.800E-06 |
| 5451 | 273 | C273L | A | 2.5e-06 | 1.5e-06 | 2.1e-06 | 2.033E-06 | 5.033E-07 | 1.500E-06 | 2.500E-06 |
| 5452 | 273 | C273K | A | 3.1e-06 | 1.6e-06 | 2.4e-06 | 2.367E-06 | 7.506E-07 | 1.600E-06 | 3.100E-06 |
| 5453 | 273 | C273M | A | 3e-06 | 3.6e-06 | 3.7e-06 | 3.433E-06 | 3.786E-07 | 3.000E-06 | 3.700E-06 |
| 5454 | 273 | C273F | A | 2.7e-06 | 3.5e-06 | 1.6e-06 | 2.600E-06 | 9.539E-07 | 1.600E-06 | 3.500E-06 |
| 5455 | 273 | C273P | A | 3.6e-06 | 2.5e-06 | 1.3e-06 | 2.467E-06 | 1.150E-06 | 1.300E-06 | 3.600E-06 |
| 5456 | 273 | C273S | A | 2.4e-06 | 1.9e-06 | 3.3e-06 | 2.533E-06 | 7.095E-07 | 1.900E-06 | 3.300E-06 |
| 5457 | 273 | C273T | A | 9.3e-07 | 9.7e-07 | 2e-06 | 1.300E-06 | 6.065E-07 | 9.300E-07 | 2.000E-06 |
| 5458 | 273 | C273W | A | 2.2e-06 | 2.8e-06 | 2.5e-06 | 2.500E-06 | 3.000E-07 | 2.200E-06 | 2.800E-06 |
| 5459 | 273 | C273Y | A | 1.3e-06 | 4.2e-06 | 2.8e-06 | 2.767E-06 | 1.450E-06 | 1.300E-06 | 4.200E-06 |
| 5460 | 273 | C273V | A | 3.6e-06 | 1.9e-06 | 2.9e-06 | 2.800E-06 | 8.544E-07 | 1.900E-06 | 3.600E-06 |
| 5461 | 274 | V274A | A | 3.2e-06 | 2.7e-06 | 2.3e-06 | 2.733E-06 | 4.509E-07 | 2.300E-06 | 3.200E-06 |
| 5462 | 274 | V274R | A | 2.7e-06 | 3.2e-06 | 2e-06 | 2.633E-06 | 6.028E-07 | 2.000E-06 | 3.200E-06 |
| 5463 | 274 | V274N | A | 2.7e-06 | 1.2e-06 | 2e-06 | 1.967E-06 | 7.506E-07 | 1.200E-06 | 2.700E-06 |
| 5464 | 274 | V274D | A | 2.5e-06 | 1.8e-06 | 2.7e-06 | 2.333E-06 | 4.726E-07 | 1.800E-06 | 2.700E-06 |
| 5465 | 274 | V274C | A | 2.4e-06 | 9.2e-07 | 2.5e-06 | 1.940E-06 | 8.848E-07 | 9.200E-07 | 2.500E-06 |
| 5466 | 274 | V274E | A | 2e-06 | 2.5e-06 | 2.1e-06 | 2.200E-06 | 2.646E-07 | 2.000E-06 | 2.500E-06 |
| 5467 | 274 | V274Q | A | 2.1e-06 | 1.9e-06 | 2.3e-06 | 2.100E-06 | 2.000E-07 | 1.900E-06 | 2.300E-06 |
| 5468 | 274 | V274G | A | 1.7e-06 | 2.3e-06 | 1.8e-06 | 1.933E-06 | 3.215E-07 | 1.700E-06 | 2.300E-06 |
| 5469 | 274 | V274H | A | 2.1e-06 | 1.9e-06 | 2.6e-06 | 2.200E-06 | 3.606E-07 | 1.900E-06 | 2.600E-06 |
| 5470 | 274 | V274I | A | 2.4e-06 | 1.7e-06 | 1.4e-06 | 1.833E-06 | 5.132E-07 | 1.400E-06 | 2.400E-06 |
| 5471 | 274 | V274L | A | 2.7e-06 | 2.3e-06 | 1.9e-06 | 2.300E-06 | 4.000E-07 | 1.900E-06 | 2.700E-06 |
| 5472 | 274 | V274K | A | 1.6e-06 | 1.7e-06 | 4.6e-06 | 2.633E-06 | 1.704E-06 | 1.600E-06 | 4.600E-06 |
| 5473 | 274 | V274M | A | 2.6e-06 | 3.3e-06 | 2.8e-06 | 2.900E-06 | 3.606E-07 | 2.600E-06 | 3.300E-06 |
| 5474 | 274 | V274F | A | 2e-06 | 2.4e-06 | 2.5e-06 | 2.300E-06 | 2.646E-07 | 2.000E-06 | 2.500E-06 |
| 5475 | 274 | V274P | A | 1.9e-06 | 1.7e-06 | 3.2e-06 | 2.267E-06 | 8.145E-07 | 1.700E-06 | 3.200E-06 |
| 5476 | 274 | V274S | A | 1.4e-06 | 3.1e-06 | 2.1e-06 | 2.200E-06 | 8.544E-07 | 1.400E-06 | 3.100E-06 |
| 5477 | 274 | V274T | A | 3e-06 | 2.7e-06 | 2.4e-06 | 2.700E-06 | 3.000E-07 | 2.400E-06 | 3.000E-06 |
| 5478 | 274 | V274W | A | 1.8e-06 | 2.3e-06 | 2.8e-06 | 2.300E-06 | 5.000E-07 | 1.800E-06 | 2.800E-06 |
| 5479 | 274 | V274Y | A | 2.5e-06 | 2.3e-06 | 2.9e-06 | 2.567E-06 | 3.055E-07 | 2.300E-06 | 2.900E-06 |
| 5480 | 274 | V274V | A | 2.3e-06 | 2.6e-06 | 2.1e-06 | 2.333E-06 | 2.517E-07 | 2.100E-06 | 2.600E-06 |
| 5481 | 275 | T275A | A | 3.5e-06 | 2.8e-06 | 1e-06 | 2.433E-06 | 1.290E-06 | 1.000E-06 | 3.500E-06 |
| 5482 | 275 | T275R | A | 2.4e-06 | 1.9e-06 | 2.4e-06 | 2.233E-06 | 2.887E-07 | 1.900E-06 | 2.400E-06 |
| 5483 | 275 | T275N | A | 1.3e-06 | 2.3e-06 | 2.9e-06 | 2.167E-06 | 8.083E-07 | 1.300E-06 | 2.900E-06 |
| 5484 | 275 | T275D | A | 2.6e-06 | 3.2e-06 | 3.4e-06 | 3.067E-06 | 4.163E-07 | 2.600E-06 | 3.400E-06 |
| 5485 | 275 | T275C | A | 2.2e-06 | 2.6e-06 | 2.7e-06 | 2.500E-06 | 2.646E-07 | 2.200E-06 | 2.700E-06 |
| 5486 | 275 | T275E | A | 1.9e-06 | 2.2e-06 | 3.1e-06 | 2.400E-06 | 6.245E-07 | 1.900E-06 | 3.100E-06 |
| 5487 | 275 | T275Q | A | 1.8e-06 | 2.7e-06 | 2.6e-06 | 2.367E-06 | 4.933E-07 | 1.800E-06 | 2.700E-06 |
| 5488 | 275 | T275G | A | 1.9e-06 | 1.8e-06 | 3.8e-06 | 2.500E-06 | 1.127E-06 | 1.800E-06 | 3.800E-06 |
| 5489 | 275 | T275H | A | 1.1e-06 | 3.3e-06 | 2.7e-06 | 2.367E-06 | 1.137E-06 | 1.100E-06 | 3.300E-06 |
| 5490 | 275 | T275I | A | 5.9e-06 | 2.3e-06 | 2.2e-06 | 3.467E-06 | 2.108E-06 | 2.200E-06 | 5.900E-06 |
| 5491 | 275 | T275L | A | 3.2e-06 | 2.3e-06 | 2.3e-06 | 2.600E-06 | 5.196E-07 | 2.300E-06 | 3.200E-06 |
| 5492 | 275 | T275K | A | 2.8e-06 | 2.9e-06 | 2e-06 | 2.567E-06 | 4.933E-07 | 2.000E-06 | 2.900E-06 |
| 5493 | 275 | T275M | A | 1.2e-06 | 2.7e-06 | 1.7e-06 | 1.867E-06 | 7.638E-07 | 1.200E-06 | 2.700E-06 |
| 5494 | 275 | T275F | A | 2.9e-06 | 3.1e-06 | 2.7e-06 | 2.900E-06 | 2.000E-07 | 2.700E-06 | 3.100E-06 |
| 5495 | 275 | T275P | A | 2e-06 | 1.9e-06 | 2.9e-06 | 2.267E-06 | 5.508E-07 | 1.900E-06 | 2.900E-06 |
| 5496 | 275 | T275S | A | 2.4e-06 | 1.4e-06 | 2.9e-06 | 2.233E-06 | 7.638E-07 | 1.400E-06 | 2.900E-06 |
| 5497 | 275 | T275T | A | 2.2e-06 | 1.2e-06 | 1.6e-06 | 1.667E-06 | 5.033E-07 | 1.200E-06 | 2.200E-06 |
| 5498 | 275 | T275W | A | 1.1e-06 | 3.9e-06 | 2.8e-06 | 2.600E-06 | 1.411E-06 | 1.100E-06 | 3.900E-06 |
| 5499 | 275 | T275Y | A | 3.4e-06 | 1.7e-06 | 1.9e-06 | 2.333E-06 | 9.292E-07 | 1.700E-06 | 3.400E-06 |
| 5500 | 275 | T275V | A | 2.4e-06 | 1.8e-06 | 2e-06 | 2.067E-06 | 3.055E-07 | 1.800E-06 | 2.400E-06 |
| 5501 | 276 | A276A | A | 1.6e-06 | 4.7e-06 | 2e-06 | 2.767E-06 | 1.686E-06 | 1.600E-06 | 4.700E-06 |
| 5502 | 276 | A276R | A | 2.2e-06 | 3.9e-06 | 3.5e-06 | 3.200E-06 | 8.888E-07 | 2.200E-06 | 3.900E-06 |
| 5503 | 276 | A276N | A | 1.9e-06 | 2.7e-06 | 2.7e-06 | 2.433E-06 | 4.619E-07 | 1.900E-06 | 2.700E-06 |
| 5504 | 276 | A276D | A | 2.4e-06 | 3e-06 | 1.7e-06 | 2.367E-06 | 6.506E-07 | 1.700E-06 | 3.000E-06 |
| 5505 | 276 | A276C | A | 1.9e-06 | 4e-06 | 2.7e-06 | 2.867E-06 | 1.060E-06 | 1.900E-06 | 4.000E-06 |
| 5506 | 276 | A276E | A | 3.7e-06 | 3.3e-06 | 3.5e-06 | 3.500E-06 | 2.000E-07 | 3.300E-06 | 3.700E-06 |
| 5507 | 276 | A276Q | A | 9.9e-07 | 3.1e-06 | 2e-06 | 2.030E-06 | 1.055E-06 | 9.900E-07 | 3.100E-06 |
| 5508 | 276 | A276G | A | 2.8e-06 | 3e-06 | 1.8e-06 | 2.533E-06 | 6.429E-07 | 1.800E-06 | 3.000E-06 |
| 5509 | 276 | A276H | A | 2.3e-06 | 2.5e-06 | 3.5e-06 | 2.767E-06 | 6.429E-07 | 2.300E-06 | 3.500E-06 |
| 5510 | 276 | A276I | A | 5.9e-06 | 2.8e-06 | 2.4e-06 | 3.700E-06 | 1.916E-06 | 2.400E-06 | 5.900E-06 |
| 5511 | 276 | A276L | A | 3.5e-06 | 2.3e-06 | 1.4e-06 | 2.400E-06 | 1.054E-06 | 1.400E-06 | 3.500E-06 |
| 5512 | 276 | A276K | A | 2.3e-06 | 1.9e-06 | 1.5e-06 | 1.900E-06 | 4.000E-07 | 1.500E-06 | 2.300E-06 |
| 5513 | 276 | A276M | A | 3.3e-06 | 1.6e-06 | 4.2e-06 | 3.033E-06 | 1.320E-06 | 1.600E-06 | 4.200E-06 |
| 5514 | 276 | A276F | A | 1.4e-06 | 3.4e-06 | 2.8e-06 | 2.533E-06 | 1.026E-06 | 1.400E-06 | 3.400E-06 |
| 5515 | 276 | A276P | A | 2.8e-06 | 1.8e-06 | 2.7e-06 | 2.433E-06 | 5.508E-07 | 1.800E-06 | 2.800E-06 |
| 5516 | 276 | A276S | A | 2.3e-06 | 1.5e-06 | 3.3e-06 | 2.367E-06 | 9.018E-07 | 1.500E-06 | 3.300E-06 |
| 5517 | 276 | A276T | A | 2.3e-06 | 1.7e-06 | 2.3e-06 | 2.100E-06 | 3.464E-07 | 1.700E-06 | 2.300E-06 |
| 5518 | 276 | A276W | A | 1.9e-06 | 2.1e-06 | 3.5e-06 | 2.500E-06 | 8.718E-07 | 1.900E-06 | 3.500E-06 |
| 5519 | 276 | A276Y | A | 2.1e-06 | 2.6e-06 | 2.3e-06 | 2.333E-06 | 2.517E-07 | 2.100E-06 | 2.600E-06 |
| 5520 | 276 | A276V | A | 1.9e-06 | 5.4e-06 | 1.7e-06 | 3.000E-06 | 2.081E-06 | 1.700E-06 | 5.400E-06 |
| 5521 | 277 | C277A | A | 2.3e-06 | 4e-06 | 2.1e-06 | 2.800E-06 | 1.044E-06 | 2.100E-06 | 4.000E-06 |
| 5522 | 277 | C277R | A | 2.1e-06 | 5.3e-06 | 1.9e-06 | 3.100E-06 | 1.908E-06 | 1.900E-06 | 5.300E-06 |
| 5523 | 277 | C277N | A | 2.3e-06 | 1.9e-06 | 2.4e-06 | 2.200E-06 | 2.646E-07 | 1.900E-06 | 2.400E-06 |
| 5524 | 277 | C277D | A | 3.6e-06 | 3e-06 | 2.7e-06 | 3.100E-06 | 4.583E-07 | 2.700E-06 | 3.600E-06 |

| | | | | | | | | | | |
|---|---|---|---|---|---|---|---|---|---|---|
| 5525 | 277 | C277C | A | 2.6e-06 | 2.6e-06 | 3e-06 | 2.733E-06 | 2.309E-07 | 2.600E-06 | 3.000E-06 |
| 5526 | 277 | C277E | A | 1.8e-06 | 3.1e-06 | 1.8e-06 | 2.233E-06 | 7.506E-07 | 1.800E-06 | 3.100E-06 |
| 5527 | 277 | C277Q | A | 1.8e-06 | 2.8e-06 | 1.5e-06 | 2.033E-06 | 6.807E-07 | 1.500E-06 | 2.800E-06 |
| 5528 | 277 | C277G | A | 2.8e-06 | 1.3e-06 | 3.1e-06 | 2.400E-06 | 9.644E-07 | 1.300E-06 | 3.100E-06 |
| 5529 | 277 | C277H | A | 2e-06 | 1.8e-06 | 2.5e-06 | 2.100E-06 | 3.606E-07 | 1.800E-06 | 2.500E-06 |
| 5530 | 277 | C277I | A | 3e-06 | 3.2e-06 | 3.3e-06 | 3.167E-06 | 1.528E-07 | 3.000E-06 | 3.300E-06 |
| 5531 | 277 | C277L | A | 4e-06 | 2.2e-06 | 3e-06 | 3.067E-06 | 9.018E-07 | 2.200E-06 | 4.000E-06 |
| 5532 | 277 | C277K | A | 1.7e-06 | 2.5e-06 | 2.4e-06 | 2.200E-06 | 4.359E-07 | 1.700E-06 | 2.500E-06 |
| 5533 | 277 | C277M | A | 2.8e-06 | 3.1e-06 | 2.3e-06 | 2.733E-06 | 4.041E-07 | 2.300E-06 | 3.100E-06 |
| 5534 | 277 | C277F | A | 1.8e-06 | 1.4e-06 | 2.2e-06 | 1.800E-06 | 4.000E-07 | 1.400E-06 | 2.200E-06 |
| 5535 | 277 | C277P | A | 2.4e-06 | 4.4e-06 | 2.2e-06 | 3.000E-06 | 1.217E-06 | 2.200E-06 | 4.400E-06 |
| 5536 | 277 | C277S | A | 3e-06 | 2.1e-06 | 5.2e-06 | 3.433E-06 | 1.595E-06 | 2.100E-06 | 5.200E-06 |
| 5537 | 277 | C277T | A | 3e-06 | 1.9e-06 | 2.2e-06 | 2.367E-06 | 5.686E-07 | 1.900E-06 | 3.000E-06 |
| 5538 | 277 | C277W | A | 2.7e-06 | 1.9e-06 | 2.6e-06 | 2.400E-06 | 4.359E-07 | 1.900E-06 | 2.700E-06 |
| 5539 | 277 | C277Y | A | 5e-06 | 3.3e-06 | 2.8e-06 | 3.700E-06 | 1.153E-06 | 2.800E-06 | 5.000E-06 |
| 5540 | 277 | C277V | A | 1.9e-06 | 2.8e-06 | 3.9e-06 | 2.867E-06 | 1.002E-06 | 1.900E-06 | 3.900E-06 |
| 5541 | 278 | P278A | A | 2.9e-06 | 3.8e-06 | 1.2e-06 | 2.633E-06 | 1.320E-06 | 1.200E-06 | 3.800E-06 |
| 5542 | 278 | P278R | A | 2.5e-06 | 1.3e-06 | 4.5e-06 | 2.767E-06 | 1.617E-06 | 1.300E-06 | 4.500E-06 |
| 5543 | 278 | P278N | A | 1.7e-06 | 2.7e-06 | 1.8e-06 | 2.067E-06 | 5.508E-07 | 1.700E-06 | 2.700E-06 |
| 5544 | 278 | P278D | A | 2.3e-06 | 3.2e-06 | 2.9e-06 | 2.800E-06 | 4.583E-07 | 2.300E-06 | 3.200E-06 |
| 5545 | 278 | P278C | A | 2.4e-06 | 3.1e-06 | 3.6e-06 | 3.033E-06 | 6.028E-07 | 2.400E-06 | 3.600E-06 |
| 5546 | 278 | P278E | A | 3.3e-06 | 2e-06 | 2e-06 | 2.433E-06 | 7.506E-07 | 2.000E-06 | 3.300E-06 |
| 5547 | 278 | P278Q | A | 2.1e-06 | 2.1e-06 | 3.2e-06 | 2.467E-06 | 6.351E-07 | 2.100E-06 | 3.200E-06 |
| 5548 | 278 | P278G | A | 2e-06 | 2.9e-06 | 1.6e-06 | 2.167E-06 | 6.658E-07 | 1.600E-06 | 2.900E-06 |
| 5549 | 278 | P278H | A | 1.6e-06 | 1.8e-06 | 2.9e-06 | 2.100E-06 | 7.000E-07 | 1.600E-06 | 2.900E-06 |
| 5550 | 278 | P278I | A | 1.6e-06 | 2.9e-06 | 2.8e-06 | 2.433E-06 | 7.234E-07 | 1.600E-06 | 2.900E-06 |
| 5551 | 278 | P278L | A | 1.2e-06 | 4e-06 | 2.5e-06 | 2.567E-06 | 1.401E-06 | 1.200E-06 | 4.000E-06 |
| 5552 | 278 | P278K | A | 3.1e-06 | 2.1e-06 | 3.5e-06 | 2.900E-06 | 7.211E-07 | 2.100E-06 | 3.500E-06 |
| 5553 | 278 | P278M | A | 1.9e-06 | 3e-06 | 1.8e-06 | 2.233E-06 | 6.658E-07 | 1.800E-06 | 3.000E-06 |
| 5554 | 278 | P278F | A | 2.7e-06 | 2e-06 | 2e-06 | 2.233E-06 | 4.041E-07 | 2.000E-06 | 2.700E-06 |
| 5555 | 278 | P278P | A | 3.8e-06 | 2.8e-06 | 2.2e-06 | 2.933E-06 | 8.083E-07 | 2.200E-06 | 3.800E-06 |
| 5556 | 278 | P278S | A | 2.7e-06 | 1.8e-06 | 2.7e-06 | 2.400E-06 | 5.196E-07 | 1.800E-06 | 2.700E-06 |
| 5557 | 278 | P278T | A | 1.5e-06 | 2.3e-06 | 1.2e-06 | 1.667E-06 | 5.686E-07 | 1.200E-06 | 2.300E-06 |
| 5558 | 278 | P278W | A | 2.7e-06 | 2.1e-06 | 2.9e-06 | 2.567E-06 | 4.163E-07 | 2.100E-06 | 2.900E-06 |
| 5559 | 278 | P278Y | A | 1.1e-06 | 1.6e-06 | 2.2e-06 | 1.633E-06 | 5.508E-07 | 1.100E-06 | 2.200E-06 |
| 5560 | 278 | P278V | A | 2.8e-06 | 2.5e-06 | 2.4e-06 | 2.567E-06 | 2.082E-07 | 2.400E-06 | 2.800E-06 |
| 5561 | 279 | Y279A | A | 2.5e-06 | 2.7e-06 | 2e-06 | 2.400E-06 | 3.606E-07 | 2.000E-06 | 2.700E-06 |
| 5562 | 279 | Y279R | A | 2.1e-06 | 1.4e-06 | 2.2e-06 | 1.900E-06 | 4.359E-07 | 1.400E-06 | 2.200E-06 |
| 5563 | 279 | Y279N | A | 1.6e-06 | 2.3e-06 | 3.6e-06 | 2.500E-06 | 1.015E-06 | 1.600E-06 | 3.600E-06 |
| 5564 | 279 | Y279D | A | 2.4e-06 | 2.8e-06 | 2.9e-06 | 2.700E-06 | 2.646E-07 | 2.400E-06 | 2.900E-06 |
| 5565 | 279 | Y279C | A | 2.7e-06 | 2.8e-06 | 2.5e-06 | 2.667E-06 | 1.528E-07 | 2.500E-06 | 2.800E-06 |
| 5566 | 279 | Y279E | A | 2.2e-06 | 3.3e-06 | 3.5e-06 | 3.000E-06 | 7.000E-07 | 2.200E-06 | 3.500E-06 |
| 5567 | 279 | Y279Q | A | 3e-06 | 1.8e-06 | 2e-06 | 2.267E-06 | 6.429E-07 | 1.800E-06 | 3.000E-06 |
| 5568 | 279 | Y279G | A | 2.5e-06 | 1.5e-06 | 2e-06 | 2.000E-06 | 5.000E-07 | 1.500E-06 | 2.500E-06 |
| 5569 | 279 | Y279H | A | 1.9e-06 | 1.8e-06 | 3.3e-06 | 2.333E-06 | 8.386E-07 | 1.800E-06 | 3.300E-06 |
| 5570 | 279 | Y279I | A | 1.7e-06 | 2.3e-06 | 2.6e-06 | 2.200E-06 | 4.583E-07 | 1.700E-06 | 2.600E-06 |
| 5571 | 279 | Y279L | A | 3e-06 | 1.7e-06 | 1.5e-06 | 2.067E-06 | 8.145E-07 | 1.500E-06 | 3.000E-06 |
| 5572 | 279 | Y279K | A | 3.2e-06 | 1.8e-06 | 2.8e-06 | 2.600E-06 | 7.211E-07 | 1.800E-06 | 3.200E-06 |
| 5573 | 279 | Y279M | A | 2.5e-06 | 3e-06 | 3e-06 | 2.833E-06 | 2.887E-07 | 2.500E-06 | 3.000E-06 |
| 5574 | 279 | Y279F | A | 3.5e-06 | 2.1e-06 | 2.7e-06 | 2.767E-06 | 7.024E-07 | 2.100E-06 | 3.500E-06 |
| 5575 | 279 | Y279P | A | 1.1e-06 | 2.7e-06 | 2.8e-06 | 2.200E-06 | 9.539E-07 | 1.100E-06 | 2.800E-06 |
| 5576 | 279 | Y279S | A | 2.9e-06 | 2.8e-06 | 2.2e-06 | 2.633E-06 | 3.786E-07 | 2.200E-06 | 2.900E-06 |
| 5577 | 279 | Y279T | A | 2.7e-06 | 1.8e-06 | 2.1e-06 | 2.200E-06 | 4.583E-07 | 1.800E-06 | 2.700E-06 |
| 5578 | 279 | Y279W | A | 3.6e-06 | 5.5e-06 | 1.4e-06 | 3.500E-06 | 2.052E-06 | 1.400E-06 | 5.500E-06 |
| 5579 | 279 | Y279Y | A | 2.7e-06 | 2.1e-06 | 2.7e-06 | 2.500E-06 | 3.464E-07 | 2.100E-06 | 2.700E-06 |
| 5580 | 279 | Y279V | A | 2.1e-06 | 1.6e-06 | 2.6e-06 | 2.100E-06 | 5.000E-07 | 1.600E-06 | 2.600E-06 |
| 5581 | 280 | N280A | A | 3.6e-06 | 2.2e-06 | 1.7e-06 | 2.500E-06 | 9.849E-07 | 1.700E-06 | 3.600E-06 |
| 5582 | 280 | N280R | A | 1.5e-06 | 2e-06 | 2.9e-06 | 2.133E-06 | 7.095E-07 | 1.500E-06 | 2.900E-06 |
| 5583 | 280 | N280N | A | 2.9e-06 | 2.5e-06 | 3.1e-06 | 2.833E-06 | 3.055E-07 | 2.500E-06 | 3.100E-06 |
| 5584 | 280 | N280D | A | 2.3e-06 | 1.9e-06 | 1.5e-06 | 1.900E-06 | 4.000E-07 | 1.500E-06 | 2.300E-06 |
| 5585 | 280 | N280C | A | 3.3e-06 | 2.8e-06 | 3.5e-06 | 3.200E-06 | 3.606E-07 | 2.800E-06 | 3.500E-06 |
| 5586 | 280 | N280E | A | 2.5e-06 | 2.6e-06 | 2.1e-06 | 2.400E-06 | 2.646E-07 | 2.100E-06 | 2.600E-06 |
| 5587 | 280 | N280Q | A | 3e-06 | 2.4e-06 | 2.4e-06 | 2.600E-06 | 3.464E-07 | 2.400E-06 | 3.000E-06 |
| 5588 | 280 | N280G | A | 2.4e-06 | 1.2e-06 | 2.2e-06 | 1.933E-06 | 6.429E-07 | 1.200E-06 | 2.400E-06 |
| 5589 | 280 | N280H | A | 2.9e-06 | 3e-06 | 1.6e-06 | 2.500E-06 | 7.810E-07 | 1.600E-06 | 3.000E-06 |
| 5590 | 280 | N280I | A | 1.9e-06 | 2.1e-06 | 2.1e-06 | 2.033E-06 | 1.155E-07 | 1.900E-06 | 2.100E-06 |
| 5591 | 280 | N280L | A | 4.3e-06 | 2.6e-06 | 1.6e-06 | 2.833E-06 | 1.365E-06 | 1.600E-06 | 4.300E-06 |
| 5592 | 280 | N280K | A | 4.4e-06 | 3.6e-06 | 2.4e-06 | 3.467E-06 | 1.007E-06 | 2.400E-06 | 4.400E-06 |
| 5593 | 280 | N280M | A | 2.7e-06 | 2.6e-06 | 1.5e-06 | 2.267E-06 | 6.658E-07 | 1.500E-06 | 2.700E-06 |
| 5594 | 280 | N280F | A | 2.3e-06 | 2.2e-06 | 4e-06 | 2.833E-06 | 1.012E-06 | 2.200E-06 | 4.000E-06 |
| 5595 | 280 | N280P | A | 3e-06 | 2.4e-06 | 2.4e-06 | 2.600E-06 | 3.464E-07 | 2.400E-06 | 3.000E-06 |
| 5596 | 280 | N280S | A | 1.2e-06 | 1.3e-06 | 2.1e-06 | 1.533E-06 | 4.933E-07 | 1.200E-06 | 2.100E-06 |
| 5597 | 280 | N280T | A | 2.8e-06 | 1.3e-06 | 2.7e-06 | 2.267E-06 | 8.386E-07 | 1.300E-06 | 2.800E-06 |
| 5598 | 280 | N280W | A | 1.9e-06 | 2.5e-06 | 2.2e-06 | 2.200E-06 | 3.000E-07 | 1.900E-06 | 2.500E-06 |
| 5599 | 280 | N280Y | A | 1.3e-06 | 3e-06 | 3.8e-06 | 2.700E-06 | 1.277E-06 | 1.300E-06 | 3.800E-06 |
| 5600 | 280 | N280V | A | 2.4e-06 | 2.3e-06 | 1.3e-06 | 2.000E-06 | 6.083E-07 | 1.300E-06 | 2.400E-06 |
| 5601 | 281 | Y281A | A | 2.3e-06 | 1.8e-06 | 2.2e-06 | 2.100E-06 | 2.646E-07 | 1.800E-06 | 2.300E-06 |
| 5602 | 281 | Y281R | A | 4.6e-06 | 1.9e-06 | 3.3e-06 | 3.267E-06 | 1.350E-06 | 1.900E-06 | 4.600E-06 |
| 5603 | 281 | Y281N | A | 2.4e-06 | 1.4e-06 | 2.3e-06 | 2.033E-06 | 5.508E-07 | 1.400E-06 | 2.400E-06 |
| 5604 | 281 | Y281D | A | 1.7e-06 | 1.5e-06 | 3.9e-06 | 2.367E-06 | 1.332E-06 | 1.500E-06 | 3.900E-06 |
| 5605 | 281 | Y281C | A | 1.5e-06 | 2.6e-06 | 2e-06 | 2.033E-06 | 5.508E-07 | 1.500E-06 | 2.600E-06 |
| 5606 | 281 | Y281E | A | 3.7e-06 | 1.6e-06 | 1.9e-06 | 2.400E-06 | 1.136E-06 | 1.600E-06 | 3.700E-06 |
| 5607 | 281 | Y281Q | A | 2.2e-06 | 1.1e-06 | 1.4e-06 | 1.567E-06 | 5.686E-07 | 1.100E-06 | 2.200E-06 |
| 5608 | 281 | Y281G | A | 2.4e-06 | 1.3e-06 | 1.7e-06 | 1.800E-06 | 5.568E-07 | 1.300E-06 | 2.400E-06 |
| 5609 | 281 | Y281H | A | 2.2e-06 | 2.8e-06 | 2.1e-06 | 2.367E-06 | 3.786E-07 | 2.100E-06 | 2.800E-06 |

| | | | | | | | | | | |
|---|---|---|---|---|---|---|---|---|---|---|
| 5610 | 281 | Y281I | A | 2.1e-06 | 3.1e-06 | 4.9e-06 | 3.367E-06 | 1.419E-06 | 2.100E-06 | 4.900E-06 |
| 5611 | 281 | Y281L | A | 2.3e-06 | 1.5e-06 | 2.2e-06 | 2.000E-06 | 4.359E-07 | 1.500E-06 | 2.300E-06 |
| 5612 | 281 | Y281K | A | 2.5e-06 | 2.5e-06 | 2e-06 | 2.333E-06 | 2.887E-07 | 2.000E-06 | 2.500E-06 |
| 5613 | 281 | Y281M | A | 1.8e-06 | 2.1e-06 | 1.6e-06 | 1.833E-06 | 2.517E-07 | 1.600E-06 | 2.100E-06 |
| 5614 | 281 | Y281F | A | 2.8e-06 | 4e-06 | 1.7e-06 | 2.833E-06 | 1.150E-06 | 1.700E-06 | 4.000E-06 |
| 5615 | 281 | Y281P | A | 2e-06 | 2.2e-06 | 3.3e-06 | 2.500E-06 | 7.000E-07 | 2.000E-06 | 3.300E-06 |
| 5616 | 281 | Y281S | A | 2.1e-06 | 1.4e-06 | 1.4e-06 | 1.633E-06 | 4.041E-07 | 1.400E-06 | 2.100E-06 |
| 5617 | 281 | Y281T | A | 2.6e-06 | 2.3e-06 | 2.4e-06 | 2.433E-06 | 1.528E-07 | 2.300E-06 | 2.600E-06 |
| 5618 | 281 | Y281W | A | 3.6e-06 | 2.4e-06 | 1.5e-06 | 2.500E-06 | 1.054E-06 | 1.500E-06 | 3.600E-06 |
| 5619 | 281 | Y281Y | A | 2.5e-06 | 1.9e-06 | 2.5e-06 | 2.300E-06 | 3.464E-07 | 1.900E-06 | 2.500E-06 |
| 5620 | 281 | Y281V | A | 2.1e-06 | 1.4e-06 | 3.3e-06 | 2.267E-06 | 9.609E-07 | 1.400E-06 | 3.300E-06 |
| 5621 | 282 | L282A | A | 2.6e-06 | 1.1e-06 | 2.4e-06 | 2.033E-06 | 8.145E-07 | 1.100E-06 | 2.600E-06 |
| 5622 | 282 | L282R | A | 1.8e-06 | 4e-06 | 3.5e-06 | 3.100E-06 | 1.153E-06 | 1.800E-06 | 4.000E-06 |
| 5623 | 282 | L282N | A | 1.1e-06 | 1.7e-06 | 2.8e-06 | 1.867E-06 | 8.622E-07 | 1.100E-06 | 2.800E-06 |
| 5624 | 282 | L282D | A | 1.5e-06 | 3.1e-06 | 1.6e-06 | 2.067E-06 | 8.963E-07 | 1.500E-06 | 3.100E-06 |
| 5625 | 282 | L282C | A | 1.9e-06 | 1.6e-06 | 2.7e-06 | 2.067E-06 | 5.686E-07 | 1.600E-06 | 2.700E-06 |
| 5626 | 282 | L282E | A | 2.4e-06 | 2.3e-06 | 1.4e-06 | 2.033E-06 | 5.508E-07 | 1.400E-06 | 2.400E-06 |
| 5627 | 282 | L282Q | A | 2.3e-06 | 2.4e-06 | 2e-06 | 2.233E-06 | 2.082E-07 | 2.000E-06 | 2.400E-06 |
| 5628 | 282 | L282G | A | 2.9e-06 | 2e-06 | 2.8e-06 | 2.567E-06 | 4.933E-07 | 2.000E-06 | 2.900E-06 |
| 5629 | 282 | L282H | A | 2.6e-06 | 2.6e-06 | 1.9e-06 | 2.367E-06 | 4.041E-07 | 1.900E-06 | 2.600E-06 |
| 5630 | 282 | L282I | A | 3.2e-06 | 1.4e-06 | 2e-06 | 2.200E-06 | 9.165E-07 | 1.400E-06 | 3.200E-06 |
| 5631 | 282 | L282L | A | 2.8e-06 | 2.5e-06 | 4.5e-06 | 3.267E-06 | 1.079E-06 | 2.500E-06 | 4.500E-06 |
| 5632 | 282 | L282K | A | 1.9e-06 | 3e-06 | 1.9e-06 | 2.267E-06 | 6.351E-07 | 1.900E-06 | 3.000E-06 |
| 5633 | 282 | L282M | A | 2.8e-06 | 2.1e-06 | 2.7e-06 | 2.533E-06 | 3.786E-07 | 2.100E-06 | 2.800E-06 |
| 5634 | 282 | L282F | A | 2.4e-06 | 2.8e-06 | 2.9e-06 | 2.700E-06 | 2.646E-07 | 2.400E-06 | 2.900E-06 |
| 5635 | 282 | L282P | A | 3.4e-06 | 2.3e-06 | 1.7e-06 | 2.467E-06 | 8.622E-07 | 1.700E-06 | 3.400E-06 |
| 5636 | 282 | L282S | A | 1.7e-06 | 3.7e-06 | 3.9e-06 | 3.100E-06 | 1.217E-06 | 1.700E-06 | 3.900E-06 |
| 5637 | 282 | L282T | A | 2.7e-06 | 3.1e-06 | 2.3e-06 | 2.700E-06 | 4.000E-07 | 2.300E-06 | 3.100E-06 |
| 5638 | 282 | L282W | A | 2.5e-06 | 2.7e-06 | 3.1e-06 | 2.767E-06 | 3.055E-07 | 2.500E-06 | 3.100E-06 |
| 5639 | 282 | L282Y | A | 1.7e-06 | 1.9e-06 | 2.5e-06 | 2.033E-06 | 4.163E-07 | 1.700E-06 | 2.500E-06 |
| 5640 | 282 | L282V | A | 3.8e-06 | 1.7e-06 | 2.3e-06 | 2.600E-06 | 1.082E-06 | 1.700E-06 | 3.800E-06 |
| 5641 | 283 | S283A | A | 3.8e-06 | 1.8e-06 | 2.3e-06 | 2.633E-06 | 1.041E-06 | 1.800E-06 | 3.800E-06 |
| 5642 | 283 | S283R | A | 3.7e-06 | 1.8e-06 | 5.1e-06 | 3.533E-06 | 1.656E-06 | 1.800E-06 | 5.100E-06 |
| 5643 | 283 | S283N | A | 1.8e-06 | 4e-06 | 2e-06 | 2.600E-06 | 1.217E-06 | 1.800E-06 | 4.000E-06 |
| 5644 | 283 | S283D | A | 1.9e-06 | 1.8e-06 | 3.1e-06 | 2.267E-06 | 7.234E-07 | 1.800E-06 | 3.100E-06 |
| 5645 | 283 | S283C | A | 2.2e-06 | 1.5e-06 | 4.7e-06 | 2.800E-06 | 1.682E-06 | 1.500E-06 | 4.700E-06 |
| 5646 | 283 | S283E | A | 2.9e-06 | 1.4e-06 | 3.9e-06 | 2.733E-06 | 1.258E-06 | 1.400E-06 | 3.900E-06 |
| 5647 | 283 | S283Q | A | 1.8e-06 | 2.8e-06 | 3e-06 | 2.533E-06 | 6.429E-07 | 1.800E-06 | 3.000E-06 |
| 5648 | 283 | S283G | A | 2.1e-06 | 2e-06 | 2.6e-06 | 2.233E-06 | 3.215E-07 | 2.000E-06 | 2.600E-06 |
| 5649 | 283 | S283H | A | 5.1e-06 | 1.8e-06 | 3.3e-06 | 3.400E-06 | 1.652E-06 | 1.800E-06 | 5.100E-06 |
| 5650 | 283 | S283I | A | 2e-06 | 2.5e-06 | 2.5e-06 | 2.333E-06 | 2.887E-07 | 2.000E-06 | 2.500E-06 |
| 5651 | 283 | S283L | A | 1.2e-06 | 1.9e-06 | 2.6e-06 | 1.900E-06 | 7.000E-07 | 1.200E-06 | 2.600E-06 |
| 5652 | 283 | S283K | A | 1.9e-06 | 1.8e-06 | 2.1e-06 | 1.933E-06 | 1.528E-07 | 1.800E-06 | 2.100E-06 |
| 5653 | 283 | S283M | A | 4.5e-06 | 1.5e-06 | 1.5e-06 | 2.500E-06 | 1.732E-06 | 1.500E-06 | 4.500E-06 |
| 5654 | 283 | S283F | A | 3.9e-06 | 1.8e-06 | 3.2e-06 | 2.967E-06 | 1.069E-06 | 1.800E-06 | 3.900E-06 |
| 5655 | 283 | S283P | A | 1.6e-06 | 3.1e-06 | 1.6e-06 | 2.100E-06 | 8.660E-07 | 1.600E-06 | 3.100E-06 |
| 5656 | 283 | S283S | A | 1.8e-06 | 1.2e-06 | 2.7e-06 | 1.900E-06 | 7.550E-07 | 1.200E-06 | 2.700E-06 |
| 5657 | 283 | S283T | A | 2.8e-06 | 2.3e-06 | 2.4e-06 | 2.500E-06 | 2.646E-07 | 2.300E-06 | 2.800E-06 |
| 5658 | 283 | S283W | A | 2e-06 | 2.3e-06 | 3e-06 | 2.433E-06 | 5.132E-07 | 2.000E-06 | 3.000E-06 |
| 5659 | 283 | S283Y | A | 1.9e-06 | 2.8e-06 | 3.2e-06 | 2.633E-06 | 6.658E-07 | 1.900E-06 | 3.200E-06 |
| 5660 | 283 | S283V | A | 6.4e-06 | 1.5e-06 | 4.1e-06 | 4.000E-06 | 2.452E-06 | 1.500E-06 | 6.400E-06 |
| 5661 | 284 | T284A | A | 1.4e-06 | 4.9e-06 | 1.8e-06 | 2.700E-06 | 1.916E-06 | 1.400E-06 | 4.900E-06 |
| 5662 | 284 | T284R | A | 2.1e-06 | 2.8e-06 | 2.3e-06 | 2.400E-06 | 3.606E-07 | 2.100E-06 | 2.800E-06 |
| 5663 | 284 | T284N | A | 2.5e-06 | 2.3e-06 | 2.1e-06 | 2.300E-06 | 2.000E-07 | 2.100E-06 | 2.500E-06 |
| 5664 | 284 | T284D | A | 2.2e-06 | 4.2e-06 | 3.2e-06 | 3.200E-06 | 1.000E-06 | 2.200E-06 | 4.200E-06 |
| 5665 | 284 | T284C | A | 1.9e-06 | 3e-06 | 2.3e-06 | 2.400E-06 | 5.568E-07 | 1.900E-06 | 3.000E-06 |
| 5666 | 284 | T284E | A | 2.2e-06 | 7.3e-06 | 3.2e-06 | 4.233E-06 | 2.702E-06 | 2.200E-06 | 7.300E-06 |
| 5667 | 284 | T284Q | A | 2.8e-06 | 2.9e-06 | 1.5e-06 | 2.400E-06 | 7.810E-07 | 1.500E-06 | 2.900E-06 |
| 5668 | 284 | T284G | A | 3e-06 | 2e-06 | 1.8e-06 | 2.267E-06 | 6.429E-07 | 1.800E-06 | 3.000E-06 |
| 5669 | 284 | T284H | A | 2.5e-06 | 2.7e-06 | 2.7e-06 | 2.633E-06 | 1.155E-07 | 2.500E-06 | 2.700E-06 |
| 5670 | 284 | T284I | A | 4.4e-06 | 2.2e-06 | 3e-06 | 3.200E-06 | 1.114E-06 | 2.200E-06 | 4.400E-06 |
| 5671 | 284 | T284L | A | 2.5e-06 | 2.9e-06 | 2.6e-06 | 2.667E-06 | 2.082E-07 | 2.500E-06 | 2.900E-06 |
| 5672 | 284 | T284K | A | 1.9e-06 | 1.8e-06 | 2.5e-06 | 2.067E-06 | 3.786E-07 | 1.800E-06 | 2.500E-06 |
| 5673 | 284 | T284M | A | 1.9e-06 | 2.8e-06 | 3.1e-06 | 2.600E-06 | 6.245E-07 | 1.900E-06 | 3.100E-06 |
| 5674 | 284 | T284F | A | 2.2e-06 | 2.9e-06 | 2.2e-06 | 2.433E-06 | 4.041E-07 | 2.200E-06 | 2.900E-06 |
| 5675 | 284 | T284P | A | 2.7e-06 | 1.7e-06 | 2.2e-06 | 2.200E-06 | 5.000E-07 | 1.700E-06 | 2.700E-06 |
| 5676 | 284 | T284S | A | 2.4e-06 | 1.4e-06 | 1.6e-06 | 1.800E-06 | 5.292E-07 | 1.400E-06 | 2.400E-06 |
| 5677 | 284 | T284T | A | 2.9e-06 | 2.8e-06 | 2e-06 | 2.567E-06 | 4.933E-07 | 2.000E-06 | 2.900E-06 |
| 5678 | 284 | T284W | A | 2.8e-06 | 3.1e-06 | 2e-06 | 2.633E-06 | 5.686E-07 | 2.000E-06 | 3.100E-06 |
| 5679 | 284 | T284Y | A | 2.9e-06 | 2.9e-06 | 2.4e-06 | 2.733E-06 | 2.887E-07 | 2.400E-06 | 2.900E-06 |
| 5680 | 284 | T284V | A | 1.4e-06 | 1.9e-06 | 1.5e-06 | 1.600E-06 | 2.646E-07 | 1.400E-06 | 1.900E-06 |
| 5681 | 285 | D285A | A | 3.6e-06 | 2.3e-06 | 2.2e-06 | 2.700E-06 | 7.810E-07 | 2.200E-06 | 3.600E-06 |
| 5682 | 285 | D285R | A | 2.9e-06 | 2.6e-06 | 2.9e-06 | 2.800E-06 | 1.732E-07 | 2.600E-06 | 2.900E-06 |
| 5683 | 285 | D285N | A | 1.9e-06 | 1.8e-06 | 2.5e-06 | 2.067E-06 | 3.786E-07 | 1.800E-06 | 2.500E-06 |
| 5684 | 285 | D285D | A | 3.4e-06 | 2.1e-06 | 2.8e-06 | 2.767E-06 | 6.506E-07 | 2.100E-06 | 3.400E-06 |
| 5685 | 285 | D285C | A | 9.4e-07 | 3.4e-06 | 1.6e-06 | 1.980E-06 | 1.273E-06 | 9.400E-07 | 3.400E-06 |
| 5686 | 285 | D285E | A | 3.1e-06 | 3.6e-06 | 1.9e-06 | 2.867E-06 | 8.737E-07 | 1.900E-06 | 3.600E-06 |
| 5687 | 285 | D285Q | A | 2.2e-06 | 3.3e-06 | 3.3e-06 | 2.933E-06 | 6.351E-07 | 2.200E-06 | 3.300E-06 |
| 5688 | 285 | D285G | A | 3.1e-06 | 2.1e-06 | 1.7e-06 | 2.300E-06 | 7.211E-07 | 1.700E-06 | 3.100E-06 |
| 5689 | 285 | D285H | A | 2.6e-06 | 2.3e-06 | 3.6e-06 | 2.833E-06 | 6.807E-07 | 2.300E-06 | 3.600E-06 |
| 5690 | 285 | D285I | A | 2.3e-06 | 1.9e-06 | 2.8e-06 | 2.333E-06 | 4.509E-07 | 1.900E-06 | 2.800E-06 |
| 5691 | 285 | D285L | A | 2.8e-06 | 2.8e-06 | 3.7e-06 | 3.100E-06 | 5.196E-07 | 2.800E-06 | 3.700E-06 |
| 5692 | 285 | D285K | A | 1.8e-06 | 2.7e-06 | 2.6e-06 | 2.367E-06 | 4.933E-07 | 1.800E-06 | 2.700E-06 |
| 5693 | 285 | D285M | A | 2.9e-06 | 2e-06 | 1.9e-06 | 2.267E-06 | 5.508E-07 | 1.900E-06 | 2.900E-06 |
| 5694 | 285 | D285F | A | 4.1e-06 | 1.8e-06 | 2.8e-06 | 2.900E-06 | 1.153E-06 | 1.800E-06 | 4.100E-06 |

| | | | | | | | | | | |
|---|---|---|---|---|---|---|---|---|---|---|
| 5695 | 285 | D285P | A | 1.7e-06 | 1.7e-06 | 1.5e-06 | 1.633E-06 | 1.155E-07 | 1.500E-06 | 1.700E-06 |
| 5696 | 285 | D285S | A | 2.7e-06 | 4.7e-06 | 1.5e-06 | 2.967E-06 | 1.617E-06 | 1.500E-06 | 4.700E-06 |
| 5697 | 285 | D285T | A | 2.1e-06 | 2.7e-06 | 5.1e-06 | 3.300E-06 | 1.587E-06 | 2.100E-06 | 5.100E-06 |
| 5698 | 285 | D285W | A | 1.6e-06 | 1.8e-06 | 2.4e-06 | 1.933E-06 | 4.163E-07 | 1.600E-06 | 2.400E-06 |
| 5699 | 285 | D285Y | A | 2.2e-06 | 2.4e-06 | 2.4e-06 | 2.333E-06 | 1.155E-07 | 2.200E-06 | 2.400E-06 |
| 5700 | 285 | D285V | A | 2.1e-06 | 2.6e-06 | 2.4e-06 | 2.367E-06 | 2.517E-07 | 2.100E-06 | 2.600E-06 |
| 5701 | 286 | V286A | A | 2.6e-06 | 2.4e-06 | 1.8e-06 | 2.267E-06 | 4.163E-07 | 1.800E-06 | 2.600E-06 |
| 5702 | 286 | V286R | A | 2.6e-06 | 1.2e-06 | 1.4e-06 | 1.733E-06 | 7.572E-07 | 1.200E-06 | 2.600E-06 |
| 5703 | 286 | V286N | A | 2.4e-06 | 2.3e-06 | 2.3e-06 | 2.333E-06 | 5.774E-08 | 2.300E-06 | 2.400E-06 |
| 5704 | 286 | V286D | A | 3.1e-06 | 1.7e-06 | 3.6e-06 | 2.800E-06 | 9.849E-07 | 1.700E-06 | 3.600E-06 |
| 5705 | 286 | V286C | A | 2.5e-06 | 2.2e-06 | 2.1e-06 | 2.267E-06 | 2.082E-07 | 2.100E-06 | 2.500E-06 |
| 5706 | 286 | V286E | A | 3.6e-06 | 4.3e-06 | 1.3e-06 | 3.067E-06 | 1.570E-06 | 1.300E-06 | 4.300E-06 |
| 5707 | 286 | V286Q | A | 2.4e-06 | 2.1e-06 | 3.8e-06 | 2.767E-06 | 9.074E-07 | 2.100E-06 | 3.800E-06 |
| 5708 | 286 | V286G | A | 2.4e-06 | 1.7e-06 | 3.6e-06 | 2.567E-06 | 9.609E-07 | 1.700E-06 | 3.600E-06 |
| 5709 | 286 | V286H | A | 2e-06 | 2.3e-06 | 2.3e-06 | 2.200E-06 | 1.732E-07 | 2.000E-06 | 2.300E-06 |
| 5710 | 286 | V286I | A | 2.7e-06 | 2.1e-06 | 2.2e-06 | 2.333E-06 | 3.215E-07 | 2.100E-06 | 2.700E-06 |
| 5711 | 286 | V286L | A | 2.3e-06 | 2.4e-06 | 2.2e-06 | 2.300E-06 | 1.000E-07 | 2.200E-06 | 2.400E-06 |
| 5712 | 286 | V286K | A | 1.2e-06 | 2.4e-06 | 2.6e-06 | 2.067E-06 | 7.572E-07 | 1.200E-06 | 2.600E-06 |
| 5713 | 286 | V286M | A | 1.8e-06 | 2.1e-06 | 1.2e-06 | 1.700E-06 | 4.583E-07 | 1.200E-06 | 2.100E-06 |
| 5714 | 286 | V286F | A | 2.8e-06 | 2.4e-06 | 1.4e-06 | 2.200E-06 | 7.211E-07 | 1.400E-06 | 2.800E-06 |
| 5715 | 286 | V286P | A | 1.5e-06 | 3.4e-06 | 2.2e-06 | 2.367E-06 | 9.609E-07 | 1.500E-06 | 3.400E-06 |
| 5716 | 286 | V286S | A | 2.6e-06 | 3.2e-06 | 2.3e-06 | 2.700E-06 | 4.583E-07 | 2.300E-06 | 3.200E-06 |
| 5717 | 286 | V286T | A | 2.7e-06 | 2e-06 | 1.9e-06 | 2.200E-06 | 4.359E-07 | 1.900E-06 | 2.700E-06 |
| 5718 | 286 | V286W | A | 4.4e-06 | 2.1e-06 | 2.7e-06 | 3.067E-06 | 1.193E-06 | 2.100E-06 | 4.400E-06 |
| 5719 | 286 | V286Y | A | 4e-06 | 2.1e-06 | 2.5e-06 | 2.867E-06 | 1.002E-06 | 2.100E-06 | 4.000E-06 |
| 5720 | 286 | V286V | A | 3.4e-06 | 1.8e-06 | 1.4e-06 | 2.200E-06 | 1.058E-06 | 1.400E-06 | 3.400E-06 |
| 5721 | 287 | G287A | A | 4.5e-06 | 2.7e-06 | 3e-06 | 3.400E-06 | 9.644E-07 | 2.700E-06 | 4.500E-06 |
| 5722 | 287 | G287R | A | 1e-06 | 1.5e-06 | 2.6e-06 | 1.700E-06 | 8.185E-07 | 1.000E-06 | 2.600E-06 |
| 5723 | 287 | G287N | A | 2.7e-06 | 2.2e-06 | 2.8e-06 | 2.567E-06 | 3.215E-07 | 2.200E-06 | 2.800E-06 |
| 5724 | 287 | G287D | A | 3.6e-06 | 2.6e-06 | 5.1e-06 | 3.767E-06 | 1.258E-06 | 2.600E-06 | 5.100E-06 |
| 5725 | 287 | G287C | A | 1.2e-06 | 4.6e-06 | 1.3e-06 | 2.367E-06 | 1.935E-06 | 1.200E-06 | 4.600E-06 |
| 5726 | 287 | G287E | A | 1.6e-06 | 2.7e-06 | 2.5e-06 | 2.267E-06 | 5.859E-07 | 1.600E-06 | 2.700E-06 |
| 5727 | 287 | G287Q | A | 1.7e-06 | 1.9e-06 | 3.1e-06 | 2.233E-06 | 7.572E-07 | 1.700E-06 | 3.100E-06 |
| 5728 | 287 | G287G | A | 2.2e-06 | 3.7e-06 | 1.4e-06 | 2.433E-06 | 1.168E-06 | 1.400E-06 | 3.700E-06 |
| 5729 | 287 | G287H | A | 1.8e-06 | 1.4e-06 | 2.5e-06 | 1.900E-06 | 5.568E-07 | 1.400E-06 | 2.500E-06 |
| 5730 | 287 | G287I | A | 2e-06 | 2.5e-06 | 2.4e-06 | 2.300E-06 | 2.646E-07 | 2.000E-06 | 2.500E-06 |
| 5731 | 287 | G287L | A | 3.5e-06 | 1.3e-06 | 2.2e-06 | 2.333E-06 | 1.106E-06 | 1.300E-06 | 3.500E-06 |
| 5732 | 287 | G287K | A | 2.8e-06 | 2.1e-06 | 3e-06 | 2.633E-06 | 4.726E-07 | 2.100E-06 | 3.000E-06 |
| 5733 | 287 | G287M | A | 2.7e-06 | 3.4e-06 | 1.8e-06 | 2.633E-06 | 8.021E-07 | 1.800E-06 | 3.400E-06 |
| 5734 | 287 | G287F | A | 1.5e-06 | 2.9e-06 | 2.4e-06 | 2.267E-06 | 7.095E-07 | 1.500E-06 | 2.900E-06 |
| 5735 | 287 | G287P | A | 2.2e-06 | 1.4e-06 | 1.9e-06 | 1.833E-06 | 4.041E-07 | 1.400E-06 | 2.200E-06 |
| 5736 | 287 | G287S | A | 2.7e-06 | 1.2e-06 | 2.7e-06 | 2.200E-06 | 8.660E-07 | 1.200E-06 | 2.700E-06 |
| 5737 | 287 | G287T | A | 4e-06 | 2.2e-06 | 2.1e-06 | 2.767E-06 | 1.069E-06 | 2.100E-06 | 4.000E-06 |
| 5738 | 287 | G287W | A | 2.1e-06 | 4.3e-06 | 3.1e-06 | 3.167E-06 | 1.102E-06 | 2.100E-06 | 4.300E-06 |
| 5739 | 287 | G287Y | A | 1e-06 | 4.4e-06 | 2.1e-06 | 2.500E-06 | 1.735E-06 | 1.000E-06 | 4.400E-06 |
| 5740 | 287 | G287V | A | 1.9e-06 | 2.7e-06 | 2.2e-06 | 2.267E-06 | 4.041E-07 | 1.900E-06 | 2.700E-06 |
| 5741 | 288 | S288A | A | 4.6e-06 | 2.5e-06 | 3.1e-06 | 3.400E-06 | 1.082E-06 | 2.500E-06 | 4.600E-06 |
| 5742 | 288 | S288R | A | 2.2e-06 | 8.5e-07 | 3e-06 | 2.017E-06 | 1.087E-06 | 8.500E-07 | 3.000E-06 |
| 5743 | 288 | S288N | A | 3.8e-06 | 2.5e-06 | 2.1e-06 | 2.800E-06 | 8.888E-07 | 2.100E-06 | 3.800E-06 |
| 5744 | 288 | S288D | A | 2.6e-06 | 3e-06 | 3.3e-06 | 2.967E-06 | 3.512E-07 | 2.600E-06 | 3.300E-06 |
| 5745 | 288 | S288C | A | 2.9e-06 | 3e-06 | 2.6e-06 | 2.833E-06 | 2.082E-07 | 2.600E-06 | 3.000E-06 |
| 5746 | 288 | S288E | A | 2.3e-06 | 1.8e-06 | 3.1e-06 | 2.400E-06 | 6.557E-07 | 1.800E-06 | 3.100E-06 |
| 5747 | 288 | S288Q | A | 1.9e-06 | 4.4e-06 | 4.1e-06 | 3.467E-06 | 1.365E-06 | 1.900E-06 | 4.400E-06 |
| 5748 | 288 | S288G | A | 4e-06 | 2.5e-06 | 2.7e-06 | 3.067E-06 | 8.145E-07 | 2.500E-06 | 4.000E-06 |
| 5749 | 288 | S288H | A | 2.2e-06 | 1.4e-06 | 2.3e-06 | 1.967E-06 | 4.933E-07 | 1.400E-06 | 2.300E-06 |
| 5750 | 288 | S288I | A | 1.6e-06 | 4.5e-06 | 1.7e-06 | 2.600E-06 | 1.646E-06 | 1.600E-06 | 4.500E-06 |
| 5751 | 288 | S288L | A | 2.3e-06 | 3e-06 | 4e-06 | 3.100E-06 | 8.544E-07 | 2.300E-06 | 4.000E-06 |
| 5752 | 288 | S288K | A | 1.9e-06 | 3.4e-06 | 3.8e-06 | 3.033E-06 | 1.002E-06 | 1.900E-06 | 3.800E-06 |
| 5753 | 288 | S288M | A | 3e-06 | 2.6e-06 | 4e-06 | 3.200E-06 | 7.211E-07 | 2.600E-06 | 4.000E-06 |
| 5754 | 288 | S288F | A | 3e-06 | 1.9e-06 | 2.4e-06 | 2.433E-06 | 5.508E-07 | 1.900E-06 | 3.000E-06 |
| 5755 | 288 | S288P | A | 4.2e-06 | 8.4e-07 | 2.7e-06 | 2.580E-06 | 1.683E-06 | 8.400E-07 | 4.200E-06 |
| 5756 | 288 | S288S | A | 5.2e-06 | 1.9e-06 | 2.1e-06 | 3.067E-06 | 1.850E-06 | 1.900E-06 | 5.200E-06 |
| 5757 | 288 | S288T | A | 1.9e-06 | 4.2e-06 | 2.9e-06 | 3.000E-06 | 1.153E-06 | 1.900E-06 | 4.200E-06 |
| 5758 | 288 | S288W | A | 1.7e-06 | 3.7e-06 | 2.3e-06 | 2.567E-06 | 1.026E-06 | 1.700E-06 | 3.700E-06 |
| 5759 | 288 | S288Y | A | 2.1e-06 | 2.1e-06 | 2.3e-06 | 2.167E-06 | 1.155E-07 | 2.100E-06 | 2.300E-06 |
| 5760 | 288 | S288V | A | 3.7e-06 | 2.4e-06 | 3.7e-06 | 3.267E-06 | 7.506E-07 | 2.400E-06 | 3.700E-06 |
| 5761 | 289 | C289A | A | 2.8e-06 | 2.4e-06 | 3e-06 | 2.733E-06 | 3.055E-07 | 2.400E-06 | 3.000E-06 |
| 5762 | 289 | C289R | A | 2.6e-06 | 7.1e-07 | 3e-06 | 2.103E-06 | 1.223E-06 | 7.100E-07 | 3.000E-06 |
| 5763 | 289 | C289N | A | 1.9e-06 | 2.5e-06 | 1.8e-06 | 2.067E-06 | 3.786E-07 | 1.800E-06 | 2.500E-06 |
| 5764 | 289 | C289D | A | 2.8e-06 | 2.2e-06 | 2.1e-06 | 2.367E-06 | 3.786E-07 | 2.100E-06 | 2.800E-06 |
| 5765 | 289 | C289C | A | 1.8e-06 | 2.7e-06 | 1.6e-06 | 2.033E-06 | 5.859E-07 | 1.600E-06 | 2.700E-06 |
| 5766 | 289 | C289E | A | 2.9e-06 | 1.9e-06 | 3.2e-06 | 2.667E-06 | 6.807E-07 | 1.900E-06 | 3.200E-06 |
| 5767 | 289 | C289Q | A | 3.1e-06 | 2.1e-06 | 1.4e-06 | 2.200E-06 | 8.544E-07 | 1.400E-06 | 3.100E-06 |
| 5768 | 289 | C289G | A | 3.7e-06 | 2.5e-06 | 1.8e-06 | 2.667E-06 | 9.609E-07 | 1.800E-06 | 3.700E-06 |
| 5769 | 289 | C289H | A | 1.3e-06 | 1.8e-06 | 1.9e-06 | 1.667E-06 | 3.215E-07 | 1.300E-06 | 1.900E-06 |
| 5770 | 289 | C289I | A | 2.5e-06 | 3.4e-06 | 2.3e-06 | 2.733E-06 | 5.859E-07 | 2.300E-06 | 3.400E-06 |
| 5771 | 289 | C289L | A | 3.7e-06 | 1.9e-06 | 3e-06 | 2.867E-06 | 9.074E-07 | 1.900E-06 | 3.700E-06 |
| 5772 | 289 | C289K | A | 2e-06 | 2.4e-06 | 1.7e-06 | 2.033E-06 | 3.512E-07 | 1.700E-06 | 2.400E-06 |
| 5773 | 289 | C289M | A | 3.9e-06 | 2.2e-06 | 2.8e-06 | 2.967E-06 | 8.622E-07 | 2.200E-06 | 3.900E-06 |
| 5774 | 289 | C289F | A | 2.9e-06 | 7.7e-06 | 2.3e-06 | 4.300E-06 | 2.960E-06 | 2.300E-06 | 7.700E-06 |
| 5775 | 289 | C289P | A | 2.4e-06 | 2.1e-06 | 2.9e-06 | 2.467E-06 | 4.041E-07 | 2.100E-06 | 2.900E-06 |
| 5776 | 289 | C289S | A | 1.9e-06 | 4.8e-06 | 1.8e-06 | 2.833E-06 | 1.704E-06 | 1.800E-06 | 4.800E-06 |
| 5777 | 289 | C289T | A | 5.4e-06 | 2.3e-06 | 2.1e-06 | 3.267E-06 | 1.850E-06 | 2.100E-06 | 5.400E-06 |
| 5778 | 289 | C289W | A | 2.4e-06 | 1.7e-06 | 2e-06 | 2.033E-06 | 3.512E-07 | 1.700E-06 | 2.400E-06 |
| 5779 | 289 | C289Y | A | 2.9e-06 | 4e-06 | 2.7e-06 | 3.200E-06 | 7.000E-07 | 2.700E-06 | 4.000E-06 |

| | | | | | | | | | | |
|---|---|---|---|---|---|---|---|---|---|---|
| 5780 | 289 | C289V | A | 2.9e-06 | 2.6e-06 | 2.6e-06 | 2.700E-06 | 1.732E-07 | 2.600E-06 | 2.900E-06 |
| 5781 | 290 | T290A | A | 1.9e-06 | 1.7e-06 | 1.7e-06 | 1.767E-06 | 1.155E-07 | 1.700E-06 | 1.900E-06 |
| 5782 | 290 | T290R | A | 2.3e-06 | 2.8e-06 | 1.4e-06 | 2.167E-06 | 7.095E-07 | 1.400E-06 | 2.800E-06 |
| 5783 | 290 | T290N | A | 3.8e-06 | 2.6e-06 | 2.2e-06 | 2.867E-06 | 8.327E-07 | 2.200E-06 | 3.800E-06 |
| 5784 | 290 | T290D | A | 2.2e-06 | 2.5e-06 | 1.4e-06 | 2.033E-06 | 5.686E-07 | 1.400E-06 | 2.500E-06 |
| 5785 | 290 | T290C | A | 3.7e-06 | 3.7e-06 | 1.2e-06 | 2.867E-06 | 1.443E-06 | 1.200E-06 | 3.700E-06 |
| 5786 | 290 | T290E | A | 1.7e-06 | 1.8e-06 | 3.4e-06 | 2.300E-06 | 9.539E-07 | 1.700E-06 | 3.400E-06 |
| 5787 | 290 | T290Q | A | 3.6e-06 | 3.1e-06 | 1.9e-06 | 2.867E-06 | 8.737E-07 | 1.900E-06 | 3.600E-06 |
| 5788 | 290 | T290G | A | 4.9e-06 | 1.5e-06 | 3.3e-06 | 3.233E-06 | 1.701E-06 | 1.500E-06 | 4.900E-06 |
| 5789 | 290 | T290H | A | 2.4e-06 | 1.2e-06 | 2.7e-06 | 2.100E-06 | 7.937E-07 | 1.200E-06 | 2.700E-06 |
| 5790 | 290 | T290I | A | 1.5e-06 | 1.8e-06 | 2.2e-06 | 1.833E-06 | 3.512E-07 | 1.500E-06 | 2.200E-06 |
| 5791 | 290 | T290L | A | 3.1e-06 | 2.8e-06 | 2.2e-06 | 2.700E-06 | 4.583E-07 | 2.200E-06 | 3.100E-06 |
| 5792 | 290 | T290K | A | 1.9e-06 | 3e-06 | 4e-06 | 2.967E-06 | 1.050E-06 | 1.900E-06 | 4.000E-06 |
| 5793 | 290 | T290M | A | 3.2e-06 | 2.4e-06 | 2.2e-06 | 2.600E-06 | 5.292E-07 | 2.200E-06 | 3.200E-06 |
| 5794 | 290 | T290F | A | 3.1e-06 | 2.4e-06 | 2.6e-06 | 2.700E-06 | 3.606E-07 | 2.400E-06 | 3.100E-06 |
| 5795 | 290 | T290P | A | 3.4e-06 | 2.9e-06 | 3.3e-06 | 3.200E-06 | 2.646E-07 | 2.900E-06 | 3.400E-06 |
| 5796 | 290 | T290S | A | 1.2e-06 | 2.3e-06 | 2.2e-06 | 1.900E-06 | 6.083E-07 | 1.200E-06 | 2.300E-06 |
| 5797 | 290 | T290T | A | 2.6e-06 | 2.4e-06 | 2.4e-06 | 2.467E-06 | 1.155E-07 | 2.400E-06 | 2.600E-06 |
| 5798 | 290 | T290W | A | 1.8e-06 | 1.9e-06 | 3.5e-06 | 2.400E-06 | 9.539E-07 | 1.800E-06 | 3.500E-06 |
| 5799 | 290 | T290Y | A | 1.5e-06 | 1.7e-06 | 1.3e-06 | 1.500E-06 | 2.000E-07 | 1.300E-06 | 1.700E-06 |
| 5800 | 290 | T290V | A | 1.7e-06 | 2.5e-06 | 2.5e-06 | 2.233E-06 | 4.619E-07 | 1.700E-06 | 2.500E-06 |
| 5801 | 291 | L291A | A | 2.2e-06 | 2.4e-06 | 1.7e-06 | 2.100E-06 | 3.606E-07 | 1.700E-06 | 2.400E-06 |
| 5802 | 291 | L291R | A | 1.5e-06 | 2.1e-06 | 2.2e-06 | 1.933E-06 | 3.786E-07 | 1.500E-06 | 2.200E-06 |
| 5803 | 291 | L291N | A | 1.9e-06 | 2.8e-06 | 2.2e-06 | 2.300E-06 | 4.583E-07 | 1.900E-06 | 2.800E-06 |
| 5804 | 291 | L291D | A | 2.3e-06 | 3.7e-06 | 2.6e-06 | 2.867E-06 | 7.371E-07 | 2.300E-06 | 3.700E-06 |
| 5805 | 291 | L291C | A | 2e-06 | 2e-06 | 1.5e-06 | 1.833E-06 | 2.887E-07 | 1.500E-06 | 2.000E-06 |
| 5806 | 291 | L291E | A | 2.2e-06 | 2.4e-06 | 1.8e-06 | 2.133E-06 | 3.055E-07 | 1.800E-06 | 2.400E-06 |
| 5807 | 291 | L291Q | A | 2e-06 | 2.1e-06 | 2.8e-06 | 2.300E-06 | 4.359E-07 | 2.000E-06 | 2.800E-06 |
| 5808 | 291 | L291G | A | 2.8e-06 | 2.1e-06 | 1.4e-06 | 2.100E-06 | 7.000E-07 | 1.400E-06 | 2.800E-06 |
| 5809 | 291 | L291H | A | 2.7e-06 | 3.6e-06 | 3.2e-06 | 3.167E-06 | 4.509E-07 | 2.700E-06 | 3.600E-06 |
| 5810 | 291 | L291I | A | 2.9e-06 | 2.3e-06 | 2.2e-06 | 2.467E-06 | 3.786E-07 | 2.200E-06 | 2.900E-06 |
| 5811 | 291 | L291L | A | 1.7e-06 | 2.9e-06 | 3.3e-06 | 2.633E-06 | 8.327E-07 | 1.700E-06 | 3.300E-06 |
| 5812 | 291 | L291K | A | 1.9e-06 | 1.8e-06 | 3.1e-06 | 2.267E-06 | 7.234E-07 | 1.800E-06 | 3.100E-06 |
| 5813 | 291 | L291M | A | 3.1e-06 | 2.3e-06 | 2.2e-06 | 2.533E-06 | 4.933E-07 | 2.200E-06 | 3.100E-06 |
| 5814 | 291 | L291F | A | 2.1e-06 | 1.6e-06 | 1.3e-06 | 1.667E-06 | 4.041E-07 | 1.300E-06 | 2.100E-06 |
| 5815 | 291 | L291P | A | 3.5e-06 | 3.5e-06 | 2e-06 | 3.000E-06 | 8.660E-07 | 2.000E-06 | 3.500E-06 |
| 5816 | 291 | L291S | A | 3.5e-06 | 2.4e-06 | 5.2e-06 | 3.700E-06 | 1.411E-06 | 2.400E-06 | 5.200E-06 |
| 5817 | 291 | L291T | A | 1.8e-06 | 2.7e-06 | 1.2e-06 | 1.900E-06 | 7.550E-07 | 1.200E-06 | 2.700E-06 |
| 5818 | 291 | L291W | A | 1.7e-06 | 1.5e-06 | 2.7e-06 | 1.967E-06 | 6.429E-07 | 1.500E-06 | 2.700E-06 |
| 5819 | 291 | L291Y | A | 1.3e-06 | 1.6e-06 | 3.1e-06 | 2.000E-06 | 9.644E-07 | 1.300E-06 | 3.100E-06 |
| 5820 | 291 | L291V | A | 2.1e-06 | 1.7e-06 | 2e-06 | 1.933E-06 | 2.082E-07 | 1.700E-06 | 2.100E-06 |
| 5821 | 292 | V292A | A | 2e-06 | 1.2e-06 | 2.2e-06 | 1.800E-06 | 5.292E-07 | 1.200E-06 | 2.200E-06 |
| 5822 | 292 | V292R | A | 3.7e-06 | 3.3e-06 | 2.4e-06 | 3.133E-06 | 6.658E-07 | 2.400E-06 | 3.700E-06 |
| 5823 | 292 | V292N | A | 1.9e-06 | 1.7e-06 | 3e-06 | 2.200E-06 | 7.000E-07 | 1.700E-06 | 3.000E-06 |
| 5824 | 292 | V292D | A | 1.5e-06 | 2e-06 | 1.9e-06 | 1.800E-06 | 2.646E-07 | 1.500E-06 | 2.000E-06 |
| 5825 | 292 | V292C | A | 5.5e-06 | 3.7e-06 | 2.1e-06 | 3.767E-06 | 1.701E-06 | 2.100E-06 | 5.500E-06 |
| 5826 | 292 | V292E | A | 1.3e-06 | 3.6e-06 | 2.7e-06 | 2.533E-06 | 1.159E-06 | 1.300E-06 | 3.600E-06 |
| 5827 | 292 | V292Q | A | 1e-06 | 1.8e-06 | 6e-06 | 2.933E-06 | 2.686E-06 | 1.000E-06 | 6.000E-06 |
| 5828 | 292 | V292G | A | 2e-06 | 3.9e-06 | 2.7e-06 | 2.867E-06 | 9.609E-07 | 2.000E-06 | 3.900E-06 |
| 5829 | 292 | V292H | A | 2.4e-06 | 4.1e-06 | 2.9e-06 | 3.133E-06 | 8.737E-07 | 2.400E-06 | 4.100E-06 |
| 5830 | 292 | V292I | A | 2.8e-06 | 2.8e-06 | 3.6e-06 | 3.067E-06 | 4.619E-07 | 2.800E-06 | 3.600E-06 |
| 5831 | 292 | V292L | A | 2.4e-06 | 2.5e-06 | 2.6e-06 | 2.500E-06 | 1.000E-07 | 2.400E-06 | 2.600E-06 |
| 5832 | 292 | V292K | A | 1.2e-06 | 1.7e-06 | 2.3e-06 | 1.733E-06 | 5.508E-07 | 1.200E-06 | 2.300E-06 |
| 5833 | 292 | V292M | A | 2.1e-06 | 2.4e-06 | 3.7e-06 | 2.733E-06 | 8.505E-07 | 2.100E-06 | 3.700E-06 |
| 5834 | 292 | V292F | A | 2.3e-06 | 2.3e-06 | 3.6e-06 | 2.733E-06 | 7.506E-07 | 2.300E-06 | 3.600E-06 |
| 5835 | 292 | V292P | A | 4.7e-06 | 1.8e-06 | 2.6e-06 | 3.033E-06 | 1.498E-06 | 1.800E-06 | 4.700E-06 |
| 5836 | 292 | V292S | A | 2.8e-06 | 2.3e-06 | 1.4e-06 | 2.167E-06 | 7.095E-07 | 1.400E-06 | 2.800E-06 |
| 5837 | 292 | V292T | A | 2.7e-06 | 1.7e-06 | 2.3e-06 | 2.233E-06 | 5.033E-07 | 1.700E-06 | 2.700E-06 |
| 5838 | 292 | V292W | A | 1.7e-06 | 2.4e-06 | 4.7e-06 | 2.933E-06 | 1.570E-06 | 1.700E-06 | 4.700E-06 |
| 5839 | 292 | V292Y | A | 2.9e-06 | 2.6e-06 | 1.8e-06 | 2.433E-06 | 5.686E-07 | 1.800E-06 | 2.900E-06 |
| 5840 | 292 | V292V | A | 1.8e-06 | 2.7e-06 | 3.7e-06 | 2.733E-06 | 9.504E-07 | 1.800E-06 | 3.700E-06 |
| 5841 | 293 | C293A | A | 3.8e-06 | 3.6e-06 | 2.8e-06 | 3.400E-06 | 5.292E-07 | 2.800E-06 | 3.800E-06 |
| 5842 | 293 | C293R | A | 3.7e-06 | 3.4e-06 | 2.2e-06 | 3.100E-06 | 7.937E-07 | 2.200E-06 | 3.700E-06 |
| 5843 | 293 | C293N | A | 1.8e-06 | 2.8e-06 | 2.4e-06 | 2.333E-06 | 5.033E-07 | 1.800E-06 | 2.800E-06 |
| 5844 | 293 | C293D | A | 1.9e-06 | 1.9e-06 | 1.3e-06 | 1.700E-06 | 3.464E-07 | 1.300E-06 | 1.900E-06 |
| 5845 | 293 | C293C | A | 1.9e-06 | 2.2e-06 | 1.7e-06 | 1.933E-06 | 2.517E-07 | 1.700E-06 | 2.200E-06 |
| 5846 | 293 | C293E | A | 2.2e-06 | 4.9e-06 | 2.8e-06 | 3.300E-06 | 1.418E-06 | 2.200E-06 | 4.900E-06 |
| 5847 | 293 | C293Q | A | 1.7e-06 | 1e-06 | 2.3e-06 | 1.667E-06 | 6.506E-07 | 1.000E-06 | 2.300E-06 |
| 5848 | 293 | C293G | A | 2.4e-06 | 2.7e-06 | 1.5e-06 | 2.200E-06 | 6.245E-07 | 1.500E-06 | 2.700E-06 |
| 5849 | 293 | C293H | A | 1.5e-06 | 2.3e-06 | 1.5e-06 | 1.767E-06 | 4.619E-07 | 1.500E-06 | 2.300E-06 |
| 5850 | 293 | C293I | A | 2.4e-06 | 2.1e-06 | 2.2e-06 | 2.233E-06 | 1.528E-07 | 2.100E-06 | 2.400E-06 |
| 5851 | 293 | C293L | A | 3.1e-06 | 2.6e-06 | 2.8e-06 | 2.833E-06 | 2.517E-07 | 2.600E-06 | 3.100E-06 |
| 5852 | 293 | C293K | A | 3.4e-06 | 2.1e-06 | 2.6e-06 | 2.700E-06 | 6.557E-07 | 2.100E-06 | 3.400E-06 |
| 5853 | 293 | C293M | A | 2.5e-06 | 2.8e-06 | 3.6e-06 | 2.967E-06 | 5.686E-07 | 2.500E-06 | 3.600E-06 |
| 5854 | 293 | C293F | A | 2.7e-06 | 2e-06 | 2e-06 | 2.233E-06 | 4.041E-07 | 2.000E-06 | 2.700E-06 |
| 5855 | 293 | C293P | A | 2.8e-06 | 2.1e-06 | 4.4e-06 | 3.100E-06 | 1.179E-06 | 2.100E-06 | 4.400E-06 |
| 5856 | 293 | C293S | A | 1.2e-06 | 2e-06 | 2.6e-06 | 1.933E-06 | 7.024E-07 | 1.200E-06 | 2.600E-06 |
| 5857 | 293 | C293T | A | 1.5e-06 | 1.5e-06 | 1.8e-06 | 1.600E-06 | 1.732E-07 | 1.500E-06 | 1.800E-06 |
| 5858 | 293 | C293W | A | 3e-06 | 3.3e-06 | 2.2e-06 | 2.833E-06 | 5.686E-07 | 2.200E-06 | 3.300E-06 |
| 5859 | 293 | C293Y | A | 3.5e-06 | 2.9e-06 | 3.2e-06 | 3.200E-06 | 3.000E-07 | 2.900E-06 | 3.500E-06 |
| 5860 | 293 | C293V | A | 1.7e-06 | 2.5e-06 | 2e-06 | 2.067E-06 | 4.041E-07 | 1.700E-06 | 2.500E-06 |
| 5861 | 294 | P294A | A | 3.7e-06 | 2.6e-06 | 2.4e-06 | 2.900E-06 | 7.000E-07 | 2.400E-06 | 3.700E-06 |
| 5862 | 294 | P294R | A | 1.9e-06 | 2e-06 | 2.4e-06 | 2.100E-06 | 2.646E-07 | 1.900E-06 | 2.400E-06 |
| 5863 | 294 | P294N | A | 4e-06 | 2.6e-06 | 2e-06 | 2.867E-06 | 1.026E-06 | 2.000E-06 | 4.000E-06 |
| 5864 | 294 | P294D | A | 3e-06 | 2.5e-06 | 2.5e-06 | 2.667E-06 | 2.887E-07 | 2.500E-06 | 3.000E-06 |

| | | | | | | | | | | |
|---|---|---|---|---|---|---|---|---|---|---|
| 5865 | 294 | P294C | A | 4.1e-06 | 3.3e-06 | 2.2e-06 | 3.200E-06 | 9.539E-07 | 2.200E-06 | 4.100E-06 |
| 5866 | 294 | P294E | A | 2.2e-06 | 2.2e-06 | 2.4e-06 | 2.267E-06 | 1.155E-07 | 2.200E-06 | 2.400E-06 |
| 5867 | 294 | P294Q | A | 2e-06 | 1.3e-06 | 2.7e-06 | 2.000E-06 | 7.000E-07 | 1.300E-06 | 2.700E-06 |
| 5868 | 294 | P294G | A | 3.4e-06 | 3.2e-06 | 1.2e-06 | 2.600E-06 | 1.217E-06 | 1.200E-06 | 3.400E-06 |
| 5869 | 294 | P294H | A | 1.4e-06 | 1.5e-06 | 2.6e-06 | 1.833E-06 | 6.658E-07 | 1.400E-06 | 2.600E-06 |
| 5870 | 294 | P294I | A | 4e-06 | 1.1e-06 | 3.2e-06 | 2.767E-06 | 1.498E-06 | 1.100E-06 | 4.000E-06 |
| 5871 | 294 | P294L | A | 2.4e-06 | 2.6e-06 | 1.9e-06 | 2.300E-06 | 3.606E-07 | 1.900E-06 | 2.600E-06 |
| 5872 | 294 | P294K | A | 2.1e-06 | 1.6e-06 | 1.7e-06 | 1.800E-06 | 2.646E-07 | 1.600E-06 | 2.100E-06 |
| 5873 | 294 | P294M | A | 2.9e-06 | 2e-06 | 1.9e-06 | 2.267E-06 | 5.508E-07 | 1.900E-06 | 2.900E-06 |
| 5874 | 294 | P294F | A | 3e-06 | 1.9e-06 | 1.9e-06 | 2.267E-06 | 6.351E-07 | 1.900E-06 | 3.000E-06 |
| 5875 | 294 | P294P | A | 3.2e-06 | 1.7e-06 | 3.1e-06 | 2.667E-06 | 8.386E-07 | 1.700E-06 | 3.200E-06 |
| 5876 | 294 | P294S | A | 3.4e-06 | 2.5e-06 | 1.6e-06 | 2.500E-06 | 9.000E-07 | 1.600E-06 | 3.400E-06 |
| 5877 | 294 | P294T | A | 2e-06 | 2.6e-06 | 3.1e-06 | 2.567E-06 | 5.508E-07 | 2.000E-06 | 3.100E-06 |
| 5878 | 294 | P294W | A | 2.8e-06 | 2.6e-06 | 2.2e-06 | 2.533E-06 | 3.055E-07 | 2.200E-06 | 2.800E-06 |
| 5879 | 294 | P294Y | A | 2.1e-06 | 3e-06 | 3e-06 | 2.700E-06 | 5.196E-07 | 2.100E-06 | 3.000E-06 |
| 5880 | 294 | P294V | A | 1.5e-06 | 4.3e-06 | 4e-06 | 3.267E-06 | 1.537E-06 | 1.500E-06 | 4.300E-06 |
| 5881 | 295 | L295A | A | 2.6e-06 | 2.9e-06 | 2.7e-06 | 2.733E-06 | 1.528E-07 | 2.600E-06 | 2.900E-06 |
| 5882 | 295 | L295R | A | 1.8e-06 | 2.4e-06 | 1.5e-06 | 1.900E-06 | 4.583E-07 | 1.500E-06 | 2.400E-06 |
| 5883 | 295 | L295N | A | 2e-06 | 4.3e-06 | 1.7e-06 | 2.667E-06 | 1.422E-06 | 1.700E-06 | 4.300E-06 |
| 5884 | 295 | L295D | A | 2.5e-06 | 3.1e-06 | 3.6e-06 | 3.067E-06 | 5.508E-07 | 2.500E-06 | 3.600E-06 |
| 5885 | 295 | L295C | A | 2e-06 | 2.8e-06 | 2e-06 | 2.267E-06 | 4.619E-07 | 2.000E-06 | 2.800E-06 |
| 5886 | 295 | L295E | A | 3e-06 | 3e-06 | 4.4e-06 | 3.467E-06 | 8.083E-07 | 3.000E-06 | 4.400E-06 |
| 5887 | 295 | L295Q | A | 2.4e-06 | 3.3e-06 | 2.3e-06 | 2.667E-06 | 5.508E-07 | 2.300E-06 | 3.300E-06 |
| 5888 | 295 | L295G | A | 2.9e-06 | 1.3e-06 | 2.6e-06 | 2.267E-06 | 8.505E-07 | 1.300E-06 | 2.900E-06 |
| 5889 | 295 | L295H | A | 2.1e-06 | 2.2e-06 | 2.6e-06 | 2.300E-06 | 2.646E-07 | 2.100E-06 | 2.600E-06 |
| 5890 | 295 | L295I | A | 1.6e-06 | 3.1e-06 | 2.9e-06 | 2.533E-06 | 8.145E-07 | 1.600E-06 | 3.100E-06 |
| 5891 | 295 | L295L | A | 1.4e-06 | 1.5e-06 | 2.1e-06 | 1.667E-06 | 3.786E-07 | 1.400E-06 | 2.100E-06 |
| 5892 | 295 | L295K | A | 2.2e-06 | 1.8e-06 | 2.5e-06 | 2.167E-06 | 3.512E-07 | 1.800E-06 | 2.500E-06 |
| 5893 | 295 | L295M | A | 3.7e-06 | 2.8e-06 | 2.5e-06 | 3.000E-06 | 6.245E-07 | 2.500E-06 | 3.700E-06 |
| 5894 | 295 | L295F | A | 1.6e-06 | 3e-06 | 1.9e-06 | 2.167E-06 | 7.371E-07 | 1.600E-06 | 3.000E-06 |
| 5895 | 295 | L295P | A | 1.3e-06 | 1.2e-06 | 2.8e-06 | 1.767E-06 | 8.963E-07 | 1.200E-06 | 2.800E-06 |
| 5896 | 295 | L295S | A | 1.7e-06 | 2.8e-06 | 3.4e-06 | 2.633E-06 | 8.622E-07 | 1.700E-06 | 3.400E-06 |
| 5897 | 295 | L295T | A | 2.9e-06 | 1.6e-06 | 2.9e-06 | 2.467E-06 | 7.506E-07 | 1.600E-06 | 2.900E-06 |
| 5898 | 295 | L295W | A | 2.7e-06 | 1.7e-06 | 4.4e-06 | 2.933E-06 | 1.365E-06 | 1.700E-06 | 4.400E-06 |
| 5899 | 295 | L295Y | A | 2.3e-06 | 3e-06 | 1.7e-06 | 2.333E-06 | 6.506E-07 | 1.700E-06 | 3.000E-06 |
| 5900 | 295 | L295V | A | 3.3e-06 | 3.9e-06 | 2.7e-06 | 3.300E-06 | 6.000E-07 | 2.700E-06 | 3.900E-06 |
| 5901 | 296 | H296A | A | 2.9e-06 | 1.5e-06 | 2.7e-06 | 2.367E-06 | 7.572E-07 | 1.500E-06 | 2.900E-06 |
| 5902 | 296 | H296R | A | 2.9e-06 | 2.2e-06 | 4.5e-06 | 3.200E-06 | 1.179E-06 | 2.200E-06 | 4.500E-06 |
| 5903 | 296 | H296N | A | 2.3e-06 | 2.6e-06 | 2.3e-06 | 2.400E-06 | 1.732E-07 | 2.300E-06 | 2.600E-06 |
| 5904 | 296 | H296D | A | 3.4e-06 | 2.7e-06 | 3.1e-06 | 3.067E-06 | 3.512E-07 | 2.700E-06 | 3.400E-06 |
| 5905 | 296 | H296C | A | 2.6e-06 | 2.4e-06 | 1.1e-06 | 2.033E-06 | 8.145E-07 | 1.100E-06 | 2.600E-06 |
| 5906 | 296 | H296E | A | 2.5e-06 | 3e-06 | 3.8e-06 | 3.100E-06 | 6.557E-07 | 2.500E-06 | 3.800E-06 |
| 5907 | 296 | H296Q | A | 2.7e-06 | 1.7e-06 | 9.1e-07 | 1.770E-06 | 8.971E-07 | 9.100E-07 | 2.700E-06 |
| 5908 | 296 | H296G | A | 2e-06 | 4.5e-06 | 1.6e-06 | 2.700E-06 | 1.572E-06 | 1.600E-06 | 4.500E-06 |
| 5909 | 296 | H296H | A | 1.5e-06 | 1.6e-06 | 2.2e-06 | 1.767E-06 | 3.786E-07 | 1.500E-06 | 2.200E-06 |
| 5910 | 296 | H296I | A | 2.7e-06 | 2.7e-06 | 1.4e-06 | 2.267E-06 | 7.506E-07 | 1.400E-06 | 2.700E-06 |
| 5911 | 296 | H296L | A | 3.1e-06 | 2.9e-06 | 2.5e-06 | 2.833E-06 | 3.055E-07 | 2.500E-06 | 3.100E-06 |
| 5912 | 296 | H296K | A | 2.3e-06 | 2.3e-06 | 2.7e-06 | 2.433E-06 | 2.309E-07 | 2.300E-06 | 2.700E-06 |
| 5913 | 296 | H296M | A | 1.4e-06 | 4.8e-06 | 1.9e-06 | 2.700E-06 | 1.836E-06 | 1.400E-06 | 4.800E-06 |
| 5914 | 296 | H296F | A | 2.8e-06 | 2.8e-06 | 2.1e-06 | 2.567E-06 | 4.041E-07 | 2.100E-06 | 2.800E-06 |
| 5915 | 296 | H296P | A | 1.4e-06 | 2.5e-06 | 2.5e-06 | 2.133E-06 | 6.351E-07 | 1.400E-06 | 2.500E-06 |
| 5916 | 296 | H296S | A | 2.2e-06 | 3e-06 | 1.9e-06 | 2.367E-06 | 5.686E-07 | 1.900E-06 | 3.000E-06 |
| 5917 | 296 | H296T | A | 2.1e-06 | 1.7e-06 | 2.5e-06 | 2.100E-06 | 4.000E-07 | 1.700E-06 | 2.500E-06 |
| 5918 | 296 | H296W | A | 3.8e-06 | 3.2e-06 | 3.4e-06 | 3.467E-06 | 3.055E-07 | 3.200E-06 | 3.800E-06 |
| 5919 | 296 | H296Y | A | 2.7e-06 | 2.7e-06 | 2.5e-06 | 2.633E-06 | 1.155E-07 | 2.500E-06 | 2.700E-06 |
| 5920 | 296 | H296V | A | 2.4e-06 | 1.8e-06 | 2.9e-06 | 2.367E-06 | 5.508E-07 | 1.800E-06 | 2.900E-06 |
| 5921 | 297 | N297A | A | 3.7e-06 | 2e-06 | 2e-06 | 2.567E-06 | 9.815E-07 | 2.000E-06 | 3.700E-06 |
| 5922 | 297 | N297R | A | 2.3e-06 | 2.6e-06 | 2.9e-06 | 2.600E-06 | 3.000E-07 | 2.300E-06 | 2.900E-06 |
| 5923 | 297 | N297N | A | 2.3e-06 | 4.6e-06 | 2.6e-06 | 3.167E-06 | 1.250E-06 | 2.300E-06 | 4.600E-06 |
| 5924 | 297 | N297D | A | 3.5e-06 | 2.5e-06 | 1.6e-06 | 2.533E-06 | 9.504E-07 | 1.600E-06 | 3.500E-06 |
| 5925 | 297 | N297C | A | 3.9e-06 | 2.1e-06 | 1.9e-06 | 2.633E-06 | 1.102E-06 | 1.900E-06 | 3.900E-06 |
| 5926 | 297 | N297E | A | 1.6e-06 | 3.9e-06 | 1.6e-06 | 2.367E-06 | 1.328E-06 | 1.600E-06 | 3.900E-06 |
| 5927 | 297 | N297Q | A | 1.2e-06 | 2.9e-06 | 1.9e-06 | 2.000E-06 | 8.544E-07 | 1.200E-06 | 2.900E-06 |
| 5928 | 297 | N297G | A | 1.7e-06 | 2.7e-06 | 3.4e-06 | 2.600E-06 | 8.544E-07 | 1.700E-06 | 3.400E-06 |
| 5929 | 297 | N297H | A | 1.4e-06 | 2.2e-06 | 3e-06 | 2.200E-06 | 8.000E-07 | 1.400E-06 | 3.000E-06 |
| 5930 | 297 | N297I | A | 1.5e-06 | 2.7e-06 | 2.8e-06 | 2.333E-06 | 7.234E-07 | 1.500E-06 | 2.800E-06 |
| 5931 | 297 | N297L | A | 1.6e-06 | 3.8e-06 | 3e-06 | 2.800E-06 | 1.114E-06 | 1.600E-06 | 3.800E-06 |
| 5932 | 297 | N297K | A | 2.7e-06 | 3.6e-06 | 4e-06 | 3.433E-06 | 6.658E-07 | 2.700E-06 | 4.000E-06 |
| 5933 | 297 | N297M | A | 2.6e-06 | 3e-06 | 3.2e-06 | 2.933E-06 | 3.055E-07 | 2.600E-06 | 3.200E-06 |
| 5934 | 297 | N297F | A | 2.5e-06 | 2e-06 | 2.8e-06 | 2.433E-06 | 4.041E-07 | 2.000E-06 | 2.800E-06 |
| 5935 | 297 | N297P | A | 3.4e-06 | 3e-06 | 2.9e-06 | 3.100E-06 | 2.646E-07 | 2.900E-06 | 3.400E-06 |
| 5936 | 297 | N297S | A | 5.1e-06 | 4.9e-06 | 1.9e-06 | 3.967E-06 | 1.793E-06 | 1.900E-06 | 5.100E-06 |
| 5937 | 297 | N297T | A | 3e-06 | 2.7e-06 | 2.4e-06 | 2.700E-06 | 3.000E-07 | 2.400E-06 | 3.000E-06 |
| 5938 | 297 | N297W | A | 2.5e-06 | 2.2e-06 | 6.8e-06 | 3.833E-06 | 2.574E-06 | 2.200E-06 | 6.800E-06 |
| 5939 | 297 | N297Y | A | 4.7e-06 | 2.2e-06 | 3.6e-06 | 3.500E-06 | 1.253E-06 | 2.200E-06 | 4.700E-06 |
| 5940 | 297 | N297V | A | 3.4e-06 | 2.8e-06 | 2.4e-06 | 2.867E-06 | 5.033E-07 | 2.400E-06 | 3.400E-06 |
| 5941 | 298 | Q298A | A | 3.5e-06 | 2.3e-06 | 2.3e-06 | 2.700E-06 | 6.928E-07 | 2.300E-06 | 3.500E-06 |
| 5942 | 298 | Q298R | A | 2.3e-06 | 6.7e-06 | 2.4e-06 | 3.800E-06 | 2.512E-06 | 2.300E-06 | 6.700E-06 |
| 5943 | 298 | Q298N | A | 2.4e-06 | 1.4e-06 | 1.4e-06 | 1.733E-06 | 5.774E-07 | 1.400E-06 | 2.400E-06 |
| 5944 | 298 | Q298D | A | 1.7e-06 | 1.9e-06 | 8.5e-07 | 1.483E-06 | 5.575E-07 | 8.500E-07 | 1.900E-06 |
| 5945 | 298 | Q298C | A | 2.7e-06 | 2.7e-06 | 2.3e-06 | 2.567E-06 | 2.309E-07 | 2.300E-06 | 2.700E-06 |
| 5946 | 298 | Q298E | A | 1.8e-06 | 2.1e-06 | 1.7e-06 | 1.867E-06 | 2.082E-07 | 1.700E-06 | 2.100E-06 |
| 5947 | 298 | Q298Q | A | 3.3e-06 | 3.4e-06 | 2.9e-06 | 3.200E-06 | 2.646E-07 | 2.900E-06 | 3.400E-06 |
| 5948 | 298 | Q298G | A | 2.7e-06 | 1.9e-06 | 3.3e-06 | 2.633E-06 | 7.024E-07 | 1.900E-06 | 3.300E-06 |
| 5949 | 298 | Q298H | A | 1.3e-06 | 2.3e-06 | 2.2e-06 | 1.933E-06 | 5.508E-07 | 1.300E-06 | 2.300E-06 |

| | | | | | | | | | | |
|---|---|---|---|---|---|---|---|---|---|---|
| 5950 | 298 | Q298I | A | 2.4e-06 | 2.4e-06 | 1.7e-06 | 2.167E-06 | 4.041E-07 | 1.700E-06 | 2.400E-06 |
| 5951 | 298 | Q298L | A | 2.4e-06 | 2.9e-06 | 2.7e-06 | 2.667E-06 | 2.517E-07 | 2.400E-06 | 2.900E-06 |
| 5952 | 298 | Q298K | A | 1.6e-06 | 2.2e-06 | 4.2e-06 | 2.667E-06 | 1.361E-06 | 1.600E-06 | 4.200E-06 |
| 5953 | 298 | Q298M | A | 1.5e-06 | 2.2e-06 | 2.8e-06 | 2.167E-06 | 6.506E-07 | 1.500E-06 | 2.800E-06 |
| 5954 | 298 | Q298F | A | 3.6e-06 | 2.5e-06 | 3.9e-06 | 3.333E-06 | 7.371E-07 | 2.500E-06 | 3.900E-06 |
| 5955 | 298 | Q298P | A | 2.3e-06 | 4.4e-06 | 2.6e-06 | 3.100E-06 | 1.136E-06 | 2.300E-06 | 4.400E-06 |
| 5956 | 298 | Q298S | A | 2.3e-06 | 2e-06 | 3.5e-06 | 2.600E-06 | 7.937E-07 | 2.000E-06 | 3.500E-06 |
| 5957 | 298 | Q298T | A | 1.6e-06 | 2.2e-06 | 1.4e-06 | 1.733E-06 | 4.163E-07 | 1.400E-06 | 2.200E-06 |
| 5958 | 298 | Q298W | A | 3.6e-06 | 1.2e-06 | 1.7e-06 | 2.167E-06 | 1.266E-06 | 1.200E-06 | 3.600E-06 |
| 5959 | 298 | Q298Y | A | 1.8e-06 | 2e-06 | 4.1e-06 | 2.633E-06 | 1.274E-06 | 1.800E-06 | 4.100E-06 |
| 5960 | 298 | Q298V | A | 2.7e-06 | 2.5e-06 | 1.1e-06 | 2.100E-06 | 8.718E-07 | 1.100E-06 | 2.700E-06 |
| 5961 | 299 | E299A | A | 1.3e-06 | 1.5e-06 | 3.3e-06 | 2.033E-06 | 1.102E-06 | 1.300E-06 | 3.300E-06 |
| 5962 | 299 | E299R | A | 3.2e-06 | 3.2e-06 | 1.5e-06 | 2.633E-06 | 9.815E-07 | 1.500E-06 | 3.200E-06 |
| 5963 | 299 | E299N | A | 2.5e-06 | 2.8e-06 | 1.3e-06 | 2.200E-06 | 7.937E-07 | 1.300E-06 | 2.800E-06 |
| 5964 | 299 | E299D | A | 1.8e-06 | 1.7e-06 | 2.5e-06 | 2.000E-06 | 4.359E-07 | 1.700E-06 | 2.500E-06 |
| 5965 | 299 | E299C | A | 4e-06 | 1.3e-06 | 2.4e-06 | 2.567E-06 | 1.358E-06 | 1.300E-06 | 4.000E-06 |
| 5966 | 299 | E299E | A | 3.5e-06 | 2.9e-06 | 2.3e-06 | 2.900E-06 | 6.000E-07 | 2.300E-06 | 3.500E-06 |
| 5967 | 299 | E299Q | A | 2.4e-06 | 2.7e-06 | 2.1e-06 | 2.400E-06 | 3.000E-07 | 2.100E-06 | 2.700E-06 |
| 5968 | 299 | E299G | A | 1.7e-06 | 2.7e-06 | 2.7e-06 | 2.367E-06 | 5.774E-07 | 1.700E-06 | 2.700E-06 |
| 5969 | 299 | E299H | A | 2.9e-06 | 3.5e-06 | 2.4e-06 | 2.933E-06 | 5.508E-07 | 2.400E-06 | 3.500E-06 |
| 5970 | 299 | E299I | A | 2.7e-06 | 2.2e-06 | 2.9e-06 | 2.600E-06 | 3.606E-07 | 2.200E-06 | 2.900E-06 |
| 5971 | 299 | E299L | A | 1.4e-06 | 1.9e-06 | 1.5e-06 | 1.600E-06 | 2.646E-07 | 1.400E-06 | 1.900E-06 |
| 5972 | 299 | E299K | A | 2.5e-06 | 3.8e-06 | 3.3e-06 | 3.200E-06 | 6.557E-07 | 2.500E-06 | 3.800E-06 |
| 5973 | 299 | E299M | A | 1.5e-06 | 2.8e-06 | 2.5e-06 | 2.267E-06 | 6.807E-07 | 1.500E-06 | 2.800E-06 |
| 5974 | 299 | E299F | A | 3.3e-06 | 3.8e-06 | 1.7e-06 | 2.933E-06 | 1.097E-06 | 1.700E-06 | 3.800E-06 |
| 5975 | 299 | E299P | A | 4.4e-06 | 2.9e-06 | 2.7e-06 | 3.333E-06 | 9.292E-07 | 2.700E-06 | 4.400E-06 |
| 5976 | 299 | E299S | A | 3.5e-06 | 2.9e-06 | 1.5e-06 | 2.633E-06 | 1.026E-06 | 1.500E-06 | 3.500E-06 |
| 5977 | 299 | E299T | A | 3e-06 | 4e-06 | 2.2e-06 | 3.067E-06 | 9.018E-07 | 2.200E-06 | 4.000E-06 |
| 5978 | 299 | E299W | A | 2.7e-06 | 3.1e-06 | 2.4e-06 | 2.733E-06 | 3.512E-07 | 2.400E-06 | 3.100E-06 |
| 5979 | 299 | E299Y | A | 2.3e-06 | 2.3e-06 | 3.6e-06 | 2.733E-06 | 7.506E-07 | 2.300E-06 | 3.600E-06 |
| 5980 | 299 | E299V | A | 1.4e-06 | 2.4e-06 | 3.3e-06 | 2.367E-06 | 9.504E-07 | 1.400E-06 | 3.300E-06 |
| 5981 | 300 | V300A | A | 3e-06 | 2.5e-06 | 2.8e-06 | 2.767E-06 | 2.517E-07 | 2.500E-06 | 3.000E-06 |
| 5982 | 300 | V300R | A | 1.4e-06 | 4.4e-06 | 2.6e-06 | 2.800E-06 | 1.510E-06 | 1.400E-06 | 4.400E-06 |
| 5983 | 300 | V300N | A | 2.9e-06 | 1.9e-06 | 1.5e-06 | 2.100E-06 | 7.211E-07 | 1.500E-06 | 2.900E-06 |
| 5984 | 300 | V300D | A | 2.1e-06 | 3.5e-06 | 2.7e-06 | 2.767E-06 | 7.024E-07 | 2.100E-06 | 3.500E-06 |
| 5985 | 300 | V300C | A | 1.9e-06 | 1.5e-06 | 2.6e-06 | 2.000E-06 | 5.568E-07 | 1.500E-06 | 2.600E-06 |
| 5986 | 300 | V300E | A | 2.5e-06 | 1.9e-06 | 2.6e-06 | 2.333E-06 | 3.786E-07 | 1.900E-06 | 2.600E-06 |
| 5987 | 300 | V300Q | A | 1.7e-06 | 3.6e-06 | 1.3e-06 | 2.200E-06 | 1.229E-06 | 1.300E-06 | 3.600E-06 |
| 5988 | 300 | V300G | A | 1.8e-06 | 1.1e-06 | 1.9e-06 | 1.600E-06 | 4.359E-07 | 1.100E-06 | 1.900E-06 |
| 5989 | 300 | V300H | A | 1.7e-06 | 2.7e-06 | 2.9e-06 | 2.433E-06 | 6.429E-07 | 1.700E-06 | 2.900E-06 |
| 5990 | 300 | V300I | A | 1.6e-06 | 4.1e-06 | 1.1e-06 | 2.267E-06 | 1.607E-06 | 1.100E-06 | 4.100E-06 |
| 5991 | 300 | V300L | A | 1.9e-06 | 1.8e-06 | 3.5e-06 | 2.400E-06 | 9.539E-07 | 1.800E-06 | 3.500E-06 |
| 5992 | 300 | V300K | A | 2.4e-06 | 2.6e-06 | 1.4e-06 | 2.133E-06 | 6.429E-07 | 1.400E-06 | 2.600E-06 |
| 5993 | 300 | V300M | A | 1.2e-06 | 1.9e-06 | 2e-06 | 1.700E-06 | 4.359E-07 | 1.200E-06 | 2.000E-06 |
| 5994 | 300 | V300F | A | 2.7e-06 | 3.1e-06 | 2.3e-06 | 2.700E-06 | 4.000E-07 | 2.300E-06 | 3.100E-06 |
| 5995 | 300 | V300P | A | 1.4e-06 | 2.7e-06 | 2.7e-06 | 2.267E-06 | 7.506E-07 | 1.400E-06 | 2.700E-06 |
| 5996 | 300 | V300S | A | 1.9e-06 | 2.8e-06 | 2.2e-06 | 2.300E-06 | 4.583E-07 | 1.900E-06 | 2.800E-06 |
| 5997 | 300 | V300T | A | 3.1e-06 | 1.8e-06 | 1.8e-06 | 2.233E-06 | 7.506E-07 | 1.800E-06 | 3.100E-06 |
| 5998 | 300 | V300W | A | 2.5e-06 | 1.9e-06 | 1.7e-06 | 2.033E-06 | 4.163E-07 | 1.700E-06 | 2.500E-06 |
| 5999 | 300 | V300Y | A | 2.3e-06 | 2e-06 | 2.7e-06 | 2.333E-06 | 3.512E-07 | 2.000E-06 | 2.700E-06 |
| 6000 | 300 | V300V | A | 2.7e-06 | 2.8e-06 | 3.3e-06 | 2.933E-06 | 3.215E-07 | 2.700E-06 | 3.300E-06 |
| 6001 | 301 | T301A | A | 2.7e-06 | 2.7e-06 | 3.5e-06 | 2.967E-06 | 4.619E-07 | 2.700E-06 | 3.500E-06 |
| 6002 | 301 | T301R | A | 2.3e-06 | 2.4e-06 | 3.7e-06 | 2.800E-06 | 7.810E-07 | 2.300E-06 | 3.700E-06 |
| 6003 | 301 | T301N | A | 3.7e-06 | 1.2e-06 | 1.8e-06 | 2.233E-06 | 1.305E-06 | 1.200E-06 | 3.700E-06 |
| 6004 | 301 | T301D | A | 5.1e-06 | 2.1e-06 | 1.8e-06 | 3.000E-06 | 1.825E-06 | 1.800E-06 | 5.100E-06 |
| 6005 | 301 | T301C | A | 2.3e-06 | 2.7e-06 | 2.8e-06 | 2.600E-06 | 2.646E-07 | 2.300E-06 | 2.800E-06 |
| 6006 | 301 | T301E | A | 2e-06 | 3.7e-06 | 2.7e-06 | 2.800E-06 | 8.544E-07 | 2.000E-06 | 3.700E-06 |
| 6007 | 301 | T301Q | A | 2.1e-06 | 2.8e-06 | 2.2e-06 | 2.367E-06 | 3.786E-07 | 2.100E-06 | 2.800E-06 |
| 6008 | 301 | T301G | A | 2.9e-06 | 1.8e-06 | 2.6e-06 | 2.433E-06 | 5.686E-07 | 1.800E-06 | 2.900E-06 |
| 6009 | 301 | T301H | A | 1.7e-06 | 2.4e-06 | 2e-06 | 2.033E-06 | 3.512E-07 | 1.700E-06 | 2.400E-06 |
| 6010 | 301 | T301I | A | 2.2e-06 | 2.6e-06 | 4e-06 | 2.933E-06 | 9.452E-07 | 2.200E-06 | 4.000E-06 |
| 6011 | 301 | T301L | A | 3.3e-06 | 3.7e-06 | 1.5e-06 | 2.833E-06 | 1.172E-06 | 1.500E-06 | 3.700E-06 |
| 6012 | 301 | T301K | A | 5e-06 | 2.3e-06 | 2.4e-06 | 3.233E-06 | 1.531E-06 | 2.300E-06 | 5.000E-06 |
| 6013 | 301 | T301M | A | 1.7e-06 | 3.2e-06 | 3e-06 | 2.633E-06 | 8.145E-07 | 1.700E-06 | 3.200E-06 |
| 6014 | 301 | T301F | A | 2e-06 | 3.1e-06 | 2e-06 | 2.367E-06 | 6.351E-07 | 2.000E-06 | 3.100E-06 |
| 6015 | 301 | T301P | A | 1.6e-06 | 2.2e-06 | 1.6e-06 | 1.800E-06 | 3.464E-07 | 1.600E-06 | 2.200E-06 |
| 6016 | 301 | T301S | A | 4.2e-06 | 1.6e-06 | 3.4e-06 | 3.067E-06 | 1.332E-06 | 1.600E-06 | 4.200E-06 |
| 6017 | 301 | T301T | A | 3e-06 | 1.1e-06 | 1.9e-06 | 2.000E-06 | 9.539E-07 | 1.100E-06 | 3.000E-06 |
| 6018 | 301 | T301W | A | 5.2e-06 | 1.6e-06 | 1.6e-06 | 2.800E-06 | 2.078E-06 | 1.600E-06 | 5.200E-06 |
| 6019 | 301 | T301Y | A | 3e-06 | 2.5e-06 | 1.6e-06 | 2.367E-06 | 7.095E-07 | 1.600E-06 | 3.000E-06 |
| 6020 | 301 | T301V | A | 2.9e-06 | 2.2e-06 | 3.8e-06 | 2.967E-06 | 8.021E-07 | 2.200E-06 | 3.800E-06 |
| 6021 | 302 | A302A | A | 2.4e-06 | 1.4e-06 | 3.6e-06 | 2.467E-06 | 1.102E-06 | 1.400E-06 | 3.600E-06 |
| 6022 | 302 | A302R | A | 2.1e-06 | 2.5e-06 | 3e-06 | 2.533E-06 | 4.509E-07 | 2.100E-06 | 3.000E-06 |
| 6023 | 302 | A302N | A | 1.1e-06 | 3.3e-06 | 3.3e-06 | 2.567E-06 | 1.270E-06 | 1.100E-06 | 3.300E-06 |
| 6024 | 302 | A302D | A | 3e-06 | 2.8e-06 | 2.2e-06 | 2.667E-06 | 4.163E-07 | 2.200E-06 | 3.000E-06 |
| 6025 | 302 | A302C | A | 1.9e-06 | 2.6e-06 | 1.9e-06 | 2.133E-06 | 4.041E-07 | 1.900E-06 | 2.600E-06 |
| 6026 | 302 | A302E | A | 2.2e-06 | 3e-06 | 1.4e-06 | 2.200E-06 | 8.000E-07 | 1.400E-06 | 3.000E-06 |
| 6027 | 302 | A302Q | A | 1.1e-06 | 2.4e-06 | 4.7e-06 | 2.733E-06 | 1.823E-06 | 1.100E-06 | 4.700E-06 |
| 6028 | 302 | A302G | A | 1.9e-06 | 2.4e-06 | 2.8e-06 | 2.367E-06 | 4.509E-07 | 1.900E-06 | 2.800E-06 |
| 6029 | 302 | A302H | A | 4e-06 | 2.9e-06 | 1.9e-06 | 2.933E-06 | 1.050E-06 | 1.900E-06 | 4.000E-06 |
| 6030 | 302 | A302I | A | 2.8e-06 | 3.2e-06 | 2.3e-06 | 2.767E-06 | 4.509E-07 | 2.300E-06 | 3.200E-06 |
| 6031 | 302 | A302L | A | 3.1e-06 | 5.1e-06 | 3.6e-06 | 3.933E-06 | 1.041E-06 | 3.100E-06 | 5.100E-06 |
| 6032 | 302 | A302K | A | 2.2e-06 | 4.2e-06 | 2.4e-06 | 2.933E-06 | 1.102E-06 | 2.200E-06 | 4.200E-06 |
| 6033 | 302 | A302M | A | 3.2e-06 | 1.4e-06 | 2e-06 | 2.200E-06 | 9.165E-07 | 1.400E-06 | 3.200E-06 |
| 6034 | 302 | A302F | A | 2.3e-06 | 3.3e-06 | 2.5e-06 | 2.700E-06 | 5.292E-07 | 2.300E-06 | 3.300E-06 |

| | | | | | | | | | | |
|---|---|---|---|---|---|---|---|---|---|---|
| 6035 | 302 | A302P | A | 2.5e-06 | 1.7e-06 | 2.6e-06 | 2.267E-06 | 4.933E-07 | 1.700E-06 | 2.600E-06 |
| 6036 | 302 | A302S | A | 2.7e-06 | 2.6e-06 | 2.8e-06 | 2.700E-06 | 1.000E-07 | 2.600E-06 | 2.800E-06 |
| 6037 | 302 | A302T | A | 1.9e-06 | 1.8e-06 | 2.5e-06 | 2.067E-06 | 3.786E-07 | 1.800E-06 | 2.500E-06 |
| 6038 | 302 | A302W | A | 9.2e-07 | 2.7e-06 | 2.8e-06 | 2.140E-06 | 1.058E-06 | 9.200E-07 | 2.800E-06 |
| 6039 | 302 | A302Y | A | 2e-06 | 2.7e-06 | 4.3e-06 | 3.000E-06 | 1.179E-06 | 2.000E-06 | 4.300E-06 |
| 6040 | 302 | A302V | A | 3.2e-06 | 4.5e-06 | 2.7e-06 | 3.467E-06 | 9.292E-07 | 2.700E-06 | 4.500E-06 |
| 6041 | 303 | E303A | A | 2.6e-06 | 2.8e-06 | 1.3e-06 | 2.233E-06 | 8.145E-07 | 1.300E-06 | 2.800E-06 |
| 6042 | 303 | E303R | A | 2.9e-06 | 1.9e-06 | 1.7e-06 | 2.167E-06 | 6.429E-07 | 1.700E-06 | 2.900E-06 |
| 6043 | 303 | E303N | A | 2.9e-06 | 2.8e-06 | 3.8e-06 | 3.167E-06 | 5.508E-07 | 2.800E-06 | 3.800E-06 |
| 6044 | 303 | E303D | A | 1.7e-06 | 1.3e-06 | 1.6e-06 | 1.533E-06 | 2.082E-07 | 1.300E-06 | 1.700E-06 |
| 6045 | 303 | E303C | A | 3e-06 | 2.2e-06 | 4.8e-06 | 3.333E-06 | 1.332E-06 | 2.200E-06 | 4.800E-06 |
| 6046 | 303 | E303E | A | 2.6e-06 | 1.6e-06 | 1.9e-06 | 2.033E-06 | 5.132E-07 | 1.600E-06 | 2.600E-06 |
| 6047 | 303 | E303Q | A | 1.5e-06 | 2.6e-06 | 2e-06 | 2.033E-06 | 5.508E-07 | 1.500E-06 | 2.600E-06 |
| 6048 | 303 | E303G | A | 2.7e-06 | 2.1e-06 | 2.1e-06 | 2.300E-06 | 3.464E-07 | 2.100E-06 | 2.700E-06 |
| 6049 | 303 | E303H | A | 2.6e-06 | 1.2e-06 | 2.7e-06 | 2.167E-06 | 8.386E-07 | 1.200E-06 | 2.700E-06 |
| 6050 | 303 | E303I | A | 3.2e-06 | 1.4e-06 | 2e-06 | 2.200E-06 | 9.165E-07 | 1.400E-06 | 3.200E-06 |
| 6051 | 303 | E303L | A | 3.8e-06 | 2.2e-06 | 2.1e-06 | 2.700E-06 | 9.539E-07 | 2.100E-06 | 3.800E-06 |
| 6052 | 303 | E303K | A | 2.3e-06 | 1.6e-06 | 2.1e-06 | 2.000E-06 | 3.606E-07 | 1.600E-06 | 2.300E-06 |
| 6053 | 303 | E303M | A | 1.9e-06 | 2.8e-06 | 1.8e-06 | 2.167E-06 | 5.508E-07 | 1.800E-06 | 2.800E-06 |
| 6054 | 303 | E303F | A | 1.8e-06 | 2.3e-06 | 2.4e-06 | 2.167E-06 | 3.215E-07 | 1.800E-06 | 2.400E-06 |
| 6055 | 303 | E303P | A | 1.9e-06 | 2.2e-06 | 9.2e-07 | 1.673E-06 | 6.694E-07 | 9.200E-07 | 2.200E-06 |
| 6056 | 303 | E303S | A | 2.7e-06 | 3.5e-06 | 1.9e-06 | 2.700E-06 | 8.000E-07 | 1.900E-06 | 3.500E-06 |
| 6057 | 303 | E303T | A | 2.9e-06 | 2.2e-06 | 2.3e-06 | 2.467E-06 | 3.786E-07 | 2.200E-06 | 2.900E-06 |
| 6058 | 303 | E303W | A | 2.5e-06 | 1.5e-06 | 2.5e-06 | 2.167E-06 | 5.774E-07 | 1.500E-06 | 2.500E-06 |
| 6059 | 303 | E303Y | A | 2.4e-06 | 1.3e-06 | 2.2e-06 | 1.967E-06 | 5.859E-07 | 1.300E-06 | 2.400E-06 |
| 6060 | 303 | E303V | A | 4e-06 | 1.9e-06 | 3.2e-06 | 3.033E-06 | 1.060E-06 | 1.900E-06 | 4.000E-06 |
| 6061 | 304 | D304A | A | 2.4e-06 | 1.4e-06 | 3e-06 | 2.267E-06 | 8.083E-07 | 1.400E-06 | 3.000E-06 |
| 6062 | 304 | D304R | A | 1.7e-06 | 1.7e-06 | 2.8e-06 | 2.067E-06 | 6.351E-07 | 1.700E-06 | 2.800E-06 |
| 6063 | 304 | D304N | A | 2.1e-06 | 6.8e-06 | 3.7e-06 | 4.200E-06 | 2.390E-06 | 2.100E-06 | 6.800E-06 |
| 6064 | 304 | D304D | A | 1.8e-06 | 3.2e-06 | 1.5e-06 | 2.167E-06 | 9.074E-07 | 1.500E-06 | 3.200E-06 |
| 6065 | 304 | D304C | A | 2.1e-06 | 1.3e-06 | 3.2e-06 | 2.200E-06 | 9.539E-07 | 1.300E-06 | 3.200E-06 |
| 6066 | 304 | D304E | A | 3.6e-06 | 1.9e-06 | 1.9e-06 | 2.467E-06 | 9.815E-07 | 1.900E-06 | 3.600E-06 |
| 6067 | 304 | D304Q | A | 2.7e-06 | 1.6e-06 | 2.6e-06 | 2.300E-06 | 6.083E-07 | 1.600E-06 | 2.700E-06 |
| 6068 | 304 | D304G | A | 1.6e-06 | 2.1e-06 | 2.4e-06 | 2.033E-06 | 4.041E-07 | 1.600E-06 | 2.400E-06 |
| 6069 | 304 | D304H | A | 2.3e-06 | 2.2e-06 | 5.2e-06 | 3.233E-06 | 1.704E-06 | 2.200E-06 | 5.200E-06 |
| 6070 | 304 | D304I | A | 3.7e-06 | 2.2e-06 | 2.2e-06 | 2.700E-06 | 8.660E-07 | 2.200E-06 | 3.700E-06 |
| 6071 | 304 | D304L | A | 2e-06 | 9.9e-07 | 4e-06 | 2.330E-06 | 1.532E-06 | 9.900E-07 | 4.000E-06 |
| 6072 | 304 | D304K | A | 2.6e-06 | 2.3e-06 | 1.5e-06 | 2.133E-06 | 5.686E-07 | 1.500E-06 | 2.600E-06 |
| 6073 | 304 | D304M | A | 5.9e-06 | 2.5e-06 | 2.3e-06 | 3.567E-06 | 2.023E-06 | 2.300E-06 | 5.900E-06 |
| 6074 | 304 | D304F | A | 8.9e-07 | 1.6e-06 | 2.3e-06 | 1.597E-06 | 7.050E-07 | 8.900E-07 | 2.300E-06 |
| 6075 | 304 | D304P | A | 1.9e-06 | 2.9e-06 | 3.1e-06 | 2.633E-06 | 6.429E-07 | 1.900E-06 | 3.100E-06 |
| 6076 | 304 | D304S | A | 3.4e-06 | 1.3e-06 | 1.9e-06 | 2.200E-06 | 1.082E-06 | 1.300E-06 | 3.400E-06 |
| 6077 | 304 | D304T | A | 2.3e-06 | 2e-06 | 2.5e-06 | 2.267E-06 | 2.517E-07 | 2.000E-06 | 2.500E-06 |
| 6078 | 304 | D304W | A | 2.4e-06 | 1.9e-06 | 1.4e-06 | 1.900E-06 | 5.000E-07 | 1.400E-06 | 2.400E-06 |
| 6079 | 304 | D304Y | A | 1.1e-06 | 4.1e-06 | 2.3e-06 | 2.500E-06 | 1.510E-06 | 1.100E-06 | 4.100E-06 |
| 6080 | 304 | D304V | A | 3.6e-06 | 2.7e-06 | 1.4e-06 | 2.567E-06 | 1.106E-06 | 1.400E-06 | 3.600E-06 |
| 6081 | 305 | G305A | A | 1.9e-06 | 2.1e-06 | 1.8e-06 | 1.933E-06 | 1.528E-07 | 1.800E-06 | 2.100E-06 |
| 6082 | 305 | G305R | A | 3.3e-06 | 4.2e-06 | 2.5e-06 | 3.333E-06 | 8.505E-07 | 2.500E-06 | 4.200E-06 |
| 6083 | 305 | G305N | A | 2.3e-06 | 1.8e-06 | 1.8e-06 | 1.967E-06 | 2.887E-07 | 1.800E-06 | 2.300E-06 |
| 6084 | 305 | G305D | A | 1.7e-06 | 2.4e-06 | 2.5e-06 | 2.200E-06 | 4.359E-07 | 1.700E-06 | 2.500E-06 |
| 6085 | 305 | G305C | A | 2.1e-06 | 1.7e-06 | 1.7e-06 | 1.833E-06 | 2.309E-07 | 1.700E-06 | 2.100E-06 |
| 6086 | 305 | G305E | A | 2.3e-06 | 4.2e-06 | 2.5e-06 | 3.000E-06 | 1.044E-06 | 2.300E-06 | 4.200E-06 |
| 6087 | 305 | G305Q | A | 2.6e-06 | 2.6e-06 | 2.7e-06 | 2.633E-06 | 5.774E-08 | 2.600E-06 | 2.700E-06 |
| 6088 | 305 | G305G | A | 2.9e-06 | 1.8e-06 | 1.1e-06 | 1.933E-06 | 9.074E-07 | 1.100E-06 | 2.900E-06 |
| 6089 | 305 | G305H | A | 3.8e-06 | 1.7e-06 | 1.8e-06 | 2.433E-06 | 1.185E-06 | 1.700E-06 | 3.800E-06 |
| 6090 | 305 | G305I | A | 1.6e-06 | 3.2e-06 | 3.2e-06 | 2.667E-06 | 9.238E-07 | 1.600E-06 | 3.200E-06 |
| 6091 | 305 | G305L | A | 2.7e-06 | 5.8e-06 | 3e-06 | 3.833E-06 | 1.710E-06 | 2.700E-06 | 5.800E-06 |
| 6092 | 305 | G305K | A | 3.5e-06 | 3.3e-06 | 1.8e-06 | 2.867E-06 | 9.292E-07 | 1.800E-06 | 3.500E-06 |
| 6093 | 305 | G305M | A | 1.2e-06 | 1.4e-06 | 3.3e-06 | 1.967E-06 | 1.159E-06 | 1.200E-06 | 3.300E-06 |
| 6094 | 305 | G305F | A | 1.7e-06 | 2.9e-06 | 1.3e-06 | 1.967E-06 | 8.327E-07 | 1.300E-06 | 2.900E-06 |
| 6095 | 305 | G305P | A | 3.3e-06 | 2e-06 | 1.8e-06 | 2.367E-06 | 8.145E-07 | 1.800E-06 | 3.300E-06 |
| 6096 | 305 | G305S | A | 2.1e-06 | 1.9e-06 | 2.1e-06 | 2.033E-06 | 1.155E-07 | 1.900E-06 | 2.100E-06 |
| 6097 | 305 | G305T | A | 3e-06 | 1.9e-06 | 1.8e-06 | 2.233E-06 | 6.658E-07 | 1.800E-06 | 3.000E-06 |
| 6098 | 305 | G305W | A | 2.1e-06 | 2e-06 | 2.5e-06 | 2.200E-06 | 2.646E-07 | 2.000E-06 | 2.500E-06 |
| 6099 | 305 | G305Y | A | 3.2e-06 | 2.7e-06 | 1.8e-06 | 2.567E-06 | 7.095E-07 | 1.800E-06 | 3.200E-06 |
| 6100 | 305 | G305V | A | 2.5e-06 | 7.8e-07 | 2.6e-06 | 1.960E-06 | 1.023E-06 | 7.800E-07 | 2.600E-06 |
| 6101 | 306 | T306A | A | 2.7e-06 | 2.7e-06 | 1.5e-06 | 2.300E-06 | 6.928E-07 | 1.500E-06 | 2.700E-06 |
| 6102 | 306 | T306R | A | 4.2e-06 | 2e-06 | 2e-06 | 2.733E-06 | 1.270E-06 | 2.000E-06 | 4.200E-06 |
| 6103 | 306 | T306N | A | 2.6e-06 | 3.8e-06 | 2.6e-06 | 3.000E-06 | 6.928E-07 | 2.600E-06 | 3.800E-06 |
| 6104 | 306 | T306D | A | 1.3e-06 | 1.7e-06 | 1.8e-06 | 1.600E-06 | 2.646E-07 | 1.300E-06 | 1.800E-06 |
| 6105 | 306 | T306C | A | 1.5e-06 | 1.8e-06 | 3e-06 | 2.100E-06 | 7.937E-07 | 1.500E-06 | 3.000E-06 |
| 6106 | 306 | T306E | A | 2.3e-06 | 2.6e-06 | 2.8e-06 | 2.567E-06 | 2.517E-07 | 2.300E-06 | 2.800E-06 |
| 6107 | 306 | T306Q | A | 2.2e-06 | 1.4e-06 | 4.1e-06 | 2.567E-06 | 1.387E-06 | 1.400E-06 | 4.100E-06 |
| 6108 | 306 | T306G | A | 2.5e-06 | 4.6e-06 | 2.8e-06 | 3.300E-06 | 1.136E-06 | 2.500E-06 | 4.600E-06 |
| 6109 | 306 | T306H | A | 3e-06 | 2.9e-06 | 3.3e-06 | 3.067E-06 | 2.082E-07 | 2.900E-06 | 3.300E-06 |
| 6110 | 306 | T306I | A | 2.3e-06 | 2.5e-06 | 2.4e-06 | 2.400E-06 | 1.000E-07 | 2.300E-06 | 2.500E-06 |
| 6111 | 306 | T306L | A | 3.7e-06 | 1.7e-06 | 1.6e-06 | 2.333E-06 | 1.185E-06 | 1.600E-06 | 3.700E-06 |
| 6112 | 306 | T306K | A | 2.9e-06 | 2.2e-06 | 2e-06 | 2.367E-06 | 4.726E-07 | 2.000E-06 | 2.900E-06 |
| 6113 | 306 | T306M | A | 2.5e-06 | 1.5e-06 | 1.8e-06 | 1.933E-06 | 5.132E-07 | 1.500E-06 | 2.500E-06 |
| 6114 | 306 | T306F | A | 4.5e-06 | 1e-06 | 2e-06 | 2.500E-06 | 1.803E-06 | 1.000E-06 | 4.500E-06 |
| 6115 | 306 | T306P | A | 4.3e-06 | 2e-06 | 2e-06 | 2.767E-06 | 1.328E-06 | 2.000E-06 | 4.300E-06 |
| 6116 | 306 | T306S | A | 1.8e-06 | 2.7e-06 | 1.8e-06 | 2.100E-06 | 5.196E-07 | 1.800E-06 | 2.700E-06 |
| 6117 | 306 | T306T | A | 1.9e-06 | 2.6e-06 | 2.2e-06 | 2.233E-06 | 3.512E-07 | 1.900E-06 | 2.600E-06 |
| 6118 | 306 | T306W | A | 1.9e-06 | 3.2e-06 | 2.1e-06 | 2.400E-06 | 7.000E-07 | 1.900E-06 | 3.200E-06 |
| 6119 | 306 | T306Y | A | 2.7e-06 | 1.6e-06 | 2.6e-06 | 2.300E-06 | 6.083E-07 | 1.600E-06 | 2.700E-06 |

| | | | | | | | | | | |
|---|---|---|---|---|---|---|---|---|---|---|
| 6120 | 306 | T306V | A | 2.5e-06 | 2e-06 | 2.3e-06 | 2.267E-06 | 2.517E-07 | 2.000E-06 | 2.500E-06 |
| 6121 | 307 | Q307A | A | 1.6e-06 | 2.1e-06 | 3.1e-06 | 2.267E-06 | 7.638E-07 | 1.600E-06 | 3.100E-06 |
| 6122 | 307 | Q307R | A | 3.6e-06 | 1.8e-06 | 2.2e-06 | 2.533E-06 | 9.452E-07 | 1.800E-06 | 3.600E-06 |
| 6123 | 307 | Q307N | A | 2.2e-06 | 1.8e-06 | 3.3e-06 | 2.433E-06 | 7.767E-07 | 1.800E-06 | 3.300E-06 |
| 6124 | 307 | Q307D | A | 1.7e-06 | 5.1e-06 | 3.3e-06 | 3.367E-06 | 1.701E-06 | 1.700E-06 | 5.100E-06 |
| 6125 | 307 | Q307C | A | 2.1e-06 | 3.4e-06 | 2.4e-06 | 2.633E-06 | 6.807E-07 | 2.100E-06 | 3.400E-06 |
| 6126 | 307 | Q307E | A | 1.3e-06 | 3e-06 | 2.7e-06 | 2.333E-06 | 9.074E-07 | 1.300E-06 | 3.000E-06 |
| 6127 | 307 | Q307Q | A | 3.6e-06 | 1.2e-06 | 2.2e-06 | 2.333E-06 | 1.206E-06 | 1.200E-06 | 3.600E-06 |
| 6128 | 307 | Q307G | A | 4.2e-06 | 3.6e-06 | 2.3e-06 | 3.367E-06 | 9.713E-07 | 2.300E-06 | 4.200E-06 |
| 6129 | 307 | Q307H | A | 2e-06 | 1.3e-06 | 4.8e-06 | 2.700E-06 | 1.852E-06 | 1.300E-06 | 4.800E-06 |
| 6130 | 307 | Q307I | A | 2.9e-06 | 4e-06 | 2.7e-06 | 3.200E-06 | 7.000E-07 | 2.700E-06 | 4.000E-06 |
| 6131 | 307 | Q307L | A | 2.2e-06 | 1.3e-06 | 2.8e-06 | 2.100E-06 | 7.550E-07 | 1.300E-06 | 2.800E-06 |
| 6132 | 307 | Q307K | A | 2.8e-06 | 2.3e-06 | 2.4e-06 | 2.500E-06 | 2.646E-07 | 2.300E-06 | 2.800E-06 |
| 6133 | 307 | Q307M | A | 2.1e-06 | 2.1e-06 | 1.7e-06 | 1.967E-06 | 2.309E-07 | 1.700E-06 | 2.100E-06 |
| 6134 | 307 | Q307F | A | 2e-06 | 1.9e-06 | 1.5e-06 | 1.800E-06 | 2.646E-07 | 1.500E-06 | 2.000E-06 |
| 6135 | 307 | Q307P | A | 2.4e-06 | 1.7e-06 | 2.9e-06 | 2.333E-06 | 6.028E-07 | 1.700E-06 | 2.900E-06 |
| 6136 | 307 | Q307S | A | 2.3e-06 | 2.8e-06 | 2.3e-06 | 2.467E-06 | 2.887E-07 | 2.300E-06 | 2.800E-06 |
| 6137 | 307 | Q307T | A | 2.5e-06 | 3.6e-06 | 2.4e-06 | 2.833E-06 | 6.658E-07 | 2.400E-06 | 3.600E-06 |
| 6138 | 307 | Q307W | A | 1.8e-06 | 2.6e-06 | 1.5e-06 | 1.967E-06 | 5.686E-07 | 1.500E-06 | 2.600E-06 |
| 6139 | 307 | Q307Y | A | 2.1e-06 | 1.7e-06 | 4.5e-06 | 2.767E-06 | 1.514E-06 | 1.700E-06 | 4.500E-06 |
| 6140 | 307 | Q307V | A | 3.3e-06 | 2e-06 | 2e-06 | 2.433E-06 | 7.506E-07 | 2.000E-06 | 3.300E-06 |
| 6141 | 308 | R308A | A | 2.7e-06 | 2.8e-06 | 2.4e-06 | 2.633E-06 | 2.082E-07 | 2.400E-06 | 2.800E-06 |
| 6142 | 308 | R308R | A | 1.6e-06 | 3.3e-06 | 3.1e-06 | 2.667E-06 | 9.292E-07 | 1.600E-06 | 3.300E-06 |
| 6143 | 308 | R308N | A | 2e-06 | 3.1e-06 | 2.6e-06 | 2.567E-06 | 5.508E-07 | 2.000E-06 | 3.100E-06 |
| 6144 | 308 | R308D | A | 2.8e-06 | 1.2e-06 | 3.4e-06 | 2.467E-06 | 1.137E-06 | 1.200E-06 | 3.400E-06 |
| 6145 | 308 | R308C | A | 2.7e-06 | 2.2e-06 | 1.5e-06 | 2.133E-06 | 6.028E-07 | 1.500E-06 | 2.700E-06 |
| 6146 | 308 | R308E | A | 1.9e-06 | 3.4e-06 | 2.4e-06 | 2.567E-06 | 7.638E-07 | 1.900E-06 | 3.400E-06 |
| 6147 | 308 | R308Q | A | 2.2e-06 | 2.4e-06 | 2.2e-06 | 2.267E-06 | 1.155E-07 | 2.200E-06 | 2.400E-06 |
| 6148 | 308 | R308G | A | 1.8e-06 | 3.3e-06 | 3.2e-06 | 2.767E-06 | 8.386E-07 | 1.800E-06 | 3.300E-06 |
| 6149 | 308 | R308H | A | 2.2e-06 | 2.6e-06 | 3.1e-06 | 2.633E-06 | 4.509E-07 | 2.200E-06 | 3.100E-06 |
| 6150 | 308 | R308I | A | 2.6e-06 | 1.9e-06 | 3e-06 | 2.500E-06 | 5.568E-07 | 1.900E-06 | 3.000E-06 |
| 6151 | 308 | R308L | A | 4.6e-06 | 2.4e-06 | 2.1e-06 | 3.033E-06 | 1.365E-06 | 2.100E-06 | 4.600E-06 |
| 6152 | 308 | R308K | A | 2.6e-06 | 1.8e-06 | 2.8e-06 | 2.400E-06 | 5.292E-07 | 1.800E-06 | 2.800E-06 |
| 6153 | 308 | R308M | A | 1.5e-06 | 2.2e-06 | 2.9e-06 | 2.200E-06 | 7.000E-07 | 1.500E-06 | 2.900E-06 |
| 6154 | 308 | R308F | A | 2.6e-06 | 1.8e-06 | 2.5e-06 | 2.300E-06 | 4.359E-07 | 1.800E-06 | 2.600E-06 |
| 6155 | 308 | R308P | A | 4.8e-06 | 2.2e-06 | 3.1e-06 | 3.367E-06 | 1.320E-06 | 2.200E-06 | 4.800E-06 |
| 6156 | 308 | R308S | A | 2.6e-06 | 2e-06 | 2.8e-06 | 2.467E-06 | 4.163E-07 | 2.000E-06 | 2.800E-06 |
| 6157 | 308 | R308T | A | 1.8e-06 | 1.6e-06 | 1.3e-06 | 1.567E-06 | 2.517E-07 | 1.300E-06 | 1.800E-06 |
| 6158 | 308 | R308W | A | 4.2e-06 | 2.7e-06 | 3.2e-06 | 3.367E-06 | 7.638E-07 | 2.700E-06 | 4.200E-06 |
| 6159 | 308 | R308Y | A | 2.2e-06 | 1.6e-06 | 2.4e-06 | 2.067E-06 | 4.163E-07 | 1.600E-06 | 2.400E-06 |
| 6160 | 308 | R308V | A | 1.8e-06 | 2.7e-06 | 3.7e-06 | 2.733E-06 | 9.504E-07 | 1.800E-06 | 3.700E-06 |
| 6161 | 309 | C309A | A | 2.3e-06 | 2.5e-06 | 3.3e-06 | 2.700E-06 | 5.292E-07 | 2.300E-06 | 3.300E-06 |
| 6162 | 309 | C309R | A | 3.2e-06 | 2.4e-06 | 2.9e-06 | 2.833E-06 | 4.041E-07 | 2.400E-06 | 3.200E-06 |
| 6163 | 309 | C309N | A | 3.7e-06 | 1.8e-06 | 3.5e-06 | 3.000E-06 | 1.044E-06 | 1.800E-06 | 3.700E-06 |
| 6164 | 309 | C309D | A | 2.2e-06 | 2.1e-06 | 2.2e-06 | 2.167E-06 | 5.774E-08 | 2.100E-06 | 2.200E-06 |
| 6165 | 309 | C309C | A | 2.5e-06 | 2.6e-06 | 2.6e-06 | 2.567E-06 | 5.774E-08 | 2.500E-06 | 2.600E-06 |
| 6166 | 309 | C309E | A | 3.4e-06 | 1.8e-06 | 1.9e-06 | 2.367E-06 | 8.963E-07 | 1.800E-06 | 3.400E-06 |
| 6167 | 309 | C309Q | A | 1.9e-06 | 3.3e-06 | 1.5e-06 | 2.233E-06 | 9.452E-07 | 1.500E-06 | 3.300E-06 |
| 6168 | 309 | C309G | A | 3e-06 | 1.7e-06 | 3.5e-06 | 2.733E-06 | 9.292E-07 | 1.700E-06 | 3.500E-06 |
| 6169 | 309 | C309H | A | 1.7e-06 | 3.1e-06 | 3.5e-06 | 2.767E-06 | 9.452E-07 | 1.700E-06 | 3.500E-06 |
| 6170 | 309 | C309I | A | 1.9e-06 | 3.5e-06 | 4.2e-06 | 3.200E-06 | 1.179E-06 | 1.900E-06 | 4.200E-06 |
| 6171 | 309 | C309L | A | 3.4e-06 | 3.4e-06 | 2e-06 | 2.933E-06 | 8.083E-07 | 2.000E-06 | 3.400E-06 |
| 6172 | 309 | C309K | A | 2.2e-06 | 3.4e-06 | 1.7e-06 | 2.433E-06 | 8.737E-07 | 1.700E-06 | 3.400E-06 |
| 6173 | 309 | C309M | A | 2.7e-06 | 2.5e-06 | 2e-06 | 2.400E-06 | 3.606E-07 | 2.000E-06 | 2.700E-06 |
| 6174 | 309 | C309F | A | 2.5e-06 | 2.1e-06 | 3.6e-06 | 2.733E-06 | 7.767E-07 | 2.100E-06 | 3.600E-06 |
| 6175 | 309 | C309P | A | 2.6e-06 | 2.3e-06 | 2.2e-06 | 2.367E-06 | 2.082E-07 | 2.200E-06 | 2.600E-06 |
| 6176 | 309 | C309S | A | 2.8e-06 | 2.5e-06 | 2.6e-06 | 2.633E-06 | 1.528E-07 | 2.500E-06 | 2.800E-06 |
| 6177 | 309 | C309T | A | 1.2e-06 | 1.7e-06 | 2.2e-06 | 1.700E-06 | 5.000E-07 | 1.200E-06 | 2.200E-06 |
| 6178 | 309 | C309W | A | 3e-06 | 2.4e-06 | 1.8e-06 | 2.400E-06 | 6.000E-07 | 1.800E-06 | 3.000E-06 |
| 6179 | 309 | C309Y | A | 2.6e-06 | 1.9e-06 | 2.3e-06 | 2.267E-06 | 3.512E-07 | 1.900E-06 | 2.600E-06 |
| 6180 | 309 | C309V | A | 2.8e-06 | 1.7e-06 | 1.7e-06 | 2.067E-06 | 6.351E-07 | 1.700E-06 | 2.800E-06 |
| 6181 | 310 | E310A | A | 2.1e-06 | 4.3e-06 | 1.8e-06 | 2.733E-06 | 1.365E-06 | 1.800E-06 | 4.300E-06 |
| 6182 | 310 | E310R | A | 2.7e-06 | 2.4e-06 | 2.3e-06 | 2.467E-06 | 2.082E-07 | 2.300E-06 | 2.700E-06 |
| 6183 | 310 | E310N | A | 1.7e-06 | 3.9e-06 | 2.3e-06 | 2.633E-06 | 1.137E-06 | 1.700E-06 | 3.900E-06 |
| 6184 | 310 | E310D | A | 1.6e-06 | 2e-06 | 2.2e-06 | 1.933E-06 | 3.055E-07 | 1.600E-06 | 2.200E-06 |
| 6185 | 310 | E310C | A | 2.1e-06 | 2.9e-06 | 2.1e-06 | 2.367E-06 | 4.619E-07 | 2.100E-06 | 2.900E-06 |
| 6186 | 310 | E310E | A | 1.8e-06 | 1.1e-06 | 1.9e-06 | 1.600E-06 | 4.359E-07 | 1.100E-06 | 1.900E-06 |
| 6187 | 310 | E310Q | A | 2.4e-06 | 2.5e-06 | 2.1e-06 | 2.333E-06 | 2.082E-07 | 2.100E-06 | 2.500E-06 |
| 6188 | 310 | E310G | A | 2e-06 | 4e-06 | 2.2e-06 | 2.733E-06 | 1.102E-06 | 2.000E-06 | 4.000E-06 |
| 6189 | 310 | E310H | A | 1.9e-06 | 1.9e-06 | 2.2e-06 | 2.000E-06 | 1.732E-07 | 1.900E-06 | 2.200E-06 |
| 6190 | 310 | E310I | A | 1.6e-06 | 3e-06 | 1.9e-06 | 2.167E-06 | 7.371E-07 | 1.600E-06 | 3.000E-06 |
| 6191 | 310 | E310L | A | 1.4e-06 | 3.8e-06 | 3.4e-06 | 2.867E-06 | 1.286E-06 | 1.400E-06 | 3.800E-06 |
| 6192 | 310 | E310K | A | 2e-06 | 1.7e-06 | 1.9e-06 | 1.867E-06 | 1.528E-07 | 1.700E-06 | 2.000E-06 |
| 6193 | 310 | E310M | A | 1.7e-06 | 2.2e-06 | 3.6e-06 | 2.500E-06 | 9.849E-07 | 1.700E-06 | 3.600E-06 |
| 6194 | 310 | E310F | A | 2.4e-06 | 2.4e-06 | 2.5e-06 | 2.433E-06 | 5.774E-08 | 2.400E-06 | 2.500E-06 |
| 6195 | 310 | E310P | A | 8.4e-07 | 3e-06 | 1.6e-06 | 1.813E-06 | 1.096E-06 | 8.400E-07 | 3.000E-06 |
| 6196 | 310 | E310S | A | 9.5e-07 | 2.5e-06 | 3e-06 | 2.150E-06 | 1.069E-06 | 9.500E-07 | 3.000E-06 |
| 6197 | 310 | E310T | A | 2.2e-06 | 2.2e-06 | 1.8e-06 | 2.067E-06 | 2.309E-07 | 1.800E-06 | 2.200E-06 |
| 6198 | 310 | E310W | A | 2.6e-06 | 3.4e-06 | 2.3e-06 | 2.767E-06 | 5.686E-07 | 2.300E-06 | 3.400E-06 |
| 6199 | 310 | E310Y | A | 1.1e-06 | 2.9e-06 | 2.8e-06 | 2.267E-06 | 1.012E-06 | 1.100E-06 | 2.900E-06 |
| 6200 | 310 | E310V | A | 2.5e-06 | 1.8e-06 | 1.9e-06 | 2.067E-06 | 3.786E-07 | 1.800E-06 | 2.500E-06 |
| 6201 | 311 | K311A | A | 9.1e-07 | 1.7e-06 | 1.8e-06 | 1.470E-06 | 4.875E-07 | 9.100E-07 | 1.800E-06 |
| 6202 | 311 | K311R | A | 3.2e-06 | 1.3e-06 | 1.6e-06 | 2.033E-06 | 1.021E-06 | 1.300E-06 | 3.200E-06 |
| 6203 | 311 | K311N | A | 1.7e-06 | 2.5e-06 | 1.9e-06 | 2.033E-06 | 4.163E-07 | 1.700E-06 | 2.500E-06 |
| 6204 | 311 | K311D | A | 3.1e-06 | 2.5e-06 | 1.7e-06 | 2.433E-06 | 7.024E-07 | 1.700E-06 | 3.100E-06 |

| | | | | | | | | | | |
|---|---|---|---|---|---|---|---|---|---|---|
| 6205 | 311 | K311C | A | 1.7e-06 | 2.1e-06 | 2.2e-06 | 2.000E-06 | 2.646E-07 | 1.700E-06 | 2.200E-06 |
| 6206 | 311 | K311E | A | 2.5e-06 | 3.5e-06 | 2.8e-06 | 2.933E-06 | 5.132E-07 | 2.500E-06 | 3.500E-06 |
| 6207 | 311 | K311Q | A | 7e-06 | 2e-06 | 1.6e-06 | 3.533E-06 | 3.009E-06 | 1.600E-06 | 7.000E-06 |
| 6208 | 311 | K311G | A | 3.1e-06 | 2.7e-06 | 2e-06 | 2.600E-06 | 5.568E-07 | 2.000E-06 | 3.100E-06 |
| 6209 | 311 | K311H | A | 1.9e-06 | 2.4e-06 | 1.7e-06 | 2.000E-06 | 3.606E-07 | 1.700E-06 | 2.400E-06 |
| 6210 | 311 | K311I | A | 1.1e-06 | 3.4e-06 | 2.1e-06 | 2.200E-06 | 1.153E-06 | 1.100E-06 | 3.400E-06 |
| 6211 | 311 | K311L | A | 1.5e-06 | 3.1e-06 | 2.1e-06 | 2.233E-06 | 8.083E-07 | 1.500E-06 | 3.100E-06 |
| 6212 | 311 | K311K | A | 2.5e-06 | 1.8e-06 | 2.2e-06 | 2.167E-06 | 3.512E-07 | 1.800E-06 | 2.500E-06 |
| 6213 | 311 | K311M | A | 3.6e-06 | 2.3e-06 | 2.8e-06 | 2.900E-06 | 6.557E-07 | 2.300E-06 | 3.600E-06 |
| 6214 | 311 | K311F | A | 2e-06 | 2.9e-06 | 2.6e-06 | 2.500E-06 | 4.583E-07 | 2.000E-06 | 2.900E-06 |
| 6215 | 311 | K311P | A | 4.1e-06 | 2.4e-06 | 2.9e-06 | 3.133E-06 | 8.737E-07 | 2.400E-06 | 4.100E-06 |
| 6216 | 311 | K311S | A | 2.6e-06 | 2.8e-06 | 2.4e-06 | 2.600E-06 | 2.000E-07 | 2.400E-06 | 2.800E-06 |
| 6217 | 311 | K311T | A | 2.9e-06 | 3.2e-06 | 1.5e-06 | 2.533E-06 | 9.074E-07 | 1.500E-06 | 3.200E-06 |
| 6218 | 311 | K311W | A | 3.2e-06 | 2.3e-06 | 2.3e-06 | 2.600E-06 | 5.196E-07 | 2.300E-06 | 3.200E-06 |
| 6219 | 311 | K311Y | A | 2.5e-06 | 3.1e-06 | 3.5e-06 | 3.033E-06 | 5.033E-07 | 2.500E-06 | 3.500E-06 |
| 6220 | 311 | K311V | A | 2.5e-06 | 2.4e-06 | 2.8e-06 | 2.567E-06 | 2.082E-07 | 2.400E-06 | 2.800E-06 |
| 6221 | 312 | C312A | A | 3.6e-06 | 2.1e-06 | 3.2e-06 | 2.967E-06 | 7.767E-07 | 2.100E-06 | 3.600E-06 |
| 6222 | 312 | C312R | A | 2.1e-06 | 3.5e-06 | 1.8e-06 | 2.467E-06 | 9.074E-07 | 1.800E-06 | 3.500E-06 |
| 6223 | 312 | C312N | A | 2.1e-06 | 2.8e-06 | 2.4e-06 | 2.433E-06 | 3.512E-07 | 2.100E-06 | 2.800E-06 |
| 6224 | 312 | C312D | A | 3.2e-06 | 1.7e-06 | 1.7e-06 | 2.200E-06 | 8.660E-07 | 1.700E-06 | 3.200E-06 |
| 6225 | 312 | C312C | A | 2.7e-06 | 1.8e-06 | 3.1e-06 | 2.533E-06 | 6.658E-07 | 1.800E-06 | 3.100E-06 |
| 6226 | 312 | C312E | A | 3e-06 | 2.9e-06 | 2.9e-06 | 2.933E-06 | 5.774E-08 | 2.900E-06 | 3.000E-06 |
| 6227 | 312 | C312Q | A | 3.1e-06 | 3.1e-06 | 2e-06 | 2.733E-06 | 6.351E-07 | 2.000E-06 | 3.100E-06 |
| 6228 | 312 | C312G | A | 2e-06 | 2.3e-06 | 2e-06 | 2.100E-06 | 1.732E-07 | 2.000E-06 | 2.300E-06 |
| 6229 | 312 | C312H | A | 1.5e-06 | 2.3e-06 | 4.7e-06 | 2.833E-06 | 1.665E-06 | 1.500E-06 | 4.700E-06 |
| 6230 | 312 | C312I | A | 2.7e-06 | 2.5e-06 | 3.2e-06 | 2.800E-06 | 3.606E-07 | 2.500E-06 | 3.200E-06 |
| 6231 | 312 | C312L | A | 2.9e-06 | 1.9e-06 | 2.2e-06 | 2.333E-06 | 5.132E-07 | 1.900E-06 | 2.900E-06 |
| 6232 | 312 | C312K | A | 1.9e-06 | 2.1e-06 | 3.3e-06 | 2.433E-06 | 7.572E-07 | 1.900E-06 | 3.300E-06 |
| 6233 | 312 | C312M | A | 2.9e-06 | 1.2e-06 | 3.8e-06 | 2.633E-06 | 1.320E-06 | 1.200E-06 | 3.800E-06 |
| 6234 | 312 | C312F | A | 2.9e-06 | 2.2e-06 | 2.4e-06 | 2.500E-06 | 3.606E-07 | 2.200E-06 | 2.900E-06 |
| 6235 | 312 | C312P | A | 3.8e-06 | 1.9e-06 | 3.5e-06 | 3.067E-06 | 1.021E-06 | 1.900E-06 | 3.800E-06 |
| 6236 | 312 | C312S | A | 3.2e-06 | 1.5e-06 | 3.1e-06 | 2.600E-06 | 9.539E-07 | 1.500E-06 | 3.200E-06 |
| 6237 | 312 | C312T | A | 1.4e-06 | 3.5e-06 | 1.9e-06 | 2.267E-06 | 1.097E-06 | 1.400E-06 | 3.500E-06 |
| 6238 | 312 | C312W | A | 2e-06 | 2.2e-06 | 2e-06 | 2.067E-06 | 1.155E-07 | 2.000E-06 | 2.200E-06 |
| 6239 | 312 | C312Y | A | 2.6e-06 | 2.1e-06 | 2.6e-06 | 2.433E-06 | 2.887E-07 | 2.100E-06 | 2.600E-06 |
| 6240 | 312 | C312V | A | 2e-06 | 3.6e-06 | 2.5e-06 | 2.700E-06 | 8.185E-07 | 2.000E-06 | 3.600E-06 |
| 6241 | 313 | S313A | A | 1.7e-06 | 1.6e-06 | 2.8e-06 | 2.033E-06 | 6.658E-07 | 1.600E-06 | 2.800E-06 |
| 6242 | 313 | S313R | A | 1.8e-06 | 2.1e-06 | 2e-06 | 1.967E-06 | 1.528E-07 | 1.800E-06 | 2.100E-06 |
| 6243 | 313 | S313N | A | 1.2e-06 | 1.5e-06 | 4e-06 | 2.233E-06 | 1.537E-06 | 1.200E-06 | 4.000E-06 |
| 6244 | 313 | S313D | A | 2.2e-06 | 1.2e-06 | 1.7e-06 | 1.700E-06 | 5.000E-07 | 1.200E-06 | 2.200E-06 |
| 6245 | 313 | S313C | A | 2.5e-06 | 3.4e-06 | 2.6e-06 | 2.833E-06 | 4.933E-07 | 2.500E-06 | 3.400E-06 |
| 6246 | 313 | S313E | A | 2.1e-06 | 6.1e-07 | 3.6e-06 | 2.103E-06 | 1.495E-06 | 6.100E-07 | 3.600E-06 |
| 6247 | 313 | S313Q | A | 2.1e-06 | 2.4e-06 | 1.9e-06 | 2.133E-06 | 2.517E-07 | 1.900E-06 | 2.400E-06 |
| 6248 | 313 | S313G | A | 4.6e-06 | 1.4e-06 | 5.3e-06 | 3.767E-06 | 2.079E-06 | 1.400E-06 | 5.300E-06 |
| 6249 | 313 | S313H | A | 2.5e-06 | 2.9e-06 | 1.6e-06 | 2.333E-06 | 6.658E-07 | 1.600E-06 | 2.900E-06 |
| 6250 | 313 | S313I | A | 1.9e-06 | 1.5e-06 | 1.5e-06 | 1.633E-06 | 2.309E-07 | 1.500E-06 | 1.900E-06 |
| 6251 | 313 | S313L | A | 4.1e-06 | 1.8e-06 | 2.8e-06 | 2.900E-06 | 1.153E-06 | 1.800E-06 | 4.100E-06 |
| 6252 | 313 | S313K | A | 1.1e-06 | 1.9e-06 | 2.9e-06 | 1.967E-06 | 9.018E-07 | 1.100E-06 | 2.900E-06 |
| 6253 | 313 | S313M | A | 3.2e-06 | 2.4e-06 | 3.3e-06 | 2.967E-06 | 4.933E-07 | 2.400E-06 | 3.300E-06 |
| 6254 | 313 | S313F | A | 1.4e-06 | 4e-06 | 2.7e-06 | 2.700E-06 | 1.300E-06 | 1.400E-06 | 4.000E-06 |
| 6255 | 313 | S313P | A | 1.9e-06 | 1.5e-06 | 2.2e-06 | 1.867E-06 | 3.512E-07 | 1.500E-06 | 2.200E-06 |
| 6256 | 313 | S313S | A | 3.4e-06 | 2.5e-06 | 1.6e-06 | 2.500E-06 | 9.000E-07 | 1.600E-06 | 3.400E-06 |
| 6257 | 313 | S313T | A | 2e-06 | 2.2e-06 | 2.4e-06 | 2.200E-06 | 2.000E-07 | 2.000E-06 | 2.400E-06 |
| 6258 | 313 | S313W | A | 2.6e-06 | 5.4e-06 | 3.1e-06 | 3.700E-06 | 1.493E-06 | 2.600E-06 | 5.400E-06 |
| 6259 | 313 | S313Y | A | 2.2e-06 | 3.1e-06 | 3e-06 | 2.767E-06 | 4.933E-07 | 2.200E-06 | 3.100E-06 |
| 6260 | 313 | S313V | A | 4.9e-06 | 2.1e-06 | 2.8e-06 | 3.267E-06 | 1.457E-06 | 2.100E-06 | 4.900E-06 |
| 6261 | 314 | K314A | A | 1.6e-06 | 3.9e-06 | 2e-06 | 2.500E-06 | 1.229E-06 | 1.600E-06 | 3.900E-06 |
| 6262 | 314 | K314R | A | 1.7e-06 | 1.6e-06 | 3.2e-06 | 2.167E-06 | 8.963E-07 | 1.600E-06 | 3.200E-06 |
| 6263 | 314 | K314N | A | 3.7e-06 | 2.5e-06 | 4.4e-06 | 3.533E-06 | 9.609E-07 | 2.500E-06 | 4.400E-06 |
| 6264 | 314 | K314D | A | 2.3e-06 | 1.5e-06 | 1.1e-06 | 1.633E-06 | 6.110E-07 | 1.100E-06 | 2.300E-06 |
| 6265 | 314 | K314C | A | 3.3e-06 | 1.8e-06 | 3.1e-06 | 2.733E-06 | 8.145E-07 | 1.800E-06 | 3.300E-06 |
| 6266 | 314 | K314E | A | 2.2e-06 | 2.6e-06 | 2e-06 | 2.267E-06 | 3.055E-07 | 2.000E-06 | 2.600E-06 |
| 6267 | 314 | K314Q | A | 2e-06 | 2.7e-06 | 1.9e-06 | 2.200E-06 | 4.359E-07 | 1.900E-06 | 2.700E-06 |
| 6268 | 314 | K314G | A | 2.6e-06 | 1.1e-06 | 2.2e-06 | 1.967E-06 | 7.767E-07 | 1.100E-06 | 2.600E-06 |
| 6269 | 314 | K314H | A | 2.9e-06 | 2.1e-06 | 2.6e-06 | 2.533E-06 | 4.041E-07 | 2.100E-06 | 2.900E-06 |
| 6270 | 314 | K314I | A | 2.2e-06 | 1.8e-06 | 4.3e-06 | 2.767E-06 | 1.343E-06 | 1.800E-06 | 4.300E-06 |
| 6271 | 314 | K314L | A | 2e-06 | 1.9e-06 | 2.5e-06 | 2.133E-06 | 3.215E-07 | 1.900E-06 | 2.500E-06 |
| 6272 | 314 | K314K | A | 2.2e-06 | 3.1e-06 | 1.7e-06 | 2.333E-06 | 7.095E-07 | 1.700E-06 | 3.100E-06 |
| 6273 | 314 | K314M | A | 2.5e-06 | 1.9e-06 | 1.5e-06 | 1.967E-06 | 5.033E-07 | 1.500E-06 | 2.500E-06 |
| 6274 | 314 | K314F | A | 1.4e-06 | 2e-06 | 3.5e-06 | 2.300E-06 | 1.082E-06 | 1.400E-06 | 3.500E-06 |
| 6275 | 314 | K314P | A | 8e-07 | 1.9e-06 | 1.6e-06 | 1.433E-06 | 5.686E-07 | 8.000E-07 | 1.900E-06 |
| 6276 | 314 | K314S | A | 2e-06 | 3e-06 | 1.7e-06 | 2.233E-06 | 6.807E-07 | 1.700E-06 | 3.000E-06 |
| 6277 | 314 | K314T | A | 2.8e-06 | 2.1e-06 | 2.6e-06 | 2.500E-06 | 3.606E-07 | 2.100E-06 | 2.800E-06 |
| 6278 | 314 | K314W | A | 1.4e-06 | 3.3e-06 | 1.1e-06 | 1.933E-06 | 1.193E-06 | 1.100E-06 | 3.300E-06 |
| 6279 | 314 | K314Y | A | 2.1e-06 | 1.9e-06 | 3.7e-06 | 2.567E-06 | 9.866E-07 | 1.900E-06 | 3.700E-06 |
| 6280 | 314 | K314V | A | 3.4e-06 | 2.7e-06 | 3.8e-06 | 3.300E-06 | 5.568E-07 | 2.700E-06 | 3.800E-06 |
| 6281 | 315 | P315A | A | 2.6e-06 | 2.4e-06 | 2.7e-06 | 2.567E-06 | 1.528E-07 | 2.400E-06 | 2.700E-06 |
| 6282 | 315 | P315R | A | 1.2e-06 | 2e-06 | 1.7e-06 | 1.633E-06 | 4.041E-07 | 1.200E-06 | 2.000E-06 |
| 6283 | 315 | P315N | A | 1.4e-06 | 1.8e-06 | 2.4e-06 | 1.867E-06 | 5.033E-07 | 1.400E-06 | 2.400E-06 |
| 6284 | 315 | P315D | A | 2.5e-06 | 1.6e-06 | 2.4e-06 | 2.167E-06 | 4.933E-07 | 1.600E-06 | 2.500E-06 |
| 6285 | 315 | P315C | A | 3.9e-06 | 2.8e-06 | 2.4e-06 | 3.033E-06 | 7.767E-07 | 2.400E-06 | 3.900E-06 |
| 6286 | 315 | P315E | A | 2.4e-06 | 2.4e-06 | 2.7e-06 | 2.500E-06 | 1.732E-07 | 2.400E-06 | 2.700E-06 |
| 6287 | 315 | P315Q | A | 1.9e-06 | 1.5e-06 | 2.7e-06 | 2.033E-06 | 6.110E-07 | 1.500E-06 | 2.700E-06 |
| 6288 | 315 | P315G | A | 2.5e-06 | 1.8e-06 | 2.5e-06 | 2.267E-06 | 4.041E-07 | 1.800E-06 | 2.500E-06 |
| 6289 | 315 | P315H | A | 2.6e-06 | 1.9e-06 | 1.4e-06 | 1.967E-06 | 6.028E-07 | 1.400E-06 | 2.600E-06 |

| | | | | | | | | | | |
|---|---|---|---|---|---|---|---|---|---|---|
| 6290 | 315 | P315I | A | 2.4e-06 | 2.1e-06 | 1.5e-06 | 2.000E-06 | 4.583E-07 | 1.500E-06 | 2.400E-06 |
| 6291 | 315 | P315L | A | 4.1e-06 | 2e-06 | 2.6e-06 | 2.900E-06 | 1.082E-06 | 2.000E-06 | 4.100E-06 |
| 6292 | 315 | P315K | A | 2.5e-06 | 2.7e-06 | 2.7e-06 | 2.633E-06 | 1.155E-07 | 2.500E-06 | 2.700E-06 |
| 6293 | 315 | P315M | A | 2.1e-06 | 4e-06 | 2.5e-06 | 2.867E-06 | 1.002E-06 | 2.100E-06 | 4.000E-06 |
| 6294 | 315 | P315F | A | 1.3e-06 | 2.8e-06 | 2.6e-06 | 2.233E-06 | 8.145E-07 | 1.300E-06 | 2.800E-06 |
| 6295 | 315 | P315P | A | 2.3e-06 | 1.9e-06 | 1.3e-06 | 1.833E-06 | 5.033E-07 | 1.300E-06 | 2.300E-06 |
| 6296 | 315 | P315S | A | 1.1e-06 | 3.8e-06 | 2.8e-06 | 2.567E-06 | 1.365E-06 | 1.100E-06 | 3.800E-06 |
| 6297 | 315 | P315T | A | 1.3e-06 | 4.2e-06 | 1.2e-06 | 2.233E-06 | 1.704E-06 | 1.200E-06 | 4.200E-06 |
| 6298 | 315 | P315W | A | 3.1e-06 | 5.1e-06 | 2.7e-06 | 3.633E-06 | 1.286E-06 | 2.700E-06 | 5.100E-06 |
| 6299 | 315 | P315Y | A | 2.6e-06 | 2.5e-06 | 1.9e-06 | 2.333E-06 | 3.786E-07 | 1.900E-06 | 2.600E-06 |
| 6300 | 315 | P315V | A | 7.9e-07 | 3e-06 | 2e-06 | 1.930E-06 | 1.107E-06 | 7.900E-07 | 3.000E-06 |
| 6301 | 316 | C316A | A | 2.9e-06 | 3e-06 | 1.2e-06 | 2.367E-06 | 1.012E-06 | 1.200E-06 | 3.000E-06 |
| 6302 | 316 | C316R | A | 2.7e-06 | 1.3e-06 | 3.7e-06 | 2.567E-06 | 1.206E-06 | 1.300E-06 | 3.700E-06 |
| 6303 | 316 | C316N | A | 2.7e-06 | 3.2e-06 | 3.4e-06 | 3.100E-06 | 3.606E-07 | 2.700E-06 | 3.400E-06 |
| 6304 | 316 | C316D | A | 1.6e-06 | 2.8e-06 | 2.7e-06 | 2.367E-06 | 6.658E-07 | 1.600E-06 | 2.800E-06 |
| 6305 | 316 | C316C | A | 3.9e-06 | 1.5e-06 | 1.7e-06 | 2.367E-06 | 1.332E-06 | 1.500E-06 | 3.900E-06 |
| 6306 | 316 | C316E | A | 1.7e-06 | 2.3e-06 | 3.3e-06 | 2.433E-06 | 8.083E-07 | 1.700E-06 | 3.300E-06 |
| 6307 | 316 | C316Q | A | 2.8e-06 | 3.3e-06 | 3.4e-06 | 3.167E-06 | 3.215E-07 | 2.800E-06 | 3.400E-06 |
| 6308 | 316 | C316G | A | 9.5e-07 | 2.2e-06 | 2.5e-06 | 1.883E-06 | 8.221E-07 | 9.500E-07 | 2.500E-06 |
| 6309 | 316 | C316H | A | 2.8e-06 | 1.6e-06 | 3e-06 | 2.467E-06 | 7.572E-07 | 1.600E-06 | 3.000E-06 |
| 6310 | 316 | C316I | A | 3e-06 | 1.5e-06 | 2.5e-06 | 2.333E-06 | 7.638E-07 | 1.500E-06 | 3.000E-06 |
| 6311 | 316 | C316L | A | 3.5e-06 | 2.4e-06 | 1.9e-06 | 2.600E-06 | 8.185E-07 | 1.900E-06 | 3.500E-06 |
| 6312 | 316 | C316K | A | 2e-06 | 2.6e-06 | 3e-06 | 2.533E-06 | 5.033E-07 | 2.000E-06 | 3.000E-06 |
| 6313 | 316 | C316M | A | 1.4e-06 | 4.2e-06 | 1.2e-06 | 2.267E-06 | 1.677E-06 | 1.200E-06 | 4.200E-06 |
| 6314 | 316 | C316F | A | 1.9e-06 | 4.1e-06 | 2.1e-06 | 2.700E-06 | 1.217E-06 | 1.900E-06 | 4.100E-06 |
| 6315 | 316 | C316P | A | 1.7e-06 | 2.7e-06 | 1.9e-06 | 2.100E-06 | 5.292E-07 | 1.700E-06 | 2.700E-06 |
| 6316 | 316 | C316S | A | 3.5e-06 | 2.1e-06 | 2e-06 | 2.533E-06 | 8.386E-07 | 2.000E-06 | 3.500E-06 |
| 6317 | 316 | C316T | A | 2.6e-06 | 1.9e-06 | 1.4e-06 | 1.967E-06 | 6.028E-07 | 1.400E-06 | 2.600E-06 |
| 6318 | 316 | C316W | A | 2.2e-06 | 2.7e-06 | 2.3e-06 | 2.400E-06 | 2.646E-07 | 2.200E-06 | 2.700E-06 |
| 6319 | 316 | C316Y | A | 1.8e-06 | 4.4e-06 | 2.5e-06 | 2.900E-06 | 1.345E-06 | 1.800E-06 | 4.400E-06 |
| 6320 | 316 | C316V | A | 2.5e-06 | 2.3e-06 | 2.9e-06 | 2.567E-06 | 3.055E-07 | 2.300E-06 | 2.900E-06 |
| 6321 | 317 | A317A | A | 2.2e-06 | 1.5e-06 | 1.3e-06 | 1.667E-06 | 4.726E-07 | 1.300E-06 | 2.200E-06 |
| 6322 | 317 | A317R | A | 3.2e-06 | 4.4e-06 | 1.5e-06 | 3.033E-06 | 1.457E-06 | 1.500E-06 | 4.400E-06 |
| 6323 | 317 | A317N | A | 1.4e-06 | 2.1e-06 | 1.4e-06 | 1.633E-06 | 4.041E-07 | 1.400E-06 | 2.100E-06 |
| 6324 | 317 | A317D | A | 3.7e-06 | 2.9e-06 | 2.7e-06 | 3.100E-06 | 5.292E-07 | 2.700E-06 | 3.700E-06 |
| 6325 | 317 | A317C | A | 1.9e-06 | 2.5e-06 | 2.6e-06 | 2.333E-06 | 3.786E-07 | 1.900E-06 | 2.600E-06 |
| 6326 | 317 | A317E | A | 2.6e-06 | 2.4e-06 | 2.4e-06 | 2.467E-06 | 1.155E-07 | 2.400E-06 | 2.600E-06 |
| 6327 | 317 | A317Q | A | 2.3e-06 | 2.2e-06 | 2.9e-06 | 2.467E-06 | 3.786E-07 | 2.200E-06 | 2.900E-06 |
| 6328 | 317 | A317G | A | 2.6e-06 | 3.4e-06 | 3.4e-06 | 3.133E-06 | 4.619E-07 | 2.600E-06 | 3.400E-06 |
| 6329 | 317 | A317H | A | 7.1e-06 | 3.4e-06 | 1.9e-06 | 4.133E-06 | 2.676E-06 | 1.900E-06 | 7.100E-06 |
| 6330 | 317 | A317I | A | 2.5e-06 | 1.9e-06 | 1.9e-06 | 2.100E-06 | 3.464E-07 | 1.900E-06 | 2.500E-06 |
| 6331 | 317 | A317L | A | 2.8e-06 | 4.1e-06 | 1.9e-06 | 2.933E-06 | 1.106E-06 | 1.900E-06 | 4.100E-06 |
| 6332 | 317 | A317K | A | 2.3e-06 | 1.9e-06 | 3.5e-06 | 2.567E-06 | 8.327E-07 | 1.900E-06 | 3.500E-06 |
| 6333 | 317 | A317M | A | 2.2e-06 | 2.8e-06 | 3e-06 | 2.667E-06 | 4.163E-07 | 2.200E-06 | 3.000E-06 |
| 6334 | 317 | A317F | A | 3.5e-06 | 2.8e-06 | 1.8e-06 | 2.700E-06 | 8.544E-07 | 1.800E-06 | 3.500E-06 |
| 6335 | 317 | A317P | A | 1.7e-06 | 3.6e-06 | 2e-06 | 2.433E-06 | 1.021E-06 | 1.700E-06 | 3.600E-06 |
| 6336 | 317 | A317S | A | 2e-06 | 3.2e-06 | 1.2e-06 | 2.133E-06 | 1.007E-06 | 1.200E-06 | 3.200E-06 |
| 6337 | 317 | A317T | A | 2.8e-06 | 1.9e-06 | 1.3e-06 | 2.000E-06 | 7.550E-07 | 1.300E-06 | 2.800E-06 |
| 6338 | 317 | A317W | A | 2.5e-06 | 1.7e-06 | 4.7e-06 | 2.967E-06 | 1.553E-06 | 1.700E-06 | 4.700E-06 |
| 6339 | 317 | A317Y | A | 3.4e-06 | 2.5e-06 | 8.5e-07 | 2.250E-06 | 1.293E-06 | 8.500E-07 | 3.400E-06 |
| 6340 | 317 | A317V | A | 2.3e-06 | 1.8e-06 | 3.9e-06 | 2.667E-06 | 1.097E-06 | 1.800E-06 | 3.900E-06 |
| 6341 | 318 | R318A | A | 3.4e-06 | 4.2e-06 | 4e-06 | 3.867E-06 | 4.163E-07 | 3.400E-06 | 4.200E-06 |
| 6342 | 318 | R318R | A | 1.9e-06 | 2.9e-06 | 3.3e-06 | 2.700E-06 | 7.211E-07 | 1.900E-06 | 3.300E-06 |
| 6343 | 318 | R318N | A | 2e-06 | 2.9e-06 | 2.7e-06 | 2.533E-06 | 4.726E-07 | 2.000E-06 | 2.900E-06 |
| 6344 | 318 | R318D | A | 2.2e-06 | 3.3e-06 | 3e-06 | 2.833E-06 | 5.686E-07 | 2.200E-06 | 3.300E-06 |
| 6345 | 318 | R318C | A | 2.4e-06 | 2.2e-06 | 3.2e-06 | 2.600E-06 | 5.292E-07 | 2.200E-06 | 3.200E-06 |
| 6346 | 318 | R318E | A | 2.1e-06 | 1e-06 | 2.9e-06 | 2.000E-06 | 9.539E-07 | 1.000E-06 | 2.900E-06 |
| 6347 | 318 | R318Q | A | 3.2e-06 | 1.8e-06 | 2e-06 | 2.333E-06 | 7.572E-07 | 1.800E-06 | 3.200E-06 |
| 6348 | 318 | R318G | A | 1.7e-06 | 2e-06 | 2e-06 | 1.900E-06 | 1.732E-07 | 1.700E-06 | 2.000E-06 |
| 6349 | 318 | R318H | A | 1.3e-06 | 2.8e-06 | 2.1e-06 | 2.067E-06 | 7.506E-07 | 1.300E-06 | 2.800E-06 |
| 6350 | 318 | R318I | A | 2.9e-06 | 1.6e-06 | 3.3e-06 | 2.600E-06 | 8.888E-07 | 1.600E-06 | 3.300E-06 |
| 6351 | 318 | R318L | A | 2.4e-06 | 3.3e-06 | 1.7e-06 | 2.467E-06 | 8.021E-07 | 1.700E-06 | 3.300E-06 |
| 6352 | 318 | R318K | A | 1.9e-06 | 2.6e-06 | 2.5e-06 | 2.333E-06 | 3.786E-07 | 1.900E-06 | 2.600E-06 |
| 6353 | 318 | R318M | A | 2.8e-06 | 7.4e-07 | 2.8e-06 | 2.113E-06 | 1.189E-06 | 7.400E-07 | 2.800E-06 |
| 6354 | 318 | R318F | A | 2.6e-06 | 2.4e-06 | 2.2e-06 | 2.400E-06 | 2.000E-07 | 2.200E-06 | 2.600E-06 |
| 6355 | 318 | R318P | A | 2.5e-06 | 2e-06 | 2.8e-06 | 2.433E-06 | 4.041E-07 | 2.000E-06 | 2.800E-06 |
| 6356 | 318 | R318S | A | 2.4e-06 | 3.3e-06 | 2e-06 | 2.567E-06 | 6.658E-07 | 2.000E-06 | 3.300E-06 |
| 6357 | 318 | R318T | A | 3e-06 | 2.3e-06 | 1.8e-06 | 2.367E-06 | 6.028E-07 | 1.800E-06 | 3.000E-06 |
| 6358 | 318 | R318W | A | 2.7e-06 | 1.6e-06 | 2.2e-06 | 2.167E-06 | 5.508E-07 | 1.600E-06 | 2.700E-06 |
| 6359 | 318 | R318Y | A | 3.4e-06 | 3e-06 | 1.4e-06 | 2.600E-06 | 1.058E-06 | 1.400E-06 | 3.400E-06 |
| 6360 | 318 | R318V | A | 4.7e-06 | 2e-06 | 3.1e-06 | 3.267E-06 | 1.358E-06 | 2.000E-06 | 4.700E-06 |
| 6361 | 319 | V319A | A | 2.2e-06 | 2.2e-06 | 2.1e-06 | 2.167E-06 | 5.774E-08 | 2.100E-06 | 2.200E-06 |
| 6362 | 319 | V319R | A | 1.3e-06 | 3.8e-06 | 4.6e-06 | 3.233E-06 | 1.721E-06 | 1.300E-06 | 4.600E-06 |
| 6363 | 319 | V319N | A | 4e-06 | 3.5e-06 | 2.6e-06 | 3.367E-06 | 7.095E-07 | 2.600E-06 | 4.000E-06 |
| 6364 | 319 | V319D | A | 2.8e-06 | 2.1e-06 | 2.1e-06 | 2.333E-06 | 4.041E-07 | 2.100E-06 | 2.800E-06 |
| 6365 | 319 | V319C | A | 3.4e-06 | 3e-06 | 3.2e-06 | 3.200E-06 | 2.000E-07 | 3.000E-06 | 3.400E-06 |
| 6366 | 319 | V319E | A | 3.3e-06 | 1.7e-06 | 9.3e-07 | 1.977E-06 | 1.209E-06 | 9.300E-07 | 3.300E-06 |
| 6367 | 319 | V319Q | A | 2.1e-06 | 2.3e-06 | 2.4e-06 | 2.267E-06 | 1.528E-07 | 2.100E-06 | 2.400E-06 |
| 6368 | 319 | V319G | A | 1.6e-06 | 1.3e-06 | 2.2e-06 | 1.700E-06 | 4.583E-07 | 1.300E-06 | 2.200E-06 |
| 6369 | 319 | V319H | A | 2e-06 | 1.7e-06 | 1.6e-06 | 1.767E-06 | 2.082E-07 | 1.600E-06 | 2.000E-06 |
| 6370 | 319 | V319I | A | 3.2e-06 | 1.7e-06 | 3.2e-06 | 2.700E-06 | 8.660E-07 | 1.700E-06 | 3.200E-06 |
| 6371 | 319 | V319L | A | 2.6e-06 | 3.4e-06 | 1.9e-06 | 2.633E-06 | 7.506E-07 | 1.900E-06 | 3.400E-06 |
| 6372 | 319 | V319K | A | 2.3e-06 | 1.3e-06 | 1.9e-06 | 1.833E-06 | 5.033E-07 | 1.300E-06 | 2.300E-06 |
| 6373 | 319 | V319M | A | 2.1e-06 | 2.3e-06 | 2.1e-06 | 2.167E-06 | 1.155E-07 | 2.100E-06 | 2.300E-06 |
| 6374 | 319 | V319F | A | 3.4e-06 | 3.3e-06 | 2.4e-06 | 3.033E-06 | 5.508E-07 | 2.400E-06 | 3.400E-06 |

| | | | | | | | | | | |
|---|---|---|---|---|---|---|---|---|---|---|
| 6375 | 319 | V319P | A | 1.9e-06 | 3.6e-06 | 1.7e-06 | 2.400E-06 | 1.044E-06 | 1.700E-06 | 3.600E-06 |
| 6376 | 319 | V319S | A | 7.4e-07 | 3.7e-06 | 2.3e-06 | 2.247E-06 | 1.481E-06 | 7.400E-07 | 3.700E-06 |
| 6377 | 319 | V319T | A | 3.8e-06 | 2.6e-06 | 1.7e-06 | 2.700E-06 | 1.054E-06 | 1.700E-06 | 3.800E-06 |
| 6378 | 319 | V319W | A | 3.8e-06 | 1.4e-06 | 1.6e-06 | 2.267E-06 | 1.332E-06 | 1.400E-06 | 3.800E-06 |
| 6379 | 319 | V319Y | A | 2.2e-06 | 2.2e-06 | 3e-06 | 2.467E-06 | 4.619E-07 | 2.200E-06 | 3.000E-06 |
| 6380 | 319 | V319V | A | 2.9e-06 | 2.1e-06 | 1.7e-06 | 2.233E-06 | 6.110E-07 | 1.700E-06 | 2.900E-06 |
| 6381 | 320 | C320A | A | 2.5e-06 | 2.7e-06 | 1.9e-06 | 2.367E-06 | 4.163E-07 | 1.900E-06 | 2.700E-06 |
| 6382 | 320 | C320R | A | 2.2e-06 | 3.4e-06 | 2.6e-06 | 2.733E-06 | 6.110E-07 | 2.200E-06 | 3.400E-06 |
| 6383 | 320 | C320N | A | 2.1e-06 | 1.4e-06 | 3.5e-06 | 2.333E-06 | 1.069E-06 | 1.400E-06 | 3.500E-06 |
| 6384 | 320 | C320D | A | 2.2e-06 | 2.7e-06 | 2.7e-06 | 2.533E-06 | 2.887E-07 | 2.200E-06 | 2.700E-06 |
| 6385 | 320 | C320C | A | 2.6e-06 | 1.3e-06 | 2.1e-06 | 2.000E-06 | 6.557E-07 | 1.300E-06 | 2.600E-06 |
| 6386 | 320 | C320E | A | 3.5e-06 | 4e-06 | 2.3e-06 | 3.267E-06 | 8.737E-07 | 2.300E-06 | 4.000E-06 |
| 6387 | 320 | C320Q | A | 2.2e-06 | 2.7e-06 | 1.9e-06 | 2.267E-06 | 4.041E-07 | 1.900E-06 | 2.700E-06 |
| 6388 | 320 | C320G | A | 2.8e-06 | 1.6e-06 | 1.7e-06 | 2.033E-06 | 6.658E-07 | 1.600E-06 | 2.800E-06 |
| 6389 | 320 | C320H | A | 4.4e-06 | 3.7e-06 | 2.4e-06 | 3.500E-06 | 1.015E-06 | 2.400E-06 | 4.400E-06 |
| 6390 | 320 | C320I | A | 2.6e-06 | 1.8e-06 | 1.7e-06 | 2.033E-06 | 4.933E-07 | 1.700E-06 | 2.600E-06 |
| 6391 | 320 | C320L | A | 2e-06 | 3.8e-06 | 3.6e-06 | 3.133E-06 | 9.866E-07 | 2.000E-06 | 3.800E-06 |
| 6392 | 320 | C320K | A | 2.2e-06 | 2.1e-06 | 2.7e-06 | 2.333E-06 | 3.215E-07 | 2.100E-06 | 2.700E-06 |
| 6393 | 320 | C320M | A | 2.8e-06 | 1.4e-06 | 3.1e-06 | 2.433E-06 | 9.074E-07 | 1.400E-06 | 3.100E-06 |
| 6394 | 320 | C320F | A | 1.9e-06 | 3.5e-06 | 2.9e-06 | 2.767E-06 | 8.083E-07 | 1.900E-06 | 3.500E-06 |
| 6395 | 320 | C320P | A | 1.7e-06 | 1.9e-06 | 2.5e-06 | 2.033E-06 | 4.163E-07 | 1.700E-06 | 2.500E-06 |
| 6396 | 320 | C320S | A | 1.4e-06 | 2.5e-06 | 2.3e-06 | 2.067E-06 | 5.859E-07 | 1.400E-06 | 2.500E-06 |
| 6397 | 320 | C320T | A | 2.6e-06 | 1.5e-06 | 2.2e-06 | 2.100E-06 | 5.568E-07 | 1.500E-06 | 2.600E-06 |
| 6398 | 320 | C320W | A | 2e-06 | 2.1e-06 | 2.6e-06 | 2.233E-06 | 3.215E-07 | 2.000E-06 | 2.600E-06 |
| 6399 | 320 | C320Y | A | 3.2e-06 | 2.1e-06 | 1.2e-06 | 2.167E-06 | 1.002E-06 | 1.200E-06 | 3.200E-06 |
| 6400 | 320 | C320V | A | 1.8e-06 | 4.1e-06 | 2.4e-06 | 2.767E-06 | 1.193E-06 | 1.800E-06 | 4.100E-06 |
| 6401 | 321 | Y321A | A | 2.2e-06 | 2.6e-06 | 2.1e-06 | 2.300E-06 | 2.646E-07 | 2.100E-06 | 2.600E-06 |
| 6402 | 321 | Y321R | A | 1.7e-06 | 8.7e-07 | 2.8e-06 | 1.790E-06 | 9.681E-07 | 8.700E-07 | 2.800E-06 |
| 6403 | 321 | Y321N | A | 1.6e-06 | 2e-06 | 3e-06 | 2.200E-06 | 7.211E-07 | 1.600E-06 | 3.000E-06 |
| 6404 | 321 | Y321D | A | 2.1e-06 | 2.3e-06 | 2.8e-06 | 2.400E-06 | 3.606E-07 | 2.100E-06 | 2.800E-06 |
| 6405 | 321 | Y321C | A | 2.4e-06 | 2.6e-06 | 2.7e-06 | 2.567E-06 | 1.528E-07 | 2.400E-06 | 2.700E-06 |
| 6406 | 321 | Y321E | A | 2.1e-06 | 2.5e-06 | 3e-06 | 2.533E-06 | 4.509E-07 | 2.100E-06 | 3.000E-06 |
| 6407 | 321 | Y321Q | A | 1.8e-06 | 2.5e-06 | 1.3e-06 | 1.867E-06 | 6.028E-07 | 1.300E-06 | 2.500E-06 |
| 6408 | 321 | Y321G | A | 3e-06 | 3.1e-06 | 2.9e-06 | 3.000E-06 | 1.000E-07 | 2.900E-06 | 3.100E-06 |
| 6409 | 321 | Y321H | A | 2.3e-06 | 2.8e-06 | 3.4e-06 | 2.833E-06 | 5.508E-07 | 2.300E-06 | 3.400E-06 |
| 6410 | 321 | Y321I | A | 1.7e-06 | 1.4e-06 | 1.8e-06 | 1.633E-06 | 2.082E-07 | 1.400E-06 | 1.800E-06 |
| 6411 | 321 | Y321L | A | 1.7e-06 | 4.3e-06 | 3.3e-06 | 3.100E-06 | 1.311E-06 | 1.700E-06 | 4.300E-06 |
| 6412 | 321 | Y321K | A | 3.6e-06 | 2.5e-06 | 2e-06 | 2.700E-06 | 8.185E-07 | 2.000E-06 | 3.600E-06 |
| 6413 | 321 | Y321M | A | 2.4e-06 | 3e-06 | 2.6e-06 | 2.667E-06 | 3.055E-07 | 2.400E-06 | 3.000E-06 |
| 6414 | 321 | Y321F | A | 2.2e-06 | 2.7e-06 | 2.2e-06 | 2.367E-06 | 2.887E-07 | 2.200E-06 | 2.700E-06 |
| 6415 | 321 | Y321P | A | 2.9e-06 | 1.5e-06 | 1.6e-06 | 2.000E-06 | 7.810E-07 | 1.500E-06 | 2.900E-06 |
| 6416 | 321 | Y321S | A | 1.9e-06 | 2.6e-06 | 2.1e-06 | 2.200E-06 | 3.606E-07 | 1.900E-06 | 2.600E-06 |
| 6417 | 321 | Y321T | A | 3e-06 | 2.1e-06 | 2.4e-06 | 2.500E-06 | 4.583E-07 | 2.100E-06 | 3.000E-06 |
| 6418 | 321 | Y321W | A | 3.6e-06 | 1.6e-06 | 2.9e-06 | 2.700E-06 | 1.015E-06 | 1.600E-06 | 3.600E-06 |
| 6419 | 321 | Y321Y | A | 3e-06 | 2.1e-06 | 2.3e-06 | 2.467E-06 | 4.726E-07 | 2.100E-06 | 3.000E-06 |
| 6420 | 321 | Y321V | A | 3e-06 | 4e-06 | 2e-06 | 3.000E-06 | 1.000E-06 | 2.000E-06 | 4.000E-06 |
| 6421 | 322 | G322A | A | 1.2e-06 | 1.8e-06 | 2.6e-06 | 1.867E-06 | 7.024E-07 | 1.200E-06 | 2.600E-06 |
| 6422 | 322 | G322R | A | 2.9e-06 | 2.6e-06 | 1.4e-06 | 2.300E-06 | 7.937E-07 | 1.400E-06 | 2.900E-06 |
| 6423 | 322 | G322N | A | 2.5e-06 | 2.6e-06 | 1.9e-06 | 2.333E-06 | 3.786E-07 | 1.900E-06 | 2.600E-06 |
| 6424 | 322 | G322D | A | 2.5e-06 | 3.1e-06 | 2.3e-06 | 2.633E-06 | 4.163E-07 | 2.300E-06 | 3.100E-06 |
| 6425 | 322 | G322C | A | 1.9e-06 | 2.3e-06 | 1.3e-06 | 1.833E-06 | 5.033E-07 | 1.300E-06 | 2.300E-06 |
| 6426 | 322 | G322E | A | 1.5e-06 | 2.9e-06 | 1.4e-06 | 1.933E-06 | 8.386E-07 | 1.400E-06 | 2.900E-06 |
| 6427 | 322 | G322Q | A | 2.5e-06 | 1.2e-06 | 1.8e-06 | 1.833E-06 | 6.506E-07 | 1.200E-06 | 2.500E-06 |
| 6428 | 322 | G322G | A | 2.5e-06 | 4.4e-06 | 1.8e-06 | 2.900E-06 | 1.345E-06 | 1.800E-06 | 4.400E-06 |
| 6429 | 322 | G322H | A | 2.3e-06 | 3.2e-06 | 3.8e-06 | 3.100E-06 | 7.550E-07 | 2.300E-06 | 3.800E-06 |
| 6430 | 322 | G322I | A | 1.9e-06 | 2.6e-06 | 1.4e-06 | 1.967E-06 | 6.028E-07 | 1.400E-06 | 2.600E-06 |
| 6431 | 322 | G322L | A | 2.6e-06 | 3.1e-06 | 2.7e-06 | 2.800E-06 | 2.646E-07 | 2.600E-06 | 3.100E-06 |
| 6432 | 322 | G322K | A | 2.7e-06 | 3e-06 | 3.8e-06 | 3.167E-06 | 5.686E-07 | 2.700E-06 | 3.800E-06 |
| 6433 | 322 | G322M | A | 3e-06 | 1.6e-06 | 1.8e-06 | 2.133E-06 | 7.572E-07 | 1.600E-06 | 3.000E-06 |
| 6434 | 322 | G322F | A | 3.9e-06 | 2.6e-06 | 1.3e-06 | 2.600E-06 | 1.300E-06 | 1.300E-06 | 3.900E-06 |
| 6435 | 322 | G322P | A | 1.4e-06 | 1.5e-06 | 1.7e-06 | 1.533E-06 | 1.528E-07 | 1.400E-06 | 1.700E-06 |
| 6436 | 322 | G322S | A | 1.7e-06 | 3.3e-06 | 2.8e-06 | 2.600E-06 | 8.185E-07 | 1.700E-06 | 3.300E-06 |
| 6437 | 322 | G322T | A | 2.8e-06 | 2.1e-06 | 1e-06 | 1.967E-06 | 9.074E-07 | 1.000E-06 | 2.800E-06 |
| 6438 | 322 | G322W | A | 1.9e-06 | 1.2e-06 | 2e-06 | 1.700E-06 | 4.359E-07 | 1.200E-06 | 2.000E-06 |
| 6439 | 322 | G322Y | A | 3.4e-06 | 3.6e-06 | 1.6e-06 | 2.867E-06 | 1.102E-06 | 1.600E-06 | 3.600E-06 |
| 6440 | 322 | G322V | A | 3e-06 | 2.4e-06 | 3e-06 | 2.800E-06 | 3.464E-07 | 2.400E-06 | 3.000E-06 |
| 6441 | 323 | L323A | A | 1.8e-06 | 2.7e-06 | 2.8e-06 | 2.433E-06 | 5.508E-07 | 1.800E-06 | 2.800E-06 |
| 6442 | 323 | L323R | A | 2.4e-06 | 2.9e-06 | 3.3e-06 | 2.867E-06 | 4.509E-07 | 2.400E-06 | 3.300E-06 |
| 6443 | 323 | L323N | A | 1.8e-06 | 3e-06 | 1.7e-06 | 2.167E-06 | 7.234E-07 | 1.700E-06 | 3.000E-06 |
| 6444 | 323 | L323D | A | 4.2e-06 | 2.3e-06 | 3e-06 | 3.167E-06 | 9.609E-07 | 2.300E-06 | 4.200E-06 |
| 6445 | 323 | L323C | A | 2.2e-06 | 3.4e-06 | 2.9e-06 | 2.833E-06 | 6.028E-07 | 2.200E-06 | 3.400E-06 |
| 6446 | 323 | L323E | A | 2.4e-06 | 1.9e-06 | 1.1e-06 | 1.800E-06 | 6.557E-07 | 1.100E-06 | 2.400E-06 |
| 6447 | 323 | L323Q | A | 2.3e-06 | 1.5e-06 | 2.3e-06 | 2.033E-06 | 4.619E-07 | 1.500E-06 | 2.300E-06 |
| 6448 | 323 | L323G | A | 1.6e-06 | 2.5e-06 | 1.1e-06 | 1.733E-06 | 7.095E-07 | 1.100E-06 | 2.500E-06 |
| 6449 | 323 | L323H | A | 2.5e-06 | 2.5e-06 | 2.3e-06 | 2.433E-06 | 1.155E-07 | 2.300E-06 | 2.500E-06 |
| 6450 | 323 | L323I | A | 2.3e-06 | 2.5e-06 | 3.4e-06 | 2.733E-06 | 5.859E-07 | 2.300E-06 | 3.400E-06 |
| 6451 | 323 | L323L | A | 3.6e-06 | 1.5e-06 | 1.8e-06 | 2.300E-06 | 1.136E-06 | 1.500E-06 | 3.600E-06 |
| 6452 | 323 | L323K | A | 2e-06 | 3.7e-06 | 2.4e-06 | 2.700E-06 | 8.888E-07 | 2.000E-06 | 3.700E-06 |
| 6453 | 323 | L323M | A | 2.6e-06 | 1.3e-06 | 1.3e-06 | 1.733E-06 | 7.506E-07 | 1.300E-06 | 2.600E-06 |
| 6454 | 323 | L323F | A | 3.3e-06 | 1.9e-06 | 2e-06 | 2.400E-06 | 7.810E-07 | 1.900E-06 | 3.300E-06 |
| 6455 | 323 | L323P | A | 2.1e-06 | 2.6e-06 | 2.1e-06 | 2.267E-06 | 2.887E-07 | 2.100E-06 | 2.600E-06 |
| 6456 | 323 | L323S | A | 1.4e-06 | 1.8e-06 | 2e-06 | 1.733E-06 | 3.055E-07 | 1.400E-06 | 2.000E-06 |
| 6457 | 323 | L323T | A | 3e-06 | 2.8e-06 | 1.4e-06 | 2.400E-06 | 8.718E-07 | 1.400E-06 | 3.000E-06 |
| 6458 | 323 | L323W | A | 2.3e-06 | 1.8e-06 | 1.8e-06 | 1.967E-06 | 2.887E-07 | 1.800E-06 | 2.300E-06 |
| 6459 | 323 | L323Y | A | 1.7e-06 | 2.7e-06 | 1.9e-06 | 2.100E-06 | 5.292E-07 | 1.700E-06 | 2.700E-06 |

| | | | | | | | | | | |
|---|---|---|---|---|---|---|---|---|---|---|
| 6460 | 323 | L323V | A | 2.5e-06 | 1.1e-06 | 2.3e-06 | 1.967E-06 | 7.572E-07 | 1.100E-06 | 2.500E-06 |
| 6461 | 324 | G324A | A | 1.6e-06 | 1.9e-06 | 2.6e-06 | 2.033E-06 | 5.132E-07 | 1.600E-06 | 2.600E-06 |
| 6462 | 324 | G324R | A | 2.7e-06 | 3.7e-06 | 3e-06 | 3.133E-06 | 5.132E-07 | 2.700E-06 | 3.700E-06 |
| 6463 | 324 | G324N | A | 1.8e-06 | 1.6e-06 | 1.6e-06 | 1.667E-06 | 1.155E-07 | 1.600E-06 | 1.800E-06 |
| 6464 | 324 | G324D | A | 2.4e-06 | 2.9e-06 | 3.6e-06 | 2.967E-06 | 6.028E-07 | 2.400E-06 | 3.600E-06 |
| 6465 | 324 | G324C | A | 2.6e-06 | 3.3e-06 | 2.4e-06 | 2.767E-06 | 4.726E-07 | 2.400E-06 | 3.300E-06 |
| 6466 | 324 | G324E | A | 2.1e-06 | 2.3e-06 | 2e-06 | 2.133E-06 | 1.528E-07 | 2.000E-06 | 2.300E-06 |
| 6467 | 324 | G324Q | A | 3.1e-06 | 2.1e-06 | 1.7e-06 | 2.300E-06 | 7.211E-07 | 1.700E-06 | 3.100E-06 |
| 6468 | 324 | G324G | A | 3.6e-06 | 1.5e-06 | 1.5e-06 | 2.200E-06 | 1.212E-06 | 1.500E-06 | 3.600E-06 |
| 6469 | 324 | G324H | A | 1.8e-06 | 1.9e-06 | 3.1e-06 | 2.267E-06 | 7.234E-07 | 1.800E-06 | 3.100E-06 |
| 6470 | 324 | G324I | A | 2.6e-06 | 2.7e-06 | 2.3e-06 | 2.533E-06 | 2.082E-07 | 2.300E-06 | 2.700E-06 |
| 6471 | 324 | G324L | A | 2.3e-06 | 1.6e-06 | 3.1e-06 | 2.333E-06 | 7.506E-07 | 1.600E-06 | 3.100E-06 |
| 6472 | 324 | G324K | A | 2.3e-06 | 1.4e-06 | 3.2e-06 | 2.300E-06 | 9.000E-07 | 1.400E-06 | 3.200E-06 |
| 6473 | 324 | G324M | A | 2.5e-06 | 1.3e-06 | 2.4e-06 | 2.067E-06 | 6.658E-07 | 1.300E-06 | 2.500E-06 |
| 6474 | 324 | G324F | A | 2.3e-06 | 1.8e-06 | 3.4e-06 | 2.500E-06 | 8.185E-07 | 1.800E-06 | 3.400E-06 |
| 6475 | 324 | G324P | A | 2.2e-06 | 1.5e-06 | 2.2e-06 | 1.967E-06 | 4.041E-07 | 1.500E-06 | 2.200E-06 |
| 6476 | 324 | G324S | A | 2.1e-06 | 1.1e-06 | 1.7e-06 | 1.633E-06 | 5.033E-07 | 1.100E-06 | 2.100E-06 |
| 6477 | 324 | G324T | A | 2e-06 | 3.5e-06 | 8.8e-07 | 2.127E-06 | 1.315E-06 | 8.800E-07 | 3.500E-06 |
| 6478 | 324 | G324W | A | 2.4e-06 | 3.2e-06 | 3.4e-06 | 3.000E-06 | 5.292E-07 | 2.400E-06 | 3.400E-06 |
| 6479 | 324 | G324Y | A | 2.3e-06 | 3e-06 | 1.9e-06 | 2.400E-06 | 5.568E-07 | 1.900E-06 | 3.000E-06 |
| 6480 | 324 | G324V | A | 2.1e-06 | 1.8e-06 | 1.4e-06 | 1.767E-06 | 3.512E-07 | 1.400E-06 | 2.100E-06 |
| 6481 | 325 | M325A | A | 1.7e-06 | 1.7e-06 | 1.9e-06 | 1.767E-06 | 1.155E-07 | 1.700E-06 | 1.900E-06 |
| 6482 | 325 | M325R | A | 2.2e-06 | 3e-06 | 3e-06 | 2.733E-06 | 4.619E-07 | 2.200E-06 | 3.000E-06 |
| 6483 | 325 | M325N | A | 2.6e-06 | 3.2e-06 | 1.5e-06 | 2.433E-06 | 8.622E-07 | 1.500E-06 | 3.200E-06 |
| 6484 | 325 | M325D | A | 2.1e-06 | 9e-07 | 9.7e-07 | 1.323E-06 | 6.735E-07 | 9.000E-07 | 2.100E-06 |
| 6485 | 325 | M325C | A | 1.2e-06 | 3.4e-06 | 2.3e-06 | 2.300E-06 | 1.100E-06 | 1.200E-06 | 3.400E-06 |
| 6486 | 325 | M325E | A | 3.2e-06 | 2.2e-06 | 2.3e-06 | 2.567E-06 | 5.508E-07 | 2.200E-06 | 3.200E-06 |
| 6487 | 325 | M325Q | A | 2.1e-06 | 1.5e-06 | 2.2e-06 | 1.933E-06 | 3.786E-07 | 1.500E-06 | 2.200E-06 |
| 6488 | 325 | M325G | A | 3.8e-06 | 3.1e-06 | 2e-06 | 2.967E-06 | 9.074E-07 | 2.000E-06 | 3.800E-06 |
| 6489 | 325 | M325H | A | 2.5e-06 | 1.7e-06 | 1.2e-06 | 1.800E-06 | 6.557E-07 | 1.200E-06 | 2.500E-06 |
| 6490 | 325 | M325I | A | 1.8e-06 | 2.2e-06 | 3.9e-06 | 2.633E-06 | 1.115E-06 | 1.800E-06 | 3.900E-06 |
| 6491 | 325 | M325L | A | 2.2e-06 | 2.2e-06 | 1.1e-06 | 1.833E-06 | 6.351E-07 | 1.100E-06 | 2.200E-06 |
| 6492 | 325 | M325K | A | 2.3e-06 | 2.1e-06 | 3.2e-06 | 2.533E-06 | 5.859E-07 | 2.100E-06 | 3.200E-06 |
| 6493 | 325 | M325M | A | 2e-06 | 3.9e-06 | 3.2e-06 | 3.033E-06 | 9.609E-07 | 2.000E-06 | 3.900E-06 |
| 6494 | 325 | M325F | A | 2.6e-06 | 1.8e-06 | 1.9e-06 | 2.100E-06 | 4.359E-07 | 1.800E-06 | 2.600E-06 |
| 6495 | 325 | M325P | A | 9.6e-07 | 1.3e-06 | 3.5e-06 | 1.920E-06 | 1.379E-06 | 9.600E-07 | 3.500E-06 |
| 6496 | 325 | M325S | A | 2.1e-06 | 2.5e-06 | 2.8e-06 | 2.467E-06 | 3.512E-07 | 2.100E-06 | 2.800E-06 |
| 6497 | 325 | M325T | A | 2.2e-06 | 2.8e-06 | 1.2e-06 | 2.067E-06 | 8.083E-07 | 1.200E-06 | 2.800E-06 |
| 6498 | 325 | M325W | A | 3e-06 | 3.1e-06 | 2.3e-06 | 2.800E-06 | 4.359E-07 | 2.300E-06 | 3.100E-06 |
| 6499 | 325 | M325Y | A | 1.6e-06 | 1.5e-06 | 1.8e-06 | 1.633E-06 | 1.528E-07 | 1.500E-06 | 1.800E-06 |
| 6500 | 325 | M325V | A | 2.1e-06 | 3.8e-06 | 2.8e-06 | 2.900E-06 | 8.544E-07 | 2.100E-06 | 3.800E-06 |
| 6501 | 326 | E326A | A | 2.4e-06 | 2.2e-06 | 1.8e-06 | 2.133E-06 | 3.055E-07 | 1.800E-06 | 2.400E-06 |
| 6502 | 326 | E326R | A | 1.4e-06 | 2.1e-06 | 3.1e-06 | 2.200E-06 | 8.544E-07 | 1.400E-06 | 3.100E-06 |
| 6503 | 326 | E326N | A | 4.3e-06 | 1.6e-06 | 1.9e-06 | 2.600E-06 | 1.480E-06 | 1.600E-06 | 4.300E-06 |
| 6504 | 326 | E326D | A | 3.6e-06 | 2.8e-06 | 2.8e-06 | 3.067E-06 | 4.619E-07 | 2.800E-06 | 3.600E-06 |
| 6505 | 326 | E326C | A | 1.6e-06 | 1.8e-06 | 2.3e-06 | 1.900E-06 | 3.606E-07 | 1.600E-06 | 2.300E-06 |
| 6506 | 326 | E326E | A | 2.7e-06 | 2.3e-06 | 3.1e-06 | 2.700E-06 | 4.000E-07 | 2.300E-06 | 3.100E-06 |
| 6507 | 326 | E326Q | A | 2.1e-06 | 2e-06 | 2.9e-06 | 2.333E-06 | 4.933E-07 | 2.000E-06 | 2.900E-06 |
| 6508 | 326 | E326G | A | 1.9e-06 | 2.5e-06 | 1.7e-06 | 2.033E-06 | 4.163E-07 | 1.700E-06 | 2.500E-06 |
| 6509 | 326 | E326H | A | 3.3e-06 | 2.2e-06 | 1.4e-06 | 2.300E-06 | 9.539E-07 | 1.400E-06 | 3.300E-06 |
| 6510 | 326 | E326I | A | 4.2e-06 | 1.3e-06 | 2.6e-06 | 2.700E-06 | 1.453E-06 | 1.300E-06 | 4.200E-06 |
| 6511 | 326 | E326L | A | 2.1e-06 | 9.6e-07 | 2.5e-06 | 1.853E-06 | 7.991E-07 | 9.600E-07 | 2.500E-06 |
| 6512 | 326 | E326K | A | 2.4e-06 | 4e-06 | 3.7e-06 | 3.367E-06 | 8.505E-07 | 2.400E-06 | 4.000E-06 |
| 6513 | 326 | E326M | A | 3.7e-06 | 3.2e-06 | 2.7e-06 | 3.200E-06 | 5.000E-07 | 2.700E-06 | 3.700E-06 |
| 6514 | 326 | E326F | A | 2.2e-06 | 2e-06 | 2.2e-06 | 2.133E-06 | 1.155E-07 | 2.000E-06 | 2.200E-06 |
| 6515 | 326 | E326P | A | 3e-06 | 2.6e-06 | 2e-06 | 2.533E-06 | 5.033E-07 | 2.000E-06 | 3.000E-06 |
| 6516 | 326 | E326S | A | 3.1e-06 | 1.6e-06 | 3.2e-06 | 2.633E-06 | 8.963E-07 | 1.600E-06 | 3.200E-06 |
| 6517 | 326 | E326T | A | 1.8e-06 | 3.1e-06 | 3.9e-06 | 2.933E-06 | 1.060E-06 | 1.800E-06 | 3.900E-06 |
| 6518 | 326 | E326W | A | 1.4e-06 | 1.4e-06 | 2.2e-06 | 1.667E-06 | 4.619E-07 | 1.400E-06 | 2.200E-06 |
| 6519 | 326 | E326Y | A | 1.8e-06 | 2.6e-06 | 3.9e-06 | 2.767E-06 | 1.060E-06 | 1.800E-06 | 3.900E-06 |
| 6520 | 326 | E326V | A | 2.1e-06 | 2.3e-06 | 3.6e-06 | 2.667E-06 | 8.145E-07 | 2.100E-06 | 3.600E-06 |
| 6521 | 327 | H327A | A | 2.5e-06 | 2.9e-06 | 2.9e-06 | 2.767E-06 | 2.309E-07 | 2.500E-06 | 2.900E-06 |
| 6522 | 327 | H327R | A | 1.5e-06 | 2.8e-06 | 3.1e-06 | 2.467E-06 | 8.505E-07 | 1.500E-06 | 3.100E-06 |
| 6523 | 327 | H327N | A | 2e-06 | 3.2e-06 | 2.9e-06 | 2.700E-06 | 6.245E-07 | 2.000E-06 | 3.200E-06 |
| 6524 | 327 | H327D | A | 1.1e-06 | 2.6e-06 | 1.9e-06 | 1.867E-06 | 7.506E-07 | 1.100E-06 | 2.600E-06 |
| 6525 | 327 | H327C | A | 1.9e-06 | 3.3e-06 | 3.3e-06 | 2.833E-06 | 8.083E-07 | 1.900E-06 | 3.300E-06 |
| 6526 | 327 | H327E | A | 2.2e-06 | 2.4e-06 | 1.7e-06 | 2.100E-06 | 3.606E-07 | 1.700E-06 | 2.400E-06 |
| 6527 | 327 | H327Q | A | 9.9e-07 | 2.5e-06 | 2e-06 | 1.830E-06 | 7.692E-07 | 9.900E-07 | 2.500E-06 |
| 6528 | 327 | H327G | A | 2e-06 | 2.6e-06 | 2.6e-06 | 2.400E-06 | 3.464E-07 | 2.000E-06 | 2.600E-06 |
| 6529 | 327 | H327H | A | 1.2e-06 | 1.2e-06 | 2.8e-06 | 1.733E-06 | 9.238E-07 | 1.200E-06 | 2.800E-06 |
| 6530 | 327 | H327I | A | 3.2e-06 | 4.1e-06 | 2.4e-06 | 3.233E-06 | 8.505E-07 | 2.400E-06 | 4.100E-06 |
| 6531 | 327 | H327L | A | 1.9e-06 | 1.7e-06 | 3.9e-06 | 2.500E-06 | 1.217E-06 | 1.700E-06 | 3.900E-06 |
| 6532 | 327 | H327K | A | 3e-06 | 2.7e-06 | 2.4e-06 | 2.700E-06 | 3.000E-07 | 2.400E-06 | 3.000E-06 |
| 6533 | 327 | H327M | A | 1.6e-06 | 2e-06 | 2.9e-06 | 2.167E-06 | 6.658E-07 | 1.600E-06 | 2.900E-06 |
| 6534 | 327 | H327F | A | 2.2e-06 | 2.2e-06 | 2.2e-06 | 2.200E-06 | 0.000E+00 | 2.200E-06 | 2.200E-06 |
| 6535 | 327 | H327P | A | 2e-06 | 3.1e-06 | 2.9e-06 | 2.667E-06 | 5.859E-07 | 2.000E-06 | 3.100E-06 |
| 6536 | 327 | H327S | A | 1.8e-06 | 2.2e-06 | 2.5e-06 | 2.167E-06 | 3.512E-07 | 1.800E-06 | 2.500E-06 |
| 6537 | 327 | H327T | A | 2.3e-06 | 2.5e-06 | 3e-06 | 2.600E-06 | 3.606E-07 | 2.300E-06 | 3.000E-06 |
| 6538 | 327 | H327W | A | 3.3e-06 | 3e-06 | 1.4e-06 | 2.567E-06 | 1.021E-06 | 1.400E-06 | 3.300E-06 |
| 6539 | 327 | H327Y | A | 1.7e-06 | 3.2e-06 | 1.5e-06 | 2.133E-06 | 9.292E-07 | 1.500E-06 | 3.200E-06 |
| 6540 | 327 | H327V | A | 2e-06 | 2.8e-06 | 3e-06 | 2.600E-06 | 5.292E-07 | 2.000E-06 | 3.000E-06 |
| 6541 | 328 | L328A | A | 1.1e-06 | 1.5e-06 | 1.9e-06 | 1.500E-06 | 4.000E-07 | 1.100E-06 | 1.900E-06 |
| 6542 | 328 | L328R | A | 3e-06 | 1.1e-06 | 2.6e-06 | 2.233E-06 | 1.002E-06 | 1.100E-06 | 3.000E-06 |
| 6543 | 328 | L328N | A | 2.1e-06 | 2.2e-06 | 1.7e-06 | 2.000E-06 | 2.646E-07 | 1.700E-06 | 2.200E-06 |
| 6544 | 328 | L328D | A | 1.7e-06 | 1.9e-06 | 3.8e-06 | 2.467E-06 | 1.159E-06 | 1.700E-06 | 3.800E-06 |

| | | | | | | | | | | |
|---|---|---|---|---|---|---|---|---|---|---|
| 6545 | 328 | L328C | A | 2.2e-06 | 4.8e-06 | 2e-06 | 3.000E-06 | 1.562E-06 | 2.000E-06 | 4.800E-06 |
| 6546 | 328 | L328E | A | 2.3e-06 | 3e-06 | 3.1e-06 | 2.800E-06 | 4.359E-07 | 2.300E-06 | 3.100E-06 |
| 6547 | 328 | L328Q | A | 2.9e-06 | 2.8e-06 | 1.2e-06 | 2.300E-06 | 9.539E-07 | 1.200E-06 | 2.900E-06 |
| 6548 | 328 | L328G | A | 2.4e-06 | 2.1e-06 | 1.9e-06 | 2.133E-06 | 2.517E-07 | 1.900E-06 | 2.400E-06 |
| 6549 | 328 | L328H | A | 2.4e-06 | 2.4e-06 | 2.4e-06 | 2.400E-06 | 0.000E+00 | 2.400E-06 | 2.400E-06 |
| 6550 | 328 | L328I | A | 2.5e-06 | 2.8e-06 | 2.4e-06 | 2.567E-06 | 2.082E-07 | 2.400E-06 | 2.800E-06 |
| 6551 | 328 | L328L | A | 1e-06 | 1.9e-06 | 1.6e-06 | 1.500E-06 | 4.583E-07 | 1.000E-06 | 1.900E-06 |
| 6552 | 328 | L328K | A | 2.4e-06 | 2.8e-06 | 2.2e-06 | 2.467E-06 | 3.055E-07 | 2.200E-06 | 2.800E-06 |
| 6553 | 328 | L328M | A | 3.5e-06 | 4.6e-06 | 2.2e-06 | 3.433E-06 | 1.201E-06 | 2.200E-06 | 4.600E-06 |
| 6554 | 328 | L328F | A | 1e-06 | 2.2e-06 | 1.6e-06 | 1.600E-06 | 6.000E-07 | 1.000E-06 | 2.200E-06 |
| 6555 | 328 | L328P | A | 1.5e-06 | 1.5e-06 | 1.2e-06 | 1.400E-06 | 1.732E-07 | 1.200E-06 | 1.500E-06 |
| 6556 | 328 | L328S | A | 2.9e-06 | 4.3e-06 | 2.8e-06 | 3.333E-06 | 8.386E-07 | 2.800E-06 | 4.300E-06 |
| 6557 | 328 | L328T | A | 2.6e-06 | 2.2e-06 | 2.2e-06 | 2.333E-06 | 2.309E-07 | 2.200E-06 | 2.600E-06 |
| 6558 | 328 | L328W | A | 2.3e-06 | 7.4e-07 | 2e-06 | 1.680E-06 | 8.278E-07 | 7.400E-07 | 2.300E-06 |
| 6559 | 328 | L328Y | A | 3.2e-06 | 2.6e-06 | 1.1e-06 | 2.300E-06 | 1.082E-06 | 1.100E-06 | 3.200E-06 |
| 6560 | 328 | L328V | A | 1.9e-06 | 1.7e-06 | 1.6e-06 | 1.733E-06 | 1.528E-07 | 1.600E-06 | 1.900E-06 |
| 6561 | 329 | R329A | A | 2.4e-06 | 1.5e-06 | 2.2e-06 | 2.033E-06 | 4.726E-07 | 1.500E-06 | 2.400E-06 |
| 6562 | 329 | R329R | A | 2.6e-06 | 2.8e-06 | 1.8e-06 | 2.400E-06 | 5.292E-07 | 1.800E-06 | 2.800E-06 |
| 6563 | 329 | R329N | A | 2.1e-06 | 2.6e-06 | 3.6e-06 | 2.767E-06 | 7.638E-07 | 2.100E-06 | 3.600E-06 |
| 6564 | 329 | R329D | A | 2e-06 | 2.4e-06 | 1.6e-06 | 2.000E-06 | 4.000E-07 | 1.600E-06 | 2.400E-06 |
| 6565 | 329 | R329C | A | 2.6e-06 | 2.6e-06 | 2.4e-06 | 2.533E-06 | 1.155E-07 | 2.400E-06 | 2.600E-06 |
| 6566 | 329 | R329E | A | 2.7e-06 | 3.4e-06 | 3.2e-06 | 3.100E-06 | 3.606E-07 | 2.700E-06 | 3.400E-06 |
| 6567 | 329 | R329Q | A | 3.1e-06 | 2.5e-06 | 1.9e-06 | 2.500E-06 | 6.000E-07 | 1.900E-06 | 3.100E-06 |
| 6568 | 329 | R329G | A | 1.9e-06 | 3.5e-06 | 3.3e-06 | 2.900E-06 | 8.718E-07 | 1.900E-06 | 3.500E-06 |
| 6569 | 329 | R329H | A | 4.3e-06 | 3.1e-06 | 2.5e-06 | 3.300E-06 | 9.165E-07 | 2.500E-06 | 4.300E-06 |
| 6570 | 329 | R329I | A | 2e-06 | 1.9e-06 | 3.8e-06 | 2.567E-06 | 1.069E-06 | 1.900E-06 | 3.800E-06 |
| 6571 | 329 | R329L | A | 4.8e-06 | 2.5e-06 | 2.6e-06 | 3.300E-06 | 1.300E-06 | 2.500E-06 | 4.800E-06 |
| 6572 | 329 | R329K | A | 1.8e-06 | 2.3e-06 | 1.7e-06 | 1.933E-06 | 3.215E-07 | 1.700E-06 | 2.300E-06 |
| 6573 | 329 | R329M | A | 2e-06 | 1.7e-06 | 1.3e-06 | 1.667E-06 | 3.512E-07 | 1.300E-06 | 2.000E-06 |
| 6574 | 329 | R329F | A | 1.8e-06 | 3.3e-06 | 2.2e-06 | 2.433E-06 | 7.767E-07 | 1.800E-06 | 3.300E-06 |
| 6575 | 329 | R329P | A | 2.7e-06 | 3.1e-06 | 2.4e-06 | 2.733E-06 | 3.512E-07 | 2.400E-06 | 3.100E-06 |
| 6576 | 329 | R329S | A | 2.4e-06 | 2e-06 | 2.1e-06 | 2.167E-06 | 2.082E-07 | 2.000E-06 | 2.400E-06 |
| 6577 | 329 | R329T | A | 2.6e-06 | 3.1e-06 | 3e-06 | 2.900E-06 | 2.646E-07 | 2.600E-06 | 3.100E-06 |
| 6578 | 329 | R329W | A | 2.1e-06 | 5.2e-06 | 2.1e-06 | 3.133E-06 | 1.790E-06 | 2.100E-06 | 5.200E-06 |
| 6579 | 329 | R329Y | A | 2.1e-06 | 1.5e-06 | 2.2e-06 | 1.933E-06 | 3.786E-07 | 1.500E-06 | 2.200E-06 |
| 6580 | 329 | R329V | A | 1.9e-06 | 2.4e-06 | 1.8e-06 | 2.033E-06 | 3.215E-07 | 1.800E-06 | 2.400E-06 |
| 6581 | 330 | E330A | A | 3.2e-06 | 2.2e-06 | 2.3e-06 | 2.567E-06 | 5.508E-07 | 2.200E-06 | 3.200E-06 |
| 6582 | 330 | E330R | A | 1.8e-06 | 2.3e-06 | 2.1e-06 | 2.067E-06 | 2.517E-07 | 1.800E-06 | 2.300E-06 |
| 6583 | 330 | E330N | A | 1.6e-06 | 1.5e-06 | 3.9e-06 | 2.333E-06 | 1.358E-06 | 1.500E-06 | 3.900E-06 |
| 6584 | 330 | E330D | A | 2.9e-06 | 1.8e-06 | 3.1e-06 | 2.600E-06 | 7.000E-07 | 1.800E-06 | 3.100E-06 |
| 6585 | 330 | E330C | A | 2e-06 | 1.9e-06 | 1.4e-06 | 1.767E-06 | 3.215E-07 | 1.400E-06 | 2.000E-06 |
| 6586 | 330 | E330E | A | 2.4e-06 | 2e-06 | 2.6e-06 | 2.333E-06 | 3.055E-07 | 2.000E-06 | 2.600E-06 |
| 6587 | 330 | E330Q | A | 3.4e-06 | 1.3e-06 | 1.4e-06 | 2.033E-06 | 1.185E-06 | 1.300E-06 | 3.400E-06 |
| 6588 | 330 | E330G | A | 2.3e-06 | 4.7e-06 | 2.3e-06 | 3.100E-06 | 1.386E-06 | 2.300E-06 | 4.700E-06 |
| 6589 | 330 | E330H | A | 2.4e-06 | 3.1e-06 | 2.7e-06 | 2.733E-06 | 3.512E-07 | 2.400E-06 | 3.100E-06 |
| 6590 | 330 | E330I | A | 4.2e-06 | 2.5e-06 | 1.1e-06 | 2.600E-06 | 1.552E-06 | 1.100E-06 | 4.200E-06 |
| 6591 | 330 | E330L | A | 3.2e-06 | 2.9e-06 | 2.7e-06 | 2.933E-06 | 2.517E-07 | 2.700E-06 | 3.200E-06 |
| 6592 | 330 | E330K | A | 2.4e-06 | 4.6e-06 | 1.5e-06 | 2.833E-06 | 1.595E-06 | 1.500E-06 | 4.600E-06 |
| 6593 | 330 | E330M | A | 3.9e-06 | 2.8e-06 | 2.6e-06 | 3.100E-06 | 7.000E-07 | 2.600E-06 | 3.900E-06 |
| 6594 | 330 | E330F | A | 1.4e-06 | 1.4e-06 | 1.5e-06 | 1.433E-06 | 5.774E-08 | 1.400E-06 | 1.500E-06 |
| 6595 | 330 | E330P | A | 2.2e-06 | 2.8e-06 | 1e-06 | 2.000E-06 | 9.165E-07 | 1.000E-06 | 2.800E-06 |
| 6596 | 330 | E330S | A | 2.6e-06 | 3.5e-06 | 2.6e-06 | 2.900E-06 | 5.196E-07 | 2.600E-06 | 3.500E-06 |
| 6597 | 330 | E330T | A | 1.5e-06 | 1.9e-06 | 2.5e-06 | 1.967E-06 | 5.033E-07 | 1.500E-06 | 2.500E-06 |
| 6598 | 330 | E330W | A | 2.8e-06 | 3.1e-06 | 2.6e-06 | 2.833E-06 | 2.517E-07 | 2.600E-06 | 3.100E-06 |
| 6599 | 330 | E330Y | A | 1.9e-06 | 2.7e-06 | 2.3e-06 | 2.300E-06 | 4.000E-07 | 1.900E-06 | 2.700E-06 |
| 6600 | 330 | E330V | A | 4.6e-06 | 2.2e-06 | 3.4e-06 | 3.400E-06 | 1.200E-06 | 2.200E-06 | 4.600E-06 |
| 6601 | 331 | V331A | A | 2.7e-06 | 1.9e-06 | 1.7e-06 | 2.100E-06 | 5.292E-07 | 1.700E-06 | 2.700E-06 |
| 6602 | 331 | V331R | A | 2.8e-06 | 1.8e-06 | 3.4e-06 | 2.667E-06 | 8.083E-07 | 1.800E-06 | 3.400E-06 |
| 6603 | 331 | V331N | A | 1.8e-06 | 2.6e-06 | 1.9e-06 | 2.100E-06 | 4.359E-07 | 1.800E-06 | 2.600E-06 |
| 6604 | 331 | V331D | A | 1.7e-06 | 1.2e-06 | 2.4e-06 | 1.767E-06 | 6.028E-07 | 1.200E-06 | 2.400E-06 |
| 6605 | 331 | V331C | A | 3.3e-06 | 1.5e-06 | 2.8e-06 | 2.533E-06 | 9.292E-07 | 1.500E-06 | 3.300E-06 |
| 6606 | 331 | V331E | A | 1.3e-06 | 1.6e-06 | 3.5e-06 | 2.133E-06 | 1.193E-06 | 1.300E-06 | 3.500E-06 |
| 6607 | 331 | V331Q | A | 3.1e-06 | 2e-06 | 1.8e-06 | 2.300E-06 | 7.000E-07 | 1.800E-06 | 3.100E-06 |
| 6608 | 331 | V331G | A | 1.7e-06 | 1.8e-06 | 1.4e-06 | 1.633E-06 | 2.082E-07 | 1.400E-06 | 1.800E-06 |
| 6609 | 331 | V331H | A | 3.6e-06 | 2.5e-06 | 3.5e-06 | 3.200E-06 | 6.083E-07 | 2.500E-06 | 3.600E-06 |
| 6610 | 331 | V331I | A | 3.1e-06 | 4.7e-06 | 2.7e-06 | 3.500E-06 | 1.058E-06 | 2.700E-06 | 4.700E-06 |
| 6611 | 331 | V331L | A | 1.3e-06 | 2.3e-06 | 2.3e-06 | 1.967E-06 | 5.774E-07 | 1.300E-06 | 2.300E-06 |
| 6612 | 331 | V331K | A | 3.2e-06 | 2.2e-06 | 2.2e-06 | 2.533E-06 | 5.774E-07 | 2.200E-06 | 3.200E-06 |
| 6613 | 331 | V331M | A | 3.6e-06 | 2.5e-06 | 2.9e-06 | 3.000E-06 | 5.568E-07 | 2.500E-06 | 3.600E-06 |
| 6614 | 331 | V331F | A | 3.2e-06 | 3.2e-06 | 1.4e-06 | 2.600E-06 | 1.039E-06 | 1.400E-06 | 3.200E-06 |
| 6615 | 331 | V331P | A | 2.4e-06 | 2.1e-06 | 2.6e-06 | 2.367E-06 | 2.517E-07 | 2.100E-06 | 2.600E-06 |
| 6616 | 331 | V331S | A | 1.7e-06 | 3.8e-06 | 2.7e-06 | 2.733E-06 | 1.050E-06 | 1.700E-06 | 3.800E-06 |
| 6617 | 331 | V331T | A | 3.8e-06 | 3.5e-06 | 2.2e-06 | 3.167E-06 | 8.505E-07 | 2.200E-06 | 3.800E-06 |
| 6618 | 331 | V331W | A | 1.3e-06 | 2.6e-06 | 1.6e-06 | 1.833E-06 | 6.807E-07 | 1.300E-06 | 2.600E-06 |
| 6619 | 331 | V331Y | A | 2.6e-06 | 2.2e-06 | 2.4e-06 | 2.400E-06 | 2.000E-07 | 2.200E-06 | 2.600E-06 |
| 6620 | 331 | V331V | A | 2.5e-06 | 2.3e-06 | 1.3e-06 | 2.033E-06 | 6.429E-07 | 1.300E-06 | 2.500E-06 |
| 6621 | 332 | R332A | A | 2.9e-06 | 2.6e-06 | 2.7e-06 | 2.733E-06 | 1.528E-07 | 2.600E-06 | 2.900E-06 |
| 6622 | 332 | R332R | A | 2.5e-06 | 2.5e-06 | 1.5e-06 | 2.167E-06 | 5.774E-07 | 1.500E-06 | 2.500E-06 |
| 6623 | 332 | R332N | A | 3.2e-06 | 2.6e-06 | 9.8e-07 | 2.260E-06 | 1.148E-06 | 9.800E-07 | 3.200E-06 |
| 6624 | 332 | R332D | A | 2.7e-06 | 2.6e-06 | 8.7e-07 | 2.057E-06 | 1.029E-06 | 8.700E-07 | 2.700E-06 |
| 6625 | 332 | R332C | A | 2.3e-06 | 1.8e-06 | 3.4e-06 | 2.500E-06 | 8.185E-07 | 1.800E-06 | 3.400E-06 |
| 6626 | 332 | R332E | A | 1.8e-06 | 3.6e-06 | 2e-06 | 2.467E-06 | 9.866E-07 | 1.800E-06 | 3.600E-06 |
| 6627 | 332 | R332Q | A | 1.9e-06 | 2e-06 | 1.4e-06 | 1.767E-06 | 3.215E-07 | 1.400E-06 | 2.000E-06 |
| 6628 | 332 | R332G | A | 2.2e-06 | 1.9e-06 | 3e-06 | 2.367E-06 | 5.686E-07 | 1.900E-06 | 3.000E-06 |
| 6629 | 332 | R332H | A | 2.5e-06 | 3.4e-06 | 1.6e-06 | 2.500E-06 | 9.000E-07 | 1.600E-06 | 3.400E-06 |

| | | | | | | | | | | |
|---|---|---|---|---|---|---|---|---|---|---|
| 6630 | 332 | R332I | A | 2.7e-06 | 2.3e-06 | 1.5e-06 | 2.167E-06 | 6.110E-07 | 1.500E-06 | 2.700E-06 |
| 6631 | 332 | R332L | A | 3.3e-06 | 1.3e-06 | 3.3e-06 | 2.633E-06 | 1.155E-06 | 1.300E-06 | 3.300E-06 |
| 6632 | 332 | R332K | A | 7.9e-07 | 2.2e-06 | 2.3e-06 | 1.763E-06 | 8.444E-07 | 7.900E-07 | 2.300E-06 |
| 6633 | 332 | R332M | A | 2e-06 | 2.3e-06 | 2.9e-06 | 2.400E-06 | 4.583E-07 | 2.000E-06 | 2.900E-06 |
| 6634 | 332 | R332F | A | 2.2e-06 | 3.7e-06 | 2.9e-06 | 2.933E-06 | 7.506E-07 | 2.200E-06 | 3.700E-06 |
| 6635 | 332 | R332P | A | 2.5e-06 | 2.2e-06 | 1.8e-06 | 2.167E-06 | 3.512E-07 | 1.800E-06 | 2.500E-06 |
| 6636 | 332 | R332S | A | 2.1e-06 | 2.6e-06 | 2.8e-06 | 2.500E-06 | 3.606E-07 | 2.100E-06 | 2.800E-06 |
| 6637 | 332 | R332T | A | 3.7e-06 | 2e-06 | 1.9e-06 | 2.533E-06 | 1.012E-06 | 1.900E-06 | 3.700E-06 |
| 6638 | 332 | R332W | A | 1.4e-06 | 2.4e-06 | 2e-06 | 1.933E-06 | 5.033E-07 | 1.400E-06 | 2.400E-06 |
| 6639 | 332 | R332Y | A | 2.6e-06 | 2.8e-06 | 2.2e-06 | 2.533E-06 | 3.055E-07 | 2.200E-06 | 2.800E-06 |
| 6640 | 332 | R332V | A | 3e-06 | 2e-06 | 1.9e-06 | 2.300E-06 | 6.083E-07 | 1.900E-06 | 3.000E-06 |
| 6641 | 333 | A333A | A | 2.3e-06 | 3.5e-06 | 2.3e-06 | 2.700E-06 | 6.928E-07 | 2.300E-06 | 3.500E-06 |
| 6642 | 333 | A333R | A | 2.1e-06 | 2.9e-06 | 1.9e-06 | 2.300E-06 | 5.292E-07 | 1.900E-06 | 2.900E-06 |
| 6643 | 333 | A333N | A | 1.7e-06 | 1.6e-06 | 8e-07 | 1.367E-06 | 4.933E-07 | 8.000E-07 | 1.700E-06 |
| 6644 | 333 | A333D | A | 4.9e-06 | 2e-06 | 2.3e-06 | 3.067E-06 | 1.595E-06 | 2.000E-06 | 4.900E-06 |
| 6645 | 333 | A333C | A | 2.8e-06 | 2.1e-06 | 2.3e-06 | 2.400E-06 | 3.606E-07 | 2.100E-06 | 2.800E-06 |
| 6646 | 333 | A333E | A | 2.4e-06 | 3.4e-06 | 1.8e-06 | 2.533E-06 | 8.083E-07 | 1.800E-06 | 3.400E-06 |
| 6647 | 333 | A333Q | A | 1.8e-06 | 3.9e-06 | 2.1e-06 | 2.600E-06 | 1.136E-06 | 1.800E-06 | 3.900E-06 |
| 6648 | 333 | A333G | A | 1.3e-06 | 3.7e-06 | 1.8e-06 | 2.267E-06 | 1.266E-06 | 1.300E-06 | 3.700E-06 |
| 6649 | 333 | A333H | A | 1.7e-06 | 2.3e-06 | 2.9e-06 | 2.300E-06 | 6.000E-07 | 1.700E-06 | 2.900E-06 |
| 6650 | 333 | A333I | A | 1.8e-06 | 1.9e-06 | 2.1e-06 | 1.933E-06 | 1.528E-07 | 1.800E-06 | 2.100E-06 |
| 6651 | 333 | A333L | A | 2.3e-06 | 1.5e-06 | 2.7e-06 | 2.167E-06 | 6.110E-07 | 1.500E-06 | 2.700E-06 |
| 6652 | 333 | A333K | A | 2.1e-06 | 2.5e-06 | 3.7e-06 | 2.767E-06 | 8.327E-07 | 2.100E-06 | 3.700E-06 |
| 6653 | 333 | A333M | A | 2.1e-06 | 2.8e-06 | 1.6e-06 | 2.167E-06 | 6.028E-07 | 1.600E-06 | 2.800E-06 |
| 6654 | 333 | A333F | A | 4.8e-06 | 2.9e-06 | 2.2e-06 | 3.300E-06 | 1.345E-06 | 2.200E-06 | 4.800E-06 |
| 6655 | 333 | A333P | A | 1.7e-06 | 4e-06 | 1.7e-06 | 2.467E-06 | 1.328E-06 | 1.700E-06 | 4.000E-06 |
| 6656 | 333 | A333S | A | 2.2e-06 | 2.8e-06 | 2.2e-06 | 2.400E-06 | 3.464E-07 | 2.200E-06 | 2.800E-06 |
| 6657 | 333 | A333T | A | 8.7e-07 | 1.6e-06 | 2.4e-06 | 1.623E-06 | 7.653E-07 | 8.700E-07 | 2.400E-06 |
| 6658 | 333 | A333W | A | 2.6e-06 | 4.2e-06 | 2.4e-06 | 3.067E-06 | 9.866E-07 | 2.400E-06 | 4.200E-06 |
| 6659 | 333 | A333Y | A | 2e-06 | 2.1e-06 | 2.8e-06 | 2.300E-06 | 4.359E-07 | 2.000E-06 | 2.800E-06 |
| 6660 | 333 | A333V | A | 1.6e-06 | 4.5e-06 | 1.6e-06 | 2.567E-06 | 1.674E-06 | 1.600E-06 | 4.500E-06 |
| 6661 | 334 | V334A | A | 3.1e-06 | 2.3e-06 | 2.2e-06 | 2.533E-06 | 4.933E-07 | 2.200E-06 | 3.100E-06 |
| 6662 | 334 | V334R | A | 3.3e-06 | 1.5e-06 | 1.8e-06 | 2.200E-06 | 9.644E-07 | 1.500E-06 | 3.300E-06 |
| 6663 | 334 | V334N | A | 1.7e-06 | 2.1e-06 | 3.5e-06 | 2.433E-06 | 9.452E-07 | 1.700E-06 | 3.500E-06 |
| 6664 | 334 | V334D | A | 1.9e-06 | 2.6e-06 | 3.1e-06 | 2.533E-06 | 6.028E-07 | 1.900E-06 | 3.100E-06 |
| 6665 | 334 | V334C | A | 2.8e-06 | 1.5e-06 | 1.9e-06 | 2.067E-06 | 6.658E-07 | 1.500E-06 | 2.800E-06 |
| 6666 | 334 | V334E | A | 1.9e-06 | 2.6e-06 | 1.7e-06 | 2.067E-06 | 4.726E-07 | 1.700E-06 | 2.600E-06 |
| 6667 | 334 | V334Q | A | 3.1e-06 | 2.4e-06 | 2.7e-06 | 2.733E-06 | 3.512E-07 | 2.400E-06 | 3.100E-06 |
| 6668 | 334 | V334G | A | 2.8e-06 | 3.7e-06 | 2.4e-06 | 2.967E-06 | 6.658E-07 | 2.400E-06 | 3.700E-06 |
| 6669 | 334 | V334H | A | 1.8e-06 | 1.3e-06 | 2e-06 | 1.700E-06 | 3.606E-07 | 1.300E-06 | 2.000E-06 |
| 6670 | 334 | V334I | A | 2.7e-06 | 3.1e-06 | 3.8e-06 | 3.200E-06 | 5.568E-07 | 2.700E-06 | 3.800E-06 |
| 6671 | 334 | V334L | A | 2.5e-06 | 2.8e-06 | 1.8e-06 | 2.367E-06 | 5.132E-07 | 1.800E-06 | 2.800E-06 |
| 6672 | 334 | V334K | A | 2.8e-06 | 2.7e-06 | 2.1e-06 | 2.533E-06 | 3.786E-07 | 2.100E-06 | 2.800E-06 |
| 6673 | 334 | V334M | A | 4.4e-06 | 2.3e-06 | 2.5e-06 | 3.067E-06 | 1.159E-06 | 2.300E-06 | 4.400E-06 |
| 6674 | 334 | V334F | A | 3e-06 | 1.6e-06 | 1.3e-06 | 1.967E-06 | 9.074E-07 | 1.300E-06 | 3.000E-06 |
| 6675 | 334 | V334P | A | 2.4e-06 | 1.2e-06 | 2.4e-06 | 2.000E-06 | 6.928E-07 | 1.200E-06 | 2.400E-06 |
| 6676 | 334 | V334S | A | 2.6e-06 | 2.1e-06 | 2.5e-06 | 2.400E-06 | 2.646E-07 | 2.100E-06 | 2.600E-06 |
| 6677 | 334 | V334T | A | 4.4e-06 | 3.5e-06 | 3.4e-06 | 3.767E-06 | 5.508E-07 | 3.400E-06 | 4.400E-06 |
| 6678 | 334 | V334W | A | 2.8e-06 | 2.2e-06 | 2.5e-06 | 2.500E-06 | 3.000E-07 | 2.200E-06 | 2.800E-06 |
| 6679 | 334 | V334Y | A | 4.3e-06 | 1.7e-06 | 2e-06 | 2.667E-06 | 1.422E-06 | 1.700E-06 | 4.300E-06 |
| 6680 | 334 | V334V | A | 3e-06 | 2.9e-06 | 2.2e-06 | 2.700E-06 | 4.359E-07 | 2.200E-06 | 3.000E-06 |
| 6681 | 335 | T335A | A | 1.9e-06 | 3e-06 | 1.4e-06 | 2.100E-06 | 8.185E-07 | 1.400E-06 | 3.000E-06 |
| 6682 | 335 | T335R | A | 1.8e-06 | 1.4e-06 | 1.6e-06 | 1.600E-06 | 2.000E-07 | 1.400E-06 | 1.800E-06 |
| 6683 | 335 | T335N | A | 2.2e-06 | 3.7e-06 | 2e-06 | 2.633E-06 | 9.292E-07 | 2.000E-06 | 3.700E-06 |
| 6684 | 335 | T335D | A | 1.4e-06 | 2.6e-06 | 2.1e-06 | 2.033E-06 | 6.028E-07 | 1.400E-06 | 2.600E-06 |
| 6685 | 335 | T335C | A | 1.9e-06 | 2.5e-06 | 2.1e-06 | 2.167E-06 | 3.055E-07 | 1.900E-06 | 2.500E-06 |
| 6686 | 335 | T335E | A | 1.3e-06 | 1.9e-06 | 1.1e-06 | 1.433E-06 | 4.163E-07 | 1.100E-06 | 1.900E-06 |
| 6687 | 335 | T335Q | A | 3.8e-06 | 2.2e-06 | 2e-06 | 2.667E-06 | 9.866E-07 | 2.000E-06 | 3.800E-06 |
| 6688 | 335 | T335G | A | 1.4e-06 | 3.6e-06 | 2.7e-06 | 2.567E-06 | 1.106E-06 | 1.400E-06 | 3.600E-06 |
| 6689 | 335 | T335H | A | 5e-06 | 2.6e-06 | 2.4e-06 | 3.333E-06 | 1.447E-06 | 2.400E-06 | 5.000E-06 |
| 6690 | 335 | T335I | A | 3.8e-06 | 2e-06 | 2e-06 | 2.600E-06 | 1.039E-06 | 2.000E-06 | 3.800E-06 |
| 6691 | 335 | T335L | A | 5.2e-06 | 3e-06 | 4.2e-06 | 4.133E-06 | 1.102E-06 | 3.000E-06 | 5.200E-06 |
| 6692 | 335 | T335K | A | 3e-06 | 3.2e-06 | 3e-06 | 3.067E-06 | 1.155E-07 | 3.000E-06 | 3.200E-06 |
| 6693 | 335 | T335M | A | 3.2e-06 | 2.7e-06 | 3.7e-06 | 3.200E-06 | 5.000E-07 | 2.700E-06 | 3.700E-06 |
| 6694 | 335 | T335F | A | 2.3e-06 | 2.7e-06 | 1.4e-06 | 2.133E-06 | 6.658E-07 | 1.400E-06 | 2.700E-06 |
| 6695 | 335 | T335P | A | 1.6e-06 | 2.5e-06 | 1.6e-06 | 1.900E-06 | 5.196E-07 | 1.600E-06 | 2.500E-06 |
| 6696 | 335 | T335S | A | 4.1e-06 | 3.7e-06 | 3.6e-06 | 3.800E-06 | 2.646E-07 | 3.600E-06 | 4.100E-06 |
| 6697 | 335 | T335T | A | 2.8e-06 | 1.8e-06 | 4.3e-06 | 2.967E-06 | 1.258E-06 | 1.800E-06 | 4.300E-06 |
| 6698 | 335 | T335W | A | 1.8e-06 | 2.7e-06 | 3.8e-06 | 2.767E-06 | 1.002E-06 | 1.800E-06 | 3.800E-06 |
| 6699 | 335 | T335Y | A | 2.6e-06 | 1.3e-06 | 2.5e-06 | 2.133E-06 | 7.234E-07 | 1.300E-06 | 2.600E-06 |
| 6700 | 335 | T335V | A | 2.5e-06 | 2e-06 | 3.1e-06 | 2.533E-06 | 5.508E-07 | 2.000E-06 | 3.100E-06 |
| 6701 | 336 | S336A | A | 2.1e-06 | 2e-06 | 2.3e-06 | 2.133E-06 | 1.528E-07 | 2.000E-06 | 2.300E-06 |
| 6702 | 336 | S336R | A | 1.8e-06 | 5e-06 | 2.1e-06 | 2.967E-06 | 1.767E-06 | 1.800E-06 | 5.000E-06 |
| 6703 | 336 | S336N | A | 1.7e-06 | 3.1e-06 | 2.9e-06 | 2.567E-06 | 7.572E-07 | 1.700E-06 | 3.100E-06 |
| 6704 | 336 | S336D | A | 2.9e-06 | 1.9e-06 | 1.9e-06 | 2.233E-06 | 5.774E-07 | 1.900E-06 | 2.900E-06 |
| 6705 | 336 | S336C | A | 1.9e-06 | 2e-06 | 2.1e-06 | 2.000E-06 | 1.000E-07 | 1.900E-06 | 2.100E-06 |
| 6706 | 336 | S336E | A | 5.9e-06 | 2.5e-06 | 2.2e-06 | 3.533E-06 | 2.055E-06 | 2.200E-06 | 5.900E-06 |
| 6707 | 336 | S336Q | A | 1.4e-06 | 2.7e-06 | 3.7e-06 | 2.600E-06 | 1.153E-06 | 1.400E-06 | 3.700E-06 |
| 6708 | 336 | S336G | A | 2.3e-06 | 2.1e-06 | 2.2e-06 | 2.200E-06 | 1.000E-07 | 2.100E-06 | 2.300E-06 |
| 6709 | 336 | S336H | A | 3e-06 | 2.6e-06 | 1e-06 | 2.200E-06 | 1.058E-06 | 1.000E-06 | 3.000E-06 |
| 6710 | 336 | S336I | A | 2.7e-06 | 1.8e-06 | 2.3e-06 | 2.267E-06 | 4.509E-07 | 1.800E-06 | 2.700E-06 |
| 6711 | 336 | S336L | A | 2.5e-06 | 2.6e-06 | 2e-06 | 2.367E-06 | 3.215E-07 | 2.000E-06 | 2.600E-06 |
| 6712 | 336 | S336K | A | 2e-06 | 2.4e-06 | 2.8e-06 | 2.400E-06 | 4.000E-07 | 2.000E-06 | 2.800E-06 |
| 6713 | 336 | S336M | A | 3.7e-06 | 3.7e-06 | 1.4e-06 | 2.933E-06 | 1.328E-06 | 1.400E-06 | 3.700E-06 |
| 6714 | 336 | S336F | A | 3.1e-06 | 1.4e-06 | 3.3e-06 | 2.600E-06 | 1.044E-06 | 1.400E-06 | 3.300E-06 |

| | | | | | | | | | | |
|---|---|---|---|---|---|---|---|---|---|---|
| 6715 | 336 | S336P | A | 3.3e-06 | 2.7e-06 | 1.3e-06 | 2.433E-06 | 1.026E-06 | 1.300E-06 | 3.300E-06 |
| 6716 | 336 | S336S | A | 2.4e-06 | 2.7e-06 | 2e-06 | 2.367E-06 | 3.512E-07 | 2.000E-06 | 2.700E-06 |
| 6717 | 336 | S336T | A | 3.9e-06 | 2.4e-06 | 2.2e-06 | 2.833E-06 | 9.292E-07 | 2.200E-06 | 3.900E-06 |
| 6718 | 336 | S336W | A | 4.7e-06 | 2e-06 | 2.3e-06 | 3.000E-06 | 1.480E-06 | 2.000E-06 | 4.700E-06 |
| 6719 | 336 | S336Y | A | 2.7e-06 | 4.2e-06 | 1.3e-06 | 2.733E-06 | 1.450E-06 | 1.300E-06 | 4.200E-06 |
| 6720 | 336 | S336V | A | 3.2e-06 | 3.7e-06 | 4.1e-06 | 3.667E-06 | 4.509E-07 | 3.200E-06 | 4.100E-06 |
| 6721 | 337 | A337A | A | 2.3e-06 | 5.9e-06 | 1.4e-06 | 3.200E-06 | 2.381E-06 | 1.400E-06 | 5.900E-06 |
| 6722 | 337 | A337R | A | 2.3e-06 | 1.4e-06 | 3.7e-06 | 2.467E-06 | 1.159E-06 | 1.400E-06 | 3.700E-06 |
| 6723 | 337 | A337N | A | 2.4e-06 | 2.1e-06 | 2.3e-06 | 2.267E-06 | 1.528E-07 | 2.100E-06 | 2.400E-06 |
| 6724 | 337 | A337D | A | 2.7e-06 | 2e-06 | 3.6e-06 | 2.767E-06 | 8.021E-07 | 2.000E-06 | 3.600E-06 |
| 6725 | 337 | A337C | A | 2.6e-06 | 3.8e-06 | 2.3e-06 | 2.900E-06 | 7.937E-07 | 2.300E-06 | 3.800E-06 |
| 6726 | 337 | A337E | A | 1.8e-06 | 4.1e-06 | 2.5e-06 | 2.800E-06 | 1.179E-06 | 1.800E-06 | 4.100E-06 |
| 6727 | 337 | A337Q | A | 3.7e-06 | 1.9e-06 | 1.2e-06 | 2.267E-06 | 1.290E-06 | 1.200E-06 | 3.700E-06 |
| 6728 | 337 | A337G | A | 1.4e-06 | 1.6e-06 | 1.1e-06 | 1.367E-06 | 2.517E-07 | 1.100E-06 | 1.600E-06 |
| 6729 | 337 | A337H | A | 3.2e-06 | 1.6e-06 | 3.6e-06 | 2.800E-06 | 1.058E-06 | 1.600E-06 | 3.600E-06 |
| 6730 | 337 | A337I | A | 2.2e-06 | 1.6e-06 | 2.7e-06 | 2.167E-06 | 5.508E-07 | 1.600E-06 | 2.700E-06 |
| 6731 | 337 | A337L | A | 2.7e-06 | 4.9e-06 | 3.2e-06 | 3.600E-06 | 1.153E-06 | 2.700E-06 | 4.900E-06 |
| 6732 | 337 | A337K | A | 2.7e-06 | 1.7e-06 | 2.2e-06 | 2.200E-06 | 5.000E-07 | 1.700E-06 | 2.700E-06 |
| 6733 | 337 | A337M | A | 2e-06 | 3.2e-06 | 2.7e-06 | 2.633E-06 | 6.028E-07 | 2.000E-06 | 3.200E-06 |
| 6734 | 337 | A337F | A | 3.4e-06 | 2.4e-06 | 2.9e-06 | 2.900E-06 | 5.000E-07 | 2.400E-06 | 3.400E-06 |
| 6735 | 337 | A337P | A | 2.5e-06 | 2.6e-06 | 3.2e-06 | 2.767E-06 | 3.786E-07 | 2.500E-06 | 3.200E-06 |
| 6736 | 337 | A337S | A | 3.7e-06 | 2.9e-06 | 3e-06 | 3.200E-06 | 4.359E-07 | 2.900E-06 | 3.700E-06 |
| 6737 | 337 | A337T | A | 1.5e-06 | 1.2e-06 | 3.1e-06 | 1.933E-06 | 1.021E-06 | 1.200E-06 | 3.100E-06 |
| 6738 | 337 | A337W | A | 1.7e-06 | 2.2e-06 | 2.7e-06 | 2.200E-06 | 5.000E-07 | 1.700E-06 | 2.700E-06 |
| 6739 | 337 | A337Y | A | 1.4e-06 | 1.1e-06 | 2.3e-06 | 1.600E-06 | 6.245E-07 | 1.100E-06 | 2.300E-06 |
| 6740 | 337 | A337V | A | 3.5e-06 | 3.4e-06 | 2.9e-06 | 3.267E-06 | 3.215E-07 | 2.900E-06 | 3.500E-06 |
| 6741 | 338 | N338A | A | 1.9e-06 | 1.8e-06 | 3.7e-06 | 2.467E-06 | 1.069E-06 | 1.800E-06 | 3.700E-06 |
| 6742 | 338 | N338R | A | 2.5e-06 | 2.1e-06 | 1.6e-06 | 2.067E-06 | 4.509E-07 | 1.600E-06 | 2.500E-06 |
| 6743 | 338 | N338N | A | 3.1e-06 | 2.7e-06 | 1.2e-06 | 2.333E-06 | 1.002E-06 | 1.200E-06 | 3.100E-06 |
| 6744 | 338 | N338D | A | 1.6e-06 | 2.6e-06 | 1.8e-06 | 2.000E-06 | 5.292E-07 | 1.600E-06 | 2.600E-06 |
| 6745 | 338 | N338C | A | 1.9e-06 | 2.4e-06 | 2.1e-06 | 2.133E-06 | 2.517E-07 | 1.900E-06 | 2.400E-06 |
| 6746 | 338 | N338E | A | 2.8e-06 | 2.6e-06 | 8.8e-07 | 2.093E-06 | 1.056E-06 | 8.800E-07 | 2.800E-06 |
| 6747 | 338 | N338Q | A | 1.7e-06 | 1.5e-06 | 1.1e-06 | 1.433E-06 | 3.055E-07 | 1.100E-06 | 1.700E-06 |
| 6748 | 338 | N338G | A | 2e-06 | 2.6e-06 | 2.2e-06 | 2.267E-06 | 3.055E-07 | 2.000E-06 | 2.600E-06 |
| 6749 | 338 | N338H | A | 2.6e-06 | 2.8e-06 | 4.5e-06 | 3.300E-06 | 1.044E-06 | 2.600E-06 | 4.500E-06 |
| 6750 | 338 | N338I | A | 2.6e-06 | 1.9e-06 | 2.8e-06 | 2.433E-06 | 4.726E-07 | 1.900E-06 | 2.800E-06 |
| 6751 | 338 | N338L | A | 2.9e-06 | 3.7e-06 | 2.6e-06 | 3.067E-06 | 5.686E-07 | 2.600E-06 | 3.700E-06 |
| 6752 | 338 | N338K | A | 1.9e-06 | 1.4e-06 | 2.3e-06 | 1.867E-06 | 4.509E-07 | 1.400E-06 | 2.300E-06 |
| 6753 | 338 | N338M | A | 1.2e-06 | 2.6e-06 | 3.3e-06 | 2.367E-06 | 1.069E-06 | 1.200E-06 | 3.300E-06 |
| 6754 | 338 | N338F | A | 3.5e-06 | 3e-06 | 3.2e-06 | 3.233E-06 | 2.517E-07 | 3.000E-06 | 3.500E-06 |
| 6755 | 338 | N338P | A | 9.6e-07 | 3.7e-06 | 2.2e-06 | 2.287E-06 | 1.372E-06 | 9.600E-07 | 3.700E-06 |
| 6756 | 338 | N338S | A | 1.5e-06 | 1.6e-06 | 2.3e-06 | 1.800E-06 | 4.359E-07 | 1.500E-06 | 2.300E-06 |
| 6757 | 338 | N338T | A | 1.2e-06 | 2.3e-06 | 2.3e-06 | 1.933E-06 | 6.351E-07 | 1.200E-06 | 2.300E-06 |
| 6758 | 338 | N338W | A | 2.5e-06 | 1.4e-06 | 2.9e-06 | 2.267E-06 | 7.767E-07 | 1.400E-06 | 2.900E-06 |
| 6759 | 338 | N338Y | A | 5.4e-06 | 2.5e-06 | 3e-06 | 3.633E-06 | 1.550E-06 | 2.500E-06 | 5.400E-06 |
| 6760 | 338 | N338V | A | 2.1e-06 | 1.6e-06 | 3e-06 | 2.233E-06 | 7.095E-07 | 1.600E-06 | 3.000E-06 |
| 6761 | 339 | I339A | A | 2.9e-06 | 3.1e-06 | 2.7e-06 | 2.900E-06 | 2.000E-07 | 2.700E-06 | 3.100E-06 |
| 6762 | 339 | I339R | A | 2.8e-06 | 2.8e-06 | 3.8e-06 | 3.133E-06 | 5.774E-07 | 2.800E-06 | 3.800E-06 |
| 6763 | 339 | I339N | A | 1.4e-06 | 2.2e-06 | 2.8e-06 | 2.133E-06 | 7.024E-07 | 1.400E-06 | 2.800E-06 |
| 6764 | 339 | I339D | A | 2.5e-06 | 1.4e-06 | 2.1e-06 | 2.000E-06 | 5.568E-07 | 1.400E-06 | 2.500E-06 |
| 6765 | 339 | I339C | A | 4.4e-06 | 2.1e-06 | 1.5e-06 | 2.667E-06 | 1.531E-06 | 1.500E-06 | 4.400E-06 |
| 6766 | 339 | I339E | A | 2.8e-06 | 2.5e-06 | 1.8e-06 | 2.367E-06 | 5.132E-07 | 1.800E-06 | 2.800E-06 |
| 6767 | 339 | I339Q | A | 2.4e-06 | 2.4e-06 | 1.5e-06 | 2.100E-06 | 5.196E-07 | 1.500E-06 | 2.400E-06 |
| 6768 | 339 | I339G | A | 1.7e-06 | 1.8e-06 | 1.6e-06 | 1.700E-06 | 1.000E-07 | 1.600E-06 | 1.800E-06 |
| 6769 | 339 | I339H | A | 2.8e-06 | 3.1e-06 | 2.5e-06 | 2.800E-06 | 3.000E-07 | 2.500E-06 | 3.100E-06 |
| 6770 | 339 | I339I | A | 3e-06 | 2.3e-06 | 3.3e-06 | 2.867E-06 | 5.132E-07 | 2.300E-06 | 3.300E-06 |
| 6771 | 339 | I339L | A | 1.5e-06 | 9.3e-07 | 2.4e-06 | 1.610E-06 | 7.411E-07 | 9.300E-07 | 2.400E-06 |
| 6772 | 339 | I339K | A | 2.8e-06 | 2e-06 | 1.6e-06 | 2.133E-06 | 6.110E-07 | 1.600E-06 | 2.800E-06 |
| 6773 | 339 | I339M | A | 2.5e-06 | 1.6e-06 | 2.9e-06 | 2.333E-06 | 6.658E-07 | 1.600E-06 | 2.900E-06 |
| 6774 | 339 | I339F | A | 3.2e-06 | 1.7e-06 | 3.7e-06 | 2.867E-06 | 1.041E-06 | 1.700E-06 | 3.700E-06 |
| 6775 | 339 | I339P | A | 1.9e-06 | 2.9e-06 | 3e-06 | 2.600E-06 | 6.083E-07 | 1.900E-06 | 3.000E-06 |
| 6776 | 339 | I339S | A | 3.1e-06 | 2.4e-06 | 1.6e-06 | 2.367E-06 | 7.506E-07 | 1.600E-06 | 3.100E-06 |
| 6777 | 339 | I339T | A | 1.7e-06 | 3.1e-06 | 2.2e-06 | 2.333E-06 | 7.095E-07 | 1.700E-06 | 3.100E-06 |
| 6778 | 339 | I339W | A | 2.5e-06 | 3.4e-06 | 2.6e-06 | 2.833E-06 | 4.933E-07 | 2.500E-06 | 3.400E-06 |
| 6779 | 339 | I339Y | A | 1.9e-06 | 1.5e-06 | 2.8e-06 | 2.067E-06 | 6.658E-07 | 1.500E-06 | 2.800E-06 |
| 6780 | 339 | I339V | A | 2e-06 | 1.5e-06 | 2.7e-06 | 2.067E-06 | 6.028E-07 | 1.500E-06 | 2.700E-06 |
| 6781 | 340 | Q340A | A | 4.3e-06 | 1.6e-06 | 2.4e-06 | 2.767E-06 | 1.387E-06 | 1.600E-06 | 4.300E-06 |
| 6782 | 340 | Q340R | A | 1.7e-06 | 2.6e-06 | 2.8e-06 | 2.367E-06 | 5.859E-07 | 1.700E-06 | 2.800E-06 |
| 6783 | 340 | Q340N | A | 3e-06 | 2.8e-06 | 2.7e-06 | 2.833E-06 | 1.528E-07 | 2.700E-06 | 3.000E-06 |
| 6784 | 340 | Q340D | A | 2.2e-06 | 2.2e-06 | 1.8e-06 | 2.067E-06 | 2.309E-07 | 1.800E-06 | 2.200E-06 |
| 6785 | 340 | Q340C | A | 2.5e-06 | 2.4e-06 | 2.5e-06 | 2.467E-06 | 5.774E-08 | 2.400E-06 | 2.500E-06 |
| 6786 | 340 | Q340E | A | 3.6e-06 | 3e-06 | 2.5e-06 | 3.033E-06 | 5.508E-07 | 2.500E-06 | 3.600E-06 |
| 6787 | 340 | Q340Q | A | 2.5e-06 | 3.5e-06 | 2.5e-06 | 2.833E-06 | 5.774E-07 | 2.500E-06 | 3.500E-06 |
| 6788 | 340 | Q340G | A | 3.3e-06 | 2.7e-06 | 2.5e-06 | 2.833E-06 | 4.163E-07 | 2.500E-06 | 3.300E-06 |
| 6789 | 340 | Q340H | A | 2.5e-06 | 2.4e-06 | 2.1e-06 | 2.333E-06 | 2.082E-07 | 2.100E-06 | 2.500E-06 |
| 6790 | 340 | Q340I | A | 2.2e-06 | 2.7e-06 | 3.4e-06 | 2.767E-06 | 6.028E-07 | 2.200E-06 | 3.400E-06 |
| 6791 | 340 | Q340L | A | 2.5e-06 | 3.5e-06 | 2.4e-06 | 2.800E-06 | 6.083E-07 | 2.400E-06 | 3.500E-06 |
| 6792 | 340 | Q340K | A | 1.7e-06 | 2.9e-06 | 2.1e-06 | 2.233E-06 | 6.110E-07 | 1.700E-06 | 2.900E-06 |
| 6793 | 340 | Q340M | A | 3.2e-06 | 2.4e-06 | 2.6e-06 | 2.733E-06 | 4.163E-07 | 2.400E-06 | 3.200E-06 |
| 6794 | 340 | Q340F | A | 2e-06 | 2.3e-06 | 1.9e-06 | 2.067E-06 | 2.082E-07 | 1.900E-06 | 2.300E-06 |
| 6795 | 340 | Q340P | A | 2.3e-06 | 2.6e-06 | 2.1e-06 | 2.333E-06 | 2.517E-07 | 2.100E-06 | 2.600E-06 |
| 6796 | 340 | Q340S | A | 3.1e-06 | 1.6e-06 | 2.9e-06 | 2.533E-06 | 8.145E-07 | 1.600E-06 | 3.100E-06 |
| 6797 | 340 | Q340T | A | 2e-06 | 2.9e-06 | 2.9e-06 | 2.600E-06 | 5.196E-07 | 2.000E-06 | 2.900E-06 |
| 6798 | 340 | Q340W | A | 3.4e-06 | 2.7e-06 | 2.7e-06 | 2.933E-06 | 4.041E-07 | 2.700E-06 | 3.400E-06 |
| 6799 | 340 | Q340Y | A | 2.6e-06 | 2.4e-06 | 1.5e-06 | 2.167E-06 | 5.859E-07 | 1.500E-06 | 2.600E-06 |

| | | | | | | | | | | |
|---|---|---|---|---|---|---|---|---|---|---|
| 6800 | 340 | Q340V | A | 1.4e-06 | 2.7e-06 | 2.1e-06 | 2.067E-06 | 6.506E-07 | 1.400E-06 | 2.700E-06 |
| 6801 | 341 | E341A | A | 3.4e-06 | 8.2e-07 | 4.5e-06 | 2.907E-06 | 1.889E-06 | 8.200E-07 | 4.500E-06 |
| 6802 | 341 | E341R | A | 4.4e-06 | 2.3e-06 | 2.6e-06 | 3.100E-06 | 1.136E-06 | 2.300E-06 | 4.400E-06 |
| 6803 | 341 | E341N | A | 2.7e-06 | 1.7e-06 | 2.8e-06 | 2.400E-06 | 6.083E-07 | 1.700E-06 | 2.800E-06 |
| 6804 | 341 | E341D | A | 2.2e-06 | 3.2e-06 | 3.6e-06 | 3.000E-06 | 7.211E-07 | 2.200E-06 | 3.600E-06 |
| 6805 | 341 | E341C | A | 2.3e-06 | 3.7e-06 | 2.1e-06 | 2.700E-06 | 8.718E-07 | 2.100E-06 | 3.700E-06 |
| 6806 | 341 | E341E | A | 1.4e-06 | 2.6e-06 | 3.1e-06 | 2.367E-06 | 8.737E-07 | 1.400E-06 | 3.100E-06 |
| 6807 | 341 | E341Q | A | 2.6e-06 | 3.9e-06 | 1.3e-06 | 2.600E-06 | 1.300E-06 | 1.300E-06 | 3.900E-06 |
| 6808 | 341 | E341G | A | 1.5e-06 | 1.9e-06 | 2.1e-06 | 1.833E-06 | 3.055E-07 | 1.500E-06 | 2.100E-06 |
| 6809 | 341 | E341H | A | 3.5e-06 | 2.1e-06 | 3e-06 | 2.867E-06 | 7.095E-07 | 2.100E-06 | 3.500E-06 |
| 6810 | 341 | E341I | A | 3.1e-06 | 2.3e-06 | 4.2e-06 | 3.200E-06 | 9.539E-07 | 2.300E-06 | 4.200E-06 |
| 6811 | 341 | E341L | A | 1.9e-06 | 2.6e-06 | 2.9e-06 | 2.467E-06 | 5.132E-07 | 1.900E-06 | 2.900E-06 |
| 6812 | 341 | E341K | A | 1.3e-06 | 2.7e-06 | 2.8e-06 | 2.267E-06 | 8.386E-07 | 1.300E-06 | 2.800E-06 |
| 6813 | 341 | E341M | A | 2.3e-06 | 2.7e-06 | 2.5e-06 | 2.500E-06 | 2.000E-07 | 2.300E-06 | 2.700E-06 |
| 6814 | 341 | E341F | A | 2.2e-06 | 2.4e-06 | 2.3e-06 | 2.300E-06 | 1.000E-07 | 2.200E-06 | 2.400E-06 |
| 6815 | 341 | E341P | A | 1.8e-06 | 1.5e-06 | 2.9e-06 | 2.067E-06 | 7.371E-07 | 1.500E-06 | 2.900E-06 |
| 6816 | 341 | E341S | A | 2.4e-06 | 1.8e-06 | 2.9e-06 | 2.367E-06 | 5.508E-07 | 1.800E-06 | 2.900E-06 |
| 6817 | 341 | E341T | A | 3e-06 | 3.2e-06 | 2.2e-06 | 2.800E-06 | 5.292E-07 | 2.200E-06 | 3.200E-06 |
| 6818 | 341 | E341W | A | 1.1e-06 | 1.4e-06 | 2.5e-06 | 1.667E-06 | 7.371E-07 | 1.100E-06 | 2.500E-06 |
| 6819 | 341 | E341Y | A | 2.1e-06 | 3.7e-06 | 2.8e-06 | 2.867E-06 | 8.021E-07 | 2.100E-06 | 3.700E-06 |
| 6820 | 341 | E341V | A | 2.7e-06 | 2.1e-06 | 2.1e-06 | 2.300E-06 | 3.464E-07 | 2.100E-06 | 2.700E-06 |
| 6821 | 342 | F342A | A | 2.7e-06 | 2.7e-06 | 3.7e-06 | 3.033E-06 | 5.774E-07 | 2.700E-06 | 3.700E-06 |
| 6822 | 342 | F342R | A | 2e-06 | 2.8e-06 | 1.6e-06 | 2.133E-06 | 6.110E-07 | 1.600E-06 | 2.800E-06 |
| 6823 | 342 | F342N | A | 2.1e-06 | 2.8e-06 | 1.8e-06 | 2.233E-06 | 5.132E-07 | 1.800E-06 | 2.800E-06 |
| 6824 | 342 | F342D | A | 2e-06 | 2.2e-06 | 2.5e-06 | 2.233E-06 | 2.517E-07 | 2.000E-06 | 2.500E-06 |
| 6825 | 342 | F342C | A | 4.7e-06 | 2.4e-06 | 1.8e-06 | 2.967E-06 | 1.531E-06 | 1.800E-06 | 4.700E-06 |
| 6826 | 342 | F342E | A | 1.9e-06 | 2.2e-06 | 1.5e-06 | 1.867E-06 | 3.512E-07 | 1.500E-06 | 2.200E-06 |
| 6827 | 342 | F342Q | A | 1.7e-06 | 4.8e-06 | 1.8e-06 | 2.767E-06 | 1.762E-06 | 1.700E-06 | 4.800E-06 |
| 6828 | 342 | F342G | A | 1.5e-06 | 2.8e-06 | 1e-06 | 1.767E-06 | 9.292E-07 | 1.000E-06 | 2.800E-06 |
| 6829 | 342 | F342H | A | 2e-06 | 2.3e-06 | 3.2e-06 | 2.500E-06 | 6.245E-07 | 2.000E-06 | 3.200E-06 |
| 6830 | 342 | F342I | A | 2.2e-06 | 4.3e-06 | 3.3e-06 | 3.267E-06 | 1.050E-06 | 2.200E-06 | 4.300E-06 |
| 6831 | 342 | F342L | A | 2.8e-06 | 2.9e-06 | 1.7e-06 | 2.467E-06 | 6.658E-07 | 1.700E-06 | 2.900E-06 |
| 6832 | 342 | F342K | A | 1.3e-06 | 3.4e-06 | 2.9e-06 | 2.533E-06 | 1.097E-06 | 1.300E-06 | 3.400E-06 |
| 6833 | 342 | F342M | A | 2e-06 | 3.3e-06 | 2.4e-06 | 2.567E-06 | 6.658E-07 | 2.000E-06 | 3.300E-06 |
| 6834 | 342 | F342F | A | 4e-06 | 1.9e-06 | 1.6e-06 | 2.500E-06 | 1.308E-06 | 1.600E-06 | 4.000E-06 |
| 6835 | 342 | F342P | A | 2.9e-06 | 2.6e-06 | 1.9e-06 | 2.467E-06 | 5.132E-07 | 1.900E-06 | 2.900E-06 |
| 6836 | 342 | F342S | A | 3.1e-06 | 2.4e-06 | 2.7e-06 | 2.733E-06 | 3.512E-07 | 2.400E-06 | 3.100E-06 |
| 6837 | 342 | F342T | A | 3.6e-06 | 1.9e-06 | 2.3e-06 | 2.600E-06 | 8.888E-07 | 1.900E-06 | 3.600E-06 |
| 6838 | 342 | F342W | A | 1.5e-06 | 3.6e-06 | 1.3e-06 | 2.133E-06 | 1.274E-06 | 1.300E-06 | 3.600E-06 |
| 6839 | 342 | F342Y | A | 2.3e-06 | 1e-06 | 2e-06 | 1.767E-06 | 6.807E-07 | 1.000E-06 | 2.300E-06 |
| 6840 | 342 | F342V | A | 2.2e-06 | 1.8e-06 | 2.9e-06 | 2.300E-06 | 5.568E-07 | 1.800E-06 | 2.900E-06 |
| 6841 | 343 | A343A | A | 2.6e-06 | 2.1e-06 | 3.8e-06 | 2.833E-06 | 8.737E-07 | 2.100E-06 | 3.800E-06 |
| 6842 | 343 | A343R | A | 3e-06 | 2e-06 | 3.4e-06 | 2.800E-06 | 7.211E-07 | 2.000E-06 | 3.400E-06 |
| 6843 | 343 | A343N | A | 1.2e-06 | 2.1e-06 | 2.9e-06 | 2.067E-06 | 8.505E-07 | 1.200E-06 | 2.900E-06 |
| 6844 | 343 | A343D | A | 2.7e-06 | 2e-06 | 2.2e-06 | 2.300E-06 | 3.606E-07 | 2.000E-06 | 2.700E-06 |
| 6845 | 343 | A343C | A | 2.9e-06 | 3.2e-06 | 2e-06 | 2.700E-06 | 6.245E-07 | 2.000E-06 | 3.200E-06 |
| 6846 | 343 | A343E | A | 2.8e-06 | 1.9e-06 | 1.4e-06 | 2.033E-06 | 7.095E-07 | 1.400E-06 | 2.800E-06 |
| 6847 | 343 | A343Q | A | 2.5e-06 | 2.4e-06 | 1.4e-06 | 2.100E-06 | 6.083E-07 | 1.400E-06 | 2.500E-06 |
| 6848 | 343 | A343G | A | 2.3e-06 | 3e-06 | 2.6e-06 | 2.633E-06 | 3.512E-07 | 2.300E-06 | 3.000E-06 |
| 6849 | 343 | A343H | A | 2.9e-06 | 2.4e-06 | 2e-06 | 2.433E-06 | 4.509E-07 | 2.000E-06 | 2.900E-06 |
| 6850 | 343 | A343I | A | 1.4e-06 | 2.9e-06 | 1.9e-06 | 2.067E-06 | 7.638E-07 | 1.400E-06 | 2.900E-06 |
| 6851 | 343 | A343L | A | 2.1e-06 | 1.1e-06 | 3.2e-06 | 2.133E-06 | 1.050E-06 | 1.100E-06 | 3.200E-06 |
| 6852 | 343 | A343K | A | 2.7e-06 | 2.1e-06 | 3.2e-06 | 2.667E-06 | 5.508E-07 | 2.100E-06 | 3.200E-06 |
| 6853 | 343 | A343M | A | 2.8e-06 | 1.7e-06 | 2.6e-06 | 2.367E-06 | 5.859E-07 | 1.700E-06 | 2.800E-06 |
| 6854 | 343 | A343F | A | 1.7e-06 | 2.7e-06 | 2e-06 | 2.133E-06 | 5.132E-07 | 1.700E-06 | 2.700E-06 |
| 6855 | 343 | A343P | A | 2.2e-06 | 5.2e-06 | 2.1e-06 | 3.167E-06 | 1.762E-06 | 2.100E-06 | 5.200E-06 |
| 6856 | 343 | A343S | A | 1.7e-06 | 2.5e-06 | 2.2e-06 | 2.133E-06 | 4.041E-07 | 1.700E-06 | 2.500E-06 |
| 6857 | 343 | A343T | A | 2e-06 | 2.9e-06 | 3.1e-06 | 2.667E-06 | 5.859E-07 | 2.000E-06 | 3.100E-06 |
| 6858 | 343 | A343W | A | 1.9e-06 | 1.4e-06 | 2.7e-06 | 2.000E-06 | 6.557E-07 | 1.400E-06 | 2.700E-06 |
| 6859 | 343 | A343Y | A | 2e-06 | 2.6e-06 | 3e-06 | 2.533E-06 | 5.033E-07 | 2.000E-06 | 3.000E-06 |
| 6860 | 343 | A343V | A | 2.8e-06 | 1.8e-06 | 1.4e-06 | 2.000E-06 | 7.211E-07 | 1.400E-06 | 2.800E-06 |
| 6861 | 344 | G344A | A | 3.8e-06 | 3.9e-06 | 2.5e-06 | 3.400E-06 | 7.810E-07 | 2.500E-06 | 3.900E-06 |
| 6862 | 344 | G344R | A | 2.8e-06 | 1.6e-06 | 3.8e-06 | 2.733E-06 | 1.102E-06 | 1.600E-06 | 3.800E-06 |
| 6863 | 344 | G344N | A | 3.4e-06 | 2.3e-06 | 2.8e-06 | 2.833E-06 | 5.508E-07 | 2.300E-06 | 3.400E-06 |
| 6864 | 344 | G344D | A | 1.3e-06 | 2.5e-06 | 2.2e-06 | 2.000E-06 | 6.245E-07 | 1.300E-06 | 2.500E-06 |
| 6865 | 344 | G344C | A | 2.7e-06 | 1.7e-06 | 1.6e-06 | 2.000E-06 | 6.083E-07 | 1.600E-06 | 2.700E-06 |
| 6866 | 344 | G344E | A | 2e-06 | 2.4e-06 | 3.4e-06 | 2.600E-06 | 7.211E-07 | 2.000E-06 | 3.400E-06 |
| 6867 | 344 | G344Q | A | 3.4e-06 | 2.7e-06 | 2.2e-06 | 2.767E-06 | 6.028E-07 | 2.200E-06 | 3.400E-06 |
| 6868 | 344 | G344G | A | 1.2e-06 | 2.8e-06 | 3.8e-06 | 2.600E-06 | 1.311E-06 | 1.200E-06 | 3.800E-06 |
| 6869 | 344 | G344H | A | 2.5e-06 | 2.2e-06 | 2.7e-06 | 2.467E-06 | 2.517E-07 | 2.200E-06 | 2.700E-06 |
| 6870 | 344 | G344I | A | 1.4e-06 | 2.6e-06 | 4e-06 | 2.667E-06 | 1.301E-06 | 1.400E-06 | 4.000E-06 |
| 6871 | 344 | G344L | A | 3e-06 | 1.7e-06 | 1.2e-06 | 1.967E-06 | 9.292E-07 | 1.200E-06 | 3.000E-06 |
| 6872 | 344 | G344K | A | 2.1e-06 | 3e-06 | 3.6e-06 | 2.900E-06 | 7.550E-07 | 2.100E-06 | 3.600E-06 |
| 6873 | 344 | G344M | A | 2.2e-06 | 2.6e-06 | 2.1e-06 | 2.300E-06 | 2.646E-07 | 2.100E-06 | 2.600E-06 |
| 6874 | 344 | G344F | A | 2.9e-06 | 1.9e-06 | 9.8e-07 | 1.927E-06 | 9.603E-07 | 9.800E-07 | 2.900E-06 |
| 6875 | 344 | G344P | A | 3.1e-06 | 1.3e-06 | 2.9e-06 | 2.433E-06 | 9.866E-07 | 1.300E-06 | 3.100E-06 |
| 6876 | 344 | G344S | A | 3.3e-06 | 2e-06 | 1.5e-06 | 2.267E-06 | 9.292E-07 | 1.500E-06 | 3.300E-06 |
| 6877 | 344 | G344T | A | 2.8e-06 | 2.1e-06 | 5.3e-06 | 3.400E-06 | 1.682E-06 | 2.100E-06 | 5.300E-06 |
| 6878 | 344 | G344W | A | 2.4e-06 | 2.5e-06 | 2.6e-06 | 2.500E-06 | 1.000E-07 | 2.400E-06 | 2.600E-06 |
| 6879 | 344 | G344Y | A | 1.9e-06 | 2.4e-06 | 2.3e-06 | 2.200E-06 | 2.646E-07 | 1.900E-06 | 2.400E-06 |
| 6880 | 344 | G344V | A | 2.7e-06 | 1.9e-06 | 1.6e-06 | 2.067E-06 | 5.686E-07 | 1.600E-06 | 2.700E-06 |
| 6881 | 345 | C345A | A | 2.3e-06 | 2.2e-06 | 3e-06 | 2.500E-06 | 4.359E-07 | 2.200E-06 | 3.000E-06 |
| 6882 | 345 | C345R | A | 1.5e-06 | 2.8e-06 | 1.9e-06 | 2.067E-06 | 6.658E-07 | 1.500E-06 | 2.800E-06 |
| 6883 | 345 | C345N | A | 2.5e-06 | 2.7e-06 | 2.2e-06 | 2.467E-06 | 2.517E-07 | 2.200E-06 | 2.700E-06 |
| 6884 | 345 | C345D | A | 2.2e-06 | 4.2e-06 | 2.9e-06 | 3.100E-06 | 1.015E-06 | 2.200E-06 | 4.200E-06 |

| | | | | | | | | | | |
|---|---|---|---|---|---|---|---|---|---|---|
| 6885 | 345 | C345C | A | 2.4e-06 | 2e-06 | 5.6e-06 | 3.333E-06 | 1.973E-06 | 2.000E-06 | 5.600E-06 |
| 6886 | 345 | C345E | A | 2e-06 | 1.3e-06 | 3.1e-06 | 2.133E-06 | 9.074E-07 | 1.300E-06 | 3.100E-06 |
| 6887 | 345 | C345Q | A | 1.5e-06 | 3.3e-06 | 2.1e-06 | 2.300E-06 | 9.165E-07 | 1.500E-06 | 3.300E-06 |
| 6888 | 345 | C345G | A | 1.2e-06 | 2.4e-06 | 2.5e-06 | 2.033E-06 | 7.234E-07 | 1.200E-06 | 2.500E-06 |
| 6889 | 345 | C345H | A | 2.2e-06 | 1.3e-06 | 1.3e-06 | 1.600E-06 | 5.196E-07 | 1.300E-06 | 2.200E-06 |
| 6890 | 345 | C345I | A | 2.5e-06 | 2.7e-06 | 2.7e-06 | 2.633E-06 | 1.155E-07 | 2.500E-06 | 2.700E-06 |
| 6891 | 345 | C345L | A | 3.2e-06 | 2.4e-06 | 2.6e-06 | 2.733E-06 | 4.163E-07 | 2.400E-06 | 3.200E-06 |
| 6892 | 345 | C345K | A | 2.8e-06 | 1.5e-06 | 1.5e-06 | 1.933E-06 | 7.506E-07 | 1.500E-06 | 2.800E-06 |
| 6893 | 345 | C345M | A | 2.7e-06 | 1.7e-06 | 3e-06 | 2.467E-06 | 6.807E-07 | 1.700E-06 | 3.000E-06 |
| 6894 | 345 | C345F | A | 1.3e-06 | 1.5e-06 | 3e-06 | 1.933E-06 | 9.292E-07 | 1.300E-06 | 3.000E-06 |
| 6895 | 345 | C345P | A | 1.5e-06 | 1.9e-06 | 2.8e-06 | 2.067E-06 | 6.658E-07 | 1.500E-06 | 2.800E-06 |
| 6896 | 345 | C345S | A | 2.6e-06 | 2e-06 | 2e-06 | 2.200E-06 | 3.464E-07 | 2.000E-06 | 2.600E-06 |
| 6897 | 345 | C345T | A | 1.6e-06 | 2.7e-06 | 2.1e-06 | 2.133E-06 | 5.508E-07 | 1.600E-06 | 2.700E-06 |
| 6898 | 345 | C345W | A | 1.8e-06 | 1.4e-06 | 3.8e-06 | 2.333E-06 | 1.286E-06 | 1.400E-06 | 3.800E-06 |
| 6899 | 345 | C345Y | A | 1.6e-06 | 2.5e-06 | 1.6e-06 | 1.900E-06 | 5.196E-07 | 1.600E-06 | 2.500E-06 |
| 6900 | 345 | C345V | A | 3.3e-06 | 2.9e-06 | 2.3e-06 | 2.833E-06 | 5.033E-07 | 2.300E-06 | 3.300E-06 |
| 6901 | 346 | K346A | A | 2.4e-06 | 2.1e-06 | 3.1e-06 | 2.533E-06 | 5.132E-07 | 2.100E-06 | 3.100E-06 |
| 6902 | 346 | K346R | A | 2.3e-06 | 2e-06 | 1.6e-06 | 1.967E-06 | 3.512E-07 | 1.600E-06 | 2.300E-06 |
| 6903 | 346 | K346N | A | 1.5e-06 | 1.8e-06 | 2.7e-06 | 2.000E-06 | 6.245E-07 | 1.500E-06 | 2.700E-06 |
| 6904 | 346 | K346D | A | 3e-06 | 1.7e-06 | 2.4e-06 | 2.367E-06 | 6.506E-07 | 1.700E-06 | 3.000E-06 |
| 6905 | 346 | K346C | A | 4.7e-06 | 2.7e-06 | 3.5e-06 | 3.633E-06 | 1.007E-06 | 2.700E-06 | 4.700E-06 |
| 6906 | 346 | K346E | A | 2e-06 | 3.2e-06 | 1.9e-06 | 2.367E-06 | 7.234E-07 | 1.900E-06 | 3.200E-06 |
| 6907 | 346 | K346Q | A | 2.4e-06 | 2.4e-06 | 1.3e-06 | 2.033E-06 | 6.351E-07 | 1.300E-06 | 2.400E-06 |
| 6908 | 346 | K346G | A | 1.7e-06 | 2e-06 | 2.9e-06 | 2.200E-06 | 6.245E-07 | 1.700E-06 | 2.900E-06 |
| 6909 | 346 | K346H | A | 2.1e-06 | 2.6e-06 | 2.5e-06 | 2.400E-06 | 2.646E-07 | 2.100E-06 | 2.600E-06 |
| 6910 | 346 | K346I | A | 3.1e-06 | 3.5e-06 | 1.8e-06 | 2.800E-06 | 8.888E-07 | 1.800E-06 | 3.500E-06 |
| 6911 | 346 | K346L | A | 2.2e-06 | 4.3e-06 | 2.6e-06 | 3.033E-06 | 1.115E-06 | 2.200E-06 | 4.300E-06 |
| 6912 | 346 | K346K | A | 1.8e-06 | 4.2e-06 | 2.6e-06 | 2.867E-06 | 1.222E-06 | 1.800E-06 | 4.200E-06 |
| 6913 | 346 | K346M | A | 4.5e-06 | 2.2e-06 | 1.7e-06 | 2.800E-06 | 1.493E-06 | 1.700E-06 | 4.500E-06 |
| 6914 | 346 | K346F | A | 2.7e-06 | 2.5e-06 | 2.8e-06 | 2.667E-06 | 1.528E-07 | 2.500E-06 | 2.800E-06 |
| 6915 | 346 | K346P | A | 1.6e-06 | 1.9e-06 | 1.6e-06 | 1.700E-06 | 1.732E-07 | 1.600E-06 | 1.900E-06 |
| 6916 | 346 | K346S | A | 1.3e-06 | 3.2e-06 | 3e-06 | 2.500E-06 | 1.044E-06 | 1.300E-06 | 3.200E-06 |
| 6917 | 346 | K346T | A | 1.3e-06 | 3.1e-06 | 3.5e-06 | 2.633E-06 | 1.172E-06 | 1.300E-06 | 3.500E-06 |
| 6918 | 346 | K346W | A | 1.3e-06 | 2.9e-06 | 1.8e-06 | 2.000E-06 | 8.185E-07 | 1.300E-06 | 2.900E-06 |
| 6919 | 346 | K346Y | A | 3.8e-06 | 2.2e-06 | 1.5e-06 | 2.500E-06 | 1.179E-06 | 1.500E-06 | 3.800E-06 |
| 6920 | 346 | K346V | A | 2.6e-06 | 3.4e-06 | 2.8e-06 | 2.933E-06 | 4.163E-07 | 2.600E-06 | 3.400E-06 |
| 6921 | 347 | K347A | A | 2.2e-06 | 2.3e-06 | 3e-06 | 2.500E-06 | 4.359E-07 | 2.200E-06 | 3.000E-06 |
| 6922 | 347 | K347R | A | 1.1e-06 | 2.3e-06 | 2.8e-06 | 2.067E-06 | 8.737E-07 | 1.100E-06 | 2.800E-06 |
| 6923 | 347 | K347N | A | 3.1e-06 | 3e-06 | 2.3e-06 | 2.800E-06 | 4.359E-07 | 2.300E-06 | 3.100E-06 |
| 6924 | 347 | K347D | A | 2.9e-06 | 2.4e-06 | 2.9e-06 | 2.733E-06 | 2.887E-07 | 2.400E-06 | 2.900E-06 |
| 6925 | 347 | K347C | A | 1.2e-06 | 2.8e-06 | 4.3e-06 | 2.767E-06 | 1.550E-06 | 1.200E-06 | 4.300E-06 |
| 6926 | 347 | K347E | A | 2e-06 | 2.7e-06 | 1.4e-06 | 2.033E-06 | 6.506E-07 | 1.400E-06 | 2.700E-06 |
| 6927 | 347 | K347Q | A | 1.8e-06 | 3.4e-06 | 2.5e-06 | 2.567E-06 | 8.021E-07 | 1.800E-06 | 3.400E-06 |
| 6928 | 347 | K347G | A | 2.6e-06 | 2.5e-06 | 1.6e-06 | 2.233E-06 | 5.508E-07 | 1.600E-06 | 2.600E-06 |
| 6929 | 347 | K347H | A | 1.6e-06 | 1.2e-06 | 3.1e-06 | 1.967E-06 | 1.002E-06 | 1.200E-06 | 3.100E-06 |
| 6930 | 347 | K347I | A | 1.6e-06 | 2.3e-06 | 2.2e-06 | 2.033E-06 | 3.786E-07 | 1.600E-06 | 2.300E-06 |
| 6931 | 347 | K347L | A | 3.1e-06 | 2.1e-06 | 2.3e-06 | 2.500E-06 | 5.292E-07 | 2.100E-06 | 3.100E-06 |
| 6932 | 347 | K347K | A | 2.6e-06 | 1.9e-06 | 2.8e-06 | 2.433E-06 | 4.726E-07 | 1.900E-06 | 2.800E-06 |
| 6933 | 347 | K347M | A | 9.2e-07 | 1.9e-06 | 2.4e-06 | 1.740E-06 | 7.529E-07 | 9.200E-07 | 2.400E-06 |
| 6934 | 347 | K347F | A | 2.6e-06 | 2.5e-06 | 2.2e-06 | 2.433E-06 | 2.082E-07 | 2.200E-06 | 2.600E-06 |
| 6935 | 347 | K347P | A | 2.4e-06 | 3.7e-06 | 1.4e-06 | 2.500E-06 | 1.153E-06 | 1.400E-06 | 3.700E-06 |
| 6936 | 347 | K347S | A | 1.2e-06 | 2.9e-06 | 1.5e-06 | 1.867E-06 | 9.074E-07 | 1.200E-06 | 2.900E-06 |
| 6937 | 347 | K347T | A | 2.8e-06 | 2.4e-06 | 2.3e-06 | 2.500E-06 | 2.646E-07 | 2.300E-06 | 2.800E-06 |
| 6938 | 347 | K347W | A | 2.8e-06 | 2.7e-06 | 1.7e-06 | 2.400E-06 | 6.083E-07 | 1.700E-06 | 2.800E-06 |
| 6939 | 347 | K347Y | A | 2e-06 | 2.9e-06 | 1.5e-06 | 2.133E-06 | 7.095E-07 | 1.500E-06 | 2.900E-06 |
| 6940 | 347 | K347V | A | 2.4e-06 | 2.5e-06 | 2e-06 | 2.300E-06 | 2.646E-07 | 2.000E-06 | 2.500E-06 |
| 6941 | 348 | I348A | A | 2.8e-06 | 2.6e-06 | 3e-06 | 2.800E-06 | 2.000E-07 | 2.600E-06 | 3.000E-06 |
| 6942 | 348 | I348R | A | 1.7e-06 | 2.3e-06 | 3e-06 | 2.333E-06 | 6.506E-07 | 1.700E-06 | 3.000E-06 |
| 6943 | 348 | I348N | A | 3.4e-06 | 2.7e-06 | 4.1e-06 | 3.400E-06 | 7.000E-07 | 2.700E-06 | 4.100E-06 |
| 6944 | 348 | I348D | A | 2e-06 | 2.6e-06 | 2.7e-06 | 2.433E-06 | 3.786E-07 | 2.000E-06 | 2.700E-06 |
| 6945 | 348 | I348C | A | 2.8e-06 | 2.5e-06 | 2.9e-06 | 2.733E-06 | 2.082E-07 | 2.500E-06 | 2.900E-06 |
| 6946 | 348 | I348E | A | 2.2e-06 | 2.9e-06 | 3.3e-06 | 2.800E-06 | 5.568E-07 | 2.200E-06 | 3.300E-06 |
| 6947 | 348 | I348Q | A | 3.2e-06 | 2.3e-06 | 2.4e-06 | 2.633E-06 | 4.933E-07 | 2.300E-06 | 3.200E-06 |
| 6948 | 348 | I348G | A | 1.8e-06 | 3.2e-06 | 2.7e-06 | 2.567E-06 | 7.095E-07 | 1.800E-06 | 3.200E-06 |
| 6949 | 348 | I348H | A | 2.2e-06 | 3.2e-06 | 1.7e-06 | 2.367E-06 | 7.638E-07 | 1.700E-06 | 3.200E-06 |
| 6950 | 348 | I348I | A | 2.9e-06 | 2.2e-06 | 3.6e-06 | 2.900E-06 | 7.000E-07 | 2.200E-06 | 3.600E-06 |
| 6951 | 348 | I348L | A | 5.2e-06 | 2.9e-06 | 2.1e-06 | 3.400E-06 | 1.609E-06 | 2.100E-06 | 5.200E-06 |
| 6952 | 348 | I348K | A | 2.2e-06 | 1.5e-06 | 3.2e-06 | 2.300E-06 | 8.544E-07 | 1.500E-06 | 3.200E-06 |
| 6953 | 348 | I348M | A | 2.3e-06 | 2.5e-06 | 4.5e-06 | 3.100E-06 | 1.217E-06 | 2.300E-06 | 4.500E-06 |
| 6954 | 348 | I348F | A | 2.4e-06 | 2.6e-06 | 3.6e-06 | 2.867E-06 | 6.429E-07 | 2.400E-06 | 3.600E-06 |
| 6955 | 348 | I348P | A | 2.3e-06 | 1.4e-06 | 2.2e-06 | 1.967E-06 | 4.933E-07 | 1.400E-06 | 2.300E-06 |
| 6956 | 348 | I348S | A | 5.1e-06 | 2.2e-06 | 2e-06 | 3.100E-06 | 1.735E-06 | 2.000E-06 | 5.100E-06 |
| 6957 | 348 | I348T | A | 2.7e-06 | 1.4e-06 | 2.7e-06 | 2.267E-06 | 7.506E-07 | 1.400E-06 | 2.700E-06 |
| 6958 | 348 | I348W | A | 3.5e-06 | 2.4e-06 | 2.4e-06 | 2.767E-06 | 6.351E-07 | 2.400E-06 | 3.500E-06 |
| 6959 | 348 | I348Y | A | 2.9e-06 | 3.3e-06 | 2.5e-06 | 2.900E-06 | 4.000E-07 | 2.500E-06 | 3.300E-06 |
| 6960 | 348 | I348V | A | 3.5e-06 | 2.5e-06 | 1.9e-06 | 2.633E-06 | 8.083E-07 | 1.900E-06 | 3.500E-06 |
| 6961 | 349 | F349A | A | 2.5e-06 | 2.5e-06 | 2.5e-06 | 2.500E-06 | 0.000E+00 | 2.500E-06 | 2.500E-06 |
| 6962 | 349 | F349R | A | 2.5e-06 | 2.3e-06 | 2.3e-06 | 2.367E-06 | 1.155E-07 | 2.300E-06 | 2.500E-06 |
| 6963 | 349 | F349N | A | 2.1e-06 | 1.5e-06 | 1.2e-06 | 1.600E-06 | 4.583E-07 | 1.200E-06 | 2.100E-06 |
| 6964 | 349 | F349D | A | 2.2e-06 | 2.7e-06 | 3.1e-06 | 2.667E-06 | 4.509E-07 | 2.200E-06 | 3.100E-06 |
| 6965 | 349 | F349C | A | 2.7e-06 | 4.3e-06 | 3.8e-06 | 3.600E-06 | 8.185E-07 | 2.700E-06 | 4.300E-06 |
| 6966 | 349 | F349E | A | 3e-06 | 2.4e-06 | 2.4e-06 | 2.600E-06 | 3.464E-07 | 2.400E-06 | 3.000E-06 |
| 6967 | 349 | F349Q | A | 3.3e-06 | 2e-06 | 1.8e-06 | 2.367E-06 | 8.145E-07 | 1.800E-06 | 3.300E-06 |
| 6968 | 349 | F349G | A | 2.9e-06 | 2.2e-06 | 2.1e-06 | 2.400E-06 | 4.359E-07 | 2.100E-06 | 2.900E-06 |
| 6969 | 349 | F349H | A | 4.6e-06 | 4.4e-06 | 1.9e-06 | 3.633E-06 | 1.504E-06 | 1.900E-06 | 4.600E-06 |

| | | | | | | | | | | |
|---|---|---|---|---|---|---|---|---|---|---|
| 6970 | 349 | F349I | A | 2.4e-06 | 1.4e-06 | 2e-06 | 1.933E-06 | 5.033E-07 | 1.400E-06 | 2.400E-06 |
| 6971 | 349 | F349L | A | 2.1e-06 | 1.9e-06 | 1.6e-06 | 1.867E-06 | 2.517E-07 | 1.600E-06 | 2.100E-06 |
| 6972 | 349 | F349K | A | 3e-06 | 2.8e-06 | 3.8e-06 | 3.200E-06 | 5.292E-07 | 2.800E-06 | 3.800E-06 |
| 6973 | 349 | F349M | A | 3e-06 | 1.9e-06 | 3.7e-06 | 2.867E-06 | 9.074E-07 | 1.900E-06 | 3.700E-06 |
| 6974 | 349 | F349F | A | 2.1e-06 | 1.8e-06 | 2.8e-06 | 2.233E-06 | 5.132E-07 | 1.800E-06 | 2.800E-06 |
| 6975 | 349 | F349P | A | 4.5e-06 | 2.9e-06 | 2.3e-06 | 3.233E-06 | 1.137E-06 | 2.300E-06 | 4.500E-06 |
| 6976 | 349 | F349S | A | 2.2e-06 | 2e-06 | 2.8e-06 | 2.333E-06 | 4.163E-07 | 2.000E-06 | 2.800E-06 |
| 6977 | 349 | F349T | A | 2.8e-06 | 2.5e-06 | 3.1e-06 | 2.800E-06 | 3.000E-07 | 2.500E-06 | 3.100E-06 |
| 6978 | 349 | F349W | A | 3.6e-06 | 1.7e-06 | 1.1e-06 | 2.133E-06 | 1.305E-06 | 1.100E-06 | 3.600E-06 |
| 6979 | 349 | F349Y | A | 1.6e-06 | 2.9e-06 | 2e-06 | 2.167E-06 | 6.658E-07 | 1.600E-06 | 2.900E-06 |
| 6980 | 349 | F349V | A | 2.4e-06 | 2.2e-06 | 3e-06 | 2.533E-06 | 4.163E-07 | 2.200E-06 | 3.000E-06 |
| 6981 | 350 | G350A | A | 4.5e-06 | 2.2e-06 | 1.8e-06 | 2.833E-06 | 1.457E-06 | 1.800E-06 | 4.500E-06 |
| 6982 | 350 | G350R | A | 3.2e-06 | 2.4e-06 | 1.6e-06 | 2.400E-06 | 8.000E-07 | 1.600E-06 | 3.200E-06 |
| 6983 | 350 | G350N | A | 1.9e-06 | 3.6e-06 | 2e-06 | 2.500E-06 | 9.539E-07 | 1.900E-06 | 3.600E-06 |
| 6984 | 350 | G350D | A | 1.3e-06 | 1.5e-06 | 1.8e-06 | 1.533E-06 | 2.517E-07 | 1.300E-06 | 1.800E-06 |
| 6985 | 350 | G350C | A | 2e-06 | 2.1e-06 | 4e-06 | 2.700E-06 | 1.127E-06 | 2.000E-06 | 4.000E-06 |
| 6986 | 350 | G350E | A | 1.2e-06 | 1.3e-06 | 2.4e-06 | 1.633E-06 | 6.658E-07 | 1.200E-06 | 2.400E-06 |
| 6987 | 350 | G350Q | A | 3.4e-06 | 2e-06 | 2e-06 | 2.467E-06 | 8.083E-07 | 2.000E-06 | 3.400E-06 |
| 6988 | 350 | G350G | A | 1.8e-06 | 1.4e-06 | 1.5e-06 | 1.567E-06 | 2.082E-07 | 1.400E-06 | 1.800E-06 |
| 6989 | 350 | G350H | A | 6e-06 | 2.3e-06 | 1.8e-06 | 3.367E-06 | 2.294E-06 | 1.800E-06 | 6.000E-06 |
| 6990 | 350 | G350I | A | 1.3e-06 | 2.4e-06 | 2.9e-06 | 2.200E-06 | 8.185E-07 | 1.300E-06 | 2.900E-06 |
| 6991 | 350 | G350L | A | 3.4e-06 | 2.4e-06 | 2.4e-06 | 2.733E-06 | 5.774E-07 | 2.400E-06 | 3.400E-06 |
| 6992 | 350 | G350K | A | 2.3e-06 | 2.4e-06 | 2.3e-06 | 2.333E-06 | 5.774E-08 | 2.300E-06 | 2.400E-06 |
| 6993 | 350 | G350M | A | 2.6e-06 | 1.6e-06 | 3.2e-06 | 2.467E-06 | 8.083E-07 | 1.600E-06 | 3.200E-06 |
| 6994 | 350 | G350F | A | 2.1e-06 | 2.9e-06 | 1.8e-06 | 2.267E-06 | 5.686E-07 | 1.800E-06 | 2.900E-06 |
| 6995 | 350 | G350P | A | 2.6e-06 | 2.5e-06 | 2.7e-06 | 2.600E-06 | 1.000E-07 | 2.500E-06 | 2.700E-06 |
| 6996 | 350 | G350S | A | 2.2e-06 | 1.6e-06 | 4e-06 | 2.600E-06 | 1.249E-06 | 1.600E-06 | 4.000E-06 |
| 6997 | 350 | G350T | A | 3.1e-06 | 2.3e-06 | 1.1e-06 | 2.167E-06 | 1.007E-06 | 1.100E-06 | 3.100E-06 |
| 6998 | 350 | G350W | A | 2.1e-06 | 2.2e-06 | 2.6e-06 | 2.300E-06 | 2.646E-07 | 2.100E-06 | 2.600E-06 |
| 6999 | 350 | G350Y | A | 2.3e-06 | 2e-06 | 3.6e-06 | 2.633E-06 | 8.505E-07 | 2.000E-06 | 3.600E-06 |
| 7000 | 350 | G350V | A | 2.6e-06 | 3.6e-06 | 2.5e-06 | 2.900E-06 | 6.083E-07 | 2.500E-06 | 3.600E-06 |
| 7001 | 351 | S351A | A | 3.4e-06 | 1.4e-06 | 9.7e-07 | 1.923E-06 | 1.297E-06 | 9.700E-07 | 3.400E-06 |
| 7002 | 351 | S351R | A | 2.1e-06 | 2.3e-06 | 2.3e-06 | 2.233E-06 | 1.155E-07 | 2.100E-06 | 2.300E-06 |
| 7003 | 351 | S351N | A | 3.4e-06 | 2.3e-06 | 2.3e-06 | 2.667E-06 | 6.351E-07 | 2.300E-06 | 3.400E-06 |
| 7004 | 351 | S351D | A | 2.3e-06 | 2.9e-06 | 2.3e-06 | 2.500E-06 | 3.464E-07 | 2.300E-06 | 2.900E-06 |
| 7005 | 351 | S351C | A | 1.5e-06 | 4.2e-06 | 1.7e-06 | 2.467E-06 | 1.504E-06 | 1.500E-06 | 4.200E-06 |
| 7006 | 351 | S351E | A | 2e-06 | 3.1e-06 | 3.2e-06 | 2.767E-06 | 6.658E-07 | 2.000E-06 | 3.200E-06 |
| 7007 | 351 | S351Q | A | 2.5e-06 | 2.5e-06 | 1.5e-06 | 2.167E-06 | 5.774E-07 | 1.500E-06 | 2.500E-06 |
| 7008 | 351 | S351G | A | 1.6e-06 | 2.6e-06 | 3.4e-06 | 2.533E-06 | 9.018E-07 | 1.600E-06 | 3.400E-06 |
| 7009 | 351 | S351H | A | 2.3e-06 | 2.3e-06 | 4.4e-06 | 3.000E-06 | 1.212E-06 | 2.300E-06 | 4.400E-06 |
| 7010 | 351 | S351I | A | 2.2e-06 | 2.4e-06 | 3.5e-06 | 2.700E-06 | 7.000E-07 | 2.200E-06 | 3.500E-06 |
| 7011 | 351 | S351L | A | 1.7e-06 | 3.5e-06 | 2.1e-06 | 2.433E-06 | 9.452E-07 | 1.700E-06 | 3.500E-06 |
| 7012 | 351 | S351K | A | 3.1e-06 | 3.6e-06 | 2.7e-06 | 3.133E-06 | 4.509E-07 | 2.700E-06 | 3.600E-06 |
| 7013 | 351 | S351M | A | 3.5e-06 | 3.3e-06 | 2.4e-06 | 3.067E-06 | 5.859E-07 | 2.400E-06 | 3.500E-06 |
| 7014 | 351 | S351F | A | 3.2e-06 | 1.4e-06 | 3e-06 | 2.533E-06 | 9.866E-07 | 1.400E-06 | 3.200E-06 |
| 7015 | 351 | S351P | A | 1.9e-06 | 1.4e-06 | 2.2e-06 | 1.833E-06 | 4.041E-07 | 1.400E-06 | 2.200E-06 |
| 7016 | 351 | S351S | A | 2.5e-06 | 2e-06 | 1.8e-06 | 2.100E-06 | 3.606E-07 | 1.800E-06 | 2.500E-06 |
| 7017 | 351 | S351T | A | 2.8e-06 | 1.8e-06 | 2e-06 | 2.200E-06 | 5.292E-07 | 1.800E-06 | 2.800E-06 |
| 7018 | 351 | S351W | A | 2.4e-06 | 2.7e-06 | 2.2e-06 | 2.433E-06 | 2.517E-07 | 2.200E-06 | 2.700E-06 |
| 7019 | 351 | S351Y | A | 3.5e-06 | 2.7e-06 | 2.2e-06 | 2.800E-06 | 6.557E-07 | 2.200E-06 | 3.500E-06 |
| 7020 | 351 | S351V | A | 2.1e-06 | 2.1e-06 | 2.8e-06 | 2.333E-06 | 4.041E-07 | 2.100E-06 | 2.800E-06 |
| 7021 | 352 | L352A | A | 3.6e-06 | 1.9e-06 | 2.9e-06 | 2.800E-06 | 8.544E-07 | 1.900E-06 | 3.600E-06 |
| 7022 | 352 | L352R | A | 3.7e-06 | 2.4e-06 | 1.7e-06 | 2.600E-06 | 1.015E-06 | 1.700E-06 | 3.700E-06 |
| 7023 | 352 | L352N | A | 1.5e-06 | 1.6e-06 | 3.2e-06 | 2.100E-06 | 9.539E-07 | 1.500E-06 | 3.200E-06 |
| 7024 | 352 | L352D | A | 2.9e-06 | 3.6e-06 | 2.7e-06 | 3.067E-06 | 4.726E-07 | 2.700E-06 | 3.600E-06 |
| 7025 | 352 | L352C | A | 2.1e-06 | 1.6e-06 | 1.9e-06 | 1.867E-06 | 2.517E-07 | 1.600E-06 | 2.100E-06 |
| 7026 | 352 | L352E | A | 2.6e-06 | 2e-06 | 4e-06 | 2.867E-06 | 1.026E-06 | 2.000E-06 | 4.000E-06 |
| 7027 | 352 | L352Q | A | 1.9e-06 | 2.9e-06 | 2.1e-06 | 2.300E-06 | 5.292E-07 | 1.900E-06 | 2.900E-06 |
| 7028 | 352 | L352G | A | 2.9e-06 | 3e-06 | 4.1e-06 | 3.333E-06 | 6.658E-07 | 2.900E-06 | 4.100E-06 |
| 7029 | 352 | L352H | A | 9.3e-07 | 1.9e-06 | 1.6e-06 | 1.477E-06 | 4.966E-07 | 9.300E-07 | 1.900E-06 |
| 7030 | 352 | L352I | A | 3.9e-06 | 2.6e-06 | 2.8e-06 | 3.100E-06 | 7.000E-07 | 2.600E-06 | 3.900E-06 |
| 7031 | 352 | L352L | A | 1.9e-06 | 2e-06 | 2.2e-06 | 2.033E-06 | 1.528E-07 | 1.900E-06 | 2.200E-06 |
| 7032 | 352 | L352K | A | 1.1e-06 | 4.7e-06 | 2.6e-06 | 2.800E-06 | 1.808E-06 | 1.100E-06 | 4.700E-06 |
| 7033 | 352 | L352M | A | 2.3e-06 | 1.9e-06 | 3.3e-06 | 2.500E-06 | 7.211E-07 | 1.900E-06 | 3.300E-06 |
| 7034 | 352 | L352F | A | 2.2e-06 | 2.5e-06 | 2.8e-06 | 2.500E-06 | 3.000E-07 | 2.200E-06 | 2.800E-06 |
| 7035 | 352 | L352P | A | 3.1e-06 | 2.9e-06 | 2.6e-06 | 2.867E-06 | 2.517E-07 | 2.600E-06 | 3.100E-06 |
| 7036 | 352 | L352S | A | 1.4e-06 | 2.8e-06 | 2.7e-06 | 2.300E-06 | 7.810E-07 | 1.400E-06 | 2.800E-06 |
| 7037 | 352 | L352T | A | 1.3e-06 | 2.1e-06 | 3.1e-06 | 2.167E-06 | 9.018E-07 | 1.300E-06 | 3.100E-06 |
| 7038 | 352 | L352W | A | 1.7e-06 | 2.1e-06 | 1.1e-06 | 1.633E-06 | 5.033E-07 | 1.100E-06 | 2.100E-06 |
| 7039 | 352 | L352Y | A | 1.4e-06 | 2.3e-06 | 1.9e-06 | 1.867E-06 | 4.509E-07 | 1.400E-06 | 2.300E-06 |
| 7040 | 352 | L352V | A | 3.4e-06 | 2.4e-06 | 2.6e-06 | 2.800E-06 | 5.292E-07 | 2.400E-06 | 3.400E-06 |
| 7041 | 353 | A353A | A | 2.7e-06 | 1.9e-06 | 1.9e-06 | 2.167E-06 | 4.619E-07 | 1.900E-06 | 2.700E-06 |
| 7042 | 353 | A353R | A | 2.7e-06 | 3.3e-06 | 2.2e-06 | 2.733E-06 | 5.508E-07 | 2.200E-06 | 3.300E-06 |
| 7043 | 353 | A353N | A | 2.6e-06 | 1.6e-06 | 3.1e-06 | 2.433E-06 | 7.638E-07 | 1.600E-06 | 3.100E-06 |
| 7044 | 353 | A353D | A | 1.9e-06 | 2.2e-06 | 3.1e-06 | 2.400E-06 | 6.245E-07 | 1.900E-06 | 3.100E-06 |
| 7045 | 353 | A353C | A | 2.8e-06 | 2.6e-06 | 1.6e-06 | 2.333E-06 | 6.429E-07 | 1.600E-06 | 2.800E-06 |
| 7046 | 353 | A353E | A | 2.2e-06 | 3.2e-06 | 2.8e-06 | 2.733E-06 | 5.033E-07 | 2.200E-06 | 3.200E-06 |
| 7047 | 353 | A353Q | A | 2e-06 | 2.2e-06 | 3.3e-06 | 2.500E-06 | 7.000E-07 | 2.000E-06 | 3.300E-06 |
| 7048 | 353 | A353G | A | 1.6e-06 | 2.5e-06 | 2.1e-06 | 2.067E-06 | 4.509E-07 | 1.600E-06 | 2.500E-06 |
| 7049 | 353 | A353H | A | 2.1e-06 | 2.5e-06 | 1.5e-06 | 2.033E-06 | 5.033E-07 | 1.500E-06 | 2.500E-06 |
| 7050 | 353 | A353I | A | 3.2e-06 | 2.2e-06 | 1.9e-06 | 2.433E-06 | 6.807E-07 | 1.900E-06 | 3.200E-06 |
| 7051 | 353 | A353L | A | 2.4e-06 | 3.2e-06 | 2.3e-06 | 2.633E-06 | 4.933E-07 | 2.300E-06 | 3.200E-06 |
| 7052 | 353 | A353K | A | 2e-06 | 4e-06 | 1.5e-06 | 2.500E-06 | 1.323E-06 | 1.500E-06 | 4.000E-06 |
| 7053 | 353 | A353M | A | 2.7e-06 | 2.2e-06 | 1.8e-06 | 2.233E-06 | 4.509E-07 | 1.800E-06 | 2.700E-06 |
| 7054 | 353 | A353F | A | 2.5e-06 | 3.7e-06 | 4.4e-06 | 3.533E-06 | 9.609E-07 | 2.500E-06 | 4.400E-06 |

| | | | | | | | | | | |
|---|---|---|---|---|---|---|---|---|---|---|
| 7055 | 353 | A353P | A | 1.4e-06 | 2.8e-06 | 3.4e-06 | 2.533E-06 | 1.026E-06 | 1.400E-06 | 3.400E-06 |
| 7056 | 353 | A353S | A | 1.3e-06 | 3.7e-06 | 2.1e-06 | 2.367E-06 | 1.222E-06 | 1.300E-06 | 3.700E-06 |
| 7057 | 353 | A353T | A | 2.8e-06 | 1.6e-06 | 2.4e-06 | 2.267E-06 | 6.110E-07 | 1.600E-06 | 2.800E-06 |
| 7058 | 353 | A353W | A | 2.1e-06 | 2.3e-06 | 2.6e-06 | 2.333E-06 | 2.517E-07 | 2.100E-06 | 2.600E-06 |
| 7059 | 353 | A353Y | A | 2.3e-06 | 2.4e-06 | 1.7e-06 | 2.133E-06 | 3.786E-07 | 1.700E-06 | 2.400E-06 |
| 7060 | 353 | A353V | A | 1.4e-06 | 1.6e-06 | 2.3e-06 | 1.767E-06 | 4.726E-07 | 1.400E-06 | 2.300E-06 |
| 7061 | 354 | F354A | A | 1.7e-06 | 2.3e-06 | 2.1e-06 | 2.033E-06 | 3.055E-07 | 1.700E-06 | 2.300E-06 |
| 7062 | 354 | F354R | A | 7.1e-06 | 2.7e-06 | 2.7e-06 | 4.167E-06 | 2.540E-06 | 2.700E-06 | 7.100E-06 |
| 7063 | 354 | F354N | A | 1.7e-06 | 8.4e-07 | 2.8e-06 | 1.780E-06 | 9.824E-07 | 8.400E-07 | 2.800E-06 |
| 7064 | 354 | F354D | A | 3.1e-06 | 5.7e-06 | 2.4e-06 | 3.733E-06 | 1.739E-06 | 2.400E-06 | 5.700E-06 |
| 7065 | 354 | F354C | A | 2.7e-06 | 3.7e-06 | 3.7e-06 | 3.367E-06 | 5.774E-07 | 2.700E-06 | 3.700E-06 |
| 7066 | 354 | F354E | A | 4.3e-06 | 1.2e-06 | 3.7e-06 | 3.067E-06 | 1.644E-06 | 1.200E-06 | 4.300E-06 |
| 7067 | 354 | F354Q | A | 2.1e-06 | 5.2e-06 | 2.3e-06 | 3.200E-06 | 1.735E-06 | 2.100E-06 | 5.200E-06 |
| 7068 | 354 | F354G | A | 4e-06 | 2.7e-06 | 4.2e-06 | 3.633E-06 | 8.145E-07 | 2.700E-06 | 4.200E-06 |
| 7069 | 354 | F354H | A | 2.7e-06 | 2.8e-06 | 2.1e-06 | 2.533E-06 | 3.786E-07 | 2.100E-06 | 2.800E-06 |
| 7070 | 354 | F354I | A | 4.6e-06 | 1.9e-06 | 1.7e-06 | 2.733E-06 | 1.620E-06 | 1.700E-06 | 4.600E-06 |
| 7071 | 354 | F354L | A | 2.7e-06 | 2.8e-06 | 3.1e-06 | 2.867E-06 | 2.082E-07 | 2.700E-06 | 3.100E-06 |
| 7072 | 354 | F354K | A | 2.6e-06 | 3e-06 | 3.2e-06 | 2.933E-06 | 3.055E-07 | 2.600E-06 | 3.200E-06 |
| 7073 | 354 | F354M | A | 2.9e-06 | 2.1e-06 | 3.3e-06 | 2.767E-06 | 6.110E-07 | 2.100E-06 | 3.300E-06 |
| 7074 | 354 | F354F | A | 1.4e-06 | 1.6e-06 | 2.3e-06 | 1.767E-06 | 4.726E-07 | 1.400E-06 | 2.300E-06 |
| 7075 | 354 | F354P | A | 2.5e-06 | 3.3e-06 | 4.3e-06 | 3.367E-06 | 9.018E-07 | 2.500E-06 | 4.300E-06 |
| 7076 | 354 | F354S | A | 2.4e-06 | 1.2e-06 | 2.5e-06 | 2.033E-06 | 7.234E-07 | 1.200E-06 | 2.500E-06 |
| 7077 | 354 | F354T | A | 1.4e-06 | 2e-06 | 3.1e-06 | 2.167E-06 | 8.622E-07 | 1.400E-06 | 3.100E-06 |
| 7078 | 354 | F354W | A | 2.1e-06 | 2.5e-06 | 1.8e-06 | 2.133E-06 | 3.512E-07 | 1.800E-06 | 2.500E-06 |
| 7079 | 354 | F354Y | A | 2.5e-06 | 2.7e-06 | 1.3e-06 | 2.167E-06 | 7.572E-07 | 1.300E-06 | 2.700E-06 |
| 7080 | 354 | F354V | A | 1.5e-06 | 3.2e-06 | 3.6e-06 | 2.767E-06 | 1.115E-06 | 1.500E-06 | 3.600E-06 |
| 7081 | 355 | L355A | A | 1.8e-06 | 1.2e-06 | 2.1e-06 | 1.700E-06 | 4.583E-07 | 1.200E-06 | 2.100E-06 |
| 7082 | 355 | L355R | A | 2.4e-06 | 1.3e-06 | 2.5e-06 | 2.067E-06 | 6.658E-07 | 1.300E-06 | 2.500E-06 |
| 7083 | 355 | L355N | A | 2.1e-06 | 1.8e-06 | 3.4e-06 | 2.433E-06 | 8.505E-07 | 1.800E-06 | 3.400E-06 |
| 7084 | 355 | L355D | A | 2.2e-06 | 2.7e-06 | 1.5e-06 | 2.133E-06 | 6.028E-07 | 1.500E-06 | 2.700E-06 |
| 7085 | 355 | L355C | A | 2.5e-06 | 1.6e-06 | 2.4e-06 | 2.167E-06 | 4.933E-07 | 1.600E-06 | 2.500E-06 |
| 7086 | 355 | L355E | A | 2e-06 | 3.5e-06 | 2.4e-06 | 2.633E-06 | 7.767E-07 | 2.000E-06 | 3.500E-06 |
| 7087 | 355 | L355Q | A | 2e-06 | 2e-06 | 9.2e-07 | 1.640E-06 | 6.235E-07 | 9.200E-07 | 2.000E-06 |
| 7088 | 355 | L355G | A | 3.1e-06 | 1.8e-06 | 1.7e-06 | 2.200E-06 | 7.810E-07 | 1.700E-06 | 3.100E-06 |
| 7089 | 355 | L355H | A | 2.5e-06 | 3.4e-06 | 2.7e-06 | 2.867E-06 | 4.726E-07 | 2.500E-06 | 3.400E-06 |
| 7090 | 355 | L355I | A | 3.4e-06 | 2.1e-06 | 1.8e-06 | 2.433E-06 | 8.505E-07 | 1.800E-06 | 3.400E-06 |
| 7091 | 355 | L355L | A | 2.1e-06 | 2.9e-06 | 3.1e-06 | 2.700E-06 | 5.292E-07 | 2.100E-06 | 3.100E-06 |
| 7092 | 355 | L355K | A | 2.2e-06 | 2.2e-06 | 2.9e-06 | 2.433E-06 | 4.041E-07 | 2.200E-06 | 2.900E-06 |
| 7093 | 355 | L355M | A | 2.5e-06 | 2e-06 | 2.3e-06 | 2.267E-06 | 2.517E-07 | 2.000E-06 | 2.500E-06 |
| 7094 | 355 | L355F | A | 2.5e-06 | 4e-06 | 1.8e-06 | 2.767E-06 | 1.124E-06 | 1.800E-06 | 4.000E-06 |
| 7095 | 355 | L355P | A | 4.4e-06 | 2.7e-06 | 2.6e-06 | 3.233E-06 | 1.012E-06 | 2.600E-06 | 4.400E-06 |
| 7096 | 355 | L355S | A | 2.1e-06 | 3.1e-06 | 1.4e-06 | 2.200E-06 | 8.544E-07 | 1.400E-06 | 3.100E-06 |
| 7097 | 355 | L355T | A | 2.3e-06 | 1.6e-06 | 3.3e-06 | 2.400E-06 | 8.544E-07 | 1.600E-06 | 3.300E-06 |
| 7098 | 355 | L355W | A | 1.1e-06 | 2.9e-06 | 1.7e-06 | 1.900E-06 | 9.165E-07 | 1.100E-06 | 2.900E-06 |
| 7099 | 355 | L355Y | A | 2.4e-06 | 1.4e-06 | 2.9e-06 | 2.233E-06 | 7.638E-07 | 1.400E-06 | 2.900E-06 |
| 7100 | 355 | L355V | A | 4.7e-06 | 1.9e-06 | 3e-06 | 3.200E-06 | 1.411E-06 | 1.900E-06 | 4.700E-06 |
| 7101 | 356 | P356A | A | 1.9e-06 | 3.1e-06 | 3.1e-06 | 2.700E-06 | 6.928E-07 | 1.900E-06 | 3.100E-06 |
| 7102 | 356 | P356R | A | 2.8e-06 | 2e-06 | 3.2e-06 | 2.667E-06 | 6.110E-07 | 2.000E-06 | 3.200E-06 |
| 7103 | 356 | P356N | A | 2.3e-06 | 2.3e-06 | 1.6e-06 | 2.067E-06 | 4.041E-07 | 1.600E-06 | 2.300E-06 |
| 7104 | 356 | P356D | A | 2.1e-06 | 2.3e-06 | 2.4e-06 | 2.267E-06 | 1.528E-07 | 2.100E-06 | 2.400E-06 |
| 7105 | 356 | P356C | A | 3.4e-06 | 2.1e-06 | 1.8e-06 | 2.433E-06 | 8.505E-07 | 1.800E-06 | 3.400E-06 |
| 7106 | 356 | P356E | A | 2.6e-06 | 2e-06 | 3.2e-06 | 2.600E-06 | 6.000E-07 | 2.000E-06 | 3.200E-06 |
| 7107 | 356 | P356Q | A | 2.8e-06 | 1.7e-06 | 2.4e-06 | 2.300E-06 | 5.568E-07 | 1.700E-06 | 2.800E-06 |
| 7108 | 356 | P356G | A | 3.4e-06 | 2e-06 | 2.4e-06 | 2.600E-06 | 7.211E-07 | 2.000E-06 | 3.400E-06 |
| 7109 | 356 | P356H | A | 2.2e-06 | 1.3e-06 | 1.6e-06 | 1.700E-06 | 4.583E-07 | 1.300E-06 | 2.200E-06 |
| 7110 | 356 | P356I | A | 2.2e-06 | 2.1e-06 | 2.7e-06 | 2.333E-06 | 3.215E-07 | 2.100E-06 | 2.700E-06 |
| 7111 | 356 | P356L | A | 2.7e-06 | 1.6e-06 | 1.5e-06 | 1.933E-06 | 6.658E-07 | 1.500E-06 | 2.700E-06 |
| 7112 | 356 | P356K | A | 1.9e-06 | 2e-06 | 2.2e-06 | 2.033E-06 | 1.528E-07 | 1.900E-06 | 2.200E-06 |
| 7113 | 356 | P356M | A | 2e-06 | 2.6e-06 | 2.6e-06 | 2.400E-06 | 3.464E-07 | 2.000E-06 | 2.600E-06 |
| 7114 | 356 | P356F | A | 2.5e-06 | 3.2e-06 | 2.3e-06 | 2.667E-06 | 4.726E-07 | 2.300E-06 | 3.200E-06 |
| 7115 | 356 | P356P | A | 1.1e-06 | 3.4e-06 | 1.4e-06 | 1.967E-06 | 1.250E-06 | 1.100E-06 | 3.400E-06 |
| 7116 | 356 | P356S | A | 2.1e-06 | 3.6e-06 | 2.5e-06 | 2.733E-06 | 7.767E-07 | 2.100E-06 | 3.600E-06 |
| 7117 | 356 | P356T | A | 2.2e-06 | 1.3e-06 | 1.9e-06 | 1.800E-06 | 4.583E-07 | 1.300E-06 | 2.200E-06 |
| 7118 | 356 | P356W | A | 2.2e-06 | 2.4e-06 | 2e-06 | 2.200E-06 | 2.000E-07 | 2.000E-06 | 2.400E-06 |
| 7119 | 356 | P356Y | A | 2e-06 | 2.6e-06 | 2.7e-06 | 2.433E-06 | 3.786E-07 | 2.000E-06 | 2.700E-06 |
| 7120 | 356 | P356V | A | 2.4e-06 | 2.3e-06 | 1.9e-06 | 2.200E-06 | 2.646E-07 | 1.900E-06 | 2.400E-06 |
| 7121 | 357 | E357A | A | 1.7e-06 | 2.5e-06 | 2.1e-06 | 2.100E-06 | 4.000E-07 | 1.700E-06 | 2.500E-06 |
| 7122 | 357 | E357R | A | 3.3e-06 | 3e-06 | 4.1e-06 | 3.467E-06 | 5.686E-07 | 3.000E-06 | 4.100E-06 |
| 7123 | 357 | E357N | A | 2.7e-06 | 2.3e-06 | 2.1e-06 | 2.367E-06 | 3.055E-07 | 2.100E-06 | 2.700E-06 |
| 7124 | 357 | E357D | A | 2e-06 | 2.3e-06 | 1.7e-06 | 2.000E-06 | 3.000E-07 | 1.700E-06 | 2.300E-06 |
| 7125 | 357 | E357C | A | 2.5e-06 | 2.2e-06 | 2.5e-06 | 2.400E-06 | 1.732E-07 | 2.200E-06 | 2.500E-06 |
| 7126 | 357 | E357E | A | 2.4e-06 | 2.1e-06 | 2.3e-06 | 2.267E-06 | 1.528E-07 | 2.100E-06 | 2.400E-06 |
| 7127 | 357 | E357Q | A | 2.5e-06 | 2.6e-06 | 2.1e-06 | 2.400E-06 | 2.646E-07 | 2.100E-06 | 2.600E-06 |
| 7128 | 357 | E357G | A | 2.9e-06 | 1.6e-06 | 1.4e-06 | 1.967E-06 | 8.145E-07 | 1.400E-06 | 2.900E-06 |
| 7129 | 357 | E357H | A | 2e-06 | 2e-06 | 2.1e-06 | 2.033E-06 | 5.774E-08 | 2.000E-06 | 2.100E-06 |
| 7130 | 357 | E357I | A | 1.3e-06 | 2.2e-06 | 2.3e-06 | 1.933E-06 | 5.508E-07 | 1.300E-06 | 2.300E-06 |
| 7131 | 357 | E357L | A | 3.7e-06 | 1.5e-06 | 2.4e-06 | 2.533E-06 | 1.106E-06 | 1.500E-06 | 3.700E-06 |
| 7132 | 357 | E357K | A | 2.4e-06 | 2.2e-06 | 2.6e-06 | 2.400E-06 | 2.000E-07 | 2.200E-06 | 2.600E-06 |
| 7133 | 357 | E357M | A | 1.3e-06 | 3e-06 | 1.4e-06 | 1.900E-06 | 9.539E-07 | 1.300E-06 | 3.000E-06 |
| 7134 | 357 | E357F | A | 2.5e-06 | 2e-06 | 2.8e-06 | 2.433E-06 | 4.041E-07 | 2.000E-06 | 2.800E-06 |
| 7135 | 357 | E357P | A | 1.5e-06 | 1.3e-06 | 1.4e-06 | 1.400E-06 | 1.000E-07 | 1.300E-06 | 1.500E-06 |
| 7136 | 357 | E357S | A | 2.5e-06 | 3.1e-06 | 2.2e-06 | 2.600E-06 | 4.583E-07 | 2.200E-06 | 3.100E-06 |
| 7137 | 357 | E357T | A | 6.4e-06 | 2.4e-06 | 3.2e-06 | 4.000E-06 | 2.117E-06 | 2.400E-06 | 6.400E-06 |
| 7138 | 357 | E357W | A | 1.8e-06 | 2.4e-06 | 1.6e-06 | 1.933E-06 | 4.163E-07 | 1.600E-06 | 2.400E-06 |
| 7139 | 357 | E357Y | A | 1.9e-06 | 2.4e-06 | 5.4e-06 | 3.233E-06 | 1.893E-06 | 1.900E-06 | 5.400E-06 |

| | | | | | | | | | | |
|---|---|---|---|---|---|---|---|---|---|---|
| 7140 | 357 | E357V | A | 2.8e-06 | 2.1e-06 | 2.5e-06 | 2.467E-06 | 3.512E-07 | 2.100E-06 | 2.800E-06 |
| 7141 | 358 | S358A | A | 4.3e-06 | 3.4e-06 | 1.7e-06 | 3.133E-06 | 1.320E-06 | 1.700E-06 | 4.300E-06 |
| 7142 | 358 | S358R | A | 1.6e-06 | 4e-06 | 2.3e-06 | 2.633E-06 | 1.234E-06 | 1.600E-06 | 4.000E-06 |
| 7143 | 358 | S358N | A | 2.1e-06 | 2.7e-06 | 2.2e-06 | 2.333E-06 | 3.215E-07 | 2.100E-06 | 2.700E-06 |
| 7144 | 358 | S358D | A | 1.2e-06 | 2.8e-06 | 2.6e-06 | 2.200E-06 | 8.718E-07 | 1.200E-06 | 2.800E-06 |
| 7145 | 358 | S358C | A | 1.1e-06 | 3.8e-06 | 1.8e-06 | 2.233E-06 | 1.401E-06 | 1.100E-06 | 3.800E-06 |
| 7146 | 358 | S358E | A | 3e-06 | 2.5e-06 | 2.1e-06 | 2.533E-06 | 4.509E-07 | 2.100E-06 | 3.000E-06 |
| 7147 | 358 | S358Q | A | 1.8e-06 | 2.8e-06 | 3.2e-06 | 2.600E-06 | 7.211E-07 | 1.800E-06 | 3.200E-06 |
| 7148 | 358 | S358G | A | 2.4e-06 | 2.6e-06 | 2.3e-06 | 2.433E-06 | 1.528E-07 | 2.300E-06 | 2.600E-06 |
| 7149 | 358 | S358H | A | 2.4e-06 | 2.9e-06 | 1.7e-06 | 2.333E-06 | 6.028E-07 | 1.700E-06 | 2.900E-06 |
| 7150 | 358 | S358I | A | 3.3e-06 | 2e-06 | 1.9e-06 | 2.400E-06 | 7.810E-07 | 1.900E-06 | 3.300E-06 |
| 7151 | 358 | S358L | A | 1.6e-06 | 2e-06 | 2.5e-06 | 2.033E-06 | 4.509E-07 | 1.600E-06 | 2.500E-06 |
| 7152 | 358 | S358K | A | 2.6e-06 | 1.3e-06 | 1.6e-06 | 1.833E-06 | 6.807E-07 | 1.300E-06 | 2.600E-06 |
| 7153 | 358 | S358M | A | 3.7e-06 | 1.8e-06 | 3.5e-06 | 3.000E-06 | 1.044E-06 | 1.800E-06 | 3.700E-06 |
| 7154 | 358 | S358F | A | 3e-06 | 2.3e-06 | 1.7e-06 | 2.333E-06 | 6.506E-07 | 1.700E-06 | 3.000E-06 |
| 7155 | 358 | S358P | A | 2.4e-06 | 2.6e-06 | 2.3e-06 | 2.433E-06 | 1.528E-07 | 2.300E-06 | 2.600E-06 |
| 7156 | 358 | S358S | A | 2.5e-06 | 1.2e-06 | 1.6e-06 | 1.767E-06 | 6.658E-07 | 1.200E-06 | 2.500E-06 |
| 7157 | 358 | S358T | A | 1e-06 | 1.4e-06 | 3.3e-06 | 1.900E-06 | 1.229E-06 | 1.000E-06 | 3.300E-06 |
| 7158 | 358 | S358W | A | 2.5e-06 | 1.6e-06 | 2.2e-06 | 2.100E-06 | 4.583E-07 | 1.600E-06 | 2.500E-06 |
| 7159 | 358 | S358Y | A | 1.9e-06 | 1.8e-06 | 1.9e-06 | 1.867E-06 | 5.774E-08 | 1.800E-06 | 1.900E-06 |
| 7160 | 358 | S358V | A | 3e-06 | 2.2e-06 | 2.5e-06 | 2.567E-06 | 4.041E-07 | 2.200E-06 | 3.000E-06 |
| 7161 | 359 | F359A | A | 2.2e-06 | 1.6e-06 | 2.6e-06 | 2.133E-06 | 5.033E-07 | 1.600E-06 | 2.600E-06 |
| 7162 | 359 | F359R | A | 2.4e-06 | 1.7e-06 | 2.2e-06 | 2.100E-06 | 3.606E-07 | 1.700E-06 | 2.400E-06 |
| 7163 | 359 | F359N | A | 2.5e-06 | 2.6e-06 | 2e-06 | 2.367E-06 | 3.215E-07 | 2.000E-06 | 2.600E-06 |
| 7164 | 359 | F359D | A | 4.4e-06 | 3.3e-06 | 1.9e-06 | 3.200E-06 | 1.253E-06 | 1.900E-06 | 4.400E-06 |
| 7165 | 359 | F359C | A | 2.7e-06 | 2.3e-06 | 1.6e-06 | 2.200E-06 | 5.568E-07 | 1.600E-06 | 2.700E-06 |
| 7166 | 359 | F359E | A | 2.5e-06 | 9.2e-07 | 3.1e-06 | 2.173E-06 | 1.126E-06 | 9.200E-07 | 3.100E-06 |
| 7167 | 359 | F359Q | A | 1.4e-06 | 2.7e-06 | 2.4e-06 | 2.167E-06 | 6.807E-07 | 1.400E-06 | 2.700E-06 |
| 7168 | 359 | F359G | A | 3.7e-06 | 2.2e-06 | 2.7e-06 | 2.867E-06 | 7.638E-07 | 2.200E-06 | 3.700E-06 |
| 7169 | 359 | F359H | A | 1.7e-06 | 3.6e-06 | 1.7e-06 | 2.333E-06 | 1.097E-06 | 1.700E-06 | 3.600E-06 |
| 7170 | 359 | F359I | A | 2.8e-06 | 5.1e-06 | 1.5e-06 | 3.133E-06 | 1.823E-06 | 1.500E-06 | 5.100E-06 |
| 7171 | 359 | F359L | A | 4.1e-06 | 3e-06 | 2.4e-06 | 3.167E-06 | 8.622E-07 | 2.400E-06 | 4.100E-06 |
| 7172 | 359 | F359K | A | 1.6e-06 | 2.2e-06 | 2.2e-06 | 2.000E-06 | 3.464E-07 | 1.600E-06 | 2.200E-06 |
| 7173 | 359 | F359M | A | 2.7e-06 | 2e-06 | 1.9e-06 | 2.200E-06 | 4.359E-07 | 1.900E-06 | 2.700E-06 |
| 7174 | 359 | F359F | A | 2.7e-06 | 2.3e-06 | 2.9e-06 | 2.633E-06 | 3.055E-07 | 2.300E-06 | 2.900E-06 |
| 7175 | 359 | F359P | A | 1.8e-06 | 9.3e-07 | 3.8e-06 | 2.177E-06 | 1.472E-06 | 9.300E-07 | 3.800E-06 |
| 7176 | 359 | F359S | A | 3.1e-06 | 2.8e-06 | 2e-06 | 2.633E-06 | 5.686E-07 | 2.000E-06 | 3.100E-06 |
| 7177 | 359 | F359T | A | 2.3e-06 | 1.7e-06 | 2.1e-06 | 2.033E-06 | 3.055E-07 | 1.700E-06 | 2.300E-06 |
| 7178 | 359 | F359W | A | 2.2e-06 | 2.5e-06 | 1.5e-06 | 2.067E-06 | 5.132E-07 | 1.500E-06 | 2.500E-06 |
| 7179 | 359 | F359Y | A | 1.5e-06 | 1.9e-06 | 3.3e-06 | 2.233E-06 | 9.452E-07 | 1.500E-06 | 3.300E-06 |
| 7180 | 359 | F359V | A | 1.7e-06 | 2.7e-06 | 2e-06 | 2.133E-06 | 5.132E-07 | 1.700E-06 | 2.700E-06 |
| 7181 | 360 | D360A | A | 2.3e-06 | 3.3e-06 | 2.4e-06 | 2.667E-06 | 5.508E-07 | 2.300E-06 | 3.300E-06 |
| 7182 | 360 | D360R | A | 2.3e-06 | 1.8e-06 | 1.3e-06 | 1.800E-06 | 5.000E-07 | 1.300E-06 | 2.300E-06 |
| 7183 | 360 | D360N | A | 2.6e-06 | 1.7e-06 | 4.2e-06 | 2.833E-06 | 1.266E-06 | 1.700E-06 | 4.200E-06 |
| 7184 | 360 | D360D | A | 1.9e-06 | 2.3e-06 | 2.9e-06 | 2.367E-06 | 5.033E-07 | 1.900E-06 | 2.900E-06 |
| 7185 | 360 | D360C | A | 1.5e-06 | 2.6e-06 | 1.2e-06 | 1.767E-06 | 7.371E-07 | 1.200E-06 | 2.600E-06 |
| 7186 | 360 | D360E | A | 3.1e-06 | 3.7e-06 | 1.8e-06 | 2.867E-06 | 9.713E-07 | 1.800E-06 | 3.700E-06 |
| 7187 | 360 | D360Q | A | 1.5e-06 | 2.7e-06 | 3.3e-06 | 2.500E-06 | 9.165E-07 | 1.500E-06 | 3.300E-06 |
| 7188 | 360 | D360G | A | 2.8e-06 | 1.8e-06 | 2.2e-06 | 2.267E-06 | 5.033E-07 | 1.800E-06 | 2.800E-06 |
| 7189 | 360 | D360H | A | 1.5e-06 | 1.8e-06 | 2e-06 | 1.767E-06 | 2.517E-07 | 1.500E-06 | 2.000E-06 |
| 7190 | 360 | D360I | A | 2.2e-06 | 1.7e-06 | 2.3e-06 | 2.067E-06 | 3.215E-07 | 1.700E-06 | 2.300E-06 |
| 7191 | 360 | D360L | A | 1.3e-06 | 2.8e-06 | 2.8e-06 | 2.300E-06 | 8.660E-07 | 1.300E-06 | 2.800E-06 |
| 7192 | 360 | D360K | A | 2.6e-06 | 3e-06 | 3.5e-06 | 3.033E-06 | 4.509E-07 | 2.600E-06 | 3.500E-06 |
| 7193 | 360 | D360M | A | 2.8e-06 | 2.4e-06 | 1.6e-06 | 2.267E-06 | 6.110E-07 | 1.600E-06 | 2.800E-06 |
| 7194 | 360 | D360F | A | 1.7e-06 | 1.1e-06 | 3.9e-06 | 2.233E-06 | 1.474E-06 | 1.100E-06 | 3.900E-06 |
| 7195 | 360 | D360P | A | 3.3e-06 | 4e-06 | 2.6e-06 | 3.300E-06 | 7.000E-07 | 2.600E-06 | 4.000E-06 |
| 7196 | 360 | D360S | A | 2.6e-06 | 1.7e-06 | 1.9e-06 | 2.067E-06 | 4.726E-07 | 1.700E-06 | 2.600E-06 |
| 7197 | 360 | D360T | A | 2.5e-06 | 4.9e-06 | 2.7e-06 | 3.367E-06 | 1.332E-06 | 2.500E-06 | 4.900E-06 |
| 7198 | 360 | D360W | A | 2.6e-06 | 1.8e-06 | 1.4e-06 | 1.933E-06 | 6.110E-07 | 1.400E-06 | 2.600E-06 |
| 7199 | 360 | D360Y | A | 2.9e-06 | 2.7e-06 | 3.8e-06 | 3.133E-06 | 5.859E-07 | 2.700E-06 | 3.800E-06 |
| 7200 | 360 | D360V | A | 2.6e-06 | 4.1e-06 | 4.4e-06 | 3.700E-06 | 9.644E-07 | 2.600E-06 | 4.400E-06 |
| 7201 | 361 | G361A | A | 3.1e-06 | 2e-06 | 3.2e-06 | 2.767E-06 | 6.658E-07 | 2.000E-06 | 3.200E-06 |
| 7202 | 361 | G361R | A | 2.6e-06 | 1.7e-06 | 2.3e-06 | 2.200E-06 | 4.583E-07 | 1.700E-06 | 2.600E-06 |
| 7203 | 361 | G361N | A | 3.9e-06 | 1.4e-06 | 1.2e-06 | 2.167E-06 | 1.504E-06 | 1.200E-06 | 3.900E-06 |
| 7204 | 361 | G361D | A | 1.9e-06 | 3e-06 | 2.5e-06 | 2.467E-06 | 5.508E-07 | 1.900E-06 | 3.000E-06 |
| 7205 | 361 | G361C | A | 1.8e-06 | 2e-06 | 1.8e-06 | 1.867E-06 | 1.155E-07 | 1.800E-06 | 2.000E-06 |
| 7206 | 361 | G361E | A | 1.7e-06 | 3.7e-06 | 1.3e-06 | 2.233E-06 | 1.286E-06 | 1.300E-06 | 3.700E-06 |
| 7207 | 361 | G361Q | A | 2.1e-06 | 1.9e-06 | 1.6e-06 | 1.867E-06 | 2.517E-07 | 1.600E-06 | 2.100E-06 |
| 7208 | 361 | G361G | A | 2.1e-06 | 2.4e-06 | 1.8e-06 | 2.100E-06 | 3.000E-07 | 1.800E-06 | 2.400E-06 |
| 7209 | 361 | G361H | A | 4.1e-06 | 2.5e-06 | 1.9e-06 | 2.833E-06 | 1.137E-06 | 1.900E-06 | 4.100E-06 |
| 7210 | 361 | G361I | A | 3.2e-06 | 2.5e-06 | 2.9e-06 | 2.867E-06 | 3.512E-07 | 2.500E-06 | 3.200E-06 |
| 7211 | 361 | G361L | A | 3e-06 | 2.4e-06 | 3.5e-06 | 2.967E-06 | 5.508E-07 | 2.400E-06 | 3.500E-06 |
| 7212 | 361 | G361K | A | 2.6e-06 | 2.8e-06 | 1.6e-06 | 2.333E-06 | 6.429E-07 | 1.600E-06 | 2.800E-06 |
| 7213 | 361 | G361M | A | 2.3e-06 | 3e-06 | 2.2e-06 | 2.500E-06 | 4.359E-07 | 2.200E-06 | 3.000E-06 |
| 7214 | 361 | G361F | A | 3e-06 | 3.3e-06 | 3.7e-06 | 3.333E-06 | 3.512E-07 | 3.000E-06 | 3.700E-06 |
| 7215 | 361 | G361P | A | 2.3e-06 | 3e-06 | 2.8e-06 | 2.700E-06 | 3.606E-07 | 2.300E-06 | 3.000E-06 |
| 7216 | 361 | G361S | A | 4.2e-06 | 5.8e-06 | 1.9e-06 | 3.967E-06 | 1.960E-06 | 1.900E-06 | 5.800E-06 |
| 7217 | 361 | G361T | A | 2.5e-06 | 3.5e-06 | 2.4e-06 | 2.800E-06 | 6.083E-07 | 2.400E-06 | 3.500E-06 |
| 7218 | 361 | G361W | A | 1.9e-06 | 2.3e-06 | 3.8e-06 | 2.667E-06 | 1.002E-06 | 1.900E-06 | 3.800E-06 |
| 7219 | 361 | G361Y | A | 2.6e-06 | 2.1e-06 | 2.7e-06 | 2.467E-06 | 3.215E-07 | 2.100E-06 | 2.700E-06 |
| 7220 | 361 | G361V | A | 2.5e-06 | 2.3e-06 | 2.1e-06 | 2.300E-06 | 2.000E-07 | 2.100E-06 | 2.500E-06 |
| 7221 | 362 | D362A | A | 2.2e-06 | 1.6e-06 | 2.6e-06 | 2.133E-06 | 5.033E-07 | 1.600E-06 | 2.600E-06 |
| 7222 | 362 | D362R | A | 1.5e-06 | 4.3e-06 | 1.9e-06 | 2.567E-06 | 1.514E-06 | 1.500E-06 | 4.300E-06 |
| 7223 | 362 | D362N | A | 1.2e-06 | 2.6e-06 | 3.7e-06 | 2.500E-06 | 1.253E-06 | 1.200E-06 | 3.700E-06 |
| 7224 | 362 | D362D | A | 1.7e-06 | 1.7e-06 | 3.2e-06 | 2.200E-06 | 8.660E-07 | 1.700E-06 | 3.200E-06 |

| | | | | | | | | | | |
|---|---|---|---|---|---|---|---|---|---|---|
| 7225 | 362 | D362C | A | 2e-06 | 2.3e-06 | 1.3e-06 | 1.867E-06 | 5.132E-07 | 1.300E-06 | 2.300E-06 |
| 7226 | 362 | D362E | A | 3.3e-06 | 2e-06 | 2.8e-06 | 2.700E-06 | 6.557E-07 | 2.000E-06 | 3.300E-06 |
| 7227 | 362 | D362Q | A | 2.1e-06 | 1.8e-06 | 2.9e-06 | 2.267E-06 | 5.686E-07 | 1.800E-06 | 2.900E-06 |
| 7228 | 362 | D362G | A | 1.2e-06 | 1.8e-06 | 3.1e-06 | 2.033E-06 | 9.713E-07 | 1.200E-06 | 3.100E-06 |
| 7229 | 362 | D362H | A | 3.5e-06 | 1.8e-06 | 1.5e-06 | 2.267E-06 | 1.079E-06 | 1.500E-06 | 3.500E-06 |
| 7230 | 362 | D362I | A | 2.7e-06 | 1.8e-06 | 2.3e-06 | 2.267E-06 | 4.509E-07 | 1.800E-06 | 2.700E-06 |
| 7231 | 362 | D362L | A | 4.1e-06 | 1.8e-06 | 2.9e-06 | 2.933E-06 | 1.150E-06 | 1.800E-06 | 4.100E-06 |
| 7232 | 362 | D362K | A | 2.2e-06 | 2e-06 | 9.6e-07 | 1.720E-06 | 6.657E-07 | 9.600E-07 | 2.200E-06 |
| 7233 | 362 | D362M | A | 2.3e-06 | 3.6e-06 | 1.9e-06 | 2.600E-06 | 8.888E-07 | 1.900E-06 | 3.600E-06 |
| 7234 | 362 | D362F | A | 1.8e-06 | 3.1e-06 | 2.3e-06 | 2.400E-06 | 6.557E-07 | 1.800E-06 | 3.100E-06 |
| 7235 | 362 | D362P | A | 1.3e-06 | 2.8e-06 | 2.4e-06 | 2.167E-06 | 7.767E-07 | 1.300E-06 | 2.800E-06 |
| 7236 | 362 | D362S | A | 1.3e-06 | 2.9e-06 | 3.2e-06 | 2.467E-06 | 1.021E-06 | 1.300E-06 | 3.200E-06 |
| 7237 | 362 | D362T | A | 2.7e-06 | 9e-07 | 2.4e-06 | 2.000E-06 | 9.644E-07 | 9.000E-07 | 2.700E-06 |
| 7238 | 362 | D362W | A | 2.4e-06 | 3.3e-06 | 2.3e-06 | 2.667E-06 | 5.508E-07 | 2.300E-06 | 3.300E-06 |
| 7239 | 362 | D362Y | A | 3e-06 | 2.2e-06 | 2.1e-06 | 2.433E-06 | 4.933E-07 | 2.100E-06 | 3.000E-06 |
| 7240 | 362 | D362V | A | 3.1e-06 | 4.6e-06 | 1.6e-06 | 3.100E-06 | 1.500E-06 | 1.600E-06 | 4.600E-06 |
| 7241 | 363 | P363A | A | 2.6e-06 | 2.4e-06 | 3.1e-06 | 2.700E-06 | 3.606E-07 | 2.400E-06 | 3.100E-06 |
| 7242 | 363 | P363R | A | 3.7e-06 | 2.3e-06 | 2.3e-06 | 2.767E-06 | 8.083E-07 | 2.300E-06 | 3.700E-06 |
| 7243 | 363 | P363N | A | 3.9e-06 | 2.7e-06 | 3.1e-06 | 3.233E-06 | 6.110E-07 | 2.700E-06 | 3.900E-06 |
| 7244 | 363 | P363D | A | 2.5e-06 | 2.1e-06 | 2.6e-06 | 2.400E-06 | 2.646E-07 | 2.100E-06 | 2.600E-06 |
| 7245 | 363 | P363C | A | 3.5e-06 | 2.2e-06 | 2.8e-06 | 2.833E-06 | 6.506E-07 | 2.200E-06 | 3.500E-06 |
| 7246 | 363 | P363E | A | 2e-06 | 2.6e-06 | 1.5e-06 | 2.033E-06 | 5.508E-07 | 1.500E-06 | 2.600E-06 |
| 7247 | 363 | P363Q | A | 1.8e-06 | 1.1e-06 | 3.6e-06 | 2.167E-06 | 1.290E-06 | 1.100E-06 | 3.600E-06 |
| 7248 | 363 | P363G | A | 1.7e-06 | 1.4e-06 | 2.8e-06 | 1.967E-06 | 7.371E-07 | 1.400E-06 | 2.800E-06 |
| 7249 | 363 | P363H | A | 2.2e-06 | 1.6e-06 | 2.4e-06 | 2.067E-06 | 4.163E-07 | 1.600E-06 | 2.400E-06 |
| 7250 | 363 | P363I | A | 2.5e-06 | 2.2e-06 | 3.2e-06 | 2.633E-06 | 5.132E-07 | 2.200E-06 | 3.200E-06 |
| 7251 | 363 | P363L | A | 2.8e-06 | 1.2e-06 | 3.3e-06 | 2.433E-06 | 1.097E-06 | 1.200E-06 | 3.300E-06 |
| 7252 | 363 | P363K | A | 3.2e-06 | 2.6e-06 | 1.5e-06 | 2.433E-06 | 8.622E-07 | 1.500E-06 | 3.200E-06 |
| 7253 | 363 | P363M | A | 3.8e-06 | 1.6e-06 | 2.4e-06 | 2.600E-06 | 1.114E-06 | 1.600E-06 | 3.800E-06 |
| 7254 | 363 | P363F | A | 1.1e-06 | 3.9e-06 | 3.3e-06 | 2.767E-06 | 1.474E-06 | 1.100E-06 | 3.900E-06 |
| 7255 | 363 | P363P | A | 6.9e-07 | 2e-06 | 2.8e-06 | 1.830E-06 | 1.065E-06 | 6.900E-07 | 2.800E-06 |
| 7256 | 363 | P363S | A | 2.4e-06 | 3.4e-06 | 1.9e-06 | 2.567E-06 | 7.638E-07 | 1.900E-06 | 3.400E-06 |
| 7257 | 363 | P363T | A | 2.9e-06 | 5.1e-06 | 3.5e-06 | 3.833E-06 | 1.137E-06 | 2.900E-06 | 5.100E-06 |
| 7258 | 363 | P363W | A | 2.2e-06 | 1.1e-06 | 2.2e-06 | 1.833E-06 | 6.351E-07 | 1.100E-06 | 2.200E-06 |
| 7259 | 363 | P363Y | A | 9.6e-07 | 5.2e-06 | 1.8e-06 | 2.653E-06 | 2.245E-06 | 9.600E-07 | 5.200E-06 |
| 7260 | 363 | P363V | A | 2.9e-06 | 3.4e-06 | 3.2e-06 | 3.167E-06 | 2.517E-07 | 2.900E-06 | 3.400E-06 |
| 7261 | 364 | A364A | A | 1.7e-06 | 2.7e-06 | 2e-06 | 2.133E-06 | 5.132E-07 | 1.700E-06 | 2.700E-06 |
| 7262 | 364 | A364R | A | 1.7e-06 | 2.1e-06 | 2.8e-06 | 2.200E-06 | 5.568E-07 | 1.700E-06 | 2.800E-06 |
| 7263 | 364 | A364N | A | 1.7e-06 | 2.4e-06 | 2.5e-06 | 2.200E-06 | 4.359E-07 | 1.700E-06 | 2.500E-06 |
| 7264 | 364 | A364D | A | 2.8e-06 | 1.3e-06 | 2.5e-06 | 2.200E-06 | 7.937E-07 | 1.300E-06 | 2.800E-06 |
| 7265 | 364 | A364C | A | 1.9e-06 | 2.4e-06 | 2.7e-06 | 2.333E-06 | 4.041E-07 | 1.900E-06 | 2.700E-06 |
| 7266 | 364 | A364E | A | 2e-06 | 3.4e-06 | 1.9e-06 | 2.433E-06 | 8.386E-07 | 1.900E-06 | 3.400E-06 |
| 7267 | 364 | A364Q | A | 1.8e-06 | 1.8e-06 | 2e-06 | 1.867E-06 | 1.155E-07 | 1.800E-06 | 2.000E-06 |
| 7268 | 364 | A364G | A | 1.9e-06 | 3.7e-06 | 1.7e-06 | 2.433E-06 | 1.102E-06 | 1.700E-06 | 3.700E-06 |
| 7269 | 364 | A364H | A | 2.1e-06 | 2.6e-06 | 3.2e-06 | 2.633E-06 | 5.508E-07 | 2.100E-06 | 3.200E-06 |
| 7270 | 364 | A364I | A | 4.9e-06 | 2e-06 | 2.8e-06 | 3.233E-06 | 1.498E-06 | 2.000E-06 | 4.900E-06 |
| 7271 | 364 | A364L | A | 3.1e-06 | 3.4e-06 | 2.4e-06 | 2.967E-06 | 5.132E-07 | 2.400E-06 | 3.400E-06 |
| 7272 | 364 | A364K | A | 3.2e-06 | 2.2e-06 | 4.7e-06 | 3.367E-06 | 1.258E-06 | 2.200E-06 | 4.700E-06 |
| 7273 | 364 | A364M | A | 3.1e-06 | 3.4e-06 | 2.2e-06 | 2.900E-06 | 6.245E-07 | 2.200E-06 | 3.400E-06 |
| 7274 | 364 | A364F | A | 3.1e-06 | 3.2e-06 | 3.6e-06 | 3.300E-06 | 2.646E-07 | 3.100E-06 | 3.600E-06 |
| 7275 | 364 | A364P | A | 2.3e-06 | 2.7e-06 | 1.6e-06 | 2.200E-06 | 5.568E-07 | 1.600E-06 | 2.700E-06 |
| 7276 | 364 | A364S | A | 2.8e-06 | 2e-06 | 2.2e-06 | 2.333E-06 | 4.163E-07 | 2.000E-06 | 2.800E-06 |
| 7277 | 364 | A364T | A | 3e-06 | 6.7e-07 | 1.9e-06 | 1.857E-06 | 1.166E-06 | 6.700E-07 | 3.000E-06 |
| 7278 | 364 | A364W | A | 1.7e-06 | 1.9e-06 | 2.7e-06 | 2.100E-06 | 5.292E-07 | 1.700E-06 | 2.700E-06 |
| 7279 | 364 | A364Y | A | 2.6e-06 | 3.2e-06 | 3.7e-06 | 3.167E-06 | 5.508E-07 | 2.600E-06 | 3.700E-06 |
| 7280 | 364 | A364V | A | 1.7e-06 | 3.3e-06 | 2e-06 | 2.333E-06 | 8.505E-07 | 1.700E-06 | 3.300E-06 |
| 7281 | 365 | S365A | A | 3.6e-06 | 1.2e-06 | 1.5e-06 | 2.100E-06 | 1.308E-06 | 1.200E-06 | 3.600E-06 |
| 7282 | 365 | S365R | A | 2.4e-06 | 3e-06 | 1.4e-06 | 2.267E-06 | 8.083E-07 | 1.400E-06 | 3.000E-06 |
| 7283 | 365 | S365N | A | 3e-06 | 2.8e-06 | 4e-06 | 3.267E-06 | 6.429E-07 | 2.800E-06 | 4.000E-06 |
| 7284 | 365 | S365D | A | 2.3e-06 | 2.2e-06 | 4.8e-06 | 3.100E-06 | 1.473E-06 | 2.200E-06 | 4.800E-06 |
| 7285 | 365 | S365C | A | 4.2e-06 | 1.7e-06 | 3.4e-06 | 3.100E-06 | 1.277E-06 | 1.700E-06 | 4.200E-06 |
| 7286 | 365 | S365E | A | 1.3e-06 | 1.6e-06 | 1.9e-06 | 1.600E-06 | 3.000E-07 | 1.300E-06 | 1.900E-06 |
| 7287 | 365 | S365Q | A | 2.6e-06 | 3.1e-06 | 2.1e-06 | 2.600E-06 | 5.000E-07 | 2.100E-06 | 3.100E-06 |
| 7288 | 365 | S365G | A | 5.3e-06 | 3.3e-06 | 2.2e-06 | 3.600E-06 | 1.572E-06 | 2.200E-06 | 5.300E-06 |
| 7289 | 365 | S365H | A | 2.8e-06 | 1.3e-06 | 3e-06 | 2.367E-06 | 9.292E-07 | 1.300E-06 | 3.000E-06 |
| 7290 | 365 | S365I | A | 2.9e-06 | 3.4e-06 | 4.2e-06 | 3.500E-06 | 6.557E-07 | 2.900E-06 | 4.200E-06 |
| 7291 | 365 | S365L | A | 2.1e-06 | 3.8e-06 | 3.1e-06 | 3.000E-06 | 8.544E-07 | 2.100E-06 | 3.800E-06 |
| 7292 | 365 | S365K | A | 2.9e-06 | 1.7e-06 | 1.9e-06 | 2.167E-06 | 6.429E-07 | 1.700E-06 | 2.900E-06 |
| 7293 | 365 | S365M | A | 2.6e-06 | 2.5e-06 | 3.1e-06 | 2.733E-06 | 3.215E-07 | 2.500E-06 | 3.100E-06 |
| 7294 | 365 | S365F | A | 1.8e-06 | 2.4e-06 | 1.9e-06 | 2.033E-06 | 3.215E-07 | 1.800E-06 | 2.400E-06 |
| 7295 | 365 | S365P | A | 2.4e-06 | 2.7e-06 | 2.4e-06 | 2.500E-06 | 1.732E-07 | 2.400E-06 | 2.700E-06 |
| 7296 | 365 | S365S | A | 1.8e-06 | 3.3e-06 | 2.6e-06 | 2.567E-06 | 7.506E-07 | 1.800E-06 | 3.300E-06 |
| 7297 | 365 | S365T | A | 2.7e-06 | 2.5e-06 | 3.1e-06 | 2.767E-06 | 3.055E-07 | 2.500E-06 | 3.100E-06 |
| 7298 | 365 | S365W | A | 2.1e-06 | 2.5e-06 | 2e-06 | 2.200E-06 | 2.646E-07 | 2.000E-06 | 2.500E-06 |
| 7299 | 365 | S365Y | A | 2.7e-06 | 2.5e-06 | 1.8e-06 | 2.333E-06 | 4.726E-07 | 1.800E-06 | 2.700E-06 |
| 7300 | 365 | S365V | A | 3.9e-06 | 2.9e-06 | 3e-06 | 3.267E-06 | 5.508E-07 | 2.900E-06 | 3.900E-06 |
| 7301 | 366 | N366A | A | 2.6e-06 | 1.8e-06 | 2.5e-06 | 2.300E-06 | 4.359E-07 | 1.800E-06 | 2.600E-06 |
| 7302 | 366 | N366R | A | 1.9e-06 | 4e-06 | 2.2e-06 | 2.700E-06 | 1.136E-06 | 1.900E-06 | 4.000E-06 |
| 7303 | 366 | N366N | A | 1.4e-06 | 3e-06 | 2.3e-06 | 2.233E-06 | 8.021E-07 | 1.400E-06 | 3.000E-06 |
| 7304 | 366 | N366D | A | 2.3e-06 | 1.4e-06 | 1.6e-06 | 1.767E-06 | 4.726E-07 | 1.400E-06 | 2.300E-06 |
| 7305 | 366 | N366C | A | 1.6e-06 | 4.1e-06 | 1.8e-06 | 2.500E-06 | 1.389E-06 | 1.600E-06 | 4.100E-06 |
| 7306 | 366 | N366E | A | 2.1e-06 | 2.4e-06 | 2.5e-06 | 2.333E-06 | 2.082E-07 | 2.100E-06 | 2.500E-06 |
| 7307 | 366 | N366Q | A | 2.4e-06 | 1.2e-06 | 2.3e-06 | 1.967E-06 | 6.658E-07 | 1.200E-06 | 2.400E-06 |
| 7308 | 366 | N366G | A | 2.5e-06 | 2.9e-06 | 3.9e-06 | 3.100E-06 | 7.211E-07 | 2.500E-06 | 3.900E-06 |
| 7309 | 366 | N366H | A | 2.6e-06 | 3.9e-06 | 2.1e-06 | 2.867E-06 | 9.292E-07 | 2.100E-06 | 3.900E-06 |

| | | | | | | | | | | |
|---|---|---|---|---|---|---|---|---|---|---|
| 7310 | 366 | N366I | A | 2.6e-06 | 3e-06 | 3.6e-06 | 3.067E-06 | 5.033E-07 | 2.600E-06 | 3.600E-06 |
| 7311 | 366 | N366L | A | 1.5e-06 | 2.6e-06 | 1.8e-06 | 1.967E-06 | 5.686E-07 | 1.500E-06 | 2.600E-06 |
| 7312 | 366 | N366K | A | 1.9e-06 | 2.1e-06 | 2e-06 | 2.000E-06 | 1.000E-07 | 1.900E-06 | 2.100E-06 |
| 7313 | 366 | N366M | A | 2.8e-06 | 1.8e-06 | 2.4e-06 | 2.333E-06 | 5.033E-07 | 1.800E-06 | 2.800E-06 |
| 7314 | 366 | N366F | A | 5.9e-06 | 5.5e-06 | 1.8e-06 | 4.400E-06 | 2.261E-06 | 1.800E-06 | 5.900E-06 |
| 7315 | 366 | N366P | A | 1.4e-06 | 4e-06 | 2.6e-06 | 2.667E-06 | 1.301E-06 | 1.400E-06 | 4.000E-06 |
| 7316 | 366 | N366S | A | 3.5e-06 | 1.8e-06 | 3.7e-06 | 3.000E-06 | 1.044E-06 | 1.800E-06 | 3.700E-06 |
| 7317 | 366 | N366T | A | 3e-06 | 1.9e-06 | 2.3e-06 | 2.400E-06 | 5.568E-07 | 1.900E-06 | 3.000E-06 |
| 7318 | 366 | N366W | A | 3.9e-06 | 1.8e-06 | 3.6e-06 | 3.100E-06 | 1.136E-06 | 1.800E-06 | 3.900E-06 |
| 7319 | 366 | N366Y | A | 3.1e-06 | 2.6e-06 | 2e-06 | 2.567E-06 | 5.508E-07 | 2.000E-06 | 3.100E-06 |
| 7320 | 366 | N366V | A | 2.8e-06 | 2.3e-06 | 2.3e-06 | 2.467E-06 | 2.887E-07 | 2.300E-06 | 2.800E-06 |
| 7321 | 367 | T367A | A | 4.4e-06 | 3.2e-06 | 2.6e-06 | 3.400E-06 | 9.165E-07 | 2.600E-06 | 4.400E-06 |
| 7322 | 367 | T367R | A | 2.4e-06 | 2e-06 | 1.4e-06 | 1.933E-06 | 5.033E-07 | 1.400E-06 | 2.400E-06 |
| 7323 | 367 | T367N | A | 2.4e-06 | 2.6e-06 | 2.7e-06 | 2.567E-06 | 1.528E-07 | 2.400E-06 | 2.700E-06 |
| 7324 | 367 | T367D | A | 2.6e-06 | 3.4e-06 | 2.7e-06 | 2.900E-06 | 4.359E-07 | 2.600E-06 | 3.400E-06 |
| 7325 | 367 | T367C | A | 2e-06 | 1.9e-06 | 4.1e-06 | 2.667E-06 | 1.242E-06 | 1.900E-06 | 4.100E-06 |
| 7326 | 367 | T367E | A | 2e-06 | 3.1e-06 | 3.2e-06 | 2.767E-06 | 6.658E-07 | 2.000E-06 | 3.200E-06 |
| 7327 | 367 | T367Q | A | 2.5e-06 | 2.7e-06 | 2.1e-06 | 2.433E-06 | 3.055E-07 | 2.100E-06 | 2.700E-06 |
| 7328 | 367 | T367G | A | 2.3e-06 | 2.4e-06 | 2.2e-06 | 2.300E-06 | 1.000E-07 | 2.200E-06 | 2.400E-06 |
| 7329 | 367 | T367H | A | 1.9e-06 | 1.4e-06 | 1.9e-06 | 1.733E-06 | 2.887E-07 | 1.400E-06 | 1.900E-06 |
| 7330 | 367 | T367I | A | 2.4e-06 | 2.7e-06 | 1.9e-06 | 2.333E-06 | 4.041E-07 | 1.900E-06 | 2.700E-06 |
| 7331 | 367 | T367L | A | 2.8e-06 | 2.9e-06 | 2.2e-06 | 2.633E-06 | 3.786E-07 | 2.200E-06 | 2.900E-06 |
| 7332 | 367 | T367K | A | 2.2e-06 | 2.7e-06 | 1.2e-06 | 2.033E-06 | 7.638E-07 | 1.200E-06 | 2.700E-06 |
| 7333 | 367 | T367M | A | 3.6e-06 | 8.3e-07 | 3.3e-06 | 2.577E-06 | 1.520E-06 | 8.300E-07 | 3.600E-06 |
| 7334 | 367 | T367F | A | 2.4e-06 | 2.9e-06 | 1.3e-06 | 2.200E-06 | 8.185E-07 | 1.300E-06 | 2.900E-06 |
| 7335 | 367 | T367P | A | 2.5e-06 | 2.2e-06 | 2.1e-06 | 2.267E-06 | 2.082E-07 | 2.100E-06 | 2.500E-06 |
| 7336 | 367 | T367S | A | 4.6e-06 | 2.6e-06 | 2.6e-06 | 3.267E-06 | 1.155E-06 | 2.600E-06 | 4.600E-06 |
| 7337 | 367 | T367T | A | 2.5e-06 | 4.7e-06 | 1.9e-06 | 3.033E-06 | 1.474E-06 | 1.900E-06 | 4.700E-06 |
| 7338 | 367 | T367W | A | 2.3e-06 | 1.9e-06 | 1.8e-06 | 2.000E-06 | 2.646E-07 | 1.800E-06 | 2.300E-06 |
| 7339 | 367 | T367Y | A | 1.4e-06 | 2.8e-06 | 1.4e-06 | 1.867E-06 | 8.083E-07 | 1.400E-06 | 2.800E-06 |
| 7340 | 367 | T367V | A | 2.7e-06 | 3.4e-06 | 1.9e-06 | 2.667E-06 | 7.506E-07 | 1.900E-06 | 3.400E-06 |
| 7341 | 368 | A368A | A | 3.3e-06 | 2.3e-06 | 2.8e-06 | 2.800E-06 | 5.000E-07 | 2.300E-06 | 3.300E-06 |
| 7342 | 368 | A368R | A | 2.3e-06 | 2.5e-06 | 2.9e-06 | 2.567E-06 | 3.055E-07 | 2.300E-06 | 2.900E-06 |
| 7343 | 368 | A368N | A | 2.1e-06 | 2.2e-06 | 2.4e-06 | 2.233E-06 | 1.528E-07 | 2.100E-06 | 2.400E-06 |
| 7344 | 368 | A368D | A | 1.7e-06 | 3e-06 | 2.9e-06 | 2.533E-06 | 7.234E-07 | 1.700E-06 | 3.000E-06 |
| 7345 | 368 | A368C | A | 1.9e-06 | 1.7e-06 | 1.6e-06 | 1.733E-06 | 1.528E-07 | 1.600E-06 | 1.900E-06 |
| 7346 | 368 | A368E | A | 2.3e-06 | 2.5e-06 | 2e-06 | 2.267E-06 | 2.517E-07 | 2.000E-06 | 2.500E-06 |
| 7347 | 368 | A368Q | A | 2.7e-06 | 4.2e-06 | 2.6e-06 | 3.167E-06 | 8.963E-07 | 2.600E-06 | 4.200E-06 |
| 7348 | 368 | A368G | A | 2e-06 | 2e-06 | 2.4e-06 | 2.133E-06 | 2.309E-07 | 2.000E-06 | 2.400E-06 |
| 7349 | 368 | A368H | A | 2.3e-06 | 1.7e-06 | 1.9e-06 | 1.967E-06 | 3.055E-07 | 1.700E-06 | 2.300E-06 |
| 7350 | 368 | A368I | A | 2e-06 | 2.1e-06 | 1.9e-06 | 2.000E-06 | 1.000E-07 | 1.900E-06 | 2.100E-06 |
| 7351 | 368 | A368L | A | 2.6e-06 | 1.8e-06 | 2.9e-06 | 2.433E-06 | 5.686E-07 | 1.800E-06 | 2.900E-06 |
| 7352 | 368 | A368K | A | 2.5e-06 | 2.2e-06 | 3.1e-06 | 2.600E-06 | 4.583E-07 | 2.200E-06 | 3.100E-06 |
| 7353 | 368 | A368M | A | 2.5e-06 | 2.3e-06 | 2.8e-06 | 2.533E-06 | 2.517E-07 | 2.300E-06 | 2.800E-06 |
| 7354 | 368 | A368F | A | 1.8e-06 | 2e-06 | 2.5e-06 | 2.100E-06 | 3.606E-07 | 1.800E-06 | 2.500E-06 |
| 7355 | 368 | A368P | A | 2.3e-06 | 1e-06 | 1.6e-06 | 1.633E-06 | 6.506E-07 | 1.000E-06 | 2.300E-06 |
| 7356 | 368 | A368S | A | 2.9e-06 | 1.5e-06 | 2.3e-06 | 2.233E-06 | 7.024E-07 | 1.500E-06 | 2.900E-06 |
| 7357 | 368 | A368T | A | 2.5e-06 | 2.7e-06 | 2.9e-06 | 2.700E-06 | 2.000E-07 | 2.500E-06 | 2.900E-06 |
| 7358 | 368 | A368W | A | 1.6e-06 | 1.5e-06 | 1.8e-06 | 1.633E-06 | 1.528E-07 | 1.500E-06 | 1.800E-06 |
| 7359 | 368 | A368Y | A | 2.2e-06 | 2.4e-06 | 2.2e-06 | 2.267E-06 | 1.155E-07 | 2.200E-06 | 2.400E-06 |
| 7360 | 368 | A368V | A | 3.2e-06 | 2.2e-06 | 3e-06 | 2.800E-06 | 5.292E-07 | 2.200E-06 | 3.200E-06 |
| 7361 | 369 | P369A | A | 2.5e-06 | 2.6e-06 | 2.6e-06 | 2.567E-06 | 5.774E-08 | 2.500E-06 | 2.600E-06 |
| 7362 | 369 | P369R | A | 2e-06 | 2.4e-06 | 1.7e-06 | 2.033E-06 | 3.512E-07 | 1.700E-06 | 2.400E-06 |
| 7363 | 369 | P369N | A | 3e-06 | 2e-06 | 2e-06 | 2.333E-06 | 5.774E-07 | 2.000E-06 | 3.000E-06 |
| 7364 | 369 | P369D | A | 3e-06 | 2.9e-06 | 4.1e-06 | 3.333E-06 | 6.658E-07 | 2.900E-06 | 4.100E-06 |
| 7365 | 369 | P369C | A | 2.3e-06 | 1.7e-06 | 2.4e-06 | 2.133E-06 | 3.786E-07 | 1.700E-06 | 2.400E-06 |
| 7366 | 369 | P369E | A | 1.7e-06 | 3.1e-06 | 2.8e-06 | 2.533E-06 | 7.371E-07 | 1.700E-06 | 3.100E-06 |
| 7367 | 369 | P369Q | A | 3.3e-06 | 1.3e-06 | 2.9e-06 | 2.500E-06 | 1.058E-06 | 1.300E-06 | 3.300E-06 |
| 7368 | 369 | P369G | A | 3.3e-06 | 2.2e-06 | 2.1e-06 | 2.533E-06 | 6.658E-07 | 2.100E-06 | 3.300E-06 |
| 7369 | 369 | P369H | A | 1.9e-06 | 1.8e-06 | 3e-06 | 2.233E-06 | 6.658E-07 | 1.800E-06 | 3.000E-06 |
| 7370 | 369 | P369I | A | 4e-06 | 1.9e-06 | 2.9e-06 | 2.933E-06 | 1.050E-06 | 1.900E-06 | 4.000E-06 |
| 7371 | 369 | P369L | A | 1.8e-06 | 2.7e-06 | 2e-06 | 2.167E-06 | 4.726E-07 | 1.800E-06 | 2.700E-06 |
| 7372 | 369 | P369K | A | 2.2e-06 | 1.3e-06 | 2.4e-06 | 1.967E-06 | 5.859E-07 | 1.300E-06 | 2.400E-06 |
| 7373 | 369 | P369M | A | 1.6e-06 | 2.6e-06 | 1.5e-06 | 1.900E-06 | 6.083E-07 | 1.500E-06 | 2.600E-06 |
| 7374 | 369 | P369F | A | 4.6e-06 | 2.8e-06 | 2.7e-06 | 3.367E-06 | 1.069E-06 | 2.700E-06 | 4.600E-06 |
| 7375 | 369 | P369P | A | 2e-06 | 3.6e-06 | 1.6e-06 | 2.400E-06 | 1.058E-06 | 1.600E-06 | 3.600E-06 |
| 7376 | 369 | P369S | A | 1.5e-06 | 2e-06 | 2.6e-06 | 2.033E-06 | 5.508E-07 | 1.500E-06 | 2.600E-06 |
| 7377 | 369 | P369T | A | 2.3e-06 | 2.9e-06 | 1.6e-06 | 2.267E-06 | 6.506E-07 | 1.600E-06 | 2.900E-06 |
| 7378 | 369 | P369W | A | 2.9e-06 | 2.2e-06 | 2.4e-06 | 2.500E-06 | 3.606E-07 | 2.200E-06 | 2.900E-06 |
| 7379 | 369 | P369Y | A | 2.8e-06 | 3e-06 | 3.5e-06 | 3.100E-06 | 3.606E-07 | 2.800E-06 | 3.500E-06 |
| 7380 | 369 | P369V | A | 2.5e-06 | 1.7e-06 | 2.8e-06 | 2.333E-06 | 5.686E-07 | 1.700E-06 | 2.800E-06 |
| 7381 | 370 | L370A | A | 1.2e-06 | 2.2e-06 | 4.1e-06 | 2.500E-06 | 1.473E-06 | 1.200E-06 | 4.100E-06 |
| 7382 | 370 | L370R | A | 2e-06 | 1.7e-06 | 3.6e-06 | 2.433E-06 | 1.021E-06 | 1.700E-06 | 3.600E-06 |
| 7383 | 370 | L370N | A | 4.6e-06 | 1.8e-06 | 2e-06 | 2.800E-06 | 1.562E-06 | 1.800E-06 | 4.600E-06 |
| 7384 | 370 | L370D | A | 2e-06 | 1.2e-06 | 4.2e-06 | 2.467E-06 | 1.553E-06 | 1.200E-06 | 4.200E-06 |
| 7385 | 370 | L370C | A | 2.1e-06 | 3.5e-06 | 1.6e-06 | 2.400E-06 | 9.849E-07 | 1.600E-06 | 3.500E-06 |
| 7386 | 370 | L370E | A | 2.9e-06 | 2.4e-06 | 2.3e-06 | 2.533E-06 | 3.215E-07 | 2.300E-06 | 2.900E-06 |
| 7387 | 370 | L370Q | A | 2.9e-06 | 2.2e-06 | 3.5e-06 | 2.867E-06 | 6.506E-07 | 2.200E-06 | 3.500E-06 |
| 7388 | 370 | L370G | A | 2.6e-06 | 3.5e-06 | 2.2e-06 | 2.767E-06 | 6.658E-07 | 2.200E-06 | 3.500E-06 |
| 7389 | 370 | L370H | A | 1.5e-06 | 1.7e-06 | 2.7e-06 | 1.967E-06 | 6.429E-07 | 1.500E-06 | 2.700E-06 |
| 7390 | 370 | L370I | A | 2.2e-06 | 1.4e-06 | 2.2e-06 | 1.933E-06 | 4.619E-07 | 1.400E-06 | 2.200E-06 |
| 7391 | 370 | L370L | A | 2.4e-06 | 2.5e-06 | 4.9e-06 | 3.267E-06 | 1.415E-06 | 2.400E-06 | 4.900E-06 |
| 7392 | 370 | L370K | A | 3.1e-06 | 3.2e-06 | 3e-06 | 3.100E-06 | 1.000E-07 | 3.000E-06 | 3.200E-06 |
| 7393 | 370 | L370M | A | 5.8e-06 | 1.9e-06 | 2.7e-06 | 3.467E-06 | 2.060E-06 | 1.900E-06 | 5.800E-06 |
| 7394 | 370 | L370F | A | 1.6e-06 | 3.1e-06 | 2.2e-06 | 2.300E-06 | 7.550E-07 | 1.600E-06 | 3.100E-06 |

| | | | | | | | | | | |
|---|---|---|---|---|---|---|---|---|---|---|
| 7395 | 370 | L370P | A | 1.7e-06 | 2.3e-06 | 2.8e-06 | 2.267E-06 | 5.508E-07 | 1.700E-06 | 2.800E-06 |
| 7396 | 370 | L370S | A | 2.4e-06 | 2.5e-06 | 1.9e-06 | 2.267E-06 | 3.215E-07 | 1.900E-06 | 2.500E-06 |
| 7397 | 370 | L370T | A | 2.7e-06 | 2.8e-06 | 3.3e-06 | 2.933E-06 | 3.215E-07 | 2.700E-06 | 3.300E-06 |
| 7398 | 370 | L370W | A | 1.5e-06 | 2.4e-06 | 2.6e-06 | 2.167E-06 | 5.859E-07 | 1.500E-06 | 2.600E-06 |
| 7399 | 370 | L370Y | A | 1.9e-06 | 1.9e-06 | 2.1e-06 | 1.967E-06 | 1.155E-07 | 1.900E-06 | 2.100E-06 |
| 7400 | 370 | L370V | A | 2.7e-06 | 3.6e-06 | 2.2e-06 | 2.833E-06 | 7.095E-07 | 2.200E-06 | 3.600E-06 |
| 7401 | 371 | Q371A | A | 2.2e-06 | 2.5e-06 | 2.8e-06 | 2.500E-06 | 3.000E-07 | 2.200E-06 | 2.800E-06 |
| 7402 | 371 | Q371R | A | 3e-06 | 2.2e-06 | 2.5e-06 | 2.567E-06 | 4.041E-07 | 2.200E-06 | 3.000E-06 |
| 7403 | 371 | Q371N | A | 1.8e-06 | 1.8e-06 | 2.3e-06 | 1.967E-06 | 2.887E-07 | 1.800E-06 | 2.300E-06 |
| 7404 | 371 | Q371D | A | 2.4e-06 | 2.1e-06 | 1.8e-06 | 2.100E-06 | 3.000E-07 | 1.800E-06 | 2.400E-06 |
| 7405 | 371 | Q371C | A | 2.5e-06 | 4.6e-06 | 3e-06 | 3.367E-06 | 1.097E-06 | 2.500E-06 | 4.600E-06 |
| 7406 | 371 | Q371E | A | 1.9e-06 | 3e-06 | 3.9e-06 | 2.933E-06 | 1.002E-06 | 1.900E-06 | 3.900E-06 |
| 7407 | 371 | Q371Q | A | 3.4e-06 | 2.1e-06 | 4.5e-06 | 3.333E-06 | 1.201E-06 | 2.100E-06 | 4.500E-06 |
| 7408 | 371 | Q371G | A | 2.3e-06 | 2.6e-06 | 1.9e-06 | 2.267E-06 | 3.512E-07 | 1.900E-06 | 2.600E-06 |
| 7409 | 371 | Q371H | A | 1.5e-06 | 2.5e-06 | 3.1e-06 | 2.367E-06 | 8.083E-07 | 1.500E-06 | 3.100E-06 |
| 7410 | 371 | Q371I | A | 1.4e-06 | 3.1e-06 | 3.8e-06 | 2.767E-06 | 1.234E-06 | 1.400E-06 | 3.800E-06 |
| 7411 | 371 | Q371L | A | 2.3e-06 | 2.8e-06 | 2.3e-06 | 2.467E-06 | 2.887E-07 | 2.300E-06 | 2.800E-06 |
| 7412 | 371 | Q371K | A | 2.7e-06 | 2.2e-06 | 3e-06 | 2.633E-06 | 4.041E-07 | 2.200E-06 | 3.000E-06 |
| 7413 | 371 | Q371M | A | 2.7e-06 | 1.7e-06 | 2.1e-06 | 2.167E-06 | 5.033E-07 | 1.700E-06 | 2.700E-06 |
| 7414 | 371 | Q371F | A | 3e-06 | 2.6e-06 | 2.4e-06 | 2.667E-06 | 3.055E-07 | 2.400E-06 | 3.000E-06 |
| 7415 | 371 | Q371P | A | 1.7e-06 | 2.6e-06 | 1.8e-06 | 2.033E-06 | 4.933E-07 | 1.700E-06 | 2.600E-06 |
| 7416 | 371 | Q371S | A | 2.1e-06 | 1.4e-06 | 3.3e-06 | 2.267E-06 | 9.609E-07 | 1.400E-06 | 3.300E-06 |
| 7417 | 371 | Q371T | A | 3.3e-06 | 2.8e-06 | 1.3e-06 | 2.467E-06 | 1.041E-06 | 1.300E-06 | 3.300E-06 |
| 7418 | 371 | Q371W | A | 2.3e-06 | 1.8e-06 | 2e-06 | 2.033E-06 | 2.517E-07 | 1.800E-06 | 2.300E-06 |
| 7419 | 371 | Q371Y | A | 3.3e-06 | 2.3e-06 | 1.4e-06 | 2.333E-06 | 9.504E-07 | 1.400E-06 | 3.300E-06 |
| 7420 | 371 | Q371V | A | 1.8e-06 | 3.5e-06 | 3.2e-06 | 2.833E-06 | 9.074E-07 | 1.800E-06 | 3.500E-06 |
| 7421 | 372 | P372A | A | 1.6e-06 | 5.4e-06 | 1.9e-06 | 2.967E-06 | 2.113E-06 | 1.600E-06 | 5.400E-06 |
| 7422 | 372 | P372R | A | 2.3e-06 | 2.7e-06 | 2.1e-06 | 2.367E-06 | 3.055E-07 | 2.100E-06 | 2.700E-06 |
| 7423 | 372 | P372N | A | 1.9e-06 | 2.3e-06 | 2.1e-06 | 2.100E-06 | 2.000E-07 | 1.900E-06 | 2.300E-06 |
| 7424 | 372 | P372D | A | 2.5e-06 | 3.1e-06 | 2.5e-06 | 2.700E-06 | 3.464E-07 | 2.500E-06 | 3.100E-06 |
| 7425 | 372 | P372C | A | 2.1e-06 | 2.4e-06 | 1.7e-06 | 2.067E-06 | 3.512E-07 | 1.700E-06 | 2.400E-06 |
| 7426 | 372 | P372E | A | 2.5e-06 | 1.1e-06 | 2.1e-06 | 1.900E-06 | 7.211E-07 | 1.100E-06 | 2.500E-06 |
| 7427 | 372 | P372Q | A | 3.6e-06 | 1.6e-06 | 1.1e-06 | 2.100E-06 | 1.323E-06 | 1.100E-06 | 3.600E-06 |
| 7428 | 372 | P372G | A | 2.6e-06 | 2.9e-06 | 3.6e-06 | 3.033E-06 | 5.132E-07 | 2.600E-06 | 3.600E-06 |
| 7429 | 372 | P372H | A | 2e-06 | 2.4e-06 | 2.2e-06 | 2.200E-06 | 2.000E-07 | 2.000E-06 | 2.400E-06 |
| 7430 | 372 | P372I | A | 4.5e-06 | 1.5e-06 | 1.7e-06 | 2.567E-06 | 1.677E-06 | 1.500E-06 | 4.500E-06 |
| 7431 | 372 | P372L | A | 2.4e-06 | 2.1e-06 | 3.2e-06 | 2.567E-06 | 5.686E-07 | 2.100E-06 | 3.200E-06 |
| 7432 | 372 | P372K | A | 2.3e-06 | 2.7e-06 | 2.2e-06 | 2.400E-06 | 2.646E-07 | 2.200E-06 | 2.700E-06 |
| 7433 | 372 | P372M | A | 3.1e-06 | 3.5e-06 | 3.4e-06 | 3.333E-06 | 2.082E-07 | 3.100E-06 | 3.500E-06 |
| 7434 | 372 | P372F | A | 2.7e-06 | 2.5e-06 | 2.5e-06 | 2.567E-06 | 1.155E-07 | 2.500E-06 | 2.700E-06 |
| 7435 | 372 | P372P | A | 1.7e-06 | 3.3e-06 | 2.8e-06 | 2.600E-06 | 8.185E-07 | 1.700E-06 | 3.300E-06 |
| 7436 | 372 | P372S | A | 2.8e-06 | 3e-06 | 1.7e-06 | 2.500E-06 | 7.000E-07 | 1.700E-06 | 3.000E-06 |
| 7437 | 372 | P372T | A | 2.8e-06 | 2.1e-06 | 1.3e-06 | 2.067E-06 | 7.506E-07 | 1.300E-06 | 2.800E-06 |
| 7438 | 372 | P372W | A | 2.5e-06 | 1.2e-06 | 2.3e-06 | 2.000E-06 | 7.000E-07 | 1.200E-06 | 2.500E-06 |
| 7439 | 372 | P372Y | A | 2.9e-06 | 2.4e-06 | 1.1e-06 | 2.133E-06 | 9.292E-07 | 1.100E-06 | 2.900E-06 |
| 7440 | 372 | P372V | A | 1.9e-06 | 2.1e-06 | 3.1e-06 | 2.367E-06 | 6.429E-07 | 1.900E-06 | 3.100E-06 |
| 7441 | 373 | E373A | A | 2.6e-06 | 2.3e-06 | 1.8e-06 | 2.233E-06 | 4.041E-07 | 1.800E-06 | 2.600E-06 |
| 7442 | 373 | E373R | A | 3e-06 | 1.7e-06 | 3e-06 | 2.567E-06 | 7.506E-07 | 1.700E-06 | 3.000E-06 |
| 7443 | 373 | E373N | A | 2e-06 | 1.8e-06 | 3.1e-06 | 2.300E-06 | 7.000E-07 | 1.800E-06 | 3.100E-06 |
| 7444 | 373 | E373D | A | 2.9e-06 | 3e-06 | 2.2e-06 | 2.700E-06 | 4.359E-07 | 2.200E-06 | 3.000E-06 |
| 7445 | 373 | E373C | A | 2.1e-06 | 2.9e-06 | 3.8e-06 | 2.933E-06 | 8.505E-07 | 2.100E-06 | 3.800E-06 |
| 7446 | 373 | E373E | A | 2.9e-06 | 1.7e-06 | 2.4e-06 | 2.333E-06 | 6.028E-07 | 1.700E-06 | 2.900E-06 |
| 7447 | 373 | E373Q | A | 2e-06 | 1.8e-06 | 2e-06 | 1.933E-06 | 1.155E-07 | 1.800E-06 | 2.000E-06 |
| 7448 | 373 | E373G | A | 2.5e-06 | 1.9e-06 | 1.4e-06 | 1.933E-06 | 5.508E-07 | 1.400E-06 | 2.500E-06 |
| 7449 | 373 | E373H | A | 2.9e-06 | 3.4e-06 | 2e-06 | 2.767E-06 | 7.095E-07 | 2.000E-06 | 3.400E-06 |
| 7450 | 373 | E373I | A | 2.5e-06 | 1.1e-06 | 3e-06 | 2.200E-06 | 9.849E-07 | 1.100E-06 | 3.000E-06 |
| 7451 | 373 | E373L | A | 3.1e-06 | 2.9e-06 | 1.8e-06 | 2.600E-06 | 7.000E-07 | 1.800E-06 | 3.100E-06 |
| 7452 | 373 | E373K | A | 2.8e-06 | 2.7e-06 | 2.2e-06 | 2.567E-06 | 3.215E-07 | 2.200E-06 | 2.800E-06 |
| 7453 | 373 | E373M | A | 2.2e-06 | 3.1e-06 | 2.8e-06 | 2.700E-06 | 4.583E-07 | 2.200E-06 | 3.100E-06 |
| 7454 | 373 | E373F | A | 2.1e-06 | 2.1e-06 | 1.4e-06 | 1.867E-06 | 4.041E-07 | 1.400E-06 | 2.100E-06 |
| 7455 | 373 | E373P | A | 2.6e-06 | 3e-06 | 2e-06 | 2.533E-06 | 5.033E-07 | 2.000E-06 | 3.000E-06 |
| 7456 | 373 | E373S | A | 4.2e-06 | 2.3e-06 | 3.2e-06 | 3.233E-06 | 9.504E-07 | 2.300E-06 | 4.200E-06 |
| 7457 | 373 | E373T | A | 1.6e-06 | 2.7e-06 | 4e-06 | 2.767E-06 | 1.201E-06 | 1.600E-06 | 4.000E-06 |
| 7458 | 373 | E373W | A | 2.8e-06 | 3.2e-06 | 2.9e-06 | 2.967E-06 | 2.082E-07 | 2.800E-06 | 3.200E-06 |
| 7459 | 373 | E373Y | A | 2.5e-06 | 2.1e-06 | 2.3e-06 | 2.300E-06 | 2.000E-07 | 2.100E-06 | 2.500E-06 |
| 7460 | 373 | E373V | A | 3.1e-06 | 2.3e-06 | 2.2e-06 | 2.533E-06 | 4.933E-07 | 2.200E-06 | 3.100E-06 |
| 7461 | 374 | Q374A | A | 1.8e-06 | 1.7e-06 | 3.7e-06 | 2.400E-06 | 1.127E-06 | 1.700E-06 | 3.700E-06 |
| 7462 | 374 | Q374R | A | 4.1e-06 | 2.4e-06 | 2.1e-06 | 2.867E-06 | 1.079E-06 | 2.100E-06 | 4.100E-06 |
| 7463 | 374 | Q374N | A | 3.1e-06 | 1.8e-06 | 2.6e-06 | 2.500E-06 | 6.557E-07 | 1.800E-06 | 3.100E-06 |
| 7464 | 374 | Q374D | A | 2.5e-06 | 2.8e-06 | 2.5e-06 | 2.600E-06 | 1.732E-07 | 2.500E-06 | 2.800E-06 |
| 7465 | 374 | Q374C | A | 2.5e-06 | 2.6e-06 | 1.8e-06 | 2.300E-06 | 4.359E-07 | 1.800E-06 | 2.600E-06 |
| 7466 | 374 | Q374E | A | 1.8e-06 | 2.4e-06 | 2.7e-06 | 2.300E-06 | 4.583E-07 | 1.800E-06 | 2.700E-06 |
| 7467 | 374 | Q374Q | A | 4.4e-06 | 1.2e-06 | 2.3e-06 | 2.633E-06 | 1.626E-06 | 1.200E-06 | 4.400E-06 |
| 7468 | 374 | Q374G | A | 3.2e-06 | 3e-06 | 3.7e-06 | 3.300E-06 | 3.606E-07 | 3.000E-06 | 3.700E-06 |
| 7469 | 374 | Q374H | A | 3.1e-06 | 2.4e-06 | 2.7e-06 | 2.733E-06 | 3.512E-07 | 2.400E-06 | 3.100E-06 |
| 7470 | 374 | Q374I | A | 3.9e-06 | 3.2e-06 | 5.1e-06 | 4.067E-06 | 9.609E-07 | 3.200E-06 | 5.100E-06 |
| 7471 | 374 | Q374L | A | 2e-06 | 2e-06 | 1.4e-06 | 1.800E-06 | 3.464E-07 | 1.400E-06 | 2.000E-06 |
| 7472 | 374 | Q374K | A | 2.4e-06 | 2.7e-06 | 3.6e-06 | 2.900E-06 | 6.245E-07 | 2.400E-06 | 3.600E-06 |
| 7473 | 374 | Q374M | A | 2.9e-06 | 1.7e-06 | 2.6e-06 | 2.400E-06 | 6.245E-07 | 1.700E-06 | 2.900E-06 |
| 7474 | 374 | Q374F | A | 2e-06 | 1.8e-06 | 3e-06 | 2.267E-06 | 6.429E-07 | 1.800E-06 | 3.000E-06 |
| 7475 | 374 | Q374P | A | 1.6e-06 | 2.3e-06 | 2.6e-06 | 2.167E-06 | 5.132E-07 | 1.600E-06 | 2.600E-06 |
| 7476 | 374 | Q374S | A | 1.9e-06 | 1.7e-06 | 2.4e-06 | 2.000E-06 | 3.606E-07 | 1.700E-06 | 2.400E-06 |
| 7477 | 374 | Q374T | A | 2.4e-06 | 1.6e-06 | 2.6e-06 | 2.200E-06 | 5.292E-07 | 1.600E-06 | 2.600E-06 |
| 7478 | 374 | Q374W | A | 1.5e-06 | 9e-07 | 2.1e-06 | 1.500E-06 | 6.000E-07 | 9.000E-07 | 2.100E-06 |
| 7479 | 374 | Q374Y | A | 2.2e-06 | 2.7e-06 | 2e-06 | 2.300E-06 | 3.606E-07 | 2.000E-06 | 2.700E-06 |

| | | | | | | | | | | |
|---|---|---|---|---|---|---|---|---|---|---|
| 7480 | 374 | Q374V | A | 5e-06 | 2.8e-06 | 2.4e-06 | 3.400E-06 | 1.400E-06 | 2.400E-06 | 5.000E-06 |
| 7481 | 375 | L375A | A | 3.3e-06 | 1.2e-06 | 2.7e-06 | 2.400E-06 | 1.082E-06 | 1.200E-06 | 3.300E-06 |
| 7482 | 375 | L375R | A | 3.1e-06 | 1.3e-06 | 1.6e-06 | 2.000E-06 | 9.644E-07 | 1.300E-06 | 3.100E-06 |
| 7483 | 375 | L375N | A | 2e-06 | 1.7e-06 | 2.1e-06 | 1.933E-06 | 2.082E-07 | 1.700E-06 | 2.100E-06 |
| 7484 | 375 | L375D | A | 3.3e-06 | 2.2e-06 | 3e-06 | 2.833E-06 | 5.686E-07 | 2.200E-06 | 3.300E-06 |
| 7485 | 375 | L375C | A | 2.1e-06 | 2.3e-06 | 2.5e-06 | 2.300E-06 | 2.000E-07 | 2.100E-06 | 2.500E-06 |
| 7486 | 375 | L375E | A | 1.9e-06 | 2.9e-06 | 2.6e-06 | 2.467E-06 | 5.132E-07 | 1.900E-06 | 2.900E-06 |
| 7487 | 375 | L375Q | A | 2.4e-06 | 8.9e-07 | 3.6e-06 | 2.297E-06 | 1.358E-06 | 8.900E-07 | 3.600E-06 |
| 7488 | 375 | L375G | A | 1.7e-06 | 1.4e-06 | 2.7e-06 | 1.933E-06 | 6.807E-07 | 1.400E-06 | 2.700E-06 |
| 7489 | 375 | L375H | A | 3e-06 | 2.4e-06 | 3.4e-06 | 2.933E-06 | 5.033E-07 | 2.400E-06 | 3.400E-06 |
| 7490 | 375 | L375I | A | 1.5e-06 | 1.9e-06 | 2e-06 | 1.800E-06 | 2.646E-07 | 1.500E-06 | 2.000E-06 |
| 7491 | 375 | L375L | A | 2.2e-06 | 1.7e-06 | 2.3e-06 | 2.067E-06 | 3.215E-07 | 1.700E-06 | 2.300E-06 |
| 7492 | 375 | L375K | A | 1.6e-06 | 1.7e-06 | 4.1e-06 | 2.467E-06 | 1.415E-06 | 1.600E-06 | 4.100E-06 |
| 7493 | 375 | L375M | A | 3e-06 | 3.3e-06 | 1.6e-06 | 2.633E-06 | 9.074E-07 | 1.600E-06 | 3.300E-06 |
| 7494 | 375 | L375F | A | 1.8e-06 | 3.8e-06 | 3.4e-06 | 3.000E-06 | 1.058E-06 | 1.800E-06 | 3.800E-06 |
| 7495 | 375 | L375P | A | 1.8e-06 | 1.7e-06 | 2.7e-06 | 2.067E-06 | 5.508E-07 | 1.700E-06 | 2.700E-06 |
| 7496 | 375 | L375S | A | 2.4e-06 | 3.1e-06 | 3.3e-06 | 2.933E-06 | 4.726E-07 | 2.400E-06 | 3.300E-06 |
| 7497 | 375 | L375T | A | 1.9e-06 | 5.4e-06 | 2.6e-06 | 3.300E-06 | 1.852E-06 | 1.900E-06 | 5.400E-06 |
| 7498 | 375 | L375W | A | 2.1e-06 | 2.9e-06 | 2.3e-06 | 2.433E-06 | 4.163E-07 | 2.100E-06 | 2.900E-06 |
| 7499 | 375 | L375Y | A | 2.5e-06 | 2.6e-06 | 2.2e-06 | 2.433E-06 | 2.082E-07 | 2.200E-06 | 2.600E-06 |
| 7500 | 375 | L375V | A | 3.3e-06 | 2.2e-06 | 3.7e-06 | 3.067E-06 | 7.767E-07 | 2.200E-06 | 3.700E-06 |
| 7501 | 376 | Q376A | A | 1.8e-06 | 2.8e-06 | 2.3e-06 | 2.300E-06 | 5.000E-07 | 1.800E-06 | 2.800E-06 |
| 7502 | 376 | Q376R | A | 1.7e-06 | 2.7e-06 | 2.9e-06 | 2.433E-06 | 6.429E-07 | 1.700E-06 | 2.900E-06 |
| 7503 | 376 | Q376N | A | 2.8e-06 | 2.3e-06 | 1.9e-06 | 2.333E-06 | 4.509E-07 | 1.900E-06 | 2.800E-06 |
| 7504 | 376 | Q376D | A | 1.8e-06 | 3.5e-06 | 2.9e-06 | 2.733E-06 | 8.622E-07 | 1.800E-06 | 3.500E-06 |
| 7505 | 376 | Q376C | A | 3e-06 | 3.3e-06 | 2.7e-06 | 3.000E-06 | 3.000E-07 | 2.700E-06 | 3.300E-06 |
| 7506 | 376 | Q376E | A | 1.6e-06 | 2.8e-06 | 2.8e-06 | 2.400E-06 | 6.928E-07 | 1.600E-06 | 2.800E-06 |
| 7507 | 376 | Q376Q | A | 1.9e-06 | 3.9e-06 | 2e-06 | 2.600E-06 | 1.127E-06 | 1.900E-06 | 3.900E-06 |
| 7508 | 376 | Q376G | A | 2.1e-06 | 1.9e-06 | 2.3e-06 | 2.100E-06 | 2.000E-07 | 1.900E-06 | 2.300E-06 |
| 7509 | 376 | Q376H | A | 1.8e-06 | 5e-06 | 2.3e-06 | 3.033E-06 | 1.721E-06 | 1.800E-06 | 5.000E-06 |
| 7510 | 376 | Q376I | A | 2.8e-06 | 2.6e-06 | 2.2e-06 | 2.533E-06 | 3.055E-07 | 2.200E-06 | 2.800E-06 |
| 7511 | 376 | Q376L | A | 4.2e-06 | 2.4e-06 | 2.2e-06 | 2.933E-06 | 1.102E-06 | 2.200E-06 | 4.200E-06 |
| 7512 | 376 | Q376K | A | 2.6e-06 | 2.4e-06 | 3.9e-06 | 2.967E-06 | 8.145E-07 | 2.400E-06 | 3.900E-06 |
| 7513 | 376 | Q376M | A | 4.2e-06 | 2.3e-06 | 2.3e-06 | 2.933E-06 | 1.097E-06 | 2.300E-06 | 4.200E-06 |
| 7514 | 376 | Q376F | A | 2.8e-06 | 2.7e-06 | 1.6e-06 | 2.367E-06 | 6.658E-07 | 1.600E-06 | 2.800E-06 |
| 7515 | 376 | Q376P | A | 4.3e-06 | 1.9e-06 | 2.5e-06 | 2.900E-06 | 1.249E-06 | 1.900E-06 | 4.300E-06 |
| 7516 | 376 | Q376S | A | 2.4e-06 | 1.9e-06 | 3e-06 | 2.433E-06 | 5.508E-07 | 1.900E-06 | 3.000E-06 |
| 7517 | 376 | Q376T | A | 3.2e-06 | 2.1e-06 | 2.8e-06 | 2.700E-06 | 5.568E-07 | 2.100E-06 | 3.200E-06 |
| 7518 | 376 | Q376W | A | 2.9e-06 | 1.3e-06 | 1.6e-06 | 1.933E-06 | 8.505E-07 | 1.300E-06 | 2.900E-06 |
| 7519 | 376 | Q376Y | A | 3.7e-06 | 2.1e-06 | 2.9e-06 | 2.900E-06 | 8.000E-07 | 2.100E-06 | 3.700E-06 |
| 7520 | 376 | Q376V | A | 2.6e-06 | 1.6e-06 | 3.1e-06 | 2.433E-06 | 7.638E-07 | 1.600E-06 | 3.100E-06 |
| 7521 | 377 | V377A | A | 3.3e-06 | 1.1e-06 | 2.9e-06 | 2.433E-06 | 1.172E-06 | 1.100E-06 | 3.300E-06 |
| 7522 | 377 | V377R | A | 1.8e-06 | 2.4e-06 | 2.7e-06 | 2.300E-06 | 4.583E-07 | 1.800E-06 | 2.700E-06 |
| 7523 | 377 | V377N | A | 2.8e-06 | 1.6e-06 | 1.6e-06 | 2.000E-06 | 6.928E-07 | 1.600E-06 | 2.800E-06 |
| 7524 | 377 | V377D | A | 2.5e-06 | 2.3e-06 | 2.9e-06 | 2.567E-06 | 3.055E-07 | 2.300E-06 | 2.900E-06 |
| 7525 | 377 | V377C | A | 2e-06 | 2e-06 | 2.6e-06 | 2.200E-06 | 3.464E-07 | 2.000E-06 | 2.600E-06 |
| 7526 | 377 | V377E | A | 1.2e-06 | 3e-06 | 2.5e-06 | 2.233E-06 | 9.292E-07 | 1.200E-06 | 3.000E-06 |
| 7527 | 377 | V377Q | A | 2.2e-06 | 4.1e-06 | 2.3e-06 | 2.867E-06 | 1.069E-06 | 2.200E-06 | 4.100E-06 |
| 7528 | 377 | V377G | A | 1.4e-06 | 2.9e-06 | 2.5e-06 | 2.267E-06 | 7.767E-07 | 1.400E-06 | 2.900E-06 |
| 7529 | 377 | V377H | A | 3e-06 | 3.1e-06 | 1.4e-06 | 2.500E-06 | 9.539E-07 | 1.400E-06 | 3.100E-06 |
| 7530 | 377 | V377I | A | 3.2e-06 | 2.8e-06 | 2.1e-06 | 2.700E-06 | 5.568E-07 | 2.100E-06 | 3.200E-06 |
| 7531 | 377 | V377L | A | 2.6e-06 | 2.2e-06 | 1.4e-06 | 2.067E-06 | 6.110E-07 | 1.400E-06 | 2.600E-06 |
| 7532 | 377 | V377K | A | 2.4e-06 | 2.2e-06 | 2.4e-06 | 2.333E-06 | 1.155E-07 | 2.200E-06 | 2.400E-06 |
| 7533 | 377 | V377M | A | 2.1e-06 | 2.8e-06 | 2.1e-06 | 2.333E-06 | 4.041E-07 | 2.100E-06 | 2.800E-06 |
| 7534 | 377 | V377F | A | 1.2e-06 | 3.3e-06 | 1.8e-06 | 2.100E-06 | 1.082E-06 | 1.200E-06 | 3.300E-06 |
| 7535 | 377 | V377P | A | 2.2e-06 | 1.9e-06 | 3.3e-06 | 2.467E-06 | 7.371E-07 | 1.900E-06 | 3.300E-06 |
| 7536 | 377 | V377S | A | 2.4e-06 | 2e-06 | 2.1e-06 | 2.167E-06 | 2.082E-07 | 2.000E-06 | 2.400E-06 |
| 7537 | 377 | V377T | A | 1.7e-06 | 2.1e-06 | 4e-06 | 2.600E-06 | 1.229E-06 | 1.700E-06 | 4.000E-06 |
| 7538 | 377 | V377W | A | 2.6e-06 | 1.7e-06 | 1e-06 | 1.767E-06 | 8.021E-07 | 1.000E-06 | 2.600E-06 |
| 7539 | 377 | V377Y | A | 2.8e-06 | 2.1e-06 | 1.6e-06 | 2.167E-06 | 6.028E-07 | 1.600E-06 | 2.800E-06 |
| 7540 | 377 | V377V | A | 1.6e-06 | 2.6e-06 | 2e-06 | 2.067E-06 | 5.033E-07 | 1.600E-06 | 2.600E-06 |
| 7541 | 378 | F378A | A | 5.4e-06 | 2.9e-06 | 3.5e-06 | 3.933E-06 | 1.305E-06 | 2.900E-06 | 5.400E-06 |
| 7542 | 378 | F378R | A | 5.7e-06 | 1.8e-06 | 3e-06 | 3.500E-06 | 1.997E-06 | 1.800E-06 | 5.700E-06 |
| 7543 | 378 | F378N | A | 3.3e-06 | 1.5e-06 | 2.1e-06 | 2.300E-06 | 9.165E-07 | 1.500E-06 | 3.300E-06 |
| 7544 | 378 | F378D | A | 1.9e-06 | 3.7e-06 | 2.7e-06 | 2.767E-06 | 9.018E-07 | 1.900E-06 | 3.700E-06 |
| 7545 | 378 | F378C | A | 3.4e-06 | 3.3e-06 | 1.7e-06 | 2.800E-06 | 9.539E-07 | 1.700E-06 | 3.400E-06 |
| 7546 | 378 | F378E | A | 2.6e-06 | 2.5e-06 | 3.3e-06 | 2.800E-06 | 4.359E-07 | 2.500E-06 | 3.300E-06 |
| 7547 | 378 | F378Q | A | 1.9e-06 | 2.9e-06 | 4.3e-06 | 3.033E-06 | 1.206E-06 | 1.900E-06 | 4.300E-06 |
| 7548 | 378 | F378G | A | 1.2e-06 | 2.9e-06 | 2e-06 | 2.033E-06 | 8.505E-07 | 1.200E-06 | 2.900E-06 |
| 7549 | 378 | F378H | A | 2.4e-06 | 2.2e-06 | 1.6e-06 | 2.067E-06 | 4.163E-07 | 1.600E-06 | 2.400E-06 |
| 7550 | 378 | F378I | A | 2.1e-06 | 2.5e-06 | 3.6e-06 | 2.733E-06 | 7.767E-07 | 2.100E-06 | 3.600E-06 |
| 7551 | 378 | F378L | A | 2e-06 | 1.3e-06 | 2.6e-06 | 1.967E-06 | 6.506E-07 | 1.300E-06 | 2.600E-06 |
| 7552 | 378 | F378K | A | 4.8e-06 | 2.4e-06 | 1.9e-06 | 3.033E-06 | 1.550E-06 | 1.900E-06 | 4.800E-06 |
| 7553 | 378 | F378M | A | 1.4e-06 | 2.3e-06 | 2.8e-06 | 2.167E-06 | 7.095E-07 | 1.400E-06 | 2.800E-06 |
| 7554 | 378 | F378F | A | 3.6e-06 | 1.8e-06 | 1.4e-06 | 2.267E-06 | 1.172E-06 | 1.400E-06 | 3.600E-06 |
| 7555 | 378 | F378P | A | 2.2e-06 | 1.9e-06 | 9.5e-07 | 1.683E-06 | 6.526E-07 | 9.500E-07 | 2.200E-06 |
| 7556 | 378 | F378S | A | 3.4e-06 | 2.2e-06 | 3.1e-06 | 2.900E-06 | 6.245E-07 | 2.200E-06 | 3.400E-06 |
| 7557 | 378 | F378T | A | 1.8e-06 | 2e-06 | 2.3e-06 | 2.033E-06 | 2.517E-07 | 1.800E-06 | 2.300E-06 |
| 7558 | 378 | F378W | A | 2.2e-06 | 1.5e-06 | 3e-06 | 2.233E-06 | 7.506E-07 | 1.500E-06 | 3.000E-06 |
| 7559 | 378 | F378Y | A | 2.1e-06 | 2.7e-06 | 2.1e-06 | 2.300E-06 | 3.464E-07 | 2.100E-06 | 2.700E-06 |
| 7560 | 378 | F378V | A | 4.2e-06 | 2.7e-06 | 1.8e-06 | 2.900E-06 | 1.212E-06 | 1.800E-06 | 4.200E-06 |
| 7561 | 379 | E379A | A | 2e-06 | 3.8e-06 | 3.5e-06 | 3.100E-06 | 9.644E-07 | 2.000E-06 | 3.800E-06 |
| 7562 | 379 | E379R | A | 1.4e-06 | 3.3e-06 | 2e-06 | 2.233E-06 | 9.713E-07 | 1.400E-06 | 3.300E-06 |
| 7563 | 379 | E379N | A | 1.9e-06 | 3.1e-06 | 4.1e-06 | 3.033E-06 | 1.102E-06 | 1.900E-06 | 4.100E-06 |
| 7564 | 379 | E379D | A | 2e-06 | 2.9e-06 | 1.9e-06 | 2.267E-06 | 5.508E-07 | 1.900E-06 | 2.900E-06 |

| | | | | | | | | | | |
|---|---|---|---|---|---|---|---|---|---|---|
| 7565 | 379 | E379C | A | 1.8e-06 | 1.8e-06 | 2.9e-06 | 2.167E-06 | 6.351E-07 | 1.800E-06 | 2.900E-06 |
| 7566 | 379 | E379E | A | 3.2e-06 | 1.4e-06 | 2e-06 | 2.200E-06 | 9.165E-07 | 1.400E-06 | 3.200E-06 |
| 7567 | 379 | E379Q | A | 3.3e-06 | 1.5e-06 | 1.3e-06 | 2.033E-06 | 1.102E-06 | 1.300E-06 | 3.300E-06 |
| 7568 | 379 | E379G | A | 2.7e-06 | 2.4e-06 | 2.5e-06 | 2.533E-06 | 1.528E-07 | 2.400E-06 | 2.700E-06 |
| 7569 | 379 | E379H | A | 1.1e-06 | 2.4e-06 | 2.2e-06 | 1.900E-06 | 7.000E-07 | 1.100E-06 | 2.400E-06 |
| 7570 | 379 | E379I | A | 2.5e-06 | 2.4e-06 | 1.5e-06 | 2.133E-06 | 5.508E-07 | 1.500E-06 | 2.500E-06 |
| 7571 | 379 | E379L | A | 2.2e-06 | 1.8e-06 | 2.4e-06 | 2.133E-06 | 3.055E-07 | 1.800E-06 | 2.400E-06 |
| 7572 | 379 | E379K | A | 2.2e-06 | 2.2e-06 | 2.4e-06 | 2.267E-06 | 1.155E-07 | 2.200E-06 | 2.400E-06 |
| 7573 | 379 | E379M | A | 2e-06 | 2.6e-06 | 2.9e-06 | 2.500E-06 | 4.583E-07 | 2.000E-06 | 2.900E-06 |
| 7574 | 379 | E379F | A | 2.5e-06 | 2e-06 | 2.6e-06 | 2.367E-06 | 3.215E-07 | 2.000E-06 | 2.600E-06 |
| 7575 | 379 | E379P | A | 3e-06 | 2.4e-06 | 2.1e-06 | 2.500E-06 | 4.583E-07 | 2.100E-06 | 3.000E-06 |
| 7576 | 379 | E379S | A | 2.7e-06 | 2.5e-06 | 3.2e-06 | 2.800E-06 | 3.606E-07 | 2.500E-06 | 3.200E-06 |
| 7577 | 379 | E379T | A | 1.8e-06 | 6.2e-06 | 2.8e-06 | 3.600E-06 | 2.307E-06 | 1.800E-06 | 6.200E-06 |
| 7578 | 379 | E379W | A | 3.3e-06 | 2.9e-06 | 2.8e-06 | 3.000E-06 | 2.646E-07 | 2.800E-06 | 3.300E-06 |
| 7579 | 379 | E379Y | A | 6.1e-06 | 2.9e-06 | 2.3e-06 | 3.767E-06 | 2.043E-06 | 2.300E-06 | 6.100E-06 |
| 7580 | 379 | E379V | A | 2.5e-06 | 3.3e-06 | 1.9e-06 | 2.567E-06 | 7.024E-07 | 1.900E-06 | 3.300E-06 |
| 7581 | 380 | T380A | A | 4.6e-06 | 2.2e-06 | 2.2e-06 | 3.000E-06 | 1.386E-06 | 2.200E-06 | 4.600E-06 |
| 7582 | 380 | T380R | A | 1.8e-06 | 2.4e-06 | 2.7e-06 | 2.300E-06 | 4.583E-07 | 1.800E-06 | 2.700E-06 |
| 7583 | 380 | T380N | A | 3.1e-06 | 1.2e-06 | 2.1e-06 | 2.133E-06 | 9.504E-07 | 1.200E-06 | 3.100E-06 |
| 7584 | 380 | T380D | A | 3e-06 | 2.4e-06 | 2.3e-06 | 2.567E-06 | 3.786E-07 | 2.300E-06 | 3.000E-06 |
| 7585 | 380 | T380C | A | 3.7e-06 | 3.5e-06 | 4.5e-06 | 3.900E-06 | 5.292E-07 | 3.500E-06 | 4.500E-06 |
| 7586 | 380 | T380E | A | 3.2e-06 | 3.9e-06 | 2.9e-06 | 3.333E-06 | 5.132E-07 | 2.900E-06 | 3.900E-06 |
| 7587 | 380 | T380Q | A | 1.8e-06 | 2.4e-06 | 2.5e-06 | 2.233E-06 | 3.786E-07 | 1.800E-06 | 2.500E-06 |
| 7588 | 380 | T380G | A | 2.6e-06 | 1.8e-06 | 1.6e-06 | 2.000E-06 | 5.292E-07 | 1.600E-06 | 2.600E-06 |
| 7589 | 380 | T380H | A | 1.8e-06 | 2e-06 | 1.7e-06 | 1.833E-06 | 1.528E-07 | 1.700E-06 | 2.000E-06 |
| 7590 | 380 | T380I | A | 2.2e-06 | 4.3e-06 | 3e-06 | 3.167E-06 | 1.060E-06 | 2.200E-06 | 4.300E-06 |
| 7591 | 380 | T380L | A | 2.6e-06 | 1.9e-06 | 2.9e-06 | 2.467E-06 | 5.132E-07 | 1.900E-06 | 2.900E-06 |
| 7592 | 380 | T380K | A | 2.7e-06 | 3.6e-06 | 1.5e-06 | 2.600E-06 | 1.054E-06 | 1.500E-06 | 3.600E-06 |
| 7593 | 380 | T380M | A | 1.8e-06 | 2.6e-06 | 3.4e-06 | 2.600E-06 | 8.000E-07 | 1.800E-06 | 3.400E-06 |
| 7594 | 380 | T380F | A | 1.9e-06 | 5.7e-06 | 2.4e-06 | 3.333E-06 | 2.065E-06 | 1.900E-06 | 5.700E-06 |
| 7595 | 380 | T380P | A | 2e-06 | 3.3e-06 | 3.9e-06 | 3.067E-06 | 9.713E-07 | 2.000E-06 | 3.900E-06 |
| 7596 | 380 | T380S | A | 2.5e-06 | 2.2e-06 | 4.6e-06 | 3.100E-06 | 1.308E-06 | 2.200E-06 | 4.600E-06 |
| 7597 | 380 | T380T | A | 3.1e-06 | 1.8e-06 | 2e-06 | 2.300E-06 | 7.000E-07 | 1.800E-06 | 3.100E-06 |
| 7598 | 380 | T380W | A | 3.3e-06 | 2.6e-06 | 3.9e-06 | 3.267E-06 | 6.506E-07 | 2.600E-06 | 3.900E-06 |
| 7599 | 380 | T380Y | A | 3.1e-06 | 2.3e-06 | 2.4e-06 | 2.600E-06 | 4.359E-07 | 2.300E-06 | 3.100E-06 |
| 7600 | 380 | T380V | A | 3.5e-06 | 2.2e-06 | 2.3e-06 | 2.667E-06 | 7.234E-07 | 2.200E-06 | 3.500E-06 |
| 7601 | 381 | L381A | A | 3.7e-06 | 2.6e-06 | 3.2e-06 | 3.167E-06 | 5.508E-07 | 2.600E-06 | 3.700E-06 |
| 7602 | 381 | L381R | A | 4.4e-06 | 1.9e-06 | 2.9e-06 | 3.067E-06 | 1.258E-06 | 1.900E-06 | 4.400E-06 |
| 7603 | 381 | L381N | A | 1.2e-06 | 4.7e-06 | 2.7e-06 | 2.867E-06 | 1.756E-06 | 1.200E-06 | 4.700E-06 |
| 7604 | 381 | L381D | A | 4.5e-06 | 1.9e-06 | 2e-06 | 2.800E-06 | 1.473E-06 | 1.900E-06 | 4.500E-06 |
| 7605 | 381 | L381C | A | 2.5e-06 | 2.2e-06 | 4.5e-06 | 3.067E-06 | 1.250E-06 | 2.200E-06 | 4.500E-06 |
| 7606 | 381 | L381E | A | 2.2e-06 | 2.8e-06 | 1.4e-06 | 2.133E-06 | 7.024E-07 | 1.400E-06 | 2.800E-06 |
| 7607 | 381 | L381Q | A | 1.4e-06 | 2.7e-06 | 1.9e-06 | 2.000E-06 | 6.557E-07 | 1.400E-06 | 2.700E-06 |
| 7608 | 381 | L381G | A | 3.1e-06 | 2.3e-06 | 2.7e-06 | 2.700E-06 | 4.000E-07 | 2.300E-06 | 3.100E-06 |
| 7609 | 381 | L381H | A | 2.2e-06 | 4.1e-06 | 1.8e-06 | 2.700E-06 | 1.229E-06 | 1.800E-06 | 4.100E-06 |
| 7610 | 381 | L381I | A | 2.5e-06 | 1.7e-06 | 2.3e-06 | 2.167E-06 | 4.163E-07 | 1.700E-06 | 2.500E-06 |
| 7611 | 381 | L381L | A | 3.2e-06 | 2.7e-06 | 1.6e-06 | 2.500E-06 | 8.185E-07 | 1.600E-06 | 3.200E-06 |
| 7612 | 381 | L381K | A | 1.7e-06 | 3.5e-06 | 2.5e-06 | 2.567E-06 | 9.018E-07 | 1.700E-06 | 3.500E-06 |
| 7613 | 381 | L381M | A | 2.2e-06 | 3.2e-06 | 1.5e-06 | 2.300E-06 | 8.544E-07 | 1.500E-06 | 3.200E-06 |
| 7614 | 381 | L381F | A | 3.7e-06 | 3.3e-06 | 2.5e-06 | 3.167E-06 | 6.110E-07 | 2.500E-06 | 3.700E-06 |
| 7615 | 381 | L381P | A | 1.6e-06 | 2.4e-06 | 3.3e-06 | 2.433E-06 | 8.505E-07 | 1.600E-06 | 3.300E-06 |
| 7616 | 381 | L381S | A | 2.8e-06 | 2.7e-06 | 2.4e-06 | 2.633E-06 | 2.082E-07 | 2.400E-06 | 2.800E-06 |
| 7617 | 381 | L381T | A | 4.6e-06 | 3.8e-06 | 3.2e-06 | 3.867E-06 | 7.024E-07 | 3.200E-06 | 4.600E-06 |
| 7618 | 381 | L381W | A | 3.4e-06 | 3.4e-06 | 2.5e-06 | 3.100E-06 | 5.196E-07 | 2.500E-06 | 3.400E-06 |
| 7619 | 381 | L381Y | A | 3.9e-06 | 2.1e-06 | 1.9e-06 | 2.633E-06 | 1.102E-06 | 1.900E-06 | 3.900E-06 |
| 7620 | 381 | L381V | A | 2.3e-06 | 1.3e-06 | 3.4e-06 | 2.333E-06 | 1.050E-06 | 1.300E-06 | 3.400E-06 |
| 7621 | 382 | E382A | A | 3.1e-06 | 2.2e-06 | 3.1e-06 | 2.800E-06 | 5.196E-07 | 2.200E-06 | 3.100E-06 |
| 7622 | 382 | E382R | A | 1.7e-06 | 3.6e-06 | 2.6e-06 | 2.633E-06 | 9.504E-07 | 1.700E-06 | 3.600E-06 |
| 7623 | 382 | E382N | A | 2.5e-06 | 2e-06 | 2.5e-06 | 2.333E-06 | 2.887E-07 | 2.000E-06 | 2.500E-06 |
| 7624 | 382 | E382D | A | 1.5e-06 | 1.6e-06 | 1.5e-06 | 1.533E-06 | 5.774E-08 | 1.500E-06 | 1.600E-06 |
| 7625 | 382 | E382C | A | 1.5e-06 | 4.4e-06 | 2.9e-06 | 2.933E-06 | 1.450E-06 | 1.500E-06 | 4.400E-06 |
| 7626 | 382 | E382E | A | 3.9e-06 | 2.3e-06 | 1.6e-06 | 2.600E-06 | 1.179E-06 | 1.600E-06 | 3.900E-06 |
| 7627 | 382 | E382Q | A | 3.5e-06 | 2.3e-06 | 2.4e-06 | 2.733E-06 | 6.658E-07 | 2.300E-06 | 3.500E-06 |
| 7628 | 382 | E382G | A | 3.7e-06 | 2.2e-06 | 3e-06 | 2.967E-06 | 7.506E-07 | 2.200E-06 | 3.700E-06 |
| 7629 | 382 | E382H | A | 6.3e-06 | 1.8e-06 | 1.3e-06 | 3.133E-06 | 2.754E-06 | 1.300E-06 | 6.300E-06 |
| 7630 | 382 | E382I | A | 2.6e-06 | 1.8e-06 | 2.2e-06 | 2.200E-06 | 4.000E-07 | 1.800E-06 | 2.600E-06 |
| 7631 | 382 | E382L | A | 3.3e-06 | 2.5e-06 | 2.9e-06 | 2.900E-06 | 4.000E-07 | 2.500E-06 | 3.300E-06 |
| 7632 | 382 | E382K | A | 3.4e-06 | 2.5e-06 | 2e-06 | 2.633E-06 | 7.095E-07 | 2.000E-06 | 3.400E-06 |
| 7633 | 382 | E382M | A | 1.3e-06 | 2.9e-06 | 2.3e-06 | 2.167E-06 | 8.083E-07 | 1.300E-06 | 2.900E-06 |
| 7634 | 382 | E382F | A | 3.5e-06 | 2.6e-06 | 3.3e-06 | 3.133E-06 | 4.726E-07 | 2.600E-06 | 3.500E-06 |
| 7635 | 382 | E382P | A | 1.8e-06 | 3.6e-06 | 2.7e-06 | 2.700E-06 | 9.000E-07 | 1.800E-06 | 3.600E-06 |
| 7636 | 382 | E382S | A | 1.8e-06 | 2.6e-06 | 2.6e-06 | 2.333E-06 | 4.619E-07 | 1.800E-06 | 2.600E-06 |
| 7637 | 382 | E382T | A | 2e-06 | 2.9e-06 | 1.3e-06 | 2.067E-06 | 8.021E-07 | 1.300E-06 | 2.900E-06 |
| 7638 | 382 | E382W | A | 1.8e-06 | 2.9e-06 | 3.3e-06 | 2.667E-06 | 7.767E-07 | 1.800E-06 | 3.300E-06 |
| 7639 | 382 | E382Y | A | 1.6e-06 | 2.3e-06 | 1.7e-06 | 1.867E-06 | 3.786E-07 | 1.600E-06 | 2.300E-06 |
| 7640 | 382 | E382V | A | 2.7e-06 | 2.6e-06 | 2.5e-06 | 2.600E-06 | 1.000E-07 | 2.500E-06 | 2.700E-06 |
| 7641 | 383 | E383A | A | 2.3e-06 | 1.6e-06 | 3.2e-06 | 2.367E-06 | 8.021E-07 | 1.600E-06 | 3.200E-06 |
| 7642 | 383 | E383R | A | 2.9e-06 | 9.6e-07 | 2e-06 | 1.953E-06 | 9.708E-07 | 9.600E-07 | 2.900E-06 |
| 7643 | 383 | E383N | A | 2.2e-06 | 1.8e-06 | 1.1e-06 | 1.700E-06 | 5.568E-07 | 1.100E-06 | 2.200E-06 |
| 7644 | 383 | E383D | A | 1.6e-06 | 2.8e-06 | 2.1e-06 | 2.167E-06 | 6.028E-07 | 1.600E-06 | 2.800E-06 |
| 7645 | 383 | E383C | A | 1.4e-06 | 2.9e-06 | 1.9e-06 | 2.067E-06 | 7.638E-07 | 1.400E-06 | 2.900E-06 |
| 7646 | 383 | E383E | A | 2.5e-06 | 4e-06 | 3e-06 | 3.167E-06 | 7.638E-07 | 2.500E-06 | 4.000E-06 |
| 7647 | 383 | E383Q | A | 1.2e-06 | 1.7e-06 | 4.1e-06 | 2.333E-06 | 1.550E-06 | 1.200E-06 | 4.100E-06 |
| 7648 | 383 | E383G | A | 2e-06 | 2e-06 | 1.8e-06 | 1.933E-06 | 1.155E-07 | 1.800E-06 | 2.000E-06 |
| 7649 | 383 | E383H | A | 4.4e-06 | 2.3e-06 | 2.6e-06 | 3.100E-06 | 1.136E-06 | 2.300E-06 | 4.400E-06 |

| | | | | | | | | | | |
|---|---|---|---|---|---|---|---|---|---|---|
| 7650 | 383 | E383I | A | 2.4e-06 | 2.4e-06 | 2.7e-06 | 2.500E-06 | 1.732E-07 | 2.400E-06 | 2.700E-06 |
| 7651 | 383 | E383L | A | 2.7e-06 | 2.8e-06 | 3.1e-06 | 2.867E-06 | 2.082E-07 | 2.700E-06 | 3.100E-06 |
| 7652 | 383 | E383K | A | 2.4e-06 | 2e-06 | 3.6e-06 | 2.667E-06 | 8.327E-07 | 2.000E-06 | 3.600E-06 |
| 7653 | 383 | E383M | A | 2.8e-06 | 2.4e-06 | 2.2e-06 | 2.467E-06 | 3.055E-07 | 2.200E-06 | 2.800E-06 |
| 7654 | 383 | E383F | A | 2.2e-06 | 1.9e-06 | 2.1e-06 | 2.067E-06 | 1.528E-07 | 1.900E-06 | 2.200E-06 |
| 7655 | 383 | E383P | A | 2.7e-06 | 4.7e-06 | 2.8e-06 | 3.400E-06 | 1.127E-06 | 2.700E-06 | 4.700E-06 |
| 7656 | 383 | E383S | A | 3.1e-06 | 3.7e-06 | 2e-06 | 2.933E-06 | 8.622E-07 | 2.000E-06 | 3.700E-06 |
| 7657 | 383 | E383T | A | 3.2e-06 | 5e-06 | 2.4e-06 | 3.533E-06 | 1.332E-06 | 2.400E-06 | 5.000E-06 |
| 7658 | 383 | E383W | A | 2.9e-06 | 1.9e-06 | 3.1e-06 | 2.633E-06 | 6.429E-07 | 1.900E-06 | 3.100E-06 |
| 7659 | 383 | E383Y | A | 2.7e-06 | 2.2e-06 | 2.4e-06 | 2.433E-06 | 2.517E-07 | 2.200E-06 | 2.700E-06 |
| 7660 | 383 | E383V | A | 1.9e-06 | 2.4e-06 | 3.6e-06 | 2.633E-06 | 8.737E-07 | 1.900E-06 | 3.600E-06 |
| 7661 | 384 | I384A | A | 3e-06 | 2.8e-06 | 2.1e-06 | 2.633E-06 | 4.726E-07 | 2.100E-06 | 3.000E-06 |
| 7662 | 384 | I384R | A | 1.9e-06 | 1.3e-06 | 1.5e-06 | 1.567E-06 | 3.055E-07 | 1.300E-06 | 1.900E-06 |
| 7663 | 384 | I384N | A | 1.7e-06 | 3e-06 | 2.6e-06 | 2.433E-06 | 6.658E-07 | 1.700E-06 | 3.000E-06 |
| 7664 | 384 | I384D | A | 1.7e-06 | 1.1e-06 | 5.2e-06 | 2.667E-06 | 2.214E-06 | 1.100E-06 | 5.200E-06 |
| 7665 | 384 | I384C | A | 2.9e-06 | 2.2e-06 | 2.5e-06 | 2.533E-06 | 3.512E-07 | 2.200E-06 | 2.900E-06 |
| 7666 | 384 | I384E | A | 2.9e-06 | 2.9e-06 | 2.8e-06 | 2.867E-06 | 5.774E-08 | 2.800E-06 | 2.900E-06 |
| 7667 | 384 | I384Q | A | 2.1e-06 | 1.6e-06 | 2e-06 | 1.900E-06 | 2.646E-07 | 1.600E-06 | 2.100E-06 |
| 7668 | 384 | I384G | A | 2.3e-06 | 2.6e-06 | 3.2e-06 | 2.700E-06 | 4.583E-07 | 2.300E-06 | 3.200E-06 |
| 7669 | 384 | I384H | A | 2e-06 | 3.9e-06 | 3.7e-06 | 3.200E-06 | 1.044E-06 | 2.000E-06 | 3.900E-06 |
| 7670 | 384 | I384I | A | 3.3e-06 | 4.5e-06 | 2.7e-06 | 3.500E-06 | 9.165E-07 | 2.700E-06 | 4.500E-06 |
| 7671 | 384 | I384L | A | 2.7e-06 | 2e-06 | 5.1e-06 | 3.267E-06 | 1.626E-06 | 2.000E-06 | 5.100E-06 |
| 7672 | 384 | I384K | A | 1.8e-06 | 1.3e-06 | 1.7e-06 | 1.600E-06 | 2.646E-07 | 1.300E-06 | 1.800E-06 |
| 7673 | 384 | I384M | A | 2.7e-06 | 2.3e-06 | 2.1e-06 | 2.367E-06 | 3.055E-07 | 2.100E-06 | 2.700E-06 |
| 7674 | 384 | I384F | A | 3.8e-06 | 2.5e-06 | 1.1e-06 | 2.467E-06 | 1.350E-06 | 1.100E-06 | 3.800E-06 |
| 7675 | 384 | I384P | A | 4e-06 | 2.2e-06 | 1.9e-06 | 2.700E-06 | 1.136E-06 | 1.900E-06 | 4.000E-06 |
| 7676 | 384 | I384S | A | 2.5e-06 | 2.6e-06 | 1.3e-06 | 2.133E-06 | 7.234E-07 | 1.300E-06 | 2.600E-06 |
| 7677 | 384 | I384T | A | 2.7e-06 | 3.6e-06 | 3.1e-06 | 3.133E-06 | 4.509E-07 | 2.700E-06 | 3.600E-06 |
| 7678 | 384 | I384W | A | 2.9e-06 | 3.3e-06 | 2.9e-06 | 3.033E-06 | 2.309E-07 | 2.900E-06 | 3.300E-06 |
| 7679 | 384 | I384Y | A | 3.9e-06 | 1.4e-06 | 2.3e-06 | 2.533E-06 | 1.266E-06 | 1.400E-06 | 3.900E-06 |
| 7680 | 384 | I384V | A | 2.5e-06 | 2.3e-06 | 2.3e-06 | 2.367E-06 | 1.155E-07 | 2.300E-06 | 2.500E-06 |
| 7681 | 385 | T385A | A | 1.4e-06 | 1.9e-06 | 2.5e-06 | 1.933E-06 | 5.508E-07 | 1.400E-06 | 2.500E-06 |
| 7682 | 385 | T385R | A | 1.8e-06 | 2.1e-06 | 2.6e-06 | 2.167E-06 | 4.041E-07 | 1.800E-06 | 2.600E-06 |
| 7683 | 385 | T385N | A | 1.9e-06 | 3.1e-06 | 1.7e-06 | 2.233E-06 | 7.572E-07 | 1.700E-06 | 3.100E-06 |
| 7684 | 385 | T385D | A | 3e-06 | 2.8e-06 | 1.8e-06 | 2.533E-06 | 6.429E-07 | 1.800E-06 | 3.000E-06 |
| 7685 | 385 | T385C | A | 2.8e-06 | 1.7e-06 | 1.9e-06 | 2.133E-06 | 5.859E-07 | 1.700E-06 | 2.800E-06 |
| 7686 | 385 | T385E | A | 2.6e-06 | 1.8e-06 | 2.3e-06 | 2.233E-06 | 4.041E-07 | 1.800E-06 | 2.600E-06 |
| 7687 | 385 | T385Q | A | 1.9e-06 | 1.4e-06 | 1.8e-06 | 1.700E-06 | 2.646E-07 | 1.400E-06 | 1.900E-06 |
| 7688 | 385 | T385G | A | 2.4e-06 | 2.4e-06 | 2.5e-06 | 2.433E-06 | 5.774E-08 | 2.400E-06 | 2.500E-06 |
| 7689 | 385 | T385H | A | 2.5e-06 | 2.1e-06 | 3.8e-06 | 2.800E-06 | 8.888E-07 | 2.100E-06 | 3.800E-06 |
| 7690 | 385 | T385I | A | 6e-06 | 2.5e-06 | 2.1e-06 | 3.533E-06 | 2.146E-06 | 2.100E-06 | 6.000E-06 |
| 7691 | 385 | T385L | A | 3.7e-06 | 2.2e-06 | 1.9e-06 | 2.600E-06 | 9.644E-07 | 1.900E-06 | 3.700E-06 |
| 7692 | 385 | T385K | A | 2.1e-06 | 2.2e-06 | 1.4e-06 | 1.900E-06 | 4.359E-07 | 1.400E-06 | 2.200E-06 |
| 7693 | 385 | T385M | A | 3.2e-06 | 2.9e-06 | 2e-06 | 2.700E-06 | 6.245E-07 | 2.000E-06 | 3.200E-06 |
| 7694 | 385 | T385F | A | 2.5e-06 | 2.9e-06 | 2.6e-06 | 2.667E-06 | 2.082E-07 | 2.500E-06 | 2.900E-06 |
| 7695 | 385 | T385P | A | 1.9e-06 | 2.3e-06 | 1.8e-06 | 2.000E-06 | 2.646E-07 | 1.800E-06 | 2.300E-06 |
| 7696 | 385 | T385S | A | 1.5e-06 | 2.2e-06 | 2.2e-06 | 1.967E-06 | 4.041E-07 | 1.500E-06 | 2.200E-06 |
| 7697 | 385 | T385T | A | 1.7e-06 | 5.2e-06 | 2.4e-06 | 3.100E-06 | 1.852E-06 | 1.700E-06 | 5.200E-06 |
| 7698 | 385 | T385W | A | 2.3e-06 | 3e-06 | 1.4e-06 | 2.233E-06 | 8.021E-07 | 1.400E-06 | 3.000E-06 |
| 7699 | 385 | T385Y | A | 2.5e-06 | 3.5e-06 | 2.7e-06 | 2.900E-06 | 5.292E-07 | 2.500E-06 | 3.500E-06 |
| 7700 | 385 | T385V | A | 4.3e-06 | 2.5e-06 | 3.7e-06 | 3.500E-06 | 9.165E-07 | 2.500E-06 | 4.300E-06 |
| 7701 | 386 | G386A | A | 2.7e-06 | 2.8e-06 | 6.7e-06 | 4.067E-06 | 2.281E-06 | 2.700E-06 | 6.700E-06 |
| 7702 | 386 | G386R | A | 1.1e-06 | 1.3e-06 | 4.2e-06 | 2.200E-06 | 1.735E-06 | 1.100E-06 | 4.200E-06 |
| 7703 | 386 | G386N | A | 1.3e-06 | 1.5e-06 | 2.3e-06 | 1.700E-06 | 5.292E-07 | 1.300E-06 | 2.300E-06 |
| 7704 | 386 | G386D | A | 1.9e-06 | 2.5e-06 | 2.8e-06 | 2.400E-06 | 4.583E-07 | 1.900E-06 | 2.800E-06 |
| 7705 | 386 | G386C | A | 1.8e-06 | 1.4e-06 | 2.4e-06 | 1.867E-06 | 5.033E-07 | 1.400E-06 | 2.400E-06 |
| 7706 | 386 | G386E | A | 2.8e-06 | 2.4e-06 | 3.7e-06 | 2.967E-06 | 6.658E-07 | 2.400E-06 | 3.700E-06 |
| 7707 | 386 | G386Q | A | 3e-06 | 3e-06 | 2.7e-06 | 2.900E-06 | 1.732E-07 | 2.700E-06 | 3.000E-06 |
| 7708 | 386 | G386G | A | 3.7e-06 | 2.1e-06 | 3.9e-06 | 3.233E-06 | 9.866E-07 | 2.100E-06 | 3.900E-06 |
| 7709 | 386 | G386H | A | 1.8e-06 | 1.3e-06 | 3.5e-06 | 2.200E-06 | 1.153E-06 | 1.300E-06 | 3.500E-06 |
| 7710 | 386 | G386I | A | 2e-06 | 2.2e-06 | 1.5e-06 | 1.900E-06 | 3.606E-07 | 1.500E-06 | 2.200E-06 |
| 7711 | 386 | G386L | A | 1.2e-06 | 2.2e-06 | 1.8e-06 | 1.733E-06 | 5.033E-07 | 1.200E-06 | 2.200E-06 |
| 7712 | 386 | G386K | A | 2.9e-06 | 1.8e-06 | 2.4e-06 | 2.367E-06 | 5.508E-07 | 1.800E-06 | 2.900E-06 |
| 7713 | 386 | G386M | A | 2.2e-06 | 2.8e-06 | 3.1e-06 | 2.700E-06 | 4.583E-07 | 2.200E-06 | 3.100E-06 |
| 7714 | 386 | G386F | A | 2e-06 | 3e-06 | 1.2e-06 | 2.067E-06 | 9.018E-07 | 1.200E-06 | 3.000E-06 |
| 7715 | 386 | G386P | A | 2.6e-06 | 3e-06 | 2.9e-06 | 2.833E-06 | 2.082E-07 | 2.600E-06 | 3.000E-06 |
| 7716 | 386 | G386S | A | 3.6e-06 | 1.9e-06 | 2e-06 | 2.500E-06 | 9.539E-07 | 1.900E-06 | 3.600E-06 |
| 7717 | 386 | G386T | A | 1.8e-06 | 4.3e-06 | 1.5e-06 | 2.533E-06 | 1.537E-06 | 1.500E-06 | 4.300E-06 |
| 7718 | 386 | G386W | A | 2.5e-06 | 3.3e-06 | 2.5e-06 | 2.767E-06 | 4.619E-07 | 2.500E-06 | 3.300E-06 |
| 7719 | 386 | G386Y | A | 2.4e-06 | 1.3e-06 | 2e-06 | 1.900E-06 | 5.568E-07 | 1.300E-06 | 2.400E-06 |
| 7720 | 386 | G386V | A | 3e-06 | 2.3e-06 | 1.5e-06 | 2.267E-06 | 7.506E-07 | 1.500E-06 | 3.000E-06 |
| 7721 | 387 | Y387A | A | 2.6e-06 | 2.7e-06 | 1.9e-06 | 2.400E-06 | 4.359E-07 | 1.900E-06 | 2.700E-06 |
| 7722 | 387 | Y387R | A | 2.8e-06 | 3.5e-06 | 1.6e-06 | 2.633E-06 | 9.609E-07 | 1.600E-06 | 3.500E-06 |
| 7723 | 387 | Y387N | A | 1.1e-06 | 2.5e-06 | 1.9e-06 | 1.833E-06 | 7.024E-07 | 1.100E-06 | 2.500E-06 |
| 7724 | 387 | Y387D | A | 1.6e-06 | 1.1e-06 | 2.3e-06 | 1.667E-06 | 6.028E-07 | 1.100E-06 | 2.300E-06 |
| 7725 | 387 | Y387C | A | 1.7e-06 | 3.3e-06 | 2.5e-06 | 2.500E-06 | 8.000E-07 | 1.700E-06 | 3.300E-06 |
| 7726 | 387 | Y387E | A | 2.6e-06 | 2e-06 | 2.1e-06 | 2.233E-06 | 3.215E-07 | 2.000E-06 | 2.600E-06 |
| 7727 | 387 | Y387Q | A | 2.9e-06 | 1.6e-06 | 3.7e-06 | 2.733E-06 | 1.060E-06 | 1.600E-06 | 3.700E-06 |
| 7728 | 387 | Y387G | A | 2.5e-06 | 2.3e-06 | 1.6e-06 | 2.133E-06 | 4.726E-07 | 1.600E-06 | 2.500E-06 |
| 7729 | 387 | Y387H | A | 2.7e-06 | 3.3e-06 | 1.5e-06 | 2.500E-06 | 9.165E-07 | 1.500E-06 | 3.300E-06 |
| 7730 | 387 | Y387I | A | 1.4e-06 | 2.6e-06 | 2.9e-06 | 2.300E-06 | 7.937E-07 | 1.400E-06 | 2.900E-06 |
| 7731 | 387 | Y387L | A | 2.1e-06 | 2.5e-06 | 1.4e-06 | 2.000E-06 | 5.568E-07 | 1.400E-06 | 2.500E-06 |
| 7732 | 387 | Y387K | A | 1.5e-06 | 2.2e-06 | 4.1e-06 | 2.600E-06 | 1.345E-06 | 1.500E-06 | 4.100E-06 |
| 7733 | 387 | Y387M | A | 2.2e-06 | 2.4e-06 | 1.7e-06 | 2.100E-06 | 3.606E-07 | 1.700E-06 | 2.400E-06 |
| 7734 | 387 | Y387F | A | 4.1e-06 | 4.1e-06 | 2.8e-06 | 3.667E-06 | 7.506E-07 | 2.800E-06 | 4.100E-06 |

| | | | | | | | | | | |
|---|---|---|---|---|---|---|---|---|---|---|
| 7735 | 387 | Y387P | A | 2.1e-06 | 2.7e-06 | 3.9e-06 | 2.900E-06 | 9.165E-07 | 2.100E-06 | 3.900E-06 |
| 7736 | 387 | Y387S | A | 1.3e-06 | 2e-06 | 2.8e-06 | 2.033E-06 | 7.506E-07 | 1.300E-06 | 2.800E-06 |
| 7737 | 387 | Y387T | A | 2.6e-06 | 2.8e-06 | 3.6e-06 | 3.000E-06 | 5.292E-07 | 2.600E-06 | 3.600E-06 |
| 7738 | 387 | Y387W | A | 1.8e-06 | 2e-06 | 2.7e-06 | 2.167E-06 | 4.726E-07 | 1.800E-06 | 2.700E-06 |
| 7739 | 387 | Y387Y | A | 1.3e-06 | 1.6e-06 | 3.2e-06 | 2.033E-06 | 1.021E-06 | 1.300E-06 | 3.200E-06 |
| 7740 | 387 | Y387V | A | 3.7e-06 | 2.8e-06 | 3.3e-06 | 3.267E-06 | 4.509E-07 | 2.800E-06 | 3.700E-06 |
| 7741 | 388 | L388A | A | 1.3e-06 | 2.4e-06 | 2e-06 | 1.900E-06 | 5.568E-07 | 1.300E-06 | 2.400E-06 |
| 7742 | 388 | L388R | A | 3.1e-06 | 1.8e-06 | 1.8e-06 | 2.233E-06 | 7.506E-07 | 1.800E-06 | 3.100E-06 |
| 7743 | 388 | L388N | A | 2.1e-06 | 2.4e-06 | 1.9e-06 | 2.133E-06 | 2.517E-07 | 1.900E-06 | 2.400E-06 |
| 7744 | 388 | L388D | A | 2.5e-06 | 2.7e-06 | 1.7e-06 | 2.300E-06 | 5.292E-07 | 1.700E-06 | 2.700E-06 |
| 7745 | 388 | L388C | A | 3.7e-06 | 2.6e-06 | 2.6e-06 | 2.967E-06 | 6.351E-07 | 2.600E-06 | 3.700E-06 |
| 7746 | 388 | L388E | A | 1.3e-06 | 2.9e-06 | 2.9e-06 | 2.367E-06 | 9.238E-07 | 1.300E-06 | 2.900E-06 |
| 7747 | 388 | L388Q | A | 1.6e-06 | 2.1e-06 | 2.2e-06 | 1.967E-06 | 3.215E-07 | 1.600E-06 | 2.200E-06 |
| 7748 | 388 | L388G | A | 4.3e-06 | 2.6e-06 | 2.9e-06 | 3.267E-06 | 9.074E-07 | 2.600E-06 | 4.300E-06 |
| 7749 | 388 | L388H | A | 1.6e-06 | 2.9e-06 | 1.6e-06 | 2.033E-06 | 7.506E-07 | 1.600E-06 | 2.900E-06 |
| 7750 | 388 | L388I | A | 2.5e-06 | 3e-06 | 4.7e-06 | 3.400E-06 | 1.153E-06 | 2.500E-06 | 4.700E-06 |
| 7751 | 388 | L388L | A | 1.2e-06 | 6e-06 | 3.9e-06 | 3.700E-06 | 2.406E-06 | 1.200E-06 | 6.000E-06 |
| 7752 | 388 | L388K | A | 2.4e-06 | 3.2e-06 | 2.1e-06 | 2.567E-06 | 5.686E-07 | 2.100E-06 | 3.200E-06 |
| 7753 | 388 | L388M | A | 2.4e-06 | 2.8e-06 | 3e-06 | 2.733E-06 | 3.055E-07 | 2.400E-06 | 3.000E-06 |
| 7754 | 388 | L388F | A | 2.8e-06 | 2.4e-06 | 3.1e-06 | 2.767E-06 | 3.512E-07 | 2.400E-06 | 3.100E-06 |
| 7755 | 388 | L388P | A | 1.5e-06 | 2.6e-06 | 1.4e-06 | 1.833E-06 | 6.658E-07 | 1.400E-06 | 2.600E-06 |
| 7756 | 388 | L388S | A | 1.7e-06 | 1.3e-06 | 2.9e-06 | 1.967E-06 | 8.327E-07 | 1.300E-06 | 2.900E-06 |
| 7757 | 388 | L388T | A | 2.7e-06 | 2.4e-06 | 3.1e-06 | 2.733E-06 | 3.512E-07 | 2.400E-06 | 3.100E-06 |
| 7758 | 388 | L388W | A | 4.1e-06 | 2.1e-06 | 4.1e-06 | 3.433E-06 | 1.155E-06 | 2.100E-06 | 4.100E-06 |
| 7759 | 388 | L388Y | A | 2e-06 | 2.9e-06 | 3.9e-06 | 2.933E-06 | 9.504E-07 | 2.000E-06 | 3.900E-06 |
| 7760 | 388 | L388V | A | 2.1e-06 | 1.9e-06 | 3.1e-06 | 2.367E-06 | 6.429E-07 | 1.900E-06 | 3.100E-06 |
| 7761 | 389 | Y389A | A | 2.1e-06 | 5.4e-06 | 4.2e-06 | 3.900E-06 | 1.670E-06 | 2.100E-06 | 5.400E-06 |
| 7762 | 389 | Y389R | A | 2e-06 | 3.7e-06 | 1.8e-06 | 2.500E-06 | 1.044E-06 | 1.800E-06 | 3.700E-06 |
| 7763 | 389 | Y389N | A | 2.9e-06 | 2.5e-06 | 2.3e-06 | 2.567E-06 | 3.055E-07 | 2.300E-06 | 2.900E-06 |
| 7764 | 389 | Y389D | A | 3.1e-06 | 3.2e-06 | 1.6e-06 | 2.633E-06 | 8.963E-07 | 1.600E-06 | 3.200E-06 |
| 7765 | 389 | Y389C | A | 3.1e-06 | 2.6e-06 | 2.1e-06 | 2.600E-06 | 5.000E-07 | 2.100E-06 | 3.100E-06 |
| 7766 | 389 | Y389E | A | 3.1e-06 | 2.1e-06 | 4.9e-06 | 3.367E-06 | 1.419E-06 | 2.100E-06 | 4.900E-06 |
| 7767 | 389 | Y389Q | A | 4.3e-06 | 2.4e-06 | 2.4e-06 | 3.033E-06 | 1.097E-06 | 2.400E-06 | 4.300E-06 |
| 7768 | 389 | Y389G | A | 1.9e-06 | 2.9e-06 | 1.9e-06 | 2.233E-06 | 5.774E-07 | 1.900E-06 | 2.900E-06 |
| 7769 | 389 | Y389H | A | 1.9e-06 | 2.5e-06 | 2.1e-06 | 2.167E-06 | 3.055E-07 | 1.900E-06 | 2.500E-06 |
| 7770 | 389 | Y389I | A | 1.7e-06 | 3.3e-06 | 2.4e-06 | 2.467E-06 | 8.021E-07 | 1.700E-06 | 3.300E-06 |
| 7771 | 389 | Y389L | A | 3.1e-06 | 3.2e-06 | 2.7e-06 | 3.000E-06 | 2.646E-07 | 2.700E-06 | 3.200E-06 |
| 7772 | 389 | Y389K | A | 2.9e-06 | 2.6e-06 | 2.2e-06 | 2.567E-06 | 3.512E-07 | 2.200E-06 | 2.900E-06 |
| 7773 | 389 | Y389M | A | 4.3e-06 | 2.2e-06 | 3.2e-06 | 3.233E-06 | 1.050E-06 | 2.200E-06 | 4.300E-06 |
| 7774 | 389 | Y389F | A | 3.6e-06 | 1.4e-06 | 1.5e-06 | 2.167E-06 | 1.242E-06 | 1.400E-06 | 3.600E-06 |
| 7775 | 389 | Y389P | A | 3.4e-06 | 3.1e-06 | 2.3e-06 | 2.933E-06 | 5.686E-07 | 2.300E-06 | 3.400E-06 |
| 7776 | 389 | Y389S | A | 4.2e-06 | 3.1e-06 | 3.4e-06 | 3.567E-06 | 5.686E-07 | 3.100E-06 | 4.200E-06 |
| 7777 | 389 | Y389T | A | 2.4e-06 | 3.3e-06 | 1.6e-06 | 2.433E-06 | 8.505E-07 | 1.600E-06 | 3.300E-06 |
| 7778 | 389 | Y389W | A | 4.1e-06 | 2.1e-06 | 2.2e-06 | 2.800E-06 | 1.127E-06 | 2.100E-06 | 4.100E-06 |
| 7779 | 389 | Y389Y | A | 3e-06 | 1.7e-06 | 2.8e-06 | 2.500E-06 | 7.000E-07 | 1.700E-06 | 3.000E-06 |
| 7780 | 389 | Y389V | A | 1.9e-06 | 2.3e-06 | 4.4e-06 | 2.867E-06 | 1.343E-06 | 1.900E-06 | 4.400E-06 |
| 7781 | 390 | I390A | A | 3.3e-06 | 3.8e-06 | 4.3e-06 | 3.800E-06 | 5.000E-07 | 3.300E-06 | 4.300E-06 |
| 7782 | 390 | I390R | A | 2.5e-06 | 3.1e-06 | 2e-06 | 2.533E-06 | 5.508E-07 | 2.000E-06 | 3.100E-06 |
| 7783 | 390 | I390N | A | 4.3e-06 | 2.6e-06 | 2e-06 | 2.967E-06 | 1.193E-06 | 2.000E-06 | 4.300E-06 |
| 7784 | 390 | I390D | A | 1.9e-06 | 3e-06 | 1.8e-06 | 2.233E-06 | 6.658E-07 | 1.800E-06 | 3.000E-06 |
| 7785 | 390 | I390C | A | 1.9e-06 | 2.2e-06 | 2.9e-06 | 2.333E-06 | 5.132E-07 | 1.900E-06 | 2.900E-06 |
| 7786 | 390 | I390E | A | 2e-06 | 1.8e-06 | 2.8e-06 | 2.200E-06 | 5.292E-07 | 1.800E-06 | 2.800E-06 |
| 7787 | 390 | I390Q | A | 2e-06 | 1.6e-06 | 2.3e-06 | 1.967E-06 | 3.512E-07 | 1.600E-06 | 2.300E-06 |
| 7788 | 390 | I390G | A | 3.5e-06 | 2e-06 | 1.9e-06 | 2.467E-06 | 8.963E-07 | 1.900E-06 | 3.500E-06 |
| 7789 | 390 | I390H | A | 2.6e-06 | 3.9e-06 | 2.9e-06 | 3.133E-06 | 6.807E-07 | 2.600E-06 | 3.900E-06 |
| 7790 | 390 | I390I | A | 2.9e-06 | 1.6e-06 | 2.3e-06 | 2.267E-06 | 6.506E-07 | 1.600E-06 | 2.900E-06 |
| 7791 | 390 | I390L | A | 4.6e-06 | 1.5e-06 | 4e-06 | 3.367E-06 | 1.644E-06 | 1.500E-06 | 4.600E-06 |
| 7792 | 390 | I390K | A | 1.5e-06 | 3.8e-06 | 2.2e-06 | 2.500E-06 | 1.179E-06 | 1.500E-06 | 3.800E-06 |
| 7793 | 390 | I390M | A | 1.5e-06 | 2e-06 | 2.3e-06 | 1.933E-06 | 4.041E-07 | 1.500E-06 | 2.300E-06 |
| 7794 | 390 | I390F | A | 1.7e-06 | 2.4e-06 | 3.6e-06 | 2.567E-06 | 9.609E-07 | 1.700E-06 | 3.600E-06 |
| 7795 | 390 | I390P | A | 3e-06 | 2.6e-06 | 1.6e-06 | 2.400E-06 | 7.211E-07 | 1.600E-06 | 3.000E-06 |
| 7796 | 390 | I390S | A | 1.9e-06 | 1.7e-06 | 1.5e-06 | 1.700E-06 | 2.000E-07 | 1.500E-06 | 1.900E-06 |
| 7797 | 390 | I390T | A | 2.6e-06 | 3.4e-06 | 2.4e-06 | 2.800E-06 | 5.292E-07 | 2.400E-06 | 3.400E-06 |
| 7798 | 390 | I390W | A | 2.9e-06 | 2.3e-06 | 1.6e-06 | 2.267E-06 | 6.506E-07 | 1.600E-06 | 2.900E-06 |
| 7799 | 390 | I390Y | A | 2.7e-06 | 3e-06 | 1.3e-06 | 2.333E-06 | 9.074E-07 | 1.300E-06 | 3.000E-06 |
| 7800 | 390 | I390V | A | 1.9e-06 | 1.7e-06 | 2e-06 | 1.867E-06 | 1.528E-07 | 1.700E-06 | 2.000E-06 |
| 7801 | 391 | S391A | A | 1.8e-06 | 2e-06 | 2.9e-06 | 2.233E-06 | 5.859E-07 | 1.800E-06 | 2.900E-06 |
| 7802 | 391 | S391R | A | 2.4e-06 | 2.9e-06 | 1.8e-06 | 2.367E-06 | 5.508E-07 | 1.800E-06 | 2.900E-06 |
| 7803 | 391 | S391N | A | 2.6e-06 | 2.2e-06 | 3.6e-06 | 2.800E-06 | 7.211E-07 | 2.200E-06 | 3.600E-06 |
| 7804 | 391 | S391D | A | 2.3e-06 | 2.5e-06 | 2.6e-06 | 2.467E-06 | 1.528E-07 | 2.300E-06 | 2.600E-06 |
| 7805 | 391 | S391C | A | 2.2e-06 | 2.3e-06 | 2.5e-06 | 2.333E-06 | 1.528E-07 | 2.200E-06 | 2.500E-06 |
| 7806 | 391 | S391E | A | 2.5e-06 | 2.4e-06 | 4e-06 | 2.967E-06 | 8.963E-07 | 2.400E-06 | 4.000E-06 |
| 7807 | 391 | S391Q | A | 1e-06 | 2.2e-06 | 2.4e-06 | 1.867E-06 | 7.572E-07 | 1.000E-06 | 2.400E-06 |
| 7808 | 391 | S391G | A | 3.9e-06 | 2.3e-06 | 2.7e-06 | 2.967E-06 | 8.327E-07 | 2.300E-06 | 3.900E-06 |
| 7809 | 391 | S391H | A | 2.4e-06 | 2.6e-06 | 2e-06 | 2.333E-06 | 3.055E-07 | 2.000E-06 | 2.600E-06 |
| 7810 | 391 | S391I | A | 2.4e-06 | 2.3e-06 | 2.2e-06 | 2.300E-06 | 1.000E-07 | 2.200E-06 | 2.400E-06 |
| 7811 | 391 | S391L | A | 4e-06 | 2.1e-06 | 1.9e-06 | 2.667E-06 | 1.159E-06 | 1.900E-06 | 4.000E-06 |
| 7812 | 391 | S391K | A | 3.8e-06 | 2.3e-06 | 2.7e-06 | 2.933E-06 | 7.767E-07 | 2.300E-06 | 3.800E-06 |
| 7813 | 391 | S391M | A | 2e-06 | 3.7e-06 | 2.6e-06 | 2.767E-06 | 8.622E-07 | 2.000E-06 | 3.700E-06 |
| 7814 | 391 | S391F | A | 1.1e-06 | 2.3e-06 | 1.4e-06 | 1.600E-06 | 6.245E-07 | 1.100E-06 | 2.300E-06 |
| 7815 | 391 | S391P | A | 2.2e-06 | 2.2e-06 | 2.3e-06 | 2.233E-06 | 5.774E-08 | 2.200E-06 | 2.300E-06 |
| 7816 | 391 | S391S | A | 3.3e-06 | 2.9e-06 | 5.5e-06 | 3.900E-06 | 1.400E-06 | 2.900E-06 | 5.500E-06 |
| 7817 | 391 | S391T | A | 1.9e-06 | 3.5e-06 | 1.9e-06 | 2.433E-06 | 9.238E-07 | 1.900E-06 | 3.500E-06 |
| 7818 | 391 | S391W | A | 3.3e-06 | 3e-06 | 2.6e-06 | 2.967E-06 | 3.512E-07 | 2.600E-06 | 3.300E-06 |
| 7819 | 391 | S391Y | A | 2.3e-06 | 2.2e-06 | 2.1e-06 | 2.200E-06 | 1.000E-07 | 2.100E-06 | 2.300E-06 |

| | | | | | | | | | | |
|---|---|---|---|---|---|---|---|---|---|---|
| 7820 | 391 | S391V | A | 2.5e-06 | 3e-06 | 4.1e-06 | 3.200E-06 | 8.185E-07 | 2.500E-06 | 4.100E-06 |
| 7821 | 392 | A392A | A | 2.7e-06 | 3.7e-06 | 2e-06 | 2.800E-06 | 8.544E-07 | 2.000E-06 | 3.700E-06 |
| 7822 | 392 | A392R | A | 1.7e-06 | 2.9e-06 | 3e-06 | 2.533E-06 | 7.234E-07 | 1.700E-06 | 3.000E-06 |
| 7823 | 392 | A392N | A | 2e-06 | 1.7e-06 | 2.9e-06 | 2.200E-06 | 6.245E-07 | 1.700E-06 | 2.900E-06 |
| 7824 | 392 | A392D | A | 1.4e-06 | 3.2e-06 | 1.2e-06 | 1.933E-06 | 1.102E-06 | 1.200E-06 | 3.200E-06 |
| 7825 | 392 | A392C | A | 1.9e-06 | 1.9e-06 | 1.5e-06 | 1.767E-06 | 2.309E-07 | 1.500E-06 | 1.900E-06 |
| 7826 | 392 | A392E | A | 2.7e-06 | 4.3e-06 | 2.5e-06 | 3.167E-06 | 9.866E-07 | 2.500E-06 | 4.300E-06 |
| 7827 | 392 | A392Q | A | 1.7e-06 | 4.4e-06 | 1.1e-06 | 2.400E-06 | 1.758E-06 | 1.100E-06 | 4.400E-06 |
| 7828 | 392 | A392G | A | 1.7e-06 | 2.2e-06 | 2.4e-06 | 2.100E-06 | 3.606E-07 | 1.700E-06 | 2.400E-06 |
| 7829 | 392 | A392H | A | 1.7e-06 | 2.5e-06 | 3e-06 | 2.400E-06 | 6.557E-07 | 1.700E-06 | 3.000E-06 |
| 7830 | 392 | A392I | A | 2.7e-06 | 2.8e-06 | 2.9e-06 | 2.800E-06 | 1.000E-07 | 2.700E-06 | 2.900E-06 |
| 7831 | 392 | A392L | A | 9e-07 | 2.7e-06 | 3.8e-06 | 2.467E-06 | 1.464E-06 | 9.000E-07 | 3.800E-06 |
| 7832 | 392 | A392K | A | 2.4e-06 | 1.7e-06 | 3.9e-06 | 2.667E-06 | 1.124E-06 | 1.700E-06 | 3.900E-06 |
| 7833 | 392 | A392M | A | 1.5e-06 | 3e-06 | 2.6e-06 | 2.367E-06 | 7.767E-07 | 1.500E-06 | 3.000E-06 |
| 7834 | 392 | A392F | A | 2.3e-06 | 1.9e-06 | 3.1e-06 | 2.433E-06 | 6.110E-07 | 1.900E-06 | 3.100E-06 |
| 7835 | 392 | A392P | A | 1.8e-06 | 2.8e-06 | 4.4e-06 | 3.000E-06 | 1.311E-06 | 1.800E-06 | 4.400E-06 |
| 7836 | 392 | A392S | A | 2.1e-06 | 1.3e-06 | 4.1e-06 | 2.500E-06 | 1.442E-06 | 1.300E-06 | 4.100E-06 |
| 7837 | 392 | A392T | A | 2.9e-06 | 2.3e-06 | 2.4e-06 | 2.533E-06 | 3.215E-07 | 2.300E-06 | 2.900E-06 |
| 7838 | 392 | A392W | A | 3e-06 | 1.2e-06 | 4e-06 | 2.733E-06 | 1.419E-06 | 1.200E-06 | 4.000E-06 |
| 7839 | 392 | A392Y | A | 1.9e-06 | 2.6e-06 | 2.8e-06 | 2.433E-06 | 4.726E-07 | 1.900E-06 | 2.800E-06 |
| 7840 | 392 | A392V | A | 1.3e-06 | 2.4e-06 | 2e-06 | 1.900E-06 | 5.568E-07 | 1.300E-06 | 2.400E-06 |
| 7841 | 393 | W393A | A | 1.7e-06 | 2.6e-06 | 2.6e-06 | 2.300E-06 | 5.196E-07 | 1.700E-06 | 2.600E-06 |
| 7842 | 393 | W393R | A | 2.5e-06 | 2.6e-06 | 4.5e-06 | 3.200E-06 | 1.127E-06 | 2.500E-06 | 4.500E-06 |
| 7843 | 393 | W393N | A | 2.3e-06 | 2e-06 | 2.8e-06 | 2.367E-06 | 4.041E-07 | 2.000E-06 | 2.800E-06 |
| 7844 | 393 | W393D | A | 3.6e-06 | 3.4e-06 | 1.3e-06 | 2.767E-06 | 1.274E-06 | 1.300E-06 | 3.600E-06 |
| 7845 | 393 | W393C | A | 4.4e-06 | 2.3e-06 | 1.9e-06 | 2.867E-06 | 1.343E-06 | 1.900E-06 | 4.400E-06 |
| 7846 | 393 | W393E | A | 3.3e-06 | 2.2e-06 | 3e-06 | 2.833E-06 | 5.686E-07 | 2.200E-06 | 3.300E-06 |
| 7847 | 393 | W393Q | A | 2.2e-06 | 4.1e-06 | 1.8e-06 | 2.700E-06 | 1.229E-06 | 1.800E-06 | 4.100E-06 |
| 7848 | 393 | W393G | A | 3.3e-06 | 2.4e-06 | 2.1e-06 | 2.600E-06 | 6.245E-07 | 2.100E-06 | 3.300E-06 |
| 7849 | 393 | W393H | A | 3.5e-06 | 2.6e-06 | 2.1e-06 | 2.733E-06 | 7.095E-07 | 2.100E-06 | 3.500E-06 |
| 7850 | 393 | W393I | A | 1.7e-06 | 3.3e-06 | 2.6e-06 | 2.533E-06 | 8.021E-07 | 1.700E-06 | 3.300E-06 |
| 7851 | 393 | W393L | A | 2.7e-06 | 1.7e-06 | 2.3e-06 | 2.233E-06 | 5.033E-07 | 1.700E-06 | 2.700E-06 |
| 7852 | 393 | W393K | A | 2.4e-06 | 5.1e-06 | 2.9e-06 | 3.467E-06 | 1.436E-06 | 2.400E-06 | 5.100E-06 |
| 7853 | 393 | W393M | A | 3e-06 | 1.9e-06 | 3.3e-06 | 2.733E-06 | 7.371E-07 | 1.900E-06 | 3.300E-06 |
| 7854 | 393 | W393F | A | 1.4e-06 | 2.6e-06 | 2e-06 | 2.000E-06 | 6.000E-07 | 1.400E-06 | 2.600E-06 |
| 7855 | 393 | W393P | A | 3.3e-06 | 1.8e-06 | 4.4e-06 | 3.167E-06 | 1.305E-06 | 1.800E-06 | 4.400E-06 |
| 7856 | 393 | W393S | A | 4e-06 | 1.8e-06 | 4e-06 | 3.267E-06 | 1.270E-06 | 1.800E-06 | 4.000E-06 |
| 7857 | 393 | W393T | A | 3.3e-06 | 4.8e-06 | 1.7e-06 | 3.267E-06 | 1.550E-06 | 1.700E-06 | 4.800E-06 |
| 7858 | 393 | W393W | A | 1.6e-06 | 4.6e-06 | 1.7e-06 | 2.633E-06 | 1.704E-06 | 1.600E-06 | 4.600E-06 |
| 7859 | 393 | W393Y | A | 2.1e-06 | 1.9e-06 | 2.7e-06 | 2.233E-06 | 4.163E-07 | 1.900E-06 | 2.700E-06 |
| 7860 | 393 | W393V | A | 3.3e-06 | 1.9e-06 | 2e-06 | 2.400E-06 | 7.810E-07 | 1.900E-06 | 3.300E-06 |
| 7861 | 394 | P394A | A | 2.8e-06 | 2.5e-06 | 2.2e-06 | 2.500E-06 | 3.000E-07 | 2.200E-06 | 2.800E-06 |
| 7862 | 394 | P394R | A | 1.8e-06 | 5.1e-06 | 2e-06 | 2.967E-06 | 1.850E-06 | 1.800E-06 | 5.100E-06 |
| 7863 | 394 | P394N | A | 1.7e-06 | 3e-06 | 1.7e-06 | 2.133E-06 | 7.506E-07 | 1.700E-06 | 3.000E-06 |
| 7864 | 394 | P394D | A | 3.9e-06 | 3.4e-06 | 2.5e-06 | 3.267E-06 | 7.095E-07 | 2.500E-06 | 3.900E-06 |
| 7865 | 394 | P394C | A | 3.6e-06 | 2.7e-06 | 2.7e-06 | 3.000E-06 | 5.196E-07 | 2.700E-06 | 3.600E-06 |
| 7866 | 394 | P394E | A | 1.9e-06 | 2.5e-06 | 1.3e-06 | 1.900E-06 | 6.000E-07 | 1.300E-06 | 2.500E-06 |
| 7867 | 394 | P394Q | A | 2.3e-06 | 2.9e-06 | 2.6e-06 | 2.600E-06 | 3.000E-07 | 2.300E-06 | 2.900E-06 |
| 7868 | 394 | P394G | A | 4.9e-06 | 3.6e-06 | 2.1e-06 | 3.533E-06 | 1.401E-06 | 2.100E-06 | 4.900E-06 |
| 7869 | 394 | P394H | A | 2.6e-06 | 2.6e-06 | 2.6e-06 | 2.600E-06 | 0.000E+00 | 2.600E-06 | 2.600E-06 |
| 7870 | 394 | P394I | A | 2.6e-06 | 1.9e-06 | 3.6e-06 | 2.700E-06 | 8.544E-07 | 1.900E-06 | 3.600E-06 |
| 7871 | 394 | P394L | A | 1.2e-06 | 2.6e-06 | 2.3e-06 | 2.033E-06 | 7.371E-07 | 1.200E-06 | 2.600E-06 |
| 7872 | 394 | P394K | A | 2.2e-06 | 1.4e-06 | 2.6e-06 | 2.067E-06 | 6.110E-07 | 1.400E-06 | 2.600E-06 |
| 7873 | 394 | P394M | A | 2.9e-06 | 3.9e-06 | 2e-06 | 2.933E-06 | 9.504E-07 | 2.000E-06 | 3.900E-06 |
| 7874 | 394 | P394F | A | 1.7e-06 | 3.3e-06 | 2.1e-06 | 2.367E-06 | 8.327E-07 | 1.700E-06 | 3.300E-06 |
| 7875 | 394 | P394P | A | 2.2e-06 | 2.2e-06 | 1.8e-06 | 2.067E-06 | 2.309E-07 | 1.800E-06 | 2.200E-06 |
| 7876 | 394 | P394S | A | 1.8e-06 | 2.9e-06 | 1.9e-06 | 2.200E-06 | 6.083E-07 | 1.800E-06 | 2.900E-06 |
| 7877 | 394 | P394T | A | 3.1e-06 | 2.8e-06 | 1.9e-06 | 2.600E-06 | 6.245E-07 | 1.900E-06 | 3.100E-06 |
| 7878 | 394 | P394W | A | 1.6e-06 | 2.6e-06 | 1.8e-06 | 2.000E-06 | 5.292E-07 | 1.600E-06 | 2.600E-06 |
| 7879 | 394 | P394Y | A | 1.1e-06 | 2.8e-06 | 2.4e-06 | 2.100E-06 | 8.888E-07 | 1.100E-06 | 2.800E-06 |
| 7880 | 394 | P394V | A | 3.4e-06 | 2.1e-06 | 3.1e-06 | 2.867E-06 | 6.807E-07 | 2.100E-06 | 3.400E-06 |
| 7881 | 395 | D395A | A | 2.2e-06 | 1.3e-06 | 1e-06 | 1.500E-06 | 6.245E-07 | 1.000E-06 | 2.200E-06 |
| 7882 | 395 | D395R | A | 3.2e-06 | 2.9e-06 | 1.7e-06 | 2.600E-06 | 7.937E-07 | 1.700E-06 | 3.200E-06 |
| 7883 | 395 | D395N | A | 1.4e-06 | 2.6e-06 | 2.1e-06 | 2.033E-06 | 6.028E-07 | 1.400E-06 | 2.600E-06 |
| 7884 | 395 | D395D | A | 2.1e-06 | 2.8e-06 | 1.8e-06 | 2.233E-06 | 5.132E-07 | 1.800E-06 | 2.800E-06 |
| 7885 | 395 | D395C | A | 2.4e-06 | 1.8e-06 | 4e-06 | 2.733E-06 | 1.137E-06 | 1.800E-06 | 4.000E-06 |
| 7886 | 395 | D395E | A | 1.7e-06 | 4.1e-06 | 2.7e-06 | 2.833E-06 | 1.206E-06 | 1.700E-06 | 4.100E-06 |
| 7887 | 395 | D395Q | A | 4.2e-06 | 2.7e-06 | 3.6e-06 | 3.500E-06 | 7.550E-07 | 2.700E-06 | 4.200E-06 |
| 7888 | 395 | D395G | A | 2.5e-06 | 2.6e-06 | 2.6e-06 | 2.567E-06 | 5.774E-08 | 2.500E-06 | 2.600E-06 |
| 7889 | 395 | D395H | A | 2.3e-06 | 2.9e-06 | 1.6e-06 | 2.267E-06 | 6.506E-07 | 1.600E-06 | 2.900E-06 |
| 7890 | 395 | D395I | A | 1.7e-06 | 3.4e-06 | 2.4e-06 | 2.500E-06 | 8.544E-07 | 1.700E-06 | 3.400E-06 |
| 7891 | 395 | D395L | A | 2.1e-06 | 2.4e-06 | 2.8e-06 | 2.433E-06 | 3.512E-07 | 2.100E-06 | 2.800E-06 |
| 7892 | 395 | D395K | A | 2.5e-06 | 2.8e-06 | 2.3e-06 | 2.533E-06 | 2.517E-07 | 2.300E-06 | 2.800E-06 |
| 7893 | 395 | D395M | A | 2.3e-06 | 3.2e-06 | 2.2e-06 | 2.567E-06 | 5.508E-07 | 2.200E-06 | 3.200E-06 |
| 7894 | 395 | D395F | A | 5.1e-06 | 3.2e-06 | 3.3e-06 | 3.867E-06 | 1.069E-06 | 3.200E-06 | 5.100E-06 |
| 7895 | 395 | D395P | A | 3.6e-06 | 3.1e-06 | 1.8e-06 | 2.833E-06 | 9.292E-07 | 1.800E-06 | 3.600E-06 |
| 7896 | 395 | D395S | A | 2.7e-06 | 3.3e-06 | 1.5e-06 | 2.500E-06 | 9.165E-07 | 1.500E-06 | 3.300E-06 |
| 7897 | 395 | D395T | A | 2.4e-06 | 3.8e-06 | 1.8e-06 | 2.667E-06 | 1.026E-06 | 1.800E-06 | 3.800E-06 |
| 7898 | 395 | D395W | A | 2.1e-06 | 2.1e-06 | 1.4e-06 | 1.867E-06 | 4.041E-07 | 1.400E-06 | 2.100E-06 |
| 7899 | 395 | D395Y | A | 4e-06 | 2e-06 | 1.7e-06 | 2.567E-06 | 1.250E-06 | 1.700E-06 | 4.000E-06 |
| 7900 | 395 | D395V | A | 4.8e-06 | 2.4e-06 | 3.4e-06 | 3.533E-06 | 1.206E-06 | 2.400E-06 | 4.800E-06 |
| 7901 | 396 | S396A | A | 2e-06 | 3.1e-06 | 3.1e-06 | 2.733E-06 | 6.351E-07 | 2.000E-06 | 3.100E-06 |
| 7902 | 396 | S396R | A | 4.3e-06 | 1.7e-06 | 2.2e-06 | 2.733E-06 | 1.380E-06 | 1.700E-06 | 4.300E-06 |
| 7903 | 396 | S396N | A | 2.2e-06 | 3.9e-06 | 2.7e-06 | 2.933E-06 | 8.737E-07 | 2.200E-06 | 3.900E-06 |
| 7904 | 396 | S396D | A | 2.7e-06 | 2.5e-06 | 1.5e-06 | 2.233E-06 | 6.429E-07 | 1.500E-06 | 2.700E-06 |

| | | | | | | | | | | |
|---|---|---|---|---|---|---|---|---|---|---|
| 7905 | 396 | S396C | A | 1.9e-06 | 2.2e-06 | 3e-06 | 2.367E-06 | 5.686E-07 | 1.900E-06 | 3.000E-06 |
| 7906 | 396 | S396E | A | 4.4e-06 | 1.6e-06 | 2e-06 | 2.667E-06 | 1.514E-06 | 1.600E-06 | 4.400E-06 |
| 7907 | 396 | S396Q | A | 1.8e-06 | 2.7e-06 | 2.4e-06 | 2.300E-06 | 4.583E-07 | 1.800E-06 | 2.700E-06 |
| 7908 | 396 | S396G | A | 3e-06 | 2.7e-06 | 2.8e-06 | 2.833E-06 | 1.528E-07 | 2.700E-06 | 3.000E-06 |
| 7909 | 396 | S396H | A | 3.4e-06 | 1.6e-06 | 2.4e-06 | 2.467E-06 | 9.018E-07 | 1.600E-06 | 3.400E-06 |
| 7910 | 396 | S396I | A | 2.2e-06 | 3.1e-06 | 2.6e-06 | 2.633E-06 | 4.509E-07 | 2.200E-06 | 3.100E-06 |
| 7911 | 396 | S396L | A | 1.8e-06 | 2.4e-06 | 1.7e-06 | 1.967E-06 | 3.786E-07 | 1.700E-06 | 2.400E-06 |
| 7912 | 396 | S396K | A | 2.3e-06 | 3e-06 | 2.3e-06 | 2.533E-06 | 4.041E-07 | 2.300E-06 | 3.000E-06 |
| 7913 | 396 | S396M | A | 2.4e-06 | 1.9e-06 | 2.8e-06 | 2.367E-06 | 4.509E-07 | 1.900E-06 | 2.800E-06 |
| 7914 | 396 | S396F | A | 1.8e-06 | 1.8e-06 | 2.2e-06 | 1.933E-06 | 2.309E-07 | 1.800E-06 | 2.200E-06 |
| 7915 | 396 | S396P | A | 3.4e-06 | 2.3e-06 | 2e-06 | 2.567E-06 | 7.371E-07 | 2.000E-06 | 3.400E-06 |
| 7916 | 396 | S396S | A | 3.7e-06 | 2.2e-06 | 3.7e-06 | 3.200E-06 | 8.660E-07 | 2.200E-06 | 3.700E-06 |
| 7917 | 396 | S396T | A | 3.3e-06 | 2.2e-06 | 1.1e-06 | 2.200E-06 | 1.100E-06 | 1.100E-06 | 3.300E-06 |
| 7918 | 396 | S396W | A | 2.1e-06 | 2.8e-06 | 1.1e-06 | 2.000E-06 | 8.544E-07 | 1.100E-06 | 2.800E-06 |
| 7919 | 396 | S396Y | A | 1.9e-06 | 1.8e-06 | 2e-06 | 1.900E-06 | 1.000E-07 | 1.800E-06 | 2.000E-06 |
| 7920 | 396 | S396V | A | 3.8e-06 | 2.2e-06 | 2.8e-06 | 2.933E-06 | 8.083E-07 | 2.200E-06 | 3.800E-06 |
| 7921 | 397 | L397A | A | 3.4e-06 | 2.7e-06 | 2.7e-06 | 2.933E-06 | 4.041E-07 | 2.700E-06 | 3.400E-06 |
| 7922 | 397 | L397R | A | 3e-06 | 3.6e-06 | 2.3e-06 | 2.967E-06 | 6.506E-07 | 2.300E-06 | 3.600E-06 |
| 7923 | 397 | L397N | A | 3.1e-06 | 3.7e-06 | 2.4e-06 | 3.067E-06 | 6.506E-07 | 2.400E-06 | 3.700E-06 |
| 7924 | 397 | L397D | A | 3.5e-06 | 4.5e-06 | 1.8e-06 | 3.267E-06 | 1.365E-06 | 1.800E-06 | 4.500E-06 |
| 7925 | 397 | L397C | A | 1.9e-06 | 1.5e-06 | 2.3e-06 | 1.900E-06 | 4.000E-07 | 1.500E-06 | 2.300E-06 |
| 7926 | 397 | L397E | A | 2.5e-06 | 1.9e-06 | 1.6e-06 | 2.000E-06 | 4.583E-07 | 1.600E-06 | 2.500E-06 |
| 7927 | 397 | L397Q | A | 2.1e-06 | 2.2e-06 | 2.3e-06 | 2.200E-06 | 1.000E-07 | 2.100E-06 | 2.300E-06 |
| 7928 | 397 | L397G | A | 1.7e-06 | 4e-06 | 2.7e-06 | 2.800E-06 | 1.153E-06 | 1.700E-06 | 4.000E-06 |
| 7929 | 397 | L397H | A | 2.1e-06 | 7.7e-07 | 2e-06 | 1.623E-06 | 7.407E-07 | 7.700E-07 | 2.100E-06 |
| 7930 | 397 | L397I | A | 2.2e-06 | 2.7e-06 | 2.2e-06 | 2.367E-06 | 2.887E-07 | 2.200E-06 | 2.700E-06 |
| 7931 | 397 | L397L | A | 3e-06 | 1.9e-06 | 2.6e-06 | 2.500E-06 | 5.568E-07 | 1.900E-06 | 3.000E-06 |
| 7932 | 397 | L397K | A | 2.8e-06 | 1.7e-06 | 2.6e-06 | 2.367E-06 | 5.859E-07 | 1.700E-06 | 2.800E-06 |
| 7933 | 397 | L397M | A | 3e-06 | 1.3e-06 | 3e-06 | 2.433E-06 | 9.815E-07 | 1.300E-06 | 3.000E-06 |
| 7934 | 397 | L397F | A | 2e-06 | 1.2e-06 | 1.9e-06 | 1.700E-06 | 4.359E-07 | 1.200E-06 | 2.000E-06 |
| 7935 | 397 | L397P | A | 2.5e-06 | 2.6e-06 | 3.8e-06 | 2.967E-06 | 7.234E-07 | 2.500E-06 | 3.800E-06 |
| 7936 | 397 | L397S | A | 3.6e-06 | 3.1e-06 | 2.1e-06 | 2.933E-06 | 7.638E-07 | 2.100E-06 | 3.600E-06 |
| 7937 | 397 | L397T | A | 2.6e-06 | 4.7e-06 | 3.6e-06 | 3.633E-06 | 1.050E-06 | 2.600E-06 | 4.700E-06 |
| 7938 | 397 | L397W | A | 2.3e-06 | 2.5e-06 | 4.1e-06 | 2.967E-06 | 9.866E-07 | 2.300E-06 | 4.100E-06 |
| 7939 | 397 | L397Y | A | 1.5e-06 | 2.1e-06 | 3.4e-06 | 2.333E-06 | 9.713E-07 | 1.500E-06 | 3.400E-06 |
| 7940 | 397 | L397V | A | 2.2e-06 | 3e-06 | 3.6e-06 | 2.933E-06 | 7.024E-07 | 2.200E-06 | 3.600E-06 |
| 7941 | 398 | P398A | A | 1.6e-06 | 1.2e-06 | 3.4e-06 | 2.067E-06 | 1.172E-06 | 1.200E-06 | 3.400E-06 |
| 7942 | 398 | P398R | A | 2.4e-06 | 3e-06 | 4.5e-06 | 3.300E-06 | 1.082E-06 | 2.400E-06 | 4.500E-06 |
| 7943 | 398 | P398N | A | 2e-06 | 1.2e-06 | 2.1e-06 | 1.767E-06 | 4.933E-07 | 1.200E-06 | 2.100E-06 |
| 7944 | 398 | P398D | A | 2.5e-06 | 2.3e-06 | 3.2e-06 | 2.667E-06 | 4.726E-07 | 2.300E-06 | 3.200E-06 |
| 7945 | 398 | P398C | A | 5.7e-06 | 1.8e-06 | 4.3e-06 | 3.933E-06 | 1.976E-06 | 1.800E-06 | 5.700E-06 |
| 7946 | 398 | P398E | A | 2.2e-06 | 1.2e-06 | 9.5e-07 | 1.450E-06 | 6.614E-07 | 9.500E-07 | 2.200E-06 |
| 7947 | 398 | P398Q | A | 2.4e-06 | 1.4e-06 | 2.4e-06 | 2.067E-06 | 5.774E-07 | 1.400E-06 | 2.400E-06 |
| 7948 | 398 | P398G | A | 2.4e-06 | 2e-06 | 1.9e-06 | 2.100E-06 | 2.646E-07 | 1.900E-06 | 2.400E-06 |
| 7949 | 398 | P398H | A | 2.6e-06 | 2e-06 | 3e-06 | 2.533E-06 | 5.033E-07 | 2.000E-06 | 3.000E-06 |
| 7950 | 398 | P398I | A | 1.5e-06 | 2.2e-06 | 1e-06 | 1.567E-06 | 6.028E-07 | 1.000E-06 | 2.200E-06 |
| 7951 | 398 | P398L | A | 2.4e-06 | 2.1e-06 | 3e-06 | 2.500E-06 | 4.583E-07 | 2.100E-06 | 3.000E-06 |
| 7952 | 398 | P398K | A | 1.4e-06 | 2.1e-06 | 2e-06 | 1.833E-06 | 3.786E-07 | 1.400E-06 | 2.100E-06 |
| 7953 | 398 | P398M | A | 2.8e-06 | 1.7e-06 | 3.2e-06 | 2.567E-06 | 7.767E-07 | 1.700E-06 | 3.200E-06 |
| 7954 | 398 | P398F | A | 2.9e-06 | 2.2e-06 | 2.3e-06 | 2.467E-06 | 3.786E-07 | 2.200E-06 | 2.900E-06 |
| 7955 | 398 | P398P | A | 1.8e-06 | 2.9e-06 | 6e-06 | 3.567E-06 | 2.178E-06 | 1.800E-06 | 6.000E-06 |
| 7956 | 398 | P398S | A | 2.8e-06 | 2.1e-06 | 1.9e-06 | 2.267E-06 | 4.726E-07 | 1.900E-06 | 2.800E-06 |
| 7957 | 398 | P398T | A | 2.4e-06 | 3.9e-06 | 1.3e-06 | 2.533E-06 | 1.305E-06 | 1.300E-06 | 3.900E-06 |
| 7958 | 398 | P398W | A | 2.8e-06 | 2.1e-06 | 3e-06 | 2.633E-06 | 4.726E-07 | 2.100E-06 | 3.000E-06 |
| 7959 | 398 | P398Y | A | 1.9e-06 | 2.7e-06 | 1.8e-06 | 2.133E-06 | 4.933E-07 | 1.800E-06 | 2.700E-06 |
| 7960 | 398 | P398V | A | 2.2e-06 | 2.9e-06 | 3e-06 | 2.700E-06 | 4.359E-07 | 2.200E-06 | 3.000E-06 |
| 7961 | 399 | D399A | A | 2e-06 | 2.8e-06 | 1.7e-06 | 2.167E-06 | 5.686E-07 | 1.700E-06 | 2.800E-06 |
| 7962 | 399 | D399R | A | 1.7e-06 | 1.9e-06 | 2e-06 | 1.867E-06 | 1.528E-07 | 1.700E-06 | 2.000E-06 |
| 7963 | 399 | D399N | A | 1.7e-06 | 3e-06 | 2.4e-06 | 2.367E-06 | 6.506E-07 | 1.700E-06 | 3.000E-06 |
| 7964 | 399 | D399D | A | 2.9e-06 | 2.7e-06 | 1.7e-06 | 2.433E-06 | 6.429E-07 | 1.700E-06 | 2.900E-06 |
| 7965 | 399 | D399C | A | 1.6e-06 | 1.5e-06 | 1.5e-06 | 1.533E-06 | 5.774E-08 | 1.500E-06 | 1.600E-06 |
| 7966 | 399 | D399E | A | 5.5e-06 | 3.7e-06 | 1.2e-06 | 3.467E-06 | 2.159E-06 | 1.200E-06 | 5.500E-06 |
| 7967 | 399 | D399Q | A | 1.6e-06 | 4.2e-06 | 1.8e-06 | 2.533E-06 | 1.447E-06 | 1.600E-06 | 4.200E-06 |
| 7968 | 399 | D399G | A | 1.4e-06 | 3e-06 | 2.8e-06 | 2.400E-06 | 8.718E-07 | 1.400E-06 | 3.000E-06 |
| 7969 | 399 | D399H | A | 2.4e-06 | 1.6e-06 | 3e-06 | 2.333E-06 | 7.024E-07 | 1.600E-06 | 3.000E-06 |
| 7970 | 399 | D399I | A | 2.3e-06 | 2.2e-06 | 1.9e-06 | 2.133E-06 | 2.082E-07 | 1.900E-06 | 2.300E-06 |
| 7971 | 399 | D399L | A | 2.3e-06 | 3.2e-06 | 2.2e-06 | 2.567E-06 | 5.508E-07 | 2.200E-06 | 3.200E-06 |
| 7972 | 399 | D399K | A | 2.8e-06 | 2.3e-06 | 1.9e-06 | 2.333E-06 | 4.509E-07 | 1.900E-06 | 2.800E-06 |
| 7973 | 399 | D399M | A | 2.5e-06 | 2.2e-06 | 1.6e-06 | 2.100E-06 | 4.583E-07 | 1.600E-06 | 2.500E-06 |
| 7974 | 399 | D399F | A | 2.5e-06 | 2.3e-06 | 1e-06 | 1.933E-06 | 8.145E-07 | 1.000E-06 | 2.500E-06 |
| 7975 | 399 | D399P | A | 2.7e-06 | 9.3e-07 | 2e-06 | 1.877E-06 | 8.914E-07 | 9.300E-07 | 2.700E-06 |
| 7976 | 399 | D399S | A | 2.6e-06 | 2.2e-06 | 3.4e-06 | 2.733E-06 | 6.110E-07 | 2.200E-06 | 3.400E-06 |
| 7977 | 399 | D399T | A | 2.9e-06 | 2.2e-06 | 2e-06 | 2.367E-06 | 4.726E-07 | 2.000E-06 | 2.900E-06 |
| 7978 | 399 | D399W | A | 1.8e-06 | 2.3e-06 | 2.7e-06 | 2.267E-06 | 4.509E-07 | 1.800E-06 | 2.700E-06 |
| 7979 | 399 | D399Y | A | 3.1e-06 | 2.9e-06 | 2.5e-06 | 2.833E-06 | 3.055E-07 | 2.500E-06 | 3.100E-06 |
| 7980 | 399 | D399V | A | 3e-06 | 2.8e-06 | 4.8e-06 | 3.533E-06 | 1.102E-06 | 2.800E-06 | 4.800E-06 |
| 7981 | 400 | L400A | A | 4.1e-06 | 2.1e-06 | 2.3e-06 | 2.833E-06 | 1.102E-06 | 2.100E-06 | 4.100E-06 |
| 7982 | 400 | L400R | A | 9.9e-07 | 1.5e-06 | 2.2e-06 | 1.563E-06 | 6.075E-07 | 9.900E-07 | 2.200E-06 |
| 7983 | 400 | L400N | A | 2.6e-06 | 2.3e-06 | 2.8e-06 | 2.567E-06 | 2.517E-07 | 2.300E-06 | 2.800E-06 |
| 7984 | 400 | L400D | A | 3.7e-06 | 4.4e-06 | 2.7e-06 | 3.600E-06 | 8.544E-07 | 2.700E-06 | 4.400E-06 |
| 7985 | 400 | L400C | A | 1.7e-06 | 3.1e-06 | 2.4e-06 | 2.400E-06 | 7.000E-07 | 1.700E-06 | 3.100E-06 |
| 7986 | 400 | L400E | A | 2.4e-06 | 2.6e-06 | 2.5e-06 | 2.500E-06 | 1.000E-07 | 2.400E-06 | 2.600E-06 |
| 7987 | 400 | L400Q | A | 1.8e-06 | 3e-06 | 2.5e-06 | 2.433E-06 | 6.028E-07 | 1.800E-06 | 3.000E-06 |
| 7988 | 400 | L400G | A | 3.7e-06 | 2.6e-06 | 4.5e-06 | 3.600E-06 | 9.539E-07 | 2.600E-06 | 4.500E-06 |
| 7989 | 400 | L400H | A | 2.1e-06 | 2.2e-06 | 3.6e-06 | 2.633E-06 | 8.386E-07 | 2.100E-06 | 3.600E-06 |

| | | | | | | | | | | |
|---|---|---|---|---|---|---|---|---|---|---|
| 7990 | 400 | L400I | A | 4.7e-06 | 1.7e-06 | 2e-06 | 2.800E-06 | 1.652E-06 | 1.700E-06 | 4.700E-06 |
| 7991 | 400 | L400L | A | 1.4e-06 | 2.9e-06 | 2e-06 | 2.100E-06 | 7.550E-07 | 1.400E-06 | 2.900E-06 |
| 7992 | 400 | L400K | A | 1.3e-06 | 1.3e-06 | 1.1e-06 | 1.233E-06 | 1.155E-07 | 1.100E-06 | 1.300E-06 |
| 7993 | 400 | L400M | A | 3.2e-06 | 2.6e-06 | 3.1e-06 | 2.967E-06 | 3.215E-07 | 2.600E-06 | 3.200E-06 |
| 7994 | 400 | L400F | A | 2.2e-06 | 2.3e-06 | 1.7e-06 | 2.067E-06 | 3.215E-07 | 1.700E-06 | 2.300E-06 |
| 7995 | 400 | L400P | A | 2.6e-06 | 2.9e-06 | 2.2e-06 | 2.567E-06 | 3.512E-07 | 2.200E-06 | 2.900E-06 |
| 7996 | 400 | L400S | A | 2.5e-06 | 3.6e-06 | 1.9e-06 | 2.667E-06 | 8.622E-07 | 1.900E-06 | 3.600E-06 |
| 7997 | 400 | L400T | A | 2.1e-06 | 5.3e-06 | 2.9e-06 | 3.433E-06 | 1.665E-06 | 2.100E-06 | 5.300E-06 |
| 7998 | 400 | L400W | A | 2.5e-06 | 2.1e-06 | 1.2e-06 | 1.933E-06 | 6.658E-07 | 1.200E-06 | 2.500E-06 |
| 7999 | 400 | L400Y | A | 2.4e-06 | 1.6e-06 | 2.9e-06 | 2.300E-06 | 6.557E-07 | 1.600E-06 | 2.900E-06 |
| 8000 | 400 | L400V | A | 2e-06 | 5.1e-06 | 1e-06 | 2.700E-06 | 2.138E-06 | 1.000E-06 | 5.100E-06 |
| 8001 | 401 | S401A | A | 4.3e-06 | 2.8e-06 | 1.6e-06 | 2.900E-06 | 1.353E-06 | 1.600E-06 | 4.300E-06 |
| 8002 | 401 | S401R | A | 4.3e-06 | 4.3e-06 | 4.5e-06 | 4.367E-06 | 1.155E-07 | 4.300E-06 | 4.500E-06 |
| 8003 | 401 | S401N | A | 1.7e-06 | 2.5e-06 | 1.4e-06 | 1.867E-06 | 5.686E-07 | 1.400E-06 | 2.500E-06 |
| 8004 | 401 | S401D | A | 4.7e-06 | 2.6e-06 | 2e-06 | 3.100E-06 | 1.418E-06 | 2.000E-06 | 4.700E-06 |
| 8005 | 401 | S401C | A | 3.1e-06 | 2.7e-06 | 1.9e-06 | 2.567E-06 | 6.110E-07 | 1.900E-06 | 3.100E-06 |
| 8006 | 401 | S401E | A | 3.4e-06 | 1.6e-06 | 3.3e-06 | 2.767E-06 | 1.012E-06 | 1.600E-06 | 3.400E-06 |
| 8007 | 401 | S401Q | A | 2.8e-06 | 2.4e-06 | 2e-06 | 2.400E-06 | 4.000E-07 | 2.000E-06 | 2.800E-06 |
| 8008 | 401 | S401G | A | 2.3e-06 | 3.7e-06 | 3.2e-06 | 3.067E-06 | 7.095E-07 | 2.300E-06 | 3.700E-06 |
| 8009 | 401 | S401H | A | 1.6e-06 | 2.3e-06 | 4e-06 | 2.633E-06 | 1.234E-06 | 1.600E-06 | 4.000E-06 |
| 8010 | 401 | S401I | A | 2.6e-06 | 1.8e-06 | 1.4e-06 | 1.933E-06 | 6.110E-07 | 1.400E-06 | 2.600E-06 |
| 8011 | 401 | S401L | A | 3.3e-06 | 2.6e-06 | 2.7e-06 | 2.867E-06 | 3.786E-07 | 2.600E-06 | 3.300E-06 |
| 8012 | 401 | S401K | A | 1.5e-06 | 2.9e-06 | 2.9e-06 | 2.433E-06 | 8.083E-07 | 1.500E-06 | 2.900E-06 |
| 8013 | 401 | S401M | A | 2.4e-06 | 1.9e-06 | 2.2e-06 | 2.167E-06 | 2.517E-07 | 1.900E-06 | 2.400E-06 |
| 8014 | 401 | S401F | A | 1.5e-06 | 1.6e-06 | 2.2e-06 | 1.767E-06 | 3.786E-07 | 1.500E-06 | 2.200E-06 |
| 8015 | 401 | S401P | A | 3.9e-06 | 2.2e-06 | 2.4e-06 | 2.833E-06 | 9.292E-07 | 2.200E-06 | 3.900E-06 |
| 8016 | 401 | S401S | A | 3.6e-06 | 3.7e-06 | 2.7e-06 | 3.333E-06 | 5.508E-07 | 2.700E-06 | 3.700E-06 |
| 8017 | 401 | S401T | A | 1.7e-06 | 2.7e-06 | 3.3e-06 | 2.567E-06 | 8.083E-07 | 1.700E-06 | 3.300E-06 |
| 8018 | 401 | S401W | A | 2.1e-06 | 1.4e-06 | 1.9e-06 | 1.800E-06 | 3.606E-07 | 1.400E-06 | 2.100E-06 |
| 8019 | 401 | S401Y | A | 2.2e-06 | 2.7e-06 | 2.1e-06 | 2.333E-06 | 3.215E-07 | 2.100E-06 | 2.700E-06 |
| 8020 | 401 | S401V | A | 1.9e-06 | 1.5e-06 | 2.7e-06 | 2.033E-06 | 6.110E-07 | 1.500E-06 | 2.700E-06 |
| 8021 | 402 | V402A | A | 2.6e-06 | 4.9e-06 | 1.1e-06 | 2.867E-06 | 1.914E-06 | 1.100E-06 | 4.900E-06 |
| 8022 | 402 | V402R | A | 3e-06 | 2.3e-06 | 4.8e-06 | 3.367E-06 | 1.290E-06 | 2.300E-06 | 4.800E-06 |
| 8023 | 402 | V402N | A | 1.9e-06 | 2.2e-06 | 4.7e-06 | 2.933E-06 | 1.537E-06 | 1.900E-06 | 4.700E-06 |
| 8024 | 402 | V402D | A | 2.3e-06 | 4e-06 | 1.4e-06 | 2.567E-06 | 1.320E-06 | 1.400E-06 | 4.000E-06 |
| 8025 | 402 | V402C | A | 2.4e-06 | 1.3e-06 | 2e-06 | 1.900E-06 | 5.568E-07 | 1.300E-06 | 2.400E-06 |
| 8026 | 402 | V402E | A | 3.3e-06 | 2.5e-06 | 4.9e-06 | 3.567E-06 | 1.222E-06 | 2.500E-06 | 4.900E-06 |
| 8027 | 402 | V402Q | A | 2.9e-06 | 5.2e-06 | 3.1e-06 | 3.733E-06 | 1.274E-06 | 2.900E-06 | 5.200E-06 |
| 8028 | 402 | V402G | A | 2.8e-06 | 3.9e-06 | 5.2e-06 | 3.967E-06 | 1.201E-06 | 2.800E-06 | 5.200E-06 |
| 8029 | 402 | V402H | A | 2.7e-06 | 3.1e-06 | 3.2e-06 | 3.000E-06 | 2.646E-07 | 2.700E-06 | 3.200E-06 |
| 8030 | 402 | V402I | A | 3.1e-06 | 3.4e-06 | 1.7e-06 | 2.733E-06 | 9.074E-07 | 1.700E-06 | 3.400E-06 |
| 8031 | 402 | V402L | A | 2.7e-06 | 2.3e-06 | 1.6e-06 | 2.200E-06 | 5.568E-07 | 1.600E-06 | 2.700E-06 |
| 8032 | 402 | V402K | A | 2.3e-06 | 2.4e-06 | 1.9e-06 | 2.200E-06 | 2.646E-07 | 1.900E-06 | 2.400E-06 |
| 8033 | 402 | V402M | A | 1.7e-06 | 4.2e-06 | 1.6e-06 | 2.500E-06 | 1.473E-06 | 1.600E-06 | 4.200E-06 |
| 8034 | 402 | V402F | A | 2.8e-06 | 4.7e-06 | 3.2e-06 | 3.567E-06 | 1.002E-06 | 2.800E-06 | 4.700E-06 |
| 8035 | 402 | V402P | A | 2.8e-06 | 2.1e-06 | 1.5e-06 | 2.133E-06 | 6.506E-07 | 1.500E-06 | 2.800E-06 |
| 8036 | 402 | V402S | A | 1.7e-06 | 1.6e-06 | 1.8e-06 | 1.700E-06 | 1.000E-07 | 1.600E-06 | 1.800E-06 |
| 8037 | 402 | V402T | A | 2.1e-06 | 3.3e-06 | 2.9e-06 | 2.767E-06 | 6.110E-07 | 2.100E-06 | 3.300E-06 |
| 8038 | 402 | V402W | A | 1.7e-06 | 1.3e-06 | 3.3e-06 | 2.100E-06 | 1.058E-06 | 1.300E-06 | 3.300E-06 |
| 8039 | 402 | V402Y | A | 2.4e-06 | 2.7e-06 | 1.4e-06 | 2.167E-06 | 6.807E-07 | 1.400E-06 | 2.700E-06 |
| 8040 | 402 | V402V | A | 2.3e-06 | 1.7e-06 | 7.5e-07 | 1.583E-06 | 7.816E-07 | 7.500E-07 | 2.300E-06 |
| 8041 | 403 | F403A | A | 5e-06 | 2.3e-06 | 2.3e-06 | 3.200E-06 | 1.559E-06 | 2.300E-06 | 5.000E-06 |
| 8042 | 403 | F403R | A | 2.2e-06 | 2.4e-06 | 1.9e-06 | 2.167E-06 | 2.517E-07 | 1.900E-06 | 2.400E-06 |
| 8043 | 403 | F403N | A | 3.4e-06 | 3.4e-06 | 2.3e-06 | 3.033E-06 | 6.351E-07 | 2.300E-06 | 3.400E-06 |
| 8044 | 403 | F403D | A | 2.4e-06 | 2.1e-06 | 3.3e-06 | 2.600E-06 | 6.245E-07 | 2.100E-06 | 3.300E-06 |
| 8045 | 403 | F403C | A | 3.8e-06 | 1.9e-06 | 2.1e-06 | 2.600E-06 | 1.044E-06 | 1.900E-06 | 3.800E-06 |
| 8046 | 403 | F403E | A | 4e-06 | 2.4e-06 | 2.4e-06 | 2.933E-06 | 9.238E-07 | 2.400E-06 | 4.000E-06 |
| 8047 | 403 | F403Q | A | 2.5e-06 | 1.6e-06 | 2.9e-06 | 2.333E-06 | 6.658E-07 | 1.600E-06 | 2.900E-06 |
| 8048 | 403 | F403G | A | 2.9e-06 | 1.8e-06 | 3.4e-06 | 2.700E-06 | 8.185E-07 | 1.800E-06 | 3.400E-06 |
| 8049 | 403 | F403H | A | 1.7e-06 | 3.8e-06 | 2e-06 | 2.500E-06 | 1.136E-06 | 1.700E-06 | 3.800E-06 |
| 8050 | 403 | F403I | A | 2.4e-06 | 1.6e-06 | 2.9e-06 | 2.300E-06 | 6.557E-07 | 1.600E-06 | 2.900E-06 |
| 8051 | 403 | F403L | A | 3e-06 | 2.6e-06 | 3.2e-06 | 2.933E-06 | 3.055E-07 | 2.600E-06 | 3.200E-06 |
| 8052 | 403 | F403K | A | 2.6e-06 | 2.9e-06 | 3.2e-06 | 2.900E-06 | 3.000E-07 | 2.600E-06 | 3.200E-06 |
| 8053 | 403 | F403M | A | 1.8e-06 | 1.7e-06 | 2.6e-06 | 2.033E-06 | 4.933E-07 | 1.700E-06 | 2.600E-06 |
| 8054 | 403 | F403F | A | 3.1e-06 | 2e-06 | 3e-06 | 2.700E-06 | 6.083E-07 | 2.000E-06 | 3.100E-06 |
| 8055 | 403 | F403P | A | 3e-06 | 2.8e-06 | 3.4e-06 | 3.067E-06 | 3.055E-07 | 2.800E-06 | 3.400E-06 |
| 8056 | 403 | F403S | A | 2.6e-06 | 4.9e-06 | 2.3e-06 | 3.267E-06 | 1.422E-06 | 2.300E-06 | 4.900E-06 |
| 8057 | 403 | F403T | A | 4e-06 | 2.2e-06 | 2.5e-06 | 2.900E-06 | 9.644E-07 | 2.200E-06 | 4.000E-06 |
| 8058 | 403 | F403W | A | 2.5e-06 | 1.4e-06 | 2.9e-06 | 2.267E-06 | 7.767E-07 | 1.400E-06 | 2.900E-06 |
| 8059 | 403 | F403Y | A | 2.9e-06 | 3.2e-06 | 4.4e-06 | 3.500E-06 | 7.937E-07 | 2.900E-06 | 4.400E-06 |
| 8060 | 403 | F403V | A | 1.5e-06 | 2.8e-06 | 3.1e-06 | 2.467E-06 | 8.505E-07 | 1.500E-06 | 3.100E-06 |
| 8061 | 404 | Q404A | A | 4.4e-06 | 2.9e-06 | 3.3e-06 | 3.533E-06 | 7.767E-07 | 2.900E-06 | 4.400E-06 |
| 8062 | 404 | Q404R | A | 3.8e-06 | 1.7e-06 | 1.9e-06 | 2.467E-06 | 1.159E-06 | 1.700E-06 | 3.800E-06 |
| 8063 | 404 | Q404N | A | 1.6e-06 | 2.7e-06 | 1.7e-06 | 2.000E-06 | 6.083E-07 | 1.600E-06 | 2.700E-06 |
| 8064 | 404 | Q404D | A | 1.9e-06 | 2.1e-06 | 2.4e-06 | 2.133E-06 | 2.517E-07 | 1.900E-06 | 2.400E-06 |
| 8065 | 404 | Q404C | A | 2.5e-06 | 2.3e-06 | 4.2e-06 | 3.000E-06 | 1.044E-06 | 2.300E-06 | 4.200E-06 |
| 8066 | 404 | Q404E | A | 2.3e-06 | 2.5e-06 | 2.3e-06 | 2.367E-06 | 1.155E-07 | 2.300E-06 | 2.500E-06 |
| 8067 | 404 | Q404Q | A | 1.5e-06 | 2.4e-06 | 2.8e-06 | 2.233E-06 | 6.658E-07 | 1.500E-06 | 2.800E-06 |
| 8068 | 404 | Q404G | A | 2.3e-06 | 2.6e-06 | 3.8e-06 | 2.900E-06 | 7.937E-07 | 2.300E-06 | 3.800E-06 |
| 8069 | 404 | Q404H | A | 2.7e-06 | 1.1e-06 | 1.4e-06 | 1.733E-06 | 8.505E-07 | 1.100E-06 | 2.700E-06 |
| 8070 | 404 | Q404I | A | 3.2e-06 | 2.1e-06 | 2e-06 | 2.433E-06 | 6.658E-07 | 2.000E-06 | 3.200E-06 |
| 8071 | 404 | Q404L | A | 3.8e-06 | 2.9e-06 | 3.5e-06 | 3.400E-06 | 4.583E-07 | 2.900E-06 | 3.800E-06 |
| 8072 | 404 | Q404K | A | 2e-06 | 2.7e-06 | 2e-06 | 2.233E-06 | 4.041E-07 | 2.000E-06 | 2.700E-06 |
| 8073 | 404 | Q404M | A | 3.5e-06 | 2.5e-06 | 1.8e-06 | 2.600E-06 | 8.544E-07 | 1.800E-06 | 3.500E-06 |
| 8074 | 404 | Q404F | A | 2.6e-06 | 3.8e-06 | 2.7e-06 | 3.033E-06 | 6.658E-07 | 2.600E-06 | 3.800E-06 |

| | | | | | | | | | | |
|---|---|---|---|---|---|---|---|---|---|---|
| 8075 | 404 | Q404P | A | 2.7e-06 | 2.8e-06 | 1.8e-06 | 2.433E-06 | 5.508E-07 | 1.800E-06 | 2.800E-06 |
| 8076 | 404 | Q404S | A | 3.2e-06 | 2.4e-06 | 2.8e-06 | 2.800E-06 | 4.000E-07 | 2.400E-06 | 3.200E-06 |
| 8077 | 404 | Q404T | A | 9.2e-07 | 1.9e-06 | 3.3e-06 | 2.040E-06 | 1.196E-06 | 9.200E-07 | 3.300E-06 |
| 8078 | 404 | Q404W | A | 2e-06 | 1.5e-06 | 1.4e-06 | 1.633E-06 | 3.215E-07 | 1.400E-06 | 2.000E-06 |
| 8079 | 404 | Q404Y | A | 2.1e-06 | 2e-06 | 2.2e-06 | 2.100E-06 | 1.000E-07 | 2.000E-06 | 2.200E-06 |
| 8080 | 404 | Q404V | A | 1.9e-06 | 2.2e-06 | 1.4e-06 | 1.833E-06 | 4.041E-07 | 1.400E-06 | 2.200E-06 |
| 8081 | 405 | N405A | A | 1.4e-06 | 2e-06 | 2e-06 | 1.800E-06 | 3.464E-07 | 1.400E-06 | 2.000E-06 |
| 8082 | 405 | N405R | A | 1.6e-06 | 2.8e-06 | 1.6e-06 | 2.000E-06 | 6.928E-07 | 1.600E-06 | 2.800E-06 |
| 8083 | 405 | N405N | A | 3.2e-06 | 3.6e-06 | 5e-06 | 3.933E-06 | 9.452E-07 | 3.200E-06 | 5.000E-06 |
| 8084 | 405 | N405D | A | 2.5e-06 | 2.6e-06 | 2.4e-06 | 2.500E-06 | 1.000E-07 | 2.400E-06 | 2.600E-06 |
| 8085 | 405 | N405C | A | 4.4e-06 | 2.3e-06 | 3.7e-06 | 3.467E-06 | 1.069E-06 | 2.300E-06 | 4.400E-06 |
| 8086 | 405 | N405E | A | 2.2e-06 | 8.3e-07 | 2.7e-06 | 1.910E-06 | 9.681E-07 | 8.300E-07 | 2.700E-06 |
| 8087 | 405 | N405Q | A | 2e-06 | 1.1e-06 | 2.9e-06 | 2.000E-06 | 9.000E-07 | 1.100E-06 | 2.900E-06 |
| 8088 | 405 | N405G | A | 2.4e-06 | 3e-06 | 2e-06 | 2.467E-06 | 5.033E-07 | 2.000E-06 | 3.000E-06 |
| 8089 | 405 | N405H | A | 1.8e-06 | 4.3e-06 | 4.8e-06 | 3.633E-06 | 1.607E-06 | 1.800E-06 | 4.800E-06 |
| 8090 | 405 | N405I | A | 3.5e-06 | 2.7e-06 | 1.8e-06 | 2.667E-06 | 8.505E-07 | 1.800E-06 | 3.500E-06 |
| 8091 | 405 | N405L | A | 2.3e-06 | 2e-06 | 3e-06 | 2.433E-06 | 5.132E-07 | 2.000E-06 | 3.000E-06 |
| 8092 | 405 | N405K | A | 1.7e-06 | 2.4e-06 | 1.8e-06 | 1.967E-06 | 3.786E-07 | 1.700E-06 | 2.400E-06 |
| 8093 | 405 | N405M | A | 1.5e-06 | 2.5e-06 | 3.1e-06 | 2.367E-06 | 8.083E-07 | 1.500E-06 | 3.100E-06 |
| 8094 | 405 | N405F | A | 3e-06 | 2.4e-06 | 1.9e-06 | 2.433E-06 | 5.508E-07 | 1.900E-06 | 3.000E-06 |
| 8095 | 405 | N405P | A | 1.8e-06 | 5.1e-06 | 2.3e-06 | 3.067E-06 | 1.779E-06 | 1.800E-06 | 5.100E-06 |
| 8096 | 405 | N405S | A | 1.7e-06 | 3.3e-06 | 1.6e-06 | 2.200E-06 | 9.539E-07 | 1.600E-06 | 3.300E-06 |
| 8097 | 405 | N405T | A | 2e-06 | 2e-06 | 2.5e-06 | 2.167E-06 | 2.887E-07 | 2.000E-06 | 2.500E-06 |
| 8098 | 405 | N405W | A | 2.6e-06 | 1.9e-06 | 2.2e-06 | 2.233E-06 | 3.512E-07 | 1.900E-06 | 2.600E-06 |
| 8099 | 405 | N405Y | A | 2.5e-06 | 2.1e-06 | 2.1e-06 | 2.233E-06 | 2.309E-07 | 2.100E-06 | 2.500E-06 |
| 8100 | 405 | N405V | A | 7.7e-06 | 3.2e-06 | 2.5e-06 | 4.467E-06 | 2.822E-06 | 2.500E-06 | 7.700E-06 |
| 8101 | 406 | L406A | A | 3.1e-06 | 2.1e-06 | 3e-06 | 2.733E-06 | 5.508E-07 | 2.100E-06 | 3.100E-06 |
| 8102 | 406 | L406R | A | 4.8e-06 | 1.9e-06 | 3.1e-06 | 3.267E-06 | 1.457E-06 | 1.900E-06 | 4.800E-06 |
| 8103 | 406 | L406N | A | 2e-06 | 1.8e-06 | 2.6e-06 | 2.133E-06 | 4.163E-07 | 1.800E-06 | 2.600E-06 |
| 8104 | 406 | L406D | A | 2.5e-06 | 3.4e-06 | 2e-06 | 2.633E-06 | 7.095E-07 | 2.000E-06 | 3.400E-06 |
| 8105 | 406 | L406C | A | 2.5e-06 | 2.4e-06 | 1.9e-06 | 2.267E-06 | 3.215E-07 | 1.900E-06 | 2.500E-06 |
| 8106 | 406 | L406E | A | 2.8e-06 | 2.8e-06 | 1.9e-06 | 2.500E-06 | 5.196E-07 | 1.900E-06 | 2.800E-06 |
| 8107 | 406 | L406Q | A | 1.6e-06 | 2.7e-06 | 1.6e-06 | 1.967E-06 | 6.351E-07 | 1.600E-06 | 2.700E-06 |
| 8108 | 406 | L406G | A | 1.8e-06 | 2.3e-06 | 2e-06 | 2.033E-06 | 2.517E-07 | 1.800E-06 | 2.300E-06 |
| 8109 | 406 | L406H | A | 3.4e-06 | 2.9e-06 | 2.7e-06 | 3.000E-06 | 3.606E-07 | 2.700E-06 | 3.400E-06 |
| 8110 | 406 | L406I | A | 1.8e-06 | 2.7e-06 | 2.8e-06 | 2.433E-06 | 5.508E-07 | 1.800E-06 | 2.800E-06 |
| 8111 | 406 | L406L | A | 1.8e-06 | 2.5e-06 | 1.7e-06 | 2.000E-06 | 4.359E-07 | 1.700E-06 | 2.500E-06 |
| 8112 | 406 | L406K | A | 8.1e-06 | 1.6e-06 | 4e-06 | 4.567E-06 | 3.287E-06 | 1.600E-06 | 8.100E-06 |
| 8113 | 406 | L406M | A | 2.3e-06 | 2.4e-06 | 2.7e-06 | 2.467E-06 | 2.082E-07 | 2.300E-06 | 2.700E-06 |
| 8114 | 406 | L406F | A | 1.7e-06 | 2.6e-06 | 2.3e-06 | 2.200E-06 | 4.583E-07 | 1.700E-06 | 2.600E-06 |
| 8115 | 406 | L406P | A | 3.2e-06 | 2.4e-06 | 2.5e-06 | 2.700E-06 | 4.359E-07 | 2.400E-06 | 3.200E-06 |
| 8116 | 406 | L406S | A | 1.9e-06 | 2.1e-06 | 1.6e-06 | 1.867E-06 | 2.517E-07 | 1.600E-06 | 2.100E-06 |
| 8117 | 406 | L406T | A | 3.4e-06 | 1.9e-06 | 2e-06 | 2.433E-06 | 8.386E-07 | 1.900E-06 | 3.400E-06 |
| 8118 | 406 | L406W | A | 3e-06 | 4.3e-06 | 1.6e-06 | 2.967E-06 | 1.350E-06 | 1.600E-06 | 4.300E-06 |
| 8119 | 406 | L406Y | A | 1.6e-06 | 2.2e-06 | 1.7e-06 | 1.833E-06 | 3.215E-07 | 1.600E-06 | 2.200E-06 |
| 8120 | 406 | L406V | A | 2.1e-06 | 2e-06 | 1.6e-06 | 1.900E-06 | 2.646E-07 | 1.600E-06 | 2.100E-06 |
| 8121 | 407 | Q407A | A | 2.3e-06 | 2.1e-06 | 2.6e-06 | 2.333E-06 | 2.517E-07 | 2.100E-06 | 2.600E-06 |
| 8122 | 407 | Q407R | A | 5.1e-06 | 1.8e-06 | 3.8e-06 | 3.567E-06 | 1.662E-06 | 1.800E-06 | 5.100E-06 |
| 8123 | 407 | Q407N | A | 4.1e-06 | 2.1e-06 | 1.7e-06 | 2.633E-06 | 1.286E-06 | 1.700E-06 | 4.100E-06 |
| 8124 | 407 | Q407D | A | 2.8e-06 | 2.7e-06 | 1.5e-06 | 2.333E-06 | 7.234E-07 | 1.500E-06 | 2.800E-06 |
| 8125 | 407 | Q407C | A | 2.8e-06 | 2.1e-06 | 1.6e-06 | 2.167E-06 | 6.028E-07 | 1.600E-06 | 2.800E-06 |
| 8126 | 407 | Q407E | A | 2.1e-06 | 1.8e-06 | 2.3e-06 | 2.067E-06 | 2.517E-07 | 1.800E-06 | 2.300E-06 |
| 8127 | 407 | Q407Q | A | 4e-06 | 3.8e-06 | 2.6e-06 | 3.467E-06 | 7.572E-07 | 2.600E-06 | 4.000E-06 |
| 8128 | 407 | Q407G | A | 3e-06 | 3e-06 | 2.2e-06 | 2.733E-06 | 4.619E-07 | 2.200E-06 | 3.000E-06 |
| 8129 | 407 | Q407H | A | 1.2e-06 | 2.1e-06 | 3.1e-06 | 2.133E-06 | 9.504E-07 | 1.200E-06 | 3.100E-06 |
| 8130 | 407 | Q407I | A | 2.2e-06 | 3e-06 | 3.7e-06 | 2.967E-06 | 7.506E-07 | 2.200E-06 | 3.700E-06 |
| 8131 | 407 | Q407L | A | 2.9e-06 | 2.1e-06 | 3.7e-06 | 2.900E-06 | 8.000E-07 | 2.100E-06 | 3.700E-06 |
| 8132 | 407 | Q407K | A | 3e-06 | 3e-06 | 2.9e-06 | 2.967E-06 | 5.774E-08 | 2.900E-06 | 3.000E-06 |
| 8133 | 407 | Q407M | A | 2.7e-06 | 1.7e-06 | 3.2e-06 | 2.533E-06 | 7.638E-07 | 1.700E-06 | 3.200E-06 |
| 8134 | 407 | Q407F | A | 1.8e-06 | 2.5e-06 | 1.8e-06 | 2.033E-06 | 4.041E-07 | 1.800E-06 | 2.500E-06 |
| 8135 | 407 | Q407P | A | 2.1e-06 | 1.9e-06 | 1.8e-06 | 1.933E-06 | 1.528E-07 | 1.800E-06 | 2.100E-06 |
| 8136 | 407 | Q407S | A | 2.5e-06 | 2.6e-06 | 2.6e-06 | 2.567E-06 | 5.774E-08 | 2.500E-06 | 2.600E-06 |
| 8137 | 407 | Q407T | A | 2.2e-06 | 1.2e-06 | 2.4e-06 | 1.933E-06 | 6.429E-07 | 1.200E-06 | 2.400E-06 |
| 8138 | 407 | Q407W | A | 4.1e-06 | 2.5e-06 | 2.4e-06 | 3.000E-06 | 9.539E-07 | 2.400E-06 | 4.100E-06 |
| 8139 | 407 | Q407Y | A | 1.9e-06 | 2.7e-06 | 2e-06 | 2.200E-06 | 4.359E-07 | 1.900E-06 | 2.700E-06 |
| 8140 | 407 | Q407V | A | 3.6e-06 | 3.6e-06 | 1.5e-06 | 2.900E-06 | 1.212E-06 | 1.500E-06 | 3.600E-06 |
| 8141 | 408 | V408A | A | 1.2e-06 | 2.9e-06 | 1.8e-06 | 1.967E-06 | 8.622E-07 | 1.200E-06 | 2.900E-06 |
| 8142 | 408 | V408R | A | 2.1e-06 | 3.9e-06 | 2.1e-06 | 2.700E-06 | 1.039E-06 | 2.100E-06 | 3.900E-06 |
| 8143 | 408 | V408N | A | 1.3e-06 | 1.2e-06 | 8e-07 | 1.100E-06 | 2.646E-07 | 8.000E-07 | 1.300E-06 |
| 8144 | 408 | V408D | A | 1.4e-06 | 2.1e-06 | 2e-06 | 1.833E-06 | 3.786E-07 | 1.400E-06 | 2.100E-06 |
| 8145 | 408 | V408C | A | 3.4e-06 | 2.7e-06 | 2.3e-06 | 2.800E-06 | 5.568E-07 | 2.300E-06 | 3.400E-06 |
| 8146 | 408 | V408E | A | 2.1e-06 | 2e-06 | 2.7e-06 | 2.267E-06 | 3.786E-07 | 2.000E-06 | 2.700E-06 |
| 8147 | 408 | V408Q | A | 1.7e-06 | 3.7e-06 | 2.7e-06 | 2.700E-06 | 1.000E-06 | 1.700E-06 | 3.700E-06 |
| 8148 | 408 | V408G | A | 1.7e-06 | 1.6e-06 | 4.3e-06 | 2.533E-06 | 1.531E-06 | 1.600E-06 | 4.300E-06 |
| 8149 | 408 | V408H | A | 2.7e-06 | 2.9e-06 | 2.3e-06 | 2.633E-06 | 3.055E-07 | 2.300E-06 | 2.900E-06 |
| 8150 | 408 | V408I | A | 2.3e-06 | 2.2e-06 | 3.9e-06 | 2.800E-06 | 9.539E-07 | 2.200E-06 | 3.900E-06 |
| 8151 | 408 | V408L | A | 3.9e-06 | 5.2e-06 | 1.7e-06 | 3.600E-06 | 1.769E-06 | 1.700E-06 | 5.200E-06 |
| 8152 | 408 | V408K | A | 1.7e-06 | 2.8e-06 | 2.3e-06 | 2.267E-06 | 5.508E-07 | 1.700E-06 | 2.800E-06 |
| 8153 | 408 | V408M | A | 1.7e-06 | 3.2e-06 | 3e-06 | 2.633E-06 | 8.145E-07 | 1.700E-06 | 3.200E-06 |
| 8154 | 408 | V408F | A | 1.5e-06 | 2.2e-06 | 1e-06 | 1.567E-06 | 6.028E-07 | 1.000E-06 | 2.200E-06 |
| 8155 | 408 | V408P | A | 1.5e-06 | 2.4e-06 | 3.1e-06 | 2.333E-06 | 8.021E-07 | 1.500E-06 | 3.100E-06 |
| 8156 | 408 | V408S | A | 2.7e-06 | 3.8e-06 | 1.9e-06 | 2.800E-06 | 9.539E-07 | 1.900E-06 | 3.800E-06 |
| 8157 | 408 | V408T | A | 1.5e-06 | 2e-06 | 1.9e-06 | 1.800E-06 | 2.646E-07 | 1.500E-06 | 2.000E-06 |
| 8158 | 408 | V408W | A | 3e-06 | 2.2e-06 | 1.4e-06 | 2.200E-06 | 8.000E-07 | 1.400E-06 | 3.000E-06 |
| 8159 | 408 | V408Y | A | 2.8e-06 | 1.4e-06 | 1.4e-06 | 1.867E-06 | 8.083E-07 | 1.400E-06 | 2.800E-06 |

| | | | | | | | | | | |
|---|---|---|---|---|---|---|---|---|---|---|
| 8160 | 408 | V408V | A | 4.6e-06 | 2.3e-06 | 2.5e-06 | 3.133E-06 | 1.274E-06 | 2.300E-06 | 4.600E-06 |
| 8161 | 409 | I409A | A | 4.6e-06 | 2.2e-06 | 2.8e-06 | 3.200E-06 | 1.249E-06 | 2.200E-06 | 4.600E-06 |
| 8162 | 409 | I409R | A | 2.4e-06 | 1.5e-06 | 1.7e-06 | 1.867E-06 | 4.726E-07 | 1.500E-06 | 2.400E-06 |
| 8163 | 409 | I409N | A | 1.9e-06 | 3e-06 | 8.8e-07 | 1.927E-06 | 1.060E-06 | 8.800E-07 | 3.000E-06 |
| 8164 | 409 | I409D | A | 3.1e-06 | 1.9e-06 | 2.7e-06 | 2.567E-06 | 6.110E-07 | 1.900E-06 | 3.100E-06 |
| 8165 | 409 | I409C | A | 3e-06 | 2e-06 | 3e-06 | 2.667E-06 | 5.774E-07 | 2.000E-06 | 3.000E-06 |
| 8166 | 409 | I409E | A | 2e-06 | 1.8e-06 | 1.6e-06 | 1.800E-06 | 2.000E-07 | 1.600E-06 | 2.000E-06 |
| 8167 | 409 | I409Q | A | 4.8e-06 | 2.3e-06 | 2.8e-06 | 3.300E-06 | 1.323E-06 | 2.300E-06 | 4.800E-06 |
| 8168 | 409 | I409G | A | 3.6e-06 | 2.5e-06 | 2e-06 | 2.700E-06 | 8.185E-07 | 2.000E-06 | 3.600E-06 |
| 8169 | 409 | I409H | A | 2e-06 | 3.3e-06 | 1.9e-06 | 2.400E-06 | 7.810E-07 | 1.900E-06 | 3.300E-06 |
| 8170 | 409 | I409I | A | 2.9e-06 | 2.8e-06 | 2e-06 | 2.567E-06 | 4.933E-07 | 2.000E-06 | 2.900E-06 |
| 8171 | 409 | I409L | A | 3e-06 | 2.9e-06 | 1.8e-06 | 2.567E-06 | 6.658E-07 | 1.800E-06 | 3.000E-06 |
| 8172 | 409 | I409K | A | 3.7e-06 | 2.1e-06 | 2.8e-06 | 2.867E-06 | 8.021E-07 | 2.100E-06 | 3.700E-06 |
| 8173 | 409 | I409M | A | 4.4e-06 | 3.9e-06 | 3.3e-06 | 3.867E-06 | 5.508E-07 | 3.300E-06 | 4.400E-06 |
| 8174 | 409 | I409F | A | 1.7e-06 | 1.9e-06 | 2.5e-06 | 2.033E-06 | 4.163E-07 | 1.700E-06 | 2.500E-06 |
| 8175 | 409 | I409P | A | 1.7e-06 | 1.2e-06 | 1.9e-06 | 1.600E-06 | 3.606E-07 | 1.200E-06 | 1.900E-06 |
| 8176 | 409 | I409S | A | 2.2e-06 | 2.8e-06 | 2.2e-06 | 2.400E-06 | 3.464E-07 | 2.200E-06 | 2.800E-06 |
| 8177 | 409 | I409T | A | 4.9e-06 | 2.9e-06 | 3.3e-06 | 3.700E-06 | 1.058E-06 | 2.900E-06 | 4.900E-06 |
| 8178 | 409 | I409W | A | 2.5e-06 | 3.9e-06 | 2.7e-06 | 3.033E-06 | 7.572E-07 | 2.500E-06 | 3.900E-06 |
| 8179 | 409 | I409Y | A | 1.6e-06 | 1.3e-06 | 2.5e-06 | 1.800E-06 | 6.245E-07 | 1.300E-06 | 2.500E-06 |
| 8180 | 409 | I409V | A | 3.5e-06 | 2.7e-06 | 1.9e-06 | 2.700E-06 | 8.000E-07 | 1.900E-06 | 3.500E-06 |
| 8181 | 410 | R410A | A | 1.9e-06 | 1.5e-06 | 2.8e-06 | 2.067E-06 | 6.658E-07 | 1.500E-06 | 2.800E-06 |
| 8182 | 410 | R410R | A | 3.2e-06 | 3.6e-06 | 2.4e-06 | 3.067E-06 | 6.110E-07 | 2.400E-06 | 3.600E-06 |
| 8183 | 410 | R410N | A | 1.6e-06 | 2.4e-06 | 2.1e-06 | 2.033E-06 | 4.041E-07 | 1.600E-06 | 2.400E-06 |
| 8184 | 410 | R410D | A | 3.7e-06 | 2.1e-06 | 3.8e-06 | 3.200E-06 | 9.539E-07 | 2.100E-06 | 3.800E-06 |
| 8185 | 410 | R410C | A | 2.5e-06 | 2.1e-06 | 2.2e-06 | 2.267E-06 | 2.082E-07 | 2.100E-06 | 2.500E-06 |
| 8186 | 410 | R410E | A | 2.9e-06 | 2.5e-06 | 2.6e-06 | 2.667E-06 | 2.082E-07 | 2.500E-06 | 2.900E-06 |
| 8187 | 410 | R410Q | A | 1.4e-06 | 2.6e-06 | 2.1e-06 | 2.033E-06 | 6.028E-07 | 1.400E-06 | 2.600E-06 |
| 8188 | 410 | R410G | A | 2.8e-06 | 3e-06 | 2.2e-06 | 2.667E-06 | 4.163E-07 | 2.200E-06 | 3.000E-06 |
| 8189 | 410 | R410H | A | 2.2e-06 | 3.7e-06 | 1.8e-06 | 2.567E-06 | 1.002E-06 | 1.800E-06 | 3.700E-06 |
| 8190 | 410 | R410I | A | 2e-06 | 2.8e-06 | 2.4e-06 | 2.400E-06 | 4.000E-07 | 2.000E-06 | 2.800E-06 |
| 8191 | 410 | R410L | A | 3.3e-06 | 3e-06 | 1.9e-06 | 2.733E-06 | 7.371E-07 | 1.900E-06 | 3.300E-06 |
| 8192 | 410 | R410K | A | 2e-06 | 2.2e-06 | 3.2e-06 | 2.467E-06 | 6.429E-07 | 2.000E-06 | 3.200E-06 |
| 8193 | 410 | R410M | A | 3e-06 | 2.2e-06 | 1.3e-06 | 2.167E-06 | 8.505E-07 | 1.300E-06 | 3.000E-06 |
| 8194 | 410 | R410F | A | 2.3e-06 | 2.1e-06 | 3.1e-06 | 2.500E-06 | 5.292E-07 | 2.100E-06 | 3.100E-06 |
| 8195 | 410 | R410P | A | 1.6e-06 | 3.6e-06 | 3e-06 | 2.733E-06 | 1.026E-06 | 1.600E-06 | 3.600E-06 |
| 8196 | 410 | R410S | A | 2.3e-06 | 2.6e-06 | 2.4e-06 | 2.433E-06 | 1.528E-07 | 2.300E-06 | 2.600E-06 |
| 8197 | 410 | R410T | A | 2.6e-06 | 1.8e-06 | 2.7e-06 | 2.367E-06 | 4.933E-07 | 1.800E-06 | 2.700E-06 |
| 8198 | 410 | R410W | A | 4.8e-06 | 2.7e-06 | 1.1e-06 | 2.867E-06 | 1.856E-06 | 1.100E-06 | 4.800E-06 |
| 8199 | 410 | R410Y | A | 3e-06 | 3.5e-06 | 3.1e-06 | 3.200E-06 | 2.646E-07 | 3.000E-06 | 3.500E-06 |
| 8200 | 410 | R410V | A | 2.7e-06 | 3.9e-06 | 1.8e-06 | 2.800E-06 | 1.054E-06 | 1.800E-06 | 3.900E-06 |
| 8201 | 411 | G411A | A | 1.9e-06 | 4e-06 | 3.7e-06 | 3.200E-06 | 1.136E-06 | 1.900E-06 | 4.000E-06 |
| 8202 | 411 | G411R | A | 3.9e-06 | 1.2e-06 | 1.9e-06 | 2.333E-06 | 1.401E-06 | 1.200E-06 | 3.900E-06 |
| 8203 | 411 | G411N | A | 2.1e-06 | 2e-06 | 1.9e-06 | 2.000E-06 | 1.000E-07 | 1.900E-06 | 2.100E-06 |
| 8204 | 411 | G411D | A | 1.9e-06 | 2.5e-06 | 2.3e-06 | 2.233E-06 | 3.055E-07 | 1.900E-06 | 2.500E-06 |
| 8205 | 411 | G411C | A | 2.9e-06 | 2.8e-06 | 2e-06 | 2.567E-06 | 4.933E-07 | 2.000E-06 | 2.900E-06 |
| 8206 | 411 | G411E | A | 3.7e-06 | 3.7e-06 | 3.4e-06 | 3.600E-06 | 1.732E-07 | 3.400E-06 | 3.700E-06 |
| 8207 | 411 | G411Q | A | 1.5e-06 | 3.9e-06 | 2.8e-06 | 2.733E-06 | 1.201E-06 | 1.500E-06 | 3.900E-06 |
| 8208 | 411 | G411G | A | 3.2e-06 | 2.7e-06 | 3.3e-06 | 3.067E-06 | 3.215E-07 | 2.700E-06 | 3.300E-06 |
| 8209 | 411 | G411H | A | 1.5e-06 | 1.9e-06 | 3.3e-06 | 2.233E-06 | 9.452E-07 | 1.500E-06 | 3.300E-06 |
| 8210 | 411 | G411I | A | 2.8e-06 | 1.9e-06 | 1.6e-06 | 2.100E-06 | 6.245E-07 | 1.600E-06 | 2.800E-06 |
| 8211 | 411 | G411L | A | 5.3e-06 | 1.6e-06 | 3.9e-06 | 3.600E-06 | 1.868E-06 | 1.600E-06 | 5.300E-06 |
| 8212 | 411 | G411K | A | 1.5e-06 | 3.3e-06 | 2.8e-06 | 2.533E-06 | 9.292E-07 | 1.500E-06 | 3.300E-06 |
| 8213 | 411 | G411M | A | 1.4e-06 | 3.3e-06 | 3.7e-06 | 2.800E-06 | 1.229E-06 | 1.400E-06 | 3.700E-06 |
| 8214 | 411 | G411F | A | 1.9e-06 | 2.1e-06 | 2.1e-06 | 2.033E-06 | 1.155E-07 | 1.900E-06 | 2.100E-06 |
| 8215 | 411 | G411P | A | 2.9e-06 | 2.3e-06 | 2.2e-06 | 2.467E-06 | 3.786E-07 | 2.200E-06 | 2.900E-06 |
| 8216 | 411 | G411S | A | 2.3e-06 | 1.6e-06 | 1.7e-06 | 1.867E-06 | 3.786E-07 | 1.600E-06 | 2.300E-06 |
| 8217 | 411 | G411T | A | 2.5e-06 | 2.4e-06 | 2.4e-06 | 2.433E-06 | 5.774E-08 | 2.400E-06 | 2.500E-06 |
| 8218 | 411 | G411W | A | 2e-06 | 3.3e-06 | 4.6e-06 | 3.300E-06 | 1.300E-06 | 2.000E-06 | 4.600E-06 |
| 8219 | 411 | G411Y | A | 3.4e-06 | 2.2e-06 | 1.7e-06 | 2.433E-06 | 8.737E-07 | 1.700E-06 | 3.400E-06 |
| 8220 | 411 | G411V | A | 2.6e-06 | 3.5e-06 | 1e-06 | 2.367E-06 | 1.266E-06 | 1.000E-06 | 3.500E-06 |
| 8221 | 412 | R412A | A | 2.7e-06 | 2.7e-06 | 1.9e-06 | 2.433E-06 | 4.619E-07 | 1.900E-06 | 2.700E-06 |
| 8222 | 412 | R412R | A | 3e-06 | 2.8e-06 | 1.3e-06 | 2.367E-06 | 9.292E-07 | 1.300E-06 | 3.000E-06 |
| 8223 | 412 | R412N | A | 1.9e-06 | 2.1e-06 | 2.7e-06 | 2.233E-06 | 4.163E-07 | 1.900E-06 | 2.700E-06 |
| 8224 | 412 | R412D | A | 3.2e-06 | 2.9e-06 | 1.9e-06 | 2.667E-06 | 6.807E-07 | 1.900E-06 | 3.200E-06 |
| 8225 | 412 | R412C | A | 3.3e-06 | 2.6e-06 | 1.5e-06 | 2.467E-06 | 9.074E-07 | 1.500E-06 | 3.300E-06 |
| 8226 | 412 | R412E | A | 1.9e-06 | 2.2e-06 | 1.9e-06 | 2.000E-06 | 1.732E-07 | 1.900E-06 | 2.200E-06 |
| 8227 | 412 | R412Q | A | 1.9e-06 | 2.5e-06 | 2.6e-06 | 2.333E-06 | 3.786E-07 | 1.900E-06 | 2.600E-06 |
| 8228 | 412 | R412G | A | 2.1e-06 | 2.5e-06 | 2e-06 | 2.200E-06 | 2.646E-07 | 2.000E-06 | 2.500E-06 |
| 8229 | 412 | R412H | A | 2.1e-06 | 2.4e-06 | 2.5e-06 | 2.333E-06 | 2.082E-07 | 2.100E-06 | 2.500E-06 |
| 8230 | 412 | R412I | A | 2.6e-06 | 3e-06 | 1.6e-06 | 2.400E-06 | 7.211E-07 | 1.600E-06 | 3.000E-06 |
| 8231 | 412 | R412L | A | 3.2e-06 | 2.2e-06 | 1.6e-06 | 2.333E-06 | 8.083E-07 | 1.600E-06 | 3.200E-06 |
| 8232 | 412 | R412K | A | 1.6e-06 | 1.8e-06 | 2.3e-06 | 1.900E-06 | 3.606E-07 | 1.600E-06 | 2.300E-06 |
| 8233 | 412 | R412M | A | 2.2e-06 | 3.2e-06 | 2.8e-06 | 2.733E-06 | 5.033E-07 | 2.200E-06 | 3.200E-06 |
| 8234 | 412 | R412F | A | 2.7e-06 | 2.4e-06 | 1.9e-06 | 2.333E-06 | 4.041E-07 | 1.900E-06 | 2.700E-06 |
| 8235 | 412 | R412P | A | 3e-06 | 3.6e-06 | 2.4e-06 | 3.000E-06 | 6.000E-07 | 2.400E-06 | 3.600E-06 |
| 8236 | 412 | R412S | A | 1.9e-06 | 1.6e-06 | 3.1e-06 | 2.200E-06 | 7.937E-07 | 1.600E-06 | 3.100E-06 |
| 8237 | 412 | R412T | A | 2.7e-06 | 2.3e-06 | 2.1e-06 | 2.367E-06 | 3.055E-07 | 2.100E-06 | 2.700E-06 |
| 8238 | 412 | R412W | A | 1.4e-06 | 2.7e-06 | 4.2e-06 | 2.767E-06 | 1.401E-06 | 1.400E-06 | 4.200E-06 |
| 8239 | 412 | R412Y | A | 3.2e-06 | 1.8e-06 | 3.4e-06 | 2.800E-06 | 8.718E-07 | 1.800E-06 | 3.400E-06 |
| 8240 | 412 | R412V | A | 2.8e-06 | 3.4e-06 | 3.9e-06 | 3.367E-06 | 5.508E-07 | 2.800E-06 | 3.900E-06 |
| 8241 | 413 | I413A | A | 4.7e-06 | 1.7e-06 | 3.7e-06 | 3.367E-06 | 1.528E-06 | 1.700E-06 | 4.700E-06 |
| 8242 | 413 | I413R | A | 2.9e-06 | 2.1e-06 | 3.4e-06 | 2.800E-06 | 6.557E-07 | 2.100E-06 | 3.400E-06 |
| 8243 | 413 | I413N | A | 2.1e-06 | 1.4e-06 | 3e-06 | 2.167E-06 | 8.021E-07 | 1.400E-06 | 3.000E-06 |
| 8244 | 413 | I413D | A | 1.4e-06 | 2.6e-06 | 2.4e-06 | 2.133E-06 | 6.429E-07 | 1.400E-06 | 2.600E-06 |

| | | | | | | | | | | |
|---|---|---|---|---|---|---|---|---|---|---|
| 8245 | 413 | I413C | A | 2.1e-06 | 2.8e-06 | 3.1e-06 | 2.667E-06 | 5.132E-07 | 2.100E-06 | 3.100E-06 |
| 8246 | 413 | I413E | A | 2e-06 | 2.2e-06 | 3.9e-06 | 2.700E-06 | 1.044E-06 | 2.000E-06 | 3.900E-06 |
| 8247 | 413 | I413Q | A | 2.4e-06 | 2.5e-06 | 4.1e-06 | 3.000E-06 | 9.539E-07 | 2.400E-06 | 4.100E-06 |
| 8248 | 413 | I413G | A | 1.4e-06 | 2.5e-06 | 1.2e-06 | 1.700E-06 | 7.000E-07 | 1.200E-06 | 2.500E-06 |
| 8249 | 413 | I413H | A | 3e-06 | 2.4e-06 | 1.6e-06 | 2.333E-06 | 7.024E-07 | 1.600E-06 | 3.000E-06 |
| 8250 | 413 | I413I | A | 3.6e-06 | 1.3e-06 | 1.7e-06 | 2.200E-06 | 1.229E-06 | 1.300E-06 | 3.600E-06 |
| 8251 | 413 | I413L | A | 2.1e-06 | 2.7e-06 | 1.6e-06 | 2.133E-06 | 5.508E-07 | 1.600E-06 | 2.700E-06 |
| 8252 | 413 | I413K | A | 2.1e-06 | 2.4e-06 | 3.1e-06 | 2.533E-06 | 5.132E-07 | 2.100E-06 | 3.100E-06 |
| 8253 | 413 | I413M | A | 3.4e-06 | 3.8e-06 | 2.6e-06 | 3.267E-06 | 6.110E-07 | 2.600E-06 | 3.800E-06 |
| 8254 | 413 | I413F | A | 2.7e-06 | 3.3e-06 | 1.7e-06 | 2.567E-06 | 8.083E-07 | 1.700E-06 | 3.300E-06 |
| 8255 | 413 | I413P | A | 2.4e-06 | 2.1e-06 | 4e-06 | 2.833E-06 | 1.021E-06 | 2.100E-06 | 4.000E-06 |
| 8256 | 413 | I413S | A | 2.4e-06 | 2.4e-06 | 4e-06 | 2.933E-06 | 9.238E-07 | 2.400E-06 | 4.000E-06 |
| 8257 | 413 | I413T | A | 2.2e-06 | 1.5e-06 | 2.6e-06 | 2.100E-06 | 5.568E-07 | 1.500E-06 | 2.600E-06 |
| 8258 | 413 | I413W | A | 2.9e-06 | 2.7e-06 | 4.2e-06 | 3.267E-06 | 8.145E-07 | 2.700E-06 | 4.200E-06 |
| 8259 | 413 | I413Y | A | 4.3e-06 | 4.1e-06 | 2.3e-06 | 3.567E-06 | 1.102E-06 | 2.300E-06 | 4.300E-06 |
| 8260 | 413 | I413V | A | 1.6e-06 | 2e-06 | 1.3e-06 | 1.633E-06 | 3.512E-07 | 1.300E-06 | 2.000E-06 |
| 8261 | 414 | L414A | A | 5.8e-06 | 2.9e-06 | 1.5e-06 | 3.400E-06 | 2.193E-06 | 1.500E-06 | 5.800E-06 |
| 8262 | 414 | L414R | A | 1.6e-06 | 3.6e-06 | 3e-06 | 2.733E-06 | 1.026E-06 | 1.600E-06 | 3.600E-06 |
| 8263 | 414 | L414N | A | 2.6e-06 | 8.8e-07 | 2.6e-06 | 2.027E-06 | 9.930E-07 | 8.800E-07 | 2.600E-06 |
| 8264 | 414 | L414D | A | 4.2e-06 | 2e-06 | 3.7e-06 | 3.300E-06 | 1.153E-06 | 2.000E-06 | 4.200E-06 |
| 8265 | 414 | L414C | A | 2.8e-06 | 2.2e-06 | 2.4e-06 | 2.467E-06 | 3.055E-07 | 2.200E-06 | 2.800E-06 |
| 8266 | 414 | L414E | A | 3.1e-06 | 2.7e-06 | 3.5e-06 | 3.100E-06 | 4.000E-07 | 2.700E-06 | 3.500E-06 |
| 8267 | 414 | L414Q | A | 2.8e-06 | 1.8e-06 | 2.5e-06 | 2.367E-06 | 5.132E-07 | 1.800E-06 | 2.800E-06 |
| 8268 | 414 | L414G | A | 1.9e-06 | 2.5e-06 | 3.3e-06 | 2.567E-06 | 7.024E-07 | 1.900E-06 | 3.300E-06 |
| 8269 | 414 | L414H | A | 3.2e-06 | 2.9e-06 | 1.9e-06 | 2.667E-06 | 6.807E-07 | 1.900E-06 | 3.200E-06 |
| 8270 | 414 | L414I | A | 6.3e-07 | 1.8e-06 | 3e-06 | 1.810E-06 | 1.185E-06 | 6.300E-07 | 3.000E-06 |
| 8271 | 414 | L414L | A | 1.1e-06 | 2.5e-06 | 1.3e-06 | 1.633E-06 | 7.572E-07 | 1.100E-06 | 2.500E-06 |
| 8272 | 414 | L414K | A | 1.7e-06 | 1.7e-06 | 2.7e-06 | 2.033E-06 | 5.774E-07 | 1.700E-06 | 2.700E-06 |
| 8273 | 414 | L414M | A | 3.8e-06 | 2.2e-06 | 2.9e-06 | 2.967E-06 | 8.021E-07 | 2.200E-06 | 3.800E-06 |
| 8274 | 414 | L414F | A | 1.8e-06 | 1.7e-06 | 2e-06 | 1.833E-06 | 1.528E-07 | 1.700E-06 | 2.000E-06 |
| 8275 | 414 | L414P | A | 1.7e-06 | 2.8e-06 | 1.3e-06 | 1.933E-06 | 7.767E-07 | 1.300E-06 | 2.800E-06 |
| 8276 | 414 | L414S | A | 3.1e-06 | 1.9e-06 | 2.2e-06 | 2.400E-06 | 6.245E-07 | 1.900E-06 | 3.100E-06 |
| 8277 | 414 | L414T | A | 1.8e-06 | 8.7e-07 | 1.5e-06 | 1.390E-06 | 4.747E-07 | 8.700E-07 | 1.800E-06 |
| 8278 | 414 | L414W | A | 1.7e-06 | 1.5e-06 | 2.5e-06 | 1.900E-06 | 5.292E-07 | 1.500E-06 | 2.500E-06 |
| 8279 | 414 | L414Y | A | 4.5e-06 | 3.4e-06 | 3.4e-06 | 3.767E-06 | 6.351E-07 | 3.400E-06 | 4.500E-06 |
| 8280 | 414 | L414V | A | 2.6e-06 | 2.8e-06 | 4.7e-06 | 3.367E-06 | 1.159E-06 | 2.600E-06 | 4.700E-06 |
| 8281 | 415 | H415A | A | 1.6e-06 | 3.4e-06 | 2.2e-06 | 2.400E-06 | 9.165E-07 | 1.600E-06 | 3.400E-06 |
| 8282 | 415 | H415R | A | 3.3e-06 | 3.4e-06 | 2e-06 | 2.900E-06 | 7.810E-07 | 2.000E-06 | 3.400E-06 |
| 8283 | 415 | H415N | A | 2.6e-06 | 2.2e-06 | 2.2e-06 | 2.333E-06 | 2.309E-07 | 2.200E-06 | 2.600E-06 |
| 8284 | 415 | H415D | A | 2.3e-06 | 3.4e-06 | 2.7e-06 | 2.800E-06 | 5.568E-07 | 2.300E-06 | 3.400E-06 |
| 8285 | 415 | H415C | A | 2.4e-06 | 2.1e-06 | 2.9e-06 | 2.467E-06 | 4.041E-07 | 2.100E-06 | 2.900E-06 |
| 8286 | 415 | H415E | A | 2e-06 | 1.7e-06 | 2.2e-06 | 1.967E-06 | 2.517E-07 | 1.700E-06 | 2.200E-06 |
| 8287 | 415 | H415Q | A | 2.3e-06 | 1e-06 | 1.5e-06 | 1.600E-06 | 6.557E-07 | 1.000E-06 | 2.300E-06 |
| 8288 | 415 | H415G | A | 4.9e-06 | 2.9e-06 | 3.3e-06 | 3.700E-06 | 1.058E-06 | 2.900E-06 | 4.900E-06 |
| 8289 | 415 | H415H | A | 3e-06 | 2.3e-06 | 3e-06 | 2.767E-06 | 4.041E-07 | 2.300E-06 | 3.000E-06 |
| 8290 | 415 | H415I | A | 2.1e-06 | 1.6e-06 | 2.4e-06 | 2.033E-06 | 4.041E-07 | 1.600E-06 | 2.400E-06 |
| 8291 | 415 | H415L | A | 4.6e-06 | 1.9e-06 | 1.6e-06 | 2.700E-06 | 1.652E-06 | 1.600E-06 | 4.600E-06 |
| 8292 | 415 | H415K | A | 3.2e-06 | 5.1e-06 | 2e-06 | 3.433E-06 | 1.563E-06 | 2.000E-06 | 5.100E-06 |
| 8293 | 415 | H415M | A | 2.3e-06 | 1.9e-06 | 1.4e-06 | 1.867E-06 | 4.509E-07 | 1.400E-06 | 2.300E-06 |
| 8294 | 415 | H415F | A | 3.5e-06 | 2.5e-06 | 3.1e-06 | 3.033E-06 | 5.033E-07 | 2.500E-06 | 3.500E-06 |
| 8295 | 415 | H415P | A | 1.3e-06 | 1.7e-06 | 2.3e-06 | 1.767E-06 | 5.033E-07 | 1.300E-06 | 2.300E-06 |
| 8296 | 415 | H415S | A | 4.1e-06 | 2.2e-06 | 2.1e-06 | 2.800E-06 | 1.127E-06 | 2.100E-06 | 4.100E-06 |
| 8297 | 415 | H415T | A | 2.6e-06 | 2e-06 | 2.3e-06 | 2.300E-06 | 3.000E-07 | 2.000E-06 | 2.600E-06 |
| 8298 | 415 | H415W | A | 3.8e-06 | 5.5e-06 | 2.2e-06 | 3.833E-06 | 1.650E-06 | 2.200E-06 | 5.500E-06 |
| 8299 | 415 | H415Y | A | 3.6e-06 | 1.9e-06 | 2.1e-06 | 2.533E-06 | 9.292E-07 | 1.900E-06 | 3.600E-06 |
| 8300 | 415 | H415V | A | 4.5e-06 | 1.3e-06 | 4.8e-06 | 3.533E-06 | 1.940E-06 | 1.300E-06 | 4.800E-06 |
| 8301 | 416 | N416A | A | 2.4e-06 | 2.5e-06 | 1.5e-06 | 2.133E-06 | 5.508E-07 | 1.500E-06 | 2.500E-06 |
| 8302 | 416 | N416R | A | 2.1e-06 | 5e-06 | 2.4e-06 | 3.167E-06 | 1.595E-06 | 2.100E-06 | 5.000E-06 |
| 8303 | 416 | N416N | A | 7.1e-06 | 2.7e-06 | 2.5e-06 | 4.100E-06 | 2.600E-06 | 2.500E-06 | 7.100E-06 |
| 8304 | 416 | N416D | A | 4.3e-06 | 1.6e-06 | 2.1e-06 | 2.667E-06 | 1.436E-06 | 1.600E-06 | 4.300E-06 |
| 8305 | 416 | N416C | A | 1.2e-06 | 2.3e-06 | 2e-06 | 1.833E-06 | 5.686E-07 | 1.200E-06 | 2.300E-06 |
| 8306 | 416 | N416E | A | 2e-06 | 4.2e-06 | 1.2e-06 | 2.467E-06 | 1.553E-06 | 1.200E-06 | 4.200E-06 |
| 8307 | 416 | N416Q | A | 3.5e-06 | 2.8e-06 | 2.5e-06 | 2.933E-06 | 5.132E-07 | 2.500E-06 | 3.500E-06 |
| 8308 | 416 | N416G | A | 1.7e-06 | 2.4e-06 | 2.3e-06 | 2.133E-06 | 3.786E-07 | 1.700E-06 | 2.400E-06 |
| 8309 | 416 | N416H | A | 2e-06 | 2.1e-06 | 1.7e-06 | 1.933E-06 | 2.082E-07 | 1.700E-06 | 2.100E-06 |
| 8310 | 416 | N416I | A | 1.7e-06 | 2.5e-06 | 2.6e-06 | 2.267E-06 | 4.933E-07 | 1.700E-06 | 2.600E-06 |
| 8311 | 416 | N416L | A | 2.3e-06 | 3.5e-06 | 2.1e-06 | 2.633E-06 | 7.572E-07 | 2.100E-06 | 3.500E-06 |
| 8312 | 416 | N416K | A | 4.5e-06 | 3.4e-06 | 2.6e-06 | 3.500E-06 | 9.539E-07 | 2.600E-06 | 4.500E-06 |
| 8313 | 416 | N416M | A | 2.7e-06 | 1.4e-06 | 1.8e-06 | 1.967E-06 | 6.658E-07 | 1.400E-06 | 2.700E-06 |
| 8314 | 416 | N416F | A | 2.7e-06 | 1.9e-06 | 1.5e-06 | 2.033E-06 | 6.110E-07 | 1.500E-06 | 2.700E-06 |
| 8315 | 416 | N416P | A | 2.5e-06 | 2.7e-06 | 2.6e-06 | 2.600E-06 | 1.000E-07 | 2.500E-06 | 2.700E-06 |
| 8316 | 416 | N416S | A | 2.3e-06 | 3.5e-06 | 4.1e-06 | 3.300E-06 | 9.165E-07 | 2.300E-06 | 4.100E-06 |
| 8317 | 416 | N416T | A | 1.8e-06 | 2.7e-06 | 1.9e-06 | 2.133E-06 | 4.933E-07 | 1.800E-06 | 2.700E-06 |
| 8318 | 416 | N416W | A | 3.2e-06 | 3.4e-06 | 3.8e-06 | 3.467E-06 | 3.055E-07 | 3.200E-06 | 3.800E-06 |
| 8319 | 416 | N416Y | A | 2.5e-06 | 2.6e-06 | 1.5e-06 | 2.200E-06 | 6.083E-07 | 1.500E-06 | 2.600E-06 |
| 8320 | 416 | N416V | A | 2.3e-06 | 2e-06 | 2.3e-06 | 2.200E-06 | 1.732E-07 | 2.000E-06 | 2.300E-06 |
| 8321 | 417 | G417A | A | 1.9e-06 | 1.9e-06 | 1.9e-06 | 1.900E-06 | 2.593E-22 | 1.900E-06 | 1.900E-06 |
| 8322 | 417 | G417R | A | 2.7e-06 | 3.5e-06 | 1.5e-06 | 2.567E-06 | 1.007E-06 | 1.500E-06 | 3.500E-06 |
| 8323 | 417 | G417N | A | 2.9e-06 | 1.6e-06 | 2.1e-06 | 2.200E-06 | 6.557E-07 | 1.600E-06 | 2.900E-06 |
| 8324 | 417 | G417D | A | 2.4e-06 | 2.5e-06 | 2.5e-06 | 2.467E-06 | 5.774E-08 | 2.400E-06 | 2.500E-06 |
| 8325 | 417 | G417C | A | 3e-06 | 2.1e-06 | 1.9e-06 | 2.333E-06 | 5.859E-07 | 1.900E-06 | 3.000E-06 |
| 8326 | 417 | G417E | A | 2.1e-06 | 2.6e-06 | 1.9e-06 | 2.200E-06 | 3.606E-07 | 1.900E-06 | 2.600E-06 |
| 8327 | 417 | G417Q | A | 1.5e-06 | 3.5e-06 | 2.8e-06 | 2.600E-06 | 1.015E-06 | 1.500E-06 | 3.500E-06 |
| 8328 | 417 | G417G | A | 2.8e-06 | 1.7e-06 | 2.3e-06 | 2.267E-06 | 5.508E-07 | 1.700E-06 | 2.800E-06 |
| 8329 | 417 | G417H | A | 2.2e-06 | 2.4e-06 | 2.8e-06 | 2.467E-06 | 3.055E-07 | 2.200E-06 | 2.800E-06 |

| | | | | | | | | | | |
|---|---|---|---|---|---|---|---|---|---|---|
| 8330 | 417 | G417I | A | 2.7e-06 | 1.3e-06 | 3.1e-06 | 2.367E-06 | 9.452E-07 | 1.300E-06 | 3.100E-06 |
| 8331 | 417 | G417L | A | 2.1e-06 | 1.4e-06 | 2e-06 | 1.833E-06 | 3.786E-07 | 1.400E-06 | 2.100E-06 |
| 8332 | 417 | G417K | A | 1.4e-06 | 2.9e-06 | 1.6e-06 | 1.967E-06 | 8.145E-07 | 1.400E-06 | 2.900E-06 |
| 8333 | 417 | G417M | A | 1.7e-06 | 2.2e-06 | 1.5e-06 | 1.800E-06 | 3.606E-07 | 1.500E-06 | 2.200E-06 |
| 8334 | 417 | G417F | A | 2e-06 | 2.2e-06 | 1.9e-06 | 2.033E-06 | 1.528E-07 | 1.900E-06 | 2.200E-06 |
| 8335 | 417 | G417P | A | 2.3e-06 | 3e-06 | 1.8e-06 | 2.367E-06 | 6.028E-07 | 1.800E-06 | 3.000E-06 |
| 8336 | 417 | G417S | A | 2.3e-06 | 2.1e-06 | 2.5e-06 | 2.300E-06 | 2.000E-07 | 2.100E-06 | 2.500E-06 |
| 8337 | 417 | G417T | A | 2.8e-06 | 2.2e-06 | 3.1e-06 | 2.700E-06 | 4.583E-07 | 2.200E-06 | 3.100E-06 |
| 8338 | 417 | G417W | A | 1.8e-06 | 2.1e-06 | 1.5e-06 | 1.800E-06 | 3.000E-07 | 1.500E-06 | 2.100E-06 |
| 8339 | 417 | G417Y | A | 2.9e-06 | 2.2e-06 | 2.7e-06 | 2.600E-06 | 3.606E-07 | 2.200E-06 | 2.900E-06 |
| 8340 | 417 | G417V | A | 2.5e-06 | 3.4e-06 | 4.9e-06 | 3.600E-06 | 1.212E-06 | 2.500E-06 | 4.900E-06 |
| 8341 | 418 | A418A | A | 3.7e-06 | 1.9e-06 | 2.7e-06 | 2.767E-06 | 9.018E-07 | 1.900E-06 | 3.700E-06 |
| 8342 | 418 | A418R | A | 2.1e-06 | 2.4e-06 | 1.3e-06 | 1.933E-06 | 5.686E-07 | 1.300E-06 | 2.400E-06 |
| 8343 | 418 | A418N | A | 1.9e-06 | 2.9e-06 | 2.8e-06 | 2.533E-06 | 5.508E-07 | 1.900E-06 | 2.900E-06 |
| 8344 | 418 | A418D | A | 2.8e-06 | 2.2e-06 | 3.4e-06 | 2.800E-06 | 6.000E-07 | 2.200E-06 | 3.400E-06 |
| 8345 | 418 | A418C | A | 1.7e-06 | 3.4e-06 | 2.6e-06 | 2.567E-06 | 8.505E-07 | 1.700E-06 | 3.400E-06 |
| 8346 | 418 | A418E | A | 2.5e-06 | 3.1e-06 | 3.1e-06 | 2.900E-06 | 3.464E-07 | 2.500E-06 | 3.100E-06 |
| 8347 | 418 | A418Q | A | 2.1e-06 | 2.7e-06 | 2.6e-06 | 2.467E-06 | 3.215E-07 | 2.100E-06 | 2.700E-06 |
| 8348 | 418 | A418G | A | 1.4e-06 | 3.5e-06 | 3.1e-06 | 2.667E-06 | 1.115E-06 | 1.400E-06 | 3.500E-06 |
| 8349 | 418 | A418H | A | 2.3e-06 | 2.1e-06 | 2.7e-06 | 2.367E-06 | 3.055E-07 | 2.100E-06 | 2.700E-06 |
| 8350 | 418 | A418I | A | 1.6e-06 | 1.8e-06 | 2.3e-06 | 1.900E-06 | 3.606E-07 | 1.600E-06 | 2.300E-06 |
| 8351 | 418 | A418L | A | 2.6e-06 | 1.6e-06 | 2.7e-06 | 2.300E-06 | 6.083E-07 | 1.600E-06 | 2.700E-06 |
| 8352 | 418 | A418K | A | 1.9e-06 | 3.5e-06 | 1.6e-06 | 2.333E-06 | 1.021E-06 | 1.600E-06 | 3.500E-06 |
| 8353 | 418 | A418M | A | 2.7e-06 | 2.7e-06 | 1.5e-06 | 2.300E-06 | 6.928E-07 | 1.500E-06 | 2.700E-06 |
| 8354 | 418 | A418F | A | 1.8e-06 | 3.3e-06 | 1.7e-06 | 2.267E-06 | 8.963E-07 | 1.700E-06 | 3.300E-06 |
| 8355 | 418 | A418P | A | 3.6e-06 | 1.5e-06 | 2.1e-06 | 2.400E-06 | 1.082E-06 | 1.500E-06 | 3.600E-06 |
| 8356 | 418 | A418S | A | 2.7e-06 | 2.6e-06 | 2.6e-06 | 2.633E-06 | 5.774E-08 | 2.600E-06 | 2.700E-06 |
| 8357 | 418 | A418T | A | 3.8e-06 | 2.6e-06 | 1.4e-06 | 2.600E-06 | 1.200E-06 | 1.400E-06 | 3.800E-06 |
| 8358 | 418 | A418W | A | 2e-06 | 3.2e-06 | 2.3e-06 | 2.500E-06 | 6.245E-07 | 2.000E-06 | 3.200E-06 |
| 8359 | 418 | A418Y | A | 1.9e-06 | 4.1e-06 | 2.8e-06 | 2.933E-06 | 1.106E-06 | 1.900E-06 | 4.100E-06 |
| 8360 | 418 | A418V | A | 1.4e-06 | 2.4e-06 | 4.5e-06 | 2.767E-06 | 1.582E-06 | 1.400E-06 | 4.500E-06 |
| 8361 | 419 | Y419A | A | 2.7e-06 | 2.7e-06 | 1.5e-06 | 2.300E-06 | 6.928E-07 | 1.500E-06 | 2.700E-06 |
| 8362 | 419 | Y419R | A | 3.2e-06 | 2.2e-06 | 3.2e-06 | 2.867E-06 | 5.774E-07 | 2.200E-06 | 3.200E-06 |
| 8363 | 419 | Y419N | A | 2.6e-06 | 2.7e-06 | 4e-06 | 3.100E-06 | 7.810E-07 | 2.600E-06 | 4.000E-06 |
| 8364 | 419 | Y419D | A | 1.7e-06 | 3.6e-06 | 1.9e-06 | 2.400E-06 | 1.044E-06 | 1.700E-06 | 3.600E-06 |
| 8365 | 419 | Y419C | A | 4.5e-06 | 2.8e-06 | 2.1e-06 | 3.133E-06 | 1.234E-06 | 2.100E-06 | 4.500E-06 |
| 8366 | 419 | Y419E | A | 3.2e-06 | 1.4e-06 | 3.5e-06 | 2.700E-06 | 1.136E-06 | 1.400E-06 | 3.500E-06 |
| 8367 | 419 | Y419Q | A | 2.9e-06 | 2.2e-06 | 2e-06 | 2.367E-06 | 4.726E-07 | 2.000E-06 | 2.900E-06 |
| 8368 | 419 | Y419G | A | 2.5e-06 | 2.7e-06 | 2.1e-06 | 2.433E-06 | 3.055E-07 | 2.100E-06 | 2.700E-06 |
| 8369 | 419 | Y419H | A | 1.8e-06 | 2e-06 | 1.5e-06 | 1.767E-06 | 2.517E-07 | 1.500E-06 | 2.000E-06 |
| 8370 | 419 | Y419I | A | 1.1e-06 | 2.2e-06 | 2.4e-06 | 1.900E-06 | 7.000E-07 | 1.100E-06 | 2.400E-06 |
| 8371 | 419 | Y419L | A | 8.7e-07 | 2e-06 | 3.9e-06 | 2.257E-06 | 1.531E-06 | 8.700E-07 | 3.900E-06 |
| 8372 | 419 | Y419K | A | 2.6e-06 | 4e-06 | 1.6e-06 | 2.733E-06 | 1.206E-06 | 1.600E-06 | 4.000E-06 |
| 8373 | 419 | Y419M | A | 3.3e-06 | 2.4e-06 | 1.6e-06 | 2.433E-06 | 8.505E-07 | 1.600E-06 | 3.300E-06 |
| 8374 | 419 | Y419F | A | 2.9e-06 | 2.8e-06 | 1.3e-06 | 2.333E-06 | 8.963E-07 | 1.300E-06 | 2.900E-06 |
| 8375 | 419 | Y419P | A | 2.3e-06 | 3.3e-06 | 2.3e-06 | 2.633E-06 | 5.774E-07 | 2.300E-06 | 3.300E-06 |
| 8376 | 419 | Y419S | A | 2.4e-06 | 2.3e-06 | 1.9e-06 | 2.200E-06 | 2.646E-07 | 1.900E-06 | 2.400E-06 |
| 8377 | 419 | Y419T | A | 3.3e-06 | 2.8e-06 | 1.7e-06 | 2.600E-06 | 8.185E-07 | 1.700E-06 | 3.300E-06 |
| 8378 | 419 | Y419W | A | 2.7e-06 | 2.3e-06 | 2.9e-06 | 2.633E-06 | 3.055E-07 | 2.300E-06 | 2.900E-06 |
| 8379 | 419 | Y419Y | A | 3.1e-06 | 2e-06 | 1.9e-06 | 2.333E-06 | 6.658E-07 | 1.900E-06 | 3.100E-06 |
| 8380 | 419 | Y419V | A | 2.3e-06 | 1.2e-06 | 1.6e-06 | 1.700E-06 | 5.568E-07 | 1.200E-06 | 2.300E-06 |
| 8381 | 420 | S420A | A | 2.9e-06 | 3.4e-06 | 2.1e-06 | 2.800E-06 | 6.557E-07 | 2.100E-06 | 3.400E-06 |
| 8382 | 420 | S420R | A | 2.6e-06 | 3.1e-06 | 2.7e-06 | 2.800E-06 | 2.646E-07 | 2.600E-06 | 3.100E-06 |
| 8383 | 420 | S420N | A | 3.4e-06 | 2e-06 | 1.9e-06 | 2.433E-06 | 8.386E-07 | 1.900E-06 | 3.400E-06 |
| 8384 | 420 | S420D | A | 2.8e-06 | 3.4e-06 | 2.9e-06 | 3.033E-06 | 3.215E-07 | 2.800E-06 | 3.400E-06 |
| 8385 | 420 | S420C | A | 2e-06 | 1.2e-06 | 2.7e-06 | 1.967E-06 | 7.506E-07 | 1.200E-06 | 2.700E-06 |
| 8386 | 420 | S420E | A | 2e-06 | 2.4e-06 | 2.9e-06 | 2.433E-06 | 4.509E-07 | 2.000E-06 | 2.900E-06 |
| 8387 | 420 | S420Q | A | 3.1e-06 | 1.6e-06 | 3.2e-06 | 2.633E-06 | 8.963E-07 | 1.600E-06 | 3.200E-06 |
| 8388 | 420 | S420G | A | 2.5e-06 | 1.7e-06 | 1.4e-06 | 1.867E-06 | 5.686E-07 | 1.400E-06 | 2.500E-06 |
| 8389 | 420 | S420H | A | 2.7e-06 | 1.5e-06 | 1.6e-06 | 1.933E-06 | 6.658E-07 | 1.500E-06 | 2.700E-06 |
| 8390 | 420 | S420I | A | 1.7e-06 | 2.1e-06 | 3.8e-06 | 2.533E-06 | 1.115E-06 | 1.700E-06 | 3.800E-06 |
| 8391 | 420 | S420L | A | 4.3e-06 | 2.5e-06 | 2.1e-06 | 2.967E-06 | 1.172E-06 | 2.100E-06 | 4.300E-06 |
| 8392 | 420 | S420K | A | 3.9e-06 | 3.2e-06 | 3e-06 | 3.367E-06 | 4.726E-07 | 3.000E-06 | 3.900E-06 |
| 8393 | 420 | S420M | A | 2.6e-06 | 1.4e-06 | 2.5e-06 | 2.167E-06 | 6.658E-07 | 1.400E-06 | 2.600E-06 |
| 8394 | 420 | S420F | A | 4.5e-06 | 2.4e-06 | 2.7e-06 | 3.200E-06 | 1.136E-06 | 2.400E-06 | 4.500E-06 |
| 8395 | 420 | S420P | A | 1.9e-06 | 2.6e-06 | 2.3e-06 | 2.267E-06 | 3.512E-07 | 1.900E-06 | 2.600E-06 |
| 8396 | 420 | S420S | A | 2.7e-06 | 1.9e-06 | 2.1e-06 | 2.233E-06 | 4.163E-07 | 1.900E-06 | 2.700E-06 |
| 8397 | 420 | S420T | A | 1.8e-06 | 2.1e-06 | 1.7e-06 | 1.867E-06 | 2.082E-07 | 1.700E-06 | 2.100E-06 |
| 8398 | 420 | S420W | A | 1.7e-06 | 2.9e-06 | 2.1e-06 | 2.233E-06 | 6.110E-07 | 1.700E-06 | 2.900E-06 |
| 8399 | 420 | S420Y | A | 1.9e-06 | 2.3e-06 | 2.3e-06 | 2.167E-06 | 2.309E-07 | 1.900E-06 | 2.300E-06 |
| 8400 | 420 | S420V | A | 2.1e-06 | 3.3e-06 | 2.2e-06 | 2.533E-06 | 6.658E-07 | 2.100E-06 | 3.300E-06 |
| 8401 | 421 | L421A | A | 1.2e-06 | 1.9e-06 | 3.5e-06 | 2.200E-06 | 1.179E-06 | 1.200E-06 | 3.500E-06 |
| 8402 | 421 | L421R | A | 3.1e-06 | 1.8e-06 | 2e-06 | 2.300E-06 | 7.000E-07 | 1.800E-06 | 3.100E-06 |
| 8403 | 421 | L421N | A | 3.2e-06 | 2.6e-06 | 2.7e-06 | 2.833E-06 | 3.215E-07 | 2.600E-06 | 3.200E-06 |
| 8404 | 421 | L421D | A | 3.5e-06 | 2.6e-06 | 3.2e-06 | 3.100E-06 | 4.583E-07 | 2.600E-06 | 3.500E-06 |
| 8405 | 421 | L421C | A | 2.6e-06 | 2.4e-06 | 3.1e-06 | 2.700E-06 | 3.606E-07 | 2.400E-06 | 3.100E-06 |
| 8406 | 421 | L421E | A | 4.8e-06 | 1.4e-06 | 1.4e-06 | 2.533E-06 | 1.963E-06 | 1.400E-06 | 4.800E-06 |
| 8407 | 421 | L421Q | A | 2.5e-06 | 2.3e-06 | 2.3e-06 | 2.367E-06 | 1.155E-07 | 2.300E-06 | 2.500E-06 |
| 8408 | 421 | L421G | A | 1.5e-06 | 2e-06 | 2.9e-06 | 2.133E-06 | 7.095E-07 | 1.500E-06 | 2.900E-06 |
| 8409 | 421 | L421H | A | 2.9e-06 | 4.1e-06 | 1.5e-06 | 2.833E-06 | 1.301E-06 | 1.500E-06 | 4.100E-06 |
| 8410 | 421 | L421I | A | 3.9e-06 | 2.7e-06 | 2.2e-06 | 2.933E-06 | 8.737E-07 | 2.200E-06 | 3.900E-06 |
| 8411 | 421 | L421L | A | 4.4e-06 | 2.9e-06 | 1.7e-06 | 3.000E-06 | 1.353E-06 | 1.700E-06 | 4.400E-06 |
| 8412 | 421 | L421K | A | 4.5e-06 | 2.7e-06 | 2.9e-06 | 3.367E-06 | 9.866E-07 | 2.700E-06 | 4.500E-06 |
| 8413 | 421 | L421M | A | 1.7e-06 | 3.5e-06 | 1.6e-06 | 2.267E-06 | 1.069E-06 | 1.600E-06 | 3.500E-06 |
| 8414 | 421 | L421F | A | 2.6e-06 | 1.9e-06 | 1.1e-06 | 1.867E-06 | 7.506E-07 | 1.100E-06 | 2.600E-06 |

| | | | | | | | | | | |
|---|---|---|---|---|---|---|---|---|---|---|
| 8415 | 421 | L421P | A | 2.8e-06 | 2.2e-06 | 3.8e-06 | 2.933E-06 | 8.083E-07 | 2.200E-06 | 3.800E-06 |
| 8416 | 421 | L421S | A | 2e-06 | 4.5e-06 | 1.7e-06 | 2.733E-06 | 1.537E-06 | 1.700E-06 | 4.500E-06 |
| 8417 | 421 | L421T | A | 3.8e-06 | 3.7e-06 | 2.6e-06 | 3.367E-06 | 6.658E-07 | 2.600E-06 | 3.800E-06 |
| 8418 | 421 | L421W | A | 2.8e-06 | 2.1e-06 | 3e-06 | 2.633E-06 | 4.726E-07 | 2.100E-06 | 3.000E-06 |
| 8419 | 421 | L421Y | A | 1.9e-06 | 3.9e-06 | 2.6e-06 | 2.800E-06 | 1.015E-06 | 1.900E-06 | 3.900E-06 |
| 8420 | 421 | L421V | A | 2.1e-06 | 2.1e-06 | 2.6e-06 | 2.267E-06 | 2.887E-07 | 2.100E-06 | 2.600E-06 |
| 8421 | 422 | T422A | A | 2.2e-06 | 2.9e-06 | 2.5e-06 | 2.533E-06 | 3.512E-07 | 2.200E-06 | 2.900E-06 |
| 8422 | 422 | T422R | A | 3.3e-06 | 2.2e-06 | 3.3e-06 | 2.933E-06 | 6.351E-07 | 2.200E-06 | 3.300E-06 |
| 8423 | 422 | T422N | A | 2.9e-06 | 2.5e-06 | 2.5e-06 | 2.633E-06 | 2.309E-07 | 2.500E-06 | 2.900E-06 |
| 8424 | 422 | T422D | A | 2.1e-06 | 8e-07 | 2.3e-06 | 1.733E-06 | 8.145E-07 | 8.000E-07 | 2.300E-06 |
| 8425 | 422 | T422C | A | 2.6e-06 | 9.9e-07 | 1.3e-06 | 1.630E-06 | 8.542E-07 | 9.900E-07 | 2.600E-06 |
| 8426 | 422 | T422E | A | 1.8e-06 | 3.8e-06 | 1.6e-06 | 2.400E-06 | 1.217E-06 | 1.600E-06 | 3.800E-06 |
| 8427 | 422 | T422Q | A | 2.5e-06 | 3.2e-06 | 2.2e-06 | 2.633E-06 | 5.132E-07 | 2.200E-06 | 3.200E-06 |
| 8428 | 422 | T422G | A | 2.9e-06 | 2.1e-06 | 2.3e-06 | 2.433E-06 | 4.163E-07 | 2.100E-06 | 2.900E-06 |
| 8429 | 422 | T422H | A | 2.3e-06 | 1.9e-06 | 3.7e-06 | 2.633E-06 | 9.452E-07 | 1.900E-06 | 3.700E-06 |
| 8430 | 422 | T422I | A | 2.6e-06 | 2.2e-06 | 2.3e-06 | 2.367E-06 | 2.082E-07 | 2.200E-06 | 2.600E-06 |
| 8431 | 422 | T422L | A | 2.5e-06 | 1.9e-06 | 2.4e-06 | 2.267E-06 | 3.215E-07 | 1.900E-06 | 2.500E-06 |
| 8432 | 422 | T422K | A | 1.7e-06 | 2e-06 | 1.5e-06 | 1.733E-06 | 2.517E-07 | 1.500E-06 | 2.000E-06 |
| 8433 | 422 | T422M | A | 3e-06 | 2.7e-06 | 1.3e-06 | 2.333E-06 | 9.074E-07 | 1.300E-06 | 3.000E-06 |
| 8434 | 422 | T422F | A | 2.1e-06 | 2.1e-06 | 4.7e-06 | 2.967E-06 | 1.501E-06 | 2.100E-06 | 4.700E-06 |
| 8435 | 422 | T422P | A | 1.5e-06 | 2.7e-06 | 1.2e-06 | 1.800E-06 | 7.937E-07 | 1.200E-06 | 2.700E-06 |
| 8436 | 422 | T422S | A | 1.9e-06 | 3e-06 | 1.8e-06 | 2.233E-06 | 6.658E-07 | 1.800E-06 | 3.000E-06 |
| 8437 | 422 | T422T | A | 2e-06 | 2.1e-06 | 2.6e-06 | 2.233E-06 | 3.215E-07 | 2.000E-06 | 2.600E-06 |
| 8438 | 422 | T422W | A | 2.7e-06 | 2.7e-06 | 2.8e-06 | 2.733E-06 | 5.774E-08 | 2.700E-06 | 2.800E-06 |
| 8439 | 422 | T422Y | A | 2.1e-06 | 3.3e-06 | 2.5e-06 | 2.633E-06 | 6.110E-07 | 2.100E-06 | 3.300E-06 |
| 8440 | 422 | T422V | A | 5.3e-06 | 3.3e-06 | 2.4e-06 | 3.667E-06 | 1.484E-06 | 2.400E-06 | 5.300E-06 |
| 8441 | 423 | L423A | A | 2.3e-06 | 2.3e-06 | 1.4e-06 | 2.000E-06 | 5.196E-07 | 1.400E-06 | 2.300E-06 |
| 8442 | 423 | L423R | A | 2.5e-06 | 3e-06 | 2.1e-06 | 2.533E-06 | 4.509E-07 | 2.100E-06 | 3.000E-06 |
| 8443 | 423 | L423N | A | 3.3e-06 | 2e-06 | 3.8e-06 | 3.033E-06 | 9.292E-07 | 2.000E-06 | 3.800E-06 |
| 8444 | 423 | L423D | A | 3e-06 | 1.2e-06 | 3e-06 | 2.400E-06 | 1.039E-06 | 1.200E-06 | 3.000E-06 |
| 8445 | 423 | L423C | A | 2.5e-06 | 1.6e-06 | 3.4e-06 | 2.500E-06 | 9.000E-07 | 1.600E-06 | 3.400E-06 |
| 8446 | 423 | L423E | A | 2.5e-06 | 2.1e-06 | 2.1e-06 | 2.233E-06 | 2.309E-07 | 2.100E-06 | 2.500E-06 |
| 8447 | 423 | L423Q | A | 2.7e-06 | 1.9e-06 | 1.9e-06 | 2.167E-06 | 4.619E-07 | 1.900E-06 | 2.700E-06 |
| 8448 | 423 | L423G | A | 3e-06 | 3.7e-06 | 1.3e-06 | 2.667E-06 | 1.234E-06 | 1.300E-06 | 3.700E-06 |
| 8449 | 423 | L423H | A | 2.6e-06 | 2.9e-06 | 1.9e-06 | 2.467E-06 | 5.132E-07 | 1.900E-06 | 2.900E-06 |
| 8450 | 423 | L423I | A | 2.5e-06 | 1.3e-06 | 2.3e-06 | 2.033E-06 | 6.429E-07 | 1.300E-06 | 2.500E-06 |
| 8451 | 423 | L423L | A | 1.2e-06 | 3.6e-06 | 2.7e-06 | 2.500E-06 | 1.212E-06 | 1.200E-06 | 3.600E-06 |
| 8452 | 423 | L423K | A | 3.7e-06 | 2.6e-06 | 2e-06 | 2.767E-06 | 8.622E-07 | 2.000E-06 | 3.700E-06 |
| 8453 | 423 | L423M | A | 3.6e-06 | 2.3e-06 | 3.5e-06 | 3.133E-06 | 7.234E-07 | 2.300E-06 | 3.600E-06 |
| 8454 | 423 | L423F | A | 2.1e-06 | 1.2e-06 | 2.6e-06 | 1.967E-06 | 7.095E-07 | 1.200E-06 | 2.600E-06 |
| 8455 | 423 | L423P | A | 2.1e-06 | 2.8e-06 | 9.4e-07 | 1.947E-06 | 9.394E-07 | 9.400E-07 | 2.800E-06 |
| 8456 | 423 | L423S | A | 2.4e-06 | 2e-06 | 1.9e-06 | 2.100E-06 | 2.646E-07 | 1.900E-06 | 2.400E-06 |
| 8457 | 423 | L423T | A | 5e-06 | 1.8e-06 | 2.5e-06 | 3.100E-06 | 1.682E-06 | 1.800E-06 | 5.000E-06 |
| 8458 | 423 | L423W | A | 2.1e-06 | 3e-06 | 1.1e-06 | 2.067E-06 | 9.504E-07 | 1.100E-06 | 3.000E-06 |
| 8459 | 423 | L423Y | A | 1.2e-06 | 2.9e-06 | 1.3e-06 | 1.800E-06 | 9.539E-07 | 1.200E-06 | 2.900E-06 |
| 8460 | 423 | L423V | A | 3.6e-06 | 1.6e-06 | 2.4e-06 | 2.533E-06 | 1.007E-06 | 1.600E-06 | 3.600E-06 |
| 8461 | 424 | Q424A | A | 1.8e-06 | 2.7e-06 | 2.2e-06 | 2.233E-06 | 4.509E-07 | 1.800E-06 | 2.700E-06 |
| 8462 | 424 | Q424R | A | 2.9e-06 | 1.6e-06 | 2.4e-06 | 2.300E-06 | 6.557E-07 | 1.600E-06 | 2.900E-06 |
| 8463 | 424 | Q424N | A | 4e-06 | 1.6e-06 | 2.6e-06 | 2.733E-06 | 1.206E-06 | 1.600E-06 | 4.000E-06 |
| 8464 | 424 | Q424D | A | 2e-06 | 4.2e-06 | 3.1e-06 | 3.100E-06 | 1.100E-06 | 2.000E-06 | 4.200E-06 |
| 8465 | 424 | Q424C | A | 2.5e-06 | 2e-06 | 3.2e-06 | 2.567E-06 | 6.028E-07 | 2.000E-06 | 3.200E-06 |
| 8466 | 424 | Q424E | A | 2.1e-06 | 3.3e-06 | 1.9e-06 | 2.433E-06 | 7.572E-07 | 1.900E-06 | 3.300E-06 |
| 8467 | 424 | Q424Q | A | 2.6e-06 | 2.5e-06 | 1.7e-06 | 2.267E-06 | 4.933E-07 | 1.700E-06 | 2.600E-06 |
| 8468 | 424 | Q424G | A | 3.1e-06 | 2.4e-06 | 2e-06 | 2.500E-06 | 5.568E-07 | 2.000E-06 | 3.100E-06 |
| 8469 | 424 | Q424H | A | 2.2e-06 | 3e-06 | 2.1e-06 | 2.433E-06 | 4.933E-07 | 2.100E-06 | 3.000E-06 |
| 8470 | 424 | Q424I | A | 2e-06 | 1.2e-06 | 1.5e-06 | 1.567E-06 | 4.041E-07 | 1.200E-06 | 2.000E-06 |
| 8471 | 424 | Q424L | A | 1.2e-06 | 1.2e-06 | 3e-06 | 1.800E-06 | 1.039E-06 | 1.200E-06 | 3.000E-06 |
| 8472 | 424 | Q424K | A | 2.8e-06 | 1.3e-06 | 2.8e-06 | 2.300E-06 | 8.660E-07 | 1.300E-06 | 2.800E-06 |
| 8473 | 424 | Q424M | A | 2.3e-06 | 2.2e-06 | 2.6e-06 | 2.367E-06 | 2.082E-07 | 2.200E-06 | 2.600E-06 |
| 8474 | 424 | Q424F | A | 1.8e-06 | 2.9e-06 | 2.5e-06 | 2.400E-06 | 5.568E-07 | 1.800E-06 | 2.900E-06 |
| 8475 | 424 | Q424P | A | 1.9e-06 | 1.9e-06 | 4.6e-06 | 2.800E-06 | 1.559E-06 | 1.900E-06 | 4.600E-06 |
| 8476 | 424 | Q424S | A | 2.5e-06 | 3.4e-06 | 2.2e-06 | 2.700E-06 | 6.245E-07 | 2.200E-06 | 3.400E-06 |
| 8477 | 424 | Q424T | A | 1.8e-06 | 1.8e-06 | 2.1e-06 | 1.900E-06 | 1.732E-07 | 1.800E-06 | 2.100E-06 |
| 8478 | 424 | Q424W | A | 2.4e-06 | 2.1e-06 | 4.2e-06 | 2.900E-06 | 1.136E-06 | 2.100E-06 | 4.200E-06 |
| 8479 | 424 | Q424Y | A | 2.3e-06 | 1.7e-06 | 2.8e-06 | 2.267E-06 | 5.508E-07 | 1.700E-06 | 2.800E-06 |
| 8480 | 424 | Q424V | A | 2.3e-06 | 3.4e-06 | 2.3e-06 | 2.667E-06 | 6.351E-07 | 2.300E-06 | 3.400E-06 |
| 8481 | 425 | G425A | A | 3.5e-06 | 1.5e-06 | 2.1e-06 | 2.367E-06 | 1.026E-06 | 1.500E-06 | 3.500E-06 |
| 8482 | 425 | G425R | A | 1.8e-06 | 2e-06 | 2e-06 | 1.933E-06 | 1.155E-07 | 1.800E-06 | 2.000E-06 |
| 8483 | 425 | G425N | A | 1.6e-06 | 2.5e-06 | 2.3e-06 | 2.133E-06 | 4.726E-07 | 1.600E-06 | 2.500E-06 |
| 8484 | 425 | G425D | A | 1.5e-06 | 3.6e-06 | 2e-06 | 2.367E-06 | 1.097E-06 | 1.500E-06 | 3.600E-06 |
| 8485 | 425 | G425C | A | 1.6e-06 | 1.2e-06 | 2.5e-06 | 1.767E-06 | 6.658E-07 | 1.200E-06 | 2.500E-06 |
| 8486 | 425 | G425E | A | 3.2e-06 | 2.3e-06 | 3.6e-06 | 3.033E-06 | 6.658E-07 | 2.300E-06 | 3.600E-06 |
| 8487 | 425 | G425Q | A | 2e-06 | 1.5e-06 | 1e-06 | 1.500E-06 | 5.000E-07 | 1.000E-06 | 2.000E-06 |
| 8488 | 425 | G425G | A | 1.7e-06 | 1.6e-06 | 2.4e-06 | 1.900E-06 | 4.359E-07 | 1.600E-06 | 2.400E-06 |
| 8489 | 425 | G425H | A | 2.4e-06 | 2.7e-06 | 1.3e-06 | 2.133E-06 | 7.371E-07 | 1.300E-06 | 2.700E-06 |
| 8490 | 425 | G425I | A | 2.6e-06 | 2.8e-06 | 1.9e-06 | 2.433E-06 | 4.726E-07 | 1.900E-06 | 2.800E-06 |
| 8491 | 425 | G425L | A | 3.8e-06 | 2e-06 | 2.4e-06 | 2.733E-06 | 9.452E-07 | 2.000E-06 | 3.800E-06 |
| 8492 | 425 | G425K | A | 1.9e-06 | 2.3e-06 | 2.4e-06 | 2.200E-06 | 2.646E-07 | 1.900E-06 | 2.400E-06 |
| 8493 | 425 | G425M | A | 1.5e-06 | 3e-06 | 2.7e-06 | 2.400E-06 | 7.937E-07 | 1.500E-06 | 3.000E-06 |
| 8494 | 425 | G425F | A | 2e-06 | 1.5e-06 | 2.1e-06 | 1.867E-06 | 3.215E-07 | 1.500E-06 | 2.100E-06 |
| 8495 | 425 | G425P | A | 1.7e-06 | 4e-06 | 1.5e-06 | 2.400E-06 | 1.389E-06 | 1.500E-06 | 4.000E-06 |
| 8496 | 425 | G425S | A | 3.9e-06 | 1.4e-06 | 2.8e-06 | 2.700E-06 | 1.253E-06 | 1.400E-06 | 3.900E-06 |
| 8497 | 425 | G425T | A | 2.1e-06 | 2.4e-06 | 7.1e-07 | 1.737E-06 | 9.017E-07 | 7.100E-07 | 2.400E-06 |
| 8498 | 425 | G425W | A | 2.2e-06 | 2.1e-06 | 1.6e-06 | 1.967E-06 | 3.215E-07 | 1.600E-06 | 2.200E-06 |
| 8499 | 425 | G425Y | A | 4e-06 | 2.3e-06 | 4.1e-06 | 3.467E-06 | 1.012E-06 | 2.300E-06 | 4.100E-06 |

| | | | | | | | | | | |
|---|---|---|---|---|---|---|---|---|---|---|
| 8500 | 425 | G425V | A | 1.6e-06 | 1.9e-06 | 1.5e-06 | 1.667E-06 | 2.082E-07 | 1.500E-06 | 1.900E-06 |
| 8501 | 426 | L426A | A | 2.1e-06 | 1.4e-06 | 2.6e-06 | 2.033E-06 | 6.028E-07 | 1.400E-06 | 2.600E-06 |
| 8502 | 426 | L426R | A | 1.5e-06 | 1.5e-06 | 2.2e-06 | 1.733E-06 | 4.041E-07 | 1.500E-06 | 2.200E-06 |
| 8503 | 426 | L426N | A | 1.4e-06 | 1.1e-06 | 2.6e-06 | 1.700E-06 | 7.937E-07 | 1.100E-06 | 2.600E-06 |
| 8504 | 426 | L426D | A | 1.9e-06 | 2.8e-06 | 3.4e-06 | 2.700E-06 | 7.550E-07 | 1.900E-06 | 3.400E-06 |
| 8505 | 426 | L426C | A | 1.7e-06 | 3.5e-06 | 5.3e-06 | 3.500E-06 | 1.800E-06 | 1.700E-06 | 5.300E-06 |
| 8506 | 426 | L426E | A | 2.4e-06 | 1.3e-06 | 2.3e-06 | 2.000E-06 | 6.083E-07 | 1.300E-06 | 2.400E-06 |
| 8507 | 426 | L426Q | A | 2.5e-06 | 1.6e-06 | 2.6e-06 | 2.233E-06 | 5.508E-07 | 1.600E-06 | 2.600E-06 |
| 8508 | 426 | L426G | A | 1.2e-06 | 2.3e-06 | 3e-06 | 2.167E-06 | 9.074E-07 | 1.200E-06 | 3.000E-06 |
| 8509 | 426 | L426H | A | 1.8e-06 | 1.8e-06 | 2.3e-06 | 1.967E-06 | 2.887E-07 | 1.800E-06 | 2.300E-06 |
| 8510 | 426 | L426I | A | 3.3e-06 | 3.2e-06 | 2.1e-06 | 2.867E-06 | 6.658E-07 | 2.100E-06 | 3.300E-06 |
| 8511 | 426 | L426L | A | 2.1e-06 | 1.7e-06 | 1.4e-06 | 1.733E-06 | 3.512E-07 | 1.400E-06 | 2.100E-06 |
| 8512 | 426 | L426K | A | 2.7e-06 | 2.3e-06 | 1.4e-06 | 2.133E-06 | 6.658E-07 | 1.400E-06 | 2.700E-06 |
| 8513 | 426 | L426M | A | 2.8e-06 | 2.5e-06 | 2.1e-06 | 2.467E-06 | 3.512E-07 | 2.100E-06 | 2.800E-06 |
| 8514 | 426 | L426F | A | 1.8e-06 | 3.1e-06 | 2.2e-06 | 2.367E-06 | 6.658E-07 | 1.800E-06 | 3.100E-06 |
| 8515 | 426 | L426P | A | 3.1e-06 | 1.9e-06 | 2.6e-06 | 2.533E-06 | 6.028E-07 | 1.900E-06 | 3.100E-06 |
| 8516 | 426 | L426S | A | 1.7e-06 | 4.7e-06 | 1.4e-06 | 2.600E-06 | 1.825E-06 | 1.400E-06 | 4.700E-06 |
| 8517 | 426 | L426T | A | 2.4e-06 | 2.1e-06 | 2.5e-06 | 2.333E-06 | 2.082E-07 | 2.100E-06 | 2.500E-06 |
| 8518 | 426 | L426W | A | 3.2e-06 | 2e-06 | 2.7e-06 | 2.633E-06 | 6.028E-07 | 2.000E-06 | 3.200E-06 |
| 8519 | 426 | L426Y | A | 5.7e-06 | 2.6e-06 | 2.9e-06 | 3.733E-06 | 1.710E-06 | 2.600E-06 | 5.700E-06 |
| 8520 | 426 | L426V | A | 1.3e-06 | 1.8e-06 | 3e-06 | 2.033E-06 | 8.737E-07 | 1.300E-06 | 3.000E-06 |
| 8521 | 427 | G427A | A | 2.6e-06 | 2.2e-06 | 1.4e-06 | 2.067E-06 | 6.110E-07 | 1.400E-06 | 2.600E-06 |
| 8522 | 427 | G427R | A | 1.6e-06 | 3.1e-06 | 2.4e-06 | 2.367E-06 | 7.506E-07 | 1.600E-06 | 3.100E-06 |
| 8523 | 427 | G427N | A | 2.1e-06 | 2.6e-06 | 2.6e-06 | 2.433E-06 | 2.887E-07 | 2.100E-06 | 2.600E-06 |
| 8524 | 427 | G427D | A | 2.5e-06 | 2.7e-06 | 1.9e-06 | 2.367E-06 | 4.163E-07 | 1.900E-06 | 2.700E-06 |
| 8525 | 427 | G427C | A | 2.3e-06 | 1.9e-06 | 1.5e-06 | 1.900E-06 | 4.000E-07 | 1.500E-06 | 2.300E-06 |
| 8526 | 427 | G427E | A | 3.4e-06 | 1.3e-06 | 5.1e-06 | 3.267E-06 | 1.904E-06 | 1.300E-06 | 5.100E-06 |
| 8527 | 427 | G427Q | A | 2.3e-06 | 2.4e-06 | 2.2e-06 | 2.300E-06 | 1.000E-07 | 2.200E-06 | 2.400E-06 |
| 8528 | 427 | G427G | A | 1.9e-06 | 1.5e-06 | 2.6e-06 | 2.000E-06 | 5.568E-07 | 1.500E-06 | 2.600E-06 |
| 8529 | 427 | G427H | A | 2.1e-06 | 3e-06 | 2.7e-06 | 2.600E-06 | 4.583E-07 | 2.100E-06 | 3.000E-06 |
| 8530 | 427 | G427I | A | 2e-06 | 2.6e-06 | 2.9e-06 | 2.500E-06 | 4.583E-07 | 2.000E-06 | 2.900E-06 |
| 8531 | 427 | G427L | A | 1.9e-06 | 2.7e-06 | 1.6e-06 | 2.067E-06 | 5.686E-07 | 1.600E-06 | 2.700E-06 |
| 8532 | 427 | G427K | A | 2.2e-06 | 3.7e-06 | 2.2e-06 | 2.700E-06 | 8.660E-07 | 2.200E-06 | 3.700E-06 |
| 8533 | 427 | G427M | A | 3.2e-06 | 1.8e-06 | 3.2e-06 | 2.733E-06 | 8.083E-07 | 1.800E-06 | 3.200E-06 |
| 8534 | 427 | G427F | A | 1.5e-06 | 1.7e-06 | 2.6e-06 | 1.933E-06 | 5.859E-07 | 1.500E-06 | 2.600E-06 |
| 8535 | 427 | G427P | A | 3e-06 | 1.9e-06 | 2.8e-06 | 2.567E-06 | 5.859E-07 | 1.900E-06 | 3.000E-06 |
| 8536 | 427 | G427S | A | 2.1e-06 | 2.6e-06 | 1.3e-06 | 2.000E-06 | 6.557E-07 | 1.300E-06 | 2.600E-06 |
| 8537 | 427 | G427T | A | 2.1e-06 | 1.8e-06 | 2.5e-06 | 2.133E-06 | 3.512E-07 | 1.800E-06 | 2.500E-06 |
| 8538 | 427 | G427W | A | 4.3e-06 | 1.6e-06 | 2.2e-06 | 2.700E-06 | 1.418E-06 | 1.600E-06 | 4.300E-06 |
| 8539 | 427 | G427Y | A | 2.9e-06 | 2.5e-06 | 3.9e-06 | 3.100E-06 | 7.211E-07 | 2.500E-06 | 3.900E-06 |
| 8540 | 427 | G427V | A | 1.9e-06 | 2e-06 | 2.7e-06 | 2.200E-06 | 4.359E-07 | 1.900E-06 | 2.700E-06 |
| 8541 | 428 | I428A | A | 3.2e-06 | 3.1e-06 | 2.5e-06 | 2.933E-06 | 3.786E-07 | 2.500E-06 | 3.200E-06 |
| 8542 | 428 | I428R | A | 2.5e-06 | 1.7e-06 | 3e-06 | 2.400E-06 | 6.557E-07 | 1.700E-06 | 3.000E-06 |
| 8543 | 428 | I428N | A | 1.9e-06 | 4.5e-06 | 2.6e-06 | 3.000E-06 | 1.345E-06 | 1.900E-06 | 4.500E-06 |
| 8544 | 428 | I428D | A | 1.3e-06 | 2.4e-06 | 4.9e-06 | 2.867E-06 | 1.845E-06 | 1.300E-06 | 4.900E-06 |
| 8545 | 428 | I428C | A | 2.1e-06 | 1.9e-06 | 3.1e-06 | 2.367E-06 | 6.429E-07 | 1.900E-06 | 3.100E-06 |
| 8546 | 428 | I428E | A | 3.4e-06 | 1.8e-06 | 2.5e-06 | 2.567E-06 | 8.021E-07 | 1.800E-06 | 3.400E-06 |
| 8547 | 428 | I428Q | A | 1.4e-06 | 2.5e-06 | 2.7e-06 | 2.200E-06 | 7.000E-07 | 1.400E-06 | 2.700E-06 |
| 8548 | 428 | I428G | A | 4.5e-06 | 1.7e-06 | 1.2e-06 | 2.467E-06 | 1.779E-06 | 1.200E-06 | 4.500E-06 |
| 8549 | 428 | I428H | A | 1.8e-06 | 1.7e-06 | 1.7e-06 | 1.733E-06 | 5.774E-08 | 1.700E-06 | 1.800E-06 |
| 8550 | 428 | I428I | A | 1.4e-06 | 2.7e-06 | 2.4e-06 | 2.167E-06 | 6.807E-07 | 1.400E-06 | 2.700E-06 |
| 8551 | 428 | I428L | A | 3.2e-06 | 2.2e-06 | 2.8e-06 | 2.733E-06 | 5.033E-07 | 2.200E-06 | 3.200E-06 |
| 8552 | 428 | I428K | A | 3.2e-06 | 2.6e-06 | 1.3e-06 | 2.367E-06 | 9.713E-07 | 1.300E-06 | 3.200E-06 |
| 8553 | 428 | I428M | A | 3.4e-06 | 2.5e-06 | 3.8e-06 | 3.233E-06 | 6.658E-07 | 2.500E-06 | 3.800E-06 |
| 8554 | 428 | I428F | A | 1.6e-06 | 4.5e-06 | 2.2e-06 | 2.767E-06 | 1.531E-06 | 1.600E-06 | 4.500E-06 |
| 8555 | 428 | I428P | A | 4.3e-06 | 2.5e-06 | 2e-06 | 2.933E-06 | 1.210E-06 | 2.000E-06 | 4.300E-06 |
| 8556 | 428 | I428S | A | 2.4e-06 | 1.7e-06 | 4.1e-06 | 2.733E-06 | 1.234E-06 | 1.700E-06 | 4.100E-06 |
| 8557 | 428 | I428T | A | 2.1e-06 | 3.7e-06 | 2.4e-06 | 2.733E-06 | 8.505E-07 | 2.100E-06 | 3.700E-06 |
| 8558 | 428 | I428W | A | 2.9e-06 | 2.4e-06 | 1.7e-06 | 2.333E-06 | 6.028E-07 | 1.700E-06 | 2.900E-06 |
| 8559 | 428 | I428Y | A | 2.8e-06 | 2.5e-06 | 2.8e-06 | 2.700E-06 | 1.732E-07 | 2.500E-06 | 2.800E-06 |
| 8560 | 428 | I428V | A | 2e-06 | 2.2e-06 | 1.9e-06 | 2.033E-06 | 1.528E-07 | 1.900E-06 | 2.200E-06 |
| 8561 | 429 | S429A | A | 2.6e-06 | 4.7e-06 | 3e-06 | 3.433E-06 | 1.115E-06 | 2.600E-06 | 4.700E-06 |
| 8562 | 429 | S429R | A | 4e-06 | 2.1e-06 | 3e-06 | 3.033E-06 | 9.504E-07 | 2.100E-06 | 4.000E-06 |
| 8563 | 429 | S429N | A | 2.4e-06 | 2.6e-06 | 3e-06 | 2.667E-06 | 3.055E-07 | 2.400E-06 | 3.000E-06 |
| 8564 | 429 | S429D | A | 2e-06 | 1.9e-06 | 2e-06 | 1.967E-06 | 5.774E-08 | 1.900E-06 | 2.000E-06 |
| 8565 | 429 | S429C | A | 1.5e-06 | 1.1e-06 | 3.1e-06 | 1.900E-06 | 1.058E-06 | 1.100E-06 | 3.100E-06 |
| 8566 | 429 | S429E | A | 3.3e-06 | 1.5e-06 | 2.4e-06 | 2.400E-06 | 9.000E-07 | 1.500E-06 | 3.300E-06 |
| 8567 | 429 | S429Q | A | 3.4e-06 | 2.7e-06 | 2.1e-06 | 2.733E-06 | 6.506E-07 | 2.100E-06 | 3.400E-06 |
| 8568 | 429 | S429G | A | 1.6e-06 | 1.9e-06 | 3.4e-06 | 2.300E-06 | 9.644E-07 | 1.600E-06 | 3.400E-06 |
| 8569 | 429 | S429H | A | 2.8e-06 | 3.3e-06 | 4e-06 | 3.367E-06 | 6.028E-07 | 2.800E-06 | 4.000E-06 |
| 8570 | 429 | S429I | A | 2.6e-06 | 2.3e-06 | 2.3e-06 | 2.400E-06 | 1.732E-07 | 2.300E-06 | 2.600E-06 |
| 8571 | 429 | S429L | A | 4.5e-06 | 3.7e-06 | 3e-06 | 3.733E-06 | 7.506E-07 | 3.000E-06 | 4.500E-06 |
| 8572 | 429 | S429K | A | 2.3e-06 | 2.2e-06 | 1.7e-06 | 2.067E-06 | 3.215E-07 | 1.700E-06 | 2.300E-06 |
| 8573 | 429 | S429M | A | 2e-06 | 4.2e-06 | 3.5e-06 | 3.233E-06 | 1.124E-06 | 2.000E-06 | 4.200E-06 |
| 8574 | 429 | S429F | A | 3.1e-06 | 1.8e-06 | 4.3e-06 | 3.067E-06 | 1.250E-06 | 1.800E-06 | 4.300E-06 |
| 8575 | 429 | S429P | A | 2.1e-06 | 2.6e-06 | 2.8e-06 | 2.500E-06 | 3.606E-07 | 2.100E-06 | 2.800E-06 |
| 8576 | 429 | S429S | A | 2e-06 | 6.2e-06 | 3e-06 | 3.733E-06 | 2.194E-06 | 2.000E-06 | 6.200E-06 |
| 8577 | 429 | S429T | A | 1.5e-06 | 1.3e-06 | 2.4e-06 | 1.733E-06 | 5.859E-07 | 1.300E-06 | 2.400E-06 |
| 8578 | 429 | S429W | A | 1.7e-06 | 3.3e-06 | 2.7e-06 | 2.567E-06 | 8.083E-07 | 1.700E-06 | 3.300E-06 |
| 8579 | 429 | S429Y | A | 2.5e-06 | 2.8e-06 | 3.2e-06 | 2.833E-06 | 3.512E-07 | 2.500E-06 | 3.200E-06 |
| 8580 | 429 | S429V | A | 1.9e-06 | 1.6e-06 | 2.6e-06 | 2.033E-06 | 5.132E-07 | 1.600E-06 | 2.600E-06 |
| 8581 | 430 | W430A | A | 2.7e-06 | 2.7e-06 | 3.1e-06 | 2.833E-06 | 2.309E-07 | 2.700E-06 | 3.100E-06 |
| 8582 | 430 | W430R | A | 1.4e-06 | 3.1e-06 | 1.9e-06 | 2.133E-06 | 8.737E-07 | 1.400E-06 | 3.100E-06 |
| 8583 | 430 | W430N | A | 2.6e-06 | 1.8e-06 | 2.7e-06 | 2.367E-06 | 4.933E-07 | 1.800E-06 | 2.700E-06 |
| 8584 | 430 | W430D | A | 2.8e-06 | 2.3e-06 | 3.1e-06 | 2.733E-06 | 4.041E-07 | 2.300E-06 | 3.100E-06 |

| | | | | | | | | | | |
|---|---|---|---|---|---|---|---|---|---|---|
| 8585 | 430 | W430C | A | 5e-06 | 4.8e-06 | 2.9e-06 | 4.233E-06 | 1.159E-06 | 2.900E-06 | 5.000E-06 |
| 8586 | 430 | W430E | A | 2.9e-06 | 4.3e-06 | 1.8e-06 | 3.000E-06 | 1.253E-06 | 1.800E-06 | 4.300E-06 |
| 8587 | 430 | W430Q | A | 1.8e-06 | 3e-06 | 2.5e-06 | 2.433E-06 | 6.028E-07 | 1.800E-06 | 3.000E-06 |
| 8588 | 430 | W430G | A | 2.3e-06 | 3.7e-06 | 1.9e-06 | 2.633E-06 | 9.452E-07 | 1.900E-06 | 3.700E-06 |
| 8589 | 430 | W430H | A | 1.9e-06 | 2.3e-06 | 1.9e-06 | 2.033E-06 | 2.309E-07 | 1.900E-06 | 2.300E-06 |
| 8590 | 430 | W430I | A | 2.6e-06 | 2.2e-06 | 2e-06 | 2.267E-06 | 3.055E-07 | 2.000E-06 | 2.600E-06 |
| 8591 | 430 | W430L | A | 3.6e-06 | 9.2e-07 | 1.8e-06 | 2.107E-06 | 1.366E-06 | 9.200E-07 | 3.600E-06 |
| 8592 | 430 | W430K | A | 2.4e-06 | 1.7e-06 | 1.4e-06 | 1.833E-06 | 5.132E-07 | 1.400E-06 | 2.400E-06 |
| 8593 | 430 | W430M | A | 1.8e-06 | 1.5e-06 | 1.9e-06 | 1.733E-06 | 2.082E-07 | 1.500E-06 | 1.900E-06 |
| 8594 | 430 | W430F | A | 2.2e-06 | 1.2e-06 | 1.6e-06 | 1.667E-06 | 5.033E-07 | 1.200E-06 | 2.200E-06 |
| 8595 | 430 | W430P | A | 2.6e-06 | 2.3e-06 | 2.1e-06 | 2.333E-06 | 2.517E-07 | 2.100E-06 | 2.600E-06 |
| 8596 | 430 | W430S | A | 1.4e-06 | 2.9e-06 | 1.9e-06 | 2.067E-06 | 7.638E-07 | 1.400E-06 | 2.900E-06 |
| 8597 | 430 | W430T | A | 2.3e-06 | 2.6e-06 | 3.4e-06 | 2.767E-06 | 5.686E-07 | 2.300E-06 | 3.400E-06 |
| 8598 | 430 | W430W | A | 2.8e-06 | 3.4e-06 | 3.1e-06 | 3.100E-06 | 3.000E-07 | 2.800E-06 | 3.400E-06 |
| 8599 | 430 | W430Y | A | 2.4e-06 | 2.1e-06 | 1.5e-06 | 2.000E-06 | 4.583E-07 | 1.500E-06 | 2.400E-06 |
| 8600 | 430 | W430V | A | 2.1e-06 | 1.7e-06 | 1.7e-06 | 1.833E-06 | 2.309E-07 | 1.700E-06 | 2.100E-06 |
| 8601 | 431 | L431A | A | 3.2e-06 | 3.6e-06 | 5.2e-06 | 4.000E-06 | 1.058E-06 | 3.200E-06 | 5.200E-06 |
| 8602 | 431 | L431R | A | 2.2e-06 | 1.1e-06 | 2.1e-06 | 1.800E-06 | 6.083E-07 | 1.100E-06 | 2.200E-06 |
| 8603 | 431 | L431N | A | 4.1e-06 | 1.7e-06 | 3.4e-06 | 3.067E-06 | 1.234E-06 | 1.700E-06 | 4.100E-06 |
| 8604 | 431 | L431D | A | 2.5e-06 | 2.1e-06 | 2.2e-06 | 2.267E-06 | 2.082E-07 | 2.100E-06 | 2.500E-06 |
| 8605 | 431 | L431C | A | 2.5e-06 | 2.6e-06 | 2e-06 | 2.367E-06 | 3.215E-07 | 2.000E-06 | 2.600E-06 |
| 8606 | 431 | L431E | A | 1.5e-06 | 2.3e-06 | 2e-06 | 1.933E-06 | 4.041E-07 | 1.500E-06 | 2.300E-06 |
| 8607 | 431 | L431Q | A | 2.2e-06 | 1.4e-06 | 2.4e-06 | 2.000E-06 | 5.292E-07 | 1.400E-06 | 2.400E-06 |
| 8608 | 431 | L431G | A | 2.2e-06 | 2.8e-06 | 2.4e-06 | 2.467E-06 | 3.055E-07 | 2.200E-06 | 2.800E-06 |
| 8609 | 431 | L431H | A | 3.3e-06 | 1.5e-06 | 3e-06 | 2.600E-06 | 9.644E-07 | 1.500E-06 | 3.300E-06 |
| 8610 | 431 | L431I | A | 3.3e-06 | 3.4e-06 | 1.9e-06 | 2.867E-06 | 8.386E-07 | 1.900E-06 | 3.400E-06 |
| 8611 | 431 | L431L | A | 2.7e-06 | 2.7e-06 | 3.7e-06 | 3.033E-06 | 5.774E-07 | 2.700E-06 | 3.700E-06 |
| 8612 | 431 | L431K | A | 2.8e-06 | 1.3e-06 | 2.7e-06 | 2.267E-06 | 8.386E-07 | 1.300E-06 | 2.800E-06 |
| 8613 | 431 | L431M | A | 2.6e-06 | 2.6e-06 | 3.5e-06 | 2.900E-06 | 5.196E-07 | 2.600E-06 | 3.500E-06 |
| 8614 | 431 | L431F | A | 2.4e-06 | 2.3e-06 | 3.5e-06 | 2.733E-06 | 6.658E-07 | 2.300E-06 | 3.500E-06 |
| 8615 | 431 | L431P | A | 3.6e-06 | 1.5e-06 | 2.7e-06 | 2.600E-06 | 1.054E-06 | 1.500E-06 | 3.600E-06 |
| 8616 | 431 | L431S | A | 2.2e-06 | 3.5e-06 | 2.9e-06 | 2.867E-06 | 6.506E-07 | 2.200E-06 | 3.500E-06 |
| 8617 | 431 | L431T | A | 1.1e-06 | 2e-06 | 1.6e-06 | 1.567E-06 | 4.509E-07 | 1.100E-06 | 2.000E-06 |
| 8618 | 431 | L431W | A | 2.5e-06 | 2.2e-06 | 3.1e-06 | 2.600E-06 | 4.583E-07 | 2.200E-06 | 3.100E-06 |
| 8619 | 431 | L431Y | A | 3.6e-06 | 1.8e-06 | 1.4e-06 | 2.267E-06 | 1.172E-06 | 1.400E-06 | 3.600E-06 |
| 8620 | 431 | L431V | A | 2.5e-06 | 2.7e-06 | 2.1e-06 | 2.433E-06 | 3.055E-07 | 2.100E-06 | 2.700E-06 |
| 8621 | 432 | G432A | A | 2.5e-06 | 1.6e-06 | 2.9e-06 | 2.333E-06 | 6.658E-07 | 1.600E-06 | 2.900E-06 |
| 8622 | 432 | G432R | A | 2e-06 | 3.1e-06 | 2.8e-06 | 2.633E-06 | 5.686E-07 | 2.000E-06 | 3.100E-06 |
| 8623 | 432 | G432N | A | 1.9e-06 | 3.4e-06 | 1.7e-06 | 2.333E-06 | 9.292E-07 | 1.700E-06 | 3.400E-06 |
| 8624 | 432 | G432D | A | 1.8e-06 | 1.7e-06 | 2.1e-06 | 1.867E-06 | 2.082E-07 | 1.700E-06 | 2.100E-06 |
| 8625 | 432 | G432C | A | 2.4e-06 | 1.7e-06 | 2.1e-06 | 2.067E-06 | 3.512E-07 | 1.700E-06 | 2.400E-06 |
| 8626 | 432 | G432E | A | 2.9e-06 | 2e-06 | 2.4e-06 | 2.433E-06 | 4.509E-07 | 2.000E-06 | 2.900E-06 |
| 8627 | 432 | G432Q | A | 3.3e-06 | 3.3e-06 | 2.3e-06 | 2.967E-06 | 5.774E-07 | 2.300E-06 | 3.300E-06 |
| 8628 | 432 | G432G | A | 2e-06 | 1.9e-06 | 3.1e-06 | 2.333E-06 | 6.658E-07 | 1.900E-06 | 3.100E-06 |
| 8629 | 432 | G432H | A | 2.3e-06 | 3.1e-06 | 2.3e-06 | 2.567E-06 | 4.619E-07 | 2.300E-06 | 3.100E-06 |
| 8630 | 432 | G432I | A | 1.6e-06 | 3.4e-06 | 1.2e-06 | 2.067E-06 | 1.172E-06 | 1.200E-06 | 3.400E-06 |
| 8631 | 432 | G432L | A | 1.2e-06 | 1.8e-06 | 1.6e-06 | 1.533E-06 | 3.055E-07 | 1.200E-06 | 1.800E-06 |
| 8632 | 432 | G432K | A | 3.1e-06 | 2.2e-06 | 3.2e-06 | 2.833E-06 | 5.508E-07 | 2.200E-06 | 3.200E-06 |
| 8633 | 432 | G432M | A | 3.4e-06 | 2.3e-06 | 1.9e-06 | 2.533E-06 | 7.767E-07 | 1.900E-06 | 3.400E-06 |
| 8634 | 432 | G432F | A | 2.6e-06 | 2.9e-06 | 2.3e-06 | 2.600E-06 | 3.000E-07 | 2.300E-06 | 2.900E-06 |
| 8635 | 432 | G432P | A | 2.4e-06 | 1.7e-06 | 2.2e-06 | 2.100E-06 | 3.606E-07 | 1.700E-06 | 2.400E-06 |
| 8636 | 432 | G432S | A | 3.5e-06 | 3.3e-06 | 1.3e-06 | 2.700E-06 | 1.217E-06 | 1.300E-06 | 3.500E-06 |
| 8637 | 432 | G432T | A | 1.5e-06 | 2.1e-06 | 1.9e-06 | 1.833E-06 | 3.055E-07 | 1.500E-06 | 2.100E-06 |
| 8638 | 432 | G432W | A | 1.8e-06 | 1.7e-06 | 1.8e-06 | 1.767E-06 | 5.774E-08 | 1.700E-06 | 1.800E-06 |
| 8639 | 432 | G432Y | A | 2.9e-06 | 2.9e-06 | 2.2e-06 | 2.667E-06 | 4.041E-07 | 2.200E-06 | 2.900E-06 |
| 8640 | 432 | G432V | A | 4.6e-06 | 3.2e-06 | 3.1e-06 | 3.633E-06 | 8.386E-07 | 3.100E-06 | 4.600E-06 |
| 8641 | 433 | L433A | A | 2.9e-06 | 5e-06 | 2.4e-06 | 3.433E-06 | 1.380E-06 | 2.400E-06 | 5.000E-06 |
| 8642 | 433 | L433R | A | 3.3e-06 | 2.2e-06 | 2.3e-06 | 2.600E-06 | 6.083E-07 | 2.200E-06 | 3.300E-06 |
| 8643 | 433 | L433N | A | 2.9e-06 | 1.8e-06 | 1.7e-06 | 2.133E-06 | 6.658E-07 | 1.700E-06 | 2.900E-06 |
| 8644 | 433 | L433D | A | 4e-06 | 2.5e-06 | 2.4e-06 | 2.967E-06 | 8.963E-07 | 2.400E-06 | 4.000E-06 |
| 8645 | 433 | L433C | A | 3.3e-06 | 2.6e-06 | 2.8e-06 | 2.900E-06 | 3.606E-07 | 2.600E-06 | 3.300E-06 |
| 8646 | 433 | L433E | A | 2.2e-06 | 2.5e-06 | 2.8e-06 | 2.500E-06 | 3.000E-07 | 2.200E-06 | 2.800E-06 |
| 8647 | 433 | L433Q | A | 2.9e-06 | 1.9e-06 | 1.9e-06 | 2.233E-06 | 5.774E-07 | 1.900E-06 | 2.900E-06 |
| 8648 | 433 | L433G | A | 2.3e-06 | 1.6e-06 | 3e-06 | 2.300E-06 | 7.000E-07 | 1.600E-06 | 3.000E-06 |
| 8649 | 433 | L433H | A | 1.9e-06 | 2e-06 | 3.9e-06 | 2.600E-06 | 1.127E-06 | 1.900E-06 | 3.900E-06 |
| 8650 | 433 | L433I | A | 2.3e-06 | 1.6e-06 | 2.2e-06 | 2.033E-06 | 3.786E-07 | 1.600E-06 | 2.300E-06 |
| 8651 | 433 | L433L | A | 1.5e-06 | 3e-06 | 2.4e-06 | 2.300E-06 | 7.550E-07 | 1.500E-06 | 3.000E-06 |
| 8652 | 433 | L433K | A | 2.8e-06 | 1.4e-06 | 2.5e-06 | 2.233E-06 | 7.371E-07 | 1.400E-06 | 2.800E-06 |
| 8653 | 433 | L433M | A | 1.2e-06 | 2.8e-06 | 1.7e-06 | 1.900E-06 | 8.185E-07 | 1.200E-06 | 2.800E-06 |
| 8654 | 433 | L433F | A | 1.1e-06 | 2.1e-06 | 2e-06 | 1.733E-06 | 5.508E-07 | 1.100E-06 | 2.100E-06 |
| 8655 | 433 | L433P | A | 3.6e-06 | 1.3e-06 | 2.6e-06 | 2.500E-06 | 1.153E-06 | 1.300E-06 | 3.600E-06 |
| 8656 | 433 | L433S | A | 1.9e-06 | 1.7e-06 | 2.1e-06 | 1.900E-06 | 2.000E-07 | 1.700E-06 | 2.100E-06 |
| 8657 | 433 | L433T | A | 1.3e-06 | 1.9e-06 | 2.2e-06 | 1.800E-06 | 4.583E-07 | 1.300E-06 | 2.200E-06 |
| 8658 | 433 | L433W | A | 2.1e-06 | 3e-06 | 3e-06 | 2.700E-06 | 5.196E-07 | 2.100E-06 | 3.000E-06 |
| 8659 | 433 | L433Y | A | 2.6e-06 | 1.4e-06 | 3.3e-06 | 2.433E-06 | 9.609E-07 | 1.400E-06 | 3.300E-06 |
| 8660 | 433 | L433V | A | 2.2e-06 | 2.4e-06 | 2.6e-06 | 2.400E-06 | 2.000E-07 | 2.200E-06 | 2.600E-06 |
| 8661 | 434 | R434A | A | 1.7e-06 | 3.3e-06 | 2e-06 | 2.333E-06 | 8.505E-07 | 1.700E-06 | 3.300E-06 |
| 8662 | 434 | R434R | A | 3.3e-06 | 2.5e-06 | 2.6e-06 | 2.800E-06 | 4.359E-07 | 2.500E-06 | 3.300E-06 |
| 8663 | 434 | R434N | A | 2.5e-06 | 2.8e-06 | 2.3e-06 | 2.533E-06 | 2.517E-07 | 2.300E-06 | 2.800E-06 |
| 8664 | 434 | R434D | A | 1.9e-06 | 2.6e-06 | 3.3e-06 | 2.600E-06 | 7.000E-07 | 1.900E-06 | 3.300E-06 |
| 8665 | 434 | R434C | A | 2.9e-06 | 1.2e-06 | 2.8e-06 | 2.300E-06 | 9.539E-07 | 1.200E-06 | 2.900E-06 |
| 8666 | 434 | R434E | A | 2.1e-06 | 2.4e-06 | 2.7e-06 | 2.400E-06 | 3.000E-07 | 2.100E-06 | 2.700E-06 |
| 8667 | 434 | R434Q | A | 2.6e-06 | 1.5e-06 | 4.3e-06 | 2.800E-06 | 1.411E-06 | 1.500E-06 | 4.300E-06 |
| 8668 | 434 | R434G | A | 1.2e-06 | 2.7e-06 | 3.7e-06 | 2.533E-06 | 1.258E-06 | 1.200E-06 | 3.700E-06 |
| 8669 | 434 | R434H | A | 1.2e-06 | 3e-06 | 2.9e-06 | 2.367E-06 | 1.012E-06 | 1.200E-06 | 3.000E-06 |

| | | | | | | | | | | |
|---|---|---|---|---|---|---|---|---|---|---|
| 8670 | 434 | R434I | A | 3.7e-06 | 1.5e-06 | 1.9e-06 | 2.367E-06 | 1.172E-06 | 1.500E-06 | 3.700E-06 |
| 8671 | 434 | R434L | A | 1.1e-06 | 2.3e-06 | 1.8e-06 | 1.733E-06 | 6.028E-07 | 1.100E-06 | 2.300E-06 |
| 8672 | 434 | R434K | A | 2.9e-06 | 2.1e-06 | 3.1e-06 | 2.700E-06 | 5.292E-07 | 2.100E-06 | 3.100E-06 |
| 8673 | 434 | R434M | A | 1.8e-06 | 1.9e-06 | 2.6e-06 | 2.100E-06 | 4.359E-07 | 1.800E-06 | 2.600E-06 |
| 8674 | 434 | R434F | A | 3.4e-06 | 4.5e-06 | 2.9e-06 | 3.600E-06 | 8.185E-07 | 2.900E-06 | 4.500E-06 |
| 8675 | 434 | R434P | A | 2e-06 | 1.8e-06 | 2.1e-06 | 1.967E-06 | 1.528E-07 | 1.800E-06 | 2.100E-06 |
| 8676 | 434 | R434S | A | 2.2e-06 | 1.5e-06 | 6.3e-06 | 3.333E-06 | 2.593E-06 | 1.500E-06 | 6.300E-06 |
| 8677 | 434 | R434T | A | 2.2e-06 | 2.2e-06 | 1.4e-06 | 1.933E-06 | 4.619E-07 | 1.400E-06 | 2.200E-06 |
| 8678 | 434 | R434W | A | 2.6e-06 | 1.7e-06 | 3e-06 | 2.433E-06 | 6.658E-07 | 1.700E-06 | 3.000E-06 |
| 8679 | 434 | R434Y | A | 2.4e-06 | 1.7e-06 | 1.9e-06 | 2.000E-06 | 3.606E-07 | 1.700E-06 | 2.400E-06 |
| 8680 | 434 | R434V | A | 1.8e-06 | 2.8e-06 | 4.7e-06 | 3.100E-06 | 1.473E-06 | 1.800E-06 | 4.700E-06 |
| 8681 | 435 | S435A | A | 2.4e-06 | 3.2e-06 | 2.1e-06 | 2.567E-06 | 5.686E-07 | 2.100E-06 | 3.200E-06 |
| 8682 | 435 | S435R | A | 1.4e-06 | 1.1e-06 | 2.3e-06 | 1.600E-06 | 6.245E-07 | 1.100E-06 | 2.300E-06 |
| 8683 | 435 | S435N | A | 4e-06 | 1.9e-06 | 2.6e-06 | 2.833E-06 | 1.069E-06 | 1.900E-06 | 4.000E-06 |
| 8684 | 435 | S435D | A | 2.2e-06 | 4e-06 | 2.4e-06 | 2.867E-06 | 9.866E-07 | 2.200E-06 | 4.000E-06 |
| 8685 | 435 | S435C | A | 2.3e-06 | 3.3e-06 | 1.1e-06 | 2.233E-06 | 1.102E-06 | 1.100E-06 | 3.300E-06 |
| 8686 | 435 | S435E | A | 2.4e-06 | 4.6e-06 | 1.5e-06 | 2.833E-06 | 1.595E-06 | 1.500E-06 | 4.600E-06 |
| 8687 | 435 | S435Q | A | 3.6e-06 | 3e-06 | 2e-06 | 2.867E-06 | 8.083E-07 | 2.000E-06 | 3.600E-06 |
| 8688 | 435 | S435G | A | 1.8e-06 | 2.9e-06 | 2e-06 | 2.233E-06 | 5.859E-07 | 1.800E-06 | 2.900E-06 |
| 8689 | 435 | S435H | A | 2.1e-06 | 2.1e-06 | 2.5e-06 | 2.233E-06 | 2.309E-07 | 2.100E-06 | 2.500E-06 |
| 8690 | 435 | S435I | A | 1.7e-06 | 1.7e-06 | 1.9e-06 | 1.767E-06 | 1.155E-07 | 1.700E-06 | 1.900E-06 |
| 8691 | 435 | S435L | A | 2.1e-06 | 1.7e-06 | 2.9e-06 | 2.233E-06 | 6.110E-07 | 1.700E-06 | 2.900E-06 |
| 8692 | 435 | S435K | A | 2.4e-06 | 1.7e-06 | 1.6e-06 | 1.900E-06 | 4.359E-07 | 1.600E-06 | 2.400E-06 |
| 8693 | 435 | S435M | A | 1.8e-06 | 2.1e-06 | 2.2e-06 | 2.033E-06 | 2.082E-07 | 1.800E-06 | 2.200E-06 |
| 8694 | 435 | S435F | A | 3.3e-06 | 2e-06 | 1.7e-06 | 2.333E-06 | 8.505E-07 | 1.700E-06 | 3.300E-06 |
| 8695 | 435 | S435P | A | 1.6e-06 | 1.6e-06 | 2.2e-06 | 1.800E-06 | 3.464E-07 | 1.600E-06 | 2.200E-06 |
| 8696 | 435 | S435S | A | 2.5e-06 | 2.6e-06 | 1.3e-06 | 2.133E-06 | 7.234E-07 | 1.300E-06 | 2.600E-06 |
| 8697 | 435 | S435T | A | 3.1e-06 | 1.1e-06 | 2.1e-06 | 2.100E-06 | 1.000E-06 | 1.100E-06 | 3.100E-06 |
| 8698 | 435 | S435W | A | 1.8e-06 | 3e-06 | 2.3e-06 | 2.367E-06 | 6.028E-07 | 1.800E-06 | 3.000E-06 |
| 8699 | 435 | S435Y | A | 2.9e-06 | 1.7e-06 | 2e-06 | 2.200E-06 | 6.245E-07 | 1.700E-06 | 2.900E-06 |
| 8700 | 435 | S435V | A | 2.6e-06 | 2.3e-06 | 3.6e-06 | 2.833E-06 | 6.807E-07 | 2.300E-06 | 3.600E-06 |
| 8701 | 436 | L436A | A | 1.7e-06 | 3.8e-06 | 2.6e-06 | 2.700E-06 | 1.054E-06 | 1.700E-06 | 3.800E-06 |
| 8702 | 436 | L436R | A | 2.3e-06 | 3e-06 | 2.6e-06 | 2.633E-06 | 3.512E-07 | 2.300E-06 | 3.000E-06 |
| 8703 | 436 | L436N | A | 2.5e-06 | 2.6e-06 | 2.9e-06 | 2.667E-06 | 2.082E-07 | 2.500E-06 | 2.900E-06 |
| 8704 | 436 | L436D | A | 4.1e-06 | 3.1e-06 | 3e-06 | 3.400E-06 | 6.083E-07 | 3.000E-06 | 4.100E-06 |
| 8705 | 436 | L436C | A | 2.2e-06 | 1.4e-06 | 3.1e-06 | 2.233E-06 | 8.505E-07 | 1.400E-06 | 3.100E-06 |
| 8706 | 436 | L436E | A | 1.5e-06 | 2.2e-06 | 1.5e-06 | 1.733E-06 | 4.041E-07 | 1.500E-06 | 2.200E-06 |
| 8707 | 436 | L436Q | A | 3.1e-06 | 2.3e-06 | 2e-06 | 2.467E-06 | 5.686E-07 | 2.000E-06 | 3.100E-06 |
| 8708 | 436 | L436G | A | 2.1e-06 | 3e-06 | 2.8e-06 | 2.633E-06 | 4.726E-07 | 2.100E-06 | 3.000E-06 |
| 8709 | 436 | L436H | A | 2.1e-06 | 3.4e-06 | 1.7e-06 | 2.400E-06 | 8.888E-07 | 1.700E-06 | 3.400E-06 |
| 8710 | 436 | L436I | A | 3.7e-06 | 2.3e-06 | 3.4e-06 | 3.133E-06 | 7.371E-07 | 2.300E-06 | 3.700E-06 |
| 8711 | 436 | L436L | A | 3.6e-06 | 1.6e-06 | 3.4e-06 | 2.867E-06 | 1.102E-06 | 1.600E-06 | 3.600E-06 |
| 8712 | 436 | L436K | A | 9.9e-07 | 1.8e-06 | 2.7e-06 | 1.830E-06 | 8.554E-07 | 9.900E-07 | 2.700E-06 |
| 8713 | 436 | L436M | A | 3.6e-06 | 1.9e-06 | 2.1e-06 | 2.533E-06 | 9.292E-07 | 1.900E-06 | 3.600E-06 |
| 8714 | 436 | L436F | A | 2.2e-06 | 3.6e-06 | 3e-06 | 2.933E-06 | 7.024E-07 | 2.200E-06 | 3.600E-06 |
| 8715 | 436 | L436P | A | 2.6e-06 | 3.2e-06 | 1.9e-06 | 2.567E-06 | 6.506E-07 | 1.900E-06 | 3.200E-06 |
| 8716 | 436 | L436S | A | 1.6e-06 | 3.2e-06 | 3.4e-06 | 2.733E-06 | 9.866E-07 | 1.600E-06 | 3.400E-06 |
| 8717 | 436 | L436T | A | 1.9e-06 | 2.3e-06 | 1.4e-06 | 1.867E-06 | 4.509E-07 | 1.400E-06 | 2.300E-06 |
| 8718 | 436 | L436W | A | 3.3e-06 | 2.2e-06 | 2.9e-06 | 2.800E-06 | 5.568E-07 | 2.200E-06 | 3.300E-06 |
| 8719 | 436 | L436Y | A | 2.9e-06 | 1.9e-06 | 1.6e-06 | 2.133E-06 | 6.807E-07 | 1.600E-06 | 2.900E-06 |
| 8720 | 436 | L436V | A | 1.3e-06 | 2.2e-06 | 1.4e-06 | 1.633E-06 | 4.933E-07 | 1.300E-06 | 2.200E-06 |
| 8721 | 437 | R437A | A | 2.1e-06 | 3.1e-06 | 1.8e-06 | 2.333E-06 | 6.807E-07 | 1.800E-06 | 3.100E-06 |
| 8722 | 437 | R437R | A | 2e-06 | 1.9e-06 | 3.8e-06 | 2.567E-06 | 1.069E-06 | 1.900E-06 | 3.800E-06 |
| 8723 | 437 | R437N | A | 2.8e-06 | 2e-06 | 1.5e-06 | 2.100E-06 | 6.557E-07 | 1.500E-06 | 2.800E-06 |
| 8724 | 437 | R437D | A | 3.2e-06 | 5.7e-06 | 1.2e-06 | 3.367E-06 | 2.255E-06 | 1.200E-06 | 5.700E-06 |
| 8725 | 437 | R437C | A | 2.1e-06 | 2.4e-06 | 2e-06 | 2.167E-06 | 2.082E-07 | 2.000E-06 | 2.400E-06 |
| 8726 | 437 | R437E | A | 2.4e-06 | 2.2e-06 | 2.8e-06 | 2.467E-06 | 3.055E-07 | 2.200E-06 | 2.800E-06 |
| 8727 | 437 | R437Q | A | 2.7e-06 | 1.5e-06 | 1.6e-06 | 1.933E-06 | 6.658E-07 | 1.500E-06 | 2.700E-06 |
| 8728 | 437 | R437G | A | 1.7e-06 | 2.6e-06 | 2.6e-06 | 2.300E-06 | 5.196E-07 | 1.700E-06 | 2.600E-06 |
| 8729 | 437 | R437H | A | 3.1e-06 | 2.5e-06 | 2.2e-06 | 2.600E-06 | 4.583E-07 | 2.200E-06 | 3.100E-06 |
| 8730 | 437 | R437I | A | 2.7e-06 | 4.3e-06 | 2e-06 | 3.000E-06 | 1.179E-06 | 2.000E-06 | 4.300E-06 |
| 8731 | 437 | R437L | A | 2.6e-06 | 1.4e-06 | 2.5e-06 | 2.167E-06 | 6.658E-07 | 1.400E-06 | 2.600E-06 |
| 8732 | 437 | R437K | A | 2.9e-06 | 8.2e-07 | 1.3e-06 | 1.673E-06 | 1.089E-06 | 8.200E-07 | 2.900E-06 |
| 8733 | 437 | R437M | A | 1.5e-06 | 1.3e-06 | 2.6e-06 | 1.800E-06 | 7.000E-07 | 1.300E-06 | 2.600E-06 |
| 8734 | 437 | R437F | A | 2.1e-06 | 2e-06 | 1.4e-06 | 1.833E-06 | 3.786E-07 | 1.400E-06 | 2.100E-06 |
| 8735 | 437 | R437P | A | 1.5e-06 | 2.1e-06 | 3e-06 | 2.200E-06 | 7.550E-07 | 1.500E-06 | 3.000E-06 |
| 8736 | 437 | R437S | A | 2.9e-06 | 1.4e-06 | 1.6e-06 | 1.967E-06 | 8.145E-07 | 1.400E-06 | 2.900E-06 |
| 8737 | 437 | R437T | A | 2.3e-06 | 1.4e-06 | 2.1e-06 | 1.933E-06 | 4.726E-07 | 1.400E-06 | 2.300E-06 |
| 8738 | 437 | R437W | A | 2.1e-06 | 2.7e-06 | 3.6e-06 | 2.800E-06 | 7.550E-07 | 2.100E-06 | 3.600E-06 |
| 8739 | 437 | R437Y | A | 2.8e-06 | 2.4e-06 | 2e-06 | 2.400E-06 | 4.000E-07 | 2.000E-06 | 2.800E-06 |
| 8740 | 437 | R437V | A | 2.8e-06 | 2.3e-06 | 1.9e-06 | 2.333E-06 | 4.509E-07 | 1.900E-06 | 2.800E-06 |
| 8741 | 438 | E438A | A | 3.6e-06 | 3e-06 | 1.9e-06 | 2.833E-06 | 8.622E-07 | 1.900E-06 | 3.600E-06 |
| 8742 | 438 | E438R | A | 1.4e-06 | 2e-06 | 5.1e-06 | 2.833E-06 | 1.986E-06 | 1.400E-06 | 5.100E-06 |
| 8743 | 438 | E438N | A | 3.3e-06 | 2.9e-06 | 2.6e-06 | 2.933E-06 | 3.512E-07 | 2.600E-06 | 3.300E-06 |
| 8744 | 438 | E438D | A | 4.1e-06 | 3.3e-06 | 2.6e-06 | 3.333E-06 | 7.506E-07 | 2.600E-06 | 4.100E-06 |
| 8745 | 438 | E438C | A | 2.8e-06 | 2e-06 | 2.3e-06 | 2.367E-06 | 4.041E-07 | 2.000E-06 | 2.800E-06 |
| 8746 | 438 | E438E | A | 4e-06 | 3.2e-06 | 2.7e-06 | 3.300E-06 | 6.557E-07 | 2.700E-06 | 4.000E-06 |
| 8747 | 438 | E438Q | A | 1.6e-06 | 3.1e-06 | 1.8e-06 | 2.167E-06 | 8.145E-07 | 1.600E-06 | 3.100E-06 |
| 8748 | 438 | E438G | A | 2.9e-06 | 2.4e-06 | 2.5e-06 | 2.600E-06 | 2.646E-07 | 2.400E-06 | 2.900E-06 |
| 8749 | 438 | E438H | A | 1.1e-06 | 1.7e-06 | 3.3e-06 | 2.033E-06 | 1.137E-06 | 1.100E-06 | 3.300E-06 |
| 8750 | 438 | E438I | A | 2.2e-06 | 2.2e-06 | 3.1e-06 | 2.500E-06 | 5.196E-07 | 2.200E-06 | 3.100E-06 |
| 8751 | 438 | E438L | A | 1.6e-06 | 2.2e-06 | 3.1e-06 | 2.300E-06 | 7.550E-07 | 1.600E-06 | 3.100E-06 |
| 8752 | 438 | E438K | A | 1.9e-06 | 2.7e-06 | 2.6e-06 | 2.400E-06 | 4.359E-07 | 1.900E-06 | 2.700E-06 |
| 8753 | 438 | E438M | A | 1.5e-06 | 1.4e-06 | 2.3e-06 | 1.733E-06 | 4.933E-07 | 1.400E-06 | 2.300E-06 |
| 8754 | 438 | E438F | A | 4.4e-06 | 2.3e-06 | 1.4e-06 | 2.700E-06 | 1.539E-06 | 1.400E-06 | 4.400E-06 |

| | | | | | | | | | | |
|---|---|---|---|---|---|---|---|---|---|---|
| 8755 | 438 | E438P | A | 3e-06 | 2.4e-06 | 3e-06 | 2.800E-06 | 3.464E-07 | 2.400E-06 | 3.000E-06 |
| 8756 | 438 | E438S | A | 1.6e-06 | 9.8e-07 | 2.2e-06 | 1.593E-06 | 6.100E-07 | 9.800E-07 | 2.200E-06 |
| 8757 | 438 | E438T | A | 2.9e-06 | 2.8e-06 | 2.1e-06 | 2.600E-06 | 4.359E-07 | 2.100E-06 | 2.900E-06 |
| 8758 | 438 | E438W | A | 1.8e-06 | 2.3e-06 | 2.3e-06 | 2.133E-06 | 2.887E-07 | 1.800E-06 | 2.300E-06 |
| 8759 | 438 | E438Y | A | 1.4e-06 | 3.5e-06 | 2.7e-06 | 2.533E-06 | 1.060E-06 | 1.400E-06 | 3.500E-06 |
| 8760 | 438 | E438V | A | 3.3e-06 | 2.4e-06 | 3.8e-06 | 3.167E-06 | 7.095E-07 | 2.400E-06 | 3.800E-06 |
| 8761 | 439 | L439A | A | 4.6e-06 | 3.8e-06 | 5e-06 | 4.467E-06 | 6.110E-07 | 3.800E-06 | 5.000E-06 |
| 8762 | 439 | L439R | A | 2.5e-06 | 2e-06 | 4e-06 | 2.833E-06 | 1.041E-06 | 2.000E-06 | 4.000E-06 |
| 8763 | 439 | L439N | A | 3.2e-06 | 1.5e-06 | 2.8e-06 | 2.500E-06 | 8.888E-07 | 1.500E-06 | 3.200E-06 |
| 8764 | 439 | L439D | A | 3.1e-06 | 3.3e-06 | 1.9e-06 | 2.767E-06 | 7.572E-07 | 1.900E-06 | 3.300E-06 |
| 8765 | 439 | L439C | A | 2.5e-06 | 1.5e-06 | 3.1e-06 | 2.367E-06 | 8.083E-07 | 1.500E-06 | 3.100E-06 |
| 8766 | 439 | L439E | A | 1.2e-06 | 9.9e-07 | 2.4e-06 | 1.530E-06 | 7.607E-07 | 9.900E-07 | 2.400E-06 |
| 8767 | 439 | L439Q | A | 1.5e-06 | 1.3e-06 | 2.7e-06 | 1.833E-06 | 7.572E-07 | 1.300E-06 | 2.700E-06 |
| 8768 | 439 | L439G | A | 2.7e-06 | 2.5e-06 | 3.5e-06 | 2.900E-06 | 5.292E-07 | 2.500E-06 | 3.500E-06 |
| 8769 | 439 | L439H | A | 2.4e-06 | 2.5e-06 | 1e-06 | 1.967E-06 | 8.386E-07 | 1.000E-06 | 2.500E-06 |
| 8770 | 439 | L439I | A | 3.2e-06 | 2.2e-06 | 2.6e-06 | 2.667E-06 | 5.033E-07 | 2.200E-06 | 3.200E-06 |
| 8771 | 439 | L439L | A | 2e-06 | 3e-06 | 1.2e-06 | 2.067E-06 | 9.018E-07 | 1.200E-06 | 3.000E-06 |
| 8772 | 439 | L439K | A | 1.5e-06 | 1.9e-06 | 2.9e-06 | 2.100E-06 | 7.211E-07 | 1.500E-06 | 2.900E-06 |
| 8773 | 439 | L439M | A | 2.7e-06 | 3.9e-06 | 2.8e-06 | 3.133E-06 | 6.658E-07 | 2.700E-06 | 3.900E-06 |
| 8774 | 439 | L439F | A | 1.8e-06 | 3.3e-06 | 3.8e-06 | 2.967E-06 | 1.041E-06 | 1.800E-06 | 3.800E-06 |
| 8775 | 439 | L439P | A | 2.5e-06 | 8.1e-07 | 1.2e-06 | 1.503E-06 | 8.849E-07 | 8.100E-07 | 2.500E-06 |
| 8776 | 439 | L439S | A | 2.8e-06 | 3.2e-06 | 2.3e-06 | 2.767E-06 | 4.509E-07 | 2.300E-06 | 3.200E-06 |
| 8777 | 439 | L439T | A | 1.9e-06 | 2.9e-06 | 2.5e-06 | 2.433E-06 | 5.033E-07 | 1.900E-06 | 2.900E-06 |
| 8778 | 439 | L439W | A | 2.5e-06 | 2.3e-06 | 2.4e-06 | 2.400E-06 | 1.000E-07 | 2.300E-06 | 2.500E-06 |
| 8779 | 439 | L439Y | A | 1.6e-06 | 4.4e-06 | 3.1e-06 | 3.033E-06 | 1.401E-06 | 1.600E-06 | 4.400E-06 |
| 8780 | 439 | L439V | A | 2.4e-06 | 1.6e-06 | 2.3e-06 | 2.100E-06 | 4.359E-07 | 1.600E-06 | 2.400E-06 |
| 8781 | 440 | G440A | A | 1.1e-06 | 2.4e-06 | 2.3e-06 | 1.933E-06 | 7.234E-07 | 1.100E-06 | 2.400E-06 |
| 8782 | 440 | G440R | A | 2.4e-06 | 2.5e-06 | 2.5e-06 | 2.467E-06 | 5.774E-08 | 2.400E-06 | 2.500E-06 |
| 8783 | 440 | G440N | A | 1e-06 | 2.1e-06 | 2.4e-06 | 1.833E-06 | 7.371E-07 | 1.000E-06 | 2.400E-06 |
| 8784 | 440 | G440D | A | 2.8e-06 | 4e-06 | 2.3e-06 | 3.033E-06 | 8.737E-07 | 2.300E-06 | 4.000E-06 |
| 8785 | 440 | G440C | A | 1.8e-06 | 1.1e-06 | 2.2e-06 | 1.700E-06 | 5.568E-07 | 1.100E-06 | 2.200E-06 |
| 8786 | 440 | G440E | A | 4.9e-06 | 1.4e-06 | 2.1e-06 | 2.800E-06 | 1.852E-06 | 1.400E-06 | 4.900E-06 |
| 8787 | 440 | G440Q | A | 3.6e-06 | 2.6e-06 | 2.2e-06 | 2.800E-06 | 7.211E-07 | 2.200E-06 | 3.600E-06 |
| 8788 | 440 | G440G | A | 1.4e-06 | 3.8e-06 | 3.2e-06 | 2.800E-06 | 1.249E-06 | 1.400E-06 | 3.800E-06 |
| 8789 | 440 | G440H | A | 1.8e-06 | 5.2e-06 | 3.6e-06 | 3.533E-06 | 1.701E-06 | 1.800E-06 | 5.200E-06 |
| 8790 | 440 | G440I | A | 1.9e-06 | 2.2e-06 | 2.9e-06 | 2.333E-06 | 5.132E-07 | 1.900E-06 | 2.900E-06 |
| 8791 | 440 | G440L | A | 4.5e-06 | 1.2e-06 | 3.6e-06 | 3.100E-06 | 1.706E-06 | 1.200E-06 | 4.500E-06 |
| 8792 | 440 | G440K | A | 2.6e-06 | 2.4e-06 | 2.2e-06 | 2.400E-06 | 2.000E-07 | 2.200E-06 | 2.600E-06 |
| 8793 | 440 | G440M | A | 2.6e-06 | 2.6e-06 | 2.6e-06 | 2.600E-06 | 0.000E+00 | 2.600E-06 | 2.600E-06 |
| 8794 | 440 | G440F | A | 1.3e-06 | 2.1e-06 | 1.7e-06 | 1.700E-06 | 4.000E-07 | 1.300E-06 | 2.100E-06 |
| 8795 | 440 | G440P | A | 1.9e-06 | 2.1e-06 | 1.8e-06 | 1.933E-06 | 1.528E-07 | 1.800E-06 | 2.100E-06 |
| 8796 | 440 | G440S | A | 2.8e-06 | 2.3e-06 | 3.9e-06 | 3.000E-06 | 8.185E-07 | 2.300E-06 | 3.900E-06 |
| 8797 | 440 | G440T | A | 1.9e-06 | 2.3e-06 | 1.6e-06 | 1.933E-06 | 3.512E-07 | 1.600E-06 | 2.300E-06 |
| 8798 | 440 | G440W | A | 2.7e-06 | 3.8e-06 | 2.7e-06 | 3.067E-06 | 6.351E-07 | 2.700E-06 | 3.800E-06 |
| 8799 | 440 | G440Y | A | 2.6e-06 | 1.3e-06 | 3.6e-06 | 2.500E-06 | 1.153E-06 | 1.300E-06 | 3.600E-06 |
| 8800 | 440 | G440V | A | 1.8e-06 | 3.2e-06 | 2.5e-06 | 2.500E-06 | 7.000E-07 | 1.800E-06 | 3.200E-06 |
| 8801 | 441 | S441A | A | 2.2e-06 | 1.9e-06 | 3.2e-06 | 2.433E-06 | 6.807E-07 | 1.900E-06 | 3.200E-06 |
| 8802 | 441 | S441R | A | 3.2e-06 | 4.4e-06 | 1.7e-06 | 3.100E-06 | 1.353E-06 | 1.700E-06 | 4.400E-06 |
| 8803 | 441 | S441N | A | 3.4e-06 | 2.3e-06 | 2.9e-06 | 2.867E-06 | 5.508E-07 | 2.300E-06 | 3.400E-06 |
| 8804 | 441 | S441D | A | 1.4e-06 | 3.8e-06 | 2.4e-06 | 2.533E-06 | 1.206E-06 | 1.400E-06 | 3.800E-06 |
| 8805 | 441 | S441C | A | 2.1e-06 | 2.6e-06 | 1.9e-06 | 2.200E-06 | 3.606E-07 | 1.900E-06 | 2.600E-06 |
| 8806 | 441 | S441E | A | 1.9e-06 | 3.2e-06 | 3.4e-06 | 2.833E-06 | 8.145E-07 | 1.900E-06 | 3.400E-06 |
| 8807 | 441 | S441Q | A | 7.4e-06 | 3.2e-06 | 1.3e-06 | 3.967E-06 | 3.121E-06 | 1.300E-06 | 7.400E-06 |
| 8808 | 441 | S441G | A | 3.6e-06 | 1.6e-06 | 1.5e-06 | 2.233E-06 | 1.185E-06 | 1.500E-06 | 3.600E-06 |
| 8809 | 441 | S441H | A | 2e-06 | 3.4e-06 | 3.5e-06 | 2.967E-06 | 8.386E-07 | 2.000E-06 | 3.500E-06 |
| 8810 | 441 | S441I | A | 2e-06 | 1.9e-06 | 2.5e-06 | 2.133E-06 | 3.215E-07 | 1.900E-06 | 2.500E-06 |
| 8811 | 441 | S441L | A | 2.6e-06 | 3.4e-06 | 2.6e-06 | 2.867E-06 | 4.619E-07 | 2.600E-06 | 3.400E-06 |
| 8812 | 441 | S441K | A | 2e-06 | 3.2e-06 | 1.8e-06 | 2.333E-06 | 7.572E-07 | 1.800E-06 | 3.200E-06 |
| 8813 | 441 | S441M | A | 2.5e-06 | 2.4e-06 | 2.2e-06 | 2.367E-06 | 1.528E-07 | 2.200E-06 | 2.500E-06 |
| 8814 | 441 | S441F | A | 2.1e-06 | 2.1e-06 | 2.4e-06 | 2.200E-06 | 1.732E-07 | 2.100E-06 | 2.400E-06 |
| 8815 | 441 | S441P | A | 1.3e-06 | 1.6e-06 | 2.7e-06 | 1.867E-06 | 7.371E-07 | 1.300E-06 | 2.700E-06 |
| 8816 | 441 | S441S | A | 2.3e-06 | 3.7e-06 | 2.6e-06 | 2.867E-06 | 7.371E-07 | 2.300E-06 | 3.700E-06 |
| 8817 | 441 | S441T | A | 3.2e-06 | 2.8e-06 | 1.9e-06 | 2.633E-06 | 6.658E-07 | 1.900E-06 | 3.200E-06 |
| 8818 | 441 | S441W | A | 1.9e-06 | 1.9e-06 | 2.3e-06 | 2.033E-06 | 2.309E-07 | 1.900E-06 | 2.300E-06 |
| 8819 | 441 | S441Y | A | 2.2e-06 | 3.2e-06 | 2.5e-06 | 2.633E-06 | 5.132E-07 | 2.200E-06 | 3.200E-06 |
| 8820 | 441 | S441V | A | 3.2e-06 | 2.9e-06 | 3.1e-06 | 3.067E-06 | 1.528E-07 | 2.900E-06 | 3.200E-06 |
| 8821 | 442 | G442A | A | 1.1e-06 | 2.1e-06 | 3e-06 | 2.067E-06 | 9.504E-07 | 1.100E-06 | 3.000E-06 |
| 8822 | 442 | G442R | A | 3.2e-06 | 2.1e-06 | 2.2e-06 | 2.500E-06 | 6.083E-07 | 2.100E-06 | 3.200E-06 |
| 8823 | 442 | G442N | A | 1.3e-06 | 3e-06 | 1.5e-06 | 1.933E-06 | 9.292E-07 | 1.300E-06 | 3.000E-06 |
| 8824 | 442 | G442D | A | 2.8e-06 | 2.6e-06 | 1.4e-06 | 2.267E-06 | 7.572E-07 | 1.400E-06 | 2.800E-06 |
| 8825 | 442 | G442C | A | 3.2e-06 | 3.3e-06 | 2.7e-06 | 3.067E-06 | 3.215E-07 | 2.700E-06 | 3.300E-06 |
| 8826 | 442 | G442E | A | 2.5e-06 | 1.7e-06 | 2.7e-06 | 2.300E-06 | 5.292E-07 | 1.700E-06 | 2.700E-06 |
| 8827 | 442 | G442Q | A | 4.7e-06 | 2.3e-06 | 3.3e-06 | 3.433E-06 | 1.206E-06 | 2.300E-06 | 4.700E-06 |
| 8828 | 442 | G442G | A | 1.4e-06 | 2.8e-06 | 2.8e-06 | 2.333E-06 | 8.083E-07 | 1.400E-06 | 2.800E-06 |
| 8829 | 442 | G442H | A | 1.8e-06 | 3.9e-06 | 1.8e-06 | 2.500E-06 | 1.212E-06 | 1.800E-06 | 3.900E-06 |
| 8830 | 442 | G442I | A | 1.2e-06 | 2.6e-06 | 2.2e-06 | 2.000E-06 | 7.211E-07 | 1.200E-06 | 2.600E-06 |
| 8831 | 442 | G442L | A | 2.3e-06 | 1.8e-06 | 2.4e-06 | 2.167E-06 | 3.215E-07 | 1.800E-06 | 2.400E-06 |
| 8832 | 442 | G442K | A | 2.7e-06 | 1.8e-06 | 2.3e-06 | 2.267E-06 | 4.509E-07 | 1.800E-06 | 2.700E-06 |
| 8833 | 442 | G442M | A | 1.3e-06 | 1.1e-06 | 1.9e-06 | 1.433E-06 | 4.163E-07 | 1.100E-06 | 1.900E-06 |
| 8834 | 442 | G442F | A | 2.2e-06 | 1.7e-06 | 2.1e-06 | 2.000E-06 | 2.646E-07 | 1.700E-06 | 2.200E-06 |
| 8835 | 442 | G442P | A | 2e-06 | 1.9e-06 | 2.9e-06 | 2.267E-06 | 5.508E-07 | 1.900E-06 | 2.900E-06 |
| 8836 | 442 | G442S | A | 1.9e-06 | 2.6e-06 | 2.8e-06 | 2.433E-06 | 4.726E-07 | 1.900E-06 | 2.800E-06 |
| 8837 | 442 | G442T | A | 2.9e-06 | 3.4e-06 | 2.5e-06 | 2.933E-06 | 4.509E-07 | 2.500E-06 | 3.400E-06 |
| 8838 | 442 | G442W | A | 1.6e-06 | 1.3e-06 | 2e-06 | 1.633E-06 | 3.512E-07 | 1.300E-06 | 2.000E-06 |
| 8839 | 442 | G442Y | A | 2.9e-06 | 1.5e-06 | 4.1e-06 | 2.833E-06 | 1.301E-06 | 1.500E-06 | 4.100E-06 |

| | | | | | | | | | | |
|---|---|---|---|---|---|---|---|---|---|---|
| 8840 | 442 | G442V | A | 1.7e-06 | 3.7e-06 | 1.8e-06 | 2.400E-06 | 1.127E-06 | 1.700E-06 | 3.700E-06 |
| 8841 | 443 | L443A | A | 2.3e-06 | 1.6e-06 | 2.4e-06 | 2.100E-06 | 4.359E-07 | 1.600E-06 | 2.400E-06 |
| 8842 | 443 | L443R | A | 3e-06 | 2.9e-06 | 2.7e-06 | 2.867E-06 | 1.528E-07 | 2.700E-06 | 3.000E-06 |
| 8843 | 443 | L443N | A | 2.4e-06 | 1.4e-06 | 2.8e-06 | 2.200E-06 | 7.211E-07 | 1.400E-06 | 2.800E-06 |
| 8844 | 443 | L443D | A | 3.5e-06 | 2.6e-06 | 1.1e-06 | 2.400E-06 | 1.212E-06 | 1.100E-06 | 3.500E-06 |
| 8845 | 443 | L443C | A | 2.5e-06 | 3.3e-06 | 1.3e-06 | 2.367E-06 | 1.007E-06 | 1.300E-06 | 3.300E-06 |
| 8846 | 443 | L443E | A | 1.2e-06 | 1.4e-06 | 2.5e-06 | 1.700E-06 | 7.000E-07 | 1.200E-06 | 2.500E-06 |
| 8847 | 443 | L443Q | A | 2.5e-06 | 3.8e-06 | 2.3e-06 | 2.867E-06 | 8.145E-07 | 2.300E-06 | 3.800E-06 |
| 8848 | 443 | L443G | A | 3.7e-06 | 2.2e-06 | 2.6e-06 | 2.833E-06 | 7.767E-07 | 2.200E-06 | 3.700E-06 |
| 8849 | 443 | L443H | A | 2.2e-06 | 2.8e-06 | 2.3e-06 | 2.433E-06 | 3.215E-07 | 2.200E-06 | 2.800E-06 |
| 8850 | 443 | L443I | A | 2.7e-06 | 2.6e-06 | 3.1e-06 | 2.800E-06 | 2.646E-07 | 2.600E-06 | 3.100E-06 |
| 8851 | 443 | L443L | A | 3.7e-06 | 2.7e-06 | 1.6e-06 | 2.667E-06 | 1.050E-06 | 1.600E-06 | 3.700E-06 |
| 8852 | 443 | L443K | A | 2.6e-06 | 2.2e-06 | 4.3e-06 | 3.033E-06 | 1.115E-06 | 2.200E-06 | 4.300E-06 |
| 8853 | 443 | L443M | A | 2.7e-06 | 2.4e-06 | 2.8e-06 | 2.633E-06 | 2.082E-07 | 2.400E-06 | 2.800E-06 |
| 8854 | 443 | L443F | A | 2.5e-06 | 2.4e-06 | 5.4e-06 | 3.433E-06 | 1.704E-06 | 2.400E-06 | 5.400E-06 |
| 8855 | 443 | L443P | A | 2.7e-06 | 1.5e-06 | 2.1e-06 | 2.100E-06 | 6.000E-07 | 1.500E-06 | 2.700E-06 |
| 8856 | 443 | L443S | A | 1.4e-06 | 2.4e-06 | 3e-06 | 2.267E-06 | 8.083E-07 | 1.400E-06 | 3.000E-06 |
| 8857 | 443 | L443T | A | 3.3e-06 | 2.7e-06 | 3.3e-06 | 3.100E-06 | 3.464E-07 | 2.700E-06 | 3.300E-06 |
| 8858 | 443 | L443W | A | 2.3e-06 | 2.9e-06 | 1.9e-06 | 2.367E-06 | 5.033E-07 | 1.900E-06 | 2.900E-06 |
| 8859 | 443 | L443Y | A | 2.5e-06 | 2.4e-06 | 1.1e-06 | 2.000E-06 | 7.810E-07 | 1.100E-06 | 2.500E-06 |
| 8860 | 443 | L443V | A | 2.3e-06 | 3.7e-06 | 1.8e-06 | 2.600E-06 | 9.849E-07 | 1.800E-06 | 3.700E-06 |
| 8861 | 444 | A444A | A | 2.5e-06 | 2.9e-06 | 2.2e-06 | 2.533E-06 | 3.512E-07 | 2.200E-06 | 2.900E-06 |
| 8862 | 444 | A444R | A | 2e-06 | 3.7e-06 | 2.2e-06 | 2.633E-06 | 9.292E-07 | 2.000E-06 | 3.700E-06 |
| 8863 | 444 | A444N | A | 1.5e-06 | 3e-06 | 2.3e-06 | 2.267E-06 | 7.506E-07 | 1.500E-06 | 3.000E-06 |
| 8864 | 444 | A444D | A | 2.2e-06 | 2.1e-06 | 2.2e-06 | 2.167E-06 | 5.774E-08 | 2.100E-06 | 2.200E-06 |
| 8865 | 444 | A444C | A | 2.8e-06 | 1.7e-06 | 2.1e-06 | 2.200E-06 | 5.568E-07 | 1.700E-06 | 2.800E-06 |
| 8866 | 444 | A444E | A | 1.9e-06 | 1.9e-06 | 2.1e-06 | 1.967E-06 | 1.155E-07 | 1.900E-06 | 2.100E-06 |
| 8867 | 444 | A444Q | A | 2.4e-06 | 2.1e-06 | 1e-06 | 1.833E-06 | 7.371E-07 | 1.000E-06 | 2.400E-06 |
| 8868 | 444 | A444G | A | 1.5e-06 | 2.1e-06 | 2.5e-06 | 2.033E-06 | 5.033E-07 | 1.500E-06 | 2.500E-06 |
| 8869 | 444 | A444H | A | 2e-06 | 1.9e-06 | 2.9e-06 | 2.267E-06 | 5.508E-07 | 1.900E-06 | 2.900E-06 |
| 8870 | 444 | A444I | A | 1.4e-06 | 1.4e-06 | 2.4e-06 | 1.733E-06 | 5.774E-07 | 1.400E-06 | 2.400E-06 |
| 8871 | 444 | A444L | A | 2.8e-06 | 2.5e-06 | 1.1e-06 | 2.133E-06 | 9.074E-07 | 1.100E-06 | 2.800E-06 |
| 8872 | 444 | A444K | A | 2.1e-06 | 1.9e-06 | 1.4e-06 | 1.800E-06 | 3.606E-07 | 1.400E-06 | 2.100E-06 |
| 8873 | 444 | A444M | A | 3.2e-06 | 4.8e-06 | 3.7e-06 | 3.900E-06 | 8.185E-07 | 3.200E-06 | 4.800E-06 |
| 8874 | 444 | A444F | A | 2.5e-06 | 2.2e-06 | 2.4e-06 | 2.367E-06 | 1.528E-07 | 2.200E-06 | 2.500E-06 |
| 8875 | 444 | A444P | A | 3.9e-06 | 2.5e-06 | 2.8e-06 | 3.067E-06 | 7.371E-07 | 2.500E-06 | 3.900E-06 |
| 8876 | 444 | A444S | A | 2e-06 | 3.1e-06 | 2.8e-06 | 2.633E-06 | 5.686E-07 | 2.000E-06 | 3.100E-06 |
| 8877 | 444 | A444T | A | 2.6e-06 | 2.2e-06 | 3.4e-06 | 2.733E-06 | 6.110E-07 | 2.200E-06 | 3.400E-06 |
| 8878 | 444 | A444W | A | 3.8e-06 | 5e-06 | 1e-06 | 3.267E-06 | 2.053E-06 | 1.000E-06 | 5.000E-06 |
| 8879 | 444 | A444Y | A | 2.7e-06 | 4.9e-06 | 2.7e-06 | 3.433E-06 | 1.270E-06 | 2.700E-06 | 4.900E-06 |
| 8880 | 444 | A444V | A | 2.4e-06 | 2.2e-06 | 4.7e-06 | 3.100E-06 | 1.389E-06 | 2.200E-06 | 4.700E-06 |
| 8881 | 445 | L445A | A | 1.8e-06 | 2.6e-06 | 2.5e-06 | 2.300E-06 | 4.359E-07 | 1.800E-06 | 2.600E-06 |
| 8882 | 445 | L445R | A | 2.3e-06 | 1.6e-06 | 2.2e-06 | 2.033E-06 | 3.786E-07 | 1.600E-06 | 2.300E-06 |
| 8883 | 445 | L445N | A | 2.5e-06 | 3.1e-06 | 3.7e-06 | 3.100E-06 | 6.000E-07 | 2.500E-06 | 3.700E-06 |
| 8884 | 445 | L445D | A | 2.7e-06 | 2.5e-06 | 1.5e-06 | 2.233E-06 | 6.429E-07 | 1.500E-06 | 2.700E-06 |
| 8885 | 445 | L445C | A | 7.4e-06 | 2.7e-06 | 2.7e-06 | 4.267E-06 | 2.714E-06 | 2.700E-06 | 7.400E-06 |
| 8886 | 445 | L445E | A | 3.5e-06 | 2.9e-06 | 2.3e-06 | 2.900E-06 | 6.000E-07 | 2.300E-06 | 3.500E-06 |
| 8887 | 445 | L445Q | A | 4.1e-06 | 1.1e-06 | 2.7e-06 | 2.633E-06 | 1.501E-06 | 1.100E-06 | 4.100E-06 |
| 8888 | 445 | L445G | A | 2.1e-06 | 2.5e-06 | 2.7e-06 | 2.433E-06 | 3.055E-07 | 2.100E-06 | 2.700E-06 |
| 8889 | 445 | L445H | A | 4e-06 | 3e-06 | 4.3e-06 | 3.767E-06 | 6.807E-07 | 3.000E-06 | 4.300E-06 |
| 8890 | 445 | L445I | A | 2e-06 | 1.9e-06 | 2.1e-06 | 2.000E-06 | 1.000E-07 | 1.900E-06 | 2.100E-06 |
| 8891 | 445 | L445L | A | 2e-06 | 2.2e-06 | 3.1e-06 | 2.433E-06 | 5.859E-07 | 2.000E-06 | 3.100E-06 |
| 8892 | 445 | L445K | A | 2.2e-06 | 3.1e-06 | 2.3e-06 | 2.533E-06 | 4.933E-07 | 2.200E-06 | 3.100E-06 |
| 8893 | 445 | L445M | A | 3.3e-06 | 1.1e-06 | 1.8e-06 | 2.067E-06 | 1.124E-06 | 1.100E-06 | 3.300E-06 |
| 8894 | 445 | L445F | A | 4.5e-06 | 3.4e-06 | 3.1e-06 | 3.667E-06 | 7.371E-07 | 3.100E-06 | 4.500E-06 |
| 8895 | 445 | L445P | A | 1.1e-06 | 2.2e-06 | 1.5e-06 | 1.600E-06 | 5.568E-07 | 1.100E-06 | 2.200E-06 |
| 8896 | 445 | L445S | A | 2e-06 | 1.8e-06 | 3.1e-06 | 2.300E-06 | 7.000E-07 | 1.800E-06 | 3.100E-06 |
| 8897 | 445 | L445T | A | 3e-06 | 1.3e-06 | 3.7e-06 | 2.667E-06 | 1.234E-06 | 1.300E-06 | 3.700E-06 |
| 8898 | 445 | L445W | A | 3.2e-06 | 3.2e-06 | 2e-06 | 2.800E-06 | 6.928E-07 | 2.000E-06 | 3.200E-06 |
| 8899 | 445 | L445Y | A | 2.2e-06 | 3.4e-06 | 1.6e-06 | 2.400E-06 | 9.165E-07 | 1.600E-06 | 3.400E-06 |
| 8900 | 445 | L445V | A | 2.5e-06 | 2.5e-06 | 2.1e-06 | 2.367E-06 | 2.309E-07 | 2.100E-06 | 2.500E-06 |
| 8901 | 446 | I446A | A | 2.5e-06 | 8.8e-07 | 2.1e-06 | 1.827E-06 | 8.439E-07 | 8.800E-07 | 2.500E-06 |
| 8902 | 446 | I446R | A | 2.3e-06 | 1.5e-06 | 3.3e-06 | 2.367E-06 | 9.018E-07 | 1.500E-06 | 3.300E-06 |
| 8903 | 446 | I446N | A | 1.9e-06 | 2.1e-06 | 2.7e-06 | 2.233E-06 | 4.163E-07 | 1.900E-06 | 2.700E-06 |
| 8904 | 446 | I446D | A | 3.7e-06 | 3e-06 | 2e-06 | 2.900E-06 | 8.544E-07 | 2.000E-06 | 3.700E-06 |
| 8905 | 446 | I446C | A | 2.3e-06 | 2.4e-06 | 1.5e-06 | 2.067E-06 | 4.933E-07 | 1.500E-06 | 2.400E-06 |
| 8906 | 446 | I446E | A | 2.7e-06 | 1.4e-06 | 1.6e-06 | 1.900E-06 | 7.000E-07 | 1.400E-06 | 2.700E-06 |
| 8907 | 446 | I446Q | A | 2.5e-06 | 3.6e-06 | 2.2e-06 | 2.767E-06 | 7.371E-07 | 2.200E-06 | 3.600E-06 |
| 8908 | 446 | I446G | A | 1.8e-06 | 2.6e-06 | 3.9e-06 | 2.767E-06 | 1.060E-06 | 1.800E-06 | 3.900E-06 |
| 8909 | 446 | I446H | A | 2.2e-06 | 2.6e-06 | 2.2e-06 | 2.333E-06 | 2.309E-07 | 2.200E-06 | 2.600E-06 |
| 8910 | 446 | I446I | A | 3.3e-06 | 2.7e-06 | 5e-06 | 3.667E-06 | 1.193E-06 | 2.700E-06 | 5.000E-06 |
| 8911 | 446 | I446L | A | 1.9e-06 | 4.9e-06 | 2.4e-06 | 3.067E-06 | 1.607E-06 | 1.900E-06 | 4.900E-06 |
| 8912 | 446 | I446K | A | 1.3e-06 | 1.6e-06 | 2.4e-06 | 1.767E-06 | 5.686E-07 | 1.300E-06 | 2.400E-06 |
| 8913 | 446 | I446M | A | 2.5e-06 | 3.8e-06 | 2.1e-06 | 2.800E-06 | 8.888E-07 | 2.100E-06 | 3.800E-06 |
| 8914 | 446 | I446F | A | 1.4e-06 | 2.3e-06 | 3.2e-06 | 2.300E-06 | 9.000E-07 | 1.400E-06 | 3.200E-06 |
| 8915 | 446 | I446P | A | 1.3e-06 | 3.3e-06 | 1.7e-06 | 2.100E-06 | 1.058E-06 | 1.300E-06 | 3.300E-06 |
| 8916 | 446 | I446S | A | 3.8e-06 | 2.2e-06 | 3.5e-06 | 3.167E-06 | 8.505E-07 | 2.200E-06 | 3.800E-06 |
| 8917 | 446 | I446T | A | 3.1e-06 | 2.2e-06 | 2.8e-06 | 2.700E-06 | 4.583E-07 | 2.200E-06 | 3.100E-06 |
| 8918 | 446 | I446W | A | 2.5e-06 | 2.9e-06 | 2.1e-06 | 2.500E-06 | 4.000E-07 | 2.100E-06 | 2.900E-06 |
| 8919 | 446 | I446Y | A | 2.3e-06 | 2.9e-06 | 3.6e-06 | 2.933E-06 | 6.506E-07 | 2.300E-06 | 3.600E-06 |
| 8920 | 446 | I446V | A | 2.7e-06 | 2.7e-06 | 1.1e-06 | 2.167E-06 | 9.238E-07 | 1.100E-06 | 2.700E-06 |
| 8921 | 447 | H447A | A | 4.1e-06 | 2.7e-06 | 2.4e-06 | 3.067E-06 | 9.074E-07 | 2.400E-06 | 4.100E-06 |
| 8922 | 447 | H447R | A | 9.9e-07 | 1.3e-06 | 1.7e-06 | 1.330E-06 | 3.559E-07 | 9.900E-07 | 1.700E-06 |
| 8923 | 447 | H447N | A | 3.1e-06 | 2e-06 | 3.2e-06 | 2.767E-06 | 6.658E-07 | 2.000E-06 | 3.200E-06 |
| 8924 | 447 | H447D | A | 4.6e-06 | 2.8e-06 | 2.4e-06 | 3.267E-06 | 1.172E-06 | 2.400E-06 | 4.600E-06 |

| | | | | | | | | | | |
|---|---|---|---|---|---|---|---|---|---|---|
| 8925 | 447 | H447C | A | 2.4e-06 | 2e-06 | 2e-06 | 2.133E-06 | 2.309E-07 | 2.000E-06 | 2.400E-06 |
| 8926 | 447 | H447E | A | 2.6e-06 | 2.1e-06 | 1.6e-06 | 2.100E-06 | 5.000E-07 | 1.600E-06 | 2.600E-06 |
| 8927 | 447 | H447Q | A | 1.9e-06 | 2e-06 | 3.9e-06 | 2.600E-06 | 1.127E-06 | 1.900E-06 | 3.900E-06 |
| 8928 | 447 | H447G | A | 2.5e-06 | 1.7e-06 | 2.7e-06 | 2.300E-06 | 5.292E-07 | 1.700E-06 | 2.700E-06 |
| 8929 | 447 | H447H | A | 4e-06 | 3e-06 | 1.6e-06 | 2.867E-06 | 1.206E-06 | 1.600E-06 | 4.000E-06 |
| 8930 | 447 | H447I | A | 2.9e-06 | 3.2e-06 | 2.9e-06 | 3.000E-06 | 1.732E-07 | 2.900E-06 | 3.200E-06 |
| 8931 | 447 | H447L | A | 2.5e-06 | 3.5e-06 | 2.6e-06 | 2.867E-06 | 5.508E-07 | 2.500E-06 | 3.500E-06 |
| 8932 | 447 | H447K | A | 2e-06 | 2e-06 | 2.5e-06 | 2.167E-06 | 2.887E-07 | 2.000E-06 | 2.500E-06 |
| 8933 | 447 | H447M | A | 3e-06 | 2e-06 | 2.3e-06 | 2.433E-06 | 5.132E-07 | 2.000E-06 | 3.000E-06 |
| 8934 | 447 | H447F | A | 1.6e-06 | 2.5e-06 | 3.7e-06 | 2.600E-06 | 1.054E-06 | 1.600E-06 | 3.700E-06 |
| 8935 | 447 | H447P | A | 1.3e-06 | 2.8e-06 | 1.7e-06 | 1.933E-06 | 7.767E-07 | 1.300E-06 | 2.800E-06 |
| 8936 | 447 | H447S | A | 3.8e-06 | 2.7e-06 | 3.4e-06 | 3.300E-06 | 5.568E-07 | 2.700E-06 | 3.800E-06 |
| 8937 | 447 | H447T | A | 3.2e-06 | 3.7e-06 | 2.2e-06 | 3.033E-06 | 7.638E-07 | 2.200E-06 | 3.700E-06 |
| 8938 | 447 | H447W | A | 3.2e-06 | 2.9e-06 | 2.2e-06 | 2.767E-06 | 5.132E-07 | 2.200E-06 | 3.200E-06 |
| 8939 | 447 | H447Y | A | 1.6e-06 | 1.9e-06 | 1.4e-06 | 1.633E-06 | 2.517E-07 | 1.400E-06 | 1.900E-06 |
| 8940 | 447 | H447V | A | 2.4e-06 | 3.6e-06 | 2e-06 | 2.667E-06 | 8.327E-07 | 2.000E-06 | 3.600E-06 |
| 8941 | 448 | H448A | A | 2.9e-06 | 1.1e-06 | 2.8e-06 | 2.267E-06 | 1.012E-06 | 1.100E-06 | 2.900E-06 |
| 8942 | 448 | H448R | A | 2.3e-06 | 3.5e-06 | 2.9e-06 | 2.900E-06 | 6.000E-07 | 2.300E-06 | 3.500E-06 |
| 8943 | 448 | H448N | A | 2.4e-06 | 2.1e-06 | 2.5e-06 | 2.333E-06 | 2.082E-07 | 2.100E-06 | 2.500E-06 |
| 8944 | 448 | H448D | A | 1.9e-06 | 3.4e-06 | 3.3e-06 | 2.867E-06 | 8.386E-07 | 1.900E-06 | 3.400E-06 |
| 8945 | 448 | H448C | A | 3.6e-06 | 1.3e-06 | 2.1e-06 | 2.333E-06 | 1.168E-06 | 1.300E-06 | 3.600E-06 |
| 8946 | 448 | H448E | A | 2.5e-06 | 3.5e-06 | 2.2e-06 | 2.733E-06 | 6.807E-07 | 2.200E-06 | 3.500E-06 |
| 8947 | 448 | H448Q | A | 1.4e-06 | 1.9e-06 | 3.1e-06 | 2.133E-06 | 8.737E-07 | 1.400E-06 | 3.100E-06 |
| 8948 | 448 | H448G | A | 2.3e-06 | 4.2e-06 | 4.2e-06 | 3.567E-06 | 1.097E-06 | 2.300E-06 | 4.200E-06 |
| 8949 | 448 | H448H | A | 1.9e-06 | 2.3e-06 | 2.7e-06 | 2.300E-06 | 4.000E-07 | 1.900E-06 | 2.700E-06 |
| 8950 | 448 | H448I | A | 1.5e-06 | 3.5e-06 | 4e-06 | 3.000E-06 | 1.323E-06 | 1.500E-06 | 4.000E-06 |
| 8951 | 448 | H448L | A | 3.9e-06 | 3e-06 | 1.2e-06 | 2.700E-06 | 1.375E-06 | 1.200E-06 | 3.900E-06 |
| 8952 | 448 | H448K | A | 1.9e-06 | 2.8e-06 | 2.3e-06 | 2.333E-06 | 4.509E-07 | 1.900E-06 | 2.800E-06 |
| 8953 | 448 | H448M | A | 2.9e-06 | 1.7e-06 | 3.5e-06 | 2.700E-06 | 9.165E-07 | 1.700E-06 | 3.500E-06 |
| 8954 | 448 | H448F | A | 2.9e-06 | 3.1e-06 | 1.6e-06 | 2.533E-06 | 8.145E-07 | 1.600E-06 | 3.100E-06 |
| 8955 | 448 | H448P | A | 2.2e-06 | 2.4e-06 | 2.4e-06 | 2.333E-06 | 1.155E-07 | 2.200E-06 | 2.400E-06 |
| 8956 | 448 | H448S | A | 2e-06 | 2e-06 | 1.8e-06 | 1.933E-06 | 1.155E-07 | 1.800E-06 | 2.000E-06 |
| 8957 | 448 | H448T | A | 1.8e-06 | 2.5e-06 | 1.7e-06 | 2.000E-06 | 4.359E-07 | 1.700E-06 | 2.500E-06 |
| 8958 | 448 | H448W | A | 2.6e-06 | 3.2e-06 | 1.6e-06 | 2.467E-06 | 8.083E-07 | 1.600E-06 | 3.200E-06 |
| 8959 | 448 | H448Y | A | 2.3e-06 | 4.4e-06 | 1.7e-06 | 2.800E-06 | 1.418E-06 | 1.700E-06 | 4.400E-06 |
| 8960 | 448 | H448V | A | 2e-06 | 3e-06 | 3.7e-06 | 2.900E-06 | 8.544E-07 | 2.000E-06 | 3.700E-06 |
| 8961 | 449 | N449A | A | 2e-06 | 1.9e-06 | 3.6e-06 | 2.500E-06 | 9.539E-07 | 1.900E-06 | 3.600E-06 |
| 8962 | 449 | N449R | A | 3.5e-06 | 1.9e-06 | 2.2e-06 | 2.533E-06 | 8.505E-07 | 1.900E-06 | 3.500E-06 |
| 8963 | 449 | N449N | A | 3.3e-06 | 3e-06 | 1.8e-06 | 2.700E-06 | 7.937E-07 | 1.800E-06 | 3.300E-06 |
| 8964 | 449 | N449D | A | 2.5e-06 | 1.5e-06 | 8.5e-07 | 1.617E-06 | 8.312E-07 | 8.500E-07 | 2.500E-06 |
| 8965 | 449 | N449C | A | 3e-06 | 4.5e-06 | 2.2e-06 | 3.233E-06 | 1.168E-06 | 2.200E-06 | 4.500E-06 |
| 8966 | 449 | N449E | A | 1.7e-06 | 1.6e-06 | 3.5e-06 | 2.267E-06 | 1.069E-06 | 1.600E-06 | 3.500E-06 |
| 8967 | 449 | N449Q | A | 2e-06 | 2.7e-06 | 2.3e-06 | 2.333E-06 | 3.512E-07 | 2.000E-06 | 2.700E-06 |
| 8968 | 449 | N449G | A | 4.7e-06 | 2.6e-06 | 1.7e-06 | 3.000E-06 | 1.539E-06 | 1.700E-06 | 4.700E-06 |
| 8969 | 449 | N449H | A | 3.1e-06 | 2.3e-06 | 3e-06 | 2.800E-06 | 4.359E-07 | 2.300E-06 | 3.100E-06 |
| 8970 | 449 | N449I | A | 3.4e-06 | 2.3e-06 | 1.8e-06 | 2.500E-06 | 8.185E-07 | 1.800E-06 | 3.400E-06 |
| 8971 | 449 | N449L | A | 2.2e-06 | 1.5e-06 | 2.2e-06 | 1.967E-06 | 4.041E-07 | 1.500E-06 | 2.200E-06 |
| 8972 | 449 | N449K | A | 1.8e-06 | 3.6e-06 | 1.8e-06 | 2.400E-06 | 1.039E-06 | 1.800E-06 | 3.600E-06 |
| 8973 | 449 | N449M | A | 1.4e-06 | 1.6e-06 | 1.6e-06 | 1.533E-06 | 1.155E-07 | 1.400E-06 | 1.600E-06 |
| 8974 | 449 | N449F | A | 2.7e-06 | 1.8e-06 | 1.8e-06 | 2.100E-06 | 5.196E-07 | 1.800E-06 | 2.700E-06 |
| 8975 | 449 | N449P | A | 2.8e-06 | 1.4e-06 | 2.2e-06 | 2.133E-06 | 7.024E-07 | 1.400E-06 | 2.800E-06 |
| 8976 | 449 | N449S | A | 1.8e-06 | 1.4e-06 | 3.9e-06 | 2.367E-06 | 1.343E-06 | 1.400E-06 | 3.900E-06 |
| 8977 | 449 | N449T | A | 3.7e-06 | 3.9e-06 | 3.8e-06 | 3.800E-06 | 1.000E-07 | 3.700E-06 | 3.900E-06 |
| 8978 | 449 | N449W | A | 2.9e-06 | 2.2e-06 | 1.9e-06 | 2.333E-06 | 5.132E-07 | 1.900E-06 | 2.900E-06 |
| 8979 | 449 | N449Y | A | 3.5e-06 | 3.4e-06 | 2.3e-06 | 3.067E-06 | 6.658E-07 | 2.300E-06 | 3.500E-06 |
| 8980 | 449 | N449V | A | 2e-06 | 1.2e-06 | 2.1e-06 | 1.767E-06 | 4.933E-07 | 1.200E-06 | 2.100E-06 |
| 8981 | 450 | T450A | A | 3.2e-06 | 2.2e-06 | 2.9e-06 | 2.767E-06 | 5.132E-07 | 2.200E-06 | 3.200E-06 |
| 8982 | 450 | T450R | A | 2.2e-06 | 2.9e-06 | 1.5e-06 | 2.200E-06 | 7.000E-07 | 1.500E-06 | 2.900E-06 |
| 8983 | 450 | T450N | A | 2.6e-06 | 3.3e-06 | 2.6e-06 | 2.833E-06 | 4.041E-07 | 2.600E-06 | 3.300E-06 |
| 8984 | 450 | T450D | A | 2.3e-06 | 3.3e-06 | 1.4e-06 | 2.333E-06 | 9.504E-07 | 1.400E-06 | 3.300E-06 |
| 8985 | 450 | T450C | A | 2.2e-06 | 1.9e-06 | 1.5e-06 | 1.867E-06 | 3.512E-07 | 1.500E-06 | 2.200E-06 |
| 8986 | 450 | T450E | A | 2.2e-06 | 2.8e-06 | 3.2e-06 | 2.733E-06 | 5.033E-07 | 2.200E-06 | 3.200E-06 |
| 8987 | 450 | T450Q | A | 2.2e-06 | 1.9e-06 | 2.7e-06 | 2.267E-06 | 4.041E-07 | 1.900E-06 | 2.700E-06 |
| 8988 | 450 | T450G | A | 3.9e-06 | 2.1e-06 | 1.7e-06 | 2.567E-06 | 1.172E-06 | 1.700E-06 | 3.900E-06 |
| 8989 | 450 | T450H | A | 2.7e-06 | 2.6e-06 | 1.8e-06 | 2.367E-06 | 4.933E-07 | 1.800E-06 | 2.700E-06 |
| 8990 | 450 | T450I | A | 2.3e-06 | 1.4e-06 | 3e-06 | 2.233E-06 | 8.021E-07 | 1.400E-06 | 3.000E-06 |
| 8991 | 450 | T450L | A | 2.9e-06 | 3.5e-06 | 1.7e-06 | 2.700E-06 | 9.165E-07 | 1.700E-06 | 3.500E-06 |
| 8992 | 450 | T450K | A | 2.7e-06 | 1.2e-06 | 2.6e-06 | 2.167E-06 | 8.386E-07 | 1.200E-06 | 2.700E-06 |
| 8993 | 450 | T450M | A | 1.7e-06 | 2.9e-06 | 3.6e-06 | 2.733E-06 | 9.609E-07 | 1.700E-06 | 3.600E-06 |
| 8994 | 450 | T450F | A | 2.3e-06 | 1.7e-06 | 1.8e-06 | 1.933E-06 | 3.215E-07 | 1.700E-06 | 2.300E-06 |
| 8995 | 450 | T450P | A | 2.5e-06 | 3.2e-06 | 3.1e-06 | 2.933E-06 | 3.786E-07 | 2.500E-06 | 3.200E-06 |
| 8996 | 450 | T450S | A | 3.6e-06 | 4.5e-06 | 2.7e-06 | 3.600E-06 | 9.000E-07 | 2.700E-06 | 4.500E-06 |
| 8997 | 450 | T450T | A | 1.4e-06 | 1.4e-06 | 3.7e-06 | 2.167E-06 | 1.328E-06 | 1.400E-06 | 3.700E-06 |
| 8998 | 450 | T450W | A | 1.7e-06 | 2e-06 | 1.6e-06 | 1.767E-06 | 2.082E-07 | 1.600E-06 | 2.000E-06 |
| 8999 | 450 | T450Y | A | 2e-06 | 2.3e-06 | 1.1e-06 | 1.800E-06 | 6.245E-07 | 1.100E-06 | 2.300E-06 |
| 9000 | 450 | T450V | A | 1.8e-06 | 2.2e-06 | 1.6e-06 | 1.867E-06 | 3.055E-07 | 1.600E-06 | 2.200E-06 |
| 9001 | 451 | H451A | A | 4.2e-06 | 2.2e-06 | 1.8e-06 | 2.733E-06 | 1.286E-06 | 1.800E-06 | 4.200E-06 |
| 9002 | 451 | H451R | A | 1.9e-06 | 1.2e-06 | 4.5e-06 | 2.533E-06 | 1.739E-06 | 1.200E-06 | 4.500E-06 |
| 9003 | 451 | H451N | A | 3e-06 | 2.3e-06 | 2.7e-06 | 2.667E-06 | 3.512E-07 | 2.300E-06 | 3.000E-06 |
| 9004 | 451 | H451D | A | 2.3e-06 | 2.2e-06 | 2.1e-06 | 2.200E-06 | 1.000E-07 | 2.100E-06 | 2.300E-06 |
| 9005 | 451 | H451C | A | 2.8e-06 | 1.8e-06 | 3.1e-06 | 2.567E-06 | 6.807E-07 | 1.800E-06 | 3.100E-06 |
| 9006 | 451 | H451E | A | 3e-06 | 3.5e-06 | 1.7e-06 | 2.733E-06 | 9.292E-07 | 1.700E-06 | 3.500E-06 |
| 9007 | 451 | H451Q | A | 4.1e-06 | 4.5e-06 | 4.1e-06 | 4.233E-06 | 2.309E-07 | 4.100E-06 | 4.500E-06 |
| 9008 | 451 | H451G | A | 1.5e-06 | 3e-06 | 1.3e-06 | 1.933E-06 | 9.292E-07 | 1.300E-06 | 3.000E-06 |
| 9009 | 451 | H451H | A | 1.5e-06 | 4e-06 | 1.9e-06 | 2.467E-06 | 1.343E-06 | 1.500E-06 | 4.000E-06 |

| | | | | | | | | | | |
|---|---|---|---|---|---|---|---|---|---|---|
| 9010 | 451 | H451I | A | 2e-06 | 1.5e-06 | 2.2e-06 | 1.900E-06 | 3.606E-07 | 1.500E-06 | 2.200E-06 |
| 9011 | 451 | H451L | A | 3.4e-06 | 2e-06 | 2.2e-06 | 2.533E-06 | 7.572E-07 | 2.000E-06 | 3.400E-06 |
| 9012 | 451 | H451K | A | 1.1e-06 | 3.6e-06 | 1.9e-06 | 2.200E-06 | 1.277E-06 | 1.100E-06 | 3.600E-06 |
| 9013 | 451 | H451M | A | 2.9e-06 | 3.1e-06 | 2.5e-06 | 2.833E-06 | 3.055E-07 | 2.500E-06 | 3.100E-06 |
| 9014 | 451 | H451F | A | 1.8e-06 | 2.3e-06 | 3.1e-06 | 2.400E-06 | 6.557E-07 | 1.800E-06 | 3.100E-06 |
| 9015 | 451 | H451P | A | 2.2e-06 | 3.7e-06 | 2.4e-06 | 2.767E-06 | 8.145E-07 | 2.200E-06 | 3.700E-06 |
| 9016 | 451 | H451S | A | 2.3e-06 | 2.6e-06 | 2.4e-06 | 2.433E-06 | 1.528E-07 | 2.300E-06 | 2.600E-06 |
| 9017 | 451 | H451T | A | 2.5e-06 | 1.8e-06 | 2.2e-06 | 2.167E-06 | 3.512E-07 | 1.800E-06 | 2.500E-06 |
| 9018 | 451 | H451W | A | 2.3e-06 | 2.3e-06 | 2.1e-06 | 2.233E-06 | 1.155E-07 | 2.100E-06 | 2.300E-06 |
| 9019 | 451 | H451Y | A | 2.7e-06 | 6e-06 | 2e-06 | 3.567E-06 | 2.136E-06 | 2.000E-06 | 6.000E-06 |
| 9020 | 451 | H451V | A | 2.2e-06 | 2.7e-06 | 2.7e-06 | 2.533E-06 | 2.887E-07 | 2.200E-06 | 2.700E-06 |
| 9021 | 452 | L452A | A | 1.7e-06 | 2.4e-06 | 2.6e-06 | 2.233E-06 | 4.726E-07 | 1.700E-06 | 2.600E-06 |
| 9022 | 452 | L452R | A | 2.5e-06 | 2.4e-06 | 1.9e-06 | 2.267E-06 | 3.215E-07 | 1.900E-06 | 2.500E-06 |
| 9023 | 452 | L452N | A | 2.1e-06 | 1.9e-06 | 3.5e-06 | 2.500E-06 | 8.718E-07 | 1.900E-06 | 3.500E-06 |
| 9024 | 452 | L452D | A | 2.1e-06 | 3e-06 | 2.8e-06 | 2.633E-06 | 4.726E-07 | 2.100E-06 | 3.000E-06 |
| 9025 | 452 | L452C | A | 2.2e-06 | 2.8e-06 | 3.1e-06 | 2.700E-06 | 4.583E-07 | 2.200E-06 | 3.100E-06 |
| 9026 | 452 | L452E | A | 2.4e-06 | 3.4e-06 | 2.7e-06 | 2.833E-06 | 5.132E-07 | 2.400E-06 | 3.400E-06 |
| 9027 | 452 | L452Q | A | 1.9e-06 | 3.1e-06 | 2e-06 | 2.333E-06 | 6.658E-07 | 1.900E-06 | 3.100E-06 |
| 9028 | 452 | L452G | A | 2.1e-06 | 2.3e-06 | 2.2e-06 | 2.200E-06 | 1.000E-07 | 2.100E-06 | 2.300E-06 |
| 9029 | 452 | L452H | A | 4.1e-06 | 1.8e-06 | 2.5e-06 | 2.800E-06 | 1.179E-06 | 1.800E-06 | 4.100E-06 |
| 9030 | 452 | L452I | A | 3.3e-06 | 1.8e-06 | 2.3e-06 | 2.467E-06 | 7.638E-07 | 1.800E-06 | 3.300E-06 |
| 9031 | 452 | L452L | A | 2.5e-06 | 2.6e-06 | 2.1e-06 | 2.400E-06 | 2.646E-07 | 2.100E-06 | 2.600E-06 |
| 9032 | 452 | L452K | A | 1.6e-06 | 2.1e-06 | 2.8e-06 | 2.167E-06 | 6.028E-07 | 1.600E-06 | 2.800E-06 |
| 9033 | 452 | L452M | A | 4.5e-06 | 1.9e-06 | 3e-06 | 3.133E-06 | 1.305E-06 | 1.900E-06 | 4.500E-06 |
| 9034 | 452 | L452F | A | 2.7e-06 | 2.7e-06 | 2.6e-06 | 2.667E-06 | 5.774E-08 | 2.600E-06 | 2.700E-06 |
| 9035 | 452 | L452P | A | 2.2e-06 | 2.4e-06 | 2.2e-06 | 2.267E-06 | 1.155E-07 | 2.200E-06 | 2.400E-06 |
| 9036 | 452 | L452S | A | 3e-06 | 3.4e-06 | 2.5e-06 | 2.967E-06 | 4.509E-07 | 2.500E-06 | 3.400E-06 |
| 9037 | 452 | L452T | A | 2.5e-06 | 3.8e-06 | 2.8e-06 | 3.033E-06 | 6.807E-07 | 2.500E-06 | 3.800E-06 |
| 9038 | 452 | L452W | A | 1.9e-06 | 3.1e-06 | 2.6e-06 | 2.533E-06 | 6.028E-07 | 1.900E-06 | 3.100E-06 |
| 9039 | 452 | L452Y | A | 2.8e-06 | 2.7e-06 | 2.7e-06 | 2.733E-06 | 5.774E-08 | 2.700E-06 | 2.800E-06 |
| 9040 | 452 | L452V | A | 1.5e-06 | 2.6e-06 | 3.2e-06 | 2.433E-06 | 8.622E-07 | 1.500E-06 | 3.200E-06 |
| 9041 | 453 | C453A | A | 2.1e-06 | 1.2e-06 | 3.5e-06 | 2.267E-06 | 1.159E-06 | 1.200E-06 | 3.500E-06 |
| 9042 | 453 | C453R | A | 2.5e-06 | 1.9e-06 | 2.2e-06 | 2.200E-06 | 3.000E-07 | 1.900E-06 | 2.500E-06 |
| 9043 | 453 | C453N | A | 3.2e-06 | 3.3e-06 | 2e-06 | 2.833E-06 | 7.234E-07 | 2.000E-06 | 3.300E-06 |
| 9044 | 453 | C453D | A | 3.2e-06 | 2.3e-06 | 2.5e-06 | 2.667E-06 | 4.726E-07 | 2.300E-06 | 3.200E-06 |
| 9045 | 453 | C453C | A | 3.1e-06 | 2e-06 | 2.5e-06 | 2.533E-06 | 5.508E-07 | 2.000E-06 | 3.100E-06 |
| 9046 | 453 | C453E | A | 2e-06 | 1.9e-06 | 2.7e-06 | 2.200E-06 | 4.359E-07 | 1.900E-06 | 2.700E-06 |
| 9047 | 453 | C453Q | A | 2.3e-06 | 1.7e-06 | 1.9e-06 | 1.967E-06 | 3.055E-07 | 1.700E-06 | 2.300E-06 |
| 9048 | 453 | C453G | A | 2.7e-06 | 3.3e-06 | 2.6e-06 | 2.867E-06 | 3.786E-07 | 2.600E-06 | 3.300E-06 |
| 9049 | 453 | C453H | A | 1.4e-06 | 1.7e-06 | 2.6e-06 | 1.900E-06 | 6.245E-07 | 1.400E-06 | 2.600E-06 |
| 9050 | 453 | C453I | A | 1.9e-06 | 1.5e-06 | 5.3e-06 | 2.900E-06 | 2.088E-06 | 1.500E-06 | 5.300E-06 |
| 9051 | 453 | C453L | A | 1.6e-06 | 2.7e-06 | 2.4e-06 | 2.233E-06 | 5.686E-07 | 1.600E-06 | 2.700E-06 |
| 9052 | 453 | C453K | A | 2.3e-06 | 2.2e-06 | 2.2e-06 | 2.233E-06 | 5.774E-08 | 2.200E-06 | 2.300E-06 |
| 9053 | 453 | C453M | A | 3.1e-06 | 2.2e-06 | 2.2e-06 | 2.500E-06 | 5.196E-07 | 2.200E-06 | 3.100E-06 |
| 9054 | 453 | C453F | A | 1.9e-06 | 6.8e-06 | 2.3e-06 | 3.667E-06 | 2.721E-06 | 1.900E-06 | 6.800E-06 |
| 9055 | 453 | C453P | A | 4.1e-06 | 3.3e-06 | 2.7e-06 | 3.367E-06 | 7.024E-07 | 2.700E-06 | 4.100E-06 |
| 9056 | 453 | C453S | A | 2.1e-06 | 3.3e-06 | 3e-06 | 2.800E-06 | 6.245E-07 | 2.100E-06 | 3.300E-06 |
| 9057 | 453 | C453T | A | 3.1e-06 | 2.5e-06 | 2e-06 | 2.533E-06 | 5.508E-07 | 2.000E-06 | 3.100E-06 |
| 9058 | 453 | C453W | A | 2.3e-06 | 1.6e-06 | 3.9e-06 | 2.600E-06 | 1.179E-06 | 1.600E-06 | 3.900E-06 |
| 9059 | 453 | C453Y | A | 2.2e-06 | 2.1e-06 | 2.4e-06 | 2.233E-06 | 1.528E-07 | 2.100E-06 | 2.400E-06 |
| 9060 | 453 | C453V | A | 2.7e-06 | 1.9e-06 | 5e-06 | 3.200E-06 | 1.609E-06 | 1.900E-06 | 5.000E-06 |
| 9061 | 454 | F454A | A | 2.1e-06 | 2.5e-06 | 1.7e-06 | 2.100E-06 | 4.000E-07 | 1.700E-06 | 2.500E-06 |
| 9062 | 454 | F454R | A | 5e-06 | 2.2e-06 | 1.8e-06 | 3.000E-06 | 1.744E-06 | 1.800E-06 | 5.000E-06 |
| 9063 | 454 | F454N | A | 3e-06 | 3.4e-06 | 3.2e-06 | 3.200E-06 | 2.000E-07 | 3.000E-06 | 3.400E-06 |
| 9064 | 454 | F454D | A | 1.8e-06 | 2.2e-06 | 3.1e-06 | 2.367E-06 | 6.658E-07 | 1.800E-06 | 3.100E-06 |
| 9065 | 454 | F454C | A | 4.3e-06 | 1.9e-06 | 2.5e-06 | 2.900E-06 | 1.249E-06 | 1.900E-06 | 4.300E-06 |
| 9066 | 454 | F454E | A | 2.4e-06 | 2.2e-06 | 3.2e-06 | 2.600E-06 | 5.292E-07 | 2.200E-06 | 3.200E-06 |
| 9067 | 454 | F454Q | A | 2.2e-06 | 2.2e-06 | 2.4e-06 | 2.267E-06 | 1.155E-07 | 2.200E-06 | 2.400E-06 |
| 9068 | 454 | F454G | A | 2.2e-06 | 1.2e-06 | 2e-06 | 1.800E-06 | 5.292E-07 | 1.200E-06 | 2.200E-06 |
| 9069 | 454 | F454H | A | 2.5e-06 | 2e-06 | 2.7e-06 | 2.400E-06 | 3.606E-07 | 2.000E-06 | 2.700E-06 |
| 9070 | 454 | F454I | A | 2.8e-06 | 2.4e-06 | 2.4e-06 | 2.533E-06 | 2.309E-07 | 2.400E-06 | 2.800E-06 |
| 9071 | 454 | F454L | A | 2.7e-06 | 2.8e-06 | 4.3e-06 | 3.267E-06 | 8.963E-07 | 2.700E-06 | 4.300E-06 |
| 9072 | 454 | F454K | A | 2e-06 | 1.9e-06 | 2.7e-06 | 2.200E-06 | 4.359E-07 | 1.900E-06 | 2.700E-06 |
| 9073 | 454 | F454M | A | 3.4e-06 | 3.1e-06 | 1e-06 | 2.500E-06 | 1.308E-06 | 1.000E-06 | 3.400E-06 |
| 9074 | 454 | F454F | A | 2e-06 | 2.3e-06 | 1.1e-06 | 1.800E-06 | 6.245E-07 | 1.100E-06 | 2.300E-06 |
| 9075 | 454 | F454P | A | 2e-06 | 2.1e-06 | 2.2e-06 | 2.100E-06 | 1.000E-07 | 2.000E-06 | 2.200E-06 |
| 9076 | 454 | F454S | A | 3.1e-06 | 2.9e-06 | 5e-06 | 3.667E-06 | 1.159E-06 | 2.900E-06 | 5.000E-06 |
| 9077 | 454 | F454T | A | 2.5e-06 | 1.9e-06 | 2.1e-06 | 2.167E-06 | 3.055E-07 | 1.900E-06 | 2.500E-06 |
| 9078 | 454 | F454W | A | 2.5e-06 | 1.8e-06 | 2.3e-06 | 2.200E-06 | 3.606E-07 | 1.800E-06 | 2.500E-06 |
| 9079 | 454 | F454Y | A | 3.2e-06 | 2e-06 | 2.4e-06 | 2.533E-06 | 6.110E-07 | 2.000E-06 | 3.200E-06 |
| 9080 | 454 | F454V | A | 2.8e-06 | 1.7e-06 | 1.7e-06 | 2.067E-06 | 6.351E-07 | 1.700E-06 | 2.800E-06 |
| 9081 | 455 | V455A | A | 2.9e-06 | 1.3e-06 | 2e-06 | 2.067E-06 | 8.021E-07 | 1.300E-06 | 2.900E-06 |
| 9082 | 455 | V455R | A | 3.7e-06 | 1.8e-06 | 2.1e-06 | 2.533E-06 | 1.021E-06 | 1.800E-06 | 3.700E-06 |
| 9083 | 455 | V455N | A | 2e-06 | 2.2e-06 | 1.9e-06 | 2.033E-06 | 1.528E-07 | 1.900E-06 | 2.200E-06 |
| 9084 | 455 | V455D | A | 1.9e-06 | 2.8e-06 | 1.8e-06 | 2.167E-06 | 5.508E-07 | 1.800E-06 | 2.800E-06 |
| 9085 | 455 | V455C | A | 3.2e-06 | 1.6e-06 | 2.6e-06 | 2.467E-06 | 8.083E-07 | 1.600E-06 | 3.200E-06 |
| 9086 | 455 | V455E | A | 1.7e-06 | 2.5e-06 | 2e-06 | 2.067E-06 | 4.041E-07 | 1.700E-06 | 2.500E-06 |
| 9087 | 455 | V455Q | A | 2.9e-06 | 2e-06 | 3.9e-06 | 2.933E-06 | 9.504E-07 | 2.000E-06 | 3.900E-06 |
| 9088 | 455 | V455G | A | 1.6e-06 | 2.1e-06 | 4.4e-06 | 2.700E-06 | 1.493E-06 | 1.600E-06 | 4.400E-06 |
| 9089 | 455 | V455H | A | 1.7e-06 | 2e-06 | 2.8e-06 | 2.167E-06 | 5.686E-07 | 1.700E-06 | 2.800E-06 |
| 9090 | 455 | V455I | A | 1.9e-06 | 4.9e-06 | 2.4e-06 | 3.067E-06 | 1.607E-06 | 1.900E-06 | 4.900E-06 |
| 9091 | 455 | V455L | A | 1.9e-06 | 2.2e-06 | 2.2e-06 | 2.100E-06 | 1.732E-07 | 1.900E-06 | 2.200E-06 |
| 9092 | 455 | V455K | A | 1.4e-06 | 3.8e-06 | 2.4e-06 | 2.533E-06 | 1.206E-06 | 1.400E-06 | 3.800E-06 |
| 9093 | 455 | V455M | A | 3e-06 | 2.5e-06 | 2.5e-06 | 2.667E-06 | 2.887E-07 | 2.500E-06 | 3.000E-06 |
| 9094 | 455 | V455F | A | 2.5e-06 | 2.2e-06 | 3.3e-06 | 2.667E-06 | 5.686E-07 | 2.200E-06 | 3.300E-06 |

| | | | | | | | | | | |
|---|---|---|---|---|---|---|---|---|---|---|
| 9095 | 455 | V455P | A | 2.6e-06 | 2.7e-06 | 2.6e-06 | 2.633E-06 | 5.774E-08 | 2.600E-06 | 2.700E-06 |
| 9096 | 455 | V455S | A | 1.8e-06 | 2.5e-06 | 4.3e-06 | 2.867E-06 | 1.290E-06 | 1.800E-06 | 4.300E-06 |
| 9097 | 455 | V455T | A | 2.2e-06 | 3.1e-06 | 2.4e-06 | 2.567E-06 | 4.726E-07 | 2.200E-06 | 3.100E-06 |
| 9098 | 455 | V455W | A | 1.4e-06 | 1.4e-06 | 2.2e-06 | 1.667E-06 | 4.619E-07 | 1.400E-06 | 2.200E-06 |
| 9099 | 455 | V455Y | A | 1e-06 | 2.1e-06 | 3.1e-06 | 2.067E-06 | 1.050E-06 | 1.000E-06 | 3.100E-06 |
| 9100 | 455 | V455V | A | 3.4e-06 | 2.1e-06 | 2.1e-06 | 2.533E-06 | 7.506E-07 | 2.100E-06 | 3.400E-06 |
| 9101 | 456 | H456A | A | 2.2e-06 | 2.7e-06 | 3.7e-06 | 2.867E-06 | 7.638E-07 | 2.200E-06 | 3.700E-06 |
| 9102 | 456 | H456R | A | 3.4e-06 | 1.7e-06 | 1.2e-06 | 2.100E-06 | 1.153E-06 | 1.200E-06 | 3.400E-06 |
| 9103 | 456 | H456N | A | 3.1e-06 | 2.6e-06 | 3.5e-06 | 3.067E-06 | 4.509E-07 | 2.600E-06 | 3.500E-06 |
| 9104 | 456 | H456D | A | 3.1e-06 | 2.2e-06 | 1.9e-06 | 2.400E-06 | 6.245E-07 | 1.900E-06 | 3.100E-06 |
| 9105 | 456 | H456C | A | 2.5e-06 | 3e-06 | 2.9e-06 | 2.800E-06 | 2.646E-07 | 2.500E-06 | 3.000E-06 |
| 9106 | 456 | H456E | A | 1.9e-06 | 2e-06 | 2.4e-06 | 2.100E-06 | 2.646E-07 | 1.900E-06 | 2.400E-06 |
| 9107 | 456 | H456Q | A | 2.5e-06 | 1.1e-06 | 3e-06 | 2.200E-06 | 9.849E-07 | 1.100E-06 | 3.000E-06 |
| 9108 | 456 | H456G | A | 3.1e-06 | 4.8e-06 | 2.2e-06 | 3.367E-06 | 1.320E-06 | 2.200E-06 | 4.800E-06 |
| 9109 | 456 | H456H | A | 1.8e-06 | 2.2e-06 | 1.3e-06 | 1.767E-06 | 4.509E-07 | 1.300E-06 | 2.200E-06 |
| 9110 | 456 | H456I | A | 3.2e-06 | 2.3e-06 | 4.4e-06 | 3.300E-06 | 1.054E-06 | 2.300E-06 | 4.400E-06 |
| 9111 | 456 | H456L | A | 2.4e-06 | 2e-06 | 1.9e-06 | 2.100E-06 | 2.646E-07 | 1.900E-06 | 2.400E-06 |
| 9112 | 456 | H456K | A | 2.3e-06 | 2.4e-06 | 4e-06 | 2.900E-06 | 9.539E-07 | 2.300E-06 | 4.000E-06 |
| 9113 | 456 | H456M | A | 2.8e-06 | 2.6e-06 | 1.5e-06 | 2.300E-06 | 7.000E-07 | 1.500E-06 | 2.800E-06 |
| 9114 | 456 | H456F | A | 4.6e-06 | 1.6e-06 | 1.5e-06 | 2.567E-06 | 1.762E-06 | 1.500E-06 | 4.600E-06 |
| 9115 | 456 | H456P | A | 2.5e-06 | 3.8e-06 | 2.6e-06 | 2.967E-06 | 7.234E-07 | 2.500E-06 | 3.800E-06 |
| 9116 | 456 | H456S | A | 2.2e-06 | 3.3e-06 | 2.9e-06 | 2.800E-06 | 5.568E-07 | 2.200E-06 | 3.300E-06 |
| 9117 | 456 | H456T | A | 2.2e-06 | 4.1e-06 | 2.8e-06 | 3.033E-06 | 9.713E-07 | 2.200E-06 | 4.100E-06 |
| 9118 | 456 | H456W | A | 3.2e-06 | 3e-06 | 1.5e-06 | 2.567E-06 | 9.292E-07 | 1.500E-06 | 3.200E-06 |
| 9119 | 456 | H456Y | A | 2.2e-06 | 2.5e-06 | 2.6e-06 | 2.433E-06 | 2.082E-07 | 2.200E-06 | 2.600E-06 |
| 9120 | 456 | H456V | A | 1.8e-06 | 3.1e-06 | 3.8e-06 | 2.900E-06 | 1.015E-06 | 1.800E-06 | 3.800E-06 |
| 9121 | 457 | T457A | A | 1.1e-06 | 3.6e-06 | 2.5e-06 | 2.400E-06 | 1.253E-06 | 1.100E-06 | 3.600E-06 |
| 9122 | 457 | T457R | A | 2.6e-06 | 3.2e-06 | 2.3e-06 | 2.700E-06 | 4.583E-07 | 2.300E-06 | 3.200E-06 |
| 9123 | 457 | T457N | A | 2e-06 | 2.2e-06 | 2.7e-06 | 2.300E-06 | 3.606E-07 | 2.000E-06 | 2.700E-06 |
| 9124 | 457 | T457D | A | 1.1e-06 | 3.7e-06 | 2e-06 | 2.267E-06 | 1.320E-06 | 1.100E-06 | 3.700E-06 |
| 9125 | 457 | T457C | A | 2.4e-06 | 2.5e-06 | 2.1e-06 | 2.333E-06 | 2.082E-07 | 2.100E-06 | 2.500E-06 |
| 9126 | 457 | T457E | A | 3.1e-06 | 3.1e-06 | 3.1e-06 | 3.100E-06 | 5.187E-22 | 3.100E-06 | 3.100E-06 |
| 9127 | 457 | T457Q | A | 2.9e-06 | 3.7e-06 | 1.9e-06 | 2.833E-06 | 9.018E-07 | 1.900E-06 | 3.700E-06 |
| 9128 | 457 | T457G | A | 3.1e-06 | 2.7e-06 | 2.7e-06 | 2.833E-06 | 2.309E-07 | 2.700E-06 | 3.100E-06 |
| 9129 | 457 | T457H | A | 2e-06 | 2.7e-06 | 2.4e-06 | 2.367E-06 | 3.512E-07 | 2.000E-06 | 2.700E-06 |
| 9130 | 457 | T457I | A | 2.9e-06 | 3.9e-06 | 1.2e-06 | 2.667E-06 | 1.365E-06 | 1.200E-06 | 3.900E-06 |
| 9131 | 457 | T457L | A | 2.2e-06 | 1.9e-06 | 1.9e-06 | 2.000E-06 | 1.732E-07 | 1.900E-06 | 2.200E-06 |
| 9132 | 457 | T457K | A | 2.1e-06 | 3e-06 | 1.9e-06 | 2.333E-06 | 5.859E-07 | 1.900E-06 | 3.000E-06 |
| 9133 | 457 | T457M | A | 3.4e-06 | 3.1e-06 | 1.4e-06 | 2.633E-06 | 1.079E-06 | 1.400E-06 | 3.400E-06 |
| 9134 | 457 | T457F | A | 3.5e-06 | 2.5e-06 | 2e-06 | 2.667E-06 | 7.638E-07 | 2.000E-06 | 3.500E-06 |
| 9135 | 457 | T457P | A | 2.7e-06 | 2.6e-06 | 1.8e-06 | 2.367E-06 | 4.933E-07 | 1.800E-06 | 2.700E-06 |
| 9136 | 457 | T457S | A | 2.3e-06 | 1.3e-06 | 2.8e-06 | 2.133E-06 | 7.638E-07 | 1.300E-06 | 2.800E-06 |
| 9137 | 457 | T457T | A | 2.8e-06 | 2.3e-06 | 3.4e-06 | 2.833E-06 | 5.508E-07 | 2.300E-06 | 3.400E-06 |
| 9138 | 457 | T457W | A | 2.8e-06 | 2.5e-06 | 1.7e-06 | 2.333E-06 | 5.686E-07 | 1.700E-06 | 2.800E-06 |
| 9139 | 457 | T457Y | A | 2.2e-06 | 1.9e-06 | 3e-06 | 2.367E-06 | 5.686E-07 | 1.900E-06 | 3.000E-06 |
| 9140 | 457 | T457V | A | 3.6e-06 | 2.3e-06 | 1.9e-06 | 2.600E-06 | 8.888E-07 | 1.900E-06 | 3.600E-06 |
| 9141 | 458 | V458A | A | 2e-06 | 3.2e-06 | 3.5e-06 | 2.900E-06 | 7.937E-07 | 2.000E-06 | 3.500E-06 |
| 9142 | 458 | V458R | A | 2.7e-06 | 2e-06 | 2.5e-06 | 2.400E-06 | 3.606E-07 | 2.000E-06 | 2.700E-06 |
| 9143 | 458 | V458N | A | 3e-06 | 2.3e-06 | 1.7e-06 | 2.333E-06 | 6.506E-07 | 1.700E-06 | 3.000E-06 |
| 9144 | 458 | V458D | A | 2.2e-06 | 1.9e-06 | 1.6e-06 | 1.900E-06 | 3.000E-07 | 1.600E-06 | 2.200E-06 |
| 9145 | 458 | V458C | A | 1.5e-06 | 3.1e-06 | 2.9e-06 | 2.500E-06 | 8.718E-07 | 1.500E-06 | 3.100E-06 |
| 9146 | 458 | V458E | A | 3.2e-06 | 1.9e-06 | 4e-06 | 3.033E-06 | 1.060E-06 | 1.900E-06 | 4.000E-06 |
| 9147 | 458 | V458Q | A | 3e-06 | 1.9e-06 | 2e-06 | 2.300E-06 | 6.083E-07 | 1.900E-06 | 3.000E-06 |
| 9148 | 458 | V458G | A | 2.2e-06 | 3.9e-06 | 3.1e-06 | 3.067E-06 | 8.505E-07 | 2.200E-06 | 3.900E-06 |
| 9149 | 458 | V458H | A | 1.8e-06 | 2.2e-06 | 2.2e-06 | 2.067E-06 | 2.309E-07 | 1.800E-06 | 2.200E-06 |
| 9150 | 458 | V458I | A | 2.6e-06 | 2.7e-06 | 3.2e-06 | 2.833E-06 | 3.215E-07 | 2.600E-06 | 3.200E-06 |
| 9151 | 458 | V458L | A | 2.7e-06 | 1.4e-06 | 2.1e-06 | 2.067E-06 | 6.506E-07 | 1.400E-06 | 2.700E-06 |
| 9152 | 458 | V458K | A | 2e-06 | 2.6e-06 | 3e-06 | 2.533E-06 | 5.033E-07 | 2.000E-06 | 3.000E-06 |
| 9153 | 458 | V458M | A | 2.8e-06 | 1.5e-06 | 3.8e-06 | 2.700E-06 | 1.153E-06 | 1.500E-06 | 3.800E-06 |
| 9154 | 458 | V458F | A | 2.7e-06 | 1.2e-06 | 2.1e-06 | 2.000E-06 | 7.550E-07 | 1.200E-06 | 2.700E-06 |
| 9155 | 458 | V458P | A | 3.4e-06 | 2.9e-06 | 2.5e-06 | 2.933E-06 | 4.509E-07 | 2.500E-06 | 3.400E-06 |
| 9156 | 458 | V458S | A | 2.7e-06 | 2.7e-06 | 3.6e-06 | 3.000E-06 | 5.196E-07 | 2.700E-06 | 3.600E-06 |
| 9157 | 458 | V458T | A | 1.7e-06 | 2.1e-06 | 2.2e-06 | 2.000E-06 | 2.646E-07 | 1.700E-06 | 2.200E-06 |
| 9158 | 458 | V458W | A | 2.4e-06 | 1.6e-06 | 2.5e-06 | 2.167E-06 | 4.933E-07 | 1.600E-06 | 2.500E-06 |
| 9159 | 458 | V458Y | A | 2.4e-06 | 2.4e-06 | 1.4e-06 | 2.067E-06 | 5.774E-07 | 1.400E-06 | 2.400E-06 |
| 9160 | 458 | V458V | A | 1.9e-06 | 2.2e-06 | 3e-06 | 2.367E-06 | 5.686E-07 | 1.900E-06 | 3.000E-06 |
| 9161 | 459 | P459A | A | 2.8e-06 | 1.6e-06 | 2.2e-06 | 2.200E-06 | 6.000E-07 | 1.600E-06 | 2.800E-06 |
| 9162 | 459 | P459R | A | 2.4e-06 | 2e-06 | 3.8e-06 | 2.733E-06 | 9.452E-07 | 2.000E-06 | 3.800E-06 |
| 9163 | 459 | P459N | A | 2.7e-06 | 2.2e-06 | 2.1e-06 | 2.333E-06 | 3.215E-07 | 2.100E-06 | 2.700E-06 |
| 9164 | 459 | P459D | A | 2.3e-06 | 1.5e-06 | 2.9e-06 | 2.233E-06 | 7.024E-07 | 1.500E-06 | 2.900E-06 |
| 9165 | 459 | P459C | A | 1.9e-06 | 1.1e-06 | 2.7e-06 | 1.900E-06 | 8.000E-07 | 1.100E-06 | 2.700E-06 |
| 9166 | 459 | P459E | A | 2.6e-06 | 1.9e-06 | 3.6e-06 | 2.700E-06 | 8.544E-07 | 1.900E-06 | 3.600E-06 |
| 9167 | 459 | P459Q | A | 2.5e-06 | 2.1e-06 | 2.6e-06 | 2.400E-06 | 2.646E-07 | 2.100E-06 | 2.600E-06 |
| 9168 | 459 | P459G | A | 2.7e-06 | 2.7e-06 | 1.5e-06 | 2.300E-06 | 6.928E-07 | 1.500E-06 | 2.700E-06 |
| 9169 | 459 | P459H | A | 2.1e-06 | 2e-06 | 2.9e-06 | 2.333E-06 | 4.933E-07 | 2.000E-06 | 2.900E-06 |
| 9170 | 459 | P459I | A | 1.4e-06 | 2e-06 | 1.9e-06 | 1.767E-06 | 3.215E-07 | 1.400E-06 | 2.000E-06 |
| 9171 | 459 | P459L | A | 2.3e-06 | 1.9e-06 | 3.8e-06 | 2.667E-06 | 1.002E-06 | 1.900E-06 | 3.800E-06 |
| 9172 | 459 | P459K | A | 2.6e-06 | 1.8e-06 | 3.1e-06 | 2.500E-06 | 6.557E-07 | 1.800E-06 | 3.100E-06 |
| 9173 | 459 | P459M | A | 4e-06 | 2.2e-06 | 2.4e-06 | 2.867E-06 | 9.866E-07 | 2.200E-06 | 4.000E-06 |
| 9174 | 459 | P459F | A | 3.4e-06 | 2.3e-06 | 1.5e-06 | 2.400E-06 | 9.539E-07 | 1.500E-06 | 3.400E-06 |
| 9175 | 459 | P459P | A | 1.6e-06 | 2.6e-06 | 2e-06 | 2.067E-06 | 5.033E-07 | 1.600E-06 | 2.600E-06 |
| 9176 | 459 | P459S | A | 4.9e-06 | 1.7e-06 | 1.8e-06 | 2.800E-06 | 1.819E-06 | 1.700E-06 | 4.900E-06 |
| 9177 | 459 | P459T | A | 3.7e-06 | 2e-06 | 2.7e-06 | 2.800E-06 | 8.544E-07 | 2.000E-06 | 3.700E-06 |
| 9178 | 459 | P459W | A | 5e-06 | 1.6e-06 | 3.2e-06 | 3.267E-06 | 1.701E-06 | 1.600E-06 | 5.000E-06 |
| 9179 | 459 | P459Y | A | 9.9e-07 | 1.9e-06 | 2.9e-06 | 1.930E-06 | 9.554E-07 | 9.900E-07 | 2.900E-06 |

| | | | | | | | | | | |
|---|---|---|---|---|---|---|---|---|---|---|
| 9180 | 459 | P459V | A | 2.1e-06 | 2.1e-06 | 2.1e-06 | 2.100E-06 | 0.000E+00 | 2.100E-06 | 2.100E-06 |
| 9181 | 460 | W460A | A | 1.8e-06 | 2.2e-06 | 2.7e-06 | 2.233E-06 | 4.509E-07 | 1.800E-06 | 2.700E-06 |
| 9182 | 460 | W460R | A | 3.6e-06 | 2.8e-06 | 1.8e-06 | 2.733E-06 | 9.018E-07 | 1.800E-06 | 3.600E-06 |
| 9183 | 460 | W460N | A | 2.5e-06 | 2.6e-06 | 3.9e-06 | 3.000E-06 | 7.810E-07 | 2.500E-06 | 3.900E-06 |
| 9184 | 460 | W460D | A | 5e-06 | 3e-06 | 9.8e-07 | 2.993E-06 | 2.010E-06 | 9.800E-07 | 5.000E-06 |
| 9185 | 460 | W460C | A | 3.7e-06 | 2.1e-06 | 3.8e-06 | 3.200E-06 | 9.539E-07 | 2.100E-06 | 3.800E-06 |
| 9186 | 460 | W460E | A | 4.6e-06 | 1.6e-06 | 2.4e-06 | 2.867E-06 | 1.553E-06 | 1.600E-06 | 4.600E-06 |
| 9187 | 460 | W460Q | A | 2.6e-06 | 3e-06 | 2.4e-06 | 2.667E-06 | 3.055E-07 | 2.400E-06 | 3.000E-06 |
| 9188 | 460 | W460G | A | 2.3e-06 | 1.4e-06 | 1.8e-06 | 1.833E-06 | 4.509E-07 | 1.400E-06 | 2.300E-06 |
| 9189 | 460 | W460H | A | 1.6e-06 | 1.9e-06 | 2.5e-06 | 2.000E-06 | 4.583E-07 | 1.600E-06 | 2.500E-06 |
| 9190 | 460 | W460I | A | 1.3e-06 | 2.9e-06 | 2.4e-06 | 2.200E-06 | 8.185E-07 | 1.300E-06 | 2.900E-06 |
| 9191 | 460 | W460L | A | 2.1e-06 | 2.8e-06 | 1.4e-06 | 2.100E-06 | 7.000E-07 | 1.400E-06 | 2.800E-06 |
| 9192 | 460 | W460K | A | 3.2e-06 | 3.4e-06 | 2.7e-06 | 3.100E-06 | 3.606E-07 | 2.700E-06 | 3.400E-06 |
| 9193 | 460 | W460M | A | 3.2e-06 | 1.3e-06 | 3.1e-06 | 2.533E-06 | 1.069E-06 | 1.300E-06 | 3.200E-06 |
| 9194 | 460 | W460F | A | 2.7e-06 | 2.2e-06 | 1.9e-06 | 2.267E-06 | 4.041E-07 | 1.900E-06 | 2.700E-06 |
| 9195 | 460 | W460P | A | 2.4e-06 | 4.6e-06 | 3.1e-06 | 3.367E-06 | 1.124E-06 | 2.400E-06 | 4.600E-06 |
| 9196 | 460 | W460S | A | 2.1e-06 | 3e-06 | 4e-06 | 3.033E-06 | 9.504E-07 | 2.100E-06 | 4.000E-06 |
| 9197 | 460 | W460T | A | 1.3e-06 | 3e-06 | 4.5e-06 | 2.933E-06 | 1.601E-06 | 1.300E-06 | 4.500E-06 |
| 9198 | 460 | W460W | A | 2.1e-06 | 1.6e-06 | 1.4e-06 | 1.700E-06 | 3.606E-07 | 1.400E-06 | 2.100E-06 |
| 9199 | 460 | W460Y | A | 3e-06 | 2.5e-06 | 1.8e-06 | 2.433E-06 | 6.028E-07 | 1.800E-06 | 3.000E-06 |
| 9200 | 460 | W460V | A | 2.4e-06 | 2.5e-06 | 2.4e-06 | 2.433E-06 | 5.774E-08 | 2.400E-06 | 2.500E-06 |
| 9201 | 461 | D461A | A | 2.5e-06 | 2.5e-06 | 2.4e-06 | 2.467E-06 | 5.774E-08 | 2.400E-06 | 2.500E-06 |
| 9202 | 461 | D461R | A | 2.6e-06 | 3.8e-06 | 2.9e-06 | 3.100E-06 | 6.245E-07 | 2.600E-06 | 3.800E-06 |
| 9203 | 461 | D461N | A | 2.7e-06 | 2e-06 | 2e-06 | 2.233E-06 | 4.041E-07 | 2.000E-06 | 2.700E-06 |
| 9204 | 461 | D461D | A | 2.7e-06 | 2.1e-06 | 2.8e-06 | 2.533E-06 | 3.786E-07 | 2.100E-06 | 2.800E-06 |
| 9205 | 461 | D461C | A | 1.5e-06 | 3.2e-06 | 2.8e-06 | 2.500E-06 | 8.888E-07 | 1.500E-06 | 3.200E-06 |
| 9206 | 461 | D461E | A | 1.5e-06 | 3.3e-06 | 1.8e-06 | 2.200E-06 | 9.644E-07 | 1.500E-06 | 3.300E-06 |
| 9207 | 461 | D461Q | A | 3e-06 | 2.8e-06 | 2.5e-06 | 2.767E-06 | 2.517E-07 | 2.500E-06 | 3.000E-06 |
| 9208 | 461 | D461G | A | 1.5e-06 | 3.7e-06 | 3.4e-06 | 2.867E-06 | 1.193E-06 | 1.500E-06 | 3.700E-06 |
| 9209 | 461 | D461H | A | 2e-06 | 2.5e-06 | 2.6e-06 | 2.367E-06 | 3.215E-07 | 2.000E-06 | 2.600E-06 |
| 9210 | 461 | D461I | A | 2.3e-06 | 1e-06 | 5.1e-06 | 2.800E-06 | 2.095E-06 | 1.000E-06 | 5.100E-06 |
| 9211 | 461 | D461L | A | 1.3e-06 | 3.7e-06 | 2.6e-06 | 2.533E-06 | 1.201E-06 | 1.300E-06 | 3.700E-06 |
| 9212 | 461 | D461K | A | 3.4e-06 | 2.4e-06 | 2.9e-06 | 2.900E-06 | 5.000E-07 | 2.400E-06 | 3.400E-06 |
| 9213 | 461 | D461M | A | 2.3e-06 | 1.9e-06 | 1.8e-06 | 2.000E-06 | 2.646E-07 | 1.800E-06 | 2.300E-06 |
| 9214 | 461 | D461F | A | 1.7e-06 | 2.6e-06 | 2e-06 | 2.100E-06 | 4.583E-07 | 1.700E-06 | 2.600E-06 |
| 9215 | 461 | D461P | A | 2.8e-06 | 4.4e-06 | 3.9e-06 | 3.700E-06 | 8.185E-07 | 2.800E-06 | 4.400E-06 |
| 9216 | 461 | D461S | A | 2.3e-06 | 3.2e-06 | 3.3e-06 | 2.933E-06 | 5.508E-07 | 2.300E-06 | 3.300E-06 |
| 9217 | 461 | D461T | A | 2.5e-06 | 1.5e-06 | 2.3e-06 | 2.100E-06 | 5.292E-07 | 1.500E-06 | 2.500E-06 |
| 9218 | 461 | D461W | A | 3.1e-06 | 1.8e-06 | 3.3e-06 | 2.733E-06 | 8.145E-07 | 1.800E-06 | 3.300E-06 |
| 9219 | 461 | D461Y | A | 1.9e-06 | 2.5e-06 | 2.5e-06 | 2.300E-06 | 3.464E-07 | 1.900E-06 | 2.500E-06 |
| 9220 | 461 | D461V | A | 2.3e-06 | 2.4e-06 | 2.2e-06 | 2.300E-06 | 1.000E-07 | 2.200E-06 | 2.400E-06 |
| 9221 | 462 | Q462A | A | 2.4e-06 | 2.2e-06 | 2.2e-06 | 2.267E-06 | 1.155E-07 | 2.200E-06 | 2.400E-06 |
| 9222 | 462 | Q462R | A | 1.8e-06 | 2.1e-06 | 3.5e-06 | 2.467E-06 | 9.074E-07 | 1.800E-06 | 3.500E-06 |
| 9223 | 462 | Q462N | A | 1.9e-06 | 1.8e-06 | 2.5e-06 | 2.067E-06 | 3.786E-07 | 1.800E-06 | 2.500E-06 |
| 9224 | 462 | Q462D | A | 2.5e-06 | 2e-06 | 2.2e-06 | 2.233E-06 | 2.517E-07 | 2.000E-06 | 2.500E-06 |
| 9225 | 462 | Q462C | A | 2.5e-06 | 2.7e-06 | 3.7e-06 | 2.967E-06 | 6.429E-07 | 2.500E-06 | 3.700E-06 |
| 9226 | 462 | Q462E | A | 2.8e-06 | 4.7e-06 | 3.3e-06 | 3.600E-06 | 9.849E-07 | 2.800E-06 | 4.700E-06 |
| 9227 | 462 | Q462Q | A | 2.5e-06 | 2.7e-06 | 2.7e-06 | 2.633E-06 | 1.155E-07 | 2.500E-06 | 2.700E-06 |
| 9228 | 462 | Q462G | A | 3e-06 | 1.6e-06 | 2.8e-06 | 2.467E-06 | 7.572E-07 | 1.600E-06 | 3.000E-06 |
| 9229 | 462 | Q462H | A | 2.2e-06 | 1.2e-06 | 1.9e-06 | 1.767E-06 | 5.132E-07 | 1.200E-06 | 2.200E-06 |
| 9230 | 462 | Q462I | A | 1.9e-06 | 1.7e-06 | 4.2e-06 | 2.600E-06 | 1.389E-06 | 1.700E-06 | 4.200E-06 |
| 9231 | 462 | Q462L | A | 2e-06 | 2.7e-06 | 2.6e-06 | 2.433E-06 | 3.786E-07 | 2.000E-06 | 2.700E-06 |
| 9232 | 462 | Q462K | A | 2.9e-06 | 1.9e-06 | 3.2e-06 | 2.667E-06 | 6.807E-07 | 1.900E-06 | 3.200E-06 |
| 9233 | 462 | Q462M | A | 1.3e-06 | 2e-06 | 3.4e-06 | 2.233E-06 | 1.069E-06 | 1.300E-06 | 3.400E-06 |
| 9234 | 462 | Q462F | A | 1.9e-06 | 2.1e-06 | 3.4e-06 | 2.467E-06 | 8.145E-07 | 1.900E-06 | 3.400E-06 |
| 9235 | 462 | Q462P | A | 2.6e-06 | 4.8e-06 | 2.1e-06 | 3.167E-06 | 1.436E-06 | 2.100E-06 | 4.800E-06 |
| 9236 | 462 | Q462S | A | 1.5e-06 | 3.4e-06 | 2.7e-06 | 2.533E-06 | 9.609E-07 | 1.500E-06 | 3.400E-06 |
| 9237 | 462 | Q462T | A | 2.7e-06 | 2.7e-06 | 1.1e-06 | 2.167E-06 | 9.238E-07 | 1.100E-06 | 2.700E-06 |
| 9238 | 462 | Q462W | A | 1.9e-06 | 1.3e-06 | 2.8e-06 | 2.000E-06 | 7.550E-07 | 1.300E-06 | 2.800E-06 |
| 9239 | 462 | Q462Y | A | 1.3e-06 | 4.6e-06 | 2.5e-06 | 2.800E-06 | 1.670E-06 | 1.300E-06 | 4.600E-06 |
| 9240 | 462 | Q462V | A | 3e-06 | 1.3e-06 | 2.1e-06 | 2.133E-06 | 8.505E-07 | 1.300E-06 | 3.000E-06 |
| 9241 | 463 | L463A | A | 2e-06 | 1.7e-06 | 2.6e-06 | 2.100E-06 | 4.583E-07 | 1.700E-06 | 2.600E-06 |
| 9242 | 463 | L463R | A | 2.6e-06 | 2.4e-06 | 2e-06 | 2.333E-06 | 3.055E-07 | 2.000E-06 | 2.600E-06 |
| 9243 | 463 | L463N | A | 2.6e-06 | 1.8e-06 | 2.1e-06 | 2.167E-06 | 4.041E-07 | 1.800E-06 | 2.600E-06 |
| 9244 | 463 | L463D | A | 3.2e-06 | 1.4e-06 | 2.3e-06 | 2.300E-06 | 9.000E-07 | 1.400E-06 | 3.200E-06 |
| 9245 | 463 | L463C | A | 2.5e-06 | 2.1e-06 | 1.9e-06 | 2.167E-06 | 3.055E-07 | 1.900E-06 | 2.500E-06 |
| 9246 | 463 | L463E | A | 3e-06 | 9.5e-07 | 3.1e-06 | 2.350E-06 | 1.213E-06 | 9.500E-07 | 3.100E-06 |
| 9247 | 463 | L463Q | A | 2.7e-06 | 2e-06 | 1.9e-06 | 2.200E-06 | 4.359E-07 | 1.900E-06 | 2.700E-06 |
| 9248 | 463 | L463G | A | 3.6e-06 | 2e-06 | 3.2e-06 | 2.933E-06 | 8.327E-07 | 2.000E-06 | 3.600E-06 |
| 9249 | 463 | L463H | A | 5.3e-06 | 2.4e-06 | 1.4e-06 | 3.033E-06 | 2.026E-06 | 1.400E-06 | 5.300E-06 |
| 9250 | 463 | L463I | A | 2.2e-06 | 2.8e-06 | 4e-06 | 3.000E-06 | 9.165E-07 | 2.200E-06 | 4.000E-06 |
| 9251 | 463 | L463L | A | 3.1e-06 | 4.4e-06 | 1.8e-06 | 3.100E-06 | 1.300E-06 | 1.800E-06 | 4.400E-06 |
| 9252 | 463 | L463K | A | 2.6e-06 | 2.1e-06 | 1.7e-06 | 2.133E-06 | 4.509E-07 | 1.700E-06 | 2.600E-06 |
| 9253 | 463 | L463M | A | 2.6e-06 | 2.3e-06 | 1.9e-06 | 2.267E-06 | 3.512E-07 | 1.900E-06 | 2.600E-06 |
| 9254 | 463 | L463F | A | 4.1e-06 | 2.5e-06 | 2.8e-06 | 3.133E-06 | 8.505E-07 | 2.500E-06 | 4.100E-06 |
| 9255 | 463 | L463P | A | 2.1e-06 | 3.2e-06 | 5e-06 | 3.433E-06 | 1.464E-06 | 2.100E-06 | 5.000E-06 |
| 9256 | 463 | L463S | A | 2.9e-06 | 2.3e-06 | 2e-06 | 2.400E-06 | 4.583E-07 | 2.000E-06 | 2.900E-06 |
| 9257 | 463 | L463T | A | 1.7e-06 | 1.9e-06 | 2.8e-06 | 2.133E-06 | 5.859E-07 | 1.700E-06 | 2.800E-06 |
| 9258 | 463 | L463W | A | 2.4e-06 | 2e-06 | 2.1e-06 | 2.167E-06 | 2.082E-07 | 2.000E-06 | 2.400E-06 |
| 9259 | 463 | L463Y | A | 2.4e-06 | 2.8e-06 | 1.9e-06 | 2.367E-06 | 4.509E-07 | 1.900E-06 | 2.800E-06 |
| 9260 | 463 | L463V | A | 3.4e-06 | 1.9e-06 | 3.7e-06 | 3.000E-06 | 9.644E-07 | 1.900E-06 | 3.700E-06 |
| 9261 | 464 | F464A | A | 2.8e-06 | 2.8e-06 | 3.3e-06 | 2.967E-06 | 2.887E-07 | 2.800E-06 | 3.300E-06 |
| 9262 | 464 | F464R | A | 2e-06 | 2.3e-06 | 1.9e-06 | 2.067E-06 | 2.082E-07 | 1.900E-06 | 2.300E-06 |
| 9263 | 464 | F464N | A | 6.1e-06 | 3.2e-06 | 2.7e-06 | 4.000E-06 | 1.836E-06 | 2.700E-06 | 6.100E-06 |
| 9264 | 464 | F464D | A | 2.6e-06 | 2.3e-06 | 2.2e-06 | 2.367E-06 | 2.082E-07 | 2.200E-06 | 2.600E-06 |

| | | | | | | | | | | |
|---|---|---|---|---|---|---|---|---|---|---|
| 9265 | 464 | F464C | A | 3.8e-06 | 2.5e-06 | 3.3e-06 | 3.200E-06 | 6.557E-07 | 2.500E-06 | 3.800E-06 |
| 9266 | 464 | F464E | A | 1.3e-06 | 2.7e-06 | 2.4e-06 | 2.133E-06 | 7.371E-07 | 1.300E-06 | 2.700E-06 |
| 9267 | 464 | F464Q | A | 2.1e-06 | 2.1e-06 | 2.8e-06 | 2.333E-06 | 4.041E-07 | 2.100E-06 | 2.800E-06 |
| 9268 | 464 | F464G | A | 1.7e-06 | 2.5e-06 | 2e-06 | 2.067E-06 | 4.041E-07 | 1.700E-06 | 2.500E-06 |
| 9269 | 464 | F464H | A | 4.5e-06 | 1.7e-06 | 2e-06 | 2.733E-06 | 1.537E-06 | 1.700E-06 | 4.500E-06 |
| 9270 | 464 | F464I | A | 1.7e-06 | 2.3e-06 | 2.3e-06 | 2.100E-06 | 3.464E-07 | 1.700E-06 | 2.300E-06 |
| 9271 | 464 | F464L | A | 1.6e-06 | 2.6e-06 | 2.3e-06 | 2.167E-06 | 5.132E-07 | 1.600E-06 | 2.600E-06 |
| 9272 | 464 | F464K | A | 2.2e-06 | 1.4e-06 | 2e-06 | 1.867E-06 | 4.163E-07 | 1.400E-06 | 2.200E-06 |
| 9273 | 464 | F464M | A | 3.4e-06 | 2.4e-06 | 2.2e-06 | 2.667E-06 | 6.429E-07 | 2.200E-06 | 3.400E-06 |
| 9274 | 464 | F464F | A | 2.3e-06 | 2.9e-06 | 2.2e-06 | 2.467E-06 | 3.786E-07 | 2.200E-06 | 2.900E-06 |
| 9275 | 464 | F464P | A | 3e-06 | 1.8e-06 | 2e-06 | 2.267E-06 | 6.429E-07 | 1.800E-06 | 3.000E-06 |
| 9276 | 464 | F464S | A | 2.6e-06 | 1.6e-06 | 3.3e-06 | 2.500E-06 | 8.544E-07 | 1.600E-06 | 3.300E-06 |
| 9277 | 464 | F464T | A | 3.9e-06 | 2e-06 | 4.3e-06 | 3.400E-06 | 1.229E-06 | 2.000E-06 | 4.300E-06 |
| 9278 | 464 | F464W | A | 2.3e-06 | 1.1e-06 | 2e-06 | 1.800E-06 | 6.245E-07 | 1.100E-06 | 2.300E-06 |
| 9279 | 464 | F464Y | A | 1.6e-06 | 2.5e-06 | 2.2e-06 | 2.100E-06 | 4.583E-07 | 1.600E-06 | 2.500E-06 |
| 9280 | 464 | F464V | A | 3.2e-06 | 3.1e-06 | 3e-06 | 3.100E-06 | 1.000E-07 | 3.000E-06 | 3.200E-06 |
| 9281 | 465 | R465A | A | 4.1e-06 | 2.3e-06 | 2.9e-06 | 3.100E-06 | 9.165E-07 | 2.300E-06 | 4.100E-06 |
| 9282 | 465 | R465R | A | 1.3e-06 | 1.2e-06 | 3.2e-06 | 1.900E-06 | 1.127E-06 | 1.200E-06 | 3.200E-06 |
| 9283 | 465 | R465N | A | 3.6e-06 | 2.2e-06 | 2e-06 | 2.600E-06 | 8.718E-07 | 2.000E-06 | 3.600E-06 |
| 9284 | 465 | R465D | A | 2.5e-06 | 3.6e-06 | 2.4e-06 | 2.833E-06 | 6.658E-07 | 2.400E-06 | 3.600E-06 |
| 9285 | 465 | R465C | A | 3e-06 | 2e-06 | 2.6e-06 | 2.533E-06 | 5.033E-07 | 2.000E-06 | 3.000E-06 |
| 9286 | 465 | R465E | A | 2.6e-06 | 2.4e-06 | 2.5e-06 | 2.500E-06 | 1.000E-07 | 2.400E-06 | 2.600E-06 |
| 9287 | 465 | R465Q | A | 2.8e-06 | 1.7e-06 | 1.5e-06 | 2.000E-06 | 7.000E-07 | 1.500E-06 | 2.800E-06 |
| 9288 | 465 | R465G | A | 3.3e-06 | 2.6e-06 | 2.1e-06 | 2.667E-06 | 6.028E-07 | 2.100E-06 | 3.300E-06 |
| 9289 | 465 | R465H | A | 2.3e-06 | 1e-06 | 2.4e-06 | 1.900E-06 | 7.810E-07 | 1.000E-06 | 2.400E-06 |
| 9290 | 465 | R465I | A | 3e-06 | 1.6e-06 | 2.7e-06 | 2.433E-06 | 7.371E-07 | 1.600E-06 | 3.000E-06 |
| 9291 | 465 | R465L | A | 2.2e-06 | 2.6e-06 | 2.9e-06 | 2.567E-06 | 3.512E-07 | 2.200E-06 | 2.900E-06 |
| 9292 | 465 | R465K | A | 4.7e-06 | 1.9e-06 | 1.2e-06 | 2.600E-06 | 1.852E-06 | 1.200E-06 | 4.700E-06 |
| 9293 | 465 | R465M | A | 2.8e-06 | 3.2e-06 | 3.2e-06 | 3.067E-06 | 2.309E-07 | 2.800E-06 | 3.200E-06 |
| 9294 | 465 | R465F | A | 1.9e-06 | 2.4e-06 | 1.8e-06 | 2.033E-06 | 3.215E-07 | 1.800E-06 | 2.400E-06 |
| 9295 | 465 | R465P | A | 5.1e-06 | 2.4e-06 | 2.2e-06 | 3.233E-06 | 1.620E-06 | 2.200E-06 | 5.100E-06 |
| 9296 | 465 | R465S | A | 2.5e-06 | 2.6e-06 | 3.5e-06 | 2.867E-06 | 5.508E-07 | 2.500E-06 | 3.500E-06 |
| 9297 | 465 | R465T | A | 2.1e-06 | 2e-06 | 2.6e-06 | 2.233E-06 | 3.215E-07 | 2.000E-06 | 2.600E-06 |
| 9298 | 465 | R465W | A | 2e-06 | 2.1e-06 | 2.6e-06 | 2.233E-06 | 3.215E-07 | 2.000E-06 | 2.600E-06 |
| 9299 | 465 | R465Y | A | 2e-06 | 4.4e-06 | 2.3e-06 | 2.900E-06 | 1.308E-06 | 2.000E-06 | 4.400E-06 |
| 9300 | 465 | R465V | A | 3.1e-06 | 2.7e-06 | 2e-06 | 2.600E-06 | 5.568E-07 | 2.000E-06 | 3.100E-06 |
| 9301 | 466 | N466A | A | 2.2e-06 | 1.5e-06 | 3.3e-06 | 2.333E-06 | 9.074E-07 | 1.500E-06 | 3.300E-06 |
| 9302 | 466 | N466R | A | 2e-06 | 3.1e-06 | 2.9e-06 | 2.667E-06 | 5.859E-07 | 2.000E-06 | 3.100E-06 |
| 9303 | 466 | N466N | A | 2e-06 | 2.8e-06 | 1.8e-06 | 2.200E-06 | 5.292E-07 | 1.800E-06 | 2.800E-06 |
| 9304 | 466 | N466D | A | 1.6e-06 | 1.7e-06 | 1.7e-06 | 1.667E-06 | 5.774E-08 | 1.600E-06 | 1.700E-06 |
| 9305 | 466 | N466C | A | 2.1e-06 | 3.1e-06 | 1.4e-06 | 2.200E-06 | 8.544E-07 | 1.400E-06 | 3.100E-06 |
| 9306 | 466 | N466E | A | 2.5e-06 | 2.8e-06 | 5e-06 | 3.433E-06 | 1.365E-06 | 2.500E-06 | 5.000E-06 |
| 9307 | 466 | N466Q | A | 4.8e-06 | 2.5e-06 | 3e-06 | 3.433E-06 | 1.210E-06 | 2.500E-06 | 4.800E-06 |
| 9308 | 466 | N466G | A | 3e-06 | 3.4e-06 | 1.6e-06 | 2.667E-06 | 9.452E-07 | 1.600E-06 | 3.400E-06 |
| 9309 | 466 | N466H | A | 2.2e-06 | 1.9e-06 | 2.1e-06 | 2.067E-06 | 1.528E-07 | 1.900E-06 | 2.200E-06 |
| 9310 | 466 | N466I | A | 2e-06 | 2.1e-06 | 4.8e-06 | 2.967E-06 | 1.589E-06 | 2.000E-06 | 4.800E-06 |
| 9311 | 466 | N466L | A | 2.2e-06 | 4.3e-06 | 2.3e-06 | 2.933E-06 | 1.185E-06 | 2.200E-06 | 4.300E-06 |
| 9312 | 466 | N466K | A | 3.3e-06 | 2.2e-06 | 2.7e-06 | 2.733E-06 | 5.508E-07 | 2.200E-06 | 3.300E-06 |
| 9313 | 466 | N466M | A | 1.2e-06 | 2.2e-06 | 3.1e-06 | 2.167E-06 | 9.504E-07 | 1.200E-06 | 3.100E-06 |
| 9314 | 466 | N466F | A | 2.5e-06 | 5.1e-06 | 2.7e-06 | 3.433E-06 | 1.447E-06 | 2.500E-06 | 5.100E-06 |
| 9315 | 466 | N466P | A | 1.6e-06 | 2.8e-06 | 2e-06 | 2.133E-06 | 6.110E-07 | 1.600E-06 | 2.800E-06 |
| 9316 | 466 | N466S | A | 2.9e-06 | 2.2e-06 | 2.4e-06 | 2.500E-06 | 3.606E-07 | 2.200E-06 | 2.900E-06 |
| 9317 | 466 | N466T | A | 1.6e-06 | 3.4e-06 | 3.1e-06 | 2.700E-06 | 9.644E-07 | 1.600E-06 | 3.400E-06 |
| 9318 | 466 | N466W | A | 2.2e-06 | 2.7e-06 | 2.5e-06 | 2.467E-06 | 2.517E-07 | 2.200E-06 | 2.700E-06 |
| 9319 | 466 | N466Y | A | 3.2e-06 | 1.1e-06 | 2.8e-06 | 2.367E-06 | 1.115E-06 | 1.100E-06 | 3.200E-06 |
| 9320 | 466 | N466V | A | 2.4e-06 | 2.6e-06 | 2.5e-06 | 2.500E-06 | 1.000E-07 | 2.400E-06 | 2.600E-06 |
| 9321 | 467 | P467A | A | 1.2e-06 | 1.5e-06 | 2.4e-06 | 1.700E-06 | 6.245E-07 | 1.200E-06 | 2.400E-06 |
| 9322 | 467 | P467R | A | 2.4e-06 | 3e-06 | 1.6e-06 | 2.333E-06 | 7.024E-07 | 1.600E-06 | 3.000E-06 |
| 9323 | 467 | P467N | A | 1.9e-06 | 2.5e-06 | 3.6e-06 | 2.667E-06 | 8.622E-07 | 1.900E-06 | 3.600E-06 |
| 9324 | 467 | P467D | A | 2e-06 | 2e-06 | 4.5e-06 | 2.833E-06 | 1.443E-06 | 2.000E-06 | 4.500E-06 |
| 9325 | 467 | P467C | A | 1.6e-06 | 2.3e-06 | 3.4e-06 | 2.433E-06 | 9.074E-07 | 1.600E-06 | 3.400E-06 |
| 9326 | 467 | P467E | A | 2.5e-06 | 1.8e-06 | 4.8e-06 | 3.033E-06 | 1.570E-06 | 1.800E-06 | 4.800E-06 |
| 9327 | 467 | P467Q | A | 1.4e-06 | 1.2e-06 | 2e-06 | 1.533E-06 | 4.163E-07 | 1.200E-06 | 2.000E-06 |
| 9328 | 467 | P467G | A | 3e-06 | 2.7e-06 | 2.3e-06 | 2.667E-06 | 3.512E-07 | 2.300E-06 | 3.000E-06 |
| 9329 | 467 | P467H | A | 2.6e-06 | 1.8e-06 | 2.1e-06 | 2.167E-06 | 4.041E-07 | 1.800E-06 | 2.600E-06 |
| 9330 | 467 | P467I | A | 2.9e-06 | 2.9e-06 | 2.2e-06 | 2.667E-06 | 4.041E-07 | 2.200E-06 | 2.900E-06 |
| 9331 | 467 | P467L | A | 2.6e-06 | 3.1e-06 | 9.5e-07 | 2.217E-06 | 1.125E-06 | 9.500E-07 | 3.100E-06 |
| 9332 | 467 | P467K | A | 2.3e-06 | 1.8e-06 | 4e-06 | 2.700E-06 | 1.153E-06 | 1.800E-06 | 4.000E-06 |
| 9333 | 467 | P467M | A | 2.1e-06 | 1.5e-06 | 2.2e-06 | 1.933E-06 | 3.786E-07 | 1.500E-06 | 2.200E-06 |
| 9334 | 467 | P467F | A | 1.8e-06 | 3.8e-06 | 2.5e-06 | 2.700E-06 | 1.015E-06 | 1.800E-06 | 3.800E-06 |
| 9335 | 467 | P467P | A | 2.7e-06 | 2.3e-06 | 9.1e-07 | 1.970E-06 | 9.395E-07 | 9.100E-07 | 2.700E-06 |
| 9336 | 467 | P467S | A | 2.2e-06 | 1.4e-06 | 1.8e-06 | 1.800E-06 | 4.000E-07 | 1.400E-06 | 2.200E-06 |
| 9337 | 467 | P467T | A | 3.9e-06 | 2e-06 | 2e-06 | 2.633E-06 | 1.097E-06 | 2.000E-06 | 3.900E-06 |
| 9338 | 467 | P467W | A | 2e-06 | 1.7e-06 | 2.3e-06 | 2.000E-06 | 3.000E-07 | 1.700E-06 | 2.300E-06 |
| 9339 | 467 | P467Y | A | 2.7e-06 | 3.6e-06 | 1.9e-06 | 2.733E-06 | 8.505E-07 | 1.900E-06 | 3.600E-06 |
| 9340 | 467 | P467V | A | 2.8e-06 | 2.6e-06 | 2.2e-06 | 2.533E-06 | 3.055E-07 | 2.200E-06 | 2.800E-06 |
| 9341 | 468 | H468A | A | 3.2e-06 | 1.6e-06 | 2.2e-06 | 2.333E-06 | 8.083E-07 | 1.600E-06 | 3.200E-06 |
| 9342 | 468 | H468R | A | 2.6e-06 | 1.1e-06 | 1.6e-06 | 1.767E-06 | 7.638E-07 | 1.100E-06 | 2.600E-06 |
| 9343 | 468 | H468N | A | 1.8e-06 | 2.2e-06 | 2.9e-06 | 2.300E-06 | 5.568E-07 | 1.800E-06 | 2.900E-06 |
| 9344 | 468 | H468D | A | 3.4e-06 | 2e-06 | 2.7e-06 | 2.700E-06 | 7.000E-07 | 2.000E-06 | 3.400E-06 |
| 9345 | 468 | H468C | A | 2.7e-06 | 5.5e-06 | 3.4e-06 | 3.867E-06 | 1.457E-06 | 2.700E-06 | 5.500E-06 |
| 9346 | 468 | H468E | A | 2.2e-06 | 4e-06 | 2.2e-06 | 2.800E-06 | 1.039E-06 | 2.200E-06 | 4.000E-06 |
| 9347 | 468 | H468Q | A | 1.7e-06 | 2.4e-06 | 2.2e-06 | 2.100E-06 | 3.606E-07 | 1.700E-06 | 2.400E-06 |
| 9348 | 468 | H468G | A | 2e-06 | 2.6e-06 | 3.1e-06 | 2.567E-06 | 5.508E-07 | 2.000E-06 | 3.100E-06 |
| 9349 | 468 | H468H | A | 2.4e-06 | 2.8e-06 | 2.7e-06 | 2.633E-06 | 2.082E-07 | 2.400E-06 | 2.800E-06 |

| | | | | | | | | | | |
|---|---|---|---|---|---|---|---|---|---|---|
| 9350 | 468 | H468I | A | 2e-06 | 3.4e-06 | 3e-06 | 2.800E-06 | 7.211E-07 | 2.000E-06 | 3.400E-06 |
| 9351 | 468 | H468L | A | 2.2e-06 | 2.5e-06 | 2.4e-06 | 2.367E-06 | 1.528E-07 | 2.200E-06 | 2.500E-06 |
| 9352 | 468 | H468K | A | 3e-06 | 3.2e-06 | 1.7e-06 | 2.633E-06 | 8.145E-07 | 1.700E-06 | 3.200E-06 |
| 9353 | 468 | H468M | A | 5.3e-06 | 1.3e-06 | 1.9e-06 | 2.833E-06 | 2.157E-06 | 1.300E-06 | 5.300E-06 |
| 9354 | 468 | H468F | A | 1.8e-06 | 2.4e-06 | 3.3e-06 | 2.500E-06 | 7.550E-07 | 1.800E-06 | 3.300E-06 |
| 9355 | 468 | H468P | A | 1.9e-06 | 1.9e-06 | 1.9e-06 | 1.900E-06 | 2.593E-22 | 1.900E-06 | 1.900E-06 |
| 9356 | 468 | H468S | A | 2.8e-06 | 2.6e-06 | 1.4e-06 | 2.267E-06 | 7.572E-07 | 1.400E-06 | 2.800E-06 |
| 9357 | 468 | H468T | A | 3.1e-06 | 4.2e-06 | 2.5e-06 | 3.267E-06 | 8.622E-07 | 2.500E-06 | 4.200E-06 |
| 9358 | 468 | H468W | A | 1.7e-06 | 2.5e-06 | 2.1e-06 | 2.100E-06 | 4.000E-07 | 1.700E-06 | 2.500E-06 |
| 9359 | 468 | H468Y | A | 2e-06 | 2e-06 | 3.1e-06 | 2.367E-06 | 6.351E-07 | 2.000E-06 | 3.100E-06 |
| 9360 | 468 | H468V | A | 3.7e-06 | 2.4e-06 | 1.4e-06 | 2.500E-06 | 1.153E-06 | 1.400E-06 | 3.700E-06 |
| 9361 | 469 | Q469A | A | 2.5e-06 | 1.8e-06 | 3e-06 | 2.433E-06 | 6.028E-07 | 1.800E-06 | 3.000E-06 |
| 9362 | 469 | Q469R | A | 3.2e-06 | 3.5e-06 | 1.5e-06 | 2.733E-06 | 1.079E-06 | 1.500E-06 | 3.500E-06 |
| 9363 | 469 | Q469N | A | 4.6e-06 | 2.2e-06 | 2e-06 | 2.933E-06 | 1.447E-06 | 2.000E-06 | 4.600E-06 |
| 9364 | 469 | Q469D | A | 3.4e-06 | 3.6e-06 | 2.8e-06 | 3.267E-06 | 4.163E-07 | 2.800E-06 | 3.600E-06 |
| 9365 | 469 | Q469C | A | 1.4e-06 | 1.1e-06 | 2.8e-06 | 1.767E-06 | 9.074E-07 | 1.100E-06 | 2.800E-06 |
| 9366 | 469 | Q469E | A | 2.1e-06 | 1.8e-06 | 3e-06 | 2.300E-06 | 6.245E-07 | 1.800E-06 | 3.000E-06 |
| 9367 | 469 | Q469Q | A | 2.5e-06 | 1.7e-06 | 2.1e-06 | 2.100E-06 | 4.000E-07 | 1.700E-06 | 2.500E-06 |
| 9368 | 469 | Q469G | A | 1.4e-06 | 2.2e-06 | 2.3e-06 | 1.967E-06 | 4.933E-07 | 1.400E-06 | 2.300E-06 |
| 9369 | 469 | Q469H | A | 2.1e-06 | 3.7e-06 | 2.5e-06 | 2.767E-06 | 8.327E-07 | 2.100E-06 | 3.700E-06 |
| 9370 | 469 | Q469I | A | 2.2e-06 | 2.5e-06 | 2.8e-06 | 2.500E-06 | 3.000E-07 | 2.200E-06 | 2.800E-06 |
| 9371 | 469 | Q469L | A | 1.5e-06 | 3.7e-06 | 2.6e-06 | 2.600E-06 | 1.100E-06 | 1.500E-06 | 3.700E-06 |
| 9372 | 469 | Q469K | A | 2.8e-06 | 2.8e-06 | 2.1e-06 | 2.567E-06 | 4.041E-07 | 2.100E-06 | 2.800E-06 |
| 9373 | 469 | Q469M | A | 2.5e-06 | 3.9e-06 | 3e-06 | 3.133E-06 | 7.095E-07 | 2.500E-06 | 3.900E-06 |
| 9374 | 469 | Q469F | A | 2.2e-06 | 2.9e-06 | 1.1e-06 | 2.067E-06 | 9.074E-07 | 1.100E-06 | 2.900E-06 |
| 9375 | 469 | Q469P | A | 2.8e-06 | 3.4e-06 | 1.3e-06 | 2.500E-06 | 1.082E-06 | 1.300E-06 | 3.400E-06 |
| 9376 | 469 | Q469S | A | 3.1e-06 | 1.1e-06 | 2.7e-06 | 2.300E-06 | 1.058E-06 | 1.100E-06 | 3.100E-06 |
| 9377 | 469 | Q469T | A | 2.4e-06 | 2.2e-06 | 2.9e-06 | 2.500E-06 | 3.606E-07 | 2.200E-06 | 2.900E-06 |
| 9378 | 469 | Q469W | A | 3.4e-06 | 1.7e-06 | 4.9e-06 | 3.333E-06 | 1.601E-06 | 1.700E-06 | 4.900E-06 |
| 9379 | 469 | Q469Y | A | 1.8e-06 | 2.5e-06 | 2.2e-06 | 2.167E-06 | 3.512E-07 | 1.800E-06 | 2.500E-06 |
| 9380 | 469 | Q469V | A | 3.7e-06 | 3e-06 | 2.9e-06 | 3.200E-06 | 4.359E-07 | 2.900E-06 | 3.700E-06 |
| 9381 | 470 | A470A | A | 2.8e-06 | 1.9e-06 | 1.9e-06 | 2.200E-06 | 5.196E-07 | 1.900E-06 | 2.800E-06 |
| 9382 | 470 | A470R | A | 3.1e-06 | 2.3e-06 | 3.2e-06 | 2.867E-06 | 4.933E-07 | 2.300E-06 | 3.200E-06 |
| 9383 | 470 | A470N | A | 1.7e-06 | 1.6e-06 | 2.4e-06 | 1.900E-06 | 4.359E-07 | 1.600E-06 | 2.400E-06 |
| 9384 | 470 | A470D | A | 2.5e-06 | 2.7e-06 | 3.2e-06 | 2.800E-06 | 3.606E-07 | 2.500E-06 | 3.200E-06 |
| 9385 | 470 | A470C | A | 1.5e-06 | 4.1e-06 | 2.2e-06 | 2.600E-06 | 1.345E-06 | 1.500E-06 | 4.100E-06 |
| 9386 | 470 | A470E | A | 2.7e-06 | 2e-06 | 3.4e-06 | 2.700E-06 | 7.000E-07 | 2.000E-06 | 3.400E-06 |
| 9387 | 470 | A470Q | A | 1.8e-06 | 2.7e-06 | 1.4e-06 | 1.967E-06 | 6.658E-07 | 1.400E-06 | 2.700E-06 |
| 9388 | 470 | A470G | A | 2.1e-06 | 3e-06 | 3.4e-06 | 2.833E-06 | 6.658E-07 | 2.100E-06 | 3.400E-06 |
| 9389 | 470 | A470H | A | 1.4e-06 | 1.9e-06 | 2.5e-06 | 1.933E-06 | 5.508E-07 | 1.400E-06 | 2.500E-06 |
| 9390 | 470 | A470I | A | 4.2e-06 | 1.9e-06 | 3.1e-06 | 3.067E-06 | 1.150E-06 | 1.900E-06 | 4.200E-06 |
| 9391 | 470 | A470L | A | 1.2e-06 | 2.2e-06 | 2.4e-06 | 1.933E-06 | 6.429E-07 | 1.200E-06 | 2.400E-06 |
| 9392 | 470 | A470K | A | 1.7e-06 | 1.2e-06 | 1.9e-06 | 1.600E-06 | 3.606E-07 | 1.200E-06 | 1.900E-06 |
| 9393 | 470 | A470M | A | 1.9e-06 | 2.8e-06 | 2.4e-06 | 2.367E-06 | 4.509E-07 | 1.900E-06 | 2.800E-06 |
| 9394 | 470 | A470F | A | 2.6e-06 | 2.6e-06 | 1.5e-06 | 2.233E-06 | 6.351E-07 | 1.500E-06 | 2.600E-06 |
| 9395 | 470 | A470P | A | 5.3e-06 | 2.1e-06 | 2.4e-06 | 3.267E-06 | 1.767E-06 | 2.100E-06 | 5.300E-06 |
| 9396 | 470 | A470S | A | 1.4e-06 | 1.6e-06 | 2.9e-06 | 1.967E-06 | 8.145E-07 | 1.400E-06 | 2.900E-06 |
| 9397 | 470 | A470T | A | 2.6e-06 | 1.8e-06 | 1.4e-06 | 1.933E-06 | 6.110E-07 | 1.400E-06 | 2.600E-06 |
| 9398 | 470 | A470W | A | 2.8e-06 | 1.9e-06 | 2.1e-06 | 2.267E-06 | 4.726E-07 | 1.900E-06 | 2.800E-06 |
| 9399 | 470 | A470Y | A | 2e-06 | 1.7e-06 | 2e-06 | 1.900E-06 | 1.732E-07 | 1.700E-06 | 2.000E-06 |
| 9400 | 470 | A470V | A | 4.2e-06 | 3.4e-06 | 1.3e-06 | 2.967E-06 | 1.498E-06 | 1.300E-06 | 4.200E-06 |
| 9401 | 471 | L471A | A | 2.1e-06 | 2.2e-06 | 1.9e-06 | 2.067E-06 | 1.528E-07 | 1.900E-06 | 2.200E-06 |
| 9402 | 471 | L471R | A | 2.1e-06 | 2e-06 | 2.7e-06 | 2.267E-06 | 3.786E-07 | 2.000E-06 | 2.700E-06 |
| 9403 | 471 | L471N | A | 2.1e-06 | 2.6e-06 | 3.4e-06 | 2.700E-06 | 6.557E-07 | 2.100E-06 | 3.400E-06 |
| 9404 | 471 | L471D | A | 2.6e-06 | 2.5e-06 | 4.1e-06 | 3.067E-06 | 8.963E-07 | 2.500E-06 | 4.100E-06 |
| 9405 | 471 | L471C | A | 2.7e-06 | 1.9e-06 | 2.6e-06 | 2.400E-06 | 4.359E-07 | 1.900E-06 | 2.700E-06 |
| 9406 | 471 | L471E | A | 1.7e-06 | 2.6e-06 | 3.1e-06 | 2.467E-06 | 7.095E-07 | 1.700E-06 | 3.100E-06 |
| 9407 | 471 | L471Q | A | 2.1e-06 | 2.4e-06 | 1.6e-06 | 2.033E-06 | 4.041E-07 | 1.600E-06 | 2.400E-06 |
| 9408 | 471 | L471G | A | 1.9e-06 | 1.1e-06 | 2.7e-06 | 1.900E-06 | 8.000E-07 | 1.100E-06 | 2.700E-06 |
| 9409 | 471 | L471H | A | 2.4e-06 | 2.3e-06 | 2e-06 | 2.233E-06 | 2.082E-07 | 2.000E-06 | 2.400E-06 |
| 9410 | 471 | L471I | A | 1.4e-06 | 2.5e-06 | 2.6e-06 | 2.167E-06 | 6.658E-07 | 1.400E-06 | 2.600E-06 |
| 9411 | 471 | L471L | A | 2.1e-06 | 2e-06 | 3.9e-06 | 2.667E-06 | 1.069E-06 | 2.000E-06 | 3.900E-06 |
| 9412 | 471 | L471K | A | 2.2e-06 | 3.2e-06 | 3.7e-06 | 3.033E-06 | 7.638E-07 | 2.200E-06 | 3.700E-06 |
| 9413 | 471 | L471M | A | 2.4e-06 | 2.6e-06 | 1.2e-06 | 2.067E-06 | 7.572E-07 | 1.200E-06 | 2.600E-06 |
| 9414 | 471 | L471F | A | 2.9e-06 | 3.1e-06 | 2.4e-06 | 2.800E-06 | 3.606E-07 | 2.400E-06 | 3.100E-06 |
| 9415 | 471 | L471P | A | 2.2e-06 | 2e-06 | 2.3e-06 | 2.167E-06 | 1.528E-07 | 2.000E-06 | 2.300E-06 |
| 9416 | 471 | L471S | A | 2.1e-06 | 2.2e-06 | 2.3e-06 | 2.200E-06 | 1.000E-07 | 2.100E-06 | 2.300E-06 |
| 9417 | 471 | L471T | A | 1.3e-06 | 2.1e-06 | 1.9e-06 | 1.767E-06 | 4.163E-07 | 1.300E-06 | 2.100E-06 |
| 9418 | 471 | L471W | A | 6.4e-06 | 2.6e-06 | 2.8e-06 | 3.933E-06 | 2.139E-06 | 2.600E-06 | 6.400E-06 |
| 9419 | 471 | L471Y | A | 4e-06 | 2.2e-06 | 3e-06 | 3.067E-06 | 9.018E-07 | 2.200E-06 | 4.000E-06 |
| 9420 | 471 | L471V | A | 3e-06 | 2.4e-06 | 2.7e-06 | 2.700E-06 | 3.000E-07 | 2.400E-06 | 3.000E-06 |
| 9421 | 472 | L472A | A | 1.8e-06 | 2.2e-06 | 2.4e-06 | 2.133E-06 | 3.055E-07 | 1.800E-06 | 2.400E-06 |
| 9422 | 472 | L472R | A | 3.1e-06 | 2.7e-06 | 2.6e-06 | 2.800E-06 | 2.646E-07 | 2.600E-06 | 3.100E-06 |
| 9423 | 472 | L472N | A | 1.8e-06 | 2e-06 | 1.2e-06 | 1.667E-06 | 4.163E-07 | 1.200E-06 | 2.000E-06 |
| 9424 | 472 | L472D | A | 2e-06 | 1.6e-06 | 2.7e-06 | 2.100E-06 | 5.568E-07 | 1.600E-06 | 2.700E-06 |
| 9425 | 472 | L472C | A | 2.2e-06 | 2.5e-06 | 1.6e-06 | 2.100E-06 | 4.583E-07 | 1.600E-06 | 2.500E-06 |
| 9426 | 472 | L472E | A | 2.5e-06 | 2e-06 | 3.3e-06 | 2.600E-06 | 6.557E-07 | 2.000E-06 | 3.300E-06 |
| 9427 | 472 | L472Q | A | 2e-06 | 1.6e-06 | 3.1e-06 | 2.233E-06 | 7.767E-07 | 1.600E-06 | 3.100E-06 |
| 9428 | 472 | L472G | A | 1.5e-06 | 1.5e-06 | 1e-06 | 1.333E-06 | 2.887E-07 | 1.000E-06 | 1.500E-06 |
| 9429 | 472 | L472H | A | 2.6e-06 | 2.5e-06 | 9.9e-07 | 2.030E-06 | 9.021E-07 | 9.900E-07 | 2.600E-06 |
| 9430 | 472 | L472I | A | 2.7e-06 | 1.8e-06 | 1.6e-06 | 2.033E-06 | 5.859E-07 | 1.600E-06 | 2.700E-06 |
| 9431 | 472 | L472L | A | 1.9e-06 | 1.2e-06 | 1.5e-06 | 1.533E-06 | 3.512E-07 | 1.200E-06 | 1.900E-06 |
| 9432 | 472 | L472K | A | 2.1e-06 | 1.8e-06 | 3.6e-06 | 2.500E-06 | 9.644E-07 | 1.800E-06 | 3.600E-06 |
| 9433 | 472 | L472M | A | 2.3e-06 | 3.4e-06 | 3.6e-06 | 3.100E-06 | 7.000E-07 | 2.300E-06 | 3.600E-06 |
| 9434 | 472 | L472F | A | 1.8e-06 | 1.6e-06 | 2.9e-06 | 2.100E-06 | 7.000E-07 | 1.600E-06 | 2.900E-06 |

| | | | | | | | | | | |
|---|---|---|---|---|---|---|---|---|---|---|
| 9435 | 472 | L472P | A | 2.2e-06 | 3.2e-06 | 4e-06 | 3.133E-06 | 9.018E-07 | 2.200E-06 | 4.000E-06 |
| 9436 | 472 | L472S | A | 2.5e-06 | 2.4e-06 | 2.3e-06 | 2.400E-06 | 1.000E-07 | 2.300E-06 | 2.500E-06 |
| 9437 | 472 | L472T | A | 2e-06 | 2e-06 | 3.7e-06 | 2.567E-06 | 9.815E-07 | 2.000E-06 | 3.700E-06 |
| 9438 | 472 | L472W | A | 2.8e-06 | 4e-06 | 2.6e-06 | 3.133E-06 | 7.572E-07 | 2.600E-06 | 4.000E-06 |
| 9439 | 472 | L472Y | A | 4.7e-06 | 2.5e-06 | 6.8e-07 | 2.627E-06 | 2.013E-06 | 6.800E-07 | 4.700E-06 |
| 9440 | 472 | L472V | A | 1.7e-06 | 3.9e-06 | 1.1e-06 | 2.233E-06 | 1.474E-06 | 1.100E-06 | 3.900E-06 |
| 9441 | 473 | H473A | A | 1.9e-06 | 1.1e-06 | 1.9e-06 | 1.633E-06 | 4.619E-07 | 1.100E-06 | 1.900E-06 |
| 9442 | 473 | H473R | A | 2.5e-06 | 2.8e-06 | 2.5e-06 | 2.600E-06 | 1.732E-07 | 2.500E-06 | 2.800E-06 |
| 9443 | 473 | H473N | A | 2.8e-06 | 2.3e-06 | 1.7e-06 | 2.267E-06 | 5.508E-07 | 1.700E-06 | 2.800E-06 |
| 9444 | 473 | H473D | A | 3e-06 | 3.8e-06 | 1.5e-06 | 2.767E-06 | 1.168E-06 | 1.500E-06 | 3.800E-06 |
| 9445 | 473 | H473C | A | 2.6e-06 | 3.4e-06 | 2.6e-06 | 2.867E-06 | 4.619E-07 | 2.600E-06 | 3.400E-06 |
| 9446 | 473 | H473E | A | 2.2e-06 | 3.6e-06 | 1.9e-06 | 2.567E-06 | 9.074E-07 | 1.900E-06 | 3.600E-06 |
| 9447 | 473 | H473Q | A | 2.6e-06 | 1.5e-06 | 1.8e-06 | 1.967E-06 | 5.686E-07 | 1.500E-06 | 2.600E-06 |
| 9448 | 473 | H473G | A | 2.7e-06 | 3e-06 | 2e-06 | 2.567E-06 | 5.132E-07 | 2.000E-06 | 3.000E-06 |
| 9449 | 473 | H473H | A | 3.3e-06 | 3.9e-06 | 2.7e-06 | 3.300E-06 | 6.000E-07 | 2.700E-06 | 3.900E-06 |
| 9450 | 473 | H473I | A | 2.7e-06 | 2.4e-06 | 2.4e-06 | 2.500E-06 | 1.732E-07 | 2.400E-06 | 2.700E-06 |
| 9451 | 473 | H473L | A | 2.5e-06 | 3.5e-06 | 3e-06 | 3.000E-06 | 5.000E-07 | 2.500E-06 | 3.500E-06 |
| 9452 | 473 | H473K | A | 2e-06 | 1.8e-06 | 2e-06 | 1.933E-06 | 1.155E-07 | 1.800E-06 | 2.000E-06 |
| 9453 | 473 | H473M | A | 4.5e-06 | 2.1e-06 | 2.7e-06 | 3.100E-06 | 1.249E-06 | 2.100E-06 | 4.500E-06 |
| 9454 | 473 | H473F | A | 2.5e-06 | 1.8e-06 | 1.7e-06 | 2.000E-06 | 4.359E-07 | 1.700E-06 | 2.500E-06 |
| 9455 | 473 | H473P | A | 1.7e-06 | 1.9e-06 | 1.1e-06 | 1.567E-06 | 4.163E-07 | 1.100E-06 | 1.900E-06 |
| 9456 | 473 | H473S | A | 2.7e-06 | 2.4e-06 | 3.1e-06 | 2.733E-06 | 3.512E-07 | 2.400E-06 | 3.100E-06 |
| 9457 | 473 | H473T | A | 1.8e-06 | 2.3e-06 | 2.1e-06 | 2.067E-06 | 2.517E-07 | 1.800E-06 | 2.300E-06 |
| 9458 | 473 | H473W | A | 3.2e-06 | 2.2e-06 | 2.5e-06 | 2.633E-06 | 5.132E-07 | 2.200E-06 | 3.200E-06 |
| 9459 | 473 | H473Y | A | 2.4e-06 | 2.5e-06 | 2e-06 | 2.300E-06 | 2.646E-07 | 2.000E-06 | 2.500E-06 |
| 9460 | 473 | H473V | A | 2e-06 | 2.3e-06 | 2.3e-06 | 2.200E-06 | 1.732E-07 | 2.000E-06 | 2.300E-06 |
| 9461 | 474 | T474A | A | 1.9e-06 | 5.4e-06 | 3.3e-06 | 3.533E-06 | 1.762E-06 | 1.900E-06 | 5.400E-06 |
| 9462 | 474 | T474R | A | 2.7e-06 | 1.6e-06 | 2.4e-06 | 2.233E-06 | 5.686E-07 | 1.600E-06 | 2.700E-06 |
| 9463 | 474 | T474N | A | 2.5e-06 | 1.8e-06 | 4.3e-06 | 2.867E-06 | 1.290E-06 | 1.800E-06 | 4.300E-06 |
| 9464 | 474 | T474D | A | 1.9e-06 | 4.7e-06 | 1.9e-06 | 2.833E-06 | 1.617E-06 | 1.900E-06 | 4.700E-06 |
| 9465 | 474 | T474C | A | 1.6e-06 | 1.8e-06 | 3.1e-06 | 2.167E-06 | 8.145E-07 | 1.600E-06 | 3.100E-06 |
| 9466 | 474 | T474E | A | 3.4e-06 | 2.8e-06 | 5e-06 | 3.733E-06 | 1.137E-06 | 2.800E-06 | 5.000E-06 |
| 9467 | 474 | T474Q | A | 3.4e-06 | 1.7e-06 | 2.7e-06 | 2.600E-06 | 8.544E-07 | 1.700E-06 | 3.400E-06 |
| 9468 | 474 | T474G | A | 3.1e-06 | 2.5e-06 | 7e-06 | 4.200E-06 | 2.443E-06 | 2.500E-06 | 7.000E-06 |
| 9469 | 474 | T474H | A | 2.5e-06 | 2.8e-06 | 6.3e-06 | 3.867E-06 | 2.113E-06 | 2.500E-06 | 6.300E-06 |
| 9470 | 474 | T474I | A | 1.5e-06 | 3.2e-06 | 2.4e-06 | 2.367E-06 | 8.505E-07 | 1.500E-06 | 3.200E-06 |
| 9471 | 474 | T474L | A | 3.7e-06 | 3.1e-06 | 2.2e-06 | 3.000E-06 | 7.550E-07 | 2.200E-06 | 3.700E-06 |
| 9472 | 474 | T474K | A | 4.5e-06 | 2.4e-06 | 1.5e-06 | 2.800E-06 | 1.539E-06 | 1.500E-06 | 4.500E-06 |
| 9473 | 474 | T474M | A | 2e-06 | 3.8e-06 | 2e-06 | 2.600E-06 | 1.039E-06 | 2.000E-06 | 3.800E-06 |
| 9474 | 474 | T474F | A | 3.4e-06 | 2.9e-06 | 2e-06 | 2.767E-06 | 7.095E-07 | 2.000E-06 | 3.400E-06 |
| 9475 | 474 | T474P | A | 2.6e-06 | 2.8e-06 | 1.6e-06 | 2.333E-06 | 6.429E-07 | 1.600E-06 | 2.800E-06 |
| 9476 | 474 | T474S | A | 1.3e-06 | 2.6e-06 | 1.5e-06 | 1.800E-06 | 7.000E-07 | 1.300E-06 | 2.600E-06 |
| 9477 | 474 | T474T | A | 2.8e-06 | 3.4e-06 | 5e-06 | 3.733E-06 | 1.137E-06 | 2.800E-06 | 5.000E-06 |
| 9478 | 474 | T474W | A | 3.1e-06 | 2.6e-06 | 1.9e-06 | 2.533E-06 | 6.028E-07 | 1.900E-06 | 3.100E-06 |
| 9479 | 474 | T474Y | A | 1.8e-06 | 3e-06 | 2.5e-06 | 2.433E-06 | 6.028E-07 | 1.800E-06 | 3.000E-06 |
| 9480 | 474 | T474V | A | 2.1e-06 | 2e-06 | 2.2e-06 | 2.100E-06 | 1.000E-07 | 2.000E-06 | 2.200E-06 |
| 9481 | 475 | A475A | A | 3.1e-06 | 1.7e-06 | 2.5e-06 | 2.433E-06 | 7.024E-07 | 1.700E-06 | 3.100E-06 |
| 9482 | 475 | A475R | A | 2e-06 | 2.8e-06 | 1.8e-06 | 2.200E-06 | 5.292E-07 | 1.800E-06 | 2.800E-06 |
| 9483 | 475 | A475N | A | 3e-06 | 1.8e-06 | 2.6e-06 | 2.467E-06 | 6.110E-07 | 1.800E-06 | 3.000E-06 |
| 9484 | 475 | A475D | A | 1.9e-06 | 2.4e-06 | 3.8e-06 | 2.700E-06 | 9.849E-07 | 1.900E-06 | 3.800E-06 |
| 9485 | 475 | A475C | A | 2.1e-06 | 1.9e-06 | 2e-06 | 2.000E-06 | 1.000E-07 | 1.900E-06 | 2.100E-06 |
| 9486 | 475 | A475E | A | 1.8e-06 | 3.2e-06 | 3.1e-06 | 2.700E-06 | 7.810E-07 | 1.800E-06 | 3.200E-06 |
| 9487 | 475 | A475Q | A | 1.8e-06 | 2.7e-06 | 2.9e-06 | 2.467E-06 | 5.859E-07 | 1.800E-06 | 2.900E-06 |
| 9488 | 475 | A475G | A | 4.7e-06 | 3.8e-06 | 1.4e-06 | 3.300E-06 | 1.706E-06 | 1.400E-06 | 4.700E-06 |
| 9489 | 475 | A475H | A | 3.6e-06 | 1.4e-06 | 3.2e-06 | 2.733E-06 | 1.172E-06 | 1.400E-06 | 3.600E-06 |
| 9490 | 475 | A475I | A | 2.4e-06 | 2.6e-06 | 2.7e-06 | 2.567E-06 | 1.528E-07 | 2.400E-06 | 2.700E-06 |
| 9491 | 475 | A475L | A | 1.4e-06 | 2.9e-06 | 3e-06 | 2.433E-06 | 8.963E-07 | 1.400E-06 | 3.000E-06 |
| 9492 | 475 | A475K | A | 4.3e-06 | 1.6e-06 | 1.8e-06 | 2.567E-06 | 1.504E-06 | 1.600E-06 | 4.300E-06 |
| 9493 | 475 | A475M | A | 2.5e-06 | 2.1e-06 | 1.5e-06 | 2.033E-06 | 5.033E-07 | 1.500E-06 | 2.500E-06 |
| 9494 | 475 | A475F | A | 3e-06 | 1.9e-06 | 3.3e-06 | 2.733E-06 | 7.371E-07 | 1.900E-06 | 3.300E-06 |
| 9495 | 475 | A475P | A | 3.1e-06 | 2.4e-06 | 1.8e-06 | 2.433E-06 | 6.506E-07 | 1.800E-06 | 3.100E-06 |
| 9496 | 475 | A475S | A | 3.1e-06 | 2.2e-06 | 2.8e-06 | 2.700E-06 | 4.583E-07 | 2.200E-06 | 3.100E-06 |
| 9497 | 475 | A475T | A | 2.2e-06 | 2.2e-06 | 3.2e-06 | 2.533E-06 | 5.774E-07 | 2.200E-06 | 3.200E-06 |
| 9498 | 475 | A475W | A | 1.6e-06 | 2.3e-06 | 1.1e-06 | 1.667E-06 | 6.028E-07 | 1.100E-06 | 2.300E-06 |
| 9499 | 475 | A475Y | A | 2.7e-06 | 2.7e-06 | 1.6e-06 | 2.333E-06 | 6.351E-07 | 1.600E-06 | 2.700E-06 |
| 9500 | 475 | A475V | A | 2.6e-06 | 2.6e-06 | 2.9e-06 | 2.700E-06 | 1.732E-07 | 2.600E-06 | 2.900E-06 |
| 9501 | 476 | N476A | A | 1.5e-06 | 2.2e-06 | 2.6e-06 | 2.100E-06 | 5.568E-07 | 1.500E-06 | 2.600E-06 |
| 9502 | 476 | N476R | A | 9.8e-07 | 2.4e-06 | 2.4e-06 | 1.927E-06 | 8.198E-07 | 9.800E-07 | 2.400E-06 |
| 9503 | 476 | N476N | A | 3.7e-06 | 9.5e-07 | 1.8e-06 | 2.150E-06 | 1.408E-06 | 9.500E-07 | 3.700E-06 |
| 9504 | 476 | N476D | A | 4e-06 | 2.6e-06 | 2.2e-06 | 2.933E-06 | 9.452E-07 | 2.200E-06 | 4.000E-06 |
| 9505 | 476 | N476C | A | 1.9e-06 | 1.3e-06 | 2.8e-06 | 2.000E-06 | 7.550E-07 | 1.300E-06 | 2.800E-06 |
| 9506 | 476 | N476E | A | 2.5e-06 | 1.6e-06 | 2.7e-06 | 2.267E-06 | 5.859E-07 | 1.600E-06 | 2.700E-06 |
| 9507 | 476 | N476Q | A | 2.1e-06 | 2.9e-06 | 2.3e-06 | 2.433E-06 | 4.163E-07 | 2.100E-06 | 2.900E-06 |
| 9508 | 476 | N476G | A | 2.7e-06 | 3.7e-06 | 2.1e-06 | 2.833E-06 | 8.083E-07 | 2.100E-06 | 3.700E-06 |
| 9509 | 476 | N476H | A | 2.5e-06 | 2e-06 | 1.8e-06 | 2.100E-06 | 3.606E-07 | 1.800E-06 | 2.500E-06 |
| 9510 | 476 | N476I | A | 2.4e-06 | 1.4e-06 | 4.2e-06 | 2.667E-06 | 1.419E-06 | 1.400E-06 | 4.200E-06 |
| 9511 | 476 | N476L | A | 1.9e-06 | 4.2e-06 | 2.2e-06 | 2.767E-06 | 1.250E-06 | 1.900E-06 | 4.200E-06 |
| 9512 | 476 | N476K | A | 2.1e-06 | 3.3e-06 | 2.9e-06 | 2.767E-06 | 6.110E-07 | 2.100E-06 | 3.300E-06 |
| 9513 | 476 | N476M | A | 3.1e-06 | 3e-06 | 2.5e-06 | 2.867E-06 | 3.215E-07 | 2.500E-06 | 3.100E-06 |
| 9514 | 476 | N476F | A | 4.3e-06 | 3.5e-06 | 1.5e-06 | 3.100E-06 | 1.442E-06 | 1.500E-06 | 4.300E-06 |
| 9515 | 476 | N476P | A | 1.8e-06 | 1.5e-06 | 2.3e-06 | 1.867E-06 | 4.041E-07 | 1.500E-06 | 2.300E-06 |
| 9516 | 476 | N476S | A | 3.7e-06 | 2.6e-06 | 2.6e-06 | 2.967E-06 | 6.351E-07 | 2.600E-06 | 3.700E-06 |
| 9517 | 476 | N476T | A | 4.7e-06 | 2.9e-06 | 2e-06 | 3.200E-06 | 1.375E-06 | 2.000E-06 | 4.700E-06 |
| 9518 | 476 | N476W | A | 4.3e-06 | 3.8e-06 | 1.8e-06 | 3.300E-06 | 1.323E-06 | 1.800E-06 | 4.300E-06 |
| 9519 | 476 | N476Y | A | 2.5e-06 | 3.3e-06 | 1.1e-06 | 2.300E-06 | 1.114E-06 | 1.100E-06 | 3.300E-06 |

| | | | | | | | | | | |
|---|---|---|---|---|---|---|---|---|---|---|
| 9520 | 476 | N476V | A | 2.5e-06 | 2.1e-06 | 1.3e-06 | 1.967E-06 | 6.110E-07 | 1.300E-06 | 2.500E-06 |
| 9521 | 477 | R477A | A | 2.6e-06 | 3e-06 | 2.2e-06 | 2.600E-06 | 4.000E-07 | 2.200E-06 | 3.000E-06 |
| 9522 | 477 | R477R | A | 1.4e-06 | 1.4e-06 | 2.9e-06 | 1.900E-06 | 8.660E-07 | 1.400E-06 | 2.900E-06 |
| 9523 | 477 | R477N | A | 1.6e-06 | 1.9e-06 | 3e-06 | 2.167E-06 | 7.371E-07 | 1.600E-06 | 3.000E-06 |
| 9524 | 477 | R477D | A | 1.9e-06 | 1.8e-06 | 2.7e-06 | 2.133E-06 | 4.933E-07 | 1.800E-06 | 2.700E-06 |
| 9525 | 477 | R477C | A | 1.5e-06 | 4.4e-06 | 1.6e-06 | 2.500E-06 | 1.646E-06 | 1.500E-06 | 4.400E-06 |
| 9526 | 477 | R477E | A | 1.9e-06 | 3.2e-06 | 2e-06 | 2.367E-06 | 7.234E-07 | 1.900E-06 | 3.200E-06 |
| 9527 | 477 | R477Q | A | 1.9e-06 | 2.8e-06 | 4.6e-06 | 3.100E-06 | 1.375E-06 | 1.900E-06 | 4.600E-06 |
| 9528 | 477 | R477G | A | 1.7e-06 | 2.1e-06 | 1.8e-06 | 1.867E-06 | 2.082E-07 | 1.700E-06 | 2.100E-06 |
| 9529 | 477 | R477H | A | 1.8e-06 | 3e-06 | 2.4e-06 | 2.400E-06 | 6.000E-07 | 1.800E-06 | 3.000E-06 |
| 9530 | 477 | R477I | A | 3e-06 | 2.9e-06 | 2.8e-06 | 2.900E-06 | 1.000E-07 | 2.800E-06 | 3.000E-06 |
| 9531 | 477 | R477L | A | 2.1e-06 | 2.1e-06 | 3.1e-06 | 2.433E-06 | 5.774E-07 | 2.100E-06 | 3.100E-06 |
| 9532 | 477 | R477K | A | 2.5e-06 | 1.9e-06 | 2.6e-06 | 2.333E-06 | 3.786E-07 | 1.900E-06 | 2.600E-06 |
| 9533 | 477 | R477M | A | 2.6e-06 | 1.9e-06 | 1.2e-06 | 1.900E-06 | 7.000E-07 | 1.200E-06 | 2.600E-06 |
| 9534 | 477 | R477F | A | 3.8e-06 | 2.7e-06 | 1.8e-06 | 2.767E-06 | 1.002E-06 | 1.800E-06 | 3.800E-06 |
| 9535 | 477 | R477P | A | 2.1e-06 | 2.7e-06 | 2.4e-06 | 2.400E-06 | 3.000E-07 | 2.100E-06 | 2.700E-06 |
| 9536 | 477 | R477S | A | 1.6e-06 | 2.7e-06 | 2.5e-06 | 2.267E-06 | 5.859E-07 | 1.600E-06 | 2.700E-06 |
| 9537 | 477 | R477T | A | 4.9e-06 | 3.5e-06 | 2.8e-06 | 3.733E-06 | 1.069E-06 | 2.800E-06 | 4.900E-06 |
| 9538 | 477 | R477W | A | 2.8e-06 | 2.2e-06 | 2.4e-06 | 2.467E-06 | 3.055E-07 | 2.200E-06 | 2.800E-06 |
| 9539 | 477 | R477Y | A | 2.7e-06 | 2.5e-06 | 2.3e-06 | 2.500E-06 | 2.000E-07 | 2.300E-06 | 2.700E-06 |
| 9540 | 477 | R477V | A | 1.8e-06 | 2.3e-06 | 3.1e-06 | 2.400E-06 | 6.557E-07 | 1.800E-06 | 3.100E-06 |
| 9541 | 478 | P478A | A | 1.8e-06 | 2.2e-06 | 1.5e-06 | 1.833E-06 | 3.512E-07 | 1.500E-06 | 2.200E-06 |
| 9542 | 478 | P478R | A | 1.6e-06 | 3e-06 | 3.1e-06 | 2.567E-06 | 8.386E-07 | 1.600E-06 | 3.100E-06 |
| 9543 | 478 | P478N | A | 2.7e-06 | 2.5e-06 | 2.8e-06 | 2.667E-06 | 1.528E-07 | 2.500E-06 | 2.800E-06 |
| 9544 | 478 | P478D | A | 2.2e-06 | 2.1e-06 | 2.4e-06 | 2.233E-06 | 1.528E-07 | 2.100E-06 | 2.400E-06 |
| 9545 | 478 | P478C | A | 2.5e-06 | 4.8e-06 | 1.9e-06 | 3.067E-06 | 1.531E-06 | 1.900E-06 | 4.800E-06 |
| 9546 | 478 | P478E | A | 2.3e-06 | 2.6e-06 | 4e-06 | 2.967E-06 | 9.074E-07 | 2.300E-06 | 4.000E-06 |
| 9547 | 478 | P478Q | A | 2.2e-06 | 2e-06 | 2.8e-06 | 2.333E-06 | 4.163E-07 | 2.000E-06 | 2.800E-06 |
| 9548 | 478 | P478G | A | 3.6e-06 | 1.2e-06 | 3.9e-06 | 2.900E-06 | 1.480E-06 | 1.200E-06 | 3.900E-06 |
| 9549 | 478 | P478H | A | 2.2e-06 | 1.6e-06 | 2.1e-06 | 1.967E-06 | 3.215E-07 | 1.600E-06 | 2.200E-06 |
| 9550 | 478 | P478I | A | 1.6e-06 | 2.6e-06 | 2.8e-06 | 2.333E-06 | 6.429E-07 | 1.600E-06 | 2.800E-06 |
| 9551 | 478 | P478L | A | 2.9e-06 | 2.7e-06 | 2.9e-06 | 2.833E-06 | 1.155E-07 | 2.700E-06 | 2.900E-06 |
| 9552 | 478 | P478K | A | 2.6e-06 | 2.5e-06 | 3.3e-06 | 2.800E-06 | 4.359E-07 | 2.500E-06 | 3.300E-06 |
| 9553 | 478 | P478M | A | 1.4e-06 | 2.4e-06 | 1.9e-06 | 1.900E-06 | 5.000E-07 | 1.400E-06 | 2.400E-06 |
| 9554 | 478 | P478F | A | 2e-06 | 2.6e-06 | 4.2e-06 | 2.933E-06 | 1.137E-06 | 2.000E-06 | 4.200E-06 |
| 9555 | 478 | P478P | A | 2.9e-06 | 4.6e-06 | 1.8e-06 | 3.100E-06 | 1.411E-06 | 1.800E-06 | 4.600E-06 |
| 9556 | 478 | P478S | A | 2.9e-06 | 1.8e-06 | 2.2e-06 | 2.300E-06 | 5.568E-07 | 1.800E-06 | 2.900E-06 |
| 9557 | 478 | P478T | A | 1.9e-06 | 1.9e-06 | 1.3e-06 | 1.700E-06 | 3.464E-07 | 1.300E-06 | 1.900E-06 |
| 9558 | 478 | P478W | A | 2.6e-06 | 8.9e-07 | 3.9e-06 | 2.463E-06 | 1.510E-06 | 8.900E-07 | 3.900E-06 |
| 9559 | 478 | P478Y | A | 1.5e-06 | 4.1e-06 | 2.6e-06 | 2.733E-06 | 1.305E-06 | 1.500E-06 | 4.100E-06 |
| 9560 | 478 | P478V | A | 1.5e-06 | 2.7e-06 | 3.3e-06 | 2.500E-06 | 9.165E-07 | 1.500E-06 | 3.300E-06 |
| 9561 | 479 | E479A | A | 2e-06 | 1.7e-06 | 2.3e-06 | 2.000E-06 | 3.000E-07 | 1.700E-06 | 2.300E-06 |
| 9562 | 479 | E479R | A | 3.3e-06 | 2e-06 | 1.8e-06 | 2.367E-06 | 8.145E-07 | 1.800E-06 | 3.300E-06 |
| 9563 | 479 | E479N | A | 1.9e-06 | 2.6e-06 | 2.8e-06 | 2.433E-06 | 4.726E-07 | 1.900E-06 | 2.800E-06 |
| 9564 | 479 | E479D | A | 8.9e-07 | 1.5e-06 | 1.9e-06 | 1.430E-06 | 5.086E-07 | 8.900E-07 | 1.900E-06 |
| 9565 | 479 | E479C | A | 3e-06 | 2e-06 | 1.3e-06 | 2.100E-06 | 8.544E-07 | 1.300E-06 | 3.000E-06 |
| 9566 | 479 | E479E | A | 1.4e-06 | 1.6e-06 | 1.7e-06 | 1.567E-06 | 1.528E-07 | 1.400E-06 | 1.700E-06 |
| 9567 | 479 | E479Q | A | 2.1e-06 | 2.2e-06 | 1.9e-06 | 2.067E-06 | 1.528E-07 | 1.900E-06 | 2.200E-06 |
| 9568 | 479 | E479G | A | 2.4e-06 | 3.9e-06 | 3.9e-06 | 3.400E-06 | 8.660E-07 | 2.400E-06 | 3.900E-06 |
| 9569 | 479 | E479H | A | 3.2e-06 | 2.3e-06 | 2.6e-06 | 2.700E-06 | 4.583E-07 | 2.300E-06 | 3.200E-06 |
| 9570 | 479 | E479I | A | 2.5e-06 | 1.4e-06 | 2.6e-06 | 2.167E-06 | 6.658E-07 | 1.400E-06 | 2.600E-06 |
| 9571 | 479 | E479L | A | 2.7e-06 | 1.8e-06 | 2.6e-06 | 2.367E-06 | 4.933E-07 | 1.800E-06 | 2.700E-06 |
| 9572 | 479 | E479K | A | 2.6e-06 | 2.4e-06 | 2e-06 | 2.333E-06 | 3.055E-07 | 2.000E-06 | 2.600E-06 |
| 9573 | 479 | E479M | A | 4.9e-06 | 1.8e-06 | 2.7e-06 | 3.133E-06 | 1.595E-06 | 1.800E-06 | 4.900E-06 |
| 9574 | 479 | E479F | A | 2.2e-06 | 2.6e-06 | 1.6e-06 | 2.133E-06 | 5.033E-07 | 1.600E-06 | 2.600E-06 |
| 9575 | 479 | E479P | A | 2.7e-06 | 1.3e-06 | 3.2e-06 | 2.400E-06 | 9.849E-07 | 1.300E-06 | 3.200E-06 |
| 9576 | 479 | E479S | A | 4.6e-06 | 2.6e-06 | 4.8e-06 | 4.000E-06 | 1.217E-06 | 2.600E-06 | 4.800E-06 |
| 9577 | 479 | E479T | A | 2.5e-06 | 2.1e-06 | 1.8e-06 | 2.133E-06 | 3.512E-07 | 1.800E-06 | 2.500E-06 |
| 9578 | 479 | E479W | A | 3.3e-06 | 2.6e-06 | 2.6e-06 | 2.833E-06 | 4.041E-07 | 2.600E-06 | 3.300E-06 |
| 9579 | 479 | E479Y | A | 4.5e-06 | 1.6e-06 | 1.5e-06 | 2.533E-06 | 1.704E-06 | 1.500E-06 | 4.500E-06 |
| 9580 | 479 | E479V | A | 4.6e-06 | 1.9e-06 | 3.3e-06 | 3.267E-06 | 1.350E-06 | 1.900E-06 | 4.600E-06 |
| 9581 | 480 | D480A | A | 2.8e-06 | 1.9e-06 | 6.4e-06 | 3.700E-06 | 2.381E-06 | 1.900E-06 | 6.400E-06 |
| 9582 | 480 | D480R | A | 2.7e-06 | 2.5e-06 | 2.5e-06 | 2.567E-06 | 1.155E-07 | 2.500E-06 | 2.700E-06 |
| 9583 | 480 | D480N | A | 1.7e-06 | 1.9e-06 | 3.8e-06 | 2.467E-06 | 1.159E-06 | 1.700E-06 | 3.800E-06 |
| 9584 | 480 | D480D | A | 1.2e-06 | 1.9e-06 | 2.8e-06 | 1.967E-06 | 8.021E-07 | 1.200E-06 | 2.800E-06 |
| 9585 | 480 | D480C | A | 2e-06 | 3.4e-06 | 1.2e-06 | 2.200E-06 | 1.114E-06 | 1.200E-06 | 3.400E-06 |
| 9586 | 480 | D480E | A | 2.8e-06 | 4.8e-06 | 1.5e-06 | 3.033E-06 | 1.662E-06 | 1.500E-06 | 4.800E-06 |
| 9587 | 480 | D480Q | A | 2.3e-06 | 3.5e-06 | 2.3e-06 | 2.700E-06 | 6.928E-07 | 2.300E-06 | 3.500E-06 |
| 9588 | 480 | D480G | A | 1.9e-06 | 3.7e-06 | 1.7e-06 | 2.433E-06 | 1.102E-06 | 1.700E-06 | 3.700E-06 |
| 9589 | 480 | D480H | A | 3e-06 | 1.6e-06 | 2.7e-06 | 2.433E-06 | 7.371E-07 | 1.600E-06 | 3.000E-06 |
| 9590 | 480 | D480I | A | 3.1e-06 | 3.8e-06 | 2.4e-06 | 3.100E-06 | 7.000E-07 | 2.400E-06 | 3.800E-06 |
| 9591 | 480 | D480L | A | 2.3e-06 | 1.2e-06 | 2.6e-06 | 2.033E-06 | 7.371E-07 | 1.200E-06 | 2.600E-06 |
| 9592 | 480 | D480K | A | 1.8e-06 | 1.6e-06 | 2.1e-06 | 1.833E-06 | 2.517E-07 | 1.600E-06 | 2.100E-06 |
| 9593 | 480 | D480M | A | 1.2e-06 | 3.9e-06 | 2.5e-06 | 2.533E-06 | 1.350E-06 | 1.200E-06 | 3.900E-06 |
| 9594 | 480 | D480F | A | 2.5e-06 | 4.1e-06 | 2.7e-06 | 3.100E-06 | 8.718E-07 | 2.500E-06 | 4.100E-06 |
| 9595 | 480 | D480P | A | 3.6e-06 | 3e-06 | 1.9e-06 | 2.833E-06 | 8.622E-07 | 1.900E-06 | 3.600E-06 |
| 9596 | 480 | D480S | A | 2.8e-06 | 1.5e-06 | 2.3e-06 | 2.200E-06 | 6.557E-07 | 1.500E-06 | 2.800E-06 |
| 9597 | 480 | D480T | A | 2.5e-06 | 1.8e-06 | 2.2e-06 | 2.167E-06 | 3.512E-07 | 1.800E-06 | 2.500E-06 |
| 9598 | 480 | D480W | A | 2.6e-06 | 2e-06 | 1.6e-06 | 2.067E-06 | 5.033E-07 | 1.600E-06 | 2.600E-06 |
| 9599 | 480 | D480Y | A | 3e-06 | 2.8e-06 | 2.4e-06 | 2.733E-06 | 3.055E-07 | 2.400E-06 | 3.000E-06 |
| 9600 | 480 | D480V | A | 2.5e-06 | 3.3e-06 | 2.1e-06 | 2.633E-06 | 6.110E-07 | 2.100E-06 | 3.300E-06 |
| 9601 | 481 | E481A | A | 3e-06 | 1.6e-06 | 2.9e-06 | 2.500E-06 | 7.810E-07 | 1.600E-06 | 3.000E-06 |
| 9602 | 481 | E481R | A | 1.9e-06 | 3.3e-06 | 2.6e-06 | 2.600E-06 | 7.000E-07 | 1.900E-06 | 3.300E-06 |
| 9603 | 481 | E481N | A | 4e-06 | 2.2e-06 | 3.3e-06 | 3.167E-06 | 9.074E-07 | 2.200E-06 | 4.000E-06 |
| 9604 | 481 | E481D | A | 3.3e-06 | 2.4e-06 | 3.4e-06 | 3.033E-06 | 5.508E-07 | 2.400E-06 | 3.400E-06 |

| | | | | | | | | | | |
|---|---|---|---|---|---|---|---|---|---|---|
| 9605 | 481 | E481C | A | 3e-06 | 3.3e-06 | 3.2e-06 | 3.167E-06 | 1.528E-07 | 3.000E-06 | 3.300E-06 |
| 9606 | 481 | E481E | A | 2.6e-06 | 2.1e-06 | 5.4e-06 | 3.367E-06 | 1.779E-06 | 2.100E-06 | 5.400E-06 |
| 9607 | 481 | E481Q | A | 1.7e-06 | 1.3e-06 | 2.7e-06 | 1.900E-06 | 7.211E-07 | 1.300E-06 | 2.700E-06 |
| 9608 | 481 | E481G | A | 2.8e-06 | 4.4e-06 | 1.8e-06 | 3.000E-06 | 1.311E-06 | 1.800E-06 | 4.400E-06 |
| 9609 | 481 | E481H | A | 1.8e-06 | 2.1e-06 | 2.4e-06 | 2.100E-06 | 3.000E-07 | 1.800E-06 | 2.400E-06 |
| 9610 | 481 | E481I | A | 2.1e-06 | 2.3e-06 | 2e-06 | 2.133E-06 | 1.528E-07 | 2.000E-06 | 2.300E-06 |
| 9611 | 481 | E481L | A | 1.7e-06 | 3.1e-06 | 2.9e-06 | 2.567E-06 | 7.572E-07 | 1.700E-06 | 3.100E-06 |
| 9612 | 481 | E481K | A | 6.6e-07 | 2.9e-06 | 1.9e-06 | 1.820E-06 | 1.122E-06 | 6.600E-07 | 2.900E-06 |
| 9613 | 481 | E481M | A | 2.3e-06 | 3.5e-06 | 3.5e-06 | 3.100E-06 | 6.928E-07 | 2.300E-06 | 3.500E-06 |
| 9614 | 481 | E481F | A | 2.1e-06 | 3e-06 | 3.6e-06 | 2.900E-06 | 7.550E-07 | 2.100E-06 | 3.600E-06 |
| 9615 | 481 | E481P | A | 1.9e-06 | 3.1e-06 | 2.7e-06 | 2.567E-06 | 6.110E-07 | 1.900E-06 | 3.100E-06 |
| 9616 | 481 | E481S | A | 2.9e-06 | 2.6e-06 | 2.7e-06 | 2.733E-06 | 1.528E-07 | 2.600E-06 | 2.900E-06 |
| 9617 | 481 | E481T | A | 2e-06 | 1.4e-06 | 3.3e-06 | 2.233E-06 | 9.713E-07 | 1.400E-06 | 3.300E-06 |
| 9618 | 481 | E481W | A | 2.1e-06 | 2e-06 | 3.7e-06 | 2.600E-06 | 9.539E-07 | 2.000E-06 | 3.700E-06 |
| 9619 | 481 | E481Y | A | 3.2e-06 | 1.7e-06 | 2.1e-06 | 2.333E-06 | 7.767E-07 | 1.700E-06 | 3.200E-06 |
| 9620 | 481 | E481V | A | 2.1e-06 | 2.6e-06 | 2.1e-06 | 2.267E-06 | 2.887E-07 | 2.100E-06 | 2.600E-06 |
| 9621 | 482 | C482A | A | 2.6e-06 | 2.3e-06 | 2.5e-06 | 2.467E-06 | 1.528E-07 | 2.300E-06 | 2.600E-06 |
| 9622 | 482 | C482R | A | 2.9e-06 | 2.1e-06 | 4.6e-06 | 3.200E-06 | 1.277E-06 | 2.100E-06 | 4.600E-06 |
| 9623 | 482 | C482N | A | 2.5e-06 | 2e-06 | 1.3e-06 | 1.933E-06 | 6.028E-07 | 1.300E-06 | 2.500E-06 |
| 9624 | 482 | C482D | A | 1.6e-06 | 1.8e-06 | 2.7e-06 | 2.033E-06 | 5.859E-07 | 1.600E-06 | 2.700E-06 |
| 9625 | 482 | C482C | A | 3.6e-06 | 2e-06 | 2.4e-06 | 2.667E-06 | 8.327E-07 | 2.000E-06 | 3.600E-06 |
| 9626 | 482 | C482E | A | 2.3e-06 | 2.5e-06 | 2.1e-06 | 2.300E-06 | 2.000E-07 | 2.100E-06 | 2.500E-06 |
| 9627 | 482 | C482Q | A | 2.8e-06 | 2.4e-06 | 1.7e-06 | 2.300E-06 | 5.568E-07 | 1.700E-06 | 2.800E-06 |
| 9628 | 482 | C482G | A | 1.9e-06 | 3.2e-06 | 4.3e-06 | 3.133E-06 | 1.201E-06 | 1.900E-06 | 4.300E-06 |
| 9629 | 482 | C482H | A | 3.2e-06 | 2.6e-06 | 2.3e-06 | 2.700E-06 | 4.583E-07 | 2.300E-06 | 3.200E-06 |
| 9630 | 482 | C482I | A | 2.6e-06 | 2.6e-06 | 3e-06 | 2.733E-06 | 2.309E-07 | 2.600E-06 | 3.000E-06 |
| 9631 | 482 | C482L | A | 3.6e-06 | 3.2e-06 | 2.4e-06 | 3.067E-06 | 6.110E-07 | 2.400E-06 | 3.600E-06 |
| 9632 | 482 | C482K | A | 5e-06 | 1.9e-06 | 1.9e-06 | 2.933E-06 | 1.790E-06 | 1.900E-06 | 5.000E-06 |
| 9633 | 482 | C482M | A | 2.9e-06 | 2.6e-06 | 1.9e-06 | 2.467E-06 | 5.132E-07 | 1.900E-06 | 2.900E-06 |
| 9634 | 482 | C482F | A | 2.3e-06 | 2.3e-06 | 2.8e-06 | 2.467E-06 | 2.887E-07 | 2.300E-06 | 2.800E-06 |
| 9635 | 482 | C482P | A | 3.3e-06 | 2.8e-06 | 4.2e-06 | 3.433E-06 | 7.095E-07 | 2.800E-06 | 4.200E-06 |
| 9636 | 482 | C482S | A | 3.4e-06 | 4.2e-06 | 3e-06 | 3.533E-06 | 6.110E-07 | 3.000E-06 | 4.200E-06 |
| 9637 | 482 | C482T | A | 2.4e-06 | 3.3e-06 | 2.2e-06 | 2.633E-06 | 5.859E-07 | 2.200E-06 | 3.300E-06 |
| 9638 | 482 | C482W | A | 2.5e-06 | 1.5e-06 | 1e-06 | 1.667E-06 | 7.638E-07 | 1.000E-06 | 2.500E-06 |
| 9639 | 482 | C482Y | A | 2.3e-06 | 1.7e-06 | 4e-06 | 2.667E-06 | 1.193E-06 | 1.700E-06 | 4.000E-06 |
| 9640 | 482 | C482V | A | 2.1e-06 | 1.4e-06 | 1.7e-06 | 1.733E-06 | 3.512E-07 | 1.400E-06 | 2.100E-06 |
| 9641 | 483 | V483A | A | 2.4e-06 | 3.9e-06 | 1.4e-06 | 2.567E-06 | 1.258E-06 | 1.400E-06 | 3.900E-06 |
| 9642 | 483 | V483R | A | 3.9e-06 | 1.7e-06 | 2.4e-06 | 2.667E-06 | 1.124E-06 | 1.700E-06 | 3.900E-06 |
| 9643 | 483 | V483N | A | 2.1e-06 | 3.2e-06 | 3.4e-06 | 2.900E-06 | 7.000E-07 | 2.100E-06 | 3.400E-06 |
| 9644 | 483 | V483D | A | 1.6e-06 | 1.1e-06 | 1.5e-06 | 1.400E-06 | 2.646E-07 | 1.100E-06 | 1.600E-06 |
| 9645 | 483 | V483C | A | 4e-06 | 2.3e-06 | 2.7e-06 | 3.000E-06 | 8.888E-07 | 2.300E-06 | 4.000E-06 |
| 9646 | 483 | V483E | A | 4.9e-06 | 3.1e-06 | 1.9e-06 | 3.300E-06 | 1.510E-06 | 1.900E-06 | 4.900E-06 |
| 9647 | 483 | V483Q | A | 1.6e-06 | 2.4e-06 | 1.9e-06 | 1.967E-06 | 4.041E-07 | 1.600E-06 | 2.400E-06 |
| 9648 | 483 | V483G | A | 3e-06 | 2.7e-06 | 1.5e-06 | 2.400E-06 | 7.937E-07 | 1.500E-06 | 3.000E-06 |
| 9649 | 483 | V483H | A | 3.2e-06 | 4e-06 | 3.2e-06 | 3.467E-06 | 4.619E-07 | 3.200E-06 | 4.000E-06 |
| 9650 | 483 | V483I | A | 3.2e-06 | 1.4e-06 | 4e-06 | 2.867E-06 | 1.332E-06 | 1.400E-06 | 4.000E-06 |
| 9651 | 483 | V483L | A | 2.7e-06 | 2e-06 | 3.5e-06 | 2.733E-06 | 7.506E-07 | 2.000E-06 | 3.500E-06 |
| 9652 | 483 | V483K | A | 3.3e-06 | 2.9e-06 | 3.7e-06 | 3.300E-06 | 4.000E-07 | 2.900E-06 | 3.700E-06 |
| 9653 | 483 | V483M | A | 2.1e-06 | 3.7e-06 | 2.7e-06 | 2.833E-06 | 8.083E-07 | 2.100E-06 | 3.700E-06 |
| 9654 | 483 | V483F | A | 2e-06 | 4.7e-06 | 1.3e-06 | 2.667E-06 | 1.795E-06 | 1.300E-06 | 4.700E-06 |
| 9655 | 483 | V483P | A | 1.6e-06 | 2.4e-06 | 3.4e-06 | 2.467E-06 | 9.018E-07 | 1.600E-06 | 3.400E-06 |
| 9656 | 483 | V483S | A | 1.6e-06 | 2.2e-06 | 2.3e-06 | 2.033E-06 | 3.786E-07 | 1.600E-06 | 2.300E-06 |
| 9657 | 483 | V483T | A | 3.8e-06 | 2e-06 | 3e-06 | 2.933E-06 | 9.018E-07 | 2.000E-06 | 3.800E-06 |
| 9658 | 483 | V483W | A | 2.7e-06 | 1.8e-06 | 3.2e-06 | 2.567E-06 | 7.095E-07 | 1.800E-06 | 3.200E-06 |
| 9659 | 483 | V483Y | A | 2.3e-06 | 2.3e-06 | 2.6e-06 | 2.400E-06 | 1.732E-07 | 2.300E-06 | 2.600E-06 |
| 9660 | 483 | V483V | A | 1.7e-06 | 3.6e-06 | 2.3e-06 | 2.533E-06 | 9.713E-07 | 1.700E-06 | 3.600E-06 |
| 9661 | 484 | G484A | A | 4e-06 | 1.2e-06 | 2e-06 | 2.400E-06 | 1.442E-06 | 1.200E-06 | 4.000E-06 |
| 9662 | 484 | G484R | A | 1.6e-06 | 1.8e-06 | 3.6e-06 | 2.333E-06 | 1.102E-06 | 1.600E-06 | 3.600E-06 |
| 9663 | 484 | G484N | A | 1.8e-06 | 1.2e-06 | 2.9e-06 | 1.967E-06 | 8.622E-07 | 1.200E-06 | 2.900E-06 |
| 9664 | 484 | G484D | A | 2.6e-06 | 2.1e-06 | 2.6e-06 | 2.433E-06 | 2.887E-07 | 2.100E-06 | 2.600E-06 |
| 9665 | 484 | G484C | A | 1.6e-06 | 1.9e-06 | 2.8e-06 | 2.100E-06 | 6.245E-07 | 1.600E-06 | 2.800E-06 |
| 9666 | 484 | G484E | A | 3.3e-06 | 3.3e-06 | 1.2e-06 | 2.600E-06 | 1.212E-06 | 1.200E-06 | 3.300E-06 |
| 9667 | 484 | G484Q | A | 2e-06 | 1.7e-06 | 1.5e-06 | 1.733E-06 | 2.517E-07 | 1.500E-06 | 2.000E-06 |
| 9668 | 484 | G484G | A | 2.6e-06 | 2.6e-06 | 1e-06 | 2.067E-06 | 9.238E-07 | 1.000E-06 | 2.600E-06 |
| 9669 | 484 | G484H | A | 4.1e-06 | 2.6e-06 | 2.2e-06 | 2.967E-06 | 1.002E-06 | 2.200E-06 | 4.100E-06 |
| 9670 | 484 | G484I | A | 2.3e-06 | 2e-06 | 2.2e-06 | 2.167E-06 | 1.528E-07 | 2.000E-06 | 2.300E-06 |
| 9671 | 484 | G484L | A | 1.7e-06 | 4.2e-06 | 2e-06 | 2.633E-06 | 1.365E-06 | 1.700E-06 | 4.200E-06 |
| 9672 | 484 | G484K | A | 2.9e-06 | 2.5e-06 | 2.4e-06 | 2.600E-06 | 2.646E-07 | 2.400E-06 | 2.900E-06 |
| 9673 | 484 | G484M | A | 2.2e-06 | 2.6e-06 | 2.8e-06 | 2.533E-06 | 3.055E-07 | 2.200E-06 | 2.800E-06 |
| 9674 | 484 | G484F | A | 3.2e-06 | 2.3e-06 | 3.2e-06 | 2.900E-06 | 5.196E-07 | 2.300E-06 | 3.200E-06 |
| 9675 | 484 | G484P | A | 4.3e-06 | 2.4e-06 | 2.1e-06 | 2.933E-06 | 1.193E-06 | 2.100E-06 | 4.300E-06 |
| 9676 | 484 | G484S | A | 3.1e-06 | 1.6e-06 | 1.7e-06 | 2.133E-06 | 8.386E-07 | 1.600E-06 | 3.100E-06 |
| 9677 | 484 | G484T | A | 2.6e-06 | 2.2e-06 | 3.7e-06 | 2.833E-06 | 7.767E-07 | 2.200E-06 | 3.700E-06 |
| 9678 | 484 | G484W | A | 2.8e-06 | 3.8e-06 | 2.4e-06 | 3.000E-06 | 7.211E-07 | 2.400E-06 | 3.800E-06 |
| 9679 | 484 | G484Y | A | 4e-06 | 4.8e-06 | 3e-06 | 3.933E-06 | 9.018E-07 | 3.000E-06 | 4.800E-06 |
| 9680 | 484 | G484V | A | 9.7e-07 | 1.5e-06 | 3.1e-06 | 1.857E-06 | 1.109E-06 | 9.700E-07 | 3.100E-06 |
| 9681 | 485 | E485A | A | 2.6e-06 | 1.7e-06 | 1e-06 | 1.767E-06 | 8.021E-07 | 1.000E-06 | 2.600E-06 |
| 9682 | 485 | E485R | A | 2.1e-06 | 2.6e-06 | 1.8e-06 | 2.167E-06 | 4.041E-07 | 1.800E-06 | 2.600E-06 |
| 9683 | 485 | E485N | A | 1.9e-06 | 2e-06 | 3.6e-06 | 2.500E-06 | 9.539E-07 | 1.900E-06 | 3.600E-06 |
| 9684 | 485 | E485D | A | 4.4e-06 | 2.5e-06 | 3.1e-06 | 3.333E-06 | 9.713E-07 | 2.500E-06 | 4.400E-06 |
| 9685 | 485 | E485C | A | 3.7e-06 | 2.9e-06 | 1.9e-06 | 2.833E-06 | 9.018E-07 | 1.900E-06 | 3.700E-06 |
| 9686 | 485 | E485E | A | 1.3e-06 | 1.4e-06 | 2.2e-06 | 1.633E-06 | 4.933E-07 | 1.300E-06 | 2.200E-06 |
| 9687 | 485 | E485Q | A | 2e-06 | 1.6e-06 | 2.3e-06 | 1.967E-06 | 3.512E-07 | 1.600E-06 | 2.300E-06 |
| 9688 | 485 | E485G | A | 1.6e-06 | 2.3e-06 | 1.7e-06 | 1.867E-06 | 3.786E-07 | 1.600E-06 | 2.300E-06 |
| 9689 | 485 | E485H | A | 3e-06 | 3.3e-06 | 2.1e-06 | 2.800E-06 | 6.245E-07 | 2.100E-06 | 3.300E-06 |

| | | | | | | | | | | |
|---|---|---|---|---|---|---|---|---|---|---|
| 9690 | 485 | E485I | A | 2.7e-06 | 4.5e-06 | 2.6e-06 | 3.267E-06 | 1.069E-06 | 2.600E-06 | 4.500E-06 |
| 9691 | 485 | E485L | A | 2.1e-06 | 4.4e-06 | 2.5e-06 | 3.000E-06 | 1.229E-06 | 2.100E-06 | 4.400E-06 |
| 9692 | 485 | E485K | A | 2.5e-06 | 1.8e-06 | 2.4e-06 | 2.233E-06 | 3.786E-07 | 1.800E-06 | 2.500E-06 |
| 9693 | 485 | E485M | A | 1.5e-06 | 2e-06 | 3.3e-06 | 2.267E-06 | 9.292E-07 | 1.500E-06 | 3.300E-06 |
| 9694 | 485 | E485F | A | 2.3e-06 | 2.1e-06 | 2.3e-06 | 2.233E-06 | 1.155E-07 | 2.100E-06 | 2.300E-06 |
| 9695 | 485 | E485P | A | 2.3e-06 | 4.2e-06 | 1.9e-06 | 2.800E-06 | 1.229E-06 | 1.900E-06 | 4.200E-06 |
| 9696 | 485 | E485S | A | 2.3e-06 | 2.6e-06 | 2e-06 | 2.300E-06 | 3.000E-07 | 2.000E-06 | 2.600E-06 |
| 9697 | 485 | E485T | A | 1.8e-06 | 2.7e-06 | 2.1e-06 | 2.200E-06 | 4.583E-07 | 1.800E-06 | 2.700E-06 |
| 9698 | 485 | E485W | A | 1.2e-06 | 1e-06 | 1.9e-06 | 1.367E-06 | 4.726E-07 | 1.000E-06 | 1.900E-06 |
| 9699 | 485 | E485Y | A | 2.1e-06 | 2.8e-06 | 2.5e-06 | 2.467E-06 | 3.512E-07 | 2.100E-06 | 2.800E-06 |
| 9700 | 485 | E485V | A | 3.6e-06 | 4.8e-06 | 2.5e-06 | 3.633E-06 | 1.150E-06 | 2.500E-06 | 4.800E-06 |
| 9701 | 486 | G486A | A | 1.4e-06 | 4.5e-06 | 2.3e-06 | 2.733E-06 | 1.595E-06 | 1.400E-06 | 4.500E-06 |
| 9702 | 486 | G486R | A | 2.5e-06 | 3e-06 | 3e-06 | 2.833E-06 | 2.887E-07 | 2.500E-06 | 3.000E-06 |
| 9703 | 486 | G486N | A | 2.9e-06 | 2.3e-06 | 2.1e-06 | 2.433E-06 | 4.163E-07 | 2.100E-06 | 2.900E-06 |
| 9704 | 486 | G486D | A | 3.2e-06 | 1.4e-06 | 3e-06 | 2.533E-06 | 9.866E-07 | 1.400E-06 | 3.200E-06 |
| 9705 | 486 | G486C | A | 2.9e-06 | 2.4e-06 | 1.3e-06 | 2.200E-06 | 8.185E-07 | 1.300E-06 | 2.900E-06 |
| 9706 | 486 | G486E | A | 1.8e-06 | 2.1e-06 | 2.9e-06 | 2.267E-06 | 5.686E-07 | 1.800E-06 | 2.900E-06 |
| 9707 | 486 | G486Q | A | 1.8e-06 | 2.1e-06 | 4.1e-06 | 2.667E-06 | 1.250E-06 | 1.800E-06 | 4.100E-06 |
| 9708 | 486 | G486G | A | 2.3e-06 | 1.8e-06 | 2.2e-06 | 2.100E-06 | 2.646E-07 | 1.800E-06 | 2.300E-06 |
| 9709 | 486 | G486H | A | 2.1e-06 | 2.6e-06 | 2.4e-06 | 2.367E-06 | 2.517E-07 | 2.100E-06 | 2.600E-06 |
| 9710 | 486 | G486I | A | 4.4e-06 | 2.5e-06 | 4.4e-06 | 3.767E-06 | 1.097E-06 | 2.500E-06 | 4.400E-06 |
| 9711 | 486 | G486L | A | 3.7e-06 | 1.4e-06 | 2.5e-06 | 2.533E-06 | 1.150E-06 | 1.400E-06 | 3.700E-06 |
| 9712 | 486 | G486K | A | 2.3e-06 | 2.2e-06 | 4.2e-06 | 2.900E-06 | 1.127E-06 | 2.200E-06 | 4.200E-06 |
| 9713 | 486 | G486M | A | 3e-06 | 2.3e-06 | 2.6e-06 | 2.633E-06 | 3.512E-07 | 2.300E-06 | 3.000E-06 |
| 9714 | 486 | G486F | A | 2.1e-06 | 1.2e-06 | 3.6e-06 | 2.300E-06 | 1.212E-06 | 1.200E-06 | 3.600E-06 |
| 9715 | 486 | G486P | A | 1.9e-06 | 1.9e-06 | 4.3e-06 | 2.700E-06 | 1.386E-06 | 1.900E-06 | 4.300E-06 |
| 9716 | 486 | G486S | A | 2.2e-06 | 2.4e-06 | 2.4e-06 | 2.333E-06 | 1.155E-07 | 2.200E-06 | 2.400E-06 |
| 9717 | 486 | G486T | A | 2.1e-06 | 2.6e-06 | 4.3e-06 | 3.000E-06 | 1.153E-06 | 2.100E-06 | 4.300E-06 |
| 9718 | 486 | G486W | A | 2.3e-06 | 1.5e-06 | 1.3e-06 | 1.700E-06 | 5.292E-07 | 1.300E-06 | 2.300E-06 |
| 9719 | 486 | G486Y | A | 2e-06 | 3.2e-06 | 2.5e-06 | 2.567E-06 | 6.028E-07 | 2.000E-06 | 3.200E-06 |
| 9720 | 486 | G486V | A | 3e-06 | 2.7e-06 | 2e-06 | 2.567E-06 | 5.132E-07 | 2.000E-06 | 3.000E-06 |
| 9721 | 487 | L487A | A | 3e-06 | 3e-06 | 1.3e-06 | 2.433E-06 | 9.815E-07 | 1.300E-06 | 3.000E-06 |
| 9722 | 487 | L487R | A | 2.3e-06 | 2.4e-06 | 2e-06 | 2.233E-06 | 2.082E-07 | 2.000E-06 | 2.400E-06 |
| 9723 | 487 | L487N | A | 8.5e-07 | 2.3e-06 | 2.6e-06 | 1.917E-06 | 9.359E-07 | 8.500E-07 | 2.600E-06 |
| 9724 | 487 | L487D | A | 2.8e-06 | 1.3e-06 | 1.3e-06 | 1.800E-06 | 8.660E-07 | 1.300E-06 | 2.800E-06 |
| 9725 | 487 | L487C | A | 2e-06 | 3.4e-06 | 1.3e-06 | 2.233E-06 | 1.069E-06 | 1.300E-06 | 3.400E-06 |
| 9726 | 487 | L487E | A | 1.6e-06 | 1.8e-06 | 2.7e-06 | 2.033E-06 | 5.859E-07 | 1.600E-06 | 2.700E-06 |
| 9727 | 487 | L487Q | A | 3.3e-06 | 3.2e-06 | 1.6e-06 | 2.700E-06 | 9.539E-07 | 1.600E-06 | 3.300E-06 |
| 9728 | 487 | L487G | A | 2.2e-06 | 1.8e-06 | 3.4e-06 | 2.467E-06 | 8.327E-07 | 1.800E-06 | 3.400E-06 |
| 9729 | 487 | L487H | A | 1.7e-06 | 2.2e-06 | 2.6e-06 | 2.167E-06 | 4.509E-07 | 1.700E-06 | 2.600E-06 |
| 9730 | 487 | L487I | A | 2.8e-06 | 2.7e-06 | 1.6e-06 | 2.367E-06 | 6.658E-07 | 1.600E-06 | 2.800E-06 |
| 9731 | 487 | L487L | A | 2.1e-06 | 3.6e-06 | 3.3e-06 | 3.000E-06 | 7.937E-07 | 2.100E-06 | 3.600E-06 |
| 9732 | 487 | L487K | A | 1.8e-06 | 3.5e-06 | 5.4e-06 | 3.567E-06 | 1.801E-06 | 1.800E-06 | 5.400E-06 |
| 9733 | 487 | L487M | A | 1.3e-06 | 3.1e-06 | 2.7e-06 | 2.367E-06 | 9.452E-07 | 1.300E-06 | 3.100E-06 |
| 9734 | 487 | L487F | A | 2.1e-06 | 2.1e-06 | 2e-06 | 2.067E-06 | 5.774E-08 | 2.000E-06 | 2.100E-06 |
| 9735 | 487 | L487P | A | 2.1e-06 | 2.3e-06 | 2.2e-06 | 2.200E-06 | 1.000E-07 | 2.100E-06 | 2.300E-06 |
| 9736 | 487 | L487S | A | 3.7e-06 | 3e-06 | 2.2e-06 | 2.967E-06 | 7.506E-07 | 2.200E-06 | 3.700E-06 |
| 9737 | 487 | L487T | A | 2.9e-06 | 3.2e-06 | 2.6e-06 | 2.900E-06 | 3.000E-07 | 2.600E-06 | 3.200E-06 |
| 9738 | 487 | L487W | A | 3e-06 | 2.8e-06 | 1.4e-06 | 2.400E-06 | 8.718E-07 | 1.400E-06 | 3.000E-06 |
| 9739 | 487 | L487Y | A | 1e-06 | 2e-06 | 2e-06 | 1.667E-06 | 5.774E-07 | 1.000E-06 | 2.000E-06 |
| 9740 | 487 | L487V | A | 1.9e-06 | 2.9e-06 | 2.8e-06 | 2.533E-06 | 5.508E-07 | 1.900E-06 | 2.900E-06 |
| 9741 | 488 | A488A | A | 2.2e-06 | 4.4e-06 | 4e-06 | 3.533E-06 | 1.172E-06 | 2.200E-06 | 4.400E-06 |
| 9742 | 488 | A488R | A | 1.7e-06 | 3.8e-06 | 1.5e-06 | 2.333E-06 | 1.274E-06 | 1.500E-06 | 3.800E-06 |
| 9743 | 488 | A488N | A | 2.8e-06 | 1.6e-06 | 2.1e-06 | 2.167E-06 | 6.028E-07 | 1.600E-06 | 2.800E-06 |
| 9744 | 488 | A488D | A | 2.2e-06 | 2.4e-06 | 2.2e-06 | 2.267E-06 | 1.155E-07 | 2.200E-06 | 2.400E-06 |
| 9745 | 488 | A488C | A | 2.2e-06 | 1.4e-06 | 2.6e-06 | 2.067E-06 | 6.110E-07 | 1.400E-06 | 2.600E-06 |
| 9746 | 488 | A488E | A | 1e-06 | 7e-06 | 3.4e-06 | 3.800E-06 | 3.020E-06 | 1.000E-06 | 7.000E-06 |
| 9747 | 488 | A488Q | A | 2.2e-06 | 2.4e-06 | 4.3e-06 | 2.967E-06 | 1.159E-06 | 2.200E-06 | 4.300E-06 |
| 9748 | 488 | A488G | A | 3e-06 | 2.3e-06 | 2.1e-06 | 2.467E-06 | 4.726E-07 | 2.100E-06 | 3.000E-06 |
| 9749 | 488 | A488H | A | 3.5e-06 | 2.3e-06 | 3.7e-06 | 3.167E-06 | 7.572E-07 | 2.300E-06 | 3.700E-06 |
| 9750 | 488 | A488I | A | 2.9e-06 | 2.6e-06 | 1.5e-06 | 2.333E-06 | 7.371E-07 | 1.500E-06 | 2.900E-06 |
| 9751 | 488 | A488L | A | 3.6e-06 | 2.6e-06 | 3.8e-06 | 3.333E-06 | 6.429E-07 | 2.600E-06 | 3.800E-06 |
| 9752 | 488 | A488K | A | 1.4e-06 | 1.5e-06 | 2.3e-06 | 1.733E-06 | 4.933E-07 | 1.400E-06 | 2.300E-06 |
| 9753 | 488 | A488M | A | 2.6e-06 | 1.3e-06 | 2.6e-06 | 2.167E-06 | 7.506E-07 | 1.300E-06 | 2.600E-06 |
| 9754 | 488 | A488F | A | 1.6e-06 | 3.8e-06 | 3.4e-06 | 2.933E-06 | 1.172E-06 | 1.600E-06 | 3.800E-06 |
| 9755 | 488 | A488P | A | 2.4e-06 | 2.1e-06 | 1.5e-06 | 2.000E-06 | 4.583E-07 | 1.500E-06 | 2.400E-06 |
| 9756 | 488 | A488S | A | 2.9e-06 | 2.3e-06 | 7.5e-07 | 1.983E-06 | 1.109E-06 | 7.500E-07 | 2.900E-06 |
| 9757 | 488 | A488T | A | 2e-06 | 1.7e-06 | 3.8e-06 | 2.500E-06 | 1.136E-06 | 1.700E-06 | 3.800E-06 |
| 9758 | 488 | A488W | A | 1.3e-06 | 2.2e-06 | 1.5e-06 | 1.667E-06 | 4.726E-07 | 1.300E-06 | 2.200E-06 |
| 9759 | 488 | A488Y | A | 3.3e-06 | 3.9e-06 | 2.7e-06 | 3.300E-06 | 6.000E-07 | 2.700E-06 | 3.900E-06 |
| 9760 | 488 | A488V | A | 2.1e-06 | 1.8e-06 | 2e-06 | 1.967E-06 | 1.528E-07 | 1.800E-06 | 2.100E-06 |
| 9761 | 489 | C489A | A | 2.6e-06 | 3.4e-06 | 2.6e-06 | 2.867E-06 | 4.619E-07 | 2.600E-06 | 3.400E-06 |
| 9762 | 489 | C489R | A | 1.8e-06 | 2.1e-06 | 2.2e-06 | 2.033E-06 | 2.082E-07 | 1.800E-06 | 2.200E-06 |
| 9763 | 489 | C489N | A | 1.5e-06 | 2.2e-06 | 1.7e-06 | 1.800E-06 | 3.606E-07 | 1.500E-06 | 2.200E-06 |
| 9764 | 489 | C489D | A | 3.6e-06 | 2.1e-06 | 1.8e-06 | 2.500E-06 | 9.644E-07 | 1.800E-06 | 3.600E-06 |
| 9765 | 489 | C489C | A | 3.3e-06 | 2.7e-06 | 1.9e-06 | 2.633E-06 | 7.024E-07 | 1.900E-06 | 3.300E-06 |
| 9766 | 489 | C489E | A | 2.4e-06 | 1.7e-06 | 1.6e-06 | 1.900E-06 | 4.359E-07 | 1.600E-06 | 2.400E-06 |
| 9767 | 489 | C489Q | A | 2.3e-06 | 3.8e-06 | 5e-06 | 3.700E-06 | 1.353E-06 | 2.300E-06 | 5.000E-06 |
| 9768 | 489 | C489G | A | 2.4e-06 | 1.5e-06 | 3.1e-06 | 2.333E-06 | 8.021E-07 | 1.500E-06 | 3.100E-06 |
| 9769 | 489 | C489H | A | 3.3e-06 | 2.7e-06 | 2.5e-06 | 2.833E-06 | 4.163E-07 | 2.500E-06 | 3.300E-06 |
| 9770 | 489 | C489I | A | 2.4e-06 | 2.3e-06 | 1.8e-06 | 2.167E-06 | 3.215E-07 | 1.800E-06 | 2.400E-06 |
| 9771 | 489 | C489L | A | 3.1e-06 | 2.7e-06 | 4e-06 | 3.267E-06 | 6.658E-07 | 2.700E-06 | 4.000E-06 |
| 9772 | 489 | C489K | A | 2.6e-06 | 2.9e-06 | 1e-06 | 2.167E-06 | 1.021E-06 | 1.000E-06 | 2.900E-06 |
| 9773 | 489 | C489M | A | 2.1e-06 | 1e-06 | 2.3e-06 | 1.800E-06 | 7.000E-07 | 1.000E-06 | 2.300E-06 |
| 9774 | 489 | C489F | A | 4.5e-06 | 3.4e-06 | 2.1e-06 | 3.333E-06 | 1.201E-06 | 2.100E-06 | 4.500E-06 |

| | | | | | | | | | | |
|---|---|---|---|---|---|---|---|---|---|---|
| 9775 | 489 | C489P | A | 3e-06 | 1.1e-06 | 2.3e-06 | 2.133E-06 | 9.609E-07 | 1.100E-06 | 3.000E-06 |
| 9776 | 489 | C489S | A | 2.2e-06 | 3e-06 | 3.8e-06 | 3.000E-06 | 8.000E-07 | 2.200E-06 | 3.800E-06 |
| 9777 | 489 | C489T | A | 2e-06 | 2.5e-06 | 3.3e-06 | 2.600E-06 | 6.557E-07 | 2.000E-06 | 3.300E-06 |
| 9778 | 489 | C489W | A | 2e-06 | 3.4e-06 | 7e-06 | 4.133E-06 | 2.579E-06 | 2.000E-06 | 7.000E-06 |
| 9779 | 489 | C489Y | A | 2.3e-06 | 3.6e-06 | 2.3e-06 | 2.733E-06 | 7.506E-07 | 2.300E-06 | 3.600E-06 |
| 9780 | 489 | C489V | A | 2.6e-06 | 2e-06 | 4.4e-06 | 3.000E-06 | 1.249E-06 | 2.000E-06 | 4.400E-06 |
| 9781 | 490 | H490A | A | 2.7e-06 | 4.7e-06 | 2.2e-06 | 3.200E-06 | 1.323E-06 | 2.200E-06 | 4.700E-06 |
| 9782 | 490 | H490R | A | 1.3e-06 | 1.6e-06 | 2.6e-06 | 1.833E-06 | 6.807E-07 | 1.300E-06 | 2.600E-06 |
| 9783 | 490 | H490N | A | 3.5e-06 | 1.6e-06 | 2.5e-06 | 2.533E-06 | 9.504E-07 | 1.600E-06 | 3.500E-06 |
| 9784 | 490 | H490D | A | 3.2e-06 | 2.5e-06 | 2.3e-06 | 2.667E-06 | 4.726E-07 | 2.300E-06 | 3.200E-06 |
| 9785 | 490 | H490C | A | 2.8e-06 | 5.3e-06 | 3.6e-06 | 3.900E-06 | 1.277E-06 | 2.800E-06 | 5.300E-06 |
| 9786 | 490 | H490E | A | 2.5e-06 | 2e-06 | 3.1e-06 | 2.533E-06 | 5.508E-07 | 2.000E-06 | 3.100E-06 |
| 9787 | 490 | H490Q | A | 2.3e-06 | 2.5e-06 | 1.7e-06 | 2.167E-06 | 4.163E-07 | 1.700E-06 | 2.500E-06 |
| 9788 | 490 | H490G | A | 1.5e-06 | 2.7e-06 | 3.4e-06 | 2.533E-06 | 9.609E-07 | 1.500E-06 | 3.400E-06 |
| 9789 | 490 | H490H | A | 5.2e-06 | 2.8e-06 | 3.1e-06 | 3.700E-06 | 1.308E-06 | 2.800E-06 | 5.200E-06 |
| 9790 | 490 | H490I | A | 1.9e-06 | 2.9e-06 | 1.7e-06 | 2.167E-06 | 6.429E-07 | 1.700E-06 | 2.900E-06 |
| 9791 | 490 | H490L | A | 1.6e-06 | 5.1e-06 | 3.3e-06 | 3.333E-06 | 1.750E-06 | 1.600E-06 | 5.100E-06 |
| 9792 | 490 | H490K | A | 2.9e-06 | 2.7e-06 | 2.7e-06 | 2.767E-06 | 1.155E-07 | 2.700E-06 | 2.900E-06 |
| 9793 | 490 | H490M | A | 2.4e-06 | 2.9e-06 | 2.6e-06 | 2.633E-06 | 2.517E-07 | 2.400E-06 | 2.900E-06 |
| 9794 | 490 | H490F | A | 2.4e-06 | 3.6e-06 | 3.9e-06 | 3.300E-06 | 7.937E-07 | 2.400E-06 | 3.900E-06 |
| 9795 | 490 | H490P | A | 1.7e-06 | 2e-06 | 3.4e-06 | 2.367E-06 | 9.074E-07 | 1.700E-06 | 3.400E-06 |
| 9796 | 490 | H490S | A | 2.1e-06 | 2.2e-06 | 1.3e-06 | 1.867E-06 | 4.933E-07 | 1.300E-06 | 2.200E-06 |
| 9797 | 490 | H490T | A | 1.8e-06 | 2.5e-06 | 2.8e-06 | 2.367E-06 | 5.132E-07 | 1.800E-06 | 2.800E-06 |
| 9798 | 490 | H490W | A | 3.2e-06 | 2.3e-06 | 2.6e-06 | 2.700E-06 | 4.583E-07 | 2.300E-06 | 3.200E-06 |
| 9799 | 490 | H490Y | A | 2.2e-06 | 2.5e-06 | 1.1e-06 | 1.933E-06 | 7.371E-07 | 1.100E-06 | 2.500E-06 |
| 9800 | 490 | H490V | A | 3e-06 | 1.6e-06 | 2e-06 | 2.200E-06 | 7.211E-07 | 1.600E-06 | 3.000E-06 |
| 9801 | 491 | Q491A | A | 1.9e-06 | 4.9e-06 | 2.2e-06 | 3.000E-06 | 1.652E-06 | 1.900E-06 | 4.900E-06 |
| 9802 | 491 | Q491R | A | 2.4e-06 | 4.3e-06 | 2.9e-06 | 3.200E-06 | 9.849E-07 | 2.400E-06 | 4.300E-06 |
| 9803 | 491 | Q491N | A | 4.9e-06 | 2.5e-06 | 2.1e-06 | 3.167E-06 | 1.514E-06 | 2.100E-06 | 4.900E-06 |
| 9804 | 491 | Q491D | A | 2.4e-06 | 1.7e-06 | 2.9e-06 | 2.333E-06 | 6.028E-07 | 1.700E-06 | 2.900E-06 |
| 9805 | 491 | Q491C | A | 4.1e-06 | 1.5e-06 | 1.6e-06 | 2.400E-06 | 1.473E-06 | 1.500E-06 | 4.100E-06 |
| 9806 | 491 | Q491E | A | 2.3e-06 | 2.3e-06 | 1.9e-06 | 2.167E-06 | 2.309E-07 | 1.900E-06 | 2.300E-06 |
| 9807 | 491 | Q491Q | A | 4.3e-06 | 3.2e-06 | 2.6e-06 | 3.367E-06 | 8.622E-07 | 2.600E-06 | 4.300E-06 |
| 9808 | 491 | Q491G | A | 1.8e-06 | 2.6e-06 | 3.2e-06 | 2.533E-06 | 7.024E-07 | 1.800E-06 | 3.200E-06 |
| 9809 | 491 | Q491H | A | 4.3e-06 | 1.6e-06 | 2.8e-06 | 2.900E-06 | 1.353E-06 | 1.600E-06 | 4.300E-06 |
| 9810 | 491 | Q491I | A | 2e-06 | 7.8e-07 | 2.7e-06 | 1.827E-06 | 9.717E-07 | 7.800E-07 | 2.700E-06 |
| 9811 | 491 | Q491L | A | 2.5e-06 | 3.2e-06 | 2.7e-06 | 2.800E-06 | 3.606E-07 | 2.500E-06 | 3.200E-06 |
| 9812 | 491 | Q491K | A | 1.7e-06 | 3.1e-06 | 2.6e-06 | 2.467E-06 | 7.095E-07 | 1.700E-06 | 3.100E-06 |
| 9813 | 491 | Q491M | A | 1.7e-06 | 2.3e-06 | 2.6e-06 | 2.200E-06 | 4.583E-07 | 1.700E-06 | 2.600E-06 |
| 9814 | 491 | Q491F | A | 4.1e-06 | 3.6e-06 | 2e-06 | 3.233E-06 | 1.097E-06 | 2.000E-06 | 4.100E-06 |
| 9815 | 491 | Q491P | A | 3.5e-06 | 3.6e-06 | 2e-06 | 3.033E-06 | 8.963E-07 | 2.000E-06 | 3.600E-06 |
| 9816 | 491 | Q491S | A | 2.7e-06 | 1.8e-06 | 2.8e-06 | 2.433E-06 | 5.508E-07 | 1.800E-06 | 2.800E-06 |
| 9817 | 491 | Q491T | A | 1.7e-06 | 2.1e-06 | 4.2e-06 | 2.667E-06 | 1.343E-06 | 1.700E-06 | 4.200E-06 |
| 9818 | 491 | Q491W | A | 2.2e-06 | 2.5e-06 | 2.1e-06 | 2.267E-06 | 2.082E-07 | 2.100E-06 | 2.500E-06 |
| 9819 | 491 | Q491Y | A | 3.2e-06 | 1.8e-06 | 3.2e-06 | 2.733E-06 | 8.083E-07 | 1.800E-06 | 3.200E-06 |
| 9820 | 491 | Q491V | A | 3.7e-06 | 2.1e-06 | 2.9e-06 | 2.900E-06 | 8.000E-07 | 2.100E-06 | 3.700E-06 |
| 9821 | 492 | L492A | A | 1.7e-06 | 1.1e-06 | 1.8e-06 | 1.533E-06 | 3.786E-07 | 1.100E-06 | 1.800E-06 |
| 9822 | 492 | L492R | A | 3e-06 | 2e-06 | 1.5e-06 | 2.167E-06 | 7.638E-07 | 1.500E-06 | 3.000E-06 |
| 9823 | 492 | L492N | A | 3.8e-06 | 2.3e-06 | 2.3e-06 | 2.800E-06 | 8.660E-07 | 2.300E-06 | 3.800E-06 |
| 9824 | 492 | L492D | A | 1.8e-06 | 1.6e-06 | 1.3e-06 | 1.567E-06 | 2.517E-07 | 1.300E-06 | 1.800E-06 |
| 9825 | 492 | L492C | A | 1.3e-06 | 3.8e-06 | 2.5e-06 | 2.533E-06 | 1.250E-06 | 1.300E-06 | 3.800E-06 |
| 9826 | 492 | L492E | A | 2.2e-06 | 2.6e-06 | 5.1e-06 | 3.300E-06 | 1.572E-06 | 2.200E-06 | 5.100E-06 |
| 9827 | 492 | L492Q | A | 4e-06 | 2.3e-06 | 2.6e-06 | 2.967E-06 | 9.074E-07 | 2.300E-06 | 4.000E-06 |
| 9828 | 492 | L492G | A | 1.8e-06 | 3e-06 | 3.1e-06 | 2.633E-06 | 7.234E-07 | 1.800E-06 | 3.100E-06 |
| 9829 | 492 | L492H | A | 1.9e-06 | 2.9e-06 | 1.5e-06 | 2.100E-06 | 7.211E-07 | 1.500E-06 | 2.900E-06 |
| 9830 | 492 | L492I | A | 3.3e-06 | 2e-06 | 2.4e-06 | 2.567E-06 | 6.658E-07 | 2.000E-06 | 3.300E-06 |
| 9831 | 492 | L492L | A | 2.2e-06 | 2.8e-06 | 4.4e-06 | 3.133E-06 | 1.137E-06 | 2.200E-06 | 4.400E-06 |
| 9832 | 492 | L492K | A | 1.8e-06 | 1.5e-06 | 2.1e-06 | 1.800E-06 | 3.000E-07 | 1.500E-06 | 2.100E-06 |
| 9833 | 492 | L492M | A | 2e-06 | 2.1e-06 | 3.2e-06 | 2.433E-06 | 6.658E-07 | 2.000E-06 | 3.200E-06 |
| 9834 | 492 | L492F | A | 2.6e-06 | 1.6e-06 | 2e-06 | 2.067E-06 | 5.033E-07 | 1.600E-06 | 2.600E-06 |
| 9835 | 492 | L492P | A | 1.2e-06 | 1.5e-06 | 2.1e-06 | 1.600E-06 | 4.583E-07 | 1.200E-06 | 2.100E-06 |
| 9836 | 492 | L492S | A | 3.4e-06 | 2.4e-06 | 2.5e-06 | 2.767E-06 | 5.508E-07 | 2.400E-06 | 3.400E-06 |
| 9837 | 492 | L492T | A | 2.2e-06 | 2.5e-06 | 1.7e-06 | 2.133E-06 | 4.041E-07 | 1.700E-06 | 2.500E-06 |
| 9838 | 492 | L492W | A | 1.6e-06 | 1.7e-06 | 4e-06 | 2.433E-06 | 1.358E-06 | 1.600E-06 | 4.000E-06 |
| 9839 | 492 | L492Y | A | 2.1e-06 | 1.8e-06 | 3.2e-06 | 2.367E-06 | 7.371E-07 | 1.800E-06 | 3.200E-06 |
| 9840 | 492 | L492V | A | 4.3e-06 | 2.2e-06 | 2.8e-06 | 3.100E-06 | 1.082E-06 | 2.200E-06 | 4.300E-06 |
| 9841 | 493 | C493A | A | 2.8e-06 | 2.5e-06 | 1.9e-06 | 2.400E-06 | 4.583E-07 | 1.900E-06 | 2.800E-06 |
| 9842 | 493 | C493R | A | 2.7e-06 | 1.7e-06 | 1.9e-06 | 2.100E-06 | 5.292E-07 | 1.700E-06 | 2.700E-06 |
| 9843 | 493 | C493N | A | 2.9e-06 | 1.9e-06 | 3.2e-06 | 2.667E-06 | 6.807E-07 | 1.900E-06 | 3.200E-06 |
| 9844 | 493 | C493D | A | 4.4e-06 | 3.3e-06 | 2.3e-06 | 3.333E-06 | 1.050E-06 | 2.300E-06 | 4.400E-06 |
| 9845 | 493 | C493C | A | 3.6e-06 | 2e-06 | 1.9e-06 | 2.500E-06 | 9.539E-07 | 1.900E-06 | 3.600E-06 |
| 9846 | 493 | C493E | A | 3.2e-06 | 1.1e-06 | 2.3e-06 | 2.200E-06 | 1.054E-06 | 1.100E-06 | 3.200E-06 |
| 9847 | 493 | C493Q | A | 1.7e-06 | 3e-06 | 3e-06 | 2.567E-06 | 7.506E-07 | 1.700E-06 | 3.000E-06 |
| 9848 | 493 | C493G | A | 3.9e-06 | 3.8e-06 | 2e-06 | 3.233E-06 | 1.069E-06 | 2.000E-06 | 3.900E-06 |
| 9849 | 493 | C493H | A | 2.2e-06 | 3.6e-06 | 4.2e-06 | 3.333E-06 | 1.026E-06 | 2.200E-06 | 4.200E-06 |
| 9850 | 493 | C493I | A | 3.7e-06 | 1.1e-06 | 5.2e-06 | 3.333E-06 | 2.074E-06 | 1.100E-06 | 5.200E-06 |
| 9851 | 493 | C493L | A | 2.1e-06 | 2.2e-06 | 2.8e-06 | 2.367E-06 | 3.786E-07 | 2.100E-06 | 2.800E-06 |
| 9852 | 493 | C493K | A | 1.8e-06 | 2.4e-06 | 2.6e-06 | 2.267E-06 | 4.163E-07 | 1.800E-06 | 2.600E-06 |
| 9853 | 493 | C493M | A | 2.2e-06 | 2.1e-06 | 1.6e-06 | 1.967E-06 | 3.215E-07 | 1.600E-06 | 2.200E-06 |
| 9854 | 493 | C493F | A | 3.1e-06 | 2.6e-06 | 2.5e-06 | 2.733E-06 | 3.215E-07 | 2.500E-06 | 3.100E-06 |
| 9855 | 493 | C493P | A | 4e-06 | 1.5e-06 | 3.7e-06 | 3.067E-06 | 1.365E-06 | 1.500E-06 | 4.000E-06 |
| 9856 | 493 | C493S | A | 5.1e-06 | 2.4e-06 | 1.4e-06 | 2.967E-06 | 1.914E-06 | 1.400E-06 | 5.100E-06 |
| 9857 | 493 | C493T | A | 3.9e-06 | 2.2e-06 | 1.4e-06 | 2.500E-06 | 1.277E-06 | 1.400E-06 | 3.900E-06 |
| 9858 | 493 | C493W | A | 3e-06 | 1.5e-06 | 1.6e-06 | 2.033E-06 | 8.386E-07 | 1.500E-06 | 3.000E-06 |
| 9859 | 493 | C493Y | A | 3.5e-06 | 2.8e-06 | 2.3e-06 | 2.867E-06 | 6.028E-07 | 2.300E-06 | 3.500E-06 |

| | | | | | | | | | | |
|---|---|---|---|---|---|---|---|---|---|---|
| 9860 | 493 | C493V | A | 2.8e-06 | 2.3e-06 | 2.4e-06 | 2.500E-06 | 2.646E-07 | 2.300E-06 | 2.800E-06 |
| 9861 | 494 | A494A | A | 2.3e-06 | 4.9e-06 | 2.6e-06 | 3.267E-06 | 1.422E-06 | 2.300E-06 | 4.900E-06 |
| 9862 | 494 | A494R | A | 2.5e-06 | 2.2e-06 | 2.4e-06 | 2.367E-06 | 1.528E-07 | 2.200E-06 | 2.500E-06 |
| 9863 | 494 | A494N | A | 9.7e-07 | 3.6e-06 | 3.3e-06 | 2.623E-06 | 1.440E-06 | 9.700E-07 | 3.600E-06 |
| 9864 | 494 | A494D | A | 2.3e-06 | 4e-06 | 1.7e-06 | 2.667E-06 | 1.193E-06 | 1.700E-06 | 4.000E-06 |
| 9865 | 494 | A494C | A | 1.6e-06 | 2.4e-06 | 2.4e-06 | 2.133E-06 | 4.619E-07 | 1.600E-06 | 2.400E-06 |
| 9866 | 494 | A494E | A | 1.7e-06 | 2.8e-06 | 5.2e-06 | 3.233E-06 | 1.790E-06 | 1.700E-06 | 5.200E-06 |
| 9867 | 494 | A494Q | A | 2.5e-06 | 1.9e-06 | 2.9e-06 | 2.433E-06 | 5.033E-07 | 1.900E-06 | 2.900E-06 |
| 9868 | 494 | A494G | A | 5.4e-06 | 5.4e-06 | 1.9e-06 | 4.233E-06 | 2.021E-06 | 1.900E-06 | 5.400E-06 |
| 9869 | 494 | A494H | A | 2e-06 | 1.4e-06 | 9e-07 | 1.433E-06 | 5.508E-07 | 9.000E-07 | 2.000E-06 |
| 9870 | 494 | A494I | A | 3.1e-06 | 2.2e-06 | 2.8e-06 | 2.700E-06 | 4.583E-07 | 2.200E-06 | 3.100E-06 |
| 9871 | 494 | A494L | A | 1.2e-06 | 2.8e-06 | 1.5e-06 | 1.833E-06 | 8.505E-07 | 1.200E-06 | 2.800E-06 |
| 9872 | 494 | A494K | A | 2.2e-06 | 3.1e-06 | 3.5e-06 | 2.933E-06 | 6.658E-07 | 2.200E-06 | 3.500E-06 |
| 9873 | 494 | A494M | A | 2.2e-06 | 3.3e-06 | 3.8e-06 | 3.100E-06 | 8.185E-07 | 2.200E-06 | 3.800E-06 |
| 9874 | 494 | A494F | A | 4.3e-06 | 3.1e-06 | 1.9e-06 | 3.100E-06 | 1.200E-06 | 1.900E-06 | 4.300E-06 |
| 9875 | 494 | A494P | A | 2.1e-06 | 2.1e-06 | 3.5e-06 | 2.567E-06 | 8.083E-07 | 2.100E-06 | 3.500E-06 |
| 9876 | 494 | A494S | A | 2.6e-06 | 2.8e-06 | 3.3e-06 | 2.900E-06 | 3.606E-07 | 2.600E-06 | 3.300E-06 |
| 9877 | 494 | A494T | A | 2.4e-06 | 2e-06 | 2.7e-06 | 2.367E-06 | 3.512E-07 | 2.000E-06 | 2.700E-06 |
| 9878 | 494 | A494W | A | 3.2e-06 | 2.7e-06 | 2.9e-06 | 2.933E-06 | 2.517E-07 | 2.700E-06 | 3.200E-06 |
| 9879 | 494 | A494Y | A | 1.8e-06 | 2.2e-06 | 1.6e-06 | 1.867E-06 | 3.055E-07 | 1.600E-06 | 2.200E-06 |
| 9880 | 494 | A494V | A | 1.4e-06 | 2.5e-06 | 1.4e-06 | 1.767E-06 | 6.351E-07 | 1.400E-06 | 2.500E-06 |
| 9881 | 495 | R495A | A | 2.4e-06 | 2.6e-06 | 4.3e-06 | 3.100E-06 | 1.044E-06 | 2.400E-06 | 4.300E-06 |
| 9882 | 495 | R495R | A | 1.8e-06 | 1.2e-06 | 2.8e-06 | 1.933E-06 | 8.083E-07 | 1.200E-06 | 2.800E-06 |
| 9883 | 495 | R495N | A | 1.7e-06 | 3e-06 | 4.3e-06 | 3.000E-06 | 1.300E-06 | 1.700E-06 | 4.300E-06 |
| 9884 | 495 | R495D | A | 3.6e-06 | 2.3e-06 | 2.7e-06 | 2.867E-06 | 6.658E-07 | 2.300E-06 | 3.600E-06 |
| 9885 | 495 | R495C | A | 1.8e-06 | 2.3e-06 | 2.6e-06 | 2.233E-06 | 4.041E-07 | 1.800E-06 | 2.600E-06 |
| 9886 | 495 | R495E | A | 1.9e-06 | 3e-06 | 2.6e-06 | 2.500E-06 | 5.568E-07 | 1.900E-06 | 3.000E-06 |
| 9887 | 495 | R495Q | A | 3.8e-06 | 2.5e-06 | 2.3e-06 | 2.867E-06 | 8.145E-07 | 2.300E-06 | 3.800E-06 |
| 9888 | 495 | R495G | A | 2.6e-06 | 2e-06 | 2.4e-06 | 2.333E-06 | 3.055E-07 | 2.000E-06 | 2.600E-06 |
| 9889 | 495 | R495H | A | 2.3e-06 | 3.1e-06 | 1.5e-06 | 2.300E-06 | 8.000E-07 | 1.500E-06 | 3.100E-06 |
| 9890 | 495 | R495I | A | 3e-06 | 2.4e-06 | 2.3e-06 | 2.567E-06 | 3.786E-07 | 2.300E-06 | 3.000E-06 |
| 9891 | 495 | R495L | A | 3.7e-06 | 3.6e-06 | 2.5e-06 | 3.267E-06 | 6.658E-07 | 2.500E-06 | 3.700E-06 |
| 9892 | 495 | R495K | A | 1.8e-06 | 1.2e-06 | 2.9e-06 | 1.967E-06 | 8.622E-07 | 1.200E-06 | 2.900E-06 |
| 9893 | 495 | R495M | A | 2.1e-06 | 3.3e-06 | 3.5e-06 | 2.967E-06 | 7.572E-07 | 2.100E-06 | 3.500E-06 |
| 9894 | 495 | R495F | A | 1.8e-06 | 3.2e-06 | 2.5e-06 | 2.500E-06 | 7.000E-07 | 1.800E-06 | 3.200E-06 |
| 9895 | 495 | R495P | A | 9e-07 | 2.3e-06 | 1.4e-06 | 1.533E-06 | 7.095E-07 | 9.000E-07 | 2.300E-06 |
| 9896 | 495 | R495S | A | 1.2e-06 | 1.7e-06 | 2.4e-06 | 1.767E-06 | 6.028E-07 | 1.200E-06 | 2.400E-06 |
| 9897 | 495 | R495T | A | 1.9e-06 | 2.4e-06 | 2.6e-06 | 2.300E-06 | 3.606E-07 | 1.900E-06 | 2.600E-06 |
| 9898 | 495 | R495W | A | 2.3e-06 | 1.4e-06 | 2.8e-06 | 2.167E-06 | 7.095E-07 | 1.400E-06 | 2.800E-06 |
| 9899 | 495 | R495Y | A | 1.2e-06 | 2.7e-06 | 4.2e-06 | 2.700E-06 | 1.500E-06 | 1.200E-06 | 4.200E-06 |
| 9900 | 495 | R495V | A | 2.4e-06 | 1.9e-06 | 1.8e-06 | 2.033E-06 | 3.215E-07 | 1.800E-06 | 2.400E-06 |
| 9901 | 496 | G496A | A | 2.2e-06 | 2.6e-06 | 1.9e-06 | 2.233E-06 | 3.512E-07 | 1.900E-06 | 2.600E-06 |
| 9902 | 496 | G496R | A | 1.6e-06 | 3.3e-06 | 1.9e-06 | 2.267E-06 | 9.074E-07 | 1.600E-06 | 3.300E-06 |
| 9903 | 496 | G496N | A | 2.3e-06 | 4e-06 | 2.3e-06 | 2.867E-06 | 9.815E-07 | 2.300E-06 | 4.000E-06 |
| 9904 | 496 | G496D | A | 4.1e-06 | 1.7e-06 | 3.4e-06 | 3.067E-06 | 1.234E-06 | 1.700E-06 | 4.100E-06 |
| 9905 | 496 | G496C | A | 2.7e-06 | 1.9e-06 | 1.2e-06 | 1.933E-06 | 7.506E-07 | 1.200E-06 | 2.700E-06 |
| 9906 | 496 | G496E | A | 1.9e-06 | 2.1e-06 | 2.3e-06 | 2.100E-06 | 2.000E-07 | 1.900E-06 | 2.300E-06 |
| 9907 | 496 | G496Q | A | 2e-06 | 2.2e-06 | 2.9e-06 | 2.367E-06 | 4.726E-07 | 2.000E-06 | 2.900E-06 |
| 9908 | 496 | G496G | A | 1.9e-06 | 2.6e-06 | 4.4e-06 | 2.967E-06 | 1.290E-06 | 1.900E-06 | 4.400E-06 |
| 9909 | 496 | G496H | A | 2.6e-06 | 2.8e-06 | 2e-06 | 2.467E-06 | 4.163E-07 | 2.000E-06 | 2.800E-06 |
| 9910 | 496 | G496I | A | 2.2e-06 | 2e-06 | 2.1e-06 | 2.100E-06 | 1.000E-07 | 2.000E-06 | 2.200E-06 |
| 9911 | 496 | G496L | A | 2.2e-06 | 2.5e-06 | 3.4e-06 | 2.700E-06 | 6.245E-07 | 2.200E-06 | 3.400E-06 |
| 9912 | 496 | G496K | A | 3e-06 | 2.1e-06 | 3e-06 | 2.700E-06 | 5.196E-07 | 2.100E-06 | 3.000E-06 |
| 9913 | 496 | G496M | A | 2.7e-06 | 2.8e-06 | 2e-06 | 2.500E-06 | 4.359E-07 | 2.000E-06 | 2.800E-06 |
| 9914 | 496 | G496F | A | 3.8e-06 | 2.9e-06 | 2.3e-06 | 3.000E-06 | 7.550E-07 | 2.300E-06 | 3.800E-06 |
| 9915 | 496 | G496P | A | 1.4e-06 | 2.9e-06 | 3.2e-06 | 2.500E-06 | 9.644E-07 | 1.400E-06 | 3.200E-06 |
| 9916 | 496 | G496S | A | 2.6e-06 | 2.1e-06 | 2.8e-06 | 2.500E-06 | 3.606E-07 | 2.100E-06 | 2.800E-06 |
| 9917 | 496 | G496T | A | 3.2e-06 | 1.3e-06 | 2.2e-06 | 2.233E-06 | 9.504E-07 | 1.300E-06 | 3.200E-06 |
| 9918 | 496 | G496W | A | 3.4e-06 | 2.4e-06 | 3.4e-06 | 3.067E-06 | 5.774E-07 | 2.400E-06 | 3.400E-06 |
| 9919 | 496 | G496Y | A | 2e-06 | 2.6e-06 | 2.2e-06 | 2.267E-06 | 3.055E-07 | 2.000E-06 | 2.600E-06 |
| 9920 | 496 | G496V | A | 3.1e-06 | 4.9e-06 | 2e-06 | 3.333E-06 | 1.464E-06 | 2.000E-06 | 4.900E-06 |
| 9921 | 497 | H497A | A | 4.6e-06 | 2e-06 | 2.2e-06 | 2.933E-06 | 1.447E-06 | 2.000E-06 | 4.600E-06 |
| 9922 | 497 | H497R | A | 2.4e-06 | 1.6e-06 | 6.4e-06 | 3.467E-06 | 2.572E-06 | 1.600E-06 | 6.400E-06 |
| 9923 | 497 | H497N | A | 2e-06 | 2.8e-06 | 2.8e-06 | 2.533E-06 | 4.619E-07 | 2.000E-06 | 2.800E-06 |
| 9924 | 497 | H497D | A | 2e-06 | 1.5e-06 | 2.9e-06 | 2.133E-06 | 7.095E-07 | 1.500E-06 | 2.900E-06 |
| 9925 | 497 | H497C | A | 3.7e-06 | 2.2e-06 | 2e-06 | 2.633E-06 | 9.292E-07 | 2.000E-06 | 3.700E-06 |
| 9926 | 497 | H497E | A | 2.5e-06 | 1.6e-06 | 3.2e-06 | 2.433E-06 | 8.021E-07 | 1.600E-06 | 3.200E-06 |
| 9927 | 497 | H497Q | A | 2.3e-06 | 2.6e-06 | 2e-06 | 2.300E-06 | 3.000E-07 | 2.000E-06 | 2.600E-06 |
| 9928 | 497 | H497G | A | 2.6e-06 | 1.5e-06 | 3e-06 | 2.367E-06 | 7.767E-07 | 1.500E-06 | 3.000E-06 |
| 9929 | 497 | H497H | A | 2e-06 | 1.9e-06 | 2.9e-06 | 2.267E-06 | 5.508E-07 | 1.900E-06 | 2.900E-06 |
| 9930 | 497 | H497I | A | 4e-06 | 2.1e-06 | 2.1e-06 | 2.733E-06 | 1.097E-06 | 2.100E-06 | 4.000E-06 |
| 9931 | 497 | H497L | A | 2.3e-06 | 2.7e-06 | 3.1e-06 | 2.700E-06 | 4.000E-07 | 2.300E-06 | 3.100E-06 |
| 9932 | 497 | H497K | A | 2.9e-06 | 3.1e-06 | 3.3e-06 | 3.100E-06 | 2.000E-07 | 2.900E-06 | 3.300E-06 |
| 9933 | 497 | H497M | A | 3.1e-06 | 2.2e-06 | 3.1e-06 | 2.800E-06 | 5.196E-07 | 2.200E-06 | 3.100E-06 |
| 9934 | 497 | H497F | A | 4.3e-06 | 2.6e-06 | 2.5e-06 | 3.133E-06 | 1.012E-06 | 2.500E-06 | 4.300E-06 |
| 9935 | 497 | H497P | A | 2.2e-06 | 2e-06 | 2e-06 | 2.067E-06 | 1.155E-07 | 2.000E-06 | 2.200E-06 |
| 9936 | 497 | H497S | A | 1.4e-06 | 1.7e-06 | 2e-06 | 1.700E-06 | 3.000E-07 | 1.400E-06 | 2.000E-06 |
| 9937 | 497 | H497T | A | 1.9e-06 | 1.9e-06 | 2.5e-06 | 2.100E-06 | 3.464E-07 | 1.900E-06 | 2.500E-06 |
| 9938 | 497 | H497W | A | 3.5e-06 | 4.2e-06 | 3.7e-06 | 3.800E-06 | 3.606E-07 | 3.500E-06 | 4.200E-06 |
| 9939 | 497 | H497Y | A | 2.1e-06 | 3.1e-06 | 1.6e-06 | 2.267E-06 | 7.638E-07 | 1.600E-06 | 3.100E-06 |
| 9940 | 497 | H497V | A | 2.8e-06 | 2e-06 | 3.1e-06 | 2.633E-06 | 5.686E-07 | 2.000E-06 | 3.100E-06 |
| 9941 | 498 | C498A | A | 2.1e-06 | 3.1e-06 | 2.8e-06 | 2.667E-06 | 5.132E-07 | 2.100E-06 | 3.100E-06 |
| 9942 | 498 | C498R | A | 1.4e-06 | 1.8e-06 | 2.8e-06 | 2.000E-06 | 7.211E-07 | 1.400E-06 | 2.800E-06 |
| 9943 | 498 | C498N | A | 3.8e-06 | 2.1e-06 | 1.6e-06 | 2.500E-06 | 1.153E-06 | 1.600E-06 | 3.800E-06 |
| 9944 | 498 | C498D | A | 1.6e-06 | 2.9e-06 | 2.6e-06 | 2.367E-06 | 6.807E-07 | 1.600E-06 | 2.900E-06 |

| | | | | | | | | | | |
|---|---|---|---|---|---|---|---|---|---|---|
| 9945 | 498 | C498C | A | 3.3e-06 | 2.7e-06 | 3.2e-06 | 3.067E-06 | 3.215E-07 | 2.700E-06 | 3.300E-06 |
| 9946 | 498 | C498E | A | 1.6e-06 | 3.1e-06 | 2.6e-06 | 2.433E-06 | 7.638E-07 | 1.600E-06 | 3.100E-06 |
| 9947 | 498 | C498Q | A | 1.7e-06 | 2.5e-06 | 2.4e-06 | 2.200E-06 | 4.359E-07 | 1.700E-06 | 2.500E-06 |
| 9948 | 498 | C498G | A | 3.1e-06 | 2.2e-06 | 2.1e-06 | 2.467E-06 | 5.508E-07 | 2.100E-06 | 3.100E-06 |
| 9949 | 498 | C498H | A | 2.1e-06 | 2.4e-06 | 2.3e-06 | 2.267E-06 | 1.528E-07 | 2.100E-06 | 2.400E-06 |
| 9950 | 498 | C498I | A | 1.3e-06 | 2.6e-06 | 2.2e-06 | 2.033E-06 | 6.658E-07 | 1.300E-06 | 2.600E-06 |
| 9951 | 498 | C498L | A | 1.8e-06 | 2.1e-06 | 1.7e-06 | 1.867E-06 | 2.082E-07 | 1.700E-06 | 2.100E-06 |
| 9952 | 498 | C498K | A | 1.7e-06 | 4.1e-06 | 1.6e-06 | 2.467E-06 | 1.415E-06 | 1.600E-06 | 4.100E-06 |
| 9953 | 498 | C498M | A | 1.3e-06 | 2.2e-06 | 2.3e-06 | 1.933E-06 | 5.508E-07 | 1.300E-06 | 2.300E-06 |
| 9954 | 498 | C498F | A | 3.2e-06 | 1.7e-06 | 1.7e-06 | 2.200E-06 | 8.660E-07 | 1.700E-06 | 3.200E-06 |
| 9955 | 498 | C498P | A | 2.2e-06 | 1.7e-06 | 2.1e-06 | 2.000E-06 | 2.646E-07 | 1.700E-06 | 2.200E-06 |
| 9956 | 498 | C498S | A | 3.6e-06 | 2.5e-06 | 2.1e-06 | 2.733E-06 | 7.767E-07 | 2.100E-06 | 3.600E-06 |
| 9957 | 498 | C498T | A | 2.9e-06 | 3.6e-06 | 1.9e-06 | 2.800E-06 | 8.544E-07 | 1.900E-06 | 3.600E-06 |
| 9958 | 498 | C498W | A | 2.7e-06 | 1.7e-06 | 3.4e-06 | 2.600E-06 | 8.544E-07 | 1.700E-06 | 3.400E-06 |
| 9959 | 498 | C498Y | A | 3.1e-06 | 1.9e-06 | 1.9e-06 | 2.300E-06 | 6.928E-07 | 1.900E-06 | 3.100E-06 |
| 9960 | 498 | C498V | A | 2.7e-06 | 2.4e-06 | 1.9e-06 | 2.333E-06 | 4.041E-07 | 1.900E-06 | 2.700E-06 |
| 9961 | 499 | W499A | A | 3.7e-06 | 1.9e-06 | 2.4e-06 | 2.667E-06 | 9.292E-07 | 1.900E-06 | 3.700E-06 |
| 9962 | 499 | W499R | A | 1.8e-06 | 2.2e-06 | 1.4e-06 | 1.800E-06 | 4.000E-07 | 1.400E-06 | 2.200E-06 |
| 9963 | 499 | W499N | A | 1.6e-06 | 2.4e-06 | 1.4e-06 | 1.800E-06 | 5.292E-07 | 1.400E-06 | 2.400E-06 |
| 9964 | 499 | W499D | A | 2.2e-06 | 3.3e-06 | 2e-06 | 2.500E-06 | 7.000E-07 | 2.000E-06 | 3.300E-06 |
| 9965 | 499 | W499C | A | 2.3e-06 | 2.2e-06 | 2.8e-06 | 2.433E-06 | 3.215E-07 | 2.200E-06 | 2.800E-06 |
| 9966 | 499 | W499E | A | 3.5e-06 | 5.2e-06 | 1.4e-06 | 3.367E-06 | 1.904E-06 | 1.400E-06 | 5.200E-06 |
| 9967 | 499 | W499Q | A | 2.6e-06 | 2.4e-06 | 2.3e-06 | 2.433E-06 | 1.528E-07 | 2.300E-06 | 2.600E-06 |
| 9968 | 499 | W499G | A | 2.2e-06 | 2.4e-06 | 2.4e-06 | 2.333E-06 | 1.155E-07 | 2.200E-06 | 2.400E-06 |
| 9969 | 499 | W499H | A | 2.7e-06 | 2.7e-06 | 3.4e-06 | 2.933E-06 | 4.041E-07 | 2.700E-06 | 3.400E-06 |
| 9970 | 499 | W499I | A | 4e-06 | 3.7e-06 | 2.7e-06 | 3.467E-06 | 6.807E-07 | 2.700E-06 | 4.000E-06 |
| 9971 | 499 | W499L | A | 1.9e-06 | 2.8e-06 | 2.7e-06 | 2.467E-06 | 4.933E-07 | 1.900E-06 | 2.800E-06 |
| 9972 | 499 | W499K | A | 1.8e-06 | 2e-06 | 2.4e-06 | 2.067E-06 | 3.055E-07 | 1.800E-06 | 2.400E-06 |
| 9973 | 499 | W499M | A | 2.3e-06 | 1.3e-06 | 2.4e-06 | 2.000E-06 | 6.083E-07 | 1.300E-06 | 2.400E-06 |
| 9974 | 499 | W499F | A | 2.9e-06 | 1.4e-06 | 3.9e-06 | 2.733E-06 | 1.258E-06 | 1.400E-06 | 3.900E-06 |
| 9975 | 499 | W499P | A | 3.9e-06 | 3.3e-06 | 3.3e-06 | 3.500E-06 | 3.464E-07 | 3.300E-06 | 3.900E-06 |
| 9976 | 499 | W499S | A | 3.3e-06 | 2.7e-06 | 4.3e-06 | 3.433E-06 | 8.083E-07 | 2.700E-06 | 4.300E-06 |
| 9977 | 499 | W499T | A | 1.9e-06 | 3e-06 | 1.8e-06 | 2.233E-06 | 6.658E-07 | 1.800E-06 | 3.000E-06 |
| 9978 | 499 | W499W | A | 1.9e-06 | 1.2e-06 | 3.6e-06 | 2.233E-06 | 1.234E-06 | 1.200E-06 | 3.600E-06 |
| 9979 | 499 | W499Y | A | 2.5e-06 | 2.1e-06 | 2e-06 | 2.200E-06 | 2.646E-07 | 2.000E-06 | 2.500E-06 |
| 9980 | 499 | W499V | A | 3.1e-06 | 2.9e-06 | 3.1e-06 | 3.033E-06 | 1.155E-07 | 2.900E-06 | 3.100E-06 |
| 9981 | 500 | G500A | A | 2.5e-06 | 1.1e-06 | 3.4e-06 | 2.333E-06 | 1.159E-06 | 1.100E-06 | 3.400E-06 |
| 9982 | 500 | G500R | A | 2.2e-06 | 2.5e-06 | 3.1e-06 | 2.600E-06 | 4.583E-07 | 2.200E-06 | 3.100E-06 |
| 9983 | 500 | G500N | A | 2.2e-06 | 2.4e-06 | 1.6e-06 | 2.067E-06 | 4.163E-07 | 1.600E-06 | 2.400E-06 |
| 9984 | 500 | G500D | A | 2.1e-06 | 2.5e-06 | 1.9e-06 | 2.167E-06 | 3.055E-07 | 1.900E-06 | 2.500E-06 |
| 9985 | 500 | G500C | A | 1.2e-06 | 2.8e-06 | 2.7e-06 | 2.233E-06 | 8.963E-07 | 1.200E-06 | 2.800E-06 |
| 9986 | 500 | G500E | A | 2e-06 | 3e-06 | 2.5e-06 | 2.500E-06 | 5.000E-07 | 2.000E-06 | 3.000E-06 |
| 9987 | 500 | G500Q | A | 2.7e-06 | 1.8e-06 | 3e-06 | 2.500E-06 | 6.245E-07 | 1.800E-06 | 3.000E-06 |
| 9988 | 500 | G500G | A | 2.6e-06 | 2.8e-06 | 2.6e-06 | 2.667E-06 | 1.155E-07 | 2.600E-06 | 2.800E-06 |
| 9989 | 500 | G500H | A | 1.8e-06 | 3.5e-06 | 1.6e-06 | 2.300E-06 | 1.044E-06 | 1.600E-06 | 3.500E-06 |
| 9990 | 500 | G500I | A | 2.6e-06 | 2.1e-06 | 2.1e-06 | 2.267E-06 | 2.887E-07 | 2.100E-06 | 2.600E-06 |
| 9991 | 500 | G500L | A | 1.5e-06 | 2.5e-06 | 5.5e-06 | 3.167E-06 | 2.082E-06 | 1.500E-06 | 5.500E-06 |
| 9992 | 500 | G500K | A | 3e-06 | 2.7e-06 | 3.3e-06 | 3.000E-06 | 3.000E-07 | 2.700E-06 | 3.300E-06 |
| 9993 | 500 | G500M | A | 2.9e-06 | 2.4e-06 | 1.6e-06 | 2.300E-06 | 6.557E-07 | 1.600E-06 | 2.900E-06 |
| 9994 | 500 | G500F | A | 3.2e-06 | 3.2e-06 | 1.9e-06 | 2.767E-06 | 7.506E-07 | 1.900E-06 | 3.200E-06 |
| 9995 | 500 | G500P | A | 3.1e-06 | 3.7e-06 | 3.9e-06 | 3.567E-06 | 4.163E-07 | 3.100E-06 | 3.900E-06 |
| 9996 | 500 | G500S | A | 2.9e-06 | 2.3e-06 | 2.6e-06 | 2.600E-06 | 3.000E-07 | 2.300E-06 | 2.900E-06 |
| 9997 | 500 | G500T | A | 2e-06 | 2e-06 | 2e-06 | 2.000E-06 | 0.000E+00 | 2.000E-06 | 2.000E-06 |
| 9998 | 500 | G500W | A | 4.2e-06 | 2.9e-06 | 1.4e-06 | 2.833E-06 | 1.401E-06 | 1.400E-06 | 4.200E-06 |
| 9999 | 500 | G500Y | A | 1.5e-06 | 3.3e-06 | 2.1e-06 | 2.300E-06 | 9.165E-07 | 1.500E-06 | 3.300E-06 |
| 10000 | 500 | G500V | A | 1.8e-06 | 2.9e-06 | 1.3e-06 | 2.000E-06 | 8.185E-07 | 1.300E-06 | 2.900E-06 |
| 10001 | 501 | P501A | A | 4.4e-06 | 1.8e-06 | 2.6e-06 | 2.933E-06 | 1.332E-06 | 1.800E-06 | 4.400E-06 |
| 10002 | 501 | P501R | A | 2.8e-06 | 1.9e-06 | 2.1e-06 | 2.267E-06 | 4.726E-07 | 1.900E-06 | 2.800E-06 |
| 10003 | 501 | P501N | A | 3.5e-06 | 2.1e-06 | 2.7e-06 | 2.767E-06 | 7.024E-07 | 2.100E-06 | 3.500E-06 |
| 10004 | 501 | P501D | A | 1.7e-06 | 2.7e-06 | 2.9e-06 | 2.433E-06 | 6.429E-07 | 1.700E-06 | 2.900E-06 |
| 10005 | 501 | P501C | A | 3.1e-06 | 2.1e-06 | 1.6e-06 | 2.267E-06 | 7.638E-07 | 1.600E-06 | 3.100E-06 |
| 10006 | 501 | P501E | A | 2.3e-06 | 2.4e-06 | 2.5e-06 | 2.400E-06 | 1.000E-07 | 2.300E-06 | 2.500E-06 |
| 10007 | 501 | P501Q | A | 2.4e-06 | 3.5e-06 | 1.2e-06 | 2.367E-06 | 1.150E-06 | 1.200E-06 | 3.500E-06 |
| 10008 | 501 | P501G | A | 2.5e-06 | 2.6e-06 | 2.8e-06 | 2.633E-06 | 1.528E-07 | 2.500E-06 | 2.800E-06 |
| 10009 | 501 | P501H | A | 1e-06 | 3.5e-06 | 1.8e-06 | 2.100E-06 | 1.277E-06 | 1.000E-06 | 3.500E-06 |
| 10010 | 501 | P501I | A | 1.9e-06 | 2.3e-06 | 2e-06 | 2.067E-06 | 2.082E-07 | 1.900E-06 | 2.300E-06 |
| 10011 | 501 | P501L | A | 1.9e-06 | 2.4e-06 | 3.1e-06 | 2.467E-06 | 6.028E-07 | 1.900E-06 | 3.100E-06 |
| 10012 | 501 | P501K | A | 2.2e-06 | 2.3e-06 | 4.2e-06 | 2.900E-06 | 1.127E-06 | 2.200E-06 | 4.200E-06 |
| 10013 | 501 | P501M | A | 1.3e-06 | 3.2e-06 | 2.4e-06 | 2.300E-06 | 9.539E-07 | 1.300E-06 | 3.200E-06 |
| 10014 | 501 | P501F | A | 1.7e-06 | 2.9e-06 | 1.8e-06 | 2.133E-06 | 6.658E-07 | 1.700E-06 | 2.900E-06 |
| 10015 | 501 | P501P | A | 2.2e-06 | 1.9e-06 | 2.2e-06 | 2.100E-06 | 1.732E-07 | 1.900E-06 | 2.200E-06 |
| 10016 | 501 | P501S | A | 2.6e-06 | 3.4e-06 | 1.3e-06 | 2.433E-06 | 1.060E-06 | 1.300E-06 | 3.400E-06 |
| 10017 | 501 | P501T | A | 1.8e-06 | 2.4e-06 | 2.8e-06 | 2.333E-06 | 5.033E-07 | 1.800E-06 | 2.800E-06 |
| 10018 | 501 | P501W | A | 2.1e-06 | 2.1e-06 | 2e-06 | 2.067E-06 | 5.774E-08 | 2.000E-06 | 2.100E-06 |
| 10019 | 501 | P501Y | A | 1.8e-06 | 1.9e-06 | 3.1e-06 | 2.267E-06 | 7.234E-07 | 1.800E-06 | 3.100E-06 |
| 10020 | 501 | P501V | A | 1.6e-06 | 2.7e-06 | 2.3e-06 | 2.200E-06 | 5.568E-07 | 1.600E-06 | 2.700E-06 |
| 10021 | 502 | G502A | A | 2.6e-06 | 1.6e-06 | 1.4e-06 | 1.867E-06 | 6.429E-07 | 1.400E-06 | 2.600E-06 |
| 10022 | 502 | G502R | A | 3.1e-06 | 9.2e-07 | 1.8e-06 | 1.940E-06 | 1.097E-06 | 9.200E-07 | 3.100E-06 |
| 10023 | 502 | G502N | A | 2.8e-06 | 3.6e-06 | 3.2e-06 | 3.200E-06 | 4.000E-07 | 2.800E-06 | 3.600E-06 |
| 10024 | 502 | G502D | A | 2.3e-06 | 1.3e-06 | 3.2e-06 | 2.267E-06 | 9.504E-07 | 1.300E-06 | 3.200E-06 |
| 10025 | 502 | G502C | A | 3.3e-06 | 1.5e-06 | 1.9e-06 | 2.233E-06 | 9.452E-07 | 1.500E-06 | 3.300E-06 |
| 10026 | 502 | G502E | A | 2.6e-06 | 3.1e-06 | 9.9e-07 | 2.230E-06 | 1.103E-06 | 9.900E-07 | 3.100E-06 |
| 10027 | 502 | G502Q | A | 1.8e-06 | 2.5e-06 | 2.9e-06 | 2.400E-06 | 5.568E-07 | 1.800E-06 | 2.900E-06 |
| 10028 | 502 | G502G | A | 1.7e-06 | 1.6e-06 | 1.9e-06 | 1.733E-06 | 1.528E-07 | 1.600E-06 | 1.900E-06 |
| 10029 | 502 | G502H | A | 2.7e-06 | 4.3e-06 | 2.1e-06 | 3.033E-06 | 1.137E-06 | 2.100E-06 | 4.300E-06 |

| | | | | | | | | | | |
|---|---|---|---|---|---|---|---|---|---|---|
| 10030 | 502 | G502I | A | 2.8e-06 | 2.5e-06 | 2.1e-06 | 2.467E-06 | 3.512E-07 | 2.100E-06 | 2.800E-06 |
| 10031 | 502 | G502L | A | 2.4e-06 | 3.2e-06 | 3.6e-06 | 3.067E-06 | 6.110E-07 | 2.400E-06 | 3.600E-06 |
| 10032 | 502 | G502K | A | 1.8e-06 | 2.8e-06 | 1.3e-06 | 1.967E-06 | 7.638E-07 | 1.300E-06 | 2.800E-06 |
| 10033 | 502 | G502M | A | 2.4e-06 | 2.7e-06 | 2.2e-06 | 2.433E-06 | 2.517E-07 | 2.200E-06 | 2.700E-06 |
| 10034 | 502 | G502F | A | 1.9e-06 | 3.2e-06 | 2.4e-06 | 2.500E-06 | 6.557E-07 | 1.900E-06 | 3.200E-06 |
| 10035 | 502 | G502P | A | 1.8e-06 | 1.6e-06 | 2.4e-06 | 1.933E-06 | 4.163E-07 | 1.600E-06 | 2.400E-06 |
| 10036 | 502 | G502S | A | 2.7e-06 | 1.8e-06 | 2.7e-06 | 2.400E-06 | 5.196E-07 | 1.800E-06 | 2.700E-06 |
| 10037 | 502 | G502T | A | 3.2e-06 | 3.8e-06 | 2.5e-06 | 3.167E-06 | 6.506E-07 | 2.500E-06 | 3.800E-06 |
| 10038 | 502 | G502W | A | 2.3e-06 | 1.9e-06 | 1.5e-06 | 1.900E-06 | 4.000E-07 | 1.500E-06 | 2.300E-06 |
| 10039 | 502 | G502Y | A | 2.8e-06 | 1.9e-06 | 1.9e-06 | 2.200E-06 | 5.196E-07 | 1.900E-06 | 2.800E-06 |
| 10040 | 502 | G502V | A | 2.3e-06 | 2.7e-06 | 2.7e-06 | 2.567E-06 | 2.309E-07 | 2.300E-06 | 2.700E-06 |
| 10041 | 503 | P503A | A | 3.3e-06 | 1.7e-06 | 1.7e-06 | 2.233E-06 | 9.238E-07 | 1.700E-06 | 3.300E-06 |
| 10042 | 503 | P503R | A | 1.6e-06 | 3e-06 | 4.5e-06 | 3.033E-06 | 1.450E-06 | 1.600E-06 | 4.500E-06 |
| 10043 | 503 | P503N | A | 1.1e-06 | 2.2e-06 | 1.4e-06 | 1.567E-06 | 5.686E-07 | 1.100E-06 | 2.200E-06 |
| 10044 | 503 | P503D | A | 1.8e-06 | 1.7e-06 | 2.3e-06 | 1.933E-06 | 3.215E-07 | 1.700E-06 | 2.300E-06 |
| 10045 | 503 | P503C | A | 2.2e-06 | 2e-06 | 2e-06 | 2.067E-06 | 1.155E-07 | 2.000E-06 | 2.200E-06 |
| 10046 | 503 | P503E | A | 2.3e-06 | 1.9e-06 | 2.2e-06 | 2.133E-06 | 2.082E-07 | 1.900E-06 | 2.300E-06 |
| 10047 | 503 | P503Q | A | 1.6e-06 | 2.3e-06 | 2.9e-06 | 2.267E-06 | 6.506E-07 | 1.600E-06 | 2.900E-06 |
| 10048 | 503 | P503G | A | 2.3e-06 | 2.8e-06 | 2.6e-06 | 2.567E-06 | 2.517E-07 | 2.300E-06 | 2.800E-06 |
| 10049 | 503 | P503H | A | 3e-06 | 2.9e-06 | 2.2e-06 | 2.700E-06 | 4.359E-07 | 2.200E-06 | 3.000E-06 |
| 10050 | 503 | P503I | A | 2.7e-06 | 1.6e-06 | 2e-06 | 2.100E-06 | 5.568E-07 | 1.600E-06 | 2.700E-06 |
| 10051 | 503 | P503L | A | 2.2e-06 | 1.9e-06 | 3.3e-06 | 2.467E-06 | 7.371E-07 | 1.900E-06 | 3.300E-06 |
| 10052 | 503 | P503K | A | 3.7e-06 | 2.7e-06 | 2.6e-06 | 3.000E-06 | 6.083E-07 | 2.600E-06 | 3.700E-06 |
| 10053 | 503 | P503M | A | 3.2e-06 | 2.4e-06 | 2.4e-06 | 2.667E-06 | 4.619E-07 | 2.400E-06 | 3.200E-06 |
| 10054 | 503 | P503F | A | 9.1e-07 | 2.9e-06 | 2.7e-06 | 2.170E-06 | 1.096E-06 | 9.100E-07 | 2.900E-06 |
| 10055 | 503 | P503P | A | 2e-06 | 2.5e-06 | 1.9e-06 | 2.133E-06 | 3.215E-07 | 1.900E-06 | 2.500E-06 |
| 10056 | 503 | P503S | A | 1.7e-06 | 2.7e-06 | 1.9e-06 | 2.100E-06 | 5.292E-07 | 1.700E-06 | 2.700E-06 |
| 10057 | 503 | P503T | A | 2.8e-06 | 2.2e-06 | 2.4e-06 | 2.467E-06 | 3.055E-07 | 2.200E-06 | 2.800E-06 |
| 10058 | 503 | P503W | A | 2.2e-06 | 2e-06 | 2.5e-06 | 2.233E-06 | 2.517E-07 | 2.000E-06 | 2.500E-06 |
| 10059 | 503 | P503Y | A | 2.9e-06 | 1.9e-06 | 2.5e-06 | 2.433E-06 | 5.033E-07 | 1.900E-06 | 2.900E-06 |
| 10060 | 503 | P503V | A | 3e-06 | 1.9e-06 | 3.1e-06 | 2.667E-06 | 6.658E-07 | 1.900E-06 | 3.100E-06 |
| 10061 | 504 | T504A | A | 2.2e-06 | 2.9e-06 | 3.4e-06 | 2.833E-06 | 6.028E-07 | 2.200E-06 | 3.400E-06 |
| 10062 | 504 | T504R | A | 3.4e-06 | 1.8e-06 | 1.9e-06 | 2.367E-06 | 8.963E-07 | 1.800E-06 | 3.400E-06 |
| 10063 | 504 | T504N | A | 1.5e-06 | 1.9e-06 | 4e-06 | 2.467E-06 | 1.343E-06 | 1.500E-06 | 4.000E-06 |
| 10064 | 504 | T504D | A | 1.9e-06 | 2.6e-06 | 2.5e-06 | 2.333E-06 | 3.786E-07 | 1.900E-06 | 2.600E-06 |
| 10065 | 504 | T504C | A | 2.8e-06 | 2.6e-06 | 1.6e-06 | 2.333E-06 | 6.429E-07 | 1.600E-06 | 2.800E-06 |
| 10066 | 504 | T504E | A | 1.6e-06 | 2.7e-06 | 2.8e-06 | 2.367E-06 | 6.658E-07 | 1.600E-06 | 2.800E-06 |
| 10067 | 504 | T504Q | A | 2.7e-06 | 2.3e-06 | 2e-06 | 2.333E-06 | 3.512E-07 | 2.000E-06 | 2.700E-06 |
| 10068 | 504 | T504G | A | 1.9e-06 | 2.4e-06 | 3.2e-06 | 2.500E-06 | 6.557E-07 | 1.900E-06 | 3.200E-06 |
| 10069 | 504 | T504H | A | 1.9e-06 | 1.9e-06 | 1.5e-06 | 1.767E-06 | 2.309E-07 | 1.500E-06 | 1.900E-06 |
| 10070 | 504 | T504I | A | 1.9e-06 | 7.2e-06 | 2.8e-06 | 3.967E-06 | 2.836E-06 | 1.900E-06 | 7.200E-06 |
| 10071 | 504 | T504L | A | 3.7e-06 | 6.1e-06 | 1.7e-06 | 3.833E-06 | 2.203E-06 | 1.700E-06 | 6.100E-06 |
| 10072 | 504 | T504K | A | 3.2e-06 | 2.5e-06 | 1.1e-06 | 2.267E-06 | 1.069E-06 | 1.100E-06 | 3.200E-06 |
| 10073 | 504 | T504M | A | 2.5e-06 | 2.6e-06 | 2.6e-06 | 2.567E-06 | 5.774E-08 | 2.500E-06 | 2.600E-06 |
| 10074 | 504 | T504F | A | 2.8e-06 | 4.4e-06 | 2.3e-06 | 3.167E-06 | 1.097E-06 | 2.300E-06 | 4.400E-06 |
| 10075 | 504 | T504P | A | 2.4e-06 | 2.5e-06 | 2.7e-06 | 2.533E-06 | 1.528E-07 | 2.400E-06 | 2.700E-06 |
| 10076 | 504 | T504S | A | 1.7e-06 | 2.3e-06 | 2.4e-06 | 2.133E-06 | 3.786E-07 | 1.700E-06 | 2.400E-06 |
| 10077 | 504 | T504T | A | 2.2e-06 | 3e-06 | 3.3e-06 | 2.833E-06 | 5.686E-07 | 2.200E-06 | 3.300E-06 |
| 10078 | 504 | T504W | A | 3.3e-06 | 1.7e-06 | 3.4e-06 | 2.800E-06 | 9.539E-07 | 1.700E-06 | 3.400E-06 |
| 10079 | 504 | T504Y | A | 3.1e-06 | 2.2e-06 | 1.8e-06 | 2.367E-06 | 6.658E-07 | 1.800E-06 | 3.100E-06 |
| 10080 | 504 | T504V | A | 1.6e-06 | 2.6e-06 | 3.7e-06 | 2.633E-06 | 1.050E-06 | 1.600E-06 | 3.700E-06 |
| 10081 | 505 | Q505A | A | 3.5e-06 | 2.1e-06 | 2.1e-06 | 2.567E-06 | 8.083E-07 | 2.100E-06 | 3.500E-06 |
| 10082 | 505 | Q505R | A | 2.6e-06 | 2.4e-06 | 2.1e-06 | 2.367E-06 | 2.517E-07 | 2.100E-06 | 2.600E-06 |
| 10083 | 505 | Q505N | A | 2.3e-06 | 2.5e-06 | 2.6e-06 | 2.467E-06 | 1.528E-07 | 2.300E-06 | 2.600E-06 |
| 10084 | 505 | Q505D | A | 4e-06 | 1.5e-06 | 1.4e-06 | 2.300E-06 | 1.473E-06 | 1.400E-06 | 4.000E-06 |
| 10085 | 505 | Q505C | A | 1.8e-06 | 2.2e-06 | 4.2e-06 | 2.733E-06 | 1.286E-06 | 1.800E-06 | 4.200E-06 |
| 10086 | 505 | Q505E | A | 2.4e-06 | 3e-06 | 4.1e-06 | 3.167E-06 | 8.622E-07 | 2.400E-06 | 4.100E-06 |
| 10087 | 505 | Q505Q | A | 2.5e-06 | 2.6e-06 | 2.7e-06 | 2.600E-06 | 1.000E-07 | 2.500E-06 | 2.700E-06 |
| 10088 | 505 | Q505G | A | 2.2e-06 | 2.1e-06 | 1.1e-06 | 1.800E-06 | 6.083E-07 | 1.100E-06 | 2.200E-06 |
| 10089 | 505 | Q505H | A | 1.7e-06 | 2.3e-06 | 2.3e-06 | 2.100E-06 | 3.464E-07 | 1.700E-06 | 2.300E-06 |
| 10090 | 505 | Q505I | A | 1.5e-06 | 2.4e-06 | 2.1e-06 | 2.000E-06 | 4.583E-07 | 1.500E-06 | 2.400E-06 |
| 10091 | 505 | Q505L | A | 2.6e-06 | 1.7e-06 | 4.5e-06 | 2.933E-06 | 1.429E-06 | 1.700E-06 | 4.500E-06 |
| 10092 | 505 | Q505K | A | 2.3e-06 | 1.7e-06 | 1.5e-06 | 1.833E-06 | 4.163E-07 | 1.500E-06 | 2.300E-06 |
| 10093 | 505 | Q505M | A | 2.8e-06 | 2.6e-06 | 4.1e-06 | 3.167E-06 | 8.145E-07 | 2.600E-06 | 4.100E-06 |
| 10094 | 505 | Q505F | A | 1.8e-06 | 2.9e-06 | 2e-06 | 2.233E-06 | 5.859E-07 | 1.800E-06 | 2.900E-06 |
| 10095 | 505 | Q505P | A | 2.2e-06 | 2.1e-06 | 1.3e-06 | 1.867E-06 | 4.933E-07 | 1.300E-06 | 2.200E-06 |
| 10096 | 505 | Q505S | A | 2.4e-06 | 1.8e-06 | 2e-06 | 2.067E-06 | 3.055E-07 | 1.800E-06 | 2.400E-06 |
| 10097 | 505 | Q505T | A | 1.8e-06 | 1.8e-06 | 1.3e-06 | 1.633E-06 | 2.887E-07 | 1.300E-06 | 1.800E-06 |
| 10098 | 505 | Q505W | A | 3.2e-06 | 1.4e-06 | 3.4e-06 | 2.667E-06 | 1.102E-06 | 1.400E-06 | 3.400E-06 |
| 10099 | 505 | Q505Y | A | 1.7e-06 | 1.6e-06 | 1.5e-06 | 1.600E-06 | 1.000E-07 | 1.500E-06 | 1.700E-06 |
| 10100 | 505 | Q505V | A | 4e-06 | 1.8e-06 | 2.5e-06 | 2.767E-06 | 1.124E-06 | 1.800E-06 | 4.000E-06 |
| 10101 | 506 | C506A | A | 1.7e-06 | 1.8e-06 | 2e-06 | 1.833E-06 | 1.528E-07 | 1.700E-06 | 2.000E-06 |
| 10102 | 506 | C506R | A | 2e-06 | 3.6e-06 | 2.2e-06 | 2.600E-06 | 8.718E-07 | 2.000E-06 | 3.600E-06 |
| 10103 | 506 | C506N | A | 2.4e-06 | 1.6e-06 | 2.1e-06 | 2.033E-06 | 4.041E-07 | 1.600E-06 | 2.400E-06 |
| 10104 | 506 | C506D | A | 2.2e-06 | 2.9e-06 | 4e-06 | 3.033E-06 | 9.074E-07 | 2.200E-06 | 4.000E-06 |
| 10105 | 506 | C506C | A | 2.8e-06 | 2.9e-06 | 2e-06 | 2.567E-06 | 4.933E-07 | 2.000E-06 | 2.900E-06 |
| 10106 | 506 | C506E | A | 4e-06 | 2.6e-06 | 2.4e-06 | 3.000E-06 | 8.718E-07 | 2.400E-06 | 4.000E-06 |
| 10107 | 506 | C506Q | A | 1.8e-06 | 2.1e-06 | 2.7e-06 | 2.200E-06 | 4.583E-07 | 1.800E-06 | 2.700E-06 |
| 10108 | 506 | C506G | A | 1.4e-06 | 2.5e-06 | 2.9e-06 | 2.267E-06 | 7.767E-07 | 1.400E-06 | 2.900E-06 |
| 10109 | 506 | C506H | A | 2e-06 | 2.5e-06 | 3.8e-06 | 2.767E-06 | 9.292E-07 | 2.000E-06 | 3.800E-06 |
| 10110 | 506 | C506I | A | 4.5e-06 | 2.4e-06 | 2.7e-06 | 3.200E-06 | 1.136E-06 | 2.400E-06 | 4.500E-06 |
| 10111 | 506 | C506L | A | 3.8e-06 | 2.2e-06 | 1.8e-06 | 2.600E-06 | 1.058E-06 | 1.800E-06 | 3.800E-06 |
| 10112 | 506 | C506K | A | 1.8e-06 | 5.4e-06 | 2.8e-06 | 3.333E-06 | 1.858E-06 | 1.800E-06 | 5.400E-06 |
| 10113 | 506 | C506M | A | 2.8e-06 | 2.3e-06 | 3.2e-06 | 2.767E-06 | 4.509E-07 | 2.300E-06 | 3.200E-06 |
| 10114 | 506 | C506F | A | 2.5e-06 | 2.8e-06 | 2.6e-06 | 2.633E-06 | 1.528E-07 | 2.500E-06 | 2.800E-06 |

| | | | | | | | | | | |
|---|---|---|---|---|---|---|---|---|---|---|
| 10115 | 506 | C506P | A | 3.1e-06 | 2e-06 | 2.9e-06 | 2.667E-06 | 5.859E-07 | 2.000E-06 | 3.100E-06 |
| 10116 | 506 | C506S | A | 4.3e-06 | 2e-06 | 3e-06 | 3.100E-06 | 1.153E-06 | 2.000E-06 | 4.300E-06 |
| 10117 | 506 | C506T | A | 1.1e-06 | 2.4e-06 | 1.5e-06 | 1.667E-06 | 6.658E-07 | 1.100E-06 | 2.400E-06 |
| 10118 | 506 | C506W | A | 2.5e-06 | 1.6e-06 | 1.5e-06 | 1.867E-06 | 5.508E-07 | 1.500E-06 | 2.500E-06 |
| 10119 | 506 | C506Y | A | 2.7e-06 | 3.5e-06 | 2.8e-06 | 3.000E-06 | 4.359E-07 | 2.700E-06 | 3.500E-06 |
| 10120 | 506 | C506V | A | 2.2e-06 | 2.7e-06 | 3e-06 | 2.633E-06 | 4.041E-07 | 2.200E-06 | 3.000E-06 |
| 10121 | 507 | V507A | A | 1.4e-06 | 1.7e-06 | 2.2e-06 | 1.767E-06 | 4.041E-07 | 1.400E-06 | 2.200E-06 |
| 10122 | 507 | V507R | A | 1.6e-06 | 2.9e-06 | 2.9e-06 | 2.467E-06 | 7.506E-07 | 1.600E-06 | 2.900E-06 |
| 10123 | 507 | V507N | A | 3.2e-06 | 2.6e-06 | 2e-06 | 2.600E-06 | 6.000E-07 | 2.000E-06 | 3.200E-06 |
| 10124 | 507 | V507D | A | 2.2e-06 | 2.5e-06 | 4e-06 | 2.900E-06 | 9.644E-07 | 2.200E-06 | 4.000E-06 |
| 10125 | 507 | V507C | A | 1.8e-06 | 2.1e-06 | 2.3e-06 | 2.067E-06 | 2.517E-07 | 1.800E-06 | 2.300E-06 |
| 10126 | 507 | V507E | A | 1.7e-06 | 2.3e-06 | 2e-06 | 2.000E-06 | 3.000E-07 | 1.700E-06 | 2.300E-06 |
| 10127 | 507 | V507Q | A | 3e-06 | 3.2e-06 | 2.2e-06 | 2.800E-06 | 5.292E-07 | 2.200E-06 | 3.200E-06 |
| 10128 | 507 | V507G | A | 1.6e-06 | 2.1e-06 | 3.3e-06 | 2.333E-06 | 8.737E-07 | 1.600E-06 | 3.300E-06 |
| 10129 | 507 | V507H | A | 4.2e-06 | 2.5e-06 | 3.6e-06 | 3.433E-06 | 8.622E-07 | 2.500E-06 | 4.200E-06 |
| 10130 | 507 | V507I | A | 2.8e-06 | 2.2e-06 | 2.5e-06 | 2.500E-06 | 3.000E-07 | 2.200E-06 | 2.800E-06 |
| 10131 | 507 | V507L | A | 3.1e-06 | 2.3e-06 | 2.8e-06 | 2.733E-06 | 4.041E-07 | 2.300E-06 | 3.100E-06 |
| 10132 | 507 | V507K | A | 3.4e-06 | 2.2e-06 | 4.3e-06 | 3.300E-06 | 1.054E-06 | 2.200E-06 | 4.300E-06 |
| 10133 | 507 | V507M | A | 1.6e-06 | 1.5e-06 | 1.9e-06 | 1.667E-06 | 2.082E-07 | 1.500E-06 | 1.900E-06 |
| 10134 | 507 | V507F | A | 2.1e-06 | 3.6e-06 | 2e-06 | 2.567E-06 | 8.963E-07 | 2.000E-06 | 3.600E-06 |
| 10135 | 507 | V507P | A | 3.6e-06 | 3.1e-06 | 3.3e-06 | 3.333E-06 | 2.517E-07 | 3.100E-06 | 3.600E-06 |
| 10136 | 507 | V507S | A | 2.3e-06 | 2.8e-06 | 2.4e-06 | 2.500E-06 | 2.646E-07 | 2.300E-06 | 2.800E-06 |
| 10137 | 507 | V507T | A | 2.6e-06 | 2e-06 | 2.5e-06 | 2.367E-06 | 3.215E-07 | 2.000E-06 | 2.600E-06 |
| 10138 | 507 | V507W | A | 4.4e-06 | 2.7e-06 | 2.2e-06 | 3.100E-06 | 1.153E-06 | 2.200E-06 | 4.400E-06 |
| 10139 | 507 | V507Y | A | 2.4e-06 | 2.1e-06 | 2.3e-06 | 2.267E-06 | 1.528E-07 | 2.100E-06 | 2.400E-06 |
| 10140 | 507 | V507V | A | 2.6e-06 | 3.7e-06 | 1.9e-06 | 2.733E-06 | 9.074E-07 | 1.900E-06 | 3.700E-06 |
| 10141 | 508 | N508A | A | 2.7e-06 | 2.8e-06 | 2.3e-06 | 2.600E-06 | 2.646E-07 | 2.300E-06 | 2.800E-06 |
| 10142 | 508 | N508R | A | 4.3e-06 | 1.8e-06 | 2.5e-06 | 2.867E-06 | 1.290E-06 | 1.800E-06 | 4.300E-06 |
| 10143 | 508 | N508N | A | 2.5e-06 | 3.7e-06 | 2.7e-06 | 2.967E-06 | 6.429E-07 | 2.500E-06 | 3.700E-06 |
| 10144 | 508 | N508D | A | 3.3e-06 | 3e-06 | 1.2e-06 | 2.500E-06 | 1.136E-06 | 1.200E-06 | 3.300E-06 |
| 10145 | 508 | N508C | A | 1.9e-06 | 2e-06 | 1.8e-06 | 1.900E-06 | 1.000E-07 | 1.800E-06 | 2.000E-06 |
| 10146 | 508 | N508E | A | 1.9e-06 | 2.9e-06 | 1.1e-06 | 1.967E-06 | 9.018E-07 | 1.100E-06 | 2.900E-06 |
| 10147 | 508 | N508Q | A | 1.7e-06 | 1.8e-06 | 1.6e-06 | 1.700E-06 | 1.000E-07 | 1.600E-06 | 1.800E-06 |
| 10148 | 508 | N508G | A | 3.2e-06 | 2.5e-06 | 3.7e-06 | 3.133E-06 | 6.028E-07 | 2.500E-06 | 3.700E-06 |
| 10149 | 508 | N508H | A | 1.6e-06 | 1e-06 | 3.7e-06 | 2.100E-06 | 1.418E-06 | 1.000E-06 | 3.700E-06 |
| 10150 | 508 | N508I | A | 3.2e-06 | 2.7e-06 | 2.4e-06 | 2.767E-06 | 4.041E-07 | 2.400E-06 | 3.200E-06 |
| 10151 | 508 | N508L | A | 1.9e-06 | 2e-06 | 8.6e-07 | 1.587E-06 | 6.313E-07 | 8.600E-07 | 2.000E-06 |
| 10152 | 508 | N508K | A | 2.2e-06 | 2.5e-06 | 2.4e-06 | 2.367E-06 | 1.528E-07 | 2.200E-06 | 2.500E-06 |
| 10153 | 508 | N508M | A | 1e-06 | 2.6e-06 | 1.9e-06 | 1.833E-06 | 8.021E-07 | 1.000E-06 | 2.600E-06 |
| 10154 | 508 | N508F | A | 1.4e-06 | 5.8e-06 | 2.1e-06 | 3.100E-06 | 2.364E-06 | 1.400E-06 | 5.800E-06 |
| 10155 | 508 | N508P | A | 2.2e-06 | 1.6e-06 | 2.9e-06 | 2.233E-06 | 6.506E-07 | 1.600E-06 | 2.900E-06 |
| 10156 | 508 | N508S | A | 2.1e-06 | 2.2e-06 | 3.1e-06 | 2.467E-06 | 5.508E-07 | 2.100E-06 | 3.100E-06 |
| 10157 | 508 | N508T | A | 2.8e-06 | 4.4e-06 | 3.2e-06 | 3.467E-06 | 8.327E-07 | 2.800E-06 | 4.400E-06 |
| 10158 | 508 | N508W | A | 2.1e-06 | 2e-06 | 1.8e-06 | 1.967E-06 | 1.528E-07 | 1.800E-06 | 2.100E-06 |
| 10159 | 508 | N508Y | A | 2.2e-06 | 2.5e-06 | 2.8e-06 | 2.500E-06 | 3.000E-07 | 2.200E-06 | 2.800E-06 |
| 10160 | 508 | N508V | A | 3.7e-06 | 5.8e-06 | 3.7e-06 | 4.400E-06 | 1.212E-06 | 3.700E-06 | 5.800E-06 |
| 10161 | 509 | C509A | A | 4.5e-06 | 6.1e-06 | 2.8e-06 | 4.467E-06 | 1.650E-06 | 2.800E-06 | 6.100E-06 |
| 10162 | 509 | C509R | A | 3.2e-06 | 1.9e-06 | 1.9e-06 | 2.333E-06 | 7.506E-07 | 1.900E-06 | 3.200E-06 |
| 10163 | 509 | C509N | A | 1.3e-06 | 2e-06 | 4.8e-06 | 2.700E-06 | 1.852E-06 | 1.300E-06 | 4.800E-06 |
| 10164 | 509 | C509D | A | 1.6e-06 | 3.2e-06 | 1.7e-06 | 2.167E-06 | 8.963E-07 | 1.600E-06 | 3.200E-06 |
| 10165 | 509 | C509C | A | 1.8e-06 | 2.2e-06 | 1.8e-06 | 1.933E-06 | 2.309E-07 | 1.800E-06 | 2.200E-06 |
| 10166 | 509 | C509E | A | 2.9e-06 | 2e-06 | 3.3e-06 | 2.733E-06 | 6.658E-07 | 2.000E-06 | 3.300E-06 |
| 10167 | 509 | C509Q | A | 3.3e-06 | 2.2e-06 | 2.7e-06 | 2.733E-06 | 5.508E-07 | 2.200E-06 | 3.300E-06 |
| 10168 | 509 | C509G | A | 2.9e-06 | 2e-06 | 3.3e-06 | 2.733E-06 | 6.658E-07 | 2.000E-06 | 3.300E-06 |
| 10169 | 509 | C509H | A | 2.5e-06 | 1.4e-06 | 2.1e-06 | 2.000E-06 | 5.568E-07 | 1.400E-06 | 2.500E-06 |
| 10170 | 509 | C509I | A | 3e-06 | 1.6e-06 | 2.1e-06 | 2.233E-06 | 7.095E-07 | 1.600E-06 | 3.000E-06 |
| 10171 | 509 | C509L | A | 2.5e-06 | 2.8e-06 | 3.1e-06 | 2.800E-06 | 3.000E-07 | 2.500E-06 | 3.100E-06 |
| 10172 | 509 | C509K | A | 1.4e-06 | 3.3e-06 | 3.7e-06 | 2.800E-06 | 1.229E-06 | 1.400E-06 | 3.700E-06 |
| 10173 | 509 | C509M | A | 3.2e-06 | 1.9e-06 | 5.5e-06 | 3.533E-06 | 1.823E-06 | 1.900E-06 | 5.500E-06 |
| 10174 | 509 | C509F | A | 5.1e-06 | 1.7e-06 | 2.6e-06 | 3.133E-06 | 1.762E-06 | 1.700E-06 | 5.100E-06 |
| 10175 | 509 | C509P | A | 2.4e-06 | 2.2e-06 | 2.4e-06 | 2.333E-06 | 1.155E-07 | 2.200E-06 | 2.400E-06 |
| 10176 | 509 | C509S | A | 3e-06 | 2e-06 | 3.6e-06 | 2.867E-06 | 8.083E-07 | 2.000E-06 | 3.600E-06 |
| 10177 | 509 | C509T | A | 2.3e-06 | 2.1e-06 | 2.3e-06 | 2.233E-06 | 1.155E-07 | 2.100E-06 | 2.300E-06 |
| 10178 | 509 | C509W | A | 2.5e-06 | 1.9e-06 | 2.5e-06 | 2.300E-06 | 3.464E-07 | 1.900E-06 | 2.500E-06 |
| 10179 | 509 | C509Y | A | 2.4e-06 | 1.5e-06 | 1.6e-06 | 1.833E-06 | 4.933E-07 | 1.500E-06 | 2.400E-06 |
| 10180 | 509 | C509V | A | 2.8e-06 | 3.2e-06 | 2.9e-06 | 2.967E-06 | 2.082E-07 | 2.800E-06 | 3.200E-06 |
| 10181 | 510 | S510A | A | 2e-06 | 1.3e-06 | 2.8e-06 | 2.033E-06 | 7.506E-07 | 1.300E-06 | 2.800E-06 |
| 10182 | 510 | S510R | A | 2e-06 | 2.3e-06 | 5.2e-06 | 3.167E-06 | 1.767E-06 | 2.000E-06 | 5.200E-06 |
| 10183 | 510 | S510N | A | 2.2e-06 | 2e-06 | 3.8e-06 | 2.667E-06 | 9.866E-07 | 2.000E-06 | 3.800E-06 |
| 10184 | 510 | S510D | A | 2.1e-06 | 2.3e-06 | 3.4e-06 | 2.600E-06 | 7.000E-07 | 2.100E-06 | 3.400E-06 |
| 10185 | 510 | S510C | A | 2.2e-06 | 2e-06 | 3.4e-06 | 2.533E-06 | 7.572E-07 | 2.000E-06 | 3.400E-06 |
| 10186 | 510 | S510E | A | 1.9e-06 | 2.6e-06 | 2e-06 | 2.167E-06 | 3.786E-07 | 1.900E-06 | 2.600E-06 |
| 10187 | 510 | S510Q | A | 1.3e-06 | 1.6e-06 | 2.7e-06 | 1.867E-06 | 7.371E-07 | 1.300E-06 | 2.700E-06 |
| 10188 | 510 | S510G | A | 2.6e-06 | 3.4e-06 | 2.7e-06 | 2.900E-06 | 4.359E-07 | 2.600E-06 | 3.400E-06 |
| 10189 | 510 | S510H | A | 2.2e-06 | 1.3e-06 | 1.3e-06 | 1.600E-06 | 5.196E-07 | 1.300E-06 | 2.200E-06 |
| 10190 | 510 | S510I | A | 2.3e-06 | 1.4e-06 | 2e-06 | 1.900E-06 | 4.583E-07 | 1.400E-06 | 2.300E-06 |
| 10191 | 510 | S510L | A | 3.8e-06 | 2.4e-06 | 1.9e-06 | 2.700E-06 | 9.849E-07 | 1.900E-06 | 3.800E-06 |
| 10192 | 510 | S510K | A | 2.3e-06 | 2.7e-06 | 1.8e-06 | 2.267E-06 | 4.509E-07 | 1.800E-06 | 2.700E-06 |
| 10193 | 510 | S510M | A | 2e-06 | 2.6e-06 | 2.1e-06 | 2.233E-06 | 3.215E-07 | 2.000E-06 | 2.600E-06 |
| 10194 | 510 | S510F | A | 3.3e-06 | 2.3e-06 | 3.2e-06 | 2.933E-06 | 5.508E-07 | 2.300E-06 | 3.300E-06 |
| 10195 | 510 | S510P | A | 2.3e-06 | 2.9e-06 | 4e-06 | 3.067E-06 | 8.622E-07 | 2.300E-06 | 4.000E-06 |
| 10196 | 510 | S510S | A | 1.8e-06 | 2.4e-06 | 2.3e-06 | 2.167E-06 | 3.215E-07 | 1.800E-06 | 2.400E-06 |
| 10197 | 510 | S510T | A | 2.5e-06 | 1.9e-06 | 2.5e-06 | 2.300E-06 | 3.464E-07 | 1.900E-06 | 2.500E-06 |
| 10198 | 510 | S510W | A | 2.6e-06 | 1.9e-06 | 2.2e-06 | 2.233E-06 | 3.512E-07 | 1.900E-06 | 2.600E-06 |
| 10199 | 510 | S510Y | A | 1.8e-06 | 2.5e-06 | 3.3e-06 | 2.533E-06 | 7.506E-07 | 1.800E-06 | 3.300E-06 |

| | | | | | | | | | | |
|---|---|---|---|---|---|---|---|---|---|---|
| 10200 | 510 | S510V | A | 2.2e-06 | 2.3e-06 | 2.2e-06 | 2.233E-06 | 5.774E-08 | 2.200E-06 | 2.300E-06 |
| 10201 | 511 | Q511A | A | 3.1e-06 | 2.9e-06 | 3.5e-06 | 3.167E-06 | 3.055E-07 | 2.900E-06 | 3.500E-06 |
| 10202 | 511 | Q511R | A | 1.5e-06 | 1.9e-06 | 3.3e-06 | 2.233E-06 | 9.452E-07 | 1.500E-06 | 3.300E-06 |
| 10203 | 511 | Q511N | A | 1.7e-06 | 1.9e-06 | 1.8e-06 | 1.800E-06 | 1.000E-07 | 1.700E-06 | 1.900E-06 |
| 10204 | 511 | Q511D | A | 3.8e-06 | 2e-06 | 2.3e-06 | 2.700E-06 | 9.644E-07 | 2.000E-06 | 3.800E-06 |
| 10205 | 511 | Q511C | A | 2e-06 | 2.4e-06 | 2.1e-06 | 2.167E-06 | 2.082E-07 | 2.000E-06 | 2.400E-06 |
| 10206 | 511 | Q511E | A | 2.2e-06 | 3e-06 | 2.6e-06 | 2.600E-06 | 4.000E-07 | 2.200E-06 | 3.000E-06 |
| 10207 | 511 | Q511Q | A | 2.9e-06 | 3.2e-06 | 1.6e-06 | 2.567E-06 | 8.505E-07 | 1.600E-06 | 3.200E-06 |
| 10208 | 511 | Q511G | A | 1.7e-06 | 2.4e-06 | 2.2e-06 | 2.100E-06 | 3.606E-07 | 1.700E-06 | 2.400E-06 |
| 10209 | 511 | Q511H | A | 2.6e-06 | 3.1e-06 | 4.8e-06 | 3.500E-06 | 1.153E-06 | 2.600E-06 | 4.800E-06 |
| 10210 | 511 | Q511I | A | 2.2e-06 | 3.6e-06 | 2.9e-06 | 2.900E-06 | 7.000E-07 | 2.200E-06 | 3.600E-06 |
| 10211 | 511 | Q511L | A | 2.3e-06 | 2.8e-06 | 2e-06 | 2.367E-06 | 4.041E-07 | 2.000E-06 | 2.800E-06 |
| 10212 | 511 | Q511K | A | 2.3e-06 | 2.8e-06 | 4e-06 | 3.033E-06 | 8.737E-07 | 2.300E-06 | 4.000E-06 |
| 10213 | 511 | Q511M | A | 3.8e-06 | 4e-06 | 5.3e-06 | 4.367E-06 | 8.145E-07 | 3.800E-06 | 5.300E-06 |
| 10214 | 511 | Q511F | A | 2e-06 | 2.1e-06 | 2.7e-06 | 2.267E-06 | 3.786E-07 | 2.000E-06 | 2.700E-06 |
| 10215 | 511 | Q511P | A | 2.8e-06 | 1.8e-06 | 2.6e-06 | 2.400E-06 | 5.292E-07 | 1.800E-06 | 2.800E-06 |
| 10216 | 511 | Q511S | A | 2.2e-06 | 3.6e-06 | 5.8e-06 | 3.867E-06 | 1.815E-06 | 2.200E-06 | 5.800E-06 |
| 10217 | 511 | Q511T | A | 2.2e-06 | 2.7e-06 | 1.8e-06 | 2.233E-06 | 4.509E-07 | 1.800E-06 | 2.700E-06 |
| 10218 | 511 | Q511W | A | 2.3e-06 | 2.2e-06 | 3.8e-06 | 2.767E-06 | 8.963E-07 | 2.200E-06 | 3.800E-06 |
| 10219 | 511 | Q511Y | A | 2.4e-06 | 2.9e-06 | 2.1e-06 | 2.467E-06 | 4.041E-07 | 2.100E-06 | 2.900E-06 |
| 10220 | 511 | Q511V | A | 5e-06 | 3e-06 | 1.4e-06 | 3.133E-06 | 1.804E-06 | 1.400E-06 | 5.000E-06 |
| 10221 | 512 | F512A | A | 2.8e-06 | 2.1e-06 | 2.3e-06 | 2.400E-06 | 3.606E-07 | 2.100E-06 | 2.800E-06 |
| 10222 | 512 | F512R | A | 2.1e-06 | 9.5e-07 | 3.7e-06 | 2.250E-06 | 1.381E-06 | 9.500E-07 | 3.700E-06 |
| 10223 | 512 | F512N | A | 2.9e-06 | 1.9e-06 | 1.9e-06 | 2.233E-06 | 5.774E-07 | 1.900E-06 | 2.900E-06 |
| 10224 | 512 | F512D | A | 2.1e-06 | 2.3e-06 | 2.3e-06 | 2.233E-06 | 1.155E-07 | 2.100E-06 | 2.300E-06 |
| 10225 | 512 | F512C | A | 2.4e-06 | 4.2e-06 | 2.6e-06 | 3.067E-06 | 9.866E-07 | 2.400E-06 | 4.200E-06 |
| 10226 | 512 | F512E | A | 2.4e-06 | 2.4e-06 | 2.9e-06 | 2.567E-06 | 2.887E-07 | 2.400E-06 | 2.900E-06 |
| 10227 | 512 | F512Q | A | 1.8e-06 | 3.4e-06 | 1.4e-06 | 2.200E-06 | 1.058E-06 | 1.400E-06 | 3.400E-06 |
| 10228 | 512 | F512G | A | 1.6e-06 | 1.7e-06 | 2.3e-06 | 1.867E-06 | 3.786E-07 | 1.600E-06 | 2.300E-06 |
| 10229 | 512 | F512H | A | 1.5e-06 | 1.4e-06 | 2.7e-06 | 1.867E-06 | 7.234E-07 | 1.400E-06 | 2.700E-06 |
| 10230 | 512 | F512I | A | 2.5e-06 | 2.3e-06 | 1.8e-06 | 2.200E-06 | 3.606E-07 | 1.800E-06 | 2.500E-06 |
| 10231 | 512 | F512L | A | 3.7e-06 | 2e-06 | 5.1e-06 | 3.600E-06 | 1.552E-06 | 2.000E-06 | 5.100E-06 |
| 10232 | 512 | F512K | A | 2.3e-06 | 3e-06 | 2.3e-06 | 2.533E-06 | 4.041E-07 | 2.300E-06 | 3.000E-06 |
| 10233 | 512 | F512M | A | 3.8e-06 | 1.8e-06 | 3.4e-06 | 3.000E-06 | 1.058E-06 | 1.800E-06 | 3.800E-06 |
| 10234 | 512 | F512F | A | 2.4e-06 | 1.7e-06 | 3e-06 | 2.367E-06 | 6.506E-07 | 1.700E-06 | 3.000E-06 |
| 10235 | 512 | F512P | A | 2.7e-06 | 2.5e-06 | 1.8e-06 | 2.333E-06 | 4.726E-07 | 1.800E-06 | 2.700E-06 |
| 10236 | 512 | F512S | A | 3e-06 | 2.4e-06 | 2.5e-06 | 2.633E-06 | 3.215E-07 | 2.400E-06 | 3.000E-06 |
| 10237 | 512 | F512T | A | 2.5e-06 | 2.2e-06 | 2e-06 | 2.233E-06 | 2.517E-07 | 2.000E-06 | 2.500E-06 |
| 10238 | 512 | F512W | A | 2.7e-06 | 1.7e-06 | 2.4e-06 | 2.267E-06 | 5.132E-07 | 1.700E-06 | 2.700E-06 |
| 10239 | 512 | F512Y | A | 4.3e-06 | 2.9e-06 | 3e-06 | 3.400E-06 | 7.810E-07 | 2.900E-06 | 4.300E-06 |
| 10240 | 512 | F512V | A | 4.5e-06 | 1.7e-06 | 2.7e-06 | 2.967E-06 | 1.419E-06 | 1.700E-06 | 4.500E-06 |
| 10241 | 513 | L513A | A | 1.9e-06 | 1.6e-06 | 2.8e-06 | 2.100E-06 | 6.245E-07 | 1.600E-06 | 2.800E-06 |
| 10242 | 513 | L513R | A | 2.2e-06 | 2.6e-06 | 2e-06 | 2.267E-06 | 3.055E-07 | 2.000E-06 | 2.600E-06 |
| 10243 | 513 | L513N | A | 1.8e-06 | 2.6e-06 | 3.6e-06 | 2.667E-06 | 9.018E-07 | 1.800E-06 | 3.600E-06 |
| 10244 | 513 | L513D | A | 1.7e-06 | 2.7e-06 | 2.1e-06 | 2.167E-06 | 5.033E-07 | 1.700E-06 | 2.700E-06 |
| 10245 | 513 | L513C | A | 2.2e-06 | 2.7e-06 | 3e-06 | 2.633E-06 | 4.041E-07 | 2.200E-06 | 3.000E-06 |
| 10246 | 513 | L513E | A | 2.6e-06 | 2.1e-06 | 2.2e-06 | 2.300E-06 | 2.646E-07 | 2.100E-06 | 2.600E-06 |
| 10247 | 513 | L513Q | A | 2.9e-06 | 2.1e-06 | 2.1e-06 | 2.367E-06 | 4.619E-07 | 2.100E-06 | 2.900E-06 |
| 10248 | 513 | L513G | A | 2.4e-06 | 2.1e-06 | 1.9e-06 | 2.133E-06 | 2.517E-07 | 1.900E-06 | 2.400E-06 |
| 10249 | 513 | L513H | A | 3e-06 | 2.3e-06 | 4.1e-06 | 3.133E-06 | 9.074E-07 | 2.300E-06 | 4.100E-06 |
| 10250 | 513 | L513I | A | 3.8e-06 | 1.8e-06 | 2.2e-06 | 2.600E-06 | 1.058E-06 | 1.800E-06 | 3.800E-06 |
| 10251 | 513 | L513L | A | 2.1e-06 | 6.3e-07 | 3.6e-06 | 2.110E-06 | 1.485E-06 | 6.300E-07 | 3.600E-06 |
| 10252 | 513 | L513K | A | 8.8e-07 | 1.2e-06 | 2.8e-06 | 1.627E-06 | 1.029E-06 | 8.800E-07 | 2.800E-06 |
| 10253 | 513 | L513M | A | 3.2e-06 | 2.2e-06 | 1.5e-06 | 2.300E-06 | 8.544E-07 | 1.500E-06 | 3.200E-06 |
| 10254 | 513 | L513F | A | 2.6e-06 | 1.9e-06 | 1.8e-06 | 2.100E-06 | 4.359E-07 | 1.800E-06 | 2.600E-06 |
| 10255 | 513 | L513P | A | 3.1e-06 | 3e-06 | 2.7e-06 | 2.933E-06 | 2.082E-07 | 2.700E-06 | 3.100E-06 |
| 10256 | 513 | L513S | A | 3.5e-06 | 1.1e-06 | 2.4e-06 | 2.333E-06 | 1.201E-06 | 1.100E-06 | 3.500E-06 |
| 10257 | 513 | L513T | A | 3.3e-06 | 3.4e-06 | 1.6e-06 | 2.767E-06 | 1.012E-06 | 1.600E-06 | 3.400E-06 |
| 10258 | 513 | L513W | A | 1.2e-06 | 2.2e-06 | 4.1e-06 | 2.500E-06 | 1.473E-06 | 1.200E-06 | 4.100E-06 |
| 10259 | 513 | L513Y | A | 2.8e-06 | 3e-06 | 3.7e-06 | 3.167E-06 | 4.726E-07 | 2.800E-06 | 3.700E-06 |
| 10260 | 513 | L513V | A | 2e-06 | 2.7e-06 | 2.5e-06 | 2.400E-06 | 3.606E-07 | 2.000E-06 | 2.700E-06 |
| 10261 | 514 | R514A | A | 2.3e-06 | 1.2e-06 | 1.9e-06 | 1.800E-06 | 5.568E-07 | 1.200E-06 | 2.300E-06 |
| 10262 | 514 | R514R | A | 2.7e-06 | 2.6e-06 | 2.8e-06 | 2.700E-06 | 1.000E-07 | 2.600E-06 | 2.800E-06 |
| 10263 | 514 | R514N | A | 1.4e-06 | 2e-06 | 4e-06 | 2.467E-06 | 1.361E-06 | 1.400E-06 | 4.000E-06 |
| 10264 | 514 | R514D | A | 2.3e-06 | 3.5e-06 | 2.1e-06 | 2.633E-06 | 7.572E-07 | 2.100E-06 | 3.500E-06 |
| 10265 | 514 | R514C | A | 2.2e-06 | 2.2e-06 | 2.5e-06 | 2.300E-06 | 1.732E-07 | 2.200E-06 | 2.500E-06 |
| 10266 | 514 | R514E | A | 3.1e-06 | 2.2e-06 | 1.8e-06 | 2.367E-06 | 6.658E-07 | 1.800E-06 | 3.100E-06 |
| 10267 | 514 | R514Q | A | 2.3e-06 | 3.6e-06 | 1.6e-06 | 2.500E-06 | 1.015E-06 | 1.600E-06 | 3.600E-06 |
| 10268 | 514 | R514G | A | 3.1e-06 | 2.7e-06 | 4.2e-06 | 3.333E-06 | 7.767E-07 | 2.700E-06 | 4.200E-06 |
| 10269 | 514 | R514H | A | 3.6e-06 | 3.4e-06 | 1.3e-06 | 2.767E-06 | 1.274E-06 | 1.300E-06 | 3.600E-06 |
| 10270 | 514 | R514I | A | 2.7e-06 | 2.6e-06 | 4.1e-06 | 3.133E-06 | 8.386E-07 | 2.600E-06 | 4.100E-06 |
| 10271 | 514 | R514L | A | 3.6e-06 | 2.2e-06 | 2.2e-06 | 2.667E-06 | 8.083E-07 | 2.200E-06 | 3.600E-06 |
| 10272 | 514 | R514K | A | 2.2e-06 | 1.9e-06 | 2.1e-06 | 2.067E-06 | 1.528E-07 | 1.900E-06 | 2.200E-06 |
| 10273 | 514 | R514M | A | 1.3e-06 | 2.5e-06 | 2.5e-06 | 2.100E-06 | 6.928E-07 | 1.300E-06 | 2.500E-06 |
| 10274 | 514 | R514F | A | 1.4e-06 | 6.2e-06 | 1.6e-06 | 3.067E-06 | 2.715E-06 | 1.400E-06 | 6.200E-06 |
| 10275 | 514 | R514P | A | 2.6e-06 | 3.1e-06 | 1.9e-06 | 2.533E-06 | 6.028E-07 | 1.900E-06 | 3.100E-06 |
| 10276 | 514 | R514S | A | 2.2e-06 | 1.7e-06 | 3.7e-06 | 2.533E-06 | 1.041E-06 | 1.700E-06 | 3.700E-06 |
| 10277 | 514 | R514T | A | 2e-06 | 1.8e-06 | 2.3e-06 | 2.033E-06 | 2.517E-07 | 1.800E-06 | 2.300E-06 |
| 10278 | 514 | R514W | A | 2.2e-06 | 1.4e-06 | 1.6e-06 | 1.733E-06 | 4.163E-07 | 1.400E-06 | 2.200E-06 |
| 10279 | 514 | R514Y | A | 1.7e-06 | 2e-06 | 2e-06 | 1.900E-06 | 1.732E-07 | 1.700E-06 | 2.000E-06 |
| 10280 | 514 | R514V | A | 1.2e-06 | 2.3e-06 | 3.2e-06 | 2.233E-06 | 1.002E-06 | 1.200E-06 | 3.200E-06 |
| 10281 | 515 | G515A | A | 2.6e-06 | 3.1e-06 | 3.8e-06 | 3.167E-06 | 6.028E-07 | 2.600E-06 | 3.800E-06 |
| 10282 | 515 | G515R | A | 3.1e-06 | 2.2e-06 | 2.7e-06 | 2.667E-06 | 4.509E-07 | 2.200E-06 | 3.100E-06 |
| 10283 | 515 | G515N | A | 1.6e-06 | 1.6e-06 | 1.3e-06 | 1.500E-06 | 1.732E-07 | 1.300E-06 | 1.600E-06 |
| 10284 | 515 | G515D | A | 3.2e-06 | 1.9e-06 | 3e-06 | 2.700E-06 | 7.000E-07 | 1.900E-06 | 3.200E-06 |

| | | | | | | | | | | |
|---|---|---|---|---|---|---|---|---|---|---|
| 10285 | 515 | G515C | A | 2.9e-06 | 2.3e-06 | 1.2e-06 | 2.133E-06 | 8.622E-07 | 1.200E-06 | 2.900E-06 |
| 10286 | 515 | G515E | A | 2.4e-06 | 2.2e-06 | 3.2e-06 | 2.600E-06 | 5.292E-07 | 2.200E-06 | 3.200E-06 |
| 10287 | 515 | G515Q | A | 2.1e-06 | 1.3e-06 | 1.6e-06 | 1.667E-06 | 4.041E-07 | 1.300E-06 | 2.100E-06 |
| 10288 | 515 | G515G | A | 3.7e-06 | 3.3e-06 | 2.3e-06 | 3.100E-06 | 7.211E-07 | 2.300E-06 | 3.700E-06 |
| 10289 | 515 | G515H | A | 2.1e-06 | 1.9e-06 | 2.3e-06 | 2.100E-06 | 2.000E-07 | 1.900E-06 | 2.300E-06 |
| 10290 | 515 | G515I | A | 1.7e-06 | 2.4e-06 | 2e-06 | 2.033E-06 | 3.512E-07 | 1.700E-06 | 2.400E-06 |
| 10291 | 515 | G515L | A | 3.5e-06 | 2.3e-06 | 1.3e-06 | 2.367E-06 | 1.102E-06 | 1.300E-06 | 3.500E-06 |
| 10292 | 515 | G515K | A | 3.3e-06 | 3.8e-06 | 3.2e-06 | 3.433E-06 | 3.215E-07 | 3.200E-06 | 3.800E-06 |
| 10293 | 515 | G515M | A | 2.9e-06 | 1.6e-06 | 3.9e-06 | 2.800E-06 | 1.153E-06 | 1.600E-06 | 3.900E-06 |
| 10294 | 515 | G515F | A | 2.4e-06 | 1.8e-06 | 4.9e-06 | 3.033E-06 | 1.644E-06 | 1.800E-06 | 4.900E-06 |
| 10295 | 515 | G515P | A | 2.3e-06 | 2.8e-06 | 3.2e-06 | 2.767E-06 | 4.509E-07 | 2.300E-06 | 3.200E-06 |
| 10296 | 515 | G515S | A | 1.8e-06 | 2.3e-06 | 1.9e-06 | 2.000E-06 | 2.646E-07 | 1.800E-06 | 2.300E-06 |
| 10297 | 515 | G515T | A | 2.8e-06 | 1.8e-06 | 1.9e-06 | 2.167E-06 | 5.508E-07 | 1.800E-06 | 2.800E-06 |
| 10298 | 515 | G515W | A | 1.5e-06 | 2.3e-06 | 4.1e-06 | 2.633E-06 | 1.332E-06 | 1.500E-06 | 4.100E-06 |
| 10299 | 515 | G515Y | A | 3.7e-06 | 1.6e-06 | 3e-06 | 2.767E-06 | 1.069E-06 | 1.600E-06 | 3.700E-06 |
| 10300 | 515 | G515V | A | 1.4e-06 | 2.1e-06 | 2.3e-06 | 1.933E-06 | 4.726E-07 | 1.400E-06 | 2.300E-06 |
| 10301 | 516 | Q516A | A | 2.7e-06 | 2.6e-06 | 2e-06 | 2.433E-06 | 3.786E-07 | 2.000E-06 | 2.700E-06 |
| 10302 | 516 | Q516R | A | 2.5e-06 | 3.4e-06 | 1.7e-06 | 2.533E-06 | 8.505E-07 | 1.700E-06 | 3.400E-06 |
| 10303 | 516 | Q516N | A | 3.9e-06 | 1.9e-06 | 2.3e-06 | 2.700E-06 | 1.058E-06 | 1.900E-06 | 3.900E-06 |
| 10304 | 516 | Q516D | A | 2.9e-06 | 2.6e-06 | 2e-06 | 2.500E-06 | 4.583E-07 | 2.000E-06 | 2.900E-06 |
| 10305 | 516 | Q516C | A | 2e-06 | 3.1e-06 | 2.3e-06 | 2.467E-06 | 5.686E-07 | 2.000E-06 | 3.100E-06 |
| 10306 | 516 | Q516E | A | 2.4e-06 | 3.1e-06 | 2.3e-06 | 2.600E-06 | 4.359E-07 | 2.300E-06 | 3.100E-06 |
| 10307 | 516 | Q516Q | A | 1.9e-06 | 3e-06 | 2.4e-06 | 2.433E-06 | 5.508E-07 | 1.900E-06 | 3.000E-06 |
| 10308 | 516 | Q516G | A | 2.6e-06 | 2.6e-06 | 2.1e-06 | 2.433E-06 | 2.887E-07 | 2.100E-06 | 2.600E-06 |
| 10309 | 516 | Q516H | A | 2e-06 | 4.4e-06 | 1.8e-06 | 2.733E-06 | 1.447E-06 | 1.800E-06 | 4.400E-06 |
| 10310 | 516 | Q516I | A | 1.4e-06 | 2.2e-06 | 2.1e-06 | 1.900E-06 | 4.359E-07 | 1.400E-06 | 2.200E-06 |
| 10311 | 516 | Q516L | A | 1.9e-06 | 2.5e-06 | 2.2e-06 | 2.200E-06 | 3.000E-07 | 1.900E-06 | 2.500E-06 |
| 10312 | 516 | Q516K | A | 2.7e-06 | 2e-06 | 3.2e-06 | 2.633E-06 | 6.028E-07 | 2.000E-06 | 3.200E-06 |
| 10313 | 516 | Q516M | A | 2e-06 | 1.3e-06 | 1.8e-06 | 1.700E-06 | 3.606E-07 | 1.300E-06 | 2.000E-06 |
| 10314 | 516 | Q516F | A | 1.9e-06 | 1.7e-06 | 1.5e-06 | 1.700E-06 | 2.000E-07 | 1.500E-06 | 1.900E-06 |
| 10315 | 516 | Q516P | A | 1.5e-06 | 2.8e-06 | 2.3e-06 | 2.200E-06 | 6.557E-07 | 1.500E-06 | 2.800E-06 |
| 10316 | 516 | Q516S | A | 2.5e-06 | 2.6e-06 | 5.9e-06 | 3.667E-06 | 1.935E-06 | 2.500E-06 | 5.900E-06 |
| 10317 | 516 | Q516T | A | 2.5e-06 | 2.7e-06 | 2.5e-06 | 2.567E-06 | 1.155E-07 | 2.500E-06 | 2.700E-06 |
| 10318 | 516 | Q516W | A | 2.8e-06 | 2.4e-06 | 2.9e-06 | 2.700E-06 | 2.646E-07 | 2.400E-06 | 2.900E-06 |
| 10319 | 516 | Q516Y | A | 2e-06 | 1.8e-06 | 4e-06 | 2.600E-06 | 1.217E-06 | 1.800E-06 | 4.000E-06 |
| 10320 | 516 | Q516V | A | 3.3e-06 | 3e-06 | 2.7e-06 | 3.000E-06 | 3.000E-07 | 2.700E-06 | 3.300E-06 |
| 10321 | 517 | E517A | A | 2.9e-06 | 2.5e-06 | 3.4e-06 | 2.933E-06 | 4.509E-07 | 2.500E-06 | 3.400E-06 |
| 10322 | 517 | E517R | A | 3.8e-06 | 2.2e-06 | 1.2e-06 | 2.400E-06 | 1.311E-06 | 1.200E-06 | 3.800E-06 |
| 10323 | 517 | E517N | A | 3.5e-06 | 2e-06 | 4.9e-06 | 3.467E-06 | 1.450E-06 | 2.000E-06 | 4.900E-06 |
| 10324 | 517 | E517D | A | 2.7e-06 | 2.7e-06 | 9.4e-07 | 2.113E-06 | 1.016E-06 | 9.400E-07 | 2.700E-06 |
| 10325 | 517 | E517C | A | 2.1e-06 | 1.8e-06 | 1.5e-06 | 1.800E-06 | 3.000E-07 | 1.500E-06 | 2.100E-06 |
| 10326 | 517 | E517E | A | 3.7e-06 | 2.2e-06 | 2e-06 | 2.633E-06 | 9.292E-07 | 2.000E-06 | 3.700E-06 |
| 10327 | 517 | E517Q | A | 1.4e-06 | 3.9e-06 | 3.5e-06 | 2.933E-06 | 1.343E-06 | 1.400E-06 | 3.900E-06 |
| 10328 | 517 | E517G | A | 2.7e-06 | 1.8e-06 | 1.7e-06 | 2.067E-06 | 5.508E-07 | 1.700E-06 | 2.700E-06 |
| 10329 | 517 | E517H | A | 2.7e-06 | 2.2e-06 | 1.7e-06 | 2.200E-06 | 5.000E-07 | 1.700E-06 | 2.700E-06 |
| 10330 | 517 | E517I | A | 4.1e-06 | 3.1e-06 | 2.9e-06 | 3.367E-06 | 6.429E-07 | 2.900E-06 | 4.100E-06 |
| 10331 | 517 | E517L | A | 3.4e-06 | 1.9e-06 | 4.5e-06 | 3.267E-06 | 1.305E-06 | 1.900E-06 | 4.500E-06 |
| 10332 | 517 | E517K | A | 2.3e-06 | 2.5e-06 | 2.7e-06 | 2.500E-06 | 2.000E-07 | 2.300E-06 | 2.700E-06 |
| 10333 | 517 | E517M | A | 3.8e-06 | 9.2e-07 | 1.8e-06 | 2.173E-06 | 1.476E-06 | 9.200E-07 | 3.800E-06 |
| 10334 | 517 | E517F | A | 2.2e-06 | 2.9e-06 | 3.7e-06 | 2.933E-06 | 7.506E-07 | 2.200E-06 | 3.700E-06 |
| 10335 | 517 | E517P | A | 1.8e-06 | 1e-06 | 2.7e-06 | 1.833E-06 | 8.505E-07 | 1.000E-06 | 2.700E-06 |
| 10336 | 517 | E517S | A | 2e-06 | 2.2e-06 | 2.5e-06 | 2.233E-06 | 2.517E-07 | 2.000E-06 | 2.500E-06 |
| 10337 | 517 | E517T | A | 2.5e-06 | 2.1e-06 | 2.8e-06 | 2.467E-06 | 3.512E-07 | 2.100E-06 | 2.800E-06 |
| 10338 | 517 | E517W | A | 1.7e-06 | 2.4e-06 | 2e-06 | 2.033E-06 | 3.512E-07 | 1.700E-06 | 2.400E-06 |
| 10339 | 517 | E517Y | A | 1.6e-06 | 2e-06 | 4.4e-06 | 2.667E-06 | 1.514E-06 | 1.600E-06 | 4.400E-06 |
| 10340 | 517 | E517V | A | 1.4e-06 | 3.3e-06 | 3.3e-06 | 2.667E-06 | 1.097E-06 | 1.400E-06 | 3.300E-06 |
| 10341 | 518 | C518A | A | 3.2e-06 | 4.3e-06 | 1.4e-06 | 2.967E-06 | 1.464E-06 | 1.400E-06 | 4.300E-06 |
| 10342 | 518 | C518R | A | 2.6e-06 | 2.1e-06 | 2.4e-06 | 2.367E-06 | 2.517E-07 | 2.100E-06 | 2.600E-06 |
| 10343 | 518 | C518N | A | 2.4e-06 | 1.9e-06 | 1.9e-06 | 2.067E-06 | 2.887E-07 | 1.900E-06 | 2.400E-06 |
| 10344 | 518 | C518D | A | 2.7e-06 | 2.6e-06 | 2.2e-06 | 2.500E-06 | 2.646E-07 | 2.200E-06 | 2.700E-06 |
| 10345 | 518 | C518C | A | 2.2e-06 | 1.9e-06 | 2.4e-06 | 2.167E-06 | 2.517E-07 | 1.900E-06 | 2.400E-06 |
| 10346 | 518 | C518E | A | 2.2e-06 | 4.4e-06 | 3.1e-06 | 3.233E-06 | 1.106E-06 | 2.200E-06 | 4.400E-06 |
| 10347 | 518 | C518Q | A | 2.5e-06 | 3e-06 | 3.6e-06 | 3.033E-06 | 5.508E-07 | 2.500E-06 | 3.600E-06 |
| 10348 | 518 | C518G | A | 2e-06 | 1.7e-06 | 1.7e-06 | 1.800E-06 | 1.732E-07 | 1.700E-06 | 2.000E-06 |
| 10349 | 518 | C518H | A | 2.7e-06 | 2.9e-06 | 1.3e-06 | 2.300E-06 | 8.718E-07 | 1.300E-06 | 2.900E-06 |
| 10350 | 518 | C518I | A | 1.1e-06 | 5.5e-06 | 2.1e-06 | 2.900E-06 | 2.307E-06 | 1.100E-06 | 5.500E-06 |
| 10351 | 518 | C518L | A | 2.5e-06 | 2.6e-06 | 2.6e-06 | 2.567E-06 | 5.774E-08 | 2.500E-06 | 2.600E-06 |
| 10352 | 518 | C518K | A | 2.6e-06 | 2.7e-06 | 3.3e-06 | 2.867E-06 | 3.786E-07 | 2.600E-06 | 3.300E-06 |
| 10353 | 518 | C518M | A | 1.9e-06 | 3.2e-06 | 3.7e-06 | 2.933E-06 | 9.292E-07 | 1.900E-06 | 3.700E-06 |
| 10354 | 518 | C518F | A | 2.4e-06 | 1.2e-06 | 2e-06 | 1.867E-06 | 6.110E-07 | 1.200E-06 | 2.400E-06 |
| 10355 | 518 | C518P | A | 2.5e-06 | 3.5e-06 | 2.3e-06 | 2.767E-06 | 6.429E-07 | 2.300E-06 | 3.500E-06 |
| 10356 | 518 | C518S | A | 1.7e-06 | 1.5e-06 | 2.4e-06 | 1.867E-06 | 4.726E-07 | 1.500E-06 | 2.400E-06 |
| 10357 | 518 | C518T | A | 2.7e-06 | 1e-06 | 2.1e-06 | 1.933E-06 | 8.622E-07 | 1.000E-06 | 2.700E-06 |
| 10358 | 518 | C518W | A | 2.9e-06 | 2.2e-06 | 2e-06 | 2.367E-06 | 4.726E-07 | 2.000E-06 | 2.900E-06 |
| 10359 | 518 | C518Y | A | 2.3e-06 | 2.9e-06 | 3e-06 | 2.733E-06 | 3.786E-07 | 2.300E-06 | 3.000E-06 |
| 10360 | 518 | C518V | A | 2.2e-06 | 2.7e-06 | 2.8e-06 | 2.567E-06 | 3.215E-07 | 2.200E-06 | 2.800E-06 |
| 10361 | 519 | V519A | A | 2.8e-06 | 1.5e-06 | 1.7e-06 | 2.000E-06 | 7.000E-07 | 1.500E-06 | 2.800E-06 |
| 10362 | 519 | V519R | A | 4.3e-06 | 1.9e-06 | 2e-06 | 2.733E-06 | 1.358E-06 | 1.900E-06 | 4.300E-06 |
| 10363 | 519 | V519N | A | 1.8e-06 | 1.8e-06 | 2e-06 | 1.867E-06 | 1.155E-07 | 1.800E-06 | 2.000E-06 |
| 10364 | 519 | V519D | A | 2.4e-06 | 1.6e-06 | 2e-06 | 2.000E-06 | 4.000E-07 | 1.600E-06 | 2.400E-06 |
| 10365 | 519 | V519C | A | 2.7e-06 | 3.1e-06 | 2e-06 | 2.600E-06 | 5.568E-07 | 2.000E-06 | 3.100E-06 |
| 10366 | 519 | V519E | A | 3.5e-06 | 2.2e-06 | 2.6e-06 | 2.767E-06 | 6.658E-07 | 2.200E-06 | 3.500E-06 |
| 10367 | 519 | V519Q | A | 2.9e-06 | 2.4e-06 | 2.2e-06 | 2.500E-06 | 3.606E-07 | 2.200E-06 | 2.900E-06 |
| 10368 | 519 | V519G | A | 2e-06 | 2.7e-06 | 1.8e-06 | 2.167E-06 | 4.726E-07 | 1.800E-06 | 2.700E-06 |
| 10369 | 519 | V519H | A | 3.2e-06 | 2e-06 | 1.5e-06 | 2.233E-06 | 8.737E-07 | 1.500E-06 | 3.200E-06 |

| | | | | | | | | | | |
|---|---|---|---|---|---|---|---|---|---|---|
| 10370 | 519 | V519I | A | 1.9e-06 | 1.3e-06 | 5.2e-06 | 2.800E-06 | 2.100E-06 | 1.300E-06 | 5.200E-06 |
| 10371 | 519 | V519L | A | 2.2e-06 | 1.5e-06 | 2.1e-06 | 1.933E-06 | 3.786E-07 | 1.500E-06 | 2.200E-06 |
| 10372 | 519 | V519K | A | 1.1e-06 | 1.9e-06 | 2.6e-06 | 1.867E-06 | 7.506E-07 | 1.100E-06 | 2.600E-06 |
| 10373 | 519 | V519M | A | 2.8e-06 | 4.1e-06 | 5.5e-06 | 4.133E-06 | 1.350E-06 | 2.800E-06 | 5.500E-06 |
| 10374 | 519 | V519F | A | 2.6e-06 | 1.7e-06 | 1.5e-06 | 1.933E-06 | 5.859E-07 | 1.500E-06 | 2.600E-06 |
| 10375 | 519 | V519P | A | 1.9e-06 | 6.6e-07 | 3.3e-06 | 1.953E-06 | 1.321E-06 | 6.600E-07 | 3.300E-06 |
| 10376 | 519 | V519S | A | 1.8e-06 | 2.1e-06 | 2.3e-06 | 2.067E-06 | 2.517E-07 | 1.800E-06 | 2.300E-06 |
| 10377 | 519 | V519T | A | 4.1e-06 | 2.5e-06 | 2e-06 | 2.867E-06 | 1.097E-06 | 2.000E-06 | 4.100E-06 |
| 10378 | 519 | V519W | A | 3.8e-06 | 4.1e-06 | 2e-06 | 3.300E-06 | 1.136E-06 | 2.000E-06 | 4.100E-06 |
| 10379 | 519 | V519Y | A | 1.6e-06 | 2.1e-06 | 2.4e-06 | 2.033E-06 | 4.041E-07 | 1.600E-06 | 2.400E-06 |
| 10380 | 519 | V519V | A | 2.4e-06 | 5e-06 | 1.4e-06 | 2.933E-06 | 1.858E-06 | 1.400E-06 | 5.000E-06 |
| 10381 | 520 | E520A | A | 2.4e-06 | 2.3e-06 | 3.9e-06 | 2.867E-06 | 8.963E-07 | 2.300E-06 | 3.900E-06 |
| 10382 | 520 | E520R | A | 2.1e-06 | 2.6e-06 | 1.4e-06 | 2.033E-06 | 6.028E-07 | 1.400E-06 | 2.600E-06 |
| 10383 | 520 | E520N | A | 3.2e-06 | 3.3e-06 | 1.6e-06 | 2.700E-06 | 9.539E-07 | 1.600E-06 | 3.300E-06 |
| 10384 | 520 | E520D | A | 2.4e-06 | 2.8e-06 | 3.1e-06 | 2.767E-06 | 3.512E-07 | 2.400E-06 | 3.100E-06 |
| 10385 | 520 | E520C | A | 1.9e-06 | 4.1e-06 | 3.9e-06 | 3.300E-06 | 1.217E-06 | 1.900E-06 | 4.100E-06 |
| 10386 | 520 | E520E | A | 2.2e-06 | 1.3e-06 | 2.3e-06 | 1.933E-06 | 5.508E-07 | 1.300E-06 | 2.300E-06 |
| 10387 | 520 | E520Q | A | 1.6e-06 | 2.3e-06 | 2.4e-06 | 2.100E-06 | 4.359E-07 | 1.600E-06 | 2.400E-06 |
| 10388 | 520 | E520G | A | 3e-06 | 5.4e-06 | 3.1e-06 | 3.833E-06 | 1.358E-06 | 3.000E-06 | 5.400E-06 |
| 10389 | 520 | E520H | A | 2.7e-06 | 3.5e-06 | 1.4e-06 | 2.533E-06 | 1.060E-06 | 1.400E-06 | 3.500E-06 |
| 10390 | 520 | E520I | A | 3.2e-06 | 3e-06 | 3.3e-06 | 3.167E-06 | 1.528E-07 | 3.000E-06 | 3.300E-06 |
| 10391 | 520 | E520L | A | 2e-06 | 2.4e-06 | 4e-06 | 2.800E-06 | 1.058E-06 | 2.000E-06 | 4.000E-06 |
| 10392 | 520 | E520K | A | 1.6e-06 | 2.4e-06 | 3.3e-06 | 2.433E-06 | 8.505E-07 | 1.600E-06 | 3.300E-06 |
| 10393 | 520 | E520M | A | 3.4e-06 | 2.2e-06 | 1.7e-06 | 2.433E-06 | 8.737E-07 | 1.700E-06 | 3.400E-06 |
| 10394 | 520 | E520F | A | 3.9e-06 | 2.6e-06 | 2.1e-06 | 2.867E-06 | 9.292E-07 | 2.100E-06 | 3.900E-06 |
| 10395 | 520 | E520P | A | 2.5e-06 | 1.6e-06 | 3.2e-06 | 2.433E-06 | 8.021E-07 | 1.600E-06 | 3.200E-06 |
| 10396 | 520 | E520S | A | 4.2e-06 | 2.3e-06 | 1.8e-06 | 2.767E-06 | 1.266E-06 | 1.800E-06 | 4.200E-06 |
| 10397 | 520 | E520T | A | 2.5e-06 | 1.6e-06 | 2.7e-06 | 2.267E-06 | 5.859E-07 | 1.600E-06 | 2.700E-06 |
| 10398 | 520 | E520W | A | 4.7e-06 | 2.8e-06 | 1.4e-06 | 2.967E-06 | 1.656E-06 | 1.400E-06 | 4.700E-06 |
| 10399 | 520 | E520Y | A | 2.6e-06 | 1.4e-06 | 1.9e-06 | 1.967E-06 | 6.028E-07 | 1.400E-06 | 2.600E-06 |
| 10400 | 520 | E520V | A | 1.7e-06 | 1.3e-06 | 2.4e-06 | 1.800E-06 | 5.568E-07 | 1.300E-06 | 2.400E-06 |
| 10401 | 521 | E521A | A | 2.8e-06 | 4.5e-06 | 3.9e-06 | 3.733E-06 | 8.622E-07 | 2.800E-06 | 4.500E-06 |
| 10402 | 521 | E521R | A | 1.7e-06 | 1.9e-06 | 2.4e-06 | 2.000E-06 | 3.606E-07 | 1.700E-06 | 2.400E-06 |
| 10403 | 521 | E521N | A | 3.2e-06 | 3.8e-06 | 2.1e-06 | 3.033E-06 | 8.622E-07 | 2.100E-06 | 3.800E-06 |
| 10404 | 521 | E521D | A | 2.9e-06 | 3.7e-06 | 2.1e-06 | 2.900E-06 | 8.000E-07 | 2.100E-06 | 3.700E-06 |
| 10405 | 521 | E521C | A | 1.9e-06 | 3.1e-06 | 3e-06 | 2.667E-06 | 6.658E-07 | 1.900E-06 | 3.100E-06 |
| 10406 | 521 | E521E | A | 2.6e-06 | 1.9e-06 | 1.5e-06 | 2.000E-06 | 5.568E-07 | 1.500E-06 | 2.600E-06 |
| 10407 | 521 | E521Q | A | 1.8e-06 | 1.5e-06 | 1.3e-06 | 1.533E-06 | 2.517E-07 | 1.300E-06 | 1.800E-06 |
| 10408 | 521 | E521G | A | 2.1e-06 | 3.3e-06 | 1.8e-06 | 2.400E-06 | 7.937E-07 | 1.800E-06 | 3.300E-06 |
| 10409 | 521 | E521H | A | 6e-06 | 2e-06 | 3.7e-06 | 3.900E-06 | 2.007E-06 | 2.000E-06 | 6.000E-06 |
| 10410 | 521 | E521I | A | 1.9e-06 | 3.2e-06 | 2.9e-06 | 2.667E-06 | 6.807E-07 | 1.900E-06 | 3.200E-06 |
| 10411 | 521 | E521L | A | 7.3e-06 | 2e-06 | 1.6e-06 | 3.633E-06 | 3.182E-06 | 1.600E-06 | 7.300E-06 |
| 10412 | 521 | E521K | A | 2.6e-06 | 1.5e-06 | 1.3e-06 | 1.800E-06 | 7.000E-07 | 1.300E-06 | 2.600E-06 |
| 10413 | 521 | E521M | A | 3e-06 | 2.9e-06 | 3.1e-06 | 3.000E-06 | 1.000E-07 | 2.900E-06 | 3.100E-06 |
| 10414 | 521 | E521F | A | 4.1e-06 | 2.3e-06 | 1.5e-06 | 2.633E-06 | 1.332E-06 | 1.500E-06 | 4.100E-06 |
| 10415 | 521 | E521P | A | 3.5e-06 | 1.5e-06 | 2.8e-06 | 2.600E-06 | 1.015E-06 | 1.500E-06 | 3.500E-06 |
| 10416 | 521 | E521S | A | 1.7e-06 | 1.9e-06 | 3.7e-06 | 2.433E-06 | 1.102E-06 | 1.700E-06 | 3.700E-06 |
| 10417 | 521 | E521T | A | 4.1e-06 | 2.5e-06 | 3.4e-06 | 3.333E-06 | 8.021E-07 | 2.500E-06 | 4.100E-06 |
| 10418 | 521 | E521W | A | 2.7e-06 | 2.6e-06 | 2.6e-06 | 2.633E-06 | 5.774E-08 | 2.600E-06 | 2.700E-06 |
| 10419 | 521 | E521Y | A | 2.4e-06 | 4.8e-06 | 3.4e-06 | 3.533E-06 | 1.206E-06 | 2.400E-06 | 4.800E-06 |
| 10420 | 521 | E521V | A | 2.6e-06 | 1.6e-06 | 2.5e-06 | 2.233E-06 | 5.508E-07 | 1.600E-06 | 2.600E-06 |
| 10421 | 522 | C522A | A | 4.7e-06 | 3.9e-06 | 2.4e-06 | 3.667E-06 | 1.168E-06 | 2.400E-06 | 4.700E-06 |
| 10422 | 522 | C522R | A | 2.3e-06 | 4.2e-06 | 3.4e-06 | 3.300E-06 | 9.539E-07 | 2.300E-06 | 4.200E-06 |
| 10423 | 522 | C522N | A | 1.1e-06 | 2.2e-06 | 2e-06 | 1.767E-06 | 5.859E-07 | 1.100E-06 | 2.200E-06 |
| 10424 | 522 | C522D | A | 2.8e-06 | 2.9e-06 | 1.3e-06 | 2.333E-06 | 8.963E-07 | 1.300E-06 | 2.900E-06 |
| 10425 | 522 | C522C | A | 1.6e-06 | 2.8e-06 | 2.8e-06 | 2.400E-06 | 6.928E-07 | 1.600E-06 | 2.800E-06 |
| 10426 | 522 | C522E | A | 2.1e-06 | 1.9e-06 | 1.9e-06 | 1.967E-06 | 1.155E-07 | 1.900E-06 | 2.100E-06 |
| 10427 | 522 | C522Q | A | 1.4e-06 | 4.3e-06 | 2.9e-06 | 2.867E-06 | 1.450E-06 | 1.400E-06 | 4.300E-06 |
| 10428 | 522 | C522G | A | 3.9e-06 | 1.2e-06 | 4.2e-06 | 3.100E-06 | 1.652E-06 | 1.200E-06 | 4.200E-06 |
| 10429 | 522 | C522H | A | 2.1e-06 | 2.8e-06 | 2.6e-06 | 2.500E-06 | 3.606E-07 | 2.100E-06 | 2.800E-06 |
| 10430 | 522 | C522I | A | 2.9e-06 | 2.6e-06 | 3.6e-06 | 3.033E-06 | 5.132E-07 | 2.600E-06 | 3.600E-06 |
| 10431 | 522 | C522L | A | 2.2e-06 | 2.3e-06 | 2.1e-06 | 2.200E-06 | 1.000E-07 | 2.100E-06 | 2.300E-06 |
| 10432 | 522 | C522K | A | 2e-06 | 1.8e-06 | 1.8e-06 | 1.867E-06 | 1.155E-07 | 1.800E-06 | 2.000E-06 |
| 10433 | 522 | C522M | A | 2.7e-06 | 3.6e-06 | 3.3e-06 | 3.200E-06 | 4.583E-07 | 2.700E-06 | 3.600E-06 |
| 10434 | 522 | C522F | A | 2.7e-06 | 2e-06 | 2.3e-06 | 2.333E-06 | 3.512E-07 | 2.000E-06 | 2.700E-06 |
| 10435 | 522 | C522P | A | 2.2e-06 | 2.7e-06 | 2.4e-06 | 2.433E-06 | 2.517E-07 | 2.200E-06 | 2.700E-06 |
| 10436 | 522 | C522S | A | 2.7e-06 | 2.3e-06 | 2.9e-06 | 2.633E-06 | 3.055E-07 | 2.300E-06 | 2.900E-06 |
| 10437 | 522 | C522T | A | 2.2e-06 | 2.2e-06 | 2.1e-06 | 2.167E-06 | 5.774E-08 | 2.100E-06 | 2.200E-06 |
| 10438 | 522 | C522W | A | 2.8e-06 | 2.3e-06 | 1.7e-06 | 2.267E-06 | 5.508E-07 | 1.700E-06 | 2.800E-06 |
| 10439 | 522 | C522Y | A | 3.9e-06 | 2.5e-06 | 3.3e-06 | 3.233E-06 | 7.024E-07 | 2.500E-06 | 3.900E-06 |
| 10440 | 522 | C522V | A | 2.6e-06 | 2.4e-06 | 3e-06 | 2.667E-06 | 3.055E-07 | 2.400E-06 | 3.000E-06 |
| 10441 | 523 | R523A | A | 1.6e-06 | 2.5e-06 | 2.7e-06 | 2.267E-06 | 5.859E-07 | 1.600E-06 | 2.700E-06 |
| 10442 | 523 | R523R | A | 1.9e-06 | 4.3e-06 | 2.2e-06 | 2.800E-06 | 1.308E-06 | 1.900E-06 | 4.300E-06 |
| 10443 | 523 | R523N | A | 2e-06 | 2e-06 | 1.5e-06 | 1.833E-06 | 2.887E-07 | 1.500E-06 | 2.000E-06 |
| 10444 | 523 | R523D | A | 2.2e-06 | 2.2e-06 | 3.7e-06 | 2.700E-06 | 8.660E-07 | 2.200E-06 | 3.700E-06 |
| 10445 | 523 | R523C | A | 3.2e-06 | 1.6e-06 | 3.1e-06 | 2.633E-06 | 8.963E-07 | 1.600E-06 | 3.200E-06 |
| 10446 | 523 | R523E | A | 1.9e-06 | 1.8e-06 | 2.7e-06 | 2.133E-06 | 4.933E-07 | 1.800E-06 | 2.700E-06 |
| 10447 | 523 | R523Q | A | 2.3e-06 | 1.7e-06 | 1.9e-06 | 1.967E-06 | 3.055E-07 | 1.700E-06 | 2.300E-06 |
| 10448 | 523 | R523G | A | 2.4e-06 | 3.4e-06 | 2e-06 | 2.600E-06 | 7.211E-07 | 2.000E-06 | 3.400E-06 |
| 10449 | 523 | R523H | A | 2.2e-06 | 1.9e-06 | 1.4e-06 | 1.833E-06 | 4.041E-07 | 1.400E-06 | 2.200E-06 |
| 10450 | 523 | R523I | A | 1.2e-06 | 2.1e-06 | 2.8e-06 | 2.033E-06 | 8.021E-07 | 1.200E-06 | 2.800E-06 |
| 10451 | 523 | R523L | A | 3.7e-06 | 2.3e-06 | 2.4e-06 | 2.800E-06 | 7.810E-07 | 2.300E-06 | 3.700E-06 |
| 10452 | 523 | R523K | A | 1.8e-06 | 3.6e-06 | 2.1e-06 | 2.500E-06 | 9.644E-07 | 1.800E-06 | 3.600E-06 |
| 10453 | 523 | R523M | A | 2.1e-06 | 1.9e-06 | 2.7e-06 | 2.233E-06 | 4.163E-07 | 1.900E-06 | 2.700E-06 |
| 10454 | 523 | R523F | A | 1.6e-06 | 1.9e-06 | 3.4e-06 | 2.300E-06 | 9.644E-07 | 1.600E-06 | 3.400E-06 |

| | | | | | | | | | | |
|---|---|---|---|---|---|---|---|---|---|---|
| 10455 | 523 | R523P | A | 2e-06 | 2.4e-06 | 2.7e-06 | 2.367E-06 | 3.512E-07 | 2.000E-06 | 2.700E-06 |
| 10456 | 523 | R523S | A | 2.5e-06 | 4.2e-06 | 3.6e-06 | 3.433E-06 | 8.622E-07 | 2.500E-06 | 4.200E-06 |
| 10457 | 523 | R523T | A | 2.4e-06 | 1.8e-06 | 3.5e-06 | 2.567E-06 | 8.622E-07 | 1.800E-06 | 3.500E-06 |
| 10458 | 523 | R523W | A | 2.3e-06 | 1.2e-06 | 2.2e-06 | 1.900E-06 | 6.083E-07 | 1.200E-06 | 2.300E-06 |
| 10459 | 523 | R523Y | A | 2.6e-06 | 2.3e-06 | 2.7e-06 | 2.533E-06 | 2.082E-07 | 2.300E-06 | 2.700E-06 |
| 10460 | 523 | R523V | A | 2.7e-06 | 2.6e-06 | 1.7e-06 | 2.333E-06 | 5.508E-07 | 1.700E-06 | 2.700E-06 |
| 10461 | 524 | V524A | A | 1.3e-06 | 1.8e-06 | 2.9e-06 | 2.000E-06 | 8.185E-07 | 1.300E-06 | 2.900E-06 |
| 10462 | 524 | V524R | A | 1.2e-06 | 1.6e-06 | 2e-06 | 1.600E-06 | 4.000E-07 | 1.200E-06 | 2.000E-06 |
| 10463 | 524 | V524N | A | 1.8e-06 | 1.6e-06 | 2.6e-06 | 2.000E-06 | 5.292E-07 | 1.600E-06 | 2.600E-06 |
| 10464 | 524 | V524D | A | 2.6e-06 | 1.6e-06 | 2.4e-06 | 2.200E-06 | 5.292E-07 | 1.600E-06 | 2.600E-06 |
| 10465 | 524 | V524C | A | 1.6e-06 | 2.3e-06 | 2e-06 | 1.967E-06 | 3.512E-07 | 1.600E-06 | 2.300E-06 |
| 10466 | 524 | V524E | A | 1.7e-06 | 2.2e-06 | 1.7e-06 | 1.867E-06 | 2.887E-07 | 1.700E-06 | 2.200E-06 |
| 10467 | 524 | V524Q | A | 2.9e-06 | 3.6e-06 | 1.6e-06 | 2.700E-06 | 1.015E-06 | 1.600E-06 | 3.600E-06 |
| 10468 | 524 | V524G | A | 1.5e-06 | 1.9e-06 | 2.3e-06 | 1.900E-06 | 4.000E-07 | 1.500E-06 | 2.300E-06 |
| 10469 | 524 | V524H | A | 2e-06 | 2.5e-06 | 1.4e-06 | 1.967E-06 | 5.508E-07 | 1.400E-06 | 2.500E-06 |
| 10470 | 524 | V524I | A | 1.8e-06 | 1.7e-06 | 5.1e-06 | 2.867E-06 | 1.935E-06 | 1.700E-06 | 5.100E-06 |
| 10471 | 524 | V524L | A | 2.8e-06 | 1.8e-06 | 3e-06 | 2.533E-06 | 6.429E-07 | 1.800E-06 | 3.000E-06 |
| 10472 | 524 | V524K | A | 1.8e-06 | 2.1e-06 | 2.5e-06 | 2.133E-06 | 3.512E-07 | 1.800E-06 | 2.500E-06 |
| 10473 | 524 | V524M | A | 3e-06 | 3.2e-06 | 4.2e-06 | 3.467E-06 | 6.429E-07 | 3.000E-06 | 4.200E-06 |
| 10474 | 524 | V524F | A | 1.5e-06 | 4.3e-06 | 3.1e-06 | 2.967E-06 | 1.405E-06 | 1.500E-06 | 4.300E-06 |
| 10475 | 524 | V524P | A | 3.2e-06 | 2.7e-06 | 1.5e-06 | 2.467E-06 | 8.737E-07 | 1.500E-06 | 3.200E-06 |
| 10476 | 524 | V524S | A | 2e-06 | 2.9e-06 | 1.7e-06 | 2.200E-06 | 6.245E-07 | 1.700E-06 | 2.900E-06 |
| 10477 | 524 | V524T | A | 2.7e-06 | 2.3e-06 | 2.2e-06 | 2.400E-06 | 2.646E-07 | 2.200E-06 | 2.700E-06 |
| 10478 | 524 | V524W | A | 2.8e-06 | 1.7e-06 | 2.4e-06 | 2.300E-06 | 5.568E-07 | 1.700E-06 | 2.800E-06 |
| 10479 | 524 | V524Y | A | 3e-06 | 1.7e-06 | 1.8e-06 | 2.167E-06 | 7.234E-07 | 1.700E-06 | 3.000E-06 |
| 10480 | 524 | V524V | A | 2.5e-06 | 2.6e-06 | 2.4e-06 | 2.500E-06 | 1.000E-07 | 2.400E-06 | 2.600E-06 |
| 10481 | 525 | L525A | A | 2e-06 | 1.6e-06 | 2.5e-06 | 2.033E-06 | 4.509E-07 | 1.600E-06 | 2.500E-06 |
| 10482 | 525 | L525R | A | 2.2e-06 | 3.7e-06 | 2.9e-06 | 2.933E-06 | 7.506E-07 | 2.200E-06 | 3.700E-06 |
| 10483 | 525 | L525N | A | 7.6e-07 | 2e-06 | 2.7e-06 | 1.820E-06 | 9.824E-07 | 7.600E-07 | 2.700E-06 |
| 10484 | 525 | L525D | A | 2.1e-06 | 1.8e-06 | 2.3e-06 | 2.067E-06 | 2.517E-07 | 1.800E-06 | 2.300E-06 |
| 10485 | 525 | L525C | A | 2.3e-06 | 2.5e-06 | 4.9e-06 | 3.233E-06 | 1.447E-06 | 2.300E-06 | 4.900E-06 |
| 10486 | 525 | L525E | A | 1.8e-06 | 7.4e-06 | 2.4e-06 | 3.867E-06 | 3.075E-06 | 1.800E-06 | 7.400E-06 |
| 10487 | 525 | L525Q | A | 1.4e-06 | 3.8e-06 | 3.6e-06 | 2.933E-06 | 1.332E-06 | 1.400E-06 | 3.800E-06 |
| 10488 | 525 | L525G | A | 3e-06 | 1.7e-06 | 6.4e-07 | 1.780E-06 | 1.182E-06 | 6.400E-07 | 3.000E-06 |
| 10489 | 525 | L525H | A | 1.8e-06 | 2.1e-06 | 1.7e-06 | 1.867E-06 | 2.082E-07 | 1.700E-06 | 2.100E-06 |
| 10490 | 525 | L525I | A | 3.1e-06 | 2.7e-06 | 2.7e-06 | 2.833E-06 | 2.309E-07 | 2.700E-06 | 3.100E-06 |
| 10491 | 525 | L525L | A | 1.3e-06 | 2e-06 | 2.8e-06 | 2.033E-06 | 7.506E-07 | 1.300E-06 | 2.800E-06 |
| 10492 | 525 | L525K | A | 2e-06 | 4.4e-06 | 2.5e-06 | 2.967E-06 | 1.266E-06 | 2.000E-06 | 4.400E-06 |
| 10493 | 525 | L525M | A | 2.2e-06 | 2.1e-06 | 2.6e-06 | 2.300E-06 | 2.646E-07 | 2.100E-06 | 2.600E-06 |
| 10494 | 525 | L525F | A | 2.1e-06 | 1.9e-06 | 1.6e-06 | 1.867E-06 | 2.517E-07 | 1.600E-06 | 2.100E-06 |
| 10495 | 525 | L525P | A | 1.8e-06 | 2.2e-06 | 1.6e-06 | 1.867E-06 | 3.055E-07 | 1.600E-06 | 2.200E-06 |
| 10496 | 525 | L525S | A | 1.8e-06 | 3.7e-06 | 2.5e-06 | 2.667E-06 | 9.609E-07 | 1.800E-06 | 3.700E-06 |
| 10497 | 525 | L525T | A | 2.3e-06 | 2e-06 | 1.6e-06 | 1.967E-06 | 3.512E-07 | 1.600E-06 | 2.300E-06 |
| 10498 | 525 | L525W | A | 3.1e-06 | 4.1e-06 | 3.5e-06 | 3.567E-06 | 5.033E-07 | 3.100E-06 | 4.100E-06 |
| 10499 | 525 | L525Y | A | 3.3e-06 | 2.1e-06 | 2.2e-06 | 2.533E-06 | 6.658E-07 | 2.100E-06 | 3.300E-06 |
| 10500 | 525 | L525V | A | 2.4e-06 | 3.2e-06 | 2.1e-06 | 2.567E-06 | 5.686E-07 | 2.100E-06 | 3.200E-06 |
| 10501 | 526 | Q526A | A | 1.4e-06 | 3.4e-06 | 3.5e-06 | 2.767E-06 | 1.185E-06 | 1.400E-06 | 3.500E-06 |
| 10502 | 526 | Q526R | A | 3e-06 | 2.3e-06 | 3.1e-06 | 2.800E-06 | 4.359E-07 | 2.300E-06 | 3.100E-06 |
| 10503 | 526 | Q526N | A | 2.1e-06 | 2.2e-06 | 4.6e-06 | 2.967E-06 | 1.415E-06 | 2.100E-06 | 4.600E-06 |
| 10504 | 526 | Q526D | A | 2.2e-06 | 2.3e-06 | 3.2e-06 | 2.567E-06 | 5.508E-07 | 2.200E-06 | 3.200E-06 |
| 10505 | 526 | Q526C | A | 3.1e-06 | 2.9e-06 | 2.8e-06 | 2.933E-06 | 1.528E-07 | 2.800E-06 | 3.100E-06 |
| 10506 | 526 | Q526E | A | 1.3e-06 | 2.3e-06 | 2e-06 | 1.867E-06 | 5.132E-07 | 1.300E-06 | 2.300E-06 |
| 10507 | 526 | Q526Q | A | 3.2e-06 | 2.7e-06 | 1.8e-06 | 2.567E-06 | 7.095E-07 | 1.800E-06 | 3.200E-06 |
| 10508 | 526 | Q526G | A | 3.2e-06 | 3.2e-06 | 3.8e-06 | 3.400E-06 | 3.464E-07 | 3.200E-06 | 3.800E-06 |
| 10509 | 526 | Q526H | A | 1.5e-06 | 1.7e-06 | 1.4e-06 | 1.533E-06 | 1.528E-07 | 1.400E-06 | 1.700E-06 |
| 10510 | 526 | Q526I | A | 5.2e-06 | 2.9e-06 | 4.1e-06 | 4.067E-06 | 1.150E-06 | 2.900E-06 | 5.200E-06 |
| 10511 | 526 | Q526L | A | 2.2e-06 | 2.5e-06 | 2.2e-06 | 2.300E-06 | 1.732E-07 | 2.200E-06 | 2.500E-06 |
| 10512 | 526 | Q526K | A | 1.6e-06 | 2.3e-06 | 3.2e-06 | 2.367E-06 | 8.021E-07 | 1.600E-06 | 3.200E-06 |
| 10513 | 526 | Q526M | A | 3e-06 | 1.3e-06 | 2.1e-06 | 2.133E-06 | 8.505E-07 | 1.300E-06 | 3.000E-06 |
| 10514 | 526 | Q526F | A | 2.2e-06 | 2.2e-06 | 8.2e-07 | 1.740E-06 | 7.967E-07 | 8.200E-07 | 2.200E-06 |
| 10515 | 526 | Q526P | A | 2.6e-06 | 2e-06 | 1.3e-06 | 1.967E-06 | 6.506E-07 | 1.300E-06 | 2.600E-06 |
| 10516 | 526 | Q526S | A | 9.2e-07 | 3.7e-06 | 1.5e-06 | 2.040E-06 | 1.467E-06 | 9.200E-07 | 3.700E-06 |
| 10517 | 526 | Q526T | A | 2.3e-06 | 2.6e-06 | 2.5e-06 | 2.467E-06 | 1.528E-07 | 2.300E-06 | 2.600E-06 |
| 10518 | 526 | Q526W | A | 2.3e-06 | 2.2e-06 | 2.3e-06 | 2.267E-06 | 5.774E-08 | 2.200E-06 | 2.300E-06 |
| 10519 | 526 | Q526Y | A | 3e-06 | 1.8e-06 | 1.6e-06 | 2.133E-06 | 7.572E-07 | 1.600E-06 | 3.000E-06 |
| 10520 | 526 | Q526V | A | 2e-06 | 4.1e-06 | 4.8e-06 | 3.633E-06 | 1.457E-06 | 2.000E-06 | 4.800E-06 |
| 10521 | 527 | G527A | A | 6.5e-06 | 2.7e-06 | 1.6e-06 | 3.600E-06 | 2.571E-06 | 1.600E-06 | 6.500E-06 |
| 10522 | 527 | G527R | A | 2.6e-06 | 2.9e-06 | 1.8e-06 | 2.433E-06 | 5.686E-07 | 1.800E-06 | 2.900E-06 |
| 10523 | 527 | G527N | A | 2.6e-06 | 2.1e-06 | 1.7e-06 | 2.133E-06 | 4.509E-07 | 1.700E-06 | 2.600E-06 |
| 10524 | 527 | G527D | A | 3e-06 | 2.2e-06 | 2.2e-06 | 2.467E-06 | 4.619E-07 | 2.200E-06 | 3.000E-06 |
| 10525 | 527 | G527C | A | 3.9e-06 | 2.1e-06 | 3.6e-06 | 3.200E-06 | 9.644E-07 | 2.100E-06 | 3.900E-06 |
| 10526 | 527 | G527E | A | 3.9e-06 | 2.7e-06 | 3.1e-06 | 3.233E-06 | 6.110E-07 | 2.700E-06 | 3.900E-06 |
| 10527 | 527 | G527Q | A | 1.4e-06 | 2.3e-06 | 2.7e-06 | 2.133E-06 | 6.658E-07 | 1.400E-06 | 2.700E-06 |
| 10528 | 527 | G527G | A | 3e-06 | 3.1e-06 | 2e-06 | 2.700E-06 | 6.083E-07 | 2.000E-06 | 3.100E-06 |
| 10529 | 527 | G527H | A | 3e-06 | 2.7e-06 | 2.1e-06 | 2.600E-06 | 4.583E-07 | 2.100E-06 | 3.000E-06 |
| 10530 | 527 | G527I | A | 2.5e-06 | 2.3e-06 | 2.4e-06 | 2.400E-06 | 1.000E-07 | 2.300E-06 | 2.500E-06 |
| 10531 | 527 | G527L | A | 2.6e-06 | 1.7e-06 | 2.4e-06 | 2.233E-06 | 4.726E-07 | 1.700E-06 | 2.600E-06 |
| 10532 | 527 | G527K | A | 3.8e-06 | 2.6e-06 | 2e-06 | 2.800E-06 | 9.165E-07 | 2.000E-06 | 3.800E-06 |
| 10533 | 527 | G527M | A | 3.1e-06 | 2.8e-06 | 2.5e-06 | 2.800E-06 | 3.000E-07 | 2.500E-06 | 3.100E-06 |
| 10534 | 527 | G527F | A | 1.3e-06 | 5.4e-06 | 1.9e-06 | 2.867E-06 | 2.214E-06 | 1.300E-06 | 5.400E-06 |
| 10535 | 527 | G527P | A | 1.6e-06 | 2.7e-06 | 2.2e-06 | 2.167E-06 | 5.508E-07 | 1.600E-06 | 2.700E-06 |
| 10536 | 527 | G527S | A | 1.5e-06 | 3e-06 | 3.1e-06 | 2.533E-06 | 8.963E-07 | 1.500E-06 | 3.100E-06 |
| 10537 | 527 | G527T | A | 3e-06 | 2.9e-06 | 2.6e-06 | 2.833E-06 | 2.082E-07 | 2.600E-06 | 3.000E-06 |
| 10538 | 527 | G527W | A | 2.3e-06 | 2e-06 | 2.1e-06 | 2.133E-06 | 1.528E-07 | 2.000E-06 | 2.300E-06 |
| 10539 | 527 | G527Y | A | 2.1e-06 | 1.8e-06 | 2.4e-06 | 2.100E-06 | 3.000E-07 | 1.800E-06 | 2.400E-06 |

| | | | | | | | | | | |
|---|---|---|---|---|---|---|---|---|---|---|
| 10540 | 527 | G527V | A | 2.7e-06 | 1.3e-06 | 2.1e-06 | 2.033E-06 | 7.024E-07 | 1.300E-06 | 2.700E-06 |
| 10541 | 528 | L528A | A | 2.5e-06 | 2.2e-06 | 3.1e-06 | 2.600E-06 | 4.583E-07 | 2.200E-06 | 3.100E-06 |
| 10542 | 528 | L528R | A | 3.8e-06 | 2.3e-06 | 1.5e-06 | 2.533E-06 | 1.168E-06 | 1.500E-06 | 3.800E-06 |
| 10543 | 528 | L528N | A | 2.6e-06 | 3.2e-06 | 1.9e-06 | 2.567E-06 | 6.506E-07 | 1.900E-06 | 3.200E-06 |
| 10544 | 528 | L528D | A | 3.7e-06 | 3.3e-06 | 3e-06 | 3.333E-06 | 3.512E-07 | 3.000E-06 | 3.700E-06 |
| 10545 | 528 | L528C | A | 2.1e-06 | 1.9e-06 | 1.5e-06 | 1.833E-06 | 3.055E-07 | 1.500E-06 | 2.100E-06 |
| 10546 | 528 | L528E | A | 2.8e-06 | 1.9e-06 | 1.4e-06 | 2.033E-06 | 7.095E-07 | 1.400E-06 | 2.800E-06 |
| 10547 | 528 | L528Q | A | 2.3e-06 | 3.5e-06 | 2.5e-06 | 2.767E-06 | 6.429E-07 | 2.300E-06 | 3.500E-06 |
| 10548 | 528 | L528G | A | 2.8e-06 | 1.6e-06 | 2e-06 | 2.133E-06 | 6.110E-07 | 1.600E-06 | 2.800E-06 |
| 10549 | 528 | L528H | A | 2.3e-06 | 3e-06 | 1.8e-06 | 2.367E-06 | 6.028E-07 | 1.800E-06 | 3.000E-06 |
| 10550 | 528 | L528I | A | 3.3e-06 | 2.8e-06 | 3.4e-06 | 3.167E-06 | 3.215E-07 | 2.800E-06 | 3.400E-06 |
| 10551 | 528 | L528L | A | 3.4e-06 | 2.9e-06 | 2.2e-06 | 2.833E-06 | 6.028E-07 | 2.200E-06 | 3.400E-06 |
| 10552 | 528 | L528K | A | 1.9e-06 | 3.4e-06 | 2.1e-06 | 2.467E-06 | 8.145E-07 | 1.900E-06 | 3.400E-06 |
| 10553 | 528 | L528M | A | 3e-06 | 1.8e-06 | 1.5e-06 | 2.100E-06 | 7.937E-07 | 1.500E-06 | 3.000E-06 |
| 10554 | 528 | L528F | A | 2.9e-06 | 2.5e-06 | 3.6e-06 | 3.000E-06 | 5.568E-07 | 2.500E-06 | 3.600E-06 |
| 10555 | 528 | L528P | A | 3.8e-06 | 3.3e-06 | 3.3e-06 | 3.467E-06 | 2.887E-07 | 3.300E-06 | 3.800E-06 |
| 10556 | 528 | L528S | A | 2.1e-06 | 2.3e-06 | 1.5e-06 | 1.967E-06 | 4.163E-07 | 1.500E-06 | 2.300E-06 |
| 10557 | 528 | L528T | A | 1.4e-06 | 3e-06 | 2e-06 | 2.133E-06 | 8.083E-07 | 1.400E-06 | 3.000E-06 |
| 10558 | 528 | L528W | A | 2.1e-06 | 1.7e-06 | 3.5e-06 | 2.433E-06 | 9.452E-07 | 1.700E-06 | 3.500E-06 |
| 10559 | 528 | L528Y | A | 1.8e-06 | 2.5e-06 | 1.6e-06 | 1.967E-06 | 4.726E-07 | 1.600E-06 | 2.500E-06 |
| 10560 | 528 | L528V | A | 2.1e-06 | 3.7e-06 | 1.6e-06 | 2.467E-06 | 1.097E-06 | 1.600E-06 | 3.700E-06 |
| 10561 | 529 | P529A | A | 1.5e-06 | 2e-06 | 2e-06 | 1.833E-06 | 2.887E-07 | 1.500E-06 | 2.000E-06 |
| 10562 | 529 | P529R | A | 2.3e-06 | 2.6e-06 | 3.5e-06 | 2.800E-06 | 6.245E-07 | 2.300E-06 | 3.500E-06 |
| 10563 | 529 | P529N | A | 2.4e-06 | 3.4e-06 | 1.5e-06 | 2.433E-06 | 9.504E-07 | 1.500E-06 | 3.400E-06 |
| 10564 | 529 | P529D | A | 3.3e-06 | 4.6e-06 | 1.8e-06 | 3.233E-06 | 1.401E-06 | 1.800E-06 | 4.600E-06 |
| 10565 | 529 | P529C | A | 2.4e-06 | 1.9e-06 | 2.8e-06 | 2.367E-06 | 4.509E-07 | 1.900E-06 | 2.800E-06 |
| 10566 | 529 | P529E | A | 2.6e-06 | 2.4e-06 | 3.3e-06 | 2.767E-06 | 4.726E-07 | 2.400E-06 | 3.300E-06 |
| 10567 | 529 | P529Q | A | 3.7e-06 | 3.2e-06 | 2.6e-06 | 3.167E-06 | 5.508E-07 | 2.600E-06 | 3.700E-06 |
| 10568 | 529 | P529G | A | 2.2e-06 | 2.3e-06 | 2.3e-06 | 2.267E-06 | 5.774E-08 | 2.200E-06 | 2.300E-06 |
| 10569 | 529 | P529H | A | 3.8e-06 | 1.7e-06 | 2.6e-06 | 2.700E-06 | 1.054E-06 | 1.700E-06 | 3.800E-06 |
| 10570 | 529 | P529I | A | 3.4e-06 | 1.8e-06 | 1.8e-06 | 2.333E-06 | 9.238E-07 | 1.800E-06 | 3.400E-06 |
| 10571 | 529 | P529L | A | 2e-06 | 1.4e-06 | 2.4e-06 | 1.933E-06 | 5.033E-07 | 1.400E-06 | 2.400E-06 |
| 10572 | 529 | P529K | A | 3e-06 | 2.2e-06 | 1.7e-06 | 2.300E-06 | 6.557E-07 | 1.700E-06 | 3.000E-06 |
| 10573 | 529 | P529M | A | 3.3e-06 | 2.3e-06 | 2.5e-06 | 2.700E-06 | 5.292E-07 | 2.300E-06 | 3.300E-06 |
| 10574 | 529 | P529F | A | 3.7e-06 | 2.6e-06 | 2.2e-06 | 2.833E-06 | 7.767E-07 | 2.200E-06 | 3.700E-06 |
| 10575 | 529 | P529P | A | 3e-06 | 2.7e-06 | 2.4e-06 | 2.700E-06 | 3.000E-07 | 2.400E-06 | 3.000E-06 |
| 10576 | 529 | P529S | A | 1.9e-06 | 1.3e-06 | 2.5e-06 | 1.900E-06 | 6.000E-07 | 1.300E-06 | 2.500E-06 |
| 10577 | 529 | P529T | A | 1.6e-06 | 2.3e-06 | 2e-06 | 1.967E-06 | 3.512E-07 | 1.600E-06 | 2.300E-06 |
| 10578 | 529 | P529W | A | 1.5e-06 | 3.6e-06 | 2.5e-06 | 2.533E-06 | 1.050E-06 | 1.500E-06 | 3.600E-06 |
| 10579 | 529 | P529Y | A | 1.7e-06 | 2.2e-06 | 2.2e-06 | 2.033E-06 | 2.887E-07 | 1.700E-06 | 2.200E-06 |
| 10580 | 529 | P529V | A | 2.2e-06 | 1.9e-06 | 2.6e-06 | 2.233E-06 | 3.512E-07 | 1.900E-06 | 2.600E-06 |
| 10581 | 530 | R530A | A | 1.6e-06 | 9.6e-07 | 2.6e-06 | 1.720E-06 | 8.266E-07 | 9.600E-07 | 2.600E-06 |
| 10582 | 530 | R530R | A | 2.8e-06 | 2.5e-06 | 1.6e-06 | 2.300E-06 | 6.245E-07 | 1.600E-06 | 2.800E-06 |
| 10583 | 530 | R530N | A | 3.4e-06 | 1.7e-06 | 3.3e-06 | 2.800E-06 | 9.539E-07 | 1.700E-06 | 3.400E-06 |
| 10584 | 530 | R530D | A | 2.4e-06 | 1.8e-06 | 2.8e-06 | 2.333E-06 | 5.033E-07 | 1.800E-06 | 2.800E-06 |
| 10585 | 530 | R530C | A | 2e-06 | 1.8e-06 | 1.3e-06 | 1.700E-06 | 3.606E-07 | 1.300E-06 | 2.000E-06 |
| 10586 | 530 | R530E | A | 1.7e-06 | 2.6e-06 | 2.7e-06 | 2.333E-06 | 5.508E-07 | 1.700E-06 | 2.700E-06 |
| 10587 | 530 | R530Q | A | 2e-06 | 2.3e-06 | 2.8e-06 | 2.367E-06 | 4.041E-07 | 2.000E-06 | 2.800E-06 |
| 10588 | 530 | R530G | A | 2.4e-06 | 3.1e-06 | 3.2e-06 | 2.900E-06 | 4.359E-07 | 2.400E-06 | 3.200E-06 |
| 10589 | 530 | R530H | A | 2.7e-06 | 1.9e-06 | 3e-06 | 2.533E-06 | 5.686E-07 | 1.900E-06 | 3.000E-06 |
| 10590 | 530 | R530I | A | 1.5e-06 | 2.1e-06 | 2.6e-06 | 2.067E-06 | 5.508E-07 | 1.500E-06 | 2.600E-06 |
| 10591 | 530 | R530L | A | 1.9e-06 | 1.6e-06 | 2.2e-06 | 1.900E-06 | 3.000E-07 | 1.600E-06 | 2.200E-06 |
| 10592 | 530 | R530K | A | 3.1e-06 | 2.6e-06 | 2.1e-06 | 2.600E-06 | 5.000E-07 | 2.100E-06 | 3.100E-06 |
| 10593 | 530 | R530M | A | 2e-06 | 2.2e-06 | 4.1e-06 | 2.767E-06 | 1.159E-06 | 2.000E-06 | 4.100E-06 |
| 10594 | 530 | R530F | A | 2.2e-06 | 2.7e-06 | 1.9e-06 | 2.267E-06 | 4.041E-07 | 1.900E-06 | 2.700E-06 |
| 10595 | 530 | R530P | A | 3.6e-06 | 2.3e-06 | 1.7e-06 | 2.533E-06 | 9.713E-07 | 1.700E-06 | 3.600E-06 |
| 10596 | 530 | R530S | A | 3.6e-06 | 1.8e-06 | 3.1e-06 | 2.833E-06 | 9.292E-07 | 1.800E-06 | 3.600E-06 |
| 10597 | 530 | R530T | A | 2.7e-06 | 2.6e-06 | 2.3e-06 | 2.533E-06 | 2.082E-07 | 2.300E-06 | 2.700E-06 |
| 10598 | 530 | R530W | A | 2.8e-06 | 2.5e-06 | 1.8e-06 | 2.367E-06 | 5.132E-07 | 1.800E-06 | 2.800E-06 |
| 10599 | 530 | R530Y | A | 2.1e-06 | 3.9e-06 | 3e-06 | 3.000E-06 | 9.000E-07 | 2.100E-06 | 3.900E-06 |
| 10600 | 530 | R530V | A | 4.3e-06 | 2e-06 | 2e-06 | 2.767E-06 | 1.328E-06 | 2.000E-06 | 4.300E-06 |
| 10601 | 531 | E531A | A | 2.9e-06 | 3.2e-06 | 1.6e-06 | 2.567E-06 | 8.505E-07 | 1.600E-06 | 3.200E-06 |
| 10602 | 531 | E531R | A | 1.8e-06 | 2.9e-06 | 1.9e-06 | 2.200E-06 | 6.083E-07 | 1.800E-06 | 2.900E-06 |
| 10603 | 531 | E531N | A | 3.8e-06 | 3e-06 | 2.5e-06 | 3.100E-06 | 6.557E-07 | 2.500E-06 | 3.800E-06 |
| 10604 | 531 | E531D | A | 3.9e-06 | 2.4e-06 | 3.8e-06 | 3.367E-06 | 8.386E-07 | 2.400E-06 | 3.900E-06 |
| 10605 | 531 | E531C | A | 3.5e-06 | 3.2e-06 | 2.2e-06 | 2.967E-06 | 6.807E-07 | 2.200E-06 | 3.500E-06 |
| 10606 | 531 | E531E | A | 2.1e-06 | 1.9e-06 | 1.6e-06 | 1.867E-06 | 2.517E-07 | 1.600E-06 | 2.100E-06 |
| 10607 | 531 | E531Q | A | 2.4e-06 | 1.9e-06 | 2e-06 | 2.100E-06 | 2.646E-07 | 1.900E-06 | 2.400E-06 |
| 10608 | 531 | E531G | A | 1.9e-06 | 2.3e-06 | 2.2e-06 | 2.133E-06 | 2.082E-07 | 1.900E-06 | 2.300E-06 |
| 10609 | 531 | E531H | A | 3.1e-06 | 2.7e-06 | 2.1e-06 | 2.633E-06 | 5.033E-07 | 2.100E-06 | 3.100E-06 |
| 10610 | 531 | E531I | A | 3.8e-06 | 2.8e-06 | 3.1e-06 | 3.233E-06 | 5.132E-07 | 2.800E-06 | 3.800E-06 |
| 10611 | 531 | E531L | A | 2.9e-06 | 1.8e-06 | 2.8e-06 | 2.500E-06 | 6.083E-07 | 1.800E-06 | 2.900E-06 |
| 10612 | 531 | E531K | A | 3.8e-06 | 3.4e-06 | 2.5e-06 | 3.233E-06 | 6.658E-07 | 2.500E-06 | 3.800E-06 |
| 10613 | 531 | E531M | A | 2.1e-06 | 1.9e-06 | 3.8e-06 | 2.600E-06 | 1.044E-06 | 1.900E-06 | 3.800E-06 |
| 10614 | 531 | E531F | A | 1.6e-06 | 3.5e-06 | 9.1e-07 | 2.003E-06 | 1.341E-06 | 9.100E-07 | 3.500E-06 |
| 10615 | 531 | E531P | A | 3.5e-06 | 2.5e-06 | 2.3e-06 | 2.767E-06 | 6.429E-07 | 2.300E-06 | 3.500E-06 |
| 10616 | 531 | E531S | A | 1.8e-06 | 2.2e-06 | 2.9e-06 | 2.300E-06 | 5.568E-07 | 1.800E-06 | 2.900E-06 |
| 10617 | 531 | E531T | A | 1.6e-06 | 1.8e-06 | 2.4e-06 | 1.933E-06 | 4.163E-07 | 1.600E-06 | 2.400E-06 |
| 10618 | 531 | E531W | A | 2.4e-06 | 2.1e-06 | 2e-06 | 2.167E-06 | 2.082E-07 | 2.000E-06 | 2.400E-06 |
| 10619 | 531 | E531Y | A | 2e-06 | 2.3e-06 | 2.8e-06 | 2.367E-06 | 4.041E-07 | 2.000E-06 | 2.800E-06 |
| 10620 | 531 | E531V | A | 1.6e-06 | 3.2e-06 | 1.8e-06 | 2.200E-06 | 8.718E-07 | 1.600E-06 | 3.200E-06 |
| 10621 | 532 | Y532A | A | 2.9e-06 | 2.2e-06 | 3.5e-06 | 2.867E-06 | 6.506E-07 | 2.200E-06 | 3.500E-06 |
| 10622 | 532 | Y532R | A | 3.2e-06 | 1.6e-06 | 2.2e-06 | 2.333E-06 | 8.083E-07 | 1.600E-06 | 3.200E-06 |
| 10623 | 532 | Y532N | A | 2e-06 | 2.7e-06 | 1.7e-06 | 2.133E-06 | 5.132E-07 | 1.700E-06 | 2.700E-06 |
| 10624 | 532 | Y532D | A | 1.4e-06 | 3.8e-06 | 2.6e-06 | 2.600E-06 | 1.200E-06 | 1.400E-06 | 3.800E-06 |

| | | | | | | | | | | |
|---|---|---|---|---|---|---|---|---|---|---|
| 10625 | 532 | Y532C | A | 2.9e-06 | 2.9e-06 | 2.6e-06 | 2.800E-06 | 1.732E-07 | 2.600E-06 | 2.900E-06 |
| 10626 | 532 | Y532E | A | 2.4e-06 | 2.2e-06 | 3e-06 | 2.533E-06 | 4.163E-07 | 2.200E-06 | 3.000E-06 |
| 10627 | 532 | Y532Q | A | 1.7e-06 | 4.4e-06 | 2.1e-06 | 2.733E-06 | 1.457E-06 | 1.700E-06 | 4.400E-06 |
| 10628 | 532 | Y532G | A | 3.3e-06 | 2.4e-06 | 3.8e-06 | 3.167E-06 | 7.095E-07 | 2.400E-06 | 3.800E-06 |
| 10629 | 532 | Y532H | A | 2.7e-06 | 1.8e-06 | 1.7e-06 | 2.067E-06 | 5.508E-07 | 1.700E-06 | 2.700E-06 |
| 10630 | 532 | Y532I | A | 2.3e-06 | 2.7e-06 | 3.5e-06 | 2.833E-06 | 6.110E-07 | 2.300E-06 | 3.500E-06 |
| 10631 | 532 | Y532L | A | 2.1e-06 | 2.4e-06 | 1.6e-06 | 2.033E-06 | 4.041E-07 | 1.600E-06 | 2.400E-06 |
| 10632 | 532 | Y532K | A | 2.3e-06 | 2.5e-06 | 2.9e-06 | 2.567E-06 | 3.055E-07 | 2.300E-06 | 2.900E-06 |
| 10633 | 532 | Y532M | A | 1.7e-06 | 2.6e-06 | 2.9e-06 | 2.400E-06 | 6.245E-07 | 1.700E-06 | 2.900E-06 |
| 10634 | 532 | Y532F | A | 3.8e-06 | 2.5e-06 | 2.4e-06 | 2.900E-06 | 7.810E-07 | 2.400E-06 | 3.800E-06 |
| 10635 | 532 | Y532P | A | 2.4e-06 | 2.5e-06 | 3.5e-06 | 2.800E-06 | 6.083E-07 | 2.400E-06 | 3.500E-06 |
| 10636 | 532 | Y532S | A | 2.7e-06 | 5.5e-06 | 2.5e-06 | 3.567E-06 | 1.677E-06 | 2.500E-06 | 5.500E-06 |
| 10637 | 532 | Y532T | A | 1.6e-06 | 4.6e-06 | 3e-06 | 3.067E-06 | 1.501E-06 | 1.600E-06 | 4.600E-06 |
| 10638 | 532 | Y532W | A | 1.9e-06 | 2.2e-06 | 1.6e-06 | 1.900E-06 | 3.000E-07 | 1.600E-06 | 2.200E-06 |
| 10639 | 532 | Y532Y | A | 1.3e-06 | 1.3e-06 | 2.7e-06 | 1.767E-06 | 8.083E-07 | 1.300E-06 | 2.700E-06 |
| 10640 | 532 | Y532V | A | 2.1e-06 | 3.3e-06 | 1.9e-06 | 2.433E-06 | 7.572E-07 | 1.900E-06 | 3.300E-06 |
| 10641 | 533 | V533A | A | 2.4e-06 | 2.6e-06 | 1.6e-06 | 2.200E-06 | 5.292E-07 | 1.600E-06 | 2.600E-06 |
| 10642 | 533 | V533R | A | 2.7e-06 | 2.3e-06 | 2.3e-06 | 2.433E-06 | 2.309E-07 | 2.300E-06 | 2.700E-06 |
| 10643 | 533 | V533N | A | 3e-06 | 1.6e-06 | 1.1e-06 | 1.900E-06 | 9.849E-07 | 1.100E-06 | 3.000E-06 |
| 10644 | 533 | V533D | A | 1.6e-06 | 1.6e-06 | 2.6e-06 | 1.933E-06 | 5.774E-07 | 1.600E-06 | 2.600E-06 |
| 10645 | 533 | V533C | A | 2.6e-06 | 3.2e-06 | 2e-06 | 2.600E-06 | 6.000E-07 | 2.000E-06 | 3.200E-06 |
| 10646 | 533 | V533E | A | 2.2e-06 | 2.3e-06 | 2.8e-06 | 2.433E-06 | 3.215E-07 | 2.200E-06 | 2.800E-06 |
| 10647 | 533 | V533Q | A | 1.1e-06 | 2.7e-06 | 2.3e-06 | 2.033E-06 | 8.327E-07 | 1.100E-06 | 2.700E-06 |
| 10648 | 533 | V533G | A | 1.2e-06 | 3.8e-06 | 1.7e-06 | 2.233E-06 | 1.380E-06 | 1.200E-06 | 3.800E-06 |
| 10649 | 533 | V533H | A | 2.2e-06 | 1.3e-06 | 1.6e-06 | 1.700E-06 | 4.583E-07 | 1.300E-06 | 2.200E-06 |
| 10650 | 533 | V533I | A | 2.1e-06 | 5e-06 | 2.1e-06 | 3.067E-06 | 1.674E-06 | 2.100E-06 | 5.000E-06 |
| 10651 | 533 | V533L | A | 2.8e-06 | 3.5e-06 | 2.3e-06 | 2.867E-06 | 6.028E-07 | 2.300E-06 | 3.500E-06 |
| 10652 | 533 | V533K | A | 2.5e-06 | 1.7e-06 | 1.6e-06 | 1.933E-06 | 4.933E-07 | 1.600E-06 | 2.500E-06 |
| 10653 | 533 | V533M | A | 3e-06 | 3.1e-06 | 1.6e-06 | 2.567E-06 | 8.386E-07 | 1.600E-06 | 3.100E-06 |
| 10654 | 533 | V533F | A | 2.3e-06 | 3.3e-06 | 3e-06 | 2.867E-06 | 5.132E-07 | 2.300E-06 | 3.300E-06 |
| 10655 | 533 | V533P | A | 2.3e-06 | 2.4e-06 | 3.7e-06 | 2.800E-06 | 7.810E-07 | 2.300E-06 | 3.700E-06 |
| 10656 | 533 | V533S | A | 2.3e-06 | 2.4e-06 | 2.4e-06 | 2.367E-06 | 5.774E-08 | 2.300E-06 | 2.400E-06 |
| 10657 | 533 | V533T | A | 2.2e-06 | 1.8e-06 | 2.2e-06 | 2.067E-06 | 2.309E-07 | 1.800E-06 | 2.200E-06 |
| 10658 | 533 | V533W | A | 3.6e-06 | 2.8e-06 | 3.6e-06 | 3.333E-06 | 4.619E-07 | 2.800E-06 | 3.600E-06 |
| 10659 | 533 | V533Y | A | 2.5e-06 | 1.3e-06 | 1.5e-06 | 1.767E-06 | 6.429E-07 | 1.300E-06 | 2.500E-06 |
| 10660 | 533 | V533V | A | 3.1e-06 | 2.1e-06 | 2.3e-06 | 2.500E-06 | 5.292E-07 | 2.100E-06 | 3.100E-06 |
| 10661 | 534 | N534A | A | 1.9e-06 | 2.1e-06 | 2.1e-06 | 2.033E-06 | 1.155E-07 | 1.900E-06 | 2.100E-06 |
| 10662 | 534 | N534R | A | 2.3e-06 | 1.5e-06 | 2.1e-06 | 1.967E-06 | 4.163E-07 | 1.500E-06 | 2.300E-06 |
| 10663 | 534 | N534N | A | 2.2e-06 | 3e-06 | 1.5e-06 | 2.233E-06 | 7.506E-07 | 1.500E-06 | 3.000E-06 |
| 10664 | 534 | N534D | A | 3.4e-06 | 1.9e-06 | 2e-06 | 2.433E-06 | 8.386E-07 | 1.900E-06 | 3.400E-06 |
| 10665 | 534 | N534C | A | 2.7e-06 | 2.2e-06 | 1.7e-06 | 2.200E-06 | 5.000E-07 | 1.700E-06 | 2.700E-06 |
| 10666 | 534 | N534E | A | 1.7e-06 | 2.3e-06 | 1.2e-06 | 1.733E-06 | 5.508E-07 | 1.200E-06 | 2.300E-06 |
| 10667 | 534 | N534Q | A | 2.1e-06 | 1.6e-06 | 2.2e-06 | 1.967E-06 | 3.215E-07 | 1.600E-06 | 2.200E-06 |
| 10668 | 534 | N534G | A | 1.9e-06 | 3.2e-06 | 2.5e-06 | 2.533E-06 | 6.506E-07 | 1.900E-06 | 3.200E-06 |
| 10669 | 534 | N534H | A | 1.4e-06 | 1.5e-06 | 2.2e-06 | 1.700E-06 | 4.359E-07 | 1.400E-06 | 2.200E-06 |
| 10670 | 534 | N534I | A | 2.3e-06 | 1.9e-06 | 2.8e-06 | 2.333E-06 | 4.509E-07 | 1.900E-06 | 2.800E-06 |
| 10671 | 534 | N534L | A | 1.8e-06 | 4e-06 | 2.5e-06 | 2.767E-06 | 1.124E-06 | 1.800E-06 | 4.000E-06 |
| 10672 | 534 | N534K | A | 2.3e-06 | 3.4e-06 | 2.9e-06 | 2.867E-06 | 5.508E-07 | 2.300E-06 | 3.400E-06 |
| 10673 | 534 | N534M | A | 3.3e-06 | 1.1e-06 | 2.2e-06 | 2.200E-06 | 1.100E-06 | 1.100E-06 | 3.300E-06 |
| 10674 | 534 | N534F | A | 3e-06 | 2.3e-06 | 4e-06 | 3.100E-06 | 8.544E-07 | 2.300E-06 | 4.000E-06 |
| 10675 | 534 | N534P | A | 1.3e-06 | 4.3e-06 | 2e-06 | 2.533E-06 | 1.570E-06 | 1.300E-06 | 4.300E-06 |
| 10676 | 534 | N534S | A | 3e-06 | 2.4e-06 | 4.2e-06 | 3.200E-06 | 9.165E-07 | 2.400E-06 | 4.200E-06 |
| 10677 | 534 | N534T | A | 1.7e-06 | 3.8e-06 | 2.9e-06 | 2.800E-06 | 1.054E-06 | 1.700E-06 | 3.800E-06 |
| 10678 | 534 | N534W | A | 2e-06 | 2.7e-06 | 2.3e-06 | 2.333E-06 | 3.512E-07 | 2.000E-06 | 2.700E-06 |
| 10679 | 534 | N534Y | A | 3.4e-06 | 2e-06 | 2.1e-06 | 2.500E-06 | 7.810E-07 | 2.000E-06 | 3.400E-06 |
| 10680 | 534 | N534V | A | 1.9e-06 | 3.9e-06 | 2.8e-06 | 2.867E-06 | 1.002E-06 | 1.900E-06 | 3.900E-06 |
| 10681 | 535 | A535A | A | 3.3e-06 | 3e-06 | 2.2e-06 | 2.833E-06 | 5.686E-07 | 2.200E-06 | 3.300E-06 |
| 10682 | 535 | A535R | A | 2.1e-06 | 3.1e-06 | 2e-06 | 2.400E-06 | 6.083E-07 | 2.000E-06 | 3.100E-06 |
| 10683 | 535 | A535N | A | 2.7e-06 | 3.2e-06 | 2.9e-06 | 2.933E-06 | 2.517E-07 | 2.700E-06 | 3.200E-06 |
| 10684 | 535 | A535D | A | 1.6e-06 | 2.7e-06 | 2.5e-06 | 2.267E-06 | 5.859E-07 | 1.600E-06 | 2.700E-06 |
| 10685 | 535 | A535C | A | 2.3e-06 | 1.8e-06 | 2.4e-06 | 2.167E-06 | 3.215E-07 | 1.800E-06 | 2.400E-06 |
| 10686 | 535 | A535E | A | 3e-06 | 2e-06 | 2.8e-06 | 2.600E-06 | 5.292E-07 | 2.000E-06 | 3.000E-06 |
| 10687 | 535 | A535Q | A | 2.9e-06 | 3.3e-06 | 3.6e-06 | 3.267E-06 | 3.512E-07 | 2.900E-06 | 3.600E-06 |
| 10688 | 535 | A535G | A | 2.1e-06 | 3.3e-06 | 1.7e-06 | 2.367E-06 | 8.327E-07 | 1.700E-06 | 3.300E-06 |
| 10689 | 535 | A535H | A | 2.5e-06 | 1.7e-06 | 1.2e-06 | 1.800E-06 | 6.557E-07 | 1.200E-06 | 2.500E-06 |
| 10690 | 535 | A535I | A | 2.5e-06 | 2.9e-06 | 4.3e-06 | 3.233E-06 | 9.452E-07 | 2.500E-06 | 4.300E-06 |
| 10691 | 535 | A535L | A | 2.7e-06 | 2.6e-06 | 2.1e-06 | 2.467E-06 | 3.215E-07 | 2.100E-06 | 2.700E-06 |
| 10692 | 535 | A535K | A | 1.5e-06 | 2.4e-06 | 3.6e-06 | 2.500E-06 | 1.054E-06 | 1.500E-06 | 3.600E-06 |
| 10693 | 535 | A535M | A | 3.3e-06 | 2.2e-06 | 2.7e-06 | 2.733E-06 | 5.508E-07 | 2.200E-06 | 3.300E-06 |
| 10694 | 535 | A535F | A | 1.7e-06 | 1.2e-06 | 1.7e-06 | 1.533E-06 | 2.887E-07 | 1.200E-06 | 1.700E-06 |
| 10695 | 535 | A535P | A | 2e-06 | 1.4e-06 | 3.8e-06 | 2.400E-06 | 1.249E-06 | 1.400E-06 | 3.800E-06 |
| 10696 | 535 | A535S | A | 1.9e-06 | 3.8e-06 | 1.5e-06 | 2.400E-06 | 1.229E-06 | 1.500E-06 | 3.800E-06 |
| 10697 | 535 | A535T | A | 1.9e-06 | 4.3e-06 | 1.7e-06 | 2.633E-06 | 1.447E-06 | 1.700E-06 | 4.300E-06 |
| 10698 | 535 | A535W | A | 1.4e-06 | 2.7e-06 | 2.2e-06 | 2.100E-06 | 6.557E-07 | 1.400E-06 | 2.700E-06 |
| 10699 | 535 | A535Y | A | 2.4e-06 | 2.8e-06 | 1.5e-06 | 2.233E-06 | 6.658E-07 | 1.500E-06 | 2.800E-06 |
| 10700 | 535 | A535V | A | 3.3e-06 | 2.2e-06 | 1.1e-06 | 2.200E-06 | 1.100E-06 | 1.100E-06 | 3.300E-06 |
| 10701 | 536 | R536A | A | 4.4e-06 | 2.2e-06 | 3.1e-06 | 3.233E-06 | 1.106E-06 | 2.200E-06 | 4.400E-06 |
| 10702 | 536 | R536R | A | 3.1e-06 | 4e-06 | 1.4e-06 | 2.833E-06 | 1.320E-06 | 1.400E-06 | 4.000E-06 |
| 10703 | 536 | R536N | A | 1.7e-06 | 5e-06 | 1.8e-06 | 2.833E-06 | 1.877E-06 | 1.700E-06 | 5.000E-06 |
| 10704 | 536 | R536D | A | 2.4e-06 | 2.2e-06 | 2.5e-06 | 2.367E-06 | 1.528E-07 | 2.200E-06 | 2.500E-06 |
| 10705 | 536 | R536C | A | 2.2e-06 | 1.7e-06 | 3.1e-06 | 2.333E-06 | 7.095E-07 | 1.700E-06 | 3.100E-06 |
| 10706 | 536 | R536E | A | 3.3e-06 | 1.8e-06 | 4.9e-06 | 3.333E-06 | 1.550E-06 | 1.800E-06 | 4.900E-06 |
| 10707 | 536 | R536Q | A | 2.6e-06 | 1.4e-06 | 3.9e-06 | 2.633E-06 | 1.250E-06 | 1.400E-06 | 3.900E-06 |
| 10708 | 536 | R536G | A | 3.5e-06 | 3.2e-06 | 3.2e-06 | 3.300E-06 | 1.732E-07 | 3.200E-06 | 3.500E-06 |
| 10709 | 536 | R536H | A | 1.8e-06 | 1.7e-06 | 1.5e-06 | 1.667E-06 | 1.528E-07 | 1.500E-06 | 1.800E-06 |

| | | | | | | | | | | |
|---|---|---|---|---|---|---|---|---|---|---|
| 10710 | 536 | R536I | A | 2.5e-06 | 3.1e-06 | 4.7e-06 | 3.433E-06 | 1.137E-06 | 2.500E-06 | 4.700E-06 |
| 10711 | 536 | R536L | A | 2.4e-06 | 1.8e-06 | 2.7e-06 | 2.300E-06 | 4.583E-07 | 1.800E-06 | 2.700E-06 |
| 10712 | 536 | R536K | A | 2.2e-06 | 2e-06 | 1.8e-06 | 2.000E-06 | 2.000E-07 | 1.800E-06 | 2.200E-06 |
| 10713 | 536 | R536M | A | 3.1e-06 | 2.1e-06 | 1.9e-06 | 2.367E-06 | 6.429E-07 | 1.900E-06 | 3.100E-06 |
| 10714 | 536 | R536F | A | 2.7e-06 | 1.1e-06 | 1.7e-06 | 1.833E-06 | 8.083E-07 | 1.100E-06 | 2.700E-06 |
| 10715 | 536 | R536P | A | 1.5e-06 | 2.3e-06 | 2e-06 | 1.933E-06 | 4.041E-07 | 1.500E-06 | 2.300E-06 |
| 10716 | 536 | R536S | A | 2.8e-06 | 2.7e-06 | 2.1e-06 | 2.533E-06 | 3.786E-07 | 2.100E-06 | 2.800E-06 |
| 10717 | 536 | R536T | A | 2.8e-06 | 4.1e-06 | 3.1e-06 | 3.333E-06 | 6.807E-07 | 2.800E-06 | 4.100E-06 |
| 10718 | 536 | R536W | A | 2e-06 | 2e-06 | 2.6e-06 | 2.200E-06 | 3.464E-07 | 2.000E-06 | 2.600E-06 |
| 10719 | 536 | R536Y | A | 2.8e-06 | 1.2e-06 | 2.3e-06 | 2.100E-06 | 8.185E-07 | 1.200E-06 | 2.800E-06 |
| 10720 | 536 | R536V | A | 2.2e-06 | 2.3e-06 | 2e-06 | 2.167E-06 | 1.528E-07 | 2.000E-06 | 2.300E-06 |
| 10721 | 537 | H537A | A | 5.6e-06 | 2.5e-06 | 3.2e-06 | 3.767E-06 | 1.626E-06 | 2.500E-06 | 5.600E-06 |
| 10722 | 537 | H537R | A | 2.4e-06 | 1.5e-06 | 1.6e-06 | 1.833E-06 | 4.933E-07 | 1.500E-06 | 2.400E-06 |
| 10723 | 537 | H537N | A | 1.3e-06 | 1.3e-06 | 1.5e-06 | 1.367E-06 | 1.155E-07 | 1.300E-06 | 1.500E-06 |
| 10724 | 537 | H537D | A | 2.7e-06 | 3.4e-06 | 3e-06 | 3.033E-06 | 3.512E-07 | 2.700E-06 | 3.400E-06 |
| 10725 | 537 | H537C | A | 2.2e-06 | 4e-06 | 1.7e-06 | 2.633E-06 | 1.210E-06 | 1.700E-06 | 4.000E-06 |
| 10726 | 537 | H537E | A | 2.2e-06 | 2e-06 | 2.9e-06 | 2.367E-06 | 4.726E-07 | 2.000E-06 | 2.900E-06 |
| 10727 | 537 | H537Q | A | 3.1e-06 | 1.6e-06 | 2.4e-06 | 2.367E-06 | 7.506E-07 | 1.600E-06 | 3.100E-06 |
| 10728 | 537 | H537G | A | 2.2e-06 | 3.9e-06 | 2.6e-06 | 2.900E-06 | 8.888E-07 | 2.200E-06 | 3.900E-06 |
| 10729 | 537 | H537H | A | 2.9e-06 | 2.6e-06 | 2.7e-06 | 2.733E-06 | 1.528E-07 | 2.600E-06 | 2.900E-06 |
| 10730 | 537 | H537I | A | 3.3e-06 | 2.2e-06 | 2.8e-06 | 2.767E-06 | 5.508E-07 | 2.200E-06 | 3.300E-06 |
| 10731 | 537 | H537L | A | 3e-06 | 2.4e-06 | 3.9e-06 | 3.100E-06 | 7.550E-07 | 2.400E-06 | 3.900E-06 |
| 10732 | 537 | H537K | A | 4.1e-06 | 3.1e-06 | 2.6e-06 | 3.267E-06 | 7.638E-07 | 2.600E-06 | 4.100E-06 |
| 10733 | 537 | H537M | A | 1.9e-06 | 2.2e-06 | 1.3e-06 | 1.800E-06 | 4.583E-07 | 1.300E-06 | 2.200E-06 |
| 10734 | 537 | H537F | A | 1.3e-06 | 2.3e-06 | 3.6e-06 | 2.400E-06 | 1.153E-06 | 1.300E-06 | 3.600E-06 |
| 10735 | 537 | H537P | A | 2.7e-06 | 4.7e-06 | 3.1e-06 | 3.500E-06 | 1.058E-06 | 2.700E-06 | 4.700E-06 |
| 10736 | 537 | H537S | A | 1.8e-06 | 4.4e-06 | 2.4e-06 | 2.867E-06 | 1.361E-06 | 1.800E-06 | 4.400E-06 |
| 10737 | 537 | H537T | A | 2.5e-06 | 2.2e-06 | 3.4e-06 | 2.700E-06 | 6.245E-07 | 2.200E-06 | 3.400E-06 |
| 10738 | 537 | H537W | A | 2.3e-06 | 2e-06 | 1.4e-06 | 1.900E-06 | 4.583E-07 | 1.400E-06 | 2.300E-06 |
| 10739 | 537 | H537Y | A | 2.2e-06 | 1.7e-06 | 2.1e-06 | 2.000E-06 | 2.646E-07 | 1.700E-06 | 2.200E-06 |
| 10740 | 537 | H537V | A | 2.9e-06 | 3.9e-06 | 1.6e-06 | 2.800E-06 | 1.153E-06 | 1.600E-06 | 3.900E-06 |
| 10741 | 538 | C538A | A | 1.3e-06 | 2.6e-06 | 1.5e-06 | 1.800E-06 | 7.000E-07 | 1.300E-06 | 2.600E-06 |
| 10742 | 538 | C538R | A | 3e-06 | 2.5e-06 | 1.9e-06 | 2.467E-06 | 5.508E-07 | 1.900E-06 | 3.000E-06 |
| 10743 | 538 | C538N | A | 1.4e-06 | 1.5e-06 | 1.6e-06 | 1.500E-06 | 1.000E-07 | 1.400E-06 | 1.600E-06 |
| 10744 | 538 | C538D | A | 2.7e-06 | 2.3e-06 | 2.2e-06 | 2.400E-06 | 2.646E-07 | 2.200E-06 | 2.700E-06 |
| 10745 | 538 | C538C | A | 3.4e-06 | 1.5e-06 | 1.3e-06 | 2.067E-06 | 1.159E-06 | 1.300E-06 | 3.400E-06 |
| 10746 | 538 | C538E | A | 3e-06 | 2.1e-06 | 3.7e-06 | 2.933E-06 | 8.021E-07 | 2.100E-06 | 3.700E-06 |
| 10747 | 538 | C538Q | A | 2.7e-06 | 3.4e-06 | 1.9e-06 | 2.667E-06 | 7.506E-07 | 1.900E-06 | 3.400E-06 |
| 10748 | 538 | C538G | A | 3.9e-06 | 1.4e-06 | 3.1e-06 | 2.800E-06 | 1.277E-06 | 1.400E-06 | 3.900E-06 |
| 10749 | 538 | C538H | A | 1.2e-06 | 1.9e-06 | 3.4e-06 | 2.167E-06 | 1.124E-06 | 1.200E-06 | 3.400E-06 |
| 10750 | 538 | C538I | A | 2.8e-06 | 2.6e-06 | 2.6e-06 | 2.667E-06 | 1.155E-07 | 2.600E-06 | 2.800E-06 |
| 10751 | 538 | C538L | A | 1.9e-06 | 1.6e-06 | 2.3e-06 | 1.933E-06 | 3.512E-07 | 1.600E-06 | 2.300E-06 |
| 10752 | 538 | C538K | A | 2.2e-06 | 2.7e-06 | 3.3e-06 | 2.733E-06 | 5.508E-07 | 2.200E-06 | 3.300E-06 |
| 10753 | 538 | C538M | A | 2.7e-06 | 3.4e-06 | 3.5e-06 | 3.200E-06 | 4.359E-07 | 2.700E-06 | 3.500E-06 |
| 10754 | 538 | C538F | A | 4.1e-06 | 2.3e-06 | 5.7e-06 | 4.033E-06 | 1.701E-06 | 2.300E-06 | 5.700E-06 |
| 10755 | 538 | C538P | A | 3.8e-06 | 1.8e-06 | 2.1e-06 | 2.567E-06 | 1.079E-06 | 1.800E-06 | 3.800E-06 |
| 10756 | 538 | C538S | A | 1.8e-06 | 2.3e-06 | 1.4e-06 | 1.833E-06 | 4.509E-07 | 1.400E-06 | 2.300E-06 |
| 10757 | 538 | C538T | A | 3e-06 | 1.8e-06 | 2.3e-06 | 2.367E-06 | 6.028E-07 | 1.800E-06 | 3.000E-06 |
| 10758 | 538 | C538W | A | 2.6e-06 | 1.8e-06 | 3.8e-06 | 2.733E-06 | 1.007E-06 | 1.800E-06 | 3.800E-06 |
| 10759 | 538 | C538Y | A | 2.8e-06 | 2.8e-06 | 1.7e-06 | 2.433E-06 | 6.351E-07 | 1.700E-06 | 2.800E-06 |
| 10760 | 538 | C538V | A | 3.8e-06 | 2.2e-06 | 2.1e-06 | 2.700E-06 | 9.539E-07 | 2.100E-06 | 3.800E-06 |
| 10761 | 539 | L539A | A | 2.3e-06 | 2.3e-06 | 1.5e-06 | 2.033E-06 | 4.619E-07 | 1.500E-06 | 2.300E-06 |
| 10762 | 539 | L539R | A | 2.4e-06 | 1.9e-06 | 1.8e-06 | 2.033E-06 | 3.215E-07 | 1.800E-06 | 2.400E-06 |
| 10763 | 539 | L539N | A | 2.8e-06 | 5.2e-06 | 2e-06 | 3.333E-06 | 1.665E-06 | 2.000E-06 | 5.200E-06 |
| 10764 | 539 | L539D | A | 2.5e-06 | 2.6e-06 | 1.4e-06 | 2.167E-06 | 6.658E-07 | 1.400E-06 | 2.600E-06 |
| 10765 | 539 | L539C | A | 1.8e-06 | 2.9e-06 | 1.9e-06 | 2.200E-06 | 6.083E-07 | 1.800E-06 | 2.900E-06 |
| 10766 | 539 | L539E | A | 1.7e-06 | 8.5e-07 | 2.2e-06 | 1.583E-06 | 6.825E-07 | 8.500E-07 | 2.200E-06 |
| 10767 | 539 | L539Q | A | 2.5e-06 | 1.4e-06 | 1.7e-06 | 1.867E-06 | 5.686E-07 | 1.400E-06 | 2.500E-06 |
| 10768 | 539 | L539G | A | 4.2e-06 | 1.6e-06 | 2.3e-06 | 2.700E-06 | 1.345E-06 | 1.600E-06 | 4.200E-06 |
| 10769 | 539 | L539H | A | 2.1e-06 | 4.3e-06 | 9e-07 | 2.433E-06 | 1.724E-06 | 9.000E-07 | 4.300E-06 |
| 10770 | 539 | L539I | A | 1.7e-06 | 1.4e-06 | 3.6e-06 | 2.233E-06 | 1.193E-06 | 1.400E-06 | 3.600E-06 |
| 10771 | 539 | L539L | A | 1.8e-06 | 1.7e-06 | 2.3e-06 | 1.933E-06 | 3.215E-07 | 1.700E-06 | 2.300E-06 |
| 10772 | 539 | L539K | A | 9.2e-07 | 2.3e-06 | 1.5e-06 | 1.573E-06 | 6.929E-07 | 9.200E-07 | 2.300E-06 |
| 10773 | 539 | L539M | A | 2.1e-06 | 2.5e-06 | 2.1e-06 | 2.233E-06 | 2.309E-07 | 2.100E-06 | 2.500E-06 |
| 10774 | 539 | L539F | A | 3.5e-06 | 2.2e-06 | 3e-06 | 2.900E-06 | 6.557E-07 | 2.200E-06 | 3.500E-06 |
| 10775 | 539 | L539P | A | 2e-06 | 3.6e-06 | 2.9e-06 | 2.833E-06 | 8.021E-07 | 2.000E-06 | 3.600E-06 |
| 10776 | 539 | L539S | A | 1.8e-06 | 1.6e-06 | 2e-06 | 1.800E-06 | 2.000E-07 | 1.600E-06 | 2.000E-06 |
| 10777 | 539 | L539T | A | 2e-06 | 1.8e-06 | 2e-06 | 1.933E-06 | 1.155E-07 | 1.800E-06 | 2.000E-06 |
| 10778 | 539 | L539W | A | 3.1e-06 | 2.1e-06 | 1.5e-06 | 2.233E-06 | 8.083E-07 | 1.500E-06 | 3.100E-06 |
| 10779 | 539 | L539Y | A | 2.2e-06 | 1.8e-06 | 2.2e-06 | 2.067E-06 | 2.309E-07 | 1.800E-06 | 2.200E-06 |
| 10780 | 539 | L539V | A | 1.9e-06 | 3e-06 | 2.5e-06 | 2.467E-06 | 5.508E-07 | 1.900E-06 | 3.000E-06 |
| 10781 | 540 | P540A | A | 3.3e-06 | 2.7e-06 | 2.9e-06 | 2.967E-06 | 3.055E-07 | 2.700E-06 | 3.300E-06 |
| 10782 | 540 | P540R | A | 1.8e-06 | 1.9e-06 | 2.1e-06 | 1.933E-06 | 1.528E-07 | 1.800E-06 | 2.100E-06 |
| 10783 | 540 | P540N | A | 1.3e-06 | 1.5e-06 | 3.6e-06 | 2.133E-06 | 1.274E-06 | 1.300E-06 | 3.600E-06 |
| 10784 | 540 | P540D | A | 4.3e-06 | 1.7e-06 | 2.8e-06 | 2.933E-06 | 1.305E-06 | 1.700E-06 | 4.300E-06 |
| 10785 | 540 | P540C | A | 1.6e-06 | 2e-06 | 1.5e-06 | 1.700E-06 | 2.646E-07 | 1.500E-06 | 2.000E-06 |
| 10786 | 540 | P540E | A | 2.5e-06 | 2.7e-06 | 1.4e-06 | 2.200E-06 | 7.000E-07 | 1.400E-06 | 2.700E-06 |
| 10787 | 540 | P540Q | A | 2e-06 | 2.1e-06 | 8e-07 | 1.633E-06 | 7.234E-07 | 8.000E-07 | 2.100E-06 |
| 10788 | 540 | P540G | A | 2.3e-06 | 5e-06 | 3e-06 | 3.433E-06 | 1.401E-06 | 2.300E-06 | 5.000E-06 |
| 10789 | 540 | P540H | A | 3.1e-06 | 3.7e-06 | 2.2e-06 | 3.000E-06 | 7.550E-07 | 2.200E-06 | 3.700E-06 |
| 10790 | 540 | P540I | A | 2.3e-06 | 2.8e-06 | 4.2e-06 | 3.100E-06 | 9.849E-07 | 2.300E-06 | 4.200E-06 |
| 10791 | 540 | P540L | A | 2.9e-06 | 3.1e-06 | 3.1e-06 | 3.033E-06 | 1.155E-07 | 2.900E-06 | 3.100E-06 |
| 10792 | 540 | P540K | A | 3.1e-06 | 2.9e-06 | 9.8e-07 | 2.327E-06 | 1.171E-06 | 9.800E-07 | 3.100E-06 |
| 10793 | 540 | P540M | A | 2.2e-06 | 2.4e-06 | 1.6e-06 | 2.067E-06 | 4.163E-07 | 1.600E-06 | 2.400E-06 |
| 10794 | 540 | P540F | A | 3.4e-06 | 3.1e-06 | 2.4e-06 | 2.967E-06 | 5.132E-07 | 2.400E-06 | 3.400E-06 |

| | | | | | | | | | | |
|---|---|---|---|---|---|---|---|---|---|---|
| 10795 | 540 | P540P | A | 3.7e-06 | 5.2e-06 | 2.3e-06 | 3.733E-06 | 1.450E-06 | 2.300E-06 | 5.200E-06 |
| 10796 | 540 | P540S | A | 3.6e-06 | 2.4e-06 | 3.3e-06 | 3.100E-06 | 6.245E-07 | 2.400E-06 | 3.600E-06 |
| 10797 | 540 | P540T | A | 1.5e-06 | 2.1e-06 | 1.7e-06 | 1.767E-06 | 3.055E-07 | 1.500E-06 | 2.100E-06 |
| 10798 | 540 | P540W | A | 2.3e-06 | 1.9e-06 | 4.2e-06 | 2.800E-06 | 1.229E-06 | 1.900E-06 | 4.200E-06 |
| 10799 | 540 | P540Y | A | 1.6e-06 | 2.2e-06 | 1.4e-06 | 1.733E-06 | 4.163E-07 | 1.400E-06 | 2.200E-06 |
| 10800 | 540 | P540V | A | 1.7e-06 | 2.5e-06 | 1.8e-06 | 2.000E-06 | 4.359E-07 | 1.700E-06 | 2.500E-06 |
| 10801 | 541 | C541A | A | 1.4e-06 | 1.6e-06 | 2.1e-06 | 1.700E-06 | 3.606E-07 | 1.400E-06 | 2.100E-06 |
| 10802 | 541 | C541R | A | 3.1e-06 | 1.9e-06 | 8.9e-07 | 1.963E-06 | 1.106E-06 | 8.900E-07 | 3.100E-06 |
| 10803 | 541 | C541N | A | 1.9e-06 | 2.9e-06 | 1.6e-06 | 2.133E-06 | 6.807E-07 | 1.600E-06 | 2.900E-06 |
| 10804 | 541 | C541D | A | 1.3e-06 | 1.6e-06 | 2.5e-06 | 1.800E-06 | 6.245E-07 | 1.300E-06 | 2.500E-06 |
| 10805 | 541 | C541C | A | 2.1e-06 | 2.2e-06 | 3e-06 | 2.433E-06 | 4.933E-07 | 2.100E-06 | 3.000E-06 |
| 10806 | 541 | C541E | A | 1.9e-06 | 1.9e-06 | 2.6e-06 | 2.133E-06 | 4.041E-07 | 1.900E-06 | 2.600E-06 |
| 10807 | 541 | C541Q | A | 2.3e-06 | 2.4e-06 | 3.1e-06 | 2.600E-06 | 4.359E-07 | 2.300E-06 | 3.100E-06 |
| 10808 | 541 | C541G | A | 2.8e-06 | 2.7e-06 | 5.2e-06 | 3.567E-06 | 1.415E-06 | 2.700E-06 | 5.200E-06 |
| 10809 | 541 | C541H | A | 1.6e-06 | 1.8e-06 | 2.1e-06 | 1.833E-06 | 2.517E-07 | 1.600E-06 | 2.100E-06 |
| 10810 | 541 | C541I | A | 3.7e-06 | 2.9e-06 | 2.7e-06 | 3.100E-06 | 5.292E-07 | 2.700E-06 | 3.700E-06 |
| 10811 | 541 | C541L | A | 3.8e-06 | 2.1e-06 | 2.5e-06 | 2.800E-06 | 8.888E-07 | 2.100E-06 | 3.800E-06 |
| 10812 | 541 | C541K | A | 1.5e-06 | 1.9e-06 | 1.6e-06 | 1.667E-06 | 2.082E-07 | 1.500E-06 | 1.900E-06 |
| 10813 | 541 | C541M | A | 3.6e-06 | 2.2e-06 | 2.8e-06 | 2.867E-06 | 7.024E-07 | 2.200E-06 | 3.600E-06 |
| 10814 | 541 | C541F | A | 2.3e-06 | 2.9e-06 | 3.1e-06 | 2.767E-06 | 4.163E-07 | 2.300E-06 | 3.100E-06 |
| 10815 | 541 | C541P | A | 3.4e-06 | 2.6e-06 | 2.2e-06 | 2.733E-06 | 6.110E-07 | 2.200E-06 | 3.400E-06 |
| 10816 | 541 | C541S | A | 2.5e-06 | 2.2e-06 | 2.3e-06 | 2.333E-06 | 1.528E-07 | 2.200E-06 | 2.500E-06 |
| 10817 | 541 | C541T | A | 2.6e-06 | 2e-06 | 1.2e-06 | 1.933E-06 | 7.024E-07 | 1.200E-06 | 2.600E-06 |
| 10818 | 541 | C541W | A | 2.8e-06 | 2.6e-06 | 2.5e-06 | 2.633E-06 | 1.528E-07 | 2.500E-06 | 2.800E-06 |
| 10819 | 541 | C541Y | A | 3.7e-06 | 1.9e-06 | 2.1e-06 | 2.567E-06 | 9.866E-07 | 1.900E-06 | 3.700E-06 |
| 10820 | 541 | C541V | A | 1.3e-06 | 2.7e-06 | 2.3e-06 | 2.100E-06 | 7.211E-07 | 1.300E-06 | 2.700E-06 |
| 10821 | 542 | H542A | A | 2.3e-06 | 2.8e-06 | 2.5e-06 | 2.533E-06 | 2.517E-07 | 2.300E-06 | 2.800E-06 |
| 10822 | 542 | H542R | A | 2.4e-06 | 2.3e-06 | 1.8e-06 | 2.167E-06 | 3.215E-07 | 1.800E-06 | 2.400E-06 |
| 10823 | 542 | H542N | A | 1.7e-06 | 2.8e-06 | 2.3e-06 | 2.267E-06 | 5.508E-07 | 1.700E-06 | 2.800E-06 |
| 10824 | 542 | H542D | A | 3.9e-06 | 2.3e-06 | 3.8e-06 | 3.333E-06 | 8.963E-07 | 2.300E-06 | 3.900E-06 |
| 10825 | 542 | H542C | A | 2.1e-06 | 3.2e-06 | 2.2e-06 | 2.500E-06 | 6.083E-07 | 2.100E-06 | 3.200E-06 |
| 10826 | 542 | H542E | A | 1.6e-06 | 1.2e-06 | 1.7e-06 | 1.500E-06 | 2.646E-07 | 1.200E-06 | 1.700E-06 |
| 10827 | 542 | H542Q | A | 2.7e-06 | 1.7e-06 | 2e-06 | 2.133E-06 | 5.132E-07 | 1.700E-06 | 2.700E-06 |
| 10828 | 542 | H542G | A | 3.6e-06 | 2.5e-06 | 4.2e-06 | 3.433E-06 | 8.622E-07 | 2.500E-06 | 4.200E-06 |
| 10829 | 542 | H542H | A | 2.1e-06 | 3.2e-06 | 1.9e-06 | 2.400E-06 | 7.000E-07 | 1.900E-06 | 3.200E-06 |
| 10830 | 542 | H542I | A | 3.1e-06 | 2.4e-06 | 2.4e-06 | 2.633E-06 | 4.041E-07 | 2.400E-06 | 3.100E-06 |
| 10831 | 542 | H542L | A | 2.3e-06 | 4.3e-06 | 1.9e-06 | 2.833E-06 | 1.286E-06 | 1.900E-06 | 4.300E-06 |
| 10832 | 542 | H542K | A | 1.8e-06 | 2.6e-06 | 2.5e-06 | 2.300E-06 | 4.359E-07 | 1.800E-06 | 2.600E-06 |
| 10833 | 542 | H542M | A | 2.8e-06 | 2.1e-06 | 6e-06 | 3.633E-06 | 2.079E-06 | 2.100E-06 | 6.000E-06 |
| 10834 | 542 | H542F | A | 2.7e-06 | 2e-06 | 1.8e-06 | 2.167E-06 | 4.726E-07 | 1.800E-06 | 2.700E-06 |
| 10835 | 542 | H542P | A | 2.3e-06 | 2.9e-06 | 3.8e-06 | 3.000E-06 | 7.550E-07 | 2.300E-06 | 3.800E-06 |
| 10836 | 542 | H542S | A | 2e-06 | 2.4e-06 | 3.2e-06 | 2.533E-06 | 6.110E-07 | 2.000E-06 | 3.200E-06 |
| 10837 | 542 | H542T | A | 2.7e-06 | 2e-06 | 2.3e-06 | 2.333E-06 | 3.512E-07 | 2.000E-06 | 2.700E-06 |
| 10838 | 542 | H542W | A | 1.6e-06 | 3.3e-06 | 1.6e-06 | 2.167E-06 | 9.815E-07 | 1.600E-06 | 3.300E-06 |
| 10839 | 542 | H542Y | A | 2.3e-06 | 3e-06 | 1.9e-06 | 2.400E-06 | 5.568E-07 | 1.900E-06 | 3.000E-06 |
| 10840 | 542 | H542V | A | 2.8e-06 | 2.9e-06 | 2.4e-06 | 2.700E-06 | 2.646E-07 | 2.400E-06 | 2.900E-06 |
| 10841 | 543 | P543A | A | 2.9e-06 | 2.4e-06 | 2.5e-06 | 2.600E-06 | 2.646E-07 | 2.400E-06 | 2.900E-06 |
| 10842 | 543 | P543R | A | 4e-06 | 3.7e-06 | 3e-06 | 3.567E-06 | 5.132E-07 | 3.000E-06 | 4.000E-06 |
| 10843 | 543 | P543N | A | 4.2e-06 | 4.1e-06 | 2e-06 | 3.433E-06 | 1.242E-06 | 2.000E-06 | 4.200E-06 |
| 10844 | 543 | P543D | A | 1.3e-06 | 1.3e-06 | 3.1e-06 | 1.900E-06 | 1.039E-06 | 1.300E-06 | 3.100E-06 |
| 10845 | 543 | P543C | A | 2.4e-06 | 1.7e-06 | 1.7e-06 | 1.933E-06 | 4.041E-07 | 1.700E-06 | 2.400E-06 |
| 10846 | 543 | P543E | A | 1.4e-06 | 2.5e-06 | 2.5e-06 | 2.133E-06 | 6.351E-07 | 1.400E-06 | 2.500E-06 |
| 10847 | 543 | P543Q | A | 2.5e-06 | 2.5e-06 | 2.2e-06 | 2.400E-06 | 1.732E-07 | 2.200E-06 | 2.500E-06 |
| 10848 | 543 | P543G | A | 2.3e-06 | 1.5e-06 | 3.8e-06 | 2.533E-06 | 1.168E-06 | 1.500E-06 | 3.800E-06 |
| 10849 | 543 | P543H | A | 2.6e-06 | 3.6e-06 | 2.7e-06 | 2.967E-06 | 5.508E-07 | 2.600E-06 | 3.600E-06 |
| 10850 | 543 | P543I | A | 2.8e-06 | 2.8e-06 | 3.8e-06 | 3.133E-06 | 5.774E-07 | 2.800E-06 | 3.800E-06 |
| 10851 | 543 | P543L | A | 1.6e-06 | 2.6e-06 | 2.1e-06 | 2.100E-06 | 5.000E-07 | 1.600E-06 | 2.600E-06 |
| 10852 | 543 | P543K | A | 1.3e-06 | 1.6e-06 | 3e-06 | 1.967E-06 | 9.074E-07 | 1.300E-06 | 3.000E-06 |
| 10853 | 543 | P543M | A | 2.4e-06 | 2e-06 | 3.3e-06 | 2.567E-06 | 6.658E-07 | 2.000E-06 | 3.300E-06 |
| 10854 | 543 | P543F | A | 2e-06 | 3.1e-06 | 2.6e-06 | 2.567E-06 | 5.508E-07 | 2.000E-06 | 3.100E-06 |
| 10855 | 543 | P543P | A | 2.3e-06 | 3.2e-06 | 2.7e-06 | 2.733E-06 | 4.509E-07 | 2.300E-06 | 3.200E-06 |
| 10856 | 543 | P543S | A | 3.2e-06 | 2.3e-06 | 2.7e-06 | 2.733E-06 | 4.509E-07 | 2.300E-06 | 3.200E-06 |
| 10857 | 543 | P543T | A | 1.6e-06 | 2e-06 | 3.2e-06 | 2.267E-06 | 8.327E-07 | 1.600E-06 | 3.200E-06 |
| 10858 | 543 | P543W | A | 2.7e-06 | 1.6e-06 | 2.9e-06 | 2.400E-06 | 7.000E-07 | 1.600E-06 | 2.900E-06 |
| 10859 | 543 | P543Y | A | 2.9e-06 | 2.2e-06 | 1e-06 | 2.033E-06 | 9.609E-07 | 1.000E-06 | 2.900E-06 |
| 10860 | 543 | P543V | A | 2.8e-06 | 1.7e-06 | 2.7e-06 | 2.400E-06 | 6.083E-07 | 1.700E-06 | 2.800E-06 |
| 10861 | 544 | E544A | A | 3.1e-06 | 3.6e-06 | 3e-06 | 3.233E-06 | 3.215E-07 | 3.000E-06 | 3.600E-06 |
| 10862 | 544 | E544R | A | 2.2e-06 | 1.5e-06 | 1.7e-06 | 1.800E-06 | 3.606E-07 | 1.500E-06 | 2.200E-06 |
| 10863 | 544 | E544N | A | 3.7e-06 | 1.3e-06 | 2.3e-06 | 2.433E-06 | 1.206E-06 | 1.300E-06 | 3.700E-06 |
| 10864 | 544 | E544D | A | 3.2e-06 | 2.3e-06 | 2.9e-06 | 2.800E-06 | 4.583E-07 | 2.300E-06 | 3.200E-06 |
| 10865 | 544 | E544C | A | 3.2e-06 | 4.2e-06 | 3.8e-06 | 3.733E-06 | 5.033E-07 | 3.200E-06 | 4.200E-06 |
| 10866 | 544 | E544E | A | 2.7e-06 | 1.8e-06 | 2.2e-06 | 2.233E-06 | 4.509E-07 | 1.800E-06 | 2.700E-06 |
| 10867 | 544 | E544Q | A | 1.5e-06 | 1e-06 | 2e-06 | 1.500E-06 | 5.000E-07 | 1.000E-06 | 2.000E-06 |
| 10868 | 544 | E544G | A | 2.1e-06 | 3.8e-06 | 1.6e-06 | 2.500E-06 | 1.153E-06 | 1.600E-06 | 3.800E-06 |
| 10869 | 544 | E544H | A | 2.2e-06 | 1.4e-06 | 2.1e-06 | 1.900E-06 | 4.359E-07 | 1.400E-06 | 2.200E-06 |
| 10870 | 544 | E544I | A | 3.4e-06 | 2.7e-06 | 2.5e-06 | 2.867E-06 | 4.726E-07 | 2.500E-06 | 3.400E-06 |
| 10871 | 544 | E544L | A | 2.6e-06 | 2.6e-06 | 2.1e-06 | 2.433E-06 | 2.887E-07 | 2.100E-06 | 2.600E-06 |
| 10872 | 544 | E544K | A | 2.2e-06 | 2e-06 | 3.9e-06 | 2.700E-06 | 1.044E-06 | 2.000E-06 | 3.900E-06 |
| 10873 | 544 | E544M | A | 2.5e-06 | 2.8e-06 | 2.3e-06 | 2.533E-06 | 2.517E-07 | 2.300E-06 | 2.800E-06 |
| 10874 | 544 | E544F | A | 3.8e-06 | 2.8e-06 | 3.6e-06 | 3.400E-06 | 5.292E-07 | 2.800E-06 | 3.800E-06 |
| 10875 | 544 | E544P | A | 3.6e-06 | 1.6e-06 | 4.4e-06 | 3.200E-06 | 1.442E-06 | 1.600E-06 | 4.400E-06 |
| 10876 | 544 | E544S | A | 2.8e-06 | 3.4e-06 | 1.6e-06 | 2.600E-06 | 9.165E-07 | 1.600E-06 | 3.400E-06 |
| 10877 | 544 | E544T | A | 2.6e-06 | 2.5e-06 | 3.2e-06 | 2.767E-06 | 3.786E-07 | 2.500E-06 | 3.200E-06 |
| 10878 | 544 | E544W | A | 2.8e-06 | 4.7e-06 | 2.2e-06 | 3.233E-06 | 1.305E-06 | 2.200E-06 | 4.700E-06 |
| 10879 | 544 | E544Y | A | 2.2e-06 | 2.6e-06 | 3.1e-06 | 2.633E-06 | 4.509E-07 | 2.200E-06 | 3.100E-06 |

| | | | | | | | | | | |
|---|---|---|---|---|---|---|---|---|---|---|
| 10880 | 544 | E544V | A | 2.4e-06 | 2.3e-06 | 4.1e-06 | 2.933E-06 | 1.012E-06 | 2.300E-06 | 4.100E-06 |
| 10881 | 545 | C545A | A | 1.7e-06 | 1.9e-06 | 3.6e-06 | 2.400E-06 | 1.044E-06 | 1.700E-06 | 3.600E-06 |
| 10882 | 545 | C545R | A | 2.2e-06 | 3.1e-06 | 6.5e-06 | 3.933E-06 | 2.268E-06 | 2.200E-06 | 6.500E-06 |
| 10883 | 545 | C545N | A | 2.4e-06 | 1.8e-06 | 2.2e-06 | 2.133E-06 | 3.055E-07 | 1.800E-06 | 2.400E-06 |
| 10884 | 545 | C545D | A | 3.3e-06 | 1.9e-06 | 2.3e-06 | 2.500E-06 | 7.211E-07 | 1.900E-06 | 3.300E-06 |
| 10885 | 545 | C545C | A | 3e-06 | 3.2e-06 | 2.1e-06 | 2.767E-06 | 5.859E-07 | 2.100E-06 | 3.200E-06 |
| 10886 | 545 | C545E | A | 3e-06 | 2.4e-06 | 1.7e-06 | 2.367E-06 | 6.506E-07 | 1.700E-06 | 3.000E-06 |
| 10887 | 545 | C545Q | A | 2.2e-06 | 1.9e-06 | 2e-06 | 2.033E-06 | 1.528E-07 | 1.900E-06 | 2.200E-06 |
| 10888 | 545 | C545G | A | 8.9e-07 | 3.2e-06 | 7.2e-07 | 1.603E-06 | 1.385E-06 | 7.200E-07 | 3.200E-06 |
| 10889 | 545 | C545H | A | 2.9e-06 | 1.4e-06 | 2.6e-06 | 2.300E-06 | 7.937E-07 | 1.400E-06 | 2.900E-06 |
| 10890 | 545 | C545I | A | 4.1e-06 | 2.9e-06 | 2.2e-06 | 3.067E-06 | 9.609E-07 | 2.200E-06 | 4.100E-06 |
| 10891 | 545 | C545L | A | 1.7e-06 | 3.3e-06 | 3.7e-06 | 2.900E-06 | 1.058E-06 | 1.700E-06 | 3.700E-06 |
| 10892 | 545 | C545K | A | 2.9e-06 | 3.1e-06 | 3.6e-06 | 3.200E-06 | 3.606E-07 | 2.900E-06 | 3.600E-06 |
| 10893 | 545 | C545M | A | 1e-06 | 2.5e-06 | 3.7e-06 | 2.400E-06 | 1.353E-06 | 1.000E-06 | 3.700E-06 |
| 10894 | 545 | C545F | A | 6e-06 | 1.8e-06 | 3e-06 | 3.600E-06 | 2.163E-06 | 1.800E-06 | 6.000E-06 |
| 10895 | 545 | C545P | A | 3.4e-06 | 2.4e-06 | 2.2e-06 | 2.667E-06 | 6.429E-07 | 2.200E-06 | 3.400E-06 |
| 10896 | 545 | C545S | A | 1.3e-06 | 2.2e-06 | 6.2e-06 | 3.233E-06 | 2.608E-06 | 1.300E-06 | 6.200E-06 |
| 10897 | 545 | C545T | A | 2.1e-06 | 3e-06 | 1.1e-06 | 2.067E-06 | 9.504E-07 | 1.100E-06 | 3.000E-06 |
| 10898 | 545 | C545W | A | 2.6e-06 | 2e-06 | 1.6e-06 | 2.067E-06 | 5.033E-07 | 1.600E-06 | 2.600E-06 |
| 10899 | 545 | C545Y | A | 2.3e-06 | 2.5e-06 | 1.8e-06 | 2.200E-06 | 3.606E-07 | 1.800E-06 | 2.500E-06 |
| 10900 | 545 | C545V | A | 2e-06 | 2.9e-06 | 3.4e-06 | 2.767E-06 | 7.095E-07 | 2.000E-06 | 3.400E-06 |
| 10901 | 546 | Q546A | A | 1.4e-06 | 2.3e-06 | 2.9e-06 | 2.200E-06 | 7.550E-07 | 1.400E-06 | 2.900E-06 |
| 10902 | 546 | Q546R | A | 1.4e-06 | 2.1e-06 | 1.5e-06 | 1.667E-06 | 3.786E-07 | 1.400E-06 | 2.100E-06 |
| 10903 | 546 | Q546N | A | 1.9e-06 | 2e-06 | 2.4e-06 | 2.100E-06 | 2.646E-07 | 1.900E-06 | 2.400E-06 |
| 10904 | 546 | Q546D | A | 1.5e-06 | 2.8e-06 | 2.4e-06 | 2.233E-06 | 6.658E-07 | 1.500E-06 | 2.800E-06 |
| 10905 | 546 | Q546C | A | 2.6e-06 | 1.1e-06 | 2e-06 | 1.900E-06 | 7.550E-07 | 1.100E-06 | 2.600E-06 |
| 10906 | 546 | Q546E | A | 4.3e-06 | 3.6e-06 | 3.3e-06 | 3.733E-06 | 5.132E-07 | 3.300E-06 | 4.300E-06 |
| 10907 | 546 | Q546Q | A | 2.5e-06 | 2.3e-06 | 1.9e-06 | 2.233E-06 | 3.055E-07 | 1.900E-06 | 2.500E-06 |
| 10908 | 546 | Q546G | A | 1.5e-06 | 2.4e-06 | 2.4e-06 | 2.100E-06 | 5.196E-07 | 1.500E-06 | 2.400E-06 |
| 10909 | 546 | Q546H | A | 2.8e-06 | 3.2e-06 | 3.1e-06 | 3.033E-06 | 2.082E-07 | 2.800E-06 | 3.200E-06 |
| 10910 | 546 | Q546I | A | 2.7e-06 | 1.6e-06 | 2.5e-06 | 2.267E-06 | 5.859E-07 | 1.600E-06 | 2.700E-06 |
| 10911 | 546 | Q546L | A | 2.7e-06 | 2.8e-06 | 4e-06 | 3.167E-06 | 7.234E-07 | 2.700E-06 | 4.000E-06 |
| 10912 | 546 | Q546K | A | 2e-06 | 2.2e-06 | 2.3e-06 | 2.167E-06 | 1.528E-07 | 2.000E-06 | 2.300E-06 |
| 10913 | 546 | Q546M | A | 3.6e-06 | 2.8e-06 | 2.1e-06 | 2.833E-06 | 7.506E-07 | 2.100E-06 | 3.600E-06 |
| 10914 | 546 | Q546F | A | 2.6e-06 | 2.1e-06 | 1.8e-06 | 2.167E-06 | 4.041E-07 | 1.800E-06 | 2.600E-06 |
| 10915 | 546 | Q546P | A | 3.1e-06 | 3.2e-06 | 2.2e-06 | 2.833E-06 | 5.508E-07 | 2.200E-06 | 3.200E-06 |
| 10916 | 546 | Q546S | A | 2.1e-06 | 3e-06 | 3.2e-06 | 2.767E-06 | 5.859E-07 | 2.100E-06 | 3.200E-06 |
| 10917 | 546 | Q546T | A | 1.5e-06 | 1.8e-06 | 3.2e-06 | 2.167E-06 | 9.074E-07 | 1.500E-06 | 3.200E-06 |
| 10918 | 546 | Q546W | A | 3.4e-06 | 1.5e-06 | 3.4e-06 | 2.767E-06 | 1.097E-06 | 1.500E-06 | 3.400E-06 |
| 10919 | 546 | Q546Y | A | 4.8e-06 | 1.9e-06 | 2.4e-06 | 3.033E-06 | 1.550E-06 | 1.900E-06 | 4.800E-06 |
| 10920 | 546 | Q546V | A | 2e-06 | 3.2e-06 | 3.9e-06 | 3.033E-06 | 9.609E-07 | 2.000E-06 | 3.900E-06 |
| 10921 | 547 | P547A | A | 2.2e-06 | 2.5e-06 | 2.3e-06 | 2.333E-06 | 1.528E-07 | 2.200E-06 | 2.500E-06 |
| 10922 | 547 | P547R | A | 3.3e-06 | 2.1e-06 | 2.7e-06 | 2.700E-06 | 6.000E-07 | 2.100E-06 | 3.300E-06 |
| 10923 | 547 | P547N | A | 2.8e-06 | 2.2e-06 | 1.4e-06 | 2.133E-06 | 7.024E-07 | 1.400E-06 | 2.800E-06 |
| 10924 | 547 | P547D | A | 1.8e-06 | 1.6e-06 | 1.3e-06 | 1.567E-06 | 2.517E-07 | 1.300E-06 | 1.800E-06 |
| 10925 | 547 | P547C | A | 3.3e-06 | 2.3e-06 | 2.1e-06 | 2.567E-06 | 6.429E-07 | 2.100E-06 | 3.300E-06 |
| 10926 | 547 | P547E | A | 1.4e-06 | 1.6e-06 | 2.7e-06 | 1.900E-06 | 7.000E-07 | 1.400E-06 | 2.700E-06 |
| 10927 | 547 | P547Q | A | 3e-06 | 3.1e-06 | 2.7e-06 | 2.933E-06 | 2.082E-07 | 2.700E-06 | 3.100E-06 |
| 10928 | 547 | P547G | A | 1.7e-06 | 8.7e-07 | 1.4e-06 | 1.323E-06 | 4.203E-07 | 8.700E-07 | 1.700E-06 |
| 10929 | 547 | P547H | A | 1.6e-06 | 2.1e-06 | 2.7e-06 | 2.133E-06 | 5.508E-07 | 1.600E-06 | 2.700E-06 |
| 10930 | 547 | P547I | A | 2.5e-06 | 3.4e-06 | 2.8e-06 | 2.900E-06 | 4.583E-07 | 2.500E-06 | 3.400E-06 |
| 10931 | 547 | P547L | A | 1.9e-06 | 1.8e-06 | 2.3e-06 | 2.000E-06 | 2.646E-07 | 1.800E-06 | 2.300E-06 |
| 10932 | 547 | P547K | A | 1.9e-06 | 1.3e-06 | 3e-06 | 2.067E-06 | 8.622E-07 | 1.300E-06 | 3.000E-06 |
| 10933 | 547 | P547M | A | 3.8e-06 | 1.7e-06 | 2.8e-06 | 2.767E-06 | 1.050E-06 | 1.700E-06 | 3.800E-06 |
| 10934 | 547 | P547F | A | 4.5e-06 | 3e-06 | 1.5e-06 | 3.000E-06 | 1.500E-06 | 1.500E-06 | 4.500E-06 |
| 10935 | 547 | P547P | A | 2.9e-06 | 2.4e-06 | 3.4e-06 | 2.900E-06 | 5.000E-07 | 2.400E-06 | 3.400E-06 |
| 10936 | 547 | P547S | A | 1.9e-06 | 3.1e-06 | 2.1e-06 | 2.367E-06 | 6.429E-07 | 1.900E-06 | 3.100E-06 |
| 10937 | 547 | P547T | A | 2.4e-06 | 1.8e-06 | 1.7e-06 | 1.967E-06 | 3.786E-07 | 1.700E-06 | 2.400E-06 |
| 10938 | 547 | P547W | A | 2.4e-06 | 3.6e-06 | 1.8e-06 | 2.600E-06 | 9.165E-07 | 1.800E-06 | 3.600E-06 |
| 10939 | 547 | P547Y | A | 2.8e-06 | 3.2e-06 | 4.6e-06 | 3.533E-06 | 9.452E-07 | 2.800E-06 | 4.600E-06 |
| 10940 | 547 | P547V | A | 3.4e-06 | 3.2e-06 | 3.5e-06 | 3.367E-06 | 1.528E-07 | 3.200E-06 | 3.500E-06 |
| 10941 | 548 | Q548A | A | 2e-06 | 3e-06 | 1.9e-06 | 2.300E-06 | 6.083E-07 | 1.900E-06 | 3.000E-06 |
| 10942 | 548 | Q548R | A | 1.8e-06 | 2.1e-06 | 1.7e-06 | 1.867E-06 | 2.082E-07 | 1.700E-06 | 2.100E-06 |
| 10943 | 548 | Q548N | A | 2e-06 | 4.1e-06 | 2.4e-06 | 2.833E-06 | 1.115E-06 | 2.000E-06 | 4.100E-06 |
| 10944 | 548 | Q548D | A | 2.3e-06 | 2.6e-06 | 1.3e-06 | 2.067E-06 | 6.807E-07 | 1.300E-06 | 2.600E-06 |
| 10945 | 548 | Q548C | A | 2.2e-06 | 2.9e-06 | 2.3e-06 | 2.467E-06 | 3.786E-07 | 2.200E-06 | 2.900E-06 |
| 10946 | 548 | Q548E | A | 1.7e-06 | 2.2e-06 | 1.9e-06 | 1.933E-06 | 2.517E-07 | 1.700E-06 | 2.200E-06 |
| 10947 | 548 | Q548Q | A | 1.9e-06 | 2.1e-06 | 2.9e-06 | 2.300E-06 | 5.292E-07 | 1.900E-06 | 2.900E-06 |
| 10948 | 548 | Q548G | A | 2.7e-06 | 2.6e-06 | 2.5e-06 | 2.600E-06 | 1.000E-07 | 2.500E-06 | 2.700E-06 |
| 10949 | 548 | Q548H | A | 1.7e-06 | 1.3e-06 | 2.5e-06 | 1.833E-06 | 6.110E-07 | 1.300E-06 | 2.500E-06 |
| 10950 | 548 | Q548I | A | 2.5e-06 | 2.3e-06 | 2.3e-06 | 2.367E-06 | 1.155E-07 | 2.300E-06 | 2.500E-06 |
| 10951 | 548 | Q548L | A | 1.2e-06 | 3.1e-06 | 3.5e-06 | 2.600E-06 | 1.229E-06 | 1.200E-06 | 3.500E-06 |
| 10952 | 548 | Q548K | A | 2.7e-06 | 1.8e-06 | 3e-06 | 2.500E-06 | 6.245E-07 | 1.800E-06 | 3.000E-06 |
| 10953 | 548 | Q548M | A | 2.9e-06 | 1.8e-06 | 1.5e-06 | 2.067E-06 | 7.371E-07 | 1.500E-06 | 2.900E-06 |
| 10954 | 548 | Q548F | A | 2.5e-06 | 2.2e-06 | 1.5e-06 | 2.067E-06 | 5.132E-07 | 1.500E-06 | 2.500E-06 |
| 10955 | 548 | Q548P | A | 2e-06 | 2.7e-06 | 2.6e-06 | 2.433E-06 | 3.786E-07 | 2.000E-06 | 2.700E-06 |
| 10956 | 548 | Q548S | A | 2.3e-06 | 1.8e-06 | 4.3e-06 | 2.800E-06 | 1.323E-06 | 1.800E-06 | 4.300E-06 |
| 10957 | 548 | Q548T | A | 1.6e-06 | 4e-06 | 4.1e-06 | 3.233E-06 | 1.415E-06 | 1.600E-06 | 4.100E-06 |
| 10958 | 548 | Q548W | A | 2.3e-06 | 4.3e-06 | 2.5e-06 | 3.033E-06 | 1.102E-06 | 2.300E-06 | 4.300E-06 |
| 10959 | 548 | Q548Y | A | 2.1e-06 | 1.2e-06 | 3e-06 | 2.100E-06 | 9.000E-07 | 1.200E-06 | 3.000E-06 |
| 10960 | 548 | Q548V | A | 3.2e-06 | 1e-06 | 2.7e-06 | 2.300E-06 | 1.153E-06 | 1.000E-06 | 3.200E-06 |
| 10961 | 549 | N549A | A | 2.7e-06 | 2e-06 | 2.8e-06 | 2.500E-06 | 4.359E-07 | 2.000E-06 | 2.800E-06 |
| 10962 | 549 | N549R | A | 2.5e-06 | 2.3e-06 | 2.8e-06 | 2.533E-06 | 2.517E-07 | 2.300E-06 | 2.800E-06 |
| 10963 | 549 | N549N | A | 1.4e-06 | 1.2e-06 | 3e-06 | 1.867E-06 | 9.866E-07 | 1.200E-06 | 3.000E-06 |
| 10964 | 549 | N549D | A | 3.6e-06 | 2.1e-06 | 4e-06 | 3.233E-06 | 1.002E-06 | 2.100E-06 | 4.000E-06 |

| | | | | | | | | | | |
|---|---|---|---|---|---|---|---|---|---|---|
| 10965 | 549 | N549C | A | 4.2e-06 | 2.3e-06 | 2.4e-06 | 2.967E-06 | 1.069E-06 | 2.300E-06 | 4.200E-06 |
| 10966 | 549 | N549E | A | 2.7e-06 | 3.1e-06 | 2.6e-06 | 2.800E-06 | 2.646E-07 | 2.600E-06 | 3.100E-06 |
| 10967 | 549 | N549Q | A | 1.6e-06 | 2.4e-06 | 2.8e-06 | 2.267E-06 | 6.110E-07 | 1.600E-06 | 2.800E-06 |
| 10968 | 549 | N549G | A | 2.4e-06 | 3.8e-06 | 3.6e-06 | 3.267E-06 | 7.572E-07 | 2.400E-06 | 3.800E-06 |
| 10969 | 549 | N549H | A | 1.9e-06 | 4.2e-06 | 1.8e-06 | 2.633E-06 | 1.358E-06 | 1.800E-06 | 4.200E-06 |
| 10970 | 549 | N549I | A | 2.9e-06 | 1.9e-06 | 2.8e-06 | 2.533E-06 | 5.508E-07 | 1.900E-06 | 2.900E-06 |
| 10971 | 549 | N549L | A | 2.3e-06 | 2.2e-06 | 2.6e-06 | 2.367E-06 | 2.082E-07 | 2.200E-06 | 2.600E-06 |
| 10972 | 549 | N549K | A | 2.2e-06 | 3e-06 | 2.6e-06 | 2.600E-06 | 4.000E-07 | 2.200E-06 | 3.000E-06 |
| 10973 | 549 | N549M | A | 1.6e-06 | 3.1e-06 | 2.8e-06 | 2.500E-06 | 7.937E-07 | 1.600E-06 | 3.100E-06 |
| 10974 | 549 | N549F | A | 2.8e-06 | 7.2e-06 | 2.2e-06 | 4.067E-06 | 2.730E-06 | 2.200E-06 | 7.200E-06 |
| 10975 | 549 | N549P | A | 3e-06 | 2.6e-06 | 1.4e-06 | 2.333E-06 | 8.327E-07 | 1.400E-06 | 3.000E-06 |
| 10976 | 549 | N549S | A | 1.5e-06 | 1.9e-06 | 1.7e-06 | 1.700E-06 | 2.000E-07 | 1.500E-06 | 1.900E-06 |
| 10977 | 549 | N549T | A | 1.8e-06 | 1.9e-06 | 2.3e-06 | 2.000E-06 | 2.646E-07 | 1.800E-06 | 2.300E-06 |
| 10978 | 549 | N549W | A | 1.7e-06 | 2.2e-06 | 2.1e-06 | 2.000E-06 | 2.646E-07 | 1.700E-06 | 2.200E-06 |
| 10979 | 549 | N549Y | A | 2.9e-06 | 1.4e-06 | 2.5e-06 | 2.267E-06 | 7.767E-07 | 1.400E-06 | 2.900E-06 |
| 10980 | 549 | N549V | A | 2.2e-06 | 2e-06 | 3.2e-06 | 2.467E-06 | 6.429E-07 | 2.000E-06 | 3.200E-06 |
| 10981 | 550 | G550A | A | 1.8e-06 | 1.7e-06 | 3.4e-06 | 2.300E-06 | 9.539E-07 | 1.700E-06 | 3.400E-06 |
| 10982 | 550 | G550R | A | 3.3e-06 | 2.2e-06 | 2e-06 | 2.500E-06 | 7.000E-07 | 2.000E-06 | 3.300E-06 |
| 10983 | 550 | G550N | A | 1.6e-06 | 2.4e-06 | 2.1e-06 | 2.033E-06 | 4.041E-07 | 1.600E-06 | 2.400E-06 |
| 10984 | 550 | G550D | A | 1.5e-06 | 3.3e-06 | 1.9e-06 | 2.233E-06 | 9.452E-07 | 1.500E-06 | 3.300E-06 |
| 10985 | 550 | G550C | A | 3.2e-06 | 1.9e-06 | 3.3e-06 | 2.800E-06 | 7.810E-07 | 1.900E-06 | 3.300E-06 |
| 10986 | 550 | G550E | A | 2.6e-06 | 1.7e-06 | 2.9e-06 | 2.400E-06 | 6.245E-07 | 1.700E-06 | 2.900E-06 |
| 10987 | 550 | G550Q | A | 3.5e-06 | 2.2e-06 | 1.3e-06 | 2.333E-06 | 1.106E-06 | 1.300E-06 | 3.500E-06 |
| 10988 | 550 | G550G | A | 1.9e-06 | 1.9e-06 | 1.3e-06 | 1.700E-06 | 3.464E-07 | 1.300E-06 | 1.900E-06 |
| 10989 | 550 | G550H | A | 2.3e-06 | 1.7e-06 | 3.3e-06 | 2.433E-06 | 8.083E-07 | 1.700E-06 | 3.300E-06 |
| 10990 | 550 | G550I | A | 4.1e-06 | 2.9e-06 | 3e-06 | 3.333E-06 | 6.658E-07 | 2.900E-06 | 4.100E-06 |
| 10991 | 550 | G550L | A | 1.2e-06 | 1.8e-06 | 1.6e-06 | 1.533E-06 | 3.055E-07 | 1.200E-06 | 1.800E-06 |
| 10992 | 550 | G550K | A | 2.1e-06 | 3.5e-06 | 2.6e-06 | 2.733E-06 | 7.095E-07 | 2.100E-06 | 3.500E-06 |
| 10993 | 550 | G550M | A | 2.5e-06 | 2e-06 | 2.6e-06 | 2.367E-06 | 3.215E-07 | 2.000E-06 | 2.600E-06 |
| 10994 | 550 | G550F | A | 1.7e-06 | 1.7e-06 | 2.6e-06 | 2.000E-06 | 5.196E-07 | 1.700E-06 | 2.600E-06 |
| 10995 | 550 | G550P | A | 1.5e-06 | 3.1e-06 | 2.1e-06 | 2.233E-06 | 8.083E-07 | 1.500E-06 | 3.100E-06 |
| 10996 | 550 | G550S | A | 2.4e-06 | 1.6e-06 | 3.1e-06 | 2.367E-06 | 7.506E-07 | 1.600E-06 | 3.100E-06 |
| 10997 | 550 | G550T | A | 2.6e-06 | 2.6e-06 | 2.1e-06 | 2.433E-06 | 2.887E-07 | 2.100E-06 | 2.600E-06 |
| 10998 | 550 | G550W | A | 1.2e-06 | 3.2e-06 | 1.2e-06 | 1.867E-06 | 1.155E-06 | 1.200E-06 | 3.200E-06 |
| 10999 | 550 | G550Y | A | 3.5e-06 | 1.6e-06 | 1.9e-06 | 2.333E-06 | 1.021E-06 | 1.600E-06 | 3.500E-06 |
| 11000 | 550 | G550V | A | 1.8e-06 | 2.9e-06 | 2.1e-06 | 2.267E-06 | 5.686E-07 | 1.800E-06 | 2.900E-06 |
| 11001 | 551 | S551A | A | 1.7e-06 | 3.5e-06 | 1.8e-06 | 2.333E-06 | 1.012E-06 | 1.700E-06 | 3.500E-06 |
| 11002 | 551 | S551R | A | 3.1e-06 | 2.8e-06 | 4.1e-06 | 3.333E-06 | 6.807E-07 | 2.800E-06 | 4.100E-06 |
| 11003 | 551 | S551N | A | 1.9e-06 | 1.9e-06 | 1.9e-06 | 1.900E-06 | 2.593E-22 | 1.900E-06 | 1.900E-06 |
| 11004 | 551 | S551D | A | 1.6e-06 | 3.1e-06 | 2.8e-06 | 2.500E-06 | 7.937E-07 | 1.600E-06 | 3.100E-06 |
| 11005 | 551 | S551C | A | 1.6e-06 | 2.7e-06 | 1.7e-06 | 2.000E-06 | 6.083E-07 | 1.600E-06 | 2.700E-06 |
| 11006 | 551 | S551E | A | 1e-06 | 3.8e-06 | 1.9e-06 | 2.233E-06 | 1.429E-06 | 1.000E-06 | 3.800E-06 |
| 11007 | 551 | S551Q | A | 3.1e-06 | 3.2e-06 | 1.6e-06 | 2.633E-06 | 8.963E-07 | 1.600E-06 | 3.200E-06 |
| 11008 | 551 | S551G | A | 4.1e-06 | 3e-06 | 2.2e-06 | 3.100E-06 | 9.539E-07 | 2.200E-06 | 4.100E-06 |
| 11009 | 551 | S551H | A | 2.8e-06 | 1.6e-06 | 2.4e-06 | 2.267E-06 | 6.110E-07 | 1.600E-06 | 2.800E-06 |
| 11010 | 551 | S551I | A | 2.2e-06 | 1.8e-06 | 2e-06 | 2.000E-06 | 2.000E-07 | 1.800E-06 | 2.200E-06 |
| 11011 | 551 | S551L | A | 2.4e-06 | 3e-06 | 2.9e-06 | 2.767E-06 | 3.215E-07 | 2.400E-06 | 3.000E-06 |
| 11012 | 551 | S551K | A | 5.1e-06 | 2.3e-06 | 2.5e-06 | 3.300E-06 | 1.562E-06 | 2.300E-06 | 5.100E-06 |
| 11013 | 551 | S551M | A | 3.9e-06 | 4.8e-06 | 2.3e-06 | 3.667E-06 | 1.266E-06 | 2.300E-06 | 4.800E-06 |
| 11014 | 551 | S551F | A | 2.6e-06 | 2.9e-06 | 2.9e-06 | 2.800E-06 | 1.732E-07 | 2.600E-06 | 2.900E-06 |
| 11015 | 551 | S551P | A | 3.1e-06 | 1.9e-06 | 2.7e-06 | 2.567E-06 | 6.110E-07 | 1.900E-06 | 3.100E-06 |
| 11016 | 551 | S551S | A | 1.8e-06 | 3.1e-06 | 2.5e-06 | 2.467E-06 | 6.506E-07 | 1.800E-06 | 3.100E-06 |
| 11017 | 551 | S551T | A | 3.5e-06 | 4.1e-06 | 2.4e-06 | 3.333E-06 | 8.622E-07 | 2.400E-06 | 4.100E-06 |
| 11018 | 551 | S551W | A | 1.7e-06 | 2.7e-06 | 1.2e-06 | 1.867E-06 | 7.638E-07 | 1.200E-06 | 2.700E-06 |
| 11019 | 551 | S551Y | A | 2.1e-06 | 2.2e-06 | 4e-06 | 2.767E-06 | 1.069E-06 | 2.100E-06 | 4.000E-06 |
| 11020 | 551 | S551V | A | 2.3e-06 | 1.8e-06 | 3.7e-06 | 2.600E-06 | 9.849E-07 | 1.800E-06 | 3.700E-06 |
| 11021 | 552 | V552A | A | 2.4e-06 | 1.4e-06 | 1.9e-06 | 1.900E-06 | 5.000E-07 | 1.400E-06 | 2.400E-06 |
| 11022 | 552 | V552R | A | 1.4e-06 | 2.7e-06 | 1.6e-06 | 1.900E-06 | 7.000E-07 | 1.400E-06 | 2.700E-06 |
| 11023 | 552 | V552N | A | 2.6e-06 | 1.8e-06 | 2.2e-06 | 2.200E-06 | 4.000E-07 | 1.800E-06 | 2.600E-06 |
| 11024 | 552 | V552D | A | 3.3e-06 | 1.6e-06 | 2.7e-06 | 2.533E-06 | 8.622E-07 | 1.600E-06 | 3.300E-06 |
| 11025 | 552 | V552C | A | 2.2e-06 | 2.4e-06 | 1.6e-06 | 2.067E-06 | 4.163E-07 | 1.600E-06 | 2.400E-06 |
| 11026 | 552 | V552E | A | 2.6e-06 | 1.6e-06 | 1.4e-06 | 1.867E-06 | 6.429E-07 | 1.400E-06 | 2.600E-06 |
| 11027 | 552 | V552Q | A | 1.5e-06 | 2.7e-06 | 1.6e-06 | 1.933E-06 | 6.658E-07 | 1.500E-06 | 2.700E-06 |
| 11028 | 552 | V552G | A | 1.9e-06 | 1.7e-06 | 3.3e-06 | 2.300E-06 | 8.718E-07 | 1.700E-06 | 3.300E-06 |
| 11029 | 552 | V552H | A | 2.7e-06 | 1.9e-06 | 2.2e-06 | 2.267E-06 | 4.041E-07 | 1.900E-06 | 2.700E-06 |
| 11030 | 552 | V552I | A | 1.9e-06 | 2.1e-06 | 2.4e-06 | 2.133E-06 | 2.517E-07 | 1.900E-06 | 2.400E-06 |
| 11031 | 552 | V552L | A | 1.4e-06 | 2.7e-06 | 3.1e-06 | 2.400E-06 | 8.888E-07 | 1.400E-06 | 3.100E-06 |
| 11032 | 552 | V552K | A | 3.8e-06 | 1.4e-06 | 1.8e-06 | 2.333E-06 | 1.286E-06 | 1.400E-06 | 3.800E-06 |
| 11033 | 552 | V552M | A | 1.3e-06 | 3.3e-06 | 2.6e-06 | 2.400E-06 | 1.015E-06 | 1.300E-06 | 3.300E-06 |
| 11034 | 552 | V552F | A | 3.7e-06 | 1.8e-06 | 2.2e-06 | 2.567E-06 | 1.002E-06 | 1.800E-06 | 3.700E-06 |
| 11035 | 552 | V552P | A | 1.4e-06 | 3.5e-06 | 3.5e-06 | 2.800E-06 | 1.212E-06 | 1.400E-06 | 3.500E-06 |
| 11036 | 552 | V552S | A | 2.1e-06 | 4e-06 | 2.2e-06 | 2.767E-06 | 1.069E-06 | 2.100E-06 | 4.000E-06 |
| 11037 | 552 | V552T | A | 3.8e-06 | 2.1e-06 | 2e-06 | 2.633E-06 | 1.012E-06 | 2.000E-06 | 3.800E-06 |
| 11038 | 552 | V552W | A | 1.6e-06 | 2.3e-06 | 1.5e-06 | 1.800E-06 | 4.359E-07 | 1.500E-06 | 2.300E-06 |
| 11039 | 552 | V552Y | A | 2.9e-06 | 2.4e-06 | 2e-06 | 2.433E-06 | 4.509E-07 | 2.000E-06 | 2.900E-06 |
| 11040 | 552 | V552V | A | 1.8e-06 | 1.7e-06 | 2.4e-06 | 1.967E-06 | 3.786E-07 | 1.700E-06 | 2.400E-06 |
| 11041 | 553 | T553A | A | 1.8e-06 | 1.2e-06 | 2.5e-06 | 1.833E-06 | 6.506E-07 | 1.200E-06 | 2.500E-06 |
| 11042 | 553 | T553R | A | 1.9e-06 | 1.7e-06 | 2.8e-06 | 2.133E-06 | 5.859E-07 | 1.700E-06 | 2.800E-06 |
| 11043 | 553 | T553N | A | 2.8e-06 | 2.4e-06 | 2.6e-06 | 2.600E-06 | 2.000E-07 | 2.400E-06 | 2.800E-06 |
| 11044 | 553 | T553D | A | 3.1e-06 | 2e-06 | 2.6e-06 | 2.567E-06 | 5.508E-07 | 2.000E-06 | 3.100E-06 |
| 11045 | 553 | T553C | A | 2e-06 | 1.7e-06 | 2.1e-06 | 1.933E-06 | 2.082E-07 | 1.700E-06 | 2.100E-06 |
| 11046 | 553 | T553E | A | 3e-06 | 2.5e-06 | 2.9e-06 | 2.800E-06 | 2.646E-07 | 2.500E-06 | 3.000E-06 |
| 11047 | 553 | T553Q | A | 1.7e-06 | 2.5e-06 | 2.6e-06 | 2.267E-06 | 4.933E-07 | 1.700E-06 | 2.600E-06 |
| 11048 | 553 | T553G | A | 4.1e-06 | 1.8e-06 | 2.3e-06 | 2.733E-06 | 1.210E-06 | 1.800E-06 | 4.100E-06 |
| 11049 | 553 | T553H | A | 2.4e-06 | 1.9e-06 | 1.6e-06 | 1.967E-06 | 4.041E-07 | 1.600E-06 | 2.400E-06 |

| | | | | | | | | | | |
|---|---|---|---|---|---|---|---|---|---|---|
| 11050 | 553 | T553I | A | 2.7e-06 | 1.8e-06 | 2.6e-06 | 2.367E-06 | 4.933E-07 | 1.800E-06 | 2.700E-06 |
| 11051 | 553 | T553L | A | 2.2e-06 | 2.5e-06 | 1.7e-06 | 2.133E-06 | 4.041E-07 | 1.700E-06 | 2.500E-06 |
| 11052 | 553 | T553K | A | 2e-06 | 2.8e-06 | 2.9e-06 | 2.567E-06 | 4.933E-07 | 2.000E-06 | 2.900E-06 |
| 11053 | 553 | T553M | A | 3.2e-06 | 7.9e-07 | 2.4e-06 | 2.130E-06 | 1.227E-06 | 7.900E-07 | 3.200E-06 |
| 11054 | 553 | T553F | A | 3.5e-06 | 2.8e-06 | 1.3e-06 | 2.533E-06 | 1.124E-06 | 1.300E-06 | 3.500E-06 |
| 11055 | 553 | T553P | A | 1.8e-06 | 2.2e-06 | 2.9e-06 | 2.300E-06 | 5.568E-07 | 1.800E-06 | 2.900E-06 |
| 11056 | 553 | T553S | A | 3.3e-06 | 1.2e-06 | 3.5e-06 | 2.667E-06 | 1.274E-06 | 1.200E-06 | 3.500E-06 |
| 11057 | 553 | T553T | A | 4.9e-06 | 2.4e-06 | 4e-06 | 3.767E-06 | 1.266E-06 | 2.400E-06 | 4.900E-06 |
| 11058 | 553 | T553W | A | 2.6e-06 | 2.7e-06 | 6.4e-06 | 3.900E-06 | 2.166E-06 | 2.600E-06 | 6.400E-06 |
| 11059 | 553 | T553Y | A | 2.8e-06 | 2.5e-06 | 1.6e-06 | 2.300E-06 | 6.245E-07 | 1.600E-06 | 2.800E-06 |
| 11060 | 553 | T553V | A | 1.5e-06 | 2.6e-06 | 2.9e-06 | 2.333E-06 | 7.371E-07 | 1.500E-06 | 2.900E-06 |
| 11061 | 554 | C554A | A | 3e-06 | 1.9e-06 | 1.8e-06 | 2.233E-06 | 6.658E-07 | 1.800E-06 | 3.000E-06 |
| 11062 | 554 | C554R | A | 2.8e-06 | 5.3e-06 | 1.2e-06 | 3.100E-06 | 2.066E-06 | 1.200E-06 | 5.300E-06 |
| 11063 | 554 | C554N | A | 1.9e-06 | 3.6e-06 | 3e-06 | 2.833E-06 | 8.622E-07 | 1.900E-06 | 3.600E-06 |
| 11064 | 554 | C554D | A | 4.8e-06 | 2e-06 | 2.9e-06 | 3.233E-06 | 1.429E-06 | 2.000E-06 | 4.800E-06 |
| 11065 | 554 | C554C | A | 1.8e-06 | 2.1e-06 | 1.3e-06 | 1.733E-06 | 4.041E-07 | 1.300E-06 | 2.100E-06 |
| 11066 | 554 | C554E | A | 1.6e-06 | 2.7e-06 | 3.2e-06 | 2.500E-06 | 8.185E-07 | 1.600E-06 | 3.200E-06 |
| 11067 | 554 | C554Q | A | 2.3e-06 | 3.5e-06 | 1.1e-06 | 2.300E-06 | 1.200E-06 | 1.100E-06 | 3.500E-06 |
| 11068 | 554 | C554G | A | 2.8e-06 | 3.3e-06 | 2.5e-06 | 2.867E-06 | 4.041E-07 | 2.500E-06 | 3.300E-06 |
| 11069 | 554 | C554H | A | 1e-06 | 3.6e-06 | 2.4e-06 | 2.333E-06 | 1.301E-06 | 1.000E-06 | 3.600E-06 |
| 11070 | 554 | C554I | A | 2.1e-06 | 2e-06 | 5.2e-06 | 3.100E-06 | 1.819E-06 | 2.000E-06 | 5.200E-06 |
| 11071 | 554 | C554L | A | 3.4e-06 | 3.5e-06 | 3.6e-06 | 3.500E-06 | 1.000E-07 | 3.400E-06 | 3.600E-06 |
| 11072 | 554 | C554K | A | 2.1e-06 | 2.8e-06 | 2.4e-06 | 2.433E-06 | 3.512E-07 | 2.100E-06 | 2.800E-06 |
| 11073 | 554 | C554M | A | 2.3e-06 | 2.1e-06 | 1.6e-06 | 2.000E-06 | 3.606E-07 | 1.600E-06 | 2.300E-06 |
| 11074 | 554 | C554F | A | 1.4e-06 | 4.1e-06 | 4.9e-06 | 3.467E-06 | 1.834E-06 | 1.400E-06 | 4.900E-06 |
| 11075 | 554 | C554P | A | 1.4e-06 | 3.2e-06 | 3.6e-06 | 2.733E-06 | 1.172E-06 | 1.400E-06 | 3.600E-06 |
| 11076 | 554 | C554S | A | 2.8e-06 | 5e-06 | 1.6e-06 | 3.133E-06 | 1.724E-06 | 1.600E-06 | 5.000E-06 |
| 11077 | 554 | C554T | A | 9.2e-07 | 3.7e-06 | 1.4e-06 | 2.007E-06 | 1.486E-06 | 9.200E-07 | 3.700E-06 |
| 11078 | 554 | C554W | A | 1.4e-06 | 3.6e-06 | 2.6e-06 | 2.533E-06 | 1.102E-06 | 1.400E-06 | 3.600E-06 |
| 11079 | 554 | C554Y | A | 2.4e-06 | 9e-07 | 1.9e-06 | 1.733E-06 | 7.638E-07 | 9.000E-07 | 2.400E-06 |
| 11080 | 554 | C554V | A | 2.2e-06 | 4.1e-06 | 1.9e-06 | 2.733E-06 | 1.193E-06 | 1.900E-06 | 4.100E-06 |
| 11081 | 555 | F555A | A | 1.8e-06 | 1.9e-06 | 1.6e-06 | 1.767E-06 | 1.528E-07 | 1.600E-06 | 1.900E-06 |
| 11082 | 555 | F555R | A | 1.7e-06 | 2.9e-06 | 4.3e-06 | 2.967E-06 | 1.301E-06 | 1.700E-06 | 4.300E-06 |
| 11083 | 555 | F555N | A | 1.8e-06 | 1.3e-06 | 2.6e-06 | 1.900E-06 | 6.557E-07 | 1.300E-06 | 2.600E-06 |
| 11084 | 555 | F555D | A | 3.5e-06 | 4.3e-06 | 2.7e-06 | 3.500E-06 | 8.000E-07 | 2.700E-06 | 4.300E-06 |
| 11085 | 555 | F555C | A | 2.2e-06 | 2.5e-06 | 4.2e-06 | 2.967E-06 | 1.079E-06 | 2.200E-06 | 4.200E-06 |
| 11086 | 555 | F555E | A | 2.7e-06 | 1.4e-06 | 2.4e-06 | 2.167E-06 | 6.807E-07 | 1.400E-06 | 2.700E-06 |
| 11087 | 555 | F555Q | A | 1.4e-06 | 3.1e-06 | 2.2e-06 | 2.233E-06 | 8.505E-07 | 1.400E-06 | 3.100E-06 |
| 11088 | 555 | F555G | A | 3.4e-06 | 2.6e-06 | 2.7e-06 | 2.900E-06 | 4.359E-07 | 2.600E-06 | 3.400E-06 |
| 11089 | 555 | F555H | A | 2.3e-06 | 3.1e-06 | 2.2e-06 | 2.533E-06 | 4.933E-07 | 2.200E-06 | 3.100E-06 |
| 11090 | 555 | F555I | A | 2.1e-06 | 1.9e-06 | 2.4e-06 | 2.133E-06 | 2.517E-07 | 1.900E-06 | 2.400E-06 |
| 11091 | 555 | F555L | A | 1.5e-06 | 1.7e-06 | 1.9e-06 | 1.700E-06 | 2.000E-07 | 1.500E-06 | 1.900E-06 |
| 11092 | 555 | F555K | A | 2.3e-06 | 2.9e-06 | 3.3e-06 | 2.833E-06 | 5.033E-07 | 2.300E-06 | 3.300E-06 |
| 11093 | 555 | F555M | A | 3.1e-06 | 3e-06 | 2.6e-06 | 2.900E-06 | 2.646E-07 | 2.600E-06 | 3.100E-06 |
| 11094 | 555 | F555F | A | 1.2e-06 | 2e-06 | 1.8e-06 | 1.667E-06 | 4.163E-07 | 1.200E-06 | 2.000E-06 |
| 11095 | 555 | F555P | A | 3e-06 | 2.5e-06 | 1.1e-06 | 2.200E-06 | 9.849E-07 | 1.100E-06 | 3.000E-06 |
| 11096 | 555 | F555S | A | 1.5e-06 | 1.8e-06 | 2.1e-06 | 1.800E-06 | 3.000E-07 | 1.500E-06 | 2.100E-06 |
| 11097 | 555 | F555T | A | 1.6e-06 | 3.8e-06 | 1.9e-06 | 2.433E-06 | 1.193E-06 | 1.600E-06 | 3.800E-06 |
| 11098 | 555 | F555W | A | 1.3e-06 | 2.4e-06 | 2.4e-06 | 2.033E-06 | 6.351E-07 | 1.300E-06 | 2.400E-06 |
| 11099 | 555 | F555Y | A | 1.5e-06 | 3.9e-06 | 1.3e-06 | 2.233E-06 | 1.447E-06 | 1.300E-06 | 3.900E-06 |
| 11100 | 555 | F555V | A | 2.2e-06 | 3.3e-06 | 2.1e-06 | 2.533E-06 | 6.658E-07 | 2.100E-06 | 3.300E-06 |
| 11101 | 556 | G556A | A | 1.9e-06 | 1.7e-06 | 1.6e-06 | 1.733E-06 | 1.528E-07 | 1.600E-06 | 1.900E-06 |
| 11102 | 556 | G556R | A | 2.4e-06 | 1.9e-06 | 2.8e-06 | 2.367E-06 | 4.509E-07 | 1.900E-06 | 2.800E-06 |
| 11103 | 556 | G556N | A | 3.6e-06 | 2.4e-06 | 3.8e-06 | 3.267E-06 | 7.572E-07 | 2.400E-06 | 3.800E-06 |
| 11104 | 556 | G556D | A | 1.5e-06 | 1.4e-06 | 2.3e-06 | 1.733E-06 | 4.933E-07 | 1.400E-06 | 2.300E-06 |
| 11105 | 556 | G556C | A | 2.3e-06 | 1.2e-06 | 1.6e-06 | 1.700E-06 | 5.568E-07 | 1.200E-06 | 2.300E-06 |
| 11106 | 556 | G556E | A | 1.7e-06 | 1.7e-06 | 1.5e-06 | 1.633E-06 | 1.155E-07 | 1.500E-06 | 1.700E-06 |
| 11107 | 556 | G556Q | A | 1.9e-06 | 1.5e-06 | 2e-06 | 1.800E-06 | 2.646E-07 | 1.500E-06 | 2.000E-06 |
| 11108 | 556 | G556G | A | 1.4e-06 | 1.8e-06 | 3.6e-06 | 2.267E-06 | 1.172E-06 | 1.400E-06 | 3.600E-06 |
| 11109 | 556 | G556H | A | 1.4e-06 | 2.2e-06 | 2.6e-06 | 2.067E-06 | 6.110E-07 | 1.400E-06 | 2.600E-06 |
| 11110 | 556 | G556I | A | 3.3e-06 | 1.9e-06 | 9.8e-07 | 2.060E-06 | 1.168E-06 | 9.800E-07 | 3.300E-06 |
| 11111 | 556 | G556L | A | 9.1e-07 | 1.1e-06 | 9.4e-07 | 9.833E-07 | 1.021E-07 | 9.100E-07 | 1.100E-06 |
| 11112 | 556 | G556K | A | 2e-06 | 1.4e-06 | 2.3e-06 | 1.900E-06 | 4.583E-07 | 1.400E-06 | 2.300E-06 |
| 11113 | 556 | G556M | A | 2e-06 | 8.9e-07 | 1e-06 | 1.297E-06 | 6.116E-07 | 8.900E-07 | 2.000E-06 |
| 11114 | 556 | G556F | A | 9.3e-07 | 2.6e-06 | 2.6e-06 | 2.043E-06 | 9.642E-07 | 9.300E-07 | 2.600E-06 |
| 11115 | 556 | G556P | A | 2.1e-06 | 3.2e-06 | 3e-06 | 2.767E-06 | 5.859E-07 | 2.100E-06 | 3.200E-06 |
| 11116 | 556 | G556S | A | 2.5e-06 | 2.2e-06 | 3e-06 | 2.567E-06 | 4.041E-07 | 2.200E-06 | 3.000E-06 |
| 11117 | 556 | G556T | A | 2e-06 | 2.4e-06 | 1.8e-06 | 2.067E-06 | 3.055E-07 | 1.800E-06 | 2.400E-06 |
| 11118 | 556 | G556W | A | 8.5e-07 | 1.6e-06 | 1.5e-06 | 1.317E-06 | 4.072E-07 | 8.500E-07 | 1.600E-06 |
| 11119 | 556 | G556Y | A | 1.3e-06 | 1.6e-06 | 1.5e-06 | 1.467E-06 | 1.528E-07 | 1.300E-06 | 1.600E-06 |
| 11120 | 556 | G556V | A | 1.9e-06 | 1.9e-06 | 9.6e-07 | 1.587E-06 | 5.427E-07 | 9.600E-07 | 1.900E-06 |
| 11121 | 557 | P557A | A | 1.8e-06 | 2.2e-06 | 2.4e-06 | 2.133E-06 | 3.055E-07 | 1.800E-06 | 2.400E-06 |
| 11122 | 557 | P557R | A | 2.7e-06 | 2.3e-06 | 1.1e-06 | 2.033E-06 | 8.327E-07 | 1.100E-06 | 2.700E-06 |
| 11123 | 557 | P557N | A | 9.3e-07 | 2.4e-06 | 2.8e-06 | 2.043E-06 | 9.847E-07 | 9.300E-07 | 2.800E-06 |
| 11124 | 557 | P557D | A | 2.2e-06 | 2.3e-06 | 2.4e-06 | 2.300E-06 | 1.000E-07 | 2.200E-06 | 2.400E-06 |
| 11125 | 557 | P557C | A | 1.5e-06 | 1.8e-06 | 2e-06 | 1.767E-06 | 2.517E-07 | 1.500E-06 | 2.000E-06 |
| 11126 | 557 | P557E | A | 1.7e-06 | 2.5e-06 | 2.3e-06 | 2.167E-06 | 4.163E-07 | 1.700E-06 | 2.500E-06 |
| 11127 | 557 | P557Q | A | 2.1e-06 | 1.5e-06 | 1.7e-06 | 1.767E-06 | 3.055E-07 | 1.500E-06 | 2.100E-06 |
| 11128 | 557 | P557G | A | 1.8e-06 | 1.7e-06 | 1.8e-06 | 1.767E-06 | 5.774E-08 | 1.700E-06 | 1.800E-06 |
| 11129 | 557 | P557H | A | 2.6e-06 | 2.6e-06 | 2.2e-06 | 2.467E-06 | 2.309E-07 | 2.200E-06 | 2.600E-06 |
| 11130 | 557 | P557I | A | 1.7e-06 | 1.8e-06 | 1.6e-06 | 1.700E-06 | 1.000E-07 | 1.600E-06 | 1.800E-06 |
| 11131 | 557 | P557L | A | 2.1e-06 | 1e-06 | 2.9e-06 | 2.000E-06 | 9.539E-07 | 1.000E-06 | 2.900E-06 |
| 11132 | 557 | P557K | A | 3.9e-06 | 2.3e-06 | 1e-06 | 2.400E-06 | 1.453E-06 | 1.000E-06 | 3.900E-06 |
| 11133 | 557 | P557M | A | 3.1e-06 | 1.4e-06 | 2.2e-06 | 2.233E-06 | 8.505E-07 | 1.400E-06 | 3.100E-06 |
| 11134 | 557 | P557F | A | 1.3e-06 | 2.1e-06 | 2.4e-06 | 1.933E-06 | 5.686E-07 | 1.300E-06 | 2.400E-06 |

| | | | | | | | | | | |
|---|---|---|---|---|---|---|---|---|---|---|
| 11135 | 557 | P557P | A | 1.8e-06 | 2.7e-06 | 2.7e-06 | 2.400E-06 | 5.196E-07 | 1.800E-06 | 2.700E-06 |
| 11136 | 557 | P557S | A | 2.3e-06 | 1.2e-06 | 1.9e-06 | 1.800E-06 | 5.568E-07 | 1.200E-06 | 2.300E-06 |
| 11137 | 557 | P557T | A | 1.2e-06 | 2.5e-06 | 1.9e-06 | 1.867E-06 | 6.506E-07 | 1.200E-06 | 2.500E-06 |
| 11138 | 557 | P557W | A | 3.1e-06 | 1.7e-06 | 1.6e-06 | 2.133E-06 | 8.386E-07 | 1.600E-06 | 3.100E-06 |
| 11139 | 557 | P557Y | A | 2.2e-06 | 2.4e-06 | 2.3e-06 | 2.300E-06 | 1.000E-07 | 2.200E-06 | 2.400E-06 |
| 11140 | 557 | P557V | A | 2.1e-06 | 2.3e-06 | 1.6e-06 | 2.000E-06 | 3.606E-07 | 1.600E-06 | 2.300E-06 |
| 11141 | 558 | E558A | A | 4.7e-06 | 3.7e-06 | 2.9e-06 | 3.767E-06 | 9.018E-07 | 2.900E-06 | 4.700E-06 |
| 11142 | 558 | E558R | A | 2.3e-06 | 2.3e-06 | 3.8e-06 | 2.800E-06 | 8.660E-07 | 2.300E-06 | 3.800E-06 |
| 11143 | 558 | E558N | A | 2.4e-06 | 1.4e-06 | 3.6e-06 | 2.467E-06 | 1.102E-06 | 1.400E-06 | 3.600E-06 |
| 11144 | 558 | E558D | A | 2.3e-06 | 3.9e-06 | 2.3e-06 | 2.833E-06 | 9.238E-07 | 2.300E-06 | 3.900E-06 |
| 11145 | 558 | E558C | A | 1.7e-06 | 2.8e-06 | 4.9e-06 | 3.133E-06 | 1.626E-06 | 1.700E-06 | 4.900E-06 |
| 11146 | 558 | E558E | A | 1.2e-06 | 1.7e-06 | 2.5e-06 | 1.800E-06 | 6.557E-07 | 1.200E-06 | 2.500E-06 |
| 11147 | 558 | E558Q | A | 1.8e-06 | 1.8e-06 | 3e-06 | 2.200E-06 | 6.928E-07 | 1.800E-06 | 3.000E-06 |
| 11148 | 558 | E558G | A | 6.9e-06 | 4.9e-06 | 5.7e-06 | 5.833E-06 | 1.007E-06 | 4.900E-06 | 6.900E-06 |
| 11149 | 558 | E558H | A | 1.2e-06 | 9.2e-07 | 1.6e-06 | 1.240E-06 | 3.418E-07 | 9.200E-07 | 1.600E-06 |
| 11150 | 558 | E558I | A | 3e-06 | 2.2e-06 | 6e-06 | 3.733E-06 | 2.003E-06 | 2.200E-06 | 6.000E-06 |
| 11151 | 558 | E558L | A | 1.7e-06 | 3.4e-06 | 2.3e-06 | 2.467E-06 | 8.622E-07 | 1.700E-06 | 3.400E-06 |
| 11152 | 558 | E558K | A | 3e-06 | 1.2e-06 | 2.2e-06 | 2.133E-06 | 9.018E-07 | 1.200E-06 | 3.000E-06 |
| 11153 | 558 | E558M | A | 1.9e-06 | 2.5e-06 | 2.9e-06 | 2.433E-06 | 5.033E-07 | 1.900E-06 | 2.900E-06 |
| 11154 | 558 | E558F | A | 1.8e-06 | 1.6e-06 | 1.9e-06 | 1.767E-06 | 1.528E-07 | 1.600E-06 | 1.900E-06 |
| 11155 | 558 | E558P | A | 3.3e-06 | 2.6e-06 | 4.3e-06 | 3.400E-06 | 8.544E-07 | 2.600E-06 | 4.300E-06 |
| 11156 | 558 | E558S | A | 4.2e-06 | 3.6e-06 | 6.2e-06 | 4.667E-06 | 1.361E-06 | 3.600E-06 | 6.200E-06 |
| 11157 | 558 | E558T | A | 1.4e-06 | 2.1e-06 | 2.3e-06 | 1.933E-06 | 4.726E-07 | 1.400E-06 | 2.300E-06 |
| 11158 | 558 | E558W | A | 2.4e-06 | 1.6e-06 | 2.2e-06 | 2.067E-06 | 4.163E-07 | 1.600E-06 | 2.400E-06 |
| 11159 | 558 | E558Y | A | 2.1e-06 | 2.7e-06 | 1.6e-06 | 2.133E-06 | 5.508E-07 | 1.600E-06 | 2.700E-06 |
| 11160 | 558 | E558V | A | 3.2e-06 | 4.8e-06 | 2.9e-06 | 3.633E-06 | 1.021E-06 | 2.900E-06 | 4.800E-06 |
| 11161 | 559 | A559A | A | 2.6e-06 | 1.6e-06 | 2.3e-06 | 2.167E-06 | 5.132E-07 | 1.600E-06 | 2.600E-06 |
| 11162 | 559 | A559R | A | 1.6e-06 | 2.2e-06 | 2.8e-06 | 2.200E-06 | 6.000E-07 | 1.600E-06 | 2.800E-06 |
| 11163 | 559 | A559N | A | 4.3e-06 | 2.2e-06 | 1.5e-06 | 2.667E-06 | 1.457E-06 | 1.500E-06 | 4.300E-06 |
| 11164 | 559 | A559D | A | 3.1e-06 | 1.4e-06 | 1.9e-06 | 2.133E-06 | 8.737E-07 | 1.400E-06 | 3.100E-06 |
| 11165 | 559 | A559C | A | 2.3e-06 | 2.6e-06 | 2.6e-06 | 2.500E-06 | 1.732E-07 | 2.300E-06 | 2.600E-06 |
| 11166 | 559 | A559E | A | 1.8e-06 | 3.7e-06 | 1.4e-06 | 2.300E-06 | 1.229E-06 | 1.400E-06 | 3.700E-06 |
| 11167 | 559 | A559Q | A | 1.3e-06 | 2e-06 | 2.3e-06 | 1.867E-06 | 5.132E-07 | 1.300E-06 | 2.300E-06 |
| 11168 | 559 | A559G | A | 2.2e-06 | 2.5e-06 | 1.6e-06 | 2.100E-06 | 4.583E-07 | 1.600E-06 | 2.500E-06 |
| 11169 | 559 | A559H | A | 2.9e-06 | 2e-06 | 6.8e-06 | 3.900E-06 | 2.551E-06 | 2.000E-06 | 6.800E-06 |
| 11170 | 559 | A559I | A | 3.7e-06 | 1.5e-06 | 2.5e-06 | 2.567E-06 | 1.102E-06 | 1.500E-06 | 3.700E-06 |
| 11171 | 559 | A559L | A | 2e-06 | 2.2e-06 | 4.4e-06 | 2.867E-06 | 1.332E-06 | 2.000E-06 | 4.400E-06 |
| 11172 | 559 | A559K | A | 1.7e-06 | 3.6e-06 | 2e-06 | 2.433E-06 | 1.021E-06 | 1.700E-06 | 3.600E-06 |
| 11173 | 559 | A559M | A | 2.7e-06 | 3.4e-06 | 3.1e-06 | 3.067E-06 | 3.512E-07 | 2.700E-06 | 3.400E-06 |
| 11174 | 559 | A559F | A | 2.2e-06 | 2.2e-06 | 2.3e-06 | 2.233E-06 | 5.774E-08 | 2.200E-06 | 2.300E-06 |
| 11175 | 559 | A559P | A | 2.3e-06 | 3.1e-06 | 2.3e-06 | 2.567E-06 | 4.619E-07 | 2.300E-06 | 3.100E-06 |
| 11176 | 559 | A559S | A | 3.3e-06 | 2e-06 | 2.8e-06 | 2.700E-06 | 6.557E-07 | 2.000E-06 | 3.300E-06 |
| 11177 | 559 | A559T | A | 2.1e-06 | 1.8e-06 | 2.7e-06 | 2.200E-06 | 4.583E-07 | 1.800E-06 | 2.700E-06 |
| 11178 | 559 | A559W | A | 2.4e-06 | 2.7e-06 | 1.5e-06 | 2.200E-06 | 6.245E-07 | 1.500E-06 | 2.700E-06 |
| 11179 | 559 | A559Y | A | 2.1e-06 | 2.2e-06 | 2.3e-06 | 2.200E-06 | 1.000E-07 | 2.100E-06 | 2.300E-06 |
| 11180 | 559 | A559V | A | 3e-06 | 2e-06 | 2.5e-06 | 2.500E-06 | 5.000E-07 | 2.000E-06 | 3.000E-06 |
| 11181 | 560 | D560A | A | 1.6e-06 | 2.3e-06 | 4e-06 | 2.633E-06 | 1.234E-06 | 1.600E-06 | 4.000E-06 |
| 11182 | 560 | D560R | A | 2.2e-06 | 1.8e-06 | 1.9e-06 | 1.967E-06 | 2.082E-07 | 1.800E-06 | 2.200E-06 |
| 11183 | 560 | D560N | A | 2.5e-06 | 1.2e-06 | 4.4e-06 | 2.700E-06 | 1.609E-06 | 1.200E-06 | 4.400E-06 |
| 11184 | 560 | D560D | A | 2.4e-06 | 2.2e-06 | 1.1e-06 | 1.900E-06 | 7.000E-07 | 1.100E-06 | 2.400E-06 |
| 11185 | 560 | D560C | A | 1.4e-06 | 2.8e-06 | 3.5e-06 | 2.567E-06 | 1.069E-06 | 1.400E-06 | 3.500E-06 |
| 11186 | 560 | D560E | A | 2.6e-06 | 2.2e-06 | 2.4e-06 | 2.400E-06 | 2.000E-07 | 2.200E-06 | 2.600E-06 |
| 11187 | 560 | D560Q | A | 3e-06 | 3.3e-06 | 3.4e-06 | 3.233E-06 | 2.082E-07 | 3.000E-06 | 3.400E-06 |
| 11188 | 560 | D560G | A | 2.4e-06 | 1.8e-06 | 8.5e-06 | 4.233E-06 | 3.707E-06 | 1.800E-06 | 8.500E-06 |
| 11189 | 560 | D560H | A | 3.9e-06 | 2.3e-06 | 3.2e-06 | 3.133E-06 | 8.021E-07 | 2.300E-06 | 3.900E-06 |
| 11190 | 560 | D560I | A | 4.1e-06 | 3.1e-06 | 2.8e-06 | 3.333E-06 | 6.807E-07 | 2.800E-06 | 4.100E-06 |
| 11191 | 560 | D560L | A | 2.3e-06 | 1.7e-06 | 1.4e-06 | 1.800E-06 | 4.583E-07 | 1.400E-06 | 2.300E-06 |
| 11192 | 560 | D560K | A | 2.3e-06 | 3.6e-06 | 3.3e-06 | 3.067E-06 | 6.807E-07 | 2.300E-06 | 3.600E-06 |
| 11193 | 560 | D560M | A | 2.8e-06 | 2e-06 | 2.2e-06 | 2.333E-06 | 4.163E-07 | 2.000E-06 | 2.800E-06 |
| 11194 | 560 | D560F | A | 2.2e-06 | 2e-06 | 2.3e-06 | 2.167E-06 | 1.528E-07 | 2.000E-06 | 2.300E-06 |
| 11195 | 560 | D560P | A | 4.1e-06 | 3.1e-06 | 3.9e-06 | 3.700E-06 | 5.292E-07 | 3.100E-06 | 4.100E-06 |
| 11196 | 560 | D560S | A | 2.6e-06 | 2.6e-06 | 5e-06 | 3.400E-06 | 1.386E-06 | 2.600E-06 | 5.000E-06 |
| 11197 | 560 | D560T | A | 2.9e-06 | 4.5e-06 | 3.5e-06 | 3.633E-06 | 8.083E-07 | 2.900E-06 | 4.500E-06 |
| 11198 | 560 | D560W | A | 2.4e-06 | 2.5e-06 | 2.9e-06 | 2.600E-06 | 2.646E-07 | 2.400E-06 | 2.900E-06 |
| 11199 | 560 | D560Y | A | 1.3e-06 | 2.7e-06 | 1.1e-06 | 1.700E-06 | 8.718E-07 | 1.100E-06 | 2.700E-06 |
| 11200 | 560 | D560V | A | 3.5e-06 | 2.1e-06 | 3e-06 | 2.867E-06 | 7.095E-07 | 2.100E-06 | 3.500E-06 |
| 11201 | 561 | Q561A | A | 3e-06 | 1.4e-06 | 2.4e-06 | 2.267E-06 | 8.083E-07 | 1.400E-06 | 3.000E-06 |
| 11202 | 561 | Q561R | A | 2.1e-06 | 1.7e-06 | 1.7e-06 | 1.833E-06 | 2.309E-07 | 1.700E-06 | 2.100E-06 |
| 11203 | 561 | Q561N | A | 1.7e-06 | 2e-06 | 3.4e-06 | 2.367E-06 | 9.074E-07 | 1.700E-06 | 3.400E-06 |
| 11204 | 561 | Q561D | A | 1.7e-06 | 1.8e-06 | 2e-06 | 1.833E-06 | 1.528E-07 | 1.700E-06 | 2.000E-06 |
| 11205 | 561 | Q561C | A | 2e-06 | 2.3e-06 | 3e-06 | 2.433E-06 | 5.132E-07 | 2.000E-06 | 3.000E-06 |
| 11206 | 561 | Q561E | A | 1.6e-06 | 1.6e-06 | 3.7e-06 | 2.300E-06 | 1.212E-06 | 1.600E-06 | 3.700E-06 |
| 11207 | 561 | Q561Q | A | 1.8e-06 | 3.1e-06 | 2.2e-06 | 2.367E-06 | 6.658E-07 | 1.800E-06 | 3.100E-06 |
| 11208 | 561 | Q561G | A | 1.9e-06 | 3.5e-06 | 3e-06 | 2.800E-06 | 8.185E-07 | 1.900E-06 | 3.500E-06 |
| 11209 | 561 | Q561H | A | 1.2e-06 | 3.3e-06 | 2e-06 | 2.167E-06 | 1.060E-06 | 1.200E-06 | 3.300E-06 |
| 11210 | 561 | Q561I | A | 2.5e-06 | 1.6e-06 | 1.3e-06 | 1.800E-06 | 6.245E-07 | 1.300E-06 | 2.500E-06 |
| 11211 | 561 | Q561L | A | 1.9e-06 | 1.1e-06 | 1.1e-06 | 1.367E-06 | 4.619E-07 | 1.100E-06 | 1.900E-06 |
| 11212 | 561 | Q561K | A | 3.1e-06 | 2.3e-06 | 4.3e-06 | 3.233E-06 | 1.007E-06 | 2.300E-06 | 4.300E-06 |
| 11213 | 561 | Q561M | A | 1.5e-06 | 1.2e-06 | 1.3e-06 | 1.333E-06 | 1.528E-07 | 1.200E-06 | 1.500E-06 |
| 11214 | 561 | Q561F | A | 1.8e-06 | 2.6e-06 | 2e-06 | 2.133E-06 | 4.163E-07 | 1.800E-06 | 2.600E-06 |
| 11215 | 561 | Q561P | A | 1.8e-06 | 2.6e-06 | 1.7e-06 | 2.033E-06 | 4.933E-07 | 1.700E-06 | 2.600E-06 |
| 11216 | 561 | Q561S | A | 1.8e-06 | 2.5e-06 | 2e-06 | 2.100E-06 | 3.606E-07 | 1.800E-06 | 2.500E-06 |
| 11217 | 561 | Q561T | A | 3.3e-06 | 4.6e-06 | 2.3e-06 | 3.400E-06 | 1.153E-06 | 2.300E-06 | 4.600E-06 |
| 11218 | 561 | Q561W | A | 8.5e-07 | 1.7e-06 | 1.3e-06 | 1.283E-06 | 4.252E-07 | 8.500E-07 | 1.700E-06 |
| 11219 | 561 | Q561Y | A | 1.5e-06 | 1.6e-06 | 2.3e-06 | 1.800E-06 | 4.359E-07 | 1.500E-06 | 2.300E-06 |

| | | | | | | | | | | |
|---|---|---|---|---|---|---|---|---|---|---|
| 11220 | 561 | Q561V | A | 2.6e-06 | 2e-06 | 2.8e-06 | 2.467E-06 | 4.163E-07 | 2.000E-06 | 2.800E-06 |
| 11221 | 562 | C562A | A | 3.5e-06 | 2.1e-06 | 1.6e-06 | 2.400E-06 | 9.849E-07 | 1.600E-06 | 3.500E-06 |
| 11222 | 562 | C562R | A | 3.3e-06 | 2.1e-06 | 2.1e-06 | 2.500E-06 | 6.928E-07 | 2.100E-06 | 3.300E-06 |
| 11223 | 562 | C562N | A | 2.8e-06 | 3.2e-06 | 2.9e-06 | 2.967E-06 | 2.082E-07 | 2.800E-06 | 3.200E-06 |
| 11224 | 562 | C562D | A | 2.4e-06 | 2.8e-06 | 1.7e-06 | 2.300E-06 | 5.568E-07 | 1.700E-06 | 2.800E-06 |
| 11225 | 562 | C562C | A | 3.5e-06 | 1.5e-06 | 2.7e-06 | 2.567E-06 | 1.007E-06 | 1.500E-06 | 3.500E-06 |
| 11226 | 562 | C562E | A | 1.6e-06 | 2.6e-06 | 2.7e-06 | 2.300E-06 | 6.083E-07 | 1.600E-06 | 2.700E-06 |
| 11227 | 562 | C562Q | A | 7.9e-07 | 2e-06 | 3.3e-06 | 2.030E-06 | 1.255E-06 | 7.900E-07 | 3.300E-06 |
| 11228 | 562 | C562G | A | 4.1e-06 | 2e-06 | 3e-06 | 3.033E-06 | 1.050E-06 | 2.000E-06 | 4.100E-06 |
| 11229 | 562 | C562H | A | 1.8e-06 | 2.9e-06 | 1.7e-06 | 2.133E-06 | 6.658E-07 | 1.700E-06 | 2.900E-06 |
| 11230 | 562 | C562I | A | 1.8e-06 | 2e-06 | 3.6e-06 | 2.467E-06 | 9.866E-07 | 1.800E-06 | 3.600E-06 |
| 11231 | 562 | C562L | A | 3.1e-06 | 2.8e-06 | 2.6e-06 | 2.833E-06 | 2.517E-07 | 2.600E-06 | 3.100E-06 |
| 11232 | 562 | C562K | A | 7.2e-07 | 2.3e-06 | 1.7e-06 | 1.573E-06 | 7.976E-07 | 7.200E-07 | 2.300E-06 |
| 11233 | 562 | C562M | A | 4e-06 | 2.5e-06 | 2.3e-06 | 2.933E-06 | 9.292E-07 | 2.300E-06 | 4.000E-06 |
| 11234 | 562 | C562F | A | 1.9e-06 | 2.9e-06 | 5.1e-06 | 3.300E-06 | 1.637E-06 | 1.900E-06 | 5.100E-06 |
| 11235 | 562 | C562P | A | 2e-06 | 2.3e-06 | 2e-06 | 2.100E-06 | 1.732E-07 | 2.000E-06 | 2.300E-06 |
| 11236 | 562 | C562S | A | 2.4e-06 | 1.7e-06 | 4.4e-06 | 2.833E-06 | 1.401E-06 | 1.700E-06 | 4.400E-06 |
| 11237 | 562 | C562T | A | 1.3e-06 | 2.5e-06 | 2.9e-06 | 2.233E-06 | 8.327E-07 | 1.300E-06 | 2.900E-06 |
| 11238 | 562 | C562W | A | 2.1e-06 | 2.1e-06 | 2.7e-06 | 2.300E-06 | 3.464E-07 | 2.100E-06 | 2.700E-06 |
| 11239 | 562 | C562Y | A | 1.3e-06 | 3.2e-06 | 1.2e-06 | 1.900E-06 | 1.127E-06 | 1.200E-06 | 3.200E-06 |
| 11240 | 562 | C562V | A | 2.9e-06 | 2.4e-06 | 1.9e-06 | 2.400E-06 | 5.000E-07 | 1.900E-06 | 2.900E-06 |
| 11241 | 563 | V563A | A | 1.5e-06 | 2.7e-06 | 2.5e-06 | 2.233E-06 | 6.429E-07 | 1.500E-06 | 2.700E-06 |
| 11242 | 563 | V563R | A | 2.6e-06 | 1.3e-06 | 8.9e-07 | 1.597E-06 | 8.928E-07 | 8.900E-07 | 2.600E-06 |
| 11243 | 563 | V563N | A | 3.3e-06 | 2.6e-06 | 2.4e-06 | 2.767E-06 | 4.726E-07 | 2.400E-06 | 3.300E-06 |
| 11244 | 563 | V563D | A | 2.4e-06 | 1.9e-06 | 3.5e-06 | 2.600E-06 | 8.185E-07 | 1.900E-06 | 3.500E-06 |
| 11245 | 563 | V563C | A | 3.7e-06 | 1.6e-06 | 3.2e-06 | 2.833E-06 | 1.097E-06 | 1.600E-06 | 3.700E-06 |
| 11246 | 563 | V563E | A | 2.2e-06 | 1.8e-06 | 2.8e-06 | 2.267E-06 | 5.033E-07 | 1.800E-06 | 2.800E-06 |
| 11247 | 563 | V563Q | A | 2.4e-06 | 1e-06 | 3.3e-06 | 2.233E-06 | 1.159E-06 | 1.000E-06 | 3.300E-06 |
| 11248 | 563 | V563G | A | 2.1e-06 | 4.4e-06 | 2.9e-06 | 3.133E-06 | 1.168E-06 | 2.100E-06 | 4.400E-06 |
| 11249 | 563 | V563H | A | 1.3e-06 | 1.9e-06 | 2.1e-06 | 1.767E-06 | 4.163E-07 | 1.300E-06 | 2.100E-06 |
| 11250 | 563 | V563I | A | 3e-06 | 3.7e-06 | 1.3e-06 | 2.667E-06 | 1.234E-06 | 1.300E-06 | 3.700E-06 |
| 11251 | 563 | V563L | A | 1.9e-06 | 1.7e-06 | 3.8e-06 | 2.467E-06 | 1.159E-06 | 1.700E-06 | 3.800E-06 |
| 11252 | 563 | V563K | A | 3.3e-06 | 2.1e-06 | 1.7e-06 | 2.367E-06 | 8.327E-07 | 1.700E-06 | 3.300E-06 |
| 11253 | 563 | V563M | A | 1.5e-06 | 2.5e-06 | 2.3e-06 | 2.100E-06 | 5.292E-07 | 1.500E-06 | 2.500E-06 |
| 11254 | 563 | V563F | A | 4.8e-06 | 2.9e-06 | 2.5e-06 | 3.400E-06 | 1.229E-06 | 2.500E-06 | 4.800E-06 |
| 11255 | 563 | V563P | A | 4e-06 | 2.6e-06 | 3.4e-06 | 3.333E-06 | 7.024E-07 | 2.600E-06 | 4.000E-06 |
| 11256 | 563 | V563S | A | 2e-06 | 6e-06 | 2.4e-06 | 3.467E-06 | 2.203E-06 | 2.000E-06 | 6.000E-06 |
| 11257 | 563 | V563T | A | 2.3e-06 | 2.1e-06 | 3.2e-06 | 2.533E-06 | 5.859E-07 | 2.100E-06 | 3.200E-06 |
| 11258 | 563 | V563W | A | 2.6e-06 | 2.3e-06 | 3.3e-06 | 2.733E-06 | 5.132E-07 | 2.300E-06 | 3.300E-06 |
| 11259 | 563 | V563Y | A | 1.9e-06 | 2.1e-06 | 1.6e-06 | 1.867E-06 | 2.517E-07 | 1.600E-06 | 2.100E-06 |
| 11260 | 563 | V563V | A | 1.9e-06 | 2.8e-06 | 1.8e-06 | 2.167E-06 | 5.508E-07 | 1.800E-06 | 2.800E-06 |
| 11261 | 564 | A564A | A | 3e-06 | 2e-06 | 1.8e-06 | 2.267E-06 | 6.429E-07 | 1.800E-06 | 3.000E-06 |
| 11262 | 564 | A564R | A | 1.4e-06 | 1.2e-06 | 2.4e-06 | 1.667E-06 | 6.429E-07 | 1.200E-06 | 2.400E-06 |
| 11263 | 564 | A564N | A | 1.5e-06 | 2.4e-06 | 2.8e-06 | 2.233E-06 | 6.658E-07 | 1.500E-06 | 2.800E-06 |
| 11264 | 564 | A564D | A | 2.7e-06 | 1.4e-06 | 1.9e-06 | 2.000E-06 | 6.557E-07 | 1.400E-06 | 2.700E-06 |
| 11265 | 564 | A564C | A | 1.7e-06 | 3.7e-06 | 2.2e-06 | 2.533E-06 | 1.041E-06 | 1.700E-06 | 3.700E-06 |
| 11266 | 564 | A564E | A | 2.7e-06 | 2.3e-06 | 2e-06 | 2.333E-06 | 3.512E-07 | 2.000E-06 | 2.700E-06 |
| 11267 | 564 | A564Q | A | 2.6e-06 | 2.2e-06 | 3.8e-06 | 2.867E-06 | 8.327E-07 | 2.200E-06 | 3.800E-06 |
| 11268 | 564 | A564G | A | 2.8e-06 | 1.3e-06 | 2.1e-06 | 2.067E-06 | 7.506E-07 | 1.300E-06 | 2.800E-06 |
| 11269 | 564 | A564H | A | 1.2e-06 | 1.6e-06 | 1.9e-06 | 1.567E-06 | 3.512E-07 | 1.200E-06 | 1.900E-06 |
| 11270 | 564 | A564I | A | 3.7e-06 | 2.6e-06 | 1.4e-06 | 2.567E-06 | 1.150E-06 | 1.400E-06 | 3.700E-06 |
| 11271 | 564 | A564L | A | 2.7e-06 | 3e-06 | 2.8e-06 | 2.833E-06 | 1.528E-07 | 2.700E-06 | 3.000E-06 |
| 11272 | 564 | A564K | A | 1.7e-06 | 1.8e-06 | 3.1e-06 | 2.200E-06 | 7.810E-07 | 1.700E-06 | 3.100E-06 |
| 11273 | 564 | A564M | A | 3.1e-06 | 2.2e-06 | 2.6e-06 | 2.633E-06 | 4.509E-07 | 2.200E-06 | 3.100E-06 |
| 11274 | 564 | A564F | A | 1.4e-06 | 2.8e-06 | 1.6e-06 | 1.933E-06 | 7.572E-07 | 1.400E-06 | 2.800E-06 |
| 11275 | 564 | A564P | A | 2.2e-06 | 1.7e-06 | 2.4e-06 | 2.100E-06 | 3.606E-07 | 1.700E-06 | 2.400E-06 |
| 11276 | 564 | A564S | A | 3.2e-06 | 1.3e-06 | 2.8e-06 | 2.433E-06 | 1.002E-06 | 1.300E-06 | 3.200E-06 |
| 11277 | 564 | A564T | A | 2.8e-06 | 2.6e-06 | 1.7e-06 | 2.367E-06 | 5.859E-07 | 1.700E-06 | 2.800E-06 |
| 11278 | 564 | A564W | A | 1.8e-06 | 2.2e-06 | 2.5e-06 | 2.167E-06 | 3.512E-07 | 1.800E-06 | 2.500E-06 |
| 11279 | 564 | A564Y | A | 1.6e-06 | 2.5e-06 | 2.6e-06 | 2.233E-06 | 5.508E-07 | 1.600E-06 | 2.600E-06 |
| 11280 | 564 | A564V | A | 1.6e-06 | 2.6e-06 | 3.1e-06 | 2.433E-06 | 7.638E-07 | 1.600E-06 | 3.100E-06 |
| 11281 | 565 | C565A | A | 6.3e-06 | 1.8e-06 | 3.7e-06 | 3.933E-06 | 2.259E-06 | 1.800E-06 | 6.300E-06 |
| 11282 | 565 | C565R | A | 3.4e-06 | 3.6e-06 | 1.8e-06 | 2.933E-06 | 9.866E-07 | 1.800E-06 | 3.600E-06 |
| 11283 | 565 | C565N | A | 5.7e-06 | 2.5e-06 | 1.7e-06 | 3.300E-06 | 2.117E-06 | 1.700E-06 | 5.700E-06 |
| 11284 | 565 | C565D | A | 3.4e-06 | 2.9e-06 | 2.1e-06 | 2.800E-06 | 6.557E-07 | 2.100E-06 | 3.400E-06 |
| 11285 | 565 | C565C | A | 2.2e-06 | 3.6e-06 | 1.8e-06 | 2.533E-06 | 9.452E-07 | 1.800E-06 | 3.600E-06 |
| 11286 | 565 | C565E | A | 2.6e-06 | 2.9e-06 | 2.9e-06 | 2.800E-06 | 1.732E-07 | 2.600E-06 | 2.900E-06 |
| 11287 | 565 | C565Q | A | 3.2e-06 | 2.5e-06 | 3.7e-06 | 3.133E-06 | 6.028E-07 | 2.500E-06 | 3.700E-06 |
| 11288 | 565 | C565G | A | 1.5e-06 | 1.4e-06 | 2.5e-06 | 1.800E-06 | 6.083E-07 | 1.400E-06 | 2.500E-06 |
| 11289 | 565 | C565H | A | 2.4e-06 | 2.2e-06 | 3.6e-06 | 2.733E-06 | 7.572E-07 | 2.200E-06 | 3.600E-06 |
| 11290 | 565 | C565I | A | 4e-06 | 1.9e-06 | 3.7e-06 | 3.200E-06 | 1.136E-06 | 1.900E-06 | 4.000E-06 |
| 11291 | 565 | C565L | A | 3.9e-06 | 1.3e-06 | 1.5e-06 | 2.233E-06 | 1.447E-06 | 1.300E-06 | 3.900E-06 |
| 11292 | 565 | C565K | A | 2.1e-06 | 2.6e-06 | 1.1e-06 | 1.933E-06 | 7.638E-07 | 1.100E-06 | 2.600E-06 |
| 11293 | 565 | C565M | A | 2.4e-06 | 2.6e-06 | 2.1e-06 | 2.367E-06 | 2.517E-07 | 2.100E-06 | 2.600E-06 |
| 11294 | 565 | C565F | A | 1.7e-06 | 2.8e-06 | 1.8e-06 | 2.100E-06 | 6.083E-07 | 1.700E-06 | 2.800E-06 |
| 11295 | 565 | C565P | A | 2.7e-06 | 3.4e-06 | 3.2e-06 | 3.100E-06 | 3.606E-07 | 2.700E-06 | 3.400E-06 |
| 11296 | 565 | C565S | A | 1.7e-06 | 1.1e-06 | 4.4e-06 | 2.400E-06 | 1.758E-06 | 1.100E-06 | 4.400E-06 |
| 11297 | 565 | C565T | A | 4.3e-06 | 3.2e-06 | 1.8e-06 | 3.100E-06 | 1.253E-06 | 1.800E-06 | 4.300E-06 |
| 11298 | 565 | C565W | A | 2e-06 | 2.3e-06 | 2.1e-06 | 2.133E-06 | 1.528E-07 | 2.000E-06 | 2.300E-06 |
| 11299 | 565 | C565Y | A | 2.1e-06 | 2.2e-06 | 3.3e-06 | 2.533E-06 | 6.658E-07 | 2.100E-06 | 3.300E-06 |
| 11300 | 565 | C565V | A | 3.1e-06 | 1.7e-06 | 1.6e-06 | 2.133E-06 | 8.386E-07 | 1.600E-06 | 3.100E-06 |
| 11301 | 566 | A566A | A | 3.9e-06 | 2.3e-06 | 2.4e-06 | 2.867E-06 | 8.963E-07 | 2.300E-06 | 3.900E-06 |
| 11302 | 566 | A566R | A | 1.9e-06 | 3e-06 | 8.5e-07 | 1.917E-06 | 1.075E-06 | 8.500E-07 | 3.000E-06 |
| 11303 | 566 | A566N | A | 1.9e-06 | 1.8e-06 | 3e-06 | 2.233E-06 | 6.658E-07 | 1.800E-06 | 3.000E-06 |
| 11304 | 566 | A566D | A | 2.2e-06 | 3.2e-06 | 2.1e-06 | 2.500E-06 | 6.083E-07 | 2.100E-06 | 3.200E-06 |

| | | | | | | | | | | |
|---|---|---|---|---|---|---|---|---|---|---|
| 11305 | 566 | A566C | A | 2e-06 | 2e-06 | 1.8e-06 | 1.933E-06 | 1.155E-07 | 1.800E-06 | 2.000E-06 |
| 11306 | 566 | A566E | A | 2.1e-06 | 2.4e-06 | 1.6e-06 | 2.033E-06 | 4.041E-07 | 1.600E-06 | 2.400E-06 |
| 11307 | 566 | A566Q | A | 4.2e-06 | 1.9e-06 | 2.7e-06 | 2.933E-06 | 1.168E-06 | 1.900E-06 | 4.200E-06 |
| 11308 | 566 | A566G | A | 2.2e-06 | 4.5e-06 | 2.4e-06 | 3.033E-06 | 1.274E-06 | 2.200E-06 | 4.500E-06 |
| 11309 | 566 | A566H | A | 2.5e-06 | 1.8e-06 | 3.6e-06 | 2.633E-06 | 9.074E-07 | 1.800E-06 | 3.600E-06 |
| 11310 | 566 | A566I | A | 2.8e-06 | 2.8e-06 | 2.3e-06 | 2.633E-06 | 2.887E-07 | 2.300E-06 | 2.800E-06 |
| 11311 | 566 | A566L | A | 1.6e-06 | 1.9e-06 | 1.6e-06 | 1.700E-06 | 1.732E-07 | 1.600E-06 | 1.900E-06 |
| 11312 | 566 | A566K | A | 2.3e-06 | 4e-06 | 2.6e-06 | 2.967E-06 | 9.074E-07 | 2.300E-06 | 4.000E-06 |
| 11313 | 566 | A566M | A | 2.4e-06 | 1.7e-06 | 2.2e-06 | 2.100E-06 | 3.606E-07 | 1.700E-06 | 2.400E-06 |
| 11314 | 566 | A566F | A | 2.5e-06 | 2.4e-06 | 2.3e-06 | 2.400E-06 | 1.000E-07 | 2.300E-06 | 2.500E-06 |
| 11315 | 566 | A566P | A | 2.4e-06 | 1.6e-06 | 1.5e-06 | 1.833E-06 | 4.933E-07 | 1.500E-06 | 2.400E-06 |
| 11316 | 566 | A566S | A | 2e-06 | 2.2e-06 | 2.4e-06 | 2.200E-06 | 2.000E-07 | 2.000E-06 | 2.400E-06 |
| 11317 | 566 | A566T | A | 2.8e-06 | 2e-06 | 3.2e-06 | 2.667E-06 | 6.110E-07 | 2.000E-06 | 3.200E-06 |
| 11318 | 566 | A566W | A | 1.9e-06 | 2.4e-06 | 2e-06 | 2.100E-06 | 2.646E-07 | 1.900E-06 | 2.400E-06 |
| 11319 | 566 | A566Y | A | 1.2e-06 | 3.2e-06 | 2.6e-06 | 2.333E-06 | 1.026E-06 | 1.200E-06 | 3.200E-06 |
| 11320 | 566 | A566V | A | 1.9e-06 | 2.2e-06 | 1.1e-06 | 1.733E-06 | 5.686E-07 | 1.100E-06 | 2.200E-06 |
| 11321 | 567 | H567A | A | 1.8e-06 | 2.2e-06 | 2.7e-06 | 2.233E-06 | 4.509E-07 | 1.800E-06 | 2.700E-06 |
| 11322 | 567 | H567R | A | 2.8e-06 | 1.5e-06 | 2e-06 | 2.100E-06 | 6.557E-07 | 1.500E-06 | 2.800E-06 |
| 11323 | 567 | H567N | A | 1.6e-06 | 1.7e-06 | 1.8e-06 | 1.700E-06 | 1.000E-07 | 1.600E-06 | 1.800E-06 |
| 11324 | 567 | H567D | A | 2.3e-06 | 2.3e-06 | 3.8e-06 | 2.800E-06 | 8.660E-07 | 2.300E-06 | 3.800E-06 |
| 11325 | 567 | H567C | A | 2.4e-06 | 2.5e-06 | 3.4e-06 | 2.767E-06 | 5.508E-07 | 2.400E-06 | 3.400E-06 |
| 11326 | 567 | H567E | A | 3e-06 | 4.2e-06 | 2.7e-06 | 3.300E-06 | 7.937E-07 | 2.700E-06 | 4.200E-06 |
| 11327 | 567 | H567Q | A | 1.9e-06 | 1.8e-06 | 1.7e-06 | 1.800E-06 | 1.000E-07 | 1.700E-06 | 1.900E-06 |
| 11328 | 567 | H567G | A | 2.5e-06 | 2.8e-06 | 2.5e-06 | 2.600E-06 | 1.732E-07 | 2.500E-06 | 2.800E-06 |
| 11329 | 567 | H567H | A | 4.6e-06 | 3.7e-06 | 2.2e-06 | 3.500E-06 | 1.212E-06 | 2.200E-06 | 4.600E-06 |
| 11330 | 567 | H567I | A | 1.7e-06 | 1.9e-06 | 1.6e-06 | 1.733E-06 | 1.528E-07 | 1.600E-06 | 1.900E-06 |
| 11331 | 567 | H567L | A | 2.8e-06 | 1.2e-06 | 4e-06 | 2.667E-06 | 1.405E-06 | 1.200E-06 | 4.000E-06 |
| 11332 | 567 | H567K | A | 2.7e-06 | 2.5e-06 | 2.7e-06 | 2.633E-06 | 1.155E-07 | 2.500E-06 | 2.700E-06 |
| 11333 | 567 | H567M | A | 3e-06 | 5.2e-06 | 2.4e-06 | 3.533E-06 | 1.474E-06 | 2.400E-06 | 5.200E-06 |
| 11334 | 567 | H567F | A | 2.1e-06 | 3.7e-06 | 2.4e-06 | 2.733E-06 | 8.505E-07 | 2.100E-06 | 3.700E-06 |
| 11335 | 567 | H567P | A | 2.1e-06 | 3.3e-06 | 3.3e-06 | 2.900E-06 | 6.928E-07 | 2.100E-06 | 3.300E-06 |
| 11336 | 567 | H567S | A | 1.5e-06 | 2.3e-06 | 3.6e-06 | 2.467E-06 | 1.060E-06 | 1.500E-06 | 3.600E-06 |
| 11337 | 567 | H567T | A | 2.8e-06 | 1.8e-06 | 1.6e-06 | 2.067E-06 | 6.429E-07 | 1.600E-06 | 2.800E-06 |
| 11338 | 567 | H567W | A | 2.9e-06 | 3e-06 | 3.3e-06 | 3.067E-06 | 2.082E-07 | 2.900E-06 | 3.300E-06 |
| 11339 | 567 | H567Y | A | 2.2e-06 | 2.7e-06 | 2.9e-06 | 2.600E-06 | 3.606E-07 | 2.200E-06 | 2.900E-06 |
| 11340 | 567 | H567V | A | 4.3e-06 | 3.1e-06 | 2.5e-06 | 3.300E-06 | 9.165E-07 | 2.500E-06 | 4.300E-06 |
| 11341 | 568 | Y568A | A | 1.8e-06 | 3.7e-06 | 2.5e-06 | 2.667E-06 | 9.609E-07 | 1.800E-06 | 3.700E-06 |
| 11342 | 568 | Y568R | A | 4e-06 | 2.3e-06 | 3e-06 | 3.100E-06 | 8.544E-07 | 2.300E-06 | 4.000E-06 |
| 11343 | 568 | Y568N | A | 2.1e-06 | 2.3e-06 | 1.9e-06 | 2.100E-06 | 2.000E-07 | 1.900E-06 | 2.300E-06 |
| 11344 | 568 | Y568D | A | 2.5e-06 | 2.4e-06 | 2.7e-06 | 2.533E-06 | 1.528E-07 | 2.400E-06 | 2.700E-06 |
| 11345 | 568 | Y568C | A | 2.9e-06 | 2.9e-06 | 3e-06 | 2.933E-06 | 5.774E-08 | 2.900E-06 | 3.000E-06 |
| 11346 | 568 | Y568E | A | 2.6e-06 | 4.4e-06 | 2.5e-06 | 3.167E-06 | 1.069E-06 | 2.500E-06 | 4.400E-06 |
| 11347 | 568 | Y568Q | A | 1.1e-06 | 2.2e-06 | 2.9e-06 | 2.067E-06 | 9.074E-07 | 1.100E-06 | 2.900E-06 |
| 11348 | 568 | Y568G | A | 2.1e-06 | 1.7e-06 | 1.3e-06 | 1.700E-06 | 4.000E-07 | 1.300E-06 | 2.100E-06 |
| 11349 | 568 | Y568H | A | 1.5e-06 | 2e-06 | 1.5e-06 | 1.667E-06 | 2.887E-07 | 1.500E-06 | 2.000E-06 |
| 11350 | 568 | Y568I | A | 2.3e-06 | 3.1e-06 | 2.4e-06 | 2.600E-06 | 4.359E-07 | 2.300E-06 | 3.100E-06 |
| 11351 | 568 | Y568L | A | 3.5e-06 | 1.3e-06 | 2.5e-06 | 2.433E-06 | 1.102E-06 | 1.300E-06 | 3.500E-06 |
| 11352 | 568 | Y568K | A | 1.3e-06 | 1.9e-06 | 2.8e-06 | 2.000E-06 | 7.550E-07 | 1.300E-06 | 2.800E-06 |
| 11353 | 568 | Y568M | A | 2.7e-06 | 1.6e-06 | 1.9e-06 | 2.067E-06 | 5.686E-07 | 1.600E-06 | 2.700E-06 |
| 11354 | 568 | Y568F | A | 2e-06 | 3e-06 | 3.4e-06 | 2.800E-06 | 7.211E-07 | 2.000E-06 | 3.400E-06 |
| 11355 | 568 | Y568P | A | 1.7e-06 | 2.9e-06 | 2.2e-06 | 2.267E-06 | 6.028E-07 | 1.700E-06 | 2.900E-06 |
| 11356 | 568 | Y568S | A | 1.1e-06 | 2.4e-06 | 3.5e-06 | 2.333E-06 | 1.201E-06 | 1.100E-06 | 3.500E-06 |
| 11357 | 568 | Y568T | A | 2.4e-06 | 2e-06 | 2.5e-06 | 2.300E-06 | 2.646E-07 | 2.000E-06 | 2.500E-06 |
| 11358 | 568 | Y568W | A | 3.5e-06 | 2.5e-06 | 1.7e-06 | 2.567E-06 | 9.018E-07 | 1.700E-06 | 3.500E-06 |
| 11359 | 568 | Y568Y | A | 2.9e-06 | 2.8e-06 | 2.8e-06 | 2.833E-06 | 5.774E-08 | 2.800E-06 | 2.900E-06 |
| 11360 | 568 | Y568V | A | 2.9e-06 | 2.7e-06 | 2.5e-06 | 2.700E-06 | 2.000E-07 | 2.500E-06 | 2.900E-06 |
| 11361 | 569 | K569A | A | 1.9e-06 | 2.8e-06 | 1e-06 | 1.900E-06 | 9.000E-07 | 1.000E-06 | 2.800E-06 |
| 11362 | 569 | K569R | A | 4.2e-06 | 2.3e-06 | 2.5e-06 | 3.000E-06 | 1.044E-06 | 2.300E-06 | 4.200E-06 |
| 11363 | 569 | K569N | A | 3.1e-06 | 1.8e-06 | 4.2e-06 | 3.033E-06 | 1.201E-06 | 1.800E-06 | 4.200E-06 |
| 11364 | 569 | K569D | A | 2.4e-06 | 4e-06 | 3e-06 | 3.133E-06 | 8.083E-07 | 2.400E-06 | 4.000E-06 |
| 11365 | 569 | K569C | A | 3e-06 | 1.4e-06 | 1.8e-06 | 2.067E-06 | 8.327E-07 | 1.400E-06 | 3.000E-06 |
| 11366 | 569 | K569E | A | 2.1e-06 | 2.2e-06 | 3.6e-06 | 2.633E-06 | 8.386E-07 | 2.100E-06 | 3.600E-06 |
| 11367 | 569 | K569Q | A | 2.1e-06 | 2.1e-06 | 9.5e-07 | 1.717E-06 | 6.640E-07 | 9.500E-07 | 2.100E-06 |
| 11368 | 569 | K569G | A | 2.2e-06 | 2e-06 | 2.9e-06 | 2.367E-06 | 4.726E-07 | 2.000E-06 | 2.900E-06 |
| 11369 | 569 | K569H | A | 2.6e-06 | 2.8e-06 | 1.7e-06 | 2.367E-06 | 5.859E-07 | 1.700E-06 | 2.800E-06 |
| 11370 | 569 | K569I | A | 1.4e-06 | 1.6e-06 | 4e-06 | 2.333E-06 | 1.447E-06 | 1.400E-06 | 4.000E-06 |
| 11371 | 569 | K569L | A | 1.5e-06 | 2.6e-06 | 3.3e-06 | 2.467E-06 | 9.074E-07 | 1.500E-06 | 3.300E-06 |
| 11372 | 569 | K569K | A | 1.9e-06 | 3.1e-06 | 2.7e-06 | 2.567E-06 | 6.110E-07 | 1.900E-06 | 3.100E-06 |
| 11373 | 569 | K569M | A | 2.4e-06 | 2.9e-06 | 2.1e-06 | 2.467E-06 | 4.041E-07 | 2.100E-06 | 2.900E-06 |
| 11374 | 569 | K569F | A | 2e-06 | 2.2e-06 | 3.7e-06 | 2.633E-06 | 9.292E-07 | 2.000E-06 | 3.700E-06 |
| 11375 | 569 | K569P | A | 2.1e-06 | 2.3e-06 | 4.2e-06 | 2.867E-06 | 1.159E-06 | 2.100E-06 | 4.200E-06 |
| 11376 | 569 | K569S | A | 2.1e-06 | 2.8e-06 | 4e-06 | 2.967E-06 | 9.609E-07 | 2.100E-06 | 4.000E-06 |
| 11377 | 569 | K569T | A | 2.1e-06 | 2.2e-06 | 1.4e-06 | 1.900E-06 | 4.359E-07 | 1.400E-06 | 2.200E-06 |
| 11378 | 569 | K569W | A | 1.7e-06 | 4.4e-06 | 2.2e-06 | 2.767E-06 | 1.436E-06 | 1.700E-06 | 4.400E-06 |
| 11379 | 569 | K569Y | A | 1.7e-06 | 2.5e-06 | 1.9e-06 | 2.033E-06 | 4.163E-07 | 1.700E-06 | 2.500E-06 |
| 11380 | 569 | K569V | A | 2.6e-06 | 2.1e-06 | 9.5e-07 | 1.883E-06 | 8.461E-07 | 9.500E-07 | 2.600E-06 |
| 11381 | 570 | D570A | A | 2.5e-06 | 2.7e-06 | 3.4e-06 | 2.867E-06 | 4.726E-07 | 2.500E-06 | 3.400E-06 |
| 11382 | 570 | D570R | A | 5.1e-06 | 5.1e-06 | 1.3e-06 | 3.833E-06 | 2.194E-06 | 1.300E-06 | 5.100E-06 |
| 11383 | 570 | D570N | A | 2.9e-06 | 1.7e-06 | 2.1e-06 | 2.233E-06 | 6.110E-07 | 1.700E-06 | 2.900E-06 |
| 11384 | 570 | D570D | A | 1.9e-06 | 3.1e-06 | 9.5e-07 | 1.983E-06 | 1.077E-06 | 9.500E-07 | 3.100E-06 |
| 11385 | 570 | D570C | A | 2e-06 | 1.1e-06 | 3.8e-06 | 2.300E-06 | 1.375E-06 | 1.100E-06 | 3.800E-06 |
| 11386 | 570 | D570E | A | 1.8e-06 | 1.2e-06 | 1.1e-06 | 1.367E-06 | 3.786E-07 | 1.100E-06 | 1.800E-06 |
| 11387 | 570 | D570Q | A | 1.6e-06 | 2.5e-06 | 2.4e-06 | 2.167E-06 | 4.933E-07 | 1.600E-06 | 2.500E-06 |
| 11388 | 570 | D570G | A | 3.1e-06 | 1.2e-06 | 6.3e-06 | 3.533E-06 | 2.577E-06 | 1.200E-06 | 6.300E-06 |
| 11389 | 570 | D570H | A | 2.6e-06 | 1.1e-06 | 1.6e-06 | 1.767E-06 | 7.638E-07 | 1.100E-06 | 2.600E-06 |

| | | | | | | | | | | |
|---|---|---|---|---|---|---|---|---|---|---|
| 11390 | 570 | D570I | A | 2.9e-06 | 5.5e-06 | 3.3e-06 | 3.900E-06 | 1.400E-06 | 2.900E-06 | 5.500E-06 |
| 11391 | 570 | D570L | A | 4.8e-06 | 2.5e-06 | 3e-06 | 3.433E-06 | 1.210E-06 | 2.500E-06 | 4.800E-06 |
| 11392 | 570 | D570K | A | 2.2e-06 | 3.5e-06 | 2.9e-06 | 2.867E-06 | 6.506E-07 | 2.200E-06 | 3.500E-06 |
| 11393 | 570 | D570M | A | 3.1e-06 | 4.9e-06 | 5.6e-06 | 4.533E-06 | 1.290E-06 | 3.100E-06 | 5.600E-06 |
| 11394 | 570 | D570F | A | 3.3e-06 | 3e-06 | 2.3e-06 | 2.867E-06 | 5.132E-07 | 2.300E-06 | 3.300E-06 |
| 11395 | 570 | D570P | A | 7.4e-06 | 5.9e-06 | 3e-06 | 5.433E-06 | 2.237E-06 | 3.000E-06 | 7.400E-06 |
| 11396 | 570 | D570S | A | 2.4e-06 | 1.7e-06 | 2.3e-06 | 2.133E-06 | 3.786E-07 | 1.700E-06 | 2.400E-06 |
| 11397 | 570 | D570T | A | 3.3e-06 | 1.5e-06 | 1.4e-06 | 2.067E-06 | 1.069E-06 | 1.400E-06 | 3.300E-06 |
| 11398 | 570 | D570W | A | 3.4e-06 | 2.5e-06 | 2.7e-06 | 2.867E-06 | 4.726E-07 | 2.500E-06 | 3.400E-06 |
| 11399 | 570 | D570Y | A | 4.4e-06 | 2.2e-06 | 6.2e-06 | 4.267E-06 | 2.003E-06 | 2.200E-06 | 6.200E-06 |
| 11400 | 570 | D570V | A | 2.7e-06 | 3.4e-06 | 2.6e-06 | 2.900E-06 | 4.359E-07 | 2.600E-06 | 3.400E-06 |
| 11401 | 571 | P571A | A | 3e-06 | 2.5e-06 | 3.6e-06 | 3.033E-06 | 5.508E-07 | 2.500E-06 | 3.600E-06 |
| 11402 | 571 | P571R | A | 1.7e-06 | 1.9e-06 | 1.9e-06 | 1.833E-06 | 1.155E-07 | 1.700E-06 | 1.900E-06 |
| 11403 | 571 | P571N | A | 1.2e-06 | 2e-06 | 1.8e-06 | 1.667E-06 | 4.163E-07 | 1.200E-06 | 2.000E-06 |
| 11404 | 571 | P571D | A | 1.8e-06 | 3.5e-06 | 1.7e-06 | 2.333E-06 | 1.012E-06 | 1.700E-06 | 3.500E-06 |
| 11405 | 571 | P571C | A | 2.5e-06 | 1.9e-06 | 2.1e-06 | 2.167E-06 | 3.055E-07 | 1.900E-06 | 2.500E-06 |
| 11406 | 571 | P571E | A | 1.5e-06 | 2.1e-06 | 2e-06 | 1.867E-06 | 3.215E-07 | 1.500E-06 | 2.100E-06 |
| 11407 | 571 | P571Q | A | 1.4e-06 | 1.5e-06 | 1.2e-06 | 1.367E-06 | 1.528E-07 | 1.200E-06 | 1.500E-06 |
| 11408 | 571 | P571G | A | 3e-06 | 2e-06 | 1.4e-06 | 2.133E-06 | 8.083E-07 | 1.400E-06 | 3.000E-06 |
| 11409 | 571 | P571H | A | 2.3e-06 | 2.2e-06 | 1.1e-06 | 1.867E-06 | 6.658E-07 | 1.100E-06 | 2.300E-06 |
| 11410 | 571 | P571I | A | 3.8e-06 | 2.2e-06 | 1.7e-06 | 2.567E-06 | 1.097E-06 | 1.700E-06 | 3.800E-06 |
| 11411 | 571 | P571L | A | 2.1e-06 | 3.5e-06 | 1.5e-06 | 2.367E-06 | 1.026E-06 | 1.500E-06 | 3.500E-06 |
| 11412 | 571 | P571K | A | 3.6e-06 | 2.2e-06 | 2e-06 | 2.600E-06 | 8.718E-07 | 2.000E-06 | 3.600E-06 |
| 11413 | 571 | P571M | A | 4e-06 | 1.9e-06 | 2.4e-06 | 2.767E-06 | 1.097E-06 | 1.900E-06 | 4.000E-06 |
| 11414 | 571 | P571F | A | 2.2e-06 | 2.9e-06 | 2.1e-06 | 2.400E-06 | 4.359E-07 | 2.100E-06 | 2.900E-06 |
| 11415 | 571 | P571P | A | 2.3e-06 | 3.5e-06 | 1.2e-06 | 2.333E-06 | 1.150E-06 | 1.200E-06 | 3.500E-06 |
| 11416 | 571 | P571S | A | 1.5e-06 | 1.6e-06 | 1.7e-06 | 1.600E-06 | 1.000E-07 | 1.500E-06 | 1.700E-06 |
| 11417 | 571 | P571T | A | 1.6e-06 | 1.7e-06 | 1.9e-06 | 1.733E-06 | 1.528E-07 | 1.600E-06 | 1.900E-06 |
| 11418 | 571 | P571W | A | 1.3e-06 | 3.2e-06 | 2.6e-06 | 2.367E-06 | 9.713E-07 | 1.300E-06 | 3.200E-06 |
| 11419 | 571 | P571Y | A | 2.7e-06 | 3.6e-06 | 3.6e-06 | 3.300E-06 | 5.196E-07 | 2.700E-06 | 3.600E-06 |
| 11420 | 571 | P571V | A | 1.6e-06 | 1.8e-06 | 3.1e-06 | 2.167E-06 | 8.145E-07 | 1.600E-06 | 3.100E-06 |
| 11421 | 572 | P572A | A | 3.5e-06 | 1.9e-06 | 3.5e-06 | 2.967E-06 | 9.238E-07 | 1.900E-06 | 3.500E-06 |
| 11422 | 572 | P572R | A | 1.4e-06 | 1.5e-06 | 6.2e-07 | 1.173E-06 | 4.818E-07 | 6.200E-07 | 1.500E-06 |
| 11423 | 572 | P572N | A | 1.7e-06 | 1.7e-06 | 1.5e-06 | 1.633E-06 | 1.155E-07 | 1.500E-06 | 1.700E-06 |
| 11424 | 572 | P572D | A | 1.3e-06 | 1.9e-06 | 1.8e-06 | 1.667E-06 | 3.215E-07 | 1.300E-06 | 1.900E-06 |
| 11425 | 572 | P572C | A | 2.2e-06 | 2.9e-06 | 3.2e-06 | 2.767E-06 | 5.132E-07 | 2.200E-06 | 3.200E-06 |
| 11426 | 572 | P572E | A | 1.9e-06 | 7.7e-07 | 1.5e-06 | 1.390E-06 | 5.730E-07 | 7.700E-07 | 1.900E-06 |
| 11427 | 572 | P572Q | A | 1.4e-06 | 1.2e-06 | 8.2e-07 | 1.140E-06 | 2.946E-07 | 8.200E-07 | 1.400E-06 |
| 11428 | 572 | P572G | A | 1.9e-06 | 2.9e-06 | 2.1e-06 | 2.300E-06 | 5.292E-07 | 1.900E-06 | 2.900E-06 |
| 11429 | 572 | P572H | A | 1.9e-06 | 1.6e-06 | 1.5e-06 | 1.667E-06 | 2.082E-07 | 1.500E-06 | 1.900E-06 |
| 11430 | 572 | P572I | A | 2.3e-06 | 1.4e-06 | 2.5e-06 | 2.067E-06 | 5.859E-07 | 1.400E-06 | 2.500E-06 |
| 11431 | 572 | P572L | A | 2.1e-06 | 5.1e-06 | 3.2e-06 | 3.467E-06 | 1.518E-06 | 2.100E-06 | 5.100E-06 |
| 11432 | 572 | P572K | A | 1.7e-06 | 1.8e-06 | 1.2e-06 | 1.567E-06 | 3.215E-07 | 1.200E-06 | 1.800E-06 |
| 11433 | 572 | P572M | A | 2.7e-06 | 3.1e-06 | 3e-06 | 2.933E-06 | 2.082E-07 | 2.700E-06 | 3.100E-06 |
| 11434 | 572 | P572F | A | 1.4e-06 | 2.4e-06 | 2.9e-06 | 2.233E-06 | 7.638E-07 | 1.400E-06 | 2.900E-06 |
| 11435 | 572 | P572P | A | 5.1e-06 | 2.2e-06 | 1.9e-06 | 3.067E-06 | 1.767E-06 | 1.900E-06 | 5.100E-06 |
| 11436 | 572 | P572S | A | 2e-06 | 1.4e-06 | 2.3e-06 | 1.900E-06 | 4.583E-07 | 1.400E-06 | 2.300E-06 |
| 11437 | 572 | P572T | A | 1.3e-06 | 2.9e-06 | 2.6e-06 | 2.267E-06 | 8.505E-07 | 1.300E-06 | 2.900E-06 |
| 11438 | 572 | P572W | A | 1.2e-06 | 1.8e-06 | 1.6e-06 | 1.533E-06 | 3.055E-07 | 1.200E-06 | 1.800E-06 |
| 11439 | 572 | P572Y | A | 1.8e-06 | 2.1e-06 | 2.6e-06 | 2.167E-06 | 4.041E-07 | 1.800E-06 | 2.600E-06 |
| 11440 | 572 | P572V | A | 2e-06 | 2.2e-06 | 1.4e-06 | 1.867E-06 | 4.163E-07 | 1.400E-06 | 2.200E-06 |
| 11441 | 573 | F573A | A | 2.1e-06 | 3e-06 | 2.3e-06 | 2.467E-06 | 4.726E-07 | 2.100E-06 | 3.000E-06 |
| 11442 | 573 | F573R | A | 1.6e-06 | 2e-06 | 1.9e-06 | 1.833E-06 | 2.082E-07 | 1.600E-06 | 2.000E-06 |
| 11443 | 573 | F573N | A | 2.2e-06 | 1.5e-06 | 1.2e-06 | 1.633E-06 | 5.132E-07 | 1.200E-06 | 2.200E-06 |
| 11444 | 573 | F573D | A | 1.7e-06 | 1.9e-06 | 1.9e-06 | 1.833E-06 | 1.155E-07 | 1.700E-06 | 1.900E-06 |
| 11445 | 573 | F573C | A | 3.2e-06 | 2.8e-06 | 2.6e-06 | 2.867E-06 | 3.055E-07 | 2.600E-06 | 3.200E-06 |
| 11446 | 573 | F573E | A | 1.9e-06 | 1.6e-06 | 2.3e-06 | 1.933E-06 | 3.512E-07 | 1.600E-06 | 2.300E-06 |
| 11447 | 573 | F573Q | A | 1.8e-06 | 1.8e-06 | 8.9e-07 | 1.497E-06 | 5.254E-07 | 8.900E-07 | 1.800E-06 |
| 11448 | 573 | F573G | A | 2.6e-06 | 3.5e-06 | 3.3e-06 | 3.133E-06 | 4.726E-07 | 2.600E-06 | 3.500E-06 |
| 11449 | 573 | F573H | A | 1.4e-06 | 1e-06 | 1.6e-06 | 1.333E-06 | 3.055E-07 | 1.000E-06 | 1.600E-06 |
| 11450 | 573 | F573I | A | 4e-06 | 3.7e-06 | 2.2e-06 | 3.300E-06 | 9.644E-07 | 2.200E-06 | 4.000E-06 |
| 11451 | 573 | F573L | A | 1.8e-06 | 2.7e-06 | 2.6e-06 | 2.367E-06 | 4.933E-07 | 1.800E-06 | 2.700E-06 |
| 11452 | 573 | F573K | A | 1.8e-06 | 1.5e-06 | 1.1e-06 | 1.467E-06 | 3.512E-07 | 1.100E-06 | 1.800E-06 |
| 11453 | 573 | F573M | A | 1.6e-06 | 2.8e-06 | 3.2e-06 | 2.533E-06 | 8.327E-07 | 1.600E-06 | 3.200E-06 |
| 11454 | 573 | F573F | A | 3.3e-06 | 2.5e-06 | 2e-06 | 2.600E-06 | 6.557E-07 | 2.000E-06 | 3.300E-06 |
| 11455 | 573 | F573P | A | 3.9e-06 | 2.6e-06 | 2.8e-06 | 3.100E-06 | 7.000E-07 | 2.600E-06 | 3.900E-06 |
| 11456 | 573 | F573S | A | 1.4e-06 | 3e-06 | 3.3e-06 | 2.567E-06 | 1.021E-06 | 1.400E-06 | 3.300E-06 |
| 11457 | 573 | F573T | A | 2.9e-06 | 2.4e-06 | 1.5e-06 | 2.267E-06 | 7.095E-07 | 1.500E-06 | 2.900E-06 |
| 11458 | 573 | F573W | A | 3.5e-06 | 1.7e-06 | 1.5e-06 | 2.233E-06 | 1.102E-06 | 1.500E-06 | 3.500E-06 |
| 11459 | 573 | F573Y | A | 1.4e-06 | 5.4e-06 | 2.3e-06 | 3.033E-06 | 2.098E-06 | 1.400E-06 | 5.400E-06 |
| 11460 | 573 | F573V | A | 2.4e-06 | 4.7e-06 | 2.3e-06 | 3.133E-06 | 1.358E-06 | 2.300E-06 | 4.700E-06 |
| 11461 | 574 | C574A | A | 3.3e-06 | 4.7e-06 | 6.1e-06 | 4.700E-06 | 1.400E-06 | 3.300E-06 | 6.100E-06 |
| 11462 | 574 | C574R | A | 3.2e-06 | 2.6e-06 | 4.2e-06 | 3.333E-06 | 8.083E-07 | 2.600E-06 | 4.200E-06 |
| 11463 | 574 | C574N | A | 1.8e-06 | 2e-06 | 1.6e-06 | 1.800E-06 | 2.000E-07 | 1.600E-06 | 2.000E-06 |
| 11464 | 574 | C574D | A | 2e-06 | 2.8e-06 | 1.4e-06 | 2.067E-06 | 7.024E-07 | 1.400E-06 | 2.800E-06 |
| 11465 | 574 | C574C | A | 2.4e-06 | 3.3e-06 | 2.5e-06 | 2.733E-06 | 4.933E-07 | 2.400E-06 | 3.300E-06 |
| 11466 | 574 | C574E | A | 2.9e-06 | 2.3e-06 | 1.9e-06 | 2.367E-06 | 5.033E-07 | 1.900E-06 | 2.900E-06 |
| 11467 | 574 | C574Q | A | 3.1e-06 | 3.8e-06 | 8.2e-07 | 2.573E-06 | 1.558E-06 | 8.200E-07 | 3.800E-06 |
| 11468 | 574 | C574G | A | 2.2e-06 | 1.9e-06 | 4.2e-06 | 2.767E-06 | 1.250E-06 | 1.900E-06 | 4.200E-06 |
| 11469 | 574 | C574H | A | 2.4e-06 | 1.9e-06 | 1.8e-06 | 2.033E-06 | 3.215E-07 | 1.800E-06 | 2.400E-06 |
| 11470 | 574 | C574I | A | 1.9e-06 | 1.9e-06 | 1.8e-06 | 1.867E-06 | 5.774E-08 | 1.800E-06 | 1.900E-06 |
| 11471 | 574 | C574L | A | 2.6e-06 | 2.2e-06 | 9.9e-07 | 1.930E-06 | 8.383E-07 | 9.900E-07 | 2.600E-06 |
| 11472 | 574 | C574K | A | 1.9e-06 | 2.9e-06 | 3.4e-06 | 2.733E-06 | 7.638E-07 | 1.900E-06 | 3.400E-06 |
| 11473 | 574 | C574M | A | 1.9e-06 | 2.2e-06 | 2.8e-06 | 2.300E-06 | 4.583E-07 | 1.900E-06 | 2.800E-06 |
| 11474 | 574 | C574F | A | 4e-06 | 2.5e-06 | 1.9e-06 | 2.800E-06 | 1.082E-06 | 1.900E-06 | 4.000E-06 |

| | | | | | | | | | | |
|---|---|---|---|---|---|---|---|---|---|---|
| 11475 | 574 | C574P | A | 4.6e-06 | 1.2e-06 | 2.9e-06 | 2.900E-06 | 1.700E-06 | 1.200E-06 | 4.600E-06 |
| 11476 | 574 | C574S | A | 3.1e-06 | 2.4e-06 | 1.9e-06 | 2.467E-06 | 6.028E-07 | 1.900E-06 | 3.100E-06 |
| 11477 | 574 | C574T | A | 2.2e-06 | 1.7e-06 | 2.4e-06 | 2.100E-06 | 3.606E-07 | 1.700E-06 | 2.400E-06 |
| 11478 | 574 | C574W | A | 2e-06 | 4.1e-06 | 1.6e-06 | 2.567E-06 | 1.343E-06 | 1.600E-06 | 4.100E-06 |
| 11479 | 574 | C574Y | A | 3.2e-06 | 2.9e-06 | 2e-06 | 2.700E-06 | 6.245E-07 | 2.000E-06 | 3.200E-06 |
| 11480 | 574 | C574V | A | 1.9e-06 | 2.6e-06 | 1.7e-06 | 2.067E-06 | 4.726E-07 | 1.700E-06 | 2.600E-06 |
| 11481 | 575 | V575A | A | 3.8e-06 | 1.7e-06 | 2.4e-06 | 2.633E-06 | 1.069E-06 | 1.700E-06 | 3.800E-06 |
| 11482 | 575 | V575R | A | 2.1e-06 | 1.7e-06 | 1.2e-06 | 1.667E-06 | 4.509E-07 | 1.200E-06 | 2.100E-06 |
| 11483 | 575 | V575N | A | 2.4e-06 | 1.6e-06 | 1.8e-06 | 1.933E-06 | 4.163E-07 | 1.600E-06 | 2.400E-06 |
| 11484 | 575 | V575D | A | 3.5e-06 | 2.8e-06 | 2.5e-06 | 2.933E-06 | 5.132E-07 | 2.500E-06 | 3.500E-06 |
| 11485 | 575 | V575C | A | 2e-06 | 1.7e-06 | 1.8e-06 | 1.833E-06 | 1.528E-07 | 1.700E-06 | 2.000E-06 |
| 11486 | 575 | V575E | A | 1.8e-06 | 1.7e-06 | 1.6e-06 | 1.700E-06 | 1.000E-07 | 1.600E-06 | 1.800E-06 |
| 11487 | 575 | V575Q | A | 1.9e-06 | 3.1e-06 | 2.4e-06 | 2.467E-06 | 6.028E-07 | 1.900E-06 | 3.100E-06 |
| 11488 | 575 | V575G | A | 2.7e-06 | 2.2e-06 | 1.5e-06 | 2.133E-06 | 6.028E-07 | 1.500E-06 | 2.700E-06 |
| 11489 | 575 | V575H | A | 2.2e-06 | 3e-06 | 2e-06 | 2.400E-06 | 5.292E-07 | 2.000E-06 | 3.000E-06 |
| 11490 | 575 | V575I | A | 1.7e-06 | 2.3e-06 | 7.9e-07 | 1.597E-06 | 7.603E-07 | 7.900E-07 | 2.300E-06 |
| 11491 | 575 | V575L | A | 2.1e-06 | 4.1e-06 | 3.3e-06 | 3.167E-06 | 1.007E-06 | 2.100E-06 | 4.100E-06 |
| 11492 | 575 | V575K | A | 1.7e-06 | 2.1e-06 | 2.3e-06 | 2.033E-06 | 3.055E-07 | 1.700E-06 | 2.300E-06 |
| 11493 | 575 | V575M | A | 3.3e-06 | 2.1e-06 | 2.6e-06 | 2.667E-06 | 6.028E-07 | 2.100E-06 | 3.300E-06 |
| 11494 | 575 | V575F | A | 3.1e-06 | 3.1e-06 | 2.6e-06 | 2.933E-06 | 2.887E-07 | 2.600E-06 | 3.100E-06 |
| 11495 | 575 | V575P | A | 2.3e-06 | 3e-06 | 1.7e-06 | 2.333E-06 | 6.506E-07 | 1.700E-06 | 3.000E-06 |
| 11496 | 575 | V575S | A | 1.4e-06 | 3.2e-06 | 2.7e-06 | 2.433E-06 | 9.292E-07 | 1.400E-06 | 3.200E-06 |
| 11497 | 575 | V575T | A | 2.2e-06 | 2.2e-06 | 1.6e-06 | 2.000E-06 | 3.464E-07 | 1.600E-06 | 2.200E-06 |
| 11498 | 575 | V575W | A | 9.2e-07 | 2.1e-06 | 1.7e-06 | 1.573E-06 | 6.001E-07 | 9.200E-07 | 2.100E-06 |
| 11499 | 575 | V575Y | A | 3.3e-06 | 2.5e-06 | 1e-06 | 2.267E-06 | 1.168E-06 | 1.000E-06 | 3.300E-06 |
| 11500 | 575 | V575V | A | 1.8e-06 | 3.9e-06 | 3.4e-06 | 3.033E-06 | 1.097E-06 | 1.800E-06 | 3.900E-06 |
| 11501 | 576 | A576A | A | 1.8e-06 | 2.2e-06 | 1.4e-06 | 1.800E-06 | 4.000E-07 | 1.400E-06 | 2.200E-06 |
| 11502 | 576 | A576R | A | 2.6e-06 | 4.1e-06 | 1.9e-06 | 2.867E-06 | 1.124E-06 | 1.900E-06 | 4.100E-06 |
| 11503 | 576 | A576N | A | 2.1e-06 | 1.8e-06 | 1.8e-06 | 1.900E-06 | 1.732E-07 | 1.800E-06 | 2.100E-06 |
| 11504 | 576 | A576D | A | 2.1e-06 | 2.1e-06 | 3.4e-06 | 2.533E-06 | 7.506E-07 | 2.100E-06 | 3.400E-06 |
| 11505 | 576 | A576C | A | 3.1e-06 | 2.4e-06 | 1.7e-06 | 2.400E-06 | 7.000E-07 | 1.700E-06 | 3.100E-06 |
| 11506 | 576 | A576E | A | 1.8e-06 | 2.2e-06 | 2.3e-06 | 2.100E-06 | 2.646E-07 | 1.800E-06 | 2.300E-06 |
| 11507 | 576 | A576Q | A | 3.1e-06 | 2.3e-06 | 3.6e-06 | 3.000E-06 | 6.557E-07 | 2.300E-06 | 3.600E-06 |
| 11508 | 576 | A576G | A | 2.2e-06 | 2.2e-06 | 2.9e-06 | 2.433E-06 | 4.041E-07 | 2.200E-06 | 2.900E-06 |
| 11509 | 576 | A576H | A | 1.2e-06 | 1.9e-06 | 2.2e-06 | 1.767E-06 | 5.132E-07 | 1.200E-06 | 2.200E-06 |
| 11510 | 576 | A576I | A | 4e-06 | 1.8e-06 | 3.9e-06 | 3.233E-06 | 1.242E-06 | 1.800E-06 | 4.000E-06 |
| 11511 | 576 | A576L | A | 2.7e-06 | 2.8e-06 | 3.1e-06 | 2.867E-06 | 2.082E-07 | 2.700E-06 | 3.100E-06 |
| 11512 | 576 | A576K | A | 1.8e-06 | 2.2e-06 | 1.9e-06 | 1.967E-06 | 2.082E-07 | 1.800E-06 | 2.200E-06 |
| 11513 | 576 | A576M | A | 1.6e-06 | 2.3e-06 | 1.8e-06 | 1.900E-06 | 3.606E-07 | 1.600E-06 | 2.300E-06 |
| 11514 | 576 | A576F | A | 2.2e-06 | 2.5e-06 | 3.3e-06 | 2.667E-06 | 5.686E-07 | 2.200E-06 | 3.300E-06 |
| 11515 | 576 | A576P | A | 2.3e-06 | 2.4e-06 | 3.2e-06 | 2.633E-06 | 4.933E-07 | 2.300E-06 | 3.200E-06 |
| 11516 | 576 | A576S | A | 2.2e-06 | 2.1e-06 | 1.1e-06 | 1.800E-06 | 6.083E-07 | 1.100E-06 | 2.200E-06 |
| 11517 | 576 | A576T | A | 2.6e-06 | 2.7e-06 | 3e-06 | 2.767E-06 | 2.082E-07 | 2.600E-06 | 3.000E-06 |
| 11518 | 576 | A576W | A | 2.2e-06 | 2.4e-06 | 2.6e-06 | 2.400E-06 | 2.000E-07 | 2.200E-06 | 2.600E-06 |
| 11519 | 576 | A576Y | A | 2.6e-06 | 3.2e-06 | 2.2e-06 | 2.667E-06 | 5.033E-07 | 2.200E-06 | 3.200E-06 |
| 11520 | 576 | A576V | A | 3e-06 | 1e-06 | 2.5e-06 | 2.167E-06 | 1.041E-06 | 1.000E-06 | 3.000E-06 |
| 11521 | 577 | R577A | A | 5.2e-06 | 2.5e-06 | 2e-06 | 3.233E-06 | 1.721E-06 | 2.000E-06 | 5.200E-06 |
| 11522 | 577 | R577R | A | 3e-06 | 2.2e-06 | 1.8e-06 | 2.333E-06 | 6.110E-07 | 1.800E-06 | 3.000E-06 |
| 11523 | 577 | R577N | A | 1.9e-06 | 1.7e-06 | 2.9e-06 | 2.167E-06 | 6.429E-07 | 1.700E-06 | 2.900E-06 |
| 11524 | 577 | R577D | A | 2.6e-06 | 5.2e-06 | 2.8e-06 | 3.533E-06 | 1.447E-06 | 2.600E-06 | 5.200E-06 |
| 11525 | 577 | R577C | A | 2.4e-06 | 1.7e-06 | 1.5e-06 | 1.867E-06 | 4.726E-07 | 1.500E-06 | 2.400E-06 |
| 11526 | 577 | R577E | A | 2.7e-06 | 2.5e-06 | 1.8e-06 | 2.333E-06 | 4.726E-07 | 1.800E-06 | 2.700E-06 |
| 11527 | 577 | R577Q | A | 2.5e-06 | 2.5e-06 | 2.3e-06 | 2.433E-06 | 1.155E-07 | 2.300E-06 | 2.500E-06 |
| 11528 | 577 | R577G | A | 1.5e-06 | 1.8e-06 | 2.4e-06 | 1.900E-06 | 4.583E-07 | 1.500E-06 | 2.400E-06 |
| 11529 | 577 | R577H | A | 2.3e-06 | 5.3e-06 | 3.2e-06 | 3.600E-06 | 1.539E-06 | 2.300E-06 | 5.300E-06 |
| 11530 | 577 | R577I | A | 1.5e-06 | 5.5e-06 | 1.3e-06 | 2.767E-06 | 2.369E-06 | 1.300E-06 | 5.500E-06 |
| 11531 | 577 | R577L | A | 4.5e-06 | 2e-06 | 2.4e-06 | 2.967E-06 | 1.343E-06 | 2.000E-06 | 4.500E-06 |
| 11532 | 577 | R577K | A | 1.5e-06 | 2.1e-06 | 3e-06 | 2.200E-06 | 7.550E-07 | 1.500E-06 | 3.000E-06 |
| 11533 | 577 | R577M | A | 2.1e-06 | 1.6e-06 | 2.3e-06 | 2.000E-06 | 3.606E-07 | 1.600E-06 | 2.300E-06 |
| 11534 | 577 | R577F | A | 2.1e-06 | 2.6e-06 | 2.9e-06 | 2.533E-06 | 4.041E-07 | 2.100E-06 | 2.900E-06 |
| 11535 | 577 | R577P | A | 2.4e-06 | 2.3e-06 | 3.3e-06 | 2.667E-06 | 5.508E-07 | 2.300E-06 | 3.300E-06 |
| 11536 | 577 | R577S | A | 2.8e-06 | 4.1e-06 | 1.9e-06 | 2.933E-06 | 1.106E-06 | 1.900E-06 | 4.100E-06 |
| 11537 | 577 | R577T | A | 1.6e-06 | 2.3e-06 | 1.5e-06 | 1.800E-06 | 4.359E-07 | 1.500E-06 | 2.300E-06 |
| 11538 | 577 | R577W | A | 2.5e-06 | 1.5e-06 | 2.4e-06 | 2.133E-06 | 5.508E-07 | 1.500E-06 | 2.500E-06 |
| 11539 | 577 | R577Y | A | 2.1e-06 | 1.3e-06 | 2e-06 | 1.800E-06 | 4.359E-07 | 1.300E-06 | 2.100E-06 |
| 11540 | 577 | R577V | A | 2.1e-06 | 2.5e-06 | 3.8e-06 | 2.800E-06 | 8.888E-07 | 2.100E-06 | 3.800E-06 |
| 11541 | 578 | C578A | A | 3.8e-06 | 3.4e-06 | 2.9e-06 | 3.367E-06 | 4.509E-07 | 2.900E-06 | 3.800E-06 |
| 11542 | 578 | C578R | A | 3.3e-06 | 3e-06 | 1.8e-06 | 2.700E-06 | 7.937E-07 | 1.800E-06 | 3.300E-06 |
| 11543 | 578 | C578N | A | 1.8e-06 | 2.3e-06 | 3.6e-06 | 2.567E-06 | 9.292E-07 | 1.800E-06 | 3.600E-06 |
| 11544 | 578 | C578D | A | 1.9e-06 | 2.3e-06 | 2.2e-06 | 2.133E-06 | 2.082E-07 | 1.900E-06 | 2.300E-06 |
| 11545 | 578 | C578C | A | 1.7e-06 | 1.7e-06 | 2.9e-06 | 2.100E-06 | 6.928E-07 | 1.700E-06 | 2.900E-06 |
| 11546 | 578 | C578E | A | 1.9e-06 | 2.2e-06 | 2.4e-06 | 2.167E-06 | 2.517E-07 | 1.900E-06 | 2.400E-06 |
| 11547 | 578 | C578Q | A | 3.7e-06 | 3.2e-06 | 2.5e-06 | 3.133E-06 | 6.028E-07 | 2.500E-06 | 3.700E-06 |
| 11548 | 578 | C578G | A | 1.7e-06 | 1.9e-06 | 2.1e-06 | 1.900E-06 | 2.000E-07 | 1.700E-06 | 2.100E-06 |
| 11549 | 578 | C578H | A | 1.4e-06 | 2.4e-06 | 1.9e-06 | 1.900E-06 | 5.000E-07 | 1.400E-06 | 2.400E-06 |
| 11550 | 578 | C578I | A | 3.7e-06 | 3.9e-06 | 2.4e-06 | 3.333E-06 | 8.145E-07 | 2.400E-06 | 3.900E-06 |
| 11551 | 578 | C578L | A | 2.3e-06 | 1.9e-06 | 1.6e-06 | 1.933E-06 | 3.512E-07 | 1.600E-06 | 2.300E-06 |
| 11552 | 578 | C578K | A | 1.9e-06 | 2.6e-06 | 1.5e-06 | 2.000E-06 | 5.568E-07 | 1.500E-06 | 2.600E-06 |
| 11553 | 578 | C578M | A | 1.6e-06 | 2.2e-06 | 1.8e-06 | 1.867E-06 | 3.055E-07 | 1.600E-06 | 2.200E-06 |
| 11554 | 578 | C578F | A | 3.3e-06 | 1.8e-06 | 3.3e-06 | 2.800E-06 | 8.660E-07 | 1.800E-06 | 3.300E-06 |
| 11555 | 578 | C578P | A | 2.3e-06 | 2.1e-06 | 5.5e-06 | 3.300E-06 | 1.908E-06 | 2.100E-06 | 5.500E-06 |
| 11556 | 578 | C578S | A | 1.5e-06 | 7.2e-07 | 9.8e-07 | 1.067E-06 | 3.972E-07 | 7.200E-07 | 1.500E-06 |
| 11557 | 578 | C578T | A | 2.7e-06 | 2.7e-06 | 2.7e-06 | 2.700E-06 | 0.000E+00 | 2.700E-06 | 2.700E-06 |
| 11558 | 578 | C578W | A | 1.6e-06 | 9e-07 | 1.9e-06 | 1.467E-06 | 5.132E-07 | 9.000E-07 | 1.900E-06 |
| 11559 | 578 | C578Y | A | 1.6e-06 | 3.1e-06 | 2.4e-06 | 2.367E-06 | 7.506E-07 | 1.600E-06 | 3.100E-06 |

| | | | | | | | | | | |
|---|---|---|---|---|---|---|---|---|---|---|
| 11560 | 578 | C578V | A | 2.1e-06 | 2.2e-06 | 2.7e-06 | 2.333E-06 | 3.215E-07 | 2.100E-06 | 2.700E-06 |
| 11561 | 579 | P579A | A | 3.1e-06 | 2.6e-06 | 2.6e-06 | 2.767E-06 | 2.887E-07 | 2.600E-06 | 3.100E-06 |
| 11562 | 579 | P579R | A | 2.2e-06 | 1.4e-06 | 1.8e-06 | 1.800E-06 | 4.000E-07 | 1.400E-06 | 2.200E-06 |
| 11563 | 579 | P579N | A | 1.4e-06 | 2.9e-06 | 5.9e-06 | 3.400E-06 | 2.291E-06 | 1.400E-06 | 5.900E-06 |
| 11564 | 579 | P579D | A | 2.1e-06 | 2.8e-06 | 2.1e-06 | 2.333E-06 | 4.041E-07 | 2.100E-06 | 2.800E-06 |
| 11565 | 579 | P579C | A | 2.3e-06 | 2.4e-06 | 1.9e-06 | 2.200E-06 | 2.646E-07 | 1.900E-06 | 2.400E-06 |
| 11566 | 579 | P579E | A | 5.5e-06 | 2.5e-06 | 1.7e-06 | 3.233E-06 | 2.003E-06 | 1.700E-06 | 5.500E-06 |
| 11567 | 579 | P579Q | A | 1.8e-06 | 1.4e-06 | 3.3e-06 | 2.167E-06 | 1.002E-06 | 1.400E-06 | 3.300E-06 |
| 11568 | 579 | P579G | A | 4.6e-06 | 3.4e-06 | 3.8e-06 | 3.933E-06 | 6.110E-07 | 3.400E-06 | 4.600E-06 |
| 11569 | 579 | P579H | A | 2.5e-06 | 2.2e-06 | 1.8e-06 | 2.167E-06 | 3.512E-07 | 1.800E-06 | 2.500E-06 |
| 11570 | 579 | P579I | A | 3.6e-06 | 1.3e-06 | 1.8e-06 | 2.233E-06 | 1.210E-06 | 1.300E-06 | 3.600E-06 |
| 11571 | 579 | P579L | A | 8e-07 | 1.1e-06 | 1.7e-06 | 1.200E-06 | 4.583E-07 | 8.000E-07 | 1.700E-06 |
| 11572 | 579 | P579K | A | 1.6e-06 | 3e-06 | 1.9e-06 | 2.167E-06 | 7.371E-07 | 1.600E-06 | 3.000E-06 |
| 11573 | 579 | P579M | A | 3.2e-06 | 2.2e-06 | 1.8e-06 | 2.400E-06 | 7.211E-07 | 1.800E-06 | 3.200E-06 |
| 11574 | 579 | P579F | A | 2.9e-06 | 2.2e-06 | 1.8e-06 | 2.300E-06 | 5.568E-07 | 1.800E-06 | 2.900E-06 |
| 11575 | 579 | P579P | A | 3e-06 | 2.5e-06 | 3.2e-06 | 2.900E-06 | 3.606E-07 | 2.500E-06 | 3.200E-06 |
| 11576 | 579 | P579S | A | 4.2e-06 | 1.2e-06 | 2e-06 | 2.467E-06 | 1.553E-06 | 1.200E-06 | 4.200E-06 |
| 11577 | 579 | P579T | A | 2.1e-06 | 2.3e-06 | 1.5e-06 | 1.967E-06 | 4.163E-07 | 1.500E-06 | 2.300E-06 |
| 11578 | 579 | P579W | A | 2.6e-06 | 1.4e-06 | 3.4e-06 | 2.467E-06 | 1.007E-06 | 1.400E-06 | 3.400E-06 |
| 11579 | 579 | P579Y | A | 2e-06 | 1.5e-06 | 1.8e-06 | 1.767E-06 | 2.517E-07 | 1.500E-06 | 2.000E-06 |
| 11580 | 579 | P579V | A | 3.2e-06 | 2.9e-06 | 1.9e-06 | 2.667E-06 | 6.807E-07 | 1.900E-06 | 3.200E-06 |
| 11581 | 580 | S580A | A | 2.2e-06 | 1.8e-06 | 2.1e-06 | 2.033E-06 | 2.082E-07 | 1.800E-06 | 2.200E-06 |
| 11582 | 580 | S580R | A | 1.5e-06 | 1e-06 | 2.4e-06 | 1.633E-06 | 7.095E-07 | 1.000E-06 | 2.400E-06 |
| 11583 | 580 | S580N | A | 3e-06 | 1.4e-06 | 2.7e-06 | 2.367E-06 | 8.505E-07 | 1.400E-06 | 3.000E-06 |
| 11584 | 580 | S580D | A | 1.6e-06 | 1.7e-06 | 2.3e-06 | 1.867E-06 | 3.786E-07 | 1.600E-06 | 2.300E-06 |
| 11585 | 580 | S580C | A | 2.1e-06 | 2.2e-06 | 2.3e-06 | 2.200E-06 | 1.000E-07 | 2.100E-06 | 2.300E-06 |
| 11586 | 580 | S580E | A | 3.1e-06 | 2.1e-06 | 1.5e-06 | 2.233E-06 | 8.083E-07 | 1.500E-06 | 3.100E-06 |
| 11587 | 580 | S580Q | A | 2.3e-06 | 2.2e-06 | 3.2e-06 | 2.567E-06 | 5.508E-07 | 2.200E-06 | 3.200E-06 |
| 11588 | 580 | S580G | A | 3e-06 | 3e-06 | 2.4e-06 | 2.800E-06 | 3.464E-07 | 2.400E-06 | 3.000E-06 |
| 11589 | 580 | S580H | A | 2.6e-06 | 2.2e-06 | 2.2e-06 | 2.333E-06 | 2.309E-07 | 2.200E-06 | 2.600E-06 |
| 11590 | 580 | S580I | A | 2.2e-06 | 2.3e-06 | 1.2e-06 | 1.900E-06 | 6.083E-07 | 1.200E-06 | 2.300E-06 |
| 11591 | 580 | S580L | A | 3.9e-06 | 3.3e-06 | 1.2e-06 | 2.800E-06 | 1.418E-06 | 1.200E-06 | 3.900E-06 |
| 11592 | 580 | S580K | A | 2.2e-06 | 2.1e-06 | 2.8e-06 | 2.367E-06 | 3.786E-07 | 2.100E-06 | 2.800E-06 |
| 11593 | 580 | S580M | A | 2.6e-06 | 3.5e-06 | 2.2e-06 | 2.767E-06 | 6.658E-07 | 2.200E-06 | 3.500E-06 |
| 11594 | 580 | S580F | A | 1.7e-06 | 1.8e-06 | 2.6e-06 | 2.033E-06 | 4.933E-07 | 1.700E-06 | 2.600E-06 |
| 11595 | 580 | S580P | A | 3.4e-06 | 3.5e-06 | 2.2e-06 | 3.033E-06 | 7.234E-07 | 2.200E-06 | 3.500E-06 |
| 11596 | 580 | S580S | A | 2.6e-06 | 2.7e-06 | 3.8e-06 | 3.033E-06 | 6.658E-07 | 2.600E-06 | 3.800E-06 |
| 11597 | 580 | S580T | A | 2e-06 | 1.9e-06 | 1.7e-06 | 1.867E-06 | 1.528E-07 | 1.700E-06 | 2.000E-06 |
| 11598 | 580 | S580W | A | 9.5e-07 | 1.7e-06 | 1.1e-06 | 1.250E-06 | 3.969E-07 | 9.500E-07 | 1.700E-06 |
| 11599 | 580 | S580Y | A | 2.4e-06 | 4e-06 | 4.5e-06 | 3.633E-06 | 1.097E-06 | 2.400E-06 | 4.500E-06 |
| 11600 | 580 | S580V | A | 4.4e-06 | 1.8e-06 | 2.8e-06 | 3.000E-06 | 1.311E-06 | 1.800E-06 | 4.400E-06 |
| 11601 | 581 | G581A | A | 2.3e-06 | 1.8e-06 | 3.3e-06 | 2.467E-06 | 7.638E-07 | 1.800E-06 | 3.300E-06 |
| 11602 | 581 | G581R | A | 2.2e-06 | 1.9e-06 | 9.9e-07 | 1.697E-06 | 6.301E-07 | 9.900E-07 | 2.200E-06 |
| 11603 | 581 | G581N | A | 1.8e-06 | 3.6e-06 | 2.2e-06 | 2.533E-06 | 9.452E-07 | 1.800E-06 | 3.600E-06 |
| 11604 | 581 | G581D | A | 3.8e-06 | 1.3e-06 | 3.7e-06 | 2.933E-06 | 1.415E-06 | 1.300E-06 | 3.800E-06 |
| 11605 | 581 | G581C | A | 8.8e-07 | 2.1e-06 | 2.2e-06 | 1.727E-06 | 7.349E-07 | 8.800E-07 | 2.200E-06 |
| 11606 | 581 | G581E | A | 1.6e-06 | 2e-06 | 1.3e-06 | 1.633E-06 | 3.512E-07 | 1.300E-06 | 2.000E-06 |
| 11607 | 581 | G581Q | A | 2.1e-06 | 3.3e-06 | 2.8e-06 | 2.733E-06 | 6.028E-07 | 2.100E-06 | 3.300E-06 |
| 11608 | 581 | G581G | A | 2.3e-06 | 1.8e-06 | 2.5e-06 | 2.200E-06 | 3.606E-07 | 1.800E-06 | 2.500E-06 |
| 11609 | 581 | G581H | A | 2.1e-06 | 1.5e-06 | 1.1e-06 | 1.567E-06 | 5.033E-07 | 1.100E-06 | 2.100E-06 |
| 11610 | 581 | G581I | A | 1.5e-06 | 1.7e-06 | 2.2e-06 | 1.800E-06 | 3.606E-07 | 1.500E-06 | 2.200E-06 |
| 11611 | 581 | G581L | A | 1.6e-06 | 1.2e-06 | 2.1e-06 | 1.633E-06 | 4.509E-07 | 1.200E-06 | 2.100E-06 |
| 11612 | 581 | G581K | A | 1.4e-06 | 1.8e-06 | 1.7e-06 | 1.633E-06 | 2.082E-07 | 1.400E-06 | 1.800E-06 |
| 11613 | 581 | G581M | A | 2e-06 | 1.5e-06 | 1.7e-06 | 1.733E-06 | 2.517E-07 | 1.500E-06 | 2.000E-06 |
| 11614 | 581 | G581F | A | 2.3e-06 | 2.1e-06 | 2.2e-06 | 2.200E-06 | 1.000E-07 | 2.100E-06 | 2.300E-06 |
| 11615 | 581 | G581P | A | 2.2e-06 | 1.9e-06 | 1.2e-06 | 1.767E-06 | 5.132E-07 | 1.200E-06 | 2.200E-06 |
| 11616 | 581 | G581S | A | 3.4e-06 | 3.5e-06 | 1.2e-06 | 2.700E-06 | 1.300E-06 | 1.200E-06 | 3.500E-06 |
| 11617 | 581 | G581T | A | 1.5e-06 | 2.4e-06 | 2.6e-06 | 2.167E-06 | 5.859E-07 | 1.500E-06 | 2.600E-06 |
| 11618 | 581 | G581W | A | 1.7e-06 | 1.2e-06 | 2.1e-06 | 1.667E-06 | 4.509E-07 | 1.200E-06 | 2.100E-06 |
| 11619 | 581 | G581Y | A | 9.3e-07 | 2.4e-06 | 1.8e-06 | 1.710E-06 | 7.391E-07 | 9.300E-07 | 2.400E-06 |
| 11620 | 581 | G581V | A | 2.2e-06 | 2e-06 | 1.7e-06 | 1.967E-06 | 2.517E-07 | 1.700E-06 | 2.200E-06 |
| 11621 | 582 | V582A | A | 3.5e-06 | 1.6e-06 | 2.5e-06 | 2.533E-06 | 9.504E-07 | 1.600E-06 | 3.500E-06 |
| 11622 | 582 | V582R | A | 1.9e-06 | 2.6e-06 | 2.7e-06 | 2.400E-06 | 4.359E-07 | 1.900E-06 | 2.700E-06 |
| 11623 | 582 | V582N | A | 2.9e-06 | 1.3e-06 | 1.7e-06 | 1.967E-06 | 8.327E-07 | 1.300E-06 | 2.900E-06 |
| 11624 | 582 | V582D | A | 2.6e-06 | 2.3e-06 | 2.1e-06 | 2.333E-06 | 2.517E-07 | 2.100E-06 | 2.600E-06 |
| 11625 | 582 | V582C | A | 3.3e-06 | 1.9e-06 | 3e-06 | 2.733E-06 | 7.371E-07 | 1.900E-06 | 3.300E-06 |
| 11626 | 582 | V582E | A | 1.7e-06 | 3.9e-06 | 1.4e-06 | 2.333E-06 | 1.365E-06 | 1.400E-06 | 3.900E-06 |
| 11627 | 582 | V582Q | A | 2.5e-06 | 1.5e-06 | 1.2e-06 | 1.733E-06 | 6.807E-07 | 1.200E-06 | 2.500E-06 |
| 11628 | 582 | V582G | A | 1.7e-06 | 2e-06 | 2.6e-06 | 2.100E-06 | 4.583E-07 | 1.700E-06 | 2.600E-06 |
| 11629 | 582 | V582H | A | 1.9e-06 | 2.7e-06 | 1.5e-06 | 2.033E-06 | 6.110E-07 | 1.500E-06 | 2.700E-06 |
| 11630 | 582 | V582I | A | 1.4e-06 | 1.8e-06 | 2.7e-06 | 1.967E-06 | 6.658E-07 | 1.400E-06 | 2.700E-06 |
| 11631 | 582 | V582L | A | 3.6e-06 | 3.6e-06 | 1.5e-06 | 2.900E-06 | 1.212E-06 | 1.500E-06 | 3.600E-06 |
| 11632 | 582 | V582K | A | 2.6e-06 | 2e-06 | 1.7e-06 | 2.100E-06 | 4.583E-07 | 1.700E-06 | 2.600E-06 |
| 11633 | 582 | V582M | A | 2.7e-06 | 2.8e-06 | 2.7e-06 | 2.733E-06 | 5.774E-08 | 2.700E-06 | 2.800E-06 |
| 11634 | 582 | V582F | A | 2e-06 | 2.2e-06 | 1.5e-06 | 1.900E-06 | 3.606E-07 | 1.500E-06 | 2.200E-06 |
| 11635 | 582 | V582P | A | 7.9e-07 | 2.6e-06 | 1.5e-06 | 1.630E-06 | 9.120E-07 | 7.900E-07 | 2.600E-06 |
| 11636 | 582 | V582S | A | 2.2e-06 | 2.9e-06 | 3.1e-06 | 2.733E-06 | 4.726E-07 | 2.200E-06 | 3.100E-06 |
| 11637 | 582 | V582T | A | 3.1e-06 | 2e-06 | 1.8e-06 | 2.300E-06 | 7.000E-07 | 1.800E-06 | 3.100E-06 |
| 11638 | 582 | V582W | A | 2.6e-06 | 2.9e-06 | 2.4e-06 | 2.633E-06 | 2.517E-07 | 2.400E-06 | 2.900E-06 |
| 11639 | 582 | V582Y | A | 2.3e-06 | 2.4e-06 | 3.2e-06 | 2.633E-06 | 4.933E-07 | 2.300E-06 | 3.200E-06 |
| 11640 | 582 | V582V | A | 2.5e-06 | 1.9e-06 | 2.3e-06 | 2.233E-06 | 3.055E-07 | 1.900E-06 | 2.500E-06 |
| 11641 | 583 | K583A | A | 1.9e-06 | 2.7e-06 | 2.7e-06 | 2.433E-06 | 4.619E-07 | 1.900E-06 | 2.700E-06 |
| 11642 | 583 | K583R | A | 2.7e-06 | 1.9e-06 | 2.3e-06 | 2.300E-06 | 4.000E-07 | 1.900E-06 | 2.700E-06 |
| 11643 | 583 | K583N | A | 1.8e-06 | 2.7e-06 | 2.2e-06 | 2.233E-06 | 4.509E-07 | 1.800E-06 | 2.700E-06 |
| 11644 | 583 | K583D | A | 4e-06 | 3.7e-06 | 2.3e-06 | 3.333E-06 | 9.074E-07 | 2.300E-06 | 4.000E-06 |

| | | | | | | | | | | |
|---|---|---|---|---|---|---|---|---|---|---|
| 11645 | 583 | K583C | A | 1.9e-06 | 1.8e-06 | 2.2e-06 | 1.967E-06 | 2.082E-07 | 1.800E-06 | 2.200E-06 |
| 11646 | 583 | K583E | A | 5.2e-06 | 2.3e-06 | 2.6e-06 | 3.367E-06 | 1.595E-06 | 2.300E-06 | 5.200E-06 |
| 11647 | 583 | K583Q | A | 4.8e-06 | 9.6e-06 | 2e-06 | 5.467E-06 | 3.844E-06 | 2.000E-06 | 9.600E-06 |
| 11648 | 583 | K583G | A | 2.2e-06 | 2.3e-06 | 2.3e-06 | 2.267E-06 | 5.774E-08 | 2.200E-06 | 2.300E-06 |
| 11649 | 583 | K583H | A | 3.8e-06 | 2.9e-06 | 2e-06 | 2.900E-06 | 9.000E-07 | 2.000E-06 | 3.800E-06 |
| 11650 | 583 | K583I | A | 1.8e-06 | 2.6e-06 | 2.9e-06 | 2.433E-06 | 5.686E-07 | 1.800E-06 | 2.900E-06 |
| 11651 | 583 | K583L | A | 2.5e-06 | 4.5e-06 | 1.9e-06 | 2.967E-06 | 1.361E-06 | 1.900E-06 | 4.500E-06 |
| 11652 | 583 | K583K | A | 3.9e-06 | 2.6e-06 | 2.1e-06 | 2.867E-06 | 9.292E-07 | 2.100E-06 | 3.900E-06 |
| 11653 | 583 | K583M | A | 3.3e-06 | 1.4e-06 | 4.3e-06 | 3.000E-06 | 1.473E-06 | 1.400E-06 | 4.300E-06 |
| 11654 | 583 | K583F | A | 1.7e-06 | 3.6e-06 | 2.3e-06 | 2.533E-06 | 9.713E-07 | 1.700E-06 | 3.600E-06 |
| 11655 | 583 | K583P | A | 3.7e-06 | 1.1e-06 | 2.9e-06 | 2.567E-06 | 1.332E-06 | 1.100E-06 | 3.700E-06 |
| 11656 | 583 | K583S | A | 2.9e-06 | 2.5e-06 | 2.1e-06 | 2.500E-06 | 4.000E-07 | 2.100E-06 | 2.900E-06 |
| 11657 | 583 | K583T | A | 3.3e-06 | 2.6e-06 | 3.2e-06 | 3.033E-06 | 3.786E-07 | 2.600E-06 | 3.300E-06 |
| 11658 | 583 | K583W | A | 4.3e-06 | 2.9e-06 | 2.3e-06 | 3.167E-06 | 1.026E-06 | 2.300E-06 | 4.300E-06 |
| 11659 | 583 | K583Y | A | 3.9e-06 | 1.5e-06 | 2.6e-06 | 2.667E-06 | 1.201E-06 | 1.500E-06 | 3.900E-06 |
| 11660 | 583 | K583V | A | 1.9e-06 | 3.3e-06 | 2e-06 | 2.400E-06 | 7.810E-07 | 1.900E-06 | 3.300E-06 |
| 11661 | 584 | P584A | A | 2.1e-06 | 1.9e-06 | 1.3e-06 | 1.767E-06 | 4.163E-07 | 1.300E-06 | 2.100E-06 |
| 11662 | 584 | P584R | A | 1.9e-06 | 2e-06 | 2.5e-06 | 2.133E-06 | 3.215E-07 | 1.900E-06 | 2.500E-06 |
| 11663 | 584 | P584N | A | 4.1e-06 | 2.7e-06 | 2e-06 | 2.933E-06 | 1.069E-06 | 2.000E-06 | 4.100E-06 |
| 11664 | 584 | P584D | A | 1.9e-06 | 3e-06 | 1.5e-06 | 2.133E-06 | 7.767E-07 | 1.500E-06 | 3.000E-06 |
| 11665 | 584 | P584C | A | 2.2e-06 | 3.3e-06 | 4.7e-06 | 3.400E-06 | 1.253E-06 | 2.200E-06 | 4.700E-06 |
| 11666 | 584 | P584E | A | 1.7e-06 | 2.3e-06 | 3.2e-06 | 2.400E-06 | 7.550E-07 | 1.700E-06 | 3.200E-06 |
| 11667 | 584 | P584Q | A | 3.4e-06 | 2.9e-06 | 2.6e-06 | 2.967E-06 | 4.041E-07 | 2.600E-06 | 3.400E-06 |
| 11668 | 584 | P584G | A | 2.1e-06 | 2.6e-06 | 3.5e-06 | 2.733E-06 | 7.095E-07 | 2.100E-06 | 3.500E-06 |
| 11669 | 584 | P584H | A | 2.5e-06 | 1.4e-06 | 3.2e-06 | 2.367E-06 | 9.074E-07 | 1.400E-06 | 3.200E-06 |
| 11670 | 584 | P584I | A | 2.5e-06 | 4.1e-06 | 1.9e-06 | 2.833E-06 | 1.137E-06 | 1.900E-06 | 4.100E-06 |
| 11671 | 584 | P584L | A | 1.7e-06 | 3.3e-06 | 2.1e-06 | 2.367E-06 | 8.327E-07 | 1.700E-06 | 3.300E-06 |
| 11672 | 584 | P584K | A | 4.9e-06 | 2e-06 | 1.3e-06 | 2.733E-06 | 1.909E-06 | 1.300E-06 | 4.900E-06 |
| 11673 | 584 | P584M | A | 2.1e-06 | 2.5e-06 | 1e-06 | 1.867E-06 | 7.767E-07 | 1.000E-06 | 2.500E-06 |
| 11674 | 584 | P584F | A | 1.5e-06 | 5e-06 | 3.1e-06 | 3.200E-06 | 1.752E-06 | 1.500E-06 | 5.000E-06 |
| 11675 | 584 | P584P | A | 1.4e-06 | 3.5e-06 | 2.7e-06 | 2.533E-06 | 1.060E-06 | 1.400E-06 | 3.500E-06 |
| 11676 | 584 | P584S | A | 2e-06 | 2.5e-06 | 1.5e-06 | 2.000E-06 | 5.000E-07 | 1.500E-06 | 2.500E-06 |
| 11677 | 584 | P584T | A | 2.6e-06 | 1.6e-06 | 2.1e-06 | 2.100E-06 | 5.000E-07 | 1.600E-06 | 2.600E-06 |
| 11678 | 584 | P584W | A | 1.4e-06 | 1.2e-06 | 1.9e-06 | 1.500E-06 | 3.606E-07 | 1.200E-06 | 1.900E-06 |
| 11679 | 584 | P584Y | A | 3.9e-06 | 3.6e-06 | 1.5e-06 | 3.000E-06 | 1.308E-06 | 1.500E-06 | 3.900E-06 |
| 11680 | 584 | P584V | A | 3.3e-06 | 1.7e-06 | 1.8e-06 | 2.267E-06 | 8.963E-07 | 1.700E-06 | 3.300E-06 |
| 11681 | 585 | D585A | A | 2.9e-06 | 2.4e-06 | 5.1e-06 | 3.467E-06 | 1.436E-06 | 2.400E-06 | 5.100E-06 |
| 11682 | 585 | D585R | A | 9.4e-07 | 1.8e-06 | 1.2e-06 | 1.313E-06 | 4.411E-07 | 9.400E-07 | 1.800E-06 |
| 11683 | 585 | D585N | A | 4.8e-06 | 2.3e-06 | 2e-06 | 3.033E-06 | 1.537E-06 | 2.000E-06 | 4.800E-06 |
| 11684 | 585 | D585D | A | 2.3e-06 | 2.5e-06 | 2.7e-06 | 2.500E-06 | 2.000E-07 | 2.300E-06 | 2.700E-06 |
| 11685 | 585 | D585C | A | 4.3e-06 | 1.5e-06 | 1.5e-06 | 2.433E-06 | 1.617E-06 | 1.500E-06 | 4.300E-06 |
| 11686 | 585 | D585E | A | 3.1e-06 | 1.2e-06 | 1.7e-06 | 2.000E-06 | 9.849E-07 | 1.200E-06 | 3.100E-06 |
| 11687 | 585 | D585Q | A | 2.4e-06 | 1.6e-06 | 3e-06 | 2.333E-06 | 7.024E-07 | 1.600E-06 | 3.000E-06 |
| 11688 | 585 | D585G | A | 3.1e-06 | 4.6e-06 | 4.1e-06 | 3.933E-06 | 7.638E-07 | 3.100E-06 | 4.600E-06 |
| 11689 | 585 | D585H | A | 2e-06 | 2.3e-06 | 1.6e-06 | 1.967E-06 | 3.512E-07 | 1.600E-06 | 2.300E-06 |
| 11690 | 585 | D585I | A | 2e-06 | 4.4e-06 | 1e-06 | 2.467E-06 | 1.747E-06 | 1.000E-06 | 4.400E-06 |
| 11691 | 585 | D585L | A | 1.6e-06 | 3.2e-06 | 3.1e-06 | 2.633E-06 | 8.963E-07 | 1.600E-06 | 3.200E-06 |
| 11692 | 585 | D585K | A | 2.5e-06 | 2.4e-06 | 2.7e-06 | 2.533E-06 | 1.528E-07 | 2.400E-06 | 2.700E-06 |
| 11693 | 585 | D585M | A | 2.7e-06 | 3.1e-06 | 2.3e-06 | 2.700E-06 | 4.000E-07 | 2.300E-06 | 3.100E-06 |
| 11694 | 585 | D585F | A | 2.1e-06 | 2.1e-06 | 1.9e-06 | 2.033E-06 | 1.155E-07 | 1.900E-06 | 2.100E-06 |
| 11695 | 585 | D585P | A | 4.1e-06 | 3.7e-06 | 3.8e-06 | 3.867E-06 | 2.082E-07 | 3.700E-06 | 4.100E-06 |
| 11696 | 585 | D585S | A | 4.2e-06 | 1.6e-06 | 3.1e-06 | 2.967E-06 | 1.305E-06 | 1.600E-06 | 4.200E-06 |
| 11697 | 585 | D585T | A | 1e-06 | 1.4e-06 | 1.7e-06 | 1.367E-06 | 3.512E-07 | 1.000E-06 | 1.700E-06 |
| 11698 | 585 | D585W | A | 2.8e-06 | 1.5e-06 | 2.2e-06 | 2.167E-06 | 6.506E-07 | 1.500E-06 | 2.800E-06 |
| 11699 | 585 | D585Y | A | 1.1e-06 | 1.8e-06 | 1.6e-06 | 1.500E-06 | 3.606E-07 | 1.100E-06 | 1.800E-06 |
| 11700 | 585 | D585V | A | 3.8e-06 | 2.2e-06 | 2.3e-06 | 2.767E-06 | 8.963E-07 | 2.200E-06 | 3.800E-06 |
| 11701 | 586 | L586A | A | 1.8e-06 | 4.2e-06 | 3.9e-06 | 3.300E-06 | 1.308E-06 | 1.800E-06 | 4.200E-06 |
| 11702 | 586 | L586R | A | 9e-07 | 1.5e-06 | 1.4e-06 | 1.267E-06 | 3.215E-07 | 9.000E-07 | 1.500E-06 |
| 11703 | 586 | L586N | A | 1.1e-06 | 1.8e-06 | 1.1e-06 | 1.333E-06 | 4.041E-07 | 1.100E-06 | 1.800E-06 |
| 11704 | 586 | L586D | A | 1.6e-06 | 2.3e-06 | 2.1e-06 | 2.000E-06 | 3.606E-07 | 1.600E-06 | 2.300E-06 |
| 11705 | 586 | L586C | A | 1.8e-06 | 3.4e-06 | 1.8e-06 | 2.333E-06 | 9.238E-07 | 1.800E-06 | 3.400E-06 |
| 11706 | 586 | L586E | A | 2.5e-06 | 1.3e-06 | 1.4e-06 | 1.733E-06 | 6.658E-07 | 1.300E-06 | 2.500E-06 |
| 11707 | 586 | L586Q | A | 8.5e-07 | 6e-07 | 1.3e-06 | 9.167E-07 | 3.547E-07 | 6.000E-07 | 1.300E-06 |
| 11708 | 586 | L586G | A | 3.6e-06 | 2.8e-06 | 1.8e-06 | 2.733E-06 | 9.018E-07 | 1.800E-06 | 3.600E-06 |
| 11709 | 586 | L586H | A | 1.8e-06 | 1.7e-06 | 1.8e-06 | 1.767E-06 | 5.774E-08 | 1.700E-06 | 1.800E-06 |
| 11710 | 586 | L586I | A | 2.1e-06 | 2.1e-06 | 2e-06 | 2.067E-06 | 5.774E-08 | 2.000E-06 | 2.100E-06 |
| 11711 | 586 | L586L | A | 2e-06 | 2.8e-06 | 2.6e-06 | 2.467E-06 | 4.163E-07 | 2.000E-06 | 2.800E-06 |
| 11712 | 586 | L586K | A | 1.6e-06 | 1.7e-06 | 9.5e-07 | 1.417E-06 | 4.072E-07 | 9.500E-07 | 1.700E-06 |
| 11713 | 586 | L586M | A | 1.3e-06 | 3.2e-06 | 2.3e-06 | 2.267E-06 | 9.504E-07 | 1.300E-06 | 3.200E-06 |
| 11714 | 586 | L586F | A | 3e-06 | 1.8e-06 | 2.1e-06 | 2.300E-06 | 6.245E-07 | 1.800E-06 | 3.000E-06 |
| 11715 | 586 | L586P | A | 2.3e-06 | 1.7e-06 | 2.8e-06 | 2.267E-06 | 5.508E-07 | 1.700E-06 | 2.800E-06 |
| 11716 | 586 | L586S | A | 2.9e-06 | 2.3e-06 | 2e-06 | 2.400E-06 | 4.583E-07 | 2.000E-06 | 2.900E-06 |
| 11717 | 586 | L586T | A | 1.5e-06 | 1.6e-06 | 1.5e-06 | 1.533E-06 | 5.774E-08 | 1.500E-06 | 1.600E-06 |
| 11718 | 586 | L586W | A | 2e-06 | 1.3e-06 | 1.7e-06 | 1.667E-06 | 3.512E-07 | 1.300E-06 | 2.000E-06 |
| 11719 | 586 | L586Y | A | 1.6e-06 | 2.3e-06 | 1.9e-06 | 1.933E-06 | 3.512E-07 | 1.600E-06 | 2.300E-06 |
| 11720 | 586 | L586V | A | 2.6e-06 | 2.1e-06 | 3.8e-06 | 2.833E-06 | 8.737E-07 | 2.100E-06 | 3.800E-06 |
| 11721 | 587 | S587A | A | 2.5e-06 | 2.9e-06 | 2.7e-06 | 2.700E-06 | 2.000E-07 | 2.500E-06 | 2.900E-06 |
| 11722 | 587 | S587R | A | 1.7e-06 | 3.7e-06 | 1.6e-06 | 2.333E-06 | 1.185E-06 | 1.600E-06 | 3.700E-06 |
| 11723 | 587 | S587N | A | 1.9e-06 | 3.1e-06 | 1.9e-06 | 2.300E-06 | 6.928E-07 | 1.900E-06 | 3.100E-06 |
| 11724 | 587 | S587D | A | 1.9e-06 | 3.2e-06 | 2.1e-06 | 2.400E-06 | 7.000E-07 | 1.900E-06 | 3.200E-06 |
| 11725 | 587 | S587C | A | 2.8e-06 | 2.1e-06 | 1.7e-06 | 2.200E-06 | 5.568E-07 | 1.700E-06 | 2.800E-06 |
| 11726 | 587 | S587E | A | 2.2e-06 | 2.9e-06 | 2.7e-06 | 2.600E-06 | 3.606E-07 | 2.200E-06 | 2.900E-06 |
| 11727 | 587 | S587Q | A | 2.2e-06 | 2.3e-06 | 2.8e-06 | 2.433E-06 | 3.215E-07 | 2.200E-06 | 2.800E-06 |
| 11728 | 587 | S587G | A | 2.3e-06 | 2.5e-06 | 3.9e-06 | 2.900E-06 | 8.718E-07 | 2.300E-06 | 3.900E-06 |
| 11729 | 587 | S587H | A | 2.1e-06 | 2.7e-06 | 3.5e-06 | 2.767E-06 | 7.024E-07 | 2.100E-06 | 3.500E-06 |

| | | | | | | | | | | |
|---|---|---|---|---|---|---|---|---|---|---|
| 11730 | 587 | S587I | A | 2.5e-06 | 2e-06 | 2.9e-06 | 2.467E-06 | 4.509E-07 | 2.000E-06 | 2.900E-06 |
| 11731 | 587 | S587L | A | 2.2e-06 | 1.7e-06 | 1.7e-06 | 1.867E-06 | 2.887E-07 | 1.700E-06 | 2.200E-06 |
| 11732 | 587 | S587K | A | 2.3e-06 | 1.7e-06 | 1.8e-06 | 1.933E-06 | 3.215E-07 | 1.700E-06 | 2.300E-06 |
| 11733 | 587 | S587M | A | 3.1e-06 | 2.4e-06 | 2.1e-06 | 2.533E-06 | 5.132E-07 | 2.100E-06 | 3.100E-06 |
| 11734 | 587 | S587F | A | 2.3e-06 | 2.2e-06 | 3.4e-06 | 2.633E-06 | 6.658E-07 | 2.200E-06 | 3.400E-06 |
| 11735 | 587 | S587P | A | 2.6e-06 | 1.7e-06 | 3.3e-06 | 2.533E-06 | 8.021E-07 | 1.700E-06 | 3.300E-06 |
| 11736 | 587 | S587S | A | 1.6e-06 | 1.8e-06 | 1.7e-06 | 1.700E-06 | 1.000E-07 | 1.600E-06 | 1.800E-06 |
| 11737 | 587 | S587T | A | 2.2e-06 | 2.1e-06 | 2.3e-06 | 2.200E-06 | 1.000E-07 | 2.100E-06 | 2.300E-06 |
| 11738 | 587 | S587W | A | 4.1e-06 | 3.1e-06 | 2.9e-06 | 3.367E-06 | 6.429E-07 | 2.900E-06 | 4.100E-06 |
| 11739 | 587 | S587Y | A | 1.9e-06 | 5.8e-06 | 3.1e-06 | 3.600E-06 | 1.997E-06 | 1.900E-06 | 5.800E-06 |
| 11740 | 587 | S587V | A | 1.6e-06 | 2.8e-06 | 1.4e-06 | 1.933E-06 | 7.572E-07 | 1.400E-06 | 2.800E-06 |
| 11741 | 588 | Y588A | A | 2.6e-06 | 2.5e-06 | 2e-06 | 2.367E-06 | 3.215E-07 | 2.000E-06 | 2.600E-06 |
| 11742 | 588 | Y588R | A | 2.4e-06 | 1.2e-06 | 2.5e-06 | 2.033E-06 | 7.234E-07 | 1.200E-06 | 2.500E-06 |
| 11743 | 588 | Y588N | A | 2e-06 | 1.5e-06 | 2.7e-06 | 2.067E-06 | 6.028E-07 | 1.500E-06 | 2.700E-06 |
| 11744 | 588 | Y588D | A | 1.3e-06 | 1.6e-06 | 3.5e-06 | 2.133E-06 | 1.193E-06 | 1.300E-06 | 3.500E-06 |
| 11745 | 588 | Y588C | A | 2.5e-06 | 2.7e-06 | 2.5e-06 | 2.567E-06 | 1.155E-07 | 2.500E-06 | 2.700E-06 |
| 11746 | 588 | Y588E | A | 2.7e-06 | 2.3e-06 | 3.7e-06 | 2.900E-06 | 7.211E-07 | 2.300E-06 | 3.700E-06 |
| 11747 | 588 | Y588Q | A | 2e-06 | 1.1e-06 | 1.2e-06 | 1.433E-06 | 4.933E-07 | 1.100E-06 | 2.000E-06 |
| 11748 | 588 | Y588G | A | 3.1e-06 | 2e-06 | 1.3e-06 | 2.133E-06 | 9.074E-07 | 1.300E-06 | 3.100E-06 |
| 11749 | 588 | Y588H | A | 3.3e-06 | 1.9e-06 | 2.3e-06 | 2.500E-06 | 7.211E-07 | 1.900E-06 | 3.300E-06 |
| 11750 | 588 | Y588I | A | 3.3e-06 | 3.6e-06 | 2.6e-06 | 3.167E-06 | 5.132E-07 | 2.600E-06 | 3.600E-06 |
| 11751 | 588 | Y588L | A | 3e-06 | 2.8e-06 | 1.2e-06 | 2.333E-06 | 9.866E-07 | 1.200E-06 | 3.000E-06 |
| 11752 | 588 | Y588K | A | 2e-06 | 2.6e-06 | 1.8e-06 | 2.133E-06 | 4.163E-07 | 1.800E-06 | 2.600E-06 |
| 11753 | 588 | Y588M | A | 2.9e-06 | 2.9e-06 | 2.1e-06 | 2.633E-06 | 4.619E-07 | 2.100E-06 | 2.900E-06 |
| 11754 | 588 | Y588F | A | 2.3e-06 | 1.6e-06 | 3.3e-06 | 2.400E-06 | 8.544E-07 | 1.600E-06 | 3.300E-06 |
| 11755 | 588 | Y588P | A | 3.2e-06 | 2.3e-06 | 3.1e-06 | 2.867E-06 | 4.933E-07 | 2.300E-06 | 3.200E-06 |
| 11756 | 588 | Y588S | A | 2.3e-06 | 1e-06 | 2.3e-06 | 1.867E-06 | 7.506E-07 | 1.000E-06 | 2.300E-06 |
| 11757 | 588 | Y588T | A | 2.5e-06 | 3.5e-06 | 4.7e-06 | 3.567E-06 | 1.102E-06 | 2.500E-06 | 4.700E-06 |
| 11758 | 588 | Y588W | A | 2.8e-06 | 5.4e-06 | 2.7e-06 | 3.633E-06 | 1.531E-06 | 2.700E-06 | 5.400E-06 |
| 11759 | 588 | Y588Y | A | 1.8e-06 | 1.2e-06 | 2.6e-06 | 1.867E-06 | 7.024E-07 | 1.200E-06 | 2.600E-06 |
| 11760 | 588 | Y588V | A | 1.7e-06 | 2.7e-06 | 1.9e-06 | 2.100E-06 | 5.292E-07 | 1.700E-06 | 2.700E-06 |
| 11761 | 589 | M589A | A | 2.2e-06 | 1.9e-06 | 3e-06 | 2.367E-06 | 5.686E-07 | 1.900E-06 | 3.000E-06 |
| 11762 | 589 | M589R | A | 3e-06 | 2.3e-06 | 2.7e-06 | 2.667E-06 | 3.512E-07 | 2.300E-06 | 3.000E-06 |
| 11763 | 589 | M589N | A | 1.5e-06 | 2.4e-06 | 2.3e-06 | 2.067E-06 | 4.933E-07 | 1.500E-06 | 2.400E-06 |
| 11764 | 589 | M589D | A | 2.4e-06 | 1.9e-06 | 2.7e-06 | 2.333E-06 | 4.041E-07 | 1.900E-06 | 2.700E-06 |
| 11765 | 589 | M589C | A | 4.2e-06 | 2.4e-06 | 3.3e-06 | 3.300E-06 | 9.000E-07 | 2.400E-06 | 4.200E-06 |
| 11766 | 589 | M589E | A | 3e-06 | 2.3e-06 | 1.3e-06 | 2.200E-06 | 8.544E-07 | 1.300E-06 | 3.000E-06 |
| 11767 | 589 | M589Q | A | 2.6e-06 | 2.2e-06 | 3.4e-06 | 2.733E-06 | 6.110E-07 | 2.200E-06 | 3.400E-06 |
| 11768 | 589 | M589G | A | 1.2e-06 | 2.1e-06 | 1.3e-06 | 1.533E-06 | 4.933E-07 | 1.200E-06 | 2.100E-06 |
| 11769 | 589 | M589H | A | 2e-06 | 2.5e-06 | 2e-06 | 2.167E-06 | 2.887E-07 | 2.000E-06 | 2.500E-06 |
| 11770 | 589 | M589I | A | 1.8e-06 | 3e-06 | 1.5e-06 | 2.100E-06 | 7.937E-07 | 1.500E-06 | 3.000E-06 |
| 11771 | 589 | M589L | A | 1.8e-06 | 3.3e-06 | 1.8e-06 | 2.300E-06 | 8.660E-07 | 1.800E-06 | 3.300E-06 |
| 11772 | 589 | M589K | A | 4e-06 | 3.4e-06 | 1.9e-06 | 3.100E-06 | 1.082E-06 | 1.900E-06 | 4.000E-06 |
| 11773 | 589 | M589M | A | 2.1e-06 | 2.6e-06 | 2.4e-06 | 2.367E-06 | 2.517E-07 | 2.100E-06 | 2.600E-06 |
| 11774 | 589 | M589F | A | 1.2e-06 | 2.6e-06 | 3.4e-06 | 2.400E-06 | 1.114E-06 | 1.200E-06 | 3.400E-06 |
| 11775 | 589 | M589P | A | 2.2e-06 | 7.1e-07 | 3e-06 | 1.970E-06 | 1.162E-06 | 7.100E-07 | 3.000E-06 |
| 11776 | 589 | M589S | A | 3.1e-06 | 2.2e-06 | 1.3e-06 | 2.200E-06 | 9.000E-07 | 1.300E-06 | 3.100E-06 |
| 11777 | 589 | M589T | A | 1.8e-06 | 2.8e-06 | 2.5e-06 | 2.367E-06 | 5.132E-07 | 1.800E-06 | 2.800E-06 |
| 11778 | 589 | M589W | A | 2.7e-06 | 1.7e-06 | 3.8e-06 | 2.733E-06 | 1.050E-06 | 1.700E-06 | 3.800E-06 |
| 11779 | 589 | M589Y | A | 2.6e-06 | 3e-06 | 2.2e-06 | 2.600E-06 | 4.000E-07 | 2.200E-06 | 3.000E-06 |
| 11780 | 589 | M589V | A | 2.1e-06 | 2.9e-06 | 2.7e-06 | 2.567E-06 | 4.163E-07 | 2.100E-06 | 2.900E-06 |
| 11781 | 590 | P590A | A | 3e-06 | 2.9e-06 | 2.2e-06 | 2.700E-06 | 4.359E-07 | 2.200E-06 | 3.000E-06 |
| 11782 | 590 | P590R | A | 2e-06 | 1.5e-06 | 1.6e-06 | 1.700E-06 | 2.646E-07 | 1.500E-06 | 2.000E-06 |
| 11783 | 590 | P590N | A | 1.6e-06 | 1.7e-06 | 7.1e-07 | 1.337E-06 | 5.450E-07 | 7.100E-07 | 1.700E-06 |
| 11784 | 590 | P590D | A | 1.8e-06 | 6.5e-07 | 1.3e-06 | 1.250E-06 | 5.766E-07 | 6.500E-07 | 1.800E-06 |
| 11785 | 590 | P590C | A | 1.3e-06 | 3e-06 | 2.3e-06 | 2.200E-06 | 8.544E-07 | 1.300E-06 | 3.000E-06 |
| 11786 | 590 | P590E | A | 1.2e-06 | 1.6e-06 | 2.1e-06 | 1.633E-06 | 4.509E-07 | 1.200E-06 | 2.100E-06 |
| 11787 | 590 | P590Q | A | 1.7e-06 | 1.5e-06 | 2.1e-06 | 1.767E-06 | 3.055E-07 | 1.500E-06 | 2.100E-06 |
| 11788 | 590 | P590G | A | 4.1e-06 | 1.3e-06 | 1.8e-06 | 2.400E-06 | 1.493E-06 | 1.300E-06 | 4.100E-06 |
| 11789 | 590 | P590H | A | 1.6e-06 | 1.2e-06 | 1.5e-06 | 1.433E-06 | 2.082E-07 | 1.200E-06 | 1.600E-06 |
| 11790 | 590 | P590I | A | 4.6e-06 | 2.6e-06 | 2.3e-06 | 3.167E-06 | 1.250E-06 | 2.300E-06 | 4.600E-06 |
| 11791 | 590 | P590L | A | 3.4e-06 | 2.4e-06 | 9e-07 | 2.233E-06 | 1.258E-06 | 9.000E-07 | 3.400E-06 |
| 11792 | 590 | P590K | A | 1.5e-06 | 2.3e-06 | 1.3e-06 | 1.700E-06 | 5.292E-07 | 1.300E-06 | 2.300E-06 |
| 11793 | 590 | P590M | A | 2.6e-06 | 2.2e-06 | 1.2e-06 | 2.000E-06 | 7.211E-07 | 1.200E-06 | 2.600E-06 |
| 11794 | 590 | P590F | A | 2.6e-06 | 1.7e-06 | 2.4e-06 | 2.233E-06 | 4.726E-07 | 1.700E-06 | 2.600E-06 |
| 11795 | 590 | P590P | A | 2e-06 | 2.9e-06 | 3.3e-06 | 2.733E-06 | 6.658E-07 | 2.000E-06 | 3.300E-06 |
| 11796 | 590 | P590S | A | 1.5e-06 | 8.4e-07 | 2e-06 | 1.447E-06 | 5.818E-07 | 8.400E-07 | 2.000E-06 |
| 11797 | 590 | P590T | A | 1.3e-06 | 1.3e-06 | 1.9e-06 | 1.500E-06 | 3.464E-07 | 1.300E-06 | 1.900E-06 |
| 11798 | 590 | P590W | A | 2.1e-06 | 1.7e-06 | 2.3e-06 | 2.033E-06 | 3.055E-07 | 1.700E-06 | 2.300E-06 |
| 11799 | 590 | P590Y | A | 2.5e-06 | 9.3e-07 | 2.2e-06 | 1.877E-06 | 8.334E-07 | 9.300E-07 | 2.500E-06 |
| 11800 | 590 | P590V | A | 2.5e-06 | 3.2e-06 | 3.3e-06 | 3.000E-06 | 4.359E-07 | 2.500E-06 | 3.300E-06 |
| 11801 | 591 | I591A | A | 2.5e-06 | 2.4e-06 | 2.3e-06 | 2.400E-06 | 1.000E-07 | 2.300E-06 | 2.500E-06 |
| 11802 | 591 | I591R | A | 3.7e-06 | 2.4e-06 | 2.2e-06 | 2.767E-06 | 8.145E-07 | 2.200E-06 | 3.700E-06 |
| 11803 | 591 | I591N | A | 1.6e-06 | 3.3e-06 | 2.8e-06 | 2.567E-06 | 8.737E-07 | 1.600E-06 | 3.300E-06 |
| 11804 | 591 | I591D | A | 2.4e-06 | 2.2e-06 | 2.5e-06 | 2.367E-06 | 1.528E-07 | 2.200E-06 | 2.500E-06 |
| 11805 | 591 | I591C | A | 3.5e-06 | 3.6e-06 | 2.4e-06 | 3.167E-06 | 6.658E-07 | 2.400E-06 | 3.600E-06 |
| 11806 | 591 | I591E | A | 1.7e-06 | 2.6e-06 | 2e-06 | 2.100E-06 | 4.583E-07 | 1.700E-06 | 2.600E-06 |
| 11807 | 591 | I591Q | A | 1.8e-06 | 4.1e-06 | 2.6e-06 | 2.833E-06 | 1.168E-06 | 1.800E-06 | 4.100E-06 |
| 11808 | 591 | I591G | A | 1.8e-06 | 2.1e-06 | 2.9e-06 | 2.267E-06 | 5.686E-07 | 1.800E-06 | 2.900E-06 |
| 11809 | 591 | I591H | A | 2.6e-06 | 1.4e-06 | 2.3e-06 | 2.100E-06 | 6.245E-07 | 1.400E-06 | 2.600E-06 |
| 11810 | 591 | I591I | A | 3.4e-06 | 2.2e-06 | 2.5e-06 | 2.700E-06 | 6.245E-07 | 2.200E-06 | 3.400E-06 |
| 11811 | 591 | I591L | A | 1.9e-06 | 3.4e-06 | 1.9e-06 | 2.400E-06 | 8.660E-07 | 1.900E-06 | 3.400E-06 |
| 11812 | 591 | I591K | A | 3.2e-06 | 1.6e-06 | 2.5e-06 | 2.433E-06 | 8.021E-07 | 1.600E-06 | 3.200E-06 |
| 11813 | 591 | I591M | A | 1.7e-06 | 3.1e-06 | 2.3e-06 | 2.367E-06 | 7.024E-07 | 1.700E-06 | 3.100E-06 |
| 11814 | 591 | I591F | A | 2.5e-06 | 3.5e-06 | 2e-06 | 2.667E-06 | 7.638E-07 | 2.000E-06 | 3.500E-06 |

| | | | | | | | | | | |
|---|---|---|---|---|---|---|---|---|---|---|
| 11815 | 591 | I591P | A | 2.4e-06 | 2.9e-06 | 1.7e-06 | 2.333E-06 | 6.028E-07 | 1.700E-06 | 2.900E-06 |
| 11816 | 591 | I591S | A | 3.4e-06 | 1.5e-06 | 2.8e-06 | 2.567E-06 | 9.713E-07 | 1.500E-06 | 3.400E-06 |
| 11817 | 591 | I591T | A | 2e-06 | 2.1e-06 | 1.5e-06 | 1.867E-06 | 3.215E-07 | 1.500E-06 | 2.100E-06 |
| 11818 | 591 | I591W | A | 2.7e-06 | 3e-06 | 2.7e-06 | 2.800E-06 | 1.732E-07 | 2.700E-06 | 3.000E-06 |
| 11819 | 591 | I591Y | A | 4.5e-06 | 1.3e-06 | 3.1e-06 | 2.967E-06 | 1.604E-06 | 1.300E-06 | 4.500E-06 |
| 11820 | 591 | I591V | A | 1.1e-06 | 2.2e-06 | 2.7e-06 | 2.000E-06 | 8.185E-07 | 1.100E-06 | 2.700E-06 |
| 11821 | 592 | W592A | A | 1.4e-06 | 2.3e-06 | 3.9e-06 | 2.533E-06 | 1.266E-06 | 1.400E-06 | 3.900E-06 |
| 11822 | 592 | W592R | A | 3.5e-06 | 2.1e-06 | 3.2e-06 | 2.933E-06 | 7.371E-07 | 2.100E-06 | 3.500E-06 |
| 11823 | 592 | W592N | A | 2.8e-06 | 2.6e-06 | 2.3e-06 | 2.567E-06 | 2.517E-07 | 2.300E-06 | 2.800E-06 |
| 11824 | 592 | W592D | A | 2.7e-06 | 3e-06 | 1.5e-06 | 2.400E-06 | 7.937E-07 | 1.500E-06 | 3.000E-06 |
| 11825 | 592 | W592C | A | 2.6e-06 | 1.3e-06 | 3.8e-06 | 2.567E-06 | 1.250E-06 | 1.300E-06 | 3.800E-06 |
| 11826 | 592 | W592E | A | 2.2e-06 | 2.2e-06 | 2.1e-06 | 2.167E-06 | 5.774E-08 | 2.100E-06 | 2.200E-06 |
| 11827 | 592 | W592Q | A | 2.4e-06 | 2.3e-06 | 3.9e-06 | 2.867E-06 | 8.963E-07 | 2.300E-06 | 3.900E-06 |
| 11828 | 592 | W592G | A | 2.3e-06 | 3.1e-06 | 3.3e-06 | 2.900E-06 | 5.292E-07 | 2.300E-06 | 3.300E-06 |
| 11829 | 592 | W592H | A | 3.5e-06 | 2.4e-06 | 3.4e-06 | 3.100E-06 | 6.083E-07 | 2.400E-06 | 3.500E-06 |
| 11830 | 592 | W592I | A | 2.9e-06 | 4.4e-06 | 3.8e-06 | 3.700E-06 | 7.550E-07 | 2.900E-06 | 4.400E-06 |
| 11831 | 592 | W592L | A | 3.3e-06 | 1.7e-06 | 2.6e-06 | 2.533E-06 | 8.021E-07 | 1.700E-06 | 3.300E-06 |
| 11832 | 592 | W592K | A | 2.2e-06 | 2.4e-06 | 2.1e-06 | 2.233E-06 | 1.528E-07 | 2.100E-06 | 2.400E-06 |
| 11833 | 592 | W592M | A | 2.4e-06 | 2.9e-06 | 3.4e-06 | 2.900E-06 | 5.000E-07 | 2.400E-06 | 3.400E-06 |
| 11834 | 592 | W592F | A | 2.5e-06 | 3.3e-06 | 4.6e-06 | 3.467E-06 | 1.060E-06 | 2.500E-06 | 4.600E-06 |
| 11835 | 592 | W592P | A | 1.8e-06 | 1.7e-06 | 2.2e-06 | 1.900E-06 | 2.646E-07 | 1.700E-06 | 2.200E-06 |
| 11836 | 592 | W592S | A | 2.3e-06 | 1.7e-06 | 2.4e-06 | 2.133E-06 | 3.786E-07 | 1.700E-06 | 2.400E-06 |
| 11837 | 592 | W592T | A | 3.5e-06 | 2.4e-06 | 3.7e-06 | 3.200E-06 | 7.000E-07 | 2.400E-06 | 3.700E-06 |
| 11838 | 592 | W592W | A | 1.2e-06 | 1.7e-06 | 2e-06 | 1.633E-06 | 4.041E-07 | 1.200E-06 | 2.000E-06 |
| 11839 | 592 | W592Y | A | 3.9e-06 | 3.2e-06 | 3.6e-06 | 3.567E-06 | 3.512E-07 | 3.200E-06 | 3.900E-06 |
| 11840 | 592 | W592V | A | 2.1e-06 | 2.1e-06 | 1.5e-06 | 1.900E-06 | 3.464E-07 | 1.500E-06 | 2.100E-06 |
| 11841 | 593 | K593A | A | 4.6e-06 | 3.7e-06 | 4.1e-06 | 4.133E-06 | 4.509E-07 | 3.700E-06 | 4.600E-06 |
| 11842 | 593 | K593R | A | 4.4e-06 | 2e-06 | 1.8e-06 | 2.733E-06 | 1.447E-06 | 1.800E-06 | 4.400E-06 |
| 11843 | 593 | K593N | A | 2.2e-06 | 2.6e-06 | 2.5e-06 | 2.433E-06 | 2.082E-07 | 2.200E-06 | 2.600E-06 |
| 11844 | 593 | K593D | A | 3.7e-06 | 2.3e-06 | 3.5e-06 | 3.167E-06 | 7.572E-07 | 2.300E-06 | 3.700E-06 |
| 11845 | 593 | K593C | A | 3e-06 | 4.3e-06 | 3.2e-06 | 3.500E-06 | 7.000E-07 | 3.000E-06 | 4.300E-06 |
| 11846 | 593 | K593E | A | 4.4e-06 | 2.7e-06 | 2.8e-06 | 3.300E-06 | 9.539E-07 | 2.700E-06 | 4.400E-06 |
| 11847 | 593 | K593Q | A | 2.7e-06 | 2e-06 | 5.6e-06 | 3.433E-06 | 1.909E-06 | 2.000E-06 | 5.600E-06 |
| 11848 | 593 | K593G | A | 7.4e-06 | 3.8e-06 | 6e-06 | 5.733E-06 | 1.815E-06 | 3.800E-06 | 7.400E-06 |
| 11849 | 593 | K593H | A | 2.3e-06 | 3.3e-06 | 1.8e-06 | 2.467E-06 | 7.638E-07 | 1.800E-06 | 3.300E-06 |
| 11850 | 593 | K593I | A | 4.6e-06 | 3.3e-06 | 4.6e-06 | 4.167E-06 | 7.506E-07 | 3.300E-06 | 4.600E-06 |
| 11851 | 593 | K593L | A | 2.5e-06 | 4.4e-06 | 4.2e-06 | 3.700E-06 | 1.044E-06 | 2.500E-06 | 4.400E-06 |
| 11852 | 593 | K593K | A | 1.8e-06 | 3.8e-06 | 3.1e-06 | 2.900E-06 | 1.015E-06 | 1.800E-06 | 3.800E-06 |
| 11853 | 593 | K593M | A | 2.9e-06 | 4.9e-06 | 3.5e-06 | 3.767E-06 | 1.026E-06 | 2.900E-06 | 4.900E-06 |
| 11854 | 593 | K593F | A | 2.8e-06 | 3.2e-06 | 5.9e-06 | 3.967E-06 | 1.686E-06 | 2.800E-06 | 5.900E-06 |
| 11855 | 593 | K593P | A | 5.1e-06 | 4.8e-06 | 3.1e-06 | 4.333E-06 | 1.079E-06 | 3.100E-06 | 5.100E-06 |
| 11856 | 593 | K593S | A | 4.9e-06 | 4.7e-06 | 4.2e-06 | 4.600E-06 | 3.606E-07 | 4.200E-06 | 4.900E-06 |
| 11857 | 593 | K593T | A | 5.3e-06 | 3.8e-06 | 4.4e-06 | 4.500E-06 | 7.550E-07 | 3.800E-06 | 5.300E-06 |
| 11858 | 593 | K593W | A | 3.4e-06 | 2.6e-06 | 2.4e-06 | 2.800E-06 | 5.292E-07 | 2.400E-06 | 3.400E-06 |
| 11859 | 593 | K593Y | A | 4.2e-06 | 5.4e-06 | 3.6e-06 | 4.400E-06 | 9.165E-07 | 3.600E-06 | 5.400E-06 |
| 11860 | 593 | K593V | A | 3.7e-06 | 2.1e-06 | 3.3e-06 | 3.033E-06 | 8.327E-07 | 2.100E-06 | 3.700E-06 |
| 11861 | 594 | F594A | A | 3.1e-06 | 3.1e-06 | 3.7e-06 | 3.300E-06 | 3.464E-07 | 3.100E-06 | 3.700E-06 |
| 11862 | 594 | F594R | A | 2e-06 | 2.6e-06 | 1.5e-06 | 2.033E-06 | 5.508E-07 | 1.500E-06 | 2.600E-06 |
| 11863 | 594 | F594N | A | 1e-06 | 5.1e-06 | 2.7e-06 | 2.933E-06 | 2.060E-06 | 1.000E-06 | 5.100E-06 |
| 11864 | 594 | F594D | A | 9.5e-07 | 2.7e-06 | 4.8e-06 | 2.817E-06 | 1.928E-06 | 9.500E-07 | 4.800E-06 |
| 11865 | 594 | F594C | A | 3.2e-06 | 1.9e-06 | 3.3e-06 | 2.800E-06 | 7.810E-07 | 1.900E-06 | 3.300E-06 |
| 11866 | 594 | F594E | A | 1.6e-06 | 3.2e-06 | 2.5e-06 | 2.433E-06 | 8.021E-07 | 1.600E-06 | 3.200E-06 |
| 11867 | 594 | F594Q | A | 1.8e-06 | 1.8e-06 | 2.4e-06 | 2.000E-06 | 3.464E-07 | 1.800E-06 | 2.400E-06 |
| 11868 | 594 | F594G | A | 1.4e-06 | 3.4e-06 | 2e-06 | 2.267E-06 | 1.026E-06 | 1.400E-06 | 3.400E-06 |
| 11869 | 594 | F594H | A | 1.6e-06 | 2.3e-06 | 3.3e-06 | 2.400E-06 | 8.544E-07 | 1.600E-06 | 3.300E-06 |
| 11870 | 594 | F594I | A | 1.9e-06 | 1.9e-06 | 2.5e-06 | 2.100E-06 | 3.464E-07 | 1.900E-06 | 2.500E-06 |
| 11871 | 594 | F594L | A | 2e-06 | 1.7e-06 | 3.2e-06 | 2.300E-06 | 7.937E-07 | 1.700E-06 | 3.200E-06 |
| 11872 | 594 | F594K | A | 2.2e-06 | 1.5e-06 | 2.8e-06 | 2.167E-06 | 6.506E-07 | 1.500E-06 | 2.800E-06 |
| 11873 | 594 | F594M | A | 2e-06 | 2.4e-06 | 2.2e-06 | 2.200E-06 | 2.000E-07 | 2.000E-06 | 2.400E-06 |
| 11874 | 594 | F594F | A | 3.1e-06 | 2e-06 | 3.1e-06 | 2.733E-06 | 6.351E-07 | 2.000E-06 | 3.100E-06 |
| 11875 | 594 | F594P | A | 2.5e-06 | 3.2e-06 | 1.6e-06 | 2.433E-06 | 8.021E-07 | 1.600E-06 | 3.200E-06 |
| 11876 | 594 | F594S | A | 1.5e-06 | 2.1e-06 | 1.1e-06 | 1.567E-06 | 5.033E-07 | 1.100E-06 | 2.100E-06 |
| 11877 | 594 | F594T | A | 3e-06 | 1.8e-06 | 3.2e-06 | 2.667E-06 | 7.572E-07 | 1.800E-06 | 3.200E-06 |
| 11878 | 594 | F594W | A | 2.3e-06 | 1.3e-06 | 1.4e-06 | 1.667E-06 | 5.508E-07 | 1.300E-06 | 2.300E-06 |
| 11879 | 594 | F594Y | A | 1.2e-06 | 4.1e-06 | 2.8e-06 | 2.700E-06 | 1.453E-06 | 1.200E-06 | 4.100E-06 |
| 11880 | 594 | F594V | A | 2.5e-06 | 4.7e-06 | 2.2e-06 | 3.133E-06 | 1.365E-06 | 2.200E-06 | 4.700E-06 |
| 11881 | 595 | P595A | A | 3.2e-06 | 3e-06 | 2e-06 | 2.733E-06 | 6.429E-07 | 2.000E-06 | 3.200E-06 |
| 11882 | 595 | P595R | A | 2.8e-06 | 2e-06 | 3.1e-06 | 2.633E-06 | 5.686E-07 | 2.000E-06 | 3.100E-06 |
| 11883 | 595 | P595N | A | 5.2e-06 | 2.4e-06 | 1.2e-06 | 2.933E-06 | 2.053E-06 | 1.200E-06 | 5.200E-06 |
| 11884 | 595 | P595D | A | 3.7e-06 | 2e-06 | 2.6e-06 | 2.767E-06 | 8.622E-07 | 2.000E-06 | 3.700E-06 |
| 11885 | 595 | P595C | A | 2.1e-06 | 1.6e-06 | 1.1e-06 | 1.600E-06 | 5.000E-07 | 1.100E-06 | 2.100E-06 |
| 11886 | 595 | P595E | A | 2.3e-06 | 1.9e-06 | 3.5e-06 | 2.567E-06 | 8.327E-07 | 1.900E-06 | 3.500E-06 |
| 11887 | 595 | P595Q | A | 2.5e-06 | 2.3e-06 | 2.8e-06 | 2.533E-06 | 2.517E-07 | 2.300E-06 | 2.800E-06 |
| 11888 | 595 | P595G | A | 3e-06 | 1.4e-06 | 3.5e-06 | 2.633E-06 | 1.097E-06 | 1.400E-06 | 3.500E-06 |
| 11889 | 595 | P595H | A | 5.1e-06 | 2.2e-06 | 2.1e-06 | 3.133E-06 | 1.704E-06 | 2.100E-06 | 5.100E-06 |
| 11890 | 595 | P595I | A | 3.9e-06 | 2.2e-06 | 2.5e-06 | 2.867E-06 | 9.074E-07 | 2.200E-06 | 3.900E-06 |
| 11891 | 595 | P595L | A | 1.6e-06 | 1.8e-06 | 1.9e-06 | 1.767E-06 | 1.528E-07 | 1.600E-06 | 1.900E-06 |
| 11892 | 595 | P595K | A | 1.9e-06 | 1.8e-06 | 3.5e-06 | 2.400E-06 | 9.539E-07 | 1.800E-06 | 3.500E-06 |
| 11893 | 595 | P595M | A | 3.4e-06 | 3.1e-06 | 2.6e-06 | 3.033E-06 | 4.041E-07 | 2.600E-06 | 3.400E-06 |
| 11894 | 595 | P595F | A | 1.8e-06 | 2.3e-06 | 3.2e-06 | 2.433E-06 | 7.095E-07 | 1.800E-06 | 3.200E-06 |
| 11895 | 595 | P595P | A | 2.2e-06 | 1.8e-06 | 3.4e-06 | 2.467E-06 | 8.327E-07 | 1.800E-06 | 3.400E-06 |
| 11896 | 595 | P595S | A | 2.4e-06 | 2.2e-06 | 4.6e-06 | 3.067E-06 | 1.332E-06 | 2.200E-06 | 4.600E-06 |
| 11897 | 595 | P595T | A | 2.1e-06 | 3.9e-06 | 1.6e-06 | 2.533E-06 | 1.210E-06 | 1.600E-06 | 3.900E-06 |
| 11898 | 595 | P595W | A | 2.3e-06 | 1.2e-06 | 1.8e-06 | 1.767E-06 | 5.508E-07 | 1.200E-06 | 2.300E-06 |
| 11899 | 595 | P595Y | A | 1.7e-06 | 3.4e-06 | 3e-06 | 2.700E-06 | 8.888E-07 | 1.700E-06 | 3.400E-06 |

| | | | | | | | | | | |
|---|---|---|---|---|---|---|---|---|---|---|
| 11900 | 595 | P595V | A | 2.7e-06 | 1.8e-06 | 1.8e-06 | 2.100E-06 | 5.196E-07 | 1.800E-06 | 2.700E-06 |
| 11901 | 596 | D596A | A | 1.9e-06 | 2.7e-06 | 2.4e-06 | 2.333E-06 | 4.041E-07 | 1.900E-06 | 2.700E-06 |
| 11902 | 596 | D596R | A | 1.8e-06 | 2.2e-06 | 3.6e-06 | 2.533E-06 | 9.452E-07 | 1.800E-06 | 3.600E-06 |
| 11903 | 596 | D596N | A | 9.6e-07 | 2.7e-06 | 3.2e-06 | 2.287E-06 | 1.176E-06 | 9.600E-07 | 3.200E-06 |
| 11904 | 596 | D596D | A | 2.5e-06 | 3.2e-06 | 4.1e-06 | 3.267E-06 | 8.021E-07 | 2.500E-06 | 4.100E-06 |
| 11905 | 596 | D596C | A | 1.8e-06 | 1.4e-06 | 1.5e-06 | 1.567E-06 | 2.082E-07 | 1.400E-06 | 1.800E-06 |
| 11906 | 596 | D596E | A | 2.9e-06 | 1.8e-06 | 2e-06 | 2.233E-06 | 5.859E-07 | 1.800E-06 | 2.900E-06 |
| 11907 | 596 | D596Q | A | 3.2e-06 | 3.1e-06 | 2.7e-06 | 3.000E-06 | 2.646E-07 | 2.700E-06 | 3.200E-06 |
| 11908 | 596 | D596G | A | 3.9e-06 | 3e-06 | 2.5e-06 | 3.133E-06 | 7.095E-07 | 2.500E-06 | 3.900E-06 |
| 11909 | 596 | D596H | A | 3.6e-06 | 3.7e-06 | 3.2e-06 | 3.500E-06 | 2.646E-07 | 3.200E-06 | 3.700E-06 |
| 11910 | 596 | D596I | A | 3.1e-06 | 4.6e-06 | 3.7e-06 | 3.800E-06 | 7.550E-07 | 3.100E-06 | 4.600E-06 |
| 11911 | 596 | D596L | A | 2.3e-06 | 4e-06 | 3.1e-06 | 3.133E-06 | 8.505E-07 | 2.300E-06 | 4.000E-06 |
| 11912 | 596 | D596K | A | 2.1e-06 | 1.6e-06 | 3.7e-06 | 2.467E-06 | 1.097E-06 | 1.600E-06 | 3.700E-06 |
| 11913 | 596 | D596M | A | 2.3e-06 | 1.5e-06 | 1.6e-06 | 1.800E-06 | 4.359E-07 | 1.500E-06 | 2.300E-06 |
| 11914 | 596 | D596F | A | 3.3e-06 | 2.1e-06 | 1.8e-06 | 2.400E-06 | 7.937E-07 | 1.800E-06 | 3.300E-06 |
| 11915 | 596 | D596P | A | 2.6e-06 | 3.7e-06 | 3.1e-06 | 3.133E-06 | 5.508E-07 | 2.600E-06 | 3.700E-06 |
| 11916 | 596 | D596S | A | 1.9e-06 | 2.3e-06 | 2e-06 | 2.067E-06 | 2.082E-07 | 1.900E-06 | 2.300E-06 |
| 11917 | 596 | D596T | A | 3.6e-06 | 2.6e-06 | 2.4e-06 | 2.867E-06 | 6.429E-07 | 2.400E-06 | 3.600E-06 |
| 11918 | 596 | D596W | A | 2.7e-06 | 3.5e-06 | 3.8e-06 | 3.333E-06 | 5.686E-07 | 2.700E-06 | 3.800E-06 |
| 11919 | 596 | D596Y | A | 2e-06 | 2.2e-06 | 1.2e-06 | 1.800E-06 | 5.292E-07 | 1.200E-06 | 2.200E-06 |
| 11920 | 596 | D596V | A | 2.2e-06 | 3.4e-06 | 2.6e-06 | 2.733E-06 | 6.110E-07 | 2.200E-06 | 3.400E-06 |
| 11921 | 597 | E597A | A | 3.3e-06 | 2.9e-06 | 2.3e-06 | 2.833E-06 | 5.033E-07 | 2.300E-06 | 3.300E-06 |
| 11922 | 597 | E597R | A | 1.9e-06 | 1.6e-06 | 2.3e-06 | 1.933E-06 | 3.512E-07 | 1.600E-06 | 2.300E-06 |
| 11923 | 597 | E597N | A | 3.4e-06 | 2.6e-06 | 4.2e-06 | 3.400E-06 | 8.000E-07 | 2.600E-06 | 4.200E-06 |
| 11924 | 597 | E597D | A | 2.7e-06 | 2.2e-06 | 2.4e-06 | 2.433E-06 | 2.517E-07 | 2.200E-06 | 2.700E-06 |
| 11925 | 597 | E597C | A | 2.4e-06 | 1.7e-06 | 3.7e-06 | 2.600E-06 | 1.015E-06 | 1.700E-06 | 3.700E-06 |
| 11926 | 597 | E597E | A | 2.4e-06 | 2.6e-06 | 1.6e-06 | 2.200E-06 | 5.292E-07 | 1.600E-06 | 2.600E-06 |
| 11927 | 597 | E597Q | A | 1.9e-06 | 2.7e-06 | 1.8e-06 | 2.133E-06 | 4.933E-07 | 1.800E-06 | 2.700E-06 |
| 11928 | 597 | E597G | A | 3.3e-06 | 2.5e-06 | 3.1e-06 | 2.967E-06 | 4.163E-07 | 2.500E-06 | 3.300E-06 |
| 11929 | 597 | E597H | A | 3.1e-06 | 2.6e-06 | 2.6e-06 | 2.767E-06 | 2.887E-07 | 2.600E-06 | 3.100E-06 |
| 11930 | 597 | E597I | A | 1.7e-06 | 1.3e-06 | 2.7e-06 | 1.900E-06 | 7.211E-07 | 1.300E-06 | 2.700E-06 |
| 11931 | 597 | E597L | A | 2.2e-06 | 1.3e-06 | 1.9e-06 | 1.800E-06 | 4.583E-07 | 1.300E-06 | 2.200E-06 |
| 11932 | 597 | E597K | A | 1.1e-06 | 1.5e-06 | 2.6e-06 | 1.733E-06 | 7.767E-07 | 1.100E-06 | 2.600E-06 |
| 11933 | 597 | E597M | A | 2.3e-06 | 2.7e-06 | 1.2e-06 | 2.067E-06 | 7.767E-07 | 1.200E-06 | 2.700E-06 |
| 11934 | 597 | E597F | A | 1.8e-06 | 1.8e-06 | 1.7e-06 | 1.767E-06 | 5.774E-08 | 1.700E-06 | 1.800E-06 |
| 11935 | 597 | E597P | A | 2.3e-06 | 4.1e-06 | 1.2e-06 | 2.533E-06 | 1.464E-06 | 1.200E-06 | 4.100E-06 |
| 11936 | 597 | E597S | A | 2.2e-06 | 2.5e-06 | 4.6e-06 | 3.100E-06 | 1.308E-06 | 2.200E-06 | 4.600E-06 |
| 11937 | 597 | E597T | A | 2.8e-06 | 1.4e-06 | 1.5e-06 | 1.900E-06 | 7.810E-07 | 1.400E-06 | 2.800E-06 |
| 11938 | 597 | E597W | A | 2.3e-06 | 2.9e-06 | 3.5e-06 | 2.900E-06 | 6.000E-07 | 2.300E-06 | 3.500E-06 |
| 11939 | 597 | E597Y | A | 1.5e-06 | 2.6e-06 | 2.1e-06 | 2.067E-06 | 5.508E-07 | 1.500E-06 | 2.600E-06 |
| 11940 | 597 | E597V | A | 1.7e-06 | 2.3e-06 | 3.4e-06 | 2.467E-06 | 8.622E-07 | 1.700E-06 | 3.400E-06 |
| 11941 | 598 | E598A | A | 2.8e-06 | 1.2e-06 | 1.6e-06 | 1.867E-06 | 8.327E-07 | 1.200E-06 | 2.800E-06 |
| 11942 | 598 | E598R | A | 2.3e-06 | 1.7e-06 | 2.2e-06 | 2.067E-06 | 3.215E-07 | 1.700E-06 | 2.300E-06 |
| 11943 | 598 | E598N | A | 1.6e-06 | 2e-06 | 3e-06 | 2.200E-06 | 7.211E-07 | 1.600E-06 | 3.000E-06 |
| 11944 | 598 | E598D | A | 1.9e-06 | 1.3e-06 | 2e-06 | 1.733E-06 | 3.786E-07 | 1.300E-06 | 2.000E-06 |
| 11945 | 598 | E598C | A | 1.8e-06 | 2.9e-06 | 1.7e-06 | 2.133E-06 | 6.658E-07 | 1.700E-06 | 2.900E-06 |
| 11946 | 598 | E598E | A | 2.7e-06 | 2.3e-06 | 2.3e-06 | 2.433E-06 | 2.309E-07 | 2.300E-06 | 2.700E-06 |
| 11947 | 598 | E598Q | A | 1.8e-06 | 1.6e-06 | 2.1e-06 | 1.833E-06 | 2.517E-07 | 1.600E-06 | 2.100E-06 |
| 11948 | 598 | E598G | A | 2.5e-06 | 1.5e-06 | 2.2e-06 | 2.067E-06 | 5.132E-07 | 1.500E-06 | 2.500E-06 |
| 11949 | 598 | E598H | A | 2.4e-06 | 2.4e-06 | 1.9e-06 | 2.233E-06 | 2.887E-07 | 1.900E-06 | 2.400E-06 |
| 11950 | 598 | E598I | A | 1.6e-06 | 4.9e-06 | 2e-06 | 2.833E-06 | 1.801E-06 | 1.600E-06 | 4.900E-06 |
| 11951 | 598 | E598L | A | 4.1e-06 | 3.1e-06 | 1.6e-06 | 2.933E-06 | 1.258E-06 | 1.600E-06 | 4.100E-06 |
| 11952 | 598 | E598K | A | 2.6e-06 | 4.5e-06 | 5e-06 | 4.033E-06 | 1.266E-06 | 2.600E-06 | 5.000E-06 |
| 11953 | 598 | E598M | A | 1.9e-06 | 3.2e-06 | 4e-06 | 3.033E-06 | 1.060E-06 | 1.900E-06 | 4.000E-06 |
| 11954 | 598 | E598F | A | 2.4e-06 | 2.6e-06 | 2.2e-06 | 2.400E-06 | 2.000E-07 | 2.200E-06 | 2.600E-06 |
| 11955 | 598 | E598P | A | 2.4e-06 | 2.6e-06 | 1.3e-06 | 2.100E-06 | 7.000E-07 | 1.300E-06 | 2.600E-06 |
| 11956 | 598 | E598S | A | 2.7e-06 | 2.4e-06 | 2.9e-06 | 2.667E-06 | 2.517E-07 | 2.400E-06 | 2.900E-06 |
| 11957 | 598 | E598T | A | 1.7e-06 | 4e-06 | 3.6e-06 | 3.100E-06 | 1.229E-06 | 1.700E-06 | 4.000E-06 |
| 11958 | 598 | E598W | A | 2.6e-06 | 3.1e-06 | 2e-06 | 2.567E-06 | 5.508E-07 | 2.000E-06 | 3.100E-06 |
| 11959 | 598 | E598Y | A | 2.7e-06 | 2.7e-06 | 4e-06 | 3.133E-06 | 7.506E-07 | 2.700E-06 | 4.000E-06 |
| 11960 | 598 | E598V | A | 3.1e-06 | 3e-06 | 3.2e-06 | 3.100E-06 | 1.000E-07 | 3.000E-06 | 3.200E-06 |
| 11961 | 599 | G599A | A | 2.8e-06 | 2.1e-06 | 2.2e-06 | 2.367E-06 | 3.786E-07 | 2.100E-06 | 2.800E-06 |
| 11962 | 599 | G599R | A | 2.8e-06 | 3e-06 | 3e-06 | 2.933E-06 | 1.155E-07 | 2.800E-06 | 3.000E-06 |
| 11963 | 599 | G599N | A | 2.5e-06 | 1.7e-06 | 1.3e-06 | 1.833E-06 | 6.110E-07 | 1.300E-06 | 2.500E-06 |
| 11964 | 599 | G599D | A | 2.7e-06 | 2.9e-06 | 2.3e-06 | 2.633E-06 | 3.055E-07 | 2.300E-06 | 2.900E-06 |
| 11965 | 599 | G599C | A | 2.9e-06 | 2.4e-06 | 1e-06 | 2.100E-06 | 9.849E-07 | 1.000E-06 | 2.900E-06 |
| 11966 | 599 | G599E | A | 1.7e-06 | 1.5e-06 | 3.6e-06 | 2.267E-06 | 1.159E-06 | 1.500E-06 | 3.600E-06 |
| 11967 | 599 | G599Q | A | 1.8e-06 | 4.4e-06 | 2.5e-06 | 2.900E-06 | 1.345E-06 | 1.800E-06 | 4.400E-06 |
| 11968 | 599 | G599G | A | 3.3e-06 | 1.6e-06 | 2.2e-06 | 2.367E-06 | 8.622E-07 | 1.600E-06 | 3.300E-06 |
| 11969 | 599 | G599H | A | 2.4e-06 | 1.9e-06 | 2.6e-06 | 2.300E-06 | 3.606E-07 | 1.900E-06 | 2.600E-06 |
| 11970 | 599 | G599I | A | 1.5e-06 | 2.2e-06 | 2.4e-06 | 2.033E-06 | 4.726E-07 | 1.500E-06 | 2.400E-06 |
| 11971 | 599 | G599L | A | 1.8e-06 | 3.2e-06 | 2.6e-06 | 2.533E-06 | 7.024E-07 | 1.800E-06 | 3.200E-06 |
| 11972 | 599 | G599K | A | 2.8e-06 | 1.6e-06 | 1.1e-06 | 1.833E-06 | 8.737E-07 | 1.100E-06 | 2.800E-06 |
| 11973 | 599 | G599M | A | 2.2e-06 | 2e-06 | 3.1e-06 | 2.433E-06 | 5.859E-07 | 2.000E-06 | 3.100E-06 |
| 11974 | 599 | G599F | A | 1.7e-06 | 2.4e-06 | 1.7e-06 | 1.933E-06 | 4.041E-07 | 1.700E-06 | 2.400E-06 |
| 11975 | 599 | G599P | A | 1.5e-06 | 1.9e-06 | 1.6e-06 | 1.667E-06 | 2.082E-07 | 1.500E-06 | 1.900E-06 |
| 11976 | 599 | G599S | A | 2.8e-06 | 2.5e-06 | 2.6e-06 | 2.633E-06 | 1.528E-07 | 2.500E-06 | 2.800E-06 |
| 11977 | 599 | G599T | A | 2.7e-06 | 4.6e-06 | 3.1e-06 | 3.467E-06 | 1.002E-06 | 2.700E-06 | 4.600E-06 |
| 11978 | 599 | G599W | A | 3.7e-06 | 1.6e-06 | 2.8e-06 | 2.700E-06 | 1.054E-06 | 1.600E-06 | 3.700E-06 |
| 11979 | 599 | G599Y | A | 2.6e-06 | 1.9e-06 | 4.1e-06 | 2.867E-06 | 1.124E-06 | 1.900E-06 | 4.100E-06 |
| 11980 | 599 | G599V | A | 2.1e-06 | 2.7e-06 | 2.2e-06 | 2.333E-06 | 3.215E-07 | 2.100E-06 | 2.700E-06 |
| 11981 | 600 | A600A | A | 2.6e-06 | 4e-06 | 2.7e-06 | 3.100E-06 | 7.810E-07 | 2.600E-06 | 4.000E-06 |
| 11982 | 600 | A600R | A | 2.7e-06 | 1.8e-06 | 2.5e-06 | 2.333E-06 | 4.726E-07 | 1.800E-06 | 2.700E-06 |
| 11983 | 600 | A600N | A | 2.3e-06 | 2e-06 | 1.7e-06 | 2.000E-06 | 3.000E-07 | 1.700E-06 | 2.300E-06 |
| 11984 | 600 | A600D | A | 3.4e-06 | 2.7e-06 | 2.2e-06 | 2.767E-06 | 6.028E-07 | 2.200E-06 | 3.400E-06 |

| | | | | | | | | | | |
|---|---|---|---|---|---|---|---|---|---|---|
| 11985 | 600 | A600C | A | 2e-06 | 3.4e-06 | 2.5e-06 | 2.633E-06 | 7.095E-07 | 2.000E-06 | 3.400E-06 |
| 11986 | 600 | A600E | A | 5.9e-06 | 2e-06 | 2.3e-06 | 3.400E-06 | 2.170E-06 | 2.000E-06 | 5.900E-06 |
| 11987 | 600 | A600Q | A | 2.4e-06 | 1.9e-06 | 2.9e-06 | 2.400E-06 | 5.000E-07 | 1.900E-06 | 2.900E-06 |
| 11988 | 600 | A600G | A | 3.2e-06 | 1.7e-06 | 1.5e-06 | 2.133E-06 | 9.292E-07 | 1.500E-06 | 3.200E-06 |
| 11989 | 600 | A600H | A | 1.7e-06 | 2.7e-06 | 1.9e-06 | 2.100E-06 | 5.292E-07 | 1.700E-06 | 2.700E-06 |
| 11990 | 600 | A600I | A | 4.5e-06 | 1.4e-06 | 2.7e-06 | 2.867E-06 | 1.557E-06 | 1.400E-06 | 4.500E-06 |
| 11991 | 600 | A600L | A | 1.4e-06 | 1.3e-06 | 2.4e-06 | 1.700E-06 | 6.083E-07 | 1.300E-06 | 2.400E-06 |
| 11992 | 600 | A600K | A | 2.2e-06 | 1.5e-06 | 2.2e-06 | 1.967E-06 | 4.041E-07 | 1.500E-06 | 2.200E-06 |
| 11993 | 600 | A600M | A | 3.1e-06 | 2.9e-06 | 3.2e-06 | 3.067E-06 | 1.528E-07 | 2.900E-06 | 3.200E-06 |
| 11994 | 600 | A600F | A | 2.9e-06 | 3e-06 | 3.8e-06 | 3.233E-06 | 4.933E-07 | 2.900E-06 | 3.800E-06 |
| 11995 | 600 | A600P | A | 2.1e-06 | 1.5e-06 | 2.5e-06 | 2.033E-06 | 5.033E-07 | 1.500E-06 | 2.500E-06 |
| 11996 | 600 | A600S | A | 2.4e-06 | 1.2e-06 | 5e-06 | 2.867E-06 | 1.943E-06 | 1.200E-06 | 5.000E-06 |
| 11997 | 600 | A600T | A | 2.7e-06 | 2.6e-06 | 1.6e-06 | 2.300E-06 | 6.083E-07 | 1.600E-06 | 2.700E-06 |
| 11998 | 600 | A600W | A | 3e-06 | 2.7e-06 | 1.3e-06 | 2.333E-06 | 9.074E-07 | 1.300E-06 | 3.000E-06 |
| 11999 | 600 | A600Y | A | 2.1e-06 | 2.7e-06 | 2.4e-06 | 2.400E-06 | 3.000E-07 | 2.100E-06 | 2.700E-06 |
| 12000 | 600 | A600V | A | 1.2e-06 | 1.8e-06 | 2e-06 | 1.667E-06 | 4.163E-07 | 1.200E-06 | 2.000E-06 |
| 12001 | 601 | C601A | A | 1.4e-06 | 1e-06 | 3.3e-06 | 1.900E-06 | 1.229E-06 | 1.000E-06 | 3.300E-06 |
| 12002 | 601 | C601R | A | 1.6e-06 | 7.9e-07 | 1.8e-06 | 1.397E-06 | 5.348E-07 | 7.900E-07 | 1.800E-06 |
| 12003 | 601 | C601N | A | 2.2e-06 | 1.5e-06 | 3.2e-06 | 2.300E-06 | 8.544E-07 | 1.500E-06 | 3.200E-06 |
| 12004 | 601 | C601D | A | 1.5e-06 | 2.6e-06 | 3.1e-06 | 2.400E-06 | 8.185E-07 | 1.500E-06 | 3.100E-06 |
| 12005 | 601 | C601C | A | 2.8e-06 | 4.9e-06 | 2.6e-06 | 3.433E-06 | 1.274E-06 | 2.600E-06 | 4.900E-06 |
| 12006 | 601 | C601E | A | 2.1e-06 | 1.4e-06 | 2.1e-06 | 1.867E-06 | 4.041E-07 | 1.400E-06 | 2.100E-06 |
| 12007 | 601 | C601Q | A | 3.6e-06 | 2e-06 | 2e-06 | 2.533E-06 | 9.238E-07 | 2.000E-06 | 3.600E-06 |
| 12008 | 601 | C601G | A | 2.4e-06 | 3.3e-06 | 1.4e-06 | 2.367E-06 | 9.504E-07 | 1.400E-06 | 3.300E-06 |
| 12009 | 601 | C601H | A | 2.1e-06 | 2.2e-06 | 1.9e-06 | 2.067E-06 | 1.528E-07 | 1.900E-06 | 2.200E-06 |
| 12010 | 601 | C601I | A | 1.5e-06 | 2.3e-06 | 2.3e-06 | 2.033E-06 | 4.619E-07 | 1.500E-06 | 2.300E-06 |
| 12011 | 601 | C601L | A | 2.4e-06 | 3.1e-06 | 2.3e-06 | 2.600E-06 | 4.359E-07 | 2.300E-06 | 3.100E-06 |
| 12012 | 601 | C601K | A | 2.4e-06 | 1.3e-06 | 2e-06 | 1.900E-06 | 5.568E-07 | 1.300E-06 | 2.400E-06 |
| 12013 | 601 | C601M | A | 1.7e-06 | 3.5e-06 | 2.4e-06 | 2.533E-06 | 9.074E-07 | 1.700E-06 | 3.500E-06 |
| 12014 | 601 | C601F | A | 1.9e-06 | 2.5e-06 | 3.6e-06 | 2.667E-06 | 8.622E-07 | 1.900E-06 | 3.600E-06 |
| 12015 | 601 | C601P | A | 2.7e-06 | 2.6e-06 | 2.3e-06 | 2.533E-06 | 2.082E-07 | 2.300E-06 | 2.700E-06 |
| 12016 | 601 | C601S | A | 2.6e-06 | 1.4e-06 | 4e-06 | 2.667E-06 | 1.301E-06 | 1.400E-06 | 4.000E-06 |
| 12017 | 601 | C601T | A | 2.9e-06 | 2.7e-06 | 2.6e-06 | 2.733E-06 | 1.528E-07 | 2.600E-06 | 2.900E-06 |
| 12018 | 601 | C601W | A | 1.8e-06 | 5e-06 | 1.8e-06 | 2.867E-06 | 1.848E-06 | 1.800E-06 | 5.000E-06 |
| 12019 | 601 | C601Y | A | 2e-06 | 3e-06 | 1.9e-06 | 2.300E-06 | 6.083E-07 | 1.900E-06 | 3.000E-06 |
| 12020 | 601 | C601V | A | 3.1e-06 | 1.4e-06 | 2.2e-06 | 2.233E-06 | 8.505E-07 | 1.400E-06 | 3.100E-06 |
| 12021 | 602 | Q602A | A | 4.4e-06 | 5.3e-06 | 2.1e-06 | 3.933E-06 | 1.650E-06 | 2.100E-06 | 5.300E-06 |
| 12022 | 602 | Q602R | A | 2.8e-06 | 4.3e-06 | 2.3e-06 | 3.133E-06 | 1.041E-06 | 2.300E-06 | 4.300E-06 |
| 12023 | 602 | Q602N | A | 2.6e-06 | 2.9e-06 | 4.2e-06 | 3.233E-06 | 8.505E-07 | 2.600E-06 | 4.200E-06 |
| 12024 | 602 | Q602D | A | 2.5e-06 | 2.5e-06 | 3.7e-06 | 2.900E-06 | 6.928E-07 | 2.500E-06 | 3.700E-06 |
| 12025 | 602 | Q602C | A | 2.4e-06 | 2.7e-06 | 3.5e-06 | 2.867E-06 | 5.686E-07 | 2.400E-06 | 3.500E-06 |
| 12026 | 602 | Q602E | A | 2.7e-06 | 2.9e-06 | 2.8e-06 | 2.800E-06 | 1.000E-07 | 2.700E-06 | 2.900E-06 |
| 12027 | 602 | Q602Q | A | 5.2e-06 | 1.8e-06 | 3e-06 | 3.333E-06 | 1.724E-06 | 1.800E-06 | 5.200E-06 |
| 12028 | 602 | Q602G | A | 2.2e-06 | 2e-06 | 2.4e-06 | 2.200E-06 | 2.000E-07 | 2.000E-06 | 2.400E-06 |
| 12029 | 602 | Q602H | A | 2.9e-06 | 2.8e-06 | 3.2e-06 | 2.967E-06 | 2.082E-07 | 2.800E-06 | 3.200E-06 |
| 12030 | 602 | Q602I | A | 1.9e-06 | 2.3e-06 | 3.9e-06 | 2.700E-06 | 1.058E-06 | 1.900E-06 | 3.900E-06 |
| 12031 | 602 | Q602L | A | 2.2e-06 | 2.6e-06 | 2.8e-06 | 2.533E-06 | 3.055E-07 | 2.200E-06 | 2.800E-06 |
| 12032 | 602 | Q602K | A | 2e-06 | 2.4e-06 | 3.4e-06 | 2.600E-06 | 7.211E-07 | 2.000E-06 | 3.400E-06 |
| 12033 | 602 | Q602M | A | 3.7e-06 | 2.1e-06 | 4.1e-06 | 3.300E-06 | 1.058E-06 | 2.100E-06 | 4.100E-06 |
| 12034 | 602 | Q602F | A | 1.3e-06 | 1.3e-06 | 2.1e-06 | 1.567E-06 | 4.619E-07 | 1.300E-06 | 2.100E-06 |
| 12035 | 602 | Q602P | A | 2.7e-06 | 3.1e-06 | 1.8e-06 | 2.533E-06 | 6.658E-07 | 1.800E-06 | 3.100E-06 |
| 12036 | 602 | Q602S | A | 2.6e-06 | 2.5e-06 | 2.3e-06 | 2.467E-06 | 1.528E-07 | 2.300E-06 | 2.600E-06 |
| 12037 | 602 | Q602T | A | 1.3e-06 | 2.2e-06 | 2.7e-06 | 2.067E-06 | 7.095E-07 | 1.300E-06 | 2.700E-06 |
| 12038 | 602 | Q602W | A | 2.9e-06 | 1e-06 | 1.6e-06 | 1.833E-06 | 9.713E-07 | 1.000E-06 | 2.900E-06 |
| 12039 | 602 | Q602Y | A | 2e-06 | 2.9e-06 | 2.6e-06 | 2.500E-06 | 4.583E-07 | 2.000E-06 | 2.900E-06 |
| 12040 | 602 | Q602V | A | 4.3e-06 | 3e-06 | 2.9e-06 | 3.400E-06 | 7.810E-07 | 2.900E-06 | 4.300E-06 |
| 12041 | 603 | P603A | A | 2.9e-06 | 1.9e-06 | 2.3e-06 | 2.367E-06 | 5.033E-07 | 1.900E-06 | 2.900E-06 |
| 12042 | 603 | P603R | A | 2.1e-06 | 1.8e-06 | 6.9e-07 | 1.530E-06 | 7.428E-07 | 6.900E-07 | 2.100E-06 |
| 12043 | 603 | P603N | A | 2.8e-06 | 1.6e-06 | 2.4e-06 | 2.267E-06 | 6.110E-07 | 1.600E-06 | 2.800E-06 |
| 12044 | 603 | P603D | A | 2.4e-06 | 2.8e-06 | 2.8e-06 | 2.667E-06 | 2.309E-07 | 2.400E-06 | 2.800E-06 |
| 12045 | 603 | P603C | A | 2.1e-06 | 1.7e-06 | 3.7e-06 | 2.500E-06 | 1.058E-06 | 1.700E-06 | 3.700E-06 |
| 12046 | 603 | P603E | A | 1.3e-06 | 2.2e-06 | 2.7e-06 | 2.067E-06 | 7.095E-07 | 1.300E-06 | 2.700E-06 |
| 12047 | 603 | P603Q | A | 3.4e-06 | 2.2e-06 | 1e-06 | 2.200E-06 | 1.200E-06 | 1.000E-06 | 3.400E-06 |
| 12048 | 603 | P603G | A | 2.1e-06 | 2.1e-06 | 4.5e-06 | 2.900E-06 | 1.386E-06 | 2.100E-06 | 4.500E-06 |
| 12049 | 603 | P603H | A | 2.5e-06 | 2.4e-06 | 3.1e-06 | 2.667E-06 | 3.786E-07 | 2.400E-06 | 3.100E-06 |
| 12050 | 603 | P603I | A | 2.2e-06 | 1.4e-06 | 2.8e-06 | 2.133E-06 | 7.024E-07 | 1.400E-06 | 2.800E-06 |
| 12051 | 603 | P603L | A | 2.3e-06 | 2.6e-06 | 2.6e-06 | 2.500E-06 | 1.732E-07 | 2.300E-06 | 2.600E-06 |
| 12052 | 603 | P603K | A | 1.6e-06 | 2.1e-06 | 2e-06 | 1.900E-06 | 2.646E-07 | 1.600E-06 | 2.100E-06 |
| 12053 | 603 | P603M | A | 2.3e-06 | 2.6e-06 | 2.1e-06 | 2.333E-06 | 2.517E-07 | 2.100E-06 | 2.600E-06 |
| 12054 | 603 | P603F | A | 1.8e-06 | 3.3e-06 | 3.2e-06 | 2.767E-06 | 8.386E-07 | 1.800E-06 | 3.300E-06 |
| 12055 | 603 | P603P | A | 1.7e-06 | 3.1e-06 | 1.8e-06 | 2.200E-06 | 7.810E-07 | 1.700E-06 | 3.100E-06 |
| 12056 | 603 | P603S | A | 2.5e-06 | 2.3e-06 | 2.8e-06 | 2.533E-06 | 2.517E-07 | 2.300E-06 | 2.800E-06 |
| 12057 | 603 | P603T | A | 1.3e-06 | 3.2e-06 | 2.5e-06 | 2.333E-06 | 9.609E-07 | 1.300E-06 | 3.200E-06 |
| 12058 | 603 | P603W | A | 2.9e-06 | 2.1e-06 | 1.7e-06 | 2.233E-06 | 6.110E-07 | 1.700E-06 | 2.900E-06 |
| 12059 | 603 | P603Y | A | 2e-06 | 1.8e-06 | 1.7e-06 | 1.833E-06 | 1.528E-07 | 1.700E-06 | 2.000E-06 |
| 12060 | 603 | P603V | A | 2.5e-06 | 2.7e-06 | 3.1e-06 | 2.767E-06 | 3.055E-07 | 2.500E-06 | 3.100E-06 |
| 12061 | 604 | C604A | A | 1.8e-06 | 2.9e-06 | 2.9e-06 | 2.533E-06 | 6.351E-07 | 1.800E-06 | 2.900E-06 |
| 12062 | 604 | C604R | A | 1.5e-06 | 4.4e-06 | 2.2e-06 | 2.700E-06 | 1.513E-06 | 1.500E-06 | 4.400E-06 |
| 12063 | 604 | C604N | A | 2.8e-06 | 2.2e-06 | 3e-06 | 2.667E-06 | 4.163E-07 | 2.200E-06 | 3.000E-06 |
| 12064 | 604 | C604D | A | 2.8e-06 | 3e-06 | 3.2e-06 | 3.000E-06 | 2.000E-07 | 2.800E-06 | 3.200E-06 |
| 12065 | 604 | C604C | A | 3.3e-06 | 2.7e-06 | 1.8e-06 | 2.600E-06 | 7.550E-07 | 1.800E-06 | 3.300E-06 |
| 12066 | 604 | C604E | A | 2e-06 | 2.7e-06 | 7.9e-07 | 1.830E-06 | 9.663E-07 | 7.900E-07 | 2.700E-06 |
| 12067 | 604 | C604Q | A | 2.5e-06 | 2.8e-06 | 2.8e-06 | 2.700E-06 | 1.732E-07 | 2.500E-06 | 2.800E-06 |
| 12068 | 604 | C604G | A | 3.2e-06 | 2.2e-06 | 2.3e-06 | 2.567E-06 | 5.508E-07 | 2.200E-06 | 3.200E-06 |
| 12069 | 604 | C604H | A | 1.6e-06 | 3.6e-06 | 2.4e-06 | 2.533E-06 | 1.007E-06 | 1.600E-06 | 3.600E-06 |

| | | | | | | | | | | |
|---|---|---|---|---|---|---|---|---|---|---|
| 12070 | 604 | C604I | A | 3e-06 | 3.2e-06 | 4.3e-06 | 3.500E-06 | 7.000E-07 | 3.000E-06 | 4.300E-06 |
| 12071 | 604 | C604L | A | 3.3e-06 | 3e-06 | 2e-06 | 2.767E-06 | 6.807E-07 | 2.000E-06 | 3.300E-06 |
| 12072 | 604 | C604K | A | 1.7e-06 | 3.9e-06 | 2.9e-06 | 2.833E-06 | 1.102E-06 | 1.700E-06 | 3.900E-06 |
| 12073 | 604 | C604M | A | 1.4e-06 | 2.9e-06 | 2.4e-06 | 2.233E-06 | 7.638E-07 | 1.400E-06 | 2.900E-06 |
| 12074 | 604 | C604F | A | 3.2e-06 | 9.4e-07 | 2.9e-06 | 2.347E-06 | 1.227E-06 | 9.400E-07 | 3.200E-06 |
| 12075 | 604 | C604P | A | 3.9e-06 | 2.1e-06 | 2.9e-06 | 2.967E-06 | 9.018E-07 | 2.100E-06 | 3.900E-06 |
| 12076 | 604 | C604S | A | 2.5e-06 | 3e-06 | 3.8e-06 | 3.100E-06 | 6.557E-07 | 2.500E-06 | 3.800E-06 |
| 12077 | 604 | C604T | A | 2e-06 | 1.5e-06 | 2.6e-06 | 2.033E-06 | 5.508E-07 | 1.500E-06 | 2.600E-06 |
| 12078 | 604 | C604W | A | 2.4e-06 | 2.4e-06 | 2.2e-06 | 2.333E-06 | 1.155E-07 | 2.200E-06 | 2.400E-06 |
| 12079 | 604 | C604Y | A | 2e-06 | 2.5e-06 | 3.2e-06 | 2.567E-06 | 6.028E-07 | 2.000E-06 | 3.200E-06 |
| 12080 | 604 | C604V | A | 2.7e-06 | 1.6e-06 | 2.8e-06 | 2.367E-06 | 6.658E-07 | 1.600E-06 | 2.800E-06 |
| 12081 | 605 | P605A | A | 1.7e-06 | 1.7e-06 | 2e-06 | 1.800E-06 | 1.732E-07 | 1.700E-06 | 2.000E-06 |
| 12082 | 605 | P605R | A | 4.4e-06 | 1.8e-06 | 1.8e-06 | 2.667E-06 | 1.501E-06 | 1.800E-06 | 4.400E-06 |
| 12083 | 605 | P605N | A | 1.7e-06 | 2.1e-06 | 3.1e-06 | 2.300E-06 | 7.211E-07 | 1.700E-06 | 3.100E-06 |
| 12084 | 605 | P605D | A | 2.2e-06 | 2.1e-06 | 2.1e-06 | 2.133E-06 | 5.774E-08 | 2.100E-06 | 2.200E-06 |
| 12085 | 605 | P605C | A | 2.3e-06 | 3.1e-06 | 2.4e-06 | 2.600E-06 | 4.359E-07 | 2.300E-06 | 3.100E-06 |
| 12086 | 605 | P605E | A | 2.2e-06 | 3.4e-06 | 2.9e-06 | 2.833E-06 | 6.028E-07 | 2.200E-06 | 3.400E-06 |
| 12087 | 605 | P605Q | A | 2.2e-06 | 1.7e-06 | 3.7e-06 | 2.533E-06 | 1.041E-06 | 1.700E-06 | 3.700E-06 |
| 12088 | 605 | P605G | A | 1.2e-06 | 1.9e-06 | 3.7e-06 | 2.267E-06 | 1.290E-06 | 1.200E-06 | 3.700E-06 |
| 12089 | 605 | P605H | A | 2.5e-06 | 3e-06 | 1.6e-06 | 2.367E-06 | 7.095E-07 | 1.600E-06 | 3.000E-06 |
| 12090 | 605 | P605I | A | 1e-06 | 2.1e-06 | 2.2e-06 | 1.767E-06 | 6.658E-07 | 1.000E-06 | 2.200E-06 |
| 12091 | 605 | P605L | A | 3.3e-06 | 3.3e-06 | 2.1e-06 | 2.900E-06 | 6.928E-07 | 2.100E-06 | 3.300E-06 |
| 12092 | 605 | P605K | A | 2e-06 | 3.7e-06 | 2.8e-06 | 2.833E-06 | 8.505E-07 | 2.000E-06 | 3.700E-06 |
| 12093 | 605 | P605M | A | 3.2e-06 | 2.7e-06 | 1.8e-06 | 2.567E-06 | 7.095E-07 | 1.800E-06 | 3.200E-06 |
| 12094 | 605 | P605F | A | 2e-06 | 2.3e-06 | 4.7e-06 | 3.000E-06 | 1.480E-06 | 2.000E-06 | 4.700E-06 |
| 12095 | 605 | P605P | A | 2.6e-06 | 2.5e-06 | 2.3e-06 | 2.467E-06 | 1.528E-07 | 2.300E-06 | 2.600E-06 |
| 12096 | 605 | P605S | A | 3e-06 | 2.6e-06 | 2.2e-06 | 2.600E-06 | 4.000E-07 | 2.200E-06 | 3.000E-06 |
| 12097 | 605 | P605T | A | 2.5e-06 | 4.3e-06 | 2e-06 | 2.933E-06 | 1.210E-06 | 2.000E-06 | 4.300E-06 |
| 12098 | 605 | P605W | A | 2.1e-06 | 2.1e-06 | 3.4e-06 | 2.533E-06 | 7.506E-07 | 2.100E-06 | 3.400E-06 |
| 12099 | 605 | P605Y | A | 2.1e-06 | 3e-06 | 1.8e-06 | 2.300E-06 | 6.245E-07 | 1.800E-06 | 3.000E-06 |
| 12100 | 605 | P605V | A | 2.2e-06 | 1.1e-06 | 2.2e-06 | 1.833E-06 | 6.351E-07 | 1.100E-06 | 2.200E-06 |
| 12101 | 606 | I606A | A | 6.7e-07 | 2.8e-06 | 1.4e-06 | 1.623E-06 | 1.082E-06 | 6.700E-07 | 2.800E-06 |
| 12102 | 606 | I606R | A | 2.7e-06 | 2.4e-06 | 1.7e-06 | 2.267E-06 | 5.132E-07 | 1.700E-06 | 2.700E-06 |
| 12103 | 606 | I606N | A | 3.5e-06 | 2.7e-06 | 2.2e-06 | 2.800E-06 | 6.557E-07 | 2.200E-06 | 3.500E-06 |
| 12104 | 606 | I606D | A | 2e-06 | 2.4e-06 | 3.2e-06 | 2.533E-06 | 6.110E-07 | 2.000E-06 | 3.200E-06 |
| 12105 | 606 | I606C | A | 3.6e-06 | 1.9e-06 | 2.6e-06 | 2.700E-06 | 8.544E-07 | 1.900E-06 | 3.600E-06 |
| 12106 | 606 | I606E | A | 2.2e-06 | 1.8e-06 | 2.1e-06 | 2.033E-06 | 2.082E-07 | 1.800E-06 | 2.200E-06 |
| 12107 | 606 | I606Q | A | 3e-06 | 1.9e-06 | 2.8e-06 | 2.567E-06 | 5.859E-07 | 1.900E-06 | 3.000E-06 |
| 12108 | 606 | I606G | A | 2.5e-06 | 2.7e-06 | 4.4e-06 | 3.200E-06 | 1.044E-06 | 2.500E-06 | 4.400E-06 |
| 12109 | 606 | I606H | A | 2.4e-06 | 1.8e-06 | 2e-06 | 2.067E-06 | 3.055E-07 | 1.800E-06 | 2.400E-06 |
| 12110 | 606 | I606I | A | 2.7e-06 | 3.2e-06 | 2e-06 | 2.633E-06 | 6.028E-07 | 2.000E-06 | 3.200E-06 |
| 12111 | 606 | I606L | A | 2.8e-06 | 2.6e-06 | 3.4e-06 | 2.933E-06 | 4.163E-07 | 2.600E-06 | 3.400E-06 |
| 12112 | 606 | I606K | A | 4.4e-06 | 2.7e-06 | 1.4e-06 | 2.833E-06 | 1.504E-06 | 1.400E-06 | 4.400E-06 |
| 12113 | 606 | I606M | A | 2.1e-06 | 2.4e-06 | 2.8e-06 | 2.433E-06 | 3.512E-07 | 2.100E-06 | 2.800E-06 |
| 12114 | 606 | I606F | A | 2.4e-06 | 3.2e-06 | 3.2e-06 | 2.933E-06 | 4.619E-07 | 2.400E-06 | 3.200E-06 |
| 12115 | 606 | I606P | A | 2.3e-06 | 3e-06 | 2.2e-06 | 2.500E-06 | 4.359E-07 | 2.200E-06 | 3.000E-06 |
| 12116 | 606 | I606S | A | 1.5e-06 | 2.2e-06 | 5.1e-06 | 2.933E-06 | 1.909E-06 | 1.500E-06 | 5.100E-06 |
| 12117 | 606 | I606T | A | 2e-06 | 1.8e-06 | 2.3e-06 | 2.033E-06 | 2.517E-07 | 1.800E-06 | 2.300E-06 |
| 12118 | 606 | I606W | A | 1.6e-06 | 2.4e-06 | 3.5e-06 | 2.500E-06 | 9.539E-07 | 1.600E-06 | 3.500E-06 |
| 12119 | 606 | I606Y | A | 2.3e-06 | 1.2e-06 | 3.1e-06 | 2.200E-06 | 9.539E-07 | 1.200E-06 | 3.100E-06 |
| 12120 | 606 | I606V | A | 2.1e-06 | 1.5e-06 | 1.4e-06 | 1.667E-06 | 3.786E-07 | 1.400E-06 | 2.100E-06 |
| 12121 | 607 | N607A | A | 3e-06 | 2.6e-06 | 2.1e-06 | 2.567E-06 | 4.509E-07 | 2.100E-06 | 3.000E-06 |
| 12122 | 607 | N607R | A | 2.3e-06 | 1.9e-06 | 2.8e-06 | 2.333E-06 | 4.509E-07 | 1.900E-06 | 2.800E-06 |
| 12123 | 607 | N607N | A | 1.8e-06 | 1.2e-06 | 3.8e-06 | 2.267E-06 | 1.361E-06 | 1.200E-06 | 3.800E-06 |
| 12124 | 607 | N607D | A | 2.7e-06 | 2.4e-06 | 2.7e-06 | 2.600E-06 | 1.732E-07 | 2.400E-06 | 2.700E-06 |
| 12125 | 607 | N607C | A | 4e-06 | 2.2e-06 | 2.5e-06 | 2.900E-06 | 9.644E-07 | 2.200E-06 | 4.000E-06 |
| 12126 | 607 | N607E | A | 2.2e-06 | 2.2e-06 | 2.7e-06 | 2.367E-06 | 2.887E-07 | 2.200E-06 | 2.700E-06 |
| 12127 | 607 | N607Q | A | 3e-06 | 1.9e-06 | 3.3e-06 | 2.733E-06 | 7.371E-07 | 1.900E-06 | 3.300E-06 |
| 12128 | 607 | N607G | A | 1.4e-06 | 1.3e-06 | 3e-06 | 1.900E-06 | 9.539E-07 | 1.300E-06 | 3.000E-06 |
| 12129 | 607 | N607H | A | 2.3e-06 | 1.5e-06 | 1.8e-06 | 1.867E-06 | 4.041E-07 | 1.500E-06 | 2.300E-06 |
| 12130 | 607 | N607I | A | 4.1e-06 | 1.6e-06 | 2.5e-06 | 2.733E-06 | 1.266E-06 | 1.600E-06 | 4.100E-06 |
| 12131 | 607 | N607L | A | 1.1e-06 | 1.1e-06 | 2.7e-06 | 1.633E-06 | 9.238E-07 | 1.100E-06 | 2.700E-06 |
| 12132 | 607 | N607K | A | 1.3e-06 | 3.6e-06 | 3.2e-06 | 2.700E-06 | 1.229E-06 | 1.300E-06 | 3.600E-06 |
| 12133 | 607 | N607M | A | 2.9e-06 | 1.6e-06 | 1.8e-06 | 2.100E-06 | 7.000E-07 | 1.600E-06 | 2.900E-06 |
| 12134 | 607 | N607F | A | 2.2e-06 | 2e-06 | 2.9e-06 | 2.367E-06 | 4.726E-07 | 2.000E-06 | 2.900E-06 |
| 12135 | 607 | N607P | A | 2.4e-06 | 1.7e-06 | 4.8e-06 | 2.967E-06 | 1.626E-06 | 1.700E-06 | 4.800E-06 |
| 12136 | 607 | N607S | A | 2.7e-06 | 3.7e-06 | 1.4e-06 | 2.600E-06 | 1.153E-06 | 1.400E-06 | 3.700E-06 |
| 12137 | 607 | N607T | A | 2.8e-06 | 4e-06 | 1.4e-06 | 2.733E-06 | 1.301E-06 | 1.400E-06 | 4.000E-06 |
| 12138 | 607 | N607W | A | 4.2e-06 | 2.6e-06 | 2.4e-06 | 3.067E-06 | 9.866E-07 | 2.400E-06 | 4.200E-06 |
| 12139 | 607 | N607Y | A | 2.1e-06 | 1.6e-06 | 2.3e-06 | 2.000E-06 | 3.606E-07 | 1.600E-06 | 2.300E-06 |
| 12140 | 607 | N607V | A | 2.3e-06 | 1.8e-06 | 2.9e-06 | 2.333E-06 | 5.508E-07 | 1.800E-06 | 2.900E-06 |
| 12141 | 608 | C608A | A | 3.7e-06 | 1.9e-06 | 2.7e-06 | 2.767E-06 | 9.018E-07 | 1.900E-06 | 3.700E-06 |
| 12142 | 608 | C608R | A | 1.4e-06 | 2.3e-06 | 1.9e-06 | 1.867E-06 | 4.509E-07 | 1.400E-06 | 2.300E-06 |
| 12143 | 608 | C608N | A | 2.3e-06 | 2.7e-06 | 2.5e-06 | 2.500E-06 | 2.000E-07 | 2.300E-06 | 2.700E-06 |
| 12144 | 608 | C608D | A | 1.4e-06 | 1.6e-06 | 4e-06 | 2.333E-06 | 1.447E-06 | 1.400E-06 | 4.000E-06 |
| 12145 | 608 | C608C | A | 2e-06 | 1.9e-06 | 2.4e-06 | 2.100E-06 | 2.646E-07 | 1.900E-06 | 2.400E-06 |
| 12146 | 608 | C608E | A | 3.4e-06 | 2.5e-06 | 2.2e-06 | 2.700E-06 | 6.245E-07 | 2.200E-06 | 3.400E-06 |
| 12147 | 608 | C608Q | A | 3e-06 | 2.7e-06 | 4.2e-06 | 3.300E-06 | 7.937E-07 | 2.700E-06 | 4.200E-06 |
| 12148 | 608 | C608G | A | 2.3e-06 | 2e-06 | 3.2e-06 | 2.500E-06 | 6.245E-07 | 2.000E-06 | 3.200E-06 |
| 12149 | 608 | C608H | A | 1.6e-06 | 1.3e-06 | 2e-06 | 1.633E-06 | 3.512E-07 | 1.300E-06 | 2.000E-06 |
| 12150 | 608 | C608I | A | 3.6e-06 | 4.1e-06 | 2.4e-06 | 3.367E-06 | 8.737E-07 | 2.400E-06 | 4.100E-06 |
| 12151 | 608 | C608L | A | 1.5e-06 | 1.8e-06 | 2.8e-06 | 2.033E-06 | 6.807E-07 | 1.500E-06 | 2.800E-06 |
| 12152 | 608 | C608K | A | 3.1e-06 | 2.7e-06 | 3e-06 | 2.933E-06 | 2.082E-07 | 2.700E-06 | 3.100E-06 |
| 12153 | 608 | C608M | A | 1.7e-06 | 2.4e-06 | 1.4e-06 | 1.833E-06 | 5.132E-07 | 1.400E-06 | 2.400E-06 |
| 12154 | 608 | C608F | A | 1.6e-06 | 3.5e-06 | 2.7e-06 | 2.600E-06 | 9.539E-07 | 1.600E-06 | 3.500E-06 |

| | | | | | | | | | | |
|---|---|---|---|---|---|---|---|---|---|---|
| 12155 | 608 | C608P | A | 2.1e-06 | 3e-06 | 1.6e-06 | 2.233E-06 | 7.095E-07 | 1.600E-06 | 3.000E-06 |
| 12156 | 608 | C608S | A | 2.3e-06 | 2.6e-06 | 1.9e-06 | 2.267E-06 | 3.512E-07 | 1.900E-06 | 2.600E-06 |
| 12157 | 608 | C608T | A | 2e-06 | 1.8e-06 | 5.1e-06 | 2.967E-06 | 1.850E-06 | 1.800E-06 | 5.100E-06 |
| 12158 | 608 | C608W | A | 1.5e-06 | 2.3e-06 | 3.9e-06 | 2.567E-06 | 1.222E-06 | 1.500E-06 | 3.900E-06 |
| 12159 | 608 | C608Y | A | 3.7e-06 | 1.2e-06 | 2.3e-06 | 2.400E-06 | 1.253E-06 | 1.200E-06 | 3.700E-06 |
| 12160 | 608 | C608V | A | 2.4e-06 | 2.3e-06 | 1.9e-06 | 2.200E-06 | 2.646E-07 | 1.900E-06 | 2.400E-06 |
| 12161 | 609 | T609A | A | 2.4e-06 | 1.5e-06 | 1.9e-06 | 1.933E-06 | 4.509E-07 | 1.500E-06 | 2.400E-06 |
| 12162 | 609 | T609R | A | 2.2e-06 | 1.3e-06 | 2.9e-06 | 2.133E-06 | 8.021E-07 | 1.300E-06 | 2.900E-06 |
| 12163 | 609 | T609N | A | 2.6e-06 | 3.6e-06 | 4.6e-06 | 3.600E-06 | 1.000E-06 | 2.600E-06 | 4.600E-06 |
| 12164 | 609 | T609D | A | 2.9e-06 | 1.8e-06 | 3.6e-06 | 2.767E-06 | 9.074E-07 | 1.800E-06 | 3.600E-06 |
| 12165 | 609 | T609C | A | 4.3e-06 | 2.3e-06 | 2e-06 | 2.867E-06 | 1.250E-06 | 2.000E-06 | 4.300E-06 |
| 12166 | 609 | T609E | A | 3.2e-06 | 1.7e-06 | 2.5e-06 | 2.467E-06 | 7.506E-07 | 1.700E-06 | 3.200E-06 |
| 12167 | 609 | T609Q | A | 2.9e-06 | 4.1e-06 | 2.9e-06 | 3.300E-06 | 6.928E-07 | 2.900E-06 | 4.100E-06 |
| 12168 | 609 | T609G | A | 2.8e-06 | 3.8e-06 | 2.9e-06 | 3.167E-06 | 5.508E-07 | 2.800E-06 | 3.800E-06 |
| 12169 | 609 | T609H | A | 2.4e-06 | 2.1e-06 | 1.8e-06 | 2.100E-06 | 3.000E-07 | 1.800E-06 | 2.400E-06 |
| 12170 | 609 | T609I | A | 4.3e-06 | 2.9e-06 | 1.2e-06 | 2.800E-06 | 1.552E-06 | 1.200E-06 | 4.300E-06 |
| 12171 | 609 | T609L | A | 2.3e-06 | 2.7e-06 | 2.3e-06 | 2.433E-06 | 2.309E-07 | 2.300E-06 | 2.700E-06 |
| 12172 | 609 | T609K | A | 2.1e-06 | 2.7e-06 | 2e-06 | 2.267E-06 | 3.786E-07 | 2.000E-06 | 2.700E-06 |
| 12173 | 609 | T609M | A | 2.1e-06 | 2.2e-06 | 2.8e-06 | 2.367E-06 | 3.786E-07 | 2.100E-06 | 2.800E-06 |
| 12174 | 609 | T609F | A | 1.3e-06 | 2.4e-06 | 1.2e-06 | 1.633E-06 | 6.658E-07 | 1.200E-06 | 2.400E-06 |
| 12175 | 609 | T609P | A | 2.6e-06 | 1.7e-06 | 3.3e-06 | 2.533E-06 | 8.021E-07 | 1.700E-06 | 3.300E-06 |
| 12176 | 609 | T609S | A | 2e-06 | 2.6e-06 | 4e-06 | 2.867E-06 | 1.026E-06 | 2.000E-06 | 4.000E-06 |
| 12177 | 609 | T609T | A | 2.2e-06 | 2.2e-06 | 2.8e-06 | 2.400E-06 | 3.464E-07 | 2.200E-06 | 2.800E-06 |
| 12178 | 609 | T609W | A | 2.8e-06 | 2.7e-06 | 3.3e-06 | 2.933E-06 | 3.215E-07 | 2.700E-06 | 3.300E-06 |
| 12179 | 609 | T609Y | A | 3.3e-06 | 2.3e-06 | 2.5e-06 | 2.700E-06 | 5.292E-07 | 2.300E-06 | 3.300E-06 |
| 12180 | 609 | T609V | A | 2.6e-06 | 2.4e-06 | 1.9e-06 | 2.300E-06 | 3.606E-07 | 1.900E-06 | 2.600E-06 |
| 12181 | 610 | H610A | A | 2.7e-06 | 3.9e-06 | 3.1e-06 | 3.233E-06 | 6.110E-07 | 2.700E-06 | 3.900E-06 |
| 12182 | 610 | H610R | A | 2.1e-06 | 2.8e-06 | 3.6e-06 | 2.833E-06 | 7.506E-07 | 2.100E-06 | 3.600E-06 |
| 12183 | 610 | H610N | A | 2.2e-06 | 2.9e-06 | 2.9e-06 | 2.667E-06 | 4.041E-07 | 2.200E-06 | 2.900E-06 |
| 12184 | 610 | H610D | A | 1.7e-06 | 1.6e-06 | 1.7e-06 | 1.667E-06 | 5.774E-08 | 1.600E-06 | 1.700E-06 |
| 12185 | 610 | H610C | A | 2.8e-06 | 2.8e-06 | 2.2e-06 | 2.600E-06 | 3.464E-07 | 2.200E-06 | 2.800E-06 |
| 12186 | 610 | H610E | A | 1.5e-06 | 1.5e-06 | 2.9e-06 | 1.967E-06 | 8.083E-07 | 1.500E-06 | 2.900E-06 |
| 12187 | 610 | H610Q | A | 2.1e-06 | 2.6e-06 | 1.6e-06 | 2.100E-06 | 5.000E-07 | 1.600E-06 | 2.600E-06 |
| 12188 | 610 | H610G | A | 2.4e-06 | 3.4e-06 | 2.7e-06 | 2.833E-06 | 5.132E-07 | 2.400E-06 | 3.400E-06 |
| 12189 | 610 | H610H | A | 2.4e-06 | 1.4e-06 | 1.6e-06 | 1.800E-06 | 5.292E-07 | 1.400E-06 | 2.400E-06 |
| 12190 | 610 | H610I | A | 2.6e-06 | 1.8e-06 | 4e-06 | 2.800E-06 | 1.114E-06 | 1.800E-06 | 4.000E-06 |
| 12191 | 610 | H610L | A | 5e-06 | 2.6e-06 | 2.1e-06 | 3.233E-06 | 1.550E-06 | 2.100E-06 | 5.000E-06 |
| 12192 | 610 | H610K | A | 1.8e-06 | 3.5e-06 | 1.7e-06 | 2.333E-06 | 1.012E-06 | 1.700E-06 | 3.500E-06 |
| 12193 | 610 | H610M | A | 1.5e-06 | 3.9e-06 | 1.8e-06 | 2.400E-06 | 1.308E-06 | 1.500E-06 | 3.900E-06 |
| 12194 | 610 | H610F | A | 2.8e-06 | 3.1e-06 | 2.9e-06 | 2.933E-06 | 1.528E-07 | 2.800E-06 | 3.100E-06 |
| 12195 | 610 | H610P | A | 1.5e-06 | 2.5e-06 | 2.3e-06 | 2.100E-06 | 5.292E-07 | 1.500E-06 | 2.500E-06 |
| 12196 | 610 | H610S | A | 2.5e-06 | 2.3e-06 | 4.5e-06 | 3.100E-06 | 1.217E-06 | 2.300E-06 | 4.500E-06 |
| 12197 | 610 | H610T | A | 3.5e-06 | 3.1e-06 | 1.9e-06 | 2.833E-06 | 8.327E-07 | 1.900E-06 | 3.500E-06 |
| 12198 | 610 | H610W | A | 2.5e-06 | 3.3e-06 | 2.4e-06 | 2.733E-06 | 4.933E-07 | 2.400E-06 | 3.300E-06 |
| 12199 | 610 | H610Y | A | 3.9e-06 | 2.1e-06 | 1.8e-06 | 2.600E-06 | 1.136E-06 | 1.800E-06 | 3.900E-06 |
| 12200 | 610 | H610V | A | 1.9e-06 | 2.1e-06 | 1.6e-06 | 1.867E-06 | 2.517E-07 | 1.600E-06 | 2.100E-06 |
| 12201 | 611 | S611A | A | 1.7e-06 | 3.1e-06 | 3.8e-06 | 2.867E-06 | 1.069E-06 | 1.700E-06 | 3.800E-06 |
| 12202 | 611 | S611R | A | 3e-06 | 2.3e-06 | 3.1e-06 | 2.800E-06 | 4.359E-07 | 2.300E-06 | 3.100E-06 |
| 12203 | 611 | S611N | A | 2.6e-06 | 1.9e-06 | 2.6e-06 | 2.367E-06 | 4.041E-07 | 1.900E-06 | 2.600E-06 |
| 12204 | 611 | S611D | A | 2.2e-06 | 2.6e-06 | 2.3e-06 | 2.367E-06 | 2.082E-07 | 2.200E-06 | 2.600E-06 |
| 12205 | 611 | S611C | A | 4e-06 | 1.3e-06 | 2.4e-06 | 2.567E-06 | 1.358E-06 | 1.300E-06 | 4.000E-06 |
| 12206 | 611 | S611E | A | 2.4e-06 | 1.5e-06 | 1.6e-06 | 1.833E-06 | 4.933E-07 | 1.500E-06 | 2.400E-06 |
| 12207 | 611 | S611Q | A | 3.1e-06 | 1.8e-06 | 2.8e-06 | 2.567E-06 | 6.807E-07 | 1.800E-06 | 3.100E-06 |
| 12208 | 611 | S611G | A | 3.1e-06 | 2.5e-06 | 2.9e-06 | 2.833E-06 | 3.055E-07 | 2.500E-06 | 3.100E-06 |
| 12209 | 611 | S611H | A | 2.6e-06 | 1.4e-06 | 2.7e-06 | 2.233E-06 | 7.234E-07 | 1.400E-06 | 2.700E-06 |
| 12210 | 611 | S611I | A | 2.2e-06 | 3.3e-06 | 1.9e-06 | 2.467E-06 | 7.371E-07 | 1.900E-06 | 3.300E-06 |
| 12211 | 611 | S611L | A | 2.5e-06 | 1.8e-06 | 3.4e-06 | 2.567E-06 | 8.021E-07 | 1.800E-06 | 3.400E-06 |
| 12212 | 611 | S611K | A | 2.6e-06 | 3.1e-06 | 1.9e-06 | 2.533E-06 | 6.028E-07 | 1.900E-06 | 3.100E-06 |
| 12213 | 611 | S611M | A | 2e-06 | 3.3e-06 | 1.5e-06 | 2.267E-06 | 9.292E-07 | 1.500E-06 | 3.300E-06 |
| 12214 | 611 | S611F | A | 3.2e-06 | 1.7e-06 | 3.3e-06 | 2.733E-06 | 8.963E-07 | 1.700E-06 | 3.300E-06 |
| 12215 | 611 | S611P | A | 2.5e-06 | 2.6e-06 | 5.6e-06 | 3.567E-06 | 1.762E-06 | 2.500E-06 | 5.600E-06 |
| 12216 | 611 | S611S | A | 1.9e-06 | 2.3e-06 | 3e-06 | 2.400E-06 | 5.568E-07 | 1.900E-06 | 3.000E-06 |
| 12217 | 611 | S611T | A | 1.4e-06 | 2.9e-06 | 2e-06 | 2.100E-06 | 7.550E-07 | 1.400E-06 | 2.900E-06 |
| 12218 | 611 | S611W | A | 1.3e-06 | 2e-06 | 2.3e-06 | 1.867E-06 | 5.132E-07 | 1.300E-06 | 2.300E-06 |
| 12219 | 611 | S611Y | A | 2e-06 | 2.8e-06 | 2.9e-06 | 2.567E-06 | 4.933E-07 | 2.000E-06 | 2.900E-06 |
| 12220 | 611 | S611V | A | 1.8e-06 | 3.9e-06 | 2e-06 | 2.567E-06 | 1.159E-06 | 1.800E-06 | 3.900E-06 |
| 12221 | 612 | C612A | A | 2.3e-06 | 3.4e-06 | 2e-06 | 2.567E-06 | 7.371E-07 | 2.000E-06 | 3.400E-06 |
| 12222 | 612 | C612R | A | 2.9e-06 | 1.9e-06 | 3e-06 | 2.600E-06 | 6.083E-07 | 1.900E-06 | 3.000E-06 |
| 12223 | 612 | C612N | A | 3.1e-06 | 4.6e-06 | 2.2e-06 | 3.300E-06 | 1.212E-06 | 2.200E-06 | 4.600E-06 |
| 12224 | 612 | C612D | A | 2.2e-06 | 4.1e-06 | 1.6e-06 | 2.633E-06 | 1.305E-06 | 1.600E-06 | 4.100E-06 |
| 12225 | 612 | C612C | A | 1.8e-06 | 2.6e-06 | 2.4e-06 | 2.267E-06 | 4.163E-07 | 1.800E-06 | 2.600E-06 |
| 12226 | 612 | C612E | A | 1.8e-06 | 3.5e-06 | 2.2e-06 | 2.500E-06 | 8.888E-07 | 1.800E-06 | 3.500E-06 |
| 12227 | 612 | C612Q | A | 3.2e-06 | 2.7e-06 | 2.6e-06 | 2.833E-06 | 3.215E-07 | 2.600E-06 | 3.200E-06 |
| 12228 | 612 | C612G | A | 2e-06 | 3.6e-06 | 3.3e-06 | 2.967E-06 | 8.505E-07 | 2.000E-06 | 3.600E-06 |
| 12229 | 612 | C612H | A | 2.4e-06 | 3.9e-06 | 3.2e-06 | 3.167E-06 | 7.506E-07 | 2.400E-06 | 3.900E-06 |
| 12230 | 612 | C612I | A | 2.7e-06 | 2.3e-06 | 2e-06 | 2.333E-06 | 3.512E-07 | 2.000E-06 | 2.700E-06 |
| 12231 | 612 | C612L | A | 1.8e-06 | 1.6e-06 | 1.7e-06 | 1.700E-06 | 1.000E-07 | 1.600E-06 | 1.800E-06 |
| 12232 | 612 | C612K | A | 3.1e-06 | 3.7e-06 | 4.1e-06 | 3.633E-06 | 5.033E-07 | 3.100E-06 | 4.100E-06 |
| 12233 | 612 | C612M | A | 2.1e-06 | 2.9e-06 | 3.2e-06 | 2.733E-06 | 5.686E-07 | 2.100E-06 | 3.200E-06 |
| 12234 | 612 | C612F | A | 1.5e-06 | 1.8e-06 | 1.4e-06 | 1.567E-06 | 2.082E-07 | 1.400E-06 | 1.800E-06 |
| 12235 | 612 | C612P | A | 3.2e-06 | 2.3e-06 | 3.4e-06 | 2.967E-06 | 5.859E-07 | 2.300E-06 | 3.400E-06 |
| 12236 | 612 | C612S | A | 1.9e-06 | 2.2e-06 | 3.9e-06 | 2.667E-06 | 1.079E-06 | 1.900E-06 | 3.900E-06 |
| 12237 | 612 | C612T | A | 3.3e-06 | 1.8e-06 | 9.7e-07 | 2.023E-06 | 1.181E-06 | 9.700E-07 | 3.300E-06 |
| 12238 | 612 | C612W | A | 2.5e-06 | 2.6e-06 | 1.5e-06 | 2.200E-06 | 6.083E-07 | 1.500E-06 | 2.600E-06 |
| 12239 | 612 | C612Y | A | 2.3e-06 | 2.1e-06 | 1.6e-06 | 2.000E-06 | 3.606E-07 | 1.600E-06 | 2.300E-06 |

| | | | | | | | | | | |
|---|---|---|---|---|---|---|---|---|---|---|
| 12240 | 612 | C612V | A | 2e-06 | 2.3e-06 | 3.6e-06 | 2.633E-06 | 8.505E-07 | 2.000E-06 | 3.600E-06 |
| 12241 | 613 | V613A | A | 2.4e-06 | 3.8e-06 | 2.4e-06 | 2.867E-06 | 8.083E-07 | 2.400E-06 | 3.800E-06 |
| 12242 | 613 | V613R | A | 1.6e-06 | 2.2e-06 | 2.7e-06 | 2.167E-06 | 5.508E-07 | 1.600E-06 | 2.700E-06 |
| 12243 | 613 | V613N | A | 1.8e-06 | 1.8e-06 | 2.4e-06 | 2.000E-06 | 3.464E-07 | 1.800E-06 | 2.400E-06 |
| 12244 | 613 | V613D | A | 4.3e-06 | 2.3e-06 | 2e-06 | 2.867E-06 | 1.250E-06 | 2.000E-06 | 4.300E-06 |
| 12245 | 613 | V613C | A | 3e-06 | 1.3e-06 | 1.5e-06 | 1.933E-06 | 9.292E-07 | 1.300E-06 | 3.000E-06 |
| 12246 | 613 | V613E | A | 2.5e-06 | 3.1e-06 | 1.8e-06 | 2.467E-06 | 6.506E-07 | 1.800E-06 | 3.100E-06 |
| 12247 | 613 | V613Q | A | 3.4e-06 | 2.8e-06 | 2.7e-06 | 2.967E-06 | 3.786E-07 | 2.700E-06 | 3.400E-06 |
| 12248 | 613 | V613G | A | 1.5e-06 | 1.4e-06 | 2.3e-06 | 1.733E-06 | 4.933E-07 | 1.400E-06 | 2.300E-06 |
| 12249 | 613 | V613H | A | 3.1e-06 | 2.3e-06 | 1.4e-06 | 2.267E-06 | 8.505E-07 | 1.400E-06 | 3.100E-06 |
| 12250 | 613 | V613I | A | 2.5e-06 | 3e-06 | 3.7e-06 | 3.067E-06 | 6.028E-07 | 2.500E-06 | 3.700E-06 |
| 12251 | 613 | V613L | A | 1.8e-06 | 2.4e-06 | 3.7e-06 | 2.633E-06 | 9.713E-07 | 1.800E-06 | 3.700E-06 |
| 12252 | 613 | V613K | A | 3.1e-06 | 2.6e-06 | 2.1e-06 | 2.600E-06 | 5.000E-07 | 2.100E-06 | 3.100E-06 |
| 12253 | 613 | V613M | A | 5.1e-06 | 1.6e-06 | 4.7e-06 | 3.800E-06 | 1.916E-06 | 1.600E-06 | 5.100E-06 |
| 12254 | 613 | V613F | A | 2.6e-06 | 3.1e-06 | 3.2e-06 | 2.967E-06 | 3.215E-07 | 2.600E-06 | 3.200E-06 |
| 12255 | 613 | V613P | A | 3.9e-06 | 2.6e-06 | 2.2e-06 | 2.900E-06 | 8.888E-07 | 2.200E-06 | 3.900E-06 |
| 12256 | 613 | V613S | A | 2.4e-06 | 3.6e-06 | 2.7e-06 | 2.900E-06 | 6.245E-07 | 2.400E-06 | 3.600E-06 |
| 12257 | 613 | V613T | A | 2.7e-06 | 1.6e-06 | 2.7e-06 | 2.333E-06 | 6.351E-07 | 1.600E-06 | 2.700E-06 |
| 12258 | 613 | V613W | A | 5.9e-06 | 5.6e-06 | 2.3e-06 | 4.600E-06 | 1.997E-06 | 2.300E-06 | 5.900E-06 |
| 12259 | 613 | V613Y | A | 2.1e-06 | 2.7e-06 | 2e-06 | 2.267E-06 | 3.786E-07 | 2.000E-06 | 2.700E-06 |
| 12260 | 613 | V613V | A | 2e-06 | 2.4e-06 | 3.5e-06 | 2.633E-06 | 7.767E-07 | 2.000E-06 | 3.500E-06 |
| 12261 | 614 | D614A | A | 1.5e-06 | 2.6e-06 | 3.9e-06 | 2.667E-06 | 1.201E-06 | 1.500E-06 | 3.900E-06 |
| 12262 | 614 | D614R | A | 2.7e-06 | 4.3e-06 | 1.3e-06 | 2.767E-06 | 1.501E-06 | 1.300E-06 | 4.300E-06 |
| 12263 | 614 | D614N | A | 2.2e-06 | 1.5e-06 | 2.7e-06 | 2.133E-06 | 6.028E-07 | 1.500E-06 | 2.700E-06 |
| 12264 | 614 | D614D | A | 1.4e-06 | 2.5e-06 | 1.3e-06 | 1.733E-06 | 6.658E-07 | 1.300E-06 | 2.500E-06 |
| 12265 | 614 | D614C | A | 2.2e-06 | 2.7e-06 | 2.3e-06 | 2.400E-06 | 2.646E-07 | 2.200E-06 | 2.700E-06 |
| 12266 | 614 | D614E | A | 1.2e-06 | 4.5e-06 | 2.4e-06 | 2.700E-06 | 1.670E-06 | 1.200E-06 | 4.500E-06 |
| 12267 | 614 | D614Q | A | 2.9e-06 | 5.9e-06 | 3.1e-06 | 3.967E-06 | 1.677E-06 | 2.900E-06 | 5.900E-06 |
| 12268 | 614 | D614G | A | 3.4e-06 | 3.2e-06 | 2e-06 | 2.867E-06 | 7.572E-07 | 2.000E-06 | 3.400E-06 |
| 12269 | 614 | D614H | A | 1.7e-06 | 2.2e-06 | 1.9e-06 | 1.933E-06 | 2.517E-07 | 1.700E-06 | 2.200E-06 |
| 12270 | 614 | D614I | A | 2.7e-06 | 1.8e-06 | 2e-06 | 2.167E-06 | 4.726E-07 | 1.800E-06 | 2.700E-06 |
| 12271 | 614 | D614L | A | 2.4e-06 | 1.7e-06 | 2.4e-06 | 2.167E-06 | 4.041E-07 | 1.700E-06 | 2.400E-06 |
| 12272 | 614 | D614K | A | 1.5e-06 | 1.6e-06 | 1.8e-06 | 1.633E-06 | 1.528E-07 | 1.500E-06 | 1.800E-06 |
| 12273 | 614 | D614M | A | 1.9e-06 | 2.1e-06 | 1.9e-06 | 1.967E-06 | 1.155E-07 | 1.900E-06 | 2.100E-06 |
| 12274 | 614 | D614F | A | 2.3e-06 | 1.1e-06 | 2.1e-06 | 1.833E-06 | 6.429E-07 | 1.100E-06 | 2.300E-06 |
| 12275 | 614 | D614P | A | 3.6e-06 | 3.2e-06 | 1.9e-06 | 2.900E-06 | 8.888E-07 | 1.900E-06 | 3.600E-06 |
| 12276 | 614 | D614S | A | 3.8e-06 | 4.3e-06 | 2.3e-06 | 3.467E-06 | 1.041E-06 | 2.300E-06 | 4.300E-06 |
| 12277 | 614 | D614T | A | 2.3e-06 | 2e-06 | 1.7e-06 | 2.000E-06 | 3.000E-07 | 1.700E-06 | 2.300E-06 |
| 12278 | 614 | D614W | A | 2e-06 | 2.2e-06 | 2.1e-06 | 2.100E-06 | 1.000E-07 | 2.000E-06 | 2.200E-06 |
| 12279 | 614 | D614Y | A | 2e-06 | 3.1e-06 | 3e-06 | 2.700E-06 | 6.083E-07 | 2.000E-06 | 3.100E-06 |
| 12280 | 614 | D614V | A | 1.6e-06 | 2.1e-06 | 1.8e-06 | 1.833E-06 | 2.517E-07 | 1.600E-06 | 2.100E-06 |
| 12281 | 615 | L615A | A | 4.4e-06 | 1.6e-06 | 1.1e-06 | 2.367E-06 | 1.779E-06 | 1.100E-06 | 4.400E-06 |
| 12282 | 615 | L615R | A | 2.2e-06 | 9.1e-07 | 2.9e-06 | 2.003E-06 | 1.009E-06 | 9.100E-07 | 2.900E-06 |
| 12283 | 615 | L615N | A | 3.5e-06 | 1.5e-06 | 2.4e-06 | 2.467E-06 | 1.002E-06 | 1.500E-06 | 3.500E-06 |
| 12284 | 615 | L615D | A | 5.9e-06 | 2.4e-06 | 3.2e-06 | 3.833E-06 | 1.834E-06 | 2.400E-06 | 5.900E-06 |
| 12285 | 615 | L615C | A | 2.3e-06 | 3.5e-06 | 1.2e-06 | 2.333E-06 | 1.150E-06 | 1.200E-06 | 3.500E-06 |
| 12286 | 615 | L615E | A | 2.8e-06 | 2.9e-06 | 3e-06 | 2.900E-06 | 1.000E-07 | 2.800E-06 | 3.000E-06 |
| 12287 | 615 | L615Q | A | 2.7e-06 | 2.4e-06 | 1.5e-06 | 2.200E-06 | 6.245E-07 | 1.500E-06 | 2.700E-06 |
| 12288 | 615 | L615G | A | 4.4e-06 | 3.1e-06 | 2.1e-06 | 3.200E-06 | 1.153E-06 | 2.100E-06 | 4.400E-06 |
| 12289 | 615 | L615H | A | 1.7e-06 | 2.1e-06 | 2.2e-06 | 2.000E-06 | 2.646E-07 | 1.700E-06 | 2.200E-06 |
| 12290 | 615 | L615I | A | 1.5e-06 | 3.8e-06 | 3e-06 | 2.767E-06 | 1.168E-06 | 1.500E-06 | 3.800E-06 |
| 12291 | 615 | L615L | A | 2.3e-06 | 1.8e-06 | 1.8e-06 | 1.967E-06 | 2.887E-07 | 1.800E-06 | 2.300E-06 |
| 12292 | 615 | L615K | A | 1.9e-06 | 1.7e-06 | 4.3e-06 | 2.633E-06 | 1.447E-06 | 1.700E-06 | 4.300E-06 |
| 12293 | 615 | L615M | A | 3.1e-06 | 1.9e-06 | 1.7e-06 | 2.233E-06 | 7.572E-07 | 1.700E-06 | 3.100E-06 |
| 12294 | 615 | L615F | A | 1.8e-06 | 2.2e-06 | 2.6e-06 | 2.200E-06 | 4.000E-07 | 1.800E-06 | 2.600E-06 |
| 12295 | 615 | L615P | A | 3.2e-06 | 2.3e-06 | 1.9e-06 | 2.467E-06 | 6.658E-07 | 1.900E-06 | 3.200E-06 |
| 12296 | 615 | L615S | A | 2.7e-06 | 1.1e-06 | 2.9e-06 | 2.233E-06 | 9.866E-07 | 1.100E-06 | 2.900E-06 |
| 12297 | 615 | L615T | A | 2.3e-06 | 3.1e-06 | 2.1e-06 | 2.500E-06 | 5.292E-07 | 2.100E-06 | 3.100E-06 |
| 12298 | 615 | L615W | A | 1.8e-06 | 2.6e-06 | 2.4e-06 | 2.267E-06 | 4.163E-07 | 1.800E-06 | 2.600E-06 |
| 12299 | 615 | L615Y | A | 3.2e-06 | 2.8e-06 | 1.5e-06 | 2.500E-06 | 8.888E-07 | 1.500E-06 | 3.200E-06 |
| 12300 | 615 | L615V | A | 3.8e-06 | 1.7e-06 | 3.4e-06 | 2.967E-06 | 1.115E-06 | 1.700E-06 | 3.800E-06 |
| 12301 | 616 | D616A | A | 2e-06 | 2.3e-06 | 1.6e-06 | 1.967E-06 | 3.512E-07 | 1.600E-06 | 2.300E-06 |
| 12302 | 616 | D616R | A | 2.7e-06 | 2.4e-06 | 3.1e-06 | 2.733E-06 | 3.512E-07 | 2.400E-06 | 3.100E-06 |
| 12303 | 616 | D616N | A | 3.3e-06 | 3e-06 | 2e-06 | 2.767E-06 | 6.807E-07 | 2.000E-06 | 3.300E-06 |
| 12304 | 616 | D616D | A | 1.2e-06 | 2.1e-06 | 4.4e-06 | 2.567E-06 | 1.650E-06 | 1.200E-06 | 4.400E-06 |
| 12305 | 616 | D616C | A | 1.6e-06 | 2.4e-06 | 1.7e-06 | 1.900E-06 | 4.359E-07 | 1.600E-06 | 2.400E-06 |
| 12306 | 616 | D616E | A | 2.6e-06 | 1.5e-06 | 1.1e-06 | 1.733E-06 | 7.767E-07 | 1.100E-06 | 2.600E-06 |
| 12307 | 616 | D616Q | A | 1.9e-06 | 1.9e-06 | 1.3e-06 | 1.700E-06 | 3.464E-07 | 1.300E-06 | 1.900E-06 |
| 12308 | 616 | D616G | A | 2.7e-06 | 2.5e-06 | 2.2e-06 | 2.467E-06 | 2.517E-07 | 2.200E-06 | 2.700E-06 |
| 12309 | 616 | D616H | A | 2.1e-06 | 2.2e-06 | 2e-06 | 2.100E-06 | 1.000E-07 | 2.000E-06 | 2.200E-06 |
| 12310 | 616 | D616I | A | 3.6e-06 | 2.7e-06 | 1.2e-06 | 2.500E-06 | 1.212E-06 | 1.200E-06 | 3.600E-06 |
| 12311 | 616 | D616L | A | 3.1e-06 | 3.2e-06 | 2.5e-06 | 2.933E-06 | 3.786E-07 | 2.500E-06 | 3.200E-06 |
| 12312 | 616 | D616K | A | 2.5e-06 | 2e-06 | 1.7e-06 | 2.067E-06 | 4.041E-07 | 1.700E-06 | 2.500E-06 |
| 12313 | 616 | D616M | A | 4.3e-06 | 1.2e-06 | 2.3e-06 | 2.600E-06 | 1.572E-06 | 1.200E-06 | 4.300E-06 |
| 12314 | 616 | D616F | A | 2.6e-06 | 2.7e-06 | 1.5e-06 | 2.267E-06 | 6.658E-07 | 1.500E-06 | 2.700E-06 |
| 12315 | 616 | D616P | A | 2.8e-06 | 2.2e-06 | 2.3e-06 | 2.433E-06 | 3.215E-07 | 2.200E-06 | 2.800E-06 |
| 12316 | 616 | D616S | A | 2e-06 | 1.1e-06 | 2.3e-06 | 1.800E-06 | 6.245E-07 | 1.100E-06 | 2.300E-06 |
| 12317 | 616 | D616T | A | 2.4e-06 | 2.2e-06 | 2.7e-06 | 2.433E-06 | 2.517E-07 | 2.200E-06 | 2.700E-06 |
| 12318 | 616 | D616W | A | 2e-06 | 2.6e-06 | 1.9e-06 | 2.167E-06 | 3.786E-07 | 1.900E-06 | 2.600E-06 |
| 12319 | 616 | D616Y | A | 3e-06 | 8.7e-07 | 3.3e-06 | 2.390E-06 | 1.325E-06 | 8.700E-07 | 3.300E-06 |
| 12320 | 616 | D616V | A | 2.1e-06 | 1.1e-06 | 2.8e-06 | 2.000E-06 | 8.544E-07 | 1.100E-06 | 2.800E-06 |
| 12321 | 617 | D617A | A | 1.8e-06 | 3e-06 | 2e-06 | 2.267E-06 | 6.429E-07 | 1.800E-06 | 3.000E-06 |
| 12322 | 617 | D617R | A | 2.4e-06 | 1.9e-06 | 3.7e-06 | 2.667E-06 | 9.292E-07 | 1.900E-06 | 3.700E-06 |
| 12323 | 617 | D617N | A | 2.4e-06 | 2.5e-06 | 1.9e-06 | 2.267E-06 | 3.215E-07 | 1.900E-06 | 2.500E-06 |
| 12324 | 617 | D617D | A | 3e-06 | 3.1e-06 | 1.8e-06 | 2.633E-06 | 7.234E-07 | 1.800E-06 | 3.100E-06 |

| | | | | | | | | | | |
|---|---|---|---|---|---|---|---|---|---|---|
| 12325 | 617 | D617C | A | 2.1e-06 | 2.2e-06 | 2.6e-06 | 2.300E-06 | 2.646E-07 | 2.100E-06 | 2.600E-06 |
| 12326 | 617 | D617E | A | 2e-06 | 2.1e-06 | 2.2e-06 | 2.100E-06 | 1.000E-07 | 2.000E-06 | 2.200E-06 |
| 12327 | 617 | D617Q | A | 2.4e-06 | 1.8e-06 | 2.6e-06 | 2.267E-06 | 4.163E-07 | 1.800E-06 | 2.600E-06 |
| 12328 | 617 | D617G | A | 2.3e-06 | 2.2e-06 | 3.4e-06 | 2.633E-06 | 6.658E-07 | 2.200E-06 | 3.400E-06 |
| 12329 | 617 | D617H | A | 2.2e-06 | 1.4e-06 | 2.4e-06 | 2.000E-06 | 5.292E-07 | 1.400E-06 | 2.400E-06 |
| 12330 | 617 | D617I | A | 2.4e-06 | 2.1e-06 | 2.8e-06 | 2.433E-06 | 3.512E-07 | 2.100E-06 | 2.800E-06 |
| 12331 | 617 | D617L | A | 3.3e-06 | 4.1e-06 | 2.2e-06 | 3.200E-06 | 9.539E-07 | 2.200E-06 | 4.100E-06 |
| 12332 | 617 | D617K | A | 2.9e-06 | 2.2e-06 | 2.8e-06 | 2.633E-06 | 3.786E-07 | 2.200E-06 | 2.900E-06 |
| 12333 | 617 | D617M | A | 2.5e-06 | 1.9e-06 | 2.6e-06 | 2.333E-06 | 3.786E-07 | 1.900E-06 | 2.600E-06 |
| 12334 | 617 | D617F | A | 2.2e-06 | 3.4e-06 | 1.6e-06 | 2.400E-06 | 9.165E-07 | 1.600E-06 | 3.400E-06 |
| 12335 | 617 | D617P | A | 3.4e-06 | 3.4e-06 | 2.2e-06 | 3.000E-06 | 6.928E-07 | 2.200E-06 | 3.400E-06 |
| 12336 | 617 | D617S | A | 2.6e-06 | 1.8e-06 | 1.5e-06 | 1.967E-06 | 5.686E-07 | 1.500E-06 | 2.600E-06 |
| 12337 | 617 | D617T | A | 3e-06 | 2.1e-06 | 3.1e-06 | 2.733E-06 | 5.508E-07 | 2.100E-06 | 3.100E-06 |
| 12338 | 617 | D617W | A | 2.8e-06 | 2.6e-06 | 2.4e-06 | 2.600E-06 | 2.000E-07 | 2.400E-06 | 2.800E-06 |
| 12339 | 617 | D617Y | A | 2.3e-06 | 2.2e-06 | 2.3e-06 | 2.267E-06 | 5.774E-08 | 2.200E-06 | 2.300E-06 |
| 12340 | 617 | D617V | A | 3.9e-06 | 3e-06 | 2.9e-06 | 3.267E-06 | 5.508E-07 | 2.900E-06 | 3.900E-06 |
| 12341 | 618 | K618A | A | 1.8e-06 | 4.1e-06 | 3.2e-06 | 3.033E-06 | 1.159E-06 | 1.800E-06 | 4.100E-06 |
| 12342 | 618 | K618R | A | 4e-06 | 1.9e-06 | 3.3e-06 | 3.067E-06 | 1.069E-06 | 1.900E-06 | 4.000E-06 |
| 12343 | 618 | K618N | A | 2.1e-06 | 2.2e-06 | 2.2e-06 | 2.167E-06 | 5.774E-08 | 2.100E-06 | 2.200E-06 |
| 12344 | 618 | K618D | A | 1.6e-06 | 1.5e-06 | 2.4e-06 | 1.833E-06 | 4.933E-07 | 1.500E-06 | 2.400E-06 |
| 12345 | 618 | K618C | A | 1.6e-06 | 1.6e-06 | 1.6e-06 | 1.600E-06 | 0.000E+00 | 1.600E-06 | 1.600E-06 |
| 12346 | 618 | K618E | A | 2.8e-06 | 4.1e-06 | 1.8e-06 | 2.900E-06 | 1.153E-06 | 1.800E-06 | 4.100E-06 |
| 12347 | 618 | K618Q | A | 2.4e-06 | 2e-06 | 2.1e-06 | 2.167E-06 | 2.082E-07 | 2.000E-06 | 2.400E-06 |
| 12348 | 618 | K618G | A | 1.7e-06 | 2.1e-06 | 3.9e-06 | 2.567E-06 | 1.172E-06 | 1.700E-06 | 3.900E-06 |
| 12349 | 618 | K618H | A | 1.9e-06 | 1.6e-06 | 1.8e-06 | 1.767E-06 | 1.528E-07 | 1.600E-06 | 1.900E-06 |
| 12350 | 618 | K618I | A | 3.1e-06 | 2.1e-06 | 1.9e-06 | 2.367E-06 | 6.429E-07 | 1.900E-06 | 3.100E-06 |
| 12351 | 618 | K618L | A | 2.2e-06 | 1.5e-06 | 2.6e-06 | 2.100E-06 | 5.568E-07 | 1.500E-06 | 2.600E-06 |
| 12352 | 618 | K618K | A | 2.6e-06 | 3.5e-06 | 2.7e-06 | 2.933E-06 | 4.933E-07 | 2.600E-06 | 3.500E-06 |
| 12353 | 618 | K618M | A | 2.5e-06 | 2.4e-06 | 3.5e-06 | 2.800E-06 | 6.083E-07 | 2.400E-06 | 3.500E-06 |
| 12354 | 618 | K618F | A | 1.5e-06 | 2.2e-06 | 3e-06 | 2.233E-06 | 7.506E-07 | 1.500E-06 | 3.000E-06 |
| 12355 | 618 | K618P | A | 1.4e-06 | 3.9e-06 | 4.1e-06 | 3.133E-06 | 1.504E-06 | 1.400E-06 | 4.100E-06 |
| 12356 | 618 | K618S | A | 2.8e-06 | 2e-06 | 2.7e-06 | 2.500E-06 | 4.359E-07 | 2.000E-06 | 2.800E-06 |
| 12357 | 618 | K618T | A | 2.5e-06 | 2.4e-06 | 1.8e-06 | 2.233E-06 | 3.786E-07 | 1.800E-06 | 2.500E-06 |
| 12358 | 618 | K618W | A | 1.8e-06 | 3.2e-06 | 2.7e-06 | 2.567E-06 | 7.095E-07 | 1.800E-06 | 3.200E-06 |
| 12359 | 618 | K618Y | A | 2.4e-06 | 2.5e-06 | 2.6e-06 | 2.500E-06 | 1.000E-07 | 2.400E-06 | 2.600E-06 |
| 12360 | 618 | K618V | A | 1.2e-06 | 1.6e-06 | 3.8e-06 | 2.200E-06 | 1.400E-06 | 1.200E-06 | 3.800E-06 |
| 12361 | 619 | G619A | A | 1.9e-06 | 2.2e-06 | 1.8e-06 | 1.967E-06 | 2.082E-07 | 1.800E-06 | 2.200E-06 |
| 12362 | 619 | G619R | A | 1.2e-06 | 2e-06 | 2.2e-06 | 1.800E-06 | 5.292E-07 | 1.200E-06 | 2.200E-06 |
| 12363 | 619 | G619N | A | 3e-06 | 2.7e-06 | 3.1e-06 | 2.933E-06 | 2.082E-07 | 2.700E-06 | 3.100E-06 |
| 12364 | 619 | G619D | A | 3.7e-06 | 2.6e-06 | 2.1e-06 | 2.800E-06 | 8.185E-07 | 2.100E-06 | 3.700E-06 |
| 12365 | 619 | G619C | A | 2e-06 | 2.7e-06 | 3.3e-06 | 2.667E-06 | 6.506E-07 | 2.000E-06 | 3.300E-06 |
| 12366 | 619 | G619E | A | 2.2e-06 | 2.7e-06 | 2.8e-06 | 2.567E-06 | 3.215E-07 | 2.200E-06 | 2.800E-06 |
| 12367 | 619 | G619Q | A | 1.2e-06 | 1.2e-06 | 2.4e-06 | 1.600E-06 | 6.928E-07 | 1.200E-06 | 2.400E-06 |
| 12368 | 619 | G619G | A | 1.9e-06 | 2.5e-06 | 2.7e-06 | 2.367E-06 | 4.163E-07 | 1.900E-06 | 2.700E-06 |
| 12369 | 619 | G619H | A | 1.8e-06 | 2.5e-06 | 3.4e-06 | 2.567E-06 | 8.021E-07 | 1.800E-06 | 3.400E-06 |
| 12370 | 619 | G619I | A | 1.8e-06 | 1.9e-06 | 2.4e-06 | 2.033E-06 | 3.215E-07 | 1.800E-06 | 2.400E-06 |
| 12371 | 619 | G619L | A | 1.6e-06 | 3.3e-06 | 2.9e-06 | 2.600E-06 | 8.888E-07 | 1.600E-06 | 3.300E-06 |
| 12372 | 619 | G619K | A | 2.4e-06 | 6.7e-07 | 1.9e-06 | 1.657E-06 | 8.903E-07 | 6.700E-07 | 2.400E-06 |
| 12373 | 619 | G619M | A | 1.7e-06 | 4.8e-06 | 1.7e-06 | 2.733E-06 | 1.790E-06 | 1.700E-06 | 4.800E-06 |
| 12374 | 619 | G619F | A | 3e-06 | 1.7e-06 | 3.2e-06 | 2.633E-06 | 8.145E-07 | 1.700E-06 | 3.200E-06 |
| 12375 | 619 | G619P | A | 2e-06 | 2.5e-06 | 1.4e-06 | 1.967E-06 | 5.508E-07 | 1.400E-06 | 2.500E-06 |
| 12376 | 619 | G619S | A | 2.1e-06 | 1.9e-06 | 2.9e-06 | 2.300E-06 | 5.292E-07 | 1.900E-06 | 2.900E-06 |
| 12377 | 619 | G619T | A | 1.6e-06 | 1.6e-06 | 2.4e-06 | 1.867E-06 | 4.619E-07 | 1.600E-06 | 2.400E-06 |
| 12378 | 619 | G619W | A | 1.9e-06 | 2.4e-06 | 2.3e-06 | 2.200E-06 | 2.646E-07 | 1.900E-06 | 2.400E-06 |
| 12379 | 619 | G619Y | A | 1.1e-06 | 2.9e-06 | 2.5e-06 | 2.167E-06 | 9.452E-07 | 1.100E-06 | 2.900E-06 |
| 12380 | 619 | G619V | A | 2.1e-06 | 2.7e-06 | 1.6e-06 | 2.133E-06 | 5.508E-07 | 1.600E-06 | 2.700E-06 |
| 12381 | 620 | C620A | A | 2.1e-06 | 2.7e-06 | 1.4e-06 | 2.067E-06 | 6.506E-07 | 1.400E-06 | 2.700E-06 |
| 12382 | 620 | C620R | A | 3.3e-06 | 2.4e-06 | 1.1e-06 | 2.267E-06 | 1.106E-06 | 1.100E-06 | 3.300E-06 |
| 12383 | 620 | C620N | A | 4.4e-06 | 3.3e-06 | 2.6e-06 | 3.433E-06 | 9.074E-07 | 2.600E-06 | 4.400E-06 |
| 12384 | 620 | C620D | A | 2e-06 | 3.1e-06 | 2.7e-06 | 2.600E-06 | 5.568E-07 | 2.000E-06 | 3.100E-06 |
| 12385 | 620 | C620C | A | 2.3e-06 | 1.5e-06 | 2.8e-06 | 2.200E-06 | 6.557E-07 | 1.500E-06 | 2.800E-06 |
| 12386 | 620 | C620E | A | 2.2e-06 | 2.6e-06 | 3e-06 | 2.600E-06 | 4.000E-07 | 2.200E-06 | 3.000E-06 |
| 12387 | 620 | C620Q | A | 2.9e-06 | 2.6e-06 | 2.4e-06 | 2.633E-06 | 2.517E-07 | 2.400E-06 | 2.900E-06 |
| 12388 | 620 | C620G | A | 3.3e-06 | 3e-06 | 2.3e-06 | 2.867E-06 | 5.132E-07 | 2.300E-06 | 3.300E-06 |
| 12389 | 620 | C620H | A | 1.9e-06 | 3e-06 | 1.9e-06 | 2.267E-06 | 6.351E-07 | 1.900E-06 | 3.000E-06 |
| 12390 | 620 | C620I | A | 2.5e-06 | 2.9e-06 | 3.1e-06 | 2.833E-06 | 3.055E-07 | 2.500E-06 | 3.100E-06 |
| 12391 | 620 | C620L | A | 3.3e-06 | 3.6e-06 | 2.7e-06 | 3.200E-06 | 4.583E-07 | 2.700E-06 | 3.600E-06 |
| 12392 | 620 | C620K | A | 3e-06 | 2.3e-06 | 2.2e-06 | 2.500E-06 | 4.359E-07 | 2.200E-06 | 3.000E-06 |
| 12393 | 620 | C620M | A | 2.4e-06 | 4.5e-06 | 1.6e-06 | 2.833E-06 | 1.498E-06 | 1.600E-06 | 4.500E-06 |
| 12394 | 620 | C620F | A | 2.9e-06 | 2.2e-06 | 2.2e-06 | 2.433E-06 | 4.041E-07 | 2.200E-06 | 2.900E-06 |
| 12395 | 620 | C620P | A | 2.4e-06 | 2.8e-06 | 4.1e-06 | 3.100E-06 | 8.888E-07 | 2.400E-06 | 4.100E-06 |
| 12396 | 620 | C620S | A | 1.9e-06 | 1.6e-06 | 4.5e-06 | 2.667E-06 | 1.595E-06 | 1.600E-06 | 4.500E-06 |
| 12397 | 620 | C620T | A | 2.2e-06 | 2e-06 | 2.8e-06 | 2.333E-06 | 4.163E-07 | 2.000E-06 | 2.800E-06 |
| 12398 | 620 | C620W | A | 3.5e-06 | 2.1e-06 | 4e-06 | 3.200E-06 | 9.849E-07 | 2.100E-06 | 4.000E-06 |
| 12399 | 620 | C620Y | A | 5.3e-06 | 1.6e-06 | 1.9e-06 | 2.933E-06 | 2.055E-06 | 1.600E-06 | 5.300E-06 |
| 12400 | 620 | C620V | A | 3.9e-06 | 2.2e-06 | 1.6e-06 | 2.567E-06 | 1.193E-06 | 1.600E-06 | 3.900E-06 |
| 12401 | 621 | P621A | A | 3.1e-06 | 2.3e-06 | 2.7e-06 | 2.700E-06 | 4.000E-07 | 2.300E-06 | 3.100E-06 |
| 12402 | 621 | P621R | A | 1.2e-06 | 1.8e-06 | 3.1e-06 | 2.033E-06 | 9.713E-07 | 1.200E-06 | 3.100E-06 |
| 12403 | 621 | P621N | A | 4.9e-06 | 3.5e-06 | 3.6e-06 | 4.000E-06 | 7.810E-07 | 3.500E-06 | 4.900E-06 |
| 12404 | 621 | P621D | A | 3.8e-06 | 1.9e-06 | 3.3e-06 | 3.000E-06 | 9.849E-07 | 1.900E-06 | 3.800E-06 |
| 12405 | 621 | P621C | A | 1.5e-06 | 1.3e-06 | 3.4e-06 | 2.067E-06 | 1.159E-06 | 1.300E-06 | 3.400E-06 |
| 12406 | 621 | P621E | A | 1.6e-06 | 2.9e-06 | 8.4e-07 | 1.780E-06 | 1.042E-06 | 8.400E-07 | 2.900E-06 |
| 12407 | 621 | P621Q | A | 3.4e-06 | 1.7e-06 | 1.2e-06 | 2.100E-06 | 1.153E-06 | 1.200E-06 | 3.400E-06 |
| 12408 | 621 | P621G | A | 2.3e-06 | 2.7e-06 | 2.4e-06 | 2.467E-06 | 2.082E-07 | 2.300E-06 | 2.700E-06 |
| 12409 | 621 | P621H | A | 2.3e-06 | 4.5e-06 | 3e-06 | 3.267E-06 | 1.124E-06 | 2.300E-06 | 4.500E-06 |

| | | | | | | | | | | |
|---|---|---|---|---|---|---|---|---|---|---|
| 12410 | 621 | P621I | A | 2.7e-06 | 1.1e-06 | 1.4e-06 | 1.733E-06 | 8.505E-07 | 1.100E-06 | 2.700E-06 |
| 12411 | 621 | P621L | A | 3.1e-06 | 2.1e-06 | 1.5e-06 | 2.233E-06 | 8.083E-07 | 1.500E-06 | 3.100E-06 |
| 12412 | 621 | P621K | A | 1.4e-06 | 2.1e-06 | 4.3e-06 | 2.600E-06 | 1.513E-06 | 1.400E-06 | 4.300E-06 |
| 12413 | 621 | P621M | A | 2.2e-06 | 1.4e-06 | 2.1e-06 | 1.900E-06 | 4.359E-07 | 1.400E-06 | 2.200E-06 |
| 12414 | 621 | P621F | A | 1.9e-06 | 3e-06 | 2.9e-06 | 2.600E-06 | 6.083E-07 | 1.900E-06 | 3.000E-06 |
| 12415 | 621 | P621P | A | 1.8e-06 | 2.4e-06 | 2.1e-06 | 2.100E-06 | 3.000E-07 | 1.800E-06 | 2.400E-06 |
| 12416 | 621 | P621S | A | 1.6e-06 | 3.3e-06 | 2.1e-06 | 2.333E-06 | 8.737E-07 | 1.600E-06 | 3.300E-06 |
| 12417 | 621 | P621T | A | 2e-06 | 2.7e-06 | 2.3e-06 | 2.333E-06 | 3.512E-07 | 2.000E-06 | 2.700E-06 |
| 12418 | 621 | P621W | A | 3.4e-06 | 1.9e-06 | 4.5e-06 | 3.267E-06 | 1.305E-06 | 1.900E-06 | 4.500E-06 |
| 12419 | 621 | P621Y | A | 2.3e-06 | 2.7e-06 | 1.8e-06 | 2.267E-06 | 4.509E-07 | 1.800E-06 | 2.700E-06 |
| 12420 | 621 | P621V | A | 3.1e-06 | 2e-06 | 5e-06 | 3.367E-06 | 1.518E-06 | 2.000E-06 | 5.000E-06 |
| 12421 | 622 | A622A | A | 2.3e-06 | 1.6e-06 | 2e-06 | 1.967E-06 | 3.512E-07 | 1.600E-06 | 2.300E-06 |
| 12422 | 622 | A622R | A | 2.2e-06 | 1.5e-06 | 2.8e-06 | 2.167E-06 | 6.506E-07 | 1.500E-06 | 2.800E-06 |
| 12423 | 622 | A622N | A | 2e-06 | 2.1e-06 | 1.3e-06 | 1.800E-06 | 4.359E-07 | 1.300E-06 | 2.100E-06 |
| 12424 | 622 | A622D | A | 1.9e-06 | 2.1e-06 | 2e-06 | 2.000E-06 | 1.000E-07 | 1.900E-06 | 2.100E-06 |
| 12425 | 622 | A622C | A | 2.6e-06 | 2.1e-06 | 1.6e-06 | 2.100E-06 | 5.000E-07 | 1.600E-06 | 2.600E-06 |
| 12426 | 622 | A622E | A | 1.1e-06 | 3.2e-06 | 1.3e-06 | 1.867E-06 | 1.159E-06 | 1.100E-06 | 3.200E-06 |
| 12427 | 622 | A622Q | A | 1.7e-06 | 2.5e-06 | 2.3e-06 | 2.167E-06 | 4.163E-07 | 1.700E-06 | 2.500E-06 |
| 12428 | 622 | A622G | A | 2e-06 | 2.1e-06 | 3e-06 | 2.367E-06 | 5.508E-07 | 2.000E-06 | 3.000E-06 |
| 12429 | 622 | A622H | A | 2.6e-06 | 3.4e-06 | 1.8e-06 | 2.600E-06 | 8.000E-07 | 1.800E-06 | 3.400E-06 |
| 12430 | 622 | A622I | A | 1.9e-06 | 1.6e-06 | 2.1e-06 | 1.867E-06 | 2.517E-07 | 1.600E-06 | 2.100E-06 |
| 12431 | 622 | A622L | A | 2.4e-06 | 1.3e-06 | 4.3e-06 | 2.667E-06 | 1.518E-06 | 1.300E-06 | 4.300E-06 |
| 12432 | 622 | A622K | A | 2.6e-06 | 1.7e-06 | 3.5e-06 | 2.600E-06 | 9.000E-07 | 1.700E-06 | 3.500E-06 |
| 12433 | 622 | A622M | A | 2.1e-06 | 2.1e-06 | 2.3e-06 | 2.167E-06 | 1.155E-07 | 2.100E-06 | 2.300E-06 |
| 12434 | 622 | A622F | A | 1.8e-06 | 1.5e-06 | 3.6e-06 | 2.300E-06 | 1.136E-06 | 1.500E-06 | 3.600E-06 |
| 12435 | 622 | A622P | A | 2.3e-06 | 1.8e-06 | 2.6e-06 | 2.233E-06 | 4.041E-07 | 1.800E-06 | 2.600E-06 |
| 12436 | 622 | A622S | A | 2.1e-06 | 2.4e-06 | 2.6e-06 | 2.367E-06 | 2.517E-07 | 2.100E-06 | 2.600E-06 |
| 12437 | 622 | A622T | A | 2.3e-06 | 3.1e-06 | 2.4e-06 | 2.600E-06 | 4.359E-07 | 2.300E-06 | 3.100E-06 |
| 12438 | 622 | A622W | A | 2.8e-06 | 1.5e-06 | 1.7e-06 | 2.000E-06 | 7.000E-07 | 1.500E-06 | 2.800E-06 |
| 12439 | 622 | A622Y | A | 3e-06 | 2e-06 | 1.8e-06 | 2.267E-06 | 6.429E-07 | 1.800E-06 | 3.000E-06 |
| 12440 | 622 | A622V | A | 1.9e-06 | 1.8e-06 | 1.8e-06 | 1.833E-06 | 5.774E-08 | 1.800E-06 | 1.900E-06 |
| 12441 | 623 | D623A | B | 3.3e-06 | 1.7e-06 | 1.5e-06 | 2.167E-06 | 9.866E-07 | 1.500E-06 | 3.300E-06 |
| 12442 | 623 | D623R | B | 2.8e-06 | 2.9e-06 | 3.6e-06 | 3.100E-06 | 4.359E-07 | 2.800E-06 | 3.600E-06 |
| 12443 | 623 | D623N | B | 2.7e-06 | 2.5e-06 | 3.7e-06 | 2.967E-06 | 6.429E-07 | 2.500E-06 | 3.700E-06 |
| 12444 | 623 | D623D | B | 3.2e-06 | 1.2e-06 | 2.1e-06 | 2.167E-06 | 1.002E-06 | 1.200E-06 | 3.200E-06 |
| 12445 | 623 | D623C | B | 3.8e-06 | 2.2e-06 | 2.3e-06 | 2.767E-06 | 8.963E-07 | 2.200E-06 | 3.800E-06 |
| 12446 | 623 | D623E | B | 1.9e-06 | 2.6e-06 | 2.2e-06 | 2.233E-06 | 3.512E-07 | 1.900E-06 | 2.600E-06 |
| 12447 | 623 | D623Q | B | 2.4e-06 | 4.2e-06 | 2.2e-06 | 2.933E-06 | 1.102E-06 | 2.200E-06 | 4.200E-06 |
| 12448 | 623 | D623G | B | 2.6e-06 | 4.4e-06 | 1.1e-06 | 2.700E-06 | 1.652E-06 | 1.100E-06 | 4.400E-06 |
| 12449 | 623 | D623H | B | 2.6e-06 | 1.8e-06 | 2.1e-06 | 2.167E-06 | 4.041E-07 | 1.800E-06 | 2.600E-06 |
| 12450 | 623 | D623I | B | 2.2e-06 | 1.2e-06 | 2.2e-06 | 1.867E-06 | 5.774E-07 | 1.200E-06 | 2.200E-06 |
| 12451 | 623 | D623L | B | 2.1e-06 | 1.2e-06 | 1.9e-06 | 1.733E-06 | 4.726E-07 | 1.200E-06 | 2.100E-06 |
| 12452 | 623 | D623K | B | 3.4e-06 | 5.9e-06 | 3.5e-06 | 4.267E-06 | 1.415E-06 | 3.400E-06 | 5.900E-06 |
| 12453 | 623 | D623M | B | 1.6e-06 | 2.6e-06 | 2.6e-06 | 2.267E-06 | 5.774E-07 | 1.600E-06 | 2.600E-06 |
| 12454 | 623 | D623F | B | 2.4e-06 | 1.9e-06 | 1.8e-06 | 2.033E-06 | 3.215E-07 | 1.800E-06 | 2.400E-06 |
| 12455 | 623 | D623P | B | 2.5e-06 | 2.6e-06 | 2.9e-06 | 2.667E-06 | 2.082E-07 | 2.500E-06 | 2.900E-06 |
| 12456 | 623 | D623S | B | 4.2e-06 | 3.5e-06 | 2.6e-06 | 3.433E-06 | 8.021E-07 | 2.600E-06 | 4.200E-06 |
| 12457 | 623 | D623T | B | 3.3e-06 | 2.1e-06 | 2.8e-06 | 2.733E-06 | 6.028E-07 | 2.100E-06 | 3.300E-06 |
| 12458 | 623 | D623W | B | 3.3e-06 | 4.1e-06 | 2.7e-06 | 3.367E-06 | 7.024E-07 | 2.700E-06 | 4.100E-06 |
| 12459 | 623 | D623Y | B | 4.2e-06 | 5e-06 | 3.2e-06 | 4.133E-06 | 9.018E-07 | 3.200E-06 | 5.000E-06 |
| 12460 | 623 | D623V | B | 2.6e-06 | 3.4e-06 | 2.4e-06 | 2.800E-06 | 5.292E-07 | 2.400E-06 | 3.400E-06 |
| 12461 | 624 | I624A | B | 1.5e-06 | 2.4e-06 | 3.2e-06 | 2.367E-06 | 8.505E-07 | 1.500E-06 | 3.200E-06 |
| 12462 | 624 | I624R | B | 2.8e-06 | 2.4e-06 | 2.3e-06 | 2.500E-06 | 2.646E-07 | 2.300E-06 | 2.800E-06 |
| 12463 | 624 | I624N | B | 2.3e-06 | 2.1e-06 | 2.1e-06 | 2.167E-06 | 1.155E-07 | 2.100E-06 | 2.300E-06 |
| 12464 | 624 | I624D | B | 2.1e-06 | 3.8e-06 | 3.4e-06 | 3.100E-06 | 8.888E-07 | 2.100E-06 | 3.800E-06 |
| 12465 | 624 | I624C | B | 1.8e-06 | 1.8e-06 | 2.3e-06 | 1.967E-06 | 2.887E-07 | 1.800E-06 | 2.300E-06 |
| 12466 | 624 | I624E | B | 1.9e-06 | 1.7e-06 | 3.4e-06 | 2.333E-06 | 9.292E-07 | 1.700E-06 | 3.400E-06 |
| 12467 | 624 | I624Q | B | 2e-06 | 3e-06 | 3.2e-06 | 2.733E-06 | 6.429E-07 | 2.000E-06 | 3.200E-06 |
| 12468 | 624 | I624G | B | 2.7e-06 | 2.1e-06 | 1.5e-06 | 2.100E-06 | 6.000E-07 | 1.500E-06 | 2.700E-06 |
| 12469 | 624 | I624H | B | 2.7e-06 | 3e-06 | 1.8e-06 | 2.500E-06 | 6.245E-07 | 1.800E-06 | 3.000E-06 |
| 12470 | 624 | I624I | B | 2.4e-06 | 3.5e-06 | 1.7e-06 | 2.533E-06 | 9.074E-07 | 1.700E-06 | 3.500E-06 |
| 12471 | 624 | I624L | B | 2.1e-06 | 2.4e-06 | 2.2e-06 | 2.233E-06 | 1.528E-07 | 2.100E-06 | 2.400E-06 |
| 12472 | 624 | I624K | B | 2.5e-06 | 1.8e-06 | 3.5e-06 | 2.600E-06 | 8.544E-07 | 1.800E-06 | 3.500E-06 |
| 12473 | 624 | I624M | B | 3.4e-06 | 1.6e-06 | 3.1e-06 | 2.700E-06 | 9.644E-07 | 1.600E-06 | 3.400E-06 |
| 12474 | 624 | I624F | B | 7.4e-07 | 2.1e-06 | 1e-06 | 1.280E-06 | 7.219E-07 | 7.400E-07 | 2.100E-06 |
| 12475 | 624 | I624P | B | 2.2e-06 | 3.1e-06 | 1.6e-06 | 2.300E-06 | 7.550E-07 | 1.600E-06 | 3.100E-06 |
| 12476 | 624 | I624S | B | 4.4e-06 | 2.8e-06 | 2e-06 | 3.067E-06 | 1.222E-06 | 2.000E-06 | 4.400E-06 |
| 12477 | 624 | I624T | B | 3.1e-06 | 1.6e-06 | 2.7e-06 | 2.467E-06 | 7.767E-07 | 1.600E-06 | 3.100E-06 |
| 12478 | 624 | I624W | B | 1.6e-06 | 1.3e-06 | 1.4e-06 | 1.433E-06 | 1.528E-07 | 1.300E-06 | 1.600E-06 |
| 12479 | 624 | I624Y | B | 2.4e-06 | 1.6e-06 | 1.9e-06 | 1.967E-06 | 4.041E-07 | 1.600E-06 | 2.400E-06 |
| 12480 | 624 | I624V | B | 1.8e-06 | 1.7e-06 | 3.1e-06 | 2.200E-06 | 7.810E-07 | 1.700E-06 | 3.100E-06 |
| 12481 | 625 | Q625A | B | 2.1e-06 | 2.1e-06 | 2.6e-06 | 2.267E-06 | 2.887E-07 | 2.100E-06 | 2.600E-06 |
| 12482 | 625 | Q625R | B | 3.8e-06 | 2.3e-06 | 3.5e-06 | 3.200E-06 | 7.937E-07 | 2.300E-06 | 3.800E-06 |
| 12483 | 625 | Q625N | B | 1.4e-06 | 2.3e-06 | 2.5e-06 | 2.067E-06 | 5.859E-07 | 1.400E-06 | 2.500E-06 |
| 12484 | 625 | Q625D | B | 3.2e-06 | 2.5e-06 | 2.4e-06 | 2.700E-06 | 4.359E-07 | 2.400E-06 | 3.200E-06 |
| 12485 | 625 | Q625C | B | 1.2e-06 | 2.6e-06 | 2.4e-06 | 2.067E-06 | 7.572E-07 | 1.200E-06 | 2.600E-06 |
| 12486 | 625 | Q625E | B | 3.9e-06 | 2.4e-06 | 3e-06 | 3.100E-06 | 7.550E-07 | 2.400E-06 | 3.900E-06 |
| 12487 | 625 | Q625Q | B | 2.5e-06 | 2.2e-06 | 2e-06 | 2.233E-06 | 2.517E-07 | 2.000E-06 | 2.500E-06 |
| 12488 | 625 | Q625G | B | 2.1e-06 | 3.1e-06 | 1.9e-06 | 2.367E-06 | 6.429E-07 | 1.900E-06 | 3.100E-06 |
| 12489 | 625 | Q625H | B | 2.2e-06 | 3.1e-06 | 3.2e-06 | 2.833E-06 | 5.508E-07 | 2.200E-06 | 3.200E-06 |
| 12490 | 625 | Q625I | B | 3e-06 | 1.6e-06 | 1.4e-06 | 2.000E-06 | 8.718E-07 | 1.400E-06 | 3.000E-06 |
| 12491 | 625 | Q625L | B | 1.9e-06 | 2.2e-06 | 2.1e-06 | 2.067E-06 | 1.528E-07 | 1.900E-06 | 2.200E-06 |
| 12492 | 625 | Q625K | B | 2.8e-06 | 4.5e-06 | 2.1e-06 | 3.133E-06 | 1.234E-06 | 2.100E-06 | 4.500E-06 |
| 12493 | 625 | Q625M | B | 2.2e-06 | 2e-06 | 2.1e-06 | 2.100E-06 | 1.000E-07 | 2.000E-06 | 2.200E-06 |
| 12494 | 625 | Q625F | B | 2.6e-06 | 2.8e-06 | 1.4e-06 | 2.267E-06 | 7.572E-07 | 1.400E-06 | 2.800E-06 |

| | | | | | | | | | | |
|---|---|---|---|---|---|---|---|---|---|---|
| 12495 | 625 | Q625P | B | 2.2e-06 | 2.9e-06 | 2.6e-06 | 2.567E-06 | 3.512E-07 | 2.200E-06 | 2.900E-06 |
| 12496 | 625 | Q625S | B | 2.3e-06 | 4e-06 | 2.9e-06 | 3.067E-06 | 8.622E-07 | 2.300E-06 | 4.000E-06 |
| 12497 | 625 | Q625T | B | 2.2e-06 | 3.4e-06 | 1.8e-06 | 2.467E-06 | 8.327E-07 | 1.800E-06 | 3.400E-06 |
| 12498 | 625 | Q625W | B | 1.5e-06 | 3.1e-06 | 1.9e-06 | 2.167E-06 | 8.327E-07 | 1.500E-06 | 3.100E-06 |
| 12499 | 625 | Q625Y | B | 4.4e-06 | 3.7e-06 | 3.3e-06 | 3.800E-06 | 5.568E-07 | 3.300E-06 | 4.400E-06 |
| 12500 | 625 | Q625V | B | 2.3e-06 | 1.3e-06 | 2.2e-06 | 1.933E-06 | 5.508E-07 | 1.300E-06 | 2.300E-06 |
| 12501 | 626 | M626A | B | 3.2e-06 | 3.6e-06 | 2.1e-06 | 2.967E-06 | 7.767E-07 | 2.100E-06 | 3.600E-06 |
| 12502 | 626 | M626R | B | 2.1e-06 | 2.2e-06 | 2e-06 | 2.100E-06 | 1.000E-07 | 2.000E-06 | 2.200E-06 |
| 12503 | 626 | M626N | B | 3.3e-06 | 2.7e-06 | 1.3e-06 | 2.433E-06 | 1.026E-06 | 1.300E-06 | 3.300E-06 |
| 12504 | 626 | M626D | B | 2.8e-06 | 2.5e-06 | 2.6e-06 | 2.633E-06 | 1.528E-07 | 2.500E-06 | 2.800E-06 |
| 12505 | 626 | M626C | B | 2.3e-06 | 2.2e-06 | 2.4e-06 | 2.300E-06 | 1.000E-07 | 2.200E-06 | 2.400E-06 |
| 12506 | 626 | M626E | B | 2.9e-06 | 2.3e-06 | 2.1e-06 | 2.433E-06 | 4.163E-07 | 2.100E-06 | 2.900E-06 |
| 12507 | 626 | M626Q | B | 2.9e-06 | 2.4e-06 | 1.9e-06 | 2.400E-06 | 5.000E-07 | 1.900E-06 | 2.900E-06 |
| 12508 | 626 | M626G | B | 2.6e-06 | 2.7e-06 | 1.9e-06 | 2.400E-06 | 4.359E-07 | 1.900E-06 | 2.700E-06 |
| 12509 | 626 | M626H | B | 4.3e-06 | 2.1e-06 | 1.6e-06 | 2.667E-06 | 1.436E-06 | 1.600E-06 | 4.300E-06 |
| 12510 | 626 | M626I | B | 3.3e-06 | 1.4e-06 | 3e-06 | 2.567E-06 | 1.021E-06 | 1.400E-06 | 3.300E-06 |
| 12511 | 626 | M626L | B | 1.3e-06 | 2.9e-06 | 2e-06 | 2.067E-06 | 8.021E-07 | 1.300E-06 | 2.900E-06 |
| 12512 | 626 | M626K | B | 2.3e-06 | 2.7e-06 | 1.3e-06 | 2.100E-06 | 7.211E-07 | 1.300E-06 | 2.700E-06 |
| 12513 | 626 | M626M | B | 3.1e-06 | 1.8e-06 | 1.8e-06 | 2.233E-06 | 7.506E-07 | 1.800E-06 | 3.100E-06 |
| 12514 | 626 | M626F | B | 2e-06 | 1.5e-06 | 1.6e-06 | 1.700E-06 | 2.646E-07 | 1.500E-06 | 2.000E-06 |
| 12515 | 626 | M626P | B | 2.9e-06 | 1.9e-06 | 1.5e-06 | 2.100E-06 | 7.211E-07 | 1.500E-06 | 2.900E-06 |
| 12516 | 626 | M626S | B | 1.7e-06 | 4.1e-06 | 3e-06 | 2.933E-06 | 1.201E-06 | 1.700E-06 | 4.100E-06 |
| 12517 | 626 | M626T | B | 2.7e-06 | 2.6e-06 | 1.8e-06 | 2.367E-06 | 4.933E-07 | 1.800E-06 | 2.700E-06 |
| 12518 | 626 | M626W | B | 3.7e-06 | 1.2e-06 | 1.2e-06 | 2.033E-06 | 1.443E-06 | 1.200E-06 | 3.700E-06 |
| 12519 | 626 | M626Y | B | 7.2e-07 | 2.1e-06 | 2.9e-06 | 1.907E-06 | 1.103E-06 | 7.200E-07 | 2.900E-06 |
| 12520 | 626 | M626V | B | 2.6e-06 | 4.7e-06 | 2.6e-06 | 3.300E-06 | 1.212E-06 | 2.600E-06 | 4.700E-06 |
| 12521 | 627 | T627A | B | 2.5e-06 | 4.4e-06 | 3.6e-06 | 3.500E-06 | 9.539E-07 | 2.500E-06 | 4.400E-06 |
| 12522 | 627 | T627R | B | 2.6e-06 | 2e-06 | 3.7e-06 | 2.767E-06 | 8.622E-07 | 2.000E-06 | 3.700E-06 |
| 12523 | 627 | T627N | B | 2.3e-06 | 2.3e-06 | 1.9e-06 | 2.167E-06 | 2.309E-07 | 1.900E-06 | 2.300E-06 |
| 12524 | 627 | T627D | B | 1.9e-06 | 3e-06 | 1e-06 | 1.967E-06 | 1.002E-06 | 1.000E-06 | 3.000E-06 |
| 12525 | 627 | T627C | B | 4.3e-06 | 2.3e-06 | 4.8e-06 | 3.800E-06 | 1.323E-06 | 2.300E-06 | 4.800E-06 |
| 12526 | 627 | T627E | B | 1.8e-06 | 2.8e-06 | 3.1e-06 | 2.567E-06 | 6.807E-07 | 1.800E-06 | 3.100E-06 |
| 12527 | 627 | T627Q | B | 1.7e-06 | 4.1e-06 | 2.8e-06 | 2.867E-06 | 1.201E-06 | 1.700E-06 | 4.100E-06 |
| 12528 | 627 | T627G | B | 4.9e-06 | 2.4e-06 | 2.1e-06 | 3.133E-06 | 1.537E-06 | 2.100E-06 | 4.900E-06 |
| 12529 | 627 | T627H | B | 2.6e-06 | 1.3e-06 | 2.7e-06 | 2.200E-06 | 7.810E-07 | 1.300E-06 | 2.700E-06 |
| 12530 | 627 | T627I | B | 2.5e-06 | 3.6e-06 | 2.5e-06 | 2.867E-06 | 6.351E-07 | 2.500E-06 | 3.600E-06 |
| 12531 | 627 | T627L | B | 1.2e-06 | 2.5e-06 | 2.6e-06 | 2.100E-06 | 7.810E-07 | 1.200E-06 | 2.600E-06 |
| 12532 | 627 | T627K | B | 3.4e-06 | 2.9e-06 | 2.3e-06 | 2.867E-06 | 5.508E-07 | 2.300E-06 | 3.400E-06 |
| 12533 | 627 | T627M | B | 2.7e-06 | 2.4e-06 | 2.2e-06 | 2.433E-06 | 2.517E-07 | 2.200E-06 | 2.700E-06 |
| 12534 | 627 | T627F | B | 3e-06 | 2.3e-06 | 3.2e-06 | 2.833E-06 | 4.726E-07 | 2.300E-06 | 3.200E-06 |
| 12535 | 627 | T627P | B | 1.2e-06 | 3.3e-06 | 3.3e-06 | 2.600E-06 | 1.212E-06 | 1.200E-06 | 3.300E-06 |
| 12536 | 627 | T627S | B | 1.8e-06 | 3.5e-06 | 2.4e-06 | 2.567E-06 | 8.622E-07 | 1.800E-06 | 3.500E-06 |
| 12537 | 627 | T627T | B | 1.7e-06 | 1.2e-06 | 2.2e-06 | 1.700E-06 | 5.000E-07 | 1.200E-06 | 2.200E-06 |
| 12538 | 627 | T627W | B | 2.6e-06 | 2e-06 | 2.4e-06 | 2.333E-06 | 3.055E-07 | 2.000E-06 | 2.600E-06 |
| 12539 | 627 | T627Y | B | 3.2e-06 | 1.1e-06 | 3.2e-06 | 2.500E-06 | 1.212E-06 | 1.100E-06 | 3.200E-06 |
| 12540 | 627 | T627V | B | 1.3e-06 | 2e-06 | 1.7e-06 | 1.667E-06 | 3.512E-07 | 1.300E-06 | 2.000E-06 |
| 12541 | 628 | Q628A | B | 2.6e-06 | 2.1e-06 | 1.7e-06 | 2.133E-06 | 4.509E-07 | 1.700E-06 | 2.600E-06 |
| 12542 | 628 | Q628R | B | 1.6e-06 | 1.9e-06 | 1.5e-06 | 1.667E-06 | 2.082E-07 | 1.500E-06 | 1.900E-06 |
| 12543 | 628 | Q628N | B | 2.6e-06 | 2.4e-06 | 4.3e-06 | 3.100E-06 | 1.044E-06 | 2.400E-06 | 4.300E-06 |
| 12544 | 628 | Q628D | B | 1.3e-06 | 2.2e-06 | 2.5e-06 | 2.000E-06 | 6.245E-07 | 1.300E-06 | 2.500E-06 |
| 12545 | 628 | Q628C | B | 4.4e-06 | 1.9e-06 | 2.5e-06 | 2.933E-06 | 1.305E-06 | 1.900E-06 | 4.400E-06 |
| 12546 | 628 | Q628E | B | 1.7e-06 | 2.9e-06 | 2.4e-06 | 2.333E-06 | 6.028E-07 | 1.700E-06 | 2.900E-06 |
| 12547 | 628 | Q628Q | B | 5.2e-06 | 3.9e-06 | 1.8e-06 | 3.633E-06 | 1.716E-06 | 1.800E-06 | 5.200E-06 |
| 12548 | 628 | Q628G | B | 2.2e-06 | 3.4e-06 | 1.6e-06 | 2.400E-06 | 9.165E-07 | 1.600E-06 | 3.400E-06 |
| 12549 | 628 | Q628H | B | 1.5e-06 | 4.1e-06 | 2.6e-06 | 2.733E-06 | 1.305E-06 | 1.500E-06 | 4.100E-06 |
| 12550 | 628 | Q628I | B | 1.5e-06 | 3.2e-06 | 2.2e-06 | 2.300E-06 | 8.544E-07 | 1.500E-06 | 3.200E-06 |
| 12551 | 628 | Q628L | B | 3.1e-06 | 2.4e-06 | 2.4e-06 | 2.633E-06 | 4.041E-07 | 2.400E-06 | 3.100E-06 |
| 12552 | 628 | Q628K | B | 1.9e-06 | 2.4e-06 | 1.8e-06 | 2.033E-06 | 3.215E-07 | 1.800E-06 | 2.400E-06 |
| 12553 | 628 | Q628M | B | 2.6e-06 | 2.1e-06 | 1.6e-06 | 2.100E-06 | 5.000E-07 | 1.600E-06 | 2.600E-06 |
| 12554 | 628 | Q628F | B | 2.6e-06 | 1.8e-06 | 2.9e-06 | 2.433E-06 | 5.686E-07 | 1.800E-06 | 2.900E-06 |
| 12555 | 628 | Q628P | B | 1.6e-06 | 1.9e-06 | 5.5e-06 | 3.000E-06 | 2.170E-06 | 1.600E-06 | 5.500E-06 |
| 12556 | 628 | Q628S | B | 2.9e-06 | 2.2e-06 | 2.5e-06 | 2.533E-06 | 3.512E-07 | 2.200E-06 | 2.900E-06 |
| 12557 | 628 | Q628T | B | 1.9e-06 | 2.3e-06 | 3.6e-06 | 2.600E-06 | 8.888E-07 | 1.900E-06 | 3.600E-06 |
| 12558 | 628 | Q628W | B | 1.7e-06 | 2.4e-06 | 2.6e-06 | 2.233E-06 | 4.726E-07 | 1.700E-06 | 2.600E-06 |
| 12559 | 628 | Q628Y | B | 2.3e-06 | 2.5e-06 | 3e-06 | 2.600E-06 | 3.606E-07 | 2.300E-06 | 3.000E-06 |
| 12560 | 628 | Q628V | B | 3e-06 | 2e-06 | 2.9e-06 | 2.633E-06 | 5.508E-07 | 2.000E-06 | 3.000E-06 |
| 12561 | 629 | S629A | B | 2.3e-06 | 2.4e-06 | 5e-06 | 3.233E-06 | 1.531E-06 | 2.300E-06 | 5.000E-06 |
| 12562 | 629 | S629R | B | 2.7e-06 | 2.9e-06 | 3e-06 | 2.867E-06 | 1.528E-07 | 2.700E-06 | 3.000E-06 |
| 12563 | 629 | S629N | B | 3.3e-06 | 2.6e-06 | 2.1e-06 | 2.667E-06 | 6.028E-07 | 2.100E-06 | 3.300E-06 |
| 12564 | 629 | S629D | B | 2.5e-06 | 2e-06 | 3e-06 | 2.500E-06 | 5.000E-07 | 2.000E-06 | 3.000E-06 |
| 12565 | 629 | S629C | B | 1.6e-06 | 1.9e-06 | 3.9e-06 | 2.467E-06 | 1.250E-06 | 1.600E-06 | 3.900E-06 |
| 12566 | 629 | S629E | B | 2.1e-06 | 2.3e-06 | 2.1e-06 | 2.167E-06 | 1.155E-07 | 2.100E-06 | 2.300E-06 |
| 12567 | 629 | S629Q | B | 2.1e-06 | 2.9e-06 | 2.1e-06 | 2.367E-06 | 4.619E-07 | 2.100E-06 | 2.900E-06 |
| 12568 | 629 | S629G | B | 3.4e-06 | 1.9e-06 | 3.6e-06 | 2.967E-06 | 9.292E-07 | 1.900E-06 | 3.600E-06 |
| 12569 | 629 | S629H | B | 1.4e-06 | 2.2e-06 | 1.3e-06 | 1.633E-06 | 4.933E-07 | 1.300E-06 | 2.200E-06 |
| 12570 | 629 | S629I | B | 2.6e-06 | 2.1e-06 | 3.9e-06 | 2.867E-06 | 9.292E-07 | 2.100E-06 | 3.900E-06 |
| 12571 | 629 | S629L | B | 2e-06 | 1.9e-06 | 2.5e-06 | 2.133E-06 | 3.215E-07 | 1.900E-06 | 2.500E-06 |
| 12572 | 629 | S629K | B | 3e-06 | 2.1e-06 | 3e-06 | 2.700E-06 | 5.196E-07 | 2.100E-06 | 3.000E-06 |
| 12573 | 629 | S629M | B | 2.8e-06 | 2e-06 | 2.6e-06 | 2.467E-06 | 4.163E-07 | 2.000E-06 | 2.800E-06 |
| 12574 | 629 | S629F | B | 2.7e-06 | 3.7e-06 | 2.8e-06 | 3.067E-06 | 5.508E-07 | 2.700E-06 | 3.700E-06 |
| 12575 | 629 | S629P | B | 2.6e-06 | 1.9e-06 | 2.4e-06 | 2.300E-06 | 3.606E-07 | 1.900E-06 | 2.600E-06 |
| 12576 | 629 | S629S | B | 4.5e-06 | 2.7e-06 | 1.8e-06 | 3.000E-06 | 1.375E-06 | 1.800E-06 | 4.500E-06 |
| 12577 | 629 | S629T | B | 1.5e-06 | 3.9e-06 | 2.2e-06 | 2.533E-06 | 1.234E-06 | 1.500E-06 | 3.900E-06 |
| 12578 | 629 | S629W | B | 2.5e-06 | 2.3e-06 | 2.1e-06 | 2.300E-06 | 2.000E-07 | 2.100E-06 | 2.500E-06 |
| 12579 | 629 | S629Y | B | 3.2e-06 | 3.3e-06 | 2.8e-06 | 3.100E-06 | 2.646E-07 | 2.800E-06 | 3.300E-06 |

| | | | | | | | | | | |
|---|---|---|---|---|---|---|---|---|---|---|
| 12580 | 629 | S629V | B | 4.1e-06 | 2.5e-06 | 3e-06 | 3.200E-06 | 8.185E-07 | 2.500E-06 | 4.100E-06 |
| 12581 | 630 | P630A | B | 1.8e-06 | 3.2e-06 | 2.3e-06 | 2.433E-06 | 7.095E-07 | 1.800E-06 | 3.200E-06 |
| 12582 | 630 | P630R | B | 2.9e-06 | 3.1e-06 | 2.4e-06 | 2.800E-06 | 3.606E-07 | 2.400E-06 | 3.100E-06 |
| 12583 | 630 | P630N | B | 3.6e-06 | 2.8e-06 | 2.2e-06 | 2.867E-06 | 7.024E-07 | 2.200E-06 | 3.600E-06 |
| 12584 | 630 | P630D | B | 1.6e-06 | 1.6e-06 | 1.9e-06 | 1.700E-06 | 1.732E-07 | 1.600E-06 | 1.900E-06 |
| 12585 | 630 | P630C | B | 2.3e-06 | 2.5e-06 | 3.1e-06 | 2.633E-06 | 4.163E-07 | 2.300E-06 | 3.100E-06 |
| 12586 | 630 | P630E | B | 3.7e-06 | 2e-06 | 2.1e-06 | 2.600E-06 | 9.539E-07 | 2.000E-06 | 3.700E-06 |
| 12587 | 630 | P630Q | B | 2.2e-06 | 2.2e-06 | 2.3e-06 | 2.233E-06 | 5.774E-08 | 2.200E-06 | 2.300E-06 |
| 12588 | 630 | P630G | B | 3.3e-06 | 1.9e-06 | 3.5e-06 | 2.900E-06 | 8.718E-07 | 1.900E-06 | 3.500E-06 |
| 12589 | 630 | P630H | B | 4.2e-06 | 3e-06 | 2.3e-06 | 3.167E-06 | 9.609E-07 | 2.300E-06 | 4.200E-06 |
| 12590 | 630 | P630I | B | 2.8e-06 | 3.1e-06 | 1.9e-06 | 2.600E-06 | 6.245E-07 | 1.900E-06 | 3.100E-06 |
| 12591 | 630 | P630L | B | 1.9e-06 | 2e-06 | 3e-06 | 2.300E-06 | 6.083E-07 | 1.900E-06 | 3.000E-06 |
| 12592 | 630 | P630K | B | 1.9e-06 | 5.2e-06 | 2.4e-06 | 3.167E-06 | 1.779E-06 | 1.900E-06 | 5.200E-06 |
| 12593 | 630 | P630M | B | 3.9e-06 | 1.9e-06 | 1.7e-06 | 2.500E-06 | 1.217E-06 | 1.700E-06 | 3.900E-06 |
| 12594 | 630 | P630F | B | 2.8e-06 | 2.2e-06 | 6.3e-07 | 1.877E-06 | 1.121E-06 | 6.300E-07 | 2.800E-06 |
| 12595 | 630 | P630P | B | 2.8e-06 | 2e-06 | 1.7e-06 | 2.167E-06 | 5.686E-07 | 1.700E-06 | 2.800E-06 |
| 12596 | 630 | P630S | B | 1.6e-06 | 3e-06 | 2.9e-06 | 2.500E-06 | 7.810E-07 | 1.600E-06 | 3.000E-06 |
| 12597 | 630 | P630T | B | 2.7e-06 | 2.1e-06 | 4.2e-06 | 3.000E-06 | 1.082E-06 | 2.100E-06 | 4.200E-06 |
| 12598 | 630 | P630W | B | 3.1e-06 | 1.5e-06 | 2.8e-06 | 2.467E-06 | 8.505E-07 | 1.500E-06 | 3.100E-06 |
| 12599 | 630 | P630Y | B | 2.4e-06 | 2.8e-06 | 3.5e-06 | 2.900E-06 | 5.568E-07 | 2.400E-06 | 3.500E-06 |
| 12600 | 630 | P630V | B | 1.6e-06 | 2.2e-06 | 1.4e-06 | 1.733E-06 | 4.163E-07 | 1.400E-06 | 2.200E-06 |
| 12601 | 631 | S631A | B | 4.7e-06 | 2.4e-06 | 3.2e-06 | 3.433E-06 | 1.168E-06 | 2.400E-06 | 4.700E-06 |
| 12602 | 631 | S631R | B | 3.7e-06 | 3.4e-06 | 2.7e-06 | 3.267E-06 | 5.132E-07 | 2.700E-06 | 3.700E-06 |
| 12603 | 631 | S631N | B | 2.3e-06 | 1.9e-06 | 1.7e-06 | 1.967E-06 | 3.055E-07 | 1.700E-06 | 2.300E-06 |
| 12604 | 631 | S631D | B | 1.8e-06 | 2.3e-06 | 2.6e-06 | 2.233E-06 | 4.041E-07 | 1.800E-06 | 2.600E-06 |
| 12605 | 631 | S631C | B | 2.6e-06 | 4.9e-06 | 2.4e-06 | 3.300E-06 | 1.389E-06 | 2.400E-06 | 4.900E-06 |
| 12606 | 631 | S631E | B | 2.3e-06 | 2e-06 | 1.6e-06 | 1.967E-06 | 3.512E-07 | 1.600E-06 | 2.300E-06 |
| 12607 | 631 | S631Q | B | 2.6e-06 | 2.3e-06 | 1.8e-06 | 2.233E-06 | 4.041E-07 | 1.800E-06 | 2.600E-06 |
| 12608 | 631 | S631G | B | 1.9e-06 | 3.3e-06 | 2.8e-06 | 2.667E-06 | 7.095E-07 | 1.900E-06 | 3.300E-06 |
| 12609 | 631 | S631H | B | 2.3e-06 | 2.1e-06 | 3.9e-06 | 2.767E-06 | 9.866E-07 | 2.100E-06 | 3.900E-06 |
| 12610 | 631 | S631I | B | 1.6e-06 | 3e-06 | 1.3e-06 | 1.967E-06 | 9.074E-07 | 1.300E-06 | 3.000E-06 |
| 12611 | 631 | S631L | B | 2.1e-06 | 2e-06 | 3.1e-06 | 2.400E-06 | 6.083E-07 | 2.000E-06 | 3.100E-06 |
| 12612 | 631 | S631K | B | 2e-06 | 2e-06 | 1.8e-06 | 1.933E-06 | 1.155E-07 | 1.800E-06 | 2.000E-06 |
| 12613 | 631 | S631M | B | 2.3e-06 | 3.8e-06 | 3e-06 | 3.033E-06 | 7.506E-07 | 2.300E-06 | 3.800E-06 |
| 12614 | 631 | S631F | B | 3.8e-06 | 2.7e-06 | 3.7e-06 | 3.400E-06 | 6.083E-07 | 2.700E-06 | 3.800E-06 |
| 12615 | 631 | S631P | B | 2.3e-06 | 2.1e-06 | 1.9e-06 | 2.100E-06 | 2.000E-07 | 1.900E-06 | 2.300E-06 |
| 12616 | 631 | S631S | B | 2.7e-06 | 2.5e-06 | 2.1e-06 | 2.433E-06 | 3.055E-07 | 2.100E-06 | 2.700E-06 |
| 12617 | 631 | S631T | B | 2.4e-06 | 4e-06 | 3e-06 | 3.133E-06 | 8.083E-07 | 2.400E-06 | 4.000E-06 |
| 12618 | 631 | S631W | B | 7.4e-07 | 2.2e-06 | 2.2e-06 | 1.713E-06 | 8.429E-07 | 7.400E-07 | 2.200E-06 |
| 12619 | 631 | S631Y | B | 2e-06 | 1.7e-06 | 1.6e-06 | 1.767E-06 | 2.082E-07 | 1.600E-06 | 2.000E-06 |
| 12620 | 631 | S631V | B | 2.3e-06 | 2.3e-06 | 4.2e-06 | 2.933E-06 | 1.097E-06 | 2.300E-06 | 4.200E-06 |
| 12621 | 632 | S632A | B | 3.2e-06 | 1.8e-06 | 1.8e-06 | 2.267E-06 | 8.083E-07 | 1.800E-06 | 3.200E-06 |
| 12622 | 632 | S632R | B | 2.1e-06 | 2.4e-06 | 2.1e-06 | 2.200E-06 | 1.732E-07 | 2.100E-06 | 2.400E-06 |
| 12623 | 632 | S632N | B | 2.6e-06 | 2.3e-06 | 2.6e-06 | 2.500E-06 | 1.732E-07 | 2.300E-06 | 2.600E-06 |
| 12624 | 632 | S632D | B | 3.4e-06 | 3.2e-06 | 3.1e-06 | 3.233E-06 | 1.528E-07 | 3.100E-06 | 3.400E-06 |
| 12625 | 632 | S632C | B | 2.2e-06 | 1.7e-06 | 2.4e-06 | 2.100E-06 | 3.606E-07 | 1.700E-06 | 2.400E-06 |
| 12626 | 632 | S632E | B | 2.6e-06 | 2.8e-06 | 1.7e-06 | 2.367E-06 | 5.859E-07 | 1.700E-06 | 2.800E-06 |
| 12627 | 632 | S632Q | B | 1.5e-06 | 2.7e-06 | 3.5e-06 | 2.567E-06 | 1.007E-06 | 1.500E-06 | 3.500E-06 |
| 12628 | 632 | S632G | B | 2.7e-06 | 2.6e-06 | 1.6e-06 | 2.300E-06 | 6.083E-07 | 1.600E-06 | 2.700E-06 |
| 12629 | 632 | S632H | B | 2.6e-06 | 3e-06 | 2e-06 | 2.533E-06 | 5.033E-07 | 2.000E-06 | 3.000E-06 |
| 12630 | 632 | S632I | B | 3.3e-06 | 2.1e-06 | 2.3e-06 | 2.567E-06 | 6.429E-07 | 2.100E-06 | 3.300E-06 |
| 12631 | 632 | S632L | B | 3.9e-06 | 2.6e-06 | 1.9e-06 | 2.800E-06 | 1.015E-06 | 1.900E-06 | 3.900E-06 |
| 12632 | 632 | S632K | B | 3.9e-06 | 1.8e-06 | 3.6e-06 | 3.100E-06 | 1.136E-06 | 1.800E-06 | 3.900E-06 |
| 12633 | 632 | S632M | B | 9e-07 | 2.7e-06 | 1.7e-06 | 1.767E-06 | 9.018E-07 | 9.000E-07 | 2.700E-06 |
| 12634 | 632 | S632F | B | 1.6e-06 | 2.6e-06 | 2e-06 | 2.067E-06 | 5.033E-07 | 1.600E-06 | 2.600E-06 |
| 12635 | 632 | S632P | B | 3.2e-06 | 3.8e-06 | 1.7e-06 | 2.900E-06 | 1.082E-06 | 1.700E-06 | 3.800E-06 |
| 12636 | 632 | S632S | B | 3.7e-06 | 2.5e-06 | 1.7e-06 | 2.633E-06 | 1.007E-06 | 1.700E-06 | 3.700E-06 |
| 12637 | 632 | S632T | B | 1.7e-06 | 3e-06 | 3.2e-06 | 2.633E-06 | 8.145E-07 | 1.700E-06 | 3.200E-06 |
| 12638 | 632 | S632W | B | 1.7e-06 | 2.5e-06 | 2.3e-06 | 2.167E-06 | 4.163E-07 | 1.700E-06 | 2.500E-06 |
| 12639 | 632 | S632Y | B | 1.9e-06 | 2.6e-06 | 1.9e-06 | 2.133E-06 | 4.041E-07 | 1.900E-06 | 2.600E-06 |
| 12640 | 632 | S632V | B | 2.9e-06 | 1.7e-06 | 2.6e-06 | 2.400E-06 | 6.245E-07 | 1.700E-06 | 2.900E-06 |
| 12641 | 633 | L633A | B | 1.8e-06 | 2.6e-06 | 2e-06 | 2.133E-06 | 4.163E-07 | 1.800E-06 | 2.600E-06 |
| 12642 | 633 | L633R | B | 1.4e-06 | 2.7e-06 | 1.7e-06 | 1.933E-06 | 6.807E-07 | 1.400E-06 | 2.700E-06 |
| 12643 | 633 | L633N | B | 2.7e-06 | 2e-06 | 1.3e-06 | 2.000E-06 | 7.000E-07 | 1.300E-06 | 2.700E-06 |
| 12644 | 633 | L633D | B | 2.9e-06 | 3e-06 | 1.8e-06 | 2.567E-06 | 6.658E-07 | 1.800E-06 | 3.000E-06 |
| 12645 | 633 | L633C | B | 1.8e-06 | 4.2e-06 | 2e-06 | 2.667E-06 | 1.332E-06 | 1.800E-06 | 4.200E-06 |
| 12646 | 633 | L633E | B | 2.5e-06 | 1.6e-06 | 4.2e-06 | 2.767E-06 | 1.320E-06 | 1.600E-06 | 4.200E-06 |
| 12647 | 633 | L633Q | B | 2.5e-06 | 3.1e-06 | 2.1e-06 | 2.567E-06 | 5.033E-07 | 2.100E-06 | 3.100E-06 |
| 12648 | 633 | L633G | B | 1.8e-06 | 2.1e-06 | 2.2e-06 | 2.033E-06 | 2.082E-07 | 1.800E-06 | 2.200E-06 |
| 12649 | 633 | L633H | B | 3e-06 | 2.3e-06 | 2.5e-06 | 2.600E-06 | 3.606E-07 | 2.300E-06 | 3.000E-06 |
| 12650 | 633 | L633I | B | 2.5e-06 | 1.6e-06 | 2.3e-06 | 2.133E-06 | 4.726E-07 | 1.600E-06 | 2.500E-06 |
| 12651 | 633 | L633L | B | 2.2e-06 | 2.9e-06 | 2.7e-06 | 2.600E-06 | 3.606E-07 | 2.200E-06 | 2.900E-06 |
| 12652 | 633 | L633K | B | 2.2e-06 | 2.2e-06 | 2.5e-06 | 2.300E-06 | 1.732E-07 | 2.200E-06 | 2.500E-06 |
| 12653 | 633 | L633M | B | 2.5e-06 | 4.2e-06 | 3.6e-06 | 3.433E-06 | 8.622E-07 | 2.500E-06 | 4.200E-06 |
| 12654 | 633 | L633F | B | 2.3e-06 | 4.7e-06 | 2.6e-06 | 3.200E-06 | 1.308E-06 | 2.300E-06 | 4.700E-06 |
| 12655 | 633 | L633P | B | 3.5e-06 | 1.5e-06 | 3.5e-06 | 2.833E-06 | 1.155E-06 | 1.500E-06 | 3.500E-06 |
| 12656 | 633 | L633S | B | 1.4e-06 | 4.2e-06 | 2.6e-06 | 2.733E-06 | 1.405E-06 | 1.400E-06 | 4.200E-06 |
| 12657 | 633 | L633T | B | 1.9e-06 | 3.2e-06 | 3.3e-06 | 2.800E-06 | 7.810E-07 | 1.900E-06 | 3.300E-06 |
| 12658 | 633 | L633W | B | 2.6e-06 | 2.8e-06 | 1.6e-06 | 2.333E-06 | 6.429E-07 | 1.600E-06 | 2.800E-06 |
| 12659 | 633 | L633Y | B | 3.5e-06 | 3.3e-06 | 2.7e-06 | 3.167E-06 | 4.163E-07 | 2.700E-06 | 3.500E-06 |
| 12660 | 633 | L633V | B | 2.7e-06 | 2.4e-06 | 1.3e-06 | 2.133E-06 | 7.371E-07 | 1.300E-06 | 2.700E-06 |
| 12661 | 634 | S634A | B | 2.4e-06 | 1.4e-06 | 2.9e-06 | 2.233E-06 | 7.638E-07 | 1.400E-06 | 2.900E-06 |
| 12662 | 634 | S634R | B | 2.9e-06 | 1.9e-06 | 1.7e-06 | 2.167E-06 | 6.429E-07 | 1.700E-06 | 2.900E-06 |
| 12663 | 634 | S634N | B | 1.2e-06 | 3.4e-06 | 2.3e-06 | 2.300E-06 | 1.100E-06 | 1.200E-06 | 3.400E-06 |
| 12664 | 634 | S634D | B | 2.4e-06 | 2.7e-06 | 1.4e-06 | 2.167E-06 | 6.807E-07 | 1.400E-06 | 2.700E-06 |

| | | | | | | | | | | |
|---|---|---|---|---|---|---|---|---|---|---|
| 12665 | 634 | S634C | B | 1.5e-06 | 1.8e-06 | 3.2e-06 | 2.167E-06 | 9.074E-07 | 1.500E-06 | 3.200E-06 |
| 12666 | 634 | S634E | B | 2.3e-06 | 3e-06 | 2.4e-06 | 2.567E-06 | 3.786E-07 | 2.300E-06 | 3.000E-06 |
| 12667 | 634 | S634Q | B | 1.7e-06 | 1.3e-06 | 2.6e-06 | 1.867E-06 | 6.658E-07 | 1.300E-06 | 2.600E-06 |
| 12668 | 634 | S634G | B | 2e-06 | 2.8e-06 | 3.8e-06 | 2.867E-06 | 9.018E-07 | 2.000E-06 | 3.800E-06 |
| 12669 | 634 | S634H | B | 3e-06 | 2.9e-06 | 2.2e-06 | 2.700E-06 | 4.359E-07 | 2.200E-06 | 3.000E-06 |
| 12670 | 634 | S634I | B | 2.2e-06 | 2.5e-06 | 3.7e-06 | 2.800E-06 | 7.937E-07 | 2.200E-06 | 3.700E-06 |
| 12671 | 634 | S634L | B | 3e-06 | 3e-06 | 3.1e-06 | 3.033E-06 | 5.774E-08 | 3.000E-06 | 3.100E-06 |
| 12672 | 634 | S634K | B | 2.6e-06 | 2.9e-06 | 3.7e-06 | 3.067E-06 | 5.686E-07 | 2.600E-06 | 3.700E-06 |
| 12673 | 634 | S634M | B | 2.2e-06 | 1.7e-06 | 1.8e-06 | 1.900E-06 | 2.646E-07 | 1.700E-06 | 2.200E-06 |
| 12674 | 634 | S634F | B | 3.2e-06 | 2.5e-06 | 2.6e-06 | 2.767E-06 | 3.786E-07 | 2.500E-06 | 3.200E-06 |
| 12675 | 634 | S634P | B | 1.7e-06 | 1.7e-06 | 3.2e-06 | 2.200E-06 | 8.660E-07 | 1.700E-06 | 3.200E-06 |
| 12676 | 634 | S634S | B | 1.5e-06 | 2e-06 | 4e-06 | 2.500E-06 | 1.323E-06 | 1.500E-06 | 4.000E-06 |
| 12677 | 634 | S634T | B | 2.8e-06 | 3.1e-06 | 2.2e-06 | 2.700E-06 | 4.583E-07 | 2.200E-06 | 3.100E-06 |
| 12678 | 634 | S634W | B | 2.6e-06 | 3.4e-06 | 2.6e-06 | 2.867E-06 | 4.619E-07 | 2.600E-06 | 3.400E-06 |
| 12679 | 634 | S634Y | B | 3.1e-06 | 2.1e-06 | 2.8e-06 | 2.667E-06 | 5.132E-07 | 2.100E-06 | 3.100E-06 |
| 12680 | 634 | S634V | B | 2.3e-06 | 1.8e-06 | 3.4e-06 | 2.500E-06 | 8.185E-07 | 1.800E-06 | 3.400E-06 |
| 12681 | 635 | A635A | B | 4.4e-06 | 1.5e-06 | 2.5e-06 | 2.800E-06 | 1.473E-06 | 1.500E-06 | 4.400E-06 |
| 12682 | 635 | A635R | B | 3.2e-06 | 3.9e-06 | 1.8e-06 | 2.967E-06 | 1.069E-06 | 1.800E-06 | 3.900E-06 |
| 12683 | 635 | A635N | B | 2.7e-06 | 3e-06 | 3.5e-06 | 3.067E-06 | 4.041E-07 | 2.700E-06 | 3.500E-06 |
| 12684 | 635 | A635D | B | 2.6e-06 | 1.8e-06 | 1.5e-06 | 1.967E-06 | 5.686E-07 | 1.500E-06 | 2.600E-06 |
| 12685 | 635 | A635C | B | 2.6e-06 | 1.8e-06 | 3.9e-06 | 2.767E-06 | 1.060E-06 | 1.800E-06 | 3.900E-06 |
| 12686 | 635 | A635E | B | 1.7e-06 | 2.9e-06 | 3e-06 | 2.533E-06 | 7.234E-07 | 1.700E-06 | 3.000E-06 |
| 12687 | 635 | A635Q | B | 2.5e-06 | 3.1e-06 | 2.1e-06 | 2.567E-06 | 5.033E-07 | 2.100E-06 | 3.100E-06 |
| 12688 | 635 | A635G | B | 2.1e-06 | 1.8e-06 | 2e-06 | 1.967E-06 | 1.528E-07 | 1.800E-06 | 2.100E-06 |
| 12689 | 635 | A635H | B | 1.8e-06 | 3.1e-06 | 2.8e-06 | 2.567E-06 | 6.807E-07 | 1.800E-06 | 3.100E-06 |
| 12690 | 635 | A635I | B | 1.4e-06 | 2.2e-06 | 3.6e-06 | 2.400E-06 | 1.114E-06 | 1.400E-06 | 3.600E-06 |
| 12691 | 635 | A635L | B | 2.1e-06 | 1.8e-06 | 3e-06 | 2.300E-06 | 6.245E-07 | 1.800E-06 | 3.000E-06 |
| 12692 | 635 | A635K | B | 3.5e-06 | 3.9e-06 | 1.9e-06 | 3.100E-06 | 1.058E-06 | 1.900E-06 | 3.900E-06 |
| 12693 | 635 | A635M | B | 2.9e-06 | 2e-06 | 2.7e-06 | 2.533E-06 | 4.726E-07 | 2.000E-06 | 2.900E-06 |
| 12694 | 635 | A635F | B | 2.1e-06 | 1.9e-06 | 2.4e-06 | 2.133E-06 | 2.517E-07 | 1.900E-06 | 2.400E-06 |
| 12695 | 635 | A635P | B | 2.3e-06 | 3.8e-06 | 1.8e-06 | 2.633E-06 | 1.041E-06 | 1.800E-06 | 3.800E-06 |
| 12696 | 635 | A635S | B | 2.3e-06 | 2.2e-06 | 2.4e-06 | 2.300E-06 | 1.000E-07 | 2.200E-06 | 2.400E-06 |
| 12697 | 635 | A635T | B | 2.7e-06 | 2.6e-06 | 2.8e-06 | 2.700E-06 | 1.000E-07 | 2.600E-06 | 2.800E-06 |
| 12698 | 635 | A635W | B | 2.3e-06 | 2.5e-06 | 5.9e-06 | 3.567E-06 | 2.023E-06 | 2.300E-06 | 5.900E-06 |
| 12699 | 635 | A635Y | B | 2.5e-06 | 2.2e-06 | 2.7e-06 | 2.467E-06 | 2.517E-07 | 2.200E-06 | 2.700E-06 |
| 12700 | 635 | A635V | B | 2.2e-06 | 3.7e-06 | 1.9e-06 | 2.600E-06 | 9.644E-07 | 1.900E-06 | 3.700E-06 |
| 12701 | 636 | S636A | B | 2.2e-06 | 3.3e-06 | 2.1e-06 | 2.533E-06 | 6.658E-07 | 2.100E-06 | 3.300E-06 |
| 12702 | 636 | S636R | B | 2.5e-06 | 3.6e-06 | 4.1e-06 | 3.400E-06 | 8.185E-07 | 2.500E-06 | 4.100E-06 |
| 12703 | 636 | S636N | B | 3e-06 | 2.7e-06 | 1.9e-06 | 2.533E-06 | 5.686E-07 | 1.900E-06 | 3.000E-06 |
| 12704 | 636 | S636D | B | 2.1e-06 | 1.5e-06 | 2.4e-06 | 2.000E-06 | 4.583E-07 | 1.500E-06 | 2.400E-06 |
| 12705 | 636 | S636C | B | 1.8e-06 | 2e-06 | 2.9e-06 | 2.233E-06 | 5.859E-07 | 1.800E-06 | 2.900E-06 |
| 12706 | 636 | S636E | B | 2.4e-06 | 2.9e-06 | 1.8e-06 | 2.367E-06 | 5.508E-07 | 1.800E-06 | 2.900E-06 |
| 12707 | 636 | S636Q | B | 2.4e-06 | 2e-06 | 2.1e-06 | 2.167E-06 | 2.082E-07 | 2.000E-06 | 2.400E-06 |
| 12708 | 636 | S636G | B | 5.6e-06 | 3.4e-06 | 2.3e-06 | 3.767E-06 | 1.680E-06 | 2.300E-06 | 5.600E-06 |
| 12709 | 636 | S636H | B | 2e-06 | 2.4e-06 | 4.2e-06 | 2.867E-06 | 1.172E-06 | 2.000E-06 | 4.200E-06 |
| 12710 | 636 | S636I | B | 3.8e-06 | 2.1e-06 | 2.9e-06 | 2.933E-06 | 8.505E-07 | 2.100E-06 | 3.800E-06 |
| 12711 | 636 | S636L | B | 2.8e-06 | 2.7e-06 | 3e-06 | 2.833E-06 | 1.528E-07 | 2.700E-06 | 3.000E-06 |
| 12712 | 636 | S636K | B | 3.9e-06 | 1.4e-06 | 2.4e-06 | 2.567E-06 | 1.258E-06 | 1.400E-06 | 3.900E-06 |
| 12713 | 636 | S636M | B | 3.1e-06 | 2e-06 | 1.9e-06 | 2.333E-06 | 6.658E-07 | 1.900E-06 | 3.100E-06 |
| 12714 | 636 | S636F | B | 2.2e-06 | 1.9e-06 | 2.5e-06 | 2.200E-06 | 3.000E-07 | 1.900E-06 | 2.500E-06 |
| 12715 | 636 | S636P | B | 3e-06 | 3.9e-06 | 2.1e-06 | 3.000E-06 | 9.000E-07 | 2.100E-06 | 3.900E-06 |
| 12716 | 636 | S636S | B | 2.7e-06 | 2.6e-06 | 1.8e-06 | 2.367E-06 | 4.933E-07 | 1.800E-06 | 2.700E-06 |
| 12717 | 636 | S636T | B | 2.8e-06 | 2.2e-06 | 2e-06 | 2.333E-06 | 4.163E-07 | 2.000E-06 | 2.800E-06 |
| 12718 | 636 | S636W | B | 2.8e-06 | 2.3e-06 | 5.3e-06 | 3.467E-06 | 1.607E-06 | 2.300E-06 | 5.300E-06 |
| 12719 | 636 | S636Y | B | 1.9e-06 | 3.1e-06 | 2.4e-06 | 2.467E-06 | 6.028E-07 | 1.900E-06 | 3.100E-06 |
| 12720 | 636 | S636V | B | 2.9e-06 | 1.1e-06 | 2.6e-06 | 2.200E-06 | 9.644E-07 | 1.100E-06 | 2.900E-06 |
| 12721 | 637 | V637A | B | 1.5e-06 | 2.8e-06 | 2.4e-06 | 2.233E-06 | 6.658E-07 | 1.500E-06 | 2.800E-06 |
| 12722 | 637 | V637R | B | 3.5e-06 | 1.6e-06 | 3.2e-06 | 2.767E-06 | 1.021E-06 | 1.600E-06 | 3.500E-06 |
| 12723 | 637 | V637N | B | 1.4e-06 | 2.2e-06 | 3e-06 | 2.200E-06 | 8.000E-07 | 1.400E-06 | 3.000E-06 |
| 12724 | 637 | V637D | B | 3.2e-06 | 1.6e-06 | 1.9e-06 | 2.233E-06 | 8.505E-07 | 1.600E-06 | 3.200E-06 |
| 12725 | 637 | V637C | B | 2.4e-06 | 2.2e-06 | 1.5e-06 | 2.033E-06 | 4.726E-07 | 1.500E-06 | 2.400E-06 |
| 12726 | 637 | V637E | B | 1.9e-06 | 3.8e-06 | 2e-06 | 2.567E-06 | 1.069E-06 | 1.900E-06 | 3.800E-06 |
| 12727 | 637 | V637Q | B | 3.5e-06 | 1.3e-06 | 2e-06 | 2.267E-06 | 1.124E-06 | 1.300E-06 | 3.500E-06 |
| 12728 | 637 | V637G | B | 3.1e-06 | 2e-06 | 2.6e-06 | 2.567E-06 | 5.508E-07 | 2.000E-06 | 3.100E-06 |
| 12729 | 637 | V637H | B | 3.2e-06 | 1.6e-06 | 2.3e-06 | 2.367E-06 | 8.021E-07 | 1.600E-06 | 3.200E-06 |
| 12730 | 637 | V637I | B | 2.2e-06 | 1.8e-06 | 2.3e-06 | 2.100E-06 | 2.646E-07 | 1.800E-06 | 2.300E-06 |
| 12731 | 637 | V637L | B | 2.7e-06 | 3.1e-06 | 1.7e-06 | 2.500E-06 | 7.211E-07 | 1.700E-06 | 3.100E-06 |
| 12732 | 637 | V637K | B | 4.5e-06 | 1.7e-06 | 2.4e-06 | 2.867E-06 | 1.457E-06 | 1.700E-06 | 4.500E-06 |
| 12733 | 637 | V637M | B | 2e-06 | 2.7e-06 | 1.7e-06 | 2.133E-06 | 5.132E-07 | 1.700E-06 | 2.700E-06 |
| 12734 | 637 | V637F | B | 2.7e-06 | 2.3e-06 | 4.2e-06 | 3.067E-06 | 1.002E-06 | 2.300E-06 | 4.200E-06 |
| 12735 | 637 | V637P | B | 3.4e-06 | 2.8e-06 | 2.7e-06 | 2.967E-06 | 3.786E-07 | 2.700E-06 | 3.400E-06 |
| 12736 | 637 | V637S | B | 1.9e-06 | 1.3e-06 | 2.1e-06 | 1.767E-06 | 4.163E-07 | 1.300E-06 | 2.100E-06 |
| 12737 | 637 | V637T | B | 3.1e-06 | 1.6e-06 | 2.5e-06 | 2.400E-06 | 7.550E-07 | 1.600E-06 | 3.100E-06 |
| 12738 | 637 | V637W | B | 2.1e-06 | 1.5e-06 | 2.7e-06 | 2.100E-06 | 6.000E-07 | 1.500E-06 | 2.700E-06 |
| 12739 | 637 | V637Y | B | 1.3e-06 | 3.1e-06 | 1.8e-06 | 2.067E-06 | 9.292E-07 | 1.300E-06 | 3.100E-06 |
| 12740 | 637 | V637V | B | 3.4e-06 | 2.8e-06 | 2.6e-06 | 2.933E-06 | 4.163E-07 | 2.600E-06 | 3.400E-06 |
| 12741 | 638 | G638A | B | 1.7e-06 | 2.3e-06 | 5.9e-06 | 3.300E-06 | 2.272E-06 | 1.700E-06 | 5.900E-06 |
| 12742 | 638 | G638R | B | 3.4e-06 | 1.3e-06 | 2.2e-06 | 2.300E-06 | 1.054E-06 | 1.300E-06 | 3.400E-06 |
| 12743 | 638 | G638N | B | 2.2e-06 | 2.2e-06 | 1.9e-06 | 2.100E-06 | 1.732E-07 | 1.900E-06 | 2.200E-06 |
| 12744 | 638 | G638D | B | 1.4e-06 | 2.5e-06 | 4e-06 | 2.633E-06 | 1.305E-06 | 1.400E-06 | 4.000E-06 |
| 12745 | 638 | G638C | B | 2e-06 | 2.1e-06 | 3e-06 | 2.367E-06 | 5.508E-07 | 2.000E-06 | 3.000E-06 |
| 12746 | 638 | G638E | B | 1.8e-06 | 2.9e-06 | 2.3e-06 | 2.333E-06 | 5.508E-07 | 1.800E-06 | 2.900E-06 |
| 12747 | 638 | G638Q | B | 2.3e-06 | 2.9e-06 | 2.1e-06 | 2.433E-06 | 4.163E-07 | 2.100E-06 | 2.900E-06 |
| 12748 | 638 | G638G | B | 2.9e-06 | 2.7e-06 | 2.4e-06 | 2.667E-06 | 2.517E-07 | 2.400E-06 | 2.900E-06 |
| 12749 | 638 | G638H | B | 2.4e-06 | 2.9e-06 | 2.2e-06 | 2.500E-06 | 3.606E-07 | 2.200E-06 | 2.900E-06 |

| | | | | | | | | | | |
|---|---|---|---|---|---|---|---|---|---|---|
| 12750 | 638 | G638I | B | 1.9e-06 | 1.1e-06 | 3.3e-06 | 2.100E-06 | 1.114E-06 | 1.100E-06 | 3.300E-06 |
| 12751 | 638 | G638L | B | 3.3e-06 | 2.2e-06 | 3.5e-06 | 3.000E-06 | 7.000E-07 | 2.200E-06 | 3.500E-06 |
| 12752 | 638 | G638K | B | 2.6e-06 | 2e-06 | 3.1e-06 | 2.567E-06 | 5.508E-07 | 2.000E-06 | 3.100E-06 |
| 12753 | 638 | G638M | B | 1.9e-06 | 2.5e-06 | 2.4e-06 | 2.267E-06 | 3.215E-07 | 1.900E-06 | 2.500E-06 |
| 12754 | 638 | G638F | B | 3.6e-06 | 2.6e-06 | 2.4e-06 | 2.867E-06 | 6.429E-07 | 2.400E-06 | 3.600E-06 |
| 12755 | 638 | G638P | B | 6.6e-06 | 1.7e-06 | 1.9e-06 | 3.400E-06 | 2.773E-06 | 1.700E-06 | 6.600E-06 |
| 12756 | 638 | G638S | B | 1.3e-06 | 2.9e-06 | 1e-06 | 1.733E-06 | 1.021E-06 | 1.000E-06 | 2.900E-06 |
| 12757 | 638 | G638T | B | 2.8e-06 | 3.1e-06 | 1.7e-06 | 2.533E-06 | 7.371E-07 | 1.700E-06 | 3.100E-06 |
| 12758 | 638 | G638W | B | 2.8e-06 | 1.2e-06 | 1.4e-06 | 1.800E-06 | 8.718E-07 | 1.200E-06 | 2.800E-06 |
| 12759 | 638 | G638Y | B | 2e-06 | 2.6e-06 | 1.8e-06 | 2.133E-06 | 4.163E-07 | 1.800E-06 | 2.600E-06 |
| 12760 | 638 | G638V | B | 2.1e-06 | 3e-06 | 3.5e-06 | 2.867E-06 | 7.095E-07 | 2.100E-06 | 3.500E-06 |
| 12761 | 639 | D639A | B | 1.7e-06 | 4.6e-06 | 3.2e-06 | 3.167E-06 | 1.450E-06 | 1.700E-06 | 4.600E-06 |
| 12762 | 639 | D639R | B | 1.3e-06 | 3.5e-06 | 3e-06 | 2.600E-06 | 1.153E-06 | 1.300E-06 | 3.500E-06 |
| 12763 | 639 | D639N | B | 1.5e-06 | 2e-06 | 1.8e-06 | 1.767E-06 | 2.517E-07 | 1.500E-06 | 2.000E-06 |
| 12764 | 639 | D639D | B | 4.4e-06 | 1.7e-06 | 2.2e-06 | 2.767E-06 | 1.436E-06 | 1.700E-06 | 4.400E-06 |
| 12765 | 639 | D639C | B | 1.2e-06 | 2.7e-06 | 2.2e-06 | 2.033E-06 | 7.638E-07 | 1.200E-06 | 2.700E-06 |
| 12766 | 639 | D639E | B | 1.7e-06 | 2.1e-06 | 2.2e-06 | 2.000E-06 | 2.646E-07 | 1.700E-06 | 2.200E-06 |
| 12767 | 639 | D639Q | B | 3.5e-06 | 1.7e-06 | 2.7e-06 | 2.633E-06 | 9.018E-07 | 1.700E-06 | 3.500E-06 |
| 12768 | 639 | D639G | B | 2.4e-06 | 2.6e-06 | 2.2e-06 | 2.400E-06 | 2.000E-07 | 2.200E-06 | 2.600E-06 |
| 12769 | 639 | D639H | B | 1.8e-06 | 2.2e-06 | 3.2e-06 | 2.400E-06 | 7.211E-07 | 1.800E-06 | 3.200E-06 |
| 12770 | 639 | D639I | B | 1.8e-06 | 2.7e-06 | 1.5e-06 | 2.000E-06 | 6.245E-07 | 1.500E-06 | 2.700E-06 |
| 12771 | 639 | D639L | B | 1.9e-06 | 2.7e-06 | 2.1e-06 | 2.233E-06 | 4.163E-07 | 1.900E-06 | 2.700E-06 |
| 12772 | 639 | D639K | B | 1.4e-06 | 3e-06 | 1.6e-06 | 2.000E-06 | 8.718E-07 | 1.400E-06 | 3.000E-06 |
| 12773 | 639 | D639M | B | 2.9e-06 | 1.6e-06 | 2.3e-06 | 2.267E-06 | 6.506E-07 | 1.600E-06 | 2.900E-06 |
| 12774 | 639 | D639F | B | 1.7e-06 | 4.1e-06 | 2.4e-06 | 2.733E-06 | 1.234E-06 | 1.700E-06 | 4.100E-06 |
| 12775 | 639 | D639P | B | 2.2e-06 | 2.9e-06 | 2.5e-06 | 2.533E-06 | 3.512E-07 | 2.200E-06 | 2.900E-06 |
| 12776 | 639 | D639S | B | 2.4e-06 | 2.5e-06 | 5.4e-06 | 3.433E-06 | 1.704E-06 | 2.400E-06 | 5.400E-06 |
| 12777 | 639 | D639T | B | 2.1e-06 | 3e-06 | 4.4e-06 | 3.167E-06 | 1.159E-06 | 2.100E-06 | 4.400E-06 |
| 12778 | 639 | D639W | B | 2.2e-06 | 2.3e-06 | 2.8e-06 | 2.433E-06 | 3.215E-07 | 2.200E-06 | 2.800E-06 |
| 12779 | 639 | D639Y | B | 2.7e-06 | 2.9e-06 | 2.6e-06 | 2.733E-06 | 1.528E-07 | 2.600E-06 | 2.900E-06 |
| 12780 | 639 | D639V | B | 1.6e-06 | 5e-06 | 4.4e-06 | 3.667E-06 | 1.815E-06 | 1.600E-06 | 5.000E-06 |
| 12781 | 640 | R640A | B | 3.2e-06 | 2.2e-06 | 3.4e-06 | 2.933E-06 | 6.429E-07 | 2.200E-06 | 3.400E-06 |
| 12782 | 640 | R640R | B | 2.7e-06 | 3.1e-06 | 2.7e-06 | 2.833E-06 | 2.309E-07 | 2.700E-06 | 3.100E-06 |
| 12783 | 640 | R640N | B | 3.4e-06 | 2.5e-06 | 3.7e-06 | 3.200E-06 | 6.245E-07 | 2.500E-06 | 3.700E-06 |
| 12784 | 640 | R640D | B | 3.1e-06 | 1e-06 | 2.8e-06 | 2.300E-06 | 1.136E-06 | 1.000E-06 | 3.100E-06 |
| 12785 | 640 | R640C | B | 2e-06 | 5.1e-06 | 2.7e-06 | 3.267E-06 | 1.626E-06 | 2.000E-06 | 5.100E-06 |
| 12786 | 640 | R640E | B | 1.9e-06 | 2.4e-06 | 2.8e-06 | 2.367E-06 | 4.509E-07 | 1.900E-06 | 2.800E-06 |
| 12787 | 640 | R640Q | B | 2.5e-06 | 2.3e-06 | 3.1e-06 | 2.633E-06 | 4.163E-07 | 2.300E-06 | 3.100E-06 |
| 12788 | 640 | R640G | B | 1.9e-06 | 2.5e-06 | 2.3e-06 | 2.233E-06 | 3.055E-07 | 1.900E-06 | 2.500E-06 |
| 12789 | 640 | R640H | B | 1.9e-06 | 3.7e-06 | 2.4e-06 | 2.667E-06 | 9.292E-07 | 1.900E-06 | 3.700E-06 |
| 12790 | 640 | R640I | B | 1.7e-06 | 2e-06 | 1.6e-06 | 1.767E-06 | 2.082E-07 | 1.600E-06 | 2.000E-06 |
| 12791 | 640 | R640L | B | 2.8e-06 | 3e-06 | 2.7e-06 | 2.833E-06 | 1.528E-07 | 2.700E-06 | 3.000E-06 |
| 12792 | 640 | R640K | B | 3.4e-06 | 2.9e-06 | 2e-06 | 2.767E-06 | 7.095E-07 | 2.000E-06 | 3.400E-06 |
| 12793 | 640 | R640M | B | 2.5e-06 | 1.9e-06 | 1.6e-06 | 2.000E-06 | 4.583E-07 | 1.600E-06 | 2.500E-06 |
| 12794 | 640 | R640F | B | 1.2e-06 | 2.3e-06 | 3.3e-06 | 2.267E-06 | 1.050E-06 | 1.200E-06 | 3.300E-06 |
| 12795 | 640 | R640P | B | 2.4e-06 | 1.4e-06 | 3.6e-06 | 2.467E-06 | 1.102E-06 | 1.400E-06 | 3.600E-06 |
| 12796 | 640 | R640S | B | 2.1e-06 | 2.5e-06 | 2.5e-06 | 2.367E-06 | 2.309E-07 | 2.100E-06 | 2.500E-06 |
| 12797 | 640 | R640T | B | 2.3e-06 | 1.2e-06 | 2.5e-06 | 2.000E-06 | 7.000E-07 | 1.200E-06 | 2.500E-06 |
| 12798 | 640 | R640W | B | 1.9e-06 | 2.9e-06 | 1.8e-06 | 2.200E-06 | 6.083E-07 | 1.800E-06 | 2.900E-06 |
| 12799 | 640 | R640Y | B | 2.5e-06 | 2.4e-06 | 1.4e-06 | 2.100E-06 | 6.083E-07 | 1.400E-06 | 2.500E-06 |
| 12800 | 640 | R640V | B | 3.6e-06 | 2.6e-06 | 3.4e-06 | 3.200E-06 | 5.292E-07 | 2.600E-06 | 3.600E-06 |
| 12801 | 641 | V641A | B | 2.2e-06 | 2.4e-06 | 2.3e-06 | 2.300E-06 | 1.000E-07 | 2.200E-06 | 2.400E-06 |
| 12802 | 641 | V641R | B | 4.6e-06 | 2.6e-06 | 3.2e-06 | 3.467E-06 | 1.026E-06 | 2.600E-06 | 4.600E-06 |
| 12803 | 641 | V641N | B | 2.1e-06 | 3.8e-06 | 3e-06 | 2.967E-06 | 8.505E-07 | 2.100E-06 | 3.800E-06 |
| 12804 | 641 | V641D | B | 1.8e-06 | 2.2e-06 | 1.8e-06 | 1.933E-06 | 2.309E-07 | 1.800E-06 | 2.200E-06 |
| 12805 | 641 | V641C | B | 3.8e-06 | 3e-06 | 4e-06 | 3.600E-06 | 5.292E-07 | 3.000E-06 | 4.000E-06 |
| 12806 | 641 | V641E | B | 2.3e-06 | 1.7e-06 | 2.3e-06 | 2.100E-06 | 3.464E-07 | 1.700E-06 | 2.300E-06 |
| 12807 | 641 | V641Q | B | 2.2e-06 | 3.1e-06 | 2.2e-06 | 2.500E-06 | 5.196E-07 | 2.200E-06 | 3.100E-06 |
| 12808 | 641 | V641G | B | 4.2e-06 | 2.5e-06 | 2.4e-06 | 3.033E-06 | 1.012E-06 | 2.400E-06 | 4.200E-06 |
| 12809 | 641 | V641H | B | 1.8e-06 | 2.7e-06 | 2.7e-06 | 2.400E-06 | 5.196E-07 | 1.800E-06 | 2.700E-06 |
| 12810 | 641 | V641I | B | 2.5e-06 | 2.2e-06 | 2e-06 | 2.233E-06 | 2.517E-07 | 2.000E-06 | 2.500E-06 |
| 12811 | 641 | V641L | B | 2.2e-06 | 3.4e-06 | 2.6e-06 | 2.733E-06 | 6.110E-07 | 2.200E-06 | 3.400E-06 |
| 12812 | 641 | V641K | B | 1.8e-06 | 2.2e-06 | 1.7e-06 | 1.900E-06 | 2.646E-07 | 1.700E-06 | 2.200E-06 |
| 12813 | 641 | V641M | B | 1.3e-06 | 2.4e-06 | 4.5e-06 | 2.733E-06 | 1.626E-06 | 1.300E-06 | 4.500E-06 |
| 12814 | 641 | V641F | B | 1.6e-06 | 1.9e-06 | 1.7e-06 | 1.733E-06 | 1.528E-07 | 1.600E-06 | 1.900E-06 |
| 12815 | 641 | V641P | B | 3.7e-06 | 2.3e-06 | 2.8e-06 | 2.933E-06 | 7.095E-07 | 2.300E-06 | 3.700E-06 |
| 12816 | 641 | V641S | B | 2.4e-06 | 2.5e-06 | 2e-06 | 2.300E-06 | 2.646E-07 | 2.000E-06 | 2.500E-06 |
| 12817 | 641 | V641T | B | 2.4e-06 | 1.5e-06 | 4.5e-06 | 2.800E-06 | 1.539E-06 | 1.500E-06 | 4.500E-06 |
| 12818 | 641 | V641W | B | 2.7e-06 | 1e-06 | 3.7e-06 | 2.467E-06 | 1.365E-06 | 1.000E-06 | 3.700E-06 |
| 12819 | 641 | V641Y | B | 2.5e-06 | 2.6e-06 | 1.4e-06 | 2.167E-06 | 6.658E-07 | 1.400E-06 | 2.600E-06 |
| 12820 | 641 | V641V | B | 1.7e-06 | 3.5e-06 | 2.4e-06 | 2.533E-06 | 9.074E-07 | 1.700E-06 | 3.500E-06 |
| 12821 | 642 | T642A | B | 1.9e-06 | 2.4e-06 | 2.5e-06 | 2.267E-06 | 3.215E-07 | 1.900E-06 | 2.500E-06 |
| 12822 | 642 | T642R | B | 3.6e-06 | 1.9e-06 | 1.9e-06 | 2.467E-06 | 9.815E-07 | 1.900E-06 | 3.600E-06 |
| 12823 | 642 | T642N | B | 2.4e-06 | 3e-06 | 2.7e-06 | 2.700E-06 | 3.000E-07 | 2.400E-06 | 3.000E-06 |
| 12824 | 642 | T642D | B | 2.3e-06 | 1.4e-06 | 2.7e-06 | 2.133E-06 | 6.658E-07 | 1.400E-06 | 2.700E-06 |
| 12825 | 642 | T642C | B | 2.2e-06 | 2.8e-06 | 2.6e-06 | 2.533E-06 | 3.055E-07 | 2.200E-06 | 2.800E-06 |
| 12826 | 642 | T642E | B | 1.5e-06 | 4.1e-06 | 2.4e-06 | 2.667E-06 | 1.320E-06 | 1.500E-06 | 4.100E-06 |
| 12827 | 642 | T642Q | B | 2.2e-06 | 2.5e-06 | 2.5e-06 | 2.400E-06 | 1.732E-07 | 2.200E-06 | 2.500E-06 |
| 12828 | 642 | T642G | B | 2.3e-06 | 2.7e-06 | 1.2e-06 | 2.067E-06 | 7.767E-07 | 1.200E-06 | 2.700E-06 |
| 12829 | 642 | T642H | B | 2.2e-06 | 2.1e-06 | 1.6e-06 | 1.967E-06 | 3.215E-07 | 1.600E-06 | 2.200E-06 |
| 12830 | 642 | T642I | B | 3.1e-06 | 3.8e-06 | 2.5e-06 | 3.133E-06 | 6.506E-07 | 2.500E-06 | 3.800E-06 |
| 12831 | 642 | T642L | B | 2.9e-06 | 2.5e-06 | 2.2e-06 | 2.533E-06 | 3.512E-07 | 2.200E-06 | 2.900E-06 |
| 12832 | 642 | T642K | B | 5e-06 | 2e-06 | 2.7e-06 | 3.233E-06 | 1.570E-06 | 2.000E-06 | 5.000E-06 |
| 12833 | 642 | T642M | B | 2.4e-06 | 1.7e-06 | 2.6e-06 | 2.233E-06 | 4.726E-07 | 1.700E-06 | 2.600E-06 |
| 12834 | 642 | T642F | B | 1.9e-06 | 2.7e-06 | 2.5e-06 | 2.367E-06 | 4.163E-07 | 1.900E-06 | 2.700E-06 |

| | | | | | | | | | | |
|---|---|---|---|---|---|---|---|---|---|---|
| 12835 | 642 | T642P | B | 2.4e-06 | 3e-06 | 2.1e-06 | 2.500E-06 | 4.583E-07 | 2.100E-06 | 3.000E-06 |
| 12836 | 642 | T642S | B | 2.3e-06 | 2.7e-06 | 3e-06 | 2.667E-06 | 3.512E-07 | 2.300E-06 | 3.000E-06 |
| 12837 | 642 | T642T | B | 2.6e-06 | 1.9e-06 | 1.7e-06 | 2.067E-06 | 4.726E-07 | 1.700E-06 | 2.600E-06 |
| 12838 | 642 | T642W | B | 8.8e-07 | 2.2e-06 | 2.6e-06 | 1.893E-06 | 9.001E-07 | 8.800E-07 | 2.600E-06 |
| 12839 | 642 | T642Y | B | 9.4e-07 | 2.6e-06 | 1.9e-06 | 1.813E-06 | 8.334E-07 | 9.400E-07 | 2.600E-06 |
| 12840 | 642 | T642V | B | 1.7e-06 | 1.1e-06 | 2.1e-06 | 1.633E-06 | 5.033E-07 | 1.100E-06 | 2.100E-06 |
| 12841 | 643 | I643A | B | 1.8e-06 | 1.8e-06 | 8e-07 | 1.467E-06 | 5.774E-07 | 8.000E-07 | 1.800E-06 |
| 12842 | 643 | I643R | B | 1.5e-06 | 2.1e-06 | 2.4e-06 | 2.000E-06 | 4.583E-07 | 1.500E-06 | 2.400E-06 |
| 12843 | 643 | I643N | B | 1.9e-06 | 2.2e-06 | 2e-06 | 2.033E-06 | 1.528E-07 | 1.900E-06 | 2.200E-06 |
| 12844 | 643 | I643D | B | 5e-06 | 1.7e-06 | 2.1e-06 | 2.933E-06 | 1.801E-06 | 1.700E-06 | 5.000E-06 |
| 12845 | 643 | I643C | B | 1.7e-06 | 5.6e-06 | 2.1e-06 | 3.133E-06 | 2.146E-06 | 1.700E-06 | 5.600E-06 |
| 12846 | 643 | I643E | B | 2.1e-06 | 2.9e-06 | 1.9e-06 | 2.300E-06 | 5.292E-07 | 1.900E-06 | 2.900E-06 |
| 12847 | 643 | I643Q | B | 1.7e-06 | 3.7e-06 | 2.1e-06 | 2.500E-06 | 1.058E-06 | 1.700E-06 | 3.700E-06 |
| 12848 | 643 | I643G | B | 1.8e-06 | 3.4e-06 | 2.5e-06 | 2.567E-06 | 8.021E-07 | 1.800E-06 | 3.400E-06 |
| 12849 | 643 | I643H | B | 2.9e-06 | 1.4e-06 | 1.4e-06 | 1.900E-06 | 8.660E-07 | 1.400E-06 | 2.900E-06 |
| 12850 | 643 | I643I | B | 2.6e-06 | 1.9e-06 | 2.6e-06 | 2.367E-06 | 4.041E-07 | 1.900E-06 | 2.600E-06 |
| 12851 | 643 | I643L | B | 1.4e-06 | 1.8e-06 | 1.5e-06 | 1.567E-06 | 2.082E-07 | 1.400E-06 | 1.800E-06 |
| 12852 | 643 | I643K | B | 2.3e-06 | 2.6e-06 | 1.5e-06 | 2.133E-06 | 5.686E-07 | 1.500E-06 | 2.600E-06 |
| 12853 | 643 | I643M | B | 1.7e-06 | 8.8e-07 | 1.5e-06 | 1.360E-06 | 4.276E-07 | 8.800E-07 | 1.700E-06 |
| 12854 | 643 | I643F | B | 2e-06 | 1.3e-06 | 2.2e-06 | 1.833E-06 | 4.726E-07 | 1.300E-06 | 2.200E-06 |
| 12855 | 643 | I643P | B | 4.3e-06 | 1.8e-06 | 3e-06 | 3.033E-06 | 1.250E-06 | 1.800E-06 | 4.300E-06 |
| 12856 | 643 | I643S | B | 2.1e-06 | 1.5e-06 | 1.8e-06 | 1.800E-06 | 3.000E-07 | 1.500E-06 | 2.100E-06 |
| 12857 | 643 | I643T | B | 1.8e-06 | 1.9e-06 | 2.3e-06 | 2.000E-06 | 2.646E-07 | 1.800E-06 | 2.300E-06 |
| 12858 | 643 | I643W | B | 1.8e-06 | 1.7e-06 | 4.2e-06 | 2.567E-06 | 1.415E-06 | 1.700E-06 | 4.200E-06 |
| 12859 | 643 | I643Y | B | 2.2e-06 | 2.3e-06 | 2.6e-06 | 2.367E-06 | 2.082E-07 | 2.200E-06 | 2.600E-06 |
| 12860 | 643 | I643V | B | 1.3e-06 | 2.8e-06 | 2.1e-06 | 2.067E-06 | 7.506E-07 | 1.300E-06 | 2.800E-06 |
| 12861 | 644 | T644A | B | 4e-06 | 3e-06 | 3.4e-06 | 3.467E-06 | 5.033E-07 | 3.000E-06 | 4.000E-06 |
| 12862 | 644 | T644R | B | 2.3e-06 | 3.7e-06 | 2.4e-06 | 2.800E-06 | 7.810E-07 | 2.300E-06 | 3.700E-06 |
| 12863 | 644 | T644N | B | 1.9e-06 | 1.7e-06 | 2.3e-06 | 1.967E-06 | 3.055E-07 | 1.700E-06 | 2.300E-06 |
| 12864 | 644 | T644D | B | 2.3e-06 | 2.6e-06 | 2.7e-06 | 2.533E-06 | 2.082E-07 | 2.300E-06 | 2.700E-06 |
| 12865 | 644 | T644C | B | 2.4e-06 | 2.2e-06 | 2.2e-06 | 2.267E-06 | 1.155E-07 | 2.200E-06 | 2.400E-06 |
| 12866 | 644 | T644E | B | 1.7e-06 | 3.2e-06 | 4.2e-06 | 3.033E-06 | 1.258E-06 | 1.700E-06 | 4.200E-06 |
| 12867 | 644 | T644Q | B | 3.1e-06 | 1.2e-06 | 3.2e-06 | 2.500E-06 | 1.127E-06 | 1.200E-06 | 3.200E-06 |
| 12868 | 644 | T644G | B | 3.5e-06 | 2.7e-06 | 2.7e-06 | 2.967E-06 | 4.619E-07 | 2.700E-06 | 3.500E-06 |
| 12869 | 644 | T644H | B | 2.1e-06 | 1.7e-06 | 1.6e-06 | 1.800E-06 | 2.646E-07 | 1.600E-06 | 2.100E-06 |
| 12870 | 644 | T644I | B | 3.6e-06 | 2e-06 | 9.6e-07 | 2.187E-06 | 1.330E-06 | 9.600E-07 | 3.600E-06 |
| 12871 | 644 | T644L | B | 2.2e-06 | 1.9e-06 | 2.6e-06 | 2.233E-06 | 3.512E-07 | 1.900E-06 | 2.600E-06 |
| 12872 | 644 | T644K | B | 2.7e-06 | 1.5e-06 | 3.6e-06 | 2.600E-06 | 1.054E-06 | 1.500E-06 | 3.600E-06 |
| 12873 | 644 | T644M | B | 3.1e-06 | 1.8e-06 | 2.2e-06 | 2.367E-06 | 6.658E-07 | 1.800E-06 | 3.100E-06 |
| 12874 | 644 | T644F | B | 1.9e-06 | 1.9e-06 | 1.8e-06 | 1.867E-06 | 5.774E-08 | 1.800E-06 | 1.900E-06 |
| 12875 | 644 | T644P | B | 3.7e-06 | 1.4e-06 | 3.2e-06 | 2.767E-06 | 1.210E-06 | 1.400E-06 | 3.700E-06 |
| 12876 | 644 | T644S | B | 2.2e-06 | 1.9e-06 | 2.7e-06 | 2.267E-06 | 4.041E-07 | 1.900E-06 | 2.700E-06 |
| 12877 | 644 | T644T | B | 4.5e-06 | 2.6e-06 | 2.6e-06 | 3.233E-06 | 1.097E-06 | 2.600E-06 | 4.500E-06 |
| 12878 | 644 | T644W | B | 1.9e-06 | 2.4e-06 | 3.7e-06 | 2.667E-06 | 9.292E-07 | 1.900E-06 | 3.700E-06 |
| 12879 | 644 | T644Y | B | 2.3e-06 | 3.4e-06 | 1.6e-06 | 2.433E-06 | 9.074E-07 | 1.600E-06 | 3.400E-06 |
| 12880 | 644 | T644V | B | 1.8e-06 | 3.4e-06 | 2.6e-06 | 2.600E-06 | 8.000E-07 | 1.800E-06 | 3.400E-06 |
| 12881 | 645 | C645A | B | 2.2e-06 | 1.4e-06 | 3e-06 | 2.200E-06 | 8.000E-07 | 1.400E-06 | 3.000E-06 |
| 12882 | 645 | C645R | B | 3e-06 | 2.3e-06 | 2.9e-06 | 2.733E-06 | 3.786E-07 | 2.300E-06 | 3.000E-06 |
| 12883 | 645 | C645N | B | 2.7e-06 | 2.1e-06 | 3.1e-06 | 2.633E-06 | 5.033E-07 | 2.100E-06 | 3.100E-06 |
| 12884 | 645 | C645D | B | 2.9e-06 | 2e-06 | 2.1e-06 | 2.333E-06 | 4.933E-07 | 2.000E-06 | 2.900E-06 |
| 12885 | 645 | C645C | B | 2.9e-06 | 2.6e-06 | 1.4e-06 | 2.300E-06 | 7.937E-07 | 1.400E-06 | 2.900E-06 |
| 12886 | 645 | C645E | B | 3e-06 | 2.8e-06 | 2.8e-06 | 2.867E-06 | 1.155E-07 | 2.800E-06 | 3.000E-06 |
| 12887 | 645 | C645Q | B | 2.6e-06 | 3.2e-06 | 3.1e-06 | 2.967E-06 | 3.215E-07 | 2.600E-06 | 3.200E-06 |
| 12888 | 645 | C645G | B | 1.9e-06 | 2.8e-06 | 3.3e-06 | 2.667E-06 | 7.095E-07 | 1.900E-06 | 3.300E-06 |
| 12889 | 645 | C645H | B | 2.2e-06 | 2.9e-06 | 2.5e-06 | 2.533E-06 | 3.512E-07 | 2.200E-06 | 2.900E-06 |
| 12890 | 645 | C645I | B | 3.5e-06 | 2.2e-06 | 2.5e-06 | 2.733E-06 | 6.807E-07 | 2.200E-06 | 3.500E-06 |
| 12891 | 645 | C645L | B | 2.1e-06 | 2.9e-06 | 3.2e-06 | 2.733E-06 | 5.686E-07 | 2.100E-06 | 3.200E-06 |
| 12892 | 645 | C645K | B | 2.7e-06 | 2.6e-06 | 2.2e-06 | 2.500E-06 | 2.646E-07 | 2.200E-06 | 2.700E-06 |
| 12893 | 645 | C645M | B | 2.6e-06 | 2.4e-06 | 4.1e-06 | 3.033E-06 | 9.292E-07 | 2.400E-06 | 4.100E-06 |
| 12894 | 645 | C645F | B | 3.5e-06 | 2.3e-06 | 2.2e-06 | 2.667E-06 | 7.234E-07 | 2.200E-06 | 3.500E-06 |
| 12895 | 645 | C645P | B | 3.1e-06 | 2.1e-06 | 3.5e-06 | 2.900E-06 | 7.211E-07 | 2.100E-06 | 3.500E-06 |
| 12896 | 645 | C645S | B | 2.7e-06 | 2.3e-06 | 1.6e-06 | 2.200E-06 | 5.568E-07 | 1.600E-06 | 2.700E-06 |
| 12897 | 645 | C645T | B | 2.3e-06 | 2.2e-06 | 1.6e-06 | 2.033E-06 | 3.786E-07 | 1.600E-06 | 2.300E-06 |
| 12898 | 645 | C645W | B | 2.1e-06 | 1.3e-06 | 2.4e-06 | 1.933E-06 | 5.686E-07 | 1.300E-06 | 2.400E-06 |
| 12899 | 645 | C645Y | B | 2.2e-06 | 2e-06 | 2.9e-06 | 2.367E-06 | 4.726E-07 | 2.000E-06 | 2.900E-06 |
| 12900 | 645 | C645V | B | 2.1e-06 | 2.6e-06 | 2.5e-06 | 2.400E-06 | 2.646E-07 | 2.100E-06 | 2.600E-06 |
| 12901 | 646 | K646A | B | 1.4e-06 | 1.7e-06 | 3.2e-06 | 2.100E-06 | 9.644E-07 | 1.400E-06 | 3.200E-06 |
| 12902 | 646 | K646R | B | 2.7e-06 | 2e-06 | 1.4e-06 | 2.033E-06 | 6.506E-07 | 1.400E-06 | 2.700E-06 |
| 12903 | 646 | K646N | B | 3.6e-06 | 2.6e-06 | 2.9e-06 | 3.033E-06 | 5.132E-07 | 2.600E-06 | 3.600E-06 |
| 12904 | 646 | K646D | B | 2e-06 | 1.4e-06 | 1.8e-06 | 1.733E-06 | 3.055E-07 | 1.400E-06 | 2.000E-06 |
| 12905 | 646 | K646C | B | 1.8e-06 | 2.7e-06 | 2.5e-06 | 2.333E-06 | 4.726E-07 | 1.800E-06 | 2.700E-06 |
| 12906 | 646 | K646E | B | 1.8e-06 | 4.4e-06 | 2.5e-06 | 2.900E-06 | 1.345E-06 | 1.800E-06 | 4.400E-06 |
| 12907 | 646 | K646Q | B | 2.5e-06 | 2.3e-06 | 3e-06 | 2.600E-06 | 3.606E-07 | 2.300E-06 | 3.000E-06 |
| 12908 | 646 | K646G | B | 2.1e-06 | 3e-06 | 2.6e-06 | 2.567E-06 | 4.509E-07 | 2.100E-06 | 3.000E-06 |
| 12909 | 646 | K646H | B | 2.3e-06 | 2.1e-06 | 3.2e-06 | 2.533E-06 | 5.859E-07 | 2.100E-06 | 3.200E-06 |
| 12910 | 646 | K646I | B | 1.7e-06 | 2.6e-06 | 3.4e-06 | 2.567E-06 | 8.505E-07 | 1.700E-06 | 3.400E-06 |
| 12911 | 646 | K646L | B | 3.3e-06 | 1.3e-06 | 2.5e-06 | 2.367E-06 | 1.007E-06 | 1.300E-06 | 3.300E-06 |
| 12912 | 646 | K646K | B | 1.8e-06 | 3.7e-06 | 5.2e-06 | 3.567E-06 | 1.704E-06 | 1.800E-06 | 5.200E-06 |
| 12913 | 646 | K646M | B | 3.6e-06 | 2.1e-06 | 3.2e-06 | 2.967E-06 | 7.767E-07 | 2.100E-06 | 3.600E-06 |
| 12914 | 646 | K646F | B | 2.5e-06 | 1.6e-06 | 1.9e-06 | 2.000E-06 | 4.583E-07 | 1.600E-06 | 2.500E-06 |
| 12915 | 646 | K646P | B | 2.5e-06 | 3e-06 | 2.6e-06 | 2.700E-06 | 2.646E-07 | 2.500E-06 | 3.000E-06 |
| 12916 | 646 | K646S | B | 1.8e-06 | 2.7e-06 | 1.8e-06 | 2.100E-06 | 5.196E-07 | 1.800E-06 | 2.700E-06 |
| 12917 | 646 | K646T | B | 1.4e-06 | 2e-06 | 2.7e-06 | 2.033E-06 | 6.506E-07 | 1.400E-06 | 2.700E-06 |
| 12918 | 646 | K646W | B | 1.9e-06 | 2.3e-06 | 2.8e-06 | 2.333E-06 | 4.509E-07 | 1.900E-06 | 2.800E-06 |
| 12919 | 646 | K646Y | B | 1.1e-06 | 3.9e-06 | 3e-06 | 2.667E-06 | 1.429E-06 | 1.100E-06 | 3.900E-06 |

| | | | | | | | | | | |
|---|---|---|---|---|---|---|---|---|---|---|
| 12920 | 646 | K646V | B | 2.4e-06 | 1.9e-06 | 3.1e-06 | 2.467E-06 | 6.028E-07 | 1.900E-06 | 3.100E-06 |
| 12921 | 647 | A647A | B | 2.2e-06 | 3.6e-06 | 3e-06 | 2.933E-06 | 7.024E-07 | 2.200E-06 | 3.600E-06 |
| 12922 | 647 | A647R | B | 3.5e-06 | 3.3e-06 | 1.6e-06 | 2.800E-06 | 1.044E-06 | 1.600E-06 | 3.500E-06 |
| 12923 | 647 | A647N | B | 2.4e-06 | 2.6e-06 | 2.3e-06 | 2.433E-06 | 1.528E-07 | 2.300E-06 | 2.600E-06 |
| 12924 | 647 | A647D | B | 2.3e-06 | 2.4e-06 | 1.3e-06 | 2.000E-06 | 6.083E-07 | 1.300E-06 | 2.400E-06 |
| 12925 | 647 | A647C | B | 2.6e-06 | 2.4e-06 | 2.4e-06 | 2.467E-06 | 1.155E-07 | 2.400E-06 | 2.600E-06 |
| 12926 | 647 | A647E | B | 2.2e-06 | 1.6e-06 | 2e-06 | 1.933E-06 | 3.055E-07 | 1.600E-06 | 2.200E-06 |
| 12927 | 647 | A647Q | B | 1.6e-06 | 3.5e-06 | 2.6e-06 | 2.567E-06 | 9.504E-07 | 1.600E-06 | 3.500E-06 |
| 12928 | 647 | A647G | B | 1.7e-06 | 2.5e-06 | 3e-06 | 2.400E-06 | 6.557E-07 | 1.700E-06 | 3.000E-06 |
| 12929 | 647 | A647H | B | 1.6e-06 | 1.5e-06 | 3.5e-06 | 2.200E-06 | 1.127E-06 | 1.500E-06 | 3.500E-06 |
| 12930 | 647 | A647I | B | 3.4e-06 | 3.2e-06 | 1.1e-06 | 2.567E-06 | 1.274E-06 | 1.100E-06 | 3.400E-06 |
| 12931 | 647 | A647L | B | 2.2e-06 | 1.8e-06 | 3.2e-06 | 2.400E-06 | 7.211E-07 | 1.800E-06 | 3.200E-06 |
| 12932 | 647 | A647K | B | 1.9e-06 | 8.7e-07 | 1.7e-06 | 1.490E-06 | 5.462E-07 | 8.700E-07 | 1.900E-06 |
| 12933 | 647 | A647M | B | 2.1e-06 | 2.2e-06 | 2e-06 | 2.100E-06 | 1.000E-07 | 2.000E-06 | 2.200E-06 |
| 12934 | 647 | A647F | B | 1.4e-06 | 1e-06 | 1.4e-06 | 1.267E-06 | 2.309E-07 | 1.000E-06 | 1.400E-06 |
| 12935 | 647 | A647P | B | 1.6e-06 | 1.4e-06 | 2.4e-06 | 1.800E-06 | 5.292E-07 | 1.400E-06 | 2.400E-06 |
| 12936 | 647 | A647S | B | 4.8e-06 | 2.5e-06 | 2.6e-06 | 3.300E-06 | 1.300E-06 | 2.500E-06 | 4.800E-06 |
| 12937 | 647 | A647T | B | 1.6e-06 | 3.2e-06 | 2.1e-06 | 2.300E-06 | 8.185E-07 | 1.600E-06 | 3.200E-06 |
| 12938 | 647 | A647W | B | 3.5e-06 | 3.2e-06 | 1.7e-06 | 2.800E-06 | 9.644E-07 | 1.700E-06 | 3.500E-06 |
| 12939 | 647 | A647Y | B | 2.1e-06 | 3.4e-06 | 2e-06 | 2.500E-06 | 7.810E-07 | 2.000E-06 | 3.400E-06 |
| 12940 | 647 | A647V | B | 2e-06 | 3.4e-06 | 2.9e-06 | 2.767E-06 | 7.095E-07 | 2.000E-06 | 3.400E-06 |
| 12941 | 648 | S648A | B | 2.4e-06 | 3.7e-06 | 2.3e-06 | 2.800E-06 | 7.810E-07 | 2.300E-06 | 3.700E-06 |
| 12942 | 648 | S648R | B | 2.8e-06 | 2.3e-06 | 1.7e-06 | 2.267E-06 | 5.508E-07 | 1.700E-06 | 2.800E-06 |
| 12943 | 648 | S648N | B | 3.4e-06 | 7.4e-06 | 2.5e-06 | 4.433E-06 | 2.608E-06 | 2.500E-06 | 7.400E-06 |
| 12944 | 648 | S648D | B | 2.2e-06 | 2.5e-06 | 1.1e-06 | 1.933E-06 | 7.371E-07 | 1.100E-06 | 2.500E-06 |
| 12945 | 648 | S648C | B | 2.4e-06 | 1.9e-06 | 5.3e-06 | 3.200E-06 | 1.836E-06 | 1.900E-06 | 5.300E-06 |
| 12946 | 648 | S648E | B | 2.1e-06 | 2.6e-06 | 2.7e-06 | 2.467E-06 | 3.215E-07 | 2.100E-06 | 2.700E-06 |
| 12947 | 648 | S648Q | B | 3.5e-06 | 1.5e-06 | 3.6e-06 | 2.867E-06 | 1.185E-06 | 1.500E-06 | 3.600E-06 |
| 12948 | 648 | S648G | B | 1.5e-06 | 3.1e-06 | 2.2e-06 | 2.267E-06 | 8.021E-07 | 1.500E-06 | 3.100E-06 |
| 12949 | 648 | S648H | B | 3.3e-06 | 3.3e-06 | 2.2e-06 | 2.933E-06 | 6.351E-07 | 2.200E-06 | 3.300E-06 |
| 12950 | 648 | S648I | B | 1.6e-06 | 1.7e-06 | 4e-06 | 2.433E-06 | 1.358E-06 | 1.600E-06 | 4.000E-06 |
| 12951 | 648 | S648L | B | 3.3e-06 | 1.3e-06 | 2e-06 | 2.200E-06 | 1.015E-06 | 1.300E-06 | 3.300E-06 |
| 12952 | 648 | S648K | B | 3.7e-06 | 1.7e-06 | 2.8e-06 | 2.733E-06 | 1.002E-06 | 1.700E-06 | 3.700E-06 |
| 12953 | 648 | S648M | B | 2.4e-06 | 2.6e-06 | 3.4e-06 | 2.800E-06 | 5.292E-07 | 2.400E-06 | 3.400E-06 |
| 12954 | 648 | S648F | B | 2.5e-06 | 2.3e-06 | 1.3e-06 | 2.033E-06 | 6.429E-07 | 1.300E-06 | 2.500E-06 |
| 12955 | 648 | S648P | B | 2.3e-06 | 3.1e-06 | 1.9e-06 | 2.433E-06 | 6.110E-07 | 1.900E-06 | 3.100E-06 |
| 12956 | 648 | S648S | B | 2.9e-06 | 3.9e-06 | 2.4e-06 | 3.067E-06 | 7.638E-07 | 2.400E-06 | 3.900E-06 |
| 12957 | 648 | S648T | B | 3.6e-06 | 2.8e-06 | 1.7e-06 | 2.700E-06 | 9.539E-07 | 1.700E-06 | 3.600E-06 |
| 12958 | 648 | S648W | B | 2.2e-06 | 2.1e-06 | 3.2e-06 | 2.500E-06 | 6.083E-07 | 2.100E-06 | 3.200E-06 |
| 12959 | 648 | S648Y | B | 2.7e-06 | 2.1e-06 | 3.2e-06 | 2.667E-06 | 5.508E-07 | 2.100E-06 | 3.200E-06 |
| 12960 | 648 | S648V | B | 2.3e-06 | 2.2e-06 | 3.3e-06 | 2.600E-06 | 6.083E-07 | 2.200E-06 | 3.300E-06 |
| 12961 | 649 | Q649A | B | 1.2e-06 | 1.8e-06 | 1.7e-06 | 1.567E-06 | 3.215E-07 | 1.200E-06 | 1.800E-06 |
| 12962 | 649 | Q649R | B | 2.4e-06 | 1.7e-06 | 4.1e-06 | 2.733E-06 | 1.234E-06 | 1.700E-06 | 4.100E-06 |
| 12963 | 649 | Q649N | B | 3.1e-06 | 1.9e-06 | 1.9e-06 | 2.300E-06 | 6.928E-07 | 1.900E-06 | 3.100E-06 |
| 12964 | 649 | Q649D | B | 2.9e-06 | 1.2e-06 | 3.9e-06 | 2.667E-06 | 1.365E-06 | 1.200E-06 | 3.900E-06 |
| 12965 | 649 | Q649C | B | 1.2e-06 | 2.1e-06 | 1.6e-06 | 1.633E-06 | 4.509E-07 | 1.200E-06 | 2.100E-06 |
| 12966 | 649 | Q649E | B | 1.5e-06 | 2.3e-06 | 3.2e-06 | 2.333E-06 | 8.505E-07 | 1.500E-06 | 3.200E-06 |
| 12967 | 649 | Q649Q | B | 3.3e-06 | 2.6e-06 | 2.2e-06 | 2.700E-06 | 5.568E-07 | 2.200E-06 | 3.300E-06 |
| 12968 | 649 | Q649G | B | 3.2e-06 | 2.8e-06 | 2.2e-06 | 2.733E-06 | 5.033E-07 | 2.200E-06 | 3.200E-06 |
| 12969 | 649 | Q649H | B | 1.9e-06 | 2.4e-06 | 3.8e-06 | 2.700E-06 | 9.849E-07 | 1.900E-06 | 3.800E-06 |
| 12970 | 649 | Q649I | B | 3.2e-06 | 1.6e-06 | 2.7e-06 | 2.500E-06 | 8.185E-07 | 1.600E-06 | 3.200E-06 |
| 12971 | 649 | Q649L | B | 2.2e-06 | 3.6e-06 | 3.2e-06 | 3.000E-06 | 7.211E-07 | 2.200E-06 | 3.600E-06 |
| 12972 | 649 | Q649K | B | 1.8e-06 | 2.8e-06 | 3.2e-06 | 2.600E-06 | 7.211E-07 | 1.800E-06 | 3.200E-06 |
| 12973 | 649 | Q649M | B | 2e-06 | 2.2e-06 | 2.5e-06 | 2.233E-06 | 2.517E-07 | 2.000E-06 | 2.500E-06 |
| 12974 | 649 | Q649F | B | 3e-06 | 3e-06 | 2.8e-06 | 2.933E-06 | 1.155E-07 | 2.800E-06 | 3.000E-06 |
| 12975 | 649 | Q649P | B | 2.3e-06 | 1.9e-06 | 2.3e-06 | 2.167E-06 | 2.309E-07 | 1.900E-06 | 2.300E-06 |
| 12976 | 649 | Q649S | B | 1.6e-06 | 1.7e-06 | 2.4e-06 | 1.900E-06 | 4.359E-07 | 1.600E-06 | 2.400E-06 |
| 12977 | 649 | Q649T | B | 3.4e-06 | 2.6e-06 | 4e-06 | 3.333E-06 | 7.024E-07 | 2.600E-06 | 4.000E-06 |
| 12978 | 649 | Q649W | B | 2.4e-06 | 2.2e-06 | 2.1e-06 | 2.233E-06 | 1.528E-07 | 2.100E-06 | 2.400E-06 |
| 12979 | 649 | Q649Y | B | 1.9e-06 | 5.3e-06 | 3.5e-06 | 3.567E-06 | 1.701E-06 | 1.900E-06 | 5.300E-06 |
| 12980 | 649 | Q649V | B | 2.1e-06 | 2.2e-06 | 1.9e-06 | 2.067E-06 | 1.528E-07 | 1.900E-06 | 2.200E-06 |
| 12981 | 650 | D650A | B | 2.9e-06 | 2.4e-06 | 2e-06 | 2.433E-06 | 4.509E-07 | 2.000E-06 | 2.900E-06 |
| 12982 | 650 | D650R | B | 2.2e-06 | 2.8e-06 | 2.8e-06 | 2.600E-06 | 3.464E-07 | 2.200E-06 | 2.800E-06 |
| 12983 | 650 | D650N | B | 1.3e-06 | 3e-06 | 2.5e-06 | 2.267E-06 | 8.737E-07 | 1.300E-06 | 3.000E-06 |
| 12984 | 650 | D650D | B | 2.3e-06 | 2.4e-06 | 2.2e-06 | 2.300E-06 | 1.000E-07 | 2.200E-06 | 2.400E-06 |
| 12985 | 650 | D650C | B | 1.6e-06 | 2.1e-06 | 2.8e-06 | 2.167E-06 | 6.028E-07 | 1.600E-06 | 2.800E-06 |
| 12986 | 650 | D650E | B | 3.7e-06 | 4.1e-06 | 2.7e-06 | 3.500E-06 | 7.211E-07 | 2.700E-06 | 4.100E-06 |
| 12987 | 650 | D650Q | B | 2.5e-06 | 1.6e-06 | 2.1e-06 | 2.067E-06 | 4.509E-07 | 1.600E-06 | 2.500E-06 |
| 12988 | 650 | D650G | B | 3.5e-06 | 2e-06 | 3.1e-06 | 2.867E-06 | 7.767E-07 | 2.000E-06 | 3.500E-06 |
| 12989 | 650 | D650H | B | 2.1e-06 | 2.3e-06 | 2.2e-06 | 2.200E-06 | 1.000E-07 | 2.100E-06 | 2.300E-06 |
| 12990 | 650 | D650I | B | 2.4e-06 | 2.2e-06 | 2e-06 | 2.200E-06 | 2.000E-07 | 2.000E-06 | 2.400E-06 |
| 12991 | 650 | D650L | B | 1.3e-06 | 6.6e-06 | 2.1e-06 | 3.333E-06 | 2.857E-06 | 1.300E-06 | 6.600E-06 |
| 12992 | 650 | D650K | B | 3.9e-06 | 1.9e-06 | 1.9e-06 | 2.567E-06 | 1.155E-06 | 1.900E-06 | 3.900E-06 |
| 12993 | 650 | D650M | B | 2.4e-06 | 3e-06 | 2.1e-06 | 2.500E-06 | 4.583E-07 | 2.100E-06 | 3.000E-06 |
| 12994 | 650 | D650F | B | 2.5e-06 | 2.2e-06 | 2.4e-06 | 2.367E-06 | 1.528E-07 | 2.200E-06 | 2.500E-06 |
| 12995 | 650 | D650P | B | 3.8e-06 | 2.4e-06 | 1.7e-06 | 2.633E-06 | 1.069E-06 | 1.700E-06 | 3.800E-06 |
| 12996 | 650 | D650S | B | 1.8e-06 | 1.2e-06 | 2.7e-06 | 1.900E-06 | 7.550E-07 | 1.200E-06 | 2.700E-06 |
| 12997 | 650 | D650T | B | 6.3e-07 | 4.9e-06 | 5.4e-06 | 3.643E-06 | 2.622E-06 | 6.300E-07 | 5.400E-06 |
| 12998 | 650 | D650W | B | 1.6e-06 | 3e-06 | 3.3e-06 | 2.633E-06 | 9.074E-07 | 1.600E-06 | 3.300E-06 |
| 12999 | 650 | D650Y | B | 2.3e-06 | 3.1e-06 | 3.2e-06 | 2.867E-06 | 4.933E-07 | 2.300E-06 | 3.200E-06 |
| 13000 | 650 | D650V | B | 1.8e-06 | 3.1e-06 | 2.4e-06 | 2.433E-06 | 6.506E-07 | 1.800E-06 | 3.100E-06 |
| 13001 | 651 | V651A | B | 3.6e-06 | 1.7e-06 | 4.6e-06 | 3.300E-06 | 1.473E-06 | 1.700E-06 | 4.600E-06 |
| 13002 | 651 | V651R | B | 1.3e-06 | 2.6e-06 | 1.4e-06 | 1.767E-06 | 7.234E-07 | 1.300E-06 | 2.600E-06 |
| 13003 | 651 | V651N | B | 1.7e-06 | 3e-06 | 2.4e-06 | 2.367E-06 | 6.506E-07 | 1.700E-06 | 3.000E-06 |
| 13004 | 651 | V651D | B | 9.7e-07 | 2.4e-06 | 2.1e-06 | 1.823E-06 | 7.541E-07 | 9.700E-07 | 2.400E-06 |

| | | | | | | | | | | |
|---|---|---|---|---|---|---|---|---|---|---|
| 13005 | 651 | V651C | B | 2.9e-06 | 2e-06 | 3e-06 | 2.633E-06 | 5.508E-07 | 2.000E-06 | 3.000E-06 |
| 13006 | 651 | V651E | B | 2.6e-06 | 1.9e-06 | 1.9e-06 | 2.133E-06 | 4.041E-07 | 1.900E-06 | 2.600E-06 |
| 13007 | 651 | V651Q | B | 2.2e-06 | 4.1e-06 | 1.6e-06 | 2.633E-06 | 1.305E-06 | 1.600E-06 | 4.100E-06 |
| 13008 | 651 | V651G | B | 2.1e-06 | 2.4e-06 | 4.1e-06 | 2.867E-06 | 1.079E-06 | 2.100E-06 | 4.100E-06 |
| 13009 | 651 | V651H | B | 1.8e-06 | 2.2e-06 | 2.5e-06 | 2.167E-06 | 3.512E-07 | 1.800E-06 | 2.500E-06 |
| 13010 | 651 | V651I | B | 3.1e-06 | 2e-06 | 3.4e-06 | 2.833E-06 | 7.371E-07 | 2.000E-06 | 3.400E-06 |
| 13011 | 651 | V651L | B | 3e-06 | 4.1e-06 | 2e-06 | 3.033E-06 | 1.050E-06 | 2.000E-06 | 4.100E-06 |
| 13012 | 651 | V651K | B | 2.7e-06 | 2.5e-06 | 2.7e-06 | 2.633E-06 | 1.155E-07 | 2.500E-06 | 2.700E-06 |
| 13013 | 651 | V651M | B | 1.6e-06 | 2.4e-06 | 3e-06 | 2.333E-06 | 7.024E-07 | 1.600E-06 | 3.000E-06 |
| 13014 | 651 | V651F | B | 2.8e-06 | 2.8e-06 | 3.1e-06 | 2.900E-06 | 1.732E-07 | 2.800E-06 | 3.100E-06 |
| 13015 | 651 | V651P | B | 2.5e-06 | 2.6e-06 | 1.3e-06 | 2.133E-06 | 7.234E-07 | 1.300E-06 | 2.600E-06 |
| 13016 | 651 | V651S | B | 2.2e-06 | 2.2e-06 | 1.6e-06 | 2.000E-06 | 3.464E-07 | 1.600E-06 | 2.200E-06 |
| 13017 | 651 | V651T | B | 2.6e-06 | 3.1e-06 | 2.6e-06 | 2.767E-06 | 2.887E-07 | 2.600E-06 | 3.100E-06 |
| 13018 | 651 | V651W | B | 2.2e-06 | 2.6e-06 | 2e-06 | 2.267E-06 | 3.055E-07 | 2.000E-06 | 2.600E-06 |
| 13019 | 651 | V651Y | B | 2.9e-06 | 1.8e-06 | 2.2e-06 | 2.300E-06 | 5.568E-07 | 1.800E-06 | 2.900E-06 |
| 13020 | 651 | V651V | B | 1.6e-06 | 2.3e-06 | 2e-06 | 1.967E-06 | 3.512E-07 | 1.600E-06 | 2.300E-06 |
| 13021 | 652 | S652A | B | 2.6e-06 | 4.6e-06 | 3.3e-06 | 3.500E-06 | 1.015E-06 | 2.600E-06 | 4.600E-06 |
| 13022 | 652 | S652R | B | 2.4e-06 | 4.5e-06 | 3.1e-06 | 3.333E-06 | 1.069E-06 | 2.400E-06 | 4.500E-06 |
| 13023 | 652 | S652N | B | 5.2e-06 | 2.5e-06 | 2.3e-06 | 3.333E-06 | 1.620E-06 | 2.300E-06 | 5.200E-06 |
| 13024 | 652 | S652D | B | 1.9e-06 | 1.9e-06 | 2.9e-06 | 2.233E-06 | 5.774E-07 | 1.900E-06 | 2.900E-06 |
| 13025 | 652 | S652C | B | 2.4e-06 | 2.3e-06 | 2.1e-06 | 2.267E-06 | 1.528E-07 | 2.100E-06 | 2.400E-06 |
| 13026 | 652 | S652E | B | 1.6e-06 | 2.3e-06 | 1.3e-06 | 1.733E-06 | 5.132E-07 | 1.300E-06 | 2.300E-06 |
| 13027 | 652 | S652Q | B | 1.7e-06 | 2.3e-06 | 2.8e-06 | 2.267E-06 | 5.508E-07 | 1.700E-06 | 2.800E-06 |
| 13028 | 652 | S652G | B | 2.5e-06 | 2.5e-06 | 3.7e-06 | 2.900E-06 | 6.928E-07 | 2.500E-06 | 3.700E-06 |
| 13029 | 652 | S652H | B | 1.6e-06 | 2.9e-06 | 2e-06 | 2.167E-06 | 6.658E-07 | 1.600E-06 | 2.900E-06 |
| 13030 | 652 | S652I | B | 1.7e-06 | 1.9e-06 | 3e-06 | 2.200E-06 | 7.000E-07 | 1.700E-06 | 3.000E-06 |
| 13031 | 652 | S652L | B | 1.7e-06 | 2.5e-06 | 2.6e-06 | 2.267E-06 | 4.933E-07 | 1.700E-06 | 2.600E-06 |
| 13032 | 652 | S652K | B | 2.2e-06 | 2.4e-06 | 2.5e-06 | 2.367E-06 | 1.528E-07 | 2.200E-06 | 2.500E-06 |
| 13033 | 652 | S652M | B | 2.2e-06 | 3e-06 | 3.1e-06 | 2.767E-06 | 4.933E-07 | 2.200E-06 | 3.100E-06 |
| 13034 | 652 | S652F | B | 1.8e-06 | 2.5e-06 | 2e-06 | 2.100E-06 | 3.606E-07 | 1.800E-06 | 2.500E-06 |
| 13035 | 652 | S652P | B | 2e-06 | 2.7e-06 | 3.5e-06 | 2.733E-06 | 7.506E-07 | 2.000E-06 | 3.500E-06 |
| 13036 | 652 | S652S | B | 2.4e-06 | 4.2e-06 | 1.2e-06 | 2.600E-06 | 1.510E-06 | 1.200E-06 | 4.200E-06 |
| 13037 | 652 | S652T | B | 2.8e-06 | 2.1e-06 | 2.4e-06 | 2.433E-06 | 3.512E-07 | 2.100E-06 | 2.800E-06 |
| 13038 | 652 | S652W | B | 1.9e-06 | 1.9e-06 | 2.7e-06 | 2.167E-06 | 4.619E-07 | 1.900E-06 | 2.700E-06 |
| 13039 | 652 | S652Y | B | 3e-06 | 2.1e-06 | 2.6e-06 | 2.567E-06 | 4.509E-07 | 2.100E-06 | 3.000E-06 |
| 13040 | 652 | S652V | B | 1.5e-06 | 4e-06 | 2.1e-06 | 2.533E-06 | 1.305E-06 | 1.500E-06 | 4.000E-06 |
| 13041 | 653 | I653A | B | 3e-06 | 2.5e-06 | 1.8e-06 | 2.433E-06 | 6.028E-07 | 1.800E-06 | 3.000E-06 |
| 13042 | 653 | I653R | B | 2.1e-06 | 1.8e-06 | 5.2e-06 | 3.033E-06 | 1.882E-06 | 1.800E-06 | 5.200E-06 |
| 13043 | 653 | I653N | B | 2e-06 | 1.1e-06 | 2.6e-06 | 1.900E-06 | 7.550E-07 | 1.100E-06 | 2.600E-06 |
| 13044 | 653 | I653D | B | 3.1e-06 | 3e-06 | 2.4e-06 | 2.833E-06 | 3.786E-07 | 2.400E-06 | 3.100E-06 |
| 13045 | 653 | I653C | B | 3.2e-06 | 1.9e-06 | 2.2e-06 | 2.433E-06 | 6.807E-07 | 1.900E-06 | 3.200E-06 |
| 13046 | 653 | I653E | B | 2.2e-06 | 3.7e-06 | 2.6e-06 | 2.833E-06 | 7.767E-07 | 2.200E-06 | 3.700E-06 |
| 13047 | 653 | I653Q | B | 2.1e-06 | 2.2e-06 | 2.7e-06 | 2.333E-06 | 3.215E-07 | 2.100E-06 | 2.700E-06 |
| 13048 | 653 | I653G | B | 2.2e-06 | 3.1e-06 | 2.9e-06 | 2.733E-06 | 4.726E-07 | 2.200E-06 | 3.100E-06 |
| 13049 | 653 | I653H | B | 2.2e-06 | 2.5e-06 | 3.2e-06 | 2.633E-06 | 5.132E-07 | 2.200E-06 | 3.200E-06 |
| 13050 | 653 | I653I | B | 2.7e-06 | 1.7e-06 | 7.5e-07 | 1.717E-06 | 9.751E-07 | 7.500E-07 | 2.700E-06 |
| 13051 | 653 | I653L | B | 2.4e-06 | 2.5e-06 | 1.4e-06 | 2.100E-06 | 6.083E-07 | 1.400E-06 | 2.500E-06 |
| 13052 | 653 | I653K | B | 3.2e-06 | 2.2e-06 | 2.3e-06 | 2.567E-06 | 5.508E-07 | 2.200E-06 | 3.200E-06 |
| 13053 | 653 | I653M | B | 2e-06 | 3.9e-06 | 4.2e-06 | 3.367E-06 | 1.193E-06 | 2.000E-06 | 4.200E-06 |
| 13054 | 653 | I653F | B | 1.1e-06 | 1.6e-06 | 2.1e-06 | 1.600E-06 | 5.000E-07 | 1.100E-06 | 2.100E-06 |
| 13055 | 653 | I653P | B | 3e-06 | 2.2e-06 | 2.9e-06 | 2.700E-06 | 4.359E-07 | 2.200E-06 | 3.000E-06 |
| 13056 | 653 | I653S | B | 2e-06 | 2e-06 | 1.8e-06 | 1.933E-06 | 1.155E-07 | 1.800E-06 | 2.000E-06 |
| 13057 | 653 | I653T | B | 2.1e-06 | 2.7e-06 | 3.3e-06 | 2.700E-06 | 6.000E-07 | 2.100E-06 | 3.300E-06 |
| 13058 | 653 | I653W | B | 2.6e-06 | 1.5e-06 | 3e-06 | 2.367E-06 | 7.767E-07 | 1.500E-06 | 3.000E-06 |
| 13059 | 653 | I653Y | B | 3.5e-06 | 1.3e-06 | 2.9e-06 | 2.567E-06 | 1.137E-06 | 1.300E-06 | 3.500E-06 |
| 13060 | 653 | I653V | B | 3.3e-06 | 2.4e-06 | 1.9e-06 | 2.533E-06 | 7.095E-07 | 1.900E-06 | 3.300E-06 |
| 13061 | 654 | G654A | B | 3.1e-06 | 3e-06 | 3.1e-06 | 3.067E-06 | 5.774E-08 | 3.000E-06 | 3.100E-06 |
| 13062 | 654 | G654R | B | 2e-06 | 3.6e-06 | 2.1e-06 | 2.567E-06 | 8.963E-07 | 2.000E-06 | 3.600E-06 |
| 13063 | 654 | G654N | B | 3e-06 | 1.4e-06 | 2.7e-06 | 2.367E-06 | 8.505E-07 | 1.400E-06 | 3.000E-06 |
| 13064 | 654 | G654D | B | 1.9e-06 | 1.3e-06 | 2.3e-06 | 1.833E-06 | 5.033E-07 | 1.300E-06 | 2.300E-06 |
| 13065 | 654 | G654C | B | 3.5e-06 | 1.9e-06 | 2.9e-06 | 2.767E-06 | 8.083E-07 | 1.900E-06 | 3.500E-06 |
| 13066 | 654 | G654E | B | 2.6e-06 | 3.2e-06 | 1.6e-06 | 2.467E-06 | 8.083E-07 | 1.600E-06 | 3.200E-06 |
| 13067 | 654 | G654Q | B | 2.6e-06 | 1.9e-06 | 3e-06 | 2.500E-06 | 5.568E-07 | 1.900E-06 | 3.000E-06 |
| 13068 | 654 | G654G | B | 7.4e-07 | 1.8e-06 | 2.1e-06 | 1.547E-06 | 7.145E-07 | 7.400E-07 | 2.100E-06 |
| 13069 | 654 | G654H | B | 1.8e-06 | 2.1e-06 | 2e-06 | 1.967E-06 | 1.528E-07 | 1.800E-06 | 2.100E-06 |
| 13070 | 654 | G654I | B | 2.7e-06 | 1.7e-06 | 4.1e-06 | 2.833E-06 | 1.206E-06 | 1.700E-06 | 4.100E-06 |
| 13071 | 654 | G654L | B | 2.9e-06 | 3.4e-06 | 3.2e-06 | 3.167E-06 | 2.517E-07 | 2.900E-06 | 3.400E-06 |
| 13072 | 654 | G654K | B | 2.7e-06 | 2e-06 | 3.3e-06 | 2.667E-06 | 6.506E-07 | 2.000E-06 | 3.300E-06 |
| 13073 | 654 | G654M | B | 2.7e-06 | 2.8e-06 | 2.7e-06 | 2.733E-06 | 5.774E-08 | 2.700E-06 | 2.800E-06 |
| 13074 | 654 | G654F | B | 3.3e-06 | 1.4e-06 | 2.3e-06 | 2.333E-06 | 9.504E-07 | 1.400E-06 | 3.300E-06 |
| 13075 | 654 | G654P | B | 2.8e-06 | 3.5e-06 | 2.3e-06 | 2.867E-06 | 6.028E-07 | 2.300E-06 | 3.500E-06 |
| 13076 | 654 | G654S | B | 2.2e-06 | 4.5e-06 | 2.2e-06 | 2.967E-06 | 1.328E-06 | 2.200E-06 | 4.500E-06 |
| 13077 | 654 | G654T | B | 2.2e-06 | 2.7e-06 | 2.3e-06 | 2.400E-06 | 2.646E-07 | 2.200E-06 | 2.700E-06 |
| 13078 | 654 | G654W | B | 2.6e-06 | 3.8e-06 | 1.9e-06 | 2.767E-06 | 9.609E-07 | 1.900E-06 | 3.800E-06 |
| 13079 | 654 | G654Y | B | 2.5e-06 | 2.8e-06 | 3.4e-06 | 2.900E-06 | 4.583E-07 | 2.500E-06 | 3.400E-06 |
| 13080 | 654 | G654V | B | 1.9e-06 | 1.5e-06 | 1.7e-06 | 1.700E-06 | 2.000E-07 | 1.500E-06 | 1.900E-06 |
| 13081 | 655 | V655A | B | 2.6e-06 | 3.1e-06 | 3.3e-06 | 3.000E-06 | 3.606E-07 | 2.600E-06 | 3.300E-06 |
| 13082 | 655 | V655R | B | 1.5e-06 | 2.5e-06 | 2.6e-06 | 2.200E-06 | 6.083E-07 | 1.500E-06 | 2.600E-06 |
| 13083 | 655 | V655N | B | 3.2e-06 | 2.4e-06 | 2e-06 | 2.533E-06 | 6.110E-07 | 2.000E-06 | 3.200E-06 |
| 13084 | 655 | V655D | B | 2.7e-06 | 2.3e-06 | 1.5e-06 | 2.167E-06 | 6.110E-07 | 1.500E-06 | 2.700E-06 |
| 13085 | 655 | V655C | B | 3e-06 | 3.3e-06 | 2.1e-06 | 2.800E-06 | 6.245E-07 | 2.100E-06 | 3.300E-06 |
| 13086 | 655 | V655E | B | 2.5e-06 | 2.5e-06 | 4.5e-06 | 3.167E-06 | 1.155E-06 | 2.500E-06 | 4.500E-06 |
| 13087 | 655 | V655Q | B | 2.6e-06 | 2.5e-06 | 1.5e-06 | 2.200E-06 | 6.083E-07 | 1.500E-06 | 2.600E-06 |
| 13088 | 655 | V655G | B | 2.6e-06 | 1.6e-06 | 2.7e-06 | 2.300E-06 | 6.083E-07 | 1.600E-06 | 2.700E-06 |
| 13089 | 655 | V655H | B | 2.7e-06 | 2.3e-06 | 2.4e-06 | 2.467E-06 | 2.082E-07 | 2.300E-06 | 2.700E-06 |

| | | | | | | | | | | |
|---|---|---|---|---|---|---|---|---|---|---|
| 13090 | 655 | V655I | B | 1.3e-06 | 2.2e-06 | 1.3e-06 | 1.600E-06 | 5.196E-07 | 1.300E-06 | 2.200E-06 |
| 13091 | 655 | V655L | B | 2.9e-06 | 3.7e-06 | 2.8e-06 | 3.133E-06 | 4.933E-07 | 2.800E-06 | 3.700E-06 |
| 13092 | 655 | V655K | B | 2.4e-06 | 2.4e-06 | 6.4e-06 | 3.733E-06 | 2.309E-06 | 2.400E-06 | 6.400E-06 |
| 13093 | 655 | V655M | B | 1.6e-06 | 2.1e-06 | 3.2e-06 | 2.300E-06 | 8.185E-07 | 1.600E-06 | 3.200E-06 |
| 13094 | 655 | V655F | B | 3.6e-06 | 2.2e-06 | 2.9e-06 | 2.900E-06 | 7.000E-07 | 2.200E-06 | 3.600E-06 |
| 13095 | 655 | V655P | B | 1.8e-06 | 2.6e-06 | 3.5e-06 | 2.633E-06 | 8.505E-07 | 1.800E-06 | 3.500E-06 |
| 13096 | 655 | V655S | B | 3e-06 | 2.3e-06 | 1.6e-06 | 2.300E-06 | 7.000E-07 | 1.600E-06 | 3.000E-06 |
| 13097 | 655 | V655T | B | 1e-06 | 2.1e-06 | 4.6e-06 | 2.567E-06 | 1.845E-06 | 1.000E-06 | 4.600E-06 |
| 13098 | 655 | V655W | B | 1.9e-06 | 5.2e-06 | 4.2e-06 | 3.767E-06 | 1.692E-06 | 1.900E-06 | 5.200E-06 |
| 13099 | 655 | V655Y | B | 2.4e-06 | 2.8e-06 | 1.4e-06 | 2.200E-06 | 7.211E-07 | 1.400E-06 | 2.800E-06 |
| 13100 | 655 | V655V | B | 2.9e-06 | 1.8e-06 | 4.8e-06 | 3.167E-06 | 1.518E-06 | 1.800E-06 | 4.800E-06 |
| 13101 | 656 | A656A | B | 3.8e-06 | 2.8e-06 | 2.9e-06 | 3.167E-06 | 5.508E-07 | 2.800E-06 | 3.800E-06 |
| 13102 | 656 | A656R | B | 1.7e-06 | 1.9e-06 | 1.6e-06 | 1.733E-06 | 1.528E-07 | 1.600E-06 | 1.900E-06 |
| 13103 | 656 | A656N | B | 2.4e-06 | 2.4e-06 | 2.3e-06 | 2.367E-06 | 5.774E-08 | 2.300E-06 | 2.400E-06 |
| 13104 | 656 | A656D | B | 1.7e-06 | 2.6e-06 | 3.9e-06 | 2.733E-06 | 1.106E-06 | 1.700E-06 | 3.900E-06 |
| 13105 | 656 | A656C | B | 2.6e-06 | 2.5e-06 | 2.9e-06 | 2.667E-06 | 2.082E-07 | 2.500E-06 | 2.900E-06 |
| 13106 | 656 | A656E | B | 2.2e-06 | 2.5e-06 | 2.1e-06 | 2.267E-06 | 2.082E-07 | 2.100E-06 | 2.500E-06 |
| 13107 | 656 | A656Q | B | 2.1e-06 | 3e-06 | 2.9e-06 | 2.667E-06 | 4.933E-07 | 2.100E-06 | 3.000E-06 |
| 13108 | 656 | A656G | B | 3.7e-06 | 1.6e-06 | 1.9e-06 | 2.400E-06 | 1.136E-06 | 1.600E-06 | 3.700E-06 |
| 13109 | 656 | A656H | B | 1.4e-06 | 3.4e-06 | 2.8e-06 | 2.533E-06 | 1.026E-06 | 1.400E-06 | 3.400E-06 |
| 13110 | 656 | A656I | B | 3.4e-06 | 1.3e-06 | 3.6e-06 | 2.767E-06 | 1.274E-06 | 1.300E-06 | 3.600E-06 |
| 13111 | 656 | A656L | B | 2.7e-06 | 2e-06 | 2e-06 | 2.233E-06 | 4.041E-07 | 2.000E-06 | 2.700E-06 |
| 13112 | 656 | A656K | B | 1.6e-06 | 2.9e-06 | 1.7e-06 | 2.067E-06 | 7.234E-07 | 1.600E-06 | 2.900E-06 |
| 13113 | 656 | A656M | B | 1.7e-06 | 2.1e-06 | 2.9e-06 | 2.233E-06 | 6.110E-07 | 1.700E-06 | 2.900E-06 |
| 13114 | 656 | A656F | B | 3.6e-06 | 2.7e-06 | 1.8e-06 | 2.700E-06 | 9.000E-07 | 1.800E-06 | 3.600E-06 |
| 13115 | 656 | A656P | B | 4.2e-06 | 2.8e-06 | 3.1e-06 | 3.367E-06 | 7.371E-07 | 2.800E-06 | 4.200E-06 |
| 13116 | 656 | A656S | B | 2.2e-06 | 2.4e-06 | 1.7e-06 | 2.100E-06 | 3.606E-07 | 1.700E-06 | 2.400E-06 |
| 13117 | 656 | A656T | B | 9.5e-07 | 4e-06 | 1.4e-06 | 2.117E-06 | 1.646E-06 | 9.500E-07 | 4.000E-06 |
| 13118 | 656 | A656W | B | 3.2e-06 | 2.6e-06 | 2.6e-06 | 2.800E-06 | 3.464E-07 | 2.600E-06 | 3.200E-06 |
| 13119 | 656 | A656Y | B | 1.5e-06 | 1.6e-06 | 3.3e-06 | 2.133E-06 | 1.012E-06 | 1.500E-06 | 3.300E-06 |
| 13120 | 656 | A656V | B | 1.9e-06 | 1.2e-06 | 2.9e-06 | 2.000E-06 | 8.544E-07 | 1.200E-06 | 2.900E-06 |
| 13121 | 657 | W657A | B | 3.9e-06 | 2.8e-06 | 2.5e-06 | 3.067E-06 | 7.371E-07 | 2.500E-06 | 3.900E-06 |
| 13122 | 657 | W657R | B | 2.6e-06 | 2e-06 | 2.6e-06 | 2.400E-06 | 3.464E-07 | 2.000E-06 | 2.600E-06 |
| 13123 | 657 | W657N | B | 1.7e-06 | 2.1e-06 | 1.9e-06 | 1.900E-06 | 2.000E-07 | 1.700E-06 | 2.100E-06 |
| 13124 | 657 | W657D | B | 1.5e-06 | 1.9e-06 | 3.6e-06 | 2.333E-06 | 1.115E-06 | 1.500E-06 | 3.600E-06 |
| 13125 | 657 | W657C | B | 1.1e-06 | 3.2e-06 | 2.3e-06 | 2.200E-06 | 1.054E-06 | 1.100E-06 | 3.200E-06 |
| 13126 | 657 | W657E | B | 2.2e-06 | 3.7e-06 | 2.6e-06 | 2.833E-06 | 7.767E-07 | 2.200E-06 | 3.700E-06 |
| 13127 | 657 | W657Q | B | 2.9e-06 | 2.7e-06 | 2.1e-06 | 2.567E-06 | 4.163E-07 | 2.100E-06 | 2.900E-06 |
| 13128 | 657 | W657G | B | 2.8e-06 | 2.6e-06 | 3.6e-06 | 3.000E-06 | 5.292E-07 | 2.600E-06 | 3.600E-06 |
| 13129 | 657 | W657H | B | 3.5e-06 | 3.7e-06 | 1.5e-06 | 2.900E-06 | 1.217E-06 | 1.500E-06 | 3.700E-06 |
| 13130 | 657 | W657I | B | 2.7e-06 | 1.7e-06 | 2.4e-06 | 2.267E-06 | 5.132E-07 | 1.700E-06 | 2.700E-06 |
| 13131 | 657 | W657L | B | 2.4e-06 | 2.4e-06 | 1.7e-06 | 2.167E-06 | 4.041E-07 | 1.700E-06 | 2.400E-06 |
| 13132 | 657 | W657K | B | 2.6e-06 | 2.1e-06 | 1.4e-06 | 2.033E-06 | 6.028E-07 | 1.400E-06 | 2.600E-06 |
| 13133 | 657 | W657M | B | 1.9e-06 | 2.4e-06 | 2.1e-06 | 2.133E-06 | 2.517E-07 | 1.900E-06 | 2.400E-06 |
| 13134 | 657 | W657F | B | 3.3e-06 | 3.9e-06 | 1.8e-06 | 3.000E-06 | 1.082E-06 | 1.800E-06 | 3.900E-06 |
| 13135 | 657 | W657P | B | 2.5e-06 | 2.7e-06 | 2.1e-06 | 2.433E-06 | 3.055E-07 | 2.100E-06 | 2.700E-06 |
| 13136 | 657 | W657S | B | 2.1e-06 | 2.5e-06 | 2.7e-06 | 2.433E-06 | 3.055E-07 | 2.100E-06 | 2.700E-06 |
| 13137 | 657 | W657T | B | 1.6e-06 | 2.2e-06 | 3.1e-06 | 2.300E-06 | 7.550E-07 | 1.600E-06 | 3.100E-06 |
| 13138 | 657 | W657W | B | 2.5e-06 | 2.5e-06 | 3e-06 | 2.667E-06 | 2.887E-07 | 2.500E-06 | 3.000E-06 |
| 13139 | 657 | W657Y | B | 2.4e-06 | 2.7e-06 | 1.6e-06 | 2.233E-06 | 5.686E-07 | 1.600E-06 | 2.700E-06 |
| 13140 | 657 | W657V | B | 2.4e-06 | 1.9e-06 | 3.1e-06 | 2.467E-06 | 6.028E-07 | 1.900E-06 | 3.100E-06 |
| 13141 | 658 | Y658A | B | 2.5e-06 | 1.8e-06 | 1.9e-06 | 2.067E-06 | 3.786E-07 | 1.800E-06 | 2.500E-06 |
| 13142 | 658 | Y658R | B | 2.6e-06 | 1.4e-06 | 3.6e-06 | 2.533E-06 | 1.102E-06 | 1.400E-06 | 3.600E-06 |
| 13143 | 658 | Y658N | B | 2.5e-06 | 2.2e-06 | 3e-06 | 2.567E-06 | 4.041E-07 | 2.200E-06 | 3.000E-06 |
| 13144 | 658 | Y658D | B | 1.7e-06 | 2.2e-06 | 2.6e-06 | 2.167E-06 | 4.509E-07 | 1.700E-06 | 2.600E-06 |
| 13145 | 658 | Y658C | B | 1.7e-06 | 2.1e-06 | 1.8e-06 | 1.867E-06 | 2.082E-07 | 1.700E-06 | 2.100E-06 |
| 13146 | 658 | Y658E | B | 1.6e-06 | 1.1e-06 | 2.9e-06 | 1.867E-06 | 9.292E-07 | 1.100E-06 | 2.900E-06 |
| 13147 | 658 | Y658Q | B | 1.8e-06 | 2.1e-06 | 2.7e-06 | 2.200E-06 | 4.583E-07 | 1.800E-06 | 2.700E-06 |
| 13148 | 658 | Y658G | B | 1.6e-06 | 2.8e-06 | 1.6e-06 | 2.000E-06 | 6.928E-07 | 1.600E-06 | 2.800E-06 |
| 13149 | 658 | Y658H | B | 2.4e-06 | 2.7e-06 | 1.7e-06 | 2.267E-06 | 5.132E-07 | 1.700E-06 | 2.700E-06 |
| 13150 | 658 | Y658I | B | 2.4e-06 | 2.4e-06 | 2.5e-06 | 2.433E-06 | 5.774E-08 | 2.400E-06 | 2.500E-06 |
| 13151 | 658 | Y658L | B | 2.5e-06 | 2.6e-06 | 2.3e-06 | 2.467E-06 | 1.528E-07 | 2.300E-06 | 2.600E-06 |
| 13152 | 658 | Y658K | B | 2.2e-06 | 1.3e-06 | 2.3e-06 | 1.933E-06 | 5.508E-07 | 1.300E-06 | 2.300E-06 |
| 13153 | 658 | Y658M | B | 1.7e-06 | 2.3e-06 | 2.3e-06 | 2.100E-06 | 3.464E-07 | 1.700E-06 | 2.300E-06 |
| 13154 | 658 | Y658F | B | 3.2e-06 | 3.7e-06 | 2.9e-06 | 3.267E-06 | 4.041E-07 | 2.900E-06 | 3.700E-06 |
| 13155 | 658 | Y658P | B | 2e-06 | 3.6e-06 | 4.1e-06 | 3.233E-06 | 1.097E-06 | 2.000E-06 | 4.100E-06 |
| 13156 | 658 | Y658S | B | 2.8e-06 | 3.7e-06 | 2.9e-06 | 3.133E-06 | 4.933E-07 | 2.800E-06 | 3.700E-06 |
| 13157 | 658 | Y658T | B | 3.3e-06 | 2.1e-06 | 2.5e-06 | 2.633E-06 | 6.110E-07 | 2.100E-06 | 3.300E-06 |
| 13158 | 658 | Y658W | B | 3.1e-06 | 1.7e-06 | 1.3e-06 | 2.033E-06 | 9.452E-07 | 1.300E-06 | 3.100E-06 |
| 13159 | 658 | Y658Y | B | 1.8e-06 | 5.5e-06 | 3.3e-06 | 3.533E-06 | 1.861E-06 | 1.800E-06 | 5.500E-06 |
| 13160 | 658 | Y658V | B | 3.6e-06 | 1.2e-06 | 2.1e-06 | 2.300E-06 | 1.212E-06 | 1.200E-06 | 3.600E-06 |
| 13161 | 659 | Q659A | B | 2.2e-06 | 1.5e-06 | 3.6e-06 | 2.433E-06 | 1.069E-06 | 1.500E-06 | 3.600E-06 |
| 13162 | 659 | Q659R | B | 2.6e-06 | 1.9e-06 | 1.8e-06 | 2.100E-06 | 4.359E-07 | 1.800E-06 | 2.600E-06 |
| 13163 | 659 | Q659N | B | 2.5e-06 | 1.1e-06 | 2.2e-06 | 1.933E-06 | 7.371E-07 | 1.100E-06 | 2.500E-06 |
| 13164 | 659 | Q659D | B | 3e-06 | 3.4e-06 | 2.3e-06 | 2.900E-06 | 5.568E-07 | 2.300E-06 | 3.400E-06 |
| 13165 | 659 | Q659C | B | 3.1e-06 | 2.5e-06 | 2.5e-06 | 2.700E-06 | 3.464E-07 | 2.500E-06 | 3.100E-06 |
| 13166 | 659 | Q659E | B | 2.6e-06 | 4.8e-06 | 3.7e-06 | 3.700E-06 | 1.100E-06 | 2.600E-06 | 4.800E-06 |
| 13167 | 659 | Q659Q | B | 4.7e-06 | 1.4e-06 | 1.5e-06 | 2.533E-06 | 1.877E-06 | 1.400E-06 | 4.700E-06 |
| 13168 | 659 | Q659G | B | 1.7e-06 | 2.5e-06 | 4.7e-06 | 2.967E-06 | 1.553E-06 | 1.700E-06 | 4.700E-06 |
| 13169 | 659 | Q659H | B | 2.4e-06 | 2.4e-06 | 1.7e-06 | 2.167E-06 | 4.041E-07 | 1.700E-06 | 2.400E-06 |
| 13170 | 659 | Q659I | B | 2.6e-06 | 4.6e-06 | 2.1e-06 | 3.100E-06 | 1.323E-06 | 2.100E-06 | 4.600E-06 |
| 13171 | 659 | Q659L | B | 3.6e-06 | 3.6e-06 | 2.1e-06 | 3.100E-06 | 8.660E-07 | 2.100E-06 | 3.600E-06 |
| 13172 | 659 | Q659K | B | 2.3e-06 | 2.5e-06 | 2.3e-06 | 2.367E-06 | 1.155E-07 | 2.300E-06 | 2.500E-06 |
| 13173 | 659 | Q659M | B | 3.1e-06 | 2.2e-06 | 1.9e-06 | 2.400E-06 | 6.245E-07 | 1.900E-06 | 3.100E-06 |
| 13174 | 659 | Q659F | B | 1.6e-06 | 1.8e-06 | 1.8e-06 | 1.733E-06 | 1.155E-07 | 1.600E-06 | 1.800E-06 |

| | | | | | | | | | | |
|---|---|---|---|---|---|---|---|---|---|---|
| 13175 | 659 | Q659P | B | 2.2e-06 | 3.1e-06 | 2e-06 | 2.433E-06 | 5.859E-07 | 2.000E-06 | 3.100E-06 |
| 13176 | 659 | Q659S | B | 3.3e-06 | 1.9e-06 | 5.5e-06 | 3.567E-06 | 1.815E-06 | 1.900E-06 | 5.500E-06 |
| 13177 | 659 | Q659T | B | 1.8e-06 | 3.1e-06 | 2.4e-06 | 2.433E-06 | 6.506E-07 | 1.800E-06 | 3.100E-06 |
| 13178 | 659 | Q659W | B | 2.6e-06 | 2.2e-06 | 3.2e-06 | 2.667E-06 | 5.033E-07 | 2.200E-06 | 3.200E-06 |
| 13179 | 659 | Q659Y | B | 1.4e-06 | 4.8e-06 | 1.3e-06 | 2.500E-06 | 1.992E-06 | 1.300E-06 | 4.800E-06 |
| 13180 | 659 | Q659V | B | 1.8e-06 | 2e-06 | 3.6e-06 | 2.467E-06 | 9.866E-07 | 1.800E-06 | 3.600E-06 |
| 13181 | 660 | Q660A | B | 1.9e-06 | 2.8e-06 | 1.8e-06 | 2.167E-06 | 5.508E-07 | 1.800E-06 | 2.800E-06 |
| 13182 | 660 | Q660R | B | 2.9e-06 | 2.6e-06 | 2.8e-06 | 2.767E-06 | 1.528E-07 | 2.600E-06 | 2.900E-06 |
| 13183 | 660 | Q660N | B | 2e-06 | 2.3e-06 | 3.3e-06 | 2.533E-06 | 6.807E-07 | 2.000E-06 | 3.300E-06 |
| 13184 | 660 | Q660D | B | 3.7e-06 | 2.2e-06 | 2.1e-06 | 2.667E-06 | 8.963E-07 | 2.100E-06 | 3.700E-06 |
| 13185 | 660 | Q660C | B | 2.7e-06 | 2.8e-06 | 3.4e-06 | 2.967E-06 | 3.786E-07 | 2.700E-06 | 3.400E-06 |
| 13186 | 660 | Q660E | B | 3.1e-06 | 3.6e-06 | 2.6e-06 | 3.100E-06 | 5.000E-07 | 2.600E-06 | 3.600E-06 |
| 13187 | 660 | Q660Q | B | 3e-06 | 2.5e-06 | 3.1e-06 | 2.867E-06 | 3.215E-07 | 2.500E-06 | 3.100E-06 |
| 13188 | 660 | Q660G | B | 2.8e-06 | 2.6e-06 | 3.3e-06 | 2.900E-06 | 3.606E-07 | 2.600E-06 | 3.300E-06 |
| 13189 | 660 | Q660H | B | 2.6e-06 | 9.8e-07 | 3e-06 | 2.193E-06 | 1.070E-06 | 9.800E-07 | 3.000E-06 |
| 13190 | 660 | Q660I | B | 1.8e-06 | 3.5e-06 | 3.6e-06 | 2.967E-06 | 1.012E-06 | 1.800E-06 | 3.600E-06 |
| 13191 | 660 | Q660L | B | 2.3e-06 | 3.1e-06 | 3e-06 | 2.800E-06 | 4.359E-07 | 2.300E-06 | 3.100E-06 |
| 13192 | 660 | Q660K | B | 2.3e-06 | 1.6e-06 | 1.8e-06 | 1.900E-06 | 3.606E-07 | 1.600E-06 | 2.300E-06 |
| 13193 | 660 | Q660M | B | 1.8e-06 | 1.6e-06 | 2.4e-06 | 1.933E-06 | 4.163E-07 | 1.600E-06 | 2.400E-06 |
| 13194 | 660 | Q660F | B | 2.6e-06 | 4.3e-06 | 3.1e-06 | 3.333E-06 | 8.737E-07 | 2.600E-06 | 4.300E-06 |
| 13195 | 660 | Q660P | B | 2.8e-06 | 3.6e-06 | 2.1e-06 | 2.833E-06 | 7.506E-07 | 2.100E-06 | 3.600E-06 |
| 13196 | 660 | Q660S | B | 1.9e-06 | 2.9e-06 | 1.8e-06 | 2.200E-06 | 6.083E-07 | 1.800E-06 | 2.900E-06 |
| 13197 | 660 | Q660T | B | 1.8e-06 | 1.8e-06 | 2.4e-06 | 2.000E-06 | 3.464E-07 | 1.800E-06 | 2.400E-06 |
| 13198 | 660 | Q660W | B | 2.4e-06 | 3.1e-06 | 1.7e-06 | 2.400E-06 | 7.000E-07 | 1.700E-06 | 3.100E-06 |
| 13199 | 660 | Q660Y | B | 2.2e-06 | 1.6e-06 | 2.5e-06 | 2.100E-06 | 4.583E-07 | 1.600E-06 | 2.500E-06 |
| 13200 | 660 | Q660V | B | 2.2e-06 | 2.7e-06 | 2.5e-06 | 2.467E-06 | 2.517E-07 | 2.200E-06 | 2.700E-06 |
| 13201 | 661 | K661A | B | 2.6e-06 | 1.5e-06 | 1.7e-06 | 1.933E-06 | 5.859E-07 | 1.500E-06 | 2.600E-06 |
| 13202 | 661 | K661R | B | 2.8e-06 | 1.6e-06 | 1.2e-06 | 1.867E-06 | 8.327E-07 | 1.200E-06 | 2.800E-06 |
| 13203 | 661 | K661N | B | 3e-06 | 2.8e-06 | 2.6e-06 | 2.800E-06 | 2.000E-07 | 2.600E-06 | 3.000E-06 |
| 13204 | 661 | K661D | B | 2.6e-06 | 2.7e-06 | 1.4e-06 | 2.233E-06 | 7.234E-07 | 1.400E-06 | 2.700E-06 |
| 13205 | 661 | K661C | B | 2.3e-06 | 3e-06 | 3.2e-06 | 2.833E-06 | 4.726E-07 | 2.300E-06 | 3.200E-06 |
| 13206 | 661 | K661E | B | 2.7e-06 | 1.4e-06 | 1.9e-06 | 2.000E-06 | 6.557E-07 | 1.400E-06 | 2.700E-06 |
| 13207 | 661 | K661Q | B | 2.8e-06 | 7.6e-06 | 2.3e-06 | 4.233E-06 | 2.926E-06 | 2.300E-06 | 7.600E-06 |
| 13208 | 661 | K661G | B | 3e-06 | 2.7e-06 | 2.1e-06 | 2.600E-06 | 4.583E-07 | 2.100E-06 | 3.000E-06 |
| 13209 | 661 | K661H | B | 2.5e-06 | 2e-06 | 4e-06 | 2.833E-06 | 1.041E-06 | 2.000E-06 | 4.000E-06 |
| 13210 | 661 | K661I | B | 1.7e-06 | 1.6e-06 | 1.6e-06 | 1.633E-06 | 5.774E-08 | 1.600E-06 | 1.700E-06 |
| 13211 | 661 | K661L | B | 2.4e-06 | 3.4e-06 | 3.2e-06 | 3.000E-06 | 5.292E-07 | 2.400E-06 | 3.400E-06 |
| 13212 | 661 | K661K | B | 2.7e-06 | 2.2e-06 | 2.3e-06 | 2.400E-06 | 2.646E-07 | 2.200E-06 | 2.700E-06 |
| 13213 | 661 | K661M | B | 4.2e-06 | 3.2e-06 | 5.1e-06 | 4.167E-06 | 9.504E-07 | 3.200E-06 | 5.100E-06 |
| 13214 | 661 | K661F | B | 2.9e-06 | 2.3e-06 | 3.1e-06 | 2.767E-06 | 4.163E-07 | 2.300E-06 | 3.100E-06 |
| 13215 | 661 | K661P | B | 2.7e-06 | 1.8e-06 | 2.4e-06 | 2.300E-06 | 4.583E-07 | 1.800E-06 | 2.700E-06 |
| 13216 | 661 | K661S | B | 1.6e-06 | 3.4e-06 | 2.5e-06 | 2.500E-06 | 9.000E-07 | 1.600E-06 | 3.400E-06 |
| 13217 | 661 | K661T | B | 3.6e-06 | 2.7e-06 | 2.9e-06 | 3.067E-06 | 4.726E-07 | 2.700E-06 | 3.600E-06 |
| 13218 | 661 | K661W | B | 3.1e-06 | 3.3e-06 | 2.5e-06 | 2.967E-06 | 4.163E-07 | 2.500E-06 | 3.300E-06 |
| 13219 | 661 | K661Y | B | 1.5e-06 | 2.4e-06 | 2.2e-06 | 2.033E-06 | 4.726E-07 | 1.500E-06 | 2.400E-06 |
| 13220 | 661 | K661V | B | 2.6e-06 | 2.2e-06 | 3.5e-06 | 2.767E-06 | 6.658E-07 | 2.200E-06 | 3.500E-06 |
| 13221 | 662 | P662A | B | 2.5e-06 | 4.9e-06 | 3.4e-06 | 3.600E-06 | 1.212E-06 | 2.500E-06 | 4.900E-06 |
| 13222 | 662 | P662R | B | 2.4e-06 | 3.7e-06 | 2.6e-06 | 2.900E-06 | 7.000E-07 | 2.400E-06 | 3.700E-06 |
| 13223 | 662 | P662N | B | 4.5e-06 | 2.2e-06 | 1.9e-06 | 2.867E-06 | 1.422E-06 | 1.900E-06 | 4.500E-06 |
| 13224 | 662 | P662D | B | 2.7e-06 | 2e-06 | 1.6e-06 | 2.100E-06 | 5.568E-07 | 1.600E-06 | 2.700E-06 |
| 13225 | 662 | P662C | B | 2.3e-06 | 2.5e-06 | 2.8e-06 | 2.533E-06 | 2.517E-07 | 2.300E-06 | 2.800E-06 |
| 13226 | 662 | P662E | B | 1.9e-06 | 3e-06 | 1.6e-06 | 2.167E-06 | 7.371E-07 | 1.600E-06 | 3.000E-06 |
| 13227 | 662 | P662Q | B | 2.1e-06 | 1.4e-06 | 1.9e-06 | 1.800E-06 | 3.606E-07 | 1.400E-06 | 2.100E-06 |
| 13228 | 662 | P662G | B | 3.1e-06 | 2.5e-06 | 2.6e-06 | 2.733E-06 | 3.215E-07 | 2.500E-06 | 3.100E-06 |
| 13229 | 662 | P662H | B | 2.4e-06 | 3.1e-06 | 2.5e-06 | 2.667E-06 | 3.786E-07 | 2.400E-06 | 3.100E-06 |
| 13230 | 662 | P662I | B | 1.2e-06 | 1.3e-06 | 1.5e-06 | 1.333E-06 | 1.528E-07 | 1.200E-06 | 1.500E-06 |
| 13231 | 662 | P662L | B | 3.1e-06 | 2.8e-06 | 2e-06 | 2.633E-06 | 5.686E-07 | 2.000E-06 | 3.100E-06 |
| 13232 | 662 | P662K | B | 2.2e-06 | 2e-06 | 2.1e-06 | 2.100E-06 | 1.000E-07 | 2.000E-06 | 2.200E-06 |
| 13233 | 662 | P662M | B | 1.9e-06 | 2.7e-06 | 4.6e-06 | 3.067E-06 | 1.387E-06 | 1.900E-06 | 4.600E-06 |
| 13234 | 662 | P662F | B | 3.7e-06 | 3.4e-06 | 2.3e-06 | 3.133E-06 | 7.371E-07 | 2.300E-06 | 3.700E-06 |
| 13235 | 662 | P662P | B | 1.5e-06 | 2.4e-06 | 2.8e-06 | 2.233E-06 | 6.658E-07 | 1.500E-06 | 2.800E-06 |
| 13236 | 662 | P662S | B | 3e-06 | 3e-06 | 2.6e-06 | 2.867E-06 | 2.309E-07 | 2.600E-06 | 3.000E-06 |
| 13237 | 662 | P662T | B | 1.8e-06 | 2.1e-06 | 3.1e-06 | 2.333E-06 | 6.807E-07 | 1.800E-06 | 3.100E-06 |
| 13238 | 662 | P662W | B | 2.8e-06 | 2.2e-06 | 2.4e-06 | 2.467E-06 | 3.055E-07 | 2.200E-06 | 2.800E-06 |
| 13239 | 662 | P662Y | B | 1.6e-06 | 3.3e-06 | 2e-06 | 2.300E-06 | 8.888E-07 | 1.600E-06 | 3.300E-06 |
| 13240 | 662 | P662V | B | 2.5e-06 | 2e-06 | 2.6e-06 | 2.367E-06 | 3.215E-07 | 2.000E-06 | 2.600E-06 |
| 13241 | 663 | G663A | B | 2.3e-06 | 2.7e-06 | 2.5e-06 | 2.500E-06 | 2.000E-07 | 2.300E-06 | 2.700E-06 |
| 13242 | 663 | G663R | B | 2.4e-06 | 2.8e-06 | 1.6e-06 | 2.267E-06 | 6.110E-07 | 1.600E-06 | 2.800E-06 |
| 13243 | 663 | G663N | B | 2.8e-06 | 2.5e-06 | 3.1e-06 | 2.800E-06 | 3.000E-07 | 2.500E-06 | 3.100E-06 |
| 13244 | 663 | G663D | B | 1.5e-06 | 2.3e-06 | 2.3e-06 | 2.033E-06 | 4.619E-07 | 1.500E-06 | 2.300E-06 |
| 13245 | 663 | G663C | B | 1.5e-06 | 2.4e-06 | 2.5e-06 | 2.133E-06 | 5.508E-07 | 1.500E-06 | 2.500E-06 |
| 13246 | 663 | G663E | B | 2.5e-06 | 3.6e-06 | 3.1e-06 | 3.067E-06 | 5.508E-07 | 2.500E-06 | 3.600E-06 |
| 13247 | 663 | G663Q | B | 1.6e-06 | 2e-06 | 2.4e-06 | 2.000E-06 | 4.000E-07 | 1.600E-06 | 2.400E-06 |
| 13248 | 663 | G663G | B | 2.3e-06 | 2.4e-06 | 2.8e-06 | 2.500E-06 | 2.646E-07 | 2.300E-06 | 2.800E-06 |
| 13249 | 663 | G663H | B | 2.5e-06 | 2.6e-06 | 1.7e-06 | 2.267E-06 | 4.933E-07 | 1.700E-06 | 2.600E-06 |
| 13250 | 663 | G663I | B | 1.5e-06 | 2.6e-06 | 2.1e-06 | 2.067E-06 | 5.508E-07 | 1.500E-06 | 2.600E-06 |
| 13251 | 663 | G663L | B | 1.5e-06 | 2.5e-06 | 4.4e-06 | 2.800E-06 | 1.473E-06 | 1.500E-06 | 4.400E-06 |
| 13252 | 663 | G663K | B | 2.9e-06 | 2.3e-06 | 2.5e-06 | 2.567E-06 | 3.055E-07 | 2.300E-06 | 2.900E-06 |
| 13253 | 663 | G663M | B | 2.9e-06 | 2.1e-06 | 3.1e-06 | 2.700E-06 | 5.292E-07 | 2.100E-06 | 3.100E-06 |
| 13254 | 663 | G663F | B | 2.6e-06 | 2.4e-06 | 1.4e-06 | 2.133E-06 | 6.429E-07 | 1.400E-06 | 2.600E-06 |
| 13255 | 663 | G663P | B | 2.2e-06 | 3.2e-06 | 3.5e-06 | 2.967E-06 | 6.807E-07 | 2.200E-06 | 3.500E-06 |
| 13256 | 663 | G663S | B | 5.5e-06 | 2.7e-06 | 2.2e-06 | 3.467E-06 | 1.779E-06 | 2.200E-06 | 5.500E-06 |
| 13257 | 663 | G663T | B | 2.1e-06 | 1.9e-06 | 2.2e-06 | 2.067E-06 | 1.528E-07 | 1.900E-06 | 2.200E-06 |
| 13258 | 663 | G663W | B | 3.4e-06 | 1.7e-06 | 2e-06 | 2.367E-06 | 9.074E-07 | 1.700E-06 | 3.400E-06 |
| 13259 | 663 | G663Y | B | 1.7e-06 | 2.8e-06 | 2.5e-06 | 2.333E-06 | 5.686E-07 | 1.700E-06 | 2.800E-06 |

| | | | | | | | | | | |
|---|---|---|---|---|---|---|---|---|---|---|
| 13260 | 663 | G663V | B | 3.2e-06 | 3.3e-06 | 2.8e-06 | 3.100E-06 | 2.646E-07 | 2.800E-06 | 3.300E-06 |
| 13261 | 664 | K664A | B | 2.4e-06 | 2.7e-06 | 1.7e-06 | 2.267E-06 | 5.132E-07 | 1.700E-06 | 2.700E-06 |
| 13262 | 664 | K664R | B | 2.7e-06 | 2.1e-06 | 1.1e-06 | 1.967E-06 | 8.083E-07 | 1.100E-06 | 2.700E-06 |
| 13263 | 664 | K664N | B | 3.1e-06 | 1.7e-06 | 3.5e-06 | 2.767E-06 | 9.452E-07 | 1.700E-06 | 3.500E-06 |
| 13264 | 664 | K664D | B | 1.5e-06 | 2.1e-06 | 1.4e-06 | 1.667E-06 | 3.786E-07 | 1.400E-06 | 2.100E-06 |
| 13265 | 664 | K664C | B | 1.6e-06 | 1.9e-06 | 1.4e-06 | 1.633E-06 | 2.517E-07 | 1.400E-06 | 1.900E-06 |
| 13266 | 664 | K664E | B | 2.9e-06 | 3.2e-06 | 1.9e-06 | 2.667E-06 | 6.807E-07 | 1.900E-06 | 3.200E-06 |
| 13267 | 664 | K664Q | B | 1.8e-06 | 1.6e-06 | 2.6e-06 | 2.000E-06 | 5.292E-07 | 1.600E-06 | 2.600E-06 |
| 13268 | 664 | K664G | B | 1.8e-06 | 2.6e-06 | 1.9e-06 | 2.100E-06 | 4.359E-07 | 1.800E-06 | 2.600E-06 |
| 13269 | 664 | K664H | B | 2.9e-06 | 2.7e-06 | 1.7e-06 | 2.433E-06 | 6.429E-07 | 1.700E-06 | 2.900E-06 |
| 13270 | 664 | K664I | B | 2.6e-06 | 2.4e-06 | 3.7e-06 | 2.900E-06 | 7.000E-07 | 2.400E-06 | 3.700E-06 |
| 13271 | 664 | K664L | B | 2.4e-06 | 2.5e-06 | 2e-06 | 2.300E-06 | 2.646E-07 | 2.000E-06 | 2.500E-06 |
| 13272 | 664 | K664K | B | 2.6e-06 | 4.7e-06 | 2.4e-06 | 3.233E-06 | 1.274E-06 | 2.400E-06 | 4.700E-06 |
| 13273 | 664 | K664M | B | 2.7e-06 | 1.9e-06 | 2.7e-06 | 2.433E-06 | 4.619E-07 | 1.900E-06 | 2.700E-06 |
| 13274 | 664 | K664F | B | 1.6e-06 | 2.3e-06 | 1.6e-06 | 1.833E-06 | 4.041E-07 | 1.600E-06 | 2.300E-06 |
| 13275 | 664 | K664P | B | 3e-06 | 1.2e-06 | 2e-06 | 2.067E-06 | 9.018E-07 | 1.200E-06 | 3.000E-06 |
| 13276 | 664 | K664S | B | 1.3e-06 | 1.9e-06 | 2.7e-06 | 1.967E-06 | 7.024E-07 | 1.300E-06 | 2.700E-06 |
| 13277 | 664 | K664T | B | 3e-06 | 2.3e-06 | 2.2e-06 | 2.500E-06 | 4.359E-07 | 2.200E-06 | 3.000E-06 |
| 13278 | 664 | K664W | B | 1.7e-06 | 1.1e-06 | 1.9e-06 | 1.567E-06 | 4.163E-07 | 1.100E-06 | 1.900E-06 |
| 13279 | 664 | K664Y | B | 3.1e-06 | 3.6e-06 | 3.5e-06 | 3.400E-06 | 2.646E-07 | 3.100E-06 | 3.600E-06 |
| 13280 | 664 | K664V | B | 3.5e-06 | 1.7e-06 | 3.5e-06 | 2.900E-06 | 1.039E-06 | 1.700E-06 | 3.500E-06 |
| 13281 | 665 | A665A | B | 1.6e-06 | 2.7e-06 | 2.4e-06 | 2.233E-06 | 5.686E-07 | 1.600E-06 | 2.700E-06 |
| 13282 | 665 | A665R | B | 4e-06 | 2.4e-06 | 1.4e-06 | 2.600E-06 | 1.311E-06 | 1.400E-06 | 4.000E-06 |
| 13283 | 665 | A665N | B | 2.4e-06 | 3.2e-06 | 2.6e-06 | 2.733E-06 | 4.163E-07 | 2.400E-06 | 3.200E-06 |
| 13284 | 665 | A665D | B | 1.8e-06 | 2.2e-06 | 2.6e-06 | 2.200E-06 | 4.000E-07 | 1.800E-06 | 2.600E-06 |
| 13285 | 665 | A665C | B | 1.4e-06 | 2.7e-06 | 2.1e-06 | 2.067E-06 | 6.506E-07 | 1.400E-06 | 2.700E-06 |
| 13286 | 665 | A665E | B | 1.8e-06 | 1.7e-06 | 2.5e-06 | 2.000E-06 | 4.359E-07 | 1.700E-06 | 2.500E-06 |
| 13287 | 665 | A665Q | B | 2.3e-06 | 1.6e-06 | 3.5e-06 | 2.467E-06 | 9.609E-07 | 1.600E-06 | 3.500E-06 |
| 13288 | 665 | A665G | B | 1.6e-06 | 6.1e-06 | 3e-06 | 3.567E-06 | 2.303E-06 | 1.600E-06 | 6.100E-06 |
| 13289 | 665 | A665H | B | 2.4e-06 | 1.8e-06 | 3.4e-06 | 2.533E-06 | 8.083E-07 | 1.800E-06 | 3.400E-06 |
| 13290 | 665 | A665I | B | 1.9e-06 | 2.1e-06 | 1.8e-06 | 1.933E-06 | 1.528E-07 | 1.800E-06 | 2.100E-06 |
| 13291 | 665 | A665L | B | 3e-06 | 1.9e-06 | 1.7e-06 | 2.200E-06 | 7.000E-07 | 1.700E-06 | 3.000E-06 |
| 13292 | 665 | A665K | B | 2.3e-06 | 3.3e-06 | 2.1e-06 | 2.567E-06 | 6.429E-07 | 2.100E-06 | 3.300E-06 |
| 13293 | 665 | A665M | B | 6.5e-06 | 1.4e-06 | 2.2e-06 | 3.367E-06 | 2.743E-06 | 1.400E-06 | 6.500E-06 |
| 13294 | 665 | A665F | B | 2.5e-06 | 1.6e-06 | 2.4e-06 | 2.167E-06 | 4.933E-07 | 1.600E-06 | 2.500E-06 |
| 13295 | 665 | A665P | B | 3.8e-06 | 2.1e-06 | 2.5e-06 | 2.800E-06 | 8.888E-07 | 2.100E-06 | 3.800E-06 |
| 13296 | 665 | A665S | B | 1.8e-06 | 1.4e-06 | 2.9e-06 | 2.033E-06 | 7.767E-07 | 1.400E-06 | 2.900E-06 |
| 13297 | 665 | A665T | B | 2.3e-06 | 2.5e-06 | 1.8e-06 | 2.200E-06 | 3.606E-07 | 1.800E-06 | 2.500E-06 |
| 13298 | 665 | A665W | B | 1.8e-06 | 1.8e-06 | 2.1e-06 | 1.900E-06 | 1.732E-07 | 1.800E-06 | 2.100E-06 |
| 13299 | 665 | A665Y | B | 3.1e-06 | 2.1e-06 | 1.9e-06 | 2.367E-06 | 6.429E-07 | 1.900E-06 | 3.100E-06 |
| 13300 | 665 | A665V | B | 2.9e-06 | 2e-06 | 2e-06 | 2.300E-06 | 5.196E-07 | 2.000E-06 | 2.900E-06 |
| 13301 | 666 | P666A | B | 3.1e-06 | 1.7e-06 | 3.2e-06 | 2.667E-06 | 8.386E-07 | 1.700E-06 | 3.200E-06 |
| 13302 | 666 | P666R | B | 2.7e-06 | 2.3e-06 | 2.1e-06 | 2.367E-06 | 3.055E-07 | 2.100E-06 | 2.700E-06 |
| 13303 | 666 | P666N | B | 2.4e-06 | 1.6e-06 | 2.1e-06 | 2.033E-06 | 4.041E-07 | 1.600E-06 | 2.400E-06 |
| 13304 | 666 | P666D | B | 3.9e-06 | 1.9e-06 | 2.2e-06 | 2.667E-06 | 1.079E-06 | 1.900E-06 | 3.900E-06 |
| 13305 | 666 | P666C | B | 2.7e-06 | 5.2e-06 | 2.6e-06 | 3.500E-06 | 1.473E-06 | 2.600E-06 | 5.200E-06 |
| 13306 | 666 | P666E | B | 2.9e-06 | 2.2e-06 | 1.9e-06 | 2.333E-06 | 5.132E-07 | 1.900E-06 | 2.900E-06 |
| 13307 | 666 | P666Q | B | 1.9e-06 | 2e-06 | 2.9e-06 | 2.267E-06 | 5.508E-07 | 1.900E-06 | 2.900E-06 |
| 13308 | 666 | P666G | B | 1.7e-06 | 2.6e-06 | 1.6e-06 | 1.967E-06 | 5.508E-07 | 1.600E-06 | 2.600E-06 |
| 13309 | 666 | P666H | B | 1.6e-06 | 4.5e-06 | 1.4e-06 | 2.500E-06 | 1.735E-06 | 1.400E-06 | 4.500E-06 |
| 13310 | 666 | P666I | B | 2.1e-06 | 4.7e-06 | 2.2e-06 | 3.000E-06 | 1.473E-06 | 2.100E-06 | 4.700E-06 |
| 13311 | 666 | P666L | B | 2.7e-06 | 1.3e-06 | 1.6e-06 | 1.867E-06 | 7.371E-07 | 1.300E-06 | 2.700E-06 |
| 13312 | 666 | P666K | B | 2.8e-06 | 2.9e-06 | 2.3e-06 | 2.667E-06 | 3.215E-07 | 2.300E-06 | 2.900E-06 |
| 13313 | 666 | P666M | B | 2.4e-06 | 1.9e-06 | 2.7e-06 | 2.333E-06 | 4.041E-07 | 1.900E-06 | 2.700E-06 |
| 13314 | 666 | P666F | B | 1.5e-06 | 2.9e-06 | 2.1e-06 | 2.167E-06 | 7.024E-07 | 1.500E-06 | 2.900E-06 |
| 13315 | 666 | P666P | B | 1.4e-06 | 2.7e-06 | 1.6e-06 | 1.900E-06 | 7.000E-07 | 1.400E-06 | 2.700E-06 |
| 13316 | 666 | P666S | B | 4.6e-06 | 2.5e-06 | 1.9e-06 | 3.000E-06 | 1.418E-06 | 1.900E-06 | 4.600E-06 |
| 13317 | 666 | P666T | B | 2.2e-06 | 3.6e-06 | 1.4e-06 | 2.400E-06 | 1.114E-06 | 1.400E-06 | 3.600E-06 |
| 13318 | 666 | P666W | B | 3.9e-06 | 2.6e-06 | 1.7e-06 | 2.733E-06 | 1.106E-06 | 1.700E-06 | 3.900E-06 |
| 13319 | 666 | P666Y | B | 1.9e-06 | 1.7e-06 | 3.2e-06 | 2.267E-06 | 8.145E-07 | 1.700E-06 | 3.200E-06 |
| 13320 | 666 | P666V | B | 3e-06 | 2.8e-06 | 1.3e-06 | 2.367E-06 | 9.292E-07 | 1.300E-06 | 3.000E-06 |
| 13321 | 667 | K667A | B | 2.7e-06 | 2.8e-06 | 1.9e-06 | 2.467E-06 | 4.933E-07 | 1.900E-06 | 2.800E-06 |
| 13322 | 667 | K667R | B | 1.9e-06 | 1.3e-06 | 1.9e-06 | 1.700E-06 | 3.464E-07 | 1.300E-06 | 1.900E-06 |
| 13323 | 667 | K667N | B | 2.5e-06 | 2.9e-06 | 2.5e-06 | 2.633E-06 | 2.309E-07 | 2.500E-06 | 2.900E-06 |
| 13324 | 667 | K667D | B | 1.8e-06 | 2.8e-06 | 2.5e-06 | 2.367E-06 | 5.132E-07 | 1.800E-06 | 2.800E-06 |
| 13325 | 667 | K667C | B | 1.6e-06 | 1.4e-06 | 3.3e-06 | 2.100E-06 | 1.044E-06 | 1.400E-06 | 3.300E-06 |
| 13326 | 667 | K667E | B | 2.8e-06 | 2.4e-06 | 2.1e-06 | 2.433E-06 | 3.512E-07 | 2.100E-06 | 2.800E-06 |
| 13327 | 667 | K667Q | B | 2.7e-06 | 2.4e-06 | 2e-06 | 2.367E-06 | 3.512E-07 | 2.000E-06 | 2.700E-06 |
| 13328 | 667 | K667G | B | 2e-06 | 2e-06 | 2.3e-06 | 2.100E-06 | 1.732E-07 | 2.000E-06 | 2.300E-06 |
| 13329 | 667 | K667H | B | 3.3e-06 | 1.9e-06 | 2.5e-06 | 2.567E-06 | 7.024E-07 | 1.900E-06 | 3.300E-06 |
| 13330 | 667 | K667I | B | 2.2e-06 | 1.1e-06 | 2.8e-06 | 2.033E-06 | 8.622E-07 | 1.100E-06 | 2.800E-06 |
| 13331 | 667 | K667L | B | 2.2e-06 | 2.7e-06 | 1.5e-06 | 2.133E-06 | 6.028E-07 | 1.500E-06 | 2.700E-06 |
| 13332 | 667 | K667K | B | 3e-06 | 2.1e-06 | 2.3e-06 | 2.467E-06 | 4.726E-07 | 2.100E-06 | 3.000E-06 |
| 13333 | 667 | K667M | B | 2.5e-06 | 2.5e-06 | 2.2e-06 | 2.400E-06 | 1.732E-07 | 2.200E-06 | 2.500E-06 |
| 13334 | 667 | K667F | B | 3.7e-06 | 3.4e-06 | 2.9e-06 | 3.333E-06 | 4.041E-07 | 2.900E-06 | 3.700E-06 |
| 13335 | 667 | K667P | B | 3.7e-06 | 2.3e-06 | 4.3e-06 | 3.433E-06 | 1.026E-06 | 2.300E-06 | 4.300E-06 |
| 13336 | 667 | K667S | B | 3.1e-06 | 3.7e-06 | 1.9e-06 | 2.900E-06 | 9.165E-07 | 1.900E-06 | 3.700E-06 |
| 13337 | 667 | K667T | B | 2.1e-06 | 2.9e-06 | 3.6e-06 | 2.867E-06 | 7.506E-07 | 2.100E-06 | 3.600E-06 |
| 13338 | 667 | K667W | B | 2.9e-06 | 1.7e-06 | 2.6e-06 | 2.400E-06 | 6.245E-07 | 1.700E-06 | 2.900E-06 |
| 13339 | 667 | K667Y | B | 3.3e-06 | 2.2e-06 | 1.8e-06 | 2.433E-06 | 7.767E-07 | 1.800E-06 | 3.300E-06 |
| 13340 | 667 | K667V | B | 2.3e-06 | 2.1e-06 | 3.1e-06 | 2.500E-06 | 5.292E-07 | 2.100E-06 | 3.100E-06 |
| 13341 | 668 | L668A | B | 2.5e-06 | 2.9e-06 | 1.2e-06 | 2.200E-06 | 8.888E-07 | 1.200E-06 | 2.900E-06 |
| 13342 | 668 | L668R | B | 2.6e-06 | 2.2e-06 | 3.5e-06 | 2.767E-06 | 6.658E-07 | 2.200E-06 | 3.500E-06 |
| 13343 | 668 | L668N | B | 1.4e-06 | 1.8e-06 | 1.3e-06 | 1.500E-06 | 2.646E-07 | 1.300E-06 | 1.800E-06 |
| 13344 | 668 | L668D | B | 4.7e-06 | 3.4e-06 | 2.4e-06 | 3.500E-06 | 1.153E-06 | 2.400E-06 | 4.700E-06 |

| | | | | | | | | | | |
|---|---|---|---|---|---|---|---|---|---|---|
| 13345 | 668 | L668C | B | 2.8e-06 | 1.8e-06 | 2.3e-06 | 2.300E-06 | 5.000E-07 | 1.800E-06 | 2.800E-06 |
| 13346 | 668 | L668E | B | 2.4e-06 | 1.7e-06 | 2.8e-06 | 2.300E-06 | 5.568E-07 | 1.700E-06 | 2.800E-06 |
| 13347 | 668 | L668Q | B | 1.6e-06 | 3.1e-06 | 3.7e-06 | 2.800E-06 | 1.082E-06 | 1.600E-06 | 3.700E-06 |
| 13348 | 668 | L668G | B | 2.7e-06 | 1.9e-06 | 2e-06 | 2.200E-06 | 4.359E-07 | 1.900E-06 | 2.700E-06 |
| 13349 | 668 | L668H | B | 2.2e-06 | 2.1e-06 | 3.1e-06 | 2.467E-06 | 5.508E-07 | 2.100E-06 | 3.100E-06 |
| 13350 | 668 | L668I | B | 1.8e-06 | 1.5e-06 | 2.3e-06 | 1.867E-06 | 4.041E-07 | 1.500E-06 | 2.300E-06 |
| 13351 | 668 | L668L | B | 2.7e-06 | 3.2e-06 | 2.6e-06 | 2.833E-06 | 3.215E-07 | 2.600E-06 | 3.200E-06 |
| 13352 | 668 | L668K | B | 9.4e-07 | 4.9e-06 | 2.8e-06 | 2.880E-06 | 1.981E-06 | 9.400E-07 | 4.900E-06 |
| 13353 | 668 | L668M | B | 2.7e-06 | 1.8e-06 | 4.9e-06 | 3.133E-06 | 1.595E-06 | 1.800E-06 | 4.900E-06 |
| 13354 | 668 | L668F | B | 1.4e-06 | 2.2e-06 | 4.2e-06 | 2.600E-06 | 1.442E-06 | 1.400E-06 | 4.200E-06 |
| 13355 | 668 | L668P | B | 2.3e-06 | 5.4e-06 | 3.7e-06 | 3.800E-06 | 1.552E-06 | 2.300E-06 | 5.400E-06 |
| 13356 | 668 | L668S | B | 2.9e-06 | 2.5e-06 | 1.3e-06 | 2.233E-06 | 8.327E-07 | 1.300E-06 | 2.900E-06 |
| 13357 | 668 | L668T | B | 3.1e-06 | 3.3e-06 | 2.3e-06 | 2.900E-06 | 5.292E-07 | 2.300E-06 | 3.300E-06 |
| 13358 | 668 | L668W | B | 1.5e-06 | 1.7e-06 | 4.1e-06 | 2.433E-06 | 1.447E-06 | 1.500E-06 | 4.100E-06 |
| 13359 | 668 | L668Y | B | 1.6e-06 | 1.1e-06 | 2.9e-06 | 1.867E-06 | 9.292E-07 | 1.100E-06 | 2.900E-06 |
| 13360 | 668 | L668V | B | 2.4e-06 | 3e-06 | 2.7e-06 | 2.700E-06 | 3.000E-07 | 2.400E-06 | 3.000E-06 |
| 13361 | 669 | L669A | B | 2.3e-06 | 1.9e-06 | 1.7e-06 | 1.967E-06 | 3.055E-07 | 1.700E-06 | 2.300E-06 |
| 13362 | 669 | L669R | B | 2e-06 | 2.4e-06 | 3.2e-06 | 2.533E-06 | 6.110E-07 | 2.000E-06 | 3.200E-06 |
| 13363 | 669 | L669N | B | 2.7e-06 | 2.3e-06 | 2.8e-06 | 2.600E-06 | 2.646E-07 | 2.300E-06 | 2.800E-06 |
| 13364 | 669 | L669D | B | 2.6e-06 | 1.4e-06 | 2.2e-06 | 2.067E-06 | 6.110E-07 | 1.400E-06 | 2.600E-06 |
| 13365 | 669 | L669C | B | 2.7e-06 | 2.1e-06 | 2.3e-06 | 2.367E-06 | 3.055E-07 | 2.100E-06 | 2.700E-06 |
| 13366 | 669 | L669E | B | 1.8e-06 | 2.2e-06 | 1.7e-06 | 1.900E-06 | 2.646E-07 | 1.700E-06 | 2.200E-06 |
| 13367 | 669 | L669Q | B | 2.1e-06 | 2.4e-06 | 1.9e-06 | 2.133E-06 | 2.517E-07 | 1.900E-06 | 2.400E-06 |
| 13368 | 669 | L669G | B | 2.3e-06 | 1.9e-06 | 3.4e-06 | 2.533E-06 | 7.767E-07 | 1.900E-06 | 3.400E-06 |
| 13369 | 669 | L669H | B | 2.6e-06 | 2.6e-06 | 2.8e-06 | 2.667E-06 | 1.155E-07 | 2.600E-06 | 2.800E-06 |
| 13370 | 669 | L669I | B | 2.7e-06 | 2.3e-06 | 2.1e-06 | 2.367E-06 | 3.055E-07 | 2.100E-06 | 2.700E-06 |
| 13371 | 669 | L669L | B | 3.9e-06 | 3.5e-06 | 2.7e-06 | 3.367E-06 | 6.110E-07 | 2.700E-06 | 3.900E-06 |
| 13372 | 669 | L669K | B | 3.1e-06 | 2e-06 | 1.9e-06 | 2.333E-06 | 6.658E-07 | 1.900E-06 | 3.100E-06 |
| 13373 | 669 | L669M | B | 3.1e-06 | 2.5e-06 | 1.9e-06 | 2.500E-06 | 6.000E-07 | 1.900E-06 | 3.100E-06 |
| 13374 | 669 | L669F | B | 2.4e-06 | 1.3e-06 | 2.4e-06 | 2.033E-06 | 6.351E-07 | 1.300E-06 | 2.400E-06 |
| 13375 | 669 | L669P | B | 3.1e-06 | 2.4e-06 | 2e-06 | 2.500E-06 | 5.568E-07 | 2.000E-06 | 3.100E-06 |
| 13376 | 669 | L669S | B | 2.3e-06 | 5.8e-06 | 1.7e-06 | 3.267E-06 | 2.214E-06 | 1.700E-06 | 5.800E-06 |
| 13377 | 669 | L669T | B | 2.8e-06 | 2.2e-06 | 2.8e-06 | 2.600E-06 | 3.464E-07 | 2.200E-06 | 2.800E-06 |
| 13378 | 669 | L669W | B | 3.2e-06 | 2.7e-06 | 3.1e-06 | 3.000E-06 | 2.646E-07 | 2.700E-06 | 3.200E-06 |
| 13379 | 669 | L669Y | B | 3.6e-06 | 2.3e-06 | 4e-06 | 3.300E-06 | 8.888E-07 | 2.300E-06 | 4.000E-06 |
| 13380 | 669 | L669V | B | 1.5e-06 | 9e-07 | 1.8e-06 | 1.400E-06 | 4.583E-07 | 9.000E-07 | 1.800E-06 |
| 13381 | 670 | I670A | B | 3e-06 | 2.6e-06 | 2.1e-06 | 2.567E-06 | 4.509E-07 | 2.100E-06 | 3.000E-06 |
| 13382 | 670 | I670R | B | 2.3e-06 | 1.7e-06 | 2.5e-06 | 2.167E-06 | 4.163E-07 | 1.700E-06 | 2.500E-06 |
| 13383 | 670 | I670N | B | 2.9e-06 | 1.9e-06 | 2.2e-06 | 2.333E-06 | 5.132E-07 | 1.900E-06 | 2.900E-06 |
| 13384 | 670 | I670D | B | 4.3e-06 | 1.6e-06 | 2.5e-06 | 2.800E-06 | 1.375E-06 | 1.600E-06 | 4.300E-06 |
| 13385 | 670 | I670C | B | 1.7e-06 | 3.3e-06 | 3.3e-06 | 2.767E-06 | 9.238E-07 | 1.700E-06 | 3.300E-06 |
| 13386 | 670 | I670E | B | 3e-06 | 3.4e-06 | 1.2e-06 | 2.533E-06 | 1.172E-06 | 1.200E-06 | 3.400E-06 |
| 13387 | 670 | I670Q | B | 2.5e-06 | 2.8e-06 | 3.5e-06 | 2.933E-06 | 5.132E-07 | 2.500E-06 | 3.500E-06 |
| 13388 | 670 | I670G | B | 2.7e-06 | 4.5e-06 | 3.7e-06 | 3.633E-06 | 9.018E-07 | 2.700E-06 | 4.500E-06 |
| 13389 | 670 | I670H | B | 3.8e-06 | 2.9e-06 | 2.7e-06 | 3.133E-06 | 5.859E-07 | 2.700E-06 | 3.800E-06 |
| 13390 | 670 | I670I | B | 2.9e-06 | 3.2e-06 | 2.3e-06 | 2.800E-06 | 4.583E-07 | 2.300E-06 | 3.200E-06 |
| 13391 | 670 | I670L | B | 4e-06 | 1.9e-06 | 2.1e-06 | 2.667E-06 | 1.159E-06 | 1.900E-06 | 4.000E-06 |
| 13392 | 670 | I670K | B | 1.9e-06 | 3.1e-06 | 2.5e-06 | 2.500E-06 | 6.000E-07 | 1.900E-06 | 3.100E-06 |
| 13393 | 670 | I670M | B | 2.2e-06 | 1.2e-06 | 2.2e-06 | 1.867E-06 | 5.774E-07 | 1.200E-06 | 2.200E-06 |
| 13394 | 670 | I670F | B | 2.1e-06 | 2.7e-06 | 2.7e-06 | 2.500E-06 | 3.464E-07 | 2.100E-06 | 2.700E-06 |
| 13395 | 670 | I670P | B | 2.5e-06 | 2.3e-06 | 4e-06 | 2.933E-06 | 9.292E-07 | 2.300E-06 | 4.000E-06 |
| 13396 | 670 | I670S | B | 3.6e-06 | 2.3e-06 | 2.6e-06 | 2.833E-06 | 6.807E-07 | 2.300E-06 | 3.600E-06 |
| 13397 | 670 | I670T | B | 2.2e-06 | 2e-06 | 2.6e-06 | 2.267E-06 | 3.055E-07 | 2.000E-06 | 2.600E-06 |
| 13398 | 670 | I670W | B | 1.6e-06 | 2.1e-06 | 2.1e-06 | 1.933E-06 | 2.887E-07 | 1.600E-06 | 2.100E-06 |
| 13399 | 670 | I670Y | B | 1.7e-06 | 2.9e-06 | 2.4e-06 | 2.333E-06 | 6.028E-07 | 1.700E-06 | 2.900E-06 |
| 13400 | 670 | I670V | B | 1.9e-06 | 2.1e-06 | 1.3e-06 | 1.767E-06 | 4.163E-07 | 1.300E-06 | 2.100E-06 |
| 13401 | 671 | Y671A | B | 2.3e-06 | 2.6e-06 | 1.2e-06 | 2.033E-06 | 7.371E-07 | 1.200E-06 | 2.600E-06 |
| 13402 | 671 | Y671R | B | 2.6e-06 | 2.6e-06 | 2.6e-06 | 2.600E-06 | 0.000E+00 | 2.600E-06 | 2.600E-06 |
| 13403 | 671 | Y671N | B | 2.7e-06 | 2.9e-06 | 2.6e-06 | 2.733E-06 | 1.528E-07 | 2.600E-06 | 2.900E-06 |
| 13404 | 671 | Y671D | B | 2e-06 | 2.6e-06 | 2.1e-06 | 2.233E-06 | 3.215E-07 | 2.000E-06 | 2.600E-06 |
| 13405 | 671 | Y671C | B | 2e-06 | 1.6e-06 | 2.5e-06 | 2.033E-06 | 4.509E-07 | 1.600E-06 | 2.500E-06 |
| 13406 | 671 | Y671E | B | 3.1e-06 | 2.2e-06 | 3.6e-06 | 2.967E-06 | 7.095E-07 | 2.200E-06 | 3.600E-06 |
| 13407 | 671 | Y671Q | B | 1.9e-06 | 1.8e-06 | 2.3e-06 | 2.000E-06 | 2.646E-07 | 1.800E-06 | 2.300E-06 |
| 13408 | 671 | Y671G | B | 1.6e-06 | 1.8e-06 | 2.1e-06 | 1.833E-06 | 2.517E-07 | 1.600E-06 | 2.100E-06 |
| 13409 | 671 | Y671H | B | 2.7e-06 | 2.4e-06 | 1.7e-06 | 2.267E-06 | 5.132E-07 | 1.700E-06 | 2.700E-06 |
| 13410 | 671 | Y671I | B | 2e-06 | 3.3e-06 | 1.8e-06 | 2.367E-06 | 8.145E-07 | 1.800E-06 | 3.300E-06 |
| 13411 | 671 | Y671L | B | 3e-06 | 2.2e-06 | 2.7e-06 | 2.633E-06 | 4.041E-07 | 2.200E-06 | 3.000E-06 |
| 13412 | 671 | Y671K | B | 3.3e-06 | 1.9e-06 | 1.5e-06 | 2.233E-06 | 9.452E-07 | 1.500E-06 | 3.300E-06 |
| 13413 | 671 | Y671M | B | 2e-06 | 2.1e-06 | 2.6e-06 | 2.233E-06 | 3.215E-07 | 2.000E-06 | 2.600E-06 |
| 13414 | 671 | Y671F | B | 2.7e-06 | 3.6e-06 | 1.3e-06 | 2.533E-06 | 1.159E-06 | 1.300E-06 | 3.600E-06 |
| 13415 | 671 | Y671P | B | 1.6e-06 | 2.4e-06 | 2.2e-06 | 2.067E-06 | 4.163E-07 | 1.600E-06 | 2.400E-06 |
| 13416 | 671 | Y671S | B | 3.7e-06 | 2.1e-06 | 1.9e-06 | 2.567E-06 | 9.866E-07 | 1.900E-06 | 3.700E-06 |
| 13417 | 671 | Y671T | B | 1.7e-06 | 1.7e-06 | 2e-06 | 1.800E-06 | 1.732E-07 | 1.700E-06 | 2.000E-06 |
| 13418 | 671 | Y671W | B | 1.7e-06 | 1.7e-06 | 2.3e-06 | 1.900E-06 | 3.464E-07 | 1.700E-06 | 2.300E-06 |
| 13419 | 671 | Y671Y | B | 2.8e-06 | 5.8e-06 | 1.8e-06 | 3.467E-06 | 2.082E-06 | 1.800E-06 | 5.800E-06 |
| 13420 | 671 | Y671V | B | 2.7e-06 | 2.2e-06 | 2.2e-06 | 2.367E-06 | 2.887E-07 | 2.200E-06 | 2.700E-06 |
| 13421 | 672 | S672A | B | 2.1e-06 | 1.6e-06 | 1.9e-06 | 1.867E-06 | 2.517E-07 | 1.600E-06 | 2.100E-06 |
| 13422 | 672 | S672R | B | 4.1e-06 | 2.6e-06 | 1.1e-06 | 2.600E-06 | 1.500E-06 | 1.100E-06 | 4.100E-06 |
| 13423 | 672 | S672N | B | 3.3e-06 | 2.3e-06 | 3.6e-06 | 3.067E-06 | 6.807E-07 | 2.300E-06 | 3.600E-06 |
| 13424 | 672 | S672D | B | 2.6e-06 | 1.9e-06 | 4.6e-06 | 3.033E-06 | 1.401E-06 | 1.900E-06 | 4.600E-06 |
| 13425 | 672 | S672C | B | 1.5e-06 | 2.8e-06 | 2.7e-06 | 2.333E-06 | 7.234E-07 | 1.500E-06 | 2.800E-06 |
| 13426 | 672 | S672E | B | 2.7e-06 | 1.9e-06 | 2.4e-06 | 2.333E-06 | 4.041E-07 | 1.900E-06 | 2.700E-06 |
| 13427 | 672 | S672Q | B | 2.7e-06 | 1.7e-06 | 1.8e-06 | 2.067E-06 | 5.508E-07 | 1.700E-06 | 2.700E-06 |
| 13428 | 672 | S672G | B | 3.2e-06 | 1.7e-06 | 1.5e-06 | 2.133E-06 | 9.292E-07 | 1.500E-06 | 3.200E-06 |
| 13429 | 672 | S672H | B | 1.4e-06 | 1.6e-06 | 1.8e-06 | 1.600E-06 | 2.000E-07 | 1.400E-06 | 1.800E-06 |

| | | | | | | | | | | |
|---|---|---|---|---|---|---|---|---|---|---|
| 13430 | 672 | S672I | B | 1.9e-06 | 3.1e-06 | 2.7e-06 | 2.567E-06 | 6.110E-07 | 1.900E-06 | 3.100E-06 |
| 13431 | 672 | S672L | B | 1.1e-06 | 3.6e-06 | 2.9e-06 | 2.533E-06 | 1.290E-06 | 1.100E-06 | 3.600E-06 |
| 13432 | 672 | S672K | B | 2.4e-06 | 2.8e-06 | 2.3e-06 | 2.500E-06 | 2.646E-07 | 2.300E-06 | 2.800E-06 |
| 13433 | 672 | S672M | B | 1.9e-06 | 2.1e-06 | 2.8e-06 | 2.267E-06 | 4.726E-07 | 1.900E-06 | 2.800E-06 |
| 13434 | 672 | S672F | B | 2.8e-06 | 2.9e-06 | 2.3e-06 | 2.667E-06 | 3.215E-07 | 2.300E-06 | 2.900E-06 |
| 13435 | 672 | S672P | B | 2e-06 | 2.6e-06 | 1.8e-06 | 2.133E-06 | 4.163E-07 | 1.800E-06 | 2.600E-06 |
| 13436 | 672 | S672S | B | 2.5e-06 | 3.1e-06 | 2.4e-06 | 2.667E-06 | 3.786E-07 | 2.400E-06 | 3.100E-06 |
| 13437 | 672 | S672T | B | 2e-06 | 8.7e-07 | 2.5e-06 | 1.790E-06 | 8.350E-07 | 8.700E-07 | 2.500E-06 |
| 13438 | 672 | S672W | B | 2.1e-06 | 2.7e-06 | 2.1e-06 | 2.300E-06 | 3.464E-07 | 2.100E-06 | 2.700E-06 |
| 13439 | 672 | S672Y | B | 2.5e-06 | 3.5e-06 | 2.7e-06 | 2.900E-06 | 5.292E-07 | 2.500E-06 | 3.500E-06 |
| 13440 | 672 | S672V | B | 2.5e-06 | 2.9e-06 | 1.8e-06 | 2.400E-06 | 5.568E-07 | 1.800E-06 | 2.900E-06 |
| 13441 | 673 | A673A | B | 3.8e-06 | 2.1e-06 | 1.8e-06 | 2.567E-06 | 1.079E-06 | 1.800E-06 | 3.800E-06 |
| 13442 | 673 | A673R | B | 1.9e-06 | 1.8e-06 | 2.9e-06 | 2.200E-06 | 6.083E-07 | 1.800E-06 | 2.900E-06 |
| 13443 | 673 | A673N | B | 2e-06 | 3.2e-06 | 2.2e-06 | 2.467E-06 | 6.429E-07 | 2.000E-06 | 3.200E-06 |
| 13444 | 673 | A673D | B | 2.3e-06 | 2e-06 | 1.6e-06 | 1.967E-06 | 3.512E-07 | 1.600E-06 | 2.300E-06 |
| 13445 | 673 | A673C | B | 2.4e-06 | 3.4e-06 | 2.6e-06 | 2.800E-06 | 5.292E-07 | 2.400E-06 | 3.400E-06 |
| 13446 | 673 | A673E | B | 2.2e-06 | 3.6e-06 | 3.1e-06 | 2.967E-06 | 7.095E-07 | 2.200E-06 | 3.600E-06 |
| 13447 | 673 | A673Q | B | 2.7e-06 | 1.3e-06 | 1.6e-06 | 1.867E-06 | 7.371E-07 | 1.300E-06 | 2.700E-06 |
| 13448 | 673 | A673G | B | 2.4e-06 | 2.7e-06 | 2.2e-06 | 2.433E-06 | 2.517E-07 | 2.200E-06 | 2.700E-06 |
| 13449 | 673 | A673H | B | 2.6e-06 | 2.9e-06 | 3.6e-06 | 3.033E-06 | 5.132E-07 | 2.600E-06 | 3.600E-06 |
| 13450 | 673 | A673I | B | 4.1e-06 | 1.8e-06 | 1.9e-06 | 2.600E-06 | 1.300E-06 | 1.800E-06 | 4.100E-06 |
| 13451 | 673 | A673L | B | 2e-06 | 1.7e-06 | 2.6e-06 | 2.100E-06 | 4.583E-07 | 1.700E-06 | 2.600E-06 |
| 13452 | 673 | A673K | B | 2.4e-06 | 1.6e-06 | 2e-06 | 2.000E-06 | 4.000E-07 | 1.600E-06 | 2.400E-06 |
| 13453 | 673 | A673M | B | 2.6e-06 | 1.9e-06 | 2.2e-06 | 2.233E-06 | 3.512E-07 | 1.900E-06 | 2.600E-06 |
| 13454 | 673 | A673F | B | 1.2e-06 | 2.9e-06 | 3.4e-06 | 2.500E-06 | 1.153E-06 | 1.200E-06 | 3.400E-06 |
| 13455 | 673 | A673P | B | 2.5e-06 | 2.5e-06 | 2.5e-06 | 2.500E-06 | 0.000E+00 | 2.500E-06 | 2.500E-06 |
| 13456 | 673 | A673S | B | 1.6e-06 | 1.7e-06 | 2.6e-06 | 1.967E-06 | 5.508E-07 | 1.600E-06 | 2.600E-06 |
| 13457 | 673 | A673T | B | 3e-06 | 1.7e-06 | 3.4e-06 | 2.700E-06 | 8.888E-07 | 1.700E-06 | 3.400E-06 |
| 13458 | 673 | A673W | B | 2.6e-06 | 2.7e-06 | 1.8e-06 | 2.367E-06 | 4.933E-07 | 1.800E-06 | 2.700E-06 |
| 13459 | 673 | A673Y | B | 2.6e-06 | 1.9e-06 | 2.9e-06 | 2.467E-06 | 5.132E-07 | 1.900E-06 | 2.900E-06 |
| 13460 | 673 | A673V | B | 2.5e-06 | 2.1e-06 | 2e-06 | 2.200E-06 | 2.646E-07 | 2.000E-06 | 2.500E-06 |
| 13461 | 674 | S674A | B | 3.2e-06 | 2.7e-06 | 2.1e-06 | 2.667E-06 | 5.508E-07 | 2.100E-06 | 3.200E-06 |
| 13462 | 674 | S674R | B | 2.4e-06 | 3.5e-06 | 2.6e-06 | 2.833E-06 | 5.859E-07 | 2.400E-06 | 3.500E-06 |
| 13463 | 674 | S674N | B | 2.3e-06 | 3e-06 | 2.2e-06 | 2.500E-06 | 4.359E-07 | 2.200E-06 | 3.000E-06 |
| 13464 | 674 | S674D | B | 5.5e-06 | 3.1e-06 | 2.9e-06 | 3.833E-06 | 1.447E-06 | 2.900E-06 | 5.500E-06 |
| 13465 | 674 | S674C | B | 1.5e-06 | 8.8e-07 | 2.2e-06 | 1.527E-06 | 6.604E-07 | 8.800E-07 | 2.200E-06 |
| 13466 | 674 | S674E | B | 3.4e-06 | 1.4e-06 | 1.9e-06 | 2.233E-06 | 1.041E-06 | 1.400E-06 | 3.400E-06 |
| 13467 | 674 | S674Q | B | 2.1e-06 | 1.8e-06 | 2e-06 | 1.967E-06 | 1.528E-07 | 1.800E-06 | 2.100E-06 |
| 13468 | 674 | S674G | B | 2.6e-06 | 1.4e-06 | 1.7e-06 | 1.900E-06 | 6.245E-07 | 1.400E-06 | 2.600E-06 |
| 13469 | 674 | S674H | B | 2.2e-06 | 1.9e-06 | 2.6e-06 | 2.233E-06 | 3.512E-07 | 1.900E-06 | 2.600E-06 |
| 13470 | 674 | S674I | B | 3.8e-06 | 2.7e-06 | 3e-06 | 3.167E-06 | 5.686E-07 | 2.700E-06 | 3.800E-06 |
| 13471 | 674 | S674L | B | 2.9e-06 | 3.7e-06 | 2.5e-06 | 3.033E-06 | 6.110E-07 | 2.500E-06 | 3.700E-06 |
| 13472 | 674 | S674K | B | 2.3e-06 | 2.3e-06 | 5.3e-06 | 3.300E-06 | 1.732E-06 | 2.300E-06 | 5.300E-06 |
| 13473 | 674 | S674M | B | 3.5e-06 | 2.1e-06 | 1.6e-06 | 2.400E-06 | 9.849E-07 | 1.600E-06 | 3.500E-06 |
| 13474 | 674 | S674F | B | 3.2e-06 | 3.7e-06 | 4.2e-06 | 3.700E-06 | 5.000E-07 | 3.200E-06 | 4.200E-06 |
| 13475 | 674 | S674P | B | 2.1e-06 | 2.1e-06 | 2.5e-06 | 2.233E-06 | 2.309E-07 | 2.100E-06 | 2.500E-06 |
| 13476 | 674 | S674S | B | 1.9e-06 | 2.9e-06 | 2e-06 | 2.267E-06 | 5.508E-07 | 1.900E-06 | 2.900E-06 |
| 13477 | 674 | S674T | B | 2.5e-06 | 1.9e-06 | 2e-06 | 2.133E-06 | 3.215E-07 | 1.900E-06 | 2.500E-06 |
| 13478 | 674 | S674W | B | 2.4e-06 | 2.6e-06 | 2.2e-06 | 2.400E-06 | 2.000E-07 | 2.200E-06 | 2.600E-06 |
| 13479 | 674 | S674Y | B | 6.3e-06 | 2.9e-06 | 1.6e-06 | 3.600E-06 | 2.427E-06 | 1.600E-06 | 6.300E-06 |
| 13480 | 674 | S674V | B | 1.9e-06 | 1.9e-06 | 2.2e-06 | 2.000E-06 | 1.732E-07 | 1.900E-06 | 2.200E-06 |
| 13481 | 675 | Y675A | B | 2.2e-06 | 1.1e-06 | 3.7e-06 | 2.333E-06 | 1.305E-06 | 1.100E-06 | 3.700E-06 |
| 13482 | 675 | Y675R | B | 3.2e-06 | 2.4e-06 | 1.8e-06 | 2.467E-06 | 7.024E-07 | 1.800E-06 | 3.200E-06 |
| 13483 | 675 | Y675N | B | 1.2e-06 | 2.2e-06 | 2e-06 | 1.800E-06 | 5.292E-07 | 1.200E-06 | 2.200E-06 |
| 13484 | 675 | Y675D | B | 1.9e-06 | 2e-06 | 3.2e-06 | 2.367E-06 | 7.234E-07 | 1.900E-06 | 3.200E-06 |
| 13485 | 675 | Y675C | B | 2e-06 | 1.7e-06 | 2.5e-06 | 2.067E-06 | 4.041E-07 | 1.700E-06 | 2.500E-06 |
| 13486 | 675 | Y675E | B | 2.3e-06 | 2.3e-06 | 2.5e-06 | 2.367E-06 | 1.155E-07 | 2.300E-06 | 2.500E-06 |
| 13487 | 675 | Y675Q | B | 3e-06 | 2.3e-06 | 3.1e-06 | 2.800E-06 | 4.359E-07 | 2.300E-06 | 3.100E-06 |
| 13488 | 675 | Y675G | B | 3.2e-06 | 2.6e-06 | 2.1e-06 | 2.633E-06 | 5.508E-07 | 2.100E-06 | 3.200E-06 |
| 13489 | 675 | Y675H | B | 2.6e-06 | 1.6e-06 | 2.7e-06 | 2.300E-06 | 6.083E-07 | 1.600E-06 | 2.700E-06 |
| 13490 | 675 | Y675I | B | 2.8e-06 | 3.8e-06 | 1.9e-06 | 2.833E-06 | 9.504E-07 | 1.900E-06 | 3.800E-06 |
| 13491 | 675 | Y675L | B | 3.4e-06 | 3.1e-06 | 2.9e-06 | 3.133E-06 | 2.517E-07 | 2.900E-06 | 3.400E-06 |
| 13492 | 675 | Y675K | B | 2.2e-06 | 2.8e-06 | 2.3e-06 | 2.433E-06 | 3.215E-07 | 2.200E-06 | 2.800E-06 |
| 13493 | 675 | Y675M | B | 1.9e-06 | 2e-06 | 1.5e-06 | 1.800E-06 | 2.646E-07 | 1.500E-06 | 2.000E-06 |
| 13494 | 675 | Y675F | B | 2.8e-06 | 1.8e-06 | 3.2e-06 | 2.600E-06 | 7.211E-07 | 1.800E-06 | 3.200E-06 |
| 13495 | 675 | Y675P | B | 1.6e-06 | 2.4e-06 | 4.8e-06 | 2.933E-06 | 1.665E-06 | 1.600E-06 | 4.800E-06 |
| 13496 | 675 | Y675S | B | 3.1e-06 | 3e-06 | 2.6e-06 | 2.900E-06 | 2.646E-07 | 2.600E-06 | 3.100E-06 |
| 13497 | 675 | Y675T | B | 2.5e-06 | 3.1e-06 | 1.2e-06 | 2.267E-06 | 9.713E-07 | 1.200E-06 | 3.100E-06 |
| 13498 | 675 | Y675W | B | 3.3e-06 | 2.4e-06 | 2.6e-06 | 2.767E-06 | 4.726E-07 | 2.400E-06 | 3.300E-06 |
| 13499 | 675 | Y675Y | B | 1.3e-06 | 1.9e-06 | 2.6e-06 | 1.933E-06 | 6.506E-07 | 1.300E-06 | 2.600E-06 |
| 13500 | 675 | Y675V | B | 2.9e-06 | 1.8e-06 | 1.1e-06 | 1.933E-06 | 9.074E-07 | 1.100E-06 | 2.900E-06 |
| 13501 | 676 | R676A | B | 2.7e-06 | 2.2e-06 | 2.4e-06 | 2.433E-06 | 2.517E-07 | 2.200E-06 | 2.700E-06 |
| 13502 | 676 | R676R | B | 2e-06 | 3.6e-06 | 2.5e-06 | 2.700E-06 | 8.185E-07 | 2.000E-06 | 3.600E-06 |
| 13503 | 676 | R676N | B | 3.1e-06 | 4.5e-06 | 1.9e-06 | 3.167E-06 | 1.301E-06 | 1.900E-06 | 4.500E-06 |
| 13504 | 676 | R676D | B | 2.5e-06 | 2.7e-06 | 2.2e-06 | 2.467E-06 | 2.517E-07 | 2.200E-06 | 2.700E-06 |
| 13505 | 676 | R676C | B | 1.3e-06 | 1.5e-06 | 2.4e-06 | 1.733E-06 | 5.859E-07 | 1.300E-06 | 2.400E-06 |
| 13506 | 676 | R676E | B | 1.4e-06 | 3e-06 | 1.4e-06 | 1.933E-06 | 9.238E-07 | 1.400E-06 | 3.000E-06 |
| 13507 | 676 | R676Q | B | 3.1e-06 | 3.1e-06 | 2.1e-06 | 2.767E-06 | 5.774E-07 | 2.100E-06 | 3.100E-06 |
| 13508 | 676 | R676G | B | 3.3e-06 | 2.2e-06 | 1.5e-06 | 2.333E-06 | 9.074E-07 | 1.500E-06 | 3.300E-06 |
| 13509 | 676 | R676H | B | 3.2e-06 | 2.9e-06 | 2.2e-06 | 2.767E-06 | 5.132E-07 | 2.200E-06 | 3.200E-06 |
| 13510 | 676 | R676I | B | 1.6e-06 | 2.2e-06 | 2.7e-06 | 2.167E-06 | 5.508E-07 | 1.600E-06 | 2.700E-06 |
| 13511 | 676 | R676L | B | 2.5e-06 | 1.9e-06 | 2.3e-06 | 2.233E-06 | 3.055E-07 | 1.900E-06 | 2.500E-06 |
| 13512 | 676 | R676K | B | 2.1e-06 | 4.7e-06 | 1.4e-06 | 2.733E-06 | 1.739E-06 | 1.400E-06 | 4.700E-06 |
| 13513 | 676 | R676M | B | 2.1e-06 | 2.5e-06 | 2.5e-06 | 2.367E-06 | 2.309E-07 | 2.100E-06 | 2.500E-06 |
| 13514 | 676 | R676F | B | 2.1e-06 | 3.4e-06 | 3.2e-06 | 2.900E-06 | 7.000E-07 | 2.100E-06 | 3.400E-06 |

| | | | | | | | | | | |
|---|---|---|---|---|---|---|---|---|---|---|
| 13515 | 676 | R676P | B | 2.1e-06 | 2.4e-06 | 3e-06 | 2.500E-06 | 4.583E-07 | 2.100E-06 | 3.000E-06 |
| 13516 | 676 | R676S | B | 4.1e-06 | 4.2e-06 | 4.2e-06 | 4.167E-06 | 5.774E-08 | 4.100E-06 | 4.200E-06 |
| 13517 | 676 | R676T | B | 3e-06 | 2.7e-06 | 2.2e-06 | 2.633E-06 | 4.041E-07 | 2.200E-06 | 3.000E-06 |
| 13518 | 676 | R676W | B | 3.4e-06 | 2.2e-06 | 2.6e-06 | 2.733E-06 | 6.110E-07 | 2.200E-06 | 3.400E-06 |
| 13519 | 676 | R676Y | B | 2.4e-06 | 2.4e-06 | 3.2e-06 | 2.667E-06 | 4.619E-07 | 2.400E-06 | 3.200E-06 |
| 13520 | 676 | R676V | B | 4e-06 | 1.2e-06 | 2.9e-06 | 2.700E-06 | 1.411E-06 | 1.200E-06 | 4.000E-06 |
| 13521 | 677 | Y677A | B | 3.8e-06 | 2e-06 | 3.1e-06 | 2.967E-06 | 9.074E-07 | 2.000E-06 | 3.800E-06 |
| 13522 | 677 | Y677R | B | 2.3e-06 | 3.9e-06 | 4.3e-06 | 3.500E-06 | 1.058E-06 | 2.300E-06 | 4.300E-06 |
| 13523 | 677 | Y677N | B | 2.4e-06 | 2.1e-06 | 2.1e-06 | 2.200E-06 | 1.732E-07 | 2.100E-06 | 2.400E-06 |
| 13524 | 677 | Y677D | B | 5.6e-06 | 3.5e-06 | 2.5e-06 | 3.867E-06 | 1.582E-06 | 2.500E-06 | 5.600E-06 |
| 13525 | 677 | Y677C | B | 2.1e-06 | 2.3e-06 | 2.3e-06 | 2.233E-06 | 1.155E-07 | 2.100E-06 | 2.300E-06 |
| 13526 | 677 | Y677E | B | 2.3e-06 | 2.3e-06 | 1e-06 | 1.867E-06 | 7.506E-07 | 1.000E-06 | 2.300E-06 |
| 13527 | 677 | Y677Q | B | 1.6e-06 | 2.3e-06 | 1.7e-06 | 1.867E-06 | 3.786E-07 | 1.600E-06 | 2.300E-06 |
| 13528 | 677 | Y677G | B | 2.3e-06 | 2.7e-06 | 2.9e-06 | 2.633E-06 | 3.055E-07 | 2.300E-06 | 2.900E-06 |
| 13529 | 677 | Y677H | B | 2.8e-06 | 2e-06 | 3.7e-06 | 2.833E-06 | 8.505E-07 | 2.000E-06 | 3.700E-06 |
| 13530 | 677 | Y677I | B | 1.3e-06 | 4.3e-06 | 8.5e-07 | 2.150E-06 | 1.875E-06 | 8.500E-07 | 4.300E-06 |
| 13531 | 677 | Y677L | B | 1.8e-06 | 1.7e-06 | 3.4e-06 | 2.300E-06 | 9.539E-07 | 1.700E-06 | 3.400E-06 |
| 13532 | 677 | Y677K | B | 3.2e-06 | 2.7e-06 | 2.7e-06 | 2.867E-06 | 2.887E-07 | 2.700E-06 | 3.200E-06 |
| 13533 | 677 | Y677M | B | 2.3e-06 | 3e-06 | 4.8e-06 | 3.367E-06 | 1.290E-06 | 2.300E-06 | 4.800E-06 |
| 13534 | 677 | Y677F | B | 1.4e-06 | 2e-06 | 1.9e-06 | 1.767E-06 | 3.215E-07 | 1.400E-06 | 2.000E-06 |
| 13535 | 677 | Y677P | B | 3.7e-06 | 2.6e-06 | 1.9e-06 | 2.733E-06 | 9.074E-07 | 1.900E-06 | 3.700E-06 |
| 13536 | 677 | Y677S | B | 1.2e-06 | 1.8e-06 | 1.8e-06 | 1.600E-06 | 3.464E-07 | 1.200E-06 | 1.800E-06 |
| 13537 | 677 | Y677T | B | 3e-06 | 2.7e-06 | 2.3e-06 | 2.667E-06 | 3.512E-07 | 2.300E-06 | 3.000E-06 |
| 13538 | 677 | Y677W | B | 3e-06 | 2.4e-06 | 2.5e-06 | 2.633E-06 | 3.215E-07 | 2.400E-06 | 3.000E-06 |
| 13539 | 677 | Y677Y | B | 2.9e-06 | 3e-06 | 2.1e-06 | 2.667E-06 | 4.933E-07 | 2.100E-06 | 3.000E-06 |
| 13540 | 677 | Y677V | B | 4e-06 | 1.9e-06 | 1.7e-06 | 2.533E-06 | 1.274E-06 | 1.700E-06 | 4.000E-06 |
| 13541 | 678 | T678A | B | 2.2e-06 | 2.2e-06 | 1.7e-06 | 2.033E-06 | 2.887E-07 | 1.700E-06 | 2.200E-06 |
| 13542 | 678 | T678R | B | 2.7e-06 | 2.5e-06 | 1.9e-06 | 2.367E-06 | 4.163E-07 | 1.900E-06 | 2.700E-06 |
| 13543 | 678 | T678N | B | 2.7e-06 | 2.1e-06 | 3e-06 | 2.600E-06 | 4.583E-07 | 2.100E-06 | 3.000E-06 |
| 13544 | 678 | T678D | B | 1.6e-06 | 2e-06 | 4.1e-06 | 2.567E-06 | 1.343E-06 | 1.600E-06 | 4.100E-06 |
| 13545 | 678 | T678C | B | 4.2e-06 | 5.9e-06 | 2e-06 | 4.033E-06 | 1.955E-06 | 2.000E-06 | 5.900E-06 |
| 13546 | 678 | T678E | B | 3.5e-06 | 1.9e-06 | 3.3e-06 | 2.900E-06 | 8.718E-07 | 1.900E-06 | 3.500E-06 |
| 13547 | 678 | T678Q | B | 3.8e-06 | 3.1e-06 | 4e-06 | 3.633E-06 | 4.726E-07 | 3.100E-06 | 4.000E-06 |
| 13548 | 678 | T678G | B | 6.2e-06 | 3.3e-06 | 2.1e-06 | 3.867E-06 | 2.108E-06 | 2.100E-06 | 6.200E-06 |
| 13549 | 678 | T678H | B | 2.6e-06 | 2.3e-06 | 2.4e-06 | 2.433E-06 | 1.528E-07 | 2.300E-06 | 2.600E-06 |
| 13550 | 678 | T678I | B | 3.3e-06 | 3.1e-06 | 2.3e-06 | 2.900E-06 | 5.292E-07 | 2.300E-06 | 3.300E-06 |
| 13551 | 678 | T678L | B | 2.8e-06 | 2.3e-06 | 1.6e-06 | 2.233E-06 | 6.028E-07 | 1.600E-06 | 2.800E-06 |
| 13552 | 678 | T678K | B | 3.3e-06 | 2.7e-06 | 1.6e-06 | 2.533E-06 | 8.622E-07 | 1.600E-06 | 3.300E-06 |
| 13553 | 678 | T678M | B | 2.1e-06 | 2.1e-06 | 4.5e-06 | 2.900E-06 | 1.386E-06 | 2.100E-06 | 4.500E-06 |
| 13554 | 678 | T678F | B | 3.6e-06 | 1.7e-06 | 8e-07 | 2.033E-06 | 1.429E-06 | 8.000E-07 | 3.600E-06 |
| 13555 | 678 | T678P | B | 2.7e-06 | 2.6e-06 | 1.7e-06 | 2.333E-06 | 5.508E-07 | 1.700E-06 | 2.700E-06 |
| 13556 | 678 | T678S | B | 1.7e-06 | 4e-06 | 3.2e-06 | 2.967E-06 | 1.168E-06 | 1.700E-06 | 4.000E-06 |
| 13557 | 678 | T678T | B | 4.9e-06 | 3.4e-06 | 1.7e-06 | 3.333E-06 | 1.601E-06 | 1.700E-06 | 4.900E-06 |
| 13558 | 678 | T678W | B | 2.4e-06 | 2.9e-06 | 2.5e-06 | 2.600E-06 | 2.646E-07 | 2.400E-06 | 2.900E-06 |
| 13559 | 678 | T678Y | B | 2.1e-06 | 1.2e-06 | 1.6e-06 | 1.633E-06 | 4.509E-07 | 1.200E-06 | 2.100E-06 |
| 13560 | 678 | T678V | B | 2.3e-06 | 1.8e-06 | 2.2e-06 | 2.100E-06 | 2.646E-07 | 1.800E-06 | 2.300E-06 |
| 13561 | 679 | G679A | B | 3.7e-06 | 2.8e-06 | 1.8e-06 | 2.767E-06 | 9.504E-07 | 1.800E-06 | 3.700E-06 |
| 13562 | 679 | G679R | B | 2.7e-06 | 2.7e-06 | 2.7e-06 | 2.700E-06 | 0.000E+00 | 2.700E-06 | 2.700E-06 |
| 13563 | 679 | G679N | B | 4.6e-06 | 2.2e-06 | 1.8e-06 | 2.867E-06 | 1.514E-06 | 1.800E-06 | 4.600E-06 |
| 13564 | 679 | G679D | B | 2.5e-06 | 1.7e-06 | 3e-06 | 2.400E-06 | 6.557E-07 | 1.700E-06 | 3.000E-06 |
| 13565 | 679 | G679C | B | 2e-06 | 2.7e-06 | 3e-06 | 2.567E-06 | 5.132E-07 | 2.000E-06 | 3.000E-06 |
| 13566 | 679 | G679E | B | 5e-06 | 1.1e-06 | 1.5e-06 | 2.533E-06 | 2.146E-06 | 1.100E-06 | 5.000E-06 |
| 13567 | 679 | G679Q | B | 2.7e-06 | 2.8e-06 | 2.1e-06 | 2.533E-06 | 3.786E-07 | 2.100E-06 | 2.800E-06 |
| 13568 | 679 | G679G | B | 1.2e-06 | 1.9e-06 | 1.4e-06 | 1.500E-06 | 3.606E-07 | 1.200E-06 | 1.900E-06 |
| 13569 | 679 | G679H | B | 3.4e-06 | 3.3e-06 | 1.1e-06 | 2.600E-06 | 1.300E-06 | 1.100E-06 | 3.400E-06 |
| 13570 | 679 | G679I | B | 1.9e-06 | 1.6e-06 | 1.9e-06 | 1.800E-06 | 1.732E-07 | 1.600E-06 | 1.900E-06 |
| 13571 | 679 | G679L | B | 2.6e-06 | 1.7e-06 | 8.5e-07 | 1.717E-06 | 8.751E-07 | 8.500E-07 | 2.600E-06 |
| 13572 | 679 | G679K | B | 2.3e-06 | 2.4e-06 | 2.1e-06 | 2.267E-06 | 1.528E-07 | 2.100E-06 | 2.400E-06 |
| 13573 | 679 | G679M | B | 1.8e-06 | 2.2e-06 | 2.3e-06 | 2.100E-06 | 2.646E-07 | 1.800E-06 | 2.300E-06 |
| 13574 | 679 | G679F | B | 2.2e-06 | 2.7e-06 | 2.8e-06 | 2.567E-06 | 3.215E-07 | 2.200E-06 | 2.800E-06 |
| 13575 | 679 | G679P | B | 1.9e-06 | 1.6e-06 | 2e-06 | 1.833E-06 | 2.082E-07 | 1.600E-06 | 2.000E-06 |
| 13576 | 679 | G679S | B | 2.7e-06 | 2.4e-06 | 2.2e-06 | 2.433E-06 | 2.517E-07 | 2.200E-06 | 2.700E-06 |
| 13577 | 679 | G679T | B | 4.7e-06 | 2.3e-06 | 1.9e-06 | 2.967E-06 | 1.514E-06 | 1.900E-06 | 4.700E-06 |
| 13578 | 679 | G679W | B | 1.5e-06 | 2.1e-06 | 1.8e-06 | 1.800E-06 | 3.000E-07 | 1.500E-06 | 2.100E-06 |
| 13579 | 679 | G679Y | B | 3.1e-06 | 2.3e-06 | 2.2e-06 | 2.533E-06 | 4.933E-07 | 2.200E-06 | 3.100E-06 |
| 13580 | 679 | G679V | B | 2.1e-06 | 1.8e-06 | 1.2e-06 | 1.700E-06 | 4.583E-07 | 1.200E-06 | 2.100E-06 |
| 13581 | 680 | V680A | B | 1.2e-06 | 2.3e-06 | 1.1e-06 | 1.533E-06 | 6.658E-07 | 1.100E-06 | 2.300E-06 |
| 13582 | 680 | V680R | B | 1.5e-06 | 2.5e-06 | 1.4e-06 | 1.800E-06 | 6.083E-07 | 1.400E-06 | 2.500E-06 |
| 13583 | 680 | V680N | B | 1.3e-06 | 1.8e-06 | 3e-06 | 2.033E-06 | 8.737E-07 | 1.300E-06 | 3.000E-06 |
| 13584 | 680 | V680D | B | 5.1e-06 | 2.2e-06 | 1.6e-06 | 2.967E-06 | 1.872E-06 | 1.600E-06 | 5.100E-06 |
| 13585 | 680 | V680C | B | 1.7e-06 | 3.3e-06 | 2.7e-06 | 2.567E-06 | 8.083E-07 | 1.700E-06 | 3.300E-06 |
| 13586 | 680 | V680E | B | 4.3e-06 | 3.3e-06 | 1.8e-06 | 3.133E-06 | 1.258E-06 | 1.800E-06 | 4.300E-06 |
| 13587 | 680 | V680Q | B | 1.5e-06 | 1.9e-06 | 2e-06 | 1.800E-06 | 2.646E-07 | 1.500E-06 | 2.000E-06 |
| 13588 | 680 | V680G | B | 2.6e-06 | 3.2e-06 | 2.8e-06 | 2.867E-06 | 3.055E-07 | 2.600E-06 | 3.200E-06 |
| 13589 | 680 | V680H | B | 2.1e-06 | 3.5e-06 | 2.2e-06 | 2.600E-06 | 7.810E-07 | 2.100E-06 | 3.500E-06 |
| 13590 | 680 | V680I | B | 3.1e-06 | 2.8e-06 | 2.1e-06 | 2.667E-06 | 5.132E-07 | 2.100E-06 | 3.100E-06 |
| 13591 | 680 | V680L | B | 2.1e-06 | 2.5e-06 | 2.8e-06 | 2.467E-06 | 3.512E-07 | 2.100E-06 | 2.800E-06 |
| 13592 | 680 | V680K | B | 2.5e-06 | 3.3e-06 | 2.5e-06 | 2.767E-06 | 4.619E-07 | 2.500E-06 | 3.300E-06 |
| 13593 | 680 | V680M | B | 2.6e-06 | 1.5e-06 | 1.2e-06 | 1.767E-06 | 7.371E-07 | 1.200E-06 | 2.600E-06 |
| 13594 | 680 | V680F | B | 2.1e-06 | 3.7e-06 | 2.9e-06 | 2.900E-06 | 8.000E-07 | 2.100E-06 | 3.700E-06 |
| 13595 | 680 | V680P | B | 1.3e-06 | 2.8e-06 | 3.2e-06 | 2.433E-06 | 1.002E-06 | 1.300E-06 | 3.200E-06 |
| 13596 | 680 | V680S | B | 1.4e-06 | 1.1e-06 | 3.7e-06 | 2.067E-06 | 1.422E-06 | 1.100E-06 | 3.700E-06 |
| 13597 | 680 | V680T | B | 1.2e-06 | 2.1e-06 | 2.5e-06 | 1.933E-06 | 6.658E-07 | 1.200E-06 | 2.500E-06 |
| 13598 | 680 | V680W | B | 3.2e-06 | 1.8e-06 | 3.1e-06 | 2.700E-06 | 7.810E-07 | 1.800E-06 | 3.200E-06 |
| 13599 | 680 | V680Y | B | 2.7e-06 | 2e-06 | 1.9e-06 | 2.200E-06 | 4.359E-07 | 1.900E-06 | 2.700E-06 |

| | | | | | | | | | | |
|---|---|---|---|---|---|---|---|---|---|---|
| 13600 | 680 | V680V | B | 3e-06 | 3.5e-06 | 2.2e-06 | 2.900E-06 | 6.557E-07 | 2.200E-06 | 3.500E-06 |
| 13601 | 681 | P681A | B | 4.3e-06 | 2.5e-06 | 3.4e-06 | 3.400E-06 | 9.000E-07 | 2.500E-06 | 4.300E-06 |
| 13602 | 681 | P681R | B | 1.3e-06 | 3.4e-06 | 2.1e-06 | 2.267E-06 | 1.060E-06 | 1.300E-06 | 3.400E-06 |
| 13603 | 681 | P681N | B | 1.1e-06 | 2.6e-06 | 2.1e-06 | 1.933E-06 | 7.638E-07 | 1.100E-06 | 2.600E-06 |
| 13604 | 681 | P681D | B | 1.5e-06 | 2.4e-06 | 1.9e-06 | 1.933E-06 | 4.509E-07 | 1.500E-06 | 2.400E-06 |
| 13605 | 681 | P681C | B | 2.6e-06 | 2.9e-06 | 1.6e-06 | 2.367E-06 | 6.807E-07 | 1.600E-06 | 2.900E-06 |
| 13606 | 681 | P681E | B | 2.7e-06 | 2.1e-06 | 2.6e-06 | 2.467E-06 | 3.215E-07 | 2.100E-06 | 2.700E-06 |
| 13607 | 681 | P681Q | B | 1.5e-06 | 3.2e-06 | 2.3e-06 | 2.333E-06 | 8.505E-07 | 1.500E-06 | 3.200E-06 |
| 13608 | 681 | P681G | B | 1.5e-06 | 2.2e-06 | 3.6e-06 | 2.433E-06 | 1.069E-06 | 1.500E-06 | 3.600E-06 |
| 13609 | 681 | P681H | B | 2.2e-06 | 3.8e-06 | 3e-06 | 3.000E-06 | 8.000E-07 | 2.200E-06 | 3.800E-06 |
| 13610 | 681 | P681I | B | 2.5e-06 | 1.1e-06 | 4.8e-06 | 2.800E-06 | 1.868E-06 | 1.100E-06 | 4.800E-06 |
| 13611 | 681 | P681L | B | 1.4e-06 | 3.4e-06 | 1.8e-06 | 2.200E-06 | 1.058E-06 | 1.400E-06 | 3.400E-06 |
| 13612 | 681 | P681K | B | 2.7e-06 | 4.1e-06 | 2.2e-06 | 3.000E-06 | 9.849E-07 | 2.200E-06 | 4.100E-06 |
| 13613 | 681 | P681M | B | 2.2e-06 | 1.8e-06 | 1.7e-06 | 1.900E-06 | 2.646E-07 | 1.700E-06 | 2.200E-06 |
| 13614 | 681 | P681F | B | 1.7e-06 | 3.1e-06 | 2.5e-06 | 2.433E-06 | 7.024E-07 | 1.700E-06 | 3.100E-06 |
| 13615 | 681 | P681P | B | 3.4e-06 | 4.3e-06 | 2e-06 | 3.233E-06 | 1.159E-06 | 2.000E-06 | 4.300E-06 |
| 13616 | 681 | P681S | B | 2.4e-06 | 1.2e-06 | 2.9e-06 | 2.167E-06 | 8.737E-07 | 1.200E-06 | 2.900E-06 |
| 13617 | 681 | P681T | B | 5.9e-06 | 1.8e-06 | 3.8e-06 | 3.833E-06 | 2.050E-06 | 1.800E-06 | 5.900E-06 |
| 13618 | 681 | P681W | B | 2.7e-06 | 2.1e-06 | 1.9e-06 | 2.233E-06 | 4.163E-07 | 1.900E-06 | 2.700E-06 |
| 13619 | 681 | P681Y | B | 2.6e-06 | 2.2e-06 | 3.6e-06 | 2.800E-06 | 7.211E-07 | 2.200E-06 | 3.600E-06 |
| 13620 | 681 | P681V | B | 3.2e-06 | 2.3e-06 | 3.6e-06 | 3.033E-06 | 6.658E-07 | 2.300E-06 | 3.600E-06 |
| 13621 | 682 | S682A | B | 1.7e-06 | 2.2e-06 | 3.2e-06 | 2.367E-06 | 7.638E-07 | 1.700E-06 | 3.200E-06 |
| 13622 | 682 | S682R | B | 2.2e-06 | 1.7e-06 | 1.3e-06 | 1.733E-06 | 4.509E-07 | 1.300E-06 | 2.200E-06 |
| 13623 | 682 | S682N | B | 2e-06 | 2.4e-06 | 2.8e-06 | 2.400E-06 | 4.000E-07 | 2.000E-06 | 2.800E-06 |
| 13624 | 682 | S682D | B | 4.3e-06 | 2.4e-06 | 3.7e-06 | 3.467E-06 | 9.713E-07 | 2.400E-06 | 4.300E-06 |
| 13625 | 682 | S682C | B | 2.9e-06 | 2.5e-06 | 5.6e-06 | 3.667E-06 | 1.686E-06 | 2.500E-06 | 5.600E-06 |
| 13626 | 682 | S682E | B | 2.4e-06 | 3.2e-06 | 1.9e-06 | 2.500E-06 | 6.557E-07 | 1.900E-06 | 3.200E-06 |
| 13627 | 682 | S682Q | B | 3.1e-06 | 2.2e-06 | 2.5e-06 | 2.600E-06 | 4.583E-07 | 2.200E-06 | 3.100E-06 |
| 13628 | 682 | S682G | B | 4.5e-06 | 2.7e-06 | 3.3e-06 | 3.500E-06 | 9.165E-07 | 2.700E-06 | 4.500E-06 |
| 13629 | 682 | S682H | B | 4.8e-06 | 2.2e-06 | 2.5e-06 | 3.167E-06 | 1.422E-06 | 2.200E-06 | 4.800E-06 |
| 13630 | 682 | S682I | B | 2.9e-06 | 3.3e-06 | 2.7e-06 | 2.967E-06 | 3.055E-07 | 2.700E-06 | 3.300E-06 |
| 13631 | 682 | S682L | B | 2.8e-06 | 1.6e-06 | 2e-06 | 2.133E-06 | 6.110E-07 | 1.600E-06 | 2.800E-06 |
| 13632 | 682 | S682K | B | 2.3e-06 | 2.5e-06 | 2.4e-06 | 2.400E-06 | 1.000E-07 | 2.300E-06 | 2.500E-06 |
| 13633 | 682 | S682M | B | 2.6e-06 | 3e-06 | 1.6e-06 | 2.400E-06 | 7.211E-07 | 1.600E-06 | 3.000E-06 |
| 13634 | 682 | S682F | B | 2.6e-06 | 2.3e-06 | 2e-06 | 2.300E-06 | 3.000E-07 | 2.000E-06 | 2.600E-06 |
| 13635 | 682 | S682P | B | 1.4e-06 | 2.8e-06 | 3.3e-06 | 2.500E-06 | 9.849E-07 | 1.400E-06 | 3.300E-06 |
| 13636 | 682 | S682S | B | 1.8e-06 | 3.5e-06 | 2.6e-06 | 2.633E-06 | 8.505E-07 | 1.800E-06 | 3.500E-06 |
| 13637 | 682 | S682T | B | 3.7e-06 | 2.1e-06 | 2.1e-06 | 2.633E-06 | 9.238E-07 | 2.100E-06 | 3.700E-06 |
| 13638 | 682 | S682W | B | 2.2e-06 | 1.2e-06 | 4e-06 | 2.467E-06 | 1.419E-06 | 1.200E-06 | 4.000E-06 |
| 13639 | 682 | S682Y | B | 1.9e-06 | 2.4e-06 | 2.5e-06 | 2.267E-06 | 3.215E-07 | 1.900E-06 | 2.500E-06 |
| 13640 | 682 | S682V | B | 1.3e-06 | 1.7e-06 | 2.2e-06 | 1.733E-06 | 4.509E-07 | 1.300E-06 | 2.200E-06 |
| 13641 | 683 | R683A | B | 2.2e-06 | 3.8e-06 | 3.3e-06 | 3.100E-06 | 8.185E-07 | 2.200E-06 | 3.800E-06 |
| 13642 | 683 | R683R | B | 2.4e-06 | 1.7e-06 | 2.6e-06 | 2.233E-06 | 4.726E-07 | 1.700E-06 | 2.600E-06 |
| 13643 | 683 | R683N | B | 2.2e-06 | 1.1e-06 | 2.7e-06 | 2.000E-06 | 8.185E-07 | 1.100E-06 | 2.700E-06 |
| 13644 | 683 | R683D | B | 4.3e-06 | 5.5e-06 | 2.6e-06 | 4.133E-06 | 1.457E-06 | 2.600E-06 | 5.500E-06 |
| 13645 | 683 | R683C | B | 3.3e-06 | 2.1e-06 | 2.2e-06 | 2.533E-06 | 6.658E-07 | 2.100E-06 | 3.300E-06 |
| 13646 | 683 | R683E | B | 2e-06 | 2.9e-06 | 3.2e-06 | 2.700E-06 | 6.245E-07 | 2.000E-06 | 3.200E-06 |
| 13647 | 683 | R683Q | B | 1.4e-06 | 2.6e-06 | 2.9e-06 | 2.300E-06 | 7.937E-07 | 1.400E-06 | 2.900E-06 |
| 13648 | 683 | R683G | B | 4e-06 | 2.3e-06 | 2.2e-06 | 2.833E-06 | 1.012E-06 | 2.200E-06 | 4.000E-06 |
| 13649 | 683 | R683H | B | 3.2e-06 | 2.8e-06 | 2.3e-06 | 2.767E-06 | 4.509E-07 | 2.300E-06 | 3.200E-06 |
| 13650 | 683 | R683I | B | 3.1e-06 | 3.8e-06 | 1.5e-06 | 2.800E-06 | 1.179E-06 | 1.500E-06 | 3.800E-06 |
| 13651 | 683 | R683L | B | 2.1e-06 | 1.9e-06 | 2.8e-06 | 2.267E-06 | 4.726E-07 | 1.900E-06 | 2.800E-06 |
| 13652 | 683 | R683K | B | 1.5e-06 | 1.6e-06 | 3.9e-06 | 2.333E-06 | 1.358E-06 | 1.500E-06 | 3.900E-06 |
| 13653 | 683 | R683M | B | 3e-06 | 1.3e-06 | 2.6e-06 | 2.300E-06 | 8.888E-07 | 1.300E-06 | 3.000E-06 |
| 13654 | 683 | R683F | B | 4.4e-06 | 1.3e-06 | 1.8e-06 | 2.500E-06 | 1.664E-06 | 1.300E-06 | 4.400E-06 |
| 13655 | 683 | R683P | B | 2.7e-06 | 1.1e-06 | 1.8e-06 | 1.867E-06 | 8.021E-07 | 1.100E-06 | 2.700E-06 |
| 13656 | 683 | R683S | B | 1.4e-06 | 2.4e-06 | 3.6e-06 | 2.467E-06 | 1.102E-06 | 1.400E-06 | 3.600E-06 |
| 13657 | 683 | R683T | B | 1.9e-06 | 2.4e-06 | 2.3e-06 | 2.200E-06 | 2.646E-07 | 1.900E-06 | 2.400E-06 |
| 13658 | 683 | R683W | B | 1.3e-06 | 3.9e-06 | 2.2e-06 | 2.467E-06 | 1.320E-06 | 1.300E-06 | 3.900E-06 |
| 13659 | 683 | R683Y | B | 3.1e-06 | 1.5e-06 | 4.3e-06 | 2.967E-06 | 1.405E-06 | 1.500E-06 | 4.300E-06 |
| 13660 | 683 | R683V | B | 3.5e-06 | 2.6e-06 | 2e-06 | 2.700E-06 | 7.550E-07 | 2.000E-06 | 3.500E-06 |
| 13661 | 684 | F684A | B | 4.8e-06 | 2.1e-06 | 2.2e-06 | 3.033E-06 | 1.531E-06 | 2.100E-06 | 4.800E-06 |
| 13662 | 684 | F684R | B | 3.4e-06 | 3e-06 | 5.2e-06 | 3.867E-06 | 1.172E-06 | 3.000E-06 | 5.200E-06 |
| 13663 | 684 | F684N | B | 2.9e-06 | 2.1e-06 | 2.8e-06 | 2.600E-06 | 4.359E-07 | 2.100E-06 | 2.900E-06 |
| 13664 | 684 | F684D | B | 2.4e-06 | 2.5e-06 | 2.4e-06 | 2.433E-06 | 5.774E-08 | 2.400E-06 | 2.500E-06 |
| 13665 | 684 | F684C | B | 2e-06 | 2e-06 | 2.3e-06 | 2.100E-06 | 1.732E-07 | 2.000E-06 | 2.300E-06 |
| 13666 | 684 | F684E | B | 3.1e-06 | 4.4e-06 | 2.1e-06 | 3.200E-06 | 1.153E-06 | 2.100E-06 | 4.400E-06 |
| 13667 | 684 | F684Q | B | 2.8e-06 | 1.9e-06 | 4.5e-06 | 3.067E-06 | 1.320E-06 | 1.900E-06 | 4.500E-06 |
| 13668 | 684 | F684G | B | 1.7e-06 | 2.2e-06 | 1.6e-06 | 1.833E-06 | 3.215E-07 | 1.600E-06 | 2.200E-06 |
| 13669 | 684 | F684H | B | 2.1e-06 | 3e-06 | 3e-06 | 2.700E-06 | 5.196E-07 | 2.100E-06 | 3.000E-06 |
| 13670 | 684 | F684I | B | 1.4e-06 | 2.1e-06 | 2.8e-06 | 2.100E-06 | 7.000E-07 | 1.400E-06 | 2.800E-06 |
| 13671 | 684 | F684L | B | 2.7e-06 | 3e-06 | 2.3e-06 | 2.667E-06 | 3.512E-07 | 2.300E-06 | 3.000E-06 |
| 13672 | 684 | F684K | B | 3.3e-06 | 2.5e-06 | 2.7e-06 | 2.833E-06 | 4.163E-07 | 2.500E-06 | 3.300E-06 |
| 13673 | 684 | F684M | B | 1.7e-06 | 1.8e-06 | 1.7e-06 | 1.733E-06 | 5.774E-08 | 1.700E-06 | 1.800E-06 |
| 13674 | 684 | F684F | B | 3.1e-06 | 3e-06 | 2.2e-06 | 2.767E-06 | 4.933E-07 | 2.200E-06 | 3.100E-06 |
| 13675 | 684 | F684P | B | 2e-06 | 3.3e-06 | 1.9e-06 | 2.400E-06 | 7.810E-07 | 1.900E-06 | 3.300E-06 |
| 13676 | 684 | F684S | B | 1.1e-06 | 2.3e-06 | 2.4e-06 | 1.933E-06 | 7.234E-07 | 1.100E-06 | 2.400E-06 |
| 13677 | 684 | F684T | B | 1.9e-06 | 4e-06 | 3.8e-06 | 3.233E-06 | 1.159E-06 | 1.900E-06 | 4.000E-06 |
| 13678 | 684 | F684W | B | 2e-06 | 1.8e-06 | 1.5e-06 | 1.767E-06 | 2.517E-07 | 1.500E-06 | 2.000E-06 |
| 13679 | 684 | F684Y | B | 8.3e-07 | 2.4e-06 | 1.4e-06 | 1.543E-06 | 7.948E-07 | 8.300E-07 | 2.400E-06 |
| 13680 | 684 | F684V | B | 5e-06 | 2.3e-06 | 1.8e-06 | 3.033E-06 | 1.721E-06 | 1.800E-06 | 5.000E-06 |
| 13681 | 685 | S685A | B | 5.4e-06 | 2.3e-06 | 2.7e-06 | 3.467E-06 | 1.686E-06 | 2.300E-06 | 5.400E-06 |
| 13682 | 685 | S685R | B | 1.7e-06 | 1.3e-06 | 2.3e-06 | 1.767E-06 | 5.033E-07 | 1.300E-06 | 2.300E-06 |
| 13683 | 685 | S685N | B | 2.2e-06 | 3e-06 | 2.6e-06 | 2.600E-06 | 4.000E-07 | 2.200E-06 | 3.000E-06 |
| 13684 | 685 | S685D | B | 2.8e-06 | 3.2e-06 | 3.1e-06 | 3.033E-06 | 2.082E-07 | 2.800E-06 | 3.200E-06 |

| | | | | | | | | | | |
|---|---|---|---|---|---|---|---|---|---|---|
| 13685 | 685 | S685C | B | 2.4e-06 | 3.1e-06 | 2.4e-06 | 2.633E-06 | 4.041E-07 | 2.400E-06 | 3.100E-06 |
| 13686 | 685 | S685E | B | 4.7e-06 | 3.1e-06 | 3.9e-06 | 3.900E-06 | 8.000E-07 | 3.100E-06 | 4.700E-06 |
| 13687 | 685 | S685Q | B | 2.7e-06 | 3.3e-06 | 2.1e-06 | 2.700E-06 | 6.000E-07 | 2.100E-06 | 3.300E-06 |
| 13688 | 685 | S685G | B | 5.7e-06 | 2.6e-06 | 2e-06 | 3.433E-06 | 1.986E-06 | 2.000E-06 | 5.700E-06 |
| 13689 | 685 | S685H | B | 2.4e-06 | 3.1e-06 | 2e-06 | 2.500E-06 | 5.568E-07 | 2.000E-06 | 3.100E-06 |
| 13690 | 685 | S685I | B | 4.3e-06 | 1.9e-06 | 1.9e-06 | 2.700E-06 | 1.386E-06 | 1.900E-06 | 4.300E-06 |
| 13691 | 685 | S685L | B | 3.2e-06 | 1.9e-06 | 3e-06 | 2.700E-06 | 7.000E-07 | 1.900E-06 | 3.200E-06 |
| 13692 | 685 | S685K | B | 4.2e-06 | 2.3e-06 | 2e-06 | 2.833E-06 | 1.193E-06 | 2.000E-06 | 4.200E-06 |
| 13693 | 685 | S685M | B | 3.1e-06 | 3.4e-06 | 4.2e-06 | 3.567E-06 | 5.686E-07 | 3.100E-06 | 4.200E-06 |
| 13694 | 685 | S685F | B | 2.7e-06 | 3.2e-06 | 2.7e-06 | 2.867E-06 | 2.887E-07 | 2.700E-06 | 3.200E-06 |
| 13695 | 685 | S685P | B | 1.7e-06 | 3.2e-06 | 3e-06 | 2.633E-06 | 8.145E-07 | 1.700E-06 | 3.200E-06 |
| 13696 | 685 | S685S | B | 2.6e-06 | 2.6e-06 | 2.5e-06 | 2.567E-06 | 5.774E-08 | 2.500E-06 | 2.600E-06 |
| 13697 | 685 | S685T | B | 2.3e-06 | 1.6e-06 | 2.2e-06 | 2.033E-06 | 3.786E-07 | 1.600E-06 | 2.300E-06 |
| 13698 | 685 | S685W | B | 1.8e-06 | 1.5e-06 | 4e-06 | 2.433E-06 | 1.365E-06 | 1.500E-06 | 4.000E-06 |
| 13699 | 685 | S685Y | B | 3.6e-06 | 3.2e-06 | 2.7e-06 | 3.167E-06 | 4.509E-07 | 2.700E-06 | 3.600E-06 |
| 13700 | 685 | S685V | B | 1.5e-06 | 2.8e-06 | 3e-06 | 2.433E-06 | 8.145E-07 | 1.500E-06 | 3.000E-06 |
| 13701 | 686 | G686A | B | 2.4e-06 | 2.9e-06 | 3.4e-06 | 2.900E-06 | 5.000E-07 | 2.400E-06 | 3.400E-06 |
| 13702 | 686 | G686R | B | 3.8e-06 | 2.8e-06 | 2.3e-06 | 2.967E-06 | 7.638E-07 | 2.300E-06 | 3.800E-06 |
| 13703 | 686 | G686N | B | 3.2e-06 | 1.4e-06 | 2.2e-06 | 2.267E-06 | 9.018E-07 | 1.400E-06 | 3.200E-06 |
| 13704 | 686 | G686D | B | 2.6e-06 | 2.6e-06 | 2.3e-06 | 2.500E-06 | 1.732E-07 | 2.300E-06 | 2.600E-06 |
| 13705 | 686 | G686C | B | 2.3e-06 | 2.6e-06 | 2e-06 | 2.300E-06 | 3.000E-07 | 2.000E-06 | 2.600E-06 |
| 13706 | 686 | G686E | B | 2.2e-06 | 2.3e-06 | 2.8e-06 | 2.433E-06 | 3.215E-07 | 2.200E-06 | 2.800E-06 |
| 13707 | 686 | G686Q | B | 2.3e-06 | 3.1e-06 | 3.2e-06 | 2.867E-06 | 4.933E-07 | 2.300E-06 | 3.200E-06 |
| 13708 | 686 | G686G | B | 3.8e-06 | 3.6e-06 | 2.3e-06 | 3.233E-06 | 8.145E-07 | 2.300E-06 | 3.800E-06 |
| 13709 | 686 | G686H | B | 1.7e-06 | 1.7e-06 | 2.6e-06 | 2.000E-06 | 5.196E-07 | 1.700E-06 | 2.600E-06 |
| 13710 | 686 | G686I | B | 3.3e-06 | 2.7e-06 | 2.9e-06 | 2.967E-06 | 3.055E-07 | 2.700E-06 | 3.300E-06 |
| 13711 | 686 | G686L | B | 2.1e-06 | 2.6e-06 | 2.3e-06 | 2.333E-06 | 2.517E-07 | 2.100E-06 | 2.600E-06 |
| 13712 | 686 | G686K | B | 3.1e-06 | 2.8e-06 | 2.1e-06 | 2.667E-06 | 5.132E-07 | 2.100E-06 | 3.100E-06 |
| 13713 | 686 | G686M | B | 2.9e-06 | 2.1e-06 | 1.3e-06 | 2.100E-06 | 8.000E-07 | 1.300E-06 | 2.900E-06 |
| 13714 | 686 | G686F | B | 3.1e-06 | 2.6e-06 | 2.3e-06 | 2.667E-06 | 4.041E-07 | 2.300E-06 | 3.100E-06 |
| 13715 | 686 | G686P | B | 1.8e-06 | 2.4e-06 | 1.5e-06 | 1.900E-06 | 4.583E-07 | 1.500E-06 | 2.400E-06 |
| 13716 | 686 | G686S | B | 2.9e-06 | 1.8e-06 | 1.6e-06 | 2.100E-06 | 7.000E-07 | 1.600E-06 | 2.900E-06 |
| 13717 | 686 | G686T | B | 2.9e-06 | 1.7e-06 | 2.9e-06 | 2.500E-06 | 6.928E-07 | 1.700E-06 | 2.900E-06 |
| 13718 | 686 | G686W | B | 2.7e-06 | 3.1e-06 | 3e-06 | 2.933E-06 | 2.082E-07 | 2.700E-06 | 3.100E-06 |
| 13719 | 686 | G686Y | B | 1.8e-06 | 1.6e-06 | 2e-06 | 1.800E-06 | 2.000E-07 | 1.600E-06 | 2.000E-06 |
| 13720 | 686 | G686V | B | 2e-06 | 3.2e-06 | 3.7e-06 | 2.967E-06 | 8.737E-07 | 2.000E-06 | 3.700E-06 |
| 13721 | 687 | S687A | B | 1.4e-06 | 1.9e-06 | 2.1e-06 | 1.800E-06 | 3.606E-07 | 1.400E-06 | 2.100E-06 |
| 13722 | 687 | S687R | B | 2.6e-06 | 2.1e-06 | 1.6e-06 | 2.100E-06 | 5.000E-07 | 1.600E-06 | 2.600E-06 |
| 13723 | 687 | S687N | B | 1.7e-06 | 2.5e-06 | 2.9e-06 | 2.367E-06 | 6.110E-07 | 1.700E-06 | 2.900E-06 |
| 13724 | 687 | S687D | B | 3.6e-06 | 1.5e-06 | 3.5e-06 | 2.867E-06 | 1.185E-06 | 1.500E-06 | 3.600E-06 |
| 13725 | 687 | S687C | B | 2.1e-06 | 2.5e-06 | 2.7e-06 | 2.433E-06 | 3.055E-07 | 2.100E-06 | 2.700E-06 |
| 13726 | 687 | S687E | B | 5.2e-06 | 4.1e-06 | 1.6e-06 | 3.633E-06 | 1.845E-06 | 1.600E-06 | 5.200E-06 |
| 13727 | 687 | S687Q | B | 1e-06 | 2.6e-06 | 1.7e-06 | 1.767E-06 | 8.021E-07 | 1.000E-06 | 2.600E-06 |
| 13728 | 687 | S687G | B | 2.6e-06 | 4.6e-06 | 2.9e-06 | 3.367E-06 | 1.079E-06 | 2.600E-06 | 4.600E-06 |
| 13729 | 687 | S687H | B | 1.6e-06 | 1.4e-06 | 1.7e-06 | 1.567E-06 | 1.528E-07 | 1.400E-06 | 1.700E-06 |
| 13730 | 687 | S687I | B | 2.2e-06 | 1.8e-06 | 2.9e-06 | 2.300E-06 | 5.568E-07 | 1.800E-06 | 2.900E-06 |
| 13731 | 687 | S687L | B | 1.2e-06 | 1.6e-06 | 2.7e-06 | 1.833E-06 | 7.767E-07 | 1.200E-06 | 2.700E-06 |
| 13732 | 687 | S687K | B | 2.2e-06 | 3.3e-06 | 1.8e-06 | 2.433E-06 | 7.767E-07 | 1.800E-06 | 3.300E-06 |
| 13733 | 687 | S687M | B | 2.6e-06 | 3e-06 | 2.3e-06 | 2.633E-06 | 3.512E-07 | 2.300E-06 | 3.000E-06 |
| 13734 | 687 | S687F | B | 2.3e-06 | 2.6e-06 | 1.8e-06 | 2.233E-06 | 4.041E-07 | 1.800E-06 | 2.600E-06 |
| 13735 | 687 | S687P | B | 3e-06 | 2.6e-06 | 1.7e-06 | 2.433E-06 | 6.658E-07 | 1.700E-06 | 3.000E-06 |
| 13736 | 687 | S687S | B | 3.1e-06 | 2.9e-06 | 1.3e-06 | 2.433E-06 | 9.866E-07 | 1.300E-06 | 3.100E-06 |
| 13737 | 687 | S687T | B | 3.5e-06 | 3.4e-06 | 2.3e-06 | 3.067E-06 | 6.658E-07 | 2.300E-06 | 3.500E-06 |
| 13738 | 687 | S687W | B | 2.5e-06 | 2.6e-06 | 2.1e-06 | 2.400E-06 | 2.646E-07 | 2.100E-06 | 2.600E-06 |
| 13739 | 687 | S687Y | B | 2.8e-06 | 2e-06 | 2.2e-06 | 2.333E-06 | 4.163E-07 | 2.000E-06 | 2.800E-06 |
| 13740 | 687 | S687V | B | 2.2e-06 | 3.2e-06 | 3.1e-06 | 2.833E-06 | 5.508E-07 | 2.200E-06 | 3.200E-06 |
| 13741 | 688 | G688A | B | 2.9e-06 | 2.3e-06 | 2.2e-06 | 2.467E-06 | 3.786E-07 | 2.200E-06 | 2.900E-06 |
| 13742 | 688 | G688R | B | 1.7e-06 | 1.5e-06 | 2.8e-06 | 2.000E-06 | 7.000E-07 | 1.500E-06 | 2.800E-06 |
| 13743 | 688 | G688N | B | 2.2e-06 | 3.3e-06 | 2.3e-06 | 2.600E-06 | 6.083E-07 | 2.200E-06 | 3.300E-06 |
| 13744 | 688 | G688D | B | 2.2e-06 | 1.8e-06 | 1.8e-06 | 1.933E-06 | 2.309E-07 | 1.800E-06 | 2.200E-06 |
| 13745 | 688 | G688C | B | 2.1e-06 | 1.5e-06 | 2.6e-06 | 2.067E-06 | 5.508E-07 | 1.500E-06 | 2.600E-06 |
| 13746 | 688 | G688E | B | 2.1e-06 | 1.3e-06 | 1.8e-06 | 1.733E-06 | 4.041E-07 | 1.300E-06 | 2.100E-06 |
| 13747 | 688 | G688Q | B | 4.2e-06 | 1.6e-06 | 1.8e-06 | 2.533E-06 | 1.447E-06 | 1.600E-06 | 4.200E-06 |
| 13748 | 688 | G688G | B | 3.6e-06 | 4.2e-06 | 3.7e-06 | 3.833E-06 | 3.215E-07 | 3.600E-06 | 4.200E-06 |
| 13749 | 688 | G688H | B | 2.4e-06 | 1.6e-06 | 2.3e-06 | 2.100E-06 | 4.359E-07 | 1.600E-06 | 2.400E-06 |
| 13750 | 688 | G688I | B | 2.6e-06 | 3e-06 | 4.3e-06 | 3.300E-06 | 8.888E-07 | 2.600E-06 | 4.300E-06 |
| 13751 | 688 | G688L | B | 8.2e-07 | 2.7e-06 | 3.4e-06 | 2.307E-06 | 1.334E-06 | 8.200E-07 | 3.400E-06 |
| 13752 | 688 | G688K | B | 3.8e-06 | 2.3e-06 | 3.3e-06 | 3.133E-06 | 7.638E-07 | 2.300E-06 | 3.800E-06 |
| 13753 | 688 | G688M | B | 2.3e-06 | 2.2e-06 | 3.1e-06 | 2.533E-06 | 4.933E-07 | 2.200E-06 | 3.100E-06 |
| 13754 | 688 | G688F | B | 3.3e-06 | 2.4e-06 | 3.2e-06 | 2.967E-06 | 4.933E-07 | 2.400E-06 | 3.300E-06 |
| 13755 | 688 | G688P | B | 3.8e-06 | 1.9e-06 | 2.6e-06 | 2.767E-06 | 9.609E-07 | 1.900E-06 | 3.800E-06 |
| 13756 | 688 | G688S | B | 1.8e-06 | 2.2e-06 | 2.1e-06 | 2.033E-06 | 2.082E-07 | 1.800E-06 | 2.200E-06 |
| 13757 | 688 | G688T | B | 3.8e-06 | 2.6e-06 | 2.3e-06 | 2.900E-06 | 7.937E-07 | 2.300E-06 | 3.800E-06 |
| 13758 | 688 | G688W | B | 2e-06 | 4.6e-06 | 1.7e-06 | 2.767E-06 | 1.595E-06 | 1.700E-06 | 4.600E-06 |
| 13759 | 688 | G688Y | B | 3.6e-06 | 2.4e-06 | 1.5e-06 | 2.500E-06 | 1.054E-06 | 1.500E-06 | 3.600E-06 |
| 13760 | 688 | G688V | B | 2.8e-06 | 2.1e-06 | 2.5e-06 | 2.467E-06 | 3.512E-07 | 2.100E-06 | 2.800E-06 |
| 13761 | 689 | S689A | B | 2.3e-06 | 3.1e-06 | 2.3e-06 | 2.567E-06 | 4.619E-07 | 2.300E-06 | 3.100E-06 |
| 13762 | 689 | S689R | B | 1.4e-06 | 2.1e-06 | 2.5e-06 | 2.000E-06 | 5.568E-07 | 1.400E-06 | 2.500E-06 |
| 13763 | 689 | S689N | B | 3.3e-06 | 2.5e-06 | 1.5e-06 | 2.433E-06 | 9.018E-07 | 1.500E-06 | 3.300E-06 |
| 13764 | 689 | S689D | B | 1.2e-06 | 1.4e-06 | 1.6e-06 | 1.400E-06 | 2.000E-07 | 1.200E-06 | 1.600E-06 |
| 13765 | 689 | S689C | B | 1.6e-06 | 4.4e-06 | 3.4e-06 | 3.133E-06 | 1.419E-06 | 1.600E-06 | 4.400E-06 |
| 13766 | 689 | S689E | B | 2.1e-06 | 1.9e-06 | 2.9e-06 | 2.300E-06 | 5.292E-07 | 1.900E-06 | 2.900E-06 |
| 13767 | 689 | S689Q | B | 2.7e-06 | 4.5e-06 | 1.8e-06 | 3.000E-06 | 1.375E-06 | 1.800E-06 | 4.500E-06 |
| 13768 | 689 | S689G | B | 3.5e-06 | 4.1e-06 | 1e-06 | 2.867E-06 | 1.644E-06 | 1.000E-06 | 4.100E-06 |
| 13769 | 689 | S689H | B | 2e-06 | 2e-06 | 1.5e-06 | 1.833E-06 | 2.887E-07 | 1.500E-06 | 2.000E-06 |

| | | | | | | | | | | |
|---|---|---|---|---|---|---|---|---|---|---|
| 13770 | 689 | S689I | B | 2.7e-06 | 2.5e-06 | 2.1e-06 | 2.433E-06 | 3.055E-07 | 2.100E-06 | 2.700E-06 |
| 13771 | 689 | S689L | B | 2.8e-06 | 2.3e-06 | 2.5e-06 | 2.533E-06 | 2.517E-07 | 2.300E-06 | 2.800E-06 |
| 13772 | 689 | S689K | B | 3.3e-06 | 1.5e-06 | 2.2e-06 | 2.333E-06 | 9.074E-07 | 1.500E-06 | 3.300E-06 |
| 13773 | 689 | S689M | B | 2.9e-06 | 4e-06 | 2.9e-06 | 3.267E-06 | 6.351E-07 | 2.900E-06 | 4.000E-06 |
| 13774 | 689 | S689F | B | 3e-06 | 3.1e-06 | 1e-06 | 2.367E-06 | 1.185E-06 | 1.000E-06 | 3.100E-06 |
| 13775 | 689 | S689P | B | 1.8e-06 | 2.4e-06 | 2.3e-06 | 2.167E-06 | 3.215E-07 | 1.800E-06 | 2.400E-06 |
| 13776 | 689 | S689S | B | 2.9e-06 | 2.3e-06 | 2.7e-06 | 2.633E-06 | 3.055E-07 | 2.300E-06 | 2.900E-06 |
| 13777 | 689 | S689T | B | 2e-06 | 2.3e-06 | 1.7e-06 | 2.000E-06 | 3.000E-07 | 1.700E-06 | 2.300E-06 |
| 13778 | 689 | S689W | B | 3.3e-06 | 2.4e-06 | 2.9e-06 | 2.867E-06 | 4.509E-07 | 2.400E-06 | 3.300E-06 |
| 13779 | 689 | S689Y | B | 2.8e-06 | 2.2e-06 | 1.7e-06 | 2.233E-06 | 5.508E-07 | 1.700E-06 | 2.800E-06 |
| 13780 | 689 | S689V | B | 5.1e-06 | 1.6e-06 | 1.9e-06 | 2.867E-06 | 1.940E-06 | 1.600E-06 | 5.100E-06 |
| 13781 | 690 | G690A | B | 1.8e-06 | 3e-06 | 2.7e-06 | 2.500E-06 | 6.245E-07 | 1.800E-06 | 3.000E-06 |
| 13782 | 690 | G690R | B | 2.4e-06 | 3e-06 | 3.6e-06 | 3.000E-06 | 6.000E-07 | 2.400E-06 | 3.600E-06 |
| 13783 | 690 | G690N | B | 2.5e-06 | 3.2e-06 | 3.3e-06 | 3.000E-06 | 4.359E-07 | 2.500E-06 | 3.300E-06 |
| 13784 | 690 | G690D | B | 3.3e-06 | 5.7e-06 | 9.8e-07 | 3.327E-06 | 2.360E-06 | 9.800E-07 | 5.700E-06 |
| 13785 | 690 | G690C | B | 2.4e-06 | 1.4e-06 | 2.1e-06 | 1.967E-06 | 5.132E-07 | 1.400E-06 | 2.400E-06 |
| 13786 | 690 | G690E | B | 2.6e-06 | 2.5e-06 | 2.6e-06 | 2.567E-06 | 5.774E-08 | 2.500E-06 | 2.600E-06 |
| 13787 | 690 | G690Q | B | 2.4e-06 | 2.5e-06 | 1.8e-06 | 2.233E-06 | 3.786E-07 | 1.800E-06 | 2.500E-06 |
| 13788 | 690 | G690G | B | 3.7e-06 | 1.8e-06 | 4.5e-06 | 3.333E-06 | 1.387E-06 | 1.800E-06 | 4.500E-06 |
| 13789 | 690 | G690H | B | 1.2e-06 | 2.5e-06 | 2.6e-06 | 2.100E-06 | 7.810E-07 | 1.200E-06 | 2.600E-06 |
| 13790 | 690 | G690I | B | 2.3e-06 | 1.7e-06 | 3.5e-06 | 2.500E-06 | 9.165E-07 | 1.700E-06 | 3.500E-06 |
| 13791 | 690 | G690L | B | 3.8e-06 | 1.9e-06 | 2.7e-06 | 2.800E-06 | 9.539E-07 | 1.900E-06 | 3.800E-06 |
| 13792 | 690 | G690K | B | 2.1e-06 | 3.9e-06 | 2.1e-06 | 2.700E-06 | 1.039E-06 | 2.100E-06 | 3.900E-06 |
| 13793 | 690 | G690M | B | 1.2e-06 | 3.4e-06 | 1.8e-06 | 2.133E-06 | 1.137E-06 | 1.200E-06 | 3.400E-06 |
| 13794 | 690 | G690F | B | 3.3e-06 | 2.1e-06 | 2.2e-06 | 2.533E-06 | 6.658E-07 | 2.100E-06 | 3.300E-06 |
| 13795 | 690 | G690P | B | 1.7e-06 | 3.9e-06 | 2.5e-06 | 2.700E-06 | 1.114E-06 | 1.700E-06 | 3.900E-06 |
| 13796 | 690 | G690S | B | 2.1e-06 | 2e-06 | 7e-07 | 1.600E-06 | 7.810E-07 | 7.000E-07 | 2.100E-06 |
| 13797 | 690 | G690T | B | 2.3e-06 | 2.1e-06 | 3.1e-06 | 2.500E-06 | 5.292E-07 | 2.100E-06 | 3.100E-06 |
| 13798 | 690 | G690W | B | 2e-06 | 3.4e-06 | 1.8e-06 | 2.400E-06 | 8.718E-07 | 1.800E-06 | 3.400E-06 |
| 13799 | 690 | G690Y | B | 2.8e-06 | 2.2e-06 | 2.6e-06 | 2.533E-06 | 3.055E-07 | 2.200E-06 | 2.800E-06 |
| 13800 | 690 | G690V | B | 2.3e-06 | 2.3e-06 | 1.3e-06 | 1.967E-06 | 5.774E-07 | 1.300E-06 | 2.300E-06 |
| 13801 | 691 | T691A | B | 2.3e-06 | 2.8e-06 | 2.5e-06 | 2.533E-06 | 2.517E-07 | 2.300E-06 | 2.800E-06 |
| 13802 | 691 | T691R | B | 4.1e-06 | 1.5e-06 | 4.1e-06 | 3.233E-06 | 1.501E-06 | 1.500E-06 | 4.100E-06 |
| 13803 | 691 | T691N | B | 2.2e-06 | 2.1e-06 | 3.2e-06 | 2.500E-06 | 6.083E-07 | 2.100E-06 | 3.200E-06 |
| 13804 | 691 | T691D | B | 2.6e-06 | 4.3e-06 | 2.8e-06 | 3.233E-06 | 9.292E-07 | 2.600E-06 | 4.300E-06 |
| 13805 | 691 | T691C | B | 2.8e-06 | 1.8e-06 | 2.3e-06 | 2.300E-06 | 5.000E-07 | 1.800E-06 | 2.800E-06 |
| 13806 | 691 | T691E | B | 1.4e-06 | 1.4e-06 | 2.4e-06 | 1.733E-06 | 5.774E-07 | 1.400E-06 | 2.400E-06 |
| 13807 | 691 | T691Q | B | 1.8e-06 | 2.3e-06 | 2e-06 | 2.033E-06 | 2.517E-07 | 1.800E-06 | 2.300E-06 |
| 13808 | 691 | T691G | B | 3e-06 | 2e-06 | 2.7e-06 | 2.567E-06 | 5.132E-07 | 2.000E-06 | 3.000E-06 |
| 13809 | 691 | T691H | B | 1.9e-06 | 2.6e-06 | 3.6e-06 | 2.700E-06 | 8.544E-07 | 1.900E-06 | 3.600E-06 |
| 13810 | 691 | T691I | B | 1.4e-06 | 2.7e-06 | 3.3e-06 | 2.467E-06 | 9.713E-07 | 1.400E-06 | 3.300E-06 |
| 13811 | 691 | T691L | B | 1.6e-06 | 2.3e-06 | 2.5e-06 | 2.133E-06 | 4.726E-07 | 1.600E-06 | 2.500E-06 |
| 13812 | 691 | T691K | B | 3.4e-06 | 2e-06 | 2.3e-06 | 2.567E-06 | 7.371E-07 | 2.000E-06 | 3.400E-06 |
| 13813 | 691 | T691M | B | 2.1e-06 | 2.5e-06 | 2.7e-06 | 2.433E-06 | 3.055E-07 | 2.100E-06 | 2.700E-06 |
| 13814 | 691 | T691F | B | 2.5e-06 | 2.7e-06 | 1.6e-06 | 2.267E-06 | 5.859E-07 | 1.600E-06 | 2.700E-06 |
| 13815 | 691 | T691P | B | 2.9e-06 | 2.3e-06 | 2.5e-06 | 2.567E-06 | 3.055E-07 | 2.300E-06 | 2.900E-06 |
| 13816 | 691 | T691S | B | 2.6e-06 | 3.2e-06 | 2.2e-06 | 2.667E-06 | 5.033E-07 | 2.200E-06 | 3.200E-06 |
| 13817 | 691 | T691T | B | 2.4e-06 | 2.1e-06 | 1.6e-06 | 2.033E-06 | 4.041E-07 | 1.600E-06 | 2.400E-06 |
| 13818 | 691 | T691W | B | 1.7e-06 | 2.3e-06 | 1.8e-06 | 1.933E-06 | 3.215E-07 | 1.700E-06 | 2.300E-06 |
| 13819 | 691 | T691Y | B | 3e-06 | 2.8e-06 | 4.8e-06 | 3.533E-06 | 1.102E-06 | 2.800E-06 | 4.800E-06 |
| 13820 | 691 | T691V | B | 3.4e-06 | 3.2e-06 | 2.6e-06 | 3.067E-06 | 4.163E-07 | 2.600E-06 | 3.400E-06 |
| 13821 | 692 | D692A | B | 3.6e-06 | 2.6e-06 | 1.8e-06 | 2.667E-06 | 9.018E-07 | 1.800E-06 | 3.600E-06 |
| 13822 | 692 | D692R | B | 3.2e-06 | 9.9e-07 | 2.2e-06 | 2.130E-06 | 1.107E-06 | 9.900E-07 | 3.200E-06 |
| 13823 | 692 | D692N | B | 2.2e-06 | 2.1e-06 | 1.9e-06 | 2.067E-06 | 1.528E-07 | 1.900E-06 | 2.200E-06 |
| 13824 | 692 | D692D | B | 2.5e-06 | 1.6e-06 | 2.5e-06 | 2.200E-06 | 5.196E-07 | 1.600E-06 | 2.500E-06 |
| 13825 | 692 | D692C | B | 2.4e-06 | 2.2e-06 | 2.9e-06 | 2.500E-06 | 3.606E-07 | 2.200E-06 | 2.900E-06 |
| 13826 | 692 | D692E | B | 2.6e-06 | 2.5e-06 | 3.9e-06 | 3.000E-06 | 7.810E-07 | 2.500E-06 | 3.900E-06 |
| 13827 | 692 | D692Q | B | 1.8e-06 | 3.2e-06 | 3e-06 | 2.667E-06 | 7.572E-07 | 1.800E-06 | 3.200E-06 |
| 13828 | 692 | D692G | B | 2.6e-06 | 3.1e-06 | 3e-06 | 2.900E-06 | 2.646E-07 | 2.600E-06 | 3.100E-06 |
| 13829 | 692 | D692H | B | 1.4e-06 | 2.7e-06 | 2e-06 | 2.033E-06 | 6.506E-07 | 1.400E-06 | 2.700E-06 |
| 13830 | 692 | D692I | B | 1.9e-06 | 2.1e-06 | 2.3e-06 | 2.100E-06 | 2.000E-07 | 1.900E-06 | 2.300E-06 |
| 13831 | 692 | D692L | B | 2.5e-06 | 2e-06 | 1.5e-06 | 2.000E-06 | 5.000E-07 | 1.500E-06 | 2.500E-06 |
| 13832 | 692 | D692K | B | 2.4e-06 | 2.8e-06 | 3.4e-06 | 2.867E-06 | 5.033E-07 | 2.400E-06 | 3.400E-06 |
| 13833 | 692 | D692M | B | 2.8e-06 | 1.3e-06 | 1.7e-06 | 1.933E-06 | 7.767E-07 | 1.300E-06 | 2.800E-06 |
| 13834 | 692 | D692F | B | 1.7e-06 | 1.8e-06 | 2.7e-06 | 2.067E-06 | 5.508E-07 | 1.700E-06 | 2.700E-06 |
| 13835 | 692 | D692P | B | 1.9e-06 | 1.8e-06 | 2.8e-06 | 2.167E-06 | 5.508E-07 | 1.800E-06 | 2.800E-06 |
| 13836 | 692 | D692S | B | 2e-06 | 3.8e-06 | 1.2e-06 | 2.333E-06 | 1.332E-06 | 1.200E-06 | 3.800E-06 |
| 13837 | 692 | D692T | B | 3.2e-06 | 2.3e-06 | 1.6e-06 | 2.367E-06 | 8.021E-07 | 1.600E-06 | 3.200E-06 |
| 13838 | 692 | D692W | B | 2.2e-06 | 2.6e-06 | 2e-06 | 2.267E-06 | 3.055E-07 | 2.000E-06 | 2.600E-06 |
| 13839 | 692 | D692Y | B | 1.8e-06 | 2.1e-06 | 3e-06 | 2.300E-06 | 6.245E-07 | 1.800E-06 | 3.000E-06 |
| 13840 | 692 | D692V | B | 2.2e-06 | 2.4e-06 | 3.9e-06 | 2.833E-06 | 9.292E-07 | 2.200E-06 | 3.900E-06 |
| 13841 | 693 | F693A | B | 2.5e-06 | 2.1e-06 | 2.3e-06 | 2.300E-06 | 2.000E-07 | 2.100E-06 | 2.500E-06 |
| 13842 | 693 | F693R | B | 1.8e-06 | 1.4e-06 | 1.9e-06 | 1.700E-06 | 2.646E-07 | 1.400E-06 | 1.900E-06 |
| 13843 | 693 | F693N | B | 2.8e-06 | 2.4e-06 | 2.7e-06 | 2.633E-06 | 2.082E-07 | 2.400E-06 | 2.800E-06 |
| 13844 | 693 | F693D | B | 2.1e-06 | 2.6e-06 | 1.9e-06 | 2.200E-06 | 3.606E-07 | 1.900E-06 | 2.600E-06 |
| 13845 | 693 | F693C | B | 4e-06 | 2.3e-06 | 2.7e-06 | 3.000E-06 | 8.888E-07 | 2.300E-06 | 4.000E-06 |
| 13846 | 693 | F693E | B | 2.3e-06 | 2.9e-06 | 3.2e-06 | 2.800E-06 | 4.583E-07 | 2.300E-06 | 3.200E-06 |
| 13847 | 693 | F693Q | B | 3.7e-06 | 2.7e-06 | 2.3e-06 | 2.900E-06 | 7.211E-07 | 2.300E-06 | 3.700E-06 |
| 13848 | 693 | F693G | B | 2.4e-06 | 2.3e-06 | 4.1e-06 | 2.933E-06 | 1.012E-06 | 2.300E-06 | 4.100E-06 |
| 13849 | 693 | F693H | B | 1.8e-06 | 1.3e-06 | 1e-06 | 1.367E-06 | 4.041E-07 | 1.000E-06 | 1.800E-06 |
| 13850 | 693 | F693I | B | 1.7e-06 | 3.6e-06 | 2.7e-06 | 2.667E-06 | 9.504E-07 | 1.700E-06 | 3.600E-06 |
| 13851 | 693 | F693L | B | 4.7e-06 | 3e-06 | 2e-06 | 3.233E-06 | 1.365E-06 | 2.000E-06 | 4.700E-06 |
| 13852 | 693 | F693K | B | 2.2e-06 | 2.3e-06 | 3.7e-06 | 2.733E-06 | 8.386E-07 | 2.200E-06 | 3.700E-06 |
| 13853 | 693 | F693M | B | 2.2e-06 | 3e-06 | 3.7e-06 | 2.967E-06 | 7.506E-07 | 2.200E-06 | 3.700E-06 |
| 13854 | 693 | F693F | B | 2.5e-06 | 3.2e-06 | 2.5e-06 | 2.733E-06 | 4.041E-07 | 2.500E-06 | 3.200E-06 |

| | | | | | | | | | | |
|---|---|---|---|---|---|---|---|---|---|---|
| 13855 | 693 | F693P | B | 2.8e-06 | 2.2e-06 | 2.7e-06 | 2.567E-06 | 3.215E-07 | 2.200E-06 | 2.800E-06 |
| 13856 | 693 | F693S | B | 3.4e-06 | 2.2e-06 | 1.7e-06 | 2.433E-06 | 8.737E-07 | 1.700E-06 | 3.400E-06 |
| 13857 | 693 | F693T | B | 2.7e-06 | 4e-06 | 1.9e-06 | 2.867E-06 | 1.060E-06 | 1.900E-06 | 4.000E-06 |
| 13858 | 693 | F693W | B | 3.4e-06 | 2.9e-06 | 3.3e-06 | 3.200E-06 | 2.646E-07 | 2.900E-06 | 3.400E-06 |
| 13859 | 693 | F693Y | B | 4.8e-06 | 2.2e-06 | 1.7e-06 | 2.900E-06 | 1.664E-06 | 1.700E-06 | 4.800E-06 |
| 13860 | 693 | F693V | B | 2.5e-06 | 1.4e-06 | 2.5e-06 | 2.133E-06 | 6.351E-07 | 1.400E-06 | 2.500E-06 |
| 13861 | 694 | T694A | B | 5.7e-06 | 2.8e-06 | 2e-06 | 3.500E-06 | 1.947E-06 | 2.000E-06 | 5.700E-06 |
| 13862 | 694 | T694R | B | 2.9e-06 | 2.5e-06 | 3.8e-06 | 3.067E-06 | 6.658E-07 | 2.500E-06 | 3.800E-06 |
| 13863 | 694 | T694N | B | 3.4e-06 | 2.3e-06 | 3.8e-06 | 3.167E-06 | 7.767E-07 | 2.300E-06 | 3.800E-06 |
| 13864 | 694 | T694D | B | 1.2e-06 | 3.1e-06 | 2.8e-06 | 2.367E-06 | 1.021E-06 | 1.200E-06 | 3.100E-06 |
| 13865 | 694 | T694C | B | 1.3e-06 | 1.9e-06 | 1.5e-06 | 1.567E-06 | 3.055E-07 | 1.300E-06 | 1.900E-06 |
| 13866 | 694 | T694E | B | 2.2e-06 | 4e-06 | 1.7e-06 | 2.633E-06 | 1.210E-06 | 1.700E-06 | 4.000E-06 |
| 13867 | 694 | T694Q | B | 2.3e-06 | 3.8e-06 | 3e-06 | 3.033E-06 | 7.506E-07 | 2.300E-06 | 3.800E-06 |
| 13868 | 694 | T694G | B | 1.7e-06 | 1.6e-06 | 3.2e-06 | 2.167E-06 | 8.963E-07 | 1.600E-06 | 3.200E-06 |
| 13869 | 694 | T694H | B | 2.6e-06 | 2.8e-06 | 2.8e-06 | 2.733E-06 | 1.155E-07 | 2.600E-06 | 2.800E-06 |
| 13870 | 694 | T694I | B | 1.6e-06 | 3.1e-06 | 2.4e-06 | 2.367E-06 | 7.506E-07 | 1.600E-06 | 3.100E-06 |
| 13871 | 694 | T694L | B | 4.4e-06 | 2.7e-06 | 3.1e-06 | 3.400E-06 | 8.888E-07 | 2.700E-06 | 4.400E-06 |
| 13872 | 694 | T694K | B | 1.8e-06 | 2.4e-06 | 2.1e-06 | 2.100E-06 | 3.000E-07 | 1.800E-06 | 2.400E-06 |
| 13873 | 694 | T694M | B | 1.9e-06 | 3.2e-06 | 2.1e-06 | 2.400E-06 | 7.000E-07 | 1.900E-06 | 3.200E-06 |
| 13874 | 694 | T694F | B | 1.6e-06 | 2.2e-06 | 2.9e-06 | 2.233E-06 | 6.506E-07 | 1.600E-06 | 2.900E-06 |
| 13875 | 694 | T694P | B | 1.4e-06 | 1.3e-06 | 1.8e-06 | 1.500E-06 | 2.646E-07 | 1.300E-06 | 1.800E-06 |
| 13876 | 694 | T694S | B | 4.2e-06 | 2.9e-06 | 1.6e-06 | 2.900E-06 | 1.300E-06 | 1.600E-06 | 4.200E-06 |
| 13877 | 694 | T694T | B | 3.4e-06 | 2.1e-06 | 1.9e-06 | 2.467E-06 | 8.145E-07 | 1.900E-06 | 3.400E-06 |
| 13878 | 694 | T694W | B | 2.3e-06 | 2e-06 | 2.6e-06 | 2.300E-06 | 3.000E-07 | 2.000E-06 | 2.600E-06 |
| 13879 | 694 | T694Y | B | 2e-06 | 3.7e-06 | 2.9e-06 | 2.867E-06 | 8.505E-07 | 2.000E-06 | 3.700E-06 |
| 13880 | 694 | T694V | B | 3e-06 | 2.8e-06 | 3.2e-06 | 3.000E-06 | 2.000E-07 | 2.800E-06 | 3.200E-06 |
| 13881 | 695 | L695A | B | 4.1e-06 | 3e-06 | 3.1e-06 | 3.400E-06 | 6.083E-07 | 3.000E-06 | 4.100E-06 |
| 13882 | 695 | L695R | B | 1.7e-06 | 1.8e-06 | 2.6e-06 | 2.033E-06 | 4.933E-07 | 1.700E-06 | 2.600E-06 |
| 13883 | 695 | L695N | B | 1.9e-06 | 2.6e-06 | 2.6e-06 | 2.367E-06 | 4.041E-07 | 1.900E-06 | 2.600E-06 |
| 13884 | 695 | L695D | B | 2.3e-06 | 1.6e-06 | 1.9e-06 | 1.933E-06 | 3.512E-07 | 1.600E-06 | 2.300E-06 |
| 13885 | 695 | L695C | B | 1.7e-06 | 2.3e-06 | 2.7e-06 | 2.233E-06 | 5.033E-07 | 1.700E-06 | 2.700E-06 |
| 13886 | 695 | L695E | B | 2.9e-06 | 2.3e-06 | 2.7e-06 | 2.633E-06 | 3.055E-07 | 2.300E-06 | 2.900E-06 |
| 13887 | 695 | L695Q | B | 3.8e-06 | 4.3e-06 | 1.5e-06 | 3.200E-06 | 1.493E-06 | 1.500E-06 | 4.300E-06 |
| 13888 | 695 | L695G | B | 1.7e-06 | 1.4e-06 | 2.1e-06 | 1.733E-06 | 3.512E-07 | 1.400E-06 | 2.100E-06 |
| 13889 | 695 | L695H | B | 2.1e-06 | 3.2e-06 | 3.3e-06 | 2.867E-06 | 6.658E-07 | 2.100E-06 | 3.300E-06 |
| 13890 | 695 | L695I | B | 1.2e-06 | 2.6e-06 | 2e-06 | 1.933E-06 | 7.024E-07 | 1.200E-06 | 2.600E-06 |
| 13891 | 695 | L695L | B | 3e-06 | 2e-06 | 2e-06 | 2.333E-06 | 5.774E-07 | 2.000E-06 | 3.000E-06 |
| 13892 | 695 | L695K | B | 3.2e-06 | 3e-06 | 2e-06 | 2.733E-06 | 6.429E-07 | 2.000E-06 | 3.200E-06 |
| 13893 | 695 | L695M | B | 1.3e-06 | 2.7e-06 | 4.1e-06 | 2.700E-06 | 1.400E-06 | 1.300E-06 | 4.100E-06 |
| 13894 | 695 | L695F | B | 3.5e-06 | 2.4e-06 | 2.7e-06 | 2.867E-06 | 5.686E-07 | 2.400E-06 | 3.500E-06 |
| 13895 | 695 | L695P | B | 1.6e-06 | 3e-06 | 2.4e-06 | 2.333E-06 | 7.024E-07 | 1.600E-06 | 3.000E-06 |
| 13896 | 695 | L695S | B | 3.3e-06 | 2.7e-06 | 2e-06 | 2.667E-06 | 6.506E-07 | 2.000E-06 | 3.300E-06 |
| 13897 | 695 | L695T | B | 5.7e-06 | 2.2e-06 | 2.3e-06 | 3.400E-06 | 1.992E-06 | 2.200E-06 | 5.700E-06 |
| 13898 | 695 | L695W | B | 2.1e-06 | 3.3e-06 | 3.1e-06 | 2.833E-06 | 6.429E-07 | 2.100E-06 | 3.300E-06 |
| 13899 | 695 | L695Y | B | 2.7e-06 | 2.7e-06 | 2.6e-06 | 2.667E-06 | 5.774E-08 | 2.600E-06 | 2.700E-06 |
| 13900 | 695 | L695V | B | 5e-06 | 2.1e-06 | 2.7e-06 | 3.267E-06 | 1.531E-06 | 2.100E-06 | 5.000E-06 |
| 13901 | 696 | T696A | B | 1.8e-06 | 1e-06 | 3.4e-06 | 2.067E-06 | 1.222E-06 | 1.000E-06 | 3.400E-06 |
| 13902 | 696 | T696R | B | 1.6e-06 | 1.4e-06 | 3e-06 | 2.000E-06 | 8.718E-07 | 1.400E-06 | 3.000E-06 |
| 13903 | 696 | T696N | B | 3.4e-06 | 3.6e-06 | 2e-06 | 3.000E-06 | 8.718E-07 | 2.000E-06 | 3.600E-06 |
| 13904 | 696 | T696D | B | 3.3e-06 | 2.7e-06 | 2.3e-06 | 2.767E-06 | 5.033E-07 | 2.300E-06 | 3.300E-06 |
| 13905 | 696 | T696C | B | 3.4e-06 | 1.9e-06 | 2.9e-06 | 2.733E-06 | 7.638E-07 | 1.900E-06 | 3.400E-06 |
| 13906 | 696 | T696E | B | 2e-06 | 1.3e-06 | 2.3e-06 | 1.867E-06 | 5.132E-07 | 1.300E-06 | 2.300E-06 |
| 13907 | 696 | T696Q | B | 2.6e-06 | 2.7e-06 | 2.7e-06 | 2.667E-06 | 5.774E-08 | 2.600E-06 | 2.700E-06 |
| 13908 | 696 | T696G | B | 1.7e-06 | 1.4e-06 | 1.9e-06 | 1.667E-06 | 2.517E-07 | 1.400E-06 | 1.900E-06 |
| 13909 | 696 | T696H | B | 2.8e-06 | 2.6e-06 | 2.5e-06 | 2.633E-06 | 1.528E-07 | 2.500E-06 | 2.800E-06 |
| 13910 | 696 | T696I | B | 1.6e-06 | 3.6e-06 | 2.9e-06 | 2.700E-06 | 1.015E-06 | 1.600E-06 | 3.600E-06 |
| 13911 | 696 | T696L | B | 4.6e-06 | 3.2e-06 | 1.2e-06 | 3.000E-06 | 1.709E-06 | 1.200E-06 | 4.600E-06 |
| 13912 | 696 | T696K | B | 3.2e-06 | 2.9e-06 | 2.2e-06 | 2.767E-06 | 5.132E-07 | 2.200E-06 | 3.200E-06 |
| 13913 | 696 | T696M | B | 3.5e-06 | 1.2e-06 | 2.1e-06 | 2.267E-06 | 1.159E-06 | 1.200E-06 | 3.500E-06 |
| 13914 | 696 | T696F | B | 2.2e-06 | 1.6e-06 | 2.2e-06 | 2.000E-06 | 3.464E-07 | 1.600E-06 | 2.200E-06 |
| 13915 | 696 | T696P | B | 3e-06 | 3.3e-06 | 1.5e-06 | 2.600E-06 | 9.644E-07 | 1.500E-06 | 3.300E-06 |
| 13916 | 696 | T696S | B | 3.5e-06 | 2.6e-06 | 1.3e-06 | 2.467E-06 | 1.106E-06 | 1.300E-06 | 3.500E-06 |
| 13917 | 696 | T696T | B | 2.6e-06 | 1.8e-06 | 2.4e-06 | 2.267E-06 | 4.163E-07 | 1.800E-06 | 2.600E-06 |
| 13918 | 696 | T696W | B | 2.1e-06 | 3e-06 | 2.1e-06 | 2.400E-06 | 5.196E-07 | 2.100E-06 | 3.000E-06 |
| 13919 | 696 | T696Y | B | 1.8e-06 | 2e-06 | 4e-06 | 2.600E-06 | 1.217E-06 | 1.800E-06 | 4.000E-06 |
| 13920 | 696 | T696V | B | 2.9e-06 | 2.8e-06 | 3.3e-06 | 3.000E-06 | 2.646E-07 | 2.800E-06 | 3.300E-06 |
| 13921 | 697 | I697A | B | 1.7e-06 | 3.4e-06 | 1.4e-06 | 2.167E-06 | 1.079E-06 | 1.400E-06 | 3.400E-06 |
| 13922 | 697 | I697R | B | 2.4e-06 | 2.6e-06 | 2.5e-06 | 2.500E-06 | 1.000E-07 | 2.400E-06 | 2.600E-06 |
| 13923 | 697 | I697N | B | 5.1e-06 | 2.7e-06 | 3.7e-06 | 3.833E-06 | 1.206E-06 | 2.700E-06 | 5.100E-06 |
| 13924 | 697 | I697D | B | 3.2e-06 | 2.5e-06 | 2.6e-06 | 2.767E-06 | 3.786E-07 | 2.500E-06 | 3.200E-06 |
| 13925 | 697 | I697C | B | 1.4e-06 | 3.3e-06 | 1.8e-06 | 2.167E-06 | 1.002E-06 | 1.400E-06 | 3.300E-06 |
| 13926 | 697 | I697E | B | 2.1e-06 | 5.1e-06 | 2.6e-06 | 3.267E-06 | 1.607E-06 | 2.100E-06 | 5.100E-06 |
| 13927 | 697 | I697Q | B | 3.6e-06 | 2.1e-06 | 2.5e-06 | 2.733E-06 | 7.767E-07 | 2.100E-06 | 3.600E-06 |
| 13928 | 697 | I697G | B | 2.5e-06 | 1.8e-06 | 3.6e-06 | 2.633E-06 | 9.074E-07 | 1.800E-06 | 3.600E-06 |
| 13929 | 697 | I697H | B | 1e-06 | 4.1e-06 | 1.9e-06 | 2.333E-06 | 1.595E-06 | 1.000E-06 | 4.100E-06 |
| 13930 | 697 | I697I | B | 7e-07 | 2.5e-06 | 2e-06 | 1.733E-06 | 9.292E-07 | 7.000E-07 | 2.500E-06 |
| 13931 | 697 | I697L | B | 2.1e-06 | 2.1e-06 | 2.3e-06 | 2.167E-06 | 1.155E-07 | 2.100E-06 | 2.300E-06 |
| 13932 | 697 | I697K | B | 1.6e-06 | 2e-06 | 3.5e-06 | 2.367E-06 | 1.002E-06 | 1.600E-06 | 3.500E-06 |
| 13933 | 697 | I697M | B | 2.8e-06 | 2.3e-06 | 2.2e-06 | 2.433E-06 | 3.215E-07 | 2.200E-06 | 2.800E-06 |
| 13934 | 697 | I697F | B | 3.1e-06 | 2.8e-06 | 1.6e-06 | 2.500E-06 | 7.937E-07 | 1.600E-06 | 3.100E-06 |
| 13935 | 697 | I697P | B | 3.9e-06 | 2.4e-06 | 2.9e-06 | 3.067E-06 | 7.638E-07 | 2.400E-06 | 3.900E-06 |
| 13936 | 697 | I697S | B | 1.5e-06 | 2.2e-06 | 2.4e-06 | 2.033E-06 | 4.726E-07 | 1.500E-06 | 2.400E-06 |
| 13937 | 697 | I697T | B | 3.1e-06 | 2.3e-06 | 3.8e-06 | 3.067E-06 | 7.506E-07 | 2.300E-06 | 3.800E-06 |
| 13938 | 697 | I697W | B | 1.2e-06 | 2.1e-06 | 2.3e-06 | 1.867E-06 | 5.859E-07 | 1.200E-06 | 2.300E-06 |
| 13939 | 697 | I697Y | B | 1.8e-06 | 1.3e-06 | 1.9e-06 | 1.667E-06 | 3.215E-07 | 1.300E-06 | 1.900E-06 |

| | | | | | | | | | | |
|---|---|---|---|---|---|---|---|---|---|---|
| 13940 | 697 | I697V | B | 2.1e-06 | 1.9e-06 | 2.5e-06 | 2.167E-06 | 3.055E-07 | 1.900E-06 | 2.500E-06 |
| 13941 | 698 | S698A | B | 2.8e-06 | 2.5e-06 | 1.8e-06 | 2.367E-06 | 5.132E-07 | 1.800E-06 | 2.800E-06 |
| 13942 | 698 | S698R | B | 4.8e-06 | 2.7e-06 | 2.2e-06 | 3.233E-06 | 1.380E-06 | 2.200E-06 | 4.800E-06 |
| 13943 | 698 | S698N | B | 2e-06 | 2.3e-06 | 1.7e-06 | 2.000E-06 | 3.000E-07 | 1.700E-06 | 2.300E-06 |
| 13944 | 698 | S698D | B | 2.7e-06 | 2.3e-06 | 3.1e-06 | 2.700E-06 | 4.000E-07 | 2.300E-06 | 3.100E-06 |
| 13945 | 698 | S698C | B | 2.3e-06 | 2.6e-06 | 1.8e-06 | 2.233E-06 | 4.041E-07 | 1.800E-06 | 2.600E-06 |
| 13946 | 698 | S698E | B | 2.2e-06 | 4.9e-06 | 2.6e-06 | 3.233E-06 | 1.457E-06 | 2.200E-06 | 4.900E-06 |
| 13947 | 698 | S698Q | B | 2.2e-06 | 2.5e-06 | 2.1e-06 | 2.267E-06 | 2.082E-07 | 2.100E-06 | 2.500E-06 |
| 13948 | 698 | S698G | B | 2.9e-06 | 1.5e-06 | 2.1e-06 | 2.167E-06 | 7.024E-07 | 1.500E-06 | 2.900E-06 |
| 13949 | 698 | S698H | B | 2.3e-06 | 1.6e-06 | 2.7e-06 | 2.200E-06 | 5.568E-07 | 1.600E-06 | 2.700E-06 |
| 13950 | 698 | S698I | B | 3.1e-06 | 2.6e-06 | 3.2e-06 | 2.967E-06 | 3.215E-07 | 2.600E-06 | 3.200E-06 |
| 13951 | 698 | S698L | B | 1.8e-06 | 2.7e-06 | 2.8e-06 | 2.433E-06 | 5.508E-07 | 1.800E-06 | 2.800E-06 |
| 13952 | 698 | S698K | B | 2.1e-06 | 1.2e-06 | 2.8e-06 | 2.033E-06 | 8.021E-07 | 1.200E-06 | 2.800E-06 |
| 13953 | 698 | S698M | B | 2.3e-06 | 2e-06 | 4.3e-06 | 2.867E-06 | 1.250E-06 | 2.000E-06 | 4.300E-06 |
| 13954 | 698 | S698F | B | 2.4e-06 | 3.9e-06 | 2.8e-06 | 3.033E-06 | 7.767E-07 | 2.400E-06 | 3.900E-06 |
| 13955 | 698 | S698P | B | 2.1e-06 | 2.7e-06 | 3e-06 | 2.600E-06 | 4.583E-07 | 2.100E-06 | 3.000E-06 |
| 13956 | 698 | S698S | B | 1.7e-06 | 2.1e-06 | 2.4e-06 | 2.067E-06 | 3.512E-07 | 1.700E-06 | 2.400E-06 |
| 13957 | 698 | S698T | B | 3.1e-06 | 3.1e-06 | 1.7e-06 | 2.633E-06 | 8.083E-07 | 1.700E-06 | 3.100E-06 |
| 13958 | 698 | S698W | B | 2.3e-06 | 2e-06 | 1.9e-06 | 2.067E-06 | 2.082E-07 | 1.900E-06 | 2.300E-06 |
| 13959 | 698 | S698Y | B | 3.1e-06 | 3.5e-06 | 2.3e-06 | 2.967E-06 | 6.110E-07 | 2.300E-06 | 3.500E-06 |
| 13960 | 698 | S698V | B | 2.6e-06 | 3.5e-06 | 2.6e-06 | 2.900E-06 | 5.196E-07 | 2.600E-06 | 3.500E-06 |
| 13961 | 699 | S699A | B | 2.9e-06 | 2.6e-06 | 2.4e-06 | 2.633E-06 | 2.517E-07 | 2.400E-06 | 2.900E-06 |
| 13962 | 699 | S699R | B | 3.8e-06 | 1.9e-06 | 3.2e-06 | 2.967E-06 | 9.713E-07 | 1.900E-06 | 3.800E-06 |
| 13963 | 699 | S699N | B | 1.4e-06 | 2.2e-06 | 2.3e-06 | 1.967E-06 | 4.933E-07 | 1.400E-06 | 2.300E-06 |
| 13964 | 699 | S699D | B | 2.6e-06 | 2.9e-06 | 3.5e-06 | 3.000E-06 | 4.583E-07 | 2.600E-06 | 3.500E-06 |
| 13965 | 699 | S699C | B | 2.5e-06 | 1.8e-06 | 2.6e-06 | 2.300E-06 | 4.359E-07 | 1.800E-06 | 2.600E-06 |
| 13966 | 699 | S699E | B | 2.2e-06 | 4.6e-06 | 2.5e-06 | 3.100E-06 | 1.308E-06 | 2.200E-06 | 4.600E-06 |
| 13967 | 699 | S699Q | B | 3e-06 | 2.5e-06 | 1.6e-06 | 2.367E-06 | 7.095E-07 | 1.600E-06 | 3.000E-06 |
| 13968 | 699 | S699G | B | 2e-06 | 1.5e-06 | 2.7e-06 | 2.067E-06 | 6.028E-07 | 1.500E-06 | 2.700E-06 |
| 13969 | 699 | S699H | B | 1.6e-06 | 2.4e-06 | 2.8e-06 | 2.267E-06 | 6.110E-07 | 1.600E-06 | 2.800E-06 |
| 13970 | 699 | S699I | B | 4.1e-06 | 5.4e-06 | 1.6e-06 | 3.700E-06 | 1.931E-06 | 1.600E-06 | 5.400E-06 |
| 13971 | 699 | S699L | B | 3.3e-06 | 3.1e-06 | 2.6e-06 | 3.000E-06 | 3.606E-07 | 2.600E-06 | 3.300E-06 |
| 13972 | 699 | S699K | B | 1.6e-06 | 2.4e-06 | 3.1e-06 | 2.367E-06 | 7.506E-07 | 1.600E-06 | 3.100E-06 |
| 13973 | 699 | S699M | B | 2.8e-06 | 2.5e-06 | 1.2e-06 | 2.167E-06 | 8.505E-07 | 1.200E-06 | 2.800E-06 |
| 13974 | 699 | S699F | B | 2.1e-06 | 4.4e-06 | 1.5e-06 | 2.667E-06 | 1.531E-06 | 1.500E-06 | 4.400E-06 |
| 13975 | 699 | S699P | B | 3.1e-06 | 4.3e-06 | 2.1e-06 | 3.167E-06 | 1.102E-06 | 2.100E-06 | 4.300E-06 |
| 13976 | 699 | S699S | B | 2.3e-06 | 2.6e-06 | 1.5e-06 | 2.133E-06 | 5.686E-07 | 1.500E-06 | 2.600E-06 |
| 13977 | 699 | S699T | B | 1.5e-06 | 2.1e-06 | 2.2e-06 | 1.933E-06 | 3.786E-07 | 1.500E-06 | 2.200E-06 |
| 13978 | 699 | S699W | B | 3e-06 | 2.4e-06 | 1.6e-06 | 2.333E-06 | 7.024E-07 | 1.600E-06 | 3.000E-06 |
| 13979 | 699 | S699Y | B | 2.4e-06 | 2.1e-06 | 2.1e-06 | 2.200E-06 | 1.732E-07 | 2.100E-06 | 2.400E-06 |
| 13980 | 699 | S699V | B | 2.7e-06 | 2.5e-06 | 2e-06 | 2.400E-06 | 3.606E-07 | 2.000E-06 | 2.700E-06 |
| 13981 | 700 | L700A | B | 3.5e-06 | 2.5e-06 | 2.4e-06 | 2.800E-06 | 6.083E-07 | 2.400E-06 | 3.500E-06 |
| 13982 | 700 | L700R | B | 2.8e-06 | 2.7e-06 | 1.4e-06 | 2.300E-06 | 7.810E-07 | 1.400E-06 | 2.800E-06 |
| 13983 | 700 | L700N | B | 5.1e-06 | 1.2e-06 | 3e-06 | 3.100E-06 | 1.952E-06 | 1.200E-06 | 5.100E-06 |
| 13984 | 700 | L700D | B | 3e-06 | 2.1e-06 | 1.2e-06 | 2.100E-06 | 9.000E-07 | 1.200E-06 | 3.000E-06 |
| 13985 | 700 | L700C | B | 2e-06 | 3.6e-06 | 1.9e-06 | 2.500E-06 | 9.539E-07 | 1.900E-06 | 3.600E-06 |
| 13986 | 700 | L700E | B | 2.5e-06 | 3.1e-06 | 1.6e-06 | 2.400E-06 | 7.550E-07 | 1.600E-06 | 3.100E-06 |
| 13987 | 700 | L700Q | B | 2.2e-06 | 2e-06 | 2e-06 | 2.067E-06 | 1.155E-07 | 2.000E-06 | 2.200E-06 |
| 13988 | 700 | L700G | B | 5.2e-06 | 2e-06 | 2e-06 | 3.067E-06 | 1.848E-06 | 2.000E-06 | 5.200E-06 |
| 13989 | 700 | L700H | B | 3.8e-06 | 3.7e-06 | 1.4e-06 | 2.967E-06 | 1.358E-06 | 1.400E-06 | 3.800E-06 |
| 13990 | 700 | L700I | B | 2.1e-06 | 1.9e-06 | 2.7e-06 | 2.233E-06 | 4.163E-07 | 1.900E-06 | 2.700E-06 |
| 13991 | 700 | L700L | B | 1.1e-06 | 2.9e-06 | 1.4e-06 | 1.800E-06 | 9.644E-07 | 1.100E-06 | 2.900E-06 |
| 13992 | 700 | L700K | B | 2.2e-06 | 1.5e-06 | 2e-06 | 1.900E-06 | 3.606E-07 | 1.500E-06 | 2.200E-06 |
| 13993 | 700 | L700M | B | 2.9e-06 | 2.6e-06 | 3.2e-06 | 2.900E-06 | 3.000E-07 | 2.600E-06 | 3.200E-06 |
| 13994 | 700 | L700F | B | 2.1e-06 | 3.6e-06 | 3e-06 | 2.900E-06 | 7.550E-07 | 2.100E-06 | 3.600E-06 |
| 13995 | 700 | L700P | B | 1.5e-06 | 2.4e-06 | 2.4e-06 | 2.100E-06 | 5.196E-07 | 1.500E-06 | 2.400E-06 |
| 13996 | 700 | L700S | B | 2.8e-06 | 2e-06 | 1.9e-06 | 2.233E-06 | 4.933E-07 | 1.900E-06 | 2.800E-06 |
| 13997 | 700 | L700T | B | 1.6e-06 | 2.2e-06 | 1.8e-06 | 1.867E-06 | 3.055E-07 | 1.600E-06 | 2.200E-06 |
| 13998 | 700 | L700W | B | 2.9e-06 | 2e-06 | 2.9e-06 | 2.600E-06 | 5.196E-07 | 2.000E-06 | 2.900E-06 |
| 13999 | 700 | L700Y | B | 2.3e-06 | 1.4e-06 | 2.6e-06 | 2.100E-06 | 6.245E-07 | 1.400E-06 | 2.600E-06 |
| 14000 | 700 | L700V | B | 3.6e-06 | 2.5e-06 | 2.4e-06 | 2.833E-06 | 6.658E-07 | 2.400E-06 | 3.600E-06 |
| 14001 | 701 | Q701A | B | 2.7e-06 | 3.2e-06 | 2.8e-06 | 2.900E-06 | 2.646E-07 | 2.700E-06 | 3.200E-06 |
| 14002 | 701 | Q701R | B | 2.6e-06 | 2.3e-06 | 3.1e-06 | 2.667E-06 | 4.041E-07 | 2.300E-06 | 3.100E-06 |
| 14003 | 701 | Q701N | B | 3.3e-06 | 2.5e-06 | 3.1e-06 | 2.967E-06 | 4.163E-07 | 2.500E-06 | 3.300E-06 |
| 14004 | 701 | Q701D | B | 2.6e-06 | 2.4e-06 | 5.8e-06 | 3.600E-06 | 1.908E-06 | 2.400E-06 | 5.800E-06 |
| 14005 | 701 | Q701C | B | 2.8e-06 | 2.2e-06 | 2.6e-06 | 2.533E-06 | 3.055E-07 | 2.200E-06 | 2.800E-06 |
| 14006 | 701 | Q701E | B | 2.6e-06 | 1.5e-06 | 4.9e-06 | 3.000E-06 | 1.735E-06 | 1.500E-06 | 4.900E-06 |
| 14007 | 701 | Q701Q | B | 2.1e-06 | 3e-06 | 2.1e-06 | 2.400E-06 | 5.196E-07 | 2.100E-06 | 3.000E-06 |
| 14008 | 701 | Q701G | B | 2.6e-06 | 2.6e-06 | 2.1e-06 | 2.433E-06 | 2.887E-07 | 2.100E-06 | 2.600E-06 |
| 14009 | 701 | Q701H | B | 2.4e-06 | 2.7e-06 | 1.8e-06 | 2.300E-06 | 4.583E-07 | 1.800E-06 | 2.700E-06 |
| 14010 | 701 | Q701I | B | 2.6e-06 | 2.5e-06 | 3e-06 | 2.700E-06 | 2.646E-07 | 2.500E-06 | 3.000E-06 |
| 14011 | 701 | Q701L | B | 1.7e-06 | 4e-06 | 1.9e-06 | 2.533E-06 | 1.274E-06 | 1.700E-06 | 4.000E-06 |
| 14012 | 701 | Q701K | B | 2e-06 | 3.5e-06 | 2.1e-06 | 2.533E-06 | 8.386E-07 | 2.000E-06 | 3.500E-06 |
| 14013 | 701 | Q701M | B | 4.3e-06 | 2.8e-06 | 2.5e-06 | 3.200E-06 | 9.644E-07 | 2.500E-06 | 4.300E-06 |
| 14014 | 701 | Q701F | B | 1.2e-06 | 3.8e-06 | 2.2e-06 | 2.400E-06 | 1.311E-06 | 1.200E-06 | 3.800E-06 |
| 14015 | 701 | Q701P | B | 1.7e-06 | 2.8e-06 | 1.3e-06 | 1.933E-06 | 7.767E-07 | 1.300E-06 | 2.800E-06 |
| 14016 | 701 | Q701S | B | 2.3e-06 | 2.6e-06 | 1.7e-06 | 2.200E-06 | 4.583E-07 | 1.700E-06 | 2.600E-06 |
| 14017 | 701 | Q701T | B | 2.1e-06 | 2.2e-06 | 3e-06 | 2.433E-06 | 4.933E-07 | 2.100E-06 | 3.000E-06 |
| 14018 | 701 | Q701W | B | 4.2e-06 | 2.6e-06 | 2.4e-06 | 3.067E-06 | 9.866E-07 | 2.400E-06 | 4.200E-06 |
| 14019 | 701 | Q701Y | B | 2.1e-06 | 2.8e-06 | 2.3e-06 | 2.400E-06 | 3.606E-07 | 2.100E-06 | 2.800E-06 |
| 14020 | 701 | Q701V | B | 3.3e-06 | 2.5e-06 | 2.2e-06 | 2.667E-06 | 5.686E-07 | 2.200E-06 | 3.300E-06 |
| 14021 | 702 | P702A | B | 2.9e-06 | 2.4e-06 | 1.7e-06 | 2.333E-06 | 6.028E-07 | 1.700E-06 | 2.900E-06 |
| 14022 | 702 | P702R | B | 1.9e-06 | 3.6e-06 | 1.1e-06 | 2.200E-06 | 1.277E-06 | 1.100E-06 | 3.600E-06 |
| 14023 | 702 | P702N | B | 3.2e-06 | 2.1e-06 | 2.4e-06 | 2.567E-06 | 5.686E-07 | 2.100E-06 | 3.200E-06 |
| 14024 | 702 | P702D | B | 3.3e-06 | 2.7e-06 | 1.9e-06 | 2.633E-06 | 7.024E-07 | 1.900E-06 | 3.300E-06 |

| | | | | | | | | | | |
|---|---|---|---|---|---|---|---|---|---|---|
| 14025 | 702 | P702C | B | 1.4e-06 | 2.3e-06 | 2.7e-06 | 2.133E-06 | 6.658E-07 | 1.400E-06 | 2.700E-06 |
| 14026 | 702 | P702E | B | 2.5e-06 | 2.7e-06 | 2.2e-06 | 2.467E-06 | 2.517E-07 | 2.200E-06 | 2.700E-06 |
| 14027 | 702 | P702Q | B | 2.2e-06 | 2.6e-06 | 9.9e-07 | 1.930E-06 | 8.383E-07 | 9.900E-07 | 2.600E-06 |
| 14028 | 702 | P702G | B | 3e-06 | 1e-06 | 2.2e-06 | 2.067E-06 | 1.007E-06 | 1.000E-06 | 3.000E-06 |
| 14029 | 702 | P702H | B | 2.5e-06 | 2.9e-06 | 2.2e-06 | 2.533E-06 | 3.512E-07 | 2.200E-06 | 2.900E-06 |
| 14030 | 702 | P702I | B | 1.2e-06 | 2.6e-06 | 2.5e-06 | 2.100E-06 | 7.810E-07 | 1.200E-06 | 2.600E-06 |
| 14031 | 702 | P702L | B | 2.2e-06 | 1.6e-06 | 1.4e-06 | 1.733E-06 | 4.163E-07 | 1.400E-06 | 2.200E-06 |
| 14032 | 702 | P702K | B | 2.7e-06 | 1.7e-06 | 3.1e-06 | 2.500E-06 | 7.211E-07 | 1.700E-06 | 3.100E-06 |
| 14033 | 702 | P702M | B | 2.8e-06 | 1.3e-06 | 2.8e-06 | 2.300E-06 | 8.660E-07 | 1.300E-06 | 2.800E-06 |
| 14034 | 702 | P702F | B | 3.6e-06 | 3.8e-06 | 1.2e-06 | 2.867E-06 | 1.447E-06 | 1.200E-06 | 3.800E-06 |
| 14035 | 702 | P702P | B | 2.3e-06 | 1.9e-06 | 1e-06 | 1.733E-06 | 6.658E-07 | 1.000E-06 | 2.300E-06 |
| 14036 | 702 | P702S | B | 1.4e-06 | 2e-06 | 1.7e-06 | 1.700E-06 | 3.000E-07 | 1.400E-06 | 2.000E-06 |
| 14037 | 702 | P702T | B | 2.5e-06 | 1.5e-06 | 1.6e-06 | 1.867E-06 | 5.508E-07 | 1.500E-06 | 2.500E-06 |
| 14038 | 702 | P702W | B | 3.2e-06 | 2.3e-06 | 2.8e-06 | 2.767E-06 | 4.509E-07 | 2.300E-06 | 3.200E-06 |
| 14039 | 702 | P702Y | B | 2.6e-06 | 2.5e-06 | 1.8e-06 | 2.300E-06 | 4.359E-07 | 1.800E-06 | 2.600E-06 |
| 14040 | 702 | P702V | B | 2.3e-06 | 1.8e-06 | 2.1e-06 | 2.067E-06 | 2.517E-07 | 1.800E-06 | 2.300E-06 |
| 14041 | 703 | E703A | B | 1.9e-06 | 2.8e-06 | 2.1e-06 | 2.267E-06 | 4.726E-07 | 1.900E-06 | 2.800E-06 |
| 14042 | 703 | E703R | B | 3.9e-06 | 2.7e-06 | 3.6e-06 | 3.400E-06 | 6.245E-07 | 2.700E-06 | 3.900E-06 |
| 14043 | 703 | E703N | B | 2.1e-06 | 2.1e-06 | 2e-06 | 2.067E-06 | 5.774E-08 | 2.000E-06 | 2.100E-06 |
| 14044 | 703 | E703D | B | 3.3e-06 | 2e-06 | 1.9e-06 | 2.400E-06 | 7.810E-07 | 1.900E-06 | 3.300E-06 |
| 14045 | 703 | E703C | B | 5.7e-06 | 3.3e-06 | 3.2e-06 | 4.067E-06 | 1.415E-06 | 3.200E-06 | 5.700E-06 |
| 14046 | 703 | E703E | B | 2.1e-06 | 3e-06 | 3.2e-06 | 2.767E-06 | 5.859E-07 | 2.100E-06 | 3.200E-06 |
| 14047 | 703 | E703Q | B | 1.8e-06 | 2.3e-06 | 2.2e-06 | 2.100E-06 | 2.646E-07 | 1.800E-06 | 2.300E-06 |
| 14048 | 703 | E703G | B | 2.2e-06 | 2.7e-06 | 1.8e-06 | 2.233E-06 | 4.509E-07 | 1.800E-06 | 2.700E-06 |
| 14049 | 703 | E703H | B | 2.8e-06 | 1.5e-06 | 2.2e-06 | 2.167E-06 | 6.506E-07 | 1.500E-06 | 2.800E-06 |
| 14050 | 703 | E703I | B | 1.7e-06 | 1.4e-06 | 2.2e-06 | 1.767E-06 | 4.041E-07 | 1.400E-06 | 2.200E-06 |
| 14051 | 703 | E703L | B | 3.6e-06 | 1.6e-06 | 2.9e-06 | 2.700E-06 | 1.015E-06 | 1.600E-06 | 3.600E-06 |
| 14052 | 703 | E703K | B | 2.1e-06 | 3.5e-06 | 2.7e-06 | 2.767E-06 | 7.024E-07 | 2.100E-06 | 3.500E-06 |
| 14053 | 703 | E703M | B | 3.1e-06 | 3.8e-06 | 3.1e-06 | 3.333E-06 | 4.041E-07 | 3.100E-06 | 3.800E-06 |
| 14054 | 703 | E703F | B | 4e-06 | 2e-06 | 1.9e-06 | 2.633E-06 | 1.185E-06 | 1.900E-06 | 4.000E-06 |
| 14055 | 703 | E703P | B | 2e-06 | 1.9e-06 | 2.7e-06 | 2.200E-06 | 4.359E-07 | 1.900E-06 | 2.700E-06 |
| 14056 | 703 | E703S | B | 3.3e-06 | 5.1e-06 | 2.7e-06 | 3.700E-06 | 1.249E-06 | 2.700E-06 | 5.100E-06 |
| 14057 | 703 | E703T | B | 2.2e-06 | 2.7e-06 | 1.4e-06 | 2.100E-06 | 6.557E-07 | 1.400E-06 | 2.700E-06 |
| 14058 | 703 | E703W | B | 2e-06 | 1.6e-06 | 5e-06 | 2.867E-06 | 1.858E-06 | 1.600E-06 | 5.000E-06 |
| 14059 | 703 | E703Y | B | 2e-06 | 2.4e-06 | 2.3e-06 | 2.233E-06 | 2.082E-07 | 2.000E-06 | 2.400E-06 |
| 14060 | 703 | E703V | B | 2.9e-06 | 2.1e-06 | 3e-06 | 2.667E-06 | 4.933E-07 | 2.100E-06 | 3.000E-06 |
| 14061 | 704 | D704A | B | 3.2e-06 | 3.6e-06 | 2.2e-06 | 3.000E-06 | 7.211E-07 | 2.200E-06 | 3.600E-06 |
| 14062 | 704 | D704R | B | 2.9e-06 | 1.3e-06 | 3.1e-06 | 2.433E-06 | 9.866E-07 | 1.300E-06 | 3.100E-06 |
| 14063 | 704 | D704N | B | 1.6e-06 | 2e-06 | 3.4e-06 | 2.333E-06 | 9.452E-07 | 1.600E-06 | 3.400E-06 |
| 14064 | 704 | D704D | B | 2.3e-06 | 3e-06 | 1.8e-06 | 2.367E-06 | 6.028E-07 | 1.800E-06 | 3.000E-06 |
| 14065 | 704 | D704C | B | 2.7e-06 | 2.4e-06 | 2e-06 | 2.367E-06 | 3.512E-07 | 2.000E-06 | 2.700E-06 |
| 14066 | 704 | D704E | B | 3e-06 | 1.8e-06 | 3.5e-06 | 2.767E-06 | 8.737E-07 | 1.800E-06 | 3.500E-06 |
| 14067 | 704 | D704Q | B | 2.3e-06 | 2.6e-06 | 3.2e-06 | 2.700E-06 | 4.583E-07 | 2.300E-06 | 3.200E-06 |
| 14068 | 704 | D704G | B | 3.5e-06 | 2.8e-06 | 2.8e-06 | 3.033E-06 | 4.041E-07 | 2.800E-06 | 3.500E-06 |
| 14069 | 704 | D704H | B | 2.3e-06 | 3.1e-06 | 2.7e-06 | 2.700E-06 | 4.000E-07 | 2.300E-06 | 3.100E-06 |
| 14070 | 704 | D704I | B | 1.5e-06 | 1.6e-06 | 2.4e-06 | 1.833E-06 | 4.933E-07 | 1.500E-06 | 2.400E-06 |
| 14071 | 704 | D704L | B | 3.1e-06 | 1.7e-06 | 1.6e-06 | 2.133E-06 | 8.386E-07 | 1.600E-06 | 3.100E-06 |
| 14072 | 704 | D704K | B | 2.7e-06 | 2.4e-06 | 3.5e-06 | 2.867E-06 | 5.686E-07 | 2.400E-06 | 3.500E-06 |
| 14073 | 704 | D704M | B | 2e-06 | 2.4e-06 | 2.9e-06 | 2.433E-06 | 4.509E-07 | 2.000E-06 | 2.900E-06 |
| 14074 | 704 | D704F | B | 2.5e-06 | 2.1e-06 | 1.9e-06 | 2.167E-06 | 3.055E-07 | 1.900E-06 | 2.500E-06 |
| 14075 | 704 | D704P | B | 2.7e-06 | 3.7e-06 | 2.7e-06 | 3.033E-06 | 5.774E-07 | 2.700E-06 | 3.700E-06 |
| 14076 | 704 | D704S | B | 3.1e-06 | 2.1e-06 | 1.8e-06 | 2.333E-06 | 6.807E-07 | 1.800E-06 | 3.100E-06 |
| 14077 | 704 | D704T | B | 2.2e-06 | 4.1e-06 | 2.4e-06 | 2.900E-06 | 1.044E-06 | 2.200E-06 | 4.100E-06 |
| 14078 | 704 | D704W | B | 2.6e-06 | 1.5e-06 | 2.1e-06 | 2.067E-06 | 5.508E-07 | 1.500E-06 | 2.600E-06 |
| 14079 | 704 | D704Y | B | 1.7e-06 | 2.2e-06 | 3.6e-06 | 2.500E-06 | 9.849E-07 | 1.700E-06 | 3.600E-06 |
| 14080 | 704 | D704V | B | 2.1e-06 | 1.9e-06 | 1.9e-06 | 1.967E-06 | 1.155E-07 | 1.900E-06 | 2.100E-06 |
| 14081 | 705 | F705A | B | 2.8e-06 | 2.7e-06 | 3e-06 | 2.833E-06 | 1.528E-07 | 2.700E-06 | 3.000E-06 |
| 14082 | 705 | F705R | B | 2.7e-06 | 2.2e-06 | 3.2e-06 | 2.700E-06 | 5.000E-07 | 2.200E-06 | 3.200E-06 |
| 14083 | 705 | F705N | B | 2.2e-06 | 2.6e-06 | 2.6e-06 | 2.467E-06 | 2.309E-07 | 2.200E-06 | 2.600E-06 |
| 14084 | 705 | F705D | B | 2.7e-06 | 3.7e-06 | 3.4e-06 | 3.267E-06 | 5.132E-07 | 2.700E-06 | 3.700E-06 |
| 14085 | 705 | F705C | B | 2.7e-06 | 1.7e-06 | 3.8e-06 | 2.733E-06 | 1.050E-06 | 1.700E-06 | 3.800E-06 |
| 14086 | 705 | F705E | B | 1.1e-06 | 3.2e-06 | 1.9e-06 | 2.067E-06 | 1.060E-06 | 1.100E-06 | 3.200E-06 |
| 14087 | 705 | F705Q | B | 2.9e-06 | 1.8e-06 | 1.7e-06 | 2.133E-06 | 6.658E-07 | 1.700E-06 | 2.900E-06 |
| 14088 | 705 | F705G | B | 2.5e-06 | 1.9e-06 | 3.3e-06 | 2.567E-06 | 7.024E-07 | 1.900E-06 | 3.300E-06 |
| 14089 | 705 | F705H | B | 3.6e-06 | 1.2e-06 | 3.6e-06 | 2.800E-06 | 1.386E-06 | 1.200E-06 | 3.600E-06 |
| 14090 | 705 | F705I | B | 1.4e-06 | 4e-06 | 1.8e-06 | 2.400E-06 | 1.400E-06 | 1.400E-06 | 4.000E-06 |
| 14091 | 705 | F705L | B | 2.3e-06 | 2.3e-06 | 1.4e-06 | 2.000E-06 | 5.196E-07 | 1.400E-06 | 2.300E-06 |
| 14092 | 705 | F705K | B | 2.3e-06 | 1.6e-06 | 5.6e-06 | 3.167E-06 | 2.136E-06 | 1.600E-06 | 5.600E-06 |
| 14093 | 705 | F705M | B | 3.3e-06 | 3e-06 | 2.2e-06 | 2.833E-06 | 5.686E-07 | 2.200E-06 | 3.300E-06 |
| 14094 | 705 | F705F | B | 2.3e-06 | 1.4e-06 | 2.4e-06 | 2.033E-06 | 5.508E-07 | 1.400E-06 | 2.400E-06 |
| 14095 | 705 | F705P | B | 1.5e-06 | 2.5e-06 | 3.6e-06 | 2.533E-06 | 1.050E-06 | 1.500E-06 | 3.600E-06 |
| 14096 | 705 | F705S | B | 2.7e-06 | 2.7e-06 | 1.9e-06 | 2.433E-06 | 4.619E-07 | 1.900E-06 | 2.700E-06 |
| 14097 | 705 | F705T | B | 2.2e-06 | 1.9e-06 | 2.5e-06 | 2.200E-06 | 3.000E-07 | 1.900E-06 | 2.500E-06 |
| 14098 | 705 | F705W | B | 1.6e-06 | 2.5e-06 | 3.9e-06 | 2.667E-06 | 1.159E-06 | 1.600E-06 | 3.900E-06 |
| 14099 | 705 | F705Y | B | 2.5e-06 | 1.6e-06 | 2.4e-06 | 2.167E-06 | 4.933E-07 | 1.600E-06 | 2.500E-06 |
| 14100 | 705 | F705V | B | 2.4e-06 | 2.1e-06 | 2.4e-06 | 2.300E-06 | 1.732E-07 | 2.100E-06 | 2.400E-06 |
| 14101 | 706 | A706A | B | 2.2e-06 | 2.4e-06 | 1.9e-06 | 2.167E-06 | 2.517E-07 | 1.900E-06 | 2.400E-06 |
| 14102 | 706 | A706R | B | 3.2e-06 | 2e-06 | 2.1e-06 | 2.433E-06 | 6.658E-07 | 2.000E-06 | 3.200E-06 |
| 14103 | 706 | A706N | B | 1.5e-06 | 1.8e-06 | 2.7e-06 | 2.000E-06 | 6.245E-07 | 1.500E-06 | 2.700E-06 |
| 14104 | 706 | A706D | B | 2.3e-06 | 3.9e-06 | 2.5e-06 | 2.900E-06 | 8.718E-07 | 2.300E-06 | 3.900E-06 |
| 14105 | 706 | A706C | B | 2.7e-06 | 2.2e-06 | 2.3e-06 | 2.400E-06 | 2.646E-07 | 2.200E-06 | 2.700E-06 |
| 14106 | 706 | A706E | B | 2.5e-06 | 1.9e-06 | 2.1e-06 | 2.167E-06 | 3.055E-07 | 1.900E-06 | 2.500E-06 |
| 14107 | 706 | A706Q | B | 3.1e-06 | 2e-06 | 2.4e-06 | 2.500E-06 | 5.568E-07 | 2.000E-06 | 3.100E-06 |
| 14108 | 706 | A706G | B | 1.6e-06 | 2.1e-06 | 2.2e-06 | 1.967E-06 | 3.215E-07 | 1.600E-06 | 2.200E-06 |
| 14109 | 706 | A706H | B | 2.2e-06 | 2.7e-06 | 1.5e-06 | 2.133E-06 | 6.028E-07 | 1.500E-06 | 2.700E-06 |

| | | | | | | | | | | |
|---|---|---|---|---|---|---|---|---|---|---|
| 14110 | 706 | A706I | B | 2.7e-06 | 2.1e-06 | 1.6e-06 | 2.133E-06 | 5.508E-07 | 1.600E-06 | 2.700E-06 |
| 14111 | 706 | A706L | B | 1.9e-06 | 2.8e-06 | 2.7e-06 | 2.467E-06 | 4.933E-07 | 1.900E-06 | 2.800E-06 |
| 14112 | 706 | A706K | B | 2.4e-06 | 2.1e-06 | 1.9e-06 | 2.133E-06 | 2.517E-07 | 1.900E-06 | 2.400E-06 |
| 14113 | 706 | A706M | B | 3.7e-06 | 2.2e-06 | 5.6e-06 | 3.833E-06 | 1.704E-06 | 2.200E-06 | 5.600E-06 |
| 14114 | 706 | A706F | B | 3.2e-06 | 1.2e-06 | 1.6e-06 | 2.000E-06 | 1.058E-06 | 1.200E-06 | 3.200E-06 |
| 14115 | 706 | A706P | B | 2.8e-06 | 8.4e-07 | 2.5e-06 | 2.047E-06 | 1.056E-06 | 8.400E-07 | 2.800E-06 |
| 14116 | 706 | A706S | B | 3.7e-06 | 1.9e-06 | 3.3e-06 | 2.967E-06 | 9.452E-07 | 1.900E-06 | 3.700E-06 |
| 14117 | 706 | A706T | B | 2.3e-06 | 2.2e-06 | 1.4e-06 | 1.967E-06 | 4.933E-07 | 1.400E-06 | 2.300E-06 |
| 14118 | 706 | A706W | B | 2.8e-06 | 1.7e-06 | 1.9e-06 | 2.133E-06 | 5.859E-07 | 1.700E-06 | 2.800E-06 |
| 14119 | 706 | A706Y | B | 2.2e-06 | 1.7e-06 | 4.4e-06 | 2.767E-06 | 1.436E-06 | 1.700E-06 | 4.400E-06 |
| 14120 | 706 | A706V | B | 2.8e-06 | 2.4e-06 | 4.4e-06 | 3.200E-06 | 1.058E-06 | 2.400E-06 | 4.400E-06 |
| 14121 | 707 | T707A | B | 2.3e-06 | 1.9e-06 | 2.1e-06 | 2.100E-06 | 2.000E-07 | 1.900E-06 | 2.300E-06 |
| 14122 | 707 | T707R | B | 2.7e-06 | 3e-06 | 2.3e-06 | 2.667E-06 | 3.512E-07 | 2.300E-06 | 3.000E-06 |
| 14123 | 707 | T707N | B | 2.5e-06 | 2.8e-06 | 2.8e-06 | 2.700E-06 | 1.732E-07 | 2.500E-06 | 2.800E-06 |
| 14124 | 707 | T707D | B | 2.7e-06 | 2.7e-06 | 2.2e-06 | 2.533E-06 | 2.887E-07 | 2.200E-06 | 2.700E-06 |
| 14125 | 707 | T707C | B | 3e-06 | 3e-06 | 7.9e-07 | 2.263E-06 | 1.276E-06 | 7.900E-07 | 3.000E-06 |
| 14126 | 707 | T707E | B | 1.6e-06 | 3.6e-06 | 3.1e-06 | 2.767E-06 | 1.041E-06 | 1.600E-06 | 3.600E-06 |
| 14127 | 707 | T707Q | B | 2.6e-06 | 2.4e-06 | 3e-06 | 2.667E-06 | 3.055E-07 | 2.400E-06 | 3.000E-06 |
| 14128 | 707 | T707G | B | 2.2e-06 | 2.4e-06 | 2.4e-06 | 2.333E-06 | 1.155E-07 | 2.200E-06 | 2.400E-06 |
| 14129 | 707 | T707H | B | 4.5e-06 | 3.2e-06 | 3.2e-06 | 3.633E-06 | 7.506E-07 | 3.200E-06 | 4.500E-06 |
| 14130 | 707 | T707I | B | 1.7e-06 | 2.6e-06 | 1.3e-06 | 1.867E-06 | 6.658E-07 | 1.300E-06 | 2.600E-06 |
| 14131 | 707 | T707L | B | 3.1e-06 | 3.5e-06 | 2.9e-06 | 3.167E-06 | 3.055E-07 | 2.900E-06 | 3.500E-06 |
| 14132 | 707 | T707K | B | 3.8e-06 | 2.2e-06 | 2.5e-06 | 2.833E-06 | 8.505E-07 | 2.200E-06 | 3.800E-06 |
| 14133 | 707 | T707M | B | 1.5e-06 | 3e-06 | 2.9e-06 | 2.467E-06 | 8.386E-07 | 1.500E-06 | 3.000E-06 |
| 14134 | 707 | T707F | B | 3.4e-06 | 3.1e-06 | 2.9e-06 | 3.133E-06 | 2.517E-07 | 2.900E-06 | 3.400E-06 |
| 14135 | 707 | T707P | B | 3.8e-06 | 4.9e-06 | 2.7e-06 | 3.800E-06 | 1.100E-06 | 2.700E-06 | 4.900E-06 |
| 14136 | 707 | T707S | B | 1.9e-06 | 2.2e-06 | 1.8e-06 | 1.967E-06 | 2.082E-07 | 1.800E-06 | 2.200E-06 |
| 14137 | 707 | T707T | B | 3e-06 | 3e-06 | 4.7e-06 | 3.567E-06 | 9.815E-07 | 3.000E-06 | 4.700E-06 |
| 14138 | 707 | T707W | B | 1.9e-06 | 2.7e-06 | 2.8e-06 | 2.467E-06 | 4.933E-07 | 1.900E-06 | 2.800E-06 |
| 14139 | 707 | T707Y | B | 4e-06 | 2.7e-06 | 1.8e-06 | 2.833E-06 | 1.106E-06 | 1.800E-06 | 4.000E-06 |
| 14140 | 707 | T707V | B | 1.6e-06 | 3.4e-06 | 3.7e-06 | 2.900E-06 | 1.136E-06 | 1.600E-06 | 3.700E-06 |
| 14141 | 708 | Y708A | B | 2.7e-06 | 1.6e-06 | 1.9e-06 | 2.067E-06 | 5.686E-07 | 1.600E-06 | 2.700E-06 |
| 14142 | 708 | Y708R | B | 2.2e-06 | 2.3e-06 | 2.7e-06 | 2.400E-06 | 2.646E-07 | 2.200E-06 | 2.700E-06 |
| 14143 | 708 | Y708N | B | 2.4e-06 | 3.5e-06 | 2.6e-06 | 2.833E-06 | 5.859E-07 | 2.400E-06 | 3.500E-06 |
| 14144 | 708 | Y708D | B | 2.8e-06 | 1.7e-06 | 2.9e-06 | 2.467E-06 | 6.658E-07 | 1.700E-06 | 2.900E-06 |
| 14145 | 708 | Y708C | B | 1.8e-06 | 2.1e-06 | 1.9e-06 | 1.933E-06 | 1.528E-07 | 1.800E-06 | 2.100E-06 |
| 14146 | 708 | Y708E | B | 2.2e-06 | 2e-06 | 3.8e-06 | 2.667E-06 | 9.866E-07 | 2.000E-06 | 3.800E-06 |
| 14147 | 708 | Y708Q | B | 2e-06 | 3e-06 | 1.9e-06 | 2.300E-06 | 6.083E-07 | 1.900E-06 | 3.000E-06 |
| 14148 | 708 | Y708G | B | 3.1e-06 | 4.1e-06 | 2.7e-06 | 3.300E-06 | 7.211E-07 | 2.700E-06 | 4.100E-06 |
| 14149 | 708 | Y708H | B | 1.9e-06 | 2.6e-06 | 2.8e-06 | 2.433E-06 | 4.726E-07 | 1.900E-06 | 2.800E-06 |
| 14150 | 708 | Y708I | B | 1.3e-06 | 1.7e-06 | 3e-06 | 2.000E-06 | 8.888E-07 | 1.300E-06 | 3.000E-06 |
| 14151 | 708 | Y708L | B | 1.9e-06 | 2.3e-06 | 3.3e-06 | 2.500E-06 | 7.211E-07 | 1.900E-06 | 3.300E-06 |
| 14152 | 708 | Y708K | B | 1.9e-06 | 1.6e-06 | 2.7e-06 | 2.067E-06 | 5.686E-07 | 1.600E-06 | 2.700E-06 |
| 14153 | 708 | Y708M | B | 2.5e-06 | 1.6e-06 | 2.3e-06 | 2.133E-06 | 4.726E-07 | 1.600E-06 | 2.500E-06 |
| 14154 | 708 | Y708F | B | 2.1e-06 | 4.5e-06 | 3.2e-06 | 3.267E-06 | 1.201E-06 | 2.100E-06 | 4.500E-06 |
| 14155 | 708 | Y708P | B | 1.7e-06 | 2.3e-06 | 1.6e-06 | 1.867E-06 | 3.786E-07 | 1.600E-06 | 2.300E-06 |
| 14156 | 708 | Y708S | B | 5.1e-06 | 3.5e-06 | 3.9e-06 | 4.167E-06 | 8.327E-07 | 3.500E-06 | 5.100E-06 |
| 14157 | 708 | Y708T | B | 2.9e-06 | 1.9e-06 | 2.5e-06 | 2.433E-06 | 5.033E-07 | 1.900E-06 | 2.900E-06 |
| 14158 | 708 | Y708W | B | 3.2e-06 | 8.5e-07 | 3.3e-06 | 2.450E-06 | 1.387E-06 | 8.500E-07 | 3.300E-06 |
| 14159 | 708 | Y708Y | B | 3.3e-06 | 2.5e-06 | 1.9e-06 | 2.567E-06 | 7.024E-07 | 1.900E-06 | 3.300E-06 |
| 14160 | 708 | Y708V | B | 4e-06 | 1.9e-06 | 1.1e-06 | 2.333E-06 | 1.498E-06 | 1.100E-06 | 4.000E-06 |
| 14161 | 709 | Y709A | B | 4.5e-06 | 3e-06 | 1.9e-06 | 3.133E-06 | 1.305E-06 | 1.900E-06 | 4.500E-06 |
| 14162 | 709 | Y709R | B | 2.4e-06 | 2.8e-06 | 2.7e-06 | 2.633E-06 | 2.082E-07 | 2.400E-06 | 2.800E-06 |
| 14163 | 709 | Y709N | B | 2.1e-06 | 3.1e-06 | 1.9e-06 | 2.367E-06 | 6.429E-07 | 1.900E-06 | 3.100E-06 |
| 14164 | 709 | Y709D | B | 3.9e-06 | 1.8e-06 | 2.5e-06 | 2.733E-06 | 1.069E-06 | 1.800E-06 | 3.900E-06 |
| 14165 | 709 | Y709C | B | 1.4e-06 | 2e-06 | 3.7e-06 | 2.367E-06 | 1.193E-06 | 1.400E-06 | 3.700E-06 |
| 14166 | 709 | Y709E | B | 1.4e-06 | 1.7e-06 | 2.2e-06 | 1.767E-06 | 4.041E-07 | 1.400E-06 | 2.200E-06 |
| 14167 | 709 | Y709Q | B | 1.6e-06 | 3.6e-06 | 2e-06 | 2.400E-06 | 1.058E-06 | 1.600E-06 | 3.600E-06 |
| 14168 | 709 | Y709G | B | 3.4e-06 | 1.5e-06 | 2.6e-06 | 2.500E-06 | 9.539E-07 | 1.500E-06 | 3.400E-06 |
| 14169 | 709 | Y709H | B | 2.8e-06 | 2.7e-06 | 2.9e-06 | 2.800E-06 | 1.000E-07 | 2.700E-06 | 2.900E-06 |
| 14170 | 709 | Y709I | B | 3e-06 | 3.8e-06 | 1.6e-06 | 2.800E-06 | 1.114E-06 | 1.600E-06 | 3.800E-06 |
| 14171 | 709 | Y709L | B | 2.9e-06 | 3.1e-06 | 1.5e-06 | 2.500E-06 | 8.718E-07 | 1.500E-06 | 3.100E-06 |
| 14172 | 709 | Y709K | B | 3e-06 | 2.4e-06 | 2.2e-06 | 2.533E-06 | 4.163E-07 | 2.200E-06 | 3.000E-06 |
| 14173 | 709 | Y709M | B | 1.2e-06 | 2e-06 | 2.1e-06 | 1.767E-06 | 4.933E-07 | 1.200E-06 | 2.100E-06 |
| 14174 | 709 | Y709F | B | 2.6e-06 | 1.7e-06 | 2.5e-06 | 2.267E-06 | 4.933E-07 | 1.700E-06 | 2.600E-06 |
| 14175 | 709 | Y709P | B | 4.4e-06 | 2.3e-06 | 5.2e-06 | 3.967E-06 | 1.498E-06 | 2.300E-06 | 5.200E-06 |
| 14176 | 709 | Y709S | B | 3.8e-06 | 2.8e-06 | 2.3e-06 | 2.967E-06 | 7.638E-07 | 2.300E-06 | 3.800E-06 |
| 14177 | 709 | Y709T | B | 2.6e-06 | 2.7e-06 | 1.9e-06 | 2.400E-06 | 4.359E-07 | 1.900E-06 | 2.700E-06 |
| 14178 | 709 | Y709W | B | 2.5e-06 | 1.6e-06 | 2.5e-06 | 2.200E-06 | 5.196E-07 | 1.600E-06 | 2.500E-06 |
| 14179 | 709 | Y709Y | B | 1.5e-06 | 3.5e-06 | 1.9e-06 | 2.300E-06 | 1.058E-06 | 1.500E-06 | 3.500E-06 |
| 14180 | 709 | Y709V | B | 2.8e-06 | 1.5e-06 | 2.2e-06 | 2.167E-06 | 6.506E-07 | 1.500E-06 | 2.800E-06 |
| 14181 | 710 | C710A | B | 2.5e-06 | 2.3e-06 | 2.5e-06 | 2.433E-06 | 1.155E-07 | 2.300E-06 | 2.500E-06 |
| 14182 | 710 | C710R | B | 2.5e-06 | 2e-06 | 4.3e-06 | 2.933E-06 | 1.210E-06 | 2.000E-06 | 4.300E-06 |
| 14183 | 710 | C710N | B | 2.9e-06 | 2.6e-06 | 2.4e-06 | 2.633E-06 | 2.517E-07 | 2.400E-06 | 2.900E-06 |
| 14184 | 710 | C710D | B | 2.6e-06 | 1.3e-06 | 1.9e-06 | 1.933E-06 | 6.506E-07 | 1.300E-06 | 2.600E-06 |
| 14185 | 710 | C710C | B | 2.3e-06 | 2.5e-06 | 2.6e-06 | 2.467E-06 | 1.528E-07 | 2.300E-06 | 2.600E-06 |
| 14186 | 710 | C710E | B | 1e-06 | 2.1e-06 | 3e-06 | 2.033E-06 | 1.002E-06 | 1.000E-06 | 3.000E-06 |
| 14187 | 710 | C710Q | B | 1.8e-06 | 2.2e-06 | 3e-06 | 2.333E-06 | 6.110E-07 | 1.800E-06 | 3.000E-06 |
| 14188 | 710 | C710G | B | 2.3e-06 | 1.5e-06 | 2e-06 | 1.933E-06 | 4.041E-07 | 1.500E-06 | 2.300E-06 |
| 14189 | 710 | C710H | B | 3.4e-06 | 2.3e-06 | 2e-06 | 2.567E-06 | 7.371E-07 | 2.000E-06 | 3.400E-06 |
| 14190 | 710 | C710I | B | 3e-06 | 2e-06 | 2.2e-06 | 2.400E-06 | 5.292E-07 | 2.000E-06 | 3.000E-06 |
| 14191 | 710 | C710L | B | 2.1e-06 | 2e-06 | 2.5e-06 | 2.200E-06 | 2.646E-07 | 2.000E-06 | 2.500E-06 |
| 14192 | 710 | C710K | B | 1.8e-06 | 2.7e-06 | 1.8e-06 | 2.100E-06 | 5.196E-07 | 1.800E-06 | 2.700E-06 |
| 14193 | 710 | C710M | B | 2.3e-06 | 2.2e-06 | 3.9e-06 | 2.800E-06 | 9.539E-07 | 2.200E-06 | 3.900E-06 |
| 14194 | 710 | C710F | B | 2.9e-06 | 1.2e-06 | 2.9e-06 | 2.333E-06 | 9.815E-07 | 1.200E-06 | 2.900E-06 |

| | | | | | | | | | | |
|---|---|---|---|---|---|---|---|---|---|---|
| 14195 | 710 | C710P | B | 2e-06 | 5.7e-06 | 3.2e-06 | 3.633E-06 | 1.888E-06 | 2.000E-06 | 5.700E-06 |
| 14196 | 710 | C710S | B | 1.4e-06 | 3e-06 | 3.1e-06 | 2.500E-06 | 9.539E-07 | 1.400E-06 | 3.100E-06 |
| 14197 | 710 | C710T | B | 2.1e-06 | 3.2e-06 | 1.6e-06 | 2.300E-06 | 8.185E-07 | 1.600E-06 | 3.200E-06 |
| 14198 | 710 | C710W | B | 1.1e-06 | 1.6e-06 | 3e-06 | 1.900E-06 | 9.849E-07 | 1.100E-06 | 3.000E-06 |
| 14199 | 710 | C710Y | B | 2.2e-06 | 2.4e-06 | 2.3e-06 | 2.300E-06 | 1.000E-07 | 2.200E-06 | 2.400E-06 |
| 14200 | 710 | C710V | B | 2.8e-06 | 4e-06 | 2.6e-06 | 3.133E-06 | 7.572E-07 | 2.600E-06 | 4.000E-06 |
| 14201 | 711 | Q711A | B | 2.9e-06 | 2.1e-06 | 2.4e-06 | 2.467E-06 | 4.041E-07 | 2.100E-06 | 2.900E-06 |
| 14202 | 711 | Q711R | B | 1.9e-06 | 3e-06 | 2.1e-06 | 2.333E-06 | 5.859E-07 | 1.900E-06 | 3.000E-06 |
| 14203 | 711 | Q711N | B | 3.8e-06 | 3.4e-06 | 3.8e-06 | 3.667E-06 | 2.309E-07 | 3.400E-06 | 3.800E-06 |
| 14204 | 711 | Q711D | B | 3.9e-06 | 2.4e-06 | 4e-06 | 3.433E-06 | 8.963E-07 | 2.400E-06 | 4.000E-06 |
| 14205 | 711 | Q711C | B | 2e-06 | 1.3e-06 | 2e-06 | 1.767E-06 | 4.041E-07 | 1.300E-06 | 2.000E-06 |
| 14206 | 711 | Q711E | B | 4.4e-06 | 1.8e-06 | 1.4e-06 | 2.533E-06 | 1.629E-06 | 1.400E-06 | 4.400E-06 |
| 14207 | 711 | Q711Q | B | 3.9e-06 | 1.4e-06 | 2.9e-06 | 2.733E-06 | 1.258E-06 | 1.400E-06 | 3.900E-06 |
| 14208 | 711 | Q711G | B | 1.8e-06 | 1.5e-06 | 1.5e-06 | 1.600E-06 | 1.732E-07 | 1.500E-06 | 1.800E-06 |
| 14209 | 711 | Q711H | B | 3.8e-06 | 3.5e-06 | 8.5e-07 | 2.717E-06 | 1.624E-06 | 8.500E-07 | 3.800E-06 |
| 14210 | 711 | Q711I | B | 2.9e-06 | 2.9e-06 | 1.9e-06 | 2.567E-06 | 5.774E-07 | 1.900E-06 | 2.900E-06 |
| 14211 | 711 | Q711L | B | 1.6e-06 | 3.5e-06 | 1.7e-06 | 2.267E-06 | 1.069E-06 | 1.600E-06 | 3.500E-06 |
| 14212 | 711 | Q711K | B | 2.1e-06 | 1.5e-06 | 2.2e-06 | 1.933E-06 | 3.786E-07 | 1.500E-06 | 2.200E-06 |
| 14213 | 711 | Q711M | B | 2.5e-06 | 4.5e-06 | 4.8e-06 | 3.933E-06 | 1.250E-06 | 2.500E-06 | 4.800E-06 |
| 14214 | 711 | Q711F | B | 2.5e-06 | 3e-06 | 1.6e-06 | 2.367E-06 | 7.095E-07 | 1.600E-06 | 3.000E-06 |
| 14215 | 711 | Q711P | B | 3.3e-06 | 2.6e-06 | 3e-06 | 2.967E-06 | 3.512E-07 | 2.600E-06 | 3.300E-06 |
| 14216 | 711 | Q711S | B | 3.1e-06 | 2.4e-06 | 2.7e-06 | 2.733E-06 | 3.512E-07 | 2.400E-06 | 3.100E-06 |
| 14217 | 711 | Q711T | B | 2.7e-06 | 2.2e-06 | 1.1e-06 | 2.000E-06 | 8.185E-07 | 1.100E-06 | 2.700E-06 |
| 14218 | 711 | Q711W | B | 2e-06 | 2.2e-06 | 3.2e-06 | 2.467E-06 | 6.429E-07 | 2.000E-06 | 3.200E-06 |
| 14219 | 711 | Q711Y | B | 1.7e-06 | 2.1e-06 | 1.6e-06 | 1.800E-06 | 2.646E-07 | 1.600E-06 | 2.100E-06 |
| 14220 | 711 | Q711V | B | 2.7e-06 | 3.2e-06 | 2e-06 | 2.633E-06 | 6.028E-07 | 2.000E-06 | 3.200E-06 |
| 14221 | 712 | Q712A | B | 4.9e-06 | 1e-06 | 3.1e-06 | 3.000E-06 | 1.952E-06 | 1.000E-06 | 4.900E-06 |
| 14222 | 712 | Q712R | B | 2.5e-06 | 2.3e-06 | 3.6e-06 | 2.800E-06 | 7.000E-07 | 2.300E-06 | 3.600E-06 |
| 14223 | 712 | Q712N | B | 2.3e-06 | 2.8e-06 | 2.8e-06 | 2.633E-06 | 2.887E-07 | 2.300E-06 | 2.800E-06 |
| 14224 | 712 | Q712D | B | 6.6e-06 | 3.3e-06 | 1.2e-06 | 3.700E-06 | 2.722E-06 | 1.200E-06 | 6.600E-06 |
| 14225 | 712 | Q712C | B | 1.8e-06 | 1.7e-06 | 2e-06 | 1.833E-06 | 1.528E-07 | 1.700E-06 | 2.000E-06 |
| 14226 | 712 | Q712E | B | 2.6e-06 | 1.1e-06 | 2.2e-06 | 1.967E-06 | 7.767E-07 | 1.100E-06 | 2.600E-06 |
| 14227 | 712 | Q712Q | B | 1.9e-06 | 2.6e-06 | 2.1e-06 | 2.200E-06 | 3.606E-07 | 1.900E-06 | 2.600E-06 |
| 14228 | 712 | Q712G | B | 1.9e-06 | 2.1e-06 | 2.4e-06 | 2.133E-06 | 2.517E-07 | 1.900E-06 | 2.400E-06 |
| 14229 | 712 | Q712H | B | 1.4e-06 | 2.3e-06 | 3.1e-06 | 2.267E-06 | 8.505E-07 | 1.400E-06 | 3.100E-06 |
| 14230 | 712 | Q712I | B | 2.8e-06 | 3.7e-06 | 1.2e-06 | 2.567E-06 | 1.266E-06 | 1.200E-06 | 3.700E-06 |
| 14231 | 712 | Q712L | B | 2.5e-06 | 2.4e-06 | 1.6e-06 | 2.167E-06 | 4.933E-07 | 1.600E-06 | 2.500E-06 |
| 14232 | 712 | Q712K | B | 4.1e-06 | 2.9e-06 | 2.8e-06 | 3.267E-06 | 7.234E-07 | 2.800E-06 | 4.100E-06 |
| 14233 | 712 | Q712M | B | 1.2e-06 | 2.7e-06 | 1.9e-06 | 1.933E-06 | 7.506E-07 | 1.200E-06 | 2.700E-06 |
| 14234 | 712 | Q712F | B | 6.6e-06 | 1.6e-06 | 2.4e-06 | 3.533E-06 | 2.686E-06 | 1.600E-06 | 6.600E-06 |
| 14235 | 712 | Q712P | B | 1.9e-06 | 2.5e-06 | 1.1e-06 | 1.833E-06 | 7.024E-07 | 1.100E-06 | 2.500E-06 |
| 14236 | 712 | Q712S | B | 1.4e-06 | 3.1e-06 | 1.5e-06 | 2.000E-06 | 9.539E-07 | 1.400E-06 | 3.100E-06 |
| 14237 | 712 | Q712T | B | 3.2e-06 | 2e-06 | 2.6e-06 | 2.600E-06 | 6.000E-07 | 2.000E-06 | 3.200E-06 |
| 14238 | 712 | Q712W | B | 3e-06 | 4.1e-06 | 2.3e-06 | 3.133E-06 | 9.074E-07 | 2.300E-06 | 4.100E-06 |
| 14239 | 712 | Q712Y | B | 2.3e-06 | 2.1e-06 | 2.1e-06 | 2.167E-06 | 1.155E-07 | 2.100E-06 | 2.300E-06 |
| 14240 | 712 | Q712V | B | 2.2e-06 | 2e-06 | 3e-06 | 2.400E-06 | 5.292E-07 | 2.000E-06 | 3.000E-06 |
| 14241 | 713 | Y713A | B | 4.2e-06 | 1.8e-06 | 2.8e-06 | 2.933E-06 | 1.206E-06 | 1.800E-06 | 4.200E-06 |
| 14242 | 713 | Y713R | B | 6.4e-06 | 3.9e-06 | 3.1e-06 | 4.467E-06 | 1.721E-06 | 3.100E-06 | 6.400E-06 |
| 14243 | 713 | Y713N | B | 4.3e-06 | 4.8e-06 | 2.5e-06 | 3.867E-06 | 1.210E-06 | 2.500E-06 | 4.800E-06 |
| 14244 | 713 | Y713D | B | 2.3e-06 | 2.8e-06 | 2.4e-06 | 2.500E-06 | 2.646E-07 | 2.300E-06 | 2.800E-06 |
| 14245 | 713 | Y713C | B | 1.7e-06 | 1.7e-06 | 2.7e-06 | 2.033E-06 | 5.774E-07 | 1.700E-06 | 2.700E-06 |
| 14246 | 713 | Y713E | B | 2.2e-06 | 2.5e-06 | 2.8e-06 | 2.500E-06 | 3.000E-07 | 2.200E-06 | 2.800E-06 |
| 14247 | 713 | Y713Q | B | 9.6e-07 | 3.6e-06 | 2.5e-06 | 2.353E-06 | 1.326E-06 | 9.600E-07 | 3.600E-06 |
| 14248 | 713 | Y713G | B | 2.3e-06 | 2.8e-06 | 1.8e-06 | 2.300E-06 | 5.000E-07 | 1.800E-06 | 2.800E-06 |
| 14249 | 713 | Y713H | B | 2.9e-06 | 1.5e-06 | 4.2e-06 | 2.867E-06 | 1.350E-06 | 1.500E-06 | 4.200E-06 |
| 14250 | 713 | Y713I | B | 2.7e-06 | 3e-06 | 2.5e-06 | 2.733E-06 | 2.517E-07 | 2.500E-06 | 3.000E-06 |
| 14251 | 713 | Y713L | B | 1.6e-06 | 1.2e-06 | 1.9e-06 | 1.567E-06 | 3.512E-07 | 1.200E-06 | 1.900E-06 |
| 14252 | 713 | Y713K | B | 2.3e-06 | 2.6e-06 | 2.5e-06 | 2.467E-06 | 1.528E-07 | 2.300E-06 | 2.600E-06 |
| 14253 | 713 | Y713M | B | 2e-06 | 2.6e-06 | 2e-06 | 2.200E-06 | 3.464E-07 | 2.000E-06 | 2.600E-06 |
| 14254 | 713 | Y713F | B | 1.7e-06 | 3.8e-06 | 2e-06 | 2.500E-06 | 1.136E-06 | 1.700E-06 | 3.800E-06 |
| 14255 | 713 | Y713P | B | 3e-06 | 2.9e-06 | 3.2e-06 | 3.033E-06 | 1.528E-07 | 2.900E-06 | 3.200E-06 |
| 14256 | 713 | Y713S | B | 4.3e-06 | 2.2e-06 | 2.5e-06 | 3.000E-06 | 1.136E-06 | 2.200E-06 | 4.300E-06 |
| 14257 | 713 | Y713T | B | 8.1e-07 | 3.8e-06 | 2e-06 | 2.203E-06 | 1.505E-06 | 8.100E-07 | 3.800E-06 |
| 14258 | 713 | Y713W | B | 3.8e-06 | 3.1e-06 | 2.8e-06 | 3.233E-06 | 5.132E-07 | 2.800E-06 | 3.800E-06 |
| 14259 | 713 | Y713Y | B | 2.4e-06 | 2.6e-06 | 1.8e-06 | 2.267E-06 | 4.163E-07 | 1.800E-06 | 2.600E-06 |
| 14260 | 713 | Y713V | B | 2.9e-06 | 2.1e-06 | 1.8e-06 | 2.267E-06 | 5.686E-07 | 1.800E-06 | 2.900E-06 |
| 14261 | 714 | Y714A | B | 2.2e-06 | 4e-06 | 3.2e-06 | 3.133E-06 | 9.018E-07 | 2.200E-06 | 4.000E-06 |
| 14262 | 714 | Y714R | B | 3e-06 | 3.2e-06 | 2.1e-06 | 2.767E-06 | 5.859E-07 | 2.100E-06 | 3.200E-06 |
| 14263 | 714 | Y714N | B | 3.4e-06 | 3.5e-06 | 2.2e-06 | 3.033E-06 | 7.234E-07 | 2.200E-06 | 3.500E-06 |
| 14264 | 714 | Y714D | B | 2.5e-06 | 2.6e-06 | 1.2e-06 | 2.100E-06 | 7.810E-07 | 1.200E-06 | 2.600E-06 |
| 14265 | 714 | Y714C | B | 2.1e-06 | 2.8e-06 | 2.4e-06 | 2.433E-06 | 3.512E-07 | 2.100E-06 | 2.800E-06 |
| 14266 | 714 | Y714E | B | 5e-06 | 2.2e-06 | 2e-06 | 3.067E-06 | 1.677E-06 | 2.000E-06 | 5.000E-06 |
| 14267 | 714 | Y714Q | B | 3.1e-06 | 3.5e-06 | 1.8e-06 | 2.800E-06 | 8.888E-07 | 1.800E-06 | 3.500E-06 |
| 14268 | 714 | Y714G | B | 2.1e-06 | 2.4e-06 | 1.3e-06 | 1.933E-06 | 5.686E-07 | 1.300E-06 | 2.400E-06 |
| 14269 | 714 | Y714H | B | 2.1e-06 | 2.9e-06 | 3.7e-06 | 2.900E-06 | 8.000E-07 | 2.100E-06 | 3.700E-06 |
| 14270 | 714 | Y714I | B | 5.3e-06 | 3e-06 | 2.8e-06 | 3.700E-06 | 1.389E-06 | 2.800E-06 | 5.300E-06 |
| 14271 | 714 | Y714L | B | 1.9e-06 | 2.3e-06 | 2.2e-06 | 2.133E-06 | 2.082E-07 | 1.900E-06 | 2.300E-06 |
| 14272 | 714 | Y714K | B | 3.1e-06 | 2.5e-06 | 2.2e-06 | 2.600E-06 | 4.583E-07 | 2.200E-06 | 3.100E-06 |
| 14273 | 714 | Y714M | B | 3.1e-06 | 2.5e-06 | 3.1e-06 | 2.900E-06 | 3.464E-07 | 2.500E-06 | 3.100E-06 |
| 14274 | 714 | Y714F | B | 4.1e-06 | 2.5e-06 | 2.6e-06 | 3.067E-06 | 8.963E-07 | 2.500E-06 | 4.100E-06 |
| 14275 | 714 | Y714P | B | 3.3e-06 | 2.2e-06 | 2.4e-06 | 2.633E-06 | 5.859E-07 | 2.200E-06 | 3.300E-06 |
| 14276 | 714 | Y714S | B | 1.1e-06 | 2.1e-06 | 2.2e-06 | 1.800E-06 | 6.083E-07 | 1.100E-06 | 2.200E-06 |
| 14277 | 714 | Y714T | B | 2.4e-06 | 1.4e-06 | 2.3e-06 | 2.033E-06 | 5.508E-07 | 1.400E-06 | 2.400E-06 |
| 14278 | 714 | Y714W | B | 1.1e-06 | 1.8e-06 | 2.9e-06 | 1.933E-06 | 9.074E-07 | 1.100E-06 | 2.900E-06 |
| 14279 | 714 | Y714Y | B | 1.6e-06 | 2.3e-06 | 1.8e-06 | 1.900E-06 | 3.606E-07 | 1.600E-06 | 2.300E-06 |

| | | | | | | | | | | |
|---|---|---|---|---|---|---|---|---|---|---|
| 14280 | 714 | Y714V | B | 1.8e-06 | 2.6e-06 | 2.1e-06 | 2.167E-06 | 4.041E-07 | 1.800E-06 | 2.600E-06 |
| 14281 | 715 | I715A | B | 3.6e-06 | 3.8e-06 | 2.7e-06 | 3.367E-06 | 5.859E-07 | 2.700E-06 | 3.800E-06 |
| 14282 | 715 | I715R | B | 1.8e-06 | 2.2e-06 | 2.3e-06 | 2.100E-06 | 2.646E-07 | 1.800E-06 | 2.300E-06 |
| 14283 | 715 | I715N | B | 2e-06 | 2.6e-06 | 2.3e-06 | 2.300E-06 | 3.000E-07 | 2.000E-06 | 2.600E-06 |
| 14284 | 715 | I715D | B | 2.6e-06 | 2e-06 | 3.6e-06 | 2.733E-06 | 8.083E-07 | 2.000E-06 | 3.600E-06 |
| 14285 | 715 | I715C | B | 4e-06 | 2.1e-06 | 2.6e-06 | 2.900E-06 | 9.849E-07 | 2.100E-06 | 4.000E-06 |
| 14286 | 715 | I715E | B | 2.3e-06 | 1.7e-06 | 1.2e-06 | 1.733E-06 | 5.508E-07 | 1.200E-06 | 2.300E-06 |
| 14287 | 715 | I715Q | B | 2.1e-06 | 1.1e-06 | 2.7e-06 | 1.967E-06 | 8.083E-07 | 1.100E-06 | 2.700E-06 |
| 14288 | 715 | I715G | B | 3.1e-06 | 4.1e-06 | 3.1e-06 | 3.433E-06 | 5.774E-07 | 3.100E-06 | 4.100E-06 |
| 14289 | 715 | I715H | B | 2.6e-06 | 2.4e-06 | 1.7e-06 | 2.233E-06 | 4.726E-07 | 1.700E-06 | 2.600E-06 |
| 14290 | 715 | I715I | B | 2.2e-06 | 1.7e-06 | 2.1e-06 | 2.000E-06 | 2.646E-07 | 1.700E-06 | 2.200E-06 |
| 14291 | 715 | I715L | B | 1.9e-06 | 4.4e-06 | 2.7e-06 | 3.000E-06 | 1.277E-06 | 1.900E-06 | 4.400E-06 |
| 14292 | 715 | I715K | B | 2.7e-06 | 1.8e-06 | 2.2e-06 | 2.233E-06 | 4.509E-07 | 1.800E-06 | 2.700E-06 |
| 14293 | 715 | I715M | B | 2.1e-06 | 1.7e-06 | 2.3e-06 | 2.033E-06 | 3.055E-07 | 1.700E-06 | 2.300E-06 |
| 14294 | 715 | I715F | B | 3e-06 | 3.1e-06 | 2.7e-06 | 2.933E-06 | 2.082E-07 | 2.700E-06 | 3.100E-06 |
| 14295 | 715 | I715P | B | 2.5e-06 | 2.9e-06 | 2.4e-06 | 2.600E-06 | 2.646E-07 | 2.400E-06 | 2.900E-06 |
| 14296 | 715 | I715S | B | 2.5e-06 | 1.3e-06 | 3.1e-06 | 2.300E-06 | 9.165E-07 | 1.300E-06 | 3.100E-06 |
| 14297 | 715 | I715T | B | 2.5e-06 | 2.5e-06 | 2.5e-06 | 2.500E-06 | 0.000E+00 | 2.500E-06 | 2.500E-06 |
| 14298 | 715 | I715W | B | 2.2e-06 | 1.3e-06 | 2.2e-06 | 1.900E-06 | 5.196E-07 | 1.300E-06 | 2.200E-06 |
| 14299 | 715 | I715Y | B | 2.6e-06 | 2.3e-06 | 2.2e-06 | 2.367E-06 | 2.082E-07 | 2.200E-06 | 2.600E-06 |
| 14300 | 715 | I715V | B | 1.9e-06 | 2e-06 | 2.4e-06 | 2.100E-06 | 2.646E-07 | 1.900E-06 | 2.400E-06 |
| 14301 | 716 | Y716A | B | 2.1e-06 | 1.8e-06 | 3.4e-06 | 2.433E-06 | 8.505E-07 | 1.800E-06 | 3.400E-06 |
| 14302 | 716 | Y716R | B | 3.2e-06 | 2.1e-06 | 3.4e-06 | 2.900E-06 | 7.000E-07 | 2.100E-06 | 3.400E-06 |
| 14303 | 716 | Y716N | B | 1.3e-06 | 5.4e-06 | 4.4e-06 | 3.700E-06 | 2.138E-06 | 1.300E-06 | 5.400E-06 |
| 14304 | 716 | Y716D | B | 2e-06 | 2.7e-06 | 1.5e-06 | 2.067E-06 | 6.028E-07 | 1.500E-06 | 2.700E-06 |
| 14305 | 716 | Y716C | B | 2.8e-06 | 2.6e-06 | 4.1e-06 | 3.167E-06 | 8.145E-07 | 2.600E-06 | 4.100E-06 |
| 14306 | 716 | Y716E | B | 2.6e-06 | 1.7e-06 | 2.7e-06 | 2.333E-06 | 5.508E-07 | 1.700E-06 | 2.700E-06 |
| 14307 | 716 | Y716Q | B | 3.5e-06 | 2.5e-06 | 3e-06 | 3.000E-06 | 5.000E-07 | 2.500E-06 | 3.500E-06 |
| 14308 | 716 | Y716G | B | 3.1e-06 | 4.2e-06 | 3.1e-06 | 3.467E-06 | 6.351E-07 | 3.100E-06 | 4.200E-06 |
| 14309 | 716 | Y716H | B | 2.3e-06 | 2.4e-06 | 2.2e-06 | 2.300E-06 | 1.000E-07 | 2.200E-06 | 2.400E-06 |
| 14310 | 716 | Y716I | B | 2.2e-06 | 2e-06 | 3.6e-06 | 2.600E-06 | 8.718E-07 | 2.000E-06 | 3.600E-06 |
| 14311 | 716 | Y716L | B | 1.8e-06 | 2.8e-06 | 1.6e-06 | 2.067E-06 | 6.429E-07 | 1.600E-06 | 2.800E-06 |
| 14312 | 716 | Y716K | B | 2.9e-06 | 2.6e-06 | 1.8e-06 | 2.433E-06 | 5.686E-07 | 1.800E-06 | 2.900E-06 |
| 14313 | 716 | Y716M | B | 2e-06 | 3.5e-06 | 1.1e-06 | 2.200E-06 | 1.212E-06 | 1.100E-06 | 3.500E-06 |
| 14314 | 716 | Y716F | B | 2.8e-06 | 1.3e-06 | 1.4e-06 | 1.833E-06 | 8.386E-07 | 1.300E-06 | 2.800E-06 |
| 14315 | 716 | Y716P | B | 1.2e-06 | 1.4e-06 | 2.3e-06 | 1.633E-06 | 5.859E-07 | 1.200E-06 | 2.300E-06 |
| 14316 | 716 | Y716S | B | 2.1e-06 | 2.2e-06 | 2.3e-06 | 2.200E-06 | 1.000E-07 | 2.100E-06 | 2.300E-06 |
| 14317 | 716 | Y716T | B | 3.2e-06 | 2.3e-06 | 1.3e-06 | 2.267E-06 | 9.504E-07 | 1.300E-06 | 3.200E-06 |
| 14318 | 716 | Y716W | B | 3.1e-06 | 5.2e-06 | 2.4e-06 | 3.567E-06 | 1.457E-06 | 2.400E-06 | 5.200E-06 |
| 14319 | 716 | Y716Y | B | 1.4e-06 | 1.6e-06 | 2.7e-06 | 1.900E-06 | 7.000E-07 | 1.400E-06 | 2.700E-06 |
| 14320 | 716 | Y716V | B | 2.6e-06 | 1.8e-06 | 2.7e-06 | 2.367E-06 | 4.933E-07 | 1.800E-06 | 2.700E-06 |
| 14321 | 717 | P717A | B | 2.5e-06 | 2.4e-06 | 3.6e-06 | 2.833E-06 | 6.658E-07 | 2.400E-06 | 3.600E-06 |
| 14322 | 717 | P717R | B | 2.4e-06 | 2.1e-06 | 1.3e-06 | 1.933E-06 | 5.686E-07 | 1.300E-06 | 2.400E-06 |
| 14323 | 717 | P717N | B | 2.3e-06 | 2e-06 | 1.5e-06 | 1.933E-06 | 4.041E-07 | 1.500E-06 | 2.300E-06 |
| 14324 | 717 | P717D | B | 2.5e-06 | 3.9e-06 | 3.1e-06 | 3.167E-06 | 7.024E-07 | 2.500E-06 | 3.900E-06 |
| 14325 | 717 | P717C | B | 2e-06 | 2e-06 | 2.4e-06 | 2.133E-06 | 2.309E-07 | 2.000E-06 | 2.400E-06 |
| 14326 | 717 | P717E | B | 1.9e-06 | 3.1e-06 | 2e-06 | 2.333E-06 | 6.658E-07 | 1.900E-06 | 3.100E-06 |
| 14327 | 717 | P717Q | B | 2.5e-06 | 3.8e-06 | 3.2e-06 | 3.167E-06 | 6.506E-07 | 2.500E-06 | 3.800E-06 |
| 14328 | 717 | P717G | B | 2.4e-06 | 2.5e-06 | 2.2e-06 | 2.367E-06 | 1.528E-07 | 2.200E-06 | 2.500E-06 |
| 14329 | 717 | P717H | B | 1.7e-06 | 3.9e-06 | 1.7e-06 | 2.433E-06 | 1.270E-06 | 1.700E-06 | 3.900E-06 |
| 14330 | 717 | P717I | B | 2.2e-06 | 1.6e-06 | 3.1e-06 | 2.300E-06 | 7.550E-07 | 1.600E-06 | 3.100E-06 |
| 14331 | 717 | P717L | B | 3.6e-06 | 2.6e-06 | 1.8e-06 | 2.667E-06 | 9.018E-07 | 1.800E-06 | 3.600E-06 |
| 14332 | 717 | P717K | B | 1.7e-06 | 4.2e-06 | 2.1e-06 | 2.667E-06 | 1.343E-06 | 1.700E-06 | 4.200E-06 |
| 14333 | 717 | P717M | B | 2.8e-06 | 3.3e-06 | 1.9e-06 | 2.667E-06 | 7.095E-07 | 1.900E-06 | 3.300E-06 |
| 14334 | 717 | P717F | B | 2.2e-06 | 2.5e-06 | 2.5e-06 | 2.400E-06 | 1.732E-07 | 2.200E-06 | 2.500E-06 |
| 14335 | 717 | P717P | B | 1.9e-06 | 2.8e-06 | 2e-06 | 2.233E-06 | 4.933E-07 | 1.900E-06 | 2.800E-06 |
| 14336 | 717 | P717S | B | 2.6e-06 | 3.2e-06 | 3.6e-06 | 3.133E-06 | 5.033E-07 | 2.600E-06 | 3.600E-06 |
| 14337 | 717 | P717T | B | 2.4e-06 | 2.7e-06 | 1.8e-06 | 2.300E-06 | 4.583E-07 | 1.800E-06 | 2.700E-06 |
| 14338 | 717 | P717W | B | 1.9e-06 | 1.4e-06 | 2.5e-06 | 1.933E-06 | 5.508E-07 | 1.400E-06 | 2.500E-06 |
| 14339 | 717 | P717Y | B | 1.4e-06 | 1.9e-06 | 3.2e-06 | 2.167E-06 | 9.292E-07 | 1.400E-06 | 3.200E-06 |
| 14340 | 717 | P717V | B | 1.4e-06 | 1.9e-06 | 3.5e-06 | 2.267E-06 | 1.097E-06 | 1.400E-06 | 3.500E-06 |
| 14341 | 718 | Y718A | B | 3.8e-06 | 3.9e-06 | 2.3e-06 | 3.333E-06 | 8.963E-07 | 2.300E-06 | 3.900E-06 |
| 14342 | 718 | Y718R | B | 2.4e-06 | 2.2e-06 | 2.1e-06 | 2.233E-06 | 1.528E-07 | 2.100E-06 | 2.400E-06 |
| 14343 | 718 | Y718N | B | 1.6e-06 | 3e-06 | 1.8e-06 | 2.133E-06 | 7.572E-07 | 1.600E-06 | 3.000E-06 |
| 14344 | 718 | Y718D | B | 4.8e-06 | 2.7e-06 | 1.7e-06 | 3.067E-06 | 1.582E-06 | 1.700E-06 | 4.800E-06 |
| 14345 | 718 | Y718C | B | 2.5e-06 | 3.4e-06 | 1.1e-06 | 2.333E-06 | 1.159E-06 | 1.100E-06 | 3.400E-06 |
| 14346 | 718 | Y718E | B | 2e-06 | 5.6e-06 | 3.7e-06 | 3.767E-06 | 1.801E-06 | 2.000E-06 | 5.600E-06 |
| 14347 | 718 | Y718Q | B | 3.6e-06 | 3.4e-06 | 1.9e-06 | 2.967E-06 | 9.292E-07 | 1.900E-06 | 3.600E-06 |
| 14348 | 718 | Y718G | B | 3e-06 | 2.2e-06 | 2.5e-06 | 2.567E-06 | 4.041E-07 | 2.200E-06 | 3.000E-06 |
| 14349 | 718 | Y718H | B | 1.2e-06 | 2.8e-06 | 1.5e-06 | 1.833E-06 | 8.505E-07 | 1.200E-06 | 2.800E-06 |
| 14350 | 718 | Y718I | B | 2.1e-06 | 3.6e-06 | 1.9e-06 | 2.533E-06 | 9.292E-07 | 1.900E-06 | 3.600E-06 |
| 14351 | 718 | Y718L | B | 4e-06 | 3.2e-06 | 2.4e-06 | 3.200E-06 | 8.000E-07 | 2.400E-06 | 4.000E-06 |
| 14352 | 718 | Y718K | B | 2.3e-06 | 9.4e-07 | 2.1e-06 | 1.780E-06 | 7.343E-07 | 9.400E-07 | 2.300E-06 |
| 14353 | 718 | Y718M | B | 2.2e-06 | 2.8e-06 | 1.9e-06 | 2.300E-06 | 4.583E-07 | 1.900E-06 | 2.800E-06 |
| 14354 | 718 | Y718F | B | 2.8e-06 | 1.6e-06 | 4e-06 | 2.800E-06 | 1.200E-06 | 1.600E-06 | 4.000E-06 |
| 14355 | 718 | Y718P | B | 1.7e-06 | 1.7e-06 | 3.5e-06 | 2.300E-06 | 1.039E-06 | 1.700E-06 | 3.500E-06 |
| 14356 | 718 | Y718S | B | 2.3e-06 | 3.9e-06 | 3e-06 | 3.067E-06 | 8.021E-07 | 2.300E-06 | 3.900E-06 |
| 14357 | 718 | Y718T | B | 2e-06 | 2.9e-06 | 1.5e-06 | 2.133E-06 | 7.095E-07 | 1.500E-06 | 2.900E-06 |
| 14358 | 718 | Y718W | B | 1.3e-06 | 3.3e-06 | 3.1e-06 | 2.567E-06 | 1.102E-06 | 1.300E-06 | 3.300E-06 |
| 14359 | 718 | Y718Y | B | 9.4e-07 | 2.5e-06 | 3e-06 | 2.147E-06 | 1.074E-06 | 9.400E-07 | 3.000E-06 |
| 14360 | 718 | Y718V | B | 3.7e-06 | 1e-06 | 2.1e-06 | 2.267E-06 | 1.358E-06 | 1.000E-06 | 3.700E-06 |
| 14361 | 719 | T719A | B | 1.9e-06 | 3.7e-06 | 3.4e-06 | 3.000E-06 | 9.644E-07 | 1.900E-06 | 3.700E-06 |
| 14362 | 719 | T719R | B | 1.5e-06 | 3e-06 | 1.9e-06 | 2.133E-06 | 7.767E-07 | 1.500E-06 | 3.000E-06 |
| 14363 | 719 | T719N | B | 1.5e-06 | 1.2e-06 | 2.3e-06 | 1.667E-06 | 5.686E-07 | 1.200E-06 | 2.300E-06 |
| 14364 | 719 | T719D | B | 2.6e-06 | 4.9e-06 | 3.3e-06 | 3.600E-06 | 1.179E-06 | 2.600E-06 | 4.900E-06 |

| | | | | | | | | | | |
|---|---|---|---|---|---|---|---|---|---|---|
| 14365 | 719 | T719C | B | 3.3e-06 | 3.2e-06 | 1.8e-06 | 2.767E-06 | 8.386E-07 | 1.800E-06 | 3.300E-06 |
| 14366 | 719 | T719E | B | 2.8e-06 | 3.4e-06 | 2.3e-06 | 2.833E-06 | 5.508E-07 | 2.300E-06 | 3.400E-06 |
| 14367 | 719 | T719Q | B | 2.3e-06 | 2.1e-06 | 2.3e-06 | 2.233E-06 | 1.155E-07 | 2.100E-06 | 2.300E-06 |
| 14368 | 719 | T719G | B | 3.9e-06 | 2.8e-06 | 1.7e-06 | 2.800E-06 | 1.100E-06 | 1.700E-06 | 3.900E-06 |
| 14369 | 719 | T719H | B | 1.9e-06 | 1.7e-06 | 1.3e-06 | 1.633E-06 | 3.055E-07 | 1.300E-06 | 1.900E-06 |
| 14370 | 719 | T719I | B | 3e-06 | 1.3e-06 | 2.4e-06 | 2.233E-06 | 8.622E-07 | 1.300E-06 | 3.000E-06 |
| 14371 | 719 | T719L | B | 2.1e-06 | 8.9e-07 | 1.7e-06 | 1.563E-06 | 6.165E-07 | 8.900E-07 | 2.100E-06 |
| 14372 | 719 | T719K | B | 2.1e-06 | 3e-06 | 3.4e-06 | 2.833E-06 | 6.658E-07 | 2.100E-06 | 3.400E-06 |
| 14373 | 719 | T719M | B | 3.8e-06 | 2.2e-06 | 1.8e-06 | 2.600E-06 | 1.058E-06 | 1.800E-06 | 3.800E-06 |
| 14374 | 719 | T719F | B | 3.9e-06 | 1.4e-06 | 2.9e-06 | 2.733E-06 | 1.258E-06 | 1.400E-06 | 3.900E-06 |
| 14375 | 719 | T719P | B | 2.8e-06 | 3.8e-06 | 1.2e-06 | 2.600E-06 | 1.311E-06 | 1.200E-06 | 3.800E-06 |
| 14376 | 719 | T719S | B | 2.3e-06 | 1.5e-06 | 2.4e-06 | 2.067E-06 | 4.933E-07 | 1.500E-06 | 2.400E-06 |
| 14377 | 719 | T719T | B | 8.9e-07 | 2.1e-06 | 3.7e-06 | 2.230E-06 | 1.410E-06 | 8.900E-07 | 3.700E-06 |
| 14378 | 719 | T719W | B | 1.4e-06 | 2.5e-06 | 2.5e-06 | 2.133E-06 | 6.351E-07 | 1.400E-06 | 2.500E-06 |
| 14379 | 719 | T719Y | B | 2.4e-06 | 1.1e-06 | 3.9e-06 | 2.467E-06 | 1.401E-06 | 1.100E-06 | 3.900E-06 |
| 14380 | 719 | T719V | B | 3.8e-06 | 2.3e-06 | 3.6e-06 | 3.233E-06 | 8.145E-07 | 2.300E-06 | 3.800E-06 |
| 14381 | 720 | F720A | B | 2.2e-06 | 2.2e-06 | 1.6e-06 | 2.000E-06 | 3.464E-07 | 1.600E-06 | 2.200E-06 |
| 14382 | 720 | F720R | B | 2.5e-06 | 4.3e-06 | 2.2e-06 | 3.000E-06 | 1.136E-06 | 2.200E-06 | 4.300E-06 |
| 14383 | 720 | F720N | B | 1.5e-06 | 1.6e-06 | 3.5e-06 | 2.200E-06 | 1.127E-06 | 1.500E-06 | 3.500E-06 |
| 14384 | 720 | F720D | B | 2.7e-06 | 2.7e-06 | 2.2e-06 | 2.533E-06 | 2.887E-07 | 2.200E-06 | 2.700E-06 |
| 14385 | 720 | F720C | B | 1.1e-06 | 1.8e-06 | 3e-06 | 1.967E-06 | 9.609E-07 | 1.100E-06 | 3.000E-06 |
| 14386 | 720 | F720E | B | 1.5e-06 | 2.7e-06 | 2.5e-06 | 2.233E-06 | 6.429E-07 | 1.500E-06 | 2.700E-06 |
| 14387 | 720 | F720Q | B | 1.9e-06 | 1.7e-06 | 2.4e-06 | 2.000E-06 | 3.606E-07 | 1.700E-06 | 2.400E-06 |
| 14388 | 720 | F720G | B | 3.1e-06 | 2.1e-06 | 2e-06 | 2.400E-06 | 6.083E-07 | 2.000E-06 | 3.100E-06 |
| 14389 | 720 | F720H | B | 2e-06 | 1.8e-06 | 1.7e-06 | 1.833E-06 | 1.528E-07 | 1.700E-06 | 2.000E-06 |
| 14390 | 720 | F720I | B | 1.9e-06 | 2.1e-06 | 2.5e-06 | 2.167E-06 | 3.055E-07 | 1.900E-06 | 2.500E-06 |
| 14391 | 720 | F720L | B | 2.4e-06 | 1.8e-06 | 2.5e-06 | 2.233E-06 | 3.786E-07 | 1.800E-06 | 2.500E-06 |
| 14392 | 720 | F720K | B | 2.3e-06 | 2.7e-06 | 1.2e-06 | 2.067E-06 | 7.767E-07 | 1.200E-06 | 2.700E-06 |
| 14393 | 720 | F720M | B | 2.8e-06 | 1.9e-06 | 2.7e-06 | 2.467E-06 | 4.933E-07 | 1.900E-06 | 2.800E-06 |
| 14394 | 720 | F720F | B | 2.9e-06 | 1.4e-06 | 1.7e-06 | 2.000E-06 | 7.937E-07 | 1.400E-06 | 2.900E-06 |
| 14395 | 720 | F720P | B | 4.1e-06 | 2e-06 | 1.6e-06 | 2.567E-06 | 1.343E-06 | 1.600E-06 | 4.100E-06 |
| 14396 | 720 | F720S | B | 1.3e-06 | 2.7e-06 | 1.5e-06 | 1.833E-06 | 7.572E-07 | 1.300E-06 | 2.700E-06 |
| 14397 | 720 | F720T | B | 2.7e-06 | 2.2e-06 | 2.1e-06 | 2.333E-06 | 3.215E-07 | 2.100E-06 | 2.700E-06 |
| 14398 | 720 | F720W | B | 3.5e-06 | 2.5e-06 | 2.6e-06 | 2.867E-06 | 5.508E-07 | 2.500E-06 | 3.500E-06 |
| 14399 | 720 | F720Y | B | 2.3e-06 | 1.6e-06 | 2.5e-06 | 2.133E-06 | 4.726E-07 | 1.600E-06 | 2.500E-06 |
| 14400 | 720 | F720V | B | 2.3e-06 | 2.3e-06 | 1.7e-06 | 2.100E-06 | 3.464E-07 | 1.700E-06 | 2.300E-06 |
| 14401 | 721 | G721A | B | 1.8e-06 | 3.1e-06 | 3e-06 | 2.633E-06 | 7.234E-07 | 1.800E-06 | 3.100E-06 |
| 14402 | 721 | G721R | B | 1.7e-06 | 2.3e-06 | 2.3e-06 | 2.100E-06 | 3.464E-07 | 1.700E-06 | 2.300E-06 |
| 14403 | 721 | G721N | B | 2.4e-06 | 1.3e-06 | 1.1e-06 | 1.600E-06 | 7.000E-07 | 1.100E-06 | 2.400E-06 |
| 14404 | 721 | G721D | B | 1.9e-06 | 1.8e-06 | 3e-06 | 2.233E-06 | 6.658E-07 | 1.800E-06 | 3.000E-06 |
| 14405 | 721 | G721C | B | 1.9e-06 | 2.6e-06 | 2.6e-06 | 2.367E-06 | 4.041E-07 | 1.900E-06 | 2.600E-06 |
| 14406 | 721 | G721E | B | 3.1e-06 | 3e-06 | 1.6e-06 | 2.567E-06 | 8.386E-07 | 1.600E-06 | 3.100E-06 |
| 14407 | 721 | G721Q | B | 2.3e-06 | 2.3e-06 | 3e-06 | 2.533E-06 | 4.041E-07 | 2.300E-06 | 3.000E-06 |
| 14408 | 721 | G721G | B | 3.3e-06 | 2e-06 | 3.4e-06 | 2.900E-06 | 7.810E-07 | 2.000E-06 | 3.400E-06 |
| 14409 | 721 | G721H | B | 3.1e-06 | 2.2e-06 | 2.5e-06 | 2.600E-06 | 4.583E-07 | 2.200E-06 | 3.100E-06 |
| 14410 | 721 | G721I | B | 2e-06 | 4.1e-06 | 1.2e-06 | 2.433E-06 | 1.498E-06 | 1.200E-06 | 4.100E-06 |
| 14411 | 721 | G721L | B | 2.7e-06 | 2.1e-06 | 2.5e-06 | 2.433E-06 | 3.055E-07 | 2.100E-06 | 2.700E-06 |
| 14412 | 721 | G721K | B | 2e-06 | 3e-06 | 1.6e-06 | 2.200E-06 | 7.211E-07 | 1.600E-06 | 3.000E-06 |
| 14413 | 721 | G721M | B | 3e-06 | 1.9e-06 | 1.9e-06 | 2.267E-06 | 6.351E-07 | 1.900E-06 | 3.000E-06 |
| 14414 | 721 | G721F | B | 2.8e-06 | 2.4e-06 | 3.6e-06 | 2.933E-06 | 6.110E-07 | 2.400E-06 | 3.600E-06 |
| 14415 | 721 | G721P | B | 3.8e-06 | 1.5e-06 | 3e-06 | 2.767E-06 | 1.168E-06 | 1.500E-06 | 3.800E-06 |
| 14416 | 721 | G721S | B | 5.1e-06 | 3.4e-06 | 2.9e-06 | 3.800E-06 | 1.153E-06 | 2.900E-06 | 5.100E-06 |
| 14417 | 721 | G721T | B | 1.3e-06 | 3.6e-06 | 1.4e-06 | 2.100E-06 | 1.300E-06 | 1.300E-06 | 3.600E-06 |
| 14418 | 721 | G721W | B | 2.1e-06 | 3e-06 | 1.9e-06 | 2.333E-06 | 5.859E-07 | 1.900E-06 | 3.000E-06 |
| 14419 | 721 | G721Y | B | 2.2e-06 | 3.3e-06 | 2.5e-06 | 2.667E-06 | 5.686E-07 | 2.200E-06 | 3.300E-06 |
| 14420 | 721 | G721V | B | 1.8e-06 | 2.9e-06 | 2.3e-06 | 2.333E-06 | 5.508E-07 | 1.800E-06 | 2.900E-06 |
| 14421 | 722 | Q722A | B | 3e-06 | 3.2e-06 | 2.5e-06 | 2.900E-06 | 3.606E-07 | 2.500E-06 | 3.200E-06 |
| 14422 | 722 | Q722R | B | 2.4e-06 | 2.7e-06 | 2e-06 | 2.367E-06 | 3.512E-07 | 2.000E-06 | 2.700E-06 |
| 14423 | 722 | Q722N | B | 2.9e-06 | 4e-06 | 3.2e-06 | 3.367E-06 | 5.686E-07 | 2.900E-06 | 4.000E-06 |
| 14424 | 722 | Q722D | B | 2.4e-06 | 1.5e-06 | 1.5e-06 | 1.800E-06 | 5.196E-07 | 1.500E-06 | 2.400E-06 |
| 14425 | 722 | Q722C | B | 2.3e-06 | 2e-06 | 1.3e-06 | 1.867E-06 | 5.132E-07 | 1.300E-06 | 2.300E-06 |
| 14426 | 722 | Q722E | B | 2.2e-06 | 2.4e-06 | 2.7e-06 | 2.433E-06 | 2.517E-07 | 2.200E-06 | 2.700E-06 |
| 14427 | 722 | Q722Q | B | 1.8e-06 | 2e-06 | 1.8e-06 | 1.867E-06 | 1.155E-07 | 1.800E-06 | 2.000E-06 |
| 14428 | 722 | Q722G | B | 3e-06 | 2e-06 | 3.2e-06 | 2.733E-06 | 6.429E-07 | 2.000E-06 | 3.200E-06 |
| 14429 | 722 | Q722H | B | 3.8e-06 | 2.6e-06 | 4e-06 | 3.467E-06 | 7.572E-07 | 2.600E-06 | 4.000E-06 |
| 14430 | 722 | Q722I | B | 3.9e-06 | 3.9e-06 | 2.1e-06 | 3.300E-06 | 1.039E-06 | 2.100E-06 | 3.900E-06 |
| 14431 | 722 | Q722L | B | 3e-06 | 2.7e-06 | 3e-06 | 2.900E-06 | 1.732E-07 | 2.700E-06 | 3.000E-06 |
| 14432 | 722 | Q722K | B | 2e-06 | 2.7e-06 | 3.2e-06 | 2.633E-06 | 6.028E-07 | 2.000E-06 | 3.200E-06 |
| 14433 | 722 | Q722M | B | 2.2e-06 | 3.1e-06 | 2.5e-06 | 2.600E-06 | 4.583E-07 | 2.200E-06 | 3.100E-06 |
| 14434 | 722 | Q722F | B | 1.4e-06 | 2.2e-06 | 3.3e-06 | 2.300E-06 | 9.539E-07 | 1.400E-06 | 3.300E-06 |
| 14435 | 722 | Q722P | B | 2.3e-06 | 2.4e-06 | 2.9e-06 | 2.533E-06 | 3.215E-07 | 2.300E-06 | 2.900E-06 |
| 14436 | 722 | Q722S | B | 2.9e-06 | 2.3e-06 | 1.5e-06 | 2.233E-06 | 7.024E-07 | 1.500E-06 | 2.900E-06 |
| 14437 | 722 | Q722T | B | 1.8e-06 | 2e-06 | 1.7e-06 | 1.833E-06 | 1.528E-07 | 1.700E-06 | 2.000E-06 |
| 14438 | 722 | Q722W | B | 4.2e-06 | 3e-06 | 3.4e-06 | 3.533E-06 | 6.110E-07 | 3.000E-06 | 4.200E-06 |
| 14439 | 722 | Q722Y | B | 2.9e-06 | 3.5e-06 | 3.1e-06 | 3.167E-06 | 3.055E-07 | 2.900E-06 | 3.500E-06 |
| 14440 | 722 | Q722V | B | 4.3e-06 | 3.4e-06 | 2.5e-06 | 3.400E-06 | 9.000E-07 | 2.500E-06 | 4.300E-06 |
| 14441 | 723 | G723A | B | 2.4e-06 | 2e-06 | 2.8e-06 | 2.400E-06 | 4.000E-07 | 2.000E-06 | 2.800E-06 |
| 14442 | 723 | G723R | B | 1e-06 | 2.2e-06 | 1.8e-06 | 1.667E-06 | 6.110E-07 | 1.000E-06 | 2.200E-06 |
| 14443 | 723 | G723N | B | 3e-06 | 3.3e-06 | 2e-06 | 2.767E-06 | 6.807E-07 | 2.000E-06 | 3.300E-06 |
| 14444 | 723 | G723D | B | 3.1e-06 | 1.2e-06 | 3.2e-06 | 2.500E-06 | 1.127E-06 | 1.200E-06 | 3.200E-06 |
| 14445 | 723 | G723C | B | 1.7e-06 | 2.1e-06 | 2.4e-06 | 2.067E-06 | 3.512E-07 | 1.700E-06 | 2.400E-06 |
| 14446 | 723 | G723E | B | 1.7e-06 | 2.7e-06 | 1.9e-06 | 2.100E-06 | 5.292E-07 | 1.700E-06 | 2.700E-06 |
| 14447 | 723 | G723Q | B | 3.6e-06 | 2.8e-06 | 2.3e-06 | 2.900E-06 | 6.557E-07 | 2.300E-06 | 3.600E-06 |
| 14448 | 723 | G723G | B | 2.3e-06 | 2.5e-06 | 2.8e-06 | 2.533E-06 | 2.517E-07 | 2.300E-06 | 2.800E-06 |
| 14449 | 723 | G723H | B | 3.5e-06 | 2.7e-06 | 2.3e-06 | 2.833E-06 | 6.110E-07 | 2.300E-06 | 3.500E-06 |

| | | | | | | | | | | |
|---|---|---|---|---|---|---|---|---|---|---|
| 14450 | 723 | G723I | B | 1.7e-06 | 1.9e-06 | 1.4e-06 | 1.667E-06 | 2.517E-07 | 1.400E-06 | 1.900E-06 |
| 14451 | 723 | G723L | B | 2e-06 | 1.8e-06 | 1.8e-06 | 1.867E-06 | 1.155E-07 | 1.800E-06 | 2.000E-06 |
| 14452 | 723 | G723K | B | 2.3e-06 | 2.4e-06 | 1.9e-06 | 2.200E-06 | 2.646E-07 | 1.900E-06 | 2.400E-06 |
| 14453 | 723 | G723M | B | 2.6e-06 | 1.7e-06 | 2.8e-06 | 2.367E-06 | 5.859E-07 | 1.700E-06 | 2.800E-06 |
| 14454 | 723 | G723F | B | 1.5e-06 | 3.4e-06 | 2.3e-06 | 2.400E-06 | 9.539E-07 | 1.500E-06 | 3.400E-06 |
| 14455 | 723 | G723P | B | 3.1e-06 | 2.5e-06 | 2.3e-06 | 2.633E-06 | 4.163E-07 | 2.300E-06 | 3.100E-06 |
| 14456 | 723 | G723S | B | 1.9e-06 | 1.4e-06 | 2e-06 | 1.767E-06 | 3.215E-07 | 1.400E-06 | 2.000E-06 |
| 14457 | 723 | G723T | B | 2.7e-06 | 1.9e-06 | 2.1e-06 | 2.233E-06 | 4.163E-07 | 1.900E-06 | 2.700E-06 |
| 14458 | 723 | G723W | B | 2.1e-06 | 2e-06 | 2.3e-06 | 2.133E-06 | 1.528E-07 | 2.000E-06 | 2.300E-06 |
| 14459 | 723 | G723Y | B | 2.1e-06 | 2.7e-06 | 2.8e-06 | 2.533E-06 | 3.786E-07 | 2.100E-06 | 2.800E-06 |
| 14460 | 723 | G723V | B | 2.3e-06 | 1.9e-06 | 2.3e-06 | 2.167E-06 | 2.309E-07 | 1.900E-06 | 2.300E-06 |
| 14461 | 724 | T724A | B | 3.8e-06 | 1.3e-06 | 3.5e-06 | 2.867E-06 | 1.365E-06 | 1.300E-06 | 3.800E-06 |
| 14462 | 724 | T724R | B | 2.6e-06 | 1.3e-06 | 2.4e-06 | 2.100E-06 | 7.000E-07 | 1.300E-06 | 2.600E-06 |
| 14463 | 724 | T724N | B | 2.5e-06 | 1.8e-06 | 1.9e-06 | 2.067E-06 | 3.786E-07 | 1.800E-06 | 2.500E-06 |
| 14464 | 724 | T724D | B | 1.4e-06 | 4.6e-06 | 4e-06 | 3.333E-06 | 1.701E-06 | 1.400E-06 | 4.600E-06 |
| 14465 | 724 | T724C | B | 1.4e-06 | 2.7e-06 | 2.6e-06 | 2.233E-06 | 7.234E-07 | 1.400E-06 | 2.700E-06 |
| 14466 | 724 | T724E | B | 3.8e-06 | 1.6e-06 | 1.1e-06 | 2.167E-06 | 1.436E-06 | 1.100E-06 | 3.800E-06 |
| 14467 | 724 | T724Q | B | 1.4e-06 | 2.4e-06 | 1.9e-06 | 1.900E-06 | 5.000E-07 | 1.400E-06 | 2.400E-06 |
| 14468 | 724 | T724G | B | 2e-06 | 2.8e-06 | 1.8e-06 | 2.200E-06 | 5.292E-07 | 1.800E-06 | 2.800E-06 |
| 14469 | 724 | T724H | B | 1.4e-06 | 1.5e-06 | 3.2e-06 | 2.033E-06 | 1.012E-06 | 1.400E-06 | 3.200E-06 |
| 14470 | 724 | T724I | B | 1.9e-06 | 2.2e-06 | 2.4e-06 | 2.167E-06 | 2.517E-07 | 1.900E-06 | 2.400E-06 |
| 14471 | 724 | T724L | B | 1.8e-06 | 2e-06 | 1.6e-06 | 1.800E-06 | 2.000E-07 | 1.600E-06 | 2.000E-06 |
| 14472 | 724 | T724K | B | 2.1e-06 | 1.8e-06 | 2.3e-06 | 2.067E-06 | 2.517E-07 | 1.800E-06 | 2.300E-06 |
| 14473 | 724 | T724M | B | 2.6e-06 | 2e-06 | 2.7e-06 | 2.433E-06 | 3.786E-07 | 2.000E-06 | 2.700E-06 |
| 14474 | 724 | T724F | B | 2.2e-06 | 3e-06 | 2.7e-06 | 2.633E-06 | 4.041E-07 | 2.200E-06 | 3.000E-06 |
| 14475 | 724 | T724P | B | 2.9e-06 | 3.4e-06 | 8.7e-07 | 2.390E-06 | 1.340E-06 | 8.700E-07 | 3.400E-06 |
| 14476 | 724 | T724S | B | 2.8e-06 | 2.8e-06 | 2.8e-06 | 2.800E-06 | 0.000E+00 | 2.800E-06 | 2.800E-06 |
| 14477 | 724 | T724T | B | 2.4e-06 | 2.4e-06 | 2.2e-06 | 2.333E-06 | 1.155E-07 | 2.200E-06 | 2.400E-06 |
| 14478 | 724 | T724W | B | 2.9e-06 | 2.6e-06 | 3.1e-06 | 2.867E-06 | 2.517E-07 | 2.600E-06 | 3.100E-06 |
| 14479 | 724 | T724Y | B | 2.3e-06 | 4.7e-06 | 2.8e-06 | 3.267E-06 | 1.266E-06 | 2.300E-06 | 4.700E-06 |
| 14480 | 724 | T724V | B | 4.1e-06 | 2.3e-06 | 2.6e-06 | 3.000E-06 | 9.644E-07 | 2.300E-06 | 4.100E-06 |
| 14481 | 725 | K725A | B | 2e-06 | 2.2e-06 | 3.2e-06 | 2.467E-06 | 6.429E-07 | 2.000E-06 | 3.200E-06 |
| 14482 | 725 | K725R | B | 2e-06 | 2.9e-06 | 2.6e-06 | 2.500E-06 | 4.583E-07 | 2.000E-06 | 2.900E-06 |
| 14483 | 725 | K725N | B | 1.6e-06 | 2.4e-06 | 2.5e-06 | 2.167E-06 | 4.933E-07 | 1.600E-06 | 2.500E-06 |
| 14484 | 725 | K725D | B | 1.4e-06 | 3.2e-06 | 3e-06 | 2.533E-06 | 9.866E-07 | 1.400E-06 | 3.200E-06 |
| 14485 | 725 | K725C | B | 1.6e-06 | 2.8e-06 | 2.2e-06 | 2.200E-06 | 6.000E-07 | 1.600E-06 | 2.800E-06 |
| 14486 | 725 | K725E | B | 2.5e-06 | 1.9e-06 | 3.1e-06 | 2.500E-06 | 6.000E-07 | 1.900E-06 | 3.100E-06 |
| 14487 | 725 | K725Q | B | 1.5e-06 | 2.1e-06 | 3.3e-06 | 2.300E-06 | 9.165E-07 | 1.500E-06 | 3.300E-06 |
| 14488 | 725 | K725G | B | 2.2e-06 | 3.6e-06 | 2.5e-06 | 2.767E-06 | 7.371E-07 | 2.200E-06 | 3.600E-06 |
| 14489 | 725 | K725H | B | 1.8e-06 | 1.1e-06 | 4.3e-06 | 2.400E-06 | 1.682E-06 | 1.100E-06 | 4.300E-06 |
| 14490 | 725 | K725I | B | 2.6e-06 | 2.9e-06 | 2.4e-06 | 2.633E-06 | 2.517E-07 | 2.400E-06 | 2.900E-06 |
| 14491 | 725 | K725L | B | 1.6e-06 | 2.7e-06 | 2.3e-06 | 2.200E-06 | 5.568E-07 | 1.600E-06 | 2.700E-06 |
| 14492 | 725 | K725K | B | 2.3e-06 | 2.1e-06 | 2.5e-06 | 2.300E-06 | 2.000E-07 | 2.100E-06 | 2.500E-06 |
| 14493 | 725 | K725M | B | 1.8e-06 | 2.9e-06 | 3.9e-06 | 2.867E-06 | 1.050E-06 | 1.800E-06 | 3.900E-06 |
| 14494 | 725 | K725F | B | 2.2e-06 | 1.7e-06 | 2e-06 | 1.967E-06 | 2.517E-07 | 1.700E-06 | 2.200E-06 |
| 14495 | 725 | K725P | B | 2e-06 | 2.2e-06 | 3.7e-06 | 2.633E-06 | 9.292E-07 | 2.000E-06 | 3.700E-06 |
| 14496 | 725 | K725S | B | 3.4e-06 | 2.1e-06 | 2.3e-06 | 2.600E-06 | 7.000E-07 | 2.100E-06 | 3.400E-06 |
| 14497 | 725 | K725T | B | 2.5e-06 | 1.8e-06 | 2.6e-06 | 2.300E-06 | 4.359E-07 | 1.800E-06 | 2.600E-06 |
| 14498 | 725 | K725W | B | 2.1e-06 | 2.2e-06 | 3.8e-06 | 2.700E-06 | 9.539E-07 | 2.100E-06 | 3.800E-06 |
| 14499 | 725 | K725Y | B | 2.7e-06 | 1.5e-06 | 3.1e-06 | 2.433E-06 | 8.327E-07 | 1.500E-06 | 3.100E-06 |
| 14500 | 725 | K725V | B | 1.3e-06 | 3.3e-06 | 2.6e-06 | 2.400E-06 | 1.015E-06 | 1.300E-06 | 3.300E-06 |
| 14501 | 726 | V726A | B | 2.9e-06 | 3e-06 | 1.5e-06 | 2.467E-06 | 8.386E-07 | 1.500E-06 | 3.000E-06 |
| 14502 | 726 | V726R | B | 1.8e-06 | 2.4e-06 | 9.8e-07 | 1.727E-06 | 7.128E-07 | 9.800E-07 | 2.400E-06 |
| 14503 | 726 | V726N | B | 2.4e-06 | 2.3e-06 | 1.4e-06 | 2.033E-06 | 5.508E-07 | 1.400E-06 | 2.400E-06 |
| 14504 | 726 | V726D | B | 2.7e-06 | 5e-06 | 1.6e-06 | 3.100E-06 | 1.735E-06 | 1.600E-06 | 5.000E-06 |
| 14505 | 726 | V726C | B | 1.9e-06 | 1.3e-06 | 2.4e-06 | 1.867E-06 | 5.508E-07 | 1.300E-06 | 2.400E-06 |
| 14506 | 726 | V726E | B | 2.5e-06 | 1.2e-06 | 2.5e-06 | 2.067E-06 | 7.506E-07 | 1.200E-06 | 2.500E-06 |
| 14507 | 726 | V726Q | B | 2.2e-06 | 1.8e-06 | 1.8e-06 | 1.933E-06 | 2.309E-07 | 1.800E-06 | 2.200E-06 |
| 14508 | 726 | V726G | B | 2.5e-06 | 1.9e-06 | 2e-06 | 2.133E-06 | 3.215E-07 | 1.900E-06 | 2.500E-06 |
| 14509 | 726 | V726H | B | 3.2e-06 | 1.7e-06 | 1.8e-06 | 2.233E-06 | 8.386E-07 | 1.700E-06 | 3.200E-06 |
| 14510 | 726 | V726I | B | 2.4e-06 | 3.7e-06 | 2.3e-06 | 2.800E-06 | 7.810E-07 | 2.300E-06 | 3.700E-06 |
| 14511 | 726 | V726L | B | 1.6e-06 | 1.8e-06 | 2.5e-06 | 1.967E-06 | 4.726E-07 | 1.600E-06 | 2.500E-06 |
| 14512 | 726 | V726K | B | 2.2e-06 | 2.6e-06 | 2.7e-06 | 2.500E-06 | 2.646E-07 | 2.200E-06 | 2.700E-06 |
| 14513 | 726 | V726M | B | 3.8e-06 | 2.1e-06 | 2e-06 | 2.633E-06 | 1.012E-06 | 2.000E-06 | 3.800E-06 |
| 14514 | 726 | V726F | B | 1.9e-06 | 2.3e-06 | 3.1e-06 | 2.433E-06 | 6.110E-07 | 1.900E-06 | 3.100E-06 |
| 14515 | 726 | V726P | B | 2.6e-06 | 1.8e-06 | 3.1e-06 | 2.500E-06 | 6.557E-07 | 1.800E-06 | 3.100E-06 |
| 14516 | 726 | V726S | B | 3.3e-06 | 2.4e-06 | 1.7e-06 | 2.467E-06 | 8.021E-07 | 1.700E-06 | 3.300E-06 |
| 14517 | 726 | V726T | B | 3e-06 | 1.5e-06 | 2.5e-06 | 2.333E-06 | 7.638E-07 | 1.500E-06 | 3.000E-06 |
| 14518 | 726 | V726W | B | 2e-06 | 5.1e-06 | 1.4e-06 | 2.833E-06 | 1.986E-06 | 1.400E-06 | 5.100E-06 |
| 14519 | 726 | V726Y | B | 2e-06 | 1.1e-06 | 1.5e-06 | 1.533E-06 | 4.509E-07 | 1.100E-06 | 2.000E-06 |
| 14520 | 726 | V726V | B | 1.7e-06 | 1.5e-06 | 1.8e-06 | 1.667E-06 | 1.528E-07 | 1.500E-06 | 1.800E-06 |
| 14521 | 727 | E727A | B | 1.6e-06 | 2.3e-06 | 2.2e-06 | 2.033E-06 | 3.786E-07 | 1.600E-06 | 2.300E-06 |
| 14522 | 727 | E727R | B | 3.6e-06 | 1.7e-06 | 1.8e-06 | 2.367E-06 | 1.069E-06 | 1.700E-06 | 3.600E-06 |
| 14523 | 727 | E727N | B | 1.5e-06 | 2.1e-06 | 2.6e-06 | 2.067E-06 | 5.508E-07 | 1.500E-06 | 2.600E-06 |
| 14524 | 727 | E727D | B | 2.6e-06 | 3.7e-06 | 3.6e-06 | 3.300E-06 | 6.083E-07 | 2.600E-06 | 3.700E-06 |
| 14525 | 727 | E727C | B | 1.6e-06 | 3.4e-06 | 2.1e-06 | 2.367E-06 | 9.292E-07 | 1.600E-06 | 3.400E-06 |
| 14526 | 727 | E727E | B | 2.4e-06 | 1.9e-06 | 3e-06 | 2.433E-06 | 5.508E-07 | 1.900E-06 | 3.000E-06 |
| 14527 | 727 | E727Q | B | 2.4e-06 | 3.9e-06 | 2.7e-06 | 3.000E-06 | 7.937E-07 | 2.400E-06 | 3.900E-06 |
| 14528 | 727 | E727G | B | 2.6e-06 | 2.2e-06 | 2.5e-06 | 2.433E-06 | 2.082E-07 | 2.200E-06 | 2.600E-06 |
| 14529 | 727 | E727H | B | 2.3e-06 | 2.7e-06 | 1.8e-06 | 2.267E-06 | 4.509E-07 | 1.800E-06 | 2.700E-06 |
| 14530 | 727 | E727I | B | 2.3e-06 | 2.4e-06 | 2.7e-06 | 2.467E-06 | 2.082E-07 | 2.300E-06 | 2.700E-06 |
| 14531 | 727 | E727L | B | 2.6e-06 | 3.6e-06 | 2.3e-06 | 2.833E-06 | 6.807E-07 | 2.300E-06 | 3.600E-06 |
| 14532 | 727 | E727K | B | 2.6e-06 | 2.1e-06 | 2e-06 | 2.233E-06 | 3.215E-07 | 2.000E-06 | 2.600E-06 |
| 14533 | 727 | E727M | B | 1.8e-06 | 3e-06 | 1.9e-06 | 2.233E-06 | 6.658E-07 | 1.800E-06 | 3.000E-06 |
| 14534 | 727 | E727F | B | 2.8e-06 | 1.3e-06 | 2.4e-06 | 2.167E-06 | 7.767E-07 | 1.300E-06 | 2.800E-06 |

| | | | | | | | | | | |
|---|---|---|---|---|---|---|---|---|---|---|
| 14535 | 727 | E727P | B | 2e-06 | 2.1e-06 | 2.6e-06 | 2.233E-06 | 3.215E-07 | 2.000E-06 | 2.600E-06 |
| 14536 | 727 | E727S | B | 1.8e-06 | 2.6e-06 | 1.9e-06 | 2.100E-06 | 4.359E-07 | 1.800E-06 | 2.600E-06 |
| 14537 | 727 | E727T | B | 2.5e-06 | 1.9e-06 | 2.1e-06 | 2.167E-06 | 3.055E-07 | 1.900E-06 | 2.500E-06 |
| 14538 | 727 | E727W | B | 3.1e-06 | 2.6e-06 | 2.9e-06 | 2.867E-06 | 2.517E-07 | 2.600E-06 | 3.100E-06 |
| 14539 | 727 | E727Y | B | 3.4e-06 | 2.6e-06 | 2.7e-06 | 2.900E-06 | 4.359E-07 | 2.600E-06 | 3.400E-06 |
| 14540 | 727 | E727V | B | 4.2e-06 | 2.1e-06 | 3e-06 | 3.100E-06 | 1.054E-06 | 2.100E-06 | 4.200E-06 |
| 14541 | 728 | I728A | B | 1.8e-06 | 2.7e-06 | 1.3e-06 | 1.933E-06 | 7.095E-07 | 1.300E-06 | 2.700E-06 |
| 14542 | 728 | I728R | B | 1.9e-06 | 1.8e-06 | 4.5e-06 | 2.733E-06 | 1.531E-06 | 1.800E-06 | 4.500E-06 |
| 14543 | 728 | I728N | B | 2.9e-06 | 2.7e-06 | 2.5e-06 | 2.700E-06 | 2.000E-07 | 2.500E-06 | 2.900E-06 |
| 14544 | 728 | I728D | B | 3.7e-06 | 3.4e-06 | 2.7e-06 | 3.267E-06 | 5.132E-07 | 2.700E-06 | 3.700E-06 |
| 14545 | 728 | I728C | B | 3e-06 | 2.6e-06 | 2.3e-06 | 2.633E-06 | 3.512E-07 | 2.300E-06 | 3.000E-06 |
| 14546 | 728 | I728E | B | 1.6e-06 | 4.3e-06 | 1.2e-06 | 2.367E-06 | 1.686E-06 | 1.200E-06 | 4.300E-06 |
| 14547 | 728 | I728Q | B | 2.7e-06 | 1.2e-06 | 2.8e-06 | 2.233E-06 | 8.963E-07 | 1.200E-06 | 2.800E-06 |
| 14548 | 728 | I728G | B | 3.5e-06 | 2.5e-06 | 1.8e-06 | 2.600E-06 | 8.544E-07 | 1.800E-06 | 3.500E-06 |
| 14549 | 728 | I728H | B | 2.2e-06 | 1.7e-06 | 1.1e-06 | 1.667E-06 | 5.508E-07 | 1.100E-06 | 2.200E-06 |
| 14550 | 728 | I728I | B | 2e-06 | 1.8e-06 | 2e-06 | 1.933E-06 | 1.155E-07 | 1.800E-06 | 2.000E-06 |
| 14551 | 728 | I728L | B | 2e-06 | 2.5e-06 | 2e-06 | 2.167E-06 | 2.887E-07 | 2.000E-06 | 2.500E-06 |
| 14552 | 728 | I728K | B | 3.4e-06 | 2.2e-06 | 3.4e-06 | 3.000E-06 | 6.928E-07 | 2.200E-06 | 3.400E-06 |
| 14553 | 728 | I728M | B | 3.4e-06 | 2.4e-06 | 2.6e-06 | 2.800E-06 | 5.292E-07 | 2.400E-06 | 3.400E-06 |
| 14554 | 728 | I728F | B | 5.1e-06 | 2.5e-06 | 1.9e-06 | 3.167E-06 | 1.701E-06 | 1.900E-06 | 5.100E-06 |
| 14555 | 728 | I728P | B | 2.8e-06 | 2e-06 | 1.6e-06 | 2.133E-06 | 6.110E-07 | 1.600E-06 | 2.800E-06 |
| 14556 | 728 | I728S | B | 2.9e-06 | 2e-06 | 2.3e-06 | 2.400E-06 | 4.583E-07 | 2.000E-06 | 2.900E-06 |
| 14557 | 728 | I728T | B | 2.4e-06 | 1.5e-06 | 2.5e-06 | 2.133E-06 | 5.508E-07 | 1.500E-06 | 2.500E-06 |
| 14558 | 728 | I728W | B | 2e-06 | 2.4e-06 | 3.6e-06 | 2.667E-06 | 8.327E-07 | 2.000E-06 | 3.600E-06 |
| 14559 | 728 | I728Y | B | 3e-06 | 2.8e-06 | 2.8e-06 | 2.867E-06 | 1.155E-07 | 2.800E-06 | 3.000E-06 |
| 14560 | 728 | I728V | B | 1.1e-06 | 3.4e-06 | 3.4e-06 | 2.633E-06 | 1.328E-06 | 1.100E-06 | 3.400E-06 |
| 14561 | 729 | K729A | B | 2.6e-06 | 1.5e-06 | 1.4e-06 | 1.833E-06 | 6.658E-07 | 1.400E-06 | 2.600E-06 |
| 14562 | 729 | K729R | B | 1.7e-06 | 1.7e-06 | 1.9e-06 | 1.767E-06 | 1.155E-07 | 1.700E-06 | 1.900E-06 |
| 14563 | 729 | K729N | B | 2.9e-06 | 4.1e-06 | 3.1e-06 | 3.367E-06 | 6.429E-07 | 2.900E-06 | 4.100E-06 |
| 14564 | 729 | K729D | B | 2.3e-06 | 2.1e-06 | 2.2e-06 | 2.200E-06 | 1.000E-07 | 2.100E-06 | 2.300E-06 |
| 14565 | 729 | K729C | B | 3e-06 | 2.2e-06 | 1.2e-06 | 2.133E-06 | 9.018E-07 | 1.200E-06 | 3.000E-06 |
| 14566 | 729 | K729E | B | 1.3e-06 | 2.1e-06 | 2.7e-06 | 2.033E-06 | 7.024E-07 | 1.300E-06 | 2.700E-06 |
| 14567 | 729 | K729Q | B | 2.2e-06 | 2.5e-06 | 3e-06 | 2.567E-06 | 4.041E-07 | 2.200E-06 | 3.000E-06 |
| 14568 | 729 | K729G | B | 9.4e-07 | 3e-06 | 1.6e-06 | 1.847E-06 | 1.052E-06 | 9.400E-07 | 3.000E-06 |
| 14569 | 729 | K729H | B | 2.9e-06 | 1.7e-06 | 1.8e-06 | 2.133E-06 | 6.658E-07 | 1.700E-06 | 2.900E-06 |
| 14570 | 729 | K729I | B | 1.8e-06 | 5.7e-06 | 3.9e-06 | 3.800E-06 | 1.952E-06 | 1.800E-06 | 5.700E-06 |
| 14571 | 729 | K729L | B | 2.2e-06 | 1.4e-06 | 2.5e-06 | 2.033E-06 | 5.686E-07 | 1.400E-06 | 2.500E-06 |
| 14572 | 729 | K729K | B | 2e-06 | 1.9e-06 | 2.3e-06 | 2.067E-06 | 2.082E-07 | 1.900E-06 | 2.300E-06 |
| 14573 | 729 | K729M | B | 2.7e-06 | 3.4e-06 | 1.8e-06 | 2.633E-06 | 8.021E-07 | 1.800E-06 | 3.400E-06 |
| 14574 | 729 | K729F | B | 1.9e-06 | 3.1e-06 | 2.8e-06 | 2.600E-06 | 6.245E-07 | 1.900E-06 | 3.100E-06 |
| 14575 | 729 | K729P | B | 2.9e-06 | 2.7e-06 | 1.8e-06 | 2.467E-06 | 5.859E-07 | 1.800E-06 | 2.900E-06 |
| 14576 | 729 | K729S | B | 3.8e-06 | 1.6e-06 | 2.7e-06 | 2.700E-06 | 1.100E-06 | 1.600E-06 | 3.800E-06 |
| 14577 | 729 | K729T | B | 2.1e-06 | 2e-06 | 3.7e-06 | 2.600E-06 | 9.539E-07 | 2.000E-06 | 3.700E-06 |
| 14578 | 729 | K729W | B | 2.1e-06 | 2.9e-06 | 1.9e-06 | 2.300E-06 | 5.292E-07 | 1.900E-06 | 2.900E-06 |
| 14579 | 729 | K729Y | B | 2.6e-06 | 3e-06 | 1.2e-06 | 2.267E-06 | 9.452E-07 | 1.200E-06 | 3.000E-06 |
| 14580 | 729 | K729V | B | 3e-06 | 1.5e-06 | 2.4e-06 | 2.300E-06 | 7.550E-07 | 1.500E-06 | 3.000E-06 |
| 14581 | 730 | R730A | B | 2.5e-06 | 2e-06 | 3.3e-06 | 2.600E-06 | 6.557E-07 | 2.000E-06 | 3.300E-06 |
| 14582 | 730 | R730R | B | 1.9e-06 | 1.5e-06 | 3.1e-06 | 2.167E-06 | 8.327E-07 | 1.500E-06 | 3.100E-06 |
| 14583 | 730 | R730N | B | 2.7e-06 | 1.3e-06 | 2.8e-06 | 2.267E-06 | 8.386E-07 | 1.300E-06 | 2.800E-06 |
| 14584 | 730 | R730D | B | 1.9e-06 | 1.4e-06 | 3.7e-06 | 2.333E-06 | 1.210E-06 | 1.400E-06 | 3.700E-06 |
| 14585 | 730 | R730C | B | 2.1e-06 | 3.4e-06 | 2e-06 | 2.500E-06 | 7.810E-07 | 2.000E-06 | 3.400E-06 |
| 14586 | 730 | R730E | B | 1.3e-06 | 1.6e-06 | 8.6e-07 | 1.253E-06 | 3.722E-07 | 8.600E-07 | 1.600E-06 |
| 14587 | 730 | R730Q | B | 2.7e-06 | 1.4e-06 | 1.2e-06 | 1.767E-06 | 8.145E-07 | 1.200E-06 | 2.700E-06 |
| 14588 | 730 | R730G | B | 2.9e-06 | 2.2e-06 | 2.2e-06 | 2.433E-06 | 4.041E-07 | 2.200E-06 | 2.900E-06 |
| 14589 | 730 | R730H | B | 1.6e-06 | 2.4e-06 | 8.1e-07 | 1.603E-06 | 7.950E-07 | 8.100E-07 | 2.400E-06 |
| 14590 | 730 | R730I | B | 2.1e-06 | 2.3e-06 | 3.2e-06 | 2.533E-06 | 5.859E-07 | 2.100E-06 | 3.200E-06 |
| 14591 | 730 | R730L | B | 3.6e-06 | 1.9e-06 | 2.6e-06 | 2.700E-06 | 8.544E-07 | 1.900E-06 | 3.600E-06 |
| 14592 | 730 | R730K | B | 1.6e-06 | 3.3e-06 | 3.4e-06 | 2.767E-06 | 1.012E-06 | 1.600E-06 | 3.400E-06 |
| 14593 | 730 | R730M | B | 1.7e-06 | 2.8e-06 | 1.8e-06 | 2.100E-06 | 6.083E-07 | 1.700E-06 | 2.800E-06 |
| 14594 | 730 | R730F | B | 2.3e-06 | 2.2e-06 | 2.3e-06 | 2.267E-06 | 5.774E-08 | 2.200E-06 | 2.300E-06 |
| 14595 | 730 | R730P | B | 1.8e-06 | 3.5e-06 | 2.9e-06 | 2.733E-06 | 8.622E-07 | 1.800E-06 | 3.500E-06 |
| 14596 | 730 | R730S | B | 1.8e-06 | 4.4e-06 | 2.7e-06 | 2.967E-06 | 1.320E-06 | 1.800E-06 | 4.400E-06 |
| 14597 | 730 | R730T | B | 4.2e-06 | 2.9e-06 | 1.8e-06 | 2.967E-06 | 1.201E-06 | 1.800E-06 | 4.200E-06 |
| 14598 | 730 | R730W | B | 2.7e-06 | 2.3e-06 | 1.7e-06 | 2.233E-06 | 5.033E-07 | 1.700E-06 | 2.700E-06 |
| 14599 | 730 | R730Y | B | 2.6e-06 | 3.7e-06 | 1.9e-06 | 2.733E-06 | 9.074E-07 | 1.900E-06 | 3.700E-06 |
| 14600 | 730 | R730V | B | 3.2e-06 | 3.4e-06 | 3.3e-06 | 3.300E-06 | 1.000E-07 | 3.200E-06 | 3.400E-06 |
| 14601 | 731 | T731A | B | 3.1e-06 | 2.5e-06 | 2.8e-06 | 2.800E-06 | 3.000E-07 | 2.500E-06 | 3.100E-06 |
| 14602 | 731 | T731R | B | 2.4e-06 | 2.5e-06 | 2.5e-06 | 2.467E-06 | 5.774E-08 | 2.400E-06 | 2.500E-06 |
| 14603 | 731 | T731N | B | 1.1e-06 | 3.1e-06 | 1.9e-06 | 2.033E-06 | 1.007E-06 | 1.100E-06 | 3.100E-06 |
| 14604 | 731 | T731D | B | 3.5e-06 | 2.4e-06 | 2.3e-06 | 2.733E-06 | 6.658E-07 | 2.300E-06 | 3.500E-06 |
| 14605 | 731 | T731C | B | 1.3e-06 | 2e-06 | 2.1e-06 | 1.800E-06 | 4.359E-07 | 1.300E-06 | 2.100E-06 |
| 14606 | 731 | T731E | B | 4.4e-06 | 3.7e-06 | 2e-06 | 3.367E-06 | 1.234E-06 | 2.000E-06 | 4.400E-06 |
| 14607 | 731 | T731Q | B | 1.9e-06 | 1.6e-06 | 2.3e-06 | 1.933E-06 | 3.512E-07 | 1.600E-06 | 2.300E-06 |
| 14608 | 731 | T731G | B | 1.3e-06 | 2.2e-06 | 2.4e-06 | 1.967E-06 | 5.859E-07 | 1.300E-06 | 2.400E-06 |
| 14609 | 731 | T731H | B | 1.9e-06 | 1.5e-06 | 1.8e-06 | 1.733E-06 | 2.082E-07 | 1.500E-06 | 1.900E-06 |
| 14610 | 731 | T731I | B | 3.3e-06 | 2.4e-06 | 2.5e-06 | 2.733E-06 | 4.933E-07 | 2.400E-06 | 3.300E-06 |
| 14611 | 731 | T731L | B | 2.2e-06 | 2.5e-06 | 1.8e-06 | 2.167E-06 | 3.512E-07 | 1.800E-06 | 2.500E-06 |
| 14612 | 731 | T731K | B | 2.5e-06 | 3.2e-06 | 4.5e-06 | 3.400E-06 | 1.015E-06 | 2.500E-06 | 4.500E-06 |
| 14613 | 731 | T731M | B | 1.8e-06 | 1.2e-06 | 1.6e-06 | 1.533E-06 | 3.055E-07 | 1.200E-06 | 1.800E-06 |
| 14614 | 731 | T731F | B | 1.5e-06 | 2.9e-06 | 4.5e-06 | 2.967E-06 | 1.501E-06 | 1.500E-06 | 4.500E-06 |
| 14615 | 731 | T731P | B | 3e-06 | 3.1e-06 | 1.8e-06 | 2.633E-06 | 7.234E-07 | 1.800E-06 | 3.100E-06 |
| 14616 | 731 | T731S | B | 1.7e-06 | 1.1e-06 | 1.9e-06 | 1.567E-06 | 4.163E-07 | 1.100E-06 | 1.900E-06 |
| 14617 | 731 | T731T | B | 1.8e-06 | 2.1e-06 | 2.5e-06 | 2.133E-06 | 3.512E-07 | 1.800E-06 | 2.500E-06 |
| 14618 | 731 | T731W | B | 2.3e-06 | 1.6e-06 | 2.1e-06 | 2.000E-06 | 3.606E-07 | 1.600E-06 | 2.300E-06 |
| 14619 | 731 | T731Y | B | 2.3e-06 | 3.8e-06 | 2.4e-06 | 2.833E-06 | 8.386E-07 | 2.300E-06 | 3.800E-06 |

| | | | | | | | | | | |
|---|---|---|---|---|---|---|---|---|---|---|
| 14620 | 731 | T731V | B | 1.3e-06 | 2.5e-06 | 3.8e-06 | 2.533E-06 | 1.250E-06 | 1.300E-06 | 3.800E-06 |
| 14621 | 732 | V732A | B | 2.1e-06 | 2.3e-06 | 2.3e-06 | 2.233E-06 | 1.155E-07 | 2.100E-06 | 2.300E-06 |
| 14622 | 732 | V732R | B | 3e-06 | 2.8e-06 | 1.9e-06 | 2.567E-06 | 5.859E-07 | 1.900E-06 | 3.000E-06 |
| 14623 | 732 | V732N | B | 2.1e-06 | 1.3e-06 | 2.8e-06 | 2.067E-06 | 7.506E-07 | 1.300E-06 | 2.800E-06 |
| 14624 | 732 | V732D | B | 2e-06 | 2e-06 | 2.9e-06 | 2.300E-06 | 5.196E-07 | 2.000E-06 | 2.900E-06 |
| 14625 | 732 | V732C | B | 2.3e-06 | 3.8e-06 | 1.9e-06 | 2.667E-06 | 1.002E-06 | 1.900E-06 | 3.800E-06 |
| 14626 | 732 | V732E | B | 4.7e-06 | 2.1e-06 | 2.4e-06 | 3.067E-06 | 1.422E-06 | 2.100E-06 | 4.700E-06 |
| 14627 | 732 | V732Q | B | 1.7e-06 | 1.8e-06 | 2.2e-06 | 1.900E-06 | 2.646E-07 | 1.700E-06 | 2.200E-06 |
| 14628 | 732 | V732G | B | 1.1e-06 | 1.6e-06 | 2.2e-06 | 1.633E-06 | 5.508E-07 | 1.100E-06 | 2.200E-06 |
| 14629 | 732 | V732H | B | 2.3e-06 | 3e-06 | 2.6e-06 | 2.633E-06 | 3.512E-07 | 2.300E-06 | 3.000E-06 |
| 14630 | 732 | V732I | B | 1.9e-06 | 3.3e-06 | 2e-06 | 2.400E-06 | 7.810E-07 | 1.900E-06 | 3.300E-06 |
| 14631 | 732 | V732L | B | 1.7e-06 | 4.3e-06 | 2.5e-06 | 2.833E-06 | 1.332E-06 | 1.700E-06 | 4.300E-06 |
| 14632 | 732 | V732K | B | 2.5e-06 | 4e-06 | 1.6e-06 | 2.700E-06 | 1.212E-06 | 1.600E-06 | 4.000E-06 |
| 14633 | 732 | V732M | B | 6.7e-06 | 3e-06 | 3.2e-06 | 4.300E-06 | 2.081E-06 | 3.000E-06 | 6.700E-06 |
| 14634 | 732 | V732F | B | 1.6e-06 | 1.9e-06 | 1.5e-06 | 1.667E-06 | 2.082E-07 | 1.500E-06 | 1.900E-06 |
| 14635 | 732 | V732P | B | 3e-06 | 2.5e-06 | 3.8e-06 | 3.100E-06 | 6.557E-07 | 2.500E-06 | 3.800E-06 |
| 14636 | 732 | V732S | B | 1.6e-06 | 2.8e-06 | 2.1e-06 | 2.167E-06 | 6.028E-07 | 1.600E-06 | 2.800E-06 |
| 14637 | 732 | V732T | B | 1.9e-06 | 3.6e-06 | 2.3e-06 | 2.600E-06 | 8.888E-07 | 1.900E-06 | 3.600E-06 |
| 14638 | 732 | V732W | B | 2e-06 | 2.3e-06 | 1.6e-06 | 1.967E-06 | 3.512E-07 | 1.600E-06 | 2.300E-06 |
| 14639 | 732 | V732Y | B | 2.9e-06 | 2.2e-06 | 2.2e-06 | 2.433E-06 | 4.041E-07 | 2.200E-06 | 2.900E-06 |
| 14640 | 732 | V732V | B | 2.6e-06 | 1.3e-06 | 3.3e-06 | 2.400E-06 | 1.015E-06 | 1.300E-06 | 3.300E-06 |
| 14641 | 733 | A733A | B | 2.3e-06 | 2.5e-06 | 2.3e-06 | 2.367E-06 | 1.155E-07 | 2.300E-06 | 2.500E-06 |
| 14642 | 733 | A733R | B | 3.4e-06 | 1.8e-06 | 2.3e-06 | 2.500E-06 | 8.185E-07 | 1.800E-06 | 3.400E-06 |
| 14643 | 733 | A733N | B | 2.6e-06 | 1.4e-06 | 2.5e-06 | 2.167E-06 | 6.658E-07 | 1.400E-06 | 2.600E-06 |
| 14644 | 733 | A733D | B | 2.3e-06 | 5.2e-06 | 2.2e-06 | 3.233E-06 | 1.704E-06 | 2.200E-06 | 5.200E-06 |
| 14645 | 733 | A733C | B | 9.7e-07 | 3e-06 | 3e-06 | 2.323E-06 | 1.172E-06 | 9.700E-07 | 3.000E-06 |
| 14646 | 733 | A733E | B | 2.2e-06 | 2.1e-06 | 2.9e-06 | 2.400E-06 | 4.359E-07 | 2.100E-06 | 2.900E-06 |
| 14647 | 733 | A733Q | B | 1.7e-06 | 1.8e-06 | 2.5e-06 | 2.000E-06 | 4.359E-07 | 1.700E-06 | 2.500E-06 |
| 14648 | 733 | A733G | B | 3.4e-06 | 3.5e-06 | 2.9e-06 | 3.267E-06 | 3.215E-07 | 2.900E-06 | 3.500E-06 |
| 14649 | 733 | A733H | B | 1.9e-06 | 1.9e-06 | 1.3e-06 | 1.700E-06 | 3.464E-07 | 1.300E-06 | 1.900E-06 |
| 14650 | 733 | A733I | B | 2.4e-06 | 1.6e-06 | 7.5e-07 | 1.583E-06 | 8.251E-07 | 7.500E-07 | 2.400E-06 |
| 14651 | 733 | A733L | B | 2.9e-06 | 5.6e-06 | 2.7e-06 | 3.733E-06 | 1.620E-06 | 2.700E-06 | 5.600E-06 |
| 14652 | 733 | A733K | B | 4.6e-06 | 1.6e-06 | 2.2e-06 | 2.800E-06 | 1.587E-06 | 1.600E-06 | 4.600E-06 |
| 14653 | 733 | A733M | B | 2.7e-06 | 1.3e-06 | 2.7e-06 | 2.233E-06 | 8.083E-07 | 1.300E-06 | 2.700E-06 |
| 14654 | 733 | A733F | B | 3e-06 | 2.7e-06 | 5.1e-06 | 3.600E-06 | 1.308E-06 | 2.700E-06 | 5.100E-06 |
| 14655 | 733 | A733P | B | 2.2e-06 | 2.2e-06 | 1.5e-06 | 1.967E-06 | 4.041E-07 | 1.500E-06 | 2.200E-06 |
| 14656 | 733 | A733S | B | 1.6e-06 | 2.6e-06 | 2.3e-06 | 2.167E-06 | 5.132E-07 | 1.600E-06 | 2.600E-06 |
| 14657 | 733 | A733T | B | 2.2e-06 | 2.8e-06 | 1.1e-06 | 2.033E-06 | 8.622E-07 | 1.100E-06 | 2.800E-06 |
| 14658 | 733 | A733W | B | 2.4e-06 | 3.8e-06 | 1.9e-06 | 2.700E-06 | 9.849E-07 | 1.900E-06 | 3.800E-06 |
| 14659 | 733 | A733Y | B | 2.6e-06 | 2.2e-06 | 3.8e-06 | 2.867E-06 | 8.327E-07 | 2.200E-06 | 3.800E-06 |
| 14660 | 733 | A733V | B | 3.1e-06 | 3.5e-06 | 3.1e-06 | 3.233E-06 | 2.309E-07 | 3.100E-06 | 3.500E-06 |
| 14661 | 734 | A734A | B | 3e-06 | 2.2e-06 | 3.3e-06 | 2.833E-06 | 5.686E-07 | 2.200E-06 | 3.300E-06 |
| 14662 | 734 | A734R | B | 2.5e-06 | 1.8e-06 | 1.7e-06 | 2.000E-06 | 4.359E-07 | 1.700E-06 | 2.500E-06 |
| 14663 | 734 | A734N | B | 3.5e-06 | 2.7e-06 | 2e-06 | 2.733E-06 | 7.506E-07 | 2.000E-06 | 3.500E-06 |
| 14664 | 734 | A734D | B | 2.5e-06 | 3.7e-06 | 2.6e-06 | 2.933E-06 | 6.658E-07 | 2.500E-06 | 3.700E-06 |
| 14665 | 734 | A734C | B | 2.8e-06 | 3.7e-06 | 2.3e-06 | 2.933E-06 | 7.095E-07 | 2.300E-06 | 3.700E-06 |
| 14666 | 734 | A734E | B | 2.8e-06 | 4.3e-06 | 1.4e-06 | 2.833E-06 | 1.450E-06 | 1.400E-06 | 4.300E-06 |
| 14667 | 734 | A734Q | B | 3.3e-06 | 2.7e-06 | 2.1e-06 | 2.700E-06 | 6.000E-07 | 2.100E-06 | 3.300E-06 |
| 14668 | 734 | A734G | B | 3.3e-06 | 3e-06 | 2.4e-06 | 2.900E-06 | 4.583E-07 | 2.400E-06 | 3.300E-06 |
| 14669 | 734 | A734H | B | 1.7e-06 | 1.6e-06 | 2.3e-06 | 1.867E-06 | 3.786E-07 | 1.600E-06 | 2.300E-06 |
| 14670 | 734 | A734I | B | 1.7e-06 | 2.8e-06 | 1.7e-06 | 2.067E-06 | 6.351E-07 | 1.700E-06 | 2.800E-06 |
| 14671 | 734 | A734L | B | 2e-06 | 1.3e-06 | 2.1e-06 | 1.800E-06 | 4.359E-07 | 1.300E-06 | 2.100E-06 |
| 14672 | 734 | A734K | B | 1.7e-06 | 2.7e-06 | 2.6e-06 | 2.333E-06 | 5.508E-07 | 1.700E-06 | 2.700E-06 |
| 14673 | 734 | A734M | B | 6.6e-07 | 2.1e-06 | 2.4e-06 | 1.720E-06 | 9.302E-07 | 6.600E-07 | 2.400E-06 |
| 14674 | 734 | A734F | B | 2.7e-06 | 2.8e-06 | 2e-06 | 2.500E-06 | 4.359E-07 | 2.000E-06 | 2.800E-06 |
| 14675 | 734 | A734P | B | 2.1e-06 | 8.9e-07 | 1.6e-06 | 1.530E-06 | 6.080E-07 | 8.900E-07 | 2.100E-06 |
| 14676 | 734 | A734S | B | 2.9e-06 | 2.6e-06 | 1.3e-06 | 2.267E-06 | 8.505E-07 | 1.300E-06 | 2.900E-06 |
| 14677 | 734 | A734T | B | 2.9e-06 | 2.5e-06 | 1.7e-06 | 2.367E-06 | 6.110E-07 | 1.700E-06 | 2.900E-06 |
| 14678 | 734 | A734W | B | 1.8e-06 | 1.4e-06 | 4.2e-06 | 2.467E-06 | 1.514E-06 | 1.400E-06 | 4.200E-06 |
| 14679 | 734 | A734Y | B | 1.6e-06 | 4.5e-06 | 3.2e-06 | 3.100E-06 | 1.453E-06 | 1.600E-06 | 4.500E-06 |
| 14680 | 734 | A734V | B | 3.6e-06 | 3.4e-06 | 2.4e-06 | 3.133E-06 | 6.429E-07 | 2.400E-06 | 3.600E-06 |
| 14681 | 735 | P735A | B | 3.8e-06 | 3.2e-06 | 2.9e-06 | 3.300E-06 | 4.583E-07 | 2.900E-06 | 3.800E-06 |
| 14682 | 735 | P735R | B | 3.5e-06 | 2.8e-06 | 2e-06 | 2.767E-06 | 7.506E-07 | 2.000E-06 | 3.500E-06 |
| 14683 | 735 | P735N | B | 2.2e-06 | 3.6e-06 | 2.3e-06 | 2.700E-06 | 7.810E-07 | 2.200E-06 | 3.600E-06 |
| 14684 | 735 | P735D | B | 3e-06 | 1.3e-06 | 2.6e-06 | 2.300E-06 | 8.888E-07 | 1.300E-06 | 3.000E-06 |
| 14685 | 735 | P735C | B | 2.1e-06 | 2.6e-06 | 1.5e-06 | 2.067E-06 | 5.508E-07 | 1.500E-06 | 2.600E-06 |
| 14686 | 735 | P735E | B | 3.2e-06 | 1.8e-06 | 2.6e-06 | 2.533E-06 | 7.024E-07 | 1.800E-06 | 3.200E-06 |
| 14687 | 735 | P735Q | B | 1.6e-06 | 1.8e-06 | 3.7e-06 | 2.367E-06 | 1.159E-06 | 1.600E-06 | 3.700E-06 |
| 14688 | 735 | P735G | B | 2.7e-06 | 3.4e-06 | 2.8e-06 | 2.967E-06 | 3.786E-07 | 2.700E-06 | 3.400E-06 |
| 14689 | 735 | P735H | B | 1.7e-06 | 2.3e-06 | 1.7e-06 | 1.900E-06 | 3.464E-07 | 1.700E-06 | 2.300E-06 |
| 14690 | 735 | P735I | B | 2.4e-06 | 3.1e-06 | 2.6e-06 | 2.700E-06 | 3.606E-07 | 2.400E-06 | 3.100E-06 |
| 14691 | 735 | P735L | B | 1.9e-06 | 3.2e-06 | 1.8e-06 | 2.300E-06 | 7.810E-07 | 1.800E-06 | 3.200E-06 |
| 14692 | 735 | P735K | B | 1.6e-06 | 5e-06 | 6.3e-06 | 4.300E-06 | 2.427E-06 | 1.600E-06 | 6.300E-06 |
| 14693 | 735 | P735M | B | 1.7e-06 | 2.4e-06 | 3.1e-06 | 2.400E-06 | 7.000E-07 | 1.700E-06 | 3.100E-06 |
| 14694 | 735 | P735F | B | 2.4e-06 | 3.8e-06 | 1.7e-06 | 2.633E-06 | 1.069E-06 | 1.700E-06 | 3.800E-06 |
| 14695 | 735 | P735P | B | 2.3e-06 | 2.5e-06 | 1.8e-06 | 2.200E-06 | 3.606E-07 | 1.800E-06 | 2.500E-06 |
| 14696 | 735 | P735S | B | 4e-06 | 2.6e-06 | 2.1e-06 | 2.900E-06 | 9.849E-07 | 2.100E-06 | 4.000E-06 |
| 14697 | 735 | P735T | B | 3.4e-06 | 2.3e-06 | 1e-06 | 2.233E-06 | 1.201E-06 | 1.000E-06 | 3.400E-06 |
| 14698 | 735 | P735W | B | 2e-06 | 1.9e-06 | 2.2e-06 | 2.033E-06 | 1.528E-07 | 1.900E-06 | 2.200E-06 |
| 14699 | 735 | P735Y | B | 4e-06 | 3.5e-06 | 2.1e-06 | 3.200E-06 | 9.849E-07 | 2.100E-06 | 4.000E-06 |
| 14700 | 735 | P735V | B | 1.9e-06 | 4.2e-06 | 2.8e-06 | 2.967E-06 | 1.159E-06 | 1.900E-06 | 4.200E-06 |
| 14701 | 736 | S736A | B | 3.1e-06 | 4.5e-06 | 2.2e-06 | 3.267E-06 | 1.159E-06 | 2.200E-06 | 4.500E-06 |
| 14702 | 736 | S736R | B | 1.8e-06 | 2.2e-06 | 2.7e-06 | 2.233E-06 | 4.509E-07 | 1.800E-06 | 2.700E-06 |
| 14703 | 736 | S736N | B | 3.3e-06 | 2.3e-06 | 2.3e-06 | 2.633E-06 | 5.774E-07 | 2.300E-06 | 3.300E-06 |
| 14704 | 736 | S736D | B | 3.6e-06 | 2.4e-06 | 2.6e-06 | 2.867E-06 | 6.429E-07 | 2.400E-06 | 3.600E-06 |

| | | | | | | | | | | |
|---|---|---|---|---|---|---|---|---|---|---|
| 14705 | 736 | S736C | B | 1.4e-06 | 1.6e-06 | 2.2e-06 | 1.733E-06 | 4.163E-07 | 1.400E-06 | 2.200E-06 |
| 14706 | 736 | S736E | B | 2.8e-06 | 2.4e-06 | 3.9e-06 | 3.033E-06 | 7.767E-07 | 2.400E-06 | 3.900E-06 |
| 14707 | 736 | S736Q | B | 1.8e-06 | 2e-06 | 2.4e-06 | 2.067E-06 | 3.055E-07 | 1.800E-06 | 2.400E-06 |
| 14708 | 736 | S736G | B | 3.7e-06 | 1.8e-06 | 3.3e-06 | 2.933E-06 | 1.002E-06 | 1.800E-06 | 3.700E-06 |
| 14709 | 736 | S736H | B | 3.2e-06 | 2.6e-06 | 2.4e-06 | 2.733E-06 | 4.163E-07 | 2.400E-06 | 3.200E-06 |
| 14710 | 736 | S736I | B | 3e-06 | 2.8e-06 | 2.6e-06 | 2.800E-06 | 2.000E-07 | 2.600E-06 | 3.000E-06 |
| 14711 | 736 | S736L | B | 3e-06 | 2.4e-06 | 3e-06 | 2.800E-06 | 3.464E-07 | 2.400E-06 | 3.000E-06 |
| 14712 | 736 | S736K | B | 1.2e-06 | 3.9e-06 | 3.1e-06 | 2.733E-06 | 1.387E-06 | 1.200E-06 | 3.900E-06 |
| 14713 | 736 | S736M | B | 2.8e-06 | 2.8e-06 | 1.9e-06 | 2.500E-06 | 5.196E-07 | 1.900E-06 | 2.800E-06 |
| 14714 | 736 | S736F | B | 3.2e-06 | 1.5e-06 | 2.7e-06 | 2.467E-06 | 8.737E-07 | 1.500E-06 | 3.200E-06 |
| 14715 | 736 | S736P | B | 2.8e-06 | 2.3e-06 | 2.2e-06 | 2.433E-06 | 3.215E-07 | 2.200E-06 | 2.800E-06 |
| 14716 | 736 | S736S | B | 2.1e-06 | 4.5e-06 | 2.4e-06 | 3.000E-06 | 1.308E-06 | 2.100E-06 | 4.500E-06 |
| 14717 | 736 | S736T | B | 3.3e-06 | 1.4e-06 | 1.1e-06 | 1.933E-06 | 1.193E-06 | 1.100E-06 | 3.300E-06 |
| 14718 | 736 | S736W | B | 1.9e-06 | 2.3e-06 | 2.9e-06 | 2.367E-06 | 5.033E-07 | 1.900E-06 | 2.900E-06 |
| 14719 | 736 | S736Y | B | 3.8e-06 | 2.6e-06 | 1.6e-06 | 2.667E-06 | 1.102E-06 | 1.600E-06 | 3.800E-06 |
| 14720 | 736 | S736V | B | 2.2e-06 | 2.9e-06 | 2.3e-06 | 2.467E-06 | 3.786E-07 | 2.200E-06 | 2.900E-06 |
| 14721 | 737 | V737A | B | 2.2e-06 | 2.8e-06 | 2.2e-06 | 2.400E-06 | 3.464E-07 | 2.200E-06 | 2.800E-06 |
| 14722 | 737 | V737R | B | 2e-06 | 1.7e-06 | 3.1e-06 | 2.267E-06 | 7.371E-07 | 1.700E-06 | 3.100E-06 |
| 14723 | 737 | V737N | B | 3e-06 | 2.6e-06 | 2.6e-06 | 2.733E-06 | 2.309E-07 | 2.600E-06 | 3.000E-06 |
| 14724 | 737 | V737D | B | 2.1e-06 | 1.8e-06 | 2.5e-06 | 2.133E-06 | 3.512E-07 | 1.800E-06 | 2.500E-06 |
| 14725 | 737 | V737C | B | 2.4e-06 | 2.8e-06 | 2.2e-06 | 2.467E-06 | 3.055E-07 | 2.200E-06 | 2.800E-06 |
| 14726 | 737 | V737E | B | 1.8e-06 | 2.2e-06 | 3.3e-06 | 2.433E-06 | 7.767E-07 | 1.800E-06 | 3.300E-06 |
| 14727 | 737 | V737Q | B | 2.3e-06 | 2.6e-06 | 2.1e-06 | 2.333E-06 | 2.517E-07 | 2.100E-06 | 2.600E-06 |
| 14728 | 737 | V737G | B | 2.6e-06 | 1.4e-06 | 2.6e-06 | 2.200E-06 | 6.928E-07 | 1.400E-06 | 2.600E-06 |
| 14729 | 737 | V737H | B | 1.5e-06 | 3.4e-06 | 1.7e-06 | 2.200E-06 | 1.044E-06 | 1.500E-06 | 3.400E-06 |
| 14730 | 737 | V737I | B | 3.6e-06 | 3e-06 | 1.5e-06 | 2.700E-06 | 1.082E-06 | 1.500E-06 | 3.600E-06 |
| 14731 | 737 | V737L | B | 2e-06 | 3e-06 | 1.9e-06 | 2.300E-06 | 6.083E-07 | 1.900E-06 | 3.000E-06 |
| 14732 | 737 | V737K | B | 2.7e-06 | 2.2e-06 | 2.6e-06 | 2.500E-06 | 2.646E-07 | 2.200E-06 | 2.700E-06 |
| 14733 | 737 | V737M | B | 3.6e-06 | 1.8e-06 | 2.2e-06 | 2.533E-06 | 9.452E-07 | 1.800E-06 | 3.600E-06 |
| 14734 | 737 | V737F | B | 2.3e-06 | 2e-06 | 2.5e-06 | 2.267E-06 | 2.517E-07 | 2.000E-06 | 2.500E-06 |
| 14735 | 737 | V737P | B | 3.6e-06 | 4.3e-06 | 2.1e-06 | 3.333E-06 | 1.124E-06 | 2.100E-06 | 4.300E-06 |
| 14736 | 737 | V737S | B | 1.3e-06 | 2.3e-06 | 1.9e-06 | 1.833E-06 | 5.033E-07 | 1.300E-06 | 2.300E-06 |
| 14737 | 737 | V737T | B | 1.9e-06 | 3.1e-06 | 3.1e-06 | 2.700E-06 | 6.928E-07 | 1.900E-06 | 3.100E-06 |
| 14738 | 737 | V737W | B | 2.6e-06 | 2.4e-06 | 1.8e-06 | 2.267E-06 | 4.163E-07 | 1.800E-06 | 2.600E-06 |
| 14739 | 737 | V737Y | B | 2.5e-06 | 2.7e-06 | 1.3e-06 | 2.167E-06 | 7.572E-07 | 1.300E-06 | 2.700E-06 |
| 14740 | 737 | V737V | B | 3e-06 | 2e-06 | 3.8e-06 | 2.933E-06 | 9.018E-07 | 2.000E-06 | 3.800E-06 |
| 14741 | 738 | F738A | B | 1.9e-06 | 1.7e-06 | 2.3e-06 | 1.967E-06 | 3.055E-07 | 1.700E-06 | 2.300E-06 |
| 14742 | 738 | F738R | B | 2.6e-06 | 1e-06 | 3.3e-06 | 2.300E-06 | 1.179E-06 | 1.000E-06 | 3.300E-06 |
| 14743 | 738 | F738N | B | 1.8e-06 | 2.1e-06 | 2.2e-06 | 2.033E-06 | 2.082E-07 | 1.800E-06 | 2.200E-06 |
| 14744 | 738 | F738D | B | 3.5e-06 | 2.4e-06 | 3.9e-06 | 3.267E-06 | 7.767E-07 | 2.400E-06 | 3.900E-06 |
| 14745 | 738 | F738C | B | 2.4e-06 | 1e-06 | 1.6e-06 | 1.667E-06 | 7.024E-07 | 1.000E-06 | 2.400E-06 |
| 14746 | 738 | F738E | B | 2.4e-06 | 2.3e-06 | 1.7e-06 | 2.133E-06 | 3.786E-07 | 1.700E-06 | 2.400E-06 |
| 14747 | 738 | F738Q | B | 1.4e-06 | 2.6e-06 | 1.4e-06 | 1.800E-06 | 6.928E-07 | 1.400E-06 | 2.600E-06 |
| 14748 | 738 | F738G | B | 2.2e-06 | 3.4e-06 | 1.6e-06 | 2.400E-06 | 9.165E-07 | 1.600E-06 | 3.400E-06 |
| 14749 | 738 | F738H | B | 1.8e-06 | 2.3e-06 | 2.1e-06 | 2.067E-06 | 2.517E-07 | 1.800E-06 | 2.300E-06 |
| 14750 | 738 | F738I | B | 2.7e-06 | 2.2e-06 | 1.8e-06 | 2.233E-06 | 4.509E-07 | 1.800E-06 | 2.700E-06 |
| 14751 | 738 | F738L | B | 2.1e-06 | 9.4e-07 | 2.3e-06 | 1.780E-06 | 7.343E-07 | 9.400E-07 | 2.300E-06 |
| 14752 | 738 | F738K | B | 2.6e-06 | 2.6e-06 | 3.3e-06 | 2.833E-06 | 4.041E-07 | 2.600E-06 | 3.300E-06 |
| 14753 | 738 | F738M | B | 1.3e-06 | 2.3e-06 | 4.3e-06 | 2.633E-06 | 1.528E-06 | 1.300E-06 | 4.300E-06 |
| 14754 | 738 | F738F | B | 2.1e-06 | 3.8e-06 | 3.4e-06 | 3.100E-06 | 8.888E-07 | 2.100E-06 | 3.800E-06 |
| 14755 | 738 | F738P | B | 4e-06 | 2.2e-06 | 1.6e-06 | 2.600E-06 | 1.249E-06 | 1.600E-06 | 4.000E-06 |
| 14756 | 738 | F738S | B | 2.3e-06 | 1.3e-06 | 1.2e-06 | 1.600E-06 | 6.083E-07 | 1.200E-06 | 2.300E-06 |
| 14757 | 738 | F738T | B | 2.2e-06 | 2.6e-06 | 1.9e-06 | 2.233E-06 | 3.512E-07 | 1.900E-06 | 2.600E-06 |
| 14758 | 738 | F738W | B | 3.8e-06 | 2.5e-06 | 5.7e-06 | 4.000E-06 | 1.609E-06 | 2.500E-06 | 5.700E-06 |
| 14759 | 738 | F738Y | B | 2.6e-06 | 2.6e-06 | 1.3e-06 | 2.167E-06 | 7.506E-07 | 1.300E-06 | 2.600E-06 |
| 14760 | 738 | F738V | B | 2.3e-06 | 3.8e-06 | 2.3e-06 | 2.800E-06 | 8.660E-07 | 2.300E-06 | 3.800E-06 |
| 14761 | 739 | I739A | B | 4.8e-06 | 2.4e-06 | 2e-06 | 3.067E-06 | 1.514E-06 | 2.000E-06 | 4.800E-06 |
| 14762 | 739 | I739R | B | 2.4e-06 | 2.6e-06 | 2.7e-06 | 2.567E-06 | 1.528E-07 | 2.400E-06 | 2.700E-06 |
| 14763 | 739 | I739N | B | 1.6e-06 | 2.5e-06 | 3.7e-06 | 2.600E-06 | 1.054E-06 | 1.600E-06 | 3.700E-06 |
| 14764 | 739 | I739D | B | 2.3e-06 | 2.6e-06 | 2.4e-06 | 2.433E-06 | 1.528E-07 | 2.300E-06 | 2.600E-06 |
| 14765 | 739 | I739C | B | 2.2e-06 | 2.2e-06 | 2.9e-06 | 2.433E-06 | 4.041E-07 | 2.200E-06 | 2.900E-06 |
| 14766 | 739 | I739E | B | 2.5e-06 | 3.1e-06 | 2e-06 | 2.533E-06 | 5.508E-07 | 2.000E-06 | 3.100E-06 |
| 14767 | 739 | I739Q | B | 3.8e-06 | 2.1e-06 | 2.2e-06 | 2.700E-06 | 9.539E-07 | 2.100E-06 | 3.800E-06 |
| 14768 | 739 | I739G | B | 3.7e-06 | 2.3e-06 | 1.6e-06 | 2.533E-06 | 1.069E-06 | 1.600E-06 | 3.700E-06 |
| 14769 | 739 | I739H | B | 2.8e-06 | 2.1e-06 | 4.1e-06 | 3.000E-06 | 1.015E-06 | 2.100E-06 | 4.100E-06 |
| 14770 | 739 | I739I | B | 1.5e-06 | 2.4e-06 | 2.2e-06 | 2.033E-06 | 4.726E-07 | 1.500E-06 | 2.400E-06 |
| 14771 | 739 | I739L | B | 2.5e-06 | 1.7e-06 | 1.6e-06 | 1.933E-06 | 4.933E-07 | 1.600E-06 | 2.500E-06 |
| 14772 | 739 | I739K | B | 3.3e-06 | 3e-06 | 2.6e-06 | 2.967E-06 | 3.512E-07 | 2.600E-06 | 3.300E-06 |
| 14773 | 739 | I739M | B | 2.3e-06 | 2.7e-06 | 4.3e-06 | 3.100E-06 | 1.058E-06 | 2.300E-06 | 4.300E-06 |
| 14774 | 739 | I739F | B | 1.9e-06 | 2e-06 | 1.6e-06 | 1.833E-06 | 2.082E-07 | 1.600E-06 | 2.000E-06 |
| 14775 | 739 | I739P | B | 4.1e-06 | 1.8e-06 | 1.4e-06 | 2.433E-06 | 1.457E-06 | 1.400E-06 | 4.100E-06 |
| 14776 | 739 | I739S | B | 1.7e-06 | 2.6e-06 | 4.5e-06 | 2.933E-06 | 1.429E-06 | 1.700E-06 | 4.500E-06 |
| 14777 | 739 | I739T | B | 2.4e-06 | 1.7e-06 | 1.7e-06 | 1.933E-06 | 4.041E-07 | 1.700E-06 | 2.400E-06 |
| 14778 | 739 | I739W | B | 1.6e-06 | 2.4e-06 | 3.4e-06 | 2.467E-06 | 9.018E-07 | 1.600E-06 | 3.400E-06 |
| 14779 | 739 | I739Y | B | 2.4e-06 | 2.1e-06 | 3.3e-06 | 2.600E-06 | 6.245E-07 | 2.100E-06 | 3.300E-06 |
| 14780 | 739 | I739V | B | 1.9e-06 | 3.7e-06 | 2.8e-06 | 2.800E-06 | 9.000E-07 | 1.900E-06 | 3.700E-06 |
| 14781 | 740 | F740A | B | 2.3e-06 | 2e-06 | 4.1e-06 | 2.800E-06 | 1.136E-06 | 2.000E-06 | 4.100E-06 |
| 14782 | 740 | F740R | B | 1.3e-06 | 1.9e-06 | 4.4e-06 | 2.533E-06 | 1.644E-06 | 1.300E-06 | 4.400E-06 |
| 14783 | 740 | F740N | B | 2.5e-06 | 2.1e-06 | 1.4e-06 | 2.000E-06 | 5.568E-07 | 1.400E-06 | 2.500E-06 |
| 14784 | 740 | F740D | B | 2.9e-06 | 1.9e-06 | 2.1e-06 | 2.300E-06 | 5.292E-07 | 1.900E-06 | 2.900E-06 |
| 14785 | 740 | F740C | B | 1.3e-06 | 3e-06 | 2.4e-06 | 2.233E-06 | 8.622E-07 | 1.300E-06 | 3.000E-06 |
| 14786 | 740 | F740E | B | 2.2e-06 | 1.2e-06 | 3.4e-06 | 2.267E-06 | 1.102E-06 | 1.200E-06 | 3.400E-06 |
| 14787 | 740 | F740Q | B | 2e-06 | 3e-06 | 3.9e-06 | 2.967E-06 | 9.504E-07 | 2.000E-06 | 3.900E-06 |
| 14788 | 740 | F740G | B | 3.9e-06 | 2.6e-06 | 2.7e-06 | 3.067E-06 | 7.234E-07 | 2.600E-06 | 3.900E-06 |
| 14789 | 740 | F740H | B | 2.1e-06 | 2.4e-06 | 2.5e-06 | 2.333E-06 | 2.082E-07 | 2.100E-06 | 2.500E-06 |

| | | | | | | | | | | |
|---|---|---|---|---|---|---|---|---|---|---|
| 14790 | 740 | F740I | B | 2.7e-06 | 2e-06 | 1.5e-06 | 2.067E-06 | 6.028E-07 | 1.500E-06 | 2.700E-06 |
| 14791 | 740 | F740L | B | 1.9e-06 | 3.1e-06 | 3e-06 | 2.667E-06 | 6.658E-07 | 1.900E-06 | 3.100E-06 |
| 14792 | 740 | F740K | B | 1.5e-06 | 2.2e-06 | 2.8e-06 | 2.167E-06 | 6.506E-07 | 1.500E-06 | 2.800E-06 |
| 14793 | 740 | F740M | B | 2.6e-06 | 1.5e-06 | 3.4e-06 | 2.500E-06 | 9.539E-07 | 1.500E-06 | 3.400E-06 |
| 14794 | 740 | F740F | B | 1.9e-06 | 2.6e-06 | 1.6e-06 | 2.033E-06 | 5.132E-07 | 1.600E-06 | 2.600E-06 |
| 14795 | 740 | F740P | B | 1.4e-06 | 2.5e-06 | 1.9e-06 | 1.933E-06 | 5.508E-07 | 1.400E-06 | 2.500E-06 |
| 14796 | 740 | F740S | B | 3.4e-06 | 2.2e-06 | 1.7e-06 | 2.433E-06 | 8.737E-07 | 1.700E-06 | 3.400E-06 |
| 14797 | 740 | F740T | B | 1.3e-06 | 2e-06 | 2.3e-06 | 1.867E-06 | 5.132E-07 | 1.300E-06 | 2.300E-06 |
| 14798 | 740 | F740W | B | 1.8e-06 | 1.9e-06 | 1.6e-06 | 1.767E-06 | 1.528E-07 | 1.600E-06 | 1.900E-06 |
| 14799 | 740 | F740Y | B | 2.2e-06 | 2.2e-06 | 1.4e-06 | 1.933E-06 | 4.619E-07 | 1.400E-06 | 2.200E-06 |
| 14800 | 740 | F740V | B | 2.4e-06 | 2.3e-06 | 1.1e-06 | 1.933E-06 | 7.234E-07 | 1.100E-06 | 2.400E-06 |
| 14801 | 741 | P741A | B | 4.5e-06 | 1.4e-06 | 1.9e-06 | 2.600E-06 | 1.664E-06 | 1.400E-06 | 4.500E-06 |
| 14802 | 741 | P741R | B | 3.3e-06 | 2.4e-06 | 1.6e-06 | 2.433E-06 | 8.505E-07 | 1.600E-06 | 3.300E-06 |
| 14803 | 741 | P741N | B | 3e-06 | 2.8e-06 | 3.3e-06 | 3.033E-06 | 2.517E-07 | 2.800E-06 | 3.300E-06 |
| 14804 | 741 | P741D | B | 2.1e-06 | 1.9e-06 | 2.4e-06 | 2.133E-06 | 2.517E-07 | 1.900E-06 | 2.400E-06 |
| 14805 | 741 | P741C | B | 1.7e-06 | 2.4e-06 | 2.2e-06 | 2.100E-06 | 3.606E-07 | 1.700E-06 | 2.400E-06 |
| 14806 | 741 | P741E | B | 3.3e-06 | 1.5e-06 | 1.9e-06 | 2.233E-06 | 9.452E-07 | 1.500E-06 | 3.300E-06 |
| 14807 | 741 | P741Q | B | 2.4e-06 | 4.1e-06 | 1.6e-06 | 2.700E-06 | 1.277E-06 | 1.600E-06 | 4.100E-06 |
| 14808 | 741 | P741G | B | 2.3e-06 | 1.7e-06 | 2.3e-06 | 2.100E-06 | 3.464E-07 | 1.700E-06 | 2.300E-06 |
| 14809 | 741 | P741H | B | 2.9e-06 | 2.8e-06 | 1.8e-06 | 2.500E-06 | 6.083E-07 | 1.800E-06 | 2.900E-06 |
| 14810 | 741 | P741I | B | 2.1e-06 | 2.6e-06 | 1.6e-06 | 2.100E-06 | 5.000E-07 | 1.600E-06 | 2.600E-06 |
| 14811 | 741 | P741L | B | 2.5e-06 | 2.1e-06 | 2e-06 | 2.200E-06 | 2.646E-07 | 2.000E-06 | 2.500E-06 |
| 14812 | 741 | P741K | B | 1.8e-06 | 1.4e-06 | 2.7e-06 | 1.967E-06 | 6.658E-07 | 1.400E-06 | 2.700E-06 |
| 14813 | 741 | P741M | B | 1.5e-06 | 4e-06 | 2.1e-06 | 2.533E-06 | 1.305E-06 | 1.500E-06 | 4.000E-06 |
| 14814 | 741 | P741F | B | 2.8e-06 | 3.8e-06 | 3.1e-06 | 3.233E-06 | 5.132E-07 | 2.800E-06 | 3.800E-06 |
| 14815 | 741 | P741P | B | 2.1e-06 | 1.7e-06 | 2.7e-06 | 2.167E-06 | 5.033E-07 | 1.700E-06 | 2.700E-06 |
| 14816 | 741 | P741S | B | 1.6e-06 | 2e-06 | 2.6e-06 | 2.067E-06 | 5.033E-07 | 1.600E-06 | 2.600E-06 |
| 14817 | 741 | P741T | B | 2e-06 | 1.8e-06 | 1.8e-06 | 1.867E-06 | 1.155E-07 | 1.800E-06 | 2.000E-06 |
| 14818 | 741 | P741W | B | 1.7e-06 | 3e-06 | 2.2e-06 | 2.300E-06 | 6.557E-07 | 1.700E-06 | 3.000E-06 |
| 14819 | 741 | P741Y | B | 1.9e-06 | 3.5e-06 | 2.2e-06 | 2.533E-06 | 8.505E-07 | 1.900E-06 | 3.500E-06 |
| 14820 | 741 | P741V | B | 2.8e-06 | 2.5e-06 | 3.5e-06 | 2.933E-06 | 5.132E-07 | 2.500E-06 | 3.500E-06 |
| 14821 | 742 | P742A | B | 1.3e-06 | 3.3e-06 | 2e-06 | 2.200E-06 | 1.015E-06 | 1.300E-06 | 3.300E-06 |
| 14822 | 742 | P742R | B | 3.6e-06 | 2e-06 | 3.5e-06 | 3.033E-06 | 8.963E-07 | 2.000E-06 | 3.600E-06 |
| 14823 | 742 | P742N | B | 2.3e-06 | 3.1e-06 | 1.9e-06 | 2.433E-06 | 6.110E-07 | 1.900E-06 | 3.100E-06 |
| 14824 | 742 | P742D | B | 4e-06 | 3.7e-06 | 1.5e-06 | 3.067E-06 | 1.365E-06 | 1.500E-06 | 4.000E-06 |
| 14825 | 742 | P742C | B | 1.3e-06 | 2.2e-06 | 2.1e-06 | 1.867E-06 | 4.933E-07 | 1.300E-06 | 2.200E-06 |
| 14826 | 742 | P742E | B | 2.2e-06 | 1.1e-06 | 2.3e-06 | 1.867E-06 | 6.658E-07 | 1.100E-06 | 2.300E-06 |
| 14827 | 742 | P742Q | B | 2.2e-06 | 2.2e-06 | 2.9e-06 | 2.433E-06 | 4.041E-07 | 2.200E-06 | 2.900E-06 |
| 14828 | 742 | P742G | B | 3e-06 | 1.2e-06 | 3.2e-06 | 2.467E-06 | 1.102E-06 | 1.200E-06 | 3.200E-06 |
| 14829 | 742 | P742H | B | 2.5e-06 | 2.4e-06 | 3.4e-06 | 2.767E-06 | 5.508E-07 | 2.400E-06 | 3.400E-06 |
| 14830 | 742 | P742I | B | 2.7e-06 | 2e-06 | 2e-06 | 2.233E-06 | 4.041E-07 | 2.000E-06 | 2.700E-06 |
| 14831 | 742 | P742L | B | 2.4e-06 | 1.8e-06 | 2.4e-06 | 2.200E-06 | 3.464E-07 | 1.800E-06 | 2.400E-06 |
| 14832 | 742 | P742K | B | 1.7e-06 | 1.6e-06 | 1.9e-06 | 1.733E-06 | 1.528E-07 | 1.600E-06 | 1.900E-06 |
| 14833 | 742 | P742M | B | 3e-06 | 1.3e-06 | 1.6e-06 | 1.967E-06 | 9.074E-07 | 1.300E-06 | 3.000E-06 |
| 14834 | 742 | P742F | B | 1.9e-06 | 3.7e-06 | 1.3e-06 | 2.300E-06 | 1.249E-06 | 1.300E-06 | 3.700E-06 |
| 14835 | 742 | P742P | B | 2.4e-06 | 1.7e-06 | 5e-06 | 3.033E-06 | 1.739E-06 | 1.700E-06 | 5.000E-06 |
| 14836 | 742 | P742S | B | 2.5e-06 | 1.3e-06 | 1.3e-06 | 1.700E-06 | 6.928E-07 | 1.300E-06 | 2.500E-06 |
| 14837 | 742 | P742T | B | 2e-06 | 2.2e-06 | 2.1e-06 | 2.100E-06 | 1.000E-07 | 2.000E-06 | 2.200E-06 |
| 14838 | 742 | P742W | B | 2.4e-06 | 1.8e-06 | 2.7e-06 | 2.300E-06 | 4.583E-07 | 1.800E-06 | 2.700E-06 |
| 14839 | 742 | P742Y | B | 1.9e-06 | 3.2e-06 | 2.2e-06 | 2.433E-06 | 6.807E-07 | 1.900E-06 | 3.200E-06 |
| 14840 | 742 | P742V | B | 4.7e-06 | 2.1e-06 | 1.8e-06 | 2.867E-06 | 1.595E-06 | 1.800E-06 | 4.700E-06 |
| 14841 | 743 | S743A | B | 3.3e-06 | 1.6e-06 | 2.8e-06 | 2.567E-06 | 8.737E-07 | 1.600E-06 | 3.300E-06 |
| 14842 | 743 | S743R | B | 2.2e-06 | 2.2e-06 | 2.9e-06 | 2.433E-06 | 4.041E-07 | 2.200E-06 | 2.900E-06 |
| 14843 | 743 | S743N | B | 2.1e-06 | 2.3e-06 | 2.5e-06 | 2.300E-06 | 2.000E-07 | 2.100E-06 | 2.500E-06 |
| 14844 | 743 | S743D | B | 2.3e-06 | 1.5e-06 | 4.2e-06 | 2.667E-06 | 1.387E-06 | 1.500E-06 | 4.200E-06 |
| 14845 | 743 | S743C | B | 1.5e-06 | 3e-06 | 2.1e-06 | 2.200E-06 | 7.550E-07 | 1.500E-06 | 3.000E-06 |
| 14846 | 743 | S743E | B | 3.7e-06 | 1.9e-06 | 3.3e-06 | 2.967E-06 | 9.452E-07 | 1.900E-06 | 3.700E-06 |
| 14847 | 743 | S743Q | B | 2.6e-06 | 2.9e-06 | 1.7e-06 | 2.400E-06 | 6.245E-07 | 1.700E-06 | 2.900E-06 |
| 14848 | 743 | S743G | B | 3.2e-06 | 2.3e-06 | 2.7e-06 | 2.733E-06 | 4.509E-07 | 2.300E-06 | 3.200E-06 |
| 14849 | 743 | S743H | B | 1.5e-06 | 1.1e-06 | 2.1e-06 | 1.567E-06 | 5.033E-07 | 1.100E-06 | 2.100E-06 |
| 14850 | 743 | S743I | B | 2.2e-06 | 2.1e-06 | 3.9e-06 | 2.733E-06 | 1.012E-06 | 2.100E-06 | 3.900E-06 |
| 14851 | 743 | S743L | B | 1.8e-06 | 4.1e-06 | 3.1e-06 | 3.000E-06 | 1.153E-06 | 1.800E-06 | 4.100E-06 |
| 14852 | 743 | S743K | B | 1.5e-06 | 2.5e-06 | 1.3e-06 | 1.767E-06 | 6.429E-07 | 1.300E-06 | 2.500E-06 |
| 14853 | 743 | S743M | B | 1.4e-06 | 2.8e-06 | 3.1e-06 | 2.433E-06 | 9.074E-07 | 1.400E-06 | 3.100E-06 |
| 14854 | 743 | S743F | B | 2.9e-06 | 2.4e-06 | 2.5e-06 | 2.600E-06 | 2.646E-07 | 2.400E-06 | 2.900E-06 |
| 14855 | 743 | S743P | B | 3.1e-06 | 1.8e-06 | 1.9e-06 | 2.267E-06 | 7.234E-07 | 1.800E-06 | 3.100E-06 |
| 14856 | 743 | S743S | B | 2.3e-06 | 2.1e-06 | 2.3e-06 | 2.233E-06 | 1.155E-07 | 2.100E-06 | 2.300E-06 |
| 14857 | 743 | S743T | B | 2.5e-06 | 1.5e-06 | 2.9e-06 | 2.300E-06 | 7.211E-07 | 1.500E-06 | 2.900E-06 |
| 14858 | 743 | S743W | B | 2.7e-06 | 2.3e-06 | 2.2e-06 | 2.400E-06 | 2.646E-07 | 2.200E-06 | 2.700E-06 |
| 14859 | 743 | S743Y | B | 2.2e-06 | 1.6e-06 | 3.5e-06 | 2.433E-06 | 9.713E-07 | 1.600E-06 | 3.500E-06 |
| 14860 | 743 | S743V | B | 2.5e-06 | 3.7e-06 | 3.1e-06 | 3.100E-06 | 6.000E-07 | 2.500E-06 | 3.700E-06 |
| 14861 | 744 | D744A | B | 1.6e-06 | 2.5e-06 | 3.3e-06 | 2.467E-06 | 8.505E-07 | 1.600E-06 | 3.300E-06 |
| 14862 | 744 | D744R | B | 2.8e-06 | 1.8e-06 | 2.9e-06 | 2.500E-06 | 6.083E-07 | 1.800E-06 | 2.900E-06 |
| 14863 | 744 | D744N | B | 1.9e-06 | 3.5e-06 | 2.2e-06 | 2.533E-06 | 8.505E-07 | 1.900E-06 | 3.500E-06 |
| 14864 | 744 | D744D | B | 3.1e-06 | 2.1e-06 | 1.6e-06 | 2.267E-06 | 7.638E-07 | 1.600E-06 | 3.100E-06 |
| 14865 | 744 | D744C | B | 2.5e-06 | 1.7e-06 | 2.5e-06 | 2.233E-06 | 4.619E-07 | 1.700E-06 | 2.500E-06 |
| 14866 | 744 | D744E | B | 2.8e-06 | 2e-06 | 1.2e-06 | 2.000E-06 | 8.000E-07 | 1.200E-06 | 2.800E-06 |
| 14867 | 744 | D744Q | B | 1.3e-06 | 2.2e-06 | 2.3e-06 | 1.933E-06 | 5.508E-07 | 1.300E-06 | 2.300E-06 |
| 14868 | 744 | D744G | B | 2.5e-06 | 2e-06 | 2.3e-06 | 2.267E-06 | 2.517E-07 | 2.000E-06 | 2.500E-06 |
| 14869 | 744 | D744H | B | 2.3e-06 | 2.6e-06 | 1.9e-06 | 2.267E-06 | 3.512E-07 | 1.900E-06 | 2.600E-06 |
| 14870 | 744 | D744I | B | 3e-06 | 2.7e-06 | 2.6e-06 | 2.767E-06 | 2.082E-07 | 2.600E-06 | 3.000E-06 |
| 14871 | 744 | D744L | B | 3.7e-06 | 1.6e-06 | 2.1e-06 | 2.467E-06 | 1.097E-06 | 1.600E-06 | 3.700E-06 |
| 14872 | 744 | D744K | B | 2.9e-06 | 2.6e-06 | 3.5e-06 | 3.000E-06 | 4.583E-07 | 2.600E-06 | 3.500E-06 |
| 14873 | 744 | D744M | B | 2.9e-06 | 2.4e-06 | 2.6e-06 | 2.633E-06 | 2.517E-07 | 2.400E-06 | 2.900E-06 |
| 14874 | 744 | D744F | B | 4.3e-06 | 1.9e-06 | 1.7e-06 | 2.633E-06 | 1.447E-06 | 1.700E-06 | 4.300E-06 |

| | | | | | | | | | | |
|---|---|---|---|---|---|---|---|---|---|---|
| 14875 | 744 | D744P | B | 1.5e-06 | 2.8e-06 | 2.3e-06 | 2.200E-06 | 6.557E-07 | 1.500E-06 | 2.800E-06 |
| 14876 | 744 | D744S | B | 2.7e-06 | 1.7e-06 | 2.5e-06 | 2.300E-06 | 5.292E-07 | 1.700E-06 | 2.700E-06 |
| 14877 | 744 | D744T | B | 1.4e-06 | 3.3e-06 | 3.2e-06 | 2.633E-06 | 1.069E-06 | 1.400E-06 | 3.300E-06 |
| 14878 | 744 | D744W | B | 1.8e-06 | 2.4e-06 | 4.2e-06 | 2.800E-06 | 1.249E-06 | 1.800E-06 | 4.200E-06 |
| 14879 | 744 | D744Y | B | 3e-06 | 1.9e-06 | 3.5e-06 | 2.800E-06 | 8.185E-07 | 1.900E-06 | 3.500E-06 |
| 14880 | 744 | D744V | B | 2.7e-06 | 2.3e-06 | 2.7e-06 | 2.567E-06 | 2.309E-07 | 2.300E-06 | 2.700E-06 |
| 14881 | 745 | E745A | B | 1.5e-06 | 2.3e-06 | 2.5e-06 | 2.100E-06 | 5.292E-07 | 1.500E-06 | 2.500E-06 |
| 14882 | 745 | E745R | B | 2.9e-06 | 2.7e-06 | 2.7e-06 | 2.767E-06 | 1.155E-07 | 2.700E-06 | 2.900E-06 |
| 14883 | 745 | E745N | B | 3e-06 | 3.8e-06 | 3.9e-06 | 3.567E-06 | 4.933E-07 | 3.000E-06 | 3.900E-06 |
| 14884 | 745 | E745D | B | 4.1e-06 | 2.3e-06 | 2.8e-06 | 3.067E-06 | 9.292E-07 | 2.300E-06 | 4.100E-06 |
| 14885 | 745 | E745C | B | 2e-06 | 3.2e-06 | 2.1e-06 | 2.433E-06 | 6.658E-07 | 2.000E-06 | 3.200E-06 |
| 14886 | 745 | E745E | B | 2.4e-06 | 2.8e-06 | 1.9e-06 | 2.367E-06 | 4.509E-07 | 1.900E-06 | 2.800E-06 |
| 14887 | 745 | E745Q | B | 2.3e-06 | 1.8e-06 | 1.4e-06 | 1.833E-06 | 4.509E-07 | 1.400E-06 | 2.300E-06 |
| 14888 | 745 | E745G | B | 2.2e-06 | 2.3e-06 | 3.6e-06 | 2.700E-06 | 7.810E-07 | 2.200E-06 | 3.600E-06 |
| 14889 | 745 | E745H | B | 2.2e-06 | 2.5e-06 | 1.9e-06 | 2.200E-06 | 3.000E-07 | 1.900E-06 | 2.500E-06 |
| 14890 | 745 | E745I | B | 3.2e-06 | 2.6e-06 | 1.7e-06 | 2.500E-06 | 7.550E-07 | 1.700E-06 | 3.200E-06 |
| 14891 | 745 | E745L | B | 2.9e-06 | 2.8e-06 | 2.3e-06 | 2.667E-06 | 3.215E-07 | 2.300E-06 | 2.900E-06 |
| 14892 | 745 | E745K | B | 8.2e-07 | 2.6e-06 | 2e-06 | 1.807E-06 | 9.056E-07 | 8.200E-07 | 2.600E-06 |
| 14893 | 745 | E745M | B | 1.4e-06 | 1.5e-06 | 4.7e-06 | 2.533E-06 | 1.877E-06 | 1.400E-06 | 4.700E-06 |
| 14894 | 745 | E745F | B | 2.1e-06 | 7.2e-06 | 3.4e-06 | 4.233E-06 | 2.650E-06 | 2.100E-06 | 7.200E-06 |
| 14895 | 745 | E745P | B | 2e-06 | 4.4e-06 | 2.2e-06 | 2.867E-06 | 1.332E-06 | 2.000E-06 | 4.400E-06 |
| 14896 | 745 | E745S | B | 2.8e-06 | 2.9e-06 | 3.7e-06 | 3.133E-06 | 4.933E-07 | 2.800E-06 | 3.700E-06 |
| 14897 | 745 | E745T | B | 2.1e-06 | 3e-06 | 2.7e-06 | 2.600E-06 | 4.583E-07 | 2.100E-06 | 3.000E-06 |
| 14898 | 745 | E745W | B | 1.4e-06 | 2.8e-06 | 1.8e-06 | 2.000E-06 | 7.211E-07 | 1.400E-06 | 2.800E-06 |
| 14899 | 745 | E745Y | B | 4.9e-06 | 2.6e-06 | 2.6e-06 | 3.367E-06 | 1.328E-06 | 2.600E-06 | 4.900E-06 |
| 14900 | 745 | E745V | B | 4.2e-06 | 2.3e-06 | 2.7e-06 | 3.067E-06 | 1.002E-06 | 2.300E-06 | 4.200E-06 |
| 14901 | 746 | Q746A | B | 3.2e-06 | 1.4e-06 | 1.6e-06 | 2.067E-06 | 9.866E-07 | 1.400E-06 | 3.200E-06 |
| 14902 | 746 | Q746R | B | 2.6e-06 | 1.9e-06 | 2.2e-06 | 2.233E-06 | 3.512E-07 | 1.900E-06 | 2.600E-06 |
| 14903 | 746 | Q746N | B | 1.5e-06 | 2e-06 | 3e-06 | 2.167E-06 | 7.638E-07 | 1.500E-06 | 3.000E-06 |
| 14904 | 746 | Q746D | B | 1.3e-06 | 1.9e-06 | 2.6e-06 | 1.933E-06 | 6.506E-07 | 1.300E-06 | 2.600E-06 |
| 14905 | 746 | Q746C | B | 2.4e-06 | 1.3e-06 | 3e-06 | 2.233E-06 | 8.622E-07 | 1.300E-06 | 3.000E-06 |
| 14906 | 746 | Q746E | B | 2e-06 | 2.4e-06 | 2.7e-06 | 2.367E-06 | 3.512E-07 | 2.000E-06 | 2.700E-06 |
| 14907 | 746 | Q746Q | B | 2.3e-06 | 2.1e-06 | 3.5e-06 | 2.633E-06 | 7.572E-07 | 2.100E-06 | 3.500E-06 |
| 14908 | 746 | Q746G | B | 1.9e-06 | 3.4e-06 | 2.9e-06 | 2.733E-06 | 7.638E-07 | 1.900E-06 | 3.400E-06 |
| 14909 | 746 | Q746H | B | 3.6e-06 | 2.2e-06 | 1.6e-06 | 2.467E-06 | 1.026E-06 | 1.600E-06 | 3.600E-06 |
| 14910 | 746 | Q746I | B | 3.4e-06 | 2.6e-06 | 2.6e-06 | 2.867E-06 | 4.619E-07 | 2.600E-06 | 3.400E-06 |
| 14911 | 746 | Q746L | B | 3.9e-06 | 1.9e-06 | 1.8e-06 | 2.533E-06 | 1.185E-06 | 1.800E-06 | 3.900E-06 |
| 14912 | 746 | Q746K | B | 2.6e-06 | 4.2e-06 | 4.5e-06 | 3.767E-06 | 1.021E-06 | 2.600E-06 | 4.500E-06 |
| 14913 | 746 | Q746M | B | 2.3e-06 | 1.7e-06 | 1.4e-06 | 1.800E-06 | 4.583E-07 | 1.400E-06 | 2.300E-06 |
| 14914 | 746 | Q746F | B | 5e-06 | 5.2e-06 | 2e-06 | 4.067E-06 | 1.793E-06 | 2.000E-06 | 5.200E-06 |
| 14915 | 746 | Q746P | B | 2.2e-06 | 3.4e-06 | 3.2e-06 | 2.933E-06 | 6.429E-07 | 2.200E-06 | 3.400E-06 |
| 14916 | 746 | Q746S | B | 2.1e-06 | 1.8e-06 | 2.3e-06 | 2.067E-06 | 2.517E-07 | 1.800E-06 | 2.300E-06 |
| 14917 | 746 | Q746T | B | 1.7e-06 | 2.9e-06 | 3e-06 | 2.533E-06 | 7.234E-07 | 1.700E-06 | 3.000E-06 |
| 14918 | 746 | Q746W | B | 2.4e-06 | 2.4e-06 | 2.4e-06 | 2.400E-06 | 0.000E+00 | 2.400E-06 | 2.400E-06 |
| 14919 | 746 | Q746Y | B | 3.6e-06 | 4.2e-06 | 3.2e-06 | 3.667E-06 | 5.033E-07 | 3.200E-06 | 4.200E-06 |
| 14920 | 746 | Q746V | B | 1.5e-06 | 2.2e-06 | 1.6e-06 | 1.767E-06 | 3.786E-07 | 1.500E-06 | 2.200E-06 |
| 14921 | 747 | L747A | B | 1.5e-06 | 1.3e-06 | 2.9e-06 | 1.900E-06 | 8.718E-07 | 1.300E-06 | 2.900E-06 |
| 14922 | 747 | L747R | B | 1.9e-06 | 1.5e-06 | 1.7e-06 | 1.700E-06 | 2.000E-07 | 1.500E-06 | 1.900E-06 |
| 14923 | 747 | L747N | B | 2.8e-06 | 2.9e-06 | 3.1e-06 | 2.933E-06 | 1.528E-07 | 2.800E-06 | 3.100E-06 |
| 14924 | 747 | L747D | B | 3.1e-06 | 1.1e-06 | 3.4e-06 | 2.533E-06 | 1.250E-06 | 1.100E-06 | 3.400E-06 |
| 14925 | 747 | L747C | B | 2.7e-06 | 2.3e-06 | 1.6e-06 | 2.200E-06 | 5.568E-07 | 1.600E-06 | 2.700E-06 |
| 14926 | 747 | L747E | B | 9.5e-07 | 2.4e-06 | 4.5e-06 | 2.617E-06 | 1.785E-06 | 9.500E-07 | 4.500E-06 |
| 14927 | 747 | L747Q | B | 1.8e-06 | 2.8e-06 | 3.4e-06 | 2.667E-06 | 8.083E-07 | 1.800E-06 | 3.400E-06 |
| 14928 | 747 | L747G | B | 4.9e-06 | 2.5e-06 | 3.5e-06 | 3.633E-06 | 1.206E-06 | 2.500E-06 | 4.900E-06 |
| 14929 | 747 | L747H | B | 2.5e-06 | 1.4e-06 | 1.4e-06 | 1.767E-06 | 6.351E-07 | 1.400E-06 | 2.500E-06 |
| 14930 | 747 | L747I | B | 1e-06 | 1.6e-06 | 3.6e-06 | 2.067E-06 | 1.361E-06 | 1.000E-06 | 3.600E-06 |
| 14931 | 747 | L747L | B | 1.6e-06 | 2.6e-06 | 2.5e-06 | 2.233E-06 | 5.508E-07 | 1.600E-06 | 2.600E-06 |
| 14932 | 747 | L747K | B | 2.2e-06 | 1.6e-06 | 2.2e-06 | 2.000E-06 | 3.464E-07 | 1.600E-06 | 2.200E-06 |
| 14933 | 747 | L747M | B | 2e-06 | 1.9e-06 | 1.8e-06 | 1.900E-06 | 1.000E-07 | 1.800E-06 | 2.000E-06 |
| 14934 | 747 | L747F | B | 2.6e-06 | 2e-06 | 2.1e-06 | 2.233E-06 | 3.215E-07 | 2.000E-06 | 2.600E-06 |
| 14935 | 747 | L747P | B | 3.6e-06 | 3e-06 | 3e-06 | 3.200E-06 | 3.464E-07 | 3.000E-06 | 3.600E-06 |
| 14936 | 747 | L747S | B | 2.2e-06 | 3.3e-06 | 2.6e-06 | 2.700E-06 | 5.568E-07 | 2.200E-06 | 3.300E-06 |
| 14937 | 747 | L747T | B | 2.9e-06 | 3.5e-06 | 1.4e-06 | 2.600E-06 | 1.082E-06 | 1.400E-06 | 3.500E-06 |
| 14938 | 747 | L747W | B | 2.2e-06 | 2.5e-06 | 1.9e-06 | 2.200E-06 | 3.000E-07 | 1.900E-06 | 2.500E-06 |
| 14939 | 747 | L747Y | B | 1.4e-06 | 2.6e-06 | 2e-06 | 2.000E-06 | 6.000E-07 | 1.400E-06 | 2.600E-06 |
| 14940 | 747 | L747V | B | 1.5e-06 | 2.6e-06 | 1.1e-06 | 1.733E-06 | 7.767E-07 | 1.100E-06 | 2.600E-06 |
| 14941 | 748 | K748A | B | 1e-06 | 3.2e-06 | 2.8e-06 | 2.333E-06 | 1.172E-06 | 1.000E-06 | 3.200E-06 |
| 14942 | 748 | K748R | B | 1.6e-06 | 1.3e-06 | 1.8e-06 | 1.567E-06 | 2.517E-07 | 1.300E-06 | 1.800E-06 |
| 14943 | 748 | K748N | B | 2.6e-06 | 9.9e-07 | 2.9e-06 | 2.163E-06 | 1.027E-06 | 9.900E-07 | 2.900E-06 |
| 14944 | 748 | K748D | B | 2.3e-06 | 2.2e-06 | 1.8e-06 | 2.100E-06 | 2.646E-07 | 1.800E-06 | 2.300E-06 |
| 14945 | 748 | K748C | B | 1.8e-06 | 2.6e-06 | 1.9e-06 | 2.100E-06 | 4.359E-07 | 1.800E-06 | 2.600E-06 |
| 14946 | 748 | K748E | B | 2.5e-06 | 1.7e-06 | 2.6e-06 | 2.267E-06 | 4.933E-07 | 1.700E-06 | 2.600E-06 |
| 14947 | 748 | K748Q | B | 2.1e-06 | 1.3e-06 | 2.7e-06 | 2.033E-06 | 7.024E-07 | 1.300E-06 | 2.700E-06 |
| 14948 | 748 | K748G | B | 2.5e-06 | 2.3e-06 | 2.3e-06 | 2.367E-06 | 1.155E-07 | 2.300E-06 | 2.500E-06 |
| 14949 | 748 | K748H | B | 1.6e-06 | 1.2e-06 | 3e-06 | 1.933E-06 | 9.452E-07 | 1.200E-06 | 3.000E-06 |
| 14950 | 748 | K748I | B | 3.2e-06 | 1.7e-06 | 1.6e-06 | 2.167E-06 | 8.963E-07 | 1.600E-06 | 3.200E-06 |
| 14951 | 748 | K748L | B | 2.9e-06 | 2.9e-06 | 2.5e-06 | 2.767E-06 | 2.309E-07 | 2.500E-06 | 2.900E-06 |
| 14952 | 748 | K748K | B | 1.6e-06 | 3.7e-06 | 2.6e-06 | 2.633E-06 | 1.050E-06 | 1.600E-06 | 3.700E-06 |
| 14953 | 748 | K748M | B | 3.4e-06 | 3.4e-06 | 2.8e-06 | 3.200E-06 | 3.464E-07 | 2.800E-06 | 3.400E-06 |
| 14954 | 748 | K748F | B | 6.2e-06 | 1.5e-06 | 4.8e-06 | 4.167E-06 | 2.413E-06 | 1.500E-06 | 6.200E-06 |
| 14955 | 748 | K748P | B | 2.9e-06 | 2.7e-06 | 2e-06 | 2.533E-06 | 4.726E-07 | 2.000E-06 | 2.900E-06 |
| 14956 | 748 | K748S | B | 3.5e-06 | 2e-06 | 2.7e-06 | 2.733E-06 | 7.506E-07 | 2.000E-06 | 3.500E-06 |
| 14957 | 748 | K748T | B | 1.5e-06 | 2.1e-06 | 1.6e-06 | 1.733E-06 | 3.215E-07 | 1.500E-06 | 2.100E-06 |
| 14958 | 748 | K748W | B | 2.5e-06 | 3.8e-06 | 2.3e-06 | 2.867E-06 | 8.145E-07 | 2.300E-06 | 3.800E-06 |
| 14959 | 748 | K748Y | B | 1.1e-06 | 2.6e-06 | 3.8e-06 | 2.500E-06 | 1.353E-06 | 1.100E-06 | 3.800E-06 |

| | | | | | | | | | | |
|---|---|---|---|---|---|---|---|---|---|---|
| 14960 | 748 | K748V | B | 1.7e-06 | 3e-06 | 2.3e-06 | 2.333E-06 | 6.506E-07 | 1.700E-06 | 3.000E-06 |
| 14961 | 749 | S749A | B | 2e-06 | 2.6e-06 | 1.9e-06 | 2.167E-06 | 3.786E-07 | 1.900E-06 | 2.600E-06 |
| 14962 | 749 | S749R | B | 1.2e-06 | 2e-06 | 1.7e-06 | 1.633E-06 | 4.041E-07 | 1.200E-06 | 2.000E-06 |
| 14963 | 749 | S749N | B | 2.2e-06 | 3.2e-06 | 2.2e-06 | 2.533E-06 | 5.774E-07 | 2.200E-06 | 3.200E-06 |
| 14964 | 749 | S749D | B | 2.4e-06 | 3.8e-06 | 2.8e-06 | 3.000E-06 | 7.211E-07 | 2.400E-06 | 3.800E-06 |
| 14965 | 749 | S749C | B | 2.4e-06 | 2.1e-06 | 3.2e-06 | 2.567E-06 | 5.686E-07 | 2.100E-06 | 3.200E-06 |
| 14966 | 749 | S749E | B | 9.8e-07 | 2.4e-06 | 1.9e-06 | 1.760E-06 | 7.203E-07 | 9.800E-07 | 2.400E-06 |
| 14967 | 749 | S749Q | B | 3.4e-06 | 2.4e-06 | 2.1e-06 | 2.633E-06 | 6.807E-07 | 2.100E-06 | 3.400E-06 |
| 14968 | 749 | S749G | B | 2.6e-06 | 2e-06 | 1e-06 | 1.867E-06 | 8.083E-07 | 1.000E-06 | 2.600E-06 |
| 14969 | 749 | S749H | B | 2e-06 | 3.3e-06 | 2.7e-06 | 2.667E-06 | 6.506E-07 | 2.000E-06 | 3.300E-06 |
| 14970 | 749 | S749I | B | 2.9e-06 | 2.4e-06 | 1.8e-06 | 2.367E-06 | 5.508E-07 | 1.800E-06 | 2.900E-06 |
| 14971 | 749 | S749L | B | 3.3e-06 | 1.7e-06 | 2.8e-06 | 2.600E-06 | 8.185E-07 | 1.700E-06 | 3.300E-06 |
| 14972 | 749 | S749K | B | 4.4e-06 | 2.9e-06 | 1.9e-06 | 3.067E-06 | 1.258E-06 | 1.900E-06 | 4.400E-06 |
| 14973 | 749 | S749M | B | 3.7e-06 | 3e-06 | 2.3e-06 | 3.000E-06 | 7.000E-07 | 2.300E-06 | 3.700E-06 |
| 14974 | 749 | S749F | B | 2.2e-06 | 2.7e-06 | 3.8e-06 | 2.900E-06 | 8.185E-07 | 2.200E-06 | 3.800E-06 |
| 14975 | 749 | S749P | B | 3e-06 | 3.1e-06 | 2e-06 | 2.700E-06 | 6.083E-07 | 2.000E-06 | 3.100E-06 |
| 14976 | 749 | S749S | B | 1.9e-06 | 3.9e-06 | 1.7e-06 | 2.500E-06 | 1.217E-06 | 1.700E-06 | 3.900E-06 |
| 14977 | 749 | S749T | B | 1.3e-06 | 2.3e-06 | 2.3e-06 | 1.967E-06 | 5.774E-07 | 1.300E-06 | 2.300E-06 |
| 14978 | 749 | S749W | B | 1.6e-06 | 1.9e-06 | 2.7e-06 | 2.067E-06 | 5.686E-07 | 1.600E-06 | 2.700E-06 |
| 14979 | 749 | S749Y | B | 3.7e-06 | 2.2e-06 | 3.8e-06 | 3.233E-06 | 8.963E-07 | 2.200E-06 | 3.800E-06 |
| 14980 | 749 | S749V | B | 2.4e-06 | 3.1e-06 | 2.4e-06 | 2.633E-06 | 4.041E-07 | 2.400E-06 | 3.100E-06 |
| 14981 | 750 | G750A | B | 2.3e-06 | 2.5e-06 | 3.8e-06 | 2.867E-06 | 8.145E-07 | 2.300E-06 | 3.800E-06 |
| 14982 | 750 | G750R | B | 2.3e-06 | 1.8e-06 | 2.5e-06 | 2.200E-06 | 3.606E-07 | 1.800E-06 | 2.500E-06 |
| 14983 | 750 | G750N | B | 3.6e-06 | 2e-06 | 3.3e-06 | 2.967E-06 | 8.505E-07 | 2.000E-06 | 3.600E-06 |
| 14984 | 750 | G750D | B | 2.2e-06 | 3e-06 | 3.3e-06 | 2.833E-06 | 5.686E-07 | 2.200E-06 | 3.300E-06 |
| 14985 | 750 | G750C | B | 1.6e-06 | 3e-06 | 3e-06 | 2.533E-06 | 8.083E-07 | 1.600E-06 | 3.000E-06 |
| 14986 | 750 | G750E | B | 2.3e-06 | 3e-06 | 2.8e-06 | 2.700E-06 | 3.606E-07 | 2.300E-06 | 3.000E-06 |
| 14987 | 750 | G750Q | B | 3.2e-06 | 2.8e-06 | 2e-06 | 2.667E-06 | 6.110E-07 | 2.000E-06 | 3.200E-06 |
| 14988 | 750 | G750G | B | 2.6e-06 | 9.6e-07 | 2.1e-06 | 1.887E-06 | 8.406E-07 | 9.600E-07 | 2.600E-06 |
| 14989 | 750 | G750H | B | 2.4e-06 | 3.2e-06 | 2.1e-06 | 2.567E-06 | 5.686E-07 | 2.100E-06 | 3.200E-06 |
| 14990 | 750 | G750I | B | 2.8e-06 | 2e-06 | 3.3e-06 | 2.700E-06 | 6.557E-07 | 2.000E-06 | 3.300E-06 |
| 14991 | 750 | G750L | B | 2.5e-06 | 1.6e-06 | 2.6e-06 | 2.233E-06 | 5.508E-07 | 1.600E-06 | 2.600E-06 |
| 14992 | 750 | G750K | B | 2.5e-06 | 2.1e-06 | 1.6e-06 | 2.067E-06 | 4.509E-07 | 1.600E-06 | 2.500E-06 |
| 14993 | 750 | G750M | B | 1.7e-06 | 2.5e-06 | 1.2e-06 | 1.800E-06 | 6.557E-07 | 1.200E-06 | 2.500E-06 |
| 14994 | 750 | G750F | B | 2.2e-06 | 1.7e-06 | 2.3e-06 | 2.067E-06 | 3.215E-07 | 1.700E-06 | 2.300E-06 |
| 14995 | 750 | G750P | B | 1.6e-06 | 3.1e-06 | 2.8e-06 | 2.500E-06 | 7.937E-07 | 1.600E-06 | 3.100E-06 |
| 14996 | 750 | G750S | B | 1.3e-06 | 3.4e-06 | 2e-06 | 2.233E-06 | 1.069E-06 | 1.300E-06 | 3.400E-06 |
| 14997 | 750 | G750T | B | 1.9e-06 | 1.5e-06 | 2.5e-06 | 1.967E-06 | 5.033E-07 | 1.500E-06 | 2.500E-06 |
| 14998 | 750 | G750W | B | 2.6e-06 | 9.4e-07 | 3.2e-06 | 2.247E-06 | 1.171E-06 | 9.400E-07 | 3.200E-06 |
| 14999 | 750 | G750Y | B | 2.3e-06 | 3.3e-06 | 3.2e-06 | 2.933E-06 | 5.508E-07 | 2.300E-06 | 3.300E-06 |
| 15000 | 750 | G750V | B | 2.2e-06 | 1.8e-06 | 3.3e-06 | 2.433E-06 | 7.767E-07 | 1.800E-06 | 3.300E-06 |
| 15001 | 751 | T751A | B | 2.1e-06 | 2.1e-06 | 2.6e-06 | 2.267E-06 | 2.887E-07 | 2.100E-06 | 2.600E-06 |
| 15002 | 751 | T751R | B | 4.8e-06 | 2.3e-06 | 1.1e-06 | 2.733E-06 | 1.888E-06 | 1.100E-06 | 4.800E-06 |
| 15003 | 751 | T751N | B | 1.4e-06 | 2.5e-06 | 2.4e-06 | 2.100E-06 | 6.083E-07 | 1.400E-06 | 2.500E-06 |
| 15004 | 751 | T751D | B | 1.9e-06 | 2.6e-06 | 2.4e-06 | 2.300E-06 | 3.606E-07 | 1.900E-06 | 2.600E-06 |
| 15005 | 751 | T751C | B | 2.2e-06 | 1.9e-06 | 2.8e-06 | 2.300E-06 | 4.583E-07 | 1.900E-06 | 2.800E-06 |
| 15006 | 751 | T751E | B | 1.9e-06 | 2.1e-06 | 2.2e-06 | 2.067E-06 | 1.528E-07 | 1.900E-06 | 2.200E-06 |
| 15007 | 751 | T751Q | B | 2.2e-06 | 4.1e-06 | 3.4e-06 | 3.233E-06 | 9.609E-07 | 2.200E-06 | 4.100E-06 |
| 15008 | 751 | T751G | B | 2.5e-06 | 3e-06 | 2.1e-06 | 2.533E-06 | 4.509E-07 | 2.100E-06 | 3.000E-06 |
| 15009 | 751 | T751H | B | 1.5e-06 | 3.9e-06 | 2.2e-06 | 2.533E-06 | 1.234E-06 | 1.500E-06 | 3.900E-06 |
| 15010 | 751 | T751I | B | 2e-06 | 3.1e-06 | 3.6e-06 | 2.900E-06 | 8.185E-07 | 2.000E-06 | 3.600E-06 |
| 15011 | 751 | T751L | B | 2.4e-06 | 2.5e-06 | 1.2e-06 | 2.033E-06 | 7.234E-07 | 1.200E-06 | 2.500E-06 |
| 15012 | 751 | T751K | B | 2.1e-06 | 1.1e-06 | 2.2e-06 | 1.800E-06 | 6.083E-07 | 1.100E-06 | 2.200E-06 |
| 15013 | 751 | T751M | B | 2.3e-06 | 2.4e-06 | 2.8e-06 | 2.500E-06 | 2.646E-07 | 2.300E-06 | 2.800E-06 |
| 15014 | 751 | T751F | B | 1.6e-06 | 4.2e-06 | 3.1e-06 | 2.967E-06 | 1.305E-06 | 1.600E-06 | 4.200E-06 |
| 15015 | 751 | T751P | B | 2.3e-06 | 2.1e-06 | 1.9e-06 | 2.100E-06 | 2.000E-07 | 1.900E-06 | 2.300E-06 |
| 15016 | 751 | T751S | B | 1.4e-06 | 2.5e-06 | 1.5e-06 | 1.800E-06 | 6.083E-07 | 1.400E-06 | 2.500E-06 |
| 15017 | 751 | T751T | B | 2.6e-06 | 2.1e-06 | 4.9e-06 | 3.200E-06 | 1.493E-06 | 2.100E-06 | 4.900E-06 |
| 15018 | 751 | T751W | B | 2e-06 | 1.7e-06 | 2.6e-06 | 2.100E-06 | 4.583E-07 | 1.700E-06 | 2.600E-06 |
| 15019 | 751 | T751Y | B | 2.3e-06 | 3.9e-06 | 2e-06 | 2.733E-06 | 1.021E-06 | 2.000E-06 | 3.900E-06 |
| 15020 | 751 | T751V | B | 1.5e-06 | 2.3e-06 | 2.5e-06 | 2.100E-06 | 5.292E-07 | 1.500E-06 | 2.500E-06 |
| 15021 | 752 | A752A | B | 1.7e-06 | 3.3e-06 | 1.4e-06 | 2.133E-06 | 1.021E-06 | 1.400E-06 | 3.300E-06 |
| 15022 | 752 | A752R | B | 3.3e-06 | 2.3e-06 | 2.9e-06 | 2.833E-06 | 5.033E-07 | 2.300E-06 | 3.300E-06 |
| 15023 | 752 | A752N | B | 2.2e-06 | 2.6e-06 | 2.3e-06 | 2.367E-06 | 2.082E-07 | 2.200E-06 | 2.600E-06 |
| 15024 | 752 | A752D | B | 2.6e-06 | 2.7e-06 | 1.5e-06 | 2.267E-06 | 6.658E-07 | 1.500E-06 | 2.700E-06 |
| 15025 | 752 | A752C | B | 2e-06 | 2.8e-06 | 2.4e-06 | 2.400E-06 | 4.000E-07 | 2.000E-06 | 2.800E-06 |
| 15026 | 752 | A752E | B | 2.1e-06 | 5e-06 | 2.3e-06 | 3.133E-06 | 1.620E-06 | 2.100E-06 | 5.000E-06 |
| 15027 | 752 | A752Q | B | 3.4e-06 | 2.3e-06 | 2.5e-06 | 2.733E-06 | 5.859E-07 | 2.300E-06 | 3.400E-06 |
| 15028 | 752 | A752G | B | 2e-06 | 3.5e-06 | 1.8e-06 | 2.433E-06 | 9.292E-07 | 1.800E-06 | 3.500E-06 |
| 15029 | 752 | A752H | B | 2.2e-06 | 2.3e-06 | 3.3e-06 | 2.600E-06 | 6.083E-07 | 2.200E-06 | 3.300E-06 |
| 15030 | 752 | A752I | B | 2.1e-06 | 2.2e-06 | 3.7e-06 | 2.667E-06 | 8.963E-07 | 2.100E-06 | 3.700E-06 |
| 15031 | 752 | A752L | B | 6.3e-07 | 1.9e-06 | 2.7e-06 | 1.743E-06 | 1.044E-06 | 6.300E-07 | 2.700E-06 |
| 15032 | 752 | A752K | B | 2.4e-06 | 2.1e-06 | 1.5e-06 | 2.000E-06 | 4.583E-07 | 1.500E-06 | 2.400E-06 |
| 15033 | 752 | A752M | B | 1.5e-06 | 2.5e-06 | 1.8e-06 | 1.933E-06 | 5.132E-07 | 1.500E-06 | 2.500E-06 |
| 15034 | 752 | A752F | B | 2.3e-06 | 2.8e-06 | 3.4e-06 | 2.833E-06 | 5.508E-07 | 2.300E-06 | 3.400E-06 |
| 15035 | 752 | A752P | B | 3.7e-06 | 2.5e-06 | 1.9e-06 | 2.700E-06 | 9.165E-07 | 1.900E-06 | 3.700E-06 |
| 15036 | 752 | A752S | B | 2.7e-06 | 1.7e-06 | 2e-06 | 2.133E-06 | 5.132E-07 | 1.700E-06 | 2.700E-06 |
| 15037 | 752 | A752T | B | 2.9e-06 | 1.7e-06 | 4.1e-06 | 2.900E-06 | 1.200E-06 | 1.700E-06 | 4.100E-06 |
| 15038 | 752 | A752W | B | 1.4e-06 | 2.1e-06 | 6e-06 | 3.167E-06 | 2.479E-06 | 1.400E-06 | 6.000E-06 |
| 15039 | 752 | A752Y | B | 2.4e-06 | 1.7e-06 | 1.9e-06 | 2.000E-06 | 3.606E-07 | 1.700E-06 | 2.400E-06 |
| 15040 | 752 | A752V | B | 1.7e-06 | 2.6e-06 | 1.7e-06 | 2.000E-06 | 5.196E-07 | 1.700E-06 | 2.600E-06 |
| 15041 | 753 | S753A | B | 2.4e-06 | 3.6e-06 | 2.1e-06 | 2.700E-06 | 7.937E-07 | 2.100E-06 | 3.600E-06 |
| 15042 | 753 | S753R | B | 1e-06 | 1.8e-06 | 3.1e-06 | 1.967E-06 | 1.060E-06 | 1.000E-06 | 3.100E-06 |
| 15043 | 753 | S753N | B | 1.6e-06 | 4.1e-06 | 2.7e-06 | 2.800E-06 | 1.253E-06 | 1.600E-06 | 4.100E-06 |
| 15044 | 753 | S753D | B | 2.6e-06 | 2.4e-06 | 1.6e-06 | 2.200E-06 | 5.292E-07 | 1.600E-06 | 2.600E-06 |

| | | | | | | | | | | |
|---|---|---|---|---|---|---|---|---|---|---|
| 15045 | 753 | S753C | B | 2.8e-06 | 2.9e-06 | 1.8e-06 | 2.500E-06 | 6.083E-07 | 1.800E-06 | 2.900E-06 |
| 15046 | 753 | S753E | B | 2.1e-06 | 3.3e-06 | 3.2e-06 | 2.867E-06 | 6.658E-07 | 2.100E-06 | 3.300E-06 |
| 15047 | 753 | S753Q | B | 2.4e-06 | 2.3e-06 | 3.3e-06 | 2.667E-06 | 5.508E-07 | 2.300E-06 | 3.300E-06 |
| 15048 | 753 | S753G | B | 3.1e-06 | 4.2e-06 | 2.2e-06 | 3.167E-06 | 1.002E-06 | 2.200E-06 | 4.200E-06 |
| 15049 | 753 | S753H | B | 2.2e-06 | 2.8e-06 | 2.8e-06 | 2.600E-06 | 3.464E-07 | 2.200E-06 | 2.800E-06 |
| 15050 | 753 | S753I | B | 2.6e-06 | 2.9e-06 | 3.9e-06 | 3.133E-06 | 6.807E-07 | 2.600E-06 | 3.900E-06 |
| 15051 | 753 | S753L | B | 2.6e-06 | 1.4e-06 | 2.2e-06 | 2.067E-06 | 6.110E-07 | 1.400E-06 | 2.600E-06 |
| 15052 | 753 | S753K | B | 2.9e-06 | 3.3e-06 | 2.4e-06 | 2.867E-06 | 4.509E-07 | 2.400E-06 | 3.300E-06 |
| 15053 | 753 | S753M | B | 2.1e-06 | 1.9e-06 | 2.1e-06 | 2.033E-06 | 1.155E-07 | 1.900E-06 | 2.100E-06 |
| 15054 | 753 | S753F | B | 2.1e-06 | 1.8e-06 | 1.6e-06 | 1.833E-06 | 2.517E-07 | 1.600E-06 | 2.100E-06 |
| 15055 | 753 | S753P | B | 2.3e-06 | 1.3e-06 | 1.9e-06 | 1.833E-06 | 5.033E-07 | 1.300E-06 | 2.300E-06 |
| 15056 | 753 | S753S | B | 3.1e-06 | 2.3e-06 | 1.7e-06 | 2.367E-06 | 7.024E-07 | 1.700E-06 | 3.100E-06 |
| 15057 | 753 | S753T | B | 2.4e-06 | 2e-06 | 3.8e-06 | 2.733E-06 | 9.452E-07 | 2.000E-06 | 3.800E-06 |
| 15058 | 753 | S753W | B | 3.3e-06 | 2e-06 | 2.8e-06 | 2.700E-06 | 6.557E-07 | 2.000E-06 | 3.300E-06 |
| 15059 | 753 | S753Y | B | 1.2e-06 | 2.5e-06 | 1.9e-06 | 1.867E-06 | 6.506E-07 | 1.200E-06 | 2.500E-06 |
| 15060 | 753 | S753V | B | 2e-06 | 1.7e-06 | 3.9e-06 | 2.533E-06 | 1.193E-06 | 1.700E-06 | 3.900E-06 |
| 15061 | 754 | V754A | B | 4.7e-06 | 7.6e-06 | 5.2e-06 | 5.833E-06 | 1.550E-06 | 4.700E-06 | 7.600E-06 |
| 15062 | 754 | V754R | B | 1.6e-06 | 2.7e-06 | 3.6e-06 | 2.633E-06 | 1.002E-06 | 1.600E-06 | 3.600E-06 |
| 15063 | 754 | V754N | B | 2.5e-06 | 9.8e-07 | 2e-06 | 1.827E-06 | 7.747E-07 | 9.800E-07 | 2.500E-06 |
| 15064 | 754 | V754D | B | 2.9e-06 | 1.8e-06 | 1.6e-06 | 2.100E-06 | 7.000E-07 | 1.600E-06 | 2.900E-06 |
| 15065 | 754 | V754C | B | 2e-06 | 2.6e-06 | 2.2e-06 | 2.267E-06 | 3.055E-07 | 2.000E-06 | 2.600E-06 |
| 15066 | 754 | V754E | B | 3.3e-06 | 2e-06 | 1.3e-06 | 2.200E-06 | 1.015E-06 | 1.300E-06 | 3.300E-06 |
| 15067 | 754 | V754Q | B | 2.3e-06 | 2.5e-06 | 1.3e-06 | 2.033E-06 | 6.429E-07 | 1.300E-06 | 2.500E-06 |
| 15068 | 754 | V754G | B | 2.8e-06 | 4.1e-06 | 2.2e-06 | 3.033E-06 | 9.713E-07 | 2.200E-06 | 4.100E-06 |
| 15069 | 754 | V754H | B | 2.8e-06 | 2.2e-06 | 2.6e-06 | 2.533E-06 | 3.055E-07 | 2.200E-06 | 2.800E-06 |
| 15070 | 754 | V754I | B | 2.4e-06 | 2.7e-06 | 2.6e-06 | 2.567E-06 | 1.528E-07 | 2.400E-06 | 2.700E-06 |
| 15071 | 754 | V754L | B | 2.9e-06 | 1.5e-06 | 1.1e-06 | 1.833E-06 | 9.452E-07 | 1.100E-06 | 2.900E-06 |
| 15072 | 754 | V754K | B | 1.3e-06 | 2.8e-06 | 2.4e-06 | 2.167E-06 | 7.767E-07 | 1.300E-06 | 2.800E-06 |
| 15073 | 754 | V754M | B | 1.1e-06 | 2.8e-06 | 4.6e-06 | 2.833E-06 | 1.750E-06 | 1.100E-06 | 4.600E-06 |
| 15074 | 754 | V754F | B | 2.3e-06 | 2.4e-06 | 1.6e-06 | 2.100E-06 | 4.359E-07 | 1.600E-06 | 2.400E-06 |
| 15075 | 754 | V754P | B | 5.1e-06 | 1.7e-06 | 2.7e-06 | 3.167E-06 | 1.747E-06 | 1.700E-06 | 5.100E-06 |
| 15076 | 754 | V754S | B | 1.9e-06 | 2.9e-06 | 1.8e-06 | 2.200E-06 | 6.083E-07 | 1.800E-06 | 2.900E-06 |
| 15077 | 754 | V754T | B | 1.6e-06 | 2.3e-06 | 1.8e-06 | 1.900E-06 | 3.606E-07 | 1.600E-06 | 2.300E-06 |
| 15078 | 754 | V754W | B | 2.8e-06 | 1.2e-06 | 2.3e-06 | 2.100E-06 | 8.185E-07 | 1.200E-06 | 2.800E-06 |
| 15079 | 754 | V754Y | B | 1.8e-06 | 2.2e-06 | 3.1e-06 | 2.367E-06 | 6.658E-07 | 1.800E-06 | 3.100E-06 |
| 15080 | 754 | V754V | B | 1.4e-06 | 1.7e-06 | 2.9e-06 | 2.000E-06 | 7.937E-07 | 1.400E-06 | 2.900E-06 |
| 15081 | 755 | V755A | B | 1.9e-06 | 1.2e-06 | 3.2e-06 | 2.100E-06 | 1.015E-06 | 1.200E-06 | 3.200E-06 |
| 15082 | 755 | V755R | B | 1.5e-06 | 2.2e-06 | 4.1e-06 | 2.600E-06 | 1.345E-06 | 1.500E-06 | 4.100E-06 |
| 15083 | 755 | V755N | B | 2.2e-06 | 2.5e-06 | 2e-06 | 2.233E-06 | 2.517E-07 | 2.000E-06 | 2.500E-06 |
| 15084 | 755 | V755D | B | 2.3e-06 | 2.2e-06 | 2.8e-06 | 2.433E-06 | 3.215E-07 | 2.200E-06 | 2.800E-06 |
| 15085 | 755 | V755C | B | 2.5e-06 | 2e-06 | 2.8e-06 | 2.433E-06 | 4.041E-07 | 2.000E-06 | 2.800E-06 |
| 15086 | 755 | V755E | B | 1.8e-06 | 2.4e-06 | 2.3e-06 | 2.167E-06 | 3.215E-07 | 1.800E-06 | 2.400E-06 |
| 15087 | 755 | V755Q | B | 1.7e-06 | 3.6e-06 | 5e-06 | 3.433E-06 | 1.656E-06 | 1.700E-06 | 5.000E-06 |
| 15088 | 755 | V755G | B | 2.7e-06 | 1.7e-06 | 3.9e-06 | 2.767E-06 | 1.102E-06 | 1.700E-06 | 3.900E-06 |
| 15089 | 755 | V755H | B | 1.8e-06 | 2.2e-06 | 3e-06 | 2.333E-06 | 6.110E-07 | 1.800E-06 | 3.000E-06 |
| 15090 | 755 | V755I | B | 1.4e-06 | 3.1e-06 | 1.6e-06 | 2.033E-06 | 9.292E-07 | 1.400E-06 | 3.100E-06 |
| 15091 | 755 | V755L | B | 1.6e-06 | 2.3e-06 | 3.7e-06 | 2.533E-06 | 1.069E-06 | 1.600E-06 | 3.700E-06 |
| 15092 | 755 | V755K | B | 2.9e-06 | 3.2e-06 | 2.8e-06 | 2.967E-06 | 2.082E-07 | 2.800E-06 | 3.200E-06 |
| 15093 | 755 | V755M | B | 2.2e-06 | 2.9e-06 | 2.3e-06 | 2.467E-06 | 3.786E-07 | 2.200E-06 | 2.900E-06 |
| 15094 | 755 | V755F | B | 2.9e-06 | 3.2e-06 | 1.9e-06 | 2.667E-06 | 6.807E-07 | 1.900E-06 | 3.200E-06 |
| 15095 | 755 | V755P | B | 2.1e-06 | 4.1e-06 | 4.2e-06 | 3.467E-06 | 1.185E-06 | 2.100E-06 | 4.200E-06 |
| 15096 | 755 | V755S | B | 1.1e-06 | 2.5e-06 | 3.2e-06 | 2.267E-06 | 1.069E-06 | 1.100E-06 | 3.200E-06 |
| 15097 | 755 | V755T | B | 2.6e-06 | 2.1e-06 | 2e-06 | 2.233E-06 | 3.215E-07 | 2.000E-06 | 2.600E-06 |
| 15098 | 755 | V755W | B | 2.8e-06 | 2.3e-06 | 2.5e-06 | 2.533E-06 | 2.517E-07 | 2.300E-06 | 2.800E-06 |
| 15099 | 755 | V755Y | B | 1.7e-06 | 1.1e-06 | 2.1e-06 | 1.633E-06 | 5.033E-07 | 1.100E-06 | 2.100E-06 |
| 15100 | 755 | V755V | B | 2.2e-06 | 3.6e-06 | 2.4e-06 | 2.733E-06 | 7.572E-07 | 2.200E-06 | 3.600E-06 |
| 15101 | 756 | C756A | B | 1.6e-06 | 2.3e-06 | 2e-06 | 1.967E-06 | 3.512E-07 | 1.600E-06 | 2.300E-06 |
| 15102 | 756 | C756R | B | 1.8e-06 | 2e-06 | 2.3e-06 | 2.033E-06 | 2.517E-07 | 1.800E-06 | 2.300E-06 |
| 15103 | 756 | C756N | B | 2.1e-06 | 3.2e-06 | 1.9e-06 | 2.400E-06 | 7.000E-07 | 1.900E-06 | 3.200E-06 |
| 15104 | 756 | C756D | B | 2.7e-06 | 3.3e-06 | 3.2e-06 | 3.067E-06 | 3.215E-07 | 2.700E-06 | 3.300E-06 |
| 15105 | 756 | C756C | B | 2.7e-06 | 5.1e-06 | 3.5e-06 | 3.767E-06 | 1.222E-06 | 2.700E-06 | 5.100E-06 |
| 15106 | 756 | C756E | B | 2e-06 | 2.6e-06 | 1.5e-06 | 2.033E-06 | 5.508E-07 | 1.500E-06 | 2.600E-06 |
| 15107 | 756 | C756Q | B | 2.1e-06 | 1.4e-06 | 1.3e-06 | 1.600E-06 | 4.359E-07 | 1.300E-06 | 2.100E-06 |
| 15108 | 756 | C756G | B | 4e-06 | 2.1e-06 | 2.7e-06 | 2.933E-06 | 9.713E-07 | 2.100E-06 | 4.000E-06 |
| 15109 | 756 | C756H | B | 3.7e-06 | 2.2e-06 | 2.7e-06 | 2.867E-06 | 7.638E-07 | 2.200E-06 | 3.700E-06 |
| 15110 | 756 | C756I | B | 2.3e-06 | 2.7e-06 | 2.2e-06 | 2.400E-06 | 2.646E-07 | 2.200E-06 | 2.700E-06 |
| 15111 | 756 | C756L | B | 1.7e-06 | 2e-06 | 2.1e-06 | 1.933E-06 | 2.082E-07 | 1.700E-06 | 2.100E-06 |
| 15112 | 756 | C756K | B | 2.6e-06 | 2.6e-06 | 2.9e-06 | 2.700E-06 | 1.732E-07 | 2.600E-06 | 2.900E-06 |
| 15113 | 756 | C756M | B | 2.9e-06 | 3.3e-06 | 3.2e-06 | 3.133E-06 | 2.082E-07 | 2.900E-06 | 3.300E-06 |
| 15114 | 756 | C756F | B | 3.2e-06 | 3e-06 | 2.9e-06 | 3.033E-06 | 1.528E-07 | 2.900E-06 | 3.200E-06 |
| 15115 | 756 | C756P | B | 2.5e-06 | 1.6e-06 | 2.4e-06 | 2.167E-06 | 4.933E-07 | 1.600E-06 | 2.500E-06 |
| 15116 | 756 | C756S | B | 2.3e-06 | 2.5e-06 | 2.3e-06 | 2.367E-06 | 1.155E-07 | 2.300E-06 | 2.500E-06 |
| 15117 | 756 | C756T | B | 3.1e-06 | 2.6e-06 | 3.6e-06 | 3.100E-06 | 5.000E-07 | 2.600E-06 | 3.600E-06 |
| 15118 | 756 | C756W | B | 3.7e-06 | 2.3e-06 | 2.6e-06 | 2.867E-06 | 7.371E-07 | 2.300E-06 | 3.700E-06 |
| 15119 | 756 | C756Y | B | 1.8e-06 | 3.4e-06 | 2.3e-06 | 2.500E-06 | 8.185E-07 | 1.800E-06 | 3.400E-06 |
| 15120 | 756 | C756V | B | 3e-06 | 2.2e-06 | 1.8e-06 | 2.333E-06 | 6.110E-07 | 1.800E-06 | 3.000E-06 |
| 15121 | 757 | L757A | B | 1.8e-06 | 1.8e-06 | 3.6e-06 | 2.400E-06 | 1.039E-06 | 1.800E-06 | 3.600E-06 |
| 15122 | 757 | L757R | B | 3.2e-06 | 2.8e-06 | 2.1e-06 | 2.700E-06 | 5.568E-07 | 2.100E-06 | 3.200E-06 |
| 15123 | 757 | L757N | B | 2.7e-06 | 2e-06 | 1.8e-06 | 2.167E-06 | 4.726E-07 | 1.800E-06 | 2.700E-06 |
| 15124 | 757 | L757D | B | 2.6e-06 | 2.2e-06 | 2.3e-06 | 2.367E-06 | 2.082E-07 | 2.200E-06 | 2.600E-06 |
| 15125 | 757 | L757C | B | 3.2e-06 | 1.4e-06 | 2.8e-06 | 2.467E-06 | 9.452E-07 | 1.400E-06 | 3.200E-06 |
| 15126 | 757 | L757E | B | 1.9e-06 | 2.4e-06 | 3.3e-06 | 2.533E-06 | 7.095E-07 | 1.900E-06 | 3.300E-06 |
| 15127 | 757 | L757Q | B | 1.4e-06 | 5e-06 | 2.3e-06 | 2.900E-06 | 1.873E-06 | 1.400E-06 | 5.000E-06 |
| 15128 | 757 | L757G | B | 2.1e-06 | 2.5e-06 | 2.7e-06 | 2.433E-06 | 3.055E-07 | 2.100E-06 | 2.700E-06 |
| 15129 | 757 | L757H | B | 1.9e-06 | 1.7e-06 | 1.6e-06 | 1.733E-06 | 1.528E-07 | 1.600E-06 | 1.900E-06 |

| | | | | | | | | | | |
|---|---|---|---|---|---|---|---|---|---|---|
| 15130 | 757 | L757I | B | 1.4e-06 | 2.2e-06 | 1.7e-06 | 1.767E-06 | 4.041E-07 | 1.400E-06 | 2.200E-06 |
| 15131 | 757 | L757L | B | 2.9e-06 | 2.3e-06 | 2.7e-06 | 2.633E-06 | 3.055E-07 | 2.300E-06 | 2.900E-06 |
| 15132 | 757 | L757K | B | 1.3e-06 | 2e-06 | 2.1e-06 | 1.800E-06 | 4.359E-07 | 1.300E-06 | 2.100E-06 |
| 15133 | 757 | L757M | B | 2.1e-06 | 3.2e-06 | 1.7e-06 | 2.333E-06 | 7.767E-07 | 1.700E-06 | 3.200E-06 |
| 15134 | 757 | L757F | B | 1.9e-06 | 3.1e-06 | 1.5e-06 | 2.167E-06 | 8.327E-07 | 1.500E-06 | 3.100E-06 |
| 15135 | 757 | L757P | B | 5.2e-06 | 2.2e-06 | 1.9e-06 | 3.100E-06 | 1.825E-06 | 1.900E-06 | 5.200E-06 |
| 15136 | 757 | L757S | B | 3.4e-06 | 3.6e-06 | 2e-06 | 3.000E-06 | 8.718E-07 | 2.000E-06 | 3.600E-06 |
| 15137 | 757 | L757T | B | 3.2e-06 | 2.1e-06 | 2.5e-06 | 2.600E-06 | 5.568E-07 | 2.100E-06 | 3.200E-06 |
| 15138 | 757 | L757W | B | 2.2e-06 | 1.9e-06 | 2.1e-06 | 2.067E-06 | 1.528E-07 | 1.900E-06 | 2.200E-06 |
| 15139 | 757 | L757Y | B | 2.4e-06 | 2.4e-06 | 2.2e-06 | 2.333E-06 | 1.155E-07 | 2.200E-06 | 2.400E-06 |
| 15140 | 757 | L757V | B | 2.8e-06 | 2.1e-06 | 1.7e-06 | 2.200E-06 | 5.568E-07 | 1.700E-06 | 2.800E-06 |
| 15141 | 758 | L758A | B | 1.4e-06 | 1.5e-06 | 2.4e-06 | 1.767E-06 | 5.508E-07 | 1.400E-06 | 2.400E-06 |
| 15142 | 758 | L758R | B | 2.3e-06 | 1.9e-06 | 2.1e-06 | 2.100E-06 | 2.000E-07 | 1.900E-06 | 2.300E-06 |
| 15143 | 758 | L758N | B | 1.7e-06 | 1.8e-06 | 4.6e-06 | 2.700E-06 | 1.646E-06 | 1.700E-06 | 4.600E-06 |
| 15144 | 758 | L758D | B | 4.9e-06 | 3.4e-06 | 2.8e-06 | 3.700E-06 | 1.082E-06 | 2.800E-06 | 4.900E-06 |
| 15145 | 758 | L758C | B | 2.1e-06 | 1.8e-06 | 3e-06 | 2.300E-06 | 6.245E-07 | 1.800E-06 | 3.000E-06 |
| 15146 | 758 | L758E | B | 2.5e-06 | 1.7e-06 | 2.5e-06 | 2.233E-06 | 4.619E-07 | 1.700E-06 | 2.500E-06 |
| 15147 | 758 | L758Q | B | 3.4e-06 | 2.6e-06 | 2.6e-06 | 2.867E-06 | 4.619E-07 | 2.600E-06 | 3.400E-06 |
| 15148 | 758 | L758G | B | 2.4e-06 | 1.8e-06 | 3.3e-06 | 2.500E-06 | 7.550E-07 | 1.800E-06 | 3.300E-06 |
| 15149 | 758 | L758H | B | 2.5e-06 | 4.1e-06 | 9.5e-07 | 2.517E-06 | 1.575E-06 | 9.500E-07 | 4.100E-06 |
| 15150 | 758 | L758I | B | 1.7e-06 | 2.5e-06 | 2.7e-06 | 2.300E-06 | 5.292E-07 | 1.700E-06 | 2.700E-06 |
| 15151 | 758 | L758L | B | 2.2e-06 | 2e-06 | 1.1e-06 | 1.767E-06 | 5.859E-07 | 1.100E-06 | 2.200E-06 |
| 15152 | 758 | L758K | B | 1.9e-06 | 2.6e-06 | 2.5e-06 | 2.333E-06 | 3.786E-07 | 1.900E-06 | 2.600E-06 |
| 15153 | 758 | L758M | B | 2.1e-06 | 2.3e-06 | 5.1e-06 | 3.167E-06 | 1.677E-06 | 2.100E-06 | 5.100E-06 |
| 15154 | 758 | L758F | B | 2.9e-06 | 2.8e-06 | 4.4e-06 | 3.367E-06 | 8.963E-07 | 2.800E-06 | 4.400E-06 |
| 15155 | 758 | L758P | B | 2.9e-06 | 3e-06 | 3.8e-06 | 3.233E-06 | 4.933E-07 | 2.900E-06 | 3.800E-06 |
| 15156 | 758 | L758S | B | 2.5e-06 | 2.1e-06 | 3.8e-06 | 2.800E-06 | 8.888E-07 | 2.100E-06 | 3.800E-06 |
| 15157 | 758 | L758T | B | 1.7e-06 | 4.6e-06 | 1.9e-06 | 2.733E-06 | 1.620E-06 | 1.700E-06 | 4.600E-06 |
| 15158 | 758 | L758W | B | 1.7e-06 | 3.3e-06 | 1.8e-06 | 2.267E-06 | 8.963E-07 | 1.700E-06 | 3.300E-06 |
| 15159 | 758 | L758Y | B | 3e-06 | 2.2e-06 | 1.6e-06 | 2.267E-06 | 7.024E-07 | 1.600E-06 | 3.000E-06 |
| 15160 | 758 | L758V | B | 4.7e-06 | 3.8e-06 | 1.4e-06 | 3.300E-06 | 1.706E-06 | 1.400E-06 | 4.700E-06 |
| 15161 | 759 | N759A | B | 2.5e-06 | 2.7e-06 | 2.5e-06 | 2.567E-06 | 1.155E-07 | 2.500E-06 | 2.700E-06 |
| 15162 | 759 | N759R | B | 1.7e-06 | 2.1e-06 | 1.8e-06 | 1.867E-06 | 2.082E-07 | 1.700E-06 | 2.100E-06 |
| 15163 | 759 | N759N | B | 4.2e-06 | 2.7e-06 | 1.5e-06 | 2.800E-06 | 1.353E-06 | 1.500E-06 | 4.200E-06 |
| 15164 | 759 | N759D | B | 2.6e-06 | 2.6e-06 | 2.3e-06 | 2.500E-06 | 1.732E-07 | 2.300E-06 | 2.600E-06 |
| 15165 | 759 | N759C | B | 2.1e-06 | 3e-06 | 2.9e-06 | 2.667E-06 | 4.933E-07 | 2.100E-06 | 3.000E-06 |
| 15166 | 759 | N759E | B | 2.8e-06 | 3.1e-06 | 2.9e-06 | 2.933E-06 | 1.528E-07 | 2.800E-06 | 3.100E-06 |
| 15167 | 759 | N759Q | B | 1.7e-06 | 2.1e-06 | 1.5e-06 | 1.767E-06 | 3.055E-07 | 1.500E-06 | 2.100E-06 |
| 15168 | 759 | N759G | B | 2.2e-06 | 2.5e-06 | 3.7e-06 | 2.800E-06 | 7.937E-07 | 2.200E-06 | 3.700E-06 |
| 15169 | 759 | N759H | B | 4e-06 | 3.8e-06 | 1.9e-06 | 3.233E-06 | 1.159E-06 | 1.900E-06 | 4.000E-06 |
| 15170 | 759 | N759I | B | 1.9e-06 | 1.9e-06 | 2.3e-06 | 2.033E-06 | 2.309E-07 | 1.900E-06 | 2.300E-06 |
| 15171 | 759 | N759L | B | 2.5e-06 | 2.7e-06 | 3.1e-06 | 2.767E-06 | 3.055E-07 | 2.500E-06 | 3.100E-06 |
| 15172 | 759 | N759K | B | 1.8e-06 | 9.9e-07 | 1.7e-06 | 1.497E-06 | 4.416E-07 | 9.900E-07 | 1.800E-06 |
| 15173 | 759 | N759M | B | 1.6e-06 | 4.4e-06 | 2e-06 | 2.667E-06 | 1.514E-06 | 1.600E-06 | 4.400E-06 |
| 15174 | 759 | N759F | B | 2.8e-06 | 2.4e-06 | 2.4e-06 | 2.533E-06 | 2.309E-07 | 2.400E-06 | 2.800E-06 |
| 15175 | 759 | N759P | B | 2.8e-06 | 2.1e-06 | 1.8e-06 | 2.233E-06 | 5.132E-07 | 1.800E-06 | 2.800E-06 |
| 15176 | 759 | N759S | B | 3.5e-06 | 2e-06 | 2.2e-06 | 2.567E-06 | 8.145E-07 | 2.000E-06 | 3.500E-06 |
| 15177 | 759 | N759T | B | 3.2e-06 | 3.3e-06 | 3e-06 | 3.167E-06 | 1.528E-07 | 3.000E-06 | 3.300E-06 |
| 15178 | 759 | N759W | B | 3.1e-06 | 1.9e-06 | 1.4e-06 | 2.133E-06 | 8.737E-07 | 1.400E-06 | 3.100E-06 |
| 15179 | 759 | N759Y | B | 3.3e-06 | 2e-06 | 1.5e-06 | 2.267E-06 | 9.292E-07 | 1.500E-06 | 3.300E-06 |
| 15180 | 759 | N759V | B | 2.1e-06 | 1.8e-06 | 1.9e-06 | 1.933E-06 | 1.528E-07 | 1.800E-06 | 2.100E-06 |
| 15181 | 760 | N760A | B | 3.8e-06 | 1.9e-06 | 3.3e-06 | 3.000E-06 | 9.849E-07 | 1.900E-06 | 3.800E-06 |
| 15182 | 760 | N760R | B | 2.7e-06 | 3.6e-06 | 2.1e-06 | 2.800E-06 | 7.550E-07 | 2.100E-06 | 3.600E-06 |
| 15183 | 760 | N760N | B | 2.4e-06 | 2e-06 | 1.8e-06 | 2.067E-06 | 3.055E-07 | 1.800E-06 | 2.400E-06 |
| 15184 | 760 | N760D | B | 1.8e-06 | 3.8e-06 | 2.4e-06 | 2.667E-06 | 1.026E-06 | 1.800E-06 | 3.800E-06 |
| 15185 | 760 | N760C | B | 3.1e-06 | 2.9e-06 | 3.5e-06 | 3.167E-06 | 3.055E-07 | 2.900E-06 | 3.500E-06 |
| 15186 | 760 | N760E | B | 1.9e-06 | 1.8e-06 | 1.5e-06 | 1.733E-06 | 2.082E-07 | 1.500E-06 | 1.900E-06 |
| 15187 | 760 | N760Q | B | 2.6e-06 | 1.4e-06 | 1.4e-06 | 1.800E-06 | 6.928E-07 | 1.400E-06 | 2.600E-06 |
| 15188 | 760 | N760G | B | 2.4e-06 | 2.9e-06 | 2.8e-06 | 2.700E-06 | 2.646E-07 | 2.400E-06 | 2.900E-06 |
| 15189 | 760 | N760H | B | 3.2e-06 | 3.1e-06 | 3e-06 | 3.100E-06 | 1.000E-07 | 3.000E-06 | 3.200E-06 |
| 15190 | 760 | N760I | B | 3.7e-06 | 3.1e-06 | 3e-06 | 3.267E-06 | 3.786E-07 | 3.000E-06 | 3.700E-06 |
| 15191 | 760 | N760L | B | 2.8e-06 | 2.4e-06 | 2.3e-06 | 2.500E-06 | 2.646E-07 | 2.300E-06 | 2.800E-06 |
| 15192 | 760 | N760K | B | 2.1e-06 | 2.4e-06 | 1.7e-06 | 2.067E-06 | 3.512E-07 | 1.700E-06 | 2.400E-06 |
| 15193 | 760 | N760M | B | 2.3e-06 | 5.1e-06 | 2.1e-06 | 3.167E-06 | 1.677E-06 | 2.100E-06 | 5.100E-06 |
| 15194 | 760 | N760F | B | 2.9e-06 | 2.9e-06 | 2.8e-06 | 2.867E-06 | 5.774E-08 | 2.800E-06 | 2.900E-06 |
| 15195 | 760 | N760P | B | 2.3e-06 | 2.5e-06 | 1.7e-06 | 2.167E-06 | 4.163E-07 | 1.700E-06 | 2.500E-06 |
| 15196 | 760 | N760S | B | 1.9e-06 | 1.9e-06 | 2e-06 | 1.933E-06 | 5.774E-08 | 1.900E-06 | 2.000E-06 |
| 15197 | 760 | N760T | B | 1.9e-06 | 2.9e-06 | 2.9e-06 | 2.567E-06 | 5.774E-07 | 1.900E-06 | 2.900E-06 |
| 15198 | 760 | N760W | B | 1.9e-06 | 3.1e-06 | 1.7e-06 | 2.233E-06 | 7.572E-07 | 1.700E-06 | 3.100E-06 |
| 15199 | 760 | N760Y | B | 2.7e-06 | 4.7e-06 | 2.4e-06 | 3.267E-06 | 1.250E-06 | 2.400E-06 | 4.700E-06 |
| 15200 | 760 | N760V | B | 2e-06 | 4.1e-06 | 2.1e-06 | 2.733E-06 | 1.185E-06 | 2.000E-06 | 4.100E-06 |
| 15201 | 761 | F761A | B | 3.1e-06 | 2e-06 | 3.7e-06 | 2.933E-06 | 8.622E-07 | 2.000E-06 | 3.700E-06 |
| 15202 | 761 | F761R | B | 2.2e-06 | 2.5e-06 | 4.3e-06 | 3.000E-06 | 1.136E-06 | 2.200E-06 | 4.300E-06 |
| 15203 | 761 | F761N | B | 3.4e-06 | 4.7e-06 | 2.5e-06 | 3.533E-06 | 1.106E-06 | 2.500E-06 | 4.700E-06 |
| 15204 | 761 | F761D | B | 1.3e-06 | 1.9e-06 | 2.8e-06 | 2.000E-06 | 7.550E-07 | 1.300E-06 | 2.800E-06 |
| 15205 | 761 | F761C | B | 1.8e-06 | 1.8e-06 | 2.4e-06 | 2.000E-06 | 3.464E-07 | 1.800E-06 | 2.400E-06 |
| 15206 | 761 | F761E | B | 2e-06 | 1.8e-06 | 1.2e-06 | 1.667E-06 | 4.163E-07 | 1.200E-06 | 2.000E-06 |
| 15207 | 761 | F761Q | B | 1.8e-06 | 2.4e-06 | 2.4e-06 | 2.200E-06 | 3.464E-07 | 1.800E-06 | 2.400E-06 |
| 15208 | 761 | F761G | B | 2.6e-06 | 2.3e-06 | 2.8e-06 | 2.567E-06 | 2.517E-07 | 2.300E-06 | 2.800E-06 |
| 15209 | 761 | F761H | B | 3.3e-06 | 2.4e-06 | 4.7e-06 | 3.467E-06 | 1.159E-06 | 2.400E-06 | 4.700E-06 |
| 15210 | 761 | F761I | B | 2.4e-06 | 1.7e-06 | 3.6e-06 | 2.567E-06 | 9.609E-07 | 1.700E-06 | 3.600E-06 |
| 15211 | 761 | F761L | B | 1.8e-06 | 1.4e-06 | 1.5e-06 | 1.567E-06 | 2.082E-07 | 1.400E-06 | 1.800E-06 |
| 15212 | 761 | F761K | B | 1.7e-06 | 2.5e-06 | 4e-06 | 2.733E-06 | 1.168E-06 | 1.700E-06 | 4.000E-06 |
| 15213 | 761 | F761M | B | 3.5e-06 | 1.7e-06 | 1.4e-06 | 2.200E-06 | 1.136E-06 | 1.400E-06 | 3.500E-06 |
| 15214 | 761 | F761F | B | 4.2e-06 | 2.2e-06 | 2.9e-06 | 3.100E-06 | 1.015E-06 | 2.200E-06 | 4.200E-06 |

| | | | | | | | | | | |
|---|---|---|---|---|---|---|---|---|---|---|
| 15215 | 761 | F761P | B | 2.9e-06 | 2.9e-06 | 2.2e-06 | 2.667E-06 | 4.041E-07 | 2.200E-06 | 2.900E-06 |
| 15216 | 761 | F761S | B | 2.3e-06 | 9.7e-07 | 2.6e-06 | 1.957E-06 | 8.675E-07 | 9.700E-07 | 2.600E-06 |
| 15217 | 761 | F761T | B | 1.1e-06 | 1.7e-06 | 2.7e-06 | 1.833E-06 | 8.083E-07 | 1.100E-06 | 2.700E-06 |
| 15218 | 761 | F761W | B | 3.1e-06 | 8.3e-07 | 2.9e-06 | 2.277E-06 | 1.257E-06 | 8.300E-07 | 3.100E-06 |
| 15219 | 761 | F761Y | B | 3.1e-06 | 3.6e-06 | 3.6e-06 | 3.433E-06 | 2.887E-07 | 3.100E-06 | 3.600E-06 |
| 15220 | 761 | F761V | B | 3.6e-06 | 1.5e-06 | 4.5e-06 | 3.200E-06 | 1.539E-06 | 1.500E-06 | 4.500E-06 |
| 15221 | 762 | Y762A | B | 1.5e-06 | 2.4e-06 | 2.5e-06 | 2.133E-06 | 5.508E-07 | 1.500E-06 | 2.500E-06 |
| 15222 | 762 | Y762R | B | 3.2e-06 | 2.3e-06 | 2.6e-06 | 2.700E-06 | 4.583E-07 | 2.300E-06 | 3.200E-06 |
| 15223 | 762 | Y762N | B | 1.6e-06 | 2.2e-06 | 2.4e-06 | 2.067E-06 | 4.163E-07 | 1.600E-06 | 2.400E-06 |
| 15224 | 762 | Y762D | B | 2.2e-06 | 5.4e-06 | 3.1e-06 | 3.567E-06 | 1.650E-06 | 2.200E-06 | 5.400E-06 |
| 15225 | 762 | Y762C | B | 1.7e-06 | 1.7e-06 | 2.3e-06 | 1.900E-06 | 3.464E-07 | 1.700E-06 | 2.300E-06 |
| 15226 | 762 | Y762E | B | 2.4e-06 | 3.7e-06 | 2.9e-06 | 3.000E-06 | 6.557E-07 | 2.400E-06 | 3.700E-06 |
| 15227 | 762 | Y762Q | B | 1.9e-06 | 3.4e-06 | 2.5e-06 | 2.600E-06 | 7.550E-07 | 1.900E-06 | 3.400E-06 |
| 15228 | 762 | Y762G | B | 2.2e-06 | 3.1e-06 | 2.1e-06 | 2.467E-06 | 5.508E-07 | 2.100E-06 | 3.100E-06 |
| 15229 | 762 | Y762H | B | 2.6e-06 | 2.2e-06 | 3.7e-06 | 2.833E-06 | 7.767E-07 | 2.200E-06 | 3.700E-06 |
| 15230 | 762 | Y762I | B | 3e-06 | 2.4e-06 | 1.9e-06 | 2.433E-06 | 5.508E-07 | 1.900E-06 | 3.000E-06 |
| 15231 | 762 | Y762L | B | 1.6e-06 | 2.8e-06 | 2.7e-06 | 2.367E-06 | 6.658E-07 | 1.600E-06 | 2.800E-06 |
| 15232 | 762 | Y762K | B | 2.7e-06 | 3.5e-06 | 3.7e-06 | 3.300E-06 | 5.292E-07 | 2.700E-06 | 3.700E-06 |
| 15233 | 762 | Y762M | B | 1.6e-06 | 1.7e-06 | 1.9e-06 | 1.733E-06 | 1.528E-07 | 1.600E-06 | 1.900E-06 |
| 15234 | 762 | Y762F | B | 3.4e-06 | 1.6e-06 | 1.9e-06 | 2.300E-06 | 9.644E-07 | 1.600E-06 | 3.400E-06 |
| 15235 | 762 | Y762P | B | 1.4e-06 | 3.7e-06 | 2e-06 | 2.367E-06 | 1.193E-06 | 1.400E-06 | 3.700E-06 |
| 15236 | 762 | Y762S | B | 2.3e-06 | 2.5e-06 | 2.6e-06 | 2.467E-06 | 1.528E-07 | 2.300E-06 | 2.600E-06 |
| 15237 | 762 | Y762T | B | 2.9e-06 | 3.2e-06 | 2.2e-06 | 2.767E-06 | 5.132E-07 | 2.200E-06 | 3.200E-06 |
| 15238 | 762 | Y762W | B | 2.3e-06 | 1.6e-06 | 2.8e-06 | 2.233E-06 | 6.028E-07 | 1.600E-06 | 2.800E-06 |
| 15239 | 762 | Y762Y | B | 2.2e-06 | 2e-06 | 3.3e-06 | 2.500E-06 | 7.000E-07 | 2.000E-06 | 3.300E-06 |
| 15240 | 762 | Y762V | B | 2.6e-06 | 2.2e-06 | 1.7e-06 | 2.167E-06 | 4.509E-07 | 1.700E-06 | 2.600E-06 |
| 15241 | 763 | P763A | B | 1.8e-06 | 1.9e-06 | 3.4e-06 | 2.367E-06 | 8.963E-07 | 1.800E-06 | 3.400E-06 |
| 15242 | 763 | P763R | B | 1.9e-06 | 2.4e-06 | 1.4e-06 | 1.900E-06 | 5.000E-07 | 1.400E-06 | 2.400E-06 |
| 15243 | 763 | P763N | B | 1.8e-06 | 2.7e-06 | 3.8e-06 | 2.767E-06 | 1.002E-06 | 1.800E-06 | 3.800E-06 |
| 15244 | 763 | P763D | B | 2.7e-06 | 2.1e-06 | 3e-06 | 2.600E-06 | 4.583E-07 | 2.100E-06 | 3.000E-06 |
| 15245 | 763 | P763C | B | 1.8e-06 | 2.3e-06 | 2.1e-06 | 2.067E-06 | 2.517E-07 | 1.800E-06 | 2.300E-06 |
| 15246 | 763 | P763E | B | 3e-06 | 1.7e-06 | 2.4e-06 | 2.367E-06 | 6.506E-07 | 1.700E-06 | 3.000E-06 |
| 15247 | 763 | P763Q | B | 2.3e-06 | 2.5e-06 | 2.3e-06 | 2.367E-06 | 1.155E-07 | 2.300E-06 | 2.500E-06 |
| 15248 | 763 | P763G | B | 2.9e-06 | 3.9e-06 | 1.6e-06 | 2.800E-06 | 1.153E-06 | 1.600E-06 | 3.900E-06 |
| 15249 | 763 | P763H | B | 2e-06 | 2.2e-06 | 5.8e-06 | 3.333E-06 | 2.139E-06 | 2.000E-06 | 5.800E-06 |
| 15250 | 763 | P763I | B | 1.9e-06 | 2.4e-06 | 1.5e-06 | 1.933E-06 | 4.509E-07 | 1.500E-06 | 2.400E-06 |
| 15251 | 763 | P763L | B | 2.5e-06 | 2.6e-06 | 3.1e-06 | 2.733E-06 | 3.215E-07 | 2.500E-06 | 3.100E-06 |
| 15252 | 763 | P763K | B | 3.3e-06 | 2.9e-06 | 2.4e-06 | 2.867E-06 | 4.509E-07 | 2.400E-06 | 3.300E-06 |
| 15253 | 763 | P763M | B | 2.5e-06 | 1.4e-06 | 3e-06 | 2.300E-06 | 8.185E-07 | 1.400E-06 | 3.000E-06 |
| 15254 | 763 | P763F | B | 1.9e-06 | 2.3e-06 | 1.9e-06 | 2.033E-06 | 2.309E-07 | 1.900E-06 | 2.300E-06 |
| 15255 | 763 | P763P | B | 1.8e-06 | 2.1e-06 | 2.1e-06 | 2.000E-06 | 1.732E-07 | 1.800E-06 | 2.100E-06 |
| 15256 | 763 | P763S | B | 2.9e-06 | 2.3e-06 | 2e-06 | 2.400E-06 | 4.583E-07 | 2.000E-06 | 2.900E-06 |
| 15257 | 763 | P763T | B | 1.3e-06 | 3.1e-06 | 3.1e-06 | 2.500E-06 | 1.039E-06 | 1.300E-06 | 3.100E-06 |
| 15258 | 763 | P763W | B | 2.8e-06 | 3e-06 | 2.1e-06 | 2.633E-06 | 4.726E-07 | 2.100E-06 | 3.000E-06 |
| 15259 | 763 | P763Y | B | 2.6e-06 | 2.6e-06 | 2.1e-06 | 2.433E-06 | 2.887E-07 | 2.100E-06 | 2.600E-06 |
| 15260 | 763 | P763V | B | 2.8e-06 | 2.3e-06 | 2.8e-06 | 2.633E-06 | 2.887E-07 | 2.300E-06 | 2.800E-06 |
| 15261 | 764 | R764A | B | 2.9e-06 | 1.6e-06 | 2.6e-06 | 2.367E-06 | 6.807E-07 | 1.600E-06 | 2.900E-06 |
| 15262 | 764 | R764R | B | 3.6e-06 | 2.6e-06 | 1.9e-06 | 2.700E-06 | 8.544E-07 | 1.900E-06 | 3.600E-06 |
| 15263 | 764 | R764N | B | 1.6e-06 | 3.4e-06 | 1.5e-06 | 2.167E-06 | 1.069E-06 | 1.500E-06 | 3.400E-06 |
| 15264 | 764 | R764D | B | 1.6e-06 | 2e-06 | 3.2e-06 | 2.267E-06 | 8.327E-07 | 1.600E-06 | 3.200E-06 |
| 15265 | 764 | R764C | B | 3.5e-06 | 9.7e-07 | 2.2e-06 | 2.223E-06 | 1.265E-06 | 9.700E-07 | 3.500E-06 |
| 15266 | 764 | R764E | B | 3.4e-06 | 1.5e-06 | 3.6e-06 | 2.833E-06 | 1.159E-06 | 1.500E-06 | 3.600E-06 |
| 15267 | 764 | R764Q | B | 1.3e-06 | 3.1e-06 | 1.8e-06 | 2.067E-06 | 9.292E-07 | 1.300E-06 | 3.100E-06 |
| 15268 | 764 | R764G | B | 2.3e-06 | 4.5e-06 | 3e-06 | 3.267E-06 | 1.124E-06 | 2.300E-06 | 4.500E-06 |
| 15269 | 764 | R764H | B | 1.8e-06 | 3.5e-06 | 2.1e-06 | 2.467E-06 | 9.074E-07 | 1.800E-06 | 3.500E-06 |
| 15270 | 764 | R764I | B | 2.7e-06 | 2.6e-06 | 4.1e-06 | 3.133E-06 | 8.386E-07 | 2.600E-06 | 4.100E-06 |
| 15271 | 764 | R764L | B | 2.7e-06 | 2.2e-06 | 2.2e-06 | 2.367E-06 | 2.887E-07 | 2.200E-06 | 2.700E-06 |
| 15272 | 764 | R764K | B | 2.8e-06 | 2.1e-06 | 2e-06 | 2.300E-06 | 4.359E-07 | 2.000E-06 | 2.800E-06 |
| 15273 | 764 | R764M | B | 2.5e-06 | 2.4e-06 | 1.5e-06 | 2.133E-06 | 5.508E-07 | 1.500E-06 | 2.500E-06 |
| 15274 | 764 | R764F | B | 3.1e-06 | 2.3e-06 | 2e-06 | 2.467E-06 | 5.686E-07 | 2.000E-06 | 3.100E-06 |
| 15275 | 764 | R764P | B | 2.6e-06 | 3.8e-06 | 2e-06 | 2.800E-06 | 9.165E-07 | 2.000E-06 | 3.800E-06 |
| 15276 | 764 | R764S | B | 2.3e-06 | 3.6e-06 | 2.8e-06 | 2.900E-06 | 6.557E-07 | 2.300E-06 | 3.600E-06 |
| 15277 | 764 | R764T | B | 2.5e-06 | 1.7e-06 | 3.7e-06 | 2.633E-06 | 1.007E-06 | 1.700E-06 | 3.700E-06 |
| 15278 | 764 | R764W | B | 2.2e-06 | 2.2e-06 | 2.5e-06 | 2.300E-06 | 1.732E-07 | 2.200E-06 | 2.500E-06 |
| 15279 | 764 | R764Y | B | 3.2e-06 | 2.4e-06 | 2.4e-06 | 2.667E-06 | 4.619E-07 | 2.400E-06 | 3.200E-06 |
| 15280 | 764 | R764V | B | 2e-06 | 2.7e-06 | 3.1e-06 | 2.600E-06 | 5.568E-07 | 2.000E-06 | 3.100E-06 |
| 15281 | 765 | E765A | B | 1.3e-06 | 1.9e-06 | 1e-06 | 1.400E-06 | 4.583E-07 | 1.000E-06 | 1.900E-06 |
| 15282 | 765 | E765R | B | 2.2e-06 | 3.1e-06 | 2.7e-06 | 2.667E-06 | 4.509E-07 | 2.200E-06 | 3.100E-06 |
| 15283 | 765 | E765N | B | 1.9e-06 | 2.7e-06 | 2.2e-06 | 2.267E-06 | 4.041E-07 | 1.900E-06 | 2.700E-06 |
| 15284 | 765 | E765D | B | 2.2e-06 | 2.2e-06 | 4.3e-06 | 2.900E-06 | 1.212E-06 | 2.200E-06 | 4.300E-06 |
| 15285 | 765 | E765C | B | 2.8e-06 | 8.7e-07 | 1.3e-06 | 1.657E-06 | 1.013E-06 | 8.700E-07 | 2.800E-06 |
| 15286 | 765 | E765E | B | 3.7e-06 | 1.8e-06 | 1.4e-06 | 2.300E-06 | 1.229E-06 | 1.400E-06 | 3.700E-06 |
| 15287 | 765 | E765Q | B | 1.7e-06 | 2e-06 | 4.8e-06 | 2.833E-06 | 1.710E-06 | 1.700E-06 | 4.800E-06 |
| 15288 | 765 | E765G | B | 2.3e-06 | 2.8e-06 | 2.4e-06 | 2.500E-06 | 2.646E-07 | 2.300E-06 | 2.800E-06 |
| 15289 | 765 | E765H | B | 1.5e-06 | 8.7e-07 | 1.3e-06 | 1.223E-06 | 3.219E-07 | 8.700E-07 | 1.500E-06 |
| 15290 | 765 | E765I | B | 2e-06 | 3.1e-06 | 1.5e-06 | 2.200E-06 | 8.185E-07 | 1.500E-06 | 3.100E-06 |
| 15291 | 765 | E765L | B | 2.4e-06 | 2.6e-06 | 2.1e-06 | 2.367E-06 | 2.517E-07 | 2.100E-06 | 2.600E-06 |
| 15292 | 765 | E765K | B | 2.2e-06 | 2e-06 | 4.3e-06 | 2.833E-06 | 1.274E-06 | 2.000E-06 | 4.300E-06 |
| 15293 | 765 | E765M | B | 2.1e-06 | 2.4e-06 | 1.8e-06 | 2.100E-06 | 3.000E-07 | 1.800E-06 | 2.400E-06 |
| 15294 | 765 | E765F | B | 1.5e-06 | 2.9e-06 | 5.1e-06 | 3.167E-06 | 1.815E-06 | 1.500E-06 | 5.100E-06 |
| 15295 | 765 | E765P | B | 2.3e-06 | 4.1e-06 | 2.1e-06 | 2.833E-06 | 1.102E-06 | 2.100E-06 | 4.100E-06 |
| 15296 | 765 | E765S | B | 1.5e-06 | 2.1e-06 | 1.6e-06 | 1.733E-06 | 3.215E-07 | 1.500E-06 | 2.100E-06 |
| 15297 | 765 | E765T | B | 2e-06 | 1.7e-06 | 2.1e-06 | 1.933E-06 | 2.082E-07 | 1.700E-06 | 2.100E-06 |
| 15298 | 765 | E765W | B | 2.2e-06 | 3e-06 | 1.9e-06 | 2.367E-06 | 5.686E-07 | 1.900E-06 | 3.000E-06 |
| 15299 | 765 | E765Y | B | 3e-06 | 1.6e-06 | 2.2e-06 | 2.267E-06 | 7.024E-07 | 1.600E-06 | 3.000E-06 |

| | | | | | | | | | | |
|---|---|---|---|---|---|---|---|---|---|---|
| 15300 | 765 | E765V | B | 2.5e-06 | 4.2e-06 | 3.1e-06 | 3.267E-06 | 8.622E-07 | 2.500E-06 | 4.200E-06 |
| 15301 | 766 | A766A | B | 3.4e-06 | 1.7e-06 | 3.2e-06 | 2.767E-06 | 9.292E-07 | 1.700E-06 | 3.400E-06 |
| 15302 | 766 | A766R | B | 2.1e-06 | 1.3e-06 | 3.9e-06 | 2.433E-06 | 1.332E-06 | 1.300E-06 | 3.900E-06 |
| 15303 | 766 | A766N | B | 3.8e-06 | 2.2e-06 | 2.9e-06 | 2.967E-06 | 8.021E-07 | 2.200E-06 | 3.800E-06 |
| 15304 | 766 | A766D | B | 2.6e-06 | 3.3e-06 | 2.9e-06 | 2.933E-06 | 3.512E-07 | 2.600E-06 | 3.300E-06 |
| 15305 | 766 | A766C | B | 1.8e-06 | 2.5e-06 | 1.9e-06 | 2.067E-06 | 3.786E-07 | 1.800E-06 | 2.500E-06 |
| 15306 | 766 | A766E | B | 2.6e-06 | 2.2e-06 | 2.7e-06 | 2.500E-06 | 2.646E-07 | 2.200E-06 | 2.700E-06 |
| 15307 | 766 | A766Q | B | 1.8e-06 | 3e-06 | 3.2e-06 | 2.667E-06 | 7.572E-07 | 1.800E-06 | 3.200E-06 |
| 15308 | 766 | A766G | B | 1.5e-06 | 3e-06 | 2.7e-06 | 2.400E-06 | 7.937E-07 | 1.500E-06 | 3.000E-06 |
| 15309 | 766 | A766H | B | 2.5e-06 | 2.5e-06 | 2e-06 | 2.333E-06 | 2.887E-07 | 2.000E-06 | 2.500E-06 |
| 15310 | 766 | A766I | B | 3.3e-06 | 2.7e-06 | 2e-06 | 2.667E-06 | 6.506E-07 | 2.000E-06 | 3.300E-06 |
| 15311 | 766 | A766L | B | 2.7e-06 | 3.4e-06 | 1.4e-06 | 2.500E-06 | 1.015E-06 | 1.400E-06 | 3.400E-06 |
| 15312 | 766 | A766K | B | 1.8e-06 | 2.3e-06 | 1.9e-06 | 2.000E-06 | 2.646E-07 | 1.800E-06 | 2.300E-06 |
| 15313 | 766 | A766M | B | 2.2e-06 | 3.3e-06 | 3.1e-06 | 2.867E-06 | 5.859E-07 | 2.200E-06 | 3.300E-06 |
| 15314 | 766 | A766F | B | 3.9e-06 | 2.8e-06 | 3.2e-06 | 3.300E-06 | 5.568E-07 | 2.800E-06 | 3.900E-06 |
| 15315 | 766 | A766P | B | 2.5e-06 | 1.8e-06 | 2.2e-06 | 2.167E-06 | 3.512E-07 | 1.800E-06 | 2.500E-06 |
| 15316 | 766 | A766S | B | 1.4e-06 | 1.2e-06 | 1.8e-06 | 1.467E-06 | 3.055E-07 | 1.200E-06 | 1.800E-06 |
| 15317 | 766 | A766T | B | 3.4e-06 | 2.5e-06 | 2.1e-06 | 2.667E-06 | 6.658E-07 | 2.100E-06 | 3.400E-06 |
| 15318 | 766 | A766W | B | 2.1e-06 | 2.5e-06 | 3e-06 | 2.533E-06 | 4.509E-07 | 2.100E-06 | 3.000E-06 |
| 15319 | 766 | A766Y | B | 1.1e-06 | 2.6e-06 | 1.1e-06 | 1.600E-06 | 8.660E-07 | 1.100E-06 | 2.600E-06 |
| 15320 | 766 | A766V | B | 2e-06 | 2.8e-06 | 2e-06 | 2.267E-06 | 4.619E-07 | 2.000E-06 | 2.800E-06 |
| 15321 | 767 | K767A | B | 3e-06 | 1.2e-06 | 2.9e-06 | 2.367E-06 | 1.012E-06 | 1.200E-06 | 3.000E-06 |
| 15322 | 767 | K767R | B | 1.7e-06 | 3.2e-06 | 3e-06 | 2.633E-06 | 8.145E-07 | 1.700E-06 | 3.200E-06 |
| 15323 | 767 | K767N | B | 3.1e-06 | 3.7e-06 | 2.2e-06 | 3.000E-06 | 7.550E-07 | 2.200E-06 | 3.700E-06 |
| 15324 | 767 | K767D | B | 2.1e-06 | 2.7e-06 | 2.9e-06 | 2.567E-06 | 4.163E-07 | 2.100E-06 | 2.900E-06 |
| 15325 | 767 | K767C | B | 3.3e-06 | 2e-06 | 1.7e-06 | 2.333E-06 | 8.505E-07 | 1.700E-06 | 3.300E-06 |
| 15326 | 767 | K767E | B | 2.6e-06 | 3.3e-06 | 3.3e-06 | 3.067E-06 | 4.041E-07 | 2.600E-06 | 3.300E-06 |
| 15327 | 767 | K767Q | B | 2e-06 | 3.2e-06 | 2.9e-06 | 2.700E-06 | 6.245E-07 | 2.000E-06 | 3.200E-06 |
| 15328 | 767 | K767G | B | 2e-06 | 1.6e-06 | 1.8e-06 | 1.800E-06 | 2.000E-07 | 1.600E-06 | 2.000E-06 |
| 15329 | 767 | K767H | B | 2.8e-06 | 4.2e-06 | 3.4e-06 | 3.467E-06 | 7.024E-07 | 2.800E-06 | 4.200E-06 |
| 15330 | 767 | K767I | B | 2.8e-06 | 3e-06 | 2.9e-06 | 2.900E-06 | 1.000E-07 | 2.800E-06 | 3.000E-06 |
| 15331 | 767 | K767L | B | 2.1e-06 | 3.2e-06 | 2.3e-06 | 2.533E-06 | 5.859E-07 | 2.100E-06 | 3.200E-06 |
| 15332 | 767 | K767K | B | 2.2e-06 | 1.9e-06 | 2.7e-06 | 2.267E-06 | 4.041E-07 | 1.900E-06 | 2.700E-06 |
| 15333 | 767 | K767M | B | 2.2e-06 | 2.1e-06 | 2.3e-06 | 2.200E-06 | 1.000E-07 | 2.100E-06 | 2.300E-06 |
| 15334 | 767 | K767F | B | 1.9e-06 | 1.4e-06 | 2.1e-06 | 1.800E-06 | 3.606E-07 | 1.400E-06 | 2.100E-06 |
| 15335 | 767 | K767P | B | 2.7e-06 | 3e-06 | 2.1e-06 | 2.600E-06 | 4.583E-07 | 2.100E-06 | 3.000E-06 |
| 15336 | 767 | K767S | B | 1.7e-06 | 4.1e-06 | 2.4e-06 | 2.733E-06 | 1.234E-06 | 1.700E-06 | 4.100E-06 |
| 15337 | 767 | K767T | B | 2.8e-06 | 1.9e-06 | 1.7e-06 | 2.133E-06 | 5.859E-07 | 1.700E-06 | 2.800E-06 |
| 15338 | 767 | K767W | B | 3.1e-06 | 2.3e-06 | 2.6e-06 | 2.667E-06 | 4.041E-07 | 2.300E-06 | 3.100E-06 |
| 15339 | 767 | K767Y | B | 1.3e-06 | 2.6e-06 | 1.4e-06 | 1.767E-06 | 7.234E-07 | 1.300E-06 | 2.600E-06 |
| 15340 | 767 | K767V | B | 3.7e-06 | 1.9e-06 | 2e-06 | 2.533E-06 | 1.012E-06 | 1.900E-06 | 3.700E-06 |
| 15341 | 768 | V768A | B | 1.4e-06 | 3.5e-06 | 1.9e-06 | 2.267E-06 | 1.097E-06 | 1.400E-06 | 3.500E-06 |
| 15342 | 768 | V768R | B | 2.5e-06 | 2.3e-06 | 1.9e-06 | 2.233E-06 | 3.055E-07 | 1.900E-06 | 2.500E-06 |
| 15343 | 768 | V768N | B | 2.1e-06 | 3.5e-06 | 3.3e-06 | 2.967E-06 | 7.572E-07 | 2.100E-06 | 3.500E-06 |
| 15344 | 768 | V768D | B | 2.4e-06 | 1.4e-06 | 2.2e-06 | 2.000E-06 | 5.292E-07 | 1.400E-06 | 2.400E-06 |
| 15345 | 768 | V768C | B | 2.6e-06 | 4.2e-06 | 1.8e-06 | 2.867E-06 | 1.222E-06 | 1.800E-06 | 4.200E-06 |
| 15346 | 768 | V768E | B | 3.3e-06 | 2.9e-06 | 3.6e-06 | 3.267E-06 | 3.512E-07 | 2.900E-06 | 3.600E-06 |
| 15347 | 768 | V768Q | B | 3.8e-06 | 1.1e-06 | 2.4e-06 | 2.433E-06 | 1.350E-06 | 1.100E-06 | 3.800E-06 |
| 15348 | 768 | V768G | B | 2.1e-06 | 2.3e-06 | 2.4e-06 | 2.267E-06 | 1.528E-07 | 2.100E-06 | 2.400E-06 |
| 15349 | 768 | V768H | B | 2e-06 | 3.3e-06 | 1.7e-06 | 2.333E-06 | 8.505E-07 | 1.700E-06 | 3.300E-06 |
| 15350 | 768 | V768I | B | 2.6e-06 | 3.6e-06 | 4e-06 | 3.400E-06 | 7.211E-07 | 2.600E-06 | 4.000E-06 |
| 15351 | 768 | V768L | B | 2.6e-06 | 2e-06 | 5.7e-06 | 3.433E-06 | 1.986E-06 | 2.000E-06 | 5.700E-06 |
| 15352 | 768 | V768K | B | 2e-06 | 1.1e-06 | 2.8e-06 | 1.967E-06 | 8.505E-07 | 1.100E-06 | 2.800E-06 |
| 15353 | 768 | V768M | B | 2.4e-06 | 2.3e-06 | 2.2e-06 | 2.300E-06 | 1.000E-07 | 2.200E-06 | 2.400E-06 |
| 15354 | 768 | V768F | B | 2.8e-06 | 2.4e-06 | 1.9e-06 | 2.367E-06 | 4.509E-07 | 1.900E-06 | 2.800E-06 |
| 15355 | 768 | V768P | B | 3.2e-06 | 1.6e-06 | 2.6e-06 | 2.467E-06 | 8.083E-07 | 1.600E-06 | 3.200E-06 |
| 15356 | 768 | V768S | B | 2.5e-06 | 2.4e-06 | 2e-06 | 2.300E-06 | 2.646E-07 | 2.000E-06 | 2.500E-06 |
| 15357 | 768 | V768T | B | 3.4e-06 | 2.2e-06 | 7.6e-07 | 2.120E-06 | 1.322E-06 | 7.600E-07 | 3.400E-06 |
| 15358 | 768 | V768W | B | 1.1e-06 | 1.6e-06 | 1.2e-06 | 1.300E-06 | 2.646E-07 | 1.100E-06 | 1.600E-06 |
| 15359 | 768 | V768Y | B | 3.1e-06 | 2e-06 | 2.1e-06 | 2.400E-06 | 6.083E-07 | 2.000E-06 | 3.100E-06 |
| 15360 | 768 | V768V | B | 1.9e-06 | 4.2e-06 | 2.4e-06 | 2.833E-06 | 1.210E-06 | 1.900E-06 | 4.200E-06 |
| 15361 | 769 | Q769A | B | 3.1e-06 | 3.1e-06 | 2.1e-06 | 2.767E-06 | 5.774E-07 | 2.100E-06 | 3.100E-06 |
| 15362 | 769 | Q769R | B | 3.3e-06 | 1.5e-06 | 2.1e-06 | 2.300E-06 | 9.165E-07 | 1.500E-06 | 3.300E-06 |
| 15363 | 769 | Q769N | B | 2.9e-06 | 2.3e-06 | 3e-06 | 2.733E-06 | 3.786E-07 | 2.300E-06 | 3.000E-06 |
| 15364 | 769 | Q769D | B | 2.9e-06 | 3.1e-06 | 2.3e-06 | 2.767E-06 | 4.163E-07 | 2.300E-06 | 3.100E-06 |
| 15365 | 769 | Q769C | B | 2.4e-06 | 2.6e-06 | 2.3e-06 | 2.433E-06 | 1.528E-07 | 2.300E-06 | 2.600E-06 |
| 15366 | 769 | Q769E | B | 2.2e-06 | 2.3e-06 | 2.4e-06 | 2.300E-06 | 1.000E-07 | 2.200E-06 | 2.400E-06 |
| 15367 | 769 | Q769Q | B | 2.1e-06 | 2.3e-06 | 1.4e-06 | 1.933E-06 | 4.726E-07 | 1.400E-06 | 2.300E-06 |
| 15368 | 769 | Q769G | B | 1.9e-06 | 4.1e-06 | 3.3e-06 | 3.100E-06 | 1.114E-06 | 1.900E-06 | 4.100E-06 |
| 15369 | 769 | Q769H | B | 5.2e-06 | 1.3e-06 | 2.8e-06 | 3.100E-06 | 1.967E-06 | 1.300E-06 | 5.200E-06 |
| 15370 | 769 | Q769I | B | 2.6e-06 | 2.9e-06 | 3.3e-06 | 2.933E-06 | 3.512E-07 | 2.600E-06 | 3.300E-06 |
| 15371 | 769 | Q769L | B | 3.1e-06 | 3.6e-06 | 2.7e-06 | 3.133E-06 | 4.509E-07 | 2.700E-06 | 3.600E-06 |
| 15372 | 769 | Q769K | B | 2.9e-06 | 2.4e-06 | 2.9e-06 | 2.733E-06 | 2.887E-07 | 2.400E-06 | 2.900E-06 |
| 15373 | 769 | Q769M | B | 4.2e-06 | 2.3e-06 | 2.7e-06 | 3.067E-06 | 1.002E-06 | 2.300E-06 | 4.200E-06 |
| 15374 | 769 | Q769F | B | 3.7e-06 | 3.1e-06 | 2.8e-06 | 3.200E-06 | 4.583E-07 | 2.800E-06 | 3.700E-06 |
| 15375 | 769 | Q769P | B | 3.2e-06 | 3.3e-06 | 1.6e-06 | 2.700E-06 | 9.539E-07 | 1.600E-06 | 3.300E-06 |
| 15376 | 769 | Q769S | B | 1.6e-06 | 1.5e-06 | 3.2e-06 | 2.100E-06 | 9.539E-07 | 1.500E-06 | 3.200E-06 |
| 15377 | 769 | Q769T | B | 1.9e-06 | 3e-06 | 3.4e-06 | 2.767E-06 | 7.767E-07 | 1.900E-06 | 3.400E-06 |
| 15378 | 769 | Q769W | B | 1.7e-06 | 1.8e-06 | 2.7e-06 | 2.067E-06 | 5.508E-07 | 1.700E-06 | 2.700E-06 |
| 15379 | 769 | Q769Y | B | 1.8e-06 | 2e-06 | 1.3e-06 | 1.700E-06 | 3.606E-07 | 1.300E-06 | 2.000E-06 |
| 15380 | 769 | Q769V | B | 1.5e-06 | 3.5e-06 | 3.2e-06 | 2.733E-06 | 1.079E-06 | 1.500E-06 | 3.500E-06 |
| 15381 | 770 | W770A | B | 3.6e-06 | 2.2e-06 | 3.7e-06 | 3.167E-06 | 8.386E-07 | 2.200E-06 | 3.700E-06 |
| 15382 | 770 | W770R | B | 2.6e-06 | 5.1e-06 | 1.9e-06 | 3.200E-06 | 1.682E-06 | 1.900E-06 | 5.100E-06 |
| 15383 | 770 | W770N | B | 2.6e-06 | 2.9e-06 | 2.8e-06 | 2.767E-06 | 1.528E-07 | 2.600E-06 | 2.900E-06 |
| 15384 | 770 | W770D | B | 2.3e-06 | 2.1e-06 | 2.5e-06 | 2.300E-06 | 2.000E-07 | 2.100E-06 | 2.500E-06 |

| | | | | | | | | | | |
|---|---|---|---|---|---|---|---|---|---|---|
| 15385 | 770 | W770C | B | 2.6e-06 | 2.5e-06 | 1.8e-06 | 2.300E-06 | 4.359E-07 | 1.800E-06 | 2.600E-06 |
| 15386 | 770 | W770E | B | 1.8e-06 | 3.2e-06 | 1.3e-06 | 2.100E-06 | 9.849E-07 | 1.300E-06 | 3.200E-06 |
| 15387 | 770 | W770Q | B | 2e-06 | 1.9e-06 | 2.6e-06 | 2.167E-06 | 3.786E-07 | 1.900E-06 | 2.600E-06 |
| 15388 | 770 | W770G | B | 2.2e-06 | 1.7e-06 | 2.1e-06 | 2.000E-06 | 2.646E-07 | 1.700E-06 | 2.200E-06 |
| 15389 | 770 | W770H | B | 2.7e-06 | 2.4e-06 | 1.6e-06 | 2.233E-06 | 5.686E-07 | 1.600E-06 | 2.700E-06 |
| 15390 | 770 | W770I | B | 3e-06 | 3.2e-06 | 2.4e-06 | 2.867E-06 | 4.163E-07 | 2.400E-06 | 3.200E-06 |
| 15391 | 770 | W770L | B | 3.2e-06 | 2.4e-06 | 2.4e-06 | 2.667E-06 | 4.619E-07 | 2.400E-06 | 3.200E-06 |
| 15392 | 770 | W770K | B | 1.4e-06 | 2.5e-06 | 2.3e-06 | 2.067E-06 | 5.859E-07 | 1.400E-06 | 2.500E-06 |
| 15393 | 770 | W770M | B | 2.5e-06 | 2.4e-06 | 2.4e-06 | 2.433E-06 | 5.774E-08 | 2.400E-06 | 2.500E-06 |
| 15394 | 770 | W770F | B | 2.7e-06 | 2.2e-06 | 3.2e-06 | 2.700E-06 | 5.000E-07 | 2.200E-06 | 3.200E-06 |
| 15395 | 770 | W770P | B | 1.9e-06 | 3e-06 | 2e-06 | 2.300E-06 | 6.083E-07 | 1.900E-06 | 3.000E-06 |
| 15396 | 770 | W770S | B | 1.7e-06 | 2.4e-06 | 3.6e-06 | 2.567E-06 | 9.609E-07 | 1.700E-06 | 3.600E-06 |
| 15397 | 770 | W770T | B | 2.9e-06 | 1.6e-06 | 3.9e-06 | 2.800E-06 | 1.153E-06 | 1.600E-06 | 3.900E-06 |
| 15398 | 770 | W770W | B | 3e-06 | 1.6e-06 | 3e-06 | 2.533E-06 | 8.083E-07 | 1.600E-06 | 3.000E-06 |
| 15399 | 770 | W770Y | B | 2.5e-06 | 1.9e-06 | 2.1e-06 | 2.167E-06 | 3.055E-07 | 1.900E-06 | 2.500E-06 |
| 15400 | 770 | W770V | B | 2.4e-06 | 2.6e-06 | 2.4e-06 | 2.467E-06 | 1.155E-07 | 2.400E-06 | 2.600E-06 |
| 15401 | 771 | K771A | B | 1.4e-06 | 2.7e-06 | 1.5e-06 | 1.867E-06 | 7.234E-07 | 1.400E-06 | 2.700E-06 |
| 15402 | 771 | K771R | B | 3.3e-06 | 2.3e-06 | 2.3e-06 | 2.633E-06 | 5.774E-07 | 2.300E-06 | 3.300E-06 |
| 15403 | 771 | K771N | B | 1.4e-06 | 2.7e-06 | 1.2e-06 | 1.767E-06 | 8.145E-07 | 1.200E-06 | 2.700E-06 |
| 15404 | 771 | K771D | B | 5.5e-06 | 2.5e-06 | 2.3e-06 | 3.433E-06 | 1.793E-06 | 2.300E-06 | 5.500E-06 |
| 15405 | 771 | K771C | B | 1.7e-06 | 2.3e-06 | 3e-06 | 2.333E-06 | 6.506E-07 | 1.700E-06 | 3.000E-06 |
| 15406 | 771 | K771E | B | 2.6e-06 | 3.7e-06 | 2.5e-06 | 2.933E-06 | 6.658E-07 | 2.500E-06 | 3.700E-06 |
| 15407 | 771 | K771Q | B | 4.1e-06 | 2.5e-06 | 2.2e-06 | 2.933E-06 | 1.021E-06 | 2.200E-06 | 4.100E-06 |
| 15408 | 771 | K771G | B | 3.3e-06 | 2.1e-06 | 2.1e-06 | 2.500E-06 | 6.928E-07 | 2.100E-06 | 3.300E-06 |
| 15409 | 771 | K771H | B | 2.2e-06 | 3e-06 | 4.7e-06 | 3.300E-06 | 1.277E-06 | 2.200E-06 | 4.700E-06 |
| 15410 | 771 | K771I | B | 2e-06 | 2.8e-06 | 1.7e-06 | 2.167E-06 | 5.686E-07 | 1.700E-06 | 2.800E-06 |
| 15411 | 771 | K771L | B | 1.4e-06 | 2.8e-06 | 1.9e-06 | 2.033E-06 | 7.095E-07 | 1.400E-06 | 2.800E-06 |
| 15412 | 771 | K771K | B | 2.4e-06 | 1.4e-06 | 3.8e-06 | 2.533E-06 | 1.206E-06 | 1.400E-06 | 3.800E-06 |
| 15413 | 771 | K771M | B | 2.4e-06 | 2.6e-06 | 1.6e-06 | 2.200E-06 | 5.292E-07 | 1.600E-06 | 2.600E-06 |
| 15414 | 771 | K771F | B | 1.8e-06 | 1.9e-06 | 2.3e-06 | 2.000E-06 | 2.646E-07 | 1.800E-06 | 2.300E-06 |
| 15415 | 771 | K771P | B | 2.8e-06 | 2.4e-06 | 2.6e-06 | 2.600E-06 | 2.000E-07 | 2.400E-06 | 2.800E-06 |
| 15416 | 771 | K771S | B | 2e-06 | 2.8e-06 | 2.5e-06 | 2.433E-06 | 4.041E-07 | 2.000E-06 | 2.800E-06 |
| 15417 | 771 | K771T | B | 3.4e-06 | 3.1e-06 | 2.9e-06 | 3.133E-06 | 2.517E-07 | 2.900E-06 | 3.400E-06 |
| 15418 | 771 | K771W | B | 1.8e-06 | 1.9e-06 | 1.9e-06 | 1.867E-06 | 5.774E-08 | 1.800E-06 | 1.900E-06 |
| 15419 | 771 | K771Y | B | 2.6e-06 | 1.8e-06 | 1.8e-06 | 2.067E-06 | 4.619E-07 | 1.800E-06 | 2.600E-06 |
| 15420 | 771 | K771V | B | 1.9e-06 | 2.2e-06 | 1.4e-06 | 1.833E-06 | 4.041E-07 | 1.400E-06 | 2.200E-06 |
| 15421 | 772 | V772A | B | 2e-06 | 1.5e-06 | 2.6e-06 | 2.033E-06 | 5.508E-07 | 1.500E-06 | 2.600E-06 |
| 15422 | 772 | V772R | B | 3.5e-06 | 2.6e-06 | 1.6e-06 | 2.567E-06 | 9.504E-07 | 1.600E-06 | 3.500E-06 |
| 15423 | 772 | V772N | B | 2.8e-06 | 3.2e-06 | 2.4e-06 | 2.800E-06 | 4.000E-07 | 2.400E-06 | 3.200E-06 |
| 15424 | 772 | V772D | B | 1.4e-06 | 7.8e-07 | 2.6e-06 | 1.593E-06 | 9.253E-07 | 7.800E-07 | 2.600E-06 |
| 15425 | 772 | V772C | B | 1.2e-06 | 1.9e-06 | 1.4e-06 | 1.500E-06 | 3.606E-07 | 1.200E-06 | 1.900E-06 |
| 15426 | 772 | V772E | B | 4.1e-06 | 1.7e-06 | 2.3e-06 | 2.700E-06 | 1.249E-06 | 1.700E-06 | 4.100E-06 |
| 15427 | 772 | V772Q | B | 3.1e-06 | 1.8e-06 | 2.3e-06 | 2.400E-06 | 6.557E-07 | 1.800E-06 | 3.100E-06 |
| 15428 | 772 | V772G | B | 2.1e-06 | 1.8e-06 | 2.9e-06 | 2.267E-06 | 5.686E-07 | 1.800E-06 | 2.900E-06 |
| 15429 | 772 | V772H | B | 2.7e-06 | 2e-06 | 3.6e-06 | 2.767E-06 | 8.021E-07 | 2.000E-06 | 3.600E-06 |
| 15430 | 772 | V772I | B | 2.6e-06 | 2.2e-06 | 1.9e-06 | 2.233E-06 | 3.512E-07 | 1.900E-06 | 2.600E-06 |
| 15431 | 772 | V772L | B | 2.2e-06 | 1.7e-06 | 2.4e-06 | 2.100E-06 | 3.606E-07 | 1.700E-06 | 2.400E-06 |
| 15432 | 772 | V772K | B | 2.5e-06 | 2.5e-06 | 1e-06 | 2.000E-06 | 8.660E-07 | 1.000E-06 | 2.500E-06 |
| 15433 | 772 | V772M | B | 2.4e-06 | 4e-06 | 2.6e-06 | 3.000E-06 | 8.718E-07 | 2.400E-06 | 4.000E-06 |
| 15434 | 772 | V772F | B | 2.7e-06 | 2.8e-06 | 2.4e-06 | 2.633E-06 | 2.082E-07 | 2.400E-06 | 2.800E-06 |
| 15435 | 772 | V772P | B | 1.9e-06 | 2.8e-06 | 2.2e-06 | 2.300E-06 | 4.583E-07 | 1.900E-06 | 2.800E-06 |
| 15436 | 772 | V772S | B | 2.9e-06 | 1.6e-06 | 2.1e-06 | 2.200E-06 | 6.557E-07 | 1.600E-06 | 2.900E-06 |
| 15437 | 772 | V772T | B | 1.4e-06 | 2.8e-06 | 3.5e-06 | 2.567E-06 | 1.069E-06 | 1.400E-06 | 3.500E-06 |
| 15438 | 772 | V772W | B | 2.2e-06 | 5.4e-06 | 2.5e-06 | 3.367E-06 | 1.767E-06 | 2.200E-06 | 5.400E-06 |
| 15439 | 772 | V772Y | B | 2.1e-06 | 2.1e-06 | 2e-06 | 2.067E-06 | 5.774E-08 | 2.000E-06 | 2.100E-06 |
| 15440 | 772 | V772V | B | 2.3e-06 | 1.6e-06 | 2.3e-06 | 2.067E-06 | 4.041E-07 | 1.600E-06 | 2.300E-06 |
| 15441 | 773 | D773A | B | 9.7e-07 | 2.3e-06 | 3.8e-06 | 2.357E-06 | 1.416E-06 | 9.700E-07 | 3.800E-06 |
| 15442 | 773 | D773R | B | 2.1e-06 | 2e-06 | 2.3e-06 | 2.133E-06 | 1.528E-07 | 2.000E-06 | 2.300E-06 |
| 15443 | 773 | D773N | B | 2.3e-06 | 5.8e-06 | 1.2e-06 | 3.100E-06 | 2.402E-06 | 1.200E-06 | 5.800E-06 |
| 15444 | 773 | D773D | B | 2.7e-06 | 1.6e-06 | 2e-06 | 2.100E-06 | 5.568E-07 | 1.600E-06 | 2.700E-06 |
| 15445 | 773 | D773C | B | 2.4e-06 | 3.9e-06 | 2.6e-06 | 2.967E-06 | 8.145E-07 | 2.400E-06 | 3.900E-06 |
| 15446 | 773 | D773E | B | 8.1e-07 | 1.6e-06 | 2.9e-06 | 1.770E-06 | 1.055E-06 | 8.100E-07 | 2.900E-06 |
| 15447 | 773 | D773Q | B | 2.1e-06 | 2.8e-06 | 2.6e-06 | 2.500E-06 | 3.606E-07 | 2.100E-06 | 2.800E-06 |
| 15448 | 773 | D773G | B | 2e-06 | 3.3e-06 | 2.4e-06 | 2.567E-06 | 6.658E-07 | 2.000E-06 | 3.300E-06 |
| 15449 | 773 | D773H | B | 2.8e-06 | 2.2e-06 | 2.1e-06 | 2.367E-06 | 3.786E-07 | 2.100E-06 | 2.800E-06 |
| 15450 | 773 | D773I | B | 2.7e-06 | 2.6e-06 | 1.8e-06 | 2.367E-06 | 4.933E-07 | 1.800E-06 | 2.700E-06 |
| 15451 | 773 | D773L | B | 3.1e-06 | 1.5e-06 | 2.3e-06 | 2.300E-06 | 8.000E-07 | 1.500E-06 | 3.100E-06 |
| 15452 | 773 | D773K | B | 2.2e-06 | 5.1e-06 | 2.7e-06 | 3.333E-06 | 1.550E-06 | 2.200E-06 | 5.100E-06 |
| 15453 | 773 | D773M | B | 2.6e-06 | 2.6e-06 | 2.2e-06 | 2.467E-06 | 2.309E-07 | 2.200E-06 | 2.600E-06 |
| 15454 | 773 | D773F | B | 1.7e-06 | 2.3e-06 | 2.9e-06 | 2.300E-06 | 6.000E-07 | 1.700E-06 | 2.900E-06 |
| 15455 | 773 | D773P | B | 3.8e-06 | 2.1e-06 | 1.7e-06 | 2.533E-06 | 1.115E-06 | 1.700E-06 | 3.800E-06 |
| 15456 | 773 | D773S | B | 2.1e-06 | 1.5e-06 | 1.7e-06 | 1.767E-06 | 3.055E-07 | 1.500E-06 | 2.100E-06 |
| 15457 | 773 | D773T | B | 2.2e-06 | 1.8e-06 | 1.9e-06 | 1.967E-06 | 2.082E-07 | 1.800E-06 | 2.200E-06 |
| 15458 | 773 | D773W | B | 2.6e-06 | 2.6e-06 | 1.2e-06 | 2.133E-06 | 8.083E-07 | 1.200E-06 | 2.600E-06 |
| 15459 | 773 | D773Y | B | 1.5e-06 | 2.4e-06 | 2.4e-06 | 2.100E-06 | 5.196E-07 | 1.500E-06 | 2.400E-06 |
| 15460 | 773 | D773V | B | 2.2e-06 | 2.6e-06 | 2.3e-06 | 2.367E-06 | 2.082E-07 | 2.200E-06 | 2.600E-06 |
| 15461 | 774 | N774A | B | 2e-06 | 3.2e-06 | 2.6e-06 | 2.600E-06 | 6.000E-07 | 2.000E-06 | 3.200E-06 |
| 15462 | 774 | N774R | B | 3.4e-06 | 3.8e-06 | 2.1e-06 | 3.100E-06 | 8.888E-07 | 2.100E-06 | 3.800E-06 |
| 15463 | 774 | N774N | B | 2.1e-06 | 3.4e-06 | 2.3e-06 | 2.600E-06 | 7.000E-07 | 2.100E-06 | 3.400E-06 |
| 15464 | 774 | N774D | B | 1.9e-06 | 2.1e-06 | 2.4e-06 | 2.133E-06 | 2.517E-07 | 1.900E-06 | 2.400E-06 |
| 15465 | 774 | N774C | B | 1.6e-06 | 2.3e-06 | 2e-06 | 1.967E-06 | 3.512E-07 | 1.600E-06 | 2.300E-06 |
| 15466 | 774 | N774E | B | 2.2e-06 | 2.2e-06 | 1.7e-06 | 2.033E-06 | 2.887E-07 | 1.700E-06 | 2.200E-06 |
| 15467 | 774 | N774Q | B | 1.8e-06 | 2.6e-06 | 2.9e-06 | 2.433E-06 | 5.686E-07 | 1.800E-06 | 2.900E-06 |
| 15468 | 774 | N774G | B | 2.8e-06 | 3.4e-06 | 2.2e-06 | 2.800E-06 | 6.000E-07 | 2.200E-06 | 3.400E-06 |
| 15469 | 774 | N774H | B | 3.1e-06 | 3.8e-06 | 2.4e-06 | 3.100E-06 | 7.000E-07 | 2.400E-06 | 3.800E-06 |

| | | | | | | | | | | |
|---|---|---|---|---|---|---|---|---|---|---|
| 15470 | 774 | N774I | B | 1.1e-06 | 2.4e-06 | 1.9e-06 | 1.800E-06 | 6.557E-07 | 1.100E-06 | 2.400E-06 |
| 15471 | 774 | N774L | B | 2.7e-06 | 1.7e-06 | 4.3e-06 | 2.900E-06 | 1.311E-06 | 1.700E-06 | 4.300E-06 |
| 15472 | 774 | N774K | B | 2.9e-06 | 2.6e-06 | 2.5e-06 | 2.667E-06 | 2.082E-07 | 2.500E-06 | 2.900E-06 |
| 15473 | 774 | N774M | B | 4e-06 | 1.9e-06 | 2.5e-06 | 2.800E-06 | 1.082E-06 | 1.900E-06 | 4.000E-06 |
| 15474 | 774 | N774F | B | 1.5e-06 | 2.7e-06 | 1.9e-06 | 2.033E-06 | 6.110E-07 | 1.500E-06 | 2.700E-06 |
| 15475 | 774 | N774P | B | 3e-06 | 1.9e-06 | 2.3e-06 | 2.400E-06 | 5.568E-07 | 1.900E-06 | 3.000E-06 |
| 15476 | 774 | N774S | B | 1.9e-06 | 1.8e-06 | 2.2e-06 | 1.967E-06 | 2.082E-07 | 1.800E-06 | 2.200E-06 |
| 15477 | 774 | N774T | B | 2.3e-06 | 1.8e-06 | 3.1e-06 | 2.400E-06 | 6.557E-07 | 1.800E-06 | 3.100E-06 |
| 15478 | 774 | N774W | B | 2.1e-06 | 2.6e-06 | 2.4e-06 | 2.367E-06 | 2.517E-07 | 2.100E-06 | 2.600E-06 |
| 15479 | 774 | N774Y | B | 2.2e-06 | 4.5e-06 | 2.1e-06 | 2.933E-06 | 1.358E-06 | 2.100E-06 | 4.500E-06 |
| 15480 | 774 | N774V | B | 3e-06 | 1.5e-06 | 1.7e-06 | 2.067E-06 | 8.145E-07 | 1.500E-06 | 3.000E-06 |
| 15481 | 775 | A775A | B | 1.8e-06 | 3.5e-06 | 4.6e-06 | 3.300E-06 | 1.411E-06 | 1.800E-06 | 4.600E-06 |
| 15482 | 775 | A775R | B | 3.4e-06 | 1.2e-06 | 2e-06 | 2.200E-06 | 1.114E-06 | 1.200E-06 | 3.400E-06 |
| 15483 | 775 | A775N | B | 1e-06 | 1.9e-06 | 2.3e-06 | 1.733E-06 | 6.658E-07 | 1.000E-06 | 2.300E-06 |
| 15484 | 775 | A775D | B | 2.9e-06 | 2.9e-06 | 2.3e-06 | 2.700E-06 | 3.464E-07 | 2.300E-06 | 2.900E-06 |
| 15485 | 775 | A775C | B | 2.3e-06 | 2.8e-06 | 4e-06 | 3.033E-06 | 8.737E-07 | 2.300E-06 | 4.000E-06 |
| 15486 | 775 | A775E | B | 2.4e-06 | 1.5e-06 | 2.1e-06 | 2.000E-06 | 4.583E-07 | 1.500E-06 | 2.400E-06 |
| 15487 | 775 | A775Q | B | 1.9e-06 | 2.6e-06 | 2.4e-06 | 2.300E-06 | 3.606E-07 | 1.900E-06 | 2.600E-06 |
| 15488 | 775 | A775G | B | 1.8e-06 | 3.8e-06 | 2.4e-06 | 2.667E-06 | 1.026E-06 | 1.800E-06 | 3.800E-06 |
| 15489 | 775 | A775H | B | 2.3e-06 | 2.2e-06 | 2.4e-06 | 2.300E-06 | 1.000E-07 | 2.200E-06 | 2.400E-06 |
| 15490 | 775 | A775I | B | 1.2e-06 | 3.5e-06 | 2.8e-06 | 2.500E-06 | 1.179E-06 | 1.200E-06 | 3.500E-06 |
| 15491 | 775 | A775L | B | 3.3e-06 | 1.8e-06 | 2.8e-06 | 2.633E-06 | 7.638E-07 | 1.800E-06 | 3.300E-06 |
| 15492 | 775 | A775K | B | 2.3e-06 | 3e-06 | 2.2e-06 | 2.500E-06 | 4.359E-07 | 2.200E-06 | 3.000E-06 |
| 15493 | 775 | A775M | B | 2.1e-06 | 2.4e-06 | 1.1e-06 | 1.867E-06 | 6.807E-07 | 1.100E-06 | 2.400E-06 |
| 15494 | 775 | A775F | B | 3.7e-06 | 2.3e-06 | 2.9e-06 | 2.967E-06 | 7.024E-07 | 2.300E-06 | 3.700E-06 |
| 15495 | 775 | A775P | B | 3.3e-06 | 2.5e-06 | 1.9e-06 | 2.567E-06 | 7.024E-07 | 1.900E-06 | 3.300E-06 |
| 15496 | 775 | A775S | B | 3e-06 | 1e-06 | 3.1e-06 | 2.367E-06 | 1.185E-06 | 1.000E-06 | 3.100E-06 |
| 15497 | 775 | A775T | B | 3.1e-06 | 2.5e-06 | 2.4e-06 | 2.667E-06 | 3.786E-07 | 2.400E-06 | 3.100E-06 |
| 15498 | 775 | A775W | B | 3.4e-06 | 3.4e-06 | 3.4e-06 | 3.400E-06 | 0.000E+00 | 3.400E-06 | 3.400E-06 |
| 15499 | 775 | A775Y | B | 1.2e-06 | 2.6e-06 | 2.6e-06 | 2.133E-06 | 8.083E-07 | 1.200E-06 | 2.600E-06 |
| 15500 | 775 | A775V | B | 1.4e-06 | 4.6e-06 | 3.3e-06 | 3.100E-06 | 1.609E-06 | 1.400E-06 | 4.600E-06 |
| 15501 | 776 | L776A | B | 1.5e-06 | 3.4e-06 | 2.4e-06 | 2.433E-06 | 9.504E-07 | 1.500E-06 | 3.400E-06 |
| 15502 | 776 | L776R | B | 5.9e-06 | 2.5e-06 | 2.6e-06 | 3.667E-06 | 1.935E-06 | 2.500E-06 | 5.900E-06 |
| 15503 | 776 | L776N | B | 3.2e-06 | 2.4e-06 | 2.5e-06 | 2.700E-06 | 4.359E-07 | 2.400E-06 | 3.200E-06 |
| 15504 | 776 | L776D | B | 4e-06 | 2.5e-06 | 1.7e-06 | 2.733E-06 | 1.168E-06 | 1.700E-06 | 4.000E-06 |
| 15505 | 776 | L776C | B | 3e-06 | 2e-06 | 2.2e-06 | 2.400E-06 | 5.292E-07 | 2.000E-06 | 3.000E-06 |
| 15506 | 776 | L776E | B | 1.8e-06 | 2.5e-06 | 1.9e-06 | 2.067E-06 | 3.786E-07 | 1.800E-06 | 2.500E-06 |
| 15507 | 776 | L776Q | B | 3.2e-06 | 1.2e-06 | 2.2e-06 | 2.200E-06 | 1.000E-06 | 1.200E-06 | 3.200E-06 |
| 15508 | 776 | L776G | B | 1.6e-06 | 3.4e-06 | 2.1e-06 | 2.367E-06 | 9.292E-07 | 1.600E-06 | 3.400E-06 |
| 15509 | 776 | L776H | B | 2.4e-06 | 2.4e-06 | 3.9e-06 | 2.900E-06 | 8.660E-07 | 2.400E-06 | 3.900E-06 |
| 15510 | 776 | L776I | B | 3e-06 | 2.3e-06 | 2.3e-06 | 2.533E-06 | 4.041E-07 | 2.300E-06 | 3.000E-06 |
| 15511 | 776 | L776L | B | 3.1e-06 | 3.1e-06 | 4e-06 | 3.400E-06 | 5.196E-07 | 3.100E-06 | 4.000E-06 |
| 15512 | 776 | L776K | B | 2.1e-06 | 3.4e-06 | 1.3e-06 | 2.267E-06 | 1.060E-06 | 1.300E-06 | 3.400E-06 |
| 15513 | 776 | L776M | B | 2.8e-06 | 2.7e-06 | 1.9e-06 | 2.467E-06 | 4.933E-07 | 1.900E-06 | 2.800E-06 |
| 15514 | 776 | L776F | B | 2.3e-06 | 4.2e-06 | 2.7e-06 | 3.067E-06 | 1.002E-06 | 2.300E-06 | 4.200E-06 |
| 15515 | 776 | L776P | B | 1.9e-06 | 3.1e-06 | 3.8e-06 | 2.933E-06 | 9.609E-07 | 1.900E-06 | 3.800E-06 |
| 15516 | 776 | L776S | B | 2.5e-06 | 2.5e-06 | 1.6e-06 | 2.200E-06 | 5.196E-07 | 1.600E-06 | 2.500E-06 |
| 15517 | 776 | L776T | B | 2e-06 | 1.7e-06 | 3e-06 | 2.233E-06 | 6.807E-07 | 1.700E-06 | 3.000E-06 |
| 15518 | 776 | L776W | B | 2.8e-06 | 1.9e-06 | 1.2e-06 | 1.967E-06 | 8.021E-07 | 1.200E-06 | 2.800E-06 |
| 15519 | 776 | L776Y | B | 2.7e-06 | 2.9e-06 | 2.7e-06 | 2.767E-06 | 1.155E-07 | 2.700E-06 | 2.900E-06 |
| 15520 | 776 | L776V | B | 3.3e-06 | 3.7e-06 | 4.3e-06 | 3.767E-06 | 5.033E-07 | 3.300E-06 | 4.300E-06 |
| 15521 | 777 | Q777A | B | 2.5e-06 | 3.2e-06 | 3.7e-06 | 3.133E-06 | 6.028E-07 | 2.500E-06 | 3.700E-06 |
| 15522 | 777 | Q777R | B | 1.3e-06 | 1.9e-06 | 2.2e-06 | 1.800E-06 | 4.583E-07 | 1.300E-06 | 2.200E-06 |
| 15523 | 777 | Q777N | B | 2.4e-06 | 2.7e-06 | 2.9e-06 | 2.667E-06 | 2.517E-07 | 2.400E-06 | 2.900E-06 |
| 15524 | 777 | Q777D | B | 1.1e-06 | 3e-06 | 1.9e-06 | 2.000E-06 | 9.539E-07 | 1.100E-06 | 3.000E-06 |
| 15525 | 777 | Q777C | B | 1.8e-06 | 3.3e-06 | 4.1e-06 | 3.067E-06 | 1.168E-06 | 1.800E-06 | 4.100E-06 |
| 15526 | 777 | Q777E | B | 2.6e-06 | 3.1e-06 | 2.6e-06 | 2.767E-06 | 2.887E-07 | 2.600E-06 | 3.100E-06 |
| 15527 | 777 | Q777Q | B | 4.3e-06 | 2.8e-06 | 3.4e-06 | 3.500E-06 | 7.550E-07 | 2.800E-06 | 4.300E-06 |
| 15528 | 777 | Q777G | B | 3.7e-06 | 3.8e-06 | 1.9e-06 | 3.133E-06 | 1.069E-06 | 1.900E-06 | 3.800E-06 |
| 15529 | 777 | Q777H | B | 2.5e-06 | 2.8e-06 | 1.7e-06 | 2.333E-06 | 5.686E-07 | 1.700E-06 | 2.800E-06 |
| 15530 | 777 | Q777I | B | 2.8e-06 | 2.6e-06 | 3e-06 | 2.800E-06 | 2.000E-07 | 2.600E-06 | 3.000E-06 |
| 15531 | 777 | Q777L | B | 2.9e-06 | 1.6e-06 | 1.9e-06 | 2.133E-06 | 6.807E-07 | 1.600E-06 | 2.900E-06 |
| 15532 | 777 | Q777K | B | 3.6e-06 | 2.6e-06 | 2.4e-06 | 2.867E-06 | 6.429E-07 | 2.400E-06 | 3.600E-06 |
| 15533 | 777 | Q777M | B | 2.7e-06 | 1.7e-06 | 2.8e-06 | 2.400E-06 | 6.083E-07 | 1.700E-06 | 2.800E-06 |
| 15534 | 777 | Q777F | B | 4.1e-06 | 3.6e-06 | 1e-06 | 2.900E-06 | 1.664E-06 | 1.000E-06 | 4.100E-06 |
| 15535 | 777 | Q777P | B | 1.7e-06 | 2.1e-06 | 2.8e-06 | 2.200E-06 | 5.568E-07 | 1.700E-06 | 2.800E-06 |
| 15536 | 777 | Q777S | B | 2.2e-06 | 2.3e-06 | 2.2e-06 | 2.233E-06 | 5.774E-08 | 2.200E-06 | 2.300E-06 |
| 15537 | 777 | Q777T | B | 2.8e-06 | 3.5e-06 | 1.5e-06 | 2.600E-06 | 1.015E-06 | 1.500E-06 | 3.500E-06 |
| 15538 | 777 | Q777W | B | 2.1e-06 | 2.5e-06 | 1.6e-06 | 2.067E-06 | 4.509E-07 | 1.600E-06 | 2.500E-06 |
| 15539 | 777 | Q777Y | B | 1.9e-06 | 1.4e-06 | 2.3e-06 | 1.867E-06 | 4.509E-07 | 1.400E-06 | 2.300E-06 |
| 15540 | 777 | Q777V | B | 3.9e-06 | 2.3e-06 | 2.6e-06 | 2.933E-06 | 8.505E-07 | 2.300E-06 | 3.900E-06 |
| 15541 | 778 | S778A | B | 3e-06 | 1.8e-06 | 2.4e-06 | 2.400E-06 | 6.000E-07 | 1.800E-06 | 3.000E-06 |
| 15542 | 778 | S778R | B | 2.2e-06 | 3.7e-06 | 1.9e-06 | 2.600E-06 | 9.644E-07 | 1.900E-06 | 3.700E-06 |
| 15543 | 778 | S778N | B | 3.7e-06 | 2.1e-06 | 2.6e-06 | 2.800E-06 | 8.185E-07 | 2.100E-06 | 3.700E-06 |
| 15544 | 778 | S778D | B | 2.4e-06 | 2.1e-06 | 1.8e-06 | 2.100E-06 | 3.000E-07 | 1.800E-06 | 2.400E-06 |
| 15545 | 778 | S778C | B | 3.1e-06 | 1.6e-06 | 3.7e-06 | 2.800E-06 | 1.082E-06 | 1.600E-06 | 3.700E-06 |
| 15546 | 778 | S778E | B | 3.5e-06 | 2.4e-06 | 4e-06 | 3.300E-06 | 8.185E-07 | 2.400E-06 | 4.000E-06 |
| 15547 | 778 | S778Q | B | 1.4e-06 | 1.7e-06 | 1.6e-06 | 1.567E-06 | 1.528E-07 | 1.400E-06 | 1.700E-06 |
| 15548 | 778 | S778G | B | 2.4e-06 | 2.3e-06 | 1.4e-06 | 2.033E-06 | 5.508E-07 | 1.400E-06 | 2.400E-06 |
| 15549 | 778 | S778H | B | 4.2e-06 | 3e-06 | 3.3e-06 | 3.500E-06 | 6.245E-07 | 3.000E-06 | 4.200E-06 |
| 15550 | 778 | S778I | B | 2.6e-06 | 1.8e-06 | 3.3e-06 | 2.567E-06 | 7.506E-07 | 1.800E-06 | 3.300E-06 |
| 15551 | 778 | S778L | B | 2.2e-06 | 2.4e-06 | 1.8e-06 | 2.133E-06 | 3.055E-07 | 1.800E-06 | 2.400E-06 |
| 15552 | 778 | S778K | B | 2.7e-06 | 3.2e-06 | 2.5e-06 | 2.800E-06 | 3.606E-07 | 2.500E-06 | 3.200E-06 |
| 15553 | 778 | S778M | B | 2.7e-06 | 3e-06 | 2.7e-06 | 2.800E-06 | 1.732E-07 | 2.700E-06 | 3.000E-06 |
| 15554 | 778 | S778F | B | 2.6e-06 | 1.6e-06 | 3.4e-06 | 2.533E-06 | 9.018E-07 | 1.600E-06 | 3.400E-06 |

| | | | | | | | | | | |
|---|---|---|---|---|---|---|---|---|---|---|
| 15555 | 778 | S778P | B | 2.1e-06 | 1.8e-06 | 1.7e-06 | 1.867E-06 | 2.082E-07 | 1.700E-06 | 2.100E-06 |
| 15556 | 778 | S778S | B | 1.5e-06 | 2.3e-06 | 3e-06 | 2.267E-06 | 7.506E-07 | 1.500E-06 | 3.000E-06 |
| 15557 | 778 | S778T | B | 2.1e-06 | 1.7e-06 | 1.3e-06 | 1.700E-06 | 4.000E-07 | 1.300E-06 | 2.100E-06 |
| 15558 | 778 | S778W | B | 1.9e-06 | 2.7e-06 | 3.4e-06 | 2.667E-06 | 7.506E-07 | 1.900E-06 | 3.400E-06 |
| 15559 | 778 | S778Y | B | 1.9e-06 | 1.6e-06 | 1.1e-06 | 1.533E-06 | 4.041E-07 | 1.100E-06 | 1.900E-06 |
| 15560 | 778 | S778V | B | 3.5e-06 | 1.9e-06 | 1.6e-06 | 2.333E-06 | 1.021E-06 | 1.600E-06 | 3.500E-06 |
| 15561 | 779 | G779A | B | 2.6e-06 | 1.3e-06 | 3.2e-06 | 2.367E-06 | 9.713E-07 | 1.300E-06 | 3.200E-06 |
| 15562 | 779 | G779R | B | 3.7e-06 | 1.5e-06 | 3.7e-06 | 2.967E-06 | 1.270E-06 | 1.500E-06 | 3.700E-06 |
| 15563 | 779 | G779N | B | 2.1e-06 | 2e-06 | 2.8e-06 | 2.300E-06 | 4.359E-07 | 2.000E-06 | 2.800E-06 |
| 15564 | 779 | G779D | B | 2.3e-06 | 2.9e-06 | 3e-06 | 2.733E-06 | 3.786E-07 | 2.300E-06 | 3.000E-06 |
| 15565 | 779 | G779C | B | 3.1e-06 | 2.9e-06 | 3.3e-06 | 3.100E-06 | 2.000E-07 | 2.900E-06 | 3.300E-06 |
| 15566 | 779 | G779E | B | 1.3e-06 | 2.1e-06 | 2.7e-06 | 2.033E-06 | 7.024E-07 | 1.300E-06 | 2.700E-06 |
| 15567 | 779 | G779Q | B | 2.3e-06 | 1.7e-06 | 3.3e-06 | 2.433E-06 | 8.083E-07 | 1.700E-06 | 3.300E-06 |
| 15568 | 779 | G779G | B | 1.4e-06 | 2.4e-06 | 2.8e-06 | 2.200E-06 | 7.211E-07 | 1.400E-06 | 2.800E-06 |
| 15569 | 779 | G779H | B | 2.2e-06 | 2.1e-06 | 4e-06 | 2.767E-06 | 1.069E-06 | 2.100E-06 | 4.000E-06 |
| 15570 | 779 | G779I | B | 1.8e-06 | 2.2e-06 | 2.9e-06 | 2.300E-06 | 5.568E-07 | 1.800E-06 | 2.900E-06 |
| 15571 | 779 | G779L | B | 4.1e-06 | 2.1e-06 | 1.3e-06 | 2.500E-06 | 1.442E-06 | 1.300E-06 | 4.100E-06 |
| 15572 | 779 | G779K | B | 3.9e-06 | 3e-06 | 1.6e-06 | 2.833E-06 | 1.159E-06 | 1.600E-06 | 3.900E-06 |
| 15573 | 779 | G779M | B | 1.5e-06 | 3.4e-06 | 2.4e-06 | 2.433E-06 | 9.504E-07 | 1.500E-06 | 3.400E-06 |
| 15574 | 779 | G779F | B | 3.2e-06 | 2.6e-06 | 3.7e-06 | 3.167E-06 | 5.508E-07 | 2.600E-06 | 3.700E-06 |
| 15575 | 779 | G779P | B | 3e-06 | 2.5e-06 | 2e-06 | 2.500E-06 | 5.000E-07 | 2.000E-06 | 3.000E-06 |
| 15576 | 779 | G779S | B | 3.6e-06 | 2.6e-06 | 2.9e-06 | 3.033E-06 | 5.132E-07 | 2.600E-06 | 3.600E-06 |
| 15577 | 779 | G779T | B | 1.7e-06 | 3e-06 | 3.1e-06 | 2.600E-06 | 7.810E-07 | 1.700E-06 | 3.100E-06 |
| 15578 | 779 | G779W | B | 2.4e-06 | 2.3e-06 | 2.1e-06 | 2.267E-06 | 1.528E-07 | 2.100E-06 | 2.400E-06 |
| 15579 | 779 | G779Y | B | 3.4e-06 | 1.5e-06 | 1.5e-06 | 2.133E-06 | 1.097E-06 | 1.500E-06 | 3.400E-06 |
| 15580 | 779 | G779V | B | 2.2e-06 | 1.9e-06 | 1.5e-06 | 1.867E-06 | 3.512E-07 | 1.500E-06 | 2.200E-06 |
| 15581 | 780 | N780A | B | 2.7e-06 | 2.1e-06 | 4.1e-06 | 2.967E-06 | 1.026E-06 | 2.100E-06 | 4.100E-06 |
| 15582 | 780 | N780R | B | 2.6e-06 | 2.5e-06 | 1.8e-06 | 2.300E-06 | 4.359E-07 | 1.800E-06 | 2.600E-06 |
| 15583 | 780 | N780N | B | 2e-06 | 2e-06 | 2.3e-06 | 2.100E-06 | 1.732E-07 | 2.000E-06 | 2.300E-06 |
| 15584 | 780 | N780D | B | 4.5e-06 | 2.4e-06 | 1.8e-06 | 2.900E-06 | 1.418E-06 | 1.800E-06 | 4.500E-06 |
| 15585 | 780 | N780C | B | 2.1e-06 | 1e-06 | 2.5e-06 | 1.867E-06 | 7.767E-07 | 1.000E-06 | 2.500E-06 |
| 15586 | 780 | N780E | B | 1.6e-06 | 2.2e-06 | 1.5e-06 | 1.767E-06 | 3.786E-07 | 1.500E-06 | 2.200E-06 |
| 15587 | 780 | N780Q | B | 2.4e-06 | 1.7e-06 | 4.6e-06 | 2.900E-06 | 1.513E-06 | 1.700E-06 | 4.600E-06 |
| 15588 | 780 | N780G | B | 2.2e-06 | 2.8e-06 | 2e-06 | 2.333E-06 | 4.163E-07 | 2.000E-06 | 2.800E-06 |
| 15589 | 780 | N780H | B | 4.1e-06 | 2.6e-06 | 2.2e-06 | 2.967E-06 | 1.002E-06 | 2.200E-06 | 4.100E-06 |
| 15590 | 780 | N780I | B | 2.7e-06 | 3.8e-06 | 2.9e-06 | 3.133E-06 | 5.859E-07 | 2.700E-06 | 3.800E-06 |
| 15591 | 780 | N780L | B | 3e-06 | 3e-06 | 2.2e-06 | 2.733E-06 | 4.619E-07 | 2.200E-06 | 3.000E-06 |
| 15592 | 780 | N780K | B | 2.5e-06 | 2e-06 | 2.3e-06 | 2.267E-06 | 2.517E-07 | 2.000E-06 | 2.500E-06 |
| 15593 | 780 | N780M | B | 3.7e-06 | 2.5e-06 | 5e-06 | 3.733E-06 | 1.250E-06 | 2.500E-06 | 5.000E-06 |
| 15594 | 780 | N780F | B | 2.9e-06 | 2.2e-06 | 3.7e-06 | 2.933E-06 | 7.506E-07 | 2.200E-06 | 3.700E-06 |
| 15595 | 780 | N780P | B | 2.1e-06 | 2.2e-06 | 2.2e-06 | 2.167E-06 | 5.774E-08 | 2.100E-06 | 2.200E-06 |
| 15596 | 780 | N780S | B | 1.8e-06 | 2.6e-06 | 3.2e-06 | 2.533E-06 | 7.024E-07 | 1.800E-06 | 3.200E-06 |
| 15597 | 780 | N780T | B | 2.5e-06 | 2.4e-06 | 1.9e-06 | 2.267E-06 | 3.215E-07 | 1.900E-06 | 2.500E-06 |
| 15598 | 780 | N780W | B | 1.9e-06 | 3.1e-06 | 1.6e-06 | 2.200E-06 | 7.937E-07 | 1.600E-06 | 3.100E-06 |
| 15599 | 780 | N780Y | B | 2.3e-06 | 2.3e-06 | 8.8e-07 | 1.827E-06 | 8.198E-07 | 8.800E-07 | 2.300E-06 |
| 15600 | 780 | N780V | B | 2.2e-06 | 3.1e-06 | 3.6e-06 | 2.967E-06 | 7.095E-07 | 2.200E-06 | 3.600E-06 |
| 15601 | 781 | S781A | B | 1.4e-06 | 2.7e-06 | 1.7e-06 | 1.933E-06 | 6.807E-07 | 1.400E-06 | 2.700E-06 |
| 15602 | 781 | S781R | B | 2.8e-06 | 4.6e-06 | 1.8e-06 | 3.067E-06 | 1.419E-06 | 1.800E-06 | 4.600E-06 |
| 15603 | 781 | S781N | B | 2.1e-06 | 1.8e-06 | 2.7e-06 | 2.200E-06 | 4.583E-07 | 1.800E-06 | 2.700E-06 |
| 15604 | 781 | S781D | B | 2.5e-06 | 2.1e-06 | 2.7e-06 | 2.433E-06 | 3.055E-07 | 2.100E-06 | 2.700E-06 |
| 15605 | 781 | S781C | B | 2.5e-06 | 3.4e-06 | 1.3e-06 | 2.400E-06 | 1.054E-06 | 1.300E-06 | 3.400E-06 |
| 15606 | 781 | S781E | B | 2.7e-06 | 2e-06 | 3.2e-06 | 2.633E-06 | 6.028E-07 | 2.000E-06 | 3.200E-06 |
| 15607 | 781 | S781Q | B | 2.1e-06 | 1.5e-06 | 1.9e-06 | 1.833E-06 | 3.055E-07 | 1.500E-06 | 2.100E-06 |
| 15608 | 781 | S781G | B | 2.6e-06 | 1.9e-06 | 1.6e-06 | 2.033E-06 | 5.132E-07 | 1.600E-06 | 2.600E-06 |
| 15609 | 781 | S781H | B | 2.4e-06 | 1.3e-06 | 2e-06 | 1.900E-06 | 5.568E-07 | 1.300E-06 | 2.400E-06 |
| 15610 | 781 | S781I | B | 5.3e-06 | 2.7e-06 | 1.5e-06 | 3.167E-06 | 1.943E-06 | 1.500E-06 | 5.300E-06 |
| 15611 | 781 | S781L | B | 3.7e-06 | 2.4e-06 | 1.7e-06 | 2.600E-06 | 1.015E-06 | 1.700E-06 | 3.700E-06 |
| 15612 | 781 | S781K | B | 3e-06 | 2.7e-06 | 1.9e-06 | 2.533E-06 | 5.686E-07 | 1.900E-06 | 3.000E-06 |
| 15613 | 781 | S781M | B | 2.4e-06 | 3.4e-06 | 1.8e-06 | 2.533E-06 | 8.083E-07 | 1.800E-06 | 3.400E-06 |
| 15614 | 781 | S781F | B | 1.6e-06 | 3.3e-06 | 2.5e-06 | 2.467E-06 | 8.505E-07 | 1.600E-06 | 3.300E-06 |
| 15615 | 781 | S781P | B | 3.5e-06 | 3.2e-06 | 2.6e-06 | 3.100E-06 | 4.583E-07 | 2.600E-06 | 3.500E-06 |
| 15616 | 781 | S781S | B | 2.2e-06 | 2.1e-06 | 3.2e-06 | 2.500E-06 | 6.083E-07 | 2.100E-06 | 3.200E-06 |
| 15617 | 781 | S781T | B | 1.3e-06 | 2e-06 | 2.5e-06 | 1.933E-06 | 6.028E-07 | 1.300E-06 | 2.500E-06 |
| 15618 | 781 | S781W | B | 2.6e-06 | 2.8e-06 | 2.1e-06 | 2.500E-06 | 3.606E-07 | 2.100E-06 | 2.800E-06 |
| 15619 | 781 | S781Y | B | 2.9e-06 | 2.9e-06 | 3.6e-06 | 3.133E-06 | 4.041E-07 | 2.900E-06 | 3.600E-06 |
| 15620 | 781 | S781V | B | 1.3e-06 | 3.6e-06 | 2.9e-06 | 2.600E-06 | 1.179E-06 | 1.300E-06 | 3.600E-06 |
| 15621 | 782 | Q782A | B | 1.5e-06 | 9.3e-07 | 2.9e-06 | 1.777E-06 | 1.014E-06 | 9.300E-07 | 2.900E-06 |
| 15622 | 782 | Q782R | B | 2.3e-06 | 1.4e-06 | 1.9e-06 | 1.867E-06 | 4.509E-07 | 1.400E-06 | 2.300E-06 |
| 15623 | 782 | Q782N | B | 1.9e-06 | 2.8e-06 | 1.6e-06 | 2.100E-06 | 6.245E-07 | 1.600E-06 | 2.800E-06 |
| 15624 | 782 | Q782D | B | 2e-06 | 2.3e-06 | 2.7e-06 | 2.333E-06 | 3.512E-07 | 2.000E-06 | 2.700E-06 |
| 15625 | 782 | Q782C | B | 1.4e-06 | 3.1e-06 | 2.7e-06 | 2.400E-06 | 8.888E-07 | 1.400E-06 | 3.100E-06 |
| 15626 | 782 | Q782E | B | 2.8e-06 | 2.2e-06 | 4e-06 | 3.000E-06 | 9.165E-07 | 2.200E-06 | 4.000E-06 |
| 15627 | 782 | Q782Q | B | 2.2e-06 | 2.2e-06 | 2e-06 | 2.133E-06 | 1.155E-07 | 2.000E-06 | 2.200E-06 |
| 15628 | 782 | Q782G | B | 2.1e-06 | 2.4e-06 | 2.2e-06 | 2.233E-06 | 1.528E-07 | 2.100E-06 | 2.400E-06 |
| 15629 | 782 | Q782H | B | 1.6e-06 | 2.9e-06 | 2.4e-06 | 2.300E-06 | 6.557E-07 | 1.600E-06 | 2.900E-06 |
| 15630 | 782 | Q782I | B | 4.1e-06 | 1.9e-06 | 3.8e-06 | 3.267E-06 | 1.193E-06 | 1.900E-06 | 4.100E-06 |
| 15631 | 782 | Q782L | B | 2.8e-06 | 2.4e-06 | 2.1e-06 | 2.433E-06 | 3.512E-07 | 2.100E-06 | 2.800E-06 |
| 15632 | 782 | Q782K | B | 1.7e-06 | 3.1e-06 | 2.6e-06 | 2.467E-06 | 7.095E-07 | 1.700E-06 | 3.100E-06 |
| 15633 | 782 | Q782M | B | 3.1e-06 | 1.9e-06 | 2.5e-06 | 2.500E-06 | 6.000E-07 | 1.900E-06 | 3.100E-06 |
| 15634 | 782 | Q782F | B | 2.6e-06 | 1.8e-06 | 3.1e-06 | 2.500E-06 | 6.557E-07 | 1.800E-06 | 3.100E-06 |
| 15635 | 782 | Q782P | B | 2.6e-06 | 3.1e-06 | 1.9e-06 | 2.533E-06 | 6.028E-07 | 1.900E-06 | 3.100E-06 |
| 15636 | 782 | Q782S | B | 1.8e-06 | 1.5e-06 | 2.6e-06 | 1.967E-06 | 5.686E-07 | 1.500E-06 | 2.600E-06 |
| 15637 | 782 | Q782T | B | 2.9e-06 | 2.8e-06 | 2.3e-06 | 2.667E-06 | 3.215E-07 | 2.300E-06 | 2.900E-06 |
| 15638 | 782 | Q782W | B | 2.8e-06 | 2.3e-06 | 3.1e-06 | 2.733E-06 | 4.041E-07 | 2.300E-06 | 3.100E-06 |
| 15639 | 782 | Q782Y | B | 2.3e-06 | 3e-06 | 2.5e-06 | 2.600E-06 | 3.606E-07 | 2.300E-06 | 3.000E-06 |

| | | | | | | | | | | |
|---|---|---|---|---|---|---|---|---|---|---|
| 15640 | 782 | Q782V | B | 2.5e-06 | 5.4e-06 | 2.7e-06 | 3.533E-06 | 1.620E-06 | 2.500E-06 | 5.400E-06 |
| 15641 | 783 | E783A | B | 3e-06 | 1.4e-06 | 3e-06 | 2.467E-06 | 9.238E-07 | 1.400E-06 | 3.000E-06 |
| 15642 | 783 | E783R | B | 2.4e-06 | 3.6e-06 | 2.9e-06 | 2.967E-06 | 6.028E-07 | 2.400E-06 | 3.600E-06 |
| 15643 | 783 | E783N | B | 2.2e-06 | 1.7e-06 | 2.8e-06 | 2.233E-06 | 5.508E-07 | 1.700E-06 | 2.800E-06 |
| 15644 | 783 | E783D | B | 5.3e-06 | 1.2e-06 | 2.1e-06 | 2.867E-06 | 2.155E-06 | 1.200E-06 | 5.300E-06 |
| 15645 | 783 | E783C | B | 2.3e-06 | 7.1e-07 | 1.4e-06 | 1.470E-06 | 7.973E-07 | 7.100E-07 | 2.300E-06 |
| 15646 | 783 | E783E | B | 1e-06 | 1.4e-06 | 1.6e-06 | 1.333E-06 | 3.055E-07 | 1.000E-06 | 1.600E-06 |
| 15647 | 783 | E783Q | B | 3.1e-06 | 2.5e-06 | 3.7e-06 | 3.100E-06 | 6.000E-07 | 2.500E-06 | 3.700E-06 |
| 15648 | 783 | E783G | B | 2.4e-06 | 2e-06 | 5.3e-06 | 3.233E-06 | 1.801E-06 | 2.000E-06 | 5.300E-06 |
| 15649 | 783 | E783H | B | 2.2e-06 | 3e-06 | 1.9e-06 | 2.367E-06 | 5.686E-07 | 1.900E-06 | 3.000E-06 |
| 15650 | 783 | E783I | B | 3e-06 | 2.5e-06 | 4.4e-06 | 3.300E-06 | 9.849E-07 | 2.500E-06 | 4.400E-06 |
| 15651 | 783 | E783L | B | 2.6e-06 | 3.8e-06 | 2.3e-06 | 2.900E-06 | 7.937E-07 | 2.300E-06 | 3.800E-06 |
| 15652 | 783 | E783K | B | 2.4e-06 | 2.7e-06 | 3.1e-06 | 2.733E-06 | 3.512E-07 | 2.400E-06 | 3.100E-06 |
| 15653 | 783 | E783M | B | 5e-06 | 1.4e-06 | 1.6e-06 | 2.667E-06 | 2.023E-06 | 1.400E-06 | 5.000E-06 |
| 15654 | 783 | E783F | B | 2.4e-06 | 3.6e-06 | 1.8e-06 | 2.600E-06 | 9.165E-07 | 1.800E-06 | 3.600E-06 |
| 15655 | 783 | E783P | B | 1.4e-06 | 2.6e-06 | 1.1e-06 | 1.700E-06 | 7.937E-07 | 1.100E-06 | 2.600E-06 |
| 15656 | 783 | E783S | B | 1.6e-06 | 2.1e-06 | 4.1e-06 | 2.600E-06 | 1.323E-06 | 1.600E-06 | 4.100E-06 |
| 15657 | 783 | E783T | B | 4e-06 | 1.1e-06 | 2.3e-06 | 2.467E-06 | 1.457E-06 | 1.100E-06 | 4.000E-06 |
| 15658 | 783 | E783W | B | 3.5e-06 | 1.5e-06 | 3.7e-06 | 2.900E-06 | 1.217E-06 | 1.500E-06 | 3.700E-06 |
| 15659 | 783 | E783Y | B | 2.1e-06 | 1.8e-06 | 2.3e-06 | 2.067E-06 | 2.517E-07 | 1.800E-06 | 2.300E-06 |
| 15660 | 783 | E783V | B | 2.1e-06 | 2.7e-06 | 3.1e-06 | 2.633E-06 | 5.033E-07 | 2.100E-06 | 3.100E-06 |
| 15661 | 784 | S784A | B | 2.6e-06 | 1.4e-06 | 3.2e-06 | 2.400E-06 | 9.165E-07 | 1.400E-06 | 3.200E-06 |
| 15662 | 784 | S784R | B | 2.5e-06 | 1.7e-06 | 2.8e-06 | 2.333E-06 | 5.686E-07 | 1.700E-06 | 2.800E-06 |
| 15663 | 784 | S784N | B | 2.6e-06 | 3.6e-06 | 1.8e-06 | 2.667E-06 | 9.018E-07 | 1.800E-06 | 3.600E-06 |
| 15664 | 784 | S784D | B | 1.7e-06 | 2.4e-06 | 4.4e-06 | 2.833E-06 | 1.401E-06 | 1.700E-06 | 4.400E-06 |
| 15665 | 784 | S784C | B | 2.4e-06 | 1.8e-06 | 2.5e-06 | 2.233E-06 | 3.786E-07 | 1.800E-06 | 2.500E-06 |
| 15666 | 784 | S784E | B | 3e-06 | 2.2e-06 | 2e-06 | 2.400E-06 | 5.292E-07 | 2.000E-06 | 3.000E-06 |
| 15667 | 784 | S784Q | B | 4e-06 | 2.6e-06 | 3.1e-06 | 3.233E-06 | 7.095E-07 | 2.600E-06 | 4.000E-06 |
| 15668 | 784 | S784G | B | 1.8e-06 | 2.1e-06 | 1.8e-06 | 1.900E-06 | 1.732E-07 | 1.800E-06 | 2.100E-06 |
| 15669 | 784 | S784H | B | 2.1e-06 | 1.9e-06 | 2.6e-06 | 2.200E-06 | 3.606E-07 | 1.900E-06 | 2.600E-06 |
| 15670 | 784 | S784I | B | 3.5e-06 | 1.7e-06 | 1.8e-06 | 2.333E-06 | 1.012E-06 | 1.700E-06 | 3.500E-06 |
| 15671 | 784 | S784L | B | 2.5e-06 | 2.4e-06 | 2.5e-06 | 2.467E-06 | 5.774E-08 | 2.400E-06 | 2.500E-06 |
| 15672 | 784 | S784K | B | 2.8e-06 | 1.7e-06 | 2.3e-06 | 2.267E-06 | 5.508E-07 | 1.700E-06 | 2.800E-06 |
| 15673 | 784 | S784M | B | 1.8e-06 | 2.8e-06 | 2.8e-06 | 2.467E-06 | 5.774E-07 | 1.800E-06 | 2.800E-06 |
| 15674 | 784 | S784F | B | 4.3e-06 | 2.9e-06 | 2e-06 | 3.067E-06 | 1.159E-06 | 2.000E-06 | 4.300E-06 |
| 15675 | 784 | S784P | B | 1.4e-06 | 3.3e-06 | 3.6e-06 | 2.767E-06 | 1.193E-06 | 1.400E-06 | 3.600E-06 |
| 15676 | 784 | S784S | B | 2.5e-06 | 4.6e-06 | 4e-06 | 3.700E-06 | 1.082E-06 | 2.500E-06 | 4.600E-06 |
| 15677 | 784 | S784T | B | 1.8e-06 | 3.5e-06 | 1.9e-06 | 2.400E-06 | 9.539E-07 | 1.800E-06 | 3.500E-06 |
| 15678 | 784 | S784W | B | 5.2e-06 | 1.9e-06 | 4.3e-06 | 3.800E-06 | 1.706E-06 | 1.900E-06 | 5.200E-06 |
| 15679 | 784 | S784Y | B | 1.4e-06 | 2.6e-06 | 1.4e-06 | 1.800E-06 | 6.928E-07 | 1.400E-06 | 2.600E-06 |
| 15680 | 784 | S784V | B | 2.6e-06 | 4.9e-06 | 2.8e-06 | 3.433E-06 | 1.274E-06 | 2.600E-06 | 4.900E-06 |
| 15681 | 785 | V785A | B | 2.3e-06 | 3.2e-06 | 1.8e-06 | 2.433E-06 | 7.095E-07 | 1.800E-06 | 3.200E-06 |
| 15682 | 785 | V785R | B | 3.3e-06 | 2.4e-06 | 2.6e-06 | 2.767E-06 | 4.726E-07 | 2.400E-06 | 3.300E-06 |
| 15683 | 785 | V785N | B | 1.8e-06 | 1.4e-06 | 1.1e-06 | 1.433E-06 | 3.512E-07 | 1.100E-06 | 1.800E-06 |
| 15684 | 785 | V785D | B | 4e-06 | 1.8e-06 | 1.4e-06 | 2.400E-06 | 1.400E-06 | 1.400E-06 | 4.000E-06 |
| 15685 | 785 | V785C | B | 4.3e-06 | 1.6e-06 | 2e-06 | 2.633E-06 | 1.457E-06 | 1.600E-06 | 4.300E-06 |
| 15686 | 785 | V785E | B | 2.6e-06 | 2.2e-06 | 2.2e-06 | 2.333E-06 | 2.309E-07 | 2.200E-06 | 2.600E-06 |
| 15687 | 785 | V785Q | B | 2e-06 | 1.1e-06 | 1.7e-06 | 1.600E-06 | 4.583E-07 | 1.100E-06 | 2.000E-06 |
| 15688 | 785 | V785G | B | 2.2e-06 | 2.1e-06 | 2.7e-06 | 2.333E-06 | 3.215E-07 | 2.100E-06 | 2.700E-06 |
| 15689 | 785 | V785H | B | 2.9e-06 | 1.8e-06 | 2.8e-06 | 2.500E-06 | 6.083E-07 | 1.800E-06 | 2.900E-06 |
| 15690 | 785 | V785I | B | 4.3e-06 | 2.8e-06 | 2.7e-06 | 3.267E-06 | 8.963E-07 | 2.700E-06 | 4.300E-06 |
| 15691 | 785 | V785L | B | 1.9e-06 | 1.6e-06 | 2.6e-06 | 2.033E-06 | 5.132E-07 | 1.600E-06 | 2.600E-06 |
| 15692 | 785 | V785K | B | 2.5e-06 | 3.5e-06 | 1.9e-06 | 2.633E-06 | 8.083E-07 | 1.900E-06 | 3.500E-06 |
| 15693 | 785 | V785M | B | 2.1e-06 | 3e-06 | 1.6e-06 | 2.233E-06 | 7.095E-07 | 1.600E-06 | 3.000E-06 |
| 15694 | 785 | V785F | B | 3e-06 | 2.4e-06 | 1.7e-06 | 2.367E-06 | 6.506E-07 | 1.700E-06 | 3.000E-06 |
| 15695 | 785 | V785P | B | 3.3e-06 | 3.1e-06 | 1.4e-06 | 2.600E-06 | 1.044E-06 | 1.400E-06 | 3.300E-06 |
| 15696 | 785 | V785S | B | 1.8e-06 | 3e-06 | 1.7e-06 | 2.167E-06 | 7.234E-07 | 1.700E-06 | 3.000E-06 |
| 15697 | 785 | V785T | B | 3.4e-06 | 2.8e-06 | 1.5e-06 | 2.567E-06 | 9.713E-07 | 1.500E-06 | 3.400E-06 |
| 15698 | 785 | V785W | B | 4.7e-06 | 2e-06 | 2.3e-06 | 3.000E-06 | 1.480E-06 | 2.000E-06 | 4.700E-06 |
| 15699 | 785 | V785Y | B | 3.9e-06 | 1.1e-06 | 2e-06 | 2.333E-06 | 1.429E-06 | 1.100E-06 | 3.900E-06 |
| 15700 | 785 | V785V | B | 1.9e-06 | 2.1e-06 | 2.2e-06 | 2.067E-06 | 1.528E-07 | 1.900E-06 | 2.200E-06 |
| 15701 | 786 | T786A | B | 2.9e-06 | 2.5e-06 | 3.7e-06 | 3.033E-06 | 6.110E-07 | 2.500E-06 | 3.700E-06 |
| 15702 | 786 | T786R | B | 1.4e-06 | 3.4e-06 | 3.6e-06 | 2.800E-06 | 1.217E-06 | 1.400E-06 | 3.600E-06 |
| 15703 | 786 | T786N | B | 2.3e-06 | 3.6e-06 | 2.7e-06 | 2.867E-06 | 6.658E-07 | 2.300E-06 | 3.600E-06 |
| 15704 | 786 | T786D | B | 2.1e-06 | 3e-06 | 1.8e-06 | 2.300E-06 | 6.245E-07 | 1.800E-06 | 3.000E-06 |
| 15705 | 786 | T786C | B | 1.5e-06 | 2e-06 | 2e-06 | 1.833E-06 | 2.887E-07 | 1.500E-06 | 2.000E-06 |
| 15706 | 786 | T786E | B | 1.3e-06 | 1.5e-06 | 2.6e-06 | 1.800E-06 | 7.000E-07 | 1.300E-06 | 2.600E-06 |
| 15707 | 786 | T786Q | B | 2e-06 | 2.1e-06 | 1.6e-06 | 1.900E-06 | 2.646E-07 | 1.600E-06 | 2.100E-06 |
| 15708 | 786 | T786G | B | 1.5e-06 | 3.2e-06 | 2.1e-06 | 2.267E-06 | 8.622E-07 | 1.500E-06 | 3.200E-06 |
| 15709 | 786 | T786H | B | 3e-06 | 1.9e-06 | 2.9e-06 | 2.600E-06 | 6.083E-07 | 1.900E-06 | 3.000E-06 |
| 15710 | 786 | T786I | B | 4.4e-06 | 1e-06 | 2.9e-06 | 2.767E-06 | 1.704E-06 | 1.000E-06 | 4.400E-06 |
| 15711 | 786 | T786L | B | 2e-06 | 5.8e-06 | 2.3e-06 | 3.367E-06 | 2.113E-06 | 2.000E-06 | 5.800E-06 |
| 15712 | 786 | T786K | B | 2.1e-06 | 3e-06 | 2.3e-06 | 2.467E-06 | 4.726E-07 | 2.100E-06 | 3.000E-06 |
| 15713 | 786 | T786M | B | 2.8e-06 | 2.1e-06 | 2.5e-06 | 2.467E-06 | 3.512E-07 | 2.100E-06 | 2.800E-06 |
| 15714 | 786 | T786F | B | 3.7e-06 | 2.8e-06 | 3.2e-06 | 3.233E-06 | 4.509E-07 | 2.800E-06 | 3.700E-06 |
| 15715 | 786 | T786P | B | 2.3e-06 | 2.2e-06 | 2.8e-06 | 2.433E-06 | 3.215E-07 | 2.200E-06 | 2.800E-06 |
| 15716 | 786 | T786S | B | 2.2e-06 | 1.9e-06 | 1.4e-06 | 1.833E-06 | 4.041E-07 | 1.400E-06 | 2.200E-06 |
| 15717 | 786 | T786T | B | 1.6e-06 | 1.6e-06 | 1.9e-06 | 1.700E-06 | 1.732E-07 | 1.600E-06 | 1.900E-06 |
| 15718 | 786 | T786W | B | 2.2e-06 | 2.7e-06 | 2.6e-06 | 2.500E-06 | 2.646E-07 | 2.200E-06 | 2.700E-06 |
| 15719 | 786 | T786Y | B | 1.8e-06 | 2.1e-06 | 1.4e-06 | 1.767E-06 | 3.512E-07 | 1.400E-06 | 2.100E-06 |
| 15720 | 786 | T786V | B | 3e-06 | 2.2e-06 | 2.2e-06 | 2.467E-06 | 4.619E-07 | 2.200E-06 | 3.000E-06 |
| 15721 | 787 | E787A | B | 1.4e-06 | 1.8e-06 | 2.3e-06 | 1.833E-06 | 4.509E-07 | 1.400E-06 | 2.300E-06 |
| 15722 | 787 | E787R | B | 3.5e-06 | 2.2e-06 | 1.2e-06 | 2.300E-06 | 1.153E-06 | 1.200E-06 | 3.500E-06 |
| 15723 | 787 | E787N | B | 3.8e-06 | 3e-06 | 2.8e-06 | 3.200E-06 | 5.292E-07 | 2.800E-06 | 3.800E-06 |
| 15724 | 787 | E787D | B | 3.3e-06 | 1.5e-06 | 3.9e-06 | 2.900E-06 | 1.249E-06 | 1.500E-06 | 3.900E-06 |

| | | | | | | | | | | |
|---|---|---|---|---|---|---|---|---|---|---|
| 15725 | 787 | E787C | B | 2.4e-06 | 3e-06 | 1.8e-06 | 2.400E-06 | 6.000E-07 | 1.800E-06 | 3.000E-06 |
| 15726 | 787 | E787E | B | 2.1e-06 | 2.1e-06 | 2.8e-06 | 2.333E-06 | 4.041E-07 | 2.100E-06 | 2.800E-06 |
| 15727 | 787 | E787Q | B | 3e-06 | 1.8e-06 | 1.8e-06 | 2.200E-06 | 6.928E-07 | 1.800E-06 | 3.000E-06 |
| 15728 | 787 | E787G | B | 3.7e-06 | 2e-06 | 2.6e-06 | 2.767E-06 | 8.622E-07 | 2.000E-06 | 3.700E-06 |
| 15729 | 787 | E787H | B | 1.6e-06 | 2e-06 | 1.6e-06 | 1.733E-06 | 2.309E-07 | 1.600E-06 | 2.000E-06 |
| 15730 | 787 | E787I | B | 1.9e-06 | 1.6e-06 | 1.6e-06 | 1.700E-06 | 1.732E-07 | 1.600E-06 | 1.900E-06 |
| 15731 | 787 | E787L | B | 3e-06 | 3.6e-06 | 2.3e-06 | 2.967E-06 | 6.506E-07 | 2.300E-06 | 3.600E-06 |
| 15732 | 787 | E787K | B | 3.1e-06 | 1.4e-06 | 1.8e-06 | 2.100E-06 | 8.888E-07 | 1.400E-06 | 3.100E-06 |
| 15733 | 787 | E787M | B | 1.3e-06 | 2.5e-06 | 2.4e-06 | 2.067E-06 | 6.658E-07 | 1.300E-06 | 2.500E-06 |
| 15734 | 787 | E787F | B | 3.8e-06 | 2.7e-06 | 1.7e-06 | 2.733E-06 | 1.050E-06 | 1.700E-06 | 3.800E-06 |
| 15735 | 787 | E787P | B | 2.4e-06 | 1.3e-06 | 1.7e-06 | 1.800E-06 | 5.568E-07 | 1.300E-06 | 2.400E-06 |
| 15736 | 787 | E787S | B | 1.5e-06 | 1.7e-06 | 2.2e-06 | 1.800E-06 | 3.606E-07 | 1.500E-06 | 2.200E-06 |
| 15737 | 787 | E787T | B | 1.5e-06 | 2.1e-06 | 3.9e-06 | 2.500E-06 | 1.249E-06 | 1.500E-06 | 3.900E-06 |
| 15738 | 787 | E787W | B | 4.3e-06 | 1.9e-06 | 2.4e-06 | 2.867E-06 | 1.266E-06 | 1.900E-06 | 4.300E-06 |
| 15739 | 787 | E787Y | B | 1.8e-06 | 2.1e-06 | 2.6e-06 | 2.167E-06 | 4.041E-07 | 1.800E-06 | 2.600E-06 |
| 15740 | 787 | E787V | B | 4.5e-06 | 1.9e-06 | 1.8e-06 | 2.733E-06 | 1.531E-06 | 1.800E-06 | 4.500E-06 |
| 15741 | 788 | Q788A | B | 2.6e-06 | 2.8e-06 | 2.3e-06 | 2.567E-06 | 2.517E-07 | 2.300E-06 | 2.800E-06 |
| 15742 | 788 | Q788R | B | 3.3e-06 | 2.4e-06 | 2.1e-06 | 2.600E-06 | 6.245E-07 | 2.100E-06 | 3.300E-06 |
| 15743 | 788 | Q788N | B | 1.9e-06 | 8.5e-07 | 1.6e-06 | 1.450E-06 | 5.408E-07 | 8.500E-07 | 1.900E-06 |
| 15744 | 788 | Q788D | B | 2.6e-06 | 2.6e-06 | 2.5e-06 | 2.567E-06 | 5.774E-08 | 2.500E-06 | 2.600E-06 |
| 15745 | 788 | Q788C | B | 2.2e-06 | 2.6e-06 | 1.6e-06 | 2.133E-06 | 5.033E-07 | 1.600E-06 | 2.600E-06 |
| 15746 | 788 | Q788E | B | 4.6e-06 | 3.2e-06 | 1e-06 | 2.933E-06 | 1.815E-06 | 1.000E-06 | 4.600E-06 |
| 15747 | 788 | Q788Q | B | 1.7e-06 | 2.5e-06 | 1.9e-06 | 2.033E-06 | 4.163E-07 | 1.700E-06 | 2.500E-06 |
| 15748 | 788 | Q788G | B | 2.4e-06 | 2.1e-06 | 2.1e-06 | 2.200E-06 | 1.732E-07 | 2.100E-06 | 2.400E-06 |
| 15749 | 788 | Q788H | B | 3.9e-06 | 2.3e-06 | 3.8e-06 | 3.333E-06 | 8.963E-07 | 2.300E-06 | 3.900E-06 |
| 15750 | 788 | Q788I | B | 3.9e-06 | 2.7e-06 | 1e-06 | 2.533E-06 | 1.457E-06 | 1.000E-06 | 3.900E-06 |
| 15751 | 788 | Q788L | B | 1.5e-06 | 3e-06 | 1.5e-06 | 2.000E-06 | 8.660E-07 | 1.500E-06 | 3.000E-06 |
| 15752 | 788 | Q788K | B | 2.8e-06 | 2.7e-06 | 2.6e-06 | 2.700E-06 | 1.000E-07 | 2.600E-06 | 2.800E-06 |
| 15753 | 788 | Q788M | B | 2.4e-06 | 3.1e-06 | 1.6e-06 | 2.367E-06 | 7.506E-07 | 1.600E-06 | 3.100E-06 |
| 15754 | 788 | Q788F | B | 3.2e-06 | 1.7e-06 | 2.3e-06 | 2.400E-06 | 7.550E-07 | 1.700E-06 | 3.200E-06 |
| 15755 | 788 | Q788P | B | 4.9e-06 | 2.8e-06 | 2.3e-06 | 3.333E-06 | 1.380E-06 | 2.300E-06 | 4.900E-06 |
| 15756 | 788 | Q788S | B | 4.4e-06 | 2.5e-06 | 2.1e-06 | 3.000E-06 | 1.229E-06 | 2.100E-06 | 4.400E-06 |
| 15757 | 788 | Q788T | B | 2.2e-06 | 1.6e-06 | 2e-06 | 1.933E-06 | 3.055E-07 | 1.600E-06 | 2.200E-06 |
| 15758 | 788 | Q788W | B | 2.2e-06 | 1.7e-06 | 2.1e-06 | 2.000E-06 | 2.646E-07 | 1.700E-06 | 2.200E-06 |
| 15759 | 788 | Q788Y | B | 3.5e-06 | 3e-06 | 3.6e-06 | 3.367E-06 | 3.215E-07 | 3.000E-06 | 3.600E-06 |
| 15760 | 788 | Q788V | B | 1.8e-06 | 1.6e-06 | 2.9e-06 | 2.100E-06 | 7.000E-07 | 1.600E-06 | 2.900E-06 |
| 15761 | 789 | D789A | B | 1.7e-06 | 2.4e-06 | 2.1e-06 | 2.067E-06 | 3.512E-07 | 1.700E-06 | 2.400E-06 |
| 15762 | 789 | D789R | B | 1.3e-06 | 3.3e-06 | 2.5e-06 | 2.367E-06 | 1.007E-06 | 1.300E-06 | 3.300E-06 |
| 15763 | 789 | D789N | B | 4e-06 | 2.3e-06 | 1.7e-06 | 2.667E-06 | 1.193E-06 | 1.700E-06 | 4.000E-06 |
| 15764 | 789 | D789D | B | 2.1e-06 | 1.4e-06 | 1.7e-06 | 1.733E-06 | 3.512E-07 | 1.400E-06 | 2.100E-06 |
| 15765 | 789 | D789C | B | 4.4e-06 | 2.5e-06 | 2.2e-06 | 3.033E-06 | 1.193E-06 | 2.200E-06 | 4.400E-06 |
| 15766 | 789 | D789E | B | 3.2e-06 | 2e-06 | 1.4e-06 | 2.200E-06 | 9.165E-07 | 1.400E-06 | 3.200E-06 |
| 15767 | 789 | D789Q | B | 2.4e-06 | 1.8e-06 | 2.8e-06 | 2.333E-06 | 5.033E-07 | 1.800E-06 | 2.800E-06 |
| 15768 | 789 | D789G | B | 2.3e-06 | 2.4e-06 | 2.9e-06 | 2.533E-06 | 3.215E-07 | 2.300E-06 | 2.900E-06 |
| 15769 | 789 | D789H | B | 2.3e-06 | 2.6e-06 | 3.7e-06 | 2.867E-06 | 7.371E-07 | 2.300E-06 | 3.700E-06 |
| 15770 | 789 | D789I | B | 3e-06 | 4e-06 | 1.7e-06 | 2.900E-06 | 1.153E-06 | 1.700E-06 | 4.000E-06 |
| 15771 | 789 | D789L | B | 2e-06 | 1.6e-06 | 2.8e-06 | 2.133E-06 | 6.110E-07 | 1.600E-06 | 2.800E-06 |
| 15772 | 789 | D789K | B | 1.8e-06 | 2.4e-06 | 2.6e-06 | 2.267E-06 | 4.163E-07 | 1.800E-06 | 2.600E-06 |
| 15773 | 789 | D789M | B | 1.7e-06 | 1.8e-06 | 9.3e-07 | 1.477E-06 | 4.761E-07 | 9.300E-07 | 1.800E-06 |
| 15774 | 789 | D789F | B | 3.2e-06 | 3e-06 | 2.9e-06 | 3.033E-06 | 1.528E-07 | 2.900E-06 | 3.200E-06 |
| 15775 | 789 | D789P | B | 2.4e-06 | 1.5e-06 | 2.4e-06 | 2.100E-06 | 5.196E-07 | 1.500E-06 | 2.400E-06 |
| 15776 | 789 | D789S | B | 2.1e-06 | 2.5e-06 | 2.6e-06 | 2.400E-06 | 2.646E-07 | 2.100E-06 | 2.600E-06 |
| 15777 | 789 | D789T | B | 2.4e-06 | 2.4e-06 | 2.1e-06 | 2.300E-06 | 1.732E-07 | 2.100E-06 | 2.400E-06 |
| 15778 | 789 | D789W | B | 1.3e-06 | 2.7e-06 | 3.1e-06 | 2.367E-06 | 9.452E-07 | 1.300E-06 | 3.100E-06 |
| 15779 | 789 | D789Y | B | 1.9e-06 | 1.5e-06 | 2.1e-06 | 1.833E-06 | 3.055E-07 | 1.500E-06 | 2.100E-06 |
| 15780 | 789 | D789V | B | 3.3e-06 | 2.1e-06 | 4.1e-06 | 3.167E-06 | 1.007E-06 | 2.100E-06 | 4.100E-06 |
| 15781 | 790 | S790A | B | 2.4e-06 | 2.4e-06 | 2.5e-06 | 2.433E-06 | 5.774E-08 | 2.400E-06 | 2.500E-06 |
| 15782 | 790 | S790R | B | 3.1e-06 | 2.4e-06 | 2e-06 | 2.500E-06 | 5.568E-07 | 2.000E-06 | 3.100E-06 |
| 15783 | 790 | S790N | B | 1.4e-06 | 1.9e-06 | 3.2e-06 | 2.167E-06 | 9.292E-07 | 1.400E-06 | 3.200E-06 |
| 15784 | 790 | S790D | B | 2.6e-06 | 4.3e-06 | 2.1e-06 | 3.000E-06 | 1.153E-06 | 2.100E-06 | 4.300E-06 |
| 15785 | 790 | S790C | B | 3.7e-06 | 1.4e-06 | 1.8e-06 | 2.300E-06 | 1.229E-06 | 1.400E-06 | 3.700E-06 |
| 15786 | 790 | S790E | B | 2.1e-06 | 3.9e-06 | 2.2e-06 | 2.733E-06 | 1.012E-06 | 2.100E-06 | 3.900E-06 |
| 15787 | 790 | S790Q | B | 2e-06 | 4e-06 | 1.8e-06 | 2.600E-06 | 1.217E-06 | 1.800E-06 | 4.000E-06 |
| 15788 | 790 | S790G | B | 1.9e-06 | 2.6e-06 | 3.3e-06 | 2.600E-06 | 7.000E-07 | 1.900E-06 | 3.300E-06 |
| 15789 | 790 | S790H | B | 2.3e-06 | 1.6e-06 | 1.8e-06 | 1.900E-06 | 3.606E-07 | 1.600E-06 | 2.300E-06 |
| 15790 | 790 | S790I | B | 1.9e-06 | 3.3e-06 | 2.6e-06 | 2.600E-06 | 7.000E-07 | 1.900E-06 | 3.300E-06 |
| 15791 | 790 | S790L | B | 4.4e-06 | 2.1e-06 | 1.8e-06 | 2.767E-06 | 1.422E-06 | 1.800E-06 | 4.400E-06 |
| 15792 | 790 | S790K | B | 2.5e-06 | 1.8e-06 | 3e-06 | 2.433E-06 | 6.028E-07 | 1.800E-06 | 3.000E-06 |
| 15793 | 790 | S790M | B | 2.1e-06 | 4.1e-06 | 2.9e-06 | 3.033E-06 | 1.007E-06 | 2.100E-06 | 4.100E-06 |
| 15794 | 790 | S790F | B | 3.9e-06 | 2.8e-06 | 1.6e-06 | 2.767E-06 | 1.150E-06 | 1.600E-06 | 3.900E-06 |
| 15795 | 790 | S790P | B | 3.2e-06 | 1.5e-06 | 3.3e-06 | 2.667E-06 | 1.012E-06 | 1.500E-06 | 3.300E-06 |
| 15796 | 790 | S790S | B | 3.6e-06 | 4.6e-06 | 3.6e-06 | 3.933E-06 | 5.774E-07 | 3.600E-06 | 4.600E-06 |
| 15797 | 790 | S790T | B | 1.6e-06 | 2.4e-06 | 1.5e-06 | 1.833E-06 | 4.933E-07 | 1.500E-06 | 2.400E-06 |
| 15798 | 790 | S790W | B | 3.6e-06 | 1.2e-06 | 3.2e-06 | 2.667E-06 | 1.286E-06 | 1.200E-06 | 3.600E-06 |
| 15799 | 790 | S790Y | B | 3.1e-06 | 1.1e-06 | 2.1e-06 | 2.100E-06 | 1.000E-06 | 1.100E-06 | 3.100E-06 |
| 15800 | 790 | S790V | B | 1.9e-06 | 2.1e-06 | 2.1e-06 | 2.033E-06 | 1.155E-07 | 1.900E-06 | 2.100E-06 |
| 15801 | 791 | K791A | B | 4e-06 | 5.6e-06 | 1.9e-06 | 3.833E-06 | 1.856E-06 | 1.900E-06 | 5.600E-06 |
| 15802 | 791 | K791R | B | 3.2e-06 | 1.7e-06 | 3.5e-06 | 2.800E-06 | 9.644E-07 | 1.700E-06 | 3.500E-06 |
| 15803 | 791 | K791N | B | 2.2e-06 | 2e-06 | 4e-06 | 2.733E-06 | 1.102E-06 | 2.000E-06 | 4.000E-06 |
| 15804 | 791 | K791D | B | 2.3e-06 | 2.5e-06 | 2.6e-06 | 2.467E-06 | 1.528E-07 | 2.300E-06 | 2.600E-06 |
| 15805 | 791 | K791C | B | 1.9e-06 | 3e-06 | 2.1e-06 | 2.333E-06 | 5.859E-07 | 1.900E-06 | 3.000E-06 |
| 15806 | 791 | K791E | B | 1.9e-06 | 2.7e-06 | 4.3e-06 | 2.967E-06 | 1.222E-06 | 1.900E-06 | 4.300E-06 |
| 15807 | 791 | K791Q | B | 2.5e-06 | 1.8e-06 | 6e-06 | 3.433E-06 | 2.250E-06 | 1.800E-06 | 6.000E-06 |
| 15808 | 791 | K791G | B | 3.3e-06 | 2.6e-06 | 3.7e-06 | 3.200E-06 | 5.568E-07 | 2.600E-06 | 3.700E-06 |
| 15809 | 791 | K791H | B | 3.1e-06 | 1.7e-06 | 2.9e-06 | 2.567E-06 | 7.572E-07 | 1.700E-06 | 3.100E-06 |

| | | | | | | | | | | |
|---|---|---|---|---|---|---|---|---|---|---|
| 15810 | 791 | K791I | B | 3.9e-06 | 2.8e-06 | 2.5e-06 | 3.067E-06 | 7.371E-07 | 2.500E-06 | 3.900E-06 |
| 15811 | 791 | K791L | B | 2.1e-06 | 5.6e-06 | 2.6e-06 | 3.433E-06 | 1.893E-06 | 2.100E-06 | 5.600E-06 |
| 15812 | 791 | K791K | B | 1.8e-06 | 2.7e-06 | 2.8e-06 | 2.433E-06 | 5.508E-07 | 1.800E-06 | 2.800E-06 |
| 15813 | 791 | K791M | B | 2.5e-06 | 1.8e-06 | 3.6e-06 | 2.633E-06 | 9.074E-07 | 1.800E-06 | 3.600E-06 |
| 15814 | 791 | K791F | B | 1.1e-06 | 2.6e-06 | 2.7e-06 | 2.133E-06 | 8.963E-07 | 1.100E-06 | 2.700E-06 |
| 15815 | 791 | K791P | B | 2.2e-06 | 2e-06 | 1.9e-06 | 2.033E-06 | 1.528E-07 | 1.900E-06 | 2.200E-06 |
| 15816 | 791 | K791S | B | 3.3e-06 | 2.3e-06 | 1.5e-06 | 2.367E-06 | 9.018E-07 | 1.500E-06 | 3.300E-06 |
| 15817 | 791 | K791T | B | 2.5e-06 | 2.5e-06 | 2.9e-06 | 2.633E-06 | 2.309E-07 | 2.500E-06 | 2.900E-06 |
| 15818 | 791 | K791W | B | 3.2e-06 | 2e-06 | 3.1e-06 | 2.767E-06 | 6.658E-07 | 2.000E-06 | 3.200E-06 |
| 15819 | 791 | K791Y | B | 2.6e-06 | 3.2e-06 | 4.3e-06 | 3.367E-06 | 8.622E-07 | 2.600E-06 | 4.300E-06 |
| 15820 | 791 | K791V | B | 1.8e-06 | 2.1e-06 | 3.5e-06 | 2.467E-06 | 9.074E-07 | 1.800E-06 | 3.500E-06 |
| 15821 | 792 | D792A | B | 5.6e-06 | 2.6e-06 | 1.6e-06 | 3.267E-06 | 2.082E-06 | 1.600E-06 | 5.600E-06 |
| 15822 | 792 | D792R | B | 1.9e-06 | 4e-06 | 2.1e-06 | 2.667E-06 | 1.159E-06 | 1.900E-06 | 4.000E-06 |
| 15823 | 792 | D792N | B | 3e-06 | 2.3e-06 | 1.9e-06 | 2.400E-06 | 5.568E-07 | 1.900E-06 | 3.000E-06 |
| 15824 | 792 | D792D | B | 5.3e-06 | 4.5e-06 | 2.3e-06 | 4.033E-06 | 1.553E-06 | 2.300E-06 | 5.300E-06 |
| 15825 | 792 | D792C | B | 2e-06 | 3e-06 | 1e-06 | 2.000E-06 | 1.000E-06 | 1.000E-06 | 3.000E-06 |
| 15826 | 792 | D792E | B | 3.8e-06 | 4.7e-06 | 1.8e-06 | 3.433E-06 | 1.484E-06 | 1.800E-06 | 4.700E-06 |
| 15827 | 792 | D792Q | B | 3.6e-06 | 1.9e-06 | 3e-06 | 2.833E-06 | 8.622E-07 | 1.900E-06 | 3.600E-06 |
| 15828 | 792 | D792G | B | 3.7e-06 | 2.4e-06 | 1.6e-06 | 2.567E-06 | 1.060E-06 | 1.600E-06 | 3.700E-06 |
| 15829 | 792 | D792H | B | 1.8e-06 | 5.1e-06 | 2.1e-06 | 3.000E-06 | 1.825E-06 | 1.800E-06 | 5.100E-06 |
| 15830 | 792 | D792I | B | 1.7e-06 | 3.5e-06 | 2.8e-06 | 2.667E-06 | 9.074E-07 | 1.700E-06 | 3.500E-06 |
| 15831 | 792 | D792L | B | 2.5e-06 | 2e-06 | 2.9e-06 | 2.467E-06 | 4.509E-07 | 2.000E-06 | 2.900E-06 |
| 15832 | 792 | D792K | B | 1.4e-06 | 1.7e-06 | 3.7e-06 | 2.267E-06 | 1.250E-06 | 1.400E-06 | 3.700E-06 |
| 15833 | 792 | D792M | B | 1.9e-06 | 1.6e-06 | 3.9e-06 | 2.467E-06 | 1.250E-06 | 1.600E-06 | 3.900E-06 |
| 15834 | 792 | D792F | B | 2.9e-06 | 1.5e-06 | 3e-06 | 2.467E-06 | 8.386E-07 | 1.500E-06 | 3.000E-06 |
| 15835 | 792 | D792P | B | 2.5e-06 | 3.4e-06 | 3.4e-06 | 3.100E-06 | 5.196E-07 | 2.500E-06 | 3.400E-06 |
| 15836 | 792 | D792S | B | 1.8e-06 | 2.9e-06 | 3.2e-06 | 2.633E-06 | 7.371E-07 | 1.800E-06 | 3.200E-06 |
| 15837 | 792 | D792T | B | 2.6e-06 | 2.1e-06 | 1.7e-06 | 2.133E-06 | 4.509E-07 | 1.700E-06 | 2.600E-06 |
| 15838 | 792 | D792W | B | 2.3e-06 | 2.2e-06 | 2.5e-06 | 2.333E-06 | 1.528E-07 | 2.200E-06 | 2.500E-06 |
| 15839 | 792 | D792Y | B | 4.6e-06 | 2.4e-06 | 3e-06 | 3.333E-06 | 1.137E-06 | 2.400E-06 | 4.600E-06 |
| 15840 | 792 | D792V | B | 1.3e-06 | 2e-06 | 1.6e-06 | 1.633E-06 | 3.512E-07 | 1.300E-06 | 2.000E-06 |
| 15841 | 793 | S793A | B | 1.9e-06 | 2.7e-06 | 2.5e-06 | 2.367E-06 | 4.163E-07 | 1.900E-06 | 2.700E-06 |
| 15842 | 793 | S793R | B | 1.9e-06 | 1.2e-06 | 1.8e-06 | 1.633E-06 | 3.786E-07 | 1.200E-06 | 1.900E-06 |
| 15843 | 793 | S793N | B | 4.7e-06 | 1.6e-06 | 1.3e-06 | 2.533E-06 | 1.882E-06 | 1.300E-06 | 4.700E-06 |
| 15844 | 793 | S793D | B | 1.3e-06 | 2.3e-06 | 2.5e-06 | 2.033E-06 | 6.429E-07 | 1.300E-06 | 2.500E-06 |
| 15845 | 793 | S793C | B | 2.2e-06 | 2.1e-06 | 2.6e-06 | 2.300E-06 | 2.646E-07 | 2.100E-06 | 2.600E-06 |
| 15846 | 793 | S793E | B | 1.4e-06 | 2.1e-06 | 2.1e-06 | 1.867E-06 | 4.041E-07 | 1.400E-06 | 2.100E-06 |
| 15847 | 793 | S793Q | B | 3.9e-06 | 2.8e-06 | 2.7e-06 | 3.133E-06 | 6.658E-07 | 2.700E-06 | 3.900E-06 |
| 15848 | 793 | S793G | B | 2.3e-06 | 2e-06 | 1.8e-06 | 2.033E-06 | 2.517E-07 | 1.800E-06 | 2.300E-06 |
| 15849 | 793 | S793H | B | 3.4e-06 | 3.5e-06 | 1.8e-06 | 2.900E-06 | 9.539E-07 | 1.800E-06 | 3.500E-06 |
| 15850 | 793 | S793I | B | 3.2e-06 | 3.4e-06 | 2.6e-06 | 3.067E-06 | 4.163E-07 | 2.600E-06 | 3.400E-06 |
| 15851 | 793 | S793L | B | 2.4e-06 | 2.5e-06 | 1.4e-06 | 2.100E-06 | 6.083E-07 | 1.400E-06 | 2.500E-06 |
| 15852 | 793 | S793K | B | 1.2e-06 | 2.5e-06 | 2.6e-06 | 2.100E-06 | 7.810E-07 | 1.200E-06 | 2.600E-06 |
| 15853 | 793 | S793M | B | 2.7e-06 | 2.1e-06 | 3.6e-06 | 2.800E-06 | 7.550E-07 | 2.100E-06 | 3.600E-06 |
| 15854 | 793 | S793F | B | 3.9e-06 | 1.8e-06 | 2.3e-06 | 2.667E-06 | 1.097E-06 | 1.800E-06 | 3.900E-06 |
| 15855 | 793 | S793P | B | 1.7e-06 | 3.1e-06 | 1.9e-06 | 2.233E-06 | 7.572E-07 | 1.700E-06 | 3.100E-06 |
| 15856 | 793 | S793S | B | 2.3e-06 | 2e-06 | 2e-06 | 2.100E-06 | 1.732E-07 | 2.000E-06 | 2.300E-06 |
| 15857 | 793 | S793T | B | 1.2e-06 | 1.6e-06 | 2.8e-06 | 1.867E-06 | 8.327E-07 | 1.200E-06 | 2.800E-06 |
| 15858 | 793 | S793W | B | 1.5e-06 | 1.8e-06 | 2.6e-06 | 1.967E-06 | 5.686E-07 | 1.500E-06 | 2.600E-06 |
| 15859 | 793 | S793Y | B | 1.5e-06 | 2.4e-06 | 3.1e-06 | 2.333E-06 | 8.021E-07 | 1.500E-06 | 3.100E-06 |
| 15860 | 793 | S793V | B | 2.5e-06 | 2.2e-06 | 1.9e-06 | 2.200E-06 | 3.000E-07 | 1.900E-06 | 2.500E-06 |
| 15861 | 794 | T794A | B | 1.9e-06 | 1.5e-06 | 3.4e-06 | 2.267E-06 | 1.002E-06 | 1.500E-06 | 3.400E-06 |
| 15862 | 794 | T794R | B | 1.7e-06 | 3.3e-06 | 3.2e-06 | 2.733E-06 | 8.963E-07 | 1.700E-06 | 3.300E-06 |
| 15863 | 794 | T794N | B | 3.4e-06 | 3e-06 | 2.6e-06 | 3.000E-06 | 4.000E-07 | 2.600E-06 | 3.400E-06 |
| 15864 | 794 | T794D | B | 2.6e-06 | 1.3e-06 | 2e-06 | 1.967E-06 | 6.506E-07 | 1.300E-06 | 2.600E-06 |
| 15865 | 794 | T794C | B | 1.8e-06 | 1.6e-06 | 1.3e-06 | 1.567E-06 | 2.517E-07 | 1.300E-06 | 1.800E-06 |
| 15866 | 794 | T794E | B | 2.4e-06 | 4e-06 | 1.8e-06 | 2.733E-06 | 1.137E-06 | 1.800E-06 | 4.000E-06 |
| 15867 | 794 | T794Q | B | 3e-06 | 3.6e-06 | 2.1e-06 | 2.900E-06 | 7.550E-07 | 2.100E-06 | 3.600E-06 |
| 15868 | 794 | T794G | B | 3e-06 | 2.1e-06 | 2.3e-06 | 2.467E-06 | 4.726E-07 | 2.100E-06 | 3.000E-06 |
| 15869 | 794 | T794H | B | 4.4e-06 | 1.4e-06 | 4.1e-06 | 3.300E-06 | 1.652E-06 | 1.400E-06 | 4.400E-06 |
| 15870 | 794 | T794I | B | 4.2e-06 | 2.2e-06 | 2.1e-06 | 2.833E-06 | 1.185E-06 | 2.100E-06 | 4.200E-06 |
| 15871 | 794 | T794L | B | 2.8e-06 | 2.4e-06 | 2.1e-06 | 2.433E-06 | 3.512E-07 | 2.100E-06 | 2.800E-06 |
| 15872 | 794 | T794K | B | 1.9e-06 | 2.1e-06 | 2.1e-06 | 2.033E-06 | 1.155E-07 | 1.900E-06 | 2.100E-06 |
| 15873 | 794 | T794M | B | 2.8e-06 | 2.7e-06 | 2.3e-06 | 2.600E-06 | 2.646E-07 | 2.300E-06 | 2.800E-06 |
| 15874 | 794 | T794F | B | 2.3e-06 | 2e-06 | 3.3e-06 | 2.533E-06 | 6.807E-07 | 2.000E-06 | 3.300E-06 |
| 15875 | 794 | T794P | B | 2.3e-06 | 2.1e-06 | 2.8e-06 | 2.400E-06 | 3.606E-07 | 2.100E-06 | 2.800E-06 |
| 15876 | 794 | T794S | B | 3.7e-06 | 3.1e-06 | 2.8e-06 | 3.200E-06 | 4.583E-07 | 2.800E-06 | 3.700E-06 |
| 15877 | 794 | T794T | B | 2.7e-06 | 1.1e-06 | 2.8e-06 | 2.200E-06 | 9.539E-07 | 1.100E-06 | 2.800E-06 |
| 15878 | 794 | T794W | B | 2e-06 | 1.4e-06 | 1.2e-06 | 1.533E-06 | 4.163E-07 | 1.200E-06 | 2.000E-06 |
| 15879 | 794 | T794Y | B | 1.7e-06 | 3.2e-06 | 2.5e-06 | 2.467E-06 | 7.506E-07 | 1.700E-06 | 3.200E-06 |
| 15880 | 794 | T794V | B | 2.2e-06 | 2.2e-06 | 4.4e-06 | 2.933E-06 | 1.270E-06 | 2.200E-06 | 4.400E-06 |
| 15881 | 795 | Y795A | B | 2.4e-06 | 1.5e-06 | 2.8e-06 | 2.233E-06 | 6.658E-07 | 1.500E-06 | 2.800E-06 |
| 15882 | 795 | Y795R | B | 3.2e-06 | 3.2e-06 | 1.5e-06 | 2.633E-06 | 9.815E-07 | 1.500E-06 | 3.200E-06 |
| 15883 | 795 | Y795N | B | 1.7e-06 | 2.5e-06 | 2.3e-06 | 2.167E-06 | 4.163E-07 | 1.700E-06 | 2.500E-06 |
| 15884 | 795 | Y795D | B | 3e-06 | 3.1e-06 | 2e-06 | 2.700E-06 | 6.083E-07 | 2.000E-06 | 3.100E-06 |
| 15885 | 795 | Y795C | B | 2.7e-06 | 2.1e-06 | 4.7e-06 | 3.167E-06 | 1.361E-06 | 2.100E-06 | 4.700E-06 |
| 15886 | 795 | Y795E | B | 1.4e-06 | 2.4e-06 | 1.5e-06 | 1.767E-06 | 5.508E-07 | 1.400E-06 | 2.400E-06 |
| 15887 | 795 | Y795Q | B | 3.1e-06 | 2.8e-06 | 2.3e-06 | 2.733E-06 | 4.041E-07 | 2.300E-06 | 3.100E-06 |
| 15888 | 795 | Y795G | B | 2.1e-06 | 2e-06 | 2.1e-06 | 2.067E-06 | 5.774E-08 | 2.000E-06 | 2.100E-06 |
| 15889 | 795 | Y795H | B | 2.5e-06 | 2.5e-06 | 2.6e-06 | 2.533E-06 | 5.774E-08 | 2.500E-06 | 2.600E-06 |
| 15890 | 795 | Y795I | B | 2.4e-06 | 2.3e-06 | 2.4e-06 | 2.367E-06 | 5.774E-08 | 2.300E-06 | 2.400E-06 |
| 15891 | 795 | Y795L | B | 2.4e-06 | 1.1e-06 | 1.9e-06 | 1.800E-06 | 6.557E-07 | 1.100E-06 | 2.400E-06 |
| 15892 | 795 | Y795K | B | 2e-06 | 1.6e-06 | 3.3e-06 | 2.300E-06 | 8.888E-07 | 1.600E-06 | 3.300E-06 |
| 15893 | 795 | Y795M | B | 3.3e-06 | 1.9e-06 | 2.5e-06 | 2.567E-06 | 7.024E-07 | 1.900E-06 | 3.300E-06 |
| 15894 | 795 | Y795F | B | 2.4e-06 | 3.3e-06 | 1.6e-06 | 2.433E-06 | 8.505E-07 | 1.600E-06 | 3.300E-06 |

| | | | | | | | | | | |
|---|---|---|---|---|---|---|---|---|---|---|
| 15895 | 795 | Y795P | B | 2e-06 | 4.1e-06 | 2.2e-06 | 2.767E-06 | 1.159E-06 | 2.000E-06 | 4.100E-06 |
| 15896 | 795 | Y795S | B | 2.3e-06 | 2.7e-06 | 2.6e-06 | 2.533E-06 | 2.082E-07 | 2.300E-06 | 2.700E-06 |
| 15897 | 795 | Y795T | B | 2.1e-06 | 2.1e-06 | 1.9e-06 | 2.033E-06 | 1.155E-07 | 1.900E-06 | 2.100E-06 |
| 15898 | 795 | Y795W | B | 2.1e-06 | 1.1e-06 | 2.3e-06 | 1.833E-06 | 6.429E-07 | 1.100E-06 | 2.300E-06 |
| 15899 | 795 | Y795Y | B | 1.2e-06 | 2.3e-06 | 1.2e-06 | 1.567E-06 | 6.351E-07 | 1.200E-06 | 2.300E-06 |
| 15900 | 795 | Y795V | B | 2.7e-06 | 3.7e-06 | 2.4e-06 | 2.933E-06 | 6.807E-07 | 2.400E-06 | 3.700E-06 |
| 15901 | 796 | S796A | B | 1.6e-06 | 1.8e-06 | 1.4e-06 | 1.600E-06 | 2.000E-07 | 1.400E-06 | 1.800E-06 |
| 15902 | 796 | S796R | B | 2.5e-06 | 1.8e-06 | 2.6e-06 | 2.300E-06 | 4.359E-07 | 1.800E-06 | 2.600E-06 |
| 15903 | 796 | S796N | B | 2.4e-06 | 2.6e-06 | 2e-06 | 2.333E-06 | 3.055E-07 | 2.000E-06 | 2.600E-06 |
| 15904 | 796 | S796D | B | 2.2e-06 | 2.2e-06 | 3.8e-06 | 2.733E-06 | 9.238E-07 | 2.200E-06 | 3.800E-06 |
| 15905 | 796 | S796C | B | 2.4e-06 | 1e-06 | 1.5e-06 | 1.633E-06 | 7.095E-07 | 1.000E-06 | 2.400E-06 |
| 15906 | 796 | S796E | B | 2.4e-06 | 2.2e-06 | 2.6e-06 | 2.400E-06 | 2.000E-07 | 2.200E-06 | 2.600E-06 |
| 15907 | 796 | S796Q | B | 1.9e-06 | 3.9e-06 | 1.8e-06 | 2.533E-06 | 1.185E-06 | 1.800E-06 | 3.900E-06 |
| 15908 | 796 | S796G | B | 2.2e-06 | 2.6e-06 | 2e-06 | 2.267E-06 | 3.055E-07 | 2.000E-06 | 2.600E-06 |
| 15909 | 796 | S796H | B | 3e-06 | 5.4e-06 | 2.8e-06 | 3.733E-06 | 1.447E-06 | 2.800E-06 | 5.400E-06 |
| 15910 | 796 | S796I | B | 1.9e-06 | 4.4e-06 | 2.9e-06 | 3.067E-06 | 1.258E-06 | 1.900E-06 | 4.400E-06 |
| 15911 | 796 | S796L | B | 3.1e-06 | 2.9e-06 | 2.6e-06 | 2.867E-06 | 2.517E-07 | 2.600E-06 | 3.100E-06 |
| 15912 | 796 | S796K | B | 1.8e-06 | 1.8e-06 | 1.9e-06 | 1.833E-06 | 5.774E-08 | 1.800E-06 | 1.900E-06 |
| 15913 | 796 | S796M | B | 2.7e-06 | 3.1e-06 | 1.2e-06 | 2.333E-06 | 1.002E-06 | 1.200E-06 | 3.100E-06 |
| 15914 | 796 | S796F | B | 2.1e-06 | 2e-06 | 2.4e-06 | 2.167E-06 | 2.082E-07 | 2.000E-06 | 2.400E-06 |
| 15915 | 796 | S796P | B | 3.4e-06 | 1.3e-06 | 2.5e-06 | 2.400E-06 | 1.054E-06 | 1.300E-06 | 3.400E-06 |
| 15916 | 796 | S796S | B | 1.9e-06 | 2.4e-06 | 2.4e-06 | 2.233E-06 | 2.887E-07 | 1.900E-06 | 2.400E-06 |
| 15917 | 796 | S796T | B | 1.2e-06 | 1.7e-06 | 3.1e-06 | 2.000E-06 | 9.849E-07 | 1.200E-06 | 3.100E-06 |
| 15918 | 796 | S796W | B | 3.8e-06 | 2.4e-06 | 2.7e-06 | 2.967E-06 | 7.371E-07 | 2.400E-06 | 3.800E-06 |
| 15919 | 796 | S796Y | B | 1.7e-06 | 6e-06 | 1.8e-06 | 3.167E-06 | 2.454E-06 | 1.700E-06 | 6.000E-06 |
| 15920 | 796 | S796V | B | 2.5e-06 | 2.5e-06 | 2.8e-06 | 2.600E-06 | 1.732E-07 | 2.500E-06 | 2.800E-06 |
| 15921 | 797 | L797A | B | 2.3e-06 | 3.3e-06 | 3.5e-06 | 3.033E-06 | 6.429E-07 | 2.300E-06 | 3.500E-06 |
| 15922 | 797 | L797R | B | 2.2e-06 | 2.5e-06 | 2.9e-06 | 2.533E-06 | 3.512E-07 | 2.200E-06 | 2.900E-06 |
| 15923 | 797 | L797N | B | 2.6e-06 | 1.5e-06 | 1.9e-06 | 2.000E-06 | 5.568E-07 | 1.500E-06 | 2.600E-06 |
| 15924 | 797 | L797D | B | 2.2e-06 | 2.5e-06 | 2.8e-06 | 2.500E-06 | 3.000E-07 | 2.200E-06 | 2.800E-06 |
| 15925 | 797 | L797C | B | 1.3e-06 | 2.6e-06 | 2.1e-06 | 2.000E-06 | 6.557E-07 | 1.300E-06 | 2.600E-06 |
| 15926 | 797 | L797E | B | 7.4e-07 | 2.7e-06 | 1.7e-06 | 1.713E-06 | 9.801E-07 | 7.400E-07 | 2.700E-06 |
| 15927 | 797 | L797Q | B | 2.8e-06 | 3.4e-06 | 1.3e-06 | 2.500E-06 | 1.082E-06 | 1.300E-06 | 3.400E-06 |
| 15928 | 797 | L797G | B | 1.6e-06 | 3.4e-06 | 2.1e-06 | 2.367E-06 | 9.292E-07 | 1.600E-06 | 3.400E-06 |
| 15929 | 797 | L797H | B | 2.2e-06 | 1e-06 | 2.2e-06 | 1.800E-06 | 6.928E-07 | 1.000E-06 | 2.200E-06 |
| 15930 | 797 | L797I | B | 2.6e-06 | 2.8e-06 | 3.1e-06 | 2.833E-06 | 2.517E-07 | 2.600E-06 | 3.100E-06 |
| 15931 | 797 | L797L | B | 2.8e-06 | 1.9e-06 | 2.7e-06 | 2.467E-06 | 4.933E-07 | 1.900E-06 | 2.800E-06 |
| 15932 | 797 | L797K | B | 1.8e-06 | 2.3e-06 | 1.5e-06 | 1.867E-06 | 4.041E-07 | 1.500E-06 | 2.300E-06 |
| 15933 | 797 | L797M | B | 2.3e-06 | 1.7e-06 | 1.9e-06 | 1.967E-06 | 3.055E-07 | 1.700E-06 | 2.300E-06 |
| 15934 | 797 | L797F | B | 2e-06 | 2.8e-06 | 3.5e-06 | 2.767E-06 | 7.506E-07 | 2.000E-06 | 3.500E-06 |
| 15935 | 797 | L797P | B | 1.8e-06 | 2.7e-06 | 3.5e-06 | 2.667E-06 | 8.505E-07 | 1.800E-06 | 3.500E-06 |
| 15936 | 797 | L797S | B | 2.1e-06 | 1.4e-06 | 1.8e-06 | 1.767E-06 | 3.512E-07 | 1.400E-06 | 2.100E-06 |
| 15937 | 797 | L797T | B | 2.9e-06 | 2e-06 | 3.2e-06 | 2.700E-06 | 6.245E-07 | 2.000E-06 | 3.200E-06 |
| 15938 | 797 | L797W | B | 1.6e-06 | 2.7e-06 | 2.1e-06 | 2.133E-06 | 5.508E-07 | 1.600E-06 | 2.700E-06 |
| 15939 | 797 | L797Y | B | 2.2e-06 | 3e-06 | 1.6e-06 | 2.267E-06 | 7.024E-07 | 1.600E-06 | 3.000E-06 |
| 15940 | 797 | L797V | B | 2.3e-06 | 1.5e-06 | 3.3e-06 | 2.367E-06 | 9.018E-07 | 1.500E-06 | 3.300E-06 |
| 15941 | 798 | S798A | B | 1.3e-06 | 3.2e-06 | 2.6e-06 | 2.367E-06 | 9.713E-07 | 1.300E-06 | 3.200E-06 |
| 15942 | 798 | S798R | B | 2.6e-06 | 2e-06 | 2.6e-06 | 2.400E-06 | 3.464E-07 | 2.000E-06 | 2.600E-06 |
| 15943 | 798 | S798N | B | 2.5e-06 | 2.3e-06 | 2.7e-06 | 2.500E-06 | 2.000E-07 | 2.300E-06 | 2.700E-06 |
| 15944 | 798 | S798D | B | 1.8e-06 | 2.5e-06 | 1.9e-06 | 2.067E-06 | 3.786E-07 | 1.800E-06 | 2.500E-06 |
| 15945 | 798 | S798C | B | 3e-06 | 2.2e-06 | 4.6e-06 | 3.267E-06 | 1.222E-06 | 2.200E-06 | 4.600E-06 |
| 15946 | 798 | S798E | B | 3e-06 | 3.1e-06 | 3.5e-06 | 3.200E-06 | 2.646E-07 | 3.000E-06 | 3.500E-06 |
| 15947 | 798 | S798Q | B | 2.8e-06 | 2.3e-06 | 2e-06 | 2.367E-06 | 4.041E-07 | 2.000E-06 | 2.800E-06 |
| 15948 | 798 | S798G | B | 2.1e-06 | 3.4e-06 | 2.7e-06 | 2.733E-06 | 6.506E-07 | 2.100E-06 | 3.400E-06 |
| 15949 | 798 | S798H | B | 2.6e-06 | 3.3e-06 | 1.5e-06 | 2.467E-06 | 9.074E-07 | 1.500E-06 | 3.300E-06 |
| 15950 | 798 | S798I | B | 2.2e-06 | 1.1e-06 | 2.1e-06 | 1.800E-06 | 6.083E-07 | 1.100E-06 | 2.200E-06 |
| 15951 | 798 | S798L | B | 3.3e-06 | 2.5e-06 | 3.1e-06 | 2.967E-06 | 4.163E-07 | 2.500E-06 | 3.300E-06 |
| 15952 | 798 | S798K | B | 1.6e-06 | 1.1e-06 | 2.4e-06 | 1.700E-06 | 6.557E-07 | 1.100E-06 | 2.400E-06 |
| 15953 | 798 | S798M | B | 1.7e-06 | 2.6e-06 | 2.6e-06 | 2.300E-06 | 5.196E-07 | 1.700E-06 | 2.600E-06 |
| 15954 | 798 | S798F | B | 2e-06 | 3.3e-06 | 3.1e-06 | 2.800E-06 | 7.000E-07 | 2.000E-06 | 3.300E-06 |
| 15955 | 798 | S798P | B | 3e-06 | 3.5e-06 | 4e-06 | 3.500E-06 | 5.000E-07 | 3.000E-06 | 4.000E-06 |
| 15956 | 798 | S798S | B | 1.9e-06 | 5.4e-06 | 1.5e-06 | 2.933E-06 | 2.146E-06 | 1.500E-06 | 5.400E-06 |
| 15957 | 798 | S798T | B | 1.8e-06 | 1.3e-06 | 2.3e-06 | 1.800E-06 | 5.000E-07 | 1.300E-06 | 2.300E-06 |
| 15958 | 798 | S798W | B | 1.6e-06 | 1.7e-06 | 2.9e-06 | 2.067E-06 | 7.234E-07 | 1.600E-06 | 2.900E-06 |
| 15959 | 798 | S798Y | B | 1.2e-06 | 2.5e-06 | 1.6e-06 | 1.767E-06 | 6.658E-07 | 1.200E-06 | 2.500E-06 |
| 15960 | 798 | S798V | B | 1.9e-06 | 2.7e-06 | 3.8e-06 | 2.800E-06 | 9.539E-07 | 1.900E-06 | 3.800E-06 |
| 15961 | 799 | S799A | B | 2.8e-06 | 1.9e-06 | 2.5e-06 | 2.400E-06 | 4.583E-07 | 1.900E-06 | 2.800E-06 |
| 15962 | 799 | S799R | B | 2.6e-06 | 2.6e-06 | 2.5e-06 | 2.567E-06 | 5.774E-08 | 2.500E-06 | 2.600E-06 |
| 15963 | 799 | S799N | B | 1.6e-06 | 2.7e-06 | 1.7e-06 | 2.000E-06 | 6.083E-07 | 1.600E-06 | 2.700E-06 |
| 15964 | 799 | S799D | B | 3.3e-06 | 2e-06 | 2.6e-06 | 2.633E-06 | 6.506E-07 | 2.000E-06 | 3.300E-06 |
| 15965 | 799 | S799C | B | 2.1e-06 | 3e-06 | 2.8e-06 | 2.633E-06 | 4.726E-07 | 2.100E-06 | 3.000E-06 |
| 15966 | 799 | S799E | B | 1.9e-06 | 2.4e-06 | 2.9e-06 | 2.400E-06 | 5.000E-07 | 1.900E-06 | 2.900E-06 |
| 15967 | 799 | S799Q | B | 4.7e-06 | 2.8e-06 | 1.5e-06 | 3.000E-06 | 1.609E-06 | 1.500E-06 | 4.700E-06 |
| 15968 | 799 | S799G | B | 1.5e-06 | 1.6e-06 | 2.3e-06 | 1.800E-06 | 4.359E-07 | 1.500E-06 | 2.300E-06 |
| 15969 | 799 | S799H | B | 1.3e-06 | 2.4e-06 | 3.7e-06 | 2.467E-06 | 1.201E-06 | 1.300E-06 | 3.700E-06 |
| 15970 | 799 | S799I | B | 2.9e-06 | 3.1e-06 | 3.4e-06 | 3.133E-06 | 2.517E-07 | 2.900E-06 | 3.400E-06 |
| 15971 | 799 | S799L | B | 2e-06 | 2e-06 | 3.8e-06 | 2.600E-06 | 1.039E-06 | 2.000E-06 | 3.800E-06 |
| 15972 | 799 | S799K | B | 3.7e-06 | 2.7e-06 | 2.3e-06 | 2.900E-06 | 7.211E-07 | 2.300E-06 | 3.700E-06 |
| 15973 | 799 | S799M | B | 2.7e-06 | 2.1e-06 | 3.1e-06 | 2.633E-06 | 5.033E-07 | 2.100E-06 | 3.100E-06 |
| 15974 | 799 | S799F | B | 2.6e-06 | 2.4e-06 | 3.2e-06 | 2.733E-06 | 4.163E-07 | 2.400E-06 | 3.200E-06 |
| 15975 | 799 | S799P | B | 3.1e-06 | 1.7e-06 | 2.6e-06 | 2.467E-06 | 7.095E-07 | 1.700E-06 | 3.100E-06 |
| 15976 | 799 | S799S | B | 1.6e-06 | 4.5e-06 | 2e-06 | 2.700E-06 | 1.572E-06 | 1.600E-06 | 4.500E-06 |
| 15977 | 799 | S799T | B | 3e-06 | 2.6e-06 | 2.2e-06 | 2.600E-06 | 4.000E-07 | 2.200E-06 | 3.000E-06 |
| 15978 | 799 | S799W | B | 2.1e-06 | 3.3e-06 | 3.2e-06 | 2.867E-06 | 6.658E-07 | 2.100E-06 | 3.300E-06 |
| 15979 | 799 | S799Y | B | 1.3e-06 | 2.4e-06 | 3e-06 | 2.233E-06 | 8.622E-07 | 1.300E-06 | 3.000E-06 |

| | | | | | | | | | | |
|---|---|---|---|---|---|---|---|---|---|---|
| 15980 | 799 | S799V | B | 1.8e-06 | 3.6e-06 | 2.6e-06 | 2.667E-06 | 9.018E-07 | 1.800E-06 | 3.600E-06 |
| 15981 | 800 | T800A | B | 1.7e-06 | 1.9e-06 | 2.5e-06 | 2.033E-06 | 4.163E-07 | 1.700E-06 | 2.500E-06 |
| 15982 | 800 | T800R | B | 1.9e-06 | 3.1e-06 | 2.2e-06 | 2.400E-06 | 6.245E-07 | 1.900E-06 | 3.100E-06 |
| 15983 | 800 | T800N | B | 2.2e-06 | 3.3e-06 | 2.1e-06 | 2.533E-06 | 6.658E-07 | 2.100E-06 | 3.300E-06 |
| 15984 | 800 | T800D | B | 1.9e-06 | 3.2e-06 | 1.4e-06 | 2.167E-06 | 9.292E-07 | 1.400E-06 | 3.200E-06 |
| 15985 | 800 | T800C | B | 1.9e-06 | 2.3e-06 | 1.5e-06 | 1.900E-06 | 4.000E-07 | 1.500E-06 | 2.300E-06 |
| 15986 | 800 | T800E | B | 2.5e-06 | 2.5e-06 | 2.9e-06 | 2.633E-06 | 2.309E-07 | 2.500E-06 | 2.900E-06 |
| 15987 | 800 | T800Q | B | 3e-06 | 3e-06 | 1.6e-06 | 2.533E-06 | 8.083E-07 | 1.600E-06 | 3.000E-06 |
| 15988 | 800 | T800G | B | 2.8e-06 | 2.8e-06 | 2.5e-06 | 2.700E-06 | 1.732E-07 | 2.500E-06 | 2.800E-06 |
| 15989 | 800 | T800H | B | 2.4e-06 | 3.2e-06 | 1.9e-06 | 2.500E-06 | 6.557E-07 | 1.900E-06 | 3.200E-06 |
| 15990 | 800 | T800I | B | 2e-06 | 2.9e-06 | 5.6e-06 | 3.500E-06 | 1.873E-06 | 2.000E-06 | 5.600E-06 |
| 15991 | 800 | T800L | B | 2.9e-06 | 1.6e-06 | 2.1e-06 | 2.200E-06 | 6.557E-07 | 1.600E-06 | 2.900E-06 |
| 15992 | 800 | T800K | B | 2.6e-06 | 5.1e-06 | 1.9e-06 | 3.200E-06 | 1.682E-06 | 1.900E-06 | 5.100E-06 |
| 15993 | 800 | T800M | B | 3.3e-06 | 1.9e-06 | 3.1e-06 | 2.767E-06 | 7.572E-07 | 1.900E-06 | 3.300E-06 |
| 15994 | 800 | T800F | B | 3.1e-06 | 2.9e-06 | 2.7e-06 | 2.900E-06 | 2.000E-07 | 2.700E-06 | 3.100E-06 |
| 15995 | 800 | T800P | B | 1.7e-06 | 1.4e-06 | 2.5e-06 | 1.867E-06 | 5.686E-07 | 1.400E-06 | 2.500E-06 |
| 15996 | 800 | T800S | B | 2.1e-06 | 3.1e-06 | 3.9e-06 | 3.033E-06 | 9.018E-07 | 2.100E-06 | 3.900E-06 |
| 15997 | 800 | T800T | B | 3.2e-06 | 2.5e-06 | 1.6e-06 | 2.433E-06 | 8.021E-07 | 1.600E-06 | 3.200E-06 |
| 15998 | 800 | T800W | B | 2.8e-06 | 1.6e-06 | 3.4e-06 | 2.600E-06 | 9.165E-07 | 1.600E-06 | 3.400E-06 |
| 15999 | 800 | T800Y | B | 1.5e-06 | 2.5e-06 | 2.4e-06 | 2.133E-06 | 5.508E-07 | 1.500E-06 | 2.500E-06 |
| 16000 | 800 | T800V | B | 2.7e-06 | 3e-06 | 1.8e-06 | 2.500E-06 | 6.245E-07 | 1.800E-06 | 3.000E-06 |
| 16001 | 801 | L801A | B | 1.8e-06 | 2.3e-06 | 1.7e-06 | 1.933E-06 | 3.215E-07 | 1.700E-06 | 2.300E-06 |
| 16002 | 801 | L801R | B | 1.6e-06 | 3.3e-06 | 1.9e-06 | 2.267E-06 | 9.074E-07 | 1.600E-06 | 3.300E-06 |
| 16003 | 801 | L801N | B | 2.8e-06 | 3.3e-06 | 1.6e-06 | 2.567E-06 | 8.737E-07 | 1.600E-06 | 3.300E-06 |
| 16004 | 801 | L801D | B | 1.8e-06 | 3.7e-06 | 2.5e-06 | 2.667E-06 | 9.609E-07 | 1.800E-06 | 3.700E-06 |
| 16005 | 801 | L801C | B | 3.3e-06 | 3e-06 | 3.1e-06 | 3.133E-06 | 1.528E-07 | 3.000E-06 | 3.300E-06 |
| 16006 | 801 | L801E | B | 2.9e-06 | 1.6e-06 | 2.5e-06 | 2.333E-06 | 6.658E-07 | 1.600E-06 | 2.900E-06 |
| 16007 | 801 | L801Q | B | 2.2e-06 | 1.3e-06 | 3.3e-06 | 2.267E-06 | 1.002E-06 | 1.300E-06 | 3.300E-06 |
| 16008 | 801 | L801G | B | 2.3e-06 | 2.1e-06 | 1.8e-06 | 2.067E-06 | 2.517E-07 | 1.800E-06 | 2.300E-06 |
| 16009 | 801 | L801H | B | 2e-06 | 3.3e-06 | 2.8e-06 | 2.700E-06 | 6.557E-07 | 2.000E-06 | 3.300E-06 |
| 16010 | 801 | L801I | B | 3.6e-06 | 2.3e-06 | 2.4e-06 | 2.767E-06 | 7.234E-07 | 2.300E-06 | 3.600E-06 |
| 16011 | 801 | L801L | B | 4.4e-06 | 3.1e-06 | 2.8e-06 | 3.433E-06 | 8.505E-07 | 2.800E-06 | 4.400E-06 |
| 16012 | 801 | L801K | B | 2.7e-06 | 1.8e-06 | 2.7e-06 | 2.400E-06 | 5.196E-07 | 1.800E-06 | 2.700E-06 |
| 16013 | 801 | L801M | B | 2.4e-06 | 1.4e-06 | 2.1e-06 | 1.967E-06 | 5.132E-07 | 1.400E-06 | 2.400E-06 |
| 16014 | 801 | L801F | B | 3.1e-06 | 3e-06 | 2.8e-06 | 2.967E-06 | 1.528E-07 | 2.800E-06 | 3.100E-06 |
| 16015 | 801 | L801P | B | 2.3e-06 | 3e-06 | 1.9e-06 | 2.400E-06 | 5.568E-07 | 1.900E-06 | 3.000E-06 |
| 16016 | 801 | L801S | B | 3.2e-06 | 3.3e-06 | 2.2e-06 | 2.900E-06 | 6.083E-07 | 2.200E-06 | 3.300E-06 |
| 16017 | 801 | L801T | B | 5.2e-06 | 3e-06 | 1.2e-06 | 3.133E-06 | 2.003E-06 | 1.200E-06 | 5.200E-06 |
| 16018 | 801 | L801W | B | 3e-06 | 2.4e-06 | 3e-06 | 2.800E-06 | 3.464E-07 | 2.400E-06 | 3.000E-06 |
| 16019 | 801 | L801Y | B | 3.2e-06 | 1.8e-06 | 1.9e-06 | 2.300E-06 | 7.810E-07 | 1.800E-06 | 3.200E-06 |
| 16020 | 801 | L801V | B | 1.3e-06 | 3.3e-06 | 1.3e-06 | 1.967E-06 | 1.155E-06 | 1.300E-06 | 3.300E-06 |
| 16021 | 802 | T802A | B | 2.3e-06 | 2e-06 | 1.4e-06 | 1.900E-06 | 4.583E-07 | 1.400E-06 | 2.300E-06 |
| 16022 | 802 | T802R | B | 5.1e-06 | 2.1e-06 | 2.5e-06 | 3.233E-06 | 1.629E-06 | 2.100E-06 | 5.100E-06 |
| 16023 | 802 | T802N | B | 1.8e-06 | 1.9e-06 | 2e-06 | 1.900E-06 | 1.000E-07 | 1.800E-06 | 2.000E-06 |
| 16024 | 802 | T802D | B | 1.7e-06 | 3.2e-06 | 1.7e-06 | 2.200E-06 | 8.660E-07 | 1.700E-06 | 3.200E-06 |
| 16025 | 802 | T802C | B | 2.3e-06 | 2.1e-06 | 2.7e-06 | 2.367E-06 | 3.055E-07 | 2.100E-06 | 2.700E-06 |
| 16026 | 802 | T802E | B | 1.8e-06 | 1.9e-06 | 1.5e-06 | 1.733E-06 | 2.082E-07 | 1.500E-06 | 1.900E-06 |
| 16027 | 802 | T802Q | B | 2.4e-06 | 1.9e-06 | 2.6e-06 | 2.300E-06 | 3.606E-07 | 1.900E-06 | 2.600E-06 |
| 16028 | 802 | T802G | B | 1.9e-06 | 4e-06 | 2.9e-06 | 2.933E-06 | 1.050E-06 | 1.900E-06 | 4.000E-06 |
| 16029 | 802 | T802H | B | 2.3e-06 | 1.8e-06 | 2.3e-06 | 2.133E-06 | 2.887E-07 | 1.800E-06 | 2.300E-06 |
| 16030 | 802 | T802I | B | 1.7e-06 | 1.9e-06 | 2.6e-06 | 2.067E-06 | 4.726E-07 | 1.700E-06 | 2.600E-06 |
| 16031 | 802 | T802L | B | 2e-06 | 2.8e-06 | 2.4e-06 | 2.400E-06 | 4.000E-07 | 2.000E-06 | 2.800E-06 |
| 16032 | 802 | T802K | B | 2.7e-06 | 2.2e-06 | 3e-06 | 2.633E-06 | 4.041E-07 | 2.200E-06 | 3.000E-06 |
| 16033 | 802 | T802M | B | 4.1e-06 | 4.7e-06 | 3.9e-06 | 4.233E-06 | 4.163E-07 | 3.900E-06 | 4.700E-06 |
| 16034 | 802 | T802F | B | 3.1e-06 | 2.7e-06 | 2.2e-06 | 2.667E-06 | 4.509E-07 | 2.200E-06 | 3.100E-06 |
| 16035 | 802 | T802P | B | 1.8e-06 | 4.5e-06 | 2e-06 | 2.767E-06 | 1.504E-06 | 1.800E-06 | 4.500E-06 |
| 16036 | 802 | T802S | B | 1.5e-06 | 3.1e-06 | 2.9e-06 | 2.500E-06 | 8.718E-07 | 1.500E-06 | 3.100E-06 |
| 16037 | 802 | T802T | B | 4.1e-06 | 9e-07 | 1.2e-06 | 2.067E-06 | 1.767E-06 | 9.000E-07 | 4.100E-06 |
| 16038 | 802 | T802W | B | 4.3e-06 | 3e-06 | 2.1e-06 | 3.133E-06 | 1.106E-06 | 2.100E-06 | 4.300E-06 |
| 16039 | 802 | T802Y | B | 2.2e-06 | 2.2e-06 | 2.1e-06 | 2.167E-06 | 5.774E-08 | 2.100E-06 | 2.200E-06 |
| 16040 | 802 | T802V | B | 2.1e-06 | 1.5e-06 | 4e-06 | 2.533E-06 | 1.305E-06 | 1.500E-06 | 4.000E-06 |
| 16041 | 803 | L803A | B | 2.7e-06 | 1.8e-06 | 3e-06 | 2.500E-06 | 6.245E-07 | 1.800E-06 | 3.000E-06 |
| 16042 | 803 | L803R | B | 2e-06 | 2.6e-06 | 3.7e-06 | 2.767E-06 | 8.622E-07 | 2.000E-06 | 3.700E-06 |
| 16043 | 803 | L803N | B | 3e-06 | 1.4e-06 | 4.7e-06 | 3.033E-06 | 1.650E-06 | 1.400E-06 | 4.700E-06 |
| 16044 | 803 | L803D | B | 1.2e-06 | 2.5e-06 | 1.8e-06 | 1.833E-06 | 6.506E-07 | 1.200E-06 | 2.500E-06 |
| 16045 | 803 | L803C | B | 2.6e-06 | 2.9e-06 | 2.7e-06 | 2.733E-06 | 1.528E-07 | 2.600E-06 | 2.900E-06 |
| 16046 | 803 | L803E | B | 2.8e-06 | 2.2e-06 | 2.4e-06 | 2.467E-06 | 3.055E-07 | 2.200E-06 | 2.800E-06 |
| 16047 | 803 | L803Q | B | 2.1e-06 | 1.4e-06 | 1.6e-06 | 1.700E-06 | 3.606E-07 | 1.400E-06 | 2.100E-06 |
| 16048 | 803 | L803G | B | 2.4e-06 | 2.2e-06 | 2.8e-06 | 2.467E-06 | 3.055E-07 | 2.200E-06 | 2.800E-06 |
| 16049 | 803 | L803H | B | 2.4e-06 | 1.4e-06 | 1.9e-06 | 1.900E-06 | 5.000E-07 | 1.400E-06 | 2.400E-06 |
| 16050 | 803 | L803I | B | 3.4e-06 | 1.5e-06 | 2.5e-06 | 2.467E-06 | 9.504E-07 | 1.500E-06 | 3.400E-06 |
| 16051 | 803 | L803L | B | 2.2e-06 | 1.9e-06 | 1.2e-06 | 1.767E-06 | 5.132E-07 | 1.200E-06 | 2.200E-06 |
| 16052 | 803 | L803K | B | 2.3e-06 | 3.3e-06 | 1.6e-06 | 2.400E-06 | 8.544E-07 | 1.600E-06 | 3.300E-06 |
| 16053 | 803 | L803M | B | 2.6e-06 | 2.4e-06 | 1.9e-06 | 2.300E-06 | 3.606E-07 | 1.900E-06 | 2.600E-06 |
| 16054 | 803 | L803F | B | 1.8e-06 | 6.4e-06 | 2.3e-06 | 3.500E-06 | 2.524E-06 | 1.800E-06 | 6.400E-06 |
| 16055 | 803 | L803P | B | 2.6e-06 | 1.4e-06 | 3.2e-06 | 2.400E-06 | 9.165E-07 | 1.400E-06 | 3.200E-06 |
| 16056 | 803 | L803S | B | 4.3e-06 | 1.7e-06 | 3.1e-06 | 3.033E-06 | 1.301E-06 | 1.700E-06 | 4.300E-06 |
| 16057 | 803 | L803T | B | 1.9e-06 | 2.4e-06 | 3e-06 | 2.433E-06 | 5.508E-07 | 1.900E-06 | 3.000E-06 |
| 16058 | 803 | L803W | B | 2.6e-06 | 3.2e-06 | 1.4e-06 | 2.400E-06 | 9.165E-07 | 1.400E-06 | 3.200E-06 |
| 16059 | 803 | L803Y | B | 2e-06 | 1.1e-06 | 1.7e-06 | 1.600E-06 | 4.583E-07 | 1.100E-06 | 2.000E-06 |
| 16060 | 803 | L803V | B | 2.8e-06 | 3.3e-06 | 1.6e-06 | 2.567E-06 | 8.737E-07 | 1.600E-06 | 3.300E-06 |
| 16061 | 804 | S804A | B | 1.4e-06 | 1.6e-06 | 1.5e-06 | 1.500E-06 | 1.000E-07 | 1.400E-06 | 1.600E-06 |
| 16062 | 804 | S804R | B | 2.3e-06 | 5e-06 | 1.7e-06 | 3.000E-06 | 1.758E-06 | 1.700E-06 | 5.000E-06 |
| 16063 | 804 | S804N | B | 1.4e-06 | 2.3e-06 | 1.6e-06 | 1.767E-06 | 4.726E-07 | 1.400E-06 | 2.300E-06 |
| 16064 | 804 | S804D | B | 1.6e-06 | 3.4e-06 | 2.2e-06 | 2.400E-06 | 9.165E-07 | 1.600E-06 | 3.400E-06 |

| | | | | | | | | | | |
|---|---|---|---|---|---|---|---|---|---|---|
| 16065 | 804 | S804C | B | 1.8e-06 | 6.1e-06 | 3.5e-06 | 3.800E-06 | 2.166E-06 | 1.800E-06 | 6.100E-06 |
| 16066 | 804 | S804E | B | 2.8e-06 | 1.7e-06 | 2.3e-06 | 2.267E-06 | 5.508E-07 | 1.700E-06 | 2.800E-06 |
| 16067 | 804 | S804Q | B | 1.6e-06 | 2.1e-06 | 2e-06 | 1.900E-06 | 2.646E-07 | 1.600E-06 | 2.100E-06 |
| 16068 | 804 | S804G | B | 2.8e-06 | 3.9e-06 | 2.2e-06 | 2.967E-06 | 8.622E-07 | 2.200E-06 | 3.900E-06 |
| 16069 | 804 | S804H | B | 3.7e-06 | 2.8e-06 | 3.1e-06 | 3.200E-06 | 4.583E-07 | 2.800E-06 | 3.700E-06 |
| 16070 | 804 | S804I | B | 2.4e-06 | 2.2e-06 | 2.8e-06 | 2.467E-06 | 3.055E-07 | 2.200E-06 | 2.800E-06 |
| 16071 | 804 | S804L | B | 1.5e-06 | 2e-06 | 2.5e-06 | 2.000E-06 | 5.000E-07 | 1.500E-06 | 2.500E-06 |
| 16072 | 804 | S804K | B | 3.7e-06 | 2.7e-06 | 3.1e-06 | 3.167E-06 | 5.033E-07 | 2.700E-06 | 3.700E-06 |
| 16073 | 804 | S804M | B | 1.4e-06 | 2.5e-06 | 2.1e-06 | 2.000E-06 | 5.568E-07 | 1.400E-06 | 2.500E-06 |
| 16074 | 804 | S804F | B | 3.1e-06 | 1.5e-06 | 2.7e-06 | 2.433E-06 | 8.327E-07 | 1.500E-06 | 3.100E-06 |
| 16075 | 804 | S804P | B | 1.4e-06 | 2.6e-06 | 1.9e-06 | 1.967E-06 | 6.028E-07 | 1.400E-06 | 2.600E-06 |
| 16076 | 804 | S804S | B | 3e-06 | 2.8e-06 | 2.6e-06 | 2.800E-06 | 2.000E-07 | 2.600E-06 | 3.000E-06 |
| 16077 | 804 | S804T | B | 3.3e-06 | 2.8e-06 | 2.8e-06 | 2.967E-06 | 2.887E-07 | 2.800E-06 | 3.300E-06 |
| 16078 | 804 | S804W | B | 2.5e-06 | 1.8e-06 | 1.7e-06 | 2.000E-06 | 4.359E-07 | 1.700E-06 | 2.500E-06 |
| 16079 | 804 | S804Y | B | 1.5e-06 | 2.4e-06 | 4.4e-06 | 2.767E-06 | 1.484E-06 | 1.500E-06 | 4.400E-06 |
| 16080 | 804 | S804V | B | 1.6e-06 | 1.8e-06 | 2.9e-06 | 2.100E-06 | 7.000E-07 | 1.600E-06 | 2.900E-06 |
| 16081 | 805 | K805A | B | 2.6e-06 | 5.6e-06 | 2.4e-06 | 3.533E-06 | 1.793E-06 | 2.400E-06 | 5.600E-06 |
| 16082 | 805 | K805R | B | 1.4e-06 | 1.8e-06 | 2.2e-06 | 1.800E-06 | 4.000E-07 | 1.400E-06 | 2.200E-06 |
| 16083 | 805 | K805N | B | 2.8e-06 | 1.7e-06 | 2.3e-06 | 2.267E-06 | 5.508E-07 | 1.700E-06 | 2.800E-06 |
| 16084 | 805 | K805D | B | 3.1e-06 | 1.7e-06 | 1.9e-06 | 2.233E-06 | 7.572E-07 | 1.700E-06 | 3.100E-06 |
| 16085 | 805 | K805C | B | 1.6e-06 | 3.1e-06 | 1.5e-06 | 2.067E-06 | 8.963E-07 | 1.500E-06 | 3.100E-06 |
| 16086 | 805 | K805E | B | 2.6e-06 | 2.5e-06 | 1.9e-06 | 2.333E-06 | 3.786E-07 | 1.900E-06 | 2.600E-06 |
| 16087 | 805 | K805Q | B | 2.2e-06 | 3.6e-06 | 2.5e-06 | 2.767E-06 | 7.371E-07 | 2.200E-06 | 3.600E-06 |
| 16088 | 805 | K805G | B | 1.9e-06 | 1.7e-06 | 1.7e-06 | 1.767E-06 | 1.155E-07 | 1.700E-06 | 1.900E-06 |
| 16089 | 805 | K805H | B | 3e-06 | 1.8e-06 | 1.8e-06 | 2.200E-06 | 6.928E-07 | 1.800E-06 | 3.000E-06 |
| 16090 | 805 | K805I | B | 1.9e-06 | 2.6e-06 | 2.4e-06 | 2.300E-06 | 3.606E-07 | 1.900E-06 | 2.600E-06 |
| 16091 | 805 | K805L | B | 1.9e-06 | 3.5e-06 | 1.2e-06 | 2.200E-06 | 1.179E-06 | 1.200E-06 | 3.500E-06 |
| 16092 | 805 | K805K | B | 1.3e-06 | 2.3e-06 | 2.4e-06 | 2.000E-06 | 6.083E-07 | 1.300E-06 | 2.400E-06 |
| 16093 | 805 | K805M | B | 2.8e-06 | 1.6e-06 | 2.6e-06 | 2.333E-06 | 6.429E-07 | 1.600E-06 | 2.800E-06 |
| 16094 | 805 | K805F | B | 1.9e-06 | 3.6e-06 | 1.5e-06 | 2.333E-06 | 1.115E-06 | 1.500E-06 | 3.600E-06 |
| 16095 | 805 | K805P | B | 3.5e-06 | 2.2e-06 | 2.2e-06 | 2.633E-06 | 7.506E-07 | 2.200E-06 | 3.500E-06 |
| 16096 | 805 | K805S | B | 1.7e-06 | 2.6e-06 | 2e-06 | 2.100E-06 | 4.583E-07 | 1.700E-06 | 2.600E-06 |
| 16097 | 805 | K805T | B | 1.5e-06 | 2.4e-06 | 3.5e-06 | 2.467E-06 | 1.002E-06 | 1.500E-06 | 3.500E-06 |
| 16098 | 805 | K805W | B | 1.8e-06 | 1.6e-06 | 5.1e-06 | 2.833E-06 | 1.966E-06 | 1.600E-06 | 5.100E-06 |
| 16099 | 805 | K805Y | B | 2.3e-06 | 3e-06 | 3.6e-06 | 2.967E-06 | 6.506E-07 | 2.300E-06 | 3.600E-06 |
| 16100 | 805 | K805V | B | 2.5e-06 | 2e-06 | 1.5e-06 | 2.000E-06 | 5.000E-07 | 1.500E-06 | 2.500E-06 |
| 16101 | 806 | A806A | B | 1.5e-06 | 2.5e-06 | 1.5e-06 | 1.833E-06 | 5.774E-07 | 1.500E-06 | 2.500E-06 |
| 16102 | 806 | A806R | B | 3.1e-06 | 2.2e-06 | 2.5e-06 | 2.600E-06 | 4.583E-07 | 2.200E-06 | 3.100E-06 |
| 16103 | 806 | A806N | B | 3.1e-06 | 3e-06 | 2.4e-06 | 2.833E-06 | 3.786E-07 | 2.400E-06 | 3.100E-06 |
| 16104 | 806 | A806D | B | 2.4e-06 | 2.6e-06 | 2.3e-06 | 2.433E-06 | 1.528E-07 | 2.300E-06 | 2.600E-06 |
| 16105 | 806 | A806C | B | 1.6e-06 | 2.2e-06 | 1.2e-06 | 1.667E-06 | 5.033E-07 | 1.200E-06 | 2.200E-06 |
| 16106 | 806 | A806E | B | 2.3e-06 | 1.4e-06 | 1.5e-06 | 1.733E-06 | 4.933E-07 | 1.400E-06 | 2.300E-06 |
| 16107 | 806 | A806Q | B | 2.4e-06 | 2.1e-06 | 2e-06 | 2.167E-06 | 2.082E-07 | 2.000E-06 | 2.400E-06 |
| 16108 | 806 | A806G | B | 3.2e-06 | 3.1e-06 | 2.1e-06 | 2.800E-06 | 6.083E-07 | 2.100E-06 | 3.200E-06 |
| 16109 | 806 | A806H | B | 2e-06 | 1.8e-06 | 1.9e-06 | 1.900E-06 | 1.000E-07 | 1.800E-06 | 2.000E-06 |
| 16110 | 806 | A806I | B | 2.4e-06 | 1.8e-06 | 1.9e-06 | 2.033E-06 | 3.215E-07 | 1.800E-06 | 2.400E-06 |
| 16111 | 806 | A806L | B | 2.9e-06 | 2.7e-06 | 2.6e-06 | 2.733E-06 | 1.528E-07 | 2.600E-06 | 2.900E-06 |
| 16112 | 806 | A806K | B | 3.2e-06 | 2.3e-06 | 2.4e-06 | 2.633E-06 | 4.933E-07 | 2.300E-06 | 3.200E-06 |
| 16113 | 806 | A806M | B | 2.3e-06 | 2.3e-06 | 1.9e-06 | 2.167E-06 | 2.309E-07 | 1.900E-06 | 2.300E-06 |
| 16114 | 806 | A806F | B | 2.8e-06 | 3.3e-06 | 2.5e-06 | 2.867E-06 | 4.041E-07 | 2.500E-06 | 3.300E-06 |
| 16115 | 806 | A806P | B | 2.7e-06 | 4.2e-06 | 3.8e-06 | 3.567E-06 | 7.767E-07 | 2.700E-06 | 4.200E-06 |
| 16116 | 806 | A806S | B | 3e-06 | 1.7e-06 | 1.9e-06 | 2.200E-06 | 7.000E-07 | 1.700E-06 | 3.000E-06 |
| 16117 | 806 | A806T | B | 4e-06 | 2e-06 | 1.8e-06 | 2.600E-06 | 1.217E-06 | 1.800E-06 | 4.000E-06 |
| 16118 | 806 | A806W | B | 2.2e-06 | 2.4e-06 | 1.1e-06 | 1.900E-06 | 7.000E-07 | 1.100E-06 | 2.400E-06 |
| 16119 | 806 | A806Y | B | 1.8e-06 | 1.5e-06 | 2.9e-06 | 2.067E-06 | 7.371E-07 | 1.500E-06 | 2.900E-06 |
| 16120 | 806 | A806V | B | 3.1e-06 | 1.6e-06 | 2.5e-06 | 2.400E-06 | 7.550E-07 | 1.600E-06 | 3.100E-06 |
| 16121 | 807 | D807A | B | 2.1e-06 | 4.1e-06 | 2.1e-06 | 2.767E-06 | 1.155E-06 | 2.100E-06 | 4.100E-06 |
| 16122 | 807 | D807R | B | 3.3e-06 | 2.8e-06 | 1.5e-06 | 2.533E-06 | 9.292E-07 | 1.500E-06 | 3.300E-06 |
| 16123 | 807 | D807N | B | 4.3e-06 | 2.8e-06 | 7.2e-06 | 4.767E-06 | 2.237E-06 | 2.800E-06 | 7.200E-06 |
| 16124 | 807 | D807D | B | 2.1e-06 | 1.8e-06 | 1.5e-06 | 1.800E-06 | 3.000E-07 | 1.500E-06 | 2.100E-06 |
| 16125 | 807 | D807C | B | 2e-06 | 1.6e-06 | 1.9e-06 | 1.833E-06 | 2.082E-07 | 1.600E-06 | 2.000E-06 |
| 16126 | 807 | D807E | B | 1.5e-06 | 3.3e-06 | 2.7e-06 | 2.500E-06 | 9.165E-07 | 1.500E-06 | 3.300E-06 |
| 16127 | 807 | D807Q | B | 2.6e-06 | 2.8e-06 | 1.8e-06 | 2.400E-06 | 5.292E-07 | 1.800E-06 | 2.800E-06 |
| 16128 | 807 | D807G | B | 3e-06 | 9.4e-07 | 2e-06 | 1.980E-06 | 1.030E-06 | 9.400E-07 | 3.000E-06 |
| 16129 | 807 | D807H | B | 3.1e-06 | 2.3e-06 | 1.8e-06 | 2.400E-06 | 6.557E-07 | 1.800E-06 | 3.100E-06 |
| 16130 | 807 | D807I | B | 3e-06 | 2.7e-06 | 2.5e-06 | 2.733E-06 | 2.517E-07 | 2.500E-06 | 3.000E-06 |
| 16131 | 807 | D807L | B | 1.6e-06 | 2.2e-06 | 2.5e-06 | 2.100E-06 | 4.583E-07 | 1.600E-06 | 2.500E-06 |
| 16132 | 807 | D807K | B | 3.1e-06 | 2.4e-06 | 1.6e-06 | 2.367E-06 | 7.506E-07 | 1.600E-06 | 3.100E-06 |
| 16133 | 807 | D807M | B | 2.3e-06 | 2.5e-06 | 1.4e-06 | 2.067E-06 | 5.859E-07 | 1.400E-06 | 2.500E-06 |
| 16134 | 807 | D807F | B | 4.3e-06 | 1.6e-06 | 3.4e-06 | 3.100E-06 | 1.375E-06 | 1.600E-06 | 4.300E-06 |
| 16135 | 807 | D807P | B | 1.9e-06 | 2.9e-06 | 1.9e-06 | 2.233E-06 | 5.774E-07 | 1.900E-06 | 2.900E-06 |
| 16136 | 807 | D807S | B | 1.8e-06 | 1.4e-06 | 2.7e-06 | 1.967E-06 | 6.658E-07 | 1.400E-06 | 2.700E-06 |
| 16137 | 807 | D807T | B | 2.5e-06 | 2e-06 | 1.9e-06 | 2.133E-06 | 3.215E-07 | 1.900E-06 | 2.500E-06 |
| 16138 | 807 | D807W | B | 2.5e-06 | 3.2e-06 | 2.5e-06 | 2.733E-06 | 4.041E-07 | 2.500E-06 | 3.200E-06 |
| 16139 | 807 | D807Y | B | 2.6e-06 | 3e-06 | 2.1e-06 | 2.567E-06 | 4.509E-07 | 2.100E-06 | 3.000E-06 |
| 16140 | 807 | D807V | B | 2.9e-06 | 2.4e-06 | 2.3e-06 | 2.533E-06 | 3.215E-07 | 2.300E-06 | 2.900E-06 |
| 16141 | 808 | Y808A | B | 2.4e-06 | 2.8e-06 | 1.5e-06 | 2.233E-06 | 6.658E-07 | 1.500E-06 | 2.800E-06 |
| 16142 | 808 | Y808R | B | 1.9e-06 | 1.7e-06 | 2.3e-06 | 1.967E-06 | 3.055E-07 | 1.700E-06 | 2.300E-06 |
| 16143 | 808 | Y808N | B | 1.4e-06 | 3.7e-06 | 1.7e-06 | 2.267E-06 | 1.250E-06 | 1.400E-06 | 3.700E-06 |
| 16144 | 808 | Y808D | B | 9.9e-07 | 3.2e-06 | 2e-06 | 2.063E-06 | 1.106E-06 | 9.900E-07 | 3.200E-06 |
| 16145 | 808 | Y808C | B | 3.2e-06 | 1.5e-06 | 2.5e-06 | 2.400E-06 | 8.544E-07 | 1.500E-06 | 3.200E-06 |
| 16146 | 808 | Y808E | B | 3.6e-06 | 1.8e-06 | 3e-06 | 2.800E-06 | 9.165E-07 | 1.800E-06 | 3.600E-06 |
| 16147 | 808 | Y808Q | B | 2.1e-06 | 3e-06 | 2.3e-06 | 2.467E-06 | 4.726E-07 | 2.100E-06 | 3.000E-06 |
| 16148 | 808 | Y808G | B | 2.8e-06 | 1.3e-06 | 3.7e-06 | 2.600E-06 | 1.212E-06 | 1.300E-06 | 3.700E-06 |
| 16149 | 808 | Y808H | B | 1.3e-06 | 3.4e-06 | 3.1e-06 | 2.600E-06 | 1.136E-06 | 1.300E-06 | 3.400E-06 |

| | | | | | | | | | | |
|---|---|---|---|---|---|---|---|---|---|---|
| 16150 | 808 | Y808I | B | 4.1e-06 | 2.4e-06 | 2.9e-06 | 3.133E-06 | 8.737E-07 | 2.400E-06 | 4.100E-06 |
| 16151 | 808 | Y808L | B | 2.4e-06 | 3.6e-06 | 2.3e-06 | 2.767E-06 | 7.234E-07 | 2.300E-06 | 3.600E-06 |
| 16152 | 808 | Y808K | B | 3.7e-06 | 4e-06 | 2.2e-06 | 3.300E-06 | 9.644E-07 | 2.200E-06 | 4.000E-06 |
| 16153 | 808 | Y808M | B | 5.2e-06 | 2.8e-06 | 2.2e-06 | 3.400E-06 | 1.587E-06 | 2.200E-06 | 5.200E-06 |
| 16154 | 808 | Y808F | B | 1.1e-06 | 1e-06 | 2.7e-06 | 1.600E-06 | 9.539E-07 | 1.000E-06 | 2.700E-06 |
| 16155 | 808 | Y808P | B | 2.8e-06 | 2.3e-06 | 2.7e-06 | 2.600E-06 | 2.646E-07 | 2.300E-06 | 2.800E-06 |
| 16156 | 808 | Y808S | B | 1.7e-06 | 1.5e-06 | 1.6e-06 | 1.600E-06 | 1.000E-07 | 1.500E-06 | 1.700E-06 |
| 16157 | 808 | Y808T | B | 2.1e-06 | 3.1e-06 | 2.2e-06 | 2.467E-06 | 5.508E-07 | 2.100E-06 | 3.100E-06 |
| 16158 | 808 | Y808W | B | 1.2e-06 | 4.9e-06 | 3.4e-06 | 3.167E-06 | 1.861E-06 | 1.200E-06 | 4.900E-06 |
| 16159 | 808 | Y808Y | B | 1.7e-06 | 1.9e-06 | 2.3e-06 | 1.967E-06 | 3.055E-07 | 1.700E-06 | 2.300E-06 |
| 16160 | 808 | Y808V | B | 2.2e-06 | 3.1e-06 | 2.2e-06 | 2.500E-06 | 5.196E-07 | 2.200E-06 | 3.100E-06 |
| 16161 | 809 | E809A | B | 2.6e-06 | 3.1e-06 | 4e-06 | 3.233E-06 | 7.095E-07 | 2.600E-06 | 4.000E-06 |
| 16162 | 809 | E809R | B | 8.1e-07 | 1.8e-06 | 1.7e-06 | 1.437E-06 | 5.450E-07 | 8.100E-07 | 1.800E-06 |
| 16163 | 809 | E809N | B | 2.7e-06 | 4.2e-06 | 1.8e-06 | 2.900E-06 | 1.212E-06 | 1.800E-06 | 4.200E-06 |
| 16164 | 809 | E809D | B | 1.9e-06 | 3.2e-06 | 2.1e-06 | 2.400E-06 | 7.000E-07 | 1.900E-06 | 3.200E-06 |
| 16165 | 809 | E809C | B | 2.7e-06 | 1.6e-06 | 3.7e-06 | 2.667E-06 | 1.050E-06 | 1.600E-06 | 3.700E-06 |
| 16166 | 809 | E809E | B | 5.8e-06 | 3.2e-06 | 2.7e-06 | 3.900E-06 | 1.664E-06 | 2.700E-06 | 5.800E-06 |
| 16167 | 809 | E809Q | B | 2.7e-06 | 1.8e-06 | 1.8e-06 | 2.100E-06 | 5.196E-07 | 1.800E-06 | 2.700E-06 |
| 16168 | 809 | E809G | B | 2.4e-06 | 3.9e-06 | 2e-06 | 2.767E-06 | 1.002E-06 | 2.000E-06 | 3.900E-06 |
| 16169 | 809 | E809H | B | 1.3e-06 | 2.7e-06 | 1.7e-06 | 1.900E-06 | 7.211E-07 | 1.300E-06 | 2.700E-06 |
| 16170 | 809 | E809I | B | 1.9e-06 | 1.9e-06 | 3.7e-06 | 2.500E-06 | 1.039E-06 | 1.900E-06 | 3.700E-06 |
| 16171 | 809 | E809L | B | 3.5e-06 | 3.2e-06 | 3.7e-06 | 3.467E-06 | 2.517E-07 | 3.200E-06 | 3.700E-06 |
| 16172 | 809 | E809K | B | 2.3e-06 | 3.2e-06 | 3.9e-06 | 3.133E-06 | 8.021E-07 | 2.300E-06 | 3.900E-06 |
| 16173 | 809 | E809M | B | 2.1e-06 | 1.4e-06 | 2.7e-06 | 2.067E-06 | 6.506E-07 | 1.400E-06 | 2.700E-06 |
| 16174 | 809 | E809F | B | 2.6e-06 | 2e-06 | 2.3e-06 | 2.300E-06 | 3.000E-07 | 2.000E-06 | 2.600E-06 |
| 16175 | 809 | E809P | B | 2.5e-06 | 6.1e-06 | 1.5e-06 | 3.367E-06 | 2.419E-06 | 1.500E-06 | 6.100E-06 |
| 16176 | 809 | E809S | B | 4.4e-06 | 2.6e-06 | 1.9e-06 | 2.967E-06 | 1.290E-06 | 1.900E-06 | 4.400E-06 |
| 16177 | 809 | E809T | B | 2.4e-06 | 2.5e-06 | 2.6e-06 | 2.500E-06 | 1.000E-07 | 2.400E-06 | 2.600E-06 |
| 16178 | 809 | E809W | B | 1.5e-06 | 2.8e-06 | 2.1e-06 | 2.133E-06 | 6.506E-07 | 1.500E-06 | 2.800E-06 |
| 16179 | 809 | E809Y | B | 1.9e-06 | 2.5e-06 | 1.9e-06 | 2.100E-06 | 3.464E-07 | 1.900E-06 | 2.500E-06 |
| 16180 | 809 | E809V | B | 4.3e-06 | 2.8e-06 | 1.4e-06 | 2.833E-06 | 1.450E-06 | 1.400E-06 | 4.300E-06 |
| 16181 | 810 | K810A | B | 2.5e-06 | 1.9e-06 | 3.4e-06 | 2.600E-06 | 7.550E-07 | 1.900E-06 | 3.400E-06 |
| 16182 | 810 | K810R | B | 2.9e-06 | 2e-06 | 1.9e-06 | 2.267E-06 | 5.508E-07 | 1.900E-06 | 2.900E-06 |
| 16183 | 810 | K810N | B | 1.3e-06 | 1.6e-06 | 3.9e-06 | 2.267E-06 | 1.422E-06 | 1.300E-06 | 3.900E-06 |
| 16184 | 810 | K810D | B | 3.3e-06 | 2.4e-06 | 1.6e-06 | 2.433E-06 | 8.505E-07 | 1.600E-06 | 3.300E-06 |
| 16185 | 810 | K810C | B | 2.3e-06 | 2.7e-06 | 2.9e-06 | 2.633E-06 | 3.055E-07 | 2.300E-06 | 2.900E-06 |
| 16186 | 810 | K810E | B | 3.4e-06 | 3e-06 | 3e-06 | 3.133E-06 | 2.309E-07 | 3.000E-06 | 3.400E-06 |
| 16187 | 810 | K810Q | B | 2.6e-06 | 5e-06 | 2.6e-06 | 3.400E-06 | 1.386E-06 | 2.600E-06 | 5.000E-06 |
| 16188 | 810 | K810G | B | 3e-06 | 1.7e-06 | 2.7e-06 | 2.467E-06 | 6.807E-07 | 1.700E-06 | 3.000E-06 |
| 16189 | 810 | K810H | B | 2.6e-06 | 1.5e-06 | 3.4e-06 | 2.500E-06 | 9.539E-07 | 1.500E-06 | 3.400E-06 |
| 16190 | 810 | K810I | B | 1.6e-06 | 2.2e-06 | 3.2e-06 | 2.333E-06 | 8.083E-07 | 1.600E-06 | 3.200E-06 |
| 16191 | 810 | K810L | B | 2.8e-06 | 2.2e-06 | 2.3e-06 | 2.433E-06 | 3.215E-07 | 2.200E-06 | 2.800E-06 |
| 16192 | 810 | K810K | B | 2.2e-06 | 2.2e-06 | 3.8e-06 | 2.733E-06 | 9.238E-07 | 2.200E-06 | 3.800E-06 |
| 16193 | 810 | K810M | B | 1.6e-06 | 2.6e-06 | 2.2e-06 | 2.133E-06 | 5.033E-07 | 1.600E-06 | 2.600E-06 |
| 16194 | 810 | K810F | B | 3.9e-06 | 1.5e-06 | 1.9e-06 | 2.433E-06 | 1.286E-06 | 1.500E-06 | 3.900E-06 |
| 16195 | 810 | K810P | B | 1.5e-06 | 1.9e-06 | 2e-06 | 1.800E-06 | 2.646E-07 | 1.500E-06 | 2.000E-06 |
| 16196 | 810 | K810S | B | 3.3e-06 | 3.6e-06 | 2.5e-06 | 3.133E-06 | 5.686E-07 | 2.500E-06 | 3.600E-06 |
| 16197 | 810 | K810T | B | 2.4e-06 | 1.7e-06 | 2.5e-06 | 2.200E-06 | 4.359E-07 | 1.700E-06 | 2.500E-06 |
| 16198 | 810 | K810W | B | 2.3e-06 | 3.4e-06 | 2.5e-06 | 2.733E-06 | 5.859E-07 | 2.300E-06 | 3.400E-06 |
| 16199 | 810 | K810Y | B | 1.7e-06 | 2e-06 | 1.6e-06 | 1.767E-06 | 2.082E-07 | 1.600E-06 | 2.000E-06 |
| 16200 | 810 | K810V | B | 3.6e-06 | 1.5e-06 | 1.1e-06 | 2.067E-06 | 1.343E-06 | 1.100E-06 | 3.600E-06 |
| 16201 | 811 | H811A | B | 2.5e-06 | 2.6e-06 | 1.8e-06 | 2.300E-06 | 4.359E-07 | 1.800E-06 | 2.600E-06 |
| 16202 | 811 | H811R | B | 9.4e-07 | 2.8e-06 | 4.3e-06 | 2.680E-06 | 1.683E-06 | 9.400E-07 | 4.300E-06 |
| 16203 | 811 | H811N | B | 3.2e-06 | 3.3e-06 | 1.7e-06 | 2.733E-06 | 8.963E-07 | 1.700E-06 | 3.300E-06 |
| 16204 | 811 | H811D | B | 3.5e-06 | 1.9e-06 | 1.7e-06 | 2.367E-06 | 9.866E-07 | 1.700E-06 | 3.500E-06 |
| 16205 | 811 | H811C | B | 1.4e-06 | 5.3e-06 | 2.4e-06 | 3.033E-06 | 2.026E-06 | 1.400E-06 | 5.300E-06 |
| 16206 | 811 | H811E | B | 1.8e-06 | 2.7e-06 | 2.9e-06 | 2.467E-06 | 5.859E-07 | 1.800E-06 | 2.900E-06 |
| 16207 | 811 | H811Q | B | 7.1e-06 | 3.6e-06 | 2.1e-06 | 4.267E-06 | 2.566E-06 | 2.100E-06 | 7.100E-06 |
| 16208 | 811 | H811G | B | 2.4e-06 | 2.9e-06 | 2.8e-06 | 2.700E-06 | 2.646E-07 | 2.400E-06 | 2.900E-06 |
| 16209 | 811 | H811H | B | 1.8e-06 | 2.2e-06 | 4.8e-06 | 2.933E-06 | 1.629E-06 | 1.800E-06 | 4.800E-06 |
| 16210 | 811 | H811I | B | 2.2e-06 | 3e-06 | 5.6e-06 | 3.600E-06 | 1.778E-06 | 2.200E-06 | 5.600E-06 |
| 16211 | 811 | H811L | B | 1.6e-06 | 1.6e-06 | 2.9e-06 | 2.033E-06 | 7.506E-07 | 1.600E-06 | 2.900E-06 |
| 16212 | 811 | H811K | B | 2.4e-06 | 3.3e-06 | 9.4e-07 | 2.213E-06 | 1.191E-06 | 9.400E-07 | 3.300E-06 |
| 16213 | 811 | H811M | B | 2.6e-06 | 2.6e-06 | 1.6e-06 | 2.267E-06 | 5.774E-07 | 1.600E-06 | 2.600E-06 |
| 16214 | 811 | H811F | B | 3e-06 | 2.1e-06 | 2.7e-06 | 2.600E-06 | 4.583E-07 | 2.100E-06 | 3.000E-06 |
| 16215 | 811 | H811P | B | 3.1e-06 | 1.6e-06 | 1.7e-06 | 2.133E-06 | 8.386E-07 | 1.600E-06 | 3.100E-06 |
| 16216 | 811 | H811S | B | 2.9e-06 | 4.5e-06 | 3.2e-06 | 3.533E-06 | 8.505E-07 | 2.900E-06 | 4.500E-06 |
| 16217 | 811 | H811T | B | 3.2e-06 | 2.6e-06 | 2.7e-06 | 2.833E-06 | 3.215E-07 | 2.600E-06 | 3.200E-06 |
| 16218 | 811 | H811W | B | 3.8e-06 | 3.1e-06 | 1.6e-06 | 2.833E-06 | 1.124E-06 | 1.600E-06 | 3.800E-06 |
| 16219 | 811 | H811Y | B | 2.8e-06 | 2.9e-06 | 2.3e-06 | 2.667E-06 | 3.215E-07 | 2.300E-06 | 2.900E-06 |
| 16220 | 811 | H811V | B | 2.8e-06 | 2e-06 | 2.5e-06 | 2.433E-06 | 4.041E-07 | 2.000E-06 | 2.800E-06 |
| 16221 | 812 | K812A | B | 1.4e-06 | 2.8e-06 | 1.7e-06 | 1.967E-06 | 7.371E-07 | 1.400E-06 | 2.800E-06 |
| 16222 | 812 | K812R | B | 2.1e-06 | 2e-06 | 3.7e-06 | 2.600E-06 | 9.539E-07 | 2.000E-06 | 3.700E-06 |
| 16223 | 812 | K812N | B | 3.1e-06 | 2.9e-06 | 1.7e-06 | 2.567E-06 | 7.572E-07 | 1.700E-06 | 3.100E-06 |
| 16224 | 812 | K812D | B | 2.6e-06 | 2.8e-06 | 2.1e-06 | 2.500E-06 | 3.606E-07 | 2.100E-06 | 2.800E-06 |
| 16225 | 812 | K812C | B | 2.5e-06 | 2.2e-06 | 2.5e-06 | 2.400E-06 | 1.732E-07 | 2.200E-06 | 2.500E-06 |
| 16226 | 812 | K812E | B | 1.8e-06 | 2.5e-06 | 2.8e-06 | 2.367E-06 | 5.132E-07 | 1.800E-06 | 2.800E-06 |
| 16227 | 812 | K812Q | B | 2.1e-06 | 2.2e-06 | 8.8e-07 | 1.727E-06 | 7.349E-07 | 8.800E-07 | 2.200E-06 |
| 16228 | 812 | K812G | B | 2.9e-06 | 3e-06 | 1.6e-06 | 2.500E-06 | 7.810E-07 | 1.600E-06 | 3.000E-06 |
| 16229 | 812 | K812H | B | 2.8e-06 | 1.9e-06 | 2.7e-06 | 2.467E-06 | 4.933E-07 | 1.900E-06 | 2.800E-06 |
| 16230 | 812 | K812I | B | 5.2e-06 | 1.8e-06 | 3.4e-06 | 3.467E-06 | 1.701E-06 | 1.800E-06 | 5.200E-06 |
| 16231 | 812 | K812L | B | 3.2e-06 | 3.2e-06 | 1.4e-06 | 2.600E-06 | 1.039E-06 | 1.400E-06 | 3.200E-06 |
| 16232 | 812 | K812K | B | 1.6e-06 | 2.8e-06 | 2.5e-06 | 2.300E-06 | 6.245E-07 | 1.600E-06 | 2.800E-06 |
| 16233 | 812 | K812M | B | 3.9e-06 | 2.2e-06 | 1.3e-06 | 2.467E-06 | 1.320E-06 | 1.300E-06 | 3.900E-06 |
| 16234 | 812 | K812F | B | 4.2e-06 | 2.6e-06 | 1.7e-06 | 2.833E-06 | 1.266E-06 | 1.700E-06 | 4.200E-06 |

| | | | | | | | | | | |
|---|---|---|---|---|---|---|---|---|---|---|
| 16235 | 812 | K812P | B | 2.9e-06 | 1.7e-06 | 2.3e-06 | 2.300E-06 | 6.000E-07 | 1.700E-06 | 2.900E-06 |
| 16236 | 812 | K812S | B | 2.8e-06 | 4.1e-06 | 1.8e-06 | 2.900E-06 | 1.153E-06 | 1.800E-06 | 4.100E-06 |
| 16237 | 812 | K812T | B | 2.4e-06 | 2.4e-06 | 1.6e-06 | 2.133E-06 | 4.619E-07 | 1.600E-06 | 2.400E-06 |
| 16238 | 812 | K812W | B | 3.2e-06 | 1.3e-06 | 2.5e-06 | 2.333E-06 | 9.609E-07 | 1.300E-06 | 3.200E-06 |
| 16239 | 812 | K812Y | B | 2.2e-06 | 2.3e-06 | 2.3e-06 | 2.267E-06 | 5.774E-08 | 2.200E-06 | 2.300E-06 |
| 16240 | 812 | K812V | B | 2.7e-06 | 3.8e-06 | 3.2e-06 | 3.233E-06 | 5.508E-07 | 2.700E-06 | 3.800E-06 |
| 16241 | 813 | V813A | B | 3.4e-06 | 2e-06 | 2.3e-06 | 2.567E-06 | 7.371E-07 | 2.000E-06 | 3.400E-06 |
| 16242 | 813 | V813R | B | 1.1e-06 | 1.8e-06 | 2.3e-06 | 1.733E-06 | 6.028E-07 | 1.100E-06 | 2.300E-06 |
| 16243 | 813 | V813N | B | 2.6e-06 | 2.1e-06 | 1.8e-06 | 2.167E-06 | 4.041E-07 | 1.800E-06 | 2.600E-06 |
| 16244 | 813 | V813D | B | 3.2e-06 | 1.6e-06 | 1.5e-06 | 2.100E-06 | 9.539E-07 | 1.500E-06 | 3.200E-06 |
| 16245 | 813 | V813C | B | 3e-06 | 2.2e-06 | 2.4e-06 | 2.533E-06 | 4.163E-07 | 2.200E-06 | 3.000E-06 |
| 16246 | 813 | V813E | B | 2.8e-06 | 1.5e-06 | 4.8e-06 | 3.033E-06 | 1.662E-06 | 1.500E-06 | 4.800E-06 |
| 16247 | 813 | V813Q | B | 5.3e-06 | 4.2e-06 | 4.5e-06 | 4.667E-06 | 5.686E-07 | 4.200E-06 | 5.300E-06 |
| 16248 | 813 | V813G | B | 2.5e-06 | 1.9e-06 | 2.5e-06 | 2.300E-06 | 3.464E-07 | 1.900E-06 | 2.500E-06 |
| 16249 | 813 | V813H | B | 2e-06 | 1.6e-06 | 2.6e-06 | 2.067E-06 | 5.033E-07 | 1.600E-06 | 2.600E-06 |
| 16250 | 813 | V813I | B | 2.7e-06 | 4.5e-06 | 2.2e-06 | 3.133E-06 | 1.210E-06 | 2.200E-06 | 4.500E-06 |
| 16251 | 813 | V813L | B | 1.6e-06 | 2.4e-06 | 2.3e-06 | 2.100E-06 | 4.359E-07 | 1.600E-06 | 2.400E-06 |
| 16252 | 813 | V813K | B | 1.8e-06 | 2.4e-06 | 4.2e-06 | 2.800E-06 | 1.249E-06 | 1.800E-06 | 4.200E-06 |
| 16253 | 813 | V813M | B | 1.4e-06 | 3.4e-06 | 3.8e-06 | 2.867E-06 | 1.286E-06 | 1.400E-06 | 3.800E-06 |
| 16254 | 813 | V813F | B | 3.8e-06 | 3e-06 | 3.7e-06 | 3.500E-06 | 4.359E-07 | 3.000E-06 | 3.800E-06 |
| 16255 | 813 | V813P | B | 2.4e-06 | 2.5e-06 | 3.2e-06 | 2.700E-06 | 4.359E-07 | 2.400E-06 | 3.200E-06 |
| 16256 | 813 | V813S | B | 3.5e-06 | 2.1e-06 | 2.1e-06 | 2.567E-06 | 8.083E-07 | 2.100E-06 | 3.500E-06 |
| 16257 | 813 | V813T | B | 2.3e-06 | 1.8e-06 | 3.5e-06 | 2.533E-06 | 8.737E-07 | 1.800E-06 | 3.500E-06 |
| 16258 | 813 | V813W | B | 2.9e-06 | 2.5e-06 | 1.5e-06 | 2.300E-06 | 7.211E-07 | 1.500E-06 | 2.900E-06 |
| 16259 | 813 | V813Y | B | 2.5e-06 | 2.5e-06 | 2.4e-06 | 2.467E-06 | 5.774E-08 | 2.400E-06 | 2.500E-06 |
| 16260 | 813 | V813V | B | 1.4e-06 | 1.9e-06 | 2.2e-06 | 1.833E-06 | 4.041E-07 | 1.400E-06 | 2.200E-06 |
| 16261 | 814 | Y814A | B | 3.1e-06 | 3e-06 | 2.2e-06 | 2.767E-06 | 4.933E-07 | 2.200E-06 | 3.100E-06 |
| 16262 | 814 | Y814R | B | 2.9e-06 | 2.7e-06 | 2.7e-06 | 2.767E-06 | 1.155E-07 | 2.700E-06 | 2.900E-06 |
| 16263 | 814 | Y814N | B | 2.7e-06 | 1.3e-06 | 3.3e-06 | 2.433E-06 | 1.026E-06 | 1.300E-06 | 3.300E-06 |
| 16264 | 814 | Y814D | B | 2.2e-06 | 1.9e-06 | 2.4e-06 | 2.167E-06 | 2.517E-07 | 1.900E-06 | 2.400E-06 |
| 16265 | 814 | Y814C | B | 2.2e-06 | 2.5e-06 | 1.3e-06 | 2.000E-06 | 6.245E-07 | 1.300E-06 | 2.500E-06 |
| 16266 | 814 | Y814E | B | 2.2e-06 | 4.1e-06 | 1.6e-06 | 2.633E-06 | 1.305E-06 | 1.600E-06 | 4.100E-06 |
| 16267 | 814 | Y814Q | B | 2e-06 | 2.1e-06 | 1.7e-06 | 1.933E-06 | 2.082E-07 | 1.700E-06 | 2.100E-06 |
| 16268 | 814 | Y814G | B | 3.8e-06 | 1.4e-06 | 1.4e-06 | 2.200E-06 | 1.386E-06 | 1.400E-06 | 3.800E-06 |
| 16269 | 814 | Y814H | B | 2.3e-06 | 2.2e-06 | 2e-06 | 2.167E-06 | 1.528E-07 | 2.000E-06 | 2.300E-06 |
| 16270 | 814 | Y814I | B | 2.3e-06 | 1.7e-06 | 1.9e-06 | 1.967E-06 | 3.055E-07 | 1.700E-06 | 2.300E-06 |
| 16271 | 814 | Y814L | B | 1.6e-06 | 3.2e-06 | 3.8e-06 | 2.867E-06 | 1.137E-06 | 1.600E-06 | 3.800E-06 |
| 16272 | 814 | Y814K | B | 2.3e-06 | 2.2e-06 | 2.1e-06 | 2.200E-06 | 1.000E-07 | 2.100E-06 | 2.300E-06 |
| 16273 | 814 | Y814M | B | 1e-06 | 2e-06 | 1.4e-06 | 1.467E-06 | 5.033E-07 | 1.000E-06 | 2.000E-06 |
| 16274 | 814 | Y814F | B | 2.1e-06 | 1.7e-06 | 2.6e-06 | 2.133E-06 | 4.509E-07 | 1.700E-06 | 2.600E-06 |
| 16275 | 814 | Y814P | B | 3.3e-06 | 3.2e-06 | 2.8e-06 | 3.100E-06 | 2.646E-07 | 2.800E-06 | 3.300E-06 |
| 16276 | 814 | Y814S | B | 2e-06 | 2e-06 | 4.6e-06 | 2.867E-06 | 1.501E-06 | 2.000E-06 | 4.600E-06 |
| 16277 | 814 | Y814T | B | 1.7e-06 | 1.9e-06 | 1.7e-06 | 1.767E-06 | 1.155E-07 | 1.700E-06 | 1.900E-06 |
| 16278 | 814 | Y814W | B | 2.3e-06 | 3.6e-06 | 2.9e-06 | 2.933E-06 | 6.506E-07 | 2.300E-06 | 3.600E-06 |
| 16279 | 814 | Y814Y | B | 1.6e-06 | 1.8e-06 | 3.2e-06 | 2.200E-06 | 8.718E-07 | 1.600E-06 | 3.200E-06 |
| 16280 | 814 | Y814V | B | 2.3e-06 | 2.2e-06 | 1.4e-06 | 1.967E-06 | 4.933E-07 | 1.400E-06 | 2.300E-06 |
| 16281 | 815 | A815A | B | 3e-06 | 1.4e-06 | 2.3e-06 | 2.233E-06 | 8.021E-07 | 1.400E-06 | 3.000E-06 |
| 16282 | 815 | A815R | B | 4.9e-06 | 2.5e-06 | 2.1e-06 | 3.167E-06 | 1.514E-06 | 2.100E-06 | 4.900E-06 |
| 16283 | 815 | A815N | B | 2.1e-06 | 1.2e-06 | 2.7e-06 | 2.000E-06 | 7.550E-07 | 1.200E-06 | 2.700E-06 |
| 16284 | 815 | A815D | B | 3.5e-06 | 2.2e-06 | 2.6e-06 | 2.767E-06 | 6.658E-07 | 2.200E-06 | 3.500E-06 |
| 16285 | 815 | A815C | B | 2.9e-06 | 5.5e-06 | 1.7e-06 | 3.367E-06 | 1.943E-06 | 1.700E-06 | 5.500E-06 |
| 16286 | 815 | A815E | B | 2.3e-06 | 2.3e-06 | 3.1e-06 | 2.567E-06 | 4.619E-07 | 2.300E-06 | 3.100E-06 |
| 16287 | 815 | A815Q | B | 3.2e-06 | 3.6e-06 | 2.7e-06 | 3.167E-06 | 4.509E-07 | 2.700E-06 | 3.600E-06 |
| 16288 | 815 | A815G | B | 3.1e-06 | 1.9e-06 | 2.4e-06 | 2.467E-06 | 6.028E-07 | 1.900E-06 | 3.100E-06 |
| 16289 | 815 | A815H | B | 3.1e-06 | 2.4e-06 | 2.5e-06 | 2.667E-06 | 3.786E-07 | 2.400E-06 | 3.100E-06 |
| 16290 | 815 | A815I | B | 2.3e-06 | 2.1e-06 | 2.7e-06 | 2.367E-06 | 3.055E-07 | 2.100E-06 | 2.700E-06 |
| 16291 | 815 | A815L | B | 2.3e-06 | 1.2e-06 | 1.6e-06 | 1.700E-06 | 5.568E-07 | 1.200E-06 | 2.300E-06 |
| 16292 | 815 | A815K | B | 2.2e-06 | 1.8e-06 | 3.4e-06 | 2.467E-06 | 8.327E-07 | 1.800E-06 | 3.400E-06 |
| 16293 | 815 | A815M | B | 3.1e-06 | 3.1e-06 | 1.4e-06 | 2.533E-06 | 9.815E-07 | 1.400E-06 | 3.100E-06 |
| 16294 | 815 | A815F | B | 1.6e-06 | 1.6e-06 | 2e-06 | 1.733E-06 | 2.309E-07 | 1.600E-06 | 2.000E-06 |
| 16295 | 815 | A815P | B | 3.1e-06 | 4.5e-06 | 1.9e-06 | 3.167E-06 | 1.301E-06 | 1.900E-06 | 4.500E-06 |
| 16296 | 815 | A815S | B | 2.8e-06 | 2.5e-06 | 2.4e-06 | 2.567E-06 | 2.082E-07 | 2.400E-06 | 2.800E-06 |
| 16297 | 815 | A815T | B | 1.9e-06 | 1.7e-06 | 2.8e-06 | 2.133E-06 | 5.859E-07 | 1.700E-06 | 2.800E-06 |
| 16298 | 815 | A815W | B | 2.5e-06 | 2.2e-06 | 2.8e-06 | 2.500E-06 | 3.000E-07 | 2.200E-06 | 2.800E-06 |
| 16299 | 815 | A815Y | B | 2e-06 | 1.7e-06 | 2.7e-06 | 2.133E-06 | 5.132E-07 | 1.700E-06 | 2.700E-06 |
| 16300 | 815 | A815V | B | 3.3e-06 | 2.6e-06 | 4.5e-06 | 3.467E-06 | 9.609E-07 | 2.600E-06 | 4.500E-06 |
| 16301 | 816 | C816A | B | 2.8e-06 | 2e-06 | 2.7e-06 | 2.500E-06 | 4.359E-07 | 2.000E-06 | 2.800E-06 |
| 16302 | 816 | C816R | B | 2.2e-06 | 1.7e-06 | 1.5e-06 | 1.800E-06 | 3.606E-07 | 1.500E-06 | 2.200E-06 |
| 16303 | 816 | C816N | B | 3.5e-06 | 2.7e-06 | 2.6e-06 | 2.933E-06 | 4.933E-07 | 2.600E-06 | 3.500E-06 |
| 16304 | 816 | C816D | B | 8.9e-07 | 1.5e-06 | 1.8e-06 | 1.397E-06 | 4.637E-07 | 8.900E-07 | 1.800E-06 |
| 16305 | 816 | C816C | B | 2.1e-06 | 2.2e-06 | 1.1e-06 | 1.800E-06 | 6.083E-07 | 1.100E-06 | 2.200E-06 |
| 16306 | 816 | C816E | B | 1.8e-06 | 2.9e-06 | 1.3e-06 | 2.000E-06 | 8.185E-07 | 1.300E-06 | 2.900E-06 |
| 16307 | 816 | C816Q | B | 3.6e-06 | 4.9e-06 | 5.8e-06 | 4.767E-06 | 1.106E-06 | 3.600E-06 | 5.800E-06 |
| 16308 | 816 | C816G | B | 1.4e-06 | 2.5e-06 | 3.3e-06 | 2.400E-06 | 9.539E-07 | 1.400E-06 | 3.300E-06 |
| 16309 | 816 | C816H | B | 1.5e-06 | 2.5e-06 | 2.5e-06 | 2.167E-06 | 5.774E-07 | 1.500E-06 | 2.500E-06 |
| 16310 | 816 | C816I | B | 3.1e-06 | 2.4e-06 | 2.3e-06 | 2.600E-06 | 4.359E-07 | 2.300E-06 | 3.100E-06 |
| 16311 | 816 | C816L | B | 2.6e-06 | 2.4e-06 | 3.4e-06 | 2.800E-06 | 5.292E-07 | 2.400E-06 | 3.400E-06 |
| 16312 | 816 | C816K | B | 1.3e-06 | 2.7e-06 | 1.6e-06 | 1.867E-06 | 7.371E-07 | 1.300E-06 | 2.700E-06 |
| 16313 | 816 | C816M | B | 1.9e-06 | 2e-06 | 1.5e-06 | 1.800E-06 | 2.646E-07 | 1.500E-06 | 2.000E-06 |
| 16314 | 816 | C816F | B | 2.8e-06 | 2.3e-06 | 1.7e-06 | 2.267E-06 | 5.508E-07 | 1.700E-06 | 2.800E-06 |
| 16315 | 816 | C816P | B | 2.4e-06 | 2.9e-06 | 2.5e-06 | 2.600E-06 | 2.646E-07 | 2.400E-06 | 2.900E-06 |
| 16316 | 816 | C816S | B | 2.9e-06 | 1.4e-06 | 1.5e-06 | 1.933E-06 | 8.386E-07 | 1.400E-06 | 2.900E-06 |
| 16317 | 816 | C816T | B | 1.4e-06 | 2.1e-06 | 3.7e-06 | 2.400E-06 | 1.179E-06 | 1.400E-06 | 3.700E-06 |
| 16318 | 816 | C816W | B | 2.5e-06 | 1.4e-06 | 1.6e-06 | 1.833E-06 | 5.859E-07 | 1.400E-06 | 2.500E-06 |
| 16319 | 816 | C816Y | B | 2.3e-06 | 3.2e-06 | 2.6e-06 | 2.700E-06 | 4.583E-07 | 2.300E-06 | 3.200E-06 |

| | | | | | | | | | | |
|---|---|---|---|---|---|---|---|---|---|---|
| 16320 | 816 | C816V | B | 1.7e-06 | 1.8e-06 | 3.8e-06 | 2.433E-06 | 1.185E-06 | 1.700E-06 | 3.800E-06 |
| 16321 | 817 | E817A | B | 2.9e-06 | 2.9e-06 | 2.9e-06 | 2.900E-06 | 5.187E-22 | 2.900E-06 | 2.900E-06 |
| 16322 | 817 | E817R | B | 1.9e-06 | 1.8e-06 | 3e-06 | 2.233E-06 | 6.658E-07 | 1.800E-06 | 3.000E-06 |
| 16323 | 817 | E817N | B | 1.1e-06 | 2.7e-06 | 2.6e-06 | 2.133E-06 | 8.963E-07 | 1.100E-06 | 2.700E-06 |
| 16324 | 817 | E817D | B | 3.5e-06 | 3.7e-06 | 2.7e-06 | 3.300E-06 | 5.292E-07 | 2.700E-06 | 3.700E-06 |
| 16325 | 817 | E817C | B | 1.2e-06 | 2e-06 | 1.8e-06 | 1.667E-06 | 4.163E-07 | 1.200E-06 | 2.000E-06 |
| 16326 | 817 | E817E | B | 4e-06 | 1.8e-06 | 1.5e-06 | 2.433E-06 | 1.365E-06 | 1.500E-06 | 4.000E-06 |
| 16327 | 817 | E817Q | B | 1.9e-06 | 3.5e-06 | 3.5e-06 | 2.967E-06 | 9.238E-07 | 1.900E-06 | 3.500E-06 |
| 16328 | 817 | E817G | B | 1.7e-06 | 2.3e-06 | 1.4e-06 | 1.800E-06 | 4.583E-07 | 1.400E-06 | 2.300E-06 |
| 16329 | 817 | E817H | B | 3.4e-06 | 2e-06 | 1.7e-06 | 2.367E-06 | 9.074E-07 | 1.700E-06 | 3.400E-06 |
| 16330 | 817 | E817I | B | 1.6e-06 | 4e-06 | 2.4e-06 | 2.667E-06 | 1.222E-06 | 1.600E-06 | 4.000E-06 |
| 16331 | 817 | E817L | B | 3.2e-06 | 1.7e-06 | 3.1e-06 | 2.667E-06 | 8.386E-07 | 1.700E-06 | 3.200E-06 |
| 16332 | 817 | E817K | B | 2.1e-06 | 2.1e-06 | 1.7e-06 | 1.967E-06 | 2.309E-07 | 1.700E-06 | 2.100E-06 |
| 16333 | 817 | E817M | B | 2.4e-06 | 1.4e-06 | 1.6e-06 | 1.800E-06 | 5.292E-07 | 1.400E-06 | 2.400E-06 |
| 16334 | 817 | E817F | B | 3.2e-06 | 4.1e-06 | 1.4e-06 | 2.900E-06 | 1.375E-06 | 1.400E-06 | 4.100E-06 |
| 16335 | 817 | E817P | B | 3.9e-06 | 2.7e-06 | 1.6e-06 | 2.733E-06 | 1.150E-06 | 1.600E-06 | 3.900E-06 |
| 16336 | 817 | E817S | B | 1.8e-06 | 2.1e-06 | 4.7e-06 | 2.867E-06 | 1.595E-06 | 1.800E-06 | 4.700E-06 |
| 16337 | 817 | E817T | B | 2.8e-06 | 1.7e-06 | 1.4e-06 | 1.967E-06 | 7.371E-07 | 1.400E-06 | 2.800E-06 |
| 16338 | 817 | E817W | B | 1.5e-06 | 1.8e-06 | 1.6e-06 | 1.633E-06 | 1.528E-07 | 1.500E-06 | 1.800E-06 |
| 16339 | 817 | E817Y | B | 2.5e-06 | 2.5e-06 | 1.3e-06 | 2.100E-06 | 6.928E-07 | 1.300E-06 | 2.500E-06 |
| 16340 | 817 | E817V | B | 3.8e-06 | 2.5e-06 | 1.9e-06 | 2.733E-06 | 9.713E-07 | 1.900E-06 | 3.800E-06 |
| 16341 | 818 | V818A | B | 1.7e-06 | 3e-06 | 5e-06 | 3.233E-06 | 1.662E-06 | 1.700E-06 | 5.000E-06 |
| 16342 | 818 | V818R | B | 2.4e-06 | 2.6e-06 | 3.7e-06 | 2.900E-06 | 7.000E-07 | 2.400E-06 | 3.700E-06 |
| 16343 | 818 | V818N | B | 2e-06 | 2.4e-06 | 2.3e-06 | 2.233E-06 | 2.082E-07 | 2.000E-06 | 2.400E-06 |
| 16344 | 818 | V818D | B | 2.1e-06 | 2.9e-06 | 1.7e-06 | 2.233E-06 | 6.110E-07 | 1.700E-06 | 2.900E-06 |
| 16345 | 818 | V818C | B | 2.3e-06 | 4e-06 | 2.2e-06 | 2.833E-06 | 1.012E-06 | 2.200E-06 | 4.000E-06 |
| 16346 | 818 | V818E | B | 1.5e-06 | 2.8e-06 | 2.2e-06 | 2.167E-06 | 6.506E-07 | 1.500E-06 | 2.800E-06 |
| 16347 | 818 | V818Q | B | 3.5e-06 | 2e-06 | 2.2e-06 | 2.567E-06 | 8.145E-07 | 2.000E-06 | 3.500E-06 |
| 16348 | 818 | V818G | B | 2.9e-06 | 2.1e-06 | 2.3e-06 | 2.433E-06 | 4.163E-07 | 2.100E-06 | 2.900E-06 |
| 16349 | 818 | V818H | B | 1.8e-06 | 3.5e-06 | 2.7e-06 | 2.667E-06 | 8.505E-07 | 1.800E-06 | 3.500E-06 |
| 16350 | 818 | V818I | B | 2e-06 | 3.7e-06 | 2.7e-06 | 2.800E-06 | 8.544E-07 | 2.000E-06 | 3.700E-06 |
| 16351 | 818 | V818L | B | 1.8e-06 | 1.5e-06 | 2.9e-06 | 2.067E-06 | 7.371E-07 | 1.500E-06 | 2.900E-06 |
| 16352 | 818 | V818K | B | 3.1e-06 | 1.9e-06 | 1.5e-06 | 2.167E-06 | 8.327E-07 | 1.500E-06 | 3.100E-06 |
| 16353 | 818 | V818M | B | 4.1e-06 | 2e-06 | 1.5e-06 | 2.533E-06 | 1.380E-06 | 1.500E-06 | 4.100E-06 |
| 16354 | 818 | V818F | B | 1.9e-06 | 3.7e-06 | 2.3e-06 | 2.633E-06 | 9.452E-07 | 1.900E-06 | 3.700E-06 |
| 16355 | 818 | V818P | B | 3.1e-06 | 3.4e-06 | 2.7e-06 | 3.067E-06 | 3.512E-07 | 2.700E-06 | 3.400E-06 |
| 16356 | 818 | V818S | B | 1.2e-06 | 2.3e-06 | 1.2e-06 | 1.567E-06 | 6.351E-07 | 1.200E-06 | 2.300E-06 |
| 16357 | 818 | V818T | B | 1.7e-06 | 3.8e-06 | 2.9e-06 | 2.800E-06 | 1.054E-06 | 1.700E-06 | 3.800E-06 |
| 16358 | 818 | V818W | B | 4.2e-06 | 2.1e-06 | 2.7e-06 | 3.000E-06 | 1.082E-06 | 2.100E-06 | 4.200E-06 |
| 16359 | 818 | V818Y | B | 1.8e-06 | 3.4e-06 | 2.6e-06 | 2.600E-06 | 8.000E-07 | 1.800E-06 | 3.400E-06 |
| 16360 | 818 | V818V | B | 1.6e-06 | 2.6e-06 | 3.2e-06 | 2.467E-06 | 8.083E-07 | 1.600E-06 | 3.200E-06 |
| 16361 | 819 | T819A | B | 2.7e-06 | 2.1e-06 | 2.4e-06 | 2.400E-06 | 3.000E-07 | 2.100E-06 | 2.700E-06 |
| 16362 | 819 | T819R | B | 1.5e-06 | 3.5e-06 | 1.9e-06 | 2.300E-06 | 1.058E-06 | 1.500E-06 | 3.500E-06 |
| 16363 | 819 | T819N | B | 2.2e-06 | 1.9e-06 | 1.6e-06 | 1.900E-06 | 3.000E-07 | 1.600E-06 | 2.200E-06 |
| 16364 | 819 | T819D | B | 2.9e-06 | 3.4e-06 | 1.9e-06 | 2.733E-06 | 7.638E-07 | 1.900E-06 | 3.400E-06 |
| 16365 | 819 | T819C | B | 2e-06 | 3.4e-06 | 2.6e-06 | 2.667E-06 | 7.024E-07 | 2.000E-06 | 3.400E-06 |
| 16366 | 819 | T819E | B | 2.1e-06 | 2.4e-06 | 2.4e-06 | 2.300E-06 | 1.732E-07 | 2.100E-06 | 2.400E-06 |
| 16367 | 819 | T819Q | B | 1.9e-06 | 1.7e-06 | 2.5e-06 | 2.033E-06 | 4.163E-07 | 1.700E-06 | 2.500E-06 |
| 16368 | 819 | T819G | B | 1.6e-06 | 1.9e-06 | 2e-06 | 1.833E-06 | 2.082E-07 | 1.600E-06 | 2.000E-06 |
| 16369 | 819 | T819H | B | 1.7e-06 | 2.9e-06 | 2.5e-06 | 2.367E-06 | 6.110E-07 | 1.700E-06 | 2.900E-06 |
| 16370 | 819 | T819I | B | 2.4e-06 | 1.8e-06 | 3.6e-06 | 2.600E-06 | 9.165E-07 | 1.800E-06 | 3.600E-06 |
| 16371 | 819 | T819L | B | 2.6e-06 | 2e-06 | 3.3e-06 | 2.633E-06 | 6.506E-07 | 2.000E-06 | 3.300E-06 |
| 16372 | 819 | T819K | B | 1.9e-06 | 2.5e-06 | 2.5e-06 | 2.300E-06 | 3.464E-07 | 1.900E-06 | 2.500E-06 |
| 16373 | 819 | T819M | B | 2.2e-06 | 2.6e-06 | 1.8e-06 | 2.200E-06 | 4.000E-07 | 1.800E-06 | 2.600E-06 |
| 16374 | 819 | T819F | B | 2.8e-06 | 1.8e-06 | 1e-06 | 1.867E-06 | 9.018E-07 | 1.000E-06 | 2.800E-06 |
| 16375 | 819 | T819P | B | 1.1e-06 | 2.2e-06 | 3e-06 | 2.100E-06 | 9.539E-07 | 1.100E-06 | 3.000E-06 |
| 16376 | 819 | T819S | B | 2.1e-06 | 1.7e-06 | 3.4e-06 | 2.400E-06 | 8.888E-07 | 1.700E-06 | 3.400E-06 |
| 16377 | 819 | T819T | B | 1.8e-06 | 3.1e-06 | 2.4e-06 | 2.433E-06 | 6.506E-07 | 1.800E-06 | 3.100E-06 |
| 16378 | 819 | T819W | B | 1.1e-06 | 2.2e-06 | 2e-06 | 1.767E-06 | 5.859E-07 | 1.100E-06 | 2.200E-06 |
| 16379 | 819 | T819Y | B | 2.6e-06 | 1.8e-06 | 4.4e-06 | 2.933E-06 | 1.332E-06 | 1.800E-06 | 4.400E-06 |
| 16380 | 819 | T819V | B | 1.5e-06 | 3.3e-06 | 4.1e-06 | 2.967E-06 | 1.332E-06 | 1.500E-06 | 4.100E-06 |
| 16381 | 820 | H820A | B | 3.2e-06 | 2.2e-06 | 2.4e-06 | 2.600E-06 | 5.292E-07 | 2.200E-06 | 3.200E-06 |
| 16382 | 820 | H820R | B | 3.5e-06 | 7.7e-07 | 2e-06 | 2.090E-06 | 1.367E-06 | 7.700E-07 | 3.500E-06 |
| 16383 | 820 | H820N | B | 1.4e-06 | 1.9e-06 | 2.5e-06 | 1.933E-06 | 5.508E-07 | 1.400E-06 | 2.500E-06 |
| 16384 | 820 | H820D | B | 4e-06 | 1.9e-06 | 2e-06 | 2.633E-06 | 1.185E-06 | 1.900E-06 | 4.000E-06 |
| 16385 | 820 | H820C | B | 2.3e-06 | 2.5e-06 | 2e-06 | 2.267E-06 | 2.517E-07 | 2.000E-06 | 2.500E-06 |
| 16386 | 820 | H820E | B | 9.6e-07 | 1.5e-06 | 1.4e-06 | 1.287E-06 | 2.873E-07 | 9.600E-07 | 1.500E-06 |
| 16387 | 820 | H820Q | B | 1.7e-06 | 2.2e-06 | 3.9e-06 | 2.600E-06 | 1.153E-06 | 1.700E-06 | 3.900E-06 |
| 16388 | 820 | H820G | B | 2.9e-06 | 3.9e-06 | 1.2e-06 | 2.667E-06 | 1.365E-06 | 1.200E-06 | 3.900E-06 |
| 16389 | 820 | H820H | B | 2.5e-06 | 1.7e-06 | 3.8e-06 | 2.667E-06 | 1.060E-06 | 1.700E-06 | 3.800E-06 |
| 16390 | 820 | H820I | B | 1.9e-06 | 1.6e-06 | 1.4e-06 | 1.633E-06 | 2.517E-07 | 1.400E-06 | 1.900E-06 |
| 16391 | 820 | H820L | B | 1.3e-06 | 1.9e-06 | 3e-06 | 2.067E-06 | 8.622E-07 | 1.300E-06 | 3.000E-06 |
| 16392 | 820 | H820K | B | 1.7e-06 | 2.5e-06 | 1.3e-06 | 1.833E-06 | 6.110E-07 | 1.300E-06 | 2.500E-06 |
| 16393 | 820 | H820M | B | 3.7e-06 | 2.5e-06 | 1.4e-06 | 2.533E-06 | 1.150E-06 | 1.400E-06 | 3.700E-06 |
| 16394 | 820 | H820F | B | 3.1e-06 | 2.7e-06 | 3.2e-06 | 3.000E-06 | 2.646E-07 | 2.700E-06 | 3.200E-06 |
| 16395 | 820 | H820P | B | 1.9e-06 | 2.3e-06 | 2.1e-06 | 2.100E-06 | 2.000E-07 | 1.900E-06 | 2.300E-06 |
| 16396 | 820 | H820S | B | 4.9e-06 | 2.6e-06 | 3.1e-06 | 3.533E-06 | 1.210E-06 | 2.600E-06 | 4.900E-06 |
| 16397 | 820 | H820T | B | 3.4e-06 | 2.2e-06 | 2.6e-06 | 2.733E-06 | 6.110E-07 | 2.200E-06 | 3.400E-06 |
| 16398 | 820 | H820W | B | 2.7e-06 | 1.8e-06 | 3e-06 | 2.500E-06 | 6.245E-07 | 1.800E-06 | 3.000E-06 |
| 16399 | 820 | H820Y | B | 1.9e-06 | 2e-06 | 9.9e-07 | 1.630E-06 | 5.565E-07 | 9.900E-07 | 2.000E-06 |
| 16400 | 820 | H820V | B | 2.2e-06 | 4.3e-06 | 3.6e-06 | 3.367E-06 | 1.069E-06 | 2.200E-06 | 4.300E-06 |
| 16401 | 821 | Q821A | B | 2.5e-06 | 2.2e-06 | 2.7e-06 | 2.467E-06 | 2.517E-07 | 2.200E-06 | 2.700E-06 |
| 16402 | 821 | Q821R | B | 3.4e-06 | 2.8e-06 | 2.2e-06 | 2.800E-06 | 6.000E-07 | 2.200E-06 | 3.400E-06 |
| 16403 | 821 | Q821N | B | 1.6e-06 | 4.4e-06 | 5.1e-06 | 3.700E-06 | 1.852E-06 | 1.600E-06 | 5.100E-06 |
| 16404 | 821 | Q821D | B | 2.1e-06 | 1.3e-06 | 3.2e-06 | 2.200E-06 | 9.539E-07 | 1.300E-06 | 3.200E-06 |

| | | | | | | | | | | |
|---|---|---|---|---|---|---|---|---|---|---|
| 16405 | 821 | Q821C | B | 3.1e-06 | 2.3e-06 | 3.2e-06 | 2.867E-06 | 4.933E-07 | 2.300E-06 | 3.200E-06 |
| 16406 | 821 | Q821E | B | 2.9e-06 | 1.4e-06 | 1.8e-06 | 2.033E-06 | 7.767E-07 | 1.400E-06 | 2.900E-06 |
| 16407 | 821 | Q821Q | B | 1.8e-06 | 2.6e-06 | 3.1e-06 | 2.500E-06 | 6.557E-07 | 1.800E-06 | 3.100E-06 |
| 16408 | 821 | Q821G | B | 3.2e-06 | 1.8e-06 | 2e-06 | 2.333E-06 | 7.572E-07 | 1.800E-06 | 3.200E-06 |
| 16409 | 821 | Q821H | B | 2.3e-06 | 2.4e-06 | 1.6e-06 | 2.100E-06 | 4.359E-07 | 1.600E-06 | 2.400E-06 |
| 16410 | 821 | Q821I | B | 1.5e-06 | 3.1e-06 | 2e-06 | 2.200E-06 | 8.185E-07 | 1.500E-06 | 3.100E-06 |
| 16411 | 821 | Q821L | B | 3.4e-06 | 2e-06 | 3.2e-06 | 2.867E-06 | 7.572E-07 | 2.000E-06 | 3.400E-06 |
| 16412 | 821 | Q821K | B | 1.4e-06 | 3.8e-06 | 3.6e-06 | 2.933E-06 | 1.332E-06 | 1.400E-06 | 3.800E-06 |
| 16413 | 821 | Q821M | B | 3.4e-06 | 5.3e-06 | 2.5e-06 | 3.733E-06 | 1.429E-06 | 2.500E-06 | 5.300E-06 |
| 16414 | 821 | Q821F | B | 2e-06 | 2.1e-06 | 2.2e-06 | 2.100E-06 | 1.000E-07 | 2.000E-06 | 2.200E-06 |
| 16415 | 821 | Q821P | B | 2.8e-06 | 1.7e-06 | 2.8e-06 | 2.433E-06 | 6.351E-07 | 1.700E-06 | 2.800E-06 |
| 16416 | 821 | Q821S | B | 2.2e-06 | 1.3e-06 | 2.1e-06 | 1.867E-06 | 4.933E-07 | 1.300E-06 | 2.200E-06 |
| 16417 | 821 | Q821T | B | 3.8e-06 | 2.2e-06 | 2.8e-06 | 2.933E-06 | 8.083E-07 | 2.200E-06 | 3.800E-06 |
| 16418 | 821 | Q821W | B | 2.5e-06 | 3e-06 | 2.3e-06 | 2.600E-06 | 3.606E-07 | 2.300E-06 | 3.000E-06 |
| 16419 | 821 | Q821Y | B | 3.7e-06 | 2.8e-06 | 4.2e-06 | 3.567E-06 | 7.095E-07 | 2.800E-06 | 4.200E-06 |
| 16420 | 821 | Q821V | B | 2.6e-06 | 2.1e-06 | 3.3e-06 | 2.667E-06 | 6.028E-07 | 2.100E-06 | 3.300E-06 |
| 16421 | 822 | G822A | B | 1.9e-06 | 2e-06 | 2.5e-06 | 2.133E-06 | 3.215E-07 | 1.900E-06 | 2.500E-06 |
| 16422 | 822 | G822R | B | 1.3e-06 | 2.5e-06 | 1.7e-06 | 1.833E-06 | 6.110E-07 | 1.300E-06 | 2.500E-06 |
| 16423 | 822 | G822N | B | 1.8e-06 | 5.8e-06 | 1.7e-06 | 3.100E-06 | 2.339E-06 | 1.700E-06 | 5.800E-06 |
| 16424 | 822 | G822D | B | 1.3e-06 | 2.4e-06 | 1.9e-06 | 1.867E-06 | 5.508E-07 | 1.300E-06 | 2.400E-06 |
| 16425 | 822 | G822C | B | 2.2e-06 | 1.4e-06 | 3.2e-06 | 2.267E-06 | 9.018E-07 | 1.400E-06 | 3.200E-06 |
| 16426 | 822 | G822E | B | 2.6e-06 | 2.6e-06 | 3.8e-06 | 3.000E-06 | 6.928E-07 | 2.600E-06 | 3.800E-06 |
| 16427 | 822 | G822Q | B | 2.5e-06 | 3e-06 | 1.2e-06 | 2.233E-06 | 9.292E-07 | 1.200E-06 | 3.000E-06 |
| 16428 | 822 | G822G | B | 4.1e-06 | 3.2e-06 | 3e-06 | 3.433E-06 | 5.859E-07 | 3.000E-06 | 4.100E-06 |
| 16429 | 822 | G822H | B | 1.9e-06 | 2.8e-06 | 2.3e-06 | 2.333E-06 | 4.509E-07 | 1.900E-06 | 2.800E-06 |
| 16430 | 822 | G822I | B | 3.2e-06 | 2.7e-06 | 2.2e-06 | 2.700E-06 | 5.000E-07 | 2.200E-06 | 3.200E-06 |
| 16431 | 822 | G822L | B | 2.6e-06 | 3.8e-06 | 1.8e-06 | 2.733E-06 | 1.007E-06 | 1.800E-06 | 3.800E-06 |
| 16432 | 822 | G822K | B | 2.8e-06 | 2.7e-06 | 2.4e-06 | 2.633E-06 | 2.082E-07 | 2.400E-06 | 2.800E-06 |
| 16433 | 822 | G822M | B | 2.9e-06 | 2.1e-06 | 2.7e-06 | 2.567E-06 | 4.163E-07 | 2.100E-06 | 2.900E-06 |
| 16434 | 822 | G822F | B | 3.1e-06 | 2.5e-06 | 1.7e-06 | 2.433E-06 | 7.024E-07 | 1.700E-06 | 3.100E-06 |
| 16435 | 822 | G822P | B | 1.6e-06 | 2.7e-06 | 2e-06 | 2.100E-06 | 5.568E-07 | 1.600E-06 | 2.700E-06 |
| 16436 | 822 | G822S | B | 2.9e-06 | 2.3e-06 | 8.5e-07 | 2.017E-06 | 1.054E-06 | 8.500E-07 | 2.900E-06 |
| 16437 | 822 | G822T | B | 2e-06 | 1.5e-06 | 3.8e-06 | 2.433E-06 | 1.210E-06 | 1.500E-06 | 3.800E-06 |
| 16438 | 822 | G822W | B | 2.3e-06 | 2.2e-06 | 2e-06 | 2.167E-06 | 1.528E-07 | 2.000E-06 | 2.300E-06 |
| 16439 | 822 | G822Y | B | 1.8e-06 | 2.3e-06 | 1.9e-06 | 2.000E-06 | 2.646E-07 | 1.800E-06 | 2.300E-06 |
| 16440 | 822 | G822V | B | 1.5e-06 | 2.8e-06 | 2.9e-06 | 2.400E-06 | 7.810E-07 | 1.500E-06 | 2.900E-06 |
| 16441 | 823 | L823A | B | 2e-06 | 2.6e-06 | 4.2e-06 | 2.933E-06 | 1.137E-06 | 2.000E-06 | 4.200E-06 |
| 16442 | 823 | L823R | B | 2e-06 | 2.3e-06 | 1.8e-06 | 2.033E-06 | 2.517E-07 | 1.800E-06 | 2.300E-06 |
| 16443 | 823 | L823N | B | 2.9e-06 | 2.8e-06 | 2.9e-06 | 2.867E-06 | 5.774E-08 | 2.800E-06 | 2.900E-06 |
| 16444 | 823 | L823D | B | 2.3e-06 | 2.8e-06 | 3e-06 | 2.700E-06 | 3.606E-07 | 2.300E-06 | 3.000E-06 |
| 16445 | 823 | L823C | B | 2.1e-06 | 3.1e-06 | 2.6e-06 | 2.600E-06 | 5.000E-07 | 2.100E-06 | 3.100E-06 |
| 16446 | 823 | L823E | B | 2.8e-06 | 2.8e-06 | 2.7e-06 | 2.767E-06 | 5.774E-08 | 2.700E-06 | 2.800E-06 |
| 16447 | 823 | L823Q | B | 2e-06 | 1.7e-06 | 2.2e-06 | 1.967E-06 | 2.517E-07 | 1.700E-06 | 2.200E-06 |
| 16448 | 823 | L823G | B | 5.4e-06 | 1.7e-06 | 2.2e-06 | 3.100E-06 | 2.007E-06 | 1.700E-06 | 5.400E-06 |
| 16449 | 823 | L823H | B | 2.2e-06 | 3e-06 | 3.4e-06 | 2.867E-06 | 6.110E-07 | 2.200E-06 | 3.400E-06 |
| 16450 | 823 | L823I | B | 2.5e-06 | 2.4e-06 | 2.7e-06 | 2.533E-06 | 1.528E-07 | 2.400E-06 | 2.700E-06 |
| 16451 | 823 | L823L | B | 3.7e-06 | 2.9e-06 | 1.3e-06 | 2.633E-06 | 1.222E-06 | 1.300E-06 | 3.700E-06 |
| 16452 | 823 | L823K | B | 2.4e-06 | 1.7e-06 | 1.9e-06 | 2.000E-06 | 3.606E-07 | 1.700E-06 | 2.400E-06 |
| 16453 | 823 | L823M | B | 2.8e-06 | 2.9e-06 | 1.2e-06 | 2.300E-06 | 9.539E-07 | 1.200E-06 | 2.900E-06 |
| 16454 | 823 | L823F | B | 1.5e-06 | 2.5e-06 | 1.7e-06 | 1.900E-06 | 5.292E-07 | 1.500E-06 | 2.500E-06 |
| 16455 | 823 | L823P | B | 4e-06 | 2.9e-06 | 2.8e-06 | 3.233E-06 | 6.658E-07 | 2.800E-06 | 4.000E-06 |
| 16456 | 823 | L823S | B | 1.6e-06 | 2.1e-06 | 2.5e-06 | 2.067E-06 | 4.509E-07 | 1.600E-06 | 2.500E-06 |
| 16457 | 823 | L823T | B | 1.4e-06 | 1.4e-06 | 2.2e-06 | 1.667E-06 | 4.619E-07 | 1.400E-06 | 2.200E-06 |
| 16458 | 823 | L823W | B | 2e-06 | 1.1e-06 | 2.4e-06 | 1.833E-06 | 6.658E-07 | 1.100E-06 | 2.400E-06 |
| 16459 | 823 | L823Y | B | 3.2e-06 | 1.9e-06 | 2.4e-06 | 2.500E-06 | 6.557E-07 | 1.900E-06 | 3.200E-06 |
| 16460 | 823 | L823V | B | 2.5e-06 | 2.8e-06 | 2.5e-06 | 2.600E-06 | 1.732E-07 | 2.500E-06 | 2.800E-06 |
| 16461 | 824 | S824A | B | 2.8e-06 | 2.3e-06 | 3.2e-06 | 2.767E-06 | 4.509E-07 | 2.300E-06 | 3.200E-06 |
| 16462 | 824 | S824R | B | 3.7e-06 | 1.9e-06 | 2e-06 | 2.533E-06 | 1.012E-06 | 1.900E-06 | 3.700E-06 |
| 16463 | 824 | S824N | B | 2.3e-06 | 2.7e-06 | 2.5e-06 | 2.500E-06 | 2.000E-07 | 2.300E-06 | 2.700E-06 |
| 16464 | 824 | S824D | B | 2.5e-06 | 2.4e-06 | 1.8e-06 | 2.233E-06 | 3.786E-07 | 1.800E-06 | 2.500E-06 |
| 16465 | 824 | S824C | B | 2.2e-06 | 2.4e-06 | 3.3e-06 | 2.633E-06 | 5.859E-07 | 2.200E-06 | 3.300E-06 |
| 16466 | 824 | S824E | B | 2.1e-06 | 3.5e-06 | 2.6e-06 | 2.733E-06 | 7.095E-07 | 2.100E-06 | 3.500E-06 |
| 16467 | 824 | S824Q | B | 1.8e-06 | 1.7e-06 | 3.7e-06 | 2.400E-06 | 1.127E-06 | 1.700E-06 | 3.700E-06 |
| 16468 | 824 | S824G | B | 2e-06 | 1.7e-06 | 2e-06 | 1.900E-06 | 1.732E-07 | 1.700E-06 | 2.000E-06 |
| 16469 | 824 | S824H | B | 4.6e-06 | 2.5e-06 | 2.9e-06 | 3.333E-06 | 1.115E-06 | 2.500E-06 | 4.600E-06 |
| 16470 | 824 | S824I | B | 2.3e-06 | 2.5e-06 | 1.1e-06 | 1.967E-06 | 7.572E-07 | 1.100E-06 | 2.500E-06 |
| 16471 | 824 | S824L | B | 3.3e-06 | 1.7e-06 | 4e-06 | 3.000E-06 | 1.179E-06 | 1.700E-06 | 4.000E-06 |
| 16472 | 824 | S824K | B | 2.9e-06 | 3.1e-06 | 2.3e-06 | 2.767E-06 | 4.163E-07 | 2.300E-06 | 3.100E-06 |
| 16473 | 824 | S824M | B | 1.9e-06 | 3e-06 | 3.7e-06 | 2.867E-06 | 9.074E-07 | 1.900E-06 | 3.700E-06 |
| 16474 | 824 | S824F | B | 2.3e-06 | 1.2e-06 | 2.6e-06 | 2.033E-06 | 7.371E-07 | 1.200E-06 | 2.600E-06 |
| 16475 | 824 | S824P | B | 2.7e-06 | 5.5e-06 | 2.1e-06 | 3.433E-06 | 1.815E-06 | 2.100E-06 | 5.500E-06 |
| 16476 | 824 | S824S | B | 1.4e-06 | 2.1e-06 | 1.9e-06 | 1.800E-06 | 3.606E-07 | 1.400E-06 | 2.100E-06 |
| 16477 | 824 | S824T | B | 2.3e-06 | 1.8e-06 | 2.2e-06 | 2.100E-06 | 2.646E-07 | 1.800E-06 | 2.300E-06 |
| 16478 | 824 | S824W | B | 3.3e-06 | 2.4e-06 | 2e-06 | 2.567E-06 | 6.658E-07 | 2.000E-06 | 3.300E-06 |
| 16479 | 824 | S824Y | B | 3.3e-06 | 3e-06 | 2.1e-06 | 2.800E-06 | 6.245E-07 | 2.100E-06 | 3.300E-06 |
| 16480 | 824 | S824V | B | 3.2e-06 | 1.8e-06 | 3.2e-06 | 2.733E-06 | 8.083E-07 | 1.800E-06 | 3.200E-06 |
| 16481 | 825 | S825A | B | 1.1e-06 | 2.9e-06 | 1.8e-06 | 1.933E-06 | 9.074E-07 | 1.100E-06 | 2.900E-06 |
| 16482 | 825 | S825R | B | 3.2e-06 | 2.1e-06 | 1.6e-06 | 2.300E-06 | 8.185E-07 | 1.600E-06 | 3.200E-06 |
| 16483 | 825 | S825N | B | 1.9e-06 | 3e-06 | 2.5e-06 | 2.467E-06 | 5.508E-07 | 1.900E-06 | 3.000E-06 |
| 16484 | 825 | S825D | B | 3.4e-06 | 2.8e-06 | 1.8e-06 | 2.667E-06 | 8.083E-07 | 1.800E-06 | 3.400E-06 |
| 16485 | 825 | S825C | B | 2.2e-06 | 2.3e-06 | 2.1e-06 | 2.200E-06 | 1.000E-07 | 2.100E-06 | 2.300E-06 |
| 16486 | 825 | S825E | B | 1.8e-06 | 1.9e-06 | 1.4e-06 | 1.700E-06 | 2.646E-07 | 1.400E-06 | 1.900E-06 |
| 16487 | 825 | S825Q | B | 2.3e-06 | 2.4e-06 | 2.9e-06 | 2.533E-06 | 3.215E-07 | 2.300E-06 | 2.900E-06 |
| 16488 | 825 | S825G | B | 1.5e-06 | 2.3e-06 | 2.8e-06 | 2.200E-06 | 6.557E-07 | 1.500E-06 | 2.800E-06 |
| 16489 | 825 | S825H | B | 1.6e-06 | 4.5e-06 | 2.5e-06 | 2.867E-06 | 1.484E-06 | 1.600E-06 | 4.500E-06 |

| | | | | | | | | | | |
|---|---|---|---|---|---|---|---|---|---|---|
| 16490 | 825 | S825I | B | 2.2e-06 | 2.1e-06 | 2.5e-06 | 2.267E-06 | 2.082E-07 | 2.100E-06 | 2.500E-06 |
| 16491 | 825 | S825L | B | 2e-06 | 2e-06 | 2.3e-06 | 2.100E-06 | 1.732E-07 | 2.000E-06 | 2.300E-06 |
| 16492 | 825 | S825K | B | 2.7e-06 | 2.4e-06 | 3.6e-06 | 2.900E-06 | 6.245E-07 | 2.400E-06 | 3.600E-06 |
| 16493 | 825 | S825M | B | 1.6e-06 | 1.5e-06 | 3.1e-06 | 2.067E-06 | 8.963E-07 | 1.500E-06 | 3.100E-06 |
| 16494 | 825 | S825F | B | 2.2e-06 | 4.3e-06 | 1.4e-06 | 2.633E-06 | 1.498E-06 | 1.400E-06 | 4.300E-06 |
| 16495 | 825 | S825P | B | 2.2e-06 | 3.7e-06 | 3e-06 | 2.967E-06 | 7.506E-07 | 2.200E-06 | 3.700E-06 |
| 16496 | 825 | S825S | B | 3.2e-06 | 1.2e-06 | 3.9e-06 | 2.767E-06 | 1.401E-06 | 1.200E-06 | 3.900E-06 |
| 16497 | 825 | S825T | B | 2.7e-06 | 3e-06 | 2.6e-06 | 2.767E-06 | 2.082E-07 | 2.600E-06 | 3.000E-06 |
| 16498 | 825 | S825W | B | 2.5e-06 | 3.2e-06 | 1.7e-06 | 2.467E-06 | 7.506E-07 | 1.700E-06 | 3.200E-06 |
| 16499 | 825 | S825Y | B | 2.2e-06 | 1.7e-06 | 1.7e-06 | 1.867E-06 | 2.887E-07 | 1.700E-06 | 2.200E-06 |
| 16500 | 825 | S825V | B | 2.9e-06 | 3.6e-06 | 3.9e-06 | 3.467E-06 | 5.132E-07 | 2.900E-06 | 3.900E-06 |
| 16501 | 826 | P826A | B | 2.9e-06 | 2.6e-06 | 2.5e-06 | 2.667E-06 | 2.082E-07 | 2.500E-06 | 2.900E-06 |
| 16502 | 826 | P826R | B | 1.8e-06 | 3e-06 | 2.4e-06 | 2.400E-06 | 6.000E-07 | 1.800E-06 | 3.000E-06 |
| 16503 | 826 | P826N | B | 1.7e-06 | 2.8e-06 | 1.7e-06 | 2.067E-06 | 6.351E-07 | 1.700E-06 | 2.800E-06 |
| 16504 | 826 | P826D | B | 3.3e-06 | 4.3e-06 | 1.9e-06 | 3.167E-06 | 1.206E-06 | 1.900E-06 | 4.300E-06 |
| 16505 | 826 | P826C | B | 2.7e-06 | 3.9e-06 | 2.9e-06 | 3.167E-06 | 6.429E-07 | 2.700E-06 | 3.900E-06 |
| 16506 | 826 | P826E | B | 2.6e-06 | 2.4e-06 | 2.9e-06 | 2.633E-06 | 2.517E-07 | 2.400E-06 | 2.900E-06 |
| 16507 | 826 | P826Q | B | 2.9e-06 | 2.3e-06 | 3.8e-06 | 3.000E-06 | 7.550E-07 | 2.300E-06 | 3.800E-06 |
| 16508 | 826 | P826G | B | 2.3e-06 | 6e-06 | 4.1e-06 | 4.133E-06 | 1.850E-06 | 2.300E-06 | 6.000E-06 |
| 16509 | 826 | P826H | B | 1.8e-06 | 1.7e-06 | 2.3e-06 | 1.933E-06 | 3.215E-07 | 1.700E-06 | 2.300E-06 |
| 16510 | 826 | P826I | B | 2.2e-06 | 3.5e-06 | 2.1e-06 | 2.600E-06 | 7.810E-07 | 2.100E-06 | 3.500E-06 |
| 16511 | 826 | P826L | B | 3.5e-06 | 1.5e-06 | 2e-06 | 2.333E-06 | 1.041E-06 | 1.500E-06 | 3.500E-06 |
| 16512 | 826 | P826K | B | 1.7e-06 | 2.7e-06 | 2.8e-06 | 2.400E-06 | 6.083E-07 | 1.700E-06 | 2.800E-06 |
| 16513 | 826 | P826M | B | 3.3e-06 | 2e-06 | 2.7e-06 | 2.667E-06 | 6.506E-07 | 2.000E-06 | 3.300E-06 |
| 16514 | 826 | P826F | B | 1.3e-06 | 2.9e-06 | 2.1e-06 | 2.100E-06 | 8.000E-07 | 1.300E-06 | 2.900E-06 |
| 16515 | 826 | P826P | B | 2.1e-06 | 9.3e-07 | 2.4e-06 | 1.810E-06 | 7.767E-07 | 9.300E-07 | 2.400E-06 |
| 16516 | 826 | P826S | B | 2.7e-06 | 2.2e-06 | 7.7e-06 | 4.200E-06 | 3.041E-06 | 2.200E-06 | 7.700E-06 |
| 16517 | 826 | P826T | B | 2.6e-06 | 2.1e-06 | 2.1e-06 | 2.267E-06 | 2.887E-07 | 2.100E-06 | 2.600E-06 |
| 16518 | 826 | P826W | B | 2.8e-06 | 1.2e-06 | 2.7e-06 | 2.233E-06 | 8.963E-07 | 1.200E-06 | 2.800E-06 |
| 16519 | 826 | P826Y | B | 2.6e-06 | 1.9e-06 | 2.6e-06 | 2.367E-06 | 4.041E-07 | 1.900E-06 | 2.600E-06 |
| 16520 | 826 | P826V | B | 3.1e-06 | 4.8e-06 | 2.1e-06 | 3.333E-06 | 1.365E-06 | 2.100E-06 | 4.800E-06 |
| 16521 | 827 | V827A | B | 4.5e-06 | 2.8e-06 | 1.7e-06 | 3.000E-06 | 1.411E-06 | 1.700E-06 | 4.500E-06 |
| 16522 | 827 | V827R | B | 1.9e-06 | 2.1e-06 | 2e-06 | 2.000E-06 | 1.000E-07 | 1.900E-06 | 2.100E-06 |
| 16523 | 827 | V827N | B | 1.5e-06 | 2e-06 | 2e-06 | 1.833E-06 | 2.887E-07 | 1.500E-06 | 2.000E-06 |
| 16524 | 827 | V827D | B | 2.1e-06 | 2.3e-06 | 2.3e-06 | 2.233E-06 | 1.155E-07 | 2.100E-06 | 2.300E-06 |
| 16525 | 827 | V827C | B | 2.3e-06 | 2.5e-06 | 2.8e-06 | 2.533E-06 | 2.517E-07 | 2.300E-06 | 2.800E-06 |
| 16526 | 827 | V827E | B | 1.6e-06 | 1.7e-06 | 2.5e-06 | 1.933E-06 | 4.933E-07 | 1.600E-06 | 2.500E-06 |
| 16527 | 827 | V827Q | B | 1.7e-06 | 2.7e-06 | 2.5e-06 | 2.300E-06 | 5.292E-07 | 1.700E-06 | 2.700E-06 |
| 16528 | 827 | V827G | B | 2.8e-06 | 1.2e-06 | 1.9e-06 | 1.967E-06 | 8.021E-07 | 1.200E-06 | 2.800E-06 |
| 16529 | 827 | V827H | B | 3e-06 | 2.4e-06 | 2.2e-06 | 2.533E-06 | 4.163E-07 | 2.200E-06 | 3.000E-06 |
| 16530 | 827 | V827I | B | 1.8e-06 | 2.9e-06 | 2.3e-06 | 2.333E-06 | 5.508E-07 | 1.800E-06 | 2.900E-06 |
| 16531 | 827 | V827L | B | 2.1e-06 | 2.6e-06 | 1.7e-06 | 2.133E-06 | 4.509E-07 | 1.700E-06 | 2.600E-06 |
| 16532 | 827 | V827K | B | 1.7e-06 | 2.6e-06 | 3.6e-06 | 2.633E-06 | 9.504E-07 | 1.700E-06 | 3.600E-06 |
| 16533 | 827 | V827M | B | 1.8e-06 | 2.7e-06 | 3.2e-06 | 2.567E-06 | 7.095E-07 | 1.800E-06 | 3.200E-06 |
| 16534 | 827 | V827F | B | 2.1e-06 | 2.2e-06 | 3.1e-06 | 2.467E-06 | 5.508E-07 | 2.100E-06 | 3.100E-06 |
| 16535 | 827 | V827P | B | 3.8e-06 | 3.2e-06 | 1.3e-06 | 2.767E-06 | 1.305E-06 | 1.300E-06 | 3.800E-06 |
| 16536 | 827 | V827S | B | 2.1e-06 | 4.1e-06 | 2.4e-06 | 2.867E-06 | 1.079E-06 | 2.100E-06 | 4.100E-06 |
| 16537 | 827 | V827T | B | 2.2e-06 | 2.4e-06 | 4.1e-06 | 2.900E-06 | 1.044E-06 | 2.200E-06 | 4.100E-06 |
| 16538 | 827 | V827W | B | 2e-06 | 1.9e-06 | 2.3e-06 | 2.067E-06 | 2.082E-07 | 1.900E-06 | 2.300E-06 |
| 16539 | 827 | V827Y | B | 1.9e-06 | 1.2e-06 | 2.2e-06 | 1.767E-06 | 5.132E-07 | 1.200E-06 | 2.200E-06 |
| 16540 | 827 | V827V | B | 3.3e-06 | 2.3e-06 | 1.3e-06 | 2.300E-06 | 1.000E-06 | 1.300E-06 | 3.300E-06 |
| 16541 | 828 | T828A | B | 3.4e-06 | 2.4e-06 | 2.1e-06 | 2.633E-06 | 6.807E-07 | 2.100E-06 | 3.400E-06 |
| 16542 | 828 | T828R | B | 3e-06 | 2.5e-06 | 1.2e-06 | 2.233E-06 | 9.292E-07 | 1.200E-06 | 3.000E-06 |
| 16543 | 828 | T828N | B | 3.4e-06 | 2e-06 | 2.2e-06 | 2.533E-06 | 7.572E-07 | 2.000E-06 | 3.400E-06 |
| 16544 | 828 | T828D | B | 2.5e-06 | 2.5e-06 | 3.2e-06 | 2.733E-06 | 4.041E-07 | 2.500E-06 | 3.200E-06 |
| 16545 | 828 | T828C | B | 2.3e-06 | 1.5e-06 | 2.2e-06 | 2.000E-06 | 4.359E-07 | 1.500E-06 | 2.300E-06 |
| 16546 | 828 | T828E | B | 2.7e-06 | 2.8e-06 | 3.2e-06 | 2.900E-06 | 2.646E-07 | 2.700E-06 | 3.200E-06 |
| 16547 | 828 | T828Q | B | 1.5e-06 | 2.8e-06 | 3.8e-06 | 2.700E-06 | 1.153E-06 | 1.500E-06 | 3.800E-06 |
| 16548 | 828 | T828G | B | 2.6e-06 | 3.6e-06 | 1.7e-06 | 2.633E-06 | 9.504E-07 | 1.700E-06 | 3.600E-06 |
| 16549 | 828 | T828H | B | 2.1e-06 | 2.8e-06 | 1.5e-06 | 2.133E-06 | 6.506E-07 | 1.500E-06 | 2.800E-06 |
| 16550 | 828 | T828I | B | 3.7e-06 | 1.8e-06 | 2.3e-06 | 2.600E-06 | 9.849E-07 | 1.800E-06 | 3.700E-06 |
| 16551 | 828 | T828L | B | 4.8e-06 | 3.2e-06 | 3e-06 | 3.667E-06 | 9.866E-07 | 3.000E-06 | 4.800E-06 |
| 16552 | 828 | T828K | B | 1.1e-06 | 2.6e-06 | 4.7e-06 | 2.800E-06 | 1.808E-06 | 1.100E-06 | 4.700E-06 |
| 16553 | 828 | T828M | B | 3.6e-06 | 1.9e-06 | 1.1e-06 | 2.200E-06 | 1.277E-06 | 1.100E-06 | 3.600E-06 |
| 16554 | 828 | T828F | B | 3.1e-06 | 2.7e-06 | 1.6e-06 | 2.467E-06 | 7.767E-07 | 1.600E-06 | 3.100E-06 |
| 16555 | 828 | T828P | B | 2.5e-06 | 4.4e-06 | 2.9e-06 | 3.267E-06 | 1.002E-06 | 2.500E-06 | 4.400E-06 |
| 16556 | 828 | T828S | B | 2e-06 | 1.6e-06 | 1.4e-06 | 1.667E-06 | 3.055E-07 | 1.400E-06 | 2.000E-06 |
| 16557 | 828 | T828T | B | 3.8e-06 | 2.7e-06 | 1.6e-06 | 2.700E-06 | 1.100E-06 | 1.600E-06 | 3.800E-06 |
| 16558 | 828 | T828W | B | 1.3e-06 | 1e-06 | 1.9e-06 | 1.400E-06 | 4.583E-07 | 1.000E-06 | 1.900E-06 |
| 16559 | 828 | T828Y | B | 2.5e-06 | 3.5e-06 | 2.5e-06 | 2.833E-06 | 5.774E-07 | 2.500E-06 | 3.500E-06 |
| 16560 | 828 | T828V | B | 2.6e-06 | 4.2e-06 | 1.9e-06 | 2.900E-06 | 1.179E-06 | 1.900E-06 | 4.200E-06 |
| 16561 | 829 | K829A | B | 3.6e-06 | 2.7e-06 | 1.6e-06 | 2.633E-06 | 1.002E-06 | 1.600E-06 | 3.600E-06 |
| 16562 | 829 | K829R | B | 3.6e-06 | 1.3e-06 | 1.9e-06 | 2.267E-06 | 1.193E-06 | 1.300E-06 | 3.600E-06 |
| 16563 | 829 | K829N | B | 2.6e-06 | 1.5e-06 | 1.3e-06 | 1.800E-06 | 7.000E-07 | 1.300E-06 | 2.600E-06 |
| 16564 | 829 | K829D | B | 1.8e-06 | 1.4e-06 | 1.9e-06 | 1.700E-06 | 2.646E-07 | 1.400E-06 | 1.900E-06 |
| 16565 | 829 | K829C | B | 2.1e-06 | 8.5e-07 | 2e-06 | 1.650E-06 | 6.946E-07 | 8.500E-07 | 2.100E-06 |
| 16566 | 829 | K829E | B | 2.4e-06 | 2e-06 | 2.8e-06 | 2.400E-06 | 4.000E-07 | 2.000E-06 | 2.800E-06 |
| 16567 | 829 | K829Q | B | 3.4e-06 | 2.8e-06 | 3e-06 | 3.067E-06 | 3.055E-07 | 2.800E-06 | 3.400E-06 |
| 16568 | 829 | K829G | B | 2.9e-06 | 2.9e-06 | 2.6e-06 | 2.800E-06 | 1.732E-07 | 2.600E-06 | 2.900E-06 |
| 16569 | 829 | K829H | B | 1.8e-06 | 2e-06 | 1.5e-06 | 1.767E-06 | 2.517E-07 | 1.500E-06 | 2.000E-06 |
| 16570 | 829 | K829I | B | 1.8e-06 | 3.1e-06 | 2.7e-06 | 2.533E-06 | 6.658E-07 | 1.800E-06 | 3.100E-06 |
| 16571 | 829 | K829L | B | 2.3e-06 | 3.5e-06 | 3.1e-06 | 2.967E-06 | 6.110E-07 | 2.300E-06 | 3.500E-06 |
| 16572 | 829 | K829K | B | 9.4e-07 | 1.8e-06 | 1.7e-06 | 1.480E-06 | 4.703E-07 | 9.400E-07 | 1.800E-06 |
| 16573 | 829 | K829M | B | 3.1e-06 | 1.7e-06 | 3.4e-06 | 2.733E-06 | 9.074E-07 | 1.700E-06 | 3.400E-06 |
| 16574 | 829 | K829F | B | 2.4e-06 | 1.2e-06 | 4.8e-06 | 2.800E-06 | 1.833E-06 | 1.200E-06 | 4.800E-06 |

| | | | | | | | | | | |
|---|---|---|---|---|---|---|---|---|---|---|
| 16575 | 829 | K829P | B | 2e-06 | 4e-06 | 1.8e-06 | 2.600E-06 | 1.217E-06 | 1.800E-06 | 4.000E-06 |
| 16576 | 829 | K829S | B | 9.8e-07 | 1.2e-06 | 1.5e-06 | 1.227E-06 | 2.610E-07 | 9.800E-07 | 1.500E-06 |
| 16577 | 829 | K829T | B | 1.7e-06 | 2.2e-06 | 3.5e-06 | 2.467E-06 | 9.292E-07 | 1.700E-06 | 3.500E-06 |
| 16578 | 829 | K829W | B | 2.7e-06 | 2.1e-06 | 1.8e-06 | 2.200E-06 | 4.583E-07 | 1.800E-06 | 2.700E-06 |
| 16579 | 829 | K829Y | B | 1.7e-06 | 3.3e-06 | 2.8e-06 | 2.600E-06 | 8.185E-07 | 1.700E-06 | 3.300E-06 |
| 16580 | 829 | K829V | B | 1.1e-06 | 2.1e-06 | 1.9e-06 | 1.700E-06 | 5.292E-07 | 1.100E-06 | 2.100E-06 |
| 16581 | 830 | S830A | B | 4.4e-06 | 3.2e-06 | 3.6e-06 | 3.733E-06 | 6.110E-07 | 3.200E-06 | 4.400E-06 |
| 16582 | 830 | S830R | B | 1.8e-06 | 2.3e-06 | 1.8e-06 | 1.967E-06 | 2.887E-07 | 1.800E-06 | 2.300E-06 |
| 16583 | 830 | S830N | B | 1.5e-06 | 3.3e-06 | 1.9e-06 | 2.233E-06 | 9.452E-07 | 1.500E-06 | 3.300E-06 |
| 16584 | 830 | S830D | B | 3.1e-06 | 3.4e-06 | 2.7e-06 | 3.067E-06 | 3.512E-07 | 2.700E-06 | 3.400E-06 |
| 16585 | 830 | S830C | B | 1.6e-06 | 3.3e-06 | 2.4e-06 | 2.433E-06 | 8.505E-07 | 1.600E-06 | 3.300E-06 |
| 16586 | 830 | S830E | B | 2.5e-06 | 1.5e-06 | 1.5e-06 | 1.833E-06 | 5.774E-07 | 1.500E-06 | 2.500E-06 |
| 16587 | 830 | S830Q | B | 3.4e-06 | 1.7e-06 | 1.9e-06 | 2.333E-06 | 9.292E-07 | 1.700E-06 | 3.400E-06 |
| 16588 | 830 | S830G | B | 1.8e-06 | 2.8e-06 | 2.9e-06 | 2.500E-06 | 6.083E-07 | 1.800E-06 | 2.900E-06 |
| 16589 | 830 | S830H | B | 1.7e-06 | 1.9e-06 | 2e-06 | 1.867E-06 | 1.528E-07 | 1.700E-06 | 2.000E-06 |
| 16590 | 830 | S830I | B | 4.8e-06 | 3.7e-06 | 1.4e-06 | 3.300E-06 | 1.735E-06 | 1.400E-06 | 4.800E-06 |
| 16591 | 830 | S830L | B | 2.5e-06 | 2.6e-06 | 2.9e-06 | 2.667E-06 | 2.082E-07 | 2.500E-06 | 2.900E-06 |
| 16592 | 830 | S830K | B | 1.8e-06 | 3.7e-06 | 2e-06 | 2.500E-06 | 1.044E-06 | 1.800E-06 | 3.700E-06 |
| 16593 | 830 | S830M | B | 2e-06 | 2.5e-06 | 2.8e-06 | 2.433E-06 | 4.041E-07 | 2.000E-06 | 2.800E-06 |
| 16594 | 830 | S830F | B | 2.5e-06 | 3.8e-06 | 2.6e-06 | 2.967E-06 | 7.234E-07 | 2.500E-06 | 3.800E-06 |
| 16595 | 830 | S830P | B | 2.5e-06 | 3.8e-06 | 1.7e-06 | 2.667E-06 | 1.060E-06 | 1.700E-06 | 3.800E-06 |
| 16596 | 830 | S830S | B | 2.6e-06 | 1.5e-06 | 3e-06 | 2.367E-06 | 7.767E-07 | 1.500E-06 | 3.000E-06 |
| 16597 | 830 | S830T | B | 2.8e-06 | 1.9e-06 | 1.5e-06 | 2.067E-06 | 6.658E-07 | 1.500E-06 | 2.800E-06 |
| 16598 | 830 | S830W | B | 1.4e-06 | 1.3e-06 | 2.9e-06 | 1.867E-06 | 8.963E-07 | 1.300E-06 | 2.900E-06 |
| 16599 | 830 | S830Y | B | 2.2e-06 | 3.3e-06 | 1.6e-06 | 2.367E-06 | 8.622E-07 | 1.600E-06 | 3.300E-06 |
| 16600 | 830 | S830V | B | 1.3e-06 | 2.2e-06 | 2e-06 | 1.833E-06 | 4.726E-07 | 1.300E-06 | 2.200E-06 |
| 16601 | 831 | F831A | B | 9.6e-07 | 3e-06 | 4.1e-06 | 2.687E-06 | 1.593E-06 | 9.600E-07 | 4.100E-06 |
| 16602 | 831 | F831R | B | 1.9e-06 | 1.4e-06 | 2.6e-06 | 1.967E-06 | 6.028E-07 | 1.400E-06 | 2.600E-06 |
| 16603 | 831 | F831N | B | 1.6e-06 | 1.6e-06 | 3e-06 | 2.067E-06 | 8.083E-07 | 1.600E-06 | 3.000E-06 |
| 16604 | 831 | F831D | B | 2.9e-06 | 3.8e-06 | 5.7e-06 | 4.133E-06 | 1.429E-06 | 2.900E-06 | 5.700E-06 |
| 16605 | 831 | F831C | B | 2.5e-06 | 2.2e-06 | 1.4e-06 | 2.033E-06 | 5.686E-07 | 1.400E-06 | 2.500E-06 |
| 16606 | 831 | F831E | B | 1.8e-06 | 3.1e-06 | 2.4e-06 | 2.433E-06 | 6.506E-07 | 1.800E-06 | 3.100E-06 |
| 16607 | 831 | F831Q | B | 2.9e-06 | 1.9e-06 | 1.8e-06 | 2.200E-06 | 6.083E-07 | 1.800E-06 | 2.900E-06 |
| 16608 | 831 | F831G | B | 4e-06 | 2e-06 | 1.8e-06 | 2.600E-06 | 1.217E-06 | 1.800E-06 | 4.000E-06 |
| 16609 | 831 | F831H | B | 2e-06 | 3.2e-06 | 1.6e-06 | 2.267E-06 | 8.327E-07 | 1.600E-06 | 3.200E-06 |
| 16610 | 831 | F831I | B | 3.1e-06 | 2.6e-06 | 2.8e-06 | 2.833E-06 | 2.517E-07 | 2.600E-06 | 3.100E-06 |
| 16611 | 831 | F831L | B | 2.5e-06 | 2.5e-06 | 1.9e-06 | 2.300E-06 | 3.464E-07 | 1.900E-06 | 2.500E-06 |
| 16612 | 831 | F831K | B | 2e-06 | 1.6e-06 | 2.2e-06 | 1.933E-06 | 3.055E-07 | 1.600E-06 | 2.200E-06 |
| 16613 | 831 | F831M | B | 3.1e-06 | 3.3e-06 | 3.8e-06 | 3.400E-06 | 3.606E-07 | 3.100E-06 | 3.800E-06 |
| 16614 | 831 | F831F | B | 2e-06 | 2.5e-06 | 2.8e-06 | 2.433E-06 | 4.041E-07 | 2.000E-06 | 2.800E-06 |
| 16615 | 831 | F831P | B | 2.3e-06 | 1.1e-06 | 1.9e-06 | 1.767E-06 | 6.110E-07 | 1.100E-06 | 2.300E-06 |
| 16616 | 831 | F831S | B | 1.4e-06 | 1.8e-06 | 1.8e-06 | 1.667E-06 | 2.309E-07 | 1.400E-06 | 1.800E-06 |
| 16617 | 831 | F831T | B | 1.6e-06 | 1.6e-06 | 2.5e-06 | 1.900E-06 | 5.196E-07 | 1.600E-06 | 2.500E-06 |
| 16618 | 831 | F831W | B | 1.4e-06 | 2.1e-06 | 1.5e-06 | 1.667E-06 | 3.786E-07 | 1.400E-06 | 2.100E-06 |
| 16619 | 831 | F831Y | B | 9.6e-07 | 4e-06 | 1.9e-06 | 2.287E-06 | 1.556E-06 | 9.600E-07 | 4.000E-06 |
| 16620 | 831 | F831V | B | 2.6e-06 | 4.6e-06 | 3.2e-06 | 3.467E-06 | 1.026E-06 | 2.600E-06 | 4.600E-06 |
| 16621 | 832 | N832A | B | 3.4e-06 | 3e-06 | 3.8e-06 | 3.400E-06 | 4.000E-07 | 3.000E-06 | 3.800E-06 |
| 16622 | 832 | N832R | B | 1.7e-06 | 3.1e-06 | 2.1e-06 | 2.300E-06 | 7.211E-07 | 1.700E-06 | 3.100E-06 |
| 16623 | 832 | N832N | B | 2.2e-06 | 2.6e-06 | 2.3e-06 | 2.367E-06 | 2.082E-07 | 2.200E-06 | 2.600E-06 |
| 16624 | 832 | N832D | B | 3.1e-06 | 3.8e-06 | 3.3e-06 | 3.400E-06 | 3.606E-07 | 3.100E-06 | 3.800E-06 |
| 16625 | 832 | N832C | B | 3.4e-06 | 2e-06 | 2.9e-06 | 2.767E-06 | 7.095E-07 | 2.000E-06 | 3.400E-06 |
| 16626 | 832 | N832E | B | 3.5e-06 | 2.4e-06 | 2.1e-06 | 2.667E-06 | 7.371E-07 | 2.100E-06 | 3.500E-06 |
| 16627 | 832 | N832Q | B | 2.1e-06 | 2.9e-06 | 1.6e-06 | 2.200E-06 | 6.557E-07 | 1.600E-06 | 2.900E-06 |
| 16628 | 832 | N832G | B | 1.5e-06 | 1.8e-06 | 3.3e-06 | 2.200E-06 | 9.644E-07 | 1.500E-06 | 3.300E-06 |
| 16629 | 832 | N832H | B | 1.5e-06 | 2.2e-06 | 3.7e-06 | 2.467E-06 | 1.124E-06 | 1.500E-06 | 3.700E-06 |
| 16630 | 832 | N832I | B | 1.4e-06 | 2.1e-06 | 4.1e-06 | 2.533E-06 | 1.401E-06 | 1.400E-06 | 4.100E-06 |
| 16631 | 832 | N832L | B | 3.7e-06 | 1.1e-06 | 1.3e-06 | 2.033E-06 | 1.447E-06 | 1.100E-06 | 3.700E-06 |
| 16632 | 832 | N832K | B | 1.5e-06 | 3.3e-06 | 2.5e-06 | 2.433E-06 | 9.018E-07 | 1.500E-06 | 3.300E-06 |
| 16633 | 832 | N832M | B | 1.8e-06 | 2.7e-06 | 1.3e-06 | 1.933E-06 | 7.095E-07 | 1.300E-06 | 2.700E-06 |
| 16634 | 832 | N832F | B | 2.1e-06 | 1.7e-06 | 2.6e-06 | 2.133E-06 | 4.509E-07 | 1.700E-06 | 2.600E-06 |
| 16635 | 832 | N832P | B | 2.6e-06 | 3.7e-06 | 2.3e-06 | 2.867E-06 | 7.371E-07 | 2.300E-06 | 3.700E-06 |
| 16636 | 832 | N832S | B | 3.1e-06 | 1.5e-06 | 2.1e-06 | 2.233E-06 | 8.083E-07 | 1.500E-06 | 3.100E-06 |
| 16637 | 832 | N832T | B | 2.9e-06 | 2.6e-06 | 4e-06 | 3.167E-06 | 7.371E-07 | 2.600E-06 | 4.000E-06 |
| 16638 | 832 | N832W | B | 2.2e-06 | 1.9e-06 | 3.6e-06 | 2.567E-06 | 9.074E-07 | 1.900E-06 | 3.600E-06 |
| 16639 | 832 | N832Y | B | 2.7e-06 | 2.6e-06 | 1.9e-06 | 2.400E-06 | 4.359E-07 | 1.900E-06 | 2.700E-06 |
| 16640 | 832 | N832V | B | 2e-06 | 1.6e-06 | 1.9e-06 | 1.833E-06 | 2.082E-07 | 1.600E-06 | 2.000E-06 |
| 16641 | 833 | R833A | B | 2.8e-06 | 1.4e-06 | 2.7e-06 | 2.300E-06 | 7.810E-07 | 1.400E-06 | 2.800E-06 |
| 16642 | 833 | R833R | B | 1.8e-06 | 1.3e-06 | 1.7e-06 | 1.600E-06 | 2.646E-07 | 1.300E-06 | 1.800E-06 |
| 16643 | 833 | R833N | B | 2.3e-06 | 2.4e-06 | 2.2e-06 | 2.300E-06 | 1.000E-07 | 2.200E-06 | 2.400E-06 |
| 16644 | 833 | R833D | B | 2.4e-06 | 2.3e-06 | 1.8e-06 | 2.167E-06 | 3.215E-07 | 1.800E-06 | 2.400E-06 |
| 16645 | 833 | R833C | B | 3.3e-06 | 2.9e-06 | 2.6e-06 | 2.933E-06 | 3.512E-07 | 2.600E-06 | 3.300E-06 |
| 16646 | 833 | R833E | B | 1.9e-06 | 1.7e-06 | 2.6e-06 | 2.067E-06 | 4.726E-07 | 1.700E-06 | 2.600E-06 |
| 16647 | 833 | R833Q | B | 3.3e-06 | 2.1e-06 | 4.1e-06 | 3.167E-06 | 1.007E-06 | 2.100E-06 | 4.100E-06 |
| 16648 | 833 | R833G | B | 1.9e-06 | 3.1e-06 | 2.2e-06 | 2.400E-06 | 6.245E-07 | 1.900E-06 | 3.100E-06 |
| 16649 | 833 | R833H | B | 3.7e-06 | 3.2e-06 | 2.5e-06 | 3.133E-06 | 6.028E-07 | 2.500E-06 | 3.700E-06 |
| 16650 | 833 | R833I | B | 1.8e-06 | 3e-06 | 2e-06 | 2.267E-06 | 6.429E-07 | 1.800E-06 | 3.000E-06 |
| 16651 | 833 | R833L | B | 2e-06 | 2.5e-06 | 3.4e-06 | 2.633E-06 | 7.095E-07 | 2.000E-06 | 3.400E-06 |
| 16652 | 833 | R833K | B | 4.4e-06 | 2.7e-06 | 1.8e-06 | 2.967E-06 | 1.320E-06 | 1.800E-06 | 4.400E-06 |
| 16653 | 833 | R833M | B | 1.4e-06 | 2.2e-06 | 1.4e-06 | 1.667E-06 | 4.619E-07 | 1.400E-06 | 2.200E-06 |
| 16654 | 833 | R833F | B | 1.5e-06 | 4.3e-06 | 1.5e-06 | 2.433E-06 | 1.617E-06 | 1.500E-06 | 4.300E-06 |
| 16655 | 833 | R833P | B | 2.9e-06 | 1.9e-06 | 2.3e-06 | 2.367E-06 | 5.033E-07 | 1.900E-06 | 2.900E-06 |
| 16656 | 833 | R833S | B | 2.4e-06 | 4.3e-06 | 2e-06 | 2.900E-06 | 1.229E-06 | 2.000E-06 | 4.300E-06 |
| 16657 | 833 | R833T | B | 2.3e-06 | 1.3e-06 | 2.4e-06 | 2.000E-06 | 6.083E-07 | 1.300E-06 | 2.400E-06 |
| 16658 | 833 | R833W | B | 1.2e-06 | 2.1e-06 | 3.1e-06 | 2.133E-06 | 9.504E-07 | 1.200E-06 | 3.100E-06 |
| 16659 | 833 | R833Y | B | 3.7e-06 | 2.5e-06 | 2.7e-06 | 2.967E-06 | 6.429E-07 | 2.500E-06 | 3.700E-06 |

| | | | | | | | | | | |
|---|---|---|---|---|---|---|---|---|---|---|
| 16660 | 833 | R833V | B | 1.9e-06 | 2.8e-06 | 4.2e-06 | 2.967E-06 | 1.159E-06 | 1.900E-06 | 4.200E-06 |
| 16661 | 834 | G834A | B | 1.2e-06 | 2e-06 | 4.4e-06 | 2.533E-06 | 1.665E-06 | 1.200E-06 | 4.400E-06 |
| 16662 | 834 | G834R | B | 4.1e-06 | 2.3e-06 | 2.2e-06 | 2.867E-06 | 1.069E-06 | 2.200E-06 | 4.100E-06 |
| 16663 | 834 | G834N | B | 2e-06 | 1.6e-06 | 2.6e-06 | 2.067E-06 | 5.033E-07 | 1.600E-06 | 2.600E-06 |
| 16664 | 834 | G834D | B | 1.7e-06 | 3.6e-06 | 2e-06 | 2.433E-06 | 1.021E-06 | 1.700E-06 | 3.600E-06 |
| 16665 | 834 | G834C | B | 2.3e-06 | 2.5e-06 | 2.7e-06 | 2.500E-06 | 2.000E-07 | 2.300E-06 | 2.700E-06 |
| 16666 | 834 | G834E | B | 3.8e-06 | 2.4e-06 | 2e-06 | 2.733E-06 | 9.452E-07 | 2.000E-06 | 3.800E-06 |
| 16667 | 834 | G834Q | B | 3.2e-06 | 1.5e-06 | 1.5e-06 | 2.067E-06 | 9.815E-07 | 1.500E-06 | 3.200E-06 |
| 16668 | 834 | G834G | B | 2.6e-06 | 2.4e-06 | 2.2e-06 | 2.400E-06 | 2.000E-07 | 2.200E-06 | 2.600E-06 |
| 16669 | 834 | G834H | B | 1.6e-06 | 4.2e-06 | 1.2e-06 | 2.333E-06 | 1.629E-06 | 1.200E-06 | 4.200E-06 |
| 16670 | 834 | G834I | B | 2e-06 | 1.4e-06 | 3.6e-06 | 2.333E-06 | 1.137E-06 | 1.400E-06 | 3.600E-06 |
| 16671 | 834 | G834L | B | 2.5e-06 | 2.4e-06 | 2.6e-06 | 2.500E-06 | 1.000E-07 | 2.400E-06 | 2.600E-06 |
| 16672 | 834 | G834K | B | 1.5e-06 | 2.7e-06 | 2.9e-06 | 2.367E-06 | 7.572E-07 | 1.500E-06 | 2.900E-06 |
| 16673 | 834 | G834M | B | 3.1e-06 | 2e-06 | 2.1e-06 | 2.400E-06 | 6.083E-07 | 2.000E-06 | 3.100E-06 |
| 16674 | 834 | G834F | B | 2.4e-06 | 2e-06 | 3.2e-06 | 2.533E-06 | 6.110E-07 | 2.000E-06 | 3.200E-06 |
| 16675 | 834 | G834P | B | 1.1e-06 | 1.7e-06 | 4.5e-06 | 2.433E-06 | 1.815E-06 | 1.100E-06 | 4.500E-06 |
| 16676 | 834 | G834S | B | 1.2e-06 | 3.2e-06 | 3.6e-06 | 2.667E-06 | 1.286E-06 | 1.200E-06 | 3.600E-06 |
| 16677 | 834 | G834T | B | 2.5e-06 | 3.2e-06 | 2.6e-06 | 2.767E-06 | 3.786E-07 | 2.500E-06 | 3.200E-06 |
| 16678 | 834 | G834W | B | 4.3e-06 | 1.2e-06 | 1.7e-06 | 2.400E-06 | 1.664E-06 | 1.200E-06 | 4.300E-06 |
| 16679 | 834 | G834Y | B | 1.4e-06 | 2.3e-06 | 3.2e-06 | 2.300E-06 | 9.000E-07 | 1.400E-06 | 3.200E-06 |
| 16680 | 834 | G834V | B | 2.1e-06 | 2.7e-06 | 1.7e-06 | 2.167E-06 | 5.033E-07 | 1.700E-06 | 2.700E-06 |
| 16681 | 835 | E835A | B | 3.4e-06 | 2.6e-06 | 1.8e-06 | 2.600E-06 | 8.000E-07 | 1.800E-06 | 3.400E-06 |
| 16682 | 835 | E835R | B | 4.6e-06 | 1.3e-06 | 1.8e-06 | 2.567E-06 | 1.779E-06 | 1.300E-06 | 4.600E-06 |
| 16683 | 835 | E835N | B | 5.4e-06 | 2.7e-06 | 4.2e-06 | 4.100E-06 | 1.353E-06 | 2.700E-06 | 5.400E-06 |
| 16684 | 835 | E835D | B | 2.1e-06 | 4.7e-06 | 2.5e-06 | 3.100E-06 | 1.400E-06 | 2.100E-06 | 4.700E-06 |
| 16685 | 835 | E835C | B | 2.5e-06 | 3.4e-06 | 3.2e-06 | 3.033E-06 | 4.726E-07 | 2.500E-06 | 3.400E-06 |
| 16686 | 835 | E835E | B | 3.1e-06 | 2.3e-06 | 2.3e-06 | 2.567E-06 | 4.619E-07 | 2.300E-06 | 3.100E-06 |
| 16687 | 835 | E835Q | B | 1.9e-06 | 2.4e-06 | 2.4e-06 | 2.233E-06 | 2.887E-07 | 1.900E-06 | 2.400E-06 |
| 16688 | 835 | E835G | B | 2e-06 | 2.3e-06 | 3.7e-06 | 2.667E-06 | 9.074E-07 | 2.000E-06 | 3.700E-06 |
| 16689 | 835 | E835H | B | 1.9e-06 | 1.2e-06 | 3e-06 | 2.033E-06 | 9.074E-07 | 1.200E-06 | 3.000E-06 |
| 16690 | 835 | E835I | B | 1.9e-06 | 2e-06 | 1.3e-06 | 1.733E-06 | 3.786E-07 | 1.300E-06 | 2.000E-06 |
| 16691 | 835 | E835L | B | 2.3e-06 | 3e-06 | 2.3e-06 | 2.533E-06 | 4.041E-07 | 2.300E-06 | 3.000E-06 |
| 16692 | 835 | E835K | B | 1.8e-06 | 2.6e-06 | 1.6e-06 | 2.000E-06 | 5.292E-07 | 1.600E-06 | 2.600E-06 |
| 16693 | 835 | E835M | B | 2.6e-06 | 1.8e-06 | 3.5e-06 | 2.633E-06 | 8.505E-07 | 1.800E-06 | 3.500E-06 |
| 16694 | 835 | E835F | B | 2.6e-06 | 1.5e-06 | 1.8e-06 | 1.967E-06 | 5.686E-07 | 1.500E-06 | 2.600E-06 |
| 16695 | 835 | E835P | B | 5.3e-06 | 3.1e-06 | 1.7e-06 | 3.367E-06 | 1.815E-06 | 1.700E-06 | 5.300E-06 |
| 16696 | 835 | E835S | B | 2e-06 | 2.2e-06 | 1.3e-06 | 1.833E-06 | 4.726E-07 | 1.300E-06 | 2.200E-06 |
| 16697 | 835 | E835T | B | 4.5e-06 | 2.4e-06 | 1.7e-06 | 2.867E-06 | 1.457E-06 | 1.700E-06 | 4.500E-06 |
| 16698 | 835 | E835W | B | 4.1e-06 | 1.5e-06 | 3.3e-06 | 2.967E-06 | 1.332E-06 | 1.500E-06 | 4.100E-06 |
| 16699 | 835 | E835Y | B | 2.1e-06 | 2.3e-06 | 2.4e-06 | 2.267E-06 | 1.528E-07 | 2.100E-06 | 2.400E-06 |
| 16700 | 835 | E835V | B | 2.5e-06 | 3.2e-06 | 1.9e-06 | 2.533E-06 | 6.506E-07 | 1.900E-06 | 3.200E-06 |
| 16701 | 836 | C836A | B | 1.6e-06 | 1.2e-06 | 2.5e-06 | 1.767E-06 | 6.658E-07 | 1.200E-06 | 2.500E-06 |
| 16702 | 836 | C836R | B | 1.3e-06 | 2.7e-06 | 2.4e-06 | 2.133E-06 | 7.371E-07 | 1.300E-06 | 2.700E-06 |
| 16703 | 836 | C836N | B | 1.9e-06 | 1.8e-06 | 2.4e-06 | 2.033E-06 | 3.215E-07 | 1.800E-06 | 2.400E-06 |
| 16704 | 836 | C836D | B | 2e-06 | 2e-06 | 3e-06 | 2.333E-06 | 5.774E-07 | 2.000E-06 | 3.000E-06 |
| 16705 | 836 | C836C | B | 2e-06 | 4.2e-06 | 2.1e-06 | 2.767E-06 | 1.242E-06 | 2.000E-06 | 4.200E-06 |
| 16706 | 836 | C836E | B | 2.3e-06 | 2.7e-06 | 3.2e-06 | 2.733E-06 | 4.509E-07 | 2.300E-06 | 3.200E-06 |
| 16707 | 836 | C836Q | B | 2.4e-06 | 3e-06 | 3.6e-06 | 3.000E-06 | 6.000E-07 | 2.400E-06 | 3.600E-06 |
| 16708 | 836 | C836G | B | 3.7e-06 | 1.6e-06 | 2.5e-06 | 2.600E-06 | 1.054E-06 | 1.600E-06 | 3.700E-06 |
| 16709 | 836 | C836H | B | 3.7e-06 | 1.5e-06 | 2.6e-06 | 2.600E-06 | 1.100E-06 | 1.500E-06 | 3.700E-06 |
| 16710 | 836 | C836I | B | 4.7e-06 | 1.4e-06 | 1.5e-06 | 2.533E-06 | 1.877E-06 | 1.400E-06 | 4.700E-06 |
| 16711 | 836 | C836L | B | 2.3e-06 | 2.5e-06 | 1.7e-06 | 2.167E-06 | 4.163E-07 | 1.700E-06 | 2.500E-06 |
| 16712 | 836 | C836K | B | 1.9e-06 | 2e-06 | 1.3e-06 | 1.733E-06 | 3.786E-07 | 1.300E-06 | 2.000E-06 |
| 16713 | 836 | C836M | B | 1.9e-06 | 1.5e-06 | 2.4e-06 | 1.933E-06 | 4.509E-07 | 1.500E-06 | 2.400E-06 |
| 16714 | 836 | C836F | B | 1.6e-06 | 1.2e-06 | 3.2e-06 | 2.000E-06 | 1.058E-06 | 1.200E-06 | 3.200E-06 |
| 16715 | 836 | C836P | B | 2.9e-06 | 2.8e-06 | 1.9e-06 | 2.533E-06 | 5.508E-07 | 1.900E-06 | 2.900E-06 |
| 16716 | 836 | C836S | B | 2.3e-06 | 2e-06 | 3.9e-06 | 2.733E-06 | 1.021E-06 | 2.000E-06 | 3.900E-06 |
| 16717 | 836 | C836T | B | 2.7e-06 | 2.7e-06 | 1.7e-06 | 2.367E-06 | 5.774E-07 | 1.700E-06 | 2.700E-06 |
| 16718 | 836 | C836W | B | 1.6e-06 | 5e-06 | 2.9e-06 | 3.167E-06 | 1.716E-06 | 1.600E-06 | 5.000E-06 |
| 16719 | 836 | C836Y | B | 3.5e-06 | 1.5e-06 | 4.1e-06 | 3.033E-06 | 1.361E-06 | 1.500E-06 | 4.100E-06 |
| 16720 | 836 | C836V | B | 3.7e-06 | 2.6e-06 | 2e-06 | 2.767E-06 | 8.622E-07 | 2.000E-06 | 3.700E-06 |
| 16721 | 837 | E837A | C | 2.3e-06 | 3.3e-06 | 2.1e-06 | 2.567E-06 | 6.429E-07 | 2.100E-06 | 3.300E-06 |
| 16722 | 837 | E837R | C | 2.7e-06 | 1.9e-06 | 1.9e-06 | 2.167E-06 | 4.619E-07 | 1.900E-06 | 2.700E-06 |
| 16723 | 837 | E837N | C | 3e-06 | 2.4e-06 | 2.8e-06 | 2.733E-06 | 3.055E-07 | 2.400E-06 | 3.000E-06 |
| 16724 | 837 | E837D | C | 2.5e-06 | 8.9e-07 | 3.8e-06 | 2.397E-06 | 1.458E-06 | 8.900E-07 | 3.800E-06 |
| 16725 | 837 | E837C | C | 2.9e-06 | 2.5e-06 | 2e-06 | 2.467E-06 | 4.509E-07 | 2.000E-06 | 2.900E-06 |
| 16726 | 837 | E837E | C | 2.7e-06 | 2e-06 | 2.3e-06 | 2.333E-06 | 3.512E-07 | 2.000E-06 | 2.700E-06 |
| 16727 | 837 | E837Q | C | 1.8e-06 | 1.3e-06 | 3.1e-06 | 2.067E-06 | 9.292E-07 | 1.300E-06 | 3.100E-06 |
| 16728 | 837 | E837G | C | 2.2e-06 | 3.2e-06 | 4.3e-06 | 3.233E-06 | 1.050E-06 | 2.200E-06 | 4.300E-06 |
| 16729 | 837 | E837H | C | 2.9e-06 | 1.2e-06 | 1.9e-06 | 2.000E-06 | 8.544E-07 | 1.200E-06 | 2.900E-06 |
| 16730 | 837 | E837I | C | 3.1e-06 | 3.8e-06 | 3.5e-06 | 3.467E-06 | 3.512E-07 | 3.100E-06 | 3.800E-06 |
| 16731 | 837 | E837L | C | 1.8e-06 | 4.5e-06 | 3.4e-06 | 3.233E-06 | 1.358E-06 | 1.800E-06 | 4.500E-06 |
| 16732 | 837 | E837K | C | 2.5e-06 | 2.5e-06 | 1.7e-06 | 2.233E-06 | 4.619E-07 | 1.700E-06 | 2.500E-06 |
| 16733 | 837 | E837M | C | 2.7e-06 | 2.4e-06 | 2.8e-06 | 2.633E-06 | 2.082E-07 | 2.400E-06 | 2.800E-06 |
| 16734 | 837 | E837F | C | 1.9e-06 | 3e-06 | 3.2e-06 | 2.700E-06 | 7.000E-07 | 1.900E-06 | 3.200E-06 |
| 16735 | 837 | E837P | C | 3.8e-06 | 2.9e-06 | 2.1e-06 | 2.933E-06 | 8.505E-07 | 2.100E-06 | 3.800E-06 |
| 16736 | 837 | E837S | C | 2.9e-06 | 2.3e-06 | 2.7e-06 | 2.633E-06 | 3.055E-07 | 2.300E-06 | 2.900E-06 |
| 16737 | 837 | E837T | C | 2.2e-06 | 2.3e-06 | 2.6e-06 | 2.367E-06 | 2.082E-07 | 2.200E-06 | 2.600E-06 |
| 16738 | 837 | E837W | C | 3.2e-06 | 1.8e-06 | 1.9e-06 | 2.300E-06 | 7.810E-07 | 1.800E-06 | 3.200E-06 |
| 16739 | 837 | E837Y | C | 1.6e-06 | 1.9e-06 | 2.9e-06 | 2.133E-06 | 6.807E-07 | 1.600E-06 | 2.900E-06 |
| 16740 | 837 | E837V | C | 2.8e-06 | 3.5e-06 | 3.1e-06 | 3.133E-06 | 3.512E-07 | 2.800E-06 | 3.500E-06 |
| 16741 | 838 | V838A | C | 2.7e-06 | 1.1e-06 | 1.8e-06 | 1.867E-06 | 8.021E-07 | 1.100E-06 | 2.700E-06 |
| 16742 | 838 | V838R | C | 2.8e-06 | 2.5e-06 | 2.9e-06 | 2.733E-06 | 2.082E-07 | 2.500E-06 | 2.900E-06 |
| 16743 | 838 | V838N | C | 3.9e-06 | 5e-06 | 1.6e-06 | 3.500E-06 | 1.735E-06 | 1.600E-06 | 5.000E-06 |
| 16744 | 838 | V838D | C | 2.7e-06 | 2.5e-06 | 3.1e-06 | 2.767E-06 | 3.055E-07 | 2.500E-06 | 3.100E-06 |

| | | | | | | | | | | |
|---|---|---|---|---|---|---|---|---|---|---|
| 16745 | 838 | V838C | C | 3.2e-06 | 2.6e-06 | 2.2e-06 | 2.667E-06 | 5.033E-07 | 2.200E-06 | 3.200E-06 |
| 16746 | 838 | V838E | C | 1.2e-06 | 1e-06 | 1.5e-06 | 1.233E-06 | 2.517E-07 | 1.000E-06 | 1.500E-06 |
| 16747 | 838 | V838Q | C | 2.1e-06 | 1.5e-06 | 2.5e-06 | 2.033E-06 | 5.033E-07 | 1.500E-06 | 2.500E-06 |
| 16748 | 838 | V838G | C | 1.1e-06 | 2.8e-06 | 1.8e-06 | 1.900E-06 | 8.544E-07 | 1.100E-06 | 2.800E-06 |
| 16749 | 838 | V838H | C | 1.8e-06 | 2.2e-06 | 3.4e-06 | 2.467E-06 | 8.327E-07 | 1.800E-06 | 3.400E-06 |
| 16750 | 838 | V838I | C | 1.2e-06 | 2.3e-06 | 1.6e-06 | 1.700E-06 | 5.568E-07 | 1.200E-06 | 2.300E-06 |
| 16751 | 838 | V838L | C | 3.8e-06 | 2.1e-06 | 1.6e-06 | 2.500E-06 | 1.153E-06 | 1.600E-06 | 3.800E-06 |
| 16752 | 838 | V838K | C | 3.8e-06 | 2.4e-06 | 3e-06 | 3.067E-06 | 7.024E-07 | 2.400E-06 | 3.800E-06 |
| 16753 | 838 | V838M | C | 2.8e-06 | 4.4e-06 | 1.4e-06 | 2.867E-06 | 1.501E-06 | 1.400E-06 | 4.400E-06 |
| 16754 | 838 | V838F | C | 1.5e-06 | 1.9e-06 | 1.8e-06 | 1.733E-06 | 2.082E-07 | 1.500E-06 | 1.900E-06 |
| 16755 | 838 | V838P | C | 5.5e-06 | 2.6e-06 | 2.2e-06 | 3.433E-06 | 1.801E-06 | 2.200E-06 | 5.500E-06 |
| 16756 | 838 | V838S | C | 2.3e-06 | 1.6e-06 | 2.5e-06 | 2.133E-06 | 4.726E-07 | 1.600E-06 | 2.500E-06 |
| 16757 | 838 | V838T | C | 2e-06 | 2.6e-06 | 2.1e-06 | 2.233E-06 | 3.215E-07 | 2.000E-06 | 2.600E-06 |
| 16758 | 838 | V838W | C | 2.7e-06 | 1.2e-06 | 3.1e-06 | 2.333E-06 | 1.002E-06 | 1.200E-06 | 3.100E-06 |
| 16759 | 838 | V838Y | C | 1.8e-06 | 2.5e-06 | 2.5e-06 | 2.267E-06 | 4.041E-07 | 1.800E-06 | 2.500E-06 |
| 16760 | 838 | V838V | C | 4.3e-06 | 4.4e-06 | 1.9e-06 | 3.533E-06 | 1.415E-06 | 1.900E-06 | 4.400E-06 |
| 16761 | 839 | Q839A | C | 3.1e-06 | 3.7e-06 | 2.2e-06 | 3.000E-06 | 7.550E-07 | 2.200E-06 | 3.700E-06 |
| 16762 | 839 | Q839R | C | 2e-06 | 2.6e-06 | 2e-06 | 2.200E-06 | 3.464E-07 | 2.000E-06 | 2.600E-06 |
| 16763 | 839 | Q839N | C | 2.8e-06 | 3e-06 | 1.9e-06 | 2.567E-06 | 5.859E-07 | 1.900E-06 | 3.000E-06 |
| 16764 | 839 | Q839D | C | 2.1e-06 | 3e-06 | 3.3e-06 | 2.800E-06 | 6.245E-07 | 2.100E-06 | 3.300E-06 |
| 16765 | 839 | Q839C | C | 1.3e-06 | 2.2e-06 | 4e-06 | 2.500E-06 | 1.375E-06 | 1.300E-06 | 4.000E-06 |
| 16766 | 839 | Q839E | C | 3.6e-06 | 1.8e-06 | 3e-06 | 2.800E-06 | 9.165E-07 | 1.800E-06 | 3.600E-06 |
| 16767 | 839 | Q839Q | C | 2.2e-06 | 3.2e-06 | 2e-06 | 2.467E-06 | 6.429E-07 | 2.000E-06 | 3.200E-06 |
| 16768 | 839 | Q839G | C | 3.3e-06 | 1.3e-06 | 3.3e-06 | 2.633E-06 | 1.155E-06 | 1.300E-06 | 3.300E-06 |
| 16769 | 839 | Q839H | C | 3e-06 | 2.3e-06 | 1.8e-06 | 2.367E-06 | 6.028E-07 | 1.800E-06 | 3.000E-06 |
| 16770 | 839 | Q839I | C | 2.3e-06 | 2.6e-06 | 1.5e-06 | 2.133E-06 | 5.686E-07 | 1.500E-06 | 2.600E-06 |
| 16771 | 839 | Q839L | C | 2.5e-06 | 1.6e-06 | 1.6e-06 | 1.900E-06 | 5.196E-07 | 1.600E-06 | 2.500E-06 |
| 16772 | 839 | Q839K | C | 3.3e-06 | 1.8e-06 | 1.5e-06 | 2.200E-06 | 9.644E-07 | 1.500E-06 | 3.300E-06 |
| 16773 | 839 | Q839M | C | 2.4e-06 | 3.3e-06 | 1.9e-06 | 2.533E-06 | 7.095E-07 | 1.900E-06 | 3.300E-06 |
| 16774 | 839 | Q839F | C | 2.1e-06 | 1.8e-06 | 4.5e-06 | 2.800E-06 | 1.480E-06 | 1.800E-06 | 4.500E-06 |
| 16775 | 839 | Q839P | C | 3.8e-06 | 2.1e-06 | 2.7e-06 | 2.867E-06 | 8.622E-07 | 2.100E-06 | 3.800E-06 |
| 16776 | 839 | Q839S | C | 2.8e-06 | 3.7e-06 | 2.3e-06 | 2.933E-06 | 7.095E-07 | 2.300E-06 | 3.700E-06 |
| 16777 | 839 | Q839T | C | 2.4e-06 | 1.7e-06 | 2.6e-06 | 2.233E-06 | 4.726E-07 | 1.700E-06 | 2.600E-06 |
| 16778 | 839 | Q839W | C | 3.8e-06 | 2.8e-06 | 2.1e-06 | 2.900E-06 | 8.544E-07 | 2.100E-06 | 3.800E-06 |
| 16779 | 839 | Q839Y | C | 2.9e-06 | 2.1e-06 | 3.4e-06 | 2.800E-06 | 6.557E-07 | 2.100E-06 | 3.400E-06 |
| 16780 | 839 | Q839V | C | 3.5e-06 | 4e-06 | 2.8e-06 | 3.433E-06 | 6.028E-07 | 2.800E-06 | 4.000E-06 |
| 16781 | 840 | L840A | C | 2.5e-06 | 2.1e-06 | 3.5e-06 | 2.700E-06 | 7.211E-07 | 2.100E-06 | 3.500E-06 |
| 16782 | 840 | L840R | C | 2.4e-06 | 2.8e-06 | 2e-06 | 2.400E-06 | 4.000E-07 | 2.000E-06 | 2.800E-06 |
| 16783 | 840 | L840N | C | 2.2e-06 | 3e-06 | 1.6e-06 | 2.267E-06 | 7.024E-07 | 1.600E-06 | 3.000E-06 |
| 16784 | 840 | L840D | C | 2.4e-06 | 2.2e-06 | 4.4e-06 | 3.000E-06 | 1.217E-06 | 2.200E-06 | 4.400E-06 |
| 16785 | 840 | L840C | C | 1.3e-06 | 2.1e-06 | 2.1e-06 | 1.833E-06 | 4.619E-07 | 1.300E-06 | 2.100E-06 |
| 16786 | 840 | L840E | C | 3.1e-06 | 2.9e-06 | 1.9e-06 | 2.633E-06 | 6.429E-07 | 1.900E-06 | 3.100E-06 |
| 16787 | 840 | L840Q | C | 2e-06 | 2.6e-06 | 2.9e-06 | 2.500E-06 | 4.583E-07 | 2.000E-06 | 2.900E-06 |
| 16788 | 840 | L840G | C | 3.3e-06 | 3.5e-06 | 1.9e-06 | 2.900E-06 | 8.718E-07 | 1.900E-06 | 3.500E-06 |
| 16789 | 840 | L840H | C | 2.8e-06 | 1.8e-06 | 2.7e-06 | 2.433E-06 | 5.508E-07 | 1.800E-06 | 2.800E-06 |
| 16790 | 840 | L840I | C | 1.9e-06 | 2.7e-06 | 1.2e-06 | 1.933E-06 | 7.506E-07 | 1.200E-06 | 2.700E-06 |
| 16791 | 840 | L840L | C | 3.3e-06 | 3.2e-06 | 2.5e-06 | 3.000E-06 | 4.359E-07 | 2.500E-06 | 3.300E-06 |
| 16792 | 840 | L840K | C | 2.5e-06 | 3.1e-06 | 1.9e-06 | 2.500E-06 | 6.000E-07 | 1.900E-06 | 3.100E-06 |
| 16793 | 840 | L840M | C | 4.5e-06 | 3.6e-06 | 2.8e-06 | 3.633E-06 | 8.505E-07 | 2.800E-06 | 4.500E-06 |
| 16794 | 840 | L840F | C | 2.9e-06 | 1.5e-06 | 3e-06 | 2.467E-06 | 8.386E-07 | 1.500E-06 | 3.000E-06 |
| 16795 | 840 | L840P | C | 3.7e-06 | 3.6e-06 | 3.1e-06 | 3.467E-06 | 3.215E-07 | 3.100E-06 | 3.700E-06 |
| 16796 | 840 | L840S | C | 2e-06 | 4.6e-06 | 1.8e-06 | 2.800E-06 | 1.562E-06 | 1.800E-06 | 4.600E-06 |
| 16797 | 840 | L840T | C | 1.8e-06 | 1.2e-06 | 2.3e-06 | 1.767E-06 | 5.508E-07 | 1.200E-06 | 2.300E-06 |
| 16798 | 840 | L840W | C | 1.7e-06 | 2.8e-06 | 2.6e-06 | 2.367E-06 | 5.859E-07 | 1.700E-06 | 2.800E-06 |
| 16799 | 840 | L840Y | C | 1.6e-06 | 4.5e-06 | 2.3e-06 | 2.800E-06 | 1.513E-06 | 1.600E-06 | 4.500E-06 |
| 16800 | 840 | L840V | C | 1.7e-06 | 2.2e-06 | 3.7e-06 | 2.533E-06 | 1.041E-06 | 1.700E-06 | 3.700E-06 |
| 16801 | 841 | V841A | C | 2.1e-06 | 2.4e-06 | 2.8e-06 | 2.433E-06 | 3.512E-07 | 2.100E-06 | 2.800E-06 |
| 16802 | 841 | V841R | C | 2.3e-06 | 2.2e-06 | 3.3e-06 | 2.600E-06 | 6.083E-07 | 2.200E-06 | 3.300E-06 |
| 16803 | 841 | V841N | C | 2.1e-06 | 3.3e-06 | 1.9e-06 | 2.433E-06 | 7.572E-07 | 1.900E-06 | 3.300E-06 |
| 16804 | 841 | V841D | C | 2.1e-06 | 2.2e-06 | 1.7e-06 | 2.000E-06 | 2.646E-07 | 1.700E-06 | 2.200E-06 |
| 16805 | 841 | V841C | C | 3.8e-06 | 2.3e-06 | 1.2e-06 | 2.433E-06 | 1.305E-06 | 1.200E-06 | 3.800E-06 |
| 16806 | 841 | V841E | C | 1.4e-06 | 2.9e-06 | 1.6e-06 | 1.967E-06 | 8.145E-07 | 1.400E-06 | 2.900E-06 |
| 16807 | 841 | V841Q | C | 1.4e-06 | 2.5e-06 | 3.1e-06 | 2.333E-06 | 8.622E-07 | 1.400E-06 | 3.100E-06 |
| 16808 | 841 | V841G | C | 1.7e-06 | 1.7e-06 | 2.3e-06 | 1.900E-06 | 3.464E-07 | 1.700E-06 | 2.300E-06 |
| 16809 | 841 | V841H | C | 2e-06 | 3.6e-06 | 9.9e-07 | 2.197E-06 | 1.316E-06 | 9.900E-07 | 3.600E-06 |
| 16810 | 841 | V841I | C | 3.2e-06 | 2.8e-06 | 2.6e-06 | 2.867E-06 | 3.055E-07 | 2.600E-06 | 3.200E-06 |
| 16811 | 841 | V841L | C | 1.9e-06 | 3.3e-06 | 2.3e-06 | 2.500E-06 | 7.211E-07 | 1.900E-06 | 3.300E-06 |
| 16812 | 841 | V841K | C | 2e-06 | 2.3e-06 | 2.1e-06 | 2.133E-06 | 1.528E-07 | 2.000E-06 | 2.300E-06 |
| 16813 | 841 | V841M | C | 3e-06 | 1.9e-06 | 2.8e-06 | 2.567E-06 | 5.859E-07 | 1.900E-06 | 3.000E-06 |
| 16814 | 841 | V841F | C | 2.2e-06 | 1.9e-06 | 4.2e-06 | 2.767E-06 | 1.250E-06 | 1.900E-06 | 4.200E-06 |
| 16815 | 841 | V841P | C | 4.9e-06 | 1.3e-06 | 2.6e-06 | 2.933E-06 | 1.823E-06 | 1.300E-06 | 4.900E-06 |
| 16816 | 841 | V841S | C | 1.9e-06 | 2.7e-06 | 1.4e-06 | 2.000E-06 | 6.557E-07 | 1.400E-06 | 2.700E-06 |
| 16817 | 841 | V841T | C | 2.5e-06 | 3e-06 | 2.2e-06 | 2.567E-06 | 4.041E-07 | 2.200E-06 | 3.000E-06 |
| 16818 | 841 | V841W | C | 1.7e-06 | 3.4e-06 | 4.1e-06 | 3.067E-06 | 1.234E-06 | 1.700E-06 | 4.100E-06 |
| 16819 | 841 | V841Y | C | 2.7e-06 | 2.3e-06 | 2.5e-06 | 2.500E-06 | 2.000E-07 | 2.300E-06 | 2.700E-06 |
| 16820 | 841 | V841V | C | 2.7e-06 | 3.7e-06 | 2e-06 | 2.800E-06 | 8.544E-07 | 2.000E-06 | 3.700E-06 |
| 16821 | 842 | E842A | C | 3.2e-06 | 3.1e-06 | 3.2e-06 | 3.167E-06 | 5.774E-08 | 3.100E-06 | 3.200E-06 |
| 16822 | 842 | E842R | C | 2e-06 | 3.1e-06 | 2.6e-06 | 2.567E-06 | 5.508E-07 | 2.000E-06 | 3.100E-06 |
| 16823 | 842 | E842N | C | 2.5e-06 | 2.2e-06 | 1.8e-06 | 2.167E-06 | 3.512E-07 | 1.800E-06 | 2.500E-06 |
| 16824 | 842 | E842D | C | 2.2e-06 | 2.5e-06 | 2.9e-06 | 2.533E-06 | 3.512E-07 | 2.200E-06 | 2.900E-06 |
| 16825 | 842 | E842C | C | 2.7e-06 | 2.3e-06 | 2.6e-06 | 2.533E-06 | 2.082E-07 | 2.300E-06 | 2.700E-06 |
| 16826 | 842 | E842E | C | 2.8e-06 | 2.3e-06 | 3.4e-06 | 2.833E-06 | 5.508E-07 | 2.300E-06 | 3.400E-06 |
| 16827 | 842 | E842Q | C | 2.3e-06 | 3.2e-06 | 1.9e-06 | 2.467E-06 | 6.658E-07 | 1.900E-06 | 3.200E-06 |
| 16828 | 842 | E842G | C | 2.2e-06 | 3.1e-06 | 2.8e-06 | 2.700E-06 | 4.583E-07 | 2.200E-06 | 3.100E-06 |
| 16829 | 842 | E842H | C | 1.8e-06 | 2.3e-06 | 1.4e-06 | 1.833E-06 | 4.509E-07 | 1.400E-06 | 2.300E-06 |

| | | | | | | | | | | |
|---|---|---|---|---|---|---|---|---|---|---|
| 16830 | 842 | E842I | C | 1.7e-06 | 1.5e-06 | 1.4e-06 | 1.533E-06 | 1.528E-07 | 1.400E-06 | 1.700E-06 |
| 16831 | 842 | E842L | C | 2.3e-06 | 2.7e-06 | 1.9e-06 | 2.300E-06 | 4.000E-07 | 1.900E-06 | 2.700E-06 |
| 16832 | 842 | E842K | C | 2.8e-06 | 1.2e-06 | 3.9e-06 | 2.633E-06 | 1.358E-06 | 1.200E-06 | 3.900E-06 |
| 16833 | 842 | E842M | C | 2.6e-06 | 4.5e-06 | 1.2e-06 | 2.767E-06 | 1.656E-06 | 1.200E-06 | 4.500E-06 |
| 16834 | 842 | E842F | C | 2.7e-06 | 1.4e-06 | 2.3e-06 | 2.133E-06 | 6.658E-07 | 1.400E-06 | 2.700E-06 |
| 16835 | 842 | E842P | C | 3.4e-06 | 3.3e-06 | 2.8e-06 | 3.167E-06 | 3.215E-07 | 2.800E-06 | 3.400E-06 |
| 16836 | 842 | E842S | C | 3e-06 | 1.9e-06 | 3.2e-06 | 2.700E-06 | 7.000E-07 | 1.900E-06 | 3.200E-06 |
| 16837 | 842 | E842T | C | 6.5e-06 | 3.4e-06 | 3.2e-06 | 4.367E-06 | 1.850E-06 | 3.200E-06 | 6.500E-06 |
| 16838 | 842 | E842W | C | 1.7e-06 | 2.6e-06 | 2e-06 | 2.100E-06 | 4.583E-07 | 1.700E-06 | 2.600E-06 |
| 16839 | 842 | E842Y | C | 2.2e-06 | 2.8e-06 | 1.4e-06 | 2.133E-06 | 7.024E-07 | 1.400E-06 | 2.800E-06 |
| 16840 | 842 | E842V | C | 1.8e-06 | 2.8e-06 | 2.6e-06 | 2.400E-06 | 5.292E-07 | 1.800E-06 | 2.800E-06 |
| 16841 | 843 | S843A | C | 2.2e-06 | 2.7e-06 | 2.4e-06 | 2.433E-06 | 2.517E-07 | 2.200E-06 | 2.700E-06 |
| 16842 | 843 | S843R | C | 2.7e-06 | 2.4e-06 | 2e-06 | 2.367E-06 | 3.512E-07 | 2.000E-06 | 2.700E-06 |
| 16843 | 843 | S843N | C | 1.3e-06 | 2.9e-06 | 2.7e-06 | 2.300E-06 | 8.718E-07 | 1.300E-06 | 2.900E-06 |
| 16844 | 843 | S843D | C | 1.8e-06 | 2e-06 | 2.9e-06 | 2.233E-06 | 5.859E-07 | 1.800E-06 | 2.900E-06 |
| 16845 | 843 | S843C | C | 1.9e-06 | 2.7e-06 | 2.9e-06 | 2.500E-06 | 5.292E-07 | 1.900E-06 | 2.900E-06 |
| 16846 | 843 | S843E | C | 2.1e-06 | 2.2e-06 | 1.9e-06 | 2.067E-06 | 1.528E-07 | 1.900E-06 | 2.200E-06 |
| 16847 | 843 | S843Q | C | 3.2e-06 | 2.6e-06 | 3.8e-06 | 3.200E-06 | 6.000E-07 | 2.600E-06 | 3.800E-06 |
| 16848 | 843 | S843G | C | 3e-06 | 1.9e-06 | 2.4e-06 | 2.433E-06 | 5.508E-07 | 1.900E-06 | 3.000E-06 |
| 16849 | 843 | S843H | C | 2.2e-06 | 9.3e-07 | 2.3e-06 | 1.810E-06 | 7.637E-07 | 9.300E-07 | 2.300E-06 |
| 16850 | 843 | S843I | C | 2.1e-06 | 6.4e-06 | 2.2e-06 | 3.567E-06 | 2.454E-06 | 2.100E-06 | 6.400E-06 |
| 16851 | 843 | S843L | C | 2.9e-06 | 2.3e-06 | 4.6e-06 | 3.267E-06 | 1.193E-06 | 2.300E-06 | 4.600E-06 |
| 16852 | 843 | S843K | C | 3.6e-06 | 1.8e-06 | 2.1e-06 | 2.500E-06 | 9.644E-07 | 1.800E-06 | 3.600E-06 |
| 16853 | 843 | S843M | C | 4.4e-06 | 3e-06 | 2.3e-06 | 3.233E-06 | 1.069E-06 | 2.300E-06 | 4.400E-06 |
| 16854 | 843 | S843F | C | 1.2e-06 | 3.1e-06 | 2.2e-06 | 2.167E-06 | 9.504E-07 | 1.200E-06 | 3.100E-06 |
| 16855 | 843 | S843P | C | 3.1e-06 | 1.4e-06 | 2.3e-06 | 2.267E-06 | 8.505E-07 | 1.400E-06 | 3.100E-06 |
| 16856 | 843 | S843S | C | 2e-06 | 1.3e-06 | 2.4e-06 | 1.900E-06 | 5.568E-07 | 1.300E-06 | 2.400E-06 |
| 16857 | 843 | S843T | C | 2.2e-06 | 2.2e-06 | 1.9e-06 | 2.100E-06 | 1.732E-07 | 1.900E-06 | 2.200E-06 |
| 16858 | 843 | S843W | C | 1.8e-06 | 3.9e-06 | 2.4e-06 | 2.700E-06 | 1.082E-06 | 1.800E-06 | 3.900E-06 |
| 16859 | 843 | S843Y | C | 3.1e-06 | 1.9e-06 | 1.6e-06 | 2.200E-06 | 7.937E-07 | 1.600E-06 | 3.100E-06 |
| 16860 | 843 | S843V | C | 2.8e-06 | 3e-06 | 3e-06 | 2.933E-06 | 1.155E-07 | 2.800E-06 | 3.000E-06 |
| 16861 | 844 | G844A | C | 9.9e-07 | 3.3e-06 | 2.7e-06 | 2.330E-06 | 1.199E-06 | 9.900E-07 | 3.300E-06 |
| 16862 | 844 | G844R | C | 2.5e-06 | 2.5e-06 | 3.1e-06 | 2.700E-06 | 3.464E-07 | 2.500E-06 | 3.100E-06 |
| 16863 | 844 | G844N | C | 1.9e-06 | 4.5e-06 | 2.5e-06 | 2.967E-06 | 1.361E-06 | 1.900E-06 | 4.500E-06 |
| 16864 | 844 | G844D | C | 1.8e-06 | 1.5e-06 | 2.2e-06 | 1.833E-06 | 3.512E-07 | 1.500E-06 | 2.200E-06 |
| 16865 | 844 | G844C | C | 1.5e-06 | 2e-06 | 1.2e-06 | 1.567E-06 | 4.041E-07 | 1.200E-06 | 2.000E-06 |
| 16866 | 844 | G844E | C | 1.3e-06 | 1.6e-06 | 1.7e-06 | 1.533E-06 | 2.082E-07 | 1.300E-06 | 1.700E-06 |
| 16867 | 844 | G844Q | C | 2.6e-06 | 2.8e-06 | 2e-06 | 2.467E-06 | 4.163E-07 | 2.000E-06 | 2.800E-06 |
| 16868 | 844 | G844G | C | 1.7e-06 | 2.9e-06 | 2.1e-06 | 2.233E-06 | 6.110E-07 | 1.700E-06 | 2.900E-06 |
| 16869 | 844 | G844H | C | 1.8e-06 | 2.3e-06 | 2.2e-06 | 2.100E-06 | 2.646E-07 | 1.800E-06 | 2.300E-06 |
| 16870 | 844 | G844I | C | 2.4e-06 | 1.5e-06 | 3.1e-06 | 2.333E-06 | 8.021E-07 | 1.500E-06 | 3.100E-06 |
| 16871 | 844 | G844L | C | 1.5e-06 | 2.5e-06 | 3.9e-06 | 2.633E-06 | 1.206E-06 | 1.500E-06 | 3.900E-06 |
| 16872 | 844 | G844K | C | 2.4e-06 | 2.2e-06 | 2.5e-06 | 2.367E-06 | 1.528E-07 | 2.200E-06 | 2.500E-06 |
| 16873 | 844 | G844M | C | 2.8e-06 | 3.7e-06 | 2.8e-06 | 3.100E-06 | 5.196E-07 | 2.800E-06 | 3.700E-06 |
| 16874 | 844 | G844F | C | 2.8e-06 | 1.5e-06 | 3.2e-06 | 2.500E-06 | 8.888E-07 | 1.500E-06 | 3.200E-06 |
| 16875 | 844 | G844P | C | 3.1e-06 | 2.4e-06 | 3.5e-06 | 3.000E-06 | 5.568E-07 | 2.400E-06 | 3.500E-06 |
| 16876 | 844 | G844S | C | 3e-06 | 2.7e-06 | 2.3e-06 | 2.667E-06 | 3.512E-07 | 2.300E-06 | 3.000E-06 |
| 16877 | 844 | G844T | C | 1.8e-06 | 2.4e-06 | 2.1e-06 | 2.100E-06 | 3.000E-07 | 1.800E-06 | 2.400E-06 |
| 16878 | 844 | G844W | C | 3.1e-06 | 2.1e-06 | 1.5e-06 | 2.233E-06 | 8.083E-07 | 1.500E-06 | 3.100E-06 |
| 16879 | 844 | G844Y | C | 2.6e-06 | 2.3e-06 | 2.9e-06 | 2.600E-06 | 3.000E-07 | 2.300E-06 | 2.900E-06 |
| 16880 | 844 | G844V | C | 1.7e-06 | 2.3e-06 | 1.8e-06 | 1.933E-06 | 3.215E-07 | 1.700E-06 | 2.300E-06 |
| 16881 | 845 | G845A | C | 2e-06 | 2.7e-06 | 2.5e-06 | 2.400E-06 | 3.606E-07 | 2.000E-06 | 2.700E-06 |
| 16882 | 845 | G845R | C | 1.5e-06 | 2e-06 | 2.7e-06 | 2.067E-06 | 6.028E-07 | 1.500E-06 | 2.700E-06 |
| 16883 | 845 | G845N | C | 2.9e-06 | 2.3e-06 | 2.2e-06 | 2.467E-06 | 3.786E-07 | 2.200E-06 | 2.900E-06 |
| 16884 | 845 | G845D | C | 2.4e-06 | 2.5e-06 | 1.3e-06 | 2.067E-06 | 6.658E-07 | 1.300E-06 | 2.500E-06 |
| 16885 | 845 | G845C | C | 2.7e-06 | 3e-06 | 4.8e-06 | 3.500E-06 | 1.136E-06 | 2.700E-06 | 4.800E-06 |
| 16886 | 845 | G845E | C | 2.2e-06 | 2e-06 | 3.2e-06 | 2.467E-06 | 6.429E-07 | 2.000E-06 | 3.200E-06 |
| 16887 | 845 | G845Q | C | 1.9e-06 | 3.1e-06 | 2.6e-06 | 2.533E-06 | 6.028E-07 | 1.900E-06 | 3.100E-06 |
| 16888 | 845 | G845G | C | 1.7e-06 | 2.5e-06 | 1.2e-06 | 1.800E-06 | 6.557E-07 | 1.200E-06 | 2.500E-06 |
| 16889 | 845 | G845H | C | 1.6e-06 | 2.2e-06 | 2.2e-06 | 2.000E-06 | 3.464E-07 | 1.600E-06 | 2.200E-06 |
| 16890 | 845 | G845I | C | 2.1e-06 | 2.1e-06 | 2e-06 | 2.067E-06 | 5.774E-08 | 2.000E-06 | 2.100E-06 |
| 16891 | 845 | G845L | C | 1.8e-06 | 3.1e-06 | 3.6e-06 | 2.833E-06 | 9.292E-07 | 1.800E-06 | 3.600E-06 |
| 16892 | 845 | G845K | C | 1.5e-06 | 2.1e-06 | 3.8e-06 | 2.467E-06 | 1.193E-06 | 1.500E-06 | 3.800E-06 |
| 16893 | 845 | G845M | C | 2.5e-06 | 2.2e-06 | 2.2e-06 | 2.300E-06 | 1.732E-07 | 2.200E-06 | 2.500E-06 |
| 16894 | 845 | G845F | C | 2.3e-06 | 1.6e-06 | 2.1e-06 | 2.000E-06 | 3.606E-07 | 1.600E-06 | 2.300E-06 |
| 16895 | 845 | G845P | C | 2.2e-06 | 1.7e-06 | 3.5e-06 | 2.467E-06 | 9.292E-07 | 1.700E-06 | 3.500E-06 |
| 16896 | 845 | G845S | C | 1.7e-06 | 3.3e-06 | 4.1e-06 | 3.033E-06 | 1.222E-06 | 1.700E-06 | 4.100E-06 |
| 16897 | 845 | G845T | C | 1.8e-06 | 1.6e-06 | 2.4e-06 | 1.933E-06 | 4.163E-07 | 1.600E-06 | 2.400E-06 |
| 16898 | 845 | G845W | C | 1.3e-06 | 1.3e-06 | 3.2e-06 | 1.933E-06 | 1.097E-06 | 1.300E-06 | 3.200E-06 |
| 16899 | 845 | G845Y | C | 2.7e-06 | 3.1e-06 | 2.5e-06 | 2.767E-06 | 3.055E-07 | 2.500E-06 | 3.100E-06 |
| 16900 | 845 | G845V | C | 4.1e-06 | 3.2e-06 | 2.1e-06 | 3.133E-06 | 1.002E-06 | 2.100E-06 | 4.100E-06 |
| 16901 | 846 | G846A | C | 2.7e-06 | 2.7e-06 | 4.4e-06 | 3.267E-06 | 9.815E-07 | 2.700E-06 | 4.400E-06 |
| 16902 | 846 | G846R | C | 2.2e-06 | 2.1e-06 | 2.6e-06 | 2.300E-06 | 2.646E-07 | 2.100E-06 | 2.600E-06 |
| 16903 | 846 | G846N | C | 2.4e-06 | 3e-06 | 3.2e-06 | 2.867E-06 | 4.163E-07 | 2.400E-06 | 3.200E-06 |
| 16904 | 846 | G846D | C | 3.8e-06 | 1.6e-06 | 2.1e-06 | 2.500E-06 | 1.153E-06 | 1.600E-06 | 3.800E-06 |
| 16905 | 846 | G846C | C | 4e-06 | 2.1e-06 | 2.5e-06 | 2.867E-06 | 1.002E-06 | 2.100E-06 | 4.000E-06 |
| 16906 | 846 | G846E | C | 1.9e-06 | 1.6e-06 | 2.3e-06 | 1.933E-06 | 3.512E-07 | 1.600E-06 | 2.300E-06 |
| 16907 | 846 | G846Q | C | 2.4e-06 | 2.1e-06 | 3.1e-06 | 2.533E-06 | 5.132E-07 | 2.100E-06 | 3.100E-06 |
| 16908 | 846 | G846G | C | 2.1e-06 | 3.2e-06 | 3.5e-06 | 2.933E-06 | 7.371E-07 | 2.100E-06 | 3.500E-06 |
| 16909 | 846 | G846H | C | 4e-06 | 1.8e-06 | 2e-06 | 2.600E-06 | 1.217E-06 | 1.800E-06 | 4.000E-06 |
| 16910 | 846 | G846I | C | 2.9e-06 | 2.9e-06 | 3.3e-06 | 3.033E-06 | 2.309E-07 | 2.900E-06 | 3.300E-06 |
| 16911 | 846 | G846L | C | 1.7e-06 | 4.8e-06 | 1.5e-06 | 2.667E-06 | 1.850E-06 | 1.500E-06 | 4.800E-06 |
| 16912 | 846 | G846K | C | 2.1e-06 | 1.9e-06 | 2.8e-06 | 2.267E-06 | 4.726E-07 | 1.900E-06 | 2.800E-06 |
| 16913 | 846 | G846M | C | 1.9e-06 | 3e-06 | 1.7e-06 | 2.200E-06 | 7.000E-07 | 1.700E-06 | 3.000E-06 |
| 16914 | 846 | G846F | C | 2.7e-06 | 2.5e-06 | 1.8e-06 | 2.333E-06 | 4.726E-07 | 1.800E-06 | 2.700E-06 |

| | | | | | | | | | | |
|---|---|---|---|---|---|---|---|---|---|---|
| 16915 | 846 | G846P | C | 2.1e-06 | 3e-06 | 1.5e-06 | 2.200E-06 | 7.550E-07 | 1.500E-06 | 3.000E-06 |
| 16916 | 846 | G846S | C | 1.6e-06 | 1.1e-06 | 3.5e-06 | 2.067E-06 | 1.266E-06 | 1.100E-06 | 3.500E-06 |
| 16917 | 846 | G846T | C | 2.6e-06 | 1.5e-06 | 2.2e-06 | 2.100E-06 | 5.568E-07 | 1.500E-06 | 2.600E-06 |
| 16918 | 846 | G846W | C | 2.2e-06 | 1.9e-06 | 3e-06 | 2.367E-06 | 5.686E-07 | 1.900E-06 | 3.000E-06 |
| 16919 | 846 | G846Y | C | 2e-06 | 2.4e-06 | 2.3e-06 | 2.233E-06 | 2.082E-07 | 2.000E-06 | 2.400E-06 |
| 16920 | 846 | G846V | C | 2.3e-06 | 2.4e-06 | 1.9e-06 | 2.200E-06 | 2.646E-07 | 1.900E-06 | 2.400E-06 |
| 16921 | 847 | L847A | C | 2.7e-06 | 3.5e-06 | 2e-06 | 2.733E-06 | 7.506E-07 | 2.000E-06 | 3.500E-06 |
| 16922 | 847 | L847R | C | 2.1e-06 | 2.6e-06 | 1.8e-06 | 2.167E-06 | 4.041E-07 | 1.800E-06 | 2.600E-06 |
| 16923 | 847 | L847N | C | 3.1e-06 | 1.9e-06 | 2.1e-06 | 2.367E-06 | 6.429E-07 | 1.900E-06 | 3.100E-06 |
| 16924 | 847 | L847D | C | 2.6e-06 | 2.1e-06 | 1.6e-06 | 2.100E-06 | 5.000E-07 | 1.600E-06 | 2.600E-06 |
| 16925 | 847 | L847C | C | 2.5e-06 | 1.3e-06 | 2.9e-06 | 2.233E-06 | 8.327E-07 | 1.300E-06 | 2.900E-06 |
| 16926 | 847 | L847E | C | 1.3e-06 | 3.1e-06 | 2.1e-06 | 2.167E-06 | 9.018E-07 | 1.300E-06 | 3.100E-06 |
| 16927 | 847 | L847Q | C | 1.7e-06 | 1.7e-06 | 2.2e-06 | 1.867E-06 | 2.887E-07 | 1.700E-06 | 2.200E-06 |
| 16928 | 847 | L847G | C | 3.4e-06 | 2.8e-06 | 1.8e-06 | 2.667E-06 | 8.083E-07 | 1.800E-06 | 3.400E-06 |
| 16929 | 847 | L847H | C | 3.2e-06 | 9.4e-07 | 3.3e-06 | 2.480E-06 | 1.335E-06 | 9.400E-07 | 3.300E-06 |
| 16930 | 847 | L847I | C | 4.5e-06 | 3.2e-06 | 3.3e-06 | 3.667E-06 | 7.234E-07 | 3.200E-06 | 4.500E-06 |
| 16931 | 847 | L847L | C | 2e-06 | 2.2e-06 | 1.8e-06 | 2.000E-06 | 2.000E-07 | 1.800E-06 | 2.200E-06 |
| 16932 | 847 | L847K | C | 1.2e-06 | 2.4e-06 | 5.6e-06 | 3.067E-06 | 2.274E-06 | 1.200E-06 | 5.600E-06 |
| 16933 | 847 | L847M | C | 2.4e-06 | 2.6e-06 | 2.6e-06 | 2.533E-06 | 1.155E-07 | 2.400E-06 | 2.600E-06 |
| 16934 | 847 | L847F | C | 2.9e-06 | 3.6e-06 | 3.7e-06 | 3.400E-06 | 4.359E-07 | 2.900E-06 | 3.700E-06 |
| 16935 | 847 | L847P | C | 1.5e-06 | 2.6e-06 | 2.9e-06 | 2.333E-06 | 7.371E-07 | 1.500E-06 | 2.900E-06 |
| 16936 | 847 | L847S | C | 2.9e-06 | 4.1e-06 | 2e-06 | 3.000E-06 | 1.054E-06 | 2.000E-06 | 4.100E-06 |
| 16937 | 847 | L847T | C | 3.5e-06 | 3e-06 | 2.3e-06 | 2.933E-06 | 6.028E-07 | 2.300E-06 | 3.500E-06 |
| 16938 | 847 | L847W | C | 2.6e-06 | 3.5e-06 | 3.3e-06 | 3.133E-06 | 4.726E-07 | 2.600E-06 | 3.500E-06 |
| 16939 | 847 | L847Y | C | 2e-06 | 3.7e-06 | 2.8e-06 | 2.833E-06 | 8.505E-07 | 2.000E-06 | 3.700E-06 |
| 16940 | 847 | L847V | C | 1.6e-06 | 3e-06 | 3.2e-06 | 2.600E-06 | 8.718E-07 | 1.600E-06 | 3.200E-06 |
| 16941 | 848 | V848A | C | 1.5e-06 | 3.3e-06 | 1.3e-06 | 2.033E-06 | 1.102E-06 | 1.300E-06 | 3.300E-06 |
| 16942 | 848 | V848R | C | 2.5e-06 | 1.8e-06 | 2.8e-06 | 2.367E-06 | 5.132E-07 | 1.800E-06 | 2.800E-06 |
| 16943 | 848 | V848N | C | 2.1e-06 | 1.3e-06 | 1.5e-06 | 1.633E-06 | 4.163E-07 | 1.300E-06 | 2.100E-06 |
| 16944 | 848 | V848D | C | 3.3e-06 | 2.6e-06 | 1.8e-06 | 2.567E-06 | 7.506E-07 | 1.800E-06 | 3.300E-06 |
| 16945 | 848 | V848C | C | 3.1e-06 | 2.3e-06 | 1.3e-06 | 2.233E-06 | 9.018E-07 | 1.300E-06 | 3.100E-06 |
| 16946 | 848 | V848E | C | 2.8e-06 | 3.4e-06 | 2.9e-06 | 3.033E-06 | 3.215E-07 | 2.800E-06 | 3.400E-06 |
| 16947 | 848 | V848Q | C | 2e-06 | 2.3e-06 | 1.8e-06 | 2.033E-06 | 2.517E-07 | 1.800E-06 | 2.300E-06 |
| 16948 | 848 | V848G | C | 3.1e-06 | 1.9e-06 | 3.1e-06 | 2.700E-06 | 6.928E-07 | 1.900E-06 | 3.100E-06 |
| 16949 | 848 | V848H | C | 7.9e-07 | 2e-06 | 2.5e-06 | 1.763E-06 | 8.792E-07 | 7.900E-07 | 2.500E-06 |
| 16950 | 848 | V848I | C | 2e-06 | 3.1e-06 | 2e-06 | 2.367E-06 | 6.351E-07 | 2.000E-06 | 3.100E-06 |
| 16951 | 848 | V848L | C | 2.1e-06 | 3e-06 | 2.2e-06 | 2.433E-06 | 4.933E-07 | 2.100E-06 | 3.000E-06 |
| 16952 | 848 | V848K | C | 3.6e-06 | 2.2e-06 | 2.9e-06 | 2.900E-06 | 7.000E-07 | 2.200E-06 | 3.600E-06 |
| 16953 | 848 | V848M | C | 2.6e-06 | 3.1e-06 | 7.5e-07 | 2.150E-06 | 1.238E-06 | 7.500E-07 | 3.100E-06 |
| 16954 | 848 | V848F | C | 1.9e-06 | 2.5e-06 | 2e-06 | 2.133E-06 | 3.215E-07 | 1.900E-06 | 2.500E-06 |
| 16955 | 848 | V848P | C | 4.4e-06 | 3.3e-06 | 3.3e-06 | 3.667E-06 | 6.351E-07 | 3.300E-06 | 4.400E-06 |
| 16956 | 848 | V848S | C | 2e-06 | 3.4e-06 | 2.8e-06 | 2.733E-06 | 7.024E-07 | 2.000E-06 | 3.400E-06 |
| 16957 | 848 | V848T | C | 5e-06 | 2.5e-06 | 1.8e-06 | 3.100E-06 | 1.682E-06 | 1.800E-06 | 5.000E-06 |
| 16958 | 848 | V848W | C | 1.6e-06 | 2e-06 | 3.1e-06 | 2.233E-06 | 7.767E-07 | 1.600E-06 | 3.100E-06 |
| 16959 | 848 | V848Y | C | 3.1e-06 | 1.9e-06 | 3.9e-06 | 2.967E-06 | 1.007E-06 | 1.900E-06 | 3.900E-06 |
| 16960 | 848 | V848V | C | 2e-06 | 2.5e-06 | 1.9e-06 | 2.133E-06 | 3.215E-07 | 1.900E-06 | 2.500E-06 |
| 16961 | 849 | Q849A | C | 3.1e-06 | 2.3e-06 | 3.5e-06 | 2.967E-06 | 6.110E-07 | 2.300E-06 | 3.500E-06 |
| 16962 | 849 | Q849R | C | 1.4e-06 | 2.8e-06 | 1.5e-06 | 1.900E-06 | 7.810E-07 | 1.400E-06 | 2.800E-06 |
| 16963 | 849 | Q849N | C | 6.9e-07 | 1.4e-06 | 1.2e-06 | 1.097E-06 | 3.661E-07 | 6.900E-07 | 1.400E-06 |
| 16964 | 849 | Q849D | C | 3.1e-06 | 2.4e-06 | 1.7e-06 | 2.400E-06 | 7.000E-07 | 1.700E-06 | 3.100E-06 |
| 16965 | 849 | Q849C | C | 3.4e-06 | 3.2e-06 | 2.1e-06 | 2.900E-06 | 7.000E-07 | 2.100E-06 | 3.400E-06 |
| 16966 | 849 | Q849E | C | 2.4e-06 | 2.7e-06 | 3.4e-06 | 2.833E-06 | 5.132E-07 | 2.400E-06 | 3.400E-06 |
| 16967 | 849 | Q849Q | C | 3.1e-06 | 2.2e-06 | 1.8e-06 | 2.367E-06 | 6.658E-07 | 1.800E-06 | 3.100E-06 |
| 16968 | 849 | Q849G | C | 4.6e-06 | 3.1e-06 | 2.2e-06 | 3.300E-06 | 1.212E-06 | 2.200E-06 | 4.600E-06 |
| 16969 | 849 | Q849H | C | 1.8e-06 | 3.2e-06 | 1.7e-06 | 2.233E-06 | 8.386E-07 | 1.700E-06 | 3.200E-06 |
| 16970 | 849 | Q849I | C | 2.3e-06 | 3.1e-06 | 3.9e-06 | 3.100E-06 | 8.000E-07 | 2.300E-06 | 3.900E-06 |
| 16971 | 849 | Q849L | C | 3e-06 | 2.8e-06 | 3.1e-06 | 2.967E-06 | 1.528E-07 | 2.800E-06 | 3.100E-06 |
| 16972 | 849 | Q849K | C | 1.5e-06 | 2.6e-06 | 2.3e-06 | 2.133E-06 | 5.686E-07 | 1.500E-06 | 2.600E-06 |
| 16973 | 849 | Q849M | C | 2e-06 | 3e-06 | 2.4e-06 | 2.467E-06 | 5.033E-07 | 2.000E-06 | 3.000E-06 |
| 16974 | 849 | Q849F | C | 1.7e-06 | 2.1e-06 | 3.3e-06 | 2.367E-06 | 8.327E-07 | 1.700E-06 | 3.300E-06 |
| 16975 | 849 | Q849P | C | 2.7e-06 | 2.4e-06 | 3.8e-06 | 2.967E-06 | 7.371E-07 | 2.400E-06 | 3.800E-06 |
| 16976 | 849 | Q849S | C | 2.3e-06 | 2.2e-06 | 2.1e-06 | 2.200E-06 | 1.000E-07 | 2.100E-06 | 2.300E-06 |
| 16977 | 849 | Q849T | C | 1.2e-06 | 1.8e-06 | 2.8e-06 | 1.933E-06 | 8.083E-07 | 1.200E-06 | 2.800E-06 |
| 16978 | 849 | Q849W | C | 2.5e-06 | 1.7e-06 | 1.8e-06 | 2.000E-06 | 4.359E-07 | 1.700E-06 | 2.500E-06 |
| 16979 | 849 | Q849Y | C | 3.2e-06 | 3.1e-06 | 2.3e-06 | 2.867E-06 | 4.933E-07 | 2.300E-06 | 3.200E-06 |
| 16980 | 849 | Q849V | C | 3e-06 | 2e-06 | 1.1e-06 | 2.033E-06 | 9.504E-07 | 1.100E-06 | 3.000E-06 |
| 16981 | 850 | P850A | C | 1.7e-06 | 2.5e-06 | 1.6e-06 | 1.933E-06 | 4.933E-07 | 1.600E-06 | 2.500E-06 |
| 16982 | 850 | P850R | C | 1.6e-06 | 1.4e-06 | 2e-06 | 1.667E-06 | 3.055E-07 | 1.400E-06 | 2.000E-06 |
| 16983 | 850 | P850N | C | 2.4e-06 | 2.1e-06 | 1.9e-06 | 2.133E-06 | 2.517E-07 | 1.900E-06 | 2.400E-06 |
| 16984 | 850 | P850D | C | 3e-06 | 2.7e-06 | 3e-06 | 2.900E-06 | 1.732E-07 | 2.700E-06 | 3.000E-06 |
| 16985 | 850 | P850C | C | 2.8e-06 | 1.1e-06 | 1.6e-06 | 1.833E-06 | 8.737E-07 | 1.100E-06 | 2.800E-06 |
| 16986 | 850 | P850E | C | 1.7e-06 | 2.8e-06 | 3.5e-06 | 2.667E-06 | 9.074E-07 | 1.700E-06 | 3.500E-06 |
| 16987 | 850 | P850Q | C | 1.3e-06 | 2.8e-06 | 1.9e-06 | 2.000E-06 | 7.550E-07 | 1.300E-06 | 2.800E-06 |
| 16988 | 850 | P850G | C | 2.3e-06 | 2.3e-06 | 2.3e-06 | 2.300E-06 | 0.000E+00 | 2.300E-06 | 2.300E-06 |
| 16989 | 850 | P850H | C | 2.6e-06 | 1.3e-06 | 2.8e-06 | 2.233E-06 | 8.145E-07 | 1.300E-06 | 2.800E-06 |
| 16990 | 850 | P850I | C | 3.4e-06 | 2.9e-06 | 3.9e-06 | 3.400E-06 | 5.000E-07 | 2.900E-06 | 3.900E-06 |
| 16991 | 850 | P850L | C | 1.6e-06 | 3.2e-06 | 3.2e-06 | 2.667E-06 | 9.238E-07 | 1.600E-06 | 3.200E-06 |
| 16992 | 850 | P850K | C | 1.3e-06 | 3.2e-06 | 1.6e-06 | 2.033E-06 | 1.021E-06 | 1.300E-06 | 3.200E-06 |
| 16993 | 850 | P850M | C | 3.1e-06 | 2.7e-06 | 1.5e-06 | 2.433E-06 | 8.327E-07 | 1.500E-06 | 3.100E-06 |
| 16994 | 850 | P850F | C | 2.1e-06 | 1.7e-06 | 2.2e-06 | 2.000E-06 | 2.646E-07 | 1.700E-06 | 2.200E-06 |
| 16995 | 850 | P850P | C | 9.7e-07 | 3e-06 | 2.4e-06 | 2.123E-06 | 1.043E-06 | 9.700E-07 | 3.000E-06 |
| 16996 | 850 | P850S | C | 1.7e-06 | 1.9e-06 | 2e-06 | 1.867E-06 | 1.528E-07 | 1.700E-06 | 2.000E-06 |
| 16997 | 850 | P850T | C | 2.3e-06 | 2.5e-06 | 2.7e-06 | 2.500E-06 | 2.000E-07 | 2.300E-06 | 2.700E-06 |
| 16998 | 850 | P850W | C | 1.4e-06 | 5.1e-06 | 3.6e-06 | 3.367E-06 | 1.861E-06 | 1.400E-06 | 5.100E-06 |
| 16999 | 850 | P850Y | C | 4.1e-06 | 1.8e-06 | 1.9e-06 | 2.600E-06 | 1.300E-06 | 1.800E-06 | 4.100E-06 |

| | | | | | | | | | | |
|---|---|---|---|---|---|---|---|---|---|---|
| 17000 | 850 | P850V | C | 2.9e-06 | 1.9e-06 | 1.7e-06 | 2.167E-06 | 6.429E-07 | 1.700E-06 | 2.900E-06 |
| 17001 | 851 | G851A | C | 1.3e-06 | 2.1e-06 | 3.3e-06 | 2.233E-06 | 1.007E-06 | 1.300E-06 | 3.300E-06 |
| 17002 | 851 | G851R | C | 6.1e-06 | 2.4e-06 | 2.1e-06 | 3.533E-06 | 2.228E-06 | 2.100E-06 | 6.100E-06 |
| 17003 | 851 | G851N | C | 1.9e-06 | 4.3e-06 | 1.5e-06 | 2.567E-06 | 1.514E-06 | 1.500E-06 | 4.300E-06 |
| 17004 | 851 | G851D | C | 1.7e-06 | 1.9e-06 | 3.1e-06 | 2.233E-06 | 7.572E-07 | 1.700E-06 | 3.100E-06 |
| 17005 | 851 | G851C | C | 2.7e-06 | 1.6e-06 | 1.9e-06 | 2.067E-06 | 5.686E-07 | 1.600E-06 | 2.700E-06 |
| 17006 | 851 | G851E | C | 3.8e-06 | 3.2e-06 | 1.7e-06 | 2.900E-06 | 1.082E-06 | 1.700E-06 | 3.800E-06 |
| 17007 | 851 | G851Q | C | 2.6e-06 | 2.7e-06 | 2.9e-06 | 2.733E-06 | 1.528E-07 | 2.600E-06 | 2.900E-06 |
| 17008 | 851 | G851G | C | 1.4e-06 | 2.5e-06 | 2.6e-06 | 2.167E-06 | 6.658E-07 | 1.400E-06 | 2.600E-06 |
| 17009 | 851 | G851H | C | 3.6e-06 | 1.4e-06 | 2.5e-06 | 2.500E-06 | 1.100E-06 | 1.400E-06 | 3.600E-06 |
| 17010 | 851 | G851I | C | 2.4e-06 | 2.3e-06 | 1.4e-06 | 2.033E-06 | 5.508E-07 | 1.400E-06 | 2.400E-06 |
| 17011 | 851 | G851L | C | 2.1e-06 | 2.3e-06 | 1.6e-06 | 2.000E-06 | 3.606E-07 | 1.600E-06 | 2.300E-06 |
| 17012 | 851 | G851K | C | 3.1e-06 | 1.2e-06 | 3.5e-06 | 2.600E-06 | 1.229E-06 | 1.200E-06 | 3.500E-06 |
| 17013 | 851 | G851M | C | 2.3e-06 | 3.1e-06 | 3.9e-06 | 3.100E-06 | 8.000E-07 | 2.300E-06 | 3.900E-06 |
| 17014 | 851 | G851F | C | 2.4e-06 | 1.8e-06 | 4.3e-06 | 2.833E-06 | 1.305E-06 | 1.800E-06 | 4.300E-06 |
| 17015 | 851 | G851P | C | 4.8e-06 | 4.4e-06 | 2.8e-06 | 4.000E-06 | 1.058E-06 | 2.800E-06 | 4.800E-06 |
| 17016 | 851 | G851S | C | 2.4e-06 | 2.3e-06 | 2.4e-06 | 2.367E-06 | 5.774E-08 | 2.300E-06 | 2.400E-06 |
| 17017 | 851 | G851T | C | 2.2e-06 | 2e-06 | 1.7e-06 | 1.967E-06 | 2.517E-07 | 1.700E-06 | 2.200E-06 |
| 17018 | 851 | G851W | C | 2.4e-06 | 2.7e-06 | 1.1e-06 | 2.067E-06 | 8.505E-07 | 1.100E-06 | 2.700E-06 |
| 17019 | 851 | G851Y | C | 3.5e-06 | 1.7e-06 | 1.1e-06 | 2.100E-06 | 1.249E-06 | 1.100E-06 | 3.500E-06 |
| 17020 | 851 | G851V | C | 2.5e-06 | 3.3e-06 | 1.6e-06 | 2.467E-06 | 8.505E-07 | 1.600E-06 | 3.300E-06 |
| 17021 | 852 | G852A | C | 3e-06 | 3.1e-06 | 3.1e-06 | 3.067E-06 | 5.774E-08 | 3.000E-06 | 3.100E-06 |
| 17022 | 852 | G852R | C | 4.4e-06 | 2.3e-06 | 1.8e-06 | 2.833E-06 | 1.380E-06 | 1.800E-06 | 4.400E-06 |
| 17023 | 852 | G852N | C | 1.3e-06 | 2.1e-06 | 1.6e-06 | 1.667E-06 | 4.041E-07 | 1.300E-06 | 2.100E-06 |
| 17024 | 852 | G852D | C | 2.3e-06 | 2.6e-06 | 2.2e-06 | 2.367E-06 | 2.082E-07 | 2.200E-06 | 2.600E-06 |
| 17025 | 852 | G852C | C | 3.4e-06 | 1.7e-06 | 2.5e-06 | 2.533E-06 | 8.505E-07 | 1.700E-06 | 3.400E-06 |
| 17026 | 852 | G852E | C | 1.6e-06 | 4.8e-06 | 2.1e-06 | 2.833E-06 | 1.721E-06 | 1.600E-06 | 4.800E-06 |
| 17027 | 852 | G852Q | C | 2.9e-06 | 2.5e-06 | 2.6e-06 | 2.667E-06 | 2.082E-07 | 2.500E-06 | 2.900E-06 |
| 17028 | 852 | G852G | C | 2.4e-06 | 2.3e-06 | 3.2e-06 | 2.633E-06 | 4.933E-07 | 2.300E-06 | 3.200E-06 |
| 17029 | 852 | G852H | C | 2.9e-06 | 3.4e-06 | 1.9e-06 | 2.733E-06 | 7.638E-07 | 1.900E-06 | 3.400E-06 |
| 17030 | 852 | G852I | C | 1.8e-06 | 2.9e-06 | 1.4e-06 | 2.033E-06 | 7.767E-07 | 1.400E-06 | 2.900E-06 |
| 17031 | 852 | G852L | C | 5e-06 | 2.8e-06 | 2.5e-06 | 3.433E-06 | 1.365E-06 | 2.500E-06 | 5.000E-06 |
| 17032 | 852 | G852K | C | 3.1e-06 | 3.5e-06 | 3.3e-06 | 3.300E-06 | 2.000E-07 | 3.100E-06 | 3.500E-06 |
| 17033 | 852 | G852M | C | 2.3e-06 | 4.1e-06 | 2.5e-06 | 2.967E-06 | 9.866E-07 | 2.300E-06 | 4.100E-06 |
| 17034 | 852 | G852F | C | 2e-06 | 2.2e-06 | 2.2e-06 | 2.133E-06 | 1.155E-07 | 2.000E-06 | 2.200E-06 |
| 17035 | 852 | G852P | C | 3e-06 | 3.1e-06 | 2.1e-06 | 2.733E-06 | 5.508E-07 | 2.100E-06 | 3.100E-06 |
| 17036 | 852 | G852S | C | 9.6e-07 | 2.8e-06 | 2.8e-06 | 2.187E-06 | 1.062E-06 | 9.600E-07 | 2.800E-06 |
| 17037 | 852 | G852T | C | 2e-06 | 3.1e-06 | 4.1e-06 | 3.067E-06 | 1.050E-06 | 2.000E-06 | 4.100E-06 |
| 17038 | 852 | G852W | C | 1.8e-06 | 1.1e-06 | 3.4e-06 | 2.100E-06 | 1.179E-06 | 1.100E-06 | 3.400E-06 |
| 17039 | 852 | G852Y | C | 1.6e-06 | 1.4e-06 | 2.7e-06 | 1.900E-06 | 7.000E-07 | 1.400E-06 | 2.700E-06 |
| 17040 | 852 | G852V | C | 2.2e-06 | 2.3e-06 | 2.1e-06 | 2.200E-06 | 1.000E-07 | 2.100E-06 | 2.300E-06 |
| 17041 | 853 | S853A | C | 4.7e-06 | 1.9e-06 | 2.3e-06 | 2.967E-06 | 1.514E-06 | 1.900E-06 | 4.700E-06 |
| 17042 | 853 | S853R | C | 2.3e-06 | 2.7e-06 | 2.2e-06 | 2.400E-06 | 2.646E-07 | 2.200E-06 | 2.700E-06 |
| 17043 | 853 | S853N | C | 2.9e-06 | 1.8e-06 | 1.5e-06 | 2.067E-06 | 7.371E-07 | 1.500E-06 | 2.900E-06 |
| 17044 | 853 | S853D | C | 3.2e-06 | 3.3e-06 | 2.3e-06 | 2.933E-06 | 5.508E-07 | 2.300E-06 | 3.300E-06 |
| 17045 | 853 | S853C | C | 4e-06 | 4.3e-06 | 3.1e-06 | 3.800E-06 | 6.245E-07 | 3.100E-06 | 4.300E-06 |
| 17046 | 853 | S853E | C | 2.4e-06 | 1.9e-06 | 1.6e-06 | 1.967E-06 | 4.041E-07 | 1.600E-06 | 2.400E-06 |
| 17047 | 853 | S853Q | C | 2.8e-06 | 3.4e-06 | 3e-06 | 3.067E-06 | 3.055E-07 | 2.800E-06 | 3.400E-06 |
| 17048 | 853 | S853G | C | 1.8e-06 | 2.8e-06 | 2.9e-06 | 2.500E-06 | 6.083E-07 | 1.800E-06 | 2.900E-06 |
| 17049 | 853 | S853H | C | 3.2e-06 | 2.5e-06 | 1.9e-06 | 2.533E-06 | 6.506E-07 | 1.900E-06 | 3.200E-06 |
| 17050 | 853 | S853I | C | 3.7e-06 | 2.2e-06 | 2.1e-06 | 2.667E-06 | 8.963E-07 | 2.100E-06 | 3.700E-06 |
| 17051 | 853 | S853L | C | 2.4e-06 | 2.2e-06 | 2.2e-06 | 2.267E-06 | 1.155E-07 | 2.200E-06 | 2.400E-06 |
| 17052 | 853 | S853K | C | 2.9e-06 | 3.2e-06 | 1.7e-06 | 2.600E-06 | 7.937E-07 | 1.700E-06 | 3.200E-06 |
| 17053 | 853 | S853M | C | 1.9e-06 | 2.3e-06 | 2.9e-06 | 2.367E-06 | 5.033E-07 | 1.900E-06 | 2.900E-06 |
| 17054 | 853 | S853F | C | 1.8e-06 | 2.3e-06 | 3.5e-06 | 2.533E-06 | 8.737E-07 | 1.800E-06 | 3.500E-06 |
| 17055 | 853 | S853P | C | 2.3e-06 | 1.9e-06 | 2.9e-06 | 2.367E-06 | 5.033E-07 | 1.900E-06 | 2.900E-06 |
| 17056 | 853 | S853S | C | 2.5e-06 | 2.5e-06 | 3.1e-06 | 2.700E-06 | 3.464E-07 | 2.500E-06 | 3.100E-06 |
| 17057 | 853 | S853T | C | 3.6e-06 | 2.4e-06 | 3e-06 | 3.000E-06 | 6.000E-07 | 2.400E-06 | 3.600E-06 |
| 17058 | 853 | S853W | C | 3.9e-06 | 2.2e-06 | 1.4e-06 | 2.500E-06 | 1.277E-06 | 1.400E-06 | 3.900E-06 |
| 17059 | 853 | S853Y | C | 1.8e-06 | 1.6e-06 | 2.1e-06 | 1.833E-06 | 2.517E-07 | 1.600E-06 | 2.100E-06 |
| 17060 | 853 | S853V | C | 2.1e-06 | 1.8e-06 | 2e-06 | 1.967E-06 | 1.528E-07 | 1.800E-06 | 2.100E-06 |
| 17061 | 854 | L854A | C | 2.1e-06 | 1.7e-06 | 3.9e-06 | 2.567E-06 | 1.172E-06 | 1.700E-06 | 3.900E-06 |
| 17062 | 854 | L854R | C | 2.2e-06 | 2.4e-06 | 3.7e-06 | 2.767E-06 | 8.145E-07 | 2.200E-06 | 3.700E-06 |
| 17063 | 854 | L854N | C | 1.1e-06 | 2.5e-06 | 1.6e-06 | 1.733E-06 | 7.095E-07 | 1.100E-06 | 2.500E-06 |
| 17064 | 854 | L854D | C | 3.2e-06 | 2.2e-06 | 3e-06 | 2.800E-06 | 5.292E-07 | 2.200E-06 | 3.200E-06 |
| 17065 | 854 | L854C | C | 2.9e-06 | 4.4e-06 | 2.9e-06 | 3.400E-06 | 8.660E-07 | 2.900E-06 | 4.400E-06 |
| 17066 | 854 | L854E | C | 3.3e-06 | 1.3e-06 | 1.5e-06 | 2.033E-06 | 1.102E-06 | 1.300E-06 | 3.300E-06 |
| 17067 | 854 | L854Q | C | 1.5e-06 | 2.2e-06 | 3.1e-06 | 2.267E-06 | 8.021E-07 | 1.500E-06 | 3.100E-06 |
| 17068 | 854 | L854G | C | 2e-06 | 1.5e-06 | 1.9e-06 | 1.800E-06 | 2.646E-07 | 1.500E-06 | 2.000E-06 |
| 17069 | 854 | L854H | C | 1.9e-06 | 2.9e-06 | 2.8e-06 | 2.533E-06 | 5.508E-07 | 1.900E-06 | 2.900E-06 |
| 17070 | 854 | L854I | C | 6.5e-06 | 3.2e-06 | 2e-06 | 3.900E-06 | 2.330E-06 | 2.000E-06 | 6.500E-06 |
| 17071 | 854 | L854L | C | 2.4e-06 | 1.5e-06 | 1.9e-06 | 1.933E-06 | 4.509E-07 | 1.500E-06 | 2.400E-06 |
| 17072 | 854 | L854K | C | 2.5e-06 | 3.1e-06 | 2e-06 | 2.533E-06 | 5.508E-07 | 2.000E-06 | 3.100E-06 |
| 17073 | 854 | L854M | C | 1.6e-06 | 2.3e-06 | 2.5e-06 | 2.133E-06 | 4.726E-07 | 1.600E-06 | 2.500E-06 |
| 17074 | 854 | L854F | C | 3.2e-06 | 2.9e-06 | 3.8e-06 | 3.300E-06 | 4.583E-07 | 2.900E-06 | 3.800E-06 |
| 17075 | 854 | L854P | C | 2.2e-06 | 3e-06 | 4.6e-06 | 3.267E-06 | 1.222E-06 | 2.200E-06 | 4.600E-06 |
| 17076 | 854 | L854S | C | 2.5e-06 | 2.8e-06 | 3.8e-06 | 3.033E-06 | 6.807E-07 | 2.500E-06 | 3.800E-06 |
| 17077 | 854 | L854T | C | 3e-06 | 2.7e-06 | 1.1e-06 | 2.267E-06 | 1.021E-06 | 1.100E-06 | 3.000E-06 |
| 17078 | 854 | L854W | C | 1.5e-06 | 1.8e-06 | 1.5e-06 | 1.600E-06 | 1.732E-07 | 1.500E-06 | 1.800E-06 |
| 17079 | 854 | L854Y | C | 2.1e-06 | 3.2e-06 | 2.1e-06 | 2.467E-06 | 6.351E-07 | 2.100E-06 | 3.200E-06 |
| 17080 | 854 | L854V | C | 2e-06 | 3e-06 | 2.3e-06 | 2.433E-06 | 5.132E-07 | 2.000E-06 | 3.000E-06 |
| 17081 | 855 | R855A | C | 3.3e-06 | 2.1e-06 | 2.5e-06 | 2.633E-06 | 6.110E-07 | 2.100E-06 | 3.300E-06 |
| 17082 | 855 | R855R | C | 2e-06 | 2.7e-06 | 2.6e-06 | 2.433E-06 | 3.786E-07 | 2.000E-06 | 2.700E-06 |
| 17083 | 855 | R855N | C | 2.4e-06 | 3.7e-06 | 2.2e-06 | 2.767E-06 | 8.145E-07 | 2.200E-06 | 3.700E-06 |
| 17084 | 855 | R855D | C | 3.1e-06 | 2.1e-06 | 2e-06 | 2.400E-06 | 6.083E-07 | 2.000E-06 | 3.100E-06 |

| | | | | | | | | | | |
|---|---|---|---|---|---|---|---|---|---|---|
| 17085 | 855 | R855C | C | 3.2e-06 | 2.7e-06 | 2.9e-06 | 2.933E-06 | 2.517E-07 | 2.700E-06 | 3.200E-06 |
| 17086 | 855 | R855E | C | 2.3e-06 | 2.2e-06 | 3.6e-06 | 2.700E-06 | 7.810E-07 | 2.200E-06 | 3.600E-06 |
| 17087 | 855 | R855Q | C | 3.5e-06 | 3.1e-06 | 3.3e-06 | 3.300E-06 | 2.000E-07 | 3.100E-06 | 3.500E-06 |
| 17088 | 855 | R855G | C | 1.3e-06 | 2e-06 | 2.1e-06 | 1.800E-06 | 4.359E-07 | 1.300E-06 | 2.100E-06 |
| 17089 | 855 | R855H | C | 3.3e-06 | 2.7e-06 | 2e-06 | 2.667E-06 | 6.506E-07 | 2.000E-06 | 3.300E-06 |
| 17090 | 855 | R855I | C | 8.9e-07 | 1.8e-06 | 2e-06 | 1.563E-06 | 5.916E-07 | 8.900E-07 | 2.000E-06 |
| 17091 | 855 | R855L | C | 2.4e-06 | 2.9e-06 | 1.8e-06 | 2.367E-06 | 5.508E-07 | 1.800E-06 | 2.900E-06 |
| 17092 | 855 | R855K | C | 2.1e-06 | 3e-06 | 3.9e-06 | 3.000E-06 | 9.000E-07 | 2.100E-06 | 3.900E-06 |
| 17093 | 855 | R855M | C | 3.1e-06 | 1.5e-06 | 2.2e-06 | 2.267E-06 | 8.021E-07 | 1.500E-06 | 3.100E-06 |
| 17094 | 855 | R855F | C | 2.9e-06 | 1.9e-06 | 1.5e-06 | 2.100E-06 | 7.211E-07 | 1.500E-06 | 2.900E-06 |
| 17095 | 855 | R855P | C | 1.8e-06 | 3e-06 | 1.4e-06 | 2.067E-06 | 8.327E-07 | 1.400E-06 | 3.000E-06 |
| 17096 | 855 | R855S | C | 3.1e-06 | 3.2e-06 | 2.9e-06 | 3.067E-06 | 1.528E-07 | 2.900E-06 | 3.200E-06 |
| 17097 | 855 | R855T | C | 2.3e-06 | 2.9e-06 | 2.2e-06 | 2.467E-06 | 3.786E-07 | 2.200E-06 | 2.900E-06 |
| 17098 | 855 | R855W | C | 1.6e-06 | 3.6e-06 | 2.9e-06 | 2.700E-06 | 1.015E-06 | 1.600E-06 | 3.600E-06 |
| 17099 | 855 | R855Y | C | 2.2e-06 | 2e-06 | 1.1e-06 | 1.767E-06 | 5.859E-07 | 1.100E-06 | 2.200E-06 |
| 17100 | 855 | R855V | C | 4.4e-06 | 2.3e-06 | 2e-06 | 2.900E-06 | 1.308E-06 | 2.000E-06 | 4.400E-06 |
| 17101 | 856 | L856A | C | 2.2e-06 | 5e-06 | 3.5e-06 | 3.567E-06 | 1.401E-06 | 2.200E-06 | 5.000E-06 |
| 17102 | 856 | L856R | C | 2.2e-06 | 2.2e-06 | 1.2e-06 | 1.867E-06 | 5.774E-07 | 1.200E-06 | 2.200E-06 |
| 17103 | 856 | L856N | C | 4.4e-06 | 2.9e-06 | 3e-06 | 3.433E-06 | 8.386E-07 | 2.900E-06 | 4.400E-06 |
| 17104 | 856 | L856D | C | 2.8e-06 | 4.3e-06 | 1.6e-06 | 2.900E-06 | 1.353E-06 | 1.600E-06 | 4.300E-06 |
| 17105 | 856 | L856C | C | 3e-06 | 1.7e-06 | 1.6e-06 | 2.100E-06 | 7.810E-07 | 1.600E-06 | 3.000E-06 |
| 17106 | 856 | L856E | C | 1.7e-06 | 4.1e-06 | 1.9e-06 | 2.567E-06 | 1.332E-06 | 1.700E-06 | 4.100E-06 |
| 17107 | 856 | L856Q | C | 2.1e-06 | 1.7e-06 | 2.9e-06 | 2.233E-06 | 6.110E-07 | 1.700E-06 | 2.900E-06 |
| 17108 | 856 | L856G | C | 4.2e-06 | 3.2e-06 | 2.5e-06 | 3.300E-06 | 8.544E-07 | 2.500E-06 | 4.200E-06 |
| 17109 | 856 | L856H | C | 3.5e-06 | 2.3e-06 | 2.7e-06 | 2.833E-06 | 6.110E-07 | 2.300E-06 | 3.500E-06 |
| 17110 | 856 | L856I | C | 2.2e-06 | 1.9e-06 | 3.2e-06 | 2.433E-06 | 6.807E-07 | 1.900E-06 | 3.200E-06 |
| 17111 | 856 | L856L | C | 2.7e-06 | 1.9e-06 | 4.2e-06 | 2.933E-06 | 1.168E-06 | 1.900E-06 | 4.200E-06 |
| 17112 | 856 | L856K | C | 1.2e-06 | 1.6e-06 | 3e-06 | 1.933E-06 | 9.452E-07 | 1.200E-06 | 3.000E-06 |
| 17113 | 856 | L856M | C | 2.8e-06 | 2.7e-06 | 1.7e-06 | 2.400E-06 | 6.083E-07 | 1.700E-06 | 2.800E-06 |
| 17114 | 856 | L856F | C | 3.2e-06 | 2.2e-06 | 2.7e-06 | 2.700E-06 | 5.000E-07 | 2.200E-06 | 3.200E-06 |
| 17115 | 856 | L856P | C | 3.2e-06 | 1.3e-06 | 2.5e-06 | 2.333E-06 | 9.609E-07 | 1.300E-06 | 3.200E-06 |
| 17116 | 856 | L856S | C | 1.7e-06 | 1.7e-06 | 1.4e-06 | 1.600E-06 | 1.732E-07 | 1.400E-06 | 1.700E-06 |
| 17117 | 856 | L856T | C | 2.8e-06 | 3.1e-06 | 4.4e-06 | 3.433E-06 | 8.505E-07 | 2.800E-06 | 4.400E-06 |
| 17118 | 856 | L856W | C | 1.9e-06 | 2.2e-06 | 2.3e-06 | 2.133E-06 | 2.082E-07 | 1.900E-06 | 2.300E-06 |
| 17119 | 856 | L856Y | C | 2.6e-06 | 1.7e-06 | 1.7e-06 | 2.000E-06 | 5.196E-07 | 1.700E-06 | 2.600E-06 |
| 17120 | 856 | L856V | C | 2.2e-06 | 1.7e-06 | 2.6e-06 | 2.167E-06 | 4.509E-07 | 1.700E-06 | 2.600E-06 |
| 17121 | 857 | S857A | C | 3.6e-06 | 2e-06 | 2.1e-06 | 2.567E-06 | 8.963E-07 | 2.000E-06 | 3.600E-06 |
| 17122 | 857 | S857R | C | 2.8e-06 | 2.8e-06 | 2.3e-06 | 2.633E-06 | 2.887E-07 | 2.300E-06 | 2.800E-06 |
| 17123 | 857 | S857N | C | 2.5e-06 | 2.7e-06 | 2.1e-06 | 2.433E-06 | 3.055E-07 | 2.100E-06 | 2.700E-06 |
| 17124 | 857 | S857D | C | 2e-06 | 1.7e-06 | 3e-06 | 2.233E-06 | 6.807E-07 | 1.700E-06 | 3.000E-06 |
| 17125 | 857 | S857C | C | 3.1e-06 | 1.9e-06 | 3e-06 | 2.667E-06 | 6.658E-07 | 1.900E-06 | 3.100E-06 |
| 17126 | 857 | S857E | C | 2.9e-06 | 2.7e-06 | 2.1e-06 | 2.567E-06 | 4.163E-07 | 2.100E-06 | 2.900E-06 |
| 17127 | 857 | S857Q | C | 1.6e-06 | 1.7e-06 | 1.6e-06 | 1.633E-06 | 5.774E-08 | 1.600E-06 | 1.700E-06 |
| 17128 | 857 | S857G | C | 3e-06 | 2.7e-06 | 1.8e-06 | 2.500E-06 | 6.245E-07 | 1.800E-06 | 3.000E-06 |
| 17129 | 857 | S857H | C | 2.9e-06 | 1.4e-06 | 2.7e-06 | 2.333E-06 | 8.145E-07 | 1.400E-06 | 2.900E-06 |
| 17130 | 857 | S857I | C | 2.3e-06 | 4.5e-06 | 4.2e-06 | 3.667E-06 | 1.193E-06 | 2.300E-06 | 4.500E-06 |
| 17131 | 857 | S857L | C | 3.6e-06 | 1.1e-06 | 2.8e-06 | 2.500E-06 | 1.277E-06 | 1.100E-06 | 3.600E-06 |
| 17132 | 857 | S857K | C | 2.4e-06 | 2.4e-06 | 3.7e-06 | 2.833E-06 | 7.506E-07 | 2.400E-06 | 3.700E-06 |
| 17133 | 857 | S857M | C | 2.6e-06 | 2.2e-06 | 2.3e-06 | 2.367E-06 | 2.082E-07 | 2.200E-06 | 2.600E-06 |
| 17134 | 857 | S857F | C | 2.9e-06 | 2.1e-06 | 2.3e-06 | 2.433E-06 | 4.163E-07 | 2.100E-06 | 2.900E-06 |
| 17135 | 857 | S857P | C | 3.4e-06 | 1.4e-06 | 3.6e-06 | 2.800E-06 | 1.217E-06 | 1.400E-06 | 3.600E-06 |
| 17136 | 857 | S857S | C | 2e-06 | 2.1e-06 | 2.9e-06 | 2.333E-06 | 4.933E-07 | 2.000E-06 | 2.900E-06 |
| 17137 | 857 | S857T | C | 1.9e-06 | 3e-06 | 2.3e-06 | 2.400E-06 | 5.568E-07 | 1.900E-06 | 3.000E-06 |
| 17138 | 857 | S857W | C | 2e-06 | 4.3e-06 | 4.8e-06 | 3.700E-06 | 1.493E-06 | 2.000E-06 | 4.800E-06 |
| 17139 | 857 | S857Y | C | 3e-06 | 2.4e-06 | 2.9e-06 | 2.767E-06 | 3.215E-07 | 2.400E-06 | 3.000E-06 |
| 17140 | 857 | S857V | C | 2.5e-06 | 2.9e-06 | 3.2e-06 | 2.867E-06 | 3.512E-07 | 2.500E-06 | 3.200E-06 |
| 17141 | 858 | C858A | C | 3.4e-06 | 3.8e-06 | 1.9e-06 | 3.033E-06 | 1.002E-06 | 1.900E-06 | 3.800E-06 |
| 17142 | 858 | C858R | C | 3e-06 | 2.8e-06 | 3.5e-06 | 3.100E-06 | 3.606E-07 | 2.800E-06 | 3.500E-06 |
| 17143 | 858 | C858N | C | 2.5e-06 | 3.1e-06 | 2.7e-06 | 2.767E-06 | 3.055E-07 | 2.500E-06 | 3.100E-06 |
| 17144 | 858 | C858D | C | 3e-06 | 2.1e-06 | 2e-06 | 2.367E-06 | 5.508E-07 | 2.000E-06 | 3.000E-06 |
| 17145 | 858 | C858C | C | 3.2e-06 | 2.5e-06 | 3.2e-06 | 2.967E-06 | 4.041E-07 | 2.500E-06 | 3.200E-06 |
| 17146 | 858 | C858E | C | 2.3e-06 | 1.8e-06 | 2.9e-06 | 2.333E-06 | 5.508E-07 | 1.800E-06 | 2.900E-06 |
| 17147 | 858 | C858Q | C | 2.7e-06 | 2.5e-06 | 3e-06 | 2.733E-06 | 2.517E-07 | 2.500E-06 | 3.000E-06 |
| 17148 | 858 | C858G | C | 1.4e-06 | 4.1e-06 | 2e-06 | 2.500E-06 | 1.418E-06 | 1.400E-06 | 4.100E-06 |
| 17149 | 858 | C858H | C | 1.6e-06 | 2.5e-06 | 1.6e-06 | 1.900E-06 | 5.196E-07 | 1.600E-06 | 2.500E-06 |
| 17150 | 858 | C858I | C | 3.3e-06 | 4.5e-06 | 1.6e-06 | 3.133E-06 | 1.457E-06 | 1.600E-06 | 4.500E-06 |
| 17151 | 858 | C858L | C | 2.5e-06 | 2.1e-06 | 1.8e-06 | 2.133E-06 | 3.512E-07 | 1.800E-06 | 2.500E-06 |
| 17152 | 858 | C858K | C | 2.4e-06 | 3.1e-06 | 1.5e-06 | 2.333E-06 | 8.021E-07 | 1.500E-06 | 3.100E-06 |
| 17153 | 858 | C858M | C | 4.4e-06 | 1.8e-06 | 2.8e-06 | 3.000E-06 | 1.311E-06 | 1.800E-06 | 4.400E-06 |
| 17154 | 858 | C858F | C | 2.4e-06 | 2.7e-06 | 3.1e-06 | 2.733E-06 | 3.512E-07 | 2.400E-06 | 3.100E-06 |
| 17155 | 858 | C858P | C | 3.2e-06 | 2.6e-06 | 1.5e-06 | 2.433E-06 | 8.622E-07 | 1.500E-06 | 3.200E-06 |
| 17156 | 858 | C858S | C | 2.4e-06 | 2.8e-06 | 2.3e-06 | 2.500E-06 | 2.646E-07 | 2.300E-06 | 2.800E-06 |
| 17157 | 858 | C858T | C | 2.6e-06 | 1e-06 | 2.6e-06 | 2.067E-06 | 9.238E-07 | 1.000E-06 | 2.600E-06 |
| 17158 | 858 | C858W | C | 2e-06 | 2.6e-06 | 3.2e-06 | 2.600E-06 | 6.000E-07 | 2.000E-06 | 3.200E-06 |
| 17159 | 858 | C858Y | C | 3.7e-06 | 2.7e-06 | 2.3e-06 | 2.900E-06 | 7.211E-07 | 2.300E-06 | 3.700E-06 |
| 17160 | 858 | C858V | C | 1.8e-06 | 2.1e-06 | 1.2e-06 | 1.700E-06 | 4.583E-07 | 1.200E-06 | 2.100E-06 |
| 17161 | 859 | A859A | C | 2.2e-06 | 1.8e-06 | 3.1e-06 | 2.367E-06 | 6.658E-07 | 1.800E-06 | 3.100E-06 |
| 17162 | 859 | A859R | C | 2.9e-06 | 3.7e-06 | 4.9e-06 | 3.833E-06 | 1.007E-06 | 2.900E-06 | 4.900E-06 |
| 17163 | 859 | A859N | C | 3.6e-06 | 2.3e-06 | 2.6e-06 | 2.833E-06 | 6.807E-07 | 2.300E-06 | 3.600E-06 |
| 17164 | 859 | A859D | C | 2.7e-06 | 2e-06 | 3.7e-06 | 2.800E-06 | 8.544E-07 | 2.000E-06 | 3.700E-06 |
| 17165 | 859 | A859C | C | 4e-06 | 3e-06 | 2.6e-06 | 3.200E-06 | 7.211E-07 | 2.600E-06 | 4.000E-06 |
| 17166 | 859 | A859E | C | 1.9e-06 | 2.6e-06 | 2.4e-06 | 2.300E-06 | 3.606E-07 | 1.900E-06 | 2.600E-06 |
| 17167 | 859 | A859Q | C | 2.5e-06 | 2.6e-06 | 1.6e-06 | 2.233E-06 | 5.508E-07 | 1.600E-06 | 2.600E-06 |
| 17168 | 859 | A859G | C | 2.6e-06 | 2.1e-06 | 1.8e-06 | 2.167E-06 | 4.041E-07 | 1.800E-06 | 2.600E-06 |
| 17169 | 859 | A859H | C | 3.5e-06 | 4.5e-06 | 3e-06 | 3.667E-06 | 7.638E-07 | 3.000E-06 | 4.500E-06 |

| | | | | | | | | | | |
|---|---|---|---|---|---|---|---|---|---|---|
| 17170 | 859 | A859I | C | 2.2e-06 | 4.2e-06 | 2.8e-06 | 3.067E-06 | 1.026E-06 | 2.200E-06 | 4.200E-06 |
| 17171 | 859 | A859L | C | 1.8e-06 | 1.8e-06 | 2.6e-06 | 2.067E-06 | 4.619E-07 | 1.800E-06 | 2.600E-06 |
| 17172 | 859 | A859K | C | 2.1e-06 | 1.7e-06 | 2.2e-06 | 2.000E-06 | 2.646E-07 | 1.700E-06 | 2.200E-06 |
| 17173 | 859 | A859M | C | 2.4e-06 | 1.3e-06 | 2.8e-06 | 2.167E-06 | 7.767E-07 | 1.300E-06 | 2.800E-06 |
| 17174 | 859 | A859F | C | 3.8e-06 | 1.4e-06 | 2.3e-06 | 2.500E-06 | 1.212E-06 | 1.400E-06 | 3.800E-06 |
| 17175 | 859 | A859P | C | 1.8e-06 | 2.5e-06 | 2.7e-06 | 2.333E-06 | 4.726E-07 | 1.800E-06 | 2.700E-06 |
| 17176 | 859 | A859S | C | 1.8e-06 | 2.5e-06 | 3.2e-06 | 2.500E-06 | 7.000E-07 | 1.800E-06 | 3.200E-06 |
| 17177 | 859 | A859T | C | 1.7e-06 | 2.6e-06 | 4.2e-06 | 2.833E-06 | 1.266E-06 | 1.700E-06 | 4.200E-06 |
| 17178 | 859 | A859W | C | 3.4e-06 | 1.5e-06 | 3e-06 | 2.633E-06 | 1.002E-06 | 1.500E-06 | 3.400E-06 |
| 17179 | 859 | A859Y | C | 2.4e-06 | 2.7e-06 | 7.1e-06 | 4.067E-06 | 2.631E-06 | 2.400E-06 | 7.100E-06 |
| 17180 | 859 | A859V | C | 2.8e-06 | 2.7e-06 | 1.9e-06 | 2.467E-06 | 4.933E-07 | 1.900E-06 | 2.800E-06 |
| 17181 | 860 | A860A | C | 3.1e-06 | 3.5e-06 | 2.8e-06 | 3.133E-06 | 3.512E-07 | 2.800E-06 | 3.500E-06 |
| 17182 | 860 | A860R | C | 3e-06 | 1.7e-06 | 2.5e-06 | 2.400E-06 | 6.557E-07 | 1.700E-06 | 3.000E-06 |
| 17183 | 860 | A860N | C | 3.1e-06 | 3e-06 | 2.2e-06 | 2.767E-06 | 4.933E-07 | 2.200E-06 | 3.100E-06 |
| 17184 | 860 | A860D | C | 2.2e-06 | 3e-06 | 4e-06 | 3.067E-06 | 9.018E-07 | 2.200E-06 | 4.000E-06 |
| 17185 | 860 | A860C | C | 2.3e-06 | 2.6e-06 | 2.2e-06 | 2.367E-06 | 2.082E-07 | 2.200E-06 | 2.600E-06 |
| 17186 | 860 | A860E | C | 2.7e-06 | 2e-06 | 3e-06 | 2.567E-06 | 5.132E-07 | 2.000E-06 | 3.000E-06 |
| 17187 | 860 | A860Q | C | 2.2e-06 | 2e-06 | 2.4e-06 | 2.200E-06 | 2.000E-07 | 2.000E-06 | 2.400E-06 |
| 17188 | 860 | A860G | C | 3.8e-06 | 2.5e-06 | 1.3e-06 | 2.533E-06 | 1.250E-06 | 1.300E-06 | 3.800E-06 |
| 17189 | 860 | A860H | C | 2.4e-06 | 1.9e-06 | 2.8e-06 | 2.367E-06 | 4.509E-07 | 1.900E-06 | 2.800E-06 |
| 17190 | 860 | A860I | C | 2e-06 | 2.9e-06 | 2.4e-06 | 2.433E-06 | 4.509E-07 | 2.000E-06 | 2.900E-06 |
| 17191 | 860 | A860L | C | 1.8e-06 | 3.2e-06 | 3e-06 | 2.667E-06 | 7.572E-07 | 1.800E-06 | 3.200E-06 |
| 17192 | 860 | A860K | C | 3e-06 | 2.8e-06 | 2.4e-06 | 2.733E-06 | 3.055E-07 | 2.400E-06 | 3.000E-06 |
| 17193 | 860 | A860M | C | 3.1e-06 | 1.4e-06 | 3.2e-06 | 2.567E-06 | 1.012E-06 | 1.400E-06 | 3.200E-06 |
| 17194 | 860 | A860F | C | 3.1e-06 | 3.6e-06 | 3.1e-06 | 3.267E-06 | 2.887E-07 | 3.100E-06 | 3.600E-06 |
| 17195 | 860 | A860P | C | 1.4e-06 | 1.7e-06 | 1.9e-06 | 1.667E-06 | 2.517E-07 | 1.400E-06 | 1.900E-06 |
| 17196 | 860 | A860S | C | 1.5e-06 | 2.6e-06 | 3.7e-06 | 2.600E-06 | 1.100E-06 | 1.500E-06 | 3.700E-06 |
| 17197 | 860 | A860T | C | 2.7e-06 | 1.7e-06 | 1.2e-06 | 1.867E-06 | 7.638E-07 | 1.200E-06 | 2.700E-06 |
| 17198 | 860 | A860W | C | 1.4e-06 | 2.8e-06 | 1.8e-06 | 2.000E-06 | 7.211E-07 | 1.400E-06 | 2.800E-06 |
| 17199 | 860 | A860Y | C | 4.5e-06 | 1.8e-06 | 3.1e-06 | 3.133E-06 | 1.350E-06 | 1.800E-06 | 4.500E-06 |
| 17200 | 860 | A860V | C | 2.7e-06 | 3.6e-06 | 4.4e-06 | 3.567E-06 | 8.505E-07 | 2.700E-06 | 4.400E-06 |
| 17201 | 861 | S861A | C | 3.3e-06 | 1.6e-06 | 2.4e-06 | 2.433E-06 | 8.505E-07 | 1.600E-06 | 3.300E-06 |
| 17202 | 861 | S861R | C | 3.6e-06 | 1.8e-06 | 2.1e-06 | 2.500E-06 | 9.644E-07 | 1.800E-06 | 3.600E-06 |
| 17203 | 861 | S861N | C | 1.1e-06 | 2.1e-06 | 3e-06 | 2.067E-06 | 9.504E-07 | 1.100E-06 | 3.000E-06 |
| 17204 | 861 | S861D | C | 3.3e-06 | 1.7e-06 | 1.8e-06 | 2.267E-06 | 8.963E-07 | 1.700E-06 | 3.300E-06 |
| 17205 | 861 | S861C | C | 1.9e-06 | 3.5e-06 | 1.3e-06 | 2.233E-06 | 1.137E-06 | 1.300E-06 | 3.500E-06 |
| 17206 | 861 | S861E | C | 2.9e-06 | 2.8e-06 | 1.8e-06 | 2.500E-06 | 6.083E-07 | 1.800E-06 | 2.900E-06 |
| 17207 | 861 | S861Q | C | 2.4e-06 | 2.2e-06 | 4.1e-06 | 2.900E-06 | 1.044E-06 | 2.200E-06 | 4.100E-06 |
| 17208 | 861 | S861G | C | 3.3e-06 | 2.2e-06 | 2.1e-06 | 2.533E-06 | 6.658E-07 | 2.100E-06 | 3.300E-06 |
| 17209 | 861 | S861H | C | 3.4e-06 | 1.4e-06 | 2.4e-06 | 2.400E-06 | 1.000E-06 | 1.400E-06 | 3.400E-06 |
| 17210 | 861 | S861I | C | 2.4e-06 | 4.2e-06 | 1.3e-06 | 2.633E-06 | 1.464E-06 | 1.300E-06 | 4.200E-06 |
| 17211 | 861 | S861L | C | 2.2e-06 | 4.4e-06 | 4.6e-06 | 3.733E-06 | 1.332E-06 | 2.200E-06 | 4.600E-06 |
| 17212 | 861 | S861K | C | 3.1e-06 | 1.6e-06 | 2.1e-06 | 2.267E-06 | 7.638E-07 | 1.600E-06 | 3.100E-06 |
| 17213 | 861 | S861M | C | 2.2e-06 | 1.8e-06 | 1.9e-06 | 1.967E-06 | 2.082E-07 | 1.800E-06 | 2.200E-06 |
| 17214 | 861 | S861F | C | 4.2e-06 | 3.1e-06 | 2.3e-06 | 3.200E-06 | 9.539E-07 | 2.300E-06 | 4.200E-06 |
| 17215 | 861 | S861P | C | 4.8e-06 | 2.3e-06 | 1.3e-06 | 2.800E-06 | 1.803E-06 | 1.300E-06 | 4.800E-06 |
| 17216 | 861 | S861S | C | 1.8e-06 | 2.2e-06 | 2.8e-06 | 2.267E-06 | 5.033E-07 | 1.800E-06 | 2.800E-06 |
| 17217 | 861 | S861T | C | 1.7e-06 | 2.3e-06 | 2.9e-06 | 2.300E-06 | 6.000E-07 | 1.700E-06 | 2.900E-06 |
| 17218 | 861 | S861W | C | 2.8e-06 | 2.8e-06 | 1.3e-06 | 2.300E-06 | 8.660E-07 | 1.300E-06 | 2.800E-06 |
| 17219 | 861 | S861Y | C | 2.4e-06 | 2.2e-06 | 2e-06 | 2.200E-06 | 2.000E-07 | 2.000E-06 | 2.400E-06 |
| 17220 | 861 | S861V | C | 5.7e-06 | 2.3e-06 | 2.2e-06 | 3.400E-06 | 1.992E-06 | 2.200E-06 | 5.700E-06 |
| 17221 | 862 | G862A | C | 1.6e-06 | 2.2e-06 | 2.6e-06 | 2.133E-06 | 5.033E-07 | 1.600E-06 | 2.600E-06 |
| 17222 | 862 | G862R | C | 1.5e-06 | 2.6e-06 | 1.6e-06 | 1.900E-06 | 6.083E-07 | 1.500E-06 | 2.600E-06 |
| 17223 | 862 | G862N | C | 1.3e-06 | 2.5e-06 | 2.7e-06 | 2.167E-06 | 7.572E-07 | 1.300E-06 | 2.700E-06 |
| 17224 | 862 | G862D | C | 1.3e-06 | 2.2e-06 | 2.5e-06 | 2.000E-06 | 6.245E-07 | 1.300E-06 | 2.500E-06 |
| 17225 | 862 | G862C | C | 2.2e-06 | 1.6e-06 | 2.9e-06 | 2.233E-06 | 6.506E-07 | 1.600E-06 | 2.900E-06 |
| 17226 | 862 | G862E | C | 4.8e-06 | 2.2e-06 | 1.4e-06 | 2.800E-06 | 1.778E-06 | 1.400E-06 | 4.800E-06 |
| 17227 | 862 | G862Q | C | 3.2e-06 | 6.8e-06 | 2.3e-06 | 4.100E-06 | 2.381E-06 | 2.300E-06 | 6.800E-06 |
| 17228 | 862 | G862G | C | 1.9e-06 | 8.8e-07 | 3e-06 | 1.927E-06 | 1.060E-06 | 8.800E-07 | 3.000E-06 |
| 17229 | 862 | G862H | C | 4.1e-06 | 2.9e-06 | 2.6e-06 | 3.200E-06 | 7.937E-07 | 2.600E-06 | 4.100E-06 |
| 17230 | 862 | G862I | C | 3.1e-06 | 2.1e-06 | 2e-06 | 2.400E-06 | 6.083E-07 | 2.000E-06 | 3.100E-06 |
| 17231 | 862 | G862L | C | 3.8e-06 | 2.3e-06 | 2.5e-06 | 2.867E-06 | 8.145E-07 | 2.300E-06 | 3.800E-06 |
| 17232 | 862 | G862K | C | 2.5e-06 | 1.3e-06 | 1.1e-06 | 1.633E-06 | 7.572E-07 | 1.100E-06 | 2.500E-06 |
| 17233 | 862 | G862M | C | 3.3e-06 | 2.1e-06 | 3.4e-06 | 2.933E-06 | 7.234E-07 | 2.100E-06 | 3.400E-06 |
| 17234 | 862 | G862F | C | 2.5e-06 | 1.8e-06 | 2.8e-06 | 2.367E-06 | 5.132E-07 | 1.800E-06 | 2.800E-06 |
| 17235 | 862 | G862P | C | 2.6e-06 | 2.3e-06 | 3.1e-06 | 2.667E-06 | 4.041E-07 | 2.300E-06 | 3.100E-06 |
| 17236 | 862 | G862S | C | 1.8e-06 | 3.2e-06 | 3.9e-06 | 2.967E-06 | 1.069E-06 | 1.800E-06 | 3.900E-06 |
| 17237 | 862 | G862T | C | 2.8e-06 | 2.3e-06 | 2.2e-06 | 2.433E-06 | 3.215E-07 | 2.200E-06 | 2.800E-06 |
| 17238 | 862 | G862W | C | 2e-06 | 2e-06 | 1.7e-06 | 1.900E-06 | 1.732E-07 | 1.700E-06 | 2.000E-06 |
| 17239 | 862 | G862Y | C | 2.4e-06 | 1.7e-06 | 2.6e-06 | 2.233E-06 | 4.726E-07 | 1.700E-06 | 2.600E-06 |
| 17240 | 862 | G862V | C | 1.6e-06 | 3.2e-06 | 1.9e-06 | 2.233E-06 | 8.505E-07 | 1.600E-06 | 3.200E-06 |
| 17241 | 863 | F863A | C | 1.7e-06 | 2.2e-06 | 3.1e-06 | 2.333E-06 | 7.095E-07 | 1.700E-06 | 3.100E-06 |
| 17242 | 863 | F863R | C | 1.6e-06 | 3.2e-06 | 4e-06 | 2.933E-06 | 1.222E-06 | 1.600E-06 | 4.000E-06 |
| 17243 | 863 | F863N | C | 3.1e-06 | 2.8e-06 | 5.1e-06 | 3.667E-06 | 1.250E-06 | 2.800E-06 | 5.100E-06 |
| 17244 | 863 | F863D | C | 2.7e-06 | 2.5e-06 | 1.9e-06 | 2.367E-06 | 4.163E-07 | 1.900E-06 | 2.700E-06 |
| 17245 | 863 | F863C | C | 1.3e-06 | 4.4e-06 | 2.2e-06 | 2.633E-06 | 1.595E-06 | 1.300E-06 | 4.400E-06 |
| 17246 | 863 | F863E | C | 2.1e-06 | 2.7e-06 | 3.2e-06 | 2.667E-06 | 5.508E-07 | 2.100E-06 | 3.200E-06 |
| 17247 | 863 | F863Q | C | 2.1e-06 | 1.6e-06 | 3.7e-06 | 2.467E-06 | 1.097E-06 | 1.600E-06 | 3.700E-06 |
| 17248 | 863 | F863G | C | 2e-06 | 2.3e-06 | 4.6e-06 | 2.967E-06 | 1.422E-06 | 2.000E-06 | 4.600E-06 |
| 17249 | 863 | F863H | C | 2.5e-06 | 8.9e-07 | 2.3e-06 | 1.897E-06 | 8.775E-07 | 8.900E-07 | 2.500E-06 |
| 17250 | 863 | F863I | C | 2.6e-06 | 2.7e-06 | 2.3e-06 | 2.533E-06 | 2.082E-07 | 2.300E-06 | 2.700E-06 |
| 17251 | 863 | F863L | C | 2.8e-06 | 1.8e-06 | 1.5e-06 | 2.033E-06 | 6.807E-07 | 1.500E-06 | 2.800E-06 |
| 17252 | 863 | F863K | C | 2e-06 | 3.1e-06 | 2.2e-06 | 2.433E-06 | 5.859E-07 | 2.000E-06 | 3.100E-06 |
| 17253 | 863 | F863M | C | 4.4e-06 | 2.2e-06 | 1.6e-06 | 2.733E-06 | 1.474E-06 | 1.600E-06 | 4.400E-06 |
| 17254 | 863 | F863F | C | 2.2e-06 | 3.2e-06 | 2.5e-06 | 2.633E-06 | 5.132E-07 | 2.200E-06 | 3.200E-06 |

| | | | | | | | | | | |
|---|---|---|---|---|---|---|---|---|---|---|
| 17255 | 863 | F863P | C | 2.4e-06 | 1.6e-06 | 3.3e-06 | 2.433E-06 | 8.505E-07 | 1.600E-06 | 3.300E-06 |
| 17256 | 863 | F863S | C | 2.2e-06 | 1.6e-06 | 2.4e-06 | 2.067E-06 | 4.163E-07 | 1.600E-06 | 2.400E-06 |
| 17257 | 863 | F863T | C | 2.4e-06 | 1.3e-06 | 2e-06 | 1.900E-06 | 5.568E-07 | 1.300E-06 | 2.400E-06 |
| 17258 | 863 | F863W | C | 4e-06 | 2.6e-06 | 1.8e-06 | 2.800E-06 | 1.114E-06 | 1.800E-06 | 4.000E-06 |
| 17259 | 863 | F863Y | C | 2.1e-06 | 2e-06 | 2e-06 | 2.033E-06 | 5.774E-08 | 2.000E-06 | 2.100E-06 |
| 17260 | 863 | F863V | C | 2.2e-06 | 3.7e-06 | 2.7e-06 | 2.867E-06 | 7.638E-07 | 2.200E-06 | 3.700E-06 |
| 17261 | 864 | T864A | C | 2.6e-06 | 2.6e-06 | 2.7e-06 | 2.633E-06 | 5.774E-08 | 2.600E-06 | 2.700E-06 |
| 17262 | 864 | T864R | C | 2.6e-06 | 2.2e-06 | 3.4e-06 | 2.733E-06 | 6.110E-07 | 2.200E-06 | 3.400E-06 |
| 17263 | 864 | T864N | C | 2.2e-06 | 3.1e-06 | 1.8e-06 | 2.367E-06 | 6.658E-07 | 1.800E-06 | 3.100E-06 |
| 17264 | 864 | T864D | C | 1.5e-06 | 3.2e-06 | 2e-06 | 2.233E-06 | 8.737E-07 | 1.500E-06 | 3.200E-06 |
| 17265 | 864 | T864C | C | 2.9e-06 | 1.4e-06 | 2.2e-06 | 2.167E-06 | 7.506E-07 | 1.400E-06 | 2.900E-06 |
| 17266 | 864 | T864E | C | 2.5e-06 | 1.7e-06 | 2.4e-06 | 2.200E-06 | 4.359E-07 | 1.700E-06 | 2.500E-06 |
| 17267 | 864 | T864Q | C | 2.4e-06 | 2.8e-06 | 2.3e-06 | 2.500E-06 | 2.646E-07 | 2.300E-06 | 2.800E-06 |
| 17268 | 864 | T864G | C | 3.6e-06 | 8.7e-07 | 2.4e-06 | 2.290E-06 | 1.368E-06 | 8.700E-07 | 3.600E-06 |
| 17269 | 864 | T864H | C | 1.1e-06 | 2.1e-06 | 1.2e-06 | 1.467E-06 | 5.508E-07 | 1.100E-06 | 2.100E-06 |
| 17270 | 864 | T864I | C | 3.2e-06 | 2.7e-06 | 3.1e-06 | 3.000E-06 | 2.646E-07 | 2.700E-06 | 3.200E-06 |
| 17271 | 864 | T864L | C | 2.6e-06 | 3.4e-06 | 2.8e-06 | 2.933E-06 | 4.163E-07 | 2.600E-06 | 3.400E-06 |
| 17272 | 864 | T864K | C | 3.5e-06 | 1.8e-06 | 2.5e-06 | 2.600E-06 | 8.544E-07 | 1.800E-06 | 3.500E-06 |
| 17273 | 864 | T864M | C | 2.2e-06 | 3.7e-06 | 3e-06 | 2.967E-06 | 7.506E-07 | 2.200E-06 | 3.700E-06 |
| 17274 | 864 | T864F | C | 2.7e-06 | 2.1e-06 | 2.4e-06 | 2.400E-06 | 3.000E-07 | 2.100E-06 | 2.700E-06 |
| 17275 | 864 | T864P | C | 2.2e-06 | 4.4e-06 | 3.2e-06 | 3.267E-06 | 1.102E-06 | 2.200E-06 | 4.400E-06 |
| 17276 | 864 | T864S | C | 1.7e-06 | 2.8e-06 | 2.4e-06 | 2.300E-06 | 5.568E-07 | 1.700E-06 | 2.800E-06 |
| 17277 | 864 | T864T | C | 1.7e-06 | 3.6e-06 | 1.7e-06 | 2.333E-06 | 1.097E-06 | 1.700E-06 | 3.600E-06 |
| 17278 | 864 | T864W | C | 3e-06 | 2e-06 | 4e-06 | 3.000E-06 | 1.000E-06 | 2.000E-06 | 4.000E-06 |
| 17279 | 864 | T864Y | C | 1.9e-06 | 3.4e-06 | 2.5e-06 | 2.600E-06 | 7.550E-07 | 1.900E-06 | 3.400E-06 |
| 17280 | 864 | T864V | C | 1.7e-06 | 3e-06 | 2.2e-06 | 2.300E-06 | 6.557E-07 | 1.700E-06 | 3.000E-06 |
| 17281 | 865 | F865A | C | 3.7e-06 | 3.5e-06 | 2.1e-06 | 3.100E-06 | 8.718E-07 | 2.100E-06 | 3.700E-06 |
| 17282 | 865 | F865R | C | 3.4e-06 | 2.7e-06 | 3.5e-06 | 3.200E-06 | 4.359E-07 | 2.700E-06 | 3.500E-06 |
| 17283 | 865 | F865N | C | 2e-06 | 1.6e-06 | 2.7e-06 | 2.100E-06 | 5.568E-07 | 1.600E-06 | 2.700E-06 |
| 17284 | 865 | F865D | C | 1.7e-06 | 2.8e-06 | 4.5e-06 | 3.000E-06 | 1.411E-06 | 1.700E-06 | 4.500E-06 |
| 17285 | 865 | F865C | C | 2.2e-06 | 2.9e-06 | 2.1e-06 | 2.400E-06 | 4.359E-07 | 2.100E-06 | 2.900E-06 |
| 17286 | 865 | F865E | C | 2.4e-06 | 1.6e-06 | 2.3e-06 | 2.100E-06 | 4.359E-07 | 1.600E-06 | 2.400E-06 |
| 17287 | 865 | F865Q | C | 1.9e-06 | 2.4e-06 | 2.3e-06 | 2.200E-06 | 2.646E-07 | 1.900E-06 | 2.400E-06 |
| 17288 | 865 | F865G | C | 3.1e-06 | 3.1e-06 | 1.9e-06 | 2.700E-06 | 6.928E-07 | 1.900E-06 | 3.100E-06 |
| 17289 | 865 | F865H | C | 4.5e-06 | 1.9e-06 | 4e-06 | 3.467E-06 | 1.380E-06 | 1.900E-06 | 4.500E-06 |
| 17290 | 865 | F865I | C | 5.3e-06 | 2.5e-06 | 1.6e-06 | 3.133E-06 | 1.930E-06 | 1.600E-06 | 5.300E-06 |
| 17291 | 865 | F865L | C | 3.1e-06 | 2.2e-06 | 3.2e-06 | 2.833E-06 | 5.508E-07 | 2.200E-06 | 3.200E-06 |
| 17292 | 865 | F865K | C | 3.2e-06 | 2.7e-06 | 2e-06 | 2.633E-06 | 6.028E-07 | 2.000E-06 | 3.200E-06 |
| 17293 | 865 | F865M | C | 2.3e-06 | 1.8e-06 | 2.4e-06 | 2.167E-06 | 3.215E-07 | 1.800E-06 | 2.400E-06 |
| 17294 | 865 | F865F | C | 3.5e-06 | 2.5e-06 | 2.1e-06 | 2.700E-06 | 7.211E-07 | 2.100E-06 | 3.500E-06 |
| 17295 | 865 | F865P | C | 2.7e-06 | 2e-06 | 1.1e-06 | 1.933E-06 | 8.021E-07 | 1.100E-06 | 2.700E-06 |
| 17296 | 865 | F865S | C | 3.6e-06 | 4.1e-06 | 1.8e-06 | 3.167E-06 | 1.210E-06 | 1.800E-06 | 4.100E-06 |
| 17297 | 865 | F865T | C | 4.2e-06 | 3e-06 | 2.2e-06 | 3.133E-06 | 1.007E-06 | 2.200E-06 | 4.200E-06 |
| 17298 | 865 | F865W | C | 2.1e-06 | 1.9e-06 | 2.3e-06 | 2.100E-06 | 2.000E-07 | 1.900E-06 | 2.300E-06 |
| 17299 | 865 | F865Y | C | 1.4e-06 | 3.1e-06 | 2.6e-06 | 2.367E-06 | 8.737E-07 | 1.400E-06 | 3.100E-06 |
| 17300 | 865 | F865V | C | 2.5e-06 | 2.5e-06 | 2.1e-06 | 2.367E-06 | 2.309E-07 | 2.100E-06 | 2.500E-06 |
| 17301 | 866 | T866A | C | 2.8e-06 | 1.8e-06 | 2.7e-06 | 2.433E-06 | 5.508E-07 | 1.800E-06 | 2.800E-06 |
| 17302 | 866 | T866R | C | 2.6e-06 | 1.2e-06 | 3.8e-06 | 2.533E-06 | 1.301E-06 | 1.200E-06 | 3.800E-06 |
| 17303 | 866 | T866N | C | 1.7e-06 | 2.8e-06 | 3.5e-06 | 2.667E-06 | 9.074E-07 | 1.700E-06 | 3.500E-06 |
| 17304 | 866 | T866D | C | 1.1e-06 | 2.1e-06 | 3e-06 | 2.067E-06 | 9.504E-07 | 1.100E-06 | 3.000E-06 |
| 17305 | 866 | T866C | C | 2e-06 | 2.7e-06 | 1.4e-06 | 2.033E-06 | 6.506E-07 | 1.400E-06 | 2.700E-06 |
| 17306 | 866 | T866E | C | 1.6e-06 | 2.6e-06 | 4.4e-06 | 2.867E-06 | 1.419E-06 | 1.600E-06 | 4.400E-06 |
| 17307 | 866 | T866Q | C | 2.2e-06 | 2.3e-06 | 3.2e-06 | 2.567E-06 | 5.508E-07 | 2.200E-06 | 3.200E-06 |
| 17308 | 866 | T866G | C | 2.8e-06 | 1.8e-06 | 2.2e-06 | 2.267E-06 | 5.033E-07 | 1.800E-06 | 2.800E-06 |
| 17309 | 866 | T866H | C | 2.4e-06 | 1.7e-06 | 3.1e-06 | 2.400E-06 | 7.000E-07 | 1.700E-06 | 3.100E-06 |
| 17310 | 866 | T866I | C | 3e-06 | 2.7e-06 | 2.6e-06 | 2.767E-06 | 2.082E-07 | 2.600E-06 | 3.000E-06 |
| 17311 | 866 | T866L | C | 3.5e-06 | 2.7e-06 | 1.5e-06 | 2.567E-06 | 1.007E-06 | 1.500E-06 | 3.500E-06 |
| 17312 | 866 | T866K | C | 2.2e-06 | 2.2e-06 | 3.4e-06 | 2.600E-06 | 6.928E-07 | 2.200E-06 | 3.400E-06 |
| 17313 | 866 | T866M | C | 2.4e-06 | 2.1e-06 | 2.9e-06 | 2.467E-06 | 4.041E-07 | 2.100E-06 | 2.900E-06 |
| 17314 | 866 | T866F | C | 3.7e-06 | 2.6e-06 | 1.4e-06 | 2.567E-06 | 1.150E-06 | 1.400E-06 | 3.700E-06 |
| 17315 | 866 | T866P | C | 3.1e-06 | 1.9e-06 | 2.9e-06 | 2.633E-06 | 6.429E-07 | 1.900E-06 | 3.100E-06 |
| 17316 | 866 | T866S | C | 2.3e-06 | 3.8e-06 | 4.1e-06 | 3.400E-06 | 9.644E-07 | 2.300E-06 | 4.100E-06 |
| 17317 | 866 | T866T | C | 2.5e-06 | 2e-06 | 2.3e-06 | 2.267E-06 | 2.517E-07 | 2.000E-06 | 2.500E-06 |
| 17318 | 866 | T866W | C | 3.2e-06 | 2.1e-06 | 1.9e-06 | 2.400E-06 | 7.000E-07 | 1.900E-06 | 3.200E-06 |
| 17319 | 866 | T866Y | C | 1.6e-06 | 2.5e-06 | 2.8e-06 | 2.300E-06 | 6.245E-07 | 1.600E-06 | 2.800E-06 |
| 17320 | 866 | T866V | C | 1.6e-06 | 2.2e-06 | 3.5e-06 | 2.433E-06 | 9.713E-07 | 1.600E-06 | 3.500E-06 |
| 17321 | 867 | D867A | C | 2.4e-06 | 1.9e-06 | 1.3e-06 | 1.867E-06 | 5.508E-07 | 1.300E-06 | 2.400E-06 |
| 17322 | 867 | D867R | C | 3.8e-06 | 2.7e-06 | 2.3e-06 | 2.933E-06 | 7.767E-07 | 2.300E-06 | 3.800E-06 |
| 17323 | 867 | D867N | C | 1.5e-06 | 3.4e-06 | 3.1e-06 | 2.667E-06 | 1.021E-06 | 1.500E-06 | 3.400E-06 |
| 17324 | 867 | D867D | C | 1.9e-06 | 3.3e-06 | 2.7e-06 | 2.633E-06 | 7.024E-07 | 1.900E-06 | 3.300E-06 |
| 17325 | 867 | D867C | C | 2.7e-06 | 2.6e-06 | 2.3e-06 | 2.533E-06 | 2.082E-07 | 2.300E-06 | 2.700E-06 |
| 17326 | 867 | D867E | C | 3.1e-06 | 2.3e-06 | 2.2e-06 | 2.533E-06 | 4.933E-07 | 2.200E-06 | 3.100E-06 |
| 17327 | 867 | D867Q | C | 2.8e-06 | 2.4e-06 | 1.4e-06 | 2.200E-06 | 7.211E-07 | 1.400E-06 | 2.800E-06 |
| 17328 | 867 | D867G | C | 3.4e-06 | 1.8e-06 | 2.2e-06 | 2.467E-06 | 8.327E-07 | 1.800E-06 | 3.400E-06 |
| 17329 | 867 | D867H | C | 2.5e-06 | 2.6e-06 | 1.8e-06 | 2.300E-06 | 4.359E-07 | 1.800E-06 | 2.600E-06 |
| 17330 | 867 | D867I | C | 1.8e-06 | 2e-06 | 1.5e-06 | 1.767E-06 | 2.517E-07 | 1.500E-06 | 2.000E-06 |
| 17331 | 867 | D867L | C | 1.7e-06 | 2.9e-06 | 2.2e-06 | 2.267E-06 | 6.028E-07 | 1.700E-06 | 2.900E-06 |
| 17332 | 867 | D867K | C | 2e-06 | 1.8e-06 | 2.4e-06 | 2.067E-06 | 3.055E-07 | 1.800E-06 | 2.400E-06 |
| 17333 | 867 | D867M | C | 2e-06 | 1.5e-06 | 3.3e-06 | 2.267E-06 | 9.292E-07 | 1.500E-06 | 3.300E-06 |
| 17334 | 867 | D867F | C | 2.8e-06 | 4.1e-06 | 2.1e-06 | 3.000E-06 | 1.015E-06 | 2.100E-06 | 4.100E-06 |
| 17335 | 867 | D867P | C | 1e-06 | 1.8e-06 | 2.1e-06 | 1.633E-06 | 5.686E-07 | 1.000E-06 | 2.100E-06 |
| 17336 | 867 | D867S | C | 2e-06 | 2.2e-06 | 1.9e-06 | 2.033E-06 | 1.528E-07 | 1.900E-06 | 2.200E-06 |
| 17337 | 867 | D867T | C | 2e-06 | 1.4e-06 | 1.9e-06 | 1.767E-06 | 3.215E-07 | 1.400E-06 | 2.000E-06 |
| 17338 | 867 | D867W | C | 2.4e-06 | 2.7e-06 | 3.5e-06 | 2.867E-06 | 5.686E-07 | 2.400E-06 | 3.500E-06 |
| 17339 | 867 | D867Y | C | 2.6e-06 | 2.4e-06 | 1.7e-06 | 2.233E-06 | 4.726E-07 | 1.700E-06 | 2.600E-06 |

| | | | | | | | | | | |
|---|---|---|---|---|---|---|---|---|---|---|
| 17340 | 867 | D867V | C | 2.2e-06 | 2.1e-06 | 4.3e-06 | 2.867E-06 | 1.242E-06 | 2.100E-06 | 4.300E-06 |
| 17341 | 868 | Y868A | C | 2.6e-06 | 1.6e-06 | 2.6e-06 | 2.267E-06 | 5.774E-07 | 1.600E-06 | 2.600E-06 |
| 17342 | 868 | Y868R | C | 4.8e-06 | 3.1e-06 | 1.4e-06 | 3.100E-06 | 1.700E-06 | 1.400E-06 | 4.800E-06 |
| 17343 | 868 | Y868N | C | 2.2e-06 | 8.7e-07 | 3.2e-06 | 2.090E-06 | 1.169E-06 | 8.700E-07 | 3.200E-06 |
| 17344 | 868 | Y868D | C | 2.1e-06 | 3.2e-06 | 1.7e-06 | 2.333E-06 | 7.767E-07 | 1.700E-06 | 3.200E-06 |
| 17345 | 868 | Y868C | C | 1.7e-06 | 2e-06 | 2.5e-06 | 2.067E-06 | 4.041E-07 | 1.700E-06 | 2.500E-06 |
| 17346 | 868 | Y868E | C | 1.7e-06 | 1.8e-06 | 2.6e-06 | 2.033E-06 | 4.933E-07 | 1.700E-06 | 2.600E-06 |
| 17347 | 868 | Y868Q | C | 2.3e-06 | 3e-06 | 1.8e-06 | 2.367E-06 | 6.028E-07 | 1.800E-06 | 3.000E-06 |
| 17348 | 868 | Y868G | C | 2.7e-06 | 2.5e-06 | 3.4e-06 | 2.867E-06 | 4.726E-07 | 2.500E-06 | 3.400E-06 |
| 17349 | 868 | Y868H | C | 1.1e-06 | 2.5e-06 | 2.7e-06 | 2.100E-06 | 8.718E-07 | 1.100E-06 | 2.700E-06 |
| 17350 | 868 | Y868I | C | 1.5e-06 | 2.2e-06 | 3.2e-06 | 2.300E-06 | 8.544E-07 | 1.500E-06 | 3.200E-06 |
| 17351 | 868 | Y868L | C | 2.3e-06 | 4e-06 | 2e-06 | 2.767E-06 | 1.079E-06 | 2.000E-06 | 4.000E-06 |
| 17352 | 868 | Y868K | C | 1.9e-06 | 2.7e-06 | 2.4e-06 | 2.333E-06 | 4.041E-07 | 1.900E-06 | 2.700E-06 |
| 17353 | 868 | Y868M | C | 3.4e-06 | 2.6e-06 | 1.5e-06 | 2.500E-06 | 9.539E-07 | 1.500E-06 | 3.400E-06 |
| 17354 | 868 | Y868F | C | 5e-06 | 2.7e-06 | 3.9e-06 | 3.867E-06 | 1.150E-06 | 2.700E-06 | 5.000E-06 |
| 17355 | 868 | Y868P | C | 2.9e-06 | 1.6e-06 | 4.2e-06 | 2.900E-06 | 1.300E-06 | 1.600E-06 | 4.200E-06 |
| 17356 | 868 | Y868S | C | 3.1e-06 | 3.5e-06 | 3e-06 | 3.200E-06 | 2.646E-07 | 3.000E-06 | 3.500E-06 |
| 17357 | 868 | Y868T | C | 2.4e-06 | 2.6e-06 | 2e-06 | 2.333E-06 | 3.055E-07 | 2.000E-06 | 2.600E-06 |
| 17358 | 868 | Y868W | C | 2.8e-06 | 3.8e-06 | 1e-06 | 2.533E-06 | 1.419E-06 | 1.000E-06 | 3.800E-06 |
| 17359 | 868 | Y868Y | C | 3.1e-06 | 2.7e-06 | 4.5e-06 | 3.433E-06 | 9.452E-07 | 2.700E-06 | 4.500E-06 |
| 17360 | 868 | Y868V | C | 2.8e-06 | 1.3e-06 | 1.6e-06 | 1.900E-06 | 7.937E-07 | 1.300E-06 | 2.800E-06 |
| 17361 | 869 | T869A | C | 2.2e-06 | 1e-06 | 3.4e-06 | 2.200E-06 | 1.200E-06 | 1.000E-06 | 3.400E-06 |
| 17362 | 869 | T869R | C | 2.3e-06 | 2.6e-06 | 4.6e-06 | 3.167E-06 | 1.250E-06 | 2.300E-06 | 4.600E-06 |
| 17363 | 869 | T869N | C | 3e-06 | 2.2e-06 | 1.6e-06 | 2.267E-06 | 7.024E-07 | 1.600E-06 | 3.000E-06 |
| 17364 | 869 | T869D | C | 2.8e-06 | 2.4e-06 | 2e-06 | 2.400E-06 | 4.000E-07 | 2.000E-06 | 2.800E-06 |
| 17365 | 869 | T869C | C | 2.5e-06 | 2.1e-06 | 1e-06 | 1.867E-06 | 7.767E-07 | 1.000E-06 | 2.500E-06 |
| 17366 | 869 | T869E | C | 2.2e-06 | 2.8e-06 | 2.1e-06 | 2.367E-06 | 3.786E-07 | 2.100E-06 | 2.800E-06 |
| 17367 | 869 | T869Q | C | 1.4e-06 | 1.9e-06 | 1.7e-06 | 1.667E-06 | 2.517E-07 | 1.400E-06 | 1.900E-06 |
| 17368 | 869 | T869G | C | 2.5e-06 | 1.9e-06 | 2.2e-06 | 2.200E-06 | 3.000E-07 | 1.900E-06 | 2.500E-06 |
| 17369 | 869 | T869H | C | 3.3e-06 | 1.3e-06 | 2.2e-06 | 2.267E-06 | 1.002E-06 | 1.300E-06 | 3.300E-06 |
| 17370 | 869 | T869I | C | 3e-06 | 2.8e-06 | 2e-06 | 2.600E-06 | 5.292E-07 | 2.000E-06 | 3.000E-06 |
| 17371 | 869 | T869L | C | 4.3e-06 | 2.5e-06 | 2.5e-06 | 3.100E-06 | 1.039E-06 | 2.500E-06 | 4.300E-06 |
| 17372 | 869 | T869K | C | 1.6e-06 | 2.3e-06 | 1.9e-06 | 1.933E-06 | 3.512E-07 | 1.600E-06 | 2.300E-06 |
| 17373 | 869 | T869M | C | 3.5e-06 | 4e-06 | 2.8e-06 | 3.433E-06 | 6.028E-07 | 2.800E-06 | 4.000E-06 |
| 17374 | 869 | T869F | C | 3.3e-06 | 1.8e-06 | 1.5e-06 | 2.200E-06 | 9.644E-07 | 1.500E-06 | 3.300E-06 |
| 17375 | 869 | T869P | C | 2.3e-06 | 2.5e-06 | 3.4e-06 | 2.733E-06 | 5.859E-07 | 2.300E-06 | 3.400E-06 |
| 17376 | 869 | T869S | C | 2.8e-06 | 1.7e-06 | 2.4e-06 | 2.300E-06 | 5.568E-07 | 1.700E-06 | 2.800E-06 |
| 17377 | 869 | T869T | C | 2.8e-06 | 6.7e-07 | 4e-06 | 2.490E-06 | 1.687E-06 | 6.700E-07 | 4.000E-06 |
| 17378 | 869 | T869W | C | 2.2e-06 | 3.7e-06 | 2.9e-06 | 2.933E-06 | 7.506E-07 | 2.200E-06 | 3.700E-06 |
| 17379 | 869 | T869Y | C | 3.4e-06 | 3.1e-06 | 1.9e-06 | 2.800E-06 | 7.937E-07 | 1.900E-06 | 3.400E-06 |
| 17380 | 869 | T869V | C | 3.4e-06 | 2.1e-06 | 2.3e-06 | 2.600E-06 | 7.000E-07 | 2.100E-06 | 3.400E-06 |
| 17381 | 870 | M870A | C | 2.7e-06 | 1.8e-06 | 2.1e-06 | 2.200E-06 | 4.583E-07 | 1.800E-06 | 2.700E-06 |
| 17382 | 870 | M870R | C | 1.5e-06 | 1.7e-06 | 1.4e-06 | 1.533E-06 | 1.528E-07 | 1.400E-06 | 1.700E-06 |
| 17383 | 870 | M870N | C | 2.7e-06 | 2.1e-06 | 3.9e-06 | 2.900E-06 | 9.165E-07 | 2.100E-06 | 3.900E-06 |
| 17384 | 870 | M870D | C | 2.4e-06 | 2e-06 | 2.7e-06 | 2.367E-06 | 3.512E-07 | 2.000E-06 | 2.700E-06 |
| 17385 | 870 | M870C | C | 2.3e-06 | 3e-06 | 2.2e-06 | 2.500E-06 | 4.359E-07 | 2.200E-06 | 3.000E-06 |
| 17386 | 870 | M870E | C | 1.4e-06 | 3.1e-06 | 1.9e-06 | 2.133E-06 | 8.737E-07 | 1.400E-06 | 3.100E-06 |
| 17387 | 870 | M870Q | C | 2e-06 | 2.6e-06 | 1.6e-06 | 2.067E-06 | 5.033E-07 | 1.600E-06 | 2.600E-06 |
| 17388 | 870 | M870G | C | 2.6e-06 | 2.3e-06 | 2e-06 | 2.300E-06 | 3.000E-07 | 2.000E-06 | 2.600E-06 |
| 17389 | 870 | M870H | C | 2.6e-06 | 2.1e-06 | 2e-06 | 2.233E-06 | 3.215E-07 | 2.000E-06 | 2.600E-06 |
| 17390 | 870 | M870I | C | 2.3e-06 | 2.1e-06 | 1.9e-06 | 2.100E-06 | 2.000E-07 | 1.900E-06 | 2.300E-06 |
| 17391 | 870 | M870L | C | 2.1e-06 | 2.4e-06 | 4.1e-06 | 2.867E-06 | 1.079E-06 | 2.100E-06 | 4.100E-06 |
| 17392 | 870 | M870K | C | 2.6e-06 | 2.2e-06 | 2.3e-06 | 2.367E-06 | 2.082E-07 | 2.200E-06 | 2.600E-06 |
| 17393 | 870 | M870M | C | 1.6e-06 | 2.4e-06 | 2.6e-06 | 2.200E-06 | 5.292E-07 | 1.600E-06 | 2.600E-06 |
| 17394 | 870 | M870F | C | 2e-06 | 1.8e-06 | 2.2e-06 | 2.000E-06 | 2.000E-07 | 1.800E-06 | 2.200E-06 |
| 17395 | 870 | M870P | C | 3.2e-06 | 3.1e-06 | 2.7e-06 | 3.000E-06 | 2.646E-07 | 2.700E-06 | 3.200E-06 |
| 17396 | 870 | M870S | C | 1.7e-06 | 2.2e-06 | 3.2e-06 | 2.367E-06 | 7.638E-07 | 1.700E-06 | 3.200E-06 |
| 17397 | 870 | M870T | C | 2.5e-06 | 2.7e-06 | 2.2e-06 | 2.467E-06 | 2.517E-07 | 2.200E-06 | 2.700E-06 |
| 17398 | 870 | M870W | C | 1.8e-06 | 2.3e-06 | 3.5e-06 | 2.533E-06 | 8.737E-07 | 1.800E-06 | 3.500E-06 |
| 17399 | 870 | M870Y | C | 2.2e-06 | 2.9e-06 | 2.4e-06 | 2.500E-06 | 3.606E-07 | 2.200E-06 | 2.900E-06 |
| 17400 | 870 | M870V | C | 2.8e-06 | 2.2e-06 | 3e-06 | 2.667E-06 | 4.163E-07 | 2.200E-06 | 3.000E-06 |
| 17401 | 871 | D871A | C | 2.1e-06 | 2.5e-06 | 3.1e-06 | 2.567E-06 | 5.033E-07 | 2.100E-06 | 3.100E-06 |
| 17402 | 871 | D871R | C | 2.8e-06 | 1.9e-06 | 2.3e-06 | 2.333E-06 | 4.509E-07 | 1.900E-06 | 2.800E-06 |
| 17403 | 871 | D871N | C | 1.6e-06 | 3.5e-06 | 2.2e-06 | 2.433E-06 | 9.713E-07 | 1.600E-06 | 3.500E-06 |
| 17404 | 871 | D871D | C | 2.7e-06 | 1.9e-06 | 3.6e-06 | 2.733E-06 | 8.505E-07 | 1.900E-06 | 3.600E-06 |
| 17405 | 871 | D871C | C | 2.5e-06 | 1.1e-06 | 1.8e-06 | 1.800E-06 | 7.000E-07 | 1.100E-06 | 2.500E-06 |
| 17406 | 871 | D871E | C | 2.5e-06 | 3.5e-06 | 1.6e-06 | 2.533E-06 | 9.504E-07 | 1.600E-06 | 3.500E-06 |
| 17407 | 871 | D871Q | C | 2.3e-06 | 1.8e-06 | 3e-06 | 2.367E-06 | 6.028E-07 | 1.800E-06 | 3.000E-06 |
| 17408 | 871 | D871G | C | 2.9e-06 | 2.7e-06 | 1.1e-06 | 2.233E-06 | 9.866E-07 | 1.100E-06 | 2.900E-06 |
| 17409 | 871 | D871H | C | 3.9e-06 | 3e-06 | 2.5e-06 | 3.133E-06 | 7.095E-07 | 2.500E-06 | 3.900E-06 |
| 17410 | 871 | D871I | C | 4.3e-06 | 3.4e-06 | 3.4e-06 | 3.700E-06 | 5.196E-07 | 3.400E-06 | 4.300E-06 |
| 17411 | 871 | D871L | C | 3.9e-06 | 2.7e-06 | 2.5e-06 | 3.033E-06 | 7.572E-07 | 2.500E-06 | 3.900E-06 |
| 17412 | 871 | D871K | C | 3.6e-06 | 2.2e-06 | 2e-06 | 2.600E-06 | 8.718E-07 | 2.000E-06 | 3.600E-06 |
| 17413 | 871 | D871M | C | 2.4e-06 | 2.4e-06 | 2e-06 | 2.267E-06 | 2.309E-07 | 2.000E-06 | 2.400E-06 |
| 17414 | 871 | D871F | C | 2.8e-06 | 2.5e-06 | 2.2e-06 | 2.500E-06 | 3.000E-07 | 2.200E-06 | 2.800E-06 |
| 17415 | 871 | D871P | C | 2.4e-06 | 1.9e-06 | 1.1e-06 | 1.800E-06 | 6.557E-07 | 1.100E-06 | 2.400E-06 |
| 17416 | 871 | D871S | C | 1.5e-06 | 3e-06 | 3.8e-06 | 2.767E-06 | 1.168E-06 | 1.500E-06 | 3.800E-06 |
| 17417 | 871 | D871T | C | 3.1e-06 | 3.6e-06 | 1.8e-06 | 2.833E-06 | 9.292E-07 | 1.800E-06 | 3.600E-06 |
| 17418 | 871 | D871W | C | 1.7e-06 | 1.7e-06 | 2.2e-06 | 1.867E-06 | 2.887E-07 | 1.700E-06 | 2.200E-06 |
| 17419 | 871 | D871Y | C | 1.8e-06 | 9.3e-07 | 3.2e-06 | 1.977E-06 | 1.145E-06 | 9.300E-07 | 3.200E-06 |
| 17420 | 871 | D871V | C | 1.8e-06 | 1.9e-06 | 2.7e-06 | 2.133E-06 | 4.933E-07 | 1.800E-06 | 2.700E-06 |
| 17421 | 872 | W872A | C | 1.8e-06 | 3.2e-06 | 1.9e-06 | 2.300E-06 | 7.810E-07 | 1.800E-06 | 3.200E-06 |
| 17422 | 872 | W872R | C | 3.5e-06 | 2.2e-06 | 2.1e-06 | 2.600E-06 | 7.810E-07 | 2.100E-06 | 3.500E-06 |
| 17423 | 872 | W872N | C | 4.3e-06 | 3.4e-06 | 2.2e-06 | 3.300E-06 | 1.054E-06 | 2.200E-06 | 4.300E-06 |
| 17424 | 872 | W872D | C | 1.7e-06 | 1.8e-06 | 1.7e-06 | 1.733E-06 | 5.774E-08 | 1.700E-06 | 1.800E-06 |

| | | | | | | | | | | |
|---|---|---|---|---|---|---|---|---|---|---|
| 17425 | 872 | W872C | C | 1.6e-06 | 2.4e-06 | 1.3e-06 | 1.767E-06 | 5.686E-07 | 1.300E-06 | 2.400E-06 |
| 17426 | 872 | W872E | C | 3.1e-06 | 2.9e-06 | 2e-06 | 2.667E-06 | 5.859E-07 | 2.000E-06 | 3.100E-06 |
| 17427 | 872 | W872Q | C | 2.3e-06 | 2.4e-06 | 2.4e-06 | 2.367E-06 | 5.774E-08 | 2.300E-06 | 2.400E-06 |
| 17428 | 872 | W872G | C | 1.8e-06 | 2.1e-06 | 2.7e-06 | 2.200E-06 | 4.583E-07 | 1.800E-06 | 2.700E-06 |
| 17429 | 872 | W872H | C | 3.3e-06 | 1.4e-06 | 3.1e-06 | 2.600E-06 | 1.044E-06 | 1.400E-06 | 3.300E-06 |
| 17430 | 872 | W872I | C | 3.8e-06 | 2.9e-06 | 2.7e-06 | 3.133E-06 | 5.859E-07 | 2.700E-06 | 3.800E-06 |
| 17431 | 872 | W872L | C | 2.3e-06 | 2.4e-06 | 1.4e-06 | 2.033E-06 | 5.508E-07 | 1.400E-06 | 2.400E-06 |
| 17432 | 872 | W872K | C | 2.4e-06 | 2e-06 | 2.6e-06 | 2.333E-06 | 3.055E-07 | 2.000E-06 | 2.600E-06 |
| 17433 | 872 | W872M | C | 1.8e-06 | 2.2e-06 | 1.7e-06 | 1.900E-06 | 2.646E-07 | 1.700E-06 | 2.200E-06 |
| 17434 | 872 | W872F | C | 2.1e-06 | 2.8e-06 | 1.7e-06 | 2.200E-06 | 5.568E-07 | 1.700E-06 | 2.800E-06 |
| 17435 | 872 | W872P | C | 2.6e-06 | 1.1e-06 | 1.2e-06 | 1.633E-06 | 8.386E-07 | 1.100E-06 | 2.600E-06 |
| 17436 | 872 | W872S | C | 2.6e-06 | 1.7e-06 | 3.1e-06 | 2.467E-06 | 7.095E-07 | 1.700E-06 | 3.100E-06 |
| 17437 | 872 | W872T | C | 2e-06 | 1.7e-06 | 3.2e-06 | 2.300E-06 | 7.937E-07 | 1.700E-06 | 3.200E-06 |
| 17438 | 872 | W872W | C | 2.4e-06 | 2.8e-06 | 2.2e-06 | 2.467E-06 | 3.055E-07 | 2.200E-06 | 2.800E-06 |
| 17439 | 872 | W872Y | C | 2.7e-06 | 1.9e-06 | 2.5e-06 | 2.367E-06 | 4.163E-07 | 1.900E-06 | 2.700E-06 |
| 17440 | 872 | W872V | C | 1.7e-06 | 2.5e-06 | 9.9e-07 | 1.730E-06 | 7.554E-07 | 9.900E-07 | 2.500E-06 |
| 17441 | 873 | V873A | C | 2.8e-06 | 1.9e-06 | 2.1e-06 | 2.267E-06 | 4.726E-07 | 1.900E-06 | 2.800E-06 |
| 17442 | 873 | V873R | C | 2.2e-06 | 2.3e-06 | 3.1e-06 | 2.533E-06 | 4.933E-07 | 2.200E-06 | 3.100E-06 |
| 17443 | 873 | V873N | C | 2.1e-06 | 2.7e-06 | 2.3e-06 | 2.367E-06 | 3.055E-07 | 2.100E-06 | 2.700E-06 |
| 17444 | 873 | V873D | C | 1.6e-06 | 2.8e-06 | 2.9e-06 | 2.433E-06 | 7.234E-07 | 1.600E-06 | 2.900E-06 |
| 17445 | 873 | V873C | C | 1.5e-06 | 2.7e-06 | 2.7e-06 | 2.300E-06 | 6.928E-07 | 1.500E-06 | 2.700E-06 |
| 17446 | 873 | V873E | C | 2.9e-06 | 1.5e-06 | 1.8e-06 | 2.067E-06 | 7.371E-07 | 1.500E-06 | 2.900E-06 |
| 17447 | 873 | V873Q | C | 1.3e-06 | 1.9e-06 | 2.4e-06 | 1.867E-06 | 5.508E-07 | 1.300E-06 | 2.400E-06 |
| 17448 | 873 | V873G | C | 2.2e-06 | 3.4e-06 | 2.7e-06 | 2.767E-06 | 6.028E-07 | 2.200E-06 | 3.400E-06 |
| 17449 | 873 | V873H | C | 2.6e-06 | 2.3e-06 | 2.5e-06 | 2.467E-06 | 1.528E-07 | 2.300E-06 | 2.600E-06 |
| 17450 | 873 | V873I | C | 3.6e-06 | 2.7e-06 | 2e-06 | 2.767E-06 | 8.021E-07 | 2.000E-06 | 3.600E-06 |
| 17451 | 873 | V873L | C | 1.7e-06 | 2.5e-06 | 1.1e-06 | 1.767E-06 | 7.024E-07 | 1.100E-06 | 2.500E-06 |
| 17452 | 873 | V873K | C | 1.8e-06 | 2.1e-06 | 1e-06 | 1.633E-06 | 5.686E-07 | 1.000E-06 | 2.100E-06 |
| 17453 | 873 | V873M | C | 3.6e-06 | 1.9e-06 | 2.8e-06 | 2.767E-06 | 8.505E-07 | 1.900E-06 | 3.600E-06 |
| 17454 | 873 | V873F | C | 1.6e-06 | 1.6e-06 | 1.9e-06 | 1.700E-06 | 1.732E-07 | 1.600E-06 | 1.900E-06 |
| 17455 | 873 | V873P | C | 1.6e-06 | 2.6e-06 | 1.7e-06 | 1.967E-06 | 5.508E-07 | 1.600E-06 | 2.600E-06 |
| 17456 | 873 | V873S | C | 3e-06 | 1.6e-06 | 1.9e-06 | 2.167E-06 | 7.371E-07 | 1.600E-06 | 3.000E-06 |
| 17457 | 873 | V873T | C | 4e-06 | 1.4e-06 | 1.3e-06 | 2.233E-06 | 1.531E-06 | 1.300E-06 | 4.000E-06 |
| 17458 | 873 | V873W | C | 3.4e-06 | 1.2e-06 | 1.8e-06 | 2.133E-06 | 1.137E-06 | 1.200E-06 | 3.400E-06 |
| 17459 | 873 | V873Y | C | 1.1e-06 | 3.8e-06 | 3.2e-06 | 2.700E-06 | 1.418E-06 | 1.100E-06 | 3.800E-06 |
| 17460 | 873 | V873V | C | 1.5e-06 | 2.2e-06 | 1.5e-06 | 1.733E-06 | 4.041E-07 | 1.500E-06 | 2.200E-06 |
| 17461 | 874 | R874A | C | 1.5e-06 | 2.2e-06 | 1.6e-06 | 1.767E-06 | 3.786E-07 | 1.500E-06 | 2.200E-06 |
| 17462 | 874 | R874R | C | 2e-06 | 2.8e-06 | 2.7e-06 | 2.500E-06 | 4.359E-07 | 2.000E-06 | 2.800E-06 |
| 17463 | 874 | R874N | C | 3.4e-06 | 1.4e-06 | 3.1e-06 | 2.633E-06 | 1.079E-06 | 1.400E-06 | 3.400E-06 |
| 17464 | 874 | R874D | C | 1.5e-06 | 3.4e-06 | 5.7e-06 | 3.533E-06 | 2.103E-06 | 1.500E-06 | 5.700E-06 |
| 17465 | 874 | R874C | C | 3.8e-06 | 3.2e-06 | 2.4e-06 | 3.133E-06 | 7.024E-07 | 2.400E-06 | 3.800E-06 |
| 17466 | 874 | R874E | C | 2.7e-06 | 1.7e-06 | 2.8e-06 | 2.400E-06 | 6.083E-07 | 1.700E-06 | 2.800E-06 |
| 17467 | 874 | R874Q | C | 1.5e-06 | 1.1e-06 | 2.9e-06 | 1.833E-06 | 9.452E-07 | 1.100E-06 | 2.900E-06 |
| 17468 | 874 | R874G | C | 2.1e-06 | 2.2e-06 | 1.6e-06 | 1.967E-06 | 3.215E-07 | 1.600E-06 | 2.200E-06 |
| 17469 | 874 | R874H | C | 3.7e-06 | 3.5e-06 | 5.6e-06 | 4.267E-06 | 1.159E-06 | 3.500E-06 | 5.600E-06 |
| 17470 | 874 | R874I | C | 3.5e-06 | 1.5e-06 | 2.6e-06 | 2.533E-06 | 1.002E-06 | 1.500E-06 | 3.500E-06 |
| 17471 | 874 | R874L | C | 2.7e-06 | 2.3e-06 | 2.5e-06 | 2.500E-06 | 2.000E-07 | 2.300E-06 | 2.700E-06 |
| 17472 | 874 | R874K | C | 3.7e-06 | 1.6e-06 | 2.5e-06 | 2.600E-06 | 1.054E-06 | 1.600E-06 | 3.700E-06 |
| 17473 | 874 | R874M | C | 1.2e-06 | 2.6e-06 | 2.2e-06 | 2.000E-06 | 7.211E-07 | 1.200E-06 | 2.600E-06 |
| 17474 | 874 | R874F | C | 2.4e-06 | 2.5e-06 | 3.5e-06 | 2.800E-06 | 6.083E-07 | 2.400E-06 | 3.500E-06 |
| 17475 | 874 | R874P | C | 2.4e-06 | 1.4e-06 | 3e-06 | 2.267E-06 | 8.083E-07 | 1.400E-06 | 3.000E-06 |
| 17476 | 874 | R874S | C | 3e-06 | 3.2e-06 | 3.1e-06 | 3.100E-06 | 1.000E-07 | 3.000E-06 | 3.200E-06 |
| 17477 | 874 | R874T | C | 2.2e-06 | 3e-06 | 4e-06 | 3.067E-06 | 9.018E-07 | 2.200E-06 | 4.000E-06 |
| 17478 | 874 | R874W | C | 1.8e-06 | 2.6e-06 | 2.1e-06 | 2.167E-06 | 4.041E-07 | 1.800E-06 | 2.600E-06 |
| 17479 | 874 | R874Y | C | 2.8e-06 | 1.5e-06 | 3.2e-06 | 2.500E-06 | 8.888E-07 | 1.500E-06 | 3.200E-06 |
| 17480 | 874 | R874V | C | 1.7e-06 | 2.7e-06 | 3.5e-06 | 2.633E-06 | 9.018E-07 | 1.700E-06 | 3.500E-06 |
| 17481 | 875 | Q875A | C | 2.7e-06 | 2.1e-06 | 1.7e-06 | 2.167E-06 | 5.033E-07 | 1.700E-06 | 2.700E-06 |
| 17482 | 875 | Q875R | C | 1.4e-06 | 3.7e-06 | 1.8e-06 | 2.300E-06 | 1.229E-06 | 1.400E-06 | 3.700E-06 |
| 17483 | 875 | Q875N | C | 3.1e-06 | 3.6e-06 | 1.6e-06 | 2.767E-06 | 1.041E-06 | 1.600E-06 | 3.600E-06 |
| 17484 | 875 | Q875D | C | 2.3e-06 | 2.7e-06 | 2.8e-06 | 2.600E-06 | 2.646E-07 | 2.300E-06 | 2.800E-06 |
| 17485 | 875 | Q875C | C | 2.4e-06 | 2.6e-06 | 1.8e-06 | 2.267E-06 | 4.163E-07 | 1.800E-06 | 2.600E-06 |
| 17486 | 875 | Q875E | C | 2.5e-06 | 2.7e-06 | 2.4e-06 | 2.533E-06 | 1.528E-07 | 2.400E-06 | 2.700E-06 |
| 17487 | 875 | Q875Q | C | 2.8e-06 | 2.3e-06 | 1.2e-06 | 2.100E-06 | 8.185E-07 | 1.200E-06 | 2.800E-06 |
| 17488 | 875 | Q875G | C | 3.4e-06 | 2.5e-06 | 2.8e-06 | 2.900E-06 | 4.583E-07 | 2.500E-06 | 3.400E-06 |
| 17489 | 875 | Q875H | C | 2e-06 | 3.2e-06 | 4e-06 | 3.067E-06 | 1.007E-06 | 2.000E-06 | 4.000E-06 |
| 17490 | 875 | Q875I | C | 2.8e-06 | 6.2e-06 | 2.2e-06 | 3.733E-06 | 2.157E-06 | 2.200E-06 | 6.200E-06 |
| 17491 | 875 | Q875L | C | 1.9e-06 | 2.9e-06 | 2e-06 | 2.267E-06 | 5.508E-07 | 1.900E-06 | 2.900E-06 |
| 17492 | 875 | Q875K | C | 2.5e-06 | 2.6e-06 | 1.2e-06 | 2.100E-06 | 7.810E-07 | 1.200E-06 | 2.600E-06 |
| 17493 | 875 | Q875M | C | 2.5e-06 | 2e-06 | 1.9e-06 | 2.133E-06 | 3.215E-07 | 1.900E-06 | 2.500E-06 |
| 17494 | 875 | Q875F | C | 2.1e-06 | 4.2e-06 | 1.8e-06 | 2.700E-06 | 1.308E-06 | 1.800E-06 | 4.200E-06 |
| 17495 | 875 | Q875P | C | 3.3e-06 | 1.6e-06 | 3.3e-06 | 2.733E-06 | 9.815E-07 | 1.600E-06 | 3.300E-06 |
| 17496 | 875 | Q875S | C | 1.8e-06 | 2.2e-06 | 3.5e-06 | 2.500E-06 | 8.888E-07 | 1.800E-06 | 3.500E-06 |
| 17497 | 875 | Q875T | C | 3.4e-06 | 2.5e-06 | 2e-06 | 2.633E-06 | 7.095E-07 | 2.000E-06 | 3.400E-06 |
| 17498 | 875 | Q875W | C | 2.6e-06 | 2.4e-06 | 2.4e-06 | 2.467E-06 | 1.155E-07 | 2.400E-06 | 2.600E-06 |
| 17499 | 875 | Q875Y | C | 2.2e-06 | 1e-06 | 1.2e-06 | 1.467E-06 | 6.429E-07 | 1.000E-06 | 2.200E-06 |
| 17500 | 875 | Q875V | C | 2.4e-06 | 3.9e-06 | 3.2e-06 | 3.167E-06 | 7.506E-07 | 2.400E-06 | 3.900E-06 |
| 17501 | 876 | A876A | C | 2.6e-06 | 3.6e-06 | 2.5e-06 | 2.900E-06 | 6.083E-07 | 2.500E-06 | 3.600E-06 |
| 17502 | 876 | A876R | C | 1.9e-06 | 2.1e-06 | 2.7e-06 | 2.233E-06 | 4.163E-07 | 1.900E-06 | 2.700E-06 |
| 17503 | 876 | A876N | C | 3e-06 | 3.4e-06 | 3.8e-06 | 3.400E-06 | 4.000E-07 | 3.000E-06 | 3.800E-06 |
| 17504 | 876 | A876D | C | 1.8e-06 | 2e-06 | 2.9e-06 | 2.233E-06 | 5.859E-07 | 1.800E-06 | 2.900E-06 |
| 17505 | 876 | A876C | C | 4.4e-06 | 2.6e-06 | 2.1e-06 | 3.033E-06 | 1.210E-06 | 2.100E-06 | 4.400E-06 |
| 17506 | 876 | A876E | C | 2e-06 | 1.7e-06 | 2.2e-06 | 1.967E-06 | 2.517E-07 | 1.700E-06 | 2.200E-06 |
| 17507 | 876 | A876Q | C | 1.8e-06 | 4e-06 | 1.4e-06 | 2.400E-06 | 1.400E-06 | 1.400E-06 | 4.000E-06 |
| 17508 | 876 | A876G | C | 1.3e-06 | 2.2e-06 | 2.9e-06 | 2.133E-06 | 8.021E-07 | 1.300E-06 | 2.900E-06 |
| 17509 | 876 | A876H | C | 2.9e-06 | 1.2e-06 | 2.6e-06 | 2.233E-06 | 9.074E-07 | 1.200E-06 | 2.900E-06 |

| | | | | | | | | | | |
|---|---|---|---|---|---|---|---|---|---|---|
| 17510 | 876 | A876I | C | 1.7e-06 | 3.3e-06 | 3.6e-06 | 2.867E-06 | 1.021E-06 | 1.700E-06 | 3.600E-06 |
| 17511 | 876 | A876L | C | 2.5e-06 | 2.9e-06 | 2.6e-06 | 2.667E-06 | 2.082E-07 | 2.500E-06 | 2.900E-06 |
| 17512 | 876 | A876K | C | 2.8e-06 | 2e-06 | 2.7e-06 | 2.500E-06 | 4.359E-07 | 2.000E-06 | 2.800E-06 |
| 17513 | 876 | A876M | C | 1.8e-06 | 2e-06 | 2.6e-06 | 2.133E-06 | 4.163E-07 | 1.800E-06 | 2.600E-06 |
| 17514 | 876 | A876F | C | 2.4e-06 | 2.4e-06 | 3.7e-06 | 2.833E-06 | 7.506E-07 | 2.400E-06 | 3.700E-06 |
| 17515 | 876 | A876P | C | 2.3e-06 | 3e-06 | 2.6e-06 | 2.633E-06 | 3.512E-07 | 2.300E-06 | 3.000E-06 |
| 17516 | 876 | A876S | C | 1.1e-06 | 2.3e-06 | 1.9e-06 | 1.767E-06 | 6.110E-07 | 1.100E-06 | 2.300E-06 |
| 17517 | 876 | A876T | C | 3e-06 | 2e-06 | 1.8e-06 | 2.267E-06 | 6.429E-07 | 1.800E-06 | 3.000E-06 |
| 17518 | 876 | A876W | C | 1.8e-06 | 9.1e-07 | 3.6e-06 | 2.103E-06 | 1.370E-06 | 9.100E-07 | 3.600E-06 |
| 17519 | 876 | A876Y | C | 2.6e-06 | 1.7e-06 | 2.7e-06 | 2.333E-06 | 5.508E-07 | 1.700E-06 | 2.700E-06 |
| 17520 | 876 | A876V | C | 3.1e-06 | 1.9e-06 | 1.8e-06 | 2.267E-06 | 7.234E-07 | 1.800E-06 | 3.100E-06 |
| 17521 | 877 | P877A | C | 2.8e-06 | 5.7e-06 | 2.3e-06 | 3.600E-06 | 1.836E-06 | 2.300E-06 | 5.700E-06 |
| 17522 | 877 | P877R | C | 2.9e-06 | 4.1e-06 | 2.5e-06 | 3.167E-06 | 8.327E-07 | 2.500E-06 | 4.100E-06 |
| 17523 | 877 | P877N | C | 2.7e-06 | 1.9e-06 | 1.6e-06 | 2.067E-06 | 5.686E-07 | 1.600E-06 | 2.700E-06 |
| 17524 | 877 | P877D | C | 3e-06 | 2.3e-06 | 3.3e-06 | 2.867E-06 | 5.132E-07 | 2.300E-06 | 3.300E-06 |
| 17525 | 877 | P877C | C | 2.5e-06 | 2e-06 | 2.1e-06 | 2.200E-06 | 2.646E-07 | 2.000E-06 | 2.500E-06 |
| 17526 | 877 | P877E | C | 2.8e-06 | 2.7e-06 | 1.7e-06 | 2.400E-06 | 6.083E-07 | 1.700E-06 | 2.800E-06 |
| 17527 | 877 | P877Q | C | 6.9e-07 | 3.3e-06 | 1.8e-06 | 1.930E-06 | 1.310E-06 | 6.900E-07 | 3.300E-06 |
| 17528 | 877 | P877G | C | 3.1e-06 | 2.8e-06 | 2.8e-06 | 2.900E-06 | 1.732E-07 | 2.800E-06 | 3.100E-06 |
| 17529 | 877 | P877H | C | 2e-06 | 2.5e-06 | 1.9e-06 | 2.133E-06 | 3.215E-07 | 1.900E-06 | 2.500E-06 |
| 17530 | 877 | P877I | C | 2.1e-06 | 2e-06 | 3e-06 | 2.367E-06 | 5.508E-07 | 2.000E-06 | 3.000E-06 |
| 17531 | 877 | P877L | C | 2.8e-06 | 2.6e-06 | 2.1e-06 | 2.500E-06 | 3.606E-07 | 2.100E-06 | 2.800E-06 |
| 17532 | 877 | P877K | C | 3.4e-06 | 1.7e-06 | 2.8e-06 | 2.633E-06 | 8.622E-07 | 1.700E-06 | 3.400E-06 |
| 17533 | 877 | P877M | C | 3.8e-06 | 2.1e-06 | 3.8e-06 | 3.233E-06 | 9.815E-07 | 2.100E-06 | 3.800E-06 |
| 17534 | 877 | P877F | C | 2.8e-06 | 2.2e-06 | 3.8e-06 | 2.933E-06 | 8.083E-07 | 2.200E-06 | 3.800E-06 |
| 17535 | 877 | P877P | C | 2.6e-06 | 1.9e-06 | 2.1e-06 | 2.200E-06 | 3.606E-07 | 1.900E-06 | 2.600E-06 |
| 17536 | 877 | P877S | C | 2.1e-06 | 3.1e-06 | 1.7e-06 | 2.300E-06 | 7.211E-07 | 1.700E-06 | 3.100E-06 |
| 17537 | 877 | P877T | C | 2.1e-06 | 1.7e-06 | 1.7e-06 | 1.833E-06 | 2.309E-07 | 1.700E-06 | 2.100E-06 |
| 17538 | 877 | P877W | C | 1.6e-06 | 1.4e-06 | 1.4e-06 | 1.467E-06 | 1.155E-07 | 1.400E-06 | 1.600E-06 |
| 17539 | 877 | P877Y | C | 2.9e-06 | 2.4e-06 | 2e-06 | 2.433E-06 | 4.509E-07 | 2.000E-06 | 2.900E-06 |
| 17540 | 877 | P877V | C | 2.6e-06 | 1.9e-06 | 2.8e-06 | 2.433E-06 | 4.726E-07 | 1.900E-06 | 2.800E-06 |
| 17541 | 878 | G878A | C | 2.3e-06 | 3.6e-06 | 1.6e-06 | 2.500E-06 | 1.015E-06 | 1.600E-06 | 3.600E-06 |
| 17542 | 878 | G878R | C | 4.7e-06 | 2.3e-06 | 2e-06 | 3.000E-06 | 1.480E-06 | 2.000E-06 | 4.700E-06 |
| 17543 | 878 | G878N | C | 1.7e-06 | 1.9e-06 | 3.1e-06 | 2.233E-06 | 7.572E-07 | 1.700E-06 | 3.100E-06 |
| 17544 | 878 | G878D | C | 3.9e-06 | 2.6e-06 | 2.4e-06 | 2.967E-06 | 8.145E-07 | 2.400E-06 | 3.900E-06 |
| 17545 | 878 | G878C | C | 1.9e-06 | 2.3e-06 | 1.9e-06 | 2.033E-06 | 2.309E-07 | 1.900E-06 | 2.300E-06 |
| 17546 | 878 | G878E | C | 3.8e-06 | 3.7e-06 | 2.7e-06 | 3.400E-06 | 6.083E-07 | 2.700E-06 | 3.800E-06 |
| 17547 | 878 | G878Q | C | 2.8e-06 | 1.4e-06 | 2.5e-06 | 2.233E-06 | 7.371E-07 | 1.400E-06 | 2.800E-06 |
| 17548 | 878 | G878G | C | 2.3e-06 | 3e-06 | 2.8e-06 | 2.700E-06 | 3.606E-07 | 2.300E-06 | 3.000E-06 |
| 17549 | 878 | G878H | C | 2.5e-06 | 1.3e-06 | 2.9e-06 | 2.233E-06 | 8.327E-07 | 1.300E-06 | 2.900E-06 |
| 17550 | 878 | G878I | C | 1.2e-06 | 2e-06 | 4.3e-06 | 2.500E-06 | 1.609E-06 | 1.200E-06 | 4.300E-06 |
| 17551 | 878 | G878L | C | 1.8e-06 | 1.9e-06 | 1.8e-06 | 1.833E-06 | 5.774E-08 | 1.800E-06 | 1.900E-06 |
| 17552 | 878 | G878K | C | 1.9e-06 | 2e-06 | 8e-07 | 1.567E-06 | 6.658E-07 | 8.000E-07 | 2.000E-06 |
| 17553 | 878 | G878M | C | 2.4e-06 | 5.6e-06 | 2.3e-06 | 3.433E-06 | 1.877E-06 | 2.300E-06 | 5.600E-06 |
| 17554 | 878 | G878F | C | 2.1e-06 | 1.4e-06 | 2.5e-06 | 2.000E-06 | 5.568E-07 | 1.400E-06 | 2.500E-06 |
| 17555 | 878 | G878P | C | 2.4e-06 | 2.4e-06 | 1.9e-06 | 2.233E-06 | 2.887E-07 | 1.900E-06 | 2.400E-06 |
| 17556 | 878 | G878S | C | 1.4e-06 | 2.3e-06 | 3.3e-06 | 2.333E-06 | 9.504E-07 | 1.400E-06 | 3.300E-06 |
| 17557 | 878 | G878T | C | 3e-06 | 1.8e-06 | 4.2e-06 | 3.000E-06 | 1.200E-06 | 1.800E-06 | 4.200E-06 |
| 17558 | 878 | G878W | C | 2.3e-06 | 1.9e-06 | 3.5e-06 | 2.567E-06 | 8.327E-07 | 1.900E-06 | 3.500E-06 |
| 17559 | 878 | G878Y | C | 3e-06 | 3.3e-06 | 2.1e-06 | 2.800E-06 | 6.245E-07 | 2.100E-06 | 3.300E-06 |
| 17560 | 878 | G878V | C | 6.6e-06 | 2.6e-06 | 1.2e-06 | 3.467E-06 | 2.802E-06 | 1.200E-06 | 6.600E-06 |
| 17561 | 879 | K879A | C | 3.3e-06 | 2.8e-06 | 2.7e-06 | 2.933E-06 | 3.215E-07 | 2.700E-06 | 3.300E-06 |
| 17562 | 879 | K879R | C | 2.9e-06 | 1.3e-06 | 2.4e-06 | 2.200E-06 | 8.185E-07 | 1.300E-06 | 2.900E-06 |
| 17563 | 879 | K879N | C | 2.7e-06 | 1.3e-06 | 1.9e-06 | 1.967E-06 | 7.024E-07 | 1.300E-06 | 2.700E-06 |
| 17564 | 879 | K879D | C | 3.6e-06 | 2.7e-06 | 2.8e-06 | 3.033E-06 | 4.933E-07 | 2.700E-06 | 3.600E-06 |
| 17565 | 879 | K879C | C | 1.9e-06 | 1.3e-06 | 3e-06 | 2.067E-06 | 8.622E-07 | 1.300E-06 | 3.000E-06 |
| 17566 | 879 | K879E | C | 4.6e-06 | 3.1e-06 | 1.9e-06 | 3.200E-06 | 1.353E-06 | 1.900E-06 | 4.600E-06 |
| 17567 | 879 | K879Q | C | 3.2e-06 | 3.9e-06 | 2.6e-06 | 3.233E-06 | 6.506E-07 | 2.600E-06 | 3.900E-06 |
| 17568 | 879 | K879G | C | 1.9e-06 | 2.9e-06 | 3.6e-06 | 2.800E-06 | 8.544E-07 | 1.900E-06 | 3.600E-06 |
| 17569 | 879 | K879H | C | 2.6e-06 | 2.3e-06 | 2.2e-06 | 2.367E-06 | 2.082E-07 | 2.200E-06 | 2.600E-06 |
| 17570 | 879 | K879I | C | 1.9e-06 | 1.8e-06 | 2.3e-06 | 2.000E-06 | 2.646E-07 | 1.800E-06 | 2.300E-06 |
| 17571 | 879 | K879L | C | 2e-06 | 1.6e-06 | 2.4e-06 | 2.000E-06 | 4.000E-07 | 1.600E-06 | 2.400E-06 |
| 17572 | 879 | K879K | C | 4.3e-06 | 2.2e-06 | 3e-06 | 3.167E-06 | 1.060E-06 | 2.200E-06 | 4.300E-06 |
| 17573 | 879 | K879M | C | 2.2e-06 | 1.4e-06 | 1.2e-06 | 1.600E-06 | 5.292E-07 | 1.200E-06 | 2.200E-06 |
| 17574 | 879 | K879F | C | 2.3e-06 | 2.5e-06 | 3.2e-06 | 2.667E-06 | 4.726E-07 | 2.300E-06 | 3.200E-06 |
| 17575 | 879 | K879P | C | 1.2e-06 | 2.3e-06 | 2.5e-06 | 2.000E-06 | 7.000E-07 | 1.200E-06 | 2.500E-06 |
| 17576 | 879 | K879S | C | 1.9e-06 | 3.1e-06 | 3.2e-06 | 2.733E-06 | 7.234E-07 | 1.900E-06 | 3.200E-06 |
| 17577 | 879 | K879T | C | 1.7e-06 | 2.2e-06 | 1.7e-06 | 1.867E-06 | 2.887E-07 | 1.700E-06 | 2.200E-06 |
| 17578 | 879 | K879W | C | 3.4e-06 | 3e-06 | 2e-06 | 2.800E-06 | 7.211E-07 | 2.000E-06 | 3.400E-06 |
| 17579 | 879 | K879Y | C | 3e-06 | 1.8e-06 | 2.9e-06 | 2.567E-06 | 6.658E-07 | 1.800E-06 | 3.000E-06 |
| 17580 | 879 | K879V | C | 1.7e-06 | 3.2e-06 | 2.1e-06 | 2.333E-06 | 7.767E-07 | 1.700E-06 | 3.200E-06 |
| 17581 | 880 | G880A | C | 1.6e-06 | 2.3e-06 | 5e-06 | 2.967E-06 | 1.795E-06 | 1.600E-06 | 5.000E-06 |
| 17582 | 880 | G880R | C | 5e-06 | 1.9e-06 | 3.3e-06 | 3.400E-06 | 1.552E-06 | 1.900E-06 | 5.000E-06 |
| 17583 | 880 | G880N | C | 2.8e-06 | 1.7e-06 | 2.4e-06 | 2.300E-06 | 5.568E-07 | 1.700E-06 | 2.800E-06 |
| 17584 | 880 | G880D | C | 2.9e-06 | 1.3e-06 | 2.4e-06 | 2.200E-06 | 8.185E-07 | 1.300E-06 | 2.900E-06 |
| 17585 | 880 | G880C | C | 3.2e-06 | 2e-06 | 3.5e-06 | 2.900E-06 | 7.937E-07 | 2.000E-06 | 3.500E-06 |
| 17586 | 880 | G880E | C | 2.1e-06 | 9.7e-07 | 8.8e-07 | 1.317E-06 | 6.799E-07 | 8.800E-07 | 2.100E-06 |
| 17587 | 880 | G880Q | C | 2.2e-06 | 2.5e-06 | 3.6e-06 | 2.767E-06 | 7.371E-07 | 2.200E-06 | 3.600E-06 |
| 17588 | 880 | G880G | C | 2.3e-06 | 1.5e-06 | 1.5e-06 | 1.767E-06 | 4.619E-07 | 1.500E-06 | 2.300E-06 |
| 17589 | 880 | G880H | C | 2.8e-06 | 2.4e-06 | 1.7e-06 | 2.300E-06 | 5.568E-07 | 1.700E-06 | 2.800E-06 |
| 17590 | 880 | G880I | C | 2.8e-06 | 5.6e-07 | 3.6e-06 | 2.320E-06 | 1.576E-06 | 5.600E-07 | 3.600E-06 |
| 17591 | 880 | G880L | C | 2.1e-06 | 1.4e-06 | 1.6e-06 | 1.700E-06 | 3.606E-07 | 1.400E-06 | 2.100E-06 |
| 17592 | 880 | G880K | C | 2.5e-06 | 3.4e-06 | 2.1e-06 | 2.667E-06 | 6.658E-07 | 2.100E-06 | 3.400E-06 |
| 17593 | 880 | G880M | C | 3.9e-06 | 2.4e-06 | 1.4e-06 | 2.567E-06 | 1.258E-06 | 1.400E-06 | 3.900E-06 |
| 17594 | 880 | G880F | C | 1.7e-06 | 1.5e-06 | 1.4e-06 | 1.533E-06 | 1.528E-07 | 1.400E-06 | 1.700E-06 |

| | | | | | | | | | | |
|---|---|---|---|---|---|---|---|---|---|---|
| 17595 | 880 | G880P | C | 5.7e-06 | 2.2e-06 | 2e-06 | 3.300E-06 | 2.081E-06 | 2.000E-06 | 5.700E-06 |
| 17596 | 880 | G880S | C | 3.1e-06 | 2e-06 | 3.1e-06 | 2.733E-06 | 6.351E-07 | 2.000E-06 | 3.100E-06 |
| 17597 | 880 | G880T | C | 1.9e-06 | 2.2e-06 | 2.3e-06 | 2.133E-06 | 2.082E-07 | 1.900E-06 | 2.300E-06 |
| 17598 | 880 | G880W | C | 1.5e-06 | 2.4e-06 | 2.5e-06 | 2.133E-06 | 5.508E-07 | 1.500E-06 | 2.500E-06 |
| 17599 | 880 | G880Y | C | 2.3e-06 | 1.4e-06 | 2.9e-06 | 2.200E-06 | 7.550E-07 | 1.400E-06 | 2.900E-06 |
| 17600 | 880 | G880V | C | 3.8e-06 | 2.5e-06 | 1.8e-06 | 2.700E-06 | 1.015E-06 | 1.800E-06 | 3.800E-06 |
| 17601 | 881 | L881A | C | 1.9e-06 | 2.9e-06 | 1.8e-06 | 2.200E-06 | 6.083E-07 | 1.800E-06 | 2.900E-06 |
| 17602 | 881 | L881R | C | 2.7e-06 | 3.5e-06 | 1.6e-06 | 2.600E-06 | 9.539E-07 | 1.600E-06 | 3.500E-06 |
| 17603 | 881 | L881N | C | 2e-06 | 1.3e-06 | 2.5e-06 | 1.933E-06 | 6.028E-07 | 1.300E-06 | 2.500E-06 |
| 17604 | 881 | L881D | C | 2e-06 | 2.7e-06 | 2e-06 | 2.233E-06 | 4.041E-07 | 2.000E-06 | 2.700E-06 |
| 17605 | 881 | L881C | C | 4.2e-06 | 2.3e-06 | 1.8e-06 | 2.767E-06 | 1.266E-06 | 1.800E-06 | 4.200E-06 |
| 17606 | 881 | L881E | C | 2.8e-06 | 2.9e-06 | 1.6e-06 | 2.433E-06 | 7.234E-07 | 1.600E-06 | 2.900E-06 |
| 17607 | 881 | L881Q | C | 2.5e-06 | 2.2e-06 | 2.4e-06 | 2.367E-06 | 1.528E-07 | 2.200E-06 | 2.500E-06 |
| 17608 | 881 | L881G | C | 2.3e-06 | 1.7e-06 | 2.8e-06 | 2.267E-06 | 5.508E-07 | 1.700E-06 | 2.800E-06 |
| 17609 | 881 | L881H | C | 2.9e-06 | 1.7e-06 | 2.2e-06 | 2.267E-06 | 6.028E-07 | 1.700E-06 | 2.900E-06 |
| 17610 | 881 | L881I | C | 2.5e-06 | 2.5e-06 | 1.8e-06 | 2.267E-06 | 4.041E-07 | 1.800E-06 | 2.500E-06 |
| 17611 | 881 | L881L | C | 3.2e-06 | 2.1e-06 | 3e-06 | 2.767E-06 | 5.859E-07 | 2.100E-06 | 3.200E-06 |
| 17612 | 881 | L881K | C | 1.9e-06 | 1.6e-06 | 2.4e-06 | 1.967E-06 | 4.041E-07 | 1.600E-06 | 2.400E-06 |
| 17613 | 881 | L881M | C | 2.1e-06 | 4.2e-06 | 2.3e-06 | 2.867E-06 | 1.159E-06 | 2.100E-06 | 4.200E-06 |
| 17614 | 881 | L881F | C | 2.2e-06 | 2.8e-06 | 3.1e-06 | 2.700E-06 | 4.583E-07 | 2.200E-06 | 3.100E-06 |
| 17615 | 881 | L881P | C | 3.3e-06 | 3.3e-06 | 2e-06 | 2.867E-06 | 7.506E-07 | 2.000E-06 | 3.300E-06 |
| 17616 | 881 | L881S | C | 2.2e-06 | 1.5e-06 | 2.9e-06 | 2.200E-06 | 7.000E-07 | 1.500E-06 | 2.900E-06 |
| 17617 | 881 | L881T | C | 2.6e-06 | 2.7e-06 | 2.2e-06 | 2.500E-06 | 2.646E-07 | 2.200E-06 | 2.700E-06 |
| 17618 | 881 | L881W | C | 3.2e-06 | 3.1e-06 | 2.7e-06 | 3.000E-06 | 2.646E-07 | 2.700E-06 | 3.200E-06 |
| 17619 | 881 | L881Y | C | 3.1e-06 | 9.9e-07 | 2.2e-06 | 2.097E-06 | 1.059E-06 | 9.900E-07 | 3.100E-06 |
| 17620 | 881 | L881V | C | 2.1e-06 | 3.2e-06 | 5.1e-06 | 3.467E-06 | 1.518E-06 | 2.100E-06 | 5.100E-06 |
| 17621 | 882 | E882A | C | 3.3e-06 | 4.6e-06 | 4.9e-06 | 4.267E-06 | 8.505E-07 | 3.300E-06 | 4.900E-06 |
| 17622 | 882 | E882R | C | 1.9e-06 | 2e-06 | 2.2e-06 | 2.033E-06 | 1.528E-07 | 1.900E-06 | 2.200E-06 |
| 17623 | 882 | E882N | C | 1.6e-06 | 3e-06 | 1.9e-06 | 2.167E-06 | 7.371E-07 | 1.600E-06 | 3.000E-06 |
| 17624 | 882 | E882D | C | 1.6e-06 | 4e-06 | 1.8e-06 | 2.467E-06 | 1.332E-06 | 1.600E-06 | 4.000E-06 |
| 17625 | 882 | E882C | C | 1.3e-06 | 2.3e-06 | 2.3e-06 | 1.967E-06 | 5.774E-07 | 1.300E-06 | 2.300E-06 |
| 17626 | 882 | E882E | C | 1.6e-06 | 2e-06 | 1.9e-06 | 1.833E-06 | 2.082E-07 | 1.600E-06 | 2.000E-06 |
| 17627 | 882 | E882Q | C | 3.9e-06 | 1.3e-06 | 2.1e-06 | 2.433E-06 | 1.332E-06 | 1.300E-06 | 3.900E-06 |
| 17628 | 882 | E882G | C | 3.7e-06 | 2.4e-06 | 2.8e-06 | 2.967E-06 | 6.658E-07 | 2.400E-06 | 3.700E-06 |
| 17629 | 882 | E882H | C | 2.1e-06 | 2.4e-06 | 2.3e-06 | 2.267E-06 | 1.528E-07 | 2.100E-06 | 2.400E-06 |
| 17630 | 882 | E882I | C | 2.4e-06 | 2.2e-06 | 4e-06 | 2.867E-06 | 9.866E-07 | 2.200E-06 | 4.000E-06 |
| 17631 | 882 | E882L | C | 1.7e-06 | 2.5e-06 | 6.2e-06 | 3.467E-06 | 2.401E-06 | 1.700E-06 | 6.200E-06 |
| 17632 | 882 | E882K | C | 3.4e-06 | 2e-06 | 2.3e-06 | 2.567E-06 | 7.371E-07 | 2.000E-06 | 3.400E-06 |
| 17633 | 882 | E882M | C | 3.1e-06 | 4.3e-06 | 2.3e-06 | 3.233E-06 | 1.007E-06 | 2.300E-06 | 4.300E-06 |
| 17634 | 882 | E882F | C | 2.7e-06 | 5.7e-06 | 1.5e-06 | 3.300E-06 | 2.163E-06 | 1.500E-06 | 5.700E-06 |
| 17635 | 882 | E882P | C | 2.3e-06 | 2e-06 | 2.5e-06 | 2.267E-06 | 2.517E-07 | 2.000E-06 | 2.500E-06 |
| 17636 | 882 | E882S | C | 2.6e-06 | 2e-06 | 2.5e-06 | 2.367E-06 | 3.215E-07 | 2.000E-06 | 2.600E-06 |
| 17637 | 882 | E882T | C | 2e-06 | 2.8e-06 | 3.1e-06 | 2.633E-06 | 5.686E-07 | 2.000E-06 | 3.100E-06 |
| 17638 | 882 | E882W | C | 3.1e-06 | 2e-06 | 3.4e-06 | 2.833E-06 | 7.371E-07 | 2.000E-06 | 3.400E-06 |
| 17639 | 882 | E882Y | C | 2.6e-06 | 2.4e-06 | 2.6e-06 | 2.533E-06 | 1.155E-07 | 2.400E-06 | 2.600E-06 |
| 17640 | 882 | E882V | C | 2.6e-06 | 1.6e-06 | 2.7e-06 | 2.300E-06 | 6.083E-07 | 1.600E-06 | 2.700E-06 |
| 17641 | 883 | W883A | C | 3.7e-06 | 2.3e-06 | 3.8e-06 | 3.267E-06 | 8.386E-07 | 2.300E-06 | 3.800E-06 |
| 17642 | 883 | W883R | C | 1.9e-06 | 2e-06 | 1.4e-06 | 1.767E-06 | 3.215E-07 | 1.400E-06 | 2.000E-06 |
| 17643 | 883 | W883N | C | 3.7e-06 | 2.2e-06 | 3.5e-06 | 3.133E-06 | 8.145E-07 | 2.200E-06 | 3.700E-06 |
| 17644 | 883 | W883D | C | 2.2e-06 | 2.3e-06 | 2.8e-06 | 2.433E-06 | 3.215E-07 | 2.200E-06 | 2.800E-06 |
| 17645 | 883 | W883C | C | 1.9e-06 | 2.3e-06 | 2.5e-06 | 2.233E-06 | 3.055E-07 | 1.900E-06 | 2.500E-06 |
| 17646 | 883 | W883E | C | 2.5e-06 | 1.4e-06 | 2.4e-06 | 2.100E-06 | 6.083E-07 | 1.400E-06 | 2.500E-06 |
| 17647 | 883 | W883Q | C | 1.6e-06 | 2.6e-06 | 1.3e-06 | 1.833E-06 | 6.807E-07 | 1.300E-06 | 2.600E-06 |
| 17648 | 883 | W883G | C | 1.9e-06 | 1.6e-06 | 1.4e-06 | 1.633E-06 | 2.517E-07 | 1.400E-06 | 1.900E-06 |
| 17649 | 883 | W883H | C | 1.5e-06 | 2.6e-06 | 3e-06 | 2.367E-06 | 7.767E-07 | 1.500E-06 | 3.000E-06 |
| 17650 | 883 | W883I | C | 3.9e-06 | 3e-06 | 2.3e-06 | 3.067E-06 | 8.021E-07 | 2.300E-06 | 3.900E-06 |
| 17651 | 883 | W883L | C | 3.2e-06 | 2.4e-06 | 1.3e-06 | 2.300E-06 | 9.539E-07 | 1.300E-06 | 3.200E-06 |
| 17652 | 883 | W883K | C | 2.7e-06 | 2.8e-06 | 3e-06 | 2.833E-06 | 1.528E-07 | 2.700E-06 | 3.000E-06 |
| 17653 | 883 | W883M | C | 3e-06 | 2.5e-06 | 2.8e-06 | 2.767E-06 | 2.517E-07 | 2.500E-06 | 3.000E-06 |
| 17654 | 883 | W883F | C | 1.7e-06 | 1.3e-06 | 2.7e-06 | 1.900E-06 | 7.211E-07 | 1.300E-06 | 2.700E-06 |
| 17655 | 883 | W883P | C | 2.4e-06 | 1.4e-06 | 2.5e-06 | 2.100E-06 | 6.083E-07 | 1.400E-06 | 2.500E-06 |
| 17656 | 883 | W883S | C | 2.4e-06 | 2e-06 | 2.8e-06 | 2.400E-06 | 4.000E-07 | 2.000E-06 | 2.800E-06 |
| 17657 | 883 | W883T | C | 2.8e-06 | 4.5e-06 | 2.7e-06 | 3.333E-06 | 1.012E-06 | 2.700E-06 | 4.500E-06 |
| 17658 | 883 | W883W | C | 1.7e-06 | 1.9e-06 | 3e-06 | 2.200E-06 | 7.000E-07 | 1.700E-06 | 3.000E-06 |
| 17659 | 883 | W883Y | C | 3.1e-06 | 2.7e-06 | 2.4e-06 | 2.733E-06 | 3.512E-07 | 2.400E-06 | 3.100E-06 |
| 17660 | 883 | W883V | C | 2.5e-06 | 2.8e-06 | 4.2e-06 | 3.167E-06 | 9.074E-07 | 2.500E-06 | 4.200E-06 |
| 17661 | 884 | V884A | C | 1.3e-06 | 2.5e-06 | 2.1e-06 | 1.967E-06 | 6.110E-07 | 1.300E-06 | 2.500E-06 |
| 17662 | 884 | V884R | C | 4e-06 | 3.8e-06 | 2.5e-06 | 3.433E-06 | 8.145E-07 | 2.500E-06 | 4.000E-06 |
| 17663 | 884 | V884N | C | 2.9e-06 | 3.6e-06 | 2.1e-06 | 2.867E-06 | 7.506E-07 | 2.100E-06 | 3.600E-06 |
| 17664 | 884 | V884D | C | 2.1e-06 | 1.8e-06 | 3e-06 | 2.300E-06 | 6.245E-07 | 1.800E-06 | 3.000E-06 |
| 17665 | 884 | V884C | C | 3.2e-06 | 2e-06 | 3.3e-06 | 2.833E-06 | 7.234E-07 | 2.000E-06 | 3.300E-06 |
| 17666 | 884 | V884E | C | 2.3e-06 | 2.9e-06 | 2.2e-06 | 2.467E-06 | 3.786E-07 | 2.200E-06 | 2.900E-06 |
| 17667 | 884 | V884Q | C | 1.8e-06 | 2.1e-06 | 2.5e-06 | 2.133E-06 | 3.512E-07 | 1.800E-06 | 2.500E-06 |
| 17668 | 884 | V884G | C | 2.8e-06 | 2.3e-06 | 2.7e-06 | 2.600E-06 | 2.646E-07 | 2.300E-06 | 2.800E-06 |
| 17669 | 884 | V884H | C | 1.9e-06 | 2.3e-06 | 1.3e-06 | 1.833E-06 | 5.033E-07 | 1.300E-06 | 2.300E-06 |
| 17670 | 884 | V884I | C | 2.6e-06 | 1.4e-06 | 1.9e-06 | 1.967E-06 | 6.028E-07 | 1.400E-06 | 2.600E-06 |
| 17671 | 884 | V884L | C | 1.9e-06 | 3e-06 | 1.2e-06 | 2.033E-06 | 9.074E-07 | 1.200E-06 | 3.000E-06 |
| 17672 | 884 | V884K | C | 3.5e-06 | 1.4e-06 | 2.9e-06 | 2.600E-06 | 1.082E-06 | 1.400E-06 | 3.500E-06 |
| 17673 | 884 | V884M | C | 2.7e-06 | 2.7e-06 | 2.1e-06 | 2.500E-06 | 3.464E-07 | 2.100E-06 | 2.700E-06 |
| 17674 | 884 | V884F | C | 2.6e-06 | 2.5e-06 | 1.6e-06 | 2.233E-06 | 5.508E-07 | 1.600E-06 | 2.600E-06 |
| 17675 | 884 | V884P | C | 1.8e-06 | 2e-06 | 1.8e-06 | 1.867E-06 | 1.155E-07 | 1.800E-06 | 2.000E-06 |
| 17676 | 884 | V884S | C | 2.2e-06 | 2.7e-06 | 2.4e-06 | 2.433E-06 | 2.517E-07 | 2.200E-06 | 2.700E-06 |
| 17677 | 884 | V884T | C | 1.8e-06 | 1.9e-06 | 1.7e-06 | 1.800E-06 | 1.000E-07 | 1.700E-06 | 1.900E-06 |
| 17678 | 884 | V884W | C | 3e-06 | 1.3e-06 | 2e-06 | 2.100E-06 | 8.544E-07 | 1.300E-06 | 3.000E-06 |
| 17679 | 884 | V884Y | C | 1.7e-06 | 1.5e-06 | 2.4e-06 | 1.867E-06 | 4.726E-07 | 1.500E-06 | 2.400E-06 |

| | | | | | | | | | | |
|---|---|---|---|---|---|---|---|---|---|---|
| 17680 | 884 | V884V | C | 1.7e-06 | 1.3e-06 | 3.4e-06 | 2.133E-06 | 1.115E-06 | 1.300E-06 | 3.400E-06 |
| 17681 | 885 | A885A | C | 4.6e-06 | 3.2e-06 | 4.7e-06 | 4.167E-06 | 8.386E-07 | 3.200E-06 | 4.700E-06 |
| 17682 | 885 | A885R | C | 1.9e-06 | 1.3e-06 | 2.2e-06 | 1.800E-06 | 4.583E-07 | 1.300E-06 | 2.200E-06 |
| 17683 | 885 | A885N | C | 1.9e-06 | 2.8e-06 | 1.6e-06 | 2.100E-06 | 6.245E-07 | 1.600E-06 | 2.800E-06 |
| 17684 | 885 | A885D | C | 2.7e-06 | 2.7e-06 | 2.7e-06 | 2.700E-06 | 0.000E+00 | 2.700E-06 | 2.700E-06 |
| 17685 | 885 | A885C | C | 3.1e-06 | 2.7e-06 | 5.4e-06 | 3.733E-06 | 1.457E-06 | 2.700E-06 | 5.400E-06 |
| 17686 | 885 | A885E | C | 1.6e-06 | 2.4e-06 | 3.4e-06 | 2.467E-06 | 9.018E-07 | 1.600E-06 | 3.400E-06 |
| 17687 | 885 | A885Q | C | 3.8e-06 | 1.8e-06 | 2.2e-06 | 2.600E-06 | 1.058E-06 | 1.800E-06 | 3.800E-06 |
| 17688 | 885 | A885G | C | 2.4e-06 | 2.3e-06 | 4.1e-06 | 2.933E-06 | 1.012E-06 | 2.300E-06 | 4.100E-06 |
| 17689 | 885 | A885H | C | 2.7e-06 | 2.6e-06 | 2.8e-06 | 2.700E-06 | 1.000E-07 | 2.600E-06 | 2.800E-06 |
| 17690 | 885 | A885I | C | 2.8e-06 | 2.5e-06 | 1.8e-06 | 2.367E-06 | 5.132E-07 | 1.800E-06 | 2.800E-06 |
| 17691 | 885 | A885L | C | 3.2e-06 | 2.7e-06 | 2.4e-06 | 2.767E-06 | 4.041E-07 | 2.400E-06 | 3.200E-06 |
| 17692 | 885 | A885K | C | 1.6e-06 | 1.7e-06 | 1.6e-06 | 1.633E-06 | 5.774E-08 | 1.600E-06 | 1.700E-06 |
| 17693 | 885 | A885M | C | 3.3e-06 | 2.3e-06 | 1.8e-06 | 2.467E-06 | 7.638E-07 | 1.800E-06 | 3.300E-06 |
| 17694 | 885 | A885F | C | 3.1e-06 | 2.3e-06 | 9.5e-07 | 2.117E-06 | 1.087E-06 | 9.500E-07 | 3.100E-06 |
| 17695 | 885 | A885P | C | 1.9e-06 | 2.3e-06 | 3.7e-06 | 2.633E-06 | 9.452E-07 | 1.900E-06 | 3.700E-06 |
| 17696 | 885 | A885S | C | 1.8e-06 | 2.6e-06 | 2.2e-06 | 2.200E-06 | 4.000E-07 | 1.800E-06 | 2.600E-06 |
| 17697 | 885 | A885T | C | 3e-06 | 3.1e-06 | 3.2e-06 | 3.100E-06 | 1.000E-07 | 3.000E-06 | 3.200E-06 |
| 17698 | 885 | A885W | C | 2.1e-06 | 1.9e-06 | 1.5e-06 | 1.833E-06 | 3.055E-07 | 1.500E-06 | 2.100E-06 |
| 17699 | 885 | A885Y | C | 2.3e-06 | 4.2e-06 | 2.6e-06 | 3.033E-06 | 1.021E-06 | 2.300E-06 | 4.200E-06 |
| 17700 | 885 | A885V | C | 2e-06 | 2.3e-06 | 1.1e-06 | 1.800E-06 | 6.245E-07 | 1.100E-06 | 2.300E-06 |
| 17701 | 886 | D886A | C | 2.1e-06 | 1.8e-06 | 1.5e-06 | 1.800E-06 | 3.000E-07 | 1.500E-06 | 2.100E-06 |
| 17702 | 886 | D886R | C | 3.1e-06 | 2e-06 | 2.1e-06 | 2.400E-06 | 6.083E-07 | 2.000E-06 | 3.100E-06 |
| 17703 | 886 | D886N | C | 2.8e-06 | 1.6e-06 | 2.9e-06 | 2.433E-06 | 7.234E-07 | 1.600E-06 | 2.900E-06 |
| 17704 | 886 | D886D | C | 2.6e-06 | 2.7e-06 | 1.5e-06 | 2.267E-06 | 6.658E-07 | 1.500E-06 | 2.700E-06 |
| 17705 | 886 | D886C | C | 2.2e-06 | 2.1e-06 | 1.7e-06 | 2.000E-06 | 2.646E-07 | 1.700E-06 | 2.200E-06 |
| 17706 | 886 | D886E | C | 2.3e-06 | 3.7e-06 | 9.7e-07 | 2.323E-06 | 1.365E-06 | 9.700E-07 | 3.700E-06 |
| 17707 | 886 | D886Q | C | 2.2e-06 | 2e-06 | 2.1e-06 | 2.100E-06 | 1.000E-07 | 2.000E-06 | 2.200E-06 |
| 17708 | 886 | D886G | C | 2.5e-06 | 1.9e-06 | 2.7e-06 | 2.367E-06 | 4.163E-07 | 1.900E-06 | 2.700E-06 |
| 17709 | 886 | D886H | C | 2.4e-06 | 3.1e-06 | 2.8e-06 | 2.767E-06 | 3.512E-07 | 2.400E-06 | 3.100E-06 |
| 17710 | 886 | D886I | C | 1.9e-06 | 2.5e-06 | 2.9e-06 | 2.433E-06 | 5.033E-07 | 1.900E-06 | 2.900E-06 |
| 17711 | 886 | D886L | C | 2.2e-06 | 4e-06 | 3.2e-06 | 3.133E-06 | 9.018E-07 | 2.200E-06 | 4.000E-06 |
| 17712 | 886 | D886K | C | 2.1e-06 | 1.6e-06 | 2.7e-06 | 2.133E-06 | 5.508E-07 | 1.600E-06 | 2.700E-06 |
| 17713 | 886 | D886M | C | 3.4e-06 | 1.9e-06 | 3e-06 | 2.767E-06 | 7.767E-07 | 1.900E-06 | 3.400E-06 |
| 17714 | 886 | D886F | C | 1.6e-06 | 3.4e-06 | 2.3e-06 | 2.433E-06 | 9.074E-07 | 1.600E-06 | 3.400E-06 |
| 17715 | 886 | D886P | C | 1.9e-06 | 3.1e-06 | 1.8e-06 | 2.267E-06 | 7.234E-07 | 1.800E-06 | 3.100E-06 |
| 17716 | 886 | D886S | C | 2.7e-06 | 3.1e-06 | 2.6e-06 | 2.800E-06 | 2.646E-07 | 2.600E-06 | 3.100E-06 |
| 17717 | 886 | D886T | C | 3e-06 | 2.8e-06 | 3.2e-06 | 3.000E-06 | 2.000E-07 | 2.800E-06 | 3.200E-06 |
| 17718 | 886 | D886W | C | 2.8e-06 | 3.2e-06 | 2.6e-06 | 2.867E-06 | 3.055E-07 | 2.600E-06 | 3.200E-06 |
| 17719 | 886 | D886Y | C | 2.2e-06 | 9.3e-07 | 1.3e-06 | 1.477E-06 | 6.532E-07 | 9.300E-07 | 2.200E-06 |
| 17720 | 886 | D886V | C | 2.4e-06 | 1.7e-06 | 3.7e-06 | 2.600E-06 | 1.015E-06 | 1.700E-06 | 3.700E-06 |
| 17721 | 887 | V887A | C | 2e-06 | 1.3e-06 | 1.5e-06 | 1.600E-06 | 3.606E-07 | 1.300E-06 | 2.000E-06 |
| 17722 | 887 | V887R | C | 1.7e-06 | 2.6e-06 | 3.5e-06 | 2.600E-06 | 9.000E-07 | 1.700E-06 | 3.500E-06 |
| 17723 | 887 | V887N | C | 3.3e-06 | 2.1e-06 | 3.9e-06 | 3.100E-06 | 9.165E-07 | 2.100E-06 | 3.900E-06 |
| 17724 | 887 | V887D | C | 1.8e-06 | 1.9e-06 | 2e-06 | 1.900E-06 | 1.000E-07 | 1.800E-06 | 2.000E-06 |
| 17725 | 887 | V887C | C | 2e-06 | 2.7e-06 | 1.3e-06 | 2.000E-06 | 7.000E-07 | 1.300E-06 | 2.700E-06 |
| 17726 | 887 | V887E | C | 1.9e-06 | 2.2e-06 | 1.7e-06 | 1.933E-06 | 2.517E-07 | 1.700E-06 | 2.200E-06 |
| 17727 | 887 | V887Q | C | 1.8e-06 | 1.7e-06 | 1.6e-06 | 1.700E-06 | 1.000E-07 | 1.600E-06 | 1.800E-06 |
| 17728 | 887 | V887G | C | 2.3e-06 | 2.2e-06 | 2.6e-06 | 2.367E-06 | 2.082E-07 | 2.200E-06 | 2.600E-06 |
| 17729 | 887 | V887H | C | 3.1e-06 | 2e-06 | 1.5e-06 | 2.200E-06 | 8.185E-07 | 1.500E-06 | 3.100E-06 |
| 17730 | 887 | V887I | C | 9.6e-07 | 2.8e-06 | 2.2e-06 | 1.987E-06 | 9.384E-07 | 9.600E-07 | 2.800E-06 |
| 17731 | 887 | V887L | C | 1.9e-06 | 2.1e-06 | 3.2e-06 | 2.400E-06 | 7.000E-07 | 1.900E-06 | 3.200E-06 |
| 17732 | 887 | V887K | C | 2.8e-06 | 2.4e-06 | 1.4e-06 | 2.200E-06 | 7.211E-07 | 1.400E-06 | 2.800E-06 |
| 17733 | 887 | V887M | C | 9.7e-07 | 3e-06 | 3.1e-06 | 2.357E-06 | 1.202E-06 | 9.700E-07 | 3.100E-06 |
| 17734 | 887 | V887F | C | 1.8e-06 | 1.8e-06 | 2.7e-06 | 2.100E-06 | 5.196E-07 | 1.800E-06 | 2.700E-06 |
| 17735 | 887 | V887P | C | 1.8e-06 | 2.2e-06 | 2.9e-06 | 2.300E-06 | 5.568E-07 | 1.800E-06 | 2.900E-06 |
| 17736 | 887 | V887S | C | 3.4e-06 | 2.4e-06 | 3.4e-06 | 3.067E-06 | 5.774E-07 | 2.400E-06 | 3.400E-06 |
| 17737 | 887 | V887T | C | 6.8e-07 | 1.1e-06 | 2.9e-06 | 1.560E-06 | 1.179E-06 | 6.800E-07 | 2.900E-06 |
| 17738 | 887 | V887W | C | 1.5e-06 | 1.5e-06 | 2.4e-06 | 1.800E-06 | 5.196E-07 | 1.500E-06 | 2.400E-06 |
| 17739 | 887 | V887Y | C | 1.7e-06 | 2.7e-06 | 4.2e-06 | 2.867E-06 | 1.258E-06 | 1.700E-06 | 4.200E-06 |
| 17740 | 887 | V887V | C | 2.5e-06 | 1.8e-06 | 1.1e-06 | 1.800E-06 | 7.000E-07 | 1.100E-06 | 2.500E-06 |
| 17741 | 888 | N888A | C | 2e-06 | 2.3e-06 | 2.5e-06 | 2.267E-06 | 2.517E-07 | 2.000E-06 | 2.500E-06 |
| 17742 | 888 | N888R | C | 4e-06 | 2.1e-06 | 1.8e-06 | 2.633E-06 | 1.193E-06 | 1.800E-06 | 4.000E-06 |
| 17743 | 888 | N888N | C | 2.5e-06 | 4.2e-06 | 2.3e-06 | 3.000E-06 | 1.044E-06 | 2.300E-06 | 4.200E-06 |
| 17744 | 888 | N888D | C | 4.1e-06 | 1.7e-06 | 3.4e-06 | 3.067E-06 | 1.234E-06 | 1.700E-06 | 4.100E-06 |
| 17745 | 888 | N888C | C | 4.4e-06 | 1.9e-06 | 2.7e-06 | 3.000E-06 | 1.277E-06 | 1.900E-06 | 4.400E-06 |
| 17746 | 888 | N888E | C | 2.3e-06 | 1.9e-06 | 1.8e-06 | 2.000E-06 | 2.646E-07 | 1.800E-06 | 2.300E-06 |
| 17747 | 888 | N888Q | C | 2.2e-06 | 2.5e-06 | 2.1e-06 | 2.267E-06 | 2.082E-07 | 2.100E-06 | 2.500E-06 |
| 17748 | 888 | N888G | C | 2.4e-06 | 2.1e-06 | 2.6e-06 | 2.367E-06 | 2.517E-07 | 2.100E-06 | 2.600E-06 |
| 17749 | 888 | N888H | C | 1.6e-06 | 2e-06 | 1.5e-06 | 1.700E-06 | 2.646E-07 | 1.500E-06 | 2.000E-06 |
| 17750 | 888 | N888I | C | 3.2e-06 | 3.2e-06 | 2.4e-06 | 2.933E-06 | 4.619E-07 | 2.400E-06 | 3.200E-06 |
| 17751 | 888 | N888L | C | 2.8e-06 | 2.3e-06 | 4.6e-06 | 3.233E-06 | 1.210E-06 | 2.300E-06 | 4.600E-06 |
| 17752 | 888 | N888K | C | 2.3e-06 | 2.7e-06 | 8e-07 | 1.933E-06 | 1.002E-06 | 8.000E-07 | 2.700E-06 |
| 17753 | 888 | N888M | C | 3.3e-06 | 4.2e-06 | 4.8e-06 | 4.100E-06 | 7.550E-07 | 3.300E-06 | 4.800E-06 |
| 17754 | 888 | N888F | C | 3e-06 | 2.3e-06 | 3e-06 | 2.767E-06 | 4.041E-07 | 2.300E-06 | 3.000E-06 |
| 17755 | 888 | N888P | C | 2.4e-06 | 2.1e-06 | 2.6e-06 | 2.367E-06 | 2.517E-07 | 2.100E-06 | 2.600E-06 |
| 17756 | 888 | N888S | C | 1.9e-06 | 1.7e-06 | 1.6e-06 | 1.733E-06 | 1.528E-07 | 1.600E-06 | 1.900E-06 |
| 17757 | 888 | N888T | C | 3.2e-06 | 6.2e-06 | 3.5e-06 | 4.300E-06 | 1.652E-06 | 3.200E-06 | 6.200E-06 |
| 17758 | 888 | N888W | C | 1.5e-06 | 3e-06 | 1.2e-06 | 1.900E-06 | 9.644E-07 | 1.200E-06 | 3.000E-06 |
| 17759 | 888 | N888Y | C | 2.3e-06 | 1.2e-06 | 1.8e-06 | 1.767E-06 | 5.508E-07 | 1.200E-06 | 2.300E-06 |
| 17760 | 888 | N888V | C | 4.3e-06 | 1.7e-06 | 3.2e-06 | 3.067E-06 | 1.305E-06 | 1.700E-06 | 4.300E-06 |
| 17761 | 889 | P889A | C | 4.3e-06 | 4.6e-06 | 2.6e-06 | 3.833E-06 | 1.079E-06 | 2.600E-06 | 4.600E-06 |
| 17762 | 889 | P889R | C | 1.5e-06 | 3.3e-06 | 2.7e-06 | 2.500E-06 | 9.165E-07 | 1.500E-06 | 3.300E-06 |
| 17763 | 889 | P889N | C | 1.1e-06 | 1.4e-06 | 2.2e-06 | 1.567E-06 | 5.686E-07 | 1.100E-06 | 2.200E-06 |
| 17764 | 889 | P889D | C | 2.5e-06 | 1.7e-06 | 3.7e-06 | 2.633E-06 | 1.007E-06 | 1.700E-06 | 3.700E-06 |

| | | | | | | | | | | |
|---|---|---|---|---|---|---|---|---|---|---|
| 17765 | 889 | P889C | C | 1.9e-06 | 1.8e-06 | 3.6e-06 | 2.433E-06 | 1.012E-06 | 1.800E-06 | 3.600E-06 |
| 17766 | 889 | P889E | C | 2.3e-06 | 2.1e-06 | 2.2e-06 | 2.200E-06 | 1.000E-07 | 2.100E-06 | 2.300E-06 |
| 17767 | 889 | P889Q | C | 2.7e-06 | 3.2e-06 | 2.5e-06 | 2.800E-06 | 3.606E-07 | 2.500E-06 | 3.200E-06 |
| 17768 | 889 | P889G | C | 2.8e-06 | 1.5e-06 | 2.1e-06 | 2.133E-06 | 6.506E-07 | 1.500E-06 | 2.800E-06 |
| 17769 | 889 | P889H | C | 3.5e-06 | 1.6e-06 | 1.7e-06 | 2.267E-06 | 1.069E-06 | 1.600E-06 | 3.500E-06 |
| 17770 | 889 | P889I | C | 1.6e-06 | 2.6e-06 | 4.6e-06 | 2.933E-06 | 1.528E-06 | 1.600E-06 | 4.600E-06 |
| 17771 | 889 | P889L | C | 2.6e-06 | 2.7e-06 | 1.9e-06 | 2.400E-06 | 4.359E-07 | 1.900E-06 | 2.700E-06 |
| 17772 | 889 | P889K | C | 2.7e-06 | 2.1e-06 | 1.3e-06 | 2.033E-06 | 7.024E-07 | 1.300E-06 | 2.700E-06 |
| 17773 | 889 | P889M | C | 2.9e-06 | 3.8e-06 | 5e-06 | 3.900E-06 | 1.054E-06 | 2.900E-06 | 5.000E-06 |
| 17774 | 889 | P889F | C | 1.2e-06 | 3.1e-06 | 2.4e-06 | 2.233E-06 | 9.609E-07 | 1.200E-06 | 3.100E-06 |
| 17775 | 889 | P889P | C | 2.3e-06 | 1.9e-06 | 1.8e-06 | 2.000E-06 | 2.646E-07 | 1.800E-06 | 2.300E-06 |
| 17776 | 889 | P889S | C | 3e-06 | 2.2e-06 | 2.4e-06 | 2.533E-06 | 4.163E-07 | 2.200E-06 | 3.000E-06 |
| 17777 | 889 | P889T | C | 1.7e-06 | 2.7e-06 | 3.2e-06 | 2.533E-06 | 7.638E-07 | 1.700E-06 | 3.200E-06 |
| 17778 | 889 | P889W | C | 1.9e-06 | 3.9e-06 | 3.2e-06 | 3.000E-06 | 1.015E-06 | 1.900E-06 | 3.900E-06 |
| 17779 | 889 | P889Y | C | 1.1e-06 | 2.2e-06 | 2.3e-06 | 1.867E-06 | 6.658E-07 | 1.100E-06 | 2.300E-06 |
| 17780 | 889 | P889V | C | 3e-06 | 4.8e-06 | 3.6e-06 | 3.800E-06 | 9.165E-07 | 3.000E-06 | 4.800E-06 |
| 17781 | 890 | N890A | C | 2.1e-06 | 2.3e-06 | 2.2e-06 | 2.200E-06 | 1.000E-07 | 2.100E-06 | 2.300E-06 |
| 17782 | 890 | N890R | C | 3.2e-06 | 2.3e-06 | 3.3e-06 | 2.933E-06 | 5.508E-07 | 2.300E-06 | 3.300E-06 |
| 17783 | 890 | N890N | C | 3.3e-06 | 3.4e-06 | 2.1e-06 | 2.933E-06 | 7.234E-07 | 2.100E-06 | 3.400E-06 |
| 17784 | 890 | N890D | C | 3.6e-06 | 1.5e-06 | 3.4e-06 | 2.833E-06 | 1.159E-06 | 1.500E-06 | 3.600E-06 |
| 17785 | 890 | N890C | C | 1.8e-06 | 1.7e-06 | 2.1e-06 | 1.867E-06 | 2.082E-07 | 1.700E-06 | 2.100E-06 |
| 17786 | 890 | N890E | C | 2.4e-06 | 2.5e-06 | 3.1e-06 | 2.667E-06 | 3.786E-07 | 2.400E-06 | 3.100E-06 |
| 17787 | 890 | N890Q | C | 3.7e-06 | 4.6e-06 | 2.3e-06 | 3.533E-06 | 1.159E-06 | 2.300E-06 | 4.600E-06 |
| 17788 | 890 | N890G | C | 1.3e-06 | 2.7e-06 | 2.5e-06 | 2.167E-06 | 7.572E-07 | 1.300E-06 | 2.700E-06 |
| 17789 | 890 | N890H | C | 2.3e-06 | 2.2e-06 | 2.5e-06 | 2.333E-06 | 1.528E-07 | 2.200E-06 | 2.500E-06 |
| 17790 | 890 | N890I | C | 2e-06 | 3.5e-06 | 1.3e-06 | 2.267E-06 | 1.124E-06 | 1.300E-06 | 3.500E-06 |
| 17791 | 890 | N890L | C | 2.8e-06 | 2.3e-06 | 1.6e-06 | 2.233E-06 | 6.028E-07 | 1.600E-06 | 2.800E-06 |
| 17792 | 890 | N890K | C | 2e-06 | 2.1e-06 | 2.7e-06 | 2.267E-06 | 3.786E-07 | 2.000E-06 | 2.700E-06 |
| 17793 | 890 | N890M | C | 2.5e-06 | 2.9e-06 | 1.4e-06 | 2.267E-06 | 7.767E-07 | 1.400E-06 | 2.900E-06 |
| 17794 | 890 | N890F | C | 2.1e-06 | 3.2e-06 | 4.7e-06 | 3.333E-06 | 1.305E-06 | 2.100E-06 | 4.700E-06 |
| 17795 | 890 | N890P | C | 1.9e-06 | 8.3e-07 | 2.9e-06 | 1.877E-06 | 1.035E-06 | 8.300E-07 | 2.900E-06 |
| 17796 | 890 | N890S | C | 3.6e-06 | 2.9e-06 | 2.5e-06 | 3.000E-06 | 5.568E-07 | 2.500E-06 | 3.600E-06 |
| 17797 | 890 | N890T | C | 2.6e-06 | 2.5e-06 | 3.2e-06 | 2.767E-06 | 3.786E-07 | 2.500E-06 | 3.200E-06 |
| 17798 | 890 | N890W | C | 1.6e-06 | 2.8e-06 | 2.2e-06 | 2.200E-06 | 6.000E-07 | 1.600E-06 | 2.800E-06 |
| 17799 | 890 | N890Y | C | 2.4e-06 | 1.5e-06 | 3.2e-06 | 2.367E-06 | 8.505E-07 | 1.500E-06 | 3.200E-06 |
| 17800 | 890 | N890V | C | 2.6e-06 | 1.9e-06 | 2.9e-06 | 2.467E-06 | 5.132E-07 | 1.900E-06 | 2.900E-06 |
| 17801 | 891 | S891A | C | 2.7e-06 | 2.5e-06 | 6e-06 | 3.733E-06 | 1.966E-06 | 2.500E-06 | 6.000E-06 |
| 17802 | 891 | S891R | C | 1.8e-06 | 2.4e-06 | 2.1e-06 | 2.100E-06 | 3.000E-07 | 1.800E-06 | 2.400E-06 |
| 17803 | 891 | S891N | C | 8.5e-07 | 2e-06 | 3.2e-06 | 2.017E-06 | 1.175E-06 | 8.500E-07 | 3.200E-06 |
| 17804 | 891 | S891D | C | 2.6e-06 | 2e-06 | 9.7e-07 | 1.857E-06 | 8.244E-07 | 9.700E-07 | 2.600E-06 |
| 17805 | 891 | S891C | C | 2.3e-06 | 2.6e-06 | 2.7e-06 | 2.533E-06 | 2.082E-07 | 2.300E-06 | 2.700E-06 |
| 17806 | 891 | S891E | C | 1.8e-06 | 3.3e-06 | 2.1e-06 | 2.400E-06 | 7.937E-07 | 1.800E-06 | 3.300E-06 |
| 17807 | 891 | S891Q | C | 2.1e-06 | 2.6e-06 | 3.8e-06 | 2.833E-06 | 8.737E-07 | 2.100E-06 | 3.800E-06 |
| 17808 | 891 | S891G | C | 3.4e-06 | 3.3e-06 | 2.6e-06 | 3.100E-06 | 4.359E-07 | 2.600E-06 | 3.400E-06 |
| 17809 | 891 | S891H | C | 1.8e-06 | 2.8e-06 | 3e-06 | 2.533E-06 | 6.429E-07 | 1.800E-06 | 3.000E-06 |
| 17810 | 891 | S891I | C | 3.4e-06 | 4.2e-06 | 1.3e-06 | 2.967E-06 | 1.498E-06 | 1.300E-06 | 4.200E-06 |
| 17811 | 891 | S891L | C | 2.1e-06 | 2.7e-06 | 1.4e-06 | 2.067E-06 | 6.506E-07 | 1.400E-06 | 2.700E-06 |
| 17812 | 891 | S891K | C | 2.7e-06 | 3.2e-06 | 1.3e-06 | 2.400E-06 | 9.849E-07 | 1.300E-06 | 3.200E-06 |
| 17813 | 891 | S891M | C | 2.8e-06 | 2.6e-06 | 2.2e-06 | 2.533E-06 | 3.055E-07 | 2.200E-06 | 2.800E-06 |
| 17814 | 891 | S891F | C | 2.3e-06 | 3.3e-06 | 2.7e-06 | 2.767E-06 | 5.033E-07 | 2.300E-06 | 3.300E-06 |
| 17815 | 891 | S891P | C | 2.5e-06 | 2.1e-06 | 1.9e-06 | 2.167E-06 | 3.055E-07 | 1.900E-06 | 2.500E-06 |
| 17816 | 891 | S891S | C | 2.3e-06 | 2.1e-06 | 2.6e-06 | 2.333E-06 | 2.517E-07 | 2.100E-06 | 2.600E-06 |
| 17817 | 891 | S891T | C | 3.1e-06 | 3.7e-06 | 1.5e-06 | 2.767E-06 | 1.137E-06 | 1.500E-06 | 3.700E-06 |
| 17818 | 891 | S891W | C | 3e-06 | 1.6e-06 | 2.6e-06 | 2.400E-06 | 7.211E-07 | 1.600E-06 | 3.000E-06 |
| 17819 | 891 | S891Y | C | 2e-06 | 1.3e-06 | 1.7e-06 | 1.667E-06 | 3.512E-07 | 1.300E-06 | 2.000E-06 |
| 17820 | 891 | S891V | C | 9.2e-07 | 1.5e-06 | 1e-06 | 1.140E-06 | 3.143E-07 | 9.200E-07 | 1.500E-06 |
| 17821 | 892 | G892A | C | 1.7e-06 | 1.6e-06 | 1.4e-06 | 1.567E-06 | 1.528E-07 | 1.400E-06 | 1.700E-06 |
| 17822 | 892 | G892R | C | 2.1e-06 | 2.3e-06 | 2.9e-06 | 2.433E-06 | 4.163E-07 | 2.100E-06 | 2.900E-06 |
| 17823 | 892 | G892N | C | 1.6e-06 | 2.4e-06 | 1.7e-06 | 1.900E-06 | 4.359E-07 | 1.600E-06 | 2.400E-06 |
| 17824 | 892 | G892D | C | 1.1e-06 | 2.2e-06 | 1.8e-06 | 1.700E-06 | 5.568E-07 | 1.100E-06 | 2.200E-06 |
| 17825 | 892 | G892C | C | 3.5e-06 | 2e-06 | 2.7e-06 | 2.733E-06 | 7.506E-07 | 2.000E-06 | 3.500E-06 |
| 17826 | 892 | G892E | C | 2.5e-06 | 2.3e-06 | 2.7e-06 | 2.500E-06 | 2.000E-07 | 2.300E-06 | 2.700E-06 |
| 17827 | 892 | G892Q | C | 1.6e-06 | 2.5e-06 | 3.8e-06 | 2.633E-06 | 1.106E-06 | 1.600E-06 | 3.800E-06 |
| 17828 | 892 | G892G | C | 1.7e-06 | 1.8e-06 | 1.4e-06 | 1.633E-06 | 2.082E-07 | 1.400E-06 | 1.800E-06 |
| 17829 | 892 | G892H | C | 3.3e-06 | 1.4e-06 | 3.7e-06 | 2.800E-06 | 1.229E-06 | 1.400E-06 | 3.700E-06 |
| 17830 | 892 | G892I | C | 2.4e-06 | 7.3e-07 | 2.1e-06 | 1.743E-06 | 8.903E-07 | 7.300E-07 | 2.400E-06 |
| 17831 | 892 | G892L | C | 3.3e-06 | 3e-06 | 1.5e-06 | 2.600E-06 | 9.644E-07 | 1.500E-06 | 3.300E-06 |
| 17832 | 892 | G892K | C | 2.6e-06 | 2.7e-06 | 3.8e-06 | 3.033E-06 | 6.658E-07 | 2.600E-06 | 3.800E-06 |
| 17833 | 892 | G892M | C | 2.5e-06 | 3.1e-06 | 2e-06 | 2.533E-06 | 5.508E-07 | 2.000E-06 | 3.100E-06 |
| 17834 | 892 | G892F | C | 1.6e-06 | 1.3e-06 | 1.9e-06 | 1.600E-06 | 3.000E-07 | 1.300E-06 | 1.900E-06 |
| 17835 | 892 | G892P | C | 2.3e-06 | 3.1e-06 | 2.9e-06 | 2.767E-06 | 4.163E-07 | 2.300E-06 | 3.100E-06 |
| 17836 | 892 | G892S | C | 2.9e-06 | 3.4e-06 | 2e-06 | 2.767E-06 | 7.095E-07 | 2.000E-06 | 3.400E-06 |
| 17837 | 892 | G892T | C | 2.9e-06 | 2.5e-06 | 7.1e-07 | 2.037E-06 | 1.166E-06 | 7.100E-07 | 2.900E-06 |
| 17838 | 892 | G892W | C | 2.7e-06 | 1.9e-06 | 3e-06 | 2.533E-06 | 5.686E-07 | 1.900E-06 | 3.000E-06 |
| 17839 | 892 | G892Y | C | 2.2e-06 | 3.5e-06 | 1.8e-06 | 2.500E-06 | 8.888E-07 | 1.800E-06 | 3.500E-06 |
| 17840 | 892 | G892V | C | 2.5e-06 | 4.3e-06 | 3.7e-06 | 3.500E-06 | 9.165E-07 | 2.500E-06 | 4.300E-06 |
| 17841 | 893 | G893A | C | 3.3e-06 | 3.8e-06 | 1.1e-06 | 2.733E-06 | 1.436E-06 | 1.100E-06 | 3.800E-06 |
| 17842 | 893 | G893R | C | 1.8e-06 | 1.6e-06 | 1.7e-06 | 1.700E-06 | 1.000E-07 | 1.600E-06 | 1.800E-06 |
| 17843 | 893 | G893N | C | 5e-06 | 3.3e-06 | 1.8e-06 | 3.367E-06 | 1.601E-06 | 1.800E-06 | 5.000E-06 |
| 17844 | 893 | G893D | C | 3.3e-06 | 2.8e-06 | 2.5e-06 | 2.867E-06 | 4.041E-07 | 2.500E-06 | 3.300E-06 |
| 17845 | 893 | G893C | C | 2e-06 | 2.9e-06 | 1.9e-06 | 2.267E-06 | 5.508E-07 | 1.900E-06 | 2.900E-06 |
| 17846 | 893 | G893E | C | 2.6e-06 | 2.7e-06 | 2.7e-06 | 2.667E-06 | 5.774E-08 | 2.600E-06 | 2.700E-06 |
| 17847 | 893 | G893Q | C | 1.5e-06 | 1.9e-06 | 5.2e-06 | 2.867E-06 | 2.031E-06 | 1.500E-06 | 5.200E-06 |
| 17848 | 893 | G893G | C | 2.1e-06 | 4.6e-06 | 3.7e-06 | 3.467E-06 | 1.266E-06 | 2.100E-06 | 4.600E-06 |
| 17849 | 893 | G893H | C | 3.4e-06 | 2.8e-06 | 2.8e-06 | 3.000E-06 | 3.464E-07 | 2.800E-06 | 3.400E-06 |

| | | | | | | | | | | |
|---|---|---|---|---|---|---|---|---|---|---|
| 17850 | 893 | G893I | C | 4e-06 | 2e-06 | 1.8e-06 | 2.600E-06 | 1.217E-06 | 1.800E-06 | 4.000E-06 |
| 17851 | 893 | G893L | C | 3.5e-06 | 3.2e-06 | 2e-06 | 2.900E-06 | 7.937E-07 | 2.000E-06 | 3.500E-06 |
| 17852 | 893 | G893K | C | 2.2e-06 | 1.6e-06 | 2.2e-06 | 2.000E-06 | 3.464E-07 | 1.600E-06 | 2.200E-06 |
| 17853 | 893 | G893M | C | 5e-06 | 2.3e-06 | 2.5e-06 | 3.267E-06 | 1.504E-06 | 2.300E-06 | 5.000E-06 |
| 17854 | 893 | G893F | C | 4.5e-06 | 2e-06 | 2.4e-06 | 2.967E-06 | 1.343E-06 | 2.000E-06 | 4.500E-06 |
| 17855 | 893 | G893P | C | 3.1e-06 | 4e-06 | 2.2e-06 | 3.100E-06 | 9.000E-07 | 2.200E-06 | 4.000E-06 |
| 17856 | 893 | G893S | C | 3.5e-06 | 3.2e-06 | 2.9e-06 | 3.200E-06 | 3.000E-07 | 2.900E-06 | 3.500E-06 |
| 17857 | 893 | G893T | C | 1.1e-06 | 1.6e-06 | 1.5e-06 | 1.400E-06 | 2.646E-07 | 1.100E-06 | 1.600E-06 |
| 17858 | 893 | G893W | C | 2.5e-06 | 1.8e-06 | 9.2e-07 | 1.740E-06 | 7.917E-07 | 9.200E-07 | 2.500E-06 |
| 17859 | 893 | G893Y | C | 2.1e-06 | 4.5e-06 | 2.6e-06 | 3.067E-06 | 1.266E-06 | 2.100E-06 | 4.500E-06 |
| 17860 | 893 | G893V | C | 2.6e-06 | 2.2e-06 | 1.1e-06 | 1.967E-06 | 7.767E-07 | 1.100E-06 | 2.600E-06 |
| 17861 | 894 | S894A | C | 2.1e-06 | 2.6e-06 | 2.4e-06 | 2.367E-06 | 2.517E-07 | 2.100E-06 | 2.600E-06 |
| 17862 | 894 | S894R | C | 2.1e-06 | 3.2e-06 | 3e-06 | 2.767E-06 | 5.859E-07 | 2.100E-06 | 3.200E-06 |
| 17863 | 894 | S894N | C | 2.3e-06 | 1.5e-06 | 1.7e-06 | 1.833E-06 | 4.163E-07 | 1.500E-06 | 2.300E-06 |
| 17864 | 894 | S894D | C | 2.2e-06 | 9e-07 | 2.9e-06 | 2.000E-06 | 1.015E-06 | 9.000E-07 | 2.900E-06 |
| 17865 | 894 | S894C | C | 2.9e-06 | 3.1e-06 | 2.6e-06 | 2.867E-06 | 2.517E-07 | 2.600E-06 | 3.100E-06 |
| 17866 | 894 | S894E | C | 2.5e-06 | 2.2e-06 | 3e-06 | 2.567E-06 | 4.041E-07 | 2.200E-06 | 3.000E-06 |
| 17867 | 894 | S894Q | C | 2.7e-06 | 1.6e-06 | 3.4e-06 | 2.567E-06 | 9.074E-07 | 1.600E-06 | 3.400E-06 |
| 17868 | 894 | S894G | C | 3.3e-06 | 1.5e-06 | 3.2e-06 | 2.667E-06 | 1.012E-06 | 1.500E-06 | 3.300E-06 |
| 17869 | 894 | S894H | C | 3.1e-06 | 1.7e-06 | 2.5e-06 | 2.433E-06 | 7.024E-07 | 1.700E-06 | 3.100E-06 |
| 17870 | 894 | S894I | C | 2.6e-06 | 3.7e-06 | 1.5e-06 | 2.600E-06 | 1.100E-06 | 1.500E-06 | 3.700E-06 |
| 17871 | 894 | S894L | C | 1.7e-06 | 2e-06 | 1.4e-06 | 1.700E-06 | 3.000E-07 | 1.400E-06 | 2.000E-06 |
| 17872 | 894 | S894K | C | 3.4e-06 | 2.2e-06 | 2.5e-06 | 2.700E-06 | 6.245E-07 | 2.200E-06 | 3.400E-06 |
| 17873 | 894 | S894M | C | 2e-06 | 3.3e-06 | 3.2e-06 | 2.833E-06 | 7.234E-07 | 2.000E-06 | 3.300E-06 |
| 17874 | 894 | S894F | C | 1.3e-06 | 2.1e-06 | 3e-06 | 2.133E-06 | 8.505E-07 | 1.300E-06 | 3.000E-06 |
| 17875 | 894 | S894P | C | 2.8e-06 | 2.1e-06 | 1.8e-06 | 2.233E-06 | 5.132E-07 | 1.800E-06 | 2.800E-06 |
| 17876 | 894 | S894S | C | 2.8e-06 | 1.3e-06 | 3.5e-06 | 2.533E-06 | 1.124E-06 | 1.300E-06 | 3.500E-06 |
| 17877 | 894 | S894T | C | 1.6e-06 | 3e-06 | 2e-06 | 2.200E-06 | 7.211E-07 | 1.600E-06 | 3.000E-06 |
| 17878 | 894 | S894W | C | 2.8e-06 | 4.2e-06 | 2.3e-06 | 3.100E-06 | 9.849E-07 | 2.300E-06 | 4.200E-06 |
| 17879 | 894 | S894Y | C | 2.3e-06 | 2.3e-06 | 1.3e-06 | 1.967E-06 | 5.774E-07 | 1.300E-06 | 2.300E-06 |
| 17880 | 894 | S894V | C | 2.4e-06 | 1.8e-06 | 2e-06 | 2.067E-06 | 3.055E-07 | 1.800E-06 | 2.400E-06 |
| 17881 | 895 | I895A | C | 2.7e-06 | 3.1e-06 | 2.5e-06 | 2.767E-06 | 3.055E-07 | 2.500E-06 | 3.100E-06 |
| 17882 | 895 | I895R | C | 1.4e-06 | 2.2e-06 | 2.1e-06 | 1.900E-06 | 4.359E-07 | 1.400E-06 | 2.200E-06 |
| 17883 | 895 | I895N | C | 3.1e-06 | 1.8e-06 | 3.3e-06 | 2.733E-06 | 8.145E-07 | 1.800E-06 | 3.300E-06 |
| 17884 | 895 | I895D | C | 3.9e-06 | 1.9e-06 | 3.2e-06 | 3.000E-06 | 1.015E-06 | 1.900E-06 | 3.900E-06 |
| 17885 | 895 | I895C | C | 1.6e-06 | 2.4e-06 | 2.3e-06 | 2.100E-06 | 4.359E-07 | 1.600E-06 | 2.400E-06 |
| 17886 | 895 | I895E | C | 3.9e-06 | 1.8e-06 | 3e-06 | 2.900E-06 | 1.054E-06 | 1.800E-06 | 3.900E-06 |
| 17887 | 895 | I895Q | C | 1.8e-06 | 2.9e-06 | 3.2e-06 | 2.633E-06 | 7.371E-07 | 1.800E-06 | 3.200E-06 |
| 17888 | 895 | I895G | C | 3.1e-06 | 1.5e-06 | 2.7e-06 | 2.433E-06 | 8.327E-07 | 1.500E-06 | 3.100E-06 |
| 17889 | 895 | I895H | C | 2.8e-06 | 2.3e-06 | 2.1e-06 | 2.400E-06 | 3.606E-07 | 2.100E-06 | 2.800E-06 |
| 17890 | 895 | I895I | C | 3.4e-06 | 2.2e-06 | 3.4e-06 | 3.000E-06 | 6.928E-07 | 2.200E-06 | 3.400E-06 |
| 17891 | 895 | I895L | C | 3.9e-06 | 2.8e-06 | 2.3e-06 | 3.000E-06 | 8.185E-07 | 2.300E-06 | 3.900E-06 |
| 17892 | 895 | I895K | C | 2.4e-06 | 2.6e-06 | 1.8e-06 | 2.267E-06 | 4.163E-07 | 1.800E-06 | 2.600E-06 |
| 17893 | 895 | I895M | C | 2.9e-06 | 1.8e-06 | 1.5e-06 | 2.067E-06 | 7.371E-07 | 1.500E-06 | 2.900E-06 |
| 17894 | 895 | I895F | C | 3.2e-06 | 2.2e-06 | 2.7e-06 | 2.700E-06 | 5.000E-07 | 2.200E-06 | 3.200E-06 |
| 17895 | 895 | I895P | C | 1.3e-06 | 2.3e-06 | 1.7e-06 | 1.767E-06 | 5.033E-07 | 1.300E-06 | 2.300E-06 |
| 17896 | 895 | I895S | C | 2.9e-06 | 2e-06 | 3e-06 | 2.633E-06 | 5.508E-07 | 2.000E-06 | 3.000E-06 |
| 17897 | 895 | I895T | C | 2.9e-06 | 3.1e-06 | 1.5e-06 | 2.500E-06 | 8.718E-07 | 1.500E-06 | 3.100E-06 |
| 17898 | 895 | I895W | C | 2.7e-06 | 3.9e-06 | 2.9e-06 | 3.167E-06 | 6.429E-07 | 2.700E-06 | 3.900E-06 |
| 17899 | 895 | I895Y | C | 1.5e-06 | 2.1e-06 | 1.5e-06 | 1.700E-06 | 3.464E-07 | 1.500E-06 | 2.100E-06 |
| 17900 | 895 | I895V | C | 4.3e-06 | 3.2e-06 | 1.1e-06 | 2.867E-06 | 1.626E-06 | 1.100E-06 | 4.300E-06 |
| 17901 | 896 | Y896A | C | 1.2e-06 | 2.3e-06 | 2.7e-06 | 2.067E-06 | 7.767E-07 | 1.200E-06 | 2.700E-06 |
| 17902 | 896 | Y896R | C | 1.5e-06 | 2.8e-06 | 1.8e-06 | 2.033E-06 | 6.807E-07 | 1.500E-06 | 2.800E-06 |
| 17903 | 896 | Y896N | C | 1.7e-06 | 2.3e-06 | 2.2e-06 | 2.067E-06 | 3.215E-07 | 1.700E-06 | 2.300E-06 |
| 17904 | 896 | Y896D | C | 2.9e-06 | 2.4e-06 | 9.7e-07 | 2.090E-06 | 1.002E-06 | 9.700E-07 | 2.900E-06 |
| 17905 | 896 | Y896C | C | 2.8e-06 | 2.3e-06 | 3.6e-06 | 2.900E-06 | 6.557E-07 | 2.300E-06 | 3.600E-06 |
| 17906 | 896 | Y896E | C | 1.9e-06 | 4.5e-06 | 2.4e-06 | 2.933E-06 | 1.380E-06 | 1.900E-06 | 4.500E-06 |
| 17907 | 896 | Y896Q | C | 3.3e-06 | 2.4e-06 | 2.2e-06 | 2.633E-06 | 5.859E-07 | 2.200E-06 | 3.300E-06 |
| 17908 | 896 | Y896G | C | 2.8e-06 | 3.6e-06 | 1.5e-06 | 2.633E-06 | 1.060E-06 | 1.500E-06 | 3.600E-06 |
| 17909 | 896 | Y896H | C | 2.3e-06 | 2.4e-06 | 2.7e-06 | 2.467E-06 | 2.082E-07 | 2.300E-06 | 2.700E-06 |
| 17910 | 896 | Y896I | C | 2.5e-06 | 3.4e-06 | 2.9e-06 | 2.933E-06 | 4.509E-07 | 2.500E-06 | 3.400E-06 |
| 17911 | 896 | Y896L | C | 2.2e-06 | 3.4e-06 | 4.8e-06 | 3.467E-06 | 1.301E-06 | 2.200E-06 | 4.800E-06 |
| 17912 | 896 | Y896K | C | 2.9e-06 | 3.4e-06 | 2e-06 | 2.767E-06 | 7.095E-07 | 2.000E-06 | 3.400E-06 |
| 17913 | 896 | Y896M | C | 4.5e-06 | 2.6e-06 | 3e-06 | 3.367E-06 | 1.002E-06 | 2.600E-06 | 4.500E-06 |
| 17914 | 896 | Y896F | C | 2.2e-06 | 3.7e-06 | 3.4e-06 | 3.100E-06 | 7.937E-07 | 2.200E-06 | 3.700E-06 |
| 17915 | 896 | Y896P | C | 9.4e-07 | 3.5e-06 | 5.6e-06 | 3.347E-06 | 2.334E-06 | 9.400E-07 | 5.600E-06 |
| 17916 | 896 | Y896S | C | 2.6e-06 | 3.1e-06 | 3.1e-06 | 2.933E-06 | 2.887E-07 | 2.600E-06 | 3.100E-06 |
| 17917 | 896 | Y896T | C | 3.4e-06 | 2.5e-06 | 2.6e-06 | 2.833E-06 | 4.933E-07 | 2.500E-06 | 3.400E-06 |
| 17918 | 896 | Y896W | C | 2.3e-06 | 2.5e-06 | 2e-06 | 2.267E-06 | 2.517E-07 | 2.000E-06 | 2.500E-06 |
| 17919 | 896 | Y896Y | C | 2.3e-06 | 1e-06 | 2.7e-06 | 2.000E-06 | 8.888E-07 | 1.000E-06 | 2.700E-06 |
| 17920 | 896 | Y896V | C | 1.2e-06 | 4.3e-06 | 2.1e-06 | 2.533E-06 | 1.595E-06 | 1.200E-06 | 4.300E-06 |
| 17921 | 897 | N897A | C | 2.4e-06 | 3.1e-06 | 2.2e-06 | 2.567E-06 | 4.726E-07 | 2.200E-06 | 3.100E-06 |
| 17922 | 897 | N897R | C | 2.3e-06 | 2.1e-06 | 2.5e-06 | 2.300E-06 | 2.000E-07 | 2.100E-06 | 2.500E-06 |
| 17923 | 897 | N897N | C | 3.5e-06 | 2.9e-06 | 1.7e-06 | 2.700E-06 | 9.165E-07 | 1.700E-06 | 3.500E-06 |
| 17924 | 897 | N897D | C | 2.3e-06 | 2.5e-06 | 2.1e-06 | 2.300E-06 | 2.000E-07 | 2.100E-06 | 2.500E-06 |
| 17925 | 897 | N897C | C | 1.9e-06 | 1.2e-06 | 1.7e-06 | 1.600E-06 | 3.606E-07 | 1.200E-06 | 1.900E-06 |
| 17926 | 897 | N897E | C | 2.4e-06 | 2.2e-06 | 4.5e-06 | 3.033E-06 | 1.274E-06 | 2.200E-06 | 4.500E-06 |
| 17927 | 897 | N897Q | C | 1.2e-06 | 1.9e-06 | 1.4e-06 | 1.500E-06 | 3.606E-07 | 1.200E-06 | 1.900E-06 |
| 17928 | 897 | N897G | C | 2.2e-06 | 1.9e-06 | 3.4e-06 | 2.500E-06 | 7.937E-07 | 1.900E-06 | 3.400E-06 |
| 17929 | 897 | N897H | C | 4.8e-06 | 2.4e-06 | 3.2e-06 | 3.467E-06 | 1.222E-06 | 2.400E-06 | 4.800E-06 |
| 17930 | 897 | N897I | C | 3.3e-06 | 2.3e-06 | 3e-06 | 2.867E-06 | 5.132E-07 | 2.300E-06 | 3.300E-06 |
| 17931 | 897 | N897L | C | 3.3e-06 | 2.1e-06 | 1.3e-06 | 2.233E-06 | 1.007E-06 | 1.300E-06 | 3.300E-06 |
| 17932 | 897 | N897K | C | 2.5e-06 | 2.4e-06 | 2.4e-06 | 2.433E-06 | 5.774E-08 | 2.400E-06 | 2.500E-06 |
| 17933 | 897 | N897M | C | 2.1e-06 | 4.2e-06 | 4.3e-06 | 3.533E-06 | 1.242E-06 | 2.100E-06 | 4.300E-06 |
| 17934 | 897 | N897F | C | 3e-06 | 2e-06 | 2.8e-06 | 2.600E-06 | 5.292E-07 | 2.000E-06 | 3.000E-06 |

| | | | | | | | | | | |
|---|---|---|---|---|---|---|---|---|---|---|
| 17935 | 897 | N897P | C | 2.1e-06 | 2.5e-06 | 3.2e-06 | 2.600E-06 | 5.568E-07 | 2.100E-06 | 3.200E-06 |
| 17936 | 897 | N897S | C | 2.1e-06 | 2e-06 | 2.4e-06 | 2.167E-06 | 2.082E-07 | 2.000E-06 | 2.400E-06 |
| 17937 | 897 | N897T | C | 2e-06 | 3.3e-06 | 1.6e-06 | 2.300E-06 | 8.888E-07 | 1.600E-06 | 3.300E-06 |
| 17938 | 897 | N897W | C | 3.2e-06 | 2.1e-06 | 2.6e-06 | 2.633E-06 | 5.508E-07 | 2.100E-06 | 3.200E-06 |
| 17939 | 897 | N897Y | C | 2.7e-06 | 2.3e-06 | 1.9e-06 | 2.300E-06 | 4.000E-07 | 1.900E-06 | 2.700E-06 |
| 17940 | 897 | N897V | C | 1.6e-06 | 1.5e-06 | 1.7e-06 | 1.600E-06 | 1.000E-07 | 1.500E-06 | 1.700E-06 |
| 17941 | 898 | Q898A | C | 1.3e-06 | 2.2e-06 | 4.7e-06 | 2.733E-06 | 1.762E-06 | 1.300E-06 | 4.700E-06 |
| 17942 | 898 | Q898R | C | 2.4e-06 | 1.9e-06 | 2.5e-06 | 2.267E-06 | 3.215E-07 | 1.900E-06 | 2.500E-06 |
| 17943 | 898 | Q898N | C | 2.3e-06 | 2.4e-06 | 2e-06 | 2.233E-06 | 2.082E-07 | 2.000E-06 | 2.400E-06 |
| 17944 | 898 | Q898D | C | 1.5e-06 | 2.3e-06 | 2.6e-06 | 2.133E-06 | 5.686E-07 | 1.500E-06 | 2.600E-06 |
| 17945 | 898 | Q898C | C | 2.9e-06 | 2.4e-06 | 1.4e-06 | 2.233E-06 | 7.638E-07 | 1.400E-06 | 2.900E-06 |
| 17946 | 898 | Q898E | C | 2.7e-06 | 2.6e-06 | 1.4e-06 | 2.233E-06 | 7.234E-07 | 1.400E-06 | 2.700E-06 |
| 17947 | 898 | Q898Q | C | 2.6e-06 | 4.3e-06 | 2.4e-06 | 3.100E-06 | 1.044E-06 | 2.400E-06 | 4.300E-06 |
| 17948 | 898 | Q898G | C | 2e-06 | 4.1e-06 | 1.3e-06 | 2.467E-06 | 1.457E-06 | 1.300E-06 | 4.100E-06 |
| 17949 | 898 | Q898H | C | 3e-06 | 2.9e-06 | 1.2e-06 | 2.367E-06 | 1.012E-06 | 1.200E-06 | 3.000E-06 |
| 17950 | 898 | Q898I | C | 1.9e-06 | 2.4e-06 | 2.8e-06 | 2.367E-06 | 4.509E-07 | 1.900E-06 | 2.800E-06 |
| 17951 | 898 | Q898L | C | 1.4e-06 | 1.8e-06 | 2.6e-06 | 1.933E-06 | 6.110E-07 | 1.400E-06 | 2.600E-06 |
| 17952 | 898 | Q898K | C | 1e-06 | 1.9e-06 | 3.1e-06 | 2.000E-06 | 1.054E-06 | 1.000E-06 | 3.100E-06 |
| 17953 | 898 | Q898M | C | 1.4e-06 | 3.4e-06 | 2.1e-06 | 2.300E-06 | 1.015E-06 | 1.400E-06 | 3.400E-06 |
| 17954 | 898 | Q898F | C | 2.2e-06 | 3.2e-06 | 2.2e-06 | 2.533E-06 | 5.774E-07 | 2.200E-06 | 3.200E-06 |
| 17955 | 898 | Q898P | C | 6.1e-06 | 2.6e-06 | 4e-06 | 4.233E-06 | 1.762E-06 | 2.600E-06 | 6.100E-06 |
| 17956 | 898 | Q898S | C | 2.5e-06 | 2.2e-06 | 3.6e-06 | 2.767E-06 | 7.371E-07 | 2.200E-06 | 3.600E-06 |
| 17957 | 898 | Q898T | C | 2.8e-06 | 2.4e-06 | 2.4e-06 | 2.533E-06 | 2.309E-07 | 2.400E-06 | 2.800E-06 |
| 17958 | 898 | Q898W | C | 3.4e-06 | 3.9e-06 | 1.7e-06 | 3.000E-06 | 1.153E-06 | 1.700E-06 | 3.900E-06 |
| 17959 | 898 | Q898Y | C | 2.2e-06 | 2.2e-06 | 2.6e-06 | 2.333E-06 | 2.309E-07 | 2.200E-06 | 2.600E-06 |
| 17960 | 898 | Q898V | C | 1.8e-06 | 1.9e-06 | 3.4e-06 | 2.367E-06 | 8.963E-07 | 1.800E-06 | 3.400E-06 |
| 17961 | 899 | R899A | C | 1.6e-06 | 1.3e-06 | 1.6e-06 | 1.500E-06 | 1.732E-07 | 1.300E-06 | 1.600E-06 |
| 17962 | 899 | R899R | C | 2.6e-06 | 4.1e-06 | 2.7e-06 | 3.133E-06 | 8.386E-07 | 2.600E-06 | 4.100E-06 |
| 17963 | 899 | R899N | C | 3e-06 | 3.7e-06 | 3.5e-06 | 3.400E-06 | 3.606E-07 | 3.000E-06 | 3.700E-06 |
| 17964 | 899 | R899D | C | 2e-06 | 2.6e-06 | 3.6e-06 | 2.733E-06 | 8.083E-07 | 2.000E-06 | 3.600E-06 |
| 17965 | 899 | R899C | C | 3.1e-06 | 3.3e-06 | 2.9e-06 | 3.100E-06 | 2.000E-07 | 2.900E-06 | 3.300E-06 |
| 17966 | 899 | R899E | C | 2.3e-06 | 4.1e-06 | 1.9e-06 | 2.767E-06 | 1.172E-06 | 1.900E-06 | 4.100E-06 |
| 17967 | 899 | R899Q | C | 1.8e-06 | 3.2e-06 | 2.2e-06 | 2.400E-06 | 7.211E-07 | 1.800E-06 | 3.200E-06 |
| 17968 | 899 | R899G | C | 3.6e-06 | 3.2e-06 | 2.4e-06 | 3.067E-06 | 6.110E-07 | 2.400E-06 | 3.600E-06 |
| 17969 | 899 | R899H | C | 2.3e-06 | 3e-06 | 3e-06 | 2.767E-06 | 4.041E-07 | 2.300E-06 | 3.000E-06 |
| 17970 | 899 | R899I | C | 3.2e-06 | 1.9e-06 | 1.8e-06 | 2.300E-06 | 7.810E-07 | 1.800E-06 | 3.200E-06 |
| 17971 | 899 | R899L | C | 2e-06 | 3.2e-06 | 3.5e-06 | 2.900E-06 | 7.937E-07 | 2.000E-06 | 3.500E-06 |
| 17972 | 899 | R899K | C | 1.8e-06 | 2.7e-06 | 2.1e-06 | 2.200E-06 | 4.583E-07 | 1.800E-06 | 2.700E-06 |
| 17973 | 899 | R899M | C | 1.8e-06 | 2.3e-06 | 1.4e-06 | 1.833E-06 | 4.509E-07 | 1.400E-06 | 2.300E-06 |
| 17974 | 899 | R899F | C | 2.8e-06 | 2.2e-06 | 1.3e-06 | 2.100E-06 | 7.550E-07 | 1.300E-06 | 2.800E-06 |
| 17975 | 899 | R899P | C | 2.1e-06 | 1.6e-06 | 2.4e-06 | 2.033E-06 | 4.041E-07 | 1.600E-06 | 2.400E-06 |
| 17976 | 899 | R899S | C | 2.4e-06 | 2.1e-06 | 1.3e-06 | 1.933E-06 | 5.686E-07 | 1.300E-06 | 2.400E-06 |
| 17977 | 899 | R899T | C | 2e-06 | 2.3e-06 | 1.8e-06 | 2.033E-06 | 2.517E-07 | 1.800E-06 | 2.300E-06 |
| 17978 | 899 | R899W | C | 1.8e-06 | 2.9e-06 | 2.1e-06 | 2.267E-06 | 5.686E-07 | 1.800E-06 | 2.900E-06 |
| 17979 | 899 | R899Y | C | 9.3e-07 | 2.8e-06 | 2.6e-06 | 2.110E-06 | 1.027E-06 | 9.300E-07 | 2.800E-06 |
| 17980 | 899 | R899V | C | 3.4e-06 | 2.6e-06 | 2e-06 | 2.667E-06 | 7.024E-07 | 2.000E-06 | 3.400E-06 |
| 17981 | 900 | F900A | C | 1.6e-06 | 1.5e-06 | 3.3e-06 | 2.133E-06 | 1.012E-06 | 1.500E-06 | 3.300E-06 |
| 17982 | 900 | F900R | C | 1.3e-06 | 4.2e-06 | 2.2e-06 | 2.567E-06 | 1.484E-06 | 1.300E-06 | 4.200E-06 |
| 17983 | 900 | F900N | C | 2.1e-06 | 2.1e-06 | 2.4e-06 | 2.200E-06 | 1.732E-07 | 2.100E-06 | 2.400E-06 |
| 17984 | 900 | F900D | C | 8.8e-07 | 1.7e-06 | 6e-06 | 2.860E-06 | 2.750E-06 | 8.800E-07 | 6.000E-06 |
| 17985 | 900 | F900C | C | 2.3e-06 | 3.3e-06 | 1.5e-06 | 2.367E-06 | 9.018E-07 | 1.500E-06 | 3.300E-06 |
| 17986 | 900 | F900E | C | 2.8e-06 | 2.3e-06 | 1.8e-06 | 2.300E-06 | 5.000E-07 | 1.800E-06 | 2.800E-06 |
| 17987 | 900 | F900Q | C | 2.2e-06 | 2.2e-06 | 2.6e-06 | 2.333E-06 | 2.309E-07 | 2.200E-06 | 2.600E-06 |
| 17988 | 900 | F900G | C | 1.6e-06 | 2.6e-06 | 3e-06 | 2.400E-06 | 7.211E-07 | 1.600E-06 | 3.000E-06 |
| 17989 | 900 | F900H | C | 3.7e-06 | 3.8e-06 | 2.5e-06 | 3.333E-06 | 7.234E-07 | 2.500E-06 | 3.800E-06 |
| 17990 | 900 | F900I | C | 2.3e-06 | 2.5e-06 | 3.2e-06 | 2.667E-06 | 4.726E-07 | 2.300E-06 | 3.200E-06 |
| 17991 | 900 | F900L | C | 3.2e-06 | 1.6e-06 | 1.2e-06 | 2.000E-06 | 1.058E-06 | 1.200E-06 | 3.200E-06 |
| 17992 | 900 | F900K | C | 3e-06 | 2.9e-06 | 2.6e-06 | 2.833E-06 | 2.082E-07 | 2.600E-06 | 3.000E-06 |
| 17993 | 900 | F900M | C | 1.8e-06 | 1.6e-06 | 1.6e-06 | 1.667E-06 | 1.155E-07 | 1.600E-06 | 1.800E-06 |
| 17994 | 900 | F900F | C | 1.6e-06 | 2.7e-06 | 1.7e-06 | 2.000E-06 | 6.083E-07 | 1.600E-06 | 2.700E-06 |
| 17995 | 900 | F900P | C | 2.1e-06 | 2.4e-06 | 1.7e-06 | 2.067E-06 | 3.512E-07 | 1.700E-06 | 2.400E-06 |
| 17996 | 900 | F900S | C | 1.7e-06 | 1.4e-06 | 2.1e-06 | 1.733E-06 | 3.512E-07 | 1.400E-06 | 2.100E-06 |
| 17997 | 900 | F900T | C | 1.8e-06 | 2.5e-06 | 3.4e-06 | 2.567E-06 | 8.021E-07 | 1.800E-06 | 3.400E-06 |
| 17998 | 900 | F900W | C | 3e-06 | 2.6e-06 | 2.5e-06 | 2.700E-06 | 2.646E-07 | 2.500E-06 | 3.000E-06 |
| 17999 | 900 | F900Y | C | 2.6e-06 | 1.8e-06 | 4.2e-06 | 2.867E-06 | 1.222E-06 | 1.800E-06 | 4.200E-06 |
| 18000 | 900 | F900V | C | 1.6e-06 | 1.5e-06 | 2.7e-06 | 1.933E-06 | 6.658E-07 | 1.500E-06 | 2.700E-06 |
| 18001 | 901 | K901A | C | 2.3e-06 | 3.1e-06 | 1.7e-06 | 2.367E-06 | 7.024E-07 | 1.700E-06 | 3.100E-06 |
| 18002 | 901 | K901R | C | 2.7e-06 | 1.5e-06 | 4.5e-06 | 2.900E-06 | 1.510E-06 | 1.500E-06 | 4.500E-06 |
| 18003 | 901 | K901N | C | 2.1e-06 | 1.9e-06 | 2.3e-06 | 2.100E-06 | 2.000E-07 | 1.900E-06 | 2.300E-06 |
| 18004 | 901 | K901D | C | 3.5e-06 | 2.5e-06 | 3.4e-06 | 3.133E-06 | 5.508E-07 | 2.500E-06 | 3.500E-06 |
| 18005 | 901 | K901C | C | 3.1e-06 | 3.8e-06 | 3.4e-06 | 3.433E-06 | 3.512E-07 | 3.100E-06 | 3.800E-06 |
| 18006 | 901 | K901E | C | 3.8e-06 | 2.9e-06 | 1.9e-06 | 2.867E-06 | 9.504E-07 | 1.900E-06 | 3.800E-06 |
| 18007 | 901 | K901Q | C | 1.1e-06 | 2.7e-06 | 3e-06 | 2.267E-06 | 1.021E-06 | 1.100E-06 | 3.000E-06 |
| 18008 | 901 | K901G | C | 3.5e-06 | 2.6e-06 | 2.8e-06 | 2.967E-06 | 4.726E-07 | 2.600E-06 | 3.500E-06 |
| 18009 | 901 | K901H | C | 1.4e-06 | 2.1e-06 | 1.5e-06 | 1.667E-06 | 3.786E-07 | 1.400E-06 | 2.100E-06 |
| 18010 | 901 | K901I | C | 2.4e-06 | 2.8e-06 | 2.9e-06 | 2.700E-06 | 2.646E-07 | 2.400E-06 | 2.900E-06 |
| 18011 | 901 | K901L | C | 3e-06 | 2.8e-06 | 2.5e-06 | 2.767E-06 | 2.517E-07 | 2.500E-06 | 3.000E-06 |
| 18012 | 901 | K901K | C | 3e-06 | 1.2e-06 | 1.8e-06 | 2.000E-06 | 9.165E-07 | 1.200E-06 | 3.000E-06 |
| 18013 | 901 | K901M | C | 2.3e-06 | 1.9e-06 | 2.4e-06 | 2.200E-06 | 2.646E-07 | 1.900E-06 | 2.400E-06 |
| 18014 | 901 | K901F | C | 1.5e-06 | 2.8e-06 | 1.8e-06 | 2.033E-06 | 6.807E-07 | 1.500E-06 | 2.800E-06 |
| 18015 | 901 | K901P | C | 1.3e-06 | 2.6e-06 | 9.6e-07 | 1.620E-06 | 8.656E-07 | 9.600E-07 | 2.600E-06 |
| 18016 | 901 | K901S | C | 1.8e-06 | 3.1e-06 | 2.6e-06 | 2.500E-06 | 6.557E-07 | 1.800E-06 | 3.100E-06 |
| 18017 | 901 | K901T | C | 2.3e-06 | 2.2e-06 | 2.2e-06 | 2.233E-06 | 5.774E-08 | 2.200E-06 | 2.300E-06 |
| 18018 | 901 | K901W | C | 1.9e-06 | 1.5e-06 | 2.6e-06 | 2.000E-06 | 5.568E-07 | 1.500E-06 | 2.600E-06 |
| 18019 | 901 | K901Y | C | 1.8e-06 | 1.8e-06 | 1.8e-06 | 1.800E-06 | 0.000E+00 | 1.800E-06 | 1.800E-06 |

| | | | | | | | | | | |
|---|---|---|---|---|---|---|---|---|---|---|
| 18020 | 901 | K901V | C | 1.3e-06 | 2.5e-06 | 2.2e-06 | 2.000E-06 | 6.245E-07 | 1.300E-06 | 2.500E-06 |
| 18021 | 902 | G902A | C | 3.1e-06 | 2.1e-06 | 3.8e-06 | 3.000E-06 | 8.544E-07 | 2.100E-06 | 3.800E-06 |
| 18022 | 902 | G902R | C | 1.9e-06 | 2.3e-06 | 1.8e-06 | 2.000E-06 | 2.646E-07 | 1.800E-06 | 2.300E-06 |
| 18023 | 902 | G902N | C | 1.1e-06 | 1.6e-06 | 3.4e-06 | 2.033E-06 | 1.210E-06 | 1.100E-06 | 3.400E-06 |
| 18024 | 902 | G902D | C | 1.4e-06 | 2.6e-06 | 2.7e-06 | 2.233E-06 | 7.234E-07 | 1.400E-06 | 2.700E-06 |
| 18025 | 902 | G902C | C | 2.7e-06 | 2.7e-06 | 1.5e-06 | 2.300E-06 | 6.928E-07 | 1.500E-06 | 2.700E-06 |
| 18026 | 902 | G902E | C | 4.5e-06 | 9.2e-07 | 2.4e-06 | 2.607E-06 | 1.799E-06 | 9.200E-07 | 4.500E-06 |
| 18027 | 902 | G902Q | C | 2e-06 | 1.6e-06 | 3.5e-06 | 2.367E-06 | 1.002E-06 | 1.600E-06 | 3.500E-06 |
| 18028 | 902 | G902G | C | 1.7e-06 | 2.1e-06 | 4e-06 | 2.600E-06 | 1.229E-06 | 1.700E-06 | 4.000E-06 |
| 18029 | 902 | G902H | C | 2e-06 | 3.4e-06 | 2.6e-06 | 2.667E-06 | 7.024E-07 | 2.000E-06 | 3.400E-06 |
| 18030 | 902 | G902I | C | 3.5e-06 | 1.6e-06 | 3e-06 | 2.700E-06 | 9.849E-07 | 1.600E-06 | 3.500E-06 |
| 18031 | 902 | G902L | C | 3.9e-06 | 3.1e-06 | 1.9e-06 | 2.967E-06 | 1.007E-06 | 1.900E-06 | 3.900E-06 |
| 18032 | 902 | G902K | C | 1.8e-06 | 1.8e-06 | 4.4e-06 | 2.667E-06 | 1.501E-06 | 1.800E-06 | 4.400E-06 |
| 18033 | 902 | G902M | C | 2.4e-06 | 2.7e-06 | 3e-06 | 2.700E-06 | 3.000E-07 | 2.400E-06 | 3.000E-06 |
| 18034 | 902 | G902F | C | 4.3e-06 | 2.9e-06 | 2e-06 | 3.067E-06 | 1.159E-06 | 2.000E-06 | 4.300E-06 |
| 18035 | 902 | G902P | C | 3e-06 | 2.8e-06 | 1.5e-06 | 2.433E-06 | 8.145E-07 | 1.500E-06 | 3.000E-06 |
| 18036 | 902 | G902S | C | 3.5e-06 | 3.6e-06 | 2.3e-06 | 3.133E-06 | 7.234E-07 | 2.300E-06 | 3.600E-06 |
| 18037 | 902 | G902T | C | 3.2e-06 | 2.8e-06 | 4e-06 | 3.333E-06 | 6.110E-07 | 2.800E-06 | 4.000E-06 |
| 18038 | 902 | G902W | C | 1.4e-06 | 2.3e-06 | 2.1e-06 | 1.933E-06 | 4.726E-07 | 1.400E-06 | 2.300E-06 |
| 18039 | 902 | G902Y | C | 2.2e-06 | 1.1e-06 | 2.5e-06 | 1.933E-06 | 7.371E-07 | 1.100E-06 | 2.500E-06 |
| 18040 | 902 | G902V | C | 2.5e-06 | 1.4e-06 | 1.5e-06 | 1.800E-06 | 6.083E-07 | 1.400E-06 | 2.500E-06 |
| 18041 | 903 | R903A | C | 2.7e-06 | 2.9e-06 | 2.1e-06 | 2.567E-06 | 4.163E-07 | 2.100E-06 | 2.900E-06 |
| 18042 | 903 | R903R | C | 1.9e-06 | 2.5e-06 | 3.2e-06 | 2.533E-06 | 6.506E-07 | 1.900E-06 | 3.200E-06 |
| 18043 | 903 | R903N | C | 1.3e-06 | 3.2e-06 | 2.6e-06 | 2.367E-06 | 9.713E-07 | 1.300E-06 | 3.200E-06 |
| 18044 | 903 | R903D | C | 1.8e-06 | 2e-06 | 2e-06 | 1.933E-06 | 1.155E-07 | 1.800E-06 | 2.000E-06 |
| 18045 | 903 | R903C | C | 1.4e-06 | 3.8e-06 | 2.6e-06 | 2.600E-06 | 1.200E-06 | 1.400E-06 | 3.800E-06 |
| 18046 | 903 | R903E | C | 2.3e-06 | 4.6e-06 | 2.5e-06 | 3.133E-06 | 1.274E-06 | 2.300E-06 | 4.600E-06 |
| 18047 | 903 | R903Q | C | 2.2e-06 | 2.4e-06 | 1.6e-06 | 2.067E-06 | 4.163E-07 | 1.600E-06 | 2.400E-06 |
| 18048 | 903 | R903G | C | 1.6e-06 | 2.6e-06 | 2.2e-06 | 2.133E-06 | 5.033E-07 | 1.600E-06 | 2.600E-06 |
| 18049 | 903 | R903H | C | 2.8e-06 | 2.7e-06 | 2.2e-06 | 2.567E-06 | 3.215E-07 | 2.200E-06 | 2.800E-06 |
| 18050 | 903 | R903I | C | 2.7e-06 | 1.7e-06 | 1.7e-06 | 2.033E-06 | 5.774E-07 | 1.700E-06 | 2.700E-06 |
| 18051 | 903 | R903L | C | 1.7e-06 | 1.8e-06 | 2.4e-06 | 1.967E-06 | 3.786E-07 | 1.700E-06 | 2.400E-06 |
| 18052 | 903 | R903K | C | 1.5e-06 | 3.3e-06 | 2.8e-06 | 2.533E-06 | 9.292E-07 | 1.500E-06 | 3.300E-06 |
| 18053 | 903 | R903M | C | 3.3e-06 | 2.8e-06 | 3.1e-06 | 3.067E-06 | 2.517E-07 | 2.800E-06 | 3.300E-06 |
| 18054 | 903 | R903F | C | 2.4e-06 | 1.4e-06 | 2.3e-06 | 2.033E-06 | 5.508E-07 | 1.400E-06 | 2.400E-06 |
| 18055 | 903 | R903P | C | 1.6e-06 | 2.6e-06 | 2.3e-06 | 2.167E-06 | 5.132E-07 | 1.600E-06 | 2.600E-06 |
| 18056 | 903 | R903S | C | 2.9e-06 | 5.2e-06 | 5.2e-06 | 4.433E-06 | 1.328E-06 | 2.900E-06 | 5.200E-06 |
| 18057 | 903 | R903T | C | 4.9e-06 | 1.7e-06 | 6.5e-06 | 4.367E-06 | 2.444E-06 | 1.700E-06 | 6.500E-06 |
| 18058 | 903 | R903W | C | 2.9e-06 | 2e-06 | 1.1e-06 | 2.000E-06 | 9.000E-07 | 1.100E-06 | 2.900E-06 |
| 18059 | 903 | R903Y | C | 2.5e-06 | 2.6e-06 | 3.6e-06 | 2.900E-06 | 6.083E-07 | 2.500E-06 | 3.600E-06 |
| 18060 | 903 | R903V | C | 2.9e-06 | 2.2e-06 | 1.7e-06 | 2.267E-06 | 6.028E-07 | 1.700E-06 | 2.900E-06 |
| 18061 | 904 | F904A | C | 3.4e-06 | 1.8e-06 | 3.1e-06 | 2.767E-06 | 8.505E-07 | 1.800E-06 | 3.400E-06 |
| 18062 | 904 | F904R | C | 3.2e-06 | 2e-06 | 1.9e-06 | 2.367E-06 | 7.234E-07 | 1.900E-06 | 3.200E-06 |
| 18063 | 904 | F904N | C | 1.8e-06 | 2e-06 | 2.2e-06 | 2.000E-06 | 2.000E-07 | 1.800E-06 | 2.200E-06 |
| 18064 | 904 | F904D | C | 1.3e-06 | 1.9e-06 | 4.3e-06 | 2.500E-06 | 1.587E-06 | 1.300E-06 | 4.300E-06 |
| 18065 | 904 | F904C | C | 2.1e-06 | 1.5e-06 | 1.8e-06 | 1.800E-06 | 3.000E-07 | 1.500E-06 | 2.100E-06 |
| 18066 | 904 | F904E | C | 2.4e-06 | 2.5e-06 | 3.7e-06 | 2.867E-06 | 7.234E-07 | 2.400E-06 | 3.700E-06 |
| 18067 | 904 | F904Q | C | 2.9e-06 | 1.5e-06 | 1.4e-06 | 1.933E-06 | 8.386E-07 | 1.400E-06 | 2.900E-06 |
| 18068 | 904 | F904G | C | 1.9e-06 | 2.3e-06 | 2.1e-06 | 2.100E-06 | 2.000E-07 | 1.900E-06 | 2.300E-06 |
| 18069 | 904 | F904H | C | 1.2e-06 | 3.4e-06 | 3.4e-06 | 2.667E-06 | 1.270E-06 | 1.200E-06 | 3.400E-06 |
| 18070 | 904 | F904I | C | 1.8e-06 | 1.8e-06 | 2.5e-06 | 2.033E-06 | 4.041E-07 | 1.800E-06 | 2.500E-06 |
| 18071 | 904 | F904L | C | 2.8e-06 | 1.7e-06 | 2.6e-06 | 2.367E-06 | 5.859E-07 | 1.700E-06 | 2.800E-06 |
| 18072 | 904 | F904K | C | 1.4e-06 | 5.2e-06 | 1.8e-06 | 2.800E-06 | 2.088E-06 | 1.400E-06 | 5.200E-06 |
| 18073 | 904 | F904M | C | 3.3e-06 | 2.7e-06 | 2.7e-06 | 2.900E-06 | 3.464E-07 | 2.700E-06 | 3.300E-06 |
| 18074 | 904 | F904F | C | 3.2e-06 | 3.5e-06 | 1.6e-06 | 2.767E-06 | 1.021E-06 | 1.600E-06 | 3.500E-06 |
| 18075 | 904 | F904P | C | 3.5e-06 | 2.5e-06 | 3.5e-06 | 3.167E-06 | 5.774E-07 | 2.500E-06 | 3.500E-06 |
| 18076 | 904 | F904S | C | 8.6e-07 | 3.4e-06 | 2.7e-06 | 2.320E-06 | 1.312E-06 | 8.600E-07 | 3.400E-06 |
| 18077 | 904 | F904T | C | 2e-06 | 2.2e-06 | 2.3e-06 | 2.167E-06 | 1.528E-07 | 2.000E-06 | 2.300E-06 |
| 18078 | 904 | F904W | C | 2.9e-06 | 1.9e-06 | 2.9e-06 | 2.567E-06 | 5.774E-07 | 1.900E-06 | 2.900E-06 |
| 18079 | 904 | F904Y | C | 2.6e-06 | 3.9e-06 | 1.6e-06 | 2.700E-06 | 1.153E-06 | 1.600E-06 | 3.900E-06 |
| 18080 | 904 | F904V | C | 5.8e-06 | 2.8e-06 | 2.4e-06 | 3.667E-06 | 1.858E-06 | 2.400E-06 | 5.800E-06 |
| 18081 | 905 | T905A | C | 2.9e-06 | 2.3e-06 | 2.7e-06 | 2.633E-06 | 3.055E-07 | 2.300E-06 | 2.900E-06 |
| 18082 | 905 | T905R | C | 1.9e-06 | 1.3e-06 | 2.4e-06 | 1.867E-06 | 5.508E-07 | 1.300E-06 | 2.400E-06 |
| 18083 | 905 | T905N | C | 2.7e-06 | 2.4e-06 | 2e-06 | 2.367E-06 | 3.512E-07 | 2.000E-06 | 2.700E-06 |
| 18084 | 905 | T905D | C | 4.4e-06 | 2.8e-06 | 2.8e-06 | 3.333E-06 | 9.238E-07 | 2.800E-06 | 4.400E-06 |
| 18085 | 905 | T905C | C | 2.7e-06 | 5.4e-06 | 2.9e-06 | 3.667E-06 | 1.504E-06 | 2.700E-06 | 5.400E-06 |
| 18086 | 905 | T905E | C | 3.2e-06 | 2.6e-06 | 1.6e-06 | 2.467E-06 | 8.083E-07 | 1.600E-06 | 3.200E-06 |
| 18087 | 905 | T905Q | C | 1e-06 | 2.9e-06 | 3.3e-06 | 2.400E-06 | 1.229E-06 | 1.000E-06 | 3.300E-06 |
| 18088 | 905 | T905G | C | 2.2e-06 | 2.3e-06 | 1.4e-06 | 1.967E-06 | 4.933E-07 | 1.400E-06 | 2.300E-06 |
| 18089 | 905 | T905H | C | 1.6e-06 | 2.2e-06 | 3.7e-06 | 2.500E-06 | 1.082E-06 | 1.600E-06 | 3.700E-06 |
| 18090 | 905 | T905I | C | 3.8e-06 | 2.2e-06 | 2e-06 | 2.667E-06 | 9.866E-07 | 2.000E-06 | 3.800E-06 |
| 18091 | 905 | T905L | C | 2.1e-06 | 1.5e-06 | 2.3e-06 | 1.967E-06 | 4.163E-07 | 1.500E-06 | 2.300E-06 |
| 18092 | 905 | T905K | C | 2.5e-06 | 2.7e-06 | 2.3e-06 | 2.500E-06 | 2.000E-07 | 2.300E-06 | 2.700E-06 |
| 18093 | 905 | T905M | C | 2.8e-06 | 2.3e-06 | 2.5e-06 | 2.533E-06 | 2.517E-07 | 2.300E-06 | 2.800E-06 |
| 18094 | 905 | T905F | C | 2.7e-06 | 2.2e-06 | 2.2e-06 | 2.367E-06 | 2.887E-07 | 2.200E-06 | 2.700E-06 |
| 18095 | 905 | T905P | C | 3.8e-06 | 2.6e-06 | 1.8e-06 | 2.733E-06 | 1.007E-06 | 1.800E-06 | 3.800E-06 |
| 18096 | 905 | T905S | C | 4.5e-06 | 1.4e-06 | 2e-06 | 2.633E-06 | 1.644E-06 | 1.400E-06 | 4.500E-06 |
| 18097 | 905 | T905T | C | 1.9e-06 | 4.6e-06 | 3.2e-06 | 3.233E-06 | 1.350E-06 | 1.900E-06 | 4.600E-06 |
| 18098 | 905 | T905W | C | 2.2e-06 | 2.2e-06 | 2.1e-06 | 2.167E-06 | 5.774E-08 | 2.100E-06 | 2.200E-06 |
| 18099 | 905 | T905Y | C | 1.2e-06 | 3.8e-06 | 1.5e-06 | 2.167E-06 | 1.422E-06 | 1.200E-06 | 3.800E-06 |
| 18100 | 905 | T905V | C | 2e-06 | 2.8e-06 | 2.5e-06 | 2.433E-06 | 4.041E-07 | 2.000E-06 | 2.800E-06 |
| 18101 | 906 | L906A | C | 2.8e-06 | 1.9e-06 | 3.9e-06 | 2.867E-06 | 1.002E-06 | 1.900E-06 | 3.900E-06 |
| 18102 | 906 | L906R | C | 4.7e-06 | 2.2e-06 | 3.2e-06 | 3.367E-06 | 1.258E-06 | 2.200E-06 | 4.700E-06 |
| 18103 | 906 | L906N | C | 2.3e-06 | 1.6e-06 | 1.5e-06 | 1.800E-06 | 4.359E-07 | 1.500E-06 | 2.300E-06 |
| 18104 | 906 | L906D | C | 1.8e-06 | 3.6e-06 | 3.6e-06 | 3.000E-06 | 1.039E-06 | 1.800E-06 | 3.600E-06 |

| | | | | | | | | | | |
|---|---|---|---|---|---|---|---|---|---|---|
| 18105 | 906 | L906C | C | 2.9e-06 | 8.5e-07 | 2e-06 | 1.917E-06 | 1.028E-06 | 8.500E-07 | 2.900E-06 |
| 18106 | 906 | L906E | C | 2.8e-06 | 2.4e-06 | 2.1e-06 | 2.433E-06 | 3.512E-07 | 2.100E-06 | 2.800E-06 |
| 18107 | 906 | L906Q | C | 2.6e-06 | 2.5e-06 | 1.8e-06 | 2.300E-06 | 4.359E-07 | 1.800E-06 | 2.600E-06 |
| 18108 | 906 | L906G | C | 2.6e-06 | 2.2e-06 | 1.8e-06 | 2.200E-06 | 4.000E-07 | 1.800E-06 | 2.600E-06 |
| 18109 | 906 | L906H | C | 1.4e-06 | 2.5e-06 | 2.1e-06 | 2.000E-06 | 5.568E-07 | 1.400E-06 | 2.500E-06 |
| 18110 | 906 | L906I | C | 2e-06 | 2.3e-06 | 2.1e-06 | 2.133E-06 | 1.528E-07 | 2.000E-06 | 2.300E-06 |
| 18111 | 906 | L906L | C | 2.4e-06 | 3.9e-06 | 2.2e-06 | 2.833E-06 | 9.292E-07 | 2.200E-06 | 3.900E-06 |
| 18112 | 906 | L906K | C | 2.7e-06 | 2.8e-06 | 2.1e-06 | 2.533E-06 | 3.786E-07 | 2.100E-06 | 2.800E-06 |
| 18113 | 906 | L906M | C | 2e-06 | 2.2e-06 | 1.1e-06 | 1.767E-06 | 5.859E-07 | 1.100E-06 | 2.200E-06 |
| 18114 | 906 | L906F | C | 3.3e-06 | 1.8e-06 | 2.1e-06 | 2.400E-06 | 7.937E-07 | 1.800E-06 | 3.300E-06 |
| 18115 | 906 | L906P | C | 1.7e-06 | 1.7e-06 | 3.4e-06 | 2.267E-06 | 9.815E-07 | 1.700E-06 | 3.400E-06 |
| 18116 | 906 | L906S | C | 2e-06 | 1.2e-06 | 2e-06 | 1.733E-06 | 4.619E-07 | 1.200E-06 | 2.000E-06 |
| 18117 | 906 | L906T | C | 4e-06 | 3.9e-06 | 2e-06 | 3.300E-06 | 1.127E-06 | 2.000E-06 | 4.000E-06 |
| 18118 | 906 | L906W | C | 5.1e-06 | 1.4e-06 | 2.5e-06 | 3.000E-06 | 1.900E-06 | 1.400E-06 | 5.100E-06 |
| 18119 | 906 | L906Y | C | 2.6e-06 | 1.5e-06 | 2.8e-06 | 2.300E-06 | 7.000E-07 | 1.500E-06 | 2.800E-06 |
| 18120 | 906 | L906V | C | 2.6e-06 | 1.7e-06 | 2.4e-06 | 2.233E-06 | 4.726E-07 | 1.700E-06 | 2.600E-06 |
| 18121 | 907 | S907A | C | 2.3e-06 | 2e-06 | 1.8e-06 | 2.033E-06 | 2.517E-07 | 1.800E-06 | 2.300E-06 |
| 18122 | 907 | S907R | C | 2.6e-06 | 1.8e-06 | 2e-06 | 2.133E-06 | 4.163E-07 | 1.800E-06 | 2.600E-06 |
| 18123 | 907 | S907N | C | 2.1e-06 | 2e-06 | 2.7e-06 | 2.267E-06 | 3.786E-07 | 2.000E-06 | 2.700E-06 |
| 18124 | 907 | S907D | C | 1.8e-06 | 2e-06 | 1.9e-06 | 1.900E-06 | 1.000E-07 | 1.800E-06 | 2.000E-06 |
| 18125 | 907 | S907C | C | 1.2e-06 | 1.5e-06 | 1.4e-06 | 1.367E-06 | 1.528E-07 | 1.200E-06 | 1.500E-06 |
| 18126 | 907 | S907E | C | 1.7e-06 | 2.7e-06 | 2.9e-06 | 2.433E-06 | 6.429E-07 | 1.700E-06 | 2.900E-06 |
| 18127 | 907 | S907Q | C | 2.2e-06 | 3.1e-06 | 3e-06 | 2.767E-06 | 4.933E-07 | 2.200E-06 | 3.100E-06 |
| 18128 | 907 | S907G | C | 3.3e-06 | 2.6e-06 | 2.7e-06 | 2.867E-06 | 3.786E-07 | 2.600E-06 | 3.300E-06 |
| 18129 | 907 | S907H | C | 2.3e-06 | 4.1e-06 | 2.1e-06 | 2.833E-06 | 1.102E-06 | 2.100E-06 | 4.100E-06 |
| 18130 | 907 | S907I | C | 2.7e-06 | 3.1e-06 | 1.3e-06 | 2.367E-06 | 9.452E-07 | 1.300E-06 | 3.100E-06 |
| 18131 | 907 | S907L | C | 2.2e-06 | 2.7e-06 | 3.7e-06 | 2.867E-06 | 7.638E-07 | 2.200E-06 | 3.700E-06 |
| 18132 | 907 | S907K | C | 2.6e-06 | 2.7e-06 | 2.2e-06 | 2.500E-06 | 2.646E-07 | 2.200E-06 | 2.700E-06 |
| 18133 | 907 | S907M | C | 2.8e-06 | 1.5e-06 | 3.4e-06 | 2.567E-06 | 9.713E-07 | 1.500E-06 | 3.400E-06 |
| 18134 | 907 | S907F | C | 2.1e-06 | 2.7e-06 | 2.5e-06 | 2.433E-06 | 3.055E-07 | 2.100E-06 | 2.700E-06 |
| 18135 | 907 | S907P | C | 2.1e-06 | 3.1e-06 | 2.6e-06 | 2.600E-06 | 5.000E-07 | 2.100E-06 | 3.100E-06 |
| 18136 | 907 | S907S | C | 1.9e-06 | 1.6e-06 | 3.2e-06 | 2.233E-06 | 8.505E-07 | 1.600E-06 | 3.200E-06 |
| 18137 | 907 | S907T | C | 1.9e-06 | 1.3e-06 | 1.6e-06 | 1.600E-06 | 3.000E-07 | 1.300E-06 | 1.900E-06 |
| 18138 | 907 | S907W | C | 2.5e-06 | 2.5e-06 | 1.8e-06 | 2.267E-06 | 4.041E-07 | 1.800E-06 | 2.500E-06 |
| 18139 | 907 | S907Y | C | 3.5e-06 | 1.6e-06 | 2.7e-06 | 2.600E-06 | 9.539E-07 | 1.600E-06 | 3.500E-06 |
| 18140 | 907 | S907V | C | 2.1e-06 | 2.3e-06 | 2.6e-06 | 2.333E-06 | 2.517E-07 | 2.100E-06 | 2.600E-06 |
| 18141 | 908 | V908A | C | 3.4e-06 | 1.6e-06 | 2.1e-06 | 2.367E-06 | 9.292E-07 | 1.600E-06 | 3.400E-06 |
| 18142 | 908 | V908R | C | 2e-06 | 1.9e-06 | 2.6e-06 | 2.167E-06 | 3.786E-07 | 1.900E-06 | 2.600E-06 |
| 18143 | 908 | V908N | C | 2.8e-06 | 3e-06 | 1.8e-06 | 2.533E-06 | 6.429E-07 | 1.800E-06 | 3.000E-06 |
| 18144 | 908 | V908D | C | 3.2e-06 | 2.5e-06 | 2.7e-06 | 2.800E-06 | 3.606E-07 | 2.500E-06 | 3.200E-06 |
| 18145 | 908 | V908C | C | 2.9e-06 | 1.6e-06 | 1.5e-06 | 2.000E-06 | 7.810E-07 | 1.500E-06 | 2.900E-06 |
| 18146 | 908 | V908E | C | 1.2e-06 | 1.7e-06 | 2.5e-06 | 1.800E-06 | 6.557E-07 | 1.200E-06 | 2.500E-06 |
| 18147 | 908 | V908Q | C | 1.9e-06 | 3.2e-06 | 2.3e-06 | 2.467E-06 | 6.658E-07 | 1.900E-06 | 3.200E-06 |
| 18148 | 908 | V908G | C | 2.6e-06 | 3e-06 | 2e-06 | 2.533E-06 | 5.033E-07 | 2.000E-06 | 3.000E-06 |
| 18149 | 908 | V908H | C | 3.4e-06 | 3.6e-06 | 8.4e-07 | 2.613E-06 | 1.539E-06 | 8.400E-07 | 3.600E-06 |
| 18150 | 908 | V908I | C | 1.6e-06 | 4.2e-06 | 1.7e-06 | 2.500E-06 | 1.473E-06 | 1.600E-06 | 4.200E-06 |
| 18151 | 908 | V908L | C | 2.3e-06 | 3.9e-06 | 2.5e-06 | 2.900E-06 | 8.718E-07 | 2.300E-06 | 3.900E-06 |
| 18152 | 908 | V908K | C | 3.3e-06 | 3e-06 | 2.8e-06 | 3.033E-06 | 2.517E-07 | 2.800E-06 | 3.300E-06 |
| 18153 | 908 | V908M | C | 8e-07 | 3.9e-06 | 2.2e-06 | 2.300E-06 | 1.552E-06 | 8.000E-07 | 3.900E-06 |
| 18154 | 908 | V908F | C | 1.3e-06 | 1.9e-06 | 5.8e-06 | 3.000E-06 | 2.443E-06 | 1.300E-06 | 5.800E-06 |
| 18155 | 908 | V908P | C | 1.6e-06 | 2.6e-06 | 1.5e-06 | 1.900E-06 | 6.083E-07 | 1.500E-06 | 2.600E-06 |
| 18156 | 908 | V908S | C | 3.1e-06 | 2.4e-06 | 1.6e-06 | 2.367E-06 | 7.506E-07 | 1.600E-06 | 3.100E-06 |
| 18157 | 908 | V908T | C | 1.4e-06 | 3.6e-06 | 1.4e-06 | 2.133E-06 | 1.270E-06 | 1.400E-06 | 3.600E-06 |
| 18158 | 908 | V908W | C | 1.9e-06 | 4.6e-06 | 2.7e-06 | 3.067E-06 | 1.387E-06 | 1.900E-06 | 4.600E-06 |
| 18159 | 908 | V908Y | C | 1.8e-06 | 2.1e-06 | 3e-06 | 2.300E-06 | 6.245E-07 | 1.800E-06 | 3.000E-06 |
| 18160 | 908 | V908V | C | 2.5e-06 | 2.5e-06 | 2.4e-06 | 2.467E-06 | 5.774E-08 | 2.400E-06 | 2.500E-06 |
| 18161 | 909 | D909A | C | 2.1e-06 | 3.2e-06 | 2.6e-06 | 2.633E-06 | 5.508E-07 | 2.100E-06 | 3.200E-06 |
| 18162 | 909 | D909R | C | 1.6e-06 | 2.1e-06 | 3.1e-06 | 2.267E-06 | 7.638E-07 | 1.600E-06 | 3.100E-06 |
| 18163 | 909 | D909N | C | 2.5e-06 | 1.6e-06 | 2.3e-06 | 2.133E-06 | 4.726E-07 | 1.600E-06 | 2.500E-06 |
| 18164 | 909 | D909D | C | 2.2e-06 | 1.3e-06 | 1.7e-06 | 1.733E-06 | 4.509E-07 | 1.300E-06 | 2.200E-06 |
| 18165 | 909 | D909C | C | 2.1e-06 | 2.8e-06 | 2.1e-06 | 2.333E-06 | 4.041E-07 | 2.100E-06 | 2.800E-06 |
| 18166 | 909 | D909E | C | 3.1e-06 | 1.8e-06 | 1.9e-06 | 2.267E-06 | 7.234E-07 | 1.800E-06 | 3.100E-06 |
| 18167 | 909 | D909Q | C | 2.5e-06 | 2.1e-06 | 2.1e-06 | 2.233E-06 | 2.309E-07 | 2.100E-06 | 2.500E-06 |
| 18168 | 909 | D909G | C | 1.9e-06 | 1.8e-06 | 2e-06 | 1.900E-06 | 1.000E-07 | 1.800E-06 | 2.000E-06 |
| 18169 | 909 | D909H | C | 2.5e-06 | 1.9e-06 | 1.1e-06 | 1.833E-06 | 7.024E-07 | 1.100E-06 | 2.500E-06 |
| 18170 | 909 | D909I | C | 1.2e-06 | 2.4e-06 | 1.6e-06 | 1.733E-06 | 6.110E-07 | 1.200E-06 | 2.400E-06 |
| 18171 | 909 | D909L | C | 2.8e-06 | 1.8e-06 | 2.7e-06 | 2.433E-06 | 5.508E-07 | 1.800E-06 | 2.800E-06 |
| 18172 | 909 | D909K | C | 1.7e-06 | 2.5e-06 | 3.5e-06 | 2.567E-06 | 9.018E-07 | 1.700E-06 | 3.500E-06 |
| 18173 | 909 | D909M | C | 2.4e-06 | 2.7e-06 | 2.8e-06 | 2.633E-06 | 2.082E-07 | 2.400E-06 | 2.800E-06 |
| 18174 | 909 | D909F | C | 3.4e-06 | 2.3e-06 | 1.7e-06 | 2.467E-06 | 8.622E-07 | 1.700E-06 | 3.400E-06 |
| 18175 | 909 | D909P | C | 2.8e-06 | 1.9e-06 | 2.7e-06 | 2.467E-06 | 4.933E-07 | 1.900E-06 | 2.800E-06 |
| 18176 | 909 | D909S | C | 2.5e-06 | 1.1e-06 | 3.6e-06 | 2.400E-06 | 1.253E-06 | 1.100E-06 | 3.600E-06 |
| 18177 | 909 | D909T | C | 1.6e-06 | 2.1e-06 | 1.2e-06 | 1.633E-06 | 4.509E-07 | 1.200E-06 | 2.100E-06 |
| 18178 | 909 | D909W | C | 1.2e-06 | 1.8e-06 | 3.4e-06 | 2.133E-06 | 1.137E-06 | 1.200E-06 | 3.400E-06 |
| 18179 | 909 | D909Y | C | 2.5e-06 | 2e-06 | 2.8e-06 | 2.433E-06 | 4.041E-07 | 2.000E-06 | 2.800E-06 |
| 18180 | 909 | D909V | C | 4.6e-06 | 1.4e-06 | 4.3e-06 | 3.433E-06 | 1.767E-06 | 1.400E-06 | 4.600E-06 |
| 18181 | 910 | R910A | C | 2.5e-06 | 1.2e-06 | 2.7e-06 | 2.133E-06 | 8.145E-07 | 1.200E-06 | 2.700E-06 |
| 18182 | 910 | R910R | C | 2.2e-06 | 2.1e-06 | 3.3e-06 | 2.533E-06 | 6.658E-07 | 2.100E-06 | 3.300E-06 |
| 18183 | 910 | R910N | C | 1.4e-06 | 3.6e-06 | 2.3e-06 | 2.433E-06 | 1.106E-06 | 1.400E-06 | 3.600E-06 |
| 18184 | 910 | R910D | C | 1.6e-06 | 3e-06 | 1.1e-06 | 1.900E-06 | 9.849E-07 | 1.100E-06 | 3.000E-06 |
| 18185 | 910 | R910C | C | 2.2e-06 | 1.3e-06 | 2.3e-06 | 1.933E-06 | 5.508E-07 | 1.300E-06 | 2.300E-06 |
| 18186 | 910 | R910E | C | 1.1e-06 | 3.5e-06 | 3.4e-06 | 2.667E-06 | 1.358E-06 | 1.100E-06 | 3.500E-06 |
| 18187 | 910 | R910Q | C | 1.6e-06 | 1.2e-06 | 3.1e-06 | 1.967E-06 | 1.002E-06 | 1.200E-06 | 3.100E-06 |
| 18188 | 910 | R910G | C | 1.6e-06 | 2.1e-06 | 1.4e-06 | 1.700E-06 | 3.606E-07 | 1.400E-06 | 2.100E-06 |
| 18189 | 910 | R910H | C | 3.1e-06 | 2.6e-06 | 2.4e-06 | 2.700E-06 | 3.606E-07 | 2.400E-06 | 3.100E-06 |

| | | | | | | | | | | |
|---|---|---|---|---|---|---|---|---|---|---|
| 18190 | 910 | R910I | C | 1.7e-06 | 2.3e-06 | 5.3e-06 | 3.100E-06 | 1.929E-06 | 1.700E-06 | 5.300E-06 |
| 18191 | 910 | R910L | C | 3.8e-06 | 3.8e-06 | 2e-06 | 3.200E-06 | 1.039E-06 | 2.000E-06 | 3.800E-06 |
| 18192 | 910 | R910K | C | 2.8e-06 | 1.7e-06 | 9.4e-07 | 1.813E-06 | 9.352E-07 | 9.400E-07 | 2.800E-06 |
| 18193 | 910 | R910M | C | 2.1e-06 | 2.8e-06 | 1.4e-06 | 2.100E-06 | 7.000E-07 | 1.400E-06 | 2.800E-06 |
| 18194 | 910 | R910F | C | 2.6e-06 | 2.3e-06 | 2.7e-06 | 2.533E-06 | 2.082E-07 | 2.300E-06 | 2.700E-06 |
| 18195 | 910 | R910P | C | 2.6e-06 | 1.6e-06 | 2.6e-06 | 2.267E-06 | 5.774E-07 | 1.600E-06 | 2.600E-06 |
| 18196 | 910 | R910S | C | 1.3e-06 | 2.2e-06 | 3.2e-06 | 2.233E-06 | 9.504E-07 | 1.300E-06 | 3.200E-06 |
| 18197 | 910 | R910T | C | 1.3e-06 | 2.2e-06 | 2.2e-06 | 1.900E-06 | 5.196E-07 | 1.300E-06 | 2.200E-06 |
| 18198 | 910 | R910W | C | 2.2e-06 | 2.8e-06 | 1.8e-06 | 2.267E-06 | 5.033E-07 | 1.800E-06 | 2.800E-06 |
| 18199 | 910 | R910Y | C | 1.1e-06 | 2.1e-06 | 3.4e-06 | 2.200E-06 | 1.153E-06 | 1.100E-06 | 3.400E-06 |
| 18200 | 910 | R910V | C | 2.1e-06 | 2.4e-06 | 3.5e-06 | 2.667E-06 | 7.371E-07 | 2.100E-06 | 3.500E-06 |
| 18201 | 911 | S911A | C | 3.6e-06 | 3e-06 | 2.5e-06 | 3.033E-06 | 5.508E-07 | 2.500E-06 | 3.600E-06 |
| 18202 | 911 | S911R | C | 2e-06 | 2.8e-06 | 2.4e-06 | 2.400E-06 | 4.000E-07 | 2.000E-06 | 2.800E-06 |
| 18203 | 911 | S911N | C | 3.5e-06 | 3.3e-06 | 2.6e-06 | 3.133E-06 | 4.726E-07 | 2.600E-06 | 3.500E-06 |
| 18204 | 911 | S911D | C | 3.4e-06 | 3.3e-06 | 5e-06 | 3.900E-06 | 9.539E-07 | 3.300E-06 | 5.000E-06 |
| 18205 | 911 | S911C | C | 2.2e-06 | 3.8e-06 | 2.7e-06 | 2.900E-06 | 8.185E-07 | 2.200E-06 | 3.800E-06 |
| 18206 | 911 | S911E | C | 3.9e-06 | 1.5e-06 | 4.2e-06 | 3.200E-06 | 1.480E-06 | 1.500E-06 | 4.200E-06 |
| 18207 | 911 | S911Q | C | 1.6e-06 | 1.4e-06 | 2.8e-06 | 1.933E-06 | 7.572E-07 | 1.400E-06 | 2.800E-06 |
| 18208 | 911 | S911G | C | 3.5e-06 | 1.7e-06 | 1.9e-06 | 2.367E-06 | 9.866E-07 | 1.700E-06 | 3.500E-06 |
| 18209 | 911 | S911H | C | 2.4e-06 | 2.7e-06 | 2.5e-06 | 2.533E-06 | 1.528E-07 | 2.400E-06 | 2.700E-06 |
| 18210 | 911 | S911I | C | 2.7e-06 | 2e-06 | 2.6e-06 | 2.433E-06 | 3.786E-07 | 2.000E-06 | 2.700E-06 |
| 18211 | 911 | S911L | C | 3.3e-06 | 2.8e-06 | 2.3e-06 | 2.800E-06 | 5.000E-07 | 2.300E-06 | 3.300E-06 |
| 18212 | 911 | S911K | C | 3.4e-06 | 2.3e-06 | 1.9e-06 | 2.533E-06 | 7.767E-07 | 1.900E-06 | 3.400E-06 |
| 18213 | 911 | S911M | C | 2.9e-06 | 2.6e-06 | 2.1e-06 | 2.533E-06 | 4.041E-07 | 2.100E-06 | 2.900E-06 |
| 18214 | 911 | S911F | C | 2.9e-06 | 1.8e-06 | 2.1e-06 | 2.267E-06 | 5.686E-07 | 1.800E-06 | 2.900E-06 |
| 18215 | 911 | S911P | C | 2.4e-06 | 2e-06 | 2.7e-06 | 2.367E-06 | 3.512E-07 | 2.000E-06 | 2.700E-06 |
| 18216 | 911 | S911S | C | 2.4e-06 | 3.6e-06 | 3.3e-06 | 3.100E-06 | 6.245E-07 | 2.400E-06 | 3.600E-06 |
| 18217 | 911 | S911T | C | 4.1e-06 | 3.3e-06 | 2.6e-06 | 3.333E-06 | 7.506E-07 | 2.600E-06 | 4.100E-06 |
| 18218 | 911 | S911W | C | 1.9e-06 | 1.9e-06 | 1.8e-06 | 1.867E-06 | 5.774E-08 | 1.800E-06 | 1.900E-06 |
| 18219 | 911 | S911Y | C | 1.8e-06 | 1.5e-06 | 3.5e-06 | 2.267E-06 | 1.079E-06 | 1.500E-06 | 3.500E-06 |
| 18220 | 911 | S911V | C | 2.4e-06 | 1.7e-06 | 2.4e-06 | 2.167E-06 | 4.041E-07 | 1.700E-06 | 2.400E-06 |
| 18221 | 912 | K912A | C | 1.1e-06 | 2.5e-06 | 2.2e-06 | 1.933E-06 | 7.371E-07 | 1.100E-06 | 2.500E-06 |
| 18222 | 912 | K912R | C | 2.6e-06 | 2.3e-06 | 2.3e-06 | 2.400E-06 | 1.732E-07 | 2.300E-06 | 2.600E-06 |
| 18223 | 912 | K912N | C | 1e-06 | 3.2e-06 | 2.2e-06 | 2.133E-06 | 1.102E-06 | 1.000E-06 | 3.200E-06 |
| 18224 | 912 | K912D | C | 1.8e-06 | 2.6e-06 | 2.3e-06 | 2.233E-06 | 4.041E-07 | 1.800E-06 | 2.600E-06 |
| 18225 | 912 | K912C | C | 3.1e-06 | 3.3e-06 | 3.1e-06 | 3.167E-06 | 1.155E-07 | 3.100E-06 | 3.300E-06 |
| 18226 | 912 | K912E | C | 1.9e-06 | 1.5e-06 | 2.7e-06 | 2.033E-06 | 6.110E-07 | 1.500E-06 | 2.700E-06 |
| 18227 | 912 | K912Q | C | 3.1e-06 | 1.9e-06 | 2.4e-06 | 2.467E-06 | 6.028E-07 | 1.900E-06 | 3.100E-06 |
| 18228 | 912 | K912G | C | 4.8e-06 | 2.5e-06 | 3.5e-06 | 3.600E-06 | 1.153E-06 | 2.500E-06 | 4.800E-06 |
| 18229 | 912 | K912H | C | 1.9e-06 | 1.5e-06 | 3.7e-06 | 2.367E-06 | 1.172E-06 | 1.500E-06 | 3.700E-06 |
| 18230 | 912 | K912I | C | 1.1e-06 | 3.7e-06 | 1.8e-06 | 2.200E-06 | 1.345E-06 | 1.100E-06 | 3.700E-06 |
| 18231 | 912 | K912L | C | 4.3e-06 | 2.3e-06 | 2.2e-06 | 2.933E-06 | 1.185E-06 | 2.200E-06 | 4.300E-06 |
| 18232 | 912 | K912K | C | 2.1e-06 | 2.4e-06 | 2.4e-06 | 2.300E-06 | 1.732E-07 | 2.100E-06 | 2.400E-06 |
| 18233 | 912 | K912M | C | 1.8e-06 | 2.2e-06 | 1.6e-06 | 1.867E-06 | 3.055E-07 | 1.600E-06 | 2.200E-06 |
| 18234 | 912 | K912F | C | 1.3e-06 | 2.7e-06 | 2.3e-06 | 2.100E-06 | 7.211E-07 | 1.300E-06 | 2.700E-06 |
| 18235 | 912 | K912P | C | 2.1e-06 | 1.9e-06 | 2.2e-06 | 2.067E-06 | 1.528E-07 | 1.900E-06 | 2.200E-06 |
| 18236 | 912 | K912S | C | 1.9e-06 | 2.2e-06 | 2e-06 | 2.033E-06 | 1.528E-07 | 1.900E-06 | 2.200E-06 |
| 18237 | 912 | K912T | C | 3e-06 | 3e-06 | 2e-06 | 2.667E-06 | 5.774E-07 | 2.000E-06 | 3.000E-06 |
| 18238 | 912 | K912W | C | 2.2e-06 | 1.9e-06 | 1.9e-06 | 2.000E-06 | 1.732E-07 | 1.900E-06 | 2.200E-06 |
| 18239 | 912 | K912Y | C | 2e-06 | 3.1e-06 | 3.9e-06 | 3.000E-06 | 9.539E-07 | 2.000E-06 | 3.900E-06 |
| 18240 | 912 | K912V | C | 1.7e-06 | 2.5e-06 | 1.5e-06 | 1.900E-06 | 5.292E-07 | 1.500E-06 | 2.500E-06 |
| 18241 | 913 | N913A | C | 1.6e-06 | 3.2e-06 | 4.1e-06 | 2.967E-06 | 1.266E-06 | 1.600E-06 | 4.100E-06 |
| 18242 | 913 | N913R | C | 2.7e-06 | 1.5e-06 | 2.3e-06 | 2.167E-06 | 6.110E-07 | 1.500E-06 | 2.700E-06 |
| 18243 | 913 | N913N | C | 2e-06 | 3.3e-06 | 2.5e-06 | 2.600E-06 | 6.557E-07 | 2.000E-06 | 3.300E-06 |
| 18244 | 913 | N913D | C | 5e-06 | 3.4e-06 | 1.8e-06 | 3.400E-06 | 1.600E-06 | 1.800E-06 | 5.000E-06 |
| 18245 | 913 | N913C | C | 2e-06 | 3e-06 | 3.1e-06 | 2.700E-06 | 6.083E-07 | 2.000E-06 | 3.100E-06 |
| 18246 | 913 | N913E | C | 2.1e-06 | 2e-06 | 2e-06 | 2.033E-06 | 5.774E-08 | 2.000E-06 | 2.100E-06 |
| 18247 | 913 | N913Q | C | 3.3e-06 | 1.7e-06 | 1.7e-06 | 2.233E-06 | 9.238E-07 | 1.700E-06 | 3.300E-06 |
| 18248 | 913 | N913G | C | 2.5e-06 | 2.6e-06 | 2.4e-06 | 2.500E-06 | 1.000E-07 | 2.400E-06 | 2.600E-06 |
| 18249 | 913 | N913H | C | 2.1e-06 | 2.5e-06 | 1.2e-06 | 1.933E-06 | 6.658E-07 | 1.200E-06 | 2.500E-06 |
| 18250 | 913 | N913I | C | 2.8e-06 | 1.9e-06 | 2.4e-06 | 2.367E-06 | 4.509E-07 | 1.900E-06 | 2.800E-06 |
| 18251 | 913 | N913L | C | 3.6e-06 | 2.3e-06 | 3.9e-06 | 3.267E-06 | 8.505E-07 | 2.300E-06 | 3.900E-06 |
| 18252 | 913 | N913K | C | 3.2e-06 | 3.2e-06 | 1.2e-06 | 2.533E-06 | 1.155E-06 | 1.200E-06 | 3.200E-06 |
| 18253 | 913 | N913M | C | 1.9e-06 | 2.2e-06 | 1.9e-06 | 2.000E-06 | 1.732E-07 | 1.900E-06 | 2.200E-06 |
| 18254 | 913 | N913F | C | 2.3e-06 | 2e-06 | 2.3e-06 | 2.200E-06 | 1.732E-07 | 2.000E-06 | 2.300E-06 |
| 18255 | 913 | N913P | C | 1.9e-06 | 2.2e-06 | 3.9e-06 | 2.667E-06 | 1.079E-06 | 1.900E-06 | 3.900E-06 |
| 18256 | 913 | N913S | C | 2.7e-06 | 2.8e-06 | 1.5e-06 | 2.333E-06 | 7.234E-07 | 1.500E-06 | 2.800E-06 |
| 18257 | 913 | N913T | C | 2.1e-06 | 2.3e-06 | 2.5e-06 | 2.300E-06 | 2.000E-07 | 2.100E-06 | 2.500E-06 |
| 18258 | 913 | N913W | C | 2.4e-06 | 2.4e-06 | 2.1e-06 | 2.300E-06 | 1.732E-07 | 2.100E-06 | 2.400E-06 |
| 18259 | 913 | N913Y | C | 2e-06 | 3.4e-06 | 2.7e-06 | 2.700E-06 | 7.000E-07 | 2.000E-06 | 3.400E-06 |
| 18260 | 913 | N913V | C | 2.4e-06 | 3.5e-06 | 2.1e-06 | 2.667E-06 | 7.371E-07 | 2.100E-06 | 3.500E-06 |
| 18261 | 914 | T914A | C | 2.1e-06 | 2.9e-06 | 2.6e-06 | 2.533E-06 | 4.041E-07 | 2.100E-06 | 2.900E-06 |
| 18262 | 914 | T914R | C | 2.4e-06 | 2.5e-06 | 2.3e-06 | 2.400E-06 | 1.000E-07 | 2.300E-06 | 2.500E-06 |
| 18263 | 914 | T914N | C | 3e-06 | 1.4e-06 | 2.6e-06 | 2.333E-06 | 8.327E-07 | 1.400E-06 | 3.000E-06 |
| 18264 | 914 | T914D | C | 2e-06 | 2e-06 | 2.6e-06 | 2.200E-06 | 3.464E-07 | 2.000E-06 | 2.600E-06 |
| 18265 | 914 | T914C | C | 1.9e-06 | 3.7e-06 | 2.1e-06 | 2.567E-06 | 9.866E-07 | 1.900E-06 | 3.700E-06 |
| 18266 | 914 | T914E | C | 2.3e-06 | 3.1e-06 | 1.8e-06 | 2.400E-06 | 6.557E-07 | 1.800E-06 | 3.100E-06 |
| 18267 | 914 | T914Q | C | 3.2e-06 | 2.5e-06 | 2.5e-06 | 2.733E-06 | 4.041E-07 | 2.500E-06 | 3.200E-06 |
| 18268 | 914 | T914G | C | 1.9e-06 | 3.6e-06 | 1.6e-06 | 2.367E-06 | 1.079E-06 | 1.600E-06 | 3.600E-06 |
| 18269 | 914 | T914H | C | 3.3e-06 | 2.7e-06 | 2.8e-06 | 2.933E-06 | 3.215E-07 | 2.700E-06 | 3.300E-06 |
| 18270 | 914 | T914I | C | 2.4e-06 | 2.6e-06 | 1.7e-06 | 2.233E-06 | 4.726E-07 | 1.700E-06 | 2.600E-06 |
| 18271 | 914 | T914L | C | 2.3e-06 | 2.6e-06 | 2e-06 | 2.300E-06 | 3.000E-07 | 2.000E-06 | 2.600E-06 |
| 18272 | 914 | T914K | C | 2.7e-06 | 3.1e-06 | 1.8e-06 | 2.533E-06 | 6.658E-07 | 1.800E-06 | 3.100E-06 |
| 18273 | 914 | T914M | C | 2e-06 | 3.5e-06 | 2.3e-06 | 2.600E-06 | 7.937E-07 | 2.000E-06 | 3.500E-06 |
| 18274 | 914 | T914F | C | 1.8e-06 | 3.2e-06 | 2.5e-06 | 2.500E-06 | 7.000E-07 | 1.800E-06 | 3.200E-06 |

| | | | | | | | | | | |
|---|---|---|---|---|---|---|---|---|---|---|
| 18275 | 914 | T914P | C | 1.3e-06 | 3e-06 | 2.6e-06 | 2.300E-06 | 8.888E-07 | 1.300E-06 | 3.000E-06 |
| 18276 | 914 | T914S | C | 2.3e-06 | 2.3e-06 | 2.7e-06 | 2.433E-06 | 2.309E-07 | 2.300E-06 | 2.700E-06 |
| 18277 | 914 | T914T | C | 2.8e-06 | 2.4e-06 | 3.9e-06 | 3.033E-06 | 7.767E-07 | 2.400E-06 | 3.900E-06 |
| 18278 | 914 | T914W | C | 2.2e-06 | 2.5e-06 | 2.1e-06 | 2.267E-06 | 2.082E-07 | 2.100E-06 | 2.500E-06 |
| 18279 | 914 | T914Y | C | 3.1e-06 | 1.8e-06 | 1.7e-06 | 2.200E-06 | 7.810E-07 | 1.700E-06 | 3.100E-06 |
| 18280 | 914 | T914V | C | 1.9e-06 | 2.3e-06 | 1.8e-06 | 2.000E-06 | 2.646E-07 | 1.800E-06 | 2.300E-06 |
| 18281 | 915 | L915A | C | 2.6e-06 | 4e-06 | 3.8e-06 | 3.467E-06 | 7.572E-07 | 2.600E-06 | 4.000E-06 |
| 18282 | 915 | L915R | C | 2.2e-06 | 1.8e-06 | 2.8e-06 | 2.267E-06 | 5.033E-07 | 1.800E-06 | 2.800E-06 |
| 18283 | 915 | L915N | C | 1.3e-06 | 4e-06 | 2.6e-06 | 2.633E-06 | 1.350E-06 | 1.300E-06 | 4.000E-06 |
| 18284 | 915 | L915D | C | 2.5e-06 | 3.5e-06 | 2.3e-06 | 2.767E-06 | 6.429E-07 | 2.300E-06 | 3.500E-06 |
| 18285 | 915 | L915C | C | 2.7e-06 | 7.8e-07 | 3.1e-06 | 2.193E-06 | 1.240E-06 | 7.800E-07 | 3.100E-06 |
| 18286 | 915 | L915E | C | 2.3e-06 | 2.5e-06 | 2.2e-06 | 2.333E-06 | 1.528E-07 | 2.200E-06 | 2.500E-06 |
| 18287 | 915 | L915Q | C | 2.3e-06 | 2.8e-06 | 1.7e-06 | 2.267E-06 | 5.508E-07 | 1.700E-06 | 2.800E-06 |
| 18288 | 915 | L915G | C | 2.3e-06 | 3.9e-06 | 3.2e-06 | 3.133E-06 | 8.021E-07 | 2.300E-06 | 3.900E-06 |
| 18289 | 915 | L915H | C | 2.9e-06 | 1.1e-06 | 4.4e-06 | 2.800E-06 | 1.652E-06 | 1.100E-06 | 4.400E-06 |
| 18290 | 915 | L915I | C | 2e-06 | 3.6e-06 | 3.3e-06 | 2.967E-06 | 8.505E-07 | 2.000E-06 | 3.600E-06 |
| 18291 | 915 | L915L | C | 2.9e-06 | 2.2e-06 | 1.6e-06 | 2.233E-06 | 6.506E-07 | 1.600E-06 | 2.900E-06 |
| 18292 | 915 | L915K | C | 1.7e-06 | 1.9e-06 | 1.2e-06 | 1.600E-06 | 3.606E-07 | 1.200E-06 | 1.900E-06 |
| 18293 | 915 | L915M | C | 1.7e-06 | 2.6e-06 | 3.2e-06 | 2.500E-06 | 7.550E-07 | 1.700E-06 | 3.200E-06 |
| 18294 | 915 | L915F | C | 6.7e-06 | 4.6e-06 | 2.9e-06 | 4.733E-06 | 1.904E-06 | 2.900E-06 | 6.700E-06 |
| 18295 | 915 | L915P | C | 2.7e-06 | 7.5e-06 | 1.9e-06 | 4.033E-06 | 3.029E-06 | 1.900E-06 | 7.500E-06 |
| 18296 | 915 | L915S | C | 2.6e-06 | 1.5e-06 | 2.6e-06 | 2.233E-06 | 6.351E-07 | 1.500E-06 | 2.600E-06 |
| 18297 | 915 | L915T | C | 3.2e-06 | 3e-06 | 1.5e-06 | 2.567E-06 | 9.292E-07 | 1.500E-06 | 3.200E-06 |
| 18298 | 915 | L915W | C | 9.7e-07 | 3.2e-06 | 1.4e-06 | 1.857E-06 | 1.183E-06 | 9.700E-07 | 3.200E-06 |
| 18299 | 915 | L915Y | C | 2.5e-06 | 1.9e-06 | 1.2e-06 | 1.867E-06 | 6.506E-07 | 1.200E-06 | 2.500E-06 |
| 18300 | 915 | L915V | C | 5.7e-06 | 2.5e-06 | 3.3e-06 | 3.833E-06 | 1.665E-06 | 2.500E-06 | 5.700E-06 |
| 18301 | 916 | Y916A | C | 1.7e-06 | 2.4e-06 | 2.5e-06 | 2.200E-06 | 4.359E-07 | 1.700E-06 | 2.500E-06 |
| 18302 | 916 | Y916R | C | 2.1e-06 | 3.2e-06 | 1.8e-06 | 2.367E-06 | 7.371E-07 | 1.800E-06 | 3.200E-06 |
| 18303 | 916 | Y916N | C | 2.1e-06 | 3.6e-06 | 2.9e-06 | 2.867E-06 | 7.506E-07 | 2.100E-06 | 3.600E-06 |
| 18304 | 916 | Y916D | C | 2e-06 | 2.7e-06 | 2.2e-06 | 2.300E-06 | 3.606E-07 | 2.000E-06 | 2.700E-06 |
| 18305 | 916 | Y916C | C | 2.6e-06 | 2.8e-06 | 2.4e-06 | 2.600E-06 | 2.000E-07 | 2.400E-06 | 2.800E-06 |
| 18306 | 916 | Y916E | C | 1.8e-06 | 3.5e-06 | 1.8e-06 | 2.367E-06 | 9.815E-07 | 1.800E-06 | 3.500E-06 |
| 18307 | 916 | Y916Q | C | 1.7e-06 | 1.7e-06 | 1.5e-06 | 1.633E-06 | 1.155E-07 | 1.500E-06 | 1.700E-06 |
| 18308 | 916 | Y916G | C | 2.3e-06 | 2e-06 | 3.6e-06 | 2.633E-06 | 8.505E-07 | 2.000E-06 | 3.600E-06 |
| 18309 | 916 | Y916H | C | 2.2e-06 | 3e-06 | 2.2e-06 | 2.467E-06 | 4.619E-07 | 2.200E-06 | 3.000E-06 |
| 18310 | 916 | Y916I | C | 2.6e-06 | 2e-06 | 4.5e-06 | 3.033E-06 | 1.305E-06 | 2.000E-06 | 4.500E-06 |
| 18311 | 916 | Y916L | C | 1.6e-06 | 3.9e-06 | 2.7e-06 | 2.733E-06 | 1.150E-06 | 1.600E-06 | 3.900E-06 |
| 18312 | 916 | Y916K | C | 1.7e-06 | 4.2e-06 | 1.4e-06 | 2.433E-06 | 1.537E-06 | 1.400E-06 | 4.200E-06 |
| 18313 | 916 | Y916M | C | 2.1e-06 | 1.5e-06 | 3.5e-06 | 2.367E-06 | 1.026E-06 | 1.500E-06 | 3.500E-06 |
| 18314 | 916 | Y916F | C | 2.9e-06 | 2.2e-06 | 2.5e-06 | 2.533E-06 | 3.512E-07 | 2.200E-06 | 2.900E-06 |
| 18315 | 916 | Y916P | C | 2.5e-06 | 2.5e-06 | 2.8e-06 | 2.600E-06 | 1.732E-07 | 2.500E-06 | 2.800E-06 |
| 18316 | 916 | Y916S | C | 2.4e-06 | 1.2e-06 | 1.6e-06 | 1.733E-06 | 6.110E-07 | 1.200E-06 | 2.400E-06 |
| 18317 | 916 | Y916T | C | 4.2e-06 | 2.9e-06 | 1.8e-06 | 2.967E-06 | 1.201E-06 | 1.800E-06 | 4.200E-06 |
| 18318 | 916 | Y916W | C | 1.6e-06 | 2.4e-06 | 2.7e-06 | 2.233E-06 | 5.686E-07 | 1.600E-06 | 2.700E-06 |
| 18319 | 916 | Y916Y | C | 2.5e-06 | 1.6e-06 | 2.6e-06 | 2.233E-06 | 5.508E-07 | 1.600E-06 | 2.600E-06 |
| 18320 | 916 | Y916V | C | 2.9e-06 | 2.3e-06 | 3.6e-06 | 2.933E-06 | 6.506E-07 | 2.300E-06 | 3.600E-06 |
| 18321 | 917 | L917A | C | 2.1e-06 | 4.2e-06 | 2e-06 | 2.767E-06 | 1.242E-06 | 2.000E-06 | 4.200E-06 |
| 18322 | 917 | L917R | C | 2.7e-06 | 4.4e-06 | 2e-06 | 3.033E-06 | 1.234E-06 | 2.000E-06 | 4.400E-06 |
| 18323 | 917 | L917N | C | 3.1e-06 | 1.5e-06 | 3.1e-06 | 2.567E-06 | 9.238E-07 | 1.500E-06 | 3.100E-06 |
| 18324 | 917 | L917D | C | 4.1e-06 | 2.2e-06 | 2.2e-06 | 2.833E-06 | 1.097E-06 | 2.200E-06 | 4.100E-06 |
| 18325 | 917 | L917C | C | 2.3e-06 | 3e-06 | 1.3e-06 | 2.200E-06 | 8.544E-07 | 1.300E-06 | 3.000E-06 |
| 18326 | 917 | L917E | C | 2.1e-06 | 2.6e-06 | 2.7e-06 | 2.467E-06 | 3.215E-07 | 2.100E-06 | 2.700E-06 |
| 18327 | 917 | L917Q | C | 1.8e-06 | 1.5e-06 | 2.1e-06 | 1.800E-06 | 3.000E-07 | 1.500E-06 | 2.100E-06 |
| 18328 | 917 | L917G | C | 3.5e-06 | 3e-06 | 2.1e-06 | 2.867E-06 | 7.095E-07 | 2.100E-06 | 3.500E-06 |
| 18329 | 917 | L917H | C | 2e-06 | 1.8e-06 | 2.7e-06 | 2.167E-06 | 4.726E-07 | 1.800E-06 | 2.700E-06 |
| 18330 | 917 | L917I | C | 2.2e-06 | 2.3e-06 | 3e-06 | 2.500E-06 | 4.359E-07 | 2.200E-06 | 3.000E-06 |
| 18331 | 917 | L917L | C | 1.9e-06 | 3.1e-06 | 3.3e-06 | 2.767E-06 | 7.572E-07 | 1.900E-06 | 3.300E-06 |
| 18332 | 917 | L917K | C | 2e-06 | 2.7e-06 | 1.7e-06 | 2.133E-06 | 5.132E-07 | 1.700E-06 | 2.700E-06 |
| 18333 | 917 | L917M | C | 1.9e-06 | 3.6e-06 | 1.4e-06 | 2.300E-06 | 1.153E-06 | 1.400E-06 | 3.600E-06 |
| 18334 | 917 | L917F | C | 3.1e-06 | 4.1e-06 | 2e-06 | 3.067E-06 | 1.050E-06 | 2.000E-06 | 4.100E-06 |
| 18335 | 917 | L917P | C | 3.5e-06 | 2.1e-06 | 1.4e-06 | 2.333E-06 | 1.069E-06 | 1.400E-06 | 3.500E-06 |
| 18336 | 917 | L917S | C | 2.8e-06 | 2.8e-06 | 3e-06 | 2.867E-06 | 1.155E-07 | 2.800E-06 | 3.000E-06 |
| 18337 | 917 | L917T | C | 2.6e-06 | 2.4e-06 | 2.3e-06 | 2.433E-06 | 1.528E-07 | 2.300E-06 | 2.600E-06 |
| 18338 | 917 | L917W | C | 1.4e-06 | 2.5e-06 | 1.5e-06 | 1.800E-06 | 6.083E-07 | 1.400E-06 | 2.500E-06 |
| 18339 | 917 | L917Y | C | 1.9e-06 | 2e-06 | 2.2e-06 | 2.033E-06 | 1.528E-07 | 1.900E-06 | 2.200E-06 |
| 18340 | 917 | L917V | C | 3.2e-06 | 2.1e-06 | 2.5e-06 | 2.600E-06 | 5.568E-07 | 2.100E-06 | 3.200E-06 |
| 18341 | 918 | Q918A | C | 3e-06 | 2e-06 | 2.3e-06 | 2.433E-06 | 5.132E-07 | 2.000E-06 | 3.000E-06 |
| 18342 | 918 | Q918R | C | 2.2e-06 | 2.5e-06 | 3.3e-06 | 2.667E-06 | 5.686E-07 | 2.200E-06 | 3.300E-06 |
| 18343 | 918 | Q918N | C | 1.7e-06 | 1.3e-06 | 2.9e-06 | 1.967E-06 | 8.327E-07 | 1.300E-06 | 2.900E-06 |
| 18344 | 918 | Q918D | C | 2e-06 | 3.1e-06 | 2.4e-06 | 2.500E-06 | 5.568E-07 | 2.000E-06 | 3.100E-06 |
| 18345 | 918 | Q918C | C | 1.6e-06 | 2.2e-06 | 2.5e-06 | 2.100E-06 | 4.583E-07 | 1.600E-06 | 2.500E-06 |
| 18346 | 918 | Q918E | C | 1.8e-06 | 2e-06 | 1.7e-06 | 1.833E-06 | 1.528E-07 | 1.700E-06 | 2.000E-06 |
| 18347 | 918 | Q918Q | C | 2.1e-06 | 2.7e-06 | 1.9e-06 | 2.233E-06 | 4.163E-07 | 1.900E-06 | 2.700E-06 |
| 18348 | 918 | Q918G | C | 1.7e-06 | 2.5e-06 | 3.2e-06 | 2.467E-06 | 7.506E-07 | 1.700E-06 | 3.200E-06 |
| 18349 | 918 | Q918H | C | 2.5e-06 | 2.9e-06 | 1.8e-06 | 2.400E-06 | 5.568E-07 | 1.800E-06 | 2.900E-06 |
| 18350 | 918 | Q918I | C | 3.4e-06 | 2.6e-06 | 3.1e-06 | 3.033E-06 | 4.041E-07 | 2.600E-06 | 3.400E-06 |
| 18351 | 918 | Q918L | C | 2.7e-06 | 2.2e-06 | 5.3e-06 | 3.400E-06 | 1.664E-06 | 2.200E-06 | 5.300E-06 |
| 18352 | 918 | Q918K | C | 2.6e-06 | 2.2e-06 | 1.9e-06 | 2.233E-06 | 3.512E-07 | 1.900E-06 | 2.600E-06 |
| 18353 | 918 | Q918M | C | 3.4e-06 | 1.8e-06 | 1.6e-06 | 2.267E-06 | 9.866E-07 | 1.600E-06 | 3.400E-06 |
| 18354 | 918 | Q918F | C | 2.2e-06 | 2.3e-06 | 4.1e-06 | 2.867E-06 | 1.069E-06 | 2.200E-06 | 4.100E-06 |
| 18355 | 918 | Q918P | C | 1.6e-06 | 2e-06 | 2.3e-06 | 1.967E-06 | 3.512E-07 | 1.600E-06 | 2.300E-06 |
| 18356 | 918 | Q918S | C | 2.8e-06 | 2e-06 | 2.8e-06 | 2.533E-06 | 4.619E-07 | 2.000E-06 | 2.800E-06 |
| 18357 | 918 | Q918T | C | 2.4e-06 | 1.6e-06 | 2.5e-06 | 2.167E-06 | 4.933E-07 | 1.600E-06 | 2.500E-06 |
| 18358 | 918 | Q918W | C | 1.9e-06 | 2.5e-06 | 2.9e-06 | 2.433E-06 | 5.033E-07 | 1.900E-06 | 2.900E-06 |
| 18359 | 918 | Q918Y | C | 4e-06 | 2.8e-06 | 2.6e-06 | 3.133E-06 | 7.572E-07 | 2.600E-06 | 4.000E-06 |

| | | | | | | | | | | |
|---|---|---|---|---|---|---|---|---|---|---|
| 18360 | 918 | Q918V | C | 2.6e-06 | 2.5e-06 | 2.5e-06 | 2.533E-06 | 5.774E-08 | 2.500E-06 | 2.600E-06 |
| 18361 | 919 | M919A | C | 2.6e-06 | 2.3e-06 | 3.9e-06 | 2.933E-06 | 8.505E-07 | 2.300E-06 | 3.900E-06 |
| 18362 | 919 | M919R | C | 1.4e-06 | 2.3e-06 | 2.5e-06 | 2.067E-06 | 5.859E-07 | 1.400E-06 | 2.500E-06 |
| 18363 | 919 | M919N | C | 1.2e-06 | 5.4e-06 | 2.4e-06 | 3.000E-06 | 2.163E-06 | 1.200E-06 | 5.400E-06 |
| 18364 | 919 | M919D | C | 4.1e-06 | 1.9e-06 | 3.1e-06 | 3.033E-06 | 1.102E-06 | 1.900E-06 | 4.100E-06 |
| 18365 | 919 | M919C | C | 2e-06 | 2.6e-06 | 2.3e-06 | 2.300E-06 | 3.000E-07 | 2.000E-06 | 2.600E-06 |
| 18366 | 919 | M919E | C | 2.7e-06 | 2.1e-06 | 1.6e-06 | 2.133E-06 | 5.508E-07 | 1.600E-06 | 2.700E-06 |
| 18367 | 919 | M919Q | C | 1.6e-06 | 2.3e-06 | 1.8e-06 | 1.900E-06 | 3.606E-07 | 1.600E-06 | 2.300E-06 |
| 18368 | 919 | M919G | C | 1.2e-06 | 5.2e-06 | 4.9e-06 | 3.767E-06 | 2.228E-06 | 1.200E-06 | 5.200E-06 |
| 18369 | 919 | M919H | C | 3.1e-06 | 1.8e-06 | 1.8e-06 | 2.233E-06 | 7.506E-07 | 1.800E-06 | 3.100E-06 |
| 18370 | 919 | M919I | C | 2.7e-06 | 1.4e-06 | 2.8e-06 | 2.300E-06 | 7.810E-07 | 1.400E-06 | 2.800E-06 |
| 18371 | 919 | M919L | C | 1.9e-06 | 2.1e-06 | 2.7e-06 | 2.233E-06 | 4.163E-07 | 1.900E-06 | 2.700E-06 |
| 18372 | 919 | M919K | C | 2.5e-06 | 1e-06 | 1.6e-06 | 1.700E-06 | 7.550E-07 | 1.000E-06 | 2.500E-06 |
| 18373 | 919 | M919M | C | 3.6e-06 | 3.3e-06 | 2.8e-06 | 3.233E-06 | 4.041E-07 | 2.800E-06 | 3.600E-06 |
| 18374 | 919 | M919F | C | 9.7e-07 | 2e-06 | 1.9e-06 | 1.623E-06 | 5.680E-07 | 9.700E-07 | 2.000E-06 |
| 18375 | 919 | M919P | C | 3.6e-06 | 2.1e-06 | 1.3e-06 | 2.333E-06 | 1.168E-06 | 1.300E-06 | 3.600E-06 |
| 18376 | 919 | M919S | C | 2.8e-06 | 2.1e-06 | 1.9e-06 | 2.267E-06 | 4.726E-07 | 1.900E-06 | 2.800E-06 |
| 18377 | 919 | M919T | C | 2.7e-06 | 3e-06 | 1.7e-06 | 2.467E-06 | 6.807E-07 | 1.700E-06 | 3.000E-06 |
| 18378 | 919 | M919W | C | 2.4e-06 | 2.2e-06 | 1.7e-06 | 2.100E-06 | 3.606E-07 | 1.700E-06 | 2.400E-06 |
| 18379 | 919 | M919Y | C | 2.8e-06 | 4e-06 | 2.7e-06 | 3.167E-06 | 7.234E-07 | 2.700E-06 | 4.000E-06 |
| 18380 | 919 | M919V | C | 1.6e-06 | 1.1e-06 | 2.9e-06 | 1.867E-06 | 9.292E-07 | 1.100E-06 | 2.900E-06 |
| 18381 | 920 | N920A | C | 1.3e-06 | 9e-07 | 2.6e-06 | 1.600E-06 | 8.888E-07 | 9.000E-07 | 2.600E-06 |
| 18382 | 920 | N920R | C | 3.8e-06 | 2.2e-06 | 2.3e-06 | 2.767E-06 | 8.963E-07 | 2.200E-06 | 3.800E-06 |
| 18383 | 920 | N920N | C | 2.4e-06 | 2.8e-06 | 1.7e-06 | 2.300E-06 | 5.568E-07 | 1.700E-06 | 2.800E-06 |
| 18384 | 920 | N920D | C | 3.2e-06 | 2.4e-06 | 3.2e-06 | 2.933E-06 | 4.619E-07 | 2.400E-06 | 3.200E-06 |
| 18385 | 920 | N920C | C | 6.1e-06 | 2.6e-06 | 2.2e-06 | 3.633E-06 | 2.146E-06 | 2.200E-06 | 6.100E-06 |
| 18386 | 920 | N920E | C | 1.5e-06 | 2.5e-06 | 2.1e-06 | 2.033E-06 | 5.033E-07 | 1.500E-06 | 2.500E-06 |
| 18387 | 920 | N920Q | C | 3.3e-06 | 2.3e-06 | 1.8e-06 | 2.467E-06 | 7.638E-07 | 1.800E-06 | 3.300E-06 |
| 18388 | 920 | N920G | C | 3.2e-06 | 3.3e-06 | 2.4e-06 | 2.967E-06 | 4.933E-07 | 2.400E-06 | 3.300E-06 |
| 18389 | 920 | N920H | C | 1.7e-06 | 2.6e-06 | 3.1e-06 | 2.467E-06 | 7.095E-07 | 1.700E-06 | 3.100E-06 |
| 18390 | 920 | N920I | C | 2.8e-06 | 3.2e-06 | 1.8e-06 | 2.600E-06 | 7.211E-07 | 1.800E-06 | 3.200E-06 |
| 18391 | 920 | N920L | C | 2.1e-06 | 2e-06 | 2.5e-06 | 2.200E-06 | 2.646E-07 | 2.000E-06 | 2.500E-06 |
| 18392 | 920 | N920K | C | 2.7e-06 | 3.5e-06 | 3.1e-06 | 3.100E-06 | 4.000E-07 | 2.700E-06 | 3.500E-06 |
| 18393 | 920 | N920M | C | 2.1e-06 | 2.5e-06 | 3.4e-06 | 2.667E-06 | 6.658E-07 | 2.100E-06 | 3.400E-06 |
| 18394 | 920 | N920F | C | 3e-06 | 2.9e-06 | 3e-06 | 2.967E-06 | 5.774E-08 | 2.900E-06 | 3.000E-06 |
| 18395 | 920 | N920P | C | 2.5e-06 | 4.2e-06 | 2.5e-06 | 3.067E-06 | 9.815E-07 | 2.500E-06 | 4.200E-06 |
| 18396 | 920 | N920S | C | 1.7e-06 | 2.2e-06 | 2.4e-06 | 2.100E-06 | 3.606E-07 | 1.700E-06 | 2.400E-06 |
| 18397 | 920 | N920T | C | 3.1e-06 | 2.1e-06 | 3.3e-06 | 2.833E-06 | 6.429E-07 | 2.100E-06 | 3.300E-06 |
| 18398 | 920 | N920W | C | 3e-06 | 1.5e-06 | 2.7e-06 | 2.400E-06 | 7.937E-07 | 1.500E-06 | 3.000E-06 |
| 18399 | 920 | N920Y | C | 2.9e-06 | 2.8e-06 | 2.4e-06 | 2.700E-06 | 2.646E-07 | 2.400E-06 | 2.900E-06 |
| 18400 | 920 | N920V | C | 1.9e-06 | 1.9e-06 | 3.5e-06 | 2.433E-06 | 9.238E-07 | 1.900E-06 | 3.500E-06 |
| 18401 | 921 | S921A | C | 2.8e-06 | 2.6e-06 | 3e-06 | 2.800E-06 | 2.000E-07 | 2.600E-06 | 3.000E-06 |
| 18402 | 921 | S921R | C | 3.6e-06 | 3.5e-06 | 1.7e-06 | 2.933E-06 | 1.069E-06 | 1.700E-06 | 3.600E-06 |
| 18403 | 921 | S921N | C | 2.6e-06 | 3.1e-06 | 1.8e-06 | 2.500E-06 | 6.557E-07 | 1.800E-06 | 3.100E-06 |
| 18404 | 921 | S921D | C | 1.6e-06 | 3.3e-06 | 3.5e-06 | 2.800E-06 | 1.044E-06 | 1.600E-06 | 3.500E-06 |
| 18405 | 921 | S921C | C | 1.2e-06 | 3e-06 | 1.1e-06 | 1.767E-06 | 1.069E-06 | 1.100E-06 | 3.000E-06 |
| 18406 | 921 | S921E | C | 3.1e-06 | 2.8e-06 | 3.3e-06 | 3.067E-06 | 2.517E-07 | 2.800E-06 | 3.300E-06 |
| 18407 | 921 | S921Q | C | 3e-06 | 1.9e-06 | 2.1e-06 | 2.333E-06 | 5.859E-07 | 1.900E-06 | 3.000E-06 |
| 18408 | 921 | S921G | C | 1.8e-06 | 3.1e-06 | 2.3e-06 | 2.400E-06 | 6.557E-07 | 1.800E-06 | 3.100E-06 |
| 18409 | 921 | S921H | C | 2.4e-06 | 3.2e-06 | 2e-06 | 2.533E-06 | 6.110E-07 | 2.000E-06 | 3.200E-06 |
| 18410 | 921 | S921I | C | 1.8e-06 | 2.4e-06 | 1.7e-06 | 1.967E-06 | 3.786E-07 | 1.700E-06 | 2.400E-06 |
| 18411 | 921 | S921L | C | 4e-06 | 2e-06 | 1.5e-06 | 2.500E-06 | 1.323E-06 | 1.500E-06 | 4.000E-06 |
| 18412 | 921 | S921K | C | 2.6e-06 | 7.1e-06 | 3e-06 | 4.233E-06 | 2.491E-06 | 2.600E-06 | 7.100E-06 |
| 18413 | 921 | S921M | C | 2.5e-06 | 2.8e-06 | 1.4e-06 | 2.233E-06 | 7.371E-07 | 1.400E-06 | 2.800E-06 |
| 18414 | 921 | S921F | C | 3.8e-06 | 2.3e-06 | 3.3e-06 | 3.133E-06 | 7.638E-07 | 2.300E-06 | 3.800E-06 |
| 18415 | 921 | S921P | C | 3e-06 | 2.1e-06 | 2.7e-06 | 2.600E-06 | 4.583E-07 | 2.100E-06 | 3.000E-06 |
| 18416 | 921 | S921S | C | 3e-06 | 1.4e-06 | 1.7e-06 | 2.033E-06 | 8.505E-07 | 1.400E-06 | 3.000E-06 |
| 18417 | 921 | S921T | C | 2.7e-06 | 1.8e-06 | 2.4e-06 | 2.300E-06 | 4.583E-07 | 1.800E-06 | 2.700E-06 |
| 18418 | 921 | S921W | C | 2.4e-06 | 2.4e-06 | 2.7e-06 | 2.500E-06 | 1.732E-07 | 2.400E-06 | 2.700E-06 |
| 18419 | 921 | S921Y | C | 1.4e-06 | 2.5e-06 | 1.2e-06 | 1.700E-06 | 7.000E-07 | 1.200E-06 | 2.500E-06 |
| 18420 | 921 | S921V | C | 2e-06 | 4e-06 | 2.5e-06 | 2.833E-06 | 1.041E-06 | 2.000E-06 | 4.000E-06 |
| 18421 | 922 | L922A | C | 1.5e-06 | 3.1e-06 | 2.4e-06 | 2.333E-06 | 8.021E-07 | 1.500E-06 | 3.100E-06 |
| 18422 | 922 | L922R | C | 1.5e-06 | 3.4e-06 | 4.7e-06 | 3.200E-06 | 1.609E-06 | 1.500E-06 | 4.700E-06 |
| 18423 | 922 | L922N | C | 2.5e-06 | 2.5e-06 | 3.1e-06 | 2.700E-06 | 3.464E-07 | 2.500E-06 | 3.100E-06 |
| 18424 | 922 | L922D | C | 3.5e-06 | 3.3e-06 | 2.1e-06 | 2.967E-06 | 7.572E-07 | 2.100E-06 | 3.500E-06 |
| 18425 | 922 | L922C | C | 1.5e-06 | 2.4e-06 | 2.3e-06 | 2.067E-06 | 4.933E-07 | 1.500E-06 | 2.400E-06 |
| 18426 | 922 | L922E | C | 1.5e-06 | 2e-06 | 2.6e-06 | 2.033E-06 | 5.508E-07 | 1.500E-06 | 2.600E-06 |
| 18427 | 922 | L922Q | C | 2.4e-06 | 3.3e-06 | 1.6e-06 | 2.433E-06 | 8.505E-07 | 1.600E-06 | 3.300E-06 |
| 18428 | 922 | L922G | C | 1.7e-06 | 3.1e-06 | 3.2e-06 | 2.667E-06 | 8.386E-07 | 1.700E-06 | 3.200E-06 |
| 18429 | 922 | L922H | C | 2e-06 | 2.2e-06 | 3.3e-06 | 2.500E-06 | 7.000E-07 | 2.000E-06 | 3.300E-06 |
| 18430 | 922 | L922I | C | 1.9e-06 | 2.5e-06 | 3.1e-06 | 2.500E-06 | 6.000E-07 | 1.900E-06 | 3.100E-06 |
| 18431 | 922 | L922L | C | 2e-06 | 1.7e-06 | 6.7e-06 | 3.467E-06 | 2.804E-06 | 1.700E-06 | 6.700E-06 |
| 18432 | 922 | L922K | C | 2.6e-06 | 3.5e-06 | 2.8e-06 | 2.967E-06 | 4.726E-07 | 2.600E-06 | 3.500E-06 |
| 18433 | 922 | L922M | C | 1.8e-06 | 3e-06 | 2.4e-06 | 2.400E-06 | 6.000E-07 | 1.800E-06 | 3.000E-06 |
| 18434 | 922 | L922F | C | 1.5e-06 | 2.5e-06 | 3.4e-06 | 2.467E-06 | 9.504E-07 | 1.500E-06 | 3.400E-06 |
| 18435 | 922 | L922P | C | 4.2e-06 | 3e-06 | 1.9e-06 | 3.033E-06 | 1.150E-06 | 1.900E-06 | 4.200E-06 |
| 18436 | 922 | L922S | C | 8.4e-07 | 2.3e-06 | 2.6e-06 | 1.913E-06 | 9.416E-07 | 8.400E-07 | 2.600E-06 |
| 18437 | 922 | L922T | C | 2.8e-06 | 2.6e-06 | 3.8e-06 | 3.067E-06 | 6.429E-07 | 2.600E-06 | 3.800E-06 |
| 18438 | 922 | L922W | C | 2.1e-06 | 2e-06 | 1.3e-06 | 1.800E-06 | 4.359E-07 | 1.300E-06 | 2.100E-06 |
| 18439 | 922 | L922Y | C | 2.1e-06 | 1.7e-06 | 1.4e-06 | 1.733E-06 | 3.512E-07 | 1.400E-06 | 2.100E-06 |
| 18440 | 922 | L922V | C | 2.4e-06 | 1.4e-06 | 1.5e-06 | 1.767E-06 | 5.508E-07 | 1.400E-06 | 2.400E-06 |
| 18441 | 923 | R923A | C | 1.8e-06 | 1.9e-06 | 1.9e-06 | 1.867E-06 | 5.774E-08 | 1.800E-06 | 1.900E-06 |
| 18442 | 923 | R923R | C | 1.2e-06 | 2.7e-06 | 2.5e-06 | 2.133E-06 | 8.145E-07 | 1.200E-06 | 2.700E-06 |
| 18443 | 923 | R923N | C | 2.3e-06 | 2.4e-06 | 1.2e-06 | 1.967E-06 | 6.658E-07 | 1.200E-06 | 2.400E-06 |
| 18444 | 923 | R923D | C | 2.6e-06 | 2.5e-06 | 1.7e-06 | 2.267E-06 | 4.933E-07 | 1.700E-06 | 2.600E-06 |

| | | | | | | | | | | |
|---|---|---|---|---|---|---|---|---|---|---|
| 18445 | 923 | R923C | C | 2.7e-06 | 2e-06 | 1.9e-06 | 2.200E-06 | 4.359E-07 | 1.900E-06 | 2.700E-06 |
| 18446 | 923 | R923E | C | 2.8e-06 | 3.2e-06 | 1.7e-06 | 2.567E-06 | 7.767E-07 | 1.700E-06 | 3.200E-06 |
| 18447 | 923 | R923Q | C | 3.3e-06 | 2.5e-06 | 2.8e-06 | 2.867E-06 | 4.041E-07 | 2.500E-06 | 3.300E-06 |
| 18448 | 923 | R923G | C | 5.1e-06 | 3.1e-06 | 2.3e-06 | 3.500E-06 | 1.442E-06 | 2.300E-06 | 5.100E-06 |
| 18449 | 923 | R923H | C | 3e-06 | 3.4e-06 | 3.1e-06 | 3.167E-06 | 2.082E-07 | 3.000E-06 | 3.400E-06 |
| 18450 | 923 | R923I | C | 1.9e-06 | 2e-06 | 2.2e-06 | 2.033E-06 | 1.528E-07 | 1.900E-06 | 2.200E-06 |
| 18451 | 923 | R923L | C | 3e-06 | 3e-06 | 1.6e-06 | 2.533E-06 | 8.083E-07 | 1.600E-06 | 3.000E-06 |
| 18452 | 923 | R923K | C | 3.1e-06 | 2.2e-06 | 2.2e-06 | 2.500E-06 | 5.196E-07 | 2.200E-06 | 3.100E-06 |
| 18453 | 923 | R923M | C | 3e-06 | 1.3e-06 | 2.5e-06 | 2.267E-06 | 8.737E-07 | 1.300E-06 | 3.000E-06 |
| 18454 | 923 | R923F | C | 1.8e-06 | 2.1e-06 | 3e-06 | 2.300E-06 | 6.245E-07 | 1.800E-06 | 3.000E-06 |
| 18455 | 923 | R923P | C | 1.8e-06 | 2.5e-06 | 1.8e-06 | 2.033E-06 | 4.041E-07 | 1.800E-06 | 2.500E-06 |
| 18456 | 923 | R923S | C | 1.1e-06 | 2.5e-06 | 2.6e-06 | 2.067E-06 | 8.386E-07 | 1.100E-06 | 2.600E-06 |
| 18457 | 923 | R923T | C | 1.1e-06 | 3.8e-06 | 2.8e-06 | 2.567E-06 | 1.365E-06 | 1.100E-06 | 3.800E-06 |
| 18458 | 923 | R923W | C | 1.6e-06 | 1.8e-06 | 2.7e-06 | 2.033E-06 | 5.859E-07 | 1.600E-06 | 2.700E-06 |
| 18459 | 923 | R923Y | C | 1.3e-06 | 2.9e-06 | 3.5e-06 | 2.567E-06 | 1.137E-06 | 1.300E-06 | 3.500E-06 |
| 18460 | 923 | R923V | C | 2.1e-06 | 2.1e-06 | 2.3e-06 | 2.167E-06 | 1.155E-07 | 2.100E-06 | 2.300E-06 |
| 18461 | 924 | A924A | C | 4.4e-06 | 2.2e-06 | 2.1e-06 | 2.900E-06 | 1.300E-06 | 2.100E-06 | 4.400E-06 |
| 18462 | 924 | A924R | C | 2.1e-06 | 2.5e-06 | 3.4e-06 | 2.667E-06 | 6.658E-07 | 2.100E-06 | 3.400E-06 |
| 18463 | 924 | A924N | C | 1.5e-06 | 9.5e-07 | 2.4e-06 | 1.617E-06 | 7.320E-07 | 9.500E-07 | 2.400E-06 |
| 18464 | 924 | A924D | C | 2.9e-06 | 4e-06 | 3e-06 | 3.300E-06 | 6.083E-07 | 2.900E-06 | 4.000E-06 |
| 18465 | 924 | A924C | C | 1.1e-06 | 2.5e-06 | 1.6e-06 | 1.733E-06 | 7.095E-07 | 1.100E-06 | 2.500E-06 |
| 18466 | 924 | A924E | C | 3.7e-06 | 1.3e-06 | 2.1e-06 | 2.367E-06 | 1.222E-06 | 1.300E-06 | 3.700E-06 |
| 18467 | 924 | A924Q | C | 2.1e-06 | 2.4e-06 | 3.2e-06 | 2.567E-06 | 5.686E-07 | 2.100E-06 | 3.200E-06 |
| 18468 | 924 | A924G | C | 1.9e-06 | 2.7e-06 | 2.3e-06 | 2.300E-06 | 4.000E-07 | 1.900E-06 | 2.700E-06 |
| 18469 | 924 | A924H | C | 2.9e-06 | 2.8e-06 | 3.1e-06 | 2.933E-06 | 1.528E-07 | 2.800E-06 | 3.100E-06 |
| 18470 | 924 | A924I | C | 1.4e-06 | 4.1e-06 | 2.1e-06 | 2.533E-06 | 1.401E-06 | 1.400E-06 | 4.100E-06 |
| 18471 | 924 | A924L | C | 2.7e-06 | 2.1e-06 | 1.9e-06 | 2.233E-06 | 4.163E-07 | 1.900E-06 | 2.700E-06 |
| 18472 | 924 | A924K | C | 2.4e-06 | 2.7e-06 | 1.9e-06 | 2.333E-06 | 4.041E-07 | 1.900E-06 | 2.700E-06 |
| 18473 | 924 | A924M | C | 2e-06 | 4.7e-06 | 2.3e-06 | 3.000E-06 | 1.480E-06 | 2.000E-06 | 4.700E-06 |
| 18474 | 924 | A924F | C | 2.8e-06 | 2.8e-06 | 2.4e-06 | 2.667E-06 | 2.309E-07 | 2.400E-06 | 2.800E-06 |
| 18475 | 924 | A924P | C | 3.4e-06 | 4.4e-06 | 2.5e-06 | 3.433E-06 | 9.504E-07 | 2.500E-06 | 4.400E-06 |
| 18476 | 924 | A924S | C | 2.1e-06 | 2e-06 | 1.5e-06 | 1.867E-06 | 3.215E-07 | 1.500E-06 | 2.100E-06 |
| 18477 | 924 | A924T | C | 2.7e-06 | 2.6e-06 | 3.9e-06 | 3.067E-06 | 7.234E-07 | 2.600E-06 | 3.900E-06 |
| 18478 | 924 | A924W | C | 3.7e-06 | 2.8e-06 | 2.5e-06 | 3.000E-06 | 6.245E-07 | 2.500E-06 | 3.700E-06 |
| 18479 | 924 | A924Y | C | 2e-06 | 1.9e-06 | 2.5e-06 | 2.133E-06 | 3.215E-07 | 1.900E-06 | 2.500E-06 |
| 18480 | 924 | A924V | C | 2.1e-06 | 2.4e-06 | 1.6e-06 | 2.033E-06 | 4.041E-07 | 1.600E-06 | 2.400E-06 |
| 18481 | 925 | E925A | C | 2.7e-06 | 1e-06 | 3.4e-06 | 2.367E-06 | 1.234E-06 | 1.000E-06 | 3.400E-06 |
| 18482 | 925 | E925R | C | 1.5e-06 | 1.2e-06 | 1.9e-06 | 1.533E-06 | 3.512E-07 | 1.200E-06 | 1.900E-06 |
| 18483 | 925 | E925N | C | 1.9e-06 | 1.8e-06 | 2.5e-06 | 2.067E-06 | 3.786E-07 | 1.800E-06 | 2.500E-06 |
| 18484 | 925 | E925D | C | 3.2e-06 | 1.8e-06 | 2.2e-06 | 2.400E-06 | 7.211E-07 | 1.800E-06 | 3.200E-06 |
| 18485 | 925 | E925C | C | 3.4e-06 | 2.6e-06 | 2.2e-06 | 2.733E-06 | 6.110E-07 | 2.200E-06 | 3.400E-06 |
| 18486 | 925 | E925E | C | 4e-06 | 2e-06 | 1.7e-06 | 2.567E-06 | 1.250E-06 | 1.700E-06 | 4.000E-06 |
| 18487 | 925 | E925Q | C | 5e-06 | 4.3e-06 | 1.6e-06 | 3.633E-06 | 1.795E-06 | 1.600E-06 | 5.000E-06 |
| 18488 | 925 | E925G | C | 2.3e-06 | 2.6e-06 | 1.1e-06 | 2.000E-06 | 7.937E-07 | 1.100E-06 | 2.600E-06 |
| 18489 | 925 | E925H | C | 3.6e-06 | 2.6e-06 | 2.7e-06 | 2.967E-06 | 5.508E-07 | 2.600E-06 | 3.600E-06 |
| 18490 | 925 | E925I | C | 1.6e-06 | 6.9e-07 | 2.9e-06 | 1.730E-06 | 1.111E-06 | 6.900E-07 | 2.900E-06 |
| 18491 | 925 | E925L | C | 2e-06 | 4.1e-06 | 2.5e-06 | 2.867E-06 | 1.097E-06 | 2.000E-06 | 4.100E-06 |
| 18492 | 925 | E925K | C | 1.4e-06 | 8.2e-07 | 2.5e-06 | 1.573E-06 | 8.533E-07 | 8.200E-07 | 2.500E-06 |
| 18493 | 925 | E925M | C | 1.5e-06 | 3e-06 | 1.4e-06 | 1.967E-06 | 8.963E-07 | 1.400E-06 | 3.000E-06 |
| 18494 | 925 | E925F | C | 2.3e-06 | 2.2e-06 | 1.9e-06 | 2.133E-06 | 2.082E-07 | 1.900E-06 | 2.300E-06 |
| 18495 | 925 | E925P | C | 5e-06 | 4.7e-06 | 1.2e-06 | 3.633E-06 | 2.113E-06 | 1.200E-06 | 5.000E-06 |
| 18496 | 925 | E925S | C | 2.2e-06 | 3.2e-06 | 1.3e-06 | 2.233E-06 | 9.504E-07 | 1.300E-06 | 3.200E-06 |
| 18497 | 925 | E925T | C | 1.2e-06 | 2.1e-06 | 2.9e-06 | 2.067E-06 | 8.505E-07 | 1.200E-06 | 2.900E-06 |
| 18498 | 925 | E925W | C | 2.1e-06 | 1.5e-06 | 1.5e-06 | 1.700E-06 | 3.464E-07 | 1.500E-06 | 2.100E-06 |
| 18499 | 925 | E925Y | C | 2.6e-06 | 3.6e-06 | 3.1e-06 | 3.100E-06 | 5.000E-07 | 2.600E-06 | 3.600E-06 |
| 18500 | 925 | E925V | C | 2.7e-06 | 2.1e-06 | 2.5e-06 | 2.433E-06 | 3.055E-07 | 2.100E-06 | 2.700E-06 |
| 18501 | 926 | D926A | C | 2e-06 | 8.1e-07 | 3.3e-06 | 2.037E-06 | 1.245E-06 | 8.100E-07 | 3.300E-06 |
| 18502 | 926 | D926R | C | 1.6e-06 | 1.8e-06 | 3.1e-06 | 2.167E-06 | 8.145E-07 | 1.600E-06 | 3.100E-06 |
| 18503 | 926 | D926N | C | 1.3e-06 | 2.4e-06 | 2.9e-06 | 2.200E-06 | 8.185E-07 | 1.300E-06 | 2.900E-06 |
| 18504 | 926 | D926D | C | 2.3e-06 | 2.6e-06 | 2.3e-06 | 2.400E-06 | 1.732E-07 | 2.300E-06 | 2.600E-06 |
| 18505 | 926 | D926C | C | 9.3e-07 | 2.1e-06 | 2.4e-06 | 1.810E-06 | 7.767E-07 | 9.300E-07 | 2.400E-06 |
| 18506 | 926 | D926E | C | 2.1e-06 | 2.6e-06 | 2.4e-06 | 2.367E-06 | 2.517E-07 | 2.100E-06 | 2.600E-06 |
| 18507 | 926 | D926Q | C | 1.9e-06 | 1.7e-06 | 2.4e-06 | 2.000E-06 | 3.606E-07 | 1.700E-06 | 2.400E-06 |
| 18508 | 926 | D926G | C | 1.8e-06 | 2.5e-06 | 1.8e-06 | 2.033E-06 | 4.041E-07 | 1.800E-06 | 2.500E-06 |
| 18509 | 926 | D926H | C | 1.6e-06 | 2.5e-06 | 3.2e-06 | 2.433E-06 | 8.021E-07 | 1.600E-06 | 3.200E-06 |
| 18510 | 926 | D926I | C | 1.1e-06 | 3.3e-06 | 4.2e-06 | 2.867E-06 | 1.595E-06 | 1.100E-06 | 4.200E-06 |
| 18511 | 926 | D926L | C | 2.1e-06 | 2.5e-06 | 1.6e-06 | 2.067E-06 | 4.509E-07 | 1.600E-06 | 2.500E-06 |
| 18512 | 926 | D926K | C | 1.6e-06 | 2.8e-06 | 1.7e-06 | 2.033E-06 | 6.658E-07 | 1.600E-06 | 2.800E-06 |
| 18513 | 926 | D926M | C | 1.9e-06 | 9.6e-07 | 1.3e-06 | 1.387E-06 | 4.760E-07 | 9.600E-07 | 1.900E-06 |
| 18514 | 926 | D926F | C | 2.8e-06 | 1.9e-06 | 2.8e-06 | 2.500E-06 | 5.196E-07 | 1.900E-06 | 2.800E-06 |
| 18515 | 926 | D926P | C | 3.8e-06 | 2.2e-06 | 3.1e-06 | 3.033E-06 | 8.021E-07 | 2.200E-06 | 3.800E-06 |
| 18516 | 926 | D926S | C | 2.3e-06 | 2.2e-06 | 1.5e-06 | 2.000E-06 | 4.359E-07 | 1.500E-06 | 2.300E-06 |
| 18517 | 926 | D926T | C | 3.2e-06 | 2.2e-06 | 3.3e-06 | 2.900E-06 | 6.083E-07 | 2.200E-06 | 3.300E-06 |
| 18518 | 926 | D926W | C | 1.2e-06 | 1.6e-06 | 1.9e-06 | 1.567E-06 | 3.512E-07 | 1.200E-06 | 1.900E-06 |
| 18519 | 926 | D926Y | C | 2.5e-06 | 3.4e-06 | 1.1e-06 | 2.333E-06 | 1.159E-06 | 1.100E-06 | 3.400E-06 |
| 18520 | 926 | D926V | C | 2.9e-06 | 2e-06 | 2e-06 | 2.300E-06 | 5.196E-07 | 2.000E-06 | 2.900E-06 |
| 18521 | 927 | T927A | C | 2.7e-06 | 3.2e-06 | 3e-06 | 2.967E-06 | 2.517E-07 | 2.700E-06 | 3.200E-06 |
| 18522 | 927 | T927R | C | 2.2e-06 | 2.1e-06 | 2.4e-06 | 2.233E-06 | 1.528E-07 | 2.100E-06 | 2.400E-06 |
| 18523 | 927 | T927N | C | 2.1e-06 | 2.8e-06 | 2.5e-06 | 2.467E-06 | 3.512E-07 | 2.100E-06 | 2.800E-06 |
| 18524 | 927 | T927D | C | 2e-06 | 4.1e-06 | 6e-06 | 4.033E-06 | 2.001E-06 | 2.000E-06 | 6.000E-06 |
| 18525 | 927 | T927C | C | 2.2e-06 | 1.9e-06 | 2.1e-06 | 2.067E-06 | 1.528E-07 | 1.900E-06 | 2.200E-06 |
| 18526 | 927 | T927E | C | 1.4e-06 | 2.3e-06 | 2.4e-06 | 2.033E-06 | 5.508E-07 | 1.400E-06 | 2.400E-06 |
| 18527 | 927 | T927Q | C | 3.3e-06 | 1.8e-06 | 2e-06 | 2.367E-06 | 8.145E-07 | 1.800E-06 | 3.300E-06 |
| 18528 | 927 | T927G | C | 2.3e-06 | 1.7e-06 | 2.1e-06 | 2.033E-06 | 3.055E-07 | 1.700E-06 | 2.300E-06 |
| 18529 | 927 | T927H | C | 2.2e-06 | 3.4e-06 | 2.1e-06 | 2.567E-06 | 7.234E-07 | 2.100E-06 | 3.400E-06 |

| | | | | | | | | | | |
|---|---|---|---|---|---|---|---|---|---|---|
| 18530 | 927 | T927I | C | 1.8e-06 | 3.3e-06 | 2.7e-06 | 2.600E-06 | 7.550E-07 | 1.800E-06 | 3.300E-06 |
| 18531 | 927 | T927L | C | 2.7e-06 | 2.5e-06 | 1e-06 | 2.067E-06 | 9.292E-07 | 1.000E-06 | 2.700E-06 |
| 18532 | 927 | T927K | C | 1.9e-06 | 1e-06 | 2.6e-06 | 1.833E-06 | 8.021E-07 | 1.000E-06 | 2.600E-06 |
| 18533 | 927 | T927M | C | 2.5e-06 | 3.4e-06 | 3.3e-06 | 3.067E-06 | 4.933E-07 | 2.500E-06 | 3.400E-06 |
| 18534 | 927 | T927F | C | 4e-06 | 1.9e-06 | 1.6e-06 | 2.500E-06 | 1.308E-06 | 1.600E-06 | 4.000E-06 |
| 18535 | 927 | T927P | C | 2.9e-06 | 2e-06 | 3.1e-06 | 2.667E-06 | 5.859E-07 | 2.000E-06 | 3.100E-06 |
| 18536 | 927 | T927S | C | 2.2e-06 | 2.6e-06 | 1.8e-06 | 2.200E-06 | 4.000E-07 | 1.800E-06 | 2.600E-06 |
| 18537 | 927 | T927T | C | 4e-06 | 2.2e-06 | 4e-06 | 3.400E-06 | 1.039E-06 | 2.200E-06 | 4.000E-06 |
| 18538 | 927 | T927W | C | 2.2e-06 | 2.3e-06 | 1.8e-06 | 2.100E-06 | 2.646E-07 | 1.800E-06 | 2.300E-06 |
| 18539 | 927 | T927Y | C | 2.8e-06 | 2.1e-06 | 2e-06 | 2.300E-06 | 4.359E-07 | 2.000E-06 | 2.800E-06 |
| 18540 | 927 | T927V | C | 1.7e-06 | 2.5e-06 | 1.9e-06 | 2.033E-06 | 4.163E-07 | 1.700E-06 | 2.500E-06 |
| 18541 | 928 | A928A | C | 1.8e-06 | 2.9e-06 | 1.1e-06 | 1.933E-06 | 9.074E-07 | 1.100E-06 | 2.900E-06 |
| 18542 | 928 | A928R | C | 1.8e-06 | 3.2e-06 | 1.7e-06 | 2.233E-06 | 8.386E-07 | 1.700E-06 | 3.200E-06 |
| 18543 | 928 | A928N | C | 2.8e-06 | 3.2e-06 | 2.3e-06 | 2.767E-06 | 4.509E-07 | 2.300E-06 | 3.200E-06 |
| 18544 | 928 | A928D | C | 2.4e-06 | 2.1e-06 | 2.3e-06 | 2.267E-06 | 1.528E-07 | 2.100E-06 | 2.400E-06 |
| 18545 | 928 | A928C | C | 2.1e-06 | 2.6e-06 | 2.5e-06 | 2.400E-06 | 2.646E-07 | 2.100E-06 | 2.600E-06 |
| 18546 | 928 | A928E | C | 3.9e-06 | 2.5e-06 | 1.6e-06 | 2.667E-06 | 1.159E-06 | 1.600E-06 | 3.900E-06 |
| 18547 | 928 | A928Q | C | 1.3e-06 | 9.9e-07 | 1.5e-06 | 1.263E-06 | 2.570E-07 | 9.900E-07 | 1.500E-06 |
| 18548 | 928 | A928G | C | 2.7e-06 | 2.1e-06 | 2.4e-06 | 2.400E-06 | 3.000E-07 | 2.100E-06 | 2.700E-06 |
| 18549 | 928 | A928H | C | 2.1e-06 | 2e-06 | 3.4e-06 | 2.500E-06 | 7.810E-07 | 2.000E-06 | 3.400E-06 |
| 18550 | 928 | A928I | C | 1.2e-06 | 1.4e-06 | 2.6e-06 | 1.733E-06 | 7.572E-07 | 1.200E-06 | 2.600E-06 |
| 18551 | 928 | A928L | C | 3e-06 | 1.6e-06 | 2.9e-06 | 2.500E-06 | 7.810E-07 | 1.600E-06 | 3.000E-06 |
| 18552 | 928 | A928K | C | 2.4e-06 | 2.2e-06 | 2.1e-06 | 2.233E-06 | 1.528E-07 | 2.100E-06 | 2.400E-06 |
| 18553 | 928 | A928M | C | 2.7e-06 | 2.6e-06 | 1.3e-06 | 2.200E-06 | 7.810E-07 | 1.300E-06 | 2.700E-06 |
| 18554 | 928 | A928F | C | 1.2e-06 | 3.1e-06 | 2e-06 | 2.100E-06 | 9.539E-07 | 1.200E-06 | 3.100E-06 |
| 18555 | 928 | A928P | C | 1.8e-06 | 2.4e-06 | 2.1e-06 | 2.100E-06 | 3.000E-07 | 1.800E-06 | 2.400E-06 |
| 18556 | 928 | A928S | C | 1.5e-06 | 2.3e-06 | 2.7e-06 | 2.167E-06 | 6.110E-07 | 1.500E-06 | 2.700E-06 |
| 18557 | 928 | A928T | C | 3.3e-06 | 1.7e-06 | 1.6e-06 | 2.200E-06 | 9.539E-07 | 1.600E-06 | 3.300E-06 |
| 18558 | 928 | A928W | C | 3.9e-06 | 3.2e-06 | 2.4e-06 | 3.167E-06 | 7.506E-07 | 2.400E-06 | 3.900E-06 |
| 18559 | 928 | A928Y | C | 2.3e-06 | 2.4e-06 | 3.3e-06 | 2.667E-06 | 5.508E-07 | 2.300E-06 | 3.300E-06 |
| 18560 | 928 | A928V | C | 2e-06 | 2e-06 | 3e-06 | 2.333E-06 | 5.774E-07 | 2.000E-06 | 3.000E-06 |
| 18561 | 929 | V929A | C | 2.2e-06 | 2.3e-06 | 1.8e-06 | 2.100E-06 | 2.646E-07 | 1.800E-06 | 2.300E-06 |
| 18562 | 929 | V929R | C | 3.3e-06 | 3.7e-06 | 1.8e-06 | 2.933E-06 | 1.002E-06 | 1.800E-06 | 3.700E-06 |
| 18563 | 929 | V929N | C | 2.3e-06 | 2e-06 | 1.6e-06 | 1.967E-06 | 3.512E-07 | 1.600E-06 | 2.300E-06 |
| 18564 | 929 | V929D | C | 1.6e-06 | 2.2e-06 | 1.7e-06 | 1.833E-06 | 3.215E-07 | 1.600E-06 | 2.200E-06 |
| 18565 | 929 | V929C | C | 1.6e-06 | 2.3e-06 | 2e-06 | 1.967E-06 | 3.512E-07 | 1.600E-06 | 2.300E-06 |
| 18566 | 929 | V929E | C | 1.9e-06 | 1.2e-06 | 2.5e-06 | 1.867E-06 | 6.506E-07 | 1.200E-06 | 2.500E-06 |
| 18567 | 929 | V929Q | C | 1.2e-06 | 1.6e-06 | 2.2e-06 | 1.667E-06 | 5.033E-07 | 1.200E-06 | 2.200E-06 |
| 18568 | 929 | V929G | C | 2.1e-06 | 2.6e-06 | 2e-06 | 2.233E-06 | 3.215E-07 | 2.000E-06 | 2.600E-06 |
| 18569 | 929 | V929H | C | 1.9e-06 | 2.7e-06 | 1.3e-06 | 1.967E-06 | 7.024E-07 | 1.300E-06 | 2.700E-06 |
| 18570 | 929 | V929I | C | 2.4e-06 | 2.7e-06 | 2.6e-06 | 2.567E-06 | 1.528E-07 | 2.400E-06 | 2.700E-06 |
| 18571 | 929 | V929L | C | 3.1e-06 | 2.5e-06 | 3.7e-06 | 3.100E-06 | 6.000E-07 | 2.500E-06 | 3.700E-06 |
| 18572 | 929 | V929K | C | 2.1e-06 | 2e-06 | 2.6e-06 | 2.233E-06 | 3.215E-07 | 2.000E-06 | 2.600E-06 |
| 18573 | 929 | V929M | C | 2.7e-06 | 4e-06 | 2.8e-06 | 3.167E-06 | 7.234E-07 | 2.700E-06 | 4.000E-06 |
| 18574 | 929 | V929F | C | 1.8e-06 | 3.1e-06 | 2.6e-06 | 2.500E-06 | 6.557E-07 | 1.800E-06 | 3.100E-06 |
| 18575 | 929 | V929P | C | 2e-06 | 2.6e-06 | 3e-06 | 2.533E-06 | 5.033E-07 | 2.000E-06 | 3.000E-06 |
| 18576 | 929 | V929S | C | 1.1e-06 | 3.4e-06 | 2.1e-06 | 2.200E-06 | 1.153E-06 | 1.100E-06 | 3.400E-06 |
| 18577 | 929 | V929T | C | 3.6e-06 | 3.7e-06 | 3.7e-06 | 3.667E-06 | 5.774E-08 | 3.600E-06 | 3.700E-06 |
| 18578 | 929 | V929W | C | 2.9e-06 | 4.4e-06 | 1.8e-06 | 3.033E-06 | 1.305E-06 | 1.800E-06 | 4.400E-06 |
| 18579 | 929 | V929Y | C | 2e-06 | 2e-06 | 2e-06 | 2.000E-06 | 0.000E+00 | 2.000E-06 | 2.000E-06 |
| 18580 | 929 | V929V | C | 1.9e-06 | 1.9e-06 | 2.2e-06 | 2.000E-06 | 1.732E-07 | 1.900E-06 | 2.200E-06 |
| 18581 | 930 | Y930A | C | 2.1e-06 | 2.5e-06 | 2.1e-06 | 2.233E-06 | 2.309E-07 | 2.100E-06 | 2.500E-06 |
| 18582 | 930 | Y930R | C | 2.5e-06 | 1.6e-06 | 2.1e-06 | 2.067E-06 | 4.509E-07 | 1.600E-06 | 2.500E-06 |
| 18583 | 930 | Y930N | C | 2.5e-06 | 2.7e-06 | 2.3e-06 | 2.500E-06 | 2.000E-07 | 2.300E-06 | 2.700E-06 |
| 18584 | 930 | Y930D | C | 1.7e-06 | 2.5e-06 | 3.5e-06 | 2.567E-06 | 9.018E-07 | 1.700E-06 | 3.500E-06 |
| 18585 | 930 | Y930C | C | 2.6e-06 | 1.9e-06 | 4e-06 | 2.833E-06 | 1.069E-06 | 1.900E-06 | 4.000E-06 |
| 18586 | 930 | Y930E | C | 2e-06 | 1.6e-06 | 2.8e-06 | 2.133E-06 | 6.110E-07 | 1.600E-06 | 2.800E-06 |
| 18587 | 930 | Y930Q | C | 1.8e-06 | 3.3e-06 | 1.6e-06 | 2.233E-06 | 9.292E-07 | 1.600E-06 | 3.300E-06 |
| 18588 | 930 | Y930G | C | 3.5e-06 | 2e-06 | 4.7e-06 | 3.400E-06 | 1.353E-06 | 2.000E-06 | 4.700E-06 |
| 18589 | 930 | Y930H | C | 3.7e-06 | 2.2e-06 | 3e-06 | 2.967E-06 | 7.506E-07 | 2.200E-06 | 3.700E-06 |
| 18590 | 930 | Y930I | C | 2.5e-06 | 2e-06 | 3e-06 | 2.500E-06 | 5.000E-07 | 2.000E-06 | 3.000E-06 |
| 18591 | 930 | Y930L | C | 2.2e-06 | 2e-06 | 1.4e-06 | 1.867E-06 | 4.163E-07 | 1.400E-06 | 2.200E-06 |
| 18592 | 930 | Y930K | C | 2.4e-06 | 2.1e-06 | 2.2e-06 | 2.233E-06 | 1.528E-07 | 2.100E-06 | 2.400E-06 |
| 18593 | 930 | Y930M | C | 3.6e-06 | 1.1e-06 | 2.7e-06 | 2.467E-06 | 1.266E-06 | 1.100E-06 | 3.600E-06 |
| 18594 | 930 | Y930F | C | 3.1e-06 | 1.6e-06 | 2e-06 | 2.233E-06 | 7.767E-07 | 1.600E-06 | 3.100E-06 |
| 18595 | 930 | Y930P | C | 3.4e-06 | 2.6e-06 | 3.4e-06 | 3.133E-06 | 4.619E-07 | 2.600E-06 | 3.400E-06 |
| 18596 | 930 | Y930S | C | 3.1e-06 | 2e-06 | 2.4e-06 | 2.500E-06 | 5.568E-07 | 2.000E-06 | 3.100E-06 |
| 18597 | 930 | Y930T | C | 2.2e-06 | 3.2e-06 | 2.2e-06 | 2.533E-06 | 5.774E-07 | 2.200E-06 | 3.200E-06 |
| 18598 | 930 | Y930W | C | 1.1e-06 | 3.7e-06 | 2.5e-06 | 2.433E-06 | 1.301E-06 | 1.100E-06 | 3.700E-06 |
| 18599 | 930 | Y930Y | C | 2.4e-06 | 1.5e-06 | 3.6e-06 | 2.500E-06 | 1.054E-06 | 1.500E-06 | 3.600E-06 |
| 18600 | 930 | Y930V | C | 1.3e-06 | 2.4e-06 | 3.2e-06 | 2.300E-06 | 9.539E-07 | 1.300E-06 | 3.200E-06 |
| 18601 | 931 | Y931A | C | 2.2e-06 | 2.6e-06 | 3.3e-06 | 2.700E-06 | 5.568E-07 | 2.200E-06 | 3.300E-06 |
| 18602 | 931 | Y931R | C | 3.6e-06 | 2.3e-06 | 2.3e-06 | 2.733E-06 | 7.506E-07 | 2.300E-06 | 3.600E-06 |
| 18603 | 931 | Y931N | C | 2.8e-06 | 3.8e-06 | 1.4e-06 | 2.667E-06 | 1.206E-06 | 1.400E-06 | 3.800E-06 |
| 18604 | 931 | Y931D | C | 2.7e-06 | 2.6e-06 | 2.4e-06 | 2.567E-06 | 1.528E-07 | 2.400E-06 | 2.700E-06 |
| 18605 | 931 | Y931C | C | 2.1e-06 | 1.4e-06 | 2.6e-06 | 2.033E-06 | 6.028E-07 | 1.400E-06 | 2.600E-06 |
| 18606 | 931 | Y931E | C | 2.8e-06 | 2.2e-06 | 2.9e-06 | 2.633E-06 | 3.786E-07 | 2.200E-06 | 2.900E-06 |
| 18607 | 931 | Y931Q | C | 2e-06 | 2.6e-06 | 2.3e-06 | 2.300E-06 | 3.000E-07 | 2.000E-06 | 2.600E-06 |
| 18608 | 931 | Y931G | C | 2.5e-06 | 1.5e-06 | 1.9e-06 | 1.967E-06 | 5.033E-07 | 1.500E-06 | 2.500E-06 |
| 18609 | 931 | Y931H | C | 2.4e-06 | 2e-06 | 2.1e-06 | 2.167E-06 | 2.082E-07 | 2.000E-06 | 2.400E-06 |
| 18610 | 931 | Y931I | C | 2e-06 | 2.2e-06 | 2.2e-06 | 2.133E-06 | 1.155E-07 | 2.000E-06 | 2.200E-06 |
| 18611 | 931 | Y931L | C | 1.2e-06 | 1.2e-06 | 2.2e-06 | 1.533E-06 | 5.774E-07 | 1.200E-06 | 2.200E-06 |
| 18612 | 931 | Y931K | C | 2.7e-06 | 2.3e-06 | 4.2e-06 | 3.067E-06 | 1.002E-06 | 2.300E-06 | 4.200E-06 |
| 18613 | 931 | Y931M | C | 1.6e-06 | 1.6e-06 | 9.5e-07 | 1.383E-06 | 3.753E-07 | 9.500E-07 | 1.600E-06 |
| 18614 | 931 | Y931F | C | 2.2e-06 | 2.5e-06 | 1.8e-06 | 2.167E-06 | 3.512E-07 | 1.800E-06 | 2.500E-06 |

| | | | | | | | | | | |
|---|---|---|---|---|---|---|---|---|---|---|
| 18615 | 931 | Y931P | C | 3.7e-06 | 3.3e-06 | 6e-06 | 4.333E-06 | 1.457E-06 | 3.300E-06 | 6.000E-06 |
| 18616 | 931 | Y931S | C | 3.8e-06 | 3.7e-06 | 3e-06 | 3.500E-06 | 4.359E-07 | 3.000E-06 | 3.800E-06 |
| 18617 | 931 | Y931T | C | 2.7e-06 | 3.2e-06 | 2.2e-06 | 2.700E-06 | 5.000E-07 | 2.200E-06 | 3.200E-06 |
| 18618 | 931 | Y931W | C | 1.8e-06 | 2.3e-06 | 2e-06 | 2.033E-06 | 2.517E-07 | 1.800E-06 | 2.300E-06 |
| 18619 | 931 | Y931Y | C | 2.5e-06 | 2e-06 | 1.5e-06 | 2.000E-06 | 5.000E-07 | 1.500E-06 | 2.500E-06 |
| 18620 | 931 | Y931V | C | 3.2e-06 | 2.3e-06 | 2.1e-06 | 2.533E-06 | 5.859E-07 | 2.100E-06 | 3.200E-06 |
| 18621 | 932 | C932A | C | 3.2e-06 | 1.8e-06 | 3e-06 | 2.667E-06 | 7.572E-07 | 1.800E-06 | 3.200E-06 |
| 18622 | 932 | C932R | C | 2.9e-06 | 2.4e-06 | 2.4e-06 | 2.567E-06 | 2.887E-07 | 2.400E-06 | 2.900E-06 |
| 18623 | 932 | C932N | C | 3e-06 | 1.9e-06 | 2.3e-06 | 2.400E-06 | 5.568E-07 | 1.900E-06 | 3.000E-06 |
| 18624 | 932 | C932D | C | 2.1e-06 | 2e-06 | 3.1e-06 | 2.400E-06 | 6.083E-07 | 2.000E-06 | 3.100E-06 |
| 18625 | 932 | C932C | C | 1.8e-06 | 1.5e-06 | 1.9e-06 | 1.733E-06 | 2.082E-07 | 1.500E-06 | 1.900E-06 |
| 18626 | 932 | C932E | C | 1.3e-06 | 2.2e-06 | 2.4e-06 | 1.967E-06 | 5.859E-07 | 1.300E-06 | 2.400E-06 |
| 18627 | 932 | C932Q | C | 1.8e-06 | 2.8e-06 | 3.8e-06 | 2.800E-06 | 1.000E-06 | 1.800E-06 | 3.800E-06 |
| 18628 | 932 | C932G | C | 1.5e-06 | 2.9e-06 | 2.1e-06 | 2.167E-06 | 7.024E-07 | 1.500E-06 | 2.900E-06 |
| 18629 | 932 | C932H | C | 2.8e-06 | 2.6e-06 | 3.7e-06 | 3.033E-06 | 5.859E-07 | 2.600E-06 | 3.700E-06 |
| 18630 | 932 | C932I | C | 2.7e-06 | 2.2e-06 | 2.4e-06 | 2.433E-06 | 2.517E-07 | 2.200E-06 | 2.700E-06 |
| 18631 | 932 | C932L | C | 2.8e-06 | 1.6e-06 | 1.7e-06 | 2.033E-06 | 6.658E-07 | 1.600E-06 | 2.800E-06 |
| 18632 | 932 | C932K | C | 2.4e-06 | 2.5e-06 | 2e-06 | 2.300E-06 | 2.646E-07 | 2.000E-06 | 2.500E-06 |
| 18633 | 932 | C932M | C | 2.2e-06 | 4e-06 | 3.1e-06 | 3.100E-06 | 9.000E-07 | 2.200E-06 | 4.000E-06 |
| 18634 | 932 | C932F | C | 2.1e-06 | 1.4e-06 | 1.6e-06 | 1.700E-06 | 3.606E-07 | 1.400E-06 | 2.100E-06 |
| 18635 | 932 | C932P | C | 3.6e-06 | 2.3e-06 | 1.4e-06 | 2.433E-06 | 1.106E-06 | 1.400E-06 | 3.600E-06 |
| 18636 | 932 | C932S | C | 2e-06 | 3.7e-06 | 1.5e-06 | 2.400E-06 | 1.153E-06 | 1.500E-06 | 3.700E-06 |
| 18637 | 932 | C932T | C | 2.5e-06 | 2.1e-06 | 1.6e-06 | 2.067E-06 | 4.509E-07 | 1.600E-06 | 2.500E-06 |
| 18638 | 932 | C932W | C | 1.1e-06 | 1.7e-06 | 3.1e-06 | 1.967E-06 | 1.026E-06 | 1.100E-06 | 3.100E-06 |
| 18639 | 932 | C932Y | C | 3.1e-06 | 2.9e-06 | 2e-06 | 2.667E-06 | 5.859E-07 | 2.000E-06 | 3.100E-06 |
| 18640 | 932 | C932V | C | 9.6e-07 | 3.3e-06 | 2.2e-06 | 2.153E-06 | 1.171E-06 | 9.600E-07 | 3.300E-06 |
| 18641 | 933 | A933A | C | 1.3e-06 | 2.5e-06 | 4.5e-06 | 2.767E-06 | 1.617E-06 | 1.300E-06 | 4.500E-06 |
| 18642 | 933 | A933R | C | 2.4e-06 | 1.6e-06 | 2.4e-06 | 2.133E-06 | 4.619E-07 | 1.600E-06 | 2.400E-06 |
| 18643 | 933 | A933N | C | 2.5e-06 | 2.8e-06 | 2.2e-06 | 2.500E-06 | 3.000E-07 | 2.200E-06 | 2.800E-06 |
| 18644 | 933 | A933D | C | 2.7e-06 | 3.7e-06 | 1.9e-06 | 2.767E-06 | 9.018E-07 | 1.900E-06 | 3.700E-06 |
| 18645 | 933 | A933C | C | 2e-06 | 1.3e-06 | 3.6e-06 | 2.300E-06 | 1.179E-06 | 1.300E-06 | 3.600E-06 |
| 18646 | 933 | A933E | C | 2.3e-06 | 1.4e-06 | 2e-06 | 1.900E-06 | 4.583E-07 | 1.400E-06 | 2.300E-06 |
| 18647 | 933 | A933Q | C | 3.5e-06 | 1.9e-06 | 1.8e-06 | 2.400E-06 | 9.539E-07 | 1.800E-06 | 3.500E-06 |
| 18648 | 933 | A933G | C | 1.7e-06 | 2.1e-06 | 3.3e-06 | 2.367E-06 | 8.327E-07 | 1.700E-06 | 3.300E-06 |
| 18649 | 933 | A933H | C | 2.2e-06 | 1.8e-06 | 2.4e-06 | 2.133E-06 | 3.055E-07 | 1.800E-06 | 2.400E-06 |
| 18650 | 933 | A933I | C | 1.5e-06 | 1.9e-06 | 1.7e-06 | 1.700E-06 | 2.000E-07 | 1.500E-06 | 1.900E-06 |
| 18651 | 933 | A933L | C | 3e-06 | 1e-06 | 1.8e-06 | 1.933E-06 | 1.007E-06 | 1.000E-06 | 3.000E-06 |
| 18652 | 933 | A933K | C | 2.1e-06 | 2.8e-06 | 3.7e-06 | 2.867E-06 | 8.021E-07 | 2.100E-06 | 3.700E-06 |
| 18653 | 933 | A933M | C | 2.9e-06 | 2.2e-06 | 2.1e-06 | 2.400E-06 | 4.359E-07 | 2.100E-06 | 2.900E-06 |
| 18654 | 933 | A933F | C | 8.9e-07 | 1.9e-06 | 4.3e-06 | 2.363E-06 | 1.752E-06 | 8.900E-07 | 4.300E-06 |
| 18655 | 933 | A933P | C | 2.7e-06 | 2.4e-06 | 1.9e-06 | 2.333E-06 | 4.041E-07 | 1.900E-06 | 2.700E-06 |
| 18656 | 933 | A933S | C | 2.7e-06 | 2.2e-06 | 2.2e-06 | 2.367E-06 | 2.887E-07 | 2.200E-06 | 2.700E-06 |
| 18657 | 933 | A933T | C | 2.9e-06 | 1.1e-06 | 1.2e-06 | 1.733E-06 | 1.012E-06 | 1.100E-06 | 2.900E-06 |
| 18658 | 933 | A933W | C | 2.9e-06 | 2.8e-06 | 2e-06 | 2.567E-06 | 4.933E-07 | 2.000E-06 | 2.900E-06 |
| 18659 | 933 | A933Y | C | 3.6e-06 | 2.2e-06 | 2.3e-06 | 2.700E-06 | 7.810E-07 | 2.200E-06 | 3.600E-06 |
| 18660 | 933 | A933V | C | 2.8e-06 | 2.3e-06 | 2.4e-06 | 2.500E-06 | 2.646E-07 | 2.300E-06 | 2.800E-06 |
| 18661 | 934 | R934A | C | 3e-06 | 1.2e-06 | 2.2e-06 | 2.133E-06 | 9.018E-07 | 1.200E-06 | 3.000E-06 |
| 18662 | 934 | R934R | C | 1.1e-06 | 2.7e-06 | 2.5e-06 | 2.100E-06 | 8.718E-07 | 1.100E-06 | 2.700E-06 |
| 18663 | 934 | R934N | C | 2.7e-06 | 3.9e-06 | 1.9e-06 | 2.833E-06 | 1.007E-06 | 1.900E-06 | 3.900E-06 |
| 18664 | 934 | R934D | C | 2e-06 | 4.3e-06 | 4.5e-06 | 3.600E-06 | 1.389E-06 | 2.000E-06 | 4.500E-06 |
| 18665 | 934 | R934C | C | 2.7e-06 | 2.1e-06 | 3.1e-06 | 2.633E-06 | 5.033E-07 | 2.100E-06 | 3.100E-06 |
| 18666 | 934 | R934E | C | 2.8e-06 | 2.2e-06 | 2.3e-06 | 2.433E-06 | 3.215E-07 | 2.200E-06 | 2.800E-06 |
| 18667 | 934 | R934Q | C | 2.6e-06 | 3.2e-06 | 2.5e-06 | 2.767E-06 | 3.786E-07 | 2.500E-06 | 3.200E-06 |
| 18668 | 934 | R934G | C | 2.5e-06 | 3.7e-06 | 2.1e-06 | 2.767E-06 | 8.327E-07 | 2.100E-06 | 3.700E-06 |
| 18669 | 934 | R934H | C | 2.7e-06 | 3.7e-06 | 3.1e-06 | 3.167E-06 | 5.033E-07 | 2.700E-06 | 3.700E-06 |
| 18670 | 934 | R934I | C | 2.2e-06 | 3e-06 | 2.5e-06 | 2.567E-06 | 4.041E-07 | 2.200E-06 | 3.000E-06 |
| 18671 | 934 | R934L | C | 1.4e-06 | 1.7e-06 | 2.3e-06 | 1.800E-06 | 4.583E-07 | 1.400E-06 | 2.300E-06 |
| 18672 | 934 | R934K | C | 2.7e-06 | 3.2e-06 | 3.7e-06 | 3.200E-06 | 5.000E-07 | 2.700E-06 | 3.700E-06 |
| 18673 | 934 | R934M | C | 2.9e-06 | 2.3e-06 | 3.7e-06 | 2.967E-06 | 7.024E-07 | 2.300E-06 | 3.700E-06 |
| 18674 | 934 | R934F | C | 2.3e-06 | 3.7e-06 | 4.3e-06 | 3.433E-06 | 1.026E-06 | 2.300E-06 | 4.300E-06 |
| 18675 | 934 | R934P | C | 3e-06 | 2.2e-06 | 3.4e-06 | 2.867E-06 | 6.110E-07 | 2.200E-06 | 3.400E-06 |
| 18676 | 934 | R934S | C | 3.1e-06 | 2.1e-06 | 2.3e-06 | 2.500E-06 | 5.292E-07 | 2.100E-06 | 3.100E-06 |
| 18677 | 934 | R934T | C | 1.8e-06 | 1.6e-06 | 1.7e-06 | 1.700E-06 | 1.000E-07 | 1.600E-06 | 1.800E-06 |
| 18678 | 934 | R934W | C | 1.7e-06 | 2.2e-06 | 2.2e-06 | 2.033E-06 | 2.887E-07 | 1.700E-06 | 2.200E-06 |
| 18679 | 934 | R934Y | C | 2.2e-06 | 4.5e-06 | 2.2e-06 | 2.967E-06 | 1.328E-06 | 2.200E-06 | 4.500E-06 |
| 18680 | 934 | R934V | C | 2.3e-06 | 2.7e-06 | 1.9e-06 | 2.300E-06 | 4.000E-07 | 1.900E-06 | 2.700E-06 |
| 18681 | 935 | N935A | C | 2.9e-06 | 3.4e-06 | 2.8e-06 | 3.033E-06 | 3.215E-07 | 2.800E-06 | 3.400E-06 |
| 18682 | 935 | N935R | C | 2.2e-06 | 1.5e-06 | 1.7e-06 | 1.800E-06 | 3.606E-07 | 1.500E-06 | 2.200E-06 |
| 18683 | 935 | N935N | C | 2.4e-06 | 2.5e-06 | 2.6e-06 | 2.500E-06 | 1.000E-07 | 2.400E-06 | 2.600E-06 |
| 18684 | 935 | N935D | C | 2.2e-06 | 3.3e-06 | 1.9e-06 | 2.467E-06 | 7.371E-07 | 1.900E-06 | 3.300E-06 |
| 18685 | 935 | N935C | C | 1.9e-06 | 2.1e-06 | 1.8e-06 | 1.933E-06 | 1.528E-07 | 1.800E-06 | 2.100E-06 |
| 18686 | 935 | N935E | C | 2.1e-06 | 3.3e-06 | 1.8e-06 | 2.400E-06 | 7.937E-07 | 1.800E-06 | 3.300E-06 |
| 18687 | 935 | N935Q | C | 3.3e-06 | 4.5e-06 | 2.1e-06 | 3.300E-06 | 1.200E-06 | 2.100E-06 | 4.500E-06 |
| 18688 | 935 | N935G | C | 1.9e-06 | 3.1e-06 | 3e-06 | 2.667E-06 | 6.658E-07 | 1.900E-06 | 3.100E-06 |
| 18689 | 935 | N935H | C | 3.5e-06 | 3.3e-06 | 2.5e-06 | 3.100E-06 | 5.292E-07 | 2.500E-06 | 3.500E-06 |
| 18690 | 935 | N935I | C | 2.8e-06 | 2.8e-06 | 3.1e-06 | 2.900E-06 | 1.732E-07 | 2.800E-06 | 3.100E-06 |
| 18691 | 935 | N935L | C | 2.3e-06 | 2.3e-06 | 2.1e-06 | 2.233E-06 | 1.155E-07 | 2.100E-06 | 2.300E-06 |
| 18692 | 935 | N935K | C | 1.5e-06 | 2.7e-06 | 1.9e-06 | 2.033E-06 | 6.110E-07 | 1.500E-06 | 2.700E-06 |
| 18693 | 935 | N935M | C | 2.7e-06 | 2.3e-06 | 4.6e-06 | 3.200E-06 | 1.229E-06 | 2.300E-06 | 4.600E-06 |
| 18694 | 935 | N935F | C | 3.3e-06 | 3.2e-06 | 2.3e-06 | 2.933E-06 | 5.508E-07 | 2.300E-06 | 3.300E-06 |
| 18695 | 935 | N935P | C | 2.5e-06 | 2.2e-06 | 2.4e-06 | 2.367E-06 | 1.528E-07 | 2.200E-06 | 2.500E-06 |
| 18696 | 935 | N935S | C | 2.9e-06 | 2.5e-06 | 2.1e-06 | 2.500E-06 | 4.000E-07 | 2.100E-06 | 2.900E-06 |
| 18697 | 935 | N935T | C | 2.2e-06 | 2.8e-06 | 2.6e-06 | 2.533E-06 | 3.055E-07 | 2.200E-06 | 2.800E-06 |
| 18698 | 935 | N935W | C | 3.6e-06 | 2.3e-06 | 3.2e-06 | 3.033E-06 | 6.658E-07 | 2.300E-06 | 3.600E-06 |
| 18699 | 935 | N935Y | C | 2.5e-06 | 2.3e-06 | 2.7e-06 | 2.500E-06 | 2.000E-07 | 2.300E-06 | 2.700E-06 |

| | | | | | | | | | | |
|---|---|---|---|---|---|---|---|---|---|---|
| 18700 | 935 | N935V | C | 1.7e-06 | 2.2e-06 | 2.3e-06 | 2.067E-06 | 3.215E-07 | 1.700E-06 | 2.300E-06 |
| 18701 | 936 | L936A | C | 1.6e-06 | 1.7e-06 | 2.2e-06 | 1.833E-06 | 3.215E-07 | 1.600E-06 | 2.200E-06 |
| 18702 | 936 | L936R | C | 2.1e-06 | 3.3e-06 | 2.7e-06 | 2.700E-06 | 6.000E-07 | 2.100E-06 | 3.300E-06 |
| 18703 | 936 | L936N | C | 1.8e-06 | 2.5e-06 | 2.7e-06 | 2.333E-06 | 4.726E-07 | 1.800E-06 | 2.700E-06 |
| 18704 | 936 | L936D | C | 2.2e-06 | 1.7e-06 | 2.2e-06 | 2.033E-06 | 2.887E-07 | 1.700E-06 | 2.200E-06 |
| 18705 | 936 | L936C | C | 2.2e-06 | 1.9e-06 | 2e-06 | 2.033E-06 | 1.528E-07 | 1.900E-06 | 2.200E-06 |
| 18706 | 936 | L936E | C | 2.8e-06 | 3.1e-06 | 2.3e-06 | 2.733E-06 | 4.041E-07 | 2.300E-06 | 3.100E-06 |
| 18707 | 936 | L936Q | C | 3.2e-06 | 2.5e-06 | 2.5e-06 | 2.733E-06 | 4.041E-07 | 2.500E-06 | 3.200E-06 |
| 18708 | 936 | L936G | C | 2e-06 | 2.4e-06 | 1.9e-06 | 2.100E-06 | 2.646E-07 | 1.900E-06 | 2.400E-06 |
| 18709 | 936 | L936H | C | 3.4e-06 | 1.6e-06 | 3.6e-06 | 2.867E-06 | 1.102E-06 | 1.600E-06 | 3.600E-06 |
| 18710 | 936 | L936I | C | 1.8e-06 | 2.7e-06 | 3.2e-06 | 2.567E-06 | 7.095E-07 | 1.800E-06 | 3.200E-06 |
| 18711 | 936 | L936L | C | 3.2e-06 | 2.2e-06 | 1.3e-06 | 2.233E-06 | 9.504E-07 | 1.300E-06 | 3.200E-06 |
| 18712 | 936 | L936K | C | 3.8e-06 | 3.2e-06 | 2.5e-06 | 3.167E-06 | 6.506E-07 | 2.500E-06 | 3.800E-06 |
| 18713 | 936 | L936M | C | 4.1e-06 | 2.2e-06 | 1.5e-06 | 2.600E-06 | 1.345E-06 | 1.500E-06 | 4.100E-06 |
| 18714 | 936 | L936F | C | 1.1e-06 | 3e-06 | 1.9e-06 | 2.000E-06 | 9.539E-07 | 1.100E-06 | 3.000E-06 |
| 18715 | 936 | L936P | C | 5.4e-06 | 2.6e-06 | 1.7e-06 | 3.233E-06 | 1.930E-06 | 1.700E-06 | 5.400E-06 |
| 18716 | 936 | L936S | C | 1.8e-06 | 2.1e-06 | 3.2e-06 | 2.367E-06 | 7.371E-07 | 1.800E-06 | 3.200E-06 |
| 18717 | 936 | L936T | C | 2.3e-06 | 2.7e-06 | 1.8e-06 | 2.267E-06 | 4.509E-07 | 1.800E-06 | 2.700E-06 |
| 18718 | 936 | L936W | C | 2e-06 | 2.5e-06 | 3.2e-06 | 2.567E-06 | 6.028E-07 | 2.000E-06 | 3.200E-06 |
| 18719 | 936 | L936Y | C | 2e-06 | 1.7e-06 | 2.3e-06 | 2.000E-06 | 3.000E-07 | 1.700E-06 | 2.300E-06 |
| 18720 | 936 | L936V | C | 2.5e-06 | 2.1e-06 | 2.4e-06 | 2.333E-06 | 2.082E-07 | 2.100E-06 | 2.500E-06 |
| 18721 | 937 | G937A | C | 2.8e-06 | 1.9e-06 | 2.3e-06 | 2.333E-06 | 4.509E-07 | 1.900E-06 | 2.800E-06 |
| 18722 | 937 | G937R | C | 1.9e-06 | 2.8e-06 | 2.7e-06 | 2.467E-06 | 4.933E-07 | 1.900E-06 | 2.800E-06 |
| 18723 | 937 | G937N | C | 2.2e-06 | 2.5e-06 | 3e-06 | 2.567E-06 | 4.041E-07 | 2.200E-06 | 3.000E-06 |
| 18724 | 937 | G937D | C | 3e-06 | 1.5e-06 | 1.6e-06 | 2.033E-06 | 8.386E-07 | 1.500E-06 | 3.000E-06 |
| 18725 | 937 | G937C | C | 3.1e-06 | 1.4e-06 | 1.1e-06 | 1.867E-06 | 1.079E-06 | 1.100E-06 | 3.100E-06 |
| 18726 | 937 | G937E | C | 2.3e-06 | 1.9e-06 | 2.4e-06 | 2.200E-06 | 2.646E-07 | 1.900E-06 | 2.400E-06 |
| 18727 | 937 | G937Q | C | 2.4e-06 | 2.3e-06 | 1.7e-06 | 2.133E-06 | 3.786E-07 | 1.700E-06 | 2.400E-06 |
| 18728 | 937 | G937G | C | 2.3e-06 | 3.3e-06 | 3.4e-06 | 3.000E-06 | 6.083E-07 | 2.300E-06 | 3.400E-06 |
| 18729 | 937 | G937H | C | 2.9e-06 | 3.9e-06 | 3.1e-06 | 3.300E-06 | 5.292E-07 | 2.900E-06 | 3.900E-06 |
| 18730 | 937 | G937I | C | 2.3e-06 | 1.8e-06 | 3e-06 | 2.367E-06 | 6.028E-07 | 1.800E-06 | 3.000E-06 |
| 18731 | 937 | G937L | C | 1.7e-06 | 3.3e-06 | 4.7e-06 | 3.233E-06 | 1.501E-06 | 1.700E-06 | 4.700E-06 |
| 18732 | 937 | G937K | C | 2.4e-06 | 2.3e-06 | 3.8e-06 | 2.833E-06 | 8.386E-07 | 2.300E-06 | 3.800E-06 |
| 18733 | 937 | G937M | C | 2e-06 | 3.3e-06 | 2.1e-06 | 2.467E-06 | 7.234E-07 | 2.000E-06 | 3.300E-06 |
| 18734 | 937 | G937F | C | 2.2e-06 | 2.4e-06 | 3.4e-06 | 2.667E-06 | 6.429E-07 | 2.200E-06 | 3.400E-06 |
| 18735 | 937 | G937P | C | 2.8e-06 | 3.4e-06 | 2.3e-06 | 2.833E-06 | 5.508E-07 | 2.300E-06 | 3.400E-06 |
| 18736 | 937 | G937S | C | 2.3e-06 | 1.8e-06 | 3.2e-06 | 2.433E-06 | 7.095E-07 | 1.800E-06 | 3.200E-06 |
| 18737 | 937 | G937T | C | 2.1e-06 | 1.8e-06 | 3.8e-06 | 2.567E-06 | 1.079E-06 | 1.800E-06 | 3.800E-06 |
| 18738 | 937 | G937W | C | 1.6e-06 | 2.3e-06 | 2e-06 | 1.967E-06 | 3.512E-07 | 1.600E-06 | 2.300E-06 |
| 18739 | 937 | G937Y | C | 2e-06 | 2.4e-06 | 1.7e-06 | 2.033E-06 | 3.512E-07 | 1.700E-06 | 2.400E-06 |
| 18740 | 937 | G937V | C | 1.7e-06 | 4.7e-06 | 2.4e-06 | 2.933E-06 | 1.570E-06 | 1.700E-06 | 4.700E-06 |
| 18741 | 938 | P938A | C | 3.2e-06 | 1.1e-06 | 2.8e-06 | 2.367E-06 | 1.115E-06 | 1.100E-06 | 3.200E-06 |
| 18742 | 938 | P938R | C | 2.3e-06 | 3.1e-06 | 3e-06 | 2.800E-06 | 4.359E-07 | 2.300E-06 | 3.100E-06 |
| 18743 | 938 | P938N | C | 2.4e-06 | 1.6e-06 | 1.8e-06 | 1.933E-06 | 4.163E-07 | 1.600E-06 | 2.400E-06 |
| 18744 | 938 | P938D | C | 1.1e-06 | 3.1e-06 | 2.4e-06 | 2.200E-06 | 1.015E-06 | 1.100E-06 | 3.100E-06 |
| 18745 | 938 | P938C | C | 1.9e-06 | 1.8e-06 | 2.1e-06 | 1.933E-06 | 1.528E-07 | 1.800E-06 | 2.100E-06 |
| 18746 | 938 | P938E | C | 3.1e-06 | 3.1e-06 | 3.3e-06 | 3.167E-06 | 1.155E-07 | 3.100E-06 | 3.300E-06 |
| 18747 | 938 | P938Q | C | 1.1e-06 | 1.5e-06 | 2.4e-06 | 1.667E-06 | 6.658E-07 | 1.100E-06 | 2.400E-06 |
| 18748 | 938 | P938G | C | 2.9e-06 | 2.2e-06 | 2.9e-06 | 2.667E-06 | 4.041E-07 | 2.200E-06 | 2.900E-06 |
| 18749 | 938 | P938H | C | 1.5e-06 | 2.8e-06 | 1.9e-06 | 2.067E-06 | 6.658E-07 | 1.500E-06 | 2.800E-06 |
| 18750 | 938 | P938I | C | 2.3e-06 | 3.2e-06 | 2.9e-06 | 2.800E-06 | 4.583E-07 | 2.300E-06 | 3.200E-06 |
| 18751 | 938 | P938L | C | 2.4e-06 | 1.9e-06 | 1.6e-06 | 1.967E-06 | 4.041E-07 | 1.600E-06 | 2.400E-06 |
| 18752 | 938 | P938K | C | 1.9e-06 | 2.2e-06 | 1.7e-06 | 1.933E-06 | 2.517E-07 | 1.700E-06 | 2.200E-06 |
| 18753 | 938 | P938M | C | 3e-06 | 2.1e-06 | 1.9e-06 | 2.333E-06 | 5.859E-07 | 1.900E-06 | 3.000E-06 |
| 18754 | 938 | P938F | C | 1.5e-06 | 1.8e-06 | 2.7e-06 | 2.000E-06 | 6.245E-07 | 1.500E-06 | 2.700E-06 |
| 18755 | 938 | P938P | C | 1.4e-06 | 3.2e-06 | 2e-06 | 2.200E-06 | 9.165E-07 | 1.400E-06 | 3.200E-06 |
| 18756 | 938 | P938S | C | 3.9e-06 | 1.5e-06 | 2.1e-06 | 2.500E-06 | 1.249E-06 | 1.500E-06 | 3.900E-06 |
| 18757 | 938 | P938T | C | 4.6e-06 | 1.8e-06 | 2.5e-06 | 2.967E-06 | 1.457E-06 | 1.800E-06 | 4.600E-06 |
| 18758 | 938 | P938W | C | 2.6e-06 | 1.2e-06 | 1.6e-06 | 1.800E-06 | 7.211E-07 | 1.200E-06 | 2.600E-06 |
| 18759 | 938 | P938Y | C | 3e-06 | 3e-06 | 1.1e-06 | 2.367E-06 | 1.097E-06 | 1.100E-06 | 3.000E-06 |
| 18760 | 938 | P938V | C | 3.8e-06 | 4.4e-06 | 2.6e-06 | 3.600E-06 | 9.165E-07 | 2.600E-06 | 4.400E-06 |
| 18761 | 939 | S939A | C | 1.4e-06 | 3e-06 | 2.4e-06 | 2.267E-06 | 8.083E-07 | 1.400E-06 | 3.000E-06 |
| 18762 | 939 | S939R | C | 1.5e-06 | 1.4e-06 | 2.4e-06 | 1.767E-06 | 5.508E-07 | 1.400E-06 | 2.400E-06 |
| 18763 | 939 | S939N | C | 1.9e-06 | 3.7e-06 | 2e-06 | 2.533E-06 | 1.012E-06 | 1.900E-06 | 3.700E-06 |
| 18764 | 939 | S939D | C | 2e-06 | 1.2e-06 | 2.7e-06 | 1.967E-06 | 7.506E-07 | 1.200E-06 | 2.700E-06 |
| 18765 | 939 | S939C | C | 2.9e-06 | 2.1e-06 | 2e-06 | 2.333E-06 | 4.933E-07 | 2.000E-06 | 2.900E-06 |
| 18766 | 939 | S939E | C | 3.7e-06 | 3.1e-06 | 3e-06 | 3.267E-06 | 3.786E-07 | 3.000E-06 | 3.700E-06 |
| 18767 | 939 | S939Q | C | 2.4e-06 | 3.1e-06 | 2.9e-06 | 2.800E-06 | 3.606E-07 | 2.400E-06 | 3.100E-06 |
| 18768 | 939 | S939G | C | 2.3e-06 | 3.2e-06 | 1.1e-06 | 2.200E-06 | 1.054E-06 | 1.100E-06 | 3.200E-06 |
| 18769 | 939 | S939H | C | 2.8e-06 | 2.4e-06 | 2.6e-06 | 2.600E-06 | 2.000E-07 | 2.400E-06 | 2.800E-06 |
| 18770 | 939 | S939I | C | 1.9e-06 | 3.7e-06 | 4.9e-06 | 3.500E-06 | 1.510E-06 | 1.900E-06 | 4.900E-06 |
| 18771 | 939 | S939L | C | 2.3e-06 | 3.6e-06 | 1.8e-06 | 2.567E-06 | 9.292E-07 | 1.800E-06 | 3.600E-06 |
| 18772 | 939 | S939K | C | 2.7e-06 | 2.5e-06 | 1.9e-06 | 2.367E-06 | 4.163E-07 | 1.900E-06 | 2.700E-06 |
| 18773 | 939 | S939M | C | 2.6e-06 | 2.9e-06 | 1.1e-06 | 2.200E-06 | 9.644E-07 | 1.100E-06 | 2.900E-06 |
| 18774 | 939 | S939F | C | 1.3e-06 | 2.2e-06 | 2.8e-06 | 2.100E-06 | 7.550E-07 | 1.300E-06 | 2.800E-06 |
| 18775 | 939 | S939P | C | 4.7e-06 | 2.5e-06 | 3.1e-06 | 3.433E-06 | 1.137E-06 | 2.500E-06 | 4.700E-06 |
| 18776 | 939 | S939S | C | 3.8e-06 | 2.1e-06 | 1.6e-06 | 2.500E-06 | 1.153E-06 | 1.600E-06 | 3.800E-06 |
| 18777 | 939 | S939T | C | 4.3e-06 | 2.8e-06 | 2.7e-06 | 3.267E-06 | 8.963E-07 | 2.700E-06 | 4.300E-06 |
| 18778 | 939 | S939W | C | 1.3e-06 | 3.4e-06 | 1.9e-06 | 2.200E-06 | 1.082E-06 | 1.300E-06 | 3.400E-06 |
| 18779 | 939 | S939Y | C | 3.2e-06 | 5.1e-06 | 3.5e-06 | 3.933E-06 | 1.021E-06 | 3.200E-06 | 5.100E-06 |
| 18780 | 939 | S939V | C | 1.8e-06 | 1.7e-06 | 1.9e-06 | 1.800E-06 | 1.000E-07 | 1.700E-06 | 1.900E-06 |
| 18781 | 940 | F940A | C | 2e-06 | 2.9e-06 | 2e-06 | 2.300E-06 | 5.196E-07 | 2.000E-06 | 2.900E-06 |
| 18782 | 940 | F940R | C | 2.1e-06 | 2.9e-06 | 1.9e-06 | 2.300E-06 | 5.292E-07 | 1.900E-06 | 2.900E-06 |
| 18783 | 940 | F940N | C | 2.2e-06 | 3.4e-06 | 2.1e-06 | 2.567E-06 | 7.234E-07 | 2.100E-06 | 3.400E-06 |
| 18784 | 940 | F940D | C | 2.4e-06 | 2.1e-06 | 2.7e-06 | 2.400E-06 | 3.000E-07 | 2.100E-06 | 2.700E-06 |

| | | | | | | | | | | |
|---|---|---|---|---|---|---|---|---|---|---|
| 18785 | 940 | F940C | C | 1.8e-06 | 2.8e-06 | 1.5e-06 | 2.033E-06 | 6.807E-07 | 1.500E-06 | 2.800E-06 |
| 18786 | 940 | F940E | C | 2.8e-06 | 2.6e-06 | 1.7e-06 | 2.367E-06 | 5.859E-07 | 1.700E-06 | 2.800E-06 |
| 18787 | 940 | F940Q | C | 2.8e-06 | 1.9e-06 | 2.3e-06 | 2.333E-06 | 4.509E-07 | 1.900E-06 | 2.800E-06 |
| 18788 | 940 | F940G | C | 3.3e-06 | 1.8e-06 | 2.3e-06 | 2.467E-06 | 7.638E-07 | 1.800E-06 | 3.300E-06 |
| 18789 | 940 | F940H | C | 8.5e-07 | 2.3e-06 | 4.3e-06 | 2.483E-06 | 1.732E-06 | 8.500E-07 | 4.300E-06 |
| 18790 | 940 | F940I | C | 2e-06 | 1.8e-06 | 2.4e-06 | 2.067E-06 | 3.055E-07 | 1.800E-06 | 2.400E-06 |
| 18791 | 940 | F940L | C | 3.5e-06 | 1.1e-06 | 4.6e-06 | 3.067E-06 | 1.790E-06 | 1.100E-06 | 4.600E-06 |
| 18792 | 940 | F940K | C | 2.2e-06 | 2.6e-06 | 4.2e-06 | 3.000E-06 | 1.058E-06 | 2.200E-06 | 4.200E-06 |
| 18793 | 940 | F940M | C | 2.6e-06 | 1.9e-06 | 2e-06 | 2.167E-06 | 3.786E-07 | 1.900E-06 | 2.600E-06 |
| 18794 | 940 | F940F | C | 2.3e-06 | 2e-06 | 2.2e-06 | 2.167E-06 | 1.528E-07 | 2.000E-06 | 2.300E-06 |
| 18795 | 940 | F940P | C | 1.1e-06 | 3.7e-06 | 2.3e-06 | 2.367E-06 | 1.301E-06 | 1.100E-06 | 3.700E-06 |
| 18796 | 940 | F940S | C | 3.1e-06 | 2.6e-06 | 1.7e-06 | 2.467E-06 | 7.095E-07 | 1.700E-06 | 3.100E-06 |
| 18797 | 940 | F940T | C | 1.9e-06 | 2.4e-06 | 1.6e-06 | 1.967E-06 | 4.041E-07 | 1.600E-06 | 2.400E-06 |
| 18798 | 940 | F940W | C | 1.8e-06 | 1.8e-06 | 1.3e-06 | 1.633E-06 | 2.887E-07 | 1.300E-06 | 1.800E-06 |
| 18799 | 940 | F940Y | C | 2.4e-06 | 2.3e-06 | 3e-06 | 2.567E-06 | 3.786E-07 | 2.300E-06 | 3.000E-06 |
| 18800 | 940 | F940V | C | 1.7e-06 | 3.8e-06 | 3.5e-06 | 3.000E-06 | 1.136E-06 | 1.700E-06 | 3.800E-06 |
| 18801 | 941 | Y941A | C | 3e-06 | 3.1e-06 | 1.7e-06 | 2.600E-06 | 7.810E-07 | 1.700E-06 | 3.100E-06 |
| 18802 | 941 | Y941R | C | 2.4e-06 | 3.1e-06 | 2e-06 | 2.500E-06 | 5.568E-07 | 2.000E-06 | 3.100E-06 |
| 18803 | 941 | Y941N | C | 2.6e-06 | 2.6e-06 | 2e-06 | 2.400E-06 | 3.464E-07 | 2.000E-06 | 2.600E-06 |
| 18804 | 941 | Y941D | C | 2.1e-06 | 2.2e-06 | 3.4e-06 | 2.567E-06 | 7.234E-07 | 2.100E-06 | 3.400E-06 |
| 18805 | 941 | Y941C | C | 2.7e-06 | 2.6e-06 | 2.7e-06 | 2.667E-06 | 5.774E-08 | 2.600E-06 | 2.700E-06 |
| 18806 | 941 | Y941E | C | 2.8e-06 | 2.8e-06 | 3.2e-06 | 2.933E-06 | 2.309E-07 | 2.800E-06 | 3.200E-06 |
| 18807 | 941 | Y941Q | C | 1.6e-06 | 3e-06 | 2.4e-06 | 2.333E-06 | 7.024E-07 | 1.600E-06 | 3.000E-06 |
| 18808 | 941 | Y941G | C | 2.7e-06 | 2.4e-06 | 1.8e-06 | 2.300E-06 | 4.583E-07 | 1.800E-06 | 2.700E-06 |
| 18809 | 941 | Y941H | C | 2.9e-06 | 3e-06 | 1.9e-06 | 2.600E-06 | 6.083E-07 | 1.900E-06 | 3.000E-06 |
| 18810 | 941 | Y941I | C | 4e-06 | 1.9e-06 | 2.1e-06 | 2.667E-06 | 1.159E-06 | 1.900E-06 | 4.000E-06 |
| 18811 | 941 | Y941L | C | 4.3e-06 | 3.1e-06 | 1.9e-06 | 3.100E-06 | 1.200E-06 | 1.900E-06 | 4.300E-06 |
| 18812 | 941 | Y941K | C | 9e-07 | 1.2e-06 | 1.7e-06 | 1.267E-06 | 4.041E-07 | 9.000E-07 | 1.700E-06 |
| 18813 | 941 | Y941M | C | 2.9e-06 | 3.6e-06 | 1.5e-06 | 2.667E-06 | 1.069E-06 | 1.500E-06 | 3.600E-06 |
| 18814 | 941 | Y941F | C | 2.7e-06 | 2.2e-06 | 3.5e-06 | 2.800E-06 | 6.557E-07 | 2.200E-06 | 3.500E-06 |
| 18815 | 941 | Y941P | C | 2.3e-06 | 2.5e-06 | 1.5e-06 | 2.100E-06 | 5.292E-07 | 1.500E-06 | 2.500E-06 |
| 18816 | 941 | Y941S | C | 1.8e-06 | 2.8e-06 | 4.8e-06 | 3.133E-06 | 1.528E-06 | 1.800E-06 | 4.800E-06 |
| 18817 | 941 | Y941T | C | 2.7e-06 | 2.9e-06 | 2.4e-06 | 2.667E-06 | 2.517E-07 | 2.400E-06 | 2.900E-06 |
| 18818 | 941 | Y941W | C | 2.3e-06 | 2.4e-06 | 1.5e-06 | 2.067E-06 | 4.933E-07 | 1.500E-06 | 2.400E-06 |
| 18819 | 941 | Y941Y | C | 4e-06 | 2.3e-06 | 4e-06 | 3.433E-06 | 9.815E-07 | 2.300E-06 | 4.000E-06 |
| 18820 | 941 | Y941V | C | 3.8e-06 | 3.1e-06 | 2.8e-06 | 3.233E-06 | 5.132E-07 | 2.800E-06 | 3.800E-06 |
| 18821 | 942 | F942A | C | 4.2e-06 | 2.2e-06 | 3.3e-06 | 3.233E-06 | 1.002E-06 | 2.200E-06 | 4.200E-06 |
| 18822 | 942 | F942R | C | 2.9e-06 | 5.6e-06 | 2.1e-06 | 3.533E-06 | 1.834E-06 | 2.100E-06 | 5.600E-06 |
| 18823 | 942 | F942N | C | 1.6e-06 | 2.8e-06 | 1.7e-06 | 2.033E-06 | 6.658E-07 | 1.600E-06 | 2.800E-06 |
| 18824 | 942 | F942D | C | 3.2e-06 | 2.3e-06 | 3.8e-06 | 3.100E-06 | 7.550E-07 | 2.300E-06 | 3.800E-06 |
| 18825 | 942 | F942C | C | 1.4e-06 | 2.1e-06 | 2e-06 | 1.833E-06 | 3.786E-07 | 1.400E-06 | 2.100E-06 |
| 18826 | 942 | F942E | C | 3.5e-06 | 3.6e-06 | 2.1e-06 | 3.067E-06 | 8.386E-07 | 2.100E-06 | 3.600E-06 |
| 18827 | 942 | F942Q | C | 2.7e-06 | 2.8e-06 | 4.1e-06 | 3.200E-06 | 7.810E-07 | 2.700E-06 | 4.100E-06 |
| 18828 | 942 | F942G | C | 2e-06 | 1.8e-06 | 2.6e-06 | 2.133E-06 | 4.163E-07 | 1.800E-06 | 2.600E-06 |
| 18829 | 942 | F942H | C | 1.8e-06 | 2.1e-06 | 1.2e-06 | 1.700E-06 | 4.583E-07 | 1.200E-06 | 2.100E-06 |
| 18830 | 942 | F942I | C | 2.3e-06 | 3.5e-06 | 2.9e-06 | 2.900E-06 | 6.000E-07 | 2.300E-06 | 3.500E-06 |
| 18831 | 942 | F942L | C | 2.6e-06 | 1.6e-06 | 1.7e-06 | 1.967E-06 | 5.508E-07 | 1.600E-06 | 2.600E-06 |
| 18832 | 942 | F942K | C | 2.1e-06 | 2e-06 | 1.9e-06 | 2.000E-06 | 1.000E-07 | 1.900E-06 | 2.100E-06 |
| 18833 | 942 | F942M | C | 1.7e-06 | 2.6e-06 | 2.7e-06 | 2.333E-06 | 5.508E-07 | 1.700E-06 | 2.700E-06 |
| 18834 | 942 | F942F | C | 9.8e-07 | 1.1e-06 | 2.5e-06 | 1.527E-06 | 8.451E-07 | 9.800E-07 | 2.500E-06 |
| 18835 | 942 | F942P | C | 2.4e-06 | 3.5e-06 | 2.5e-06 | 2.800E-06 | 6.083E-07 | 2.400E-06 | 3.500E-06 |
| 18836 | 942 | F942S | C | 1.7e-06 | 1.6e-06 | 1.6e-06 | 1.633E-06 | 5.774E-08 | 1.600E-06 | 1.700E-06 |
| 18837 | 942 | F942T | C | 1.9e-06 | 1.4e-06 | 2.7e-06 | 2.000E-06 | 6.557E-07 | 1.400E-06 | 2.700E-06 |
| 18838 | 942 | F942W | C | 1.3e-06 | 3.1e-06 | 2.3e-06 | 2.233E-06 | 9.018E-07 | 1.300E-06 | 3.100E-06 |
| 18839 | 942 | F942Y | C | 1.9e-06 | 2.5e-06 | 1.8e-06 | 2.067E-06 | 3.786E-07 | 1.800E-06 | 2.500E-06 |
| 18840 | 942 | F942V | C | 3.1e-06 | 1.8e-06 | 1.4e-06 | 2.100E-06 | 8.888E-07 | 1.400E-06 | 3.100E-06 |
| 18841 | 943 | D943A | C | 3.4e-06 | 2.6e-06 | 1.6e-06 | 2.533E-06 | 9.018E-07 | 1.600E-06 | 3.400E-06 |
| 18842 | 943 | D943R | C | 3.6e-06 | 2.7e-06 | 2.2e-06 | 2.833E-06 | 7.095E-07 | 2.200E-06 | 3.600E-06 |
| 18843 | 943 | D943N | C | 2.2e-06 | 4.1e-06 | 3.3e-06 | 3.200E-06 | 9.539E-07 | 2.200E-06 | 4.100E-06 |
| 18844 | 943 | D943D | C | 1.9e-06 | 1.6e-06 | 2e-06 | 1.833E-06 | 2.082E-07 | 1.600E-06 | 2.000E-06 |
| 18845 | 943 | D943C | C | 2.6e-06 | 2.2e-06 | 2e-06 | 2.267E-06 | 3.055E-07 | 2.000E-06 | 2.600E-06 |
| 18846 | 943 | D943E | C | 1.8e-06 | 1.7e-06 | 2.8e-06 | 2.100E-06 | 6.083E-07 | 1.700E-06 | 2.800E-06 |
| 18847 | 943 | D943Q | C | 2.1e-06 | 2.4e-06 | 3.5e-06 | 2.667E-06 | 7.371E-07 | 2.100E-06 | 3.500E-06 |
| 18848 | 943 | D943G | C | 4.3e-06 | 1.8e-06 | 2.6e-06 | 2.900E-06 | 1.277E-06 | 1.800E-06 | 4.300E-06 |
| 18849 | 943 | D943H | C | 2.9e-06 | 1.8e-06 | 2.1e-06 | 2.267E-06 | 5.686E-07 | 1.800E-06 | 2.900E-06 |
| 18850 | 943 | D943I | C | 1.6e-06 | 1.6e-06 | 3.8e-06 | 2.333E-06 | 1.270E-06 | 1.600E-06 | 3.800E-06 |
| 18851 | 943 | D943L | C | 2.7e-06 | 3.3e-06 | 1.1e-06 | 2.367E-06 | 1.137E-06 | 1.100E-06 | 3.300E-06 |
| 18852 | 943 | D943K | C | 1.8e-06 | 1.6e-06 | 2.1e-06 | 1.833E-06 | 2.517E-07 | 1.600E-06 | 2.100E-06 |
| 18853 | 943 | D943M | C | 3.1e-06 | 2.8e-06 | 2.2e-06 | 2.700E-06 | 4.583E-07 | 2.200E-06 | 3.100E-06 |
| 18854 | 943 | D943F | C | 2e-06 | 2.1e-06 | 3.6e-06 | 2.567E-06 | 8.963E-07 | 2.000E-06 | 3.600E-06 |
| 18855 | 943 | D943P | C | 2.8e-06 | 1.5e-06 | 2.5e-06 | 2.267E-06 | 6.807E-07 | 1.500E-06 | 2.800E-06 |
| 18856 | 943 | D943S | C | 1.6e-06 | 1.8e-06 | 2.4e-06 | 1.933E-06 | 4.163E-07 | 1.600E-06 | 2.400E-06 |
| 18857 | 943 | D943T | C | 3.6e-06 | 3.8e-06 | 1.8e-06 | 3.067E-06 | 1.102E-06 | 1.800E-06 | 3.800E-06 |
| 18858 | 943 | D943W | C | 2.5e-06 | 2.7e-06 | 2.5e-06 | 2.567E-06 | 1.155E-07 | 2.500E-06 | 2.700E-06 |
| 18859 | 943 | D943Y | C | 2.7e-06 | 2.8e-06 | 3.4e-06 | 2.967E-06 | 3.786E-07 | 2.700E-06 | 3.400E-06 |
| 18860 | 943 | D943V | C | 3.9e-06 | 3.2e-06 | 2e-06 | 3.033E-06 | 9.609E-07 | 2.000E-06 | 3.900E-06 |
| 18861 | 944 | Y944A | C | 2.4e-06 | 1.7e-06 | 2.2e-06 | 2.100E-06 | 3.606E-07 | 1.700E-06 | 2.400E-06 |
| 18862 | 944 | Y944R | C | 1.6e-06 | 3.5e-06 | 3.1e-06 | 2.733E-06 | 1.002E-06 | 1.600E-06 | 3.500E-06 |
| 18863 | 944 | Y944N | C | 1.8e-06 | 3.2e-06 | 2.4e-06 | 2.467E-06 | 7.024E-07 | 1.800E-06 | 3.200E-06 |
| 18864 | 944 | Y944D | C | 2e-06 | 2.3e-06 | 1.3e-06 | 1.867E-06 | 5.132E-07 | 1.300E-06 | 2.300E-06 |
| 18865 | 944 | Y944C | C | 1.7e-06 | 1.7e-06 | 1.6e-06 | 1.667E-06 | 5.774E-08 | 1.600E-06 | 1.700E-06 |
| 18866 | 944 | Y944E | C | 3.6e-06 | 2.6e-06 | 2.6e-06 | 2.933E-06 | 5.774E-07 | 2.600E-06 | 3.600E-06 |
| 18867 | 944 | Y944Q | C | 2.5e-06 | 1.9e-06 | 1.9e-06 | 2.100E-06 | 3.464E-07 | 1.900E-06 | 2.500E-06 |
| 18868 | 944 | Y944G | C | 1.9e-06 | 2e-06 | 2.1e-06 | 2.000E-06 | 1.000E-07 | 1.900E-06 | 2.100E-06 |
| 18869 | 944 | Y944H | C | 1.2e-06 | 4e-06 | 1.8e-06 | 2.333E-06 | 1.474E-06 | 1.200E-06 | 4.000E-06 |

| | | | | | | | | | | |
|---|---|---|---|---|---|---|---|---|---|---|
| 18870 | 944 | Y944I | C | 4.4e-06 | 3.2e-06 | 3.2e-06 | 3.600E-06 | 6.928E-07 | 3.200E-06 | 4.400E-06 |
| 18871 | 944 | Y944L | C | 1.7e-06 | 2.2e-06 | 1.7e-06 | 1.867E-06 | 2.887E-07 | 1.700E-06 | 2.200E-06 |
| 18872 | 944 | Y944K | C | 2.2e-06 | 3e-06 | 2e-06 | 2.400E-06 | 5.292E-07 | 2.000E-06 | 3.000E-06 |
| 18873 | 944 | Y944M | C | 2.9e-06 | 9e-07 | 3.2e-06 | 2.333E-06 | 1.250E-06 | 9.000E-07 | 3.200E-06 |
| 18874 | 944 | Y944F | C | 2.2e-06 | 3.3e-06 | 1.6e-06 | 2.367E-06 | 8.622E-07 | 1.600E-06 | 3.300E-06 |
| 18875 | 944 | Y944P | C | 1.9e-06 | 2.9e-06 | 3.7e-06 | 2.833E-06 | 9.018E-07 | 1.900E-06 | 3.700E-06 |
| 18876 | 944 | Y944S | C | 2.3e-06 | 1.6e-06 | 1.7e-06 | 1.867E-06 | 3.786E-07 | 1.600E-06 | 2.300E-06 |
| 18877 | 944 | Y944T | C | 2.2e-06 | 3.1e-06 | 2.9e-06 | 2.733E-06 | 4.726E-07 | 2.200E-06 | 3.100E-06 |
| 18878 | 944 | Y944W | C | 2e-06 | 2.3e-06 | 1.5e-06 | 1.933E-06 | 4.041E-07 | 1.500E-06 | 2.300E-06 |
| 18879 | 944 | Y944Y | C | 5.4e-06 | 1.6e-06 | 2.9e-06 | 3.300E-06 | 1.931E-06 | 1.600E-06 | 5.400E-06 |
| 18880 | 944 | Y944V | C | 2.4e-06 | 3e-06 | 2.7e-06 | 2.700E-06 | 3.000E-07 | 2.400E-06 | 3.000E-06 |
| 18881 | 945 | W945A | C | 3e-06 | 2.8e-06 | 1.8e-06 | 2.533E-06 | 6.429E-07 | 1.800E-06 | 3.000E-06 |
| 18882 | 945 | W945R | C | 1.9e-06 | 4.1e-06 | 2.3e-06 | 2.767E-06 | 1.172E-06 | 1.900E-06 | 4.100E-06 |
| 18883 | 945 | W945N | C | 3.6e-06 | 2.5e-06 | 2e-06 | 2.700E-06 | 8.185E-07 | 2.000E-06 | 3.600E-06 |
| 18884 | 945 | W945D | C | 2e-06 | 2.7e-06 | 1.5e-06 | 2.067E-06 | 6.028E-07 | 1.500E-06 | 2.700E-06 |
| 18885 | 945 | W945C | C | 2.1e-06 | 4.5e-06 | 1.8e-06 | 2.800E-06 | 1.480E-06 | 1.800E-06 | 4.500E-06 |
| 18886 | 945 | W945E | C | 4.1e-06 | 4.5e-06 | 3.2e-06 | 3.933E-06 | 6.658E-07 | 3.200E-06 | 4.500E-06 |
| 18887 | 945 | W945Q | C | 1.5e-06 | 1.7e-06 | 1.1e-06 | 1.433E-06 | 3.055E-07 | 1.100E-06 | 1.700E-06 |
| 18888 | 945 | W945G | C | 1.9e-06 | 4.8e-06 | 1.8e-06 | 2.833E-06 | 1.704E-06 | 1.800E-06 | 4.800E-06 |
| 18889 | 945 | W945H | C | 3e-06 | 1.9e-06 | 1.8e-06 | 2.233E-06 | 6.658E-07 | 1.800E-06 | 3.000E-06 |
| 18890 | 945 | W945I | C | 2.5e-06 | 2.7e-06 | 1.9e-06 | 2.367E-06 | 4.163E-07 | 1.900E-06 | 2.700E-06 |
| 18891 | 945 | W945L | C | 1.3e-06 | 2.3e-06 | 3.9e-06 | 2.500E-06 | 1.311E-06 | 1.300E-06 | 3.900E-06 |
| 18892 | 945 | W945K | C | 1.5e-06 | 1.9e-06 | 1.6e-06 | 1.667E-06 | 2.082E-07 | 1.500E-06 | 1.900E-06 |
| 18893 | 945 | W945M | C | 3.7e-06 | 2.2e-06 | 2.2e-06 | 2.700E-06 | 8.660E-07 | 2.200E-06 | 3.700E-06 |
| 18894 | 945 | W945F | C | 1.1e-06 | 2e-06 | 4.7e-06 | 2.600E-06 | 1.873E-06 | 1.100E-06 | 4.700E-06 |
| 18895 | 945 | W945P | C | 1.4e-06 | 4.5e-06 | 2.6e-06 | 2.833E-06 | 1.563E-06 | 1.400E-06 | 4.500E-06 |
| 18896 | 945 | W945S | C | 2.6e-06 | 3.3e-06 | 2.3e-06 | 2.733E-06 | 5.132E-07 | 2.300E-06 | 3.300E-06 |
| 18897 | 945 | W945T | C | 2.3e-06 | 2.8e-06 | 9.5e-07 | 2.017E-06 | 9.570E-07 | 9.500E-07 | 2.800E-06 |
| 18898 | 945 | W945W | C | 2e-06 | 2.8e-06 | 1.7e-06 | 2.167E-06 | 5.686E-07 | 1.700E-06 | 2.800E-06 |
| 18899 | 945 | W945Y | C | 2.9e-06 | 1.6e-06 | 3e-06 | 2.500E-06 | 7.810E-07 | 1.600E-06 | 3.000E-06 |
| 18900 | 945 | W945V | C | 3.1e-06 | 3.2e-06 | 2.5e-06 | 2.933E-06 | 3.786E-07 | 2.500E-06 | 3.200E-06 |
| 18901 | 946 | G946A | C | 2.1e-06 | 1.1e-06 | 1.5e-06 | 1.567E-06 | 5.033E-07 | 1.100E-06 | 2.100E-06 |
| 18902 | 946 | G946R | C | 2.6e-06 | 2.3e-06 | 2.8e-06 | 2.567E-06 | 2.517E-07 | 2.300E-06 | 2.800E-06 |
| 18903 | 946 | G946N | C | 2.5e-06 | 2.7e-06 | 2.3e-06 | 2.500E-06 | 2.000E-07 | 2.300E-06 | 2.700E-06 |
| 18904 | 946 | G946D | C | 2.6e-06 | 2.6e-06 | 2e-06 | 2.400E-06 | 3.464E-07 | 2.000E-06 | 2.600E-06 |
| 18905 | 946 | G946C | C | 2e-06 | 1.9e-06 | 2.8e-06 | 2.233E-06 | 4.933E-07 | 1.900E-06 | 2.800E-06 |
| 18906 | 946 | G946E | C | 2.5e-06 | 1.9e-06 | 2.2e-06 | 2.200E-06 | 3.000E-07 | 1.900E-06 | 2.500E-06 |
| 18907 | 946 | G946Q | C | 1.4e-06 | 2.1e-06 | 1.7e-06 | 1.733E-06 | 3.512E-07 | 1.400E-06 | 2.100E-06 |
| 18908 | 946 | G946G | C | 3.2e-06 | 1.9e-06 | 2e-06 | 2.367E-06 | 7.234E-07 | 1.900E-06 | 3.200E-06 |
| 18909 | 946 | G946H | C | 2.3e-06 | 2.5e-06 | 3.3e-06 | 2.700E-06 | 5.292E-07 | 2.300E-06 | 3.300E-06 |
| 18910 | 946 | G946I | C | 1.7e-06 | 1.7e-06 | 2.4e-06 | 1.933E-06 | 4.041E-07 | 1.700E-06 | 2.400E-06 |
| 18911 | 946 | G946L | C | 2.6e-06 | 2.4e-06 | 2.4e-06 | 2.467E-06 | 1.155E-07 | 2.400E-06 | 2.600E-06 |
| 18912 | 946 | G946K | C | 2.3e-06 | 2.1e-06 | 3.4e-06 | 2.600E-06 | 7.000E-07 | 2.100E-06 | 3.400E-06 |
| 18913 | 946 | G946M | C | 2.3e-06 | 2e-06 | 2.4e-06 | 2.233E-06 | 2.082E-07 | 2.000E-06 | 2.400E-06 |
| 18914 | 946 | G946F | C | 1.7e-06 | 2.2e-06 | 2.9e-06 | 2.267E-06 | 6.028E-07 | 1.700E-06 | 2.900E-06 |
| 18915 | 946 | G946P | C | 2.3e-06 | 2.1e-06 | 1.5e-06 | 1.967E-06 | 4.163E-07 | 1.500E-06 | 2.300E-06 |
| 18916 | 946 | G946S | C | 2.5e-06 | 2.6e-06 | 2.2e-06 | 2.433E-06 | 2.082E-07 | 2.200E-06 | 2.600E-06 |
| 18917 | 946 | G946T | C | 2.7e-06 | 2.4e-06 | 1.9e-06 | 2.333E-06 | 4.041E-07 | 1.900E-06 | 2.700E-06 |
| 18918 | 946 | G946W | C | 2.2e-06 | 5.2e-06 | 1.9e-06 | 3.100E-06 | 1.825E-06 | 1.900E-06 | 5.200E-06 |
| 18919 | 946 | G946Y | C | 1.8e-06 | 2.4e-06 | 2.6e-06 | 2.267E-06 | 4.163E-07 | 1.800E-06 | 2.600E-06 |
| 18920 | 946 | G946V | C | 2.8e-06 | 1.7e-06 | 2.1e-06 | 2.200E-06 | 5.568E-07 | 1.700E-06 | 2.800E-06 |
| 18921 | 947 | Q947A | C | 2.3e-06 | 3.4e-06 | 2.5e-06 | 2.733E-06 | 5.859E-07 | 2.300E-06 | 3.400E-06 |
| 18922 | 947 | Q947R | C | 2.8e-06 | 3.4e-06 | 2.1e-06 | 2.767E-06 | 6.506E-07 | 2.100E-06 | 3.400E-06 |
| 18923 | 947 | Q947N | C | 2.6e-06 | 2.3e-06 | 3.2e-06 | 2.700E-06 | 4.583E-07 | 2.300E-06 | 3.200E-06 |
| 18924 | 947 | Q947D | C | 1.9e-06 | 2e-06 | 2.7e-06 | 2.200E-06 | 4.359E-07 | 1.900E-06 | 2.700E-06 |
| 18925 | 947 | Q947C | C | 4.4e-06 | 1.4e-06 | 2e-06 | 2.600E-06 | 1.587E-06 | 1.400E-06 | 4.400E-06 |
| 18926 | 947 | Q947E | C | 1.5e-06 | 2.3e-06 | 2.4e-06 | 2.067E-06 | 4.933E-07 | 1.500E-06 | 2.400E-06 |
| 18927 | 947 | Q947Q | C | 3.1e-06 | 1.5e-06 | 4.3e-06 | 2.967E-06 | 1.405E-06 | 1.500E-06 | 4.300E-06 |
| 18928 | 947 | Q947G | C | 2.3e-06 | 2.4e-06 | 1.8e-06 | 2.167E-06 | 3.215E-07 | 1.800E-06 | 2.400E-06 |
| 18929 | 947 | Q947H | C | 2.1e-06 | 2e-06 | 2.9e-06 | 2.333E-06 | 4.933E-07 | 2.000E-06 | 2.900E-06 |
| 18930 | 947 | Q947I | C | 4e-06 | 1.7e-06 | 2.2e-06 | 2.633E-06 | 1.210E-06 | 1.700E-06 | 4.000E-06 |
| 18931 | 947 | Q947L | C | 3.2e-06 | 4e-06 | 2.1e-06 | 3.100E-06 | 9.539E-07 | 2.100E-06 | 4.000E-06 |
| 18932 | 947 | Q947K | C | 1.8e-06 | 1.7e-06 | 1.6e-06 | 1.700E-06 | 1.000E-07 | 1.600E-06 | 1.800E-06 |
| 18933 | 947 | Q947M | C | 2.1e-06 | 3.7e-06 | 3.9e-06 | 3.233E-06 | 9.866E-07 | 2.100E-06 | 3.900E-06 |
| 18934 | 947 | Q947F | C | 2.1e-06 | 3.3e-06 | 1.6e-06 | 2.333E-06 | 8.737E-07 | 1.600E-06 | 3.300E-06 |
| 18935 | 947 | Q947P | C | 3.2e-06 | 1.6e-06 | 2.5e-06 | 2.433E-06 | 8.021E-07 | 1.600E-06 | 3.200E-06 |
| 18936 | 947 | Q947S | C | 2.7e-06 | 2.2e-06 | 2.3e-06 | 2.400E-06 | 2.646E-07 | 2.200E-06 | 2.700E-06 |
| 18937 | 947 | Q947T | C | 1.9e-06 | 3.4e-06 | 4.1e-06 | 3.133E-06 | 1.124E-06 | 1.900E-06 | 4.100E-06 |
| 18938 | 947 | Q947W | C | 1.9e-06 | 3e-06 | 2e-06 | 2.300E-06 | 6.083E-07 | 1.900E-06 | 3.000E-06 |
| 18939 | 947 | Q947Y | C | 2.3e-06 | 2.2e-06 | 3.1e-06 | 2.533E-06 | 4.933E-07 | 2.200E-06 | 3.100E-06 |
| 18940 | 947 | Q947V | C | 2.8e-06 | 3e-06 | 2.3e-06 | 2.700E-06 | 3.606E-07 | 2.300E-06 | 3.000E-06 |
| 18941 | 948 | G948A | C | 3.8e-06 | 1.9e-06 | 2.9e-06 | 2.867E-06 | 9.504E-07 | 1.900E-06 | 3.800E-06 |
| 18942 | 948 | G948R | C | 1.4e-06 | 1.5e-06 | 2.4e-06 | 1.767E-06 | 5.508E-07 | 1.400E-06 | 2.400E-06 |
| 18943 | 948 | G948N | C | 3.5e-06 | 1.7e-06 | 2.9e-06 | 2.700E-06 | 9.165E-07 | 1.700E-06 | 3.500E-06 |
| 18944 | 948 | G948D | C | 5.7e-06 | 2.3e-06 | 1.4e-06 | 3.133E-06 | 2.268E-06 | 1.400E-06 | 5.700E-06 |
| 18945 | 948 | G948C | C | 2.5e-06 | 2e-06 | 2.4e-06 | 2.300E-06 | 2.646E-07 | 2.000E-06 | 2.500E-06 |
| 18946 | 948 | G948E | C | 1.8e-06 | 1.8e-06 | 3.2e-06 | 2.267E-06 | 8.083E-07 | 1.800E-06 | 3.200E-06 |
| 18947 | 948 | G948Q | C | 3.6e-06 | 6.5e-06 | 1.3e-06 | 3.800E-06 | 2.606E-06 | 1.300E-06 | 6.500E-06 |
| 18948 | 948 | G948G | C | 2.2e-06 | 2e-06 | 1.9e-06 | 2.033E-06 | 1.528E-07 | 1.900E-06 | 2.200E-06 |
| 18949 | 948 | G948H | C | 4.3e-06 | 2.2e-06 | 1.8e-06 | 2.767E-06 | 1.343E-06 | 1.800E-06 | 4.300E-06 |
| 18950 | 948 | G948I | C | 3.4e-06 | 3.4e-06 | 2e-06 | 2.933E-06 | 8.083E-07 | 2.000E-06 | 3.400E-06 |
| 18951 | 948 | G948L | C | 4.4e-06 | 2.4e-06 | 3.8e-06 | 3.533E-06 | 1.026E-06 | 2.400E-06 | 4.400E-06 |
| 18952 | 948 | G948K | C | 1.2e-06 | 1.1e-06 | 1.5e-06 | 1.267E-06 | 2.082E-07 | 1.100E-06 | 1.500E-06 |
| 18953 | 948 | G948M | C | 2.4e-06 | 2.6e-06 | 4e-06 | 3.000E-06 | 8.718E-07 | 2.400E-06 | 4.000E-06 |
| 18954 | 948 | G948F | C | 9.5e-07 | 2.3e-06 | 1.7e-06 | 1.650E-06 | 6.764E-07 | 9.500E-07 | 2.300E-06 |

| | | | | | | | | | | |
|---|---|---|---|---|---|---|---|---|---|---|
| 18955 | 948 | G948P | C | 1.9e-06 | 1.6e-06 | 2e-06 | 1.833E-06 | 2.082E-07 | 1.600E-06 | 2.000E-06 |
| 18956 | 948 | G948S | C | 3e-06 | 2e-06 | 2.5e-06 | 2.500E-06 | 5.000E-07 | 2.000E-06 | 3.000E-06 |
| 18957 | 948 | G948T | C | 1.3e-06 | 2.9e-06 | 3.9e-06 | 2.700E-06 | 1.311E-06 | 1.300E-06 | 3.900E-06 |
| 18958 | 948 | G948W | C | 1.4e-06 | 9.6e-07 | 2.9e-06 | 1.753E-06 | 1.017E-06 | 9.600E-07 | 2.900E-06 |
| 18959 | 948 | G948Y | C | 3.1e-06 | 1.6e-06 | 2.3e-06 | 2.333E-06 | 7.506E-07 | 1.600E-06 | 3.100E-06 |
| 18960 | 948 | G948V | C | 2.5e-06 | 2.7e-06 | 2.6e-06 | 2.600E-06 | 1.000E-07 | 2.500E-06 | 2.700E-06 |
| 18961 | 949 | T949A | C | 3.8e-06 | 2.1e-06 | 2.1e-06 | 2.667E-06 | 9.815E-07 | 2.100E-06 | 3.800E-06 |
| 18962 | 949 | T949R | C | 1.7e-06 | 2.7e-06 | 2.5e-06 | 2.300E-06 | 5.292E-07 | 1.700E-06 | 2.700E-06 |
| 18963 | 949 | T949N | C | 2.4e-06 | 3.1e-06 | 2.6e-06 | 2.700E-06 | 3.606E-07 | 2.400E-06 | 3.100E-06 |
| 18964 | 949 | T949D | C | 2.6e-06 | 3.9e-06 | 2.6e-06 | 3.033E-06 | 7.506E-07 | 2.600E-06 | 3.900E-06 |
| 18965 | 949 | T949C | C | 2e-06 | 2.9e-06 | 1.6e-06 | 2.167E-06 | 6.658E-07 | 1.600E-06 | 2.900E-06 |
| 18966 | 949 | T949E | C | 3.5e-06 | 3e-06 | 1.9e-06 | 2.800E-06 | 8.185E-07 | 1.900E-06 | 3.500E-06 |
| 18967 | 949 | T949Q | C | 4.2e-06 | 5e-06 | 3.6e-06 | 4.267E-06 | 7.024E-07 | 3.600E-06 | 5.000E-06 |
| 18968 | 949 | T949G | C | 2.2e-06 | 1.8e-06 | 2.8e-06 | 2.267E-06 | 5.033E-07 | 1.800E-06 | 2.800E-06 |
| 18969 | 949 | T949H | C | 3.6e-06 | 2.1e-06 | 2.3e-06 | 2.667E-06 | 8.145E-07 | 2.100E-06 | 3.600E-06 |
| 18970 | 949 | T949I | C | 2.1e-06 | 1.6e-06 | 2e-06 | 1.900E-06 | 2.646E-07 | 1.600E-06 | 2.100E-06 |
| 18971 | 949 | T949L | C | 2.5e-06 | 2e-06 | 1.8e-06 | 2.100E-06 | 3.606E-07 | 1.800E-06 | 2.500E-06 |
| 18972 | 949 | T949K | C | 3.5e-06 | 3.5e-06 | 4.1e-06 | 3.700E-06 | 3.464E-07 | 3.500E-06 | 4.100E-06 |
| 18973 | 949 | T949M | C | 2.4e-06 | 2e-06 | 2.3e-06 | 2.233E-06 | 2.082E-07 | 2.000E-06 | 2.400E-06 |
| 18974 | 949 | T949F | C | 3.1e-06 | 1.2e-06 | 3e-06 | 2.433E-06 | 1.069E-06 | 1.200E-06 | 3.100E-06 |
| 18975 | 949 | T949P | C | 2.4e-06 | 2.3e-06 | 2.8e-06 | 2.500E-06 | 2.646E-07 | 2.300E-06 | 2.800E-06 |
| 18976 | 949 | T949S | C | 2.8e-06 | 1.8e-06 | 4.2e-06 | 2.933E-06 | 1.206E-06 | 1.800E-06 | 4.200E-06 |
| 18977 | 949 | T949T | C | 2e-06 | 2.1e-06 | 1.4e-06 | 1.833E-06 | 3.786E-07 | 1.400E-06 | 2.100E-06 |
| 18978 | 949 | T949W | C | 2.6e-06 | 1.4e-06 | 1.6e-06 | 1.867E-06 | 6.429E-07 | 1.400E-06 | 2.600E-06 |
| 18979 | 949 | T949Y | C | 2e-06 | 2.6e-06 | 6.1e-06 | 3.567E-06 | 2.214E-06 | 2.000E-06 | 6.100E-06 |
| 18980 | 949 | T949V | C | 2.1e-06 | 2.5e-06 | 2.7e-06 | 2.433E-06 | 3.055E-07 | 2.100E-06 | 2.700E-06 |
| 18981 | 950 | L950A | C | 2e-06 | 1.5e-06 | 2.7e-06 | 2.067E-06 | 6.028E-07 | 1.500E-06 | 2.700E-06 |
| 18982 | 950 | L950R | C | 1.9e-06 | 2.1e-06 | 2.8e-06 | 2.267E-06 | 4.726E-07 | 1.900E-06 | 2.800E-06 |
| 18983 | 950 | L950N | C | 2.9e-06 | 2.3e-06 | 1.6e-06 | 2.267E-06 | 6.506E-07 | 1.600E-06 | 2.900E-06 |
| 18984 | 950 | L950D | C | 2.4e-06 | 2.3e-06 | 2.9e-06 | 2.533E-06 | 3.215E-07 | 2.300E-06 | 2.900E-06 |
| 18985 | 950 | L950C | C | 1.1e-06 | 2.7e-06 | 2.2e-06 | 2.000E-06 | 8.185E-07 | 1.100E-06 | 2.700E-06 |
| 18986 | 950 | L950E | C | 2.2e-06 | 2.8e-06 | 1.2e-06 | 2.067E-06 | 8.083E-07 | 1.200E-06 | 2.800E-06 |
| 18987 | 950 | L950Q | C | 2.4e-06 | 2e-06 | 2e-06 | 2.133E-06 | 2.309E-07 | 2.000E-06 | 2.400E-06 |
| 18988 | 950 | L950G | C | 3.2e-06 | 1.9e-06 | 2.6e-06 | 2.567E-06 | 6.506E-07 | 1.900E-06 | 3.200E-06 |
| 18989 | 950 | L950H | C | 2.7e-06 | 2.4e-06 | 2.3e-06 | 2.467E-06 | 2.082E-07 | 2.300E-06 | 2.700E-06 |
| 18990 | 950 | L950I | C | 1.8e-06 | 2.1e-06 | 2e-06 | 1.967E-06 | 1.528E-07 | 1.800E-06 | 2.100E-06 |
| 18991 | 950 | L950L | C | 4e-06 | 1.8e-06 | 2.8e-06 | 2.867E-06 | 1.102E-06 | 1.800E-06 | 4.000E-06 |
| 18992 | 950 | L950K | C | 7.4e-06 | 3e-06 | 5.2e-06 | 5.200E-06 | 2.200E-06 | 3.000E-06 | 7.400E-06 |
| 18993 | 950 | L950M | C | 1.3e-06 | 2e-06 | 1.3e-06 | 1.533E-06 | 4.041E-07 | 1.300E-06 | 2.000E-06 |
| 18994 | 950 | L950F | C | 5.6e-06 | 3.8e-06 | 3.2e-06 | 4.200E-06 | 1.249E-06 | 3.200E-06 | 5.600E-06 |
| 18995 | 950 | L950P | C | 3e-06 | 2.6e-06 | 2.9e-06 | 2.833E-06 | 2.082E-07 | 2.600E-06 | 3.000E-06 |
| 18996 | 950 | L950S | C | 4.6e-06 | 1e-06 | 2.6e-06 | 2.733E-06 | 1.804E-06 | 1.000E-06 | 4.600E-06 |
| 18997 | 950 | L950T | C | 2.8e-06 | 1.9e-06 | 1.9e-06 | 2.200E-06 | 5.196E-07 | 1.900E-06 | 2.800E-06 |
| 18998 | 950 | L950W | C | 2.5e-06 | 3.3e-06 | 2.3e-06 | 2.700E-06 | 5.292E-07 | 2.300E-06 | 3.300E-06 |
| 18999 | 950 | L950Y | C | 1.9e-06 | 2.6e-06 | 3.1e-06 | 2.533E-06 | 6.028E-07 | 1.900E-06 | 3.100E-06 |
| 19000 | 950 | L950V | C | 9.7e-07 | 2.7e-06 | 5.7e-06 | 3.123E-06 | 2.393E-06 | 9.700E-07 | 5.700E-06 |
| 19001 | 951 | V951A | C | 2.4e-06 | 2.6e-06 | 3.6e-06 | 2.867E-06 | 6.429E-07 | 2.400E-06 | 3.600E-06 |
| 19002 | 951 | V951R | C | 1.3e-06 | 2.8e-06 | 4.5e-06 | 2.867E-06 | 1.601E-06 | 1.300E-06 | 4.500E-06 |
| 19003 | 951 | V951N | C | 3.1e-06 | 2.7e-06 | 2.5e-06 | 2.767E-06 | 3.055E-07 | 2.500E-06 | 3.100E-06 |
| 19004 | 951 | V951D | C | 2.3e-06 | 2.4e-06 | 2.7e-06 | 2.467E-06 | 2.082E-07 | 2.300E-06 | 2.700E-06 |
| 19005 | 951 | V951C | C | 2.6e-06 | 3.1e-06 | 2.3e-06 | 2.667E-06 | 4.041E-07 | 2.300E-06 | 3.100E-06 |
| 19006 | 951 | V951E | C | 1.8e-06 | 3.4e-06 | 2.3e-06 | 2.500E-06 | 8.185E-07 | 1.800E-06 | 3.400E-06 |
| 19007 | 951 | V951Q | C | 1.1e-06 | 2.6e-06 | 8.4e-07 | 1.513E-06 | 9.500E-07 | 8.400E-07 | 2.600E-06 |
| 19008 | 951 | V951G | C | 2.3e-06 | 1.2e-06 | 2.5e-06 | 2.000E-06 | 7.000E-07 | 1.200E-06 | 2.500E-06 |
| 19009 | 951 | V951H | C | 2.7e-06 | 3.2e-06 | 2.4e-06 | 2.767E-06 | 4.041E-07 | 2.400E-06 | 3.200E-06 |
| 19010 | 951 | V951I | C | 4.6e-06 | 1.9e-06 | 3.2e-06 | 3.233E-06 | 1.350E-06 | 1.900E-06 | 4.600E-06 |
| 19011 | 951 | V951L | C | 2.3e-06 | 3.3e-06 | 1.9e-06 | 2.500E-06 | 7.211E-07 | 1.900E-06 | 3.300E-06 |
| 19012 | 951 | V951K | C | 1.8e-06 | 2.2e-06 | 2.4e-06 | 2.133E-06 | 3.055E-07 | 1.800E-06 | 2.400E-06 |
| 19013 | 951 | V951M | C | 1.9e-06 | 1.7e-06 | 3.2e-06 | 2.267E-06 | 8.145E-07 | 1.700E-06 | 3.200E-06 |
| 19014 | 951 | V951F | C | 2.3e-06 | 2.6e-06 | 4.3e-06 | 3.067E-06 | 1.079E-06 | 2.300E-06 | 4.300E-06 |
| 19015 | 951 | V951P | C | 1.9e-06 | 1.5e-06 | 2.4e-06 | 1.933E-06 | 4.509E-07 | 1.500E-06 | 2.400E-06 |
| 19016 | 951 | V951S | C | 2.4e-06 | 2.7e-06 | 2.5e-06 | 2.533E-06 | 1.528E-07 | 2.400E-06 | 2.700E-06 |
| 19017 | 951 | V951T | C | 1.9e-06 | 2.3e-06 | 1.9e-06 | 2.033E-06 | 2.309E-07 | 1.900E-06 | 2.300E-06 |
| 19018 | 951 | V951W | C | 1.8e-06 | 2e-06 | 5.3e-06 | 3.033E-06 | 1.966E-06 | 1.800E-06 | 5.300E-06 |
| 19019 | 951 | V951Y | C | 2.6e-06 | 2.8e-06 | 5.6e-06 | 3.667E-06 | 1.677E-06 | 2.600E-06 | 5.600E-06 |
| 19020 | 951 | V951V | C | 2.8e-06 | 2.7e-06 | 2.9e-06 | 2.800E-06 | 1.000E-07 | 2.700E-06 | 2.900E-06 |
| 19021 | 952 | T952A | C | 2.4e-06 | 1.7e-06 | 3.1e-06 | 2.400E-06 | 7.000E-07 | 1.700E-06 | 3.100E-06 |
| 19022 | 952 | T952R | C | 3.2e-06 | 1.9e-06 | 1.7e-06 | 2.267E-06 | 8.145E-07 | 1.700E-06 | 3.200E-06 |
| 19023 | 952 | T952N | C | 3.2e-06 | 2.4e-06 | 3.1e-06 | 2.900E-06 | 4.359E-07 | 2.400E-06 | 3.200E-06 |
| 19024 | 952 | T952D | C | 3.1e-06 | 3.1e-06 | 2.3e-06 | 2.833E-06 | 4.619E-07 | 2.300E-06 | 3.100E-06 |
| 19025 | 952 | T952C | C | 2.8e-06 | 1.9e-06 | 2.6e-06 | 2.433E-06 | 4.726E-07 | 1.900E-06 | 2.800E-06 |
| 19026 | 952 | T952E | C | 3e-06 | 5.3e-06 | 2.2e-06 | 3.500E-06 | 1.609E-06 | 2.200E-06 | 5.300E-06 |
| 19027 | 952 | T952Q | C | 3.2e-06 | 1.4e-06 | 2.7e-06 | 2.433E-06 | 9.292E-07 | 1.400E-06 | 3.200E-06 |
| 19028 | 952 | T952G | C | 2.7e-06 | 2.4e-06 | 2.8e-06 | 2.633E-06 | 2.082E-07 | 2.400E-06 | 2.800E-06 |
| 19029 | 952 | T952H | C | 1.7e-06 | 3e-06 | 3.1e-06 | 2.600E-06 | 7.810E-07 | 1.700E-06 | 3.100E-06 |
| 19030 | 952 | T952I | C | 3.4e-06 | 2.3e-06 | 3.3e-06 | 3.000E-06 | 6.083E-07 | 2.300E-06 | 3.400E-06 |
| 19031 | 952 | T952L | C | 2.3e-06 | 3.5e-06 | 1.7e-06 | 2.500E-06 | 9.165E-07 | 1.700E-06 | 3.500E-06 |
| 19032 | 952 | T952K | C | 3.6e-06 | 2.4e-06 | 3.3e-06 | 3.100E-06 | 6.245E-07 | 2.400E-06 | 3.600E-06 |
| 19033 | 952 | T952M | C | 1.8e-06 | 2.3e-06 | 2.8e-06 | 2.300E-06 | 5.000E-07 | 1.800E-06 | 2.800E-06 |
| 19034 | 952 | T952F | C | 2.5e-06 | 2.4e-06 | 3e-06 | 2.633E-06 | 3.215E-07 | 2.400E-06 | 3.000E-06 |
| 19035 | 952 | T952P | C | 1.8e-06 | 1.9e-06 | 2.4e-06 | 2.033E-06 | 3.215E-07 | 1.800E-06 | 2.400E-06 |
| 19036 | 952 | T952S | C | 1.5e-06 | 1.1e-06 | 3e-06 | 1.867E-06 | 1.002E-06 | 1.100E-06 | 3.000E-06 |
| 19037 | 952 | T952T | C | 3.6e-06 | 2.4e-06 | 3.2e-06 | 3.067E-06 | 6.110E-07 | 2.400E-06 | 3.600E-06 |
| 19038 | 952 | T952W | C | 2.4e-06 | 1.6e-06 | 1.3e-06 | 1.767E-06 | 5.686E-07 | 1.300E-06 | 2.400E-06 |
| 19039 | 952 | T952Y | C | 3.1e-06 | 2.8e-06 | 4.9e-06 | 3.600E-06 | 1.136E-06 | 2.800E-06 | 4.900E-06 |

| | | | | | | | | | | |
|---|---|---|---|---|---|---|---|---|---|---|
| 19040 | 952 | T952V | C | 3e-06 | 3.2e-06 | 2.4e-06 | 2.867E-06 | 4.163E-07 | 2.400E-06 | 3.200E-06 |
| 19041 | 953 | V953A | C | 1.4e-06 | 1.8e-06 | 2.6e-06 | 1.933E-06 | 6.110E-07 | 1.400E-06 | 2.600E-06 |
| 19042 | 953 | V953R | C | 1.8e-06 | 2.2e-06 | 4.1e-06 | 2.700E-06 | 1.229E-06 | 1.800E-06 | 4.100E-06 |
| 19043 | 953 | V953N | C | 2.2e-06 | 2e-06 | 3.6e-06 | 2.600E-06 | 8.718E-07 | 2.000E-06 | 3.600E-06 |
| 19044 | 953 | V953D | C | 1.9e-06 | 2.9e-06 | 3.2e-06 | 2.667E-06 | 6.807E-07 | 1.900E-06 | 3.200E-06 |
| 19045 | 953 | V953C | C | 2e-06 | 2.5e-06 | 2.8e-06 | 2.433E-06 | 4.041E-07 | 2.000E-06 | 2.800E-06 |
| 19046 | 953 | V953E | C | 3.4e-06 | 1.7e-06 | 3.4e-06 | 2.833E-06 | 9.815E-07 | 1.700E-06 | 3.400E-06 |
| 19047 | 953 | V953Q | C | 2.1e-06 | 2.8e-06 | 2.1e-06 | 2.333E-06 | 4.041E-07 | 2.100E-06 | 2.800E-06 |
| 19048 | 953 | V953G | C | 2e-06 | 3.5e-06 | 1.7e-06 | 2.400E-06 | 9.644E-07 | 1.700E-06 | 3.500E-06 |
| 19049 | 953 | V953H | C | 3.5e-06 | 2.9e-06 | 2.5e-06 | 2.967E-06 | 5.033E-07 | 2.500E-06 | 3.500E-06 |
| 19050 | 953 | V953I | C | 3.3e-06 | 2.5e-06 | 2.4e-06 | 2.733E-06 | 4.933E-07 | 2.400E-06 | 3.300E-06 |
| 19051 | 953 | V953L | C | 1.1e-06 | 1.4e-06 | 2.8e-06 | 1.767E-06 | 9.074E-07 | 1.100E-06 | 2.800E-06 |
| 19052 | 953 | V953K | C | 9.7e-07 | 1.3e-06 | 2.4e-06 | 1.557E-06 | 7.488E-07 | 9.700E-07 | 2.400E-06 |
| 19053 | 953 | V953M | C | 2.6e-06 | 3e-06 | 4.1e-06 | 3.233E-06 | 7.767E-07 | 2.600E-06 | 4.100E-06 |
| 19054 | 953 | V953F | C | 1.8e-06 | 1.5e-06 | 2.7e-06 | 2.000E-06 | 6.245E-07 | 1.500E-06 | 2.700E-06 |
| 19055 | 953 | V953P | C | 1.2e-06 | 1.5e-06 | 1.9e-06 | 1.533E-06 | 3.512E-07 | 1.200E-06 | 1.900E-06 |
| 19056 | 953 | V953S | C | 2e-06 | 2.3e-06 | 2.2e-06 | 2.167E-06 | 1.528E-07 | 2.000E-06 | 2.300E-06 |
| 19057 | 953 | V953T | C | 1.7e-06 | 1.9e-06 | 2e-06 | 1.867E-06 | 1.528E-07 | 1.700E-06 | 2.000E-06 |
| 19058 | 953 | V953W | C | 2.5e-06 | 2e-06 | 2.1e-06 | 2.200E-06 | 2.646E-07 | 2.000E-06 | 2.500E-06 |
| 19059 | 953 | V953Y | C | 1.5e-06 | 2.5e-06 | 2.1e-06 | 2.033E-06 | 5.033E-07 | 1.500E-06 | 2.500E-06 |
| 19060 | 953 | V953V | C | 3e-06 | 4.1e-06 | 2.8e-06 | 3.300E-06 | 7.000E-07 | 2.800E-06 | 4.100E-06 |
| 19061 | 954 | S954A | C | 1.5e-06 | 3.4e-06 | 1.2e-06 | 2.033E-06 | 1.193E-06 | 1.200E-06 | 3.400E-06 |
| 19062 | 954 | S954R | C | 3.2e-06 | 2.9e-06 | 2.6e-06 | 2.900E-06 | 3.000E-07 | 2.600E-06 | 3.200E-06 |
| 19063 | 954 | S954N | C | 1.7e-06 | 2.3e-06 | 2.1e-06 | 2.033E-06 | 3.055E-07 | 1.700E-06 | 2.300E-06 |
| 19064 | 954 | S954D | C | 1.7e-06 | 2.3e-06 | 3.5e-06 | 2.500E-06 | 9.165E-07 | 1.700E-06 | 3.500E-06 |
| 19065 | 954 | S954C | C | 1.2e-06 | 2.9e-06 | 2.6e-06 | 2.233E-06 | 9.074E-07 | 1.200E-06 | 2.900E-06 |
| 19066 | 954 | S954E | C | 2.5e-06 | 3.9e-06 | 2.7e-06 | 3.033E-06 | 7.572E-07 | 2.500E-06 | 3.900E-06 |
| 19067 | 954 | S954Q | C | 2.8e-06 | 2.7e-06 | 2.1e-06 | 2.533E-06 | 3.786E-07 | 2.100E-06 | 2.800E-06 |
| 19068 | 954 | S954G | C | 1.8e-06 | 2.6e-06 | 1.7e-06 | 2.033E-06 | 4.933E-07 | 1.700E-06 | 2.600E-06 |
| 19069 | 954 | S954H | C | 2.3e-06 | 2.1e-06 | 1.2e-06 | 1.867E-06 | 5.859E-07 | 1.200E-06 | 2.300E-06 |
| 19070 | 954 | S954I | C | 1.8e-06 | 2.6e-06 | 1.5e-06 | 1.967E-06 | 5.686E-07 | 1.500E-06 | 2.600E-06 |
| 19071 | 954 | S954L | C | 2e-06 | 2.4e-06 | 1.7e-06 | 2.033E-06 | 3.512E-07 | 1.700E-06 | 2.400E-06 |
| 19072 | 954 | S954K | C | 2.1e-06 | 2.4e-06 | 2e-06 | 2.167E-06 | 2.082E-07 | 2.000E-06 | 2.400E-06 |
| 19073 | 954 | S954M | C | 3.5e-06 | 3e-06 | 1.9e-06 | 2.800E-06 | 8.185E-07 | 1.900E-06 | 3.500E-06 |
| 19074 | 954 | S954F | C | 2.3e-06 | 1.4e-06 | 4.4e-06 | 2.700E-06 | 1.539E-06 | 1.400E-06 | 4.400E-06 |
| 19075 | 954 | S954P | C | 3.8e-06 | 1.8e-06 | 4.9e-06 | 3.500E-06 | 1.572E-06 | 1.800E-06 | 4.900E-06 |
| 19076 | 954 | S954S | C | 2.4e-06 | 3.2e-06 | 1.6e-06 | 2.400E-06 | 8.000E-07 | 1.600E-06 | 3.200E-06 |
| 19077 | 954 | S954T | C | 2.4e-06 | 2.1e-06 | 2.2e-06 | 2.233E-06 | 1.528E-07 | 2.100E-06 | 2.400E-06 |
| 19078 | 954 | S954W | C | 2e-06 | 1.1e-06 | 3e-06 | 2.033E-06 | 9.504E-07 | 1.100E-06 | 3.000E-06 |
| 19079 | 954 | S954Y | C | 2.5e-06 | 2.7e-06 | 3.3e-06 | 2.833E-06 | 4.163E-07 | 2.500E-06 | 3.300E-06 |
| 19080 | 954 | S954V | C | 2.9e-06 | 2.8e-06 | 4.2e-06 | 3.300E-06 | 7.810E-07 | 2.800E-06 | 4.200E-06 |
| 19081 | 955 | S955A | C | 1e-06 | 2.8e-06 | 2.2e-06 | 2.000E-06 | 9.165E-07 | 1.000E-06 | 2.800E-06 |
| 19082 | 955 | S955R | C | 9.7e-07 | 2.3e-06 | 2.6e-06 | 1.957E-06 | 8.675E-07 | 9.700E-07 | 2.600E-06 |
| 19083 | 955 | S955N | C | 2.2e-06 | 1.8e-06 | 2.8e-06 | 2.267E-06 | 5.033E-07 | 1.800E-06 | 2.800E-06 |
| 19084 | 955 | S955D | C | 1.6e-06 | 4.5e-06 | 2e-06 | 2.700E-06 | 1.572E-06 | 1.600E-06 | 4.500E-06 |
| 19085 | 955 | S955C | C | 1.5e-06 | 2.4e-06 | 2.1e-06 | 2.000E-06 | 4.583E-07 | 1.500E-06 | 2.400E-06 |
| 19086 | 955 | S955E | C | 1.4e-06 | 2.8e-06 | 2.1e-06 | 2.100E-06 | 7.000E-07 | 1.400E-06 | 2.800E-06 |
| 19087 | 955 | S955Q | C | 2.1e-06 | 1.5e-06 | 2.8e-06 | 2.133E-06 | 6.506E-07 | 1.500E-06 | 2.800E-06 |
| 19088 | 955 | S955G | C | 2.3e-06 | 1.7e-06 | 2.3e-06 | 2.100E-06 | 3.464E-07 | 1.700E-06 | 2.300E-06 |
| 19089 | 955 | S955H | C | 1.9e-06 | 3.6e-06 | 3.1e-06 | 2.867E-06 | 8.737E-07 | 1.900E-06 | 3.600E-06 |
| 19090 | 955 | S955I | C | 2.5e-06 | 2.9e-06 | 2.2e-06 | 2.533E-06 | 3.512E-07 | 2.200E-06 | 2.900E-06 |
| 19091 | 955 | S955L | C | 1.4e-06 | 3.4e-06 | 2.8e-06 | 2.533E-06 | 1.026E-06 | 1.400E-06 | 3.400E-06 |
| 19092 | 955 | S955K | C | 2.4e-06 | 2.8e-06 | 2.2e-06 | 2.467E-06 | 3.055E-07 | 2.200E-06 | 2.800E-06 |
| 19093 | 955 | S955M | C | 1.9e-06 | 3.4e-06 | 8.6e-07 | 2.053E-06 | 1.277E-06 | 8.600E-07 | 3.400E-06 |
| 19094 | 955 | S955F | C | 1.6e-06 | 2.6e-06 | 2.5e-06 | 2.233E-06 | 5.508E-07 | 1.600E-06 | 2.600E-06 |
| 19095 | 955 | S955P | C | 2.3e-06 | 1.9e-06 | 2e-06 | 2.067E-06 | 2.082E-07 | 1.900E-06 | 2.300E-06 |
| 19096 | 955 | S955S | C | 2.3e-06 | 2.4e-06 | 1.8e-06 | 2.167E-06 | 3.215E-07 | 1.800E-06 | 2.400E-06 |
| 19097 | 955 | S955T | C | 1.8e-06 | 1.8e-06 | 1.3e-06 | 1.633E-06 | 2.887E-07 | 1.300E-06 | 1.800E-06 |
| 19098 | 955 | S955W | C | 1.7e-06 | 2.6e-06 | 3.1e-06 | 2.467E-06 | 7.095E-07 | 1.700E-06 | 3.100E-06 |
| 19099 | 955 | S955Y | C | 2.8e-06 | 2.2e-06 | 4.1e-06 | 3.033E-06 | 9.713E-07 | 2.200E-06 | 4.100E-06 |
| 19100 | 955 | S955V | C | 1.5e-06 | 2.7e-06 | 2.7e-06 | 2.300E-06 | 6.928E-07 | 1.500E-06 | 2.700E-06 |
| 19101 | 956 | A956A | C | 2.1e-06 | 2.3e-06 | 2.6e-06 | 2.333E-06 | 2.517E-07 | 2.100E-06 | 2.600E-06 |
| 19102 | 956 | A956R | C | 2.3e-06 | 2.4e-06 | 2.7e-06 | 2.467E-06 | 2.082E-07 | 2.300E-06 | 2.700E-06 |
| 19103 | 956 | A956N | C | 2.2e-06 | 2.6e-06 | 1.4e-06 | 2.067E-06 | 6.110E-07 | 1.400E-06 | 2.600E-06 |
| 19104 | 956 | A956D | C | 2.7e-06 | 2.5e-06 | 1.8e-06 | 2.333E-06 | 4.726E-07 | 1.800E-06 | 2.700E-06 |
| 19105 | 956 | A956C | C | 2e-06 | 1.6e-06 | 2.8e-06 | 2.133E-06 | 6.110E-07 | 1.600E-06 | 2.800E-06 |
| 19106 | 956 | A956E | C | 1.9e-06 | 2.6e-06 | 1.5e-06 | 2.000E-06 | 5.568E-07 | 1.500E-06 | 2.600E-06 |
| 19107 | 956 | A956Q | C | 1.5e-06 | 3.2e-06 | 2.9e-06 | 2.533E-06 | 9.074E-07 | 1.500E-06 | 3.200E-06 |
| 19108 | 956 | A956G | C | 3.3e-06 | 3e-06 | 3.1e-06 | 3.133E-06 | 1.528E-07 | 3.000E-06 | 3.300E-06 |
| 19109 | 956 | A956H | C | 3.1e-06 | 1.9e-06 | 1.6e-06 | 2.200E-06 | 7.937E-07 | 1.600E-06 | 3.100E-06 |
| 19110 | 956 | A956I | C | 2.6e-06 | 2.1e-06 | 2.4e-06 | 2.367E-06 | 2.517E-07 | 2.100E-06 | 2.600E-06 |
| 19111 | 956 | A956L | C | 2.8e-06 | 1.9e-06 | 2.4e-06 | 2.367E-06 | 4.509E-07 | 1.900E-06 | 2.800E-06 |
| 19112 | 956 | A956K | C | 3.1e-06 | 3.1e-06 | 2.7e-06 | 2.967E-06 | 2.309E-07 | 2.700E-06 | 3.100E-06 |
| 19113 | 956 | A956M | C | 1.8e-06 | 4.2e-06 | 1.8e-06 | 2.600E-06 | 1.386E-06 | 1.800E-06 | 4.200E-06 |
| 19114 | 956 | A956F | C | 3.7e-06 | 3e-06 | 2.7e-06 | 3.133E-06 | 5.132E-07 | 2.700E-06 | 3.700E-06 |
| 19115 | 956 | A956P | C | 3.9e-06 | 1.7e-06 | 3.5e-06 | 3.033E-06 | 1.172E-06 | 1.700E-06 | 3.900E-06 |
| 19116 | 956 | A956S | C | 2.5e-06 | 2.4e-06 | 3.2e-06 | 2.700E-06 | 4.359E-07 | 2.400E-06 | 3.200E-06 |
| 19117 | 956 | A956T | C | 3.2e-06 | 1.8e-06 | 1.1e-06 | 2.033E-06 | 1.069E-06 | 1.100E-06 | 3.200E-06 |
| 19118 | 956 | A956W | C | 1.3e-06 | 3.4e-06 | 1.7e-06 | 2.133E-06 | 1.115E-06 | 1.300E-06 | 3.400E-06 |
| 19119 | 956 | A956Y | C | 1.7e-06 | 3.6e-06 | 3.2e-06 | 2.833E-06 | 1.002E-06 | 1.700E-06 | 3.600E-06 |
| 19120 | 956 | A956V | C | 2.7e-06 | 2e-06 | 2.6e-06 | 2.433E-06 | 3.786E-07 | 2.000E-06 | 2.700E-06 |
| 19121 | 957 | S957A | C | 2.6e-06 | 3.4e-06 | 3e-06 | 3.000E-06 | 4.000E-07 | 2.600E-06 | 3.400E-06 |
| 19122 | 957 | S957R | C | 1.3e-06 | 3e-06 | 1.7e-06 | 2.000E-06 | 8.888E-07 | 1.300E-06 | 3.000E-06 |
| 19123 | 957 | S957N | C | 3.1e-06 | 2.8e-06 | 2.2e-06 | 2.700E-06 | 4.583E-07 | 2.200E-06 | 3.100E-06 |
| 19124 | 957 | S957D | C | 3.7e-06 | 3.6e-06 | 2.3e-06 | 3.200E-06 | 7.810E-07 | 2.300E-06 | 3.700E-06 |

| | | | | | | | | | | |
|---|---|---|---|---|---|---|---|---|---|---|
| 19125 | 957 | S957C | C | 2.7e-06 | 2.5e-06 | 1.7e-06 | 2.300E-06 | 5.292E-07 | 1.700E-06 | 2.700E-06 |
| 19126 | 957 | S957E | C | 2.3e-06 | 2.7e-06 | 2.2e-06 | 2.400E-06 | 2.646E-07 | 2.200E-06 | 2.700E-06 |
| 19127 | 957 | S957Q | C | 1.3e-06 | 1.5e-06 | 1.5e-06 | 1.433E-06 | 1.155E-07 | 1.300E-06 | 1.500E-06 |
| 19128 | 957 | S957G | C | 2.2e-06 | 5e-06 | 2.5e-06 | 3.233E-06 | 1.537E-06 | 2.200E-06 | 5.000E-06 |
| 19129 | 957 | S957H | C | 2.8e-06 | 3.1e-06 | 2.4e-06 | 2.767E-06 | 3.512E-07 | 2.400E-06 | 3.100E-06 |
| 19130 | 957 | S957I | C | 2.3e-06 | 4.5e-06 | 2.4e-06 | 3.067E-06 | 1.242E-06 | 2.300E-06 | 4.500E-06 |
| 19131 | 957 | S957L | C | 3e-06 | 2.7e-06 | 2.2e-06 | 2.633E-06 | 4.041E-07 | 2.200E-06 | 3.000E-06 |
| 19132 | 957 | S957K | C | 3.9e-06 | 5e-06 | 2.4e-06 | 3.767E-06 | 1.305E-06 | 2.400E-06 | 5.000E-06 |
| 19133 | 957 | S957M | C | 5.4e-06 | 3.2e-06 | 2e-06 | 3.533E-06 | 1.724E-06 | 2.000E-06 | 5.400E-06 |
| 19134 | 957 | S957F | C | 2.7e-06 | 3e-06 | 2.6e-06 | 2.767E-06 | 2.082E-07 | 2.600E-06 | 3.000E-06 |
| 19135 | 957 | S957P | C | 1.6e-06 | 3.2e-06 | 2.7e-06 | 2.500E-06 | 8.185E-07 | 1.600E-06 | 3.200E-06 |
| 19136 | 957 | S957S | C | 3.3e-06 | 1.1e-06 | 2.5e-06 | 2.300E-06 | 1.114E-06 | 1.100E-06 | 3.300E-06 |
| 19137 | 957 | S957T | C | 3e-06 | 3.1e-06 | 2.2e-06 | 2.767E-06 | 4.933E-07 | 2.200E-06 | 3.100E-06 |
| 19138 | 957 | S957W | C | 2e-06 | 2.2e-06 | 1.3e-06 | 1.833E-06 | 4.726E-07 | 1.300E-06 | 2.200E-06 |
| 19139 | 957 | S957Y | C | 1.7e-06 | 1.9e-06 | 1.3e-06 | 1.633E-06 | 3.055E-07 | 1.300E-06 | 1.900E-06 |
| 19140 | 957 | S957V | C | 1.9e-06 | 3.4e-06 | 3.2e-06 | 2.833E-06 | 8.145E-07 | 1.900E-06 | 3.400E-06 |
| 19141 | 958 | T958A | C | 2e-06 | 1.8e-06 | 1.3e-06 | 1.700E-06 | 3.606E-07 | 1.300E-06 | 2.000E-06 |
| 19142 | 958 | T958R | C | 1.5e-06 | 3.3e-06 | 3.1e-06 | 2.633E-06 | 9.866E-07 | 1.500E-06 | 3.300E-06 |
| 19143 | 958 | T958N | C | 2.4e-06 | 1.6e-06 | 4e-06 | 2.667E-06 | 1.222E-06 | 1.600E-06 | 4.000E-06 |
| 19144 | 958 | T958D | C | 3.6e-06 | 3.4e-06 | 1.8e-06 | 2.933E-06 | 9.866E-07 | 1.800E-06 | 3.600E-06 |
| 19145 | 958 | T958C | C | 2.6e-06 | 1.8e-06 | 1.2e-06 | 1.867E-06 | 7.024E-07 | 1.200E-06 | 2.600E-06 |
| 19146 | 958 | T958E | C | 3.6e-06 | 3.8e-06 | 1.9e-06 | 3.100E-06 | 1.044E-06 | 1.900E-06 | 3.800E-06 |
| 19147 | 958 | T958Q | C | 4e-06 | 2.6e-06 | 2.3e-06 | 2.967E-06 | 9.074E-07 | 2.300E-06 | 4.000E-06 |
| 19148 | 958 | T958G | C | 2.9e-06 | 2.6e-06 | 2.1e-06 | 2.533E-06 | 4.041E-07 | 2.100E-06 | 2.900E-06 |
| 19149 | 958 | T958H | C | 2.8e-06 | 2.3e-06 | 2.9e-06 | 2.667E-06 | 3.215E-07 | 2.300E-06 | 2.900E-06 |
| 19150 | 958 | T958I | C | 2.8e-06 | 2e-06 | 3.2e-06 | 2.667E-06 | 6.110E-07 | 2.000E-06 | 3.200E-06 |
| 19151 | 958 | T958L | C | 1.8e-06 | 2.5e-06 | 2.3e-06 | 2.200E-06 | 3.606E-07 | 1.800E-06 | 2.500E-06 |
| 19152 | 958 | T958K | C | 1.2e-06 | 3.2e-06 | 1.4e-06 | 1.933E-06 | 1.102E-06 | 1.200E-06 | 3.200E-06 |
| 19153 | 958 | T958M | C | 1.4e-06 | 1.8e-06 | 1e-06 | 1.400E-06 | 4.000E-07 | 1.000E-06 | 1.800E-06 |
| 19154 | 958 | T958F | C | 3e-06 | 2.3e-06 | 2.3e-06 | 2.533E-06 | 4.041E-07 | 2.300E-06 | 3.000E-06 |
| 19155 | 958 | T958P | C | 2.4e-06 | 1.9e-06 | 2.6e-06 | 2.300E-06 | 3.606E-07 | 1.900E-06 | 2.600E-06 |
| 19156 | 958 | T958S | C | 2.5e-06 | 1.5e-06 | 3.1e-06 | 2.367E-06 | 8.083E-07 | 1.500E-06 | 3.100E-06 |
| 19157 | 958 | T958T | C | 2.6e-06 | 2.6e-06 | 1.7e-06 | 2.300E-06 | 5.196E-07 | 1.700E-06 | 2.600E-06 |
| 19158 | 958 | T958W | C | 4e-06 | 1.8e-06 | 2.7e-06 | 2.833E-06 | 1.106E-06 | 1.800E-06 | 4.000E-06 |
| 19159 | 958 | T958Y | C | 2.1e-06 | 2.1e-06 | 2.3e-06 | 2.167E-06 | 1.155E-07 | 2.100E-06 | 2.300E-06 |
| 19160 | 958 | T958V | C | 4.1e-06 | 1.5e-06 | 2.3e-06 | 2.633E-06 | 1.332E-06 | 1.500E-06 | 4.100E-06 |
| 19161 | 959 | K959A | C | 2.1e-06 | 2.8e-06 | 1.6e-06 | 2.167E-06 | 6.028E-07 | 1.600E-06 | 2.800E-06 |
| 19162 | 959 | K959R | C | 2.9e-06 | 3.9e-06 | 2.5e-06 | 3.100E-06 | 7.211E-07 | 2.500E-06 | 3.900E-06 |
| 19163 | 959 | K959N | C | 4.1e-06 | 2.7e-06 | 2.1e-06 | 2.967E-06 | 1.026E-06 | 2.100E-06 | 4.100E-06 |
| 19164 | 959 | K959D | C | 1.9e-06 | 3.1e-06 | 1.7e-06 | 2.233E-06 | 7.572E-07 | 1.700E-06 | 3.100E-06 |
| 19165 | 959 | K959C | C | 2.7e-06 | 2.7e-06 | 3.1e-06 | 2.833E-06 | 2.309E-07 | 2.700E-06 | 3.100E-06 |
| 19166 | 959 | K959E | C | 4.4e-06 | 2.7e-06 | 1.8e-06 | 2.967E-06 | 1.320E-06 | 1.800E-06 | 4.400E-06 |
| 19167 | 959 | K959Q | C | 2.9e-06 | 1.8e-06 | 3e-06 | 2.567E-06 | 6.658E-07 | 1.800E-06 | 3.000E-06 |
| 19168 | 959 | K959G | C | 3.8e-06 | 1.8e-06 | 2.3e-06 | 2.633E-06 | 1.041E-06 | 1.800E-06 | 3.800E-06 |
| 19169 | 959 | K959H | C | 1.7e-06 | 1.8e-06 | 2.5e-06 | 2.000E-06 | 4.359E-07 | 1.700E-06 | 2.500E-06 |
| 19170 | 959 | K959I | C | 1.6e-06 | 2.4e-06 | 2.6e-06 | 2.200E-06 | 5.292E-07 | 1.600E-06 | 2.600E-06 |
| 19171 | 959 | K959L | C | 2.7e-06 | 2.3e-06 | 2.9e-06 | 2.633E-06 | 3.055E-07 | 2.300E-06 | 2.900E-06 |
| 19172 | 959 | K959K | C | 1.1e-06 | 2.8e-06 | 1.8e-06 | 1.900E-06 | 8.544E-07 | 1.100E-06 | 2.800E-06 |
| 19173 | 959 | K959M | C | 1.5e-06 | 2.5e-06 | 6.5e-06 | 3.500E-06 | 2.646E-06 | 1.500E-06 | 6.500E-06 |
| 19174 | 959 | K959F | C | 1.8e-06 | 3.4e-06 | 1.9e-06 | 2.367E-06 | 8.963E-07 | 1.800E-06 | 3.400E-06 |
| 19175 | 959 | K959P | C | 3.5e-06 | 2.3e-06 | 3e-06 | 2.933E-06 | 6.028E-07 | 2.300E-06 | 3.500E-06 |
| 19176 | 959 | K959S | C | 2e-06 | 1.6e-06 | 2.2e-06 | 1.933E-06 | 3.055E-07 | 1.600E-06 | 2.200E-06 |
| 19177 | 959 | K959T | C | 2.6e-06 | 2.6e-06 | 2.5e-06 | 2.567E-06 | 5.774E-08 | 2.500E-06 | 2.600E-06 |
| 19178 | 959 | K959W | C | 3.9e-06 | 2.8e-06 | 2.1e-06 | 2.933E-06 | 9.074E-07 | 2.100E-06 | 3.900E-06 |
| 19179 | 959 | K959Y | C | 4.2e-06 | 2.2e-06 | 2.2e-06 | 2.867E-06 | 1.155E-06 | 2.200E-06 | 4.200E-06 |
| 19180 | 959 | K959V | C | 3.4e-06 | 1.8e-06 | 2e-06 | 2.400E-06 | 8.718E-07 | 1.800E-06 | 3.400E-06 |
| 19181 | 960 | G960A | C | 1.8e-06 | 2.2e-06 | 2.9e-06 | 2.300E-06 | 5.568E-07 | 1.800E-06 | 2.900E-06 |
| 19182 | 960 | G960R | C | 1.3e-06 | 1.8e-06 | 1.5e-06 | 1.533E-06 | 2.517E-07 | 1.300E-06 | 1.800E-06 |
| 19183 | 960 | G960N | C | 2.8e-06 | 5.4e-06 | 2.8e-06 | 3.667E-06 | 1.501E-06 | 2.800E-06 | 5.400E-06 |
| 19184 | 960 | G960D | C | 2.2e-06 | 2.7e-06 | 2.6e-06 | 2.500E-06 | 2.646E-07 | 2.200E-06 | 2.700E-06 |
| 19185 | 960 | G960C | C | 1.5e-06 | 9.7e-07 | 3.2e-06 | 1.890E-06 | 1.165E-06 | 9.700E-07 | 3.200E-06 |
| 19186 | 960 | G960E | C | 3.8e-06 | 2e-06 | 1.7e-06 | 2.500E-06 | 1.136E-06 | 1.700E-06 | 3.800E-06 |
| 19187 | 960 | G960Q | C | 1.4e-06 | 2.4e-06 | 2.6e-06 | 2.133E-06 | 6.429E-07 | 1.400E-06 | 2.600E-06 |
| 19188 | 960 | G960G | C | 1.7e-06 | 3.7e-06 | 2.5e-06 | 2.633E-06 | 1.007E-06 | 1.700E-06 | 3.700E-06 |
| 19189 | 960 | G960H | C | 1.9e-06 | 2.4e-06 | 2.2e-06 | 2.167E-06 | 2.517E-07 | 1.900E-06 | 2.400E-06 |
| 19190 | 960 | G960I | C | 2.9e-06 | 2.4e-06 | 2.8e-06 | 2.700E-06 | 2.646E-07 | 2.400E-06 | 2.900E-06 |
| 19191 | 960 | G960L | C | 1.7e-06 | 1.3e-06 | 3.1e-06 | 2.033E-06 | 9.452E-07 | 1.300E-06 | 3.100E-06 |
| 19192 | 960 | G960K | C | 2e-06 | 3.1e-06 | 1.7e-06 | 2.267E-06 | 7.371E-07 | 1.700E-06 | 3.100E-06 |
| 19193 | 960 | G960M | C | 5.9e-06 | 2.4e-06 | 3.8e-06 | 4.033E-06 | 1.762E-06 | 2.400E-06 | 5.900E-06 |
| 19194 | 960 | G960F | C | 4.3e-06 | 1.5e-06 | 2.6e-06 | 2.800E-06 | 1.411E-06 | 1.500E-06 | 4.300E-06 |
| 19195 | 960 | G960P | C | 3.4e-06 | 2.4e-06 | 1.7e-06 | 2.500E-06 | 8.544E-07 | 1.700E-06 | 3.400E-06 |
| 19196 | 960 | G960S | C | 3.7e-06 | 2.6e-06 | 3.4e-06 | 3.233E-06 | 5.686E-07 | 2.600E-06 | 3.700E-06 |
| 19197 | 960 | G960T | C | 1.1e-06 | 2.1e-06 | 2.9e-06 | 2.033E-06 | 9.018E-07 | 1.100E-06 | 2.900E-06 |
| 19198 | 960 | G960W | C | 3e-06 | 2.4e-06 | 2.5e-06 | 2.633E-06 | 3.215E-07 | 2.400E-06 | 3.000E-06 |
| 19199 | 960 | G960Y | C | 1.6e-06 | 2.6e-06 | 1.5e-06 | 1.900E-06 | 6.083E-07 | 1.500E-06 | 2.600E-06 |
| 19200 | 960 | G960V | C | 2.2e-06 | 1.8e-06 | 1.9e-06 | 1.967E-06 | 2.082E-07 | 1.800E-06 | 2.200E-06 |
| 19201 | 961 | P961A | C | 1.9e-06 | 3.8e-06 | 2.9e-06 | 2.867E-06 | 9.504E-07 | 1.900E-06 | 3.800E-06 |
| 19202 | 961 | P961R | C | 2.2e-06 | 5e-06 | 1.8e-06 | 3.000E-06 | 1.744E-06 | 1.800E-06 | 5.000E-06 |
| 19203 | 961 | P961N | C | 3.1e-06 | 1.5e-06 | 3.6e-06 | 2.733E-06 | 1.097E-06 | 1.500E-06 | 3.600E-06 |
| 19204 | 961 | P961D | C | 2.5e-06 | 2.4e-06 | 2.3e-06 | 2.400E-06 | 1.000E-07 | 2.300E-06 | 2.500E-06 |
| 19205 | 961 | P961C | C | 2e-06 | 2.9e-06 | 1.6e-06 | 2.167E-06 | 6.658E-07 | 1.600E-06 | 2.900E-06 |
| 19206 | 961 | P961E | C | 2.7e-06 | 3.4e-06 | 1.5e-06 | 2.533E-06 | 9.609E-07 | 1.500E-06 | 3.400E-06 |
| 19207 | 961 | P961Q | C | 1.8e-06 | 2.2e-06 | 2.3e-06 | 2.100E-06 | 2.646E-07 | 1.800E-06 | 2.300E-06 |
| 19208 | 961 | P961G | C | 1.8e-06 | 2.1e-06 | 2.1e-06 | 2.000E-06 | 1.732E-07 | 1.800E-06 | 2.100E-06 |
| 19209 | 961 | P961H | C | 7.9e-07 | 3.5e-06 | 3.9e-06 | 2.730E-06 | 1.692E-06 | 7.900E-07 | 3.900E-06 |

| | | | | | | | | | | |
|---|---|---|---|---|---|---|---|---|---|---|
| 19210 | 961 | P961I | C | 1.5e-06 | 2.2e-06 | 2e-06 | 1.900E-06 | 3.606E-07 | 1.500E-06 | 2.200E-06 |
| 19211 | 961 | P961L | C | 2.5e-06 | 3.9e-06 | 1.8e-06 | 2.733E-06 | 1.069E-06 | 1.800E-06 | 3.900E-06 |
| 19212 | 961 | P961K | C | 2.2e-06 | 2.8e-06 | 3.8e-06 | 2.933E-06 | 8.083E-07 | 2.200E-06 | 3.800E-06 |
| 19213 | 961 | P961M | C | 2.7e-06 | 1.5e-06 | 3.6e-06 | 2.600E-06 | 1.054E-06 | 1.500E-06 | 3.600E-06 |
| 19214 | 961 | P961F | C | 1.8e-06 | 2.9e-06 | 3.2e-06 | 2.633E-06 | 7.371E-07 | 1.800E-06 | 3.200E-06 |
| 19215 | 961 | P961P | C | 1.7e-06 | 4.4e-06 | 2e-06 | 2.700E-06 | 1.480E-06 | 1.700E-06 | 4.400E-06 |
| 19216 | 961 | P961S | C | 3e-06 | 2.2e-06 | 3e-06 | 2.733E-06 | 4.619E-07 | 2.200E-06 | 3.000E-06 |
| 19217 | 961 | P961T | C | 3.6e-06 | 3.1e-06 | 2.3e-06 | 3.000E-06 | 6.557E-07 | 2.300E-06 | 3.600E-06 |
| 19218 | 961 | P961W | C | 1.3e-06 | 3e-06 | 2.2e-06 | 2.167E-06 | 8.505E-07 | 1.300E-06 | 3.000E-06 |
| 19219 | 961 | P961Y | C | 3.6e-06 | 1.9e-06 | 2.6e-06 | 2.700E-06 | 8.544E-07 | 1.900E-06 | 3.600E-06 |
| 19220 | 961 | P961V | C | 7.4e-07 | 4.1e-06 | 9.4e-07 | 1.927E-06 | 1.885E-06 | 7.400E-07 | 4.100E-06 |
| 19221 | 962 | S962A | C | 2.7e-06 | 2.8e-06 | 2.8e-06 | 2.767E-06 | 5.774E-08 | 2.700E-06 | 2.800E-06 |
| 19222 | 962 | S962R | C | 2.3e-06 | 1.4e-06 | 3.8e-06 | 2.500E-06 | 1.212E-06 | 1.400E-06 | 3.800E-06 |
| 19223 | 962 | S962N | C | 3e-06 | 1.9e-06 | 2.8e-06 | 2.567E-06 | 5.859E-07 | 1.900E-06 | 3.000E-06 |
| 19224 | 962 | S962D | C | 3.3e-06 | 3.1e-06 | 2.7e-06 | 3.033E-06 | 3.055E-07 | 2.700E-06 | 3.300E-06 |
| 19225 | 962 | S962C | C | 2.5e-06 | 1.4e-06 | 1.5e-06 | 1.800E-06 | 6.083E-07 | 1.400E-06 | 2.500E-06 |
| 19226 | 962 | S962E | C | 1.1e-06 | 3.2e-06 | 2.1e-06 | 2.133E-06 | 1.050E-06 | 1.100E-06 | 3.200E-06 |
| 19227 | 962 | S962Q | C | 1e-06 | 4.3e-06 | 2.3e-06 | 2.533E-06 | 1.662E-06 | 1.000E-06 | 4.300E-06 |
| 19228 | 962 | S962G | C | 2.6e-06 | 2.1e-06 | 4.6e-06 | 3.100E-06 | 1.323E-06 | 2.100E-06 | 4.600E-06 |
| 19229 | 962 | S962H | C | 3e-06 | 2.2e-06 | 2.6e-06 | 2.600E-06 | 4.000E-07 | 2.200E-06 | 3.000E-06 |
| 19230 | 962 | S962I | C | 2.2e-06 | 4.3e-06 | 2.5e-06 | 3.000E-06 | 1.136E-06 | 2.200E-06 | 4.300E-06 |
| 19231 | 962 | S962L | C | 3.1e-06 | 2.2e-06 | 2e-06 | 2.433E-06 | 5.859E-07 | 2.000E-06 | 3.100E-06 |
| 19232 | 962 | S962K | C | 3.2e-06 | 1.8e-06 | 1.9e-06 | 2.300E-06 | 7.810E-07 | 1.800E-06 | 3.200E-06 |
| 19233 | 962 | S962M | C | 5e-06 | 1.9e-06 | 2.6e-06 | 3.167E-06 | 1.626E-06 | 1.900E-06 | 5.000E-06 |
| 19234 | 962 | S962F | C | 1.9e-06 | 2.9e-06 | 2.7e-06 | 2.500E-06 | 5.292E-07 | 1.900E-06 | 2.900E-06 |
| 19235 | 962 | S962P | C | 1.4e-06 | 1.8e-06 | 1.3e-06 | 1.500E-06 | 2.646E-07 | 1.300E-06 | 1.800E-06 |
| 19236 | 962 | S962S | C | 2.1e-06 | 1.1e-06 | 3.9e-06 | 2.367E-06 | 1.419E-06 | 1.100E-06 | 3.900E-06 |
| 19237 | 962 | S962T | C | 3.4e-06 | 2.8e-06 | 2.6e-06 | 2.933E-06 | 4.163E-07 | 2.600E-06 | 3.400E-06 |
| 19238 | 962 | S962W | C | 1.7e-06 | 4.4e-06 | 1.6e-06 | 2.567E-06 | 1.589E-06 | 1.600E-06 | 4.400E-06 |
| 19239 | 962 | S962Y | C | 1.8e-06 | 2.5e-06 | 4.2e-06 | 2.833E-06 | 1.234E-06 | 1.800E-06 | 4.200E-06 |
| 19240 | 962 | S962V | C | 2.3e-06 | 2.1e-06 | 2.6e-06 | 2.333E-06 | 2.517E-07 | 2.100E-06 | 2.600E-06 |
| 19241 | 963 | V963A | C | 2.3e-06 | 2.1e-06 | 2.9e-06 | 2.433E-06 | 4.163E-07 | 2.100E-06 | 2.900E-06 |
| 19242 | 963 | V963R | C | 1.8e-06 | 3.7e-06 | 2.3e-06 | 2.600E-06 | 9.849E-07 | 1.800E-06 | 3.700E-06 |
| 19243 | 963 | V963N | C | 2.1e-06 | 2.3e-06 | 2.3e-06 | 2.233E-06 | 1.155E-07 | 2.100E-06 | 2.300E-06 |
| 19244 | 963 | V963D | C | 2.8e-06 | 3.8e-06 | 1.8e-06 | 2.800E-06 | 1.000E-06 | 1.800E-06 | 3.800E-06 |
| 19245 | 963 | V963C | C | 1.8e-06 | 2.9e-06 | 1.8e-06 | 2.167E-06 | 6.351E-07 | 1.800E-06 | 2.900E-06 |
| 19246 | 963 | V963E | C | 2.3e-06 | 3.7e-06 | 2.3e-06 | 2.767E-06 | 8.083E-07 | 2.300E-06 | 3.700E-06 |
| 19247 | 963 | V963Q | C | 4.2e-06 | 1.5e-06 | 2.4e-06 | 2.700E-06 | 1.375E-06 | 1.500E-06 | 4.200E-06 |
| 19248 | 963 | V963G | C | 1.4e-06 | 3.2e-06 | 4.5e-06 | 3.033E-06 | 1.557E-06 | 1.400E-06 | 4.500E-06 |
| 19249 | 963 | V963H | C | 2.4e-06 | 2.3e-06 | 2.5e-06 | 2.400E-06 | 1.000E-07 | 2.300E-06 | 2.500E-06 |
| 19250 | 963 | V963I | C | 4e-06 | 1.2e-06 | 2.2e-06 | 2.467E-06 | 1.419E-06 | 1.200E-06 | 4.000E-06 |
| 19251 | 963 | V963L | C | 3.2e-06 | 1.7e-06 | 2.3e-06 | 2.400E-06 | 7.550E-07 | 1.700E-06 | 3.200E-06 |
| 19252 | 963 | V963K | C | 1.2e-06 | 1.9e-06 | 2e-06 | 1.700E-06 | 4.359E-07 | 1.200E-06 | 2.000E-06 |
| 19253 | 963 | V963M | C | 2.5e-06 | 2.6e-06 | 1.4e-06 | 2.167E-06 | 6.658E-07 | 1.400E-06 | 2.600E-06 |
| 19254 | 963 | V963F | C | 2.2e-06 | 4.2e-06 | 1.6e-06 | 2.667E-06 | 1.361E-06 | 1.600E-06 | 4.200E-06 |
| 19255 | 963 | V963P | C | 3.1e-06 | 2.5e-06 | 1.6e-06 | 2.400E-06 | 7.550E-07 | 1.600E-06 | 3.100E-06 |
| 19256 | 963 | V963S | C | 1.7e-06 | 3.5e-06 | 2.7e-06 | 2.633E-06 | 9.018E-07 | 1.700E-06 | 3.500E-06 |
| 19257 | 963 | V963T | C | 2.5e-06 | 2e-06 | 2.7e-06 | 2.400E-06 | 3.606E-07 | 2.000E-06 | 2.700E-06 |
| 19258 | 963 | V963W | C | 1.8e-06 | 2.3e-06 | 4.2e-06 | 2.767E-06 | 1.266E-06 | 1.800E-06 | 4.200E-06 |
| 19259 | 963 | V963Y | C | 1.5e-06 | 3.2e-06 | 2e-06 | 2.233E-06 | 8.737E-07 | 1.500E-06 | 3.200E-06 |
| 19260 | 963 | V963V | C | 1.4e-06 | 2.9e-06 | 2.3e-06 | 2.200E-06 | 7.550E-07 | 1.400E-06 | 2.900E-06 |
| 19261 | 964 | F964A | C | 2.4e-06 | 3.7e-06 | 3.2e-06 | 3.100E-06 | 6.557E-07 | 2.400E-06 | 3.700E-06 |
| 19262 | 964 | F964R | C | 5.8e-06 | 1.9e-06 | 4.7e-06 | 4.133E-06 | 2.011E-06 | 1.900E-06 | 5.800E-06 |
| 19263 | 964 | F964N | C | 1.3e-06 | 4.3e-06 | 2.4e-06 | 2.667E-06 | 1.518E-06 | 1.300E-06 | 4.300E-06 |
| 19264 | 964 | F964D | C | 1.9e-06 | 2.1e-06 | 2.2e-06 | 2.067E-06 | 1.528E-07 | 1.900E-06 | 2.200E-06 |
| 19265 | 964 | F964C | C | 2.5e-06 | 2.1e-06 | 3.1e-06 | 2.567E-06 | 5.033E-07 | 2.100E-06 | 3.100E-06 |
| 19266 | 964 | F964E | C | 2.3e-06 | 1.8e-06 | 3e-06 | 2.367E-06 | 6.028E-07 | 1.800E-06 | 3.000E-06 |
| 19267 | 964 | F964Q | C | 4.8e-06 | 2.6e-06 | 1.8e-06 | 3.067E-06 | 1.553E-06 | 1.800E-06 | 4.800E-06 |
| 19268 | 964 | F964G | C | 1.9e-06 | 2.6e-06 | 2.2e-06 | 2.233E-06 | 3.512E-07 | 1.900E-06 | 2.600E-06 |
| 19269 | 964 | F964H | C | 4.9e-06 | 2.9e-06 | 2.2e-06 | 3.333E-06 | 1.401E-06 | 2.200E-06 | 4.900E-06 |
| 19270 | 964 | F964I | C | 2.5e-06 | 4.3e-06 | 1.5e-06 | 2.767E-06 | 1.419E-06 | 1.500E-06 | 4.300E-06 |
| 19271 | 964 | F964L | C | 2.5e-06 | 2.3e-06 | 1.5e-06 | 2.100E-06 | 5.292E-07 | 1.500E-06 | 2.500E-06 |
| 19272 | 964 | F964K | C | 2.1e-06 | 3.2e-06 | 3.6e-06 | 2.967E-06 | 7.767E-07 | 2.100E-06 | 3.600E-06 |
| 19273 | 964 | F964M | C | 4.4e-06 | 1.9e-06 | 3.5e-06 | 3.267E-06 | 1.266E-06 | 1.900E-06 | 4.400E-06 |
| 19274 | 964 | F964F | C | 2.7e-06 | 2.8e-06 | 1.4e-06 | 2.300E-06 | 7.810E-07 | 1.400E-06 | 2.800E-06 |
| 19275 | 964 | F964P | C | 2.1e-06 | 2.5e-06 | 2e-06 | 2.200E-06 | 2.646E-07 | 2.000E-06 | 2.500E-06 |
| 19276 | 964 | F964S | C | 1.7e-06 | 2.1e-06 | 1.9e-06 | 1.900E-06 | 2.000E-07 | 1.700E-06 | 2.100E-06 |
| 19277 | 964 | F964T | C | 2.9e-06 | 3e-06 | 1.6e-06 | 2.500E-06 | 7.810E-07 | 1.600E-06 | 3.000E-06 |
| 19278 | 964 | F964W | C | 2.3e-06 | 1.4e-06 | 2.4e-06 | 2.033E-06 | 5.508E-07 | 1.400E-06 | 2.400E-06 |
| 19279 | 964 | F964Y | C | 2.2e-06 | 2.4e-06 | 3.5e-06 | 2.700E-06 | 7.000E-07 | 2.200E-06 | 3.500E-06 |
| 19280 | 964 | F964V | C | 1.6e-06 | 2.2e-06 | 2.6e-06 | 2.133E-06 | 5.033E-07 | 1.600E-06 | 2.600E-06 |
| 19281 | 965 | P965A | C | 2.4e-06 | 1.7e-06 | 2.4e-06 | 2.167E-06 | 4.041E-07 | 1.700E-06 | 2.400E-06 |
| 19282 | 965 | P965R | C | 2.5e-06 | 1.5e-06 | 1e-06 | 1.667E-06 | 7.638E-07 | 1.000E-06 | 2.500E-06 |
| 19283 | 965 | P965N | C | 1.9e-06 | 2.4e-06 | 2.2e-06 | 2.167E-06 | 2.517E-07 | 1.900E-06 | 2.400E-06 |
| 19284 | 965 | P965D | C | 1.8e-06 | 2.7e-06 | 3.1e-06 | 2.533E-06 | 6.658E-07 | 1.800E-06 | 3.100E-06 |
| 19285 | 965 | P965C | C | 2.3e-06 | 2.7e-06 | 3e-06 | 2.667E-06 | 3.512E-07 | 2.300E-06 | 3.000E-06 |
| 19286 | 965 | P965E | C | 2.4e-06 | 4.3e-06 | 2.6e-06 | 3.100E-06 | 1.044E-06 | 2.400E-06 | 4.300E-06 |
| 19287 | 965 | P965Q | C | 3.4e-06 | 1.6e-06 | 2.3e-06 | 2.433E-06 | 9.074E-07 | 1.600E-06 | 3.400E-06 |
| 19288 | 965 | P965G | C | 1.2e-06 | 4.5e-06 | 3.3e-06 | 3.000E-06 | 1.670E-06 | 1.200E-06 | 4.500E-06 |
| 19289 | 965 | P965H | C | 3.3e-06 | 2.5e-06 | 1.5e-06 | 2.433E-06 | 9.018E-07 | 1.500E-06 | 3.300E-06 |
| 19290 | 965 | P965I | C | 2e-06 | 2.9e-06 | 3.2e-06 | 2.700E-06 | 6.245E-07 | 2.000E-06 | 3.200E-06 |
| 19291 | 965 | P965L | C | 3.5e-06 | 2.2e-06 | 1.7e-06 | 2.467E-06 | 9.292E-07 | 1.700E-06 | 3.500E-06 |
| 19292 | 965 | P965K | C | 2.4e-06 | 1.3e-06 | 4e-06 | 2.567E-06 | 1.358E-06 | 1.300E-06 | 4.000E-06 |
| 19293 | 965 | P965M | C | 3.8e-06 | 1.5e-06 | 4.5e-06 | 3.267E-06 | 1.570E-06 | 1.500E-06 | 4.500E-06 |
| 19294 | 965 | P965F | C | 1.3e-06 | 1.6e-06 | 1.9e-06 | 1.600E-06 | 3.000E-07 | 1.300E-06 | 1.900E-06 |

| | | | | | | | | | | |
|---|---|---|---|---|---|---|---|---|---|---|
| 19295 | 965 | P965P | C | 9.6e-07 | 2.1e-06 | 2.4e-06 | 1.820E-06 | 7.597E-07 | 9.600E-07 | 2.400E-06 |
| 19296 | 965 | P965S | C | 2.3e-06 | 2.7e-06 | 2.2e-06 | 2.400E-06 | 2.646E-07 | 2.200E-06 | 2.700E-06 |
| 19297 | 965 | P965T | C | 1.7e-06 | 2e-06 | 1.7e-06 | 1.800E-06 | 1.732E-07 | 1.700E-06 | 2.000E-06 |
| 19298 | 965 | P965W | C | 1.7e-06 | 2.5e-06 | 3.7e-06 | 2.633E-06 | 1.007E-06 | 1.700E-06 | 3.700E-06 |
| 19299 | 965 | P965Y | C | 1.6e-06 | 1.8e-06 | 1.2e-06 | 1.533E-06 | 3.055E-07 | 1.200E-06 | 1.800E-06 |
| 19300 | 965 | P965V | C | 1.5e-06 | 1.8e-06 | 3.2e-06 | 2.167E-06 | 9.074E-07 | 1.500E-06 | 3.200E-06 |
| 19301 | 966 | L966A | C | 3.8e-06 | 2.6e-06 | 3.8e-06 | 3.400E-06 | 6.928E-07 | 2.600E-06 | 3.800E-06 |
| 19302 | 966 | L966R | C | 3.1e-06 | 2e-06 | 2.3e-06 | 2.467E-06 | 5.686E-07 | 2.000E-06 | 3.100E-06 |
| 19303 | 966 | L966N | C | 1.8e-06 | 3.6e-06 | 3.8e-06 | 3.067E-06 | 1.102E-06 | 1.800E-06 | 3.800E-06 |
| 19304 | 966 | L966D | C | 3.5e-06 | 2.5e-06 | 2.6e-06 | 2.867E-06 | 5.508E-07 | 2.500E-06 | 3.500E-06 |
| 19305 | 966 | L966C | C | 3.4e-06 | 1.8e-06 | 2.6e-06 | 2.600E-06 | 8.000E-07 | 1.800E-06 | 3.400E-06 |
| 19306 | 966 | L966E | C | 2.6e-06 | 2.8e-06 | 2.3e-06 | 2.567E-06 | 2.517E-07 | 2.300E-06 | 2.800E-06 |
| 19307 | 966 | L966Q | C | 1.9e-06 | 2.7e-06 | 2.2e-06 | 2.267E-06 | 4.041E-07 | 1.900E-06 | 2.700E-06 |
| 19308 | 966 | L966G | C | 2.3e-06 | 2e-06 | 2.7e-06 | 2.333E-06 | 3.512E-07 | 2.000E-06 | 2.700E-06 |
| 19309 | 966 | L966H | C | 2.1e-06 | 2.4e-06 | 2.3e-06 | 2.267E-06 | 1.528E-07 | 2.100E-06 | 2.400E-06 |
| 19310 | 966 | L966I | C | 2.8e-06 | 2.3e-06 | 2.6e-06 | 2.567E-06 | 2.517E-07 | 2.300E-06 | 2.800E-06 |
| 19311 | 966 | L966L | C | 2e-06 | 1.7e-06 | 1.7e-06 | 1.800E-06 | 1.732E-07 | 1.700E-06 | 2.000E-06 |
| 19312 | 966 | L966K | C | 1.9e-06 | 3.4e-06 | 3.6e-06 | 2.967E-06 | 9.292E-07 | 1.900E-06 | 3.600E-06 |
| 19313 | 966 | L966M | C | 2.7e-06 | 4.6e-06 | 1.9e-06 | 3.067E-06 | 1.387E-06 | 1.900E-06 | 4.600E-06 |
| 19314 | 966 | L966F | C | 2.6e-06 | 3e-06 | 2.5e-06 | 2.700E-06 | 2.646E-07 | 2.500E-06 | 3.000E-06 |
| 19315 | 966 | L966P | C | 2.2e-06 | 3.6e-06 | 1.1e-06 | 2.300E-06 | 1.253E-06 | 1.100E-06 | 3.600E-06 |
| 19316 | 966 | L966S | C | 2.1e-06 | 1.6e-06 | 3.9e-06 | 2.533E-06 | 1.210E-06 | 1.600E-06 | 3.900E-06 |
| 19317 | 966 | L966T | C | 2.7e-06 | 2.3e-06 | 1.6e-06 | 2.200E-06 | 5.568E-07 | 1.600E-06 | 2.700E-06 |
| 19318 | 966 | L966W | C | 1.8e-06 | 2.1e-06 | 2.7e-06 | 2.200E-06 | 4.583E-07 | 1.800E-06 | 2.700E-06 |
| 19319 | 966 | L966Y | C | 1.4e-06 | 1.2e-06 | 3e-06 | 1.867E-06 | 9.866E-07 | 1.200E-06 | 3.000E-06 |
| 19320 | 966 | L966V | C | 1.4e-06 | 1.3e-06 | 2.4e-06 | 1.700E-06 | 6.083E-07 | 1.300E-06 | 2.400E-06 |
| 19321 | 967 | A967A | C | 5.9e-06 | 2.7e-06 | 3e-06 | 3.867E-06 | 1.767E-06 | 2.700E-06 | 5.900E-06 |
| 19322 | 967 | A967R | C | 1.8e-06 | 1.7e-06 | 1.7e-06 | 1.733E-06 | 5.774E-08 | 1.700E-06 | 1.800E-06 |
| 19323 | 967 | A967N | C | 2.2e-06 | 2.1e-06 | 3.1e-06 | 2.467E-06 | 5.508E-07 | 2.100E-06 | 3.100E-06 |
| 19324 | 967 | A967D | C | 2.5e-06 | 2.2e-06 | 3.4e-06 | 2.700E-06 | 6.245E-07 | 2.200E-06 | 3.400E-06 |
| 19325 | 967 | A967C | C | 2.5e-06 | 3.9e-06 | 2.3e-06 | 2.900E-06 | 8.718E-07 | 2.300E-06 | 3.900E-06 |
| 19326 | 967 | A967E | C | 2.9e-06 | 3.1e-06 | 2e-06 | 2.667E-06 | 5.859E-07 | 2.000E-06 | 3.100E-06 |
| 19327 | 967 | A967Q | C | 1.8e-06 | 3.3e-06 | 1.6e-06 | 2.233E-06 | 9.292E-07 | 1.600E-06 | 3.300E-06 |
| 19328 | 967 | A967G | C | 2.2e-06 | 4.1e-06 | 2.6e-06 | 2.967E-06 | 1.002E-06 | 2.200E-06 | 4.100E-06 |
| 19329 | 967 | A967H | C | 1.9e-06 | 1.9e-06 | 1.3e-06 | 1.700E-06 | 3.464E-07 | 1.300E-06 | 1.900E-06 |
| 19330 | 967 | A967I | C | 1.8e-06 | 1.3e-06 | 1.9e-06 | 1.667E-06 | 3.215E-07 | 1.300E-06 | 1.900E-06 |
| 19331 | 967 | A967L | C | 1.1e-06 | 3.8e-06 | 3.1e-06 | 2.667E-06 | 1.401E-06 | 1.100E-06 | 3.800E-06 |
| 19332 | 967 | A967K | C | 1.5e-06 | 3.6e-06 | 1.5e-06 | 2.200E-06 | 1.212E-06 | 1.500E-06 | 3.600E-06 |
| 19333 | 967 | A967M | C | 2e-06 | 4.7e-06 | 1.7e-06 | 2.800E-06 | 1.652E-06 | 1.700E-06 | 4.700E-06 |
| 19334 | 967 | A967F | C | 3.1e-06 | 7.1e-07 | 2.8e-06 | 2.203E-06 | 1.302E-06 | 7.100E-07 | 3.100E-06 |
| 19335 | 967 | A967P | C | 2.2e-06 | 2.7e-06 | 2.1e-06 | 2.333E-06 | 3.215E-07 | 2.100E-06 | 2.700E-06 |
| 19336 | 967 | A967S | C | 3e-06 | 2.2e-06 | 1.9e-06 | 2.367E-06 | 5.686E-07 | 1.900E-06 | 3.000E-06 |
| 19337 | 967 | A967T | C | 2.2e-06 | 2.3e-06 | 2e-06 | 2.167E-06 | 1.528E-07 | 2.000E-06 | 2.300E-06 |
| 19338 | 967 | A967W | C | 2.9e-06 | 2.9e-06 | 3.3e-06 | 3.033E-06 | 2.309E-07 | 2.900E-06 | 3.300E-06 |
| 19339 | 967 | A967Y | C | 2.6e-06 | 2.1e-06 | 2.2e-06 | 2.300E-06 | 2.646E-07 | 2.100E-06 | 2.600E-06 |
| 19340 | 967 | A967V | C | 2.2e-06 | 2.5e-06 | 2.1e-06 | 2.267E-06 | 2.082E-07 | 2.100E-06 | 2.500E-06 |
| 19341 | 968 | P968A | C | 2.7e-06 | 1.5e-06 | 1.9e-06 | 2.033E-06 | 6.110E-07 | 1.500E-06 | 2.700E-06 |
| 19342 | 968 | P968R | C | 2.6e-06 | 3.6e-06 | 3.4e-06 | 3.200E-06 | 5.292E-07 | 2.600E-06 | 3.600E-06 |
| 19343 | 968 | P968N | C | 2.6e-06 | 3.4e-06 | 1.1e-06 | 2.367E-06 | 1.168E-06 | 1.100E-06 | 3.400E-06 |
| 19344 | 968 | P968D | C | 1.7e-06 | 2e-06 | 2.6e-06 | 2.100E-06 | 4.583E-07 | 1.700E-06 | 2.600E-06 |
| 19345 | 968 | P968C | C | 2.1e-06 | 2.4e-06 | 2.6e-06 | 2.367E-06 | 2.517E-07 | 2.100E-06 | 2.600E-06 |
| 19346 | 968 | P968E | C | 2.3e-06 | 2.1e-06 | 1.2e-06 | 1.867E-06 | 5.859E-07 | 1.200E-06 | 2.300E-06 |
| 19347 | 968 | P968Q | C | 2.5e-06 | 3.1e-06 | 2.7e-06 | 2.767E-06 | 3.055E-07 | 2.500E-06 | 3.100E-06 |
| 19348 | 968 | P968G | C | 2.2e-06 | 1.9e-06 | 2.2e-06 | 2.100E-06 | 1.732E-07 | 1.900E-06 | 2.200E-06 |
| 19349 | 968 | P968H | C | 2e-06 | 3.2e-06 | 3.4e-06 | 2.867E-06 | 7.572E-07 | 2.000E-06 | 3.400E-06 |
| 19350 | 968 | P968I | C | 1.7e-06 | 2.3e-06 | 2.2e-06 | 2.067E-06 | 3.215E-07 | 1.700E-06 | 2.300E-06 |
| 19351 | 968 | P968L | C | 2.9e-06 | 2.2e-06 | 1.5e-06 | 2.200E-06 | 7.000E-07 | 1.500E-06 | 2.900E-06 |
| 19352 | 968 | P968K | C | 2.8e-06 | 2.9e-06 | 1.6e-06 | 2.433E-06 | 7.234E-07 | 1.600E-06 | 2.900E-06 |
| 19353 | 968 | P968M | C | 2.5e-06 | 1.6e-06 | 2.1e-06 | 2.067E-06 | 4.509E-07 | 1.600E-06 | 2.500E-06 |
| 19354 | 968 | P968F | C | 2.3e-06 | 1.7e-06 | 2.2e-06 | 2.067E-06 | 3.215E-07 | 1.700E-06 | 2.300E-06 |
| 19355 | 968 | P968P | C | 1.6e-06 | 2.8e-06 | 2.1e-06 | 2.167E-06 | 6.028E-07 | 1.600E-06 | 2.800E-06 |
| 19356 | 968 | P968S | C | 3.1e-06 | 2e-06 | 1.7e-06 | 2.267E-06 | 7.371E-07 | 1.700E-06 | 3.100E-06 |
| 19357 | 968 | P968T | C | 1.3e-06 | 1.2e-06 | 1.1e-06 | 1.200E-06 | 1.000E-07 | 1.100E-06 | 1.300E-06 |
| 19358 | 968 | P968W | C | 1.6e-06 | 1.5e-06 | 1.7e-06 | 1.600E-06 | 1.000E-07 | 1.500E-06 | 1.700E-06 |
| 19359 | 968 | P968Y | C | 1.8e-06 | 2.7e-06 | 1.8e-06 | 2.100E-06 | 5.196E-07 | 1.800E-06 | 2.700E-06 |
| 19360 | 968 | P968V | C | 2.1e-06 | 1.7e-06 | 2.2e-06 | 2.000E-06 | 2.646E-07 | 1.700E-06 | 2.200E-06 |
| 19361 | 969 | S969A | C | 2.5e-06 | 5.3e-06 | 2.3e-06 | 3.367E-06 | 1.677E-06 | 2.300E-06 | 5.300E-06 |
| 19362 | 969 | S969R | C | 2.3e-06 | 1.8e-06 | 1.4e-06 | 1.833E-06 | 4.509E-07 | 1.400E-06 | 2.300E-06 |
| 19363 | 969 | S969N | C | 3.7e-06 | 2e-06 | 3.7e-06 | 3.133E-06 | 9.815E-07 | 2.000E-06 | 3.700E-06 |
| 19364 | 969 | S969D | C | 9.9e-07 | 2.8e-06 | 2.7e-06 | 2.163E-06 | 1.017E-06 | 9.900E-07 | 2.800E-06 |
| 19365 | 969 | S969C | C | 3.2e-06 | 2.7e-06 | 2.1e-06 | 2.667E-06 | 5.508E-07 | 2.100E-06 | 3.200E-06 |
| 19366 | 969 | S969E | C | 9.9e-07 | 1.1e-06 | 3.9e-06 | 1.997E-06 | 1.649E-06 | 9.900E-07 | 3.900E-06 |
| 19367 | 969 | S969Q | C | 1.1e-06 | 1.6e-06 | 2.9e-06 | 1.867E-06 | 9.292E-07 | 1.100E-06 | 2.900E-06 |
| 19368 | 969 | S969G | C | 2e-06 | 1.7e-06 | 1.8e-06 | 1.833E-06 | 1.528E-07 | 1.700E-06 | 2.000E-06 |
| 19369 | 969 | S969H | C | 1.3e-06 | 1.7e-06 | 3.9e-06 | 2.300E-06 | 1.400E-06 | 1.300E-06 | 3.900E-06 |
| 19370 | 969 | S969I | C | 2.3e-06 | 3.4e-06 | 2.4e-06 | 2.700E-06 | 6.083E-07 | 2.300E-06 | 3.400E-06 |
| 19371 | 969 | S969L | C | 2.2e-06 | 2.8e-06 | 1.7e-06 | 2.233E-06 | 5.508E-07 | 1.700E-06 | 2.800E-06 |
| 19372 | 969 | S969K | C | 2.6e-06 | 2.1e-06 | 1.4e-06 | 2.033E-06 | 6.028E-07 | 1.400E-06 | 2.600E-06 |
| 19373 | 969 | S969M | C | 2.4e-06 | 2e-06 | 2.5e-06 | 2.300E-06 | 2.646E-07 | 2.000E-06 | 2.500E-06 |
| 19374 | 969 | S969F | C | 2e-06 | 2.7e-06 | 2.9e-06 | 2.533E-06 | 4.726E-07 | 2.000E-06 | 2.900E-06 |
| 19375 | 969 | S969P | C | 2.6e-06 | 1.1e-06 | 1.2e-06 | 1.633E-06 | 8.386E-07 | 1.100E-06 | 2.600E-06 |
| 19376 | 969 | S969S | C | 1.9e-06 | 2.3e-06 | 2.7e-06 | 2.300E-06 | 4.000E-07 | 1.900E-06 | 2.700E-06 |
| 19377 | 969 | S969T | C | 2.8e-06 | 4.3e-06 | 1.9e-06 | 3.000E-06 | 1.212E-06 | 1.900E-06 | 4.300E-06 |
| 19378 | 969 | S969W | C | 1.4e-06 | 2.5e-06 | 2.4e-06 | 2.100E-06 | 6.083E-07 | 1.400E-06 | 2.500E-06 |
| 19379 | 969 | S969Y | C | 1.2e-06 | 2.5e-06 | 3.7e-06 | 2.467E-06 | 1.250E-06 | 1.200E-06 | 3.700E-06 |

| | | | | | | | | | | |
|---|---|---|---|---|---|---|---|---|---|---|
| 19380 | 969 | S969V | C | 2.4e-06 | 2.6e-06 | 3.9e-06 | 2.967E-06 | 8.145E-07 | 2.400E-06 | 3.900E-06 |
| 19381 | 970 | S970A | C | 2.6e-06 | 1.9e-06 | 1.9e-06 | 2.133E-06 | 4.041E-07 | 1.900E-06 | 2.600E-06 |
| 19382 | 970 | S970R | C | 3.7e-06 | 4.3e-06 | 3.2e-06 | 3.733E-06 | 5.508E-07 | 3.200E-06 | 4.300E-06 |
| 19383 | 970 | S970N | C | 3.7e-06 | 2.2e-06 | 2.4e-06 | 2.767E-06 | 8.145E-07 | 2.200E-06 | 3.700E-06 |
| 19384 | 970 | S970D | C | 4.8e-06 | 1.9e-06 | 1.5e-06 | 2.733E-06 | 1.801E-06 | 1.500E-06 | 4.800E-06 |
| 19385 | 970 | S970C | C | 1.5e-06 | 2.4e-06 | 3.5e-06 | 2.467E-06 | 1.002E-06 | 1.500E-06 | 3.500E-06 |
| 19386 | 970 | S970E | C | 3.5e-06 | 2.9e-06 | 2.6e-06 | 3.000E-06 | 4.583E-07 | 2.600E-06 | 3.500E-06 |
| 19387 | 970 | S970Q | C | 2.5e-06 | 2.8e-06 | 3.9e-06 | 3.067E-06 | 7.371E-07 | 2.500E-06 | 3.900E-06 |
| 19388 | 970 | S970G | C | 2.2e-06 | 1.3e-06 | 2.3e-06 | 1.933E-06 | 5.508E-07 | 1.300E-06 | 2.300E-06 |
| 19389 | 970 | S970H | C | 2.1e-06 | 2.4e-06 | 2.2e-06 | 2.233E-06 | 1.528E-07 | 2.100E-06 | 2.400E-06 |
| 19390 | 970 | S970I | C | 1.3e-06 | 2.8e-06 | 2.1e-06 | 2.067E-06 | 7.506E-07 | 1.300E-06 | 2.800E-06 |
| 19391 | 970 | S970L | C | 1.5e-06 | 4.7e-06 | 8.5e-06 | 4.900E-06 | 3.504E-06 | 1.500E-06 | 8.500E-06 |
| 19392 | 970 | S970K | C | 3.8e-06 | 2.8e-06 | 1.7e-06 | 2.767E-06 | 1.050E-06 | 1.700E-06 | 3.800E-06 |
| 19393 | 970 | S970M | C | 1.4e-06 | 3.3e-06 | 2.5e-06 | 2.400E-06 | 9.539E-07 | 1.400E-06 | 3.300E-06 |
| 19394 | 970 | S970F | C | 4.5e-06 | 4.3e-06 | 2.9e-06 | 3.900E-06 | 8.718E-07 | 2.900E-06 | 4.500E-06 |
| 19395 | 970 | S970P | C | 1.5e-06 | 2.2e-06 | 4.5e-06 | 2.733E-06 | 1.570E-06 | 1.500E-06 | 4.500E-06 |
| 19396 | 970 | S970S | C | 2.8e-06 | 4.7e-06 | 1.8e-06 | 3.100E-06 | 1.473E-06 | 1.800E-06 | 4.700E-06 |
| 19397 | 970 | S970T | C | 2.2e-06 | 2.3e-06 | 2.5e-06 | 2.333E-06 | 1.528E-07 | 2.200E-06 | 2.500E-06 |
| 19398 | 970 | S970W | C | 4.4e-06 | 2.1e-06 | 1.1e-06 | 2.533E-06 | 1.692E-06 | 1.100E-06 | 4.400E-06 |
| 19399 | 970 | S970Y | C | 3.2e-06 | 3.8e-06 | 7.5e-07 | 2.583E-06 | 1.616E-06 | 7.500E-07 | 3.800E-06 |
| 19400 | 970 | S970V | C | 2.1e-06 | 2.1e-06 | 1.5e-06 | 1.900E-06 | 3.464E-07 | 1.500E-06 | 2.100E-06 |
| 19401 | 971 | K971A | C | 4.7e-06 | 2.6e-06 | 2.6e-06 | 3.300E-06 | 1.212E-06 | 2.600E-06 | 4.700E-06 |
| 19402 | 971 | K971R | C | 1.4e-06 | 3.2e-06 | 3.1e-06 | 2.567E-06 | 1.012E-06 | 1.400E-06 | 3.200E-06 |
| 19403 | 971 | K971N | C | 1.5e-06 | 2.4e-06 | 1.9e-06 | 1.933E-06 | 4.509E-07 | 1.500E-06 | 2.400E-06 |
| 19404 | 971 | K971D | C | 2e-06 | 2.2e-06 | 2.5e-06 | 2.233E-06 | 2.517E-07 | 2.000E-06 | 2.500E-06 |
| 19405 | 971 | K971C | C | 4e-06 | 2.1e-06 | 3.8e-06 | 3.300E-06 | 1.044E-06 | 2.100E-06 | 4.000E-06 |
| 19406 | 971 | K971E | C | 2.8e-06 | 1.6e-06 | 2.3e-06 | 2.233E-06 | 6.028E-07 | 1.600E-06 | 2.800E-06 |
| 19407 | 971 | K971Q | C | 2.2e-06 | 2.4e-06 | 1.9e-06 | 2.167E-06 | 2.517E-07 | 1.900E-06 | 2.400E-06 |
| 19408 | 971 | K971G | C | 2.1e-06 | 2.6e-06 | 2.9e-06 | 2.533E-06 | 4.041E-07 | 2.100E-06 | 2.900E-06 |
| 19409 | 971 | K971H | C | 3.2e-06 | 2.9e-06 | 1.5e-06 | 2.533E-06 | 9.074E-07 | 1.500E-06 | 3.200E-06 |
| 19410 | 971 | K971I | C | 2.5e-06 | 2.2e-06 | 1.2e-06 | 1.967E-06 | 6.807E-07 | 1.200E-06 | 2.500E-06 |
| 19411 | 971 | K971L | C | 6.8e-06 | 3.3e-06 | 2.5e-06 | 4.200E-06 | 2.287E-06 | 2.500E-06 | 6.800E-06 |
| 19412 | 971 | K971K | C | 2.7e-06 | 1.9e-06 | 2.4e-06 | 2.333E-06 | 4.041E-07 | 1.900E-06 | 2.700E-06 |
| 19413 | 971 | K971M | C | 3.7e-06 | 1.5e-06 | 2.4e-06 | 2.533E-06 | 1.106E-06 | 1.500E-06 | 3.700E-06 |
| 19414 | 971 | K971F | C | 3.7e-06 | 2.7e-06 | 1.5e-06 | 2.633E-06 | 1.102E-06 | 1.500E-06 | 3.700E-06 |
| 19415 | 971 | K971P | C | 4.8e-06 | 4.1e-06 | 1.8e-06 | 3.567E-06 | 1.570E-06 | 1.800E-06 | 4.800E-06 |
| 19416 | 971 | K971S | C | 1.6e-06 | 1.7e-06 | 2.3e-06 | 1.867E-06 | 3.786E-07 | 1.600E-06 | 2.300E-06 |
| 19417 | 971 | K971T | C | 1.3e-06 | 1.9e-06 | 4.5e-06 | 2.567E-06 | 1.701E-06 | 1.300E-06 | 4.500E-06 |
| 19418 | 971 | K971W | C | 2.5e-06 | 1.4e-06 | 1.9e-06 | 1.933E-06 | 5.508E-07 | 1.400E-06 | 2.500E-06 |
| 19419 | 971 | K971Y | C | 2.6e-06 | 4.7e-06 | 3.1e-06 | 3.467E-06 | 1.097E-06 | 2.600E-06 | 4.700E-06 |
| 19420 | 971 | K971V | C | 3.7e-06 | 2.4e-06 | 2.2e-06 | 2.767E-06 | 8.145E-07 | 2.200E-06 | 3.700E-06 |
| 19421 | 972 | S972A | C | 3.3e-06 | 1.8e-06 | 2.2e-06 | 2.433E-06 | 7.767E-07 | 1.800E-06 | 3.300E-06 |
| 19422 | 972 | S972R | C | 1.7e-06 | 1.8e-06 | 1.8e-06 | 1.767E-06 | 5.774E-08 | 1.700E-06 | 1.800E-06 |
| 19423 | 972 | S972N | C | 2.5e-06 | 1.6e-06 | 3.5e-06 | 2.533E-06 | 9.504E-07 | 1.600E-06 | 3.500E-06 |
| 19424 | 972 | S972D | C | 2.2e-06 | 2.1e-06 | 2.9e-06 | 2.400E-06 | 4.359E-07 | 2.100E-06 | 2.900E-06 |
| 19425 | 972 | S972C | C | 1.2e-06 | 3.1e-06 | 2e-06 | 2.100E-06 | 9.539E-07 | 1.200E-06 | 3.100E-06 |
| 19426 | 972 | S972E | C | 1.6e-06 | 2.5e-06 | 3.7e-06 | 2.600E-06 | 1.054E-06 | 1.600E-06 | 3.700E-06 |
| 19427 | 972 | S972Q | C | 1.6e-06 | 1.8e-06 | 2.2e-06 | 1.867E-06 | 3.055E-07 | 1.600E-06 | 2.200E-06 |
| 19428 | 972 | S972G | C | 3.2e-06 | 2e-06 | 3e-06 | 2.733E-06 | 6.429E-07 | 2.000E-06 | 3.200E-06 |
| 19429 | 972 | S972H | C | 2.2e-06 | 2.4e-06 | 1.8e-06 | 2.133E-06 | 3.055E-07 | 1.800E-06 | 2.400E-06 |
| 19430 | 972 | S972I | C | 2.1e-06 | 2.8e-06 | 2.7e-06 | 2.533E-06 | 3.786E-07 | 2.100E-06 | 2.800E-06 |
| 19431 | 972 | S972L | C | 1.6e-06 | 1.9e-06 | 1.9e-06 | 1.800E-06 | 1.732E-07 | 1.600E-06 | 1.900E-06 |
| 19432 | 972 | S972K | C | 1.5e-06 | 2e-06 | 1.8e-06 | 1.767E-06 | 2.517E-07 | 1.500E-06 | 2.000E-06 |
| 19433 | 972 | S972M | C | 5.2e-06 | 3.3e-06 | 1.9e-06 | 3.467E-06 | 1.656E-06 | 1.900E-06 | 5.200E-06 |
| 19434 | 972 | S972F | C | 2.6e-06 | 2.8e-06 | 2.9e-06 | 2.767E-06 | 1.528E-07 | 2.600E-06 | 2.900E-06 |
| 19435 | 972 | S972P | C | 3.1e-06 | 2.2e-06 | 1.7e-06 | 2.333E-06 | 7.095E-07 | 1.700E-06 | 3.100E-06 |
| 19436 | 972 | S972S | C | 2.7e-06 | 1.9e-06 | 1.7e-06 | 2.100E-06 | 5.292E-07 | 1.700E-06 | 2.700E-06 |
| 19437 | 972 | S972T | C | 3.1e-06 | 2.5e-06 | 2e-06 | 2.533E-06 | 5.508E-07 | 2.000E-06 | 3.100E-06 |
| 19438 | 972 | S972W | C | 3.4e-06 | 6.3e-06 | 3.1e-06 | 4.267E-06 | 1.767E-06 | 3.100E-06 | 6.300E-06 |
| 19439 | 972 | S972Y | C | 1.9e-06 | 1.9e-06 | 3.1e-06 | 2.300E-06 | 6.928E-07 | 1.900E-06 | 3.100E-06 |
| 19440 | 972 | S972V | C | 2.7e-06 | 2e-06 | 2.1e-06 | 2.267E-06 | 3.786E-07 | 2.000E-06 | 2.700E-06 |
| 19441 | 973 | T973A | C | 2.7e-06 | 1.9e-06 | 2.4e-06 | 2.333E-06 | 4.041E-07 | 1.900E-06 | 2.700E-06 |
| 19442 | 973 | T973R | C | 1.7e-06 | 2.8e-06 | 2.4e-06 | 2.300E-06 | 5.568E-07 | 1.700E-06 | 2.800E-06 |
| 19443 | 973 | T973N | C | 2.6e-06 | 1.8e-06 | 3.2e-06 | 2.533E-06 | 7.024E-07 | 1.800E-06 | 3.200E-06 |
| 19444 | 973 | T973D | C | 3.1e-06 | 3.4e-06 | 2.9e-06 | 3.133E-06 | 2.517E-07 | 2.900E-06 | 3.400E-06 |
| 19445 | 973 | T973C | C | 3.3e-06 | 3.4e-06 | 2.6e-06 | 3.100E-06 | 4.359E-07 | 2.600E-06 | 3.400E-06 |
| 19446 | 973 | T973E | C | 2.1e-06 | 3.8e-06 | 2.1e-06 | 2.667E-06 | 9.815E-07 | 2.100E-06 | 3.800E-06 |
| 19447 | 973 | T973Q | C | 3.1e-06 | 1.9e-06 | 1.8e-06 | 2.267E-06 | 7.234E-07 | 1.800E-06 | 3.100E-06 |
| 19448 | 973 | T973G | C | 1.5e-06 | 2.9e-06 | 3.1e-06 | 2.500E-06 | 8.718E-07 | 1.500E-06 | 3.100E-06 |
| 19449 | 973 | T973H | C | 3e-06 | 3.2e-06 | 1.4e-06 | 2.533E-06 | 9.866E-07 | 1.400E-06 | 3.200E-06 |
| 19450 | 973 | T973I | C | 1.7e-06 | 1.9e-06 | 2.7e-06 | 2.100E-06 | 5.292E-07 | 1.700E-06 | 2.700E-06 |
| 19451 | 973 | T973L | C | 2.2e-06 | 3.3e-06 | 2.3e-06 | 2.600E-06 | 6.083E-07 | 2.200E-06 | 3.300E-06 |
| 19452 | 973 | T973K | C | 2.9e-06 | 4.1e-06 | 2.5e-06 | 3.167E-06 | 8.327E-07 | 2.500E-06 | 4.100E-06 |
| 19453 | 973 | T973M | C | 2e-06 | 2.1e-06 | 2.4e-06 | 2.167E-06 | 2.082E-07 | 2.000E-06 | 2.400E-06 |
| 19454 | 973 | T973F | C | 4.6e-06 | 3.3e-06 | 3.3e-06 | 3.733E-06 | 7.506E-07 | 3.300E-06 | 4.600E-06 |
| 19455 | 973 | T973P | C | 1.6e-06 | 2.6e-06 | 2.1e-06 | 2.100E-06 | 5.000E-07 | 1.600E-06 | 2.600E-06 |
| 19456 | 973 | T973S | C | 5e-06 | 1.9e-06 | 3.2e-06 | 3.367E-06 | 1.557E-06 | 1.900E-06 | 5.000E-06 |
| 19457 | 973 | T973T | C | 1.7e-06 | 2.8e-06 | 2e-06 | 2.167E-06 | 5.686E-07 | 1.700E-06 | 2.800E-06 |
| 19458 | 973 | T973W | C | 2.2e-06 | 3e-06 | 1.6e-06 | 2.267E-06 | 7.024E-07 | 1.600E-06 | 3.000E-06 |
| 19459 | 973 | T973Y | C | 2.6e-06 | 9.8e-07 | 1.3e-06 | 1.627E-06 | 8.580E-07 | 9.800E-07 | 2.600E-06 |
| 19460 | 973 | T973V | C | 3.1e-06 | 2.7e-06 | 2.7e-06 | 2.833E-06 | 2.309E-07 | 2.700E-06 | 3.100E-06 |
| 19461 | 974 | S974A | C | 1.5e-06 | 3e-06 | 2.5e-06 | 2.333E-06 | 7.638E-07 | 1.500E-06 | 3.000E-06 |
| 19462 | 974 | S974R | C | 1.8e-06 | 3.2e-06 | 2.8e-06 | 2.600E-06 | 7.211E-07 | 1.800E-06 | 3.200E-06 |
| 19463 | 974 | S974N | C | 3.8e-06 | 2e-06 | 1.7e-06 | 2.500E-06 | 1.136E-06 | 1.700E-06 | 3.800E-06 |
| 19464 | 974 | S974D | C | 2.1e-06 | 1.5e-06 | 2e-06 | 1.867E-06 | 3.215E-07 | 1.500E-06 | 2.100E-06 |

| | | | | | | | | | | |
|---|---|---|---|---|---|---|---|---|---|---|
| 19465 | 974 | S974C | C | 1.9e-06 | 1.9e-06 | 2.7e-06 | 2.167E-06 | 4.619E-07 | 1.900E-06 | 2.700E-06 |
| 19466 | 974 | S974E | C | 3.4e-06 | 4.4e-06 | 1.4e-06 | 3.067E-06 | 1.528E-06 | 1.400E-06 | 4.400E-06 |
| 19467 | 974 | S974Q | C | 2.7e-06 | 3e-06 | 2.1e-06 | 2.600E-06 | 4.583E-07 | 2.100E-06 | 3.000E-06 |
| 19468 | 974 | S974G | C | 2.7e-06 | 1.4e-06 | 1.6e-06 | 1.900E-06 | 7.000E-07 | 1.400E-06 | 2.700E-06 |
| 19469 | 974 | S974H | C | 1.4e-06 | 4.3e-06 | 1.9e-06 | 2.533E-06 | 1.550E-06 | 1.400E-06 | 4.300E-06 |
| 19470 | 974 | S974I | C | 7.5e-07 | 2.3e-06 | 3.9e-06 | 2.317E-06 | 1.575E-06 | 7.500E-07 | 3.900E-06 |
| 19471 | 974 | S974L | C | 2.7e-06 | 2.5e-06 | 3.2e-06 | 2.800E-06 | 3.606E-07 | 2.500E-06 | 3.200E-06 |
| 19472 | 974 | S974K | C | 3.2e-06 | 2.6e-06 | 3e-06 | 2.933E-06 | 3.055E-07 | 2.600E-06 | 3.200E-06 |
| 19473 | 974 | S974M | C | 1.4e-06 | 2.5e-06 | 4.2e-06 | 2.700E-06 | 1.411E-06 | 1.400E-06 | 4.200E-06 |
| 19474 | 974 | S974F | C | 3.7e-06 | 4e-06 | 2.9e-06 | 3.533E-06 | 5.686E-07 | 2.900E-06 | 4.000E-06 |
| 19475 | 974 | S974P | C | 3.3e-06 | 1.3e-06 | 2.2e-06 | 2.267E-06 | 1.002E-06 | 1.300E-06 | 3.300E-06 |
| 19476 | 974 | S974S | C | 1.8e-06 | 3.2e-06 | 2.8e-06 | 2.600E-06 | 7.211E-07 | 1.800E-06 | 3.200E-06 |
| 19477 | 974 | S974T | C | 7.5e-07 | 2e-06 | 2.2e-06 | 1.650E-06 | 7.858E-07 | 7.500E-07 | 2.200E-06 |
| 19478 | 974 | S974W | C | 1.6e-06 | 3.4e-06 | 1.5e-06 | 2.167E-06 | 1.069E-06 | 1.500E-06 | 3.400E-06 |
| 19479 | 974 | S974Y | C | 2.5e-06 | 2.1e-06 | 2e-06 | 2.200E-06 | 2.646E-07 | 2.000E-06 | 2.500E-06 |
| 19480 | 974 | S974V | C | 3e-06 | 2.6e-06 | 3.5e-06 | 3.033E-06 | 4.509E-07 | 2.600E-06 | 3.500E-06 |
| 19481 | 975 | G975A | C | 2.6e-06 | 1.9e-06 | 2.9e-06 | 2.467E-06 | 5.132E-07 | 1.900E-06 | 2.900E-06 |
| 19482 | 975 | G975R | C | 3.7e-06 | 1.3e-06 | 2.7e-06 | 2.567E-06 | 1.206E-06 | 1.300E-06 | 3.700E-06 |
| 19483 | 975 | G975N | C | 2.6e-06 | 3.6e-06 | 2.7e-06 | 2.967E-06 | 5.508E-07 | 2.600E-06 | 3.600E-06 |
| 19484 | 975 | G975D | C | 1.8e-06 | 1.4e-06 | 2.6e-06 | 1.933E-06 | 6.110E-07 | 1.400E-06 | 2.600E-06 |
| 19485 | 975 | G975C | C | 1.8e-06 | 1.8e-06 | 1.8e-06 | 1.800E-06 | 0.000E+00 | 1.800E-06 | 1.800E-06 |
| 19486 | 975 | G975E | C | 2.3e-06 | 1.9e-06 | 1.6e-06 | 1.933E-06 | 3.512E-07 | 1.600E-06 | 2.300E-06 |
| 19487 | 975 | G975Q | C | 2.4e-06 | 2.6e-06 | 2.2e-06 | 2.400E-06 | 2.000E-07 | 2.200E-06 | 2.600E-06 |
| 19488 | 975 | G975G | C | 1.8e-06 | 2.6e-06 | 1.4e-06 | 1.933E-06 | 6.110E-07 | 1.400E-06 | 2.600E-06 |
| 19489 | 975 | G975H | C | 2.3e-06 | 1.5e-06 | 2.8e-06 | 2.200E-06 | 6.557E-07 | 1.500E-06 | 2.800E-06 |
| 19490 | 975 | G975I | C | 3.7e-06 | 2.2e-06 | 3.6e-06 | 3.167E-06 | 8.386E-07 | 2.200E-06 | 3.700E-06 |
| 19491 | 975 | G975L | C | 2.3e-06 | 2.1e-06 | 2.4e-06 | 2.267E-06 | 1.528E-07 | 2.100E-06 | 2.400E-06 |
| 19492 | 975 | G975K | C | 2.9e-06 | 3.5e-06 | 3.1e-06 | 3.167E-06 | 3.055E-07 | 2.900E-06 | 3.500E-06 |
| 19493 | 975 | G975M | C | 4e-06 | 4e-06 | 2.3e-06 | 3.433E-06 | 9.815E-07 | 2.300E-06 | 4.000E-06 |
| 19494 | 975 | G975F | C | 2.6e-06 | 2.6e-06 | 1.5e-06 | 2.233E-06 | 6.351E-07 | 1.500E-06 | 2.600E-06 |
| 19495 | 975 | G975P | C | 3.4e-06 | 2.4e-06 | 3.5e-06 | 3.100E-06 | 6.083E-07 | 2.400E-06 | 3.500E-06 |
| 19496 | 975 | G975S | C | 6.2e-06 | 1.8e-06 | 3.3e-06 | 3.767E-06 | 2.237E-06 | 1.800E-06 | 6.200E-06 |
| 19497 | 975 | G975T | C | 3.1e-06 | 3e-06 | 2.2e-06 | 2.767E-06 | 4.933E-07 | 2.200E-06 | 3.100E-06 |
| 19498 | 975 | G975W | C | 3e-06 | 1.8e-06 | 2e-06 | 2.267E-06 | 6.429E-07 | 1.800E-06 | 3.000E-06 |
| 19499 | 975 | G975Y | C | 3.7e-06 | 1.1e-06 | 2.4e-06 | 2.400E-06 | 1.300E-06 | 1.100E-06 | 3.700E-06 |
| 19500 | 975 | G975V | C | 2.7e-06 | 2.3e-06 | 3.2e-06 | 2.733E-06 | 4.509E-07 | 2.300E-06 | 3.200E-06 |
| 19501 | 976 | G976A | C | 3e-06 | 9.1e-07 | 9.9e-07 | 1.633E-06 | 1.184E-06 | 9.100E-07 | 3.000E-06 |
| 19502 | 976 | G976R | C | 1.5e-06 | 2.6e-06 | 1.4e-06 | 1.833E-06 | 6.658E-07 | 1.400E-06 | 2.600E-06 |
| 19503 | 976 | G976N | C | 1.5e-06 | 2.7e-06 | 3.4e-06 | 2.533E-06 | 9.609E-07 | 1.500E-06 | 3.400E-06 |
| 19504 | 976 | G976D | C | 2.3e-06 | 1.7e-06 | 2e-06 | 2.000E-06 | 3.000E-07 | 1.700E-06 | 2.300E-06 |
| 19505 | 976 | G976C | C | 1.6e-06 | 2.3e-06 | 4.5e-06 | 2.800E-06 | 1.513E-06 | 1.600E-06 | 4.500E-06 |
| 19506 | 976 | G976E | C | 2.3e-06 | 2.3e-06 | 1.6e-06 | 2.067E-06 | 4.041E-07 | 1.600E-06 | 2.300E-06 |
| 19507 | 976 | G976Q | C | 2e-06 | 1.5e-06 | 1.4e-06 | 1.633E-06 | 3.215E-07 | 1.400E-06 | 2.000E-06 |
| 19508 | 976 | G976G | C | 2.9e-06 | 1.4e-06 | 2.2e-06 | 2.167E-06 | 7.506E-07 | 1.400E-06 | 2.900E-06 |
| 19509 | 976 | G976H | C | 2.1e-06 | 2e-06 | 2.8e-06 | 2.300E-06 | 4.359E-07 | 2.000E-06 | 2.800E-06 |
| 19510 | 976 | G976I | C | 2.3e-06 | 2.1e-06 | 2.2e-06 | 2.200E-06 | 1.000E-07 | 2.100E-06 | 2.300E-06 |
| 19511 | 976 | G976L | C | 1.6e-06 | 2.5e-06 | 2.1e-06 | 2.067E-06 | 4.509E-07 | 1.600E-06 | 2.500E-06 |
| 19512 | 976 | G976K | C | 5.6e-06 | 2.6e-06 | 3.1e-06 | 3.767E-06 | 1.607E-06 | 2.600E-06 | 5.600E-06 |
| 19513 | 976 | G976M | C | 2.5e-06 | 1.8e-06 | 1.7e-06 | 2.000E-06 | 4.359E-07 | 1.700E-06 | 2.500E-06 |
| 19514 | 976 | G976F | C | 1.8e-06 | 3.2e-06 | 2.4e-06 | 2.467E-06 | 7.024E-07 | 1.800E-06 | 3.200E-06 |
| 19515 | 976 | G976P | C | 1.9e-06 | 1.4e-06 | 1.9e-06 | 1.733E-06 | 2.887E-07 | 1.400E-06 | 1.900E-06 |
| 19516 | 976 | G976S | C | 1.7e-06 | 4.9e-06 | 1.8e-06 | 2.800E-06 | 1.819E-06 | 1.700E-06 | 4.900E-06 |
| 19517 | 976 | G976T | C | 3.3e-06 | 2.9e-06 | 1.3e-06 | 2.500E-06 | 1.058E-06 | 1.300E-06 | 3.300E-06 |
| 19518 | 976 | G976W | C | 2.8e-06 | 2.7e-06 | 2.3e-06 | 2.600E-06 | 2.646E-07 | 2.300E-06 | 2.800E-06 |
| 19519 | 976 | G976Y | C | 2.8e-06 | 3.8e-06 | 2.4e-06 | 3.000E-06 | 7.211E-07 | 2.400E-06 | 3.800E-06 |
| 19520 | 976 | G976V | C | 1.5e-06 | 3.9e-06 | 1.9e-06 | 2.433E-06 | 1.286E-06 | 1.500E-06 | 3.900E-06 |
| 19521 | 977 | T977A | C | 3.7e-06 | 4.3e-06 | 2.5e-06 | 3.500E-06 | 9.165E-07 | 2.500E-06 | 4.300E-06 |
| 19522 | 977 | T977R | C | 3.2e-06 | 1.5e-06 | 1.9e-06 | 2.200E-06 | 8.888E-07 | 1.500E-06 | 3.200E-06 |
| 19523 | 977 | T977N | C | 2.3e-06 | 3.9e-06 | 1.9e-06 | 2.700E-06 | 1.058E-06 | 1.900E-06 | 3.900E-06 |
| 19524 | 977 | T977D | C | 2.7e-06 | 2.7e-06 | 1.9e-06 | 2.433E-06 | 4.619E-07 | 1.900E-06 | 2.700E-06 |
| 19525 | 977 | T977C | C | 1.9e-06 | 2.1e-06 | 2.5e-06 | 2.167E-06 | 3.055E-07 | 1.900E-06 | 2.500E-06 |
| 19526 | 977 | T977E | C | 2.4e-06 | 2.6e-06 | 1.9e-06 | 2.300E-06 | 3.606E-07 | 1.900E-06 | 2.600E-06 |
| 19527 | 977 | T977Q | C | 2.8e-06 | 1.2e-06 | 1.8e-06 | 1.933E-06 | 8.083E-07 | 1.200E-06 | 2.800E-06 |
| 19528 | 977 | T977G | C | 3.1e-06 | 3.1e-06 | 2.2e-06 | 2.800E-06 | 5.196E-07 | 2.200E-06 | 3.100E-06 |
| 19529 | 977 | T977H | C | 2.4e-06 | 2.9e-06 | 2.8e-06 | 2.700E-06 | 2.646E-07 | 2.400E-06 | 2.900E-06 |
| 19530 | 977 | T977I | C | 1.9e-06 | 2.4e-06 | 2.7e-06 | 2.333E-06 | 4.041E-07 | 1.900E-06 | 2.700E-06 |
| 19531 | 977 | T977L | C | 2.7e-06 | 1.5e-06 | 2.7e-06 | 2.300E-06 | 6.928E-07 | 1.500E-06 | 2.700E-06 |
| 19532 | 977 | T977K | C | 3.2e-06 | 4.2e-06 | 4e-06 | 3.800E-06 | 5.292E-07 | 3.200E-06 | 4.200E-06 |
| 19533 | 977 | T977M | C | 1.6e-06 | 1.2e-06 | 2.2e-06 | 1.667E-06 | 5.033E-07 | 1.200E-06 | 2.200E-06 |
| 19534 | 977 | T977F | C | 2.8e-06 | 2e-06 | 3.3e-06 | 2.700E-06 | 6.557E-07 | 2.000E-06 | 3.300E-06 |
| 19535 | 977 | T977P | C | 2.4e-06 | 3.9e-06 | 2e-06 | 2.767E-06 | 1.002E-06 | 2.000E-06 | 3.900E-06 |
| 19536 | 977 | T977S | C | 2.1e-06 | 3.6e-06 | 2e-06 | 2.567E-06 | 8.963E-07 | 2.000E-06 | 3.600E-06 |
| 19537 | 977 | T977T | C | 2.2e-06 | 2.4e-06 | 2.6e-06 | 2.400E-06 | 2.000E-07 | 2.200E-06 | 2.600E-06 |
| 19538 | 977 | T977W | C | 3.2e-06 | 2e-06 | 3.6e-06 | 2.933E-06 | 8.327E-07 | 2.000E-06 | 3.600E-06 |
| 19539 | 977 | T977Y | C | 2.5e-06 | 2e-06 | 3.1e-06 | 2.533E-06 | 5.508E-07 | 2.000E-06 | 3.100E-06 |
| 19540 | 977 | T977V | C | 2.6e-06 | 2.5e-06 | 2.3e-06 | 2.467E-06 | 1.528E-07 | 2.300E-06 | 2.600E-06 |
| 19541 | 978 | A978A | C | 2.9e-06 | 2.8e-06 | 2.3e-06 | 2.667E-06 | 3.215E-07 | 2.300E-06 | 2.900E-06 |
| 19542 | 978 | A978R | C | 2.4e-06 | 1.2e-06 | 2.6e-06 | 2.067E-06 | 7.572E-07 | 1.200E-06 | 2.600E-06 |
| 19543 | 978 | A978N | C | 5.2e-06 | 3.1e-06 | 2e-06 | 3.433E-06 | 1.626E-06 | 2.000E-06 | 5.200E-06 |
| 19544 | 978 | A978D | C | 3.3e-06 | 3.8e-06 | 2.4e-06 | 3.167E-06 | 7.095E-07 | 2.400E-06 | 3.800E-06 |
| 19545 | 978 | A978C | C | 2.9e-06 | 2e-06 | 2.5e-06 | 2.467E-06 | 4.509E-07 | 2.000E-06 | 2.900E-06 |
| 19546 | 978 | A978E | C | 2.7e-06 | 2.4e-06 | 2.1e-06 | 2.400E-06 | 3.000E-07 | 2.100E-06 | 2.700E-06 |
| 19547 | 978 | A978Q | C | 4.4e-06 | 2e-06 | 4.5e-06 | 3.633E-06 | 1.415E-06 | 2.000E-06 | 4.500E-06 |
| 19548 | 978 | A978G | C | 1.8e-06 | 3.3e-06 | 3.8e-06 | 2.967E-06 | 1.041E-06 | 1.800E-06 | 3.800E-06 |
| 19549 | 978 | A978H | C | 2.4e-06 | 2.9e-06 | 2.3e-06 | 2.533E-06 | 3.215E-07 | 2.300E-06 | 2.900E-06 |

| | | | | | | | | | | |
|---|---|---|---|---|---|---|---|---|---|---|
| 19550 | 978 | A978I | C | 1.8e-06 | 1.9e-06 | 1.6e-06 | 1.767E-06 | 1.528E-07 | 1.600E-06 | 1.900E-06 |
| 19551 | 978 | A978L | C | 3.1e-06 | 2.4e-06 | 2.6e-06 | 2.700E-06 | 3.606E-07 | 2.400E-06 | 3.100E-06 |
| 19552 | 978 | A978K | C | 1.6e-06 | 1.7e-06 | 2.6e-06 | 1.967E-06 | 5.508E-07 | 1.600E-06 | 2.600E-06 |
| 19553 | 978 | A978M | C | 2.4e-06 | 1.9e-06 | 3.1e-06 | 2.467E-06 | 6.028E-07 | 1.900E-06 | 3.100E-06 |
| 19554 | 978 | A978F | C | 3.2e-06 | 2e-06 | 7.5e-07 | 1.983E-06 | 1.225E-06 | 7.500E-07 | 3.200E-06 |
| 19555 | 978 | A978P | C | 2.3e-06 | 1.2e-06 | 3e-06 | 2.167E-06 | 9.074E-07 | 1.200E-06 | 3.000E-06 |
| 19556 | 978 | A978S | C | 2.5e-06 | 4e-06 | 2.1e-06 | 2.867E-06 | 1.002E-06 | 2.100E-06 | 4.000E-06 |
| 19557 | 978 | A978T | C | 3.1e-06 | 2.4e-06 | 2.3e-06 | 2.600E-06 | 4.359E-07 | 2.300E-06 | 3.100E-06 |
| 19558 | 978 | A978W | C | 2.6e-06 | 1.3e-06 | 1.9e-06 | 1.933E-06 | 6.506E-07 | 1.300E-06 | 2.600E-06 |
| 19559 | 978 | A978Y | C | 2e-06 | 1.7e-06 | 2.4e-06 | 2.033E-06 | 3.512E-07 | 1.700E-06 | 2.400E-06 |
| 19560 | 978 | A978V | C | 3e-06 | 2.8e-06 | 2.1e-06 | 2.633E-06 | 4.726E-07 | 2.100E-06 | 3.000E-06 |
| 19561 | 979 | A979A | C | 2.3e-06 | 1.2e-06 | 2.2e-06 | 1.900E-06 | 6.083E-07 | 1.200E-06 | 2.300E-06 |
| 19562 | 979 | A979R | C | 1.9e-06 | 4.1e-06 | 1.7e-06 | 2.567E-06 | 1.332E-06 | 1.700E-06 | 4.100E-06 |
| 19563 | 979 | A979N | C | 3.1e-06 | 3.1e-06 | 2.4e-06 | 2.867E-06 | 4.041E-07 | 2.400E-06 | 3.100E-06 |
| 19564 | 979 | A979D | C | 4.8e-06 | 2.6e-06 | 3.6e-06 | 3.667E-06 | 1.102E-06 | 2.600E-06 | 4.800E-06 |
| 19565 | 979 | A979C | C | 2.1e-06 | 1.7e-06 | 2.5e-06 | 2.100E-06 | 4.000E-07 | 1.700E-06 | 2.500E-06 |
| 19566 | 979 | A979E | C | 2.4e-06 | 2.2e-06 | 1.6e-06 | 2.067E-06 | 4.163E-07 | 1.600E-06 | 2.400E-06 |
| 19567 | 979 | A979Q | C | 1.3e-06 | 2.2e-06 | 2.4e-06 | 1.967E-06 | 5.859E-07 | 1.300E-06 | 2.400E-06 |
| 19568 | 979 | A979G | C | 1.5e-06 | 1.4e-06 | 2.4e-06 | 1.767E-06 | 5.508E-07 | 1.400E-06 | 2.400E-06 |
| 19569 | 979 | A979H | C | 2.7e-06 | 1.7e-06 | 2.5e-06 | 2.300E-06 | 5.292E-07 | 1.700E-06 | 2.700E-06 |
| 19570 | 979 | A979I | C | 2.3e-06 | 1.8e-06 | 3.3e-06 | 2.467E-06 | 7.638E-07 | 1.800E-06 | 3.300E-06 |
| 19571 | 979 | A979L | C | 3.9e-06 | 2.1e-06 | 2.1e-06 | 2.700E-06 | 1.039E-06 | 2.100E-06 | 3.900E-06 |
| 19572 | 979 | A979K | C | 2.9e-06 | 1.5e-06 | 2.5e-06 | 2.300E-06 | 7.211E-07 | 1.500E-06 | 2.900E-06 |
| 19573 | 979 | A979M | C | 2.2e-06 | 4e-06 | 1.2e-06 | 2.467E-06 | 1.419E-06 | 1.200E-06 | 4.000E-06 |
| 19574 | 979 | A979F | C | 7.6e-07 | 2.7e-06 | 3.6e-06 | 2.353E-06 | 1.451E-06 | 7.600E-07 | 3.600E-06 |
| 19575 | 979 | A979P | C | 3.1e-06 | 2.1e-06 | 2.3e-06 | 2.500E-06 | 5.292E-07 | 2.100E-06 | 3.100E-06 |
| 19576 | 979 | A979S | C | 1.4e-06 | 1.1e-06 | 3.6e-06 | 2.033E-06 | 1.365E-06 | 1.100E-06 | 3.600E-06 |
| 19577 | 979 | A979T | C | 2.2e-06 | 2.7e-06 | 2.1e-06 | 2.333E-06 | 3.215E-07 | 2.100E-06 | 2.700E-06 |
| 19578 | 979 | A979W | C | 2e-06 | 3.5e-06 | 2.5e-06 | 2.667E-06 | 7.638E-07 | 2.000E-06 | 3.500E-06 |
| 19579 | 979 | A979Y | C | 2.7e-06 | 3.7e-06 | 2.4e-06 | 2.933E-06 | 6.807E-07 | 2.400E-06 | 3.700E-06 |
| 19580 | 979 | A979V | C | 1.9e-06 | 2.2e-06 | 3.4e-06 | 2.500E-06 | 7.937E-07 | 1.900E-06 | 3.400E-06 |
| 19581 | 980 | L980A | C | 2.5e-06 | 1.9e-06 | 2.6e-06 | 2.333E-06 | 3.786E-07 | 1.900E-06 | 2.600E-06 |
| 19582 | 980 | L980R | C | 3.5e-06 | 2.1e-06 | 2.5e-06 | 2.700E-06 | 7.211E-07 | 2.100E-06 | 3.500E-06 |
| 19583 | 980 | L980N | C | 2.3e-06 | 2.1e-06 | 2.2e-06 | 2.200E-06 | 1.000E-07 | 2.100E-06 | 2.300E-06 |
| 19584 | 980 | L980D | C | 9e-07 | 3.1e-06 | 1.9e-06 | 1.967E-06 | 1.102E-06 | 9.000E-07 | 3.100E-06 |
| 19585 | 980 | L980C | C | 2.4e-06 | 2.4e-06 | 2e-06 | 2.267E-06 | 2.309E-07 | 2.000E-06 | 2.400E-06 |
| 19586 | 980 | L980E | C | 3.4e-06 | 2e-06 | 4.1e-06 | 3.167E-06 | 1.069E-06 | 2.000E-06 | 4.100E-06 |
| 19587 | 980 | L980Q | C | 3.7e-06 | 1.4e-06 | 2.4e-06 | 2.500E-06 | 1.153E-06 | 1.400E-06 | 3.700E-06 |
| 19588 | 980 | L980G | C | 4.4e-06 | 2e-06 | 2.6e-06 | 3.000E-06 | 1.249E-06 | 2.000E-06 | 4.400E-06 |
| 19589 | 980 | L980H | C | 2.2e-06 | 1.9e-06 | 3.1e-06 | 2.400E-06 | 6.245E-07 | 1.900E-06 | 3.100E-06 |
| 19590 | 980 | L980I | C | 1.6e-06 | 2e-06 | 3.2e-06 | 2.267E-06 | 8.327E-07 | 1.600E-06 | 3.200E-06 |
| 19591 | 980 | L980L | C | 1.8e-06 | 1.9e-06 | 1.8e-06 | 1.833E-06 | 5.774E-08 | 1.800E-06 | 1.900E-06 |
| 19592 | 980 | L980K | C | 2.6e-06 | 2.7e-06 | 4.3e-06 | 3.200E-06 | 9.539E-07 | 2.600E-06 | 4.300E-06 |
| 19593 | 980 | L980M | C | 2.8e-06 | 9.6e-07 | 3.5e-06 | 2.420E-06 | 1.312E-06 | 9.600E-07 | 3.500E-06 |
| 19594 | 980 | L980F | C | 8e-07 | 1.4e-06 | 4.6e-06 | 2.267E-06 | 2.043E-06 | 8.000E-07 | 4.600E-06 |
| 19595 | 980 | L980P | C | 1.7e-06 | 2.1e-06 | 2.4e-06 | 2.067E-06 | 3.512E-07 | 1.700E-06 | 2.400E-06 |
| 19596 | 980 | L980S | C | 2.9e-06 | 2.9e-06 | 2.6e-06 | 2.800E-06 | 1.732E-07 | 2.600E-06 | 2.900E-06 |
| 19597 | 980 | L980T | C | 2.9e-06 | 2.9e-06 | 3.1e-06 | 2.967E-06 | 1.155E-07 | 2.900E-06 | 3.100E-06 |
| 19598 | 980 | L980W | C | 2.8e-06 | 2.6e-06 | 2.8e-06 | 2.733E-06 | 1.155E-07 | 2.600E-06 | 2.800E-06 |
| 19599 | 980 | L980Y | C | 2.3e-06 | 1.7e-06 | 1.9e-06 | 1.967E-06 | 3.055E-07 | 1.700E-06 | 2.300E-06 |
| 19600 | 980 | L980V | C | 2.4e-06 | 2.3e-06 | 2.4e-06 | 2.367E-06 | 5.774E-08 | 2.300E-06 | 2.400E-06 |
| 19601 | 981 | G981A | C | 2.8e-06 | 3.1e-06 | 4.1e-06 | 3.333E-06 | 6.807E-07 | 2.800E-06 | 4.100E-06 |
| 19602 | 981 | G981R | C | 1.8e-06 | 3.7e-06 | 3.4e-06 | 2.967E-06 | 1.021E-06 | 1.800E-06 | 3.700E-06 |
| 19603 | 981 | G981N | C | 3e-06 | 2.5e-06 | 1.4e-06 | 2.300E-06 | 8.185E-07 | 1.400E-06 | 3.000E-06 |
| 19604 | 981 | G981D | C | 2.1e-06 | 2.7e-06 | 2e-06 | 2.267E-06 | 3.786E-07 | 2.000E-06 | 2.700E-06 |
| 19605 | 981 | G981C | C | 4.4e-06 | 2.6e-06 | 1.1e-06 | 2.700E-06 | 1.652E-06 | 1.100E-06 | 4.400E-06 |
| 19606 | 981 | G981E | C | 4.8e-06 | 2.8e-06 | 3.1e-06 | 3.567E-06 | 1.079E-06 | 2.800E-06 | 4.800E-06 |
| 19607 | 981 | G981Q | C | 1.7e-06 | 2.2e-06 | 2.3e-06 | 2.067E-06 | 3.215E-07 | 1.700E-06 | 2.300E-06 |
| 19608 | 981 | G981G | C | 2.8e-06 | 3.7e-06 | 2.1e-06 | 2.867E-06 | 8.021E-07 | 2.100E-06 | 3.700E-06 |
| 19609 | 981 | G981H | C | 3.3e-06 | 2.2e-06 | 3.4e-06 | 2.967E-06 | 6.658E-07 | 2.200E-06 | 3.400E-06 |
| 19610 | 981 | G981I | C | 2.6e-06 | 1.4e-06 | 1.7e-06 | 1.900E-06 | 6.245E-07 | 1.400E-06 | 2.600E-06 |
| 19611 | 981 | G981L | C | 2.9e-06 | 3.1e-06 | 3.1e-06 | 3.033E-06 | 1.155E-07 | 2.900E-06 | 3.100E-06 |
| 19612 | 981 | G981K | C | 1.6e-06 | 2.9e-06 | 2.3e-06 | 2.267E-06 | 6.506E-07 | 1.600E-06 | 2.900E-06 |
| 19613 | 981 | G981M | C | 3.3e-06 | 1.2e-06 | 2.3e-06 | 2.267E-06 | 1.050E-06 | 1.200E-06 | 3.300E-06 |
| 19614 | 981 | G981F | C | 1.8e-06 | 1.2e-06 | 2.8e-06 | 1.933E-06 | 8.083E-07 | 1.200E-06 | 2.800E-06 |
| 19615 | 981 | G981P | C | 1.3e-06 | 1.3e-06 | 2.2e-06 | 1.600E-06 | 5.196E-07 | 1.300E-06 | 2.200E-06 |
| 19616 | 981 | G981S | C | 3.6e-06 | 3e-06 | 2.7e-06 | 3.100E-06 | 4.583E-07 | 2.700E-06 | 3.600E-06 |
| 19617 | 981 | G981T | C | 2.6e-06 | 2.3e-06 | 2.8e-06 | 2.567E-06 | 2.517E-07 | 2.300E-06 | 2.800E-06 |
| 19618 | 981 | G981W | C | 1.8e-06 | 3e-06 | 2.1e-06 | 2.300E-06 | 6.245E-07 | 1.800E-06 | 3.000E-06 |
| 19619 | 981 | G981Y | C | 2.1e-06 | 3.2e-06 | 2.7e-06 | 2.667E-06 | 5.508E-07 | 2.100E-06 | 3.200E-06 |
| 19620 | 981 | G981V | C | 1.7e-06 | 2e-06 | 2.3e-06 | 2.000E-06 | 3.000E-07 | 1.700E-06 | 2.300E-06 |
| 19621 | 982 | C982A | C | 2.2e-06 | 1.9e-06 | 5.2e-06 | 3.100E-06 | 1.825E-06 | 1.900E-06 | 5.200E-06 |
| 19622 | 982 | C982R | C | 2.6e-06 | 2.8e-06 | 2.4e-06 | 2.600E-06 | 2.000E-07 | 2.400E-06 | 2.800E-06 |
| 19623 | 982 | C982N | C | 1.4e-06 | 2e-06 | 2.8e-06 | 2.067E-06 | 7.024E-07 | 1.400E-06 | 2.800E-06 |
| 19624 | 982 | C982D | C | 2.7e-06 | 3.5e-06 | 1.4e-06 | 2.533E-06 | 1.060E-06 | 1.400E-06 | 3.500E-06 |
| 19625 | 982 | C982C | C | 3.1e-06 | 1.4e-06 | 1.8e-06 | 2.100E-06 | 8.888E-07 | 1.400E-06 | 3.100E-06 |
| 19626 | 982 | C982E | C | 1.9e-06 | 2.9e-06 | 3.8e-06 | 2.867E-06 | 9.504E-07 | 1.900E-06 | 3.800E-06 |
| 19627 | 982 | C982Q | C | 1.8e-06 | 3.6e-06 | 1.8e-06 | 2.400E-06 | 1.039E-06 | 1.800E-06 | 3.600E-06 |
| 19628 | 982 | C982G | C | 3.2e-06 | 2.7e-06 | 1.4e-06 | 2.433E-06 | 9.292E-07 | 1.400E-06 | 3.200E-06 |
| 19629 | 982 | C982H | C | 2e-06 | 2.5e-06 | 1.8e-06 | 2.100E-06 | 3.606E-07 | 1.800E-06 | 2.500E-06 |
| 19630 | 982 | C982I | C | 2.1e-06 | 2.3e-06 | 1.5e-06 | 1.967E-06 | 4.163E-07 | 1.500E-06 | 2.300E-06 |
| 19631 | 982 | C982L | C | 2.7e-06 | 3e-06 | 2.3e-06 | 2.667E-06 | 3.512E-07 | 2.300E-06 | 3.000E-06 |
| 19632 | 982 | C982K | C | 1.4e-06 | 1.2e-06 | 2.1e-06 | 1.567E-06 | 4.726E-07 | 1.200E-06 | 2.100E-06 |
| 19633 | 982 | C982M | C | 2.9e-06 | 3.1e-06 | 2.9e-06 | 2.967E-06 | 1.155E-07 | 2.900E-06 | 3.100E-06 |
| 19634 | 982 | C982F | C | 2e-06 | 2.6e-06 | 1.9e-06 | 2.167E-06 | 3.786E-07 | 1.900E-06 | 2.600E-06 |

| | | | | | | | | | | |
|---|---|---|---|---|---|---|---|---|---|---|
| 19635 | 982 | C982P | C | 2.2e-06 | 1.2e-06 | 2.6e-06 | 2.000E-06 | 7.211E-07 | 1.200E-06 | 2.600E-06 |
| 19636 | 982 | C982S | C | 2.3e-06 | 4.7e-06 | 3.1e-06 | 3.367E-06 | 1.222E-06 | 2.300E-06 | 4.700E-06 |
| 19637 | 982 | C982T | C | 2.9e-06 | 2e-06 | 2.6e-06 | 2.500E-06 | 4.583E-07 | 2.000E-06 | 2.900E-06 |
| 19638 | 982 | C982W | C | 2.6e-06 | 1.5e-06 | 5.2e-06 | 3.100E-06 | 1.900E-06 | 1.500E-06 | 5.200E-06 |
| 19639 | 982 | C982Y | C | 1.8e-06 | 2.2e-06 | 3.9e-06 | 2.633E-06 | 1.115E-06 | 1.800E-06 | 3.900E-06 |
| 19640 | 982 | C982V | C | 3.6e-06 | 2.4e-06 | 2.1e-06 | 2.700E-06 | 7.937E-07 | 2.100E-06 | 3.600E-06 |
| 19641 | 983 | L983A | C | 3.6e-06 | 2.8e-06 | 2.2e-06 | 2.867E-06 | 7.024E-07 | 2.200E-06 | 3.600E-06 |
| 19642 | 983 | L983R | C | 2.4e-06 | 2.5e-06 | 1.4e-06 | 2.100E-06 | 6.083E-07 | 1.400E-06 | 2.500E-06 |
| 19643 | 983 | L983N | C | 3.6e-06 | 1.6e-06 | 3.6e-06 | 2.933E-06 | 1.155E-06 | 1.600E-06 | 3.600E-06 |
| 19644 | 983 | L983D | C | 2.6e-06 | 2.3e-06 | 2.4e-06 | 2.433E-06 | 1.528E-07 | 2.300E-06 | 2.600E-06 |
| 19645 | 983 | L983C | C | 2.5e-06 | 2e-06 | 3.2e-06 | 2.567E-06 | 6.028E-07 | 2.000E-06 | 3.200E-06 |
| 19646 | 983 | L983E | C | 3.6e-06 | 9.4e-07 | 1.5e-06 | 2.013E-06 | 1.402E-06 | 9.400E-07 | 3.600E-06 |
| 19647 | 983 | L983Q | C | 1.9e-06 | 1.6e-06 | 2.1e-06 | 1.867E-06 | 2.517E-07 | 1.600E-06 | 2.100E-06 |
| 19648 | 983 | L983G | C | 3.7e-06 | 3.6e-06 | 2e-06 | 3.100E-06 | 9.539E-07 | 2.000E-06 | 3.700E-06 |
| 19649 | 983 | L983H | C | 1.1e-06 | 2.8e-06 | 2.2e-06 | 2.033E-06 | 8.622E-07 | 1.100E-06 | 2.800E-06 |
| 19650 | 983 | L983I | C | 2e-06 | 3.5e-06 | 2.3e-06 | 2.600E-06 | 7.937E-07 | 2.000E-06 | 3.500E-06 |
| 19651 | 983 | L983L | C | 1.6e-06 | 2.8e-06 | 1.9e-06 | 2.100E-06 | 6.245E-07 | 1.600E-06 | 2.800E-06 |
| 19652 | 983 | L983K | C | 2.6e-06 | 2.1e-06 | 3.5e-06 | 2.733E-06 | 7.095E-07 | 2.100E-06 | 3.500E-06 |
| 19653 | 983 | L983M | C | 3.9e-06 | 1.7e-06 | 4.1e-06 | 3.233E-06 | 1.332E-06 | 1.700E-06 | 4.100E-06 |
| 19654 | 983 | L983F | C | 2.9e-06 | 4e-06 | 2.6e-06 | 3.167E-06 | 7.371E-07 | 2.600E-06 | 4.000E-06 |
| 19655 | 983 | L983P | C | 2.8e-06 | 1.2e-06 | 2.1e-06 | 2.033E-06 | 8.021E-07 | 1.200E-06 | 2.800E-06 |
| 19656 | 983 | L983S | C | 2.1e-06 | 2.4e-06 | 2.5e-06 | 2.333E-06 | 2.082E-07 | 2.100E-06 | 2.500E-06 |
| 19657 | 983 | L983T | C | 3.1e-06 | 1.9e-06 | 2.4e-06 | 2.467E-06 | 6.028E-07 | 1.900E-06 | 3.100E-06 |
| 19658 | 983 | L983W | C | 2.5e-06 | 2.3e-06 | 3.3e-06 | 2.700E-06 | 5.292E-07 | 2.300E-06 | 3.300E-06 |
| 19659 | 983 | L983Y | C | 4.1e-06 | 1.8e-06 | 2.3e-06 | 2.733E-06 | 1.210E-06 | 1.800E-06 | 4.100E-06 |
| 19660 | 983 | L983V | C | 2e-06 | 2.5e-06 | 3.9e-06 | 2.800E-06 | 9.849E-07 | 2.000E-06 | 3.900E-06 |
| 19661 | 984 | V984A | C | 4.5e-06 | 3.5e-06 | 2.6e-06 | 3.533E-06 | 9.504E-07 | 2.600E-06 | 4.500E-06 |
| 19662 | 984 | V984R | C | 1.3e-06 | 2.8e-06 | 1.9e-06 | 2.000E-06 | 7.550E-07 | 1.300E-06 | 2.800E-06 |
| 19663 | 984 | V984N | C | 3e-06 | 1.5e-06 | 1.6e-06 | 2.033E-06 | 8.386E-07 | 1.500E-06 | 3.000E-06 |
| 19664 | 984 | V984D | C | 3.4e-06 | 2.3e-06 | 2.6e-06 | 2.767E-06 | 5.686E-07 | 2.300E-06 | 3.400E-06 |
| 19665 | 984 | V984C | C | 9.5e-07 | 2.6e-06 | 1.4e-06 | 1.650E-06 | 8.529E-07 | 9.500E-07 | 2.600E-06 |
| 19666 | 984 | V984E | C | 2.4e-06 | 3.7e-06 | 1.8e-06 | 2.633E-06 | 9.713E-07 | 1.800E-06 | 3.700E-06 |
| 19667 | 984 | V984Q | C | 2e-06 | 2e-06 | 4e-06 | 2.667E-06 | 1.155E-06 | 2.000E-06 | 4.000E-06 |
| 19668 | 984 | V984G | C | 2.1e-06 | 3.3e-06 | 3.1e-06 | 2.833E-06 | 6.429E-07 | 2.100E-06 | 3.300E-06 |
| 19669 | 984 | V984H | C | 3e-06 | 1.6e-06 | 2.4e-06 | 2.333E-06 | 7.024E-07 | 1.600E-06 | 3.000E-06 |
| 19670 | 984 | V984I | C | 1.7e-06 | 2.8e-06 | 2.8e-06 | 2.433E-06 | 6.351E-07 | 1.700E-06 | 2.800E-06 |
| 19671 | 984 | V984L | C | 3.4e-06 | 1.5e-06 | 2.2e-06 | 2.367E-06 | 9.609E-07 | 1.500E-06 | 3.400E-06 |
| 19672 | 984 | V984K | C | 1.6e-06 | 3.4e-06 | 1.6e-06 | 2.200E-06 | 1.039E-06 | 1.600E-06 | 3.400E-06 |
| 19673 | 984 | V984M | C | 1.6e-06 | 2.7e-06 | 2.3e-06 | 2.200E-06 | 5.568E-07 | 1.600E-06 | 2.700E-06 |
| 19674 | 984 | V984F | C | 2.7e-06 | 2.7e-06 | 1.6e-06 | 2.333E-06 | 6.351E-07 | 1.600E-06 | 2.700E-06 |
| 19675 | 984 | V984P | C | 2.5e-06 | 6.2e-07 | 2.5e-06 | 1.873E-06 | 1.085E-06 | 6.200E-07 | 2.500E-06 |
| 19676 | 984 | V984S | C | 2e-06 | 1.4e-06 | 9.5e-07 | 1.450E-06 | 5.268E-07 | 9.500E-07 | 2.000E-06 |
| 19677 | 984 | V984T | C | 2.6e-06 | 3e-06 | 3.3e-06 | 2.967E-06 | 3.512E-07 | 2.600E-06 | 3.300E-06 |
| 19678 | 984 | V984W | C | 1.9e-06 | 2.7e-06 | 1.6e-06 | 2.067E-06 | 5.686E-07 | 1.600E-06 | 2.700E-06 |
| 19679 | 984 | V984Y | C | 3.2e-06 | 1.6e-06 | 1.9e-06 | 2.233E-06 | 8.505E-07 | 1.600E-06 | 3.200E-06 |
| 19680 | 984 | V984V | C | 3.3e-06 | 2.6e-06 | 1.9e-06 | 2.600E-06 | 7.000E-07 | 1.900E-06 | 3.300E-06 |
| 19681 | 985 | K985A | C | 2.2e-06 | 2.2e-06 | 1.9e-06 | 2.100E-06 | 1.732E-07 | 1.900E-06 | 2.200E-06 |
| 19682 | 985 | K985R | C | 4e-06 | 2.1e-06 | 2.7e-06 | 2.933E-06 | 9.713E-07 | 2.100E-06 | 4.000E-06 |
| 19683 | 985 | K985N | C | 1.5e-06 | 2.6e-06 | 2.2e-06 | 2.100E-06 | 5.568E-07 | 1.500E-06 | 2.600E-06 |
| 19684 | 985 | K985D | C | 2.2e-06 | 3.8e-06 | 1.9e-06 | 2.633E-06 | 1.021E-06 | 1.900E-06 | 3.800E-06 |
| 19685 | 985 | K985C | C | 2.1e-06 | 4.6e-06 | 2.4e-06 | 3.033E-06 | 1.365E-06 | 2.100E-06 | 4.600E-06 |
| 19686 | 985 | K985E | C | 3e-06 | 2.9e-06 | 2.4e-06 | 2.767E-06 | 3.215E-07 | 2.400E-06 | 3.000E-06 |
| 19687 | 985 | K985Q | C | 4.7e-06 | 2.2e-06 | 1.9e-06 | 2.933E-06 | 1.537E-06 | 1.900E-06 | 4.700E-06 |
| 19688 | 985 | K985G | C | 2.4e-06 | 3.3e-06 | 1.7e-06 | 2.467E-06 | 8.021E-07 | 1.700E-06 | 3.300E-06 |
| 19689 | 985 | K985H | C | 2.6e-06 | 3.3e-06 | 3.2e-06 | 3.033E-06 | 3.786E-07 | 2.600E-06 | 3.300E-06 |
| 19690 | 985 | K985I | C | 3e-06 | 1.8e-06 | 2.7e-06 | 2.500E-06 | 6.245E-07 | 1.800E-06 | 3.000E-06 |
| 19691 | 985 | K985L | C | 1.7e-06 | 3.3e-06 | 3.4e-06 | 2.800E-06 | 9.539E-07 | 1.700E-06 | 3.400E-06 |
| 19692 | 985 | K985K | C | 2.7e-06 | 2.4e-06 | 2e-06 | 2.367E-06 | 3.512E-07 | 2.000E-06 | 2.700E-06 |
| 19693 | 985 | K985M | C | 2.5e-06 | 2.4e-06 | 4.3e-06 | 3.067E-06 | 1.069E-06 | 2.400E-06 | 4.300E-06 |
| 19694 | 985 | K985F | C | 2.6e-06 | 2e-06 | 2.2e-06 | 2.267E-06 | 3.055E-07 | 2.000E-06 | 2.600E-06 |
| 19695 | 985 | K985P | C | 3.7e-06 | 3.1e-06 | 2.4e-06 | 3.067E-06 | 6.506E-07 | 2.400E-06 | 3.700E-06 |
| 19696 | 985 | K985S | C | 1.4e-06 | 2.8e-06 | 2.6e-06 | 2.267E-06 | 7.572E-07 | 1.400E-06 | 2.800E-06 |
| 19697 | 985 | K985T | C | 2.2e-06 | 2.7e-06 | 8.2e-07 | 1.907E-06 | 9.737E-07 | 8.200E-07 | 2.700E-06 |
| 19698 | 985 | K985W | C | 2.6e-06 | 2.3e-06 | 1.5e-06 | 2.133E-06 | 5.686E-07 | 1.500E-06 | 2.600E-06 |
| 19699 | 985 | K985Y | C | 2.1e-06 | 3.9e-06 | 1.9e-06 | 2.633E-06 | 1.102E-06 | 1.900E-06 | 3.900E-06 |
| 19700 | 985 | K985V | C | 1.8e-06 | 1.9e-06 | 3e-06 | 2.233E-06 | 6.658E-07 | 1.800E-06 | 3.000E-06 |
| 19701 | 986 | D986A | C | 2.3e-06 | 3.8e-06 | 1.2e-06 | 2.433E-06 | 1.305E-06 | 1.200E-06 | 3.800E-06 |
| 19702 | 986 | D986R | C | 1.6e-06 | 2.4e-06 | 1.6e-06 | 1.867E-06 | 4.619E-07 | 1.600E-06 | 2.400E-06 |
| 19703 | 986 | D986N | C | 2.1e-06 | 3.6e-06 | 1.4e-06 | 2.367E-06 | 1.124E-06 | 1.400E-06 | 3.600E-06 |
| 19704 | 986 | D986D | C | 1.9e-06 | 3.1e-06 | 3.2e-06 | 2.733E-06 | 7.234E-07 | 1.900E-06 | 3.200E-06 |
| 19705 | 986 | D986C | C | 2.3e-06 | 2.2e-06 | 2.1e-06 | 2.200E-06 | 1.000E-07 | 2.100E-06 | 2.300E-06 |
| 19706 | 986 | D986E | C | 1.9e-06 | 3.3e-06 | 1.8e-06 | 2.333E-06 | 8.386E-07 | 1.800E-06 | 3.300E-06 |
| 19707 | 986 | D986Q | C | 1.4e-06 | 2.2e-06 | 2.8e-06 | 2.133E-06 | 7.024E-07 | 1.400E-06 | 2.800E-06 |
| 19708 | 986 | D986G | C | 3.4e-06 | 1.1e-06 | 1.8e-06 | 2.100E-06 | 1.179E-06 | 1.100E-06 | 3.400E-06 |
| 19709 | 986 | D986H | C | 2.6e-06 | 2.6e-06 | 2.5e-06 | 2.567E-06 | 5.774E-08 | 2.500E-06 | 2.600E-06 |
| 19710 | 986 | D986I | C | 1.7e-06 | 2.4e-06 | 1.9e-06 | 2.000E-06 | 3.606E-07 | 1.700E-06 | 2.400E-06 |
| 19711 | 986 | D986L | C | 2.6e-06 | 2.2e-06 | 2.2e-06 | 2.333E-06 | 2.309E-07 | 2.200E-06 | 2.600E-06 |
| 19712 | 986 | D986K | C | 2.5e-06 | 3.6e-06 | 2.8e-06 | 2.967E-06 | 5.686E-07 | 2.500E-06 | 3.600E-06 |
| 19713 | 986 | D986M | C | 2.5e-06 | 3.5e-06 | 2e-06 | 2.667E-06 | 7.638E-07 | 2.000E-06 | 3.500E-06 |
| 19714 | 986 | D986F | C | 1.9e-06 | 2.6e-06 | 2.6e-06 | 2.367E-06 | 4.041E-07 | 1.900E-06 | 2.600E-06 |
| 19715 | 986 | D986P | C | 1.9e-06 | 2.8e-06 | 2.5e-06 | 2.400E-06 | 4.583E-07 | 1.900E-06 | 2.800E-06 |
| 19716 | 986 | D986S | C | 2e-06 | 3.1e-06 | 2e-06 | 2.367E-06 | 6.351E-07 | 2.000E-06 | 3.100E-06 |
| 19717 | 986 | D986T | C | 1.7e-06 | 1.8e-06 | 2e-06 | 1.833E-06 | 1.528E-07 | 1.700E-06 | 2.000E-06 |
| 19718 | 986 | D986W | C | 2.4e-06 | 2.4e-06 | 2e-06 | 2.267E-06 | 2.309E-07 | 2.000E-06 | 2.400E-06 |
| 19719 | 986 | D986Y | C | 2.6e-06 | 2.1e-06 | 3.3e-06 | 2.667E-06 | 6.028E-07 | 2.100E-06 | 3.300E-06 |

| | | | | | | | | | | |
|---|---|---|---|---|---|---|---|---|---|---|
| 19720 | 986 | D986V | C | 2.6e-06 | 2.7e-06 | 3.8e-06 | 3.033E-06 | 6.658E-07 | 2.600E-06 | 3.800E-06 |
| 19721 | 987 | Y987A | C | 3e-06 | 2.7e-06 | 8.9e-07 | 2.197E-06 | 1.142E-06 | 8.900E-07 | 3.000E-06 |
| 19722 | 987 | Y987R | C | 1.9e-06 | 3.2e-06 | 2.6e-06 | 2.567E-06 | 6.506E-07 | 1.900E-06 | 3.200E-06 |
| 19723 | 987 | Y987N | C | 5e-06 | 3.1e-06 | 3e-06 | 3.700E-06 | 1.127E-06 | 3.000E-06 | 5.000E-06 |
| 19724 | 987 | Y987D | C | 1.9e-06 | 3e-06 | 2.3e-06 | 2.400E-06 | 5.568E-07 | 1.900E-06 | 3.000E-06 |
| 19725 | 987 | Y987C | C | 2.5e-06 | 2e-06 | 2.7e-06 | 2.400E-06 | 3.606E-07 | 2.000E-06 | 2.700E-06 |
| 19726 | 987 | Y987E | C | 4.4e-06 | 3.7e-06 | 3e-06 | 3.700E-06 | 7.000E-07 | 3.000E-06 | 4.400E-06 |
| 19727 | 987 | Y987Q | C | 3.7e-06 | 2.8e-06 | 2.9e-06 | 3.133E-06 | 4.933E-07 | 2.800E-06 | 3.700E-06 |
| 19728 | 987 | Y987G | C | 3.3e-06 | 2.5e-06 | 1.9e-06 | 2.567E-06 | 7.024E-07 | 1.900E-06 | 3.300E-06 |
| 19729 | 987 | Y987H | C | 3.5e-06 | 3.1e-06 | 1.4e-06 | 2.667E-06 | 1.115E-06 | 1.400E-06 | 3.500E-06 |
| 19730 | 987 | Y987I | C | 2.1e-06 | 2.6e-06 | 2.1e-06 | 2.267E-06 | 2.887E-07 | 2.100E-06 | 2.600E-06 |
| 19731 | 987 | Y987L | C | 1.5e-06 | 3e-06 | 3e-06 | 2.500E-06 | 8.660E-07 | 1.500E-06 | 3.000E-06 |
| 19732 | 987 | Y987K | C | 2e-06 | 2e-06 | 1.7e-06 | 1.900E-06 | 1.732E-07 | 1.700E-06 | 2.000E-06 |
| 19733 | 987 | Y987M | C | 1.7e-06 | 3.1e-06 | 3.7e-06 | 2.833E-06 | 1.026E-06 | 1.700E-06 | 3.700E-06 |
| 19734 | 987 | Y987F | C | 3.3e-06 | 2.9e-06 | 2.6e-06 | 2.933E-06 | 3.512E-07 | 2.600E-06 | 3.300E-06 |
| 19735 | 987 | Y987P | C | 2.6e-06 | 2.4e-06 | 2.9e-06 | 2.633E-06 | 2.517E-07 | 2.400E-06 | 2.900E-06 |
| 19736 | 987 | Y987S | C | 1.9e-06 | 2.9e-06 | 1.3e-06 | 2.033E-06 | 8.083E-07 | 1.300E-06 | 2.900E-06 |
| 19737 | 987 | Y987T | C | 2.8e-06 | 2e-06 | 3.1e-06 | 2.633E-06 | 5.686E-07 | 2.000E-06 | 3.100E-06 |
| 19738 | 987 | Y987W | C | 3.9e-06 | 2.6e-06 | 3.1e-06 | 3.200E-06 | 6.557E-07 | 2.600E-06 | 3.900E-06 |
| 19739 | 987 | Y987Y | C | 2.7e-06 | 3e-06 | 2.1e-06 | 2.600E-06 | 4.583E-07 | 2.100E-06 | 3.000E-06 |
| 19740 | 987 | Y987V | C | 3.1e-06 | 2.2e-06 | 2.1e-06 | 2.467E-06 | 5.508E-07 | 2.100E-06 | 3.100E-06 |
| 19741 | 988 | F988A | C | 2.7e-06 | 2.5e-06 | 2.5e-06 | 2.567E-06 | 1.155E-07 | 2.500E-06 | 2.700E-06 |
| 19742 | 988 | F988R | C | 3e-06 | 2e-06 | 2.5e-06 | 2.500E-06 | 5.000E-07 | 2.000E-06 | 3.000E-06 |
| 19743 | 988 | F988N | C | 2.9e-06 | 2.4e-06 | 2.5e-06 | 2.600E-06 | 2.646E-07 | 2.400E-06 | 2.900E-06 |
| 19744 | 988 | F988D | C | 1.9e-06 | 3.5e-06 | 5.2e-06 | 3.533E-06 | 1.650E-06 | 1.900E-06 | 5.200E-06 |
| 19745 | 988 | F988C | C | 1.4e-06 | 4e-06 | 1.2e-06 | 2.200E-06 | 1.562E-06 | 1.200E-06 | 4.000E-06 |
| 19746 | 988 | F988E | C | 2.8e-06 | 9.5e-07 | 2.5e-06 | 2.083E-06 | 9.929E-07 | 9.500E-07 | 2.800E-06 |
| 19747 | 988 | F988Q | C | 1.6e-06 | 2.9e-06 | 5.7e-06 | 3.400E-06 | 2.095E-06 | 1.600E-06 | 5.700E-06 |
| 19748 | 988 | F988G | C | 1.7e-06 | 1.9e-06 | 4.9e-06 | 2.833E-06 | 1.793E-06 | 1.700E-06 | 4.900E-06 |
| 19749 | 988 | F988H | C | 2.4e-06 | 1.2e-06 | 5.3e-06 | 2.967E-06 | 2.108E-06 | 1.200E-06 | 5.300E-06 |
| 19750 | 988 | F988I | C | 4.1e-06 | 2.1e-06 | 1.7e-06 | 2.633E-06 | 1.286E-06 | 1.700E-06 | 4.100E-06 |
| 19751 | 988 | F988L | C | 2.9e-06 | 3.9e-06 | 2.6e-06 | 3.133E-06 | 6.807E-07 | 2.600E-06 | 3.900E-06 |
| 19752 | 988 | F988K | C | 2.8e-06 | 2.7e-06 | 1.8e-06 | 2.433E-06 | 5.508E-07 | 1.800E-06 | 2.800E-06 |
| 19753 | 988 | F988M | C | 3.3e-06 | 1.1e-06 | 1.9e-06 | 2.100E-06 | 1.114E-06 | 1.100E-06 | 3.300E-06 |
| 19754 | 988 | F988F | C | 2.5e-06 | 2.1e-06 | 2.8e-06 | 2.467E-06 | 3.512E-07 | 2.100E-06 | 2.800E-06 |
| 19755 | 988 | F988P | C | 2.4e-06 | 1.8e-06 | 3.1e-06 | 2.433E-06 | 6.506E-07 | 1.800E-06 | 3.100E-06 |
| 19756 | 988 | F988S | C | 2.8e-06 | 2.5e-06 | 2.7e-06 | 2.667E-06 | 1.528E-07 | 2.500E-06 | 2.800E-06 |
| 19757 | 988 | F988T | C | 1.5e-06 | 2.3e-06 | 3.9e-06 | 2.567E-06 | 1.222E-06 | 1.500E-06 | 3.900E-06 |
| 19758 | 988 | F988W | C | 2.4e-06 | 3.1e-06 | 1.7e-06 | 2.400E-06 | 7.000E-07 | 1.700E-06 | 3.100E-06 |
| 19759 | 988 | F988Y | C | 2.2e-06 | 2.9e-06 | 2.5e-06 | 2.533E-06 | 3.512E-07 | 2.200E-06 | 2.900E-06 |
| 19760 | 988 | F988V | C | 2e-06 | 2.3e-06 | 1.5e-06 | 1.933E-06 | 4.041E-07 | 1.500E-06 | 2.300E-06 |
| 19761 | 989 | P989A | C | 4.2e-06 | 3.2e-06 | 2.4e-06 | 3.267E-06 | 9.018E-07 | 2.400E-06 | 4.200E-06 |
| 19762 | 989 | P989R | C | 3.4e-06 | 3.9e-06 | 2.4e-06 | 3.233E-06 | 7.638E-07 | 2.400E-06 | 3.900E-06 |
| 19763 | 989 | P989N | C | 2.9e-06 | 2.2e-06 | 2.6e-06 | 2.567E-06 | 3.512E-07 | 2.200E-06 | 2.900E-06 |
| 19764 | 989 | P989D | C | 3.4e-06 | 2.5e-06 | 1.9e-06 | 2.600E-06 | 7.550E-07 | 1.900E-06 | 3.400E-06 |
| 19765 | 989 | P989C | C | 1.5e-06 | 1.2e-06 | 1.9e-06 | 1.533E-06 | 3.512E-07 | 1.200E-06 | 1.900E-06 |
| 19766 | 989 | P989E | C | 2.8e-06 | 1.8e-06 | 3.6e-06 | 2.733E-06 | 9.018E-07 | 1.800E-06 | 3.600E-06 |
| 19767 | 989 | P989Q | C | 1.7e-06 | 1.8e-06 | 2.2e-06 | 1.900E-06 | 2.646E-07 | 1.700E-06 | 2.200E-06 |
| 19768 | 989 | P989G | C | 2.6e-06 | 3.1e-06 | 1.6e-06 | 2.433E-06 | 7.638E-07 | 1.600E-06 | 3.100E-06 |
| 19769 | 989 | P989H | C | 2.6e-06 | 1.8e-06 | 1.3e-06 | 1.900E-06 | 6.557E-07 | 1.300E-06 | 2.600E-06 |
| 19770 | 989 | P989I | C | 2.8e-06 | 2.9e-06 | 2e-06 | 2.567E-06 | 4.933E-07 | 2.000E-06 | 2.900E-06 |
| 19771 | 989 | P989L | C | 2e-06 | 2e-06 | 1.2e-06 | 1.733E-06 | 4.619E-07 | 1.200E-06 | 2.000E-06 |
| 19772 | 989 | P989K | C | 3.1e-06 | 2.3e-06 | 2.3e-06 | 2.567E-06 | 4.619E-07 | 2.300E-06 | 3.100E-06 |
| 19773 | 989 | P989M | C | 2.2e-06 | 2.1e-06 | 4.4e-06 | 2.900E-06 | 1.300E-06 | 2.100E-06 | 4.400E-06 |
| 19774 | 989 | P989F | C | 1.9e-06 | 1.8e-06 | 2.6e-06 | 2.100E-06 | 4.359E-07 | 1.800E-06 | 2.600E-06 |
| 19775 | 989 | P989P | C | 2.8e-06 | 1.6e-06 | 2e-06 | 2.133E-06 | 6.110E-07 | 1.600E-06 | 2.800E-06 |
| 19776 | 989 | P989S | C | 3.6e-06 | 3.6e-06 | 2.3e-06 | 3.167E-06 | 7.506E-07 | 2.300E-06 | 3.600E-06 |
| 19777 | 989 | P989T | C | 3.3e-06 | 2.5e-06 | 1.9e-06 | 2.567E-06 | 7.024E-07 | 1.900E-06 | 3.300E-06 |
| 19778 | 989 | P989W | C | 2.7e-06 | 2.9e-06 | 2.5e-06 | 2.700E-06 | 2.000E-07 | 2.500E-06 | 2.900E-06 |
| 19779 | 989 | P989Y | C | 1.9e-06 | 2.3e-06 | 2.8e-06 | 2.333E-06 | 4.509E-07 | 1.900E-06 | 2.800E-06 |
| 19780 | 989 | P989V | C | 3.2e-06 | 3.2e-06 | 2.3e-06 | 2.900E-06 | 5.196E-07 | 2.300E-06 | 3.200E-06 |
| 19781 | 990 | E990A | C | 3.8e-06 | 2.3e-06 | 1.8e-06 | 2.633E-06 | 1.041E-06 | 1.800E-06 | 3.800E-06 |
| 19782 | 990 | E990R | C | 1.9e-06 | 4e-06 | 2.6e-06 | 2.833E-06 | 1.069E-06 | 1.900E-06 | 4.000E-06 |
| 19783 | 990 | E990N | C | 1.6e-06 | 2.3e-06 | 2e-06 | 1.967E-06 | 3.512E-07 | 1.600E-06 | 2.300E-06 |
| 19784 | 990 | E990D | C | 2.9e-06 | 2.3e-06 | 2.3e-06 | 2.500E-06 | 3.464E-07 | 2.300E-06 | 2.900E-06 |
| 19785 | 990 | E990C | C | 2.8e-06 | 1.3e-06 | 3.8e-06 | 2.633E-06 | 1.258E-06 | 1.300E-06 | 3.800E-06 |
| 19786 | 990 | E990E | C | 4.5e-06 | 2.1e-06 | 2.8e-06 | 3.133E-06 | 1.234E-06 | 2.100E-06 | 4.500E-06 |
| 19787 | 990 | E990Q | C | 1.1e-06 | 1.4e-06 | 2e-06 | 1.500E-06 | 4.583E-07 | 1.100E-06 | 2.000E-06 |
| 19788 | 990 | E990G | C | 1.3e-06 | 2.3e-06 | 2.6e-06 | 2.067E-06 | 6.807E-07 | 1.300E-06 | 2.600E-06 |
| 19789 | 990 | E990H | C | 3.3e-06 | 1.6e-06 | 1e-06 | 1.967E-06 | 1.193E-06 | 1.000E-06 | 3.300E-06 |
| 19790 | 990 | E990I | C | 1.8e-06 | 2.8e-06 | 2.3e-06 | 2.300E-06 | 5.000E-07 | 1.800E-06 | 2.800E-06 |
| 19791 | 990 | E990L | C | 3e-06 | 3.6e-06 | 3.6e-06 | 3.400E-06 | 3.464E-07 | 3.000E-06 | 3.600E-06 |
| 19792 | 990 | E990K | C | 2.7e-06 | 1.7e-06 | 1.7e-06 | 2.033E-06 | 5.774E-07 | 1.700E-06 | 2.700E-06 |
| 19793 | 990 | E990M | C | 2.2e-06 | 3e-06 | 1.9e-06 | 2.367E-06 | 5.686E-07 | 1.900E-06 | 3.000E-06 |
| 19794 | 990 | E990F | C | 2.1e-06 | 3.3e-06 | 1.8e-06 | 2.400E-06 | 7.937E-07 | 1.800E-06 | 3.300E-06 |
| 19795 | 990 | E990P | C | 1.1e-06 | 4e-06 | 2.9e-06 | 2.667E-06 | 1.464E-06 | 1.100E-06 | 4.000E-06 |
| 19796 | 990 | E990S | C | 3e-06 | 3.2e-06 | 2.4e-06 | 2.867E-06 | 4.163E-07 | 2.400E-06 | 3.200E-06 |
| 19797 | 990 | E990T | C | 1.3e-06 | 1.2e-06 | 2.4e-06 | 1.633E-06 | 6.658E-07 | 1.200E-06 | 2.400E-06 |
| 19798 | 990 | E990W | C | 2.6e-06 | 1.5e-06 | 3e-06 | 2.367E-06 | 7.767E-07 | 1.500E-06 | 3.000E-06 |
| 19799 | 990 | E990Y | C | 2.7e-06 | 2.2e-06 | 1.1e-06 | 2.000E-06 | 8.185E-07 | 1.100E-06 | 2.700E-06 |
| 19800 | 990 | E990V | C | 1.6e-06 | 2.9e-06 | 3.5e-06 | 2.667E-06 | 9.713E-07 | 1.600E-06 | 3.500E-06 |
| 19801 | 991 | P991A | C | 2.1e-06 | 1.7e-06 | 3.3e-06 | 2.367E-06 | 8.327E-07 | 1.700E-06 | 3.300E-06 |
| 19802 | 991 | P991R | C | 2.7e-06 | 2.1e-06 | 5.2e-06 | 3.333E-06 | 1.644E-06 | 2.100E-06 | 5.200E-06 |
| 19803 | 991 | P991N | C | 2.8e-06 | 2.8e-06 | 2.7e-06 | 2.767E-06 | 5.774E-08 | 2.700E-06 | 2.800E-06 |
| 19804 | 991 | P991D | C | 2.4e-06 | 3e-06 | 2.7e-06 | 2.700E-06 | 3.000E-07 | 2.400E-06 | 3.000E-06 |

| | | | | | | | | | | |
|---|---|---|---|---|---|---|---|---|---|---|
| 19805 | 991 | P991C | C | 1.5e-06 | 1.3e-06 | 2.7e-06 | 1.833E-06 | 7.572E-07 | 1.300E-06 | 2.700E-06 |
| 19806 | 991 | P991E | C | 2e-06 | 2.5e-06 | 2e-06 | 2.167E-06 | 2.887E-07 | 2.000E-06 | 2.500E-06 |
| 19807 | 991 | P991Q | C | 2.8e-06 | 3.1e-06 | 2.3e-06 | 2.733E-06 | 4.041E-07 | 2.300E-06 | 3.100E-06 |
| 19808 | 991 | P991G | C | 2.3e-06 | 2.3e-06 | 2.5e-06 | 2.367E-06 | 1.155E-07 | 2.300E-06 | 2.500E-06 |
| 19809 | 991 | P991H | C | 3e-06 | 1.7e-06 | 1.2e-06 | 1.967E-06 | 9.292E-07 | 1.200E-06 | 3.000E-06 |
| 19810 | 991 | P991I | C | 9.8e-07 | 2.3e-06 | 3.7e-06 | 2.327E-06 | 1.360E-06 | 9.800E-07 | 3.700E-06 |
| 19811 | 991 | P991L | C | 1.9e-06 | 2.4e-06 | 3.5e-06 | 2.600E-06 | 8.185E-07 | 1.900E-06 | 3.500E-06 |
| 19812 | 991 | P991K | C | 1.7e-06 | 2.5e-06 | 2.5e-06 | 2.233E-06 | 4.619E-07 | 1.700E-06 | 2.500E-06 |
| 19813 | 991 | P991M | C | 5.4e-06 | 2.5e-06 | 2.4e-06 | 3.433E-06 | 1.704E-06 | 2.400E-06 | 5.400E-06 |
| 19814 | 991 | P991F | C | 1.5e-06 | 2.6e-06 | 3e-06 | 2.367E-06 | 7.767E-07 | 1.500E-06 | 3.000E-06 |
| 19815 | 991 | P991P | C | 2.4e-06 | 2.1e-06 | 1.5e-06 | 2.000E-06 | 4.583E-07 | 1.500E-06 | 2.400E-06 |
| 19816 | 991 | P991S | C | 1.8e-06 | 2.3e-06 | 2.7e-06 | 2.267E-06 | 4.509E-07 | 1.800E-06 | 2.700E-06 |
| 19817 | 991 | P991T | C | 3.5e-06 | 2.6e-06 | 3.9e-06 | 3.333E-06 | 6.658E-07 | 2.600E-06 | 3.900E-06 |
| 19818 | 991 | P991W | C | 1.7e-06 | 1.8e-06 | 2.1e-06 | 1.867E-06 | 2.082E-07 | 1.700E-06 | 2.100E-06 |
| 19819 | 991 | P991Y | C | 2.4e-06 | 1.5e-06 | 1.9e-06 | 1.933E-06 | 4.509E-07 | 1.500E-06 | 2.400E-06 |
| 19820 | 991 | P991V | C | 2e-06 | 2.8e-06 | 4.3e-06 | 3.033E-06 | 1.168E-06 | 2.000E-06 | 4.300E-06 |
| 19821 | 992 | V992A | C | 2.1e-06 | 2.6e-06 | 3.2e-06 | 2.633E-06 | 5.508E-07 | 2.100E-06 | 3.200E-06 |
| 19822 | 992 | V992R | C | 1.1e-06 | 1.3e-06 | 2.6e-06 | 1.667E-06 | 8.145E-07 | 1.100E-06 | 2.600E-06 |
| 19823 | 992 | V992N | C | 2.7e-06 | 2.2e-06 | 2.4e-06 | 2.433E-06 | 2.517E-07 | 2.200E-06 | 2.700E-06 |
| 19824 | 992 | V992D | C | 4.9e-06 | 2.5e-06 | 2.1e-06 | 3.167E-06 | 1.514E-06 | 2.100E-06 | 4.900E-06 |
| 19825 | 992 | V992C | C | 2.7e-06 | 2.5e-06 | 2.2e-06 | 2.467E-06 | 2.517E-07 | 2.200E-06 | 2.700E-06 |
| 19826 | 992 | V992E | C | 2.3e-06 | 2.7e-06 | 1.5e-06 | 2.167E-06 | 6.110E-07 | 1.500E-06 | 2.700E-06 |
| 19827 | 992 | V992Q | C | 1.6e-06 | 2.7e-06 | 2.1e-06 | 2.133E-06 | 5.508E-07 | 1.600E-06 | 2.700E-06 |
| 19828 | 992 | V992G | C | 1.2e-06 | 2.6e-06 | 4.2e-06 | 2.667E-06 | 1.501E-06 | 1.200E-06 | 4.200E-06 |
| 19829 | 992 | V992H | C | 2.1e-06 | 2.2e-06 | 2.1e-06 | 2.133E-06 | 5.774E-08 | 2.100E-06 | 2.200E-06 |
| 19830 | 992 | V992I | C | 2.8e-06 | 1e-06 | 3e-06 | 2.267E-06 | 1.102E-06 | 1.000E-06 | 3.000E-06 |
| 19831 | 992 | V992L | C | 2e-06 | 2e-06 | 2.4e-06 | 2.133E-06 | 2.309E-07 | 2.000E-06 | 2.400E-06 |
| 19832 | 992 | V992K | C | 2.3e-06 | 2.6e-06 | 3.4e-06 | 2.767E-06 | 5.686E-07 | 2.300E-06 | 3.400E-06 |
| 19833 | 992 | V992M | C | 2.6e-06 | 2e-06 | 2.9e-06 | 2.500E-06 | 4.583E-07 | 2.000E-06 | 2.900E-06 |
| 19834 | 992 | V992F | C | 3.4e-06 | 2.3e-06 | 2.1e-06 | 2.600E-06 | 7.000E-07 | 2.100E-06 | 3.400E-06 |
| 19835 | 992 | V992P | C | 1.8e-06 | 3.3e-06 | 1.5e-06 | 2.200E-06 | 9.644E-07 | 1.500E-06 | 3.300E-06 |
| 19836 | 992 | V992S | C | 2.4e-06 | 2.8e-06 | 1.9e-06 | 2.367E-06 | 4.509E-07 | 1.900E-06 | 2.800E-06 |
| 19837 | 992 | V992T | C | 1.6e-06 | 2.1e-06 | 2e-06 | 1.900E-06 | 2.646E-07 | 1.600E-06 | 2.100E-06 |
| 19838 | 992 | V992W | C | 2.9e-06 | 2.9e-06 | 1.9e-06 | 2.567E-06 | 5.774E-07 | 1.900E-06 | 2.900E-06 |
| 19839 | 992 | V992Y | C | 2e-06 | 4.1e-06 | 3e-06 | 3.033E-06 | 1.050E-06 | 2.000E-06 | 4.100E-06 |
| 19840 | 992 | V992V | C | 2.1e-06 | 3.1e-06 | 3.1e-06 | 2.767E-06 | 5.774E-07 | 2.100E-06 | 3.100E-06 |
| 19841 | 993 | T993A | C | 2.6e-06 | 3.2e-06 | 2.4e-06 | 2.733E-06 | 4.163E-07 | 2.400E-06 | 3.200E-06 |
| 19842 | 993 | T993R | C | 2e-06 | 2.4e-06 | 3.4e-06 | 2.600E-06 | 7.211E-07 | 2.000E-06 | 3.400E-06 |
| 19843 | 993 | T993N | C | 2.6e-06 | 1.9e-06 | 2.8e-06 | 2.433E-06 | 4.726E-07 | 1.900E-06 | 2.800E-06 |
| 19844 | 993 | T993D | C | 2.9e-06 | 3.1e-06 | 2.4e-06 | 2.800E-06 | 3.606E-07 | 2.400E-06 | 3.100E-06 |
| 19845 | 993 | T993C | C | 2.1e-06 | 1.7e-06 | 3.1e-06 | 2.300E-06 | 7.211E-07 | 1.700E-06 | 3.100E-06 |
| 19846 | 993 | T993E | C | 1.6e-06 | 1.9e-06 | 3.7e-06 | 2.400E-06 | 1.136E-06 | 1.600E-06 | 3.700E-06 |
| 19847 | 993 | T993Q | C | 2.1e-06 | 1.8e-06 | 1.3e-06 | 1.733E-06 | 4.041E-07 | 1.300E-06 | 2.100E-06 |
| 19848 | 993 | T993G | C | 2.1e-06 | 1.7e-06 | 1.6e-06 | 1.800E-06 | 2.646E-07 | 1.600E-06 | 2.100E-06 |
| 19849 | 993 | T993H | C | 2.5e-06 | 1.3e-06 | 1.9e-06 | 1.900E-06 | 6.000E-07 | 1.300E-06 | 2.500E-06 |
| 19850 | 993 | T993I | C | 2.2e-06 | 1.8e-06 | 2.8e-06 | 2.267E-06 | 5.033E-07 | 1.800E-06 | 2.800E-06 |
| 19851 | 993 | T993L | C | 2.1e-06 | 2.9e-06 | 2.8e-06 | 2.600E-06 | 4.359E-07 | 2.100E-06 | 2.900E-06 |
| 19852 | 993 | T993K | C | 2.5e-06 | 3.1e-06 | 2.8e-06 | 2.800E-06 | 3.000E-07 | 2.500E-06 | 3.100E-06 |
| 19853 | 993 | T993M | C | 3.8e-06 | 3.8e-06 | 3.1e-06 | 3.567E-06 | 4.041E-07 | 3.100E-06 | 3.800E-06 |
| 19854 | 993 | T993F | C | 2.9e-06 | 1.2e-06 | 2.9e-06 | 2.333E-06 | 9.815E-07 | 1.200E-06 | 2.900E-06 |
| 19855 | 993 | T993P | C | 3.5e-06 | 2.9e-06 | 1.8e-06 | 2.733E-06 | 8.622E-07 | 1.800E-06 | 3.500E-06 |
| 19856 | 993 | T993S | C | 2.2e-06 | 2e-06 | 1.7e-06 | 1.967E-06 | 2.517E-07 | 1.700E-06 | 2.200E-06 |
| 19857 | 993 | T993T | C | 2.3e-06 | 3.1e-06 | 2.1e-06 | 2.500E-06 | 5.292E-07 | 2.100E-06 | 3.100E-06 |
| 19858 | 993 | T993W | C | 2.5e-06 | 2.2e-06 | 2.2e-06 | 2.300E-06 | 1.732E-07 | 2.200E-06 | 2.500E-06 |
| 19859 | 993 | T993Y | C | 1.7e-06 | 2e-06 | 1.9e-06 | 1.867E-06 | 1.528E-07 | 1.700E-06 | 2.000E-06 |
| 19860 | 993 | T993V | C | 2e-06 | 3.5e-06 | 1.8e-06 | 2.433E-06 | 9.292E-07 | 1.800E-06 | 3.500E-06 |
| 19861 | 994 | V994A | C | 1.8e-06 | 2.9e-06 | 2.6e-06 | 2.433E-06 | 5.686E-07 | 1.800E-06 | 2.900E-06 |
| 19862 | 994 | V994R | C | 2.8e-06 | 2.3e-06 | 2.2e-06 | 2.433E-06 | 3.215E-07 | 2.200E-06 | 2.800E-06 |
| 19863 | 994 | V994N | C | 1.4e-06 | 1.9e-06 | 1.4e-06 | 1.567E-06 | 2.887E-07 | 1.400E-06 | 1.900E-06 |
| 19864 | 994 | V994D | C | 2.7e-06 | 1.1e-06 | 2e-06 | 1.933E-06 | 8.021E-07 | 1.100E-06 | 2.700E-06 |
| 19865 | 994 | V994C | C | 1.5e-06 | 2.1e-06 | 3e-06 | 2.200E-06 | 7.550E-07 | 1.500E-06 | 3.000E-06 |
| 19866 | 994 | V994E | C | 2.7e-06 | 2.6e-06 | 2.4e-06 | 2.567E-06 | 1.528E-07 | 2.400E-06 | 2.700E-06 |
| 19867 | 994 | V994Q | C | 2.4e-06 | 2.1e-06 | 2.8e-06 | 2.433E-06 | 3.512E-07 | 2.100E-06 | 2.800E-06 |
| 19868 | 994 | V994G | C | 2.1e-06 | 4.4e-06 | 2e-06 | 2.833E-06 | 1.358E-06 | 2.000E-06 | 4.400E-06 |
| 19869 | 994 | V994H | C | 1.4e-06 | 1.6e-06 | 3.2e-06 | 2.067E-06 | 9.866E-07 | 1.400E-06 | 3.200E-06 |
| 19870 | 994 | V994I | C | 2.1e-06 | 2.1e-06 | 3.3e-06 | 2.500E-06 | 6.928E-07 | 2.100E-06 | 3.300E-06 |
| 19871 | 994 | V994L | C | 2.4e-06 | 1.9e-06 | 1.6e-06 | 1.967E-06 | 4.041E-07 | 1.600E-06 | 2.400E-06 |
| 19872 | 994 | V994K | C | 2.3e-06 | 2.3e-06 | 1.3e-06 | 1.967E-06 | 5.774E-07 | 1.300E-06 | 2.300E-06 |
| 19873 | 994 | V994M | C | 2e-06 | 2.6e-06 | 2.3e-06 | 2.300E-06 | 3.000E-07 | 2.000E-06 | 2.600E-06 |
| 19874 | 994 | V994F | C | 3.3e-06 | 2.6e-06 | 2.2e-06 | 2.700E-06 | 5.568E-07 | 2.200E-06 | 3.300E-06 |
| 19875 | 994 | V994P | C | 1.5e-06 | 2.2e-06 | 1.7e-06 | 1.800E-06 | 3.606E-07 | 1.500E-06 | 2.200E-06 |
| 19876 | 994 | V994S | C | 1.8e-06 | 1.7e-06 | 2.7e-06 | 2.067E-06 | 5.508E-07 | 1.700E-06 | 2.700E-06 |
| 19877 | 994 | V994T | C | 1.4e-06 | 1.2e-06 | 2.6e-06 | 1.733E-06 | 7.572E-07 | 1.200E-06 | 2.600E-06 |
| 19878 | 994 | V994W | C | 2.6e-06 | 3.3e-06 | 2.6e-06 | 2.833E-06 | 4.041E-07 | 2.600E-06 | 3.300E-06 |
| 19879 | 994 | V994Y | C | 2.3e-06 | 2.1e-06 | 2.4e-06 | 2.267E-06 | 1.528E-07 | 2.100E-06 | 2.400E-06 |
| 19880 | 994 | V994V | C | 2.4e-06 | 2.4e-06 | 2.8e-06 | 2.533E-06 | 2.309E-07 | 2.400E-06 | 2.800E-06 |
| 19881 | 995 | S995A | C | 2.7e-06 | 2.5e-06 | 2.2e-06 | 2.467E-06 | 2.517E-07 | 2.200E-06 | 2.700E-06 |
| 19882 | 995 | S995R | C | 3.5e-06 | 2.5e-06 | 2.5e-06 | 2.833E-06 | 5.774E-07 | 2.500E-06 | 3.500E-06 |
| 19883 | 995 | S995N | C | 1.5e-06 | 2.5e-06 | 2.5e-06 | 2.167E-06 | 5.774E-07 | 1.500E-06 | 2.500E-06 |
| 19884 | 995 | S995D | C | 2.6e-06 | 2.2e-06 | 2.2e-06 | 2.333E-06 | 2.309E-07 | 2.200E-06 | 2.600E-06 |
| 19885 | 995 | S995C | C | 2.3e-06 | 2.7e-06 | 2.7e-06 | 2.567E-06 | 2.309E-07 | 2.300E-06 | 2.700E-06 |
| 19886 | 995 | S995E | C | 2.1e-06 | 3.3e-06 | 1.9e-06 | 2.433E-06 | 7.572E-07 | 1.900E-06 | 3.300E-06 |
| 19887 | 995 | S995Q | C | 1.8e-06 | 2.3e-06 | 1.6e-06 | 1.900E-06 | 3.606E-07 | 1.600E-06 | 2.300E-06 |
| 19888 | 995 | S995G | C | 2.3e-06 | 2.5e-06 | 2.3e-06 | 2.367E-06 | 1.155E-07 | 2.300E-06 | 2.500E-06 |
| 19889 | 995 | S995H | C | 2.1e-06 | 3.4e-06 | 2.3e-06 | 2.600E-06 | 7.000E-07 | 2.100E-06 | 3.400E-06 |

| | | | | | | | | | | |
|---|---|---|---|---|---|---|---|---|---|---|
| 19890 | 995 | S995I | C | 3.2e-06 | 1.5e-06 | 2.9e-06 | 2.533E-06 | 9.074E-07 | 1.500E-06 | 3.200E-06 |
| 19891 | 995 | S995L | C | 3.6e-06 | 3.6e-06 | 1.8e-06 | 3.000E-06 | 1.039E-06 | 1.800E-06 | 3.600E-06 |
| 19892 | 995 | S995K | C | 1.6e-06 | 2.4e-06 | 1.2e-06 | 1.733E-06 | 6.110E-07 | 1.200E-06 | 2.400E-06 |
| 19893 | 995 | S995M | C | 3.1e-06 | 2.6e-06 | 1.2e-06 | 2.300E-06 | 9.849E-07 | 1.200E-06 | 3.100E-06 |
| 19894 | 995 | S995F | C | 2.8e-06 | 1.8e-06 | 2.3e-06 | 2.300E-06 | 5.000E-07 | 1.800E-06 | 2.800E-06 |
| 19895 | 995 | S995P | C | 3.1e-06 | 2.8e-06 | 3.4e-06 | 3.100E-06 | 3.000E-07 | 2.800E-06 | 3.400E-06 |
| 19896 | 995 | S995S | C | 2.1e-06 | 2.4e-06 | 2e-06 | 2.167E-06 | 2.082E-07 | 2.000E-06 | 2.400E-06 |
| 19897 | 995 | S995T | C | 1.4e-06 | 1.6e-06 | 3.9e-06 | 2.300E-06 | 1.389E-06 | 1.400E-06 | 3.900E-06 |
| 19898 | 995 | S995W | C | 1.9e-06 | 1.8e-06 | 2.6e-06 | 2.100E-06 | 4.359E-07 | 1.800E-06 | 2.600E-06 |
| 19899 | 995 | S995Y | C | 3e-06 | 3.4e-06 | 2.3e-06 | 2.900E-06 | 5.568E-07 | 2.300E-06 | 3.400E-06 |
| 19900 | 995 | S995V | C | 1.5e-06 | 2.8e-06 | 2.3e-06 | 2.200E-06 | 6.557E-07 | 1.500E-06 | 2.800E-06 |
| 19901 | 996 | W996A | C | 3.2e-06 | 1.9e-06 | 2.5e-06 | 2.533E-06 | 6.506E-07 | 1.900E-06 | 3.200E-06 |
| 19902 | 996 | W996R | C | 1.6e-06 | 1.8e-06 | 2.8e-06 | 2.067E-06 | 6.429E-07 | 1.600E-06 | 2.800E-06 |
| 19903 | 996 | W996N | C | 2e-06 | 1.7e-06 | 2.1e-06 | 1.933E-06 | 2.082E-07 | 1.700E-06 | 2.100E-06 |
| 19904 | 996 | W996D | C | 1.8e-06 | 1.8e-06 | 1.1e-06 | 1.567E-06 | 4.041E-07 | 1.100E-06 | 1.800E-06 |
| 19905 | 996 | W996C | C | 3.9e-06 | 3.6e-06 | 2.6e-06 | 3.367E-06 | 6.807E-07 | 2.600E-06 | 3.900E-06 |
| 19906 | 996 | W996E | C | 2.2e-06 | 2.8e-06 | 2.9e-06 | 2.633E-06 | 3.786E-07 | 2.200E-06 | 2.900E-06 |
| 19907 | 996 | W996Q | C | 1.4e-06 | 1.9e-06 | 3.5e-06 | 2.267E-06 | 1.097E-06 | 1.400E-06 | 3.500E-06 |
| 19908 | 996 | W996G | C | 2.5e-06 | 2e-06 | 2.9e-06 | 2.467E-06 | 4.509E-07 | 2.000E-06 | 2.900E-06 |
| 19909 | 996 | W996H | C | 1.1e-06 | 1.6e-06 | 1.6e-06 | 1.433E-06 | 2.887E-07 | 1.100E-06 | 1.600E-06 |
| 19910 | 996 | W996I | C | 3.3e-06 | 1.6e-06 | 1.7e-06 | 2.200E-06 | 9.539E-07 | 1.600E-06 | 3.300E-06 |
| 19911 | 996 | W996L | C | 1.5e-06 | 1.7e-06 | 2.3e-06 | 1.833E-06 | 4.163E-07 | 1.500E-06 | 2.300E-06 |
| 19912 | 996 | W996K | C | 3e-06 | 1.9e-06 | 1.5e-06 | 2.133E-06 | 7.767E-07 | 1.500E-06 | 3.000E-06 |
| 19913 | 996 | W996M | C | 3.4e-06 | 2.8e-06 | 2.2e-06 | 2.800E-06 | 6.000E-07 | 2.200E-06 | 3.400E-06 |
| 19914 | 996 | W996F | C | 1.4e-06 | 1.3e-06 | 3.3e-06 | 2.000E-06 | 1.127E-06 | 1.300E-06 | 3.300E-06 |
| 19915 | 996 | W996P | C | 2.4e-06 | 3.7e-06 | 6.5e-06 | 4.200E-06 | 2.095E-06 | 2.400E-06 | 6.500E-06 |
| 19916 | 996 | W996S | C | 3.5e-06 | 2.9e-06 | 2e-06 | 2.800E-06 | 7.550E-07 | 2.000E-06 | 3.500E-06 |
| 19917 | 996 | W996T | C | 2.3e-06 | 2.4e-06 | 2.1e-06 | 2.267E-06 | 1.528E-07 | 2.100E-06 | 2.400E-06 |
| 19918 | 996 | W996W | C | 3.2e-06 | 2.6e-06 | 2.3e-06 | 2.700E-06 | 4.583E-07 | 2.300E-06 | 3.200E-06 |
| 19919 | 996 | W996Y | C | 1.8e-06 | 2.5e-06 | 3.8e-06 | 2.700E-06 | 1.015E-06 | 1.800E-06 | 3.800E-06 |
| 19920 | 996 | W996V | C | 2e-06 | 1.2e-06 | 2.2e-06 | 1.800E-06 | 5.292E-07 | 1.200E-06 | 2.200E-06 |
| 19921 | 997 | N997A | C | 2.3e-06 | 2e-06 | 9.8e-07 | 1.760E-06 | 6.920E-07 | 9.800E-07 | 2.300E-06 |
| 19922 | 997 | N997R | C | 1.5e-06 | 2e-06 | 2.4e-06 | 1.967E-06 | 4.509E-07 | 1.500E-06 | 2.400E-06 |
| 19923 | 997 | N997N | C | 2.7e-06 | 2.3e-06 | 3.1e-06 | 2.700E-06 | 4.000E-07 | 2.300E-06 | 3.100E-06 |
| 19924 | 997 | N997D | C | 2e-06 | 1.5e-06 | 2.1e-06 | 1.867E-06 | 3.215E-07 | 1.500E-06 | 2.100E-06 |
| 19925 | 997 | N997C | C | 1.5e-06 | 2.9e-06 | 3.2e-06 | 2.533E-06 | 9.074E-07 | 1.500E-06 | 3.200E-06 |
| 19926 | 997 | N997E | C | 2.1e-06 | 2.1e-06 | 2.9e-06 | 2.367E-06 | 4.619E-07 | 2.100E-06 | 2.900E-06 |
| 19927 | 997 | N997Q | C | 2.2e-06 | 9.2e-07 | 1.4e-06 | 1.507E-06 | 6.466E-07 | 9.200E-07 | 2.200E-06 |
| 19928 | 997 | N997G | C | 1.1e-06 | 2.7e-06 | 2.3e-06 | 2.033E-06 | 8.327E-07 | 1.100E-06 | 2.700E-06 |
| 19929 | 997 | N997H | C | 4e-06 | 1.9e-06 | 2.8e-06 | 2.900E-06 | 1.054E-06 | 1.900E-06 | 4.000E-06 |
| 19930 | 997 | N997I | C | 3.5e-06 | 2.3e-06 | 1.8e-06 | 2.533E-06 | 8.737E-07 | 1.800E-06 | 3.500E-06 |
| 19931 | 997 | N997L | C | 1e-06 | 3.6e-06 | 3.1e-06 | 2.567E-06 | 1.380E-06 | 1.000E-06 | 3.600E-06 |
| 19932 | 997 | N997K | C | 2.4e-06 | 3.5e-06 | 3.1e-06 | 3.000E-06 | 5.568E-07 | 2.400E-06 | 3.500E-06 |
| 19933 | 997 | N997M | C | 2.7e-06 | 2.9e-06 | 1.1e-06 | 2.233E-06 | 9.866E-07 | 1.100E-06 | 2.900E-06 |
| 19934 | 997 | N997F | C | 2.3e-06 | 1.8e-06 | 2.9e-06 | 2.333E-06 | 5.508E-07 | 1.800E-06 | 2.900E-06 |
| 19935 | 997 | N997P | C | 2.8e-06 | 2.2e-06 | 1.8e-06 | 2.267E-06 | 5.033E-07 | 1.800E-06 | 2.800E-06 |
| 19936 | 997 | N997S | C | 2.2e-06 | 2.7e-06 | 3.2e-06 | 2.700E-06 | 5.000E-07 | 2.200E-06 | 3.200E-06 |
| 19937 | 997 | N997T | C | 2.2e-06 | 2.8e-06 | 2.6e-06 | 2.533E-06 | 3.055E-07 | 2.200E-06 | 2.800E-06 |
| 19938 | 997 | N997W | C | 2.8e-06 | 2.1e-06 | 3.5e-06 | 2.800E-06 | 7.000E-07 | 2.100E-06 | 3.500E-06 |
| 19939 | 997 | N997Y | C | 4.2e-06 | 2e-06 | 2.5e-06 | 2.900E-06 | 1.153E-06 | 2.000E-06 | 4.200E-06 |
| 19940 | 997 | N997V | C | 1.7e-06 | 2.3e-06 | 3.3e-06 | 2.433E-06 | 8.083E-07 | 1.700E-06 | 3.300E-06 |
| 19941 | 998 | S998A | C | 2.3e-06 | 3e-06 | 6.7e-07 | 1.990E-06 | 1.196E-06 | 6.700E-07 | 3.000E-06 |
| 19942 | 998 | S998R | C | 3.2e-06 | 2.9e-06 | 2.8e-06 | 2.967E-06 | 2.082E-07 | 2.800E-06 | 3.200E-06 |
| 19943 | 998 | S998N | C | 4.7e-06 | 2.3e-06 | 3.3e-06 | 3.433E-06 | 1.206E-06 | 2.300E-06 | 4.700E-06 |
| 19944 | 998 | S998D | C | 2.5e-06 | 2.6e-06 | 3.2e-06 | 2.767E-06 | 3.786E-07 | 2.500E-06 | 3.200E-06 |
| 19945 | 998 | S998C | C | 2.3e-06 | 1.2e-06 | 1.8e-06 | 1.767E-06 | 5.508E-07 | 1.200E-06 | 2.300E-06 |
| 19946 | 998 | S998E | C | 1.3e-06 | 1.6e-06 | 2.9e-06 | 1.933E-06 | 8.505E-07 | 1.300E-06 | 2.900E-06 |
| 19947 | 998 | S998Q | C | 2.5e-06 | 2.6e-06 | 1.6e-06 | 2.233E-06 | 5.508E-07 | 1.600E-06 | 2.600E-06 |
| 19948 | 998 | S998G | C | 3.6e-06 | 1.4e-06 | 1.5e-06 | 2.167E-06 | 1.242E-06 | 1.400E-06 | 3.600E-06 |
| 19949 | 998 | S998H | C | 1.8e-06 | 9.5e-07 | 4.1e-06 | 2.283E-06 | 1.630E-06 | 9.500E-07 | 4.100E-06 |
| 19950 | 998 | S998I | C | 3.7e-06 | 2.9e-06 | 2.4e-06 | 3.000E-06 | 6.557E-07 | 2.400E-06 | 3.700E-06 |
| 19951 | 998 | S998L | C | 2.6e-06 | 2.4e-06 | 2.6e-06 | 2.533E-06 | 1.155E-07 | 2.400E-06 | 2.600E-06 |
| 19952 | 998 | S998K | C | 3.4e-06 | 1.6e-06 | 1.6e-06 | 2.200E-06 | 1.039E-06 | 1.600E-06 | 3.400E-06 |
| 19953 | 998 | S998M | C | 2.8e-06 | 2e-06 | 2.5e-06 | 2.433E-06 | 4.041E-07 | 2.000E-06 | 2.800E-06 |
| 19954 | 998 | S998F | C | 2.3e-06 | 3e-06 | 3.6e-06 | 2.967E-06 | 6.506E-07 | 2.300E-06 | 3.600E-06 |
| 19955 | 998 | S998P | C | 2e-06 | 2.1e-06 | 1.6e-06 | 1.900E-06 | 2.646E-07 | 1.600E-06 | 2.100E-06 |
| 19956 | 998 | S998S | C | 1.8e-06 | 2.8e-06 | 2.8e-06 | 2.467E-06 | 5.774E-07 | 1.800E-06 | 2.800E-06 |
| 19957 | 998 | S998T | C | 2.7e-06 | 2.5e-06 | 1.1e-06 | 2.100E-06 | 8.718E-07 | 1.100E-06 | 2.700E-06 |
| 19958 | 998 | S998W | C | 3.4e-06 | 5.6e-06 | 1.8e-06 | 3.600E-06 | 1.908E-06 | 1.800E-06 | 5.600E-06 |
| 19959 | 998 | S998Y | C | 3.1e-06 | 2.2e-06 | 2.4e-06 | 2.567E-06 | 4.726E-07 | 2.200E-06 | 3.100E-06 |
| 19960 | 998 | S998V | C | 3.6e-06 | 1.9e-06 | 1.7e-06 | 2.400E-06 | 1.044E-06 | 1.700E-06 | 3.600E-06 |
| 19961 | 999 | G999A | C | 2.5e-06 | 3.7e-06 | 2.4e-06 | 2.867E-06 | 7.234E-07 | 2.400E-06 | 3.700E-06 |
| 19962 | 999 | G999R | C | 3.2e-06 | 2.7e-06 | 2e-06 | 2.633E-06 | 6.028E-07 | 2.000E-06 | 3.200E-06 |
| 19963 | 999 | G999N | C | 2.1e-06 | 1.7e-06 | 2e-06 | 1.933E-06 | 2.082E-07 | 1.700E-06 | 2.100E-06 |
| 19964 | 999 | G999D | C | 2e-06 | 2.2e-06 | 2.1e-06 | 2.100E-06 | 1.000E-07 | 2.000E-06 | 2.200E-06 |
| 19965 | 999 | G999C | C | 2.9e-06 | 1.5e-06 | 3.2e-06 | 2.533E-06 | 9.074E-07 | 1.500E-06 | 3.200E-06 |
| 19966 | 999 | G999E | C | 1.9e-06 | 2.6e-06 | 2.5e-06 | 2.333E-06 | 3.786E-07 | 1.900E-06 | 2.600E-06 |
| 19967 | 999 | G999Q | C | 2.2e-06 | 2e-06 | 5.7e-06 | 3.300E-06 | 2.081E-06 | 2.000E-06 | 5.700E-06 |
| 19968 | 999 | G999G | C | 3.1e-06 | 2.8e-06 | 2.6e-06 | 2.833E-06 | 2.517E-07 | 2.600E-06 | 3.100E-06 |
| 19969 | 999 | G999H | C | 2e-06 | 2.3e-06 | 2.2e-06 | 2.167E-06 | 1.528E-07 | 2.000E-06 | 2.300E-06 |
| 19970 | 999 | G999I | C | 2.2e-06 | 4.3e-06 | 7.1e-07 | 2.403E-06 | 1.804E-06 | 7.100E-07 | 4.300E-06 |
| 19971 | 999 | G999L | C | 2.5e-06 | 2.3e-06 | 3.3e-06 | 2.700E-06 | 5.292E-07 | 2.300E-06 | 3.300E-06 |
| 19972 | 999 | G999K | C | 2.2e-06 | 2.6e-06 | 1.7e-06 | 2.167E-06 | 4.509E-07 | 1.700E-06 | 2.600E-06 |
| 19973 | 999 | G999M | C | 3e-06 | 1.8e-06 | 2.3e-06 | 2.367E-06 | 6.028E-07 | 1.800E-06 | 3.000E-06 |
| 19974 | 999 | G999F | C | 2.3e-06 | 2.7e-06 | 1.5e-06 | 2.167E-06 | 6.110E-07 | 1.500E-06 | 2.700E-06 |

| | | | | | | | | | | |
|---|---|---|---|---|---|---|---|---|---|---|
| 19975 | 999 | G999P | C | 1.9e-06 | 1.4e-06 | 2e-06 | 1.767E-06 | 3.215E-07 | 1.400E-06 | 2.000E-06 |
| 19976 | 999 | G999S | C | 3.1e-06 | 2.5e-06 | 1.4e-06 | 2.333E-06 | 8.622E-07 | 1.400E-06 | 3.100E-06 |
| 19977 | 999 | G999T | C | 3.3e-06 | 2.8e-06 | 2.6e-06 | 2.900E-06 | 3.606E-07 | 2.600E-06 | 3.300E-06 |
| 19978 | 999 | G999W | C | 2.4e-06 | 4.4e-06 | 2.5e-06 | 3.100E-06 | 1.127E-06 | 2.400E-06 | 4.400E-06 |
| 19979 | 999 | G999Y | C | 2.1e-06 | 2.9e-06 | 2.9e-06 | 2.633E-06 | 4.619E-07 | 2.100E-06 | 2.900E-06 |
| 19980 | 999 | G999V | C | 3.5e-06 | 2.8e-06 | 1.7e-06 | 2.667E-06 | 9.074E-07 | 1.700E-06 | 3.500E-06 |
| 19981 | 1000 | A1000A | C | 1.6e-06 | 2.5e-06 | 2.5e-06 | 2.200E-06 | 5.196E-07 | 1.600E-06 | 2.500E-06 |
| 19982 | 1000 | A1000R | C | 2.8e-06 | 2.7e-06 | 2.9e-06 | 2.800E-06 | 1.000E-07 | 2.700E-06 | 2.900E-06 |
| 19983 | 1000 | A1000N | C | 2.6e-06 | 2.4e-06 | 2.7e-06 | 2.567E-06 | 1.528E-07 | 2.400E-06 | 2.700E-06 |
| 19984 | 1000 | A1000D | C | 3.2e-06 | 4.1e-06 | 2.5e-06 | 3.267E-06 | 8.021E-07 | 2.500E-06 | 4.100E-06 |
| 19985 | 1000 | A1000C | C | 2.4e-06 | 2.1e-06 | 2.8e-06 | 2.433E-06 | 3.512E-07 | 2.100E-06 | 2.800E-06 |
| 19986 | 1000 | A1000E | C | 3.2e-06 | 1.4e-06 | 2.7e-06 | 2.433E-06 | 9.292E-07 | 1.400E-06 | 3.200E-06 |
| 19987 | 1000 | A1000Q | C | 2.9e-06 | 1.8e-06 | 2e-06 | 2.233E-06 | 5.859E-07 | 1.800E-06 | 2.900E-06 |
| 19988 | 1000 | A1000G | C | 4.7e-06 | 3.7e-06 | 1.7e-06 | 3.367E-06 | 1.528E-06 | 1.700E-06 | 4.700E-06 |
| 19989 | 1000 | A1000H | C | 1.9e-06 | 2.5e-06 | 3.7e-06 | 2.700E-06 | 9.165E-07 | 1.900E-06 | 3.700E-06 |
| 19990 | 1000 | A1000I | C | 2.8e-06 | 2.3e-06 | 3.1e-06 | 2.733E-06 | 4.041E-07 | 2.300E-06 | 3.100E-06 |
| 19991 | 1000 | A1000L | C | 2.7e-06 | 2.4e-06 | 1.8e-06 | 2.300E-06 | 4.583E-07 | 1.800E-06 | 2.700E-06 |
| 19992 | 1000 | A1000K | C | 1.3e-06 | 2.3e-06 | 2.6e-06 | 2.067E-06 | 6.807E-07 | 1.300E-06 | 2.600E-06 |
| 19993 | 1000 | A1000M | C | 2.1e-06 | 2.2e-06 | 2e-06 | 2.100E-06 | 1.000E-07 | 2.000E-06 | 2.200E-06 |
| 19994 | 1000 | A1000F | C | 2.8e-06 | 2.4e-06 | 8e-07 | 2.000E-06 | 1.058E-06 | 8.000E-07 | 2.800E-06 |
| 19995 | 1000 | A1000P | C | 2.9e-06 | 1.2e-06 | 3.6e-06 | 2.567E-06 | 1.234E-06 | 1.200E-06 | 3.600E-06 |
| 19996 | 1000 | A1000S | C | 2.4e-06 | 4.8e-06 | 2e-06 | 3.067E-06 | 1.514E-06 | 2.000E-06 | 4.800E-06 |
| 19997 | 1000 | A1000T | C | 2e-06 | 1.6e-06 | 2.7e-06 | 2.100E-06 | 5.568E-07 | 1.600E-06 | 2.700E-06 |
| 19998 | 1000 | A1000W | C | 1.5e-06 | 2e-06 | 1.7e-06 | 1.733E-06 | 2.517E-07 | 1.500E-06 | 2.000E-06 |
| 19999 | 1000 | A1000Y | C | 1.4e-06 | 1.6e-06 | 2.2e-06 | 1.733E-06 | 4.163E-07 | 1.400E-06 | 2.200E-06 |
| 20000 | 1000 | A1000V | C | 2.7e-06 | 2.1e-06 | 2.1e-06 | 2.300E-06 | 3.464E-07 | 2.100E-06 | 2.700E-06 |
| 20001 | 1001 | L1001A | C | 2.4e-06 | 1.5e-06 | 1.7e-06 | 1.867E-06 | 4.726E-07 | 1.500E-06 | 2.400E-06 |
| 20002 | 1001 | L1001R | C | 2.4e-06 | 2.8e-06 | 1.5e-06 | 2.233E-06 | 6.658E-07 | 1.500E-06 | 2.800E-06 |
| 20003 | 1001 | L1001N | C | 2.6e-06 | 2.3e-06 | 2e-06 | 2.300E-06 | 3.000E-07 | 2.000E-06 | 2.600E-06 |
| 20004 | 1001 | L1001D | C | 2.7e-06 | 8.3e-07 | 3.2e-06 | 2.243E-06 | 1.249E-06 | 8.300E-07 | 3.200E-06 |
| 20005 | 1001 | L1001C | C | 2.2e-06 | 2.5e-06 | 3.1e-06 | 2.600E-06 | 4.583E-07 | 2.200E-06 | 3.100E-06 |
| 20006 | 1001 | L1001E | C | 1.1e-06 | 1.8e-06 | 2.8e-06 | 1.900E-06 | 8.544E-07 | 1.100E-06 | 2.800E-06 |
| 20007 | 1001 | L1001Q | C | 3.7e-06 | 2.3e-06 | 3.3e-06 | 3.100E-06 | 7.211E-07 | 2.300E-06 | 3.700E-06 |
| 20008 | 1001 | L1001G | C | 5.5e-06 | 1.5e-06 | 1.2e-06 | 2.733E-06 | 2.401E-06 | 1.200E-06 | 5.500E-06 |
| 20009 | 1001 | L1001H | C | 2.9e-06 | 1.3e-06 | 1.3e-06 | 1.833E-06 | 9.238E-07 | 1.300E-06 | 2.900E-06 |
| 20010 | 1001 | L1001I | C | 2.4e-06 | 2.4e-06 | 1.5e-06 | 2.100E-06 | 5.196E-07 | 1.500E-06 | 2.400E-06 |
| 20011 | 1001 | L1001L | C | 1e-06 | 2.7e-06 | 2e-06 | 1.900E-06 | 8.544E-07 | 1.000E-06 | 2.700E-06 |
| 20012 | 1001 | L1001K | C | 2.2e-06 | 2.4e-06 | 3.3e-06 | 2.633E-06 | 5.859E-07 | 2.200E-06 | 3.300E-06 |
| 20013 | 1001 | L1001M | C | 1.3e-06 | 1.5e-06 | 2.2e-06 | 1.667E-06 | 4.726E-07 | 1.300E-06 | 2.200E-06 |
| 20014 | 1001 | L1001F | C | 2.5e-06 | 2.8e-06 | 2.5e-06 | 2.600E-06 | 1.732E-07 | 2.500E-06 | 2.800E-06 |
| 20015 | 1001 | L1001P | C | 1.7e-06 | 1.7e-06 | 3.2e-06 | 2.200E-06 | 8.660E-07 | 1.700E-06 | 3.200E-06 |
| 20016 | 1001 | L1001S | C | 3e-06 | 3.9e-06 | 2.7e-06 | 3.200E-06 | 6.245E-07 | 2.700E-06 | 3.900E-06 |
| 20017 | 1001 | L1001T | C | 1.6e-06 | 1.9e-06 | 2.5e-06 | 2.000E-06 | 4.583E-07 | 1.600E-06 | 2.500E-06 |
| 20018 | 1001 | L1001W | C | 2.9e-06 | 1.1e-06 | 1.9e-06 | 1.967E-06 | 9.018E-07 | 1.100E-06 | 2.900E-06 |
| 20019 | 1001 | L1001Y | C | 1.8e-06 | 2.1e-06 | 2.2e-06 | 2.033E-06 | 2.082E-07 | 1.800E-06 | 2.200E-06 |
| 20020 | 1001 | L1001V | C | 2.3e-06 | 3.2e-06 | 3.2e-06 | 2.900E-06 | 5.196E-07 | 2.300E-06 | 3.200E-06 |
| 20021 | 1002 | T1002A | C | 2.3e-06 | 1.3e-06 | 1.9e-06 | 1.833E-06 | 5.033E-07 | 1.300E-06 | 2.300E-06 |
| 20022 | 1002 | T1002R | C | 2.6e-06 | 2.8e-06 | 2.6e-06 | 2.667E-06 | 1.155E-07 | 2.600E-06 | 2.800E-06 |
| 20023 | 1002 | T1002N | C | 2e-06 | 3.7e-06 | 1.5e-06 | 2.400E-06 | 1.153E-06 | 1.500E-06 | 3.700E-06 |
| 20024 | 1002 | T1002D | C | 2e-06 | 2.4e-06 | 2.1e-06 | 2.167E-06 | 2.082E-07 | 2.000E-06 | 2.400E-06 |
| 20025 | 1002 | T1002C | C | 3.2e-06 | 1.4e-06 | 5.4e-06 | 3.333E-06 | 2.003E-06 | 1.400E-06 | 5.400E-06 |
| 20026 | 1002 | T1002E | C | 2e-06 | 2.1e-06 | 2.5e-06 | 2.200E-06 | 2.646E-07 | 2.000E-06 | 2.500E-06 |
| 20027 | 1002 | T1002Q | C | 1.7e-06 | 2.9e-06 | 2.6e-06 | 2.400E-06 | 6.245E-07 | 1.700E-06 | 2.900E-06 |
| 20028 | 1002 | T1002G | C | 2.3e-06 | 3e-06 | 2e-06 | 2.433E-06 | 5.132E-07 | 2.000E-06 | 3.000E-06 |
| 20029 | 1002 | T1002H | C | 3.6e-06 | 2.7e-06 | 3.1e-06 | 3.133E-06 | 4.509E-07 | 2.700E-06 | 3.600E-06 |
| 20030 | 1002 | T1002I | C | 3.6e-06 | 4.3e-06 | 2.7e-06 | 3.533E-06 | 8.021E-07 | 2.700E-06 | 4.300E-06 |
| 20031 | 1002 | T1002L | C | 3.6e-06 | 2.1e-06 | 2.2e-06 | 2.633E-06 | 8.386E-07 | 2.100E-06 | 3.600E-06 |
| 20032 | 1002 | T1002K | C | 2e-06 | 3.8e-06 | 2.2e-06 | 2.667E-06 | 9.866E-07 | 2.000E-06 | 3.800E-06 |
| 20033 | 1002 | T1002M | C | 2.7e-06 | 3.1e-06 | 2.9e-06 | 2.900E-06 | 2.000E-07 | 2.700E-06 | 3.100E-06 |
| 20034 | 1002 | T1002F | C | 2.9e-06 | 2.3e-06 | 3.4e-06 | 2.867E-06 | 5.508E-07 | 2.300E-06 | 3.400E-06 |
| 20035 | 1002 | T1002P | C | 3.7e-06 | 2.5e-06 | 1.9e-06 | 2.700E-06 | 9.165E-07 | 1.900E-06 | 3.700E-06 |
| 20036 | 1002 | T1002S | C | 1.3e-06 | 2.4e-06 | 2.9e-06 | 2.200E-06 | 8.185E-07 | 1.300E-06 | 2.900E-06 |
| 20037 | 1002 | T1002T | C | 2.2e-06 | 2.1e-06 | 1.8e-06 | 2.033E-06 | 2.082E-07 | 1.800E-06 | 2.200E-06 |
| 20038 | 1002 | T1002W | C | 5.4e-06 | 1.4e-06 | 2.6e-06 | 3.133E-06 | 2.053E-06 | 1.400E-06 | 5.400E-06 |
| 20039 | 1002 | T1002Y | C | 2.8e-06 | 5.3e-06 | 2.8e-06 | 3.633E-06 | 1.443E-06 | 2.800E-06 | 5.300E-06 |
| 20040 | 1002 | T1002V | C | 2.6e-06 | 3.2e-06 | 1.8e-06 | 2.533E-06 | 7.024E-07 | 1.800E-06 | 3.200E-06 |
| 20041 | 1003 | S1003A | C | 2.1e-06 | 2.6e-06 | 3.8e-06 | 2.833E-06 | 8.737E-07 | 2.100E-06 | 3.800E-06 |
| 20042 | 1003 | S1003R | C | 2.6e-06 | 3.7e-06 | 1.3e-06 | 2.533E-06 | 1.201E-06 | 1.300E-06 | 3.700E-06 |
| 20043 | 1003 | S1003N | C | 2.3e-06 | 2.4e-06 | 3.7e-06 | 2.800E-06 | 7.810E-07 | 2.300E-06 | 3.700E-06 |
| 20044 | 1003 | S1003D | C | 5.5e-06 | 1.6e-06 | 2.8e-06 | 3.300E-06 | 1.997E-06 | 1.600E-06 | 5.500E-06 |
| 20045 | 1003 | S1003C | C | 1.7e-06 | 2e-06 | 2.6e-06 | 2.100E-06 | 4.583E-07 | 1.700E-06 | 2.600E-06 |
| 20046 | 1003 | S1003E | C | 2.4e-06 | 1.9e-06 | 4.8e-06 | 3.033E-06 | 1.550E-06 | 1.900E-06 | 4.800E-06 |
| 20047 | 1003 | S1003Q | C | 1.9e-06 | 3.1e-06 | 2e-06 | 2.333E-06 | 6.658E-07 | 1.900E-06 | 3.100E-06 |
| 20048 | 1003 | S1003G | C | 2.3e-06 | 4e-06 | 2.4e-06 | 2.900E-06 | 9.539E-07 | 2.300E-06 | 4.000E-06 |
| 20049 | 1003 | S1003H | C | 2.2e-06 | 2.5e-06 | 3.4e-06 | 2.700E-06 | 6.245E-07 | 2.200E-06 | 3.400E-06 |
| 20050 | 1003 | S1003I | C | 2.4e-06 | 3.5e-06 | 5.6e-06 | 3.833E-06 | 1.626E-06 | 2.400E-06 | 5.600E-06 |
| 20051 | 1003 | S1003L | C | 1.7e-06 | 2.1e-06 | 2.6e-06 | 2.133E-06 | 4.509E-07 | 1.700E-06 | 2.600E-06 |
| 20052 | 1003 | S1003K | C | 1.6e-06 | 1.2e-06 | 2e-06 | 1.600E-06 | 4.000E-07 | 1.200E-06 | 2.000E-06 |
| 20053 | 1003 | S1003M | C | 3.8e-06 | 1.6e-06 | 3.1e-06 | 2.833E-06 | 1.124E-06 | 1.600E-06 | 3.800E-06 |
| 20054 | 1003 | S1003F | C | 2e-06 | 2.4e-06 | 3.2e-06 | 2.533E-06 | 6.110E-07 | 2.000E-06 | 3.200E-06 |
| 20055 | 1003 | S1003P | C | 3.6e-06 | 8.7e-07 | 2.3e-06 | 2.257E-06 | 1.366E-06 | 8.700E-07 | 3.600E-06 |
| 20056 | 1003 | S1003S | C | 2.5e-06 | 3.3e-06 | 2.8e-06 | 2.867E-06 | 4.041E-07 | 2.500E-06 | 3.300E-06 |
| 20057 | 1003 | S1003T | C | 4.9e-06 | 2.1e-06 | 1.7e-06 | 2.900E-06 | 1.744E-06 | 1.700E-06 | 4.900E-06 |
| 20058 | 1003 | S1003W | C | 1.3e-06 | 2e-06 | 2e-06 | 1.767E-06 | 4.041E-07 | 1.300E-06 | 2.000E-06 |
| 20059 | 1003 | S1003Y | C | 1.7e-06 | 4.5e-06 | 5.2e-06 | 3.800E-06 | 1.852E-06 | 1.700E-06 | 5.200E-06 |

| | | | | | | | | | | |
|---|---|---|---|---|---|---|---|---|---|---|
| 20060 | 1003 | S1003V | C | 3e-06 | 1.7e-06 | 2.3e-06 | 2.333E-06 | 6.506E-07 | 1.700E-06 | 3.000E-06 |
| 20061 | 1004 | G1004A | C | 1.9e-06 | 2.2e-06 | 1.9e-06 | 2.000E-06 | 1.732E-07 | 1.900E-06 | 2.200E-06 |
| 20062 | 1004 | G1004R | C | 8.7e-07 | 2.6e-06 | 2.7e-06 | 2.057E-06 | 1.029E-06 | 8.700E-07 | 2.700E-06 |
| 20063 | 1004 | G1004N | C | 2.4e-06 | 2.4e-06 | 3.5e-06 | 2.767E-06 | 6.351E-07 | 2.400E-06 | 3.500E-06 |
| 20064 | 1004 | G1004D | C | 2.2e-06 | 2.6e-06 | 2.9e-06 | 2.567E-06 | 3.512E-07 | 2.200E-06 | 2.900E-06 |
| 20065 | 1004 | G1004C | C | 3e-06 | 1.5e-06 | 2.5e-06 | 2.333E-06 | 7.638E-07 | 1.500E-06 | 3.000E-06 |
| 20066 | 1004 | G1004E | C | 2.5e-06 | 2.4e-06 | 1.9e-06 | 2.267E-06 | 3.215E-07 | 1.900E-06 | 2.500E-06 |
| 20067 | 1004 | G1004Q | C | 2.9e-06 | 1.3e-06 | 2.6e-06 | 2.267E-06 | 8.505E-07 | 1.300E-06 | 2.900E-06 |
| 20068 | 1004 | G1004G | C | 2.4e-06 | 2.2e-06 | 2.1e-06 | 2.233E-06 | 1.528E-07 | 2.100E-06 | 2.400E-06 |
| 20069 | 1004 | G1004H | C | 1.6e-06 | 1e-06 | 2.2e-06 | 1.600E-06 | 6.000E-07 | 1.000E-06 | 2.200E-06 |
| 20070 | 1004 | G1004I | C | 1.1e-06 | 3.2e-06 | 2.7e-06 | 2.333E-06 | 1.097E-06 | 1.100E-06 | 3.200E-06 |
| 20071 | 1004 | G1004L | C | 2.3e-06 | 2.1e-06 | 1.9e-06 | 2.100E-06 | 2.000E-07 | 1.900E-06 | 2.300E-06 |
| 20072 | 1004 | G1004K | C | 2.3e-06 | 2.2e-06 | 3.7e-06 | 2.733E-06 | 8.386E-07 | 2.200E-06 | 3.700E-06 |
| 20073 | 1004 | G1004M | C | 4.1e-06 | 2.4e-06 | 1.3e-06 | 2.600E-06 | 1.411E-06 | 1.300E-06 | 4.100E-06 |
| 20074 | 1004 | G1004F | C | 2.3e-06 | 2e-06 | 2.2e-06 | 2.167E-06 | 1.528E-07 | 2.000E-06 | 2.300E-06 |
| 20075 | 1004 | G1004P | C | 1.8e-06 | 2.3e-06 | 4.6e-06 | 2.900E-06 | 1.493E-06 | 1.800E-06 | 4.600E-06 |
| 20076 | 1004 | G1004S | C | 2.5e-06 | 2.8e-06 | 1.9e-06 | 2.400E-06 | 4.583E-07 | 1.900E-06 | 2.800E-06 |
| 20077 | 1004 | G1004T | C | 3.1e-06 | 1.9e-06 | 1.1e-06 | 2.033E-06 | 1.007E-06 | 1.100E-06 | 3.100E-06 |
| 20078 | 1004 | G1004W | C | 2.3e-06 | 2.7e-06 | 2.6e-06 | 2.533E-06 | 2.082E-07 | 2.300E-06 | 2.700E-06 |
| 20079 | 1004 | G1004Y | C | 1.6e-06 | 2.2e-06 | 2.3e-06 | 2.033E-06 | 3.786E-07 | 1.600E-06 | 2.300E-06 |
| 20080 | 1004 | G1004V | C | 2.1e-06 | 1.9e-06 | 2.4e-06 | 2.133E-06 | 2.517E-07 | 1.900E-06 | 2.400E-06 |
| 20081 | 1005 | V1005A | C | 2.9e-06 | 2.3e-06 | 2.1e-06 | 2.433E-06 | 4.163E-07 | 2.100E-06 | 2.900E-06 |
| 20082 | 1005 | V1005R | C | 2.2e-06 | 3.4e-06 | 2.2e-06 | 2.600E-06 | 6.928E-07 | 2.200E-06 | 3.400E-06 |
| 20083 | 1005 | V1005N | C | 1.9e-06 | 1.4e-06 | 2.3e-06 | 1.867E-06 | 4.509E-07 | 1.400E-06 | 2.300E-06 |
| 20084 | 1005 | V1005D | C | 2.5e-06 | 1.9e-06 | 1.6e-06 | 2.000E-06 | 4.583E-07 | 1.600E-06 | 2.500E-06 |
| 20085 | 1005 | V1005C | C | 1.5e-06 | 1.9e-06 | 2.3e-06 | 1.900E-06 | 4.000E-07 | 1.500E-06 | 2.300E-06 |
| 20086 | 1005 | V1005E | C | 3.3e-06 | 1.7e-06 | 1.9e-06 | 2.300E-06 | 8.718E-07 | 1.700E-06 | 3.300E-06 |
| 20087 | 1005 | V1005Q | C | 2.1e-06 | 2.2e-06 | 2.1e-06 | 2.133E-06 | 5.774E-08 | 2.100E-06 | 2.200E-06 |
| 20088 | 1005 | V1005G | C | 2.3e-06 | 2e-06 | 4.1e-06 | 2.800E-06 | 1.136E-06 | 2.000E-06 | 4.100E-06 |
| 20089 | 1005 | V1005H | C | 2.4e-06 | 2.4e-06 | 1.9e-06 | 2.233E-06 | 2.887E-07 | 1.900E-06 | 2.400E-06 |
| 20090 | 1005 | V1005I | C | 1.8e-06 | 2e-06 | 4.3e-06 | 2.700E-06 | 1.389E-06 | 1.800E-06 | 4.300E-06 |
| 20091 | 1005 | V1005L | C | 2.7e-06 | 4.1e-06 | 2.6e-06 | 3.133E-06 | 8.386E-07 | 2.600E-06 | 4.100E-06 |
| 20092 | 1005 | V1005K | C | 1.5e-06 | 1.3e-06 | 2.6e-06 | 1.800E-06 | 7.000E-07 | 1.300E-06 | 2.600E-06 |
| 20093 | 1005 | V1005M | C | 1.7e-06 | 1.3e-06 | 2.4e-06 | 1.800E-06 | 5.568E-07 | 1.300E-06 | 2.400E-06 |
| 20094 | 1005 | V1005F | C | 1.7e-06 | 3.1e-06 | 2.5e-06 | 2.433E-06 | 7.024E-07 | 1.700E-06 | 3.100E-06 |
| 20095 | 1005 | V1005P | C | 3.8e-06 | 3.3e-06 | 2.8e-06 | 3.300E-06 | 5.000E-07 | 2.800E-06 | 3.800E-06 |
| 20096 | 1005 | V1005S | C | 2.6e-06 | 1.4e-06 | 2.1e-06 | 2.033E-06 | 6.028E-07 | 1.400E-06 | 2.600E-06 |
| 20097 | 1005 | V1005T | C | 3.5e-06 | 2.1e-06 | 2.8e-06 | 2.800E-06 | 7.000E-07 | 2.100E-06 | 3.500E-06 |
| 20098 | 1005 | V1005W | C | 1.4e-06 | 2.8e-06 | 2.9e-06 | 2.367E-06 | 8.386E-07 | 1.400E-06 | 2.900E-06 |
| 20099 | 1005 | V1005Y | C | 3.1e-06 | 3.5e-06 | 2.5e-06 | 3.033E-06 | 5.033E-07 | 2.500E-06 | 3.500E-06 |
| 20100 | 1005 | V1005V | C | 3e-06 | 3.7e-06 | 3.9e-06 | 3.533E-06 | 4.726E-07 | 3.000E-06 | 3.900E-06 |
| 20101 | 1006 | H1006A | C | 2.7e-06 | 2.1e-06 | 3.1e-06 | 2.633E-06 | 5.033E-07 | 2.100E-06 | 3.100E-06 |
| 20102 | 1006 | H1006R | C | 2.1e-06 | 2.7e-06 | 2e-06 | 2.267E-06 | 3.786E-07 | 2.000E-06 | 2.700E-06 |
| 20103 | 1006 | H1006N | C | 7.9e-07 | 2.5e-06 | 1.6e-06 | 1.630E-06 | 8.554E-07 | 7.900E-07 | 2.500E-06 |
| 20104 | 1006 | H1006D | C | 2.7e-06 | 2.8e-06 | 1.7e-06 | 2.400E-06 | 6.083E-07 | 1.700E-06 | 2.800E-06 |
| 20105 | 1006 | H1006C | C | 1.6e-06 | 3.4e-06 | 8.9e-07 | 1.963E-06 | 1.294E-06 | 8.900E-07 | 3.400E-06 |
| 20106 | 1006 | H1006E | C | 1.6e-06 | 2.3e-06 | 2.6e-06 | 2.167E-06 | 5.132E-07 | 1.600E-06 | 2.600E-06 |
| 20107 | 1006 | H1006Q | C | 3e-06 | 1.9e-06 | 2.6e-06 | 2.500E-06 | 5.568E-07 | 1.900E-06 | 3.000E-06 |
| 20108 | 1006 | H1006G | C | 3e-06 | 1.7e-06 | 1.9e-06 | 2.200E-06 | 7.000E-07 | 1.700E-06 | 3.000E-06 |
| 20109 | 1006 | H1006H | C | 3.7e-06 | 1.3e-06 | 2.3e-06 | 2.433E-06 | 1.206E-06 | 1.300E-06 | 3.700E-06 |
| 20110 | 1006 | H1006I | C | 2.1e-06 | 2.4e-06 | 2.8e-06 | 2.433E-06 | 3.512E-07 | 2.100E-06 | 2.800E-06 |
| 20111 | 1006 | H1006L | C | 2.8e-06 | 3.9e-06 | 3.8e-06 | 3.500E-06 | 6.083E-07 | 2.800E-06 | 3.900E-06 |
| 20112 | 1006 | H1006K | C | 3.3e-06 | 3.1e-06 | 1.9e-06 | 2.767E-06 | 7.572E-07 | 1.900E-06 | 3.300E-06 |
| 20113 | 1006 | H1006M | C | 2.2e-06 | 1.8e-06 | 3.1e-06 | 2.367E-06 | 6.658E-07 | 1.800E-06 | 3.100E-06 |
| 20114 | 1006 | H1006F | C | 2.2e-06 | 2.5e-06 | 1.9e-06 | 2.200E-06 | 3.000E-07 | 1.900E-06 | 2.500E-06 |
| 20115 | 1006 | H1006P | C | 1.7e-06 | 3.1e-06 | 2.4e-06 | 2.400E-06 | 7.000E-07 | 1.700E-06 | 3.100E-06 |
| 20116 | 1006 | H1006S | C | 2.1e-06 | 2.2e-06 | 2.7e-06 | 2.333E-06 | 3.215E-07 | 2.100E-06 | 2.700E-06 |
| 20117 | 1006 | H1006T | C | 2.4e-06 | 2.1e-06 | 1.5e-06 | 2.000E-06 | 4.583E-07 | 1.500E-06 | 2.400E-06 |
| 20118 | 1006 | H1006W | C | 3.2e-06 | 3.6e-06 | 2.3e-06 | 3.033E-06 | 6.658E-07 | 2.300E-06 | 3.600E-06 |
| 20119 | 1006 | H1006Y | C | 1.8e-06 | 2.2e-06 | 1.9e-06 | 1.967E-06 | 2.082E-07 | 1.800E-06 | 2.200E-06 |
| 20120 | 1006 | H1006V | C | 2.4e-06 | 3.2e-06 | 1.9e-06 | 2.500E-06 | 6.557E-07 | 1.900E-06 | 3.200E-06 |
| 20121 | 1007 | T1007A | C | 5.1e-06 | 2e-06 | 2.5e-06 | 3.200E-06 | 1.664E-06 | 2.000E-06 | 5.100E-06 |
| 20122 | 1007 | T1007R | C | 4.4e-06 | 2.2e-06 | 3.1e-06 | 3.233E-06 | 1.106E-06 | 2.200E-06 | 4.400E-06 |
| 20123 | 1007 | T1007N | C | 2.4e-06 | 1.5e-06 | 4e-06 | 2.633E-06 | 1.266E-06 | 1.500E-06 | 4.000E-06 |
| 20124 | 1007 | T1007D | C | 2.7e-06 | 2.4e-06 | 2.4e-06 | 2.500E-06 | 1.732E-07 | 2.400E-06 | 2.700E-06 |
| 20125 | 1007 | T1007C | C | 1.6e-06 | 1.6e-06 | 2.6e-06 | 1.933E-06 | 5.774E-07 | 1.600E-06 | 2.600E-06 |
| 20126 | 1007 | T1007E | C | 3.1e-06 | 2.2e-06 | 2.1e-06 | 2.467E-06 | 5.508E-07 | 2.100E-06 | 3.100E-06 |
| 20127 | 1007 | T1007Q | C | 1.5e-06 | 2.3e-06 | 4.2e-06 | 2.667E-06 | 1.387E-06 | 1.500E-06 | 4.200E-06 |
| 20128 | 1007 | T1007G | C | 1.7e-06 | 2.6e-06 | 3.5e-06 | 2.600E-06 | 9.000E-07 | 1.700E-06 | 3.500E-06 |
| 20129 | 1007 | T1007H | C | 2e-06 | 1.8e-06 | 1.7e-06 | 1.833E-06 | 1.528E-07 | 1.700E-06 | 2.000E-06 |
| 20130 | 1007 | T1007I | C | 2.4e-06 | 1.6e-06 | 2.4e-06 | 2.133E-06 | 4.619E-07 | 1.600E-06 | 2.400E-06 |
| 20131 | 1007 | T1007L | C | 2.8e-06 | 1e-06 | 2.5e-06 | 2.100E-06 | 9.644E-07 | 1.000E-06 | 2.800E-06 |
| 20132 | 1007 | T1007K | C | 2.1e-06 | 2.5e-06 | 1.7e-06 | 2.100E-06 | 4.000E-07 | 1.700E-06 | 2.500E-06 |
| 20133 | 1007 | T1007M | C | 1e-06 | 2.5e-06 | 1.5e-06 | 1.667E-06 | 7.638E-07 | 1.000E-06 | 2.500E-06 |
| 20134 | 1007 | T1007F | C | 2.6e-06 | 3.2e-06 | 1.2e-06 | 2.333E-06 | 1.026E-06 | 1.200E-06 | 3.200E-06 |
| 20135 | 1007 | T1007P | C | 3.7e-06 | 3.4e-06 | 3.7e-06 | 3.600E-06 | 1.732E-07 | 3.400E-06 | 3.700E-06 |
| 20136 | 1007 | T1007S | C | 2.2e-06 | 2.7e-06 | 2.9e-06 | 2.600E-06 | 3.606E-07 | 2.200E-06 | 2.900E-06 |
| 20137 | 1007 | T1007T | C | 2e-06 | 2.3e-06 | 2e-06 | 2.100E-06 | 1.732E-07 | 2.000E-06 | 2.300E-06 |
| 20138 | 1007 | T1007W | C | 4e-06 | 2.3e-06 | 2.7e-06 | 3.000E-06 | 8.888E-07 | 2.300E-06 | 4.000E-06 |
| 20139 | 1007 | T1007Y | C | 1.6e-06 | 1.4e-06 | 1.7e-06 | 1.567E-06 | 1.528E-07 | 1.400E-06 | 1.700E-06 |
| 20140 | 1007 | T1007V | C | 2.5e-06 | 2.3e-06 | 3.1e-06 | 2.633E-06 | 4.163E-07 | 2.300E-06 | 3.100E-06 |
| 20141 | 1008 | F1008A | C | 2.6e-06 | 4.6e-06 | 4.1e-06 | 3.767E-06 | 1.041E-06 | 2.600E-06 | 4.600E-06 |
| 20142 | 1008 | F1008R | C | 2.2e-06 | 2.7e-06 | 1.9e-06 | 2.267E-06 | 4.041E-07 | 1.900E-06 | 2.700E-06 |
| 20143 | 1008 | F1008N | C | 3.1e-06 | 3.7e-06 | 1.7e-06 | 2.833E-06 | 1.026E-06 | 1.700E-06 | 3.700E-06 |
| 20144 | 1008 | F1008D | C | 2.6e-06 | 3.2e-06 | 3.1e-06 | 2.967E-06 | 3.215E-07 | 2.600E-06 | 3.200E-06 |

| | | | | | | | | | | |
|---|---|---|---|---|---|---|---|---|---|---|
| 20145 | 1008 | F1008C | C | 1.4e-06 | 1.4e-06 | 3.4e-06 | 2.067E-06 | 1.155E-06 | 1.400E-06 | 3.400E-06 |
| 20146 | 1008 | F1008E | C | 2.7e-06 | 2.7e-06 | 2.3e-06 | 2.567E-06 | 2.309E-07 | 2.300E-06 | 2.700E-06 |
| 20147 | 1008 | F1008Q | C | 2.5e-06 | 3.2e-06 | 4.6e-06 | 3.433E-06 | 1.069E-06 | 2.500E-06 | 4.600E-06 |
| 20148 | 1008 | F1008G | C | 1.9e-06 | 2.3e-06 | 3e-06 | 2.400E-06 | 5.568E-07 | 1.900E-06 | 3.000E-06 |
| 20149 | 1008 | F1008H | C | 1.6e-06 | 3e-06 | 2e-06 | 2.200E-06 | 7.211E-07 | 1.600E-06 | 3.000E-06 |
| 20150 | 1008 | F1008I | C | 2.3e-06 | 2.5e-06 | 2.2e-06 | 2.333E-06 | 1.528E-07 | 2.200E-06 | 2.500E-06 |
| 20151 | 1008 | F1008L | C | 9.3e-07 | 2.6e-06 | 2.1e-06 | 1.877E-06 | 8.571E-07 | 9.300E-07 | 2.600E-06 |
| 20152 | 1008 | F1008K | C | 2.7e-06 | 2.2e-06 | 2.6e-06 | 2.500E-06 | 2.646E-07 | 2.200E-06 | 2.700E-06 |
| 20153 | 1008 | F1008M | C | 2.8e-06 | 2.4e-06 | 2.6e-06 | 2.600E-06 | 2.000E-07 | 2.400E-06 | 2.800E-06 |
| 20154 | 1008 | F1008F | C | 1.8e-06 | 2.2e-06 | 1.7e-06 | 1.900E-06 | 2.646E-07 | 1.700E-06 | 2.200E-06 |
| 20155 | 1008 | F1008P | C | 1.1e-06 | 2.1e-06 | 2.9e-06 | 2.033E-06 | 9.018E-07 | 1.100E-06 | 2.900E-06 |
| 20156 | 1008 | F1008S | C | 2.6e-06 | 1.7e-06 | 2.1e-06 | 2.133E-06 | 4.509E-07 | 1.700E-06 | 2.600E-06 |
| 20157 | 1008 | F1008T | C | 2.1e-06 | 3.9e-06 | 1.5e-06 | 2.500E-06 | 1.249E-06 | 1.500E-06 | 3.900E-06 |
| 20158 | 1008 | F1008W | C | 2.6e-06 | 2.1e-06 | 2.1e-06 | 2.267E-06 | 2.887E-07 | 2.100E-06 | 2.600E-06 |
| 20159 | 1008 | F1008Y | C | 2e-06 | 1.9e-06 | 1.7e-06 | 1.867E-06 | 1.528E-07 | 1.700E-06 | 2.000E-06 |
| 20160 | 1008 | F1008V | C | 1.9e-06 | 2.9e-06 | 3e-06 | 2.600E-06 | 6.083E-07 | 1.900E-06 | 3.000E-06 |
| 20161 | 1009 | P1009A | C | 2.5e-06 | 2e-06 | 1.7e-06 | 2.067E-06 | 4.041E-07 | 1.700E-06 | 2.500E-06 |
| 20162 | 1009 | P1009R | C | 2.6e-06 | 2.1e-06 | 3e-06 | 2.567E-06 | 4.509E-07 | 2.100E-06 | 3.000E-06 |
| 20163 | 1009 | P1009N | C | 2.3e-06 | 1.1e-06 | 4.5e-06 | 2.633E-06 | 1.724E-06 | 1.100E-06 | 4.500E-06 |
| 20164 | 1009 | P1009D | C | 3.1e-06 | 2.6e-06 | 9.2e-07 | 2.207E-06 | 1.142E-06 | 9.200E-07 | 3.100E-06 |
| 20165 | 1009 | P1009C | C | 1.7e-06 | 2.2e-06 | 1.4e-06 | 1.767E-06 | 4.041E-07 | 1.400E-06 | 2.200E-06 |
| 20166 | 1009 | P1009E | C | 2.3e-06 | 1.5e-06 | 2.1e-06 | 1.967E-06 | 4.163E-07 | 1.500E-06 | 2.300E-06 |
| 20167 | 1009 | P1009Q | C | 2.6e-06 | 2.9e-06 | 3.4e-06 | 2.967E-06 | 4.041E-07 | 2.600E-06 | 3.400E-06 |
| 20168 | 1009 | P1009G | C | 3.6e-06 | 1.1e-06 | 2.3e-06 | 2.333E-06 | 1.250E-06 | 1.100E-06 | 3.600E-06 |
| 20169 | 1009 | P1009H | C | 2.2e-06 | 2.1e-06 | 3.2e-06 | 2.500E-06 | 6.083E-07 | 2.100E-06 | 3.200E-06 |
| 20170 | 1009 | P1009I | C | 1.5e-06 | 2.2e-06 | 1.5e-06 | 1.733E-06 | 4.041E-07 | 1.500E-06 | 2.200E-06 |
| 20171 | 1009 | P1009L | C | 1.9e-06 | 2.1e-06 | 2.4e-06 | 2.133E-06 | 2.517E-07 | 1.900E-06 | 2.400E-06 |
| 20172 | 1009 | P1009K | C | 3e-06 | 3.2e-06 | 2.1e-06 | 2.767E-06 | 5.859E-07 | 2.100E-06 | 3.200E-06 |
| 20173 | 1009 | P1009M | C | 2.7e-06 | 1.5e-06 | 3.7e-06 | 2.633E-06 | 1.102E-06 | 1.500E-06 | 3.700E-06 |
| 20174 | 1009 | P1009F | C | 1.6e-06 | 2.3e-06 | 2.6e-06 | 2.167E-06 | 5.132E-07 | 1.600E-06 | 2.600E-06 |
| 20175 | 1009 | P1009P | C | 1.8e-06 | 2.5e-06 | 2.3e-06 | 2.200E-06 | 3.606E-07 | 1.800E-06 | 2.500E-06 |
| 20176 | 1009 | P1009S | C | 1.7e-06 | 3e-06 | 2.4e-06 | 2.367E-06 | 6.506E-07 | 1.700E-06 | 3.000E-06 |
| 20177 | 1009 | P1009T | C | 2.5e-06 | 2.7e-06 | 2.1e-06 | 2.433E-06 | 3.055E-07 | 2.100E-06 | 2.700E-06 |
| 20178 | 1009 | P1009W | C | 2.1e-06 | 2e-06 | 1.8e-06 | 1.967E-06 | 1.528E-07 | 1.800E-06 | 2.100E-06 |
| 20179 | 1009 | P1009Y | C | 1.4e-06 | 1.4e-06 | 1.6e-06 | 1.467E-06 | 1.155E-07 | 1.400E-06 | 1.600E-06 |
| 20180 | 1009 | P1009V | C | 2.2e-06 | 1.8e-06 | 1.4e-06 | 1.800E-06 | 4.000E-07 | 1.400E-06 | 2.200E-06 |
| 20181 | 1010 | A1010A | C | 1.5e-06 | 1.9e-06 | 3.4e-06 | 2.267E-06 | 1.002E-06 | 1.500E-06 | 3.400E-06 |
| 20182 | 1010 | A1010R | C | 1.9e-06 | 2.1e-06 | 2.5e-06 | 2.167E-06 | 3.055E-07 | 1.900E-06 | 2.500E-06 |
| 20183 | 1010 | A1010N | C | 2.8e-06 | 1.8e-06 | 1.9e-06 | 2.167E-06 | 5.508E-07 | 1.800E-06 | 2.800E-06 |
| 20184 | 1010 | A1010D | C | 2e-06 | 3.4e-06 | 8.1e-07 | 2.070E-06 | 1.296E-06 | 8.100E-07 | 3.400E-06 |
| 20185 | 1010 | A1010C | C | 2.7e-06 | 1.8e-06 | 2.7e-06 | 2.400E-06 | 5.196E-07 | 1.800E-06 | 2.700E-06 |
| 20186 | 1010 | A1010E | C | 2.3e-06 | 2.4e-06 | 2e-06 | 2.233E-06 | 2.082E-07 | 2.000E-06 | 2.400E-06 |
| 20187 | 1010 | A1010Q | C | 1.8e-06 | 2.5e-06 | 2.9e-06 | 2.400E-06 | 5.568E-07 | 1.800E-06 | 2.900E-06 |
| 20188 | 1010 | A1010G | C | 1.9e-06 | 2e-06 | 2.5e-06 | 2.133E-06 | 3.215E-07 | 1.900E-06 | 2.500E-06 |
| 20189 | 1010 | A1010H | C | 1.8e-06 | 1.4e-06 | 1.7e-06 | 1.633E-06 | 2.082E-07 | 1.400E-06 | 1.800E-06 |
| 20190 | 1010 | A1010I | C | 3e-06 | 2.7e-06 | 3.6e-06 | 3.100E-06 | 4.583E-07 | 2.700E-06 | 3.600E-06 |
| 20191 | 1010 | A1010L | C | 3.2e-06 | 2.2e-06 | 1.1e-06 | 2.167E-06 | 1.050E-06 | 1.100E-06 | 3.200E-06 |
| 20192 | 1010 | A1010K | C | 2.6e-06 | 3.5e-06 | 2.1e-06 | 2.733E-06 | 7.095E-07 | 2.100E-06 | 3.500E-06 |
| 20193 | 1010 | A1010M | C | 1.7e-06 | 2.6e-06 | 1.8e-06 | 2.033E-06 | 4.933E-07 | 1.700E-06 | 2.600E-06 |
| 20194 | 1010 | A1010F | C | 1.8e-06 | 2.1e-06 | 2.9e-06 | 2.267E-06 | 5.686E-07 | 1.800E-06 | 2.900E-06 |
| 20195 | 1010 | A1010P | C | 2.4e-06 | 1.8e-06 | 2.6e-06 | 2.267E-06 | 4.163E-07 | 1.800E-06 | 2.600E-06 |
| 20196 | 1010 | A1010S | C | 3.3e-06 | 3e-06 | 3.1e-06 | 3.133E-06 | 1.528E-07 | 3.000E-06 | 3.300E-06 |
| 20197 | 1010 | A1010T | C | 2.7e-06 | 3.2e-06 | 6.1e-06 | 4.000E-06 | 1.836E-06 | 2.700E-06 | 6.100E-06 |
| 20198 | 1010 | A1010W | C | 2.6e-06 | 2.9e-06 | 3.6e-06 | 3.033E-06 | 5.132E-07 | 2.600E-06 | 3.600E-06 |
| 20199 | 1010 | A1010Y | C | 2.2e-06 | 1.7e-06 | 3.5e-06 | 2.467E-06 | 9.292E-07 | 1.700E-06 | 3.500E-06 |
| 20200 | 1010 | A1010V | C | 3.3e-06 | 2.8e-06 | 2.5e-06 | 2.867E-06 | 4.041E-07 | 2.500E-06 | 3.300E-06 |
| 20201 | 1011 | V1011A | C | 2.1e-06 | 1.9e-06 | 3.4e-06 | 2.467E-06 | 8.145E-07 | 1.900E-06 | 3.400E-06 |
| 20202 | 1011 | V1011R | C | 1.8e-06 | 3.2e-06 | 1.4e-06 | 2.133E-06 | 9.452E-07 | 1.400E-06 | 3.200E-06 |
| 20203 | 1011 | V1011N | C | 2.9e-06 | 2.2e-06 | 2.3e-06 | 2.467E-06 | 3.786E-07 | 2.200E-06 | 2.900E-06 |
| 20204 | 1011 | V1011D | C | 2.3e-06 | 3.8e-06 | 2.3e-06 | 2.800E-06 | 8.660E-07 | 2.300E-06 | 3.800E-06 |
| 20205 | 1011 | V1011C | C | 2.9e-06 | 1.8e-06 | 1.6e-06 | 2.100E-06 | 7.000E-07 | 1.600E-06 | 2.900E-06 |
| 20206 | 1011 | V1011E | C | 3e-06 | 1.9e-06 | 2.3e-06 | 2.400E-06 | 5.568E-07 | 1.900E-06 | 3.000E-06 |
| 20207 | 1011 | V1011Q | C | 3.4e-06 | 3.9e-06 | 2.6e-06 | 3.300E-06 | 6.557E-07 | 2.600E-06 | 3.900E-06 |
| 20208 | 1011 | V1011G | C | 2.7e-06 | 2.4e-06 | 2.7e-06 | 2.600E-06 | 1.732E-07 | 2.400E-06 | 2.700E-06 |
| 20209 | 1011 | V1011H | C | 1.6e-06 | 3.6e-06 | 1.8e-06 | 2.333E-06 | 1.102E-06 | 1.600E-06 | 3.600E-06 |
| 20210 | 1011 | V1011I | C | 1.7e-06 | 1.9e-06 | 1.8e-06 | 1.800E-06 | 1.000E-07 | 1.700E-06 | 1.900E-06 |
| 20211 | 1011 | V1011L | C | 2.6e-06 | 1.9e-06 | 3.2e-06 | 2.567E-06 | 6.506E-07 | 1.900E-06 | 3.200E-06 |
| 20212 | 1011 | V1011K | C | 1.4e-06 | 1.1e-06 | 4e-06 | 2.167E-06 | 1.595E-06 | 1.100E-06 | 4.000E-06 |
| 20213 | 1011 | V1011M | C | 3.6e-06 | 2.8e-06 | 8.8e-07 | 2.427E-06 | 1.398E-06 | 8.800E-07 | 3.600E-06 |
| 20214 | 1011 | V1011F | C | 2e-06 | 4e-06 | 2.8e-06 | 2.933E-06 | 1.007E-06 | 2.000E-06 | 4.000E-06 |
| 20215 | 1011 | V1011P | C | 2.8e-06 | 3.1e-06 | 1.1e-06 | 2.333E-06 | 1.079E-06 | 1.100E-06 | 3.100E-06 |
| 20216 | 1011 | V1011S | C | 2.8e-06 | 2.3e-06 | 3e-06 | 2.700E-06 | 3.606E-07 | 2.300E-06 | 3.000E-06 |
| 20217 | 1011 | V1011T | C | 3.2e-06 | 2.9e-06 | 1.3e-06 | 2.467E-06 | 1.021E-06 | 1.300E-06 | 3.200E-06 |
| 20218 | 1011 | V1011W | C | 2.8e-06 | 2.9e-06 | 6.1e-06 | 3.933E-06 | 1.877E-06 | 2.800E-06 | 6.100E-06 |
| 20219 | 1011 | V1011Y | C | 2.3e-06 | 3.7e-06 | 1.9e-06 | 2.633E-06 | 9.452E-07 | 1.900E-06 | 3.700E-06 |
| 20220 | 1011 | V1011V | C | 1.9e-06 | 2.3e-06 | 2.2e-06 | 2.133E-06 | 2.082E-07 | 1.900E-06 | 2.300E-06 |
| 20221 | 1012 | L1012A | C | 3.6e-06 | 2e-06 | 1.6e-06 | 2.400E-06 | 1.058E-06 | 1.600E-06 | 3.600E-06 |
| 20222 | 1012 | L1012R | C | 2.4e-06 | 1.9e-06 | 3.2e-06 | 2.500E-06 | 6.557E-07 | 1.900E-06 | 3.200E-06 |
| 20223 | 1012 | L1012N | C | 2.4e-06 | 2.6e-06 | 2.7e-06 | 2.567E-06 | 1.528E-07 | 2.400E-06 | 2.700E-06 |
| 20224 | 1012 | L1012D | C | 2.8e-06 | 3e-06 | 2.1e-06 | 2.633E-06 | 4.726E-07 | 2.100E-06 | 3.000E-06 |
| 20225 | 1012 | L1012C | C | 3.1e-06 | 2.6e-06 | 3.1e-06 | 2.933E-06 | 2.887E-07 | 2.600E-06 | 3.100E-06 |
| 20226 | 1012 | L1012E | C | 2.8e-06 | 3.6e-06 | 2.4e-06 | 2.933E-06 | 6.110E-07 | 2.400E-06 | 3.600E-06 |
| 20227 | 1012 | L1012Q | C | 3.6e-06 | 2.2e-06 | 3.6e-06 | 3.133E-06 | 8.083E-07 | 2.200E-06 | 3.600E-06 |
| 20228 | 1012 | L1012G | C | 3.4e-06 | 2.4e-06 | 3.6e-06 | 3.133E-06 | 6.429E-07 | 2.400E-06 | 3.600E-06 |
| 20229 | 1012 | L1012H | C | 1.4e-06 | 3.2e-06 | 2.1e-06 | 2.233E-06 | 9.074E-07 | 1.400E-06 | 3.200E-06 |

| | | | | | | | | | | |
|---|---|---|---|---|---|---|---|---|---|---|
| 20230 | 1012 | L1012I | C | 2.6e-06 | 2e-06 | 2.2e-06 | 2.267E-06 | 3.055E-07 | 2.000E-06 | 2.600E-06 |
| 20231 | 1012 | L1012L | C | 2.2e-06 | 1.6e-06 | 3.1e-06 | 2.300E-06 | 7.550E-07 | 1.600E-06 | 3.100E-06 |
| 20232 | 1012 | L1012K | C | 1.7e-06 | 3.5e-06 | 1.3e-06 | 2.167E-06 | 1.172E-06 | 1.300E-06 | 3.500E-06 |
| 20233 | 1012 | L1012M | C | 2.6e-06 | 2.2e-06 | 2.6e-06 | 2.467E-06 | 2.309E-07 | 2.200E-06 | 2.600E-06 |
| 20234 | 1012 | L1012F | C | 1.7e-06 | 2.2e-06 | 1.6e-06 | 1.833E-06 | 3.215E-07 | 1.600E-06 | 2.200E-06 |
| 20235 | 1012 | L1012P | C | 1.9e-06 | 1.3e-06 | 2.4e-06 | 1.867E-06 | 5.508E-07 | 1.300E-06 | 2.400E-06 |
| 20236 | 1012 | L1012S | C | 3.3e-06 | 1.9e-06 | 1.7e-06 | 2.300E-06 | 8.718E-07 | 1.700E-06 | 3.300E-06 |
| 20237 | 1012 | L1012T | C | 1.5e-06 | 1.7e-06 | 2.3e-06 | 1.833E-06 | 4.163E-07 | 1.500E-06 | 2.300E-06 |
| 20238 | 1012 | L1012W | C | 3.8e-06 | 2.8e-06 | 1.8e-06 | 2.800E-06 | 1.000E-06 | 1.800E-06 | 3.800E-06 |
| 20239 | 1012 | L1012Y | C | 4.3e-06 | 2.7e-06 | 2.6e-06 | 3.200E-06 | 9.539E-07 | 2.600E-06 | 4.300E-06 |
| 20240 | 1012 | L1012V | C | 2.3e-06 | 1.6e-06 | 2.6e-06 | 2.167E-06 | 5.132E-07 | 1.600E-06 | 2.600E-06 |
| 20241 | 1013 | Q1013A | C | 1.6e-06 | 3.1e-06 | 2.7e-06 | 2.467E-06 | 7.767E-07 | 1.600E-06 | 3.100E-06 |
| 20242 | 1013 | Q1013R | C | 2.5e-06 | 1.7e-06 | 1.9e-06 | 2.033E-06 | 4.163E-07 | 1.700E-06 | 2.500E-06 |
| 20243 | 1013 | Q1013N | C | 3.2e-06 | 2.1e-06 | 3.2e-06 | 2.833E-06 | 6.351E-07 | 2.100E-06 | 3.200E-06 |
| 20244 | 1013 | Q1013D | C | 2.8e-06 | 2.5e-06 | 2e-06 | 2.433E-06 | 4.041E-07 | 2.000E-06 | 2.800E-06 |
| 20245 | 1013 | Q1013C | C | 1.2e-06 | 2.8e-06 | 2.6e-06 | 2.200E-06 | 8.718E-07 | 1.200E-06 | 2.800E-06 |
| 20246 | 1013 | Q1013E | C | 2.1e-06 | 1.6e-06 | 1.9e-06 | 1.867E-06 | 2.517E-07 | 1.600E-06 | 2.100E-06 |
| 20247 | 1013 | Q1013Q | C | 1.1e-06 | 2.8e-06 | 1.8e-06 | 1.900E-06 | 8.544E-07 | 1.100E-06 | 2.800E-06 |
| 20248 | 1013 | Q1013G | C | 1.9e-06 | 2.9e-06 | 2.5e-06 | 2.433E-06 | 5.033E-07 | 1.900E-06 | 2.900E-06 |
| 20249 | 1013 | Q1013H | C | 1.3e-06 | 2.3e-06 | 1.3e-06 | 1.633E-06 | 5.774E-07 | 1.300E-06 | 2.300E-06 |
| 20250 | 1013 | Q1013I | C | 2.3e-06 | 1.9e-06 | 1.8e-06 | 2.000E-06 | 2.646E-07 | 1.800E-06 | 2.300E-06 |
| 20251 | 1013 | Q1013L | C | 1.6e-06 | 2.9e-06 | 1.7e-06 | 2.067E-06 | 7.234E-07 | 1.600E-06 | 2.900E-06 |
| 20252 | 1013 | Q1013K | C | 2.2e-06 | 2.4e-06 | 3.1e-06 | 2.567E-06 | 4.726E-07 | 2.200E-06 | 3.100E-06 |
| 20253 | 1013 | Q1013M | C | 2.5e-06 | 2.2e-06 | 2.1e-06 | 2.267E-06 | 2.082E-07 | 2.100E-06 | 2.500E-06 |
| 20254 | 1013 | Q1013F | C | 1.7e-06 | 2.2e-06 | 2e-06 | 1.967E-06 | 2.517E-07 | 1.700E-06 | 2.200E-06 |
| 20255 | 1013 | Q1013P | C | 1.1e-06 | 2.1e-06 | 2.4e-06 | 1.867E-06 | 6.807E-07 | 1.100E-06 | 2.400E-06 |
| 20256 | 1013 | Q1013S | C | 2.1e-06 | 2.6e-06 | 2.3e-06 | 2.333E-06 | 2.517E-07 | 2.100E-06 | 2.600E-06 |
| 20257 | 1013 | Q1013T | C | 4.7e-06 | 1.9e-06 | 2.6e-06 | 3.067E-06 | 1.457E-06 | 1.900E-06 | 4.700E-06 |
| 20258 | 1013 | Q1013W | C | 1.3e-06 | 2.9e-06 | 3.1e-06 | 2.433E-06 | 9.866E-07 | 1.300E-06 | 3.100E-06 |
| 20259 | 1013 | Q1013Y | C | 3.3e-06 | 1.9e-06 | 2.8e-06 | 2.667E-06 | 7.095E-07 | 1.900E-06 | 3.300E-06 |
| 20260 | 1013 | Q1013V | C | 2e-06 | 2.2e-06 | 2.3e-06 | 2.167E-06 | 1.528E-07 | 2.000E-06 | 2.300E-06 |
| 20261 | 1014 | S1014A | C | 3.4e-06 | 1.7e-06 | 2.3e-06 | 2.467E-06 | 8.622E-07 | 1.700E-06 | 3.400E-06 |
| 20262 | 1014 | S1014R | C | 1.9e-06 | 1.7e-06 | 4.4e-06 | 2.667E-06 | 1.504E-06 | 1.700E-06 | 4.400E-06 |
| 20263 | 1014 | S1014N | C | 3.1e-06 | 2.7e-06 | 2.4e-06 | 2.733E-06 | 3.512E-07 | 2.400E-06 | 3.100E-06 |
| 20264 | 1014 | S1014D | C | 3.1e-06 | 2.3e-06 | 2.4e-06 | 2.600E-06 | 4.359E-07 | 2.300E-06 | 3.100E-06 |
| 20265 | 1014 | S1014C | C | 1.3e-06 | 3.1e-06 | 2e-06 | 2.133E-06 | 9.074E-07 | 1.300E-06 | 3.100E-06 |
| 20266 | 1014 | S1014E | C | 2.3e-06 | 2.3e-06 | 3.1e-06 | 2.567E-06 | 4.619E-07 | 2.300E-06 | 3.100E-06 |
| 20267 | 1014 | S1014Q | C | 1.2e-06 | 2e-06 | 1.9e-06 | 1.700E-06 | 4.359E-07 | 1.200E-06 | 2.000E-06 |
| 20268 | 1014 | S1014G | C | 3.4e-06 | 2.3e-06 | 1.7e-06 | 2.467E-06 | 8.622E-07 | 1.700E-06 | 3.400E-06 |
| 20269 | 1014 | S1014H | C | 2.6e-06 | 2.2e-06 | 1.5e-06 | 2.100E-06 | 5.568E-07 | 1.500E-06 | 2.600E-06 |
| 20270 | 1014 | S1014I | C | 2.2e-06 | 2.8e-06 | 3e-06 | 2.667E-06 | 4.163E-07 | 2.200E-06 | 3.000E-06 |
| 20271 | 1014 | S1014L | C | 2.9e-06 | 4e-06 | 3.6e-06 | 3.500E-06 | 5.568E-07 | 2.900E-06 | 4.000E-06 |
| 20272 | 1014 | S1014K | C | 1.8e-06 | 2.5e-06 | 2.6e-06 | 2.300E-06 | 4.359E-07 | 1.800E-06 | 2.600E-06 |
| 20273 | 1014 | S1014M | C | 1.6e-06 | 4.8e-06 | 2.9e-06 | 3.100E-06 | 1.609E-06 | 1.600E-06 | 4.800E-06 |
| 20274 | 1014 | S1014F | C | 9.7e-07 | 3.8e-06 | 4.3e-06 | 3.023E-06 | 1.796E-06 | 9.700E-07 | 4.300E-06 |
| 20275 | 1014 | S1014P | C | 2.7e-06 | 2e-06 | 2.3e-06 | 2.333E-06 | 3.512E-07 | 2.000E-06 | 2.700E-06 |
| 20276 | 1014 | S1014S | C | 2e-06 | 2.4e-06 | 2.9e-06 | 2.433E-06 | 4.509E-07 | 2.000E-06 | 2.900E-06 |
| 20277 | 1014 | S1014T | C | 2.2e-06 | 2.6e-06 | 3.8e-06 | 2.867E-06 | 8.327E-07 | 2.200E-06 | 3.800E-06 |
| 20278 | 1014 | S1014W | C | 1.9e-06 | 2.2e-06 | 2.5e-06 | 2.200E-06 | 3.000E-07 | 1.900E-06 | 2.500E-06 |
| 20279 | 1014 | S1014Y | C | 2.5e-06 | 3.9e-06 | 2.6e-06 | 3.000E-06 | 7.810E-07 | 2.500E-06 | 3.900E-06 |
| 20280 | 1014 | S1014V | C | 2.2e-06 | 3e-06 | 2.4e-06 | 2.533E-06 | 4.163E-07 | 2.200E-06 | 3.000E-06 |
| 20281 | 1015 | S1015A | C | 2.4e-06 | 1.8e-06 | 2.6e-06 | 2.267E-06 | 4.163E-07 | 1.800E-06 | 2.600E-06 |
| 20282 | 1015 | S1015R | C | 2.7e-06 | 2.3e-06 | 2.4e-06 | 2.467E-06 | 2.082E-07 | 2.300E-06 | 2.700E-06 |
| 20283 | 1015 | S1015N | C | 3.1e-06 | 4.4e-06 | 2.5e-06 | 3.333E-06 | 9.713E-07 | 2.500E-06 | 4.400E-06 |
| 20284 | 1015 | S1015D | C | 1.9e-06 | 1.4e-06 | 1.8e-06 | 1.700E-06 | 2.646E-07 | 1.400E-06 | 1.900E-06 |
| 20285 | 1015 | S1015C | C | 2e-06 | 1.8e-06 | 1.7e-06 | 1.833E-06 | 1.528E-07 | 1.700E-06 | 2.000E-06 |
| 20286 | 1015 | S1015E | C | 3.3e-06 | 1.8e-06 | 2.4e-06 | 2.500E-06 | 7.550E-07 | 1.800E-06 | 3.300E-06 |
| 20287 | 1015 | S1015Q | C | 2.2e-06 | 1.9e-06 | 2.8e-06 | 2.300E-06 | 4.583E-07 | 1.900E-06 | 2.800E-06 |
| 20288 | 1015 | S1015G | C | 2.6e-06 | 1.7e-06 | 3e-06 | 2.433E-06 | 6.658E-07 | 1.700E-06 | 3.000E-06 |
| 20289 | 1015 | S1015H | C | 1.7e-06 | 2.8e-06 | 3e-06 | 2.500E-06 | 7.000E-07 | 1.700E-06 | 3.000E-06 |
| 20290 | 1015 | S1015I | C | 2.8e-06 | 3.6e-06 | 3e-06 | 3.133E-06 | 4.163E-07 | 2.800E-06 | 3.600E-06 |
| 20291 | 1015 | S1015L | C | 1.6e-06 | 2.2e-06 | 2.5e-06 | 2.100E-06 | 4.583E-07 | 1.600E-06 | 2.500E-06 |
| 20292 | 1015 | S1015K | C | 2.5e-06 | 2e-06 | 2.3e-06 | 2.267E-06 | 2.517E-07 | 2.000E-06 | 2.500E-06 |
| 20293 | 1015 | S1015M | C | 1.4e-06 | 1.9e-06 | 2.2e-06 | 1.833E-06 | 4.041E-07 | 1.400E-06 | 2.200E-06 |
| 20294 | 1015 | S1015F | C | 2.3e-06 | 1.2e-06 | 1.4e-06 | 1.633E-06 | 5.859E-07 | 1.200E-06 | 2.300E-06 |
| 20295 | 1015 | S1015P | C | 2.4e-06 | 2.6e-06 | 1.6e-06 | 2.200E-06 | 5.292E-07 | 1.600E-06 | 2.600E-06 |
| 20296 | 1015 | S1015S | C | 3.1e-06 | 2.6e-06 | 2.6e-06 | 2.767E-06 | 2.887E-07 | 2.600E-06 | 3.100E-06 |
| 20297 | 1015 | S1015T | C | 1.9e-06 | 2.6e-06 | 2.1e-06 | 2.200E-06 | 3.606E-07 | 1.900E-06 | 2.600E-06 |
| 20298 | 1015 | S1015W | C | 3.2e-06 | 2.9e-06 | 1.9e-06 | 2.667E-06 | 6.807E-07 | 1.900E-06 | 3.200E-06 |
| 20299 | 1015 | S1015Y | C | 3.8e-06 | 2.6e-06 | 3.9e-06 | 3.433E-06 | 7.234E-07 | 2.600E-06 | 3.900E-06 |
| 20300 | 1015 | S1015V | C | 1.7e-06 | 2.5e-06 | 2.7e-06 | 2.300E-06 | 5.292E-07 | 1.700E-06 | 2.700E-06 |
| 20301 | 1016 | G1016A | C | 3.1e-06 | 2.6e-06 | 2.2e-06 | 2.633E-06 | 4.509E-07 | 2.200E-06 | 3.100E-06 |
| 20302 | 1016 | G1016R | C | 2.6e-06 | 2.3e-06 | 3.1e-06 | 2.667E-06 | 4.041E-07 | 2.300E-06 | 3.100E-06 |
| 20303 | 1016 | G1016N | C | 3.7e-06 | 3.1e-06 | 2.2e-06 | 3.000E-06 | 7.550E-07 | 2.200E-06 | 3.700E-06 |
| 20304 | 1016 | G1016D | C | 1.9e-06 | 3e-06 | 3e-06 | 2.633E-06 | 6.351E-07 | 1.900E-06 | 3.000E-06 |
| 20305 | 1016 | G1016C | C | 2.4e-06 | 1.4e-06 | 3e-06 | 2.267E-06 | 8.083E-07 | 1.400E-06 | 3.000E-06 |
| 20306 | 1016 | G1016E | C | 1.3e-06 | 3.7e-06 | 2.2e-06 | 2.400E-06 | 1.212E-06 | 1.300E-06 | 3.700E-06 |
| 20307 | 1016 | G1016Q | C | 1.9e-06 | 1.5e-06 | 2.2e-06 | 1.867E-06 | 3.512E-07 | 1.500E-06 | 2.200E-06 |
| 20308 | 1016 | G1016G | C | 1.5e-06 | 2.6e-06 | 2.6e-06 | 2.233E-06 | 6.351E-07 | 1.500E-06 | 2.600E-06 |
| 20309 | 1016 | G1016H | C | 2.5e-06 | 3e-06 | 3.3e-06 | 2.933E-06 | 4.041E-07 | 2.500E-06 | 3.300E-06 |
| 20310 | 1016 | G1016I | C | 1.9e-06 | 2.4e-06 | 2e-06 | 2.100E-06 | 2.646E-07 | 1.900E-06 | 2.400E-06 |
| 20311 | 1016 | G1016L | C | 2.7e-06 | 1.5e-06 | 3e-06 | 2.400E-06 | 7.937E-07 | 1.500E-06 | 3.000E-06 |
| 20312 | 1016 | G1016K | C | 2.7e-06 | 2e-06 | 1.9e-06 | 2.200E-06 | 4.359E-07 | 1.900E-06 | 2.700E-06 |
| 20313 | 1016 | G1016M | C | 2.5e-06 | 1.3e-06 | 1.3e-06 | 1.700E-06 | 6.928E-07 | 1.300E-06 | 2.500E-06 |
| 20314 | 1016 | G1016F | C | 3e-06 | 3.9e-06 | 2.8e-06 | 3.233E-06 | 5.859E-07 | 2.800E-06 | 3.900E-06 |

| | | | | | | | | | | |
|---|---|---|---|---|---|---|---|---|---|---|
| 20315 | 1016 | G1016P | C | 2.1e-06 | 2e-06 | 2e-06 | 2.033E-06 | 5.774E-08 | 2.000E-06 | 2.100E-06 |
| 20316 | 1016 | G1016S | C | 2.9e-06 | 2.4e-06 | 2.4e-06 | 2.567E-06 | 2.887E-07 | 2.400E-06 | 2.900E-06 |
| 20317 | 1016 | G1016T | C | 2.5e-06 | 3.4e-06 | 3.1e-06 | 3.000E-06 | 4.583E-07 | 2.500E-06 | 3.400E-06 |
| 20318 | 1016 | G1016W | C | 2.3e-06 | 3.5e-06 | 2e-06 | 2.600E-06 | 7.937E-07 | 2.000E-06 | 3.500E-06 |
| 20319 | 1016 | G1016Y | C | 2.9e-06 | 2.6e-06 | 2.7e-06 | 2.733E-06 | 1.528E-07 | 2.600E-06 | 2.900E-06 |
| 20320 | 1016 | G1016V | C | 4.2e-06 | 3.2e-06 | 2.9e-06 | 3.433E-06 | 6.807E-07 | 2.900E-06 | 4.200E-06 |
| 20321 | 1017 | L1017A | C | 4.3e-06 | 2.8e-06 | 2e-06 | 3.033E-06 | 1.168E-06 | 2.000E-06 | 4.300E-06 |
| 20322 | 1017 | L1017R | C | 5.6e-06 | 3.1e-06 | 1.9e-06 | 3.533E-06 | 1.888E-06 | 1.900E-06 | 5.600E-06 |
| 20323 | 1017 | L1017N | C | 2.1e-06 | 2.4e-06 | 2.1e-06 | 2.200E-06 | 1.732E-07 | 2.100E-06 | 2.400E-06 |
| 20324 | 1017 | L1017D | C | 3.4e-06 | 3.1e-06 | 2.3e-06 | 2.933E-06 | 5.686E-07 | 2.300E-06 | 3.400E-06 |
| 20325 | 1017 | L1017C | C | 1.6e-06 | 3.1e-06 | 2.7e-06 | 2.467E-06 | 7.767E-07 | 1.600E-06 | 3.100E-06 |
| 20326 | 1017 | L1017E | C | 2.3e-06 | 3.5e-06 | 1.3e-06 | 2.367E-06 | 1.102E-06 | 1.300E-06 | 3.500E-06 |
| 20327 | 1017 | L1017Q | C | 3.9e-06 | 3.5e-06 | 2.3e-06 | 3.233E-06 | 8.327E-07 | 2.300E-06 | 3.900E-06 |
| 20328 | 1017 | L1017G | C | 2.6e-06 | 1.6e-06 | 3.1e-06 | 2.433E-06 | 7.638E-07 | 1.600E-06 | 3.100E-06 |
| 20329 | 1017 | L1017H | C | 3.5e-06 | 2.5e-06 | 2.6e-06 | 2.867E-06 | 5.508E-07 | 2.500E-06 | 3.500E-06 |
| 20330 | 1017 | L1017I | C | 1.8e-06 | 1.9e-06 | 5.4e-06 | 3.033E-06 | 2.050E-06 | 1.800E-06 | 5.400E-06 |
| 20331 | 1017 | L1017L | C | 1.6e-06 | 2.8e-06 | 2.8e-06 | 2.400E-06 | 6.928E-07 | 1.600E-06 | 2.800E-06 |
| 20332 | 1017 | L1017K | C | 3.6e-06 | 2.7e-06 | 1.5e-06 | 2.600E-06 | 1.054E-06 | 1.500E-06 | 3.600E-06 |
| 20333 | 1017 | L1017M | C | 2.1e-06 | 2.5e-06 | 2.2e-06 | 2.267E-06 | 2.082E-07 | 2.100E-06 | 2.500E-06 |
| 20334 | 1017 | L1017F | C | 2.2e-06 | 3.6e-06 | 2.4e-06 | 2.733E-06 | 7.572E-07 | 2.200E-06 | 3.600E-06 |
| 20335 | 1017 | L1017P | C | 1.5e-06 | 2e-06 | 2.4e-06 | 1.967E-06 | 4.509E-07 | 1.500E-06 | 2.400E-06 |
| 20336 | 1017 | L1017S | C | 1.4e-06 | 2.3e-06 | 2.5e-06 | 2.067E-06 | 5.859E-07 | 1.400E-06 | 2.500E-06 |
| 20337 | 1017 | L1017T | C | 9.7e-07 | 2.1e-06 | 1.6e-06 | 1.557E-06 | 5.662E-07 | 9.700E-07 | 2.100E-06 |
| 20338 | 1017 | L1017W | C | 2.3e-06 | 1.9e-06 | 3.3e-06 | 2.500E-06 | 7.211E-07 | 1.900E-06 | 3.300E-06 |
| 20339 | 1017 | L1017Y | C | 1.9e-06 | 1.9e-06 | 2.6e-06 | 2.133E-06 | 4.041E-07 | 1.900E-06 | 2.600E-06 |
| 20340 | 1017 | L1017V | C | 3e-06 | 2.8e-06 | 2.7e-06 | 2.833E-06 | 1.528E-07 | 2.700E-06 | 3.000E-06 |
| 20341 | 1018 | Y1018A | C | 1.7e-06 | 2.5e-06 | 2e-06 | 2.067E-06 | 4.041E-07 | 1.700E-06 | 2.500E-06 |
| 20342 | 1018 | Y1018R | C | 1.8e-06 | 4.4e-06 | 2.2e-06 | 2.800E-06 | 1.400E-06 | 1.800E-06 | 4.400E-06 |
| 20343 | 1018 | Y1018N | C | 2.3e-06 | 2.4e-06 | 2.2e-06 | 2.300E-06 | 1.000E-07 | 2.200E-06 | 2.400E-06 |
| 20344 | 1018 | Y1018D | C | 2e-06 | 3.7e-06 | 1.7e-06 | 2.467E-06 | 1.079E-06 | 1.700E-06 | 3.700E-06 |
| 20345 | 1018 | Y1018C | C | 3e-06 | 1.3e-06 | 2.3e-06 | 2.200E-06 | 8.544E-07 | 1.300E-06 | 3.000E-06 |
| 20346 | 1018 | Y1018E | C | 2.4e-06 | 3.3e-06 | 4.7e-06 | 3.467E-06 | 1.159E-06 | 2.400E-06 | 4.700E-06 |
| 20347 | 1018 | Y1018Q | C | 1.8e-06 | 1.9e-06 | 2.2e-06 | 1.967E-06 | 2.082E-07 | 1.800E-06 | 2.200E-06 |
| 20348 | 1018 | Y1018G | C | 2.3e-06 | 3.1e-06 | 4.4e-06 | 3.267E-06 | 1.060E-06 | 2.300E-06 | 4.400E-06 |
| 20349 | 1018 | Y1018H | C | 2.3e-06 | 1.1e-06 | 2.2e-06 | 1.867E-06 | 6.658E-07 | 1.100E-06 | 2.300E-06 |
| 20350 | 1018 | Y1018I | C | 2.7e-06 | 2.2e-06 | 2.4e-06 | 2.433E-06 | 2.517E-07 | 2.200E-06 | 2.700E-06 |
| 20351 | 1018 | Y1018L | C | 1.5e-06 | 4.5e-06 | 2e-06 | 2.667E-06 | 1.607E-06 | 1.500E-06 | 4.500E-06 |
| 20352 | 1018 | Y1018K | C | 1.6e-06 | 2.3e-06 | 1.7e-06 | 1.867E-06 | 3.786E-07 | 1.600E-06 | 2.300E-06 |
| 20353 | 1018 | Y1018M | C | 2.1e-06 | 3.3e-06 | 1.3e-06 | 2.233E-06 | 1.007E-06 | 1.300E-06 | 3.300E-06 |
| 20354 | 1018 | Y1018F | C | 1.5e-06 | 1.9e-06 | 9.6e-07 | 1.453E-06 | 4.717E-07 | 9.600E-07 | 1.900E-06 |
| 20355 | 1018 | Y1018P | C | 3e-06 | 4e-06 | 3.1e-06 | 3.367E-06 | 5.508E-07 | 3.000E-06 | 4.000E-06 |
| 20356 | 1018 | Y1018S | C | 2.7e-06 | 3.3e-06 | 2.2e-06 | 2.733E-06 | 5.508E-07 | 2.200E-06 | 3.300E-06 |
| 20357 | 1018 | Y1018T | C | 2.6e-06 | 1.6e-06 | 1.4e-06 | 1.867E-06 | 6.429E-07 | 1.400E-06 | 2.600E-06 |
| 20358 | 1018 | Y1018W | C | 1.5e-06 | 1.4e-06 | 2.3e-06 | 1.733E-06 | 4.933E-07 | 1.400E-06 | 2.300E-06 |
| 20359 | 1018 | Y1018Y | C | 3e-06 | 2.2e-06 | 4e-06 | 3.067E-06 | 9.018E-07 | 2.200E-06 | 4.000E-06 |
| 20360 | 1018 | Y1018V | C | 1.4e-06 | 2.6e-06 | 2.2e-06 | 2.067E-06 | 6.110E-07 | 1.400E-06 | 2.600E-06 |
| 20361 | 1019 | S1019A | C | 2.5e-06 | 4e-06 | 2.2e-06 | 2.900E-06 | 9.644E-07 | 2.200E-06 | 4.000E-06 |
| 20362 | 1019 | S1019R | C | 2.1e-06 | 3.9e-06 | 3.2e-06 | 3.067E-06 | 9.074E-07 | 2.100E-06 | 3.900E-06 |
| 20363 | 1019 | S1019N | C | 2.8e-06 | 2.3e-06 | 3.1e-06 | 2.733E-06 | 4.041E-07 | 2.300E-06 | 3.100E-06 |
| 20364 | 1019 | S1019D | C | 1.4e-06 | 3.6e-06 | 2.1e-06 | 2.367E-06 | 1.124E-06 | 1.400E-06 | 3.600E-06 |
| 20365 | 1019 | S1019C | C | 1.7e-06 | 2.2e-06 | 2.5e-06 | 2.133E-06 | 4.041E-07 | 1.700E-06 | 2.500E-06 |
| 20366 | 1019 | S1019E | C | 2.8e-06 | 2.8e-06 | 3.5e-06 | 3.033E-06 | 4.041E-07 | 2.800E-06 | 3.500E-06 |
| 20367 | 1019 | S1019Q | C | 2.6e-06 | 3.1e-06 | 1.8e-06 | 2.500E-06 | 6.557E-07 | 1.800E-06 | 3.100E-06 |
| 20368 | 1019 | S1019G | C | 2.6e-06 | 2.5e-06 | 2.8e-06 | 2.633E-06 | 1.528E-07 | 2.500E-06 | 2.800E-06 |
| 20369 | 1019 | S1019H | C | 3.2e-06 | 2.9e-06 | 2.1e-06 | 2.733E-06 | 5.686E-07 | 2.100E-06 | 3.200E-06 |
| 20370 | 1019 | S1019I | C | 2.2e-06 | 2.8e-06 | 2.8e-06 | 2.600E-06 | 3.464E-07 | 2.200E-06 | 2.800E-06 |
| 20371 | 1019 | S1019L | C | 2e-06 | 3.7e-06 | 4.1e-06 | 3.267E-06 | 1.115E-06 | 2.000E-06 | 4.100E-06 |
| 20372 | 1019 | S1019K | C | 3.3e-06 | 2.2e-06 | 2.4e-06 | 2.633E-06 | 5.859E-07 | 2.200E-06 | 3.300E-06 |
| 20373 | 1019 | S1019M | C | 2.9e-06 | 2.2e-06 | 1.7e-06 | 2.267E-06 | 6.028E-07 | 1.700E-06 | 2.900E-06 |
| 20374 | 1019 | S1019F | C | 3.5e-06 | 1.4e-06 | 2e-06 | 2.300E-06 | 1.082E-06 | 1.400E-06 | 3.500E-06 |
| 20375 | 1019 | S1019P | C | 1.4e-06 | 2.1e-06 | 4.3e-06 | 2.600E-06 | 1.513E-06 | 1.400E-06 | 4.300E-06 |
| 20376 | 1019 | S1019S | C | 2.8e-06 | 2e-06 | 1.8e-06 | 2.200E-06 | 5.292E-07 | 1.800E-06 | 2.800E-06 |
| 20377 | 1019 | S1019T | C | 3.1e-06 | 2e-06 | 2.5e-06 | 2.533E-06 | 5.508E-07 | 2.000E-06 | 3.100E-06 |
| 20378 | 1019 | S1019W | C | 2.5e-06 | 3.2e-06 | 1.9e-06 | 2.533E-06 | 6.506E-07 | 1.900E-06 | 3.200E-06 |
| 20379 | 1019 | S1019Y | C | 4.1e-06 | 2.6e-06 | 2.8e-06 | 3.167E-06 | 8.145E-07 | 2.600E-06 | 4.100E-06 |
| 20380 | 1019 | S1019V | C | 4.9e-06 | 2.4e-06 | 2.2e-06 | 3.167E-06 | 1.504E-06 | 2.200E-06 | 4.900E-06 |
| 20381 | 1020 | L1020A | C | 1.5e-06 | 2.6e-06 | 2.2e-06 | 2.100E-06 | 5.568E-07 | 1.500E-06 | 2.600E-06 |
| 20382 | 1020 | L1020R | C | 7.5e-07 | 1.8e-06 | 1.2e-06 | 1.250E-06 | 5.268E-07 | 7.500E-07 | 1.800E-06 |
| 20383 | 1020 | L1020N | C | 2.5e-06 | 2.4e-06 | 3.3e-06 | 2.733E-06 | 4.933E-07 | 2.400E-06 | 3.300E-06 |
| 20384 | 1020 | L1020D | C | 2.5e-06 | 2.2e-06 | 2.2e-06 | 2.300E-06 | 1.732E-07 | 2.200E-06 | 2.500E-06 |
| 20385 | 1020 | L1020C | C | 3.5e-06 | 1.3e-06 | 2.8e-06 | 2.533E-06 | 1.124E-06 | 1.300E-06 | 3.500E-06 |
| 20386 | 1020 | L1020E | C | 2e-06 | 2.3e-06 | 3.6e-06 | 2.633E-06 | 8.505E-07 | 2.000E-06 | 3.600E-06 |
| 20387 | 1020 | L1020Q | C | 1.6e-06 | 2.6e-06 | 2.8e-06 | 2.333E-06 | 6.429E-07 | 1.600E-06 | 2.800E-06 |
| 20388 | 1020 | L1020G | C | 2.5e-06 | 2e-06 | 2.8e-06 | 2.433E-06 | 4.041E-07 | 2.000E-06 | 2.800E-06 |
| 20389 | 1020 | L1020H | C | 5.1e-06 | 1.9e-06 | 3.6e-06 | 3.533E-06 | 1.601E-06 | 1.900E-06 | 5.100E-06 |
| 20390 | 1020 | L1020I | C | 2e-06 | 2.7e-06 | 1.8e-06 | 2.167E-06 | 4.726E-07 | 1.800E-06 | 2.700E-06 |
| 20391 | 1020 | L1020L | C | 1.6e-06 | 1.9e-06 | 1.8e-06 | 1.767E-06 | 1.528E-07 | 1.600E-06 | 1.900E-06 |
| 20392 | 1020 | L1020K | C | 1.3e-06 | 1.8e-06 | 4.5e-06 | 2.533E-06 | 1.721E-06 | 1.300E-06 | 4.500E-06 |
| 20393 | 1020 | L1020M | C | 3.1e-06 | 3e-06 | 1.6e-06 | 2.567E-06 | 8.386E-07 | 1.600E-06 | 3.100E-06 |
| 20394 | 1020 | L1020F | C | 3e-06 | 2.8e-06 | 1.9e-06 | 2.567E-06 | 5.859E-07 | 1.900E-06 | 3.000E-06 |
| 20395 | 1020 | L1020P | C | 3.1e-06 | 4e-06 | 2.8e-06 | 3.300E-06 | 6.245E-07 | 2.800E-06 | 4.000E-06 |
| 20396 | 1020 | L1020S | C | 3.9e-06 | 2.6e-06 | 2.3e-06 | 2.933E-06 | 8.505E-07 | 2.300E-06 | 3.900E-06 |
| 20397 | 1020 | L1020T | C | 1.2e-06 | 2.1e-06 | 4e-06 | 2.433E-06 | 1.429E-06 | 1.200E-06 | 4.000E-06 |
| 20398 | 1020 | L1020W | C | 4.8e-06 | 2.6e-06 | 2.5e-06 | 3.300E-06 | 1.300E-06 | 2.500E-06 | 4.800E-06 |
| 20399 | 1020 | L1020Y | C | 1.3e-06 | 2e-06 | 1.9e-06 | 1.733E-06 | 3.786E-07 | 1.300E-06 | 2.000E-06 |

| | | | | | | | | | | |
|---|---|---|---|---|---|---|---|---|---|---|
| 20400 | 1020 | L1020V | C | 3.1e-06 | 2.4e-06 | 2.1e-06 | 2.533E-06 | 5.132E-07 | 2.100E-06 | 3.100E-06 |
| 20401 | 1021 | S1021A | C | 1.9e-06 | 1.9e-06 | 2.3e-06 | 2.033E-06 | 2.309E-07 | 1.900E-06 | 2.300E-06 |
| 20402 | 1021 | S1021R | C | 3.3e-06 | 4.4e-06 | 1.9e-06 | 3.200E-06 | 1.253E-06 | 1.900E-06 | 4.400E-06 |
| 20403 | 1021 | S1021N | C | 3.5e-06 | 3.6e-06 | 2.2e-06 | 3.100E-06 | 7.810E-07 | 2.200E-06 | 3.600E-06 |
| 20404 | 1021 | S1021D | C | 4.2e-06 | 1.3e-06 | 1.6e-06 | 2.367E-06 | 1.595E-06 | 1.300E-06 | 4.200E-06 |
| 20405 | 1021 | S1021C | C | 1.3e-06 | 3.5e-06 | 4.7e-06 | 3.167E-06 | 1.724E-06 | 1.300E-06 | 4.700E-06 |
| 20406 | 1021 | S1021E | C | 2.4e-06 | 2.3e-06 | 1.8e-06 | 2.167E-06 | 3.215E-07 | 1.800E-06 | 2.400E-06 |
| 20407 | 1021 | S1021Q | C | 1.8e-06 | 2.1e-06 | 2e-06 | 1.967E-06 | 1.528E-07 | 1.800E-06 | 2.100E-06 |
| 20408 | 1021 | S1021G | C | 3.8e-06 | 2.1e-06 | 3.7e-06 | 3.200E-06 | 9.539E-07 | 2.100E-06 | 3.800E-06 |
| 20409 | 1021 | S1021H | C | 2.6e-06 | 1.8e-06 | 1.9e-06 | 2.100E-06 | 4.359E-07 | 1.800E-06 | 2.600E-06 |
| 20410 | 1021 | S1021I | C | 2.3e-06 | 3.8e-06 | 2.9e-06 | 3.000E-06 | 7.550E-07 | 2.300E-06 | 3.800E-06 |
| 20411 | 1021 | S1021L | C | 2.7e-06 | 2.4e-06 | 2.6e-06 | 2.567E-06 | 1.528E-07 | 2.400E-06 | 2.700E-06 |
| 20412 | 1021 | S1021K | C | 2e-06 | 1.7e-06 | 2.3e-06 | 2.000E-06 | 3.000E-07 | 1.700E-06 | 2.300E-06 |
| 20413 | 1021 | S1021M | C | 3.3e-06 | 3e-06 | 3.1e-06 | 3.133E-06 | 1.528E-07 | 3.000E-06 | 3.300E-06 |
| 20414 | 1021 | S1021F | C | 2.2e-06 | 3e-06 | 2.3e-06 | 2.500E-06 | 4.359E-07 | 2.200E-06 | 3.000E-06 |
| 20415 | 1021 | S1021P | C | 2.4e-06 | 1.6e-06 | 2.9e-06 | 2.300E-06 | 6.557E-07 | 1.600E-06 | 2.900E-06 |
| 20416 | 1021 | S1021S | C | 2e-06 | 4.6e-06 | 8.4e-07 | 2.480E-06 | 1.925E-06 | 8.400E-07 | 4.600E-06 |
| 20417 | 1021 | S1021T | C | 2e-06 | 2.4e-06 | 3.2e-06 | 2.533E-06 | 6.110E-07 | 2.000E-06 | 3.200E-06 |
| 20418 | 1021 | S1021W | C | 2.3e-06 | 3.3e-06 | 3.3e-06 | 2.967E-06 | 5.774E-07 | 2.300E-06 | 3.300E-06 |
| 20419 | 1021 | S1021Y | C | 1.5e-06 | 3e-06 | 4.4e-06 | 2.967E-06 | 1.450E-06 | 1.500E-06 | 4.400E-06 |
| 20420 | 1021 | S1021V | C | 2.2e-06 | 2.8e-06 | 2.4e-06 | 2.467E-06 | 3.055E-07 | 2.200E-06 | 2.800E-06 |
| 20421 | 1022 | S1022A | C | 2.1e-06 | 2.6e-06 | 2.7e-06 | 2.467E-06 | 3.215E-07 | 2.100E-06 | 2.700E-06 |
| 20422 | 1022 | S1022R | C | 1.1e-06 | 2.2e-06 | 2e-06 | 1.767E-06 | 5.859E-07 | 1.100E-06 | 2.200E-06 |
| 20423 | 1022 | S1022N | C | 2.1e-06 | 1.9e-06 | 1.8e-06 | 1.933E-06 | 1.528E-07 | 1.800E-06 | 2.100E-06 |
| 20424 | 1022 | S1022D | C | 2.1e-06 | 3.6e-06 | 1.6e-06 | 2.433E-06 | 1.041E-06 | 1.600E-06 | 3.600E-06 |
| 20425 | 1022 | S1022C | C | 2.3e-06 | 3.3e-06 | 1.7e-06 | 2.433E-06 | 8.083E-07 | 1.700E-06 | 3.300E-06 |
| 20426 | 1022 | S1022E | C | 5.7e-06 | 2.5e-06 | 2.1e-06 | 3.433E-06 | 1.973E-06 | 2.100E-06 | 5.700E-06 |
| 20427 | 1022 | S1022Q | C | 2.2e-06 | 3.3e-06 | 3e-06 | 2.833E-06 | 5.686E-07 | 2.200E-06 | 3.300E-06 |
| 20428 | 1022 | S1022G | C | 2.8e-06 | 3.9e-06 | 1.8e-06 | 2.833E-06 | 1.050E-06 | 1.800E-06 | 3.900E-06 |
| 20429 | 1022 | S1022H | C | 2e-06 | 3.5e-06 | 2.6e-06 | 2.700E-06 | 7.550E-07 | 2.000E-06 | 3.500E-06 |
| 20430 | 1022 | S1022I | C | 2.7e-06 | 1.4e-06 | 1.7e-06 | 1.933E-06 | 6.807E-07 | 1.400E-06 | 2.700E-06 |
| 20431 | 1022 | S1022L | C | 3.5e-06 | 2.5e-06 | 1.3e-06 | 2.433E-06 | 1.102E-06 | 1.300E-06 | 3.500E-06 |
| 20432 | 1022 | S1022K | C | 2.3e-06 | 3.2e-06 | 3.7e-06 | 3.067E-06 | 7.095E-07 | 2.300E-06 | 3.700E-06 |
| 20433 | 1022 | S1022M | C | 4e-06 | 1.5e-06 | 1.8e-06 | 2.433E-06 | 1.365E-06 | 1.500E-06 | 4.000E-06 |
| 20434 | 1022 | S1022F | C | 2.9e-06 | 3.8e-06 | 2.2e-06 | 2.967E-06 | 8.021E-07 | 2.200E-06 | 3.800E-06 |
| 20435 | 1022 | S1022P | C | 1.9e-06 | 1.7e-06 | 2.5e-06 | 2.033E-06 | 4.163E-07 | 1.700E-06 | 2.500E-06 |
| 20436 | 1022 | S1022S | C | 4.4e-06 | 1.6e-06 | 3.8e-06 | 3.267E-06 | 1.474E-06 | 1.600E-06 | 4.400E-06 |
| 20437 | 1022 | S1022T | C | 4.2e-06 | 1.4e-06 | 3.7e-06 | 3.100E-06 | 1.493E-06 | 1.400E-06 | 4.200E-06 |
| 20438 | 1022 | S1022W | C | 3.5e-06 | 2.3e-06 | 2.2e-06 | 2.667E-06 | 7.234E-07 | 2.200E-06 | 3.500E-06 |
| 20439 | 1022 | S1022Y | C | 1.9e-06 | 1.7e-06 | 2e-06 | 1.867E-06 | 1.528E-07 | 1.700E-06 | 2.000E-06 |
| 20440 | 1022 | S1022V | C | 2.2e-06 | 3.8e-06 | 2.4e-06 | 2.800E-06 | 8.718E-07 | 2.200E-06 | 3.800E-06 |
| 20441 | 1023 | V1023A | C | 3.1e-06 | 2.4e-06 | 2.4e-06 | 2.633E-06 | 4.041E-07 | 2.400E-06 | 3.100E-06 |
| 20442 | 1023 | V1023R | C | 4e-06 | 4.1e-06 | 2.3e-06 | 3.467E-06 | 1.012E-06 | 2.300E-06 | 4.100E-06 |
| 20443 | 1023 | V1023N | C | 1.5e-06 | 2.8e-06 | 3.8e-06 | 2.700E-06 | 1.153E-06 | 1.500E-06 | 3.800E-06 |
| 20444 | 1023 | V1023D | C | 1.9e-06 | 1.8e-06 | 1.9e-06 | 1.867E-06 | 5.774E-08 | 1.800E-06 | 1.900E-06 |
| 20445 | 1023 | V1023C | C | 2e-06 | 1.9e-06 | 2.4e-06 | 2.100E-06 | 2.646E-07 | 1.900E-06 | 2.400E-06 |
| 20446 | 1023 | V1023E | C | 1.1e-06 | 3e-06 | 2.7e-06 | 2.267E-06 | 1.021E-06 | 1.100E-06 | 3.000E-06 |
| 20447 | 1023 | V1023Q | C | 1.9e-06 | 1.1e-06 | 3.2e-06 | 2.067E-06 | 1.060E-06 | 1.100E-06 | 3.200E-06 |
| 20448 | 1023 | V1023G | C | 3.7e-06 | 2.4e-06 | 2e-06 | 2.700E-06 | 8.888E-07 | 2.000E-06 | 3.700E-06 |
| 20449 | 1023 | V1023H | C | 2.7e-06 | 2.7e-06 | 3.1e-06 | 2.833E-06 | 2.309E-07 | 2.700E-06 | 3.100E-06 |
| 20450 | 1023 | V1023I | C | 4.2e-06 | 2.5e-06 | 3.1e-06 | 3.267E-06 | 8.622E-07 | 2.500E-06 | 4.200E-06 |
| 20451 | 1023 | V1023L | C | 2.5e-06 | 1.6e-06 | 2.8e-06 | 2.300E-06 | 6.245E-07 | 1.600E-06 | 2.800E-06 |
| 20452 | 1023 | V1023K | C | 2e-06 | 3.6e-06 | 1.9e-06 | 2.500E-06 | 9.539E-07 | 1.900E-06 | 3.600E-06 |
| 20453 | 1023 | V1023M | C | 1.5e-06 | 1.8e-06 | 3.2e-06 | 2.167E-06 | 9.074E-07 | 1.500E-06 | 3.200E-06 |
| 20454 | 1023 | V1023F | C | 2.6e-06 | 2.8e-06 | 1.9e-06 | 2.433E-06 | 4.726E-07 | 1.900E-06 | 2.800E-06 |
| 20455 | 1023 | V1023P | C | 1.6e-06 | 2.4e-06 | 2.2e-06 | 2.067E-06 | 4.163E-07 | 1.600E-06 | 2.400E-06 |
| 20456 | 1023 | V1023S | C | 3.9e-06 | 2.7e-06 | 1.9e-06 | 2.833E-06 | 1.007E-06 | 1.900E-06 | 3.900E-06 |
| 20457 | 1023 | V1023T | C | 2.9e-06 | 1.7e-06 | 1.4e-06 | 2.000E-06 | 7.937E-07 | 1.400E-06 | 2.900E-06 |
| 20458 | 1023 | V1023W | C | 1.8e-06 | 2.2e-06 | 2.4e-06 | 2.133E-06 | 3.055E-07 | 1.800E-06 | 2.400E-06 |
| 20459 | 1023 | V1023Y | C | 3.1e-06 | 1.3e-06 | 1.5e-06 | 1.967E-06 | 9.866E-07 | 1.300E-06 | 3.100E-06 |
| 20460 | 1023 | V1023V | C | 2.9e-06 | 1.7e-06 | 2.1e-06 | 2.233E-06 | 6.110E-07 | 1.700E-06 | 2.900E-06 |
| 20461 | 1024 | V1024A | C | 2e-06 | 3e-06 | 1.5e-06 | 2.167E-06 | 7.638E-07 | 1.500E-06 | 3.000E-06 |
| 20462 | 1024 | V1024R | C | 2.2e-06 | 3.4e-06 | 1.5e-06 | 2.367E-06 | 9.609E-07 | 1.500E-06 | 3.400E-06 |
| 20463 | 1024 | V1024N | C | 4.4e-06 | 3.6e-06 | 2.3e-06 | 3.433E-06 | 1.060E-06 | 2.300E-06 | 4.400E-06 |
| 20464 | 1024 | V1024D | C | 2.1e-06 | 3.6e-06 | 1.7e-06 | 2.467E-06 | 1.002E-06 | 1.700E-06 | 3.600E-06 |
| 20465 | 1024 | V1024C | C | 1.5e-06 | 1.3e-06 | 3e-06 | 1.933E-06 | 9.292E-07 | 1.300E-06 | 3.000E-06 |
| 20466 | 1024 | V1024E | C | 3.1e-06 | 2.4e-06 | 2.7e-06 | 2.733E-06 | 3.512E-07 | 2.400E-06 | 3.100E-06 |
| 20467 | 1024 | V1024Q | C | 2.4e-06 | 3.9e-06 | 2.3e-06 | 2.867E-06 | 8.963E-07 | 2.300E-06 | 3.900E-06 |
| 20468 | 1024 | V1024G | C | 1.3e-06 | 2e-06 | 3.1e-06 | 2.133E-06 | 9.074E-07 | 1.300E-06 | 3.100E-06 |
| 20469 | 1024 | V1024H | C | 2e-06 | 1.2e-06 | 5.3e-06 | 2.833E-06 | 2.173E-06 | 1.200E-06 | 5.300E-06 |
| 20470 | 1024 | V1024I | C | 1.5e-06 | 2.9e-06 | 2e-06 | 2.133E-06 | 7.095E-07 | 1.500E-06 | 2.900E-06 |
| 20471 | 1024 | V1024L | C | 1.9e-06 | 2.1e-06 | 1.6e-06 | 1.867E-06 | 2.517E-07 | 1.600E-06 | 2.100E-06 |
| 20472 | 1024 | V1024K | C | 2.6e-06 | 1.2e-06 | 4.6e-06 | 2.800E-06 | 1.709E-06 | 1.200E-06 | 4.600E-06 |
| 20473 | 1024 | V1024M | C | 3.2e-06 | 3.6e-06 | 2.1e-06 | 2.967E-06 | 7.767E-07 | 2.100E-06 | 3.600E-06 |
| 20474 | 1024 | V1024F | C | 2.6e-06 | 1.8e-06 | 2.2e-06 | 2.200E-06 | 4.000E-07 | 1.800E-06 | 2.600E-06 |
| 20475 | 1024 | V1024P | C | 2e-06 | 2.1e-06 | 2.1e-06 | 2.067E-06 | 5.774E-08 | 2.000E-06 | 2.100E-06 |
| 20476 | 1024 | V1024S | C | 1.4e-06 | 1.5e-06 | 2.9e-06 | 1.933E-06 | 8.386E-07 | 1.400E-06 | 2.900E-06 |
| 20477 | 1024 | V1024T | C | 3.8e-06 | 2.2e-06 | 2.1e-06 | 2.700E-06 | 9.539E-07 | 2.100E-06 | 3.800E-06 |
| 20478 | 1024 | V1024W | C | 2.6e-06 | 2.5e-06 | 3.3e-06 | 2.800E-06 | 4.359E-07 | 2.500E-06 | 3.300E-06 |
| 20479 | 1024 | V1024Y | C | 1.9e-06 | 3.2e-06 | 2.2e-06 | 2.433E-06 | 6.807E-07 | 1.900E-06 | 3.200E-06 |
| 20480 | 1024 | V1024V | C | 1.3e-06 | 2.5e-06 | 3e-06 | 2.267E-06 | 8.737E-07 | 1.300E-06 | 3.000E-06 |
| 20481 | 1025 | T1025A | C | 1.3e-06 | 2.7e-06 | 3.6e-06 | 2.533E-06 | 1.159E-06 | 1.300E-06 | 3.600E-06 |
| 20482 | 1025 | T1025R | C | 1.5e-06 | 3.3e-06 | 2.1e-06 | 2.300E-06 | 9.165E-07 | 1.500E-06 | 3.300E-06 |
| 20483 | 1025 | T1025N | C | 2.1e-06 | 2.8e-06 | 2e-06 | 2.300E-06 | 4.359E-07 | 2.000E-06 | 2.800E-06 |
| 20484 | 1025 | T1025D | C | 2.9e-06 | 1.7e-06 | 1.7e-06 | 2.100E-06 | 6.928E-07 | 1.700E-06 | 2.900E-06 |

| | | | | | | | | | | |
|---|---|---|---|---|---|---|---|---|---|---|
| 20485 | 1025 | T1025C | C | 5e-06 | 1.4e-06 | 1.4e-06 | 2.600E-06 | 2.078E-06 | 1.400E-06 | 5.000E-06 |
| 20486 | 1025 | T1025E | C | 2.3e-06 | 1.5e-06 | 3.9e-06 | 2.567E-06 | 1.222E-06 | 1.500E-06 | 3.900E-06 |
| 20487 | 1025 | T1025Q | C | 3.3e-06 | 1.7e-06 | 3.3e-06 | 2.767E-06 | 9.238E-07 | 1.700E-06 | 3.300E-06 |
| 20488 | 1025 | T1025G | C | 1.7e-06 | 2.5e-06 | 3.4e-06 | 2.533E-06 | 8.505E-07 | 1.700E-06 | 3.400E-06 |
| 20489 | 1025 | T1025H | C | 2.8e-06 | 1.5e-06 | 2.8e-06 | 2.367E-06 | 7.506E-07 | 1.500E-06 | 2.800E-06 |
| 20490 | 1025 | T1025I | C | 1.2e-06 | 4.4e-06 | 2.7e-06 | 2.767E-06 | 1.601E-06 | 1.200E-06 | 4.400E-06 |
| 20491 | 1025 | T1025L | C | 2.1e-06 | 2e-06 | 3.5e-06 | 2.533E-06 | 8.386E-07 | 2.000E-06 | 3.500E-06 |
| 20492 | 1025 | T1025K | C | 1.8e-06 | 2.6e-06 | 2.2e-06 | 2.200E-06 | 4.000E-07 | 1.800E-06 | 2.600E-06 |
| 20493 | 1025 | T1025M | C | 2.4e-06 | 2.9e-06 | 2.2e-06 | 2.500E-06 | 3.606E-07 | 2.200E-06 | 2.900E-06 |
| 20494 | 1025 | T1025F | C | 1.9e-06 | 2.3e-06 | 3.3e-06 | 2.500E-06 | 7.211E-07 | 1.900E-06 | 3.300E-06 |
| 20495 | 1025 | T1025P | C | 1.7e-06 | 1.8e-06 | 1.6e-06 | 1.700E-06 | 1.000E-07 | 1.600E-06 | 1.800E-06 |
| 20496 | 1025 | T1025S | C | 2.1e-06 | 2.3e-06 | 1.6e-06 | 2.000E-06 | 3.606E-07 | 1.600E-06 | 2.300E-06 |
| 20497 | 1025 | T1025T | C | 1.4e-06 | 4.2e-06 | 2.5e-06 | 2.700E-06 | 1.411E-06 | 1.400E-06 | 4.200E-06 |
| 20498 | 1025 | T1025W | C | 2e-06 | 2.5e-06 | 3.1e-06 | 2.533E-06 | 5.508E-07 | 2.000E-06 | 3.100E-06 |
| 20499 | 1025 | T1025Y | C | 1.9e-06 | 4e-06 | 2.2e-06 | 2.700E-06 | 1.136E-06 | 1.900E-06 | 4.000E-06 |
| 20500 | 1025 | T1025V | C | 1.7e-06 | 1.3e-06 | 1.9e-06 | 1.633E-06 | 3.055E-07 | 1.300E-06 | 1.900E-06 |
| 20501 | 1026 | V1026A | C | 1.8e-06 | 1.8e-06 | 1.1e-06 | 1.567E-06 | 4.041E-07 | 1.100E-06 | 1.800E-06 |
| 20502 | 1026 | V1026R | C | 1.7e-06 | 2e-06 | 2.6e-06 | 2.100E-06 | 4.583E-07 | 1.700E-06 | 2.600E-06 |
| 20503 | 1026 | V1026N | C | 3.5e-06 | 1.7e-06 | 1.3e-06 | 2.167E-06 | 1.172E-06 | 1.300E-06 | 3.500E-06 |
| 20504 | 1026 | V1026D | C | 4.3e-06 | 2.3e-06 | 2.8e-06 | 3.133E-06 | 1.041E-06 | 2.300E-06 | 4.300E-06 |
| 20505 | 1026 | V1026C | C | 3e-06 | 2.8e-06 | 1.8e-06 | 2.533E-06 | 6.429E-07 | 1.800E-06 | 3.000E-06 |
| 20506 | 1026 | V1026E | C | 4.1e-06 | 3.6e-06 | 3.8e-06 | 3.833E-06 | 2.517E-07 | 3.600E-06 | 4.100E-06 |
| 20507 | 1026 | V1026Q | C | 1.4e-06 | 3.6e-06 | 1.9e-06 | 2.300E-06 | 1.153E-06 | 1.400E-06 | 3.600E-06 |
| 20508 | 1026 | V1026G | C | 2.2e-06 | 2.8e-06 | 3.2e-06 | 2.733E-06 | 5.033E-07 | 2.200E-06 | 3.200E-06 |
| 20509 | 1026 | V1026H | C | 2.5e-06 | 3.2e-06 | 2.4e-06 | 2.700E-06 | 4.359E-07 | 2.400E-06 | 3.200E-06 |
| 20510 | 1026 | V1026I | C | 1.9e-06 | 1.7e-06 | 2.3e-06 | 1.967E-06 | 3.055E-07 | 1.700E-06 | 2.300E-06 |
| 20511 | 1026 | V1026L | C | 2.1e-06 | 2.6e-06 | 1.9e-06 | 2.200E-06 | 3.606E-07 | 1.900E-06 | 2.600E-06 |
| 20512 | 1026 | V1026K | C | 3.8e-06 | 2.4e-06 | 2.3e-06 | 2.833E-06 | 8.386E-07 | 2.300E-06 | 3.800E-06 |
| 20513 | 1026 | V1026M | C | 2.6e-06 | 1.9e-06 | 2.7e-06 | 2.400E-06 | 4.359E-07 | 1.900E-06 | 2.700E-06 |
| 20514 | 1026 | V1026F | C | 2.7e-06 | 3.5e-06 | 3.4e-06 | 3.200E-06 | 4.359E-07 | 2.700E-06 | 3.500E-06 |
| 20515 | 1026 | V1026P | C | 4.7e-06 | 1.9e-06 | 4.6e-06 | 3.733E-06 | 1.589E-06 | 1.900E-06 | 4.700E-06 |
| 20516 | 1026 | V1026S | C | 2.4e-06 | 1.6e-06 | 2e-06 | 2.000E-06 | 4.000E-07 | 1.600E-06 | 2.400E-06 |
| 20517 | 1026 | V1026T | C | 2.6e-06 | 2.3e-06 | 3e-06 | 2.633E-06 | 3.512E-07 | 2.300E-06 | 3.000E-06 |
| 20518 | 1026 | V1026W | C | 2.3e-06 | 9.6e-07 | 2e-06 | 1.753E-06 | 7.032E-07 | 9.600E-07 | 2.300E-06 |
| 20519 | 1026 | V1026Y | C | 2.1e-06 | 3.4e-06 | 1.9e-06 | 2.467E-06 | 8.145E-07 | 1.900E-06 | 3.400E-06 |
| 20520 | 1026 | V1026V | C | 3.2e-06 | 2.6e-06 | 2.3e-06 | 2.700E-06 | 4.583E-07 | 2.300E-06 | 3.200E-06 |
| 20521 | 1027 | P1027A | C | 2.5e-06 | 2.3e-06 | 3.3e-06 | 2.700E-06 | 5.292E-07 | 2.300E-06 | 3.300E-06 |
| 20522 | 1027 | P1027R | C | 3.1e-06 | 2.4e-06 | 2.9e-06 | 2.800E-06 | 3.606E-07 | 2.400E-06 | 3.100E-06 |
| 20523 | 1027 | P1027N | C | 3.3e-06 | 3.1e-06 | 1.9e-06 | 2.767E-06 | 7.572E-07 | 1.900E-06 | 3.300E-06 |
| 20524 | 1027 | P1027D | C | 2.1e-06 | 1.2e-06 | 9.9e-07 | 1.430E-06 | 5.897E-07 | 9.900E-07 | 2.100E-06 |
| 20525 | 1027 | P1027C | C | 4.1e-06 | 1.7e-06 | 4.9e-06 | 3.567E-06 | 1.665E-06 | 1.700E-06 | 4.900E-06 |
| 20526 | 1027 | P1027E | C | 2.6e-06 | 2.2e-06 | 2.7e-06 | 2.500E-06 | 2.646E-07 | 2.200E-06 | 2.700E-06 |
| 20527 | 1027 | P1027Q | C | 1.7e-06 | 1.6e-06 | 3.2e-06 | 2.167E-06 | 8.963E-07 | 1.600E-06 | 3.200E-06 |
| 20528 | 1027 | P1027G | C | 1.8e-06 | 3.5e-06 | 3.8e-06 | 3.033E-06 | 1.079E-06 | 1.800E-06 | 3.800E-06 |
| 20529 | 1027 | P1027H | C | 2.6e-06 | 2.8e-06 | 2e-06 | 2.467E-06 | 4.163E-07 | 2.000E-06 | 2.800E-06 |
| 20530 | 1027 | P1027I | C | 3.7e-06 | 2.4e-06 | 2.7e-06 | 2.933E-06 | 6.807E-07 | 2.400E-06 | 3.700E-06 |
| 20531 | 1027 | P1027L | C | 2.3e-06 | 2.8e-06 | 3.2e-06 | 2.767E-06 | 4.509E-07 | 2.300E-06 | 3.200E-06 |
| 20532 | 1027 | P1027K | C | 1.9e-06 | 2e-06 | 4.1e-06 | 2.667E-06 | 1.242E-06 | 1.900E-06 | 4.100E-06 |
| 20533 | 1027 | P1027M | C | 1.3e-06 | 2.2e-06 | 2.3e-06 | 1.933E-06 | 5.508E-07 | 1.300E-06 | 2.300E-06 |
| 20534 | 1027 | P1027F | C | 2.2e-06 | 2.6e-06 | 2.2e-06 | 2.333E-06 | 2.309E-07 | 2.200E-06 | 2.600E-06 |
| 20535 | 1027 | P1027P | C | 1.8e-06 | 1.6e-06 | 3.8e-06 | 2.400E-06 | 1.217E-06 | 1.600E-06 | 3.800E-06 |
| 20536 | 1027 | P1027S | C | 3.4e-06 | 2.2e-06 | 2.9e-06 | 2.833E-06 | 6.028E-07 | 2.200E-06 | 3.400E-06 |
| 20537 | 1027 | P1027T | C | 3.1e-06 | 2.4e-06 | 3.2e-06 | 2.900E-06 | 4.359E-07 | 2.400E-06 | 3.200E-06 |
| 20538 | 1027 | P1027W | C | 3.3e-06 | 1.8e-06 | 3e-06 | 2.700E-06 | 7.937E-07 | 1.800E-06 | 3.300E-06 |
| 20539 | 1027 | P1027Y | C | 2e-06 | 2.3e-06 | 2.4e-06 | 2.233E-06 | 2.082E-07 | 2.000E-06 | 2.400E-06 |
| 20540 | 1027 | P1027V | C | 2.5e-06 | 4.3e-06 | 2.4e-06 | 3.067E-06 | 1.069E-06 | 2.400E-06 | 4.300E-06 |
| 20541 | 1028 | S1028A | C | 2e-06 | 1.9e-06 | 1.4e-06 | 1.767E-06 | 3.215E-07 | 1.400E-06 | 2.000E-06 |
| 20542 | 1028 | S1028R | C | 2.4e-06 | 2.5e-06 | 4.6e-06 | 3.167E-06 | 1.242E-06 | 2.400E-06 | 4.600E-06 |
| 20543 | 1028 | S1028N | C | 1.7e-06 | 3.4e-06 | 3.1e-06 | 2.733E-06 | 9.074E-07 | 1.700E-06 | 3.400E-06 |
| 20544 | 1028 | S1028D | C | 2.8e-06 | 4.5e-06 | 1.9e-06 | 3.067E-06 | 1.320E-06 | 1.900E-06 | 4.500E-06 |
| 20545 | 1028 | S1028C | C | 2.4e-06 | 2.1e-06 | 2.1e-06 | 2.200E-06 | 1.732E-07 | 2.100E-06 | 2.400E-06 |
| 20546 | 1028 | S1028E | C | 3.6e-06 | 1.7e-06 | 4.1e-06 | 3.133E-06 | 1.266E-06 | 1.700E-06 | 4.100E-06 |
| 20547 | 1028 | S1028Q | C | 1.3e-06 | 3.6e-06 | 1.2e-06 | 2.033E-06 | 1.358E-06 | 1.200E-06 | 3.600E-06 |
| 20548 | 1028 | S1028G | C | 3.3e-06 | 3.3e-06 | 2.5e-06 | 3.033E-06 | 4.619E-07 | 2.500E-06 | 3.300E-06 |
| 20549 | 1028 | S1028H | C | 1.8e-06 | 2.4e-06 | 2.6e-06 | 2.267E-06 | 4.163E-07 | 1.800E-06 | 2.600E-06 |
| 20550 | 1028 | S1028I | C | 2.7e-06 | 2.1e-06 | 3.3e-06 | 2.700E-06 | 6.000E-07 | 2.100E-06 | 3.300E-06 |
| 20551 | 1028 | S1028L | C | 2.9e-06 | 2.3e-06 | 1.7e-06 | 2.300E-06 | 6.000E-07 | 1.700E-06 | 2.900E-06 |
| 20552 | 1028 | S1028K | C | 1.9e-06 | 2.4e-06 | 4.3e-06 | 2.867E-06 | 1.266E-06 | 1.900E-06 | 4.300E-06 |
| 20553 | 1028 | S1028M | C | 2.6e-06 | 2.8e-06 | 3.3e-06 | 2.900E-06 | 3.606E-07 | 2.600E-06 | 3.300E-06 |
| 20554 | 1028 | S1028F | C | 1.8e-06 | 3.5e-06 | 3.4e-06 | 2.900E-06 | 9.539E-07 | 1.800E-06 | 3.500E-06 |
| 20555 | 1028 | S1028P | C | 3.7e-06 | 2.3e-06 | 6.4e-06 | 4.133E-06 | 2.084E-06 | 2.300E-06 | 6.400E-06 |
| 20556 | 1028 | S1028S | C | 2.4e-06 | 2.1e-06 | 1.2e-06 | 1.900E-06 | 6.245E-07 | 1.200E-06 | 2.400E-06 |
| 20557 | 1028 | S1028T | C | 1.9e-06 | 1.8e-06 | 2.6e-06 | 2.100E-06 | 4.359E-07 | 1.800E-06 | 2.600E-06 |
| 20558 | 1028 | S1028W | C | 2.7e-06 | 1.9e-06 | 3.2e-06 | 2.600E-06 | 6.557E-07 | 1.900E-06 | 3.200E-06 |
| 20559 | 1028 | S1028Y | C | 1.8e-06 | 2.8e-06 | 4e-06 | 2.867E-06 | 1.102E-06 | 1.800E-06 | 4.000E-06 |
| 20560 | 1028 | S1028V | C | 2.5e-06 | 1.7e-06 | 1.8e-06 | 2.000E-06 | 4.359E-07 | 1.700E-06 | 2.500E-06 |
| 20561 | 1029 | S1029A | C | 1.6e-06 | 2.6e-06 | 2.6e-06 | 2.267E-06 | 5.774E-07 | 1.600E-06 | 2.600E-06 |
| 20562 | 1029 | S1029R | C | 1.6e-06 | 2.8e-06 | 2.9e-06 | 2.433E-06 | 7.234E-07 | 1.600E-06 | 2.900E-06 |
| 20563 | 1029 | S1029N | C | 1.9e-06 | 3.4e-06 | 1.6e-06 | 2.300E-06 | 9.644E-07 | 1.600E-06 | 3.400E-06 |
| 20564 | 1029 | S1029D | C | 2.2e-06 | 2.5e-06 | 3.2e-06 | 2.633E-06 | 5.132E-07 | 2.200E-06 | 3.200E-06 |
| 20565 | 1029 | S1029C | C | 1.6e-06 | 2.9e-06 | 3.3e-06 | 2.600E-06 | 8.888E-07 | 1.600E-06 | 3.300E-06 |
| 20566 | 1029 | S1029E | C | 2.4e-06 | 2.1e-06 | 2.2e-06 | 2.233E-06 | 1.528E-07 | 2.100E-06 | 2.400E-06 |
| 20567 | 1029 | S1029Q | C | 2.2e-06 | 2.8e-06 | 2.1e-06 | 2.367E-06 | 3.786E-07 | 2.100E-06 | 2.800E-06 |
| 20568 | 1029 | S1029G | C | 2.3e-06 | 3.5e-06 | 1.7e-06 | 2.500E-06 | 9.165E-07 | 1.700E-06 | 3.500E-06 |
| 20569 | 1029 | S1029H | C | 4.4e-06 | 4.2e-06 | 3.9e-06 | 4.167E-06 | 2.517E-07 | 3.900E-06 | 4.400E-06 |

| | | | | | | | | | | |
|---|---|---|---|---|---|---|---|---|---|---|
| 20570 | 1029 | S1029I | C | 2.4e-06 | 4.9e-06 | 2.1e-06 | 3.133E-06 | 1.537E-06 | 2.100E-06 | 4.900E-06 |
| 20571 | 1029 | S1029L | C | 2.6e-06 | 3.7e-06 | 2.1e-06 | 2.800E-06 | 8.185E-07 | 2.100E-06 | 3.700E-06 |
| 20572 | 1029 | S1029K | C | 2.6e-06 | 2.2e-06 | 1.9e-06 | 2.233E-06 | 3.512E-07 | 1.900E-06 | 2.600E-06 |
| 20573 | 1029 | S1029M | C | 4.5e-06 | 2.9e-06 | 2.3e-06 | 3.233E-06 | 1.137E-06 | 2.300E-06 | 4.500E-06 |
| 20574 | 1029 | S1029F | C | 2.8e-06 | 1.4e-06 | 4e-06 | 2.733E-06 | 1.301E-06 | 1.400E-06 | 4.000E-06 |
| 20575 | 1029 | S1029P | C | 2.8e-06 | 2.4e-06 | 2.8e-06 | 2.667E-06 | 2.309E-07 | 2.400E-06 | 2.800E-06 |
| 20576 | 1029 | S1029S | C | 2.7e-06 | 2.6e-06 | 3.6e-06 | 2.967E-06 | 5.508E-07 | 2.600E-06 | 3.600E-06 |
| 20577 | 1029 | S1029T | C | 1.7e-06 | 3.4e-06 | 2.6e-06 | 2.567E-06 | 8.505E-07 | 1.700E-06 | 3.400E-06 |
| 20578 | 1029 | S1029W | C | 2.7e-06 | 2.1e-06 | 2.4e-06 | 2.400E-06 | 3.000E-07 | 2.100E-06 | 2.700E-06 |
| 20579 | 1029 | S1029Y | C | 3.2e-06 | 9.8e-07 | 2e-06 | 2.060E-06 | 1.111E-06 | 9.800E-07 | 3.200E-06 |
| 20580 | 1029 | S1029V | C | 2.2e-06 | 2.1e-06 | 2.9e-06 | 2.400E-06 | 4.359E-07 | 2.100E-06 | 2.900E-06 |
| 20581 | 1030 | S1030A | C | 1.7e-06 | 2.9e-06 | 1.6e-06 | 2.067E-06 | 7.234E-07 | 1.600E-06 | 2.900E-06 |
| 20582 | 1030 | S1030R | C | 1.7e-06 | 1.9e-06 | 3.7e-06 | 2.433E-06 | 1.102E-06 | 1.700E-06 | 3.700E-06 |
| 20583 | 1030 | S1030N | C | 2.9e-06 | 1.9e-06 | 2.7e-06 | 2.500E-06 | 5.292E-07 | 1.900E-06 | 2.900E-06 |
| 20584 | 1030 | S1030D | C | 5e-06 | 2.5e-06 | 2.7e-06 | 3.400E-06 | 1.389E-06 | 2.500E-06 | 5.000E-06 |
| 20585 | 1030 | S1030C | C | 2.1e-06 | 2.8e-06 | 1.5e-06 | 2.133E-06 | 6.506E-07 | 1.500E-06 | 2.800E-06 |
| 20586 | 1030 | S1030E | C | 2.7e-06 | 3.2e-06 | 3.5e-06 | 3.133E-06 | 4.041E-07 | 2.700E-06 | 3.500E-06 |
| 20587 | 1030 | S1030Q | C | 2.3e-06 | 2.6e-06 | 2e-06 | 2.300E-06 | 3.000E-07 | 2.000E-06 | 2.600E-06 |
| 20588 | 1030 | S1030G | C | 1.3e-06 | 3.3e-06 | 2.4e-06 | 2.333E-06 | 1.002E-06 | 1.300E-06 | 3.300E-06 |
| 20589 | 1030 | S1030H | C | 2.7e-06 | 2.9e-06 | 2.6e-06 | 2.733E-06 | 1.528E-07 | 2.600E-06 | 2.900E-06 |
| 20590 | 1030 | S1030I | C | 2.1e-06 | 1.7e-06 | 2.4e-06 | 2.067E-06 | 3.512E-07 | 1.700E-06 | 2.400E-06 |
| 20591 | 1030 | S1030L | C | 2.1e-06 | 3.8e-06 | 2.6e-06 | 2.833E-06 | 8.737E-07 | 2.100E-06 | 3.800E-06 |
| 20592 | 1030 | S1030K | C | 2e-06 | 2.2e-06 | 2.9e-06 | 2.367E-06 | 4.726E-07 | 2.000E-06 | 2.900E-06 |
| 20593 | 1030 | S1030M | C | 2.8e-06 | 2.7e-06 | 3.8e-06 | 3.100E-06 | 6.083E-07 | 2.700E-06 | 3.800E-06 |
| 20594 | 1030 | S1030F | C | 2.5e-06 | 3.9e-06 | 3.1e-06 | 3.167E-06 | 7.024E-07 | 2.500E-06 | 3.900E-06 |
| 20595 | 1030 | S1030P | C | 3.6e-06 | 4.9e-06 | 3.5e-06 | 4.000E-06 | 7.810E-07 | 3.500E-06 | 4.900E-06 |
| 20596 | 1030 | S1030S | C | 1.9e-06 | 2.1e-06 | 1.5e-06 | 1.833E-06 | 3.055E-07 | 1.500E-06 | 2.100E-06 |
| 20597 | 1030 | S1030T | C | 1.3e-06 | 2.8e-06 | 2.5e-06 | 2.200E-06 | 7.937E-07 | 1.300E-06 | 2.800E-06 |
| 20598 | 1030 | S1030W | C | 2.5e-06 | 2.3e-06 | 2.4e-06 | 2.400E-06 | 1.000E-07 | 2.300E-06 | 2.500E-06 |
| 20599 | 1030 | S1030Y | C | 2.3e-06 | 3.6e-06 | 2.8e-06 | 2.900E-06 | 6.557E-07 | 2.300E-06 | 3.600E-06 |
| 20600 | 1030 | S1030V | C | 2.2e-06 | 3e-06 | 2.2e-06 | 2.467E-06 | 4.619E-07 | 2.200E-06 | 3.000E-06 |
| 20601 | 1031 | L1031A | C | 4.3e-06 | 2.6e-06 | 2.5e-06 | 3.133E-06 | 1.012E-06 | 2.500E-06 | 4.300E-06 |
| 20602 | 1031 | L1031R | C | 1.4e-06 | 2.1e-06 | 2.3e-06 | 1.933E-06 | 4.726E-07 | 1.400E-06 | 2.300E-06 |
| 20603 | 1031 | L1031N | C | 2.1e-06 | 2.1e-06 | 2.5e-06 | 2.233E-06 | 2.309E-07 | 2.100E-06 | 2.500E-06 |
| 20604 | 1031 | L1031D | C | 2e-06 | 1.6e-06 | 2.8e-06 | 2.133E-06 | 6.110E-07 | 1.600E-06 | 2.800E-06 |
| 20605 | 1031 | L1031C | C | 1.7e-06 | 2.8e-06 | 2.4e-06 | 2.300E-06 | 5.568E-07 | 1.700E-06 | 2.800E-06 |
| 20606 | 1031 | L1031E | C | 1.4e-06 | 1.8e-06 | 2.5e-06 | 1.900E-06 | 5.568E-07 | 1.400E-06 | 2.500E-06 |
| 20607 | 1031 | L1031Q | C | 2.4e-06 | 2.9e-06 | 1.9e-06 | 2.400E-06 | 5.000E-07 | 1.900E-06 | 2.900E-06 |
| 20608 | 1031 | L1031G | C | 2.6e-06 | 7.1e-07 | 2.1e-06 | 1.803E-06 | 9.793E-07 | 7.100E-07 | 2.600E-06 |
| 20609 | 1031 | L1031H | C | 1.8e-06 | 2.3e-06 | 2.6e-06 | 2.233E-06 | 4.041E-07 | 1.800E-06 | 2.600E-06 |
| 20610 | 1031 | L1031I | C | 2.5e-06 | 1.9e-06 | 2.4e-06 | 2.267E-06 | 3.215E-07 | 1.900E-06 | 2.500E-06 |
| 20611 | 1031 | L1031L | C | 1.1e-06 | 2.3e-06 | 2.9e-06 | 2.100E-06 | 9.165E-07 | 1.100E-06 | 2.900E-06 |
| 20612 | 1031 | L1031K | C | 2.2e-06 | 2.3e-06 | 2e-06 | 2.167E-06 | 1.528E-07 | 2.000E-06 | 2.300E-06 |
| 20613 | 1031 | L1031M | C | 2.8e-06 | 2.3e-06 | 2.6e-06 | 2.567E-06 | 2.517E-07 | 2.300E-06 | 2.800E-06 |
| 20614 | 1031 | L1031F | C | 1.5e-06 | 1.4e-06 | 1.8e-06 | 1.567E-06 | 2.082E-07 | 1.400E-06 | 1.800E-06 |
| 20615 | 1031 | L1031P | C | 2e-06 | 1.9e-06 | 6.9e-06 | 3.600E-06 | 2.858E-06 | 1.900E-06 | 6.900E-06 |
| 20616 | 1031 | L1031S | C | 1.2e-06 | 2.2e-06 | 2.7e-06 | 2.033E-06 | 7.638E-07 | 1.200E-06 | 2.700E-06 |
| 20617 | 1031 | L1031T | C | 2.1e-06 | 2.5e-06 | 2.6e-06 | 2.400E-06 | 2.646E-07 | 2.100E-06 | 2.600E-06 |
| 20618 | 1031 | L1031W | C | 3.1e-06 | 1.3e-06 | 2.2e-06 | 2.200E-06 | 9.000E-07 | 1.300E-06 | 3.100E-06 |
| 20619 | 1031 | L1031Y | C | 4.2e-06 | 2.7e-06 | 2.6e-06 | 3.167E-06 | 8.963E-07 | 2.600E-06 | 4.200E-06 |
| 20620 | 1031 | L1031V | C | 2.3e-06 | 1.4e-06 | 2e-06 | 1.900E-06 | 4.583E-07 | 1.400E-06 | 2.300E-06 |
| 20621 | 1032 | G1032A | C | 3.7e-06 | 2.5e-06 | 5.4e-06 | 3.867E-06 | 1.457E-06 | 2.500E-06 | 5.400E-06 |
| 20622 | 1032 | G1032R | C | 1.9e-06 | 2.7e-06 | 3.6e-06 | 2.733E-06 | 8.505E-07 | 1.900E-06 | 3.600E-06 |
| 20623 | 1032 | G1032N | C | 2.2e-06 | 2.1e-06 | 2.4e-06 | 2.233E-06 | 1.528E-07 | 2.100E-06 | 2.400E-06 |
| 20624 | 1032 | G1032D | C | 2.7e-06 | 1.8e-06 | 2.3e-06 | 2.267E-06 | 4.509E-07 | 1.800E-06 | 2.700E-06 |
| 20625 | 1032 | G1032C | C | 2.7e-06 | 1.5e-06 | 2.1e-06 | 2.100E-06 | 6.000E-07 | 1.500E-06 | 2.700E-06 |
| 20626 | 1032 | G1032E | C | 1.8e-06 | 2.6e-06 | 3e-06 | 2.467E-06 | 6.110E-07 | 1.800E-06 | 3.000E-06 |
| 20627 | 1032 | G1032Q | C | 1.9e-06 | 2.7e-06 | 2e-06 | 2.200E-06 | 4.359E-07 | 1.900E-06 | 2.700E-06 |
| 20628 | 1032 | G1032G | C | 2e-06 | 3.6e-06 | 2.7e-06 | 2.767E-06 | 8.021E-07 | 2.000E-06 | 3.600E-06 |
| 20629 | 1032 | G1032H | C | 3e-06 | 2.6e-06 | 1.3e-06 | 2.300E-06 | 8.888E-07 | 1.300E-06 | 3.000E-06 |
| 20630 | 1032 | G1032I | C | 3e-06 | 1.4e-06 | 2.4e-06 | 2.267E-06 | 8.083E-07 | 1.400E-06 | 3.000E-06 |
| 20631 | 1032 | G1032L | C | 2e-06 | 2.1e-06 | 2.9e-06 | 2.333E-06 | 4.933E-07 | 2.000E-06 | 2.900E-06 |
| 20632 | 1032 | G1032K | C | 1.7e-06 | 1.5e-06 | 2.9e-06 | 2.033E-06 | 7.572E-07 | 1.500E-06 | 2.900E-06 |
| 20633 | 1032 | G1032M | C | 1.8e-06 | 1.4e-06 | 2e-06 | 1.733E-06 | 3.055E-07 | 1.400E-06 | 2.000E-06 |
| 20634 | 1032 | G1032F | C | 1.8e-06 | 1.6e-06 | 2.9e-06 | 2.100E-06 | 7.000E-07 | 1.600E-06 | 2.900E-06 |
| 20635 | 1032 | G1032P | C | 1.1e-06 | 2.3e-06 | 4.1e-06 | 2.500E-06 | 1.510E-06 | 1.100E-06 | 4.100E-06 |
| 20636 | 1032 | G1032S | C | 3.4e-06 | 2e-06 | 2e-06 | 2.467E-06 | 8.083E-07 | 2.000E-06 | 3.400E-06 |
| 20637 | 1032 | G1032T | C | 1.7e-06 | 2.9e-06 | 3e-06 | 2.533E-06 | 7.234E-07 | 1.700E-06 | 3.000E-06 |
| 20638 | 1032 | G1032W | C | 2.1e-06 | 2.9e-06 | 1.9e-06 | 2.300E-06 | 5.292E-07 | 1.900E-06 | 2.900E-06 |
| 20639 | 1032 | G1032Y | C | 1.1e-06 | 1.6e-06 | 2e-06 | 1.567E-06 | 4.509E-07 | 1.100E-06 | 2.000E-06 |
| 20640 | 1032 | G1032V | C | 3.6e-06 | 1.4e-06 | 1e-06 | 2.000E-06 | 1.400E-06 | 1.000E-06 | 3.600E-06 |
| 20641 | 1033 | T1033A | C | 2.2e-06 | 1.6e-06 | 2.5e-06 | 2.100E-06 | 4.583E-07 | 1.600E-06 | 2.500E-06 |
| 20642 | 1033 | T1033R | C | 1.9e-06 | 1.9e-06 | 2.7e-06 | 2.167E-06 | 4.619E-07 | 1.900E-06 | 2.700E-06 |
| 20643 | 1033 | T1033N | C | 1.4e-06 | 1.9e-06 | 2.7e-06 | 2.000E-06 | 6.557E-07 | 1.400E-06 | 2.700E-06 |
| 20644 | 1033 | T1033D | C | 1.2e-06 | 2.2e-06 | 4e-06 | 2.467E-06 | 1.419E-06 | 1.200E-06 | 4.000E-06 |
| 20645 | 1033 | T1033C | C | 4.5e-06 | 1.9e-06 | 2.6e-06 | 3.000E-06 | 1.345E-06 | 1.900E-06 | 4.500E-06 |
| 20646 | 1033 | T1033E | C | 1.7e-06 | 1.7e-06 | 3.7e-06 | 2.367E-06 | 1.155E-06 | 1.700E-06 | 3.700E-06 |
| 20647 | 1033 | T1033Q | C | 1e-06 | 2e-06 | 3e-06 | 2.000E-06 | 1.000E-06 | 1.000E-06 | 3.000E-06 |
| 20648 | 1033 | T1033G | C | 3.1e-06 | 1.9e-06 | 2.7e-06 | 2.567E-06 | 6.110E-07 | 1.900E-06 | 3.100E-06 |
| 20649 | 1033 | T1033H | C | 3e-06 | 3e-06 | 3e-06 | 3.000E-06 | 0.000E+00 | 3.000E-06 | 3.000E-06 |
| 20650 | 1033 | T1033I | C | 5.2e-06 | 1.2e-06 | 2e-06 | 2.800E-06 | 2.117E-06 | 1.200E-06 | 5.200E-06 |
| 20651 | 1033 | T1033L | C | 2.3e-06 | 1.9e-06 | 2.8e-06 | 2.333E-06 | 4.509E-07 | 1.900E-06 | 2.800E-06 |
| 20652 | 1033 | T1033K | C | 1.6e-06 | 1.9e-06 | 1.2e-06 | 1.567E-06 | 3.512E-07 | 1.200E-06 | 1.900E-06 |
| 20653 | 1033 | T1033M | C | 2.4e-06 | 2.9e-06 | 2.7e-06 | 2.667E-06 | 2.517E-07 | 2.400E-06 | 2.900E-06 |
| 20654 | 1033 | T1033F | C | 2.6e-06 | 3.2e-06 | 1.8e-06 | 2.533E-06 | 7.024E-07 | 1.800E-06 | 3.200E-06 |

| | | | | | | | | | | |
|---|---|---|---|---|---|---|---|---|---|---|
| 20655 | 1033 | T1033P | C | 2.7e-06 | 5.1e-06 | 2.4e-06 | 3.400E-06 | 1.480E-06 | 2.400E-06 | 5.100E-06 |
| 20656 | 1033 | T1033S | C | 3.1e-06 | 8.4e-06 | 2.8e-06 | 4.767E-06 | 3.150E-06 | 2.800E-06 | 8.400E-06 |
| 20657 | 1033 | T1033T | C | 3.7e-06 | 2.6e-06 | 3.6e-06 | 3.300E-06 | 6.083E-07 | 2.600E-06 | 3.700E-06 |
| 20658 | 1033 | T1033W | C | 2.6e-06 | 2.3e-06 | 2.5e-06 | 2.467E-06 | 1.528E-07 | 2.300E-06 | 2.600E-06 |
| 20659 | 1033 | T1033Y | C | 1.9e-06 | 3e-06 | 2.2e-06 | 2.367E-06 | 5.686E-07 | 1.900E-06 | 3.000E-06 |
| 20660 | 1033 | T1033V | C | 2.5e-06 | 3.5e-06 | 1.2e-06 | 2.400E-06 | 1.153E-06 | 1.200E-06 | 3.500E-06 |
| 20661 | 1034 | Q1034A | C | 3e-06 | 2.5e-06 | 1.5e-06 | 2.333E-06 | 7.638E-07 | 1.500E-06 | 3.000E-06 |
| 20662 | 1034 | Q1034R | C | 4.1e-06 | 1.9e-06 | 1.6e-06 | 2.533E-06 | 1.365E-06 | 1.600E-06 | 4.100E-06 |
| 20663 | 1034 | Q1034N | C | 2.4e-06 | 1.8e-06 | 2.4e-06 | 2.200E-06 | 3.464E-07 | 1.800E-06 | 2.400E-06 |
| 20664 | 1034 | Q1034D | C | 2.3e-06 | 2.5e-06 | 2.1e-06 | 2.300E-06 | 2.000E-07 | 2.100E-06 | 2.500E-06 |
| 20665 | 1034 | Q1034C | C | 3.1e-06 | 2.8e-06 | 1.4e-06 | 2.433E-06 | 9.074E-07 | 1.400E-06 | 3.100E-06 |
| 20666 | 1034 | Q1034E | C | 2.6e-06 | 2.4e-06 | 1.8e-06 | 2.267E-06 | 4.163E-07 | 1.800E-06 | 2.600E-06 |
| 20667 | 1034 | Q1034Q | C | 1.6e-06 | 3.4e-06 | 3.1e-06 | 2.700E-06 | 9.644E-07 | 1.600E-06 | 3.400E-06 |
| 20668 | 1034 | Q1034G | C | 2.1e-06 | 3e-06 | 2.5e-06 | 2.533E-06 | 4.509E-07 | 2.100E-06 | 3.000E-06 |
| 20669 | 1034 | Q1034H | C | 1.6e-06 | 2.3e-06 | 2.1e-06 | 2.000E-06 | 3.606E-07 | 1.600E-06 | 2.300E-06 |
| 20670 | 1034 | Q1034I | C | 2.1e-06 | 3.6e-06 | 3.1e-06 | 2.933E-06 | 7.638E-07 | 2.100E-06 | 3.600E-06 |
| 20671 | 1034 | Q1034L | C | 2.4e-06 | 2e-06 | 3e-06 | 2.467E-06 | 5.033E-07 | 2.000E-06 | 3.000E-06 |
| 20672 | 1034 | Q1034K | C | 2.6e-06 | 1.9e-06 | 2.2e-06 | 2.233E-06 | 3.512E-07 | 1.900E-06 | 2.600E-06 |
| 20673 | 1034 | Q1034M | C | 2.4e-06 | 2.5e-06 | 3.9e-06 | 2.933E-06 | 8.386E-07 | 2.400E-06 | 3.900E-06 |
| 20674 | 1034 | Q1034F | C | 2.3e-06 | 6.1e-06 | 3.4e-06 | 3.933E-06 | 1.955E-06 | 2.300E-06 | 6.100E-06 |
| 20675 | 1034 | Q1034P | C | 1.4e-06 | 2.8e-06 | 3.2e-06 | 2.467E-06 | 9.452E-07 | 1.400E-06 | 3.200E-06 |
| 20676 | 1034 | Q1034S | C | 1.5e-06 | 2e-06 | 2.4e-06 | 1.967E-06 | 4.509E-07 | 1.500E-06 | 2.400E-06 |
| 20677 | 1034 | Q1034T | C | 2.2e-06 | 4.4e-06 | 2e-06 | 2.867E-06 | 1.332E-06 | 2.000E-06 | 4.400E-06 |
| 20678 | 1034 | Q1034W | C | 2.8e-06 | 1.6e-06 | 1.9e-06 | 2.100E-06 | 6.245E-07 | 1.600E-06 | 2.800E-06 |
| 20679 | 1034 | Q1034Y | C | 3.8e-06 | 3.8e-06 | 2.1e-06 | 3.233E-06 | 9.815E-07 | 2.100E-06 | 3.800E-06 |
| 20680 | 1034 | Q1034V | C | 2.5e-06 | 2.4e-06 | 3.6e-06 | 2.833E-06 | 6.658E-07 | 2.400E-06 | 3.600E-06 |
| 20681 | 1035 | T1035A | C | 2.6e-06 | 1.9e-06 | 1.6e-06 | 2.033E-06 | 5.132E-07 | 1.600E-06 | 2.600E-06 |
| 20682 | 1035 | T1035R | C | 1.9e-06 | 2.8e-06 | 4.4e-06 | 3.033E-06 | 1.266E-06 | 1.900E-06 | 4.400E-06 |
| 20683 | 1035 | T1035N | C | 1.5e-06 | 1.8e-06 | 1.6e-06 | 1.633E-06 | 1.528E-07 | 1.500E-06 | 1.800E-06 |
| 20684 | 1035 | T1035D | C | 2.6e-06 | 2.7e-06 | 3e-06 | 2.767E-06 | 2.082E-07 | 2.600E-06 | 3.000E-06 |
| 20685 | 1035 | T1035C | C | 3.9e-06 | 3.2e-06 | 5.4e-06 | 4.167E-06 | 1.124E-06 | 3.200E-06 | 5.400E-06 |
| 20686 | 1035 | T1035E | C | 2.4e-06 | 2e-06 | 2.2e-06 | 2.200E-06 | 2.000E-07 | 2.000E-06 | 2.400E-06 |
| 20687 | 1035 | T1035Q | C | 1.9e-06 | 2e-06 | 2.7e-06 | 2.200E-06 | 4.359E-07 | 1.900E-06 | 2.700E-06 |
| 20688 | 1035 | T1035G | C | 2.7e-06 | 2.1e-06 | 3.3e-06 | 2.700E-06 | 6.000E-07 | 2.100E-06 | 3.300E-06 |
| 20689 | 1035 | T1035H | C | 2e-06 | 2.2e-06 | 3e-06 | 2.400E-06 | 5.292E-07 | 2.000E-06 | 3.000E-06 |
| 20690 | 1035 | T1035I | C | 3.7e-06 | 2.3e-06 | 3.4e-06 | 3.133E-06 | 7.371E-07 | 2.300E-06 | 3.700E-06 |
| 20691 | 1035 | T1035L | C | 2.2e-06 | 2.4e-06 | 3.4e-06 | 2.667E-06 | 6.429E-07 | 2.200E-06 | 3.400E-06 |
| 20692 | 1035 | T1035K | C | 1.8e-06 | 1.3e-06 | 3.1e-06 | 2.067E-06 | 9.292E-07 | 1.300E-06 | 3.100E-06 |
| 20693 | 1035 | T1035M | C | 2.1e-06 | 2.2e-06 | 2.1e-06 | 2.133E-06 | 5.774E-08 | 2.100E-06 | 2.200E-06 |
| 20694 | 1035 | T1035F | C | 1.9e-06 | 2.5e-06 | 2.7e-06 | 2.367E-06 | 4.163E-07 | 1.900E-06 | 2.700E-06 |
| 20695 | 1035 | T1035P | C | 4.4e-06 | 1.9e-06 | 2.4e-06 | 2.900E-06 | 1.323E-06 | 1.900E-06 | 4.400E-06 |
| 20696 | 1035 | T1035S | C | 1.6e-06 | 1.2e-06 | 2.6e-06 | 1.800E-06 | 7.211E-07 | 1.200E-06 | 2.600E-06 |
| 20697 | 1035 | T1035T | C | 5.3e-06 | 1.4e-06 | 2e-06 | 2.900E-06 | 2.100E-06 | 1.400E-06 | 5.300E-06 |
| 20698 | 1035 | T1035W | C | 1.8e-06 | 1.9e-06 | 2.9e-06 | 2.200E-06 | 6.083E-07 | 1.800E-06 | 2.900E-06 |
| 20699 | 1035 | T1035Y | C | 2.8e-06 | 2.2e-06 | 1.8e-06 | 2.267E-06 | 5.033E-07 | 1.800E-06 | 2.800E-06 |
| 20700 | 1035 | T1035V | C | 2.8e-06 | 2.9e-06 | 4e-06 | 3.233E-06 | 6.658E-07 | 2.800E-06 | 4.000E-06 |
| 20701 | 1036 | Y1036A | C | 2.6e-06 | 1.8e-06 | 3.1e-06 | 2.500E-06 | 6.557E-07 | 1.800E-06 | 3.100E-06 |
| 20702 | 1036 | Y1036R | C | 4.8e-06 | 2.3e-06 | 3.4e-06 | 3.500E-06 | 1.253E-06 | 2.300E-06 | 4.800E-06 |
| 20703 | 1036 | Y1036N | C | 3e-06 | 2.1e-06 | 2.2e-06 | 2.433E-06 | 4.933E-07 | 2.100E-06 | 3.000E-06 |
| 20704 | 1036 | Y1036D | C | 1.5e-06 | 1.9e-06 | 1.7e-06 | 1.700E-06 | 2.000E-07 | 1.500E-06 | 1.900E-06 |
| 20705 | 1036 | Y1036C | C | 2.2e-06 | 2.7e-06 | 3.3e-06 | 2.733E-06 | 5.508E-07 | 2.200E-06 | 3.300E-06 |
| 20706 | 1036 | Y1036E | C | 1.9e-06 | 2.3e-06 | 1.1e-06 | 1.767E-06 | 6.110E-07 | 1.100E-06 | 2.300E-06 |
| 20707 | 1036 | Y1036Q | C | 2.5e-06 | 3.2e-06 | 2e-06 | 2.567E-06 | 6.028E-07 | 2.000E-06 | 3.200E-06 |
| 20708 | 1036 | Y1036G | C | 2.7e-06 | 3.1e-06 | 2.9e-06 | 2.900E-06 | 2.000E-07 | 2.700E-06 | 3.100E-06 |
| 20709 | 1036 | Y1036H | C | 2.1e-06 | 2e-06 | 2.4e-06 | 2.167E-06 | 2.082E-07 | 2.000E-06 | 2.400E-06 |
| 20710 | 1036 | Y1036I | C | 3.5e-06 | 3.7e-06 | 1.9e-06 | 3.033E-06 | 9.866E-07 | 1.900E-06 | 3.700E-06 |
| 20711 | 1036 | Y1036L | C | 1.4e-06 | 2.6e-06 | 2.4e-06 | 2.133E-06 | 6.429E-07 | 1.400E-06 | 2.600E-06 |
| 20712 | 1036 | Y1036K | C | 4.6e-06 | 1.9e-06 | 3.3e-06 | 3.267E-06 | 1.350E-06 | 1.900E-06 | 4.600E-06 |
| 20713 | 1036 | Y1036M | C | 1.2e-06 | 2.8e-06 | 2.5e-06 | 2.167E-06 | 8.505E-07 | 1.200E-06 | 2.800E-06 |
| 20714 | 1036 | Y1036F | C | 1.6e-06 | 1.6e-06 | 2.7e-06 | 1.967E-06 | 6.351E-07 | 1.600E-06 | 2.700E-06 |
| 20715 | 1036 | Y1036P | C | 2.3e-06 | 3.2e-06 | 2.9e-06 | 2.800E-06 | 4.583E-07 | 2.300E-06 | 3.200E-06 |
| 20716 | 1036 | Y1036S | C | 1.7e-06 | 2.8e-06 | 2e-06 | 2.167E-06 | 5.686E-07 | 1.700E-06 | 2.800E-06 |
| 20717 | 1036 | Y1036T | C | 2.2e-06 | 2.7e-06 | 2.1e-06 | 2.333E-06 | 3.215E-07 | 2.100E-06 | 2.700E-06 |
| 20718 | 1036 | Y1036W | C | 1.9e-06 | 1.4e-06 | 2.2e-06 | 1.833E-06 | 4.041E-07 | 1.400E-06 | 2.200E-06 |
| 20719 | 1036 | Y1036Y | C | 4.1e-06 | 1.7e-06 | 3.1e-06 | 2.967E-06 | 1.206E-06 | 1.700E-06 | 4.100E-06 |
| 20720 | 1036 | Y1036V | C | 1.8e-06 | 1.9e-06 | 2.9e-06 | 2.200E-06 | 6.083E-07 | 1.800E-06 | 2.900E-06 |
| 20721 | 1037 | I1037A | C | 2.6e-06 | 2.5e-06 | 2.9e-06 | 2.667E-06 | 2.082E-07 | 2.500E-06 | 2.900E-06 |
| 20722 | 1037 | I1037R | C | 3.9e-06 | 1.8e-06 | 2e-06 | 2.567E-06 | 1.159E-06 | 1.800E-06 | 3.900E-06 |
| 20723 | 1037 | I1037N | C | 1.8e-06 | 3.1e-06 | 1.5e-06 | 2.133E-06 | 8.505E-07 | 1.500E-06 | 3.100E-06 |
| 20724 | 1037 | I1037D | C | 2.4e-06 | 3.2e-06 | 2e-06 | 2.533E-06 | 6.110E-07 | 2.000E-06 | 3.200E-06 |
| 20725 | 1037 | I1037C | C | 2.8e-06 | 2.3e-06 | 1.8e-06 | 2.300E-06 | 5.000E-07 | 1.800E-06 | 2.800E-06 |
| 20726 | 1037 | I1037E | C | 2.7e-06 | 2.7e-06 | 2.8e-06 | 2.733E-06 | 5.774E-08 | 2.700E-06 | 2.800E-06 |
| 20727 | 1037 | I1037Q | C | 5.3e-06 | 2.3e-06 | 2.1e-06 | 3.233E-06 | 1.793E-06 | 2.100E-06 | 5.300E-06 |
| 20728 | 1037 | I1037G | C | 5.6e-06 | 2.3e-06 | 2.4e-06 | 3.433E-06 | 1.877E-06 | 2.300E-06 | 5.600E-06 |
| 20729 | 1037 | I1037H | C | 2.8e-06 | 1.7e-06 | 3.5e-06 | 2.667E-06 | 9.074E-07 | 1.700E-06 | 3.500E-06 |
| 20730 | 1037 | I1037I | C | 1.6e-06 | 2.7e-06 | 1.6e-06 | 1.967E-06 | 6.351E-07 | 1.600E-06 | 2.700E-06 |
| 20731 | 1037 | I1037L | C | 2.3e-06 | 1.8e-06 | 1.4e-06 | 1.833E-06 | 4.509E-07 | 1.400E-06 | 2.300E-06 |
| 20732 | 1037 | I1037K | C | 2.1e-06 | 3.5e-06 | 4.4e-06 | 3.333E-06 | 1.159E-06 | 2.100E-06 | 4.400E-06 |
| 20733 | 1037 | I1037M | C | 1.8e-06 | 1.7e-06 | 3.7e-06 | 2.400E-06 | 1.127E-06 | 1.700E-06 | 3.700E-06 |
| 20734 | 1037 | I1037F | C | 1.4e-06 | 3.4e-06 | 1.2e-06 | 2.000E-06 | 1.217E-06 | 1.200E-06 | 3.400E-06 |
| 20735 | 1037 | I1037P | C | 1.2e-06 | 2.7e-06 | 2.2e-06 | 2.033E-06 | 7.638E-07 | 1.200E-06 | 2.700E-06 |
| 20736 | 1037 | I1037S | C | 1.6e-06 | 2.2e-06 | 2.7e-06 | 2.167E-06 | 5.508E-07 | 1.600E-06 | 2.700E-06 |
| 20737 | 1037 | I1037T | C | 2.6e-06 | 2.4e-06 | 2.7e-06 | 2.567E-06 | 1.528E-07 | 2.400E-06 | 2.700E-06 |
| 20738 | 1037 | I1037W | C | 1.6e-06 | 2.5e-06 | 1.1e-06 | 1.733E-06 | 7.095E-07 | 1.100E-06 | 2.500E-06 |
| 20739 | 1037 | I1037Y | C | 2.5e-06 | 3.1e-06 | 1.9e-06 | 2.500E-06 | 6.000E-07 | 1.900E-06 | 3.100E-06 |

| | | | | | | | | | | |
|---|---|---|---|---|---|---|---|---|---|---|
| 20740 | 1037 | I1037V | C | 2.8e-06 | 2.1e-06 | 2.5e-06 | 2.467E-06 | 3.512E-07 | 2.100E-06 | 2.800E-06 |
| 20741 | 1038 | C1038A | C | 2.5e-06 | 3.1e-06 | 2.9e-06 | 2.833E-06 | 3.055E-07 | 2.500E-06 | 3.100E-06 |
| 20742 | 1038 | C1038R | C | 3.1e-06 | 2e-06 | 2.9e-06 | 2.667E-06 | 5.859E-07 | 2.000E-06 | 3.100E-06 |
| 20743 | 1038 | C1038N | C | 2.2e-06 | 3.3e-06 | 2.4e-06 | 2.633E-06 | 5.859E-07 | 2.200E-06 | 3.300E-06 |
| 20744 | 1038 | C1038D | C | 1.6e-06 | 3.1e-06 | 1.8e-06 | 2.167E-06 | 8.145E-07 | 1.600E-06 | 3.100E-06 |
| 20745 | 1038 | C1038C | C | 1.8e-06 | 1.4e-06 | 2.2e-06 | 1.800E-06 | 4.000E-07 | 1.400E-06 | 2.200E-06 |
| 20746 | 1038 | C1038E | C | 1.4e-06 | 1.9e-06 | 1.8e-06 | 1.700E-06 | 2.646E-07 | 1.400E-06 | 1.900E-06 |
| 20747 | 1038 | C1038Q | C | 1.7e-06 | 2.6e-06 | 2.7e-06 | 2.333E-06 | 5.508E-07 | 1.700E-06 | 2.700E-06 |
| 20748 | 1038 | C1038G | C | 2e-06 | 4.4e-06 | 2.9e-06 | 3.100E-06 | 1.212E-06 | 2.000E-06 | 4.400E-06 |
| 20749 | 1038 | C1038H | C | 2.8e-06 | 3.9e-06 | 1.3e-06 | 2.667E-06 | 1.305E-06 | 1.300E-06 | 3.900E-06 |
| 20750 | 1038 | C1038I | C | 3e-06 | 2.8e-06 | 1.2e-06 | 2.333E-06 | 9.866E-07 | 1.200E-06 | 3.000E-06 |
| 20751 | 1038 | C1038L | C | 2.1e-06 | 4e-06 | 1.8e-06 | 2.633E-06 | 1.193E-06 | 1.800E-06 | 4.000E-06 |
| 20752 | 1038 | C1038K | C | 1.9e-06 | 2.4e-06 | 1.2e-06 | 1.833E-06 | 6.028E-07 | 1.200E-06 | 2.400E-06 |
| 20753 | 1038 | C1038M | C | 2e-06 | 1.6e-06 | 2.9e-06 | 2.167E-06 | 6.658E-07 | 1.600E-06 | 2.900E-06 |
| 20754 | 1038 | C1038F | C | 4.4e-06 | 3.3e-06 | 4e-06 | 3.900E-06 | 5.568E-07 | 3.300E-06 | 4.400E-06 |
| 20755 | 1038 | C1038P | C | 4e-06 | 1.8e-06 | 1.9e-06 | 2.567E-06 | 1.242E-06 | 1.800E-06 | 4.000E-06 |
| 20756 | 1038 | C1038S | C | 2.7e-06 | 1.6e-06 | 2.8e-06 | 2.367E-06 | 6.658E-07 | 1.600E-06 | 2.800E-06 |
| 20757 | 1038 | C1038T | C | 2.7e-06 | 2.6e-06 | 3.8e-06 | 3.033E-06 | 6.658E-07 | 2.600E-06 | 3.800E-06 |
| 20758 | 1038 | C1038W | C | 2.5e-06 | 1.4e-06 | 2.1e-06 | 2.000E-06 | 5.568E-07 | 1.400E-06 | 2.500E-06 |
| 20759 | 1038 | C1038Y | C | 3.2e-06 | 1.9e-06 | 2.2e-06 | 2.433E-06 | 6.807E-07 | 1.900E-06 | 3.200E-06 |
| 20760 | 1038 | C1038V | C | 3.6e-06 | 3.5e-06 | 2.7e-06 | 3.267E-06 | 4.933E-07 | 2.700E-06 | 3.600E-06 |
| 20761 | 1039 | N1039A | C | 2.3e-06 | 1.6e-06 | 2.2e-06 | 2.033E-06 | 3.786E-07 | 1.600E-06 | 2.300E-06 |
| 20762 | 1039 | N1039R | C | 2.7e-06 | 1.5e-06 | 2e-06 | 2.067E-06 | 6.028E-07 | 1.500E-06 | 2.700E-06 |
| 20763 | 1039 | N1039N | C | 3.4e-06 | 3.1e-06 | 2e-06 | 2.833E-06 | 7.371E-07 | 2.000E-06 | 3.400E-06 |
| 20764 | 1039 | N1039D | C | 2e-06 | 2.7e-06 | 3.5e-06 | 2.733E-06 | 7.506E-07 | 2.000E-06 | 3.500E-06 |
| 20765 | 1039 | N1039C | C | 3e-06 | 2.5e-06 | 2.2e-06 | 2.567E-06 | 4.041E-07 | 2.200E-06 | 3.000E-06 |
| 20766 | 1039 | N1039E | C | 2.2e-06 | 2.5e-06 | 2.3e-06 | 2.333E-06 | 1.528E-07 | 2.200E-06 | 2.500E-06 |
| 20767 | 1039 | N1039Q | C | 1.7e-06 | 2.9e-06 | 2.6e-06 | 2.400E-06 | 6.245E-07 | 1.700E-06 | 2.900E-06 |
| 20768 | 1039 | N1039G | C | 2.8e-06 | 2.2e-06 | 1.9e-06 | 2.300E-06 | 4.583E-07 | 1.900E-06 | 2.800E-06 |
| 20769 | 1039 | N1039H | C | 1.2e-06 | 5e-06 | 1.7e-06 | 2.633E-06 | 2.065E-06 | 1.200E-06 | 5.000E-06 |
| 20770 | 1039 | N1039I | C | 1.6e-06 | 2e-06 | 1.9e-06 | 1.833E-06 | 2.082E-07 | 1.600E-06 | 2.000E-06 |
| 20771 | 1039 | N1039L | C | 2.1e-06 | 1.5e-06 | 3.1e-06 | 2.233E-06 | 8.083E-07 | 1.500E-06 | 3.100E-06 |
| 20772 | 1039 | N1039K | C | 2.9e-06 | 2.1e-06 | 2e-06 | 2.333E-06 | 4.933E-07 | 2.000E-06 | 2.900E-06 |
| 20773 | 1039 | N1039M | C | 3.5e-06 | 2.1e-06 | 2.1e-06 | 2.567E-06 | 8.083E-07 | 2.100E-06 | 3.500E-06 |
| 20774 | 1039 | N1039F | C | 1.7e-06 | 3.3e-06 | 2.3e-06 | 2.433E-06 | 8.083E-07 | 1.700E-06 | 3.300E-06 |
| 20775 | 1039 | N1039P | C | 2.6e-06 | 4.3e-06 | 1.7e-06 | 2.867E-06 | 1.320E-06 | 1.700E-06 | 4.300E-06 |
| 20776 | 1039 | N1039S | C | 1.6e-06 | 2.1e-06 | 1.6e-06 | 1.767E-06 | 2.887E-07 | 1.600E-06 | 2.100E-06 |
| 20777 | 1039 | N1039T | C | 2.5e-06 | 2.9e-06 | 3.1e-06 | 2.833E-06 | 3.055E-07 | 2.500E-06 | 3.100E-06 |
| 20778 | 1039 | N1039W | C | 2e-06 | 2.1e-06 | 2.3e-06 | 2.133E-06 | 1.528E-07 | 2.000E-06 | 2.300E-06 |
| 20779 | 1039 | N1039Y | C | 2.3e-06 | 2.7e-06 | 2.7e-06 | 2.567E-06 | 2.309E-07 | 2.300E-06 | 2.700E-06 |
| 20780 | 1039 | N1039V | C | 1.9e-06 | 1.8e-06 | 2.6e-06 | 2.100E-06 | 4.359E-07 | 1.800E-06 | 2.600E-06 |
| 20781 | 1040 | V1040A | C | 2.1e-06 | 2.3e-06 | 1.5e-06 | 1.967E-06 | 4.163E-07 | 1.500E-06 | 2.300E-06 |
| 20782 | 1040 | V1040R | C | 2.2e-06 | 2.9e-06 | 3.1e-06 | 2.733E-06 | 4.726E-07 | 2.200E-06 | 3.100E-06 |
| 20783 | 1040 | V1040N | C | 2.7e-06 | 2.8e-06 | 2.4e-06 | 2.633E-06 | 2.082E-07 | 2.400E-06 | 2.800E-06 |
| 20784 | 1040 | V1040D | C | 2.2e-06 | 1.9e-06 | 1.5e-06 | 1.867E-06 | 3.512E-07 | 1.500E-06 | 2.200E-06 |
| 20785 | 1040 | V1040C | C | 2.6e-06 | 4.9e-06 | 1.9e-06 | 3.133E-06 | 1.570E-06 | 1.900E-06 | 4.900E-06 |
| 20786 | 1040 | V1040E | C | 2e-06 | 2.2e-06 | 2.5e-06 | 2.233E-06 | 2.517E-07 | 2.000E-06 | 2.500E-06 |
| 20787 | 1040 | V1040Q | C | 2.5e-06 | 1.7e-06 | 2.3e-06 | 2.167E-06 | 4.163E-07 | 1.700E-06 | 2.500E-06 |
| 20788 | 1040 | V1040G | C | 3.6e-06 | 4.5e-06 | 1.8e-06 | 3.300E-06 | 1.375E-06 | 1.800E-06 | 4.500E-06 |
| 20789 | 1040 | V1040H | C | 3.5e-06 | 2.1e-06 | 3e-06 | 2.867E-06 | 7.095E-07 | 2.100E-06 | 3.500E-06 |
| 20790 | 1040 | V1040I | C | 4e-06 | 3e-06 | 2e-06 | 3.000E-06 | 1.000E-06 | 2.000E-06 | 4.000E-06 |
| 20791 | 1040 | V1040L | C | 1.7e-06 | 4.2e-06 | 2.1e-06 | 2.667E-06 | 1.343E-06 | 1.700E-06 | 4.200E-06 |
| 20792 | 1040 | V1040K | C | 2.5e-06 | 2.9e-06 | 3.1e-06 | 2.833E-06 | 3.055E-07 | 2.500E-06 | 3.100E-06 |
| 20793 | 1040 | V1040M | C | 2.6e-06 | 2.1e-06 | 2.8e-06 | 2.500E-06 | 3.606E-07 | 2.100E-06 | 2.800E-06 |
| 20794 | 1040 | V1040F | C | 3.9e-06 | 2.4e-06 | 3.5e-06 | 3.267E-06 | 7.767E-07 | 2.400E-06 | 3.900E-06 |
| 20795 | 1040 | V1040P | C | 1.5e-06 | 2e-06 | 2.1e-06 | 1.867E-06 | 3.215E-07 | 1.500E-06 | 2.100E-06 |
| 20796 | 1040 | V1040S | C | 2e-06 | 3.2e-06 | 3.2e-06 | 2.800E-06 | 6.928E-07 | 2.000E-06 | 3.200E-06 |
| 20797 | 1040 | V1040T | C | 2e-06 | 1.6e-06 | 2.3e-06 | 1.967E-06 | 3.512E-07 | 1.600E-06 | 2.300E-06 |
| 20798 | 1040 | V1040W | C | 2.7e-06 | 1.2e-06 | 3.8e-06 | 2.567E-06 | 1.305E-06 | 1.200E-06 | 3.800E-06 |
| 20799 | 1040 | V1040Y | C | 5.1e-06 | 2.5e-06 | 2.3e-06 | 3.300E-06 | 1.562E-06 | 2.300E-06 | 5.100E-06 |
| 20800 | 1040 | V1040V | C | 5.4e-06 | 2.1e-06 | 1.2e-06 | 2.900E-06 | 2.211E-06 | 1.200E-06 | 5.400E-06 |
| 20801 | 1041 | N1041A | C | 4.6e-06 | 1.9e-06 | 1.7e-06 | 2.733E-06 | 1.620E-06 | 1.700E-06 | 4.600E-06 |
| 20802 | 1041 | N1041R | C | 3.4e-06 | 2.4e-06 | 2.1e-06 | 2.633E-06 | 6.807E-07 | 2.100E-06 | 3.400E-06 |
| 20803 | 1041 | N1041N | C | 1.8e-06 | 1.8e-06 | 2.8e-06 | 2.133E-06 | 5.774E-07 | 1.800E-06 | 2.800E-06 |
| 20804 | 1041 | N1041D | C | 1.9e-06 | 2.6e-06 | 3.1e-06 | 2.533E-06 | 6.028E-07 | 1.900E-06 | 3.100E-06 |
| 20805 | 1041 | N1041C | C | 2e-06 | 4.3e-06 | 3e-06 | 3.100E-06 | 1.153E-06 | 2.000E-06 | 4.300E-06 |
| 20806 | 1041 | N1041E | C | 2.4e-06 | 1.2e-06 | 2.9e-06 | 2.167E-06 | 8.737E-07 | 1.200E-06 | 2.900E-06 |
| 20807 | 1041 | N1041Q | C | 1.6e-06 | 3.2e-06 | 3.1e-06 | 2.633E-06 | 8.963E-07 | 1.600E-06 | 3.200E-06 |
| 20808 | 1041 | N1041G | C | 3.2e-06 | 2.6e-06 | 1.2e-06 | 2.333E-06 | 1.026E-06 | 1.200E-06 | 3.200E-06 |
| 20809 | 1041 | N1041H | C | 1.3e-06 | 2.3e-06 | 2.7e-06 | 2.100E-06 | 7.211E-07 | 1.300E-06 | 2.700E-06 |
| 20810 | 1041 | N1041I | C | 1.7e-06 | 2.8e-06 | 1.2e-06 | 1.900E-06 | 8.185E-07 | 1.200E-06 | 2.800E-06 |
| 20811 | 1041 | N1041L | C | 2e-06 | 2.9e-06 | 2.3e-06 | 2.400E-06 | 4.583E-07 | 2.000E-06 | 2.900E-06 |
| 20812 | 1041 | N1041K | C | 2.6e-06 | 2.1e-06 | 3.2e-06 | 2.633E-06 | 5.508E-07 | 2.100E-06 | 3.200E-06 |
| 20813 | 1041 | N1041M | C | 3.1e-06 | 2e-06 | 3.7e-06 | 2.933E-06 | 8.622E-07 | 2.000E-06 | 3.700E-06 |
| 20814 | 1041 | N1041F | C | 1.4e-06 | 2.2e-06 | 4.1e-06 | 2.567E-06 | 1.387E-06 | 1.400E-06 | 4.100E-06 |
| 20815 | 1041 | N1041P | C | 2.1e-06 | 3.2e-06 | 1.7e-06 | 2.333E-06 | 7.767E-07 | 1.700E-06 | 3.200E-06 |
| 20816 | 1041 | N1041S | C | 1.8e-06 | 2e-06 | 1.8e-06 | 1.867E-06 | 1.155E-07 | 1.800E-06 | 2.000E-06 |
| 20817 | 1041 | N1041T | C | 2.5e-06 | 3.2e-06 | 2.1e-06 | 2.600E-06 | 5.568E-07 | 2.100E-06 | 3.200E-06 |
| 20818 | 1041 | N1041W | C | 1.6e-06 | 4.4e-06 | 1.8e-06 | 2.600E-06 | 1.562E-06 | 1.600E-06 | 4.400E-06 |
| 20819 | 1041 | N1041Y | C | 1.9e-06 | 2e-06 | 1.8e-06 | 1.900E-06 | 1.000E-07 | 1.800E-06 | 2.000E-06 |
| 20820 | 1041 | N1041V | C | 2.4e-06 | 4.4e-06 | 2.7e-06 | 3.167E-06 | 1.079E-06 | 2.400E-06 | 4.400E-06 |
| 20821 | 1042 | H1042A | C | 4.6e-06 | 2.4e-06 | 3.2e-06 | 3.400E-06 | 1.114E-06 | 2.400E-06 | 4.600E-06 |
| 20822 | 1042 | H1042R | C | 5.8e-06 | 6.4e-06 | 3.1e-06 | 5.100E-06 | 1.758E-06 | 3.100E-06 | 6.400E-06 |
| 20823 | 1042 | H1042N | C | 2.5e-06 | 2.4e-06 | 2.9e-06 | 2.600E-06 | 2.646E-07 | 2.400E-06 | 2.900E-06 |
| 20824 | 1042 | H1042D | C | 2e-06 | 2e-06 | 1.4e-06 | 1.800E-06 | 3.464E-07 | 1.400E-06 | 2.000E-06 |

| | | | | | | | | | | |
|---|---|---|---|---|---|---|---|---|---|---|
| 20825 | 1042 | H1042C | C | 1.5e-06 | 2e-06 | 7.4e-07 | 1.413E-06 | 6.345E-07 | 7.400E-07 | 2.000E-06 |
| 20826 | 1042 | H1042E | C | 2.8e-06 | 2.1e-06 | 1e-06 | 1.967E-06 | 9.074E-07 | 1.000E-06 | 2.800E-06 |
| 20827 | 1042 | H1042Q | C | 1.6e-06 | 2.7e-06 | 2.6e-06 | 2.300E-06 | 6.083E-07 | 1.600E-06 | 2.700E-06 |
| 20828 | 1042 | H1042G | C | 3.7e-06 | 2.7e-06 | 3.7e-06 | 3.367E-06 | 5.774E-07 | 2.700E-06 | 3.700E-06 |
| 20829 | 1042 | H1042H | C | 2.9e-06 | 3.8e-06 | 2.6e-06 | 3.100E-06 | 6.245E-07 | 2.600E-06 | 3.800E-06 |
| 20830 | 1042 | H1042I | C | 2.1e-06 | 2.3e-06 | 2.4e-06 | 2.267E-06 | 1.528E-07 | 2.100E-06 | 2.400E-06 |
| 20831 | 1042 | H1042L | C | 1.7e-06 | 2.4e-06 | 2.4e-06 | 2.167E-06 | 4.041E-07 | 1.700E-06 | 2.400E-06 |
| 20832 | 1042 | H1042K | C | 1.6e-06 | 2.9e-06 | 1.9e-06 | 2.133E-06 | 6.807E-07 | 1.600E-06 | 2.900E-06 |
| 20833 | 1042 | H1042M | C | 1.5e-06 | 4.8e-06 | 2.8e-06 | 3.033E-06 | 1.662E-06 | 1.500E-06 | 4.800E-06 |
| 20834 | 1042 | H1042F | C | 1.9e-06 | 1.9e-06 | 1.8e-06 | 1.867E-06 | 5.774E-08 | 1.800E-06 | 1.900E-06 |
| 20835 | 1042 | H1042P | C | 4.2e-06 | 2.5e-06 | 2.5e-06 | 3.067E-06 | 9.815E-07 | 2.500E-06 | 4.200E-06 |
| 20836 | 1042 | H1042S | C | 2.4e-06 | 3.2e-06 | 3e-06 | 2.867E-06 | 4.163E-07 | 2.400E-06 | 3.200E-06 |
| 20837 | 1042 | H1042T | C | 2.1e-06 | 2.4e-06 | 2.9e-06 | 2.467E-06 | 4.041E-07 | 2.100E-06 | 2.900E-06 |
| 20838 | 1042 | H1042W | C | 2.9e-06 | 9.3e-07 | 1.8e-06 | 1.877E-06 | 9.872E-07 | 9.300E-07 | 2.900E-06 |
| 20839 | 1042 | H1042Y | C | 3.6e-06 | 2e-06 | 4.6e-06 | 3.400E-06 | 1.311E-06 | 2.000E-06 | 4.600E-06 |
| 20840 | 1042 | H1042V | C | 2.5e-06 | 1.9e-06 | 2.2e-06 | 2.200E-06 | 3.000E-07 | 1.900E-06 | 2.500E-06 |
| 20841 | 1043 | K1043A | C | 1.9e-06 | 2e-06 | 4.1e-06 | 2.667E-06 | 1.242E-06 | 1.900E-06 | 4.100E-06 |
| 20842 | 1043 | K1043R | C | 1.7e-06 | 1.7e-06 | 2.3e-06 | 1.900E-06 | 3.464E-07 | 1.700E-06 | 2.300E-06 |
| 20843 | 1043 | K1043N | C | 2.1e-06 | 9.5e-07 | 1.9e-06 | 1.650E-06 | 6.144E-07 | 9.500E-07 | 2.100E-06 |
| 20844 | 1043 | K1043D | C | 1.5e-06 | 1.8e-06 | 5.6e-06 | 2.967E-06 | 2.285E-06 | 1.500E-06 | 5.600E-06 |
| 20845 | 1043 | K1043C | C | 2.4e-06 | 2.2e-06 | 3e-06 | 2.533E-06 | 4.163E-07 | 2.200E-06 | 3.000E-06 |
| 20846 | 1043 | K1043E | C | 3.8e-06 | 2.5e-06 | 2.7e-06 | 3.000E-06 | 7.000E-07 | 2.500E-06 | 3.800E-06 |
| 20847 | 1043 | K1043Q | C | 2.3e-06 | 1.9e-06 | 2.5e-06 | 2.233E-06 | 3.055E-07 | 1.900E-06 | 2.500E-06 |
| 20848 | 1043 | K1043G | C | 1.4e-06 | 1.7e-06 | 3e-06 | 2.033E-06 | 8.505E-07 | 1.400E-06 | 3.000E-06 |
| 20849 | 1043 | K1043H | C | 3e-06 | 1.5e-06 | 2.2e-06 | 2.233E-06 | 7.506E-07 | 1.500E-06 | 3.000E-06 |
| 20850 | 1043 | K1043I | C | 2.2e-06 | 4.3e-06 | 2.7e-06 | 3.067E-06 | 1.097E-06 | 2.200E-06 | 4.300E-06 |
| 20851 | 1043 | K1043L | C | 2e-06 | 1.6e-06 | 3.3e-06 | 2.300E-06 | 8.888E-07 | 1.600E-06 | 3.300E-06 |
| 20852 | 1043 | K1043K | C | 2e-06 | 2.5e-06 | 1.8e-06 | 2.100E-06 | 3.606E-07 | 1.800E-06 | 2.500E-06 |
| 20853 | 1043 | K1043M | C | 2.8e-06 | 2.3e-06 | 3.4e-06 | 2.833E-06 | 5.508E-07 | 2.300E-06 | 3.400E-06 |
| 20854 | 1043 | K1043F | C | 1.8e-06 | 3.1e-06 | 1.4e-06 | 2.100E-06 | 8.888E-07 | 1.400E-06 | 3.100E-06 |
| 20855 | 1043 | K1043P | C | 2.3e-06 | 2.8e-06 | 1.6e-06 | 2.233E-06 | 6.028E-07 | 1.600E-06 | 2.800E-06 |
| 20856 | 1043 | K1043S | C | 2.4e-06 | 3.6e-06 | 1.5e-06 | 2.500E-06 | 1.054E-06 | 1.500E-06 | 3.600E-06 |
| 20857 | 1043 | K1043T | C | 2.3e-06 | 3.8e-06 | 2.3e-06 | 2.800E-06 | 8.660E-07 | 2.300E-06 | 3.800E-06 |
| 20858 | 1043 | K1043W | C | 2.1e-06 | 2.5e-06 | 1.4e-06 | 2.000E-06 | 5.568E-07 | 1.400E-06 | 2.500E-06 |
| 20859 | 1043 | K1043Y | C | 1.2e-06 | 1.9e-06 | 2.6e-06 | 1.900E-06 | 7.000E-07 | 1.200E-06 | 2.600E-06 |
| 20860 | 1043 | K1043V | C | 3.6e-06 | 2.4e-06 | 2.7e-06 | 2.900E-06 | 6.245E-07 | 2.400E-06 | 3.600E-06 |
| 20861 | 1044 | P1044A | C | 3.6e-06 | 2.5e-06 | 5.9e-06 | 4.000E-06 | 1.735E-06 | 2.500E-06 | 5.900E-06 |
| 20862 | 1044 | P1044R | C | 2.3e-06 | 3.1e-06 | 2e-06 | 2.467E-06 | 5.686E-07 | 2.000E-06 | 3.100E-06 |
| 20863 | 1044 | P1044N | C | 2.2e-06 | 2.9e-06 | 2.4e-06 | 2.500E-06 | 3.606E-07 | 2.200E-06 | 2.900E-06 |
| 20864 | 1044 | P1044D | C | 3.4e-06 | 1.8e-06 | 2.8e-06 | 2.667E-06 | 8.083E-07 | 1.800E-06 | 3.400E-06 |
| 20865 | 1044 | P1044C | C | 3.2e-06 | 2.5e-06 | 2.1e-06 | 2.600E-06 | 5.568E-07 | 2.100E-06 | 3.200E-06 |
| 20866 | 1044 | P1044E | C | 2.1e-06 | 3.1e-06 | 1.8e-06 | 2.333E-06 | 6.807E-07 | 1.800E-06 | 3.100E-06 |
| 20867 | 1044 | P1044Q | C | 2.1e-06 | 3.9e-06 | 3.1e-06 | 3.033E-06 | 9.018E-07 | 2.100E-06 | 3.900E-06 |
| 20868 | 1044 | P1044G | C | 2.2e-06 | 2.3e-06 | 2.9e-06 | 2.467E-06 | 3.786E-07 | 2.200E-06 | 2.900E-06 |
| 20869 | 1044 | P1044H | C | 3.7e-06 | 2.6e-06 | 3e-06 | 3.100E-06 | 5.568E-07 | 2.600E-06 | 3.700E-06 |
| 20870 | 1044 | P1044I | C | 1.6e-06 | 2.6e-06 | 2.5e-06 | 2.233E-06 | 5.508E-07 | 1.600E-06 | 2.600E-06 |
| 20871 | 1044 | P1044L | C | 1.3e-06 | 2.7e-06 | 1.1e-06 | 1.700E-06 | 8.718E-07 | 1.100E-06 | 2.700E-06 |
| 20872 | 1044 | P1044K | C | 2.5e-06 | 2.5e-06 | 2.2e-06 | 2.400E-06 | 1.732E-07 | 2.200E-06 | 2.500E-06 |
| 20873 | 1044 | P1044M | C | 2.7e-06 | 4.2e-06 | 2.2e-06 | 3.033E-06 | 1.041E-06 | 2.200E-06 | 4.200E-06 |
| 20874 | 1044 | P1044F | C | 1.2e-06 | 1.7e-06 | 5.3e-06 | 2.733E-06 | 2.237E-06 | 1.200E-06 | 5.300E-06 |
| 20875 | 1044 | P1044P | C | 1.9e-06 | 2.2e-06 | 3.5e-06 | 2.533E-06 | 8.505E-07 | 1.900E-06 | 3.500E-06 |
| 20876 | 1044 | P1044S | C | 3.3e-06 | 2.1e-06 | 2.4e-06 | 2.600E-06 | 6.245E-07 | 2.100E-06 | 3.300E-06 |
| 20877 | 1044 | P1044T | C | 2.5e-06 | 2.7e-06 | 2.1e-06 | 2.433E-06 | 3.055E-07 | 2.100E-06 | 2.700E-06 |
| 20878 | 1044 | P1044W | C | 1.7e-06 | 1.6e-06 | 1.9e-06 | 1.733E-06 | 1.528E-07 | 1.600E-06 | 1.900E-06 |
| 20879 | 1044 | P1044Y | C | 2.2e-06 | 2e-06 | 1.4e-06 | 1.867E-06 | 4.163E-07 | 1.400E-06 | 2.200E-06 |
| 20880 | 1044 | P1044V | C | 3.4e-06 | 1.7e-06 | 3.3e-06 | 2.800E-06 | 9.539E-07 | 1.700E-06 | 3.400E-06 |
| 20881 | 1045 | S1045A | C | 1.9e-06 | 2e-06 | 3.2e-06 | 2.367E-06 | 7.234E-07 | 1.900E-06 | 3.200E-06 |
| 20882 | 1045 | S1045R | C | 2.4e-06 | 2.9e-06 | 4.4e-06 | 3.233E-06 | 1.041E-06 | 2.400E-06 | 4.400E-06 |
| 20883 | 1045 | S1045N | C | 4.1e-06 | 1.1e-06 | 1.9e-06 | 2.367E-06 | 1.553E-06 | 1.100E-06 | 4.100E-06 |
| 20884 | 1045 | S1045D | C | 3.6e-06 | 2e-06 | 2.6e-06 | 2.733E-06 | 8.083E-07 | 2.000E-06 | 3.600E-06 |
| 20885 | 1045 | S1045C | C | 1e-06 | 3.1e-06 | 1.6e-06 | 1.900E-06 | 1.082E-06 | 1.000E-06 | 3.100E-06 |
| 20886 | 1045 | S1045E | C | 3.6e-06 | 2.9e-06 | 1.7e-06 | 2.733E-06 | 9.609E-07 | 1.700E-06 | 3.600E-06 |
| 20887 | 1045 | S1045Q | C | 4.8e-06 | 2.2e-06 | 2e-06 | 3.000E-06 | 1.562E-06 | 2.000E-06 | 4.800E-06 |
| 20888 | 1045 | S1045G | C | 2.8e-06 | 2e-06 | 4.1e-06 | 2.967E-06 | 1.060E-06 | 2.000E-06 | 4.100E-06 |
| 20889 | 1045 | S1045H | C | 2.8e-06 | 2.1e-06 | 2.7e-06 | 2.533E-06 | 3.786E-07 | 2.100E-06 | 2.800E-06 |
| 20890 | 1045 | S1045I | C | 2.7e-06 | 3.3e-06 | 2.3e-06 | 2.767E-06 | 5.033E-07 | 2.300E-06 | 3.300E-06 |
| 20891 | 1045 | S1045L | C | 2.6e-06 | 3e-06 | 4.1e-06 | 3.233E-06 | 7.767E-07 | 2.600E-06 | 4.100E-06 |
| 20892 | 1045 | S1045K | C | 1.5e-06 | 2.5e-06 | 1.8e-06 | 1.933E-06 | 5.132E-07 | 1.500E-06 | 2.500E-06 |
| 20893 | 1045 | S1045M | C | 2.9e-06 | 2.2e-06 | 2.1e-06 | 2.400E-06 | 4.359E-07 | 2.100E-06 | 2.900E-06 |
| 20894 | 1045 | S1045F | C | 2.7e-06 | 1.3e-06 | 2.3e-06 | 2.100E-06 | 7.211E-07 | 1.300E-06 | 2.700E-06 |
| 20895 | 1045 | S1045P | C | 1.2e-06 | 1.9e-06 | 1.5e-06 | 1.533E-06 | 3.512E-07 | 1.200E-06 | 1.900E-06 |
| 20896 | 1045 | S1045S | C | 4.6e-06 | 3.1e-06 | 2.6e-06 | 3.433E-06 | 1.041E-06 | 2.600E-06 | 4.600E-06 |
| 20897 | 1045 | S1045T | C | 3.3e-06 | 3.1e-06 | 2.4e-06 | 2.933E-06 | 4.726E-07 | 2.400E-06 | 3.300E-06 |
| 20898 | 1045 | S1045W | C | 1.5e-06 | 3.3e-06 | 2.1e-06 | 2.300E-06 | 9.165E-07 | 1.500E-06 | 3.300E-06 |
| 20899 | 1045 | S1045Y | C | 2.7e-06 | 1.5e-06 | 3.3e-06 | 2.500E-06 | 9.165E-07 | 1.500E-06 | 3.300E-06 |
| 20900 | 1045 | S1045V | C | 3.1e-06 | 1.5e-06 | 2.2e-06 | 2.267E-06 | 8.021E-07 | 1.500E-06 | 3.100E-06 |
| 20901 | 1046 | N1046A | C | 2.7e-06 | 6.9e-06 | 2.4e-06 | 4.000E-06 | 2.516E-06 | 2.400E-06 | 6.900E-06 |
| 20902 | 1046 | N1046R | C | 3.6e-06 | 3.2e-06 | 1.9e-06 | 2.900E-06 | 8.888E-07 | 1.900E-06 | 3.600E-06 |
| 20903 | 1046 | N1046N | C | 2.2e-06 | 1.8e-06 | 2.6e-06 | 2.200E-06 | 4.000E-07 | 1.800E-06 | 2.600E-06 |
| 20904 | 1046 | N1046D | C | 3.7e-06 | 2.7e-06 | 1.3e-06 | 2.567E-06 | 1.206E-06 | 1.300E-06 | 3.700E-06 |
| 20905 | 1046 | N1046C | C | 3.3e-06 | 3.2e-06 | 2e-06 | 2.833E-06 | 7.234E-07 | 2.000E-06 | 3.300E-06 |
| 20906 | 1046 | N1046E | C | 2.7e-06 | 4.5e-06 | 4e-06 | 3.733E-06 | 9.292E-07 | 2.700E-06 | 4.500E-06 |
| 20907 | 1046 | N1046Q | C | 1.6e-06 | 2e-06 | 1.5e-06 | 1.700E-06 | 2.646E-07 | 1.500E-06 | 2.000E-06 |
| 20908 | 1046 | N1046G | C | 1.4e-06 | 3.9e-06 | 3.9e-06 | 3.067E-06 | 1.443E-06 | 1.400E-06 | 3.900E-06 |
| 20909 | 1046 | N1046H | C | 1.4e-06 | 3.9e-06 | 2e-06 | 2.433E-06 | 1.305E-06 | 1.400E-06 | 3.900E-06 |

| | | | | | | | | | | |
|---|---|---|---|---|---|---|---|---|---|---|
| 20910 | 1046 | N1046I | C | 1.3e-06 | 1.9e-06 | 2.3e-06 | 1.833E-06 | 5.033E-07 | 1.300E-06 | 2.300E-06 |
| 20911 | 1046 | N1046L | C | 3.1e-06 | 2.4e-06 | 2.8e-06 | 2.767E-06 | 3.512E-07 | 2.400E-06 | 3.100E-06 |
| 20912 | 1046 | N1046K | C | 2.1e-06 | 2.5e-06 | 1.8e-06 | 2.133E-06 | 3.512E-07 | 1.800E-06 | 2.500E-06 |
| 20913 | 1046 | N1046M | C | 2.3e-06 | 2.1e-06 | 5.5e-06 | 3.300E-06 | 1.908E-06 | 2.100E-06 | 5.500E-06 |
| 20914 | 1046 | N1046F | C | 2.2e-06 | 1.8e-06 | 2.4e-06 | 2.133E-06 | 3.055E-07 | 1.800E-06 | 2.400E-06 |
| 20915 | 1046 | N1046P | C | 2e-06 | 2.4e-06 | 2.4e-06 | 2.267E-06 | 2.309E-07 | 2.000E-06 | 2.400E-06 |
| 20916 | 1046 | N1046S | C | 3.1e-06 | 2.8e-06 | 2.4e-06 | 2.767E-06 | 3.512E-07 | 2.400E-06 | 3.100E-06 |
| 20917 | 1046 | N1046T | C | 1.8e-06 | 2e-06 | 7.6e-07 | 1.520E-06 | 6.657E-07 | 7.600E-07 | 2.000E-06 |
| 20918 | 1046 | N1046W | C | 2.9e-06 | 1.5e-06 | 3.2e-06 | 2.533E-06 | 9.074E-07 | 1.500E-06 | 3.200E-06 |
| 20919 | 1046 | N1046Y | C | 1.8e-06 | 2.4e-06 | 2.9e-06 | 2.367E-06 | 5.508E-07 | 1.800E-06 | 2.900E-06 |
| 20920 | 1046 | N1046V | C | 2.1e-06 | 2.7e-06 | 1.6e-06 | 2.133E-06 | 5.508E-07 | 1.600E-06 | 2.700E-06 |
| 20921 | 1047 | T1047A | C | 2e-06 | 2.8e-06 | 2.3e-06 | 2.367E-06 | 4.041E-07 | 2.000E-06 | 2.800E-06 |
| 20922 | 1047 | T1047R | C | 4e-06 | 2.5e-06 | 2.4e-06 | 2.967E-06 | 8.963E-07 | 2.400E-06 | 4.000E-06 |
| 20923 | 1047 | T1047N | C | 1.1e-06 | 2.4e-06 | 2.6e-06 | 2.033E-06 | 8.145E-07 | 1.100E-06 | 2.600E-06 |
| 20924 | 1047 | T1047D | C | 2e-06 | 4.5e-06 | 3.7e-06 | 3.400E-06 | 1.277E-06 | 2.000E-06 | 4.500E-06 |
| 20925 | 1047 | T1047C | C | 2.2e-06 | 2.6e-06 | 1.4e-06 | 2.067E-06 | 6.110E-07 | 1.400E-06 | 2.600E-06 |
| 20926 | 1047 | T1047E | C | 2.9e-06 | 2.9e-06 | 3.9e-06 | 3.233E-06 | 5.774E-07 | 2.900E-06 | 3.900E-06 |
| 20927 | 1047 | T1047Q | C | 2.6e-06 | 3.9e-06 | 1.9e-06 | 2.800E-06 | 1.015E-06 | 1.900E-06 | 3.900E-06 |
| 20928 | 1047 | T1047G | C | 1.5e-06 | 1.3e-06 | 2.1e-06 | 1.633E-06 | 4.163E-07 | 1.300E-06 | 2.100E-06 |
| 20929 | 1047 | T1047H | C | 2.3e-06 | 1e-06 | 2.6e-06 | 1.967E-06 | 8.505E-07 | 1.000E-06 | 2.600E-06 |
| 20930 | 1047 | T1047I | C | 2.4e-06 | 1.5e-06 | 2.4e-06 | 2.100E-06 | 5.196E-07 | 1.500E-06 | 2.400E-06 |
| 20931 | 1047 | T1047L | C | 2.6e-06 | 1.5e-06 | 3.5e-06 | 2.533E-06 | 1.002E-06 | 1.500E-06 | 3.500E-06 |
| 20932 | 1047 | T1047K | C | 1.1e-06 | 1.9e-06 | 3.2e-06 | 2.067E-06 | 1.060E-06 | 1.100E-06 | 3.200E-06 |
| 20933 | 1047 | T1047M | C | 2.4e-06 | 3.3e-06 | 2.3e-06 | 2.667E-06 | 5.508E-07 | 2.300E-06 | 3.300E-06 |
| 20934 | 1047 | T1047F | C | 2.8e-06 | 3.3e-06 | 2e-06 | 2.700E-06 | 6.557E-07 | 2.000E-06 | 3.300E-06 |
| 20935 | 1047 | T1047P | C | 2.3e-06 | 2.4e-06 | 2.8e-06 | 2.500E-06 | 2.646E-07 | 2.300E-06 | 2.800E-06 |
| 20936 | 1047 | T1047S | C | 3.7e-06 | 4.8e-06 | 2.8e-06 | 3.767E-06 | 1.002E-06 | 2.800E-06 | 4.800E-06 |
| 20937 | 1047 | T1047T | C | 1.9e-06 | 1.9e-06 | 1.1e-06 | 1.633E-06 | 4.619E-07 | 1.100E-06 | 1.900E-06 |
| 20938 | 1047 | T1047W | C | 1.9e-06 | 2.6e-06 | 2.4e-06 | 2.300E-06 | 3.606E-07 | 1.900E-06 | 2.600E-06 |
| 20939 | 1047 | T1047Y | C | 1.7e-06 | 1.7e-06 | 2.6e-06 | 2.000E-06 | 5.196E-07 | 1.700E-06 | 2.600E-06 |
| 20940 | 1047 | T1047V | C | 3.6e-06 | 3e-06 | 1.6e-06 | 2.733E-06 | 1.026E-06 | 1.600E-06 | 3.600E-06 |
| 20941 | 1048 | K1048A | C | 2.7e-06 | 3.2e-06 | 2.2e-06 | 2.700E-06 | 5.000E-07 | 2.200E-06 | 3.200E-06 |
| 20942 | 1048 | K1048R | C | 1.8e-06 | 3.2e-06 | 2.9e-06 | 2.633E-06 | 7.371E-07 | 1.800E-06 | 3.200E-06 |
| 20943 | 1048 | K1048N | C | 3.4e-06 | 1.6e-06 | 4e-06 | 3.000E-06 | 1.249E-06 | 1.600E-06 | 4.000E-06 |
| 20944 | 1048 | K1048D | C | 1.6e-06 | 2.7e-06 | 2.3e-06 | 2.200E-06 | 5.568E-07 | 1.600E-06 | 2.700E-06 |
| 20945 | 1048 | K1048C | C | 2.8e-06 | 1.7e-06 | 2.3e-06 | 2.267E-06 | 5.508E-07 | 1.700E-06 | 2.800E-06 |
| 20946 | 1048 | K1048E | C | 2e-06 | 2e-06 | 2.6e-06 | 2.200E-06 | 3.464E-07 | 2.000E-06 | 2.600E-06 |
| 20947 | 1048 | K1048Q | C | 1.4e-06 | 2.1e-06 | 4.7e-06 | 2.733E-06 | 1.739E-06 | 1.400E-06 | 4.700E-06 |
| 20948 | 1048 | K1048G | C | 2.4e-06 | 2.6e-06 | 3.1e-06 | 2.700E-06 | 3.606E-07 | 2.400E-06 | 3.100E-06 |
| 20949 | 1048 | K1048H | C | 1.5e-06 | 1.9e-06 | 2.1e-06 | 1.833E-06 | 3.055E-07 | 1.500E-06 | 2.100E-06 |
| 20950 | 1048 | K1048I | C | 4.3e-06 | 2.5e-06 | 2.7e-06 | 3.167E-06 | 9.866E-07 | 2.500E-06 | 4.300E-06 |
| 20951 | 1048 | K1048L | C | 2.1e-06 | 2.6e-06 | 2.1e-06 | 2.267E-06 | 2.887E-07 | 2.100E-06 | 2.600E-06 |
| 20952 | 1048 | K1048K | C | 3.4e-06 | 1.8e-06 | 2.7e-06 | 2.633E-06 | 8.021E-07 | 1.800E-06 | 3.400E-06 |
| 20953 | 1048 | K1048M | C | 3.8e-06 | 2.4e-06 | 2.1e-06 | 2.767E-06 | 9.074E-07 | 2.100E-06 | 3.800E-06 |
| 20954 | 1048 | K1048F | C | 1.7e-06 | 2.8e-06 | 3.1e-06 | 2.533E-06 | 7.371E-07 | 1.700E-06 | 3.100E-06 |
| 20955 | 1048 | K1048P | C | 5.3e-06 | 2e-06 | 3e-06 | 3.433E-06 | 1.692E-06 | 2.000E-06 | 5.300E-06 |
| 20956 | 1048 | K1048S | C | 1.8e-06 | 2.5e-06 | 3.9e-06 | 2.733E-06 | 1.069E-06 | 1.800E-06 | 3.900E-06 |
| 20957 | 1048 | K1048T | C | 2.1e-06 | 1.7e-06 | 5.4e-06 | 3.067E-06 | 2.031E-06 | 1.700E-06 | 5.400E-06 |
| 20958 | 1048 | K1048W | C | 8.6e-07 | 2.2e-06 | 2.1e-06 | 1.720E-06 | 7.465E-07 | 8.600E-07 | 2.200E-06 |
| 20959 | 1048 | K1048Y | C | 2.9e-06 | 3.1e-06 | 2e-06 | 2.667E-06 | 5.859E-07 | 2.000E-06 | 3.100E-06 |
| 20960 | 1048 | K1048V | C | 2.1e-06 | 1.7e-06 | 1.6e-06 | 1.800E-06 | 2.646E-07 | 1.600E-06 | 2.100E-06 |
| 20961 | 1049 | V1049A | C | 1.3e-06 | 2.9e-06 | 2.4e-06 | 2.200E-06 | 8.185E-07 | 1.300E-06 | 2.900E-06 |
| 20962 | 1049 | V1049R | C | 3.3e-06 | 2.5e-06 | 3e-06 | 2.933E-06 | 4.041E-07 | 2.500E-06 | 3.300E-06 |
| 20963 | 1049 | V1049N | C | 2.5e-06 | 1.7e-06 | 3.1e-06 | 2.433E-06 | 7.024E-07 | 1.700E-06 | 3.100E-06 |
| 20964 | 1049 | V1049D | C | 3.8e-06 | 1.3e-06 | 1.7e-06 | 2.267E-06 | 1.343E-06 | 1.300E-06 | 3.800E-06 |
| 20965 | 1049 | V1049C | C | 1.5e-06 | 2.1e-06 | 3e-06 | 2.200E-06 | 7.550E-07 | 1.500E-06 | 3.000E-06 |
| 20966 | 1049 | V1049E | C | 2.5e-06 | 1.5e-06 | 2.7e-06 | 2.233E-06 | 6.429E-07 | 1.500E-06 | 2.700E-06 |
| 20967 | 1049 | V1049Q | C | 1.6e-06 | 1.8e-06 | 1.6e-06 | 1.667E-06 | 1.155E-07 | 1.600E-06 | 1.800E-06 |
| 20968 | 1049 | V1049G | C | 3.4e-06 | 3.3e-06 | 1.2e-06 | 2.633E-06 | 1.242E-06 | 1.200E-06 | 3.400E-06 |
| 20969 | 1049 | V1049H | C | 3e-06 | 2.2e-06 | 2.6e-06 | 2.600E-06 | 4.000E-07 | 2.200E-06 | 3.000E-06 |
| 20970 | 1049 | V1049I | C | 1.6e-06 | 3.1e-06 | 2.1e-06 | 2.267E-06 | 7.638E-07 | 1.600E-06 | 3.100E-06 |
| 20971 | 1049 | V1049L | C | 2.6e-06 | 2.5e-06 | 3e-06 | 2.700E-06 | 2.646E-07 | 2.500E-06 | 3.000E-06 |
| 20972 | 1049 | V1049K | C | 3.1e-06 | 3e-06 | 1.5e-06 | 2.533E-06 | 8.963E-07 | 1.500E-06 | 3.100E-06 |
| 20973 | 1049 | V1049M | C | 2.1e-06 | 2.4e-06 | 1.6e-06 | 2.033E-06 | 4.041E-07 | 1.600E-06 | 2.400E-06 |
| 20974 | 1049 | V1049F | C | 3.5e-06 | 1.6e-06 | 3.2e-06 | 2.767E-06 | 1.021E-06 | 1.600E-06 | 3.500E-06 |
| 20975 | 1049 | V1049P | C | 2.5e-06 | 2.2e-06 | 2.5e-06 | 2.400E-06 | 1.732E-07 | 2.200E-06 | 2.500E-06 |
| 20976 | 1049 | V1049S | C | 2.9e-06 | 3.2e-06 | 2.3e-06 | 2.800E-06 | 4.583E-07 | 2.300E-06 | 3.200E-06 |
| 20977 | 1049 | V1049T | C | 1.4e-06 | 1.9e-06 | 2.6e-06 | 1.967E-06 | 6.028E-07 | 1.400E-06 | 2.600E-06 |
| 20978 | 1049 | V1049W | C | 2.7e-06 | 1.7e-06 | 1.9e-06 | 2.100E-06 | 5.292E-07 | 1.700E-06 | 2.700E-06 |
| 20979 | 1049 | V1049Y | C | 2.8e-06 | 1.9e-06 | 2.8e-06 | 2.500E-06 | 5.196E-07 | 1.900E-06 | 2.800E-06 |
| 20980 | 1049 | V1049V | C | 2.1e-06 | 1.6e-06 | 9.7e-07 | 1.557E-06 | 5.662E-07 | 9.700E-07 | 2.100E-06 |
| 20981 | 1050 | D1050A | C | 3.7e-06 | 2.8e-06 | 3e-06 | 3.167E-06 | 4.726E-07 | 2.800E-06 | 3.700E-06 |
| 20982 | 1050 | D1050R | C | 3.2e-06 | 2.7e-06 | 2e-06 | 2.633E-06 | 6.028E-07 | 2.000E-06 | 3.200E-06 |
| 20983 | 1050 | D1050N | C | 1.7e-06 | 2.4e-06 | 2.6e-06 | 2.233E-06 | 4.726E-07 | 1.700E-06 | 2.600E-06 |
| 20984 | 1050 | D1050D | C | 3.5e-06 | 4.6e-06 | 2.3e-06 | 3.467E-06 | 1.150E-06 | 2.300E-06 | 4.600E-06 |
| 20985 | 1050 | D1050C | C | 3.8e-06 | 2.1e-06 | 3.8e-06 | 3.233E-06 | 9.815E-07 | 2.100E-06 | 3.800E-06 |
| 20986 | 1050 | D1050E | C | 1.5e-06 | 2.1e-06 | 2.1e-06 | 1.900E-06 | 3.464E-07 | 1.500E-06 | 2.100E-06 |
| 20987 | 1050 | D1050Q | C | 2.3e-06 | 2.8e-06 | 2.9e-06 | 2.667E-06 | 3.215E-07 | 2.300E-06 | 2.900E-06 |
| 20988 | 1050 | D1050G | C | 2.3e-06 | 1.7e-06 | 2.4e-06 | 2.133E-06 | 3.786E-07 | 1.700E-06 | 2.400E-06 |
| 20989 | 1050 | D1050H | C | 2.8e-06 | 2.6e-06 | 2.5e-06 | 2.633E-06 | 1.528E-07 | 2.500E-06 | 2.800E-06 |
| 20990 | 1050 | D1050I | C | 2.9e-06 | 3e-06 | 6.2e-06 | 4.033E-06 | 1.877E-06 | 2.900E-06 | 6.200E-06 |
| 20991 | 1050 | D1050L | C | 2e-06 | 2.2e-06 | 1.7e-06 | 1.967E-06 | 2.517E-07 | 1.700E-06 | 2.200E-06 |
| 20992 | 1050 | D1050K | C | 2.8e-06 | 2.2e-06 | 3.2e-06 | 2.733E-06 | 5.033E-07 | 2.200E-06 | 3.200E-06 |
| 20993 | 1050 | D1050M | C | 3e-06 | 1.3e-06 | 1.9e-06 | 2.067E-06 | 8.622E-07 | 1.300E-06 | 3.000E-06 |
| 20994 | 1050 | D1050F | C | 2.2e-06 | 1.6e-06 | 3.2e-06 | 2.333E-06 | 8.083E-07 | 1.600E-06 | 3.200E-06 |

| | | | | | | | | | | |
|---|---|---|---|---|---|---|---|---|---|---|
| 20995 | 1050 | D1050P | C | 2.9e-06 | 2.2e-06 | 1.9e-06 | 2.333E-06 | 5.132E-07 | 1.900E-06 | 2.900E-06 |
| 20996 | 1050 | D1050S | C | 2.1e-06 | 1.7e-06 | 2.3e-06 | 2.033E-06 | 3.055E-07 | 1.700E-06 | 2.300E-06 |
| 20997 | 1050 | D1050T | C | 3.4e-06 | 2.2e-06 | 3.1e-06 | 2.900E-06 | 6.245E-07 | 2.200E-06 | 3.400E-06 |
| 20998 | 1050 | D1050W | C | 8.1e-07 | 3.6e-06 | 3.4e-06 | 2.603E-06 | 1.556E-06 | 8.100E-07 | 3.600E-06 |
| 20999 | 1050 | D1050Y | C | 1.3e-06 | 2.3e-06 | 2.3e-06 | 1.967E-06 | 5.774E-07 | 1.300E-06 | 2.300E-06 |
| 21000 | 1050 | D1050V | C | 3.4e-06 | 2.6e-06 | 2.4e-06 | 2.800E-06 | 5.292E-07 | 2.400E-06 | 3.400E-06 |
| 21001 | 1051 | K1051A | C | 2e-06 | 1.9e-06 | 2.3e-06 | 2.067E-06 | 2.082E-07 | 1.900E-06 | 2.300E-06 |
| 21002 | 1051 | K1051R | C | 3.3e-06 | 2.4e-06 | 2.5e-06 | 2.733E-06 | 4.933E-07 | 2.400E-06 | 3.300E-06 |
| 21003 | 1051 | K1051N | C | 1.8e-06 | 1.8e-06 | 3.1e-06 | 2.233E-06 | 7.506E-07 | 1.800E-06 | 3.100E-06 |
| 21004 | 1051 | K1051D | C | 3.6e-06 | 3.7e-06 | 1.4e-06 | 2.900E-06 | 1.300E-06 | 1.400E-06 | 3.700E-06 |
| 21005 | 1051 | K1051C | C | 1.5e-06 | 2.1e-06 | 2.7e-06 | 2.100E-06 | 6.000E-07 | 1.500E-06 | 2.700E-06 |
| 21006 | 1051 | K1051E | C | 2.2e-06 | 3.7e-06 | 1.4e-06 | 2.433E-06 | 1.168E-06 | 1.400E-06 | 3.700E-06 |
| 21007 | 1051 | K1051Q | C | 1.1e-06 | 1.8e-06 | 1.5e-06 | 1.467E-06 | 3.512E-07 | 1.100E-06 | 1.800E-06 |
| 21008 | 1051 | K1051G | C | 1.9e-06 | 2.4e-06 | 2.8e-06 | 2.367E-06 | 4.509E-07 | 1.900E-06 | 2.800E-06 |
| 21009 | 1051 | K1051H | C | 2.2e-06 | 1.1e-06 | 1.5e-06 | 1.600E-06 | 5.568E-07 | 1.100E-06 | 2.200E-06 |
| 21010 | 1051 | K1051I | C | 2.7e-06 | 2e-06 | 3.8e-06 | 2.833E-06 | 9.074E-07 | 2.000E-06 | 3.800E-06 |
| 21011 | 1051 | K1051L | C | 3.5e-06 | 2.2e-06 | 2.3e-06 | 2.667E-06 | 7.234E-07 | 2.200E-06 | 3.500E-06 |
| 21012 | 1051 | K1051K | C | 2.8e-06 | 3e-06 | 2.1e-06 | 2.633E-06 | 4.726E-07 | 2.100E-06 | 3.000E-06 |
| 21013 | 1051 | K1051M | C | 2.2e-06 | 2.1e-06 | 2.6e-06 | 2.300E-06 | 2.646E-07 | 2.100E-06 | 2.600E-06 |
| 21014 | 1051 | K1051F | C | 2.4e-06 | 2.6e-06 | 1.8e-06 | 2.267E-06 | 4.163E-07 | 1.800E-06 | 2.600E-06 |
| 21015 | 1051 | K1051P | C | 2.3e-06 | 3.6e-06 | 3e-06 | 2.967E-06 | 6.506E-07 | 2.300E-06 | 3.600E-06 |
| 21016 | 1051 | K1051S | C | 1.1e-06 | 3.2e-06 | 2.3e-06 | 2.200E-06 | 1.054E-06 | 1.100E-06 | 3.200E-06 |
| 21017 | 1051 | K1051T | C | 4.6e-06 | 1.9e-06 | 8.7e-07 | 2.457E-06 | 1.926E-06 | 8.700E-07 | 4.600E-06 |
| 21018 | 1051 | K1051W | C | 2.3e-06 | 2.1e-06 | 1.9e-06 | 2.100E-06 | 2.000E-07 | 1.900E-06 | 2.300E-06 |
| 21019 | 1051 | K1051Y | C | 2e-06 | 2.8e-06 | 2.4e-06 | 2.400E-06 | 4.000E-07 | 2.000E-06 | 2.800E-06 |
| 21020 | 1051 | K1051V | C | 2.8e-06 | 2.5e-06 | 3.2e-06 | 2.833E-06 | 3.512E-07 | 2.500E-06 | 3.200E-06 |
| 21021 | 1052 | K1052A | C | 3.6e-06 | 3e-06 | 1.8e-06 | 2.800E-06 | 9.165E-07 | 1.800E-06 | 3.600E-06 |
| 21022 | 1052 | K1052R | C | 1.8e-06 | 2.2e-06 | 3.1e-06 | 2.367E-06 | 6.658E-07 | 1.800E-06 | 3.100E-06 |
| 21023 | 1052 | K1052N | C | 3e-06 | 1.3e-06 | 1.3e-06 | 1.867E-06 | 9.815E-07 | 1.300E-06 | 3.000E-06 |
| 21024 | 1052 | K1052D | C | 2.2e-06 | 1.6e-06 | 4e-06 | 2.600E-06 | 1.249E-06 | 1.600E-06 | 4.000E-06 |
| 21025 | 1052 | K1052C | C | 2.9e-06 | 3.1e-06 | 4.3e-06 | 3.433E-06 | 7.572E-07 | 2.900E-06 | 4.300E-06 |
| 21026 | 1052 | K1052E | C | 4.2e-06 | 2.3e-06 | 2.2e-06 | 2.900E-06 | 1.127E-06 | 2.200E-06 | 4.200E-06 |
| 21027 | 1052 | K1052Q | C | 4.1e-06 | 1.7e-06 | 1.8e-06 | 2.533E-06 | 1.358E-06 | 1.700E-06 | 4.100E-06 |
| 21028 | 1052 | K1052G | C | 1.2e-06 | 3.1e-06 | 2.2e-06 | 2.167E-06 | 9.504E-07 | 1.200E-06 | 3.100E-06 |
| 21029 | 1052 | K1052H | C | 1.6e-06 | 2.7e-06 | 1.4e-06 | 1.900E-06 | 7.000E-07 | 1.400E-06 | 2.700E-06 |
| 21030 | 1052 | K1052I | C | 1.2e-06 | 1.5e-06 | 3.6e-06 | 2.100E-06 | 1.308E-06 | 1.200E-06 | 3.600E-06 |
| 21031 | 1052 | K1052L | C | 2.4e-06 | 3.2e-06 | 2.3e-06 | 2.633E-06 | 4.933E-07 | 2.300E-06 | 3.200E-06 |
| 21032 | 1052 | K1052K | C | 3.2e-06 | 3e-06 | 2.4e-06 | 2.867E-06 | 4.163E-07 | 2.400E-06 | 3.200E-06 |
| 21033 | 1052 | K1052M | C | 1.8e-06 | 4.5e-06 | 3.1e-06 | 3.133E-06 | 1.350E-06 | 1.800E-06 | 4.500E-06 |
| 21034 | 1052 | K1052F | C | 2.3e-06 | 2e-06 | 1.4e-06 | 1.900E-06 | 4.583E-07 | 1.400E-06 | 2.300E-06 |
| 21035 | 1052 | K1052P | C | 2e-06 | 2.7e-06 | 1.9e-06 | 2.200E-06 | 4.359E-07 | 1.900E-06 | 2.700E-06 |
| 21036 | 1052 | K1052S | C | 2.6e-06 | 1.9e-06 | 2.6e-06 | 2.367E-06 | 4.041E-07 | 1.900E-06 | 2.600E-06 |
| 21037 | 1052 | K1052T | C | 2e-06 | 3.8e-06 | 2.1e-06 | 2.633E-06 | 1.012E-06 | 2.000E-06 | 3.800E-06 |
| 21038 | 1052 | K1052W | C | 3.1e-06 | 3.9e-06 | 1.9e-06 | 2.967E-06 | 1.007E-06 | 1.900E-06 | 3.900E-06 |
| 21039 | 1052 | K1052Y | C | 1.6e-06 | 2e-06 | 3.4e-06 | 2.333E-06 | 9.452E-07 | 1.600E-06 | 3.400E-06 |
| 21040 | 1052 | K1052V | C | 2e-06 | 1.6e-06 | 3.2e-06 | 2.267E-06 | 8.327E-07 | 1.600E-06 | 3.200E-06 |
| 21041 | 1053 | V1053A | C | 2.3e-06 | 2.9e-06 | 2.9e-06 | 2.700E-06 | 3.464E-07 | 2.300E-06 | 2.900E-06 |
| 21042 | 1053 | V1053R | C | 1.9e-06 | 2.6e-06 | 3.6e-06 | 2.700E-06 | 8.544E-07 | 1.900E-06 | 3.600E-06 |
| 21043 | 1053 | V1053N | C | 3.2e-06 | 2.6e-06 | 3.1e-06 | 2.967E-06 | 3.215E-07 | 2.600E-06 | 3.200E-06 |
| 21044 | 1053 | V1053D | C | 2.5e-06 | 2.8e-06 | 1.3e-06 | 2.200E-06 | 7.937E-07 | 1.300E-06 | 2.800E-06 |
| 21045 | 1053 | V1053C | C | 3.3e-06 | 1.9e-06 | 2.6e-06 | 2.600E-06 | 7.000E-07 | 1.900E-06 | 3.300E-06 |
| 21046 | 1053 | V1053E | C | 2.4e-06 | 1.5e-06 | 2.4e-06 | 2.100E-06 | 5.196E-07 | 1.500E-06 | 2.400E-06 |
| 21047 | 1053 | V1053Q | C | 2.4e-06 | 1.1e-06 | 2.6e-06 | 2.033E-06 | 8.145E-07 | 1.100E-06 | 2.600E-06 |
| 21048 | 1053 | V1053G | C | 2.1e-06 | 2.8e-06 | 5e-06 | 3.300E-06 | 1.513E-06 | 2.100E-06 | 5.000E-06 |
| 21049 | 1053 | V1053H | C | 1.8e-06 | 2e-06 | 2.5e-06 | 2.100E-06 | 3.606E-07 | 1.800E-06 | 2.500E-06 |
| 21050 | 1053 | V1053I | C | 2.2e-06 | 3.2e-06 | 2.1e-06 | 2.500E-06 | 6.083E-07 | 2.100E-06 | 3.200E-06 |
| 21051 | 1053 | V1053L | C | 3.5e-06 | 1.6e-06 | 3.2e-06 | 2.767E-06 | 1.021E-06 | 1.600E-06 | 3.500E-06 |
| 21052 | 1053 | V1053K | C | 2e-06 | 2.1e-06 | 1.3e-06 | 1.800E-06 | 4.359E-07 | 1.300E-06 | 2.100E-06 |
| 21053 | 1053 | V1053M | C | 1.9e-06 | 6.7e-06 | 2.3e-06 | 3.633E-06 | 2.663E-06 | 1.900E-06 | 6.700E-06 |
| 21054 | 1053 | V1053F | C | 2.7e-06 | 1.7e-06 | 1.7e-06 | 2.033E-06 | 5.774E-07 | 1.700E-06 | 2.700E-06 |
| 21055 | 1053 | V1053P | C | 3.5e-06 | 1.9e-06 | 3.4e-06 | 2.933E-06 | 8.963E-07 | 1.900E-06 | 3.500E-06 |
| 21056 | 1053 | V1053S | C | 2.3e-06 | 1.8e-06 | 2.1e-06 | 2.067E-06 | 2.517E-07 | 1.800E-06 | 2.300E-06 |
| 21057 | 1053 | V1053T | C | 1.8e-06 | 3.9e-06 | 2.5e-06 | 2.733E-06 | 1.069E-06 | 1.800E-06 | 3.900E-06 |
| 21058 | 1053 | V1053W | C | 1.2e-06 | 1.9e-06 | 2.1e-06 | 1.733E-06 | 4.726E-07 | 1.200E-06 | 2.100E-06 |
| 21059 | 1053 | V1053Y | C | 2.3e-06 | 1.8e-06 | 2e-06 | 2.033E-06 | 2.517E-07 | 1.800E-06 | 2.300E-06 |
| 21060 | 1053 | V1053V | C | 3.5e-06 | 2.3e-06 | 2.2e-06 | 2.667E-06 | 7.234E-07 | 2.200E-06 | 3.500E-06 |
| 21061 | 1054 | E1054A | C | 3.1e-06 | 2.2e-06 | 3.3e-06 | 2.867E-06 | 5.859E-07 | 2.200E-06 | 3.300E-06 |
| 21062 | 1054 | E1054R | C | 2.2e-06 | 2.2e-06 | 3.2e-06 | 2.533E-06 | 5.774E-07 | 2.200E-06 | 3.200E-06 |
| 21063 | 1054 | E1054N | C | 1.7e-06 | 2e-06 | 1.9e-06 | 1.867E-06 | 1.528E-07 | 1.700E-06 | 2.000E-06 |
| 21064 | 1054 | E1054D | C | 2.5e-06 | 1.5e-06 | 2.1e-06 | 2.033E-06 | 5.033E-07 | 1.500E-06 | 2.500E-06 |
| 21065 | 1054 | E1054C | C | 2.2e-06 | 1.9e-06 | 1.6e-06 | 1.900E-06 | 3.000E-07 | 1.600E-06 | 2.200E-06 |
| 21066 | 1054 | E1054E | C | 2.8e-06 | 2e-06 | 2.2e-06 | 2.333E-06 | 4.163E-07 | 2.000E-06 | 2.800E-06 |
| 21067 | 1054 | E1054Q | C | 1.8e-06 | 1.8e-06 | 3.1e-06 | 2.233E-06 | 7.506E-07 | 1.800E-06 | 3.100E-06 |
| 21068 | 1054 | E1054G | C | 1.9e-06 | 1.6e-06 | 3.1e-06 | 2.200E-06 | 7.937E-07 | 1.600E-06 | 3.100E-06 |
| 21069 | 1054 | E1054H | C | 2.4e-06 | 1.6e-06 | 1.6e-06 | 1.867E-06 | 4.619E-07 | 1.600E-06 | 2.400E-06 |
| 21070 | 1054 | E1054I | C | 2.4e-06 | 2.5e-06 | 3.3e-06 | 2.733E-06 | 4.933E-07 | 2.400E-06 | 3.300E-06 |
| 21071 | 1054 | E1054L | C | 2.6e-06 | 1.2e-06 | 3.6e-06 | 2.467E-06 | 1.206E-06 | 1.200E-06 | 3.600E-06 |
| 21072 | 1054 | E1054K | C | 4.4e-06 | 2.8e-06 | 1.5e-06 | 2.900E-06 | 1.453E-06 | 1.500E-06 | 4.400E-06 |
| 21073 | 1054 | E1054M | C | 3e-06 | 2.9e-06 | 2.6e-06 | 2.833E-06 | 2.082E-07 | 2.600E-06 | 3.000E-06 |
| 21074 | 1054 | E1054F | C | 2.4e-06 | 2e-06 | 3.1e-06 | 2.500E-06 | 5.568E-07 | 2.000E-06 | 3.100E-06 |
| 21075 | 1054 | E1054P | C | 4.1e-06 | 2.5e-06 | 1.2e-06 | 2.600E-06 | 1.453E-06 | 1.200E-06 | 4.100E-06 |
| 21076 | 1054 | E1054S | C | 2.5e-06 | 3.3e-06 | 2.9e-06 | 2.900E-06 | 4.000E-07 | 2.500E-06 | 3.300E-06 |
| 21077 | 1054 | E1054T | C | 1.8e-06 | 3.7e-06 | 2.7e-06 | 2.733E-06 | 9.504E-07 | 1.800E-06 | 3.700E-06 |
| 21078 | 1054 | E1054W | C | 3.9e-06 | 2.9e-06 | 2.1e-06 | 2.967E-06 | 9.018E-07 | 2.100E-06 | 3.900E-06 |
| 21079 | 1054 | E1054Y | C | 3.3e-06 | 2e-06 | 2.7e-06 | 2.667E-06 | 6.506E-07 | 2.000E-06 | 3.300E-06 |

| | | | | | | | | | | |
|---|---|---|---|---|---|---|---|---|---|---|
| 21080 | 1054 | E1054V | C | 2.3e-06 | 1.7e-06 | 1.9e-06 | 1.967E-06 | 3.055E-07 | 1.700E-06 | 2.300E-06 |
| 21081 | 1055 | P1055A | C | 1.8e-06 | 1e-06 | 2.6e-06 | 1.800E-06 | 8.000E-07 | 1.000E-06 | 2.600E-06 |
| 21082 | 1055 | P1055R | C | 2.1e-06 | 1.4e-06 | 2.6e-06 | 2.033E-06 | 6.028E-07 | 1.400E-06 | 2.600E-06 |
| 21083 | 1055 | P1055N | C | 2.2e-06 | 1.6e-06 | 1.4e-06 | 1.733E-06 | 4.163E-07 | 1.400E-06 | 2.200E-06 |
| 21084 | 1055 | P1055D | C | 1.8e-06 | 2.6e-06 | 2.4e-06 | 2.267E-06 | 4.163E-07 | 1.800E-06 | 2.600E-06 |
| 21085 | 1055 | P1055C | C | 2.1e-06 | 2.6e-06 | 2.3e-06 | 2.333E-06 | 2.517E-07 | 2.100E-06 | 2.600E-06 |
| 21086 | 1055 | P1055E | C | 9.2e-07 | 2.3e-06 | 2.8e-06 | 2.007E-06 | 9.737E-07 | 9.200E-07 | 2.800E-06 |
| 21087 | 1055 | P1055Q | C | 2.6e-06 | 2.7e-06 | 2.5e-06 | 2.600E-06 | 1.000E-07 | 2.500E-06 | 2.700E-06 |
| 21088 | 1055 | P1055G | C | 1.6e-06 | 3.2e-06 | 2.3e-06 | 2.367E-06 | 8.021E-07 | 1.600E-06 | 3.200E-06 |
| 21089 | 1055 | P1055H | C | 2.1e-06 | 2.8e-06 | 2.5e-06 | 2.467E-06 | 3.512E-07 | 2.100E-06 | 2.800E-06 |
| 21090 | 1055 | P1055I | C | 3.1e-06 | 1.4e-06 | 1.6e-06 | 2.033E-06 | 9.292E-07 | 1.400E-06 | 3.100E-06 |
| 21091 | 1055 | P1055L | C | 1.1e-06 | 2.1e-06 | 2.3e-06 | 1.833E-06 | 6.429E-07 | 1.100E-06 | 2.300E-06 |
| 21092 | 1055 | P1055K | C | 2.4e-06 | 2.8e-06 | 2.2e-06 | 2.467E-06 | 3.055E-07 | 2.200E-06 | 2.800E-06 |
| 21093 | 1055 | P1055M | C | 2.1e-06 | 1.9e-06 | 3.1e-06 | 2.367E-06 | 6.429E-07 | 1.900E-06 | 3.100E-06 |
| 21094 | 1055 | P1055F | C | 1.6e-06 | 2.2e-06 | 2.2e-06 | 2.000E-06 | 3.464E-07 | 1.600E-06 | 2.200E-06 |
| 21095 | 1055 | P1055P | C | 1.9e-06 | 2.9e-06 | 3e-06 | 2.600E-06 | 6.083E-07 | 1.900E-06 | 3.000E-06 |
| 21096 | 1055 | P1055S | C | 1.1e-06 | 3.5e-06 | 2.5e-06 | 2.367E-06 | 1.206E-06 | 1.100E-06 | 3.500E-06 |
| 21097 | 1055 | P1055T | C | 2.3e-06 | 4.2e-06 | 2.7e-06 | 3.067E-06 | 1.002E-06 | 2.300E-06 | 4.200E-06 |
| 21098 | 1055 | P1055W | C | 2.2e-06 | 1.8e-06 | 3.1e-06 | 2.367E-06 | 6.658E-07 | 1.800E-06 | 3.100E-06 |
| 21099 | 1055 | P1055Y | C | 1.6e-06 | 3.5e-06 | 2e-06 | 2.367E-06 | 1.002E-06 | 1.600E-06 | 3.500E-06 |
| 21100 | 1055 | P1055V | C | 3.7e-06 | 2.7e-06 | 2.5e-06 | 2.967E-06 | 6.429E-07 | 2.500E-06 | 3.700E-06 |
| 21101 | 1056 | K1056A | C | 2.7e-06 | 2.4e-06 | 2.4e-06 | 2.500E-06 | 1.732E-07 | 2.400E-06 | 2.700E-06 |
| 21102 | 1056 | K1056R | C | 3.7e-06 | 2.8e-06 | 2.7e-06 | 3.067E-06 | 5.508E-07 | 2.700E-06 | 3.700E-06 |
| 21103 | 1056 | K1056N | C | 2.8e-06 | 2.3e-06 | 2.3e-06 | 2.467E-06 | 2.887E-07 | 2.300E-06 | 2.800E-06 |
| 21104 | 1056 | K1056D | C | 3.3e-06 | 2e-06 | 2.3e-06 | 2.533E-06 | 6.807E-07 | 2.000E-06 | 3.300E-06 |
| 21105 | 1056 | K1056C | C | 1.6e-06 | 2.9e-06 | 1.8e-06 | 2.100E-06 | 7.000E-07 | 1.600E-06 | 2.900E-06 |
| 21106 | 1056 | K1056E | C | 4.3e-06 | 8.5e-07 | 1.9e-06 | 2.350E-06 | 1.768E-06 | 8.500E-07 | 4.300E-06 |
| 21107 | 1056 | K1056Q | C | 2.9e-06 | 2.5e-06 | 3.2e-06 | 2.867E-06 | 3.512E-07 | 2.500E-06 | 3.200E-06 |
| 21108 | 1056 | K1056G | C | 2.2e-06 | 2.8e-06 | 2.2e-06 | 2.400E-06 | 3.464E-07 | 2.200E-06 | 2.800E-06 |
| 21109 | 1056 | K1056H | C | 2.6e-06 | 3.1e-06 | 3e-06 | 2.900E-06 | 2.646E-07 | 2.600E-06 | 3.100E-06 |
| 21110 | 1056 | K1056I | C | 3.6e-06 | 2.2e-06 | 2.6e-06 | 2.800E-06 | 7.211E-07 | 2.200E-06 | 3.600E-06 |
| 21111 | 1056 | K1056L | C | 2.7e-06 | 9.6e-07 | 2e-06 | 1.887E-06 | 8.755E-07 | 9.600E-07 | 2.700E-06 |
| 21112 | 1056 | K1056K | C | 2.3e-06 | 1.5e-06 | 1.4e-06 | 1.733E-06 | 4.933E-07 | 1.400E-06 | 2.300E-06 |
| 21113 | 1056 | K1056M | C | 1.7e-06 | 2.3e-06 | 1e-06 | 1.667E-06 | 6.506E-07 | 1.000E-06 | 2.300E-06 |
| 21114 | 1056 | K1056F | C | 3.5e-06 | 3.1e-06 | 2.1e-06 | 2.900E-06 | 7.211E-07 | 2.100E-06 | 3.500E-06 |
| 21115 | 1056 | K1056P | C | 2.8e-06 | 2.6e-06 | 2.7e-06 | 2.700E-06 | 1.000E-07 | 2.600E-06 | 2.800E-06 |
| 21116 | 1056 | K1056S | C | 3.4e-06 | 1.6e-06 | 1.3e-06 | 2.100E-06 | 1.136E-06 | 1.300E-06 | 3.400E-06 |
| 21117 | 1056 | K1056T | C | 1.7e-06 | 3.2e-06 | 1.9e-06 | 2.267E-06 | 8.145E-07 | 1.700E-06 | 3.200E-06 |
| 21118 | 1056 | K1056W | C | 1.6e-06 | 2.3e-06 | 3e-06 | 2.300E-06 | 7.000E-07 | 1.600E-06 | 3.000E-06 |
| 21119 | 1056 | K1056Y | C | 2.2e-06 | 3.2e-06 | 1.5e-06 | 2.300E-06 | 8.544E-07 | 1.500E-06 | 3.200E-06 |
| 21120 | 1056 | K1056V | C | 2.4e-06 | 2.4e-06 | 3.1e-06 | 2.633E-06 | 4.041E-07 | 2.400E-06 | 3.100E-06 |
| 21121 | 1057 | S1057A | C | 1.2e-06 | 1.7e-06 | 3.1e-06 | 2.000E-06 | 9.849E-07 | 1.200E-06 | 3.100E-06 |
| 21122 | 1057 | S1057R | C | 2.2e-06 | 2.5e-06 | 2.9e-06 | 2.533E-06 | 3.512E-07 | 2.200E-06 | 2.900E-06 |
| 21123 | 1057 | S1057N | C | 1.7e-06 | 2.4e-06 | 2.7e-06 | 2.267E-06 | 5.132E-07 | 1.700E-06 | 2.700E-06 |
| 21124 | 1057 | S1057D | C | 2.5e-06 | 2.1e-06 | 3.8e-06 | 2.800E-06 | 8.888E-07 | 2.100E-06 | 3.800E-06 |
| 21125 | 1057 | S1057C | C | 1.2e-06 | 4.2e-06 | 2.2e-06 | 2.533E-06 | 1.528E-06 | 1.200E-06 | 4.200E-06 |
| 21126 | 1057 | S1057E | C | 1.5e-06 | 1.9e-06 | 2e-06 | 1.800E-06 | 2.646E-07 | 1.500E-06 | 2.000E-06 |
| 21127 | 1057 | S1057Q | C | 3.3e-06 | 1.5e-06 | 3.1e-06 | 2.633E-06 | 9.866E-07 | 1.500E-06 | 3.300E-06 |
| 21128 | 1057 | S1057G | C | 3e-06 | 1.4e-06 | 3.9e-06 | 2.767E-06 | 1.266E-06 | 1.400E-06 | 3.900E-06 |
| 21129 | 1057 | S1057H | C | 3.2e-06 | 1.8e-06 | 3.5e-06 | 2.833E-06 | 9.074E-07 | 1.800E-06 | 3.500E-06 |
| 21130 | 1057 | S1057I | C | 1.6e-06 | 2.5e-06 | 2.1e-06 | 2.067E-06 | 4.509E-07 | 1.600E-06 | 2.500E-06 |
| 21131 | 1057 | S1057L | C | 2.3e-06 | 2.5e-06 | 2.2e-06 | 2.333E-06 | 1.528E-07 | 2.200E-06 | 2.500E-06 |
| 21132 | 1057 | S1057K | C | 3.2e-06 | 2.1e-06 | 2.3e-06 | 2.533E-06 | 5.859E-07 | 2.100E-06 | 3.200E-06 |
| 21133 | 1057 | S1057M | C | 3.6e-06 | 1.4e-06 | 3.5e-06 | 2.833E-06 | 1.242E-06 | 1.400E-06 | 3.600E-06 |
| 21134 | 1057 | S1057F | C | 1.9e-06 | 2.3e-06 | 3.3e-06 | 2.500E-06 | 7.211E-07 | 1.900E-06 | 3.300E-06 |
| 21135 | 1057 | S1057P | C | 1.2e-06 | 3.3e-06 | 4e-06 | 2.833E-06 | 1.457E-06 | 1.200E-06 | 4.000E-06 |
| 21136 | 1057 | S1057S | C | 1.6e-06 | 2.4e-06 | 2.5e-06 | 2.167E-06 | 4.933E-07 | 1.600E-06 | 2.500E-06 |
| 21137 | 1057 | S1057T | C | 2.8e-06 | 2.5e-06 | 1.7e-06 | 2.333E-06 | 5.686E-07 | 1.700E-06 | 2.800E-06 |
| 21138 | 1057 | S1057W | C | 2e-06 | 1.1e-06 | 2.5e-06 | 1.867E-06 | 7.095E-07 | 1.100E-06 | 2.500E-06 |
| 21139 | 1057 | S1057Y | C | 9.6e-07 | 3.4e-06 | 1.3e-06 | 1.887E-06 | 1.322E-06 | 9.600E-07 | 3.400E-06 |
| 21140 | 1057 | S1057V | C | 2e-06 | 2.2e-06 | 5e-06 | 3.067E-06 | 1.677E-06 | 2.000E-06 | 5.000E-06 |
| 21141 | 1058 | C1058A | C | 2.9e-06 | 2.3e-06 | 2.8e-06 | 2.667E-06 | 3.215E-07 | 2.300E-06 | 2.900E-06 |
| 21142 | 1058 | C1058R | C | 2.4e-06 | 3.5e-06 | 2.4e-06 | 2.767E-06 | 6.351E-07 | 2.400E-06 | 3.500E-06 |
| 21143 | 1058 | C1058N | C | 3.7e-06 | 3.1e-06 | 2.4e-06 | 3.067E-06 | 6.506E-07 | 2.400E-06 | 3.700E-06 |
| 21144 | 1058 | C1058D | C | 1.8e-06 | 2.8e-06 | 2.4e-06 | 2.333E-06 | 5.033E-07 | 1.800E-06 | 2.800E-06 |
| 21145 | 1058 | C1058C | C | 3.1e-06 | 2.1e-06 | 2.1e-06 | 2.433E-06 | 5.774E-07 | 2.100E-06 | 3.100E-06 |
| 21146 | 1058 | C1058E | C | 2.7e-06 | 1.9e-06 | 3.7e-06 | 2.767E-06 | 9.018E-07 | 1.900E-06 | 3.700E-06 |
| 21147 | 1058 | C1058Q | C | 1.9e-06 | 1.9e-06 | 3.7e-06 | 2.500E-06 | 1.039E-06 | 1.900E-06 | 3.700E-06 |
| 21148 | 1058 | C1058G | C | 3.4e-06 | 3.6e-06 | 3.1e-06 | 3.367E-06 | 2.517E-07 | 3.100E-06 | 3.600E-06 |
| 21149 | 1058 | C1058H | C | 2.1e-06 | 2.5e-06 | 3e-06 | 2.533E-06 | 4.509E-07 | 2.100E-06 | 3.000E-06 |
| 21150 | 1058 | C1058I | C | 4.6e-06 | 2.5e-06 | 2.7e-06 | 3.267E-06 | 1.159E-06 | 2.500E-06 | 4.600E-06 |
| 21151 | 1058 | C1058L | C | 2.3e-06 | 3.6e-06 | 2.5e-06 | 2.800E-06 | 7.000E-07 | 2.300E-06 | 3.600E-06 |
| 21152 | 1058 | C1058K | C | 2.8e-06 | 2.5e-06 | 4.4e-06 | 3.233E-06 | 1.021E-06 | 2.500E-06 | 4.400E-06 |
| 21153 | 1058 | C1058M | C | 2.7e-06 | 3.1e-06 | 2.8e-06 | 2.867E-06 | 2.082E-07 | 2.700E-06 | 3.100E-06 |
| 21154 | 1058 | C1058F | C | 2.5e-06 | 2e-06 | 3.3e-06 | 2.600E-06 | 6.557E-07 | 2.000E-06 | 3.300E-06 |
| 21155 | 1058 | C1058P | C | 2.5e-06 | 2.1e-06 | 4.1e-06 | 2.900E-06 | 1.058E-06 | 2.100E-06 | 4.100E-06 |
| 21156 | 1058 | C1058S | C | 3e-06 | 2.7e-06 | 1.2e-06 | 2.300E-06 | 9.644E-07 | 1.200E-06 | 3.000E-06 |
| 21157 | 1058 | C1058T | C | 3e-06 | 1.9e-06 | 3.9e-06 | 2.933E-06 | 1.002E-06 | 1.900E-06 | 3.900E-06 |
| 21158 | 1058 | C1058W | C | 1.5e-06 | 3.5e-06 | 1.6e-06 | 2.200E-06 | 1.127E-06 | 1.500E-06 | 3.500E-06 |
| 21159 | 1058 | C1058Y | C | 2e-06 | 2.5e-06 | 2.9e-06 | 2.467E-06 | 4.509E-07 | 2.000E-06 | 2.900E-06 |
| 21160 | 1058 | C1058V | C | 1.9e-06 | 1.9e-06 | 3e-06 | 2.267E-06 | 6.351E-07 | 1.900E-06 | 3.000E-06 |
| 21161 | 1059 | D1059A | D | 2.5e-06 | 2.6e-06 | 2.3e-06 | 2.467E-06 | 1.528E-07 | 2.300E-06 | 2.600E-06 |
| 21162 | 1059 | D1059R | D | 1.4e-06 | 1.4e-06 | 2.5e-06 | 1.767E-06 | 6.351E-07 | 1.400E-06 | 2.500E-06 |
| 21163 | 1059 | D1059N | D | 3.4e-06 | 3.2e-06 | 1.9e-06 | 2.833E-06 | 8.145E-07 | 1.900E-06 | 3.400E-06 |
| 21164 | 1059 | D1059D | D | 9.1e-07 | 3.2e-06 | 4.6e-06 | 2.903E-06 | 1.863E-06 | 9.100E-07 | 4.600E-06 |

| | | | | | | | | | | |
|---|---|---|---|---|---|---|---|---|---|---|
| 21165 | 1059 | D1059C | D | 2.9e-06 | 2e-06 | 2e-06 | 2.300E-06 | 5.196E-07 | 2.000E-06 | 2.900E-06 |
| 21166 | 1059 | D1059E | D | 2.1e-06 | 1.8e-06 | 3.5e-06 | 2.467E-06 | 9.074E-07 | 1.800E-06 | 3.500E-06 |
| 21167 | 1059 | D1059Q | D | 1.6e-06 | 1.8e-06 | 1.8e-06 | 1.733E-06 | 1.155E-07 | 1.600E-06 | 1.800E-06 |
| 21168 | 1059 | D1059G | D | 3e-06 | 2.7e-06 | 1.9e-06 | 2.533E-06 | 5.686E-07 | 1.900E-06 | 3.000E-06 |
| 21169 | 1059 | D1059H | D | 1.6e-06 | 3.7e-06 | 3.3e-06 | 2.867E-06 | 1.115E-06 | 1.600E-06 | 3.700E-06 |
| 21170 | 1059 | D1059I | D | 3.1e-06 | 2.2e-06 | 2.1e-06 | 2.467E-06 | 5.508E-07 | 2.100E-06 | 3.100E-06 |
| 21171 | 1059 | D1059L | D | 2.6e-06 | 2.5e-06 | 1.9e-06 | 2.333E-06 | 3.786E-07 | 1.900E-06 | 2.600E-06 |
| 21172 | 1059 | D1059K | D | 1.5e-06 | 2.3e-06 | 1.9e-06 | 1.900E-06 | 4.000E-07 | 1.500E-06 | 2.300E-06 |
| 21173 | 1059 | D1059M | D | 2.6e-06 | 3e-06 | 2.1e-06 | 2.567E-06 | 4.509E-07 | 2.100E-06 | 3.000E-06 |
| 21174 | 1059 | D1059F | D | 3.2e-06 | 2.4e-06 | 2.2e-06 | 2.600E-06 | 5.292E-07 | 2.200E-06 | 3.200E-06 |
| 21175 | 1059 | D1059P | D | 2.8e-06 | 2.2e-06 | 2.9e-06 | 2.633E-06 | 3.786E-07 | 2.200E-06 | 2.900E-06 |
| 21176 | 1059 | D1059S | D | 3.5e-06 | 2.4e-06 | 3.1e-06 | 3.000E-06 | 5.568E-07 | 2.400E-06 | 3.500E-06 |
| 21177 | 1059 | D1059T | D | 3.8e-06 | 2.5e-06 | 1.9e-06 | 2.733E-06 | 9.713E-07 | 1.900E-06 | 3.800E-06 |
| 21178 | 1059 | D1059W | D | 1.9e-06 | 1.7e-06 | 4.4e-06 | 2.667E-06 | 1.504E-06 | 1.700E-06 | 4.400E-06 |
| 21179 | 1059 | D1059Y | D | 2.2e-06 | 2e-06 | 2.1e-06 | 2.100E-06 | 1.000E-07 | 2.000E-06 | 2.200E-06 |
| 21180 | 1059 | D1059V | D | 1.7e-06 | 2.5e-06 | 3.6e-06 | 2.600E-06 | 9.539E-07 | 1.700E-06 | 3.600E-06 |
| 21181 | 1060 | I1060A | D | 2.5e-06 | 1.3e-06 | 1.9e-06 | 1.900E-06 | 6.000E-07 | 1.300E-06 | 2.500E-06 |
| 21182 | 1060 | I1060R | D | 2.7e-06 | 2.3e-06 | 2.7e-06 | 2.567E-06 | 2.309E-07 | 2.300E-06 | 2.700E-06 |
| 21183 | 1060 | I1060N | D | 2e-06 | 1.7e-06 | 2.7e-06 | 2.133E-06 | 5.132E-07 | 1.700E-06 | 2.700E-06 |
| 21184 | 1060 | I1060D | D | 3e-06 | 1.5e-06 | 2.3e-06 | 2.267E-06 | 7.506E-07 | 1.500E-06 | 3.000E-06 |
| 21185 | 1060 | I1060C | D | 2.8e-06 | 2.9e-06 | 1.6e-06 | 2.433E-06 | 7.234E-07 | 1.600E-06 | 2.900E-06 |
| 21186 | 1060 | I1060E | D | 3e-06 | 1.4e-06 | 2.2e-06 | 2.200E-06 | 8.000E-07 | 1.400E-06 | 3.000E-06 |
| 21187 | 1060 | I1060Q | D | 1.6e-06 | 2.2e-06 | 1.9e-06 | 1.900E-06 | 3.000E-07 | 1.600E-06 | 2.200E-06 |
| 21188 | 1060 | I1060G | D | 2.8e-06 | 1.8e-06 | 2.5e-06 | 2.367E-06 | 5.132E-07 | 1.800E-06 | 2.800E-06 |
| 21189 | 1060 | I1060H | D | 2.3e-06 | 3.7e-06 | 2.2e-06 | 2.733E-06 | 8.386E-07 | 2.200E-06 | 3.700E-06 |
| 21190 | 1060 | I1060I | D | 3.4e-06 | 1.2e-06 | 2.2e-06 | 2.267E-06 | 1.102E-06 | 1.200E-06 | 3.400E-06 |
| 21191 | 1060 | I1060L | D | 2.5e-06 | 1.3e-06 | 2.2e-06 | 2.000E-06 | 6.245E-07 | 1.300E-06 | 2.500E-06 |
| 21192 | 1060 | I1060K | D | 1.7e-06 | 3.1e-06 | 1.7e-06 | 2.167E-06 | 8.083E-07 | 1.700E-06 | 3.100E-06 |
| 21193 | 1060 | I1060M | D | 1.8e-06 | 2.9e-06 | 2.3e-06 | 2.333E-06 | 5.508E-07 | 1.800E-06 | 2.900E-06 |
| 21194 | 1060 | I1060F | D | 3.1e-06 | 2.1e-06 | 3.3e-06 | 2.833E-06 | 6.429E-07 | 2.100E-06 | 3.300E-06 |
| 21195 | 1060 | I1060P | D | 2.1e-06 | 2.8e-06 | 3.3e-06 | 2.733E-06 | 6.028E-07 | 2.100E-06 | 3.300E-06 |
| 21196 | 1060 | I1060S | D | 1.9e-06 | 5.4e-06 | 3.5e-06 | 3.600E-06 | 1.752E-06 | 1.900E-06 | 5.400E-06 |
| 21197 | 1060 | I1060T | D | 4.2e-06 | 2.4e-06 | 3e-06 | 3.200E-06 | 9.165E-07 | 2.400E-06 | 4.200E-06 |
| 21198 | 1060 | I1060W | D | 2.8e-06 | 3.4e-06 | 3.6e-06 | 3.267E-06 | 4.163E-07 | 2.800E-06 | 3.600E-06 |
| 21199 | 1060 | I1060Y | D | 4.5e-06 | 2.5e-06 | 1.4e-06 | 2.800E-06 | 1.572E-06 | 1.400E-06 | 4.500E-06 |
| 21200 | 1060 | I1060V | D | 3e-06 | 2.5e-06 | 2.4e-06 | 2.633E-06 | 3.215E-07 | 2.400E-06 | 3.000E-06 |
| 21201 | 1061 | Q1061A | D | 1.8e-06 | 5e-06 | 2.6e-06 | 3.133E-06 | 1.665E-06 | 1.800E-06 | 5.000E-06 |
| 21202 | 1061 | Q1061R | D | 2.6e-06 | 2.1e-06 | 2.8e-06 | 2.500E-06 | 3.606E-07 | 2.100E-06 | 2.800E-06 |
| 21203 | 1061 | Q1061N | D | 2.7e-06 | 1.6e-06 | 1.8e-06 | 2.033E-06 | 5.859E-07 | 1.600E-06 | 2.700E-06 |
| 21204 | 1061 | Q1061D | D | 2.6e-06 | 3.1e-06 | 3.4e-06 | 3.033E-06 | 4.041E-07 | 2.600E-06 | 3.400E-06 |
| 21205 | 1061 | Q1061C | D | 3.1e-06 | 2.1e-06 | 2.2e-06 | 2.467E-06 | 5.508E-07 | 2.100E-06 | 3.100E-06 |
| 21206 | 1061 | Q1061E | D | 1.8e-06 | 2.5e-06 | 3.2e-06 | 2.500E-06 | 7.000E-07 | 1.800E-06 | 3.200E-06 |
| 21207 | 1061 | Q1061Q | D | 3.8e-06 | 3.5e-06 | 2.3e-06 | 3.200E-06 | 7.937E-07 | 2.300E-06 | 3.800E-06 |
| 21208 | 1061 | Q1061G | D | 2.7e-06 | 1.5e-06 | 2.6e-06 | 2.267E-06 | 6.658E-07 | 1.500E-06 | 2.700E-06 |
| 21209 | 1061 | Q1061H | D | 3.4e-06 | 2.1e-06 | 1.6e-06 | 2.367E-06 | 9.292E-07 | 1.600E-06 | 3.400E-06 |
| 21210 | 1061 | Q1061I | D | 2.8e-06 | 1.7e-06 | 2.9e-06 | 2.467E-06 | 6.658E-07 | 1.700E-06 | 2.900E-06 |
| 21211 | 1061 | Q1061L | D | 1.5e-06 | 2.2e-06 | 2.6e-06 | 2.100E-06 | 5.568E-07 | 1.500E-06 | 2.600E-06 |
| 21212 | 1061 | Q1061K | D | 8.4e-07 | 2.1e-06 | 1.8e-06 | 1.580E-06 | 6.582E-07 | 8.400E-07 | 2.100E-06 |
| 21213 | 1061 | Q1061M | D | 2.2e-06 | 3.2e-06 | 4.8e-06 | 3.400E-06 | 1.311E-06 | 2.200E-06 | 4.800E-06 |
| 21214 | 1061 | Q1061F | D | 1.6e-06 | 3e-06 | 2.3e-06 | 2.300E-06 | 7.000E-07 | 1.600E-06 | 3.000E-06 |
| 21215 | 1061 | Q1061P | D | 2.2e-06 | 2.5e-06 | 2.5e-06 | 2.400E-06 | 1.732E-07 | 2.200E-06 | 2.500E-06 |
| 21216 | 1061 | Q1061S | D | 4e-06 | 2.3e-06 | 2e-06 | 2.767E-06 | 1.079E-06 | 2.000E-06 | 4.000E-06 |
| 21217 | 1061 | Q1061T | D | 2.6e-06 | 3.2e-06 | 5.5e-06 | 3.767E-06 | 1.531E-06 | 2.600E-06 | 5.500E-06 |
| 21218 | 1061 | Q1061W | D | 3.7e-06 | 3.7e-06 | 1.6e-06 | 3.000E-06 | 1.212E-06 | 1.600E-06 | 3.700E-06 |
| 21219 | 1061 | Q1061Y | D | 4.1e-06 | 5.8e-06 | 1.4e-06 | 3.767E-06 | 2.219E-06 | 1.400E-06 | 5.800E-06 |
| 21220 | 1061 | Q1061V | D | 2.1e-06 | 8.9e-07 | 6.2e-06 | 3.063E-06 | 2.783E-06 | 8.900E-07 | 6.200E-06 |
| 21221 | 1062 | M1062A | D | 2.8e-06 | 2.2e-06 | 1.8e-06 | 2.267E-06 | 5.033E-07 | 1.800E-06 | 2.800E-06 |
| 21222 | 1062 | M1062R | D | 3.6e-06 | 1.7e-06 | 1.7e-06 | 2.333E-06 | 1.097E-06 | 1.700E-06 | 3.600E-06 |
| 21223 | 1062 | M1062N | D | 4e-06 | 1.9e-06 | 3.1e-06 | 3.000E-06 | 1.054E-06 | 1.900E-06 | 4.000E-06 |
| 21224 | 1062 | M1062D | D | 2.2e-06 | 2.7e-06 | 2e-06 | 2.300E-06 | 3.606E-07 | 2.000E-06 | 2.700E-06 |
| 21225 | 1062 | M1062C | D | 2e-06 | 1.4e-06 | 1.4e-06 | 1.600E-06 | 3.464E-07 | 1.400E-06 | 2.000E-06 |
| 21226 | 1062 | M1062E | D | 1.9e-06 | 2.6e-06 | 2e-06 | 2.167E-06 | 3.786E-07 | 1.900E-06 | 2.600E-06 |
| 21227 | 1062 | M1062Q | D | 2.1e-06 | 2.2e-06 | 2.8e-06 | 2.367E-06 | 3.786E-07 | 2.100E-06 | 2.800E-06 |
| 21228 | 1062 | M1062G | D | 4.4e-06 | 9.9e-07 | 1.6e-06 | 2.330E-06 | 1.818E-06 | 9.900E-07 | 4.400E-06 |
| 21229 | 1062 | M1062H | D | 1.5e-06 | 2.2e-06 | 2.2e-06 | 1.967E-06 | 4.041E-07 | 1.500E-06 | 2.200E-06 |
| 21230 | 1062 | M1062I | D | 2.4e-06 | 2.2e-06 | 3.4e-06 | 2.667E-06 | 6.429E-07 | 2.200E-06 | 3.400E-06 |
| 21231 | 1062 | M1062L | D | 1.7e-06 | 2.6e-06 | 1.7e-06 | 2.000E-06 | 5.196E-07 | 1.700E-06 | 2.600E-06 |
| 21232 | 1062 | M1062K | D | 1.6e-06 | 3.5e-06 | 2.6e-06 | 2.567E-06 | 9.504E-07 | 1.600E-06 | 3.500E-06 |
| 21233 | 1062 | M1062M | D | 4.8e-06 | 2.4e-06 | 1.3e-06 | 2.833E-06 | 1.790E-06 | 1.300E-06 | 4.800E-06 |
| 21234 | 1062 | M1062F | D | 1.6e-06 | 2e-06 | 4.4e-06 | 2.667E-06 | 1.514E-06 | 1.600E-06 | 4.400E-06 |
| 21235 | 1062 | M1062P | D | 4.4e-06 | 2.8e-06 | 2e-06 | 3.067E-06 | 1.222E-06 | 2.000E-06 | 4.400E-06 |
| 21236 | 1062 | M1062S | D | 3.1e-06 | 2.3e-06 | 1.7e-06 | 2.367E-06 | 7.024E-07 | 1.700E-06 | 3.100E-06 |
| 21237 | 1062 | M1062T | D | 1.6e-06 | 2e-06 | 2.3e-06 | 1.967E-06 | 3.512E-07 | 1.600E-06 | 2.300E-06 |
| 21238 | 1062 | M1062W | D | 1.4e-06 | 1.9e-06 | 2e-06 | 1.767E-06 | 3.215E-07 | 1.400E-06 | 2.000E-06 |
| 21239 | 1062 | M1062Y | D | 2.8e-06 | 2.6e-06 | 2.1e-06 | 2.500E-06 | 3.606E-07 | 2.100E-06 | 2.800E-06 |
| 21240 | 1062 | M1062V | D | 2.6e-06 | 2.6e-06 | 3.4e-06 | 2.867E-06 | 4.619E-07 | 2.600E-06 | 3.400E-06 |
| 21241 | 1063 | T1063A | D | 3.5e-06 | 1.7e-06 | 2.2e-06 | 2.467E-06 | 9.292E-07 | 1.700E-06 | 3.500E-06 |
| 21242 | 1063 | T1063R | D | 2.3e-06 | 2e-06 | 4e-06 | 2.767E-06 | 1.079E-06 | 2.000E-06 | 4.000E-06 |
| 21243 | 1063 | T1063N | D | 4e-06 | 1.9e-06 | 2.5e-06 | 2.800E-06 | 1.082E-06 | 1.900E-06 | 4.000E-06 |
| 21244 | 1063 | T1063D | D | 1.5e-06 | 1.5e-06 | 2.3e-06 | 1.767E-06 | 4.619E-07 | 1.500E-06 | 2.300E-06 |
| 21245 | 1063 | T1063C | D | 1.4e-06 | 1.6e-06 | 2.8e-06 | 1.933E-06 | 7.572E-07 | 1.400E-06 | 2.800E-06 |
| 21246 | 1063 | T1063E | D | 4.7e-06 | 2.5e-06 | 2.8e-06 | 3.333E-06 | 1.193E-06 | 2.500E-06 | 4.700E-06 |
| 21247 | 1063 | T1063Q | D | 3.3e-06 | 2.7e-06 | 2.3e-06 | 2.767E-06 | 5.033E-07 | 2.300E-06 | 3.300E-06 |
| 21248 | 1063 | T1063G | D | 2.5e-06 | 2.6e-06 | 2.8e-06 | 2.633E-06 | 1.528E-07 | 2.500E-06 | 2.800E-06 |
| 21249 | 1063 | T1063H | D | 2.2e-06 | 1.7e-06 | 3.1e-06 | 2.333E-06 | 7.095E-07 | 1.700E-06 | 3.100E-06 |

| | | | | | | | | | | |
|---|---|---|---|---|---|---|---|---|---|---|
| 21250 | 1063 | T1063I | D | 2.3e-06 | 3e-06 | 8.2e-07 | 2.040E-06 | 1.113E-06 | 8.200E-07 | 3.000E-06 |
| 21251 | 1063 | T1063L | D | 3.6e-06 | 1.8e-06 | 3.2e-06 | 2.867E-06 | 9.452E-07 | 1.800E-06 | 3.600E-06 |
| 21252 | 1063 | T1063K | D | 2.1e-06 | 2e-06 | 2.8e-06 | 2.300E-06 | 4.359E-07 | 2.000E-06 | 2.800E-06 |
| 21253 | 1063 | T1063M | D | 2.5e-06 | 3.8e-06 | 2.2e-06 | 2.833E-06 | 8.505E-07 | 2.200E-06 | 3.800E-06 |
| 21254 | 1063 | T1063F | D | 4.4e-06 | 1.1e-06 | 2.4e-06 | 2.633E-06 | 1.662E-06 | 1.100E-06 | 4.400E-06 |
| 21255 | 1063 | T1063P | D | 3.3e-06 | 1.5e-06 | 2.1e-06 | 2.300E-06 | 9.165E-07 | 1.500E-06 | 3.300E-06 |
| 21256 | 1063 | T1063S | D | 4.7e-06 | 2.2e-06 | 2.5e-06 | 3.133E-06 | 1.365E-06 | 2.200E-06 | 4.700E-06 |
| 21257 | 1063 | T1063T | D | 1.9e-06 | 2.7e-06 | 2.7e-06 | 2.433E-06 | 4.619E-07 | 1.900E-06 | 2.700E-06 |
| 21258 | 1063 | T1063W | D | 1.8e-06 | 2.6e-06 | 1.9e-06 | 2.100E-06 | 4.359E-07 | 1.800E-06 | 2.600E-06 |
| 21259 | 1063 | T1063Y | D | 2.2e-06 | 2.1e-06 | 1.7e-06 | 2.000E-06 | 2.646E-07 | 1.700E-06 | 2.200E-06 |
| 21260 | 1063 | T1063V | D | 3e-06 | 1.7e-06 | 2.5e-06 | 2.400E-06 | 6.557E-07 | 1.700E-06 | 3.000E-06 |
| 21261 | 1064 | Q1064A | D | 3.6e-06 | 3.1e-06 | 3.6e-06 | 3.433E-06 | 2.887E-07 | 3.100E-06 | 3.600E-06 |
| 21262 | 1064 | Q1064R | D | 2.2e-06 | 3.7e-06 | 3.8e-06 | 3.233E-06 | 8.963E-07 | 2.200E-06 | 3.800E-06 |
| 21263 | 1064 | Q1064N | D | 2.2e-06 | 2.9e-06 | 2.5e-06 | 2.533E-06 | 3.512E-07 | 2.200E-06 | 2.900E-06 |
| 21264 | 1064 | Q1064D | D | 3.5e-06 | 3.3e-06 | 2.7e-06 | 3.167E-06 | 4.163E-07 | 2.700E-06 | 3.500E-06 |
| 21265 | 1064 | Q1064C | D | 4.2e-06 | 2.2e-06 | 1.6e-06 | 2.667E-06 | 1.361E-06 | 1.600E-06 | 4.200E-06 |
| 21266 | 1064 | Q1064E | D | 4e-06 | 2.1e-06 | 1.8e-06 | 2.633E-06 | 1.193E-06 | 1.800E-06 | 4.000E-06 |
| 21267 | 1064 | Q1064Q | D | 2e-06 | 2.4e-06 | 3e-06 | 2.467E-06 | 5.033E-07 | 2.000E-06 | 3.000E-06 |
| 21268 | 1064 | Q1064G | D | 2.8e-06 | 2.4e-06 | 2.5e-06 | 2.567E-06 | 2.082E-07 | 2.400E-06 | 2.800E-06 |
| 21269 | 1064 | Q1064H | D | 2.5e-06 | 1.8e-06 | 1.6e-06 | 1.967E-06 | 4.726E-07 | 1.600E-06 | 2.500E-06 |
| 21270 | 1064 | Q1064I | D | 2.1e-06 | 2.2e-06 | 1.6e-06 | 1.967E-06 | 3.215E-07 | 1.600E-06 | 2.200E-06 |
| 21271 | 1064 | Q1064L | D | 1.3e-06 | 2.1e-06 | 3.9e-06 | 2.433E-06 | 1.332E-06 | 1.300E-06 | 3.900E-06 |
| 21272 | 1064 | Q1064K | D | 2.2e-06 | 2.3e-06 | 2e-06 | 2.167E-06 | 1.528E-07 | 2.000E-06 | 2.300E-06 |
| 21273 | 1064 | Q1064M | D | 2.4e-06 | 3.4e-06 | 2.3e-06 | 2.700E-06 | 6.083E-07 | 2.300E-06 | 3.400E-06 |
| 21274 | 1064 | Q1064F | D | 4.7e-06 | 1.3e-06 | 3e-06 | 3.000E-06 | 1.700E-06 | 1.300E-06 | 4.700E-06 |
| 21275 | 1064 | Q1064P | D | 2.2e-06 | 2.5e-06 | 3e-06 | 2.567E-06 | 4.041E-07 | 2.200E-06 | 3.000E-06 |
| 21276 | 1064 | Q1064S | D | 1.3e-06 | 1.5e-06 | 3.2e-06 | 2.000E-06 | 1.044E-06 | 1.300E-06 | 3.200E-06 |
| 21277 | 1064 | Q1064T | D | 2.2e-06 | 4.9e-06 | 2.3e-06 | 3.133E-06 | 1.531E-06 | 2.200E-06 | 4.900E-06 |
| 21278 | 1064 | Q1064W | D | 2.1e-06 | 1.8e-06 | 3.2e-06 | 2.367E-06 | 7.371E-07 | 1.800E-06 | 3.200E-06 |
| 21279 | 1064 | Q1064Y | D | 2.7e-06 | 2.3e-06 | 2.3e-06 | 2.433E-06 | 2.309E-07 | 2.300E-06 | 2.700E-06 |
| 21280 | 1064 | Q1064V | D | 2.5e-06 | 5e-06 | 3.3e-06 | 3.600E-06 | 1.277E-06 | 2.500E-06 | 5.000E-06 |
| 21281 | 1065 | S1065A | D | 2.8e-06 | 2.5e-06 | 2e-06 | 2.433E-06 | 4.041E-07 | 2.000E-06 | 2.800E-06 |
| 21282 | 1065 | S1065R | D | 2.4e-06 | 1.3e-06 | 2.5e-06 | 2.067E-06 | 6.658E-07 | 1.300E-06 | 2.500E-06 |
| 21283 | 1065 | S1065N | D | 1.7e-06 | 2.9e-06 | 2.8e-06 | 2.467E-06 | 6.658E-07 | 1.700E-06 | 2.900E-06 |
| 21284 | 1065 | S1065D | D | 2.2e-06 | 1.9e-06 | 3.8e-06 | 2.633E-06 | 1.021E-06 | 1.900E-06 | 3.800E-06 |
| 21285 | 1065 | S1065C | D | 2.9e-06 | 3.6e-06 | 2.3e-06 | 2.933E-06 | 6.506E-07 | 2.300E-06 | 3.600E-06 |
| 21286 | 1065 | S1065E | D | 3.4e-06 | 2.2e-06 | 1.9e-06 | 2.500E-06 | 7.937E-07 | 1.900E-06 | 3.400E-06 |
| 21287 | 1065 | S1065Q | D | 1.5e-06 | 2e-06 | 2e-06 | 1.833E-06 | 2.887E-07 | 1.500E-06 | 2.000E-06 |
| 21288 | 1065 | S1065G | D | 1.3e-06 | 3.7e-06 | 3.1e-06 | 2.700E-06 | 1.249E-06 | 1.300E-06 | 3.700E-06 |
| 21289 | 1065 | S1065H | D | 2.1e-06 | 2.3e-06 | 2.4e-06 | 2.267E-06 | 1.528E-07 | 2.100E-06 | 2.400E-06 |
| 21290 | 1065 | S1065I | D | 3.1e-06 | 2.4e-06 | 2.6e-06 | 2.700E-06 | 3.606E-07 | 2.400E-06 | 3.100E-06 |
| 21291 | 1065 | S1065L | D | 3.2e-06 | 2.3e-06 | 2.1e-06 | 2.533E-06 | 5.859E-07 | 2.100E-06 | 3.200E-06 |
| 21292 | 1065 | S1065K | D | 1.7e-06 | 1.4e-06 | 3.5e-06 | 2.200E-06 | 1.136E-06 | 1.400E-06 | 3.500E-06 |
| 21293 | 1065 | S1065M | D | 2.5e-06 | 3.3e-06 | 1.8e-06 | 2.533E-06 | 7.506E-07 | 1.800E-06 | 3.300E-06 |
| 21294 | 1065 | S1065F | D | 3.1e-06 | 3.7e-06 | 2.6e-06 | 3.133E-06 | 5.508E-07 | 2.600E-06 | 3.700E-06 |
| 21295 | 1065 | S1065P | D | 1.4e-06 | 2.5e-06 | 7.2e-06 | 3.700E-06 | 3.081E-06 | 1.400E-06 | 7.200E-06 |
| 21296 | 1065 | S1065S | D | 1.2e-06 | 4e-06 | 2.6e-06 | 2.600E-06 | 1.400E-06 | 1.200E-06 | 4.000E-06 |
| 21297 | 1065 | S1065T | D | 3.6e-06 | 1.2e-06 | 2.2e-06 | 2.333E-06 | 1.206E-06 | 1.200E-06 | 3.600E-06 |
| 21298 | 1065 | S1065W | D | 1.3e-06 | 3.6e-06 | 2.3e-06 | 2.400E-06 | 1.153E-06 | 1.300E-06 | 3.600E-06 |
| 21299 | 1065 | S1065Y | D | 1.1e-06 | 2.9e-06 | 1.6e-06 | 1.867E-06 | 9.292E-07 | 1.100E-06 | 2.900E-06 |
| 21300 | 1065 | S1065V | D | 4e-06 | 3.5e-06 | 2.9e-06 | 3.467E-06 | 5.508E-07 | 2.900E-06 | 4.000E-06 |
| 21301 | 1066 | P1066A | D | 2e-06 | 2.2e-06 | 2.3e-06 | 2.167E-06 | 1.528E-07 | 2.000E-06 | 2.300E-06 |
| 21302 | 1066 | P1066R | D | 2.8e-06 | 2.3e-06 | 1.9e-06 | 2.333E-06 | 4.509E-07 | 1.900E-06 | 2.800E-06 |
| 21303 | 1066 | P1066N | D | 2.5e-06 | 1.2e-06 | 1.7e-06 | 1.800E-06 | 6.557E-07 | 1.200E-06 | 2.500E-06 |
| 21304 | 1066 | P1066D | D | 3.2e-06 | 3e-06 | 4.9e-06 | 3.700E-06 | 1.044E-06 | 3.000E-06 | 4.900E-06 |
| 21305 | 1066 | P1066C | D | 3.4e-06 | 2.5e-06 | 2.1e-06 | 2.667E-06 | 6.658E-07 | 2.100E-06 | 3.400E-06 |
| 21306 | 1066 | P1066E | D | 3e-06 | 2.2e-06 | 2.6e-06 | 2.600E-06 | 4.000E-07 | 2.200E-06 | 3.000E-06 |
| 21307 | 1066 | P1066Q | D | 1.7e-06 | 1.3e-06 | 2.6e-06 | 1.867E-06 | 6.658E-07 | 1.300E-06 | 2.600E-06 |
| 21308 | 1066 | P1066G | D | 5.3e-06 | 2.8e-06 | 2.4e-06 | 3.500E-06 | 1.572E-06 | 2.400E-06 | 5.300E-06 |
| 21309 | 1066 | P1066H | D | 1.3e-06 | 1.7e-06 | 3.4e-06 | 2.133E-06 | 1.115E-06 | 1.300E-06 | 3.400E-06 |
| 21310 | 1066 | P1066I | D | 1.8e-06 | 1.8e-06 | 3.5e-06 | 2.367E-06 | 9.815E-07 | 1.800E-06 | 3.500E-06 |
| 21311 | 1066 | P1066L | D | 1.9e-06 | 3.7e-06 | 2.2e-06 | 2.600E-06 | 9.644E-07 | 1.900E-06 | 3.700E-06 |
| 21312 | 1066 | P1066K | D | 1.7e-06 | 2.6e-06 | 2.3e-06 | 2.200E-06 | 4.583E-07 | 1.700E-06 | 2.600E-06 |
| 21313 | 1066 | P1066M | D | 2.7e-06 | 2e-06 | 1.7e-06 | 2.133E-06 | 5.132E-07 | 1.700E-06 | 2.700E-06 |
| 21314 | 1066 | P1066F | D | 2.3e-06 | 2.4e-06 | 2e-06 | 2.233E-06 | 2.082E-07 | 2.000E-06 | 2.400E-06 |
| 21315 | 1066 | P1066P | D | 1.6e-06 | 2.8e-06 | 2.4e-06 | 2.267E-06 | 6.110E-07 | 1.600E-06 | 2.800E-06 |
| 21316 | 1066 | P1066S | D | 2.4e-06 | 2.7e-06 | 2.2e-06 | 2.433E-06 | 2.517E-07 | 2.200E-06 | 2.700E-06 |
| 21317 | 1066 | P1066T | D | 4.9e-06 | 1.2e-06 | 1.9e-06 | 2.667E-06 | 1.966E-06 | 1.200E-06 | 4.900E-06 |
| 21318 | 1066 | P1066W | D | 3.1e-06 | 2.2e-06 | 2.4e-06 | 2.567E-06 | 4.726E-07 | 2.200E-06 | 3.100E-06 |
| 21319 | 1066 | P1066Y | D | 2.7e-06 | 3.7e-06 | 2.4e-06 | 2.933E-06 | 6.807E-07 | 2.400E-06 | 3.700E-06 |
| 21320 | 1066 | P1066V | D | 2e-06 | 2.1e-06 | 3.5e-06 | 2.533E-06 | 8.386E-07 | 2.000E-06 | 3.500E-06 |
| 21321 | 1067 | S1067A | D | 2.4e-06 | 4.4e-06 | 3.6e-06 | 3.467E-06 | 1.007E-06 | 2.400E-06 | 4.400E-06 |
| 21322 | 1067 | S1067R | D | 3.4e-06 | 3e-06 | 2.3e-06 | 2.900E-06 | 5.568E-07 | 2.300E-06 | 3.400E-06 |
| 21323 | 1067 | S1067N | D | 1.1e-06 | 2.1e-06 | 1.5e-06 | 1.567E-06 | 5.033E-07 | 1.100E-06 | 2.100E-06 |
| 21324 | 1067 | S1067D | D | 2.2e-06 | 2.6e-06 | 3.2e-06 | 2.667E-06 | 5.033E-07 | 2.200E-06 | 3.200E-06 |
| 21325 | 1067 | S1067C | D | 4.2e-06 | 3.2e-06 | 2.3e-06 | 3.233E-06 | 9.504E-07 | 2.300E-06 | 4.200E-06 |
| 21326 | 1067 | S1067E | D | 4.7e-06 | 2.2e-06 | 2.9e-06 | 3.267E-06 | 1.290E-06 | 2.200E-06 | 4.700E-06 |
| 21327 | 1067 | S1067Q | D | 2.6e-06 | 2.2e-06 | 1.7e-06 | 2.167E-06 | 4.509E-07 | 1.700E-06 | 2.600E-06 |
| 21328 | 1067 | S1067G | D | 2.7e-06 | 3.8e-06 | 2.1e-06 | 2.867E-06 | 8.622E-07 | 2.100E-06 | 3.800E-06 |
| 21329 | 1067 | S1067H | D | 1.7e-06 | 1.9e-06 | 2.8e-06 | 2.133E-06 | 5.859E-07 | 1.700E-06 | 2.800E-06 |
| 21330 | 1067 | S1067I | D | 1.7e-06 | 3.2e-06 | 2.7e-06 | 2.533E-06 | 7.638E-07 | 1.700E-06 | 3.200E-06 |
| 21331 | 1067 | S1067L | D | 1.3e-06 | 9.2e-07 | 1.9e-06 | 1.373E-06 | 4.941E-07 | 9.200E-07 | 1.900E-06 |
| 21332 | 1067 | S1067K | D | 2e-06 | 2e-06 | 1.3e-06 | 1.767E-06 | 4.041E-07 | 1.300E-06 | 2.000E-06 |
| 21333 | 1067 | S1067M | D | 2e-06 | 1.5e-06 | 4e-06 | 2.500E-06 | 1.323E-06 | 1.500E-06 | 4.000E-06 |
| 21334 | 1067 | S1067F | D | 2.8e-06 | 3e-06 | 1.6e-06 | 2.467E-06 | 7.572E-07 | 1.600E-06 | 3.000E-06 |

| | | | | | | | | | | |
|---|---|---|---|---|---|---|---|---|---|---|
| 21335 | 1067 | S1067P | D | 2.1e-06 | 2.6e-06 | 1.1e-06 | 1.933E-06 | 7.638E-07 | 1.100E-06 | 2.600E-06 |
| 21336 | 1067 | S1067S | D | 1.6e-06 | 2.2e-06 | 1.8e-06 | 1.867E-06 | 3.055E-07 | 1.600E-06 | 2.200E-06 |
| 21337 | 1067 | S1067T | D | 3.3e-06 | 3.1e-06 | 2.4e-06 | 2.933E-06 | 4.726E-07 | 2.400E-06 | 3.300E-06 |
| 21338 | 1067 | S1067W | D | 3.1e-06 | 3.2e-06 | 5.3e-06 | 3.867E-06 | 1.242E-06 | 3.100E-06 | 5.300E-06 |
| 21339 | 1067 | S1067Y | D | 2.9e-06 | 2.6e-06 | 3.2e-06 | 2.900E-06 | 3.000E-07 | 2.600E-06 | 3.200E-06 |
| 21340 | 1067 | S1067V | D | 3e-06 | 2.3e-06 | 2e-06 | 2.433E-06 | 5.132E-07 | 2.000E-06 | 3.000E-06 |
| 21341 | 1068 | S1068A | D | 1.6e-06 | 2.1e-06 | 1.8e-06 | 1.833E-06 | 2.517E-07 | 1.600E-06 | 2.100E-06 |
| 21342 | 1068 | S1068R | D | 6.1e-06 | 2e-06 | 1.5e-06 | 3.200E-06 | 2.524E-06 | 1.500E-06 | 6.100E-06 |
| 21343 | 1068 | S1068N | D | 2.6e-06 | 1.4e-06 | 2.9e-06 | 2.300E-06 | 7.937E-07 | 1.400E-06 | 2.900E-06 |
| 21344 | 1068 | S1068D | D | 2.9e-06 | 2.9e-06 | 3.9e-06 | 3.233E-06 | 5.774E-07 | 2.900E-06 | 3.900E-06 |
| 21345 | 1068 | S1068C | D | 1.5e-06 | 1.3e-06 | 4e-06 | 2.267E-06 | 1.504E-06 | 1.300E-06 | 4.000E-06 |
| 21346 | 1068 | S1068E | D | 2.2e-06 | 4.4e-06 | 3.8e-06 | 3.467E-06 | 1.137E-06 | 2.200E-06 | 4.400E-06 |
| 21347 | 1068 | S1068Q | D | 2.3e-06 | 1.8e-06 | 2.2e-06 | 2.100E-06 | 2.646E-07 | 1.800E-06 | 2.300E-06 |
| 21348 | 1068 | S1068G | D | 3e-06 | 3.2e-06 | 2.1e-06 | 2.767E-06 | 5.859E-07 | 2.100E-06 | 3.200E-06 |
| 21349 | 1068 | S1068H | D | 3e-06 | 1.7e-06 | 2.7e-06 | 2.467E-06 | 6.807E-07 | 1.700E-06 | 3.000E-06 |
| 21350 | 1068 | S1068I | D | 2.8e-06 | 1.5e-06 | 2.4e-06 | 2.233E-06 | 6.658E-07 | 1.500E-06 | 2.800E-06 |
| 21351 | 1068 | S1068L | D | 1.6e-06 | 2.4e-06 | 2e-06 | 2.000E-06 | 4.000E-07 | 1.600E-06 | 2.400E-06 |
| 21352 | 1068 | S1068K | D | 2.3e-06 | 2.1e-06 | 5.5e-06 | 3.300E-06 | 1.908E-06 | 2.100E-06 | 5.500E-06 |
| 21353 | 1068 | S1068M | D | 3.2e-06 | 1.5e-06 | 1.9e-06 | 2.200E-06 | 8.888E-07 | 1.500E-06 | 3.200E-06 |
| 21354 | 1068 | S1068F | D | 2.1e-06 | 2.6e-06 | 4.3e-06 | 3.000E-06 | 1.153E-06 | 2.100E-06 | 4.300E-06 |
| 21355 | 1068 | S1068P | D | 1.6e-06 | 4.2e-06 | 1.4e-06 | 2.400E-06 | 1.562E-06 | 1.400E-06 | 4.200E-06 |
| 21356 | 1068 | S1068S | D | 1.9e-06 | 2.3e-06 | 5.2e-06 | 3.133E-06 | 1.801E-06 | 1.900E-06 | 5.200E-06 |
| 21357 | 1068 | S1068T | D | 1.2e-06 | 2.6e-06 | 2.3e-06 | 2.033E-06 | 7.371E-07 | 1.200E-06 | 2.600E-06 |
| 21358 | 1068 | S1068W | D | 3e-06 | 2.4e-06 | 2.5e-06 | 2.633E-06 | 3.215E-07 | 2.400E-06 | 3.000E-06 |
| 21359 | 1068 | S1068Y | D | 2.5e-06 | 1.5e-06 | 2.6e-06 | 2.200E-06 | 6.083E-07 | 1.500E-06 | 2.600E-06 |
| 21360 | 1068 | S1068V | D | 1.8e-06 | 1.5e-06 | 1.8e-06 | 1.700E-06 | 1.732E-07 | 1.500E-06 | 1.800E-06 |
| 21361 | 1069 | L1069A | D | 1.8e-06 | 2.8e-06 | 1.5e-06 | 2.033E-06 | 6.807E-07 | 1.500E-06 | 2.800E-06 |
| 21362 | 1069 | L1069R | D | 3e-06 | 1.9e-06 | 1.9e-06 | 2.267E-06 | 6.351E-07 | 1.900E-06 | 3.000E-06 |
| 21363 | 1069 | L1069N | D | 1.6e-06 | 2.6e-06 | 2.3e-06 | 2.167E-06 | 5.132E-07 | 1.600E-06 | 2.600E-06 |
| 21364 | 1069 | L1069D | D | 1.7e-06 | 2.6e-06 | 2.9e-06 | 2.400E-06 | 6.245E-07 | 1.700E-06 | 2.900E-06 |
| 21365 | 1069 | L1069C | D | 3.2e-06 | 1.7e-06 | 1.8e-06 | 2.233E-06 | 8.386E-07 | 1.700E-06 | 3.200E-06 |
| 21366 | 1069 | L1069E | D | 1.3e-06 | 3e-06 | 1.3e-06 | 1.867E-06 | 9.815E-07 | 1.300E-06 | 3.000E-06 |
| 21367 | 1069 | L1069Q | D | 3e-06 | 1.8e-06 | 3e-06 | 2.600E-06 | 6.928E-07 | 1.800E-06 | 3.000E-06 |
| 21368 | 1069 | L1069G | D | 3.7e-06 | 2.4e-06 | 5e-06 | 3.700E-06 | 1.300E-06 | 2.400E-06 | 5.000E-06 |
| 21369 | 1069 | L1069H | D | 2.1e-06 | 1.9e-06 | 2.2e-06 | 2.067E-06 | 1.528E-07 | 1.900E-06 | 2.200E-06 |
| 21370 | 1069 | L1069I | D | 2.1e-06 | 2.9e-06 | 2.3e-06 | 2.433E-06 | 4.163E-07 | 2.100E-06 | 2.900E-06 |
| 21371 | 1069 | L1069L | D | 1.9e-06 | 2.6e-06 | 3.1e-06 | 2.533E-06 | 6.028E-07 | 1.900E-06 | 3.100E-06 |
| 21372 | 1069 | L1069K | D | 6.8e-06 | 2.2e-06 | 2.1e-06 | 3.700E-06 | 2.685E-06 | 2.100E-06 | 6.800E-06 |
| 21373 | 1069 | L1069M | D | 1.8e-06 | 2.5e-06 | 1.8e-06 | 2.033E-06 | 4.041E-07 | 1.800E-06 | 2.500E-06 |
| 21374 | 1069 | L1069F | D | 1.1e-06 | 4.9e-06 | 2.3e-06 | 2.767E-06 | 1.943E-06 | 1.100E-06 | 4.900E-06 |
| 21375 | 1069 | L1069P | D | 3.2e-06 | 3.1e-06 | 2.1e-06 | 2.800E-06 | 6.083E-07 | 2.100E-06 | 3.200E-06 |
| 21376 | 1069 | L1069S | D | 2.3e-06 | 2.7e-06 | 3.1e-06 | 2.700E-06 | 4.000E-07 | 2.300E-06 | 3.100E-06 |
| 21377 | 1069 | L1069T | D | 2.4e-06 | 2.2e-06 | 1.2e-06 | 1.933E-06 | 6.429E-07 | 1.200E-06 | 2.400E-06 |
| 21378 | 1069 | L1069W | D | 2.4e-06 | 1.6e-06 | 3.1e-06 | 2.367E-06 | 7.506E-07 | 1.600E-06 | 3.100E-06 |
| 21379 | 1069 | L1069Y | D | 3.8e-06 | 2.8e-06 | 1.9e-06 | 2.833E-06 | 9.504E-07 | 1.900E-06 | 3.800E-06 |
| 21380 | 1069 | L1069V | D | 5e-06 | 2.3e-06 | 1.9e-06 | 3.067E-06 | 1.686E-06 | 1.900E-06 | 5.000E-06 |
| 21381 | 1070 | S1070A | D | 3.4e-06 | 1.9e-06 | 3.2e-06 | 2.833E-06 | 8.145E-07 | 1.900E-06 | 3.400E-06 |
| 21382 | 1070 | S1070R | D | 3.5e-06 | 3e-06 | 3e-06 | 3.167E-06 | 2.887E-07 | 3.000E-06 | 3.500E-06 |
| 21383 | 1070 | S1070N | D | 2e-06 | 2.5e-06 | 4e-06 | 2.833E-06 | 1.041E-06 | 2.000E-06 | 4.000E-06 |
| 21384 | 1070 | S1070D | D | 1.6e-06 | 2.2e-06 | 2e-06 | 1.933E-06 | 3.055E-07 | 1.600E-06 | 2.200E-06 |
| 21385 | 1070 | S1070C | D | 3.6e-06 | 1.7e-06 | 4.1e-06 | 3.133E-06 | 1.266E-06 | 1.700E-06 | 4.100E-06 |
| 21386 | 1070 | S1070E | D | 3.4e-06 | 6.3e-06 | 2e-06 | 3.900E-06 | 2.193E-06 | 2.000E-06 | 6.300E-06 |
| 21387 | 1070 | S1070Q | D | 2.4e-06 | 2.2e-06 | 2.6e-06 | 2.400E-06 | 2.000E-07 | 2.200E-06 | 2.600E-06 |
| 21388 | 1070 | S1070G | D | 1.7e-06 | 1.9e-06 | 3.2e-06 | 2.267E-06 | 8.145E-07 | 1.700E-06 | 3.200E-06 |
| 21389 | 1070 | S1070H | D | 2.2e-06 | 1.9e-06 | 1.9e-06 | 2.000E-06 | 1.732E-07 | 1.900E-06 | 2.200E-06 |
| 21390 | 1070 | S1070I | D | 1e-06 | 8.6e-07 | 2.6e-06 | 1.487E-06 | 9.667E-07 | 8.600E-07 | 2.600E-06 |
| 21391 | 1070 | S1070L | D | 6.5e-06 | 1.2e-06 | 6.5e-07 | 2.783E-06 | 3.230E-06 | 6.500E-07 | 6.500E-06 |
| 21392 | 1070 | S1070K | D | 2.9e-06 | 1.8e-06 | 3.8e-06 | 2.833E-06 | 1.002E-06 | 1.800E-06 | 3.800E-06 |
| 21393 | 1070 | S1070M | D | 2.5e-06 | 2.2e-06 | 2e-06 | 2.233E-06 | 2.517E-07 | 2.000E-06 | 2.500E-06 |
| 21394 | 1070 | S1070F | D | 2.2e-06 | 2.9e-06 | 3.9e-06 | 3.000E-06 | 8.544E-07 | 2.200E-06 | 3.900E-06 |
| 21395 | 1070 | S1070P | D | 1.4e-06 | 1.4e-06 | 3.6e-06 | 2.133E-06 | 1.270E-06 | 1.400E-06 | 3.600E-06 |
| 21396 | 1070 | S1070S | D | 2.4e-06 | 2.3e-06 | 1.9e-06 | 2.200E-06 | 2.646E-07 | 1.900E-06 | 2.400E-06 |
| 21397 | 1070 | S1070T | D | 2.2e-06 | 3.7e-06 | 2e-06 | 2.633E-06 | 9.292E-07 | 2.000E-06 | 3.700E-06 |
| 21398 | 1070 | S1070W | D | 2.7e-06 | 2.7e-06 | 1.7e-06 | 2.367E-06 | 5.774E-07 | 1.700E-06 | 2.700E-06 |
| 21399 | 1070 | S1070Y | D | 2.5e-06 | 2.5e-06 | 1.7e-06 | 2.233E-06 | 4.619E-07 | 1.700E-06 | 2.500E-06 |
| 21400 | 1070 | S1070V | D | 1.7e-06 | 2.4e-06 | 1.2e-06 | 1.767E-06 | 6.028E-07 | 1.200E-06 | 2.400E-06 |
| 21401 | 1071 | A1071A | D | 1.6e-06 | 1.9e-06 | 2.5e-06 | 2.000E-06 | 4.583E-07 | 1.600E-06 | 2.500E-06 |
| 21402 | 1071 | A1071R | D | 3.2e-06 | 2.1e-06 | 1.7e-06 | 2.333E-06 | 7.767E-07 | 1.700E-06 | 3.200E-06 |
| 21403 | 1071 | A1071N | D | 2.9e-06 | 2.3e-06 | 1.5e-06 | 2.233E-06 | 7.024E-07 | 1.500E-06 | 2.900E-06 |
| 21404 | 1071 | A1071D | D | 1.3e-06 | 2.8e-06 | 3e-06 | 2.367E-06 | 9.292E-07 | 1.300E-06 | 3.000E-06 |
| 21405 | 1071 | A1071C | D | 2.9e-06 | 4.7e-06 | 2.4e-06 | 3.333E-06 | 1.210E-06 | 2.400E-06 | 4.700E-06 |
| 21406 | 1071 | A1071E | D | 1.1e-06 | 1.5e-06 | 3.2e-06 | 1.933E-06 | 1.115E-06 | 1.100E-06 | 3.200E-06 |
| 21407 | 1071 | A1071Q | D | 2.4e-06 | 2.7e-06 | 2.4e-06 | 2.500E-06 | 1.732E-07 | 2.400E-06 | 2.700E-06 |
| 21408 | 1071 | A1071G | D | 2.5e-06 | 1.4e-06 | 3.3e-06 | 2.400E-06 | 9.539E-07 | 1.400E-06 | 3.300E-06 |
| 21409 | 1071 | A1071H | D | 2.1e-06 | 2.2e-06 | 2.4e-06 | 2.233E-06 | 1.528E-07 | 2.100E-06 | 2.400E-06 |
| 21410 | 1071 | A1071I | D | 1.6e-06 | 4e-06 | 3.4e-06 | 3.000E-06 | 1.249E-06 | 1.600E-06 | 4.000E-06 |
| 21411 | 1071 | A1071L | D | 2.7e-06 | 2.7e-06 | 3.2e-06 | 2.867E-06 | 2.887E-07 | 2.700E-06 | 3.200E-06 |
| 21412 | 1071 | A1071K | D | 2.2e-06 | 2e-06 | 1.1e-06 | 1.767E-06 | 5.859E-07 | 1.100E-06 | 2.200E-06 |
| 21413 | 1071 | A1071M | D | 2.3e-06 | 2.3e-06 | 2.9e-06 | 2.500E-06 | 3.464E-07 | 2.300E-06 | 2.900E-06 |
| 21414 | 1071 | A1071F | D | 2.7e-06 | 2.4e-06 | 2.4e-06 | 2.500E-06 | 1.732E-07 | 2.400E-06 | 2.700E-06 |
| 21415 | 1071 | A1071P | D | 2.4e-06 | 2.4e-06 | 1.8e-06 | 2.200E-06 | 3.464E-07 | 1.800E-06 | 2.400E-06 |
| 21416 | 1071 | A1071S | D | 2.8e-06 | 4.7e-06 | 2.8e-06 | 3.433E-06 | 1.097E-06 | 2.800E-06 | 4.700E-06 |
| 21417 | 1071 | A1071T | D | 1.7e-06 | 5e-06 | 2.3e-06 | 3.000E-06 | 1.758E-06 | 1.700E-06 | 5.000E-06 |
| 21418 | 1071 | A1071W | D | 1.4e-06 | 2.8e-06 | 2.3e-06 | 2.167E-06 | 7.095E-07 | 1.400E-06 | 2.800E-06 |
| 21419 | 1071 | A1071Y | D | 2.9e-06 | 2.5e-06 | 2.7e-06 | 2.700E-06 | 2.000E-07 | 2.500E-06 | 2.900E-06 |

| | | | | | | | | | | |
|---|---|---|---|---|---|---|---|---|---|---|
| 21420 | 1071 | A1071V | D | 2.8e-06 | 3.3e-06 | 1.1e-06 | 2.400E-06 | 1.153E-06 | 1.100E-06 | 3.300E-06 |
| 21421 | 1072 | S1072A | D | 2.3e-06 | 3.6e-06 | 2.2e-06 | 2.700E-06 | 7.810E-07 | 2.200E-06 | 3.600E-06 |
| 21422 | 1072 | S1072R | D | 1.1e-06 | 1.7e-06 | 3.3e-06 | 2.033E-06 | 1.137E-06 | 1.100E-06 | 3.300E-06 |
| 21423 | 1072 | S1072N | D | 1.3e-06 | 2.7e-06 | 2.3e-06 | 2.100E-06 | 7.211E-07 | 1.300E-06 | 2.700E-06 |
| 21424 | 1072 | S1072D | D | 1.4e-06 | 2.5e-06 | 1.2e-06 | 1.700E-06 | 7.000E-07 | 1.200E-06 | 2.500E-06 |
| 21425 | 1072 | S1072C | D | 2.3e-06 | 2.6e-06 | 2.3e-06 | 2.400E-06 | 1.732E-07 | 2.300E-06 | 2.600E-06 |
| 21426 | 1072 | S1072E | D | 2.4e-06 | 3.2e-06 | 2.3e-06 | 2.633E-06 | 4.933E-07 | 2.300E-06 | 3.200E-06 |
| 21427 | 1072 | S1072Q | D | 2.1e-06 | 2.4e-06 | 1.5e-06 | 2.000E-06 | 4.583E-07 | 1.500E-06 | 2.400E-06 |
| 21428 | 1072 | S1072G | D | 2.7e-06 | 2e-06 | 3e-06 | 2.567E-06 | 5.132E-07 | 2.000E-06 | 3.000E-06 |
| 21429 | 1072 | S1072H | D | 3.1e-06 | 2.7e-06 | 2.5e-06 | 2.767E-06 | 3.055E-07 | 2.500E-06 | 3.100E-06 |
| 21430 | 1072 | S1072I | D | 1.5e-06 | 1.5e-06 | 2.4e-06 | 1.800E-06 | 5.196E-07 | 1.500E-06 | 2.400E-06 |
| 21431 | 1072 | S1072L | D | 2.4e-06 | 2.7e-06 | 3.9e-06 | 3.000E-06 | 7.937E-07 | 2.400E-06 | 3.900E-06 |
| 21432 | 1072 | S1072K | D | 2.6e-06 | 2.9e-06 | 1.3e-06 | 2.267E-06 | 8.505E-07 | 1.300E-06 | 2.900E-06 |
| 21433 | 1072 | S1072M | D | 2.4e-06 | 2.6e-06 | 2.1e-06 | 2.367E-06 | 2.517E-07 | 2.100E-06 | 2.600E-06 |
| 21434 | 1072 | S1072F | D | 3e-06 | 1.5e-06 | 3e-06 | 2.500E-06 | 8.660E-07 | 1.500E-06 | 3.000E-06 |
| 21435 | 1072 | S1072P | D | 3e-06 | 2e-06 | 2.1e-06 | 2.367E-06 | 5.508E-07 | 2.000E-06 | 3.000E-06 |
| 21436 | 1072 | S1072S | D | 3.5e-06 | 1.9e-06 | 2.4e-06 | 2.600E-06 | 8.185E-07 | 1.900E-06 | 3.500E-06 |
| 21437 | 1072 | S1072T | D | 2.5e-06 | 2.1e-06 | 3.3e-06 | 2.633E-06 | 6.110E-07 | 2.100E-06 | 3.300E-06 |
| 21438 | 1072 | S1072W | D | 3.8e-06 | 2.4e-06 | 1.9e-06 | 2.700E-06 | 9.849E-07 | 1.900E-06 | 3.800E-06 |
| 21439 | 1072 | S1072Y | D | 3.8e-06 | 2e-06 | 2.2e-06 | 2.667E-06 | 9.866E-07 | 2.000E-06 | 3.800E-06 |
| 21440 | 1072 | S1072V | D | 1.6e-06 | 2.3e-06 | 2.6e-06 | 2.167E-06 | 5.132E-07 | 1.600E-06 | 2.600E-06 |
| 21441 | 1073 | V1073A | D | 4.3e-06 | 2.2e-06 | 2.1e-06 | 2.867E-06 | 1.242E-06 | 2.100E-06 | 4.300E-06 |
| 21442 | 1073 | V1073R | D | 3.3e-06 | 4.5e-06 | 2.4e-06 | 3.400E-06 | 1.054E-06 | 2.400E-06 | 4.500E-06 |
| 21443 | 1073 | V1073N | D | 1.8e-06 | 2.2e-06 | 1.6e-06 | 1.867E-06 | 3.055E-07 | 1.600E-06 | 2.200E-06 |
| 21444 | 1073 | V1073D | D | 2.7e-06 | 1.7e-06 | 3.1e-06 | 2.500E-06 | 7.211E-07 | 1.700E-06 | 3.100E-06 |
| 21445 | 1073 | V1073C | D | 1.4e-06 | 4e-06 | 2e-06 | 2.467E-06 | 1.361E-06 | 1.400E-06 | 4.000E-06 |
| 21446 | 1073 | V1073E | D | 1.4e-06 | 2.5e-06 | 1.5e-06 | 1.800E-06 | 6.083E-07 | 1.400E-06 | 2.500E-06 |
| 21447 | 1073 | V1073Q | D | 1.5e-06 | 2.7e-06 | 1.7e-06 | 1.967E-06 | 6.429E-07 | 1.500E-06 | 2.700E-06 |
| 21448 | 1073 | V1073G | D | 2.3e-06 | 4.2e-06 | 3e-06 | 3.167E-06 | 9.609E-07 | 2.300E-06 | 4.200E-06 |
| 21449 | 1073 | V1073H | D | 2.5e-06 | 3.2e-06 | 1.5e-06 | 2.400E-06 | 8.544E-07 | 1.500E-06 | 3.200E-06 |
| 21450 | 1073 | V1073I | D | 3e-06 | 2.6e-06 | 1.5e-06 | 2.367E-06 | 7.767E-07 | 1.500E-06 | 3.000E-06 |
| 21451 | 1073 | V1073L | D | 3.1e-06 | 1.4e-06 | 1.8e-06 | 2.100E-06 | 8.888E-07 | 1.400E-06 | 3.100E-06 |
| 21452 | 1073 | V1073K | D | 2.2e-06 | 1.8e-06 | 1.6e-06 | 1.867E-06 | 3.055E-07 | 1.600E-06 | 2.200E-06 |
| 21453 | 1073 | V1073M | D | 2.3e-06 | 2.5e-06 | 2.8e-06 | 2.533E-06 | 2.517E-07 | 2.300E-06 | 2.800E-06 |
| 21454 | 1073 | V1073F | D | 2.4e-06 | 1.2e-06 | 2.2e-06 | 1.933E-06 | 6.429E-07 | 1.200E-06 | 2.400E-06 |
| 21455 | 1073 | V1073P | D | 9.8e-07 | 4.2e-06 | 1.5e-06 | 2.227E-06 | 1.729E-06 | 9.800E-07 | 4.200E-06 |
| 21456 | 1073 | V1073S | D | 1.7e-06 | 3.5e-06 | 4e-06 | 3.067E-06 | 1.210E-06 | 1.700E-06 | 4.000E-06 |
| 21457 | 1073 | V1073T | D | 1.9e-06 | 2.3e-06 | 2.4e-06 | 2.200E-06 | 2.646E-07 | 1.900E-06 | 2.400E-06 |
| 21458 | 1073 | V1073W | D | 1.5e-06 | 2.4e-06 | 1.7e-06 | 1.867E-06 | 4.726E-07 | 1.500E-06 | 2.400E-06 |
| 21459 | 1073 | V1073Y | D | 2.8e-06 | 1.5e-06 | 1.2e-06 | 1.833E-06 | 8.505E-07 | 1.200E-06 | 2.800E-06 |
| 21460 | 1073 | V1073V | D | 3.1e-06 | 1.6e-06 | 1.7e-06 | 2.133E-06 | 8.386E-07 | 1.600E-06 | 3.100E-06 |
| 21461 | 1074 | G1074A | D | 2.7e-06 | 1.6e-06 | 2.1e-06 | 2.133E-06 | 5.508E-07 | 1.600E-06 | 2.700E-06 |
| 21462 | 1074 | G1074R | D | 2.9e-06 | 3.2e-06 | 1.9e-06 | 2.667E-06 | 6.807E-07 | 1.900E-06 | 3.200E-06 |
| 21463 | 1074 | G1074N | D | 2.7e-06 | 2.9e-06 | 2.4e-06 | 2.667E-06 | 2.517E-07 | 2.400E-06 | 2.900E-06 |
| 21464 | 1074 | G1074D | D | 2.1e-06 | 2.8e-06 | 2.2e-06 | 2.367E-06 | 3.786E-07 | 2.100E-06 | 2.800E-06 |
| 21465 | 1074 | G1074C | D | 4.6e-06 | 2.2e-06 | 1.7e-06 | 2.833E-06 | 1.550E-06 | 1.700E-06 | 4.600E-06 |
| 21466 | 1074 | G1074E | D | 2.2e-06 | 2.6e-06 | 3.3e-06 | 2.700E-06 | 5.568E-07 | 2.200E-06 | 3.300E-06 |
| 21467 | 1074 | G1074Q | D | 2.4e-06 | 2.2e-06 | 2.7e-06 | 2.433E-06 | 2.517E-07 | 2.200E-06 | 2.700E-06 |
| 21468 | 1074 | G1074G | D | 2.2e-06 | 2.6e-06 | 2.3e-06 | 2.367E-06 | 2.082E-07 | 2.200E-06 | 2.600E-06 |
| 21469 | 1074 | G1074H | D | 2.7e-06 | 3.6e-06 | 1.6e-06 | 2.633E-06 | 1.002E-06 | 1.600E-06 | 3.600E-06 |
| 21470 | 1074 | G1074I | D | 1.9e-06 | 2.8e-06 | 3.6e-06 | 2.767E-06 | 8.505E-07 | 1.900E-06 | 3.600E-06 |
| 21471 | 1074 | G1074L | D | 3.8e-06 | 2.3e-06 | 2.2e-06 | 2.767E-06 | 8.963E-07 | 2.200E-06 | 3.800E-06 |
| 21472 | 1074 | G1074K | D | 1.9e-06 | 2e-06 | 1.4e-06 | 1.767E-06 | 3.215E-07 | 1.400E-06 | 2.000E-06 |
| 21473 | 1074 | G1074M | D | 1.7e-06 | 3.2e-06 | 2.9e-06 | 2.600E-06 | 7.937E-07 | 1.700E-06 | 3.200E-06 |
| 21474 | 1074 | G1074F | D | 1.4e-06 | 2.5e-06 | 2.9e-06 | 2.267E-06 | 7.767E-07 | 1.400E-06 | 2.900E-06 |
| 21475 | 1074 | G1074P | D | 2.1e-06 | 2.5e-06 | 2.9e-06 | 2.500E-06 | 4.000E-07 | 2.100E-06 | 2.900E-06 |
| 21476 | 1074 | G1074S | D | 5.1e-06 | 2.3e-06 | 3e-06 | 3.467E-06 | 1.457E-06 | 2.300E-06 | 5.100E-06 |
| 21477 | 1074 | G1074T | D | 2.8e-06 | 2.1e-06 | 1.6e-06 | 2.167E-06 | 6.028E-07 | 1.600E-06 | 2.800E-06 |
| 21478 | 1074 | G1074W | D | 2.1e-06 | 2.4e-06 | 1.9e-06 | 2.133E-06 | 2.517E-07 | 1.900E-06 | 2.400E-06 |
| 21479 | 1074 | G1074Y | D | 2.2e-06 | 2.7e-06 | 2.3e-06 | 2.400E-06 | 2.646E-07 | 2.200E-06 | 2.700E-06 |
| 21480 | 1074 | G1074V | D | 1.9e-06 | 2.1e-06 | 3.7e-06 | 2.567E-06 | 9.866E-07 | 1.900E-06 | 3.700E-06 |
| 21481 | 1075 | D1075A | D | 2.3e-06 | 1.8e-06 | 2.2e-06 | 2.100E-06 | 2.646E-07 | 1.800E-06 | 2.300E-06 |
| 21482 | 1075 | D1075R | D | 3.5e-06 | 1.8e-06 | 1.8e-06 | 2.367E-06 | 9.815E-07 | 1.800E-06 | 3.500E-06 |
| 21483 | 1075 | D1075N | D | 2.3e-06 | 4.6e-06 | 2.3e-06 | 3.067E-06 | 1.328E-06 | 2.300E-06 | 4.600E-06 |
| 21484 | 1075 | D1075D | D | 4.1e-06 | 3.4e-06 | 1.7e-06 | 3.067E-06 | 1.234E-06 | 1.700E-06 | 4.100E-06 |
| 21485 | 1075 | D1075C | D | 2.1e-06 | 1.4e-06 | 4.3e-06 | 2.600E-06 | 1.513E-06 | 1.400E-06 | 4.300E-06 |
| 21486 | 1075 | D1075E | D | 1.8e-06 | 1.9e-06 | 4.4e-06 | 2.700E-06 | 1.473E-06 | 1.800E-06 | 4.400E-06 |
| 21487 | 1075 | D1075Q | D | 2.4e-06 | 2.6e-06 | 2.3e-06 | 2.433E-06 | 1.528E-07 | 2.300E-06 | 2.600E-06 |
| 21488 | 1075 | D1075G | D | 2.2e-06 | 2.2e-06 | 2.1e-06 | 2.167E-06 | 5.774E-08 | 2.100E-06 | 2.200E-06 |
| 21489 | 1075 | D1075H | D | 3.2e-06 | 3.1e-06 | 2e-06 | 2.767E-06 | 6.658E-07 | 2.000E-06 | 3.200E-06 |
| 21490 | 1075 | D1075I | D | 2.2e-06 | 2.3e-06 | 8.8e-07 | 1.793E-06 | 7.925E-07 | 8.800E-07 | 2.300E-06 |
| 21491 | 1075 | D1075L | D | 2.5e-06 | 1.7e-06 | 2.8e-06 | 2.333E-06 | 5.686E-07 | 1.700E-06 | 2.800E-06 |
| 21492 | 1075 | D1075K | D | 2.6e-06 | 2.1e-06 | 2.1e-06 | 2.267E-06 | 2.887E-07 | 2.100E-06 | 2.600E-06 |
| 21493 | 1075 | D1075M | D | 1.8e-06 | 2.9e-06 | 1.4e-06 | 2.033E-06 | 7.767E-07 | 1.400E-06 | 2.900E-06 |
| 21494 | 1075 | D1075F | D | 1.1e-06 | 2e-06 | 3.6e-06 | 2.233E-06 | 1.266E-06 | 1.100E-06 | 3.600E-06 |
| 21495 | 1075 | D1075P | D | 2.9e-06 | 1.9e-06 | 1.6e-06 | 2.133E-06 | 6.807E-07 | 1.600E-06 | 2.900E-06 |
| 21496 | 1075 | D1075S | D | 2.6e-06 | 2.1e-06 | 1.9e-06 | 2.200E-06 | 3.606E-07 | 1.900E-06 | 2.600E-06 |
| 21497 | 1075 | D1075T | D | 1.3e-06 | 2.2e-06 | 2.7e-06 | 2.067E-06 | 7.095E-07 | 1.300E-06 | 2.700E-06 |
| 21498 | 1075 | D1075W | D | 1.8e-06 | 2.5e-06 | 7.5e-07 | 1.683E-06 | 8.808E-07 | 7.500E-07 | 2.500E-06 |
| 21499 | 1075 | D1075Y | D | 1.9e-06 | 9.6e-07 | 2.8e-06 | 1.887E-06 | 9.201E-07 | 9.600E-07 | 2.800E-06 |
| 21500 | 1075 | D1075V | D | 2.1e-06 | 2.9e-06 | 2.6e-06 | 2.533E-06 | 4.041E-07 | 2.100E-06 | 2.900E-06 |
| 21501 | 1076 | R1076A | D | 1.7e-06 | 2.4e-06 | 2.3e-06 | 2.133E-06 | 3.786E-07 | 1.700E-06 | 2.400E-06 |
| 21502 | 1076 | R1076R | D | 2e-06 | 2.5e-06 | 2.3e-06 | 2.267E-06 | 2.517E-07 | 2.000E-06 | 2.500E-06 |
| 21503 | 1076 | R1076N | D | 1.8e-06 | 1.5e-06 | 2.9e-06 | 2.067E-06 | 7.371E-07 | 1.500E-06 | 2.900E-06 |
| 21504 | 1076 | R1076D | D | 1.2e-06 | 3.2e-06 | 2.2e-06 | 2.200E-06 | 1.000E-06 | 1.200E-06 | 3.200E-06 |

| | | | | | | | | | | |
|---|---|---|---|---|---|---|---|---|---|---|
| 21505 | 1076 | R1076C | D | 1.5e-06 | 2.4e-06 | 3.2e-06 | 2.367E-06 | 8.505E-07 | 1.500E-06 | 3.200E-06 |
| 21506 | 1076 | R1076E | D | 1.4e-06 | 1.5e-06 | 1.8e-06 | 1.567E-06 | 2.082E-07 | 1.400E-06 | 1.800E-06 |
| 21507 | 1076 | R1076Q | D | 1.5e-06 | 2.1e-06 | 2.1e-06 | 1.900E-06 | 3.464E-07 | 1.500E-06 | 2.100E-06 |
| 21508 | 1076 | R1076G | D | 2.1e-06 | 3.1e-06 | 2.2e-06 | 2.467E-06 | 5.508E-07 | 2.100E-06 | 3.100E-06 |
| 21509 | 1076 | R1076H | D | 2.1e-06 | 2.1e-06 | 3.8e-06 | 2.667E-06 | 9.815E-07 | 2.100E-06 | 3.800E-06 |
| 21510 | 1076 | R1076I | D | 3.9e-06 | 1.8e-06 | 1.8e-06 | 2.500E-06 | 1.212E-06 | 1.800E-06 | 3.900E-06 |
| 21511 | 1076 | R1076L | D | 1.9e-06 | 1.8e-06 | 2.1e-06 | 1.933E-06 | 1.528E-07 | 1.800E-06 | 2.100E-06 |
| 21512 | 1076 | R1076K | D | 2.1e-06 | 2.2e-06 | 2.9e-06 | 2.400E-06 | 4.359E-07 | 2.100E-06 | 2.900E-06 |
| 21513 | 1076 | R1076M | D | 4.9e-06 | 2.5e-06 | 2.3e-06 | 3.233E-06 | 1.447E-06 | 2.300E-06 | 4.900E-06 |
| 21514 | 1076 | R1076F | D | 3.9e-06 | 2.1e-06 | 2.3e-06 | 2.767E-06 | 9.866E-07 | 2.100E-06 | 3.900E-06 |
| 21515 | 1076 | R1076P | D | 2.9e-06 | 1.6e-06 | 3.6e-06 | 2.700E-06 | 1.015E-06 | 1.600E-06 | 3.600E-06 |
| 21516 | 1076 | R1076S | D | 3e-06 | 1.7e-06 | 1.8e-06 | 2.167E-06 | 7.234E-07 | 1.700E-06 | 3.000E-06 |
| 21517 | 1076 | R1076T | D | 2.8e-06 | 2.2e-06 | 4.9e-06 | 3.300E-06 | 1.418E-06 | 2.200E-06 | 4.900E-06 |
| 21518 | 1076 | R1076W | D | 1.7e-06 | 2.7e-06 | 2.5e-06 | 2.300E-06 | 5.292E-07 | 1.700E-06 | 2.700E-06 |
| 21519 | 1076 | R1076Y | D | 3.9e-06 | 2.5e-06 | 2e-06 | 2.800E-06 | 9.849E-07 | 2.000E-06 | 3.900E-06 |
| 21520 | 1076 | R1076V | D | 2e-06 | 2.5e-06 | 3.1e-06 | 2.533E-06 | 5.508E-07 | 2.000E-06 | 3.100E-06 |
| 21521 | 1077 | V1077A | D | 2.1e-06 | 1.9e-06 | 2.3e-06 | 2.100E-06 | 2.000E-07 | 1.900E-06 | 2.300E-06 |
| 21522 | 1077 | V1077R | D | 3.5e-06 | 1.6e-06 | 1.8e-06 | 2.300E-06 | 1.044E-06 | 1.600E-06 | 3.500E-06 |
| 21523 | 1077 | V1077N | D | 2e-06 | 2.7e-06 | 4e-06 | 2.900E-06 | 1.015E-06 | 2.000E-06 | 4.000E-06 |
| 21524 | 1077 | V1077D | D | 1.5e-06 | 3.7e-06 | 2.7e-06 | 2.633E-06 | 1.102E-06 | 1.500E-06 | 3.700E-06 |
| 21525 | 1077 | V1077C | D | 1.8e-06 | 3.6e-06 | 1.9e-06 | 2.433E-06 | 1.012E-06 | 1.800E-06 | 3.600E-06 |
| 21526 | 1077 | V1077E | D | 2.4e-06 | 2e-06 | 2.7e-06 | 2.367E-06 | 3.512E-07 | 2.000E-06 | 2.700E-06 |
| 21527 | 1077 | V1077Q | D | 2.3e-06 | 2.7e-06 | 1.4e-06 | 2.133E-06 | 6.658E-07 | 1.400E-06 | 2.700E-06 |
| 21528 | 1077 | V1077G | D | 1.8e-06 | 2.5e-06 | 1.8e-06 | 2.033E-06 | 4.041E-07 | 1.800E-06 | 2.500E-06 |
| 21529 | 1077 | V1077H | D | 2.3e-06 | 3.7e-06 | 3e-06 | 3.000E-06 | 7.000E-07 | 2.300E-06 | 3.700E-06 |
| 21530 | 1077 | V1077I | D | 2.8e-06 | 2.2e-06 | 2.7e-06 | 2.567E-06 | 3.215E-07 | 2.200E-06 | 2.800E-06 |
| 21531 | 1077 | V1077L | D | 2.4e-06 | 3e-06 | 3e-06 | 2.800E-06 | 3.464E-07 | 2.400E-06 | 3.000E-06 |
| 21532 | 1077 | V1077K | D | 3.8e-06 | 2.4e-06 | 3.2e-06 | 3.133E-06 | 7.024E-07 | 2.400E-06 | 3.800E-06 |
| 21533 | 1077 | V1077M | D | 2.8e-06 | 3.2e-06 | 2.3e-06 | 2.767E-06 | 4.509E-07 | 2.300E-06 | 3.200E-06 |
| 21534 | 1077 | V1077F | D | 2.5e-06 | 2.2e-06 | 2.3e-06 | 2.333E-06 | 1.528E-07 | 2.200E-06 | 2.500E-06 |
| 21535 | 1077 | V1077P | D | 2.9e-06 | 3.2e-06 | 2.5e-06 | 2.867E-06 | 3.512E-07 | 2.500E-06 | 3.200E-06 |
| 21536 | 1077 | V1077S | D | 1.4e-06 | 1.8e-06 | 8.2e-07 | 1.340E-06 | 4.927E-07 | 8.200E-07 | 1.800E-06 |
| 21537 | 1077 | V1077T | D | 2.8e-06 | 2.7e-06 | 1.5e-06 | 2.333E-06 | 7.234E-07 | 1.500E-06 | 2.800E-06 |
| 21538 | 1077 | V1077W | D | 1.5e-06 | 1.3e-06 | 2.2e-06 | 1.667E-06 | 4.726E-07 | 1.300E-06 | 2.200E-06 |
| 21539 | 1077 | V1077Y | D | 1.7e-06 | 2e-06 | 3.1e-06 | 2.267E-06 | 7.371E-07 | 1.700E-06 | 3.100E-06 |
| 21540 | 1077 | V1077V | D | 2.3e-06 | 2.9e-06 | 3.6e-06 | 2.933E-06 | 6.506E-07 | 2.300E-06 | 3.600E-06 |
| 21541 | 1078 | T1078A | D | 2.8e-06 | 2.8e-06 | 2.5e-06 | 2.700E-06 | 1.732E-07 | 2.500E-06 | 2.800E-06 |
| 21542 | 1078 | T1078R | D | 3.5e-06 | 2e-06 | 1.4e-06 | 2.300E-06 | 1.082E-06 | 1.400E-06 | 3.500E-06 |
| 21543 | 1078 | T1078N | D | 3.1e-06 | 2.5e-06 | 2.4e-06 | 2.667E-06 | 3.786E-07 | 2.400E-06 | 3.100E-06 |
| 21544 | 1078 | T1078D | D | 3e-06 | 2.4e-06 | 3.3e-06 | 2.900E-06 | 4.583E-07 | 2.400E-06 | 3.300E-06 |
| 21545 | 1078 | T1078C | D | 2e-06 | 1.7e-06 | 2.7e-06 | 2.133E-06 | 5.132E-07 | 1.700E-06 | 2.700E-06 |
| 21546 | 1078 | T1078E | D | 1.3e-06 | 2.1e-06 | 1.9e-06 | 1.767E-06 | 4.163E-07 | 1.300E-06 | 2.100E-06 |
| 21547 | 1078 | T1078Q | D | 2.8e-06 | 3.3e-06 | 2.3e-06 | 2.800E-06 | 5.000E-07 | 2.300E-06 | 3.300E-06 |
| 21548 | 1078 | T1078G | D | 2.5e-06 | 2.6e-06 | 2.7e-06 | 2.600E-06 | 1.000E-07 | 2.500E-06 | 2.700E-06 |
| 21549 | 1078 | T1078H | D | 2.2e-06 | 1.5e-06 | 2.6e-06 | 2.100E-06 | 5.568E-07 | 1.500E-06 | 2.600E-06 |
| 21550 | 1078 | T1078I | D | 2.5e-06 | 2.6e-06 | 1.3e-06 | 2.133E-06 | 7.234E-07 | 1.300E-06 | 2.600E-06 |
| 21551 | 1078 | T1078L | D | 2.2e-06 | 2.1e-06 | 4.3e-06 | 2.867E-06 | 1.242E-06 | 2.100E-06 | 4.300E-06 |
| 21552 | 1078 | T1078K | D | 1.5e-06 | 2.7e-06 | 2.2e-06 | 2.133E-06 | 6.028E-07 | 1.500E-06 | 2.700E-06 |
| 21553 | 1078 | T1078M | D | 2.6e-06 | 3.9e-06 | 3.5e-06 | 3.333E-06 | 6.658E-07 | 2.600E-06 | 3.900E-06 |
| 21554 | 1078 | T1078F | D | 1.3e-06 | 4.3e-06 | 2.3e-06 | 2.633E-06 | 1.528E-06 | 1.300E-06 | 4.300E-06 |
| 21555 | 1078 | T1078P | D | 3.3e-06 | 2.3e-06 | 2.6e-06 | 2.733E-06 | 5.132E-07 | 2.300E-06 | 3.300E-06 |
| 21556 | 1078 | T1078S | D | 1.7e-06 | 3.7e-06 | 2.5e-06 | 2.633E-06 | 1.007E-06 | 1.700E-06 | 3.700E-06 |
| 21557 | 1078 | T1078T | D | 2e-06 | 1.9e-06 | 2.6e-06 | 2.167E-06 | 3.786E-07 | 1.900E-06 | 2.600E-06 |
| 21558 | 1078 | T1078W | D | 2.6e-06 | 3e-06 | 1.6e-06 | 2.400E-06 | 7.211E-07 | 1.600E-06 | 3.000E-06 |
| 21559 | 1078 | T1078Y | D | 1.7e-06 | 2.6e-06 | 2.4e-06 | 2.233E-06 | 4.726E-07 | 1.700E-06 | 2.600E-06 |
| 21560 | 1078 | T1078V | D | 2.4e-06 | 1.6e-06 | 3e-06 | 2.333E-06 | 7.024E-07 | 1.600E-06 | 3.000E-06 |
| 21561 | 1079 | I1079A | D | 1.4e-06 | 4.7e-06 | 2.6e-06 | 2.900E-06 | 1.670E-06 | 1.400E-06 | 4.700E-06 |
| 21562 | 1079 | I1079R | D | 1.8e-06 | 2.6e-06 | 2e-06 | 2.133E-06 | 4.163E-07 | 1.800E-06 | 2.600E-06 |
| 21563 | 1079 | I1079N | D | 2e-06 | 1.9e-06 | 1.4e-06 | 1.767E-06 | 3.215E-07 | 1.400E-06 | 2.000E-06 |
| 21564 | 1079 | I1079D | D | 2.1e-06 | 2e-06 | 2.5e-06 | 2.200E-06 | 2.646E-07 | 2.000E-06 | 2.500E-06 |
| 21565 | 1079 | I1079C | D | 1.8e-06 | 2.8e-06 | 2.2e-06 | 2.267E-06 | 5.033E-07 | 1.800E-06 | 2.800E-06 |
| 21566 | 1079 | I1079E | D | 3e-06 | 3.2e-06 | 3.3e-06 | 3.167E-06 | 1.528E-07 | 3.000E-06 | 3.300E-06 |
| 21567 | 1079 | I1079Q | D | 2.7e-06 | 1.4e-06 | 3.8e-06 | 2.633E-06 | 1.201E-06 | 1.400E-06 | 3.800E-06 |
| 21568 | 1079 | I1079G | D | 1.8e-06 | 3.6e-06 | 1.2e-06 | 2.200E-06 | 1.249E-06 | 1.200E-06 | 3.600E-06 |
| 21569 | 1079 | I1079H | D | 2e-06 | 4e-06 | 2.6e-06 | 2.867E-06 | 1.026E-06 | 2.000E-06 | 4.000E-06 |
| 21570 | 1079 | I1079I | D | 2.5e-06 | 3.1e-06 | 1.8e-06 | 2.467E-06 | 6.506E-07 | 1.800E-06 | 3.100E-06 |
| 21571 | 1079 | I1079L | D | 4.7e-06 | 2.8e-06 | 1.6e-06 | 3.033E-06 | 1.563E-06 | 1.600E-06 | 4.700E-06 |
| 21572 | 1079 | I1079K | D | 3.8e-06 | 4.1e-06 | 5.4e-06 | 4.433E-06 | 8.505E-07 | 3.800E-06 | 5.400E-06 |
| 21573 | 1079 | I1079M | D | 1.4e-06 | 1.5e-06 | 1.1e-06 | 1.333E-06 | 2.082E-07 | 1.100E-06 | 1.500E-06 |
| 21574 | 1079 | I1079F | D | 2.1e-06 | 4e-06 | 6.4e-06 | 4.167E-06 | 2.155E-06 | 2.100E-06 | 6.400E-06 |
| 21575 | 1079 | I1079P | D | 2.1e-06 | 4.3e-06 | 2.3e-06 | 2.900E-06 | 1.217E-06 | 2.100E-06 | 4.300E-06 |
| 21576 | 1079 | I1079S | D | 3.3e-06 | 2.8e-06 | 2e-06 | 2.700E-06 | 6.557E-07 | 2.000E-06 | 3.300E-06 |
| 21577 | 1079 | I1079T | D | 1.8e-06 | 4e-06 | 2.3e-06 | 2.700E-06 | 1.153E-06 | 1.800E-06 | 4.000E-06 |
| 21578 | 1079 | I1079W | D | 1.9e-06 | 2.5e-06 | 2.8e-06 | 2.400E-06 | 4.583E-07 | 1.900E-06 | 2.800E-06 |
| 21579 | 1079 | I1079Y | D | 2.7e-06 | 2.3e-06 | 2e-06 | 2.333E-06 | 3.512E-07 | 2.000E-06 | 2.700E-06 |
| 21580 | 1079 | I1079V | D | 1.8e-06 | 2.1e-06 | 2.3e-06 | 2.067E-06 | 2.517E-07 | 1.800E-06 | 2.300E-06 |
| 21581 | 1080 | T1080A | D | 3.9e-06 | 4.2e-06 | 2.7e-06 | 3.600E-06 | 7.937E-07 | 2.700E-06 | 4.200E-06 |
| 21582 | 1080 | T1080R | D | 2.5e-06 | 1.6e-06 | 2.6e-06 | 2.233E-06 | 5.508E-07 | 1.600E-06 | 2.600E-06 |
| 21583 | 1080 | T1080N | D | 3.7e-06 | 5.3e-06 | 2.3e-06 | 3.767E-06 | 1.501E-06 | 2.300E-06 | 5.300E-06 |
| 21584 | 1080 | T1080D | D | 3.3e-06 | 1.6e-06 | 2e-06 | 2.300E-06 | 8.888E-07 | 1.600E-06 | 3.300E-06 |
| 21585 | 1080 | T1080C | D | 2.3e-06 | 3.2e-06 | 3e-06 | 2.833E-06 | 4.726E-07 | 2.300E-06 | 3.200E-06 |
| 21586 | 1080 | T1080E | D | 2.2e-06 | 2e-06 | 3e-06 | 2.400E-06 | 5.292E-07 | 2.000E-06 | 3.000E-06 |
| 21587 | 1080 | T1080Q | D | 9.9e-07 | 1.2e-06 | 2.8e-06 | 1.663E-06 | 9.900E-07 | 9.900E-07 | 2.800E-06 |
| 21588 | 1080 | T1080G | D | 2.4e-06 | 3.1e-06 | 2.5e-06 | 2.667E-06 | 3.786E-07 | 2.400E-06 | 3.100E-06 |
| 21589 | 1080 | T1080H | D | 2.7e-06 | 1.6e-06 | 2.6e-06 | 2.300E-06 | 6.083E-07 | 1.600E-06 | 2.700E-06 |

| | | | | | | | | | | |
|---|---|---|---|---|---|---|---|---|---|---|
| 21590 | 1080 | T1080I | D | 1.9e-06 | 2.3e-06 | 1.3e-06 | 1.833E-06 | 5.033E-07 | 1.300E-06 | 2.300E-06 |
| 21591 | 1080 | T1080L | D | 4e-06 | 2e-06 | 1.7e-06 | 2.567E-06 | 1.250E-06 | 1.700E-06 | 4.000E-06 |
| 21592 | 1080 | T1080K | D | 3.1e-06 | 2.2e-06 | 2.4e-06 | 2.567E-06 | 4.726E-07 | 2.200E-06 | 3.100E-06 |
| 21593 | 1080 | T1080M | D | 1.7e-06 | 2.5e-06 | 3.3e-06 | 2.500E-06 | 8.000E-07 | 1.700E-06 | 3.300E-06 |
| 21594 | 1080 | T1080F | D | 3.3e-06 | 1.4e-06 | 3.3e-06 | 2.667E-06 | 1.097E-06 | 1.400E-06 | 3.300E-06 |
| 21595 | 1080 | T1080P | D | 2.6e-06 | 3.3e-06 | 2.6e-06 | 2.833E-06 | 4.041E-07 | 2.600E-06 | 3.300E-06 |
| 21596 | 1080 | T1080S | D | 2.9e-06 | 1.5e-06 | 2.2e-06 | 2.200E-06 | 7.000E-07 | 1.500E-06 | 2.900E-06 |
| 21597 | 1080 | T1080T | D | 2.3e-06 | 2.4e-06 | 2.7e-06 | 2.467E-06 | 2.082E-07 | 2.300E-06 | 2.700E-06 |
| 21598 | 1080 | T1080W | D | 3.3e-06 | 3.7e-06 | 1.5e-06 | 2.833E-06 | 1.172E-06 | 1.500E-06 | 3.700E-06 |
| 21599 | 1080 | T1080Y | D | 2.8e-06 | 2.5e-06 | 1.9e-06 | 2.400E-06 | 4.583E-07 | 1.900E-06 | 2.800E-06 |
| 21600 | 1080 | T1080V | D | 1.8e-06 | 2.7e-06 | 4.2e-06 | 2.900E-06 | 1.212E-06 | 1.800E-06 | 4.200E-06 |
| 21601 | 1081 | C1081A | D | 3e-06 | 2.3e-06 | 1.6e-06 | 2.300E-06 | 7.000E-07 | 1.600E-06 | 3.000E-06 |
| 21602 | 1081 | C1081R | D | 2.2e-06 | 2.3e-06 | 2.5e-06 | 2.333E-06 | 1.528E-07 | 2.200E-06 | 2.500E-06 |
| 21603 | 1081 | C1081N | D | 2.5e-06 | 3.6e-06 | 2.4e-06 | 2.833E-06 | 6.658E-07 | 2.400E-06 | 3.600E-06 |
| 21604 | 1081 | C1081D | D | 3.3e-06 | 2.8e-06 | 2.3e-06 | 2.800E-06 | 5.000E-07 | 2.300E-06 | 3.300E-06 |
| 21605 | 1081 | C1081C | D | 2e-06 | 2.5e-06 | 2.3e-06 | 2.267E-06 | 2.517E-07 | 2.000E-06 | 2.500E-06 |
| 21606 | 1081 | C1081E | D | 2.9e-06 | 2.7e-06 | 1.9e-06 | 2.500E-06 | 5.292E-07 | 1.900E-06 | 2.900E-06 |
| 21607 | 1081 | C1081Q | D | 1.8e-06 | 3.1e-06 | 2.4e-06 | 2.433E-06 | 6.506E-07 | 1.800E-06 | 3.100E-06 |
| 21608 | 1081 | C1081G | D | 1.7e-06 | 2.7e-06 | 3.7e-06 | 2.700E-06 | 1.000E-06 | 1.700E-06 | 3.700E-06 |
| 21609 | 1081 | C1081H | D | 2.7e-06 | 2.6e-06 | 4.1e-06 | 3.133E-06 | 8.386E-07 | 2.600E-06 | 4.100E-06 |
| 21610 | 1081 | C1081I | D | 1.5e-06 | 3.1e-06 | 1.8e-06 | 2.133E-06 | 8.505E-07 | 1.500E-06 | 3.100E-06 |
| 21611 | 1081 | C1081L | D | 1.6e-06 | 3.2e-06 | 2e-06 | 2.267E-06 | 8.327E-07 | 1.600E-06 | 3.200E-06 |
| 21612 | 1081 | C1081K | D | 2.2e-06 | 7.6e-07 | 2.6e-06 | 1.853E-06 | 9.677E-07 | 7.600E-07 | 2.600E-06 |
| 21613 | 1081 | C1081M | D | 2.2e-06 | 1.9e-06 | 3.7e-06 | 2.600E-06 | 9.644E-07 | 1.900E-06 | 3.700E-06 |
| 21614 | 1081 | C1081F | D | 2.8e-06 | 2e-06 | 1.8e-06 | 2.200E-06 | 5.292E-07 | 1.800E-06 | 2.800E-06 |
| 21615 | 1081 | C1081P | D | 2.7e-06 | 2.5e-06 | 2e-06 | 2.400E-06 | 3.606E-07 | 2.000E-06 | 2.700E-06 |
| 21616 | 1081 | C1081S | D | 1e-06 | 3e-06 | 2e-06 | 2.000E-06 | 1.000E-06 | 1.000E-06 | 3.000E-06 |
| 21617 | 1081 | C1081T | D | 2.4e-06 | 3.1e-06 | 1.6e-06 | 2.367E-06 | 7.506E-07 | 1.600E-06 | 3.100E-06 |
| 21618 | 1081 | C1081W | D | 3.5e-06 | 2e-06 | 3.3e-06 | 2.933E-06 | 8.145E-07 | 2.000E-06 | 3.500E-06 |
| 21619 | 1081 | C1081Y | D | 2.4e-06 | 1.3e-06 | 2.5e-06 | 2.067E-06 | 6.658E-07 | 1.300E-06 | 2.500E-06 |
| 21620 | 1081 | C1081V | D | 1.6e-06 | 1.6e-06 | 1.9e-06 | 1.700E-06 | 1.732E-07 | 1.600E-06 | 1.900E-06 |
| 21621 | 1082 | R1082A | D | 3.3e-06 | 2.1e-06 | 2.2e-06 | 2.533E-06 | 6.658E-07 | 2.100E-06 | 3.300E-06 |
| 21622 | 1082 | R1082R | D | 2.4e-06 | 1.3e-06 | 3e-06 | 2.233E-06 | 8.622E-07 | 1.300E-06 | 3.000E-06 |
| 21623 | 1082 | R1082N | D | 3.6e-06 | 4.6e-06 | 2.2e-06 | 3.467E-06 | 1.206E-06 | 2.200E-06 | 4.600E-06 |
| 21624 | 1082 | R1082D | D | 4.3e-06 | 2.3e-06 | 3.6e-06 | 3.400E-06 | 1.015E-06 | 2.300E-06 | 4.300E-06 |
| 21625 | 1082 | R1082C | D | 1.4e-06 | 3.9e-06 | 3.8e-06 | 3.033E-06 | 1.415E-06 | 1.400E-06 | 3.900E-06 |
| 21626 | 1082 | R1082E | D | 1.7e-06 | 3.4e-06 | 2.1e-06 | 2.400E-06 | 8.888E-07 | 1.700E-06 | 3.400E-06 |
| 21627 | 1082 | R1082Q | D | 2.6e-06 | 2.8e-06 | 4e-06 | 3.133E-06 | 7.572E-07 | 2.600E-06 | 4.000E-06 |
| 21628 | 1082 | R1082G | D | 2.1e-06 | 2.1e-06 | 2.7e-06 | 2.300E-06 | 3.464E-07 | 2.100E-06 | 2.700E-06 |
| 21629 | 1082 | R1082H | D | 1.5e-06 | 1.8e-06 | 1.4e-06 | 1.567E-06 | 2.082E-07 | 1.400E-06 | 1.800E-06 |
| 21630 | 1082 | R1082I | D | 1.5e-06 | 1.3e-06 | 2.7e-06 | 1.833E-06 | 7.572E-07 | 1.300E-06 | 2.700E-06 |
| 21631 | 1082 | R1082L | D | 3.7e-06 | 3.3e-06 | 5.3e-06 | 4.100E-06 | 1.058E-06 | 3.300E-06 | 5.300E-06 |
| 21632 | 1082 | R1082K | D | 2.1e-06 | 3.8e-06 | 2.8e-06 | 2.900E-06 | 8.544E-07 | 2.100E-06 | 3.800E-06 |
| 21633 | 1082 | R1082M | D | 1.6e-06 | 2.2e-06 | 1.2e-06 | 1.667E-06 | 5.033E-07 | 1.200E-06 | 2.200E-06 |
| 21634 | 1082 | R1082F | D | 2.6e-06 | 1.3e-06 | 2.3e-06 | 2.067E-06 | 6.807E-07 | 1.300E-06 | 2.600E-06 |
| 21635 | 1082 | R1082P | D | 1.5e-06 | 3.9e-06 | 4.3e-06 | 3.233E-06 | 1.514E-06 | 1.500E-06 | 4.300E-06 |
| 21636 | 1082 | R1082S | D | 2.5e-06 | 3.4e-06 | 3e-06 | 2.967E-06 | 4.509E-07 | 2.500E-06 | 3.400E-06 |
| 21637 | 1082 | R1082T | D | 3e-06 | 2.7e-06 | 2.8e-06 | 2.833E-06 | 1.528E-07 | 2.700E-06 | 3.000E-06 |
| 21638 | 1082 | R1082W | D | 2.5e-06 | 1.3e-06 | 1.6e-06 | 1.800E-06 | 6.245E-07 | 1.300E-06 | 2.500E-06 |
| 21639 | 1082 | R1082Y | D | 2.3e-06 | 1.8e-06 | 2.3e-06 | 2.133E-06 | 2.887E-07 | 1.800E-06 | 2.300E-06 |
| 21640 | 1082 | R1082V | D | 2.4e-06 | 2.4e-06 | 1.5e-06 | 2.100E-06 | 5.196E-07 | 1.500E-06 | 2.400E-06 |
| 21641 | 1083 | A1083A | D | 2.1e-06 | 3.5e-06 | 3.1e-06 | 2.900E-06 | 7.211E-07 | 2.100E-06 | 3.500E-06 |
| 21642 | 1083 | A1083R | D | 1.3e-06 | 2.4e-06 | 3.3e-06 | 2.333E-06 | 1.002E-06 | 1.300E-06 | 3.300E-06 |
| 21643 | 1083 | A1083N | D | 1.2e-06 | 2.5e-06 | 3.4e-06 | 2.367E-06 | 1.106E-06 | 1.200E-06 | 3.400E-06 |
| 21644 | 1083 | A1083D | D | 2.8e-06 | 2.7e-06 | 1.5e-06 | 2.333E-06 | 7.234E-07 | 1.500E-06 | 2.800E-06 |
| 21645 | 1083 | A1083C | D | 2.6e-06 | 2.8e-06 | 2.2e-06 | 2.533E-06 | 3.055E-07 | 2.200E-06 | 2.800E-06 |
| 21646 | 1083 | A1083E | D | 3e-06 | 2.5e-06 | 1.4e-06 | 2.300E-06 | 8.185E-07 | 1.400E-06 | 3.000E-06 |
| 21647 | 1083 | A1083Q | D | 1.8e-06 | 2.3e-06 | 1.5e-06 | 1.867E-06 | 4.041E-07 | 1.500E-06 | 2.300E-06 |
| 21648 | 1083 | A1083G | D | 1.1e-06 | 2e-06 | 2.3e-06 | 1.800E-06 | 6.245E-07 | 1.100E-06 | 2.300E-06 |
| 21649 | 1083 | A1083H | D | 2.6e-06 | 3e-06 | 2e-06 | 2.533E-06 | 5.033E-07 | 2.000E-06 | 3.000E-06 |
| 21650 | 1083 | A1083I | D | 2.9e-06 | 3.1e-06 | 1.7e-06 | 2.567E-06 | 7.572E-07 | 1.700E-06 | 3.100E-06 |
| 21651 | 1083 | A1083L | D | 2.1e-06 | 3e-06 | 2.6e-06 | 2.567E-06 | 4.509E-07 | 2.100E-06 | 3.000E-06 |
| 21652 | 1083 | A1083K | D | 2.5e-06 | 1.7e-06 | 1.7e-06 | 1.967E-06 | 4.619E-07 | 1.700E-06 | 2.500E-06 |
| 21653 | 1083 | A1083M | D | 2.9e-06 | 2.3e-06 | 2.3e-06 | 2.500E-06 | 3.464E-07 | 2.300E-06 | 2.900E-06 |
| 21654 | 1083 | A1083F | D | 2.7e-06 | 2.2e-06 | 1.7e-06 | 2.200E-06 | 5.000E-07 | 1.700E-06 | 2.700E-06 |
| 21655 | 1083 | A1083P | D | 2.1e-06 | 2.9e-06 | 2.7e-06 | 2.567E-06 | 4.163E-07 | 2.100E-06 | 2.900E-06 |
| 21656 | 1083 | A1083S | D | 2.2e-06 | 1.5e-06 | 2.2e-06 | 1.967E-06 | 4.041E-07 | 1.500E-06 | 2.200E-06 |
| 21657 | 1083 | A1083T | D | 2.2e-06 | 2.6e-06 | 2e-06 | 2.267E-06 | 3.055E-07 | 2.000E-06 | 2.600E-06 |
| 21658 | 1083 | A1083W | D | 1.4e-06 | 2.8e-06 | 3.1e-06 | 2.433E-06 | 9.074E-07 | 1.400E-06 | 3.100E-06 |
| 21659 | 1083 | A1083Y | D | 2.9e-06 | 2.3e-06 | 1.6e-06 | 2.267E-06 | 6.506E-07 | 1.600E-06 | 2.900E-06 |
| 21660 | 1083 | A1083V | D | 2.8e-06 | 1.1e-06 | 3.9e-06 | 2.600E-06 | 1.411E-06 | 1.100E-06 | 3.900E-06 |
| 21661 | 1084 | S1084A | D | 1.9e-06 | 3e-06 | 3.3e-06 | 2.733E-06 | 7.371E-07 | 1.900E-06 | 3.300E-06 |
| 21662 | 1084 | S1084R | D | 1.9e-06 | 3.6e-06 | 3.2e-06 | 2.900E-06 | 8.888E-07 | 1.900E-06 | 3.600E-06 |
| 21663 | 1084 | S1084N | D | 4.3e-06 | 2.5e-06 | 2.5e-06 | 3.100E-06 | 1.039E-06 | 2.500E-06 | 4.300E-06 |
| 21664 | 1084 | S1084D | D | 2.9e-06 | 1.8e-06 | 2.4e-06 | 2.367E-06 | 5.508E-07 | 1.800E-06 | 2.900E-06 |
| 21665 | 1084 | S1084C | D | 3e-06 | 3.9e-06 | 3.6e-06 | 3.500E-06 | 4.583E-07 | 3.000E-06 | 3.900E-06 |
| 21666 | 1084 | S1084E | D | 2.7e-06 | 2.9e-06 | 1.4e-06 | 2.333E-06 | 8.145E-07 | 1.400E-06 | 2.900E-06 |
| 21667 | 1084 | S1084Q | D | 1.9e-06 | 1.9e-06 | 3.4e-06 | 2.400E-06 | 8.660E-07 | 1.900E-06 | 3.400E-06 |
| 21668 | 1084 | S1084G | D | 2.9e-06 | 3.7e-06 | 2e-06 | 2.867E-06 | 8.505E-07 | 2.000E-06 | 3.700E-06 |
| 21669 | 1084 | S1084H | D | 2.9e-06 | 2.3e-06 | 3.7e-06 | 2.967E-06 | 7.024E-07 | 2.300E-06 | 3.700E-06 |
| 21670 | 1084 | S1084I | D | 2.2e-06 | 4e-06 | 2.7e-06 | 2.967E-06 | 9.292E-07 | 2.200E-06 | 4.000E-06 |
| 21671 | 1084 | S1084L | D | 1e-06 | 2.3e-06 | 2.8e-06 | 2.033E-06 | 9.292E-07 | 1.000E-06 | 2.800E-06 |
| 21672 | 1084 | S1084K | D | 1.8e-06 | 2.2e-06 | 2.2e-06 | 2.067E-06 | 2.309E-07 | 1.800E-06 | 2.200E-06 |
| 21673 | 1084 | S1084M | D | 2.2e-06 | 3.7e-06 | 5.2e-06 | 3.700E-06 | 1.500E-06 | 2.200E-06 | 5.200E-06 |
| 21674 | 1084 | S1084F | D | 1.1e-06 | 3e-06 | 1.3e-06 | 1.800E-06 | 1.044E-06 | 1.100E-06 | 3.000E-06 |

| | | | | | | | | | | |
|---|---|---|---|---|---|---|---|---|---|---|
| 21675 | 1084 | S1084P | D | 1.9e-06 | 1.7e-06 | 2e-06 | 1.867E-06 | 1.528E-07 | 1.700E-06 | 2.000E-06 |
| 21676 | 1084 | S1084S | D | 5.5e-06 | 1.8e-06 | 1.9e-06 | 3.067E-06 | 2.108E-06 | 1.800E-06 | 5.500E-06 |
| 21677 | 1084 | S1084T | D | 2.4e-06 | 2.7e-06 | 4.8e-06 | 3.300E-06 | 1.308E-06 | 2.400E-06 | 4.800E-06 |
| 21678 | 1084 | S1084W | D | 1.4e-06 | 3e-06 | 3e-06 | 2.467E-06 | 9.238E-07 | 1.400E-06 | 3.000E-06 |
| 21679 | 1084 | S1084Y | D | 3.1e-06 | 3.6e-06 | 1.9e-06 | 2.867E-06 | 8.737E-07 | 1.900E-06 | 3.600E-06 |
| 21680 | 1084 | S1084V | D | 3e-06 | 2e-06 | 2.8e-06 | 2.600E-06 | 5.292E-07 | 2.000E-06 | 3.000E-06 |
| 21681 | 1085 | Q1085A | D | 2.9e-06 | 2.1e-06 | 1e-06 | 2.000E-06 | 9.539E-07 | 1.000E-06 | 2.900E-06 |
| 21682 | 1085 | Q1085R | D | 1.2e-06 | 1.9e-06 | 1.3e-06 | 1.467E-06 | 3.786E-07 | 1.200E-06 | 1.900E-06 |
| 21683 | 1085 | Q1085N | D | 4e-06 | 2.4e-06 | 1.7e-06 | 2.700E-06 | 1.179E-06 | 1.700E-06 | 4.000E-06 |
| 21684 | 1085 | Q1085D | D | 3.6e-06 | 3.8e-06 | 2.1e-06 | 3.167E-06 | 9.292E-07 | 2.100E-06 | 3.800E-06 |
| 21685 | 1085 | Q1085C | D | 3.1e-06 | 3.8e-06 | 2.6e-06 | 3.167E-06 | 6.028E-07 | 2.600E-06 | 3.800E-06 |
| 21686 | 1085 | Q1085E | D | 3.8e-06 | 1.5e-06 | 3.7e-06 | 3.000E-06 | 1.300E-06 | 1.500E-06 | 3.800E-06 |
| 21687 | 1085 | Q1085Q | D | 4.4e-06 | 2.5e-06 | 2.3e-06 | 3.067E-06 | 1.159E-06 | 2.300E-06 | 4.400E-06 |
| 21688 | 1085 | Q1085G | D | 2.4e-06 | 2.5e-06 | 2.1e-06 | 2.333E-06 | 2.082E-07 | 2.100E-06 | 2.500E-06 |
| 21689 | 1085 | Q1085H | D | 1.9e-06 | 4.3e-06 | 3.4e-06 | 3.200E-06 | 1.212E-06 | 1.900E-06 | 4.300E-06 |
| 21690 | 1085 | Q1085I | D | 2e-06 | 3.4e-06 | 1.8e-06 | 2.400E-06 | 8.718E-07 | 1.800E-06 | 3.400E-06 |
| 21691 | 1085 | Q1085L | D | 1.6e-06 | 1.7e-06 | 1.7e-06 | 1.667E-06 | 5.774E-08 | 1.600E-06 | 1.700E-06 |
| 21692 | 1085 | Q1085K | D | 4.9e-06 | 2.2e-06 | 1.2e-06 | 2.767E-06 | 1.914E-06 | 1.200E-06 | 4.900E-06 |
| 21693 | 1085 | Q1085M | D | 2.3e-06 | 3.7e-06 | 4.8e-06 | 3.600E-06 | 1.253E-06 | 2.300E-06 | 4.800E-06 |
| 21694 | 1085 | Q1085F | D | 1.9e-06 | 3.3e-06 | 2.3e-06 | 2.500E-06 | 7.211E-07 | 1.900E-06 | 3.300E-06 |
| 21695 | 1085 | Q1085P | D | 3.1e-06 | 2.6e-06 | 2.1e-06 | 2.600E-06 | 5.000E-07 | 2.100E-06 | 3.100E-06 |
| 21696 | 1085 | Q1085S | D | 3.4e-06 | 2e-06 | 2.5e-06 | 2.633E-06 | 7.095E-07 | 2.000E-06 | 3.400E-06 |
| 21697 | 1085 | Q1085T | D | 2.8e-06 | 3e-06 | 2e-06 | 2.600E-06 | 5.292E-07 | 2.000E-06 | 3.000E-06 |
| 21698 | 1085 | Q1085W | D | 2.2e-06 | 2.4e-06 | 2.3e-06 | 2.300E-06 | 1.000E-07 | 2.200E-06 | 2.400E-06 |
| 21699 | 1085 | Q1085Y | D | 2.8e-06 | 2e-06 | 2e-06 | 2.267E-06 | 4.619E-07 | 2.000E-06 | 2.800E-06 |
| 21700 | 1085 | Q1085V | D | 2.9e-06 | 4e-06 | 3.2e-06 | 3.367E-06 | 5.686E-07 | 2.900E-06 | 4.000E-06 |
| 21701 | 1086 | D1086A | D | 3.3e-06 | 2.1e-06 | 1.1e-06 | 2.167E-06 | 1.102E-06 | 1.100E-06 | 3.300E-06 |
| 21702 | 1086 | D1086R | D | 1.8e-06 | 5.3e-06 | 1.5e-06 | 2.867E-06 | 2.113E-06 | 1.500E-06 | 5.300E-06 |
| 21703 | 1086 | D1086N | D | 2e-06 | 2.6e-06 | 2.8e-06 | 2.467E-06 | 4.163E-07 | 2.000E-06 | 2.800E-06 |
| 21704 | 1086 | D1086D | D | 1.7e-06 | 1.3e-06 | 2.3e-06 | 1.767E-06 | 5.033E-07 | 1.300E-06 | 2.300E-06 |
| 21705 | 1086 | D1086C | D | 2.7e-06 | 2.1e-06 | 2.7e-06 | 2.500E-06 | 3.464E-07 | 2.100E-06 | 2.700E-06 |
| 21706 | 1086 | D1086E | D | 2e-06 | 2e-06 | 2.2e-06 | 2.067E-06 | 1.155E-07 | 2.000E-06 | 2.200E-06 |
| 21707 | 1086 | D1086Q | D | 2e-06 | 1.6e-06 | 2.6e-06 | 2.067E-06 | 5.033E-07 | 1.600E-06 | 2.600E-06 |
| 21708 | 1086 | D1086G | D | 2.5e-06 | 2.3e-06 | 3.1e-06 | 2.633E-06 | 4.163E-07 | 2.300E-06 | 3.100E-06 |
| 21709 | 1086 | D1086H | D | 2.1e-06 | 2e-06 | 3.8e-06 | 2.633E-06 | 1.012E-06 | 2.000E-06 | 3.800E-06 |
| 21710 | 1086 | D1086I | D | 2.7e-06 | 2e-06 | 3e-06 | 2.567E-06 | 5.132E-07 | 2.000E-06 | 3.000E-06 |
| 21711 | 1086 | D1086L | D | 5.7e-06 | 2.9e-06 | 2.7e-06 | 3.767E-06 | 1.677E-06 | 2.700E-06 | 5.700E-06 |
| 21712 | 1086 | D1086K | D | 2.1e-06 | 1.6e-06 | 2.7e-06 | 2.133E-06 | 5.508E-07 | 1.600E-06 | 2.700E-06 |
| 21713 | 1086 | D1086M | D | 3.6e-06 | 1.6e-06 | 1.8e-06 | 2.333E-06 | 1.102E-06 | 1.600E-06 | 3.600E-06 |
| 21714 | 1086 | D1086F | D | 3.5e-06 | 2.5e-06 | 1.5e-06 | 2.500E-06 | 1.000E-06 | 1.500E-06 | 3.500E-06 |
| 21715 | 1086 | D1086P | D | 2.4e-06 | 3.2e-06 | 2e-06 | 2.533E-06 | 6.110E-07 | 2.000E-06 | 3.200E-06 |
| 21716 | 1086 | D1086S | D | 3.3e-06 | 1.8e-06 | 9.3e-07 | 2.010E-06 | 1.199E-06 | 9.300E-07 | 3.300E-06 |
| 21717 | 1086 | D1086T | D | 2e-06 | 1.9e-06 | 3e-06 | 2.300E-06 | 6.083E-07 | 1.900E-06 | 3.000E-06 |
| 21718 | 1086 | D1086W | D | 1.6e-06 | 1.6e-06 | 2.9e-06 | 2.033E-06 | 7.506E-07 | 1.600E-06 | 2.900E-06 |
| 21719 | 1086 | D1086Y | D | 3.8e-06 | 2.7e-06 | 2.4e-06 | 2.967E-06 | 7.371E-07 | 2.400E-06 | 3.800E-06 |
| 21720 | 1086 | D1086V | D | 2.8e-06 | 4e-06 | 2.1e-06 | 2.967E-06 | 9.609E-07 | 2.100E-06 | 4.000E-06 |
| 21721 | 1087 | V1087A | D | 1.5e-06 | 2.9e-06 | 2.3e-06 | 2.233E-06 | 7.024E-07 | 1.500E-06 | 2.900E-06 |
| 21722 | 1087 | V1087R | D | 1.5e-06 | 2e-06 | 2.2e-06 | 1.900E-06 | 3.606E-07 | 1.500E-06 | 2.200E-06 |
| 21723 | 1087 | V1087N | D | 1.2e-06 | 1.9e-06 | 2.2e-06 | 1.767E-06 | 5.132E-07 | 1.200E-06 | 2.200E-06 |
| 21724 | 1087 | V1087D | D | 3e-06 | 9.7e-07 | 2.7e-06 | 2.223E-06 | 1.096E-06 | 9.700E-07 | 3.000E-06 |
| 21725 | 1087 | V1087C | D | 3.4e-06 | 1.5e-06 | 3.1e-06 | 2.667E-06 | 1.021E-06 | 1.500E-06 | 3.400E-06 |
| 21726 | 1087 | V1087E | D | 2.1e-06 | 4.3e-06 | 2.2e-06 | 2.867E-06 | 1.242E-06 | 2.100E-06 | 4.300E-06 |
| 21727 | 1087 | V1087Q | D | 2.5e-06 | 2.3e-06 | 2.8e-06 | 2.533E-06 | 2.517E-07 | 2.300E-06 | 2.800E-06 |
| 21728 | 1087 | V1087G | D | 1.3e-06 | 1.7e-06 | 3.1e-06 | 2.033E-06 | 9.452E-07 | 1.300E-06 | 3.100E-06 |
| 21729 | 1087 | V1087H | D | 2.8e-06 | 1.8e-06 | 2.5e-06 | 2.367E-06 | 5.132E-07 | 1.800E-06 | 2.800E-06 |
| 21730 | 1087 | V1087I | D | 2.5e-06 | 3.2e-06 | 3.8e-06 | 3.167E-06 | 6.506E-07 | 2.500E-06 | 3.800E-06 |
| 21731 | 1087 | V1087L | D | 1.9e-06 | 2e-06 | 2.9e-06 | 2.267E-06 | 5.508E-07 | 1.900E-06 | 2.900E-06 |
| 21732 | 1087 | V1087K | D | 1.2e-06 | 2e-06 | 2.9e-06 | 2.033E-06 | 8.505E-07 | 1.200E-06 | 2.900E-06 |
| 21733 | 1087 | V1087M | D | 4.2e-06 | 2.5e-06 | 2.6e-06 | 3.100E-06 | 9.539E-07 | 2.500E-06 | 4.200E-06 |
| 21734 | 1087 | V1087F | D | 2.1e-06 | 2.5e-06 | 2.2e-06 | 2.267E-06 | 2.082E-07 | 2.100E-06 | 2.500E-06 |
| 21735 | 1087 | V1087P | D | 1.2e-06 | 1.7e-06 | 2.1e-06 | 1.667E-06 | 4.509E-07 | 1.200E-06 | 2.100E-06 |
| 21736 | 1087 | V1087S | D | 2.5e-06 | 1.7e-06 | 3.1e-06 | 2.433E-06 | 7.024E-07 | 1.700E-06 | 3.100E-06 |
| 21737 | 1087 | V1087T | D | 2.1e-06 | 1.7e-06 | 2.2e-06 | 2.000E-06 | 2.646E-07 | 1.700E-06 | 2.200E-06 |
| 21738 | 1087 | V1087W | D | 3.9e-06 | 3.5e-06 | 2.8e-06 | 3.400E-06 | 5.568E-07 | 2.800E-06 | 3.900E-06 |
| 21739 | 1087 | V1087Y | D | 2.3e-06 | 2.7e-06 | 1.8e-06 | 2.267E-06 | 4.509E-07 | 1.800E-06 | 2.700E-06 |
| 21740 | 1087 | V1087V | D | 1.8e-06 | 2.7e-06 | 1.2e-06 | 1.900E-06 | 7.550E-07 | 1.200E-06 | 2.700E-06 |
| 21741 | 1088 | N1088A | D | 3.7e-06 | 2.4e-06 | 2.3e-06 | 2.800E-06 | 7.810E-07 | 2.300E-06 | 3.700E-06 |
| 21742 | 1088 | N1088R | D | 2.5e-06 | 3e-06 | 3.8e-06 | 3.100E-06 | 6.557E-07 | 2.500E-06 | 3.800E-06 |
| 21743 | 1088 | N1088N | D | 2.6e-06 | 2.4e-06 | 2.5e-06 | 2.500E-06 | 1.000E-07 | 2.400E-06 | 2.600E-06 |
| 21744 | 1088 | N1088D | D | 2.1e-06 | 1.9e-06 | 3.2e-06 | 2.400E-06 | 7.000E-07 | 1.900E-06 | 3.200E-06 |
| 21745 | 1088 | N1088C | D | 1.9e-06 | 2.7e-06 | 1.7e-06 | 2.100E-06 | 5.292E-07 | 1.700E-06 | 2.700E-06 |
| 21746 | 1088 | N1088E | D | 1.9e-06 | 2e-06 | 4.2e-06 | 2.700E-06 | 1.300E-06 | 1.900E-06 | 4.200E-06 |
| 21747 | 1088 | N1088Q | D | 2e-06 | 2.5e-06 | 3.2e-06 | 2.567E-06 | 6.028E-07 | 2.000E-06 | 3.200E-06 |
| 21748 | 1088 | N1088G | D | 2.3e-06 | 2.5e-06 | 4.8e-06 | 3.200E-06 | 1.389E-06 | 2.300E-06 | 4.800E-06 |
| 21749 | 1088 | N1088H | D | 2.2e-06 | 2.2e-06 | 3.3e-06 | 2.567E-06 | 6.351E-07 | 2.200E-06 | 3.300E-06 |
| 21750 | 1088 | N1088I | D | 3.3e-06 | 1.5e-06 | 2.8e-06 | 2.533E-06 | 9.292E-07 | 1.500E-06 | 3.300E-06 |
| 21751 | 1088 | N1088L | D | 2.6e-06 | 2.2e-06 | 3.6e-06 | 2.800E-06 | 7.211E-07 | 2.200E-06 | 3.600E-06 |
| 21752 | 1088 | N1088K | D | 3.1e-06 | 1.9e-06 | 2.5e-06 | 2.500E-06 | 6.000E-07 | 1.900E-06 | 3.100E-06 |
| 21753 | 1088 | N1088M | D | 2.7e-06 | 2e-06 | 2e-06 | 2.233E-06 | 4.041E-07 | 2.000E-06 | 2.700E-06 |
| 21754 | 1088 | N1088F | D | 1.8e-06 | 2e-06 | 1.6e-06 | 1.800E-06 | 2.000E-07 | 1.600E-06 | 2.000E-06 |
| 21755 | 1088 | N1088P | D | 2.4e-06 | 2e-06 | 2.8e-06 | 2.400E-06 | 4.000E-07 | 2.000E-06 | 2.800E-06 |
| 21756 | 1088 | N1088S | D | 3.2e-06 | 3.3e-06 | 1.9e-06 | 2.800E-06 | 7.810E-07 | 1.900E-06 | 3.300E-06 |
| 21757 | 1088 | N1088T | D | 3.1e-06 | 2.3e-06 | 3.1e-06 | 2.833E-06 | 4.619E-07 | 2.300E-06 | 3.100E-06 |
| 21758 | 1088 | N1088W | D | 1.2e-06 | 2.5e-06 | 2.7e-06 | 2.133E-06 | 8.145E-07 | 1.200E-06 | 2.700E-06 |
| 21759 | 1088 | N1088Y | D | 3.5e-06 | 3.9e-06 | 4.3e-06 | 3.900E-06 | 4.000E-07 | 3.500E-06 | 4.300E-06 |

| | | | | | | | | | | |
|---|---|---|---|---|---|---|---|---|---|---|
| 21760 | 1088 | N1088V | D | 4.5e-06 | 3.2e-06 | 3.4e-06 | 3.700E-06 | 7.000E-07 | 3.200E-06 | 4.500E-06 |
| 21761 | 1089 | T1089A | D | 2.4e-06 | 3.6e-06 | 2.6e-06 | 2.867E-06 | 6.429E-07 | 2.400E-06 | 3.600E-06 |
| 21762 | 1089 | T1089R | D | 1.6e-06 | 2.8e-06 | 3.3e-06 | 2.567E-06 | 8.737E-07 | 1.600E-06 | 3.300E-06 |
| 21763 | 1089 | T1089N | D | 1.8e-06 | 1.4e-06 | 2.1e-06 | 1.767E-06 | 3.512E-07 | 1.400E-06 | 2.100E-06 |
| 21764 | 1089 | T1089D | D | 2.4e-06 | 3.2e-06 | 5.3e-06 | 3.633E-06 | 1.498E-06 | 2.400E-06 | 5.300E-06 |
| 21765 | 1089 | T1089C | D | 2.4e-06 | 2e-06 | 1.7e-06 | 2.033E-06 | 3.512E-07 | 1.700E-06 | 2.400E-06 |
| 21766 | 1089 | T1089E | D | 2.4e-06 | 2.2e-06 | 1.9e-06 | 2.167E-06 | 2.517E-07 | 1.900E-06 | 2.400E-06 |
| 21767 | 1089 | T1089Q | D | 3.7e-06 | 1.4e-06 | 2.9e-06 | 2.667E-06 | 1.168E-06 | 1.400E-06 | 3.700E-06 |
| 21768 | 1089 | T1089G | D | 5.5e-06 | 3.3e-06 | 1.3e-06 | 3.367E-06 | 2.101E-06 | 1.300E-06 | 5.500E-06 |
| 21769 | 1089 | T1089H | D | 2.5e-06 | 8.7e-07 | 8.8e-07 | 1.417E-06 | 9.382E-07 | 8.700E-07 | 2.500E-06 |
| 21770 | 1089 | T1089I | D | 2.4e-06 | 1.1e-06 | 3e-06 | 2.167E-06 | 9.713E-07 | 1.100E-06 | 3.000E-06 |
| 21771 | 1089 | T1089L | D | 2.9e-06 | 1.9e-06 | 1.5e-06 | 2.100E-06 | 7.211E-07 | 1.500E-06 | 2.900E-06 |
| 21772 | 1089 | T1089K | D | 1.2e-06 | 3.5e-06 | 2.5e-06 | 2.400E-06 | 1.153E-06 | 1.200E-06 | 3.500E-06 |
| 21773 | 1089 | T1089M | D | 2.3e-06 | 2.8e-06 | 2.3e-06 | 2.467E-06 | 2.887E-07 | 2.300E-06 | 2.800E-06 |
| 21774 | 1089 | T1089F | D | 3.9e-06 | 2.5e-06 | 1.3e-06 | 2.567E-06 | 1.301E-06 | 1.300E-06 | 3.900E-06 |
| 21775 | 1089 | T1089P | D | 2.7e-06 | 1.4e-06 | 2.3e-06 | 2.133E-06 | 6.658E-07 | 1.400E-06 | 2.700E-06 |
| 21776 | 1089 | T1089S | D | 4.9e-06 | 1.5e-06 | 2.5e-06 | 2.967E-06 | 1.747E-06 | 1.500E-06 | 4.900E-06 |
| 21777 | 1089 | T1089T | D | 2.4e-06 | 2e-06 | 1.3e-06 | 1.900E-06 | 5.568E-07 | 1.300E-06 | 2.400E-06 |
| 21778 | 1089 | T1089W | D | 1.7e-06 | 2.9e-06 | 3.6e-06 | 2.733E-06 | 9.609E-07 | 1.700E-06 | 3.600E-06 |
| 21779 | 1089 | T1089Y | D | 2.7e-06 | 2.9e-06 | 1.5e-06 | 2.367E-06 | 7.572E-07 | 1.500E-06 | 2.900E-06 |
| 21780 | 1089 | T1089V | D | 3.3e-06 | 2.3e-06 | 2.4e-06 | 2.667E-06 | 5.508E-07 | 2.300E-06 | 3.300E-06 |
| 21781 | 1090 | A1090A | D | 3.9e-06 | 1.3e-06 | 1.9e-06 | 2.367E-06 | 1.361E-06 | 1.300E-06 | 3.900E-06 |
| 21782 | 1090 | A1090R | D | 2.5e-06 | 1.6e-06 | 2e-06 | 2.033E-06 | 4.509E-07 | 1.600E-06 | 2.500E-06 |
| 21783 | 1090 | A1090N | D | 2.1e-06 | 2.1e-06 | 1.4e-06 | 1.867E-06 | 4.041E-07 | 1.400E-06 | 2.100E-06 |
| 21784 | 1090 | A1090D | D | 2.1e-06 | 2.1e-06 | 2.1e-06 | 2.100E-06 | 0.000E+00 | 2.100E-06 | 2.100E-06 |
| 21785 | 1090 | A1090C | D | 2.3e-06 | 2.7e-06 | 3.8e-06 | 2.933E-06 | 7.767E-07 | 2.300E-06 | 3.800E-06 |
| 21786 | 1090 | A1090E | D | 1.9e-06 | 1.8e-06 | 3.7e-06 | 2.467E-06 | 1.069E-06 | 1.800E-06 | 3.700E-06 |
| 21787 | 1090 | A1090Q | D | 2.9e-06 | 1.4e-06 | 2.3e-06 | 2.200E-06 | 7.550E-07 | 1.400E-06 | 2.900E-06 |
| 21788 | 1090 | A1090G | D | 2.4e-06 | 2.2e-06 | 2.2e-06 | 2.267E-06 | 1.155E-07 | 2.200E-06 | 2.400E-06 |
| 21789 | 1090 | A1090H | D | 2.1e-06 | 2.1e-06 | 2.6e-06 | 2.267E-06 | 2.887E-07 | 2.100E-06 | 2.600E-06 |
| 21790 | 1090 | A1090I | D | 2.1e-06 | 2.6e-06 | 1.3e-06 | 2.000E-06 | 6.557E-07 | 1.300E-06 | 2.600E-06 |
| 21791 | 1090 | A1090L | D | 1.3e-06 | 1.6e-06 | 1.9e-06 | 1.600E-06 | 3.000E-07 | 1.300E-06 | 1.900E-06 |
| 21792 | 1090 | A1090K | D | 2.7e-06 | 2.5e-06 | 2.7e-06 | 2.633E-06 | 1.155E-07 | 2.500E-06 | 2.700E-06 |
| 21793 | 1090 | A1090M | D | 2.5e-06 | 2.7e-06 | 2.8e-06 | 2.667E-06 | 1.528E-07 | 2.500E-06 | 2.800E-06 |
| 21794 | 1090 | A1090F | D | 2e-06 | 3.1e-06 | 2.5e-06 | 2.533E-06 | 5.508E-07 | 2.000E-06 | 3.100E-06 |
| 21795 | 1090 | A1090P | D | 4.2e-06 | 1.2e-06 | 2.7e-06 | 2.700E-06 | 1.500E-06 | 1.200E-06 | 4.200E-06 |
| 21796 | 1090 | A1090S | D | 3.2e-06 | 1.6e-06 | 2.4e-06 | 2.400E-06 | 8.000E-07 | 1.600E-06 | 3.200E-06 |
| 21797 | 1090 | A1090T | D | 2.7e-06 | 1.8e-06 | 2.3e-06 | 2.267E-06 | 4.509E-07 | 1.800E-06 | 2.700E-06 |
| 21798 | 1090 | A1090W | D | 2e-06 | 3.8e-06 | 3.7e-06 | 3.167E-06 | 1.012E-06 | 2.000E-06 | 3.800E-06 |
| 21799 | 1090 | A1090Y | D | 1.2e-06 | 2.8e-06 | 3.8e-06 | 2.600E-06 | 1.311E-06 | 1.200E-06 | 3.800E-06 |
| 21800 | 1090 | A1090V | D | 2.4e-06 | 1.2e-06 | 2.1e-06 | 1.900E-06 | 6.245E-07 | 1.200E-06 | 2.400E-06 |
| 21801 | 1091 | V1091A | D | 2.7e-06 | 2.7e-06 | 2.7e-06 | 2.700E-06 | 0.000E+00 | 2.700E-06 | 2.700E-06 |
| 21802 | 1091 | V1091R | D | 2.2e-06 | 3.8e-06 | 2.4e-06 | 2.800E-06 | 8.718E-07 | 2.200E-06 | 3.800E-06 |
| 21803 | 1091 | V1091N | D | 1.6e-06 | 3.4e-06 | 2.2e-06 | 2.400E-06 | 9.165E-07 | 1.600E-06 | 3.400E-06 |
| 21804 | 1091 | V1091D | D | 2.3e-06 | 3.6e-06 | 1.8e-06 | 2.567E-06 | 9.292E-07 | 1.800E-06 | 3.600E-06 |
| 21805 | 1091 | V1091C | D | 2.7e-06 | 2.1e-06 | 2.4e-06 | 2.400E-06 | 3.000E-07 | 2.100E-06 | 2.700E-06 |
| 21806 | 1091 | V1091E | D | 1.4e-06 | 1.7e-06 | 1.9e-06 | 1.667E-06 | 2.517E-07 | 1.400E-06 | 1.900E-06 |
| 21807 | 1091 | V1091Q | D | 1.5e-06 | 1.5e-06 | 2.8e-06 | 1.933E-06 | 7.506E-07 | 1.500E-06 | 2.800E-06 |
| 21808 | 1091 | V1091G | D | 3.1e-06 | 3.4e-06 | 1.7e-06 | 2.733E-06 | 9.074E-07 | 1.700E-06 | 3.400E-06 |
| 21809 | 1091 | V1091H | D | 2.1e-06 | 3.1e-06 | 3.4e-06 | 2.867E-06 | 6.807E-07 | 2.100E-06 | 3.400E-06 |
| 21810 | 1091 | V1091I | D | 2.1e-06 | 1.9e-06 | 2.1e-06 | 2.033E-06 | 1.155E-07 | 1.900E-06 | 2.100E-06 |
| 21811 | 1091 | V1091L | D | 3.5e-06 | 1.4e-06 | 2.1e-06 | 2.333E-06 | 1.069E-06 | 1.400E-06 | 3.500E-06 |
| 21812 | 1091 | V1091K | D | 1.6e-06 | 1.9e-06 | 2.5e-06 | 2.000E-06 | 4.583E-07 | 1.600E-06 | 2.500E-06 |
| 21813 | 1091 | V1091M | D | 2.2e-06 | 2.3e-06 | 2.7e-06 | 2.400E-06 | 2.646E-07 | 2.200E-06 | 2.700E-06 |
| 21814 | 1091 | V1091F | D | 3.2e-06 | 1.6e-06 | 2.3e-06 | 2.367E-06 | 8.021E-07 | 1.600E-06 | 3.200E-06 |
| 21815 | 1091 | V1091P | D | 3e-06 | 4e-06 | 2.5e-06 | 3.167E-06 | 7.638E-07 | 2.500E-06 | 4.000E-06 |
| 21816 | 1091 | V1091S | D | 3.3e-06 | 2e-06 | 1.5e-06 | 2.267E-06 | 9.292E-07 | 1.500E-06 | 3.300E-06 |
| 21817 | 1091 | V1091T | D | 2.9e-06 | 3e-06 | 2.6e-06 | 2.833E-06 | 2.082E-07 | 2.600E-06 | 3.000E-06 |
| 21818 | 1091 | V1091W | D | 1.9e-06 | 2.1e-06 | 3.8e-06 | 2.600E-06 | 1.044E-06 | 1.900E-06 | 3.800E-06 |
| 21819 | 1091 | V1091Y | D | 1.4e-06 | 1.7e-06 | 1.7e-06 | 1.600E-06 | 1.732E-07 | 1.400E-06 | 1.700E-06 |
| 21820 | 1091 | V1091V | D | 2.5e-06 | 9.7e-07 | 1.9e-06 | 1.790E-06 | 7.709E-07 | 9.700E-07 | 2.500E-06 |
| 21821 | 1092 | A1092A | D | 1.6e-06 | 1.7e-06 | 2.5e-06 | 1.933E-06 | 4.933E-07 | 1.600E-06 | 2.500E-06 |
| 21822 | 1092 | A1092R | D | 1.8e-06 | 9.8e-07 | 3.2e-06 | 1.993E-06 | 1.123E-06 | 9.800E-07 | 3.200E-06 |
| 21823 | 1092 | A1092N | D | 2.6e-06 | 1.9e-06 | 4.6e-06 | 3.033E-06 | 1.401E-06 | 1.900E-06 | 4.600E-06 |
| 21824 | 1092 | A1092D | D | 2.1e-06 | 3.8e-06 | 4.2e-06 | 3.367E-06 | 1.115E-06 | 2.100E-06 | 4.200E-06 |
| 21825 | 1092 | A1092C | D | 3.4e-06 | 3.4e-06 | 1.9e-06 | 2.900E-06 | 8.660E-07 | 1.900E-06 | 3.400E-06 |
| 21826 | 1092 | A1092E | D | 4e-06 | 2.9e-06 | 2.8e-06 | 3.233E-06 | 6.658E-07 | 2.800E-06 | 4.000E-06 |
| 21827 | 1092 | A1092Q | D | 2.6e-06 | 3.1e-06 | 1.2e-06 | 2.300E-06 | 9.849E-07 | 1.200E-06 | 3.100E-06 |
| 21828 | 1092 | A1092G | D | 2.7e-06 | 4.3e-06 | 1.1e-06 | 2.700E-06 | 1.600E-06 | 1.100E-06 | 4.300E-06 |
| 21829 | 1092 | A1092H | D | 3.5e-06 | 3.6e-06 | 2e-06 | 3.033E-06 | 8.963E-07 | 2.000E-06 | 3.600E-06 |
| 21830 | 1092 | A1092I | D | 3e-06 | 2.4e-06 | 1.8e-06 | 2.400E-06 | 6.000E-07 | 1.800E-06 | 3.000E-06 |
| 21831 | 1092 | A1092L | D | 2e-06 | 2.5e-06 | 1.8e-06 | 2.100E-06 | 3.606E-07 | 1.800E-06 | 2.500E-06 |
| 21832 | 1092 | A1092K | D | 3.2e-06 | 2.2e-06 | 2.1e-06 | 2.500E-06 | 6.083E-07 | 2.100E-06 | 3.200E-06 |
| 21833 | 1092 | A1092M | D | 2.2e-06 | 1.8e-06 | 2.4e-06 | 2.133E-06 | 3.055E-07 | 1.800E-06 | 2.400E-06 |
| 21834 | 1092 | A1092F | D | 2.8e-06 | 2.8e-06 | 1.8e-06 | 2.467E-06 | 5.774E-07 | 1.800E-06 | 2.800E-06 |
| 21835 | 1092 | A1092P | D | 1.5e-06 | 2.4e-06 | 1.9e-06 | 1.933E-06 | 4.509E-07 | 1.500E-06 | 2.400E-06 |
| 21836 | 1092 | A1092S | D | 2.1e-06 | 1.8e-06 | 4.3e-06 | 2.733E-06 | 1.365E-06 | 1.800E-06 | 4.300E-06 |
| 21837 | 1092 | A1092T | D | 2.4e-06 | 2.6e-06 | 2.6e-06 | 2.533E-06 | 1.155E-07 | 2.400E-06 | 2.600E-06 |
| 21838 | 1092 | A1092W | D | 2.5e-06 | 1.8e-06 | 2.1e-06 | 2.133E-06 | 3.512E-07 | 1.800E-06 | 2.500E-06 |
| 21839 | 1092 | A1092Y | D | 2.1e-06 | 4.2e-06 | 2.8e-06 | 3.033E-06 | 1.069E-06 | 2.100E-06 | 4.200E-06 |
| 21840 | 1092 | A1092V | D | 3.2e-06 | 2.2e-06 | 3.4e-06 | 2.933E-06 | 6.429E-07 | 2.200E-06 | 3.400E-06 |
| 21841 | 1093 | W1093A | D | 2.3e-06 | 2.4e-06 | 2.4e-06 | 2.367E-06 | 5.774E-08 | 2.300E-06 | 2.400E-06 |
| 21842 | 1093 | W1093R | D | 2.7e-06 | 4.3e-06 | 2.3e-06 | 3.100E-06 | 1.058E-06 | 2.300E-06 | 4.300E-06 |
| 21843 | 1093 | W1093N | D | 1.8e-06 | 3.2e-06 | 1.9e-06 | 2.300E-06 | 7.810E-07 | 1.800E-06 | 3.200E-06 |
| 21844 | 1093 | W1093D | D | 2e-06 | 2.2e-06 | 2.4e-06 | 2.200E-06 | 2.000E-07 | 2.000E-06 | 2.400E-06 |

| | | | | | | | | | | |
|---|---|---|---|---|---|---|---|---|---|---|
| 21845 | 1093 | W1093C | D | 2.1e-06 | 3.6e-06 | 4.1e-06 | 3.267E-06 | 1.041E-06 | 2.100E-06 | 4.100E-06 |
| 21846 | 1093 | W1093E | D | 2e-06 | 2e-06 | 2.5e-06 | 2.167E-06 | 2.887E-07 | 2.000E-06 | 2.500E-06 |
| 21847 | 1093 | W1093Q | D | 2e-06 | 3.2e-06 | 2.4e-06 | 2.533E-06 | 6.110E-07 | 2.000E-06 | 3.200E-06 |
| 21848 | 1093 | W1093G | D | 3.2e-06 | 2.4e-06 | 3.6e-06 | 3.067E-06 | 6.110E-07 | 2.400E-06 | 3.600E-06 |
| 21849 | 1093 | W1093H | D | 3.7e-06 | 2.7e-06 | 5e-06 | 3.800E-06 | 1.153E-06 | 2.700E-06 | 5.000E-06 |
| 21850 | 1093 | W1093I | D | 2.3e-06 | 1.9e-06 | 2.4e-06 | 2.200E-06 | 2.646E-07 | 1.900E-06 | 2.400E-06 |
| 21851 | 1093 | W1093L | D | 1.9e-06 | 9.5e-07 | 1.7e-06 | 1.517E-06 | 5.008E-07 | 9.500E-07 | 1.900E-06 |
| 21852 | 1093 | W1093K | D | 1.5e-06 | 3.7e-06 | 1.9e-06 | 2.367E-06 | 1.172E-06 | 1.500E-06 | 3.700E-06 |
| 21853 | 1093 | W1093M | D | 3.2e-06 | 1.8e-06 | 3.9e-06 | 2.967E-06 | 1.069E-06 | 1.800E-06 | 3.900E-06 |
| 21854 | 1093 | W1093F | D | 2.2e-06 | 1.6e-06 | 2.6e-06 | 2.133E-06 | 5.033E-07 | 1.600E-06 | 2.600E-06 |
| 21855 | 1093 | W1093P | D | 1.6e-06 | 2.7e-06 | 2.2e-06 | 2.167E-06 | 5.508E-07 | 1.600E-06 | 2.700E-06 |
| 21856 | 1093 | W1093S | D | 2.3e-06 | 2.8e-06 | 2.5e-06 | 2.533E-06 | 2.517E-07 | 2.300E-06 | 2.800E-06 |
| 21857 | 1093 | W1093T | D | 2.6e-06 | 1.9e-06 | 1.9e-06 | 2.133E-06 | 4.041E-07 | 1.900E-06 | 2.600E-06 |
| 21858 | 1093 | W1093W | D | 1.4e-06 | 1.4e-06 | 1.9e-06 | 1.567E-06 | 2.887E-07 | 1.400E-06 | 1.900E-06 |
| 21859 | 1093 | W1093Y | D | 2.8e-06 | 1.8e-06 | 3.2e-06 | 2.600E-06 | 7.211E-07 | 1.800E-06 | 3.200E-06 |
| 21860 | 1093 | W1093V | D | 3.6e-06 | 1.2e-06 | 4e-06 | 2.933E-06 | 1.514E-06 | 1.200E-06 | 4.000E-06 |
| 21861 | 1094 | Y1094A | D | 4.9e-06 | 2.7e-06 | 1.5e-06 | 3.033E-06 | 1.724E-06 | 1.500E-06 | 4.900E-06 |
| 21862 | 1094 | Y1094R | D | 2.5e-06 | 1.8e-06 | 1.8e-06 | 2.033E-06 | 4.041E-07 | 1.800E-06 | 2.500E-06 |
| 21863 | 1094 | Y1094N | D | 2e-06 | 5e-06 | 3.1e-06 | 3.367E-06 | 1.518E-06 | 2.000E-06 | 5.000E-06 |
| 21864 | 1094 | Y1094D | D | 2.1e-06 | 2.4e-06 | 1.9e-06 | 2.133E-06 | 2.517E-07 | 1.900E-06 | 2.400E-06 |
| 21865 | 1094 | Y1094C | D | 2.2e-06 | 1.9e-06 | 1.4e-06 | 1.833E-06 | 4.041E-07 | 1.400E-06 | 2.200E-06 |
| 21866 | 1094 | Y1094E | D | 1.7e-06 | 2.8e-06 | 2e-06 | 2.167E-06 | 5.686E-07 | 1.700E-06 | 2.800E-06 |
| 21867 | 1094 | Y1094Q | D | 1.8e-06 | 4.6e-06 | 2.6e-06 | 3.000E-06 | 1.442E-06 | 1.800E-06 | 4.600E-06 |
| 21868 | 1094 | Y1094G | D | 2e-06 | 2.3e-06 | 8.6e-07 | 1.720E-06 | 7.597E-07 | 8.600E-07 | 2.300E-06 |
| 21869 | 1094 | Y1094H | D | 1.9e-06 | 3.3e-06 | 2.2e-06 | 2.467E-06 | 7.371E-07 | 1.900E-06 | 3.300E-06 |
| 21870 | 1094 | Y1094I | D | 1.6e-06 | 3e-06 | 3.2e-06 | 2.600E-06 | 8.718E-07 | 1.600E-06 | 3.200E-06 |
| 21871 | 1094 | Y1094L | D | 3.2e-06 | 2.7e-06 | 2.2e-06 | 2.700E-06 | 5.000E-07 | 2.200E-06 | 3.200E-06 |
| 21872 | 1094 | Y1094K | D | 3.2e-06 | 2.3e-06 | 3.3e-06 | 2.933E-06 | 5.508E-07 | 2.300E-06 | 3.300E-06 |
| 21873 | 1094 | Y1094M | D | 2.1e-06 | 3.2e-06 | 2.5e-06 | 2.600E-06 | 5.568E-07 | 2.100E-06 | 3.200E-06 |
| 21874 | 1094 | Y1094F | D | 2.1e-06 | 1.6e-06 | 1.8e-06 | 1.833E-06 | 2.517E-07 | 1.600E-06 | 2.100E-06 |
| 21875 | 1094 | Y1094P | D | 2.2e-06 | 1.9e-06 | 1.6e-06 | 1.900E-06 | 3.000E-07 | 1.600E-06 | 2.200E-06 |
| 21876 | 1094 | Y1094S | D | 2.5e-06 | 2.3e-06 | 3.3e-06 | 2.700E-06 | 5.292E-07 | 2.300E-06 | 3.300E-06 |
| 21877 | 1094 | Y1094T | D | 1.7e-06 | 2.2e-06 | 1.7e-06 | 1.867E-06 | 2.887E-07 | 1.700E-06 | 2.200E-06 |
| 21878 | 1094 | Y1094W | D | 2.2e-06 | 9.8e-07 | 3.8e-06 | 2.327E-06 | 1.414E-06 | 9.800E-07 | 3.800E-06 |
| 21879 | 1094 | Y1094Y | D | 1.7e-06 | 1.3e-06 | 2.7e-06 | 1.900E-06 | 7.211E-07 | 1.300E-06 | 2.700E-06 |
| 21880 | 1094 | Y1094V | D | 2.9e-06 | 2.3e-06 | 2.4e-06 | 2.533E-06 | 3.215E-07 | 2.300E-06 | 2.900E-06 |
| 21881 | 1095 | Q1095A | D | 1.8e-06 | 1.7e-06 | 3e-06 | 2.167E-06 | 7.234E-07 | 1.700E-06 | 3.000E-06 |
| 21882 | 1095 | Q1095R | D | 1.7e-06 | 2.5e-06 | 2.7e-06 | 2.300E-06 | 5.292E-07 | 1.700E-06 | 2.700E-06 |
| 21883 | 1095 | Q1095N | D | 2.9e-06 | 1.2e-06 | 2.4e-06 | 2.167E-06 | 8.737E-07 | 1.200E-06 | 2.900E-06 |
| 21884 | 1095 | Q1095D | D | 1.7e-06 | 2.8e-06 | 2.1e-06 | 2.200E-06 | 5.568E-07 | 1.700E-06 | 2.800E-06 |
| 21885 | 1095 | Q1095C | D | 3e-06 | 2.3e-06 | 1.9e-06 | 2.400E-06 | 5.568E-07 | 1.900E-06 | 3.000E-06 |
| 21886 | 1095 | Q1095E | D | 3.6e-06 | 1.8e-06 | 2.6e-06 | 2.667E-06 | 9.018E-07 | 1.800E-06 | 3.600E-06 |
| 21887 | 1095 | Q1095Q | D | 3.8e-06 | 2.4e-06 | 2.6e-06 | 2.933E-06 | 7.572E-07 | 2.400E-06 | 3.800E-06 |
| 21888 | 1095 | Q1095G | D | 2.5e-06 | 3.6e-06 | 2.5e-06 | 2.867E-06 | 6.351E-07 | 2.500E-06 | 3.600E-06 |
| 21889 | 1095 | Q1095H | D | 1.9e-06 | 3e-06 | 3.7e-06 | 2.867E-06 | 9.074E-07 | 1.900E-06 | 3.700E-06 |
| 21890 | 1095 | Q1095I | D | 2.7e-06 | 2.3e-06 | 3.6e-06 | 2.867E-06 | 6.658E-07 | 2.300E-06 | 3.600E-06 |
| 21891 | 1095 | Q1095L | D | 2.4e-06 | 1.8e-06 | 2.2e-06 | 2.133E-06 | 3.055E-07 | 1.800E-06 | 2.400E-06 |
| 21892 | 1095 | Q1095K | D | 1.5e-06 | 2.4e-06 | 1.5e-06 | 1.800E-06 | 5.196E-07 | 1.500E-06 | 2.400E-06 |
| 21893 | 1095 | Q1095M | D | 2.9e-06 | 2.8e-06 | 2.2e-06 | 2.633E-06 | 3.786E-07 | 2.200E-06 | 2.900E-06 |
| 21894 | 1095 | Q1095F | D | 1.7e-06 | 2.3e-06 | 3.5e-06 | 2.500E-06 | 9.165E-07 | 1.700E-06 | 3.500E-06 |
| 21895 | 1095 | Q1095P | D | 3.2e-06 | 1.3e-06 | 3.2e-06 | 2.567E-06 | 1.097E-06 | 1.300E-06 | 3.200E-06 |
| 21896 | 1095 | Q1095S | D | 2.1e-06 | 2.4e-06 | 8.1e-06 | 4.200E-06 | 3.381E-06 | 2.100E-06 | 8.100E-06 |
| 21897 | 1095 | Q1095T | D | 2.4e-06 | 1.1e-06 | 2e-06 | 1.833E-06 | 6.658E-07 | 1.100E-06 | 2.400E-06 |
| 21898 | 1095 | Q1095W | D | 2.7e-06 | 3e-06 | 2.3e-06 | 2.667E-06 | 3.512E-07 | 2.300E-06 | 3.000E-06 |
| 21899 | 1095 | Q1095Y | D | 1.7e-06 | 1.5e-06 | 3.3e-06 | 2.167E-06 | 9.866E-07 | 1.500E-06 | 3.300E-06 |
| 21900 | 1095 | Q1095V | D | 3.8e-06 | 2.1e-06 | 1.8e-06 | 2.567E-06 | 1.079E-06 | 1.800E-06 | 3.800E-06 |
| 21901 | 1096 | Q1096A | D | 1.4e-06 | 3.7e-06 | 4e-06 | 3.033E-06 | 1.422E-06 | 1.400E-06 | 4.000E-06 |
| 21902 | 1096 | Q1096R | D | 2.7e-06 | 3.1e-06 | 2e-06 | 2.600E-06 | 5.568E-07 | 2.000E-06 | 3.100E-06 |
| 21903 | 1096 | Q1096N | D | 2e-06 | 1.2e-06 | 2.1e-06 | 1.767E-06 | 4.933E-07 | 1.200E-06 | 2.100E-06 |
| 21904 | 1096 | Q1096D | D | 1e-06 | 4.4e-06 | 3.2e-06 | 2.867E-06 | 1.724E-06 | 1.000E-06 | 4.400E-06 |
| 21905 | 1096 | Q1096C | D | 4.1e-06 | 1.8e-06 | 6.4e-07 | 2.180E-06 | 1.761E-06 | 6.400E-07 | 4.100E-06 |
| 21906 | 1096 | Q1096E | D | 1.9e-06 | 2.4e-06 | 4.2e-06 | 2.833E-06 | 1.210E-06 | 1.900E-06 | 4.200E-06 |
| 21907 | 1096 | Q1096Q | D | 1.1e-06 | 1.8e-06 | 1.7e-06 | 1.533E-06 | 3.786E-07 | 1.100E-06 | 1.800E-06 |
| 21908 | 1096 | Q1096G | D | 1.7e-06 | 2.1e-06 | 2.9e-06 | 2.233E-06 | 6.110E-07 | 1.700E-06 | 2.900E-06 |
| 21909 | 1096 | Q1096H | D | 1.3e-06 | 1.4e-06 | 3e-06 | 1.900E-06 | 9.539E-07 | 1.300E-06 | 3.000E-06 |
| 21910 | 1096 | Q1096I | D | 1.7e-06 | 1.7e-06 | 2.8e-06 | 2.067E-06 | 6.351E-07 | 1.700E-06 | 2.800E-06 |
| 21911 | 1096 | Q1096L | D | 4.7e-06 | 3e-06 | 2.4e-06 | 3.367E-06 | 1.193E-06 | 2.400E-06 | 4.700E-06 |
| 21912 | 1096 | Q1096K | D | 3.4e-06 | 3.4e-06 | 1.6e-06 | 2.800E-06 | 1.039E-06 | 1.600E-06 | 3.400E-06 |
| 21913 | 1096 | Q1096M | D | 4e-06 | 4.8e-06 | 2.5e-06 | 3.767E-06 | 1.168E-06 | 2.500E-06 | 4.800E-06 |
| 21914 | 1096 | Q1096F | D | 1.6e-06 | 2.3e-06 | 2.9e-06 | 2.267E-06 | 6.506E-07 | 1.600E-06 | 2.900E-06 |
| 21915 | 1096 | Q1096P | D | 3.7e-06 | 2.7e-06 | 2.2e-06 | 2.867E-06 | 7.638E-07 | 2.200E-06 | 3.700E-06 |
| 21916 | 1096 | Q1096S | D | 2e-06 | 3.1e-06 | 2.3e-06 | 2.467E-06 | 5.686E-07 | 2.000E-06 | 3.100E-06 |
| 21917 | 1096 | Q1096T | D | 2e-06 | 1.6e-06 | 3.3e-06 | 2.300E-06 | 8.888E-07 | 1.600E-06 | 3.300E-06 |
| 21918 | 1096 | Q1096W | D | 2.5e-06 | 2.2e-06 | 1.1e-06 | 1.933E-06 | 7.371E-07 | 1.100E-06 | 2.500E-06 |
| 21919 | 1096 | Q1096Y | D | 3e-06 | 2.8e-06 | 3.4e-06 | 3.067E-06 | 3.055E-07 | 2.800E-06 | 3.400E-06 |
| 21920 | 1096 | Q1096V | D | 3.1e-06 | 1.9e-06 | 8.3e-07 | 1.943E-06 | 1.136E-06 | 8.300E-07 | 3.100E-06 |
| 21921 | 1097 | K1097A | D | 3e-06 | 4e-06 | 1.5e-06 | 2.833E-06 | 1.258E-06 | 1.500E-06 | 4.000E-06 |
| 21922 | 1097 | K1097R | D | 2.9e-06 | 3.4e-06 | 2.8e-06 | 3.033E-06 | 3.215E-07 | 2.800E-06 | 3.400E-06 |
| 21923 | 1097 | K1097N | D | 3.3e-06 | 2e-06 | 2e-06 | 2.433E-06 | 7.506E-07 | 2.000E-06 | 3.300E-06 |
| 21924 | 1097 | K1097D | D | 1.8e-06 | 3.1e-06 | 2e-06 | 2.300E-06 | 7.000E-07 | 1.800E-06 | 3.100E-06 |
| 21925 | 1097 | K1097C | D | 1.8e-06 | 1.8e-06 | 1.9e-06 | 1.833E-06 | 5.774E-08 | 1.800E-06 | 1.900E-06 |
| 21926 | 1097 | K1097E | D | 3.6e-06 | 1.6e-06 | 9.7e-07 | 2.057E-06 | 1.373E-06 | 9.700E-07 | 3.600E-06 |
| 21927 | 1097 | K1097Q | D | 2.2e-06 | 2.7e-06 | 2.1e-06 | 2.333E-06 | 3.215E-07 | 2.100E-06 | 2.700E-06 |
| 21928 | 1097 | K1097G | D | 1.9e-06 | 3.8e-06 | 2.3e-06 | 2.667E-06 | 1.002E-06 | 1.900E-06 | 3.800E-06 |
| 21929 | 1097 | K1097H | D | 1.9e-06 | 3.3e-06 | 2.4e-06 | 2.533E-06 | 7.095E-07 | 1.900E-06 | 3.300E-06 |

| | | | | | | | | | | |
|---|---|---|---|---|---|---|---|---|---|---|
| 21930 | 1097 | K1097I | D | 4.9e-06 | 2.8e-06 | 2.1e-06 | 3.267E-06 | 1.457E-06 | 2.100E-06 | 4.900E-06 |
| 21931 | 1097 | K1097L | D | 2.1e-06 | 4.7e-06 | 1.4e-06 | 2.733E-06 | 1.739E-06 | 1.400E-06 | 4.700E-06 |
| 21932 | 1097 | K1097K | D | 2.5e-06 | 1.6e-06 | 1.9e-06 | 2.000E-06 | 4.583E-07 | 1.600E-06 | 2.500E-06 |
| 21933 | 1097 | K1097M | D | 7.1e-07 | 2.1e-06 | 2.5e-06 | 1.770E-06 | 9.395E-07 | 7.100E-07 | 2.500E-06 |
| 21934 | 1097 | K1097F | D | 1.9e-06 | 2.6e-06 | 2.6e-06 | 2.367E-06 | 4.041E-07 | 1.900E-06 | 2.600E-06 |
| 21935 | 1097 | K1097P | D | 1.6e-06 | 2.3e-06 | 2.7e-06 | 2.200E-06 | 5.568E-07 | 1.600E-06 | 2.700E-06 |
| 21936 | 1097 | K1097S | D | 4.3e-06 | 3.5e-06 | 3.9e-06 | 3.900E-06 | 4.000E-07 | 3.500E-06 | 4.300E-06 |
| 21937 | 1097 | K1097T | D | 2.5e-06 | 4.4e-06 | 2.2e-06 | 3.033E-06 | 1.193E-06 | 2.200E-06 | 4.400E-06 |
| 21938 | 1097 | K1097W | D | 2.4e-06 | 1.8e-06 | 1.8e-06 | 2.000E-06 | 3.464E-07 | 1.800E-06 | 2.400E-06 |
| 21939 | 1097 | K1097Y | D | 2.8e-06 | 2.1e-06 | 2.2e-06 | 2.367E-06 | 3.786E-07 | 2.100E-06 | 2.800E-06 |
| 21940 | 1097 | K1097V | D | 3.4e-06 | 2.7e-06 | 2.6e-06 | 2.900E-06 | 4.359E-07 | 2.600E-06 | 3.400E-06 |
| 21941 | 1098 | P1098A | D | 4.3e-06 | 3.8e-06 | 2.1e-06 | 3.400E-06 | 1.153E-06 | 2.100E-06 | 4.300E-06 |
| 21942 | 1098 | P1098R | D | 3.9e-06 | 3e-06 | 2.2e-06 | 3.033E-06 | 8.505E-07 | 2.200E-06 | 3.900E-06 |
| 21943 | 1098 | P1098N | D | 2.2e-06 | 1.3e-06 | 1.8e-06 | 1.767E-06 | 4.509E-07 | 1.300E-06 | 2.200E-06 |
| 21944 | 1098 | P1098D | D | 2.5e-06 | 3.5e-06 | 1.4e-06 | 2.467E-06 | 1.050E-06 | 1.400E-06 | 3.500E-06 |
| 21945 | 1098 | P1098C | D | 3.1e-06 | 1.6e-06 | 3e-06 | 2.567E-06 | 8.386E-07 | 1.600E-06 | 3.100E-06 |
| 21946 | 1098 | P1098E | D | 1.4e-06 | 2.9e-06 | 2.7e-06 | 2.333E-06 | 8.145E-07 | 1.400E-06 | 2.900E-06 |
| 21947 | 1098 | P1098Q | D | 2.1e-06 | 3e-06 | 3.2e-06 | 2.767E-06 | 5.859E-07 | 2.100E-06 | 3.200E-06 |
| 21948 | 1098 | P1098G | D | 2.2e-06 | 3.9e-06 | 2.8e-06 | 2.967E-06 | 8.622E-07 | 2.200E-06 | 3.900E-06 |
| 21949 | 1098 | P1098H | D | 2.1e-06 | 2.3e-06 | 5.1e-06 | 3.167E-06 | 1.677E-06 | 2.100E-06 | 5.100E-06 |
| 21950 | 1098 | P1098I | D | 2e-06 | 2.8e-06 | 4.4e-06 | 3.067E-06 | 1.222E-06 | 2.000E-06 | 4.400E-06 |
| 21951 | 1098 | P1098L | D | 1.8e-06 | 2e-06 | 1.8e-06 | 1.867E-06 | 1.155E-07 | 1.800E-06 | 2.000E-06 |
| 21952 | 1098 | P1098K | D | 5.2e-06 | 2.3e-06 | 2.6e-06 | 3.367E-06 | 1.595E-06 | 2.300E-06 | 5.200E-06 |
| 21953 | 1098 | P1098M | D | 1.8e-06 | 2.1e-06 | 1.7e-06 | 1.867E-06 | 2.082E-07 | 1.700E-06 | 2.100E-06 |
| 21954 | 1098 | P1098F | D | 2.5e-06 | 3e-06 | 3e-06 | 2.833E-06 | 2.887E-07 | 2.500E-06 | 3.000E-06 |
| 21955 | 1098 | P1098P | D | 2.7e-06 | 2.2e-06 | 4e-06 | 2.967E-06 | 9.292E-07 | 2.200E-06 | 4.000E-06 |
| 21956 | 1098 | P1098S | D | 3.6e-06 | 3.1e-06 | 2.6e-06 | 3.100E-06 | 5.000E-07 | 2.600E-06 | 3.600E-06 |
| 21957 | 1098 | P1098T | D | 1.7e-06 | 2.1e-06 | 1.3e-06 | 1.700E-06 | 4.000E-07 | 1.300E-06 | 2.100E-06 |
| 21958 | 1098 | P1098W | D | 1.5e-06 | 2.4e-06 | 2.1e-06 | 2.000E-06 | 4.583E-07 | 1.500E-06 | 2.400E-06 |
| 21959 | 1098 | P1098Y | D | 2.2e-06 | 4.4e-06 | 1.8e-06 | 2.800E-06 | 1.400E-06 | 1.800E-06 | 4.400E-06 |
| 21960 | 1098 | P1098V | D | 2.8e-06 | 5.7e-06 | 2e-06 | 3.500E-06 | 1.947E-06 | 2.000E-06 | 5.700E-06 |
| 21961 | 1099 | G1099A | D | 3.8e-06 | 1.9e-06 | 3.4e-06 | 3.033E-06 | 1.002E-06 | 1.900E-06 | 3.800E-06 |
| 21962 | 1099 | G1099R | D | 2.6e-06 | 1.4e-06 | 2.4e-06 | 2.133E-06 | 6.429E-07 | 1.400E-06 | 2.600E-06 |
| 21963 | 1099 | G1099N | D | 2.8e-06 | 4.8e-06 | 2.7e-06 | 3.433E-06 | 1.185E-06 | 2.700E-06 | 4.800E-06 |
| 21964 | 1099 | G1099D | D | 2.5e-06 | 2.8e-06 | 2.3e-06 | 2.533E-06 | 2.517E-07 | 2.300E-06 | 2.800E-06 |
| 21965 | 1099 | G1099C | D | 2.2e-06 | 2.4e-06 | 1.9e-06 | 2.167E-06 | 2.517E-07 | 1.900E-06 | 2.400E-06 |
| 21966 | 1099 | G1099E | D | 2.3e-06 | 1.1e-06 | 1.8e-06 | 1.733E-06 | 6.028E-07 | 1.100E-06 | 2.300E-06 |
| 21967 | 1099 | G1099Q | D | 2.3e-06 | 2.2e-06 | 3.6e-06 | 2.700E-06 | 7.810E-07 | 2.200E-06 | 3.600E-06 |
| 21968 | 1099 | G1099G | D | 2.7e-06 | 2.7e-06 | 2.5e-06 | 2.633E-06 | 1.155E-07 | 2.500E-06 | 2.700E-06 |
| 21969 | 1099 | G1099H | D | 2.8e-06 | 2.8e-06 | 2.6e-06 | 2.733E-06 | 1.155E-07 | 2.600E-06 | 2.800E-06 |
| 21970 | 1099 | G1099I | D | 2.6e-06 | 3.8e-06 | 2.5e-06 | 2.967E-06 | 7.234E-07 | 2.500E-06 | 3.800E-06 |
| 21971 | 1099 | G1099L | D | 2.5e-06 | 1.8e-06 | 3.1e-06 | 2.467E-06 | 6.506E-07 | 1.800E-06 | 3.100E-06 |
| 21972 | 1099 | G1099K | D | 2.5e-06 | 2.4e-06 | 1.5e-06 | 2.133E-06 | 5.508E-07 | 1.500E-06 | 2.500E-06 |
| 21973 | 1099 | G1099M | D | 1.7e-06 | 1.9e-06 | 2.7e-06 | 2.100E-06 | 5.292E-07 | 1.700E-06 | 2.700E-06 |
| 21974 | 1099 | G1099F | D | 2.9e-06 | 3.1e-06 | 4.3e-06 | 3.433E-06 | 7.572E-07 | 2.900E-06 | 4.300E-06 |
| 21975 | 1099 | G1099P | D | 2.1e-06 | 4.3e-06 | 1.7e-06 | 2.700E-06 | 1.400E-06 | 1.700E-06 | 4.300E-06 |
| 21976 | 1099 | G1099S | D | 2.1e-06 | 2.1e-06 | 2.3e-06 | 2.167E-06 | 1.155E-07 | 2.100E-06 | 2.300E-06 |
| 21977 | 1099 | G1099T | D | 1.3e-06 | 3.3e-06 | 1.7e-06 | 2.100E-06 | 1.058E-06 | 1.300E-06 | 3.300E-06 |
| 21978 | 1099 | G1099W | D | 1.9e-06 | 3e-06 | 2.4e-06 | 2.433E-06 | 5.508E-07 | 1.900E-06 | 3.000E-06 |
| 21979 | 1099 | G1099Y | D | 2e-06 | 2.3e-06 | 2.6e-06 | 2.300E-06 | 3.000E-07 | 2.000E-06 | 2.600E-06 |
| 21980 | 1099 | G1099V | D | 2.3e-06 | 2.4e-06 | 2.6e-06 | 2.433E-06 | 1.528E-07 | 2.300E-06 | 2.600E-06 |
| 21981 | 1100 | K1100A | D | 3e-06 | 2.2e-06 | 1.6e-06 | 2.267E-06 | 7.024E-07 | 1.600E-06 | 3.000E-06 |
| 21982 | 1100 | K1100R | D | 1.8e-06 | 2.2e-06 | 1.9e-06 | 1.967E-06 | 2.082E-07 | 1.800E-06 | 2.200E-06 |
| 21983 | 1100 | K1100N | D | 2e-06 | 1.8e-06 | 2.3e-06 | 2.033E-06 | 2.517E-07 | 1.800E-06 | 2.300E-06 |
| 21984 | 1100 | K1100D | D | 4.7e-06 | 2.5e-06 | 1.8e-06 | 3.000E-06 | 1.513E-06 | 1.800E-06 | 4.700E-06 |
| 21985 | 1100 | K1100C | D | 2.1e-06 | 2.3e-06 | 3.6e-06 | 2.667E-06 | 8.145E-07 | 2.100E-06 | 3.600E-06 |
| 21986 | 1100 | K1100E | D | 2.2e-06 | 1.8e-06 | 2.3e-06 | 2.100E-06 | 2.646E-07 | 1.800E-06 | 2.300E-06 |
| 21987 | 1100 | K1100Q | D | 1.4e-06 | 3.5e-06 | 2.9e-06 | 2.600E-06 | 1.082E-06 | 1.400E-06 | 3.500E-06 |
| 21988 | 1100 | K1100G | D | 1.8e-06 | 1.7e-06 | 2e-06 | 1.833E-06 | 1.528E-07 | 1.700E-06 | 2.000E-06 |
| 21989 | 1100 | K1100H | D | 1.8e-06 | 2.5e-06 | 4.8e-06 | 3.033E-06 | 1.570E-06 | 1.800E-06 | 4.800E-06 |
| 21990 | 1100 | K1100I | D | 2.6e-06 | 2.2e-06 | 1.6e-06 | 2.133E-06 | 5.033E-07 | 1.600E-06 | 2.600E-06 |
| 21991 | 1100 | K1100L | D | 2.5e-06 | 3.3e-06 | 3.1e-06 | 2.967E-06 | 4.163E-07 | 2.500E-06 | 3.300E-06 |
| 21992 | 1100 | K1100K | D | 3.4e-06 | 3.1e-06 | 2e-06 | 2.833E-06 | 7.371E-07 | 2.000E-06 | 3.400E-06 |
| 21993 | 1100 | K1100M | D | 2.3e-06 | 1.7e-06 | 2.1e-06 | 2.033E-06 | 3.055E-07 | 1.700E-06 | 2.300E-06 |
| 21994 | 1100 | K1100F | D | 2.2e-06 | 1.2e-06 | 1.5e-06 | 1.633E-06 | 5.132E-07 | 1.200E-06 | 2.200E-06 |
| 21995 | 1100 | K1100P | D | 2.6e-06 | 2.5e-06 | 1.8e-06 | 2.300E-06 | 4.359E-07 | 1.800E-06 | 2.600E-06 |
| 21996 | 1100 | K1100S | D | 2.3e-06 | 1.4e-06 | 3.2e-06 | 2.300E-06 | 9.000E-07 | 1.400E-06 | 3.200E-06 |
| 21997 | 1100 | K1100T | D | 2.5e-06 | 2.3e-06 | 2.3e-06 | 2.367E-06 | 1.155E-07 | 2.300E-06 | 2.500E-06 |
| 21998 | 1100 | K1100W | D | 1e-06 | 2.9e-06 | 2.9e-06 | 2.267E-06 | 1.097E-06 | 1.000E-06 | 2.900E-06 |
| 21999 | 1100 | K1100Y | D | 3.2e-06 | 1.6e-06 | 3.2e-06 | 2.667E-06 | 9.238E-07 | 1.600E-06 | 3.200E-06 |
| 22000 | 1100 | K1100V | D | 2.7e-06 | 9e-07 | 3.1e-06 | 2.233E-06 | 1.172E-06 | 9.000E-07 | 3.100E-06 |
| 22001 | 1101 | A1101A | D | 1.5e-06 | 3.2e-06 | 1.9e-06 | 2.200E-06 | 8.888E-07 | 1.500E-06 | 3.200E-06 |
| 22002 | 1101 | A1101R | D | 3.7e-06 | 2.6e-06 | 3e-06 | 3.100E-06 | 5.568E-07 | 2.600E-06 | 3.700E-06 |
| 22003 | 1101 | A1101N | D | 1.1e-06 | 3.1e-06 | 1.4e-06 | 1.867E-06 | 1.079E-06 | 1.100E-06 | 3.100E-06 |
| 22004 | 1101 | A1101D | D | 3.2e-06 | 2.3e-06 | 1.9e-06 | 2.467E-06 | 6.658E-07 | 1.900E-06 | 3.200E-06 |
| 22005 | 1101 | A1101C | D | 2.9e-06 | 2.5e-06 | 1.4e-06 | 2.267E-06 | 7.767E-07 | 1.400E-06 | 2.900E-06 |
| 22006 | 1101 | A1101E | D | 2.7e-06 | 1.9e-06 | 2.9e-06 | 2.500E-06 | 5.292E-07 | 1.900E-06 | 2.900E-06 |
| 22007 | 1101 | A1101Q | D | 3e-06 | 3.4e-06 | 2.9e-06 | 3.100E-06 | 2.646E-07 | 2.900E-06 | 3.400E-06 |
| 22008 | 1101 | A1101G | D | 2.2e-06 | 2.3e-06 | 1.3e-06 | 1.933E-06 | 5.508E-07 | 1.300E-06 | 2.300E-06 |
| 22009 | 1101 | A1101H | D | 2.7e-06 | 1.4e-06 | 2.6e-06 | 2.233E-06 | 7.234E-07 | 1.400E-06 | 2.700E-06 |
| 22010 | 1101 | A1101I | D | 2.8e-06 | 2.1e-06 | 2.3e-06 | 2.400E-06 | 3.606E-07 | 2.100E-06 | 2.800E-06 |
| 22011 | 1101 | A1101L | D | 1.9e-06 | 1.9e-06 | 2.8e-06 | 2.200E-06 | 5.196E-07 | 1.900E-06 | 2.800E-06 |
| 22012 | 1101 | A1101K | D | 3.4e-06 | 2.5e-06 | 3.6e-06 | 3.167E-06 | 5.859E-07 | 2.500E-06 | 3.600E-06 |
| 22013 | 1101 | A1101M | D | 1.7e-06 | 2.5e-06 | 1.8e-06 | 2.000E-06 | 4.359E-07 | 1.700E-06 | 2.500E-06 |
| 22014 | 1101 | A1101F | D | 2.6e-06 | 2.5e-06 | 2.3e-06 | 2.467E-06 | 1.528E-07 | 2.300E-06 | 2.600E-06 |

| | | | | | | | | | | |
|---|---|---|---|---|---|---|---|---|---|---|
| 22015 | 1101 | A1101P | D | 3.2e-06 | 2.4e-06 | 3.2e-06 | 2.933E-06 | 4.619E-07 | 2.400E-06 | 3.200E-06 |
| 22016 | 1101 | A1101S | D | 2.1e-06 | 1.7e-06 | 2.3e-06 | 2.033E-06 | 3.055E-07 | 1.700E-06 | 2.300E-06 |
| 22017 | 1101 | A1101T | D | 2.2e-06 | 2.8e-06 | 2.7e-06 | 2.567E-06 | 3.215E-07 | 2.200E-06 | 2.800E-06 |
| 22018 | 1101 | A1101W | D | 2.2e-06 | 2.9e-06 | 2.5e-06 | 2.533E-06 | 3.512E-07 | 2.200E-06 | 2.900E-06 |
| 22019 | 1101 | A1101Y | D | 2.2e-06 | 3.3e-06 | 2.4e-06 | 2.633E-06 | 5.859E-07 | 2.200E-06 | 3.300E-06 |
| 22020 | 1101 | A1101V | D | 1.9e-06 | 3.9e-06 | 2.2e-06 | 2.667E-06 | 1.079E-06 | 1.900E-06 | 3.900E-06 |
| 22021 | 1102 | P1102A | D | 2.7e-06 | 2e-06 | 1.3e-06 | 2.000E-06 | 7.000E-07 | 1.300E-06 | 2.700E-06 |
| 22022 | 1102 | P1102R | D | 3.1e-06 | 1.9e-06 | 2.1e-06 | 2.367E-06 | 6.429E-07 | 1.900E-06 | 3.100E-06 |
| 22023 | 1102 | P1102N | D | 1.4e-06 | 2.4e-06 | 3.1e-06 | 2.300E-06 | 8.544E-07 | 1.400E-06 | 3.100E-06 |
| 22024 | 1102 | P1102D | D | 2.1e-06 | 1.2e-06 | 2e-06 | 1.767E-06 | 4.933E-07 | 1.200E-06 | 2.100E-06 |
| 22025 | 1102 | P1102C | D | 2.3e-06 | 3.5e-06 | 3e-06 | 2.933E-06 | 6.028E-07 | 2.300E-06 | 3.500E-06 |
| 22026 | 1102 | P1102E | D | 2.3e-06 | 2.2e-06 | 2.2e-06 | 2.233E-06 | 5.774E-08 | 2.200E-06 | 2.300E-06 |
| 22027 | 1102 | P1102Q | D | 3.3e-06 | 1.1e-06 | 2.1e-06 | 2.167E-06 | 1.102E-06 | 1.100E-06 | 3.300E-06 |
| 22028 | 1102 | P1102G | D | 3.8e-06 | 2.6e-06 | 2.3e-06 | 2.900E-06 | 7.937E-07 | 2.300E-06 | 3.800E-06 |
| 22029 | 1102 | P1102H | D | 1.5e-06 | 2.9e-06 | 4.3e-06 | 2.900E-06 | 1.400E-06 | 1.500E-06 | 4.300E-06 |
| 22030 | 1102 | P1102I | D | 2.6e-06 | 4.8e-06 | 2.1e-06 | 3.167E-06 | 1.436E-06 | 2.100E-06 | 4.800E-06 |
| 22031 | 1102 | P1102L | D | 2.6e-06 | 2.6e-06 | 2.1e-06 | 2.433E-06 | 2.887E-07 | 2.100E-06 | 2.600E-06 |
| 22032 | 1102 | P1102K | D | 3.7e-06 | 3.6e-06 | 2.3e-06 | 3.200E-06 | 7.810E-07 | 2.300E-06 | 3.700E-06 |
| 22033 | 1102 | P1102M | D | 4.2e-06 | 2.8e-06 | 3.5e-06 | 3.500E-06 | 7.000E-07 | 2.800E-06 | 4.200E-06 |
| 22034 | 1102 | P1102F | D | 1.3e-06 | 2.3e-06 | 2e-06 | 1.867E-06 | 5.132E-07 | 1.300E-06 | 2.300E-06 |
| 22035 | 1102 | P1102P | D | 4.1e-06 | 1.5e-06 | 1e-06 | 2.200E-06 | 1.664E-06 | 1.000E-06 | 4.100E-06 |
| 22036 | 1102 | P1102S | D | 2e-06 | 1.6e-06 | 1.7e-06 | 1.767E-06 | 2.082E-07 | 1.600E-06 | 2.000E-06 |
| 22037 | 1102 | P1102T | D | 2e-06 | 1.6e-06 | 2.5e-06 | 2.033E-06 | 4.509E-07 | 1.600E-06 | 2.500E-06 |
| 22038 | 1102 | P1102W | D | 2.9e-06 | 2.8e-06 | 2e-06 | 2.567E-06 | 4.933E-07 | 2.000E-06 | 2.900E-06 |
| 22039 | 1102 | P1102Y | D | 3.1e-06 | 2.9e-06 | 1.7e-06 | 2.567E-06 | 7.572E-07 | 1.700E-06 | 3.100E-06 |
| 22040 | 1102 | P1102V | D | 2.1e-06 | 1.2e-06 | 8.5e-07 | 1.383E-06 | 6.449E-07 | 8.500E-07 | 2.100E-06 |
| 22041 | 1103 | K1103A | D | 2.6e-06 | 4.4e-06 | 2.6e-06 | 3.200E-06 | 1.039E-06 | 2.600E-06 | 4.400E-06 |
| 22042 | 1103 | K1103R | D | 2.2e-06 | 2e-06 | 1.6e-06 | 1.933E-06 | 3.055E-07 | 1.600E-06 | 2.200E-06 |
| 22043 | 1103 | K1103N | D | 2.9e-06 | 2.5e-06 | 3.2e-06 | 2.867E-06 | 3.512E-07 | 2.500E-06 | 3.200E-06 |
| 22044 | 1103 | K1103D | D | 2.1e-06 | 9.5e-07 | 3.1e-06 | 2.050E-06 | 1.076E-06 | 9.500E-07 | 3.100E-06 |
| 22045 | 1103 | K1103C | D | 2.1e-06 | 1.2e-06 | 1.8e-06 | 1.700E-06 | 4.583E-07 | 1.200E-06 | 2.100E-06 |
| 22046 | 1103 | K1103E | D | 1.4e-06 | 2.2e-06 | 1.4e-06 | 1.667E-06 | 4.619E-07 | 1.400E-06 | 2.200E-06 |
| 22047 | 1103 | K1103Q | D | 3.5e-06 | 1.9e-06 | 2.4e-06 | 2.600E-06 | 8.185E-07 | 1.900E-06 | 3.500E-06 |
| 22048 | 1103 | K1103G | D | 1.4e-06 | 2e-06 | 2.8e-06 | 2.067E-06 | 7.024E-07 | 1.400E-06 | 2.800E-06 |
| 22049 | 1103 | K1103H | D | 4e-06 | 3e-06 | 1.5e-06 | 2.833E-06 | 1.258E-06 | 1.500E-06 | 4.000E-06 |
| 22050 | 1103 | K1103I | D | 2.9e-06 | 2.3e-06 | 2.6e-06 | 2.600E-06 | 3.000E-07 | 2.300E-06 | 2.900E-06 |
| 22051 | 1103 | K1103L | D | 2.9e-06 | 2.3e-06 | 3e-06 | 2.733E-06 | 3.786E-07 | 2.300E-06 | 3.000E-06 |
| 22052 | 1103 | K1103K | D | 2.9e-06 | 4e-06 | 9.9e-07 | 2.630E-06 | 1.523E-06 | 9.900E-07 | 4.000E-06 |
| 22053 | 1103 | K1103M | D | 2.7e-06 | 1.6e-06 | 2e-06 | 2.100E-06 | 5.568E-07 | 1.600E-06 | 2.700E-06 |
| 22054 | 1103 | K1103F | D | 1.5e-06 | 2.8e-06 | 2.3e-06 | 2.200E-06 | 6.557E-07 | 1.500E-06 | 2.800E-06 |
| 22055 | 1103 | K1103P | D | 2.3e-06 | 1.4e-06 | 3e-06 | 2.233E-06 | 8.021E-07 | 1.400E-06 | 3.000E-06 |
| 22056 | 1103 | K1103S | D | 3.2e-06 | 2e-06 | 2.1e-06 | 2.433E-06 | 6.658E-07 | 2.000E-06 | 3.200E-06 |
| 22057 | 1103 | K1103T | D | 1.5e-06 | 1.6e-06 | 2.6e-06 | 1.900E-06 | 6.083E-07 | 1.500E-06 | 2.600E-06 |
| 22058 | 1103 | K1103W | D | 4.8e-06 | 1.5e-06 | 4e-06 | 3.433E-06 | 1.721E-06 | 1.500E-06 | 4.800E-06 |
| 22059 | 1103 | K1103Y | D | 2.3e-06 | 4.2e-06 | 8.4e-07 | 2.447E-06 | 1.685E-06 | 8.400E-07 | 4.200E-06 |
| 22060 | 1103 | K1103V | D | 1.4e-06 | 2.7e-06 | 2.3e-06 | 2.133E-06 | 6.658E-07 | 1.400E-06 | 2.700E-06 |
| 22061 | 1104 | L1104A | D | 2.2e-06 | 2.8e-06 | 1.7e-06 | 2.233E-06 | 5.508E-07 | 1.700E-06 | 2.800E-06 |
| 22062 | 1104 | L1104R | D | 1.3e-06 | 2.1e-06 | 2.6e-06 | 2.000E-06 | 6.557E-07 | 1.300E-06 | 2.600E-06 |
| 22063 | 1104 | L1104N | D | 2.9e-06 | 3.4e-06 | 2.2e-06 | 2.833E-06 | 6.028E-07 | 2.200E-06 | 3.400E-06 |
| 22064 | 1104 | L1104D | D | 3.6e-06 | 2.3e-06 | 2.1e-06 | 2.667E-06 | 8.145E-07 | 2.100E-06 | 3.600E-06 |
| 22065 | 1104 | L1104C | D | 3e-06 | 1.9e-06 | 2.9e-06 | 2.600E-06 | 6.083E-07 | 1.900E-06 | 3.000E-06 |
| 22066 | 1104 | L1104E | D | 2.3e-06 | 2.5e-06 | 1e-06 | 1.933E-06 | 8.145E-07 | 1.000E-06 | 2.500E-06 |
| 22067 | 1104 | L1104Q | D | 2.1e-06 | 1.2e-06 | 1.8e-06 | 1.700E-06 | 4.583E-07 | 1.200E-06 | 2.100E-06 |
| 22068 | 1104 | L1104G | D | 2.5e-06 | 2.9e-06 | 1.7e-06 | 2.367E-06 | 6.110E-07 | 1.700E-06 | 2.900E-06 |
| 22069 | 1104 | L1104H | D | 1.7e-06 | 3.5e-06 | 2.9e-06 | 2.700E-06 | 9.165E-07 | 1.700E-06 | 3.500E-06 |
| 22070 | 1104 | L1104I | D | 3.1e-06 | 3.1e-06 | 1.9e-06 | 2.700E-06 | 6.928E-07 | 1.900E-06 | 3.100E-06 |
| 22071 | 1104 | L1104L | D | 3.9e-06 | 2.6e-06 | 8.4e-07 | 2.447E-06 | 1.536E-06 | 8.400E-07 | 3.900E-06 |
| 22072 | 1104 | L1104K | D | 3.3e-06 | 2.3e-06 | 2e-06 | 2.533E-06 | 6.807E-07 | 2.000E-06 | 3.300E-06 |
| 22073 | 1104 | L1104M | D | 2e-06 | 1.3e-06 | 3.1e-06 | 2.133E-06 | 9.074E-07 | 1.300E-06 | 3.100E-06 |
| 22074 | 1104 | L1104F | D | 2.5e-06 | 2.2e-06 | 3.6e-06 | 2.767E-06 | 7.371E-07 | 2.200E-06 | 3.600E-06 |
| 22075 | 1104 | L1104P | D | 1.6e-06 | 4.4e-06 | 3.2e-06 | 3.067E-06 | 1.405E-06 | 1.600E-06 | 4.400E-06 |
| 22076 | 1104 | L1104S | D | 2.2e-06 | 2.9e-06 | 2.4e-06 | 2.500E-06 | 3.606E-07 | 2.200E-06 | 2.900E-06 |
| 22077 | 1104 | L1104T | D | 4.1e-06 | 8.3e-07 | 1.1e-06 | 2.010E-06 | 1.815E-06 | 8.300E-07 | 4.100E-06 |
| 22078 | 1104 | L1104W | D | 1.9e-06 | 2.5e-06 | 2.3e-06 | 2.233E-06 | 3.055E-07 | 1.900E-06 | 2.500E-06 |
| 22079 | 1104 | L1104Y | D | 1.7e-06 | 4e-06 | 2.3e-06 | 2.667E-06 | 1.193E-06 | 1.700E-06 | 4.000E-06 |
| 22080 | 1104 | L1104V | D | 2e-06 | 1.7e-06 | 2.1e-06 | 1.933E-06 | 2.082E-07 | 1.700E-06 | 2.100E-06 |
| 22081 | 1105 | L1105A | D | 1.9e-06 | 3.8e-06 | 3.2e-06 | 2.967E-06 | 9.713E-07 | 1.900E-06 | 3.800E-06 |
| 22082 | 1105 | L1105R | D | 2.3e-06 | 3.4e-06 | 3.1e-06 | 2.933E-06 | 5.686E-07 | 2.300E-06 | 3.400E-06 |
| 22083 | 1105 | L1105N | D | 5.9e-07 | 3e-06 | 2.6e-06 | 2.063E-06 | 1.292E-06 | 5.900E-07 | 3.000E-06 |
| 22084 | 1105 | L1105D | D | 1.8e-06 | 8.5e-07 | 2.1e-06 | 1.583E-06 | 6.526E-07 | 8.500E-07 | 2.100E-06 |
| 22085 | 1105 | L1105C | D | 3.4e-06 | 3.6e-06 | 1.8e-06 | 2.933E-06 | 9.866E-07 | 1.800E-06 | 3.600E-06 |
| 22086 | 1105 | L1105E | D | 2.4e-06 | 1.4e-06 | 7.6e-06 | 3.800E-06 | 3.329E-06 | 1.400E-06 | 7.600E-06 |
| 22087 | 1105 | L1105Q | D | 2e-06 | 1.8e-06 | 1.7e-06 | 1.833E-06 | 1.528E-07 | 1.700E-06 | 2.000E-06 |
| 22088 | 1105 | L1105G | D | 3.3e-06 | 4.4e-06 | 3.3e-06 | 3.667E-06 | 6.351E-07 | 3.300E-06 | 4.400E-06 |
| 22089 | 1105 | L1105H | D | 2.3e-06 | 2.8e-06 | 2.4e-06 | 2.500E-06 | 2.646E-07 | 2.300E-06 | 2.800E-06 |
| 22090 | 1105 | L1105I | D | 3.7e-06 | 2.3e-06 | 3.2e-06 | 3.067E-06 | 7.095E-07 | 2.300E-06 | 3.700E-06 |
| 22091 | 1105 | L1105L | D | 2.4e-06 | 1.7e-06 | 2.5e-06 | 2.200E-06 | 4.359E-07 | 1.700E-06 | 2.500E-06 |
| 22092 | 1105 | L1105K | D | 2.2e-06 | 3.8e-06 | 2e-06 | 2.667E-06 | 9.866E-07 | 2.000E-06 | 3.800E-06 |
| 22093 | 1105 | L1105M | D | 4.5e-06 | 1.3e-06 | 3.3e-06 | 3.033E-06 | 1.617E-06 | 1.300E-06 | 4.500E-06 |
| 22094 | 1105 | L1105F | D | 2.1e-06 | 2.4e-06 | 2e-06 | 2.167E-06 | 2.082E-07 | 2.000E-06 | 2.400E-06 |
| 22095 | 1105 | L1105P | D | 3.1e-06 | 3.5e-06 | 3.2e-06 | 3.267E-06 | 2.082E-07 | 3.100E-06 | 3.500E-06 |
| 22096 | 1105 | L1105S | D | 1.8e-06 | 1.9e-06 | 2.9e-06 | 2.200E-06 | 6.083E-07 | 1.800E-06 | 2.900E-06 |
| 22097 | 1105 | L1105T | D | 2.7e-06 | 3.9e-06 | 3.6e-06 | 3.400E-06 | 6.245E-07 | 2.700E-06 | 3.900E-06 |
| 22098 | 1105 | L1105W | D | 2.8e-06 | 2.4e-06 | 2.3e-06 | 2.500E-06 | 2.646E-07 | 2.300E-06 | 2.800E-06 |
| 22099 | 1105 | L1105Y | D | 2.7e-06 | 2.6e-06 | 2.1e-06 | 2.467E-06 | 3.215E-07 | 2.100E-06 | 2.700E-06 |

| | | | | | | | | | | |
|---|---|---|---|---|---|---|---|---|---|---|
| 22100 | 1105 | L1105V | D | 2e-06 | 2.3e-06 | 7e-06 | 3.767E-06 | 2.804E-06 | 2.000E-06 | 7.000E-06 |
| 22101 | 1106 | I1106A | D | 5.7e-06 | 1.4e-06 | 1.1e-06 | 2.733E-06 | 2.574E-06 | 1.100E-06 | 5.700E-06 |
| 22102 | 1106 | I1106R | D | 5e-06 | 3.8e-06 | 1.6e-06 | 3.467E-06 | 1.724E-06 | 1.600E-06 | 5.000E-06 |
| 22103 | 1106 | I1106N | D | 3.3e-06 | 2.9e-06 | 1.1e-06 | 2.433E-06 | 1.172E-06 | 1.100E-06 | 3.300E-06 |
| 22104 | 1106 | I1106D | D | 1.4e-06 | 2.1e-06 | 4.1e-06 | 2.533E-06 | 1.401E-06 | 1.400E-06 | 4.100E-06 |
| 22105 | 1106 | I1106C | D | 5.2e-06 | 3.5e-06 | 2.5e-06 | 3.733E-06 | 1.365E-06 | 2.500E-06 | 5.200E-06 |
| 22106 | 1106 | I1106E | D | 1.5e-06 | 1.8e-06 | 2.2e-06 | 1.833E-06 | 3.512E-07 | 1.500E-06 | 2.200E-06 |
| 22107 | 1106 | I1106Q | D | 2.9e-06 | 1.5e-06 | 3.3e-06 | 2.567E-06 | 9.452E-07 | 1.500E-06 | 3.300E-06 |
| 22108 | 1106 | I1106G | D | 2.9e-06 | 2.1e-06 | 3.2e-06 | 2.733E-06 | 5.686E-07 | 2.100E-06 | 3.200E-06 |
| 22109 | 1106 | I1106H | D | 3.6e-06 | 2.9e-06 | 2.6e-06 | 3.033E-06 | 5.132E-07 | 2.600E-06 | 3.600E-06 |
| 22110 | 1106 | I1106I | D | 3.1e-06 | 2.9e-06 | 3.4e-06 | 3.133E-06 | 2.517E-07 | 2.900E-06 | 3.400E-06 |
| 22111 | 1106 | I1106L | D | 2.4e-06 | 2.2e-06 | 3.5e-06 | 2.700E-06 | 7.000E-07 | 2.200E-06 | 3.500E-06 |
| 22112 | 1106 | I1106K | D | 3e-06 | 2.8e-06 | 2.4e-06 | 2.733E-06 | 3.055E-07 | 2.400E-06 | 3.000E-06 |
| 22113 | 1106 | I1106M | D | 1e-06 | 2.6e-06 | 3e-06 | 2.200E-06 | 1.058E-06 | 1.000E-06 | 3.000E-06 |
| 22114 | 1106 | I1106F | D | 2.6e-06 | 1.3e-06 | 1.8e-06 | 1.900E-06 | 6.557E-07 | 1.300E-06 | 2.600E-06 |
| 22115 | 1106 | I1106P | D | 2e-06 | 2.7e-06 | 1.5e-06 | 2.067E-06 | 6.028E-07 | 1.500E-06 | 2.700E-06 |
| 22116 | 1106 | I1106S | D | 1.5e-06 | 4.2e-06 | 2.3e-06 | 2.667E-06 | 1.387E-06 | 1.500E-06 | 4.200E-06 |
| 22117 | 1106 | I1106T | D | 1.2e-06 | 2.9e-06 | 2.7e-06 | 2.267E-06 | 9.292E-07 | 1.200E-06 | 2.900E-06 |
| 22118 | 1106 | I1106W | D | 2.5e-06 | 2.6e-06 | 1.1e-06 | 2.067E-06 | 8.386E-07 | 1.100E-06 | 2.600E-06 |
| 22119 | 1106 | I1106Y | D | 2e-06 | 2.1e-06 | 2.2e-06 | 2.100E-06 | 1.000E-07 | 2.000E-06 | 2.200E-06 |
| 22120 | 1106 | I1106V | D | 2.8e-06 | 2.7e-06 | 2.7e-06 | 2.733E-06 | 5.774E-08 | 2.700E-06 | 2.800E-06 |
| 22121 | 1107 | Y1107A | D | 1.2e-06 | 1.6e-06 | 1.9e-06 | 1.567E-06 | 3.512E-07 | 1.200E-06 | 1.900E-06 |
| 22122 | 1107 | Y1107R | D | 4.8e-06 | 2e-06 | 3.3e-06 | 3.367E-06 | 1.401E-06 | 2.000E-06 | 4.800E-06 |
| 22123 | 1107 | Y1107N | D | 2.6e-06 | 2.7e-06 | 1.6e-06 | 2.300E-06 | 6.083E-07 | 1.600E-06 | 2.700E-06 |
| 22124 | 1107 | Y1107D | D | 4e-06 | 2.5e-06 | 2e-06 | 2.833E-06 | 1.041E-06 | 2.000E-06 | 4.000E-06 |
| 22125 | 1107 | Y1107C | D | 2.5e-06 | 1.2e-06 | 2.7e-06 | 2.133E-06 | 8.145E-07 | 1.200E-06 | 2.700E-06 |
| 22126 | 1107 | Y1107E | D | 1.5e-06 | 2.8e-06 | 3.1e-06 | 2.467E-06 | 8.505E-07 | 1.500E-06 | 3.100E-06 |
| 22127 | 1107 | Y1107Q | D | 1.5e-06 | 1.6e-06 | 2.3e-06 | 1.800E-06 | 4.359E-07 | 1.500E-06 | 2.300E-06 |
| 22128 | 1107 | Y1107G | D | 3.2e-06 | 2.8e-06 | 2.8e-06 | 2.933E-06 | 2.309E-07 | 2.800E-06 | 3.200E-06 |
| 22129 | 1107 | Y1107H | D | 2.3e-06 | 2.5e-06 | 2.9e-06 | 2.567E-06 | 3.055E-07 | 2.300E-06 | 2.900E-06 |
| 22130 | 1107 | Y1107I | D | 3.3e-06 | 1.6e-06 | 1.2e-06 | 2.033E-06 | 1.115E-06 | 1.200E-06 | 3.300E-06 |
| 22131 | 1107 | Y1107L | D | 4e-06 | 1.6e-06 | 3.2e-06 | 2.933E-06 | 1.222E-06 | 1.600E-06 | 4.000E-06 |
| 22132 | 1107 | Y1107K | D | 1.6e-06 | 2.6e-06 | 1.9e-06 | 2.033E-06 | 5.132E-07 | 1.600E-06 | 2.600E-06 |
| 22133 | 1107 | Y1107M | D | 2.1e-06 | 2.3e-06 | 2.8e-06 | 2.400E-06 | 3.606E-07 | 2.100E-06 | 2.800E-06 |
| 22134 | 1107 | Y1107F | D | 4.5e-06 | 2e-06 | 3e-06 | 3.167E-06 | 1.258E-06 | 2.000E-06 | 4.500E-06 |
| 22135 | 1107 | Y1107P | D | 2.8e-06 | 5.4e-06 | 2.1e-06 | 3.433E-06 | 1.739E-06 | 2.100E-06 | 5.400E-06 |
| 22136 | 1107 | Y1107S | D | 2.6e-06 | 1.5e-06 | 2e-06 | 2.033E-06 | 5.508E-07 | 1.500E-06 | 2.600E-06 |
| 22137 | 1107 | Y1107T | D | 5.5e-06 | 3.2e-06 | 3.1e-06 | 3.933E-06 | 1.358E-06 | 3.100E-06 | 5.500E-06 |
| 22138 | 1107 | Y1107W | D | 2.7e-06 | 3.3e-06 | 2.5e-06 | 2.833E-06 | 4.163E-07 | 2.500E-06 | 3.300E-06 |
| 22139 | 1107 | Y1107Y | D | 2.1e-06 | 4.1e-06 | 2.3e-06 | 2.833E-06 | 1.102E-06 | 2.100E-06 | 4.100E-06 |
| 22140 | 1107 | Y1107V | D | 1.7e-06 | 2.6e-06 | 1.4e-06 | 1.900E-06 | 6.245E-07 | 1.400E-06 | 2.600E-06 |
| 22141 | 1108 | S1108A | D | 6.5e-06 | 1.7e-06 | 1.5e-06 | 3.233E-06 | 2.831E-06 | 1.500E-06 | 6.500E-06 |
| 22142 | 1108 | S1108R | D | 1.8e-06 | 2.5e-06 | 2.1e-06 | 2.133E-06 | 3.512E-07 | 1.800E-06 | 2.500E-06 |
| 22143 | 1108 | S1108N | D | 2.6e-06 | 2.1e-06 | 2.4e-06 | 2.367E-06 | 2.517E-07 | 2.100E-06 | 2.600E-06 |
| 22144 | 1108 | S1108D | D | 2.7e-06 | 2e-06 | 2.7e-06 | 2.467E-06 | 4.041E-07 | 2.000E-06 | 2.700E-06 |
| 22145 | 1108 | S1108C | D | 2.4e-06 | 1.5e-06 | 1.9e-06 | 1.933E-06 | 4.509E-07 | 1.500E-06 | 2.400E-06 |
| 22146 | 1108 | S1108E | D | 2.8e-06 | 8.7e-07 | 2.1e-06 | 1.923E-06 | 9.771E-07 | 8.700E-07 | 2.800E-06 |
| 22147 | 1108 | S1108Q | D | 2.1e-06 | 1.4e-06 | 1.7e-06 | 1.733E-06 | 3.512E-07 | 1.400E-06 | 2.100E-06 |
| 22148 | 1108 | S1108G | D | 1.5e-06 | 2e-06 | 1.5e-06 | 1.667E-06 | 2.887E-07 | 1.500E-06 | 2.000E-06 |
| 22149 | 1108 | S1108H | D | 1e-06 | 2.1e-06 | 3.1e-06 | 2.067E-06 | 1.050E-06 | 1.000E-06 | 3.100E-06 |
| 22150 | 1108 | S1108I | D | 1.9e-06 | 2.5e-06 | 1.4e-06 | 1.933E-06 | 5.508E-07 | 1.400E-06 | 2.500E-06 |
| 22151 | 1108 | S1108L | D | 2.4e-06 | 2.9e-06 | 2.6e-06 | 2.633E-06 | 2.517E-07 | 2.400E-06 | 2.900E-06 |
| 22152 | 1108 | S1108K | D | 1.5e-06 | 2.6e-06 | 2.8e-06 | 2.300E-06 | 7.000E-07 | 1.500E-06 | 2.800E-06 |
| 22153 | 1108 | S1108M | D | 2.5e-06 | 4.3e-06 | 2.3e-06 | 3.033E-06 | 1.102E-06 | 2.300E-06 | 4.300E-06 |
| 22154 | 1108 | S1108F | D | 3e-06 | 2.4e-06 | 1.7e-06 | 2.367E-06 | 6.506E-07 | 1.700E-06 | 3.000E-06 |
| 22155 | 1108 | S1108P | D | 1.9e-06 | 2e-06 | 2.7e-06 | 2.200E-06 | 4.359E-07 | 1.900E-06 | 2.700E-06 |
| 22156 | 1108 | S1108S | D | 1.8e-06 | 2.7e-06 | 3.2e-06 | 2.567E-06 | 7.095E-07 | 1.800E-06 | 3.200E-06 |
| 22157 | 1108 | S1108T | D | 1.4e-06 | 2.3e-06 | 4.6e-06 | 2.767E-06 | 1.650E-06 | 1.400E-06 | 4.600E-06 |
| 22158 | 1108 | S1108W | D | 2.2e-06 | 1.7e-06 | 3.5e-06 | 2.467E-06 | 9.292E-07 | 1.700E-06 | 3.500E-06 |
| 22159 | 1108 | S1108Y | D | 1.9e-06 | 3.1e-06 | 3e-06 | 2.667E-06 | 6.658E-07 | 1.900E-06 | 3.100E-06 |
| 22160 | 1108 | S1108V | D | 3.2e-06 | 2.3e-06 | 2.3e-06 | 2.600E-06 | 5.196E-07 | 2.300E-06 | 3.200E-06 |
| 22161 | 1109 | A1109A | D | 3.3e-06 | 2.1e-06 | 2.7e-06 | 2.700E-06 | 6.000E-07 | 2.100E-06 | 3.300E-06 |
| 22162 | 1109 | A1109R | D | 3.1e-06 | 2.3e-06 | 2e-06 | 2.467E-06 | 5.686E-07 | 2.000E-06 | 3.100E-06 |
| 22163 | 1109 | A1109N | D | 4.2e-06 | 3.8e-06 | 2.5e-06 | 3.500E-06 | 8.888E-07 | 2.500E-06 | 4.200E-06 |
| 22164 | 1109 | A1109D | D | 1.5e-06 | 1.1e-06 | 1.6e-06 | 1.400E-06 | 2.646E-07 | 1.100E-06 | 1.600E-06 |
| 22165 | 1109 | A1109C | D | 3.1e-06 | 2.3e-06 | 1.8e-06 | 2.400E-06 | 6.557E-07 | 1.800E-06 | 3.100E-06 |
| 22166 | 1109 | A1109E | D | 2.3e-06 | 2.6e-06 | 2.4e-06 | 2.433E-06 | 1.528E-07 | 2.300E-06 | 2.600E-06 |
| 22167 | 1109 | A1109Q | D | 1.9e-06 | 1.8e-06 | 3.2e-06 | 2.300E-06 | 7.810E-07 | 1.800E-06 | 3.200E-06 |
| 22168 | 1109 | A1109G | D | 1.7e-06 | 2.8e-06 | 2.7e-06 | 2.400E-06 | 6.083E-07 | 1.700E-06 | 2.800E-06 |
| 22169 | 1109 | A1109H | D | 9.7e-07 | 2.6e-06 | 3e-06 | 2.190E-06 | 1.075E-06 | 9.700E-07 | 3.000E-06 |
| 22170 | 1109 | A1109I | D | 3e-06 | 3.2e-06 | 2.9e-06 | 3.033E-06 | 1.528E-07 | 2.900E-06 | 3.200E-06 |
| 22171 | 1109 | A1109L | D | 1.3e-06 | 2.3e-06 | 1.9e-06 | 1.833E-06 | 5.033E-07 | 1.300E-06 | 2.300E-06 |
| 22172 | 1109 | A1109K | D | 1.2e-06 | 1.7e-06 | 1.1e-06 | 1.333E-06 | 3.215E-07 | 1.100E-06 | 1.700E-06 |
| 22173 | 1109 | A1109M | D | 2.7e-06 | 2.7e-06 | 2.5e-06 | 2.633E-06 | 1.155E-07 | 2.500E-06 | 2.700E-06 |
| 22174 | 1109 | A1109F | D | 1.5e-06 | 2e-06 | 2.4e-06 | 1.967E-06 | 4.509E-07 | 1.500E-06 | 2.400E-06 |
| 22175 | 1109 | A1109P | D | 2.8e-06 | 3.5e-06 | 2.8e-06 | 3.033E-06 | 4.041E-07 | 2.800E-06 | 3.500E-06 |
| 22176 | 1109 | A1109S | D | 2.3e-06 | 3e-06 | 2.4e-06 | 2.567E-06 | 3.786E-07 | 2.300E-06 | 3.000E-06 |
| 22177 | 1109 | A1109T | D | 1.6e-06 | 3e-06 | 2.2e-06 | 2.267E-06 | 7.024E-07 | 1.600E-06 | 3.000E-06 |
| 22178 | 1109 | A1109W | D | 3.1e-06 | 2.1e-06 | 8.8e-07 | 2.027E-06 | 1.112E-06 | 8.800E-07 | 3.100E-06 |
| 22179 | 1109 | A1109Y | D | 2.9e-06 | 1.8e-06 | 1.8e-06 | 2.167E-06 | 6.351E-07 | 1.800E-06 | 2.900E-06 |
| 22180 | 1109 | A1109V | D | 2e-06 | 3.6e-06 | 2.6e-06 | 2.733E-06 | 8.083E-07 | 2.000E-06 | 3.600E-06 |
| 22181 | 1110 | S1110A | D | 2.4e-06 | 2.2e-06 | 1.5e-06 | 2.033E-06 | 4.726E-07 | 1.500E-06 | 2.400E-06 |
| 22182 | 1110 | S1110R | D | 2.6e-06 | 2.3e-06 | 3.7e-06 | 2.867E-06 | 7.371E-07 | 2.300E-06 | 3.700E-06 |
| 22183 | 1110 | S1110N | D | 2.4e-06 | 2.9e-06 | 1.3e-06 | 2.200E-06 | 8.185E-07 | 1.300E-06 | 2.900E-06 |
| 22184 | 1110 | S1110D | D | 3.7e-06 | 2e-06 | 2.1e-06 | 2.600E-06 | 9.539E-07 | 2.000E-06 | 3.700E-06 |

| | | | | | | | | | | |
|---|---|---|---|---|---|---|---|---|---|---|
| 22185 | 1110 | S1110C | D | 2.8e-06 | 1.8e-06 | 1.8e-06 | 2.133E-06 | 5.774E-07 | 1.800E-06 | 2.800E-06 |
| 22186 | 1110 | S1110E | D | 1.4e-06 | 1.6e-06 | 4.3e-06 | 2.433E-06 | 1.620E-06 | 1.400E-06 | 4.300E-06 |
| 22187 | 1110 | S1110Q | D | 2.6e-06 | 1.4e-06 | 2.1e-06 | 2.033E-06 | 6.028E-07 | 1.400E-06 | 2.600E-06 |
| 22188 | 1110 | S1110G | D | 3.7e-06 | 1.6e-06 | 2.9e-06 | 2.733E-06 | 1.060E-06 | 1.600E-06 | 3.700E-06 |
| 22189 | 1110 | S1110H | D | 1.8e-06 | 3.3e-06 | 1.4e-06 | 2.167E-06 | 1.002E-06 | 1.400E-06 | 3.300E-06 |
| 22190 | 1110 | S1110I | D | 2.5e-06 | 3.1e-06 | 1.8e-06 | 2.467E-06 | 6.506E-07 | 1.800E-06 | 3.100E-06 |
| 22191 | 1110 | S1110L | D | 1.3e-06 | 3.9e-06 | 1.9e-06 | 2.367E-06 | 1.361E-06 | 1.300E-06 | 3.900E-06 |
| 22192 | 1110 | S1110K | D | 3.1e-06 | 3.8e-06 | 1.1e-06 | 2.667E-06 | 1.401E-06 | 1.100E-06 | 3.800E-06 |
| 22193 | 1110 | S1110M | D | 3e-06 | 2.1e-06 | 1.7e-06 | 2.267E-06 | 6.658E-07 | 1.700E-06 | 3.000E-06 |
| 22194 | 1110 | S1110F | D | 2e-06 | 2e-06 | 2.3e-06 | 2.100E-06 | 1.732E-07 | 2.000E-06 | 2.300E-06 |
| 22195 | 1110 | S1110P | D | 3.5e-06 | 1.9e-06 | 3e-06 | 2.800E-06 | 8.185E-07 | 1.900E-06 | 3.500E-06 |
| 22196 | 1110 | S1110S | D | 2.5e-06 | 2.8e-06 | 3.2e-06 | 2.833E-06 | 3.512E-07 | 2.500E-06 | 3.200E-06 |
| 22197 | 1110 | S1110T | D | 2.3e-06 | 3.5e-06 | 3.4e-06 | 3.067E-06 | 6.658E-07 | 2.300E-06 | 3.500E-06 |
| 22198 | 1110 | S1110W | D | 2e-06 | 4.8e-06 | 1.1e-06 | 2.633E-06 | 1.930E-06 | 1.100E-06 | 4.800E-06 |
| 22199 | 1110 | S1110Y | D | 1.3e-06 | 9.2e-07 | 2.7e-06 | 1.640E-06 | 9.374E-07 | 9.200E-07 | 2.700E-06 |
| 22200 | 1110 | S1110V | D | 3.4e-06 | 4.1e-06 | 1.8e-06 | 3.100E-06 | 1.179E-06 | 1.800E-06 | 4.100E-06 |
| 22201 | 1111 | F1111A | D | 1.9e-06 | 1e-06 | 9.8e-07 | 1.293E-06 | 5.255E-07 | 9.800E-07 | 1.900E-06 |
| 22202 | 1111 | F1111R | D | 2.5e-06 | 9.1e-07 | 2.3e-06 | 1.903E-06 | 8.660E-07 | 9.100E-07 | 2.500E-06 |
| 22203 | 1111 | F1111N | D | 3.8e-06 | 2.9e-06 | 5e-06 | 3.900E-06 | 1.054E-06 | 2.900E-06 | 5.000E-06 |
| 22204 | 1111 | F1111D | D | 1.3e-06 | 2e-06 | 2.6e-06 | 1.967E-06 | 6.506E-07 | 1.300E-06 | 2.600E-06 |
| 22205 | 1111 | F1111C | D | 2.6e-06 | 2.9e-06 | 2.6e-06 | 2.700E-06 | 1.732E-07 | 2.600E-06 | 2.900E-06 |
| 22206 | 1111 | F1111E | D | 1.5e-06 | 1.6e-06 | 2.4e-06 | 1.833E-06 | 4.933E-07 | 1.500E-06 | 2.400E-06 |
| 22207 | 1111 | F1111Q | D | 1.8e-06 | 2.1e-06 | 2.7e-06 | 2.200E-06 | 4.583E-07 | 1.800E-06 | 2.700E-06 |
| 22208 | 1111 | F1111G | D | 1.4e-06 | 1.7e-06 | 1.1e-06 | 1.400E-06 | 3.000E-07 | 1.100E-06 | 1.700E-06 |
| 22209 | 1111 | F1111H | D | 2.9e-06 | 1.4e-06 | 3.6e-06 | 2.633E-06 | 1.124E-06 | 1.400E-06 | 3.600E-06 |
| 22210 | 1111 | F1111I | D | 1.4e-06 | 2.6e-06 | 2e-06 | 2.000E-06 | 6.000E-07 | 1.400E-06 | 2.600E-06 |
| 22211 | 1111 | F1111L | D | 3.9e-06 | 1.5e-06 | 1.7e-06 | 2.367E-06 | 1.332E-06 | 1.500E-06 | 3.900E-06 |
| 22212 | 1111 | F1111K | D | 4.6e-06 | 1.8e-06 | 2.4e-06 | 2.933E-06 | 1.474E-06 | 1.800E-06 | 4.600E-06 |
| 22213 | 1111 | F1111M | D | 3.1e-06 | 2.3e-06 | 2.1e-06 | 2.500E-06 | 5.292E-07 | 2.100E-06 | 3.100E-06 |
| 22214 | 1111 | F1111F | D | 2.4e-06 | 2.2e-06 | 3e-06 | 2.533E-06 | 4.163E-07 | 2.200E-06 | 3.000E-06 |
| 22215 | 1111 | F1111P | D | 2.2e-06 | 3e-06 | 1.7e-06 | 2.300E-06 | 6.557E-07 | 1.700E-06 | 3.000E-06 |
| 22216 | 1111 | F1111S | D | 2.3e-06 | 1.7e-06 | 2.6e-06 | 2.200E-06 | 4.583E-07 | 1.700E-06 | 2.600E-06 |
| 22217 | 1111 | F1111T | D | 2.9e-06 | 3.1e-06 | 2e-06 | 2.667E-06 | 5.859E-07 | 2.000E-06 | 3.100E-06 |
| 22218 | 1111 | F1111W | D | 1.8e-06 | 1.8e-06 | 2.8e-06 | 2.133E-06 | 5.774E-07 | 1.800E-06 | 2.800E-06 |
| 22219 | 1111 | F1111Y | D | 3.2e-06 | 1.7e-06 | 2.3e-06 | 2.400E-06 | 7.550E-07 | 1.700E-06 | 3.200E-06 |
| 22220 | 1111 | F1111V | D | 1.7e-06 | 2.3e-06 | 1.9e-06 | 1.967E-06 | 3.055E-07 | 1.700E-06 | 2.300E-06 |
| 22221 | 1112 | L1112A | D | 3.1e-06 | 1.6e-06 | 2.3e-06 | 2.333E-06 | 7.506E-07 | 1.600E-06 | 3.100E-06 |
| 22222 | 1112 | L1112R | D | 2.2e-06 | 4e-06 | 1.8e-06 | 2.667E-06 | 1.172E-06 | 1.800E-06 | 4.000E-06 |
| 22223 | 1112 | L1112N | D | 2.6e-06 | 1.6e-06 | 1.5e-06 | 1.900E-06 | 6.083E-07 | 1.500E-06 | 2.600E-06 |
| 22224 | 1112 | L1112D | D | 2.4e-06 | 2.4e-06 | 1.9e-06 | 2.233E-06 | 2.887E-07 | 1.900E-06 | 2.400E-06 |
| 22225 | 1112 | L1112C | D | 2.2e-06 | 1.2e-06 | 2.6e-06 | 2.000E-06 | 7.211E-07 | 1.200E-06 | 2.600E-06 |
| 22226 | 1112 | L1112E | D | 2.3e-06 | 3e-06 | 2.1e-06 | 2.467E-06 | 4.726E-07 | 2.100E-06 | 3.000E-06 |
| 22227 | 1112 | L1112Q | D | 2.6e-06 | 2.4e-06 | 2.7e-06 | 2.567E-06 | 1.528E-07 | 2.400E-06 | 2.700E-06 |
| 22228 | 1112 | L1112G | D | 2.2e-06 | 3.2e-06 | 3.2e-06 | 2.867E-06 | 5.774E-07 | 2.200E-06 | 3.200E-06 |
| 22229 | 1112 | L1112H | D | 2.5e-06 | 1.5e-06 | 1.7e-06 | 1.900E-06 | 5.292E-07 | 1.500E-06 | 2.500E-06 |
| 22230 | 1112 | L1112I | D | 3e-06 | 3.1e-06 | 1.3e-06 | 2.467E-06 | 1.012E-06 | 1.300E-06 | 3.100E-06 |
| 22231 | 1112 | L1112L | D | 2.2e-06 | 2.1e-06 | 3e-06 | 2.433E-06 | 4.933E-07 | 2.100E-06 | 3.000E-06 |
| 22232 | 1112 | L1112K | D | 2.7e-06 | 2.5e-06 | 2.5e-06 | 2.567E-06 | 1.155E-07 | 2.500E-06 | 2.700E-06 |
| 22233 | 1112 | L1112M | D | 2.3e-06 | 2.1e-06 | 3e-06 | 2.467E-06 | 4.726E-07 | 2.100E-06 | 3.000E-06 |
| 22234 | 1112 | L1112F | D | 3.4e-06 | 3.9e-06 | 3e-06 | 3.433E-06 | 4.509E-07 | 3.000E-06 | 3.900E-06 |
| 22235 | 1112 | L1112P | D | 2.9e-06 | 2.2e-06 | 1.6e-06 | 2.233E-06 | 6.506E-07 | 1.600E-06 | 2.900E-06 |
| 22236 | 1112 | L1112S | D | 2.3e-06 | 1.7e-06 | 2.2e-06 | 2.067E-06 | 3.215E-07 | 1.700E-06 | 2.300E-06 |
| 22237 | 1112 | L1112T | D | 3.7e-06 | 2.8e-06 | 2.3e-06 | 2.933E-06 | 7.095E-07 | 2.300E-06 | 3.700E-06 |
| 22238 | 1112 | L1112W | D | 1.4e-06 | 2.2e-06 | 2e-06 | 1.867E-06 | 4.163E-07 | 1.400E-06 | 2.200E-06 |
| 22239 | 1112 | L1112Y | D | 2.5e-06 | 2e-06 | 3.2e-06 | 2.567E-06 | 6.028E-07 | 2.000E-06 | 3.200E-06 |
| 22240 | 1112 | L1112V | D | 2.4e-06 | 2.5e-06 | 2.5e-06 | 2.467E-06 | 5.774E-08 | 2.400E-06 | 2.500E-06 |
| 22241 | 1113 | Y1113A | D | 1.9e-06 | 2.2e-06 | 1.5e-06 | 1.867E-06 | 3.512E-07 | 1.500E-06 | 2.200E-06 |
| 22242 | 1113 | Y1113R | D | 1.3e-06 | 3.2e-06 | 2.9e-06 | 2.467E-06 | 1.021E-06 | 1.300E-06 | 3.200E-06 |
| 22243 | 1113 | Y1113N | D | 1.5e-06 | 2.9e-06 | 2e-06 | 2.133E-06 | 7.095E-07 | 1.500E-06 | 2.900E-06 |
| 22244 | 1113 | Y1113D | D | 3.4e-06 | 2.8e-06 | 1.9e-06 | 2.700E-06 | 7.550E-07 | 1.900E-06 | 3.400E-06 |
| 22245 | 1113 | Y1113C | D | 2e-06 | 2.9e-06 | 1e-06 | 1.967E-06 | 9.504E-07 | 1.000E-06 | 2.900E-06 |
| 22246 | 1113 | Y1113E | D | 2.8e-06 | 2.2e-06 | 1.5e-06 | 2.167E-06 | 6.506E-07 | 1.500E-06 | 2.800E-06 |
| 22247 | 1113 | Y1113Q | D | 2.1e-06 | 1.6e-06 | 2.6e-06 | 2.100E-06 | 5.000E-07 | 1.600E-06 | 2.600E-06 |
| 22248 | 1113 | Y1113G | D | 3.7e-06 | 2.1e-06 | 2.3e-06 | 2.700E-06 | 8.718E-07 | 2.100E-06 | 3.700E-06 |
| 22249 | 1113 | Y1113H | D | 2.5e-06 | 3.5e-06 | 1.3e-06 | 2.433E-06 | 1.102E-06 | 1.300E-06 | 3.500E-06 |
| 22250 | 1113 | Y1113I | D | 2.3e-06 | 2.5e-06 | 2.5e-06 | 2.433E-06 | 1.155E-07 | 2.300E-06 | 2.500E-06 |
| 22251 | 1113 | Y1113L | D | 2.2e-06 | 2.4e-06 | 2.1e-06 | 2.233E-06 | 1.528E-07 | 2.100E-06 | 2.400E-06 |
| 22252 | 1113 | Y1113K | D | 4.8e-06 | 2.7e-06 | 3.2e-06 | 3.567E-06 | 1.097E-06 | 2.700E-06 | 4.800E-06 |
| 22253 | 1113 | Y1113M | D | 1.1e-06 | 5.2e-06 | 2.4e-06 | 2.900E-06 | 2.095E-06 | 1.100E-06 | 5.200E-06 |
| 22254 | 1113 | Y1113F | D | 1.9e-06 | 2.5e-06 | 3.2e-06 | 2.533E-06 | 6.506E-07 | 1.900E-06 | 3.200E-06 |
| 22255 | 1113 | Y1113P | D | 3e-06 | 2.1e-06 | 2.1e-06 | 2.400E-06 | 5.196E-07 | 2.100E-06 | 3.000E-06 |
| 22256 | 1113 | Y1113S | D | 2.5e-06 | 2e-06 | 3.2e-06 | 2.567E-06 | 6.028E-07 | 2.000E-06 | 3.200E-06 |
| 22257 | 1113 | Y1113T | D | 5e-06 | 1.3e-06 | 2.6e-06 | 2.967E-06 | 1.877E-06 | 1.300E-06 | 5.000E-06 |
| 22258 | 1113 | Y1113W | D | 1.4e-06 | 2.2e-06 | 2.5e-06 | 2.033E-06 | 5.686E-07 | 1.400E-06 | 2.500E-06 |
| 22259 | 1113 | Y1113Y | D | 2.9e-06 | 3.4e-06 | 2.8e-06 | 3.033E-06 | 3.215E-07 | 2.800E-06 | 3.400E-06 |
| 22260 | 1113 | Y1113V | D | 2.2e-06 | 2.2e-06 | 5e-06 | 3.133E-06 | 1.617E-06 | 2.200E-06 | 5.000E-06 |
| 22261 | 1114 | S1114A | D | 2.4e-06 | 3.2e-06 | 1.1e-06 | 2.233E-06 | 1.060E-06 | 1.100E-06 | 3.200E-06 |
| 22262 | 1114 | S1114R | D | 2.5e-06 | 1.1e-06 | 2.8e-06 | 2.133E-06 | 9.074E-07 | 1.100E-06 | 2.800E-06 |
| 22263 | 1114 | S1114N | D | 2.7e-06 | 2.5e-06 | 1.9e-06 | 2.367E-06 | 4.163E-07 | 1.900E-06 | 2.700E-06 |
| 22264 | 1114 | S1114D | D | 3.4e-06 | 1.6e-06 | 2.9e-06 | 2.633E-06 | 9.292E-07 | 1.600E-06 | 3.400E-06 |
| 22265 | 1114 | S1114C | D | 2.9e-06 | 1.3e-06 | 2.9e-06 | 2.367E-06 | 9.238E-07 | 1.300E-06 | 2.900E-06 |
| 22266 | 1114 | S1114E | D | 2.2e-06 | 2.2e-06 | 2.6e-06 | 2.333E-06 | 2.309E-07 | 2.200E-06 | 2.600E-06 |
| 22267 | 1114 | S1114Q | D | 3e-06 | 1.1e-06 | 2.5e-06 | 2.200E-06 | 9.849E-07 | 1.100E-06 | 3.000E-06 |
| 22268 | 1114 | S1114G | D | 2.4e-06 | 1.8e-06 | 3.4e-06 | 2.533E-06 | 8.083E-07 | 1.800E-06 | 3.400E-06 |
| 22269 | 1114 | S1114H | D | 2e-06 | 1.7e-06 | 2.1e-06 | 1.933E-06 | 2.082E-07 | 1.700E-06 | 2.100E-06 |

| | | | | | | | | | | |
|---|---|---|---|---|---|---|---|---|---|---|
| 22270 | 1114 | S1114I | D | 1.6e-06 | 2.4e-06 | 2.4e-06 | 2.133E-06 | 4.619E-07 | 1.600E-06 | 2.400E-06 |
| 22271 | 1114 | S1114L | D | 1.4e-06 | 3.2e-06 | 2e-06 | 2.200E-06 | 9.165E-07 | 1.400E-06 | 3.200E-06 |
| 22272 | 1114 | S1114K | D | 2.8e-06 | 3.5e-06 | 3e-06 | 3.100E-06 | 3.606E-07 | 2.800E-06 | 3.500E-06 |
| 22273 | 1114 | S1114M | D | 2.2e-06 | 2.1e-06 | 3.8e-06 | 2.700E-06 | 9.539E-07 | 2.100E-06 | 3.800E-06 |
| 22274 | 1114 | S1114F | D | 2.2e-06 | 2.2e-06 | 2.7e-06 | 2.367E-06 | 2.887E-07 | 2.200E-06 | 2.700E-06 |
| 22275 | 1114 | S1114P | D | 2.4e-06 | 2.3e-06 | 1.9e-06 | 2.200E-06 | 2.646E-07 | 1.900E-06 | 2.400E-06 |
| 22276 | 1114 | S1114S | D | 1.9e-06 | 2.2e-06 | 3.2e-06 | 2.433E-06 | 6.807E-07 | 1.900E-06 | 3.200E-06 |
| 22277 | 1114 | S1114T | D | 2e-06 | 1.8e-06 | 8.4e-07 | 1.547E-06 | 6.201E-07 | 8.400E-07 | 2.000E-06 |
| 22278 | 1114 | S1114W | D | 5.2e-06 | 3e-06 | 3e-06 | 3.733E-06 | 1.270E-06 | 3.000E-06 | 5.200E-06 |
| 22279 | 1114 | S1114Y | D | 1.8e-06 | 1.8e-06 | 2.4e-06 | 2.000E-06 | 3.464E-07 | 1.800E-06 | 2.400E-06 |
| 22280 | 1114 | S1114V | D | 2.3e-06 | 2e-06 | 1.9e-06 | 2.067E-06 | 2.082E-07 | 1.900E-06 | 2.300E-06 |
| 22281 | 1115 | G1115A | D | 2.5e-06 | 1.8e-06 | 1.7e-06 | 2.000E-06 | 4.359E-07 | 1.700E-06 | 2.500E-06 |
| 22282 | 1115 | G1115R | D | 2.4e-06 | 2.2e-06 | 2e-06 | 2.200E-06 | 2.000E-07 | 2.000E-06 | 2.400E-06 |
| 22283 | 1115 | G1115N | D | 1.3e-06 | 1.4e-06 | 1.5e-06 | 1.400E-06 | 1.000E-07 | 1.300E-06 | 1.500E-06 |
| 22284 | 1115 | G1115D | D | 3.3e-06 | 4.1e-06 | 2e-06 | 3.133E-06 | 1.060E-06 | 2.000E-06 | 4.100E-06 |
| 22285 | 1115 | G1115C | D | 2.1e-06 | 1.8e-06 | 1.4e-06 | 1.767E-06 | 3.512E-07 | 1.400E-06 | 2.100E-06 |
| 22286 | 1115 | G1115E | D | 1.9e-06 | 3.4e-06 | 2.5e-06 | 2.600E-06 | 7.550E-07 | 1.900E-06 | 3.400E-06 |
| 22287 | 1115 | G1115Q | D | 2.4e-06 | 2.7e-06 | 1.9e-06 | 2.333E-06 | 4.041E-07 | 1.900E-06 | 2.700E-06 |
| 22288 | 1115 | G1115G | D | 4.1e-06 | 1.4e-06 | 2.9e-06 | 2.800E-06 | 1.353E-06 | 1.400E-06 | 4.100E-06 |
| 22289 | 1115 | G1115H | D | 1.8e-06 | 3e-06 | 2.4e-06 | 2.400E-06 | 6.000E-07 | 1.800E-06 | 3.000E-06 |
| 22290 | 1115 | G1115I | D | 2.2e-06 | 2.6e-06 | 2.6e-06 | 2.467E-06 | 2.309E-07 | 2.200E-06 | 2.600E-06 |
| 22291 | 1115 | G1115L | D | 1.5e-06 | 3.3e-06 | 3e-06 | 2.600E-06 | 9.644E-07 | 1.500E-06 | 3.300E-06 |
| 22292 | 1115 | G1115K | D | 5.3e-06 | 1.3e-06 | 2.7e-06 | 3.100E-06 | 2.030E-06 | 1.300E-06 | 5.300E-06 |
| 22293 | 1115 | G1115M | D | 1.6e-06 | 2e-06 | 2e-06 | 1.867E-06 | 2.309E-07 | 1.600E-06 | 2.000E-06 |
| 22294 | 1115 | G1115F | D | 1.2e-06 | 9.3e-07 | 2e-06 | 1.377E-06 | 5.564E-07 | 9.300E-07 | 2.000E-06 |
| 22295 | 1115 | G1115P | D | 2.3e-06 | 3.1e-06 | 2.8e-06 | 2.733E-06 | 4.041E-07 | 2.300E-06 | 3.100E-06 |
| 22296 | 1115 | G1115S | D | 2.8e-06 | 2.3e-06 | 2.8e-06 | 2.633E-06 | 2.887E-07 | 2.300E-06 | 2.800E-06 |
| 22297 | 1115 | G1115T | D | 2.1e-06 | 2.2e-06 | 2.3e-06 | 2.200E-06 | 1.000E-07 | 2.100E-06 | 2.300E-06 |
| 22298 | 1115 | G1115W | D | 2.3e-06 | 3.6e-06 | 1.5e-06 | 2.467E-06 | 1.060E-06 | 1.500E-06 | 3.600E-06 |
| 22299 | 1115 | G1115Y | D | 1.8e-06 | 3e-06 | 2.1e-06 | 2.300E-06 | 6.245E-07 | 1.800E-06 | 3.000E-06 |
| 22300 | 1115 | G1115V | D | 1.7e-06 | 3.5e-06 | 2.7e-06 | 2.633E-06 | 9.018E-07 | 1.700E-06 | 3.500E-06 |
| 22301 | 1116 | V1116A | D | 3.1e-06 | 2.1e-06 | 2.8e-06 | 2.667E-06 | 5.132E-07 | 2.100E-06 | 3.100E-06 |
| 22302 | 1116 | V1116R | D | 2.7e-06 | 2.2e-06 | 2.4e-06 | 2.433E-06 | 2.517E-07 | 2.200E-06 | 2.700E-06 |
| 22303 | 1116 | V1116N | D | 1.4e-06 | 2.2e-06 | 2.5e-06 | 2.033E-06 | 5.686E-07 | 1.400E-06 | 2.500E-06 |
| 22304 | 1116 | V1116D | D | 2e-06 | 2.8e-06 | 1.9e-06 | 2.233E-06 | 4.933E-07 | 1.900E-06 | 2.800E-06 |
| 22305 | 1116 | V1116C | D | 1.5e-06 | 1.2e-06 | 3.1e-06 | 1.933E-06 | 1.021E-06 | 1.200E-06 | 3.100E-06 |
| 22306 | 1116 | V1116E | D | 2.2e-06 | 2.8e-06 | 3.3e-06 | 2.767E-06 | 5.508E-07 | 2.200E-06 | 3.300E-06 |
| 22307 | 1116 | V1116Q | D | 2.8e-06 | 3.2e-06 | 2.5e-06 | 2.833E-06 | 3.512E-07 | 2.500E-06 | 3.200E-06 |
| 22308 | 1116 | V1116G | D | 3.5e-06 | 1.7e-06 | 6.6e-06 | 3.933E-06 | 2.479E-06 | 1.700E-06 | 6.600E-06 |
| 22309 | 1116 | V1116H | D | 2e-06 | 2.5e-06 | 3.7e-06 | 2.733E-06 | 8.737E-07 | 2.000E-06 | 3.700E-06 |
| 22310 | 1116 | V1116I | D | 3.9e-06 | 2.8e-06 | 1.8e-06 | 2.833E-06 | 1.050E-06 | 1.800E-06 | 3.900E-06 |
| 22311 | 1116 | V1116L | D | 3e-06 | 1.9e-06 | 1.4e-06 | 2.100E-06 | 8.185E-07 | 1.400E-06 | 3.000E-06 |
| 22312 | 1116 | V1116K | D | 1.8e-06 | 2e-06 | 2.7e-06 | 2.167E-06 | 4.726E-07 | 1.800E-06 | 2.700E-06 |
| 22313 | 1116 | V1116M | D | 2e-06 | 1.4e-06 | 3.4e-06 | 2.267E-06 | 1.026E-06 | 1.400E-06 | 3.400E-06 |
| 22314 | 1116 | V1116F | D | 1.3e-06 | 2.8e-06 | 2.3e-06 | 2.133E-06 | 7.638E-07 | 1.300E-06 | 2.800E-06 |
| 22315 | 1116 | V1116P | D | 3.3e-06 | 8e-07 | 2.9e-06 | 2.333E-06 | 1.343E-06 | 8.000E-07 | 3.300E-06 |
| 22316 | 1116 | V1116S | D | 2.1e-06 | 2.6e-06 | 2.6e-06 | 2.433E-06 | 2.887E-07 | 2.100E-06 | 2.600E-06 |
| 22317 | 1116 | V1116T | D | 2.2e-06 | 1.8e-06 | 2.5e-06 | 2.167E-06 | 3.512E-07 | 1.800E-06 | 2.500E-06 |
| 22318 | 1116 | V1116W | D | 2.6e-06 | 2.9e-06 | 1.9e-06 | 2.467E-06 | 5.132E-07 | 1.900E-06 | 2.900E-06 |
| 22319 | 1116 | V1116Y | D | 3e-06 | 2.4e-06 | 2e-06 | 2.467E-06 | 5.033E-07 | 2.000E-06 | 3.000E-06 |
| 22320 | 1116 | V1116V | D | 2.1e-06 | 2.3e-06 | 5.3e-06 | 3.233E-06 | 1.793E-06 | 2.100E-06 | 5.300E-06 |
| 22321 | 1117 | P1117A | D | 3.7e-06 | 2.7e-06 | 1.7e-06 | 2.700E-06 | 1.000E-06 | 1.700E-06 | 3.700E-06 |
| 22322 | 1117 | P1117R | D | 1.8e-06 | 2.9e-06 | 3.8e-06 | 2.833E-06 | 1.002E-06 | 1.800E-06 | 3.800E-06 |
| 22323 | 1117 | P1117N | D | 1.6e-06 | 1.6e-06 | 2.4e-06 | 1.867E-06 | 4.619E-07 | 1.600E-06 | 2.400E-06 |
| 22324 | 1117 | P1117D | D | 3.3e-06 | 1.8e-06 | 2.1e-06 | 2.400E-06 | 7.937E-07 | 1.800E-06 | 3.300E-06 |
| 22325 | 1117 | P1117C | D | 2.7e-06 | 3.8e-06 | 4.2e-06 | 3.567E-06 | 7.767E-07 | 2.700E-06 | 4.200E-06 |
| 22326 | 1117 | P1117E | D | 1.8e-06 | 3.5e-06 | 1.7e-06 | 2.333E-06 | 1.012E-06 | 1.700E-06 | 3.500E-06 |
| 22327 | 1117 | P1117Q | D | 2.4e-06 | 2.4e-06 | 2.5e-06 | 2.433E-06 | 5.774E-08 | 2.400E-06 | 2.500E-06 |
| 22328 | 1117 | P1117G | D | 1.7e-06 | 2.7e-06 | 2.7e-06 | 2.367E-06 | 5.774E-07 | 1.700E-06 | 2.700E-06 |
| 22329 | 1117 | P1117H | D | 2.7e-06 | 2e-06 | 2.7e-06 | 2.467E-06 | 4.041E-07 | 2.000E-06 | 2.700E-06 |
| 22330 | 1117 | P1117I | D | 1.8e-06 | 3.5e-06 | 2.3e-06 | 2.533E-06 | 8.737E-07 | 1.800E-06 | 3.500E-06 |
| 22331 | 1117 | P1117L | D | 1.2e-06 | 3.7e-06 | 1.6e-06 | 2.167E-06 | 1.343E-06 | 1.200E-06 | 3.700E-06 |
| 22332 | 1117 | P1117K | D | 2.1e-06 | 2.9e-06 | 3e-06 | 2.667E-06 | 4.933E-07 | 2.100E-06 | 3.000E-06 |
| 22333 | 1117 | P1117M | D | 2.8e-06 | 1.3e-06 | 2.2e-06 | 2.100E-06 | 7.550E-07 | 1.300E-06 | 2.800E-06 |
| 22334 | 1117 | P1117F | D | 2.4e-06 | 1.2e-06 | 2.3e-06 | 1.967E-06 | 6.658E-07 | 1.200E-06 | 2.400E-06 |
| 22335 | 1117 | P1117P | D | 2.4e-06 | 1.7e-06 | 4e-06 | 2.700E-06 | 1.179E-06 | 1.700E-06 | 4.000E-06 |
| 22336 | 1117 | P1117S | D | 3.5e-06 | 2.7e-06 | 5e-06 | 3.733E-06 | 1.168E-06 | 2.700E-06 | 5.000E-06 |
| 22337 | 1117 | P1117T | D | 1.8e-06 | 1e-06 | 9.9e-07 | 1.263E-06 | 4.648E-07 | 9.900E-07 | 1.800E-06 |
| 22338 | 1117 | P1117W | D | 1.3e-06 | 1.9e-06 | 3.2e-06 | 2.133E-06 | 9.713E-07 | 1.300E-06 | 3.200E-06 |
| 22339 | 1117 | P1117Y | D | 2.5e-06 | 9.7e-07 | 3.5e-06 | 2.323E-06 | 1.274E-06 | 9.700E-07 | 3.500E-06 |
| 22340 | 1117 | P1117V | D | 2.9e-06 | 2.2e-06 | 5.4e-06 | 3.500E-06 | 1.682E-06 | 2.200E-06 | 5.400E-06 |
| 22341 | 1118 | S1118A | D | 2.3e-06 | 1.2e-06 | 2.9e-06 | 2.133E-06 | 8.622E-07 | 1.200E-06 | 2.900E-06 |
| 22342 | 1118 | S1118R | D | 9.3e-07 | 3.6e-06 | 2.3e-06 | 2.277E-06 | 1.335E-06 | 9.300E-07 | 3.600E-06 |
| 22343 | 1118 | S1118N | D | 1.4e-06 | 2.7e-06 | 2.4e-06 | 2.167E-06 | 6.807E-07 | 1.400E-06 | 2.700E-06 |
| 22344 | 1118 | S1118D | D | 1.7e-06 | 2.3e-06 | 1.7e-06 | 1.900E-06 | 3.464E-07 | 1.700E-06 | 2.300E-06 |
| 22345 | 1118 | S1118C | D | 2.2e-06 | 1.6e-06 | 1.5e-06 | 1.767E-06 | 3.786E-07 | 1.500E-06 | 2.200E-06 |
| 22346 | 1118 | S1118E | D | 1.8e-06 | 2.3e-06 | 1.6e-06 | 1.900E-06 | 3.606E-07 | 1.600E-06 | 2.300E-06 |
| 22347 | 1118 | S1118Q | D | 1.7e-06 | 3.2e-06 | 2.8e-06 | 2.567E-06 | 7.767E-07 | 1.700E-06 | 3.200E-06 |
| 22348 | 1118 | S1118G | D | 2.8e-06 | 2.3e-06 | 2.3e-06 | 2.467E-06 | 2.887E-07 | 2.300E-06 | 2.800E-06 |
| 22349 | 1118 | S1118H | D | 2.6e-06 | 1.4e-06 | 3.4e-06 | 2.467E-06 | 1.007E-06 | 1.400E-06 | 3.400E-06 |
| 22350 | 1118 | S1118I | D | 3.4e-06 | 2.6e-06 | 2.6e-06 | 2.867E-06 | 4.619E-07 | 2.600E-06 | 3.400E-06 |
| 22351 | 1118 | S1118L | D | 1.4e-06 | 2.5e-06 | 3.2e-06 | 2.367E-06 | 9.074E-07 | 1.400E-06 | 3.200E-06 |
| 22352 | 1118 | S1118K | D | 4.2e-06 | 1.9e-06 | 2.4e-06 | 2.833E-06 | 1.210E-06 | 1.900E-06 | 4.200E-06 |
| 22353 | 1118 | S1118M | D | 8.3e-07 | 1.9e-06 | 3.3e-06 | 2.010E-06 | 1.239E-06 | 8.300E-07 | 3.300E-06 |
| 22354 | 1118 | S1118F | D | 2.5e-06 | 2.6e-06 | 2e-06 | 2.367E-06 | 3.215E-07 | 2.000E-06 | 2.600E-06 |

| | | | | | | | | | | |
|---|---|---|---|---|---|---|---|---|---|---|
| 22355 | 1118 | S1118P | D | 1.7e-06 | 3.5e-06 | 3.1e-06 | 2.767E-06 | 9.452E-07 | 1.700E-06 | 3.500E-06 |
| 22356 | 1118 | S1118S | D | 1.9e-06 | 1.7e-06 | 3e-06 | 2.200E-06 | 7.000E-07 | 1.700E-06 | 3.000E-06 |
| 22357 | 1118 | S1118T | D | 2.4e-06 | 2.4e-06 | 3.1e-06 | 2.633E-06 | 4.041E-07 | 2.400E-06 | 3.100E-06 |
| 22358 | 1118 | S1118W | D | 5.1e-06 | 2.1e-06 | 2e-06 | 3.067E-06 | 1.762E-06 | 2.000E-06 | 5.100E-06 |
| 22359 | 1118 | S1118Y | D | 2.2e-06 | 2.8e-06 | 2.3e-06 | 2.433E-06 | 3.215E-07 | 2.200E-06 | 2.800E-06 |
| 22360 | 1118 | S1118V | D | 2.2e-06 | 2.4e-06 | 2.5e-06 | 2.367E-06 | 1.528E-07 | 2.200E-06 | 2.500E-06 |
| 22361 | 1119 | R1119A | D | 2.3e-06 | 1.8e-06 | 2e-06 | 2.033E-06 | 2.517E-07 | 1.800E-06 | 2.300E-06 |
| 22362 | 1119 | R1119R | D | 2.7e-06 | 2e-06 | 1.7e-06 | 2.133E-06 | 5.132E-07 | 1.700E-06 | 2.700E-06 |
| 22363 | 1119 | R1119N | D | 1.8e-06 | 2.2e-06 | 2e-06 | 2.000E-06 | 2.000E-07 | 1.800E-06 | 2.200E-06 |
| 22364 | 1119 | R1119D | D | 2.8e-06 | 3.2e-06 | 3.7e-06 | 3.233E-06 | 4.509E-07 | 2.800E-06 | 3.700E-06 |
| 22365 | 1119 | R1119C | D | 2.5e-06 | 1.9e-06 | 2.9e-06 | 2.433E-06 | 5.033E-07 | 1.900E-06 | 2.900E-06 |
| 22366 | 1119 | R1119E | D | 3.1e-06 | 1.9e-06 | 1.9e-06 | 2.300E-06 | 6.928E-07 | 1.900E-06 | 3.100E-06 |
| 22367 | 1119 | R1119Q | D | 1.8e-06 | 3e-06 | 1.8e-06 | 2.200E-06 | 6.928E-07 | 1.800E-06 | 3.000E-06 |
| 22368 | 1119 | R1119G | D | 2.2e-06 | 4.4e-06 | 2e-06 | 2.867E-06 | 1.332E-06 | 2.000E-06 | 4.400E-06 |
| 22369 | 1119 | R1119H | D | 2.5e-06 | 2.2e-06 | 7.8e-07 | 1.827E-06 | 9.188E-07 | 7.800E-07 | 2.500E-06 |
| 22370 | 1119 | R1119I | D | 3.7e-06 | 3.5e-06 | 1.2e-06 | 2.800E-06 | 1.389E-06 | 1.200E-06 | 3.700E-06 |
| 22371 | 1119 | R1119L | D | 2.1e-06 | 2.5e-06 | 3.8e-06 | 2.800E-06 | 8.888E-07 | 2.100E-06 | 3.800E-06 |
| 22372 | 1119 | R1119K | D | 2.6e-06 | 2.8e-06 | 2.2e-06 | 2.533E-06 | 3.055E-07 | 2.200E-06 | 2.800E-06 |
| 22373 | 1119 | R1119M | D | 2.3e-06 | 1.2e-06 | 1.8e-06 | 1.767E-06 | 5.508E-07 | 1.200E-06 | 2.300E-06 |
| 22374 | 1119 | R1119F | D | 2.4e-06 | 3e-06 | 2.6e-06 | 2.667E-06 | 3.055E-07 | 2.400E-06 | 3.000E-06 |
| 22375 | 1119 | R1119P | D | 1.8e-06 | 2.3e-06 | 2.1e-06 | 2.067E-06 | 2.517E-07 | 1.800E-06 | 2.300E-06 |
| 22376 | 1119 | R1119S | D | 2.3e-06 | 2.2e-06 | 1.7e-06 | 2.067E-06 | 3.215E-07 | 1.700E-06 | 2.300E-06 |
| 22377 | 1119 | R1119T | D | 2.6e-06 | 2.6e-06 | 1.7e-06 | 2.300E-06 | 5.196E-07 | 1.700E-06 | 2.600E-06 |
| 22378 | 1119 | R1119W | D | 1.5e-06 | 2.5e-06 | 4.8e-06 | 2.933E-06 | 1.692E-06 | 1.500E-06 | 4.800E-06 |
| 22379 | 1119 | R1119Y | D | 1.6e-06 | 2.6e-06 | 1.7e-06 | 1.967E-06 | 5.508E-07 | 1.600E-06 | 2.600E-06 |
| 22380 | 1119 | R1119V | D | 2.6e-06 | 2.4e-06 | 2.2e-06 | 2.400E-06 | 2.000E-07 | 2.200E-06 | 2.600E-06 |
| 22381 | 1120 | F1120A | D | 3e-06 | 1.8e-06 | 2.1e-06 | 2.300E-06 | 6.245E-07 | 1.800E-06 | 3.000E-06 |
| 22382 | 1120 | F1120R | D | 4.1e-06 | 2.3e-06 | 3.2e-06 | 3.200E-06 | 9.000E-07 | 2.300E-06 | 4.100E-06 |
| 22383 | 1120 | F1120N | D | 1.7e-06 | 2.6e-06 | 1.9e-06 | 2.067E-06 | 4.726E-07 | 1.700E-06 | 2.600E-06 |
| 22384 | 1120 | F1120D | D | 3.2e-06 | 1.9e-06 | 1.6e-06 | 2.233E-06 | 8.505E-07 | 1.600E-06 | 3.200E-06 |
| 22385 | 1120 | F1120C | D | 1.3e-06 | 3.7e-06 | 3.5e-06 | 2.833E-06 | 1.332E-06 | 1.300E-06 | 3.700E-06 |
| 22386 | 1120 | F1120E | D | 1.6e-06 | 2.8e-06 | 2.2e-06 | 2.200E-06 | 6.000E-07 | 1.600E-06 | 2.800E-06 |
| 22387 | 1120 | F1120Q | D | 3.4e-06 | 2.9e-06 | 2.4e-06 | 2.900E-06 | 5.000E-07 | 2.400E-06 | 3.400E-06 |
| 22388 | 1120 | F1120G | D | 2.2e-06 | 3.9e-06 | 2.6e-06 | 2.900E-06 | 8.888E-07 | 2.200E-06 | 3.900E-06 |
| 22389 | 1120 | F1120H | D | 1.1e-06 | 2.6e-06 | 2.6e-06 | 2.100E-06 | 8.660E-07 | 1.100E-06 | 2.600E-06 |
| 22390 | 1120 | F1120I | D | 2.7e-06 | 3e-06 | 5.5e-06 | 3.733E-06 | 1.537E-06 | 2.700E-06 | 5.500E-06 |
| 22391 | 1120 | F1120L | D | 1.7e-06 | 2.2e-06 | 3.4e-06 | 2.433E-06 | 8.737E-07 | 1.700E-06 | 3.400E-06 |
| 22392 | 1120 | F1120K | D | 2.3e-06 | 1.6e-06 | 2.7e-06 | 2.200E-06 | 5.568E-07 | 1.600E-06 | 2.700E-06 |
| 22393 | 1120 | F1120M | D | 2.1e-06 | 2.1e-06 | 1.7e-06 | 1.967E-06 | 2.309E-07 | 1.700E-06 | 2.100E-06 |
| 22394 | 1120 | F1120F | D | 2.2e-06 | 2.1e-06 | 1.7e-06 | 2.000E-06 | 2.646E-07 | 1.700E-06 | 2.200E-06 |
| 22395 | 1120 | F1120P | D | 2.3e-06 | 2.2e-06 | 4.4e-06 | 2.967E-06 | 1.242E-06 | 2.200E-06 | 4.400E-06 |
| 22396 | 1120 | F1120S | D | 2e-06 | 2.5e-06 | 2.3e-06 | 2.267E-06 | 2.517E-07 | 2.000E-06 | 2.500E-06 |
| 22397 | 1120 | F1120T | D | 3e-06 | 2.7e-06 | 1.6e-06 | 2.433E-06 | 7.371E-07 | 1.600E-06 | 3.000E-06 |
| 22398 | 1120 | F1120W | D | 1.7e-06 | 2e-06 | 7.8e-07 | 1.493E-06 | 6.357E-07 | 7.800E-07 | 2.000E-06 |
| 22399 | 1120 | F1120Y | D | 2.8e-06 | 4.2e-06 | 2e-06 | 3.000E-06 | 1.114E-06 | 2.000E-06 | 4.200E-06 |
| 22400 | 1120 | F1120V | D | 3.6e-06 | 2e-06 | 2.3e-06 | 2.633E-06 | 8.505E-07 | 2.000E-06 | 3.600E-06 |
| 22401 | 1121 | S1121A | D | 3.1e-06 | 2.5e-06 | 5e-06 | 3.533E-06 | 1.305E-06 | 2.500E-06 | 5.000E-06 |
| 22402 | 1121 | S1121R | D | 2.3e-06 | 2.3e-06 | 1.6e-06 | 2.067E-06 | 4.041E-07 | 1.600E-06 | 2.300E-06 |
| 22403 | 1121 | S1121N | D | 2.9e-06 | 1.8e-06 | 2.3e-06 | 2.333E-06 | 5.508E-07 | 1.800E-06 | 2.900E-06 |
| 22404 | 1121 | S1121D | D | 1.1e-06 | 2.3e-06 | 1.8e-06 | 1.733E-06 | 6.028E-07 | 1.100E-06 | 2.300E-06 |
| 22405 | 1121 | S1121C | D | 2.2e-06 | 2e-06 | 1.6e-06 | 1.933E-06 | 3.055E-07 | 1.600E-06 | 2.200E-06 |
| 22406 | 1121 | S1121E | D | 3.4e-06 | 3.8e-06 | 1e-06 | 2.733E-06 | 1.514E-06 | 1.000E-06 | 3.800E-06 |
| 22407 | 1121 | S1121Q | D | 1.8e-06 | 3.6e-06 | 3.5e-06 | 2.967E-06 | 1.012E-06 | 1.800E-06 | 3.600E-06 |
| 22408 | 1121 | S1121G | D | 1.5e-06 | 2.9e-06 | 1e-06 | 1.800E-06 | 9.849E-07 | 1.000E-06 | 2.900E-06 |
| 22409 | 1121 | S1121H | D | 1.7e-06 | 2.7e-06 | 1.5e-06 | 1.967E-06 | 6.429E-07 | 1.500E-06 | 2.700E-06 |
| 22410 | 1121 | S1121I | D | 2e-06 | 2.3e-06 | 1.8e-06 | 2.033E-06 | 2.517E-07 | 1.800E-06 | 2.300E-06 |
| 22411 | 1121 | S1121L | D | 2.1e-06 | 2.6e-06 | 2.8e-06 | 2.500E-06 | 3.606E-07 | 2.100E-06 | 2.800E-06 |
| 22412 | 1121 | S1121K | D | 2.3e-06 | 2.3e-06 | 2.6e-06 | 2.400E-06 | 1.732E-07 | 2.300E-06 | 2.600E-06 |
| 22413 | 1121 | S1121M | D | 1.4e-06 | 3.2e-06 | 3.1e-06 | 2.567E-06 | 1.012E-06 | 1.400E-06 | 3.200E-06 |
| 22414 | 1121 | S1121F | D | 2.3e-06 | 2.2e-06 | 2.7e-06 | 2.400E-06 | 2.646E-07 | 2.200E-06 | 2.700E-06 |
| 22415 | 1121 | S1121P | D | 3e-06 | 1.9e-06 | 2e-06 | 2.300E-06 | 6.083E-07 | 1.900E-06 | 3.000E-06 |
| 22416 | 1121 | S1121S | D | 1.9e-06 | 3.8e-06 | 2.9e-06 | 2.867E-06 | 9.504E-07 | 1.900E-06 | 3.800E-06 |
| 22417 | 1121 | S1121T | D | 2e-06 | 1.8e-06 | 2.1e-06 | 1.967E-06 | 1.528E-07 | 1.800E-06 | 2.100E-06 |
| 22418 | 1121 | S1121W | D | 1.7e-06 | 3.7e-06 | 3.1e-06 | 2.833E-06 | 1.026E-06 | 1.700E-06 | 3.700E-06 |
| 22419 | 1121 | S1121Y | D | 2.7e-06 | 1.8e-06 | 3.7e-06 | 2.733E-06 | 9.504E-07 | 1.800E-06 | 3.700E-06 |
| 22420 | 1121 | S1121V | D | 2.2e-06 | 1.7e-06 | 5.1e-06 | 3.000E-06 | 1.836E-06 | 1.700E-06 | 5.100E-06 |
| 22421 | 1122 | G1122A | D | 2.1e-06 | 2.9e-06 | 3.2e-06 | 2.733E-06 | 5.686E-07 | 2.100E-06 | 3.200E-06 |
| 22422 | 1122 | G1122R | D | 2.2e-06 | 3.1e-06 | 3.1e-06 | 2.800E-06 | 5.196E-07 | 2.200E-06 | 3.100E-06 |
| 22423 | 1122 | G1122N | D | 1.4e-06 | 2e-06 | 3.1e-06 | 2.167E-06 | 8.622E-07 | 1.400E-06 | 3.100E-06 |
| 22424 | 1122 | G1122D | D | 2e-06 | 2.3e-06 | 1.6e-06 | 1.967E-06 | 3.512E-07 | 1.600E-06 | 2.300E-06 |
| 22425 | 1122 | G1122C | D | 2.4e-06 | 3.1e-06 | 2.8e-06 | 2.767E-06 | 3.512E-07 | 2.400E-06 | 3.100E-06 |
| 22426 | 1122 | G1122E | D | 1.9e-06 | 3.1e-06 | 4e-06 | 3.000E-06 | 1.054E-06 | 1.900E-06 | 4.000E-06 |
| 22427 | 1122 | G1122Q | D | 2.4e-06 | 1.7e-06 | 2.6e-06 | 2.233E-06 | 4.726E-07 | 1.700E-06 | 2.600E-06 |
| 22428 | 1122 | G1122G | D | 2.2e-06 | 3.5e-06 | 1.8e-06 | 2.500E-06 | 8.888E-07 | 1.800E-06 | 3.500E-06 |
| 22429 | 1122 | G1122H | D | 2.2e-06 | 2.1e-06 | 2.2e-06 | 2.167E-06 | 5.774E-08 | 2.100E-06 | 2.200E-06 |
| 22430 | 1122 | G1122I | D | 2e-06 | 1.8e-06 | 2.8e-06 | 2.200E-06 | 5.292E-07 | 1.800E-06 | 2.800E-06 |
| 22431 | 1122 | G1122L | D | 2.7e-06 | 1.9e-06 | 2.2e-06 | 2.267E-06 | 4.041E-07 | 1.900E-06 | 2.700E-06 |
| 22432 | 1122 | G1122K | D | 4e-06 | 2e-06 | 3.4e-06 | 3.133E-06 | 1.026E-06 | 2.000E-06 | 4.000E-06 |
| 22433 | 1122 | G1122M | D | 2.6e-06 | 6.6e-06 | 2e-06 | 3.733E-06 | 2.501E-06 | 2.000E-06 | 6.600E-06 |
| 22434 | 1122 | G1122F | D | 1.5e-06 | 3.1e-06 | 1.2e-06 | 1.933E-06 | 1.021E-06 | 1.200E-06 | 3.100E-06 |
| 22435 | 1122 | G1122P | D | 1.4e-06 | 2.5e-06 | 3.2e-06 | 2.367E-06 | 9.074E-07 | 1.400E-06 | 3.200E-06 |
| 22436 | 1122 | G1122S | D | 1.9e-06 | 2.4e-06 | 2.6e-06 | 2.300E-06 | 3.606E-07 | 1.900E-06 | 2.600E-06 |
| 22437 | 1122 | G1122T | D | 1.7e-06 | 2.1e-06 | 2.1e-06 | 1.967E-06 | 2.309E-07 | 1.700E-06 | 2.100E-06 |
| 22438 | 1122 | G1122W | D | 2.8e-06 | 2.4e-06 | 2.1e-06 | 2.433E-06 | 3.512E-07 | 2.100E-06 | 2.800E-06 |
| 22439 | 1122 | G1122Y | D | 1.6e-06 | 3.9e-06 | 2.3e-06 | 2.600E-06 | 1.179E-06 | 1.600E-06 | 3.900E-06 |

| | | | | | | | | | | |
|---|---|---|---|---|---|---|---|---|---|---|
| 22440 | 1122 | G1122V | D | 1.6e-06 | 3.1e-06 | 4.7e-06 | 3.133E-06 | 1.550E-06 | 1.600E-06 | 4.700E-06 |
| 22441 | 1123 | S1123A | D | 3.4e-06 | 2.4e-06 | 1.9e-06 | 2.567E-06 | 7.638E-07 | 1.900E-06 | 3.400E-06 |
| 22442 | 1123 | S1123R | D | 2.1e-06 | 1.1e-06 | 3.2e-06 | 2.133E-06 | 1.050E-06 | 1.100E-06 | 3.200E-06 |
| 22443 | 1123 | S1123N | D | 3.1e-06 | 2.1e-06 | 3.4e-06 | 2.867E-06 | 6.807E-07 | 2.100E-06 | 3.400E-06 |
| 22444 | 1123 | S1123D | D | 2.1e-06 | 1.9e-06 | 2.4e-06 | 2.133E-06 | 2.517E-07 | 1.900E-06 | 2.400E-06 |
| 22445 | 1123 | S1123C | D | 1.8e-06 | 4.1e-06 | 2.9e-06 | 2.933E-06 | 1.150E-06 | 1.800E-06 | 4.100E-06 |
| 22446 | 1123 | S1123E | D | 3.4e-06 | 2.4e-06 | 1.9e-06 | 2.567E-06 | 7.638E-07 | 1.900E-06 | 3.400E-06 |
| 22447 | 1123 | S1123Q | D | 2.2e-06 | 2.5e-06 | 1.9e-06 | 2.200E-06 | 3.000E-07 | 1.900E-06 | 2.500E-06 |
| 22448 | 1123 | S1123G | D | 2.9e-06 | 3.2e-06 | 2e-06 | 2.700E-06 | 6.245E-07 | 2.000E-06 | 3.200E-06 |
| 22449 | 1123 | S1123H | D | 3e-06 | 2.9e-06 | 1.1e-06 | 2.333E-06 | 1.069E-06 | 1.100E-06 | 3.000E-06 |
| 22450 | 1123 | S1123I | D | 1.2e-06 | 2.1e-06 | 2.5e-06 | 1.933E-06 | 6.658E-07 | 1.200E-06 | 2.500E-06 |
| 22451 | 1123 | S1123L | D | 2.1e-06 | 3e-06 | 2.3e-06 | 2.467E-06 | 4.726E-07 | 2.100E-06 | 3.000E-06 |
| 22452 | 1123 | S1123K | D | 2e-06 | 2.2e-06 | 1.8e-06 | 2.000E-06 | 2.000E-07 | 1.800E-06 | 2.200E-06 |
| 22453 | 1123 | S1123M | D | 2.9e-06 | 2.4e-06 | 2.1e-06 | 2.467E-06 | 4.041E-07 | 2.100E-06 | 2.900E-06 |
| 22454 | 1123 | S1123F | D | 2.7e-06 | 2.7e-06 | 2.6e-06 | 2.667E-06 | 5.774E-08 | 2.600E-06 | 2.700E-06 |
| 22455 | 1123 | S1123P | D | 2.9e-06 | 1.8e-06 | 4.1e-06 | 2.933E-06 | 1.150E-06 | 1.800E-06 | 4.100E-06 |
| 22456 | 1123 | S1123S | D | 1.9e-06 | 3.3e-06 | 1.9e-06 | 2.367E-06 | 8.083E-07 | 1.900E-06 | 3.300E-06 |
| 22457 | 1123 | S1123T | D | 2.5e-06 | 2.3e-06 | 2.8e-06 | 2.533E-06 | 2.517E-07 | 2.300E-06 | 2.800E-06 |
| 22458 | 1123 | S1123W | D | 4.1e-06 | 1.7e-06 | 2.9e-06 | 2.900E-06 | 1.200E-06 | 1.700E-06 | 4.100E-06 |
| 22459 | 1123 | S1123Y | D | 2.7e-06 | 1.4e-06 | 3.2e-06 | 2.433E-06 | 9.292E-07 | 1.400E-06 | 3.200E-06 |
| 22460 | 1123 | S1123V | D | 3.3e-06 | 2.1e-06 | 4.2e-06 | 3.200E-06 | 1.054E-06 | 2.100E-06 | 4.200E-06 |
| 22461 | 1124 | R1124A | D | 2.7e-06 | 3.2e-06 | 2.4e-06 | 2.767E-06 | 4.041E-07 | 2.400E-06 | 3.200E-06 |
| 22462 | 1124 | R1124R | D | 3.5e-06 | 2.9e-06 | 1.7e-06 | 2.700E-06 | 9.165E-07 | 1.700E-06 | 3.500E-06 |
| 22463 | 1124 | R1124N | D | 2.7e-06 | 1.4e-06 | 2.5e-06 | 2.200E-06 | 7.000E-07 | 1.400E-06 | 2.700E-06 |
| 22464 | 1124 | R1124D | D | 2.2e-06 | 1.9e-06 | 2.7e-06 | 2.267E-06 | 4.041E-07 | 1.900E-06 | 2.700E-06 |
| 22465 | 1124 | R1124C | D | 1.3e-06 | 4.6e-06 | 2.9e-06 | 2.933E-06 | 1.650E-06 | 1.300E-06 | 4.600E-06 |
| 22466 | 1124 | R1124E | D | 5.4e-06 | 4.3e-06 | 2.1e-06 | 3.933E-06 | 1.680E-06 | 2.100E-06 | 5.400E-06 |
| 22467 | 1124 | R1124Q | D | 2.2e-06 | 5.8e-06 | 4.3e-06 | 4.100E-06 | 1.808E-06 | 2.200E-06 | 5.800E-06 |
| 22468 | 1124 | R1124G | D | 2.6e-06 | 1.9e-06 | 2.3e-06 | 2.267E-06 | 3.512E-07 | 1.900E-06 | 2.600E-06 |
| 22469 | 1124 | R1124H | D | 3.9e-06 | 2.7e-06 | 1.6e-06 | 2.733E-06 | 1.150E-06 | 1.600E-06 | 3.900E-06 |
| 22470 | 1124 | R1124I | D | 2.3e-06 | 4.6e-06 | 2.2e-06 | 3.033E-06 | 1.358E-06 | 2.200E-06 | 4.600E-06 |
| 22471 | 1124 | R1124L | D | 2.9e-06 | 1.6e-06 | 4.6e-06 | 3.033E-06 | 1.504E-06 | 1.600E-06 | 4.600E-06 |
| 22472 | 1124 | R1124K | D | 1.7e-06 | 2.7e-06 | 3.8e-06 | 2.733E-06 | 1.050E-06 | 1.700E-06 | 3.800E-06 |
| 22473 | 1124 | R1124M | D | 2.8e-06 | 1.4e-06 | 1.3e-06 | 1.833E-06 | 8.386E-07 | 1.300E-06 | 2.800E-06 |
| 22474 | 1124 | R1124F | D | 3.5e-06 | 2e-06 | 2.1e-06 | 2.533E-06 | 8.386E-07 | 2.000E-06 | 3.500E-06 |
| 22475 | 1124 | R1124P | D | 1.7e-06 | 2.2e-06 | 3.2e-06 | 2.367E-06 | 7.638E-07 | 1.700E-06 | 3.200E-06 |
| 22476 | 1124 | R1124S | D | 1.8e-06 | 2.5e-06 | 2.4e-06 | 2.233E-06 | 3.786E-07 | 1.800E-06 | 2.500E-06 |
| 22477 | 1124 | R1124T | D | 2.2e-06 | 3e-06 | 2.3e-06 | 2.500E-06 | 4.359E-07 | 2.200E-06 | 3.000E-06 |
| 22478 | 1124 | R1124W | D | 2.8e-06 | 2.4e-06 | 2.5e-06 | 2.567E-06 | 2.082E-07 | 2.400E-06 | 2.800E-06 |
| 22479 | 1124 | R1124Y | D | 2.8e-06 | 3.1e-06 | 2e-06 | 2.633E-06 | 5.686E-07 | 2.000E-06 | 3.100E-06 |
| 22480 | 1124 | R1124V | D | 2.6e-06 | 4.3e-06 | 2.7e-06 | 3.200E-06 | 9.539E-07 | 2.600E-06 | 4.300E-06 |
| 22481 | 1125 | S1125A | D | 2e-06 | 2.2e-06 | 1.9e-06 | 2.033E-06 | 1.528E-07 | 1.900E-06 | 2.200E-06 |
| 22482 | 1125 | S1125R | D | 2.7e-06 | 3.4e-06 | 3e-06 | 3.033E-06 | 3.512E-07 | 2.700E-06 | 3.400E-06 |
| 22483 | 1125 | S1125N | D | 3.1e-06 | 1.2e-06 | 2.1e-06 | 2.133E-06 | 9.504E-07 | 1.200E-06 | 3.100E-06 |
| 22484 | 1125 | S1125D | D | 2e-06 | 3.4e-06 | 1.6e-06 | 2.333E-06 | 9.452E-07 | 1.600E-06 | 3.400E-06 |
| 22485 | 1125 | S1125C | D | 2.8e-06 | 2.2e-06 | 1.8e-06 | 2.267E-06 | 5.033E-07 | 1.800E-06 | 2.800E-06 |
| 22486 | 1125 | S1125E | D | 4.3e-06 | 1.6e-06 | 2.2e-06 | 2.700E-06 | 1.418E-06 | 1.600E-06 | 4.300E-06 |
| 22487 | 1125 | S1125Q | D | 2.2e-06 | 2.5e-06 | 3.3e-06 | 2.667E-06 | 5.686E-07 | 2.200E-06 | 3.300E-06 |
| 22488 | 1125 | S1125G | D | 4e-06 | 1.9e-06 | 3.4e-06 | 3.100E-06 | 1.082E-06 | 1.900E-06 | 4.000E-06 |
| 22489 | 1125 | S1125H | D | 3.4e-06 | 2.1e-06 | 1.9e-06 | 2.467E-06 | 8.145E-07 | 1.900E-06 | 3.400E-06 |
| 22490 | 1125 | S1125I | D | 3.2e-06 | 3.5e-06 | 1e-06 | 2.567E-06 | 1.365E-06 | 1.000E-06 | 3.500E-06 |
| 22491 | 1125 | S1125L | D | 3.2e-06 | 3.4e-06 | 1.5e-06 | 2.700E-06 | 1.044E-06 | 1.500E-06 | 3.400E-06 |
| 22492 | 1125 | S1125K | D | 2e-06 | 2.5e-06 | 2.3e-06 | 2.267E-06 | 2.517E-07 | 2.000E-06 | 2.500E-06 |
| 22493 | 1125 | S1125M | D | 1.8e-06 | 2.3e-06 | 3.3e-06 | 2.467E-06 | 7.638E-07 | 1.800E-06 | 3.300E-06 |
| 22494 | 1125 | S1125F | D | 2.2e-06 | 1.8e-06 | 3.2e-06 | 2.400E-06 | 7.211E-07 | 1.800E-06 | 3.200E-06 |
| 22495 | 1125 | S1125P | D | 1.7e-06 | 2.3e-06 | 4.2e-06 | 2.733E-06 | 1.305E-06 | 1.700E-06 | 4.200E-06 |
| 22496 | 1125 | S1125S | D | 2.5e-06 | 6.2e-06 | 3e-06 | 3.900E-06 | 2.007E-06 | 2.500E-06 | 6.200E-06 |
| 22497 | 1125 | S1125T | D | 3.1e-06 | 4.2e-06 | 1.9e-06 | 3.067E-06 | 1.150E-06 | 1.900E-06 | 4.200E-06 |
| 22498 | 1125 | S1125W | D | 2.5e-06 | 2.2e-06 | 1.9e-06 | 2.200E-06 | 3.000E-07 | 1.900E-06 | 2.500E-06 |
| 22499 | 1125 | S1125Y | D | 3e-06 | 3.8e-06 | 2.3e-06 | 3.033E-06 | 7.506E-07 | 2.300E-06 | 3.800E-06 |
| 22500 | 1125 | S1125V | D | 3.9e-06 | 1.8e-06 | 2.8e-06 | 2.833E-06 | 1.050E-06 | 1.800E-06 | 3.900E-06 |
| 22501 | 1126 | G1126A | D | 2.5e-06 | 1.7e-06 | 2.6e-06 | 2.267E-06 | 4.933E-07 | 1.700E-06 | 2.600E-06 |
| 22502 | 1126 | G1126R | D | 2e-06 | 3.7e-06 | 2.9e-06 | 2.867E-06 | 8.505E-07 | 2.000E-06 | 3.700E-06 |
| 22503 | 1126 | G1126N | D | 2.3e-06 | 2.7e-06 | 2.4e-06 | 2.467E-06 | 2.082E-07 | 2.300E-06 | 2.700E-06 |
| 22504 | 1126 | G1126D | D | 3.4e-06 | 3e-06 | 1.2e-06 | 2.533E-06 | 1.172E-06 | 1.200E-06 | 3.400E-06 |
| 22505 | 1126 | G1126C | D | 3.3e-06 | 2.6e-06 | 3.1e-06 | 3.000E-06 | 3.606E-07 | 2.600E-06 | 3.300E-06 |
| 22506 | 1126 | G1126E | D | 2.1e-06 | 1.9e-06 | 1.9e-06 | 1.967E-06 | 1.155E-07 | 1.900E-06 | 2.100E-06 |
| 22507 | 1126 | G1126Q | D | 2.4e-06 | 2.1e-06 | 1.4e-06 | 1.967E-06 | 5.132E-07 | 1.400E-06 | 2.400E-06 |
| 22508 | 1126 | G1126G | D | 2.7e-06 | 2.4e-06 | 2.5e-06 | 2.533E-06 | 1.528E-07 | 2.400E-06 | 2.700E-06 |
| 22509 | 1126 | G1126H | D | 2.2e-06 | 2.8e-06 | 1.4e-06 | 2.133E-06 | 7.024E-07 | 1.400E-06 | 2.800E-06 |
| 22510 | 1126 | G1126I | D | 2.3e-06 | 2.3e-06 | 3e-06 | 2.533E-06 | 4.041E-07 | 2.300E-06 | 3.000E-06 |
| 22511 | 1126 | G1126L | D | 2.9e-06 | 2e-06 | 2.9e-06 | 2.600E-06 | 5.196E-07 | 2.000E-06 | 2.900E-06 |
| 22512 | 1126 | G1126K | D | 2.1e-06 | 2.8e-06 | 1.6e-06 | 2.167E-06 | 6.028E-07 | 1.600E-06 | 2.800E-06 |
| 22513 | 1126 | G1126M | D | 3.3e-06 | 1.8e-06 | 3.3e-06 | 2.800E-06 | 8.660E-07 | 1.800E-06 | 3.300E-06 |
| 22514 | 1126 | G1126F | D | 2.9e-06 | 2.1e-06 | 2.9e-06 | 2.633E-06 | 4.619E-07 | 2.100E-06 | 2.900E-06 |
| 22515 | 1126 | G1126P | D | 2.1e-06 | 2e-06 | 2.6e-06 | 2.233E-06 | 3.215E-07 | 2.000E-06 | 2.600E-06 |
| 22516 | 1126 | G1126S | D | 1.7e-06 | 9.8e-07 | 1.9e-06 | 1.527E-06 | 4.839E-07 | 9.800E-07 | 1.900E-06 |
| 22517 | 1126 | G1126T | D | 2.7e-06 | 2.5e-06 | 1.3e-06 | 2.167E-06 | 7.572E-07 | 1.300E-06 | 2.700E-06 |
| 22518 | 1126 | G1126W | D | 2.2e-06 | 2e-06 | 2.5e-06 | 2.233E-06 | 2.517E-07 | 2.000E-06 | 2.500E-06 |
| 22519 | 1126 | G1126Y | D | 2.7e-06 | 4.9e-06 | 2.1e-06 | 3.233E-06 | 1.474E-06 | 2.100E-06 | 4.900E-06 |
| 22520 | 1126 | G1126V | D | 2.3e-06 | 2.5e-06 | 3.3e-06 | 2.700E-06 | 5.292E-07 | 2.300E-06 | 3.300E-06 |
| 22521 | 1127 | T1127A | D | 2.6e-06 | 2.9e-06 | 3.2e-06 | 2.900E-06 | 3.000E-07 | 2.600E-06 | 3.200E-06 |
| 22522 | 1127 | T1127R | D | 2.6e-06 | 2.4e-06 | 2.7e-06 | 2.567E-06 | 1.528E-07 | 2.400E-06 | 2.700E-06 |
| 22523 | 1127 | T1127N | D | 2.8e-06 | 1.2e-06 | 3.1e-06 | 2.367E-06 | 1.021E-06 | 1.200E-06 | 3.100E-06 |
| 22524 | 1127 | T1127D | D | 4.7e-06 | 2.1e-06 | 2.2e-06 | 3.000E-06 | 1.473E-06 | 2.100E-06 | 4.700E-06 |

| | | | | | | | | | | |
|---|---|---|---|---|---|---|---|---|---|---|
| 22525 | 1127 | T1127C | D | 2.8e-06 | 1.3e-06 | 2.6e-06 | 2.233E-06 | 8.145E-07 | 1.300E-06 | 2.800E-06 |
| 22526 | 1127 | T1127E | D | 1.9e-06 | 4.2e-06 | 2.4e-06 | 2.833E-06 | 1.210E-06 | 1.900E-06 | 4.200E-06 |
| 22527 | 1127 | T1127Q | D | 2.4e-06 | 1.9e-06 | 2.9e-06 | 2.400E-06 | 5.000E-07 | 1.900E-06 | 2.900E-06 |
| 22528 | 1127 | T1127G | D | 3.5e-06 | 1.8e-06 | 3.4e-06 | 2.900E-06 | 9.539E-07 | 1.800E-06 | 3.500E-06 |
| 22529 | 1127 | T1127H | D | 8.8e-07 | 4e-06 | 1.6e-06 | 2.160E-06 | 1.634E-06 | 8.800E-07 | 4.000E-06 |
| 22530 | 1127 | T1127I | D | 4e-06 | 3.6e-06 | 2.4e-06 | 3.333E-06 | 8.327E-07 | 2.400E-06 | 4.000E-06 |
| 22531 | 1127 | T1127L | D | 2.6e-06 | 2.9e-06 | 2.9e-06 | 2.800E-06 | 1.732E-07 | 2.600E-06 | 2.900E-06 |
| 22532 | 1127 | T1127K | D | 2.6e-06 | 2.7e-06 | 2e-06 | 2.433E-06 | 3.786E-07 | 2.000E-06 | 2.700E-06 |
| 22533 | 1127 | T1127M | D | 2.8e-06 | 1.5e-06 | 2.3e-06 | 2.200E-06 | 6.557E-07 | 1.500E-06 | 2.800E-06 |
| 22534 | 1127 | T1127F | D | 3.9e-06 | 1.9e-06 | 2.5e-06 | 2.767E-06 | 1.026E-06 | 1.900E-06 | 3.900E-06 |
| 22535 | 1127 | T1127P | D | 2.5e-06 | 2.2e-06 | 2.4e-06 | 2.367E-06 | 1.528E-07 | 2.200E-06 | 2.500E-06 |
| 22536 | 1127 | T1127S | D | 1.2e-06 | 1.9e-06 | 3.4e-06 | 2.167E-06 | 1.124E-06 | 1.200E-06 | 3.400E-06 |
| 22537 | 1127 | T1127T | D | 2.9e-06 | 4.2e-06 | 1.5e-06 | 2.867E-06 | 1.350E-06 | 1.500E-06 | 4.200E-06 |
| 22538 | 1127 | T1127W | D | 3.5e-06 | 3.7e-06 | 2.2e-06 | 3.133E-06 | 8.145E-07 | 2.200E-06 | 3.700E-06 |
| 22539 | 1127 | T1127Y | D | 2e-06 | 1.8e-06 | 2.8e-06 | 2.200E-06 | 5.292E-07 | 1.800E-06 | 2.800E-06 |
| 22540 | 1127 | T1127V | D | 2.1e-06 | 1.7e-06 | 2.7e-06 | 2.167E-06 | 5.033E-07 | 1.700E-06 | 2.700E-06 |
| 22541 | 1128 | D1128A | D | 1.9e-06 | 2.2e-06 | 2.1e-06 | 2.067E-06 | 1.528E-07 | 1.900E-06 | 2.200E-06 |
| 22542 | 1128 | D1128R | D | 1.3e-06 | 2.8e-06 | 2.1e-06 | 2.067E-06 | 7.506E-07 | 1.300E-06 | 2.800E-06 |
| 22543 | 1128 | D1128N | D | 2.8e-06 | 2.6e-06 | 2.1e-06 | 2.500E-06 | 3.606E-07 | 2.100E-06 | 2.800E-06 |
| 22544 | 1128 | D1128D | D | 4.1e-06 | 1.9e-06 | 1.5e-06 | 2.500E-06 | 1.400E-06 | 1.500E-06 | 4.100E-06 |
| 22545 | 1128 | D1128C | D | 2.3e-06 | 3.5e-06 | 2.4e-06 | 2.733E-06 | 6.658E-07 | 2.300E-06 | 3.500E-06 |
| 22546 | 1128 | D1128E | D | 3.4e-06 | 2.3e-06 | 2e-06 | 2.567E-06 | 7.371E-07 | 2.000E-06 | 3.400E-06 |
| 22547 | 1128 | D1128Q | D | 2.2e-06 | 3e-06 | 1.5e-06 | 2.233E-06 | 7.506E-07 | 1.500E-06 | 3.000E-06 |
| 22548 | 1128 | D1128G | D | 2e-06 | 1.8e-06 | 4.3e-06 | 2.700E-06 | 1.389E-06 | 1.800E-06 | 4.300E-06 |
| 22549 | 1128 | D1128H | D | 1.5e-06 | 1.5e-06 | 2.5e-06 | 1.833E-06 | 5.774E-07 | 1.500E-06 | 2.500E-06 |
| 22550 | 1128 | D1128I | D | 2.2e-06 | 3.6e-06 | 3e-06 | 2.933E-06 | 7.024E-07 | 2.200E-06 | 3.600E-06 |
| 22551 | 1128 | D1128L | D | 3.3e-06 | 2.3e-06 | 2.7e-06 | 2.767E-06 | 5.033E-07 | 2.300E-06 | 3.300E-06 |
| 22552 | 1128 | D1128K | D | 2.6e-06 | 1.3e-06 | 2.3e-06 | 2.067E-06 | 6.807E-07 | 1.300E-06 | 2.600E-06 |
| 22553 | 1128 | D1128M | D | 1.9e-06 | 3.6e-06 | 2.4e-06 | 2.633E-06 | 8.737E-07 | 1.900E-06 | 3.600E-06 |
| 22554 | 1128 | D1128F | D | 2.1e-06 | 3.1e-06 | 8.4e-07 | 2.013E-06 | 1.132E-06 | 8.400E-07 | 3.100E-06 |
| 22555 | 1128 | D1128P | D | 2.4e-06 | 2.8e-06 | 2.8e-06 | 2.667E-06 | 2.309E-07 | 2.400E-06 | 2.800E-06 |
| 22556 | 1128 | D1128S | D | 2.8e-06 | 1.4e-06 | 2.2e-06 | 2.133E-06 | 7.024E-07 | 1.400E-06 | 2.800E-06 |
| 22557 | 1128 | D1128T | D | 3.4e-06 | 3.2e-06 | 2.8e-06 | 3.133E-06 | 3.055E-07 | 2.800E-06 | 3.400E-06 |
| 22558 | 1128 | D1128W | D | 2e-06 | 2.6e-06 | 1.9e-06 | 2.167E-06 | 3.786E-07 | 1.900E-06 | 2.600E-06 |
| 22559 | 1128 | D1128Y | D | 3e-06 | 2.5e-06 | 2.4e-06 | 2.633E-06 | 3.215E-07 | 2.400E-06 | 3.000E-06 |
| 22560 | 1128 | D1128V | D | 3e-06 | 2.5e-06 | 2.1e-06 | 2.533E-06 | 4.509E-07 | 2.100E-06 | 3.000E-06 |
| 22561 | 1129 | F1129A | D | 1.6e-06 | 3.4e-06 | 2.8e-06 | 2.600E-06 | 9.165E-07 | 1.600E-06 | 3.400E-06 |
| 22562 | 1129 | F1129R | D | 1.7e-06 | 1.7e-06 | 2.7e-06 | 2.033E-06 | 5.774E-07 | 1.700E-06 | 2.700E-06 |
| 22563 | 1129 | F1129N | D | 1.6e-06 | 4.3e-06 | 2e-06 | 2.633E-06 | 1.457E-06 | 1.600E-06 | 4.300E-06 |
| 22564 | 1129 | F1129D | D | 2.3e-06 | 2.7e-06 | 3.4e-06 | 2.800E-06 | 5.568E-07 | 2.300E-06 | 3.400E-06 |
| 22565 | 1129 | F1129C | D | 3.4e-06 | 1.5e-06 | 2.2e-06 | 2.367E-06 | 9.609E-07 | 1.500E-06 | 3.400E-06 |
| 22566 | 1129 | F1129E | D | 3e-06 | 1.6e-06 | 2.4e-06 | 2.333E-06 | 7.024E-07 | 1.600E-06 | 3.000E-06 |
| 22567 | 1129 | F1129Q | D | 2.1e-06 | 2.4e-06 | 3.6e-06 | 2.700E-06 | 7.937E-07 | 2.100E-06 | 3.600E-06 |
| 22568 | 1129 | F1129G | D | 2.5e-06 | 2.8e-06 | 3.3e-06 | 2.867E-06 | 4.041E-07 | 2.500E-06 | 3.300E-06 |
| 22569 | 1129 | F1129H | D | 1.8e-06 | 3.5e-06 | 2.3e-06 | 2.533E-06 | 8.737E-07 | 1.800E-06 | 3.500E-06 |
| 22570 | 1129 | F1129I | D | 3.7e-06 | 2e-06 | 3.2e-06 | 2.967E-06 | 8.737E-07 | 2.000E-06 | 3.700E-06 |
| 22571 | 1129 | F1129L | D | 2.9e-06 | 2.6e-06 | 2.9e-06 | 2.800E-06 | 1.732E-07 | 2.600E-06 | 2.900E-06 |
| 22572 | 1129 | F1129K | D | 2.2e-06 | 2.1e-06 | 4e-06 | 2.767E-06 | 1.069E-06 | 2.100E-06 | 4.000E-06 |
| 22573 | 1129 | F1129M | D | 1.8e-06 | 1.9e-06 | 2.1e-06 | 1.933E-06 | 1.528E-07 | 1.800E-06 | 2.100E-06 |
| 22574 | 1129 | F1129F | D | 3.1e-06 | 4.7e-06 | 1.5e-06 | 3.100E-06 | 1.600E-06 | 1.500E-06 | 4.700E-06 |
| 22575 | 1129 | F1129P | D | 2.6e-06 | 1.3e-06 | 2.3e-06 | 2.067E-06 | 6.807E-07 | 1.300E-06 | 2.600E-06 |
| 22576 | 1129 | F1129S | D | 3.7e-06 | 2.5e-06 | 1.4e-06 | 2.533E-06 | 1.150E-06 | 1.400E-06 | 3.700E-06 |
| 22577 | 1129 | F1129T | D | 2.3e-06 | 2.4e-06 | 2.2e-06 | 2.300E-06 | 1.000E-07 | 2.200E-06 | 2.400E-06 |
| 22578 | 1129 | F1129W | D | 2.5e-06 | 1.3e-06 | 3e-06 | 2.267E-06 | 8.737E-07 | 1.300E-06 | 3.000E-06 |
| 22579 | 1129 | F1129Y | D | 1.8e-06 | 1.9e-06 | 2.1e-06 | 1.933E-06 | 1.528E-07 | 1.800E-06 | 2.100E-06 |
| 22580 | 1129 | F1129V | D | 1.9e-06 | 1.8e-06 | 2.1e-06 | 1.933E-06 | 1.528E-07 | 1.800E-06 | 2.100E-06 |
| 22581 | 1130 | T1130A | D | 2.7e-06 | 3.6e-06 | 2.3e-06 | 2.867E-06 | 6.658E-07 | 2.300E-06 | 3.600E-06 |
| 22582 | 1130 | T1130R | D | 1.7e-06 | 3.7e-06 | 2.2e-06 | 2.533E-06 | 1.041E-06 | 1.700E-06 | 3.700E-06 |
| 22583 | 1130 | T1130N | D | 2e-06 | 2.6e-06 | 3.5e-06 | 2.700E-06 | 7.550E-07 | 2.000E-06 | 3.500E-06 |
| 22584 | 1130 | T1130D | D | 2e-06 | 3.2e-06 | 2.2e-06 | 2.467E-06 | 6.429E-07 | 2.000E-06 | 3.200E-06 |
| 22585 | 1130 | T1130C | D | 2.2e-06 | 2.2e-06 | 2.3e-06 | 2.233E-06 | 5.774E-08 | 2.200E-06 | 2.300E-06 |
| 22586 | 1130 | T1130E | D | 1.4e-06 | 2.2e-06 | 1.8e-06 | 1.800E-06 | 4.000E-07 | 1.400E-06 | 2.200E-06 |
| 22587 | 1130 | T1130Q | D | 2.8e-06 | 1.7e-06 | 2e-06 | 2.167E-06 | 5.686E-07 | 1.700E-06 | 2.800E-06 |
| 22588 | 1130 | T1130G | D | 2.4e-06 | 2.4e-06 | 2.8e-06 | 2.533E-06 | 2.309E-07 | 2.400E-06 | 2.800E-06 |
| 22589 | 1130 | T1130H | D | 1.9e-06 | 3.3e-06 | 2.9e-06 | 2.700E-06 | 7.211E-07 | 1.900E-06 | 3.300E-06 |
| 22590 | 1130 | T1130I | D | 1.8e-06 | 1.4e-06 | 2.3e-06 | 1.833E-06 | 4.509E-07 | 1.400E-06 | 2.300E-06 |
| 22591 | 1130 | T1130L | D | 1.6e-06 | 2.3e-06 | 2e-06 | 1.967E-06 | 3.512E-07 | 1.600E-06 | 2.300E-06 |
| 22592 | 1130 | T1130K | D | 2.7e-06 | 3.1e-06 | 2.6e-06 | 2.800E-06 | 2.646E-07 | 2.600E-06 | 3.100E-06 |
| 22593 | 1130 | T1130M | D | 1.9e-06 | 2.8e-06 | 1.7e-06 | 2.133E-06 | 5.859E-07 | 1.700E-06 | 2.800E-06 |
| 22594 | 1130 | T1130F | D | 2.4e-06 | 1.9e-06 | 2.8e-06 | 2.367E-06 | 4.509E-07 | 1.900E-06 | 2.800E-06 |
| 22595 | 1130 | T1130P | D | 3.3e-06 | 2.3e-06 | 2.9e-06 | 2.833E-06 | 5.033E-07 | 2.300E-06 | 3.300E-06 |
| 22596 | 1130 | T1130S | D | 1.9e-06 | 1.7e-06 | 3.6e-06 | 2.400E-06 | 1.044E-06 | 1.700E-06 | 3.600E-06 |
| 22597 | 1130 | T1130T | D | 4.8e-06 | 1.4e-06 | 3e-06 | 3.067E-06 | 1.701E-06 | 1.400E-06 | 4.800E-06 |
| 22598 | 1130 | T1130W | D | 2.1e-06 | 2.4e-06 | 2.6e-06 | 2.367E-06 | 2.517E-07 | 2.100E-06 | 2.600E-06 |
| 22599 | 1130 | T1130Y | D | 3.5e-06 | 3.3e-06 | 2.9e-06 | 3.233E-06 | 3.055E-07 | 2.900E-06 | 3.500E-06 |
| 22600 | 1130 | T1130V | D | 3.2e-06 | 4.7e-06 | 2.9e-06 | 3.600E-06 | 9.644E-07 | 2.900E-06 | 4.700E-06 |
| 22601 | 1131 | L1131A | D | 1.5e-06 | 1.3e-06 | 3.5e-06 | 2.100E-06 | 1.217E-06 | 1.300E-06 | 3.500E-06 |
| 22602 | 1131 | L1131R | D | 2.1e-06 | 3.9e-06 | 2e-06 | 2.667E-06 | 1.069E-06 | 2.000E-06 | 3.900E-06 |
| 22603 | 1131 | L1131N | D | 2.2e-06 | 2e-06 | 2.3e-06 | 2.167E-06 | 1.528E-07 | 2.000E-06 | 2.300E-06 |
| 22604 | 1131 | L1131D | D | 2.8e-06 | 3.1e-06 | 2.2e-06 | 2.700E-06 | 4.583E-07 | 2.200E-06 | 3.100E-06 |
| 22605 | 1131 | L1131C | D | 3e-06 | 1.9e-06 | 2.8e-06 | 2.567E-06 | 5.859E-07 | 1.900E-06 | 3.000E-06 |
| 22606 | 1131 | L1131E | D | 3.1e-06 | 3.6e-06 | 2.3e-06 | 3.000E-06 | 6.557E-07 | 2.300E-06 | 3.600E-06 |
| 22607 | 1131 | L1131Q | D | 2.9e-06 | 2.4e-06 | 1.5e-06 | 2.267E-06 | 7.095E-07 | 1.500E-06 | 2.900E-06 |
| 22608 | 1131 | L1131G | D | 3.1e-06 | 1.7e-06 | 2.3e-06 | 2.367E-06 | 7.024E-07 | 1.700E-06 | 3.100E-06 |
| 22609 | 1131 | L1131H | D | 3.4e-06 | 2.6e-06 | 2.5e-06 | 2.833E-06 | 4.933E-07 | 2.500E-06 | 3.400E-06 |

| | | | | | | | | | | |
|---|---|---|---|---|---|---|---|---|---|---|
| 22610 | 1131 | L1131I | D | 2.4e-06 | 3.4e-06 | 3.8e-06 | 3.200E-06 | 7.211E-07 | 2.400E-06 | 3.800E-06 |
| 22611 | 1131 | L1131L | D | 1.8e-06 | 1.6e-06 | 2.5e-06 | 1.967E-06 | 4.726E-07 | 1.600E-06 | 2.500E-06 |
| 22612 | 1131 | L1131K | D | 2.2e-06 | 3.5e-06 | 3e-06 | 2.900E-06 | 6.557E-07 | 2.200E-06 | 3.500E-06 |
| 22613 | 1131 | L1131M | D | 1.7e-06 | 2e-06 | 2.1e-06 | 1.933E-06 | 2.082E-07 | 1.700E-06 | 2.100E-06 |
| 22614 | 1131 | L1131F | D | 2.6e-06 | 3.3e-06 | 2e-06 | 2.633E-06 | 6.506E-07 | 2.000E-06 | 3.300E-06 |
| 22615 | 1131 | L1131P | D | 2.5e-06 | 2e-06 | 1.9e-06 | 2.133E-06 | 3.215E-07 | 1.900E-06 | 2.500E-06 |
| 22616 | 1131 | L1131S | D | 1.4e-06 | 2.1e-06 | 2.3e-06 | 1.933E-06 | 4.726E-07 | 1.400E-06 | 2.300E-06 |
| 22617 | 1131 | L1131T | D | 1.9e-06 | 2.9e-06 | 3.7e-06 | 2.833E-06 | 9.018E-07 | 1.900E-06 | 3.700E-06 |
| 22618 | 1131 | L1131W | D | 2e-06 | 2.8e-06 | 2.4e-06 | 2.400E-06 | 4.000E-07 | 2.000E-06 | 2.800E-06 |
| 22619 | 1131 | L1131Y | D | 2.4e-06 | 2.2e-06 | 3.1e-06 | 2.567E-06 | 4.726E-07 | 2.200E-06 | 3.100E-06 |
| 22620 | 1131 | L1131V | D | 1.9e-06 | 1.8e-06 | 1.5e-06 | 1.733E-06 | 2.082E-07 | 1.500E-06 | 1.900E-06 |
| 22621 | 1132 | T1132A | D | 4.1e-06 | 2.7e-06 | 1.7e-06 | 2.833E-06 | 1.206E-06 | 1.700E-06 | 4.100E-06 |
| 22622 | 1132 | T1132R | D | 3.2e-06 | 2.7e-06 | 3e-06 | 2.967E-06 | 2.517E-07 | 2.700E-06 | 3.200E-06 |
| 22623 | 1132 | T1132N | D | 2.3e-06 | 1.5e-06 | 1.8e-06 | 1.867E-06 | 4.041E-07 | 1.500E-06 | 2.300E-06 |
| 22624 | 1132 | T1132D | D | 2.9e-06 | 3.8e-06 | 2.9e-06 | 3.200E-06 | 5.196E-07 | 2.900E-06 | 3.800E-06 |
| 22625 | 1132 | T1132C | D | 2.3e-06 | 2.8e-06 | 3.7e-06 | 2.933E-06 | 7.095E-07 | 2.300E-06 | 3.700E-06 |
| 22626 | 1132 | T1132E | D | 3.2e-06 | 2.3e-06 | 3.4e-06 | 2.967E-06 | 5.859E-07 | 2.300E-06 | 3.400E-06 |
| 22627 | 1132 | T1132Q | D | 3.1e-06 | 2.7e-06 | 9.9e-07 | 2.263E-06 | 1.121E-06 | 9.900E-07 | 3.100E-06 |
| 22628 | 1132 | T1132G | D | 1.9e-06 | 2.1e-06 | 2.7e-06 | 2.233E-06 | 4.163E-07 | 1.900E-06 | 2.700E-06 |
| 22629 | 1132 | T1132H | D | 3.3e-06 | 1.5e-06 | 2e-06 | 2.267E-06 | 9.292E-07 | 1.500E-06 | 3.300E-06 |
| 22630 | 1132 | T1132I | D | 1.8e-06 | 2.7e-06 | 3.5e-06 | 2.667E-06 | 8.505E-07 | 1.800E-06 | 3.500E-06 |
| 22631 | 1132 | T1132L | D | 2.4e-06 | 1.4e-06 | 1.1e-06 | 1.633E-06 | 6.807E-07 | 1.100E-06 | 2.400E-06 |
| 22632 | 1132 | T1132K | D | 2.2e-06 | 1.8e-06 | 2.4e-06 | 2.133E-06 | 3.055E-07 | 1.800E-06 | 2.400E-06 |
| 22633 | 1132 | T1132M | D | 2e-06 | 2.4e-06 | 2.3e-06 | 2.233E-06 | 2.082E-07 | 2.000E-06 | 2.400E-06 |
| 22634 | 1132 | T1132F | D | 1.5e-06 | 3.1e-06 | 2.8e-06 | 2.467E-06 | 8.505E-07 | 1.500E-06 | 3.100E-06 |
| 22635 | 1132 | T1132P | D | 2e-06 | 2.5e-06 | 2.2e-06 | 2.233E-06 | 2.517E-07 | 2.000E-06 | 2.500E-06 |
| 22636 | 1132 | T1132S | D | 1.3e-06 | 2.2e-06 | 2.4e-06 | 1.967E-06 | 5.859E-07 | 1.300E-06 | 2.400E-06 |
| 22637 | 1132 | T1132T | D | 2.6e-06 | 2.2e-06 | 2.3e-06 | 2.367E-06 | 2.082E-07 | 2.200E-06 | 2.600E-06 |
| 22638 | 1132 | T1132W | D | 2.5e-06 | 2.6e-06 | 1.9e-06 | 2.333E-06 | 3.786E-07 | 1.900E-06 | 2.600E-06 |
| 22639 | 1132 | T1132Y | D | 2.4e-06 | 1.3e-06 | 3e-06 | 2.233E-06 | 8.622E-07 | 1.300E-06 | 3.000E-06 |
| 22640 | 1132 | T1132V | D | 2e-06 | 1.7e-06 | 4.1e-06 | 2.600E-06 | 1.308E-06 | 1.700E-06 | 4.100E-06 |
| 22641 | 1133 | I1133A | D | 1.6e-06 | 2.3e-06 | 3.4e-06 | 2.433E-06 | 9.074E-07 | 1.600E-06 | 3.400E-06 |
| 22642 | 1133 | I1133R | D | 2.1e-06 | 1.8e-06 | 2e-06 | 1.967E-06 | 1.528E-07 | 1.800E-06 | 2.100E-06 |
| 22643 | 1133 | I1133N | D | 1e-06 | 3.2e-06 | 1.6e-06 | 1.933E-06 | 1.137E-06 | 1.000E-06 | 3.200E-06 |
| 22644 | 1133 | I1133D | D | 2.2e-06 | 3.2e-06 | 1.3e-06 | 2.233E-06 | 9.504E-07 | 1.300E-06 | 3.200E-06 |
| 22645 | 1133 | I1133C | D | 4.7e-06 | 3.1e-06 | 2.6e-06 | 3.467E-06 | 1.097E-06 | 2.600E-06 | 4.700E-06 |
| 22646 | 1133 | I1133E | D | 1.8e-06 | 2e-06 | 1.8e-06 | 1.867E-06 | 1.155E-07 | 1.800E-06 | 2.000E-06 |
| 22647 | 1133 | I1133Q | D | 2.4e-06 | 2.7e-06 | 1.5e-06 | 2.200E-06 | 6.245E-07 | 1.500E-06 | 2.700E-06 |
| 22648 | 1133 | I1133G | D | 2.1e-06 | 1.3e-06 | 4.2e-06 | 2.533E-06 | 1.498E-06 | 1.300E-06 | 4.200E-06 |
| 22649 | 1133 | I1133H | D | 1.3e-06 | 2.6e-06 | 1.7e-06 | 1.867E-06 | 6.658E-07 | 1.300E-06 | 2.600E-06 |
| 22650 | 1133 | I1133I | D | 1.1e-06 | 2.2e-06 | 3e-06 | 2.100E-06 | 9.539E-07 | 1.100E-06 | 3.000E-06 |
| 22651 | 1133 | I1133L | D | 3.3e-06 | 2.4e-06 | 2.5e-06 | 2.733E-06 | 4.933E-07 | 2.400E-06 | 3.300E-06 |
| 22652 | 1133 | I1133K | D | 2.5e-06 | 4.4e-06 | 2.3e-06 | 3.067E-06 | 1.159E-06 | 2.300E-06 | 4.400E-06 |
| 22653 | 1133 | I1133M | D | 2.5e-06 | 2.5e-06 | 2.2e-06 | 2.400E-06 | 1.732E-07 | 2.200E-06 | 2.500E-06 |
| 22654 | 1133 | I1133F | D | 2.5e-06 | 1.5e-06 | 1.4e-06 | 1.800E-06 | 6.083E-07 | 1.400E-06 | 2.500E-06 |
| 22655 | 1133 | I1133P | D | 2.5e-06 | 2.4e-06 | 2e-06 | 2.300E-06 | 2.646E-07 | 2.000E-06 | 2.500E-06 |
| 22656 | 1133 | I1133S | D | 2.2e-06 | 1.8e-06 | 1.1e-06 | 1.700E-06 | 5.568E-07 | 1.100E-06 | 2.200E-06 |
| 22657 | 1133 | I1133T | D | 3.6e-06 | 2.7e-06 | 1.6e-06 | 2.633E-06 | 1.002E-06 | 1.600E-06 | 3.600E-06 |
| 22658 | 1133 | I1133W | D | 2e-06 | 1.5e-06 | 3.4e-06 | 2.300E-06 | 9.849E-07 | 1.500E-06 | 3.400E-06 |
| 22659 | 1133 | I1133Y | D | 2.5e-06 | 2.1e-06 | 2.6e-06 | 2.400E-06 | 2.646E-07 | 2.100E-06 | 2.600E-06 |
| 22660 | 1133 | I1133V | D | 1.7e-06 | 2.4e-06 | 2.7e-06 | 2.267E-06 | 5.132E-07 | 1.700E-06 | 2.700E-06 |
| 22661 | 1134 | S1134A | D | 2.8e-06 | 2.2e-06 | 1.7e-06 | 2.233E-06 | 5.508E-07 | 1.700E-06 | 2.800E-06 |
| 22662 | 1134 | S1134R | D | 3e-06 | 2.1e-06 | 3.4e-06 | 2.833E-06 | 6.658E-07 | 2.100E-06 | 3.400E-06 |
| 22663 | 1134 | S1134N | D | 2.4e-06 | 2.7e-06 | 2.5e-06 | 2.533E-06 | 1.528E-07 | 2.400E-06 | 2.700E-06 |
| 22664 | 1134 | S1134D | D | 3.3e-06 | 2.8e-06 | 3.6e-06 | 3.233E-06 | 4.041E-07 | 2.800E-06 | 3.600E-06 |
| 22665 | 1134 | S1134C | D | 2.5e-06 | 2.3e-06 | 2.9e-06 | 2.567E-06 | 3.055E-07 | 2.300E-06 | 2.900E-06 |
| 22666 | 1134 | S1134E | D | 2.2e-06 | 1.8e-06 | 1.6e-06 | 1.867E-06 | 3.055E-07 | 1.600E-06 | 2.200E-06 |
| 22667 | 1134 | S1134Q | D | 2.2e-06 | 2e-06 | 2.7e-06 | 2.300E-06 | 3.606E-07 | 2.000E-06 | 2.700E-06 |
| 22668 | 1134 | S1134G | D | 4.5e-06 | 2.6e-06 | 1.9e-06 | 3.000E-06 | 1.345E-06 | 1.900E-06 | 4.500E-06 |
| 22669 | 1134 | S1134H | D | 2.8e-06 | 1.7e-06 | 3.7e-06 | 2.733E-06 | 1.002E-06 | 1.700E-06 | 3.700E-06 |
| 22670 | 1134 | S1134I | D | 2.8e-06 | 4.7e-06 | 2.3e-06 | 3.267E-06 | 1.266E-06 | 2.300E-06 | 4.700E-06 |
| 22671 | 1134 | S1134L | D | 3.7e-06 | 2.2e-06 | 1.7e-06 | 2.533E-06 | 1.041E-06 | 1.700E-06 | 3.700E-06 |
| 22672 | 1134 | S1134K | D | 1.8e-06 | 3.2e-06 | 2.2e-06 | 2.400E-06 | 7.211E-07 | 1.800E-06 | 3.200E-06 |
| 22673 | 1134 | S1134M | D | 3.5e-06 | 1.7e-06 | 3.9e-06 | 3.033E-06 | 1.172E-06 | 1.700E-06 | 3.900E-06 |
| 22674 | 1134 | S1134F | D | 2.8e-06 | 3.2e-06 | 1e-06 | 2.333E-06 | 1.172E-06 | 1.000E-06 | 3.200E-06 |
| 22675 | 1134 | S1134P | D | 2.7e-06 | 2.1e-06 | 2.1e-06 | 2.300E-06 | 3.464E-07 | 2.100E-06 | 2.700E-06 |
| 22676 | 1134 | S1134S | D | 2.5e-06 | 2e-06 | 2.1e-06 | 2.200E-06 | 2.646E-07 | 2.000E-06 | 2.500E-06 |
| 22677 | 1134 | S1134T | D | 2.7e-06 | 1.9e-06 | 2.4e-06 | 2.333E-06 | 4.041E-07 | 1.900E-06 | 2.700E-06 |
| 22678 | 1134 | S1134W | D | 4.9e-06 | 2.5e-06 | 2.2e-06 | 3.200E-06 | 1.480E-06 | 2.200E-06 | 4.900E-06 |
| 22679 | 1134 | S1134Y | D | 2.9e-06 | 2.5e-06 | 1.7e-06 | 2.367E-06 | 6.110E-07 | 1.700E-06 | 2.900E-06 |
| 22680 | 1134 | S1134V | D | 1.7e-06 | 1.4e-06 | 2.4e-06 | 1.833E-06 | 5.132E-07 | 1.400E-06 | 2.400E-06 |
| 22681 | 1135 | S1135A | D | 1.9e-06 | 2.9e-06 | 2.8e-06 | 2.533E-06 | 5.508E-07 | 1.900E-06 | 2.900E-06 |
| 22682 | 1135 | S1135R | D | 2.5e-06 | 1.3e-06 | 3.2e-06 | 2.333E-06 | 9.609E-07 | 1.300E-06 | 3.200E-06 |
| 22683 | 1135 | S1135N | D | 2.5e-06 | 1.9e-06 | 3.5e-06 | 2.633E-06 | 8.083E-07 | 1.900E-06 | 3.500E-06 |
| 22684 | 1135 | S1135D | D | 2.7e-06 | 2.3e-06 | 3.5e-06 | 2.833E-06 | 6.110E-07 | 2.300E-06 | 3.500E-06 |
| 22685 | 1135 | S1135C | D | 3e-06 | 2.3e-06 | 2.3e-06 | 2.533E-06 | 4.041E-07 | 2.300E-06 | 3.000E-06 |
| 22686 | 1135 | S1135E | D | 2.6e-06 | 1.6e-06 | 3.8e-06 | 2.667E-06 | 1.102E-06 | 1.600E-06 | 3.800E-06 |
| 22687 | 1135 | S1135Q | D | 3.5e-06 | 2.4e-06 | 2e-06 | 2.633E-06 | 7.767E-07 | 2.000E-06 | 3.500E-06 |
| 22688 | 1135 | S1135G | D | 2e-06 | 3.1e-06 | 1.6e-06 | 2.233E-06 | 7.767E-07 | 1.600E-06 | 3.100E-06 |
| 22689 | 1135 | S1135H | D | 3.9e-06 | 2.7e-06 | 1.8e-06 | 2.800E-06 | 1.054E-06 | 1.800E-06 | 3.900E-06 |
| 22690 | 1135 | S1135I | D | 3e-06 | 2.5e-06 | 2e-06 | 2.500E-06 | 5.000E-07 | 2.000E-06 | 3.000E-06 |
| 22691 | 1135 | S1135L | D | 1.5e-06 | 4.4e-06 | 2.4e-06 | 2.767E-06 | 1.484E-06 | 1.500E-06 | 4.400E-06 |
| 22692 | 1135 | S1135K | D | 2.7e-06 | 2.6e-06 | 2.4e-06 | 2.567E-06 | 1.528E-07 | 2.400E-06 | 2.700E-06 |
| 22693 | 1135 | S1135M | D | 2.8e-06 | 1.9e-06 | 3.6e-06 | 2.767E-06 | 8.505E-07 | 1.900E-06 | 3.600E-06 |
| 22694 | 1135 | S1135F | D | 2.5e-06 | 3e-06 | 2.2e-06 | 2.567E-06 | 4.041E-07 | 2.200E-06 | 3.000E-06 |

| | | | | | | | | | | |
|---|---|---|---|---|---|---|---|---|---|---|
| 22695 | 1135 | S1135P | D | 2.1e-06 | 1.4e-06 | 1.7e-06 | 1.733E-06 | 3.512E-07 | 1.400E-06 | 2.100E-06 |
| 22696 | 1135 | S1135S | D | 2.1e-06 | 2.2e-06 | 1e-06 | 1.767E-06 | 6.658E-07 | 1.000E-06 | 2.200E-06 |
| 22697 | 1135 | S1135T | D | 2e-06 | 2.3e-06 | 3.2e-06 | 2.500E-06 | 6.245E-07 | 2.000E-06 | 3.200E-06 |
| 22698 | 1135 | S1135W | D | 3.9e-06 | 1.6e-06 | 2e-06 | 2.500E-06 | 1.229E-06 | 1.600E-06 | 3.900E-06 |
| 22699 | 1135 | S1135Y | D | 2.7e-06 | 3e-06 | 1.9e-06 | 2.533E-06 | 5.686E-07 | 1.900E-06 | 3.000E-06 |
| 22700 | 1135 | S1135V | D | 2.8e-06 | 1.6e-06 | 1.5e-06 | 1.967E-06 | 7.234E-07 | 1.500E-06 | 2.800E-06 |
| 22701 | 1136 | L1136A | D | 2.2e-06 | 2.6e-06 | 2.8e-06 | 2.533E-06 | 3.055E-07 | 2.200E-06 | 2.800E-06 |
| 22702 | 1136 | L1136R | D | 2.4e-06 | 2.4e-06 | 2.3e-06 | 2.367E-06 | 5.774E-08 | 2.300E-06 | 2.400E-06 |
| 22703 | 1136 | L1136N | D | 3.6e-06 | 3.6e-06 | 2.8e-06 | 3.333E-06 | 4.619E-07 | 2.800E-06 | 3.600E-06 |
| 22704 | 1136 | L1136D | D | 3.6e-06 | 2.9e-06 | 2.6e-06 | 3.033E-06 | 5.132E-07 | 2.600E-06 | 3.600E-06 |
| 22705 | 1136 | L1136C | D | 2.3e-06 | 1.7e-06 | 2e-06 | 2.000E-06 | 3.000E-07 | 1.700E-06 | 2.300E-06 |
| 22706 | 1136 | L1136E | D | 2.8e-06 | 2.4e-06 | 2.9e-06 | 2.700E-06 | 2.646E-07 | 2.400E-06 | 2.900E-06 |
| 22707 | 1136 | L1136Q | D | 1.5e-06 | 2.3e-06 | 2.9e-06 | 2.233E-06 | 7.024E-07 | 1.500E-06 | 2.900E-06 |
| 22708 | 1136 | L1136G | D | 2.1e-06 | 2.1e-06 | 1.8e-06 | 2.000E-06 | 1.732E-07 | 1.800E-06 | 2.100E-06 |
| 22709 | 1136 | L1136H | D | 4.8e-06 | 3.1e-06 | 2.1e-06 | 3.333E-06 | 1.365E-06 | 2.100E-06 | 4.800E-06 |
| 22710 | 1136 | L1136I | D | 3e-06 | 2.5e-06 | 2.2e-06 | 2.567E-06 | 4.041E-07 | 2.200E-06 | 3.000E-06 |
| 22711 | 1136 | L1136L | D | 3e-06 | 2.3e-06 | 3.9e-06 | 3.067E-06 | 8.021E-07 | 2.300E-06 | 3.900E-06 |
| 22712 | 1136 | L1136K | D | 3.4e-06 | 2.4e-06 | 2.5e-06 | 2.767E-06 | 5.508E-07 | 2.400E-06 | 3.400E-06 |
| 22713 | 1136 | L1136M | D | 2.4e-06 | 2.4e-06 | 2.5e-06 | 2.433E-06 | 5.774E-08 | 2.400E-06 | 2.500E-06 |
| 22714 | 1136 | L1136F | D | 2.5e-06 | 4e-06 | 2.6e-06 | 3.033E-06 | 8.386E-07 | 2.500E-06 | 4.000E-06 |
| 22715 | 1136 | L1136P | D | 4.6e-06 | 2.3e-06 | 2.8e-06 | 3.233E-06 | 1.210E-06 | 2.300E-06 | 4.600E-06 |
| 22716 | 1136 | L1136S | D | 2.2e-06 | 2.8e-06 | 1.4e-06 | 2.133E-06 | 7.024E-07 | 1.400E-06 | 2.800E-06 |
| 22717 | 1136 | L1136T | D | 1.4e-06 | 3.1e-06 | 1.9e-06 | 2.133E-06 | 8.737E-07 | 1.400E-06 | 3.100E-06 |
| 22718 | 1136 | L1136W | D | 2.8e-06 | 2.6e-06 | 1.9e-06 | 2.433E-06 | 4.726E-07 | 1.900E-06 | 2.800E-06 |
| 22719 | 1136 | L1136Y | D | 2e-06 | 2e-06 | 2.8e-06 | 2.267E-06 | 4.619E-07 | 2.000E-06 | 2.800E-06 |
| 22720 | 1136 | L1136V | D | 1.9e-06 | 1.5e-06 | 1e-06 | 1.467E-06 | 4.509E-07 | 1.000E-06 | 1.900E-06 |
| 22721 | 1137 | Q1137A | D | 1.7e-06 | 2.9e-06 | 1.6e-06 | 2.067E-06 | 7.234E-07 | 1.600E-06 | 2.900E-06 |
| 22722 | 1137 | Q1137R | D | 2e-06 | 1.2e-06 | 4.6e-06 | 2.600E-06 | 1.778E-06 | 1.200E-06 | 4.600E-06 |
| 22723 | 1137 | Q1137N | D | 1.6e-06 | 2.5e-06 | 3.3e-06 | 2.467E-06 | 8.505E-07 | 1.600E-06 | 3.300E-06 |
| 22724 | 1137 | Q1137D | D | 3.3e-06 | 1.5e-06 | 3.7e-06 | 2.833E-06 | 1.172E-06 | 1.500E-06 | 3.700E-06 |
| 22725 | 1137 | Q1137C | D | 2.3e-06 | 1.6e-06 | 3.1e-06 | 2.333E-06 | 7.506E-07 | 1.600E-06 | 3.100E-06 |
| 22726 | 1137 | Q1137E | D | 3.7e-06 | 3.2e-06 | 3.9e-06 | 3.600E-06 | 3.606E-07 | 3.200E-06 | 3.900E-06 |
| 22727 | 1137 | Q1137Q | D | 2.5e-06 | 3.3e-06 | 2.6e-06 | 2.800E-06 | 4.359E-07 | 2.500E-06 | 3.300E-06 |
| 22728 | 1137 | Q1137G | D | 2.3e-06 | 9.5e-07 | 2.9e-06 | 2.050E-06 | 9.987E-07 | 9.500E-07 | 2.900E-06 |
| 22729 | 1137 | Q1137H | D | 2.3e-06 | 2.2e-06 | 1.7e-06 | 2.067E-06 | 3.215E-07 | 1.700E-06 | 2.300E-06 |
| 22730 | 1137 | Q1137I | D | 4.5e-06 | 1.8e-06 | 2.4e-06 | 2.900E-06 | 1.418E-06 | 1.800E-06 | 4.500E-06 |
| 22731 | 1137 | Q1137L | D | 4.1e-06 | 1.3e-06 | 2.4e-06 | 2.600E-06 | 1.411E-06 | 1.300E-06 | 4.100E-06 |
| 22732 | 1137 | Q1137K | D | 1.6e-06 | 3.4e-06 | 1.4e-06 | 2.133E-06 | 1.102E-06 | 1.400E-06 | 3.400E-06 |
| 22733 | 1137 | Q1137M | D | 2.5e-06 | 2e-06 | 3.2e-06 | 2.567E-06 | 6.028E-07 | 2.000E-06 | 3.200E-06 |
| 22734 | 1137 | Q1137F | D | 2.2e-06 | 2.2e-06 | 1.4e-06 | 1.933E-06 | 4.619E-07 | 1.400E-06 | 2.200E-06 |
| 22735 | 1137 | Q1137P | D | 2.1e-06 | 1.8e-06 | 2.3e-06 | 2.067E-06 | 2.517E-07 | 1.800E-06 | 2.300E-06 |
| 22736 | 1137 | Q1137S | D | 3.4e-06 | 2.1e-06 | 2.5e-06 | 2.667E-06 | 6.658E-07 | 2.100E-06 | 3.400E-06 |
| 22737 | 1137 | Q1137T | D | 2e-06 | 2.4e-06 | 1.5e-06 | 1.967E-06 | 4.509E-07 | 1.500E-06 | 2.400E-06 |
| 22738 | 1137 | Q1137W | D | 2.7e-06 | 2.6e-06 | 2.3e-06 | 2.533E-06 | 2.082E-07 | 2.300E-06 | 2.700E-06 |
| 22739 | 1137 | Q1137Y | D | 2.5e-06 | 1.7e-06 | 3.3e-06 | 2.500E-06 | 8.000E-07 | 1.700E-06 | 3.300E-06 |
| 22740 | 1137 | Q1137V | D | 2.7e-06 | 2.1e-06 | 1.8e-06 | 2.200E-06 | 4.583E-07 | 1.800E-06 | 2.700E-06 |
| 22741 | 1138 | P1138A | D | 2.8e-06 | 1.8e-06 | 1.6e-06 | 2.067E-06 | 6.429E-07 | 1.600E-06 | 2.800E-06 |
| 22742 | 1138 | P1138R | D | 1.4e-06 | 2.6e-06 | 1.9e-06 | 1.967E-06 | 6.028E-07 | 1.400E-06 | 2.600E-06 |
| 22743 | 1138 | P1138N | D | 3.2e-06 | 2.8e-06 | 2.1e-06 | 2.700E-06 | 5.568E-07 | 2.100E-06 | 3.200E-06 |
| 22744 | 1138 | P1138D | D | 5e-06 | 3e-06 | 2.7e-06 | 3.567E-06 | 1.250E-06 | 2.700E-06 | 5.000E-06 |
| 22745 | 1138 | P1138C | D | 2.5e-06 | 1.4e-06 | 2.1e-06 | 2.000E-06 | 5.568E-07 | 1.400E-06 | 2.500E-06 |
| 22746 | 1138 | P1138E | D | 3.6e-06 | 1.8e-06 | 2.4e-06 | 2.600E-06 | 9.165E-07 | 1.800E-06 | 3.600E-06 |
| 22747 | 1138 | P1138Q | D | 2e-06 | 5.4e-06 | 2.1e-06 | 3.167E-06 | 1.935E-06 | 2.000E-06 | 5.400E-06 |
| 22748 | 1138 | P1138G | D | 2.5e-06 | 2.6e-06 | 2.9e-06 | 2.667E-06 | 2.082E-07 | 2.500E-06 | 2.900E-06 |
| 22749 | 1138 | P1138H | D | 2.2e-06 | 2.3e-06 | 2.8e-06 | 2.433E-06 | 3.215E-07 | 2.200E-06 | 2.800E-06 |
| 22750 | 1138 | P1138I | D | 2.1e-06 | 2.7e-06 | 2.6e-06 | 2.467E-06 | 3.215E-07 | 2.100E-06 | 2.700E-06 |
| 22751 | 1138 | P1138L | D | 1.5e-06 | 3.2e-06 | 1.8e-06 | 2.167E-06 | 9.074E-07 | 1.500E-06 | 3.200E-06 |
| 22752 | 1138 | P1138K | D | 1.7e-06 | 2.4e-06 | 3e-06 | 2.367E-06 | 6.506E-07 | 1.700E-06 | 3.000E-06 |
| 22753 | 1138 | P1138M | D | 3.1e-06 | 2e-06 | 1.4e-06 | 2.167E-06 | 8.622E-07 | 1.400E-06 | 3.100E-06 |
| 22754 | 1138 | P1138F | D | 2.3e-06 | 2.1e-06 | 2.2e-06 | 2.200E-06 | 1.000E-07 | 2.100E-06 | 2.300E-06 |
| 22755 | 1138 | P1138P | D | 2.7e-06 | 2.4e-06 | 1.6e-06 | 2.233E-06 | 5.686E-07 | 1.600E-06 | 2.700E-06 |
| 22756 | 1138 | P1138S | D | 2e-06 | 2e-06 | 2.2e-06 | 2.067E-06 | 1.155E-07 | 2.000E-06 | 2.200E-06 |
| 22757 | 1138 | P1138T | D | 2.4e-06 | 2.2e-06 | 3.7e-06 | 2.767E-06 | 8.145E-07 | 2.200E-06 | 3.700E-06 |
| 22758 | 1138 | P1138W | D | 2.1e-06 | 2.4e-06 | 1.7e-06 | 2.067E-06 | 3.512E-07 | 1.700E-06 | 2.400E-06 |
| 22759 | 1138 | P1138Y | D | 2.1e-06 | 2.8e-06 | 2.5e-06 | 2.467E-06 | 3.512E-07 | 2.100E-06 | 2.800E-06 |
| 22760 | 1138 | P1138V | D | 1.7e-06 | 1.9e-06 | 1.3e-06 | 1.633E-06 | 3.055E-07 | 1.300E-06 | 1.900E-06 |
| 22761 | 1139 | E1139A | D | 1.7e-06 | 3.2e-06 | 5.3e-06 | 3.400E-06 | 1.808E-06 | 1.700E-06 | 5.300E-06 |
| 22762 | 1139 | E1139R | D | 2.4e-06 | 1.8e-06 | 7e-06 | 3.733E-06 | 2.845E-06 | 1.800E-06 | 7.000E-06 |
| 22763 | 1139 | E1139N | D | 3e-06 | 1.5e-06 | 2.4e-06 | 2.300E-06 | 7.550E-07 | 1.500E-06 | 3.000E-06 |
| 22764 | 1139 | E1139D | D | 2.6e-06 | 4.1e-06 | 3.5e-06 | 3.400E-06 | 7.550E-07 | 2.600E-06 | 4.100E-06 |
| 22765 | 1139 | E1139C | D | 2.5e-06 | 2.3e-06 | 2.8e-06 | 2.533E-06 | 2.517E-07 | 2.300E-06 | 2.800E-06 |
| 22766 | 1139 | E1139E | D | 2.2e-06 | 2.6e-06 | 4.9e-06 | 3.233E-06 | 1.457E-06 | 2.200E-06 | 4.900E-06 |
| 22767 | 1139 | E1139Q | D | 2e-06 | 2.6e-06 | 1.4e-06 | 2.000E-06 | 6.000E-07 | 1.400E-06 | 2.600E-06 |
| 22768 | 1139 | E1139G | D | 2.6e-06 | 5.5e-06 | 2.2e-06 | 3.433E-06 | 1.801E-06 | 2.200E-06 | 5.500E-06 |
| 22769 | 1139 | E1139H | D | 2.5e-06 | 2.4e-06 | 1.8e-06 | 2.233E-06 | 3.786E-07 | 1.800E-06 | 2.500E-06 |
| 22770 | 1139 | E1139I | D | 2.7e-06 | 4.6e-06 | 2.8e-06 | 3.367E-06 | 1.069E-06 | 2.700E-06 | 4.600E-06 |
| 22771 | 1139 | E1139L | D | 3.1e-06 | 2.7e-06 | 3.7e-06 | 3.167E-06 | 5.033E-07 | 2.700E-06 | 3.700E-06 |
| 22772 | 1139 | E1139K | D | 1.5e-06 | 1.5e-06 | 2.6e-06 | 1.867E-06 | 6.351E-07 | 1.500E-06 | 2.600E-06 |
| 22773 | 1139 | E1139M | D | 2e-06 | 2.5e-06 | 2.6e-06 | 2.367E-06 | 3.215E-07 | 2.000E-06 | 2.600E-06 |
| 22774 | 1139 | E1139F | D | 3.6e-06 | 2.5e-06 | 1.6e-06 | 2.567E-06 | 1.002E-06 | 1.600E-06 | 3.600E-06 |
| 22775 | 1139 | E1139P | D | 2.3e-06 | 1.8e-06 | 2.5e-06 | 2.200E-06 | 3.606E-07 | 1.800E-06 | 2.500E-06 |
| 22776 | 1139 | E1139S | D | 3.2e-06 | 2.7e-06 | 2.7e-06 | 2.867E-06 | 2.887E-07 | 2.700E-06 | 3.200E-06 |
| 22777 | 1139 | E1139T | D | 1.4e-06 | 2.6e-06 | 2.1e-06 | 2.033E-06 | 6.028E-07 | 1.400E-06 | 2.600E-06 |
| 22778 | 1139 | E1139W | D | 2.4e-06 | 1.2e-06 | 3.7e-06 | 2.433E-06 | 1.250E-06 | 1.200E-06 | 3.700E-06 |
| 22779 | 1139 | E1139Y | D | 1.7e-06 | 1.7e-06 | 3.3e-06 | 2.233E-06 | 9.238E-07 | 1.700E-06 | 3.300E-06 |

| | | | | | | | | | | |
|---|---|---|---|---|---|---|---|---|---|---|
| 22780 | 1139 | E1139V | D | 3e-06 | 2.5e-06 | 1.9e-06 | 2.467E-06 | 5.508E-07 | 1.900E-06 | 3.000E-06 |
| 22781 | 1140 | D1140A | D | 1.7e-06 | 3.5e-06 | 2.8e-06 | 2.667E-06 | 9.074E-07 | 1.700E-06 | 3.500E-06 |
| 22782 | 1140 | D1140R | D | 2e-06 | 2.5e-06 | 2.5e-06 | 2.333E-06 | 2.887E-07 | 2.000E-06 | 2.500E-06 |
| 22783 | 1140 | D1140N | D | 2.1e-06 | 3.2e-06 | 2.7e-06 | 2.667E-06 | 5.508E-07 | 2.100E-06 | 3.200E-06 |
| 22784 | 1140 | D1140D | D | 3.9e-06 | 2.6e-06 | 2.4e-06 | 2.967E-06 | 8.145E-07 | 2.400E-06 | 3.900E-06 |
| 22785 | 1140 | D1140C | D | 3.4e-06 | 2.9e-06 | 2.4e-06 | 2.900E-06 | 5.000E-07 | 2.400E-06 | 3.400E-06 |
| 22786 | 1140 | D1140E | D | 2.9e-06 | 2e-06 | 2.5e-06 | 2.467E-06 | 4.509E-07 | 2.000E-06 | 2.900E-06 |
| 22787 | 1140 | D1140Q | D | 3.3e-06 | 1.8e-06 | 2.6e-06 | 2.567E-06 | 7.506E-07 | 1.800E-06 | 3.300E-06 |
| 22788 | 1140 | D1140G | D | 1.5e-06 | 2.4e-06 | 2.3e-06 | 2.067E-06 | 4.933E-07 | 1.500E-06 | 2.400E-06 |
| 22789 | 1140 | D1140H | D | 1.7e-06 | 3.7e-06 | 3.1e-06 | 2.833E-06 | 1.026E-06 | 1.700E-06 | 3.700E-06 |
| 22790 | 1140 | D1140I | D | 1.7e-06 | 3.3e-06 | 1.9e-06 | 2.300E-06 | 8.718E-07 | 1.700E-06 | 3.300E-06 |
| 22791 | 1140 | D1140L | D | 3.9e-06 | 2.5e-06 | 2.5e-06 | 2.967E-06 | 8.083E-07 | 2.500E-06 | 3.900E-06 |
| 22792 | 1140 | D1140K | D | 3.6e-06 | 1.2e-06 | 1.7e-06 | 2.167E-06 | 1.266E-06 | 1.200E-06 | 3.600E-06 |
| 22793 | 1140 | D1140M | D | 2.1e-06 | 2.1e-06 | 1.8e-06 | 2.000E-06 | 1.732E-07 | 1.800E-06 | 2.100E-06 |
| 22794 | 1140 | D1140F | D | 4.6e-06 | 2e-06 | 2.2e-06 | 2.933E-06 | 1.447E-06 | 2.000E-06 | 4.600E-06 |
| 22795 | 1140 | D1140P | D | 3.1e-06 | 2.5e-06 | 2.6e-06 | 2.733E-06 | 3.215E-07 | 2.500E-06 | 3.100E-06 |
| 22796 | 1140 | D1140S | D | 2.6e-06 | 2.3e-06 | 3.8e-06 | 2.900E-06 | 7.937E-07 | 2.300E-06 | 3.800E-06 |
| 22797 | 1140 | D1140T | D | 2e-06 | 2.9e-06 | 2.7e-06 | 2.533E-06 | 4.726E-07 | 2.000E-06 | 2.900E-06 |
| 22798 | 1140 | D1140W | D | 2.8e-06 | 1.2e-06 | 1.8e-06 | 1.933E-06 | 8.083E-07 | 1.200E-06 | 2.800E-06 |
| 22799 | 1140 | D1140Y | D | 3.3e-06 | 1.8e-06 | 3.4e-06 | 2.833E-06 | 8.963E-07 | 1.800E-06 | 3.400E-06 |
| 22800 | 1140 | D1140V | D | 2.6e-06 | 2.3e-06 | 1.7e-06 | 2.200E-06 | 4.583E-07 | 1.700E-06 | 2.600E-06 |
| 22801 | 1141 | F1141A | D | 3.1e-06 | 2.5e-06 | 1.5e-06 | 2.367E-06 | 8.083E-07 | 1.500E-06 | 3.100E-06 |
| 22802 | 1141 | F1141R | D | 1.5e-06 | 3e-06 | 5.1e-06 | 3.200E-06 | 1.808E-06 | 1.500E-06 | 5.100E-06 |
| 22803 | 1141 | F1141N | D | 2.4e-06 | 1.4e-06 | 2.6e-06 | 2.133E-06 | 6.429E-07 | 1.400E-06 | 2.600E-06 |
| 22804 | 1141 | F1141D | D | 2.2e-06 | 2.9e-06 | 2.8e-06 | 2.633E-06 | 3.786E-07 | 2.200E-06 | 2.900E-06 |
| 22805 | 1141 | F1141C | D | 2.1e-06 | 2.3e-06 | 2.9e-06 | 2.433E-06 | 4.163E-07 | 2.100E-06 | 2.900E-06 |
| 22806 | 1141 | F1141E | D | 2.6e-06 | 2.5e-06 | 2.6e-06 | 2.567E-06 | 5.774E-08 | 2.500E-06 | 2.600E-06 |
| 22807 | 1141 | F1141Q | D | 1.3e-06 | 3.6e-06 | 1.9e-06 | 2.267E-06 | 1.193E-06 | 1.300E-06 | 3.600E-06 |
| 22808 | 1141 | F1141G | D | 1.5e-06 | 2.2e-06 | 1.4e-06 | 1.700E-06 | 4.359E-07 | 1.400E-06 | 2.200E-06 |
| 22809 | 1141 | F1141H | D | 3e-06 | 3.8e-06 | 2.3e-06 | 3.033E-06 | 7.506E-07 | 2.300E-06 | 3.800E-06 |
| 22810 | 1141 | F1141I | D | 2.6e-06 | 1.9e-06 | 2.2e-06 | 2.233E-06 | 3.512E-07 | 1.900E-06 | 2.600E-06 |
| 22811 | 1141 | F1141L | D | 1.3e-06 | 2.3e-06 | 3.8e-06 | 2.467E-06 | 1.258E-06 | 1.300E-06 | 3.800E-06 |
| 22812 | 1141 | F1141K | D | 1.6e-06 | 2.8e-06 | 2.9e-06 | 2.433E-06 | 7.234E-07 | 1.600E-06 | 2.900E-06 |
| 22813 | 1141 | F1141M | D | 5.7e-06 | 2e-06 | 2e-06 | 3.233E-06 | 2.136E-06 | 2.000E-06 | 5.700E-06 |
| 22814 | 1141 | F1141F | D | 3.5e-06 | 2.6e-06 | 2.4e-06 | 2.833E-06 | 5.859E-07 | 2.400E-06 | 3.500E-06 |
| 22815 | 1141 | F1141P | D | 2.9e-06 | 2e-06 | 2.2e-06 | 2.367E-06 | 4.726E-07 | 2.000E-06 | 2.900E-06 |
| 22816 | 1141 | F1141S | D | 1.8e-06 | 3e-06 | 2.5e-06 | 2.433E-06 | 6.028E-07 | 1.800E-06 | 3.000E-06 |
| 22817 | 1141 | F1141T | D | 2.7e-06 | 1.8e-06 | 2.7e-06 | 2.400E-06 | 5.196E-07 | 1.800E-06 | 2.700E-06 |
| 22818 | 1141 | F1141W | D | 2.7e-06 | 2.2e-06 | 2e-06 | 2.300E-06 | 3.606E-07 | 2.000E-06 | 2.700E-06 |
| 22819 | 1141 | F1141Y | D | 3.1e-06 | 4.8e-06 | 3e-06 | 3.633E-06 | 1.012E-06 | 3.000E-06 | 4.800E-06 |
| 22820 | 1141 | F1141V | D | 2.1e-06 | 2e-06 | 3.4e-06 | 2.500E-06 | 7.810E-07 | 2.000E-06 | 3.400E-06 |
| 22821 | 1142 | A1142A | D | 2.9e-06 | 3.4e-06 | 2.2e-06 | 2.833E-06 | 6.028E-07 | 2.200E-06 | 3.400E-06 |
| 22822 | 1142 | A1142R | D | 3.4e-06 | 3.2e-06 | 2.6e-06 | 3.067E-06 | 4.163E-07 | 2.600E-06 | 3.400E-06 |
| 22823 | 1142 | A1142N | D | 4.1e-06 | 1.7e-06 | 1.3e-06 | 2.367E-06 | 1.514E-06 | 1.300E-06 | 4.100E-06 |
| 22824 | 1142 | A1142D | D | 2.4e-06 | 2.3e-06 | 2.2e-06 | 2.300E-06 | 1.000E-07 | 2.200E-06 | 2.400E-06 |
| 22825 | 1142 | A1142C | D | 2.6e-06 | 2.3e-06 | 1.9e-06 | 2.267E-06 | 3.512E-07 | 1.900E-06 | 2.600E-06 |
| 22826 | 1142 | A1142E | D | 2e-06 | 2.1e-06 | 2.9e-06 | 2.333E-06 | 4.933E-07 | 2.000E-06 | 2.900E-06 |
| 22827 | 1142 | A1142Q | D | 2.2e-06 | 1.9e-06 | 2.2e-06 | 2.100E-06 | 1.732E-07 | 1.900E-06 | 2.200E-06 |
| 22828 | 1142 | A1142G | D | 2.1e-06 | 9.6e-07 | 1.7e-06 | 1.587E-06 | 5.784E-07 | 9.600E-07 | 2.100E-06 |
| 22829 | 1142 | A1142H | D | 1.7e-06 | 1.9e-06 | 2.6e-06 | 2.067E-06 | 4.726E-07 | 1.700E-06 | 2.600E-06 |
| 22830 | 1142 | A1142I | D | 1.4e-06 | 2.2e-06 | 2.5e-06 | 2.033E-06 | 5.686E-07 | 1.400E-06 | 2.500E-06 |
| 22831 | 1142 | A1142L | D | 3.2e-06 | 1.7e-06 | 2.8e-06 | 2.567E-06 | 7.767E-07 | 1.700E-06 | 3.200E-06 |
| 22832 | 1142 | A1142K | D | 1.9e-06 | 3.2e-06 | 2.7e-06 | 2.600E-06 | 6.557E-07 | 1.900E-06 | 3.200E-06 |
| 22833 | 1142 | A1142M | D | 3.8e-06 | 2.5e-06 | 2.8e-06 | 3.033E-06 | 6.807E-07 | 2.500E-06 | 3.800E-06 |
| 22834 | 1142 | A1142F | D | 2e-06 | 3.3e-06 | 1.8e-06 | 2.367E-06 | 8.145E-07 | 1.800E-06 | 3.300E-06 |
| 22835 | 1142 | A1142P | D | 1.7e-06 | 3.4e-06 | 2.1e-06 | 2.400E-06 | 8.888E-07 | 1.700E-06 | 3.400E-06 |
| 22836 | 1142 | A1142S | D | 1.5e-06 | 1.9e-06 | 3.1e-06 | 2.167E-06 | 8.327E-07 | 1.500E-06 | 3.100E-06 |
| 22837 | 1142 | A1142T | D | 2.2e-06 | 1.7e-06 | 3.5e-06 | 2.467E-06 | 9.292E-07 | 1.700E-06 | 3.500E-06 |
| 22838 | 1142 | A1142W | D | 9.2e-07 | 2.8e-06 | 3e-06 | 2.240E-06 | 1.148E-06 | 9.200E-07 | 3.000E-06 |
| 22839 | 1142 | A1142Y | D | 2.9e-06 | 5e-06 | 1.6e-06 | 3.167E-06 | 1.716E-06 | 1.600E-06 | 5.000E-06 |
| 22840 | 1142 | A1142V | D | 2.4e-06 | 2.6e-06 | 2.1e-06 | 2.367E-06 | 2.517E-07 | 2.100E-06 | 2.600E-06 |
| 22841 | 1143 | T1143A | D | 2.8e-06 | 2.1e-06 | 2.2e-06 | 2.367E-06 | 3.786E-07 | 2.100E-06 | 2.800E-06 |
| 22842 | 1143 | T1143R | D | 2.7e-06 | 1.9e-06 | 1.4e-06 | 2.000E-06 | 6.557E-07 | 1.400E-06 | 2.700E-06 |
| 22843 | 1143 | T1143N | D | 3.6e-06 | 3.2e-06 | 2.4e-06 | 3.067E-06 | 6.110E-07 | 2.400E-06 | 3.600E-06 |
| 22844 | 1143 | T1143D | D | 2.4e-06 | 1.7e-06 | 2.9e-06 | 2.333E-06 | 6.028E-07 | 1.700E-06 | 2.900E-06 |
| 22845 | 1143 | T1143C | D | 3.5e-06 | 3.6e-06 | 3.5e-06 | 3.533E-06 | 5.774E-08 | 3.500E-06 | 3.600E-06 |
| 22846 | 1143 | T1143E | D | 2.1e-06 | 4.9e-06 | 1.8e-06 | 2.933E-06 | 1.710E-06 | 1.800E-06 | 4.900E-06 |
| 22847 | 1143 | T1143Q | D | 2.9e-06 | 2.3e-06 | 1.7e-06 | 2.300E-06 | 6.000E-07 | 1.700E-06 | 2.900E-06 |
| 22848 | 1143 | T1143G | D | 3.1e-06 | 1.8e-06 | 1.6e-06 | 2.167E-06 | 8.145E-07 | 1.600E-06 | 3.100E-06 |
| 22849 | 1143 | T1143H | D | 2.7e-06 | 5.4e-06 | 3.5e-06 | 3.867E-06 | 1.387E-06 | 2.700E-06 | 5.400E-06 |
| 22850 | 1143 | T1143I | D | 1.5e-06 | 3e-06 | 2.6e-06 | 2.367E-06 | 7.767E-07 | 1.500E-06 | 3.000E-06 |
| 22851 | 1143 | T1143L | D | 2.4e-06 | 2.7e-06 | 3.3e-06 | 2.800E-06 | 4.583E-07 | 2.400E-06 | 3.300E-06 |
| 22852 | 1143 | T1143K | D | 2.2e-06 | 3.4e-06 | 2.5e-06 | 2.700E-06 | 6.245E-07 | 2.200E-06 | 3.400E-06 |
| 22853 | 1143 | T1143M | D | 3.2e-06 | 1.4e-06 | 2.6e-06 | 2.400E-06 | 9.165E-07 | 1.400E-06 | 3.200E-06 |
| 22854 | 1143 | T1143F | D | 2.6e-06 | 3.5e-06 | 2.2e-06 | 2.767E-06 | 6.658E-07 | 2.200E-06 | 3.500E-06 |
| 22855 | 1143 | T1143P | D | 6.4e-06 | 2.5e-06 | 3e-06 | 3.967E-06 | 2.122E-06 | 2.500E-06 | 6.400E-06 |
| 22856 | 1143 | T1143S | D | 3.2e-06 | 2.6e-06 | 3.7e-06 | 3.167E-06 | 5.508E-07 | 2.600E-06 | 3.700E-06 |
| 22857 | 1143 | T1143T | D | 2.2e-06 | 2.7e-06 | 3.2e-06 | 2.700E-06 | 5.000E-07 | 2.200E-06 | 3.200E-06 |
| 22858 | 1143 | T1143W | D | 2.9e-06 | 4.1e-06 | 1.3e-06 | 2.767E-06 | 1.405E-06 | 1.300E-06 | 4.100E-06 |
| 22859 | 1143 | T1143Y | D | 1.8e-06 | 2.3e-06 | 1.7e-06 | 1.933E-06 | 3.215E-07 | 1.700E-06 | 2.300E-06 |
| 22860 | 1143 | T1143V | D | 1.8e-06 | 1.3e-06 | 2.1e-06 | 1.733E-06 | 4.041E-07 | 1.300E-06 | 2.100E-06 |
| 22861 | 1144 | Y1144A | D | 2.5e-06 | 3.9e-06 | 2.1e-06 | 2.833E-06 | 9.452E-07 | 2.100E-06 | 3.900E-06 |
| 22862 | 1144 | Y1144R | D | 2.1e-06 | 2.7e-06 | 1.3e-06 | 2.033E-06 | 7.024E-07 | 1.300E-06 | 2.700E-06 |
| 22863 | 1144 | Y1144N | D | 2.1e-06 | 2.7e-06 | 2.3e-06 | 2.367E-06 | 3.055E-07 | 2.100E-06 | 2.700E-06 |
| 22864 | 1144 | Y1144D | D | 2.3e-06 | 2.4e-06 | 2.9e-06 | 2.533E-06 | 3.215E-07 | 2.300E-06 | 2.900E-06 |

| | | | | | | | | | | |
|---|---|---|---|---|---|---|---|---|---|---|
| 22865 | 1144 | Y1144C | D | 1.6e-06 | 1.9e-06 | 2.4e-06 | 1.967E-06 | 4.041E-07 | 1.600E-06 | 2.400E-06 |
| 22866 | 1144 | Y1144E | D | 5.2e-06 | 1.4e-06 | 4.5e-06 | 3.700E-06 | 2.022E-06 | 1.400E-06 | 5.200E-06 |
| 22867 | 1144 | Y1144Q | D | 3.1e-06 | 3.1e-06 | 4.3e-06 | 3.500E-06 | 6.928E-07 | 3.100E-06 | 4.300E-06 |
| 22868 | 1144 | Y1144G | D | 2.6e-06 | 3e-06 | 2.3e-06 | 2.633E-06 | 3.512E-07 | 2.300E-06 | 3.000E-06 |
| 22869 | 1144 | Y1144H | D | 3.7e-06 | 1.6e-06 | 1.7e-06 | 2.333E-06 | 1.185E-06 | 1.600E-06 | 3.700E-06 |
| 22870 | 1144 | Y1144I | D | 2.6e-06 | 2.3e-06 | 2.4e-06 | 2.433E-06 | 1.528E-07 | 2.300E-06 | 2.600E-06 |
| 22871 | 1144 | Y1144L | D | 3.2e-06 | 1.5e-06 | 1e-06 | 1.900E-06 | 1.153E-06 | 1.000E-06 | 3.200E-06 |
| 22872 | 1144 | Y1144K | D | 2.5e-06 | 2.6e-06 | 4.3e-06 | 3.133E-06 | 1.012E-06 | 2.500E-06 | 4.300E-06 |
| 22873 | 1144 | Y1144M | D | 2.2e-06 | 1.2e-06 | 2.1e-06 | 1.833E-06 | 5.508E-07 | 1.200E-06 | 2.200E-06 |
| 22874 | 1144 | Y1144F | D | 1.8e-06 | 1.6e-06 | 3.2e-06 | 2.200E-06 | 8.718E-07 | 1.600E-06 | 3.200E-06 |
| 22875 | 1144 | Y1144P | D | 3.1e-06 | 3.2e-06 | 3.3e-06 | 3.200E-06 | 1.000E-07 | 3.100E-06 | 3.300E-06 |
| 22876 | 1144 | Y1144S | D | 2.9e-06 | 4e-06 | 1.2e-06 | 2.700E-06 | 1.411E-06 | 1.200E-06 | 4.000E-06 |
| 22877 | 1144 | Y1144T | D | 2.8e-06 | 2.2e-06 | 2.4e-06 | 2.467E-06 | 3.055E-07 | 2.200E-06 | 2.800E-06 |
| 22878 | 1144 | Y1144W | D | 2.9e-06 | 2.6e-06 | 3e-06 | 2.833E-06 | 2.082E-07 | 2.600E-06 | 3.000E-06 |
| 22879 | 1144 | Y1144Y | D | 2.6e-06 | 2.5e-06 | 1.4e-06 | 2.167E-06 | 6.658E-07 | 1.400E-06 | 2.600E-06 |
| 22880 | 1144 | Y1144V | D | 1.2e-06 | 3e-06 | 1.6e-06 | 1.933E-06 | 9.452E-07 | 1.200E-06 | 3.000E-06 |
| 22881 | 1145 | Y1145A | D | 2.5e-06 | 2.4e-06 | 2.2e-06 | 2.367E-06 | 1.528E-07 | 2.200E-06 | 2.500E-06 |
| 22882 | 1145 | Y1145R | D | 2.5e-06 | 1.4e-06 | 2.2e-06 | 2.033E-06 | 5.686E-07 | 1.400E-06 | 2.500E-06 |
| 22883 | 1145 | Y1145N | D | 1.6e-06 | 1.6e-06 | 3.2e-06 | 2.133E-06 | 9.238E-07 | 1.600E-06 | 3.200E-06 |
| 22884 | 1145 | Y1145D | D | 3.4e-06 | 2.6e-06 | 3.1e-06 | 3.033E-06 | 4.041E-07 | 2.600E-06 | 3.400E-06 |
| 22885 | 1145 | Y1145C | D | 2.3e-06 | 2.9e-06 | 3.9e-06 | 3.033E-06 | 8.083E-07 | 2.300E-06 | 3.900E-06 |
| 22886 | 1145 | Y1145E | D | 2.1e-06 | 3.2e-06 | 1.8e-06 | 2.367E-06 | 7.371E-07 | 1.800E-06 | 3.200E-06 |
| 22887 | 1145 | Y1145Q | D | 2.4e-06 | 2.3e-06 | 2e-06 | 2.233E-06 | 2.082E-07 | 2.000E-06 | 2.400E-06 |
| 22888 | 1145 | Y1145G | D | 2.3e-06 | 1.8e-06 | 2.4e-06 | 2.167E-06 | 3.215E-07 | 1.800E-06 | 2.400E-06 |
| 22889 | 1145 | Y1145H | D | 2.3e-06 | 1.7e-06 | 4.3e-06 | 2.767E-06 | 1.361E-06 | 1.700E-06 | 4.300E-06 |
| 22890 | 1145 | Y1145I | D | 2.5e-06 | 2.8e-06 | 2.2e-06 | 2.500E-06 | 3.000E-07 | 2.200E-06 | 2.800E-06 |
| 22891 | 1145 | Y1145L | D | 1.6e-06 | 2e-06 | 5.1e-06 | 2.900E-06 | 1.916E-06 | 1.600E-06 | 5.100E-06 |
| 22892 | 1145 | Y1145K | D | 3e-06 | 1.9e-06 | 2.9e-06 | 2.600E-06 | 6.083E-07 | 1.900E-06 | 3.000E-06 |
| 22893 | 1145 | Y1145M | D | 3.7e-06 | 1.8e-06 | 2.5e-06 | 2.667E-06 | 9.609E-07 | 1.800E-06 | 3.700E-06 |
| 22894 | 1145 | Y1145F | D | 2.7e-06 | 2.7e-06 | 2e-06 | 2.467E-06 | 4.041E-07 | 2.000E-06 | 2.700E-06 |
| 22895 | 1145 | Y1145P | D | 2.5e-06 | 3.7e-06 | 2.6e-06 | 2.933E-06 | 6.658E-07 | 2.500E-06 | 3.700E-06 |
| 22896 | 1145 | Y1145S | D | 2.3e-06 | 5.4e-06 | 2e-06 | 3.233E-06 | 1.882E-06 | 2.000E-06 | 5.400E-06 |
| 22897 | 1145 | Y1145T | D | 4.8e-06 | 2.3e-06 | 2.8e-06 | 3.300E-06 | 1.323E-06 | 2.300E-06 | 4.800E-06 |
| 22898 | 1145 | Y1145W | D | 1.8e-06 | 2.7e-06 | 3.1e-06 | 2.533E-06 | 6.658E-07 | 1.800E-06 | 3.100E-06 |
| 22899 | 1145 | Y1145Y | D | 2.2e-06 | 1.5e-06 | 2.4e-06 | 2.033E-06 | 4.726E-07 | 1.500E-06 | 2.400E-06 |
| 22900 | 1145 | Y1145V | D | 1.8e-06 | 3.6e-06 | 2.9e-06 | 2.767E-06 | 9.074E-07 | 1.800E-06 | 3.600E-06 |
| 22901 | 1146 | C1146A | D | 3.9e-06 | 1.5e-06 | 3.2e-06 | 2.867E-06 | 1.234E-06 | 1.500E-06 | 3.900E-06 |
| 22902 | 1146 | C1146R | D | 1.8e-06 | 1.8e-06 | 2.4e-06 | 2.000E-06 | 3.464E-07 | 1.800E-06 | 2.400E-06 |
| 22903 | 1146 | C1146N | D | 2.8e-06 | 3.7e-06 | 3.9e-06 | 3.467E-06 | 5.859E-07 | 2.800E-06 | 3.900E-06 |
| 22904 | 1146 | C1146D | D | 1.8e-06 | 2e-06 | 3.7e-06 | 2.500E-06 | 1.044E-06 | 1.800E-06 | 3.700E-06 |
| 22905 | 1146 | C1146C | D | 2.3e-06 | 1.8e-06 | 2.3e-06 | 2.133E-06 | 2.887E-07 | 1.800E-06 | 2.300E-06 |
| 22906 | 1146 | C1146E | D | 5.4e-06 | 1.4e-06 | 2.1e-06 | 2.967E-06 | 2.136E-06 | 1.400E-06 | 5.400E-06 |
| 22907 | 1146 | C1146Q | D | 2e-06 | 1.2e-06 | 5.1e-06 | 2.767E-06 | 2.060E-06 | 1.200E-06 | 5.100E-06 |
| 22908 | 1146 | C1146G | D | 3.4e-06 | 3e-06 | 2.6e-06 | 3.000E-06 | 4.000E-07 | 2.600E-06 | 3.400E-06 |
| 22909 | 1146 | C1146H | D | 2.2e-06 | 1.7e-06 | 2.9e-06 | 2.267E-06 | 6.028E-07 | 1.700E-06 | 2.900E-06 |
| 22910 | 1146 | C1146I | D | 2.5e-06 | 2.5e-06 | 2.6e-06 | 2.533E-06 | 5.774E-08 | 2.500E-06 | 2.600E-06 |
| 22911 | 1146 | C1146L | D | 2.8e-06 | 2.5e-06 | 2.9e-06 | 2.733E-06 | 2.082E-07 | 2.500E-06 | 2.900E-06 |
| 22912 | 1146 | C1146K | D | 1e-06 | 2e-06 | 3.5e-06 | 2.167E-06 | 1.258E-06 | 1.000E-06 | 3.500E-06 |
| 22913 | 1146 | C1146M | D | 1.6e-06 | 3.4e-06 | 1.7e-06 | 2.233E-06 | 1.012E-06 | 1.600E-06 | 3.400E-06 |
| 22914 | 1146 | C1146F | D | 2.5e-06 | 2.3e-06 | 3.3e-06 | 2.700E-06 | 5.292E-07 | 2.300E-06 | 3.300E-06 |
| 22915 | 1146 | C1146P | D | 1.7e-06 | 2.1e-06 | 2.2e-06 | 2.000E-06 | 2.646E-07 | 1.700E-06 | 2.200E-06 |
| 22916 | 1146 | C1146S | D | 2.6e-06 | 4.3e-06 | 3.3e-06 | 3.400E-06 | 8.544E-07 | 2.600E-06 | 4.300E-06 |
| 22917 | 1146 | C1146T | D | 1.4e-06 | 3.4e-06 | 4e-06 | 2.933E-06 | 1.361E-06 | 1.400E-06 | 4.000E-06 |
| 22918 | 1146 | C1146W | D | 2.9e-06 | 1.1e-06 | 3.7e-06 | 2.567E-06 | 1.332E-06 | 1.100E-06 | 3.700E-06 |
| 22919 | 1146 | C1146Y | D | 2.1e-06 | 1.5e-06 | 2.5e-06 | 2.033E-06 | 5.033E-07 | 1.500E-06 | 2.500E-06 |
| 22920 | 1146 | C1146V | D | 2.6e-06 | 3.5e-06 | 2.2e-06 | 2.767E-06 | 6.658E-07 | 2.200E-06 | 3.500E-06 |
| 22921 | 1147 | Q1147A | D | 1.8e-06 | 3.6e-06 | 2.6e-06 | 2.667E-06 | 9.018E-07 | 1.800E-06 | 3.600E-06 |
| 22922 | 1147 | Q1147R | D | 1.9e-06 | 3.5e-06 | 2.9e-06 | 2.767E-06 | 8.083E-07 | 1.900E-06 | 3.500E-06 |
| 22923 | 1147 | Q1147N | D | 4.9e-06 | 1.7e-06 | 3.4e-06 | 3.333E-06 | 1.601E-06 | 1.700E-06 | 4.900E-06 |
| 22924 | 1147 | Q1147D | D | 2.2e-06 | 3.3e-06 | 2.6e-06 | 2.700E-06 | 5.568E-07 | 2.200E-06 | 3.300E-06 |
| 22925 | 1147 | Q1147C | D | 3e-06 | 4e-06 | 2.2e-06 | 3.067E-06 | 9.018E-07 | 2.200E-06 | 4.000E-06 |
| 22926 | 1147 | Q1147E | D | 2.9e-06 | 2.9e-06 | 1.2e-06 | 2.333E-06 | 9.815E-07 | 1.200E-06 | 2.900E-06 |
| 22927 | 1147 | Q1147Q | D | 2.8e-06 | 1.8e-06 | 1.4e-06 | 2.000E-06 | 7.211E-07 | 1.400E-06 | 2.800E-06 |
| 22928 | 1147 | Q1147G | D | 4e-06 | 4e-06 | 2.2e-06 | 3.400E-06 | 1.039E-06 | 2.200E-06 | 4.000E-06 |
| 22929 | 1147 | Q1147H | D | 1.4e-06 | 2.3e-06 | 1.3e-06 | 1.667E-06 | 5.508E-07 | 1.300E-06 | 2.300E-06 |
| 22930 | 1147 | Q1147I | D | 1.8e-06 | 2.3e-06 | 1.7e-06 | 1.933E-06 | 3.215E-07 | 1.700E-06 | 2.300E-06 |
| 22931 | 1147 | Q1147L | D | 3.5e-06 | 4.7e-06 | 2.4e-06 | 3.533E-06 | 1.150E-06 | 2.400E-06 | 4.700E-06 |
| 22932 | 1147 | Q1147K | D | 5e-06 | 2.5e-06 | 3.5e-06 | 3.667E-06 | 1.258E-06 | 2.500E-06 | 5.000E-06 |
| 22933 | 1147 | Q1147M | D | 3.9e-06 | 2.2e-06 | 1.5e-06 | 2.533E-06 | 1.234E-06 | 1.500E-06 | 3.900E-06 |
| 22934 | 1147 | Q1147F | D | 2.6e-06 | 2.2e-06 | 2.2e-06 | 2.333E-06 | 2.309E-07 | 2.200E-06 | 2.600E-06 |
| 22935 | 1147 | Q1147P | D | 3.2e-06 | 1.6e-06 | 3.1e-06 | 2.633E-06 | 8.963E-07 | 1.600E-06 | 3.200E-06 |
| 22936 | 1147 | Q1147S | D | 2.2e-06 | 1.8e-06 | 3.5e-06 | 2.500E-06 | 8.888E-07 | 1.800E-06 | 3.500E-06 |
| 22937 | 1147 | Q1147T | D | 2.2e-06 | 1.1e-06 | 1.7e-06 | 1.667E-06 | 5.508E-07 | 1.100E-06 | 2.200E-06 |
| 22938 | 1147 | Q1147W | D | 2.8e-06 | 2.2e-06 | 1.4e-06 | 2.133E-06 | 7.024E-07 | 1.400E-06 | 2.800E-06 |
| 22939 | 1147 | Q1147Y | D | 1e-06 | 3.6e-06 | 3.9e-06 | 2.833E-06 | 1.595E-06 | 1.000E-06 | 3.900E-06 |
| 22940 | 1147 | Q1147V | D | 2.4e-06 | 2.7e-06 | 9.5e-07 | 2.017E-06 | 9.359E-07 | 9.500E-07 | 2.700E-06 |
| 22941 | 1148 | Q1148A | D | 2.1e-06 | 2.2e-06 | 1.8e-06 | 2.033E-06 | 2.082E-07 | 1.800E-06 | 2.200E-06 |
| 22942 | 1148 | Q1148R | D | 3.2e-06 | 1.8e-06 | 2.3e-06 | 2.433E-06 | 7.095E-07 | 1.800E-06 | 3.200E-06 |
| 22943 | 1148 | Q1148N | D | 2.2e-06 | 2.3e-06 | 1.9e-06 | 2.133E-06 | 2.082E-07 | 1.900E-06 | 2.300E-06 |
| 22944 | 1148 | Q1148D | D | 2.6e-06 | 2.7e-06 | 2.3e-06 | 2.533E-06 | 2.082E-07 | 2.300E-06 | 2.700E-06 |
| 22945 | 1148 | Q1148C | D | 2.5e-06 | 9.3e-07 | 3.8e-06 | 2.410E-06 | 1.437E-06 | 9.300E-07 | 3.800E-06 |
| 22946 | 1148 | Q1148E | D | 1.9e-06 | 2.3e-06 | 1.8e-06 | 2.000E-06 | 2.646E-07 | 1.800E-06 | 2.300E-06 |
| 22947 | 1148 | Q1148Q | D | 1.9e-06 | 2.6e-06 | 2.6e-06 | 2.367E-06 | 4.041E-07 | 1.900E-06 | 2.600E-06 |
| 22948 | 1148 | Q1148G | D | 2.6e-06 | 1.6e-06 | 1.4e-06 | 1.867E-06 | 6.429E-07 | 1.400E-06 | 2.600E-06 |
| 22949 | 1148 | Q1148H | D | 1.8e-06 | 1.8e-06 | 1.2e-06 | 1.600E-06 | 3.464E-07 | 1.200E-06 | 1.800E-06 |

| | | | | | | | | | | |
|---|---|---|---|---|---|---|---|---|---|---|
| 22950 | 1148 | Q1148I | D | 1.3e-06 | 2e-06 | 1.6e-06 | 1.633E-06 | 3.512E-07 | 1.300E-06 | 2.000E-06 |
| 22951 | 1148 | Q1148L | D | 3.3e-06 | 1.6e-06 | 1.8e-06 | 2.233E-06 | 9.292E-07 | 1.600E-06 | 3.300E-06 |
| 22952 | 1148 | Q1148K | D | 2e-06 | 2.2e-06 | 2e-06 | 2.067E-06 | 1.155E-07 | 2.000E-06 | 2.200E-06 |
| 22953 | 1148 | Q1148M | D | 2.6e-06 | 2.4e-06 | 1.7e-06 | 2.233E-06 | 4.726E-07 | 1.700E-06 | 2.600E-06 |
| 22954 | 1148 | Q1148F | D | 1.9e-06 | 3.2e-06 | 3e-06 | 2.700E-06 | 7.000E-07 | 1.900E-06 | 3.200E-06 |
| 22955 | 1148 | Q1148P | D | 1.1e-06 | 2.8e-06 | 2.9e-06 | 2.267E-06 | 1.012E-06 | 1.100E-06 | 2.900E-06 |
| 22956 | 1148 | Q1148S | D | 2.1e-06 | 2.2e-06 | 2.5e-06 | 2.267E-06 | 2.082E-07 | 2.100E-06 | 2.500E-06 |
| 22957 | 1148 | Q1148T | D | 2.5e-06 | 1.9e-06 | 1.1e-06 | 1.833E-06 | 7.024E-07 | 1.100E-06 | 2.500E-06 |
| 22958 | 1148 | Q1148W | D | 3e-06 | 3.8e-06 | 3.3e-06 | 3.367E-06 | 4.041E-07 | 3.000E-06 | 3.800E-06 |
| 22959 | 1148 | Q1148Y | D | 3.5e-06 | 2.4e-06 | 2.9e-06 | 2.933E-06 | 5.508E-07 | 2.400E-06 | 3.500E-06 |
| 22960 | 1148 | Q1148V | D | 3e-06 | 1.6e-06 | 1.1e-06 | 1.900E-06 | 9.849E-07 | 1.100E-06 | 3.000E-06 |
| 22961 | 1149 | H1149A | D | 1.6e-06 | 3.9e-06 | 1.7e-06 | 2.400E-06 | 1.300E-06 | 1.600E-06 | 3.900E-06 |
| 22962 | 1149 | H1149R | D | 3.1e-06 | 2.3e-06 | 2e-06 | 2.467E-06 | 5.686E-07 | 2.000E-06 | 3.100E-06 |
| 22963 | 1149 | H1149N | D | 2.5e-06 | 3.5e-06 | 2.1e-06 | 2.700E-06 | 7.211E-07 | 2.100E-06 | 3.500E-06 |
| 22964 | 1149 | H1149D | D | 3.2e-06 | 1.8e-06 | 2.4e-06 | 2.467E-06 | 7.024E-07 | 1.800E-06 | 3.200E-06 |
| 22965 | 1149 | H1149C | D | 9.9e-07 | 1.9e-06 | 1.9e-06 | 1.597E-06 | 5.254E-07 | 9.900E-07 | 1.900E-06 |
| 22966 | 1149 | H1149E | D | 1.5e-06 | 2.1e-06 | 2e-06 | 1.867E-06 | 3.215E-07 | 1.500E-06 | 2.100E-06 |
| 22967 | 1149 | H1149Q | D | 2.5e-06 | 1.2e-06 | 3.1e-06 | 2.267E-06 | 9.713E-07 | 1.200E-06 | 3.100E-06 |
| 22968 | 1149 | H1149G | D | 3.9e-06 | 2e-06 | 3.5e-06 | 3.133E-06 | 1.002E-06 | 2.000E-06 | 3.900E-06 |
| 22969 | 1149 | H1149H | D | 1.9e-06 | 2.9e-06 | 3.7e-06 | 2.833E-06 | 9.018E-07 | 1.900E-06 | 3.700E-06 |
| 22970 | 1149 | H1149I | D | 1.4e-06 | 4.6e-06 | 2.5e-06 | 2.833E-06 | 1.626E-06 | 1.400E-06 | 4.600E-06 |
| 22971 | 1149 | H1149L | D | 2.6e-06 | 1.4e-06 | 3.3e-06 | 2.433E-06 | 9.609E-07 | 1.400E-06 | 3.300E-06 |
| 22972 | 1149 | H1149K | D | 2.8e-06 | 3e-06 | 2.5e-06 | 2.767E-06 | 2.517E-07 | 2.500E-06 | 3.000E-06 |
| 22973 | 1149 | H1149M | D | 5.5e-06 | 2.7e-06 | 2.4e-06 | 3.533E-06 | 1.710E-06 | 2.400E-06 | 5.500E-06 |
| 22974 | 1149 | H1149F | D | 4.5e-06 | 1.6e-06 | 5e-06 | 3.700E-06 | 1.836E-06 | 1.600E-06 | 5.000E-06 |
| 22975 | 1149 | H1149P | D | 3.5e-06 | 3.6e-06 | 3.9e-06 | 3.667E-06 | 2.082E-07 | 3.500E-06 | 3.900E-06 |
| 22976 | 1149 | H1149S | D | 2.7e-06 | 2e-06 | 2.6e-06 | 2.433E-06 | 3.786E-07 | 2.000E-06 | 2.700E-06 |
| 22977 | 1149 | H1149T | D | 1.5e-06 | 2.6e-06 | 2.7e-06 | 2.267E-06 | 6.658E-07 | 1.500E-06 | 2.700E-06 |
| 22978 | 1149 | H1149W | D | 3.3e-06 | 1.3e-06 | 2.1e-06 | 2.233E-06 | 1.007E-06 | 1.300E-06 | 3.300E-06 |
| 22979 | 1149 | H1149Y | D | 5e-06 | 2.4e-06 | 2.5e-06 | 3.300E-06 | 1.473E-06 | 2.400E-06 | 5.000E-06 |
| 22980 | 1149 | H1149V | D | 3.3e-06 | 2.8e-06 | 1.9e-06 | 2.667E-06 | 7.095E-07 | 1.900E-06 | 3.300E-06 |
| 22981 | 1150 | Y1150A | D | 2.7e-06 | 2.3e-06 | 2.9e-06 | 2.633E-06 | 3.055E-07 | 2.300E-06 | 2.900E-06 |
| 22982 | 1150 | Y1150R | D | 1.8e-06 | 2.8e-06 | 2.9e-06 | 2.500E-06 | 6.083E-07 | 1.800E-06 | 2.900E-06 |
| 22983 | 1150 | Y1150N | D | 2.5e-06 | 3.3e-06 | 2e-06 | 2.600E-06 | 6.557E-07 | 2.000E-06 | 3.300E-06 |
| 22984 | 1150 | Y1150D | D | 3.7e-06 | 4.2e-06 | 1.4e-06 | 3.100E-06 | 1.493E-06 | 1.400E-06 | 4.200E-06 |
| 22985 | 1150 | Y1150C | D | 2e-06 | 2.8e-06 | 3.6e-06 | 2.800E-06 | 8.000E-07 | 2.000E-06 | 3.600E-06 |
| 22986 | 1150 | Y1150E | D | 2.1e-06 | 2e-06 | 1.8e-06 | 1.967E-06 | 1.528E-07 | 1.800E-06 | 2.100E-06 |
| 22987 | 1150 | Y1150Q | D | 2.3e-06 | 3.3e-06 | 2.6e-06 | 2.733E-06 | 5.132E-07 | 2.300E-06 | 3.300E-06 |
| 22988 | 1150 | Y1150G | D | 2.9e-06 | 4.5e-06 | 2.2e-06 | 3.200E-06 | 1.179E-06 | 2.200E-06 | 4.500E-06 |
| 22989 | 1150 | Y1150H | D | 2.8e-06 | 3.4e-06 | 2.6e-06 | 2.933E-06 | 4.163E-07 | 2.600E-06 | 3.400E-06 |
| 22990 | 1150 | Y1150I | D | 3.8e-06 | 2.1e-06 | 3e-06 | 2.967E-06 | 8.505E-07 | 2.100E-06 | 3.800E-06 |
| 22991 | 1150 | Y1150L | D | 1.9e-06 | 2e-06 | 2.7e-06 | 2.200E-06 | 4.359E-07 | 1.900E-06 | 2.700E-06 |
| 22992 | 1150 | Y1150K | D | 2.9e-06 | 3.4e-06 | 3e-06 | 3.100E-06 | 2.646E-07 | 2.900E-06 | 3.400E-06 |
| 22993 | 1150 | Y1150M | D | 3.6e-06 | 3.2e-06 | 1.7e-06 | 2.833E-06 | 1.002E-06 | 1.700E-06 | 3.600E-06 |
| 22994 | 1150 | Y1150F | D | 2.7e-06 | 3.2e-06 | 2.4e-06 | 2.767E-06 | 4.041E-07 | 2.400E-06 | 3.200E-06 |
| 22995 | 1150 | Y1150P | D | 1.2e-06 | 2.7e-06 | 2.9e-06 | 2.267E-06 | 9.292E-07 | 1.200E-06 | 2.900E-06 |
| 22996 | 1150 | Y1150S | D | 2.7e-06 | 2.6e-06 | 2.1e-06 | 2.467E-06 | 3.215E-07 | 2.100E-06 | 2.700E-06 |
| 22997 | 1150 | Y1150T | D | 1.9e-06 | 3.4e-06 | 2.6e-06 | 2.633E-06 | 7.506E-07 | 1.900E-06 | 3.400E-06 |
| 22998 | 1150 | Y1150W | D | 2e-06 | 3.2e-06 | 1.3e-06 | 2.167E-06 | 9.609E-07 | 1.300E-06 | 3.200E-06 |
| 22999 | 1150 | Y1150Y | D | 1.4e-06 | 3e-06 | 2.8e-06 | 2.400E-06 | 8.718E-07 | 1.400E-06 | 3.000E-06 |
| 23000 | 1150 | Y1150V | D | 3.4e-06 | 2e-06 | 1.7e-06 | 2.367E-06 | 9.074E-07 | 1.700E-06 | 3.400E-06 |
| 23001 | 1151 | T1151A | D | 1.6e-06 | 1.8e-06 | 2.7e-06 | 2.033E-06 | 5.859E-07 | 1.600E-06 | 2.700E-06 |
| 23002 | 1151 | T1151R | D | 2.4e-06 | 2.4e-06 | 2.5e-06 | 2.433E-06 | 5.774E-08 | 2.400E-06 | 2.500E-06 |
| 23003 | 1151 | T1151N | D | 9.2e-07 | 1.6e-06 | 2.1e-06 | 1.540E-06 | 5.923E-07 | 9.200E-07 | 2.100E-06 |
| 23004 | 1151 | T1151D | D | 1.9e-06 | 1.8e-06 | 3.8e-06 | 2.500E-06 | 1.127E-06 | 1.800E-06 | 3.800E-06 |
| 23005 | 1151 | T1151C | D | 3.3e-06 | 3.3e-06 | 2.8e-06 | 3.133E-06 | 2.887E-07 | 2.800E-06 | 3.300E-06 |
| 23006 | 1151 | T1151E | D | 3.5e-06 | 1.6e-06 | 2.1e-06 | 2.400E-06 | 9.849E-07 | 1.600E-06 | 3.500E-06 |
| 23007 | 1151 | T1151Q | D | 2.9e-06 | 2.2e-06 | 2.7e-06 | 2.600E-06 | 3.606E-07 | 2.200E-06 | 2.900E-06 |
| 23008 | 1151 | T1151G | D | 2.3e-06 | 9.4e-07 | 3e-06 | 2.080E-06 | 1.047E-06 | 9.400E-07 | 3.000E-06 |
| 23009 | 1151 | T1151H | D | 2e-06 | 2.4e-06 | 1.9e-06 | 2.100E-06 | 2.646E-07 | 1.900E-06 | 2.400E-06 |
| 23010 | 1151 | T1151I | D | 2e-06 | 3.3e-06 | 3.8e-06 | 3.033E-06 | 9.292E-07 | 2.000E-06 | 3.800E-06 |
| 23011 | 1151 | T1151L | D | 2.3e-06 | 2.7e-06 | 1.5e-06 | 2.167E-06 | 6.110E-07 | 1.500E-06 | 2.700E-06 |
| 23012 | 1151 | T1151K | D | 2.4e-06 | 2.7e-06 | 3.1e-06 | 2.733E-06 | 3.512E-07 | 2.400E-06 | 3.100E-06 |
| 23013 | 1151 | T1151M | D | 2.2e-06 | 1.8e-06 | 1.7e-06 | 1.900E-06 | 2.646E-07 | 1.700E-06 | 2.200E-06 |
| 23014 | 1151 | T1151F | D | 1.8e-06 | 2.5e-06 | 2.1e-06 | 2.133E-06 | 3.512E-07 | 1.800E-06 | 2.500E-06 |
| 23015 | 1151 | T1151P | D | 3.8e-06 | 3.2e-06 | 2.7e-06 | 3.233E-06 | 5.508E-07 | 2.700E-06 | 3.800E-06 |
| 23016 | 1151 | T1151S | D | 2.1e-06 | 1.8e-06 | 2e-06 | 1.967E-06 | 1.528E-07 | 1.800E-06 | 2.100E-06 |
| 23017 | 1151 | T1151T | D | 3.2e-06 | 2.7e-06 | 1.4e-06 | 2.433E-06 | 9.292E-07 | 1.400E-06 | 3.200E-06 |
| 23018 | 1151 | T1151W | D | 2.6e-06 | 2.1e-06 | 2.6e-06 | 2.433E-06 | 2.887E-07 | 2.100E-06 | 2.600E-06 |
| 23019 | 1151 | T1151Y | D | 3.4e-06 | 2.8e-06 | 3e-06 | 3.067E-06 | 3.055E-07 | 2.800E-06 | 3.400E-06 |
| 23020 | 1151 | T1151V | D | 2e-06 | 1.3e-06 | 2.8e-06 | 2.033E-06 | 7.506E-07 | 1.300E-06 | 2.800E-06 |
| 23021 | 1152 | T1152A | D | 2.1e-06 | 2.1e-06 | 4.8e-06 | 3.000E-06 | 1.559E-06 | 2.100E-06 | 4.800E-06 |
| 23022 | 1152 | T1152R | D | 2.8e-06 | 1.2e-06 | 2.9e-06 | 2.300E-06 | 9.539E-07 | 1.200E-06 | 2.900E-06 |
| 23023 | 1152 | T1152N | D | 2.5e-06 | 1.6e-06 | 2.4e-06 | 2.167E-06 | 4.933E-07 | 1.600E-06 | 2.500E-06 |
| 23024 | 1152 | T1152D | D | 1.2e-06 | 2.2e-06 | 1.4e-06 | 1.600E-06 | 5.292E-07 | 1.200E-06 | 2.200E-06 |
| 23025 | 1152 | T1152C | D | 3.5e-06 | 3.8e-06 | 2.7e-06 | 3.333E-06 | 5.686E-07 | 2.700E-06 | 3.800E-06 |
| 23026 | 1152 | T1152E | D | 3.8e-06 | 3.1e-06 | 1.7e-06 | 2.867E-06 | 1.069E-06 | 1.700E-06 | 3.800E-06 |
| 23027 | 1152 | T1152Q | D | 2.3e-06 | 1.7e-06 | 1e-06 | 1.667E-06 | 6.506E-07 | 1.000E-06 | 2.300E-06 |
| 23028 | 1152 | T1152G | D | 3.4e-06 | 2.8e-06 | 2.4e-06 | 2.867E-06 | 5.033E-07 | 2.400E-06 | 3.400E-06 |
| 23029 | 1152 | T1152H | D | 1.4e-06 | 2.2e-06 | 2.3e-06 | 1.967E-06 | 4.933E-07 | 1.400E-06 | 2.300E-06 |
| 23030 | 1152 | T1152I | D | 1.2e-06 | 3.1e-06 | 2.7e-06 | 2.333E-06 | 1.002E-06 | 1.200E-06 | 3.100E-06 |
| 23031 | 1152 | T1152L | D | 3.7e-06 | 1.4e-06 | 2.8e-06 | 2.633E-06 | 1.159E-06 | 1.400E-06 | 3.700E-06 |
| 23032 | 1152 | T1152K | D | 2e-06 | 2.1e-06 | 2.1e-06 | 2.067E-06 | 5.774E-08 | 2.000E-06 | 2.100E-06 |
| 23033 | 1152 | T1152M | D | 3e-06 | 2.6e-06 | 1.7e-06 | 2.433E-06 | 6.658E-07 | 1.700E-06 | 3.000E-06 |
| 23034 | 1152 | T1152F | D | 2.6e-06 | 4.5e-06 | 2.4e-06 | 3.167E-06 | 1.159E-06 | 2.400E-06 | 4.500E-06 |

| | | | | | | | | | | |
|---|---|---|---|---|---|---|---|---|---|---|
| 23035 | 1152 | T1152P | D | 1.7e-06 | 1.9e-06 | 2.9e-06 | 2.167E-06 | 6.429E-07 | 1.700E-06 | 2.900E-06 |
| 23036 | 1152 | T1152S | D | 2.5e-06 | 2.1e-06 | 3.1e-06 | 2.567E-06 | 5.033E-07 | 2.100E-06 | 3.100E-06 |
| 23037 | 1152 | T1152T | D | 2.2e-06 | 3.6e-06 | 4e-06 | 3.267E-06 | 9.452E-07 | 2.200E-06 | 4.000E-06 |
| 23038 | 1152 | T1152W | D | 2.7e-06 | 2.2e-06 | 1.3e-06 | 2.067E-06 | 7.095E-07 | 1.300E-06 | 2.700E-06 |
| 23039 | 1152 | T1152Y | D | 1e-06 | 3.5e-06 | 2.5e-06 | 2.333E-06 | 1.258E-06 | 1.000E-06 | 3.500E-06 |
| 23040 | 1152 | T1152V | D | 1.5e-06 | 2.2e-06 | 2.5e-06 | 2.067E-06 | 5.132E-07 | 1.500E-06 | 2.500E-06 |
| 23041 | 1153 | P1153A | D | 4.4e-06 | 5.6e-06 | 3.7e-06 | 4.567E-06 | 9.609E-07 | 3.700E-06 | 5.600E-06 |
| 23042 | 1153 | P1153R | D | 3.2e-06 | 1.8e-06 | 2.9e-06 | 2.633E-06 | 7.371E-07 | 1.800E-06 | 3.200E-06 |
| 23043 | 1153 | P1153N | D | 1.2e-06 | 3.6e-06 | 3.3e-06 | 2.700E-06 | 1.308E-06 | 1.200E-06 | 3.600E-06 |
| 23044 | 1153 | P1153D | D | 2.3e-06 | 2.9e-06 | 2.9e-06 | 2.700E-06 | 3.464E-07 | 2.300E-06 | 2.900E-06 |
| 23045 | 1153 | P1153C | D | 1.2e-06 | 2.3e-06 | 1.9e-06 | 1.800E-06 | 5.568E-07 | 1.200E-06 | 2.300E-06 |
| 23046 | 1153 | P1153E | D | 3.2e-06 | 1.7e-06 | 3e-06 | 2.633E-06 | 8.145E-07 | 1.700E-06 | 3.200E-06 |
| 23047 | 1153 | P1153Q | D | 3e-06 | 1.2e-06 | 1.7e-06 | 1.967E-06 | 9.292E-07 | 1.200E-06 | 3.000E-06 |
| 23048 | 1153 | P1153G | D | 2.1e-06 | 1.5e-06 | 2.7e-06 | 2.100E-06 | 6.000E-07 | 1.500E-06 | 2.700E-06 |
| 23049 | 1153 | P1153H | D | 2.5e-06 | 4.3e-06 | 2.4e-06 | 3.067E-06 | 1.069E-06 | 2.400E-06 | 4.300E-06 |
| 23050 | 1153 | P1153I | D | 3.2e-06 | 2.2e-06 | 2.5e-06 | 2.633E-06 | 5.132E-07 | 2.200E-06 | 3.200E-06 |
| 23051 | 1153 | P1153L | D | 1.8e-06 | 2e-06 | 1.2e-06 | 1.667E-06 | 4.163E-07 | 1.200E-06 | 2.000E-06 |
| 23052 | 1153 | P1153K | D | 3.4e-06 | 2.5e-06 | 2.3e-06 | 2.733E-06 | 5.859E-07 | 2.300E-06 | 3.400E-06 |
| 23053 | 1153 | P1153M | D | 2.8e-06 | 1.8e-06 | 2.4e-06 | 2.333E-06 | 5.033E-07 | 1.800E-06 | 2.800E-06 |
| 23054 | 1153 | P1153F | D | 2.6e-06 | 3e-06 | 2.7e-06 | 2.767E-06 | 2.082E-07 | 2.600E-06 | 3.000E-06 |
| 23055 | 1153 | P1153P | D | 2.8e-06 | 1.7e-06 | 2.3e-06 | 2.267E-06 | 5.508E-07 | 1.700E-06 | 2.800E-06 |
| 23056 | 1153 | P1153S | D | 2.7e-06 | 2.9e-06 | 2.3e-06 | 2.633E-06 | 3.055E-07 | 2.300E-06 | 2.900E-06 |
| 23057 | 1153 | P1153T | D | 1.5e-06 | 2.1e-06 | 1.4e-06 | 1.667E-06 | 3.786E-07 | 1.400E-06 | 2.100E-06 |
| 23058 | 1153 | P1153W | D | 1.8e-06 | 1.8e-06 | 2.8e-06 | 2.133E-06 | 5.774E-07 | 1.800E-06 | 2.800E-06 |
| 23059 | 1153 | P1153Y | D | 4.2e-06 | 1.8e-06 | 1.3e-06 | 2.433E-06 | 1.550E-06 | 1.300E-06 | 4.200E-06 |
| 23060 | 1153 | P1153V | D | 9e-07 | 3.2e-06 | 2.6e-06 | 2.233E-06 | 1.193E-06 | 9.000E-07 | 3.200E-06 |
| 23061 | 1154 | P1154A | D | 3.2e-06 | 1e-06 | 2.5e-06 | 2.233E-06 | 1.124E-06 | 1.000E-06 | 3.200E-06 |
| 23062 | 1154 | P1154R | D | 1.3e-06 | 2.6e-06 | 3.7e-06 | 2.533E-06 | 1.201E-06 | 1.300E-06 | 3.700E-06 |
| 23063 | 1154 | P1154N | D | 2.6e-06 | 3.2e-06 | 1.7e-06 | 2.500E-06 | 7.550E-07 | 1.700E-06 | 3.200E-06 |
| 23064 | 1154 | P1154D | D | 2.4e-06 | 2.2e-06 | 3e-06 | 2.533E-06 | 4.163E-07 | 2.200E-06 | 3.000E-06 |
| 23065 | 1154 | P1154C | D | 4.7e-06 | 2.7e-06 | 2.2e-06 | 3.200E-06 | 1.323E-06 | 2.200E-06 | 4.700E-06 |
| 23066 | 1154 | P1154E | D | 3.1e-06 | 2.6e-06 | 3.8e-06 | 3.167E-06 | 6.028E-07 | 2.600E-06 | 3.800E-06 |
| 23067 | 1154 | P1154Q | D | 2.1e-06 | 1.9e-06 | 4.7e-06 | 2.900E-06 | 1.562E-06 | 1.900E-06 | 4.700E-06 |
| 23068 | 1154 | P1154G | D | 1.5e-06 | 2.6e-06 | 1.3e-06 | 1.800E-06 | 7.000E-07 | 1.300E-06 | 2.600E-06 |
| 23069 | 1154 | P1154H | D | 2.1e-06 | 2.1e-06 | 2.4e-06 | 2.200E-06 | 1.732E-07 | 2.100E-06 | 2.400E-06 |
| 23070 | 1154 | P1154I | D | 2.5e-06 | 2.4e-06 | 1.2e-06 | 2.033E-06 | 7.234E-07 | 1.200E-06 | 2.500E-06 |
| 23071 | 1154 | P1154L | D | 2.2e-06 | 2.9e-06 | 3.4e-06 | 2.833E-06 | 6.028E-07 | 2.200E-06 | 3.400E-06 |
| 23072 | 1154 | P1154K | D | 2.8e-06 | 2e-06 | 1.6e-06 | 2.133E-06 | 6.110E-07 | 1.600E-06 | 2.800E-06 |
| 23073 | 1154 | P1154M | D | 1.7e-06 | 1.1e-06 | 2e-06 | 1.600E-06 | 4.583E-07 | 1.100E-06 | 2.000E-06 |
| 23074 | 1154 | P1154F | D | 3.2e-06 | 2.3e-06 | 1.9e-06 | 2.467E-06 | 6.658E-07 | 1.900E-06 | 3.200E-06 |
| 23075 | 1154 | P1154P | D | 4.9e-06 | 1.6e-06 | 1.6e-06 | 2.700E-06 | 1.905E-06 | 1.600E-06 | 4.900E-06 |
| 23076 | 1154 | P1154S | D | 2.3e-06 | 1.5e-06 | 2.5e-06 | 2.100E-06 | 5.292E-07 | 1.500E-06 | 2.500E-06 |
| 23077 | 1154 | P1154T | D | 2.6e-06 | 2.6e-06 | 2.3e-06 | 2.500E-06 | 1.732E-07 | 2.300E-06 | 2.600E-06 |
| 23078 | 1154 | P1154W | D | 2.8e-06 | 2.3e-06 | 2.6e-06 | 2.567E-06 | 2.517E-07 | 2.300E-06 | 2.800E-06 |
| 23079 | 1154 | P1154Y | D | 2.2e-06 | 4.9e-06 | 1.9e-06 | 3.000E-06 | 1.652E-06 | 1.900E-06 | 4.900E-06 |
| 23080 | 1154 | P1154V | D | 2.5e-06 | 2.4e-06 | 1.8e-06 | 2.233E-06 | 3.786E-07 | 1.800E-06 | 2.500E-06 |
| 23081 | 1155 | T1155A | D | 1.8e-06 | 4.7e-06 | 2.6e-06 | 3.033E-06 | 1.498E-06 | 1.800E-06 | 4.700E-06 |
| 23082 | 1155 | T1155R | D | 1.8e-06 | 2e-06 | 2.2e-06 | 2.000E-06 | 2.000E-07 | 1.800E-06 | 2.200E-06 |
| 23083 | 1155 | T1155N | D | 1.9e-06 | 3.5e-06 | 1.8e-06 | 2.400E-06 | 9.539E-07 | 1.800E-06 | 3.500E-06 |
| 23084 | 1155 | T1155D | D | 2.3e-06 | 1.9e-06 | 2.6e-06 | 2.267E-06 | 3.512E-07 | 1.900E-06 | 2.600E-06 |
| 23085 | 1155 | T1155C | D | 1.5e-06 | 2e-06 | 2.6e-06 | 2.033E-06 | 5.508E-07 | 1.500E-06 | 2.600E-06 |
| 23086 | 1155 | T1155E | D | 2.5e-06 | 1.3e-06 | 2e-06 | 1.933E-06 | 6.028E-07 | 1.300E-06 | 2.500E-06 |
| 23087 | 1155 | T1155Q | D | 1.4e-06 | 2.4e-06 | 2.1e-06 | 1.967E-06 | 5.132E-07 | 1.400E-06 | 2.400E-06 |
| 23088 | 1155 | T1155G | D | 2.4e-06 | 4.8e-06 | 2.1e-06 | 3.100E-06 | 1.480E-06 | 2.100E-06 | 4.800E-06 |
| 23089 | 1155 | T1155H | D | 2.2e-06 | 3.1e-06 | 2.7e-06 | 2.667E-06 | 4.509E-07 | 2.200E-06 | 3.100E-06 |
| 23090 | 1155 | T1155I | D | 2.2e-06 | 2.8e-06 | 3.3e-06 | 2.767E-06 | 5.508E-07 | 2.200E-06 | 3.300E-06 |
| 23091 | 1155 | T1155L | D | 3.1e-06 | 2.8e-06 | 2.1e-06 | 2.667E-06 | 5.132E-07 | 2.100E-06 | 3.100E-06 |
| 23092 | 1155 | T1155K | D | 3.5e-06 | 2.2e-06 | 3.6e-06 | 3.100E-06 | 7.810E-07 | 2.200E-06 | 3.600E-06 |
| 23093 | 1155 | T1155M | D | 3.3e-06 | 3.6e-06 | 2e-06 | 2.967E-06 | 8.505E-07 | 2.000E-06 | 3.600E-06 |
| 23094 | 1155 | T1155F | D | 3.7e-06 | 2.2e-06 | 2.7e-06 | 2.867E-06 | 7.638E-07 | 2.200E-06 | 3.700E-06 |
| 23095 | 1155 | T1155P | D | 2.6e-06 | 2e-06 | 3.1e-06 | 2.567E-06 | 5.508E-07 | 2.000E-06 | 3.100E-06 |
| 23096 | 1155 | T1155S | D | 1.8e-06 | 2.4e-06 | 1.9e-06 | 2.033E-06 | 3.215E-07 | 1.800E-06 | 2.400E-06 |
| 23097 | 1155 | T1155T | D | 2.2e-06 | 2.3e-06 | 2.7e-06 | 2.400E-06 | 2.646E-07 | 2.200E-06 | 2.700E-06 |
| 23098 | 1155 | T1155W | D | 2.5e-06 | 2.7e-06 | 1.8e-06 | 2.333E-06 | 4.726E-07 | 1.800E-06 | 2.700E-06 |
| 23099 | 1155 | T1155Y | D | 3.4e-06 | 2.9e-06 | 1.2e-06 | 2.500E-06 | 1.153E-06 | 1.200E-06 | 3.400E-06 |
| 23100 | 1155 | T1155V | D | 4.3e-06 | 1.1e-06 | 2.2e-06 | 2.533E-06 | 1.626E-06 | 1.100E-06 | 4.300E-06 |
| 23101 | 1156 | F1156A | D | 2.4e-06 | 2.8e-06 | 2.6e-06 | 2.600E-06 | 2.000E-07 | 2.400E-06 | 2.800E-06 |
| 23102 | 1156 | F1156R | D | 3.7e-06 | 2.8e-06 | 1.3e-06 | 2.600E-06 | 1.212E-06 | 1.300E-06 | 3.700E-06 |
| 23103 | 1156 | F1156N | D | 2.6e-06 | 2.3e-06 | 3.1e-06 | 2.667E-06 | 4.041E-07 | 2.300E-06 | 3.100E-06 |
| 23104 | 1156 | F1156D | D | 1.4e-06 | 4.5e-06 | 1.8e-06 | 2.567E-06 | 1.686E-06 | 1.400E-06 | 4.500E-06 |
| 23105 | 1156 | F1156C | D | 1.7e-06 | 2.9e-06 | 1.6e-06 | 2.067E-06 | 7.234E-07 | 1.600E-06 | 2.900E-06 |
| 23106 | 1156 | F1156E | D | 3.5e-06 | 2.8e-06 | 3.4e-06 | 3.233E-06 | 3.786E-07 | 2.800E-06 | 3.500E-06 |
| 23107 | 1156 | F1156Q | D | 2.1e-06 | 1.9e-06 | 1.4e-06 | 1.800E-06 | 3.606E-07 | 1.400E-06 | 2.100E-06 |
| 23108 | 1156 | F1156G | D | 1.8e-06 | 1.8e-06 | 2.5e-06 | 2.033E-06 | 4.041E-07 | 1.800E-06 | 2.500E-06 |
| 23109 | 1156 | F1156H | D | 2.9e-06 | 1.9e-06 | 1.6e-06 | 2.133E-06 | 6.807E-07 | 1.600E-06 | 2.900E-06 |
| 23110 | 1156 | F1156I | D | 2.7e-06 | 1.8e-06 | 2.2e-06 | 2.233E-06 | 4.509E-07 | 1.800E-06 | 2.700E-06 |
| 23111 | 1156 | F1156L | D | 2.6e-06 | 1.7e-06 | 1.1e-06 | 1.800E-06 | 7.550E-07 | 1.100E-06 | 2.600E-06 |
| 23112 | 1156 | F1156K | D | 1.5e-06 | 2.1e-06 | 3.3e-06 | 2.300E-06 | 9.165E-07 | 1.500E-06 | 3.300E-06 |
| 23113 | 1156 | F1156M | D | 2.8e-06 | 4.1e-06 | 2.3e-06 | 3.067E-06 | 9.292E-07 | 2.300E-06 | 4.100E-06 |
| 23114 | 1156 | F1156F | D | 3.5e-06 | 6e-06 | 1.7e-06 | 3.733E-06 | 2.159E-06 | 1.700E-06 | 6.000E-06 |
| 23115 | 1156 | F1156P | D | 3e-06 | 2.2e-06 | 1.8e-06 | 2.333E-06 | 6.110E-07 | 1.800E-06 | 3.000E-06 |
| 23116 | 1156 | F1156S | D | 3.6e-06 | 2.2e-06 | 2.4e-06 | 2.733E-06 | 7.572E-07 | 2.200E-06 | 3.600E-06 |
| 23117 | 1156 | F1156T | D | 1.7e-06 | 3.2e-06 | 4.1e-06 | 3.000E-06 | 1.212E-06 | 1.700E-06 | 4.100E-06 |
| 23118 | 1156 | F1156W | D | 2.3e-06 | 2.5e-06 | 3.3e-06 | 2.700E-06 | 5.292E-07 | 2.300E-06 | 3.300E-06 |
| 23119 | 1156 | F1156Y | D | 3.1e-06 | 2.6e-06 | 3.9e-06 | 3.200E-06 | 6.557E-07 | 2.600E-06 | 3.900E-06 |

| | | | | | | | | | | |
|---|---|---|---|---|---|---|---|---|---|---|
| 23120 | 1156 | F1156V | D | 1.9e-06 | 2.2e-06 | 2.3e-06 | 2.133E-06 | 2.082E-07 | 1.900E-06 | 2.300E-06 |
| 23121 | 1157 | G1157A | D | 2.2e-06 | 2e-06 | 2.8e-06 | 2.333E-06 | 4.163E-07 | 2.000E-06 | 2.800E-06 |
| 23122 | 1157 | G1157R | D | 1.5e-06 | 1.7e-06 | 2.1e-06 | 1.767E-06 | 3.055E-07 | 1.500E-06 | 2.100E-06 |
| 23123 | 1157 | G1157N | D | 1.3e-06 | 2.8e-06 | 2.5e-06 | 2.200E-06 | 7.937E-07 | 1.300E-06 | 2.800E-06 |
| 23124 | 1157 | G1157D | D | 1.5e-06 | 2.5e-06 | 3.1e-06 | 2.367E-06 | 8.083E-07 | 1.500E-06 | 3.100E-06 |
| 23125 | 1157 | G1157C | D | 2.2e-06 | 1.9e-06 | 2.5e-06 | 2.200E-06 | 3.000E-07 | 1.900E-06 | 2.500E-06 |
| 23126 | 1157 | G1157E | D | 1.1e-06 | 3.3e-06 | 2.7e-06 | 2.367E-06 | 1.137E-06 | 1.100E-06 | 3.300E-06 |
| 23127 | 1157 | G1157Q | D | 2.3e-06 | 1.7e-06 | 3e-06 | 2.333E-06 | 6.506E-07 | 1.700E-06 | 3.000E-06 |
| 23128 | 1157 | G1157G | D | 4.5e-06 | 2.2e-06 | 3.6e-06 | 3.433E-06 | 1.159E-06 | 2.200E-06 | 4.500E-06 |
| 23129 | 1157 | G1157H | D | 2.9e-06 | 2.6e-06 | 2.1e-06 | 2.533E-06 | 4.041E-07 | 2.100E-06 | 2.900E-06 |
| 23130 | 1157 | G1157I | D | 2.2e-06 | 2.3e-06 | 2.9e-06 | 2.467E-06 | 3.786E-07 | 2.200E-06 | 2.900E-06 |
| 23131 | 1157 | G1157L | D | 2.4e-06 | 3.2e-06 | 2.9e-06 | 2.833E-06 | 4.041E-07 | 2.400E-06 | 3.200E-06 |
| 23132 | 1157 | G1157K | D | 1.6e-06 | 1.8e-06 | 9.1e-07 | 1.437E-06 | 4.669E-07 | 9.100E-07 | 1.800E-06 |
| 23133 | 1157 | G1157M | D | 2.6e-06 | 2.5e-06 | 2.6e-06 | 2.567E-06 | 5.774E-08 | 2.500E-06 | 2.600E-06 |
| 23134 | 1157 | G1157F | D | 1.9e-06 | 3.6e-06 | 2.1e-06 | 2.533E-06 | 9.292E-07 | 1.900E-06 | 3.600E-06 |
| 23135 | 1157 | G1157P | D | 3.6e-06 | 2.3e-06 | 1.7e-06 | 2.533E-06 | 9.713E-07 | 1.700E-06 | 3.600E-06 |
| 23136 | 1157 | G1157S | D | 3.5e-06 | 2.4e-06 | 1.3e-06 | 2.400E-06 | 1.100E-06 | 1.300E-06 | 3.500E-06 |
| 23137 | 1157 | G1157T | D | 4.1e-06 | 2.2e-06 | 1.3e-06 | 2.533E-06 | 1.429E-06 | 1.300E-06 | 4.100E-06 |
| 23138 | 1157 | G1157W | D | 2.7e-06 | 2.3e-06 | 4.9e-06 | 3.300E-06 | 1.400E-06 | 2.300E-06 | 4.900E-06 |
| 23139 | 1157 | G1157Y | D | 2.2e-06 | 1.5e-06 | 2.4e-06 | 2.033E-06 | 4.726E-07 | 1.500E-06 | 2.400E-06 |
| 23140 | 1157 | G1157V | D | 2.7e-06 | 2.7e-06 | 2.5e-06 | 2.633E-06 | 1.155E-07 | 2.500E-06 | 2.700E-06 |
| 23141 | 1158 | Q1158A | D | 2.2e-06 | 4.4e-06 | 2.4e-06 | 3.000E-06 | 1.217E-06 | 2.200E-06 | 4.400E-06 |
| 23142 | 1158 | Q1158R | D | 2.7e-06 | 1.5e-06 | 1.9e-06 | 2.033E-06 | 6.110E-07 | 1.500E-06 | 2.700E-06 |
| 23143 | 1158 | Q1158N | D | 2.6e-06 | 1.5e-06 | 1.8e-06 | 1.967E-06 | 5.686E-07 | 1.500E-06 | 2.600E-06 |
| 23144 | 1158 | Q1158D | D | 1.3e-06 | 2.8e-06 | 3.8e-06 | 2.633E-06 | 1.258E-06 | 1.300E-06 | 3.800E-06 |
| 23145 | 1158 | Q1158C | D | 3.1e-06 | 2e-06 | 1.7e-06 | 2.267E-06 | 7.371E-07 | 1.700E-06 | 3.100E-06 |
| 23146 | 1158 | Q1158E | D | 2e-06 | 1.8e-06 | 1.7e-06 | 1.833E-06 | 1.528E-07 | 1.700E-06 | 2.000E-06 |
| 23147 | 1158 | Q1158Q | D | 2.4e-06 | 1.9e-06 | 2.6e-06 | 2.300E-06 | 3.606E-07 | 1.900E-06 | 2.600E-06 |
| 23148 | 1158 | Q1158G | D | 2.7e-06 | 2.1e-06 | 3.1e-06 | 2.633E-06 | 5.033E-07 | 2.100E-06 | 3.100E-06 |
| 23149 | 1158 | Q1158H | D | 2.4e-06 | 2.1e-06 | 4e-06 | 2.833E-06 | 1.021E-06 | 2.100E-06 | 4.000E-06 |
| 23150 | 1158 | Q1158I | D | 3.2e-06 | 3.9e-06 | 1.7e-06 | 2.933E-06 | 1.124E-06 | 1.700E-06 | 3.900E-06 |
| 23151 | 1158 | Q1158L | D | 2.6e-06 | 2.3e-06 | 1.4e-06 | 2.100E-06 | 6.245E-07 | 1.400E-06 | 2.600E-06 |
| 23152 | 1158 | Q1158K | D | 1.4e-06 | 2.4e-06 | 2.3e-06 | 2.033E-06 | 5.508E-07 | 1.400E-06 | 2.400E-06 |
| 23153 | 1158 | Q1158M | D | 2.3e-06 | 1.5e-06 | 2.5e-06 | 2.100E-06 | 5.292E-07 | 1.500E-06 | 2.500E-06 |
| 23154 | 1158 | Q1158F | D | 2.6e-06 | 2.5e-06 | 2.9e-06 | 2.667E-06 | 2.082E-07 | 2.500E-06 | 2.900E-06 |
| 23155 | 1158 | Q1158P | D | 2.9e-06 | 2.1e-06 | 1.9e-06 | 2.300E-06 | 5.292E-07 | 1.900E-06 | 2.900E-06 |
| 23156 | 1158 | Q1158S | D | 2.1e-06 | 2.1e-06 | 2.4e-06 | 2.200E-06 | 1.732E-07 | 2.100E-06 | 2.400E-06 |
| 23157 | 1158 | Q1158T | D | 2.7e-06 | 1.9e-06 | 4.4e-06 | 3.000E-06 | 1.277E-06 | 1.900E-06 | 4.400E-06 |
| 23158 | 1158 | Q1158W | D | 3.7e-06 | 1.5e-06 | 1.9e-06 | 2.367E-06 | 1.172E-06 | 1.500E-06 | 3.700E-06 |
| 23159 | 1158 | Q1158Y | D | 2.4e-06 | 2.4e-06 | 2e-06 | 2.267E-06 | 2.309E-07 | 2.000E-06 | 2.400E-06 |
| 23160 | 1158 | Q1158V | D | 1.5e-06 | 1.2e-06 | 2.1e-06 | 1.600E-06 | 4.583E-07 | 1.200E-06 | 2.100E-06 |
| 23161 | 1159 | G1159A | D | 2.9e-06 | 1.3e-06 | 3.4e-06 | 2.533E-06 | 1.097E-06 | 1.300E-06 | 3.400E-06 |
| 23162 | 1159 | G1159R | D | 1e-06 | 3.5e-06 | 1.9e-06 | 2.133E-06 | 1.266E-06 | 1.000E-06 | 3.500E-06 |
| 23163 | 1159 | G1159N | D | 1.6e-06 | 1.5e-06 | 5.8e-06 | 2.967E-06 | 2.454E-06 | 1.500E-06 | 5.800E-06 |
| 23164 | 1159 | G1159D | D | 1.8e-06 | 3.4e-06 | 1.6e-06 | 2.267E-06 | 9.866E-07 | 1.600E-06 | 3.400E-06 |
| 23165 | 1159 | G1159C | D | 2.2e-06 | 1.9e-06 | 3.6e-06 | 2.567E-06 | 9.074E-07 | 1.900E-06 | 3.600E-06 |
| 23166 | 1159 | G1159E | D | 3e-06 | 2.5e-06 | 3.8e-06 | 3.100E-06 | 6.557E-07 | 2.500E-06 | 3.800E-06 |
| 23167 | 1159 | G1159Q | D | 6.6e-06 | 1.1e-06 | 2.1e-06 | 3.267E-06 | 2.930E-06 | 1.100E-06 | 6.600E-06 |
| 23168 | 1159 | G1159G | D | 3.9e-06 | 1.8e-06 | 3.2e-06 | 2.967E-06 | 1.069E-06 | 1.800E-06 | 3.900E-06 |
| 23169 | 1159 | G1159H | D | 3.1e-06 | 3e-06 | 3.1e-06 | 3.067E-06 | 5.774E-08 | 3.000E-06 | 3.100E-06 |
| 23170 | 1159 | G1159I | D | 3.3e-06 | 1.3e-06 | 3.3e-06 | 2.633E-06 | 1.155E-06 | 1.300E-06 | 3.300E-06 |
| 23171 | 1159 | G1159L | D | 1.5e-06 | 3e-06 | 2.3e-06 | 2.267E-06 | 7.506E-07 | 1.500E-06 | 3.000E-06 |
| 23172 | 1159 | G1159K | D | 2.8e-06 | 1.7e-06 | 2.3e-06 | 2.267E-06 | 5.508E-07 | 1.700E-06 | 2.800E-06 |
| 23173 | 1159 | G1159M | D | 2.1e-06 | 2e-06 | 3.7e-06 | 2.600E-06 | 9.539E-07 | 2.000E-06 | 3.700E-06 |
| 23174 | 1159 | G1159F | D | 4.2e-06 | 3.2e-06 | 1.6e-06 | 3.000E-06 | 1.311E-06 | 1.600E-06 | 4.200E-06 |
| 23175 | 1159 | G1159P | D | 2.2e-06 | 3.3e-06 | 2.6e-06 | 2.700E-06 | 5.568E-07 | 2.200E-06 | 3.300E-06 |
| 23176 | 1159 | G1159S | D | 2.1e-06 | 2.7e-06 | 1.4e-06 | 2.067E-06 | 6.506E-07 | 1.400E-06 | 2.700E-06 |
| 23177 | 1159 | G1159T | D | 2.4e-06 | 2.2e-06 | 2e-06 | 2.200E-06 | 2.000E-07 | 2.000E-06 | 2.400E-06 |
| 23178 | 1159 | G1159W | D | 2e-06 | 2.9e-06 | 1.6e-06 | 2.167E-06 | 6.658E-07 | 1.600E-06 | 2.900E-06 |
| 23179 | 1159 | G1159Y | D | 1.7e-06 | 1.2e-06 | 2.5e-06 | 1.800E-06 | 6.557E-07 | 1.200E-06 | 2.500E-06 |
| 23180 | 1159 | G1159V | D | 3.2e-06 | 1.9e-06 | 2.2e-06 | 2.433E-06 | 6.807E-07 | 1.900E-06 | 3.200E-06 |
| 23181 | 1160 | T1160A | D | 1.3e-06 | 4.5e-06 | 1.9e-06 | 2.567E-06 | 1.701E-06 | 1.300E-06 | 4.500E-06 |
| 23182 | 1160 | T1160R | D | 2.4e-06 | 2.8e-06 | 2.1e-06 | 2.433E-06 | 3.512E-07 | 2.100E-06 | 2.800E-06 |
| 23183 | 1160 | T1160N | D | 2.3e-06 | 3.8e-06 | 3e-06 | 3.033E-06 | 7.506E-07 | 2.300E-06 | 3.800E-06 |
| 23184 | 1160 | T1160D | D | 3.3e-06 | 3.4e-06 | 4.9e-06 | 3.867E-06 | 8.963E-07 | 3.300E-06 | 4.900E-06 |
| 23185 | 1160 | T1160C | D | 2.1e-06 | 2.5e-06 | 2.5e-06 | 2.367E-06 | 2.309E-07 | 2.100E-06 | 2.500E-06 |
| 23186 | 1160 | T1160E | D | 2.2e-06 | 2.6e-06 | 2.1e-06 | 2.300E-06 | 2.646E-07 | 2.100E-06 | 2.600E-06 |
| 23187 | 1160 | T1160Q | D | 4.4e-06 | 4.8e-06 | 3.2e-06 | 4.133E-06 | 8.327E-07 | 3.200E-06 | 4.800E-06 |
| 23188 | 1160 | T1160G | D | 4.7e-06 | 2.8e-06 | 2.2e-06 | 3.233E-06 | 1.305E-06 | 2.200E-06 | 4.700E-06 |
| 23189 | 1160 | T1160H | D | 2.3e-06 | 2.2e-06 | 2.6e-06 | 2.367E-06 | 2.082E-07 | 2.200E-06 | 2.600E-06 |
| 23190 | 1160 | T1160I | D | 2.6e-06 | 3.2e-06 | 2.2e-06 | 2.667E-06 | 5.033E-07 | 2.200E-06 | 3.200E-06 |
| 23191 | 1160 | T1160L | D | 2.5e-06 | 2.1e-06 | 2e-06 | 2.200E-06 | 2.646E-07 | 2.000E-06 | 2.500E-06 |
| 23192 | 1160 | T1160K | D | 3.3e-06 | 2.2e-06 | 1.7e-06 | 2.400E-06 | 8.185E-07 | 1.700E-06 | 3.300E-06 |
| 23193 | 1160 | T1160M | D | 2.6e-06 | 3.2e-06 | 1.2e-06 | 2.333E-06 | 1.026E-06 | 1.200E-06 | 3.200E-06 |
| 23194 | 1160 | T1160F | D | 2.8e-06 | 3.1e-06 | 1.3e-06 | 2.400E-06 | 9.644E-07 | 1.300E-06 | 3.100E-06 |
| 23195 | 1160 | T1160P | D | 1.9e-06 | 1.9e-06 | 2.6e-06 | 2.133E-06 | 4.041E-07 | 1.900E-06 | 2.600E-06 |
| 23196 | 1160 | T1160S | D | 3e-06 | 2.4e-06 | 2.3e-06 | 2.567E-06 | 3.786E-07 | 2.300E-06 | 3.000E-06 |
| 23197 | 1160 | T1160T | D | 2.4e-06 | 1.7e-06 | 3.9e-06 | 2.667E-06 | 1.124E-06 | 1.700E-06 | 3.900E-06 |
| 23198 | 1160 | T1160W | D | 1.4e-06 | 2.6e-06 | 1.6e-06 | 1.867E-06 | 6.429E-07 | 1.400E-06 | 2.600E-06 |
| 23199 | 1160 | T1160Y | D | 2.6e-06 | 1.3e-06 | 3.8e-06 | 2.567E-06 | 1.250E-06 | 1.300E-06 | 3.800E-06 |
| 23200 | 1160 | T1160V | D | 2.1e-06 | 1e-06 | 3.5e-06 | 2.200E-06 | 1.253E-06 | 1.000E-06 | 3.500E-06 |
| 23201 | 1161 | K1161A | D | 2e-06 | 2.2e-06 | 4e-06 | 2.733E-06 | 1.102E-06 | 2.000E-06 | 4.000E-06 |
| 23202 | 1161 | K1161R | D | 2e-06 | 1.9e-06 | 1.8e-06 | 1.900E-06 | 1.000E-07 | 1.800E-06 | 2.000E-06 |
| 23203 | 1161 | K1161N | D | 1.9e-06 | 1.5e-06 | 2.2e-06 | 1.867E-06 | 3.512E-07 | 1.500E-06 | 2.200E-06 |
| 23204 | 1161 | K1161D | D | 1.9e-06 | 4.3e-06 | 3.9e-06 | 3.367E-06 | 1.286E-06 | 1.900E-06 | 4.300E-06 |

| | | | | | | | | | | |
|---|---|---|---|---|---|---|---|---|---|---|
| 23205 | 1161 | K1161C | D | 1.8e-06 | 6.2e-06 | 1.4e-06 | 3.133E-06 | 2.663E-06 | 1.400E-06 | 6.200E-06 |
| 23206 | 1161 | K1161E | D | 3.2e-06 | 3.1e-06 | 4.5e-06 | 3.600E-06 | 7.810E-07 | 3.100E-06 | 4.500E-06 |
| 23207 | 1161 | K1161Q | D | 2.4e-06 | 2e-06 | 2.9e-06 | 2.433E-06 | 4.509E-07 | 2.000E-06 | 2.900E-06 |
| 23208 | 1161 | K1161G | D | 2.8e-06 | 2.2e-06 | 2.8e-06 | 2.600E-06 | 3.464E-07 | 2.200E-06 | 2.800E-06 |
| 23209 | 1161 | K1161H | D | 1.3e-06 | 1.2e-06 | 3.4e-06 | 1.967E-06 | 1.242E-06 | 1.200E-06 | 3.400E-06 |
| 23210 | 1161 | K1161I | D | 2.2e-06 | 3.1e-06 | 1.6e-06 | 2.300E-06 | 7.550E-07 | 1.600E-06 | 3.100E-06 |
| 23211 | 1161 | K1161L | D | 3.1e-06 | 1.8e-06 | 1e-06 | 1.967E-06 | 1.060E-06 | 1.000E-06 | 3.100E-06 |
| 23212 | 1161 | K1161K | D | 2.4e-06 | 4.7e-06 | 1.7e-06 | 2.933E-06 | 1.570E-06 | 1.700E-06 | 4.700E-06 |
| 23213 | 1161 | K1161M | D | 3.2e-06 | 2.8e-06 | 2.4e-06 | 2.800E-06 | 4.000E-07 | 2.400E-06 | 3.200E-06 |
| 23214 | 1161 | K1161F | D | 4.6e-06 | 2.4e-06 | 3.3e-06 | 3.433E-06 | 1.106E-06 | 2.400E-06 | 4.600E-06 |
| 23215 | 1161 | K1161P | D | 2.3e-06 | 2.6e-06 | 2e-06 | 2.300E-06 | 3.000E-07 | 2.000E-06 | 2.600E-06 |
| 23216 | 1161 | K1161S | D | 2.5e-06 | 1.5e-06 | 2e-06 | 2.000E-06 | 5.000E-07 | 1.500E-06 | 2.500E-06 |
| 23217 | 1161 | K1161T | D | 2.1e-06 | 2.1e-06 | 2.3e-06 | 2.167E-06 | 1.155E-07 | 2.100E-06 | 2.300E-06 |
| 23218 | 1161 | K1161W | D | 3.2e-06 | 2.9e-06 | 2.9e-06 | 3.000E-06 | 1.732E-07 | 2.900E-06 | 3.200E-06 |
| 23219 | 1161 | K1161Y | D | 2.1e-06 | 4.7e-06 | 1.6e-06 | 2.800E-06 | 1.664E-06 | 1.600E-06 | 4.700E-06 |
| 23220 | 1161 | K1161V | D | 3.7e-06 | 2.4e-06 | 2e-06 | 2.700E-06 | 8.888E-07 | 2.000E-06 | 3.700E-06 |
| 23221 | 1162 | V1162A | D | 2.3e-06 | 2.9e-06 | 1.7e-06 | 2.300E-06 | 6.000E-07 | 1.700E-06 | 2.900E-06 |
| 23222 | 1162 | V1162R | D | 3.8e-06 | 1.5e-06 | 2.4e-06 | 2.567E-06 | 1.159E-06 | 1.500E-06 | 3.800E-06 |
| 23223 | 1162 | V1162N | D | 2.6e-06 | 2.6e-06 | 4.3e-06 | 3.167E-06 | 9.815E-07 | 2.600E-06 | 4.300E-06 |
| 23224 | 1162 | V1162D | D | 3.4e-06 | 2.6e-06 | 3.6e-06 | 3.200E-06 | 5.292E-07 | 2.600E-06 | 3.600E-06 |
| 23225 | 1162 | V1162C | D | 3.3e-06 | 2.5e-06 | 1.1e-06 | 2.300E-06 | 1.114E-06 | 1.100E-06 | 3.300E-06 |
| 23226 | 1162 | V1162E | D | 1.1e-06 | 2.3e-06 | 3.3e-06 | 2.233E-06 | 1.102E-06 | 1.100E-06 | 3.300E-06 |
| 23227 | 1162 | V1162Q | D | 2.3e-06 | 1.6e-06 | 1.7e-06 | 1.867E-06 | 3.786E-07 | 1.600E-06 | 2.300E-06 |
| 23228 | 1162 | V1162G | D | 2.7e-06 | 2.1e-06 | 1.6e-06 | 2.133E-06 | 5.508E-07 | 1.600E-06 | 2.700E-06 |
| 23229 | 1162 | V1162H | D | 1.9e-06 | 1.9e-06 | 2.1e-06 | 1.967E-06 | 1.155E-07 | 1.900E-06 | 2.100E-06 |
| 23230 | 1162 | V1162I | D | 4.4e-06 | 1.4e-06 | 3.1e-06 | 2.967E-06 | 1.504E-06 | 1.400E-06 | 4.400E-06 |
| 23231 | 1162 | V1162L | D | 1.9e-06 | 3.5e-06 | 3.5e-06 | 2.967E-06 | 9.238E-07 | 1.900E-06 | 3.500E-06 |
| 23232 | 1162 | V1162K | D | 4e-06 | 2.2e-06 | 2.7e-06 | 2.967E-06 | 9.292E-07 | 2.200E-06 | 4.000E-06 |
| 23233 | 1162 | V1162M | D | 3.7e-06 | 2.4e-06 | 1.1e-06 | 2.400E-06 | 1.300E-06 | 1.100E-06 | 3.700E-06 |
| 23234 | 1162 | V1162F | D | 1.7e-06 | 1.7e-06 | 2.1e-06 | 1.833E-06 | 2.309E-07 | 1.700E-06 | 2.100E-06 |
| 23235 | 1162 | V1162P | D | 3e-06 | 2.1e-06 | 1.7e-06 | 2.267E-06 | 6.658E-07 | 1.700E-06 | 3.000E-06 |
| 23236 | 1162 | V1162S | D | 2e-06 | 2.7e-06 | 1.8e-06 | 2.167E-06 | 4.726E-07 | 1.800E-06 | 2.700E-06 |
| 23237 | 1162 | V1162T | D | 1.6e-06 | 3.5e-06 | 3.3e-06 | 2.800E-06 | 1.044E-06 | 1.600E-06 | 3.500E-06 |
| 23238 | 1162 | V1162W | D | 4.3e-06 | 1.5e-06 | 4.9e-06 | 3.567E-06 | 1.815E-06 | 1.500E-06 | 4.900E-06 |
| 23239 | 1162 | V1162Y | D | 2.9e-06 | 1.6e-06 | 3e-06 | 2.500E-06 | 7.810E-07 | 1.600E-06 | 3.000E-06 |
| 23240 | 1162 | V1162V | D | 2.1e-06 | 1.8e-06 | 1.7e-06 | 1.867E-06 | 2.082E-07 | 1.700E-06 | 2.100E-06 |
| 23241 | 1163 | E1163A | D | 1.5e-06 | 2.6e-06 | 2e-06 | 2.033E-06 | 5.508E-07 | 1.500E-06 | 2.600E-06 |
| 23242 | 1163 | E1163R | D | 2.1e-06 | 3e-06 | 2e-06 | 2.367E-06 | 5.508E-07 | 2.000E-06 | 3.000E-06 |
| 23243 | 1163 | E1163N | D | 2.3e-06 | 5.5e-06 | 1.8e-06 | 3.200E-06 | 2.007E-06 | 1.800E-06 | 5.500E-06 |
| 23244 | 1163 | E1163D | D | 2.1e-06 | 2.2e-06 | 2.5e-06 | 2.267E-06 | 2.082E-07 | 2.100E-06 | 2.500E-06 |
| 23245 | 1163 | E1163C | D | 2.3e-06 | 3.2e-06 | 1.9e-06 | 2.467E-06 | 6.658E-07 | 1.900E-06 | 3.200E-06 |
| 23246 | 1163 | E1163E | D | 1.6e-06 | 2.2e-06 | 2.2e-06 | 2.000E-06 | 3.464E-07 | 1.600E-06 | 2.200E-06 |
| 23247 | 1163 | E1163Q | D | 1.9e-06 | 2.6e-06 | 1.3e-06 | 1.933E-06 | 6.506E-07 | 1.300E-06 | 2.600E-06 |
| 23248 | 1163 | E1163G | D | 2.2e-06 | 2.2e-06 | 2.1e-06 | 2.167E-06 | 5.774E-08 | 2.100E-06 | 2.200E-06 |
| 23249 | 1163 | E1163H | D | 1.7e-06 | 3.5e-06 | 1.7e-06 | 2.300E-06 | 1.039E-06 | 1.700E-06 | 3.500E-06 |
| 23250 | 1163 | E1163I | D | 2e-06 | 2.2e-06 | 5.1e-06 | 3.100E-06 | 1.735E-06 | 2.000E-06 | 5.100E-06 |
| 23251 | 1163 | E1163L | D | 1.7e-06 | 2.6e-06 | 2.1e-06 | 2.133E-06 | 4.509E-07 | 1.700E-06 | 2.600E-06 |
| 23252 | 1163 | E1163K | D | 1.7e-06 | 1.2e-06 | 3e-06 | 1.967E-06 | 9.292E-07 | 1.200E-06 | 3.000E-06 |
| 23253 | 1163 | E1163M | D | 2.8e-06 | 2.3e-06 | 2.9e-06 | 2.667E-06 | 3.215E-07 | 2.300E-06 | 2.900E-06 |
| 23254 | 1163 | E1163F | D | 4.1e-06 | 2.9e-06 | 3.7e-06 | 3.567E-06 | 6.110E-07 | 2.900E-06 | 4.100E-06 |
| 23255 | 1163 | E1163P | D | 1.8e-06 | 1.1e-06 | 2.4e-06 | 1.767E-06 | 6.506E-07 | 1.100E-06 | 2.400E-06 |
| 23256 | 1163 | E1163S | D | 3e-06 | 2.7e-06 | 1.6e-06 | 2.433E-06 | 7.371E-07 | 1.600E-06 | 3.000E-06 |
| 23257 | 1163 | E1163T | D | 1.7e-06 | 4e-06 | 1.4e-06 | 2.367E-06 | 1.422E-06 | 1.400E-06 | 4.000E-06 |
| 23258 | 1163 | E1163W | D | 2e-06 | 1.6e-06 | 2.2e-06 | 1.933E-06 | 3.055E-07 | 1.600E-06 | 2.200E-06 |
| 23259 | 1163 | E1163Y | D | 2e-06 | 2.6e-06 | 1.7e-06 | 2.100E-06 | 4.583E-07 | 1.700E-06 | 2.600E-06 |
| 23260 | 1163 | E1163V | D | 1.4e-06 | 1.9e-06 | 2.5e-06 | 1.933E-06 | 5.508E-07 | 1.400E-06 | 2.500E-06 |
| 23261 | 1164 | I1164A | D | 1.5e-06 | 1.9e-06 | 1.6e-06 | 1.667E-06 | 2.082E-07 | 1.500E-06 | 1.900E-06 |
| 23262 | 1164 | I1164R | D | 1.6e-06 | 3.3e-06 | 1.6e-06 | 2.167E-06 | 9.815E-07 | 1.600E-06 | 3.300E-06 |
| 23263 | 1164 | I1164N | D | 2.8e-06 | 2.1e-06 | 2.3e-06 | 2.400E-06 | 3.606E-07 | 2.100E-06 | 2.800E-06 |
| 23264 | 1164 | I1164D | D | 1.6e-06 | 3.2e-06 | 2.2e-06 | 2.333E-06 | 8.083E-07 | 1.600E-06 | 3.200E-06 |
| 23265 | 1164 | I1164C | D | 3.5e-06 | 3e-06 | 2.6e-06 | 3.033E-06 | 4.509E-07 | 2.600E-06 | 3.500E-06 |
| 23266 | 1164 | I1164E | D | 2.4e-06 | 2.4e-06 | 2.3e-06 | 2.367E-06 | 5.774E-08 | 2.300E-06 | 2.400E-06 |
| 23267 | 1164 | I1164Q | D | 1.4e-06 | 5.3e-06 | 1.9e-06 | 2.867E-06 | 2.122E-06 | 1.400E-06 | 5.300E-06 |
| 23268 | 1164 | I1164G | D | 4.5e-06 | 1.8e-06 | 3.9e-06 | 3.400E-06 | 1.418E-06 | 1.800E-06 | 4.500E-06 |
| 23269 | 1164 | I1164H | D | 3e-06 | 1.7e-06 | 3.2e-06 | 2.633E-06 | 8.145E-07 | 1.700E-06 | 3.200E-06 |
| 23270 | 1164 | I1164I | D | 1.9e-06 | 2e-06 | 2.2e-06 | 2.033E-06 | 1.528E-07 | 1.900E-06 | 2.200E-06 |
| 23271 | 1164 | I1164L | D | 2.2e-06 | 2.9e-06 | 2.5e-06 | 2.533E-06 | 3.512E-07 | 2.200E-06 | 2.900E-06 |
| 23272 | 1164 | I1164K | D | 1.1e-06 | 2.1e-06 | 2.1e-06 | 1.767E-06 | 5.774E-07 | 1.100E-06 | 2.100E-06 |
| 23273 | 1164 | I1164M | D | 3.6e-06 | 1.7e-06 | 1.4e-06 | 2.233E-06 | 1.193E-06 | 1.400E-06 | 3.600E-06 |
| 23274 | 1164 | I1164F | D | 3e-06 | 2.7e-06 | 2e-06 | 2.567E-06 | 5.132E-07 | 2.000E-06 | 3.000E-06 |
| 23275 | 1164 | I1164P | D | 2.5e-06 | 2.5e-06 | 4.3e-06 | 3.100E-06 | 1.039E-06 | 2.500E-06 | 4.300E-06 |
| 23276 | 1164 | I1164S | D | 1.7e-06 | 3e-06 | 9.2e-07 | 1.873E-06 | 1.051E-06 | 9.200E-07 | 3.000E-06 |
| 23277 | 1164 | I1164T | D | 3.4e-06 | 2.3e-06 | 2.4e-06 | 2.700E-06 | 6.083E-07 | 2.300E-06 | 3.400E-06 |
| 23278 | 1164 | I1164W | D | 3.6e-06 | 2e-06 | 2.5e-06 | 2.700E-06 | 8.185E-07 | 2.000E-06 | 3.600E-06 |
| 23279 | 1164 | I1164Y | D | 1.9e-06 | 2e-06 | 1.2e-06 | 1.700E-06 | 4.359E-07 | 1.200E-06 | 2.000E-06 |
| 23280 | 1164 | I1164V | D | 2.2e-06 | 2.3e-06 | 1.7e-06 | 2.067E-06 | 3.215E-07 | 1.700E-06 | 2.300E-06 |
| 23281 | 1165 | K1165A | D | 2e-06 | 2.2e-06 | 4e-06 | 2.733E-06 | 1.102E-06 | 2.000E-06 | 4.000E-06 |
| 23282 | 1165 | K1165R | D | 2e-06 | 2.5e-06 | 2.8e-06 | 2.433E-06 | 4.041E-07 | 2.000E-06 | 2.800E-06 |
| 23283 | 1165 | K1165N | D | 9.4e-07 | 2.3e-06 | 2.5e-06 | 1.913E-06 | 8.488E-07 | 9.400E-07 | 2.500E-06 |
| 23284 | 1165 | K1165D | D | 2.6e-06 | 2.9e-06 | 1.4e-06 | 2.300E-06 | 7.937E-07 | 1.400E-06 | 2.900E-06 |
| 23285 | 1165 | K1165C | D | 1.2e-06 | 1.9e-06 | 1.9e-06 | 1.667E-06 | 4.041E-07 | 1.200E-06 | 1.900E-06 |
| 23286 | 1165 | K1165E | D | 2.2e-06 | 2.2e-06 | 2.2e-06 | 2.200E-06 | 0.000E+00 | 2.200E-06 | 2.200E-06 |
| 23287 | 1165 | K1165Q | D | 2.1e-06 | 8.5e-07 | 2.6e-06 | 1.850E-06 | 9.014E-07 | 8.500E-07 | 2.600E-06 |
| 23288 | 1165 | K1165G | D | 1.4e-06 | 2.3e-06 | 4.3e-06 | 2.667E-06 | 1.484E-06 | 1.400E-06 | 4.300E-06 |
| 23289 | 1165 | K1165H | D | 3e-06 | 2.1e-06 | 1.9e-06 | 2.333E-06 | 5.859E-07 | 1.900E-06 | 3.000E-06 |

| | | | | | | | | | | |
|---|---|---|---|---|---|---|---|---|---|---|
| 23290 | 1165 | K1165I | D | 2.2e-06 | 2.7e-06 | 3.8e-06 | 2.900E-06 | 8.185E-07 | 2.200E-06 | 3.800E-06 |
| 23291 | 1165 | K1165L | D | 3.2e-06 | 4e-06 | 2.3e-06 | 3.167E-06 | 8.505E-07 | 2.300E-06 | 4.000E-06 |
| 23292 | 1165 | K1165K | D | 3.8e-06 | 3.3e-06 | 3e-06 | 3.367E-06 | 4.041E-07 | 3.000E-06 | 3.800E-06 |
| 23293 | 1165 | K1165M | D | 1.7e-06 | 3.8e-06 | 2.8e-06 | 2.767E-06 | 1.050E-06 | 1.700E-06 | 3.800E-06 |
| 23294 | 1165 | K1165F | D | 2.6e-06 | 2.2e-06 | 3.3e-06 | 2.700E-06 | 5.568E-07 | 2.200E-06 | 3.300E-06 |
| 23295 | 1165 | K1165P | D | 2.5e-06 | 5e-06 | 3.4e-06 | 3.633E-06 | 1.266E-06 | 2.500E-06 | 5.000E-06 |
| 23296 | 1165 | K1165S | D | 3.1e-06 | 2.2e-06 | 2.6e-06 | 2.633E-06 | 4.509E-07 | 2.200E-06 | 3.100E-06 |
| 23297 | 1165 | K1165T | D | 9.8e-07 | 2.7e-06 | 1.7e-06 | 1.793E-06 | 8.638E-07 | 9.800E-07 | 2.700E-06 |
| 23298 | 1165 | K1165W | D | 3.1e-06 | 2.7e-06 | 2.4e-06 | 2.733E-06 | 3.512E-07 | 2.400E-06 | 3.100E-06 |
| 23299 | 1165 | K1165Y | D | 1.8e-06 | 2.6e-06 | 2.3e-06 | 2.233E-06 | 4.041E-07 | 1.800E-06 | 2.600E-06 |
| 23300 | 1165 | K1165V | D | 1.6e-06 | 1.6e-06 | 4.1e-06 | 2.433E-06 | 1.443E-06 | 1.600E-06 | 4.100E-06 |
| 23301 | 1166 | R1166A | D | 3.2e-06 | 2.6e-06 | 2.3e-06 | 2.700E-06 | 4.583E-07 | 2.300E-06 | 3.200E-06 |
| 23302 | 1166 | R1166R | D | 1.6e-06 | 1.9e-06 | 1.2e-06 | 1.567E-06 | 3.512E-07 | 1.200E-06 | 1.900E-06 |
| 23303 | 1166 | R1166N | D | 3e-06 | 1.6e-06 | 3.1e-06 | 2.567E-06 | 8.386E-07 | 1.600E-06 | 3.100E-06 |
| 23304 | 1166 | R1166D | D | 1.8e-06 | 2e-06 | 1.9e-06 | 1.900E-06 | 1.000E-07 | 1.800E-06 | 2.000E-06 |
| 23305 | 1166 | R1166C | D | 3.1e-06 | 1.7e-06 | 3e-06 | 2.600E-06 | 7.810E-07 | 1.700E-06 | 3.100E-06 |
| 23306 | 1166 | R1166E | D | 1.8e-06 | 3.5e-06 | 2.7e-06 | 2.667E-06 | 8.505E-07 | 1.800E-06 | 3.500E-06 |
| 23307 | 1166 | R1166Q | D | 3.2e-06 | 2.5e-06 | 3e-06 | 2.900E-06 | 3.606E-07 | 2.500E-06 | 3.200E-06 |
| 23308 | 1166 | R1166G | D | 3.1e-06 | 3e-06 | 1.8e-06 | 2.633E-06 | 7.234E-07 | 1.800E-06 | 3.100E-06 |
| 23309 | 1166 | R1166H | D | 2.1e-06 | 1.6e-06 | 2.6e-06 | 2.100E-06 | 5.000E-07 | 1.600E-06 | 2.600E-06 |
| 23310 | 1166 | R1166I | D | 2e-06 | 2.3e-06 | 2.4e-06 | 2.233E-06 | 2.082E-07 | 2.000E-06 | 2.400E-06 |
| 23311 | 1166 | R1166L | D | 2.4e-06 | 2.6e-06 | 4.4e-06 | 3.133E-06 | 1.102E-06 | 2.400E-06 | 4.400E-06 |
| 23312 | 1166 | R1166K | D | 2.2e-06 | 1.7e-06 | 3.2e-06 | 2.367E-06 | 7.638E-07 | 1.700E-06 | 3.200E-06 |
| 23313 | 1166 | R1166M | D | 2.2e-06 | 2.5e-06 | 2.7e-06 | 2.467E-06 | 2.517E-07 | 2.200E-06 | 2.700E-06 |
| 23314 | 1166 | R1166F | D | 1.9e-06 | 2.2e-06 | 1.8e-06 | 1.967E-06 | 2.082E-07 | 1.800E-06 | 2.200E-06 |
| 23315 | 1166 | R1166P | D | 1.8e-06 | 1.2e-06 | 1.3e-06 | 1.433E-06 | 3.215E-07 | 1.200E-06 | 1.800E-06 |
| 23316 | 1166 | R1166S | D | 2.4e-06 | 9.6e-07 | 2.7e-06 | 2.020E-06 | 9.302E-07 | 9.600E-07 | 2.700E-06 |
| 23317 | 1166 | R1166T | D | 1.4e-06 | 2.6e-06 | 2.7e-06 | 2.233E-06 | 7.234E-07 | 1.400E-06 | 2.700E-06 |
| 23318 | 1166 | R1166W | D | 2.1e-06 | 1.6e-06 | 2.1e-06 | 1.933E-06 | 2.887E-07 | 1.600E-06 | 2.100E-06 |
| 23319 | 1166 | R1166Y | D | 2.7e-06 | 2.3e-06 | 3e-06 | 2.667E-06 | 3.512E-07 | 2.300E-06 | 3.000E-06 |
| 23320 | 1166 | R1166V | D | 2.8e-06 | 3.3e-06 | 3.3e-06 | 3.133E-06 | 2.887E-07 | 2.800E-06 | 3.300E-06 |
| 23321 | 1167 | T1167A | D | 1.8e-06 | 2.6e-06 | 3.2e-06 | 2.533E-06 | 7.024E-07 | 1.800E-06 | 3.200E-06 |
| 23322 | 1167 | T1167R | D | 3.2e-06 | 2.4e-06 | 3.6e-06 | 3.067E-06 | 6.110E-07 | 2.400E-06 | 3.600E-06 |
| 23323 | 1167 | T1167N | D | 1.5e-06 | 2.3e-06 | 2.3e-06 | 2.033E-06 | 4.619E-07 | 1.500E-06 | 2.300E-06 |
| 23324 | 1167 | T1167D | D | 2e-06 | 2.3e-06 | 2.2e-06 | 2.167E-06 | 1.528E-07 | 2.000E-06 | 2.300E-06 |
| 23325 | 1167 | T1167C | D | 1.8e-06 | 3e-06 | 2e-06 | 2.267E-06 | 6.429E-07 | 1.800E-06 | 3.000E-06 |
| 23326 | 1167 | T1167E | D | 2.4e-06 | 2.7e-06 | 1.7e-06 | 2.267E-06 | 5.132E-07 | 1.700E-06 | 2.700E-06 |
| 23327 | 1167 | T1167Q | D | 1.6e-06 | 2.2e-06 | 2.9e-06 | 2.233E-06 | 6.506E-07 | 1.600E-06 | 2.900E-06 |
| 23328 | 1167 | T1167G | D | 1.8e-06 | 2.9e-06 | 2.4e-06 | 2.367E-06 | 5.508E-07 | 1.800E-06 | 2.900E-06 |
| 23329 | 1167 | T1167H | D | 2.9e-06 | 2.5e-06 | 1.7e-06 | 2.367E-06 | 6.110E-07 | 1.700E-06 | 2.900E-06 |
| 23330 | 1167 | T1167I | D | 2.8e-06 | 2.2e-06 | 2.6e-06 | 2.533E-06 | 3.055E-07 | 2.200E-06 | 2.800E-06 |
| 23331 | 1167 | T1167L | D | 2.3e-06 | 1.8e-06 | 2.6e-06 | 2.233E-06 | 4.041E-07 | 1.800E-06 | 2.600E-06 |
| 23332 | 1167 | T1167K | D | 3.1e-06 | 2.1e-06 | 1.9e-06 | 2.367E-06 | 6.429E-07 | 1.900E-06 | 3.100E-06 |
| 23333 | 1167 | T1167M | D | 1.7e-06 | 2.3e-06 | 4.1e-06 | 2.700E-06 | 1.249E-06 | 1.700E-06 | 4.100E-06 |
| 23334 | 1167 | T1167F | D | 2.2e-06 | 1.7e-06 | 4e-06 | 2.633E-06 | 1.210E-06 | 1.700E-06 | 4.000E-06 |
| 23335 | 1167 | T1167P | D | 3.1e-06 | 3.4e-06 | 2.1e-06 | 2.867E-06 | 6.807E-07 | 2.100E-06 | 3.400E-06 |
| 23336 | 1167 | T1167S | D | 1.6e-06 | 2.8e-06 | 2.1e-06 | 2.167E-06 | 6.028E-07 | 1.600E-06 | 2.800E-06 |
| 23337 | 1167 | T1167T | D | 2.2e-06 | 1.9e-06 | 2.1e-06 | 2.067E-06 | 1.528E-07 | 1.900E-06 | 2.200E-06 |
| 23338 | 1167 | T1167W | D | 3.8e-06 | 2.1e-06 | 2.6e-06 | 2.833E-06 | 8.737E-07 | 2.100E-06 | 3.800E-06 |
| 23339 | 1167 | T1167Y | D | 2.6e-06 | 1.8e-06 | 2.4e-06 | 2.267E-06 | 4.163E-07 | 1.800E-06 | 2.600E-06 |
| 23340 | 1167 | T1167V | D | 3.9e-06 | 1.9e-06 | 3.9e-06 | 3.233E-06 | 1.155E-06 | 1.900E-06 | 3.900E-06 |
| 23341 | 1168 | V1168A | D | 1.4e-06 | 2.3e-06 | 2.9e-06 | 2.200E-06 | 7.550E-07 | 1.400E-06 | 2.900E-06 |
| 23342 | 1168 | V1168R | D | 2.8e-06 | 2.3e-06 | 4.4e-06 | 3.167E-06 | 1.097E-06 | 2.300E-06 | 4.400E-06 |
| 23343 | 1168 | V1168N | D | 3.3e-06 | 2.5e-06 | 2.1e-06 | 2.633E-06 | 6.110E-07 | 2.100E-06 | 3.300E-06 |
| 23344 | 1168 | V1168D | D | 2.5e-06 | 2e-06 | 1.3e-06 | 1.933E-06 | 6.028E-07 | 1.300E-06 | 2.500E-06 |
| 23345 | 1168 | V1168C | D | 1.2e-06 | 1.9e-06 | 2.7e-06 | 1.933E-06 | 7.506E-07 | 1.200E-06 | 2.700E-06 |
| 23346 | 1168 | V1168E | D | 2.8e-06 | 2e-06 | 2.5e-06 | 2.433E-06 | 4.041E-07 | 2.000E-06 | 2.800E-06 |
| 23347 | 1168 | V1168Q | D | 1.8e-06 | 1.7e-06 | 2e-06 | 1.833E-06 | 1.528E-07 | 1.700E-06 | 2.000E-06 |
| 23348 | 1168 | V1168G | D | 3.3e-06 | 1.9e-06 | 3.3e-06 | 2.833E-06 | 8.083E-07 | 1.900E-06 | 3.300E-06 |
| 23349 | 1168 | V1168H | D | 2.6e-06 | 2.1e-06 | 2.3e-06 | 2.333E-06 | 2.517E-07 | 2.100E-06 | 2.600E-06 |
| 23350 | 1168 | V1168I | D | 3.6e-06 | 1.2e-06 | 2.8e-06 | 2.533E-06 | 1.222E-06 | 1.200E-06 | 3.600E-06 |
| 23351 | 1168 | V1168L | D | 1.7e-06 | 1.6e-06 | 3e-06 | 2.100E-06 | 7.810E-07 | 1.600E-06 | 3.000E-06 |
| 23352 | 1168 | V1168K | D | 1.9e-06 | 2.1e-06 | 2.8e-06 | 2.267E-06 | 4.726E-07 | 1.900E-06 | 2.800E-06 |
| 23353 | 1168 | V1168M | D | 2.4e-06 | 2.2e-06 | 3e-06 | 2.533E-06 | 4.163E-07 | 2.200E-06 | 3.000E-06 |
| 23354 | 1168 | V1168F | D | 3.6e-06 | 1.9e-06 | 2.6e-06 | 2.700E-06 | 8.544E-07 | 1.900E-06 | 3.600E-06 |
| 23355 | 1168 | V1168P | D | 3.1e-06 | 2.8e-06 | 2.5e-06 | 2.800E-06 | 3.000E-07 | 2.500E-06 | 3.100E-06 |
| 23356 | 1168 | V1168S | D | 2.6e-06 | 2.7e-06 | 2e-06 | 2.433E-06 | 3.786E-07 | 2.000E-06 | 2.700E-06 |
| 23357 | 1168 | V1168T | D | 2.4e-06 | 2e-06 | 4.2e-06 | 2.867E-06 | 1.172E-06 | 2.000E-06 | 4.200E-06 |
| 23358 | 1168 | V1168W | D | 1.2e-06 | 3.1e-06 | 2.3e-06 | 2.200E-06 | 9.539E-07 | 1.200E-06 | 3.100E-06 |
| 23359 | 1168 | V1168Y | D | 3.1e-06 | 1.5e-06 | 2.3e-06 | 2.300E-06 | 8.000E-07 | 1.500E-06 | 3.100E-06 |
| 23360 | 1168 | V1168V | D | 3.3e-06 | 3e-06 | 2.1e-06 | 2.800E-06 | 6.245E-07 | 2.100E-06 | 3.300E-06 |
| 23361 | 1169 | A1169A | D | 2.2e-06 | 2.5e-06 | 2.6e-06 | 2.433E-06 | 2.082E-07 | 2.200E-06 | 2.600E-06 |
| 23362 | 1169 | A1169R | D | 2.2e-06 | 2.9e-06 | 1.8e-06 | 2.300E-06 | 5.568E-07 | 1.800E-06 | 2.900E-06 |
| 23363 | 1169 | A1169N | D | 1.3e-06 | 1.3e-06 | 2.4e-06 | 1.667E-06 | 6.351E-07 | 1.300E-06 | 2.400E-06 |
| 23364 | 1169 | A1169D | D | 1.8e-06 | 3.5e-06 | 2.9e-06 | 2.733E-06 | 8.622E-07 | 1.800E-06 | 3.500E-06 |
| 23365 | 1169 | A1169C | D | 3e-06 | 1.3e-06 | 2.8e-06 | 2.367E-06 | 9.292E-07 | 1.300E-06 | 3.000E-06 |
| 23366 | 1169 | A1169E | D | 2.2e-06 | 1.7e-06 | 2.3e-06 | 2.067E-06 | 3.215E-07 | 1.700E-06 | 2.300E-06 |
| 23367 | 1169 | A1169Q | D | 1.8e-06 | 2.1e-06 | 4.5e-06 | 2.800E-06 | 1.480E-06 | 1.800E-06 | 4.500E-06 |
| 23368 | 1169 | A1169G | D | 2.7e-06 | 1.8e-06 | 2.2e-06 | 2.233E-06 | 4.509E-07 | 1.800E-06 | 2.700E-06 |
| 23369 | 1169 | A1169H | D | 1.5e-06 | 1.9e-06 | 2.2e-06 | 1.867E-06 | 3.512E-07 | 1.500E-06 | 2.200E-06 |
| 23370 | 1169 | A1169I | D | 1.4e-06 | 2.7e-06 | 3.3e-06 | 2.467E-06 | 9.713E-07 | 1.400E-06 | 3.300E-06 |
| 23371 | 1169 | A1169L | D | 2.3e-06 | 2.5e-06 | 1.6e-06 | 2.133E-06 | 4.726E-07 | 1.600E-06 | 2.500E-06 |
| 23372 | 1169 | A1169K | D | 1.6e-06 | 2.2e-06 | 2.4e-06 | 2.067E-06 | 4.163E-07 | 1.600E-06 | 2.400E-06 |
| 23373 | 1169 | A1169M | D | 2.9e-06 | 2.3e-06 | 1.8e-06 | 2.333E-06 | 5.508E-07 | 1.800E-06 | 2.900E-06 |
| 23374 | 1169 | A1169F | D | 1.3e-06 | 2.8e-06 | 2.4e-06 | 2.167E-06 | 7.767E-07 | 1.300E-06 | 2.800E-06 |

| | | | | | | | | | | |
|---|---|---|---|---|---|---|---|---|---|---|
| 23375 | 1169 | A1169P | D | 2.6e-06 | 3.9e-06 | 3.5e-06 | 3.333E-06 | 6.658E-07 | 2.600E-06 | 3.900E-06 |
| 23376 | 1169 | A1169S | D | 2.3e-06 | 1.6e-06 | 2.6e-06 | 2.167E-06 | 5.132E-07 | 1.600E-06 | 2.600E-06 |
| 23377 | 1169 | A1169T | D | 2.5e-06 | 2.3e-06 | 2.3e-06 | 2.367E-06 | 1.155E-07 | 2.300E-06 | 2.500E-06 |
| 23378 | 1169 | A1169W | D | 1.9e-06 | 2.4e-06 | 2.3e-06 | 2.200E-06 | 2.646E-07 | 1.900E-06 | 2.400E-06 |
| 23379 | 1169 | A1169Y | D | 2.3e-06 | 1.7e-06 | 1.8e-06 | 1.933E-06 | 3.215E-07 | 1.700E-06 | 2.300E-06 |
| 23380 | 1169 | A1169V | D | 3.4e-06 | 3.2e-06 | 3.8e-06 | 3.467E-06 | 3.055E-07 | 3.200E-06 | 3.800E-06 |
| 23381 | 1170 | A1170A | D | 3.4e-06 | 1.3e-06 | 2.5e-06 | 2.400E-06 | 1.054E-06 | 1.300E-06 | 3.400E-06 |
| 23382 | 1170 | A1170R | D | 3e-06 | 2.1e-06 | 2.2e-06 | 2.433E-06 | 4.933E-07 | 2.100E-06 | 3.000E-06 |
| 23383 | 1170 | A1170N | D | 3.1e-06 | 3.2e-06 | 1.8e-06 | 2.700E-06 | 7.810E-07 | 1.800E-06 | 3.200E-06 |
| 23384 | 1170 | A1170D | D | 2e-06 | 2.4e-06 | 3.3e-06 | 2.567E-06 | 6.658E-07 | 2.000E-06 | 3.300E-06 |
| 23385 | 1170 | A1170C | D | 2.2e-06 | 3.5e-06 | 2.5e-06 | 2.733E-06 | 6.807E-07 | 2.200E-06 | 3.500E-06 |
| 23386 | 1170 | A1170E | D | 2.3e-06 | 2.3e-06 | 1.9e-06 | 2.167E-06 | 2.309E-07 | 1.900E-06 | 2.300E-06 |
| 23387 | 1170 | A1170Q | D | 2.6e-06 | 2.1e-06 | 1.6e-06 | 2.100E-06 | 5.000E-07 | 1.600E-06 | 2.600E-06 |
| 23388 | 1170 | A1170G | D | 1.7e-06 | 1.5e-06 | 2.2e-06 | 1.800E-06 | 3.606E-07 | 1.500E-06 | 2.200E-06 |
| 23389 | 1170 | A1170H | D | 1.9e-06 | 3.1e-06 | 3.2e-06 | 2.733E-06 | 7.234E-07 | 1.900E-06 | 3.200E-06 |
| 23390 | 1170 | A1170I | D | 2.4e-06 | 2.5e-06 | 3.5e-06 | 2.800E-06 | 6.083E-07 | 2.400E-06 | 3.500E-06 |
| 23391 | 1170 | A1170L | D | 2.1e-06 | 1.8e-06 | 2.6e-06 | 2.167E-06 | 4.041E-07 | 1.800E-06 | 2.600E-06 |
| 23392 | 1170 | A1170K | D | 1.9e-06 | 1.7e-06 | 2.7e-06 | 2.100E-06 | 5.292E-07 | 1.700E-06 | 2.700E-06 |
| 23393 | 1170 | A1170M | D | 2.1e-06 | 2.7e-06 | 2.1e-06 | 2.300E-06 | 3.464E-07 | 2.100E-06 | 2.700E-06 |
| 23394 | 1170 | A1170F | D | 2e-06 | 1.9e-06 | 2.2e-06 | 2.033E-06 | 1.528E-07 | 1.900E-06 | 2.200E-06 |
| 23395 | 1170 | A1170P | D | 3.9e-06 | 3.1e-06 | 3.1e-06 | 3.367E-06 | 4.619E-07 | 3.100E-06 | 3.900E-06 |
| 23396 | 1170 | A1170S | D | 2e-06 | 4.5e-06 | 1.5e-06 | 2.667E-06 | 1.607E-06 | 1.500E-06 | 4.500E-06 |
| 23397 | 1170 | A1170T | D | 2.2e-06 | 1.7e-06 | 2.2e-06 | 2.033E-06 | 2.887E-07 | 1.700E-06 | 2.200E-06 |
| 23398 | 1170 | A1170W | D | 1.8e-06 | 1.7e-06 | 1.1e-06 | 1.533E-06 | 3.786E-07 | 1.100E-06 | 1.800E-06 |
| 23399 | 1170 | A1170Y | D | 2.9e-06 | 1.8e-06 | 3.4e-06 | 2.700E-06 | 8.185E-07 | 1.800E-06 | 3.400E-06 |
| 23400 | 1170 | A1170V | D | 3.3e-06 | 2.6e-06 | 4e-06 | 3.300E-06 | 7.000E-07 | 2.600E-06 | 4.000E-06 |
| 23401 | 1171 | P1171A | D | 2.6e-06 | 3.2e-06 | 1.6e-06 | 2.467E-06 | 8.083E-07 | 1.600E-06 | 3.200E-06 |
| 23402 | 1171 | P1171R | D | 1.5e-06 | 2.4e-06 | 1.9e-06 | 1.933E-06 | 4.509E-07 | 1.500E-06 | 2.400E-06 |
| 23403 | 1171 | P1171N | D | 2.1e-06 | 4e-06 | 2.6e-06 | 2.900E-06 | 9.849E-07 | 2.100E-06 | 4.000E-06 |
| 23404 | 1171 | P1171D | D | 2.2e-06 | 2.7e-06 | 5.2e-06 | 3.367E-06 | 1.607E-06 | 2.200E-06 | 5.200E-06 |
| 23405 | 1171 | P1171C | D | 3.7e-06 | 3e-06 | 2.9e-06 | 3.200E-06 | 4.359E-07 | 2.900E-06 | 3.700E-06 |
| 23406 | 1171 | P1171E | D | 2e-06 | 2.2e-06 | 2.4e-06 | 2.200E-06 | 2.000E-07 | 2.000E-06 | 2.400E-06 |
| 23407 | 1171 | P1171Q | D | 2e-06 | 1.3e-06 | 2.5e-06 | 1.933E-06 | 6.028E-07 | 1.300E-06 | 2.500E-06 |
| 23408 | 1171 | P1171G | D | 2.6e-06 | 2.2e-06 | 1.3e-06 | 2.033E-06 | 6.658E-07 | 1.300E-06 | 2.600E-06 |
| 23409 | 1171 | P1171H | D | 1.6e-06 | 1.8e-06 | 2.1e-06 | 1.833E-06 | 2.517E-07 | 1.600E-06 | 2.100E-06 |
| 23410 | 1171 | P1171I | D | 2.3e-06 | 2.7e-06 | 3.3e-06 | 2.767E-06 | 5.033E-07 | 2.300E-06 | 3.300E-06 |
| 23411 | 1171 | P1171L | D | 3.7e-06 | 2.1e-06 | 3.2e-06 | 3.000E-06 | 8.185E-07 | 2.100E-06 | 3.700E-06 |
| 23412 | 1171 | P1171K | D | 3.1e-06 | 4e-06 | 2.1e-06 | 3.067E-06 | 9.504E-07 | 2.100E-06 | 4.000E-06 |
| 23413 | 1171 | P1171M | D | 1.7e-06 | 1.4e-06 | 2.1e-06 | 1.733E-06 | 3.512E-07 | 1.400E-06 | 2.100E-06 |
| 23414 | 1171 | P1171F | D | 2.2e-06 | 2.2e-06 | 1.4e-06 | 1.933E-06 | 4.619E-07 | 1.400E-06 | 2.200E-06 |
| 23415 | 1171 | P1171P | D | 4e-06 | 2e-06 | 3.8e-06 | 3.267E-06 | 1.102E-06 | 2.000E-06 | 4.000E-06 |
| 23416 | 1171 | P1171S | D | 1.8e-06 | 1.5e-06 | 2e-06 | 1.767E-06 | 2.517E-07 | 1.500E-06 | 2.000E-06 |
| 23417 | 1171 | P1171T | D | 2.3e-06 | 2.4e-06 | 2.4e-06 | 2.367E-06 | 5.774E-08 | 2.300E-06 | 2.400E-06 |
| 23418 | 1171 | P1171W | D | 3.5e-06 | 1.9e-06 | 2e-06 | 2.467E-06 | 8.963E-07 | 1.900E-06 | 3.500E-06 |
| 23419 | 1171 | P1171Y | D | 1.9e-06 | 2.3e-06 | 2e-06 | 2.067E-06 | 2.082E-07 | 1.900E-06 | 2.300E-06 |
| 23420 | 1171 | P1171V | D | 3.1e-06 | 3.1e-06 | 4.2e-06 | 3.467E-06 | 6.351E-07 | 3.100E-06 | 4.200E-06 |
| 23421 | 1172 | S1172A | D | 2.9e-06 | 2e-06 | 1.7e-06 | 2.200E-06 | 6.245E-07 | 1.700E-06 | 2.900E-06 |
| 23422 | 1172 | S1172R | D | 1.4e-06 | 2.9e-06 | 2.6e-06 | 2.300E-06 | 7.937E-07 | 1.400E-06 | 2.900E-06 |
| 23423 | 1172 | S1172N | D | 2.1e-06 | 2.5e-06 | 2.2e-06 | 2.267E-06 | 2.082E-07 | 2.100E-06 | 2.500E-06 |
| 23424 | 1172 | S1172D | D | 2.9e-06 | 2.6e-06 | 2.4e-06 | 2.633E-06 | 2.517E-07 | 2.400E-06 | 2.900E-06 |
| 23425 | 1172 | S1172C | D | 3.5e-06 | 2.7e-06 | 1.3e-06 | 2.500E-06 | 1.114E-06 | 1.300E-06 | 3.500E-06 |
| 23426 | 1172 | S1172E | D | 2.4e-06 | 2.8e-06 | 2.5e-06 | 2.567E-06 | 2.082E-07 | 2.400E-06 | 2.800E-06 |
| 23427 | 1172 | S1172Q | D | 1.4e-06 | 2.3e-06 | 4.3e-06 | 2.667E-06 | 1.484E-06 | 1.400E-06 | 4.300E-06 |
| 23428 | 1172 | S1172G | D | 3.2e-06 | 1.4e-06 | 2.7e-06 | 2.433E-06 | 9.292E-07 | 1.400E-06 | 3.200E-06 |
| 23429 | 1172 | S1172H | D | 2e-06 | 1.7e-06 | 2.8e-06 | 2.167E-06 | 5.686E-07 | 1.700E-06 | 2.800E-06 |
| 23430 | 1172 | S1172I | D | 9.5e-07 | 3.3e-06 | 3.1e-06 | 2.450E-06 | 1.303E-06 | 9.500E-07 | 3.300E-06 |
| 23431 | 1172 | S1172L | D | 2.1e-06 | 2.5e-06 | 2.1e-06 | 2.233E-06 | 2.309E-07 | 2.100E-06 | 2.500E-06 |
| 23432 | 1172 | S1172K | D | 3.1e-06 | 2.1e-06 | 2.3e-06 | 2.500E-06 | 5.292E-07 | 2.100E-06 | 3.100E-06 |
| 23433 | 1172 | S1172M | D | 3.5e-06 | 1.3e-06 | 2.5e-06 | 2.433E-06 | 1.102E-06 | 1.300E-06 | 3.500E-06 |
| 23434 | 1172 | S1172F | D | 1.5e-06 | 2e-06 | 2e-06 | 1.833E-06 | 2.887E-07 | 1.500E-06 | 2.000E-06 |
| 23435 | 1172 | S1172P | D | 1.5e-06 | 3.4e-06 | 2e-06 | 2.300E-06 | 9.849E-07 | 1.500E-06 | 3.400E-06 |
| 23436 | 1172 | S1172S | D | 1.1e-06 | 2.1e-06 | 3.5e-06 | 2.233E-06 | 1.206E-06 | 1.100E-06 | 3.500E-06 |
| 23437 | 1172 | S1172T | D | 2.1e-06 | 1.4e-06 | 4.7e-06 | 2.733E-06 | 1.739E-06 | 1.400E-06 | 4.700E-06 |
| 23438 | 1172 | S1172W | D | 2.2e-06 | 1.3e-06 | 3.6e-06 | 2.367E-06 | 1.159E-06 | 1.300E-06 | 3.600E-06 |
| 23439 | 1172 | S1172Y | D | 3e-06 | 1.5e-06 | 2e-06 | 2.167E-06 | 7.638E-07 | 1.500E-06 | 3.000E-06 |
| 23440 | 1172 | S1172V | D | 2.9e-06 | 2.6e-06 | 2.3e-06 | 2.600E-06 | 3.000E-07 | 2.300E-06 | 2.900E-06 |
| 23441 | 1173 | V1173A | D | 2.4e-06 | 1.8e-06 | 1.6e-06 | 1.933E-06 | 4.163E-07 | 1.600E-06 | 2.400E-06 |
| 23442 | 1173 | V1173R | D | 2.3e-06 | 2.1e-06 | 2.6e-06 | 2.333E-06 | 2.517E-07 | 2.100E-06 | 2.600E-06 |
| 23443 | 1173 | V1173N | D | 3.1e-06 | 1.6e-06 | 2.3e-06 | 2.333E-06 | 7.506E-07 | 1.600E-06 | 3.100E-06 |
| 23444 | 1173 | V1173D | D | 1.8e-06 | 2.9e-06 | 3.5e-06 | 2.733E-06 | 8.622E-07 | 1.800E-06 | 3.500E-06 |
| 23445 | 1173 | V1173C | D | 2.9e-06 | 1.8e-06 | 3.1e-06 | 2.600E-06 | 7.000E-07 | 1.800E-06 | 3.100E-06 |
| 23446 | 1173 | V1173E | D | 2.7e-06 | 2.3e-06 | 2.3e-06 | 2.433E-06 | 2.309E-07 | 2.300E-06 | 2.700E-06 |
| 23447 | 1173 | V1173Q | D | 2.5e-06 | 2.7e-06 | 2.2e-06 | 2.467E-06 | 2.517E-07 | 2.200E-06 | 2.700E-06 |
| 23448 | 1173 | V1173G | D | 2.5e-06 | 1.2e-06 | 1.8e-06 | 1.833E-06 | 6.506E-07 | 1.200E-06 | 2.500E-06 |
| 23449 | 1173 | V1173H | D | 1.7e-06 | 3e-06 | 2.2e-06 | 2.300E-06 | 6.557E-07 | 1.700E-06 | 3.000E-06 |
| 23450 | 1173 | V1173I | D | 2.5e-06 | 2.8e-06 | 2.5e-06 | 2.600E-06 | 1.732E-07 | 2.500E-06 | 2.800E-06 |
| 23451 | 1173 | V1173L | D | 3.9e-06 | 2.2e-06 | 2.7e-06 | 2.933E-06 | 8.737E-07 | 2.200E-06 | 3.900E-06 |
| 23452 | 1173 | V1173K | D | 4.2e-06 | 1.3e-06 | 2.4e-06 | 2.633E-06 | 1.464E-06 | 1.300E-06 | 4.200E-06 |
| 23453 | 1173 | V1173M | D | 1.9e-06 | 2e-06 | 2.5e-06 | 2.133E-06 | 3.215E-07 | 1.900E-06 | 2.500E-06 |
| 23454 | 1173 | V1173F | D | 2.3e-06 | 2.7e-06 | 4e-06 | 3.000E-06 | 8.888E-07 | 2.300E-06 | 4.000E-06 |
| 23455 | 1173 | V1173P | D | 2.5e-06 | 2.1e-06 | 1.7e-06 | 2.100E-06 | 4.000E-07 | 1.700E-06 | 2.500E-06 |
| 23456 | 1173 | V1173S | D | 2e-06 | 1.7e-06 | 1.6e-06 | 1.767E-06 | 2.082E-07 | 1.600E-06 | 2.000E-06 |
| 23457 | 1173 | V1173T | D | 3.6e-06 | 2.5e-06 | 1.8e-06 | 2.633E-06 | 9.074E-07 | 1.800E-06 | 3.600E-06 |
| 23458 | 1173 | V1173W | D | 3.1e-06 | 2.4e-06 | 2e-06 | 2.500E-06 | 5.568E-07 | 2.000E-06 | 3.100E-06 |
| 23459 | 1173 | V1173Y | D | 4.5e-06 | 2.5e-06 | 1.8e-06 | 2.933E-06 | 1.401E-06 | 1.800E-06 | 4.500E-06 |

| | | | | | | | | | | |
|---|---|---|---|---|---|---|---|---|---|---|
| 23460 | 1173 | V1173V | D | 3.8e-06 | 3.3e-06 | 3e-06 | 3.367E-06 | 4.041E-07 | 3.000E-06 | 3.800E-06 |
| 23461 | 1174 | F1174A | D | 1.3e-06 | 2.3e-06 | 2.7e-06 | 2.100E-06 | 7.211E-07 | 1.300E-06 | 2.700E-06 |
| 23462 | 1174 | F1174R | D | 3.2e-06 | 2.3e-06 | 2.5e-06 | 2.667E-06 | 4.726E-07 | 2.300E-06 | 3.200E-06 |
| 23463 | 1174 | F1174N | D | 2e-06 | 1.9e-06 | 3.2e-06 | 2.367E-06 | 7.234E-07 | 1.900E-06 | 3.200E-06 |
| 23464 | 1174 | F1174D | D | 2.7e-06 | 2.2e-06 | 1.7e-06 | 2.200E-06 | 5.000E-07 | 1.700E-06 | 2.700E-06 |
| 23465 | 1174 | F1174C | D | 2.6e-06 | 1.9e-06 | 9.8e-07 | 1.827E-06 | 8.125E-07 | 9.800E-07 | 2.600E-06 |
| 23466 | 1174 | F1174E | D | 2.6e-06 | 3.4e-06 | 4.3e-06 | 3.433E-06 | 8.505E-07 | 2.600E-06 | 4.300E-06 |
| 23467 | 1174 | F1174Q | D | 2.4e-06 | 2.3e-06 | 3e-06 | 2.567E-06 | 3.786E-07 | 2.300E-06 | 3.000E-06 |
| 23468 | 1174 | F1174G | D | 1.3e-06 | 3.1e-06 | 2e-06 | 2.133E-06 | 9.074E-07 | 1.300E-06 | 3.100E-06 |
| 23469 | 1174 | F1174H | D | 2.8e-06 | 2.5e-06 | 1.6e-06 | 2.300E-06 | 6.245E-07 | 1.600E-06 | 2.800E-06 |
| 23470 | 1174 | F1174I | D | 3.5e-06 | 4e-06 | 2.3e-06 | 3.267E-06 | 8.737E-07 | 2.300E-06 | 4.000E-06 |
| 23471 | 1174 | F1174L | D | 2e-06 | 1.7e-06 | 2.9e-06 | 2.200E-06 | 6.245E-07 | 1.700E-06 | 2.900E-06 |
| 23472 | 1174 | F1174K | D | 2.5e-06 | 3.2e-06 | 2.5e-06 | 2.733E-06 | 4.041E-07 | 2.500E-06 | 3.200E-06 |
| 23473 | 1174 | F1174M | D | 5e-06 | 2.4e-06 | 1.7e-06 | 3.033E-06 | 1.739E-06 | 1.700E-06 | 5.000E-06 |
| 23474 | 1174 | F1174F | D | 1.5e-06 | 1.5e-06 | 3.1e-06 | 2.033E-06 | 9.238E-07 | 1.500E-06 | 3.100E-06 |
| 23475 | 1174 | F1174P | D | 2.3e-06 | 3.9e-06 | 2.2e-06 | 2.800E-06 | 9.539E-07 | 2.200E-06 | 3.900E-06 |
| 23476 | 1174 | F1174S | D | 2.3e-06 | 1.5e-06 | 2.8e-06 | 2.200E-06 | 6.557E-07 | 1.500E-06 | 2.800E-06 |
| 23477 | 1174 | F1174T | D | 1.8e-06 | 2.5e-06 | 3.4e-06 | 2.567E-06 | 8.021E-07 | 1.800E-06 | 3.400E-06 |
| 23478 | 1174 | F1174W | D | 2.5e-06 | 2.3e-06 | 2.3e-06 | 2.367E-06 | 1.155E-07 | 2.300E-06 | 2.500E-06 |
| 23479 | 1174 | F1174Y | D | 2.7e-06 | 2.1e-06 | 2e-06 | 2.267E-06 | 3.786E-07 | 2.000E-06 | 2.700E-06 |
| 23480 | 1174 | F1174V | D | 3.1e-06 | 2.9e-06 | 2.6e-06 | 2.867E-06 | 2.517E-07 | 2.600E-06 | 3.100E-06 |
| 23481 | 1175 | I1175A | D | 1.4e-06 | 2.6e-06 | 2.1e-06 | 2.033E-06 | 6.028E-07 | 1.400E-06 | 2.600E-06 |
| 23482 | 1175 | I1175R | D | 2.2e-06 | 3.1e-06 | 2.4e-06 | 2.567E-06 | 4.726E-07 | 2.200E-06 | 3.100E-06 |
| 23483 | 1175 | I1175N | D | 3.6e-06 | 3.1e-06 | 9.8e-07 | 2.560E-06 | 1.391E-06 | 9.800E-07 | 3.600E-06 |
| 23484 | 1175 | I1175D | D | 2.2e-06 | 2.3e-06 | 2e-06 | 2.167E-06 | 1.528E-07 | 2.000E-06 | 2.300E-06 |
| 23485 | 1175 | I1175C | D | 2.8e-06 | 3.9e-06 | 1.9e-06 | 2.867E-06 | 1.002E-06 | 1.900E-06 | 3.900E-06 |
| 23486 | 1175 | I1175E | D | 3.1e-06 | 2e-06 | 2.2e-06 | 2.433E-06 | 5.859E-07 | 2.000E-06 | 3.100E-06 |
| 23487 | 1175 | I1175Q | D | 1.5e-06 | 2e-06 | 2.1e-06 | 1.867E-06 | 3.215E-07 | 1.500E-06 | 2.100E-06 |
| 23488 | 1175 | I1175G | D | 2.4e-06 | 1.8e-06 | 3.1e-06 | 2.433E-06 | 6.506E-07 | 1.800E-06 | 3.100E-06 |
| 23489 | 1175 | I1175H | D | 5e-06 | 2.7e-06 | 1.7e-06 | 3.133E-06 | 1.692E-06 | 1.700E-06 | 5.000E-06 |
| 23490 | 1175 | I1175I | D | 2.8e-06 | 2.3e-06 | 2.7e-06 | 2.600E-06 | 2.646E-07 | 2.300E-06 | 2.800E-06 |
| 23491 | 1175 | I1175L | D | 2e-06 | 3.5e-06 | 1.8e-06 | 2.433E-06 | 9.292E-07 | 1.800E-06 | 3.500E-06 |
| 23492 | 1175 | I1175K | D | 2.7e-06 | 1.3e-06 | 1.6e-06 | 1.867E-06 | 7.371E-07 | 1.300E-06 | 2.700E-06 |
| 23493 | 1175 | I1175M | D | 1.3e-06 | 2.7e-06 | 2.6e-06 | 2.200E-06 | 7.810E-07 | 1.300E-06 | 2.700E-06 |
| 23494 | 1175 | I1175F | D | 1.4e-06 | 2.5e-06 | 9.1e-07 | 1.603E-06 | 8.143E-07 | 9.100E-07 | 2.500E-06 |
| 23495 | 1175 | I1175P | D | 2e-06 | 3.3e-06 | 1.9e-06 | 2.400E-06 | 7.810E-07 | 1.900E-06 | 3.300E-06 |
| 23496 | 1175 | I1175S | D | 2.2e-06 | 2.1e-06 | 4.2e-06 | 2.833E-06 | 1.185E-06 | 2.100E-06 | 4.200E-06 |
| 23497 | 1175 | I1175T | D | 3.9e-06 | 1.7e-06 | 2.5e-06 | 2.700E-06 | 1.114E-06 | 1.700E-06 | 3.900E-06 |
| 23498 | 1175 | I1175W | D | 2e-06 | 2.3e-06 | 3e-06 | 2.433E-06 | 5.132E-07 | 2.000E-06 | 3.000E-06 |
| 23499 | 1175 | I1175Y | D | 2.6e-06 | 1.7e-06 | 3.5e-06 | 2.600E-06 | 9.000E-07 | 1.700E-06 | 3.500E-06 |
| 23500 | 1175 | I1175V | D | 2.7e-06 | 5e-06 | 1.9e-06 | 3.200E-06 | 1.609E-06 | 1.900E-06 | 5.000E-06 |
| 23501 | 1176 | F1176A | D | 2.5e-06 | 2.8e-06 | 2.2e-06 | 2.500E-06 | 3.000E-07 | 2.200E-06 | 2.800E-06 |
| 23502 | 1176 | F1176R | D | 2.4e-06 | 3.9e-06 | 3.3e-06 | 3.200E-06 | 7.550E-07 | 2.400E-06 | 3.900E-06 |
| 23503 | 1176 | F1176N | D | 2.2e-06 | 1.8e-06 | 2.2e-06 | 2.067E-06 | 2.309E-07 | 1.800E-06 | 2.200E-06 |
| 23504 | 1176 | F1176D | D | 9.3e-07 | 3.2e-06 | 1.1e-06 | 1.743E-06 | 1.264E-06 | 9.300E-07 | 3.200E-06 |
| 23505 | 1176 | F1176C | D | 4.2e-06 | 2.3e-06 | 2.6e-06 | 3.033E-06 | 1.021E-06 | 2.300E-06 | 4.200E-06 |
| 23506 | 1176 | F1176E | D | 2.6e-06 | 2.6e-06 | 3.6e-06 | 2.933E-06 | 5.774E-07 | 2.600E-06 | 3.600E-06 |
| 23507 | 1176 | F1176Q | D | 3e-06 | 1.9e-06 | 2.4e-06 | 2.433E-06 | 5.508E-07 | 1.900E-06 | 3.000E-06 |
| 23508 | 1176 | F1176G | D | 4.1e-06 | 2.8e-06 | 2.2e-06 | 3.033E-06 | 9.713E-07 | 2.200E-06 | 4.100E-06 |
| 23509 | 1176 | F1176H | D | 2.9e-06 | 2.1e-06 | 2.6e-06 | 2.533E-06 | 4.041E-07 | 2.100E-06 | 2.900E-06 |
| 23510 | 1176 | F1176I | D | 3.6e-06 | 3.1e-06 | 2.7e-06 | 3.133E-06 | 4.509E-07 | 2.700E-06 | 3.600E-06 |
| 23511 | 1176 | F1176L | D | 2.2e-06 | 2.8e-06 | 2.9e-06 | 2.633E-06 | 3.786E-07 | 2.200E-06 | 2.900E-06 |
| 23512 | 1176 | F1176K | D | 2.5e-06 | 3.4e-06 | 2.1e-06 | 2.667E-06 | 6.658E-07 | 2.100E-06 | 3.400E-06 |
| 23513 | 1176 | F1176M | D | 3e-06 | 2.4e-06 | 2.8e-06 | 2.733E-06 | 3.055E-07 | 2.400E-06 | 3.000E-06 |
| 23514 | 1176 | F1176F | D | 3e-06 | 9.6e-07 | 2e-06 | 1.987E-06 | 1.020E-06 | 9.600E-07 | 3.000E-06 |
| 23515 | 1176 | F1176P | D | 4.8e-06 | 2.1e-06 | 2.5e-06 | 3.133E-06 | 1.457E-06 | 2.100E-06 | 4.800E-06 |
| 23516 | 1176 | F1176S | D | 9.8e-07 | 2.9e-06 | 2.8e-06 | 2.227E-06 | 1.081E-06 | 9.800E-07 | 2.900E-06 |
| 23517 | 1176 | F1176T | D | 3.1e-06 | 2.9e-06 | 2e-06 | 2.667E-06 | 5.859E-07 | 2.000E-06 | 3.100E-06 |
| 23518 | 1176 | F1176W | D | 2.9e-06 | 1.6e-06 | 4.1e-06 | 2.867E-06 | 1.250E-06 | 1.600E-06 | 4.100E-06 |
| 23519 | 1176 | F1176Y | D | 3.7e-06 | 1.3e-06 | 2.6e-06 | 2.533E-06 | 1.201E-06 | 1.300E-06 | 3.700E-06 |
| 23520 | 1176 | F1176V | D | 3.7e-06 | 2.8e-06 | 1.5e-06 | 2.667E-06 | 1.106E-06 | 1.500E-06 | 3.700E-06 |
| 23521 | 1177 | P1177A | D | 1.1e-06 | 2.7e-06 | 3.1e-06 | 2.300E-06 | 1.058E-06 | 1.100E-06 | 3.100E-06 |
| 23522 | 1177 | P1177R | D | 1.9e-06 | 1.5e-06 | 1.6e-06 | 1.667E-06 | 2.082E-07 | 1.500E-06 | 1.900E-06 |
| 23523 | 1177 | P1177N | D | 2.9e-06 | 2.7e-06 | 3.3e-06 | 2.967E-06 | 3.055E-07 | 2.700E-06 | 3.300E-06 |
| 23524 | 1177 | P1177D | D | 3.1e-06 | 4e-06 | 3.7e-06 | 3.600E-06 | 4.583E-07 | 3.100E-06 | 4.000E-06 |
| 23525 | 1177 | P1177C | D | 1.5e-06 | 3e-06 | 3.8e-06 | 2.767E-06 | 1.168E-06 | 1.500E-06 | 3.800E-06 |
| 23526 | 1177 | P1177E | D | 8.5e-07 | 1.8e-06 | 2.8e-06 | 1.817E-06 | 9.751E-07 | 8.500E-07 | 2.800E-06 |
| 23527 | 1177 | P1177Q | D | 1.1e-06 | 2.4e-06 | 1.7e-06 | 1.733E-06 | 6.506E-07 | 1.100E-06 | 2.400E-06 |
| 23528 | 1177 | P1177G | D | 1.1e-06 | 2.6e-06 | 1.5e-06 | 1.733E-06 | 7.767E-07 | 1.100E-06 | 2.600E-06 |
| 23529 | 1177 | P1177H | D | 1.1e-06 | 2.1e-06 | 3.7e-06 | 2.300E-06 | 1.311E-06 | 1.100E-06 | 3.700E-06 |
| 23530 | 1177 | P1177I | D | 1.5e-06 | 2.7e-06 | 2.3e-06 | 2.167E-06 | 6.110E-07 | 1.500E-06 | 2.700E-06 |
| 23531 | 1177 | P1177L | D | 2.6e-06 | 1.5e-06 | 2.1e-06 | 2.067E-06 | 5.508E-07 | 1.500E-06 | 2.600E-06 |
| 23532 | 1177 | P1177K | D | 2.1e-06 | 2.3e-06 | 2.1e-06 | 2.167E-06 | 1.155E-07 | 2.100E-06 | 2.300E-06 |
| 23533 | 1177 | P1177M | D | 3.1e-06 | 4.4e-06 | 3.3e-06 | 3.600E-06 | 7.000E-07 | 3.100E-06 | 4.400E-06 |
| 23534 | 1177 | P1177F | D | 1.6e-06 | 2.8e-06 | 2.1e-06 | 2.167E-06 | 6.028E-07 | 1.600E-06 | 2.800E-06 |
| 23535 | 1177 | P1177P | D | 3.6e-06 | 3.7e-06 | 1.7e-06 | 3.000E-06 | 1.127E-06 | 1.700E-06 | 3.700E-06 |
| 23536 | 1177 | P1177S | D | 2.3e-06 | 2.6e-06 | 1.7e-06 | 2.200E-06 | 4.583E-07 | 1.700E-06 | 2.600E-06 |
| 23537 | 1177 | P1177T | D | 3.6e-06 | 4.4e-06 | 3.6e-06 | 3.867E-06 | 4.619E-07 | 3.600E-06 | 4.400E-06 |
| 23538 | 1177 | P1177W | D | 2.1e-06 | 2.4e-06 | 2.3e-06 | 2.267E-06 | 1.528E-07 | 2.100E-06 | 2.400E-06 |
| 23539 | 1177 | P1177Y | D | 2.1e-06 | 1.1e-06 | 2.4e-06 | 1.867E-06 | 6.807E-07 | 1.100E-06 | 2.400E-06 |
| 23540 | 1177 | P1177V | D | 1.9e-06 | 1.9e-06 | 2.9e-06 | 2.233E-06 | 5.774E-07 | 1.900E-06 | 2.900E-06 |
| 23541 | 1178 | P1178A | D | 1.3e-06 | 3.6e-06 | 2.3e-06 | 2.400E-06 | 1.153E-06 | 1.300E-06 | 3.600E-06 |
| 23542 | 1178 | P1178R | D | 3e-06 | 2.7e-06 | 2.7e-06 | 2.800E-06 | 1.732E-07 | 2.700E-06 | 3.000E-06 |
| 23543 | 1178 | P1178N | D | 2.3e-06 | 1.6e-06 | 4e-06 | 2.633E-06 | 1.234E-06 | 1.600E-06 | 4.000E-06 |
| 23544 | 1178 | P1178D | D | 3.5e-06 | 2.8e-06 | 3.1e-06 | 3.133E-06 | 3.512E-07 | 2.800E-06 | 3.500E-06 |

| | | | | | | | | | | |
|---|---|---|---|---|---|---|---|---|---|---|
| 23545 | 1178 | P1178C | D | 2.4e-06 | 1.8e-06 | 3.8e-06 | 2.667E-06 | 1.026E-06 | 1.800E-06 | 3.800E-06 |
| 23546 | 1178 | P1178E | D | 1.2e-06 | 3e-06 | 1.9e-06 | 2.033E-06 | 9.074E-07 | 1.200E-06 | 3.000E-06 |
| 23547 | 1178 | P1178Q | D | 2.5e-06 | 3.5e-06 | 2.2e-06 | 2.733E-06 | 6.807E-07 | 2.200E-06 | 3.500E-06 |
| 23548 | 1178 | P1178G | D | 2.4e-06 | 2.6e-06 | 2.8e-06 | 2.600E-06 | 2.000E-07 | 2.400E-06 | 2.800E-06 |
| 23549 | 1178 | P1178H | D | 2.6e-06 | 1.3e-06 | 2.6e-06 | 2.167E-06 | 7.506E-07 | 1.300E-06 | 2.600E-06 |
| 23550 | 1178 | P1178I | D | 2.5e-06 | 2.2e-06 | 2.9e-06 | 2.533E-06 | 3.512E-07 | 2.200E-06 | 2.900E-06 |
| 23551 | 1178 | P1178L | D | 1.8e-06 | 3.4e-06 | 1.7e-06 | 2.300E-06 | 9.539E-07 | 1.700E-06 | 3.400E-06 |
| 23552 | 1178 | P1178K | D | 1.6e-06 | 2.9e-06 | 3.3e-06 | 2.600E-06 | 8.888E-07 | 1.600E-06 | 3.300E-06 |
| 23553 | 1178 | P1178M | D | 2.6e-06 | 2.8e-06 | 2.6e-06 | 2.667E-06 | 1.155E-07 | 2.600E-06 | 2.800E-06 |
| 23554 | 1178 | P1178F | D | 2.7e-06 | 2.2e-06 | 2.2e-06 | 2.367E-06 | 2.887E-07 | 2.200E-06 | 2.700E-06 |
| 23555 | 1178 | P1178P | D | 2.7e-06 | 1.9e-06 | 3.3e-06 | 2.633E-06 | 7.024E-07 | 1.900E-06 | 3.300E-06 |
| 23556 | 1178 | P1178S | D | 2.8e-06 | 1.2e-06 | 2.1e-06 | 2.033E-06 | 8.021E-07 | 1.200E-06 | 2.800E-06 |
| 23557 | 1178 | P1178T | D | 2.8e-06 | 1.3e-06 | 2.6e-06 | 2.233E-06 | 8.145E-07 | 1.300E-06 | 2.800E-06 |
| 23558 | 1178 | P1178W | D | 1.7e-06 | 2.5e-06 | 2.9e-06 | 2.367E-06 | 6.110E-07 | 1.700E-06 | 2.900E-06 |
| 23559 | 1178 | P1178Y | D | 3.1e-06 | 2.1e-06 | 2.6e-06 | 2.600E-06 | 5.000E-07 | 2.100E-06 | 3.100E-06 |
| 23560 | 1178 | P1178V | D | 4e-06 | 3.9e-06 | 1.8e-06 | 3.233E-06 | 1.242E-06 | 1.800E-06 | 4.000E-06 |
| 23561 | 1179 | S1179A | D | 4.6e-06 | 2.8e-06 | 1.2e-06 | 2.867E-06 | 1.701E-06 | 1.200E-06 | 4.600E-06 |
| 23562 | 1179 | S1179R | D | 1.6e-06 | 2.3e-06 | 2.8e-06 | 2.233E-06 | 6.028E-07 | 1.600E-06 | 2.800E-06 |
| 23563 | 1179 | S1179N | D | 1.3e-06 | 2.7e-06 | 3.5e-06 | 2.500E-06 | 1.114E-06 | 1.300E-06 | 3.500E-06 |
| 23564 | 1179 | S1179D | D | 5.8e-06 | 2.5e-06 | 2.9e-06 | 3.733E-06 | 1.801E-06 | 2.500E-06 | 5.800E-06 |
| 23565 | 1179 | S1179C | D | 1.7e-06 | 2.3e-06 | 1.8e-06 | 1.933E-06 | 3.215E-07 | 1.700E-06 | 2.300E-06 |
| 23566 | 1179 | S1179E | D | 3.8e-06 | 1.8e-06 | 4.3e-06 | 3.300E-06 | 1.323E-06 | 1.800E-06 | 4.300E-06 |
| 23567 | 1179 | S1179Q | D | 2.5e-06 | 1.9e-06 | 3.3e-06 | 2.567E-06 | 7.024E-07 | 1.900E-06 | 3.300E-06 |
| 23568 | 1179 | S1179G | D | 3.2e-06 | 1.6e-06 | 2.7e-06 | 2.500E-06 | 8.185E-07 | 1.600E-06 | 3.200E-06 |
| 23569 | 1179 | S1179H | D | 2.3e-06 | 3.2e-06 | 4e-06 | 3.167E-06 | 8.505E-07 | 2.300E-06 | 4.000E-06 |
| 23570 | 1179 | S1179I | D | 2.2e-06 | 2.6e-06 | 2.1e-06 | 2.300E-06 | 2.646E-07 | 2.100E-06 | 2.600E-06 |
| 23571 | 1179 | S1179L | D | 2.2e-06 | 4e-06 | 1.4e-06 | 2.533E-06 | 1.332E-06 | 1.400E-06 | 4.000E-06 |
| 23572 | 1179 | S1179K | D | 3.1e-06 | 1.2e-06 | 2.8e-06 | 2.367E-06 | 1.021E-06 | 1.200E-06 | 3.100E-06 |
| 23573 | 1179 | S1179M | D | 2.6e-06 | 2e-06 | 1.6e-06 | 2.067E-06 | 5.033E-07 | 1.600E-06 | 2.600E-06 |
| 23574 | 1179 | S1179F | D | 1.8e-06 | 1.9e-06 | 2.7e-06 | 2.133E-06 | 4.933E-07 | 1.800E-06 | 2.700E-06 |
| 23575 | 1179 | S1179P | D | 3.3e-06 | 1.9e-06 | 2.7e-06 | 2.633E-06 | 7.024E-07 | 1.900E-06 | 3.300E-06 |
| 23576 | 1179 | S1179S | D | 2.9e-06 | 2.6e-06 | 2.1e-06 | 2.533E-06 | 4.041E-07 | 2.100E-06 | 2.900E-06 |
| 23577 | 1179 | S1179T | D | 2.2e-06 | 2.7e-06 | 1.9e-06 | 2.267E-06 | 4.041E-07 | 1.900E-06 | 2.700E-06 |
| 23578 | 1179 | S1179W | D | 2.6e-06 | 3.6e-06 | 2.4e-06 | 2.867E-06 | 6.429E-07 | 2.400E-06 | 3.600E-06 |
| 23579 | 1179 | S1179Y | D | 2e-06 | 1.5e-06 | 1.8e-06 | 1.767E-06 | 2.517E-07 | 1.500E-06 | 2.000E-06 |
| 23580 | 1179 | S1179V | D | 1.6e-06 | 2.8e-06 | 2.7e-06 | 2.367E-06 | 6.658E-07 | 1.600E-06 | 2.800E-06 |
| 23581 | 1180 | D1180A | D | 1.2e-06 | 2.1e-06 | 2.5e-06 | 1.933E-06 | 6.658E-07 | 1.200E-06 | 2.500E-06 |
| 23582 | 1180 | D1180R | D | 2.6e-06 | 1.5e-06 | 2.8e-06 | 2.300E-06 | 7.000E-07 | 1.500E-06 | 2.800E-06 |
| 23583 | 1180 | D1180N | D | 2.2e-06 | 2.5e-06 | 1.9e-06 | 2.200E-06 | 3.000E-07 | 1.900E-06 | 2.500E-06 |
| 23584 | 1180 | D1180D | D | 1.7e-06 | 2.5e-06 | 3.1e-06 | 2.433E-06 | 7.024E-07 | 1.700E-06 | 3.100E-06 |
| 23585 | 1180 | D1180C | D | 3.1e-06 | 2.7e-06 | 2.5e-06 | 2.767E-06 | 3.055E-07 | 2.500E-06 | 3.100E-06 |
| 23586 | 1180 | D1180E | D | 3.1e-06 | 2.6e-06 | 2.1e-06 | 2.600E-06 | 5.000E-07 | 2.100E-06 | 3.100E-06 |
| 23587 | 1180 | D1180Q | D | 2.6e-06 | 2.2e-06 | 1.9e-06 | 2.233E-06 | 3.512E-07 | 1.900E-06 | 2.600E-06 |
| 23588 | 1180 | D1180G | D | 2.8e-06 | 2e-06 | 1.9e-06 | 2.233E-06 | 4.933E-07 | 1.900E-06 | 2.800E-06 |
| 23589 | 1180 | D1180H | D | 2.2e-06 | 2.9e-06 | 2.1e-06 | 2.400E-06 | 4.359E-07 | 2.100E-06 | 2.900E-06 |
| 23590 | 1180 | D1180I | D | 2.4e-06 | 2.3e-06 | 2.5e-06 | 2.400E-06 | 1.000E-07 | 2.300E-06 | 2.500E-06 |
| 23591 | 1180 | D1180L | D | 2e-06 | 5.3e-06 | 2e-06 | 3.100E-06 | 1.905E-06 | 2.000E-06 | 5.300E-06 |
| 23592 | 1180 | D1180K | D | 2.7e-06 | 3.1e-06 | 2.2e-06 | 2.667E-06 | 4.509E-07 | 2.200E-06 | 3.100E-06 |
| 23593 | 1180 | D1180M | D | 2.6e-06 | 2.6e-06 | 4e-06 | 3.067E-06 | 8.083E-07 | 2.600E-06 | 4.000E-06 |
| 23594 | 1180 | D1180F | D | 2.1e-06 | 1.6e-06 | 2.7e-06 | 2.133E-06 | 5.508E-07 | 1.600E-06 | 2.700E-06 |
| 23595 | 1180 | D1180P | D | 2.4e-06 | 2e-06 | 2e-06 | 2.133E-06 | 2.309E-07 | 2.000E-06 | 2.400E-06 |
| 23596 | 1180 | D1180S | D | 2.3e-06 | 2.6e-06 | 2.8e-06 | 2.567E-06 | 2.517E-07 | 2.300E-06 | 2.800E-06 |
| 23597 | 1180 | D1180T | D | 2.8e-06 | 2.1e-06 | 2.5e-06 | 2.467E-06 | 3.512E-07 | 2.100E-06 | 2.800E-06 |
| 23598 | 1180 | D1180W | D | 1.6e-06 | 3.6e-06 | 1.8e-06 | 2.333E-06 | 1.102E-06 | 1.600E-06 | 3.600E-06 |
| 23599 | 1180 | D1180Y | D | 1.5e-06 | 3e-06 | 3.8e-06 | 2.767E-06 | 1.168E-06 | 1.500E-06 | 3.800E-06 |
| 23600 | 1180 | D1180V | D | 5e-06 | 3.1e-06 | 2e-06 | 3.367E-06 | 1.518E-06 | 2.000E-06 | 5.000E-06 |
| 23601 | 1181 | E1181A | D | 2.5e-06 | 2.2e-06 | 2.2e-06 | 2.300E-06 | 1.732E-07 | 2.200E-06 | 2.500E-06 |
| 23602 | 1181 | E1181R | D | 2.4e-06 | 2.8e-06 | 2.6e-06 | 2.600E-06 | 2.000E-07 | 2.400E-06 | 2.800E-06 |
| 23603 | 1181 | E1181N | D | 2.9e-06 | 3.1e-06 | 1.8e-06 | 2.600E-06 | 7.000E-07 | 1.800E-06 | 3.100E-06 |
| 23604 | 1181 | E1181D | D | 2.5e-06 | 1.9e-06 | 2.3e-06 | 2.233E-06 | 3.055E-07 | 1.900E-06 | 2.500E-06 |
| 23605 | 1181 | E1181C | D | 2.1e-06 | 3.5e-06 | 1.3e-06 | 2.300E-06 | 1.114E-06 | 1.300E-06 | 3.500E-06 |
| 23606 | 1181 | E1181E | D | 1.8e-06 | 2.5e-06 | 3.4e-06 | 2.567E-06 | 8.021E-07 | 1.800E-06 | 3.400E-06 |
| 23607 | 1181 | E1181Q | D | 1.4e-06 | 4.4e-06 | 2.2e-06 | 2.667E-06 | 1.553E-06 | 1.400E-06 | 4.400E-06 |
| 23608 | 1181 | E1181G | D | 3e-06 | 2.5e-06 | 2.2e-06 | 2.567E-06 | 4.041E-07 | 2.200E-06 | 3.000E-06 |
| 23609 | 1181 | E1181H | D | 2.2e-06 | 1.4e-06 | 2.5e-06 | 2.033E-06 | 5.686E-07 | 1.400E-06 | 2.500E-06 |
| 23610 | 1181 | E1181I | D | 2e-06 | 2.6e-06 | 2e-06 | 2.200E-06 | 3.464E-07 | 2.000E-06 | 2.600E-06 |
| 23611 | 1181 | E1181L | D | 2.8e-06 | 2.9e-06 | 2e-06 | 2.567E-06 | 4.933E-07 | 2.000E-06 | 2.900E-06 |
| 23612 | 1181 | E1181K | D | 1.7e-06 | 1.4e-06 | 1.9e-06 | 1.667E-06 | 2.517E-07 | 1.400E-06 | 1.900E-06 |
| 23613 | 1181 | E1181M | D | 2.3e-06 | 2e-06 | 2.5e-06 | 2.267E-06 | 2.517E-07 | 2.000E-06 | 2.500E-06 |
| 23614 | 1181 | E1181F | D | 3.1e-06 | 2.7e-06 | 3.9e-06 | 3.233E-06 | 6.110E-07 | 2.700E-06 | 3.900E-06 |
| 23615 | 1181 | E1181P | D | 1.6e-06 | 1.6e-06 | 2.8e-06 | 2.000E-06 | 6.928E-07 | 1.600E-06 | 2.800E-06 |
| 23616 | 1181 | E1181S | D | 1.7e-06 | 2.1e-06 | 1.6e-06 | 1.800E-06 | 2.646E-07 | 1.600E-06 | 2.100E-06 |
| 23617 | 1181 | E1181T | D | 2.7e-06 | 2e-06 | 2e-06 | 2.233E-06 | 4.041E-07 | 2.000E-06 | 2.700E-06 |
| 23618 | 1181 | E1181W | D | 1.3e-06 | 2.7e-06 | 2.1e-06 | 2.033E-06 | 7.024E-07 | 1.300E-06 | 2.700E-06 |
| 23619 | 1181 | E1181Y | D | 3.7e-06 | 1.5e-06 | 2.9e-06 | 2.700E-06 | 1.114E-06 | 1.500E-06 | 3.700E-06 |
| 23620 | 1181 | E1181V | D | 1.6e-06 | 3.1e-06 | 2e-06 | 2.233E-06 | 7.767E-07 | 1.600E-06 | 3.100E-06 |
| 23621 | 1182 | Q1182A | D | 3.7e-06 | 3.6e-06 | 4.6e-06 | 3.967E-06 | 5.508E-07 | 3.600E-06 | 4.600E-06 |
| 23622 | 1182 | Q1182R | D | 2.5e-06 | 2.6e-06 | 8.9e-07 | 1.997E-06 | 9.597E-07 | 8.900E-07 | 2.600E-06 |
| 23623 | 1182 | Q1182N | D | 2.2e-06 | 1.2e-06 | 1.7e-06 | 1.700E-06 | 5.000E-07 | 1.200E-06 | 2.200E-06 |
| 23624 | 1182 | Q1182D | D | 1.6e-06 | 2.6e-06 | 4.3e-06 | 2.833E-06 | 1.365E-06 | 1.600E-06 | 4.300E-06 |
| 23625 | 1182 | Q1182C | D | 2.5e-06 | 2.6e-06 | 2.4e-06 | 2.500E-06 | 1.000E-07 | 2.400E-06 | 2.600E-06 |
| 23626 | 1182 | Q1182E | D | 2.5e-06 | 4.3e-06 | 3.2e-06 | 3.333E-06 | 9.074E-07 | 2.500E-06 | 4.300E-06 |
| 23627 | 1182 | Q1182Q | D | 2.5e-06 | 2e-06 | 3.5e-06 | 2.667E-06 | 7.638E-07 | 2.000E-06 | 3.500E-06 |
| 23628 | 1182 | Q1182G | D | 2.1e-06 | 1.8e-06 | 4.2e-06 | 2.700E-06 | 1.308E-06 | 1.800E-06 | 4.200E-06 |
| 23629 | 1182 | Q1182H | D | 2.2e-06 | 2e-06 | 1.4e-06 | 1.867E-06 | 4.163E-07 | 1.400E-06 | 2.200E-06 |

| | | | | | | | | | | |
|---|---|---|---|---|---|---|---|---|---|---|
| 23630 | 1182 | Q1182I | D | 1.9e-06 | 3e-06 | 3.4e-06 | 2.767E-06 | 7.767E-07 | 1.900E-06 | 3.400E-06 |
| 23631 | 1182 | Q1182L | D | 3.6e-06 | 3.2e-06 | 2.1e-06 | 2.967E-06 | 7.767E-07 | 2.100E-06 | 3.600E-06 |
| 23632 | 1182 | Q1182K | D | 3.9e-06 | 2.3e-06 | 3.4e-06 | 3.200E-06 | 8.185E-07 | 2.300E-06 | 3.900E-06 |
| 23633 | 1182 | Q1182M | D | 2.6e-06 | 1.6e-06 | 2.2e-06 | 2.133E-06 | 5.033E-07 | 1.600E-06 | 2.600E-06 |
| 23634 | 1182 | Q1182F | D | 3.4e-06 | 1.7e-06 | 2.1e-06 | 2.400E-06 | 8.888E-07 | 1.700E-06 | 3.400E-06 |
| 23635 | 1182 | Q1182P | D | 1.7e-06 | 1.8e-06 | 2.3e-06 | 1.933E-06 | 3.215E-07 | 1.700E-06 | 2.300E-06 |
| 23636 | 1182 | Q1182S | D | 1.4e-06 | 2e-06 | 2.6e-06 | 2.000E-06 | 6.000E-07 | 1.400E-06 | 2.600E-06 |
| 23637 | 1182 | Q1182T | D | 2.7e-06 | 2.9e-06 | 3.3e-06 | 2.967E-06 | 3.055E-07 | 2.700E-06 | 3.300E-06 |
| 23638 | 1182 | Q1182W | D | 1.9e-06 | 2.2e-06 | 3.6e-06 | 2.567E-06 | 9.074E-07 | 1.900E-06 | 3.600E-06 |
| 23639 | 1182 | Q1182Y | D | 2.6e-06 | 2.1e-06 | 3.1e-06 | 2.600E-06 | 5.000E-07 | 2.100E-06 | 3.100E-06 |
| 23640 | 1182 | Q1182V | D | 2.4e-06 | 1.7e-06 | 2.6e-06 | 2.233E-06 | 4.726E-07 | 1.700E-06 | 2.600E-06 |
| 23641 | 1183 | L1183A | D | 1.3e-06 | 2.5e-06 | 2.4e-06 | 2.067E-06 | 6.658E-07 | 1.300E-06 | 2.500E-06 |
| 23642 | 1183 | L1183R | D | 2.9e-06 | 1.6e-06 | 2.2e-06 | 2.233E-06 | 6.506E-07 | 1.600E-06 | 2.900E-06 |
| 23643 | 1183 | L1183N | D | 1.4e-06 | 2e-06 | 1.8e-06 | 1.733E-06 | 3.055E-07 | 1.400E-06 | 2.000E-06 |
| 23644 | 1183 | L1183D | D | 4.4e-06 | 3.2e-06 | 2.5e-06 | 3.367E-06 | 9.609E-07 | 2.500E-06 | 4.400E-06 |
| 23645 | 1183 | L1183C | D | 1.7e-06 | 1.8e-06 | 1.8e-06 | 1.767E-06 | 5.774E-08 | 1.700E-06 | 1.800E-06 |
| 23646 | 1183 | L1183E | D | 2.1e-06 | 3.4e-06 | 2e-06 | 2.500E-06 | 7.810E-07 | 2.000E-06 | 3.400E-06 |
| 23647 | 1183 | L1183Q | D | 3.1e-06 | 2.4e-06 | 2e-06 | 2.500E-06 | 5.568E-07 | 2.000E-06 | 3.100E-06 |
| 23648 | 1183 | L1183G | D | 2.9e-06 | 3e-06 | 1.6e-06 | 2.500E-06 | 7.810E-07 | 1.600E-06 | 3.000E-06 |
| 23649 | 1183 | L1183H | D | 1.7e-06 | 3.5e-06 | 2.9e-06 | 2.700E-06 | 9.165E-07 | 1.700E-06 | 3.500E-06 |
| 23650 | 1183 | L1183I | D | 3.4e-06 | 2.6e-06 | 2.8e-06 | 2.933E-06 | 4.163E-07 | 2.600E-06 | 3.400E-06 |
| 23651 | 1183 | L1183L | D | 1.4e-06 | 2.1e-06 | 2.9e-06 | 2.133E-06 | 7.506E-07 | 1.400E-06 | 2.900E-06 |
| 23652 | 1183 | L1183K | D | 1.8e-06 | 1.5e-06 | 2.1e-06 | 1.800E-06 | 3.000E-07 | 1.500E-06 | 2.100E-06 |
| 23653 | 1183 | L1183M | D | 2.9e-06 | 1.8e-06 | 4.7e-06 | 3.133E-06 | 1.464E-06 | 1.800E-06 | 4.700E-06 |
| 23654 | 1183 | L1183F | D | 1.5e-06 | 3.7e-06 | 1.4e-06 | 2.200E-06 | 1.300E-06 | 1.400E-06 | 3.700E-06 |
| 23655 | 1183 | L1183P | D | 3.4e-06 | 2.4e-06 | 2.7e-06 | 2.833E-06 | 5.132E-07 | 2.400E-06 | 3.400E-06 |
| 23656 | 1183 | L1183S | D | 2.2e-06 | 1.1e-06 | 1.8e-06 | 1.700E-06 | 5.568E-07 | 1.100E-06 | 2.200E-06 |
| 23657 | 1183 | L1183T | D | 1.6e-06 | 2.7e-06 | 1.9e-06 | 2.067E-06 | 5.686E-07 | 1.600E-06 | 2.700E-06 |
| 23658 | 1183 | L1183W | D | 1.7e-06 | 3e-06 | 2.1e-06 | 2.267E-06 | 6.658E-07 | 1.700E-06 | 3.000E-06 |
| 23659 | 1183 | L1183Y | D | 1.4e-06 | 1.9e-06 | 1.8e-06 | 1.700E-06 | 2.646E-07 | 1.400E-06 | 1.900E-06 |
| 23660 | 1183 | L1183V | D | 1.9e-06 | 2e-06 | 2.3e-06 | 2.067E-06 | 2.082E-07 | 1.900E-06 | 2.300E-06 |
| 23661 | 1184 | K1184A | D | 1.2e-06 | 2.8e-06 | 1.4e-06 | 1.800E-06 | 8.718E-07 | 1.200E-06 | 2.800E-06 |
| 23662 | 1184 | K1184R | D | 2.2e-06 | 2.9e-06 | 4.3e-06 | 3.133E-06 | 1.069E-06 | 2.200E-06 | 4.300E-06 |
| 23663 | 1184 | K1184N | D | 1.8e-06 | 1.3e-06 | 1.7e-06 | 1.600E-06 | 2.646E-07 | 1.300E-06 | 1.800E-06 |
| 23664 | 1184 | K1184D | D | 1.9e-06 | 1.6e-06 | 1.4e-06 | 1.633E-06 | 2.517E-07 | 1.400E-06 | 1.900E-06 |
| 23665 | 1184 | K1184C | D | 4e-06 | 1.7e-06 | 3e-06 | 2.900E-06 | 1.153E-06 | 1.700E-06 | 4.000E-06 |
| 23666 | 1184 | K1184E | D | 2.2e-06 | 2.4e-06 | 1.1e-06 | 1.900E-06 | 7.000E-07 | 1.100E-06 | 2.400E-06 |
| 23667 | 1184 | K1184Q | D | 2.4e-06 | 2.1e-06 | 1.2e-06 | 1.900E-06 | 6.245E-07 | 1.200E-06 | 2.400E-06 |
| 23668 | 1184 | K1184G | D | 1.9e-06 | 3.2e-06 | 2.9e-06 | 2.667E-06 | 6.807E-07 | 1.900E-06 | 3.200E-06 |
| 23669 | 1184 | K1184H | D | 1.8e-06 | 1.9e-06 | 3.9e-06 | 2.533E-06 | 1.185E-06 | 1.800E-06 | 3.900E-06 |
| 23670 | 1184 | K1184I | D | 3.4e-06 | 2.7e-06 | 2.1e-06 | 2.733E-06 | 6.506E-07 | 2.100E-06 | 3.400E-06 |
| 23671 | 1184 | K1184L | D | 1.6e-06 | 1.9e-06 | 3.5e-06 | 2.333E-06 | 1.021E-06 | 1.600E-06 | 3.500E-06 |
| 23672 | 1184 | K1184K | D | 2.2e-06 | 1.9e-06 | 3.6e-06 | 2.567E-06 | 9.074E-07 | 1.900E-06 | 3.600E-06 |
| 23673 | 1184 | K1184M | D | 1.8e-06 | 2e-06 | 2.9e-06 | 2.233E-06 | 5.859E-07 | 1.800E-06 | 2.900E-06 |
| 23674 | 1184 | K1184F | D | 2.8e-06 | 3e-06 | 3.6e-06 | 3.133E-06 | 4.163E-07 | 2.800E-06 | 3.600E-06 |
| 23675 | 1184 | K1184P | D | 2.5e-06 | 2.7e-06 | 1.6e-06 | 2.267E-06 | 5.859E-07 | 1.600E-06 | 2.700E-06 |
| 23676 | 1184 | K1184S | D | 4.5e-06 | 2.2e-06 | 1.9e-06 | 2.867E-06 | 1.422E-06 | 1.900E-06 | 4.500E-06 |
| 23677 | 1184 | K1184T | D | 1.9e-06 | 3.2e-06 | 1.8e-06 | 2.300E-06 | 7.810E-07 | 1.800E-06 | 3.200E-06 |
| 23678 | 1184 | K1184W | D | 2.4e-06 | 3.4e-06 | 1.3e-06 | 2.367E-06 | 1.050E-06 | 1.300E-06 | 3.400E-06 |
| 23679 | 1184 | K1184Y | D | 4.9e-06 | 1.5e-06 | 3.7e-06 | 3.367E-06 | 1.724E-06 | 1.500E-06 | 4.900E-06 |
| 23680 | 1184 | K1184V | D | 2.5e-06 | 1.9e-06 | 2.5e-06 | 2.300E-06 | 3.464E-07 | 1.900E-06 | 2.500E-06 |
| 23681 | 1185 | S1185A | D | 2.1e-06 | 1.9e-06 | 4.3e-06 | 2.767E-06 | 1.332E-06 | 1.900E-06 | 4.300E-06 |
| 23682 | 1185 | S1185R | D | 2.7e-06 | 3.1e-06 | 1.4e-06 | 2.400E-06 | 8.888E-07 | 1.400E-06 | 3.100E-06 |
| 23683 | 1185 | S1185N | D | 2.2e-06 | 2.1e-06 | 2.7e-06 | 2.333E-06 | 3.215E-07 | 2.100E-06 | 2.700E-06 |
| 23684 | 1185 | S1185D | D | 1.4e-06 | 2.1e-06 | 1.6e-06 | 1.700E-06 | 3.606E-07 | 1.400E-06 | 2.100E-06 |
| 23685 | 1185 | S1185C | D | 2.3e-06 | 4e-06 | 2.5e-06 | 2.933E-06 | 9.292E-07 | 2.300E-06 | 4.000E-06 |
| 23686 | 1185 | S1185E | D | 2.6e-06 | 2.5e-06 | 1.6e-06 | 2.233E-06 | 5.508E-07 | 1.600E-06 | 2.600E-06 |
| 23687 | 1185 | S1185Q | D | 1.9e-06 | 2.3e-06 | 1.7e-06 | 1.967E-06 | 3.055E-07 | 1.700E-06 | 2.300E-06 |
| 23688 | 1185 | S1185G | D | 2.3e-06 | 2.6e-06 | 8.8e-07 | 1.927E-06 | 9.188E-07 | 8.800E-07 | 2.600E-06 |
| 23689 | 1185 | S1185H | D | 1.7e-06 | 2.5e-06 | 2.3e-06 | 2.167E-06 | 4.163E-07 | 1.700E-06 | 2.500E-06 |
| 23690 | 1185 | S1185I | D | 2.7e-06 | 1.8e-06 | 2.7e-06 | 2.400E-06 | 5.196E-07 | 1.800E-06 | 2.700E-06 |
| 23691 | 1185 | S1185L | D | 1.8e-06 | 3.6e-06 | 2.9e-06 | 2.767E-06 | 9.074E-07 | 1.800E-06 | 3.600E-06 |
| 23692 | 1185 | S1185K | D | 1.8e-06 | 2.6e-06 | 2.2e-06 | 2.200E-06 | 4.000E-07 | 1.800E-06 | 2.600E-06 |
| 23693 | 1185 | S1185M | D | 1.3e-06 | 3.3e-06 | 1.9e-06 | 2.167E-06 | 1.026E-06 | 1.300E-06 | 3.300E-06 |
| 23694 | 1185 | S1185F | D | 2.2e-06 | 2.2e-06 | 1.5e-06 | 1.967E-06 | 4.041E-07 | 1.500E-06 | 2.200E-06 |
| 23695 | 1185 | S1185P | D | 2.7e-06 | 2e-06 | 2.1e-06 | 2.267E-06 | 3.786E-07 | 2.000E-06 | 2.700E-06 |
| 23696 | 1185 | S1185S | D | 1.8e-06 | 3.2e-06 | 2.9e-06 | 2.633E-06 | 7.371E-07 | 1.800E-06 | 3.200E-06 |
| 23697 | 1185 | S1185T | D | 3e-06 | 9.4e-07 | 2e-06 | 1.980E-06 | 1.030E-06 | 9.400E-07 | 3.000E-06 |
| 23698 | 1185 | S1185W | D | 3.5e-06 | 3.4e-06 | 3.1e-06 | 3.333E-06 | 2.082E-07 | 3.100E-06 | 3.500E-06 |
| 23699 | 1185 | S1185Y | D | 2.5e-06 | 2.1e-06 | 2.4e-06 | 2.333E-06 | 2.082E-07 | 2.100E-06 | 2.500E-06 |
| 23700 | 1185 | S1185V | D | 2.5e-06 | 2.1e-06 | 2e-06 | 2.200E-06 | 2.646E-07 | 2.000E-06 | 2.500E-06 |
| 23701 | 1186 | G1186A | D | 2.4e-06 | 2e-06 | 2.4e-06 | 2.267E-06 | 2.309E-07 | 2.000E-06 | 2.400E-06 |
| 23702 | 1186 | G1186R | D | 2.4e-06 | 2.6e-06 | 1.1e-06 | 2.033E-06 | 8.145E-07 | 1.100E-06 | 2.600E-06 |
| 23703 | 1186 | G1186N | D | 1.6e-06 | 2.3e-06 | 2.1e-06 | 2.000E-06 | 3.606E-07 | 1.600E-06 | 2.300E-06 |
| 23704 | 1186 | G1186D | D | 3.2e-06 | 3e-06 | 1.9e-06 | 2.700E-06 | 7.000E-07 | 1.900E-06 | 3.200E-06 |
| 23705 | 1186 | G1186C | D | 2.4e-06 | 1.8e-06 | 1.9e-06 | 2.033E-06 | 3.215E-07 | 1.800E-06 | 2.400E-06 |
| 23706 | 1186 | G1186E | D | 2.7e-06 | 2.6e-06 | 2.4e-06 | 2.567E-06 | 1.528E-07 | 2.400E-06 | 2.700E-06 |
| 23707 | 1186 | G1186Q | D | 2.1e-06 | 1.5e-06 | 2.5e-06 | 2.033E-06 | 5.033E-07 | 1.500E-06 | 2.500E-06 |
| 23708 | 1186 | G1186G | D | 1.9e-06 | 2.6e-06 | 2.5e-06 | 2.333E-06 | 3.786E-07 | 1.900E-06 | 2.600E-06 |
| 23709 | 1186 | G1186H | D | 2.7e-06 | 3.8e-06 | 3.5e-06 | 3.333E-06 | 5.686E-07 | 2.700E-06 | 3.800E-06 |
| 23710 | 1186 | G1186I | D | 1.9e-06 | 3.1e-06 | 1.9e-06 | 2.300E-06 | 6.928E-07 | 1.900E-06 | 3.100E-06 |
| 23711 | 1186 | G1186L | D | 1.8e-06 | 2.2e-06 | 3e-06 | 2.333E-06 | 6.110E-07 | 1.800E-06 | 3.000E-06 |
| 23712 | 1186 | G1186K | D | 2.2e-06 | 2e-06 | 1.9e-06 | 2.033E-06 | 1.528E-07 | 1.900E-06 | 2.200E-06 |
| 23713 | 1186 | G1186M | D | 2.4e-06 | 2.1e-06 | 1.6e-06 | 2.033E-06 | 4.041E-07 | 1.600E-06 | 2.400E-06 |
| 23714 | 1186 | G1186F | D | 3.7e-06 | 2.4e-06 | 2.3e-06 | 2.800E-06 | 7.810E-07 | 2.300E-06 | 3.700E-06 |

| | | | | | | | | | | |
|---|---|---|---|---|---|---|---|---|---|---|
| 23715 | 1186 | G1186P | D | 1.4e-06 | 2.1e-06 | 3.2e-06 | 2.233E-06 | 9.074E-07 | 1.400E-06 | 3.200E-06 |
| 23716 | 1186 | G1186S | D | 1.9e-06 | 2.2e-06 | 2.1e-06 | 2.067E-06 | 1.528E-07 | 1.900E-06 | 2.200E-06 |
| 23717 | 1186 | G1186T | D | 2.2e-06 | 1.8e-06 | 2.1e-06 | 2.033E-06 | 2.082E-07 | 1.800E-06 | 2.200E-06 |
| 23718 | 1186 | G1186W | D | 3.5e-06 | 2.8e-06 | 1.8e-06 | 2.700E-06 | 8.544E-07 | 1.800E-06 | 3.500E-06 |
| 23719 | 1186 | G1186Y | D | 1.7e-06 | 2.1e-06 | 1.7e-06 | 1.833E-06 | 2.309E-07 | 1.700E-06 | 2.100E-06 |
| 23720 | 1186 | G1186V | D | 3.4e-06 | 2.8e-06 | 2.2e-06 | 2.800E-06 | 6.000E-07 | 2.200E-06 | 3.400E-06 |
| 23721 | 1187 | T1187A | D | 4.5e-06 | 2.3e-06 | 9.6e-07 | 2.587E-06 | 1.787E-06 | 9.600E-07 | 4.500E-06 |
| 23722 | 1187 | T1187R | D | 1.8e-06 | 2.1e-06 | 1.6e-06 | 1.833E-06 | 2.517E-07 | 1.600E-06 | 2.100E-06 |
| 23723 | 1187 | T1187N | D | 1.7e-06 | 1.9e-06 | 3.6e-06 | 2.400E-06 | 1.044E-06 | 1.700E-06 | 3.600E-06 |
| 23724 | 1187 | T1187D | D | 1.2e-06 | 3.1e-06 | 3.1e-06 | 2.467E-06 | 1.097E-06 | 1.200E-06 | 3.100E-06 |
| 23725 | 1187 | T1187C | D | 1.9e-06 | 3e-06 | 2.7e-06 | 2.533E-06 | 5.686E-07 | 1.900E-06 | 3.000E-06 |
| 23726 | 1187 | T1187E | D | 3e-06 | 1.6e-06 | 1.6e-06 | 2.067E-06 | 8.083E-07 | 1.600E-06 | 3.000E-06 |
| 23727 | 1187 | T1187Q | D | 2.3e-06 | 4.3e-06 | 1.3e-06 | 2.633E-06 | 1.528E-06 | 1.300E-06 | 4.300E-06 |
| 23728 | 1187 | T1187G | D | 2.3e-06 | 2.3e-06 | 4e-06 | 2.867E-06 | 9.815E-07 | 2.300E-06 | 4.000E-06 |
| 23729 | 1187 | T1187H | D | 2.3e-06 | 1.5e-06 | 3.6e-06 | 2.467E-06 | 1.060E-06 | 1.500E-06 | 3.600E-06 |
| 23730 | 1187 | T1187I | D | 1.8e-06 | 4.2e-06 | 2.3e-06 | 2.767E-06 | 1.266E-06 | 1.800E-06 | 4.200E-06 |
| 23731 | 1187 | T1187L | D | 2.9e-06 | 1.3e-06 | 1.6e-06 | 1.933E-06 | 8.505E-07 | 1.300E-06 | 2.900E-06 |
| 23732 | 1187 | T1187K | D | 2.1e-06 | 3.3e-06 | 1.7e-06 | 2.367E-06 | 8.327E-07 | 1.700E-06 | 3.300E-06 |
| 23733 | 1187 | T1187M | D | 3.2e-06 | 3e-06 | 2.3e-06 | 2.833E-06 | 4.726E-07 | 2.300E-06 | 3.200E-06 |
| 23734 | 1187 | T1187F | D | 2.8e-06 | 4.1e-06 | 2.4e-06 | 3.100E-06 | 8.888E-07 | 2.400E-06 | 4.100E-06 |
| 23735 | 1187 | T1187P | D | 1.6e-06 | 2.5e-06 | 1.6e-06 | 1.900E-06 | 5.196E-07 | 1.600E-06 | 2.500E-06 |
| 23736 | 1187 | T1187S | D | 2.1e-06 | 1.6e-06 | 2.2e-06 | 1.967E-06 | 3.215E-07 | 1.600E-06 | 2.200E-06 |
| 23737 | 1187 | T1187T | D | 2e-06 | 1.4e-06 | 2.8e-06 | 2.067E-06 | 7.024E-07 | 1.400E-06 | 2.800E-06 |
| 23738 | 1187 | T1187W | D | 2.4e-06 | 2.8e-06 | 2.6e-06 | 2.600E-06 | 2.000E-07 | 2.400E-06 | 2.800E-06 |
| 23739 | 1187 | T1187Y | D | 2.7e-06 | 1.2e-06 | 1.8e-06 | 1.900E-06 | 7.550E-07 | 1.200E-06 | 2.700E-06 |
| 23740 | 1187 | T1187V | D | 1.8e-06 | 3.1e-06 | 1.4e-06 | 2.100E-06 | 8.888E-07 | 1.400E-06 | 3.100E-06 |
| 23741 | 1188 | A1188A | D | 2.6e-06 | 2.3e-06 | 2.4e-06 | 2.433E-06 | 1.528E-07 | 2.300E-06 | 2.600E-06 |
| 23742 | 1188 | A1188R | D | 2.7e-06 | 4.7e-06 | 1.8e-06 | 3.067E-06 | 1.484E-06 | 1.800E-06 | 4.700E-06 |
| 23743 | 1188 | A1188N | D | 3.5e-06 | 3.3e-06 | 2e-06 | 2.933E-06 | 8.145E-07 | 2.000E-06 | 3.500E-06 |
| 23744 | 1188 | A1188D | D | 4e-06 | 1.5e-06 | 1.6e-06 | 2.367E-06 | 1.415E-06 | 1.500E-06 | 4.000E-06 |
| 23745 | 1188 | A1188C | D | 2.2e-06 | 1.9e-06 | 2.1e-06 | 2.067E-06 | 1.528E-07 | 1.900E-06 | 2.200E-06 |
| 23746 | 1188 | A1188E | D | 2.7e-06 | 2.5e-06 | 2.1e-06 | 2.433E-06 | 3.055E-07 | 2.100E-06 | 2.700E-06 |
| 23747 | 1188 | A1188Q | D | 1.4e-06 | 4.1e-06 | 2.2e-06 | 2.567E-06 | 1.387E-06 | 1.400E-06 | 4.100E-06 |
| 23748 | 1188 | A1188G | D | 2.4e-06 | 3.1e-06 | 1.2e-06 | 2.233E-06 | 9.609E-07 | 1.200E-06 | 3.100E-06 |
| 23749 | 1188 | A1188H | D | 3.6e-06 | 3e-06 | 2.3e-06 | 2.967E-06 | 6.506E-07 | 2.300E-06 | 3.600E-06 |
| 23750 | 1188 | A1188I | D | 4.3e-06 | 4e-06 | 1.6e-06 | 3.300E-06 | 1.480E-06 | 1.600E-06 | 4.300E-06 |
| 23751 | 1188 | A1188L | D | 1.7e-06 | 2.6e-06 | 2.1e-06 | 2.133E-06 | 4.509E-07 | 1.700E-06 | 2.600E-06 |
| 23752 | 1188 | A1188K | D | 2.1e-06 | 1.6e-06 | 2e-06 | 1.900E-06 | 2.646E-07 | 1.600E-06 | 2.100E-06 |
| 23753 | 1188 | A1188M | D | 2.6e-06 | 2.7e-06 | 2.5e-06 | 2.600E-06 | 1.000E-07 | 2.500E-06 | 2.700E-06 |
| 23754 | 1188 | A1188F | D | 4.4e-06 | 1.7e-06 | 1.9e-06 | 2.667E-06 | 1.504E-06 | 1.700E-06 | 4.400E-06 |
| 23755 | 1188 | A1188P | D | 3e-06 | 2.2e-06 | 2.3e-06 | 2.500E-06 | 4.359E-07 | 2.200E-06 | 3.000E-06 |
| 23756 | 1188 | A1188S | D | 1e-06 | 2.3e-06 | 4.5e-06 | 2.600E-06 | 1.769E-06 | 1.000E-06 | 4.500E-06 |
| 23757 | 1188 | A1188T | D | 2.6e-06 | 2.8e-06 | 3.9e-06 | 3.100E-06 | 7.000E-07 | 2.600E-06 | 3.900E-06 |
| 23758 | 1188 | A1188W | D | 2e-06 | 1.6e-06 | 1.5e-06 | 1.700E-06 | 2.646E-07 | 1.500E-06 | 2.000E-06 |
| 23759 | 1188 | A1188Y | D | 2.7e-06 | 2.5e-06 | 1.5e-06 | 2.233E-06 | 6.429E-07 | 1.500E-06 | 2.700E-06 |
| 23760 | 1188 | A1188V | D | 3.7e-06 | 1.7e-06 | 3.1e-06 | 2.833E-06 | 1.026E-06 | 1.700E-06 | 3.700E-06 |
| 23761 | 1189 | S1189A | D | 2.8e-06 | 2.9e-06 | 2.5e-06 | 2.733E-06 | 2.082E-07 | 2.500E-06 | 2.900E-06 |
| 23762 | 1189 | S1189R | D | 2.7e-06 | 1.3e-06 | 2.6e-06 | 2.200E-06 | 7.810E-07 | 1.300E-06 | 2.700E-06 |
| 23763 | 1189 | S1189N | D | 1.6e-06 | 2.8e-06 | 1.8e-06 | 2.067E-06 | 6.429E-07 | 1.600E-06 | 2.800E-06 |
| 23764 | 1189 | S1189D | D | 2.8e-06 | 4.2e-06 | 2.4e-06 | 3.133E-06 | 9.452E-07 | 2.400E-06 | 4.200E-06 |
| 23765 | 1189 | S1189C | D | 3.2e-06 | 2.9e-06 | 3.1e-06 | 3.067E-06 | 1.528E-07 | 2.900E-06 | 3.200E-06 |
| 23766 | 1189 | S1189E | D | 1.9e-06 | 2.3e-06 | 1.7e-06 | 1.967E-06 | 3.055E-07 | 1.700E-06 | 2.300E-06 |
| 23767 | 1189 | S1189Q | D | 3.3e-06 | 3.5e-06 | 2.1e-06 | 2.967E-06 | 7.572E-07 | 2.100E-06 | 3.500E-06 |
| 23768 | 1189 | S1189G | D | 2.8e-06 | 2.3e-06 | 2.5e-06 | 2.533E-06 | 2.517E-07 | 2.300E-06 | 2.800E-06 |
| 23769 | 1189 | S1189H | D | 2.5e-06 | 1.6e-06 | 1.4e-06 | 1.833E-06 | 5.859E-07 | 1.400E-06 | 2.500E-06 |
| 23770 | 1189 | S1189I | D | 2.2e-06 | 2.6e-06 | 2.9e-06 | 2.567E-06 | 3.512E-07 | 2.200E-06 | 2.900E-06 |
| 23771 | 1189 | S1189L | D | 1.9e-06 | 1.7e-06 | 2.7e-06 | 2.100E-06 | 5.292E-07 | 1.700E-06 | 2.700E-06 |
| 23772 | 1189 | S1189K | D | 1.8e-06 | 2.1e-06 | 3.7e-06 | 2.533E-06 | 1.021E-06 | 1.800E-06 | 3.700E-06 |
| 23773 | 1189 | S1189M | D | 2.5e-06 | 2.7e-06 | 2.4e-06 | 2.533E-06 | 1.528E-07 | 2.400E-06 | 2.700E-06 |
| 23774 | 1189 | S1189F | D | 3.1e-06 | 2.1e-06 | 3.1e-06 | 2.767E-06 | 5.774E-07 | 2.100E-06 | 3.100E-06 |
| 23775 | 1189 | S1189P | D | 2.2e-06 | 2e-06 | 1.8e-06 | 2.000E-06 | 2.000E-07 | 1.800E-06 | 2.200E-06 |
| 23776 | 1189 | S1189S | D | 5.3e-06 | 1.8e-06 | 3e-06 | 3.367E-06 | 1.779E-06 | 1.800E-06 | 5.300E-06 |
| 23777 | 1189 | S1189T | D | 1.3e-06 | 2.1e-06 | 2.6e-06 | 2.000E-06 | 6.557E-07 | 1.300E-06 | 2.600E-06 |
| 23778 | 1189 | S1189W | D | 2.4e-06 | 2.9e-06 | 3.3e-06 | 2.867E-06 | 4.509E-07 | 2.400E-06 | 3.300E-06 |
| 23779 | 1189 | S1189Y | D | 2.3e-06 | 2.6e-06 | 1.4e-06 | 2.100E-06 | 6.245E-07 | 1.400E-06 | 2.600E-06 |
| 23780 | 1189 | S1189V | D | 1.9e-06 | 2.6e-06 | 2.2e-06 | 2.233E-06 | 3.512E-07 | 1.900E-06 | 2.600E-06 |
| 23781 | 1190 | V1190A | D | 2.1e-06 | 1.5e-06 | 4.3e-06 | 2.633E-06 | 1.474E-06 | 1.500E-06 | 4.300E-06 |
| 23782 | 1190 | V1190R | D | 8.6e-07 | 3.4e-06 | 1.9e-06 | 2.053E-06 | 1.277E-06 | 8.600E-07 | 3.400E-06 |
| 23783 | 1190 | V1190N | D | 4.2e-06 | 2.1e-06 | 3.7e-06 | 3.333E-06 | 1.097E-06 | 2.100E-06 | 4.200E-06 |
| 23784 | 1190 | V1190D | D | 2.5e-06 | 2.2e-06 | 1.8e-06 | 2.167E-06 | 3.512E-07 | 1.800E-06 | 2.500E-06 |
| 23785 | 1190 | V1190C | D | 2.9e-06 | 2.4e-06 | 2.9e-06 | 2.733E-06 | 2.887E-07 | 2.400E-06 | 2.900E-06 |
| 23786 | 1190 | V1190E | D | 1.9e-06 | 2.4e-06 | 3.6e-06 | 2.633E-06 | 8.737E-07 | 1.900E-06 | 3.600E-06 |
| 23787 | 1190 | V1190Q | D | 1.5e-06 | 1.9e-06 | 2.4e-06 | 1.933E-06 | 4.509E-07 | 1.500E-06 | 2.400E-06 |
| 23788 | 1190 | V1190G | D | 2.6e-06 | 2.4e-06 | 2.4e-06 | 2.467E-06 | 1.155E-07 | 2.400E-06 | 2.600E-06 |
| 23789 | 1190 | V1190H | D | 2.8e-06 | 2.1e-06 | 1.4e-06 | 2.100E-06 | 7.000E-07 | 1.400E-06 | 2.800E-06 |
| 23790 | 1190 | V1190I | D | 3.4e-06 | 2.2e-06 | 2.1e-06 | 2.567E-06 | 7.234E-07 | 2.100E-06 | 3.400E-06 |
| 23791 | 1190 | V1190L | D | 2.8e-06 | 2.4e-06 | 1.8e-06 | 2.333E-06 | 5.033E-07 | 1.800E-06 | 2.800E-06 |
| 23792 | 1190 | V1190K | D | 2.4e-06 | 2.6e-06 | 1.9e-06 | 2.300E-06 | 3.606E-07 | 1.900E-06 | 2.600E-06 |
| 23793 | 1190 | V1190M | D | 2.2e-06 | 1.6e-06 | 1.9e-06 | 1.900E-06 | 3.000E-07 | 1.600E-06 | 2.200E-06 |
| 23794 | 1190 | V1190F | D | 1.3e-06 | 2.4e-06 | 1.2e-06 | 1.633E-06 | 6.658E-07 | 1.200E-06 | 2.400E-06 |
| 23795 | 1190 | V1190P | D | 4.9e-06 | 2.2e-06 | 3.2e-06 | 3.433E-06 | 1.365E-06 | 2.200E-06 | 4.900E-06 |
| 23796 | 1190 | V1190S | D | 3.5e-06 | 1e-06 | 2.7e-06 | 2.400E-06 | 1.277E-06 | 1.000E-06 | 3.500E-06 |
| 23797 | 1190 | V1190T | D | 2.5e-06 | 3.1e-06 | 2.8e-06 | 2.800E-06 | 3.000E-07 | 2.500E-06 | 3.100E-06 |
| 23798 | 1190 | V1190W | D | 2.5e-06 | 3.6e-06 | 2.7e-06 | 2.933E-06 | 5.859E-07 | 2.500E-06 | 3.600E-06 |
| 23799 | 1190 | V1190Y | D | 1.7e-06 | 2.2e-06 | 2e-06 | 1.967E-06 | 2.517E-07 | 1.700E-06 | 2.200E-06 |

| | | | | | | | | | | |
|---|---|---|---|---|---|---|---|---|---|---|
| 23800 | 1190 | V1190V | D | 2.8e-06 | 2.6e-06 | 1.1e-06 | 2.167E-06 | 9.292E-07 | 1.100E-06 | 2.800E-06 |
| 23801 | 1191 | V1191A | D | 2.3e-06 | 1.7e-06 | 2.4e-06 | 2.133E-06 | 3.786E-07 | 1.700E-06 | 2.400E-06 |
| 23802 | 1191 | V1191R | D | 2e-06 | 2.1e-06 | 2.1e-06 | 2.067E-06 | 5.774E-08 | 2.000E-06 | 2.100E-06 |
| 23803 | 1191 | V1191N | D | 2.6e-06 | 2.1e-06 | 4.2e-06 | 2.967E-06 | 1.097E-06 | 2.100E-06 | 4.200E-06 |
| 23804 | 1191 | V1191D | D | 1.4e-06 | 1.9e-06 | 4e-06 | 2.433E-06 | 1.380E-06 | 1.400E-06 | 4.000E-06 |
| 23805 | 1191 | V1191C | D | 2.5e-06 | 2.8e-06 | 1.6e-06 | 2.300E-06 | 6.245E-07 | 1.600E-06 | 2.800E-06 |
| 23806 | 1191 | V1191E | D | 1.5e-06 | 1.5e-06 | 2.4e-06 | 1.800E-06 | 5.196E-07 | 1.500E-06 | 2.400E-06 |
| 23807 | 1191 | V1191Q | D | 2.2e-06 | 2.5e-06 | 2e-06 | 2.233E-06 | 2.517E-07 | 2.000E-06 | 2.500E-06 |
| 23808 | 1191 | V1191G | D | 2.4e-06 | 2.1e-06 | 1.9e-06 | 2.133E-06 | 2.517E-07 | 1.900E-06 | 2.400E-06 |
| 23809 | 1191 | V1191H | D | 1e-06 | 2.2e-06 | 3.1e-06 | 2.100E-06 | 1.054E-06 | 1.000E-06 | 3.100E-06 |
| 23810 | 1191 | V1191I | D | 2e-06 | 1.6e-06 | 1.7e-06 | 1.767E-06 | 2.082E-07 | 1.600E-06 | 2.000E-06 |
| 23811 | 1191 | V1191L | D | 1.7e-06 | 2.7e-06 | 4.9e-06 | 3.100E-06 | 1.637E-06 | 1.700E-06 | 4.900E-06 |
| 23812 | 1191 | V1191K | D | 2.2e-06 | 1.8e-06 | 3.1e-06 | 2.367E-06 | 6.658E-07 | 1.800E-06 | 3.100E-06 |
| 23813 | 1191 | V1191M | D | 2.6e-06 | 2.1e-06 | 3e-06 | 2.567E-06 | 4.509E-07 | 2.100E-06 | 3.000E-06 |
| 23814 | 1191 | V1191F | D | 2.1e-06 | 2.2e-06 | 3e-06 | 2.433E-06 | 4.933E-07 | 2.100E-06 | 3.000E-06 |
| 23815 | 1191 | V1191P | D | 1.8e-06 | 2.3e-06 | 3.4e-06 | 2.500E-06 | 8.185E-07 | 1.800E-06 | 3.400E-06 |
| 23816 | 1191 | V1191S | D | 3.2e-06 | 1.9e-06 | 2.1e-06 | 2.400E-06 | 7.000E-07 | 1.900E-06 | 3.200E-06 |
| 23817 | 1191 | V1191T | D | 1.7e-06 | 2.2e-06 | 2.7e-06 | 2.200E-06 | 5.000E-07 | 1.700E-06 | 2.700E-06 |
| 23818 | 1191 | V1191W | D | 2.2e-06 | 2.2e-06 | 2.4e-06 | 2.267E-06 | 1.155E-07 | 2.200E-06 | 2.400E-06 |
| 23819 | 1191 | V1191Y | D | 2.8e-06 | 2.6e-06 | 2.4e-06 | 2.600E-06 | 2.000E-07 | 2.400E-06 | 2.800E-06 |
| 23820 | 1191 | V1191V | D | 3.5e-06 | 3e-06 | 2.3e-06 | 2.933E-06 | 6.028E-07 | 2.300E-06 | 3.500E-06 |
| 23821 | 1192 | C1192A | D | 2.2e-06 | 3.8e-06 | 1.9e-06 | 2.633E-06 | 1.021E-06 | 1.900E-06 | 3.800E-06 |
| 23822 | 1192 | C1192R | D | 1.9e-06 | 1.5e-06 | 2e-06 | 1.800E-06 | 2.646E-07 | 1.500E-06 | 2.000E-06 |
| 23823 | 1192 | C1192N | D | 2.5e-06 | 2.5e-06 | 3e-06 | 2.667E-06 | 2.887E-07 | 2.500E-06 | 3.000E-06 |
| 23824 | 1192 | C1192D | D | 3e-06 | 2.7e-06 | 5.5e-06 | 3.733E-06 | 1.537E-06 | 2.700E-06 | 5.500E-06 |
| 23825 | 1192 | C1192C | D | 3.4e-06 | 2.7e-06 | 1.9e-06 | 2.667E-06 | 7.506E-07 | 1.900E-06 | 3.400E-06 |
| 23826 | 1192 | C1192E | D | 2.8e-06 | 1.4e-06 | 3.5e-06 | 2.567E-06 | 1.069E-06 | 1.400E-06 | 3.500E-06 |
| 23827 | 1192 | C1192Q | D | 1.8e-06 | 2.9e-06 | 7.9e-07 | 1.830E-06 | 1.055E-06 | 7.900E-07 | 2.900E-06 |
| 23828 | 1192 | C1192G | D | 2.3e-06 | 3e-06 | 2e-06 | 2.433E-06 | 5.132E-07 | 2.000E-06 | 3.000E-06 |
| 23829 | 1192 | C1192H | D | 3.2e-06 | 3.6e-06 | 4.1e-06 | 3.633E-06 | 4.509E-07 | 3.200E-06 | 4.100E-06 |
| 23830 | 1192 | C1192I | D | 4.6e-06 | 1.5e-06 | 9.2e-07 | 2.340E-06 | 1.979E-06 | 9.200E-07 | 4.600E-06 |
| 23831 | 1192 | C1192L | D | 1.2e-06 | 1.3e-06 | 1.3e-06 | 1.267E-06 | 5.774E-08 | 1.200E-06 | 1.300E-06 |
| 23832 | 1192 | C1192K | D | 1.8e-06 | 2.2e-06 | 2.5e-06 | 2.167E-06 | 3.512E-07 | 1.800E-06 | 2.500E-06 |
| 23833 | 1192 | C1192M | D | 2.7e-06 | 2.1e-06 | 2.2e-06 | 2.333E-06 | 3.215E-07 | 2.100E-06 | 2.700E-06 |
| 23834 | 1192 | C1192F | D | 2e-06 | 1.4e-06 | 3e-06 | 2.133E-06 | 8.083E-07 | 1.400E-06 | 3.000E-06 |
| 23835 | 1192 | C1192P | D | 3.6e-06 | 2.5e-06 | 2.4e-06 | 2.833E-06 | 6.658E-07 | 2.400E-06 | 3.600E-06 |
| 23836 | 1192 | C1192S | D | 2.4e-06 | 4.3e-06 | 6.8e-06 | 4.500E-06 | 2.207E-06 | 2.400E-06 | 6.800E-06 |
| 23837 | 1192 | C1192T | D | 2.1e-06 | 2.1e-06 | 3.9e-06 | 2.700E-06 | 1.039E-06 | 2.100E-06 | 3.900E-06 |
| 23838 | 1192 | C1192W | D | 1.8e-06 | 2.8e-06 | 2e-06 | 2.200E-06 | 5.292E-07 | 1.800E-06 | 2.800E-06 |
| 23839 | 1192 | C1192Y | D | 1.6e-06 | 1.8e-06 | 2.7e-06 | 2.033E-06 | 5.859E-07 | 1.600E-06 | 2.700E-06 |
| 23840 | 1192 | C1192V | D | 2.4e-06 | 1.9e-06 | 4.6e-06 | 2.967E-06 | 1.436E-06 | 1.900E-06 | 4.600E-06 |
| 23841 | 1193 | L1193A | D | 2.2e-06 | 2.3e-06 | 2.1e-06 | 2.200E-06 | 1.000E-07 | 2.100E-06 | 2.300E-06 |
| 23842 | 1193 | L1193R | D | 2.1e-06 | 2.1e-06 | 2.1e-06 | 2.100E-06 | 0.000E+00 | 2.100E-06 | 2.100E-06 |
| 23843 | 1193 | L1193N | D | 3.2e-06 | 1.5e-06 | 2e-06 | 2.233E-06 | 8.737E-07 | 1.500E-06 | 3.200E-06 |
| 23844 | 1193 | L1193D | D | 2.9e-06 | 2.6e-06 | 2e-06 | 2.500E-06 | 4.583E-07 | 2.000E-06 | 2.900E-06 |
| 23845 | 1193 | L1193C | D | 2.3e-06 | 2.1e-06 | 1.3e-06 | 1.900E-06 | 5.292E-07 | 1.300E-06 | 2.300E-06 |
| 23846 | 1193 | L1193E | D | 4.6e-06 | 2.5e-06 | 1.8e-06 | 2.967E-06 | 1.457E-06 | 1.800E-06 | 4.600E-06 |
| 23847 | 1193 | L1193Q | D | 2.6e-06 | 1.6e-06 | 2.1e-06 | 2.100E-06 | 5.000E-07 | 1.600E-06 | 2.600E-06 |
| 23848 | 1193 | L1193G | D | 2.9e-06 | 2.5e-06 | 2.8e-06 | 2.733E-06 | 2.082E-07 | 2.500E-06 | 2.900E-06 |
| 23849 | 1193 | L1193H | D | 2.1e-06 | 1.6e-06 | 2.1e-06 | 1.933E-06 | 2.887E-07 | 1.600E-06 | 2.100E-06 |
| 23850 | 1193 | L1193I | D | 4.9e-06 | 2.8e-06 | 2.9e-06 | 3.533E-06 | 1.185E-06 | 2.800E-06 | 4.900E-06 |
| 23851 | 1193 | L1193L | D | 3.7e-06 | 2.5e-06 | 3.8e-06 | 3.333E-06 | 7.234E-07 | 2.500E-06 | 3.800E-06 |
| 23852 | 1193 | L1193K | D | 2.4e-06 | 1.9e-06 | 1.9e-06 | 2.067E-06 | 2.887E-07 | 1.900E-06 | 2.400E-06 |
| 23853 | 1193 | L1193M | D | 3.4e-06 | 2e-06 | 2.8e-06 | 2.733E-06 | 7.024E-07 | 2.000E-06 | 3.400E-06 |
| 23854 | 1193 | L1193F | D | 1.5e-06 | 2.1e-06 | 2.6e-06 | 2.067E-06 | 5.508E-07 | 1.500E-06 | 2.600E-06 |
| 23855 | 1193 | L1193P | D | 2.7e-06 | 3.1e-06 | 3.1e-06 | 2.967E-06 | 2.309E-07 | 2.700E-06 | 3.100E-06 |
| 23856 | 1193 | L1193S | D | 1.6e-06 | 2.6e-06 | 2.4e-06 | 2.200E-06 | 5.292E-07 | 1.600E-06 | 2.600E-06 |
| 23857 | 1193 | L1193T | D | 3.9e-06 | 3.8e-06 | 3.4e-06 | 3.700E-06 | 2.646E-07 | 3.400E-06 | 3.900E-06 |
| 23858 | 1193 | L1193W | D | 3.2e-06 | 2.3e-06 | 2.3e-06 | 2.600E-06 | 5.196E-07 | 2.300E-06 | 3.200E-06 |
| 23859 | 1193 | L1193Y | D | 3.1e-06 | 2.1e-06 | 2.3e-06 | 2.500E-06 | 5.292E-07 | 2.100E-06 | 3.100E-06 |
| 23860 | 1193 | L1193V | D | 2.3e-06 | 2.4e-06 | 2e-06 | 2.233E-06 | 2.082E-07 | 2.000E-06 | 2.400E-06 |
| 23861 | 1194 | L1194A | D | 3.4e-06 | 2.2e-06 | 3.1e-06 | 2.900E-06 | 6.245E-07 | 2.200E-06 | 3.400E-06 |
| 23862 | 1194 | L1194R | D | 2.6e-06 | 3.5e-06 | 5.1e-06 | 3.733E-06 | 1.266E-06 | 2.600E-06 | 5.100E-06 |
| 23863 | 1194 | L1194N | D | 2.3e-06 | 2.6e-06 | 2.5e-06 | 2.467E-06 | 1.528E-07 | 2.300E-06 | 2.600E-06 |
| 23864 | 1194 | L1194D | D | 2.1e-06 | 2.5e-06 | 1.1e-06 | 1.900E-06 | 7.211E-07 | 1.100E-06 | 2.500E-06 |
| 23865 | 1194 | L1194C | D | 3e-06 | 2.2e-06 | 1.3e-06 | 2.167E-06 | 8.505E-07 | 1.300E-06 | 3.000E-06 |
| 23866 | 1194 | L1194E | D | 2.6e-06 | 1.4e-06 | 1.8e-06 | 1.933E-06 | 6.110E-07 | 1.400E-06 | 2.600E-06 |
| 23867 | 1194 | L1194Q | D | 2e-06 | 2.1e-06 | 1.9e-06 | 2.000E-06 | 1.000E-07 | 1.900E-06 | 2.100E-06 |
| 23868 | 1194 | L1194G | D | 1.6e-06 | 2.5e-06 | 1.8e-06 | 1.967E-06 | 4.726E-07 | 1.600E-06 | 2.500E-06 |
| 23869 | 1194 | L1194H | D | 2e-06 | 1.9e-06 | 3.9e-06 | 2.600E-06 | 1.127E-06 | 1.900E-06 | 3.900E-06 |
| 23870 | 1194 | L1194I | D | 2.8e-06 | 2.4e-06 | 3.6e-06 | 2.933E-06 | 6.110E-07 | 2.400E-06 | 3.600E-06 |
| 23871 | 1194 | L1194L | D | 1.5e-06 | 1.5e-06 | 1.8e-06 | 1.600E-06 | 1.732E-07 | 1.500E-06 | 1.800E-06 |
| 23872 | 1194 | L1194K | D | 2.4e-06 | 2.7e-06 | 2e-06 | 2.367E-06 | 3.512E-07 | 2.000E-06 | 2.700E-06 |
| 23873 | 1194 | L1194M | D | 2.7e-06 | 3.5e-06 | 4.9e-06 | 3.700E-06 | 1.114E-06 | 2.700E-06 | 4.900E-06 |
| 23874 | 1194 | L1194F | D | 2.4e-06 | 1.4e-06 | 2.6e-06 | 2.133E-06 | 6.429E-07 | 1.400E-06 | 2.600E-06 |
| 23875 | 1194 | L1194P | D | 2.4e-06 | 2.2e-06 | 3.1e-06 | 2.567E-06 | 4.726E-07 | 2.200E-06 | 3.100E-06 |
| 23876 | 1194 | L1194S | D | 2.2e-06 | 3.5e-06 | 2e-06 | 2.567E-06 | 8.145E-07 | 2.000E-06 | 3.500E-06 |
| 23877 | 1194 | L1194T | D | 1.8e-06 | 2.5e-06 | 1.8e-06 | 2.033E-06 | 4.041E-07 | 1.800E-06 | 2.500E-06 |
| 23878 | 1194 | L1194W | D | 1.9e-06 | 3.1e-06 | 1.5e-06 | 2.167E-06 | 8.327E-07 | 1.500E-06 | 3.100E-06 |
| 23879 | 1194 | L1194Y | D | 1.9e-06 | 2.6e-06 | 1.5e-06 | 2.000E-06 | 5.568E-07 | 1.500E-06 | 2.600E-06 |
| 23880 | 1194 | L1194V | D | 3e-06 | 1.7e-06 | 2.1e-06 | 2.267E-06 | 6.658E-07 | 1.700E-06 | 3.000E-06 |
| 23881 | 1195 | N1195A | D | 2e-06 | 3e-06 | 3.7e-06 | 2.900E-06 | 8.544E-07 | 2.000E-06 | 3.700E-06 |
| 23882 | 1195 | N1195R | D | 2.3e-06 | 3e-06 | 2e-06 | 2.433E-06 | 5.132E-07 | 2.000E-06 | 3.000E-06 |
| 23883 | 1195 | N1195N | D | 4.6e-06 | 3.2e-06 | 2.2e-06 | 3.333E-06 | 1.206E-06 | 2.200E-06 | 4.600E-06 |
| 23884 | 1195 | N1195D | D | 2.1e-06 | 5.1e-06 | 2.5e-06 | 3.233E-06 | 1.629E-06 | 2.100E-06 | 5.100E-06 |

| | | | | | | | | | | |
|---|---|---|---|---|---|---|---|---|---|---|
| 23885 | 1195 | N1195C | D | 1.6e-06 | 1.7e-06 | 2.5e-06 | 1.933E-06 | 4.933E-07 | 1.600E-06 | 2.500E-06 |
| 23886 | 1195 | N1195E | D | 2.1e-06 | 1.6e-06 | 1.8e-06 | 1.833E-06 | 2.517E-07 | 1.600E-06 | 2.100E-06 |
| 23887 | 1195 | N1195Q | D | 2.4e-06 | 2.2e-06 | 2.4e-06 | 2.333E-06 | 1.155E-07 | 2.200E-06 | 2.400E-06 |
| 23888 | 1195 | N1195G | D | 2.4e-06 | 1.5e-06 | 2.6e-06 | 2.167E-06 | 5.859E-07 | 1.500E-06 | 2.600E-06 |
| 23889 | 1195 | N1195H | D | 2.4e-06 | 2e-06 | 4.5e-06 | 2.967E-06 | 1.343E-06 | 2.000E-06 | 4.500E-06 |
| 23890 | 1195 | N1195I | D | 2e-06 | 1.8e-06 | 2.9e-06 | 2.233E-06 | 5.859E-07 | 1.800E-06 | 2.900E-06 |
| 23891 | 1195 | N1195L | D | 2.2e-06 | 2.3e-06 | 1.3e-06 | 1.933E-06 | 5.508E-07 | 1.300E-06 | 2.300E-06 |
| 23892 | 1195 | N1195K | D | 2.7e-06 | 2.5e-06 | 2.1e-06 | 2.433E-06 | 3.055E-07 | 2.100E-06 | 2.700E-06 |
| 23893 | 1195 | N1195M | D | 3.6e-06 | 2.5e-06 | 5.1e-06 | 3.733E-06 | 1.305E-06 | 2.500E-06 | 5.100E-06 |
| 23894 | 1195 | N1195F | D | 1.7e-06 | 2e-06 | 3.5e-06 | 2.400E-06 | 9.644E-07 | 1.700E-06 | 3.500E-06 |
| 23895 | 1195 | N1195P | D | 3.1e-06 | 1.4e-06 | 2.6e-06 | 2.367E-06 | 8.737E-07 | 1.400E-06 | 3.100E-06 |
| 23896 | 1195 | N1195S | D | 3e-06 | 2.2e-06 | 1.7e-06 | 2.300E-06 | 6.557E-07 | 1.700E-06 | 3.000E-06 |
| 23897 | 1195 | N1195T | D | 2.4e-06 | 1.8e-06 | 1.7e-06 | 1.967E-06 | 3.786E-07 | 1.700E-06 | 2.400E-06 |
| 23898 | 1195 | N1195W | D | 1.7e-06 | 1.5e-06 | 1.5e-06 | 1.567E-06 | 1.155E-07 | 1.500E-06 | 1.700E-06 |
| 23899 | 1195 | N1195Y | D | 1.3e-06 | 2.3e-06 | 3.7e-06 | 2.433E-06 | 1.206E-06 | 1.300E-06 | 3.700E-06 |
| 23900 | 1195 | N1195V | D | 2.6e-06 | 1.7e-06 | 1.2e-06 | 1.833E-06 | 7.095E-07 | 1.200E-06 | 2.600E-06 |
| 23901 | 1196 | N1196A | D | 1.8e-06 | 1.6e-06 | 7.7e-07 | 1.390E-06 | 5.462E-07 | 7.700E-07 | 1.800E-06 |
| 23902 | 1196 | N1196R | D | 1.6e-06 | 1.9e-06 | 3.4e-06 | 2.300E-06 | 9.644E-07 | 1.600E-06 | 3.400E-06 |
| 23903 | 1196 | N1196N | D | 2.1e-06 | 3e-06 | 2.4e-06 | 2.500E-06 | 4.583E-07 | 2.100E-06 | 3.000E-06 |
| 23904 | 1196 | N1196D | D | 1.7e-06 | 2.7e-06 | 1e-06 | 1.800E-06 | 8.544E-07 | 1.000E-06 | 2.700E-06 |
| 23905 | 1196 | N1196C | D | 2.3e-06 | 2e-06 | 2.2e-06 | 2.167E-06 | 1.528E-07 | 2.000E-06 | 2.300E-06 |
| 23906 | 1196 | N1196E | D | 2.1e-06 | 2.1e-06 | 1e-06 | 1.733E-06 | 6.351E-07 | 1.000E-06 | 2.100E-06 |
| 23907 | 1196 | N1196Q | D | 2.6e-06 | 3e-06 | 2.7e-06 | 2.767E-06 | 2.082E-07 | 2.600E-06 | 3.000E-06 |
| 23908 | 1196 | N1196G | D | 1.1e-06 | 1.3e-06 | 1.3e-06 | 1.233E-06 | 1.155E-07 | 1.100E-06 | 1.300E-06 |
| 23909 | 1196 | N1196H | D | 2.8e-06 | 2.1e-06 | 2.4e-06 | 2.433E-06 | 3.512E-07 | 2.100E-06 | 2.800E-06 |
| 23910 | 1196 | N1196I | D | 2.8e-06 | 2.4e-06 | 1.5e-06 | 2.233E-06 | 6.658E-07 | 1.500E-06 | 2.800E-06 |
| 23911 | 1196 | N1196L | D | 1.6e-06 | 2.3e-06 | 2.5e-06 | 2.133E-06 | 4.726E-07 | 1.600E-06 | 2.500E-06 |
| 23912 | 1196 | N1196K | D | 2.4e-06 | 1.4e-06 | 2e-06 | 1.933E-06 | 5.033E-07 | 1.400E-06 | 2.400E-06 |
| 23913 | 1196 | N1196M | D | 2.9e-06 | 2.5e-06 | 3.2e-06 | 2.867E-06 | 3.512E-07 | 2.500E-06 | 3.200E-06 |
| 23914 | 1196 | N1196F | D | 1.7e-06 | 2.7e-06 | 2.9e-06 | 2.433E-06 | 6.429E-07 | 1.700E-06 | 2.900E-06 |
| 23915 | 1196 | N1196P | D | 1.2e-06 | 2.3e-06 | 1.6e-06 | 1.700E-06 | 5.568E-07 | 1.200E-06 | 2.300E-06 |
| 23916 | 1196 | N1196S | D | 2.3e-06 | 2.6e-06 | 1.9e-06 | 2.267E-06 | 3.512E-07 | 1.900E-06 | 2.600E-06 |
| 23917 | 1196 | N1196T | D | 2.6e-06 | 2.9e-06 | 1.7e-06 | 2.400E-06 | 6.245E-07 | 1.700E-06 | 2.900E-06 |
| 23918 | 1196 | N1196W | D | 1.9e-06 | 2e-06 | 1.9e-06 | 1.933E-06 | 5.774E-08 | 1.900E-06 | 2.000E-06 |
| 23919 | 1196 | N1196Y | D | 3.5e-06 | 2.2e-06 | 1.2e-06 | 2.300E-06 | 1.153E-06 | 1.200E-06 | 3.500E-06 |
| 23920 | 1196 | N1196V | D | 1.8e-06 | 3.3e-06 | 2.4e-06 | 2.500E-06 | 7.550E-07 | 1.800E-06 | 3.300E-06 |
| 23921 | 1197 | F1197A | D | 2.4e-06 | 3.3e-06 | 2.8e-06 | 2.833E-06 | 4.509E-07 | 2.400E-06 | 3.300E-06 |
| 23922 | 1197 | F1197R | D | 2.8e-06 | 3.4e-06 | 2.8e-06 | 3.000E-06 | 3.464E-07 | 2.800E-06 | 3.400E-06 |
| 23923 | 1197 | F1197N | D | 1.6e-06 | 1.8e-06 | 1.2e-06 | 1.533E-06 | 3.055E-07 | 1.200E-06 | 1.800E-06 |
| 23924 | 1197 | F1197D | D | 2.3e-06 | 2.2e-06 | 3.7e-06 | 2.733E-06 | 8.386E-07 | 2.200E-06 | 3.700E-06 |
| 23925 | 1197 | F1197C | D | 3.9e-06 | 3.1e-06 | 1.8e-06 | 2.933E-06 | 1.060E-06 | 1.800E-06 | 3.900E-06 |
| 23926 | 1197 | F1197E | D | 1.7e-06 | 2.5e-06 | 2.4e-06 | 2.200E-06 | 4.359E-07 | 1.700E-06 | 2.500E-06 |
| 23927 | 1197 | F1197Q | D | 1.2e-06 | 3.2e-06 | 2e-06 | 2.133E-06 | 1.007E-06 | 1.200E-06 | 3.200E-06 |
| 23928 | 1197 | F1197G | D | 3.4e-06 | 2.9e-06 | 1.7e-06 | 2.667E-06 | 8.737E-07 | 1.700E-06 | 3.400E-06 |
| 23929 | 1197 | F1197H | D | 1.8e-06 | 4.6e-06 | 3.9e-06 | 3.433E-06 | 1.457E-06 | 1.800E-06 | 4.600E-06 |
| 23930 | 1197 | F1197I | D | 2.8e-06 | 1.9e-06 | 2.6e-06 | 2.433E-06 | 4.726E-07 | 1.900E-06 | 2.800E-06 |
| 23931 | 1197 | F1197L | D | 1.8e-06 | 1.9e-06 | 2.6e-06 | 2.100E-06 | 4.359E-07 | 1.800E-06 | 2.600E-06 |
| 23932 | 1197 | F1197K | D | 1.8e-06 | 7.9e-07 | 1.3e-06 | 1.297E-06 | 5.050E-07 | 7.900E-07 | 1.800E-06 |
| 23933 | 1197 | F1197M | D | 6.5e-06 | 3e-06 | 1.8e-06 | 3.767E-06 | 2.442E-06 | 1.800E-06 | 6.500E-06 |
| 23934 | 1197 | F1197F | D | 1.6e-06 | 4.4e-06 | 3.4e-06 | 3.133E-06 | 1.419E-06 | 1.600E-06 | 4.400E-06 |
| 23935 | 1197 | F1197P | D | 2.9e-06 | 2.7e-06 | 1.8e-06 | 2.467E-06 | 5.859E-07 | 1.800E-06 | 2.900E-06 |
| 23936 | 1197 | F1197S | D | 2.2e-06 | 1.7e-06 | 2.6e-06 | 2.167E-06 | 4.509E-07 | 1.700E-06 | 2.600E-06 |
| 23937 | 1197 | F1197T | D | 2.2e-06 | 2.4e-06 | 2.2e-06 | 2.267E-06 | 1.155E-07 | 2.200E-06 | 2.400E-06 |
| 23938 | 1197 | F1197W | D | 3.5e-06 | 3.6e-06 | 2.3e-06 | 3.133E-06 | 7.234E-07 | 2.300E-06 | 3.600E-06 |
| 23939 | 1197 | F1197Y | D | 3.5e-06 | 4e-06 | 1.9e-06 | 3.133E-06 | 1.097E-06 | 1.900E-06 | 4.000E-06 |
| 23940 | 1197 | F1197V | D | 1.8e-06 | 2.7e-06 | 1.2e-06 | 1.900E-06 | 7.550E-07 | 1.200E-06 | 2.700E-06 |
| 23941 | 1198 | Y1198A | D | 1.9e-06 | 3.2e-06 | 3.6e-06 | 2.900E-06 | 8.888E-07 | 1.900E-06 | 3.600E-06 |
| 23942 | 1198 | Y1198R | D | 2.9e-06 | 1.7e-06 | 2.1e-06 | 2.233E-06 | 6.110E-07 | 1.700E-06 | 2.900E-06 |
| 23943 | 1198 | Y1198N | D | 5.8e-06 | 2.1e-06 | 4.3e-06 | 4.067E-06 | 1.861E-06 | 2.100E-06 | 5.800E-06 |
| 23944 | 1198 | Y1198D | D | 3.1e-06 | 1.5e-06 | 3e-06 | 2.533E-06 | 8.963E-07 | 1.500E-06 | 3.100E-06 |
| 23945 | 1198 | Y1198C | D | 2.8e-06 | 3e-06 | 3e-06 | 2.933E-06 | 1.155E-07 | 2.800E-06 | 3.000E-06 |
| 23946 | 1198 | Y1198E | D | 2.1e-06 | 2e-06 | 2.1e-06 | 2.067E-06 | 5.774E-08 | 2.000E-06 | 2.100E-06 |
| 23947 | 1198 | Y1198Q | D | 2.1e-06 | 2.3e-06 | 9.7e-07 | 1.790E-06 | 7.171E-07 | 9.700E-07 | 2.300E-06 |
| 23948 | 1198 | Y1198G | D | 2.2e-06 | 1.3e-06 | 1.7e-06 | 1.733E-06 | 4.509E-07 | 1.300E-06 | 2.200E-06 |
| 23949 | 1198 | Y1198H | D | 3.1e-06 | 2.6e-06 | 2.9e-06 | 2.867E-06 | 2.517E-07 | 2.600E-06 | 3.100E-06 |
| 23950 | 1198 | Y1198I | D | 2e-06 | 3e-06 | 2.9e-06 | 2.633E-06 | 5.508E-07 | 2.000E-06 | 3.000E-06 |
| 23951 | 1198 | Y1198L | D | 2.2e-06 | 1e-06 | 1.9e-06 | 1.700E-06 | 6.245E-07 | 1.000E-06 | 2.200E-06 |
| 23952 | 1198 | Y1198K | D | 2.9e-06 | 2.6e-06 | 1.1e-06 | 2.200E-06 | 9.644E-07 | 1.100E-06 | 2.900E-06 |
| 23953 | 1198 | Y1198M | D | 2.4e-06 | 1.6e-06 | 2.9e-06 | 2.300E-06 | 6.557E-07 | 1.600E-06 | 2.900E-06 |
| 23954 | 1198 | Y1198F | D | 2.2e-06 | 3.4e-06 | 2.4e-06 | 2.667E-06 | 6.429E-07 | 2.200E-06 | 3.400E-06 |
| 23955 | 1198 | Y1198P | D | 1.8e-06 | 1.7e-06 | 2.3e-06 | 1.933E-06 | 3.215E-07 | 1.700E-06 | 2.300E-06 |
| 23956 | 1198 | Y1198S | D | 2.7e-06 | 4.1e-06 | 1.2e-06 | 2.667E-06 | 1.450E-06 | 1.200E-06 | 4.100E-06 |
| 23957 | 1198 | Y1198T | D | 2.2e-06 | 3.2e-06 | 2.2e-06 | 2.533E-06 | 5.774E-07 | 2.200E-06 | 3.200E-06 |
| 23958 | 1198 | Y1198W | D | 1.9e-06 | 3.9e-06 | 2.2e-06 | 2.667E-06 | 1.079E-06 | 1.900E-06 | 3.900E-06 |
| 23959 | 1198 | Y1198Y | D | 4.2e-06 | 2e-06 | 2.7e-06 | 2.967E-06 | 1.124E-06 | 2.000E-06 | 4.200E-06 |
| 23960 | 1198 | Y1198V | D | 3.4e-06 | 2.8e-06 | 1.6e-06 | 2.600E-06 | 9.165E-07 | 1.600E-06 | 3.400E-06 |
| 23961 | 1199 | P1199A | D | 2.1e-06 | 4.9e-06 | 1.9e-06 | 2.967E-06 | 1.677E-06 | 1.900E-06 | 4.900E-06 |
| 23962 | 1199 | P1199R | D | 1.6e-06 | 1.8e-06 | 2.6e-06 | 2.000E-06 | 5.292E-07 | 1.600E-06 | 2.600E-06 |
| 23963 | 1199 | P1199N | D | 1.9e-06 | 2.9e-06 | 1.6e-06 | 2.133E-06 | 6.807E-07 | 1.600E-06 | 2.900E-06 |
| 23964 | 1199 | P1199D | D | 3.2e-06 | 3e-06 | 2.5e-06 | 2.900E-06 | 3.606E-07 | 2.500E-06 | 3.200E-06 |
| 23965 | 1199 | P1199C | D | 6.7e-07 | 2e-06 | 2.6e-06 | 1.757E-06 | 9.877E-07 | 6.700E-07 | 2.600E-06 |
| 23966 | 1199 | P1199E | D | 2.6e-06 | 5.1e-06 | 2.5e-06 | 3.400E-06 | 1.473E-06 | 2.500E-06 | 5.100E-06 |
| 23967 | 1199 | P1199Q | D | 2.2e-06 | 2.1e-06 | 4.4e-06 | 2.900E-06 | 1.300E-06 | 2.100E-06 | 4.400E-06 |
| 23968 | 1199 | P1199G | D | 2.5e-06 | 3e-06 | 3.3e-06 | 2.933E-06 | 4.041E-07 | 2.500E-06 | 3.300E-06 |
| 23969 | 1199 | P1199H | D | 3.4e-06 | 2.1e-06 | 1.5e-06 | 2.333E-06 | 9.713E-07 | 1.500E-06 | 3.400E-06 |

| | | | | | | | | | | |
|---|---|---|---|---|---|---|---|---|---|---|
| 23970 | 1199 | P1199I | D | 2.9e-06 | 2.7e-06 | 2.6e-06 | 2.733E-06 | 1.528E-07 | 2.600E-06 | 2.900E-06 |
| 23971 | 1199 | P1199L | D | 3.4e-06 | 2.3e-06 | 1.5e-06 | 2.400E-06 | 9.539E-07 | 1.500E-06 | 3.400E-06 |
| 23972 | 1199 | P1199K | D | 3.7e-06 | 2.7e-06 | 1.7e-06 | 2.700E-06 | 1.000E-06 | 1.700E-06 | 3.700E-06 |
| 23973 | 1199 | P1199M | D | 3.9e-06 | 1.9e-06 | 2.2e-06 | 2.667E-06 | 1.079E-06 | 1.900E-06 | 3.900E-06 |
| 23974 | 1199 | P1199F | D | 2e-06 | 1.2e-06 | 2e-06 | 1.733E-06 | 4.619E-07 | 1.200E-06 | 2.000E-06 |
| 23975 | 1199 | P1199P | D | 2e-06 | 3.9e-06 | 1.8e-06 | 2.567E-06 | 1.159E-06 | 1.800E-06 | 3.900E-06 |
| 23976 | 1199 | P1199S | D | 2.1e-06 | 2.3e-06 | 3.2e-06 | 2.533E-06 | 5.859E-07 | 2.100E-06 | 3.200E-06 |
| 23977 | 1199 | P1199T | D | 2.9e-06 | 2.2e-06 | 2.5e-06 | 2.533E-06 | 3.512E-07 | 2.200E-06 | 2.900E-06 |
| 23978 | 1199 | P1199W | D | 2e-06 | 3.1e-06 | 2e-06 | 2.367E-06 | 6.351E-07 | 2.000E-06 | 3.100E-06 |
| 23979 | 1199 | P1199Y | D | 1.6e-06 | 1.6e-06 | 2.3e-06 | 1.833E-06 | 4.041E-07 | 1.600E-06 | 2.300E-06 |
| 23980 | 1199 | P1199V | D | 2.4e-06 | 1.9e-06 | 1.4e-06 | 1.900E-06 | 5.000E-07 | 1.400E-06 | 2.400E-06 |
| 23981 | 1200 | R1200A | D | 2.5e-06 | 2.5e-06 | 2.5e-06 | 2.500E-06 | 0.000E+00 | 2.500E-06 | 2.500E-06 |
| 23982 | 1200 | R1200R | D | 1.4e-06 | 3e-06 | 1.6e-06 | 2.000E-06 | 8.718E-07 | 1.400E-06 | 3.000E-06 |
| 23983 | 1200 | R1200N | D | 2e-06 | 3.1e-06 | 2.9e-06 | 2.667E-06 | 5.859E-07 | 2.000E-06 | 3.100E-06 |
| 23984 | 1200 | R1200D | D | 4.9e-06 | 3.3e-06 | 2.5e-06 | 3.567E-06 | 1.222E-06 | 2.500E-06 | 4.900E-06 |
| 23985 | 1200 | R1200C | D | 2.2e-06 | 1.3e-06 | 2.3e-06 | 1.933E-06 | 5.508E-07 | 1.300E-06 | 2.300E-06 |
| 23986 | 1200 | R1200E | D | 1.9e-06 | 1.7e-06 | 3.9e-06 | 2.500E-06 | 1.217E-06 | 1.700E-06 | 3.900E-06 |
| 23987 | 1200 | R1200Q | D | 9.8e-07 | 3.3e-06 | 2.3e-06 | 2.193E-06 | 1.164E-06 | 9.800E-07 | 3.300E-06 |
| 23988 | 1200 | R1200G | D | 1.6e-06 | 3.1e-06 | 1.7e-06 | 2.133E-06 | 8.386E-07 | 1.600E-06 | 3.100E-06 |
| 23989 | 1200 | R1200H | D | 2.5e-06 | 2.4e-06 | 2.6e-06 | 2.500E-06 | 1.000E-07 | 2.400E-06 | 2.600E-06 |
| 23990 | 1200 | R1200I | D | 3.3e-06 | 2.8e-06 | 2.3e-06 | 2.800E-06 | 5.000E-07 | 2.300E-06 | 3.300E-06 |
| 23991 | 1200 | R1200L | D | 2.7e-06 | 3.9e-06 | 1.1e-06 | 2.567E-06 | 1.405E-06 | 1.100E-06 | 3.900E-06 |
| 23992 | 1200 | R1200K | D | 1.7e-06 | 1.6e-06 | 1.4e-06 | 1.567E-06 | 1.528E-07 | 1.400E-06 | 1.700E-06 |
| 23993 | 1200 | R1200M | D | 2.6e-06 | 2.1e-06 | 2e-06 | 2.233E-06 | 3.215E-07 | 2.000E-06 | 2.600E-06 |
| 23994 | 1200 | R1200F | D | 2.1e-06 | 2.7e-06 | 2.2e-06 | 2.333E-06 | 3.215E-07 | 2.100E-06 | 2.700E-06 |
| 23995 | 1200 | R1200P | D | 2.7e-06 | 3e-06 | 1.7e-06 | 2.467E-06 | 6.807E-07 | 1.700E-06 | 3.000E-06 |
| 23996 | 1200 | R1200S | D | 1.9e-06 | 1.7e-06 | 3.4e-06 | 2.333E-06 | 9.292E-07 | 1.700E-06 | 3.400E-06 |
| 23997 | 1200 | R1200T | D | 3e-06 | 1.9e-06 | 2.9e-06 | 2.600E-06 | 6.083E-07 | 1.900E-06 | 3.000E-06 |
| 23998 | 1200 | R1200W | D | 2e-06 | 1.7e-06 | 1.7e-06 | 1.800E-06 | 1.732E-07 | 1.700E-06 | 2.000E-06 |
| 23999 | 1200 | R1200Y | D | 4.9e-06 | 2.2e-06 | 2.8e-06 | 3.300E-06 | 1.418E-06 | 2.200E-06 | 4.900E-06 |
| 24000 | 1200 | R1200V | D | 4.3e-06 | 3.9e-06 | 1.6e-06 | 3.267E-06 | 1.457E-06 | 1.600E-06 | 4.300E-06 |
| 24001 | 1201 | E1201A | D | 2.4e-06 | 2.8e-06 | 1.7e-06 | 2.300E-06 | 5.568E-07 | 1.700E-06 | 2.800E-06 |
| 24002 | 1201 | E1201R | D | 2.9e-06 | 2.3e-06 | 2.4e-06 | 2.533E-06 | 3.215E-07 | 2.300E-06 | 2.900E-06 |
| 24003 | 1201 | E1201N | D | 1.6e-06 | 5.1e-06 | 2.1e-06 | 2.933E-06 | 1.893E-06 | 1.600E-06 | 5.100E-06 |
| 24004 | 1201 | E1201D | D | 2.7e-06 | 1.2e-06 | 2.6e-06 | 2.167E-06 | 8.386E-07 | 1.200E-06 | 2.700E-06 |
| 24005 | 1201 | E1201C | D | 5e-06 | 1.1e-06 | 1.7e-06 | 2.600E-06 | 2.100E-06 | 1.100E-06 | 5.000E-06 |
| 24006 | 1201 | E1201E | D | 2.8e-06 | 2.1e-06 | 3e-06 | 2.633E-06 | 4.726E-07 | 2.100E-06 | 3.000E-06 |
| 24007 | 1201 | E1201Q | D | 2.1e-06 | 2e-06 | 3.3e-06 | 2.467E-06 | 7.234E-07 | 2.000E-06 | 3.300E-06 |
| 24008 | 1201 | E1201G | D | 2.6e-06 | 2.3e-06 | 2e-06 | 2.300E-06 | 3.000E-07 | 2.000E-06 | 2.600E-06 |
| 24009 | 1201 | E1201H | D | 3.3e-06 | 2.2e-06 | 2.8e-06 | 2.767E-06 | 5.508E-07 | 2.200E-06 | 3.300E-06 |
| 24010 | 1201 | E1201I | D | 2.2e-06 | 3.1e-06 | 2.8e-06 | 2.700E-06 | 4.583E-07 | 2.200E-06 | 3.100E-06 |
| 24011 | 1201 | E1201L | D | 2.4e-06 | 1.9e-06 | 2e-06 | 2.100E-06 | 2.646E-07 | 1.900E-06 | 2.400E-06 |
| 24012 | 1201 | E1201K | D | 4.5e-06 | 2.8e-06 | 2.6e-06 | 3.300E-06 | 1.044E-06 | 2.600E-06 | 4.500E-06 |
| 24013 | 1201 | E1201M | D | 1.3e-06 | 1.8e-06 | 1.6e-06 | 1.567E-06 | 2.517E-07 | 1.300E-06 | 1.800E-06 |
| 24014 | 1201 | E1201F | D | 2.6e-06 | 1.9e-06 | 4.5e-06 | 3.000E-06 | 1.345E-06 | 1.900E-06 | 4.500E-06 |
| 24015 | 1201 | E1201P | D | 1.8e-06 | 2.1e-06 | 3.9e-06 | 2.600E-06 | 1.136E-06 | 1.800E-06 | 3.900E-06 |
| 24016 | 1201 | E1201S | D | 4.2e-06 | 2.1e-06 | 3.3e-06 | 3.200E-06 | 1.054E-06 | 2.100E-06 | 4.200E-06 |
| 24017 | 1201 | E1201T | D | 2.2e-06 | 1.6e-06 | 1.7e-06 | 1.833E-06 | 3.215E-07 | 1.600E-06 | 2.200E-06 |
| 24018 | 1201 | E1201W | D | 2.5e-06 | 1.3e-06 | 1.5e-06 | 1.767E-06 | 6.429E-07 | 1.300E-06 | 2.500E-06 |
| 24019 | 1201 | E1201Y | D | 3.9e-06 | 2.2e-06 | 1.9e-06 | 2.667E-06 | 1.079E-06 | 1.900E-06 | 3.900E-06 |
| 24020 | 1201 | E1201V | D | 2.9e-06 | 4.7e-06 | 1.9e-06 | 3.167E-06 | 1.419E-06 | 1.900E-06 | 4.700E-06 |
| 24021 | 1202 | A1202A | D | 2.3e-06 | 2e-06 | 2.3e-06 | 2.200E-06 | 1.732E-07 | 2.000E-06 | 2.300E-06 |
| 24022 | 1202 | A1202R | D | 2.2e-06 | 1.1e-06 | 2.1e-06 | 1.800E-06 | 6.083E-07 | 1.100E-06 | 2.200E-06 |
| 24023 | 1202 | A1202N | D | 2.5e-06 | 3.9e-06 | 3.6e-06 | 3.333E-06 | 7.371E-07 | 2.500E-06 | 3.900E-06 |
| 24024 | 1202 | A1202D | D | 2.5e-06 | 3.7e-06 | 2e-06 | 2.733E-06 | 8.737E-07 | 2.000E-06 | 3.700E-06 |
| 24025 | 1202 | A1202C | D | 2e-06 | 2.9e-06 | 2.8e-06 | 2.567E-06 | 4.933E-07 | 2.000E-06 | 2.900E-06 |
| 24026 | 1202 | A1202E | D | 3.1e-06 | 2.7e-06 | 3.3e-06 | 3.033E-06 | 3.055E-07 | 2.700E-06 | 3.300E-06 |
| 24027 | 1202 | A1202Q | D | 2.4e-06 | 3.5e-06 | 1.4e-06 | 2.433E-06 | 1.050E-06 | 1.400E-06 | 3.500E-06 |
| 24028 | 1202 | A1202G | D | 1.5e-06 | 3.9e-06 | 1.7e-06 | 2.367E-06 | 1.332E-06 | 1.500E-06 | 3.900E-06 |
| 24029 | 1202 | A1202H | D | 2.9e-06 | 1.7e-06 | 1.8e-06 | 2.133E-06 | 6.658E-07 | 1.700E-06 | 2.900E-06 |
| 24030 | 1202 | A1202I | D | 2.8e-06 | 2.2e-06 | 2.8e-06 | 2.600E-06 | 3.464E-07 | 2.200E-06 | 2.800E-06 |
| 24031 | 1202 | A1202L | D | 3.4e-06 | 3.4e-06 | 3.2e-06 | 3.333E-06 | 1.155E-07 | 3.200E-06 | 3.400E-06 |
| 24032 | 1202 | A1202K | D | 2.7e-06 | 2.6e-06 | 2.2e-06 | 2.500E-06 | 2.646E-07 | 2.200E-06 | 2.700E-06 |
| 24033 | 1202 | A1202M | D | 5.1e-06 | 2.3e-06 | 1.2e-06 | 2.867E-06 | 2.011E-06 | 1.200E-06 | 5.100E-06 |
| 24034 | 1202 | A1202F | D | 3.2e-06 | 3.4e-06 | 1.4e-06 | 2.667E-06 | 1.102E-06 | 1.400E-06 | 3.400E-06 |
| 24035 | 1202 | A1202P | D | 5.2e-06 | 3.8e-06 | 4.5e-06 | 4.500E-06 | 7.000E-07 | 3.800E-06 | 5.200E-06 |
| 24036 | 1202 | A1202S | D | 2.6e-06 | 2e-06 | 1.7e-06 | 2.100E-06 | 4.583E-07 | 1.700E-06 | 2.600E-06 |
| 24037 | 1202 | A1202T | D | 2.6e-06 | 1.5e-06 | 2.5e-06 | 2.200E-06 | 6.083E-07 | 1.500E-06 | 2.600E-06 |
| 24038 | 1202 | A1202W | D | 1.4e-06 | 3e-06 | 2e-06 | 2.133E-06 | 8.083E-07 | 1.400E-06 | 3.000E-06 |
| 24039 | 1202 | A1202Y | D | 1.8e-06 | 2.3e-06 | 1.8e-06 | 1.967E-06 | 2.887E-07 | 1.800E-06 | 2.300E-06 |
| 24040 | 1202 | A1202V | D | 3.1e-06 | 3.6e-06 | 1.8e-06 | 2.833E-06 | 9.292E-07 | 1.800E-06 | 3.600E-06 |
| 24041 | 1203 | K1203A | D | 3.8e-06 | 1.8e-06 | 2.3e-06 | 2.633E-06 | 1.041E-06 | 1.800E-06 | 3.800E-06 |
| 24042 | 1203 | K1203R | D | 1.2e-06 | 2.3e-06 | 1.1e-06 | 1.533E-06 | 6.658E-07 | 1.100E-06 | 2.300E-06 |
| 24043 | 1203 | K1203N | D | 2.7e-06 | 2.7e-06 | 1e-06 | 2.133E-06 | 9.815E-07 | 1.000E-06 | 2.700E-06 |
| 24044 | 1203 | K1203D | D | 2.5e-06 | 2.7e-06 | 4e-06 | 3.067E-06 | 8.145E-07 | 2.500E-06 | 4.000E-06 |
| 24045 | 1203 | K1203C | D | 3.6e-06 | 1.8e-06 | 2.9e-06 | 2.767E-06 | 9.074E-07 | 1.800E-06 | 3.600E-06 |
| 24046 | 1203 | K1203E | D | 7.1e-06 | 2.4e-06 | 2.2e-06 | 3.900E-06 | 2.773E-06 | 2.200E-06 | 7.100E-06 |
| 24047 | 1203 | K1203Q | D | 2.3e-06 | 1.3e-06 | 1.5e-06 | 1.700E-06 | 5.292E-07 | 1.300E-06 | 2.300E-06 |
| 24048 | 1203 | K1203G | D | 2.1e-06 | 3.1e-06 | 2.1e-06 | 2.433E-06 | 5.774E-07 | 2.100E-06 | 3.100E-06 |
| 24049 | 1203 | K1203H | D | 3e-06 | 2.9e-06 | 1.3e-06 | 2.400E-06 | 9.539E-07 | 1.300E-06 | 3.000E-06 |
| 24050 | 1203 | K1203I | D | 2.2e-06 | 1.8e-06 | 3e-06 | 2.333E-06 | 6.110E-07 | 1.800E-06 | 3.000E-06 |
| 24051 | 1203 | K1203L | D | 2.5e-06 | 2.3e-06 | 1.9e-06 | 2.233E-06 | 3.055E-07 | 1.900E-06 | 2.500E-06 |
| 24052 | 1203 | K1203K | D | 1.5e-06 | 2.7e-06 | 2.9e-06 | 2.367E-06 | 7.572E-07 | 1.500E-06 | 2.900E-06 |
| 24053 | 1203 | K1203M | D | 2.3e-06 | 2e-06 | 2.6e-06 | 2.300E-06 | 3.000E-07 | 2.000E-06 | 2.600E-06 |
| 24054 | 1203 | K1203F | D | 2.7e-06 | 1.3e-06 | 2.1e-06 | 2.033E-06 | 7.024E-07 | 1.300E-06 | 2.700E-06 |

| | | | | | | | | | | |
|---|---|---|---|---|---|---|---|---|---|---|
| 24055 | 1203 | K1203P | D | 3.6e-06 | 2.8e-06 | 2e-06 | 2.800E-06 | 8.000E-07 | 2.000E-06 | 3.600E-06 |
| 24056 | 1203 | K1203S | D | 3.5e-06 | 2.8e-06 | 1.7e-06 | 2.667E-06 | 9.074E-07 | 1.700E-06 | 3.500E-06 |
| 24057 | 1203 | K1203T | D | 2.8e-06 | 2.3e-06 | 2e-06 | 2.367E-06 | 4.041E-07 | 2.000E-06 | 2.800E-06 |
| 24058 | 1203 | K1203W | D | 1.6e-06 | 1.8e-06 | 3.7e-06 | 2.367E-06 | 1.159E-06 | 1.600E-06 | 3.700E-06 |
| 24059 | 1203 | K1203Y | D | 1.5e-06 | 1.6e-06 | 1.2e-06 | 1.433E-06 | 2.082E-07 | 1.200E-06 | 1.600E-06 |
| 24060 | 1203 | K1203V | D | 3.1e-06 | 2.8e-06 | 2.9e-06 | 2.933E-06 | 1.528E-07 | 2.800E-06 | 3.100E-06 |
| 24061 | 1204 | V1204A | D | 1.7e-06 | 1.4e-06 | 3.8e-06 | 2.300E-06 | 1.308E-06 | 1.400E-06 | 3.800E-06 |
| 24062 | 1204 | V1204R | D | 3.1e-06 | 1.8e-06 | 1.4e-06 | 2.100E-06 | 8.888E-07 | 1.400E-06 | 3.100E-06 |
| 24063 | 1204 | V1204N | D | 2.1e-06 | 2.4e-06 | 3.3e-06 | 2.600E-06 | 6.245E-07 | 2.100E-06 | 3.300E-06 |
| 24064 | 1204 | V1204D | D | 1.6e-06 | 2.7e-06 | 3.4e-06 | 2.567E-06 | 9.074E-07 | 1.600E-06 | 3.400E-06 |
| 24065 | 1204 | V1204C | D | 2e-06 | 1.8e-06 | 1.7e-06 | 1.833E-06 | 1.528E-07 | 1.700E-06 | 2.000E-06 |
| 24066 | 1204 | V1204E | D | 4.1e-06 | 2.4e-06 | 1.7e-06 | 2.733E-06 | 1.234E-06 | 1.700E-06 | 4.100E-06 |
| 24067 | 1204 | V1204Q | D | 2.2e-06 | 2.7e-06 | 3.1e-06 | 2.667E-06 | 4.509E-07 | 2.200E-06 | 3.100E-06 |
| 24068 | 1204 | V1204G | D | 1.7e-06 | 4.4e-06 | 1.4e-06 | 2.500E-06 | 1.652E-06 | 1.400E-06 | 4.400E-06 |
| 24069 | 1204 | V1204H | D | 2.9e-06 | 3.8e-06 | 2e-06 | 2.900E-06 | 9.000E-07 | 2.000E-06 | 3.800E-06 |
| 24070 | 1204 | V1204I | D | 1.3e-06 | 3.8e-06 | 4.1e-06 | 3.067E-06 | 1.537E-06 | 1.300E-06 | 4.100E-06 |
| 24071 | 1204 | V1204L | D | 3.6e-06 | 2e-06 | 2.1e-06 | 2.567E-06 | 8.963E-07 | 2.000E-06 | 3.600E-06 |
| 24072 | 1204 | V1204K | D | 2.5e-06 | 1.9e-06 | 3.6e-06 | 2.667E-06 | 8.622E-07 | 1.900E-06 | 3.600E-06 |
| 24073 | 1204 | V1204M | D | 2.1e-06 | 4.5e-06 | 3.7e-06 | 3.433E-06 | 1.222E-06 | 2.100E-06 | 4.500E-06 |
| 24074 | 1204 | V1204F | D | 2.6e-06 | 2.9e-06 | 2.8e-06 | 2.767E-06 | 1.528E-07 | 2.600E-06 | 2.900E-06 |
| 24075 | 1204 | V1204P | D | 1.4e-06 | 1.5e-06 | 4.7e-06 | 2.533E-06 | 1.877E-06 | 1.400E-06 | 4.700E-06 |
| 24076 | 1204 | V1204S | D | 1.9e-06 | 3.1e-06 | 2.8e-06 | 2.600E-06 | 6.245E-07 | 1.900E-06 | 3.100E-06 |
| 24077 | 1204 | V1204T | D | 2.5e-06 | 2.3e-06 | 2.1e-06 | 2.300E-06 | 2.000E-07 | 2.100E-06 | 2.500E-06 |
| 24078 | 1204 | V1204W | D | 1.7e-06 | 3.1e-06 | 2e-06 | 2.267E-06 | 7.371E-07 | 1.700E-06 | 3.100E-06 |
| 24079 | 1204 | V1204Y | D | 1.8e-06 | 9.6e-07 | 1.9e-06 | 1.553E-06 | 5.163E-07 | 9.600E-07 | 1.900E-06 |
| 24080 | 1204 | V1204V | D | 1.4e-06 | 3.4e-06 | 2.8e-06 | 2.533E-06 | 1.026E-06 | 1.400E-06 | 3.400E-06 |
| 24081 | 1205 | Q1205A | D | 2.9e-06 | 2.9e-06 | 2.4e-06 | 2.733E-06 | 2.887E-07 | 2.400E-06 | 2.900E-06 |
| 24082 | 1205 | Q1205R | D | 1.7e-06 | 2.6e-06 | 1.7e-06 | 2.000E-06 | 5.196E-07 | 1.700E-06 | 2.600E-06 |
| 24083 | 1205 | Q1205N | D | 2.2e-06 | 3.6e-06 | 2e-06 | 2.600E-06 | 8.718E-07 | 2.000E-06 | 3.600E-06 |
| 24084 | 1205 | Q1205D | D | 2.5e-06 | 2.1e-06 | 2.4e-06 | 2.333E-06 | 2.082E-07 | 2.100E-06 | 2.500E-06 |
| 24085 | 1205 | Q1205C | D | 1.9e-06 | 1.3e-06 | 2.1e-06 | 1.767E-06 | 4.163E-07 | 1.300E-06 | 2.100E-06 |
| 24086 | 1205 | Q1205E | D | 3.2e-06 | 3.2e-06 | 1.4e-06 | 2.600E-06 | 1.039E-06 | 1.400E-06 | 3.200E-06 |
| 24087 | 1205 | Q1205Q | D | 2.8e-06 | 1.4e-06 | 3e-06 | 2.400E-06 | 8.718E-07 | 1.400E-06 | 3.000E-06 |
| 24088 | 1205 | Q1205G | D | 2.3e-06 | 3.1e-06 | 3.2e-06 | 2.867E-06 | 4.933E-07 | 2.300E-06 | 3.200E-06 |
| 24089 | 1205 | Q1205H | D | 1.9e-06 | 1.5e-06 | 1.8e-06 | 1.733E-06 | 2.082E-07 | 1.500E-06 | 1.900E-06 |
| 24090 | 1205 | Q1205I | D | 3e-06 | 1.4e-06 | 2.2e-06 | 2.200E-06 | 8.000E-07 | 1.400E-06 | 3.000E-06 |
| 24091 | 1205 | Q1205L | D | 2.1e-06 | 3.1e-06 | 2.5e-06 | 2.567E-06 | 5.033E-07 | 2.100E-06 | 3.100E-06 |
| 24092 | 1205 | Q1205K | D | 2.3e-06 | 2.5e-06 | 2.5e-06 | 2.433E-06 | 1.155E-07 | 2.300E-06 | 2.500E-06 |
| 24093 | 1205 | Q1205M | D | 2.6e-06 | 2.1e-06 | 3.1e-06 | 2.600E-06 | 5.000E-07 | 2.100E-06 | 3.100E-06 |
| 24094 | 1205 | Q1205F | D | 4.4e-06 | 3e-06 | 1.8e-06 | 3.067E-06 | 1.301E-06 | 1.800E-06 | 4.400E-06 |
| 24095 | 1205 | Q1205P | D | 3e-06 | 3e-06 | 2.6e-06 | 2.867E-06 | 2.309E-07 | 2.600E-06 | 3.000E-06 |
| 24096 | 1205 | Q1205S | D | 2.3e-06 | 2.2e-06 | 2.4e-06 | 2.300E-06 | 1.000E-07 | 2.200E-06 | 2.400E-06 |
| 24097 | 1205 | Q1205T | D | 2.1e-06 | 1.4e-06 | 1.8e-06 | 1.767E-06 | 3.512E-07 | 1.400E-06 | 2.100E-06 |
| 24098 | 1205 | Q1205W | D | 1.4e-06 | 3.4e-06 | 2.2e-06 | 2.333E-06 | 1.007E-06 | 1.400E-06 | 3.400E-06 |
| 24099 | 1205 | Q1205Y | D | 2.6e-06 | 2.3e-06 | 3.8e-06 | 2.900E-06 | 7.937E-07 | 2.300E-06 | 3.800E-06 |
| 24100 | 1205 | Q1205V | D | 2.3e-06 | 2.3e-06 | 3.2e-06 | 2.600E-06 | 5.196E-07 | 2.300E-06 | 3.200E-06 |
| 24101 | 1206 | W1206A | D | 3.6e-06 | 1.1e-06 | 2e-06 | 2.233E-06 | 1.266E-06 | 1.100E-06 | 3.600E-06 |
| 24102 | 1206 | W1206R | D | 1.9e-06 | 3.4e-06 | 2.7e-06 | 2.667E-06 | 7.506E-07 | 1.900E-06 | 3.400E-06 |
| 24103 | 1206 | W1206N | D | 3.6e-06 | 2.8e-06 | 2.2e-06 | 2.867E-06 | 7.024E-07 | 2.200E-06 | 3.600E-06 |
| 24104 | 1206 | W1206D | D | 2.3e-06 | 1.9e-06 | 2.7e-06 | 2.300E-06 | 4.000E-07 | 1.900E-06 | 2.700E-06 |
| 24105 | 1206 | W1206C | D | 3e-06 | 4.7e-06 | 3e-06 | 3.567E-06 | 9.815E-07 | 3.000E-06 | 4.700E-06 |
| 24106 | 1206 | W1206E | D | 2.7e-06 | 1.7e-06 | 2.4e-06 | 2.267E-06 | 5.132E-07 | 1.700E-06 | 2.700E-06 |
| 24107 | 1206 | W1206Q | D | 3.5e-06 | 1.4e-06 | 1.8e-06 | 2.233E-06 | 1.115E-06 | 1.400E-06 | 3.500E-06 |
| 24108 | 1206 | W1206G | D | 2.3e-06 | 2.9e-06 | 2.8e-06 | 2.667E-06 | 3.215E-07 | 2.300E-06 | 2.900E-06 |
| 24109 | 1206 | W1206H | D | 2.5e-06 | 3.5e-06 | 1.4e-06 | 2.467E-06 | 1.050E-06 | 1.400E-06 | 3.500E-06 |
| 24110 | 1206 | W1206I | D | 2e-06 | 2.1e-06 | 2.6e-06 | 2.233E-06 | 3.215E-07 | 2.000E-06 | 2.600E-06 |
| 24111 | 1206 | W1206L | D | 2.7e-06 | 2.1e-06 | 1.8e-06 | 2.200E-06 | 4.583E-07 | 1.800E-06 | 2.700E-06 |
| 24112 | 1206 | W1206K | D | 2.3e-06 | 1.8e-06 | 1.7e-06 | 1.933E-06 | 3.215E-07 | 1.700E-06 | 2.300E-06 |
| 24113 | 1206 | W1206M | D | 4.3e-06 | 1.3e-06 | 2.8e-06 | 2.800E-06 | 1.500E-06 | 1.300E-06 | 4.300E-06 |
| 24114 | 1206 | W1206F | D | 1.8e-06 | 2.1e-06 | 3.1e-06 | 2.333E-06 | 6.807E-07 | 1.800E-06 | 3.100E-06 |
| 24115 | 1206 | W1206P | D | 1.8e-06 | 3.5e-06 | 2.1e-06 | 2.467E-06 | 9.074E-07 | 1.800E-06 | 3.500E-06 |
| 24116 | 1206 | W1206S | D | 2.8e-06 | 1.5e-06 | 3e-06 | 2.433E-06 | 8.145E-07 | 1.500E-06 | 3.000E-06 |
| 24117 | 1206 | W1206T | D | 2.1e-06 | 2.8e-06 | 2.5e-06 | 2.467E-06 | 3.512E-07 | 2.100E-06 | 2.800E-06 |
| 24118 | 1206 | W1206W | D | 1.9e-06 | 2.2e-06 | 2.6e-06 | 2.233E-06 | 3.512E-07 | 1.900E-06 | 2.600E-06 |
| 24119 | 1206 | W1206Y | D | 3e-06 | 3.6e-06 | 1.7e-06 | 2.767E-06 | 9.713E-07 | 1.700E-06 | 3.600E-06 |
| 24120 | 1206 | W1206V | D | 3.6e-06 | 2.8e-06 | 2.8e-06 | 3.067E-06 | 4.619E-07 | 2.800E-06 | 3.600E-06 |
| 24121 | 1207 | K1207A | D | 3e-06 | 2.5e-06 | 1.3e-06 | 2.267E-06 | 8.737E-07 | 1.300E-06 | 3.000E-06 |
| 24122 | 1207 | K1207R | D | 2e-06 | 2.8e-06 | 3e-06 | 2.600E-06 | 5.292E-07 | 2.000E-06 | 3.000E-06 |
| 24123 | 1207 | K1207N | D | 2e-06 | 1.5e-06 | 4.4e-06 | 2.633E-06 | 1.550E-06 | 1.500E-06 | 4.400E-06 |
| 24124 | 1207 | K1207D | D | 2.6e-06 | 1.7e-06 | 3.4e-06 | 2.567E-06 | 8.505E-07 | 1.700E-06 | 3.400E-06 |
| 24125 | 1207 | K1207C | D | 1.8e-06 | 2e-06 | 2.6e-06 | 2.133E-06 | 4.163E-07 | 1.800E-06 | 2.600E-06 |
| 24126 | 1207 | K1207E | D | 3.6e-06 | 2.6e-06 | 3e-06 | 3.067E-06 | 5.033E-07 | 2.600E-06 | 3.600E-06 |
| 24127 | 1207 | K1207Q | D | 2.9e-06 | 2.7e-06 | 1.6e-06 | 2.400E-06 | 7.000E-07 | 1.600E-06 | 2.900E-06 |
| 24128 | 1207 | K1207G | D | 2e-06 | 1.4e-06 | 1.8e-06 | 1.733E-06 | 3.055E-07 | 1.400E-06 | 2.000E-06 |
| 24129 | 1207 | K1207H | D | 2.2e-06 | 1.9e-06 | 3.2e-06 | 2.433E-06 | 6.807E-07 | 1.900E-06 | 3.200E-06 |
| 24130 | 1207 | K1207I | D | 2.1e-06 | 1.4e-06 | 1.5e-06 | 1.667E-06 | 3.786E-07 | 1.400E-06 | 2.100E-06 |
| 24131 | 1207 | K1207L | D | 3e-06 | 3.2e-06 | 1.8e-06 | 2.667E-06 | 7.572E-07 | 1.800E-06 | 3.200E-06 |
| 24132 | 1207 | K1207K | D | 2e-06 | 1.5e-06 | 2.1e-06 | 1.867E-06 | 3.215E-07 | 1.500E-06 | 2.100E-06 |
| 24133 | 1207 | K1207M | D | 1e-06 | 1.8e-06 | 1.9e-06 | 1.567E-06 | 4.933E-07 | 1.000E-06 | 1.900E-06 |
| 24134 | 1207 | K1207F | D | 2e-06 | 2.5e-06 | 3.7e-06 | 2.733E-06 | 8.737E-07 | 2.000E-06 | 3.700E-06 |
| 24135 | 1207 | K1207P | D | 2.6e-06 | 1.3e-06 | 2.4e-06 | 2.100E-06 | 7.000E-07 | 1.300E-06 | 2.600E-06 |
| 24136 | 1207 | K1207S | D | 1.9e-06 | 1.8e-06 | 2.1e-06 | 1.933E-06 | 1.528E-07 | 1.800E-06 | 2.100E-06 |
| 24137 | 1207 | K1207T | D | 2.5e-06 | 1.9e-06 | 2.6e-06 | 2.333E-06 | 3.786E-07 | 1.900E-06 | 2.600E-06 |
| 24138 | 1207 | K1207W | D | 1.2e-06 | 2.4e-06 | 3.3e-06 | 2.300E-06 | 1.054E-06 | 1.200E-06 | 3.300E-06 |
| 24139 | 1207 | K1207Y | D | 3.3e-06 | 1.8e-06 | 1.5e-06 | 2.200E-06 | 9.644E-07 | 1.500E-06 | 3.300E-06 |

| | | | | | | | | | | |
|---|---|---|---|---|---|---|---|---|---|---|
| 24140 | 1207 | K1207V | D | 2e-06 | 2.1e-06 | 3.5e-06 | 2.533E-06 | 8.386E-07 | 2.000E-06 | 3.500E-06 |
| 24141 | 1208 | V1208A | D | 2.3e-06 | 2.1e-06 | 2.7e-06 | 2.367E-06 | 3.055E-07 | 2.100E-06 | 2.700E-06 |
| 24142 | 1208 | V1208R | D | 2e-06 | 2.1e-06 | 2.1e-06 | 2.067E-06 | 5.774E-08 | 2.000E-06 | 2.100E-06 |
| 24143 | 1208 | V1208N | D | 2.8e-06 | 3e-06 | 2.1e-06 | 2.633E-06 | 4.726E-07 | 2.100E-06 | 3.000E-06 |
| 24144 | 1208 | V1208D | D | 3.8e-06 | 1.5e-06 | 2.9e-06 | 2.733E-06 | 1.159E-06 | 1.500E-06 | 3.800E-06 |
| 24145 | 1208 | V1208C | D | 3.6e-06 | 2e-06 | 2.4e-06 | 2.667E-06 | 8.327E-07 | 2.000E-06 | 3.600E-06 |
| 24146 | 1208 | V1208E | D | 2.2e-06 | 2.6e-06 | 2.2e-06 | 2.333E-06 | 2.309E-07 | 2.200E-06 | 2.600E-06 |
| 24147 | 1208 | V1208Q | D | 4e-06 | 2.3e-06 | 3e-06 | 3.100E-06 | 8.544E-07 | 2.300E-06 | 4.000E-06 |
| 24148 | 1208 | V1208G | D | 2.3e-06 | 1.5e-06 | 3e-06 | 2.267E-06 | 7.506E-07 | 1.500E-06 | 3.000E-06 |
| 24149 | 1208 | V1208H | D | 2.4e-06 | 3.7e-06 | 1.6e-06 | 2.567E-06 | 1.060E-06 | 1.600E-06 | 3.700E-06 |
| 24150 | 1208 | V1208I | D | 2.4e-06 | 4.2e-06 | 2.5e-06 | 3.033E-06 | 1.012E-06 | 2.400E-06 | 4.200E-06 |
| 24151 | 1208 | V1208L | D | 3.7e-06 | 2.2e-06 | 2.6e-06 | 2.833E-06 | 7.767E-07 | 2.200E-06 | 3.700E-06 |
| 24152 | 1208 | V1208K | D | 2.3e-06 | 2.8e-06 | 4.5e-06 | 3.200E-06 | 1.153E-06 | 2.300E-06 | 4.500E-06 |
| 24153 | 1208 | V1208M | D | 2.7e-06 | 1.9e-06 | 1.9e-06 | 2.167E-06 | 4.619E-07 | 1.900E-06 | 2.700E-06 |
| 24154 | 1208 | V1208F | D | 2.1e-06 | 1.5e-06 | 1.6e-06 | 1.733E-06 | 3.215E-07 | 1.500E-06 | 2.100E-06 |
| 24155 | 1208 | V1208P | D | 2.5e-06 | 2.4e-06 | 1.9e-06 | 2.267E-06 | 3.215E-07 | 1.900E-06 | 2.500E-06 |
| 24156 | 1208 | V1208S | D | 2.2e-06 | 3e-06 | 4.4e-06 | 3.200E-06 | 1.114E-06 | 2.200E-06 | 4.400E-06 |
| 24157 | 1208 | V1208T | D | 1.8e-06 | 1.8e-06 | 1.5e-06 | 1.700E-06 | 1.732E-07 | 1.500E-06 | 1.800E-06 |
| 24158 | 1208 | V1208W | D | 1.8e-06 | 2.9e-06 | 1.8e-06 | 2.167E-06 | 6.351E-07 | 1.800E-06 | 2.900E-06 |
| 24159 | 1208 | V1208Y | D | 3.9e-06 | 2.3e-06 | 2.7e-06 | 2.967E-06 | 8.327E-07 | 2.300E-06 | 3.900E-06 |
| 24160 | 1208 | V1208V | D | 2.6e-06 | 2e-06 | 1.9e-06 | 2.167E-06 | 3.786E-07 | 1.900E-06 | 2.600E-06 |
| 24161 | 1209 | D1209A | D | 2.8e-06 | 2.5e-06 | 1.2e-06 | 2.167E-06 | 8.505E-07 | 1.200E-06 | 2.800E-06 |
| 24162 | 1209 | D1209R | D | 3.1e-06 | 3.1e-06 | 3.2e-06 | 3.133E-06 | 5.774E-08 | 3.100E-06 | 3.200E-06 |
| 24163 | 1209 | D1209N | D | 3.1e-06 | 1.4e-06 | 1.8e-06 | 2.100E-06 | 8.888E-07 | 1.400E-06 | 3.100E-06 |
| 24164 | 1209 | D1209D | D | 2.6e-06 | 3.6e-06 | 4.2e-06 | 3.467E-06 | 8.083E-07 | 2.600E-06 | 4.200E-06 |
| 24165 | 1209 | D1209C | D | 5.3e-06 | 1.8e-06 | 3.4e-06 | 3.500E-06 | 1.752E-06 | 1.800E-06 | 5.300E-06 |
| 24166 | 1209 | D1209E | D | 2.2e-06 | 2.6e-06 | 2.2e-06 | 2.333E-06 | 2.309E-07 | 2.200E-06 | 2.600E-06 |
| 24167 | 1209 | D1209Q | D | 2.3e-06 | 3.1e-06 | 2.4e-06 | 2.600E-06 | 4.359E-07 | 2.300E-06 | 3.100E-06 |
| 24168 | 1209 | D1209G | D | 2.7e-06 | 9.7e-07 | 1.8e-06 | 1.823E-06 | 8.652E-07 | 9.700E-07 | 2.700E-06 |
| 24169 | 1209 | D1209H | D | 7.2e-07 | 1.9e-06 | 3.4e-06 | 2.007E-06 | 1.343E-06 | 7.200E-07 | 3.400E-06 |
| 24170 | 1209 | D1209I | D | 3.5e-06 | 1.9e-06 | 1.9e-06 | 2.433E-06 | 9.238E-07 | 1.900E-06 | 3.500E-06 |
| 24171 | 1209 | D1209L | D | 1.9e-06 | 2.6e-06 | 2.1e-06 | 2.200E-06 | 3.606E-07 | 1.900E-06 | 2.600E-06 |
| 24172 | 1209 | D1209K | D | 9.8e-07 | 2.3e-06 | 1.2e-06 | 1.493E-06 | 7.072E-07 | 9.800E-07 | 2.300E-06 |
| 24173 | 1209 | D1209M | D | 2e-06 | 3.6e-06 | 2.8e-06 | 2.800E-06 | 8.000E-07 | 2.000E-06 | 3.600E-06 |
| 24174 | 1209 | D1209F | D | 2.1e-06 | 1.8e-06 | 2.6e-06 | 2.167E-06 | 4.041E-07 | 1.800E-06 | 2.600E-06 |
| 24175 | 1209 | D1209P | D | 1.8e-06 | 1.8e-06 | 3.8e-06 | 2.467E-06 | 1.155E-06 | 1.800E-06 | 3.800E-06 |
| 24176 | 1209 | D1209S | D | 2.5e-06 | 2.8e-06 | 1.1e-06 | 2.133E-06 | 9.074E-07 | 1.100E-06 | 2.800E-06 |
| 24177 | 1209 | D1209T | D | 2.8e-06 | 1.5e-06 | 2.3e-06 | 2.200E-06 | 6.557E-07 | 1.500E-06 | 2.800E-06 |
| 24178 | 1209 | D1209W | D | 2.8e-06 | 1.8e-06 | 2.4e-06 | 2.333E-06 | 5.033E-07 | 1.800E-06 | 2.800E-06 |
| 24179 | 1209 | D1209Y | D | 2.1e-06 | 3.5e-06 | 1.4e-06 | 2.333E-06 | 1.069E-06 | 1.400E-06 | 3.500E-06 |
| 24180 | 1209 | D1209V | D | 2.6e-06 | 2.2e-06 | 3.8e-06 | 2.867E-06 | 8.327E-07 | 2.200E-06 | 3.800E-06 |
| 24181 | 1210 | N1210A | D | 2.3e-06 | 1.3e-06 | 3.8e-06 | 2.467E-06 | 1.258E-06 | 1.300E-06 | 3.800E-06 |
| 24182 | 1210 | N1210R | D | 2.4e-06 | 1.4e-06 | 2.8e-06 | 2.200E-06 | 7.211E-07 | 1.400E-06 | 2.800E-06 |
| 24183 | 1210 | N1210N | D | 1.9e-06 | 2.9e-06 | 4.7e-06 | 3.167E-06 | 1.419E-06 | 1.900E-06 | 4.700E-06 |
| 24184 | 1210 | N1210D | D | 1.5e-06 | 1.6e-06 | 2.3e-06 | 1.800E-06 | 4.359E-07 | 1.500E-06 | 2.300E-06 |
| 24185 | 1210 | N1210C | D | 3.1e-06 | 2.6e-06 | 2.9e-06 | 2.867E-06 | 2.517E-07 | 2.600E-06 | 3.100E-06 |
| 24186 | 1210 | N1210E | D | 1.2e-06 | 1.8e-06 | 1.4e-06 | 1.467E-06 | 3.055E-07 | 1.200E-06 | 1.800E-06 |
| 24187 | 1210 | N1210Q | D | 3.3e-06 | 2.7e-06 | 2.3e-06 | 2.767E-06 | 5.033E-07 | 2.300E-06 | 3.300E-06 |
| 24188 | 1210 | N1210G | D | 2.6e-06 | 3.8e-06 | 1.1e-06 | 2.500E-06 | 1.353E-06 | 1.100E-06 | 3.800E-06 |
| 24189 | 1210 | N1210H | D | 2.2e-06 | 3.4e-06 | 1.7e-06 | 2.433E-06 | 8.737E-07 | 1.700E-06 | 3.400E-06 |
| 24190 | 1210 | N1210I | D | 2.7e-06 | 3.1e-06 | 2.2e-06 | 2.667E-06 | 4.509E-07 | 2.200E-06 | 3.100E-06 |
| 24191 | 1210 | N1210L | D | 2.7e-06 | 3e-06 | 2.7e-06 | 2.800E-06 | 1.732E-07 | 2.700E-06 | 3.000E-06 |
| 24192 | 1210 | N1210K | D | 2.7e-06 | 2.1e-06 | 4.3e-06 | 3.033E-06 | 1.137E-06 | 2.100E-06 | 4.300E-06 |
| 24193 | 1210 | N1210M | D | 3.1e-06 | 4.3e-06 | 1.9e-06 | 3.100E-06 | 1.200E-06 | 1.900E-06 | 4.300E-06 |
| 24194 | 1210 | N1210F | D | 2e-06 | 1.6e-06 | 3.4e-06 | 2.333E-06 | 9.452E-07 | 1.600E-06 | 3.400E-06 |
| 24195 | 1210 | N1210P | D | 1.8e-06 | 1.9e-06 | 2e-06 | 1.900E-06 | 1.000E-07 | 1.800E-06 | 2.000E-06 |
| 24196 | 1210 | N1210S | D | 3.3e-06 | 2.7e-06 | 2.9e-06 | 2.967E-06 | 3.055E-07 | 2.700E-06 | 3.300E-06 |
| 24197 | 1210 | N1210T | D | 3.1e-06 | 1.7e-06 | 2.9e-06 | 2.567E-06 | 7.572E-07 | 1.700E-06 | 3.100E-06 |
| 24198 | 1210 | N1210W | D | 2.3e-06 | 2.4e-06 | 2.3e-06 | 2.333E-06 | 5.774E-08 | 2.300E-06 | 2.400E-06 |
| 24199 | 1210 | N1210Y | D | 3.3e-06 | 1.6e-06 | 4e-06 | 2.967E-06 | 1.234E-06 | 1.600E-06 | 4.000E-06 |
| 24200 | 1210 | N1210V | D | 3.3e-06 | 4.4e-06 | 3.6e-06 | 3.767E-06 | 5.686E-07 | 3.300E-06 | 4.400E-06 |
| 24201 | 1211 | A1211A | D | 1.8e-06 | 3.6e-06 | 3.5e-06 | 2.967E-06 | 1.012E-06 | 1.800E-06 | 3.600E-06 |
| 24202 | 1211 | A1211R | D | 1.3e-06 | 1.4e-06 | 2.9e-06 | 1.867E-06 | 8.963E-07 | 1.300E-06 | 2.900E-06 |
| 24203 | 1211 | A1211N | D | 1.8e-06 | 2.3e-06 | 2e-06 | 2.033E-06 | 2.517E-07 | 1.800E-06 | 2.300E-06 |
| 24204 | 1211 | A1211D | D | 2.4e-06 | 1.8e-06 | 2.8e-06 | 2.333E-06 | 5.033E-07 | 1.800E-06 | 2.800E-06 |
| 24205 | 1211 | A1211C | D | 3.6e-06 | 1.2e-06 | 3.4e-06 | 2.733E-06 | 1.332E-06 | 1.200E-06 | 3.600E-06 |
| 24206 | 1211 | A1211E | D | 1.7e-06 | 2e-06 | 1.6e-06 | 1.767E-06 | 2.082E-07 | 1.600E-06 | 2.000E-06 |
| 24207 | 1211 | A1211Q | D | 1.8e-06 | 3.4e-06 | 2.4e-06 | 2.533E-06 | 8.083E-07 | 1.800E-06 | 3.400E-06 |
| 24208 | 1211 | A1211G | D | 1.5e-06 | 2.8e-06 | 4.5e-06 | 2.933E-06 | 1.504E-06 | 1.500E-06 | 4.500E-06 |
| 24209 | 1211 | A1211H | D | 2.8e-06 | 2.9e-06 | 3.3e-06 | 3.000E-06 | 2.646E-07 | 2.800E-06 | 3.300E-06 |
| 24210 | 1211 | A1211I | D | 1.9e-06 | 3.1e-06 | 2.2e-06 | 2.400E-06 | 6.245E-07 | 1.900E-06 | 3.100E-06 |
| 24211 | 1211 | A1211L | D | 3.1e-06 | 9.7e-07 | 3.3e-06 | 2.457E-06 | 1.291E-06 | 9.700E-07 | 3.300E-06 |
| 24212 | 1211 | A1211K | D | 3.9e-06 | 1.6e-06 | 1.9e-06 | 2.467E-06 | 1.250E-06 | 1.600E-06 | 3.900E-06 |
| 24213 | 1211 | A1211M | D | 1.4e-06 | 2.7e-06 | 2.4e-06 | 2.167E-06 | 6.807E-07 | 1.400E-06 | 2.700E-06 |
| 24214 | 1211 | A1211F | D | 8.5e-07 | 3.1e-06 | 1.7e-06 | 1.883E-06 | 1.136E-06 | 8.500E-07 | 3.100E-06 |
| 24215 | 1211 | A1211P | D | 1.7e-06 | 2.2e-06 | 2.7e-06 | 2.200E-06 | 5.000E-07 | 1.700E-06 | 2.700E-06 |
| 24216 | 1211 | A1211S | D | 2e-06 | 2.5e-06 | 3.2e-06 | 2.567E-06 | 6.028E-07 | 2.000E-06 | 3.200E-06 |
| 24217 | 1211 | A1211T | D | 2.1e-06 | 2.1e-06 | 1.6e-06 | 1.933E-06 | 2.887E-07 | 1.600E-06 | 2.100E-06 |
| 24218 | 1211 | A1211W | D | 1.6e-06 | 2.5e-06 | 2.1e-06 | 2.067E-06 | 4.509E-07 | 1.600E-06 | 2.500E-06 |
| 24219 | 1211 | A1211Y | D | 2.1e-06 | 2.2e-06 | 1.7e-06 | 2.000E-06 | 2.646E-07 | 1.700E-06 | 2.200E-06 |
| 24220 | 1211 | A1211V | D | 9.5e-07 | 9.9e-07 | 1.8e-06 | 1.247E-06 | 4.796E-07 | 9.500E-07 | 1.800E-06 |
| 24221 | 1212 | L1212A | D | 2.1e-06 | 2.2e-06 | 2.2e-06 | 2.167E-06 | 5.774E-08 | 2.100E-06 | 2.200E-06 |
| 24222 | 1212 | L1212R | D | 2.4e-06 | 1.3e-06 | 2.5e-06 | 2.067E-06 | 6.658E-07 | 1.300E-06 | 2.500E-06 |
| 24223 | 1212 | L1212N | D | 2.8e-06 | 2.9e-06 | 3.1e-06 | 2.933E-06 | 1.528E-07 | 2.800E-06 | 3.100E-06 |
| 24224 | 1212 | L1212D | D | 2.3e-06 | 2.7e-06 | 2.3e-06 | 2.433E-06 | 2.309E-07 | 2.300E-06 | 2.700E-06 |

| | | | | | | | | | | |
|---|---|---|---|---|---|---|---|---|---|---|
| 24225 | 1212 | L1212C | D | 2.2e-06 | 2.8e-06 | 2.9e-06 | 2.633E-06 | 3.786E-07 | 2.200E-06 | 2.900E-06 |
| 24226 | 1212 | L1212E | D | 2.2e-06 | 1.6e-06 | 1.9e-06 | 1.900E-06 | 3.000E-07 | 1.600E-06 | 2.200E-06 |
| 24227 | 1212 | L1212Q | D | 1.9e-06 | 2.2e-06 | 2.7e-06 | 2.267E-06 | 4.041E-07 | 1.900E-06 | 2.700E-06 |
| 24228 | 1212 | L1212G | D | 2.1e-06 | 2.9e-06 | 1.7e-06 | 2.233E-06 | 6.110E-07 | 1.700E-06 | 2.900E-06 |
| 24229 | 1212 | L1212H | D | 2.8e-06 | 6.1e-06 | 2.1e-06 | 3.667E-06 | 2.136E-06 | 2.100E-06 | 6.100E-06 |
| 24230 | 1212 | L1212I | D | 1.3e-06 | 2.4e-06 | 2.2e-06 | 1.967E-06 | 5.859E-07 | 1.300E-06 | 2.400E-06 |
| 24231 | 1212 | L1212L | D | 2.7e-06 | 1.5e-06 | 1.8e-06 | 2.000E-06 | 6.245E-07 | 1.500E-06 | 2.700E-06 |
| 24232 | 1212 | L1212K | D | 2.3e-06 | 3.1e-06 | 1.5e-06 | 2.300E-06 | 8.000E-07 | 1.500E-06 | 3.100E-06 |
| 24233 | 1212 | L1212M | D | 1.6e-06 | 1.5e-06 | 3e-06 | 2.033E-06 | 8.386E-07 | 1.500E-06 | 3.000E-06 |
| 24234 | 1212 | L1212F | D | 2.3e-06 | 2e-06 | 1.4e-06 | 1.900E-06 | 4.583E-07 | 1.400E-06 | 2.300E-06 |
| 24235 | 1212 | L1212P | D | 1.8e-06 | 2.4e-06 | 1.8e-06 | 2.000E-06 | 3.464E-07 | 1.800E-06 | 2.400E-06 |
| 24236 | 1212 | L1212S | D | 2.7e-06 | 4.1e-06 | 3e-06 | 3.267E-06 | 7.371E-07 | 2.700E-06 | 4.100E-06 |
| 24237 | 1212 | L1212T | D | 3.5e-06 | 2.5e-06 | 3.4e-06 | 3.133E-06 | 5.508E-07 | 2.500E-06 | 3.500E-06 |
| 24238 | 1212 | L1212W | D | 1.4e-06 | 2.7e-06 | 2.3e-06 | 2.133E-06 | 6.658E-07 | 1.400E-06 | 2.700E-06 |
| 24239 | 1212 | L1212Y | D | 2.5e-06 | 2.4e-06 | 1.9e-06 | 2.267E-06 | 3.215E-07 | 1.900E-06 | 2.500E-06 |
| 24240 | 1212 | L1212V | D | 2.9e-06 | 3.6e-06 | 2.7e-06 | 3.067E-06 | 4.726E-07 | 2.700E-06 | 3.600E-06 |
| 24241 | 1213 | Q1213A | D | 3.3e-06 | 1.3e-06 | 3.3e-06 | 2.633E-06 | 1.155E-06 | 1.300E-06 | 3.300E-06 |
| 24242 | 1213 | Q1213R | D | 2.2e-06 | 2.5e-06 | 1.8e-06 | 2.167E-06 | 3.512E-07 | 1.800E-06 | 2.500E-06 |
| 24243 | 1213 | Q1213N | D | 2e-06 | 3.8e-06 | 1.8e-06 | 2.533E-06 | 1.102E-06 | 1.800E-06 | 3.800E-06 |
| 24244 | 1213 | Q1213D | D | 3.2e-06 | 2.6e-06 | 3e-06 | 2.933E-06 | 3.055E-07 | 2.600E-06 | 3.200E-06 |
| 24245 | 1213 | Q1213C | D | 1.9e-06 | 2.8e-06 | 2.1e-06 | 2.267E-06 | 4.726E-07 | 1.900E-06 | 2.800E-06 |
| 24246 | 1213 | Q1213E | D | 1.3e-06 | 2.7e-06 | 4e-06 | 2.667E-06 | 1.350E-06 | 1.300E-06 | 4.000E-06 |
| 24247 | 1213 | Q1213Q | D | 2.8e-06 | 1.7e-06 | 3.2e-06 | 2.567E-06 | 7.767E-07 | 1.700E-06 | 3.200E-06 |
| 24248 | 1213 | Q1213G | D | 1.8e-06 | 1.6e-06 | 6.9e-06 | 3.433E-06 | 3.004E-06 | 1.600E-06 | 6.900E-06 |
| 24249 | 1213 | Q1213H | D | 2.4e-06 | 2.5e-06 | 2e-06 | 2.300E-06 | 2.646E-07 | 2.000E-06 | 2.500E-06 |
| 24250 | 1213 | Q1213I | D | 3e-06 | 2.3e-06 | 2.1e-06 | 2.467E-06 | 4.726E-07 | 2.100E-06 | 3.000E-06 |
| 24251 | 1213 | Q1213L | D | 2.9e-06 | 1.7e-06 | 4e-06 | 2.867E-06 | 1.150E-06 | 1.700E-06 | 4.000E-06 |
| 24252 | 1213 | Q1213K | D | 1.1e-06 | 3.2e-06 | 2.8e-06 | 2.367E-06 | 1.115E-06 | 1.100E-06 | 3.200E-06 |
| 24253 | 1213 | Q1213M | D | 3.7e-06 | 2.7e-06 | 2.3e-06 | 2.900E-06 | 7.211E-07 | 2.300E-06 | 3.700E-06 |
| 24254 | 1213 | Q1213F | D | 1.8e-06 | 1.8e-06 | 2.1e-06 | 1.900E-06 | 1.732E-07 | 1.800E-06 | 2.100E-06 |
| 24255 | 1213 | Q1213P | D | 1.6e-06 | 3.4e-06 | 1.9e-06 | 2.300E-06 | 9.644E-07 | 1.600E-06 | 3.400E-06 |
| 24256 | 1213 | Q1213S | D | 2.1e-06 | 4.4e-06 | 2.4e-06 | 2.967E-06 | 1.250E-06 | 2.100E-06 | 4.400E-06 |
| 24257 | 1213 | Q1213T | D | 3.3e-06 | 1.9e-06 | 3e-06 | 2.733E-06 | 7.371E-07 | 1.900E-06 | 3.300E-06 |
| 24258 | 1213 | Q1213W | D | 9.1e-07 | 2.9e-06 | 2.2e-06 | 2.003E-06 | 1.009E-06 | 9.100E-07 | 2.900E-06 |
| 24259 | 1213 | Q1213Y | D | 2.2e-06 | 2.5e-06 | 3.2e-06 | 2.633E-06 | 5.132E-07 | 2.200E-06 | 3.200E-06 |
| 24260 | 1213 | Q1213V | D | 1.7e-06 | 2.7e-06 | 3.2e-06 | 2.533E-06 | 7.638E-07 | 1.700E-06 | 3.200E-06 |
| 24261 | 1214 | S1214A | D | 9.9e-07 | 4.1e-06 | 2.8e-06 | 2.630E-06 | 1.562E-06 | 9.900E-07 | 4.100E-06 |
| 24262 | 1214 | S1214R | D | 2.3e-06 | 2.8e-06 | 1.6e-06 | 2.233E-06 | 6.028E-07 | 1.600E-06 | 2.800E-06 |
| 24263 | 1214 | S1214N | D | 1e-06 | 1.8e-06 | 2.7e-06 | 1.833E-06 | 8.505E-07 | 1.000E-06 | 2.700E-06 |
| 24264 | 1214 | S1214D | D | 3.2e-06 | 1.8e-06 | 2e-06 | 2.333E-06 | 7.572E-07 | 1.800E-06 | 3.200E-06 |
| 24265 | 1214 | S1214C | D | 2.6e-06 | 3.5e-06 | 2.3e-06 | 2.800E-06 | 6.245E-07 | 2.300E-06 | 3.500E-06 |
| 24266 | 1214 | S1214E | D | 2.6e-06 | 1.8e-06 | 3.3e-06 | 2.567E-06 | 7.506E-07 | 1.800E-06 | 3.300E-06 |
| 24267 | 1214 | S1214Q | D | 2e-06 | 3.1e-06 | 2.8e-06 | 2.633E-06 | 5.686E-07 | 2.000E-06 | 3.100E-06 |
| 24268 | 1214 | S1214G | D | 2.2e-06 | 2.7e-06 | 1.8e-06 | 2.233E-06 | 4.509E-07 | 1.800E-06 | 2.700E-06 |
| 24269 | 1214 | S1214H | D | 2.8e-06 | 1.5e-06 | 1.9e-06 | 2.067E-06 | 6.658E-07 | 1.500E-06 | 2.800E-06 |
| 24270 | 1214 | S1214I | D | 2.3e-06 | 2.6e-06 | 2.4e-06 | 2.433E-06 | 1.528E-07 | 2.300E-06 | 2.600E-06 |
| 24271 | 1214 | S1214L | D | 2.3e-06 | 2.4e-06 | 3.9e-06 | 2.867E-06 | 8.963E-07 | 2.300E-06 | 3.900E-06 |
| 24272 | 1214 | S1214K | D | 4.6e-06 | 2e-06 | 1.7e-06 | 2.767E-06 | 1.595E-06 | 1.700E-06 | 4.600E-06 |
| 24273 | 1214 | S1214M | D | 3.5e-06 | 5.9e-06 | 1.6e-06 | 3.667E-06 | 2.155E-06 | 1.600E-06 | 5.900E-06 |
| 24274 | 1214 | S1214F | D | 2.6e-06 | 2.2e-06 | 2.9e-06 | 2.567E-06 | 3.512E-07 | 2.200E-06 | 2.900E-06 |
| 24275 | 1214 | S1214P | D | 1.7e-06 | 2.2e-06 | 1.4e-06 | 1.767E-06 | 4.041E-07 | 1.400E-06 | 2.200E-06 |
| 24276 | 1214 | S1214S | D | 1.1e-06 | 2.9e-06 | 2.5e-06 | 2.167E-06 | 9.452E-07 | 1.100E-06 | 2.900E-06 |
| 24277 | 1214 | S1214T | D | 3.7e-06 | 1.8e-06 | 2.3e-06 | 2.600E-06 | 9.849E-07 | 1.800E-06 | 3.700E-06 |
| 24278 | 1214 | S1214W | D | 3e-06 | 1e-06 | 2.9e-06 | 2.300E-06 | 1.127E-06 | 1.000E-06 | 3.000E-06 |
| 24279 | 1214 | S1214Y | D | 1.2e-06 | 2.1e-06 | 1.8e-06 | 1.700E-06 | 4.583E-07 | 1.200E-06 | 2.100E-06 |
| 24280 | 1214 | S1214V | D | 2.7e-06 | 2.9e-06 | 2.2e-06 | 2.600E-06 | 3.606E-07 | 2.200E-06 | 2.900E-06 |
| 24281 | 1215 | G1215A | D | 1.4e-06 | 3.1e-06 | 2.4e-06 | 2.300E-06 | 8.544E-07 | 1.400E-06 | 3.100E-06 |
| 24282 | 1215 | G1215R | D | 5e-06 | 2.3e-06 | 1.8e-06 | 3.033E-06 | 1.721E-06 | 1.800E-06 | 5.000E-06 |
| 24283 | 1215 | G1215N | D | 1.7e-06 | 1.5e-06 | 1.4e-06 | 1.533E-06 | 1.528E-07 | 1.400E-06 | 1.700E-06 |
| 24284 | 1215 | G1215D | D | 2.9e-06 | 1.4e-06 | 2.2e-06 | 2.167E-06 | 7.506E-07 | 1.400E-06 | 2.900E-06 |
| 24285 | 1215 | G1215C | D | 2.3e-06 | 2.7e-06 | 2.7e-06 | 2.567E-06 | 2.309E-07 | 2.300E-06 | 2.700E-06 |
| 24286 | 1215 | G1215E | D | 3.4e-06 | 3.1e-06 | 3.4e-06 | 3.300E-06 | 1.732E-07 | 3.100E-06 | 3.400E-06 |
| 24287 | 1215 | G1215Q | D | 2.9e-06 | 2.1e-06 | 3.3e-06 | 2.767E-06 | 6.110E-07 | 2.100E-06 | 3.300E-06 |
| 24288 | 1215 | G1215G | D | 4.4e-06 | 2e-06 | 1.6e-06 | 2.667E-06 | 1.514E-06 | 1.600E-06 | 4.400E-06 |
| 24289 | 1215 | G1215H | D | 1.6e-06 | 2.1e-06 | 1.7e-06 | 1.800E-06 | 2.646E-07 | 1.600E-06 | 2.100E-06 |
| 24290 | 1215 | G1215I | D | 1.6e-06 | 2.6e-06 | 2.2e-06 | 2.133E-06 | 5.033E-07 | 1.600E-06 | 2.600E-06 |
| 24291 | 1215 | G1215L | D | 1.7e-06 | 2e-06 | 3.1e-06 | 2.267E-06 | 7.371E-07 | 1.700E-06 | 3.100E-06 |
| 24292 | 1215 | G1215K | D | 2.8e-06 | 3.1e-06 | 1.7e-06 | 2.533E-06 | 7.371E-07 | 1.700E-06 | 3.100E-06 |
| 24293 | 1215 | G1215M | D | 2.1e-06 | 3.2e-06 | 1.5e-06 | 2.267E-06 | 8.622E-07 | 1.500E-06 | 3.200E-06 |
| 24294 | 1215 | G1215F | D | 2.6e-06 | 1.2e-06 | 2.3e-06 | 2.033E-06 | 7.371E-07 | 1.200E-06 | 2.600E-06 |
| 24295 | 1215 | G1215P | D | 2.3e-06 | 3.2e-06 | 2.4e-06 | 2.633E-06 | 4.933E-07 | 2.300E-06 | 3.200E-06 |
| 24296 | 1215 | G1215S | D | 2.3e-06 | 2.4e-06 | 2.4e-06 | 2.367E-06 | 5.774E-08 | 2.300E-06 | 2.400E-06 |
| 24297 | 1215 | G1215T | D | 1.7e-06 | 2e-06 | 2.4e-06 | 2.033E-06 | 3.512E-07 | 1.700E-06 | 2.400E-06 |
| 24298 | 1215 | G1215W | D | 3e-06 | 1.8e-06 | 2.1e-06 | 2.300E-06 | 6.245E-07 | 1.800E-06 | 3.000E-06 |
| 24299 | 1215 | G1215Y | D | 1.7e-06 | 2.2e-06 | 1.9e-06 | 1.933E-06 | 2.517E-07 | 1.700E-06 | 2.200E-06 |
| 24300 | 1215 | G1215V | D | 1.8e-06 | 1.7e-06 | 2.2e-06 | 1.900E-06 | 2.646E-07 | 1.700E-06 | 2.200E-06 |
| 24301 | 1216 | N1216A | D | 1.8e-06 | 1.8e-06 | 3.1e-06 | 2.233E-06 | 7.506E-07 | 1.800E-06 | 3.100E-06 |
| 24302 | 1216 | N1216R | D | 2.1e-06 | 9.7e-07 | 3e-06 | 2.023E-06 | 1.017E-06 | 9.700E-07 | 3.000E-06 |
| 24303 | 1216 | N1216N | D | 1.6e-06 | 2.6e-06 | 2.7e-06 | 2.300E-06 | 6.083E-07 | 1.600E-06 | 2.700E-06 |
| 24304 | 1216 | N1216D | D | 2.6e-06 | 1.8e-06 | 1.5e-06 | 1.967E-06 | 5.686E-07 | 1.500E-06 | 2.600E-06 |
| 24305 | 1216 | N1216C | D | 3.3e-06 | 2.6e-06 | 1.7e-06 | 2.533E-06 | 8.021E-07 | 1.700E-06 | 3.300E-06 |
| 24306 | 1216 | N1216E | D | 2.1e-06 | 2.5e-06 | 2.5e-06 | 2.367E-06 | 2.309E-07 | 2.100E-06 | 2.500E-06 |
| 24307 | 1216 | N1216Q | D | 2.7e-06 | 2.9e-06 | 3.6e-06 | 3.067E-06 | 4.726E-07 | 2.700E-06 | 3.600E-06 |
| 24308 | 1216 | N1216G | D | 1.7e-06 | 2.8e-06 | 2.2e-06 | 2.233E-06 | 5.508E-07 | 1.700E-06 | 2.800E-06 |
| 24309 | 1216 | N1216H | D | 2.4e-06 | 2.7e-06 | 1.5e-06 | 2.200E-06 | 6.245E-07 | 1.500E-06 | 2.700E-06 |

| | | | | | | | | | | |
|---|---|---|---|---|---|---|---|---|---|---|
| 24310 | 1216 | N1216I | D | 3e-06 | 2.9e-06 | 1.2e-06 | 2.367E-06 | 1.012E-06 | 1.200E-06 | 3.000E-06 |
| 24311 | 1216 | N1216L | D | 2.3e-06 | 1.8e-06 | 2.6e-06 | 2.233E-06 | 4.041E-07 | 1.800E-06 | 2.600E-06 |
| 24312 | 1216 | N1216K | D | 3e-06 | 1.7e-06 | 1.8e-06 | 2.167E-06 | 7.234E-07 | 1.700E-06 | 3.000E-06 |
| 24313 | 1216 | N1216M | D | 2e-06 | 1.2e-06 | 1.5e-06 | 1.567E-06 | 4.041E-07 | 1.200E-06 | 2.000E-06 |
| 24314 | 1216 | N1216F | D | 2.9e-06 | 3.1e-06 | 2.4e-06 | 2.800E-06 | 3.606E-07 | 2.400E-06 | 3.100E-06 |
| 24315 | 1216 | N1216P | D | 2.1e-06 | 1.3e-06 | 1.6e-06 | 1.667E-06 | 4.041E-07 | 1.300E-06 | 2.100E-06 |
| 24316 | 1216 | N1216S | D | 3e-06 | 2.3e-06 | 1.3e-06 | 2.200E-06 | 8.544E-07 | 1.300E-06 | 3.000E-06 |
| 24317 | 1216 | N1216T | D | 2e-06 | 2.6e-06 | 2.3e-06 | 2.300E-06 | 3.000E-07 | 2.000E-06 | 2.600E-06 |
| 24318 | 1216 | N1216W | D | 2.1e-06 | 2.3e-06 | 1.9e-06 | 2.100E-06 | 2.000E-07 | 1.900E-06 | 2.300E-06 |
| 24319 | 1216 | N1216Y | D | 2.9e-06 | 2.4e-06 | 2.8e-06 | 2.700E-06 | 2.646E-07 | 2.400E-06 | 2.900E-06 |
| 24320 | 1216 | N1216V | D | 3.1e-06 | 2.6e-06 | 2.4e-06 | 2.700E-06 | 3.606E-07 | 2.400E-06 | 3.100E-06 |
| 24321 | 1217 | S1217A | D | 1.7e-06 | 2.5e-06 | 1.5e-06 | 1.900E-06 | 5.292E-07 | 1.500E-06 | 2.500E-06 |
| 24322 | 1217 | S1217R | D | 2.4e-06 | 3.6e-06 | 1.2e-06 | 2.400E-06 | 1.200E-06 | 1.200E-06 | 3.600E-06 |
| 24323 | 1217 | S1217N | D | 2.6e-06 | 2.8e-06 | 1.6e-06 | 2.333E-06 | 6.429E-07 | 1.600E-06 | 2.800E-06 |
| 24324 | 1217 | S1217D | D | 1.5e-06 | 4.1e-06 | 2.5e-06 | 2.700E-06 | 1.311E-06 | 1.500E-06 | 4.100E-06 |
| 24325 | 1217 | S1217C | D | 1.6e-06 | 3.1e-06 | 1.6e-06 | 2.100E-06 | 8.660E-07 | 1.600E-06 | 3.100E-06 |
| 24326 | 1217 | S1217E | D | 3.2e-06 | 3.1e-06 | 2.1e-06 | 2.800E-06 | 6.083E-07 | 2.100E-06 | 3.200E-06 |
| 24327 | 1217 | S1217Q | D | 2.4e-06 | 2.3e-06 | 2.3e-06 | 2.333E-06 | 5.774E-08 | 2.300E-06 | 2.400E-06 |
| 24328 | 1217 | S1217G | D | 2.6e-06 | 3.3e-06 | 2e-06 | 2.633E-06 | 6.506E-07 | 2.000E-06 | 3.300E-06 |
| 24329 | 1217 | S1217H | D | 2.8e-06 | 3.1e-06 | 1.7e-06 | 2.533E-06 | 7.371E-07 | 1.700E-06 | 3.100E-06 |
| 24330 | 1217 | S1217I | D | 1.8e-06 | 2.6e-06 | 1.2e-06 | 1.867E-06 | 7.024E-07 | 1.200E-06 | 2.600E-06 |
| 24331 | 1217 | S1217L | D | 2.8e-06 | 3.4e-06 | 3e-06 | 3.067E-06 | 3.055E-07 | 2.800E-06 | 3.400E-06 |
| 24332 | 1217 | S1217K | D | 3.3e-06 | 2.8e-06 | 1.9e-06 | 2.667E-06 | 7.095E-07 | 1.900E-06 | 3.300E-06 |
| 24333 | 1217 | S1217M | D | 1.5e-06 | 2.3e-06 | 2.3e-06 | 2.033E-06 | 4.619E-07 | 1.500E-06 | 2.300E-06 |
| 24334 | 1217 | S1217F | D | 2.8e-06 | 1.7e-06 | 2.8e-06 | 2.433E-06 | 6.351E-07 | 1.700E-06 | 2.800E-06 |
| 24335 | 1217 | S1217P | D | 1.6e-06 | 3.1e-06 | 2e-06 | 2.233E-06 | 7.767E-07 | 1.600E-06 | 3.100E-06 |
| 24336 | 1217 | S1217S | D | 3.6e-06 | 1.8e-06 | 1.4e-06 | 2.267E-06 | 1.172E-06 | 1.400E-06 | 3.600E-06 |
| 24337 | 1217 | S1217T | D | 3.1e-06 | 3.6e-06 | 1.3e-06 | 2.667E-06 | 1.210E-06 | 1.300E-06 | 3.600E-06 |
| 24338 | 1217 | S1217W | D | 3.8e-06 | 1.8e-06 | 1.9e-06 | 2.500E-06 | 1.127E-06 | 1.800E-06 | 3.800E-06 |
| 24339 | 1217 | S1217Y | D | 2.2e-06 | 2.6e-06 | 2.4e-06 | 2.400E-06 | 2.000E-07 | 2.200E-06 | 2.600E-06 |
| 24340 | 1217 | S1217V | D | 3.8e-06 | 3.3e-06 | 1.8e-06 | 2.967E-06 | 1.041E-06 | 1.800E-06 | 3.800E-06 |
| 24341 | 1218 | Q1218A | D | 2.5e-06 | 2.2e-06 | 1.5e-06 | 2.067E-06 | 5.132E-07 | 1.500E-06 | 2.500E-06 |
| 24342 | 1218 | Q1218R | D | 1.9e-06 | 2.8e-06 | 3.7e-06 | 2.800E-06 | 9.000E-07 | 1.900E-06 | 3.700E-06 |
| 24343 | 1218 | Q1218N | D | 1.5e-06 | 1.8e-06 | 4e-06 | 2.433E-06 | 1.365E-06 | 1.500E-06 | 4.000E-06 |
| 24344 | 1218 | Q1218D | D | 3.2e-06 | 1.4e-06 | 2.7e-06 | 2.433E-06 | 9.292E-07 | 1.400E-06 | 3.200E-06 |
| 24345 | 1218 | Q1218C | D | 3.2e-06 | 3.9e-06 | 3.2e-06 | 3.433E-06 | 4.041E-07 | 3.200E-06 | 3.900E-06 |
| 24346 | 1218 | Q1218E | D | 3.3e-06 | 2.9e-06 | 3.1e-06 | 3.100E-06 | 2.000E-07 | 2.900E-06 | 3.300E-06 |
| 24347 | 1218 | Q1218Q | D | 1.9e-06 | 2.6e-06 | 2.5e-06 | 2.333E-06 | 3.786E-07 | 1.900E-06 | 2.600E-06 |
| 24348 | 1218 | Q1218G | D | 1.6e-06 | 2e-06 | 2e-06 | 1.867E-06 | 2.309E-07 | 1.600E-06 | 2.000E-06 |
| 24349 | 1218 | Q1218H | D | 3.2e-06 | 3.7e-06 | 3.9e-06 | 3.600E-06 | 3.606E-07 | 3.200E-06 | 3.900E-06 |
| 24350 | 1218 | Q1218I | D | 1.3e-06 | 2.3e-06 | 2.7e-06 | 2.100E-06 | 7.211E-07 | 1.300E-06 | 2.700E-06 |
| 24351 | 1218 | Q1218L | D | 2e-06 | 3.1e-06 | 2.4e-06 | 2.500E-06 | 5.568E-07 | 2.000E-06 | 3.100E-06 |
| 24352 | 1218 | Q1218K | D | 3.6e-06 | 2e-06 | 1.9e-06 | 2.500E-06 | 9.539E-07 | 1.900E-06 | 3.600E-06 |
| 24353 | 1218 | Q1218M | D | 1.6e-06 | 2.2e-06 | 2.3e-06 | 2.033E-06 | 3.786E-07 | 1.600E-06 | 2.300E-06 |
| 24354 | 1218 | Q1218F | D | 1.4e-06 | 3.2e-06 | 1.6e-06 | 2.067E-06 | 9.866E-07 | 1.400E-06 | 3.200E-06 |
| 24355 | 1218 | Q1218P | D | 2.9e-06 | 1.4e-06 | 2.6e-06 | 2.300E-06 | 7.937E-07 | 1.400E-06 | 2.900E-06 |
| 24356 | 1218 | Q1218S | D | 1.4e-06 | 3.2e-06 | 2.6e-06 | 2.400E-06 | 9.165E-07 | 1.400E-06 | 3.200E-06 |
| 24357 | 1218 | Q1218T | D | 1.9e-06 | 1.5e-06 | 3.2e-06 | 2.200E-06 | 8.888E-07 | 1.500E-06 | 3.200E-06 |
| 24358 | 1218 | Q1218W | D | 2.4e-06 | 2.6e-06 | 3.1e-06 | 2.700E-06 | 3.606E-07 | 2.400E-06 | 3.100E-06 |
| 24359 | 1218 | Q1218Y | D | 2.4e-06 | 1.7e-06 | 1.9e-06 | 2.000E-06 | 3.606E-07 | 1.700E-06 | 2.400E-06 |
| 24360 | 1218 | Q1218V | D | 2.5e-06 | 2.8e-06 | 2.3e-06 | 2.533E-06 | 2.517E-07 | 2.300E-06 | 2.800E-06 |
| 24361 | 1219 | E1219A | D | 2.7e-06 | 2e-06 | 2.5e-06 | 2.400E-06 | 3.606E-07 | 2.000E-06 | 2.700E-06 |
| 24362 | 1219 | E1219R | D | 2.2e-06 | 1.9e-06 | 3.1e-06 | 2.400E-06 | 6.245E-07 | 1.900E-06 | 3.100E-06 |
| 24363 | 1219 | E1219N | D | 2.9e-06 | 1.7e-06 | 2.5e-06 | 2.367E-06 | 6.110E-07 | 1.700E-06 | 2.900E-06 |
| 24364 | 1219 | E1219D | D | 1.1e-06 | 3e-06 | 8.3e-07 | 1.643E-06 | 1.183E-06 | 8.300E-07 | 3.000E-06 |
| 24365 | 1219 | E1219C | D | 2e-06 | 3.8e-06 | 2.4e-06 | 2.733E-06 | 9.452E-07 | 2.000E-06 | 3.800E-06 |
| 24366 | 1219 | E1219E | D | 2e-06 | 2.2e-06 | 1e-06 | 1.733E-06 | 6.429E-07 | 1.000E-06 | 2.200E-06 |
| 24367 | 1219 | E1219Q | D | 1.8e-06 | 2.7e-06 | 1.9e-06 | 2.133E-06 | 4.933E-07 | 1.800E-06 | 2.700E-06 |
| 24368 | 1219 | E1219G | D | 2.7e-06 | 3.3e-06 | 2.3e-06 | 2.767E-06 | 5.033E-07 | 2.300E-06 | 3.300E-06 |
| 24369 | 1219 | E1219H | D | 1.9e-06 | 1.7e-06 | 2.6e-06 | 2.067E-06 | 4.726E-07 | 1.700E-06 | 2.600E-06 |
| 24370 | 1219 | E1219I | D | 2e-06 | 3.1e-06 | 2.3e-06 | 2.467E-06 | 5.686E-07 | 2.000E-06 | 3.100E-06 |
| 24371 | 1219 | E1219L | D | 2.1e-06 | 2.2e-06 | 1.9e-06 | 2.067E-06 | 1.528E-07 | 1.900E-06 | 2.200E-06 |
| 24372 | 1219 | E1219K | D | 1.3e-06 | 2e-06 | 1.4e-06 | 1.567E-06 | 3.786E-07 | 1.300E-06 | 2.000E-06 |
| 24373 | 1219 | E1219M | D | 4.1e-06 | 3e-06 | 3.1e-06 | 3.400E-06 | 6.083E-07 | 3.000E-06 | 4.100E-06 |
| 24374 | 1219 | E1219F | D | 2.4e-06 | 1.9e-06 | 2e-06 | 2.100E-06 | 2.646E-07 | 1.900E-06 | 2.400E-06 |
| 24375 | 1219 | E1219P | D | 2.6e-06 | 2.6e-06 | 1.6e-06 | 2.267E-06 | 5.774E-07 | 1.600E-06 | 2.600E-06 |
| 24376 | 1219 | E1219S | D | 2.6e-06 | 2.1e-06 | 1.8e-06 | 2.167E-06 | 4.041E-07 | 1.800E-06 | 2.600E-06 |
| 24377 | 1219 | E1219T | D | 3.1e-06 | 2.1e-06 | 1.8e-06 | 2.333E-06 | 6.807E-07 | 1.800E-06 | 3.100E-06 |
| 24378 | 1219 | E1219W | D | 2.2e-06 | 2.2e-06 | 2.3e-06 | 2.233E-06 | 5.774E-08 | 2.200E-06 | 2.300E-06 |
| 24379 | 1219 | E1219Y | D | 1.8e-06 | 3.6e-06 | 2.6e-06 | 2.667E-06 | 9.018E-07 | 1.800E-06 | 3.600E-06 |
| 24380 | 1219 | E1219V | D | 2.4e-06 | 2.3e-06 | 1.4e-06 | 2.033E-06 | 5.508E-07 | 1.400E-06 | 2.400E-06 |
| 24381 | 1220 | S1220A | D | 2.2e-06 | 3.5e-06 | 1.7e-06 | 2.467E-06 | 9.292E-07 | 1.700E-06 | 3.500E-06 |
| 24382 | 1220 | S1220R | D | 3.7e-06 | 2.8e-06 | 2.5e-06 | 3.000E-06 | 6.245E-07 | 2.500E-06 | 3.700E-06 |
| 24383 | 1220 | S1220N | D | 2.9e-06 | 2.3e-06 | 1.8e-06 | 2.333E-06 | 5.508E-07 | 1.800E-06 | 2.900E-06 |
| 24384 | 1220 | S1220D | D | 3.1e-06 | 1.3e-06 | 3.3e-06 | 2.567E-06 | 1.102E-06 | 1.300E-06 | 3.300E-06 |
| 24385 | 1220 | S1220C | D | 1.3e-06 | 2.5e-06 | 1.9e-06 | 1.900E-06 | 6.000E-07 | 1.300E-06 | 2.500E-06 |
| 24386 | 1220 | S1220E | D | 8.9e-07 | 2.4e-06 | 2.2e-06 | 1.830E-06 | 8.202E-07 | 8.900E-07 | 2.400E-06 |
| 24387 | 1220 | S1220Q | D | 1.4e-06 | 3.1e-06 | 4.1e-06 | 2.867E-06 | 1.365E-06 | 1.400E-06 | 4.100E-06 |
| 24388 | 1220 | S1220G | D | 4.1e-06 | 2.5e-06 | 3.1e-06 | 3.233E-06 | 8.083E-07 | 2.500E-06 | 4.100E-06 |
| 24389 | 1220 | S1220H | D | 3.1e-06 | 1.8e-06 | 2e-06 | 2.300E-06 | 7.000E-07 | 1.800E-06 | 3.100E-06 |
| 24390 | 1220 | S1220I | D | 2.6e-06 | 3.1e-06 | 2.5e-06 | 2.733E-06 | 3.215E-07 | 2.500E-06 | 3.100E-06 |
| 24391 | 1220 | S1220L | D | 1.9e-06 | 1.9e-06 | 1.2e-06 | 1.667E-06 | 4.041E-07 | 1.200E-06 | 1.900E-06 |
| 24392 | 1220 | S1220K | D | 3.7e-06 | 1.3e-06 | 2.5e-06 | 2.500E-06 | 1.200E-06 | 1.300E-06 | 3.700E-06 |
| 24393 | 1220 | S1220M | D | 1.6e-06 | 2.7e-06 | 1.7e-06 | 2.000E-06 | 6.083E-07 | 1.600E-06 | 2.700E-06 |
| 24394 | 1220 | S1220F | D | 3.3e-06 | 2.6e-06 | 3.3e-06 | 3.067E-06 | 4.041E-07 | 2.600E-06 | 3.300E-06 |

| | | | | | | | | | | |
|---|---|---|---|---|---|---|---|---|---|---|
| 24395 | 1220 | S1220P | D | 9.8e-07 | 3e-06 | 1.3e-06 | 1.760E-06 | 1.086E-06 | 9.800E-07 | 3.000E-06 |
| 24396 | 1220 | S1220S | D | 1.7e-06 | 1.8e-06 | 3.4e-06 | 2.300E-06 | 9.539E-07 | 1.700E-06 | 3.400E-06 |
| 24397 | 1220 | S1220T | D | 2.1e-06 | 2.9e-06 | 1.7e-06 | 2.233E-06 | 6.110E-07 | 1.700E-06 | 2.900E-06 |
| 24398 | 1220 | S1220W | D | 2.3e-06 | 2e-06 | 2.5e-06 | 2.267E-06 | 2.517E-07 | 2.000E-06 | 2.500E-06 |
| 24399 | 1220 | S1220Y | D | 2.4e-06 | 1e-06 | 3.2e-06 | 2.200E-06 | 1.114E-06 | 1.000E-06 | 3.200E-06 |
| 24400 | 1220 | S1220V | D | 2.3e-06 | 2e-06 | 1.9e-06 | 2.067E-06 | 2.082E-07 | 1.900E-06 | 2.300E-06 |
| 24401 | 1221 | V1221A | D | 1.8e-06 | 2e-06 | 1.6e-06 | 1.800E-06 | 2.000E-07 | 1.600E-06 | 2.000E-06 |
| 24402 | 1221 | V1221R | D | 3.8e-06 | 9e-07 | 3e-06 | 2.567E-06 | 1.498E-06 | 9.000E-07 | 3.800E-06 |
| 24403 | 1221 | V1221N | D | 3.8e-06 | 2.7e-06 | 2.7e-06 | 3.067E-06 | 6.351E-07 | 2.700E-06 | 3.800E-06 |
| 24404 | 1221 | V1221D | D | 1.9e-06 | 1.9e-06 | 1.9e-06 | 1.900E-06 | 2.593E-22 | 1.900E-06 | 1.900E-06 |
| 24405 | 1221 | V1221C | D | 1.7e-06 | 2.4e-06 | 2.2e-06 | 2.100E-06 | 3.606E-07 | 1.700E-06 | 2.400E-06 |
| 24406 | 1221 | V1221E | D | 1.3e-06 | 2e-06 | 2.9e-06 | 2.067E-06 | 8.021E-07 | 1.300E-06 | 2.900E-06 |
| 24407 | 1221 | V1221Q | D | 3.4e-06 | 3e-06 | 2.5e-06 | 2.967E-06 | 4.509E-07 | 2.500E-06 | 3.400E-06 |
| 24408 | 1221 | V1221G | D | 2e-06 | 2.8e-06 | 1.3e-06 | 2.033E-06 | 7.506E-07 | 1.300E-06 | 2.800E-06 |
| 24409 | 1221 | V1221H | D | 3.3e-06 | 2e-06 | 1.5e-06 | 2.267E-06 | 9.292E-07 | 1.500E-06 | 3.300E-06 |
| 24410 | 1221 | V1221I | D | 2.7e-06 | 2.4e-06 | 2.5e-06 | 2.533E-06 | 1.528E-07 | 2.400E-06 | 2.700E-06 |
| 24411 | 1221 | V1221L | D | 2.4e-06 | 2.3e-06 | 1.8e-06 | 2.167E-06 | 3.215E-07 | 1.800E-06 | 2.400E-06 |
| 24412 | 1221 | V1221K | D | 3e-06 | 3.1e-06 | 1.6e-06 | 2.567E-06 | 8.386E-07 | 1.600E-06 | 3.100E-06 |
| 24413 | 1221 | V1221M | D | 2.2e-06 | 3e-06 | 2.9e-06 | 2.700E-06 | 4.359E-07 | 2.200E-06 | 3.000E-06 |
| 24414 | 1221 | V1221F | D | 2.7e-06 | 1.9e-06 | 2.5e-06 | 2.367E-06 | 4.163E-07 | 1.900E-06 | 2.700E-06 |
| 24415 | 1221 | V1221P | D | 4e-06 | 1.9e-06 | 3.5e-06 | 3.133E-06 | 1.097E-06 | 1.900E-06 | 4.000E-06 |
| 24416 | 1221 | V1221S | D | 3.3e-06 | 3e-06 | 3e-06 | 3.100E-06 | 1.732E-07 | 3.000E-06 | 3.300E-06 |
| 24417 | 1221 | V1221T | D | 5.8e-06 | 4.1e-06 | 2.1e-06 | 4.000E-06 | 1.852E-06 | 2.100E-06 | 5.800E-06 |
| 24418 | 1221 | V1221W | D | 1.8e-06 | 1.4e-06 | 3e-06 | 2.067E-06 | 8.327E-07 | 1.400E-06 | 3.000E-06 |
| 24419 | 1221 | V1221Y | D | 3e-06 | 4.1e-06 | 2.2e-06 | 3.100E-06 | 9.539E-07 | 2.200E-06 | 4.100E-06 |
| 24420 | 1221 | V1221V | D | 1.3e-06 | 2e-06 | 2.4e-06 | 1.900E-06 | 5.568E-07 | 1.300E-06 | 2.400E-06 |
| 24421 | 1222 | T1222A | D | 2e-06 | 2.1e-06 | 1.8e-06 | 1.967E-06 | 1.528E-07 | 1.800E-06 | 2.100E-06 |
| 24422 | 1222 | T1222R | D | 2.3e-06 | 2.4e-06 | 4.3e-06 | 3.000E-06 | 1.127E-06 | 2.300E-06 | 4.300E-06 |
| 24423 | 1222 | T1222N | D | 3.4e-06 | 1.8e-06 | 3.6e-06 | 2.933E-06 | 9.866E-07 | 1.800E-06 | 3.600E-06 |
| 24424 | 1222 | T1222D | D | 1.5e-06 | 2e-06 | 2.4e-06 | 1.967E-06 | 4.509E-07 | 1.500E-06 | 2.400E-06 |
| 24425 | 1222 | T1222C | D | 3.2e-06 | 2.1e-06 | 2.2e-06 | 2.500E-06 | 6.083E-07 | 2.100E-06 | 3.200E-06 |
| 24426 | 1222 | T1222E | D | 3.4e-06 | 2.3e-06 | 4e-06 | 3.233E-06 | 8.622E-07 | 2.300E-06 | 4.000E-06 |
| 24427 | 1222 | T1222Q | D | 9.2e-07 | 2.5e-06 | 2.1e-06 | 1.840E-06 | 8.215E-07 | 9.200E-07 | 2.500E-06 |
| 24428 | 1222 | T1222G | D | 1.9e-06 | 3e-06 | 2.6e-06 | 2.500E-06 | 5.568E-07 | 1.900E-06 | 3.000E-06 |
| 24429 | 1222 | T1222H | D | 1.8e-06 | 2.6e-06 | 1.8e-06 | 2.067E-06 | 4.619E-07 | 1.800E-06 | 2.600E-06 |
| 24430 | 1222 | T1222I | D | 3.3e-06 | 2.5e-06 | 2.6e-06 | 2.800E-06 | 4.359E-07 | 2.500E-06 | 3.300E-06 |
| 24431 | 1222 | T1222L | D | 3e-06 | 1.9e-06 | 9.9e-07 | 1.963E-06 | 1.006E-06 | 9.900E-07 | 3.000E-06 |
| 24432 | 1222 | T1222K | D | 2.9e-06 | 2.5e-06 | 2e-06 | 2.467E-06 | 4.509E-07 | 2.000E-06 | 2.900E-06 |
| 24433 | 1222 | T1222M | D | 2.2e-06 | 2.2e-06 | 2.8e-06 | 2.400E-06 | 3.464E-07 | 2.200E-06 | 2.800E-06 |
| 24434 | 1222 | T1222F | D | 3.2e-06 | 2.6e-06 | 3.4e-06 | 3.067E-06 | 4.163E-07 | 2.600E-06 | 3.400E-06 |
| 24435 | 1222 | T1222P | D | 2.5e-06 | 2.1e-06 | 2.5e-06 | 2.367E-06 | 2.309E-07 | 2.100E-06 | 2.500E-06 |
| 24436 | 1222 | T1222S | D | 1.9e-06 | 2e-06 | 1.7e-06 | 1.867E-06 | 1.528E-07 | 1.700E-06 | 2.000E-06 |
| 24437 | 1222 | T1222T | D | 2.4e-06 | 2.5e-06 | 1.4e-06 | 2.100E-06 | 6.083E-07 | 1.400E-06 | 2.500E-06 |
| 24438 | 1222 | T1222W | D | 2.7e-06 | 2.8e-06 | 3.3e-06 | 2.933E-06 | 3.215E-07 | 2.700E-06 | 3.300E-06 |
| 24439 | 1222 | T1222Y | D | 1.9e-06 | 4.2e-06 | 1.4e-06 | 2.500E-06 | 1.493E-06 | 1.400E-06 | 4.200E-06 |
| 24440 | 1222 | T1222V | D | 3.1e-06 | 9.6e-07 | 3.1e-06 | 2.387E-06 | 1.236E-06 | 9.600E-07 | 3.100E-06 |
| 24441 | 1223 | E1223A | D | 2.2e-06 | 2.7e-06 | 2.4e-06 | 2.433E-06 | 2.517E-07 | 2.200E-06 | 2.700E-06 |
| 24442 | 1223 | E1223R | D | 3.6e-06 | 2.2e-06 | 2.6e-06 | 2.800E-06 | 7.211E-07 | 2.200E-06 | 3.600E-06 |
| 24443 | 1223 | E1223N | D | 2.1e-06 | 3.2e-06 | 1.3e-06 | 2.200E-06 | 9.539E-07 | 1.300E-06 | 3.200E-06 |
| 24444 | 1223 | E1223D | D | 3e-06 | 2.2e-06 | 4e-06 | 3.067E-06 | 9.018E-07 | 2.200E-06 | 4.000E-06 |
| 24445 | 1223 | E1223C | D | 1.3e-06 | 2e-06 | 3.3e-06 | 2.200E-06 | 1.015E-06 | 1.300E-06 | 3.300E-06 |
| 24446 | 1223 | E1223E | D | 2.9e-06 | 1.9e-06 | 3.1e-06 | 2.633E-06 | 6.429E-07 | 1.900E-06 | 3.100E-06 |
| 24447 | 1223 | E1223Q | D | 2e-06 | 1.9e-06 | 3.7e-06 | 2.533E-06 | 1.012E-06 | 1.900E-06 | 3.700E-06 |
| 24448 | 1223 | E1223G | D | 2.6e-06 | 3.1e-06 | 2.7e-06 | 2.800E-06 | 2.646E-07 | 2.600E-06 | 3.100E-06 |
| 24449 | 1223 | E1223H | D | 2.8e-06 | 1.8e-06 | 2.6e-06 | 2.400E-06 | 5.292E-07 | 1.800E-06 | 2.800E-06 |
| 24450 | 1223 | E1223I | D | 2.1e-06 | 3.5e-06 | 2.6e-06 | 2.733E-06 | 7.095E-07 | 2.100E-06 | 3.500E-06 |
| 24451 | 1223 | E1223L | D | 4.6e-06 | 1.6e-06 | 3.8e-06 | 3.333E-06 | 1.553E-06 | 1.600E-06 | 4.600E-06 |
| 24452 | 1223 | E1223K | D | 2.2e-06 | 2.4e-06 | 2e-06 | 2.200E-06 | 2.000E-07 | 2.000E-06 | 2.400E-06 |
| 24453 | 1223 | E1223M | D | 2.4e-06 | 2.7e-06 | 3.3e-06 | 2.800E-06 | 4.583E-07 | 2.400E-06 | 3.300E-06 |
| 24454 | 1223 | E1223F | D | 1.9e-06 | 3.2e-06 | 2.7e-06 | 2.600E-06 | 6.557E-07 | 1.900E-06 | 3.200E-06 |
| 24455 | 1223 | E1223P | D | 1.9e-06 | 2.4e-06 | 3.2e-06 | 2.500E-06 | 6.557E-07 | 1.900E-06 | 3.200E-06 |
| 24456 | 1223 | E1223S | D | 2.7e-06 | 2.8e-06 | 2.1e-06 | 2.533E-06 | 3.786E-07 | 2.100E-06 | 2.800E-06 |
| 24457 | 1223 | E1223T | D | 2.3e-06 | 1.4e-06 | 2.3e-06 | 2.000E-06 | 5.196E-07 | 1.400E-06 | 2.300E-06 |
| 24458 | 1223 | E1223W | D | 4.3e-06 | 3.3e-06 | 3.1e-06 | 3.567E-06 | 6.429E-07 | 3.100E-06 | 4.300E-06 |
| 24459 | 1223 | E1223Y | D | 3.6e-06 | 3.4e-06 | 4.8e-06 | 3.933E-06 | 7.572E-07 | 3.400E-06 | 4.800E-06 |
| 24460 | 1223 | E1223V | D | 4.4e-06 | 2e-06 | 2.2e-06 | 2.867E-06 | 1.332E-06 | 2.000E-06 | 4.400E-06 |
| 24461 | 1224 | Q1224A | D | 2.9e-06 | 3.3e-06 | 2.8e-06 | 3.000E-06 | 2.646E-07 | 2.800E-06 | 3.300E-06 |
| 24462 | 1224 | Q1224R | D | 2.7e-06 | 2.1e-06 | 3.6e-06 | 2.800E-06 | 7.550E-07 | 2.100E-06 | 3.600E-06 |
| 24463 | 1224 | Q1224N | D | 2.2e-06 | 2e-06 | 2.3e-06 | 2.167E-06 | 1.528E-07 | 2.000E-06 | 2.300E-06 |
| 24464 | 1224 | Q1224D | D | 2.7e-06 | 1.9e-06 | 1.3e-06 | 1.967E-06 | 7.024E-07 | 1.300E-06 | 2.700E-06 |
| 24465 | 1224 | Q1224C | D | 2.9e-06 | 2.2e-06 | 2.2e-06 | 2.433E-06 | 4.041E-07 | 2.200E-06 | 2.900E-06 |
| 24466 | 1224 | Q1224E | D | 1.1e-06 | 3.8e-06 | 2.9e-06 | 2.600E-06 | 1.375E-06 | 1.100E-06 | 3.800E-06 |
| 24467 | 1224 | Q1224Q | D | 2.4e-06 | 2.3e-06 | 3.6e-06 | 2.767E-06 | 7.234E-07 | 2.300E-06 | 3.600E-06 |
| 24468 | 1224 | Q1224G | D | 2.1e-06 | 1.5e-06 | 3.1e-06 | 2.233E-06 | 8.083E-07 | 1.500E-06 | 3.100E-06 |
| 24469 | 1224 | Q1224H | D | 1.9e-06 | 2.1e-06 | 2.8e-06 | 2.267E-06 | 4.726E-07 | 1.900E-06 | 2.800E-06 |
| 24470 | 1224 | Q1224I | D | 2.6e-06 | 3.3e-06 | 3.2e-06 | 3.033E-06 | 3.786E-07 | 2.600E-06 | 3.300E-06 |
| 24471 | 1224 | Q1224L | D | 2e-06 | 2.5e-06 | 2.4e-06 | 2.300E-06 | 2.646E-07 | 2.000E-06 | 2.500E-06 |
| 24472 | 1224 | Q1224K | D | 3.1e-06 | 2.7e-06 | 2e-06 | 2.600E-06 | 5.568E-07 | 2.000E-06 | 3.100E-06 |
| 24473 | 1224 | Q1224M | D | 1.4e-06 | 2e-06 | 3.6e-06 | 2.333E-06 | 1.137E-06 | 1.400E-06 | 3.600E-06 |
| 24474 | 1224 | Q1224F | D | 2.2e-06 | 1.1e-06 | 2.7e-06 | 2.000E-06 | 8.185E-07 | 1.100E-06 | 2.700E-06 |
| 24475 | 1224 | Q1224P | D | 3.7e-06 | 2.4e-06 | 2e-06 | 2.700E-06 | 8.888E-07 | 2.000E-06 | 3.700E-06 |
| 24476 | 1224 | Q1224S | D | 1.5e-06 | 3.4e-06 | 1.8e-06 | 2.233E-06 | 1.021E-06 | 1.500E-06 | 3.400E-06 |
| 24477 | 1224 | Q1224T | D | 1.9e-06 | 2.2e-06 | 1.8e-06 | 1.967E-06 | 2.082E-07 | 1.800E-06 | 2.200E-06 |
| 24478 | 1224 | Q1224W | D | 2.3e-06 | 2e-06 | 2.6e-06 | 2.300E-06 | 3.000E-07 | 2.000E-06 | 2.600E-06 |
| 24479 | 1224 | Q1224Y | D | 3.6e-06 | 2.1e-06 | 2.5e-06 | 2.733E-06 | 7.767E-07 | 2.100E-06 | 3.600E-06 |

| | | | | | | | | | | |
|---|---|---|---|---|---|---|---|---|---|---|
| 24480 | 1224 | Q1224V | D | 2.3e-06 | 3.4e-06 | 2e-06 | 2.567E-06 | 7.371E-07 | 2.000E-06 | 3.400E-06 |
| 24481 | 1225 | D1225A | D | 3e-06 | 1.2e-06 | 2.7e-06 | 2.300E-06 | 9.644E-07 | 1.200E-06 | 3.000E-06 |
| 24482 | 1225 | D1225R | D | 2.3e-06 | 2.2e-06 | 3.3e-06 | 2.600E-06 | 6.083E-07 | 2.200E-06 | 3.300E-06 |
| 24483 | 1225 | D1225N | D | 3.1e-06 | 1.5e-06 | 1.8e-06 | 2.133E-06 | 8.505E-07 | 1.500E-06 | 3.100E-06 |
| 24484 | 1225 | D1225D | D | 2.1e-06 | 2.7e-06 | 2.7e-06 | 2.500E-06 | 3.464E-07 | 2.100E-06 | 2.700E-06 |
| 24485 | 1225 | D1225C | D | 2e-06 | 1.6e-06 | 2e-06 | 1.867E-06 | 2.309E-07 | 1.600E-06 | 2.000E-06 |
| 24486 | 1225 | D1225E | D | 2.2e-06 | 2.7e-06 | 2.3e-06 | 2.400E-06 | 2.646E-07 | 2.200E-06 | 2.700E-06 |
| 24487 | 1225 | D1225Q | D | 4.6e-06 | 3.4e-06 | 1.8e-06 | 3.267E-06 | 1.405E-06 | 1.800E-06 | 4.600E-06 |
| 24488 | 1225 | D1225G | D | 2.3e-06 | 2.1e-06 | 1.3e-06 | 1.900E-06 | 5.292E-07 | 1.300E-06 | 2.300E-06 |
| 24489 | 1225 | D1225H | D | 2.2e-06 | 2.5e-06 | 2.3e-06 | 2.333E-06 | 1.528E-07 | 2.200E-06 | 2.500E-06 |
| 24490 | 1225 | D1225I | D | 2.1e-06 | 3.3e-06 | 3.1e-06 | 2.833E-06 | 6.429E-07 | 2.100E-06 | 3.300E-06 |
| 24491 | 1225 | D1225L | D | 2.4e-06 | 3e-06 | 2.6e-06 | 2.667E-06 | 3.055E-07 | 2.400E-06 | 3.000E-06 |
| 24492 | 1225 | D1225K | D | 2.5e-06 | 2.3e-06 | 3.8e-06 | 2.867E-06 | 8.145E-07 | 2.300E-06 | 3.800E-06 |
| 24493 | 1225 | D1225M | D | 2.3e-06 | 1.9e-06 | 2.1e-06 | 2.100E-06 | 2.000E-07 | 1.900E-06 | 2.300E-06 |
| 24494 | 1225 | D1225F | D | 3.1e-06 | 3.4e-06 | 2.2e-06 | 2.900E-06 | 6.245E-07 | 2.200E-06 | 3.400E-06 |
| 24495 | 1225 | D1225P | D | 2.3e-06 | 3.1e-06 | 2.1e-06 | 2.500E-06 | 5.292E-07 | 2.100E-06 | 3.100E-06 |
| 24496 | 1225 | D1225S | D | 2.6e-06 | 2.1e-06 | 2.2e-06 | 2.300E-06 | 2.646E-07 | 2.100E-06 | 2.600E-06 |
| 24497 | 1225 | D1225T | D | 2.4e-06 | 2.4e-06 | 1.6e-06 | 2.133E-06 | 4.619E-07 | 1.600E-06 | 2.400E-06 |
| 24498 | 1225 | D1225W | D | 2.7e-06 | 2.5e-06 | 2.8e-06 | 2.667E-06 | 1.528E-07 | 2.500E-06 | 2.800E-06 |
| 24499 | 1225 | D1225Y | D | 3.4e-06 | 1.1e-06 | 1.4e-06 | 1.967E-06 | 1.250E-06 | 1.100E-06 | 3.400E-06 |
| 24500 | 1225 | D1225V | D | 2.1e-06 | 2.5e-06 | 1.7e-06 | 2.100E-06 | 4.000E-07 | 1.700E-06 | 2.500E-06 |
| 24501 | 1226 | S1226A | D | 2e-06 | 2.6e-06 | 2.3e-06 | 2.300E-06 | 3.000E-07 | 2.000E-06 | 2.600E-06 |
| 24502 | 1226 | S1226R | D | 1.8e-06 | 2.3e-06 | 2.1e-06 | 2.067E-06 | 2.517E-07 | 1.800E-06 | 2.300E-06 |
| 24503 | 1226 | S1226N | D | 1.9e-06 | 2.3e-06 | 2e-06 | 2.067E-06 | 2.082E-07 | 1.900E-06 | 2.300E-06 |
| 24504 | 1226 | S1226D | D | 2.5e-06 | 1.6e-06 | 2.5e-06 | 2.200E-06 | 5.196E-07 | 1.600E-06 | 2.500E-06 |
| 24505 | 1226 | S1226C | D | 1.6e-06 | 2.3e-06 | 2.4e-06 | 2.100E-06 | 4.359E-07 | 1.600E-06 | 2.400E-06 |
| 24506 | 1226 | S1226E | D | 2.8e-06 | 3e-06 | 1.8e-06 | 2.533E-06 | 6.429E-07 | 1.800E-06 | 3.000E-06 |
| 24507 | 1226 | S1226Q | D | 1.4e-06 | 1e-06 | 2.1e-06 | 1.500E-06 | 5.568E-07 | 1.000E-06 | 2.100E-06 |
| 24508 | 1226 | S1226G | D | 2.9e-06 | 1.8e-06 | 1.8e-06 | 2.167E-06 | 6.351E-07 | 1.800E-06 | 2.900E-06 |
| 24509 | 1226 | S1226H | D | 2.4e-06 | 1.8e-06 | 3.5e-06 | 2.567E-06 | 8.622E-07 | 1.800E-06 | 3.500E-06 |
| 24510 | 1226 | S1226I | D | 2.8e-06 | 3.7e-06 | 2.5e-06 | 3.000E-06 | 6.245E-07 | 2.500E-06 | 3.700E-06 |
| 24511 | 1226 | S1226L | D | 2.1e-06 | 2.8e-06 | 1.7e-06 | 2.200E-06 | 5.568E-07 | 1.700E-06 | 2.800E-06 |
| 24512 | 1226 | S1226K | D | 2.1e-06 | 3.1e-06 | 2.1e-06 | 2.433E-06 | 5.774E-07 | 2.100E-06 | 3.100E-06 |
| 24513 | 1226 | S1226M | D | 2.4e-06 | 2.4e-06 | 2.3e-06 | 2.367E-06 | 5.774E-08 | 2.300E-06 | 2.400E-06 |
| 24514 | 1226 | S1226F | D | 2.6e-06 | 2.4e-06 | 2.6e-06 | 2.533E-06 | 1.155E-07 | 2.400E-06 | 2.600E-06 |
| 24515 | 1226 | S1226P | D | 2e-06 | 2e-06 | 2e-06 | 2.000E-06 | 0.000E+00 | 2.000E-06 | 2.000E-06 |
| 24516 | 1226 | S1226S | D | 3e-06 | 2.2e-06 | 2.1e-06 | 2.433E-06 | 4.933E-07 | 2.100E-06 | 3.000E-06 |
| 24517 | 1226 | S1226T | D | 2.3e-06 | 2.4e-06 | 1.6e-06 | 2.100E-06 | 4.359E-07 | 1.600E-06 | 2.400E-06 |
| 24518 | 1226 | S1226W | D | 4.6e-06 | 2.6e-06 | 1.6e-06 | 2.933E-06 | 1.528E-06 | 1.600E-06 | 4.600E-06 |
| 24519 | 1226 | S1226Y | D | 1.2e-06 | 1.5e-06 | 3e-06 | 1.900E-06 | 9.644E-07 | 1.200E-06 | 3.000E-06 |
| 24520 | 1226 | S1226V | D | 3.3e-06 | 1.5e-06 | 2.6e-06 | 2.467E-06 | 9.074E-07 | 1.500E-06 | 3.300E-06 |
| 24521 | 1227 | K1227A | D | 1.8e-06 | 3.3e-06 | 2.1e-06 | 2.400E-06 | 7.937E-07 | 1.800E-06 | 3.300E-06 |
| 24522 | 1227 | K1227R | D | 1.9e-06 | 2.3e-06 | 2.1e-06 | 2.100E-06 | 2.000E-07 | 1.900E-06 | 2.300E-06 |
| 24523 | 1227 | K1227N | D | 1.7e-06 | 2.5e-06 | 1.5e-06 | 1.900E-06 | 5.292E-07 | 1.500E-06 | 2.500E-06 |
| 24524 | 1227 | K1227D | D | 1e-06 | 3.2e-06 | 1.8e-06 | 2.000E-06 | 1.114E-06 | 1.000E-06 | 3.200E-06 |
| 24525 | 1227 | K1227C | D | 1.9e-06 | 2.4e-06 | 1.7e-06 | 2.000E-06 | 3.606E-07 | 1.700E-06 | 2.400E-06 |
| 24526 | 1227 | K1227E | D | 2.8e-06 | 2.1e-06 | 2.8e-06 | 2.567E-06 | 4.041E-07 | 2.100E-06 | 2.800E-06 |
| 24527 | 1227 | K1227Q | D | 1.7e-06 | 1.7e-06 | 1.6e-06 | 1.667E-06 | 5.774E-08 | 1.600E-06 | 1.700E-06 |
| 24528 | 1227 | K1227G | D | 2.3e-06 | 1.1e-06 | 2.3e-06 | 1.900E-06 | 6.928E-07 | 1.100E-06 | 2.300E-06 |
| 24529 | 1227 | K1227H | D | 1.4e-06 | 2.2e-06 | 3.4e-06 | 2.333E-06 | 1.007E-06 | 1.400E-06 | 3.400E-06 |
| 24530 | 1227 | K1227I | D | 1.4e-06 | 3e-06 | 2.4e-06 | 2.267E-06 | 8.083E-07 | 1.400E-06 | 3.000E-06 |
| 24531 | 1227 | K1227L | D | 4.3e-06 | 2.3e-06 | 3.3e-06 | 3.300E-06 | 1.000E-06 | 2.300E-06 | 4.300E-06 |
| 24532 | 1227 | K1227K | D | 3.7e-06 | 2.3e-06 | 1.9e-06 | 2.633E-06 | 9.452E-07 | 1.900E-06 | 3.700E-06 |
| 24533 | 1227 | K1227M | D | 2.7e-06 | 2.9e-06 | 2.4e-06 | 2.667E-06 | 2.517E-07 | 2.400E-06 | 2.900E-06 |
| 24534 | 1227 | K1227F | D | 1.7e-06 | 3.3e-06 | 1.4e-06 | 2.133E-06 | 1.021E-06 | 1.400E-06 | 3.300E-06 |
| 24535 | 1227 | K1227P | D | 3.9e-06 | 3e-06 | 3e-06 | 3.300E-06 | 5.196E-07 | 3.000E-06 | 3.900E-06 |
| 24536 | 1227 | K1227S | D | 2.4e-06 | 1.7e-06 | 3.5e-06 | 2.533E-06 | 9.074E-07 | 1.700E-06 | 3.500E-06 |
| 24537 | 1227 | K1227T | D | 3.1e-06 | 5.5e-06 | 2.8e-06 | 3.800E-06 | 1.480E-06 | 2.800E-06 | 5.500E-06 |
| 24538 | 1227 | K1227W | D | 1.3e-06 | 2e-06 | 2.5e-06 | 1.933E-06 | 6.028E-07 | 1.300E-06 | 2.500E-06 |
| 24539 | 1227 | K1227Y | D | 2.7e-06 | 2.2e-06 | 1.2e-06 | 2.033E-06 | 7.638E-07 | 1.200E-06 | 2.700E-06 |
| 24540 | 1227 | K1227V | D | 2.1e-06 | 3.1e-06 | 2.1e-06 | 2.433E-06 | 5.774E-07 | 2.100E-06 | 3.100E-06 |
| 24541 | 1228 | D1228A | D | 2e-06 | 3.6e-06 | 2.3e-06 | 2.633E-06 | 8.505E-07 | 2.000E-06 | 3.600E-06 |
| 24542 | 1228 | D1228R | D | 2.7e-06 | 2e-06 | 3.5e-06 | 2.733E-06 | 7.506E-07 | 2.000E-06 | 3.500E-06 |
| 24543 | 1228 | D1228N | D | 1.9e-06 | 1.1e-06 | 3.7e-06 | 2.233E-06 | 1.332E-06 | 1.100E-06 | 3.700E-06 |
| 24544 | 1228 | D1228D | D | 3.1e-06 | 2.9e-06 | 2.6e-06 | 2.867E-06 | 2.517E-07 | 2.600E-06 | 3.100E-06 |
| 24545 | 1228 | D1228C | D | 3.7e-06 | 1.7e-06 | 3e-06 | 2.800E-06 | 1.015E-06 | 1.700E-06 | 3.700E-06 |
| 24546 | 1228 | D1228E | D | 3.4e-06 | 3.4e-06 | 2.5e-06 | 3.100E-06 | 5.196E-07 | 2.500E-06 | 3.400E-06 |
| 24547 | 1228 | D1228Q | D | 3.2e-06 | 2.9e-06 | 4.8e-06 | 3.633E-06 | 1.021E-06 | 2.900E-06 | 4.800E-06 |
| 24548 | 1228 | D1228G | D | 1.3e-06 | 3.2e-06 | 2.7e-06 | 2.400E-06 | 9.849E-07 | 1.300E-06 | 3.200E-06 |
| 24549 | 1228 | D1228H | D | 3.7e-06 | 3.5e-06 | 2.7e-06 | 3.300E-06 | 5.292E-07 | 2.700E-06 | 3.700E-06 |
| 24550 | 1228 | D1228I | D | 1.5e-06 | 2.5e-06 | 5.3e-06 | 3.100E-06 | 1.970E-06 | 1.500E-06 | 5.300E-06 |
| 24551 | 1228 | D1228L | D | 2.5e-06 | 3.3e-06 | 2.2e-06 | 2.667E-06 | 5.686E-07 | 2.200E-06 | 3.300E-06 |
| 24552 | 1228 | D1228K | D | 2.6e-06 | 3.3e-06 | 1.7e-06 | 2.533E-06 | 8.021E-07 | 1.700E-06 | 3.300E-06 |
| 24553 | 1228 | D1228M | D | 2.3e-06 | 2.1e-06 | 1.6e-06 | 2.000E-06 | 3.606E-07 | 1.600E-06 | 2.300E-06 |
| 24554 | 1228 | D1228F | D | 1.5e-06 | 2.5e-06 | 1.4e-06 | 1.800E-06 | 6.083E-07 | 1.400E-06 | 2.500E-06 |
| 24555 | 1228 | D1228P | D | 3.7e-06 | 2.9e-06 | 2.7e-06 | 3.100E-06 | 5.292E-07 | 2.700E-06 | 3.700E-06 |
| 24556 | 1228 | D1228S | D | 2.5e-06 | 1.3e-06 | 1.2e-06 | 1.667E-06 | 7.234E-07 | 1.200E-06 | 2.500E-06 |
| 24557 | 1228 | D1228T | D | 1.8e-06 | 2e-06 | 2.6e-06 | 2.133E-06 | 4.163E-07 | 1.800E-06 | 2.600E-06 |
| 24558 | 1228 | D1228W | D | 2.9e-06 | 2.1e-06 | 4e-06 | 3.000E-06 | 9.539E-07 | 2.100E-06 | 4.000E-06 |
| 24559 | 1228 | D1228Y | D | 3.2e-06 | 2.6e-06 | 2.2e-06 | 2.667E-06 | 5.033E-07 | 2.200E-06 | 3.200E-06 |
| 24560 | 1228 | D1228V | D | 2.6e-06 | 1.8e-06 | 3.3e-06 | 2.567E-06 | 7.506E-07 | 1.800E-06 | 3.300E-06 |
| 24561 | 1229 | S1229A | D | 2.1e-06 | 1.8e-06 | 1.2e-06 | 1.700E-06 | 4.583E-07 | 1.200E-06 | 2.100E-06 |
| 24562 | 1229 | S1229R | D | 2.4e-06 | 2.5e-06 | 3.3e-06 | 2.733E-06 | 4.933E-07 | 2.400E-06 | 3.300E-06 |
| 24563 | 1229 | S1229N | D | 3e-06 | 1.5e-06 | 1.7e-06 | 2.067E-06 | 8.145E-07 | 1.500E-06 | 3.000E-06 |
| 24564 | 1229 | S1229D | D | 3.2e-06 | 2.2e-06 | 1.5e-06 | 2.300E-06 | 8.544E-07 | 1.500E-06 | 3.200E-06 |

| | | | | | | | | | | |
|---|---|---|---|---|---|---|---|---|---|---|
| 24565 | 1229 | S1229C | D | 1.7e-06 | 1.3e-06 | 1.8e-06 | 1.600E-06 | 2.646E-07 | 1.300E-06 | 1.800E-06 |
| 24566 | 1229 | S1229E | D | 2.4e-06 | 2e-06 | 1.7e-06 | 2.033E-06 | 3.512E-07 | 1.700E-06 | 2.400E-06 |
| 24567 | 1229 | S1229Q | D | 1.7e-06 | 2.9e-06 | 2.2e-06 | 2.267E-06 | 6.028E-07 | 1.700E-06 | 2.900E-06 |
| 24568 | 1229 | S1229G | D | 1.9e-06 | 3.4e-06 | 2.5e-06 | 2.600E-06 | 7.550E-07 | 1.900E-06 | 3.400E-06 |
| 24569 | 1229 | S1229H | D | 3.5e-06 | 3.4e-06 | 2e-06 | 2.967E-06 | 8.386E-07 | 2.000E-06 | 3.500E-06 |
| 24570 | 1229 | S1229I | D | 2e-06 | 1.6e-06 | 1.9e-06 | 1.833E-06 | 2.082E-07 | 1.600E-06 | 2.000E-06 |
| 24571 | 1229 | S1229L | D | 2.5e-06 | 1.5e-06 | 1.9e-06 | 1.967E-06 | 5.033E-07 | 1.500E-06 | 2.500E-06 |
| 24572 | 1229 | S1229K | D | 2.4e-06 | 2e-06 | 2.7e-06 | 2.367E-06 | 3.512E-07 | 2.000E-06 | 2.700E-06 |
| 24573 | 1229 | S1229M | D | 4.4e-06 | 2.3e-06 | 2.2e-06 | 2.967E-06 | 1.242E-06 | 2.200E-06 | 4.400E-06 |
| 24574 | 1229 | S1229F | D | 2.5e-06 | 2.4e-06 | 2.8e-06 | 2.567E-06 | 2.082E-07 | 2.400E-06 | 2.800E-06 |
| 24575 | 1229 | S1229P | D | 2.7e-06 | 2e-06 | 1.6e-06 | 2.100E-06 | 5.568E-07 | 1.600E-06 | 2.700E-06 |
| 24576 | 1229 | S1229S | D | 4.6e-06 | 2e-06 | 2.9e-06 | 3.167E-06 | 1.320E-06 | 2.000E-06 | 4.600E-06 |
| 24577 | 1229 | S1229T | D | 3.5e-06 | 9.5e-07 | 2.2e-06 | 2.217E-06 | 1.275E-06 | 9.500E-07 | 3.500E-06 |
| 24578 | 1229 | S1229W | D | 2.5e-06 | 2.8e-06 | 2.5e-06 | 2.600E-06 | 1.732E-07 | 2.500E-06 | 2.800E-06 |
| 24579 | 1229 | S1229Y | D | 2.1e-06 | 2.9e-06 | 2.7e-06 | 2.567E-06 | 4.163E-07 | 2.100E-06 | 2.900E-06 |
| 24580 | 1229 | S1229V | D | 2.6e-06 | 2e-06 | 3.5e-06 | 2.700E-06 | 7.550E-07 | 2.000E-06 | 3.500E-06 |
| 24581 | 1230 | T1230A | D | 2.2e-06 | 3e-06 | 2.1e-06 | 2.433E-06 | 4.933E-07 | 2.100E-06 | 3.000E-06 |
| 24582 | 1230 | T1230R | D | 3.3e-06 | 3.2e-06 | 2.3e-06 | 2.933E-06 | 5.508E-07 | 2.300E-06 | 3.300E-06 |
| 24583 | 1230 | T1230N | D | 1.7e-06 | 3.6e-06 | 2.4e-06 | 2.567E-06 | 9.609E-07 | 1.700E-06 | 3.600E-06 |
| 24584 | 1230 | T1230D | D | 2.6e-06 | 2.1e-06 | 1.5e-06 | 2.067E-06 | 5.508E-07 | 1.500E-06 | 2.600E-06 |
| 24585 | 1230 | T1230C | D | 4.6e-06 | 4.8e-06 | 3.8e-06 | 4.400E-06 | 5.292E-07 | 3.800E-06 | 4.800E-06 |
| 24586 | 1230 | T1230E | D | 2.2e-06 | 2.5e-06 | 2.2e-06 | 2.300E-06 | 1.732E-07 | 2.200E-06 | 2.500E-06 |
| 24587 | 1230 | T1230Q | D | 1.2e-06 | 2.3e-06 | 2.9e-06 | 2.133E-06 | 8.622E-07 | 1.200E-06 | 2.900E-06 |
| 24588 | 1230 | T1230G | D | 2.3e-06 | 2e-06 | 3.4e-06 | 2.567E-06 | 7.371E-07 | 2.000E-06 | 3.400E-06 |
| 24589 | 1230 | T1230H | D | 2.4e-06 | 3.1e-06 | 2.3e-06 | 2.600E-06 | 4.359E-07 | 2.300E-06 | 3.100E-06 |
| 24590 | 1230 | T1230I | D | 2.7e-06 | 2.7e-06 | 3.5e-06 | 2.967E-06 | 4.619E-07 | 2.700E-06 | 3.500E-06 |
| 24591 | 1230 | T1230L | D | 1.8e-06 | 2.7e-06 | 1.7e-06 | 2.067E-06 | 5.508E-07 | 1.700E-06 | 2.700E-06 |
| 24592 | 1230 | T1230K | D | 2.2e-06 | 1.4e-06 | 1.2e-06 | 1.600E-06 | 5.292E-07 | 1.200E-06 | 2.200E-06 |
| 24593 | 1230 | T1230M | D | 1.9e-06 | 1.9e-06 | 1.8e-06 | 1.867E-06 | 5.774E-08 | 1.800E-06 | 1.900E-06 |
| 24594 | 1230 | T1230F | D | 3.2e-06 | 1.3e-06 | 2.4e-06 | 2.300E-06 | 9.539E-07 | 1.300E-06 | 3.200E-06 |
| 24595 | 1230 | T1230P | D | 1.6e-06 | 2.4e-06 | 2.3e-06 | 2.100E-06 | 4.359E-07 | 1.600E-06 | 2.400E-06 |
| 24596 | 1230 | T1230S | D | 2.6e-06 | 2e-06 | 1.4e-06 | 2.000E-06 | 6.000E-07 | 1.400E-06 | 2.600E-06 |
| 24597 | 1230 | T1230T | D | 2.9e-06 | 3.2e-06 | 2e-06 | 2.700E-06 | 6.245E-07 | 2.000E-06 | 3.200E-06 |
| 24598 | 1230 | T1230W | D | 2.1e-06 | 2.6e-06 | 2.8e-06 | 2.500E-06 | 3.606E-07 | 2.100E-06 | 2.800E-06 |
| 24599 | 1230 | T1230Y | D | 1.4e-06 | 1.8e-06 | 3.3e-06 | 2.167E-06 | 1.002E-06 | 1.400E-06 | 3.300E-06 |
| 24600 | 1230 | T1230V | D | 2.3e-06 | 1.7e-06 | 3.1e-06 | 2.367E-06 | 7.024E-07 | 1.700E-06 | 3.100E-06 |
| 24601 | 1231 | Y1231A | D | 2e-06 | 1.6e-06 | 2.5e-06 | 2.033E-06 | 4.509E-07 | 1.600E-06 | 2.500E-06 |
| 24602 | 1231 | Y1231R | D | 3.2e-06 | 2.6e-06 | 1.5e-06 | 2.433E-06 | 8.622E-07 | 1.500E-06 | 3.200E-06 |
| 24603 | 1231 | Y1231N | D | 1.4e-06 | 2.2e-06 | 3.1e-06 | 2.233E-06 | 8.505E-07 | 1.400E-06 | 3.100E-06 |
| 24604 | 1231 | Y1231D | D | 3.2e-06 | 2.5e-06 | 2.7e-06 | 2.800E-06 | 3.606E-07 | 2.500E-06 | 3.200E-06 |
| 24605 | 1231 | Y1231C | D | 2.9e-06 | 1.4e-06 | 2.8e-06 | 2.367E-06 | 8.386E-07 | 1.400E-06 | 2.900E-06 |
| 24606 | 1231 | Y1231E | D | 3.8e-06 | 2.1e-06 | 1.7e-06 | 2.533E-06 | 1.115E-06 | 1.700E-06 | 3.800E-06 |
| 24607 | 1231 | Y1231Q | D | 2.6e-06 | 1.9e-06 | 1.2e-06 | 1.900E-06 | 7.000E-07 | 1.200E-06 | 2.600E-06 |
| 24608 | 1231 | Y1231G | D | 2.9e-06 | 1.6e-06 | 3.6e-06 | 2.700E-06 | 1.015E-06 | 1.600E-06 | 3.600E-06 |
| 24609 | 1231 | Y1231H | D | 2.3e-06 | 1.9e-06 | 1.7e-06 | 1.967E-06 | 3.055E-07 | 1.700E-06 | 2.300E-06 |
| 24610 | 1231 | Y1231I | D | 2.6e-06 | 1.1e-06 | 2.1e-06 | 1.933E-06 | 7.638E-07 | 1.100E-06 | 2.600E-06 |
| 24611 | 1231 | Y1231L | D | 1.1e-06 | 3.5e-06 | 2.1e-06 | 2.233E-06 | 1.206E-06 | 1.100E-06 | 3.500E-06 |
| 24612 | 1231 | Y1231K | D | 3.1e-06 | 2.4e-06 | 3.6e-06 | 3.033E-06 | 6.028E-07 | 2.400E-06 | 3.600E-06 |
| 24613 | 1231 | Y1231M | D | 3.3e-06 | 2.2e-06 | 3.2e-06 | 2.900E-06 | 6.083E-07 | 2.200E-06 | 3.300E-06 |
| 24614 | 1231 | Y1231F | D | 3.1e-06 | 2.2e-06 | 2e-06 | 2.433E-06 | 5.859E-07 | 2.000E-06 | 3.100E-06 |
| 24615 | 1231 | Y1231P | D | 2.6e-06 | 2.7e-06 | 3.3e-06 | 2.867E-06 | 3.786E-07 | 2.600E-06 | 3.300E-06 |
| 24616 | 1231 | Y1231S | D | 2.2e-06 | 2.4e-06 | 2.2e-06 | 2.267E-06 | 1.155E-07 | 2.200E-06 | 2.400E-06 |
| 24617 | 1231 | Y1231T | D | 1.9e-06 | 1.9e-06 | 3.5e-06 | 2.433E-06 | 9.238E-07 | 1.900E-06 | 3.500E-06 |
| 24618 | 1231 | Y1231W | D | 3.2e-06 | 3.5e-06 | 2.3e-06 | 3.000E-06 | 6.245E-07 | 2.300E-06 | 3.500E-06 |
| 24619 | 1231 | Y1231Y | D | 3.9e-06 | 2.6e-06 | 3.4e-06 | 3.300E-06 | 6.557E-07 | 2.600E-06 | 3.900E-06 |
| 24620 | 1231 | Y1231V | D | 2.2e-06 | 2.3e-06 | 2.4e-06 | 2.300E-06 | 1.000E-07 | 2.200E-06 | 2.400E-06 |
| 24621 | 1232 | S1232A | D | 3e-06 | 2.2e-06 | 3.3e-06 | 2.833E-06 | 5.686E-07 | 2.200E-06 | 3.300E-06 |
| 24622 | 1232 | S1232R | D | 1.9e-06 | 1.7e-06 | 1.9e-06 | 1.833E-06 | 1.155E-07 | 1.700E-06 | 1.900E-06 |
| 24623 | 1232 | S1232N | D | 2.4e-06 | 2.6e-06 | 1.4e-06 | 2.133E-06 | 6.429E-07 | 1.400E-06 | 2.600E-06 |
| 24624 | 1232 | S1232D | D | 2.8e-06 | 3.2e-06 | 2e-06 | 2.667E-06 | 6.110E-07 | 2.000E-06 | 3.200E-06 |
| 24625 | 1232 | S1232C | D | 1.3e-06 | 2.8e-06 | 2.6e-06 | 2.233E-06 | 8.145E-07 | 1.300E-06 | 2.800E-06 |
| 24626 | 1232 | S1232E | D | 3.6e-06 | 2.2e-06 | 2e-06 | 2.600E-06 | 8.718E-07 | 2.000E-06 | 3.600E-06 |
| 24627 | 1232 | S1232Q | D | 2.4e-06 | 2e-06 | 1.9e-06 | 2.100E-06 | 2.646E-07 | 1.900E-06 | 2.400E-06 |
| 24628 | 1232 | S1232G | D | 1.6e-06 | 1.9e-06 | 1.7e-06 | 1.733E-06 | 1.528E-07 | 1.600E-06 | 1.900E-06 |
| 24629 | 1232 | S1232H | D | 2.3e-06 | 3.1e-06 | 1.5e-06 | 2.300E-06 | 8.000E-07 | 1.500E-06 | 3.100E-06 |
| 24630 | 1232 | S1232I | D | 2.2e-06 | 2e-06 | 3.5e-06 | 2.567E-06 | 8.145E-07 | 2.000E-06 | 3.500E-06 |
| 24631 | 1232 | S1232L | D | 2e-06 | 2.4e-06 | 2e-06 | 2.133E-06 | 2.309E-07 | 2.000E-06 | 2.400E-06 |
| 24632 | 1232 | S1232K | D | 3.2e-06 | 2.3e-06 | 2e-06 | 2.500E-06 | 6.245E-07 | 2.000E-06 | 3.200E-06 |
| 24633 | 1232 | S1232M | D | 1.5e-06 | 3.5e-06 | 1.3e-06 | 2.100E-06 | 1.217E-06 | 1.300E-06 | 3.500E-06 |
| 24634 | 1232 | S1232F | D | 2.2e-06 | 3e-06 | 3.2e-06 | 2.800E-06 | 5.292E-07 | 2.200E-06 | 3.200E-06 |
| 24635 | 1232 | S1232P | D | 2.8e-06 | 1.7e-06 | 2.2e-06 | 2.233E-06 | 5.508E-07 | 1.700E-06 | 2.800E-06 |
| 24636 | 1232 | S1232S | D | 2.6e-06 | 2.1e-06 | 1.5e-06 | 2.067E-06 | 5.508E-07 | 1.500E-06 | 2.600E-06 |
| 24637 | 1232 | S1232T | D | 3.2e-06 | 3.1e-06 | 1.1e-06 | 2.467E-06 | 1.185E-06 | 1.100E-06 | 3.200E-06 |
| 24638 | 1232 | S1232W | D | 4e-06 | 1.5e-06 | 2.2e-06 | 2.567E-06 | 1.290E-06 | 1.500E-06 | 4.000E-06 |
| 24639 | 1232 | S1232Y | D | 2.6e-06 | 1.9e-06 | 2.7e-06 | 2.400E-06 | 4.359E-07 | 1.900E-06 | 2.700E-06 |
| 24640 | 1232 | S1232V | D | 1.8e-06 | 1.5e-06 | 2.8e-06 | 2.033E-06 | 6.807E-07 | 1.500E-06 | 2.800E-06 |
| 24641 | 1233 | L1233A | D | 1.8e-06 | 3e-06 | 1.1e-06 | 1.967E-06 | 9.609E-07 | 1.100E-06 | 3.000E-06 |
| 24642 | 1233 | L1233R | D | 2e-06 | 2.8e-06 | 3.7e-06 | 2.833E-06 | 8.505E-07 | 2.000E-06 | 3.700E-06 |
| 24643 | 1233 | L1233N | D | 1.8e-06 | 1.7e-06 | 3.4e-06 | 2.300E-06 | 9.539E-07 | 1.700E-06 | 3.400E-06 |
| 24644 | 1233 | L1233D | D | 3.6e-06 | 2.5e-06 | 3e-06 | 3.033E-06 | 5.508E-07 | 2.500E-06 | 3.600E-06 |
| 24645 | 1233 | L1233C | D | 1.5e-06 | 2.5e-06 | 2.2e-06 | 2.067E-06 | 5.132E-07 | 1.500E-06 | 2.500E-06 |
| 24646 | 1233 | L1233E | D | 1.1e-06 | 4.7e-06 | 3.1e-06 | 2.967E-06 | 1.804E-06 | 1.100E-06 | 4.700E-06 |
| 24647 | 1233 | L1233Q | D | 3.4e-06 | 3.1e-06 | 3.8e-06 | 3.433E-06 | 3.512E-07 | 3.100E-06 | 3.800E-06 |
| 24648 | 1233 | L1233G | D | 3.2e-06 | 2.3e-06 | 2.1e-06 | 2.533E-06 | 5.859E-07 | 2.100E-06 | 3.200E-06 |
| 24649 | 1233 | L1233H | D | 1.4e-06 | 1.7e-06 | 1.5e-06 | 1.533E-06 | 1.528E-07 | 1.400E-06 | 1.700E-06 |

| | | | | | | | | | | |
|---|---|---|---|---|---|---|---|---|---|---|
| 24650 | 1233 | L1233I | D | 1.6e-06 | 2.4e-06 | 2.7e-06 | 2.233E-06 | 5.686E-07 | 1.600E-06 | 2.700E-06 |
| 24651 | 1233 | L1233L | D | 1.4e-06 | 5.1e-06 | 4.4e-06 | 3.633E-06 | 1.966E-06 | 1.400E-06 | 5.100E-06 |
| 24652 | 1233 | L1233K | D | 2.7e-06 | 2.1e-06 | 2e-06 | 2.267E-06 | 3.786E-07 | 2.000E-06 | 2.700E-06 |
| 24653 | 1233 | L1233M | D | 2.5e-06 | 1.3e-06 | 2.8e-06 | 2.200E-06 | 7.937E-07 | 1.300E-06 | 2.800E-06 |
| 24654 | 1233 | L1233F | D | 2.3e-06 | 1.4e-06 | 2.5e-06 | 2.067E-06 | 5.859E-07 | 1.400E-06 | 2.500E-06 |
| 24655 | 1233 | L1233P | D | 2.2e-06 | 3e-06 | 2.4e-06 | 2.533E-06 | 4.163E-07 | 2.200E-06 | 3.000E-06 |
| 24656 | 1233 | L1233S | D | 1.2e-06 | 1.7e-06 | 1.9e-06 | 1.600E-06 | 3.606E-07 | 1.200E-06 | 1.900E-06 |
| 24657 | 1233 | L1233T | D | 2.7e-06 | 2e-06 | 3.3e-06 | 2.667E-06 | 6.506E-07 | 2.000E-06 | 3.300E-06 |
| 24658 | 1233 | L1233W | D | 2.7e-06 | 2.5e-06 | 2.1e-06 | 2.433E-06 | 3.055E-07 | 2.100E-06 | 2.700E-06 |
| 24659 | 1233 | L1233Y | D | 4.3e-06 | 1.7e-06 | 2.9e-06 | 2.967E-06 | 1.301E-06 | 1.700E-06 | 4.300E-06 |
| 24660 | 1233 | L1233V | D | 1.8e-06 | 2.5e-06 | 3.8e-06 | 2.700E-06 | 1.015E-06 | 1.800E-06 | 3.800E-06 |
| 24661 | 1234 | S1234A | D | 3.3e-06 | 3.4e-06 | 2.1e-06 | 2.933E-06 | 7.234E-07 | 2.100E-06 | 3.400E-06 |
| 24662 | 1234 | S1234R | D | 3.1e-06 | 1.3e-06 | 3e-06 | 2.467E-06 | 1.012E-06 | 1.300E-06 | 3.100E-06 |
| 24663 | 1234 | S1234N | D | 2e-06 | 1.9e-06 | 2.3e-06 | 2.067E-06 | 2.082E-07 | 1.900E-06 | 2.300E-06 |
| 24664 | 1234 | S1234D | D | 2.2e-06 | 2.8e-06 | 2.1e-06 | 2.367E-06 | 3.786E-07 | 2.100E-06 | 2.800E-06 |
| 24665 | 1234 | S1234C | D | 2.2e-06 | 3.4e-06 | 3.1e-06 | 2.900E-06 | 6.245E-07 | 2.200E-06 | 3.400E-06 |
| 24666 | 1234 | S1234E | D | 2.2e-06 | 1.6e-06 | 3.5e-06 | 2.433E-06 | 9.713E-07 | 1.600E-06 | 3.500E-06 |
| 24667 | 1234 | S1234Q | D | 3.1e-06 | 3.2e-06 | 2.3e-06 | 2.867E-06 | 4.933E-07 | 2.300E-06 | 3.200E-06 |
| 24668 | 1234 | S1234G | D | 2.3e-06 | 1.7e-06 | 2.3e-06 | 2.100E-06 | 3.464E-07 | 1.700E-06 | 2.300E-06 |
| 24669 | 1234 | S1234H | D | 1.6e-06 | 3.5e-06 | 4.1e-06 | 3.067E-06 | 1.305E-06 | 1.600E-06 | 4.100E-06 |
| 24670 | 1234 | S1234I | D | 1.6e-06 | 2.5e-06 | 2.6e-06 | 2.233E-06 | 5.508E-07 | 1.600E-06 | 2.600E-06 |
| 24671 | 1234 | S1234L | D | 2.7e-06 | 2.3e-06 | 2.2e-06 | 2.400E-06 | 2.646E-07 | 2.200E-06 | 2.700E-06 |
| 24672 | 1234 | S1234K | D | 2.5e-06 | 1.5e-06 | 4.9e-06 | 2.967E-06 | 1.747E-06 | 1.500E-06 | 4.900E-06 |
| 24673 | 1234 | S1234M | D | 2.2e-06 | 1.4e-06 | 1.8e-06 | 1.800E-06 | 4.000E-07 | 1.400E-06 | 2.200E-06 |
| 24674 | 1234 | S1234F | D | 3.5e-06 | 2.7e-06 | 3.1e-06 | 3.100E-06 | 4.000E-07 | 2.700E-06 | 3.500E-06 |
| 24675 | 1234 | S1234P | D | 2.9e-06 | 2.2e-06 | 3.1e-06 | 2.733E-06 | 4.726E-07 | 2.200E-06 | 3.100E-06 |
| 24676 | 1234 | S1234S | D | 3.7e-06 | 2.1e-06 | 1.7e-06 | 2.500E-06 | 1.058E-06 | 1.700E-06 | 3.700E-06 |
| 24677 | 1234 | S1234T | D | 2.5e-06 | 3.6e-06 | 2.2e-06 | 2.767E-06 | 7.371E-07 | 2.200E-06 | 3.600E-06 |
| 24678 | 1234 | S1234W | D | 2.3e-06 | 3.1e-06 | 1.5e-06 | 2.300E-06 | 8.000E-07 | 1.500E-06 | 3.100E-06 |
| 24679 | 1234 | S1234Y | D | 2.7e-06 | 2.3e-06 | 3.6e-06 | 2.867E-06 | 6.658E-07 | 2.300E-06 | 3.600E-06 |
| 24680 | 1234 | S1234V | D | 2.5e-06 | 2.6e-06 | 2e-06 | 2.367E-06 | 3.215E-07 | 2.000E-06 | 2.600E-06 |
| 24681 | 1235 | S1235A | D | 2.3e-06 | 1.1e-06 | 3.3e-06 | 2.233E-06 | 1.102E-06 | 1.100E-06 | 3.300E-06 |
| 24682 | 1235 | S1235R | D | 2.2e-06 | 3.2e-06 | 2.2e-06 | 2.533E-06 | 5.774E-07 | 2.200E-06 | 3.200E-06 |
| 24683 | 1235 | S1235N | D | 1.5e-06 | 2.3e-06 | 1.9e-06 | 1.900E-06 | 4.000E-07 | 1.500E-06 | 2.300E-06 |
| 24684 | 1235 | S1235D | D | 2.4e-06 | 2.6e-06 | 2.4e-06 | 2.467E-06 | 1.155E-07 | 2.400E-06 | 2.600E-06 |
| 24685 | 1235 | S1235C | D | 3.1e-06 | 3.1e-06 | 2.4e-06 | 2.867E-06 | 4.041E-07 | 2.400E-06 | 3.100E-06 |
| 24686 | 1235 | S1235E | D | 2.5e-06 | 1.7e-06 | 4.7e-06 | 2.967E-06 | 1.553E-06 | 1.700E-06 | 4.700E-06 |
| 24687 | 1235 | S1235Q | D | 2.4e-06 | 3.5e-06 | 2.3e-06 | 2.733E-06 | 6.658E-07 | 2.300E-06 | 3.500E-06 |
| 24688 | 1235 | S1235G | D | 1.9e-06 | 2.6e-06 | 2.9e-06 | 2.467E-06 | 5.132E-07 | 1.900E-06 | 2.900E-06 |
| 24689 | 1235 | S1235H | D | 2.3e-06 | 2.9e-06 | 8.3e-07 | 2.010E-06 | 1.065E-06 | 8.300E-07 | 2.900E-06 |
| 24690 | 1235 | S1235I | D | 4e-06 | 3.6e-06 | 2.5e-06 | 3.367E-06 | 7.767E-07 | 2.500E-06 | 4.000E-06 |
| 24691 | 1235 | S1235L | D | 1.8e-06 | 1.8e-06 | 8.6e-07 | 1.487E-06 | 5.427E-07 | 8.600E-07 | 1.800E-06 |
| 24692 | 1235 | S1235K | D | 2.6e-06 | 2.7e-06 | 1.8e-06 | 2.367E-06 | 4.933E-07 | 1.800E-06 | 2.700E-06 |
| 24693 | 1235 | S1235M | D | 7.9e-07 | 2.8e-06 | 2.7e-06 | 2.097E-06 | 1.133E-06 | 7.900E-07 | 2.800E-06 |
| 24694 | 1235 | S1235F | D | 3.3e-06 | 1.5e-06 | 2.6e-06 | 2.467E-06 | 9.074E-07 | 1.500E-06 | 3.300E-06 |
| 24695 | 1235 | S1235P | D | 3.3e-06 | 3.2e-06 | 2.6e-06 | 3.033E-06 | 3.786E-07 | 2.600E-06 | 3.300E-06 |
| 24696 | 1235 | S1235S | D | 2.5e-06 | 2.9e-06 | 3.8e-06 | 3.067E-06 | 6.658E-07 | 2.500E-06 | 3.800E-06 |
| 24697 | 1235 | S1235T | D | 2.5e-06 | 4.2e-06 | 2.7e-06 | 3.133E-06 | 9.292E-07 | 2.500E-06 | 4.200E-06 |
| 24698 | 1235 | S1235W | D | 3.1e-06 | 3.8e-06 | 2.5e-06 | 3.133E-06 | 6.506E-07 | 2.500E-06 | 3.800E-06 |
| 24699 | 1235 | S1235Y | D | 3.9e-06 | 2.2e-06 | 3.1e-06 | 3.067E-06 | 8.505E-07 | 2.200E-06 | 3.900E-06 |
| 24700 | 1235 | S1235V | D | 2.6e-06 | 3.5e-06 | 2.5e-06 | 2.867E-06 | 5.508E-07 | 2.500E-06 | 3.500E-06 |
| 24701 | 1236 | T1236A | D | 2.6e-06 | 2e-06 | 2.5e-06 | 2.367E-06 | 3.215E-07 | 2.000E-06 | 2.600E-06 |
| 24702 | 1236 | T1236R | D | 1.4e-06 | 1.8e-06 | 2.5e-06 | 1.900E-06 | 5.568E-07 | 1.400E-06 | 2.500E-06 |
| 24703 | 1236 | T1236N | D | 1.4e-06 | 2.7e-06 | 1.6e-06 | 1.900E-06 | 7.000E-07 | 1.400E-06 | 2.700E-06 |
| 24704 | 1236 | T1236D | D | 5.1e-06 | 2.2e-06 | 1.7e-06 | 3.000E-06 | 1.836E-06 | 1.700E-06 | 5.100E-06 |
| 24705 | 1236 | T1236C | D | 1.7e-06 | 2.2e-06 | 3e-06 | 2.300E-06 | 6.557E-07 | 1.700E-06 | 3.000E-06 |
| 24706 | 1236 | T1236E | D | 1.5e-06 | 2.3e-06 | 1.5e-06 | 1.767E-06 | 4.619E-07 | 1.500E-06 | 2.300E-06 |
| 24707 | 1236 | T1236Q | D | 3.7e-06 | 3.8e-06 | 2.1e-06 | 3.200E-06 | 9.539E-07 | 2.100E-06 | 3.800E-06 |
| 24708 | 1236 | T1236G | D | 1.9e-06 | 3.3e-06 | 2e-06 | 2.400E-06 | 7.810E-07 | 1.900E-06 | 3.300E-06 |
| 24709 | 1236 | T1236H | D | 2.2e-06 | 1.6e-06 | 2.4e-06 | 2.067E-06 | 4.163E-07 | 1.600E-06 | 2.400E-06 |
| 24710 | 1236 | T1236I | D | 3.8e-06 | 3.8e-06 | 2e-06 | 3.200E-06 | 1.039E-06 | 2.000E-06 | 3.800E-06 |
| 24711 | 1236 | T1236L | D | 2.8e-06 | 2.1e-06 | 2e-06 | 2.300E-06 | 4.359E-07 | 2.000E-06 | 2.800E-06 |
| 24712 | 1236 | T1236K | D | 2.4e-06 | 4e-06 | 2.3e-06 | 2.900E-06 | 9.539E-07 | 2.300E-06 | 4.000E-06 |
| 24713 | 1236 | T1236M | D | 3.4e-06 | 2.6e-06 | 2.5e-06 | 2.833E-06 | 4.933E-07 | 2.500E-06 | 3.400E-06 |
| 24714 | 1236 | T1236F | D | 3e-06 | 1.5e-06 | 3.3e-06 | 2.600E-06 | 9.644E-07 | 1.500E-06 | 3.300E-06 |
| 24715 | 1236 | T1236P | D | 3e-06 | 2.6e-06 | 2.7e-06 | 2.767E-06 | 2.082E-07 | 2.600E-06 | 3.000E-06 |
| 24716 | 1236 | T1236S | D | 1.6e-06 | 2e-06 | 5.7e-06 | 3.100E-06 | 2.261E-06 | 1.600E-06 | 5.700E-06 |
| 24717 | 1236 | T1236T | D | 1.7e-06 | 1.6e-06 | 3.4e-06 | 2.233E-06 | 1.012E-06 | 1.600E-06 | 3.400E-06 |
| 24718 | 1236 | T1236W | D | 2.6e-06 | 4.2e-06 | 1.4e-06 | 2.733E-06 | 1.405E-06 | 1.400E-06 | 4.200E-06 |
| 24719 | 1236 | T1236Y | D | 3.1e-06 | 1.5e-06 | 1.8e-06 | 2.133E-06 | 8.505E-07 | 1.500E-06 | 3.100E-06 |
| 24720 | 1236 | T1236V | D | 2.8e-06 | 2.8e-06 | 3e-06 | 2.867E-06 | 1.155E-07 | 2.800E-06 | 3.000E-06 |
| 24721 | 1237 | L1237A | D | 2.5e-06 | 3.8e-06 | 1.8e-06 | 2.700E-06 | 1.015E-06 | 1.800E-06 | 3.800E-06 |
| 24722 | 1237 | L1237R | D | 4e-06 | 2.9e-06 | 3.8e-06 | 3.567E-06 | 5.859E-07 | 2.900E-06 | 4.000E-06 |
| 24723 | 1237 | L1237N | D | 5.4e-06 | 1.9e-06 | 3.1e-06 | 3.467E-06 | 1.779E-06 | 1.900E-06 | 5.400E-06 |
| 24724 | 1237 | L1237D | D | 1.8e-06 | 1.5e-06 | 1.3e-06 | 1.533E-06 | 2.517E-07 | 1.300E-06 | 1.800E-06 |
| 24725 | 1237 | L1237C | D | 1.7e-06 | 2.4e-06 | 1.8e-06 | 1.967E-06 | 3.786E-07 | 1.700E-06 | 2.400E-06 |
| 24726 | 1237 | L1237E | D | 4.6e-06 | 2.8e-06 | 2.1e-06 | 3.167E-06 | 1.290E-06 | 2.100E-06 | 4.600E-06 |
| 24727 | 1237 | L1237Q | D | 1.5e-06 | 2.3e-06 | 2.1e-06 | 1.967E-06 | 4.163E-07 | 1.500E-06 | 2.300E-06 |
| 24728 | 1237 | L1237G | D | 2.6e-06 | 2.6e-06 | 3.4e-06 | 2.867E-06 | 4.619E-07 | 2.600E-06 | 3.400E-06 |
| 24729 | 1237 | L1237H | D | 1.8e-06 | 2.9e-06 | 1.3e-06 | 2.000E-06 | 8.185E-07 | 1.300E-06 | 2.900E-06 |
| 24730 | 1237 | L1237I | D | 2.6e-06 | 1.6e-06 | 3.1e-06 | 2.433E-06 | 7.638E-07 | 1.600E-06 | 3.100E-06 |
| 24731 | 1237 | L1237L | D | 2.4e-06 | 2.3e-06 | 2.8e-06 | 2.500E-06 | 2.646E-07 | 2.300E-06 | 2.800E-06 |
| 24732 | 1237 | L1237K | D | 1.7e-06 | 1.5e-06 | 3e-06 | 2.067E-06 | 8.145E-07 | 1.500E-06 | 3.000E-06 |
| 24733 | 1237 | L1237M | D | 2.5e-06 | 2.5e-06 | 2.7e-06 | 2.567E-06 | 1.155E-07 | 2.500E-06 | 2.700E-06 |
| 24734 | 1237 | L1237F | D | 3.3e-06 | 2e-06 | 2.3e-06 | 2.533E-06 | 6.807E-07 | 2.000E-06 | 3.300E-06 |

| | | | | | | | | | | |
|---|---|---|---|---|---|---|---|---|---|---|
| 24735 | 1237 | L1237P | D | 1.9e-06 | 2.4e-06 | 3.3e-06 | 2.533E-06 | 7.095E-07 | 1.900E-06 | 3.300E-06 |
| 24736 | 1237 | L1237S | D | 1.7e-06 | 3.6e-06 | 3.2e-06 | 2.833E-06 | 1.002E-06 | 1.700E-06 | 3.600E-06 |
| 24737 | 1237 | L1237T | D | 3.2e-06 | 3.3e-06 | 2.8e-06 | 3.100E-06 | 2.646E-07 | 2.800E-06 | 3.300E-06 |
| 24738 | 1237 | L1237W | D | 3.9e-06 | 3.2e-06 | 3.6e-06 | 3.567E-06 | 3.512E-07 | 3.200E-06 | 3.900E-06 |
| 24739 | 1237 | L1237Y | D | 3.3e-06 | 3.4e-06 | 2.9e-06 | 3.200E-06 | 2.646E-07 | 2.900E-06 | 3.400E-06 |
| 24740 | 1237 | L1237V | D | 2.2e-06 | 2.3e-06 | 1.1e-06 | 1.867E-06 | 6.658E-07 | 1.100E-06 | 2.300E-06 |
| 24741 | 1238 | T1238A | D | 1.8e-06 | 4.4e-06 | 2.8e-06 | 3.000E-06 | 1.311E-06 | 1.800E-06 | 4.400E-06 |
| 24742 | 1238 | T1238R | D | 2.4e-06 | 2.7e-06 | 3.6e-06 | 2.900E-06 | 6.245E-07 | 2.400E-06 | 3.600E-06 |
| 24743 | 1238 | T1238N | D | 3.4e-06 | 4.3e-06 | 4.7e-06 | 4.133E-06 | 6.658E-07 | 3.400E-06 | 4.700E-06 |
| 24744 | 1238 | T1238D | D | 3.8e-06 | 1.3e-06 | 2e-06 | 2.367E-06 | 1.290E-06 | 1.300E-06 | 3.800E-06 |
| 24745 | 1238 | T1238C | D | 2.1e-06 | 1.6e-06 | 3.3e-06 | 2.333E-06 | 8.737E-07 | 1.600E-06 | 3.300E-06 |
| 24746 | 1238 | T1238E | D | 3.2e-06 | 2.6e-06 | 2.7e-06 | 2.833E-06 | 3.215E-07 | 2.600E-06 | 3.200E-06 |
| 24747 | 1238 | T1238Q | D | 3.9e-06 | 2.8e-06 | 2.9e-06 | 3.200E-06 | 6.083E-07 | 2.800E-06 | 3.900E-06 |
| 24748 | 1238 | T1238G | D | 3.4e-06 | 2.3e-06 | 1.8e-06 | 2.500E-06 | 8.185E-07 | 1.800E-06 | 3.400E-06 |
| 24749 | 1238 | T1238H | D | 2e-06 | 2.4e-06 | 1.9e-06 | 2.100E-06 | 2.646E-07 | 1.900E-06 | 2.400E-06 |
| 24750 | 1238 | T1238I | D | 4.1e-06 | 2.3e-06 | 3.2e-06 | 3.200E-06 | 9.000E-07 | 2.300E-06 | 4.100E-06 |
| 24751 | 1238 | T1238L | D | 2.2e-06 | 3.4e-06 | 3.7e-06 | 3.100E-06 | 7.937E-07 | 2.200E-06 | 3.700E-06 |
| 24752 | 1238 | T1238K | D | 3.3e-06 | 1.4e-06 | 2.7e-06 | 2.467E-06 | 9.713E-07 | 1.400E-06 | 3.300E-06 |
| 24753 | 1238 | T1238M | D | 2.6e-06 | 2.8e-06 | 3.2e-06 | 2.867E-06 | 3.055E-07 | 2.600E-06 | 3.200E-06 |
| 24754 | 1238 | T1238F | D | 1.7e-06 | 3.6e-06 | 1.9e-06 | 2.400E-06 | 1.044E-06 | 1.700E-06 | 3.600E-06 |
| 24755 | 1238 | T1238P | D | 3.8e-06 | 2.8e-06 | 2.5e-06 | 3.033E-06 | 6.807E-07 | 2.500E-06 | 3.800E-06 |
| 24756 | 1238 | T1238S | D | 1.5e-06 | 2.4e-06 | 3.1e-06 | 2.333E-06 | 8.021E-07 | 1.500E-06 | 3.100E-06 |
| 24757 | 1238 | T1238T | D | 4.1e-06 | 2.7e-06 | 1.6e-06 | 2.800E-06 | 1.253E-06 | 1.600E-06 | 4.100E-06 |
| 24758 | 1238 | T1238W | D | 2.8e-06 | 1.9e-06 | 1.6e-06 | 2.100E-06 | 6.245E-07 | 1.600E-06 | 2.800E-06 |
| 24759 | 1238 | T1238Y | D | 2.4e-06 | 4.8e-06 | 2.8e-06 | 3.333E-06 | 1.286E-06 | 2.400E-06 | 4.800E-06 |
| 24760 | 1238 | T1238V | D | 2.5e-06 | 2.3e-06 | 2.8e-06 | 2.533E-06 | 2.517E-07 | 2.300E-06 | 2.800E-06 |
| 24761 | 1239 | L1239A | D | 4.3e-06 | 2.8e-06 | 1.7e-06 | 2.933E-06 | 1.305E-06 | 1.700E-06 | 4.300E-06 |
| 24762 | 1239 | L1239R | D | 2.2e-06 | 3.8e-06 | 3e-06 | 3.000E-06 | 8.000E-07 | 2.200E-06 | 3.800E-06 |
| 24763 | 1239 | L1239N | D | 2e-06 | 2.2e-06 | 3.6e-06 | 2.600E-06 | 8.718E-07 | 2.000E-06 | 3.600E-06 |
| 24764 | 1239 | L1239D | D | 3e-06 | 2.8e-06 | 2.3e-06 | 2.700E-06 | 3.606E-07 | 2.300E-06 | 3.000E-06 |
| 24765 | 1239 | L1239C | D | 2.5e-06 | 2.6e-06 | 2.6e-06 | 2.567E-06 | 5.774E-08 | 2.500E-06 | 2.600E-06 |
| 24766 | 1239 | L1239E | D | 2.4e-06 | 3.4e-06 | 2.9e-06 | 2.900E-06 | 5.000E-07 | 2.400E-06 | 3.400E-06 |
| 24767 | 1239 | L1239Q | D | 2.1e-06 | 2.6e-06 | 1.5e-06 | 2.067E-06 | 5.508E-07 | 1.500E-06 | 2.600E-06 |
| 24768 | 1239 | L1239G | D | 4.8e-06 | 2.3e-06 | 4.4e-06 | 3.833E-06 | 1.343E-06 | 2.300E-06 | 4.800E-06 |
| 24769 | 1239 | L1239H | D | 3.6e-06 | 3.2e-06 | 2.2e-06 | 3.000E-06 | 7.211E-07 | 2.200E-06 | 3.600E-06 |
| 24770 | 1239 | L1239I | D | 2.9e-06 | 4.7e-06 | 2.3e-06 | 3.300E-06 | 1.249E-06 | 2.300E-06 | 4.700E-06 |
| 24771 | 1239 | L1239L | D | 1.7e-06 | 2.3e-06 | 2.1e-06 | 2.033E-06 | 3.055E-07 | 1.700E-06 | 2.300E-06 |
| 24772 | 1239 | L1239K | D | 3e-06 | 1.2e-06 | 1.3e-06 | 1.833E-06 | 1.012E-06 | 1.200E-06 | 3.000E-06 |
| 24773 | 1239 | L1239M | D | 2.2e-06 | 2.2e-06 | 2.6e-06 | 2.333E-06 | 2.309E-07 | 2.200E-06 | 2.600E-06 |
| 24774 | 1239 | L1239F | D | 2.5e-06 | 3.4e-06 | 2.6e-06 | 2.833E-06 | 4.933E-07 | 2.500E-06 | 3.400E-06 |
| 24775 | 1239 | L1239P | D | 2.3e-06 | 2.7e-06 | 2e-06 | 2.333E-06 | 3.512E-07 | 2.000E-06 | 2.700E-06 |
| 24776 | 1239 | L1239S | D | 5e-06 | 2.8e-06 | 1.7e-06 | 3.167E-06 | 1.680E-06 | 1.700E-06 | 5.000E-06 |
| 24777 | 1239 | L1239T | D | 1.9e-06 | 2.3e-06 | 2.3e-06 | 2.167E-06 | 2.309E-07 | 1.900E-06 | 2.300E-06 |
| 24778 | 1239 | L1239W | D | 2.2e-06 | 2.8e-06 | 1.7e-06 | 2.233E-06 | 5.508E-07 | 1.700E-06 | 2.800E-06 |
| 24779 | 1239 | L1239Y | D | 1.4e-06 | 3.5e-06 | 3.3e-06 | 2.733E-06 | 1.159E-06 | 1.400E-06 | 3.500E-06 |
| 24780 | 1239 | L1239V | D | 1.9e-06 | 2.8e-06 | 2.1e-06 | 2.267E-06 | 4.726E-07 | 1.900E-06 | 2.800E-06 |
| 24781 | 1240 | S1240A | D | 2.4e-06 | 2.7e-06 | 2.5e-06 | 2.533E-06 | 1.528E-07 | 2.400E-06 | 2.700E-06 |
| 24782 | 1240 | S1240R | D | 2.7e-06 | 4.3e-06 | 2.7e-06 | 3.233E-06 | 9.238E-07 | 2.700E-06 | 4.300E-06 |
| 24783 | 1240 | S1240N | D | 2.1e-06 | 2.2e-06 | 1.2e-06 | 1.833E-06 | 5.508E-07 | 1.200E-06 | 2.200E-06 |
| 24784 | 1240 | S1240D | D | 2.4e-06 | 3e-06 | 2.1e-06 | 2.500E-06 | 4.583E-07 | 2.100E-06 | 3.000E-06 |
| 24785 | 1240 | S1240C | D | 1.3e-06 | 3.2e-06 | 2.8e-06 | 2.433E-06 | 1.002E-06 | 1.300E-06 | 3.200E-06 |
| 24786 | 1240 | S1240E | D | 3.3e-06 | 2.7e-06 | 2.8e-06 | 2.933E-06 | 3.215E-07 | 2.700E-06 | 3.300E-06 |
| 24787 | 1240 | S1240Q | D | 2.3e-06 | 1.9e-06 | 1.7e-06 | 1.967E-06 | 3.055E-07 | 1.700E-06 | 2.300E-06 |
| 24788 | 1240 | S1240G | D | 1.8e-06 | 1.8e-06 | 2.4e-06 | 2.000E-06 | 3.464E-07 | 1.800E-06 | 2.400E-06 |
| 24789 | 1240 | S1240H | D | 1.6e-06 | 4.6e-06 | 2.1e-06 | 2.767E-06 | 1.607E-06 | 1.600E-06 | 4.600E-06 |
| 24790 | 1240 | S1240I | D | 2.4e-06 | 3.1e-06 | 1.4e-06 | 2.300E-06 | 8.544E-07 | 1.400E-06 | 3.100E-06 |
| 24791 | 1240 | S1240L | D | 3.3e-06 | 2.5e-06 | 2.1e-06 | 2.633E-06 | 6.110E-07 | 2.100E-06 | 3.300E-06 |
| 24792 | 1240 | S1240K | D | 2.6e-06 | 3.1e-06 | 2.4e-06 | 2.700E-06 | 3.606E-07 | 2.400E-06 | 3.100E-06 |
| 24793 | 1240 | S1240M | D | 2.2e-06 | 2.4e-06 | 1.6e-06 | 2.067E-06 | 4.163E-07 | 1.600E-06 | 2.400E-06 |
| 24794 | 1240 | S1240F | D | 2e-06 | 2.5e-06 | 1.7e-06 | 2.067E-06 | 4.041E-07 | 1.700E-06 | 2.500E-06 |
| 24795 | 1240 | S1240P | D | 2.5e-06 | 1.8e-06 | 3.5e-06 | 2.600E-06 | 8.544E-07 | 1.800E-06 | 3.500E-06 |
| 24796 | 1240 | S1240S | D | 1.8e-06 | 2.4e-06 | 2.6e-06 | 2.267E-06 | 4.163E-07 | 1.800E-06 | 2.600E-06 |
| 24797 | 1240 | S1240T | D | 2.5e-06 | 2e-06 | 1.6e-06 | 2.033E-06 | 4.509E-07 | 1.600E-06 | 2.500E-06 |
| 24798 | 1240 | S1240W | D | 2.7e-06 | 2.5e-06 | 2.5e-06 | 2.567E-06 | 1.155E-07 | 2.500E-06 | 2.700E-06 |
| 24799 | 1240 | S1240Y | D | 2e-06 | 2e-06 | 2.5e-06 | 2.167E-06 | 2.887E-07 | 2.000E-06 | 2.500E-06 |
| 24800 | 1240 | S1240V | D | 3.2e-06 | 4.2e-06 | 2.3e-06 | 3.233E-06 | 9.504E-07 | 2.300E-06 | 4.200E-06 |
| 24801 | 1241 | K1241A | D | 2.1e-06 | 2.4e-06 | 1.5e-06 | 2.000E-06 | 4.583E-07 | 1.500E-06 | 2.400E-06 |
| 24802 | 1241 | K1241R | D | 1.5e-06 | 2e-06 | 2e-06 | 1.833E-06 | 2.887E-07 | 1.500E-06 | 2.000E-06 |
| 24803 | 1241 | K1241N | D | 3e-06 | 2e-06 | 1.2e-06 | 2.067E-06 | 9.018E-07 | 1.200E-06 | 3.000E-06 |
| 24804 | 1241 | K1241D | D | 1.7e-06 | 2.2e-06 | 3.6e-06 | 2.500E-06 | 9.849E-07 | 1.700E-06 | 3.600E-06 |
| 24805 | 1241 | K1241C | D | 2.5e-06 | 3.1e-06 | 3e-06 | 2.867E-06 | 3.215E-07 | 2.500E-06 | 3.100E-06 |
| 24806 | 1241 | K1241E | D | 3e-06 | 2.5e-06 | 2.9e-06 | 2.800E-06 | 2.646E-07 | 2.500E-06 | 3.000E-06 |
| 24807 | 1241 | K1241Q | D | 1.6e-06 | 2.9e-06 | 1.3e-06 | 1.933E-06 | 8.505E-07 | 1.300E-06 | 2.900E-06 |
| 24808 | 1241 | K1241G | D | 2.8e-06 | 2.9e-06 | 2.1e-06 | 2.600E-06 | 4.359E-07 | 2.100E-06 | 2.900E-06 |
| 24809 | 1241 | K1241H | D | 1.5e-06 | 1.4e-06 | 2.8e-06 | 1.900E-06 | 7.810E-07 | 1.400E-06 | 2.800E-06 |
| 24810 | 1241 | K1241I | D | 2.2e-06 | 3.1e-06 | 2.5e-06 | 2.600E-06 | 4.583E-07 | 2.200E-06 | 3.100E-06 |
| 24811 | 1241 | K1241L | D | 1.7e-06 | 2e-06 | 2.5e-06 | 2.067E-06 | 4.041E-07 | 1.700E-06 | 2.500E-06 |
| 24812 | 1241 | K1241K | D | 1.3e-06 | 5e-06 | 2.6e-06 | 2.967E-06 | 1.877E-06 | 1.300E-06 | 5.000E-06 |
| 24813 | 1241 | K1241M | D | 1.8e-06 | 2.2e-06 | 2.6e-06 | 2.200E-06 | 4.000E-07 | 1.800E-06 | 2.600E-06 |
| 24814 | 1241 | K1241F | D | 2.1e-06 | 1.1e-06 | 1.5e-06 | 1.567E-06 | 5.033E-07 | 1.100E-06 | 2.100E-06 |
| 24815 | 1241 | K1241P | D | 2.4e-06 | 2.1e-06 | 2.3e-06 | 2.267E-06 | 1.528E-07 | 2.100E-06 | 2.400E-06 |
| 24816 | 1241 | K1241S | D | 3e-06 | 2.1e-06 | 1.5e-06 | 2.200E-06 | 7.550E-07 | 1.500E-06 | 3.000E-06 |
| 24817 | 1241 | K1241T | D | 2.4e-06 | 4.2e-06 | 2.7e-06 | 3.100E-06 | 9.644E-07 | 2.400E-06 | 4.200E-06 |
| 24818 | 1241 | K1241W | D | 2.4e-06 | 2.3e-06 | 2.6e-06 | 2.433E-06 | 1.528E-07 | 2.300E-06 | 2.600E-06 |
| 24819 | 1241 | K1241Y | D | 2.4e-06 | 2.3e-06 | 9.9e-07 | 1.897E-06 | 7.868E-07 | 9.900E-07 | 2.400E-06 |

| | | | | | | | | | | |
|---|---|---|---|---|---|---|---|---|---|---|
| 24820 | 1241 | K1241V | D | 2.6e-06 | 1.7e-06 | 2.9e-06 | 2.400E-06 | 6.245E-07 | 1.700E-06 | 2.900E-06 |
| 24821 | 1242 | A1242A | D | 1.9e-06 | 3.3e-06 | 3.1e-06 | 2.767E-06 | 7.572E-07 | 1.900E-06 | 3.300E-06 |
| 24822 | 1242 | A1242R | D | 2.5e-06 | 2.2e-06 | 1.7e-06 | 2.133E-06 | 4.041E-07 | 1.700E-06 | 2.500E-06 |
| 24823 | 1242 | A1242N | D | 1.9e-06 | 2.8e-06 | 1.3e-06 | 2.000E-06 | 7.550E-07 | 1.300E-06 | 2.800E-06 |
| 24824 | 1242 | A1242D | D | 2e-06 | 2e-06 | 2.6e-06 | 2.200E-06 | 3.464E-07 | 2.000E-06 | 2.600E-06 |
| 24825 | 1242 | A1242C | D | 1.6e-06 | 3.8e-06 | 2.9e-06 | 2.767E-06 | 1.106E-06 | 1.600E-06 | 3.800E-06 |
| 24826 | 1242 | A1242E | D | 1.8e-06 | 2.9e-06 | 2.9e-06 | 2.533E-06 | 6.351E-07 | 1.800E-06 | 2.900E-06 |
| 24827 | 1242 | A1242Q | D | 1.5e-06 | 2e-06 | 2.1e-06 | 1.867E-06 | 3.215E-07 | 1.500E-06 | 2.100E-06 |
| 24828 | 1242 | A1242G | D | 2.1e-06 | 4.8e-06 | 1.6e-06 | 2.833E-06 | 1.721E-06 | 1.600E-06 | 4.800E-06 |
| 24829 | 1242 | A1242H | D | 1.7e-06 | 1.2e-06 | 2.6e-06 | 1.833E-06 | 7.095E-07 | 1.200E-06 | 2.600E-06 |
| 24830 | 1242 | A1242I | D | 5e-06 | 3.4e-06 | 2.7e-06 | 3.700E-06 | 1.179E-06 | 2.700E-06 | 5.000E-06 |
| 24831 | 1242 | A1242L | D | 2.5e-06 | 2.4e-06 | 2.3e-06 | 2.400E-06 | 1.000E-07 | 2.300E-06 | 2.500E-06 |
| 24832 | 1242 | A1242K | D | 2.5e-06 | 3.3e-06 | 2.2e-06 | 2.667E-06 | 5.686E-07 | 2.200E-06 | 3.300E-06 |
| 24833 | 1242 | A1242M | D | 2.6e-06 | 1.7e-06 | 2.2e-06 | 2.167E-06 | 4.509E-07 | 1.700E-06 | 2.600E-06 |
| 24834 | 1242 | A1242F | D | 1.8e-06 | 2.2e-06 | 3.3e-06 | 2.433E-06 | 7.767E-07 | 1.800E-06 | 3.300E-06 |
| 24835 | 1242 | A1242P | D | 2.8e-06 | 2.8e-06 | 2.1e-06 | 2.567E-06 | 4.041E-07 | 2.100E-06 | 2.800E-06 |
| 24836 | 1242 | A1242S | D | 1.8e-06 | 1.7e-06 | 3.2e-06 | 2.233E-06 | 8.386E-07 | 1.700E-06 | 3.200E-06 |
| 24837 | 1242 | A1242T | D | 1.3e-06 | 2.4e-06 | 2.1e-06 | 1.933E-06 | 5.686E-07 | 1.300E-06 | 2.400E-06 |
| 24838 | 1242 | A1242W | D | 4.5e-06 | 2.6e-06 | 3.5e-06 | 3.533E-06 | 9.504E-07 | 2.600E-06 | 4.500E-06 |
| 24839 | 1242 | A1242Y | D | 1.3e-06 | 1.9e-06 | 3.6e-06 | 2.267E-06 | 1.193E-06 | 1.300E-06 | 3.600E-06 |
| 24840 | 1242 | A1242V | D | 3.2e-06 | 6.6e-06 | 3.1e-06 | 4.300E-06 | 1.992E-06 | 3.100E-06 | 6.600E-06 |
| 24841 | 1243 | D1243A | D | 2.4e-06 | 1.8e-06 | 3.1e-06 | 2.433E-06 | 6.506E-07 | 1.800E-06 | 3.100E-06 |
| 24842 | 1243 | D1243R | D | 1.7e-06 | 2.8e-06 | 2.3e-06 | 2.267E-06 | 5.508E-07 | 1.700E-06 | 2.800E-06 |
| 24843 | 1243 | D1243N | D | 2e-06 | 4.1e-06 | 1.8e-06 | 2.633E-06 | 1.274E-06 | 1.800E-06 | 4.100E-06 |
| 24844 | 1243 | D1243D | D | 2.8e-06 | 2e-06 | 1.6e-06 | 2.133E-06 | 6.110E-07 | 1.600E-06 | 2.800E-06 |
| 24845 | 1243 | D1243C | D | 3.1e-06 | 2.7e-06 | 2.8e-06 | 2.867E-06 | 2.082E-07 | 2.700E-06 | 3.100E-06 |
| 24846 | 1243 | D1243E | D | 2.1e-06 | 2.9e-06 | 1.1e-06 | 2.033E-06 | 9.018E-07 | 1.100E-06 | 2.900E-06 |
| 24847 | 1243 | D1243Q | D | 2.2e-06 | 1.9e-06 | 2.9e-06 | 2.333E-06 | 5.132E-07 | 1.900E-06 | 2.900E-06 |
| 24848 | 1243 | D1243G | D | 2.1e-06 | 1.4e-06 | 3.1e-06 | 2.200E-06 | 8.544E-07 | 1.400E-06 | 3.100E-06 |
| 24849 | 1243 | D1243H | D | 1.8e-06 | 2.2e-06 | 2.7e-06 | 2.233E-06 | 4.509E-07 | 1.800E-06 | 2.700E-06 |
| 24850 | 1243 | D1243I | D | 1.7e-06 | 1.5e-06 | 1.6e-06 | 1.600E-06 | 1.000E-07 | 1.500E-06 | 1.700E-06 |
| 24851 | 1243 | D1243L | D | 2.2e-06 | 3.4e-06 | 2.1e-06 | 2.567E-06 | 7.234E-07 | 2.100E-06 | 3.400E-06 |
| 24852 | 1243 | D1243K | D | 1.6e-06 | 1.8e-06 | 2.9e-06 | 2.100E-06 | 7.000E-07 | 1.600E-06 | 2.900E-06 |
| 24853 | 1243 | D1243M | D | 3.6e-06 | 2.5e-06 | 1.8e-06 | 2.633E-06 | 9.074E-07 | 1.800E-06 | 3.600E-06 |
| 24854 | 1243 | D1243F | D | 3.3e-06 | 1.9e-06 | 2.5e-06 | 2.567E-06 | 7.024E-07 | 1.900E-06 | 3.300E-06 |
| 24855 | 1243 | D1243P | D | 2.4e-06 | 1.9e-06 | 2.2e-06 | 2.167E-06 | 2.517E-07 | 1.900E-06 | 2.400E-06 |
| 24856 | 1243 | D1243S | D | 3.6e-06 | 1.6e-06 | 2.4e-06 | 2.533E-06 | 1.007E-06 | 1.600E-06 | 3.600E-06 |
| 24857 | 1243 | D1243T | D | 2.4e-06 | 3.8e-06 | 2.3e-06 | 2.833E-06 | 8.386E-07 | 2.300E-06 | 3.800E-06 |
| 24858 | 1243 | D1243W | D | 1.1e-06 | 3.1e-06 | 2.6e-06 | 2.267E-06 | 1.041E-06 | 1.100E-06 | 3.100E-06 |
| 24859 | 1243 | D1243Y | D | 2.1e-06 | 2e-06 | 1.7e-06 | 1.933E-06 | 2.082E-07 | 1.700E-06 | 2.100E-06 |
| 24860 | 1243 | D1243V | D | 2e-06 | 2e-06 | 1.8e-06 | 1.933E-06 | 1.155E-07 | 1.800E-06 | 2.000E-06 |
| 24861 | 1244 | Y1244A | D | 1.4e-06 | 2.4e-06 | 3.7e-06 | 2.500E-06 | 1.153E-06 | 1.400E-06 | 3.700E-06 |
| 24862 | 1244 | Y1244R | D | 2.2e-06 | 1.6e-06 | 3e-06 | 2.267E-06 | 7.024E-07 | 1.600E-06 | 3.000E-06 |
| 24863 | 1244 | Y1244N | D | 3.1e-06 | 3.2e-06 | 3e-06 | 3.100E-06 | 1.000E-07 | 3.000E-06 | 3.200E-06 |
| 24864 | 1244 | Y1244D | D | 1.9e-06 | 2.6e-06 | 2e-06 | 2.167E-06 | 3.786E-07 | 1.900E-06 | 2.600E-06 |
| 24865 | 1244 | Y1244C | D | 1.6e-06 | 3.1e-06 | 1.1e-06 | 1.933E-06 | 1.041E-06 | 1.100E-06 | 3.100E-06 |
| 24866 | 1244 | Y1244E | D | 1.2e-06 | 1.4e-06 | 2.8e-06 | 1.800E-06 | 8.718E-07 | 1.200E-06 | 2.800E-06 |
| 24867 | 1244 | Y1244Q | D | 2e-06 | 3.2e-06 | 4.6e-06 | 3.267E-06 | 1.301E-06 | 2.000E-06 | 4.600E-06 |
| 24868 | 1244 | Y1244G | D | 2.6e-06 | 2.1e-06 | 2.9e-06 | 2.533E-06 | 4.041E-07 | 2.100E-06 | 2.900E-06 |
| 24869 | 1244 | Y1244H | D | 3.4e-06 | 3.8e-06 | 2.4e-06 | 3.200E-06 | 7.211E-07 | 2.400E-06 | 3.800E-06 |
| 24870 | 1244 | Y1244I | D | 2.2e-06 | 4e-06 | 2.4e-06 | 2.867E-06 | 9.866E-07 | 2.200E-06 | 4.000E-06 |
| 24871 | 1244 | Y1244L | D | 1.7e-06 | 1.9e-06 | 2.3e-06 | 1.967E-06 | 3.055E-07 | 1.700E-06 | 2.300E-06 |
| 24872 | 1244 | Y1244K | D | 2.9e-06 | 1.8e-06 | 2.2e-06 | 2.300E-06 | 5.568E-07 | 1.800E-06 | 2.900E-06 |
| 24873 | 1244 | Y1244M | D | 3.2e-06 | 2.4e-06 | 2.4e-06 | 2.667E-06 | 4.619E-07 | 2.400E-06 | 3.200E-06 |
| 24874 | 1244 | Y1244F | D | 3.8e-06 | 1.8e-06 | 1.9e-06 | 2.500E-06 | 1.127E-06 | 1.800E-06 | 3.800E-06 |
| 24875 | 1244 | Y1244P | D | 2.5e-06 | 2.5e-06 | 2.4e-06 | 2.467E-06 | 5.774E-08 | 2.400E-06 | 2.500E-06 |
| 24876 | 1244 | Y1244S | D | 2.6e-06 | 3.6e-06 | 1.5e-06 | 2.567E-06 | 1.050E-06 | 1.500E-06 | 3.600E-06 |
| 24877 | 1244 | Y1244T | D | 2.5e-06 | 2.7e-06 | 2.6e-06 | 2.600E-06 | 1.000E-07 | 2.500E-06 | 2.700E-06 |
| 24878 | 1244 | Y1244W | D | 3.7e-06 | 1.6e-06 | 3.2e-06 | 2.833E-06 | 1.097E-06 | 1.600E-06 | 3.700E-06 |
| 24879 | 1244 | Y1244Y | D | 3.1e-06 | 3.1e-06 | 2.1e-06 | 2.767E-06 | 5.774E-07 | 2.100E-06 | 3.100E-06 |
| 24880 | 1244 | Y1244V | D | 3.6e-06 | 1.3e-06 | 3.3e-06 | 2.733E-06 | 1.250E-06 | 1.300E-06 | 3.600E-06 |
| 24881 | 1245 | E1245A | D | 2.2e-06 | 1.2e-06 | 2.6e-06 | 2.000E-06 | 7.211E-07 | 1.200E-06 | 2.600E-06 |
| 24882 | 1245 | E1245R | D | 2e-06 | 3.1e-06 | 2.9e-06 | 2.667E-06 | 5.859E-07 | 2.000E-06 | 3.100E-06 |
| 24883 | 1245 | E1245N | D | 2.8e-06 | 2.7e-06 | 2.7e-06 | 2.733E-06 | 5.774E-08 | 2.700E-06 | 2.800E-06 |
| 24884 | 1245 | E1245D | D | 2.9e-06 | 2.1e-06 | 2.8e-06 | 2.600E-06 | 4.359E-07 | 2.100E-06 | 2.900E-06 |
| 24885 | 1245 | E1245C | D | 2.5e-06 | 1.9e-06 | 2.3e-06 | 2.233E-06 | 3.055E-07 | 1.900E-06 | 2.500E-06 |
| 24886 | 1245 | E1245E | D | 2.8e-06 | 1.8e-06 | 1.6e-06 | 2.067E-06 | 6.429E-07 | 1.600E-06 | 2.800E-06 |
| 24887 | 1245 | E1245Q | D | 2.8e-06 | 3.1e-06 | 1.9e-06 | 2.600E-06 | 6.245E-07 | 1.900E-06 | 3.100E-06 |
| 24888 | 1245 | E1245G | D | 2.4e-06 | 1.9e-06 | 3.3e-06 | 2.533E-06 | 7.095E-07 | 1.900E-06 | 3.300E-06 |
| 24889 | 1245 | E1245H | D | 5.4e-06 | 2.5e-06 | 2.7e-06 | 3.533E-06 | 1.620E-06 | 2.500E-06 | 5.400E-06 |
| 24890 | 1245 | E1245I | D | 2.9e-06 | 1.8e-06 | 2.2e-06 | 2.300E-06 | 5.568E-07 | 1.800E-06 | 2.900E-06 |
| 24891 | 1245 | E1245L | D | 3.9e-06 | 1.5e-06 | 3.2e-06 | 2.867E-06 | 1.234E-06 | 1.500E-06 | 3.900E-06 |
| 24892 | 1245 | E1245K | D | 3.1e-06 | 2.5e-06 | 2.8e-06 | 2.800E-06 | 3.000E-07 | 2.500E-06 | 3.100E-06 |
| 24893 | 1245 | E1245M | D | 2.8e-06 | 2.4e-06 | 2.7e-06 | 2.633E-06 | 2.082E-07 | 2.400E-06 | 2.800E-06 |
| 24894 | 1245 | E1245F | D | 3.1e-06 | 8e-07 | 3e-06 | 2.300E-06 | 1.300E-06 | 8.000E-07 | 3.100E-06 |
| 24895 | 1245 | E1245P | D | 1.9e-06 | 1.3e-06 | 3.2e-06 | 2.133E-06 | 9.713E-07 | 1.300E-06 | 3.200E-06 |
| 24896 | 1245 | E1245S | D | 4.2e-06 | 3.2e-06 | 2.5e-06 | 3.300E-06 | 8.544E-07 | 2.500E-06 | 4.200E-06 |
| 24897 | 1245 | E1245T | D | 2.7e-06 | 2.2e-06 | 3.4e-06 | 2.767E-06 | 6.028E-07 | 2.200E-06 | 3.400E-06 |
| 24898 | 1245 | E1245W | D | 1.9e-06 | 2.8e-06 | 2e-06 | 2.233E-06 | 4.933E-07 | 1.900E-06 | 2.800E-06 |
| 24899 | 1245 | E1245Y | D | 1e-06 | 4.1e-06 | 2.4e-06 | 2.500E-06 | 1.552E-06 | 1.000E-06 | 4.100E-06 |
| 24900 | 1245 | E1245V | D | 1.5e-06 | 3.5e-06 | 3.2e-06 | 2.733E-06 | 1.079E-06 | 1.500E-06 | 3.500E-06 |
| 24901 | 1246 | K1246A | D | 1.8e-06 | 1.9e-06 | 2.3e-06 | 2.000E-06 | 2.646E-07 | 1.800E-06 | 2.300E-06 |
| 24902 | 1246 | K1246R | D | 2.7e-06 | 2.4e-06 | 1.6e-06 | 2.233E-06 | 5.686E-07 | 1.600E-06 | 2.700E-06 |
| 24903 | 1246 | K1246N | D | 1.5e-06 | 1.9e-06 | 1.9e-06 | 1.767E-06 | 2.309E-07 | 1.500E-06 | 1.900E-06 |
| 24904 | 1246 | K1246D | D | 1.9e-06 | 3e-06 | 2.6e-06 | 2.500E-06 | 5.568E-07 | 1.900E-06 | 3.000E-06 |

| | | | | | | | | | | |
|---|---|---|---|---|---|---|---|---|---|---|
| 24905 | 1246 | K1246C | D | 2.1e-06 | 3.2e-06 | 2.7e-06 | 2.667E-06 | 5.508E-07 | 2.100E-06 | 3.200E-06 |
| 24906 | 1246 | K1246E | D | 3.4e-06 | 1.7e-06 | 1.3e-06 | 2.133E-06 | 1.115E-06 | 1.300E-06 | 3.400E-06 |
| 24907 | 1246 | K1246Q | D | 2.1e-06 | 3e-06 | 5.1e-06 | 3.400E-06 | 1.539E-06 | 2.100E-06 | 5.100E-06 |
| 24908 | 1246 | K1246G | D | 1.4e-06 | 2.2e-06 | 2e-06 | 1.867E-06 | 4.163E-07 | 1.400E-06 | 2.200E-06 |
| 24909 | 1246 | K1246H | D | 3.1e-06 | 2.5e-06 | 1.9e-06 | 2.500E-06 | 6.000E-07 | 1.900E-06 | 3.100E-06 |
| 24910 | 1246 | K1246I | D | 3.1e-06 | 2.7e-06 | 2.6e-06 | 2.800E-06 | 2.646E-07 | 2.600E-06 | 3.100E-06 |
| 24911 | 1246 | K1246L | D | 3e-06 | 2.5e-06 | 3.4e-06 | 2.967E-06 | 4.509E-07 | 2.500E-06 | 3.400E-06 |
| 24912 | 1246 | K1246K | D | 2.1e-06 | 4.5e-06 | 1.7e-06 | 2.767E-06 | 1.514E-06 | 1.700E-06 | 4.500E-06 |
| 24913 | 1246 | K1246M | D | 2.5e-06 | 2.9e-06 | 3.2e-06 | 2.867E-06 | 3.512E-07 | 2.500E-06 | 3.200E-06 |
| 24914 | 1246 | K1246F | D | 1.8e-06 | 2.9e-06 | 2.6e-06 | 2.433E-06 | 5.686E-07 | 1.800E-06 | 2.900E-06 |
| 24915 | 1246 | K1246P | D | 2.7e-06 | 4.4e-06 | 2.2e-06 | 3.100E-06 | 1.153E-06 | 2.200E-06 | 4.400E-06 |
| 24916 | 1246 | K1246S | D | 2.8e-06 | 2.2e-06 | 1.5e-06 | 2.167E-06 | 6.506E-07 | 1.500E-06 | 2.800E-06 |
| 24917 | 1246 | K1246T | D | 3.6e-06 | 1.6e-06 | 3.5e-06 | 2.900E-06 | 1.127E-06 | 1.600E-06 | 3.600E-06 |
| 24918 | 1246 | K1246W | D | 3e-06 | 1.9e-06 | 1.4e-06 | 2.100E-06 | 8.185E-07 | 1.400E-06 | 3.000E-06 |
| 24919 | 1246 | K1246Y | D | 2.7e-06 | 2.7e-06 | 4e-06 | 3.133E-06 | 7.506E-07 | 2.700E-06 | 4.000E-06 |
| 24920 | 1246 | K1246V | D | 2.2e-06 | 3.1e-06 | 2.3e-06 | 2.533E-06 | 4.933E-07 | 2.200E-06 | 3.100E-06 |
| 24921 | 1247 | H1247A | D | 1.6e-06 | 2.2e-06 | 1.6e-06 | 1.800E-06 | 3.464E-07 | 1.600E-06 | 2.200E-06 |
| 24922 | 1247 | H1247R | D | 2.6e-06 | 2.5e-06 | 3.4e-06 | 2.833E-06 | 4.933E-07 | 2.500E-06 | 3.400E-06 |
| 24923 | 1247 | H1247N | D | 3e-06 | 2.3e-06 | 2.3e-06 | 2.533E-06 | 4.041E-07 | 2.300E-06 | 3.000E-06 |
| 24924 | 1247 | H1247D | D | 1.6e-06 | 3.2e-06 | 1.8e-06 | 2.200E-06 | 8.718E-07 | 1.600E-06 | 3.200E-06 |
| 24925 | 1247 | H1247C | D | 3e-06 | 2.5e-06 | 2.5e-06 | 2.667E-06 | 2.887E-07 | 2.500E-06 | 3.000E-06 |
| 24926 | 1247 | H1247E | D | 2.7e-06 | 2.4e-06 | 2.2e-06 | 2.433E-06 | 2.517E-07 | 2.200E-06 | 2.700E-06 |
| 24927 | 1247 | H1247Q | D | 1.7e-06 | 1.2e-06 | 1.8e-06 | 1.567E-06 | 3.215E-07 | 1.200E-06 | 1.800E-06 |
| 24928 | 1247 | H1247G | D | 3.6e-06 | 3.7e-06 | 2.7e-06 | 3.333E-06 | 5.508E-07 | 2.700E-06 | 3.700E-06 |
| 24929 | 1247 | H1247H | D | 1.8e-06 | 1.6e-06 | 1.6e-06 | 1.667E-06 | 1.155E-07 | 1.600E-06 | 1.800E-06 |
| 24930 | 1247 | H1247I | D | 2.7e-06 | 3.3e-06 | 2.4e-06 | 2.800E-06 | 4.583E-07 | 2.400E-06 | 3.300E-06 |
| 24931 | 1247 | H1247L | D | 3.4e-06 | 3.7e-06 | 1.4e-06 | 2.833E-06 | 1.250E-06 | 1.400E-06 | 3.700E-06 |
| 24932 | 1247 | H1247K | D | 2.1e-06 | 2.4e-06 | 1.7e-06 | 2.067E-06 | 3.512E-07 | 1.700E-06 | 2.400E-06 |
| 24933 | 1247 | H1247M | D | 3.6e-06 | 1.9e-06 | 3e-06 | 2.833E-06 | 8.622E-07 | 1.900E-06 | 3.600E-06 |
| 24934 | 1247 | H1247F | D | 1.6e-06 | 3.8e-06 | 4.8e-06 | 3.400E-06 | 1.637E-06 | 1.600E-06 | 4.800E-06 |
| 24935 | 1247 | H1247P | D | 1.9e-06 | 2.2e-06 | 1.8e-06 | 1.967E-06 | 2.082E-07 | 1.800E-06 | 2.200E-06 |
| 24936 | 1247 | H1247S | D | 4.6e-06 | 5e-06 | 3.1e-06 | 4.233E-06 | 1.002E-06 | 3.100E-06 | 5.000E-06 |
| 24937 | 1247 | H1247T | D | 2.2e-06 | 2e-06 | 1.4e-06 | 1.867E-06 | 4.163E-07 | 1.400E-06 | 2.200E-06 |
| 24938 | 1247 | H1247W | D | 2.1e-06 | 2.7e-06 | 2.3e-06 | 2.367E-06 | 3.055E-07 | 2.100E-06 | 2.700E-06 |
| 24939 | 1247 | H1247Y | D | 1.9e-06 | 1.2e-06 | 2.1e-06 | 1.733E-06 | 4.726E-07 | 1.200E-06 | 2.100E-06 |
| 24940 | 1247 | H1247V | D | 2.2e-06 | 2.2e-06 | 1.7e-06 | 2.033E-06 | 2.887E-07 | 1.700E-06 | 2.200E-06 |
| 24941 | 1248 | K1248A | D | 2.2e-06 | 4.1e-06 | 2.6e-06 | 2.967E-06 | 1.002E-06 | 2.200E-06 | 4.100E-06 |
| 24942 | 1248 | K1248R | D | 1.6e-06 | 2.6e-06 | 3.4e-06 | 2.533E-06 | 9.018E-07 | 1.600E-06 | 3.400E-06 |
| 24943 | 1248 | K1248N | D | 2e-06 | 1.8e-06 | 2.1e-06 | 1.967E-06 | 1.528E-07 | 1.800E-06 | 2.100E-06 |
| 24944 | 1248 | K1248D | D | 2.5e-06 | 1.6e-06 | 2.5e-06 | 2.200E-06 | 5.196E-07 | 1.600E-06 | 2.500E-06 |
| 24945 | 1248 | K1248C | D | 2.4e-06 | 3.1e-06 | 2e-06 | 2.500E-06 | 5.568E-07 | 2.000E-06 | 3.100E-06 |
| 24946 | 1248 | K1248E | D | 4.8e-06 | 4.6e-06 | 1.3e-06 | 3.567E-06 | 1.966E-06 | 1.300E-06 | 4.800E-06 |
| 24947 | 1248 | K1248Q | D | 2e-06 | 2.3e-06 | 3.8e-06 | 2.700E-06 | 9.644E-07 | 2.000E-06 | 3.800E-06 |
| 24948 | 1248 | K1248G | D | 2.5e-06 | 2.3e-06 | 2.6e-06 | 2.467E-06 | 1.528E-07 | 2.300E-06 | 2.600E-06 |
| 24949 | 1248 | K1248H | D | 2.9e-06 | 2.2e-06 | 1.8e-06 | 2.300E-06 | 5.568E-07 | 1.800E-06 | 2.900E-06 |
| 24950 | 1248 | K1248I | D | 3e-06 | 2.7e-06 | 2.7e-06 | 2.800E-06 | 1.732E-07 | 2.700E-06 | 3.000E-06 |
| 24951 | 1248 | K1248L | D | 2.1e-06 | 2.2e-06 | 3.1e-06 | 2.467E-06 | 5.508E-07 | 2.100E-06 | 3.100E-06 |
| 24952 | 1248 | K1248K | D | 4.2e-06 | 2.9e-06 | 3.5e-06 | 3.533E-06 | 6.506E-07 | 2.900E-06 | 4.200E-06 |
| 24953 | 1248 | K1248M | D | 2.7e-06 | 1e-06 | 2.2e-06 | 1.967E-06 | 8.737E-07 | 1.000E-06 | 2.700E-06 |
| 24954 | 1248 | K1248F | D | 2.7e-06 | 2.6e-06 | 1.6e-06 | 2.300E-06 | 6.083E-07 | 1.600E-06 | 2.700E-06 |
| 24955 | 1248 | K1248P | D | 1.1e-06 | 3.5e-06 | 3.3e-06 | 2.633E-06 | 1.332E-06 | 1.100E-06 | 3.500E-06 |
| 24956 | 1248 | K1248S | D | 2.3e-06 | 2.1e-06 | 2.9e-06 | 2.433E-06 | 4.163E-07 | 2.100E-06 | 2.900E-06 |
| 24957 | 1248 | K1248T | D | 1.6e-06 | 2.1e-06 | 2e-06 | 1.900E-06 | 2.646E-07 | 1.600E-06 | 2.100E-06 |
| 24958 | 1248 | K1248W | D | 2.5e-06 | 3.4e-06 | 2.6e-06 | 2.833E-06 | 4.933E-07 | 2.500E-06 | 3.400E-06 |
| 24959 | 1248 | K1248Y | D | 3.4e-06 | 4e-06 | 2.9e-06 | 3.433E-06 | 5.508E-07 | 2.900E-06 | 4.000E-06 |
| 24960 | 1248 | K1248V | D | 2.4e-06 | 3.7e-06 | 2.3e-06 | 2.800E-06 | 7.810E-07 | 2.300E-06 | 3.700E-06 |
| 24961 | 1249 | V1249A | D | 1.8e-06 | 2.8e-06 | 1.8e-06 | 2.133E-06 | 5.774E-07 | 1.800E-06 | 2.800E-06 |
| 24962 | 1249 | V1249R | D | 1.3e-06 | 2.2e-06 | 2.5e-06 | 2.000E-06 | 6.245E-07 | 1.300E-06 | 2.500E-06 |
| 24963 | 1249 | V1249N | D | 1.9e-06 | 4.3e-06 | 9.5e-07 | 2.383E-06 | 1.727E-06 | 9.500E-07 | 4.300E-06 |
| 24964 | 1249 | V1249D | D | 1.5e-06 | 1.6e-06 | 1.7e-06 | 1.600E-06 | 1.000E-07 | 1.500E-06 | 1.700E-06 |
| 24965 | 1249 | V1249C | D | 2.1e-06 | 1.8e-06 | 2.8e-06 | 2.233E-06 | 5.132E-07 | 1.800E-06 | 2.800E-06 |
| 24966 | 1249 | V1249E | D | 1.1e-06 | 2.7e-06 | 3.1e-06 | 2.300E-06 | 1.058E-06 | 1.100E-06 | 3.100E-06 |
| 24967 | 1249 | V1249Q | D | 1.4e-06 | 1.5e-06 | 3.1e-06 | 2.000E-06 | 9.539E-07 | 1.400E-06 | 3.100E-06 |
| 24968 | 1249 | V1249G | D | 2.9e-06 | 1.9e-06 | 1.1e-06 | 1.967E-06 | 9.018E-07 | 1.100E-06 | 2.900E-06 |
| 24969 | 1249 | V1249H | D | 2.4e-06 | 2.4e-06 | 3.1e-06 | 2.633E-06 | 4.041E-07 | 2.400E-06 | 3.100E-06 |
| 24970 | 1249 | V1249I | D | 1.8e-06 | 2.4e-06 | 1.3e-06 | 1.833E-06 | 5.508E-07 | 1.300E-06 | 2.400E-06 |
| 24971 | 1249 | V1249L | D | 2e-06 | 2.9e-06 | 1.5e-06 | 2.133E-06 | 7.095E-07 | 1.500E-06 | 2.900E-06 |
| 24972 | 1249 | V1249K | D | 3.3e-06 | 3.7e-06 | 1.4e-06 | 2.800E-06 | 1.229E-06 | 1.400E-06 | 3.700E-06 |
| 24973 | 1249 | V1249M | D | 2.4e-06 | 1.6e-06 | 3.2e-06 | 2.400E-06 | 8.000E-07 | 1.600E-06 | 3.200E-06 |
| 24974 | 1249 | V1249F | D | 2.7e-06 | 1.8e-06 | 1.8e-06 | 2.100E-06 | 5.196E-07 | 1.800E-06 | 2.700E-06 |
| 24975 | 1249 | V1249P | D | 3.3e-06 | 4.4e-06 | 3.2e-06 | 3.633E-06 | 6.658E-07 | 3.200E-06 | 4.400E-06 |
| 24976 | 1249 | V1249S | D | 1.3e-06 | 1.4e-06 | 2.6e-06 | 1.767E-06 | 7.234E-07 | 1.300E-06 | 2.600E-06 |
| 24977 | 1249 | V1249T | D | 3.4e-06 | 1.6e-06 | 2.4e-06 | 2.467E-06 | 9.018E-07 | 1.600E-06 | 3.400E-06 |
| 24978 | 1249 | V1249W | D | 1.8e-06 | 1.7e-06 | 2e-06 | 1.833E-06 | 1.528E-07 | 1.700E-06 | 2.000E-06 |
| 24979 | 1249 | V1249Y | D | 1.5e-06 | 2e-06 | 1.9e-06 | 1.800E-06 | 2.646E-07 | 1.500E-06 | 2.000E-06 |
| 24980 | 1249 | V1249V | D | 1.5e-06 | 3.2e-06 | 1.6e-06 | 2.100E-06 | 9.539E-07 | 1.500E-06 | 3.200E-06 |
| 24981 | 1250 | Y1250A | D | 1.7e-06 | 3.6e-06 | 1.9e-06 | 2.400E-06 | 1.044E-06 | 1.700E-06 | 3.600E-06 |
| 24982 | 1250 | Y1250R | D | 1e-06 | 2.5e-06 | 1.2e-06 | 1.567E-06 | 8.145E-07 | 1.000E-06 | 2.500E-06 |
| 24983 | 1250 | Y1250N | D | 4.2e-06 | 1.9e-06 | 3.7e-06 | 3.267E-06 | 1.210E-06 | 1.900E-06 | 4.200E-06 |
| 24984 | 1250 | Y1250D | D | 2.4e-06 | 2.3e-06 | 2.7e-06 | 2.467E-06 | 2.082E-07 | 2.300E-06 | 2.700E-06 |
| 24985 | 1250 | Y1250C | D | 3.4e-06 | 2.6e-06 | 2.5e-06 | 2.833E-06 | 4.933E-07 | 2.500E-06 | 3.400E-06 |
| 24986 | 1250 | Y1250E | D | 2.4e-06 | 2.5e-06 | 2e-06 | 2.300E-06 | 2.646E-07 | 2.000E-06 | 2.500E-06 |
| 24987 | 1250 | Y1250Q | D | 1.9e-06 | 2e-06 | 2e-06 | 1.967E-06 | 5.774E-08 | 1.900E-06 | 2.000E-06 |
| 24988 | 1250 | Y1250G | D | 4.1e-06 | 2.1e-06 | 2.4e-06 | 2.867E-06 | 1.079E-06 | 2.100E-06 | 4.100E-06 |
| 24989 | 1250 | Y1250H | D | 3.9e-06 | 1.3e-06 | 2.2e-06 | 2.467E-06 | 1.320E-06 | 1.300E-06 | 3.900E-06 |

| | | | | | | | | | | |
|---|---|---|---|---|---|---|---|---|---|---|
| 24990 | 1250 | Y1250I | D | 2.2e-06 | 2.4e-06 | 2.5e-06 | 2.367E-06 | 1.528E-07 | 2.200E-06 | 2.500E-06 |
| 24991 | 1250 | Y1250L | D | 2.9e-06 | 3.3e-06 | 2.9e-06 | 3.033E-06 | 2.309E-07 | 2.900E-06 | 3.300E-06 |
| 24992 | 1250 | Y1250K | D | 2.6e-06 | 1.8e-06 | 2.8e-06 | 2.400E-06 | 5.292E-07 | 1.800E-06 | 2.800E-06 |
| 24993 | 1250 | Y1250M | D | 2.5e-06 | 2.1e-06 | 2.2e-06 | 2.267E-06 | 2.082E-07 | 2.100E-06 | 2.500E-06 |
| 24994 | 1250 | Y1250F | D | 2e-06 | 1.7e-06 | 3e-06 | 2.233E-06 | 6.807E-07 | 1.700E-06 | 3.000E-06 |
| 24995 | 1250 | Y1250P | D | 2.1e-06 | 1.6e-06 | 2.6e-06 | 2.100E-06 | 5.000E-07 | 1.600E-06 | 2.600E-06 |
| 24996 | 1250 | Y1250S | D | 4.6e-06 | 1.6e-06 | 2.7e-06 | 2.967E-06 | 1.518E-06 | 1.600E-06 | 4.600E-06 |
| 24997 | 1250 | Y1250T | D | 1.6e-06 | 3.5e-06 | 2.5e-06 | 2.533E-06 | 9.504E-07 | 1.600E-06 | 3.500E-06 |
| 24998 | 1250 | Y1250W | D | 2.6e-06 | 3.4e-06 | 2.3e-06 | 2.767E-06 | 5.686E-07 | 2.300E-06 | 3.400E-06 |
| 24999 | 1250 | Y1250Y | D | 2.6e-06 | 2.6e-06 | 2.5e-06 | 2.567E-06 | 5.774E-08 | 2.500E-06 | 2.600E-06 |
| 25000 | 1250 | Y1250V | D | 1.3e-06 | 2.2e-06 | 3.6e-06 | 2.367E-06 | 1.159E-06 | 1.300E-06 | 3.600E-06 |
| 25001 | 1251 | A1251A | D | 1.6e-06 | 3.4e-06 | 1.9e-06 | 2.300E-06 | 9.644E-07 | 1.600E-06 | 3.400E-06 |
| 25002 | 1251 | A1251R | D | 4.3e-06 | 2.2e-06 | 5.5e-06 | 4.000E-06 | 1.670E-06 | 2.200E-06 | 5.500E-06 |
| 25003 | 1251 | A1251N | D | 2.2e-06 | 2.7e-06 | 1.9e-06 | 2.267E-06 | 4.041E-07 | 1.900E-06 | 2.700E-06 |
| 25004 | 1251 | A1251D | D | 2.3e-06 | 2.3e-06 | 2.9e-06 | 2.500E-06 | 3.464E-07 | 2.300E-06 | 2.900E-06 |
| 25005 | 1251 | A1251C | D | 1.6e-06 | 2e-06 | 2.8e-06 | 2.133E-06 | 6.110E-07 | 1.600E-06 | 2.800E-06 |
| 25006 | 1251 | A1251E | D | 2.1e-06 | 2.8e-06 | 4.2e-06 | 3.033E-06 | 1.069E-06 | 2.100E-06 | 4.200E-06 |
| 25007 | 1251 | A1251Q | D | 1.7e-06 | 2.6e-06 | 2.6e-06 | 2.300E-06 | 5.196E-07 | 1.700E-06 | 2.600E-06 |
| 25008 | 1251 | A1251G | D | 2.5e-06 | 4.4e-06 | 3.2e-06 | 3.367E-06 | 9.609E-07 | 2.500E-06 | 4.400E-06 |
| 25009 | 1251 | A1251H | D | 1.9e-06 | 3e-06 | 3.2e-06 | 2.700E-06 | 7.000E-07 | 1.900E-06 | 3.200E-06 |
| 25010 | 1251 | A1251I | D | 2.5e-06 | 3.3e-06 | 2.4e-06 | 2.733E-06 | 4.933E-07 | 2.400E-06 | 3.300E-06 |
| 25011 | 1251 | A1251L | D | 3.5e-06 | 2.4e-06 | 3.8e-06 | 3.233E-06 | 7.371E-07 | 2.400E-06 | 3.800E-06 |
| 25012 | 1251 | A1251K | D | 1.5e-06 | 1.5e-06 | 2.6e-06 | 1.867E-06 | 6.351E-07 | 1.500E-06 | 2.600E-06 |
| 25013 | 1251 | A1251M | D | 1.2e-06 | 2e-06 | 2.5e-06 | 1.900E-06 | 6.557E-07 | 1.200E-06 | 2.500E-06 |
| 25014 | 1251 | A1251F | D | 1.7e-06 | 1.5e-06 | 2.4e-06 | 1.867E-06 | 4.726E-07 | 1.500E-06 | 2.400E-06 |
| 25015 | 1251 | A1251P | D | 8.5e-07 | 3.1e-06 | 1.1e-06 | 1.683E-06 | 1.233E-06 | 8.500E-07 | 3.100E-06 |
| 25016 | 1251 | A1251S | D | 2.2e-06 | 2.5e-06 | 2.3e-06 | 2.333E-06 | 1.528E-07 | 2.200E-06 | 2.500E-06 |
| 25017 | 1251 | A1251T | D | 1.6e-06 | 2.8e-06 | 2.7e-06 | 2.367E-06 | 6.658E-07 | 1.600E-06 | 2.800E-06 |
| 25018 | 1251 | A1251W | D | 1.6e-06 | 2.8e-06 | 1.3e-06 | 1.900E-06 | 7.937E-07 | 1.300E-06 | 2.800E-06 |
| 25019 | 1251 | A1251Y | D | 2.6e-06 | 2.9e-06 | 2.4e-06 | 2.633E-06 | 2.517E-07 | 2.400E-06 | 2.900E-06 |
| 25020 | 1251 | A1251V | D | 2.1e-06 | 2.5e-06 | 2.5e-06 | 2.367E-06 | 2.309E-07 | 2.100E-06 | 2.500E-06 |
| 25021 | 1252 | C1252A | D | 3.5e-06 | 2e-06 | 1.9e-06 | 2.467E-06 | 8.963E-07 | 1.900E-06 | 3.500E-06 |
| 25022 | 1252 | C1252R | D | 3.1e-06 | 2.5e-06 | 3.9e-06 | 3.167E-06 | 7.024E-07 | 2.500E-06 | 3.900E-06 |
| 25023 | 1252 | C1252N | D | 2.1e-06 | 2.6e-06 | 2.1e-06 | 2.267E-06 | 2.887E-07 | 2.100E-06 | 2.600E-06 |
| 25024 | 1252 | C1252D | D | 2.3e-06 | 2.1e-06 | 2.4e-06 | 2.267E-06 | 1.528E-07 | 2.100E-06 | 2.400E-06 |
| 25025 | 1252 | C1252C | D | 2.4e-06 | 2.4e-06 | 1.6e-06 | 2.133E-06 | 4.619E-07 | 1.600E-06 | 2.400E-06 |
| 25026 | 1252 | C1252E | D | 4.3e-06 | 1.9e-06 | 2.4e-06 | 2.867E-06 | 1.266E-06 | 1.900E-06 | 4.300E-06 |
| 25027 | 1252 | C1252Q | D | 2e-06 | 3.7e-06 | 2.8e-06 | 2.833E-06 | 8.505E-07 | 2.000E-06 | 3.700E-06 |
| 25028 | 1252 | C1252G | D | 2.3e-06 | 3.9e-06 | 1.5e-06 | 2.567E-06 | 1.222E-06 | 1.500E-06 | 3.900E-06 |
| 25029 | 1252 | C1252H | D | 2.6e-06 | 2.6e-06 | 2.7e-06 | 2.633E-06 | 5.774E-08 | 2.600E-06 | 2.700E-06 |
| 25030 | 1252 | C1252I | D | 1.8e-06 | 2e-06 | 1.2e-06 | 1.667E-06 | 4.163E-07 | 1.200E-06 | 2.000E-06 |
| 25031 | 1252 | C1252L | D | 2.7e-06 | 2.4e-06 | 4e-06 | 3.033E-06 | 8.505E-07 | 2.400E-06 | 4.000E-06 |
| 25032 | 1252 | C1252K | D | 2.9e-06 | 5.3e-06 | 3.4e-06 | 3.867E-06 | 1.266E-06 | 2.900E-06 | 5.300E-06 |
| 25033 | 1252 | C1252M | D | 2.6e-06 | 2.3e-06 | 1.6e-06 | 2.167E-06 | 5.132E-07 | 1.600E-06 | 2.600E-06 |
| 25034 | 1252 | C1252F | D | 1.2e-06 | 2.3e-06 | 3e-06 | 2.167E-06 | 9.074E-07 | 1.200E-06 | 3.000E-06 |
| 25035 | 1252 | C1252P | D | 3.6e-06 | 2.1e-06 | 2.2e-06 | 2.633E-06 | 8.386E-07 | 2.100E-06 | 3.600E-06 |
| 25036 | 1252 | C1252S | D | 1.4e-06 | 2.5e-06 | 3.7e-06 | 2.533E-06 | 1.150E-06 | 1.400E-06 | 3.700E-06 |
| 25037 | 1252 | C1252T | D | 2.3e-06 | 2.8e-06 | 1.6e-06 | 2.233E-06 | 6.028E-07 | 1.600E-06 | 2.800E-06 |
| 25038 | 1252 | C1252W | D | 1.9e-06 | 3.5e-06 | 1.5e-06 | 2.300E-06 | 1.058E-06 | 1.500E-06 | 3.500E-06 |
| 25039 | 1252 | C1252Y | D | 1.8e-06 | 1.8e-06 | 2e-06 | 1.867E-06 | 1.155E-07 | 1.800E-06 | 2.000E-06 |
| 25040 | 1252 | C1252V | D | 3.1e-06 | 2.7e-06 | 3.3e-06 | 3.033E-06 | 3.055E-07 | 2.700E-06 | 3.300E-06 |
| 25041 | 1253 | E1253A | D | 1.9e-06 | 2.1e-06 | 2.5e-06 | 2.167E-06 | 3.055E-07 | 1.900E-06 | 2.500E-06 |
| 25042 | 1253 | E1253R | D | 3.7e-06 | 4.4e-06 | 3.1e-06 | 3.733E-06 | 6.506E-07 | 3.100E-06 | 4.400E-06 |
| 25043 | 1253 | E1253N | D | 2e-06 | 1.5e-06 | 1.9e-06 | 1.800E-06 | 2.646E-07 | 1.500E-06 | 2.000E-06 |
| 25044 | 1253 | E1253D | D | 2.4e-06 | 1.4e-06 | 2.3e-06 | 2.033E-06 | 5.508E-07 | 1.400E-06 | 2.400E-06 |
| 25045 | 1253 | E1253C | D | 2.2e-06 | 1.6e-06 | 1.9e-06 | 1.900E-06 | 3.000E-07 | 1.600E-06 | 2.200E-06 |
| 25046 | 1253 | E1253E | D | 3.2e-06 | 3e-06 | 1.7e-06 | 2.633E-06 | 8.145E-07 | 1.700E-06 | 3.200E-06 |
| 25047 | 1253 | E1253Q | D | 3.4e-06 | 1.6e-06 | 2.3e-06 | 2.433E-06 | 9.074E-07 | 1.600E-06 | 3.400E-06 |
| 25048 | 1253 | E1253G | D | 3.1e-06 | 2e-06 | 2.9e-06 | 2.667E-06 | 5.859E-07 | 2.000E-06 | 3.100E-06 |
| 25049 | 1253 | E1253H | D | 2.2e-06 | 2e-06 | 2.4e-06 | 2.200E-06 | 2.000E-07 | 2.000E-06 | 2.400E-06 |
| 25050 | 1253 | E1253I | D | 2.8e-06 | 2.5e-06 | 2.3e-06 | 2.533E-06 | 2.517E-07 | 2.300E-06 | 2.800E-06 |
| 25051 | 1253 | E1253L | D | 1.9e-06 | 2.1e-06 | 2.9e-06 | 2.300E-06 | 5.292E-07 | 1.900E-06 | 2.900E-06 |
| 25052 | 1253 | E1253K | D | 2.4e-06 | 2.3e-06 | 1.9e-06 | 2.200E-06 | 2.646E-07 | 1.900E-06 | 2.400E-06 |
| 25053 | 1253 | E1253M | D | 1.6e-06 | 2.1e-06 | 1.8e-06 | 1.833E-06 | 2.517E-07 | 1.600E-06 | 2.100E-06 |
| 25054 | 1253 | E1253F | D | 1.9e-06 | 2.5e-06 | 1.9e-06 | 2.100E-06 | 3.464E-07 | 1.900E-06 | 2.500E-06 |
| 25055 | 1253 | E1253P | D | 2.4e-06 | 2.5e-06 | 2.6e-06 | 2.500E-06 | 1.000E-07 | 2.400E-06 | 2.600E-06 |
| 25056 | 1253 | E1253S | D | 3.1e-06 | 1.6e-06 | 2.2e-06 | 2.300E-06 | 7.550E-07 | 1.600E-06 | 3.100E-06 |
| 25057 | 1253 | E1253T | D | 2.3e-06 | 2.1e-06 | 1.5e-06 | 1.967E-06 | 4.163E-07 | 1.500E-06 | 2.300E-06 |
| 25058 | 1253 | E1253W | D | 2.6e-06 | 1.6e-06 | 1.7e-06 | 1.967E-06 | 5.508E-07 | 1.600E-06 | 2.600E-06 |
| 25059 | 1253 | E1253Y | D | 2.1e-06 | 2.1e-06 | 2.2e-06 | 2.133E-06 | 5.774E-08 | 2.100E-06 | 2.200E-06 |
| 25060 | 1253 | E1253V | D | 2.7e-06 | 2.9e-06 | 3.1e-06 | 2.900E-06 | 2.000E-07 | 2.700E-06 | 3.100E-06 |
| 25061 | 1254 | V1254A | D | 1.3e-06 | 3.2e-06 | 1.7e-06 | 2.067E-06 | 1.002E-06 | 1.300E-06 | 3.200E-06 |
| 25062 | 1254 | V1254R | D | 3.2e-06 | 1.9e-06 | 5e-06 | 3.367E-06 | 1.557E-06 | 1.900E-06 | 5.000E-06 |
| 25063 | 1254 | V1254N | D | 2.9e-06 | 2.4e-06 | 2.5e-06 | 2.600E-06 | 2.646E-07 | 2.400E-06 | 2.900E-06 |
| 25064 | 1254 | V1254D | D | 2.1e-06 | 3.2e-06 | 1.5e-06 | 2.267E-06 | 8.622E-07 | 1.500E-06 | 3.200E-06 |
| 25065 | 1254 | V1254C | D | 2.2e-06 | 2e-06 | 2.2e-06 | 2.133E-06 | 1.155E-07 | 2.000E-06 | 2.200E-06 |
| 25066 | 1254 | V1254E | D | 1.9e-06 | 2.9e-06 | 2.4e-06 | 2.400E-06 | 5.000E-07 | 1.900E-06 | 2.900E-06 |
| 25067 | 1254 | V1254Q | D | 3.7e-06 | 2.4e-06 | 1.5e-06 | 2.533E-06 | 1.106E-06 | 1.500E-06 | 3.700E-06 |
| 25068 | 1254 | V1254G | D | 3.1e-06 | 3.2e-06 | 1.5e-06 | 2.600E-06 | 9.539E-07 | 1.500E-06 | 3.200E-06 |
| 25069 | 1254 | V1254H | D | 3.8e-06 | 1.7e-06 | 1.5e-06 | 2.333E-06 | 1.274E-06 | 1.500E-06 | 3.800E-06 |
| 25070 | 1254 | V1254I | D | 2.4e-06 | 2.4e-06 | 1.6e-06 | 2.133E-06 | 4.619E-07 | 1.600E-06 | 2.400E-06 |
| 25071 | 1254 | V1254L | D | 2.9e-06 | 2.4e-06 | 2.3e-06 | 2.533E-06 | 3.215E-07 | 2.300E-06 | 2.900E-06 |
| 25072 | 1254 | V1254K | D | 1.7e-06 | 1.2e-06 | 1.3e-06 | 1.400E-06 | 2.646E-07 | 1.200E-06 | 1.700E-06 |
| 25073 | 1254 | V1254M | D | 1.9e-06 | 2.3e-06 | 1.7e-06 | 1.967E-06 | 3.055E-07 | 1.700E-06 | 2.300E-06 |
| 25074 | 1254 | V1254F | D | 1.4e-06 | 1.7e-06 | 5.4e-06 | 2.833E-06 | 2.228E-06 | 1.400E-06 | 5.400E-06 |

| | | | | | | | | | | |
|---|---|---|---|---|---|---|---|---|---|---|
| 25075 | 1254 | V1254P | D | 1.8e-06 | 2.2e-06 | 2.1e-06 | 2.033E-06 | 2.082E-07 | 1.800E-06 | 2.200E-06 |
| 25076 | 1254 | V1254S | D | 2.3e-06 | 1.9e-06 | 2.4e-06 | 2.200E-06 | 2.646E-07 | 1.900E-06 | 2.400E-06 |
| 25077 | 1254 | V1254T | D | 2.8e-06 | 1.9e-06 | 2e-06 | 2.233E-06 | 4.933E-07 | 1.900E-06 | 2.800E-06 |
| 25078 | 1254 | V1254W | D | 2e-06 | 3.6e-06 | 2.4e-06 | 2.667E-06 | 8.327E-07 | 2.000E-06 | 3.600E-06 |
| 25079 | 1254 | V1254Y | D | 2.5e-06 | 1.2e-06 | 2.1e-06 | 1.933E-06 | 6.658E-07 | 1.200E-06 | 2.500E-06 |
| 25080 | 1254 | V1254V | D | 2e-06 | 1.9e-06 | 1.5e-06 | 1.800E-06 | 2.646E-07 | 1.500E-06 | 2.000E-06 |
| 25081 | 1255 | T1255A | D | 2.9e-06 | 3.2e-06 | 2.6e-06 | 2.900E-06 | 3.000E-07 | 2.600E-06 | 3.200E-06 |
| 25082 | 1255 | T1255R | D | 3.3e-06 | 1.3e-06 | 3.3e-06 | 2.633E-06 | 1.155E-06 | 1.300E-06 | 3.300E-06 |
| 25083 | 1255 | T1255N | D | 3.4e-06 | 2.7e-06 | 1.8e-06 | 2.633E-06 | 8.021E-07 | 1.800E-06 | 3.400E-06 |
| 25084 | 1255 | T1255D | D | 2.6e-06 | 2.6e-06 | 3e-06 | 2.733E-06 | 2.309E-07 | 2.600E-06 | 3.000E-06 |
| 25085 | 1255 | T1255C | D | 3.1e-06 | 2.6e-06 | 2.1e-06 | 2.600E-06 | 5.000E-07 | 2.100E-06 | 3.100E-06 |
| 25086 | 1255 | T1255E | D | 2.4e-06 | 3.1e-06 | 2.6e-06 | 2.700E-06 | 3.606E-07 | 2.400E-06 | 3.100E-06 |
| 25087 | 1255 | T1255Q | D | 2.3e-06 | 2.6e-06 | 2.8e-06 | 2.567E-06 | 2.517E-07 | 2.300E-06 | 2.800E-06 |
| 25088 | 1255 | T1255G | D | 2.7e-06 | 1.1e-06 | 2.6e-06 | 2.133E-06 | 8.963E-07 | 1.100E-06 | 2.700E-06 |
| 25089 | 1255 | T1255H | D | 2.6e-06 | 3.3e-06 | 1.4e-06 | 2.433E-06 | 9.609E-07 | 1.400E-06 | 3.300E-06 |
| 25090 | 1255 | T1255I | D | 2.2e-06 | 2e-06 | 2.3e-06 | 2.167E-06 | 1.528E-07 | 2.000E-06 | 2.300E-06 |
| 25091 | 1255 | T1255L | D | 3.2e-06 | 3.9e-06 | 4.2e-06 | 3.767E-06 | 5.132E-07 | 3.200E-06 | 4.200E-06 |
| 25092 | 1255 | T1255K | D | 1.4e-06 | 2e-06 | 1.6e-06 | 1.667E-06 | 3.055E-07 | 1.400E-06 | 2.000E-06 |
| 25093 | 1255 | T1255M | D | 2e-06 | 2.8e-06 | 2.9e-06 | 2.567E-06 | 4.933E-07 | 2.000E-06 | 2.900E-06 |
| 25094 | 1255 | T1255F | D | 2.6e-06 | 3.4e-06 | 2.7e-06 | 2.900E-06 | 4.359E-07 | 2.600E-06 | 3.400E-06 |
| 25095 | 1255 | T1255P | D | 1.5e-06 | 2.7e-06 | 1.3e-06 | 1.833E-06 | 7.572E-07 | 1.300E-06 | 2.700E-06 |
| 25096 | 1255 | T1255S | D | 1.3e-06 | 1.9e-06 | 2.5e-06 | 1.900E-06 | 6.000E-07 | 1.300E-06 | 2.500E-06 |
| 25097 | 1255 | T1255T | D | 2.5e-06 | 1.3e-06 | 2.2e-06 | 2.000E-06 | 6.245E-07 | 1.300E-06 | 2.500E-06 |
| 25098 | 1255 | T1255W | D | 3.1e-06 | 3.1e-06 | 2.7e-06 | 2.967E-06 | 2.309E-07 | 2.700E-06 | 3.100E-06 |
| 25099 | 1255 | T1255Y | D | 3.2e-06 | 1.6e-06 | 1.5e-06 | 2.100E-06 | 9.539E-07 | 1.500E-06 | 3.200E-06 |
| 25100 | 1255 | T1255V | D | 3.1e-06 | 1.7e-06 | 1.8e-06 | 2.200E-06 | 7.810E-07 | 1.700E-06 | 3.100E-06 |
| 25101 | 1256 | H1256A | D | 2.2e-06 | 2.7e-06 | 2.3e-06 | 2.400E-06 | 2.646E-07 | 2.200E-06 | 2.700E-06 |
| 25102 | 1256 | H1256R | D | 2.2e-06 | 1.8e-06 | 2.8e-06 | 2.267E-06 | 5.033E-07 | 1.800E-06 | 2.800E-06 |
| 25103 | 1256 | H1256N | D | 1.9e-06 | 1.3e-06 | 2.2e-06 | 1.800E-06 | 4.583E-07 | 1.300E-06 | 2.200E-06 |
| 25104 | 1256 | H1256D | D | 3e-06 | 1.4e-06 | 2.2e-06 | 2.200E-06 | 8.000E-07 | 1.400E-06 | 3.000E-06 |
| 25105 | 1256 | H1256C | D | 3.2e-06 | 1.7e-06 | 3.3e-06 | 2.733E-06 | 8.963E-07 | 1.700E-06 | 3.300E-06 |
| 25106 | 1256 | H1256E | D | 2e-06 | 2.7e-06 | 1.9e-06 | 2.200E-06 | 4.359E-07 | 1.900E-06 | 2.700E-06 |
| 25107 | 1256 | H1256Q | D | 2.1e-06 | 2.6e-06 | 2.6e-06 | 2.433E-06 | 2.887E-07 | 2.100E-06 | 2.600E-06 |
| 25108 | 1256 | H1256G | D | 2.5e-06 | 1.8e-06 | 4.5e-06 | 2.933E-06 | 1.401E-06 | 1.800E-06 | 4.500E-06 |
| 25109 | 1256 | H1256H | D | 2.2e-06 | 1.1e-06 | 2.3e-06 | 1.867E-06 | 6.658E-07 | 1.100E-06 | 2.300E-06 |
| 25110 | 1256 | H1256I | D | 1.8e-06 | 1.7e-06 | 2.3e-06 | 1.933E-06 | 3.215E-07 | 1.700E-06 | 2.300E-06 |
| 25111 | 1256 | H1256L | D | 1.9e-06 | 2.1e-06 | 3.9e-06 | 2.633E-06 | 1.102E-06 | 1.900E-06 | 3.900E-06 |
| 25112 | 1256 | H1256K | D | 1.1e-06 | 2.3e-06 | 3.4e-06 | 2.267E-06 | 1.150E-06 | 1.100E-06 | 3.400E-06 |
| 25113 | 1256 | H1256M | D | 2e-06 | 2e-06 | 1.6e-06 | 1.867E-06 | 2.309E-07 | 1.600E-06 | 2.000E-06 |
| 25114 | 1256 | H1256F | D | 2.6e-06 | 2.7e-06 | 2.9e-06 | 2.733E-06 | 1.528E-07 | 2.600E-06 | 2.900E-06 |
| 25115 | 1256 | H1256P | D | 2e-06 | 2.2e-06 | 2.7e-06 | 2.300E-06 | 3.606E-07 | 2.000E-06 | 2.700E-06 |
| 25116 | 1256 | H1256S | D | 1.2e-06 | 2.4e-06 | 2.8e-06 | 2.133E-06 | 8.327E-07 | 1.200E-06 | 2.800E-06 |
| 25117 | 1256 | H1256T | D | 2.6e-06 | 2.3e-06 | 2.9e-06 | 2.600E-06 | 3.000E-07 | 2.300E-06 | 2.900E-06 |
| 25118 | 1256 | H1256W | D | 1.6e-06 | 2.9e-06 | 3.3e-06 | 2.600E-06 | 8.888E-07 | 1.600E-06 | 3.300E-06 |
| 25119 | 1256 | H1256Y | D | 3.9e-06 | 1.4e-06 | 3.3e-06 | 2.867E-06 | 1.305E-06 | 1.400E-06 | 3.900E-06 |
| 25120 | 1256 | H1256V | D | 3e-06 | 2.5e-06 | 2.6e-06 | 2.700E-06 | 2.646E-07 | 2.500E-06 | 3.000E-06 |
| 25121 | 1257 | Q1257A | D | 4.1e-06 | 3.1e-06 | 2.8e-06 | 3.333E-06 | 6.807E-07 | 2.800E-06 | 4.100E-06 |
| 25122 | 1257 | Q1257R | D | 4.8e-06 | 3.1e-06 | 1.9e-06 | 3.267E-06 | 1.457E-06 | 1.900E-06 | 4.800E-06 |
| 25123 | 1257 | Q1257N | D | 1.5e-06 | 2.7e-06 | 2.3e-06 | 2.167E-06 | 6.110E-07 | 1.500E-06 | 2.700E-06 |
| 25124 | 1257 | Q1257D | D | 2.3e-06 | 2.5e-06 | 1.9e-06 | 2.233E-06 | 3.055E-07 | 1.900E-06 | 2.500E-06 |
| 25125 | 1257 | Q1257C | D | 2.1e-06 | 1.9e-06 | 2e-06 | 2.000E-06 | 1.000E-07 | 1.900E-06 | 2.100E-06 |
| 25126 | 1257 | Q1257E | D | 4.1e-06 | 4.1e-06 | 1.9e-06 | 3.367E-06 | 1.270E-06 | 1.900E-06 | 4.100E-06 |
| 25127 | 1257 | Q1257Q | D | 2.7e-06 | 3.1e-06 | 4.2e-06 | 3.333E-06 | 7.767E-07 | 2.700E-06 | 4.200E-06 |
| 25128 | 1257 | Q1257G | D | 3e-06 | 3.2e-06 | 2.6e-06 | 2.933E-06 | 3.055E-07 | 2.600E-06 | 3.200E-06 |
| 25129 | 1257 | Q1257H | D | 2.2e-06 | 1.2e-06 | 2.7e-06 | 2.033E-06 | 7.638E-07 | 1.200E-06 | 2.700E-06 |
| 25130 | 1257 | Q1257I | D | 2.4e-06 | 1.8e-06 | 3.6e-06 | 2.600E-06 | 9.165E-07 | 1.800E-06 | 3.600E-06 |
| 25131 | 1257 | Q1257L | D | 1.8e-06 | 1.9e-06 | 3.3e-06 | 2.333E-06 | 8.386E-07 | 1.800E-06 | 3.300E-06 |
| 25132 | 1257 | Q1257K | D | 1.5e-06 | 2e-06 | 3.3e-06 | 2.267E-06 | 9.292E-07 | 1.500E-06 | 3.300E-06 |
| 25133 | 1257 | Q1257M | D | 2.8e-06 | 1.2e-06 | 3.1e-06 | 2.367E-06 | 1.021E-06 | 1.200E-06 | 3.100E-06 |
| 25134 | 1257 | Q1257F | D | 4.2e-06 | 3.1e-06 | 1.3e-06 | 2.867E-06 | 1.464E-06 | 1.300E-06 | 4.200E-06 |
| 25135 | 1257 | Q1257P | D | 3e-06 | 2.4e-06 | 1.3e-06 | 2.233E-06 | 8.622E-07 | 1.300E-06 | 3.000E-06 |
| 25136 | 1257 | Q1257S | D | 3.9e-06 | 2.4e-06 | 2e-06 | 2.767E-06 | 1.002E-06 | 2.000E-06 | 3.900E-06 |
| 25137 | 1257 | Q1257T | D | 3.3e-06 | 3.1e-06 | 2.2e-06 | 2.867E-06 | 5.859E-07 | 2.200E-06 | 3.300E-06 |
| 25138 | 1257 | Q1257W | D | 5e-06 | 1.2e-06 | 3.8e-06 | 3.333E-06 | 1.943E-06 | 1.200E-06 | 5.000E-06 |
| 25139 | 1257 | Q1257Y | D | 1.6e-06 | 1.9e-06 | 2.9e-06 | 2.133E-06 | 6.807E-07 | 1.600E-06 | 2.900E-06 |
| 25140 | 1257 | Q1257V | D | 3.8e-06 | 2.2e-06 | 2.5e-06 | 2.833E-06 | 8.505E-07 | 2.200E-06 | 3.800E-06 |
| 25141 | 1258 | G1258A | D | 2.3e-06 | 1.2e-06 | 2.4e-06 | 1.967E-06 | 6.658E-07 | 1.200E-06 | 2.400E-06 |
| 25142 | 1258 | G1258R | D | 1.3e-06 | 2.8e-06 | 1.8e-06 | 1.967E-06 | 7.638E-07 | 1.300E-06 | 2.800E-06 |
| 25143 | 1258 | G1258N | D | 2.1e-06 | 1.5e-06 | 1.6e-06 | 1.733E-06 | 3.215E-07 | 1.500E-06 | 2.100E-06 |
| 25144 | 1258 | G1258D | D | 1.8e-06 | 2.3e-06 | 1.6e-06 | 1.900E-06 | 3.606E-07 | 1.600E-06 | 2.300E-06 |
| 25145 | 1258 | G1258C | D | 2e-06 | 2e-06 | 1.8e-06 | 1.933E-06 | 1.155E-07 | 1.800E-06 | 2.000E-06 |
| 25146 | 1258 | G1258E | D | 3.3e-06 | 2e-06 | 1.8e-06 | 2.367E-06 | 8.145E-07 | 1.800E-06 | 3.300E-06 |
| 25147 | 1258 | G1258Q | D | 2.2e-06 | 2.9e-06 | 2.6e-06 | 2.567E-06 | 3.512E-07 | 2.200E-06 | 2.900E-06 |
| 25148 | 1258 | G1258G | D | 7.5e-07 | 2.5e-06 | 3.1e-06 | 2.117E-06 | 1.221E-06 | 7.500E-07 | 3.100E-06 |
| 25149 | 1258 | G1258H | D | 8.8e-07 | 2.2e-06 | 3.1e-06 | 2.060E-06 | 1.117E-06 | 8.800E-07 | 3.100E-06 |
| 25150 | 1258 | G1258I | D | 1.8e-06 | 2.2e-06 | 4.2e-06 | 2.733E-06 | 1.286E-06 | 1.800E-06 | 4.200E-06 |
| 25151 | 1258 | G1258L | D | 3.2e-06 | 1.9e-06 | 2.4e-06 | 2.500E-06 | 6.557E-07 | 1.900E-06 | 3.200E-06 |
| 25152 | 1258 | G1258K | D | 4.4e-06 | 2.5e-06 | 1.9e-06 | 2.933E-06 | 1.305E-06 | 1.900E-06 | 4.400E-06 |
| 25153 | 1258 | G1258M | D | 2.4e-06 | 2.3e-06 | 2.8e-06 | 2.500E-06 | 2.646E-07 | 2.300E-06 | 2.800E-06 |
| 25154 | 1258 | G1258F | D | 2.1e-06 | 2.4e-06 | 1.2e-06 | 1.900E-06 | 6.245E-07 | 1.200E-06 | 2.400E-06 |
| 25155 | 1258 | G1258P | D | 2.2e-06 | 2.6e-06 | 2.4e-06 | 2.400E-06 | 2.000E-07 | 2.200E-06 | 2.600E-06 |
| 25156 | 1258 | G1258S | D | 2.2e-06 | 1.7e-06 | 2.6e-06 | 2.167E-06 | 4.509E-07 | 1.700E-06 | 2.600E-06 |
| 25157 | 1258 | G1258T | D | 1.1e-06 | 2.2e-06 | 2.2e-06 | 1.833E-06 | 6.351E-07 | 1.100E-06 | 2.200E-06 |
| 25158 | 1258 | G1258W | D | 2e-06 | 3.9e-06 | 3.2e-06 | 3.033E-06 | 9.609E-07 | 2.000E-06 | 3.900E-06 |
| 25159 | 1258 | G1258Y | D | 2.9e-06 | 2.5e-06 | 1.6e-06 | 2.333E-06 | 6.658E-07 | 1.600E-06 | 2.900E-06 |

| | | | | | | | | | | |
|---|---|---|---|---|---|---|---|---|---|---|
| 25160 | 1258 | G1258V | D | 3.3e-06 | 2.5e-06 | 2.7e-06 | 2.833E-06 | 4.163E-07 | 2.500E-06 | 3.300E-06 |
| 25161 | 1259 | L1259A | D | 1.8e-06 | 2.4e-06 | 3.3e-06 | 2.500E-06 | 7.550E-07 | 1.800E-06 | 3.300E-06 |
| 25162 | 1259 | L1259R | D | 1.7e-06 | 2.6e-06 | 1.9e-06 | 2.067E-06 | 4.726E-07 | 1.700E-06 | 2.600E-06 |
| 25163 | 1259 | L1259N | D | 2.6e-06 | 2.2e-06 | 3.2e-06 | 2.667E-06 | 5.033E-07 | 2.200E-06 | 3.200E-06 |
| 25164 | 1259 | L1259D | D | 1.1e-06 | 2.6e-06 | 2.8e-06 | 2.167E-06 | 9.292E-07 | 1.100E-06 | 2.800E-06 |
| 25165 | 1259 | L1259C | D | 1.3e-06 | 3.7e-06 | 3.7e-06 | 2.900E-06 | 1.386E-06 | 1.300E-06 | 3.700E-06 |
| 25166 | 1259 | L1259E | D | 1.2e-06 | 4.3e-06 | 2.3e-06 | 2.600E-06 | 1.572E-06 | 1.200E-06 | 4.300E-06 |
| 25167 | 1259 | L1259Q | D | 2.4e-06 | 2.2e-06 | 2e-06 | 2.200E-06 | 2.000E-07 | 2.000E-06 | 2.400E-06 |
| 25168 | 1259 | L1259G | D | 1.6e-06 | 3e-06 | 2.4e-06 | 2.333E-06 | 7.024E-07 | 1.600E-06 | 3.000E-06 |
| 25169 | 1259 | L1259H | D | 1.8e-06 | 1e-06 | 1.8e-06 | 1.533E-06 | 4.619E-07 | 1.000E-06 | 1.800E-06 |
| 25170 | 1259 | L1259I | D | 4.2e-06 | 2.1e-06 | 3e-06 | 3.100E-06 | 1.054E-06 | 2.100E-06 | 4.200E-06 |
| 25171 | 1259 | L1259L | D | 4.2e-06 | 2.8e-06 | 3.6e-06 | 3.533E-06 | 7.024E-07 | 2.800E-06 | 4.200E-06 |
| 25172 | 1259 | L1259K | D | 2.9e-06 | 2.3e-06 | 2.2e-06 | 2.467E-06 | 3.786E-07 | 2.200E-06 | 2.900E-06 |
| 25173 | 1259 | L1259M | D | 2.2e-06 | 1.6e-06 | 1.2e-06 | 1.667E-06 | 5.033E-07 | 1.200E-06 | 2.200E-06 |
| 25174 | 1259 | L1259F | D | 2e-06 | 3.7e-06 | 1.2e-06 | 2.300E-06 | 1.277E-06 | 1.200E-06 | 3.700E-06 |
| 25175 | 1259 | L1259P | D | 2.2e-06 | 2.3e-06 | 2.2e-06 | 2.233E-06 | 5.774E-08 | 2.200E-06 | 2.300E-06 |
| 25176 | 1259 | L1259S | D | 7.1e-06 | 2.3e-06 | 2.6e-06 | 4.000E-06 | 2.689E-06 | 2.300E-06 | 7.100E-06 |
| 25177 | 1259 | L1259T | D | 1.3e-06 | 2.3e-06 | 2.4e-06 | 2.000E-06 | 6.083E-07 | 1.300E-06 | 2.400E-06 |
| 25178 | 1259 | L1259W | D | 2.3e-06 | 9.9e-07 | 2.3e-06 | 1.863E-06 | 7.563E-07 | 9.900E-07 | 2.300E-06 |
| 25179 | 1259 | L1259Y | D | 2.3e-06 | 1.6e-06 | 2.2e-06 | 2.033E-06 | 3.786E-07 | 1.600E-06 | 2.300E-06 |
| 25180 | 1259 | L1259V | D | 3.3e-06 | 2.9e-06 | 2.6e-06 | 2.933E-06 | 3.512E-07 | 2.600E-06 | 3.300E-06 |
| 25181 | 1260 | S1260A | D | 1.9e-06 | 2.3e-06 | 2.1e-06 | 2.100E-06 | 2.000E-07 | 1.900E-06 | 2.300E-06 |
| 25182 | 1260 | S1260R | D | 1.8e-06 | 1.8e-06 | 2.4e-06 | 2.000E-06 | 3.464E-07 | 1.800E-06 | 2.400E-06 |
| 25183 | 1260 | S1260N | D | 2e-06 | 2.5e-06 | 2.2e-06 | 2.233E-06 | 2.517E-07 | 2.000E-06 | 2.500E-06 |
| 25184 | 1260 | S1260D | D | 2.1e-06 | 3e-06 | 3e-06 | 2.700E-06 | 5.196E-07 | 2.100E-06 | 3.000E-06 |
| 25185 | 1260 | S1260C | D | 1.9e-06 | 8.5e-07 | 1.6e-06 | 1.450E-06 | 5.408E-07 | 8.500E-07 | 1.900E-06 |
| 25186 | 1260 | S1260E | D | 2e-06 | 4.4e-06 | 2.9e-06 | 3.100E-06 | 1.212E-06 | 2.000E-06 | 4.400E-06 |
| 25187 | 1260 | S1260Q | D | 2.6e-06 | 1.4e-06 | 1.9e-06 | 1.967E-06 | 6.028E-07 | 1.400E-06 | 2.600E-06 |
| 25188 | 1260 | S1260G | D | 1.4e-06 | 2.3e-06 | 2e-06 | 1.900E-06 | 4.583E-07 | 1.400E-06 | 2.300E-06 |
| 25189 | 1260 | S1260H | D | 2.2e-06 | 1.8e-06 | 1.2e-06 | 1.733E-06 | 5.033E-07 | 1.200E-06 | 2.200E-06 |
| 25190 | 1260 | S1260I | D | 2.3e-06 | 1.4e-06 | 1.9e-06 | 1.867E-06 | 4.509E-07 | 1.400E-06 | 2.300E-06 |
| 25191 | 1260 | S1260L | D | 2.1e-06 | 3.8e-06 | 3.1e-06 | 3.000E-06 | 8.544E-07 | 2.100E-06 | 3.800E-06 |
| 25192 | 1260 | S1260K | D | 2.8e-06 | 4.4e-06 | 3.2e-06 | 3.467E-06 | 8.327E-07 | 2.800E-06 | 4.400E-06 |
| 25193 | 1260 | S1260M | D | 1.3e-06 | 2.2e-06 | 2.5e-06 | 2.000E-06 | 6.245E-07 | 1.300E-06 | 2.500E-06 |
| 25194 | 1260 | S1260F | D | 2.3e-06 | 2.8e-06 | 2.5e-06 | 2.533E-06 | 2.517E-07 | 2.300E-06 | 2.800E-06 |
| 25195 | 1260 | S1260P | D | 1.2e-06 | 3.1e-06 | 3.7e-06 | 2.667E-06 | 1.305E-06 | 1.200E-06 | 3.700E-06 |
| 25196 | 1260 | S1260S | D | 3e-06 | 3e-06 | 3.5e-06 | 3.167E-06 | 2.887E-07 | 3.000E-06 | 3.500E-06 |
| 25197 | 1260 | S1260T | D | 1.9e-06 | 2.2e-06 | 1.7e-06 | 1.933E-06 | 2.517E-07 | 1.700E-06 | 2.200E-06 |
| 25198 | 1260 | S1260W | D | 3.9e-06 | 2.3e-06 | 3.7e-06 | 3.300E-06 | 8.718E-07 | 2.300E-06 | 3.900E-06 |
| 25199 | 1260 | S1260Y | D | 2.2e-06 | 2.6e-06 | 1.8e-06 | 2.200E-06 | 4.000E-07 | 1.800E-06 | 2.600E-06 |
| 25200 | 1260 | S1260V | D | 2.9e-06 | 1.7e-06 | 2.9e-06 | 2.500E-06 | 6.928E-07 | 1.700E-06 | 2.900E-06 |
| 25201 | 1261 | S1261A | D | 3.6e-06 | 2.6e-06 | 2e-06 | 2.733E-06 | 8.083E-07 | 2.000E-06 | 3.600E-06 |
| 25202 | 1261 | S1261R | D | 3.1e-06 | 2.3e-06 | 4.4e-06 | 3.267E-06 | 1.060E-06 | 2.300E-06 | 4.400E-06 |
| 25203 | 1261 | S1261N | D | 2.5e-06 | 2e-06 | 2.8e-06 | 2.433E-06 | 4.041E-07 | 2.000E-06 | 2.800E-06 |
| 25204 | 1261 | S1261D | D | 2.3e-06 | 2.1e-06 | 1.8e-06 | 2.067E-06 | 2.517E-07 | 1.800E-06 | 2.300E-06 |
| 25205 | 1261 | S1261C | D | 2.1e-06 | 3.8e-06 | 3e-06 | 2.967E-06 | 8.505E-07 | 2.100E-06 | 3.800E-06 |
| 25206 | 1261 | S1261E | D | 2.8e-06 | 2.1e-06 | 3.5e-06 | 2.800E-06 | 7.000E-07 | 2.100E-06 | 3.500E-06 |
| 25207 | 1261 | S1261Q | D | 2.9e-06 | 2.1e-06 | 1.8e-06 | 2.267E-06 | 5.686E-07 | 1.800E-06 | 2.900E-06 |
| 25208 | 1261 | S1261G | D | 2.7e-06 | 2.7e-06 | 2.2e-06 | 2.533E-06 | 2.887E-07 | 2.200E-06 | 2.700E-06 |
| 25209 | 1261 | S1261H | D | 8.7e-07 | 1.8e-06 | 2.9e-06 | 1.857E-06 | 1.016E-06 | 8.700E-07 | 2.900E-06 |
| 25210 | 1261 | S1261I | D | 3.3e-06 | 1.9e-06 | 2.5e-06 | 2.567E-06 | 7.024E-07 | 1.900E-06 | 3.300E-06 |
| 25211 | 1261 | S1261L | D | 2.6e-06 | 4.1e-06 | 3e-06 | 3.233E-06 | 7.767E-07 | 2.600E-06 | 4.100E-06 |
| 25212 | 1261 | S1261K | D | 3.1e-06 | 2.9e-06 | 1e-06 | 2.333E-06 | 1.159E-06 | 1.000E-06 | 3.100E-06 |
| 25213 | 1261 | S1261M | D | 3.6e-06 | 2.8e-06 | 2.3e-06 | 2.900E-06 | 6.557E-07 | 2.300E-06 | 3.600E-06 |
| 25214 | 1261 | S1261F | D | 2.9e-06 | 2.6e-06 | 2e-06 | 2.500E-06 | 4.583E-07 | 2.000E-06 | 2.900E-06 |
| 25215 | 1261 | S1261P | D | 3.3e-06 | 3.6e-06 | 4.4e-06 | 3.767E-06 | 5.686E-07 | 3.300E-06 | 4.400E-06 |
| 25216 | 1261 | S1261S | D | 1.2e-06 | 4.3e-06 | 2.3e-06 | 2.600E-06 | 1.572E-06 | 1.200E-06 | 4.300E-06 |
| 25217 | 1261 | S1261T | D | 1.7e-06 | 1.5e-06 | 1.1e-06 | 1.433E-06 | 3.055E-07 | 1.100E-06 | 1.700E-06 |
| 25218 | 1261 | S1261W | D | 2.3e-06 | 3.5e-06 | 1.8e-06 | 2.533E-06 | 8.737E-07 | 1.800E-06 | 3.500E-06 |
| 25219 | 1261 | S1261Y | D | 1.9e-06 | 2.2e-06 | 2.4e-06 | 2.167E-06 | 2.517E-07 | 1.900E-06 | 2.400E-06 |
| 25220 | 1261 | S1261V | D | 2.4e-06 | 1.8e-06 | 2.7e-06 | 2.300E-06 | 4.583E-07 | 1.800E-06 | 2.700E-06 |
| 25221 | 1262 | P1262A | D | 1.7e-06 | 2.2e-06 | 1.7e-06 | 1.867E-06 | 2.887E-07 | 1.700E-06 | 2.200E-06 |
| 25222 | 1262 | P1262R | D | 3e-06 | 2.5e-06 | 2.3e-06 | 2.600E-06 | 3.606E-07 | 2.300E-06 | 3.000E-06 |
| 25223 | 1262 | P1262N | D | 2.9e-06 | 2.1e-06 | 2.5e-06 | 2.500E-06 | 4.000E-07 | 2.100E-06 | 2.900E-06 |
| 25224 | 1262 | P1262D | D | 2e-06 | 1.8e-06 | 2.2e-06 | 2.000E-06 | 2.000E-07 | 1.800E-06 | 2.200E-06 |
| 25225 | 1262 | P1262C | D | 1.9e-06 | 2.5e-06 | 2.9e-06 | 2.433E-06 | 5.033E-07 | 1.900E-06 | 2.900E-06 |
| 25226 | 1262 | P1262E | D | 3.6e-06 | 4.7e-06 | 1.2e-06 | 3.167E-06 | 1.790E-06 | 1.200E-06 | 4.700E-06 |
| 25227 | 1262 | P1262Q | D | 2.8e-06 | 2.2e-06 | 1.7e-06 | 2.233E-06 | 5.508E-07 | 1.700E-06 | 2.800E-06 |
| 25228 | 1262 | P1262G | D | 6.4e-06 | 2.1e-06 | 2.3e-06 | 3.600E-06 | 2.427E-06 | 2.100E-06 | 6.400E-06 |
| 25229 | 1262 | P1262H | D | 2.4e-06 | 1.5e-06 | 1.6e-06 | 1.833E-06 | 4.933E-07 | 1.500E-06 | 2.400E-06 |
| 25230 | 1262 | P1262I | D | 2.4e-06 | 2.2e-06 | 3.3e-06 | 2.633E-06 | 5.859E-07 | 2.200E-06 | 3.300E-06 |
| 25231 | 1262 | P1262L | D | 1.4e-06 | 3.1e-06 | 2.4e-06 | 2.300E-06 | 8.544E-07 | 1.400E-06 | 3.100E-06 |
| 25232 | 1262 | P1262K | D | 2e-06 | 4.4e-06 | 1.9e-06 | 2.767E-06 | 1.415E-06 | 1.900E-06 | 4.400E-06 |
| 25233 | 1262 | P1262M | D | 3.4e-06 | 2.5e-06 | 2.1e-06 | 2.667E-06 | 6.658E-07 | 2.100E-06 | 3.400E-06 |
| 25234 | 1262 | P1262F | D | 3.1e-06 | 2.5e-06 | 2.9e-06 | 2.833E-06 | 3.055E-07 | 2.500E-06 | 3.100E-06 |
| 25235 | 1262 | P1262P | D | 2.1e-06 | 2e-06 | 3.7e-06 | 2.600E-06 | 9.539E-07 | 2.000E-06 | 3.700E-06 |
| 25236 | 1262 | P1262S | D | 2.3e-06 | 4.9e-06 | 1.6e-06 | 2.933E-06 | 1.739E-06 | 1.600E-06 | 4.900E-06 |
| 25237 | 1262 | P1262T | D | 2.8e-06 | 3.2e-06 | 2.2e-06 | 2.733E-06 | 5.033E-07 | 2.200E-06 | 3.200E-06 |
| 25238 | 1262 | P1262W | D | 2.5e-06 | 3.4e-06 | 3e-06 | 2.967E-06 | 4.509E-07 | 2.500E-06 | 3.400E-06 |
| 25239 | 1262 | P1262Y | D | 2.8e-06 | 1.2e-06 | 2.9e-06 | 2.300E-06 | 9.539E-07 | 1.200E-06 | 2.900E-06 |
| 25240 | 1262 | P1262V | D | 2.5e-06 | 2.3e-06 | 3e-06 | 2.600E-06 | 3.606E-07 | 2.300E-06 | 3.000E-06 |
| 25241 | 1263 | V1263A | D | 2.2e-06 | 2.2e-06 | 3.3e-06 | 2.567E-06 | 6.351E-07 | 2.200E-06 | 3.300E-06 |
| 25242 | 1263 | V1263R | D | 2.8e-06 | 2.1e-06 | 2e-06 | 2.300E-06 | 4.359E-07 | 2.000E-06 | 2.800E-06 |
| 25243 | 1263 | V1263N | D | 1.9e-06 | 9.6e-07 | 2.4e-06 | 1.753E-06 | 7.311E-07 | 9.600E-07 | 2.400E-06 |
| 25244 | 1263 | V1263D | D | 2.6e-06 | 1.6e-06 | 1.6e-06 | 1.933E-06 | 5.774E-07 | 1.600E-06 | 2.600E-06 |

| | | | | | | | | | | |
|---|---|---|---|---|---|---|---|---|---|---|
| 25245 | 1263 | V1263C | D | 1.4e-06 | 2.6e-06 | 2.7e-06 | 2.233E-06 | 7.234E-07 | 1.400E-06 | 2.700E-06 |
| 25246 | 1263 | V1263E | D | 2.7e-06 | 2.6e-06 | 2.1e-06 | 2.467E-06 | 3.215E-07 | 2.100E-06 | 2.700E-06 |
| 25247 | 1263 | V1263Q | D | 1.8e-06 | 3.1e-06 | 2.3e-06 | 2.400E-06 | 6.557E-07 | 1.800E-06 | 3.100E-06 |
| 25248 | 1263 | V1263G | D | 3.1e-06 | 1.5e-06 | 1.7e-06 | 2.100E-06 | 8.718E-07 | 1.500E-06 | 3.100E-06 |
| 25249 | 1263 | V1263H | D | 2.6e-06 | 5e-06 | 2.8e-06 | 3.467E-06 | 1.332E-06 | 2.600E-06 | 5.000E-06 |
| 25250 | 1263 | V1263I | D | 2.1e-06 | 2.6e-06 | 1.7e-06 | 2.133E-06 | 4.509E-07 | 1.700E-06 | 2.600E-06 |
| 25251 | 1263 | V1263L | D | 3e-06 | 2.8e-06 | 3.1e-06 | 2.967E-06 | 1.528E-07 | 2.800E-06 | 3.100E-06 |
| 25252 | 1263 | V1263K | D | 3.7e-06 | 2.2e-06 | 2.4e-06 | 2.767E-06 | 8.145E-07 | 2.200E-06 | 3.700E-06 |
| 25253 | 1263 | V1263M | D | 1.5e-06 | 9.5e-07 | 1.6e-06 | 1.350E-06 | 3.500E-07 | 9.500E-07 | 1.600E-06 |
| 25254 | 1263 | V1263F | D | 3.1e-06 | 2.1e-06 | 1.9e-06 | 2.367E-06 | 6.429E-07 | 1.900E-06 | 3.100E-06 |
| 25255 | 1263 | V1263P | D | 2.1e-06 | 2.5e-06 | 2.5e-06 | 2.367E-06 | 2.309E-07 | 2.100E-06 | 2.500E-06 |
| 25256 | 1263 | V1263S | D | 3e-06 | 2.6e-06 | 2.2e-06 | 2.600E-06 | 4.000E-07 | 2.200E-06 | 3.000E-06 |
| 25257 | 1263 | V1263T | D | 1.4e-06 | 2e-06 | 3.6e-06 | 2.333E-06 | 1.137E-06 | 1.400E-06 | 3.600E-06 |
| 25258 | 1263 | V1263W | D | 4e-06 | 2e-06 | 2.2e-06 | 2.733E-06 | 1.102E-06 | 2.000E-06 | 4.000E-06 |
| 25259 | 1263 | V1263Y | D | 2.1e-06 | 1.8e-06 | 1.9e-06 | 1.933E-06 | 1.528E-07 | 1.800E-06 | 2.100E-06 |
| 25260 | 1263 | V1263V | D | 2.4e-06 | 3.1e-06 | 1.9e-06 | 2.467E-06 | 6.028E-07 | 1.900E-06 | 3.100E-06 |
| 25261 | 1264 | T1264A | D | 3e-06 | 2.9e-06 | 4.5e-06 | 3.467E-06 | 8.963E-07 | 2.900E-06 | 4.500E-06 |
| 25262 | 1264 | T1264R | D | 2e-06 | 2.2e-06 | 1.5e-06 | 1.900E-06 | 3.606E-07 | 1.500E-06 | 2.200E-06 |
| 25263 | 1264 | T1264N | D | 2.1e-06 | 2.8e-06 | 1.7e-06 | 2.200E-06 | 5.568E-07 | 1.700E-06 | 2.800E-06 |
| 25264 | 1264 | T1264D | D | 2.8e-06 | 1.6e-06 | 3.6e-06 | 2.667E-06 | 1.007E-06 | 1.600E-06 | 3.600E-06 |
| 25265 | 1264 | T1264C | D | 1e-06 | 3.1e-06 | 4.3e-06 | 2.800E-06 | 1.670E-06 | 1.000E-06 | 4.300E-06 |
| 25266 | 1264 | T1264E | D | 1.6e-06 | 1.7e-06 | 2.1e-06 | 1.800E-06 | 2.646E-07 | 1.600E-06 | 2.100E-06 |
| 25267 | 1264 | T1264Q | D | 2.1e-06 | 4.2e-06 | 2.4e-06 | 2.900E-06 | 1.136E-06 | 2.100E-06 | 4.200E-06 |
| 25268 | 1264 | T1264G | D | 2.8e-06 | 1.8e-06 | 2.3e-06 | 2.300E-06 | 5.000E-07 | 1.800E-06 | 2.800E-06 |
| 25269 | 1264 | T1264H | D | 1.9e-06 | 3e-06 | 2.7e-06 | 2.533E-06 | 5.686E-07 | 1.900E-06 | 3.000E-06 |
| 25270 | 1264 | T1264I | D | 2.9e-06 | 3e-06 | 2e-06 | 2.633E-06 | 5.508E-07 | 2.000E-06 | 3.000E-06 |
| 25271 | 1264 | T1264L | D | 4.5e-06 | 2.1e-06 | 2.3e-06 | 2.967E-06 | 1.332E-06 | 2.100E-06 | 4.500E-06 |
| 25272 | 1264 | T1264K | D | 1.7e-06 | 3.3e-06 | 2.2e-06 | 2.400E-06 | 8.185E-07 | 1.700E-06 | 3.300E-06 |
| 25273 | 1264 | T1264M | D | 2e-06 | 3.3e-06 | 2e-06 | 2.433E-06 | 7.506E-07 | 2.000E-06 | 3.300E-06 |
| 25274 | 1264 | T1264F | D | 1.9e-06 | 2.7e-06 | 3.3e-06 | 2.633E-06 | 7.024E-07 | 1.900E-06 | 3.300E-06 |
| 25275 | 1264 | T1264P | D | 2.5e-06 | 1.5e-06 | 2.8e-06 | 2.267E-06 | 6.807E-07 | 1.500E-06 | 2.800E-06 |
| 25276 | 1264 | T1264S | D | 3.9e-06 | 3.9e-06 | 3e-06 | 3.600E-06 | 5.196E-07 | 3.000E-06 | 3.900E-06 |
| 25277 | 1264 | T1264T | D | 3.6e-06 | 3.5e-06 | 3.2e-06 | 3.433E-06 | 2.082E-07 | 3.200E-06 | 3.600E-06 |
| 25278 | 1264 | T1264W | D | 6.3e-07 | 1.9e-06 | 4.6e-06 | 2.377E-06 | 2.027E-06 | 6.300E-07 | 4.600E-06 |
| 25279 | 1264 | T1264Y | D | 3.1e-06 | 2.3e-06 | 2.2e-06 | 2.533E-06 | 4.933E-07 | 2.200E-06 | 3.100E-06 |
| 25280 | 1264 | T1264V | D | 2.2e-06 | 2.3e-06 | 2.5e-06 | 2.333E-06 | 1.528E-07 | 2.200E-06 | 2.500E-06 |
| 25281 | 1265 | K1265A | D | 1.9e-06 | 2.4e-06 | 2.3e-06 | 2.200E-06 | 2.646E-07 | 1.900E-06 | 2.400E-06 |
| 25282 | 1265 | K1265R | D | 2e-06 | 2.6e-06 | 1.7e-06 | 2.100E-06 | 4.583E-07 | 1.700E-06 | 2.600E-06 |
| 25283 | 1265 | K1265N | D | 1.8e-06 | 1.5e-06 | 3.4e-06 | 2.233E-06 | 1.021E-06 | 1.500E-06 | 3.400E-06 |
| 25284 | 1265 | K1265D | D | 4.1e-06 | 2e-06 | 1.8e-06 | 2.633E-06 | 1.274E-06 | 1.800E-06 | 4.100E-06 |
| 25285 | 1265 | K1265C | D | 3.6e-06 | 4.9e-06 | 2.9e-06 | 3.800E-06 | 1.015E-06 | 2.900E-06 | 4.900E-06 |
| 25286 | 1265 | K1265E | D | 3e-06 | 2e-06 | 2.9e-06 | 2.633E-06 | 5.508E-07 | 2.000E-06 | 3.000E-06 |
| 25287 | 1265 | K1265Q | D | 3e-06 | 1.4e-06 | 2.4e-06 | 2.267E-06 | 8.083E-07 | 1.400E-06 | 3.000E-06 |
| 25288 | 1265 | K1265G | D | 2.7e-06 | 1.8e-06 | 2.4e-06 | 2.300E-06 | 4.583E-07 | 1.800E-06 | 2.700E-06 |
| 25289 | 1265 | K1265H | D | 2.4e-06 | 2.4e-06 | 2.2e-06 | 2.333E-06 | 1.155E-07 | 2.200E-06 | 2.400E-06 |
| 25290 | 1265 | K1265I | D | 2.9e-06 | 1.8e-06 | 1.2e-06 | 1.967E-06 | 8.622E-07 | 1.200E-06 | 2.900E-06 |
| 25291 | 1265 | K1265L | D | 2.8e-06 | 3.1e-06 | 3e-06 | 2.967E-06 | 1.528E-07 | 2.800E-06 | 3.100E-06 |
| 25292 | 1265 | K1265K | D | 2e-06 | 1.9e-06 | 2.3e-06 | 2.067E-06 | 2.082E-07 | 1.900E-06 | 2.300E-06 |
| 25293 | 1265 | K1265M | D | 3.2e-06 | 2.7e-06 | 1.7e-06 | 2.533E-06 | 7.638E-07 | 1.700E-06 | 3.200E-06 |
| 25294 | 1265 | K1265F | D | 2.8e-06 | 2e-06 | 2.1e-06 | 2.300E-06 | 4.359E-07 | 2.000E-06 | 2.800E-06 |
| 25295 | 1265 | K1265P | D | 8.9e-07 | 2.3e-06 | 1.2e-06 | 1.463E-06 | 7.410E-07 | 8.900E-07 | 2.300E-06 |
| 25296 | 1265 | K1265S | D | 3e-06 | 2.4e-06 | 3.8e-06 | 3.067E-06 | 7.024E-07 | 2.400E-06 | 3.800E-06 |
| 25297 | 1265 | K1265T | D | 2.6e-06 | 5e-06 | 1.6e-06 | 3.067E-06 | 1.747E-06 | 1.600E-06 | 5.000E-06 |
| 25298 | 1265 | K1265W | D | 3e-06 | 3.6e-06 | 2.7e-06 | 3.100E-06 | 4.583E-07 | 2.700E-06 | 3.600E-06 |
| 25299 | 1265 | K1265Y | D | 3.1e-06 | 2.6e-06 | 2.7e-06 | 2.800E-06 | 2.646E-07 | 2.600E-06 | 3.100E-06 |
| 25300 | 1265 | K1265V | D | 2.7e-06 | 2.2e-06 | 1.5e-06 | 2.133E-06 | 6.028E-07 | 1.500E-06 | 2.700E-06 |
| 25301 | 1266 | S1266A | D | 3.1e-06 | 2.9e-06 | 1.2e-06 | 2.400E-06 | 1.044E-06 | 1.200E-06 | 3.100E-06 |
| 25302 | 1266 | S1266R | D | 2.3e-06 | 2.2e-06 | 2.1e-06 | 2.200E-06 | 1.000E-07 | 2.100E-06 | 2.300E-06 |
| 25303 | 1266 | S1266N | D | 3.1e-06 | 2.2e-06 | 1.7e-06 | 2.333E-06 | 7.095E-07 | 1.700E-06 | 3.100E-06 |
| 25304 | 1266 | S1266D | D | 1.8e-06 | 2.5e-06 | 2.8e-06 | 2.367E-06 | 5.132E-07 | 1.800E-06 | 2.800E-06 |
| 25305 | 1266 | S1266C | D | 2.3e-06 | 2.2e-06 | 2.5e-06 | 2.333E-06 | 1.528E-07 | 2.200E-06 | 2.500E-06 |
| 25306 | 1266 | S1266E | D | 2.2e-06 | 2.1e-06 | 3.2e-06 | 2.500E-06 | 6.083E-07 | 2.100E-06 | 3.200E-06 |
| 25307 | 1266 | S1266Q | D | 2.9e-06 | 2.3e-06 | 2.7e-06 | 2.633E-06 | 3.055E-07 | 2.300E-06 | 2.900E-06 |
| 25308 | 1266 | S1266G | D | 2e-06 | 4.7e-06 | 3.6e-06 | 3.433E-06 | 1.358E-06 | 2.000E-06 | 4.700E-06 |
| 25309 | 1266 | S1266H | D | 2.2e-06 | 1.8e-06 | 1.9e-06 | 1.967E-06 | 2.082E-07 | 1.800E-06 | 2.200E-06 |
| 25310 | 1266 | S1266I | D | 6.6e-06 | 2.9e-06 | 3.5e-06 | 4.333E-06 | 1.986E-06 | 2.900E-06 | 6.600E-06 |
| 25311 | 1266 | S1266L | D | 2.6e-06 | 2.2e-06 | 3.1e-06 | 2.633E-06 | 4.509E-07 | 2.200E-06 | 3.100E-06 |
| 25312 | 1266 | S1266K | D | 3.8e-06 | 2.4e-06 | 1.3e-06 | 2.500E-06 | 1.253E-06 | 1.300E-06 | 3.800E-06 |
| 25313 | 1266 | S1266M | D | 2e-06 | 2.5e-06 | 2.4e-06 | 2.300E-06 | 2.646E-07 | 2.000E-06 | 2.500E-06 |
| 25314 | 1266 | S1266F | D | 2.3e-06 | 3.2e-06 | 2.9e-06 | 2.800E-06 | 4.583E-07 | 2.300E-06 | 3.200E-06 |
| 25315 | 1266 | S1266P | D | 3.8e-06 | 2.7e-06 | 2.1e-06 | 2.867E-06 | 8.622E-07 | 2.100E-06 | 3.800E-06 |
| 25316 | 1266 | S1266S | D | 2e-06 | 2e-06 | 2.8e-06 | 2.267E-06 | 4.619E-07 | 2.000E-06 | 2.800E-06 |
| 25317 | 1266 | S1266T | D | 3.1e-06 | 2.2e-06 | 1.8e-06 | 2.367E-06 | 6.658E-07 | 1.800E-06 | 3.100E-06 |
| 25318 | 1266 | S1266W | D | 2e-06 | 2.3e-06 | 2.7e-06 | 2.333E-06 | 3.512E-07 | 2.000E-06 | 2.700E-06 |
| 25319 | 1266 | S1266Y | D | 2.1e-06 | 1.9e-06 | 1.7e-06 | 1.900E-06 | 2.000E-07 | 1.700E-06 | 2.100E-06 |
| 25320 | 1266 | S1266V | D | 2.5e-06 | 3.9e-06 | 2.6e-06 | 3.000E-06 | 7.810E-07 | 2.500E-06 | 3.900E-06 |
| 25321 | 1267 | F1267A | D | 5.3e-06 | 3.1e-06 | 3.7e-06 | 4.033E-06 | 1.137E-06 | 3.100E-06 | 5.300E-06 |
| 25322 | 1267 | F1267R | D | 4.2e-06 | 3.5e-06 | 2.2e-06 | 3.300E-06 | 1.015E-06 | 2.200E-06 | 4.200E-06 |
| 25323 | 1267 | F1267N | D | 1.6e-06 | 3.8e-06 | 3.6e-06 | 3.000E-06 | 1.217E-06 | 1.600E-06 | 3.800E-06 |
| 25324 | 1267 | F1267D | D | 1.8e-06 | 2.6e-06 | 2.2e-06 | 2.200E-06 | 4.000E-07 | 1.800E-06 | 2.600E-06 |
| 25325 | 1267 | F1267C | D | 2.1e-06 | 1.7e-06 | 2.7e-06 | 2.167E-06 | 5.033E-07 | 1.700E-06 | 2.700E-06 |
| 25326 | 1267 | F1267E | D | 1.9e-06 | 2.7e-06 | 3.1e-06 | 2.567E-06 | 6.110E-07 | 1.900E-06 | 3.100E-06 |
| 25327 | 1267 | F1267Q | D | 1.5e-06 | 5.5e-06 | 2.7e-06 | 3.233E-06 | 2.053E-06 | 1.500E-06 | 5.500E-06 |
| 25328 | 1267 | F1267G | D | 9.9e-07 | 2.2e-06 | 3.6e-06 | 2.263E-06 | 1.306E-06 | 9.900E-07 | 3.600E-06 |
| 25329 | 1267 | F1267H | D | 2.9e-06 | 2.5e-06 | 3.1e-06 | 2.833E-06 | 3.055E-07 | 2.500E-06 | 3.100E-06 |

| | | | | | | | | | | |
|---|---|---|---|---|---|---|---|---|---|---|
| 25330 | 1267 | F1267I | D | 2.7e-06 | 1.3e-06 | 2.7e-06 | 2.233E-06 | 8.083E-07 | 1.300E-06 | 2.700E-06 |
| 25331 | 1267 | F1267L | D | 2.3e-06 | 1.8e-06 | 1.6e-06 | 1.900E-06 | 3.606E-07 | 1.600E-06 | 2.300E-06 |
| 25332 | 1267 | F1267K | D | 3.3e-06 | 2.5e-06 | 5.3e-06 | 3.700E-06 | 1.442E-06 | 2.500E-06 | 5.300E-06 |
| 25333 | 1267 | F1267M | D | 1.3e-06 | 1.7e-06 | 3.2e-06 | 2.067E-06 | 1.002E-06 | 1.300E-06 | 3.200E-06 |
| 25334 | 1267 | F1267F | D | 2.2e-06 | 2.1e-06 | 2.7e-06 | 2.333E-06 | 3.215E-07 | 2.100E-06 | 2.700E-06 |
| 25335 | 1267 | F1267P | D | 2.1e-06 | 3.1e-06 | 4.4e-06 | 3.200E-06 | 1.153E-06 | 2.100E-06 | 4.400E-06 |
| 25336 | 1267 | F1267S | D | 2.4e-06 | 2.4e-06 | 3.5e-06 | 2.767E-06 | 6.351E-07 | 2.400E-06 | 3.500E-06 |
| 25337 | 1267 | F1267T | D | 1.7e-06 | 2.5e-06 | 4.5e-06 | 2.900E-06 | 1.442E-06 | 1.700E-06 | 4.500E-06 |
| 25338 | 1267 | F1267W | D | 2.9e-06 | 2.6e-06 | 2e-06 | 2.500E-06 | 4.583E-07 | 2.000E-06 | 2.900E-06 |
| 25339 | 1267 | F1267Y | D | 1.4e-06 | 1.6e-06 | 3.1e-06 | 2.033E-06 | 9.292E-07 | 1.400E-06 | 3.100E-06 |
| 25340 | 1267 | F1267V | D | 4e-06 | 2.9e-06 | 1.9e-06 | 2.933E-06 | 1.050E-06 | 1.900E-06 | 4.000E-06 |
| 25341 | 1268 | N1268A | D | 2.9e-06 | 2.4e-06 | 2.5e-06 | 2.600E-06 | 2.646E-07 | 2.400E-06 | 2.900E-06 |
| 25342 | 1268 | N1268R | D | 2.3e-06 | 9.7e-07 | 2.6e-06 | 1.957E-06 | 8.675E-07 | 9.700E-07 | 2.600E-06 |
| 25343 | 1268 | N1268N | D | 1.2e-06 | 2e-06 | 4.8e-06 | 2.667E-06 | 1.890E-06 | 1.200E-06 | 4.800E-06 |
| 25344 | 1268 | N1268D | D | 2.6e-06 | 2.2e-06 | 3.2e-06 | 2.667E-06 | 5.033E-07 | 2.200E-06 | 3.200E-06 |
| 25345 | 1268 | N1268C | D | 2.4e-06 | 2e-06 | 2.4e-06 | 2.267E-06 | 2.309E-07 | 2.000E-06 | 2.400E-06 |
| 25346 | 1268 | N1268E | D | 1.4e-06 | 2.8e-06 | 1.6e-06 | 1.933E-06 | 7.572E-07 | 1.400E-06 | 2.800E-06 |
| 25347 | 1268 | N1268Q | D | 2.9e-06 | 2.7e-06 | 2.5e-06 | 2.700E-06 | 2.000E-07 | 2.500E-06 | 2.900E-06 |
| 25348 | 1268 | N1268G | D | 2.2e-06 | 1.9e-06 | 3.2e-06 | 2.433E-06 | 6.807E-07 | 1.900E-06 | 3.200E-06 |
| 25349 | 1268 | N1268H | D | 2.3e-06 | 2.2e-06 | 2.6e-06 | 2.367E-06 | 2.082E-07 | 2.200E-06 | 2.600E-06 |
| 25350 | 1268 | N1268I | D | 1.4e-06 | 4.2e-06 | 2.8e-06 | 2.800E-06 | 1.400E-06 | 1.400E-06 | 4.200E-06 |
| 25351 | 1268 | N1268L | D | 2.8e-06 | 2.8e-06 | 1.2e-06 | 2.267E-06 | 9.238E-07 | 1.200E-06 | 2.800E-06 |
| 25352 | 1268 | N1268K | D | 2.8e-06 | 2.8e-06 | 1.4e-06 | 2.333E-06 | 8.083E-07 | 1.400E-06 | 2.800E-06 |
| 25353 | 1268 | N1268M | D | 2.4e-06 | 2.1e-06 | 2.4e-06 | 2.300E-06 | 1.732E-07 | 2.100E-06 | 2.400E-06 |
| 25354 | 1268 | N1268F | D | 3.1e-06 | 3.2e-06 | 2.4e-06 | 2.900E-06 | 4.359E-07 | 2.400E-06 | 3.200E-06 |
| 25355 | 1268 | N1268P | D | 1.9e-06 | 2.3e-06 | 1.7e-06 | 1.967E-06 | 3.055E-07 | 1.700E-06 | 2.300E-06 |
| 25356 | 1268 | N1268S | D | 2e-06 | 3.6e-06 | 2.7e-06 | 2.767E-06 | 8.021E-07 | 2.000E-06 | 3.600E-06 |
| 25357 | 1268 | N1268T | D | 1.2e-06 | 2.1e-06 | 1.8e-06 | 1.700E-06 | 4.583E-07 | 1.200E-06 | 2.100E-06 |
| 25358 | 1268 | N1268W | D | 3.3e-06 | 1.3e-06 | 3.9e-06 | 2.833E-06 | 1.361E-06 | 1.300E-06 | 3.900E-06 |
| 25359 | 1268 | N1268Y | D | 1.6e-06 | 3.5e-06 | 1.7e-06 | 2.267E-06 | 1.069E-06 | 1.600E-06 | 3.500E-06 |
| 25360 | 1268 | N1268V | D | 2.1e-06 | 1.5e-06 | 2.9e-06 | 2.167E-06 | 7.024E-07 | 1.500E-06 | 2.900E-06 |
| 25361 | 1269 | R1269A | D | 2.5e-06 | 2.1e-06 | 2e-06 | 2.200E-06 | 2.646E-07 | 2.000E-06 | 2.500E-06 |
| 25362 | 1269 | R1269R | D | 5.3e-06 | 1.6e-06 | 2.6e-06 | 3.167E-06 | 1.914E-06 | 1.600E-06 | 5.300E-06 |
| 25363 | 1269 | R1269N | D | 3e-06 | 1.8e-06 | 2.1e-06 | 2.300E-06 | 6.245E-07 | 1.800E-06 | 3.000E-06 |
| 25364 | 1269 | R1269D | D | 3e-06 | 2.7e-06 | 1.6e-06 | 2.433E-06 | 7.371E-07 | 1.600E-06 | 3.000E-06 |
| 25365 | 1269 | R1269C | D | 3.2e-06 | 2.6e-06 | 1.1e-06 | 2.300E-06 | 1.082E-06 | 1.100E-06 | 3.200E-06 |
| 25366 | 1269 | R1269E | D | 1.4e-06 | 2.9e-06 | 2.9e-06 | 2.400E-06 | 8.660E-07 | 1.400E-06 | 2.900E-06 |
| 25367 | 1269 | R1269Q | D | 1.5e-06 | 3.7e-06 | 1.7e-06 | 2.300E-06 | 1.217E-06 | 1.500E-06 | 3.700E-06 |
| 25368 | 1269 | R1269G | D | 2e-06 | 2.4e-06 | 1.9e-06 | 2.100E-06 | 2.646E-07 | 1.900E-06 | 2.400E-06 |
| 25369 | 1269 | R1269H | D | 4.5e-06 | 2.4e-06 | 2.8e-06 | 3.233E-06 | 1.115E-06 | 2.400E-06 | 4.500E-06 |
| 25370 | 1269 | R1269I | D | 2.5e-06 | 1.3e-06 | 1.4e-06 | 1.733E-06 | 6.658E-07 | 1.300E-06 | 2.500E-06 |
| 25371 | 1269 | R1269L | D | 3.6e-06 | 3.4e-06 | 1.9e-06 | 2.967E-06 | 9.292E-07 | 1.900E-06 | 3.600E-06 |
| 25372 | 1269 | R1269K | D | 1.9e-06 | 2.1e-06 | 3e-06 | 2.333E-06 | 5.859E-07 | 1.900E-06 | 3.000E-06 |
| 25373 | 1269 | R1269M | D | 2.1e-06 | 2.5e-06 | 3.7e-06 | 2.767E-06 | 8.327E-07 | 2.100E-06 | 3.700E-06 |
| 25374 | 1269 | R1269F | D | 1.8e-06 | 2.3e-06 | 2.6e-06 | 2.233E-06 | 4.041E-07 | 1.800E-06 | 2.600E-06 |
| 25375 | 1269 | R1269P | D | 3.1e-06 | 1.6e-06 | 2.5e-06 | 2.400E-06 | 7.550E-07 | 1.600E-06 | 3.100E-06 |
| 25376 | 1269 | R1269S | D | 2.5e-06 | 2.9e-06 | 3e-06 | 2.800E-06 | 2.646E-07 | 2.500E-06 | 3.000E-06 |
| 25377 | 1269 | R1269T | D | 2.2e-06 | 1.7e-06 | 2.3e-06 | 2.067E-06 | 3.215E-07 | 1.700E-06 | 2.300E-06 |
| 25378 | 1269 | R1269W | D | 1.7e-06 | 3e-06 | 1.4e-06 | 2.033E-06 | 8.505E-07 | 1.400E-06 | 3.000E-06 |
| 25379 | 1269 | R1269Y | D | 2e-06 | 3e-06 | 1.7e-06 | 2.233E-06 | 6.807E-07 | 1.700E-06 | 3.000E-06 |
| 25380 | 1269 | R1269V | D | 3.4e-06 | 1.1e-06 | 1.8e-06 | 2.100E-06 | 1.179E-06 | 1.100E-06 | 3.400E-06 |
| 25381 | 1270 | G1270A | D | 1e-06 | 2.1e-06 | 3.2e-06 | 2.100E-06 | 1.100E-06 | 1.000E-06 | 3.200E-06 |
| 25382 | 1270 | G1270R | D | 2e-06 | 6.1e-06 | 2.2e-06 | 3.433E-06 | 2.312E-06 | 2.000E-06 | 6.100E-06 |
| 25383 | 1270 | G1270N | D | 3e-06 | 6.7e-06 | 2.5e-06 | 4.067E-06 | 2.294E-06 | 2.500E-06 | 6.700E-06 |
| 25384 | 1270 | G1270D | D | 1.8e-06 | 2.8e-06 | 1.2e-06 | 1.933E-06 | 8.083E-07 | 1.200E-06 | 2.800E-06 |
| 25385 | 1270 | G1270C | D | 2.2e-06 | 3.5e-06 | 2.3e-06 | 2.667E-06 | 7.234E-07 | 2.200E-06 | 3.500E-06 |
| 25386 | 1270 | G1270E | D | 1.4e-06 | 4e-06 | 3.4e-06 | 2.933E-06 | 1.361E-06 | 1.400E-06 | 4.000E-06 |
| 25387 | 1270 | G1270Q | D | 3.2e-06 | 2.6e-06 | 9.7e-07 | 2.257E-06 | 1.154E-06 | 9.700E-07 | 3.200E-06 |
| 25388 | 1270 | G1270G | D | 3e-06 | 2.7e-06 | 4.1e-06 | 3.267E-06 | 7.371E-07 | 2.700E-06 | 4.100E-06 |
| 25389 | 1270 | G1270H | D | 1.9e-06 | 2.1e-06 | 1.7e-06 | 1.900E-06 | 2.000E-07 | 1.700E-06 | 2.100E-06 |
| 25390 | 1270 | G1270I | D | 2.9e-06 | 1.9e-06 | 1.9e-06 | 2.233E-06 | 5.774E-07 | 1.900E-06 | 2.900E-06 |
| 25391 | 1270 | G1270L | D | 2.6e-06 | 2e-06 | 2.2e-06 | 2.267E-06 | 3.055E-07 | 2.000E-06 | 2.600E-06 |
| 25392 | 1270 | G1270K | D | 1.4e-06 | 1.6e-06 | 2.5e-06 | 1.833E-06 | 5.859E-07 | 1.400E-06 | 2.500E-06 |
| 25393 | 1270 | G1270M | D | 3.7e-06 | 2.8e-06 | 1.8e-06 | 2.767E-06 | 9.504E-07 | 1.800E-06 | 3.700E-06 |
| 25394 | 1270 | G1270F | D | 3.3e-06 | 2.6e-06 | 2.3e-06 | 2.733E-06 | 5.132E-07 | 2.300E-06 | 3.300E-06 |
| 25395 | 1270 | G1270P | D | 3.5e-06 | 1.8e-06 | 1.7e-06 | 2.333E-06 | 1.012E-06 | 1.700E-06 | 3.500E-06 |
| 25396 | 1270 | G1270S | D | 2e-06 | 1.9e-06 | 2.4e-06 | 2.100E-06 | 2.646E-07 | 1.900E-06 | 2.400E-06 |
| 25397 | 1270 | G1270T | D | 3.1e-06 | 3.1e-06 | 1.1e-06 | 2.433E-06 | 1.155E-06 | 1.100E-06 | 3.100E-06 |
| 25398 | 1270 | G1270W | D | 2.4e-06 | 4.1e-06 | 1.9e-06 | 2.800E-06 | 1.153E-06 | 1.900E-06 | 4.100E-06 |
| 25399 | 1270 | G1270Y | D | 1.9e-06 | 2.1e-06 | 3.3e-06 | 2.433E-06 | 7.572E-07 | 1.900E-06 | 3.300E-06 |
| 25400 | 1270 | G1270V | D | 4.1e-06 | 3.1e-06 | 2.4e-06 | 3.200E-06 | 8.544E-07 | 2.400E-06 | 4.100E-06 |
| 25401 | 1271 | E1271A | D | 3.3e-06 | 2.1e-06 | 2.7e-06 | 2.700E-06 | 6.000E-07 | 2.100E-06 | 3.300E-06 |
| 25402 | 1271 | E1271R | D | 2.7e-06 | 3.7e-06 | 2.4e-06 | 2.933E-06 | 6.807E-07 | 2.400E-06 | 3.700E-06 |
| 25403 | 1271 | E1271N | D | 2.7e-06 | 1.8e-06 | 3.5e-06 | 2.667E-06 | 8.505E-07 | 1.800E-06 | 3.500E-06 |
| 25404 | 1271 | E1271D | D | 3.6e-06 | 2.6e-06 | 2.3e-06 | 2.833E-06 | 6.807E-07 | 2.300E-06 | 3.600E-06 |
| 25405 | 1271 | E1271C | D | 1.5e-06 | 1.9e-06 | 2.1e-06 | 1.833E-06 | 3.055E-07 | 1.500E-06 | 2.100E-06 |
| 25406 | 1271 | E1271E | D | 1.8e-06 | 4.1e-06 | 2.6e-06 | 2.833E-06 | 1.168E-06 | 1.800E-06 | 4.100E-06 |
| 25407 | 1271 | E1271Q | D | 1.2e-06 | 2.7e-06 | 2e-06 | 1.967E-06 | 7.506E-07 | 1.200E-06 | 2.700E-06 |
| 25408 | 1271 | E1271G | D | 2.7e-06 | 3.6e-06 | 3.3e-06 | 3.200E-06 | 4.583E-07 | 2.700E-06 | 3.600E-06 |
| 25409 | 1271 | E1271H | D | 2.4e-06 | 3.1e-06 | 5.2e-06 | 3.567E-06 | 1.457E-06 | 2.400E-06 | 5.200E-06 |
| 25410 | 1271 | E1271I | D | 2e-06 | 1.8e-06 | 3.6e-06 | 2.467E-06 | 9.866E-07 | 1.800E-06 | 3.600E-06 |
| 25411 | 1271 | E1271L | D | 1.6e-06 | 2.7e-06 | 2e-06 | 2.100E-06 | 5.568E-07 | 1.600E-06 | 2.700E-06 |
| 25412 | 1271 | E1271K | D | 1.6e-06 | 1.8e-06 | 2.7e-06 | 2.033E-06 | 5.859E-07 | 1.600E-06 | 2.700E-06 |
| 25413 | 1271 | E1271M | D | 2.5e-06 | 2.4e-06 | 2.4e-06 | 2.433E-06 | 5.774E-08 | 2.400E-06 | 2.500E-06 |
| 25414 | 1271 | E1271F | D | 4.2e-06 | 3.6e-06 | 2.2e-06 | 3.333E-06 | 1.026E-06 | 2.200E-06 | 4.200E-06 |

| | | | | | | | | | | |
|---|---|---|---|---|---|---|---|---|---|---|
| 25415 | 1271 | E1271P | D | 2.4e-06 | 2e-06 | 2.6e-06 | 2.333E-06 | 3.055E-07 | 2.000E-06 | 2.600E-06 |
| 25416 | 1271 | E1271S | D | 2.1e-06 | 3e-06 | 2.2e-06 | 2.433E-06 | 4.933E-07 | 2.100E-06 | 3.000E-06 |
| 25417 | 1271 | E1271T | D | 3e-06 | 2.5e-06 | 2.5e-06 | 2.667E-06 | 2.887E-07 | 2.500E-06 | 3.000E-06 |
| 25418 | 1271 | E1271W | D | 2e-06 | 3.2e-06 | 3.6e-06 | 2.933E-06 | 8.327E-07 | 2.000E-06 | 3.600E-06 |
| 25419 | 1271 | E1271Y | D | 1.9e-06 | 2.5e-06 | 2.1e-06 | 2.167E-06 | 3.055E-07 | 1.900E-06 | 2.500E-06 |
| 25420 | 1271 | E1271V | D | 2.2e-06 | 2.7e-06 | 2.6e-06 | 2.500E-06 | 2.646E-07 | 2.200E-06 | 2.700E-06 |
| 25421 | 1272 | C1272A | D | 2.5e-06 | 2e-06 | 1.1e-06 | 1.867E-06 | 7.095E-07 | 1.100E-06 | 2.500E-06 |
| 25422 | 1272 | C1272R | D | 1.5e-06 | 2.3e-06 | 2.1e-06 | 1.967E-06 | 4.163E-07 | 1.500E-06 | 2.300E-06 |
| 25423 | 1272 | C1272N | D | 2.7e-06 | 2.4e-06 | 2.2e-06 | 2.433E-06 | 2.517E-07 | 2.200E-06 | 2.700E-06 |
| 25424 | 1272 | C1272D | D | 3e-06 | 2.2e-06 | 4e-06 | 3.067E-06 | 9.018E-07 | 2.200E-06 | 4.000E-06 |
| 25425 | 1272 | C1272C | D | 1.8e-06 | 2e-06 | 2.5e-06 | 2.100E-06 | 3.606E-07 | 1.800E-06 | 2.500E-06 |
| 25426 | 1272 | C1272E | D | 2.2e-06 | 2.2e-06 | 2.7e-06 | 2.367E-06 | 2.887E-07 | 2.200E-06 | 2.700E-06 |
| 25427 | 1272 | C1272Q | D | 2.6e-06 | 3.5e-06 | 1.5e-06 | 2.533E-06 | 1.002E-06 | 1.500E-06 | 3.500E-06 |
| 25428 | 1272 | C1272G | D | 2e-06 | 2.3e-06 | 1.1e-06 | 1.800E-06 | 6.245E-07 | 1.100E-06 | 2.300E-06 |
| 25429 | 1272 | C1272H | D | 2.7e-06 | 2.1e-06 | 2e-06 | 2.267E-06 | 3.786E-07 | 2.000E-06 | 2.700E-06 |
| 25430 | 1272 | C1272I | D | 2.1e-06 | 1.8e-06 | 3.1e-06 | 2.333E-06 | 6.807E-07 | 1.800E-06 | 3.100E-06 |
| 25431 | 1272 | C1272L | D | 2.4e-06 | 3.6e-06 | 2.4e-06 | 2.800E-06 | 6.928E-07 | 2.400E-06 | 3.600E-06 |
| 25432 | 1272 | C1272K | D | 1.3e-06 | 2.6e-06 | 3.1e-06 | 2.333E-06 | 9.292E-07 | 1.300E-06 | 3.100E-06 |
| 25433 | 1272 | C1272M | D | 1.8e-06 | 2.5e-06 | 2.1e-06 | 2.133E-06 | 3.512E-07 | 1.800E-06 | 2.500E-06 |
| 25434 | 1272 | C1272F | D | 2.7e-06 | 3.5e-06 | 2.8e-06 | 3.000E-06 | 4.359E-07 | 2.700E-06 | 3.500E-06 |
| 25435 | 1272 | C1272P | D | 5.2e-06 | 2.7e-06 | 3.2e-06 | 3.700E-06 | 1.323E-06 | 2.700E-06 | 5.200E-06 |
| 25436 | 1272 | C1272S | D | 2.7e-06 | 1.3e-06 | 1.9e-06 | 1.967E-06 | 7.024E-07 | 1.300E-06 | 2.700E-06 |
| 25437 | 1272 | C1272T | D | 2.8e-06 | 9.9e-07 | 1.7e-06 | 1.830E-06 | 9.120E-07 | 9.900E-07 | 2.800E-06 |
| 25438 | 1272 | C1272W | D | 3.1e-06 | 1.5e-06 | 2.2e-06 | 2.267E-06 | 8.021E-07 | 1.500E-06 | 3.100E-06 |
| 25439 | 1272 | C1272Y | D | 2e-06 | 2.5e-06 | 3.1e-06 | 2.533E-06 | 5.508E-07 | 2.000E-06 | 3.100E-06 |
| 25440 | 1272 | C1272V | D | 2.3e-06 | 2.6e-06 | 4.3e-06 | 3.067E-06 | 1.079E-06 | 2.300E-06 | 4.300E-06 |
| 25441 | 1273 | E1273A | E | 3.4e-06 | 1.8e-06 | 2.6e-06 | 2.600E-06 | 8.000E-07 | 1.800E-06 | 3.400E-06 |
| 25442 | 1273 | E1273R | E | 2.1e-06 | 3.6e-06 | 2.7e-06 | 2.800E-06 | 7.550E-07 | 2.100E-06 | 3.600E-06 |
| 25443 | 1273 | E1273N | E | 1.4e-06 | 1.8e-06 | 1.1e-06 | 1.433E-06 | 3.512E-07 | 1.100E-06 | 1.800E-06 |
| 25444 | 1273 | E1273D | E | 4.1e-06 | 4.1e-06 | 2.6e-06 | 3.600E-06 | 8.660E-07 | 2.600E-06 | 4.100E-06 |
| 25445 | 1273 | E1273C | E | 2e-06 | 2.5e-06 | 2.8e-06 | 2.433E-06 | 4.041E-07 | 2.000E-06 | 2.800E-06 |
| 25446 | 1273 | E1273E | E | 2.7e-06 | 2.3e-06 | 3.2e-06 | 2.733E-06 | 4.509E-07 | 2.300E-06 | 3.200E-06 |
| 25447 | 1273 | E1273Q | E | 1.8e-06 | 1.6e-06 | 2.5e-06 | 1.967E-06 | 4.726E-07 | 1.600E-06 | 2.500E-06 |
| 25448 | 1273 | E1273G | E | 1.6e-06 | 3.3e-06 | 3.2e-06 | 2.700E-06 | 9.539E-07 | 1.600E-06 | 3.300E-06 |
| 25449 | 1273 | E1273H | E | 2.9e-06 | 2.7e-06 | 2.6e-06 | 2.733E-06 | 1.528E-07 | 2.600E-06 | 2.900E-06 |
| 25450 | 1273 | E1273I | E | 2.8e-06 | 3.8e-06 | 2.9e-06 | 3.167E-06 | 5.508E-07 | 2.800E-06 | 3.800E-06 |
| 25451 | 1273 | E1273L | E | 3.2e-06 | 2.9e-06 | 1.4e-06 | 2.500E-06 | 9.644E-07 | 1.400E-06 | 3.200E-06 |
| 25452 | 1273 | E1273K | E | 2.2e-06 | 3.3e-06 | 2.1e-06 | 2.533E-06 | 6.658E-07 | 2.100E-06 | 3.300E-06 |
| 25453 | 1273 | E1273M | E | 2.8e-06 | 2.5e-06 | 2e-06 | 2.433E-06 | 4.041E-07 | 2.000E-06 | 2.800E-06 |
| 25454 | 1273 | E1273F | E | 1.5e-06 | 1.9e-06 | 2.5e-06 | 1.967E-06 | 5.033E-07 | 1.500E-06 | 2.500E-06 |
| 25455 | 1273 | E1273P | E | 2.9e-06 | 2.1e-06 | 4.6e-06 | 3.200E-06 | 1.277E-06 | 2.100E-06 | 4.600E-06 |
| 25456 | 1273 | E1273S | E | 1.2e-06 | 1.8e-06 | 1.4e-06 | 1.467E-06 | 3.055E-07 | 1.200E-06 | 1.800E-06 |
| 25457 | 1273 | E1273T | E | 3.6e-06 | 2.3e-06 | 2.2e-06 | 2.700E-06 | 7.810E-07 | 2.200E-06 | 3.600E-06 |
| 25458 | 1273 | E1273W | E | 2.6e-06 | 1.7e-06 | 3e-06 | 2.433E-06 | 6.658E-07 | 1.700E-06 | 3.000E-06 |
| 25459 | 1273 | E1273Y | E | 2.2e-06 | 2.2e-06 | 1.8e-06 | 2.067E-06 | 2.309E-07 | 1.800E-06 | 2.200E-06 |
| 25460 | 1273 | E1273V | E | 3.9e-06 | 2.8e-06 | 2.5e-06 | 3.067E-06 | 7.371E-07 | 2.500E-06 | 3.900E-06 |
| 25461 | 1274 | V1274A | E | 2.3e-06 | 2.2e-06 | 2.2e-06 | 2.233E-06 | 5.774E-08 | 2.200E-06 | 2.300E-06 |
| 25462 | 1274 | V1274R | E | 2e-06 | 2.6e-06 | 1.7e-06 | 2.100E-06 | 4.583E-07 | 1.700E-06 | 2.600E-06 |
| 25463 | 1274 | V1274N | E | 1.1e-06 | 1.3e-06 | 1.1e-06 | 1.167E-06 | 1.155E-07 | 1.100E-06 | 1.300E-06 |
| 25464 | 1274 | V1274D | E | 1.5e-06 | 1e-06 | 1.7e-06 | 1.400E-06 | 3.606E-07 | 1.000E-06 | 1.700E-06 |
| 25465 | 1274 | V1274C | E | 2.3e-06 | 3.1e-06 | 1.3e-06 | 2.233E-06 | 9.018E-07 | 1.300E-06 | 3.100E-06 |
| 25466 | 1274 | V1274E | E | 1.5e-06 | 1.2e-06 | 1.2e-06 | 1.300E-06 | 1.732E-07 | 1.200E-06 | 1.500E-06 |
| 25467 | 1274 | V1274Q | E | 2.3e-06 | 2.4e-06 | 8.3e-07 | 1.843E-06 | 8.790E-07 | 8.300E-07 | 2.400E-06 |
| 25468 | 1274 | V1274G | E | 2.8e-06 | 4.5e-06 | 1.7e-06 | 3.000E-06 | 1.411E-06 | 1.700E-06 | 4.500E-06 |
| 25469 | 1274 | V1274H | E | 1.9e-06 | 2e-06 | 1.7e-06 | 1.867E-06 | 1.528E-07 | 1.700E-06 | 2.000E-06 |
| 25470 | 1274 | V1274I | E | 2.5e-06 | 1.6e-06 | 2.3e-06 | 2.133E-06 | 4.726E-07 | 1.600E-06 | 2.500E-06 |
| 25471 | 1274 | V1274L | E | 2.7e-06 | 2.9e-06 | 8e-07 | 2.133E-06 | 1.159E-06 | 8.000E-07 | 2.900E-06 |
| 25472 | 1274 | V1274K | E | 1.1e-06 | 3.2e-06 | 2.3e-06 | 2.200E-06 | 1.054E-06 | 1.100E-06 | 3.200E-06 |
| 25473 | 1274 | V1274M | E | 1.9e-06 | 1.6e-06 | 3e-06 | 2.167E-06 | 7.371E-07 | 1.600E-06 | 3.000E-06 |
| 25474 | 1274 | V1274F | E | 1.9e-06 | 2.2e-06 | 2.6e-06 | 2.233E-06 | 3.512E-07 | 1.900E-06 | 2.600E-06 |
| 25475 | 1274 | V1274P | E | 2.3e-06 | 2.4e-06 | 2.4e-06 | 2.367E-06 | 5.774E-08 | 2.300E-06 | 2.400E-06 |
| 25476 | 1274 | V1274S | E | 1.6e-06 | 1.1e-06 | 2.9e-06 | 1.867E-06 | 9.292E-07 | 1.100E-06 | 2.900E-06 |
| 25477 | 1274 | V1274T | E | 2e-06 | 2.6e-06 | 1.5e-06 | 2.033E-06 | 5.508E-07 | 1.500E-06 | 2.600E-06 |
| 25478 | 1274 | V1274W | E | 2.6e-06 | 2.7e-06 | 2e-06 | 2.433E-06 | 3.786E-07 | 2.000E-06 | 2.700E-06 |
| 25479 | 1274 | V1274Y | E | 2.6e-06 | 1.7e-06 | 3e-06 | 2.433E-06 | 6.658E-07 | 1.700E-06 | 3.000E-06 |
| 25480 | 1274 | V1274V | E | 1.5e-06 | 4.3e-06 | 2.2e-06 | 2.667E-06 | 1.457E-06 | 1.500E-06 | 4.300E-06 |
| 25481 | 1275 | Q1275A | E | 3.4e-06 | 1.9e-06 | 2.8e-06 | 2.700E-06 | 7.550E-07 | 1.900E-06 | 3.400E-06 |
| 25482 | 1275 | Q1275R | E | 2.5e-06 | 2.3e-06 | 3.2e-06 | 2.667E-06 | 4.726E-07 | 2.300E-06 | 3.200E-06 |
| 25483 | 1275 | Q1275N | E | 1.7e-06 | 3.7e-06 | 1.4e-06 | 2.267E-06 | 1.250E-06 | 1.400E-06 | 3.700E-06 |
| 25484 | 1275 | Q1275D | E | 2.6e-06 | 3.1e-06 | 2.3e-06 | 2.667E-06 | 4.041E-07 | 2.300E-06 | 3.100E-06 |
| 25485 | 1275 | Q1275C | E | 4e-06 | 2.5e-06 | 2.1e-06 | 2.867E-06 | 1.002E-06 | 2.100E-06 | 4.000E-06 |
| 25486 | 1275 | Q1275E | E | 1.4e-06 | 4.7e-06 | 3e-06 | 3.033E-06 | 1.650E-06 | 1.400E-06 | 4.700E-06 |
| 25487 | 1275 | Q1275Q | E | 1.8e-06 | 2.1e-06 | 2.7e-06 | 2.200E-06 | 4.583E-07 | 1.800E-06 | 2.700E-06 |
| 25488 | 1275 | Q1275G | E | 4e-06 | 1.7e-06 | 2.3e-06 | 2.667E-06 | 1.193E-06 | 1.700E-06 | 4.000E-06 |
| 25489 | 1275 | Q1275H | E | 2.7e-06 | 1.2e-06 | 3.5e-06 | 2.467E-06 | 1.168E-06 | 1.200E-06 | 3.500E-06 |
| 25490 | 1275 | Q1275I | E | 2.8e-06 | 3.2e-06 | 2.6e-06 | 2.867E-06 | 3.055E-07 | 2.600E-06 | 3.200E-06 |
| 25491 | 1275 | Q1275L | E | 1.4e-06 | 2.8e-06 | 1.9e-06 | 2.033E-06 | 7.095E-07 | 1.400E-06 | 2.800E-06 |
| 25492 | 1275 | Q1275K | E | 3.2e-06 | 2.3e-06 | 2.1e-06 | 2.533E-06 | 5.859E-07 | 2.100E-06 | 3.200E-06 |
| 25493 | 1275 | Q1275M | E | 4.6e-06 | 2.5e-06 | 2e-06 | 3.033E-06 | 1.380E-06 | 2.000E-06 | 4.600E-06 |
| 25494 | 1275 | Q1275F | E | 1.7e-06 | 2.9e-06 | 2.8e-06 | 2.467E-06 | 6.658E-07 | 1.700E-06 | 2.900E-06 |
| 25495 | 1275 | Q1275P | E | 1.9e-06 | 2.2e-06 | 4e-06 | 2.700E-06 | 1.136E-06 | 1.900E-06 | 4.000E-06 |
| 25496 | 1275 | Q1275S | E | 1.6e-06 | 3e-06 | 4.2e-06 | 2.933E-06 | 1.301E-06 | 1.600E-06 | 4.200E-06 |
| 25497 | 1275 | Q1275T | E | 2.3e-06 | 2.7e-06 | 3.4e-06 | 2.800E-06 | 5.568E-07 | 2.300E-06 | 3.400E-06 |
| 25498 | 1275 | Q1275W | E | 2.2e-06 | 4.4e-06 | 1.1e-06 | 2.567E-06 | 1.680E-06 | 1.100E-06 | 4.400E-06 |
| 25499 | 1275 | Q1275Y | E | 2.9e-06 | 1.5e-06 | 3.1e-06 | 2.500E-06 | 8.718E-07 | 1.500E-06 | 3.100E-06 |

| | | | | | | | | | | |
|---|---|---|---|---|---|---|---|---|---|---|
| 25500 | 1275 | Q1275V | E | 9.9e-07 | 2.5e-06 | 2.2e-06 | 1.897E-06 | 7.994E-07 | 9.900E-07 | 2.500E-06 |
| 25501 | 1276 | L1276A | E | 2.8e-06 | 3.4e-06 | 2.8e-06 | 3.000E-06 | 3.464E-07 | 2.800E-06 | 3.400E-06 |
| 25502 | 1276 | L1276R | E | 2.5e-06 | 2.4e-06 | 3.9e-06 | 2.933E-06 | 8.386E-07 | 2.400E-06 | 3.900E-06 |
| 25503 | 1276 | L1276N | E | 2.5e-06 | 2e-06 | 2.9e-06 | 2.467E-06 | 4.509E-07 | 2.000E-06 | 2.900E-06 |
| 25504 | 1276 | L1276D | E | 4.4e-06 | 1e-06 | 2.3e-06 | 2.567E-06 | 1.716E-06 | 1.000E-06 | 4.400E-06 |
| 25505 | 1276 | L1276C | E | 2.1e-06 | 2.4e-06 | 3e-06 | 2.500E-06 | 4.583E-07 | 2.100E-06 | 3.000E-06 |
| 25506 | 1276 | L1276E | E | 1.8e-06 | 3e-06 | 2e-06 | 2.267E-06 | 6.429E-07 | 1.800E-06 | 3.000E-06 |
| 25507 | 1276 | L1276Q | E | 3e-06 | 2.9e-06 | 2.9e-06 | 2.933E-06 | 5.774E-08 | 2.900E-06 | 3.000E-06 |
| 25508 | 1276 | L1276G | E | 2.4e-06 | 3.8e-06 | 2.1e-06 | 2.767E-06 | 9.074E-07 | 2.100E-06 | 3.800E-06 |
| 25509 | 1276 | L1276H | E | 1.7e-06 | 3.7e-06 | 4.2e-06 | 3.200E-06 | 1.323E-06 | 1.700E-06 | 4.200E-06 |
| 25510 | 1276 | L1276I | E | 1.8e-06 | 2.8e-06 | 2.7e-06 | 2.433E-06 | 5.508E-07 | 1.800E-06 | 2.800E-06 |
| 25511 | 1276 | L1276L | E | 2.6e-06 | 1.8e-06 | 5.5e-06 | 3.300E-06 | 1.947E-06 | 1.800E-06 | 5.500E-06 |
| 25512 | 1276 | L1276K | E | 3.5e-06 | 5.2e-06 | 2e-06 | 3.567E-06 | 1.601E-06 | 2.000E-06 | 5.200E-06 |
| 25513 | 1276 | L1276M | E | 4e-06 | 2.6e-06 | 1.7e-06 | 2.767E-06 | 1.159E-06 | 1.700E-06 | 4.000E-06 |
| 25514 | 1276 | L1276F | E | 1.9e-06 | 2.2e-06 | 1e-06 | 1.700E-06 | 6.245E-07 | 1.000E-06 | 2.200E-06 |
| 25515 | 1276 | L1276P | E | 1.9e-06 | 2.5e-06 | 2.2e-06 | 2.200E-06 | 3.000E-07 | 1.900E-06 | 2.500E-06 |
| 25516 | 1276 | L1276S | E | 2.7e-06 | 2.5e-06 | 1.7e-06 | 2.300E-06 | 5.292E-07 | 1.700E-06 | 2.700E-06 |
| 25517 | 1276 | L1276T | E | 2.4e-06 | 2.9e-06 | 2.7e-06 | 2.667E-06 | 2.517E-07 | 2.400E-06 | 2.900E-06 |
| 25518 | 1276 | L1276W | E | 5.6e-06 | 1.8e-06 | 2.8e-06 | 3.400E-06 | 1.970E-06 | 1.800E-06 | 5.600E-06 |
| 25519 | 1276 | L1276Y | E | 3.2e-06 | 2.3e-06 | 2.1e-06 | 2.533E-06 | 5.859E-07 | 2.100E-06 | 3.200E-06 |
| 25520 | 1276 | L1276V | E | 2.6e-06 | 2.1e-06 | 3e-06 | 2.567E-06 | 4.509E-07 | 2.100E-06 | 3.000E-06 |
| 25521 | 1277 | V1277A | E | 2.2e-06 | 2.7e-06 | 2.2e-06 | 2.367E-06 | 2.887E-07 | 2.200E-06 | 2.700E-06 |
| 25522 | 1277 | V1277R | E | 2.3e-06 | 3e-06 | 2.5e-06 | 2.600E-06 | 3.606E-07 | 2.300E-06 | 3.000E-06 |
| 25523 | 1277 | V1277N | E | 2e-06 | 1.6e-06 | 2.4e-06 | 2.000E-06 | 4.000E-07 | 1.600E-06 | 2.400E-06 |
| 25524 | 1277 | V1277D | E | 2.1e-06 | 2.1e-06 | 2.1e-06 | 2.100E-06 | 0.000E+00 | 2.100E-06 | 2.100E-06 |
| 25525 | 1277 | V1277C | E | 3.3e-06 | 3.5e-06 | 7.6e-07 | 2.520E-06 | 1.527E-06 | 7.600E-07 | 3.500E-06 |
| 25526 | 1277 | V1277E | E | 2e-06 | 1.9e-06 | 2e-06 | 1.967E-06 | 5.774E-08 | 1.900E-06 | 2.000E-06 |
| 25527 | 1277 | V1277Q | E | 2.6e-06 | 3e-06 | 5.9e-06 | 3.833E-06 | 1.801E-06 | 2.600E-06 | 5.900E-06 |
| 25528 | 1277 | V1277G | E | 3e-06 | 2.4e-06 | 4.2e-06 | 3.200E-06 | 9.165E-07 | 2.400E-06 | 4.200E-06 |
| 25529 | 1277 | V1277H | E | 2.5e-06 | 2.4e-06 | 1.7e-06 | 2.200E-06 | 4.359E-07 | 1.700E-06 | 2.500E-06 |
| 25530 | 1277 | V1277I | E | 2.1e-06 | 2.4e-06 | 1.1e-06 | 1.867E-06 | 6.807E-07 | 1.100E-06 | 2.400E-06 |
| 25531 | 1277 | V1277L | E | 3.3e-06 | 3.5e-06 | 2.6e-06 | 3.133E-06 | 4.726E-07 | 2.600E-06 | 3.500E-06 |
| 25532 | 1277 | V1277K | E | 4.6e-06 | 2.8e-06 | 2.3e-06 | 3.233E-06 | 1.210E-06 | 2.300E-06 | 4.600E-06 |
| 25533 | 1277 | V1277M | E | 3.9e-06 | 1.6e-06 | 6.5e-06 | 4.000E-06 | 2.452E-06 | 1.600E-06 | 6.500E-06 |
| 25534 | 1277 | V1277F | E | 2.2e-06 | 3e-06 | 1.7e-06 | 2.300E-06 | 6.557E-07 | 1.700E-06 | 3.000E-06 |
| 25535 | 1277 | V1277P | E | 2.9e-06 | 2.1e-06 | 1.5e-06 | 2.167E-06 | 7.024E-07 | 1.500E-06 | 2.900E-06 |
| 25536 | 1277 | V1277S | E | 3.8e-06 | 2.2e-06 | 2.1e-06 | 2.700E-06 | 9.539E-07 | 2.100E-06 | 3.800E-06 |
| 25537 | 1277 | V1277T | E | 4.9e-06 | 1.7e-06 | 3.7e-06 | 3.433E-06 | 1.617E-06 | 1.700E-06 | 4.900E-06 |
| 25538 | 1277 | V1277W | E | 2e-06 | 2.4e-06 | 2.8e-06 | 2.400E-06 | 4.000E-07 | 2.000E-06 | 2.800E-06 |
| 25539 | 1277 | V1277Y | E | 9e-07 | 1.9e-06 | 1.6e-06 | 1.467E-06 | 5.132E-07 | 9.000E-07 | 1.900E-06 |
| 25540 | 1277 | V1277V | E | 2.1e-06 | 6.9e-07 | 2.5e-06 | 1.763E-06 | 9.508E-07 | 6.900E-07 | 2.500E-06 |
| 25541 | 1278 | E1278A | E | 2e-06 | 2.8e-06 | 1.8e-06 | 2.200E-06 | 5.292E-07 | 1.800E-06 | 2.800E-06 |
| 25542 | 1278 | E1278R | E | 1.9e-06 | 1.9e-06 | 3.2e-06 | 2.333E-06 | 7.506E-07 | 1.900E-06 | 3.200E-06 |
| 25543 | 1278 | E1278N | E | 4.5e-06 | 1.6e-06 | 3.2e-06 | 3.100E-06 | 1.453E-06 | 1.600E-06 | 4.500E-06 |
| 25544 | 1278 | E1278D | E | 1.5e-06 | 3.8e-06 | 1.5e-06 | 2.267E-06 | 1.328E-06 | 1.500E-06 | 3.800E-06 |
| 25545 | 1278 | E1278C | E | 2.1e-06 | 2e-06 | 1.6e-06 | 1.900E-06 | 2.646E-07 | 1.600E-06 | 2.100E-06 |
| 25546 | 1278 | E1278E | E | 2.1e-06 | 3.2e-06 | 2.6e-06 | 2.633E-06 | 5.508E-07 | 2.100E-06 | 3.200E-06 |
| 25547 | 1278 | E1278Q | E | 3.1e-06 | 2.1e-06 | 2.6e-06 | 2.600E-06 | 5.000E-07 | 2.100E-06 | 3.100E-06 |
| 25548 | 1278 | E1278G | E | 5.4e-06 | 4.7e-06 | 9.6e-07 | 3.687E-06 | 2.387E-06 | 9.600E-07 | 5.400E-06 |
| 25549 | 1278 | E1278H | E | 2e-06 | 3.3e-06 | 1.7e-06 | 2.333E-06 | 8.505E-07 | 1.700E-06 | 3.300E-06 |
| 25550 | 1278 | E1278I | E | 1.9e-06 | 2.6e-06 | 2.8e-06 | 2.433E-06 | 4.726E-07 | 1.900E-06 | 2.800E-06 |
| 25551 | 1278 | E1278L | E | 3.7e-06 | 2.3e-06 | 3e-06 | 3.000E-06 | 7.000E-07 | 2.300E-06 | 3.700E-06 |
| 25552 | 1278 | E1278K | E | 2.4e-06 | 4.7e-06 | 2e-06 | 3.033E-06 | 1.457E-06 | 2.000E-06 | 4.700E-06 |
| 25553 | 1278 | E1278M | E | 1.8e-06 | 1.9e-06 | 3.1e-06 | 2.267E-06 | 7.234E-07 | 1.800E-06 | 3.100E-06 |
| 25554 | 1278 | E1278F | E | 3.7e-06 | 2.6e-06 | 1.2e-06 | 2.500E-06 | 1.253E-06 | 1.200E-06 | 3.700E-06 |
| 25555 | 1278 | E1278P | E | 2.8e-06 | 3.3e-06 | 2e-06 | 2.700E-06 | 6.557E-07 | 2.000E-06 | 3.300E-06 |
| 25556 | 1278 | E1278S | E | 2.6e-06 | 3.9e-06 | 3e-06 | 3.167E-06 | 6.658E-07 | 2.600E-06 | 3.900E-06 |
| 25557 | 1278 | E1278T | E | 2.3e-06 | 3.5e-06 | 2.3e-06 | 2.700E-06 | 6.928E-07 | 2.300E-06 | 3.500E-06 |
| 25558 | 1278 | E1278W | E | 1.4e-06 | 4.4e-06 | 4.7e-06 | 3.500E-06 | 1.825E-06 | 1.400E-06 | 4.700E-06 |
| 25559 | 1278 | E1278Y | E | 3e-06 | 2.9e-06 | 3.2e-06 | 3.033E-06 | 1.528E-07 | 2.900E-06 | 3.200E-06 |
| 25560 | 1278 | E1278V | E | 3.2e-06 | 2.3e-06 | 2.2e-06 | 2.567E-06 | 5.508E-07 | 2.200E-06 | 3.200E-06 |
| 25561 | 1279 | S1279A | E | 2.5e-06 | 5.3e-06 | 1.2e-06 | 3.000E-06 | 2.095E-06 | 1.200E-06 | 5.300E-06 |
| 25562 | 1279 | S1279R | E | 9.5e-07 | 2.8e-06 | 3.6e-06 | 2.450E-06 | 1.359E-06 | 9.500E-07 | 3.600E-06 |
| 25563 | 1279 | S1279N | E | 3.7e-06 | 4.2e-06 | 1.9e-06 | 3.267E-06 | 1.210E-06 | 1.900E-06 | 4.200E-06 |
| 25564 | 1279 | S1279D | E | 2.9e-06 | 2.7e-06 | 1.6e-06 | 2.400E-06 | 7.000E-07 | 1.600E-06 | 2.900E-06 |
| 25565 | 1279 | S1279C | E | 1.8e-06 | 1.5e-06 | 1.6e-06 | 1.633E-06 | 1.528E-07 | 1.500E-06 | 1.800E-06 |
| 25566 | 1279 | S1279E | E | 3.5e-06 | 2.4e-06 | 3.7e-06 | 3.200E-06 | 7.000E-07 | 2.400E-06 | 3.700E-06 |
| 25567 | 1279 | S1279Q | E | 2.8e-06 | 1.7e-06 | 1.5e-06 | 2.000E-06 | 7.000E-07 | 1.500E-06 | 2.800E-06 |
| 25568 | 1279 | S1279G | E | 1.3e-06 | 2.3e-06 | 2.4e-06 | 2.000E-06 | 6.083E-07 | 1.300E-06 | 2.400E-06 |
| 25569 | 1279 | S1279H | E | 3.7e-06 | 1.2e-06 | 3.4e-06 | 2.767E-06 | 1.365E-06 | 1.200E-06 | 3.700E-06 |
| 25570 | 1279 | S1279I | E | 2.6e-06 | 2.2e-06 | 1.9e-06 | 2.233E-06 | 3.512E-07 | 1.900E-06 | 2.600E-06 |
| 25571 | 1279 | S1279L | E | 5.5e-06 | 2.3e-06 | 3.6e-06 | 3.800E-06 | 1.609E-06 | 2.300E-06 | 5.500E-06 |
| 25572 | 1279 | S1279K | E | 3.7e-06 | 1.8e-06 | 1.8e-06 | 2.433E-06 | 1.097E-06 | 1.800E-06 | 3.700E-06 |
| 25573 | 1279 | S1279M | E | 1.3e-06 | 2.1e-06 | 3.5e-06 | 2.300E-06 | 1.114E-06 | 1.300E-06 | 3.500E-06 |
| 25574 | 1279 | S1279F | E | 4.8e-06 | 1e-06 | 1.8e-06 | 2.533E-06 | 2.003E-06 | 1.000E-06 | 4.800E-06 |
| 25575 | 1279 | S1279P | E | 1.8e-06 | 1.6e-06 | 3e-06 | 2.133E-06 | 7.572E-07 | 1.600E-06 | 3.000E-06 |
| 25576 | 1279 | S1279S | E | 3e-06 | 2.8e-06 | 3.3e-06 | 3.033E-06 | 2.517E-07 | 2.800E-06 | 3.300E-06 |
| 25577 | 1279 | S1279T | E | 2.2e-06 | 1.5e-06 | 3.8e-06 | 2.500E-06 | 1.179E-06 | 1.500E-06 | 3.800E-06 |
| 25578 | 1279 | S1279W | E | 1.7e-06 | 1.5e-06 | 3.5e-06 | 2.233E-06 | 1.102E-06 | 1.500E-06 | 3.500E-06 |
| 25579 | 1279 | S1279Y | E | 2.1e-06 | 3.3e-06 | 1.7e-06 | 2.367E-06 | 8.327E-07 | 1.700E-06 | 3.300E-06 |
| 25580 | 1279 | S1279V | E | 2.9e-06 | 2.4e-06 | 2.3e-06 | 2.533E-06 | 3.215E-07 | 2.300E-06 | 2.900E-06 |
| 25581 | 1280 | G1280A | E | 1.8e-06 | 2e-06 | 1.5e-06 | 1.767E-06 | 2.517E-07 | 1.500E-06 | 2.000E-06 |
| 25582 | 1280 | G1280R | E | 1.1e-06 | 1.9e-06 | 2.4e-06 | 1.800E-06 | 6.557E-07 | 1.100E-06 | 2.400E-06 |
| 25583 | 1280 | G1280N | E | 2.5e-06 | 2.4e-06 | 4.4e-06 | 3.100E-06 | 1.127E-06 | 2.400E-06 | 4.400E-06 |
| 25584 | 1280 | G1280D | E | 1.9e-06 | 1.8e-06 | 1.7e-06 | 1.800E-06 | 1.000E-07 | 1.700E-06 | 1.900E-06 |

| | | | | | | | | | | |
|---|---|---|---|---|---|---|---|---|---|---|
| 25585 | 1280 | G1280C | E | 2.2e-06 | 3.1e-06 | 2e-06 | 2.433E-06 | 5.859E-07 | 2.000E-06 | 3.100E-06 |
| 25586 | 1280 | G1280E | E | 1.5e-06 | 3e-06 | 2.2e-06 | 2.233E-06 | 7.506E-07 | 1.500E-06 | 3.000E-06 |
| 25587 | 1280 | G1280Q | E | 2.4e-06 | 2.7e-06 | 1.6e-06 | 2.233E-06 | 5.686E-07 | 1.600E-06 | 2.700E-06 |
| 25588 | 1280 | G1280G | E | 3.1e-06 | 3.3e-06 | 1.3e-06 | 2.567E-06 | 1.102E-06 | 1.300E-06 | 3.300E-06 |
| 25589 | 1280 | G1280H | E | 2.8e-06 | 1.1e-06 | 2.2e-06 | 2.033E-06 | 8.622E-07 | 1.100E-06 | 2.800E-06 |
| 25590 | 1280 | G1280I | E | 4e-06 | 2.5e-06 | 3.4e-06 | 3.300E-06 | 7.550E-07 | 2.500E-06 | 4.000E-06 |
| 25591 | 1280 | G1280L | E | 3.5e-06 | 2.8e-06 | 2.6e-06 | 2.967E-06 | 4.726E-07 | 2.600E-06 | 3.500E-06 |
| 25592 | 1280 | G1280K | E | 2.5e-06 | 2.1e-06 | 2e-06 | 2.200E-06 | 2.646E-07 | 2.000E-06 | 2.500E-06 |
| 25593 | 1280 | G1280M | E | 3e-06 | 2.4e-06 | 2e-06 | 2.467E-06 | 5.033E-07 | 2.000E-06 | 3.000E-06 |
| 25594 | 1280 | G1280F | E | 4.3e-06 | 1.3e-06 | 2e-06 | 2.533E-06 | 1.570E-06 | 1.300E-06 | 4.300E-06 |
| 25595 | 1280 | G1280P | E | 3.8e-06 | 2.2e-06 | 2.1e-06 | 2.700E-06 | 9.539E-07 | 2.100E-06 | 3.800E-06 |
| 25596 | 1280 | G1280S | E | 2.2e-06 | 2e-06 | 2.7e-06 | 2.300E-06 | 3.606E-07 | 2.000E-06 | 2.700E-06 |
| 25597 | 1280 | G1280T | E | 4.2e-06 | 2.8e-06 | 2.4e-06 | 3.133E-06 | 9.452E-07 | 2.400E-06 | 4.200E-06 |
| 25598 | 1280 | G1280W | E | 2.5e-06 | 2.3e-06 | 2.2e-06 | 2.333E-06 | 1.528E-07 | 2.200E-06 | 2.500E-06 |
| 25599 | 1280 | G1280Y | E | 2.3e-06 | 2.4e-06 | 3.1e-06 | 2.600E-06 | 4.359E-07 | 2.300E-06 | 3.100E-06 |
| 25600 | 1280 | G1280V | E | 1.7e-06 | 2.9e-06 | 3.7e-06 | 2.767E-06 | 1.007E-06 | 1.700E-06 | 3.700E-06 |
| 25601 | 1281 | G1281A | E | 1.3e-06 | 2.6e-06 | 2.8e-06 | 2.233E-06 | 8.145E-07 | 1.300E-06 | 2.800E-06 |
| 25602 | 1281 | G1281R | E | 1.6e-06 | 2.4e-06 | 2.1e-06 | 2.033E-06 | 4.041E-07 | 1.600E-06 | 2.400E-06 |
| 25603 | 1281 | G1281N | E | 1.2e-06 | 2.5e-06 | 1.8e-06 | 1.833E-06 | 6.506E-07 | 1.200E-06 | 2.500E-06 |
| 25604 | 1281 | G1281D | E | 2.9e-06 | 1.9e-06 | 1.4e-06 | 2.067E-06 | 7.638E-07 | 1.400E-06 | 2.900E-06 |
| 25605 | 1281 | G1281C | E | 2.1e-06 | 2.4e-06 | 2.5e-06 | 2.333E-06 | 2.082E-07 | 2.100E-06 | 2.500E-06 |
| 25606 | 1281 | G1281E | E | 1.5e-06 | 2.1e-06 | 2.2e-06 | 1.933E-06 | 3.786E-07 | 1.500E-06 | 2.200E-06 |
| 25607 | 1281 | G1281Q | E | 2.2e-06 | 1.8e-06 | 1.5e-06 | 1.833E-06 | 3.512E-07 | 1.500E-06 | 2.200E-06 |
| 25608 | 1281 | G1281G | E | 1.1e-06 | 4e-06 | 4.7e-06 | 3.267E-06 | 1.909E-06 | 1.100E-06 | 4.700E-06 |
| 25609 | 1281 | G1281H | E | 1.5e-06 | 1.7e-06 | 2.4e-06 | 1.867E-06 | 4.726E-07 | 1.500E-06 | 2.400E-06 |
| 25610 | 1281 | G1281I | E | 1.1e-06 | 3e-06 | 2.7e-06 | 2.267E-06 | 1.021E-06 | 1.100E-06 | 3.000E-06 |
| 25611 | 1281 | G1281L | E | 1.4e-06 | 3.2e-06 | 3.5e-06 | 2.700E-06 | 1.136E-06 | 1.400E-06 | 3.500E-06 |
| 25612 | 1281 | G1281K | E | 2.5e-06 | 2.7e-06 | 3.3e-06 | 2.833E-06 | 4.163E-07 | 2.500E-06 | 3.300E-06 |
| 25613 | 1281 | G1281M | E | 2.2e-06 | 5e-06 | 2.2e-06 | 3.133E-06 | 1.617E-06 | 2.200E-06 | 5.000E-06 |
| 25614 | 1281 | G1281F | E | 3.9e-06 | 2e-06 | 1.2e-06 | 2.367E-06 | 1.387E-06 | 1.200E-06 | 3.900E-06 |
| 25615 | 1281 | G1281P | E | 2.1e-06 | 2.2e-06 | 2.3e-06 | 2.200E-06 | 1.000E-07 | 2.100E-06 | 2.300E-06 |
| 25616 | 1281 | G1281S | E | 2.8e-06 | 2.8e-06 | 3.2e-06 | 2.933E-06 | 2.309E-07 | 2.800E-06 | 3.200E-06 |
| 25617 | 1281 | G1281T | E | 2.5e-06 | 1.2e-06 | 1.9e-06 | 1.867E-06 | 6.506E-07 | 1.200E-06 | 2.500E-06 |
| 25618 | 1281 | G1281W | E | 2e-06 | 2.8e-06 | 2.5e-06 | 2.433E-06 | 4.041E-07 | 2.000E-06 | 2.800E-06 |
| 25619 | 1281 | G1281Y | E | 3.2e-06 | 2.7e-06 | 1.6e-06 | 2.500E-06 | 8.185E-07 | 1.600E-06 | 3.200E-06 |
| 25620 | 1281 | G1281V | E | 2.2e-06 | 1.9e-06 | 1.2e-06 | 1.767E-06 | 5.132E-07 | 1.200E-06 | 2.200E-06 |
| 25621 | 1282 | G1282A | E | 2.9e-06 | 2.7e-06 | 2.1e-06 | 2.567E-06 | 4.163E-07 | 2.100E-06 | 2.900E-06 |
| 25622 | 1282 | G1282R | E | 2.5e-06 | 3.2e-06 | 4.1e-06 | 3.267E-06 | 8.021E-07 | 2.500E-06 | 4.100E-06 |
| 25623 | 1282 | G1282N | E | 1.5e-06 | 2.4e-06 | 2.2e-06 | 2.033E-06 | 4.726E-07 | 1.500E-06 | 2.400E-06 |
| 25624 | 1282 | G1282D | E | 2.4e-06 | 2.2e-06 | 2.9e-06 | 2.500E-06 | 3.606E-07 | 2.200E-06 | 2.900E-06 |
| 25625 | 1282 | G1282C | E | 1.5e-06 | 2.2e-06 | 1.5e-06 | 1.733E-06 | 4.041E-07 | 1.500E-06 | 2.200E-06 |
| 25626 | 1282 | G1282E | E | 3.6e-06 | 1.4e-06 | 1.8e-06 | 2.267E-06 | 1.172E-06 | 1.400E-06 | 3.600E-06 |
| 25627 | 1282 | G1282Q | E | 2.6e-06 | 1.5e-06 | 1.3e-06 | 1.800E-06 | 7.000E-07 | 1.300E-06 | 2.600E-06 |
| 25628 | 1282 | G1282G | E | 2e-06 | 1.7e-06 | 2.5e-06 | 2.067E-06 | 4.041E-07 | 1.700E-06 | 2.500E-06 |
| 25629 | 1282 | G1282H | E | 1.2e-06 | 3.6e-06 | 1.7e-06 | 2.167E-06 | 1.266E-06 | 1.200E-06 | 3.600E-06 |
| 25630 | 1282 | G1282I | E | 1.9e-06 | 3.4e-06 | 2.8e-06 | 2.700E-06 | 7.550E-07 | 1.900E-06 | 3.400E-06 |
| 25631 | 1282 | G1282L | E | 1.6e-06 | 3e-06 | 3e-06 | 2.533E-06 | 8.083E-07 | 1.600E-06 | 3.000E-06 |
| 25632 | 1282 | G1282K | E | 2.8e-06 | 3.1e-06 | 3.3e-06 | 3.067E-06 | 2.517E-07 | 2.800E-06 | 3.300E-06 |
| 25633 | 1282 | G1282M | E | 2.2e-06 | 2.5e-06 | 1.3e-06 | 2.000E-06 | 6.245E-07 | 1.300E-06 | 2.500E-06 |
| 25634 | 1282 | G1282F | E | 2.9e-06 | 1.9e-06 | 4e-06 | 2.933E-06 | 1.050E-06 | 1.900E-06 | 4.000E-06 |
| 25635 | 1282 | G1282P | E | 4.2e-06 | 3.8e-06 | 2.2e-06 | 3.400E-06 | 1.058E-06 | 2.200E-06 | 4.200E-06 |
| 25636 | 1282 | G1282S | E | 3e-06 | 1.3e-06 | 2.8e-06 | 2.367E-06 | 9.292E-07 | 1.300E-06 | 3.000E-06 |
| 25637 | 1282 | G1282T | E | 3e-06 | 1.8e-06 | 2.2e-06 | 2.333E-06 | 6.110E-07 | 1.800E-06 | 3.000E-06 |
| 25638 | 1282 | G1282W | E | 3.4e-06 | 1.9e-06 | 1.9e-06 | 2.400E-06 | 8.660E-07 | 1.900E-06 | 3.400E-06 |
| 25639 | 1282 | G1282Y | E | 2.9e-06 | 1.9e-06 | 3.8e-06 | 2.867E-06 | 9.504E-07 | 1.900E-06 | 3.800E-06 |
| 25640 | 1282 | G1282V | E | 2.7e-06 | 2.5e-06 | 2.3e-06 | 2.500E-06 | 2.000E-07 | 2.300E-06 | 2.700E-06 |
| 25641 | 1283 | L1283A | E | 2.4e-06 | 2.7e-06 | 2.5e-06 | 2.533E-06 | 1.528E-07 | 2.400E-06 | 2.700E-06 |
| 25642 | 1283 | L1283R | E | 2.7e-06 | 1.8e-06 | 3.1e-06 | 2.533E-06 | 6.658E-07 | 1.800E-06 | 3.100E-06 |
| 25643 | 1283 | L1283N | E | 2.3e-06 | 3e-06 | 3.6e-06 | 2.967E-06 | 6.506E-07 | 2.300E-06 | 3.600E-06 |
| 25644 | 1283 | L1283D | E | 1.7e-06 | 2.4e-06 | 2.1e-06 | 2.067E-06 | 3.512E-07 | 1.700E-06 | 2.400E-06 |
| 25645 | 1283 | L1283C | E | 1.2e-06 | 2e-06 | 1.7e-06 | 1.633E-06 | 4.041E-07 | 1.200E-06 | 2.000E-06 |
| 25646 | 1283 | L1283E | E | 1.5e-06 | 2.2e-06 | 1.2e-06 | 1.633E-06 | 5.132E-07 | 1.200E-06 | 2.200E-06 |
| 25647 | 1283 | L1283Q | E | 2.2e-06 | 2.1e-06 | 2e-06 | 2.100E-06 | 1.000E-07 | 2.000E-06 | 2.200E-06 |
| 25648 | 1283 | L1283G | E | 2.7e-06 | 2.2e-06 | 3e-06 | 2.633E-06 | 4.041E-07 | 2.200E-06 | 3.000E-06 |
| 25649 | 1283 | L1283H | E | 2.2e-06 | 4e-06 | 1.8e-06 | 2.667E-06 | 1.172E-06 | 1.800E-06 | 4.000E-06 |
| 25650 | 1283 | L1283I | E | 2.4e-06 | 1.7e-06 | 4.4e-06 | 2.833E-06 | 1.401E-06 | 1.700E-06 | 4.400E-06 |
| 25651 | 1283 | L1283L | E | 1.5e-06 | 2.5e-06 | 1.7e-06 | 1.900E-06 | 5.292E-07 | 1.500E-06 | 2.500E-06 |
| 25652 | 1283 | L1283K | E | 2.1e-06 | 2.3e-06 | 2.2e-06 | 2.200E-06 | 1.000E-07 | 2.100E-06 | 2.300E-06 |
| 25653 | 1283 | L1283M | E | 2.4e-06 | 4.7e-06 | 2.2e-06 | 3.100E-06 | 1.389E-06 | 2.200E-06 | 4.700E-06 |
| 25654 | 1283 | L1283F | E | 1.8e-06 | 3e-06 | 2.1e-06 | 2.300E-06 | 6.245E-07 | 1.800E-06 | 3.000E-06 |
| 25655 | 1283 | L1283P | E | 2e-06 | 2.6e-06 | 3.3e-06 | 2.633E-06 | 6.506E-07 | 2.000E-06 | 3.300E-06 |
| 25656 | 1283 | L1283S | E | 8.4e-07 | 2.8e-06 | 5.6e-06 | 3.080E-06 | 2.392E-06 | 8.400E-07 | 5.600E-06 |
| 25657 | 1283 | L1283T | E | 2.2e-06 | 3.1e-06 | 2e-06 | 2.433E-06 | 5.859E-07 | 2.000E-06 | 3.100E-06 |
| 25658 | 1283 | L1283W | E | 2.2e-06 | 2.9e-06 | 2e-06 | 2.367E-06 | 4.726E-07 | 2.000E-06 | 2.900E-06 |
| 25659 | 1283 | L1283Y | E | 4.8e-06 | 1.8e-06 | 3.2e-06 | 3.267E-06 | 1.501E-06 | 1.800E-06 | 4.800E-06 |
| 25660 | 1283 | L1283V | E | 1.4e-06 | 2e-06 | 1.3e-06 | 1.567E-06 | 3.786E-07 | 1.300E-06 | 2.000E-06 |
| 25661 | 1284 | V1284A | E | 2.6e-06 | 4.1e-06 | 2.3e-06 | 3.000E-06 | 9.644E-07 | 2.300E-06 | 4.100E-06 |
| 25662 | 1284 | V1284R | E | 1.1e-06 | 2.4e-06 | 3.2e-06 | 2.233E-06 | 1.060E-06 | 1.100E-06 | 3.200E-06 |
| 25663 | 1284 | V1284N | E | 3e-06 | 2.1e-06 | 1.7e-06 | 2.267E-06 | 6.658E-07 | 1.700E-06 | 3.000E-06 |
| 25664 | 1284 | V1284D | E | 2.9e-06 | 2.8e-06 | 3.4e-06 | 3.033E-06 | 3.215E-07 | 2.800E-06 | 3.400E-06 |
| 25665 | 1284 | V1284C | E | 2.2e-06 | 2.8e-06 | 2.9e-06 | 2.633E-06 | 3.786E-07 | 2.200E-06 | 2.900E-06 |
| 25666 | 1284 | V1284E | E | 3.5e-06 | 2.8e-06 | 3.7e-06 | 3.333E-06 | 4.726E-07 | 2.800E-06 | 3.700E-06 |
| 25667 | 1284 | V1284Q | E | 2.1e-06 | 2.5e-06 | 1.7e-06 | 2.100E-06 | 4.000E-07 | 1.700E-06 | 2.500E-06 |
| 25668 | 1284 | V1284G | E | 1.4e-06 | 2.6e-06 | 3.7e-06 | 2.567E-06 | 1.150E-06 | 1.400E-06 | 3.700E-06 |
| 25669 | 1284 | V1284H | E | 1.9e-06 | 3e-06 | 3.6e-06 | 2.833E-06 | 8.622E-07 | 1.900E-06 | 3.600E-06 |

| | | | | | | | | | | |
|---|---|---|---|---|---|---|---|---|---|---|
| 25670 | 1284 | V1284I | E | 3.3e-06 | 2.6e-06 | 2.6e-06 | 2.833E-06 | 4.041E-07 | 2.600E-06 | 3.300E-06 |
| 25671 | 1284 | V1284L | E | 1.6e-06 | 2e-06 | 2.1e-06 | 1.900E-06 | 2.646E-07 | 1.600E-06 | 2.100E-06 |
| 25672 | 1284 | V1284K | E | 2.9e-06 | 2.3e-06 | 1.9e-06 | 2.367E-06 | 5.033E-07 | 1.900E-06 | 2.900E-06 |
| 25673 | 1284 | V1284M | E | 2.2e-06 | 3.5e-06 | 1.7e-06 | 2.467E-06 | 9.292E-07 | 1.700E-06 | 3.500E-06 |
| 25674 | 1284 | V1284F | E | 6.4e-07 | 1.1e-06 | 2.1e-06 | 1.280E-06 | 7.465E-07 | 6.400E-07 | 2.100E-06 |
| 25675 | 1284 | V1284P | E | 3.6e-06 | 3e-06 | 2.1e-06 | 2.900E-06 | 7.550E-07 | 2.100E-06 | 3.600E-06 |
| 25676 | 1284 | V1284S | E | 1.1e-06 | 4.2e-06 | 1.9e-06 | 2.400E-06 | 1.609E-06 | 1.100E-06 | 4.200E-06 |
| 25677 | 1284 | V1284T | E | 2.2e-06 | 2.7e-06 | 2.8e-06 | 2.567E-06 | 3.215E-07 | 2.200E-06 | 2.800E-06 |
| 25678 | 1284 | V1284W | E | 2.2e-06 | 2.4e-06 | 3.1e-06 | 2.567E-06 | 4.726E-07 | 2.200E-06 | 3.100E-06 |
| 25679 | 1284 | V1284Y | E | 1.9e-06 | 1.2e-06 | 2.3e-06 | 1.800E-06 | 5.568E-07 | 1.200E-06 | 2.300E-06 |
| 25680 | 1284 | V1284V | E | 2e-06 | 2.2e-06 | 2.4e-06 | 2.200E-06 | 2.000E-07 | 2.000E-06 | 2.400E-06 |
| 25681 | 1285 | Q1285A | E | 1.7e-06 | 3.2e-06 | 1.5e-06 | 2.133E-06 | 9.292E-07 | 1.500E-06 | 3.200E-06 |
| 25682 | 1285 | Q1285R | E | 2.7e-06 | 1.5e-06 | 2.3e-06 | 2.167E-06 | 6.110E-07 | 1.500E-06 | 2.700E-06 |
| 25683 | 1285 | Q1285N | E | 1.4e-06 | 2.1e-06 | 1.1e-06 | 1.533E-06 | 5.132E-07 | 1.100E-06 | 2.100E-06 |
| 25684 | 1285 | Q1285D | E | 2.3e-06 | 2.1e-06 | 2.1e-06 | 2.167E-06 | 1.155E-07 | 2.100E-06 | 2.300E-06 |
| 25685 | 1285 | Q1285C | E | 2.9e-06 | 2e-06 | 2e-06 | 2.300E-06 | 5.196E-07 | 2.000E-06 | 2.900E-06 |
| 25686 | 1285 | Q1285E | E | 1.4e-06 | 2.9e-06 | 2.7e-06 | 2.333E-06 | 8.145E-07 | 1.400E-06 | 2.900E-06 |
| 25687 | 1285 | Q1285Q | E | 3.4e-06 | 2.6e-06 | 2e-06 | 2.667E-06 | 7.024E-07 | 2.000E-06 | 3.400E-06 |
| 25688 | 1285 | Q1285G | E | 1.6e-06 | 2.3e-06 | 1.9e-06 | 1.933E-06 | 3.512E-07 | 1.600E-06 | 2.300E-06 |
| 25689 | 1285 | Q1285H | E | 1.9e-06 | 2.8e-06 | 2.6e-06 | 2.433E-06 | 4.726E-07 | 1.900E-06 | 2.800E-06 |
| 25690 | 1285 | Q1285I | E | 2.5e-06 | 2.9e-06 | 2.9e-06 | 2.767E-06 | 2.309E-07 | 2.500E-06 | 2.900E-06 |
| 25691 | 1285 | Q1285L | E | 2.5e-06 | 1.9e-06 | 2.3e-06 | 2.233E-06 | 3.055E-07 | 1.900E-06 | 2.500E-06 |
| 25692 | 1285 | Q1285K | E | 3.2e-06 | 1.9e-06 | 1.6e-06 | 2.233E-06 | 8.505E-07 | 1.600E-06 | 3.200E-06 |
| 25693 | 1285 | Q1285M | E | 4e-06 | 4.6e-06 | 1.8e-06 | 3.467E-06 | 1.474E-06 | 1.800E-06 | 4.600E-06 |
| 25694 | 1285 | Q1285F | E | 2.6e-06 | 2.1e-06 | 1.7e-06 | 2.133E-06 | 4.509E-07 | 1.700E-06 | 2.600E-06 |
| 25695 | 1285 | Q1285P | E | 2.2e-06 | 1.3e-06 | 3.3e-06 | 2.267E-06 | 1.002E-06 | 1.300E-06 | 3.300E-06 |
| 25696 | 1285 | Q1285S | E | 2e-06 | 2.1e-06 | 2.9e-06 | 2.333E-06 | 4.933E-07 | 2.000E-06 | 2.900E-06 |
| 25697 | 1285 | Q1285T | E | 2.7e-06 | 4.9e-06 | 3.5e-06 | 3.700E-06 | 1.114E-06 | 2.700E-06 | 4.900E-06 |
| 25698 | 1285 | Q1285W | E | 3.2e-06 | 2e-06 | 3.5e-06 | 2.900E-06 | 7.937E-07 | 2.000E-06 | 3.500E-06 |
| 25699 | 1285 | Q1285Y | E | 2.1e-06 | 2.7e-06 | 3e-06 | 2.600E-06 | 4.583E-07 | 2.100E-06 | 3.000E-06 |
| 25700 | 1285 | Q1285V | E | 3.1e-06 | 4.4e-06 | 2e-06 | 3.167E-06 | 1.201E-06 | 2.000E-06 | 4.400E-06 |
| 25701 | 1286 | P1286A | E | 2.1e-06 | 2.6e-06 | 2.5e-06 | 2.400E-06 | 2.646E-07 | 2.100E-06 | 2.600E-06 |
| 25702 | 1286 | P1286R | E | 2.1e-06 | 1.9e-06 | 2.2e-06 | 2.067E-06 | 1.528E-07 | 1.900E-06 | 2.200E-06 |
| 25703 | 1286 | P1286N | E | 3e-06 | 4e-06 | 1.6e-06 | 2.867E-06 | 1.206E-06 | 1.600E-06 | 4.000E-06 |
| 25704 | 1286 | P1286D | E | 4.6e-06 | 2e-06 | 2.6e-06 | 3.067E-06 | 1.361E-06 | 2.000E-06 | 4.600E-06 |
| 25705 | 1286 | P1286C | E | 1.6e-06 | 2.2e-06 | 1.3e-06 | 1.700E-06 | 4.583E-07 | 1.300E-06 | 2.200E-06 |
| 25706 | 1286 | P1286E | E | 2.9e-06 | 2e-06 | 2.8e-06 | 2.567E-06 | 4.933E-07 | 2.000E-06 | 2.900E-06 |
| 25707 | 1286 | P1286Q | E | 2.9e-06 | 1.6e-06 | 2.3e-06 | 2.267E-06 | 6.506E-07 | 1.600E-06 | 2.900E-06 |
| 25708 | 1286 | P1286G | E | 2.2e-06 | 1.6e-06 | 3e-06 | 2.267E-06 | 7.024E-07 | 1.600E-06 | 3.000E-06 |
| 25709 | 1286 | P1286H | E | 2.2e-06 | 2.2e-06 | 3e-06 | 2.467E-06 | 4.619E-07 | 2.200E-06 | 3.000E-06 |
| 25710 | 1286 | P1286I | E | 2.2e-06 | 5.6e-06 | 2e-06 | 3.267E-06 | 2.023E-06 | 2.000E-06 | 5.600E-06 |
| 25711 | 1286 | P1286L | E | 1.6e-06 | 1.6e-06 | 2.2e-06 | 1.800E-06 | 3.464E-07 | 1.600E-06 | 2.200E-06 |
| 25712 | 1286 | P1286K | E | 1.6e-06 | 2.6e-06 | 2.4e-06 | 2.200E-06 | 5.292E-07 | 1.600E-06 | 2.600E-06 |
| 25713 | 1286 | P1286M | E | 2.8e-06 | 2.6e-06 | 3.6e-06 | 3.000E-06 | 5.292E-07 | 2.600E-06 | 3.600E-06 |
| 25714 | 1286 | P1286F | E | 2.8e-06 | 2.2e-06 | 2.7e-06 | 2.567E-06 | 3.215E-07 | 2.200E-06 | 2.800E-06 |
| 25715 | 1286 | P1286P | E | 3e-06 | 1.8e-06 | 2.7e-06 | 2.500E-06 | 6.245E-07 | 1.800E-06 | 3.000E-06 |
| 25716 | 1286 | P1286S | E | 2.4e-06 | 1.2e-06 | 1.6e-06 | 1.733E-06 | 6.110E-07 | 1.200E-06 | 2.400E-06 |
| 25717 | 1286 | P1286T | E | 1.2e-06 | 2.8e-06 | 2.7e-06 | 2.233E-06 | 8.963E-07 | 1.200E-06 | 2.800E-06 |
| 25718 | 1286 | P1286W | E | 2.1e-06 | 1.8e-06 | 3.5e-06 | 2.467E-06 | 9.074E-07 | 1.800E-06 | 3.500E-06 |
| 25719 | 1286 | P1286Y | E | 2.9e-06 | 3.4e-06 | 1.9e-06 | 2.733E-06 | 7.638E-07 | 1.900E-06 | 3.400E-06 |
| 25720 | 1286 | P1286V | E | 3e-06 | 3.5e-06 | 6.9e-07 | 2.397E-06 | 1.499E-06 | 6.900E-07 | 3.500E-06 |
| 25721 | 1287 | G1287A | E | 2e-06 | 1.7e-06 | 2.2e-06 | 1.967E-06 | 2.517E-07 | 1.700E-06 | 2.200E-06 |
| 25722 | 1287 | G1287R | E | 1.6e-06 | 3.4e-06 | 2.6e-06 | 2.533E-06 | 9.018E-07 | 1.600E-06 | 3.400E-06 |
| 25723 | 1287 | G1287N | E | 3.3e-06 | 2.8e-06 | 2.2e-06 | 2.767E-06 | 5.508E-07 | 2.200E-06 | 3.300E-06 |
| 25724 | 1287 | G1287D | E | 2.1e-06 | 2e-06 | 2.5e-06 | 2.200E-06 | 2.646E-07 | 2.000E-06 | 2.500E-06 |
| 25725 | 1287 | G1287C | E | 1.6e-06 | 2e-06 | 4.4e-06 | 2.667E-06 | 1.514E-06 | 1.600E-06 | 4.400E-06 |
| 25726 | 1287 | G1287E | E | 3e-06 | 2.2e-06 | 2.9e-06 | 2.700E-06 | 4.359E-07 | 2.200E-06 | 3.000E-06 |
| 25727 | 1287 | G1287Q | E | 2e-06 | 1.8e-06 | 2.3e-06 | 2.033E-06 | 2.517E-07 | 1.800E-06 | 2.300E-06 |
| 25728 | 1287 | G1287G | E | 2.2e-06 | 2.3e-06 | 2.7e-06 | 2.400E-06 | 2.646E-07 | 2.200E-06 | 2.700E-06 |
| 25729 | 1287 | G1287H | E | 1.8e-06 | 2.8e-06 | 3.7e-06 | 2.767E-06 | 9.504E-07 | 1.800E-06 | 3.700E-06 |
| 25730 | 1287 | G1287I | E | 2.3e-06 | 1.7e-06 | 2.9e-06 | 2.300E-06 | 6.000E-07 | 1.700E-06 | 2.900E-06 |
| 25731 | 1287 | G1287L | E | 2.7e-06 | 3.3e-06 | 2.2e-06 | 2.733E-06 | 5.508E-07 | 2.200E-06 | 3.300E-06 |
| 25732 | 1287 | G1287K | E | 2.2e-06 | 1.1e-06 | 2.7e-06 | 2.000E-06 | 8.185E-07 | 1.100E-06 | 2.700E-06 |
| 25733 | 1287 | G1287M | E | 2.5e-06 | 2.1e-06 | 2.5e-06 | 2.367E-06 | 2.309E-07 | 2.100E-06 | 2.500E-06 |
| 25734 | 1287 | G1287F | E | 2.5e-06 | 3.1e-06 | 1.8e-06 | 2.467E-06 | 6.506E-07 | 1.800E-06 | 3.100E-06 |
| 25735 | 1287 | G1287P | E | 2.3e-06 | 3.4e-06 | 2.4e-06 | 2.700E-06 | 6.083E-07 | 2.300E-06 | 3.400E-06 |
| 25736 | 1287 | G1287S | E | 3.1e-06 | 2.7e-06 | 2.6e-06 | 2.800E-06 | 2.646E-07 | 2.600E-06 | 3.100E-06 |
| 25737 | 1287 | G1287T | E | 2.7e-06 | 2e-06 | 2.3e-06 | 2.333E-06 | 3.512E-07 | 2.000E-06 | 2.700E-06 |
| 25738 | 1287 | G1287W | E | 2.9e-06 | 2.5e-06 | 3.4e-06 | 2.933E-06 | 4.509E-07 | 2.500E-06 | 3.400E-06 |
| 25739 | 1287 | G1287Y | E | 1.5e-06 | 3.3e-06 | 2.2e-06 | 2.333E-06 | 9.074E-07 | 1.500E-06 | 3.300E-06 |
| 25740 | 1287 | G1287V | E | 2.2e-06 | 4.4e-06 | 2e-06 | 2.867E-06 | 1.332E-06 | 2.000E-06 | 4.400E-06 |
| 25741 | 1288 | G1288A | E | 2.6e-06 | 2.5e-06 | 1.4e-06 | 2.167E-06 | 6.658E-07 | 1.400E-06 | 2.600E-06 |
| 25742 | 1288 | G1288R | E | 2.8e-06 | 2.7e-06 | 1.9e-06 | 2.467E-06 | 4.933E-07 | 1.900E-06 | 2.800E-06 |
| 25743 | 1288 | G1288N | E | 1.8e-06 | 1.4e-06 | 2.5e-06 | 1.900E-06 | 5.568E-07 | 1.400E-06 | 2.500E-06 |
| 25744 | 1288 | G1288D | E | 3.2e-06 | 3.1e-06 | 2.3e-06 | 2.867E-06 | 4.933E-07 | 2.300E-06 | 3.200E-06 |
| 25745 | 1288 | G1288C | E | 1.6e-06 | 1.5e-06 | 1.6e-06 | 1.567E-06 | 5.774E-08 | 1.500E-06 | 1.600E-06 |
| 25746 | 1288 | G1288E | E | 1.5e-06 | 2.3e-06 | 2.1e-06 | 1.967E-06 | 4.163E-07 | 1.500E-06 | 2.300E-06 |
| 25747 | 1288 | G1288Q | E | 1.8e-06 | 3e-06 | 1.9e-06 | 2.233E-06 | 6.658E-07 | 1.800E-06 | 3.000E-06 |
| 25748 | 1288 | G1288G | E | 3.3e-06 | 1.9e-06 | 2.1e-06 | 2.433E-06 | 7.572E-07 | 1.900E-06 | 3.300E-06 |
| 25749 | 1288 | G1288H | E | 2.4e-06 | 2.2e-06 | 3.8e-06 | 2.800E-06 | 8.718E-07 | 2.200E-06 | 3.800E-06 |
| 25750 | 1288 | G1288I | E | 2.6e-06 | 4.4e-06 | 2e-06 | 3.000E-06 | 1.249E-06 | 2.000E-06 | 4.400E-06 |
| 25751 | 1288 | G1288L | E | 2.6e-06 | 2.4e-06 | 2.4e-06 | 2.467E-06 | 1.155E-07 | 2.400E-06 | 2.600E-06 |
| 25752 | 1288 | G1288K | E | 1.5e-06 | 3.2e-06 | 2.3e-06 | 2.333E-06 | 8.505E-07 | 1.500E-06 | 3.200E-06 |
| 25753 | 1288 | G1288M | E | 1.6e-06 | 2e-06 | 2.2e-06 | 1.933E-06 | 3.055E-07 | 1.600E-06 | 2.200E-06 |
| 25754 | 1288 | G1288F | E | 1.5e-06 | 1.7e-06 | 2.7e-06 | 1.967E-06 | 6.429E-07 | 1.500E-06 | 2.700E-06 |

| | | | | | | | | | | |
|---|---|---|---|---|---|---|---|---|---|---|
| 25755 | 1288 | G1288P | E | 2.6e-06 | 3.3e-06 | 1.7e-06 | 2.533E-06 | 8.021E-07 | 1.700E-06 | 3.300E-06 |
| 25756 | 1288 | G1288S | E | 2.8e-06 | 2.4e-06 | 2.8e-06 | 2.667E-06 | 2.309E-07 | 2.400E-06 | 2.800E-06 |
| 25757 | 1288 | G1288T | E | 2.2e-06 | 2.4e-06 | 3.1e-06 | 2.567E-06 | 4.726E-07 | 2.200E-06 | 3.100E-06 |
| 25758 | 1288 | G1288W | E | 2.2e-06 | 2.5e-06 | 2e-06 | 2.233E-06 | 2.517E-07 | 2.000E-06 | 2.500E-06 |
| 25759 | 1288 | G1288Y | E | 2.5e-06 | 2.5e-06 | 2.9e-06 | 2.633E-06 | 2.309E-07 | 2.500E-06 | 2.900E-06 |
| 25760 | 1288 | G1288V | E | 2.2e-06 | 2.2e-06 | 2.3e-06 | 2.233E-06 | 5.774E-08 | 2.200E-06 | 2.300E-06 |
| 25761 | 1289 | S1289A | E | 3.6e-06 | 2.2e-06 | 1.8e-06 | 2.533E-06 | 9.452E-07 | 1.800E-06 | 3.600E-06 |
| 25762 | 1289 | S1289R | E | 3.5e-06 | 3.2e-06 | 1.1e-06 | 2.600E-06 | 1.308E-06 | 1.100E-06 | 3.500E-06 |
| 25763 | 1289 | S1289N | E | 2.1e-06 | 5.3e-06 | 1.8e-06 | 3.067E-06 | 1.940E-06 | 1.800E-06 | 5.300E-06 |
| 25764 | 1289 | S1289D | E | 2.8e-06 | 2e-06 | 2.6e-06 | 2.467E-06 | 4.163E-07 | 2.000E-06 | 2.800E-06 |
| 25765 | 1289 | S1289C | E | 1.8e-06 | 2e-06 | 1.5e-06 | 1.767E-06 | 2.517E-07 | 1.500E-06 | 2.000E-06 |
| 25766 | 1289 | S1289E | E | 2.8e-06 | 1.9e-06 | 4e-06 | 2.900E-06 | 1.054E-06 | 1.900E-06 | 4.000E-06 |
| 25767 | 1289 | S1289Q | E | 3e-06 | 2.7e-06 | 3e-06 | 2.900E-06 | 1.732E-07 | 2.700E-06 | 3.000E-06 |
| 25768 | 1289 | S1289G | E | 2e-06 | 1.8e-06 | 3.6e-06 | 2.467E-06 | 9.866E-07 | 1.800E-06 | 3.600E-06 |
| 25769 | 1289 | S1289H | E | 2.4e-06 | 1.3e-06 | 1.7e-06 | 1.800E-06 | 5.568E-07 | 1.300E-06 | 2.400E-06 |
| 25770 | 1289 | S1289I | E | 2.7e-06 | 1.4e-06 | 3.4e-06 | 2.500E-06 | 1.015E-06 | 1.400E-06 | 3.400E-06 |
| 25771 | 1289 | S1289L | E | 2.5e-06 | 2.2e-06 | 2e-06 | 2.233E-06 | 2.517E-07 | 2.000E-06 | 2.500E-06 |
| 25772 | 1289 | S1289K | E | 3.2e-06 | 2.8e-06 | 4.2e-06 | 3.400E-06 | 7.211E-07 | 2.800E-06 | 4.200E-06 |
| 25773 | 1289 | S1289M | E | 3.3e-06 | 3.2e-06 | 2e-06 | 2.833E-06 | 7.234E-07 | 2.000E-06 | 3.300E-06 |
| 25774 | 1289 | S1289F | E | 3.7e-06 | 2.1e-06 | 2e-06 | 2.600E-06 | 9.539E-07 | 2.000E-06 | 3.700E-06 |
| 25775 | 1289 | S1289P | E | 2.8e-06 | 2.3e-06 | 2e-06 | 2.367E-06 | 4.041E-07 | 2.000E-06 | 2.800E-06 |
| 25776 | 1289 | S1289S | E | 1.5e-06 | 3e-06 | 5.7e-06 | 3.400E-06 | 2.128E-06 | 1.500E-06 | 5.700E-06 |
| 25777 | 1289 | S1289T | E | 3e-06 | 2.4e-06 | 2.1e-06 | 2.500E-06 | 4.583E-07 | 2.100E-06 | 3.000E-06 |
| 25778 | 1289 | S1289W | E | 4.2e-06 | 2.8e-06 | 2.3e-06 | 3.100E-06 | 9.849E-07 | 2.300E-06 | 4.200E-06 |
| 25779 | 1289 | S1289Y | E | 2.3e-06 | 2.5e-06 | 1.4e-06 | 2.067E-06 | 5.859E-07 | 1.400E-06 | 2.500E-06 |
| 25780 | 1289 | S1289V | E | 2.6e-06 | 3.2e-06 | 3.9e-06 | 3.233E-06 | 6.506E-07 | 2.600E-06 | 3.900E-06 |
| 25781 | 1290 | L1290A | E | 1.3e-06 | 2.3e-06 | 3e-06 | 2.200E-06 | 8.544E-07 | 1.300E-06 | 3.000E-06 |
| 25782 | 1290 | L1290R | E | 2.2e-06 | 1.8e-06 | 2.9e-06 | 2.300E-06 | 5.568E-07 | 1.800E-06 | 2.900E-06 |
| 25783 | 1290 | L1290N | E | 2.3e-06 | 2.5e-06 | 2.3e-06 | 2.367E-06 | 1.155E-07 | 2.300E-06 | 2.500E-06 |
| 25784 | 1290 | L1290D | E | 1.8e-06 | 2.3e-06 | 2.4e-06 | 2.167E-06 | 3.215E-07 | 1.800E-06 | 2.400E-06 |
| 25785 | 1290 | L1290C | E | 1.5e-06 | 1.9e-06 | 2.1e-06 | 1.833E-06 | 3.055E-07 | 1.500E-06 | 2.100E-06 |
| 25786 | 1290 | L1290E | E | 3.8e-06 | 3.1e-06 | 1.8e-06 | 2.900E-06 | 1.015E-06 | 1.800E-06 | 3.800E-06 |
| 25787 | 1290 | L1290Q | E | 1.4e-06 | 2e-06 | 3.9e-06 | 2.433E-06 | 1.305E-06 | 1.400E-06 | 3.900E-06 |
| 25788 | 1290 | L1290G | E | 2.3e-06 | 3.1e-06 | 2.7e-06 | 2.700E-06 | 4.000E-07 | 2.300E-06 | 3.100E-06 |
| 25789 | 1290 | L1290H | E | 2.1e-06 | 3.5e-06 | 3.1e-06 | 2.900E-06 | 7.211E-07 | 2.100E-06 | 3.500E-06 |
| 25790 | 1290 | L1290I | E | 3.1e-06 | 6.4e-06 | 1.4e-06 | 3.633E-06 | 2.542E-06 | 1.400E-06 | 6.400E-06 |
| 25791 | 1290 | L1290L | E | 1.5e-06 | 2.1e-06 | 1.4e-06 | 1.667E-06 | 3.786E-07 | 1.400E-06 | 2.100E-06 |
| 25792 | 1290 | L1290K | E | 1.9e-06 | 2.4e-06 | 3.5e-06 | 2.600E-06 | 8.185E-07 | 1.900E-06 | 3.500E-06 |
| 25793 | 1290 | L1290M | E | 2.4e-06 | 1.6e-06 | 3.1e-06 | 2.367E-06 | 7.506E-07 | 1.600E-06 | 3.100E-06 |
| 25794 | 1290 | L1290F | E | 1.9e-06 | 2.9e-06 | 1.2e-06 | 2.000E-06 | 8.544E-07 | 1.200E-06 | 2.900E-06 |
| 25795 | 1290 | L1290P | E | 1.5e-06 | 2.2e-06 | 2.4e-06 | 2.033E-06 | 4.726E-07 | 1.500E-06 | 2.400E-06 |
| 25796 | 1290 | L1290S | E | 2.1e-06 | 1.4e-06 | 2.2e-06 | 1.900E-06 | 4.359E-07 | 1.400E-06 | 2.200E-06 |
| 25797 | 1290 | L1290T | E | 8.4e-07 | 1e-06 | 1.9e-06 | 1.247E-06 | 5.714E-07 | 8.400E-07 | 1.900E-06 |
| 25798 | 1290 | L1290W | E | 3.2e-06 | 2e-06 | 1.8e-06 | 2.333E-06 | 7.572E-07 | 1.800E-06 | 3.200E-06 |
| 25799 | 1290 | L1290Y | E | 1.3e-06 | 1.6e-06 | 2.9e-06 | 1.933E-06 | 8.505E-07 | 1.300E-06 | 2.900E-06 |
| 25800 | 1290 | L1290V | E | 3.3e-06 | 2.4e-06 | 2.1e-06 | 2.600E-06 | 6.245E-07 | 2.100E-06 | 3.300E-06 |
| 25801 | 1291 | R1291A | E | 4.4e-06 | 2e-06 | 2e-06 | 2.800E-06 | 1.386E-06 | 2.000E-06 | 4.400E-06 |
| 25802 | 1291 | R1291R | E | 2.7e-06 | 4.3e-06 | 3.6e-06 | 3.533E-06 | 8.021E-07 | 2.700E-06 | 4.300E-06 |
| 25803 | 1291 | R1291N | E | 3.2e-06 | 2.5e-06 | 2.8e-06 | 2.833E-06 | 3.512E-07 | 2.500E-06 | 3.200E-06 |
| 25804 | 1291 | R1291D | E | 1.5e-06 | 2.3e-06 | 1.9e-06 | 1.900E-06 | 4.000E-07 | 1.500E-06 | 2.300E-06 |
| 25805 | 1291 | R1291C | E | 2.7e-06 | 1.6e-06 | 2.2e-06 | 2.167E-06 | 5.508E-07 | 1.600E-06 | 2.700E-06 |
| 25806 | 1291 | R1291E | E | 2.1e-06 | 2.6e-06 | 2.1e-06 | 2.267E-06 | 2.887E-07 | 2.100E-06 | 2.600E-06 |
| 25807 | 1291 | R1291Q | E | 1e-06 | 2.5e-06 | 2.7e-06 | 2.067E-06 | 9.292E-07 | 1.000E-06 | 2.700E-06 |
| 25808 | 1291 | R1291G | E | 2.5e-06 | 2.1e-06 | 1.4e-06 | 2.000E-06 | 5.568E-07 | 1.400E-06 | 2.500E-06 |
| 25809 | 1291 | R1291H | E | 2.6e-06 | 2.8e-06 | 1.3e-06 | 2.233E-06 | 8.145E-07 | 1.300E-06 | 2.800E-06 |
| 25810 | 1291 | R1291I | E | 2.4e-06 | 5.8e-06 | 2.2e-06 | 3.467E-06 | 2.023E-06 | 2.200E-06 | 5.800E-06 |
| 25811 | 1291 | R1291L | E | 2.2e-06 | 2.6e-06 | 1.7e-06 | 2.167E-06 | 4.509E-07 | 1.700E-06 | 2.600E-06 |
| 25812 | 1291 | R1291K | E | 2.1e-06 | 2.5e-06 | 3.8e-06 | 2.800E-06 | 8.888E-07 | 2.100E-06 | 3.800E-06 |
| 25813 | 1291 | R1291M | E | 2.4e-06 | 3.4e-06 | 3.5e-06 | 3.100E-06 | 6.083E-07 | 2.400E-06 | 3.500E-06 |
| 25814 | 1291 | R1291F | E | 2.2e-06 | 3.7e-06 | 2.7e-06 | 2.867E-06 | 7.638E-07 | 2.200E-06 | 3.700E-06 |
| 25815 | 1291 | R1291P | E | 2.4e-06 | 1.6e-06 | 1.1e-06 | 1.700E-06 | 6.557E-07 | 1.100E-06 | 2.400E-06 |
| 25816 | 1291 | R1291S | E | 2.2e-06 | 2.3e-06 | 3.1e-06 | 2.533E-06 | 4.933E-07 | 2.200E-06 | 3.100E-06 |
| 25817 | 1291 | R1291T | E | 2.2e-06 | 2.3e-06 | 2e-06 | 2.167E-06 | 1.528E-07 | 2.000E-06 | 2.300E-06 |
| 25818 | 1291 | R1291W | E | 2.5e-06 | 2.1e-06 | 2.8e-06 | 2.467E-06 | 3.512E-07 | 2.100E-06 | 2.800E-06 |
| 25819 | 1291 | R1291Y | E | 1.7e-06 | 2.4e-06 | 3.8e-06 | 2.633E-06 | 1.069E-06 | 1.700E-06 | 3.800E-06 |
| 25820 | 1291 | R1291V | E | 1.9e-06 | 2.4e-06 | 2.9e-06 | 2.400E-06 | 5.000E-07 | 1.900E-06 | 2.900E-06 |
| 25821 | 1292 | L1292A | E | 2.6e-06 | 2.2e-06 | 2.1e-06 | 2.300E-06 | 2.646E-07 | 2.100E-06 | 2.600E-06 |
| 25822 | 1292 | L1292R | E | 2e-06 | 2.6e-06 | 2.4e-06 | 2.333E-06 | 3.055E-07 | 2.000E-06 | 2.600E-06 |
| 25823 | 1292 | L1292N | E | 2.4e-06 | 2.2e-06 | 2.3e-06 | 2.300E-06 | 1.000E-07 | 2.200E-06 | 2.400E-06 |
| 25824 | 1292 | L1292D | E | 2.6e-06 | 2.4e-06 | 2.5e-06 | 2.500E-06 | 1.000E-07 | 2.400E-06 | 2.600E-06 |
| 25825 | 1292 | L1292C | E | 2.8e-06 | 3.1e-06 | 2.6e-06 | 2.833E-06 | 2.517E-07 | 2.600E-06 | 3.100E-06 |
| 25826 | 1292 | L1292E | E | 1.9e-06 | 2.9e-06 | 2.9e-06 | 2.567E-06 | 5.774E-07 | 1.900E-06 | 2.900E-06 |
| 25827 | 1292 | L1292Q | E | 4.6e-06 | 3.9e-06 | 2.6e-06 | 3.700E-06 | 1.015E-06 | 2.600E-06 | 4.600E-06 |
| 25828 | 1292 | L1292G | E | 3.3e-06 | 1.4e-06 | 2.2e-06 | 2.300E-06 | 9.539E-07 | 1.400E-06 | 3.300E-06 |
| 25829 | 1292 | L1292H | E | 1.9e-06 | 2.7e-06 | 2.8e-06 | 2.467E-06 | 4.933E-07 | 1.900E-06 | 2.800E-06 |
| 25830 | 1292 | L1292I | E | 1.3e-06 | 1.9e-06 | 4.3e-06 | 2.500E-06 | 1.587E-06 | 1.300E-06 | 4.300E-06 |
| 25831 | 1292 | L1292L | E | 1.7e-06 | 2.4e-06 | 2.2e-06 | 2.100E-06 | 3.606E-07 | 1.700E-06 | 2.400E-06 |
| 25832 | 1292 | L1292K | E | 2.6e-06 | 2.1e-06 | 1.9e-06 | 2.200E-06 | 3.606E-07 | 1.900E-06 | 2.600E-06 |
| 25833 | 1292 | L1292M | E | 1.8e-06 | 1.8e-06 | 2.5e-06 | 2.033E-06 | 4.041E-07 | 1.800E-06 | 2.500E-06 |
| 25834 | 1292 | L1292F | E | 1.6e-06 | 2e-06 | 4.2e-06 | 2.600E-06 | 1.400E-06 | 1.600E-06 | 4.200E-06 |
| 25835 | 1292 | L1292P | E | 2.6e-06 | 2.6e-06 | 2.3e-06 | 2.500E-06 | 1.732E-07 | 2.300E-06 | 2.600E-06 |
| 25836 | 1292 | L1292S | E | 2e-06 | 2.5e-06 | 2.7e-06 | 2.400E-06 | 3.606E-07 | 2.000E-06 | 2.700E-06 |
| 25837 | 1292 | L1292T | E | 3.8e-06 | 3e-06 | 2.5e-06 | 3.100E-06 | 6.557E-07 | 2.500E-06 | 3.800E-06 |
| 25838 | 1292 | L1292W | E | 2e-06 | 3.3e-06 | 2.5e-06 | 2.600E-06 | 6.557E-07 | 2.000E-06 | 3.300E-06 |
| 25839 | 1292 | L1292Y | E | 2.7e-06 | 2.1e-06 | 2e-06 | 2.267E-06 | 3.786E-07 | 2.000E-06 | 2.700E-06 |

| | | | | | | | | | | |
|---|---|---|---|---|---|---|---|---|---|---|
| 25840 | 1292 | L1292V | E | 1.8e-06 | 2.7e-06 | 2.9e-06 | 2.467E-06 | 5.859E-07 | 1.800E-06 | 2.900E-06 |
| 25841 | 1293 | S1293A | E | 2.3e-06 | 1.7e-06 | 2.1e-06 | 2.033E-06 | 3.055E-07 | 1.700E-06 | 2.300E-06 |
| 25842 | 1293 | S1293R | E | 1.8e-06 | 3.1e-06 | 2.3e-06 | 2.400E-06 | 6.557E-07 | 1.800E-06 | 3.100E-06 |
| 25843 | 1293 | S1293N | E | 2.4e-06 | 2.5e-06 | 4e-06 | 2.967E-06 | 8.963E-07 | 2.400E-06 | 4.000E-06 |
| 25844 | 1293 | S1293D | E | 2.1e-06 | 2.1e-06 | 2.5e-06 | 2.233E-06 | 2.309E-07 | 2.100E-06 | 2.500E-06 |
| 25845 | 1293 | S1293C | E | 1.2e-06 | 1.8e-06 | 1.4e-06 | 1.467E-06 | 3.055E-07 | 1.200E-06 | 1.800E-06 |
| 25846 | 1293 | S1293E | E | 2.2e-06 | 2e-06 | 1.1e-06 | 1.767E-06 | 5.859E-07 | 1.100E-06 | 2.200E-06 |
| 25847 | 1293 | S1293Q | E | 2.2e-06 | 1.1e-06 | 2.4e-06 | 1.900E-06 | 7.000E-07 | 1.100E-06 | 2.400E-06 |
| 25848 | 1293 | S1293G | E | 4.1e-06 | 1.3e-06 | 2.3e-06 | 2.567E-06 | 1.419E-06 | 1.300E-06 | 4.100E-06 |
| 25849 | 1293 | S1293H | E | 7.5e-07 | 1.1e-06 | 3.2e-06 | 1.683E-06 | 1.325E-06 | 7.500E-07 | 3.200E-06 |
| 25850 | 1293 | S1293I | E | 2.5e-06 | 2.7e-06 | 1.2e-06 | 2.133E-06 | 8.145E-07 | 1.200E-06 | 2.700E-06 |
| 25851 | 1293 | S1293L | E | 2.1e-06 | 3.1e-06 | 1.8e-06 | 2.333E-06 | 6.807E-07 | 1.800E-06 | 3.100E-06 |
| 25852 | 1293 | S1293K | E | 2.8e-06 | 2.5e-06 | 2.8e-06 | 2.700E-06 | 1.732E-07 | 2.500E-06 | 2.800E-06 |
| 25853 | 1293 | S1293M | E | 3.9e-06 | 2.1e-06 | 3.4e-06 | 3.133E-06 | 9.292E-07 | 2.100E-06 | 3.900E-06 |
| 25854 | 1293 | S1293F | E | 3.9e-06 | 2.1e-06 | 2.3e-06 | 2.767E-06 | 9.866E-07 | 2.100E-06 | 3.900E-06 |
| 25855 | 1293 | S1293P | E | 1.1e-06 | 2.3e-06 | 2.3e-06 | 1.900E-06 | 6.928E-07 | 1.100E-06 | 2.300E-06 |
| 25856 | 1293 | S1293S | E | 2.8e-06 | 2.3e-06 | 2.1e-06 | 2.400E-06 | 3.606E-07 | 2.100E-06 | 2.800E-06 |
| 25857 | 1293 | S1293T | E | 4.1e-06 | 2.5e-06 | 3.9e-06 | 3.500E-06 | 8.718E-07 | 2.500E-06 | 4.100E-06 |
| 25858 | 1293 | S1293W | E | 2.7e-06 | 2.8e-06 | 1.7e-06 | 2.400E-06 | 6.083E-07 | 1.700E-06 | 2.800E-06 |
| 25859 | 1293 | S1293Y | E | 2.4e-06 | 2.2e-06 | 3.2e-06 | 2.600E-06 | 5.292E-07 | 2.200E-06 | 3.200E-06 |
| 25860 | 1293 | S1293V | E | 2.4e-06 | 3.6e-06 | 1.3e-06 | 2.433E-06 | 1.150E-06 | 1.300E-06 | 3.600E-06 |
| 25861 | 1294 | C1294A | E | 3.3e-06 | 2.4e-06 | 1.4e-06 | 2.367E-06 | 9.504E-07 | 1.400E-06 | 3.300E-06 |
| 25862 | 1294 | C1294R | E | 1.7e-06 | 1.1e-06 | 1e-06 | 1.267E-06 | 3.786E-07 | 1.000E-06 | 1.700E-06 |
| 25863 | 1294 | C1294N | E | 2.2e-06 | 2e-06 | 2.3e-06 | 2.167E-06 | 1.528E-07 | 2.000E-06 | 2.300E-06 |
| 25864 | 1294 | C1294D | E | 2e-06 | 3e-06 | 2.2e-06 | 2.400E-06 | 5.292E-07 | 2.000E-06 | 3.000E-06 |
| 25865 | 1294 | C1294C | E | 3.9e-06 | 2.9e-06 | 2.3e-06 | 3.033E-06 | 8.083E-07 | 2.300E-06 | 3.900E-06 |
| 25866 | 1294 | C1294E | E | 2e-06 | 2.4e-06 | 2.6e-06 | 2.333E-06 | 3.055E-07 | 2.000E-06 | 2.600E-06 |
| 25867 | 1294 | C1294Q | E | 3.7e-06 | 2.5e-06 | 2.3e-06 | 2.833E-06 | 7.572E-07 | 2.300E-06 | 3.700E-06 |
| 25868 | 1294 | C1294G | E | 2e-06 | 2.9e-06 | 2.7e-06 | 2.533E-06 | 4.726E-07 | 2.000E-06 | 2.900E-06 |
| 25869 | 1294 | C1294H | E | 6.9e-07 | 1.5e-06 | 3.4e-06 | 1.863E-06 | 1.391E-06 | 6.900E-07 | 3.400E-06 |
| 25870 | 1294 | C1294I | E | 2.5e-06 | 2.9e-06 | 1.7e-06 | 2.367E-06 | 6.110E-07 | 1.700E-06 | 2.900E-06 |
| 25871 | 1294 | C1294L | E | 2.5e-06 | 2.1e-06 | 1.5e-06 | 2.033E-06 | 5.033E-07 | 1.500E-06 | 2.500E-06 |
| 25872 | 1294 | C1294K | E | 2e-06 | 2.1e-06 | 2.5e-06 | 2.200E-06 | 2.646E-07 | 2.000E-06 | 2.500E-06 |
| 25873 | 1294 | C1294M | E | 1.9e-06 | 2.7e-06 | 3.5e-06 | 2.700E-06 | 8.000E-07 | 1.900E-06 | 3.500E-06 |
| 25874 | 1294 | C1294F | E | 2.1e-06 | 1.2e-06 | 9.7e-07 | 1.423E-06 | 5.972E-07 | 9.700E-07 | 2.100E-06 |
| 25875 | 1294 | C1294P | E | 3.1e-06 | 3.3e-06 | 2.7e-06 | 3.033E-06 | 3.055E-07 | 2.700E-06 | 3.300E-06 |
| 25876 | 1294 | C1294S | E | 2.3e-06 | 2.1e-06 | 2e-06 | 2.133E-06 | 1.528E-07 | 2.000E-06 | 2.300E-06 |
| 25877 | 1294 | C1294T | E | 1.5e-06 | 3.2e-06 | 3.6e-06 | 2.767E-06 | 1.115E-06 | 1.500E-06 | 3.600E-06 |
| 25878 | 1294 | C1294W | E | 2.9e-06 | 3.2e-06 | 1.9e-06 | 2.667E-06 | 6.807E-07 | 1.900E-06 | 3.200E-06 |
| 25879 | 1294 | C1294Y | E | 1.3e-06 | 3.4e-06 | 2.6e-06 | 2.433E-06 | 1.060E-06 | 1.300E-06 | 3.400E-06 |
| 25880 | 1294 | C1294V | E | 3.4e-06 | 3e-06 | 1.1e-06 | 2.500E-06 | 1.229E-06 | 1.100E-06 | 3.400E-06 |
| 25881 | 1295 | A1295A | E | 1.7e-06 | 2e-06 | 3e-06 | 2.233E-06 | 6.807E-07 | 1.700E-06 | 3.000E-06 |
| 25882 | 1295 | A1295R | E | 2.1e-06 | 3e-06 | 1.7e-06 | 2.267E-06 | 6.658E-07 | 1.700E-06 | 3.000E-06 |
| 25883 | 1295 | A1295N | E | 2.1e-06 | 2.9e-06 | 3.2e-06 | 2.733E-06 | 5.686E-07 | 2.100E-06 | 3.200E-06 |
| 25884 | 1295 | A1295D | E | 2.7e-06 | 2.7e-06 | 2.2e-06 | 2.533E-06 | 2.887E-07 | 2.200E-06 | 2.700E-06 |
| 25885 | 1295 | A1295C | E | 3.1e-06 | 1.7e-06 | 2.1e-06 | 2.300E-06 | 7.211E-07 | 1.700E-06 | 3.100E-06 |
| 25886 | 1295 | A1295E | E | 2.1e-06 | 1.6e-06 | 1.8e-06 | 1.833E-06 | 2.517E-07 | 1.600E-06 | 2.100E-06 |
| 25887 | 1295 | A1295Q | E | 9.6e-07 | 2.7e-06 | 2.6e-06 | 2.087E-06 | 9.770E-07 | 9.600E-07 | 2.700E-06 |
| 25888 | 1295 | A1295G | E | 4.1e-06 | 4.1e-06 | 1.4e-06 | 3.200E-06 | 1.559E-06 | 1.400E-06 | 4.100E-06 |
| 25889 | 1295 | A1295H | E | 5.1e-06 | 2.5e-06 | 1.7e-06 | 3.100E-06 | 1.778E-06 | 1.700E-06 | 5.100E-06 |
| 25890 | 1295 | A1295I | E | 4.2e-06 | 2.9e-06 | 1.7e-06 | 2.933E-06 | 1.250E-06 | 1.700E-06 | 4.200E-06 |
| 25891 | 1295 | A1295L | E | 2.6e-06 | 2.7e-06 | 1.5e-06 | 2.267E-06 | 6.658E-07 | 1.500E-06 | 2.700E-06 |
| 25892 | 1295 | A1295K | E | 3.4e-06 | 1.2e-06 | 2.5e-06 | 2.367E-06 | 1.106E-06 | 1.200E-06 | 3.400E-06 |
| 25893 | 1295 | A1295M | E | 2e-06 | 3.3e-06 | 3.7e-06 | 3.000E-06 | 8.888E-07 | 2.000E-06 | 3.700E-06 |
| 25894 | 1295 | A1295F | E | 1.7e-06 | 2e-06 | 1.7e-06 | 1.800E-06 | 1.732E-07 | 1.700E-06 | 2.000E-06 |
| 25895 | 1295 | A1295P | E | 4.2e-06 | 1.7e-06 | 3.9e-06 | 3.267E-06 | 1.365E-06 | 1.700E-06 | 4.200E-06 |
| 25896 | 1295 | A1295S | E | 2.7e-06 | 2.2e-06 | 2.7e-06 | 2.533E-06 | 2.887E-07 | 2.200E-06 | 2.700E-06 |
| 25897 | 1295 | A1295T | E | 2.7e-06 | 3e-06 | 1.1e-06 | 2.267E-06 | 1.021E-06 | 1.100E-06 | 3.000E-06 |
| 25898 | 1295 | A1295W | E | 4.2e-06 | 2.4e-06 | 2.6e-06 | 3.067E-06 | 9.866E-07 | 2.400E-06 | 4.200E-06 |
| 25899 | 1295 | A1295Y | E | 1.8e-06 | 3.5e-06 | 1.9e-06 | 2.400E-06 | 9.539E-07 | 1.800E-06 | 3.500E-06 |
| 25900 | 1295 | A1295V | E | 2.1e-06 | 2e-06 | 3e-06 | 2.367E-06 | 5.508E-07 | 2.000E-06 | 3.000E-06 |
| 25901 | 1296 | A1296A | E | 3.3e-06 | 1.9e-06 | 2e-06 | 2.400E-06 | 7.810E-07 | 1.900E-06 | 3.300E-06 |
| 25902 | 1296 | A1296R | E | 2.2e-06 | 2.5e-06 | 1.9e-06 | 2.200E-06 | 3.000E-07 | 1.900E-06 | 2.500E-06 |
| 25903 | 1296 | A1296N | E | 2.6e-06 | 2.6e-06 | 1.9e-06 | 2.367E-06 | 4.041E-07 | 1.900E-06 | 2.600E-06 |
| 25904 | 1296 | A1296D | E | 2.1e-06 | 1.8e-06 | 1.6e-06 | 1.833E-06 | 2.517E-07 | 1.600E-06 | 2.100E-06 |
| 25905 | 1296 | A1296C | E | 2.2e-06 | 2.6e-06 | 2.1e-06 | 2.300E-06 | 2.646E-07 | 2.100E-06 | 2.600E-06 |
| 25906 | 1296 | A1296E | E | 1.7e-06 | 2.3e-06 | 3.9e-06 | 2.633E-06 | 1.137E-06 | 1.700E-06 | 3.900E-06 |
| 25907 | 1296 | A1296Q | E | 1.7e-06 | 1.4e-06 | 2.3e-06 | 1.800E-06 | 4.583E-07 | 1.400E-06 | 2.300E-06 |
| 25908 | 1296 | A1296G | E | 2.9e-06 | 1.8e-06 | 2.1e-06 | 2.267E-06 | 5.686E-07 | 1.800E-06 | 2.900E-06 |
| 25909 | 1296 | A1296H | E | 1.6e-06 | 1.8e-06 | 2.2e-06 | 1.867E-06 | 3.055E-07 | 1.600E-06 | 2.200E-06 |
| 25910 | 1296 | A1296I | E | 2.1e-06 | 2.9e-06 | 3.8e-06 | 2.933E-06 | 8.505E-07 | 2.100E-06 | 3.800E-06 |
| 25911 | 1296 | A1296L | E | 4.4e-06 | 2.6e-06 | 1.9e-06 | 2.967E-06 | 1.290E-06 | 1.900E-06 | 4.400E-06 |
| 25912 | 1296 | A1296K | E | 2.1e-06 | 1.9e-06 | 2.5e-06 | 2.167E-06 | 3.055E-07 | 1.900E-06 | 2.500E-06 |
| 25913 | 1296 | A1296M | E | 1.1e-06 | 2.1e-06 | 2.5e-06 | 1.900E-06 | 7.211E-07 | 1.100E-06 | 2.500E-06 |
| 25914 | 1296 | A1296F | E | 1.7e-06 | 2.8e-06 | 1.8e-06 | 2.100E-06 | 6.083E-07 | 1.700E-06 | 2.800E-06 |
| 25915 | 1296 | A1296P | E | 2.2e-06 | 2.1e-06 | 2.7e-06 | 2.333E-06 | 3.215E-07 | 2.100E-06 | 2.700E-06 |
| 25916 | 1296 | A1296S | E | 2.5e-06 | 1.9e-06 | 3.2e-06 | 2.533E-06 | 6.506E-07 | 1.900E-06 | 3.200E-06 |
| 25917 | 1296 | A1296T | E | 2.1e-06 | 3.8e-06 | 2.9e-06 | 2.933E-06 | 8.505E-07 | 2.100E-06 | 3.800E-06 |
| 25918 | 1296 | A1296W | E | 3.3e-06 | 1.5e-06 | 2e-06 | 2.267E-06 | 9.292E-07 | 1.500E-06 | 3.300E-06 |
| 25919 | 1296 | A1296Y | E | 8.5e-07 | 2.7e-06 | 2.8e-06 | 2.117E-06 | 1.098E-06 | 8.500E-07 | 2.800E-06 |
| 25920 | 1296 | A1296V | E | 1.8e-06 | 3.1e-06 | 2.7e-06 | 2.533E-06 | 6.658E-07 | 1.800E-06 | 3.100E-06 |
| 25921 | 1297 | S1297A | E | 2.7e-06 | 2.5e-06 | 2.7e-06 | 2.633E-06 | 1.155E-07 | 2.500E-06 | 2.700E-06 |
| 25922 | 1297 | S1297R | E | 2.9e-06 | 4.5e-06 | 3.1e-06 | 3.500E-06 | 8.718E-07 | 2.900E-06 | 4.500E-06 |
| 25923 | 1297 | S1297N | E | 3e-06 | 2.1e-06 | 2.1e-06 | 2.400E-06 | 5.196E-07 | 2.100E-06 | 3.000E-06 |
| 25924 | 1297 | S1297D | E | 2.4e-06 | 2.7e-06 | 3.5e-06 | 2.867E-06 | 5.686E-07 | 2.400E-06 | 3.500E-06 |

| | | | | | | | | | | |
|---|---|---|---|---|---|---|---|---|---|---|
| 25925 | 1297 | S1297C | E | 4e-06 | 3.4e-06 | 4e-06 | 3.800E-06 | 3.464E-07 | 3.400E-06 | 4.000E-06 |
| 25926 | 1297 | S1297E | E | 2.3e-06 | 2.4e-06 | 2.2e-06 | 2.300E-06 | 1.000E-07 | 2.200E-06 | 2.400E-06 |
| 25927 | 1297 | S1297Q | E | 1.7e-06 | 2.1e-06 | 1.8e-06 | 1.867E-06 | 2.082E-07 | 1.700E-06 | 2.100E-06 |
| 25928 | 1297 | S1297G | E | 2.7e-06 | 2.7e-06 | 3.4e-06 | 2.933E-06 | 4.041E-07 | 2.700E-06 | 3.400E-06 |
| 25929 | 1297 | S1297H | E | 1.9e-06 | 2.9e-06 | 2.4e-06 | 2.400E-06 | 5.000E-07 | 1.900E-06 | 2.900E-06 |
| 25930 | 1297 | S1297I | E | 1.7e-06 | 2.3e-06 | 2.5e-06 | 2.167E-06 | 4.163E-07 | 1.700E-06 | 2.500E-06 |
| 25931 | 1297 | S1297L | E | 2.2e-06 | 4.1e-06 | 2.9e-06 | 3.067E-06 | 9.609E-07 | 2.200E-06 | 4.100E-06 |
| 25932 | 1297 | S1297K | E | 1.8e-06 | 1.3e-06 | 1.7e-06 | 1.600E-06 | 2.646E-07 | 1.300E-06 | 1.800E-06 |
| 25933 | 1297 | S1297M | E | 2.3e-06 | 3.1e-06 | 2.6e-06 | 2.667E-06 | 4.041E-07 | 2.300E-06 | 3.100E-06 |
| 25934 | 1297 | S1297F | E | 2.4e-06 | 2.9e-06 | 3.4e-06 | 2.900E-06 | 5.000E-07 | 2.400E-06 | 3.400E-06 |
| 25935 | 1297 | S1297P | E | 2.9e-06 | 1.6e-06 | 2.6e-06 | 2.367E-06 | 6.807E-07 | 1.600E-06 | 2.900E-06 |
| 25936 | 1297 | S1297S | E | 4e-06 | 4.9e-06 | 2.8e-06 | 3.900E-06 | 1.054E-06 | 2.800E-06 | 4.900E-06 |
| 25937 | 1297 | S1297T | E | 2.6e-06 | 2.2e-06 | 1.6e-06 | 2.133E-06 | 5.033E-07 | 1.600E-06 | 2.600E-06 |
| 25938 | 1297 | S1297W | E | 2.8e-06 | 2.1e-06 | 2.9e-06 | 2.600E-06 | 4.359E-07 | 2.100E-06 | 2.900E-06 |
| 25939 | 1297 | S1297Y | E | 1e-06 | 2.5e-06 | 2.1e-06 | 1.867E-06 | 7.767E-07 | 1.000E-06 | 2.500E-06 |
| 25940 | 1297 | S1297V | E | 2.6e-06 | 2.3e-06 | 2e-06 | 2.300E-06 | 3.000E-07 | 2.000E-06 | 2.600E-06 |
| 25941 | 1298 | G1298A | E | 1.1e-06 | 3.3e-06 | 2.2e-06 | 2.200E-06 | 1.100E-06 | 1.100E-06 | 3.300E-06 |
| 25942 | 1298 | G1298R | E | 4e-06 | 2e-06 | 3.2e-06 | 3.067E-06 | 1.007E-06 | 2.000E-06 | 4.000E-06 |
| 25943 | 1298 | G1298N | E | 2.4e-06 | 3e-06 | 1.8e-06 | 2.400E-06 | 6.000E-07 | 1.800E-06 | 3.000E-06 |
| 25944 | 1298 | G1298D | E | 1.2e-06 | 1.8e-06 | 2.6e-06 | 1.867E-06 | 7.024E-07 | 1.200E-06 | 2.600E-06 |
| 25945 | 1298 | G1298C | E | 2.6e-06 | 2.1e-06 | 1.6e-06 | 2.100E-06 | 5.000E-07 | 1.600E-06 | 2.600E-06 |
| 25946 | 1298 | G1298E | E | 2.4e-06 | 2.3e-06 | 5.1e-06 | 3.267E-06 | 1.589E-06 | 2.300E-06 | 5.100E-06 |
| 25947 | 1298 | G1298Q | E | 2.9e-06 | 3.9e-06 | 1.8e-06 | 2.867E-06 | 1.050E-06 | 1.800E-06 | 3.900E-06 |
| 25948 | 1298 | G1298G | E | 3.8e-06 | 2.4e-06 | 1.2e-06 | 2.467E-06 | 1.301E-06 | 1.200E-06 | 3.800E-06 |
| 25949 | 1298 | G1298H | E | 2.5e-06 | 1.4e-06 | 1.6e-06 | 1.833E-06 | 5.859E-07 | 1.400E-06 | 2.500E-06 |
| 25950 | 1298 | G1298I | E | 2.6e-06 | 2.5e-06 | 2.2e-06 | 2.433E-06 | 2.082E-07 | 2.200E-06 | 2.600E-06 |
| 25951 | 1298 | G1298L | E | 2e-06 | 3.5e-06 | 2.4e-06 | 2.633E-06 | 7.767E-07 | 2.000E-06 | 3.500E-06 |
| 25952 | 1298 | G1298K | E | 2.4e-06 | 2.4e-06 | 3.2e-06 | 2.667E-06 | 4.619E-07 | 2.400E-06 | 3.200E-06 |
| 25953 | 1298 | G1298M | E | 2.4e-06 | 1.8e-06 | 2.6e-06 | 2.267E-06 | 4.163E-07 | 1.800E-06 | 2.600E-06 |
| 25954 | 1298 | G1298F | E | 2.4e-06 | 3.5e-06 | 2.2e-06 | 2.700E-06 | 7.000E-07 | 2.200E-06 | 3.500E-06 |
| 25955 | 1298 | G1298P | E | 1.7e-06 | 2.8e-06 | 2.7e-06 | 2.400E-06 | 6.083E-07 | 1.700E-06 | 2.800E-06 |
| 25956 | 1298 | G1298S | E | 1.6e-06 | 2.9e-06 | 1.8e-06 | 2.100E-06 | 7.000E-07 | 1.600E-06 | 2.900E-06 |
| 25957 | 1298 | G1298T | E | 4.6e-06 | 3.2e-06 | 1.5e-06 | 3.100E-06 | 1.552E-06 | 1.500E-06 | 4.600E-06 |
| 25958 | 1298 | G1298W | E | 2.9e-06 | 2.2e-06 | 2.3e-06 | 2.467E-06 | 3.786E-07 | 2.200E-06 | 2.900E-06 |
| 25959 | 1298 | G1298Y | E | 2.4e-06 | 2.9e-06 | 2.5e-06 | 2.600E-06 | 2.646E-07 | 2.400E-06 | 2.900E-06 |
| 25960 | 1298 | G1298V | E | 2.9e-06 | 2.2e-06 | 2.1e-06 | 2.400E-06 | 4.359E-07 | 2.100E-06 | 2.900E-06 |
| 25961 | 1299 | F1299A | E | 1.5e-06 | 2.6e-06 | 2.4e-06 | 2.167E-06 | 5.859E-07 | 1.500E-06 | 2.600E-06 |
| 25962 | 1299 | F1299R | E | 2.2e-06 | 1.1e-06 | 2.2e-06 | 1.833E-06 | 6.351E-07 | 1.100E-06 | 2.200E-06 |
| 25963 | 1299 | F1299N | E | 2.9e-06 | 2.9e-06 | 3.4e-06 | 3.067E-06 | 2.887E-07 | 2.900E-06 | 3.400E-06 |
| 25964 | 1299 | F1299D | E | 2.5e-06 | 2e-06 | 3.6e-06 | 2.700E-06 | 8.185E-07 | 2.000E-06 | 3.600E-06 |
| 25965 | 1299 | F1299C | E | 1.6e-06 | 2.7e-06 | 2.1e-06 | 2.133E-06 | 5.508E-07 | 1.600E-06 | 2.700E-06 |
| 25966 | 1299 | F1299E | E | 2.1e-06 | 3.1e-06 | 1.2e-06 | 2.133E-06 | 9.504E-07 | 1.200E-06 | 3.100E-06 |
| 25967 | 1299 | F1299Q | E | 1.2e-06 | 1.8e-06 | 2.7e-06 | 1.900E-06 | 7.550E-07 | 1.200E-06 | 2.700E-06 |
| 25968 | 1299 | F1299G | E | 4.3e-06 | 3.9e-06 | 3.1e-06 | 3.767E-06 | 6.110E-07 | 3.100E-06 | 4.300E-06 |
| 25969 | 1299 | F1299H | E | 2.9e-06 | 2.2e-06 | 4.3e-06 | 3.133E-06 | 1.069E-06 | 2.200E-06 | 4.300E-06 |
| 25970 | 1299 | F1299I | E | 3.5e-06 | 4.6e-06 | 3.1e-06 | 3.733E-06 | 7.767E-07 | 3.100E-06 | 4.600E-06 |
| 25971 | 1299 | F1299L | E | 2e-06 | 1.9e-06 | 3.1e-06 | 2.333E-06 | 6.658E-07 | 1.900E-06 | 3.100E-06 |
| 25972 | 1299 | F1299K | E | 2.9e-06 | 1.6e-06 | 9.5e-07 | 1.817E-06 | 9.929E-07 | 9.500E-07 | 2.900E-06 |
| 25973 | 1299 | F1299M | E | 1.6e-06 | 2.7e-06 | 2.3e-06 | 2.200E-06 | 5.568E-07 | 1.600E-06 | 2.700E-06 |
| 25974 | 1299 | F1299F | E | 2.3e-06 | 1.9e-06 | 2.9e-06 | 2.367E-06 | 5.033E-07 | 1.900E-06 | 2.900E-06 |
| 25975 | 1299 | F1299P | E | 2.3e-06 | 3.2e-06 | 2.2e-06 | 2.567E-06 | 5.508E-07 | 2.200E-06 | 3.200E-06 |
| 25976 | 1299 | F1299S | E | 3e-06 | 4.3e-06 | 1.3e-06 | 2.867E-06 | 1.504E-06 | 1.300E-06 | 4.300E-06 |
| 25977 | 1299 | F1299T | E | 1.9e-06 | 1.2e-06 | 5e-06 | 2.700E-06 | 2.022E-06 | 1.200E-06 | 5.000E-06 |
| 25978 | 1299 | F1299W | E | 2e-06 | 1.9e-06 | 3e-06 | 2.300E-06 | 6.083E-07 | 1.900E-06 | 3.000E-06 |
| 25979 | 1299 | F1299Y | E | 9.4e-07 | 2.9e-06 | 1.2e-06 | 1.680E-06 | 1.065E-06 | 9.400E-07 | 2.900E-06 |
| 25980 | 1299 | F1299V | E | 2.2e-06 | 2.4e-06 | 1.5e-06 | 2.033E-06 | 4.726E-07 | 1.500E-06 | 2.400E-06 |
| 25981 | 1300 | N1300A | E | 1.8e-06 | 2.3e-06 | 1.7e-06 | 1.933E-06 | 3.215E-07 | 1.700E-06 | 2.300E-06 |
| 25982 | 1300 | N1300R | E | 8.6e-07 | 2e-06 | 1.5e-06 | 1.453E-06 | 5.714E-07 | 8.600E-07 | 2.000E-06 |
| 25983 | 1300 | N1300N | E | 2.2e-06 | 3.4e-06 | 2e-06 | 2.533E-06 | 7.572E-07 | 2.000E-06 | 3.400E-06 |
| 25984 | 1300 | N1300D | E | 2.5e-06 | 2.6e-06 | 1.6e-06 | 2.233E-06 | 5.508E-07 | 1.600E-06 | 2.600E-06 |
| 25985 | 1300 | N1300C | E | 2.2e-06 | 1.8e-06 | 2.1e-06 | 2.033E-06 | 2.082E-07 | 1.800E-06 | 2.200E-06 |
| 25986 | 1300 | N1300E | E | 2.3e-06 | 3e-06 | 1.8e-06 | 2.367E-06 | 6.028E-07 | 1.800E-06 | 3.000E-06 |
| 25987 | 1300 | N1300Q | E | 2.2e-06 | 2.4e-06 | 1.5e-06 | 2.033E-06 | 4.726E-07 | 1.500E-06 | 2.400E-06 |
| 25988 | 1300 | N1300G | E | 1.3e-06 | 2.7e-06 | 1.8e-06 | 1.933E-06 | 7.095E-07 | 1.300E-06 | 2.700E-06 |
| 25989 | 1300 | N1300H | E | 2e-06 | 1.6e-06 | 1.3e-06 | 1.633E-06 | 3.512E-07 | 1.300E-06 | 2.000E-06 |
| 25990 | 1300 | N1300I | E | 1.5e-06 | 1.9e-06 | 2.2e-06 | 1.867E-06 | 3.512E-07 | 1.500E-06 | 2.200E-06 |
| 25991 | 1300 | N1300L | E | 1.4e-06 | 1.8e-06 | 2.2e-06 | 1.800E-06 | 4.000E-07 | 1.400E-06 | 2.200E-06 |
| 25992 | 1300 | N1300K | E | 1.1e-06 | 2.8e-06 | 1.6e-06 | 1.833E-06 | 8.737E-07 | 1.100E-06 | 2.800E-06 |
| 25993 | 1300 | N1300M | E | 9.8e-07 | 1.5e-06 | 2.7e-06 | 1.727E-06 | 8.821E-07 | 9.800E-07 | 2.700E-06 |
| 25994 | 1300 | N1300F | E | 2.3e-06 | 1.4e-06 | 1.2e-06 | 1.633E-06 | 5.859E-07 | 1.200E-06 | 2.300E-06 |
| 25995 | 1300 | N1300P | E | 1.2e-06 | 1.7e-06 | 1.2e-06 | 1.367E-06 | 2.887E-07 | 1.200E-06 | 1.700E-06 |
| 25996 | 1300 | N1300S | E | 1.7e-06 | 3.1e-06 | 1.9e-06 | 2.233E-06 | 7.572E-07 | 1.700E-06 | 3.100E-06 |
| 25997 | 1300 | N1300T | E | 1.7e-06 | 2.4e-06 | 3.7e-06 | 2.600E-06 | 1.015E-06 | 1.700E-06 | 3.700E-06 |
| 25998 | 1300 | N1300W | E | 2.1e-06 | 1.9e-06 | 2.5e-06 | 2.167E-06 | 3.055E-07 | 1.900E-06 | 2.500E-06 |
| 25999 | 1300 | N1300Y | E | 1.8e-06 | 3.7e-06 | 1.9e-06 | 2.467E-06 | 1.069E-06 | 1.800E-06 | 3.700E-06 |
| 26000 | 1300 | N1300V | E | 1.1e-06 | 1.8e-06 | 2.7e-06 | 1.867E-06 | 8.021E-07 | 1.100E-06 | 2.700E-06 |
| 26001 | 1301 | I1301A | E | 3.1e-06 | 2.8e-06 | 3.3e-06 | 3.067E-06 | 2.517E-07 | 2.800E-06 | 3.300E-06 |
| 26002 | 1301 | I1301R | E | 2e-06 | 2.9e-06 | 2.1e-06 | 2.333E-06 | 4.933E-07 | 2.000E-06 | 2.900E-06 |
| 26003 | 1301 | I1301N | E | 2.5e-06 | 2.4e-06 | 1.8e-06 | 2.233E-06 | 3.786E-07 | 1.800E-06 | 2.500E-06 |
| 26004 | 1301 | I1301D | E | 1.5e-06 | 1.6e-06 | 2.2e-06 | 1.767E-06 | 3.786E-07 | 1.500E-06 | 2.200E-06 |
| 26005 | 1301 | I1301C | E | 2.8e-06 | 2.4e-06 | 2.4e-06 | 2.533E-06 | 2.309E-07 | 2.400E-06 | 2.800E-06 |
| 26006 | 1301 | I1301E | E | 1.9e-06 | 2.2e-06 | 2.7e-06 | 2.267E-06 | 4.041E-07 | 1.900E-06 | 2.700E-06 |
| 26007 | 1301 | I1301Q | E | 2.4e-06 | 3.1e-06 | 3.8e-06 | 3.100E-06 | 7.000E-07 | 2.400E-06 | 3.800E-06 |
| 26008 | 1301 | I1301G | E | 1.2e-06 | 1.8e-06 | 2e-06 | 1.667E-06 | 4.163E-07 | 1.200E-06 | 2.000E-06 |
| 26009 | 1301 | I1301H | E | 2.3e-06 | 2.6e-06 | 2.1e-06 | 2.333E-06 | 2.517E-07 | 2.100E-06 | 2.600E-06 |

| | | | | | | | | | | |
|---|---|---|---|---|---|---|---|---|---|---|
| 26010 | 1301 | I1301I | E | 2e-06 | 2.1e-06 | 2.3e-06 | 2.133E-06 | 1.528E-07 | 2.000E-06 | 2.300E-06 |
| 26011 | 1301 | I1301L | E | 2.9e-06 | 2.2e-06 | 1.8e-06 | 2.300E-06 | 5.568E-07 | 1.800E-06 | 2.900E-06 |
| 26012 | 1301 | I1301K | E | 2.4e-06 | 2.1e-06 | 1.9e-06 | 2.133E-06 | 2.517E-07 | 1.900E-06 | 2.400E-06 |
| 26013 | 1301 | I1301M | E | 1.3e-06 | 2.9e-06 | 3e-06 | 2.400E-06 | 9.539E-07 | 1.300E-06 | 3.000E-06 |
| 26014 | 1301 | I1301F | E | 1.8e-06 | 2.8e-06 | 3.1e-06 | 2.567E-06 | 6.807E-07 | 1.800E-06 | 3.100E-06 |
| 26015 | 1301 | I1301P | E | 2.8e-06 | 2.8e-06 | 3.1e-06 | 2.900E-06 | 1.732E-07 | 2.800E-06 | 3.100E-06 |
| 26016 | 1301 | I1301S | E | 3.2e-06 | 2.8e-06 | 2.7e-06 | 2.900E-06 | 2.646E-07 | 2.700E-06 | 3.200E-06 |
| 26017 | 1301 | I1301T | E | 2e-06 | 2.3e-06 | 2.9e-06 | 2.400E-06 | 4.583E-07 | 2.000E-06 | 2.900E-06 |
| 26018 | 1301 | I1301W | E | 1.6e-06 | 3e-06 | 4.4e-06 | 3.000E-06 | 1.400E-06 | 1.600E-06 | 4.400E-06 |
| 26019 | 1301 | I1301Y | E | 3.9e-06 | 3.4e-06 | 2.3e-06 | 3.200E-06 | 8.185E-07 | 2.300E-06 | 3.900E-06 |
| 26020 | 1301 | I1301V | E | 2.3e-06 | 3.8e-06 | 3.7e-06 | 3.267E-06 | 8.386E-07 | 2.300E-06 | 3.800E-06 |
| 26021 | 1302 | K1302A | E | 3e-06 | 2.1e-06 | 2.6e-06 | 2.567E-06 | 4.509E-07 | 2.100E-06 | 3.000E-06 |
| 26022 | 1302 | K1302R | E | 2.7e-06 | 2.2e-06 | 3.1e-06 | 2.667E-06 | 4.509E-07 | 2.200E-06 | 3.100E-06 |
| 26023 | 1302 | K1302N | E | 2.4e-06 | 1.1e-06 | 2.7e-06 | 2.067E-06 | 8.505E-07 | 1.100E-06 | 2.700E-06 |
| 26024 | 1302 | K1302D | E | 2.8e-06 | 2.2e-06 | 2.8e-06 | 2.600E-06 | 3.464E-07 | 2.200E-06 | 2.800E-06 |
| 26025 | 1302 | K1302C | E | 1.9e-06 | 2.3e-06 | 2.9e-06 | 2.367E-06 | 5.033E-07 | 1.900E-06 | 2.900E-06 |
| 26026 | 1302 | K1302E | E | 2.4e-06 | 3e-06 | 1.6e-06 | 2.333E-06 | 7.024E-07 | 1.600E-06 | 3.000E-06 |
| 26027 | 1302 | K1302Q | E | 2.8e-06 | 3e-06 | 1.8e-06 | 2.533E-06 | 6.429E-07 | 1.800E-06 | 3.000E-06 |
| 26028 | 1302 | K1302G | E | 2.8e-06 | 1.9e-06 | 2.1e-06 | 2.267E-06 | 4.726E-07 | 1.900E-06 | 2.800E-06 |
| 26029 | 1302 | K1302H | E | 3.7e-06 | 2.2e-06 | 3.2e-06 | 3.033E-06 | 7.638E-07 | 2.200E-06 | 3.700E-06 |
| 26030 | 1302 | K1302I | E | 2.4e-06 | 3.1e-06 | 1.6e-06 | 2.367E-06 | 7.506E-07 | 1.600E-06 | 3.100E-06 |
| 26031 | 1302 | K1302L | E | 2.1e-06 | 4.9e-06 | 1.4e-06 | 2.800E-06 | 1.852E-06 | 1.400E-06 | 4.900E-06 |
| 26032 | 1302 | K1302K | E | 1.9e-06 | 2.4e-06 | 3.1e-06 | 2.467E-06 | 6.028E-07 | 1.900E-06 | 3.100E-06 |
| 26033 | 1302 | K1302M | E | 3.1e-06 | 3.1e-06 | 2.9e-06 | 3.033E-06 | 1.155E-07 | 2.900E-06 | 3.100E-06 |
| 26034 | 1302 | K1302F | E | 2.1e-06 | 1.9e-06 | 2.8e-06 | 2.267E-06 | 4.726E-07 | 1.900E-06 | 2.800E-06 |
| 26035 | 1302 | K1302P | E | 1.1e-06 | 2e-06 | 1.8e-06 | 1.633E-06 | 4.726E-07 | 1.100E-06 | 2.000E-06 |
| 26036 | 1302 | K1302S | E | 1.8e-06 | 3.6e-06 | 2.1e-06 | 2.500E-06 | 9.644E-07 | 1.800E-06 | 3.600E-06 |
| 26037 | 1302 | K1302T | E | 2.3e-06 | 2.8e-06 | 2.7e-06 | 2.600E-06 | 2.646E-07 | 2.300E-06 | 2.800E-06 |
| 26038 | 1302 | K1302W | E | 2.9e-06 | 2.2e-06 | 2.2e-06 | 2.433E-06 | 4.041E-07 | 2.200E-06 | 2.900E-06 |
| 26039 | 1302 | K1302Y | E | 3.9e-06 | 3.6e-06 | 3.8e-06 | 3.767E-06 | 1.528E-07 | 3.600E-06 | 3.900E-06 |
| 26040 | 1302 | K1302V | E | 2.5e-06 | 1.5e-06 | 2.7e-06 | 2.233E-06 | 6.429E-07 | 1.500E-06 | 2.700E-06 |
| 26041 | 1303 | D1303A | E | 1.9e-06 | 3.1e-06 | 1.8e-06 | 2.267E-06 | 7.234E-07 | 1.800E-06 | 3.100E-06 |
| 26042 | 1303 | D1303R | E | 1.7e-06 | 1.6e-06 | 2.1e-06 | 1.800E-06 | 2.646E-07 | 1.600E-06 | 2.100E-06 |
| 26043 | 1303 | D1303N | E | 1.8e-06 | 1.4e-06 | 3e-06 | 2.067E-06 | 8.327E-07 | 1.400E-06 | 3.000E-06 |
| 26044 | 1303 | D1303D | E | 1.5e-06 | 2.7e-06 | 1.4e-06 | 1.867E-06 | 7.234E-07 | 1.400E-06 | 2.700E-06 |
| 26045 | 1303 | D1303C | E | 2e-06 | 3.2e-06 | 2.5e-06 | 2.567E-06 | 6.028E-07 | 2.000E-06 | 3.200E-06 |
| 26046 | 1303 | D1303E | E | 1.8e-06 | 1.7e-06 | 1.6e-06 | 1.700E-06 | 1.000E-07 | 1.600E-06 | 1.800E-06 |
| 26047 | 1303 | D1303Q | E | 1.8e-06 | 1.4e-06 | 3.5e-06 | 2.233E-06 | 1.115E-06 | 1.400E-06 | 3.500E-06 |
| 26048 | 1303 | D1303G | E | 2.5e-06 | 2.5e-06 | 3.5e-06 | 2.833E-06 | 5.774E-07 | 2.500E-06 | 3.500E-06 |
| 26049 | 1303 | D1303H | E | 8.1e-07 | 1.8e-06 | 2e-06 | 1.537E-06 | 6.372E-07 | 8.100E-07 | 2.000E-06 |
| 26050 | 1303 | D1303I | E | 2.5e-06 | 2e-06 | 2e-06 | 2.167E-06 | 2.887E-07 | 2.000E-06 | 2.500E-06 |
| 26051 | 1303 | D1303L | E | 1.8e-06 | 1.2e-06 | 1.8e-06 | 1.600E-06 | 3.464E-07 | 1.200E-06 | 1.800E-06 |
| 26052 | 1303 | D1303K | E | 1.2e-06 | 3.3e-06 | 1.5e-06 | 2.000E-06 | 1.136E-06 | 1.200E-06 | 3.300E-06 |
| 26053 | 1303 | D1303M | E | 1.9e-06 | 2.5e-06 | 1.9e-06 | 2.100E-06 | 3.464E-07 | 1.900E-06 | 2.500E-06 |
| 26054 | 1303 | D1303F | E | 2e-06 | 2e-06 | 4.8e-06 | 2.933E-06 | 1.617E-06 | 2.000E-06 | 4.800E-06 |
| 26055 | 1303 | D1303P | E | 2.7e-06 | 2.8e-06 | 2.4e-06 | 2.633E-06 | 2.082E-07 | 2.400E-06 | 2.800E-06 |
| 26056 | 1303 | D1303S | E | 1.2e-06 | 3e-06 | 2.5e-06 | 2.233E-06 | 9.292E-07 | 1.200E-06 | 3.000E-06 |
| 26057 | 1303 | D1303T | E | 2.3e-06 | 3.9e-06 | 2.4e-06 | 2.867E-06 | 8.963E-07 | 2.300E-06 | 3.900E-06 |
| 26058 | 1303 | D1303W | E | 3e-06 | 3.5e-06 | 1.8e-06 | 2.767E-06 | 8.737E-07 | 1.800E-06 | 3.500E-06 |
| 26059 | 1303 | D1303Y | E | 3.4e-06 | 1.7e-06 | 2.8e-06 | 2.633E-06 | 8.622E-07 | 1.700E-06 | 3.400E-06 |
| 26060 | 1303 | D1303V | E | 1.7e-06 | 2.5e-06 | 1.9e-06 | 2.033E-06 | 4.163E-07 | 1.700E-06 | 2.500E-06 |
| 26061 | 1304 | T1304A | E | 3e-06 | 2.6e-06 | 2e-06 | 2.533E-06 | 5.033E-07 | 2.000E-06 | 3.000E-06 |
| 26062 | 1304 | T1304R | E | 2.1e-06 | 2.3e-06 | 1.3e-06 | 1.900E-06 | 5.292E-07 | 1.300E-06 | 2.300E-06 |
| 26063 | 1304 | T1304N | E | 3.1e-06 | 2.4e-06 | 2.4e-06 | 2.633E-06 | 4.041E-07 | 2.400E-06 | 3.100E-06 |
| 26064 | 1304 | T1304D | E | 3.2e-06 | 1.8e-06 | 1.7e-06 | 2.233E-06 | 8.386E-07 | 1.700E-06 | 3.200E-06 |
| 26065 | 1304 | T1304C | E | 1.5e-06 | 2.8e-06 | 2.9e-06 | 2.400E-06 | 7.810E-07 | 1.500E-06 | 2.900E-06 |
| 26066 | 1304 | T1304E | E | 1.5e-06 | 3.6e-06 | 1.9e-06 | 2.333E-06 | 1.115E-06 | 1.500E-06 | 3.600E-06 |
| 26067 | 1304 | T1304Q | E | 2.3e-06 | 2.6e-06 | 1.6e-06 | 2.167E-06 | 5.132E-07 | 1.600E-06 | 2.600E-06 |
| 26068 | 1304 | T1304G | E | 2.8e-06 | 2.3e-06 | 4.5e-06 | 3.200E-06 | 1.153E-06 | 2.300E-06 | 4.500E-06 |
| 26069 | 1304 | T1304H | E | 4.3e-06 | 2.6e-06 | 2e-06 | 2.967E-06 | 1.193E-06 | 2.000E-06 | 4.300E-06 |
| 26070 | 1304 | T1304I | E | 3.9e-06 | 1.4e-06 | 3.6e-06 | 2.967E-06 | 1.365E-06 | 1.400E-06 | 3.900E-06 |
| 26071 | 1304 | T1304L | E | 2.2e-06 | 2.4e-06 | 2.2e-06 | 2.267E-06 | 1.155E-07 | 2.200E-06 | 2.400E-06 |
| 26072 | 1304 | T1304K | E | 2.7e-06 | 1.3e-06 | 1.6e-06 | 1.867E-06 | 7.371E-07 | 1.300E-06 | 2.700E-06 |
| 26073 | 1304 | T1304M | E | 3.1e-06 | 2.6e-06 | 1.5e-06 | 2.400E-06 | 8.185E-07 | 1.500E-06 | 3.100E-06 |
| 26074 | 1304 | T1304F | E | 2.2e-06 | 2.8e-06 | 2.8e-06 | 2.600E-06 | 3.464E-07 | 2.200E-06 | 2.800E-06 |
| 26075 | 1304 | T1304P | E | 1.9e-06 | 2.2e-06 | 2.3e-06 | 2.133E-06 | 2.082E-07 | 1.900E-06 | 2.300E-06 |
| 26076 | 1304 | T1304S | E | 3.7e-06 | 4.3e-06 | 1.5e-06 | 3.167E-06 | 1.474E-06 | 1.500E-06 | 4.300E-06 |
| 26077 | 1304 | T1304T | E | 3.1e-06 | 2.3e-06 | 1.6e-06 | 2.333E-06 | 7.506E-07 | 1.600E-06 | 3.100E-06 |
| 26078 | 1304 | T1304W | E | 2.5e-06 | 2.4e-06 | 3.1e-06 | 2.667E-06 | 3.786E-07 | 2.400E-06 | 3.100E-06 |
| 26079 | 1304 | T1304Y | E | 2.7e-06 | 4.8e-06 | 2.9e-06 | 3.467E-06 | 1.159E-06 | 2.700E-06 | 4.800E-06 |
| 26080 | 1304 | T1304V | E | 1.4e-06 | 1.5e-06 | 1.3e-06 | 1.400E-06 | 1.000E-07 | 1.300E-06 | 1.500E-06 |
| 26081 | 1305 | Y1305A | E | 2.5e-06 | 3.6e-06 | 2.2e-06 | 2.767E-06 | 7.371E-07 | 2.200E-06 | 3.600E-06 |
| 26082 | 1305 | Y1305R | E | 2.8e-06 | 1.5e-06 | 1.6e-06 | 1.967E-06 | 7.234E-07 | 1.500E-06 | 2.800E-06 |
| 26083 | 1305 | Y1305N | E | 9.2e-07 | 1.9e-06 | 4e-06 | 2.273E-06 | 1.574E-06 | 9.200E-07 | 4.000E-06 |
| 26084 | 1305 | Y1305D | E | 2.7e-06 | 1.2e-06 | 4.2e-06 | 2.700E-06 | 1.500E-06 | 1.200E-06 | 4.200E-06 |
| 26085 | 1305 | Y1305C | E | 3.8e-06 | 2.1e-06 | 2.8e-06 | 2.900E-06 | 8.544E-07 | 2.100E-06 | 3.800E-06 |
| 26086 | 1305 | Y1305E | E | 3.2e-06 | 1.8e-06 | 4.3e-06 | 3.100E-06 | 1.253E-06 | 1.800E-06 | 4.300E-06 |
| 26087 | 1305 | Y1305Q | E | 2.2e-06 | 2.5e-06 | 2.1e-06 | 2.267E-06 | 2.082E-07 | 2.100E-06 | 2.500E-06 |
| 26088 | 1305 | Y1305G | E | 3.1e-06 | 3e-06 | 1.6e-06 | 2.567E-06 | 8.386E-07 | 1.600E-06 | 3.100E-06 |
| 26089 | 1305 | Y1305H | E | 3.1e-06 | 4.1e-06 | 2.4e-06 | 3.200E-06 | 8.544E-07 | 2.400E-06 | 4.100E-06 |
| 26090 | 1305 | Y1305I | E | 3.7e-06 | 2e-06 | 2.6e-06 | 2.767E-06 | 8.622E-07 | 2.000E-06 | 3.700E-06 |
| 26091 | 1305 | Y1305L | E | 2.6e-06 | 3.2e-06 | 3.1e-06 | 2.967E-06 | 3.215E-07 | 2.600E-06 | 3.200E-06 |
| 26092 | 1305 | Y1305K | E | 1.8e-06 | 2.2e-06 | 2.2e-06 | 2.067E-06 | 2.309E-07 | 1.800E-06 | 2.200E-06 |
| 26093 | 1305 | Y1305M | E | 2.3e-06 | 2.4e-06 | 2.3e-06 | 2.333E-06 | 5.774E-08 | 2.300E-06 | 2.400E-06 |
| 26094 | 1305 | Y1305F | E | 2.9e-06 | 1.7e-06 | 3.1e-06 | 2.567E-06 | 7.572E-07 | 1.700E-06 | 3.100E-06 |

| | | | | | | | | | | |
|---|---|---|---|---|---|---|---|---|---|---|
| 26095 | 1305 | Y1305P | E | 3.6e-06 | 1.6e-06 | 3.6e-06 | 2.933E-06 | 1.155E-06 | 1.600E-06 | 3.600E-06 |
| 26096 | 1305 | Y1305S | E | 2.3e-06 | 2.5e-06 | 2.5e-06 | 2.433E-06 | 1.155E-07 | 2.300E-06 | 2.500E-06 |
| 26097 | 1305 | Y1305T | E | 2.3e-06 | 4e-06 | 2.4e-06 | 2.900E-06 | 9.539E-07 | 2.300E-06 | 4.000E-06 |
| 26098 | 1305 | Y1305W | E | 2.3e-06 | 2.7e-06 | 2.6e-06 | 2.533E-06 | 2.082E-07 | 2.300E-06 | 2.700E-06 |
| 26099 | 1305 | Y1305Y | E | 2.9e-06 | 2e-06 | 1.2e-06 | 2.033E-06 | 8.505E-07 | 1.200E-06 | 2.900E-06 |
| 26100 | 1305 | Y1305V | E | 5.8e-06 | 3.2e-06 | 2.6e-06 | 3.867E-06 | 1.701E-06 | 2.600E-06 | 5.800E-06 |
| 26101 | 1306 | I1306A | E | 3.7e-06 | 2.3e-06 | 2.7e-06 | 2.900E-06 | 7.211E-07 | 2.300E-06 | 3.700E-06 |
| 26102 | 1306 | I1306R | E | 2.8e-06 | 1.9e-06 | 2.4e-06 | 2.367E-06 | 4.509E-07 | 1.900E-06 | 2.800E-06 |
| 26103 | 1306 | I1306N | E | 2.7e-06 | 2.5e-06 | 3e-06 | 2.733E-06 | 2.517E-07 | 2.500E-06 | 3.000E-06 |
| 26104 | 1306 | I1306D | E | 1.9e-06 | 2.6e-06 | 2.4e-06 | 2.300E-06 | 3.606E-07 | 1.900E-06 | 2.600E-06 |
| 26105 | 1306 | I1306C | E | 1.5e-06 | 1.5e-06 | 2.3e-06 | 1.767E-06 | 4.619E-07 | 1.500E-06 | 2.300E-06 |
| 26106 | 1306 | I1306E | E | 4.3e-06 | 1.4e-06 | 3.5e-06 | 3.067E-06 | 1.498E-06 | 1.400E-06 | 4.300E-06 |
| 26107 | 1306 | I1306Q | E | 2.9e-06 | 2.7e-06 | 2.2e-06 | 2.600E-06 | 3.606E-07 | 2.200E-06 | 2.900E-06 |
| 26108 | 1306 | I1306G | E | 1.9e-06 | 2.1e-06 | 2.3e-06 | 2.100E-06 | 2.000E-07 | 1.900E-06 | 2.300E-06 |
| 26109 | 1306 | I1306H | E | 1.7e-06 | 1.1e-06 | 2.8e-06 | 1.867E-06 | 8.622E-07 | 1.100E-06 | 2.800E-06 |
| 26110 | 1306 | I1306I | E | 1.6e-06 | 1.8e-06 | 1.1e-06 | 1.500E-06 | 3.606E-07 | 1.100E-06 | 1.800E-06 |
| 26111 | 1306 | I1306L | E | 2.3e-06 | 2.7e-06 | 2.2e-06 | 2.400E-06 | 2.646E-07 | 2.200E-06 | 2.700E-06 |
| 26112 | 1306 | I1306K | E | 3.2e-06 | 1.9e-06 | 4.5e-06 | 3.200E-06 | 1.300E-06 | 1.900E-06 | 4.500E-06 |
| 26113 | 1306 | I1306M | E | 2.3e-06 | 2.7e-06 | 2.4e-06 | 2.467E-06 | 2.082E-07 | 2.300E-06 | 2.700E-06 |
| 26114 | 1306 | I1306F | E | 3.7e-06 | 2.1e-06 | 1.3e-06 | 2.367E-06 | 1.222E-06 | 1.300E-06 | 3.700E-06 |
| 26115 | 1306 | I1306P | E | 2.6e-06 | 3.4e-06 | 2.4e-06 | 2.800E-06 | 5.292E-07 | 2.400E-06 | 3.400E-06 |
| 26116 | 1306 | I1306S | E | 2.5e-06 | 1.2e-06 | 2e-06 | 1.900E-06 | 6.557E-07 | 1.200E-06 | 2.500E-06 |
| 26117 | 1306 | I1306T | E | 2.3e-06 | 1.8e-06 | 1.9e-06 | 2.000E-06 | 2.646E-07 | 1.800E-06 | 2.300E-06 |
| 26118 | 1306 | I1306W | E | 6.7e-06 | 1.4e-06 | 1.7e-06 | 3.267E-06 | 2.977E-06 | 1.400E-06 | 6.700E-06 |
| 26119 | 1306 | I1306Y | E | 2.3e-06 | 3.8e-06 | 1.9e-06 | 2.667E-06 | 1.002E-06 | 1.900E-06 | 3.800E-06 |
| 26120 | 1306 | I1306V | E | 2.3e-06 | 1.7e-06 | 2.9e-06 | 2.300E-06 | 6.000E-07 | 1.700E-06 | 2.900E-06 |
| 26121 | 1307 | H1307A | E | 1.9e-06 | 2.3e-06 | 1.7e-06 | 1.967E-06 | 3.055E-07 | 1.700E-06 | 2.300E-06 |
| 26122 | 1307 | H1307R | E | 1.9e-06 | 2.6e-06 | 2.2e-06 | 2.233E-06 | 3.512E-07 | 1.900E-06 | 2.600E-06 |
| 26123 | 1307 | H1307N | E | 3.5e-06 | 3.9e-06 | 3.1e-06 | 3.500E-06 | 4.000E-07 | 3.100E-06 | 3.900E-06 |
| 26124 | 1307 | H1307D | E | 2e-06 | 2.9e-06 | 2.8e-06 | 2.567E-06 | 4.933E-07 | 2.000E-06 | 2.900E-06 |
| 26125 | 1307 | H1307C | E | 2.5e-06 | 2.2e-06 | 2.9e-06 | 2.533E-06 | 3.512E-07 | 2.200E-06 | 2.900E-06 |
| 26126 | 1307 | H1307E | E | 2.7e-06 | 2.4e-06 | 2.7e-06 | 2.600E-06 | 1.732E-07 | 2.400E-06 | 2.700E-06 |
| 26127 | 1307 | H1307Q | E | 2.9e-06 | 1.9e-06 | 3.3e-06 | 2.700E-06 | 7.211E-07 | 1.900E-06 | 3.300E-06 |
| 26128 | 1307 | H1307G | E | 1.8e-06 | 1.7e-06 | 2.5e-06 | 2.000E-06 | 4.359E-07 | 1.700E-06 | 2.500E-06 |
| 26129 | 1307 | H1307H | E | 1.9e-06 | 2.3e-06 | 2.6e-06 | 2.267E-06 | 3.512E-07 | 1.900E-06 | 2.600E-06 |
| 26130 | 1307 | H1307I | E | 3.2e-06 | 1.4e-06 | 1.1e-06 | 1.900E-06 | 1.136E-06 | 1.100E-06 | 3.200E-06 |
| 26131 | 1307 | H1307L | E | 2.2e-06 | 3.1e-06 | 3.4e-06 | 2.900E-06 | 6.245E-07 | 2.200E-06 | 3.400E-06 |
| 26132 | 1307 | H1307K | E | 1.6e-06 | 1.6e-06 | 1.9e-06 | 1.700E-06 | 1.732E-07 | 1.600E-06 | 1.900E-06 |
| 26133 | 1307 | H1307M | E | 2.4e-06 | 4.1e-06 | 2.7e-06 | 3.067E-06 | 9.074E-07 | 2.400E-06 | 4.100E-06 |
| 26134 | 1307 | H1307F | E | 2.1e-06 | 2.8e-06 | 4.6e-06 | 3.167E-06 | 1.290E-06 | 2.100E-06 | 4.600E-06 |
| 26135 | 1307 | H1307P | E | 2.7e-06 | 2e-06 | 1.2e-06 | 1.967E-06 | 7.506E-07 | 1.200E-06 | 2.700E-06 |
| 26136 | 1307 | H1307S | E | 3.6e-06 | 4.2e-06 | 1.6e-06 | 3.133E-06 | 1.361E-06 | 1.600E-06 | 4.200E-06 |
| 26137 | 1307 | H1307T | E | 2e-06 | 3.4e-06 | 3.1e-06 | 2.833E-06 | 7.371E-07 | 2.000E-06 | 3.400E-06 |
| 26138 | 1307 | H1307W | E | 1.4e-06 | 2.8e-06 | 8.3e-07 | 1.677E-06 | 1.014E-06 | 8.300E-07 | 2.800E-06 |
| 26139 | 1307 | H1307Y | E | 1.9e-06 | 1.7e-06 | 4.3e-06 | 2.633E-06 | 1.447E-06 | 1.700E-06 | 4.300E-06 |
| 26140 | 1307 | H1307V | E | 1.4e-06 | 1.7e-06 | 2.7e-06 | 1.933E-06 | 6.807E-07 | 1.400E-06 | 2.700E-06 |
| 26141 | 1308 | W1308A | E | 2.8e-06 | 2e-06 | 4.1e-06 | 2.967E-06 | 1.060E-06 | 2.000E-06 | 4.100E-06 |
| 26142 | 1308 | W1308R | E | 2.5e-06 | 9.3e-07 | 3.2e-06 | 2.210E-06 | 1.162E-06 | 9.300E-07 | 3.200E-06 |
| 26143 | 1308 | W1308N | E | 1.8e-06 | 3e-06 | 3.8e-06 | 2.867E-06 | 1.007E-06 | 1.800E-06 | 3.800E-06 |
| 26144 | 1308 | W1308D | E | 2.1e-06 | 2.8e-06 | 1.7e-06 | 2.200E-06 | 5.568E-07 | 1.700E-06 | 2.800E-06 |
| 26145 | 1308 | W1308C | E | 2.2e-06 | 2.2e-06 | 2.7e-06 | 2.367E-06 | 2.887E-07 | 2.200E-06 | 2.700E-06 |
| 26146 | 1308 | W1308E | E | 2.1e-06 | 2.3e-06 | 2.7e-06 | 2.367E-06 | 3.055E-07 | 2.100E-06 | 2.700E-06 |
| 26147 | 1308 | W1308Q | E | 1.3e-06 | 3.8e-06 | 2.3e-06 | 2.467E-06 | 1.258E-06 | 1.300E-06 | 3.800E-06 |
| 26148 | 1308 | W1308G | E | 1.7e-06 | 3.1e-06 | 4e-06 | 2.933E-06 | 1.159E-06 | 1.700E-06 | 4.000E-06 |
| 26149 | 1308 | W1308H | E | 2.4e-06 | 1.3e-06 | 2.4e-06 | 2.033E-06 | 6.351E-07 | 1.300E-06 | 2.400E-06 |
| 26150 | 1308 | W1308I | E | 2.3e-06 | 4.3e-06 | 2.9e-06 | 3.167E-06 | 1.026E-06 | 2.300E-06 | 4.300E-06 |
| 26151 | 1308 | W1308L | E | 1.6e-06 | 2.3e-06 | 1.8e-06 | 1.900E-06 | 3.606E-07 | 1.600E-06 | 2.300E-06 |
| 26152 | 1308 | W1308K | E | 2e-06 | 1.8e-06 | 2.5e-06 | 2.100E-06 | 3.606E-07 | 1.800E-06 | 2.500E-06 |
| 26153 | 1308 | W1308M | E | 2.4e-06 | 2.8e-06 | 2.5e-06 | 2.567E-06 | 2.082E-07 | 2.400E-06 | 2.800E-06 |
| 26154 | 1308 | W1308F | E | 2.7e-06 | 2.9e-06 | 2.2e-06 | 2.600E-06 | 3.606E-07 | 2.200E-06 | 2.900E-06 |
| 26155 | 1308 | W1308P | E | 2.7e-06 | 1.6e-06 | 3.1e-06 | 2.467E-06 | 7.767E-07 | 1.600E-06 | 3.100E-06 |
| 26156 | 1308 | W1308S | E | 2.6e-06 | 1.3e-06 | 2.4e-06 | 2.100E-06 | 7.000E-07 | 1.300E-06 | 2.600E-06 |
| 26157 | 1308 | W1308T | E | 1.3e-06 | 3.4e-06 | 1.7e-06 | 2.133E-06 | 1.115E-06 | 1.300E-06 | 3.400E-06 |
| 26158 | 1308 | W1308W | E | 7.3e-07 | 3.4e-06 | 3.1e-06 | 2.410E-06 | 1.463E-06 | 7.300E-07 | 3.400E-06 |
| 26159 | 1308 | W1308Y | E | 1.7e-06 | 3.6e-06 | 2.8e-06 | 2.700E-06 | 9.539E-07 | 1.700E-06 | 3.600E-06 |
| 26160 | 1308 | W1308V | E | 2e-06 | 2.1e-06 | 3.2e-06 | 2.433E-06 | 6.658E-07 | 2.000E-06 | 3.200E-06 |
| 26161 | 1309 | V1309A | E | 2.1e-06 | 1.4e-06 | 4.6e-06 | 2.700E-06 | 1.682E-06 | 1.400E-06 | 4.600E-06 |
| 26162 | 1309 | V1309R | E | 2.5e-06 | 3.6e-06 | 4.1e-06 | 3.400E-06 | 8.185E-07 | 2.500E-06 | 4.100E-06 |
| 26163 | 1309 | V1309N | E | 3e-06 | 2e-06 | 2.1e-06 | 2.367E-06 | 5.508E-07 | 2.000E-06 | 3.000E-06 |
| 26164 | 1309 | V1309D | E | 1.5e-06 | 4e-06 | 3.4e-06 | 2.967E-06 | 1.305E-06 | 1.500E-06 | 4.000E-06 |
| 26165 | 1309 | V1309C | E | 3e-06 | 1.5e-06 | 2.3e-06 | 2.267E-06 | 7.506E-07 | 1.500E-06 | 3.000E-06 |
| 26166 | 1309 | V1309E | E | 1.5e-06 | 4.7e-06 | 4e-06 | 3.400E-06 | 1.682E-06 | 1.500E-06 | 4.700E-06 |
| 26167 | 1309 | V1309Q | E | 2.9e-06 | 2.7e-06 | 1.7e-06 | 2.433E-06 | 6.429E-07 | 1.700E-06 | 2.900E-06 |
| 26168 | 1309 | V1309G | E | 1.5e-06 | 2.8e-06 | 3.3e-06 | 2.533E-06 | 9.292E-07 | 1.500E-06 | 3.300E-06 |
| 26169 | 1309 | V1309H | E | 4e-06 | 2.6e-06 | 2.4e-06 | 3.000E-06 | 8.718E-07 | 2.400E-06 | 4.000E-06 |
| 26170 | 1309 | V1309I | E | 1.8e-06 | 2.5e-06 | 1.9e-06 | 2.067E-06 | 3.786E-07 | 1.800E-06 | 2.500E-06 |
| 26171 | 1309 | V1309L | E | 3.4e-06 | 3e-06 | 1.3e-06 | 2.567E-06 | 1.115E-06 | 1.300E-06 | 3.400E-06 |
| 26172 | 1309 | V1309K | E | 1.6e-06 | 2.6e-06 | 2.5e-06 | 2.233E-06 | 5.508E-07 | 1.600E-06 | 2.600E-06 |
| 26173 | 1309 | V1309M | E | 1.4e-06 | 1.3e-06 | 2.9e-06 | 1.867E-06 | 8.963E-07 | 1.300E-06 | 2.900E-06 |
| 26174 | 1309 | V1309F | E | 3.3e-06 | 1.5e-06 | 1.7e-06 | 2.167E-06 | 9.866E-07 | 1.500E-06 | 3.300E-06 |
| 26175 | 1309 | V1309P | E | 2.3e-06 | 2.4e-06 | 3.9e-06 | 2.867E-06 | 8.963E-07 | 2.300E-06 | 3.900E-06 |
| 26176 | 1309 | V1309S | E | 3.9e-06 | 5.9e-06 | 1.8e-06 | 3.867E-06 | 2.050E-06 | 1.800E-06 | 5.900E-06 |
| 26177 | 1309 | V1309T | E | 2e-06 | 1.9e-06 | 2.8e-06 | 2.233E-06 | 4.933E-07 | 1.900E-06 | 2.800E-06 |
| 26178 | 1309 | V1309W | E | 2.3e-06 | 1.6e-06 | 1.9e-06 | 1.933E-06 | 3.512E-07 | 1.600E-06 | 2.300E-06 |
| 26179 | 1309 | V1309Y | E | 2.5e-06 | 3e-06 | 2e-06 | 2.500E-06 | 5.000E-07 | 2.000E-06 | 3.000E-06 |

| | | | | | | | | | | |
|---|---|---|---|---|---|---|---|---|---|---|
| 26180 | 1309 | V1309V | E | 2.7e-06 | 1.8e-06 | 2.7e-06 | 2.400E-06 | 5.196E-07 | 1.800E-06 | 2.700E-06 |
| 26181 | 1310 | R1310A | E | 2.2e-06 | 1.6e-06 | 2.3e-06 | 2.033E-06 | 3.786E-07 | 1.600E-06 | 2.300E-06 |
| 26182 | 1310 | R1310R | E | 2.8e-06 | 3.2e-06 | 1.6e-06 | 2.533E-06 | 8.327E-07 | 1.600E-06 | 3.200E-06 |
| 26183 | 1310 | R1310N | E | 1.5e-06 | 2.1e-06 | 1.4e-06 | 1.667E-06 | 3.786E-07 | 1.400E-06 | 2.100E-06 |
| 26184 | 1310 | R1310D | E | 2.1e-06 | 4.7e-06 | 2.2e-06 | 3.000E-06 | 1.473E-06 | 2.100E-06 | 4.700E-06 |
| 26185 | 1310 | R1310C | E | 2e-06 | 2.3e-06 | 2.6e-06 | 2.300E-06 | 3.000E-07 | 2.000E-06 | 2.600E-06 |
| 26186 | 1310 | R1310E | E | 2.6e-06 | 2.3e-06 | 4.2e-06 | 3.033E-06 | 1.021E-06 | 2.300E-06 | 4.200E-06 |
| 26187 | 1310 | R1310Q | E | 3.2e-06 | 2.4e-06 | 1.6e-06 | 2.400E-06 | 8.000E-07 | 1.600E-06 | 3.200E-06 |
| 26188 | 1310 | R1310G | E | 2.5e-06 | 2.5e-06 | 3.3e-06 | 2.767E-06 | 4.619E-07 | 2.500E-06 | 3.300E-06 |
| 26189 | 1310 | R1310H | E | 2.2e-06 | 2.1e-06 | 2.7e-06 | 2.333E-06 | 3.215E-07 | 2.100E-06 | 2.700E-06 |
| 26190 | 1310 | R1310I | E | 1.8e-06 | 2.9e-06 | 2.7e-06 | 2.467E-06 | 5.859E-07 | 1.800E-06 | 2.900E-06 |
| 26191 | 1310 | R1310L | E | 2.2e-06 | 4.4e-06 | 2.2e-06 | 2.933E-06 | 1.270E-06 | 2.200E-06 | 4.400E-06 |
| 26192 | 1310 | R1310K | E | 1.1e-06 | 3.1e-06 | 3.2e-06 | 2.467E-06 | 1.185E-06 | 1.100E-06 | 3.200E-06 |
| 26193 | 1310 | R1310M | E | 2.5e-06 | 2e-06 | 2.6e-06 | 2.367E-06 | 3.215E-07 | 2.000E-06 | 2.600E-06 |
| 26194 | 1310 | R1310F | E | 1.8e-06 | 1.6e-06 | 5.3e-06 | 2.900E-06 | 2.081E-06 | 1.600E-06 | 5.300E-06 |
| 26195 | 1310 | R1310P | E | 1.3e-06 | 3.7e-06 | 1.3e-06 | 2.100E-06 | 1.386E-06 | 1.300E-06 | 3.700E-06 |
| 26196 | 1310 | R1310S | E | 3.1e-06 | 3e-06 | 1.7e-06 | 2.600E-06 | 7.810E-07 | 1.700E-06 | 3.100E-06 |
| 26197 | 1310 | R1310T | E | 3.1e-06 | 3.5e-06 | 1.8e-06 | 2.800E-06 | 8.888E-07 | 1.800E-06 | 3.500E-06 |
| 26198 | 1310 | R1310W | E | 2.1e-06 | 4.4e-06 | 3e-06 | 3.167E-06 | 1.159E-06 | 2.100E-06 | 4.400E-06 |
| 26199 | 1310 | R1310Y | E | 1.7e-06 | 2e-06 | 2.3e-06 | 2.000E-06 | 3.000E-07 | 1.700E-06 | 2.300E-06 |
| 26200 | 1310 | R1310V | E | 3.5e-06 | 1.6e-06 | 2.8e-06 | 2.633E-06 | 9.609E-07 | 1.600E-06 | 3.500E-06 |
| 26201 | 1311 | Q1311A | E | 2.2e-06 | 3.4e-06 | 2.3e-06 | 2.633E-06 | 6.658E-07 | 2.200E-06 | 3.400E-06 |
| 26202 | 1311 | Q1311R | E | 2.5e-06 | 1.5e-06 | 1.6e-06 | 1.867E-06 | 5.508E-07 | 1.500E-06 | 2.500E-06 |
| 26203 | 1311 | Q1311N | E | 3.4e-06 | 1.8e-06 | 2.8e-06 | 2.667E-06 | 8.083E-07 | 1.800E-06 | 3.400E-06 |
| 26204 | 1311 | Q1311D | E | 1.7e-06 | 2.5e-06 | 2e-06 | 2.067E-06 | 4.041E-07 | 1.700E-06 | 2.500E-06 |
| 26205 | 1311 | Q1311C | E | 1.2e-06 | 2.9e-06 | 2.3e-06 | 2.133E-06 | 8.622E-07 | 1.200E-06 | 2.900E-06 |
| 26206 | 1311 | Q1311E | E | 1.2e-06 | 1.8e-06 | 1.2e-06 | 1.400E-06 | 3.464E-07 | 1.200E-06 | 1.800E-06 |
| 26207 | 1311 | Q1311Q | E | 2.5e-06 | 2.3e-06 | 1.5e-06 | 2.100E-06 | 5.292E-07 | 1.500E-06 | 2.500E-06 |
| 26208 | 1311 | Q1311G | E | 2.4e-06 | 2.4e-06 | 2e-06 | 2.267E-06 | 2.309E-07 | 2.000E-06 | 2.400E-06 |
| 26209 | 1311 | Q1311H | E | 2.3e-06 | 2.4e-06 | 2.5e-06 | 2.400E-06 | 1.000E-07 | 2.300E-06 | 2.500E-06 |
| 26210 | 1311 | Q1311I | E | 2.2e-06 | 2.7e-06 | 2.5e-06 | 2.467E-06 | 2.517E-07 | 2.200E-06 | 2.700E-06 |
| 26211 | 1311 | Q1311L | E | 3.4e-06 | 3.2e-06 | 9.4e-07 | 2.513E-06 | 1.366E-06 | 9.400E-07 | 3.400E-06 |
| 26212 | 1311 | Q1311K | E | 1.4e-06 | 2e-06 | 3.2e-06 | 2.200E-06 | 9.165E-07 | 1.400E-06 | 3.200E-06 |
| 26213 | 1311 | Q1311M | E | 2.7e-06 | 2.6e-06 | 3.2e-06 | 2.833E-06 | 3.215E-07 | 2.600E-06 | 3.200E-06 |
| 26214 | 1311 | Q1311F | E | 2.7e-06 | 3e-06 | 1.3e-06 | 2.333E-06 | 9.074E-07 | 1.300E-06 | 3.000E-06 |
| 26215 | 1311 | Q1311P | E | 2.2e-06 | 2.8e-06 | 2.6e-06 | 2.533E-06 | 3.055E-07 | 2.200E-06 | 2.800E-06 |
| 26216 | 1311 | Q1311S | E | 3.2e-06 | 8.8e-07 | 1.6e-06 | 1.893E-06 | 1.187E-06 | 8.800E-07 | 3.200E-06 |
| 26217 | 1311 | Q1311T | E | 4.1e-06 | 2.2e-06 | 2.1e-06 | 2.800E-06 | 1.127E-06 | 2.100E-06 | 4.100E-06 |
| 26218 | 1311 | Q1311W | E | 1.9e-06 | 2.7e-06 | 1.9e-06 | 2.167E-06 | 4.619E-07 | 1.900E-06 | 2.700E-06 |
| 26219 | 1311 | Q1311Y | E | 2.7e-06 | 1.6e-06 | 1.8e-06 | 2.033E-06 | 5.859E-07 | 1.600E-06 | 2.700E-06 |
| 26220 | 1311 | Q1311V | E | 1.9e-06 | 1.8e-06 | 2.5e-06 | 2.067E-06 | 3.786E-07 | 1.800E-06 | 2.500E-06 |
| 26221 | 1312 | A1312A | E | 2.9e-06 | 3e-06 | 2.2e-06 | 2.700E-06 | 4.359E-07 | 2.200E-06 | 3.000E-06 |
| 26222 | 1312 | A1312R | E | 3e-06 | 1.7e-06 | 2e-06 | 2.233E-06 | 6.807E-07 | 1.700E-06 | 3.000E-06 |
| 26223 | 1312 | A1312N | E | 2e-06 | 1.8e-06 | 2.3e-06 | 2.033E-06 | 2.517E-07 | 1.800E-06 | 2.300E-06 |
| 26224 | 1312 | A1312D | E | 1.9e-06 | 2.2e-06 | 1.6e-06 | 1.900E-06 | 3.000E-07 | 1.600E-06 | 2.200E-06 |
| 26225 | 1312 | A1312C | E | 1e-06 | 1.7e-06 | 2.1e-06 | 1.600E-06 | 5.568E-07 | 1.000E-06 | 2.100E-06 |
| 26226 | 1312 | A1312E | E | 3.2e-06 | 4.9e-06 | 1.7e-06 | 3.267E-06 | 1.601E-06 | 1.700E-06 | 4.900E-06 |
| 26227 | 1312 | A1312Q | E | 2.3e-06 | 1.8e-06 | 2.3e-06 | 2.133E-06 | 2.887E-07 | 1.800E-06 | 2.300E-06 |
| 26228 | 1312 | A1312G | E | 1.7e-06 | 2.2e-06 | 2.6e-06 | 2.167E-06 | 4.509E-07 | 1.700E-06 | 2.600E-06 |
| 26229 | 1312 | A1312H | E | 2.5e-06 | 2.8e-06 | 2.3e-06 | 2.533E-06 | 2.517E-07 | 2.300E-06 | 2.800E-06 |
| 26230 | 1312 | A1312I | E | 3.2e-06 | 2.1e-06 | 2.2e-06 | 2.500E-06 | 6.083E-07 | 2.100E-06 | 3.200E-06 |
| 26231 | 1312 | A1312L | E | 3.7e-06 | 2.5e-06 | 2.5e-06 | 2.900E-06 | 6.928E-07 | 2.500E-06 | 3.700E-06 |
| 26232 | 1312 | A1312K | E | 2.8e-06 | 3.6e-06 | 3.7e-06 | 3.367E-06 | 4.933E-07 | 2.800E-06 | 3.700E-06 |
| 26233 | 1312 | A1312M | E | 2.5e-06 | 1.9e-06 | 1.8e-06 | 2.067E-06 | 3.786E-07 | 1.800E-06 | 2.500E-06 |
| 26234 | 1312 | A1312F | E | 2.4e-06 | 2.6e-06 | 2.3e-06 | 2.433E-06 | 1.528E-07 | 2.300E-06 | 2.600E-06 |
| 26235 | 1312 | A1312P | E | 2e-06 | 2.7e-06 | 2.6e-06 | 2.433E-06 | 3.786E-07 | 2.000E-06 | 2.700E-06 |
| 26236 | 1312 | A1312S | E | 2.6e-06 | 2.6e-06 | 1.9e-06 | 2.367E-06 | 4.041E-07 | 1.900E-06 | 2.600E-06 |
| 26237 | 1312 | A1312T | E | 2.6e-06 | 2.1e-06 | 1.9e-06 | 2.200E-06 | 3.606E-07 | 1.900E-06 | 2.600E-06 |
| 26238 | 1312 | A1312W | E | 4.9e-06 | 1.8e-06 | 1.9e-06 | 2.867E-06 | 1.762E-06 | 1.800E-06 | 4.900E-06 |
| 26239 | 1312 | A1312Y | E | 2.2e-06 | 6.4e-06 | 1.8e-06 | 3.467E-06 | 2.548E-06 | 1.800E-06 | 6.400E-06 |
| 26240 | 1312 | A1312V | E | 2.6e-06 | 2.2e-06 | 4.7e-06 | 3.167E-06 | 1.343E-06 | 2.200E-06 | 4.700E-06 |
| 26241 | 1313 | P1313A | E | 2.1e-06 | 3.7e-06 | 1.6e-06 | 2.467E-06 | 1.097E-06 | 1.600E-06 | 3.700E-06 |
| 26242 | 1313 | P1313R | E | 3.5e-06 | 1.8e-06 | 2.2e-06 | 2.500E-06 | 8.888E-07 | 1.800E-06 | 3.500E-06 |
| 26243 | 1313 | P1313N | E | 3.6e-06 | 1.5e-06 | 2.9e-06 | 2.667E-06 | 1.069E-06 | 1.500E-06 | 3.600E-06 |
| 26244 | 1313 | P1313D | E | 2.4e-06 | 2e-06 | 2.8e-06 | 2.400E-06 | 4.000E-07 | 2.000E-06 | 2.800E-06 |
| 26245 | 1313 | P1313C | E | 2.9e-06 | 2.1e-06 | 3e-06 | 2.667E-06 | 4.933E-07 | 2.100E-06 | 3.000E-06 |
| 26246 | 1313 | P1313E | E | 1.8e-06 | 2.1e-06 | 2.9e-06 | 2.267E-06 | 5.686E-07 | 1.800E-06 | 2.900E-06 |
| 26247 | 1313 | P1313Q | E | 2.3e-06 | 2.5e-06 | 2.5e-06 | 2.433E-06 | 1.155E-07 | 2.300E-06 | 2.500E-06 |
| 26248 | 1313 | P1313G | E | 1.5e-06 | 1.8e-06 | 2e-06 | 1.767E-06 | 2.517E-07 | 1.500E-06 | 2.000E-06 |
| 26249 | 1313 | P1313H | E | 2.4e-06 | 1.2e-06 | 4.6e-06 | 2.733E-06 | 1.724E-06 | 1.200E-06 | 4.600E-06 |
| 26250 | 1313 | P1313I | E | 3.9e-06 | 2.6e-06 | 4.4e-06 | 3.633E-06 | 9.292E-07 | 2.600E-06 | 4.400E-06 |
| 26251 | 1313 | P1313L | E | 2.7e-06 | 1.7e-06 | 2.7e-06 | 2.367E-06 | 5.774E-07 | 1.700E-06 | 2.700E-06 |
| 26252 | 1313 | P1313K | E | 9.8e-07 | 2.8e-06 | 2.6e-06 | 2.127E-06 | 9.981E-07 | 9.800E-07 | 2.800E-06 |
| 26253 | 1313 | P1313M | E | 1.9e-06 | 3.4e-06 | 2.5e-06 | 2.600E-06 | 7.550E-07 | 1.900E-06 | 3.400E-06 |
| 26254 | 1313 | P1313F | E | 2.8e-06 | 2.2e-06 | 1.9e-06 | 2.300E-06 | 4.583E-07 | 1.900E-06 | 2.800E-06 |
| 26255 | 1313 | P1313P | E | 4.1e-06 | 2.2e-06 | 3.1e-06 | 3.133E-06 | 9.504E-07 | 2.200E-06 | 4.100E-06 |
| 26256 | 1313 | P1313S | E | 3.7e-06 | 1.7e-06 | 3.3e-06 | 2.900E-06 | 1.058E-06 | 1.700E-06 | 3.700E-06 |
| 26257 | 1313 | P1313T | E | 2.9e-06 | 1.4e-06 | 2.9e-06 | 2.400E-06 | 8.660E-07 | 1.400E-06 | 2.900E-06 |
| 26258 | 1313 | P1313W | E | 3.7e-06 | 2.6e-06 | 2.7e-06 | 3.000E-06 | 6.083E-07 | 2.600E-06 | 3.700E-06 |
| 26259 | 1313 | P1313Y | E | 3.2e-06 | 2.8e-06 | 4.1e-06 | 3.367E-06 | 6.658E-07 | 2.800E-06 | 4.100E-06 |
| 26260 | 1313 | P1313V | E | 1.8e-06 | 3.4e-06 | 1.7e-06 | 2.300E-06 | 9.539E-07 | 1.700E-06 | 3.400E-06 |
| 26261 | 1314 | G1314A | E | 1.5e-06 | 2e-06 | 6.6e-06 | 3.367E-06 | 2.811E-06 | 1.500E-06 | 6.600E-06 |
| 26262 | 1314 | G1314R | E | 1.9e-06 | 2.5e-06 | 2e-06 | 2.133E-06 | 3.215E-07 | 1.900E-06 | 2.500E-06 |
| 26263 | 1314 | G1314N | E | 2.1e-06 | 3.1e-06 | 2.1e-06 | 2.433E-06 | 5.774E-07 | 2.100E-06 | 3.100E-06 |
| 26264 | 1314 | G1314D | E | 5.1e-06 | 3.1e-06 | 2.6e-06 | 3.600E-06 | 1.323E-06 | 2.600E-06 | 5.100E-06 |

| | | | | | | | | | | |
|---|---|---|---|---|---|---|---|---|---|---|
| 26265 | 1314 | G1314C | E | 2.1e-06 | 1.2e-06 | 2.4e-06 | 1.900E-06 | 6.245E-07 | 1.200E-06 | 2.400E-06 |
| 26266 | 1314 | G1314E | E | 3.5e-06 | 2.6e-06 | 1.3e-06 | 2.467E-06 | 1.106E-06 | 1.300E-06 | 3.500E-06 |
| 26267 | 1314 | G1314Q | E | 1.3e-06 | 2.5e-06 | 1.8e-06 | 1.867E-06 | 6.028E-07 | 1.300E-06 | 2.500E-06 |
| 26268 | 1314 | G1314G | E | 3e-06 | 2.6e-06 | 1.5e-06 | 2.367E-06 | 7.767E-07 | 1.500E-06 | 3.000E-06 |
| 26269 | 1314 | G1314H | E | 2.9e-06 | 2.8e-06 | 4.5e-06 | 3.400E-06 | 9.539E-07 | 2.800E-06 | 4.500E-06 |
| 26270 | 1314 | G1314I | E | 2.6e-06 | 1.3e-06 | 2.5e-06 | 2.133E-06 | 7.234E-07 | 1.300E-06 | 2.600E-06 |
| 26271 | 1314 | G1314L | E | 2.1e-06 | 2.4e-06 | 3e-06 | 2.500E-06 | 4.583E-07 | 2.100E-06 | 3.000E-06 |
| 26272 | 1314 | G1314K | E | 1.9e-06 | 2.3e-06 | 5.8e-06 | 3.333E-06 | 2.146E-06 | 1.900E-06 | 5.800E-06 |
| 26273 | 1314 | G1314M | E | 2.6e-06 | 2.8e-06 | 1.6e-06 | 2.333E-06 | 6.429E-07 | 1.600E-06 | 2.800E-06 |
| 26274 | 1314 | G1314F | E | 3.7e-06 | 1.5e-06 | 2.1e-06 | 2.433E-06 | 1.137E-06 | 1.500E-06 | 3.700E-06 |
| 26275 | 1314 | G1314P | E | 2.2e-06 | 2.2e-06 | 2.5e-06 | 2.300E-06 | 1.732E-07 | 2.200E-06 | 2.500E-06 |
| 26276 | 1314 | G1314S | E | 1.7e-06 | 2.6e-06 | 2.3e-06 | 2.200E-06 | 4.583E-07 | 1.700E-06 | 2.600E-06 |
| 26277 | 1314 | G1314T | E | 1.8e-06 | 2.2e-06 | 2.6e-06 | 2.200E-06 | 4.000E-07 | 1.800E-06 | 2.600E-06 |
| 26278 | 1314 | G1314W | E | 3.6e-06 | 2.2e-06 | 3.8e-06 | 3.200E-06 | 8.718E-07 | 2.200E-06 | 3.800E-06 |
| 26279 | 1314 | G1314Y | E | 1.2e-06 | 2.1e-06 | 2.1e-06 | 1.800E-06 | 5.196E-07 | 1.200E-06 | 2.100E-06 |
| 26280 | 1314 | G1314V | E | 2.2e-06 | 2.5e-06 | 2.3e-06 | 2.333E-06 | 1.528E-07 | 2.200E-06 | 2.500E-06 |
| 26281 | 1315 | K1315A | E | 1.6e-06 | 1.9e-06 | 2e-06 | 1.833E-06 | 2.082E-07 | 1.600E-06 | 2.000E-06 |
| 26282 | 1315 | K1315R | E | 1.9e-06 | 2.5e-06 | 3.4e-06 | 2.600E-06 | 7.550E-07 | 1.900E-06 | 3.400E-06 |
| 26283 | 1315 | K1315N | E | 1.6e-06 | 2.5e-06 | 3.3e-06 | 2.467E-06 | 8.505E-07 | 1.600E-06 | 3.300E-06 |
| 26284 | 1315 | K1315D | E | 2.4e-06 | 3.2e-06 | 2.4e-06 | 2.667E-06 | 4.619E-07 | 2.400E-06 | 3.200E-06 |
| 26285 | 1315 | K1315C | E | 2.7e-06 | 2e-06 | 2.5e-06 | 2.400E-06 | 3.606E-07 | 2.000E-06 | 2.700E-06 |
| 26286 | 1315 | K1315E | E | 1.6e-06 | 2.8e-06 | 1.6e-06 | 2.000E-06 | 6.928E-07 | 1.600E-06 | 2.800E-06 |
| 26287 | 1315 | K1315Q | E | 2.9e-06 | 1.6e-06 | 2.2e-06 | 2.233E-06 | 6.506E-07 | 1.600E-06 | 2.900E-06 |
| 26288 | 1315 | K1315G | E | 2.5e-06 | 3.2e-06 | 2.4e-06 | 2.700E-06 | 4.359E-07 | 2.400E-06 | 3.200E-06 |
| 26289 | 1315 | K1315H | E | 1.5e-06 | 1.7e-06 | 1.8e-06 | 1.667E-06 | 1.528E-07 | 1.500E-06 | 1.800E-06 |
| 26290 | 1315 | K1315I | E | 2.1e-06 | 2.2e-06 | 2.4e-06 | 2.233E-06 | 1.528E-07 | 2.100E-06 | 2.400E-06 |
| 26291 | 1315 | K1315L | E | 3.8e-06 | 2e-06 | 1.9e-06 | 2.567E-06 | 1.069E-06 | 1.900E-06 | 3.800E-06 |
| 26292 | 1315 | K1315K | E | 1.5e-06 | 1.4e-06 | 1.9e-06 | 1.600E-06 | 2.646E-07 | 1.400E-06 | 1.900E-06 |
| 26293 | 1315 | K1315M | E | 3.8e-06 | 1.1e-06 | 2e-06 | 2.300E-06 | 1.375E-06 | 1.100E-06 | 3.800E-06 |
| 26294 | 1315 | K1315F | E | 1.8e-06 | 1.6e-06 | 2.3e-06 | 1.900E-06 | 3.606E-07 | 1.600E-06 | 2.300E-06 |
| 26295 | 1315 | K1315P | E | 1.5e-06 | 2.7e-06 | 1.4e-06 | 1.867E-06 | 7.234E-07 | 1.400E-06 | 2.700E-06 |
| 26296 | 1315 | K1315S | E | 4e-06 | 3.3e-06 | 1.6e-06 | 2.967E-06 | 1.234E-06 | 1.600E-06 | 4.000E-06 |
| 26297 | 1315 | K1315T | E | 1.3e-06 | 3.6e-06 | 3.1e-06 | 2.667E-06 | 1.210E-06 | 1.300E-06 | 3.600E-06 |
| 26298 | 1315 | K1315W | E | 1.5e-06 | 1.7e-06 | 1.9e-06 | 1.700E-06 | 2.000E-07 | 1.500E-06 | 1.900E-06 |
| 26299 | 1315 | K1315Y | E | 1.8e-06 | 3.4e-06 | 1.9e-06 | 2.367E-06 | 8.963E-07 | 1.800E-06 | 3.400E-06 |
| 26300 | 1315 | K1315V | E | 3.3e-06 | 3e-06 | 1.7e-06 | 2.667E-06 | 8.505E-07 | 1.700E-06 | 3.300E-06 |
| 26301 | 1316 | G1316A | E | 3.4e-06 | 1.4e-06 | 2.7e-06 | 2.500E-06 | 1.015E-06 | 1.400E-06 | 3.400E-06 |
| 26302 | 1316 | G1316R | E | 1.9e-06 | 2.2e-06 | 2.5e-06 | 2.200E-06 | 3.000E-07 | 1.900E-06 | 2.500E-06 |
| 26303 | 1316 | G1316N | E | 2.9e-06 | 2.3e-06 | 2.1e-06 | 2.433E-06 | 4.163E-07 | 2.100E-06 | 2.900E-06 |
| 26304 | 1316 | G1316D | E | 1.3e-06 | 3.1e-06 | 2.4e-06 | 2.267E-06 | 9.074E-07 | 1.300E-06 | 3.100E-06 |
| 26305 | 1316 | G1316C | E | 2.3e-06 | 1.6e-06 | 2.2e-06 | 2.033E-06 | 3.786E-07 | 1.600E-06 | 2.300E-06 |
| 26306 | 1316 | G1316E | E | 1.5e-06 | 2.7e-06 | 2.5e-06 | 2.233E-06 | 6.429E-07 | 1.500E-06 | 2.700E-06 |
| 26307 | 1316 | G1316Q | E | 1.7e-06 | 1.4e-06 | 1.5e-06 | 1.533E-06 | 1.528E-07 | 1.400E-06 | 1.700E-06 |
| 26308 | 1316 | G1316G | E | 1.9e-06 | 1.5e-06 | 1.8e-06 | 1.733E-06 | 2.082E-07 | 1.500E-06 | 1.900E-06 |
| 26309 | 1316 | G1316H | E | 3e-06 | 1.8e-06 | 2.2e-06 | 2.333E-06 | 6.110E-07 | 1.800E-06 | 3.000E-06 |
| 26310 | 1316 | G1316I | E | 2.2e-06 | 2.5e-06 | 1.8e-06 | 2.167E-06 | 3.512E-07 | 1.800E-06 | 2.500E-06 |
| 26311 | 1316 | G1316L | E | 1.9e-06 | 2.9e-06 | 1.8e-06 | 2.200E-06 | 6.083E-07 | 1.800E-06 | 2.900E-06 |
| 26312 | 1316 | G1316K | E | 2.5e-06 | 1.7e-06 | 2.5e-06 | 2.233E-06 | 4.619E-07 | 1.700E-06 | 2.500E-06 |
| 26313 | 1316 | G1316M | E | 1.9e-06 | 2.5e-06 | 2.3e-06 | 2.233E-06 | 3.055E-07 | 1.900E-06 | 2.500E-06 |
| 26314 | 1316 | G1316F | E | 2.7e-06 | 2.6e-06 | 2.6e-06 | 2.633E-06 | 5.774E-08 | 2.600E-06 | 2.700E-06 |
| 26315 | 1316 | G1316P | E | 1.5e-06 | 2.8e-06 | 1.7e-06 | 2.000E-06 | 7.000E-07 | 1.500E-06 | 2.800E-06 |
| 26316 | 1316 | G1316S | E | 2.3e-06 | 1.6e-06 | 4.9e-06 | 2.933E-06 | 1.739E-06 | 1.600E-06 | 4.900E-06 |
| 26317 | 1316 | G1316T | E | 2.7e-06 | 2.4e-06 | 2.3e-06 | 2.467E-06 | 2.082E-07 | 2.300E-06 | 2.700E-06 |
| 26318 | 1316 | G1316W | E | 1.7e-06 | 1.8e-06 | 2.5e-06 | 2.000E-06 | 4.359E-07 | 1.700E-06 | 2.500E-06 |
| 26319 | 1316 | G1316Y | E | 3.1e-06 | 2.7e-06 | 2.8e-06 | 2.867E-06 | 2.082E-07 | 2.700E-06 | 3.100E-06 |
| 26320 | 1316 | G1316V | E | 2.1e-06 | 2.5e-06 | 1.3e-06 | 1.967E-06 | 6.110E-07 | 1.300E-06 | 2.500E-06 |
| 26321 | 1317 | L1317A | E | 1.7e-06 | 2.4e-06 | 2.6e-06 | 2.233E-06 | 4.726E-07 | 1.700E-06 | 2.600E-06 |
| 26322 | 1317 | L1317R | E | 2.3e-06 | 2.4e-06 | 1.6e-06 | 2.100E-06 | 4.359E-07 | 1.600E-06 | 2.400E-06 |
| 26323 | 1317 | L1317N | E | 2.3e-06 | 1.8e-06 | 1.9e-06 | 2.000E-06 | 2.646E-07 | 1.800E-06 | 2.300E-06 |
| 26324 | 1317 | L1317D | E | 2e-06 | 2.6e-06 | 3.3e-06 | 2.633E-06 | 6.506E-07 | 2.000E-06 | 3.300E-06 |
| 26325 | 1317 | L1317C | E | 3.4e-06 | 5.2e-06 | 2.3e-06 | 3.633E-06 | 1.464E-06 | 2.300E-06 | 5.200E-06 |
| 26326 | 1317 | L1317E | E | 1.9e-06 | 1.8e-06 | 2.5e-06 | 2.067E-06 | 3.786E-07 | 1.800E-06 | 2.500E-06 |
| 26327 | 1317 | L1317Q | E | 1.5e-06 | 2.1e-06 | 2.4e-06 | 2.000E-06 | 4.583E-07 | 1.500E-06 | 2.400E-06 |
| 26328 | 1317 | L1317G | E | 2.3e-06 | 2.5e-06 | 1.7e-06 | 2.167E-06 | 4.163E-07 | 1.700E-06 | 2.500E-06 |
| 26329 | 1317 | L1317H | E | 2e-06 | 3e-06 | 2.3e-06 | 2.433E-06 | 5.132E-07 | 2.000E-06 | 3.000E-06 |
| 26330 | 1317 | L1317I | E | 2.6e-06 | 1.9e-06 | 1.1e-06 | 1.867E-06 | 7.506E-07 | 1.100E-06 | 2.600E-06 |
| 26331 | 1317 | L1317L | E | 2.3e-06 | 2.7e-06 | 3.6e-06 | 2.867E-06 | 6.658E-07 | 2.300E-06 | 3.600E-06 |
| 26332 | 1317 | L1317K | E | 1.5e-06 | 1.6e-06 | 2.3e-06 | 1.800E-06 | 4.359E-07 | 1.500E-06 | 2.300E-06 |
| 26333 | 1317 | L1317M | E | 2.6e-06 | 2.6e-06 | 3.4e-06 | 2.867E-06 | 4.619E-07 | 2.600E-06 | 3.400E-06 |
| 26334 | 1317 | L1317F | E | 3e-06 | 2.9e-06 | 2.6e-06 | 2.833E-06 | 2.082E-07 | 2.600E-06 | 3.000E-06 |
| 26335 | 1317 | L1317P | E | 1.2e-06 | 2.3e-06 | 2.4e-06 | 1.967E-06 | 6.658E-07 | 1.200E-06 | 2.400E-06 |
| 26336 | 1317 | L1317S | E | 2.6e-06 | 2e-06 | 2.1e-06 | 2.233E-06 | 3.215E-07 | 2.000E-06 | 2.600E-06 |
| 26337 | 1317 | L1317T | E | 1.5e-06 | 1.9e-06 | 2.2e-06 | 1.867E-06 | 3.512E-07 | 1.500E-06 | 2.200E-06 |
| 26338 | 1317 | L1317W | E | 3.2e-06 | 2.2e-06 | 1.9e-06 | 2.433E-06 | 6.807E-07 | 1.900E-06 | 3.200E-06 |
| 26339 | 1317 | L1317Y | E | 3.6e-06 | 2.9e-06 | 1.6e-06 | 2.700E-06 | 1.015E-06 | 1.600E-06 | 3.600E-06 |
| 26340 | 1317 | L1317V | E | 3.4e-06 | 1.8e-06 | 1.8e-06 | 2.333E-06 | 9.238E-07 | 1.800E-06 | 3.400E-06 |
| 26341 | 1318 | E1318A | E | 3.8e-06 | 2.1e-06 | 2.9e-06 | 2.933E-06 | 8.505E-07 | 2.100E-06 | 3.800E-06 |
| 26342 | 1318 | E1318R | E | 2.7e-06 | 2.9e-06 | 2.6e-06 | 2.733E-06 | 1.528E-07 | 2.600E-06 | 2.900E-06 |
| 26343 | 1318 | E1318N | E | 2e-06 | 1.6e-06 | 2.6e-06 | 2.067E-06 | 5.033E-07 | 1.600E-06 | 2.600E-06 |
| 26344 | 1318 | E1318D | E | 2.8e-06 | 2.7e-06 | 3e-06 | 2.833E-06 | 1.528E-07 | 2.700E-06 | 3.000E-06 |
| 26345 | 1318 | E1318C | E | 2.1e-06 | 2.3e-06 | 2.9e-06 | 2.433E-06 | 4.163E-07 | 2.100E-06 | 2.900E-06 |
| 26346 | 1318 | E1318E | E | 2.5e-06 | 1.4e-06 | 2e-06 | 1.967E-06 | 5.508E-07 | 1.400E-06 | 2.500E-06 |
| 26347 | 1318 | E1318Q | E | 2.7e-06 | 2e-06 | 1.6e-06 | 2.100E-06 | 5.568E-07 | 1.600E-06 | 2.700E-06 |
| 26348 | 1318 | E1318G | E | 1.9e-06 | 3.5e-06 | 2.4e-06 | 2.600E-06 | 8.185E-07 | 1.900E-06 | 3.500E-06 |
| 26349 | 1318 | E1318H | E | 2.5e-06 | 1.1e-06 | 2.2e-06 | 1.933E-06 | 7.371E-07 | 1.100E-06 | 2.500E-06 |

| | | | | | | | | | | |
|---|---|---|---|---|---|---|---|---|---|---|
| 26350 | 1318 | E1318I | E | 2.4e-06 | 1.1e-06 | 3.4e-06 | 2.300E-06 | 1.153E-06 | 1.100E-06 | 3.400E-06 |
| 26351 | 1318 | E1318L | E | 3.3e-06 | 2.3e-06 | 1.9e-06 | 2.500E-06 | 7.211E-07 | 1.900E-06 | 3.300E-06 |
| 26352 | 1318 | E1318K | E | 3.5e-06 | 2.1e-06 | 2.9e-06 | 2.833E-06 | 7.024E-07 | 2.100E-06 | 3.500E-06 |
| 26353 | 1318 | E1318M | E | 2e-06 | 2.4e-06 | 3e-06 | 2.467E-06 | 5.033E-07 | 2.000E-06 | 3.000E-06 |
| 26354 | 1318 | E1318F | E | 2.2e-06 | 2.7e-06 | 2.6e-06 | 2.500E-06 | 2.646E-07 | 2.200E-06 | 2.700E-06 |
| 26355 | 1318 | E1318P | E | 1.9e-06 | 2.8e-06 | 1.5e-06 | 2.067E-06 | 6.658E-07 | 1.500E-06 | 2.800E-06 |
| 26356 | 1318 | E1318S | E | 1.3e-06 | 2.3e-06 | 2.3e-06 | 1.967E-06 | 5.774E-07 | 1.300E-06 | 2.300E-06 |
| 26357 | 1318 | E1318T | E | 4.7e-06 | 1.8e-06 | 1.5e-06 | 2.667E-06 | 1.767E-06 | 1.500E-06 | 4.700E-06 |
| 26358 | 1318 | E1318W | E | 2.2e-06 | 2.8e-06 | 2.1e-06 | 2.367E-06 | 3.786E-07 | 2.100E-06 | 2.800E-06 |
| 26359 | 1318 | E1318Y | E | 3.6e-06 | 2.7e-06 | 2.7e-06 | 3.000E-06 | 5.196E-07 | 2.700E-06 | 3.600E-06 |
| 26360 | 1318 | E1318V | E | 1.2e-06 | 2.6e-06 | 1.6e-06 | 1.800E-06 | 7.211E-07 | 1.200E-06 | 2.600E-06 |
| 26361 | 1319 | W1319A | E | 2.4e-06 | 2e-06 | 2.5e-06 | 2.300E-06 | 2.646E-07 | 2.000E-06 | 2.500E-06 |
| 26362 | 1319 | W1319R | E | 1.9e-06 | 2.3e-06 | 2.5e-06 | 2.233E-06 | 3.055E-07 | 1.900E-06 | 2.500E-06 |
| 26363 | 1319 | W1319N | E | 1.8e-06 | 2.1e-06 | 2.1e-06 | 2.000E-06 | 1.732E-07 | 1.800E-06 | 2.100E-06 |
| 26364 | 1319 | W1319D | E | 3.5e-06 | 2e-06 | 3e-06 | 2.833E-06 | 7.638E-07 | 2.000E-06 | 3.500E-06 |
| 26365 | 1319 | W1319C | E | 2.2e-06 | 2.5e-06 | 2.2e-06 | 2.300E-06 | 1.732E-07 | 2.200E-06 | 2.500E-06 |
| 26366 | 1319 | W1319E | E | 3.9e-06 | 3.9e-06 | 4.3e-06 | 4.033E-06 | 2.309E-07 | 3.900E-06 | 4.300E-06 |
| 26367 | 1319 | W1319Q | E | 2.6e-06 | 1.6e-06 | 1.3e-06 | 1.833E-06 | 6.807E-07 | 1.300E-06 | 2.600E-06 |
| 26368 | 1319 | W1319G | E | 1.9e-06 | 2.6e-06 | 1.4e-06 | 1.967E-06 | 6.028E-07 | 1.400E-06 | 2.600E-06 |
| 26369 | 1319 | W1319H | E | 2.5e-06 | 2.1e-06 | 2.6e-06 | 2.400E-06 | 2.646E-07 | 2.100E-06 | 2.600E-06 |
| 26370 | 1319 | W1319I | E | 1.8e-06 | 2.7e-06 | 2e-06 | 2.167E-06 | 4.726E-07 | 1.800E-06 | 2.700E-06 |
| 26371 | 1319 | W1319L | E | 1.6e-06 | 3.5e-06 | 2.7e-06 | 2.600E-06 | 9.539E-07 | 1.600E-06 | 3.500E-06 |
| 26372 | 1319 | W1319K | E | 3.1e-06 | 2.4e-06 | 3.7e-06 | 3.067E-06 | 6.506E-07 | 2.400E-06 | 3.700E-06 |
| 26373 | 1319 | W1319M | E | 2.9e-06 | 1.8e-06 | 2.3e-06 | 2.333E-06 | 5.508E-07 | 1.800E-06 | 2.900E-06 |
| 26374 | 1319 | W1319F | E | 1.5e-06 | 1.5e-06 | 1.4e-06 | 1.467E-06 | 5.774E-08 | 1.400E-06 | 1.500E-06 |
| 26375 | 1319 | W1319P | E | 1.4e-06 | 3.4e-06 | 2.5e-06 | 2.433E-06 | 1.002E-06 | 1.400E-06 | 3.400E-06 |
| 26376 | 1319 | W1319S | E | 1.5e-06 | 2e-06 | 2.6e-06 | 2.033E-06 | 5.508E-07 | 1.500E-06 | 2.600E-06 |
| 26377 | 1319 | W1319T | E | 3.1e-06 | 2.2e-06 | 1.9e-06 | 2.400E-06 | 6.245E-07 | 1.900E-06 | 3.100E-06 |
| 26378 | 1319 | W1319W | E | 2.7e-06 | 3.6e-06 | 1.8e-06 | 2.700E-06 | 9.000E-07 | 1.800E-06 | 3.600E-06 |
| 26379 | 1319 | W1319Y | E | 2e-06 | 4.7e-06 | 1.3e-06 | 2.667E-06 | 1.795E-06 | 1.300E-06 | 4.700E-06 |
| 26380 | 1319 | W1319V | E | 3.3e-06 | 3.9e-06 | 2.3e-06 | 3.167E-06 | 8.083E-07 | 2.300E-06 | 3.900E-06 |
| 26381 | 1320 | V1320A | E | 3.3e-06 | 2.4e-06 | 1.6e-06 | 2.433E-06 | 8.505E-07 | 1.600E-06 | 3.300E-06 |
| 26382 | 1320 | V1320R | E | 3.8e-06 | 2.2e-06 | 3.5e-06 | 3.167E-06 | 8.505E-07 | 2.200E-06 | 3.800E-06 |
| 26383 | 1320 | V1320N | E | 2.8e-06 | 1.7e-06 | 2.1e-06 | 2.200E-06 | 5.568E-07 | 1.700E-06 | 2.800E-06 |
| 26384 | 1320 | V1320D | E | 1.9e-06 | 2.4e-06 | 3.9e-06 | 2.733E-06 | 1.041E-06 | 1.900E-06 | 3.900E-06 |
| 26385 | 1320 | V1320C | E | 2.9e-06 | 2.5e-06 | 2.6e-06 | 2.667E-06 | 2.082E-07 | 2.500E-06 | 2.900E-06 |
| 26386 | 1320 | V1320E | E | 1.9e-06 | 1.8e-06 | 3.3e-06 | 2.333E-06 | 8.386E-07 | 1.800E-06 | 3.300E-06 |
| 26387 | 1320 | V1320Q | E | 1.6e-06 | 3.8e-06 | 1.7e-06 | 2.367E-06 | 1.242E-06 | 1.600E-06 | 3.800E-06 |
| 26388 | 1320 | V1320G | E | 1.5e-06 | 1.6e-06 | 2.7e-06 | 1.933E-06 | 6.658E-07 | 1.500E-06 | 2.700E-06 |
| 26389 | 1320 | V1320H | E | 1.4e-06 | 2.3e-06 | 2.4e-06 | 2.033E-06 | 5.508E-07 | 1.400E-06 | 2.400E-06 |
| 26390 | 1320 | V1320I | E | 3e-06 | 2.2e-06 | 1.6e-06 | 2.267E-06 | 7.024E-07 | 1.600E-06 | 3.000E-06 |
| 26391 | 1320 | V1320L | E | 3.7e-06 | 2.8e-06 | 3.3e-06 | 3.267E-06 | 4.509E-07 | 2.800E-06 | 3.700E-06 |
| 26392 | 1320 | V1320K | E | 2.4e-06 | 1.7e-06 | 4e-06 | 2.700E-06 | 1.179E-06 | 1.700E-06 | 4.000E-06 |
| 26393 | 1320 | V1320M | E | 2.3e-06 | 3e-06 | 2.1e-06 | 2.467E-06 | 4.726E-07 | 2.100E-06 | 3.000E-06 |
| 26394 | 1320 | V1320F | E | 1.8e-06 | 3.7e-06 | 2.2e-06 | 2.567E-06 | 1.002E-06 | 1.800E-06 | 3.700E-06 |
| 26395 | 1320 | V1320P | E | 2.2e-06 | 2.7e-06 | 2.2e-06 | 2.367E-06 | 2.887E-07 | 2.200E-06 | 2.700E-06 |
| 26396 | 1320 | V1320S | E | 1.8e-06 | 2.7e-06 | 1.9e-06 | 2.133E-06 | 4.933E-07 | 1.800E-06 | 2.700E-06 |
| 26397 | 1320 | V1320T | E | 3.7e-06 | 2.4e-06 | 1.7e-06 | 2.600E-06 | 1.015E-06 | 1.700E-06 | 3.700E-06 |
| 26398 | 1320 | V1320W | E | 1.4e-06 | 3.3e-06 | 3.8e-06 | 2.833E-06 | 1.266E-06 | 1.400E-06 | 3.800E-06 |
| 26399 | 1320 | V1320Y | E | 2.8e-06 | 1.7e-06 | 1.6e-06 | 2.033E-06 | 6.658E-07 | 1.600E-06 | 2.800E-06 |
| 26400 | 1320 | V1320V | E | 3.4e-06 | 2.1e-06 | 2.5e-06 | 2.667E-06 | 6.658E-07 | 2.100E-06 | 3.400E-06 |
| 26401 | 1321 | A1321A | E | 2.5e-06 | 1.8e-06 | 2.3e-06 | 2.200E-06 | 3.606E-07 | 1.800E-06 | 2.500E-06 |
| 26402 | 1321 | A1321R | E | 1.8e-06 | 1.9e-06 | 2.8e-06 | 2.167E-06 | 5.508E-07 | 1.800E-06 | 2.800E-06 |
| 26403 | 1321 | A1321N | E | 2.5e-06 | 3.2e-06 | 3.4e-06 | 3.033E-06 | 4.726E-07 | 2.500E-06 | 3.400E-06 |
| 26404 | 1321 | A1321D | E | 1.4e-06 | 2e-06 | 2.2e-06 | 1.867E-06 | 4.163E-07 | 1.400E-06 | 2.200E-06 |
| 26405 | 1321 | A1321C | E | 2.5e-06 | 1.7e-06 | 2.8e-06 | 2.333E-06 | 5.686E-07 | 1.700E-06 | 2.800E-06 |
| 26406 | 1321 | A1321E | E | 2.5e-06 | 2.7e-06 | 2e-06 | 2.400E-06 | 3.606E-07 | 2.000E-06 | 2.700E-06 |
| 26407 | 1321 | A1321Q | E | 3e-06 | 3.2e-06 | 2.7e-06 | 2.967E-06 | 2.517E-07 | 2.700E-06 | 3.200E-06 |
| 26408 | 1321 | A1321G | E | 2.2e-06 | 2.6e-06 | 3.2e-06 | 2.667E-06 | 5.033E-07 | 2.200E-06 | 3.200E-06 |
| 26409 | 1321 | A1321H | E | 3.2e-06 | 1.7e-06 | 1.6e-06 | 2.167E-06 | 8.963E-07 | 1.600E-06 | 3.200E-06 |
| 26410 | 1321 | A1321I | E | 1.9e-06 | 2.6e-06 | 1.1e-06 | 1.867E-06 | 7.506E-07 | 1.100E-06 | 2.600E-06 |
| 26411 | 1321 | A1321L | E | 1.7e-06 | 2.5e-06 | 1.8e-06 | 2.000E-06 | 4.359E-07 | 1.700E-06 | 2.500E-06 |
| 26412 | 1321 | A1321K | E | 2.5e-06 | 3e-06 | 2.8e-06 | 2.767E-06 | 2.517E-07 | 2.500E-06 | 3.000E-06 |
| 26413 | 1321 | A1321M | E | 1.6e-06 | 2.5e-06 | 1.9e-06 | 2.000E-06 | 4.583E-07 | 1.600E-06 | 2.500E-06 |
| 26414 | 1321 | A1321F | E | 3e-06 | 4.5e-06 | 1.6e-06 | 3.033E-06 | 1.450E-06 | 1.600E-06 | 4.500E-06 |
| 26415 | 1321 | A1321P | E | 1.5e-06 | 2.9e-06 | 4e-06 | 2.800E-06 | 1.253E-06 | 1.500E-06 | 4.000E-06 |
| 26416 | 1321 | A1321S | E | 3.1e-06 | 2.5e-06 | 1.6e-06 | 2.400E-06 | 7.550E-07 | 1.600E-06 | 3.100E-06 |
| 26417 | 1321 | A1321T | E | 2e-06 | 1.7e-06 | 6.6e-07 | 1.453E-06 | 7.032E-07 | 6.600E-07 | 2.000E-06 |
| 26418 | 1321 | A1321W | E | 2e-06 | 2.3e-06 | 2.4e-06 | 2.233E-06 | 2.082E-07 | 2.000E-06 | 2.400E-06 |
| 26419 | 1321 | A1321Y | E | 1.1e-06 | 2.2e-06 | 2.4e-06 | 1.900E-06 | 7.000E-07 | 1.100E-06 | 2.400E-06 |
| 26420 | 1321 | A1321V | E | 2.8e-06 | 5.1e-06 | 1.4e-06 | 3.100E-06 | 1.868E-06 | 1.400E-06 | 5.100E-06 |
| 26421 | 1322 | R1322A | E | 2e-06 | 3e-06 | 1.9e-06 | 2.300E-06 | 6.083E-07 | 1.900E-06 | 3.000E-06 |
| 26422 | 1322 | R1322R | E | 3.6e-06 | 4.2e-06 | 3.3e-06 | 3.700E-06 | 4.583E-07 | 3.300E-06 | 4.200E-06 |
| 26423 | 1322 | R1322N | E | 4.4e-06 | 3.9e-06 | 5.6e-06 | 4.633E-06 | 8.737E-07 | 3.900E-06 | 5.600E-06 |
| 26424 | 1322 | R1322D | E | 3.1e-06 | 4.3e-06 | 3.9e-06 | 3.767E-06 | 6.110E-07 | 3.100E-06 | 4.300E-06 |
| 26425 | 1322 | R1322C | E | 2.9e-06 | 3.9e-06 | 3.3e-06 | 3.367E-06 | 5.033E-07 | 2.900E-06 | 3.900E-06 |
| 26426 | 1322 | R1322E | E | 3e-06 | 2.8e-06 | 2.3e-06 | 2.700E-06 | 3.606E-07 | 2.300E-06 | 3.000E-06 |
| 26427 | 1322 | R1322Q | E | 2.2e-06 | 3.9e-06 | 3.8e-06 | 3.300E-06 | 9.539E-07 | 2.200E-06 | 3.900E-06 |
| 26428 | 1322 | R1322G | E | 6.6e-06 | 3.8e-06 | 6.6e-06 | 5.667E-06 | 1.617E-06 | 3.800E-06 | 6.600E-06 |
| 26429 | 1322 | R1322H | E | 4.1e-06 | 4.6e-06 | 5e-06 | 4.567E-06 | 4.509E-07 | 4.100E-06 | 5.000E-06 |
| 26430 | 1322 | R1322I | E | 4e-06 | 3.2e-06 | 2e-06 | 3.067E-06 | 1.007E-06 | 2.000E-06 | 4.000E-06 |
| 26431 | 1322 | R1322L | E | 2.6e-06 | 3.5e-06 | 4.2e-06 | 3.433E-06 | 8.021E-07 | 2.600E-06 | 4.200E-06 |
| 26432 | 1322 | R1322K | E | 3.6e-06 | 2.6e-06 | 2.1e-06 | 2.767E-06 | 7.638E-07 | 2.100E-06 | 3.600E-06 |
| 26433 | 1322 | R1322M | E | 2.5e-06 | 7.7e-06 | 2.7e-06 | 4.300E-06 | 2.946E-06 | 2.500E-06 | 7.700E-06 |
| 26434 | 1322 | R1322F | E | 3.1e-06 | 2.2e-06 | 3.7e-06 | 3.000E-06 | 7.550E-07 | 2.200E-06 | 3.700E-06 |

| | | | | | | | | | | |
|---|---|---|---|---|---|---|---|---|---|---|
| 26435 | 1322 | R1322P | E | 3.3e-06 | 5.1e-06 | 3.3e-06 | 3.900E-06 | 1.039E-06 | 3.300E-06 | 5.100E-06 |
| 26436 | 1322 | R1322S | E | 4.7e-06 | 5.1e-06 | 6.8e-06 | 5.533E-06 | 1.115E-06 | 4.700E-06 | 6.800E-06 |
| 26437 | 1322 | R1322T | E | 3.1e-06 | 5.3e-06 | 3.6e-06 | 4.000E-06 | 1.153E-06 | 3.100E-06 | 5.300E-06 |
| 26438 | 1322 | R1322W | E | 3e-06 | 4.1e-06 | 5.4e-06 | 4.167E-06 | 1.201E-06 | 3.000E-06 | 5.400E-06 |
| 26439 | 1322 | R1322Y | E | 5e-06 | 2.9e-06 | 1.4e-06 | 3.100E-06 | 1.808E-06 | 1.400E-06 | 5.000E-06 |
| 26440 | 1322 | R1322V | E | 2.9e-06 | 4.9e-06 | 5e-06 | 4.267E-06 | 1.185E-06 | 2.900E-06 | 5.000E-06 |
| 26441 | 1323 | I1323A | E | 3.4e-06 | 2.7e-06 | 3.3e-06 | 3.133E-06 | 3.786E-07 | 2.700E-06 | 3.400E-06 |
| 26442 | 1323 | I1323R | E | 2.4e-06 | 3.9e-06 | 2.4e-06 | 2.900E-06 | 8.660E-07 | 2.400E-06 | 3.900E-06 |
| 26443 | 1323 | I1323N | E | 3e-06 | 3.5e-06 | 2.2e-06 | 2.900E-06 | 6.557E-07 | 2.200E-06 | 3.500E-06 |
| 26444 | 1323 | I1323D | E | 2.5e-06 | 1.6e-06 | 2.2e-06 | 2.100E-06 | 4.583E-07 | 1.600E-06 | 2.500E-06 |
| 26445 | 1323 | I1323C | E | 1.8e-06 | 2.2e-06 | 1.4e-06 | 1.800E-06 | 4.000E-07 | 1.400E-06 | 2.200E-06 |
| 26446 | 1323 | I1323E | E | 1.9e-06 | 1.3e-06 | 2.7e-06 | 1.967E-06 | 7.024E-07 | 1.300E-06 | 2.700E-06 |
| 26447 | 1323 | I1323Q | E | 1.9e-06 | 2.5e-06 | 3.2e-06 | 2.533E-06 | 6.506E-07 | 1.900E-06 | 3.200E-06 |
| 26448 | 1323 | I1323G | E | 2e-06 | 1.7e-06 | 1.8e-06 | 1.833E-06 | 1.528E-07 | 1.700E-06 | 2.000E-06 |
| 26449 | 1323 | I1323H | E | 2.4e-06 | 1.6e-06 | 4.2e-06 | 2.733E-06 | 1.332E-06 | 1.600E-06 | 4.200E-06 |
| 26450 | 1323 | I1323I | E | 3e-06 | 1.6e-06 | 1.7e-06 | 2.100E-06 | 7.810E-07 | 1.600E-06 | 3.000E-06 |
| 26451 | 1323 | I1323L | E | 3.1e-06 | 4.5e-06 | 2.1e-06 | 3.233E-06 | 1.206E-06 | 2.100E-06 | 4.500E-06 |
| 26452 | 1323 | I1323K | E | 3.4e-06 | 2.6e-06 | 3.4e-06 | 3.133E-06 | 4.619E-07 | 2.600E-06 | 3.400E-06 |
| 26453 | 1323 | I1323M | E | 1.6e-06 | 2.9e-06 | 2.8e-06 | 2.433E-06 | 7.234E-07 | 1.600E-06 | 2.900E-06 |
| 26454 | 1323 | I1323F | E | 2.3e-06 | 3.6e-06 | 1.4e-06 | 2.433E-06 | 1.106E-06 | 1.400E-06 | 3.600E-06 |
| 26455 | 1323 | I1323P | E | 2.3e-06 | 2.2e-06 | 2.1e-06 | 2.200E-06 | 1.000E-07 | 2.100E-06 | 2.300E-06 |
| 26456 | 1323 | I1323S | E | 3.3e-06 | 4.9e-06 | 2.8e-06 | 3.667E-06 | 1.097E-06 | 2.800E-06 | 4.900E-06 |
| 26457 | 1323 | I1323T | E | 3.3e-06 | 3.4e-06 | 2.2e-06 | 2.967E-06 | 6.658E-07 | 2.200E-06 | 3.400E-06 |
| 26458 | 1323 | I1323W | E | 3.9e-06 | 2.1e-06 | 2.4e-06 | 2.800E-06 | 9.644E-07 | 2.100E-06 | 3.900E-06 |
| 26459 | 1323 | I1323Y | E | 2e-06 | 3.3e-06 | 4.7e-06 | 3.333E-06 | 1.350E-06 | 2.000E-06 | 4.700E-06 |
| 26460 | 1323 | I1323V | E | 2e-06 | 2.1e-06 | 1e-06 | 1.700E-06 | 6.083E-07 | 1.000E-06 | 2.100E-06 |
| 26461 | 1324 | Y1324A | E | 2e-06 | 2.7e-06 | 2.7e-06 | 2.467E-06 | 4.041E-07 | 2.000E-06 | 2.700E-06 |
| 26462 | 1324 | Y1324R | E | 1.3e-06 | 1.5e-06 | 2.2e-06 | 1.667E-06 | 4.726E-07 | 1.300E-06 | 2.200E-06 |
| 26463 | 1324 | Y1324N | E | 2.1e-06 | 2.2e-06 | 4.7e-06 | 3.000E-06 | 1.473E-06 | 2.100E-06 | 4.700E-06 |
| 26464 | 1324 | Y1324D | E | 1.9e-06 | 2.6e-06 | 2.5e-06 | 2.333E-06 | 3.786E-07 | 1.900E-06 | 2.600E-06 |
| 26465 | 1324 | Y1324C | E | 1.6e-06 | 2.8e-06 | 2.3e-06 | 2.233E-06 | 6.028E-07 | 1.600E-06 | 2.800E-06 |
| 26466 | 1324 | Y1324E | E | 2.4e-06 | 2.7e-06 | 1e-06 | 2.033E-06 | 9.074E-07 | 1.000E-06 | 2.700E-06 |
| 26467 | 1324 | Y1324Q | E | 4.1e-06 | 2e-06 | 2e-06 | 2.700E-06 | 1.212E-06 | 2.000E-06 | 4.100E-06 |
| 26468 | 1324 | Y1324G | E | 2.5e-06 | 3.1e-06 | 3.1e-06 | 2.900E-06 | 3.464E-07 | 2.500E-06 | 3.100E-06 |
| 26469 | 1324 | Y1324H | E | 1.6e-06 | 2.1e-06 | 3.1e-06 | 2.267E-06 | 7.638E-07 | 1.600E-06 | 3.100E-06 |
| 26470 | 1324 | Y1324I | E | 4.7e-06 | 2.4e-06 | 2.7e-06 | 3.267E-06 | 1.250E-06 | 2.400E-06 | 4.700E-06 |
| 26471 | 1324 | Y1324L | E | 3.9e-06 | 3e-06 | 3.3e-06 | 3.400E-06 | 4.583E-07 | 3.000E-06 | 3.900E-06 |
| 26472 | 1324 | Y1324K | E | 2.8e-06 | 2e-06 | 2.5e-06 | 2.433E-06 | 4.041E-07 | 2.000E-06 | 2.800E-06 |
| 26473 | 1324 | Y1324M | E | 2.9e-06 | 3.4e-06 | 1.5e-06 | 2.600E-06 | 9.849E-07 | 1.500E-06 | 3.400E-06 |
| 26474 | 1324 | Y1324F | E | 3e-06 | 1.6e-06 | 1.6e-06 | 2.067E-06 | 8.083E-07 | 1.600E-06 | 3.000E-06 |
| 26475 | 1324 | Y1324P | E | 2.1e-06 | 2.9e-06 | 2.8e-06 | 2.600E-06 | 4.359E-07 | 2.100E-06 | 2.900E-06 |
| 26476 | 1324 | Y1324S | E | 2.8e-06 | 2.7e-06 | 2.6e-06 | 2.700E-06 | 1.000E-07 | 2.600E-06 | 2.800E-06 |
| 26477 | 1324 | Y1324T | E | 5.4e-06 | 2.3e-06 | 2.6e-06 | 3.433E-06 | 1.710E-06 | 2.300E-06 | 5.400E-06 |
| 26478 | 1324 | Y1324W | E | 4.2e-06 | 1.4e-06 | 2.6e-06 | 2.733E-06 | 1.405E-06 | 1.400E-06 | 4.200E-06 |
| 26479 | 1324 | Y1324Y | E | 2.1e-06 | 2.5e-06 | 2.1e-06 | 2.233E-06 | 2.309E-07 | 2.100E-06 | 2.500E-06 |
| 26480 | 1324 | Y1324V | E | 3.1e-06 | 1.2e-06 | 1.4e-06 | 1.900E-06 | 1.044E-06 | 1.200E-06 | 3.100E-06 |
| 26481 | 1325 | P1325A | E | 3e-06 | 1.7e-06 | 2.4e-06 | 2.367E-06 | 6.506E-07 | 1.700E-06 | 3.000E-06 |
| 26482 | 1325 | P1325R | E | 1.7e-06 | 1.6e-06 | 2.3e-06 | 1.867E-06 | 3.786E-07 | 1.600E-06 | 2.300E-06 |
| 26483 | 1325 | P1325N | E | 2.3e-06 | 2.7e-06 | 2.4e-06 | 2.467E-06 | 2.082E-07 | 2.300E-06 | 2.700E-06 |
| 26484 | 1325 | P1325D | E | 2.1e-06 | 2e-06 | 2.6e-06 | 2.233E-06 | 3.215E-07 | 2.000E-06 | 2.600E-06 |
| 26485 | 1325 | P1325C | E | 2.1e-06 | 1.4e-06 | 2.2e-06 | 1.900E-06 | 4.359E-07 | 1.400E-06 | 2.200E-06 |
| 26486 | 1325 | P1325E | E | 3.4e-06 | 3.2e-06 | 2.6e-06 | 3.067E-06 | 4.163E-07 | 2.600E-06 | 3.400E-06 |
| 26487 | 1325 | P1325Q | E | 2e-06 | 3e-06 | 2.6e-06 | 2.533E-06 | 5.033E-07 | 2.000E-06 | 3.000E-06 |
| 26488 | 1325 | P1325G | E | 2.1e-06 | 1.3e-06 | 4.7e-06 | 2.700E-06 | 1.778E-06 | 1.300E-06 | 4.700E-06 |
| 26489 | 1325 | P1325H | E | 2.7e-06 | 2.9e-06 | 2.6e-06 | 2.733E-06 | 1.528E-07 | 2.600E-06 | 2.900E-06 |
| 26490 | 1325 | P1325I | E | 1.5e-06 | 2.3e-06 | 1.1e-06 | 1.633E-06 | 6.110E-07 | 1.100E-06 | 2.300E-06 |
| 26491 | 1325 | P1325L | E | 1.3e-06 | 3.3e-06 | 4e-06 | 2.867E-06 | 1.401E-06 | 1.300E-06 | 4.000E-06 |
| 26492 | 1325 | P1325K | E | 2.7e-06 | 4.4e-06 | 2.8e-06 | 3.300E-06 | 9.539E-07 | 2.700E-06 | 4.400E-06 |
| 26493 | 1325 | P1325M | E | 4.5e-06 | 2.5e-06 | 2.5e-06 | 3.167E-06 | 1.155E-06 | 2.500E-06 | 4.500E-06 |
| 26494 | 1325 | P1325F | E | 2.7e-06 | 3.8e-06 | 3.4e-06 | 3.300E-06 | 5.568E-07 | 2.700E-06 | 3.800E-06 |
| 26495 | 1325 | P1325P | E | 4.1e-06 | 2e-06 | 3e-06 | 3.033E-06 | 1.050E-06 | 2.000E-06 | 4.100E-06 |
| 26496 | 1325 | P1325S | E | 2.7e-06 | 2.3e-06 | 3.7e-06 | 2.900E-06 | 7.211E-07 | 2.300E-06 | 3.700E-06 |
| 26497 | 1325 | P1325T | E | 2.2e-06 | 2.1e-06 | 2.8e-06 | 2.367E-06 | 3.786E-07 | 2.100E-06 | 2.800E-06 |
| 26498 | 1325 | P1325W | E | 2e-06 | 1.9e-06 | 2.4e-06 | 2.100E-06 | 2.646E-07 | 1.900E-06 | 2.400E-06 |
| 26499 | 1325 | P1325Y | E | 2.5e-06 | 2.7e-06 | 2.7e-06 | 2.633E-06 | 1.155E-07 | 2.500E-06 | 2.700E-06 |
| 26500 | 1325 | P1325V | E | 2.6e-06 | 2.6e-06 | 3.1e-06 | 2.767E-06 | 2.887E-07 | 2.600E-06 | 3.100E-06 |
| 26501 | 1326 | T1326A | E | 3.1e-06 | 1.9e-06 | 5e-06 | 3.333E-06 | 1.563E-06 | 1.900E-06 | 5.000E-06 |
| 26502 | 1326 | T1326R | E | 3.1e-06 | 4e-06 | 2.3e-06 | 3.133E-06 | 8.505E-07 | 2.300E-06 | 4.000E-06 |
| 26503 | 1326 | T1326N | E | 2.5e-06 | 2.4e-06 | 3.2e-06 | 2.700E-06 | 4.359E-07 | 2.400E-06 | 3.200E-06 |
| 26504 | 1326 | T1326D | E | 1.9e-06 | 3.3e-06 | 4e-06 | 3.067E-06 | 1.069E-06 | 1.900E-06 | 4.000E-06 |
| 26505 | 1326 | T1326C | E | 2.6e-06 | 3.2e-06 | 3.8e-06 | 3.200E-06 | 6.000E-07 | 2.600E-06 | 3.800E-06 |
| 26506 | 1326 | T1326E | E | 3.4e-06 | 2.6e-06 | 2.5e-06 | 2.833E-06 | 4.933E-07 | 2.500E-06 | 3.400E-06 |
| 26507 | 1326 | T1326Q | E | 3.2e-06 | 1.6e-06 | 2e-06 | 2.267E-06 | 8.327E-07 | 1.600E-06 | 3.200E-06 |
| 26508 | 1326 | T1326G | E | 4e-06 | 2.8e-06 | 1.9e-06 | 2.900E-06 | 1.054E-06 | 1.900E-06 | 4.000E-06 |
| 26509 | 1326 | T1326H | E | 1.7e-06 | 1.4e-06 | 1.7e-06 | 1.600E-06 | 1.732E-07 | 1.400E-06 | 1.700E-06 |
| 26510 | 1326 | T1326I | E | 2.4e-06 | 1.8e-06 | 2.9e-06 | 2.367E-06 | 5.508E-07 | 1.800E-06 | 2.900E-06 |
| 26511 | 1326 | T1326L | E | 2.2e-06 | 3.6e-06 | 2.8e-06 | 2.867E-06 | 7.024E-07 | 2.200E-06 | 3.600E-06 |
| 26512 | 1326 | T1326K | E | 2.3e-06 | 2.5e-06 | 3.5e-06 | 2.767E-06 | 6.429E-07 | 2.300E-06 | 3.500E-06 |
| 26513 | 1326 | T1326M | E | 3.6e-06 | 3e-06 | 2.5e-06 | 3.033E-06 | 5.508E-07 | 2.500E-06 | 3.600E-06 |
| 26514 | 1326 | T1326F | E | 1.8e-06 | 2.2e-06 | 1.7e-06 | 1.900E-06 | 2.646E-07 | 1.700E-06 | 2.200E-06 |
| 26515 | 1326 | T1326P | E | 3.3e-06 | 1.8e-06 | 3.9e-06 | 3.000E-06 | 1.082E-06 | 1.800E-06 | 3.900E-06 |
| 26516 | 1326 | T1326S | E | 3.4e-06 | 2.2e-06 | 2.1e-06 | 2.567E-06 | 7.234E-07 | 2.100E-06 | 3.400E-06 |
| 26517 | 1326 | T1326T | E | 3.1e-06 | 1.8e-06 | 1.9e-06 | 2.267E-06 | 7.234E-07 | 1.800E-06 | 3.100E-06 |
| 26518 | 1326 | T1326W | E | 1.2e-06 | 4.8e-06 | 2.5e-06 | 2.833E-06 | 1.823E-06 | 1.200E-06 | 4.800E-06 |
| 26519 | 1326 | T1326Y | E | 2.4e-06 | 2.1e-06 | 3.1e-06 | 2.533E-06 | 5.132E-07 | 2.100E-06 | 3.100E-06 |

| | | | | | | | | | | |
|---|---|---|---|---|---|---|---|---|---|---|
| 26520 | 1326 | T1326V | E | 2.1e-06 | 2.6e-06 | 3.5e-06 | 2.733E-06 | 7.095E-07 | 2.100E-06 | 3.500E-06 |
| 26521 | 1327 | N1327A | E | 3e-06 | 3.1e-06 | 1.9e-06 | 2.667E-06 | 6.658E-07 | 1.900E-06 | 3.100E-06 |
| 26522 | 1327 | N1327R | E | 1.7e-06 | 2.8e-06 | 1.2e-06 | 1.900E-06 | 8.185E-07 | 1.200E-06 | 2.800E-06 |
| 26523 | 1327 | N1327N | E | 3.3e-06 | 3.2e-06 | 2.2e-06 | 2.900E-06 | 6.083E-07 | 2.200E-06 | 3.300E-06 |
| 26524 | 1327 | N1327D | E | 3e-06 | 2.6e-06 | 3.3e-06 | 2.967E-06 | 3.512E-07 | 2.600E-06 | 3.300E-06 |
| 26525 | 1327 | N1327C | E | 1.5e-06 | 1.9e-06 | 1.9e-06 | 1.767E-06 | 2.309E-07 | 1.500E-06 | 1.900E-06 |
| 26526 | 1327 | N1327E | E | 1.9e-06 | 1.4e-06 | 2.8e-06 | 2.033E-06 | 7.095E-07 | 1.400E-06 | 2.800E-06 |
| 26527 | 1327 | N1327Q | E | 2.2e-06 | 2.9e-06 | 1.7e-06 | 2.267E-06 | 6.028E-07 | 1.700E-06 | 2.900E-06 |
| 26528 | 1327 | N1327G | E | 3.3e-06 | 1.6e-06 | 2.4e-06 | 2.433E-06 | 8.505E-07 | 1.600E-06 | 3.300E-06 |
| 26529 | 1327 | N1327H | E | 1.9e-06 | 1.9e-06 | 2.5e-06 | 2.100E-06 | 3.464E-07 | 1.900E-06 | 2.500E-06 |
| 26530 | 1327 | N1327I | E | 1.4e-06 | 2.2e-06 | 2.2e-06 | 1.933E-06 | 4.619E-07 | 1.400E-06 | 2.200E-06 |
| 26531 | 1327 | N1327L | E | 7.9e-07 | 2.4e-06 | 1.9e-06 | 1.697E-06 | 8.240E-07 | 7.900E-07 | 2.400E-06 |
| 26532 | 1327 | N1327K | E | 2.5e-06 | 2e-06 | 2.5e-06 | 2.333E-06 | 2.887E-07 | 2.000E-06 | 2.500E-06 |
| 26533 | 1327 | N1327M | E | 2.3e-06 | 3.4e-06 | 2.2e-06 | 2.633E-06 | 6.658E-07 | 2.200E-06 | 3.400E-06 |
| 26534 | 1327 | N1327F | E | 1.8e-06 | 2.6e-06 | 2.4e-06 | 2.267E-06 | 4.163E-07 | 1.800E-06 | 2.600E-06 |
| 26535 | 1327 | N1327P | E | 1e-06 | 2.2e-06 | 2.6e-06 | 1.933E-06 | 8.327E-07 | 1.000E-06 | 2.600E-06 |
| 26536 | 1327 | N1327S | E | 4.1e-06 | 2.7e-06 | 5.1e-06 | 3.967E-06 | 1.206E-06 | 2.700E-06 | 5.100E-06 |
| 26537 | 1327 | N1327T | E | 2.8e-06 | 1.8e-06 | 3.1e-06 | 2.567E-06 | 6.807E-07 | 1.800E-06 | 3.100E-06 |
| 26538 | 1327 | N1327W | E | 4.3e-06 | 2.2e-06 | 1.3e-06 | 2.600E-06 | 1.539E-06 | 1.300E-06 | 4.300E-06 |
| 26539 | 1327 | N1327Y | E | 2.2e-06 | 2.4e-06 | 2.9e-06 | 2.500E-06 | 3.606E-07 | 2.200E-06 | 2.900E-06 |
| 26540 | 1327 | N1327V | E | 1.8e-06 | 3.2e-06 | 1.9e-06 | 2.300E-06 | 7.810E-07 | 1.800E-06 | 3.200E-06 |
| 26541 | 1328 | G1328A | E | 3.2e-06 | 1.8e-06 | 2.3e-06 | 2.433E-06 | 7.095E-07 | 1.800E-06 | 3.200E-06 |
| 26542 | 1328 | G1328R | E | 2.6e-06 | 2.4e-06 | 2.5e-06 | 2.500E-06 | 1.000E-07 | 2.400E-06 | 2.600E-06 |
| 26543 | 1328 | G1328N | E | 4e-06 | 1.8e-06 | 3e-06 | 2.933E-06 | 1.102E-06 | 1.800E-06 | 4.000E-06 |
| 26544 | 1328 | G1328D | E | 2.2e-06 | 2.2e-06 | 2.6e-06 | 2.333E-06 | 2.309E-07 | 2.200E-06 | 2.600E-06 |
| 26545 | 1328 | G1328C | E | 4.1e-06 | 3e-06 | 1.7e-06 | 2.933E-06 | 1.201E-06 | 1.700E-06 | 4.100E-06 |
| 26546 | 1328 | G1328E | E | 2.8e-06 | 2e-06 | 1.6e-06 | 2.133E-06 | 6.110E-07 | 1.600E-06 | 2.800E-06 |
| 26547 | 1328 | G1328Q | E | 2.7e-06 | 3e-06 | 2.2e-06 | 2.633E-06 | 4.041E-07 | 2.200E-06 | 3.000E-06 |
| 26548 | 1328 | G1328G | E | 2.9e-06 | 1.4e-06 | 1.8e-06 | 2.033E-06 | 7.767E-07 | 1.400E-06 | 2.900E-06 |
| 26549 | 1328 | G1328H | E | 2.7e-06 | 3.3e-06 | 2.2e-06 | 2.733E-06 | 5.508E-07 | 2.200E-06 | 3.300E-06 |
| 26550 | 1328 | G1328I | E | 2.1e-06 | 3.3e-06 | 2.8e-06 | 2.733E-06 | 6.028E-07 | 2.100E-06 | 3.300E-06 |
| 26551 | 1328 | G1328L | E | 4.2e-06 | 3e-06 | 3.9e-06 | 3.700E-06 | 6.245E-07 | 3.000E-06 | 4.200E-06 |
| 26552 | 1328 | G1328K | E | 3.6e-06 | 2.3e-06 | 2.1e-06 | 2.667E-06 | 8.145E-07 | 2.100E-06 | 3.600E-06 |
| 26553 | 1328 | G1328M | E | 1.1e-06 | 3.4e-06 | 1.8e-06 | 2.100E-06 | 1.179E-06 | 1.100E-06 | 3.400E-06 |
| 26554 | 1328 | G1328F | E | 2.1e-06 | 1.8e-06 | 2.1e-06 | 2.000E-06 | 1.732E-07 | 1.800E-06 | 2.100E-06 |
| 26555 | 1328 | G1328P | E | 2.6e-06 | 2.1e-06 | 1.4e-06 | 2.033E-06 | 6.028E-07 | 1.400E-06 | 2.600E-06 |
| 26556 | 1328 | G1328S | E | 8.8e-07 | 2.9e-06 | 3e-06 | 2.260E-06 | 1.196E-06 | 8.800E-07 | 3.000E-06 |
| 26557 | 1328 | G1328T | E | 2e-06 | 3.5e-06 | 2.3e-06 | 2.600E-06 | 7.937E-07 | 2.000E-06 | 3.500E-06 |
| 26558 | 1328 | G1328W | E | 1.9e-06 | 2.4e-06 | 2.9e-06 | 2.400E-06 | 5.000E-07 | 1.900E-06 | 2.900E-06 |
| 26559 | 1328 | G1328Y | E | 3.1e-06 | 9.9e-07 | 2.2e-06 | 2.097E-06 | 1.059E-06 | 9.900E-07 | 3.100E-06 |
| 26560 | 1328 | G1328V | E | 2.2e-06 | 2.4e-06 | 2.3e-06 | 2.300E-06 | 1.000E-07 | 2.200E-06 | 2.400E-06 |
| 26561 | 1329 | Y1329A | E | 1.7e-06 | 1.8e-06 | 1.8e-06 | 1.767E-06 | 5.774E-08 | 1.700E-06 | 1.800E-06 |
| 26562 | 1329 | Y1329R | E | 1.1e-06 | 1.8e-06 | 1.7e-06 | 1.533E-06 | 3.786E-07 | 1.100E-06 | 1.800E-06 |
| 26563 | 1329 | Y1329N | E | 3.6e-06 | 2.7e-06 | 8.8e-07 | 2.393E-06 | 1.386E-06 | 8.800E-07 | 3.600E-06 |
| 26564 | 1329 | Y1329D | E | 2.5e-06 | 1.2e-06 | 2.6e-06 | 2.100E-06 | 7.810E-07 | 1.200E-06 | 2.600E-06 |
| 26565 | 1329 | Y1329C | E | 1.8e-06 | 2.5e-06 | 2.6e-06 | 2.300E-06 | 4.359E-07 | 1.800E-06 | 2.600E-06 |
| 26566 | 1329 | Y1329E | E | 2.1e-06 | 9.6e-07 | 1.7e-06 | 1.587E-06 | 5.784E-07 | 9.600E-07 | 2.100E-06 |
| 26567 | 1329 | Y1329Q | E | 1.2e-06 | 1.8e-06 | 8.3e-07 | 1.277E-06 | 4.895E-07 | 8.300E-07 | 1.800E-06 |
| 26568 | 1329 | Y1329G | E | 3.1e-06 | 4e-06 | 9.6e-07 | 2.687E-06 | 1.562E-06 | 9.600E-07 | 4.000E-06 |
| 26569 | 1329 | Y1329H | E | 1.4e-06 | 1.6e-06 | 2.1e-06 | 1.700E-06 | 3.606E-07 | 1.400E-06 | 2.100E-06 |
| 26570 | 1329 | Y1329I | E | 2.6e-06 | 1.3e-06 | 1.7e-06 | 1.867E-06 | 6.658E-07 | 1.300E-06 | 2.600E-06 |
| 26571 | 1329 | Y1329L | E | 3e-06 | 3.3e-06 | 3.2e-06 | 3.167E-06 | 1.528E-07 | 3.000E-06 | 3.300E-06 |
| 26572 | 1329 | Y1329K | E | 1.6e-06 | 1.8e-06 | 2.6e-06 | 2.000E-06 | 5.292E-07 | 1.600E-06 | 2.600E-06 |
| 26573 | 1329 | Y1329M | E | 2.9e-06 | 3.6e-06 | 1.2e-06 | 2.567E-06 | 1.234E-06 | 1.200E-06 | 3.600E-06 |
| 26574 | 1329 | Y1329F | E | 1.8e-06 | 3.8e-06 | 1.6e-06 | 2.400E-06 | 1.217E-06 | 1.600E-06 | 3.800E-06 |
| 26575 | 1329 | Y1329P | E | 1.9e-06 | 2.5e-06 | 2.4e-06 | 2.267E-06 | 3.215E-07 | 1.900E-06 | 2.500E-06 |
| 26576 | 1329 | Y1329S | E | 1.9e-06 | 1e-06 | 1.4e-06 | 1.433E-06 | 4.509E-07 | 1.000E-06 | 1.900E-06 |
| 26577 | 1329 | Y1329T | E | 2.7e-06 | 1.1e-06 | 1.8e-06 | 1.867E-06 | 8.021E-07 | 1.100E-06 | 2.700E-06 |
| 26578 | 1329 | Y1329W | E | 1.7e-06 | 2.2e-06 | 1.9e-06 | 1.933E-06 | 2.517E-07 | 1.700E-06 | 2.200E-06 |
| 26579 | 1329 | Y1329Y | E | 2.9e-06 | 1.9e-06 | 2.1e-06 | 2.300E-06 | 5.292E-07 | 1.900E-06 | 2.900E-06 |
| 26580 | 1329 | Y1329V | E | 1.9e-06 | 2.6e-06 | 2e-06 | 2.167E-06 | 3.786E-07 | 1.900E-06 | 2.600E-06 |
| 26581 | 1330 | T1330A | E | 2.6e-06 | 1.9e-06 | 1.5e-06 | 2.000E-06 | 5.568E-07 | 1.500E-06 | 2.600E-06 |
| 26582 | 1330 | T1330R | E | 2.5e-06 | 1.8e-06 | 2e-06 | 2.100E-06 | 3.606E-07 | 1.800E-06 | 2.500E-06 |
| 26583 | 1330 | T1330N | E | 1.9e-06 | 3.9e-06 | 2.6e-06 | 2.800E-06 | 1.015E-06 | 1.900E-06 | 3.900E-06 |
| 26584 | 1330 | T1330D | E | 2.3e-06 | 1.4e-06 | 1.3e-06 | 1.667E-06 | 5.508E-07 | 1.300E-06 | 2.300E-06 |
| 26585 | 1330 | T1330C | E | 1.9e-06 | 2.7e-06 | 1.8e-06 | 2.133E-06 | 4.933E-07 | 1.800E-06 | 2.700E-06 |
| 26586 | 1330 | T1330E | E | 1.1e-06 | 1.9e-06 | 1.8e-06 | 1.600E-06 | 4.359E-07 | 1.100E-06 | 1.900E-06 |
| 26587 | 1330 | T1330Q | E | 1.9e-06 | 2.4e-06 | 2.5e-06 | 2.267E-06 | 3.215E-07 | 1.900E-06 | 2.500E-06 |
| 26588 | 1330 | T1330G | E | 2.8e-06 | 1.7e-06 | 1.8e-06 | 2.100E-06 | 6.083E-07 | 1.700E-06 | 2.800E-06 |
| 26589 | 1330 | T1330H | E | 2.7e-06 | 1.6e-06 | 2.5e-06 | 2.267E-06 | 5.859E-07 | 1.600E-06 | 2.700E-06 |
| 26590 | 1330 | T1330I | E | 2.3e-06 | 1.4e-06 | 2.2e-06 | 1.967E-06 | 4.933E-07 | 1.400E-06 | 2.300E-06 |
| 26591 | 1330 | T1330L | E | 2.5e-06 | 2.7e-06 | 1.9e-06 | 2.367E-06 | 4.163E-07 | 1.900E-06 | 2.700E-06 |
| 26592 | 1330 | T1330K | E | 2.3e-06 | 2.2e-06 | 2.7e-06 | 2.400E-06 | 2.646E-07 | 2.200E-06 | 2.700E-06 |
| 26593 | 1330 | T1330M | E | 1.9e-06 | 1.9e-06 | 1.4e-06 | 1.733E-06 | 2.887E-07 | 1.400E-06 | 1.900E-06 |
| 26594 | 1330 | T1330F | E | 2.8e-06 | 2.8e-06 | 2.2e-06 | 2.600E-06 | 3.464E-07 | 2.200E-06 | 2.800E-06 |
| 26595 | 1330 | T1330P | E | 1.6e-06 | 1.3e-06 | 2.2e-06 | 1.700E-06 | 4.583E-07 | 1.300E-06 | 2.200E-06 |
| 26596 | 1330 | T1330S | E | 2.1e-06 | 4.3e-06 | 3.4e-06 | 3.267E-06 | 1.106E-06 | 2.100E-06 | 4.300E-06 |
| 26597 | 1330 | T1330T | E | 2.7e-06 | 2.1e-06 | 1.7e-06 | 2.167E-06 | 5.033E-07 | 1.700E-06 | 2.700E-06 |
| 26598 | 1330 | T1330W | E | 1.8e-06 | 3.8e-06 | 1.7e-06 | 2.433E-06 | 1.185E-06 | 1.700E-06 | 3.800E-06 |
| 26599 | 1330 | T1330Y | E | 1.9e-06 | 1.7e-06 | 2.7e-06 | 2.100E-06 | 5.292E-07 | 1.700E-06 | 2.700E-06 |
| 26600 | 1330 | T1330V | E | 1.3e-06 | 2.3e-06 | 2.2e-06 | 1.933E-06 | 5.508E-07 | 1.300E-06 | 2.300E-06 |
| 26601 | 1331 | R1331A | E | 2.7e-06 | 2.5e-06 | 3.3e-06 | 2.833E-06 | 4.163E-07 | 2.500E-06 | 3.300E-06 |
| 26602 | 1331 | R1331R | E | 2.3e-06 | 1.8e-06 | 2.5e-06 | 2.200E-06 | 3.606E-07 | 1.800E-06 | 2.500E-06 |
| 26603 | 1331 | R1331N | E | 3.8e-06 | 3e-06 | 3.8e-06 | 3.533E-06 | 4.619E-07 | 3.000E-06 | 3.800E-06 |
| 26604 | 1331 | R1331D | E | 3e-06 | 2.8e-06 | 4.3e-06 | 3.367E-06 | 8.145E-07 | 2.800E-06 | 4.300E-06 |

| | | | | | | | | | | |
|---|---|---|---|---|---|---|---|---|---|---|
| 26605 | 1331 | R1331C | E | 2.1e-06 | 2.1e-06 | 1.8e-06 | 2.000E-06 | 1.732E-07 | 1.800E-06 | 2.100E-06 |
| 26606 | 1331 | R1331E | E | 3.4e-06 | 3.8e-06 | 3.6e-06 | 3.600E-06 | 2.000E-07 | 3.400E-06 | 3.800E-06 |
| 26607 | 1331 | R1331Q | E | 5e-06 | 4.5e-06 | 3.8e-06 | 4.433E-06 | 6.028E-07 | 3.800E-06 | 5.000E-06 |
| 26608 | 1331 | R1331G | E | 3e-06 | 3.1e-06 | 2.2e-06 | 2.767E-06 | 4.933E-07 | 2.200E-06 | 3.100E-06 |
| 26609 | 1331 | R1331H | E | 2.1e-06 | 1.9e-06 | 2.9e-06 | 2.300E-06 | 5.292E-07 | 1.900E-06 | 2.900E-06 |
| 26610 | 1331 | R1331I | E | 3e-06 | 1.7e-06 | 2.6e-06 | 2.433E-06 | 6.658E-07 | 1.700E-06 | 3.000E-06 |
| 26611 | 1331 | R1331L | E | 2.9e-06 | 2.2e-06 | 2.7e-06 | 2.600E-06 | 3.606E-07 | 2.200E-06 | 2.900E-06 |
| 26612 | 1331 | R1331K | E | 2.5e-06 | 1.9e-06 | 2.6e-06 | 2.333E-06 | 3.786E-07 | 1.900E-06 | 2.600E-06 |
| 26613 | 1331 | R1331M | E | 2.7e-06 | 1.4e-06 | 1.8e-06 | 1.967E-06 | 6.658E-07 | 1.400E-06 | 2.700E-06 |
| 26614 | 1331 | R1331F | E | 1.5e-06 | 2.3e-06 | 2.1e-06 | 1.967E-06 | 4.163E-07 | 1.500E-06 | 2.300E-06 |
| 26615 | 1331 | R1331P | E | 3.3e-06 | 2.7e-06 | 2.2e-06 | 2.733E-06 | 5.508E-07 | 2.200E-06 | 3.300E-06 |
| 26616 | 1331 | R1331S | E | 6e-06 | 2.9e-06 | 4.6e-06 | 4.500E-06 | 1.552E-06 | 2.900E-06 | 6.000E-06 |
| 26617 | 1331 | R1331T | E | 3.3e-06 | 5.1e-06 | 6.7e-06 | 5.033E-06 | 1.701E-06 | 3.300E-06 | 6.700E-06 |
| 26618 | 1331 | R1331W | E | 9.2e-07 | 3.8e-06 | 3e-06 | 2.573E-06 | 1.487E-06 | 9.200E-07 | 3.800E-06 |
| 26619 | 1331 | R1331Y | E | 9.1e-07 | 1.5e-06 | 2.5e-06 | 1.637E-06 | 8.038E-07 | 9.100E-07 | 2.500E-06 |
| 26620 | 1331 | R1331V | E | 2.4e-06 | 4.1e-06 | 1.7e-06 | 2.733E-06 | 1.234E-06 | 1.700E-06 | 4.100E-06 |
| 26621 | 1332 | Y1332A | E | 1.8e-06 | 2.1e-06 | 1.7e-06 | 1.867E-06 | 2.082E-07 | 1.700E-06 | 2.100E-06 |
| 26622 | 1332 | Y1332R | E | 2.7e-06 | 3.1e-06 | 1.3e-06 | 2.367E-06 | 9.452E-07 | 1.300E-06 | 3.100E-06 |
| 26623 | 1332 | Y1332N | E | 2.6e-06 | 2.2e-06 | 5e-06 | 3.267E-06 | 1.514E-06 | 2.200E-06 | 5.000E-06 |
| 26624 | 1332 | Y1332D | E | 4.7e-06 | 2.2e-06 | 2.3e-06 | 3.067E-06 | 1.415E-06 | 2.200E-06 | 4.700E-06 |
| 26625 | 1332 | Y1332C | E | 2.2e-06 | 3.2e-06 | 1.7e-06 | 2.367E-06 | 7.638E-07 | 1.700E-06 | 3.200E-06 |
| 26626 | 1332 | Y1332E | E | 3.1e-06 | 2e-06 | 2.4e-06 | 2.500E-06 | 5.568E-07 | 2.000E-06 | 3.100E-06 |
| 26627 | 1332 | Y1332Q | E | 2.8e-06 | 3.4e-06 | 2.6e-06 | 2.933E-06 | 4.163E-07 | 2.600E-06 | 3.400E-06 |
| 26628 | 1332 | Y1332G | E | 4e-06 | 2e-06 | 1.6e-06 | 2.533E-06 | 1.286E-06 | 1.600E-06 | 4.000E-06 |
| 26629 | 1332 | Y1332H | E | 2.7e-06 | 2.7e-06 | 2.1e-06 | 2.500E-06 | 3.464E-07 | 2.100E-06 | 2.700E-06 |
| 26630 | 1332 | Y1332I | E | 2.6e-06 | 2.2e-06 | 2.3e-06 | 2.367E-06 | 2.082E-07 | 2.200E-06 | 2.600E-06 |
| 26631 | 1332 | Y1332L | E | 2.2e-06 | 2.3e-06 | 3.2e-06 | 2.567E-06 | 5.508E-07 | 2.200E-06 | 3.200E-06 |
| 26632 | 1332 | Y1332K | E | 3.7e-06 | 3.1e-06 | 3.2e-06 | 3.333E-06 | 3.215E-07 | 3.100E-06 | 3.700E-06 |
| 26633 | 1332 | Y1332M | E | 1.7e-06 | 1.2e-06 | 3.8e-06 | 2.233E-06 | 1.380E-06 | 1.200E-06 | 3.800E-06 |
| 26634 | 1332 | Y1332F | E | 1.9e-06 | 3.1e-06 | 1.2e-06 | 2.067E-06 | 9.609E-07 | 1.200E-06 | 3.100E-06 |
| 26635 | 1332 | Y1332P | E | 3.3e-06 | 1.7e-06 | 3e-06 | 2.667E-06 | 8.505E-07 | 1.700E-06 | 3.300E-06 |
| 26636 | 1332 | Y1332S | E | 2.2e-06 | 1.7e-06 | 2.2e-06 | 2.033E-06 | 2.887E-07 | 1.700E-06 | 2.200E-06 |
| 26637 | 1332 | Y1332T | E | 2.2e-06 | 1.8e-06 | 3.6e-06 | 2.533E-06 | 9.452E-07 | 1.800E-06 | 3.600E-06 |
| 26638 | 1332 | Y1332W | E | 3.2e-06 | 2.6e-06 | 1.9e-06 | 2.567E-06 | 6.506E-07 | 1.900E-06 | 3.200E-06 |
| 26639 | 1332 | Y1332Y | E | 2.8e-06 | 3.4e-06 | 1.3e-06 | 2.500E-06 | 1.082E-06 | 1.300E-06 | 3.400E-06 |
| 26640 | 1332 | Y1332V | E | 3.9e-06 | 1.8e-06 | 1e-06 | 2.233E-06 | 1.498E-06 | 1.000E-06 | 3.900E-06 |
| 26641 | 1333 | A1333A | E | 2.6e-06 | 2.1e-06 | 2.6e-06 | 2.433E-06 | 2.887E-07 | 2.100E-06 | 2.600E-06 |
| 26642 | 1333 | A1333R | E | 2e-06 | 1.9e-06 | 2.1e-06 | 2.000E-06 | 1.000E-07 | 1.900E-06 | 2.100E-06 |
| 26643 | 1333 | A1333N | E | 2e-06 | 2.3e-06 | 2.3e-06 | 2.200E-06 | 1.732E-07 | 2.000E-06 | 2.300E-06 |
| 26644 | 1333 | A1333D | E | 1.5e-06 | 1.5e-06 | 3.9e-06 | 2.300E-06 | 1.386E-06 | 1.500E-06 | 3.900E-06 |
| 26645 | 1333 | A1333C | E | 2.3e-06 | 2.5e-06 | 2.6e-06 | 2.467E-06 | 1.528E-07 | 2.300E-06 | 2.600E-06 |
| 26646 | 1333 | A1333E | E | 3.8e-06 | 3.7e-06 | 2.2e-06 | 3.233E-06 | 8.963E-07 | 2.200E-06 | 3.800E-06 |
| 26647 | 1333 | A1333Q | E | 2.4e-06 | 1.3e-06 | 1.7e-06 | 1.800E-06 | 5.568E-07 | 1.300E-06 | 2.400E-06 |
| 26648 | 1333 | A1333G | E | 4e-06 | 1.8e-06 | 2.1e-06 | 2.633E-06 | 1.193E-06 | 1.800E-06 | 4.000E-06 |
| 26649 | 1333 | A1333H | E | 1.3e-06 | 3e-06 | 2.5e-06 | 2.267E-06 | 8.737E-07 | 1.300E-06 | 3.000E-06 |
| 26650 | 1333 | A1333I | E | 1.4e-06 | 1.9e-06 | 3.7e-06 | 2.333E-06 | 1.210E-06 | 1.400E-06 | 3.700E-06 |
| 26651 | 1333 | A1333L | E | 2.5e-06 | 1.6e-06 | 3.3e-06 | 2.467E-06 | 8.505E-07 | 1.600E-06 | 3.300E-06 |
| 26652 | 1333 | A1333K | E | 2.6e-06 | 2.1e-06 | 1.4e-06 | 2.033E-06 | 6.028E-07 | 1.400E-06 | 2.600E-06 |
| 26653 | 1333 | A1333M | E | 3.5e-06 | 1.5e-06 | 2.4e-06 | 2.467E-06 | 1.002E-06 | 1.500E-06 | 3.500E-06 |
| 26654 | 1333 | A1333F | E | 1.9e-06 | 1.7e-06 | 3.4e-06 | 2.333E-06 | 9.292E-07 | 1.700E-06 | 3.400E-06 |
| 26655 | 1333 | A1333P | E | 2.5e-06 | 3.1e-06 | 2.2e-06 | 2.600E-06 | 4.583E-07 | 2.200E-06 | 3.100E-06 |
| 26656 | 1333 | A1333S | E | 2.2e-06 | 2e-06 | 2.3e-06 | 2.167E-06 | 1.528E-07 | 2.000E-06 | 2.300E-06 |
| 26657 | 1333 | A1333T | E | 2.9e-06 | 2.6e-06 | 2e-06 | 2.500E-06 | 4.583E-07 | 2.000E-06 | 2.900E-06 |
| 26658 | 1333 | A1333W | E | 1.6e-06 | 1.4e-06 | 2.1e-06 | 1.700E-06 | 3.606E-07 | 1.400E-06 | 2.100E-06 |
| 26659 | 1333 | A1333Y | E | 2.8e-06 | 2.3e-06 | 3.4e-06 | 2.833E-06 | 5.508E-07 | 2.300E-06 | 3.400E-06 |
| 26660 | 1333 | A1333V | E | 4.5e-06 | 2.8e-06 | 2.1e-06 | 3.133E-06 | 1.234E-06 | 2.100E-06 | 4.500E-06 |
| 26661 | 1334 | D1334A | E | 2.1e-06 | 1.3e-06 | 2.5e-06 | 1.967E-06 | 6.110E-07 | 1.300E-06 | 2.500E-06 |
| 26662 | 1334 | D1334R | E | 3.5e-06 | 2.6e-06 | 2.6e-06 | 2.900E-06 | 5.196E-07 | 2.600E-06 | 3.500E-06 |
| 26663 | 1334 | D1334N | E | 2.7e-06 | 2.6e-06 | 1.7e-06 | 2.333E-06 | 5.508E-07 | 1.700E-06 | 2.700E-06 |
| 26664 | 1334 | D1334D | E | 2.2e-06 | 2.5e-06 | 2.8e-06 | 2.500E-06 | 3.000E-07 | 2.200E-06 | 2.800E-06 |
| 26665 | 1334 | D1334C | E | 2.9e-06 | 2.4e-06 | 1.4e-06 | 2.233E-06 | 7.638E-07 | 1.400E-06 | 2.900E-06 |
| 26666 | 1334 | D1334E | E | 2.2e-06 | 2e-06 | 3.8e-06 | 2.667E-06 | 9.866E-07 | 2.000E-06 | 3.800E-06 |
| 26667 | 1334 | D1334Q | E | 2.8e-06 | 2.9e-06 | 1.7e-06 | 2.467E-06 | 6.658E-07 | 1.700E-06 | 2.900E-06 |
| 26668 | 1334 | D1334G | E | 2.5e-06 | 2.3e-06 | 1.6e-06 | 2.133E-06 | 4.726E-07 | 1.600E-06 | 2.500E-06 |
| 26669 | 1334 | D1334H | E | 2.8e-06 | 2.2e-06 | 2.6e-06 | 2.533E-06 | 3.055E-07 | 2.200E-06 | 2.800E-06 |
| 26670 | 1334 | D1334I | E | 2.5e-06 | 2.3e-06 | 1.4e-06 | 2.067E-06 | 5.859E-07 | 1.400E-06 | 2.500E-06 |
| 26671 | 1334 | D1334L | E | 2.6e-06 | 2.9e-06 | 3.6e-06 | 3.033E-06 | 5.132E-07 | 2.600E-06 | 3.600E-06 |
| 26672 | 1334 | D1334K | E | 2.3e-06 | 3.9e-06 | 2.7e-06 | 2.967E-06 | 8.327E-07 | 2.300E-06 | 3.900E-06 |
| 26673 | 1334 | D1334M | E | 2.2e-06 | 4.8e-06 | 1.6e-06 | 2.867E-06 | 1.701E-06 | 1.600E-06 | 4.800E-06 |
| 26674 | 1334 | D1334F | E | 3.1e-06 | 2.8e-06 | 1.3e-06 | 2.400E-06 | 9.644E-07 | 1.300E-06 | 3.100E-06 |
| 26675 | 1334 | D1334P | E | 2.5e-06 | 3.1e-06 | 3.6e-06 | 3.067E-06 | 5.508E-07 | 2.500E-06 | 3.600E-06 |
| 26676 | 1334 | D1334S | E | 2.4e-06 | 2.2e-06 | 1.5e-06 | 2.033E-06 | 4.726E-07 | 1.500E-06 | 2.400E-06 |
| 26677 | 1334 | D1334T | E | 2.1e-06 | 2.1e-06 | 2.4e-06 | 2.200E-06 | 1.732E-07 | 2.100E-06 | 2.400E-06 |
| 26678 | 1334 | D1334W | E | 4.9e-06 | 3.7e-06 | 3.3e-06 | 3.967E-06 | 8.327E-07 | 3.300E-06 | 4.900E-06 |
| 26679 | 1334 | D1334Y | E | 2.6e-06 | 2.1e-06 | 4.1e-06 | 2.933E-06 | 1.041E-06 | 2.100E-06 | 4.100E-06 |
| 26680 | 1334 | D1334V | E | 2.8e-06 | 2e-06 | 2.1e-06 | 2.300E-06 | 4.359E-07 | 2.000E-06 | 2.800E-06 |
| 26681 | 1335 | S1335A | E | 2.2e-06 | 2.2e-06 | 1.6e-06 | 2.000E-06 | 3.464E-07 | 1.600E-06 | 2.200E-06 |
| 26682 | 1335 | S1335R | E | 3.6e-06 | 1.9e-06 | 1.2e-06 | 2.233E-06 | 1.234E-06 | 1.200E-06 | 3.600E-06 |
| 26683 | 1335 | S1335N | E | 2.7e-06 | 2.5e-06 | 2.2e-06 | 2.467E-06 | 2.517E-07 | 2.200E-06 | 2.700E-06 |
| 26684 | 1335 | S1335D | E | 2e-06 | 2.8e-06 | 3.1e-06 | 2.633E-06 | 5.686E-07 | 2.000E-06 | 3.100E-06 |
| 26685 | 1335 | S1335C | E | 1.7e-06 | 2e-06 | 4.4e-06 | 2.700E-06 | 1.480E-06 | 1.700E-06 | 4.400E-06 |
| 26686 | 1335 | S1335E | E | 1.5e-06 | 2.2e-06 | 2.3e-06 | 2.000E-06 | 4.359E-07 | 1.500E-06 | 2.300E-06 |
| 26687 | 1335 | S1335Q | E | 1.4e-06 | 3.9e-06 | 3.5e-06 | 2.933E-06 | 1.343E-06 | 1.400E-06 | 3.900E-06 |
| 26688 | 1335 | S1335G | E | 8.8e-07 | 2.1e-06 | 1.5e-06 | 1.493E-06 | 6.100E-07 | 8.800E-07 | 2.100E-06 |
| 26689 | 1335 | S1335H | E | 1.1e-06 | 3.8e-06 | 2.2e-06 | 2.367E-06 | 1.358E-06 | 1.100E-06 | 3.800E-06 |

| | | | | | | | | | | |
|---|---|---|---|---|---|---|---|---|---|---|
| 26690 | 1335 | S1335I | E | 2.2e-06 | 2.1e-06 | 5.7e-06 | 3.333E-06 | 2.050E-06 | 2.100E-06 | 5.700E-06 |
| 26691 | 1335 | S1335L | E | 3.6e-06 | 1.7e-06 | 2e-06 | 2.433E-06 | 1.021E-06 | 1.700E-06 | 3.600E-06 |
| 26692 | 1335 | S1335K | E | 2.7e-06 | 2.7e-06 | 4.3e-06 | 3.233E-06 | 9.238E-07 | 2.700E-06 | 4.300E-06 |
| 26693 | 1335 | S1335M | E | 2.4e-06 | 3e-06 | 2.3e-06 | 2.567E-06 | 3.786E-07 | 2.300E-06 | 3.000E-06 |
| 26694 | 1335 | S1335F | E | 3.1e-06 | 2.1e-06 | 3.5e-06 | 2.900E-06 | 7.211E-07 | 2.100E-06 | 3.500E-06 |
| 26695 | 1335 | S1335P | E | 3.6e-06 | 2.5e-06 | 3.3e-06 | 3.133E-06 | 5.686E-07 | 2.500E-06 | 3.600E-06 |
| 26696 | 1335 | S1335S | E | 2.2e-06 | 3.5e-06 | 3.3e-06 | 3.000E-06 | 7.000E-07 | 2.200E-06 | 3.500E-06 |
| 26697 | 1335 | S1335T | E | 1.3e-06 | 1.2e-06 | 1.8e-06 | 1.433E-06 | 3.215E-07 | 1.200E-06 | 1.800E-06 |
| 26698 | 1335 | S1335W | E | 2.3e-06 | 2.5e-06 | 9.2e-07 | 1.907E-06 | 8.603E-07 | 9.200E-07 | 2.500E-06 |
| 26699 | 1335 | S1335Y | E | 2.6e-06 | 5.4e-06 | 2.4e-06 | 3.467E-06 | 1.677E-06 | 2.400E-06 | 5.400E-06 |
| 26700 | 1335 | S1335V | E | 2.9e-06 | 3.1e-06 | 1.6e-06 | 2.533E-06 | 8.145E-07 | 1.600E-06 | 3.100E-06 |
| 26701 | 1336 | V1336A | E | 1.2e-06 | 2.5e-06 | 2.2e-06 | 1.967E-06 | 6.807E-07 | 1.200E-06 | 2.500E-06 |
| 26702 | 1336 | V1336R | E | 1.8e-06 | 1.9e-06 | 4.4e-06 | 2.700E-06 | 1.473E-06 | 1.800E-06 | 4.400E-06 |
| 26703 | 1336 | V1336N | E | 1.3e-06 | 1.8e-06 | 1.5e-06 | 1.533E-06 | 2.517E-07 | 1.300E-06 | 1.800E-06 |
| 26704 | 1336 | V1336D | E | 2.4e-06 | 1.9e-06 | 4.2e-06 | 2.833E-06 | 1.210E-06 | 1.900E-06 | 4.200E-06 |
| 26705 | 1336 | V1336C | E | 2.7e-06 | 3.4e-06 | 1.9e-06 | 2.667E-06 | 7.506E-07 | 1.900E-06 | 3.400E-06 |
| 26706 | 1336 | V1336E | E | 2.8e-06 | 3.2e-06 | 2.3e-06 | 2.767E-06 | 4.509E-07 | 2.300E-06 | 3.200E-06 |
| 26707 | 1336 | V1336Q | E | 1.9e-06 | 1.5e-06 | 2.8e-06 | 2.067E-06 | 6.658E-07 | 1.500E-06 | 2.800E-06 |
| 26708 | 1336 | V1336G | E | 1.8e-06 | 3.7e-06 | 2.2e-06 | 2.567E-06 | 1.002E-06 | 1.800E-06 | 3.700E-06 |
| 26709 | 1336 | V1336H | E | 2.9e-06 | 1.9e-06 | 2.6e-06 | 2.467E-06 | 5.132E-07 | 1.900E-06 | 2.900E-06 |
| 26710 | 1336 | V1336I | E | 1.8e-06 | 3.5e-06 | 1.2e-06 | 2.167E-06 | 1.193E-06 | 1.200E-06 | 3.500E-06 |
| 26711 | 1336 | V1336L | E | 2.2e-06 | 2.8e-06 | 2.2e-06 | 2.400E-06 | 3.464E-07 | 2.200E-06 | 2.800E-06 |
| 26712 | 1336 | V1336K | E | 2.5e-06 | 2.6e-06 | 2.4e-06 | 2.500E-06 | 1.000E-07 | 2.400E-06 | 2.600E-06 |
| 26713 | 1336 | V1336M | E | 2.5e-06 | 2.4e-06 | 2.4e-06 | 2.433E-06 | 5.774E-08 | 2.400E-06 | 2.500E-06 |
| 26714 | 1336 | V1336F | E | 2e-06 | 1.8e-06 | 2.6e-06 | 2.133E-06 | 4.163E-07 | 1.800E-06 | 2.600E-06 |
| 26715 | 1336 | V1336P | E | 1.5e-06 | 1.9e-06 | 4.7e-06 | 2.700E-06 | 1.744E-06 | 1.500E-06 | 4.700E-06 |
| 26716 | 1336 | V1336S | E | 2.5e-06 | 2.3e-06 | 2.8e-06 | 2.533E-06 | 2.517E-07 | 2.300E-06 | 2.800E-06 |
| 26717 | 1336 | V1336T | E | 3.8e-06 | 2.5e-06 | 2.1e-06 | 2.800E-06 | 8.888E-07 | 2.100E-06 | 3.800E-06 |
| 26718 | 1336 | V1336W | E | 2.3e-06 | 1.6e-06 | 2.3e-06 | 2.067E-06 | 4.041E-07 | 1.600E-06 | 2.300E-06 |
| 26719 | 1336 | V1336Y | E | 1.9e-06 | 1.3e-06 | 2.1e-06 | 1.767E-06 | 4.163E-07 | 1.300E-06 | 2.100E-06 |
| 26720 | 1336 | V1336V | E | 2.3e-06 | 2.3e-06 | 2.5e-06 | 2.367E-06 | 1.155E-07 | 2.300E-06 | 2.500E-06 |
| 26721 | 1337 | K1337A | E | 2.5e-06 | 2.9e-06 | 2e-06 | 2.467E-06 | 4.509E-07 | 2.000E-06 | 2.900E-06 |
| 26722 | 1337 | K1337R | E | 1.5e-06 | 2.6e-06 | 1.7e-06 | 1.933E-06 | 5.859E-07 | 1.500E-06 | 2.600E-06 |
| 26723 | 1337 | K1337N | E | 2.9e-06 | 1.6e-06 | 3.2e-06 | 2.567E-06 | 8.505E-07 | 1.600E-06 | 3.200E-06 |
| 26724 | 1337 | K1337D | E | 1.3e-06 | 2.9e-06 | 2.7e-06 | 2.300E-06 | 8.718E-07 | 1.300E-06 | 2.900E-06 |
| 26725 | 1337 | K1337C | E | 2.1e-06 | 1.4e-06 | 3.9e-06 | 2.467E-06 | 1.290E-06 | 1.400E-06 | 3.900E-06 |
| 26726 | 1337 | K1337E | E | 1.2e-06 | 2.6e-06 | 2.7e-06 | 2.167E-06 | 8.386E-07 | 1.200E-06 | 2.700E-06 |
| 26727 | 1337 | K1337Q | E | 2.3e-06 | 2.4e-06 | 3.6e-06 | 2.767E-06 | 7.234E-07 | 2.300E-06 | 3.600E-06 |
| 26728 | 1337 | K1337G | E | 3.8e-06 | 2.2e-06 | 2.4e-06 | 2.800E-06 | 8.718E-07 | 2.200E-06 | 3.800E-06 |
| 26729 | 1337 | K1337H | E | 2.3e-06 | 2.8e-06 | 1.8e-06 | 2.300E-06 | 5.000E-07 | 1.800E-06 | 2.800E-06 |
| 26730 | 1337 | K1337I | E | 1.3e-06 | 2e-06 | 3.9e-06 | 2.400E-06 | 1.345E-06 | 1.300E-06 | 3.900E-06 |
| 26731 | 1337 | K1337L | E | 3.2e-06 | 1.9e-06 | 2e-06 | 2.367E-06 | 7.234E-07 | 1.900E-06 | 3.200E-06 |
| 26732 | 1337 | K1337K | E | 1.8e-06 | 1.7e-06 | 4.4e-06 | 2.633E-06 | 1.531E-06 | 1.700E-06 | 4.400E-06 |
| 26733 | 1337 | K1337M | E | 3.4e-06 | 1.9e-06 | 3e-06 | 2.767E-06 | 7.767E-07 | 1.900E-06 | 3.400E-06 |
| 26734 | 1337 | K1337F | E | 2.4e-06 | 2.5e-06 | 2.7e-06 | 2.533E-06 | 1.528E-07 | 2.400E-06 | 2.700E-06 |
| 26735 | 1337 | K1337P | E | 3e-06 | 2.6e-06 | 1.7e-06 | 2.433E-06 | 6.658E-07 | 1.700E-06 | 3.000E-06 |
| 26736 | 1337 | K1337S | E | 2.5e-06 | 1.6e-06 | 2.5e-06 | 2.200E-06 | 5.196E-07 | 1.600E-06 | 2.500E-06 |
| 26737 | 1337 | K1337T | E | 3.2e-06 | 2.3e-06 | 2.6e-06 | 2.700E-06 | 4.583E-07 | 2.300E-06 | 3.200E-06 |
| 26738 | 1337 | K1337W | E | 1.4e-06 | 3e-06 | 2.4e-06 | 2.267E-06 | 8.083E-07 | 1.400E-06 | 3.000E-06 |
| 26739 | 1337 | K1337Y | E | 2.5e-06 | 2e-06 | 2.2e-06 | 2.233E-06 | 2.517E-07 | 2.000E-06 | 2.500E-06 |
| 26740 | 1337 | K1337V | E | 2.1e-06 | 3e-06 | 2.1e-06 | 2.400E-06 | 5.196E-07 | 2.100E-06 | 3.000E-06 |
| 26741 | 1338 | G1338A | E | 1.5e-06 | 2.4e-06 | 5.7e-06 | 3.200E-06 | 2.211E-06 | 1.500E-06 | 5.700E-06 |
| 26742 | 1338 | G1338R | E | 1.6e-06 | 2.6e-06 | 2.3e-06 | 2.167E-06 | 5.132E-07 | 1.600E-06 | 2.600E-06 |
| 26743 | 1338 | G1338N | E | 2.4e-06 | 3e-06 | 1.3e-06 | 2.233E-06 | 8.622E-07 | 1.300E-06 | 3.000E-06 |
| 26744 | 1338 | G1338D | E | 2.3e-06 | 1.9e-06 | 2.4e-06 | 2.200E-06 | 2.646E-07 | 1.900E-06 | 2.400E-06 |
| 26745 | 1338 | G1338C | E | 3.2e-06 | 2.7e-06 | 1.2e-06 | 2.367E-06 | 1.041E-06 | 1.200E-06 | 3.200E-06 |
| 26746 | 1338 | G1338E | E | 2.3e-06 | 2.5e-06 | 2.4e-06 | 2.400E-06 | 1.000E-07 | 2.300E-06 | 2.500E-06 |
| 26747 | 1338 | G1338Q | E | 2.2e-06 | 3e-06 | 2.7e-06 | 2.633E-06 | 4.041E-07 | 2.200E-06 | 3.000E-06 |
| 26748 | 1338 | G1338G | E | 3.2e-06 | 1.7e-06 | 2e-06 | 2.300E-06 | 7.937E-07 | 1.700E-06 | 3.200E-06 |
| 26749 | 1338 | G1338H | E | 2.7e-06 | 3.7e-06 | 2.4e-06 | 2.933E-06 | 6.807E-07 | 2.400E-06 | 3.700E-06 |
| 26750 | 1338 | G1338I | E | 1.8e-06 | 2.7e-06 | 2.3e-06 | 2.267E-06 | 4.509E-07 | 1.800E-06 | 2.700E-06 |
| 26751 | 1338 | G1338L | E | 2.4e-06 | 1.9e-06 | 2.8e-06 | 2.367E-06 | 4.509E-07 | 1.900E-06 | 2.800E-06 |
| 26752 | 1338 | G1338K | E | 2.2e-06 | 2.5e-06 | 1.5e-06 | 2.067E-06 | 5.132E-07 | 1.500E-06 | 2.500E-06 |
| 26753 | 1338 | G1338M | E | 1.4e-06 | 2.2e-06 | 2.6e-06 | 2.067E-06 | 6.110E-07 | 1.400E-06 | 2.600E-06 |
| 26754 | 1338 | G1338F | E | 3.7e-06 | 3e-06 | 2.3e-06 | 3.000E-06 | 7.000E-07 | 2.300E-06 | 3.700E-06 |
| 26755 | 1338 | G1338P | E | 1.1e-06 | 2.6e-06 | 3.5e-06 | 2.400E-06 | 1.212E-06 | 1.100E-06 | 3.500E-06 |
| 26756 | 1338 | G1338S | E | 3e-06 | 3.1e-06 | 1.9e-06 | 2.667E-06 | 6.658E-07 | 1.900E-06 | 3.100E-06 |
| 26757 | 1338 | G1338T | E | 2.4e-06 | 3e-06 | 1.9e-06 | 2.433E-06 | 5.508E-07 | 1.900E-06 | 3.000E-06 |
| 26758 | 1338 | G1338W | E | 2.5e-06 | 1.6e-06 | 3.7e-06 | 2.600E-06 | 1.054E-06 | 1.600E-06 | 3.700E-06 |
| 26759 | 1338 | G1338Y | E | 2.5e-06 | 2.9e-06 | 1.9e-06 | 2.433E-06 | 5.033E-07 | 1.900E-06 | 2.900E-06 |
| 26760 | 1338 | G1338V | E | 2.7e-06 | 5.3e-06 | 1.9e-06 | 3.300E-06 | 1.778E-06 | 1.900E-06 | 5.300E-06 |
| 26761 | 1339 | R1339A | E | 3.5e-06 | 2.2e-06 | 1.9e-06 | 2.533E-06 | 8.505E-07 | 1.900E-06 | 3.500E-06 |
| 26762 | 1339 | R1339R | E | 2.8e-06 | 2.4e-06 | 2.9e-06 | 2.700E-06 | 2.646E-07 | 2.400E-06 | 2.900E-06 |
| 26763 | 1339 | R1339N | E | 2.5e-06 | 1.2e-06 | 2.4e-06 | 2.033E-06 | 7.234E-07 | 1.200E-06 | 2.500E-06 |
| 26764 | 1339 | R1339D | E | 1.5e-06 | 3e-06 | 2.4e-06 | 2.300E-06 | 7.550E-07 | 1.500E-06 | 3.000E-06 |
| 26765 | 1339 | R1339C | E | 2.1e-06 | 5.3e-06 | 2.2e-06 | 3.200E-06 | 1.819E-06 | 2.100E-06 | 5.300E-06 |
| 26766 | 1339 | R1339E | E | 1.2e-06 | 2.3e-06 | 1.6e-06 | 1.700E-06 | 5.568E-07 | 1.200E-06 | 2.300E-06 |
| 26767 | 1339 | R1339Q | E | 1.5e-06 | 3.7e-06 | 2.8e-06 | 2.667E-06 | 1.106E-06 | 1.500E-06 | 3.700E-06 |
| 26768 | 1339 | R1339G | E | 2.2e-06 | 3.6e-06 | 1.6e-06 | 2.467E-06 | 1.026E-06 | 1.600E-06 | 3.600E-06 |
| 26769 | 1339 | R1339H | E | 2.3e-06 | 2.2e-06 | 2.2e-06 | 2.233E-06 | 5.774E-08 | 2.200E-06 | 2.300E-06 |
| 26770 | 1339 | R1339I | E | 1.6e-06 | 3.1e-06 | 1.9e-06 | 2.200E-06 | 7.937E-07 | 1.600E-06 | 3.100E-06 |
| 26771 | 1339 | R1339L | E | 2.9e-06 | 1.7e-06 | 2.3e-06 | 2.300E-06 | 6.000E-07 | 1.700E-06 | 2.900E-06 |
| 26772 | 1339 | R1339K | E | 1.4e-06 | 2.2e-06 | 2.6e-06 | 2.067E-06 | 6.110E-07 | 1.400E-06 | 2.600E-06 |
| 26773 | 1339 | R1339M | E | 3.3e-06 | 2.4e-06 | 1.6e-06 | 2.433E-06 | 8.505E-07 | 1.600E-06 | 3.300E-06 |
| 26774 | 1339 | R1339F | E | 1.7e-06 | 3.9e-06 | 1.9e-06 | 2.500E-06 | 1.217E-06 | 1.700E-06 | 3.900E-06 |

| | | | | | | | | | | |
|---|---|---|---|---|---|---|---|---|---|---|
| 26775 | 1339 | R1339P | E | 1.7e-06 | 1.3e-06 | 2.6e-06 | 1.867E-06 | 6.658E-07 | 1.300E-06 | 2.600E-06 |
| 26776 | 1339 | R1339S | E | 1.4e-06 | 3.1e-06 | 2.7e-06 | 2.400E-06 | 8.888E-07 | 1.400E-06 | 3.100E-06 |
| 26777 | 1339 | R1339T | E | 2.5e-06 | 2.4e-06 | 2.2e-06 | 2.367E-06 | 1.528E-07 | 2.200E-06 | 2.500E-06 |
| 26778 | 1339 | R1339W | E | 2.6e-06 | 2.8e-06 | 2.7e-06 | 2.700E-06 | 1.000E-07 | 2.600E-06 | 2.800E-06 |
| 26779 | 1339 | R1339Y | E | 2.6e-06 | 2.5e-06 | 3.4e-06 | 2.833E-06 | 4.933E-07 | 2.500E-06 | 3.400E-06 |
| 26780 | 1339 | R1339V | E | 2.8e-06 | 2.2e-06 | 2.3e-06 | 2.433E-06 | 3.215E-07 | 2.200E-06 | 2.800E-06 |
| 26781 | 1340 | F1340A | E | 3.4e-06 | 2.6e-06 | 3.3e-06 | 3.100E-06 | 4.359E-07 | 2.600E-06 | 3.400E-06 |
| 26782 | 1340 | F1340R | E | 2.4e-06 | 3.8e-06 | 1.8e-06 | 2.667E-06 | 1.026E-06 | 1.800E-06 | 3.800E-06 |
| 26783 | 1340 | F1340N | E | 3.2e-06 | 3.4e-06 | 3e-06 | 3.200E-06 | 2.000E-07 | 3.000E-06 | 3.400E-06 |
| 26784 | 1340 | F1340D | E | 2.2e-06 | 2.8e-06 | 3.6e-06 | 2.867E-06 | 7.024E-07 | 2.200E-06 | 3.600E-06 |
| 26785 | 1340 | F1340C | E | 1.2e-06 | 2.5e-06 | 1.5e-06 | 1.733E-06 | 6.807E-07 | 1.200E-06 | 2.500E-06 |
| 26786 | 1340 | F1340E | E | 3e-06 | 2.7e-06 | 2.9e-06 | 2.867E-06 | 1.528E-07 | 2.700E-06 | 3.000E-06 |
| 26787 | 1340 | F1340Q | E | 2.4e-06 | 2.3e-06 | 1e-06 | 1.900E-06 | 7.810E-07 | 1.000E-06 | 2.400E-06 |
| 26788 | 1340 | F1340G | E | 1.6e-06 | 2.1e-06 | 2.1e-06 | 1.933E-06 | 2.887E-07 | 1.600E-06 | 2.100E-06 |
| 26789 | 1340 | F1340H | E | 2.2e-06 | 2.7e-06 | 1.6e-06 | 2.167E-06 | 5.508E-07 | 1.600E-06 | 2.700E-06 |
| 26790 | 1340 | F1340I | E | 1.2e-06 | 2.8e-06 | 4.5e-06 | 2.833E-06 | 1.650E-06 | 1.200E-06 | 4.500E-06 |
| 26791 | 1340 | F1340L | E | 1.5e-06 | 2.8e-06 | 1.7e-06 | 2.000E-06 | 7.000E-07 | 1.500E-06 | 2.800E-06 |
| 26792 | 1340 | F1340K | E | 1.9e-06 | 2.6e-06 | 1.4e-06 | 1.967E-06 | 6.028E-07 | 1.400E-06 | 2.600E-06 |
| 26793 | 1340 | F1340M | E | 2.1e-06 | 1.3e-06 | 2.3e-06 | 1.900E-06 | 5.292E-07 | 1.300E-06 | 2.300E-06 |
| 26794 | 1340 | F1340F | E | 2.2e-06 | 2.3e-06 | 1.9e-06 | 2.133E-06 | 2.082E-07 | 1.900E-06 | 2.300E-06 |
| 26795 | 1340 | F1340P | E | 3.3e-06 | 1.4e-06 | 1.6e-06 | 2.100E-06 | 1.044E-06 | 1.400E-06 | 3.300E-06 |
| 26796 | 1340 | F1340S | E | 2.9e-06 | 2.1e-06 | 2.5e-06 | 2.500E-06 | 4.000E-07 | 2.100E-06 | 2.900E-06 |
| 26797 | 1340 | F1340T | E | 2.7e-06 | 2.4e-06 | 2.5e-06 | 2.533E-06 | 1.528E-07 | 2.400E-06 | 2.700E-06 |
| 26798 | 1340 | F1340W | E | 2.3e-06 | 4.5e-06 | 3e-06 | 3.267E-06 | 1.124E-06 | 2.300E-06 | 4.500E-06 |
| 26799 | 1340 | F1340Y | E | 1.8e-06 | 3.2e-06 | 2.2e-06 | 2.400E-06 | 7.211E-07 | 1.800E-06 | 3.200E-06 |
| 26800 | 1340 | F1340V | E | 2.1e-06 | 1.5e-06 | 2.3e-06 | 1.967E-06 | 4.163E-07 | 1.500E-06 | 2.300E-06 |
| 26801 | 1341 | T1341A | E | 2.7e-06 | 3.1e-06 | 3.5e-06 | 3.100E-06 | 4.000E-07 | 2.700E-06 | 3.500E-06 |
| 26802 | 1341 | T1341R | E | 2.6e-06 | 2.1e-06 | 1.6e-06 | 2.100E-06 | 5.000E-07 | 1.600E-06 | 2.600E-06 |
| 26803 | 1341 | T1341N | E | 3.1e-06 | 6.8e-07 | 1.7e-06 | 1.827E-06 | 1.215E-06 | 6.800E-07 | 3.100E-06 |
| 26804 | 1341 | T1341D | E | 2.2e-06 | 1.7e-06 | 1.5e-06 | 1.800E-06 | 3.606E-07 | 1.500E-06 | 2.200E-06 |
| 26805 | 1341 | T1341C | E | 1.2e-06 | 1.8e-06 | 2.4e-06 | 1.800E-06 | 6.000E-07 | 1.200E-06 | 2.400E-06 |
| 26806 | 1341 | T1341E | E | 3e-06 | 1.8e-06 | 1.2e-06 | 2.000E-06 | 9.165E-07 | 1.200E-06 | 3.000E-06 |
| 26807 | 1341 | T1341Q | E | 2e-06 | 3.2e-06 | 1.6e-06 | 2.267E-06 | 8.327E-07 | 1.600E-06 | 3.200E-06 |
| 26808 | 1341 | T1341G | E | 2e-06 | 2.1e-06 | 3.8e-06 | 2.633E-06 | 1.012E-06 | 2.000E-06 | 3.800E-06 |
| 26809 | 1341 | T1341H | E | 3.4e-06 | 2.8e-06 | 2.3e-06 | 2.833E-06 | 5.508E-07 | 2.300E-06 | 3.400E-06 |
| 26810 | 1341 | T1341I | E | 2.4e-06 | 2.8e-06 | 2.2e-06 | 2.467E-06 | 3.055E-07 | 2.200E-06 | 2.800E-06 |
| 26811 | 1341 | T1341L | E | 2e-06 | 4.4e-06 | 2.3e-06 | 2.900E-06 | 1.308E-06 | 2.000E-06 | 4.400E-06 |
| 26812 | 1341 | T1341K | E | 3.2e-06 | 1.9e-06 | 2.9e-06 | 2.667E-06 | 6.807E-07 | 1.900E-06 | 3.200E-06 |
| 26813 | 1341 | T1341M | E | 1.2e-06 | 3.3e-06 | 1.9e-06 | 2.133E-06 | 1.069E-06 | 1.200E-06 | 3.300E-06 |
| 26814 | 1341 | T1341F | E | 2.2e-06 | 1.7e-06 | 2.7e-06 | 2.200E-06 | 5.000E-07 | 1.700E-06 | 2.700E-06 |
| 26815 | 1341 | T1341P | E | 2.2e-06 | 3.5e-06 | 1.2e-06 | 2.300E-06 | 1.153E-06 | 1.200E-06 | 3.500E-06 |
| 26816 | 1341 | T1341S | E | 2e-06 | 3.7e-06 | 9.7e-07 | 2.223E-06 | 1.379E-06 | 9.700E-07 | 3.700E-06 |
| 26817 | 1341 | T1341T | E | 2.5e-06 | 3.1e-06 | 2.8e-06 | 2.800E-06 | 3.000E-07 | 2.500E-06 | 3.100E-06 |
| 26818 | 1341 | T1341W | E | 2.1e-06 | 2.5e-06 | 2.5e-06 | 2.367E-06 | 2.309E-07 | 2.100E-06 | 2.500E-06 |
| 26819 | 1341 | T1341Y | E | 1.4e-06 | 1.5e-06 | 1.8e-06 | 1.567E-06 | 2.082E-07 | 1.400E-06 | 1.800E-06 |
| 26820 | 1341 | T1341V | E | 3.2e-06 | 3.3e-06 | 2.8e-06 | 3.100E-06 | 2.646E-07 | 2.800E-06 | 3.300E-06 |
| 26821 | 1342 | I1342A | E | 2.2e-06 | 2.2e-06 | 2.3e-06 | 2.233E-06 | 5.774E-08 | 2.200E-06 | 2.300E-06 |
| 26822 | 1342 | I1342R | E | 2.5e-06 | 1.2e-06 | 3.2e-06 | 2.300E-06 | 1.015E-06 | 1.200E-06 | 3.200E-06 |
| 26823 | 1342 | I1342N | E | 2.2e-06 | 1.6e-06 | 2.8e-06 | 2.200E-06 | 6.000E-07 | 1.600E-06 | 2.800E-06 |
| 26824 | 1342 | I1342D | E | 2.4e-06 | 3.5e-06 | 2.3e-06 | 2.733E-06 | 6.658E-07 | 2.300E-06 | 3.500E-06 |
| 26825 | 1342 | I1342C | E | 3.9e-06 | 2.9e-06 | 2.6e-06 | 3.133E-06 | 6.807E-07 | 2.600E-06 | 3.900E-06 |
| 26826 | 1342 | I1342E | E | 3.2e-06 | 2.2e-06 | 3.9e-06 | 3.100E-06 | 8.544E-07 | 2.200E-06 | 3.900E-06 |
| 26827 | 1342 | I1342Q | E | 1.8e-06 | 2.2e-06 | 2e-06 | 2.000E-06 | 2.000E-07 | 1.800E-06 | 2.200E-06 |
| 26828 | 1342 | I1342G | E | 2.7e-06 | 2.2e-06 | 2e-06 | 2.300E-06 | 3.606E-07 | 2.000E-06 | 2.700E-06 |
| 26829 | 1342 | I1342H | E | 1.6e-06 | 8.8e-07 | 1.5e-06 | 1.327E-06 | 3.900E-07 | 8.800E-07 | 1.600E-06 |
| 26830 | 1342 | I1342I | E | 2.5e-06 | 1.3e-06 | 1.9e-06 | 1.900E-06 | 6.000E-07 | 1.300E-06 | 2.500E-06 |
| 26831 | 1342 | I1342L | E | 1.7e-06 | 2.6e-06 | 2e-06 | 2.100E-06 | 4.583E-07 | 1.700E-06 | 2.600E-06 |
| 26832 | 1342 | I1342K | E | 2e-06 | 2.2e-06 | 1.9e-06 | 2.033E-06 | 1.528E-07 | 1.900E-06 | 2.200E-06 |
| 26833 | 1342 | I1342M | E | 1.7e-06 | 4.9e-06 | 2.5e-06 | 3.033E-06 | 1.665E-06 | 1.700E-06 | 4.900E-06 |
| 26834 | 1342 | I1342F | E | 4.8e-06 | 3.2e-06 | 2.2e-06 | 3.400E-06 | 1.311E-06 | 2.200E-06 | 4.800E-06 |
| 26835 | 1342 | I1342P | E | 4e-06 | 2.4e-06 | 1.4e-06 | 2.600E-06 | 1.311E-06 | 1.400E-06 | 4.000E-06 |
| 26836 | 1342 | I1342S | E | 2e-06 | 1.8e-06 | 2.5e-06 | 2.100E-06 | 3.606E-07 | 1.800E-06 | 2.500E-06 |
| 26837 | 1342 | I1342T | E | 3e-06 | 1.8e-06 | 2.4e-06 | 2.400E-06 | 6.000E-07 | 1.800E-06 | 3.000E-06 |
| 26838 | 1342 | I1342W | E | 1.1e-06 | 9.3e-07 | 2.7e-06 | 1.577E-06 | 9.765E-07 | 9.300E-07 | 2.700E-06 |
| 26839 | 1342 | I1342Y | E | 1.5e-06 | 1.7e-06 | 2.9e-06 | 2.033E-06 | 7.572E-07 | 1.500E-06 | 2.900E-06 |
| 26840 | 1342 | I1342V | E | 3.2e-06 | 2.3e-06 | 2.6e-06 | 2.700E-06 | 4.583E-07 | 2.300E-06 | 3.200E-06 |
| 26841 | 1343 | S1343A | E | 1.5e-06 | 2.3e-06 | 3.5e-06 | 2.433E-06 | 1.007E-06 | 1.500E-06 | 3.500E-06 |
| 26842 | 1343 | S1343R | E | 1.6e-06 | 2.3e-06 | 2.6e-06 | 2.167E-06 | 5.132E-07 | 1.600E-06 | 2.600E-06 |
| 26843 | 1343 | S1343N | E | 2.7e-06 | 2.5e-06 | 2e-06 | 2.400E-06 | 3.606E-07 | 2.000E-06 | 2.700E-06 |
| 26844 | 1343 | S1343D | E | 2.4e-06 | 2e-06 | 4.3e-06 | 2.900E-06 | 1.229E-06 | 2.000E-06 | 4.300E-06 |
| 26845 | 1343 | S1343C | E | 2.4e-06 | 9.8e-07 | 1.6e-06 | 1.660E-06 | 7.119E-07 | 9.800E-07 | 2.400E-06 |
| 26846 | 1343 | S1343E | E | 2.2e-06 | 1.7e-06 | 4.2e-06 | 2.700E-06 | 1.323E-06 | 1.700E-06 | 4.200E-06 |
| 26847 | 1343 | S1343Q | E | 2.9e-06 | 3.2e-06 | 4.2e-06 | 3.433E-06 | 6.807E-07 | 2.900E-06 | 4.200E-06 |
| 26848 | 1343 | S1343G | E | 2.2e-06 | 2.8e-06 | 1.9e-06 | 2.300E-06 | 4.583E-07 | 1.900E-06 | 2.800E-06 |
| 26849 | 1343 | S1343H | E | 2e-06 | 1.8e-06 | 3.2e-06 | 2.333E-06 | 7.572E-07 | 1.800E-06 | 3.200E-06 |
| 26850 | 1343 | S1343I | E | 2.6e-06 | 3.9e-06 | 3.6e-06 | 3.367E-06 | 6.807E-07 | 2.600E-06 | 3.900E-06 |
| 26851 | 1343 | S1343L | E | 4.2e-06 | 4.5e-06 | 3.5e-06 | 4.067E-06 | 5.132E-07 | 3.500E-06 | 4.500E-06 |
| 26852 | 1343 | S1343K | E | 3.4e-06 | 1.6e-06 | 2e-06 | 2.333E-06 | 9.452E-07 | 1.600E-06 | 3.400E-06 |
| 26853 | 1343 | S1343M | E | 2.8e-06 | 1.6e-06 | 2.4e-06 | 2.267E-06 | 6.110E-07 | 1.600E-06 | 2.800E-06 |
| 26854 | 1343 | S1343F | E | 2.2e-06 | 1.9e-06 | 2.9e-06 | 2.333E-06 | 5.132E-07 | 1.900E-06 | 2.900E-06 |
| 26855 | 1343 | S1343P | E | 2.2e-06 | 4.1e-06 | 2e-06 | 2.767E-06 | 1.159E-06 | 2.000E-06 | 4.100E-06 |
| 26856 | 1343 | S1343S | E | 2.4e-06 | 2.3e-06 | 4.8e-06 | 3.167E-06 | 1.415E-06 | 2.300E-06 | 4.800E-06 |
| 26857 | 1343 | S1343T | E | 3.4e-06 | 2.6e-06 | 3.9e-06 | 3.300E-06 | 6.557E-07 | 2.600E-06 | 3.900E-06 |
| 26858 | 1343 | S1343W | E | 1.2e-06 | 3.1e-06 | 3.2e-06 | 2.500E-06 | 1.127E-06 | 1.200E-06 | 3.200E-06 |
| 26859 | 1343 | S1343Y | E | 2.8e-06 | 2.5e-06 | 2.2e-06 | 2.500E-06 | 3.000E-07 | 2.200E-06 | 2.800E-06 |

| | | | | | | | | | | |
|---|---|---|---|---|---|---|---|---|---|---|
| 26860 | 1343 | S1343V | E | 2.8e-06 | 2.7e-06 | 2.4e-06 | 2.633E-06 | 2.082E-07 | 2.400E-06 | 2.800E-06 |
| 26861 | 1344 | A1344A | E | 1.6e-06 | 2.1e-06 | 2e-06 | 1.900E-06 | 2.646E-07 | 1.600E-06 | 2.100E-06 |
| 26862 | 1344 | A1344R | E | 2.6e-06 | 2.2e-06 | 3.6e-06 | 2.800E-06 | 7.211E-07 | 2.200E-06 | 3.600E-06 |
| 26863 | 1344 | A1344N | E | 3.1e-06 | 2e-06 | 1.6e-06 | 2.233E-06 | 7.767E-07 | 1.600E-06 | 3.100E-06 |
| 26864 | 1344 | A1344D | E | 2.7e-06 | 2.3e-06 | 3.1e-06 | 2.700E-06 | 4.000E-07 | 2.300E-06 | 3.100E-06 |
| 26865 | 1344 | A1344C | E | 3e-06 | 1.6e-06 | 2e-06 | 2.200E-06 | 7.211E-07 | 1.600E-06 | 3.000E-06 |
| 26866 | 1344 | A1344E | E | 2.2e-06 | 2.3e-06 | 3e-06 | 2.500E-06 | 4.359E-07 | 2.200E-06 | 3.000E-06 |
| 26867 | 1344 | A1344Q | E | 2.4e-06 | 1.9e-06 | 2.1e-06 | 2.133E-06 | 2.517E-07 | 1.900E-06 | 2.400E-06 |
| 26868 | 1344 | A1344G | E | 2.3e-06 | 2.7e-06 | 3.2e-06 | 2.733E-06 | 4.509E-07 | 2.300E-06 | 3.200E-06 |
| 26869 | 1344 | A1344H | E | 2.3e-06 | 1.8e-06 | 2.4e-06 | 2.167E-06 | 3.215E-07 | 1.800E-06 | 2.400E-06 |
| 26870 | 1344 | A1344I | E | 2e-06 | 1.3e-06 | 2.9e-06 | 2.067E-06 | 8.021E-07 | 1.300E-06 | 2.900E-06 |
| 26871 | 1344 | A1344L | E | 2e-06 | 3.8e-06 | 2.6e-06 | 2.800E-06 | 9.165E-07 | 2.000E-06 | 3.800E-06 |
| 26872 | 1344 | A1344K | E | 2.2e-06 | 2.1e-06 | 2e-06 | 2.100E-06 | 1.000E-07 | 2.000E-06 | 2.200E-06 |
| 26873 | 1344 | A1344M | E | 2.1e-06 | 3.1e-06 | 2.4e-06 | 2.533E-06 | 5.132E-07 | 2.100E-06 | 3.100E-06 |
| 26874 | 1344 | A1344F | E | 1.4e-06 | 2.7e-06 | 1.3e-06 | 1.800E-06 | 7.810E-07 | 1.300E-06 | 2.700E-06 |
| 26875 | 1344 | A1344P | E | 2.7e-06 | 2e-06 | 2.6e-06 | 2.433E-06 | 3.786E-07 | 2.000E-06 | 2.700E-06 |
| 26876 | 1344 | A1344S | E | 1.3e-06 | 2.4e-06 | 2e-06 | 1.900E-06 | 5.568E-07 | 1.300E-06 | 2.400E-06 |
| 26877 | 1344 | A1344T | E | 9.6e-07 | 2.4e-06 | 8.8e-07 | 1.413E-06 | 8.554E-07 | 8.800E-07 | 2.400E-06 |
| 26878 | 1344 | A1344W | E | 2e-06 | 1.9e-06 | 2.2e-06 | 2.033E-06 | 1.528E-07 | 1.900E-06 | 2.200E-06 |
| 26879 | 1344 | A1344Y | E | 3.1e-06 | 1.3e-06 | 2.2e-06 | 2.200E-06 | 9.000E-07 | 1.300E-06 | 3.100E-06 |
| 26880 | 1344 | A1344V | E | 1.8e-06 | 1.2e-06 | 2.5e-06 | 1.833E-06 | 6.506E-07 | 1.200E-06 | 2.500E-06 |
| 26881 | 1345 | D1345A | E | 1.6e-06 | 3.2e-06 | 2e-06 | 2.267E-06 | 8.327E-07 | 1.600E-06 | 3.200E-06 |
| 26882 | 1345 | D1345R | E | 2.3e-06 | 2.1e-06 | 1.7e-06 | 2.033E-06 | 3.055E-07 | 1.700E-06 | 2.300E-06 |
| 26883 | 1345 | D1345N | E | 1.9e-06 | 2.9e-06 | 1.9e-06 | 2.233E-06 | 5.774E-07 | 1.900E-06 | 2.900E-06 |
| 26884 | 1345 | D1345D | E | 2.2e-06 | 1.5e-06 | 1.8e-06 | 1.833E-06 | 3.512E-07 | 1.500E-06 | 2.200E-06 |
| 26885 | 1345 | D1345C | E | 4.2e-06 | 1.8e-06 | 1.6e-06 | 2.533E-06 | 1.447E-06 | 1.600E-06 | 4.200E-06 |
| 26886 | 1345 | D1345E | E | 3.1e-06 | 1.5e-06 | 3.2e-06 | 2.600E-06 | 9.539E-07 | 1.500E-06 | 3.200E-06 |
| 26887 | 1345 | D1345Q | E | 2.6e-06 | 1.7e-06 | 3.1e-06 | 2.467E-06 | 7.095E-07 | 1.700E-06 | 3.100E-06 |
| 26888 | 1345 | D1345G | E | 2.3e-06 | 3.3e-06 | 2.6e-06 | 2.733E-06 | 5.132E-07 | 2.300E-06 | 3.300E-06 |
| 26889 | 1345 | D1345H | E | 1.6e-06 | 2.2e-06 | 2e-06 | 1.933E-06 | 3.055E-07 | 1.600E-06 | 2.200E-06 |
| 26890 | 1345 | D1345I | E | 1.7e-06 | 2.1e-06 | 2.3e-06 | 2.033E-06 | 3.055E-07 | 1.700E-06 | 2.300E-06 |
| 26891 | 1345 | D1345L | E | 3.7e-06 | 4.4e-06 | 2.7e-06 | 3.600E-06 | 8.544E-07 | 2.700E-06 | 4.400E-06 |
| 26892 | 1345 | D1345K | E | 2.8e-06 | 2.6e-06 | 2.5e-06 | 2.633E-06 | 1.528E-07 | 2.500E-06 | 2.800E-06 |
| 26893 | 1345 | D1345M | E | 2e-06 | 1.8e-06 | 1.6e-06 | 1.800E-06 | 2.000E-07 | 1.600E-06 | 2.000E-06 |
| 26894 | 1345 | D1345F | E | 2.2e-06 | 2.2e-06 | 2.3e-06 | 2.233E-06 | 5.774E-08 | 2.200E-06 | 2.300E-06 |
| 26895 | 1345 | D1345P | E | 4.9e-06 | 1.9e-06 | 2.6e-06 | 3.133E-06 | 1.570E-06 | 1.900E-06 | 4.900E-06 |
| 26896 | 1345 | D1345S | E | 2.5e-06 | 2e-06 | 2.5e-06 | 2.333E-06 | 2.887E-07 | 2.000E-06 | 2.500E-06 |
| 26897 | 1345 | D1345T | E | 1.6e-06 | 2.1e-06 | 1.8e-06 | 1.833E-06 | 2.517E-07 | 1.600E-06 | 2.100E-06 |
| 26898 | 1345 | D1345W | E | 1.4e-06 | 2.1e-06 | 2.2e-06 | 1.900E-06 | 4.359E-07 | 1.400E-06 | 2.200E-06 |
| 26899 | 1345 | D1345Y | E | 1.4e-06 | 1.9e-06 | 2.1e-06 | 1.800E-06 | 3.606E-07 | 1.400E-06 | 2.100E-06 |
| 26900 | 1345 | D1345V | E | 2.3e-06 | 2.9e-06 | 2e-06 | 2.400E-06 | 4.583E-07 | 2.000E-06 | 2.900E-06 |
| 26901 | 1346 | T1346A | E | 1.9e-06 | 2.7e-06 | 2.2e-06 | 2.267E-06 | 4.041E-07 | 1.900E-06 | 2.700E-06 |
| 26902 | 1346 | T1346R | E | 2.5e-06 | 2e-06 | 2e-06 | 2.167E-06 | 2.887E-07 | 2.000E-06 | 2.500E-06 |
| 26903 | 1346 | T1346N | E | 2.5e-06 | 2.4e-06 | 1.6e-06 | 2.167E-06 | 4.933E-07 | 1.600E-06 | 2.500E-06 |
| 26904 | 1346 | T1346D | E | 2.5e-06 | 3.5e-06 | 1.5e-06 | 2.500E-06 | 1.000E-06 | 1.500E-06 | 3.500E-06 |
| 26905 | 1346 | T1346C | E | 1e-06 | 1.9e-06 | 3.5e-06 | 2.133E-06 | 1.266E-06 | 1.000E-06 | 3.500E-06 |
| 26906 | 1346 | T1346E | E | 3.4e-06 | 2.4e-06 | 2.4e-06 | 2.733E-06 | 5.774E-07 | 2.400E-06 | 3.400E-06 |
| 26907 | 1346 | T1346Q | E | 2.2e-06 | 2.8e-06 | 2.3e-06 | 2.433E-06 | 3.215E-07 | 2.200E-06 | 2.800E-06 |
| 26908 | 1346 | T1346G | E | 3.8e-06 | 2.6e-06 | 2.7e-06 | 3.033E-06 | 6.658E-07 | 2.600E-06 | 3.800E-06 |
| 26909 | 1346 | T1346H | E | 2.4e-06 | 1.9e-06 | 3.2e-06 | 2.500E-06 | 6.557E-07 | 1.900E-06 | 3.200E-06 |
| 26910 | 1346 | T1346I | E | 2.5e-06 | 2.7e-06 | 2.2e-06 | 2.467E-06 | 2.517E-07 | 2.200E-06 | 2.700E-06 |
| 26911 | 1346 | T1346L | E | 2e-06 | 2.9e-06 | 2.9e-06 | 2.600E-06 | 5.196E-07 | 2.000E-06 | 2.900E-06 |
| 26912 | 1346 | T1346K | E | 2e-06 | 1.6e-06 | 1.9e-06 | 1.833E-06 | 2.082E-07 | 1.600E-06 | 2.000E-06 |
| 26913 | 1346 | T1346M | E | 4.2e-06 | 3.1e-06 | 2e-06 | 3.100E-06 | 1.100E-06 | 2.000E-06 | 4.200E-06 |
| 26914 | 1346 | T1346F | E | 2.5e-06 | 2.5e-06 | 1.3e-06 | 2.100E-06 | 6.928E-07 | 1.300E-06 | 2.500E-06 |
| 26915 | 1346 | T1346P | E | 3.2e-06 | 2.6e-06 | 1.6e-06 | 2.467E-06 | 8.083E-07 | 1.600E-06 | 3.200E-06 |
| 26916 | 1346 | T1346S | E | 2.5e-06 | 1.8e-06 | 3.8e-06 | 2.700E-06 | 1.015E-06 | 1.800E-06 | 3.800E-06 |
| 26917 | 1346 | T1346T | E | 2.3e-06 | 1.4e-06 | 2.7e-06 | 2.133E-06 | 6.658E-07 | 1.400E-06 | 2.700E-06 |
| 26918 | 1346 | T1346W | E | 2.7e-06 | 4.3e-06 | 5.2e-06 | 4.067E-06 | 1.266E-06 | 2.700E-06 | 5.200E-06 |
| 26919 | 1346 | T1346Y | E | 2.9e-06 | 2.2e-06 | 3.3e-06 | 2.800E-06 | 5.568E-07 | 2.200E-06 | 3.300E-06 |
| 26920 | 1346 | T1346V | E | 4.6e-06 | 1.3e-06 | 5e-06 | 3.633E-06 | 2.031E-06 | 1.300E-06 | 5.000E-06 |
| 26921 | 1347 | S1347A | E | 3.4e-06 | 2.8e-06 | 4.3e-06 | 3.500E-06 | 7.550E-07 | 2.800E-06 | 4.300E-06 |
| 26922 | 1347 | S1347R | E | 2.3e-06 | 3.1e-06 | 2.3e-06 | 2.567E-06 | 4.619E-07 | 2.300E-06 | 3.100E-06 |
| 26923 | 1347 | S1347N | E | 2e-06 | 2.5e-06 | 4.1e-06 | 2.867E-06 | 1.097E-06 | 2.000E-06 | 4.100E-06 |
| 26924 | 1347 | S1347D | E | 3.7e-06 | 2.3e-06 | 1.8e-06 | 2.600E-06 | 9.849E-07 | 1.800E-06 | 3.700E-06 |
| 26925 | 1347 | S1347C | E | 3.3e-06 | 2.6e-06 | 2e-06 | 2.633E-06 | 6.506E-07 | 2.000E-06 | 3.300E-06 |
| 26926 | 1347 | S1347E | E | 2.1e-06 | 2e-06 | 2.2e-06 | 2.100E-06 | 1.000E-07 | 2.000E-06 | 2.200E-06 |
| 26927 | 1347 | S1347Q | E | 2.4e-06 | 1.5e-06 | 2.5e-06 | 2.133E-06 | 5.508E-07 | 1.500E-06 | 2.500E-06 |
| 26928 | 1347 | S1347G | E | 2.3e-06 | 3.8e-06 | 3.1e-06 | 3.067E-06 | 7.506E-07 | 2.300E-06 | 3.800E-06 |
| 26929 | 1347 | S1347H | E | 3.2e-06 | 2.2e-06 | 3.3e-06 | 2.900E-06 | 6.083E-07 | 2.200E-06 | 3.300E-06 |
| 26930 | 1347 | S1347I | E | 3.7e-06 | 1.7e-06 | 2.2e-06 | 2.533E-06 | 1.041E-06 | 1.700E-06 | 3.700E-06 |
| 26931 | 1347 | S1347L | E | 2.9e-06 | 1.8e-06 | 2.5e-06 | 2.400E-06 | 5.568E-07 | 1.800E-06 | 2.900E-06 |
| 26932 | 1347 | S1347K | E | 2.3e-06 | 1.7e-06 | 3.4e-06 | 2.467E-06 | 8.622E-07 | 1.700E-06 | 3.400E-06 |
| 26933 | 1347 | S1347M | E | 2.2e-06 | 2.8e-06 | 4.6e-06 | 3.200E-06 | 1.249E-06 | 2.200E-06 | 4.600E-06 |
| 26934 | 1347 | S1347F | E | 3e-06 | 1.9e-06 | 3.2e-06 | 2.700E-06 | 7.000E-07 | 1.900E-06 | 3.200E-06 |
| 26935 | 1347 | S1347P | E | 2.2e-06 | 5.6e-06 | 1.8e-06 | 3.200E-06 | 2.088E-06 | 1.800E-06 | 5.600E-06 |
| 26936 | 1347 | S1347S | E | 2.3e-06 | 2.1e-06 | 2.9e-06 | 2.433E-06 | 4.163E-07 | 2.100E-06 | 2.900E-06 |
| 26937 | 1347 | S1347T | E | 1.8e-06 | 2.5e-06 | 1e-06 | 1.767E-06 | 7.506E-07 | 1.000E-06 | 2.500E-06 |
| 26938 | 1347 | S1347W | E | 2.3e-06 | 1.3e-06 | 3.2e-06 | 2.267E-06 | 9.504E-07 | 1.300E-06 | 3.200E-06 |
| 26939 | 1347 | S1347Y | E | 3.2e-06 | 2e-06 | 3.1e-06 | 2.767E-06 | 6.658E-07 | 2.000E-06 | 3.200E-06 |
| 26940 | 1347 | S1347V | E | 1.9e-06 | 1e-06 | 2.2e-06 | 1.700E-06 | 6.245E-07 | 1.000E-06 | 2.200E-06 |
| 26941 | 1348 | K1348A | E | 3.2e-06 | 2.9e-06 | 2.7e-06 | 2.933E-06 | 2.517E-07 | 2.700E-06 | 3.200E-06 |
| 26942 | 1348 | K1348R | E | 2.3e-06 | 3.6e-06 | 3.6e-06 | 3.167E-06 | 7.506E-07 | 2.300E-06 | 3.600E-06 |
| 26943 | 1348 | K1348N | E | 2.3e-06 | 1.8e-06 | 1.3e-06 | 1.800E-06 | 5.000E-07 | 1.300E-06 | 2.300E-06 |
| 26944 | 1348 | K1348D | E | 2.6e-06 | 1.8e-06 | 2e-06 | 2.133E-06 | 4.163E-07 | 1.800E-06 | 2.600E-06 |

| | | | | | | | | | | |
|---|---|---|---|---|---|---|---|---|---|---|
| 26945 | 1348 | K1348C | E | 1.7e-06 | 1.3e-06 | 2.6e-06 | 1.867E-06 | 6.658E-07 | 1.300E-06 | 2.600E-06 |
| 26946 | 1348 | K1348E | E | 4.3e-06 | 3.2e-06 | 3e-06 | 3.500E-06 | 7.000E-07 | 3.000E-06 | 4.300E-06 |
| 26947 | 1348 | K1348Q | E | 2.4e-06 | 1.7e-06 | 1.8e-06 | 1.967E-06 | 3.786E-07 | 1.700E-06 | 2.400E-06 |
| 26948 | 1348 | K1348G | E | 4.4e-06 | 3e-06 | 2.4e-06 | 3.267E-06 | 1.026E-06 | 2.400E-06 | 4.400E-06 |
| 26949 | 1348 | K1348H | E | 1.8e-06 | 1.5e-06 | 1.9e-06 | 1.733E-06 | 2.082E-07 | 1.500E-06 | 1.900E-06 |
| 26950 | 1348 | K1348I | E | 1.7e-06 | 1.2e-06 | 1.7e-06 | 1.533E-06 | 2.887E-07 | 1.200E-06 | 1.700E-06 |
| 26951 | 1348 | K1348L | E | 2.5e-06 | 2.5e-06 | 3.6e-06 | 2.867E-06 | 6.351E-07 | 2.500E-06 | 3.600E-06 |
| 26952 | 1348 | K1348K | E | 2.1e-06 | 3.6e-06 | 3e-06 | 2.900E-06 | 7.550E-07 | 2.100E-06 | 3.600E-06 |
| 26953 | 1348 | K1348M | E | 3.5e-06 | 2e-06 | 1.8e-06 | 2.433E-06 | 9.292E-07 | 1.800E-06 | 3.500E-06 |
| 26954 | 1348 | K1348F | E | 1.7e-06 | 2.8e-06 | 2e-06 | 2.167E-06 | 5.686E-07 | 1.700E-06 | 2.800E-06 |
| 26955 | 1348 | K1348P | E | 3e-06 | 2.4e-06 | 1.9e-06 | 2.433E-06 | 5.508E-07 | 1.900E-06 | 3.000E-06 |
| 26956 | 1348 | K1348S | E | 2.1e-06 | 4.4e-06 | 2.1e-06 | 2.867E-06 | 1.328E-06 | 2.100E-06 | 4.400E-06 |
| 26957 | 1348 | K1348T | E | 2.3e-06 | 2.6e-06 | 2e-06 | 2.300E-06 | 3.000E-07 | 2.000E-06 | 2.600E-06 |
| 26958 | 1348 | K1348W | E | 1.5e-06 | 2.6e-06 | 4.7e-06 | 2.933E-06 | 1.626E-06 | 1.500E-06 | 4.700E-06 |
| 26959 | 1348 | K1348Y | E | 2.6e-06 | 3.2e-06 | 2.9e-06 | 2.900E-06 | 3.000E-07 | 2.600E-06 | 3.200E-06 |
| 26960 | 1348 | K1348V | E | 2.3e-06 | 3.3e-06 | 3.2e-06 | 2.933E-06 | 5.508E-07 | 2.300E-06 | 3.300E-06 |
| 26961 | 1349 | N1349A | E | 2e-06 | 1.6e-06 | 2.9e-06 | 2.167E-06 | 6.658E-07 | 1.600E-06 | 2.900E-06 |
| 26962 | 1349 | N1349R | E | 2.1e-06 | 3.4e-06 | 2.6e-06 | 2.700E-06 | 6.557E-07 | 2.100E-06 | 3.400E-06 |
| 26963 | 1349 | N1349N | E | 3e-06 | 1.3e-06 | 9.3e-07 | 1.743E-06 | 1.104E-06 | 9.300E-07 | 3.000E-06 |
| 26964 | 1349 | N1349D | E | 2e-06 | 3.1e-06 | 1.7e-06 | 2.267E-06 | 7.371E-07 | 1.700E-06 | 3.100E-06 |
| 26965 | 1349 | N1349C | E | 2.2e-06 | 2.4e-06 | 2.5e-06 | 2.367E-06 | 1.528E-07 | 2.200E-06 | 2.500E-06 |
| 26966 | 1349 | N1349E | E | 2.1e-06 | 2.3e-06 | 2e-06 | 2.133E-06 | 1.528E-07 | 2.000E-06 | 2.300E-06 |
| 26967 | 1349 | N1349Q | E | 1.5e-06 | 2.6e-06 | 2.4e-06 | 2.167E-06 | 5.859E-07 | 1.500E-06 | 2.600E-06 |
| 26968 | 1349 | N1349G | E | 3.6e-06 | 2.6e-06 | 1.1e-06 | 2.433E-06 | 1.258E-06 | 1.100E-06 | 3.600E-06 |
| 26969 | 1349 | N1349H | E | 2.4e-06 | 2.9e-06 | 2.4e-06 | 2.567E-06 | 2.887E-07 | 2.400E-06 | 2.900E-06 |
| 26970 | 1349 | N1349I | E | 3.6e-06 | 5.1e-06 | 1.8e-06 | 3.500E-06 | 1.652E-06 | 1.800E-06 | 5.100E-06 |
| 26971 | 1349 | N1349L | E | 3.7e-06 | 1.8e-06 | 3.7e-06 | 3.067E-06 | 1.097E-06 | 1.800E-06 | 3.700E-06 |
| 26972 | 1349 | N1349K | E | 1.7e-06 | 2.6e-06 | 1.9e-06 | 2.067E-06 | 4.726E-07 | 1.700E-06 | 2.600E-06 |
| 26973 | 1349 | N1349M | E | 2.7e-06 | 3e-06 | 1.1e-06 | 2.267E-06 | 1.021E-06 | 1.100E-06 | 3.000E-06 |
| 26974 | 1349 | N1349F | E | 2.2e-06 | 2.9e-06 | 1.4e-06 | 2.167E-06 | 7.506E-07 | 1.400E-06 | 2.900E-06 |
| 26975 | 1349 | N1349P | E | 3.2e-06 | 1.7e-06 | 2.3e-06 | 2.400E-06 | 7.550E-07 | 1.700E-06 | 3.200E-06 |
| 26976 | 1349 | N1349S | E | 2.4e-06 | 1.3e-06 | 2.7e-06 | 2.133E-06 | 7.371E-07 | 1.300E-06 | 2.700E-06 |
| 26977 | 1349 | N1349T | E | 2.1e-06 | 4e-06 | 2.4e-06 | 2.833E-06 | 1.021E-06 | 2.100E-06 | 4.000E-06 |
| 26978 | 1349 | N1349W | E | 3.3e-06 | 4e-06 | 2.4e-06 | 3.233E-06 | 8.021E-07 | 2.400E-06 | 4.000E-06 |
| 26979 | 1349 | N1349Y | E | 2e-06 | 2.9e-06 | 2.9e-06 | 2.600E-06 | 5.196E-07 | 2.000E-06 | 2.900E-06 |
| 26980 | 1349 | N1349V | E | 2e-06 | 7.5e-07 | 2.1e-06 | 1.617E-06 | 7.522E-07 | 7.500E-07 | 2.100E-06 |
| 26981 | 1350 | T1350A | E | 2.1e-06 | 3.1e-06 | 4.4e-06 | 3.200E-06 | 1.153E-06 | 2.100E-06 | 4.400E-06 |
| 26982 | 1350 | T1350R | E | 2.3e-06 | 1.9e-06 | 2.4e-06 | 2.200E-06 | 2.646E-07 | 1.900E-06 | 2.400E-06 |
| 26983 | 1350 | T1350N | E | 2.6e-06 | 1.7e-06 | 2.9e-06 | 2.400E-06 | 6.245E-07 | 1.700E-06 | 2.900E-06 |
| 26984 | 1350 | T1350D | E | 2.2e-06 | 2.8e-06 | 2.2e-06 | 2.400E-06 | 3.464E-07 | 2.200E-06 | 2.800E-06 |
| 26985 | 1350 | T1350C | E | 3.9e-06 | 2.4e-06 | 1.3e-06 | 2.533E-06 | 1.305E-06 | 1.300E-06 | 3.900E-06 |
| 26986 | 1350 | T1350E | E | 1.7e-06 | 1.9e-06 | 3.7e-06 | 2.433E-06 | 1.102E-06 | 1.700E-06 | 3.700E-06 |
| 26987 | 1350 | T1350Q | E | 1.8e-06 | 2.2e-06 | 2e-06 | 2.000E-06 | 2.000E-07 | 1.800E-06 | 2.200E-06 |
| 26988 | 1350 | T1350G | E | 3.1e-06 | 3.4e-06 | 1.9e-06 | 2.800E-06 | 7.937E-07 | 1.900E-06 | 3.400E-06 |
| 26989 | 1350 | T1350H | E | 2e-06 | 2.3e-06 | 3e-06 | 2.433E-06 | 5.132E-07 | 2.000E-06 | 3.000E-06 |
| 26990 | 1350 | T1350I | E | 1.5e-06 | 1.2e-06 | 1.4e-06 | 1.367E-06 | 1.528E-07 | 1.200E-06 | 1.500E-06 |
| 26991 | 1350 | T1350L | E | 3.1e-06 | 2.2e-06 | 1.3e-06 | 2.200E-06 | 9.000E-07 | 1.300E-06 | 3.100E-06 |
| 26992 | 1350 | T1350K | E | 2.2e-06 | 3e-06 | 1.6e-06 | 2.267E-06 | 7.024E-07 | 1.600E-06 | 3.000E-06 |
| 26993 | 1350 | T1350M | E | 2.1e-06 | 2.8e-06 | 2.9e-06 | 2.600E-06 | 4.359E-07 | 2.100E-06 | 2.900E-06 |
| 26994 | 1350 | T1350F | E | 3e-06 | 1.3e-06 | 3.4e-06 | 2.567E-06 | 1.115E-06 | 1.300E-06 | 3.400E-06 |
| 26995 | 1350 | T1350P | E | 1.9e-06 | 2.2e-06 | 1.6e-06 | 1.900E-06 | 3.000E-07 | 1.600E-06 | 2.200E-06 |
| 26996 | 1350 | T1350S | E | 3e-06 | 3.1e-06 | 1.3e-06 | 2.467E-06 | 1.012E-06 | 1.300E-06 | 3.100E-06 |
| 26997 | 1350 | T1350T | E | 2e-06 | 2.5e-06 | 1.9e-06 | 2.133E-06 | 3.215E-07 | 1.900E-06 | 2.500E-06 |
| 26998 | 1350 | T1350W | E | 2.7e-06 | 3.4e-06 | 2e-06 | 2.700E-06 | 7.000E-07 | 2.000E-06 | 3.400E-06 |
| 26999 | 1350 | T1350Y | E | 1.9e-06 | 2.5e-06 | 1.9e-06 | 2.100E-06 | 3.464E-07 | 1.900E-06 | 2.500E-06 |
| 27000 | 1350 | T1350V | E | 2.8e-06 | 2.4e-06 | 2.5e-06 | 2.567E-06 | 2.082E-07 | 2.400E-06 | 2.800E-06 |
| 27001 | 1351 | A1351A | E | 2.6e-06 | 2.3e-06 | 2.7e-06 | 2.533E-06 | 2.082E-07 | 2.300E-06 | 2.700E-06 |
| 27002 | 1351 | A1351R | E | 2e-06 | 2.9e-06 | 3.1e-06 | 2.667E-06 | 5.859E-07 | 2.000E-06 | 3.100E-06 |
| 27003 | 1351 | A1351N | E | 1.4e-06 | 2.2e-06 | 2.7e-06 | 2.100E-06 | 6.557E-07 | 1.400E-06 | 2.700E-06 |
| 27004 | 1351 | A1351D | E | 2.4e-06 | 3.9e-06 | 3e-06 | 3.100E-06 | 7.550E-07 | 2.400E-06 | 3.900E-06 |
| 27005 | 1351 | A1351C | E | 2.3e-06 | 1.8e-06 | 2.5e-06 | 2.200E-06 | 3.606E-07 | 1.800E-06 | 2.500E-06 |
| 27006 | 1351 | A1351E | E | 3.1e-06 | 2.1e-06 | 4.5e-06 | 3.233E-06 | 1.206E-06 | 2.100E-06 | 4.500E-06 |
| 27007 | 1351 | A1351Q | E | 2e-06 | 5.7e-06 | 1.8e-06 | 3.167E-06 | 2.196E-06 | 1.800E-06 | 5.700E-06 |
| 27008 | 1351 | A1351G | E | 4e-06 | 1.5e-06 | 3e-06 | 2.833E-06 | 1.258E-06 | 1.500E-06 | 4.000E-06 |
| 27009 | 1351 | A1351H | E | 2.4e-06 | 2.3e-06 | 1.9e-06 | 2.200E-06 | 2.646E-07 | 1.900E-06 | 2.400E-06 |
| 27010 | 1351 | A1351I | E | 2.1e-06 | 2.2e-06 | 1.9e-06 | 2.067E-06 | 1.528E-07 | 1.900E-06 | 2.200E-06 |
| 27011 | 1351 | A1351L | E | 2.7e-06 | 3.1e-06 | 2.4e-06 | 2.733E-06 | 3.512E-07 | 2.400E-06 | 3.100E-06 |
| 27012 | 1351 | A1351K | E | 1.3e-06 | 2.5e-06 | 2.4e-06 | 2.067E-06 | 6.658E-07 | 1.300E-06 | 2.500E-06 |
| 27013 | 1351 | A1351M | E | 2.1e-06 | 2.7e-06 | 2.2e-06 | 2.333E-06 | 3.215E-07 | 2.100E-06 | 2.700E-06 |
| 27014 | 1351 | A1351F | E | 3e-06 | 2e-06 | 2.9e-06 | 2.633E-06 | 5.508E-07 | 2.000E-06 | 3.000E-06 |
| 27015 | 1351 | A1351P | E | 2.6e-06 | 2.3e-06 | 2.4e-06 | 2.433E-06 | 1.528E-07 | 2.300E-06 | 2.600E-06 |
| 27016 | 1351 | A1351S | E | 3.6e-06 | 1.8e-06 | 3.4e-06 | 2.933E-06 | 9.866E-07 | 1.800E-06 | 3.600E-06 |
| 27017 | 1351 | A1351T | E | 2.4e-06 | 1.7e-06 | 3.6e-06 | 2.567E-06 | 9.609E-07 | 1.700E-06 | 3.600E-06 |
| 27018 | 1351 | A1351W | E | 2.6e-06 | 2.5e-06 | 2.1e-06 | 2.400E-06 | 2.646E-07 | 2.100E-06 | 2.600E-06 |
| 27019 | 1351 | A1351Y | E | 9.7e-07 | 2.1e-06 | 1.7e-06 | 1.590E-06 | 5.730E-07 | 9.700E-07 | 2.100E-06 |
| 27020 | 1351 | A1351V | E | 2.8e-06 | 2.3e-06 | 4.7e-06 | 3.267E-06 | 1.266E-06 | 2.300E-06 | 4.700E-06 |
| 27021 | 1352 | Y1352A | E | 2.3e-06 | 2.1e-06 | 3e-06 | 2.467E-06 | 4.726E-07 | 2.100E-06 | 3.000E-06 |
| 27022 | 1352 | Y1352R | E | 1.8e-06 | 2.5e-06 | 2.2e-06 | 2.167E-06 | 3.512E-07 | 1.800E-06 | 2.500E-06 |
| 27023 | 1352 | Y1352N | E | 3.7e-06 | 4.2e-06 | 2e-06 | 3.300E-06 | 1.153E-06 | 2.000E-06 | 4.200E-06 |
| 27024 | 1352 | Y1352D | E | 1.8e-06 | 1.1e-06 | 2.4e-06 | 1.767E-06 | 6.506E-07 | 1.100E-06 | 2.400E-06 |
| 27025 | 1352 | Y1352C | E | 1.5e-06 | 1.8e-06 | 3e-06 | 2.100E-06 | 7.937E-07 | 1.500E-06 | 3.000E-06 |
| 27026 | 1352 | Y1352E | E | 1.6e-06 | 2.8e-06 | 2.7e-06 | 2.367E-06 | 6.658E-07 | 1.600E-06 | 2.800E-06 |
| 27027 | 1352 | Y1352Q | E | 2.7e-06 | 3.4e-06 | 2.3e-06 | 2.800E-06 | 5.568E-07 | 2.300E-06 | 3.400E-06 |
| 27028 | 1352 | Y1352G | E | 1.4e-06 | 2e-06 | 1.7e-06 | 1.700E-06 | 3.000E-07 | 1.400E-06 | 2.000E-06 |
| 27029 | 1352 | Y1352H | E | 3.5e-06 | 1.9e-06 | 2.2e-06 | 2.533E-06 | 8.505E-07 | 1.900E-06 | 3.500E-06 |

| | | | | | | | | | | |
|---|---|---|---|---|---|---|---|---|---|---|
| 27030 | 1352 | Y1352I | E | 2.5e-06 | 2.3e-06 | 2e-06 | 2.267E-06 | 2.517E-07 | 2.000E-06 | 2.500E-06 |
| 27031 | 1352 | Y1352L | E | 2.8e-06 | 2.2e-06 | 1.8e-06 | 2.267E-06 | 5.033E-07 | 1.800E-06 | 2.800E-06 |
| 27032 | 1352 | Y1352K | E | 2e-06 | 2e-06 | 2.4e-06 | 2.133E-06 | 2.309E-07 | 2.000E-06 | 2.400E-06 |
| 27033 | 1352 | Y1352M | E | 3.8e-06 | 3.1e-06 | 1.7e-06 | 2.867E-06 | 1.069E-06 | 1.700E-06 | 3.800E-06 |
| 27034 | 1352 | Y1352F | E | 4.6e-06 | 2.8e-06 | 5.1e-06 | 4.167E-06 | 1.210E-06 | 2.800E-06 | 5.100E-06 |
| 27035 | 1352 | Y1352P | E | 1.4e-06 | 2.5e-06 | 1.1e-06 | 1.667E-06 | 7.371E-07 | 1.100E-06 | 2.500E-06 |
| 27036 | 1352 | Y1352S | E | 2.5e-06 | 1.1e-06 | 2.9e-06 | 2.167E-06 | 9.452E-07 | 1.100E-06 | 2.900E-06 |
| 27037 | 1352 | Y1352T | E | 3.6e-06 | 2.4e-06 | 2.5e-06 | 2.833E-06 | 6.658E-07 | 2.400E-06 | 3.600E-06 |
| 27038 | 1352 | Y1352W | E | 4.7e-06 | 2.4e-06 | 2.2e-06 | 3.100E-06 | 1.389E-06 | 2.200E-06 | 4.700E-06 |
| 27039 | 1352 | Y1352Y | E | 3.2e-06 | 2.8e-06 | 3e-06 | 3.000E-06 | 2.000E-07 | 2.800E-06 | 3.200E-06 |
| 27040 | 1352 | Y1352V | E | 4.6e-06 | 1.9e-06 | 1.9e-06 | 2.800E-06 | 1.559E-06 | 1.900E-06 | 4.600E-06 |
| 27041 | 1353 | L1353A | E | 2.7e-06 | 1.9e-06 | 2.7e-06 | 2.433E-06 | 4.619E-07 | 1.900E-06 | 2.700E-06 |
| 27042 | 1353 | L1353R | E | 4.7e-06 | 3.4e-06 | 2e-06 | 3.367E-06 | 1.350E-06 | 2.000E-06 | 4.700E-06 |
| 27043 | 1353 | L1353N | E | 2.9e-06 | 2.2e-06 | 2.5e-06 | 2.533E-06 | 3.512E-07 | 2.200E-06 | 2.900E-06 |
| 27044 | 1353 | L1353D | E | 3.8e-06 | 2.6e-06 | 3e-06 | 3.133E-06 | 6.110E-07 | 2.600E-06 | 3.800E-06 |
| 27045 | 1353 | L1353C | E | 1.8e-06 | 4e-06 | 3.3e-06 | 3.033E-06 | 1.124E-06 | 1.800E-06 | 4.000E-06 |
| 27046 | 1353 | L1353E | E | 1.7e-06 | 2e-06 | 2e-06 | 1.900E-06 | 1.732E-07 | 1.700E-06 | 2.000E-06 |
| 27047 | 1353 | L1353Q | E | 3.2e-06 | 2.3e-06 | 2.3e-06 | 2.600E-06 | 5.196E-07 | 2.300E-06 | 3.200E-06 |
| 27048 | 1353 | L1353G | E | 2.5e-06 | 2.6e-06 | 1e-06 | 2.033E-06 | 8.963E-07 | 1.000E-06 | 2.600E-06 |
| 27049 | 1353 | L1353H | E | 2.5e-06 | 2.7e-06 | 2.4e-06 | 2.533E-06 | 1.528E-07 | 2.400E-06 | 2.700E-06 |
| 27050 | 1353 | L1353I | E | 3.3e-06 | 2.7e-06 | 3.1e-06 | 3.033E-06 | 3.055E-07 | 2.700E-06 | 3.300E-06 |
| 27051 | 1353 | L1353L | E | 1.5e-06 | 1.4e-06 | 3.1e-06 | 2.000E-06 | 9.539E-07 | 1.400E-06 | 3.100E-06 |
| 27052 | 1353 | L1353K | E | 2.5e-06 | 2e-06 | 2.9e-06 | 2.467E-06 | 4.509E-07 | 2.000E-06 | 2.900E-06 |
| 27053 | 1353 | L1353M | E | 2.5e-06 | 2.9e-06 | 1.2e-06 | 2.200E-06 | 8.888E-07 | 1.200E-06 | 2.900E-06 |
| 27054 | 1353 | L1353F | E | 2.4e-06 | 2.2e-06 | 3.1e-06 | 2.567E-06 | 4.726E-07 | 2.200E-06 | 3.100E-06 |
| 27055 | 1353 | L1353P | E | 2.3e-06 | 2.4e-06 | 1.7e-06 | 2.133E-06 | 3.786E-07 | 1.700E-06 | 2.400E-06 |
| 27056 | 1353 | L1353S | E | 1.6e-06 | 3e-06 | 1.2e-06 | 1.933E-06 | 9.452E-07 | 1.200E-06 | 3.000E-06 |
| 27057 | 1353 | L1353T | E | 3.3e-06 | 2.6e-06 | 2.1e-06 | 2.667E-06 | 6.028E-07 | 2.100E-06 | 3.300E-06 |
| 27058 | 1353 | L1353W | E | 2.7e-06 | 1.2e-06 | 2.3e-06 | 2.067E-06 | 7.767E-07 | 1.200E-06 | 2.700E-06 |
| 27059 | 1353 | L1353Y | E | 3.3e-06 | 1.7e-06 | 2.4e-06 | 2.467E-06 | 8.021E-07 | 1.700E-06 | 3.300E-06 |
| 27060 | 1353 | L1353V | E | 2.4e-06 | 3.2e-06 | 3.2e-06 | 2.933E-06 | 4.619E-07 | 2.400E-06 | 3.200E-06 |
| 27061 | 1354 | Q1354A | E | 2.9e-06 | 1.1e-06 | 2e-06 | 2.000E-06 | 9.000E-07 | 1.100E-06 | 2.900E-06 |
| 27062 | 1354 | Q1354R | E | 1.3e-06 | 2.8e-06 | 2.2e-06 | 2.100E-06 | 7.550E-07 | 1.300E-06 | 2.800E-06 |
| 27063 | 1354 | Q1354N | E | 2.2e-06 | 3e-06 | 2.7e-06 | 2.633E-06 | 4.041E-07 | 2.200E-06 | 3.000E-06 |
| 27064 | 1354 | Q1354D | E | 4.4e-06 | 1.9e-06 | 2.2e-06 | 2.833E-06 | 1.365E-06 | 1.900E-06 | 4.400E-06 |
| 27065 | 1354 | Q1354C | E | 2.3e-06 | 1.8e-06 | 2.8e-06 | 2.300E-06 | 5.000E-07 | 1.800E-06 | 2.800E-06 |
| 27066 | 1354 | Q1354E | E | 3e-06 | 2.2e-06 | 1.3e-06 | 2.167E-06 | 8.505E-07 | 1.300E-06 | 3.000E-06 |
| 27067 | 1354 | Q1354Q | E | 2.6e-06 | 3.8e-06 | 1.5e-06 | 2.633E-06 | 1.150E-06 | 1.500E-06 | 3.800E-06 |
| 27068 | 1354 | Q1354G | E | 2.9e-06 | 1.6e-06 | 2.1e-06 | 2.200E-06 | 6.557E-07 | 1.600E-06 | 2.900E-06 |
| 27069 | 1354 | Q1354H | E | 2.8e-06 | 2.7e-06 | 1.4e-06 | 2.300E-06 | 7.810E-07 | 1.400E-06 | 2.800E-06 |
| 27070 | 1354 | Q1354I | E | 4.8e-06 | 3.3e-06 | 4.4e-06 | 4.167E-06 | 7.767E-07 | 3.300E-06 | 4.800E-06 |
| 27071 | 1354 | Q1354L | E | 2.2e-06 | 2.2e-06 | 1.2e-06 | 1.867E-06 | 5.774E-07 | 1.200E-06 | 2.200E-06 |
| 27072 | 1354 | Q1354K | E | 2.5e-06 | 2.2e-06 | 1.5e-06 | 2.067E-06 | 5.132E-07 | 1.500E-06 | 2.500E-06 |
| 27073 | 1354 | Q1354M | E | 1.7e-06 | 2.5e-06 | 3.9e-06 | 2.700E-06 | 1.114E-06 | 1.700E-06 | 3.900E-06 |
| 27074 | 1354 | Q1354F | E | 2.5e-06 | 3.6e-06 | 2.4e-06 | 2.833E-06 | 6.658E-07 | 2.400E-06 | 3.600E-06 |
| 27075 | 1354 | Q1354P | E | 2e-06 | 2.8e-06 | 3.7e-06 | 2.833E-06 | 8.505E-07 | 2.000E-06 | 3.700E-06 |
| 27076 | 1354 | Q1354S | E | 2.7e-06 | 2.2e-06 | 1.8e-06 | 2.233E-06 | 4.509E-07 | 1.800E-06 | 2.700E-06 |
| 27077 | 1354 | Q1354T | E | 2e-06 | 1.3e-06 | 1.8e-06 | 1.700E-06 | 3.606E-07 | 1.300E-06 | 2.000E-06 |
| 27078 | 1354 | Q1354W | E | 2.2e-06 | 1.3e-06 | 1.8e-06 | 1.767E-06 | 4.509E-07 | 1.300E-06 | 2.200E-06 |
| 27079 | 1354 | Q1354Y | E | 1.2e-06 | 1.9e-06 | 1.8e-06 | 1.633E-06 | 3.786E-07 | 1.200E-06 | 1.900E-06 |
| 27080 | 1354 | Q1354V | E | 3.3e-06 | 2.2e-06 | 4.1e-06 | 3.200E-06 | 9.539E-07 | 2.200E-06 | 4.100E-06 |
| 27081 | 1355 | M1355A | E | 1.6e-06 | 1.7e-06 | 2.2e-06 | 1.833E-06 | 3.215E-07 | 1.600E-06 | 2.200E-06 |
| 27082 | 1355 | M1355R | E | 2.3e-06 | 2.1e-06 | 9.1e-07 | 1.770E-06 | 7.515E-07 | 9.100E-07 | 2.300E-06 |
| 27083 | 1355 | M1355N | E | 2.1e-06 | 2.7e-06 | 1.1e-06 | 1.967E-06 | 8.083E-07 | 1.100E-06 | 2.700E-06 |
| 27084 | 1355 | M1355D | E | 2.1e-06 | 3.4e-06 | 4.3e-06 | 3.267E-06 | 1.106E-06 | 2.100E-06 | 4.300E-06 |
| 27085 | 1355 | M1355C | E | 3.6e-06 | 2.7e-06 | 4.2e-06 | 3.500E-06 | 7.550E-07 | 2.700E-06 | 4.200E-06 |
| 27086 | 1355 | M1355E | E | 2.6e-06 | 2.9e-06 | 2.1e-06 | 2.533E-06 | 4.041E-07 | 2.100E-06 | 2.900E-06 |
| 27087 | 1355 | M1355Q | E | 2.4e-06 | 3.3e-06 | 2.4e-06 | 2.700E-06 | 5.196E-07 | 2.400E-06 | 3.300E-06 |
| 27088 | 1355 | M1355G | E | 2.1e-06 | 2.1e-06 | 3.2e-06 | 2.467E-06 | 6.351E-07 | 2.100E-06 | 3.200E-06 |
| 27089 | 1355 | M1355H | E | 2.3e-06 | 4.5e-06 | 1.7e-06 | 2.833E-06 | 1.474E-06 | 1.700E-06 | 4.500E-06 |
| 27090 | 1355 | M1355I | E | 2.3e-06 | 2.5e-06 | 3.7e-06 | 2.833E-06 | 7.572E-07 | 2.300E-06 | 3.700E-06 |
| 27091 | 1355 | M1355L | E | 1.8e-06 | 2.3e-06 | 2.7e-06 | 2.267E-06 | 4.509E-07 | 1.800E-06 | 2.700E-06 |
| 27092 | 1355 | M1355K | E | 2.5e-06 | 2.3e-06 | 1.5e-06 | 2.100E-06 | 5.292E-07 | 1.500E-06 | 2.500E-06 |
| 27093 | 1355 | M1355M | E | 2.2e-06 | 2.1e-06 | 3.1e-06 | 2.467E-06 | 5.508E-07 | 2.100E-06 | 3.100E-06 |
| 27094 | 1355 | M1355F | E | 2.8e-06 | 2.4e-06 | 2.5e-06 | 2.567E-06 | 2.082E-07 | 2.400E-06 | 2.800E-06 |
| 27095 | 1355 | M1355P | E | 2.3e-06 | 2.6e-06 | 4.5e-06 | 3.133E-06 | 1.193E-06 | 2.300E-06 | 4.500E-06 |
| 27096 | 1355 | M1355S | E | 2.2e-06 | 1.9e-06 | 2.2e-06 | 2.100E-06 | 1.732E-07 | 1.900E-06 | 2.200E-06 |
| 27097 | 1355 | M1355T | E | 2.3e-06 | 5.2e-06 | 2.1e-06 | 3.200E-06 | 1.735E-06 | 2.100E-06 | 5.200E-06 |
| 27098 | 1355 | M1355W | E | 2.5e-06 | 3.2e-06 | 2.1e-06 | 2.600E-06 | 5.568E-07 | 2.100E-06 | 3.200E-06 |
| 27099 | 1355 | M1355Y | E | 2.6e-06 | 1.2e-06 | 3.9e-06 | 2.567E-06 | 1.350E-06 | 1.200E-06 | 3.900E-06 |
| 27100 | 1355 | M1355V | E | 3e-06 | 3.1e-06 | 3.9e-06 | 3.333E-06 | 4.933E-07 | 3.000E-06 | 3.900E-06 |
| 27101 | 1356 | N1356A | E | 1.7e-06 | 3.1e-06 | 1.4e-06 | 2.067E-06 | 9.074E-07 | 1.400E-06 | 3.100E-06 |
| 27102 | 1356 | N1356R | E | 5.8e-06 | 2.6e-06 | 3.4e-06 | 3.933E-06 | 1.665E-06 | 2.600E-06 | 5.800E-06 |
| 27103 | 1356 | N1356N | E | 1.7e-06 | 1.4e-06 | 1.5e-06 | 1.533E-06 | 1.528E-07 | 1.400E-06 | 1.700E-06 |
| 27104 | 1356 | N1356D | E | 2.8e-06 | 2.1e-06 | 2.8e-06 | 2.567E-06 | 4.041E-07 | 2.100E-06 | 2.800E-06 |
| 27105 | 1356 | N1356C | E | 1e-06 | 3.3e-06 | 2e-06 | 2.100E-06 | 1.153E-06 | 1.000E-06 | 3.300E-06 |
| 27106 | 1356 | N1356E | E | 2.6e-06 | 2e-06 | 2.9e-06 | 2.500E-06 | 4.583E-07 | 2.000E-06 | 2.900E-06 |
| 27107 | 1356 | N1356Q | E | 2.5e-06 | 1.3e-06 | 2.8e-06 | 2.200E-06 | 7.937E-07 | 1.300E-06 | 2.800E-06 |
| 27108 | 1356 | N1356G | E | 3.3e-06 | 2.8e-06 | 4.3e-06 | 3.467E-06 | 7.638E-07 | 2.800E-06 | 4.300E-06 |
| 27109 | 1356 | N1356H | E | 2.2e-06 | 7.7e-07 | 3.5e-06 | 2.157E-06 | 1.366E-06 | 7.700E-07 | 3.500E-06 |
| 27110 | 1356 | N1356I | E | 2.3e-06 | 4.3e-06 | 1.8e-06 | 2.800E-06 | 1.323E-06 | 1.800E-06 | 4.300E-06 |
| 27111 | 1356 | N1356L | E | 2.1e-06 | 2.3e-06 | 1.9e-06 | 2.100E-06 | 2.000E-07 | 1.900E-06 | 2.300E-06 |
| 27112 | 1356 | N1356K | E | 3.9e-06 | 2.3e-06 | 2.1e-06 | 2.767E-06 | 9.866E-07 | 2.100E-06 | 3.900E-06 |
| 27113 | 1356 | N1356M | E | 2.9e-06 | 2.8e-06 | 2.1e-06 | 2.600E-06 | 4.359E-07 | 2.100E-06 | 2.900E-06 |
| 27114 | 1356 | N1356F | E | 2.6e-06 | 1.8e-06 | 1.9e-06 | 2.100E-06 | 4.359E-07 | 1.800E-06 | 2.600E-06 |

| | | | | | | | | | | |
|---|---|---|---|---|---|---|---|---|---|---|
| 27115 | 1356 | N1356P | E | 2.9e-06 | 1.8e-06 | 3.2e-06 | 2.633E-06 | 7.371E-07 | 1.800E-06 | 3.200E-06 |
| 27116 | 1356 | N1356S | E | 2.5e-06 | 2.3e-06 | 3.9e-06 | 2.900E-06 | 8.718E-07 | 2.300E-06 | 3.900E-06 |
| 27117 | 1356 | N1356T | E | 3.2e-06 | 1.2e-06 | 2e-06 | 2.133E-06 | 1.007E-06 | 1.200E-06 | 3.200E-06 |
| 27118 | 1356 | N1356W | E | 2.9e-06 | 3.5e-06 | 2.7e-06 | 3.033E-06 | 4.163E-07 | 2.700E-06 | 3.500E-06 |
| 27119 | 1356 | N1356Y | E | 1.7e-06 | 2.5e-06 | 3e-06 | 2.400E-06 | 6.557E-07 | 1.700E-06 | 3.000E-06 |
| 27120 | 1356 | N1356V | E | 2e-06 | 2.1e-06 | 7.4e-07 | 1.613E-06 | 7.580E-07 | 7.400E-07 | 2.100E-06 |
| 27121 | 1357 | S1357A | E | 2e-06 | 2.1e-06 | 1.9e-06 | 2.000E-06 | 1.000E-07 | 1.900E-06 | 2.100E-06 |
| 27122 | 1357 | S1357R | E | 1.3e-06 | 1.5e-06 | 1.5e-06 | 1.433E-06 | 1.155E-07 | 1.300E-06 | 1.500E-06 |
| 27123 | 1357 | S1357N | E | 1.3e-06 | 2.5e-06 | 3.8e-06 | 2.533E-06 | 1.250E-06 | 1.300E-06 | 3.800E-06 |
| 27124 | 1357 | S1357D | E | 2.6e-06 | 1.9e-06 | 2.5e-06 | 2.333E-06 | 3.786E-07 | 1.900E-06 | 2.600E-06 |
| 27125 | 1357 | S1357C | E | 2.1e-06 | 1.7e-06 | 3.3e-06 | 2.367E-06 | 8.327E-07 | 1.700E-06 | 3.300E-06 |
| 27126 | 1357 | S1357E | E | 3.9e-06 | 2e-06 | 2.8e-06 | 2.900E-06 | 9.539E-07 | 2.000E-06 | 3.900E-06 |
| 27127 | 1357 | S1357Q | E | 2.7e-06 | 2.6e-06 | 2.2e-06 | 2.500E-06 | 2.646E-07 | 2.200E-06 | 2.700E-06 |
| 27128 | 1357 | S1357G | E | 2.8e-06 | 3e-06 | 2.5e-06 | 2.767E-06 | 2.517E-07 | 2.500E-06 | 3.000E-06 |
| 27129 | 1357 | S1357H | E | 2.5e-06 | 2.7e-06 | 5.3e-06 | 3.500E-06 | 1.562E-06 | 2.500E-06 | 5.300E-06 |
| 27130 | 1357 | S1357I | E | 3.6e-06 | 2.8e-06 | 2.1e-06 | 2.833E-06 | 7.506E-07 | 2.100E-06 | 3.600E-06 |
| 27131 | 1357 | S1357L | E | 3.9e-06 | 3.2e-06 | 2.9e-06 | 3.333E-06 | 5.132E-07 | 2.900E-06 | 3.900E-06 |
| 27132 | 1357 | S1357K | E | 2.2e-06 | 2.2e-06 | 2.3e-06 | 2.233E-06 | 5.774E-08 | 2.200E-06 | 2.300E-06 |
| 27133 | 1357 | S1357M | E | 1.6e-06 | 3.2e-06 | 2e-06 | 2.267E-06 | 8.327E-07 | 1.600E-06 | 3.200E-06 |
| 27134 | 1357 | S1357F | E | 1.3e-06 | 1.1e-06 | 1.6e-06 | 1.333E-06 | 2.517E-07 | 1.100E-06 | 1.600E-06 |
| 27135 | 1357 | S1357P | E | 1.6e-06 | 1.9e-06 | 2.1e-06 | 1.867E-06 | 2.517E-07 | 1.600E-06 | 2.100E-06 |
| 27136 | 1357 | S1357S | E | 2.5e-06 | 2.4e-06 | 3.7e-06 | 2.867E-06 | 7.234E-07 | 2.400E-06 | 3.700E-06 |
| 27137 | 1357 | S1357T | E | 4.9e-06 | 1.9e-06 | 2.7e-06 | 3.167E-06 | 1.553E-06 | 1.900E-06 | 4.900E-06 |
| 27138 | 1357 | S1357W | E | 3.2e-06 | 2.7e-06 | 3.5e-06 | 3.133E-06 | 4.041E-07 | 2.700E-06 | 3.500E-06 |
| 27139 | 1357 | S1357Y | E | 2.1e-06 | 3.1e-06 | 2.8e-06 | 2.667E-06 | 5.132E-07 | 2.100E-06 | 3.100E-06 |
| 27140 | 1357 | S1357V | E | 1.6e-06 | 1.5e-06 | 3.2e-06 | 2.100E-06 | 9.539E-07 | 1.500E-06 | 3.200E-06 |
| 27141 | 1358 | L1358A | E | 2.7e-06 | 2.3e-06 | 2.4e-06 | 2.467E-06 | 2.082E-07 | 2.300E-06 | 2.700E-06 |
| 27142 | 1358 | L1358R | E | 1e-06 | 2.1e-06 | 2.1e-06 | 1.733E-06 | 6.351E-07 | 1.000E-06 | 2.100E-06 |
| 27143 | 1358 | L1358N | E | 2.3e-06 | 1.7e-06 | 2e-06 | 2.000E-06 | 3.000E-07 | 1.700E-06 | 2.300E-06 |
| 27144 | 1358 | L1358D | E | 2.6e-06 | 3.1e-06 | 2.4e-06 | 2.700E-06 | 3.606E-07 | 2.400E-06 | 3.100E-06 |
| 27145 | 1358 | L1358C | E | 1.3e-06 | 1.2e-06 | 2.3e-06 | 1.600E-06 | 6.083E-07 | 1.200E-06 | 2.300E-06 |
| 27146 | 1358 | L1358E | E | 2.3e-06 | 3.7e-06 | 3.2e-06 | 3.067E-06 | 7.095E-07 | 2.300E-06 | 3.700E-06 |
| 27147 | 1358 | L1358Q | E | 2.4e-06 | 2.3e-06 | 1.9e-06 | 2.200E-06 | 2.646E-07 | 1.900E-06 | 2.400E-06 |
| 27148 | 1358 | L1358G | E | 1.9e-06 | 2.3e-06 | 2.2e-06 | 2.133E-06 | 2.082E-07 | 1.900E-06 | 2.300E-06 |
| 27149 | 1358 | L1358H | E | 2.4e-06 | 1.8e-06 | 2e-06 | 2.067E-06 | 3.055E-07 | 1.800E-06 | 2.400E-06 |
| 27150 | 1358 | L1358I | E | 2.7e-06 | 2.3e-06 | 3.3e-06 | 2.767E-06 | 5.033E-07 | 2.300E-06 | 3.300E-06 |
| 27151 | 1358 | L1358L | E | 1.9e-06 | 2.9e-06 | 2.6e-06 | 2.467E-06 | 5.132E-07 | 1.900E-06 | 2.900E-06 |
| 27152 | 1358 | L1358K | E | 2.4e-06 | 2.3e-06 | 2.6e-06 | 2.433E-06 | 1.528E-07 | 2.300E-06 | 2.600E-06 |
| 27153 | 1358 | L1358M | E | 3.2e-06 | 1.1e-06 | 2.2e-06 | 2.167E-06 | 1.050E-06 | 1.100E-06 | 3.200E-06 |
| 27154 | 1358 | L1358F | E | 1.5e-06 | 1.8e-06 | 3.4e-06 | 2.233E-06 | 1.021E-06 | 1.500E-06 | 3.400E-06 |
| 27155 | 1358 | L1358P | E | 3.5e-06 | 2.6e-06 | 4.1e-06 | 3.400E-06 | 7.550E-07 | 2.600E-06 | 4.100E-06 |
| 27156 | 1358 | L1358S | E | 1.9e-06 | 3.3e-06 | 2.5e-06 | 2.567E-06 | 7.024E-07 | 1.900E-06 | 3.300E-06 |
| 27157 | 1358 | L1358T | E | 1.3e-06 | 1.7e-06 | 5.3e-06 | 2.767E-06 | 2.203E-06 | 1.300E-06 | 5.300E-06 |
| 27158 | 1358 | L1358W | E | 2e-06 | 2e-06 | 1.9e-06 | 1.967E-06 | 5.774E-08 | 1.900E-06 | 2.000E-06 |
| 27159 | 1358 | L1358Y | E | 3.8e-06 | 2.6e-06 | 5.3e-06 | 3.900E-06 | 1.353E-06 | 2.600E-06 | 5.300E-06 |
| 27160 | 1358 | L1358V | E | 4.3e-06 | 2.5e-06 | 2.8e-06 | 3.200E-06 | 9.644E-07 | 2.500E-06 | 4.300E-06 |
| 27161 | 1359 | R1359A | E | 1.1e-06 | 2.1e-06 | 2.2e-06 | 1.800E-06 | 6.083E-07 | 1.100E-06 | 2.200E-06 |
| 27162 | 1359 | R1359R | E | 3.1e-06 | 2.4e-06 | 1.7e-06 | 2.400E-06 | 7.000E-07 | 1.700E-06 | 3.100E-06 |
| 27163 | 1359 | R1359N | E | 3.2e-06 | 3.2e-06 | 4.6e-06 | 3.667E-06 | 8.083E-07 | 3.200E-06 | 4.600E-06 |
| 27164 | 1359 | R1359D | E | 2.7e-06 | 2.1e-06 | 1.9e-06 | 2.233E-06 | 4.163E-07 | 1.900E-06 | 2.700E-06 |
| 27165 | 1359 | R1359C | E | 2.1e-06 | 2.1e-06 | 3.1e-06 | 2.433E-06 | 5.774E-07 | 2.100E-06 | 3.100E-06 |
| 27166 | 1359 | R1359E | E | 2.4e-06 | 2.4e-06 | 2.9e-06 | 2.567E-06 | 2.887E-07 | 2.400E-06 | 2.900E-06 |
| 27167 | 1359 | R1359Q | E | 2.8e-06 | 2.4e-06 | 1.8e-06 | 2.333E-06 | 5.033E-07 | 1.800E-06 | 2.800E-06 |
| 27168 | 1359 | R1359G | E | 2.5e-06 | 2.5e-06 | 5e-06 | 3.333E-06 | 1.443E-06 | 2.500E-06 | 5.000E-06 |
| 27169 | 1359 | R1359H | E | 1.7e-06 | 2.1e-06 | 1e-06 | 1.600E-06 | 5.568E-07 | 1.000E-06 | 2.100E-06 |
| 27170 | 1359 | R1359I | E | 2.3e-06 | 1.6e-06 | 2.7e-06 | 2.200E-06 | 5.568E-07 | 1.600E-06 | 2.700E-06 |
| 27171 | 1359 | R1359L | E | 3e-06 | 2.2e-06 | 2.3e-06 | 2.500E-06 | 4.359E-07 | 2.200E-06 | 3.000E-06 |
| 27172 | 1359 | R1359K | E | 3.8e-06 | 1.4e-06 | 2.2e-06 | 2.467E-06 | 1.222E-06 | 1.400E-06 | 3.800E-06 |
| 27173 | 1359 | R1359M | E | 1.7e-06 | 1.9e-06 | 1.9e-06 | 1.833E-06 | 1.155E-07 | 1.700E-06 | 1.900E-06 |
| 27174 | 1359 | R1359F | E | 3e-06 | 2.6e-06 | 5.2e-06 | 3.600E-06 | 1.400E-06 | 2.600E-06 | 5.200E-06 |
| 27175 | 1359 | R1359P | E | 2e-06 | 1.7e-06 | 2.2e-06 | 1.967E-06 | 2.517E-07 | 1.700E-06 | 2.200E-06 |
| 27176 | 1359 | R1359S | E | 3.4e-06 | 4.7e-06 | 3.6e-06 | 3.900E-06 | 7.000E-07 | 3.400E-06 | 4.700E-06 |
| 27177 | 1359 | R1359T | E | 1.9e-06 | 2.9e-06 | 2.4e-06 | 2.400E-06 | 5.000E-07 | 1.900E-06 | 2.900E-06 |
| 27178 | 1359 | R1359W | E | 3e-06 | 2.3e-06 | 2.6e-06 | 2.633E-06 | 3.512E-07 | 2.300E-06 | 3.000E-06 |
| 27179 | 1359 | R1359Y | E | 3.1e-06 | 2.5e-06 | 1.9e-06 | 2.500E-06 | 6.000E-07 | 1.900E-06 | 3.100E-06 |
| 27180 | 1359 | R1359V | E | 1.1e-06 | 3.2e-06 | 2e-06 | 2.100E-06 | 1.054E-06 | 1.100E-06 | 3.200E-06 |
| 27181 | 1360 | A1360A | E | 1.2e-06 | 5.7e-06 | 2.5e-06 | 3.133E-06 | 2.316E-06 | 1.200E-06 | 5.700E-06 |
| 27182 | 1360 | A1360R | E | 1.8e-06 | 2e-06 | 1.7e-06 | 1.833E-06 | 1.528E-07 | 1.700E-06 | 2.000E-06 |
| 27183 | 1360 | A1360N | E | 1.4e-06 | 2.6e-06 | 2.8e-06 | 2.267E-06 | 7.572E-07 | 1.400E-06 | 2.800E-06 |
| 27184 | 1360 | A1360D | E | 2.1e-06 | 1.1e-06 | 2.2e-06 | 1.800E-06 | 6.083E-07 | 1.100E-06 | 2.200E-06 |
| 27185 | 1360 | A1360C | E | 2e-06 | 2.7e-06 | 2.6e-06 | 2.433E-06 | 3.786E-07 | 2.000E-06 | 2.700E-06 |
| 27186 | 1360 | A1360E | E | 1.5e-06 | 3.6e-06 | 1.9e-06 | 2.333E-06 | 1.115E-06 | 1.500E-06 | 3.600E-06 |
| 27187 | 1360 | A1360Q | E | 3.1e-06 | 1.9e-06 | 2.3e-06 | 2.433E-06 | 6.110E-07 | 1.900E-06 | 3.100E-06 |
| 27188 | 1360 | A1360G | E | 3.4e-06 | 2e-06 | 2.9e-06 | 2.767E-06 | 7.095E-07 | 2.000E-06 | 3.400E-06 |
| 27189 | 1360 | A1360H | E | 1e-06 | 4.1e-06 | 2.6e-06 | 2.567E-06 | 1.550E-06 | 1.000E-06 | 4.100E-06 |
| 27190 | 1360 | A1360I | E | 2.5e-06 | 2.3e-06 | 2.1e-06 | 2.300E-06 | 2.000E-07 | 2.100E-06 | 2.500E-06 |
| 27191 | 1360 | A1360L | E | 3.5e-06 | 3.6e-06 | 2.5e-06 | 3.200E-06 | 6.083E-07 | 2.500E-06 | 3.600E-06 |
| 27192 | 1360 | A1360K | E | 2.2e-06 | 3e-06 | 1.9e-06 | 2.367E-06 | 5.686E-07 | 1.900E-06 | 3.000E-06 |
| 27193 | 1360 | A1360M | E | 2e-06 | 2.1e-06 | 2.5e-06 | 2.200E-06 | 2.646E-07 | 2.000E-06 | 2.500E-06 |
| 27194 | 1360 | A1360F | E | 3.6e-06 | 2.4e-06 | 1.6e-06 | 2.533E-06 | 1.007E-06 | 1.600E-06 | 3.600E-06 |
| 27195 | 1360 | A1360P | E | 2.1e-06 | 3.2e-06 | 1.9e-06 | 2.400E-06 | 7.000E-07 | 1.900E-06 | 3.200E-06 |
| 27196 | 1360 | A1360S | E | 2.4e-06 | 1.5e-06 | 1.8e-06 | 1.900E-06 | 4.583E-07 | 1.500E-06 | 2.400E-06 |
| 27197 | 1360 | A1360T | E | 3.4e-06 | 2.2e-06 | 1.6e-06 | 2.400E-06 | 9.165E-07 | 1.600E-06 | 3.400E-06 |
| 27198 | 1360 | A1360W | E | 1.1e-06 | 2.4e-06 | 4.7e-06 | 2.733E-06 | 1.823E-06 | 1.100E-06 | 4.700E-06 |
| 27199 | 1360 | A1360Y | E | 3.3e-06 | 1.7e-06 | 3e-06 | 2.667E-06 | 8.505E-07 | 1.700E-06 | 3.300E-06 |

| | | | | | | | | | | |
|---|---|---|---|---|---|---|---|---|---|---|
| 27200 | 1360 | A1360V | E | 2.4e-06 | 4e-06 | 3.3e-06 | 3.233E-06 | 8.021E-07 | 2.400E-06 | 4.000E-06 |
| 27201 | 1361 | E1361A | E | 1.6e-06 | 2.2e-06 | 1.8e-06 | 1.867E-06 | 3.055E-07 | 1.600E-06 | 2.200E-06 |
| 27202 | 1361 | E1361R | E | 1.2e-06 | 3e-06 | 2.2e-06 | 2.133E-06 | 9.018E-07 | 1.200E-06 | 3.000E-06 |
| 27203 | 1361 | E1361N | E | 2.1e-06 | 2.9e-06 | 1.6e-06 | 2.200E-06 | 6.557E-07 | 1.600E-06 | 2.900E-06 |
| 27204 | 1361 | E1361D | E | 1.6e-06 | 2.7e-06 | 2.5e-06 | 2.267E-06 | 5.859E-07 | 1.600E-06 | 2.700E-06 |
| 27205 | 1361 | E1361C | E | 2.3e-06 | 4.3e-06 | 1.5e-06 | 2.700E-06 | 1.442E-06 | 1.500E-06 | 4.300E-06 |
| 27206 | 1361 | E1361E | E | 2.5e-06 | 1.8e-06 | 8e-07 | 1.700E-06 | 8.544E-07 | 8.000E-07 | 2.500E-06 |
| 27207 | 1361 | E1361Q | E | 2e-06 | 2.9e-06 | 3.6e-06 | 2.833E-06 | 8.021E-07 | 2.000E-06 | 3.600E-06 |
| 27208 | 1361 | E1361G | E | 3.4e-06 | 2.1e-06 | 4.1e-06 | 3.200E-06 | 1.015E-06 | 2.100E-06 | 4.100E-06 |
| 27209 | 1361 | E1361H | E | 5.3e-06 | 8.7e-07 | 3.2e-06 | 3.123E-06 | 2.216E-06 | 8.700E-07 | 5.300E-06 |
| 27210 | 1361 | E1361I | E | 3.5e-06 | 2.8e-06 | 1.6e-06 | 2.633E-06 | 9.609E-07 | 1.600E-06 | 3.500E-06 |
| 27211 | 1361 | E1361L | E | 3.8e-06 | 2.6e-06 | 2.3e-06 | 2.900E-06 | 7.937E-07 | 2.300E-06 | 3.800E-06 |
| 27212 | 1361 | E1361K | E | 3.3e-06 | 2.5e-06 | 2.6e-06 | 2.800E-06 | 4.359E-07 | 2.500E-06 | 3.300E-06 |
| 27213 | 1361 | E1361M | E | 2.3e-06 | 3.2e-06 | 3.2e-06 | 2.900E-06 | 5.196E-07 | 2.300E-06 | 3.200E-06 |
| 27214 | 1361 | E1361F | E | 3.7e-06 | 2.4e-06 | 1.5e-06 | 2.533E-06 | 1.106E-06 | 1.500E-06 | 3.700E-06 |
| 27215 | 1361 | E1361P | E | 5e-06 | 4.6e-06 | 1.4e-06 | 3.667E-06 | 1.973E-06 | 1.400E-06 | 5.000E-06 |
| 27216 | 1361 | E1361S | E | 2.1e-06 | 2.2e-06 | 2.4e-06 | 2.233E-06 | 1.528E-07 | 2.100E-06 | 2.400E-06 |
| 27217 | 1361 | E1361T | E | 4.9e-06 | 2.5e-06 | 2e-06 | 3.133E-06 | 1.550E-06 | 2.000E-06 | 4.900E-06 |
| 27218 | 1361 | E1361W | E | 2.3e-06 | 2.6e-06 | 1.9e-06 | 2.267E-06 | 3.512E-07 | 1.900E-06 | 2.600E-06 |
| 27219 | 1361 | E1361Y | E | 4.4e-06 | 2.7e-06 | 2.7e-06 | 3.267E-06 | 9.815E-07 | 2.700E-06 | 4.400E-06 |
| 27220 | 1361 | E1361V | E | 1.8e-06 | 2.6e-06 | 1.6e-06 | 2.000E-06 | 5.292E-07 | 1.600E-06 | 2.600E-06 |
| 27221 | 1362 | D1362A | E | 2.9e-06 | 2.6e-06 | 2.1e-06 | 2.533E-06 | 4.041E-07 | 2.100E-06 | 2.900E-06 |
| 27222 | 1362 | D1362R | E | 2.1e-06 | 1.6e-06 | 2.6e-06 | 2.100E-06 | 5.000E-07 | 1.600E-06 | 2.600E-06 |
| 27223 | 1362 | D1362N | E | 1.6e-06 | 3.2e-06 | 4.8e-06 | 3.200E-06 | 1.600E-06 | 1.600E-06 | 4.800E-06 |
| 27224 | 1362 | D1362D | E | 1.6e-06 | 1.9e-06 | 4.6e-06 | 2.700E-06 | 1.652E-06 | 1.600E-06 | 4.600E-06 |
| 27225 | 1362 | D1362C | E | 3.7e-06 | 3.5e-06 | 3.6e-06 | 3.600E-06 | 1.000E-07 | 3.500E-06 | 3.700E-06 |
| 27226 | 1362 | D1362E | E | 3.7e-06 | 2.4e-06 | 2.3e-06 | 2.800E-06 | 7.810E-07 | 2.300E-06 | 3.700E-06 |
| 27227 | 1362 | D1362Q | E | 2.4e-06 | 2.5e-06 | 1.9e-06 | 2.267E-06 | 3.215E-07 | 1.900E-06 | 2.500E-06 |
| 27228 | 1362 | D1362G | E | 3.3e-06 | 3.7e-06 | 1.8e-06 | 2.933E-06 | 1.002E-06 | 1.800E-06 | 3.700E-06 |
| 27229 | 1362 | D1362H | E | 2.8e-06 | 3e-06 | 2.4e-06 | 2.733E-06 | 3.055E-07 | 2.400E-06 | 3.000E-06 |
| 27230 | 1362 | D1362I | E | 1.5e-06 | 1.8e-06 | 3.2e-06 | 2.167E-06 | 9.074E-07 | 1.500E-06 | 3.200E-06 |
| 27231 | 1362 | D1362L | E | 4.3e-06 | 2.9e-06 | 2.9e-06 | 3.367E-06 | 8.083E-07 | 2.900E-06 | 4.300E-06 |
| 27232 | 1362 | D1362K | E | 2.7e-06 | 2.3e-06 | 2.9e-06 | 2.633E-06 | 3.055E-07 | 2.300E-06 | 2.900E-06 |
| 27233 | 1362 | D1362M | E | 2.3e-06 | 2.3e-06 | 1.9e-06 | 2.167E-06 | 2.309E-07 | 1.900E-06 | 2.300E-06 |
| 27234 | 1362 | D1362F | E | 1.7e-06 | 2.4e-06 | 3.3e-06 | 2.467E-06 | 8.021E-07 | 1.700E-06 | 3.300E-06 |
| 27235 | 1362 | D1362P | E | 2e-06 | 2.1e-06 | 2e-06 | 2.033E-06 | 5.774E-08 | 2.000E-06 | 2.100E-06 |
| 27236 | 1362 | D1362S | E | 1.6e-06 | 2e-06 | 1.9e-06 | 1.833E-06 | 2.082E-07 | 1.600E-06 | 2.000E-06 |
| 27237 | 1362 | D1362T | E | 1.3e-06 | 4.5e-06 | 2.5e-06 | 2.767E-06 | 1.617E-06 | 1.300E-06 | 4.500E-06 |
| 27238 | 1362 | D1362W | E | 1.9e-06 | 2.7e-06 | 2.5e-06 | 2.367E-06 | 4.163E-07 | 1.900E-06 | 2.700E-06 |
| 27239 | 1362 | D1362Y | E | 1.8e-06 | 1.7e-06 | 2.1e-06 | 1.867E-06 | 2.082E-07 | 1.700E-06 | 2.100E-06 |
| 27240 | 1362 | D1362V | E | 2.3e-06 | 2.7e-06 | 2.9e-06 | 2.633E-06 | 3.055E-07 | 2.300E-06 | 2.900E-06 |
| 27241 | 1363 | T1363A | E | 2.7e-06 | 2.2e-06 | 3.6e-06 | 2.833E-06 | 7.095E-07 | 2.200E-06 | 3.600E-06 |
| 27242 | 1363 | T1363R | E | 2.4e-06 | 3.8e-06 | 2.3e-06 | 2.833E-06 | 8.386E-07 | 2.300E-06 | 3.800E-06 |
| 27243 | 1363 | T1363N | E | 2.6e-06 | 1.5e-06 | 2.9e-06 | 2.333E-06 | 7.371E-07 | 1.500E-06 | 2.900E-06 |
| 27244 | 1363 | T1363D | E | 2.9e-06 | 3.1e-06 | 2.6e-06 | 2.867E-06 | 2.517E-07 | 2.600E-06 | 3.100E-06 |
| 27245 | 1363 | T1363C | E | 3.1e-06 | 2.1e-06 | 2.1e-06 | 2.433E-06 | 5.774E-07 | 2.100E-06 | 3.100E-06 |
| 27246 | 1363 | T1363E | E | 2.3e-06 | 3.2e-06 | 2.5e-06 | 2.667E-06 | 4.726E-07 | 2.300E-06 | 3.200E-06 |
| 27247 | 1363 | T1363Q | E | 1.6e-06 | 1.6e-06 | 3.5e-06 | 2.233E-06 | 1.097E-06 | 1.600E-06 | 3.500E-06 |
| 27248 | 1363 | T1363G | E | 1.7e-06 | 3.1e-06 | 1.5e-06 | 2.100E-06 | 8.718E-07 | 1.500E-06 | 3.100E-06 |
| 27249 | 1363 | T1363H | E | 2.3e-06 | 1.6e-06 | 3.1e-06 | 2.333E-06 | 7.506E-07 | 1.600E-06 | 3.100E-06 |
| 27250 | 1363 | T1363I | E | 2.4e-06 | 2.2e-06 | 2e-06 | 2.200E-06 | 2.000E-07 | 2.000E-06 | 2.400E-06 |
| 27251 | 1363 | T1363L | E | 1.8e-06 | 2e-06 | 1.9e-06 | 1.900E-06 | 1.000E-07 | 1.800E-06 | 2.000E-06 |
| 27252 | 1363 | T1363K | E | 1.3e-06 | 2.2e-06 | 2.8e-06 | 2.100E-06 | 7.550E-07 | 1.300E-06 | 2.800E-06 |
| 27253 | 1363 | T1363M | E | 2.2e-06 | 2.9e-06 | 2.3e-06 | 2.467E-06 | 3.786E-07 | 2.200E-06 | 2.900E-06 |
| 27254 | 1363 | T1363F | E | 1.9e-06 | 1.9e-06 | 2.5e-06 | 2.100E-06 | 3.464E-07 | 1.900E-06 | 2.500E-06 |
| 27255 | 1363 | T1363P | E | 2.8e-06 | 1.8e-06 | 2.2e-06 | 2.267E-06 | 5.033E-07 | 1.800E-06 | 2.800E-06 |
| 27256 | 1363 | T1363S | E | 3e-06 | 2.3e-06 | 2.4e-06 | 2.567E-06 | 3.786E-07 | 2.300E-06 | 3.000E-06 |
| 27257 | 1363 | T1363T | E | 1.8e-06 | 3.5e-06 | 3.9e-06 | 3.067E-06 | 1.115E-06 | 1.800E-06 | 3.900E-06 |
| 27258 | 1363 | T1363W | E | 1.4e-06 | 2.3e-06 | 2.7e-06 | 2.133E-06 | 6.658E-07 | 1.400E-06 | 2.700E-06 |
| 27259 | 1363 | T1363Y | E | 2.8e-06 | 2.2e-06 | 2.2e-06 | 2.400E-06 | 3.464E-07 | 2.200E-06 | 2.800E-06 |
| 27260 | 1363 | T1363V | E | 2.7e-06 | 3.4e-06 | 2.5e-06 | 2.867E-06 | 4.726E-07 | 2.500E-06 | 3.400E-06 |
| 27261 | 1364 | A1364A | E | 2.6e-06 | 3.1e-06 | 2.2e-06 | 2.633E-06 | 4.509E-07 | 2.200E-06 | 3.100E-06 |
| 27262 | 1364 | A1364R | E | 2.5e-06 | 2e-06 | 2e-06 | 2.167E-06 | 2.887E-07 | 2.000E-06 | 2.500E-06 |
| 27263 | 1364 | A1364N | E | 2.3e-06 | 2.3e-06 | 3.1e-06 | 2.567E-06 | 4.619E-07 | 2.300E-06 | 3.100E-06 |
| 27264 | 1364 | A1364D | E | 1.4e-06 | 1.9e-06 | 2.4e-06 | 1.900E-06 | 5.000E-07 | 1.400E-06 | 2.400E-06 |
| 27265 | 1364 | A1364C | E | 1.8e-06 | 2.3e-06 | 2.2e-06 | 2.100E-06 | 2.646E-07 | 1.800E-06 | 2.300E-06 |
| 27266 | 1364 | A1364E | E | 3.6e-06 | 1.9e-06 | 2.2e-06 | 2.567E-06 | 9.074E-07 | 1.900E-06 | 3.600E-06 |
| 27267 | 1364 | A1364Q | E | 3.2e-06 | 2.2e-06 | 2.1e-06 | 2.500E-06 | 6.083E-07 | 2.100E-06 | 3.200E-06 |
| 27268 | 1364 | A1364G | E | 2.9e-06 | 1.8e-06 | 1.7e-06 | 2.133E-06 | 6.658E-07 | 1.700E-06 | 2.900E-06 |
| 27269 | 1364 | A1364H | E | 3.5e-06 | 4.2e-06 | 1.9e-06 | 3.200E-06 | 1.179E-06 | 1.900E-06 | 4.200E-06 |
| 27270 | 1364 | A1364I | E | 2.3e-06 | 9e-07 | 1.1e-06 | 1.433E-06 | 7.572E-07 | 9.000E-07 | 2.300E-06 |
| 27271 | 1364 | A1364L | E | 3.4e-06 | 2.2e-06 | 2.7e-06 | 2.767E-06 | 6.028E-07 | 2.200E-06 | 3.400E-06 |
| 27272 | 1364 | A1364K | E | 2.3e-06 | 2.2e-06 | 2.1e-06 | 2.200E-06 | 1.000E-07 | 2.100E-06 | 2.300E-06 |
| 27273 | 1364 | A1364M | E | 2.7e-06 | 2.7e-06 | 1.8e-06 | 2.400E-06 | 5.196E-07 | 1.800E-06 | 2.700E-06 |
| 27274 | 1364 | A1364F | E | 2.9e-06 | 1.2e-06 | 2.2e-06 | 2.100E-06 | 8.544E-07 | 1.200E-06 | 2.900E-06 |
| 27275 | 1364 | A1364P | E | 3.8e-06 | 3.3e-06 | 2.4e-06 | 3.167E-06 | 7.095E-07 | 2.400E-06 | 3.800E-06 |
| 27276 | 1364 | A1364S | E | 2.4e-06 | 2.6e-06 | 1.6e-06 | 2.200E-06 | 5.292E-07 | 1.600E-06 | 2.600E-06 |
| 27277 | 1364 | A1364T | E | 2.3e-06 | 2.9e-06 | 2.1e-06 | 2.433E-06 | 4.163E-07 | 2.100E-06 | 2.900E-06 |
| 27278 | 1364 | A1364W | E | 2.1e-06 | 3.4e-06 | 1.4e-06 | 2.300E-06 | 1.015E-06 | 1.400E-06 | 3.400E-06 |
| 27279 | 1364 | A1364Y | E | 3.2e-06 | 1.8e-06 | 3.1e-06 | 2.700E-06 | 7.810E-07 | 1.800E-06 | 3.200E-06 |
| 27280 | 1364 | A1364V | E | 3e-06 | 1.9e-06 | 2.1e-06 | 2.333E-06 | 5.859E-07 | 1.900E-06 | 3.000E-06 |
| 27281 | 1365 | V1365A | E | 2e-06 | 1.3e-06 | 3.1e-06 | 2.133E-06 | 9.074E-07 | 1.300E-06 | 3.100E-06 |
| 27282 | 1365 | V1365R | E | 3.5e-06 | 3.4e-06 | 4.2e-06 | 3.700E-06 | 4.359E-07 | 3.400E-06 | 4.200E-06 |
| 27283 | 1365 | V1365N | E | 1.5e-06 | 1.7e-06 | 1.9e-06 | 1.700E-06 | 2.000E-07 | 1.500E-06 | 1.900E-06 |
| 27284 | 1365 | V1365D | E | 3.8e-06 | 1.8e-06 | 2.8e-06 | 2.800E-06 | 1.000E-06 | 1.800E-06 | 3.800E-06 |

| | | | | | | | | | | |
|---|---|---|---|---|---|---|---|---|---|---|
| 27285 | 1365 | V1365C | E | 1.4e-06 | 2.7e-06 | 3.4e-06 | 2.500E-06 | 1.015E-06 | 1.400E-06 | 3.400E-06 |
| 27286 | 1365 | V1365E | E | 2.2e-06 | 2.5e-06 | 2e-06 | 2.233E-06 | 2.517E-07 | 2.000E-06 | 2.500E-06 |
| 27287 | 1365 | V1365Q | E | 2.1e-06 | 1.5e-06 | 2.5e-06 | 2.033E-06 | 5.033E-07 | 1.500E-06 | 2.500E-06 |
| 27288 | 1365 | V1365G | E | 1.8e-06 | 2e-06 | 1.7e-06 | 1.833E-06 | 1.528E-07 | 1.700E-06 | 2.000E-06 |
| 27289 | 1365 | V1365H | E | 1.1e-06 | 1.4e-06 | 1.4e-06 | 1.300E-06 | 1.732E-07 | 1.100E-06 | 1.400E-06 |
| 27290 | 1365 | V1365I | E | 1.2e-06 | 1.8e-06 | 2.6e-06 | 1.867E-06 | 7.024E-07 | 1.200E-06 | 2.600E-06 |
| 27291 | 1365 | V1365L | E | 2.8e-06 | 2.2e-06 | 1.2e-06 | 2.067E-06 | 8.083E-07 | 1.200E-06 | 2.800E-06 |
| 27292 | 1365 | V1365K | E | 3.5e-06 | 3.7e-06 | 2.2e-06 | 3.133E-06 | 8.145E-07 | 2.200E-06 | 3.700E-06 |
| 27293 | 1365 | V1365M | E | 2.6e-06 | 2.1e-06 | 1.2e-06 | 1.967E-06 | 7.095E-07 | 1.200E-06 | 2.600E-06 |
| 27294 | 1365 | V1365F | E | 2.6e-06 | 4.3e-06 | 2.3e-06 | 3.067E-06 | 1.079E-06 | 2.300E-06 | 4.300E-06 |
| 27295 | 1365 | V1365P | E | 2.5e-06 | 2.1e-06 | 1.7e-06 | 2.100E-06 | 4.000E-07 | 1.700E-06 | 2.500E-06 |
| 27296 | 1365 | V1365S | E | 3.1e-06 | 2.1e-06 | 2e-06 | 2.400E-06 | 6.083E-07 | 2.000E-06 | 3.100E-06 |
| 27297 | 1365 | V1365T | E | 2.5e-06 | 2.1e-06 | 1.5e-06 | 2.033E-06 | 5.033E-07 | 1.500E-06 | 2.500E-06 |
| 27298 | 1365 | V1365W | E | 1.4e-06 | 2.5e-06 | 2.3e-06 | 2.067E-06 | 5.859E-07 | 1.400E-06 | 2.500E-06 |
| 27299 | 1365 | V1365Y | E | 3e-06 | 1.8e-06 | 1.4e-06 | 2.067E-06 | 8.327E-07 | 1.400E-06 | 3.000E-06 |
| 27300 | 1365 | V1365V | E | 1.9e-06 | 1.8e-06 | 2.6e-06 | 2.100E-06 | 4.359E-07 | 1.800E-06 | 2.600E-06 |
| 27301 | 1366 | Y1366A | E | 3.4e-06 | 3.1e-06 | 1.5e-06 | 2.667E-06 | 1.021E-06 | 1.500E-06 | 3.400E-06 |
| 27302 | 1366 | Y1366R | E | 2.1e-06 | 3.3e-06 | 1.9e-06 | 2.433E-06 | 7.572E-07 | 1.900E-06 | 3.300E-06 |
| 27303 | 1366 | Y1366N | E | 1.2e-06 | 2.1e-06 | 2.7e-06 | 2.000E-06 | 7.550E-07 | 1.200E-06 | 2.700E-06 |
| 27304 | 1366 | Y1366D | E | 2.3e-06 | 3.4e-06 | 2.5e-06 | 2.733E-06 | 5.859E-07 | 2.300E-06 | 3.400E-06 |
| 27305 | 1366 | Y1366C | E | 2.4e-06 | 2.9e-06 | 2.8e-06 | 2.700E-06 | 2.646E-07 | 2.400E-06 | 2.900E-06 |
| 27306 | 1366 | Y1366E | E | 2.2e-06 | 4.2e-06 | 1.9e-06 | 2.767E-06 | 1.250E-06 | 1.900E-06 | 4.200E-06 |
| 27307 | 1366 | Y1366Q | E | 2.6e-06 | 3.9e-06 | 2.3e-06 | 2.933E-06 | 8.505E-07 | 2.300E-06 | 3.900E-06 |
| 27308 | 1366 | Y1366G | E | 3.4e-06 | 2.7e-06 | 3e-06 | 3.033E-06 | 3.512E-07 | 2.700E-06 | 3.400E-06 |
| 27309 | 1366 | Y1366H | E | 2.1e-06 | 2e-06 | 2.5e-06 | 2.200E-06 | 2.646E-07 | 2.000E-06 | 2.500E-06 |
| 27310 | 1366 | Y1366I | E | 3.3e-06 | 4.6e-06 | 2.5e-06 | 3.467E-06 | 1.060E-06 | 2.500E-06 | 4.600E-06 |
| 27311 | 1366 | Y1366L | E | 3.1e-06 | 1.9e-06 | 3.2e-06 | 2.733E-06 | 7.234E-07 | 1.900E-06 | 3.200E-06 |
| 27312 | 1366 | Y1366K | E | 2.4e-06 | 1.6e-06 | 3.9e-06 | 2.633E-06 | 1.168E-06 | 1.600E-06 | 3.900E-06 |
| 27313 | 1366 | Y1366M | E | 1.5e-06 | 3.5e-06 | 2.5e-06 | 2.500E-06 | 1.000E-06 | 1.500E-06 | 3.500E-06 |
| 27314 | 1366 | Y1366F | E | 3.6e-06 | 2.3e-06 | 1.1e-06 | 2.333E-06 | 1.250E-06 | 1.100E-06 | 3.600E-06 |
| 27315 | 1366 | Y1366P | E | 2.7e-06 | 3.7e-06 | 2.9e-06 | 3.100E-06 | 5.292E-07 | 2.700E-06 | 3.700E-06 |
| 27316 | 1366 | Y1366S | E | 2.4e-06 | 3.4e-06 | 2.4e-06 | 2.733E-06 | 5.774E-07 | 2.400E-06 | 3.400E-06 |
| 27317 | 1366 | Y1366T | E | 2.8e-06 | 2.3e-06 | 3.3e-06 | 2.800E-06 | 5.000E-07 | 2.300E-06 | 3.300E-06 |
| 27318 | 1366 | Y1366W | E | 1.6e-06 | 2.1e-06 | 2e-06 | 1.900E-06 | 2.646E-07 | 1.600E-06 | 2.100E-06 |
| 27319 | 1366 | Y1366Y | E | 1.6e-06 | 1.7e-06 | 3.8e-06 | 2.367E-06 | 1.242E-06 | 1.600E-06 | 3.800E-06 |
| 27320 | 1366 | Y1366V | E | 2.4e-06 | 5.3e-06 | 2.7e-06 | 3.467E-06 | 1.595E-06 | 2.400E-06 | 5.300E-06 |
| 27321 | 1367 | Y1367A | E | 1.9e-06 | 2.6e-06 | 1.5e-06 | 2.000E-06 | 5.568E-07 | 1.500E-06 | 2.600E-06 |
| 27322 | 1367 | Y1367R | E | 2.2e-06 | 2.2e-06 | 2.4e-06 | 2.267E-06 | 1.155E-07 | 2.200E-06 | 2.400E-06 |
| 27323 | 1367 | Y1367N | E | 2.2e-06 | 1.9e-06 | 2.6e-06 | 2.233E-06 | 3.512E-07 | 1.900E-06 | 2.600E-06 |
| 27324 | 1367 | Y1367D | E | 2.4e-06 | 2.3e-06 | 2.4e-06 | 2.367E-06 | 5.774E-08 | 2.300E-06 | 2.400E-06 |
| 27325 | 1367 | Y1367C | E | 2.2e-06 | 4e-06 | 3e-06 | 3.067E-06 | 9.018E-07 | 2.200E-06 | 4.000E-06 |
| 27326 | 1367 | Y1367E | E | 2.7e-06 | 1.9e-06 | 1.6e-06 | 2.067E-06 | 5.686E-07 | 1.600E-06 | 2.700E-06 |
| 27327 | 1367 | Y1367Q | E | 2.3e-06 | 1.8e-06 | 2e-06 | 2.033E-06 | 2.517E-07 | 1.800E-06 | 2.300E-06 |
| 27328 | 1367 | Y1367G | E | 2.2e-06 | 2.3e-06 | 1.6e-06 | 2.033E-06 | 3.786E-07 | 1.600E-06 | 2.300E-06 |
| 27329 | 1367 | Y1367H | E | 2.5e-06 | 2.1e-06 | 5.2e-06 | 3.267E-06 | 1.686E-06 | 2.100E-06 | 5.200E-06 |
| 27330 | 1367 | Y1367I | E | 1.4e-06 | 4.6e-06 | 3e-06 | 3.000E-06 | 1.600E-06 | 1.400E-06 | 4.600E-06 |
| 27331 | 1367 | Y1367L | E | 1.5e-06 | 2.8e-06 | 1.6e-06 | 1.967E-06 | 7.234E-07 | 1.500E-06 | 2.800E-06 |
| 27332 | 1367 | Y1367K | E | 1.4e-06 | 2.9e-06 | 4.4e-06 | 2.900E-06 | 1.500E-06 | 1.400E-06 | 4.400E-06 |
| 27333 | 1367 | Y1367M | E | 3.2e-06 | 1.3e-06 | 3.8e-06 | 2.767E-06 | 1.305E-06 | 1.300E-06 | 3.800E-06 |
| 27334 | 1367 | Y1367F | E | 2.5e-06 | 3.2e-06 | 2.6e-06 | 2.767E-06 | 3.786E-07 | 2.500E-06 | 3.200E-06 |
| 27335 | 1367 | Y1367P | E | 2e-06 | 2.6e-06 | 2.2e-06 | 2.267E-06 | 3.055E-07 | 2.000E-06 | 2.600E-06 |
| 27336 | 1367 | Y1367S | E | 2.9e-06 | 2.9e-06 | 2e-06 | 2.600E-06 | 5.196E-07 | 2.000E-06 | 2.900E-06 |
| 27337 | 1367 | Y1367T | E | 2.1e-06 | 2.9e-06 | 2.2e-06 | 2.400E-06 | 4.359E-07 | 2.100E-06 | 2.900E-06 |
| 27338 | 1367 | Y1367W | E | 3.4e-06 | 2.4e-06 | 2.3e-06 | 2.700E-06 | 6.083E-07 | 2.300E-06 | 3.400E-06 |
| 27339 | 1367 | Y1367Y | E | 2.3e-06 | 2.8e-06 | 1.8e-06 | 2.300E-06 | 5.000E-07 | 1.800E-06 | 2.800E-06 |
| 27340 | 1367 | Y1367V | E | 1.7e-06 | 2.9e-06 | 2e-06 | 2.200E-06 | 6.245E-07 | 1.700E-06 | 2.900E-06 |
| 27341 | 1368 | C1368A | E | 2.6e-06 | 2.2e-06 | 3.1e-06 | 2.633E-06 | 4.509E-07 | 2.200E-06 | 3.100E-06 |
| 27342 | 1368 | C1368R | E | 2.1e-06 | 1.3e-06 | 2.6e-06 | 2.000E-06 | 6.557E-07 | 1.300E-06 | 2.600E-06 |
| 27343 | 1368 | C1368N | E | 3.5e-06 | 2.1e-06 | 2.5e-06 | 2.700E-06 | 7.211E-07 | 2.100E-06 | 3.500E-06 |
| 27344 | 1368 | C1368D | E | 2e-06 | 2.6e-06 | 2.5e-06 | 2.367E-06 | 3.215E-07 | 2.000E-06 | 2.600E-06 |
| 27345 | 1368 | C1368C | E | 1.7e-06 | 3.9e-06 | 2.7e-06 | 2.767E-06 | 1.102E-06 | 1.700E-06 | 3.900E-06 |
| 27346 | 1368 | C1368E | E | 3.1e-06 | 2.9e-06 | 3.2e-06 | 3.067E-06 | 1.528E-07 | 2.900E-06 | 3.200E-06 |
| 27347 | 1368 | C1368Q | E | 1.5e-06 | 2.2e-06 | 2.9e-06 | 2.200E-06 | 7.000E-07 | 1.500E-06 | 2.900E-06 |
| 27348 | 1368 | C1368G | E | 2.1e-06 | 2.2e-06 | 2.5e-06 | 2.267E-06 | 2.082E-07 | 2.100E-06 | 2.500E-06 |
| 27349 | 1368 | C1368H | E | 4.5e-06 | 3.2e-06 | 2.2e-06 | 3.300E-06 | 1.153E-06 | 2.200E-06 | 4.500E-06 |
| 27350 | 1368 | C1368I | E | 2.2e-06 | 2.4e-06 | 2.6e-06 | 2.400E-06 | 2.000E-07 | 2.200E-06 | 2.600E-06 |
| 27351 | 1368 | C1368L | E | 3.3e-06 | 2.6e-06 | 1.9e-06 | 2.600E-06 | 7.000E-07 | 1.900E-06 | 3.300E-06 |
| 27352 | 1368 | C1368K | E | 3e-06 | 4e-06 | 2.3e-06 | 3.100E-06 | 8.544E-07 | 2.300E-06 | 4.000E-06 |
| 27353 | 1368 | C1368M | E | 1.6e-06 | 2.6e-06 | 3e-06 | 2.400E-06 | 7.211E-07 | 1.600E-06 | 3.000E-06 |
| 27354 | 1368 | C1368F | E | 2.5e-06 | 2e-06 | 2.6e-06 | 2.367E-06 | 3.215E-07 | 2.000E-06 | 2.600E-06 |
| 27355 | 1368 | C1368P | E | 3.8e-06 | 1.8e-06 | 2.6e-06 | 2.733E-06 | 1.007E-06 | 1.800E-06 | 3.800E-06 |
| 27356 | 1368 | C1368S | E | 1.6e-06 | 4.2e-06 | 2e-06 | 2.600E-06 | 1.400E-06 | 1.600E-06 | 4.200E-06 |
| 27357 | 1368 | C1368T | E | 1.9e-06 | 2.9e-06 | 1.7e-06 | 2.167E-06 | 6.429E-07 | 1.700E-06 | 2.900E-06 |
| 27358 | 1368 | C1368W | E | 1.9e-06 | 2.4e-06 | 2.2e-06 | 2.167E-06 | 2.517E-07 | 1.900E-06 | 2.400E-06 |
| 27359 | 1368 | C1368Y | E | 1.6e-06 | 2.2e-06 | 2.6e-06 | 2.133E-06 | 5.033E-07 | 1.600E-06 | 2.600E-06 |
| 27360 | 1368 | C1368V | E | 1.8e-06 | 2.5e-06 | 3.1e-06 | 2.467E-06 | 6.506E-07 | 1.800E-06 | 3.100E-06 |
| 27361 | 1369 | S1369A | E | 2.3e-06 | 2.1e-06 | 1.4e-06 | 1.933E-06 | 4.726E-07 | 1.400E-06 | 2.300E-06 |
| 27362 | 1369 | S1369R | E | 2.8e-06 | 3.9e-06 | 2.2e-06 | 2.967E-06 | 8.622E-07 | 2.200E-06 | 3.900E-06 |
| 27363 | 1369 | S1369N | E | 3.5e-06 | 1.9e-06 | 2.7e-06 | 2.700E-06 | 8.000E-07 | 1.900E-06 | 3.500E-06 |
| 27364 | 1369 | S1369D | E | 2.5e-06 | 2.1e-06 | 1.4e-06 | 2.000E-06 | 5.568E-07 | 1.400E-06 | 2.500E-06 |
| 27365 | 1369 | S1369C | E | 2.2e-06 | 1.9e-06 | 2.3e-06 | 2.133E-06 | 2.082E-07 | 1.900E-06 | 2.300E-06 |
| 27366 | 1369 | S1369E | E | 3.3e-06 | 1.5e-06 | 1.7e-06 | 2.167E-06 | 9.866E-07 | 1.500E-06 | 3.300E-06 |
| 27367 | 1369 | S1369Q | E | 2.4e-06 | 2e-06 | 1.8e-06 | 2.067E-06 | 3.055E-07 | 1.800E-06 | 2.400E-06 |
| 27368 | 1369 | S1369G | E | 3.2e-06 | 1.3e-06 | 2.7e-06 | 2.400E-06 | 9.849E-07 | 1.300E-06 | 3.200E-06 |
| 27369 | 1369 | S1369H | E | 2e-06 | 2.6e-06 | 3e-06 | 2.533E-06 | 5.033E-07 | 2.000E-06 | 3.000E-06 |

| | | | | | | | | | | |
|---|---|---|---|---|---|---|---|---|---|---|
| 27370 | 1369 | S1369I | E | 3.7e-06 | 1.9e-06 | 2.2e-06 | 2.600E-06 | 9.644E-07 | 1.900E-06 | 3.700E-06 |
| 27371 | 1369 | S1369L | E | 2.6e-06 | 2.6e-06 | 3e-06 | 2.733E-06 | 2.309E-07 | 2.600E-06 | 3.000E-06 |
| 27372 | 1369 | S1369K | E | 1.7e-06 | 1.9e-06 | 2.6e-06 | 2.067E-06 | 4.726E-07 | 1.700E-06 | 2.600E-06 |
| 27373 | 1369 | S1369M | E | 1.7e-06 | 1.7e-06 | 2.1e-06 | 1.833E-06 | 2.309E-07 | 1.700E-06 | 2.100E-06 |
| 27374 | 1369 | S1369F | E | 3.7e-06 | 3e-06 | 9.4e-07 | 2.547E-06 | 1.435E-06 | 9.400E-07 | 3.700E-06 |
| 27375 | 1369 | S1369P | E | 1.2e-06 | 1.9e-06 | 1.8e-06 | 1.633E-06 | 3.786E-07 | 1.200E-06 | 1.900E-06 |
| 27376 | 1369 | S1369S | E | 2.2e-06 | 2.2e-06 | 1.5e-06 | 1.967E-06 | 4.041E-07 | 1.500E-06 | 2.200E-06 |
| 27377 | 1369 | S1369T | E | 2.4e-06 | 1.8e-06 | 2.2e-06 | 2.133E-06 | 3.055E-07 | 1.800E-06 | 2.400E-06 |
| 27378 | 1369 | S1369W | E | 2.4e-06 | 1.9e-06 | 2e-06 | 2.100E-06 | 2.646E-07 | 1.900E-06 | 2.400E-06 |
| 27379 | 1369 | S1369Y | E | 2.8e-06 | 1.6e-06 | 1.4e-06 | 1.933E-06 | 7.572E-07 | 1.400E-06 | 2.800E-06 |
| 27380 | 1369 | S1369V | E | 2.1e-06 | 2e-06 | 2.8e-06 | 2.300E-06 | 4.359E-07 | 2.000E-06 | 2.800E-06 |
| 27381 | 1370 | R1370A | E | 2.9e-06 | 3.8e-06 | 3.6e-06 | 3.433E-06 | 4.726E-07 | 2.900E-06 | 3.800E-06 |
| 27382 | 1370 | R1370R | E | 2.6e-06 | 2.9e-06 | 3.4e-06 | 2.967E-06 | 4.041E-07 | 2.600E-06 | 3.400E-06 |
| 27383 | 1370 | R1370N | E | 2.1e-06 | 3.5e-06 | 3.7e-06 | 3.100E-06 | 8.718E-07 | 2.100E-06 | 3.700E-06 |
| 27384 | 1370 | R1370D | E | 4.3e-06 | 2.9e-06 | 6e-06 | 4.400E-06 | 1.552E-06 | 2.900E-06 | 6.000E-06 |
| 27385 | 1370 | R1370C | E | 2.2e-06 | 3.8e-06 | 4.3e-06 | 3.433E-06 | 1.097E-06 | 2.200E-06 | 4.300E-06 |
| 27386 | 1370 | R1370E | E | 3.7e-06 | 1.4e-06 | 2.5e-06 | 2.533E-06 | 1.150E-06 | 1.400E-06 | 3.700E-06 |
| 27387 | 1370 | R1370Q | E | 2.5e-06 | 1.3e-06 | 2e-06 | 1.933E-06 | 6.028E-07 | 1.300E-06 | 2.500E-06 |
| 27388 | 1370 | R1370G | E | 2.3e-06 | 4.8e-06 | 2.8e-06 | 3.300E-06 | 1.323E-06 | 2.300E-06 | 4.800E-06 |
| 27389 | 1370 | R1370H | E | 3.7e-06 | 1.6e-06 | 4e-06 | 3.100E-06 | 1.308E-06 | 1.600E-06 | 4.000E-06 |
| 27390 | 1370 | R1370I | E | 3.4e-06 | 2.9e-06 | 3.3e-06 | 3.200E-06 | 2.646E-07 | 2.900E-06 | 3.400E-06 |
| 27391 | 1370 | R1370L | E | 2.3e-06 | 3.1e-06 | 5.4e-06 | 3.600E-06 | 1.609E-06 | 2.300E-06 | 5.400E-06 |
| 27392 | 1370 | R1370K | E | 9.9e-07 | 3.2e-06 | 2.4e-06 | 2.197E-06 | 1.119E-06 | 9.900E-07 | 3.200E-06 |
| 27393 | 1370 | R1370M | E | 2.4e-06 | 2.4e-06 | 3.9e-06 | 2.900E-06 | 8.660E-07 | 2.400E-06 | 3.900E-06 |
| 27394 | 1370 | R1370F | E | 2.8e-06 | 1.1e-06 | 2.7e-06 | 2.200E-06 | 9.539E-07 | 1.100E-06 | 2.800E-06 |
| 27395 | 1370 | R1370P | E | 6.6e-06 | 2.8e-06 | 3.2e-06 | 4.200E-06 | 2.088E-06 | 2.800E-06 | 6.600E-06 |
| 27396 | 1370 | R1370S | E | 3e-06 | 3.1e-06 | 2.5e-06 | 2.867E-06 | 3.215E-07 | 2.500E-06 | 3.100E-06 |
| 27397 | 1370 | R1370T | E | 4.6e-06 | 3.5e-06 | 3.4e-06 | 3.833E-06 | 6.658E-07 | 3.400E-06 | 4.600E-06 |
| 27398 | 1370 | R1370W | E | 2.8e-06 | 1.5e-06 | 1.9e-06 | 2.067E-06 | 6.658E-07 | 1.500E-06 | 2.800E-06 |
| 27399 | 1370 | R1370Y | E | 2.6e-06 | 2.7e-06 | 2.5e-06 | 2.600E-06 | 1.000E-07 | 2.500E-06 | 2.700E-06 |
| 27400 | 1370 | R1370V | E | 2.8e-06 | 4.6e-06 | 1.5e-06 | 2.967E-06 | 1.557E-06 | 1.500E-06 | 4.600E-06 |
| 27401 | 1371 | W1371A | E | 2.9e-06 | 3.4e-06 | 2e-06 | 2.767E-06 | 7.095E-07 | 2.000E-06 | 3.400E-06 |
| 27402 | 1371 | W1371R | E | 8.6e-07 | 1.9e-06 | 3.2e-06 | 1.987E-06 | 1.172E-06 | 8.600E-07 | 3.200E-06 |
| 27403 | 1371 | W1371N | E | 1.4e-06 | 1.7e-06 | 3.8e-06 | 2.300E-06 | 1.308E-06 | 1.400E-06 | 3.800E-06 |
| 27404 | 1371 | W1371D | E | 2.6e-06 | 1.6e-06 | 2.4e-06 | 2.200E-06 | 5.292E-07 | 1.600E-06 | 2.600E-06 |
| 27405 | 1371 | W1371C | E | 2.5e-06 | 2.6e-06 | 2.6e-06 | 2.567E-06 | 5.774E-08 | 2.500E-06 | 2.600E-06 |
| 27406 | 1371 | W1371E | E | 1.8e-06 | 2.6e-06 | 1.5e-06 | 1.967E-06 | 5.686E-07 | 1.500E-06 | 2.600E-06 |
| 27407 | 1371 | W1371Q | E | 2.7e-06 | 1.3e-06 | 1.8e-06 | 1.933E-06 | 7.095E-07 | 1.300E-06 | 2.700E-06 |
| 27408 | 1371 | W1371G | E | 2.6e-06 | 3.3e-06 | 3e-06 | 2.967E-06 | 3.512E-07 | 2.600E-06 | 3.300E-06 |
| 27409 | 1371 | W1371H | E | 2.4e-06 | 2e-06 | 3.6e-06 | 2.667E-06 | 8.327E-07 | 2.000E-06 | 3.600E-06 |
| 27410 | 1371 | W1371I | E | 2.7e-06 | 2.1e-06 | 2e-06 | 2.267E-06 | 3.786E-07 | 2.000E-06 | 2.700E-06 |
| 27411 | 1371 | W1371L | E | 1.5e-06 | 2.8e-06 | 2.7e-06 | 2.333E-06 | 7.234E-07 | 1.500E-06 | 2.800E-06 |
| 27412 | 1371 | W1371K | E | 2.2e-06 | 2e-06 | 1.9e-06 | 2.033E-06 | 1.528E-07 | 1.900E-06 | 2.200E-06 |
| 27413 | 1371 | W1371M | E | 2.8e-06 | 2.8e-06 | 2.3e-06 | 2.633E-06 | 2.887E-07 | 2.300E-06 | 2.800E-06 |
| 27414 | 1371 | W1371F | E | 4.2e-06 | 2.2e-06 | 3e-06 | 3.133E-06 | 1.007E-06 | 2.200E-06 | 4.200E-06 |
| 27415 | 1371 | W1371P | E | 2e-06 | 2.8e-06 | 1.8e-06 | 2.200E-06 | 5.292E-07 | 1.800E-06 | 2.800E-06 |
| 27416 | 1371 | W1371S | E | 2.3e-06 | 2.5e-06 | 3.6e-06 | 2.800E-06 | 7.000E-07 | 2.300E-06 | 3.600E-06 |
| 27417 | 1371 | W1371T | E | 1.8e-06 | 3.3e-06 | 1.3e-06 | 2.133E-06 | 1.041E-06 | 1.300E-06 | 3.300E-06 |
| 27418 | 1371 | W1371W | E | 1.7e-06 | 1.7e-06 | 2.7e-06 | 2.033E-06 | 5.774E-07 | 1.700E-06 | 2.700E-06 |
| 27419 | 1371 | W1371Y | E | 2.6e-06 | 3.9e-06 | 2.6e-06 | 3.033E-06 | 7.506E-07 | 2.600E-06 | 3.900E-06 |
| 27420 | 1371 | W1371V | E | 2.3e-06 | 2.9e-06 | 2.3e-06 | 2.500E-06 | 3.464E-07 | 2.300E-06 | 2.900E-06 |
| 27421 | 1372 | G1372A | E | 4.4e-06 | 3e-06 | 1.2e-06 | 2.867E-06 | 1.604E-06 | 1.200E-06 | 4.400E-06 |
| 27422 | 1372 | G1372R | E | 1.7e-06 | 2.3e-06 | 3.3e-06 | 2.433E-06 | 8.083E-07 | 1.700E-06 | 3.300E-06 |
| 27423 | 1372 | G1372N | E | 2e-06 | 1.3e-06 | 2.1e-06 | 1.800E-06 | 4.359E-07 | 1.300E-06 | 2.100E-06 |
| 27424 | 1372 | G1372D | E | 1.1e-06 | 1.3e-06 | 3.4e-06 | 1.933E-06 | 1.274E-06 | 1.100E-06 | 3.400E-06 |
| 27425 | 1372 | G1372C | E | 1.8e-06 | 2.5e-06 | 2.8e-06 | 2.367E-06 | 5.132E-07 | 1.800E-06 | 2.800E-06 |
| 27426 | 1372 | G1372E | E | 1.6e-06 | 1.8e-06 | 1.5e-06 | 1.633E-06 | 1.528E-07 | 1.500E-06 | 1.800E-06 |
| 27427 | 1372 | G1372Q | E | 1e-06 | 1.4e-06 | 1.3e-06 | 1.233E-06 | 2.082E-07 | 1.000E-06 | 1.400E-06 |
| 27428 | 1372 | G1372G | E | 1.6e-06 | 5.6e-06 | 4.1e-06 | 3.767E-06 | 2.021E-06 | 1.600E-06 | 5.600E-06 |
| 27429 | 1372 | G1372H | E | 1.1e-06 | 2e-06 | 1.2e-06 | 1.433E-06 | 4.933E-07 | 1.100E-06 | 2.000E-06 |
| 27430 | 1372 | G1372I | E | 1.9e-06 | 3.6e-06 | 2.1e-06 | 2.533E-06 | 9.292E-07 | 1.900E-06 | 3.600E-06 |
| 27431 | 1372 | G1372L | E | 2.3e-06 | 2.8e-06 | 3.6e-06 | 2.900E-06 | 6.557E-07 | 2.300E-06 | 3.600E-06 |
| 27432 | 1372 | G1372K | E | 2.4e-06 | 2.2e-06 | 1.6e-06 | 2.067E-06 | 4.163E-07 | 1.600E-06 | 2.400E-06 |
| 27433 | 1372 | G1372M | E | 1.7e-06 | 2.7e-06 | 2.3e-06 | 2.233E-06 | 5.033E-07 | 1.700E-06 | 2.700E-06 |
| 27434 | 1372 | G1372F | E | 1.8e-06 | 2.2e-06 | 1.9e-06 | 1.967E-06 | 2.082E-07 | 1.800E-06 | 2.200E-06 |
| 27435 | 1372 | G1372P | E | 2.1e-06 | 9.5e-07 | 1.4e-06 | 1.483E-06 | 5.795E-07 | 9.500E-07 | 2.100E-06 |
| 27436 | 1372 | G1372S | E | 1.6e-06 | 1.2e-06 | 2.1e-06 | 1.633E-06 | 4.509E-07 | 1.200E-06 | 2.100E-06 |
| 27437 | 1372 | G1372T | E | 8.6e-07 | 1.3e-06 | 1.8e-06 | 1.320E-06 | 4.703E-07 | 8.600E-07 | 1.800E-06 |
| 27438 | 1372 | G1372W | E | 9.5e-07 | 2.5e-06 | 2.8e-06 | 2.083E-06 | 9.929E-07 | 9.500E-07 | 2.800E-06 |
| 27439 | 1372 | G1372Y | E | 1.5e-06 | 1.9e-06 | 1.5e-06 | 1.633E-06 | 2.309E-07 | 1.500E-06 | 1.900E-06 |
| 27440 | 1372 | G1372V | E | 2.6e-06 | 2.5e-06 | 2.2e-06 | 2.433E-06 | 2.082E-07 | 2.200E-06 | 2.600E-06 |
| 27441 | 1373 | G1373A | E | 3.1e-06 | 2.7e-06 | 3.9e-06 | 3.233E-06 | 6.110E-07 | 2.700E-06 | 3.900E-06 |
| 27442 | 1373 | G1373R | E | 1.4e-06 | 1.1e-06 | 1.6e-06 | 1.367E-06 | 2.517E-07 | 1.100E-06 | 1.600E-06 |
| 27443 | 1373 | G1373N | E | 6.9e-07 | 5.8e-07 | 8.4e-07 | 7.033E-07 | 1.305E-07 | 5.800E-07 | 8.400E-07 |
| 27444 | 1373 | G1373D | E | 9.2e-07 | 8.3e-07 | 2.5e-06 | 1.417E-06 | 9.393E-07 | 8.300E-07 | 2.500E-06 |
| 27445 | 1373 | G1373C | E | 2.2e-06 | 3.1e-06 | 2.5e-06 | 2.600E-06 | 4.583E-07 | 2.200E-06 | 3.100E-06 |
| 27446 | 1373 | G1373E | E | 1.4e-06 | 8.7e-07 | 1.1e-06 | 1.123E-06 | 2.658E-07 | 8.700E-07 | 1.400E-06 |
| 27447 | 1373 | G1373Q | E | 6.2e-07 | 4.3e-07 | 3.7e-07 | 4.733E-07 | 1.305E-07 | 3.700E-07 | 6.200E-07 |
| 27448 | 1373 | G1373G | E | 1.7e-06 | 2.2e-06 | 2e-06 | 1.967E-06 | 2.517E-07 | 1.700E-06 | 2.200E-06 |
| 27449 | 1373 | G1373H | E | 7.8e-07 | 7.3e-07 | 7.2e-07 | 7.433E-07 | 3.215E-08 | 7.200E-07 | 7.800E-07 |
| 27450 | 1373 | G1373I | E | 2.6e-06 | 5.5e-06 | 1.9e-06 | 3.333E-06 | 1.909E-06 | 1.900E-06 | 5.500E-06 |
| 27451 | 1373 | G1373L | E | 8.9e-07 | 2.8e-06 | 1.7e-06 | 1.797E-06 | 9.587E-07 | 8.900E-07 | 2.800E-06 |
| 27452 | 1373 | G1373K | E | 1.2e-06 | 1.1e-06 | 2.5e-06 | 1.600E-06 | 7.810E-07 | 1.100E-06 | 2.500E-06 |
| 27453 | 1373 | G1373M | E | 2.6e-06 | 2.2e-06 | 1.8e-06 | 2.200E-06 | 4.000E-07 | 1.800E-06 | 2.600E-06 |
| 27454 | 1373 | G1373F | E | 1.1e-06 | 2.5e-06 | 9.4e-07 | 1.513E-06 | 8.582E-07 | 9.400E-07 | 2.500E-06 |

| | | | | | | | | | | |
|---|---|---|---|---|---|---|---|---|---|---|
| 27455 | 1373 | G1373P | E | 2.5e-06 | 2.6e-06 | 2.2e-06 | 2.433E-06 | 2.082E-07 | 2.200E-06 | 2.600E-06 |
| 27456 | 1373 | G1373S | E | 4.8e-07 | 9e-07 | 6.9e-07 | 6.900E-07 | 2.100E-07 | 4.800E-07 | 9.000E-07 |
| 27457 | 1373 | G1373T | E | 6.3e-07 | 2.6e-07 | 1e-06 | 6.300E-07 | 3.700E-07 | 2.600E-07 | 1.000E-06 |
| 27458 | 1373 | G1373W | E | 1.7e-06 | 1.2e-06 | 1.3e-06 | 1.400E-06 | 2.646E-07 | 1.200E-06 | 1.700E-06 |
| 27459 | 1373 | G1373Y | E | 3e-06 | 1.4e-06 | 1.1e-06 | 1.833E-06 | 1.021E-06 | 1.100E-06 | 3.000E-06 |
| 27460 | 1373 | G1373V | E | 2.6e-06 | 1.8e-06 | 1.5e-06 | 1.967E-06 | 5.686E-07 | 1.500E-06 | 2.600E-06 |
| 27461 | 1374 | D1374A | E | 2.1e-06 | 5.8e-06 | 2.9e-06 | 3.600E-06 | 1.947E-06 | 2.100E-06 | 5.800E-06 |
| 27462 | 1374 | D1374R | E | 8.4e-07 | 2.5e-06 | 1.3e-06 | 1.547E-06 | 8.570E-07 | 8.400E-07 | 2.500E-06 |
| 27463 | 1374 | D1374N | E | 3.5e-06 | 2.7e-06 | 3.2e-06 | 3.133E-06 | 4.041E-07 | 2.700E-06 | 3.500E-06 |
| 27464 | 1374 | D1374D | E | 1.8e-06 | 2.4e-06 | 2.6e-06 | 2.267E-06 | 4.163E-07 | 1.800E-06 | 2.600E-06 |
| 27465 | 1374 | D1374C | E | 2.1e-06 | 2.1e-06 | 2.8e-06 | 2.333E-06 | 4.041E-07 | 2.100E-06 | 2.800E-06 |
| 27466 | 1374 | D1374E | E | 2.6e-06 | 1.3e-06 | 1.8e-06 | 1.900E-06 | 6.557E-07 | 1.300E-06 | 2.600E-06 |
| 27467 | 1374 | D1374Q | E | 1.1e-06 | 1.1e-06 | 1.4e-06 | 1.200E-06 | 1.732E-07 | 1.100E-06 | 1.400E-06 |
| 27468 | 1374 | D1374G | E | 2.6e-06 | 4.7e-06 | 1.9e-06 | 3.067E-06 | 1.457E-06 | 1.900E-06 | 4.700E-06 |
| 27469 | 1374 | D1374H | E | 1.1e-06 | 2.9e-06 | 2.3e-06 | 2.100E-06 | 9.165E-07 | 1.100E-06 | 2.900E-06 |
| 27470 | 1374 | D1374I | E | 4.4e-06 | 3.7e-06 | 4e-06 | 4.033E-06 | 3.512E-07 | 3.700E-06 | 4.400E-06 |
| 27471 | 1374 | D1374L | E | 2.5e-06 | 2.6e-06 | 5.3e-06 | 3.467E-06 | 1.589E-06 | 2.500E-06 | 5.300E-06 |
| 27472 | 1374 | D1374K | E | 2.8e-06 | 2.6e-06 | 1.2e-06 | 2.200E-06 | 8.718E-07 | 1.200E-06 | 2.800E-06 |
| 27473 | 1374 | D1374M | E | 1.9e-06 | 3e-06 | 3.2e-06 | 2.700E-06 | 7.000E-07 | 1.900E-06 | 3.200E-06 |
| 27474 | 1374 | D1374F | E | 2e-06 | 4.3e-06 | 2.4e-06 | 2.900E-06 | 1.229E-06 | 2.000E-06 | 4.300E-06 |
| 27475 | 1374 | D1374P | E | 1.2e-06 | 3.9e-06 | 3.5e-06 | 2.867E-06 | 1.457E-06 | 1.200E-06 | 3.900E-06 |
| 27476 | 1374 | D1374S | E | 2.3e-06 | 2.2e-06 | 2.7e-06 | 2.400E-06 | 2.646E-07 | 2.200E-06 | 2.700E-06 |
| 27477 | 1374 | D1374T | E | 2e-06 | 2.2e-06 | 2.6e-06 | 2.267E-06 | 3.055E-07 | 2.000E-06 | 2.600E-06 |
| 27478 | 1374 | D1374W | E | 2.1e-06 | 3.3e-06 | 2.7e-06 | 2.700E-06 | 6.000E-07 | 2.100E-06 | 3.300E-06 |
| 27479 | 1374 | D1374Y | E | 3.7e-06 | 7.4e-06 | 2e-06 | 4.367E-06 | 2.761E-06 | 2.000E-06 | 7.400E-06 |
| 27480 | 1374 | D1374V | E | 3.2e-06 | 4e-06 | 3.7e-06 | 3.633E-06 | 4.041E-07 | 3.200E-06 | 4.000E-06 |
| 27481 | 1375 | G1375A | E | 2.4e-06 | 2.8e-06 | 3e-06 | 2.733E-06 | 3.055E-07 | 2.400E-06 | 3.000E-06 |
| 27482 | 1375 | G1375R | E | 1.4e-06 | 8.9e-07 | 1e-06 | 1.097E-06 | 2.684E-07 | 8.900E-07 | 1.400E-06 |
| 27483 | 1375 | G1375N | E | 5.3e-07 | 1.3e-06 | 9.9e-07 | 9.400E-07 | 3.874E-07 | 5.300E-07 | 1.300E-06 |
| 27484 | 1375 | G1375D | E | 2.7e-06 | 1.9e-06 | 1.6e-06 | 2.067E-06 | 5.686E-07 | 1.600E-06 | 2.700E-06 |
| 27485 | 1375 | G1375C | E | 1.1e-06 | 1.3e-06 | 3.6e-06 | 2.000E-06 | 1.389E-06 | 1.100E-06 | 3.600E-06 |
| 27486 | 1375 | G1375E | E | 1.7e-06 | 1.1e-06 | 1.5e-06 | 1.433E-06 | 3.055E-07 | 1.100E-06 | 1.700E-06 |
| 27487 | 1375 | G1375Q | E | 6.9e-07 | 3.7e-07 | 7.9e-07 | 6.167E-07 | 2.194E-07 | 3.700E-07 | 7.900E-07 |
| 27488 | 1375 | G1375G | E | 1.1e-06 | 1.9e-06 | 1.8e-06 | 1.600E-06 | 4.359E-07 | 1.100E-06 | 1.900E-06 |
| 27489 | 1375 | G1375H | E | 9.6e-07 | 1.3e-06 | 8.3e-07 | 1.030E-06 | 2.427E-07 | 8.300E-07 | 1.300E-06 |
| 27490 | 1375 | G1375I | E | 1.6e-06 | 2.5e-06 | 2.6e-06 | 2.233E-06 | 5.508E-07 | 1.600E-06 | 2.600E-06 |
| 27491 | 1375 | G1375L | E | 2.3e-06 | 4.2e-06 | 1.4e-06 | 2.633E-06 | 1.429E-06 | 1.400E-06 | 4.200E-06 |
| 27492 | 1375 | G1375K | E | 1.7e-06 | 1.4e-06 | 8.6e-07 | 1.320E-06 | 4.257E-07 | 8.600E-07 | 1.700E-06 |
| 27493 | 1375 | G1375M | E | 3.3e-06 | 2.6e-06 | 1.8e-06 | 2.567E-06 | 7.506E-07 | 1.800E-06 | 3.300E-06 |
| 27494 | 1375 | G1375F | E | 2e-06 | 1e-06 | 1.2e-06 | 1.400E-06 | 5.292E-07 | 1.000E-06 | 2.000E-06 |
| 27495 | 1375 | G1375P | E | 4.3e-06 | 4.2e-06 | 2.6e-06 | 3.700E-06 | 9.539E-07 | 2.600E-06 | 4.300E-06 |
| 27496 | 1375 | G1375S | E | 7.7e-07 | 7.8e-07 | 6.4e-07 | 7.300E-07 | 7.810E-08 | 6.400E-07 | 7.800E-07 |
| 27497 | 1375 | G1375T | E | 8.3e-07 | 8.6e-07 | 5.9e-07 | 7.600E-07 | 1.480E-07 | 5.900E-07 | 8.600E-07 |
| 27498 | 1375 | G1375W | E | 2e-06 | 3.6e-06 | 2.8e-06 | 2.800E-06 | 8.000E-07 | 2.000E-06 | 3.600E-06 |
| 27499 | 1375 | G1375Y | E | 2.1e-06 | 3.1e-06 | 2.7e-06 | 2.633E-06 | 5.033E-07 | 2.100E-06 | 3.100E-06 |
| 27500 | 1375 | G1375V | E | 2.6e-06 | 3.6e-06 | 2.6e-06 | 2.933E-06 | 5.774E-07 | 2.600E-06 | 3.600E-06 |
| 27501 | 1376 | F1376A | E | 1.9e-06 | 2e-06 | 3.3e-06 | 2.400E-06 | 7.810E-07 | 1.900E-06 | 3.300E-06 |
| 27502 | 1376 | F1376R | E | 1.7e-06 | 2.6e-06 | 1.6e-06 | 1.967E-06 | 5.508E-07 | 1.600E-06 | 2.600E-06 |
| 27503 | 1376 | F1376N | E | 1.9e-06 | 1.8e-06 | 1.4e-06 | 1.700E-06 | 2.646E-07 | 1.400E-06 | 1.900E-06 |
| 27504 | 1376 | F1376D | E | 2e-06 | 4e-06 | 4.2e-06 | 3.400E-06 | 1.217E-06 | 2.000E-06 | 4.200E-06 |
| 27505 | 1376 | F1376C | E | 2.3e-06 | 2.2e-06 | 3.9e-06 | 2.800E-06 | 9.539E-07 | 2.200E-06 | 3.900E-06 |
| 27506 | 1376 | F1376E | E | 2.7e-06 | 2.1e-06 | 1.5e-06 | 2.100E-06 | 6.000E-07 | 1.500E-06 | 2.700E-06 |
| 27507 | 1376 | F1376Q | E | 1.1e-06 | 6.8e-07 | 1.6e-06 | 1.127E-06 | 4.606E-07 | 6.800E-07 | 1.600E-06 |
| 27508 | 1376 | F1376G | E | 2e-06 | 1.1e-06 | 2.1e-06 | 1.733E-06 | 5.508E-07 | 1.100E-06 | 2.100E-06 |
| 27509 | 1376 | F1376H | E | 2.7e-06 | 2.1e-06 | 2.4e-06 | 2.400E-06 | 3.000E-07 | 2.100E-06 | 2.700E-06 |
| 27510 | 1376 | F1376I | E | 2.8e-06 | 2.6e-06 | 2.5e-06 | 2.633E-06 | 1.528E-07 | 2.500E-06 | 2.800E-06 |
| 27511 | 1376 | F1376L | E | 1.4e-06 | 2e-06 | 2.8e-06 | 2.067E-06 | 7.024E-07 | 1.400E-06 | 2.800E-06 |
| 27512 | 1376 | F1376K | E | 1.6e-06 | 1.4e-06 | 2.3e-06 | 1.767E-06 | 4.726E-07 | 1.400E-06 | 2.300E-06 |
| 27513 | 1376 | F1376M | E | 2.8e-06 | 4.1e-06 | 1.5e-06 | 2.800E-06 | 1.300E-06 | 1.500E-06 | 4.100E-06 |
| 27514 | 1376 | F1376F | E | 4.5e-06 | 1.6e-06 | 2.7e-06 | 2.933E-06 | 1.464E-06 | 1.600E-06 | 4.500E-06 |
| 27515 | 1376 | F1376P | E | 2.2e-06 | 2e-06 | 1.6e-06 | 1.933E-06 | 3.055E-07 | 1.600E-06 | 2.200E-06 |
| 27516 | 1376 | F1376S | E | 2.7e-06 | 1.3e-06 | 2.1e-06 | 2.033E-06 | 7.024E-07 | 1.300E-06 | 2.700E-06 |
| 27517 | 1376 | F1376T | E | 3.9e-06 | 2.1e-06 | 4.7e-06 | 3.567E-06 | 1.332E-06 | 2.100E-06 | 4.700E-06 |
| 27518 | 1376 | F1376W | E | 2.5e-06 | 2.5e-06 | 2.4e-06 | 2.467E-06 | 5.774E-08 | 2.400E-06 | 2.500E-06 |
| 27519 | 1376 | F1376Y | E | 2.2e-06 | 2.5e-06 | 2.8e-06 | 2.500E-06 | 3.000E-07 | 2.200E-06 | 2.800E-06 |
| 27520 | 1376 | F1376V | E | 3.2e-06 | 2.5e-06 | 2.6e-06 | 2.767E-06 | 3.786E-07 | 2.500E-06 | 3.200E-06 |
| 27521 | 1377 | Y1377A | E | 2.6e-06 | 3.9e-06 | 2e-06 | 2.833E-06 | 9.713E-07 | 2.000E-06 | 3.900E-06 |
| 27522 | 1377 | Y1377R | E | 1.7e-06 | 1.3e-06 | 1.4e-06 | 1.467E-06 | 2.082E-07 | 1.300E-06 | 1.700E-06 |
| 27523 | 1377 | Y1377N | E | 1.8e-06 | 9.9e-07 | 9e-07 | 1.230E-06 | 4.957E-07 | 9.000E-07 | 1.800E-06 |
| 27524 | 1377 | Y1377D | E | 1.9e-06 | 2.2e-06 | 2.8e-06 | 2.300E-06 | 4.583E-07 | 1.900E-06 | 2.800E-06 |
| 27525 | 1377 | Y1377C | E | 1.9e-06 | 2e-06 | 1.8e-06 | 1.900E-06 | 1.000E-07 | 1.800E-06 | 2.000E-06 |
| 27526 | 1377 | Y1377E | E | 2.7e-06 | 2.2e-06 | 1e-06 | 1.967E-06 | 8.737E-07 | 1.000E-06 | 2.700E-06 |
| 27527 | 1377 | Y1377Q | E | 1.6e-06 | 1.3e-06 | 1.6e-06 | 1.500E-06 | 1.732E-07 | 1.300E-06 | 1.600E-06 |
| 27528 | 1377 | Y1377G | E | 3.5e-06 | 2.5e-06 | 4.9e-06 | 3.633E-06 | 1.206E-06 | 2.500E-06 | 4.900E-06 |
| 27529 | 1377 | Y1377H | E | 1.1e-06 | 9.1e-07 | 1.4e-06 | 1.137E-06 | 2.470E-07 | 9.100E-07 | 1.400E-06 |
| 27530 | 1377 | Y1377I | E | 1e-06 | 2.2e-06 | 1.8e-06 | 1.667E-06 | 6.110E-07 | 1.000E-06 | 2.200E-06 |
| 27531 | 1377 | Y1377L | E | 1.9e-06 | 3.8e-06 | 2.9e-06 | 2.867E-06 | 9.504E-07 | 1.900E-06 | 3.800E-06 |
| 27532 | 1377 | Y1377K | E | 2.2e-06 | 1.1e-06 | 1.4e-06 | 1.567E-06 | 5.686E-07 | 1.100E-06 | 2.200E-06 |
| 27533 | 1377 | Y1377M | E | 2.7e-06 | 3e-06 | 2.7e-06 | 2.800E-06 | 1.732E-07 | 2.700E-06 | 3.000E-06 |
| 27534 | 1377 | Y1377F | E | 2.5e-06 | 2.5e-06 | 3.1e-06 | 2.700E-06 | 3.464E-07 | 2.500E-06 | 3.100E-06 |
| 27535 | 1377 | Y1377P | E | 1.7e-06 | 4.1e-06 | 2.5e-06 | 2.767E-06 | 1.222E-06 | 1.700E-06 | 4.100E-06 |
| 27536 | 1377 | Y1377S | E | 1.7e-06 | 8e-07 | 1.5e-06 | 1.333E-06 | 4.726E-07 | 8.000E-07 | 1.700E-06 |
| 27537 | 1377 | Y1377T | E | 6e-07 | 1.3e-06 | 1e-06 | 9.667E-07 | 3.512E-07 | 6.000E-07 | 1.300E-06 |
| 27538 | 1377 | Y1377W | E | 1.9e-06 | 3.2e-06 | 9.4e-07 | 2.013E-06 | 1.134E-06 | 9.400E-07 | 3.200E-06 |
| 27539 | 1377 | Y1377Y | E | 2.4e-06 | 3.8e-06 | 2.5e-06 | 2.900E-06 | 7.810E-07 | 2.400E-06 | 3.800E-06 |

| | | | | | | | | | | |
|---|---|---|---|---|---|---|---|---|---|---|
| 27540 | 1377 | Y1377V | E | 3.7e-06 | 2.8e-06 | 1.7e-06 | 2.733E-06 | 1.002E-06 | 1.700E-06 | 3.700E-06 |
| 27541 | 1378 | A1378A | E | 1.6e-06 | 3.6e-06 | 2.3e-06 | 2.500E-06 | 1.015E-06 | 1.600E-06 | 3.600E-06 |
| 27542 | 1378 | A1378R | E | 3.8e-06 | 2.8e-06 | 2.3e-06 | 2.967E-06 | 7.638E-07 | 2.300E-06 | 3.800E-06 |
| 27543 | 1378 | A1378N | E | 2.5e-06 | 2.9e-06 | 4.5e-06 | 3.300E-06 | 1.058E-06 | 2.500E-06 | 4.500E-06 |
| 27544 | 1378 | A1378D | E | 3e-06 | 2.6e-06 | 2.4e-06 | 2.667E-06 | 3.055E-07 | 2.400E-06 | 3.000E-06 |
| 27545 | 1378 | A1378C | E | 2.2e-06 | 3.8e-06 | 2e-06 | 2.667E-06 | 9.866E-07 | 2.000E-06 | 3.800E-06 |
| 27546 | 1378 | A1378E | E | 1.7e-06 | 2.5e-06 | 1.3e-06 | 1.833E-06 | 6.110E-07 | 1.300E-06 | 2.500E-06 |
| 27547 | 1378 | A1378Q | E | 2.3e-06 | 2.2e-06 | 2.4e-06 | 2.300E-06 | 1.000E-07 | 2.200E-06 | 2.400E-06 |
| 27548 | 1378 | A1378G | E | 2.7e-06 | 2.9e-06 | 2.4e-06 | 2.667E-06 | 2.517E-07 | 2.400E-06 | 2.900E-06 |
| 27549 | 1378 | A1378H | E | 2.5e-06 | 1.6e-06 | 2.1e-06 | 2.067E-06 | 4.509E-07 | 1.600E-06 | 2.500E-06 |
| 27550 | 1378 | A1378I | E | 2.6e-06 | 3.7e-06 | 3.5e-06 | 3.267E-06 | 5.859E-07 | 2.600E-06 | 3.700E-06 |
| 27551 | 1378 | A1378L | E | 2.1e-06 | 2.6e-06 | 2.7e-06 | 2.467E-06 | 3.215E-07 | 2.100E-06 | 2.700E-06 |
| 27552 | 1378 | A1378K | E | 2.5e-06 | 4.2e-06 | 1.8e-06 | 2.833E-06 | 1.234E-06 | 1.800E-06 | 4.200E-06 |
| 27553 | 1378 | A1378M | E | 1.5e-06 | 2e-06 | 1.8e-06 | 1.767E-06 | 2.517E-07 | 1.500E-06 | 2.000E-06 |
| 27554 | 1378 | A1378F | E | 3.8e-06 | 2e-06 | 1.7e-06 | 2.500E-06 | 1.136E-06 | 1.700E-06 | 3.800E-06 |
| 27555 | 1378 | A1378P | E | 2.3e-06 | 3e-06 | 2.1e-06 | 2.467E-06 | 4.726E-07 | 2.100E-06 | 3.000E-06 |
| 27556 | 1378 | A1378S | E | 2.6e-06 | 2.6e-06 | 1.6e-06 | 2.267E-06 | 5.774E-07 | 1.600E-06 | 2.600E-06 |
| 27557 | 1378 | A1378T | E | 2.3e-06 | 1.7e-06 | 1.8e-06 | 1.933E-06 | 3.215E-07 | 1.700E-06 | 2.300E-06 |
| 27558 | 1378 | A1378W | E | 2.5e-06 | 1.7e-06 | 5.9e-06 | 3.367E-06 | 2.230E-06 | 1.700E-06 | 5.900E-06 |
| 27559 | 1378 | A1378Y | E | 3.2e-06 | 2.9e-06 | 2e-06 | 2.700E-06 | 6.245E-07 | 2.000E-06 | 3.200E-06 |
| 27560 | 1378 | A1378V | E | 1.7e-06 | 2.7e-06 | 1.9e-06 | 2.100E-06 | 5.292E-07 | 1.700E-06 | 2.700E-06 |
| 27561 | 1379 | M1379A | E | 3.4e-06 | 3e-06 | 1.6e-06 | 2.667E-06 | 9.452E-07 | 1.600E-06 | 3.400E-06 |
| 27562 | 1379 | M1379R | E | 1.9e-06 | 1.9e-06 | 3.4e-06 | 2.400E-06 | 8.660E-07 | 1.900E-06 | 3.400E-06 |
| 27563 | 1379 | M1379N | E | 2.3e-06 | 2.4e-06 | 3.3e-06 | 2.667E-06 | 5.508E-07 | 2.300E-06 | 3.300E-06 |
| 27564 | 1379 | M1379D | E | 2.4e-06 | 2.6e-06 | 1.9e-06 | 2.300E-06 | 3.606E-07 | 1.900E-06 | 2.600E-06 |
| 27565 | 1379 | M1379C | E | 1.9e-06 | 2e-06 | 2.3e-06 | 2.067E-06 | 2.082E-07 | 1.900E-06 | 2.300E-06 |
| 27566 | 1379 | M1379E | E | 2.1e-06 | 3.3e-06 | 4.2e-06 | 3.200E-06 | 1.054E-06 | 2.100E-06 | 4.200E-06 |
| 27567 | 1379 | M1379Q | E | 1.8e-06 | 2.3e-06 | 3.2e-06 | 2.433E-06 | 7.095E-07 | 1.800E-06 | 3.200E-06 |
| 27568 | 1379 | M1379G | E | 2.1e-06 | 2.3e-06 | 1.8e-06 | 2.067E-06 | 2.517E-07 | 1.800E-06 | 2.300E-06 |
| 27569 | 1379 | M1379H | E | 2.8e-06 | 3e-06 | 3.4e-06 | 3.067E-06 | 3.055E-07 | 2.800E-06 | 3.400E-06 |
| 27570 | 1379 | M1379I | E | 2e-06 | 3.3e-06 | 2.4e-06 | 2.567E-06 | 6.658E-07 | 2.000E-06 | 3.300E-06 |
| 27571 | 1379 | M1379L | E | 1.6e-06 | 3.5e-06 | 2.3e-06 | 2.467E-06 | 9.609E-07 | 1.600E-06 | 3.500E-06 |
| 27572 | 1379 | M1379K | E | 1.7e-06 | 3.4e-06 | 3.5e-06 | 2.867E-06 | 1.012E-06 | 1.700E-06 | 3.500E-06 |
| 27573 | 1379 | M1379M | E | 2.6e-06 | 3.1e-06 | 2.3e-06 | 2.667E-06 | 4.041E-07 | 2.300E-06 | 3.100E-06 |
| 27574 | 1379 | M1379F | E | 1.9e-06 | 3.5e-06 | 2.6e-06 | 2.667E-06 | 8.021E-07 | 1.900E-06 | 3.500E-06 |
| 27575 | 1379 | M1379P | E | 2.4e-06 | 2e-06 | 2.6e-06 | 2.333E-06 | 3.055E-07 | 2.000E-06 | 2.600E-06 |
| 27576 | 1379 | M1379S | E | 3.7e-06 | 1.9e-06 | 1.7e-06 | 2.433E-06 | 1.102E-06 | 1.700E-06 | 3.700E-06 |
| 27577 | 1379 | M1379T | E | 2.1e-06 | 1.7e-06 | 2.6e-06 | 2.133E-06 | 4.509E-07 | 1.700E-06 | 2.600E-06 |
| 27578 | 1379 | M1379W | E | 4.1e-06 | 3.2e-06 | 2.8e-06 | 3.367E-06 | 6.658E-07 | 2.800E-06 | 4.100E-06 |
| 27579 | 1379 | M1379Y | E | 2.3e-06 | 1.8e-06 | 1.4e-06 | 1.833E-06 | 4.509E-07 | 1.400E-06 | 2.300E-06 |
| 27580 | 1379 | M1379V | E | 1.7e-06 | 2.1e-06 | 3.4e-06 | 2.400E-06 | 8.888E-07 | 1.700E-06 | 3.400E-06 |
| 27581 | 1380 | D1380A | E | 3.6e-06 | 1.6e-06 | 9.9e-07 | 2.063E-06 | 1.365E-06 | 9.900E-07 | 3.600E-06 |
| 27582 | 1380 | D1380R | E | 1e-06 | 2.9e-06 | 1.6e-06 | 1.833E-06 | 9.713E-07 | 1.000E-06 | 2.900E-06 |
| 27583 | 1380 | D1380N | E | 2e-06 | 1.7e-06 | 2.8e-06 | 2.167E-06 | 5.686E-07 | 1.700E-06 | 2.800E-06 |
| 27584 | 1380 | D1380D | E | 3.1e-06 | 2.3e-06 | 2.3e-06 | 2.567E-06 | 4.619E-07 | 2.300E-06 | 3.100E-06 |
| 27585 | 1380 | D1380C | E | 1.7e-06 | 2.3e-06 | 2.2e-06 | 2.067E-06 | 3.215E-07 | 1.700E-06 | 2.300E-06 |
| 27586 | 1380 | D1380E | E | 1.7e-06 | 3.1e-06 | 2.4e-06 | 2.400E-06 | 7.000E-07 | 1.700E-06 | 3.100E-06 |
| 27587 | 1380 | D1380Q | E | 1.9e-06 | 2.5e-06 | 1.7e-06 | 2.033E-06 | 4.163E-07 | 1.700E-06 | 2.500E-06 |
| 27588 | 1380 | D1380G | E | 1.2e-06 | 4.4e-06 | 2e-06 | 2.533E-06 | 1.665E-06 | 1.200E-06 | 4.400E-06 |
| 27589 | 1380 | D1380H | E | 2.2e-06 | 3.3e-06 | 1.8e-06 | 2.433E-06 | 7.767E-07 | 1.800E-06 | 3.300E-06 |
| 27590 | 1380 | D1380I | E | 4.1e-06 | 3e-06 | 1.9e-06 | 3.000E-06 | 1.100E-06 | 1.900E-06 | 4.100E-06 |
| 27591 | 1380 | D1380L | E | 2.9e-06 | 2.4e-06 | 1.6e-06 | 2.300E-06 | 6.557E-07 | 1.600E-06 | 2.900E-06 |
| 27592 | 1380 | D1380K | E | 2.7e-06 | 2.2e-06 | 1.6e-06 | 2.167E-06 | 5.508E-07 | 1.600E-06 | 2.700E-06 |
| 27593 | 1380 | D1380M | E | 1.9e-06 | 2.7e-06 | 2.9e-06 | 2.500E-06 | 5.292E-07 | 1.900E-06 | 2.900E-06 |
| 27594 | 1380 | D1380F | E | 3.4e-06 | 2.6e-06 | 2.3e-06 | 2.767E-06 | 5.686E-07 | 2.300E-06 | 3.400E-06 |
| 27595 | 1380 | D1380P | E | 1.9e-06 | 1.8e-06 | 2.2e-06 | 1.967E-06 | 2.082E-07 | 1.800E-06 | 2.200E-06 |
| 27596 | 1380 | D1380S | E | 1.7e-06 | 2.4e-06 | 2.6e-06 | 2.233E-06 | 4.726E-07 | 1.700E-06 | 2.600E-06 |
| 27597 | 1380 | D1380T | E | 2.6e-06 | 2e-06 | 3e-06 | 2.533E-06 | 5.033E-07 | 2.000E-06 | 3.000E-06 |
| 27598 | 1380 | D1380W | E | 4.3e-06 | 3.2e-06 | 4e-06 | 3.833E-06 | 5.686E-07 | 3.200E-06 | 4.300E-06 |
| 27599 | 1380 | D1380Y | E | 1.4e-06 | 1.6e-06 | 3.3e-06 | 2.100E-06 | 1.044E-06 | 1.400E-06 | 3.300E-06 |
| 27600 | 1380 | D1380V | E | 2.5e-06 | 2.9e-06 | 2.6e-06 | 2.667E-06 | 2.082E-07 | 2.500E-06 | 2.900E-06 |
| 27601 | 1381 | Y1381A | E | 1.9e-06 | 2.8e-06 | 1.4e-06 | 2.033E-06 | 7.095E-07 | 1.400E-06 | 2.800E-06 |
| 27602 | 1381 | Y1381R | E | 1.4e-06 | 2.4e-06 | 2.7e-06 | 2.167E-06 | 6.807E-07 | 1.400E-06 | 2.700E-06 |
| 27603 | 1381 | Y1381N | E | 2.3e-06 | 2.4e-06 | 1.9e-06 | 2.200E-06 | 2.646E-07 | 1.900E-06 | 2.400E-06 |
| 27604 | 1381 | Y1381D | E | 2.4e-06 | 2.9e-06 | 3.6e-06 | 2.967E-06 | 6.028E-07 | 2.400E-06 | 3.600E-06 |
| 27605 | 1381 | Y1381C | E | 2.3e-06 | 2.8e-06 | 3e-06 | 2.700E-06 | 3.606E-07 | 2.300E-06 | 3.000E-06 |
| 27606 | 1381 | Y1381E | E | 3.3e-06 | 4.6e-06 | 3.5e-06 | 3.800E-06 | 7.000E-07 | 3.300E-06 | 4.600E-06 |
| 27607 | 1381 | Y1381Q | E | 3.2e-06 | 2.6e-06 | 2.3e-06 | 2.700E-06 | 4.583E-07 | 2.300E-06 | 3.200E-06 |
| 27608 | 1381 | Y1381G | E | 1.8e-06 | 3.2e-06 | 2.7e-06 | 2.567E-06 | 7.095E-07 | 1.800E-06 | 3.200E-06 |
| 27609 | 1381 | Y1381H | E | 1.9e-06 | 1.6e-06 | 2.2e-06 | 1.900E-06 | 3.000E-07 | 1.600E-06 | 2.200E-06 |
| 27610 | 1381 | Y1381I | E | 1.7e-06 | 3.3e-06 | 2.4e-06 | 2.467E-06 | 8.021E-07 | 1.700E-06 | 3.300E-06 |
| 27611 | 1381 | Y1381L | E | 1.1e-06 | 1.9e-06 | 1.2e-06 | 1.400E-06 | 4.359E-07 | 1.100E-06 | 1.900E-06 |
| 27612 | 1381 | Y1381K | E | 1.7e-06 | 2.6e-06 | 2.3e-06 | 2.200E-06 | 4.583E-07 | 1.700E-06 | 2.600E-06 |
| 27613 | 1381 | Y1381M | E | 3.4e-06 | 2.8e-06 | 3.1e-06 | 3.100E-06 | 3.000E-07 | 2.800E-06 | 3.400E-06 |
| 27614 | 1381 | Y1381F | E | 2.9e-06 | 4.2e-06 | 2.2e-06 | 3.100E-06 | 1.015E-06 | 2.200E-06 | 4.200E-06 |
| 27615 | 1381 | Y1381P | E | 1.5e-06 | 2.1e-06 | 1.7e-06 | 1.767E-06 | 3.055E-07 | 1.500E-06 | 2.100E-06 |
| 27616 | 1381 | Y1381S | E | 2.7e-06 | 1.6e-06 | 1.9e-06 | 2.067E-06 | 5.686E-07 | 1.600E-06 | 2.700E-06 |
| 27617 | 1381 | Y1381T | E | 2.9e-06 | 2.1e-06 | 1.7e-06 | 2.233E-06 | 6.110E-07 | 1.700E-06 | 2.900E-06 |
| 27618 | 1381 | Y1381W | E | 2e-06 | 2.9e-06 | 3.2e-06 | 2.700E-06 | 6.245E-07 | 2.000E-06 | 3.200E-06 |
| 27619 | 1381 | Y1381Y | E | 2.8e-06 | 2.5e-06 | 3.3e-06 | 2.867E-06 | 4.041E-07 | 2.500E-06 | 3.300E-06 |
| 27620 | 1381 | Y1381V | E | 3.4e-06 | 2.4e-06 | 2.3e-06 | 2.700E-06 | 6.083E-07 | 2.300E-06 | 3.400E-06 |
| 27621 | 1382 | W1382A | E | 2.5e-06 | 2e-06 | 1.8e-06 | 2.100E-06 | 3.606E-07 | 1.800E-06 | 2.500E-06 |
| 27622 | 1382 | W1382R | E | 1.1e-06 | 2e-06 | 2.4e-06 | 1.833E-06 | 6.658E-07 | 1.100E-06 | 2.400E-06 |
| 27623 | 1382 | W1382N | E | 2.1e-06 | 2.7e-06 | 2.8e-06 | 2.533E-06 | 3.786E-07 | 2.100E-06 | 2.800E-06 |
| 27624 | 1382 | W1382D | E | 1.8e-06 | 3.6e-06 | 2.2e-06 | 2.533E-06 | 9.452E-07 | 1.800E-06 | 3.600E-06 |

| | | | | | | | | | | |
|---|---|---|---|---|---|---|---|---|---|---|
| 27625 | 1382 | W1382C | E | 1.3e-06 | 2.1e-06 | 2.3e-06 | 1.900E-06 | 5.292E-07 | 1.300E-06 | 2.300E-06 |
| 27626 | 1382 | W1382E | E | 1.7e-06 | 2.4e-06 | 2e-06 | 2.033E-06 | 3.512E-07 | 1.700E-06 | 2.400E-06 |
| 27627 | 1382 | W1382Q | E | 1.4e-06 | 1.9e-06 | 1.6e-06 | 1.633E-06 | 2.517E-07 | 1.400E-06 | 1.900E-06 |
| 27628 | 1382 | W1382G | E | 2.6e-06 | 1.7e-06 | 1.8e-06 | 2.033E-06 | 4.933E-07 | 1.700E-06 | 2.600E-06 |
| 27629 | 1382 | W1382H | E | 2.7e-06 | 1.7e-06 | 2.3e-06 | 2.233E-06 | 5.033E-07 | 1.700E-06 | 2.700E-06 |
| 27630 | 1382 | W1382I | E | 3.3e-06 | 2.8e-06 | 3.9e-06 | 3.333E-06 | 5.508E-07 | 2.800E-06 | 3.900E-06 |
| 27631 | 1382 | W1382L | E | 1.1e-06 | 1.7e-06 | 1.9e-06 | 1.567E-06 | 4.163E-07 | 1.100E-06 | 1.900E-06 |
| 27632 | 1382 | W1382K | E | 1.5e-06 | 1.6e-06 | 1.1e-06 | 1.400E-06 | 2.646E-07 | 1.100E-06 | 1.600E-06 |
| 27633 | 1382 | W1382M | E | 2.1e-06 | 5.1e-06 | 2.2e-06 | 3.133E-06 | 1.704E-06 | 2.100E-06 | 5.100E-06 |
| 27634 | 1382 | W1382F | E | 2.4e-06 | 2.8e-06 | 2.3e-06 | 2.500E-06 | 2.646E-07 | 2.300E-06 | 2.800E-06 |
| 27635 | 1382 | W1382P | E | 2.1e-06 | 2.6e-06 | 2.1e-06 | 2.267E-06 | 2.887E-07 | 2.100E-06 | 2.600E-06 |
| 27636 | 1382 | W1382S | E | 4.8e-06 | 2e-06 | 3.4e-06 | 3.400E-06 | 1.400E-06 | 2.000E-06 | 4.800E-06 |
| 27637 | 1382 | W1382T | E | 1.9e-06 | 1.3e-06 | 1.7e-06 | 1.633E-06 | 3.055E-07 | 1.300E-06 | 1.900E-06 |
| 27638 | 1382 | W1382W | E | 1.8e-06 | 1.6e-06 | 2.2e-06 | 1.867E-06 | 3.055E-07 | 1.600E-06 | 2.200E-06 |
| 27639 | 1382 | W1382Y | E | 2.4e-06 | 2.8e-06 | 2.2e-06 | 2.467E-06 | 3.055E-07 | 2.200E-06 | 2.800E-06 |
| 27640 | 1382 | W1382V | E | 3.6e-06 | 1.9e-06 | 2.9e-06 | 2.800E-06 | 8.544E-07 | 1.900E-06 | 3.600E-06 |
| 27641 | 1383 | G1383A | E | 1.9e-06 | 1.8e-06 | 2.2e-06 | 1.967E-06 | 2.082E-07 | 1.800E-06 | 2.200E-06 |
| 27642 | 1383 | G1383R | E | 1.9e-06 | 2.2e-06 | 1.8e-06 | 1.967E-06 | 2.082E-07 | 1.800E-06 | 2.200E-06 |
| 27643 | 1383 | G1383N | E | 2.7e-06 | 2.7e-06 | 2.2e-06 | 2.533E-06 | 2.887E-07 | 2.200E-06 | 2.700E-06 |
| 27644 | 1383 | G1383D | E | 1.8e-06 | 2e-06 | 3.7e-06 | 2.500E-06 | 1.044E-06 | 1.800E-06 | 3.700E-06 |
| 27645 | 1383 | G1383C | E | 2.6e-06 | 1.3e-06 | 1.6e-06 | 1.833E-06 | 6.807E-07 | 1.300E-06 | 2.600E-06 |
| 27646 | 1383 | G1383E | E | 3.1e-06 | 1.5e-06 | 3.4e-06 | 2.667E-06 | 1.021E-06 | 1.500E-06 | 3.400E-06 |
| 27647 | 1383 | G1383Q | E | 3.8e-06 | 1.5e-06 | 2.9e-06 | 2.733E-06 | 1.159E-06 | 1.500E-06 | 3.800E-06 |
| 27648 | 1383 | G1383G | E | 2.3e-06 | 3.8e-06 | 1.8e-06 | 2.633E-06 | 1.041E-06 | 1.800E-06 | 3.800E-06 |
| 27649 | 1383 | G1383H | E | 1.7e-06 | 3.1e-06 | 1.7e-06 | 2.167E-06 | 8.083E-07 | 1.700E-06 | 3.100E-06 |
| 27650 | 1383 | G1383I | E | 9.1e-07 | 4.2e-06 | 1.9e-06 | 2.337E-06 | 1.688E-06 | 9.100E-07 | 4.200E-06 |
| 27651 | 1383 | G1383L | E | 2e-06 | 2.8e-06 | 3.6e-06 | 2.800E-06 | 8.000E-07 | 2.000E-06 | 3.600E-06 |
| 27652 | 1383 | G1383K | E | 2.6e-06 | 2e-06 | 2.4e-06 | 2.333E-06 | 3.055E-07 | 2.000E-06 | 2.600E-06 |
| 27653 | 1383 | G1383M | E | 2.9e-06 | 2.7e-06 | 3.4e-06 | 3.000E-06 | 3.606E-07 | 2.700E-06 | 3.400E-06 |
| 27654 | 1383 | G1383F | E | 2.2e-06 | 2.3e-06 | 3.4e-06 | 2.633E-06 | 6.658E-07 | 2.200E-06 | 3.400E-06 |
| 27655 | 1383 | G1383P | E | 8.5e-07 | 1.7e-06 | 2.2e-06 | 1.583E-06 | 6.825E-07 | 8.500E-07 | 2.200E-06 |
| 27656 | 1383 | G1383S | E | 3.4e-06 | 1.7e-06 | 2.7e-06 | 2.600E-06 | 8.544E-07 | 1.700E-06 | 3.400E-06 |
| 27657 | 1383 | G1383T | E | 3e-06 | 2.4e-06 | 2.4e-06 | 2.600E-06 | 3.464E-07 | 2.400E-06 | 3.000E-06 |
| 27658 | 1383 | G1383W | E | 2.2e-06 | 2.2e-06 | 2.2e-06 | 2.200E-06 | 0.000E+00 | 2.200E-06 | 2.200E-06 |
| 27659 | 1383 | G1383Y | E | 3.4e-06 | 3.9e-06 | 3.7e-06 | 3.667E-06 | 2.517E-07 | 3.400E-06 | 3.900E-06 |
| 27660 | 1383 | G1383V | E | 2.8e-06 | 2.3e-06 | 3.3e-06 | 2.800E-06 | 5.000E-07 | 2.300E-06 | 3.300E-06 |
| 27661 | 1384 | Q1384A | E | 2.9e-06 | 2.5e-06 | 2.1e-06 | 2.500E-06 | 4.000E-07 | 2.100E-06 | 2.900E-06 |
| 27662 | 1384 | Q1384R | E | 2.5e-06 | 1.3e-06 | 2e-06 | 1.933E-06 | 6.028E-07 | 1.300E-06 | 2.500E-06 |
| 27663 | 1384 | Q1384N | E | 1.6e-06 | 2.1e-06 | 3.2e-06 | 2.300E-06 | 8.185E-07 | 1.600E-06 | 3.200E-06 |
| 27664 | 1384 | Q1384D | E | 2.3e-06 | 3.2e-06 | 3e-06 | 2.833E-06 | 4.726E-07 | 2.300E-06 | 3.200E-06 |
| 27665 | 1384 | Q1384C | E | 2.7e-06 | 2.5e-06 | 2.4e-06 | 2.533E-06 | 1.528E-07 | 2.400E-06 | 2.700E-06 |
| 27666 | 1384 | Q1384E | E | 2.6e-06 | 1.1e-06 | 2.7e-06 | 2.133E-06 | 8.963E-07 | 1.100E-06 | 2.700E-06 |
| 27667 | 1384 | Q1384Q | E | 2.6e-06 | 2.4e-06 | 2.3e-06 | 2.433E-06 | 1.528E-07 | 2.300E-06 | 2.600E-06 |
| 27668 | 1384 | Q1384G | E | 2.9e-06 | 3.1e-06 | 2.3e-06 | 2.767E-06 | 4.163E-07 | 2.300E-06 | 3.100E-06 |
| 27669 | 1384 | Q1384H | E | 2.6e-06 | 2.3e-06 | 3.1e-06 | 2.667E-06 | 4.041E-07 | 2.300E-06 | 3.100E-06 |
| 27670 | 1384 | Q1384I | E | 3.1e-06 | 2.5e-06 | 2.8e-06 | 2.800E-06 | 3.000E-07 | 2.500E-06 | 3.100E-06 |
| 27671 | 1384 | Q1384L | E | 4.4e-06 | 2.8e-06 | 2.6e-06 | 3.267E-06 | 9.866E-07 | 2.600E-06 | 4.400E-06 |
| 27672 | 1384 | Q1384K | E | 1.8e-06 | 3.8e-06 | 3.2e-06 | 2.933E-06 | 1.026E-06 | 1.800E-06 | 3.800E-06 |
| 27673 | 1384 | Q1384M | E | 2.7e-06 | 2.6e-06 | 1.7e-06 | 2.333E-06 | 5.508E-07 | 1.700E-06 | 2.700E-06 |
| 27674 | 1384 | Q1384F | E | 2.3e-06 | 2.4e-06 | 2.7e-06 | 2.467E-06 | 2.082E-07 | 2.300E-06 | 2.700E-06 |
| 27675 | 1384 | Q1384P | E | 1.5e-06 | 2.6e-06 | 2.9e-06 | 2.333E-06 | 7.371E-07 | 1.500E-06 | 2.900E-06 |
| 27676 | 1384 | Q1384S | E | 2.1e-06 | 2.1e-06 | 2.7e-06 | 2.300E-06 | 3.464E-07 | 2.100E-06 | 2.700E-06 |
| 27677 | 1384 | Q1384T | E | 2e-06 | 1.1e-06 | 2.3e-06 | 1.800E-06 | 6.245E-07 | 1.100E-06 | 2.300E-06 |
| 27678 | 1384 | Q1384W | E | 2.6e-06 | 2.5e-06 | 2.7e-06 | 2.600E-06 | 1.000E-07 | 2.500E-06 | 2.700E-06 |
| 27679 | 1384 | Q1384Y | E | 3.1e-06 | 3e-06 | 2.1e-06 | 2.733E-06 | 5.508E-07 | 2.100E-06 | 3.100E-06 |
| 27680 | 1384 | Q1384V | E | 5.8e-06 | 2.8e-06 | 2.3e-06 | 3.633E-06 | 1.893E-06 | 2.300E-06 | 5.800E-06 |
| 27681 | 1385 | G1385A | E | 2e-06 | 2.2e-06 | 3.7e-06 | 2.633E-06 | 9.292E-07 | 2.000E-06 | 3.700E-06 |
| 27682 | 1385 | G1385R | E | 3.3e-06 | 2.7e-06 | 5.4e-06 | 3.800E-06 | 1.418E-06 | 2.700E-06 | 5.400E-06 |
| 27683 | 1385 | G1385N | E | 2.7e-06 | 2.1e-06 | 1.9e-06 | 2.233E-06 | 4.163E-07 | 1.900E-06 | 2.700E-06 |
| 27684 | 1385 | G1385D | E | 2.2e-06 | 2.3e-06 | 2e-06 | 2.167E-06 | 1.528E-07 | 2.000E-06 | 2.300E-06 |
| 27685 | 1385 | G1385C | E | 2.7e-06 | 1.3e-06 | 3.3e-06 | 2.433E-06 | 1.026E-06 | 1.300E-06 | 3.300E-06 |
| 27686 | 1385 | G1385E | E | 2.4e-06 | 2.3e-06 | 1.8e-06 | 2.167E-06 | 3.215E-07 | 1.800E-06 | 2.400E-06 |
| 27687 | 1385 | G1385Q | E | 3.3e-06 | 1.9e-06 | 1.9e-06 | 2.367E-06 | 8.083E-07 | 1.900E-06 | 3.300E-06 |
| 27688 | 1385 | G1385G | E | 2.2e-06 | 3.2e-06 | 2.9e-06 | 2.767E-06 | 5.132E-07 | 2.200E-06 | 3.200E-06 |
| 27689 | 1385 | G1385H | E | 1.9e-06 | 2.6e-06 | 2e-06 | 2.167E-06 | 3.786E-07 | 1.900E-06 | 2.600E-06 |
| 27690 | 1385 | G1385I | E | 3.5e-06 | 2.5e-06 | 1.4e-06 | 2.467E-06 | 1.050E-06 | 1.400E-06 | 3.500E-06 |
| 27691 | 1385 | G1385L | E | 2.2e-06 | 2.3e-06 | 3.2e-06 | 2.567E-06 | 5.508E-07 | 2.200E-06 | 3.200E-06 |
| 27692 | 1385 | G1385K | E | 4e-06 | 2.2e-06 | 2.6e-06 | 2.933E-06 | 9.452E-07 | 2.200E-06 | 4.000E-06 |
| 27693 | 1385 | G1385M | E | 2.1e-06 | 2.4e-06 | 1.9e-06 | 2.133E-06 | 2.517E-07 | 1.900E-06 | 2.400E-06 |
| 27694 | 1385 | G1385F | E | 2.6e-06 | 2.5e-06 | 2.4e-06 | 2.500E-06 | 1.000E-07 | 2.400E-06 | 2.600E-06 |
| 27695 | 1385 | G1385P | E | 2.4e-06 | 1.6e-06 | 2e-06 | 2.000E-06 | 4.000E-07 | 1.600E-06 | 2.400E-06 |
| 27696 | 1385 | G1385S | E | 4e-06 | 2.5e-06 | 2.7e-06 | 3.067E-06 | 8.145E-07 | 2.500E-06 | 4.000E-06 |
| 27697 | 1385 | G1385T | E | 2e-06 | 1.8e-06 | 3e-06 | 2.267E-06 | 6.429E-07 | 1.800E-06 | 3.000E-06 |
| 27698 | 1385 | G1385W | E | 1.3e-06 | 3.6e-06 | 3e-06 | 2.633E-06 | 1.193E-06 | 1.300E-06 | 3.600E-06 |
| 27699 | 1385 | G1385Y | E | 2.1e-06 | 3.8e-06 | 2.7e-06 | 2.867E-06 | 8.622E-07 | 2.100E-06 | 3.800E-06 |
| 27700 | 1385 | G1385V | E | 1.9e-06 | 1.8e-06 | 2.8e-06 | 2.167E-06 | 5.508E-07 | 1.800E-06 | 2.800E-06 |
| 27701 | 1386 | T1386A | E | 3.5e-06 | 2.4e-06 | 3.1e-06 | 3.000E-06 | 5.568E-07 | 2.400E-06 | 3.500E-06 |
| 27702 | 1386 | T1386R | E | 2.4e-06 | 3e-06 | 2.5e-06 | 2.633E-06 | 3.215E-07 | 2.400E-06 | 3.000E-06 |
| 27703 | 1386 | T1386N | E | 2.2e-06 | 7.2e-07 | 2.7e-06 | 1.873E-06 | 1.030E-06 | 7.200E-07 | 2.700E-06 |
| 27704 | 1386 | T1386D | E | 2e-06 | 3.2e-06 | 2.5e-06 | 2.567E-06 | 6.028E-07 | 2.000E-06 | 3.200E-06 |
| 27705 | 1386 | T1386C | E | 2.4e-06 | 2e-06 | 3.6e-06 | 2.667E-06 | 8.327E-07 | 2.000E-06 | 3.600E-06 |
| 27706 | 1386 | T1386E | E | 1.9e-06 | 1.4e-06 | 3.4e-06 | 2.233E-06 | 1.041E-06 | 1.400E-06 | 3.400E-06 |
| 27707 | 1386 | T1386Q | E | 4.1e-06 | 2.4e-06 | 5.6e-06 | 4.033E-06 | 1.601E-06 | 2.400E-06 | 5.600E-06 |
| 27708 | 1386 | T1386G | E | 3.4e-06 | 3.1e-06 | 1.3e-06 | 2.600E-06 | 1.136E-06 | 1.300E-06 | 3.400E-06 |
| 27709 | 1386 | T1386H | E | 2.7e-06 | 2.5e-06 | 3.3e-06 | 2.833E-06 | 4.163E-07 | 2.500E-06 | 3.300E-06 |

| | | | | | | | | | | |
|---|---|---|---|---|---|---|---|---|---|---|
| 27710 | 1386 | T1386I | E | 1.8e-06 | 2.9e-06 | 2.3e-06 | 2.333E-06 | 5.508E-07 | 1.800E-06 | 2.900E-06 |
| 27711 | 1386 | T1386L | E | 3e-06 | 2.3e-06 | 1.5e-06 | 2.267E-06 | 7.506E-07 | 1.500E-06 | 3.000E-06 |
| 27712 | 1386 | T1386K | E | 2.6e-06 | 1.5e-06 | 1.9e-06 | 2.000E-06 | 5.568E-07 | 1.500E-06 | 2.600E-06 |
| 27713 | 1386 | T1386M | E | 2.3e-06 | 2.5e-06 | 2.4e-06 | 2.400E-06 | 1.000E-07 | 2.300E-06 | 2.500E-06 |
| 27714 | 1386 | T1386F | E | 2.4e-06 | 4.3e-06 | 2.7e-06 | 3.133E-06 | 1.021E-06 | 2.400E-06 | 4.300E-06 |
| 27715 | 1386 | T1386P | E | 1.6e-06 | 1.9e-06 | 1.2e-06 | 1.567E-06 | 3.512E-07 | 1.200E-06 | 1.900E-06 |
| 27716 | 1386 | T1386S | E | 4.5e-06 | 2.7e-06 | 3e-06 | 3.400E-06 | 9.644E-07 | 2.700E-06 | 4.500E-06 |
| 27717 | 1386 | T1386T | E | 2.1e-06 | 2.1e-06 | 2.7e-06 | 2.300E-06 | 3.464E-07 | 2.100E-06 | 2.700E-06 |
| 27718 | 1386 | T1386W | E | 2e-06 | 4.6e-06 | 2.3e-06 | 2.967E-06 | 1.422E-06 | 2.000E-06 | 4.600E-06 |
| 27719 | 1386 | T1386Y | E | 2.6e-06 | 2.7e-06 | 2e-06 | 2.433E-06 | 3.786E-07 | 2.000E-06 | 2.700E-06 |
| 27720 | 1386 | T1386V | E | 2.4e-06 | 2.1e-06 | 2.1e-06 | 2.200E-06 | 1.732E-07 | 2.100E-06 | 2.400E-06 |
| 27721 | 1387 | L1387A | E | 2.2e-06 | 1.9e-06 | 2.5e-06 | 2.200E-06 | 3.000E-07 | 1.900E-06 | 2.500E-06 |
| 27722 | 1387 | L1387R | E | 2.5e-06 | 9.6e-07 | 1.7e-06 | 1.720E-06 | 7.702E-07 | 9.600E-07 | 2.500E-06 |
| 27723 | 1387 | L1387N | E | 2.3e-06 | 2.5e-06 | 3e-06 | 2.600E-06 | 3.606E-07 | 2.300E-06 | 3.000E-06 |
| 27724 | 1387 | L1387D | E | 2.1e-06 | 2.1e-06 | 3.6e-06 | 2.600E-06 | 8.660E-07 | 2.100E-06 | 3.600E-06 |
| 27725 | 1387 | L1387C | E | 2.1e-06 | 1.7e-06 | 2.4e-06 | 2.067E-06 | 3.512E-07 | 1.700E-06 | 2.400E-06 |
| 27726 | 1387 | L1387E | E | 1.9e-06 | 2.1e-06 | 2.2e-06 | 2.067E-06 | 1.528E-07 | 1.900E-06 | 2.200E-06 |
| 27727 | 1387 | L1387Q | E | 2.9e-06 | 2.5e-06 | 2.6e-06 | 2.667E-06 | 2.082E-07 | 2.500E-06 | 2.900E-06 |
| 27728 | 1387 | L1387G | E | 3.3e-06 | 2.5e-06 | 1.4e-06 | 2.400E-06 | 9.539E-07 | 1.400E-06 | 3.300E-06 |
| 27729 | 1387 | L1387H | E | 3.4e-06 | 2.7e-06 | 2e-06 | 2.700E-06 | 7.000E-07 | 2.000E-06 | 3.400E-06 |
| 27730 | 1387 | L1387I | E | 2.1e-06 | 2.7e-06 | 1.6e-06 | 2.133E-06 | 5.508E-07 | 1.600E-06 | 2.700E-06 |
| 27731 | 1387 | L1387L | E | 2.6e-06 | 2.1e-06 | 2.8e-06 | 2.500E-06 | 3.606E-07 | 2.100E-06 | 2.800E-06 |
| 27732 | 1387 | L1387K | E | 3e-06 | 2e-06 | 1.4e-06 | 2.133E-06 | 8.083E-07 | 1.400E-06 | 3.000E-06 |
| 27733 | 1387 | L1387M | E | 2.3e-06 | 3.1e-06 | 5.2e-06 | 3.533E-06 | 1.498E-06 | 2.300E-06 | 5.200E-06 |
| 27734 | 1387 | L1387F | E | 1.8e-06 | 1.5e-06 | 1.7e-06 | 1.667E-06 | 1.528E-07 | 1.500E-06 | 1.800E-06 |
| 27735 | 1387 | L1387P | E | 3.2e-06 | 1.8e-06 | 2e-06 | 2.333E-06 | 7.572E-07 | 1.800E-06 | 3.200E-06 |
| 27736 | 1387 | L1387S | E | 2.2e-06 | 2.7e-06 | 2.1e-06 | 2.333E-06 | 3.215E-07 | 2.100E-06 | 2.700E-06 |
| 27737 | 1387 | L1387T | E | 2.2e-06 | 2.9e-06 | 2.1e-06 | 2.400E-06 | 4.359E-07 | 2.100E-06 | 2.900E-06 |
| 27738 | 1387 | L1387W | E | 1.9e-06 | 3e-06 | 2.6e-06 | 2.500E-06 | 5.568E-07 | 1.900E-06 | 3.000E-06 |
| 27739 | 1387 | L1387Y | E | 2.5e-06 | 1e-06 | 2.1e-06 | 1.867E-06 | 7.767E-07 | 1.000E-06 | 2.500E-06 |
| 27740 | 1387 | L1387V | E | 2.6e-06 | 2.3e-06 | 1.7e-06 | 2.200E-06 | 4.583E-07 | 1.700E-06 | 2.600E-06 |
| 27741 | 1388 | V1388A | E | 3.6e-06 | 2.6e-06 | 1.2e-06 | 2.467E-06 | 1.206E-06 | 1.200E-06 | 3.600E-06 |
| 27742 | 1388 | V1388R | E | 2.8e-06 | 2.5e-06 | 2.5e-06 | 2.600E-06 | 1.732E-07 | 2.500E-06 | 2.800E-06 |
| 27743 | 1388 | V1388N | E | 1.4e-06 | 3e-06 | 2.2e-06 | 2.200E-06 | 8.000E-07 | 1.400E-06 | 3.000E-06 |
| 27744 | 1388 | V1388D | E | 3.1e-06 | 1.4e-06 | 1.9e-06 | 2.133E-06 | 8.737E-07 | 1.400E-06 | 3.100E-06 |
| 27745 | 1388 | V1388C | E | 1.8e-06 | 2.2e-06 | 2e-06 | 2.000E-06 | 2.000E-07 | 1.800E-06 | 2.200E-06 |
| 27746 | 1388 | V1388E | E | 1.6e-06 | 2e-06 | 2.3e-06 | 1.967E-06 | 3.512E-07 | 1.600E-06 | 2.300E-06 |
| 27747 | 1388 | V1388Q | E | 2.8e-06 | 2.8e-06 | 2.7e-06 | 2.767E-06 | 5.774E-08 | 2.700E-06 | 2.800E-06 |
| 27748 | 1388 | V1388G | E | 1.9e-06 | 2.6e-06 | 1.7e-06 | 2.067E-06 | 4.726E-07 | 1.700E-06 | 2.600E-06 |
| 27749 | 1388 | V1388H | E | 2.5e-06 | 2.5e-06 | 3.2e-06 | 2.733E-06 | 4.041E-07 | 2.500E-06 | 3.200E-06 |
| 27750 | 1388 | V1388I | E | 3.2e-06 | 2.3e-06 | 2e-06 | 2.500E-06 | 6.245E-07 | 2.000E-06 | 3.200E-06 |
| 27751 | 1388 | V1388L | E | 2.5e-06 | 1.8e-06 | 1.2e-06 | 1.833E-06 | 6.506E-07 | 1.200E-06 | 2.500E-06 |
| 27752 | 1388 | V1388K | E | 2e-06 | 1.9e-06 | 3e-06 | 2.300E-06 | 6.083E-07 | 1.900E-06 | 3.000E-06 |
| 27753 | 1388 | V1388M | E | 2.3e-06 | 1.5e-06 | 2.2e-06 | 2.000E-06 | 4.359E-07 | 1.500E-06 | 2.300E-06 |
| 27754 | 1388 | V1388F | E | 1.8e-06 | 1.6e-06 | 2.3e-06 | 1.900E-06 | 3.606E-07 | 1.600E-06 | 2.300E-06 |
| 27755 | 1388 | V1388P | E | 1.7e-06 | 1.8e-06 | 2.7e-06 | 2.067E-06 | 5.508E-07 | 1.700E-06 | 2.700E-06 |
| 27756 | 1388 | V1388S | E | 2.4e-06 | 2.2e-06 | 2.5e-06 | 2.367E-06 | 1.528E-07 | 2.200E-06 | 2.500E-06 |
| 27757 | 1388 | V1388T | E | 2e-06 | 2.4e-06 | 2.3e-06 | 2.233E-06 | 2.082E-07 | 2.000E-06 | 2.400E-06 |
| 27758 | 1388 | V1388W | E | 3e-06 | 1.8e-06 | 3e-06 | 2.600E-06 | 6.928E-07 | 1.800E-06 | 3.000E-06 |
| 27759 | 1388 | V1388Y | E | 4.7e-06 | 1e-06 | 2.3e-06 | 2.667E-06 | 1.877E-06 | 1.000E-06 | 4.700E-06 |
| 27760 | 1388 | V1388V | E | 2e-06 | 1.8e-06 | 2e-06 | 1.933E-06 | 1.155E-07 | 1.800E-06 | 2.000E-06 |
| 27761 | 1389 | T1389A | E | 2.4e-06 | 2.2e-06 | 1.9e-06 | 2.167E-06 | 2.517E-07 | 1.900E-06 | 2.400E-06 |
| 27762 | 1389 | T1389R | E | 2.4e-06 | 1.7e-06 | 1.3e-06 | 1.800E-06 | 5.568E-07 | 1.300E-06 | 2.400E-06 |
| 27763 | 1389 | T1389N | E | 1.7e-06 | 2.9e-06 | 2.3e-06 | 2.300E-06 | 6.000E-07 | 1.700E-06 | 2.900E-06 |
| 27764 | 1389 | T1389D | E | 2.1e-06 | 1.9e-06 | 3.9e-06 | 2.633E-06 | 1.102E-06 | 1.900E-06 | 3.900E-06 |
| 27765 | 1389 | T1389C | E | 2.7e-06 | 2.6e-06 | 2.1e-06 | 2.467E-06 | 3.215E-07 | 2.100E-06 | 2.700E-06 |
| 27766 | 1389 | T1389E | E | 1.5e-06 | 1.6e-06 | 1.7e-06 | 1.600E-06 | 1.000E-07 | 1.500E-06 | 1.700E-06 |
| 27767 | 1389 | T1389Q | E | 2.1e-06 | 2.7e-06 | 2.9e-06 | 2.567E-06 | 4.163E-07 | 2.100E-06 | 2.900E-06 |
| 27768 | 1389 | T1389G | E | 2.5e-06 | 3.4e-06 | 1.9e-06 | 2.600E-06 | 7.550E-07 | 1.900E-06 | 3.400E-06 |
| 27769 | 1389 | T1389H | E | 2e-06 | 1.9e-06 | 4.5e-06 | 2.800E-06 | 1.473E-06 | 1.900E-06 | 4.500E-06 |
| 27770 | 1389 | T1389I | E | 1.7e-06 | 2.1e-06 | 4.2e-06 | 2.667E-06 | 1.343E-06 | 1.700E-06 | 4.200E-06 |
| 27771 | 1389 | T1389L | E | 2.8e-06 | 1.8e-06 | 1.9e-06 | 2.167E-06 | 5.508E-07 | 1.800E-06 | 2.800E-06 |
| 27772 | 1389 | T1389K | E | 2.4e-06 | 1.5e-06 | 3e-06 | 2.300E-06 | 7.550E-07 | 1.500E-06 | 3.000E-06 |
| 27773 | 1389 | T1389M | E | 1.2e-06 | 1.6e-06 | 3.8e-06 | 2.200E-06 | 1.400E-06 | 1.200E-06 | 3.800E-06 |
| 27774 | 1389 | T1389F | E | 1.5e-06 | 2.9e-06 | 7.4e-07 | 1.713E-06 | 1.096E-06 | 7.400E-07 | 2.900E-06 |
| 27775 | 1389 | T1389P | E | 1.7e-06 | 1.6e-06 | 2.4e-06 | 1.900E-06 | 4.359E-07 | 1.600E-06 | 2.400E-06 |
| 27776 | 1389 | T1389S | E | 3e-06 | 1.7e-06 | 2.2e-06 | 2.300E-06 | 6.557E-07 | 1.700E-06 | 3.000E-06 |
| 27777 | 1389 | T1389T | E | 2.8e-06 | 1.9e-06 | 3.6e-06 | 2.767E-06 | 8.505E-07 | 1.900E-06 | 3.600E-06 |
| 27778 | 1389 | T1389W | E | 1.6e-06 | 1.9e-06 | 2.5e-06 | 2.000E-06 | 4.583E-07 | 1.600E-06 | 2.500E-06 |
| 27779 | 1389 | T1389Y | E | 1.4e-06 | 2.1e-06 | 4.1e-06 | 2.533E-06 | 1.401E-06 | 1.400E-06 | 4.100E-06 |
| 27780 | 1389 | T1389V | E | 1.7e-06 | 3.1e-06 | 2.1e-06 | 2.300E-06 | 7.211E-07 | 1.700E-06 | 3.100E-06 |
| 27781 | 1390 | V1390A | E | 1.5e-06 | 2.5e-06 | 3e-06 | 2.333E-06 | 7.638E-07 | 1.500E-06 | 3.000E-06 |
| 27782 | 1390 | V1390R | E | 2e-06 | 1.9e-06 | 2.1e-06 | 2.000E-06 | 1.000E-07 | 1.900E-06 | 2.100E-06 |
| 27783 | 1390 | V1390N | E | 1.7e-06 | 2.6e-06 | 2.8e-06 | 2.367E-06 | 5.859E-07 | 1.700E-06 | 2.800E-06 |
| 27784 | 1390 | V1390D | E | 1.9e-06 | 2.7e-06 | 2.6e-06 | 2.400E-06 | 4.359E-07 | 1.900E-06 | 2.700E-06 |
| 27785 | 1390 | V1390C | E | 1.9e-06 | 9.8e-07 | 3.3e-06 | 2.060E-06 | 1.168E-06 | 9.800E-07 | 3.300E-06 |
| 27786 | 1390 | V1390E | E | 2.1e-06 | 1.8e-06 | 2.6e-06 | 2.167E-06 | 4.041E-07 | 1.800E-06 | 2.600E-06 |
| 27787 | 1390 | V1390Q | E | 1.9e-06 | 1.4e-06 | 2e-06 | 1.767E-06 | 3.215E-07 | 1.400E-06 | 2.000E-06 |
| 27788 | 1390 | V1390G | E | 2e-06 | 2.6e-06 | 2.1e-06 | 2.233E-06 | 3.215E-07 | 2.000E-06 | 2.600E-06 |
| 27789 | 1390 | V1390H | E | 3.5e-06 | 2.3e-06 | 1.7e-06 | 2.500E-06 | 9.165E-07 | 1.700E-06 | 3.500E-06 |
| 27790 | 1390 | V1390I | E | 2.5e-06 | 3.3e-06 | 2.5e-06 | 2.767E-06 | 4.619E-07 | 2.500E-06 | 3.300E-06 |
| 27791 | 1390 | V1390L | E | 3.3e-06 | 4e-06 | 3.2e-06 | 3.500E-06 | 4.359E-07 | 3.200E-06 | 4.000E-06 |
| 27792 | 1390 | V1390K | E | 2.3e-06 | 2.8e-06 | 2.6e-06 | 2.567E-06 | 2.517E-07 | 2.300E-06 | 2.800E-06 |
| 27793 | 1390 | V1390M | E | 2.7e-06 | 2.9e-06 | 2.5e-06 | 2.700E-06 | 2.000E-07 | 2.500E-06 | 2.900E-06 |
| 27794 | 1390 | V1390F | E | 1.5e-06 | 1.9e-06 | 3e-06 | 2.133E-06 | 7.767E-07 | 1.500E-06 | 3.000E-06 |

| | | | | | | | | | | |
|---|---|---|---|---|---|---|---|---|---|---|
| 27795 | 1390 | V1390P | E | 2.1e-06 | 2e-06 | 2.5e-06 | 2.200E-06 | 2.646E-07 | 2.000E-06 | 2.500E-06 |
| 27796 | 1390 | V1390S | E | 2.1e-06 | 3.8e-06 | 1.4e-06 | 2.433E-06 | 1.234E-06 | 1.400E-06 | 3.800E-06 |
| 27797 | 1390 | V1390T | E | 1.5e-06 | 2.1e-06 | 2e-06 | 1.867E-06 | 3.215E-07 | 1.500E-06 | 2.100E-06 |
| 27798 | 1390 | V1390W | E | 2.5e-06 | 2.1e-06 | 2e-06 | 2.200E-06 | 2.646E-07 | 2.000E-06 | 2.500E-06 |
| 27799 | 1390 | V1390Y | E | 3.5e-06 | 3.3e-06 | 1.8e-06 | 2.867E-06 | 9.292E-07 | 1.800E-06 | 3.500E-06 |
| 27800 | 1390 | V1390V | E | 1.4e-06 | 2e-06 | 2.9e-06 | 2.100E-06 | 7.550E-07 | 1.400E-06 | 2.900E-06 |
| 27801 | 1391 | S1391A | E | 1.5e-06 | 1.7e-06 | 4.3e-06 | 2.500E-06 | 1.562E-06 | 1.500E-06 | 4.300E-06 |
| 27802 | 1391 | S1391R | E | 1.9e-06 | 2e-06 | 1.7e-06 | 1.867E-06 | 1.528E-07 | 1.700E-06 | 2.000E-06 |
| 27803 | 1391 | S1391N | E | 3.5e-06 | 2.4e-06 | 2.1e-06 | 2.667E-06 | 7.371E-07 | 2.100E-06 | 3.500E-06 |
| 27804 | 1391 | S1391D | E | 2.3e-06 | 1.9e-06 | 2.7e-06 | 2.300E-06 | 4.000E-07 | 1.900E-06 | 2.700E-06 |
| 27805 | 1391 | S1391C | E | 4.1e-06 | 4.3e-06 | 1.8e-06 | 3.400E-06 | 1.389E-06 | 1.800E-06 | 4.300E-06 |
| 27806 | 1391 | S1391E | E | 2.2e-06 | 2.7e-06 | 2.9e-06 | 2.600E-06 | 3.606E-07 | 2.200E-06 | 2.900E-06 |
| 27807 | 1391 | S1391Q | E | 1.3e-06 | 1.6e-06 | 2.2e-06 | 1.700E-06 | 4.583E-07 | 1.300E-06 | 2.200E-06 |
| 27808 | 1391 | S1391G | E | 1.5e-06 | 1.7e-06 | 2.8e-06 | 2.000E-06 | 7.000E-07 | 1.500E-06 | 2.800E-06 |
| 27809 | 1391 | S1391H | E | 2.3e-06 | 2.6e-06 | 3.5e-06 | 2.800E-06 | 6.245E-07 | 2.300E-06 | 3.500E-06 |
| 27810 | 1391 | S1391I | E | 2.5e-06 | 2.6e-06 | 3e-06 | 2.700E-06 | 2.646E-07 | 2.500E-06 | 3.000E-06 |
| 27811 | 1391 | S1391L | E | 2e-06 | 2e-06 | 4.7e-06 | 2.900E-06 | 1.559E-06 | 2.000E-06 | 4.700E-06 |
| 27812 | 1391 | S1391K | E | 2.1e-06 | 3.9e-06 | 3.5e-06 | 3.167E-06 | 9.452E-07 | 2.100E-06 | 3.900E-06 |
| 27813 | 1391 | S1391M | E | 2.4e-06 | 2.9e-06 | 3.9e-06 | 3.067E-06 | 7.638E-07 | 2.400E-06 | 3.900E-06 |
| 27814 | 1391 | S1391F | E | 1.5e-06 | 3.1e-06 | 2.4e-06 | 2.333E-06 | 8.021E-07 | 1.500E-06 | 3.100E-06 |
| 27815 | 1391 | S1391P | E | 2.9e-06 | 3.9e-06 | 2.4e-06 | 3.067E-06 | 7.638E-07 | 2.400E-06 | 3.900E-06 |
| 27816 | 1391 | S1391S | E | 1.9e-06 | 1.5e-06 | 2e-06 | 1.800E-06 | 2.646E-07 | 1.500E-06 | 2.000E-06 |
| 27817 | 1391 | S1391T | E | 2.8e-06 | 2.3e-06 | 2.3e-06 | 2.467E-06 | 2.887E-07 | 2.300E-06 | 2.800E-06 |
| 27818 | 1391 | S1391W | E | 3e-06 | 2.3e-06 | 1.7e-06 | 2.333E-06 | 6.506E-07 | 1.700E-06 | 3.000E-06 |
| 27819 | 1391 | S1391Y | E | 1.8e-06 | 1.8e-06 | 4.1e-06 | 2.567E-06 | 1.328E-06 | 1.800E-06 | 4.100E-06 |
| 27820 | 1391 | S1391V | E | 2.6e-06 | 2.8e-06 | 2.4e-06 | 2.600E-06 | 2.000E-07 | 2.400E-06 | 2.800E-06 |
| 27821 | 1392 | S1392A | E | 3.8e-06 | 2.6e-06 | 3.3e-06 | 3.233E-06 | 6.028E-07 | 2.600E-06 | 3.800E-06 |
| 27822 | 1392 | S1392R | E | 4e-06 | 2.1e-06 | 1.2e-06 | 2.433E-06 | 1.429E-06 | 1.200E-06 | 4.000E-06 |
| 27823 | 1392 | S1392N | E | 2.8e-06 | 2.5e-06 | 2.7e-06 | 2.667E-06 | 1.528E-07 | 2.500E-06 | 2.800E-06 |
| 27824 | 1392 | S1392D | E | 2.3e-06 | 2.3e-06 | 3e-06 | 2.533E-06 | 4.041E-07 | 2.300E-06 | 3.000E-06 |
| 27825 | 1392 | S1392C | E | 1.9e-06 | 3.1e-06 | 1.2e-06 | 2.067E-06 | 9.609E-07 | 1.200E-06 | 3.100E-06 |
| 27826 | 1392 | S1392E | E | 2.3e-06 | 2.4e-06 | 1.7e-06 | 2.133E-06 | 3.786E-07 | 1.700E-06 | 2.400E-06 |
| 27827 | 1392 | S1392Q | E | 1.7e-06 | 1.5e-06 | 3.5e-06 | 2.233E-06 | 1.102E-06 | 1.500E-06 | 3.500E-06 |
| 27828 | 1392 | S1392G | E | 2.8e-06 | 3.2e-06 | 2.5e-06 | 2.833E-06 | 3.512E-07 | 2.500E-06 | 3.200E-06 |
| 27829 | 1392 | S1392H | E | 2.9e-06 | 2e-06 | 1.5e-06 | 2.133E-06 | 7.095E-07 | 1.500E-06 | 2.900E-06 |
| 27830 | 1392 | S1392I | E | 2.1e-06 | 4.1e-06 | 3.3e-06 | 3.167E-06 | 1.007E-06 | 2.100E-06 | 4.100E-06 |
| 27831 | 1392 | S1392L | E | 1.2e-06 | 1.4e-06 | 1.8e-06 | 1.467E-06 | 3.055E-07 | 1.200E-06 | 1.800E-06 |
| 27832 | 1392 | S1392K | E | 2.4e-06 | 2.3e-06 | 2.6e-06 | 2.433E-06 | 1.528E-07 | 2.300E-06 | 2.600E-06 |
| 27833 | 1392 | S1392M | E | 1.8e-06 | 2.7e-06 | 1.8e-06 | 2.100E-06 | 5.196E-07 | 1.800E-06 | 2.700E-06 |
| 27834 | 1392 | S1392F | E | 1.4e-06 | 2.3e-06 | 4.1e-06 | 2.600E-06 | 1.375E-06 | 1.400E-06 | 4.100E-06 |
| 27835 | 1392 | S1392P | E | 1.7e-06 | 1.7e-06 | 2.4e-06 | 1.933E-06 | 4.041E-07 | 1.700E-06 | 2.400E-06 |
| 27836 | 1392 | S1392S | E | 2.6e-06 | 1.4e-06 | 2.7e-06 | 2.233E-06 | 7.234E-07 | 1.400E-06 | 2.700E-06 |
| 27837 | 1392 | S1392T | E | 3.9e-06 | 2.6e-06 | 2.6e-06 | 3.033E-06 | 7.506E-07 | 2.600E-06 | 3.900E-06 |
| 27838 | 1392 | S1392W | E | 3.4e-06 | 2.2e-06 | 1.7e-06 | 2.433E-06 | 8.737E-07 | 1.700E-06 | 3.400E-06 |
| 27839 | 1392 | S1392Y | E | 1.5e-06 | 1.2e-06 | 3.3e-06 | 2.000E-06 | 1.136E-06 | 1.200E-06 | 3.300E-06 |
| 27840 | 1392 | S1392V | E | 1.3e-06 | 3.1e-06 | 2.7e-06 | 2.367E-06 | 9.452E-07 | 1.300E-06 | 3.100E-06 |
| 27841 | 1393 | A1393A | E | 1.8e-06 | 1.8e-06 | 2.9e-06 | 2.167E-06 | 6.351E-07 | 1.800E-06 | 2.900E-06 |
| 27842 | 1393 | A1393R | E | 2.6e-06 | 1.4e-06 | 1.3e-06 | 1.767E-06 | 7.234E-07 | 1.300E-06 | 2.600E-06 |
| 27843 | 1393 | A1393N | E | 2.1e-06 | 2e-06 | 2.7e-06 | 2.267E-06 | 3.786E-07 | 2.000E-06 | 2.700E-06 |
| 27844 | 1393 | A1393D | E | 1.8e-06 | 3.5e-06 | 1.7e-06 | 2.333E-06 | 1.012E-06 | 1.700E-06 | 3.500E-06 |
| 27845 | 1393 | A1393C | E | 2.5e-06 | 5.2e-06 | 2.2e-06 | 3.300E-06 | 1.652E-06 | 2.200E-06 | 5.200E-06 |
| 27846 | 1393 | A1393E | E | 4.6e-06 | 5.8e-06 | 1.5e-06 | 3.967E-06 | 2.219E-06 | 1.500E-06 | 5.800E-06 |
| 27847 | 1393 | A1393Q | E | 1.2e-06 | 2.3e-06 | 2.2e-06 | 1.900E-06 | 6.083E-07 | 1.200E-06 | 2.300E-06 |
| 27848 | 1393 | A1393G | E | 2.8e-06 | 2.1e-06 | 3.8e-06 | 2.900E-06 | 8.544E-07 | 2.100E-06 | 3.800E-06 |
| 27849 | 1393 | A1393H | E | 1.9e-06 | 2.3e-06 | 1.5e-06 | 1.900E-06 | 4.000E-07 | 1.500E-06 | 2.300E-06 |
| 27850 | 1393 | A1393I | E | 2.6e-06 | 3.7e-06 | 1.4e-06 | 2.567E-06 | 1.150E-06 | 1.400E-06 | 3.700E-06 |
| 27851 | 1393 | A1393L | E | 2e-06 | 1.9e-06 | 3.8e-06 | 2.567E-06 | 1.069E-06 | 1.900E-06 | 3.800E-06 |
| 27852 | 1393 | A1393K | E | 4.7e-06 | 3.3e-06 | 3.7e-06 | 3.900E-06 | 7.211E-07 | 3.300E-06 | 4.700E-06 |
| 27853 | 1393 | A1393M | E | 2.2e-06 | 1.8e-06 | 2.3e-06 | 2.100E-06 | 2.646E-07 | 1.800E-06 | 2.300E-06 |
| 27854 | 1393 | A1393F | E | 1.9e-06 | 1.5e-06 | 2e-06 | 1.800E-06 | 2.646E-07 | 1.500E-06 | 2.000E-06 |
| 27855 | 1393 | A1393P | E | 3e-06 | 2.6e-06 | 4.1e-06 | 3.233E-06 | 7.767E-07 | 2.600E-06 | 4.100E-06 |
| 27856 | 1393 | A1393S | E | 2.9e-06 | 2.2e-06 | 2.6e-06 | 2.567E-06 | 3.512E-07 | 2.200E-06 | 2.900E-06 |
| 27857 | 1393 | A1393T | E | 2.6e-06 | 2.9e-06 | 3.7e-06 | 3.067E-06 | 5.686E-07 | 2.600E-06 | 3.700E-06 |
| 27858 | 1393 | A1393W | E | 2.3e-06 | 1.4e-06 | 2.9e-06 | 2.200E-06 | 7.550E-07 | 1.400E-06 | 2.900E-06 |
| 27859 | 1393 | A1393Y | E | 3e-06 | 2.6e-06 | 2.8e-06 | 2.800E-06 | 2.000E-07 | 2.600E-06 | 3.000E-06 |
| 27860 | 1393 | A1393V | E | 2.3e-06 | 2.3e-06 | 1.9e-06 | 2.167E-06 | 2.309E-07 | 1.900E-06 | 2.300E-06 |
| 27861 | 1394 | S1394A | E | 3e-06 | 3.7e-06 | 2.4e-06 | 3.033E-06 | 6.506E-07 | 2.400E-06 | 3.700E-06 |
| 27862 | 1394 | S1394R | E | 2.8e-06 | 1.5e-06 | 2.5e-06 | 2.267E-06 | 6.807E-07 | 1.500E-06 | 2.800E-06 |
| 27863 | 1394 | S1394N | E | 1.4e-06 | 3.7e-06 | 3.6e-06 | 2.900E-06 | 1.300E-06 | 1.400E-06 | 3.700E-06 |
| 27864 | 1394 | S1394D | E | 1.9e-06 | 1.1e-06 | 2.7e-06 | 1.900E-06 | 8.000E-07 | 1.100E-06 | 2.700E-06 |
| 27865 | 1394 | S1394C | E | 3.3e-06 | 1.7e-06 | 2e-06 | 2.333E-06 | 8.505E-07 | 1.700E-06 | 3.300E-06 |
| 27866 | 1394 | S1394E | E | 2.3e-06 | 2.2e-06 | 1.9e-06 | 2.133E-06 | 2.082E-07 | 1.900E-06 | 2.300E-06 |
| 27867 | 1394 | S1394Q | E | 1.3e-06 | 1.4e-06 | 3e-06 | 1.900E-06 | 9.539E-07 | 1.300E-06 | 3.000E-06 |
| 27868 | 1394 | S1394G | E | 4.6e-06 | 2.4e-06 | 3e-06 | 3.333E-06 | 1.137E-06 | 2.400E-06 | 4.600E-06 |
| 27869 | 1394 | S1394H | E | 2.6e-06 | 2.7e-06 | 2.6e-06 | 2.633E-06 | 5.774E-08 | 2.600E-06 | 2.700E-06 |
| 27870 | 1394 | S1394I | E | 2.5e-06 | 3.2e-06 | 2.2e-06 | 2.633E-06 | 5.132E-07 | 2.200E-06 | 3.200E-06 |
| 27871 | 1394 | S1394L | E | 2.1e-06 | 1.9e-06 | 3.1e-06 | 2.367E-06 | 6.429E-07 | 1.900E-06 | 3.100E-06 |
| 27872 | 1394 | S1394K | E | 1.7e-06 | 3e-06 | 3.2e-06 | 2.633E-06 | 8.145E-07 | 1.700E-06 | 3.200E-06 |
| 27873 | 1394 | S1394M | E | 2e-06 | 2.7e-06 | 1.5e-06 | 2.067E-06 | 6.028E-07 | 1.500E-06 | 2.700E-06 |
| 27874 | 1394 | S1394F | E | 2.1e-06 | 2e-06 | 2.5e-06 | 2.200E-06 | 2.646E-07 | 2.000E-06 | 2.500E-06 |
| 27875 | 1394 | S1394P | E | 2.7e-06 | 2e-06 | 2.4e-06 | 2.367E-06 | 3.512E-07 | 2.000E-06 | 2.700E-06 |
| 27876 | 1394 | S1394S | E | 2.1e-06 | 1.6e-06 | 2.4e-06 | 2.033E-06 | 4.041E-07 | 1.600E-06 | 2.400E-06 |
| 27877 | 1394 | S1394T | E | 1.7e-06 | 2.5e-06 | 2.1e-06 | 2.100E-06 | 4.000E-07 | 1.700E-06 | 2.500E-06 |
| 27878 | 1394 | S1394W | E | 3.3e-06 | 1.8e-06 | 2e-06 | 2.367E-06 | 8.145E-07 | 1.800E-06 | 3.300E-06 |
| 27879 | 1394 | S1394Y | E | 1.1e-06 | 1.6e-06 | 2.2e-06 | 1.633E-06 | 5.508E-07 | 1.100E-06 | 2.200E-06 |

| | | | | | | | | | | |
|---|---|---|---|---|---|---|---|---|---|---|
| 27880 | 1394 | S1394V | E | 2.3e-06 | 2e-06 | 2.1e-06 | 2.133E-06 | 1.528E-07 | 2.000E-06 | 2.300E-06 |
| 27881 | 1395 | T1395A | E | 3.3e-06 | 2.3e-06 | 1.8e-06 | 2.467E-06 | 7.638E-07 | 1.800E-06 | 3.300E-06 |
| 27882 | 1395 | T1395R | E | 2.4e-06 | 3.6e-06 | 2.8e-06 | 2.933E-06 | 6.110E-07 | 2.400E-06 | 3.600E-06 |
| 27883 | 1395 | T1395N | E | 3.7e-06 | 2.7e-06 | 3.1e-06 | 3.167E-06 | 5.033E-07 | 2.700E-06 | 3.700E-06 |
| 27884 | 1395 | T1395D | E | 2e-06 | 2.9e-06 | 1.7e-06 | 2.200E-06 | 6.245E-07 | 1.700E-06 | 2.900E-06 |
| 27885 | 1395 | T1395C | E | 2.2e-06 | 1.9e-06 | 2.5e-06 | 2.200E-06 | 3.000E-07 | 1.900E-06 | 2.500E-06 |
| 27886 | 1395 | T1395E | E | 2.5e-06 | 2.5e-06 | 2.2e-06 | 2.400E-06 | 1.732E-07 | 2.200E-06 | 2.500E-06 |
| 27887 | 1395 | T1395Q | E | 1.4e-06 | 2.5e-06 | 2.6e-06 | 2.167E-06 | 6.658E-07 | 1.400E-06 | 2.600E-06 |
| 27888 | 1395 | T1395G | E | 2.7e-06 | 1.3e-06 | 2.7e-06 | 2.233E-06 | 8.083E-07 | 1.300E-06 | 2.700E-06 |
| 27889 | 1395 | T1395H | E | 3.4e-06 | 2.6e-06 | 2.3e-06 | 2.767E-06 | 5.686E-07 | 2.300E-06 | 3.400E-06 |
| 27890 | 1395 | T1395I | E | 1.6e-06 | 2.8e-06 | 3.6e-06 | 2.667E-06 | 1.007E-06 | 1.600E-06 | 3.600E-06 |
| 27891 | 1395 | T1395L | E | 3e-06 | 2.7e-06 | 2.6e-06 | 2.767E-06 | 2.082E-07 | 2.600E-06 | 3.000E-06 |
| 27892 | 1395 | T1395K | E | 2.9e-06 | 4e-06 | 4.3e-06 | 3.733E-06 | 7.371E-07 | 2.900E-06 | 4.300E-06 |
| 27893 | 1395 | T1395M | E | 1.9e-06 | 4e-06 | 2.2e-06 | 2.700E-06 | 1.136E-06 | 1.900E-06 | 4.000E-06 |
| 27894 | 1395 | T1395F | E | 2.4e-06 | 2.7e-06 | 3.5e-06 | 2.867E-06 | 5.686E-07 | 2.400E-06 | 3.500E-06 |
| 27895 | 1395 | T1395P | E | 4.9e-06 | 2.3e-06 | 2.6e-06 | 3.267E-06 | 1.422E-06 | 2.300E-06 | 4.900E-06 |
| 27896 | 1395 | T1395S | E | 1.8e-06 | 3.2e-06 | 4.3e-06 | 3.100E-06 | 1.253E-06 | 1.800E-06 | 4.300E-06 |
| 27897 | 1395 | T1395T | E | 3e-06 | 2.6e-06 | 1.4e-06 | 2.333E-06 | 8.327E-07 | 1.400E-06 | 3.000E-06 |
| 27898 | 1395 | T1395W | E | 1.2e-06 | 1.6e-06 | 3.1e-06 | 1.967E-06 | 1.002E-06 | 1.200E-06 | 3.100E-06 |
| 27899 | 1395 | T1395Y | E | 3.5e-06 | 3.9e-06 | 3.5e-06 | 3.633E-06 | 2.309E-07 | 3.500E-06 | 3.900E-06 |
| 27900 | 1395 | T1395V | E | 1.6e-06 | 2.3e-06 | 9.2e-07 | 1.607E-06 | 6.900E-07 | 9.200E-07 | 2.300E-06 |
| 27901 | 1396 | K1396A | E | 3.1e-06 | 3e-06 | 2.4e-06 | 2.833E-06 | 3.786E-07 | 2.400E-06 | 3.100E-06 |
| 27902 | 1396 | K1396R | E | 6.2e-06 | 2.6e-06 | 1.8e-06 | 3.533E-06 | 2.344E-06 | 1.800E-06 | 6.200E-06 |
| 27903 | 1396 | K1396N | E | 2.7e-06 | 1.9e-06 | 1.6e-06 | 2.067E-06 | 5.686E-07 | 1.600E-06 | 2.700E-06 |
| 27904 | 1396 | K1396D | E | 1.3e-06 | 2.5e-06 | 2.5e-06 | 2.100E-06 | 6.928E-07 | 1.300E-06 | 2.500E-06 |
| 27905 | 1396 | K1396C | E | 2.2e-06 | 2.8e-06 | 2.8e-06 | 2.600E-06 | 3.464E-07 | 2.200E-06 | 2.800E-06 |
| 27906 | 1396 | K1396E | E | 2.6e-06 | 2.9e-06 | 1.9e-06 | 2.467E-06 | 5.132E-07 | 1.900E-06 | 2.900E-06 |
| 27907 | 1396 | K1396Q | E | 2.9e-06 | 1.6e-06 | 3.2e-06 | 2.567E-06 | 8.505E-07 | 1.600E-06 | 3.200E-06 |
| 27908 | 1396 | K1396G | E | 2.3e-06 | 1.6e-06 | 1.9e-06 | 1.933E-06 | 3.512E-07 | 1.600E-06 | 2.300E-06 |
| 27909 | 1396 | K1396H | E | 2.8e-06 | 3.6e-06 | 1.7e-06 | 2.700E-06 | 9.539E-07 | 1.700E-06 | 3.600E-06 |
| 27910 | 1396 | K1396I | E | 3.7e-06 | 3.7e-06 | 2.2e-06 | 3.200E-06 | 8.660E-07 | 2.200E-06 | 3.700E-06 |
| 27911 | 1396 | K1396L | E | 2.1e-06 | 3.3e-06 | 3.4e-06 | 2.933E-06 | 7.234E-07 | 2.100E-06 | 3.400E-06 |
| 27912 | 1396 | K1396K | E | 2.4e-06 | 2e-06 | 2e-06 | 2.133E-06 | 2.309E-07 | 2.000E-06 | 2.400E-06 |
| 27913 | 1396 | K1396M | E | 3.6e-06 | 3e-06 | 2.1e-06 | 2.900E-06 | 7.550E-07 | 2.100E-06 | 3.600E-06 |
| 27914 | 1396 | K1396F | E | 2.3e-06 | 2.2e-06 | 1.8e-06 | 2.100E-06 | 2.646E-07 | 1.800E-06 | 2.300E-06 |
| 27915 | 1396 | K1396P | E | 2.8e-06 | 3.7e-06 | 3.2e-06 | 3.233E-06 | 4.509E-07 | 2.800E-06 | 3.700E-06 |
| 27916 | 1396 | K1396S | E | 1.8e-06 | 1.9e-06 | 1.5e-06 | 1.733E-06 | 2.082E-07 | 1.500E-06 | 1.900E-06 |
| 27917 | 1396 | K1396T | E | 1.8e-06 | 1.6e-06 | 2.6e-06 | 2.000E-06 | 5.292E-07 | 1.600E-06 | 2.600E-06 |
| 27918 | 1396 | K1396W | E | 2.1e-06 | 2.5e-06 | 1.7e-06 | 2.100E-06 | 4.000E-07 | 1.700E-06 | 2.500E-06 |
| 27919 | 1396 | K1396Y | E | 2.7e-06 | 5.8e-06 | 1.9e-06 | 3.467E-06 | 2.060E-06 | 1.900E-06 | 5.800E-06 |
| 27920 | 1396 | K1396V | E | 1.8e-06 | 1.4e-06 | 1.7e-06 | 1.633E-06 | 2.082E-07 | 1.400E-06 | 1.800E-06 |
| 27921 | 1397 | G1397A | E | 1.8e-06 | 2e-06 | 3e-06 | 2.267E-06 | 6.429E-07 | 1.800E-06 | 3.000E-06 |
| 27922 | 1397 | G1397R | E | 2.7e-06 | 2.5e-06 | 1.6e-06 | 2.267E-06 | 5.859E-07 | 1.600E-06 | 2.700E-06 |
| 27923 | 1397 | G1397N | E | 1.5e-06 | 2.6e-06 | 2.6e-06 | 2.233E-06 | 6.351E-07 | 1.500E-06 | 2.600E-06 |
| 27924 | 1397 | G1397D | E | 4e-06 | 2e-06 | 3.9e-06 | 3.300E-06 | 1.127E-06 | 2.000E-06 | 4.000E-06 |
| 27925 | 1397 | G1397C | E | 2.6e-06 | 3.1e-06 | 3.3e-06 | 3.000E-06 | 3.606E-07 | 2.600E-06 | 3.300E-06 |
| 27926 | 1397 | G1397E | E | 2.2e-06 | 9.8e-07 | 8.5e-07 | 1.343E-06 | 7.447E-07 | 8.500E-07 | 2.200E-06 |
| 27927 | 1397 | G1397Q | E | 2.2e-06 | 2.4e-06 | 2.6e-06 | 2.400E-06 | 2.000E-07 | 2.200E-06 | 2.600E-06 |
| 27928 | 1397 | G1397G | E | 1.8e-06 | 1.9e-06 | 3.7e-06 | 2.467E-06 | 1.069E-06 | 1.800E-06 | 3.700E-06 |
| 27929 | 1397 | G1397H | E | 2.8e-06 | 2.8e-06 | 1.9e-06 | 2.500E-06 | 5.196E-07 | 1.900E-06 | 2.800E-06 |
| 27930 | 1397 | G1397I | E | 2.8e-06 | 2.7e-06 | 2e-06 | 2.500E-06 | 4.359E-07 | 2.000E-06 | 2.800E-06 |
| 27931 | 1397 | G1397L | E | 3e-06 | 2.5e-06 | 1.2e-06 | 2.233E-06 | 9.292E-07 | 1.200E-06 | 3.000E-06 |
| 27932 | 1397 | G1397K | E | 1e-06 | 2.4e-06 | 1.9e-06 | 1.767E-06 | 7.095E-07 | 1.000E-06 | 2.400E-06 |
| 27933 | 1397 | G1397M | E | 2.2e-06 | 2.4e-06 | 2.4e-06 | 2.333E-06 | 1.155E-07 | 2.200E-06 | 2.400E-06 |
| 27934 | 1397 | G1397F | E | 2.3e-06 | 3.9e-06 | 2.5e-06 | 2.900E-06 | 8.718E-07 | 2.300E-06 | 3.900E-06 |
| 27935 | 1397 | G1397P | E | 2.4e-06 | 2.5e-06 | 1.5e-06 | 2.133E-06 | 5.508E-07 | 1.500E-06 | 2.500E-06 |
| 27936 | 1397 | G1397S | E | 1.6e-06 | 2.2e-06 | 2.4e-06 | 2.067E-06 | 4.163E-07 | 1.600E-06 | 2.400E-06 |
| 27937 | 1397 | G1397T | E | 1.9e-06 | 1.8e-06 | 1.6e-06 | 1.767E-06 | 1.528E-07 | 1.600E-06 | 1.900E-06 |
| 27938 | 1397 | G1397W | E | 2e-06 | 2.9e-06 | 1.7e-06 | 2.200E-06 | 6.245E-07 | 1.700E-06 | 2.900E-06 |
| 27939 | 1397 | G1397Y | E | 2.8e-06 | 2.2e-06 | 4.4e-06 | 3.133E-06 | 1.137E-06 | 2.200E-06 | 4.400E-06 |
| 27940 | 1397 | G1397V | E | 2.3e-06 | 2.1e-06 | 3.6e-06 | 2.667E-06 | 8.145E-07 | 2.100E-06 | 3.600E-06 |
| 27941 | 1398 | P1398A | E | 1.6e-06 | 2.5e-06 | 2e-06 | 2.033E-06 | 4.509E-07 | 1.600E-06 | 2.500E-06 |
| 27942 | 1398 | P1398R | E | 2e-06 | 1.3e-06 | 2.6e-06 | 1.967E-06 | 6.506E-07 | 1.300E-06 | 2.600E-06 |
| 27943 | 1398 | P1398N | E | 3.1e-06 | 2.8e-06 | 2.3e-06 | 2.733E-06 | 4.041E-07 | 2.300E-06 | 3.100E-06 |
| 27944 | 1398 | P1398D | E | 3.6e-06 | 2e-06 | 2.3e-06 | 2.633E-06 | 8.505E-07 | 2.000E-06 | 3.600E-06 |
| 27945 | 1398 | P1398C | E | 3.1e-06 | 1.1e-06 | 3e-06 | 2.400E-06 | 1.127E-06 | 1.100E-06 | 3.100E-06 |
| 27946 | 1398 | P1398E | E | 3.2e-06 | 2.9e-06 | 2.4e-06 | 2.833E-06 | 4.041E-07 | 2.400E-06 | 3.200E-06 |
| 27947 | 1398 | P1398Q | E | 2.6e-06 | 2.7e-06 | 3.5e-06 | 2.933E-06 | 4.933E-07 | 2.600E-06 | 3.500E-06 |
| 27948 | 1398 | P1398G | E | 2.4e-06 | 2.2e-06 | 2e-06 | 2.200E-06 | 2.000E-07 | 2.000E-06 | 2.400E-06 |
| 27949 | 1398 | P1398H | E | 2.2e-06 | 3.7e-06 | 2.3e-06 | 2.733E-06 | 8.386E-07 | 2.200E-06 | 3.700E-06 |
| 27950 | 1398 | P1398I | E | 2.2e-06 | 2.1e-06 | 2.4e-06 | 2.233E-06 | 1.528E-07 | 2.100E-06 | 2.400E-06 |
| 27951 | 1398 | P1398L | E | 3.9e-06 | 1.9e-06 | 2e-06 | 2.600E-06 | 1.127E-06 | 1.900E-06 | 3.900E-06 |
| 27952 | 1398 | P1398K | E | 2.3e-06 | 2.6e-06 | 1.7e-06 | 2.200E-06 | 4.583E-07 | 1.700E-06 | 2.600E-06 |
| 27953 | 1398 | P1398M | E | 2.3e-06 | 1.5e-06 | 1.3e-06 | 1.700E-06 | 5.292E-07 | 1.300E-06 | 2.300E-06 |
| 27954 | 1398 | P1398F | E | 2.9e-06 | 2.4e-06 | 2.5e-06 | 2.600E-06 | 2.646E-07 | 2.400E-06 | 2.900E-06 |
| 27955 | 1398 | P1398P | E | 3.1e-06 | 2.6e-06 | 2.3e-06 | 2.667E-06 | 4.041E-07 | 2.300E-06 | 3.100E-06 |
| 27956 | 1398 | P1398S | E | 2.9e-06 | 1.5e-06 | 3.2e-06 | 2.533E-06 | 9.074E-07 | 1.500E-06 | 3.200E-06 |
| 27957 | 1398 | P1398T | E | 1.9e-06 | 4.1e-06 | 1.9e-06 | 2.633E-06 | 1.270E-06 | 1.900E-06 | 4.100E-06 |
| 27958 | 1398 | P1398W | E | 3.1e-06 | 2e-06 | 2.8e-06 | 2.633E-06 | 5.686E-07 | 2.000E-06 | 3.100E-06 |
| 27959 | 1398 | P1398Y | E | 2.7e-06 | 2.6e-06 | 1.1e-06 | 2.133E-06 | 8.963E-07 | 1.100E-06 | 2.700E-06 |
| 27960 | 1398 | P1398V | E | 2.9e-06 | 1.7e-06 | 2.3e-06 | 2.300E-06 | 6.000E-07 | 1.700E-06 | 2.900E-06 |
| 27961 | 1399 | S1399A | E | 2.4e-06 | 3.4e-06 | 4.5e-06 | 3.433E-06 | 1.050E-06 | 2.400E-06 | 4.500E-06 |
| 27962 | 1399 | S1399R | E | 2.3e-06 | 2.7e-06 | 2.1e-06 | 2.367E-06 | 3.055E-07 | 2.100E-06 | 2.700E-06 |
| 27963 | 1399 | S1399N | E | 4.6e-06 | 2.2e-06 | 1.9e-06 | 2.900E-06 | 1.480E-06 | 1.900E-06 | 4.600E-06 |
| 27964 | 1399 | S1399D | E | 4e-06 | 4.2e-06 | 2.5e-06 | 3.567E-06 | 9.292E-07 | 2.500E-06 | 4.200E-06 |

| | | | | | | | | | | |
|---|---|---|---|---|---|---|---|---|---|---|
| 27965 | 1399 | S1399C | E | 2.8e-06 | 2e-06 | 1.5e-06 | 2.100E-06 | 6.557E-07 | 1.500E-06 | 2.800E-06 |
| 27966 | 1399 | S1399E | E | 2.6e-06 | 5.2e-06 | 3.6e-06 | 3.800E-06 | 1.311E-06 | 2.600E-06 | 5.200E-06 |
| 27967 | 1399 | S1399Q | E | 2.2e-06 | 2.6e-06 | 2.5e-06 | 2.433E-06 | 2.082E-07 | 2.200E-06 | 2.600E-06 |
| 27968 | 1399 | S1399G | E | 1.4e-06 | 2.2e-06 | 2.5e-06 | 2.033E-06 | 5.686E-07 | 1.400E-06 | 2.500E-06 |
| 27969 | 1399 | S1399H | E | 1.4e-06 | 2.4e-06 | 2e-06 | 1.933E-06 | 5.033E-07 | 1.400E-06 | 2.400E-06 |
| 27970 | 1399 | S1399I | E | 1.8e-06 | 5.4e-06 | 2.6e-06 | 3.267E-06 | 1.890E-06 | 1.800E-06 | 5.400E-06 |
| 27971 | 1399 | S1399L | E | 2.7e-06 | 2e-06 | 1.4e-06 | 2.033E-06 | 6.506E-07 | 1.400E-06 | 2.700E-06 |
| 27972 | 1399 | S1399K | E | 3.7e-06 | 1.8e-06 | 2.7e-06 | 2.733E-06 | 9.504E-07 | 1.800E-06 | 3.700E-06 |
| 27973 | 1399 | S1399M | E | 2.7e-06 | 2.6e-06 | 2.3e-06 | 2.533E-06 | 2.082E-07 | 2.300E-06 | 2.700E-06 |
| 27974 | 1399 | S1399F | E | 2.4e-06 | 1.7e-06 | 3.2e-06 | 2.433E-06 | 7.506E-07 | 1.700E-06 | 3.200E-06 |
| 27975 | 1399 | S1399P | E | 2.2e-06 | 3.2e-06 | 2.6e-06 | 2.667E-06 | 5.033E-07 | 2.200E-06 | 3.200E-06 |
| 27976 | 1399 | S1399S | E | 1.4e-06 | 1.2e-06 | 2.5e-06 | 1.700E-06 | 7.000E-07 | 1.200E-06 | 2.500E-06 |
| 27977 | 1399 | S1399T | E | 2.6e-06 | 2.2e-06 | 1.1e-06 | 1.967E-06 | 7.767E-07 | 1.100E-06 | 2.600E-06 |
| 27978 | 1399 | S1399W | E | 2.4e-06 | 1.5e-06 | 2e-06 | 1.967E-06 | 4.509E-07 | 1.500E-06 | 2.400E-06 |
| 27979 | 1399 | S1399Y | E | 2.7e-06 | 2e-06 | 1.8e-06 | 2.167E-06 | 4.726E-07 | 1.800E-06 | 2.700E-06 |
| 27980 | 1399 | S1399V | E | 2e-06 | 3.8e-06 | 3.4e-06 | 3.067E-06 | 9.452E-07 | 2.000E-06 | 3.800E-06 |
| 27981 | 1400 | V1400A | E | 2.1e-06 | 2.3e-06 | 1.9e-06 | 2.100E-06 | 2.000E-07 | 1.900E-06 | 2.300E-06 |
| 27982 | 1400 | V1400R | E | 2.3e-06 | 1.7e-06 | 2.2e-06 | 2.067E-06 | 3.215E-07 | 1.700E-06 | 2.300E-06 |
| 27983 | 1400 | V1400N | E | 2.4e-06 | 2.7e-06 | 2.6e-06 | 2.567E-06 | 1.528E-07 | 2.400E-06 | 2.700E-06 |
| 27984 | 1400 | V1400D | E | 2e-06 | 2.8e-06 | 1.2e-06 | 2.000E-06 | 8.000E-07 | 1.200E-06 | 2.800E-06 |
| 27985 | 1400 | V1400C | E | 1.8e-06 | 1.6e-06 | 3e-06 | 2.133E-06 | 7.572E-07 | 1.600E-06 | 3.000E-06 |
| 27986 | 1400 | V1400E | E | 2.8e-06 | 2.2e-06 | 2.9e-06 | 2.633E-06 | 3.786E-07 | 2.200E-06 | 2.900E-06 |
| 27987 | 1400 | V1400Q | E | 2.5e-06 | 2.3e-06 | 2.6e-06 | 2.467E-06 | 1.528E-07 | 2.300E-06 | 2.600E-06 |
| 27988 | 1400 | V1400G | E | 3.3e-06 | 2.9e-06 | 1.7e-06 | 2.633E-06 | 8.327E-07 | 1.700E-06 | 3.300E-06 |
| 27989 | 1400 | V1400H | E | 3.2e-06 | 1.7e-06 | 2.7e-06 | 2.533E-06 | 7.638E-07 | 1.700E-06 | 3.200E-06 |
| 27990 | 1400 | V1400I | E | 2.3e-06 | 2.4e-06 | 2.2e-06 | 2.300E-06 | 1.000E-07 | 2.200E-06 | 2.400E-06 |
| 27991 | 1400 | V1400L | E | 3.6e-06 | 3.8e-06 | 1.3e-06 | 2.900E-06 | 1.389E-06 | 1.300E-06 | 3.800E-06 |
| 27992 | 1400 | V1400K | E | 2e-06 | 2e-06 | 3.7e-06 | 2.567E-06 | 9.815E-07 | 2.000E-06 | 3.700E-06 |
| 27993 | 1400 | V1400M | E | 1.8e-06 | 2.2e-06 | 1.8e-06 | 1.933E-06 | 2.309E-07 | 1.800E-06 | 2.200E-06 |
| 27994 | 1400 | V1400F | E | 2e-06 | 2e-06 | 2.9e-06 | 2.300E-06 | 5.196E-07 | 2.000E-06 | 2.900E-06 |
| 27995 | 1400 | V1400P | E | 2.7e-06 | 2.5e-06 | 3.3e-06 | 2.833E-06 | 4.163E-07 | 2.500E-06 | 3.300E-06 |
| 27996 | 1400 | V1400S | E | 1.9e-06 | 3.9e-06 | 2.9e-06 | 2.900E-06 | 1.000E-06 | 1.900E-06 | 3.900E-06 |
| 27997 | 1400 | V1400T | E | 3.6e-06 | 2.6e-06 | 2e-06 | 2.733E-06 | 8.083E-07 | 2.000E-06 | 3.600E-06 |
| 27998 | 1400 | V1400W | E | 1.2e-06 | 1.7e-06 | 3.6e-06 | 2.167E-06 | 1.266E-06 | 1.200E-06 | 3.600E-06 |
| 27999 | 1400 | V1400Y | E | 1.8e-06 | 2.2e-06 | 2.3e-06 | 2.100E-06 | 2.646E-07 | 1.800E-06 | 2.300E-06 |
| 28000 | 1400 | V1400V | E | 3.3e-06 | 9.7e-07 | 4e-06 | 2.757E-06 | 1.586E-06 | 9.700E-07 | 4.000E-06 |
| 28001 | 1401 | F1401A | E | 1.3e-06 | 3.1e-06 | 2.2e-06 | 2.200E-06 | 9.000E-07 | 1.300E-06 | 3.100E-06 |
| 28002 | 1401 | F1401R | E | 3.1e-06 | 2.5e-06 | 1.6e-06 | 2.400E-06 | 7.550E-07 | 1.600E-06 | 3.100E-06 |
| 28003 | 1401 | F1401N | E | 2.3e-06 | 2e-06 | 2.4e-06 | 2.233E-06 | 2.082E-07 | 2.000E-06 | 2.400E-06 |
| 28004 | 1401 | F1401D | E | 3.1e-06 | 1.9e-06 | 2.9e-06 | 2.633E-06 | 6.429E-07 | 1.900E-06 | 3.100E-06 |
| 28005 | 1401 | F1401C | E | 2.6e-06 | 2.2e-06 | 1.8e-06 | 2.200E-06 | 4.000E-07 | 1.800E-06 | 2.600E-06 |
| 28006 | 1401 | F1401E | E | 2e-06 | 3.2e-06 | 1.9e-06 | 2.367E-06 | 7.234E-07 | 1.900E-06 | 3.200E-06 |
| 28007 | 1401 | F1401Q | E | 1.4e-06 | 2.6e-06 | 2.1e-06 | 2.033E-06 | 6.028E-07 | 1.400E-06 | 2.600E-06 |
| 28008 | 1401 | F1401G | E | 1.3e-06 | 2.1e-06 | 2.8e-06 | 2.067E-06 | 7.506E-07 | 1.300E-06 | 2.800E-06 |
| 28009 | 1401 | F1401H | E | 2.6e-06 | 2.2e-06 | 1.9e-06 | 2.233E-06 | 3.512E-07 | 1.900E-06 | 2.600E-06 |
| 28010 | 1401 | F1401I | E | 3.2e-06 | 2.7e-06 | 1.1e-06 | 2.333E-06 | 1.097E-06 | 1.100E-06 | 3.200E-06 |
| 28011 | 1401 | F1401L | E | 3.6e-06 | 2.7e-06 | 2.9e-06 | 3.067E-06 | 4.726E-07 | 2.700E-06 | 3.600E-06 |
| 28012 | 1401 | F1401K | E | 3.2e-06 | 2.5e-06 | 1.7e-06 | 2.467E-06 | 7.506E-07 | 1.700E-06 | 3.200E-06 |
| 28013 | 1401 | F1401M | E | 5.2e-06 | 1.4e-06 | 1.8e-06 | 2.800E-06 | 2.088E-06 | 1.400E-06 | 5.200E-06 |
| 28014 | 1401 | F1401F | E | 2.8e-06 | 3.7e-06 | 1.5e-06 | 2.667E-06 | 1.106E-06 | 1.500E-06 | 3.700E-06 |
| 28015 | 1401 | F1401P | E | 2.1e-06 | 1.9e-06 | 2.7e-06 | 2.233E-06 | 4.163E-07 | 1.900E-06 | 2.700E-06 |
| 28016 | 1401 | F1401S | E | 2e-06 | 4.1e-06 | 4.3e-06 | 3.467E-06 | 1.274E-06 | 2.000E-06 | 4.300E-06 |
| 28017 | 1401 | F1401T | E | 1.9e-06 | 2.5e-06 | 2.4e-06 | 2.267E-06 | 3.215E-07 | 1.900E-06 | 2.500E-06 |
| 28018 | 1401 | F1401W | E | 2e-06 | 1.1e-06 | 2.3e-06 | 1.800E-06 | 6.245E-07 | 1.100E-06 | 2.300E-06 |
| 28019 | 1401 | F1401Y | E | 3.7e-06 | 2.2e-06 | 3.6e-06 | 3.167E-06 | 8.386E-07 | 2.200E-06 | 3.700E-06 |
| 28020 | 1401 | F1401V | E | 2.4e-06 | 3.2e-06 | 2.7e-06 | 2.767E-06 | 4.041E-07 | 2.400E-06 | 3.200E-06 |
| 28021 | 1402 | P1402A | E | 2e-06 | 2.8e-06 | 1.5e-06 | 2.100E-06 | 6.557E-07 | 1.500E-06 | 2.800E-06 |
| 28022 | 1402 | P1402R | E | 2.5e-06 | 1.5e-06 | 4.9e-06 | 2.967E-06 | 1.747E-06 | 1.500E-06 | 4.900E-06 |
| 28023 | 1402 | P1402N | E | 3.7e-06 | 2.5e-06 | 3.5e-06 | 3.233E-06 | 6.429E-07 | 2.500E-06 | 3.700E-06 |
| 28024 | 1402 | P1402D | E | 2.5e-06 | 3.2e-06 | 2.1e-06 | 2.600E-06 | 5.568E-07 | 2.100E-06 | 3.200E-06 |
| 28025 | 1402 | P1402C | E | 1.7e-06 | 3e-06 | 3e-06 | 2.567E-06 | 7.506E-07 | 1.700E-06 | 3.000E-06 |
| 28026 | 1402 | P1402E | E | 2.5e-06 | 2.3e-06 | 3e-06 | 2.600E-06 | 3.606E-07 | 2.300E-06 | 3.000E-06 |
| 28027 | 1402 | P1402Q | E | 1e-06 | 2.5e-06 | 3.3e-06 | 2.267E-06 | 1.168E-06 | 1.000E-06 | 3.300E-06 |
| 28028 | 1402 | P1402G | E | 3e-06 | 4e-06 | 1.4e-06 | 2.800E-06 | 1.311E-06 | 1.400E-06 | 4.000E-06 |
| 28029 | 1402 | P1402H | E | 2.5e-06 | 1.9e-06 | 2.2e-06 | 2.200E-06 | 3.000E-07 | 1.900E-06 | 2.500E-06 |
| 28030 | 1402 | P1402I | E | 2.9e-06 | 2.7e-06 | 2.6e-06 | 2.733E-06 | 1.528E-07 | 2.600E-06 | 2.900E-06 |
| 28031 | 1402 | P1402L | E | 1.2e-06 | 1.4e-06 | 3e-06 | 1.867E-06 | 9.866E-07 | 1.200E-06 | 3.000E-06 |
| 28032 | 1402 | P1402K | E | 2.8e-06 | 1.8e-06 | 2.7e-06 | 2.433E-06 | 5.508E-07 | 1.800E-06 | 2.800E-06 |
| 28033 | 1402 | P1402M | E | 2.4e-06 | 2.9e-06 | 2.3e-06 | 2.533E-06 | 3.215E-07 | 2.300E-06 | 2.900E-06 |
| 28034 | 1402 | P1402F | E | 1e-06 | 2e-06 | 1.9e-06 | 1.633E-06 | 5.508E-07 | 1.000E-06 | 2.000E-06 |
| 28035 | 1402 | P1402P | E | 2.6e-06 | 3.3e-06 | 2e-06 | 2.633E-06 | 6.506E-07 | 2.000E-06 | 3.300E-06 |
| 28036 | 1402 | P1402S | E | 2.7e-06 | 2.5e-06 | 2.2e-06 | 2.467E-06 | 2.517E-07 | 2.200E-06 | 2.700E-06 |
| 28037 | 1402 | P1402T | E | 1.8e-06 | 3.1e-06 | 2.4e-06 | 2.433E-06 | 6.506E-07 | 1.800E-06 | 3.100E-06 |
| 28038 | 1402 | P1402W | E | 6.5e-06 | 2e-06 | 2.3e-06 | 3.600E-06 | 2.516E-06 | 2.000E-06 | 6.500E-06 |
| 28039 | 1402 | P1402Y | E | 1.5e-06 | 3.7e-06 | 2.1e-06 | 2.433E-06 | 1.137E-06 | 1.500E-06 | 3.700E-06 |
| 28040 | 1402 | P1402V | E | 2.7e-06 | 2.4e-06 | 2.2e-06 | 2.433E-06 | 2.517E-07 | 2.200E-06 | 2.700E-06 |
| 28041 | 1403 | L1403A | E | 1.9e-06 | 2.6e-06 | 1.9e-06 | 2.133E-06 | 4.041E-07 | 1.900E-06 | 2.600E-06 |
| 28042 | 1403 | L1403R | E | 2.6e-06 | 2.1e-06 | 2.3e-06 | 2.333E-06 | 2.517E-07 | 2.100E-06 | 2.600E-06 |
| 28043 | 1403 | L1403N | E | 2e-06 | 2e-06 | 2.7e-06 | 2.233E-06 | 4.041E-07 | 2.000E-06 | 2.700E-06 |
| 28044 | 1403 | L1403D | E | 2.3e-06 | 2e-06 | 2.9e-06 | 2.400E-06 | 4.583E-07 | 2.000E-06 | 2.900E-06 |
| 28045 | 1403 | L1403C | E | 1.5e-06 | 3.6e-06 | 2.3e-06 | 2.467E-06 | 1.060E-06 | 1.500E-06 | 3.600E-06 |
| 28046 | 1403 | L1403E | E | 2.6e-06 | 1.9e-06 | 2.1e-06 | 2.200E-06 | 3.606E-07 | 1.900E-06 | 2.600E-06 |
| 28047 | 1403 | L1403Q | E | 2.2e-06 | 3e-06 | 3e-06 | 2.733E-06 | 4.619E-07 | 2.200E-06 | 3.000E-06 |
| 28048 | 1403 | L1403G | E | 3.2e-06 | 2.6e-06 | 2e-06 | 2.600E-06 | 6.000E-07 | 2.000E-06 | 3.200E-06 |
| 28049 | 1403 | L1403H | E | 3.9e-06 | 5.4e-06 | 3.2e-06 | 4.167E-06 | 1.124E-06 | 3.200E-06 | 5.400E-06 |

| | | | | | | | | | | |
|---|---|---|---|---|---|---|---|---|---|---|
| 28050 | 1403 | L1403I | E | 2.1e-06 | 3.2e-06 | 1.9e-06 | 2.400E-06 | 7.000E-07 | 1.900E-06 | 3.200E-06 |
| 28051 | 1403 | L1403L | E | 3.1e-06 | 3.1e-06 | 2.7e-06 | 2.967E-06 | 2.309E-07 | 2.700E-06 | 3.100E-06 |
| 28052 | 1403 | L1403K | E | 1e-06 | 3.9e-06 | 3.2e-06 | 2.700E-06 | 1.513E-06 | 1.000E-06 | 3.900E-06 |
| 28053 | 1403 | L1403M | E | 3e-06 | 2.4e-06 | 2.5e-06 | 2.633E-06 | 3.215E-07 | 2.400E-06 | 3.000E-06 |
| 28054 | 1403 | L1403F | E | 2.5e-06 | 2.4e-06 | 3.4e-06 | 2.767E-06 | 5.508E-07 | 2.400E-06 | 3.400E-06 |
| 28055 | 1403 | L1403P | E | 1.3e-06 | 1.9e-06 | 2.5e-06 | 1.900E-06 | 6.000E-07 | 1.300E-06 | 2.500E-06 |
| 28056 | 1403 | L1403S | E | 1.7e-06 | 2.3e-06 | 2.9e-06 | 2.300E-06 | 6.000E-07 | 1.700E-06 | 2.900E-06 |
| 28057 | 1403 | L1403T | E | 2.6e-06 | 2.5e-06 | 1.6e-06 | 2.233E-06 | 5.508E-07 | 1.600E-06 | 2.600E-06 |
| 28058 | 1403 | L1403W | E | 5.1e-06 | 3.2e-06 | 4e-06 | 4.100E-06 | 9.539E-07 | 3.200E-06 | 5.100E-06 |
| 28059 | 1403 | L1403Y | E | 2.2e-06 | 2.6e-06 | 1.3e-06 | 2.033E-06 | 6.658E-07 | 1.300E-06 | 2.600E-06 |
| 28060 | 1403 | L1403V | E | 2.6e-06 | 3.5e-06 | 2.1e-06 | 2.733E-06 | 7.095E-07 | 2.100E-06 | 3.500E-06 |
| 28061 | 1404 | A1404A | E | 3.5e-06 | 3.5e-06 | 1.6e-06 | 2.867E-06 | 1.097E-06 | 1.600E-06 | 3.500E-06 |
| 28062 | 1404 | A1404R | E | 1.6e-06 | 2.2e-06 | 2.6e-06 | 2.133E-06 | 5.033E-07 | 1.600E-06 | 2.600E-06 |
| 28063 | 1404 | A1404N | E | 2.2e-06 | 3e-06 | 1.3e-06 | 2.167E-06 | 8.505E-07 | 1.300E-06 | 3.000E-06 |
| 28064 | 1404 | A1404D | E | 3.6e-06 | 2.7e-06 | 4.7e-06 | 3.667E-06 | 1.002E-06 | 2.700E-06 | 4.700E-06 |
| 28065 | 1404 | A1404C | E | 4.1e-06 | 2.7e-06 | 3.1e-06 | 3.300E-06 | 7.211E-07 | 2.700E-06 | 4.100E-06 |
| 28066 | 1404 | A1404E | E | 3.4e-06 | 3.1e-06 | 2.5e-06 | 3.000E-06 | 4.583E-07 | 2.500E-06 | 3.400E-06 |
| 28067 | 1404 | A1404Q | E | 2.1e-06 | 3.4e-06 | 2.2e-06 | 2.567E-06 | 7.234E-07 | 2.100E-06 | 3.400E-06 |
| 28068 | 1404 | A1404G | E | 2.6e-06 | 3.1e-06 | 1.8e-06 | 2.500E-06 | 6.557E-07 | 1.800E-06 | 3.100E-06 |
| 28069 | 1404 | A1404H | E | 1.2e-06 | 1.8e-06 | 1.4e-06 | 1.467E-06 | 3.055E-07 | 1.200E-06 | 1.800E-06 |
| 28070 | 1404 | A1404I | E | 2.4e-06 | 2.2e-06 | 2.5e-06 | 2.367E-06 | 1.528E-07 | 2.200E-06 | 2.500E-06 |
| 28071 | 1404 | A1404L | E | 3e-06 | 2.3e-06 | 3.2e-06 | 2.833E-06 | 4.726E-07 | 2.300E-06 | 3.200E-06 |
| 28072 | 1404 | A1404K | E | 1.7e-06 | 3.9e-06 | 2.6e-06 | 2.733E-06 | 1.106E-06 | 1.700E-06 | 3.900E-06 |
| 28073 | 1404 | A1404M | E | 2.7e-06 | 2.1e-06 | 2.6e-06 | 2.467E-06 | 3.215E-07 | 2.100E-06 | 2.700E-06 |
| 28074 | 1404 | A1404F | E | 2.5e-06 | 1.7e-06 | 3.8e-06 | 2.667E-06 | 1.060E-06 | 1.700E-06 | 3.800E-06 |
| 28075 | 1404 | A1404P | E | 3.6e-06 | 2.9e-06 | 1.8e-06 | 2.767E-06 | 9.074E-07 | 1.800E-06 | 3.600E-06 |
| 28076 | 1404 | A1404S | E | 2.6e-06 | 2e-06 | 2.6e-06 | 2.400E-06 | 3.464E-07 | 2.000E-06 | 2.600E-06 |
| 28077 | 1404 | A1404T | E | 9.5e-07 | 2.6e-06 | 3.6e-06 | 2.383E-06 | 1.338E-06 | 9.500E-07 | 3.600E-06 |
| 28078 | 1404 | A1404W | E | 1.4e-06 | 1.8e-06 | 2.1e-06 | 1.767E-06 | 3.512E-07 | 1.400E-06 | 2.100E-06 |
| 28079 | 1404 | A1404Y | E | 2.3e-06 | 1.4e-06 | 3.7e-06 | 2.467E-06 | 1.159E-06 | 1.400E-06 | 3.700E-06 |
| 28080 | 1404 | A1404V | E | 3.1e-06 | 2.1e-06 | 2.8e-06 | 2.667E-06 | 5.132E-07 | 2.100E-06 | 3.100E-06 |
| 28081 | 1405 | P1405A | E | 3.2e-06 | 2.2e-06 | 1.2e-06 | 2.200E-06 | 1.000E-06 | 1.200E-06 | 3.200E-06 |
| 28082 | 1405 | P1405R | E | 2.5e-06 | 2.8e-06 | 3.1e-06 | 2.800E-06 | 3.000E-07 | 2.500E-06 | 3.100E-06 |
| 28083 | 1405 | P1405N | E | 9.1e-07 | 1.7e-06 | 3.7e-06 | 2.103E-06 | 1.438E-06 | 9.100E-07 | 3.700E-06 |
| 28084 | 1405 | P1405D | E | 2.5e-06 | 1.8e-06 | 3.7e-06 | 2.667E-06 | 9.609E-07 | 1.800E-06 | 3.700E-06 |
| 28085 | 1405 | P1405C | E | 1.5e-06 | 3.7e-06 | 2.3e-06 | 2.500E-06 | 1.114E-06 | 1.500E-06 | 3.700E-06 |
| 28086 | 1405 | P1405E | E | 2.4e-06 | 3e-06 | 3.5e-06 | 2.967E-06 | 5.508E-07 | 2.400E-06 | 3.500E-06 |
| 28087 | 1405 | P1405Q | E | 1.9e-06 | 2.1e-06 | 3.1e-06 | 2.367E-06 | 6.429E-07 | 1.900E-06 | 3.100E-06 |
| 28088 | 1405 | P1405G | E | 4.6e-06 | 1.3e-06 | 2.3e-06 | 2.733E-06 | 1.692E-06 | 1.300E-06 | 4.600E-06 |
| 28089 | 1405 | P1405H | E | 2.3e-06 | 1.8e-06 | 2.7e-06 | 2.267E-06 | 4.509E-07 | 1.800E-06 | 2.700E-06 |
| 28090 | 1405 | P1405I | E | 1.7e-06 | 1.4e-06 | 2.3e-06 | 1.800E-06 | 4.583E-07 | 1.400E-06 | 2.300E-06 |
| 28091 | 1405 | P1405L | E | 2.2e-06 | 2.6e-06 | 2.1e-06 | 2.300E-06 | 2.646E-07 | 2.100E-06 | 2.600E-06 |
| 28092 | 1405 | P1405K | E | 1.4e-06 | 2e-06 | 2e-06 | 1.800E-06 | 3.464E-07 | 1.400E-06 | 2.000E-06 |
| 28093 | 1405 | P1405M | E | 2.8e-06 | 2.8e-06 | 2.4e-06 | 2.667E-06 | 2.309E-07 | 2.400E-06 | 2.800E-06 |
| 28094 | 1405 | P1405F | E | 2.7e-06 | 2.5e-06 | 2.4e-06 | 2.533E-06 | 1.528E-07 | 2.400E-06 | 2.700E-06 |
| 28095 | 1405 | P1405P | E | 1.3e-06 | 3.6e-06 | 2.2e-06 | 2.367E-06 | 1.159E-06 | 1.300E-06 | 3.600E-06 |
| 28096 | 1405 | P1405S | E | 3.1e-06 | 9.3e-07 | 2.3e-06 | 2.110E-06 | 1.097E-06 | 9.300E-07 | 3.100E-06 |
| 28097 | 1405 | P1405T | E | 2.6e-06 | 1.4e-06 | 2.9e-06 | 2.300E-06 | 7.937E-07 | 1.400E-06 | 2.900E-06 |
| 28098 | 1405 | P1405W | E | 1.7e-06 | 2.3e-06 | 2.3e-06 | 2.100E-06 | 3.464E-07 | 1.700E-06 | 2.300E-06 |
| 28099 | 1405 | P1405Y | E | 3.1e-06 | 1.9e-06 | 2.3e-06 | 2.433E-06 | 6.110E-07 | 1.900E-06 | 3.100E-06 |
| 28100 | 1405 | P1405V | E | 2.5e-06 | 3.4e-06 | 2.7e-06 | 2.867E-06 | 4.726E-07 | 2.500E-06 | 3.400E-06 |
| 28101 | 1406 | S1406A | E | 2.7e-06 | 1.5e-06 | 2.1e-06 | 2.100E-06 | 6.000E-07 | 1.500E-06 | 2.700E-06 |
| 28102 | 1406 | S1406R | E | 1.9e-06 | 2.2e-06 | 2.7e-06 | 2.267E-06 | 4.041E-07 | 1.900E-06 | 2.700E-06 |
| 28103 | 1406 | S1406N | E | 3.2e-06 | 3.8e-06 | 2.6e-06 | 3.200E-06 | 6.000E-07 | 2.600E-06 | 3.800E-06 |
| 28104 | 1406 | S1406D | E | 2.8e-06 | 3.9e-06 | 2.3e-06 | 3.000E-06 | 8.185E-07 | 2.300E-06 | 3.900E-06 |
| 28105 | 1406 | S1406C | E | 2.4e-06 | 1.6e-06 | 1.8e-06 | 1.933E-06 | 4.163E-07 | 1.600E-06 | 2.400E-06 |
| 28106 | 1406 | S1406E | E | 2.3e-06 | 2.8e-06 | 2.2e-06 | 2.433E-06 | 3.215E-07 | 2.200E-06 | 2.800E-06 |
| 28107 | 1406 | S1406Q | E | 2.6e-06 | 2.2e-06 | 3e-06 | 2.600E-06 | 4.000E-07 | 2.200E-06 | 3.000E-06 |
| 28108 | 1406 | S1406G | E | 2.9e-06 | 1.2e-06 | 3.1e-06 | 2.400E-06 | 1.044E-06 | 1.200E-06 | 3.100E-06 |
| 28109 | 1406 | S1406H | E | 2.8e-06 | 3.1e-06 | 1.7e-06 | 2.533E-06 | 7.371E-07 | 1.700E-06 | 3.100E-06 |
| 28110 | 1406 | S1406I | E | 2.6e-06 | 2.1e-06 | 2.6e-06 | 2.433E-06 | 2.887E-07 | 2.100E-06 | 2.600E-06 |
| 28111 | 1406 | S1406L | E | 2e-06 | 2.5e-06 | 1.6e-06 | 2.033E-06 | 4.509E-07 | 1.600E-06 | 2.500E-06 |
| 28112 | 1406 | S1406K | E | 3.2e-06 | 1e-06 | 3e-06 | 2.400E-06 | 1.217E-06 | 1.000E-06 | 3.200E-06 |
| 28113 | 1406 | S1406M | E | 2e-06 | 3.2e-06 | 2.8e-06 | 2.667E-06 | 6.110E-07 | 2.000E-06 | 3.200E-06 |
| 28114 | 1406 | S1406F | E | 2.7e-06 | 3.3e-06 | 1.7e-06 | 2.567E-06 | 8.083E-07 | 1.700E-06 | 3.300E-06 |
| 28115 | 1406 | S1406P | E | 1.1e-06 | 1.4e-06 | 4.2e-06 | 2.233E-06 | 1.710E-06 | 1.100E-06 | 4.200E-06 |
| 28116 | 1406 | S1406S | E | 1.6e-06 | 2.7e-06 | 1.9e-06 | 2.067E-06 | 5.686E-07 | 1.600E-06 | 2.700E-06 |
| 28117 | 1406 | S1406T | E | 1.4e-06 | 2.9e-06 | 3.3e-06 | 2.533E-06 | 1.002E-06 | 1.400E-06 | 3.300E-06 |
| 28118 | 1406 | S1406W | E | 2.6e-06 | 1e-06 | 2.4e-06 | 2.000E-06 | 8.718E-07 | 1.000E-06 | 2.600E-06 |
| 28119 | 1406 | S1406Y | E | 2e-06 | 2.2e-06 | 2e-06 | 2.067E-06 | 1.155E-07 | 2.000E-06 | 2.200E-06 |
| 28120 | 1406 | S1406V | E | 2.6e-06 | 1.8e-06 | 2.7e-06 | 2.367E-06 | 4.933E-07 | 1.800E-06 | 2.700E-06 |
| 28121 | 1407 | S1407A | E | 2.3e-06 | 2.7e-06 | 2.1e-06 | 2.367E-06 | 3.055E-07 | 2.100E-06 | 2.700E-06 |
| 28122 | 1407 | S1407R | E | 3e-06 | 1.7e-06 | 3.9e-06 | 2.867E-06 | 1.106E-06 | 1.700E-06 | 3.900E-06 |
| 28123 | 1407 | S1407N | E | 2.7e-06 | 6.4e-06 | 2.8e-06 | 3.967E-06 | 2.108E-06 | 2.700E-06 | 6.400E-06 |
| 28124 | 1407 | S1407D | E | 2.2e-06 | 3e-06 | 2.9e-06 | 2.700E-06 | 4.359E-07 | 2.200E-06 | 3.000E-06 |
| 28125 | 1407 | S1407C | E | 2.7e-06 | 3.4e-06 | 3.3e-06 | 3.133E-06 | 3.786E-07 | 2.700E-06 | 3.400E-06 |
| 28126 | 1407 | S1407E | E | 2.6e-06 | 1.4e-06 | 3.2e-06 | 2.400E-06 | 9.165E-07 | 1.400E-06 | 3.200E-06 |
| 28127 | 1407 | S1407Q | E | 2.6e-06 | 2.1e-06 | 2.4e-06 | 2.367E-06 | 2.517E-07 | 2.100E-06 | 2.600E-06 |
| 28128 | 1407 | S1407G | E | 3.1e-06 | 2.7e-06 | 2.2e-06 | 2.667E-06 | 4.509E-07 | 2.200E-06 | 3.100E-06 |
| 28129 | 1407 | S1407H | E | 2.4e-06 | 2.1e-06 | 2.1e-06 | 2.200E-06 | 1.732E-07 | 2.100E-06 | 2.400E-06 |
| 28130 | 1407 | S1407I | E | 1.9e-06 | 2.8e-06 | 1.8e-06 | 2.167E-06 | 5.508E-07 | 1.800E-06 | 2.800E-06 |
| 28131 | 1407 | S1407L | E | 2.1e-06 | 1.8e-06 | 1.5e-06 | 1.800E-06 | 3.000E-07 | 1.500E-06 | 2.100E-06 |
| 28132 | 1407 | S1407K | E | 2.1e-06 | 4e-06 | 3e-06 | 3.033E-06 | 9.504E-07 | 2.100E-06 | 4.000E-06 |
| 28133 | 1407 | S1407M | E | 2.5e-06 | 4.7e-06 | 2.2e-06 | 3.133E-06 | 1.365E-06 | 2.200E-06 | 4.700E-06 |
| 28134 | 1407 | S1407F | E | 2.6e-06 | 1.3e-06 | 3e-06 | 2.300E-06 | 8.888E-07 | 1.300E-06 | 3.000E-06 |

| | | | | | | | | | | |
|---|---|---|---|---|---|---|---|---|---|---|
| 28135 | 1407 | S1407P | E | 1.8e-06 | 2.4e-06 | 1.7e-06 | 1.967E-06 | 3.786E-07 | 1.700E-06 | 2.400E-06 |
| 28136 | 1407 | S1407S | E | 2e-06 | 3.7e-06 | 1.9e-06 | 2.533E-06 | 1.012E-06 | 1.900E-06 | 3.700E-06 |
| 28137 | 1407 | S1407T | E | 2.9e-06 | 2.2e-06 | 2.4e-06 | 2.500E-06 | 3.606E-07 | 2.200E-06 | 2.900E-06 |
| 28138 | 1407 | S1407W | E | 1.6e-06 | 3.8e-06 | 1.8e-06 | 2.400E-06 | 1.217E-06 | 1.600E-06 | 3.800E-06 |
| 28139 | 1407 | S1407Y | E | 4.1e-06 | 2.5e-06 | 2.9e-06 | 3.167E-06 | 8.327E-07 | 2.500E-06 | 4.100E-06 |
| 28140 | 1407 | S1407V | E | 1.7e-06 | 2.6e-06 | 3.5e-06 | 2.600E-06 | 9.000E-07 | 1.700E-06 | 3.500E-06 |
| 28141 | 1408 | K1408A | E | 1.8e-06 | 9.4e-07 | 1.2e-06 | 1.313E-06 | 4.411E-07 | 9.400E-07 | 1.800E-06 |
| 28142 | 1408 | K1408R | E | 2.5e-06 | 1.5e-06 | 2.9e-06 | 2.300E-06 | 7.211E-07 | 1.500E-06 | 2.900E-06 |
| 28143 | 1408 | K1408N | E | 2e-06 | 1.9e-06 | 1.1e-06 | 1.667E-06 | 4.933E-07 | 1.100E-06 | 2.000E-06 |
| 28144 | 1408 | K1408D | E | 3.4e-06 | 3.1e-06 | 3.1e-06 | 3.200E-06 | 1.732E-07 | 3.100E-06 | 3.400E-06 |
| 28145 | 1408 | K1408C | E | 3.3e-06 | 2.2e-06 | 2.6e-06 | 2.700E-06 | 5.568E-07 | 2.200E-06 | 3.300E-06 |
| 28146 | 1408 | K1408E | E | 2.4e-06 | 4.7e-06 | 1.7e-06 | 2.933E-06 | 1.570E-06 | 1.700E-06 | 4.700E-06 |
| 28147 | 1408 | K1408Q | E | 2.4e-06 | 1.6e-06 | 2.9e-06 | 2.300E-06 | 6.557E-07 | 1.600E-06 | 2.900E-06 |
| 28148 | 1408 | K1408G | E | 3.2e-06 | 3.7e-06 | 2.8e-06 | 3.233E-06 | 4.509E-07 | 2.800E-06 | 3.700E-06 |
| 28149 | 1408 | K1408H | E | 3.2e-06 | 3.1e-06 | 2.7e-06 | 3.000E-06 | 2.646E-07 | 2.700E-06 | 3.200E-06 |
| 28150 | 1408 | K1408I | E | 3.3e-06 | 3e-06 | 1.7e-06 | 2.667E-06 | 8.505E-07 | 1.700E-06 | 3.300E-06 |
| 28151 | 1408 | K1408L | E | 2.4e-06 | 1.2e-06 | 2.6e-06 | 2.067E-06 | 7.572E-07 | 1.200E-06 | 2.600E-06 |
| 28152 | 1408 | K1408K | E | 3.1e-06 | 2.1e-06 | 1.7e-06 | 2.300E-06 | 7.211E-07 | 1.700E-06 | 3.100E-06 |
| 28153 | 1408 | K1408M | E | 1.8e-06 | 2.2e-06 | 1.9e-06 | 1.967E-06 | 2.082E-07 | 1.800E-06 | 2.200E-06 |
| 28154 | 1408 | K1408F | E | 3.4e-06 | 2.3e-06 | 3.6e-06 | 3.100E-06 | 7.000E-07 | 2.300E-06 | 3.600E-06 |
| 28155 | 1408 | K1408P | E | 2.6e-06 | 1.6e-06 | 2.7e-06 | 2.300E-06 | 6.083E-07 | 1.600E-06 | 2.700E-06 |
| 28156 | 1408 | K1408S | E | 2.3e-06 | 2.1e-06 | 2.5e-06 | 2.300E-06 | 2.000E-07 | 2.100E-06 | 2.500E-06 |
| 28157 | 1408 | K1408T | E | 2.2e-06 | 3.6e-06 | 1.7e-06 | 2.500E-06 | 9.849E-07 | 1.700E-06 | 3.600E-06 |
| 28158 | 1408 | K1408W | E | 4.6e-06 | 2.1e-06 | 3.7e-06 | 3.467E-06 | 1.266E-06 | 2.100E-06 | 4.600E-06 |
| 28159 | 1408 | K1408Y | E | 1.8e-06 | 2e-06 | 1.9e-06 | 1.900E-06 | 1.000E-07 | 1.800E-06 | 2.000E-06 |
| 28160 | 1408 | K1408V | E | 2.6e-06 | 2.1e-06 | 2.3e-06 | 2.333E-06 | 2.517E-07 | 2.100E-06 | 2.600E-06 |
| 28161 | 1409 | S1409A | E | 1.9e-06 | 2.2e-06 | 2.7e-06 | 2.267E-06 | 4.041E-07 | 1.900E-06 | 2.700E-06 |
| 28162 | 1409 | S1409R | E | 4e-06 | 2.8e-06 | 2.5e-06 | 3.100E-06 | 7.937E-07 | 2.500E-06 | 4.000E-06 |
| 28163 | 1409 | S1409N | E | 2e-06 | 2.6e-06 | 2.3e-06 | 2.300E-06 | 3.000E-07 | 2.000E-06 | 2.600E-06 |
| 28164 | 1409 | S1409D | E | 2.5e-06 | 2.4e-06 | 1.8e-06 | 2.233E-06 | 3.786E-07 | 1.800E-06 | 2.500E-06 |
| 28165 | 1409 | S1409C | E | 2e-06 | 2.1e-06 | 1.7e-06 | 1.933E-06 | 2.082E-07 | 1.700E-06 | 2.100E-06 |
| 28166 | 1409 | S1409E | E | 1.7e-06 | 2.5e-06 | 3.1e-06 | 2.433E-06 | 7.024E-07 | 1.700E-06 | 3.100E-06 |
| 28167 | 1409 | S1409Q | E | 2.6e-06 | 3.5e-06 | 2.4e-06 | 2.833E-06 | 5.859E-07 | 2.400E-06 | 3.500E-06 |
| 28168 | 1409 | S1409G | E | 3.1e-06 | 2e-06 | 2.5e-06 | 2.533E-06 | 5.508E-07 | 2.000E-06 | 3.100E-06 |
| 28169 | 1409 | S1409H | E | 1.5e-06 | 2.3e-06 | 2.4e-06 | 2.067E-06 | 4.933E-07 | 1.500E-06 | 2.400E-06 |
| 28170 | 1409 | S1409I | E | 1.6e-06 | 1.7e-06 | 2.2e-06 | 1.833E-06 | 3.215E-07 | 1.600E-06 | 2.200E-06 |
| 28171 | 1409 | S1409L | E | 3.6e-06 | 2.9e-06 | 1.3e-06 | 2.600E-06 | 1.179E-06 | 1.300E-06 | 3.600E-06 |
| 28172 | 1409 | S1409K | E | 2.4e-06 | 2.2e-06 | 4.7e-06 | 3.100E-06 | 1.389E-06 | 2.200E-06 | 4.700E-06 |
| 28173 | 1409 | S1409M | E | 3.1e-06 | 2.7e-06 | 2e-06 | 2.600E-06 | 5.568E-07 | 2.000E-06 | 3.100E-06 |
| 28174 | 1409 | S1409F | E | 1.6e-06 | 2.6e-06 | 1.5e-06 | 1.900E-06 | 6.083E-07 | 1.500E-06 | 2.600E-06 |
| 28175 | 1409 | S1409P | E | 2.3e-06 | 2.3e-06 | 2.3e-06 | 2.300E-06 | 0.000E+00 | 2.300E-06 | 2.300E-06 |
| 28176 | 1409 | S1409S | E | 2.6e-06 | 2.6e-06 | 1.5e-06 | 2.233E-06 | 6.351E-07 | 1.500E-06 | 2.600E-06 |
| 28177 | 1409 | S1409T | E | 2.3e-06 | 2.4e-06 | 1.9e-06 | 2.200E-06 | 2.646E-07 | 1.900E-06 | 2.400E-06 |
| 28178 | 1409 | S1409W | E | 3.4e-06 | 2.9e-06 | 3.6e-06 | 3.300E-06 | 3.606E-07 | 2.900E-06 | 3.600E-06 |
| 28179 | 1409 | S1409Y | E | 2.5e-06 | 1.7e-06 | 1e-06 | 1.733E-06 | 7.506E-07 | 1.000E-06 | 2.500E-06 |
| 28180 | 1409 | S1409V | E | 1.5e-06 | 2.1e-06 | 2.7e-06 | 2.100E-06 | 6.000E-07 | 1.500E-06 | 2.700E-06 |
| 28181 | 1410 | T1410A | E | 2.9e-06 | 1.9e-06 | 2.2e-06 | 2.333E-06 | 5.132E-07 | 1.900E-06 | 2.900E-06 |
| 28182 | 1410 | T1410R | E | 2.7e-06 | 8.5e-07 | 2.2e-06 | 1.917E-06 | 9.570E-07 | 8.500E-07 | 2.700E-06 |
| 28183 | 1410 | T1410N | E | 2.7e-06 | 1.4e-06 | 2e-06 | 2.033E-06 | 6.506E-07 | 1.400E-06 | 2.700E-06 |
| 28184 | 1410 | T1410D | E | 2.9e-06 | 3.5e-06 | 2.7e-06 | 3.033E-06 | 4.163E-07 | 2.700E-06 | 3.500E-06 |
| 28185 | 1410 | T1410C | E | 2.9e-06 | 1.9e-06 | 2e-06 | 2.267E-06 | 5.508E-07 | 1.900E-06 | 2.900E-06 |
| 28186 | 1410 | T1410E | E | 3.1e-06 | 2.5e-06 | 1.5e-06 | 2.367E-06 | 8.083E-07 | 1.500E-06 | 3.100E-06 |
| 28187 | 1410 | T1410Q | E | 3.7e-06 | 2.5e-06 | 2.9e-06 | 3.033E-06 | 6.110E-07 | 2.500E-06 | 3.700E-06 |
| 28188 | 1410 | T1410G | E | 2e-06 | 2.5e-06 | 2.5e-06 | 2.333E-06 | 2.887E-07 | 2.000E-06 | 2.500E-06 |
| 28189 | 1410 | T1410H | E | 2.4e-06 | 1.8e-06 | 3.2e-06 | 2.467E-06 | 7.024E-07 | 1.800E-06 | 3.200E-06 |
| 28190 | 1410 | T1410I | E | 2.6e-06 | 3.3e-06 | 2e-06 | 2.633E-06 | 6.506E-07 | 2.000E-06 | 3.300E-06 |
| 28191 | 1410 | T1410L | E | 2.3e-06 | 3.1e-06 | 3.1e-06 | 2.833E-06 | 4.619E-07 | 2.300E-06 | 3.100E-06 |
| 28192 | 1410 | T1410K | E | 3.2e-06 | 2.1e-06 | 2.4e-06 | 2.567E-06 | 5.686E-07 | 2.100E-06 | 3.200E-06 |
| 28193 | 1410 | T1410M | E | 1.4e-06 | 1.8e-06 | 4.1e-06 | 2.433E-06 | 1.457E-06 | 1.400E-06 | 4.100E-06 |
| 28194 | 1410 | T1410F | E | 1.6e-06 | 2.7e-06 | 2.2e-06 | 2.167E-06 | 5.508E-07 | 1.600E-06 | 2.700E-06 |
| 28195 | 1410 | T1410P | E | 2.7e-06 | 3.2e-06 | 2.3e-06 | 2.733E-06 | 4.509E-07 | 2.300E-06 | 3.200E-06 |
| 28196 | 1410 | T1410S | E | 1.9e-06 | 2e-06 | 2.8e-06 | 2.233E-06 | 4.933E-07 | 1.900E-06 | 2.800E-06 |
| 28197 | 1410 | T1410T | E | 2.2e-06 | 2.6e-06 | 2e-06 | 2.267E-06 | 3.055E-07 | 2.000E-06 | 2.600E-06 |
| 28198 | 1410 | T1410W | E | 3.1e-06 | 1.2e-06 | 9.7e-07 | 1.757E-06 | 1.169E-06 | 9.700E-07 | 3.100E-06 |
| 28199 | 1410 | T1410Y | E | 2.8e-06 | 2.7e-06 | 2.2e-06 | 2.567E-06 | 3.215E-07 | 2.200E-06 | 2.800E-06 |
| 28200 | 1410 | T1410V | E | 2.3e-06 | 5e-06 | 1.9e-06 | 3.067E-06 | 1.686E-06 | 1.900E-06 | 5.000E-06 |
| 28201 | 1411 | S1411A | E | 2.9e-06 | 2.4e-06 | 3.7e-06 | 3.000E-06 | 6.557E-07 | 2.400E-06 | 3.700E-06 |
| 28202 | 1411 | S1411R | E | 2.4e-06 | 5.1e-06 | 2.1e-06 | 3.200E-06 | 1.652E-06 | 2.100E-06 | 5.100E-06 |
| 28203 | 1411 | S1411N | E | 1.3e-06 | 3.9e-06 | 2.5e-06 | 2.567E-06 | 1.301E-06 | 1.300E-06 | 3.900E-06 |
| 28204 | 1411 | S1411D | E | 2.8e-06 | 1.7e-06 | 1.8e-06 | 2.100E-06 | 6.083E-07 | 1.700E-06 | 2.800E-06 |
| 28205 | 1411 | S1411C | E | 4e-06 | 2e-06 | 2.8e-06 | 2.933E-06 | 1.007E-06 | 2.000E-06 | 4.000E-06 |
| 28206 | 1411 | S1411E | E | 3.6e-06 | 2e-06 | 4.1e-06 | 3.233E-06 | 1.097E-06 | 2.000E-06 | 4.100E-06 |
| 28207 | 1411 | S1411Q | E | 2.1e-06 | 1.8e-06 | 1.8e-06 | 1.900E-06 | 1.732E-07 | 1.800E-06 | 2.100E-06 |
| 28208 | 1411 | S1411G | E | 2.7e-06 | 2.5e-06 | 2.9e-06 | 2.700E-06 | 2.000E-07 | 2.500E-06 | 2.900E-06 |
| 28209 | 1411 | S1411H | E | 3.1e-06 | 3.9e-06 | 2.6e-06 | 3.200E-06 | 6.557E-07 | 2.600E-06 | 3.900E-06 |
| 28210 | 1411 | S1411I | E | 2.4e-06 | 1.8e-06 | 2.2e-06 | 2.133E-06 | 3.055E-07 | 1.800E-06 | 2.400E-06 |
| 28211 | 1411 | S1411L | E | 2.2e-06 | 1.5e-06 | 1.7e-06 | 1.800E-06 | 3.606E-07 | 1.500E-06 | 2.200E-06 |
| 28212 | 1411 | S1411K | E | 3.9e-06 | 4.8e-06 | 3.7e-06 | 4.133E-06 | 5.859E-07 | 3.700E-06 | 4.800E-06 |
| 28213 | 1411 | S1411M | E | 2.7e-06 | 2.1e-06 | 2.5e-06 | 2.433E-06 | 3.055E-07 | 2.100E-06 | 2.700E-06 |
| 28214 | 1411 | S1411F | E | 3.7e-06 | 3.7e-06 | 2.1e-06 | 3.167E-06 | 9.238E-07 | 2.100E-06 | 3.700E-06 |
| 28215 | 1411 | S1411P | E | 3.8e-06 | 2.1e-06 | 1.4e-06 | 2.433E-06 | 1.234E-06 | 1.400E-06 | 3.800E-06 |
| 28216 | 1411 | S1411S | E | 2.7e-06 | 3e-06 | 2.8e-06 | 2.833E-06 | 1.528E-07 | 2.700E-06 | 3.000E-06 |
| 28217 | 1411 | S1411T | E | 1.9e-06 | 2.6e-06 | 1.1e-06 | 1.867E-06 | 7.506E-07 | 1.100E-06 | 2.600E-06 |
| 28218 | 1411 | S1411W | E | 2.2e-06 | 1.7e-06 | 4e-06 | 2.633E-06 | 1.210E-06 | 1.700E-06 | 4.000E-06 |
| 28219 | 1411 | S1411Y | E | 2.3e-06 | 3.1e-06 | 2.9e-06 | 2.767E-06 | 4.163E-07 | 2.300E-06 | 3.100E-06 |

| | | | | | | | | | | |
|---|---|---|---|---|---|---|---|---|---|---|
| 28220 | 1411 | S1411V | E | 3.7e-06 | 3.1e-06 | 2.5e-06 | 3.100E-06 | 6.000E-07 | 2.500E-06 | 3.700E-06 |
| 28221 | 1412 | G1412A | E | 1.8e-06 | 2.7e-06 | 6.2e-06 | 3.567E-06 | 2.325E-06 | 1.800E-06 | 6.200E-06 |
| 28222 | 1412 | G1412R | E | 2.3e-06 | 4.2e-06 | 1.9e-06 | 2.800E-06 | 1.229E-06 | 1.900E-06 | 4.200E-06 |
| 28223 | 1412 | G1412N | E | 1.6e-06 | 2.9e-06 | 3.3e-06 | 2.600E-06 | 8.888E-07 | 1.600E-06 | 3.300E-06 |
| 28224 | 1412 | G1412D | E | 1.9e-06 | 2.3e-06 | 2.2e-06 | 2.133E-06 | 2.082E-07 | 1.900E-06 | 2.300E-06 |
| 28225 | 1412 | G1412C | E | 2.8e-06 | 3.7e-06 | 2.3e-06 | 2.933E-06 | 7.095E-07 | 2.300E-06 | 3.700E-06 |
| 28226 | 1412 | G1412E | E | 2.1e-06 | 2.5e-06 | 2e-06 | 2.200E-06 | 2.646E-07 | 2.000E-06 | 2.500E-06 |
| 28227 | 1412 | G1412Q | E | 2.6e-06 | 3.6e-06 | 2.8e-06 | 3.000E-06 | 5.292E-07 | 2.600E-06 | 3.600E-06 |
| 28228 | 1412 | G1412G | E | 1.8e-06 | 2.2e-06 | 2e-06 | 2.000E-06 | 2.000E-07 | 1.800E-06 | 2.200E-06 |
| 28229 | 1412 | G1412H | E | 2.2e-06 | 2.5e-06 | 1.5e-06 | 2.067E-06 | 5.132E-07 | 1.500E-06 | 2.500E-06 |
| 28230 | 1412 | G1412I | E | 1.6e-06 | 2.9e-06 | 3.4e-06 | 2.633E-06 | 9.292E-07 | 1.600E-06 | 3.400E-06 |
| 28231 | 1412 | G1412L | E | 3.7e-06 | 1.7e-06 | 4.9e-06 | 3.433E-06 | 1.617E-06 | 1.700E-06 | 4.900E-06 |
| 28232 | 1412 | G1412K | E | 3.8e-06 | 2.6e-06 | 2.1e-06 | 2.833E-06 | 8.737E-07 | 2.100E-06 | 3.800E-06 |
| 28233 | 1412 | G1412M | E | 2.3e-06 | 2.2e-06 | 2e-06 | 2.167E-06 | 1.528E-07 | 2.000E-06 | 2.300E-06 |
| 28234 | 1412 | G1412F | E | 1.8e-06 | 2.1e-06 | 2e-06 | 1.967E-06 | 1.528E-07 | 1.800E-06 | 2.100E-06 |
| 28235 | 1412 | G1412P | E | 2.2e-06 | 2.5e-06 | 2.6e-06 | 2.433E-06 | 2.082E-07 | 2.200E-06 | 2.600E-06 |
| 28236 | 1412 | G1412S | E | 2.8e-06 | 2.6e-06 | 1.5e-06 | 2.300E-06 | 7.000E-07 | 1.500E-06 | 2.800E-06 |
| 28237 | 1412 | G1412T | E | 1.9e-06 | 2.3e-06 | 2.5e-06 | 2.233E-06 | 3.055E-07 | 1.900E-06 | 2.500E-06 |
| 28238 | 1412 | G1412W | E | 2.2e-06 | 3.7e-06 | 1.9e-06 | 2.600E-06 | 9.644E-07 | 1.900E-06 | 3.700E-06 |
| 28239 | 1412 | G1412Y | E | 1.6e-06 | 2.9e-06 | 4.7e-06 | 3.067E-06 | 1.557E-06 | 1.600E-06 | 4.700E-06 |
| 28240 | 1412 | G1412V | E | 1.8e-06 | 1.2e-06 | 2e-06 | 1.667E-06 | 4.163E-07 | 1.200E-06 | 2.000E-06 |
| 28241 | 1413 | G1413A | E | 2.9e-06 | 2e-06 | 2.2e-06 | 2.367E-06 | 4.726E-07 | 2.000E-06 | 2.900E-06 |
| 28242 | 1413 | G1413R | E | 4.2e-06 | 2.4e-06 | 4.1e-06 | 3.567E-06 | 1.012E-06 | 2.400E-06 | 4.200E-06 |
| 28243 | 1413 | G1413N | E | 2.8e-06 | 4.2e-06 | 3.3e-06 | 3.433E-06 | 7.095E-07 | 2.800E-06 | 4.200E-06 |
| 28244 | 1413 | G1413D | E | 3.7e-06 | 3.9e-06 | 3.1e-06 | 3.567E-06 | 4.163E-07 | 3.100E-06 | 3.900E-06 |
| 28245 | 1413 | G1413C | E | 2.2e-06 | 1.3e-06 | 1.8e-06 | 1.767E-06 | 4.509E-07 | 1.300E-06 | 2.200E-06 |
| 28246 | 1413 | G1413E | E | 3e-06 | 1.6e-06 | 1.7e-06 | 2.100E-06 | 7.810E-07 | 1.600E-06 | 3.000E-06 |
| 28247 | 1413 | G1413Q | E | 2.6e-06 | 1.3e-06 | 2.2e-06 | 2.033E-06 | 6.658E-07 | 1.300E-06 | 2.600E-06 |
| 28248 | 1413 | G1413G | E | 9.3e-07 | 3e-06 | 3.1e-06 | 2.343E-06 | 1.225E-06 | 9.300E-07 | 3.100E-06 |
| 28249 | 1413 | G1413H | E | 1.7e-06 | 2e-06 | 1.9e-06 | 1.867E-06 | 1.528E-07 | 1.700E-06 | 2.000E-06 |
| 28250 | 1413 | G1413I | E | 1.8e-06 | 1.4e-06 | 2.3e-06 | 1.833E-06 | 4.509E-07 | 1.400E-06 | 2.300E-06 |
| 28251 | 1413 | G1413L | E | 4.1e-06 | 1.9e-06 | 2.2e-06 | 2.733E-06 | 1.193E-06 | 1.900E-06 | 4.100E-06 |
| 28252 | 1413 | G1413K | E | 1.7e-06 | 2.2e-06 | 1.8e-06 | 1.900E-06 | 2.646E-07 | 1.700E-06 | 2.200E-06 |
| 28253 | 1413 | G1413M | E | 2.3e-06 | 1.9e-06 | 2.8e-06 | 2.333E-06 | 4.509E-07 | 1.900E-06 | 2.800E-06 |
| 28254 | 1413 | G1413F | E | 2.7e-06 | 3.4e-06 | 1e-06 | 2.367E-06 | 1.234E-06 | 1.000E-06 | 3.400E-06 |
| 28255 | 1413 | G1413P | E | 7.4e-07 | 3.4e-06 | 1.6e-06 | 1.913E-06 | 1.357E-06 | 7.400E-07 | 3.400E-06 |
| 28256 | 1413 | G1413S | E | 2e-06 | 1.6e-06 | 1.8e-06 | 1.800E-06 | 2.000E-07 | 1.600E-06 | 2.000E-06 |
| 28257 | 1413 | G1413T | E | 2.2e-06 | 2.9e-06 | 2.8e-06 | 2.633E-06 | 3.786E-07 | 2.200E-06 | 2.900E-06 |
| 28258 | 1413 | G1413W | E | 2.2e-06 | 2.3e-06 | 2e-06 | 2.167E-06 | 1.528E-07 | 2.000E-06 | 2.300E-06 |
| 28259 | 1413 | G1413Y | E | 2.2e-06 | 2.9e-06 | 2.5e-06 | 2.533E-06 | 3.512E-07 | 2.200E-06 | 2.900E-06 |
| 28260 | 1413 | G1413V | E | 2.6e-06 | 3.1e-06 | 1.5e-06 | 2.400E-06 | 8.185E-07 | 1.500E-06 | 3.100E-06 |
| 28261 | 1414 | T1414A | E | 2.7e-06 | 1.6e-06 | 1.1e-06 | 1.800E-06 | 8.185E-07 | 1.100E-06 | 2.700E-06 |
| 28262 | 1414 | T1414R | E | 2.6e-06 | 3.6e-06 | 2.4e-06 | 2.867E-06 | 6.429E-07 | 2.400E-06 | 3.600E-06 |
| 28263 | 1414 | T1414N | E | 2.1e-06 | 2.6e-06 | 3.9e-06 | 2.867E-06 | 9.292E-07 | 2.100E-06 | 3.900E-06 |
| 28264 | 1414 | T1414D | E | 2.5e-06 | 2.8e-06 | 3.8e-06 | 3.033E-06 | 6.807E-07 | 2.500E-06 | 3.800E-06 |
| 28265 | 1414 | T1414C | E | 3.8e-06 | 2.3e-06 | 3.4e-06 | 3.167E-06 | 7.767E-07 | 2.300E-06 | 3.800E-06 |
| 28266 | 1414 | T1414E | E | 1.9e-06 | 2.5e-06 | 3e-06 | 2.467E-06 | 5.508E-07 | 1.900E-06 | 3.000E-06 |
| 28267 | 1414 | T1414Q | E | 1.7e-06 | 2.1e-06 | 1.8e-06 | 1.867E-06 | 2.082E-07 | 1.700E-06 | 2.100E-06 |
| 28268 | 1414 | T1414G | E | 2.7e-06 | 3e-06 | 1.7e-06 | 2.467E-06 | 6.807E-07 | 1.700E-06 | 3.000E-06 |
| 28269 | 1414 | T1414H | E | 2.6e-06 | 3.5e-06 | 2.4e-06 | 2.833E-06 | 5.859E-07 | 2.400E-06 | 3.500E-06 |
| 28270 | 1414 | T1414I | E | 3.8e-06 | 1.2e-06 | 2.7e-06 | 2.567E-06 | 1.305E-06 | 1.200E-06 | 3.800E-06 |
| 28271 | 1414 | T1414L | E | 2.2e-06 | 1.5e-06 | 2.6e-06 | 2.100E-06 | 5.568E-07 | 1.500E-06 | 2.600E-06 |
| 28272 | 1414 | T1414K | E | 2.9e-06 | 3.1e-06 | 4.4e-06 | 3.467E-06 | 8.145E-07 | 2.900E-06 | 4.400E-06 |
| 28273 | 1414 | T1414M | E | 1.7e-06 | 3.1e-06 | 2.4e-06 | 2.400E-06 | 7.000E-07 | 1.700E-06 | 3.100E-06 |
| 28274 | 1414 | T1414F | E | 5e-06 | 6e-06 | 2.2e-06 | 4.400E-06 | 1.970E-06 | 2.200E-06 | 6.000E-06 |
| 28275 | 1414 | T1414P | E | 1.5e-06 | 6.1e-06 | 2.4e-06 | 3.333E-06 | 2.438E-06 | 1.500E-06 | 6.100E-06 |
| 28276 | 1414 | T1414S | E | 1.8e-06 | 1.9e-06 | 2.2e-06 | 1.967E-06 | 2.082E-07 | 1.800E-06 | 2.200E-06 |
| 28277 | 1414 | T1414T | E | 2.1e-06 | 2.7e-06 | 3e-06 | 2.600E-06 | 4.583E-07 | 2.100E-06 | 3.000E-06 |
| 28278 | 1414 | T1414W | E | 2.7e-06 | 2.2e-06 | 2.9e-06 | 2.600E-06 | 3.606E-07 | 2.200E-06 | 2.900E-06 |
| 28279 | 1414 | T1414Y | E | 2.7e-06 | 1.7e-06 | 1.7e-06 | 2.033E-06 | 5.774E-07 | 1.700E-06 | 2.700E-06 |
| 28280 | 1414 | T1414V | E | 4e-06 | 3.2e-06 | 2e-06 | 3.067E-06 | 1.007E-06 | 2.000E-06 | 4.000E-06 |
| 28281 | 1415 | A1415A | E | 2.5e-06 | 8.4e-07 | 1.9e-06 | 1.747E-06 | 8.406E-07 | 8.400E-07 | 2.500E-06 |
| 28282 | 1415 | A1415R | E | 2.1e-06 | 3.2e-06 | 1.3e-06 | 2.200E-06 | 9.539E-07 | 1.300E-06 | 3.200E-06 |
| 28283 | 1415 | A1415N | E | 2.6e-06 | 1.3e-06 | 2.8e-06 | 2.233E-06 | 8.145E-07 | 1.300E-06 | 2.800E-06 |
| 28284 | 1415 | A1415D | E | 2.3e-06 | 2.2e-06 | 2.2e-06 | 2.233E-06 | 5.774E-08 | 2.200E-06 | 2.300E-06 |
| 28285 | 1415 | A1415C | E | 2.9e-06 | 3.2e-06 | 3.3e-06 | 3.133E-06 | 2.082E-07 | 2.900E-06 | 3.300E-06 |
| 28286 | 1415 | A1415E | E | 1.3e-06 | 2.9e-06 | 1e-06 | 1.733E-06 | 1.021E-06 | 1.000E-06 | 2.900E-06 |
| 28287 | 1415 | A1415Q | E | 2.1e-06 | 1.8e-06 | 2.4e-06 | 2.100E-06 | 3.000E-07 | 1.800E-06 | 2.400E-06 |
| 28288 | 1415 | A1415G | E | 2.3e-06 | 1.5e-06 | 3.8e-06 | 2.533E-06 | 1.168E-06 | 1.500E-06 | 3.800E-06 |
| 28289 | 1415 | A1415H | E | 2.5e-06 | 2.4e-06 | 1.9e-06 | 2.267E-06 | 3.215E-07 | 1.900E-06 | 2.500E-06 |
| 28290 | 1415 | A1415I | E | 2.4e-06 | 2.4e-06 | 2.2e-06 | 2.333E-06 | 1.155E-07 | 2.200E-06 | 2.400E-06 |
| 28291 | 1415 | A1415L | E | 2.3e-06 | 3.1e-06 | 2e-06 | 2.467E-06 | 5.686E-07 | 2.000E-06 | 3.100E-06 |
| 28292 | 1415 | A1415K | E | 1.4e-06 | 3.3e-06 | 2.2e-06 | 2.300E-06 | 9.539E-07 | 1.400E-06 | 3.300E-06 |
| 28293 | 1415 | A1415M | E | 1.3e-06 | 2.6e-06 | 2.8e-06 | 2.233E-06 | 8.145E-07 | 1.300E-06 | 2.800E-06 |
| 28294 | 1415 | A1415F | E | 3.2e-06 | 1.9e-06 | 2.2e-06 | 2.433E-06 | 6.807E-07 | 1.900E-06 | 3.200E-06 |
| 28295 | 1415 | A1415P | E | 2.7e-06 | 1.3e-06 | 2.2e-06 | 2.067E-06 | 7.095E-07 | 1.300E-06 | 2.700E-06 |
| 28296 | 1415 | A1415S | E | 1.5e-06 | 2.7e-06 | 3.5e-06 | 2.567E-06 | 1.007E-06 | 1.500E-06 | 3.500E-06 |
| 28297 | 1415 | A1415T | E | 5.8e-07 | 1.8e-06 | 3e-06 | 1.793E-06 | 1.210E-06 | 5.800E-07 | 3.000E-06 |
| 28298 | 1415 | A1415W | E | 3e-06 | 2.8e-06 | 3.1e-06 | 2.967E-06 | 1.528E-07 | 2.800E-06 | 3.100E-06 |
| 28299 | 1415 | A1415Y | E | 2.3e-06 | 2.6e-06 | 3.4e-06 | 2.767E-06 | 5.686E-07 | 2.300E-06 | 3.400E-06 |
| 28300 | 1415 | A1415V | E | 1.7e-06 | 2.6e-06 | 2e-06 | 2.100E-06 | 4.583E-07 | 1.700E-06 | 2.600E-06 |
| 28301 | 1416 | A1416A | E | 3.2e-06 | 2.7e-06 | 1.7e-06 | 2.533E-06 | 7.638E-07 | 1.700E-06 | 3.200E-06 |
| 28302 | 1416 | A1416R | E | 3.7e-06 | 1.2e-06 | 2.2e-06 | 2.367E-06 | 1.258E-06 | 1.200E-06 | 3.700E-06 |
| 28303 | 1416 | A1416N | E | 2.1e-06 | 4.6e-06 | 2.5e-06 | 3.067E-06 | 1.343E-06 | 2.100E-06 | 4.600E-06 |
| 28304 | 1416 | A1416D | E | 2.4e-06 | 3.2e-06 | 3.4e-06 | 3.000E-06 | 5.292E-07 | 2.400E-06 | 3.400E-06 |

| | | | | | | | | | | |
|---|---|---|---|---|---|---|---|---|---|---|
| 28305 | 1416 | A1416C | E | 2e-06 | 1.9e-06 | 1.7e-06 | 1.867E-06 | 1.528E-07 | 1.700E-06 | 2.000E-06 |
| 28306 | 1416 | A1416E | E | 2.9e-06 | 1.7e-06 | 2.3e-06 | 2.300E-06 | 6.000E-07 | 1.700E-06 | 2.900E-06 |
| 28307 | 1416 | A1416Q | E | 1.5e-06 | 2.7e-06 | 2.4e-06 | 2.200E-06 | 6.245E-07 | 1.500E-06 | 2.700E-06 |
| 28308 | 1416 | A1416G | E | 2.7e-06 | 1.9e-06 | 3.7e-06 | 2.767E-06 | 9.018E-07 | 1.900E-06 | 3.700E-06 |
| 28309 | 1416 | A1416H | E | 2.5e-06 | 3.3e-06 | 1.7e-06 | 2.500E-06 | 8.000E-07 | 1.700E-06 | 3.300E-06 |
| 28310 | 1416 | A1416I | E | 2.3e-06 | 9.8e-07 | 2.7e-06 | 1.993E-06 | 9.001E-07 | 9.800E-07 | 2.700E-06 |
| 28311 | 1416 | A1416L | E | 1.9e-06 | 4.4e-06 | 2.6e-06 | 2.967E-06 | 1.290E-06 | 1.900E-06 | 4.400E-06 |
| 28312 | 1416 | A1416K | E | 1.2e-06 | 1.6e-06 | 1.7e-06 | 1.500E-06 | 2.646E-07 | 1.200E-06 | 1.700E-06 |
| 28313 | 1416 | A1416M | E | 1.8e-06 | 2.4e-06 | 1.5e-06 | 1.900E-06 | 4.583E-07 | 1.500E-06 | 2.400E-06 |
| 28314 | 1416 | A1416F | E | 2.1e-06 | 2e-06 | 2.8e-06 | 2.300E-06 | 4.359E-07 | 2.000E-06 | 2.800E-06 |
| 28315 | 1416 | A1416P | E | 1.6e-06 | 2.5e-06 | 2.5e-06 | 2.200E-06 | 5.196E-07 | 1.600E-06 | 2.500E-06 |
| 28316 | 1416 | A1416S | E | 1.1e-06 | 4.1e-06 | 1.6e-06 | 2.267E-06 | 1.607E-06 | 1.100E-06 | 4.100E-06 |
| 28317 | 1416 | A1416T | E | 2.9e-06 | 2.4e-06 | 2.1e-06 | 2.467E-06 | 4.041E-07 | 2.100E-06 | 2.900E-06 |
| 28318 | 1416 | A1416W | E | 9.6e-07 | 6e-06 | 1.3e-06 | 2.753E-06 | 2.817E-06 | 9.600E-07 | 6.000E-06 |
| 28319 | 1416 | A1416Y | E | 2.4e-06 | 2.2e-06 | 2.6e-06 | 2.400E-06 | 2.000E-07 | 2.200E-06 | 2.600E-06 |
| 28320 | 1416 | A1416V | E | 1.1e-06 | 2.4e-06 | 2.1e-06 | 1.867E-06 | 6.807E-07 | 1.100E-06 | 2.400E-06 |
| 28321 | 1417 | L1417A | E | 2.9e-06 | 2.2e-06 | 2e-06 | 2.367E-06 | 4.726E-07 | 2.000E-06 | 2.900E-06 |
| 28322 | 1417 | L1417R | E | 2e-06 | 1.7e-06 | 2.6e-06 | 2.100E-06 | 4.583E-07 | 1.700E-06 | 2.600E-06 |
| 28323 | 1417 | L1417N | E | 2.3e-06 | 1.6e-06 | 1.3e-06 | 1.733E-06 | 5.132E-07 | 1.300E-06 | 2.300E-06 |
| 28324 | 1417 | L1417D | E | 2.5e-06 | 3.3e-06 | 2.7e-06 | 2.833E-06 | 4.163E-07 | 2.500E-06 | 3.300E-06 |
| 28325 | 1417 | L1417C | E | 2.5e-06 | 2.4e-06 | 2.6e-06 | 2.500E-06 | 1.000E-07 | 2.400E-06 | 2.600E-06 |
| 28326 | 1417 | L1417E | E | 2.3e-06 | 3.5e-06 | 3e-06 | 2.933E-06 | 6.028E-07 | 2.300E-06 | 3.500E-06 |
| 28327 | 1417 | L1417Q | E | 2.2e-06 | 4.3e-06 | 1.9e-06 | 2.800E-06 | 1.308E-06 | 1.900E-06 | 4.300E-06 |
| 28328 | 1417 | L1417G | E | 2.8e-06 | 1.8e-06 | 2.1e-06 | 2.233E-06 | 5.132E-07 | 1.800E-06 | 2.800E-06 |
| 28329 | 1417 | L1417H | E | 4.5e-06 | 5.1e-06 | 1.5e-06 | 3.700E-06 | 1.929E-06 | 1.500E-06 | 5.100E-06 |
| 28330 | 1417 | L1417I | E | 2.3e-06 | 1.3e-06 | 2.2e-06 | 1.933E-06 | 5.508E-07 | 1.300E-06 | 2.300E-06 |
| 28331 | 1417 | L1417L | E | 3e-06 | 2.3e-06 | 1.4e-06 | 2.233E-06 | 8.021E-07 | 1.400E-06 | 3.000E-06 |
| 28332 | 1417 | L1417K | E | 3.2e-06 | 2.1e-06 | 1.2e-06 | 2.167E-06 | 1.002E-06 | 1.200E-06 | 3.200E-06 |
| 28333 | 1417 | L1417M | E | 2.4e-06 | 2.7e-06 | 1.8e-06 | 2.300E-06 | 4.583E-07 | 1.800E-06 | 2.700E-06 |
| 28334 | 1417 | L1417F | E | 2.6e-06 | 2.6e-06 | 2.2e-06 | 2.467E-06 | 2.309E-07 | 2.200E-06 | 2.600E-06 |
| 28335 | 1417 | L1417P | E | 2.2e-06 | 1.9e-06 | 2.5e-06 | 2.200E-06 | 3.000E-07 | 1.900E-06 | 2.500E-06 |
| 28336 | 1417 | L1417S | E | 1.4e-06 | 3e-06 | 1.6e-06 | 2.000E-06 | 8.718E-07 | 1.400E-06 | 3.000E-06 |
| 28337 | 1417 | L1417T | E | 2.2e-06 | 2e-06 | 2.7e-06 | 2.300E-06 | 3.606E-07 | 2.000E-06 | 2.700E-06 |
| 28338 | 1417 | L1417W | E | 2.1e-06 | 3.3e-06 | 2.1e-06 | 2.500E-06 | 6.928E-07 | 2.100E-06 | 3.300E-06 |
| 28339 | 1417 | L1417Y | E | 1.7e-06 | 1.7e-06 | 3.3e-06 | 2.233E-06 | 9.238E-07 | 1.700E-06 | 3.300E-06 |
| 28340 | 1417 | L1417V | E | 4e-06 | 2.2e-06 | 1.4e-06 | 2.533E-06 | 1.332E-06 | 1.400E-06 | 4.000E-06 |
| 28341 | 1418 | G1418A | E | 2.1e-06 | 1.9e-06 | 2.4e-06 | 2.133E-06 | 2.517E-07 | 1.900E-06 | 2.400E-06 |
| 28342 | 1418 | G1418R | E | 2.1e-06 | 2e-06 | 2.6e-06 | 2.233E-06 | 3.215E-07 | 2.000E-06 | 2.600E-06 |
| 28343 | 1418 | G1418N | E | 1.9e-06 | 2.2e-06 | 3.5e-06 | 2.533E-06 | 8.505E-07 | 1.900E-06 | 3.500E-06 |
| 28344 | 1418 | G1418D | E | 1.9e-06 | 1.8e-06 | 1.9e-06 | 1.867E-06 | 5.774E-08 | 1.800E-06 | 1.900E-06 |
| 28345 | 1418 | G1418C | E | 2.3e-06 | 1.6e-06 | 2.1e-06 | 2.000E-06 | 3.606E-07 | 1.600E-06 | 2.300E-06 |
| 28346 | 1418 | G1418E | E | 1.9e-06 | 2.5e-06 | 2.1e-06 | 2.167E-06 | 3.055E-07 | 1.900E-06 | 2.500E-06 |
| 28347 | 1418 | G1418Q | E | 1.9e-06 | 2.2e-06 | 4e-06 | 2.700E-06 | 1.136E-06 | 1.900E-06 | 4.000E-06 |
| 28348 | 1418 | G1418G | E | 2e-06 | 2.1e-06 | 2.2e-06 | 2.100E-06 | 1.000E-07 | 2.000E-06 | 2.200E-06 |
| 28349 | 1418 | G1418H | E | 2.6e-06 | 3.4e-06 | 2.7e-06 | 2.900E-06 | 4.359E-07 | 2.600E-06 | 3.400E-06 |
| 28350 | 1418 | G1418I | E | 1.7e-06 | 2.4e-06 | 2.4e-06 | 2.167E-06 | 4.041E-07 | 1.700E-06 | 2.400E-06 |
| 28351 | 1418 | G1418L | E | 4.5e-06 | 3.3e-06 | 1.8e-06 | 3.200E-06 | 1.353E-06 | 1.800E-06 | 4.500E-06 |
| 28352 | 1418 | G1418K | E | 2.7e-06 | 3.2e-06 | 2e-06 | 2.633E-06 | 6.028E-07 | 2.000E-06 | 3.200E-06 |
| 28353 | 1418 | G1418M | E | 3.5e-06 | 3.5e-06 | 2.4e-06 | 3.133E-06 | 6.351E-07 | 2.400E-06 | 3.500E-06 |
| 28354 | 1418 | G1418F | E | 2e-06 | 1.7e-06 | 1.3e-06 | 1.667E-06 | 3.512E-07 | 1.300E-06 | 2.000E-06 |
| 28355 | 1418 | G1418P | E | 2.5e-06 | 1.5e-06 | 2.7e-06 | 2.233E-06 | 6.429E-07 | 1.500E-06 | 2.700E-06 |
| 28356 | 1418 | G1418S | E | 1e-06 | 2.4e-06 | 2.3e-06 | 1.900E-06 | 7.810E-07 | 1.000E-06 | 2.400E-06 |
| 28357 | 1418 | G1418T | E | 3e-06 | 1e-06 | 1.6e-06 | 1.867E-06 | 1.026E-06 | 1.000E-06 | 3.000E-06 |
| 28358 | 1418 | G1418W | E | 1.5e-06 | 2.3e-06 | 1.9e-06 | 1.900E-06 | 4.000E-07 | 1.500E-06 | 2.300E-06 |
| 28359 | 1418 | G1418Y | E | 1.9e-06 | 2.8e-06 | 2e-06 | 2.233E-06 | 4.933E-07 | 1.900E-06 | 2.800E-06 |
| 28360 | 1418 | G1418V | E | 4.4e-06 | 3.2e-06 | 1.1e-06 | 2.900E-06 | 1.670E-06 | 1.100E-06 | 4.400E-06 |
| 28361 | 1419 | C1419A | E | 1.6e-06 | 2.3e-06 | 1.9e-06 | 1.933E-06 | 3.512E-07 | 1.600E-06 | 2.300E-06 |
| 28362 | 1419 | C1419R | E | 2e-06 | 2.7e-06 | 3.8e-06 | 2.833E-06 | 9.074E-07 | 2.000E-06 | 3.800E-06 |
| 28363 | 1419 | C1419N | E | 2.6e-06 | 3.8e-06 | 2.5e-06 | 2.967E-06 | 7.234E-07 | 2.500E-06 | 3.800E-06 |
| 28364 | 1419 | C1419D | E | 2.5e-06 | 3.2e-06 | 1.6e-06 | 2.433E-06 | 8.021E-07 | 1.600E-06 | 3.200E-06 |
| 28365 | 1419 | C1419C | E | 2.3e-06 | 1.5e-06 | 2.8e-06 | 2.200E-06 | 6.557E-07 | 1.500E-06 | 2.800E-06 |
| 28366 | 1419 | C1419E | E | 2.4e-06 | 2.7e-06 | 2.4e-06 | 2.500E-06 | 1.732E-07 | 2.400E-06 | 2.700E-06 |
| 28367 | 1419 | C1419Q | E | 3.7e-06 | 3.9e-06 | 1.4e-06 | 3.000E-06 | 1.389E-06 | 1.400E-06 | 3.900E-06 |
| 28368 | 1419 | C1419G | E | 1.2e-06 | 2.1e-06 | 3.8e-06 | 2.367E-06 | 1.320E-06 | 1.200E-06 | 3.800E-06 |
| 28369 | 1419 | C1419H | E | 2.9e-06 | 5e-06 | 2.9e-06 | 3.600E-06 | 1.212E-06 | 2.900E-06 | 5.000E-06 |
| 28370 | 1419 | C1419I | E | 2.8e-06 | 3.9e-06 | 3.8e-06 | 3.500E-06 | 6.083E-07 | 2.800E-06 | 3.900E-06 |
| 28371 | 1419 | C1419L | E | 2.3e-06 | 2.4e-06 | 2.1e-06 | 2.267E-06 | 1.528E-07 | 2.100E-06 | 2.400E-06 |
| 28372 | 1419 | C1419K | E | 2.3e-06 | 1.7e-06 | 1.6e-06 | 1.867E-06 | 3.786E-07 | 1.600E-06 | 2.300E-06 |
| 28373 | 1419 | C1419M | E | 3.5e-06 | 3.8e-06 | 2.1e-06 | 3.133E-06 | 9.074E-07 | 2.100E-06 | 3.800E-06 |
| 28374 | 1419 | C1419F | E | 2.9e-06 | 1.7e-06 | 2.6e-06 | 2.400E-06 | 6.245E-07 | 1.700E-06 | 2.900E-06 |
| 28375 | 1419 | C1419P | E | 2.7e-06 | 2.6e-06 | 2e-06 | 2.433E-06 | 3.786E-07 | 2.000E-06 | 2.700E-06 |
| 28376 | 1419 | C1419S | E | 2.4e-06 | 2.4e-06 | 1.2e-06 | 2.000E-06 | 6.928E-07 | 1.200E-06 | 2.400E-06 |
| 28377 | 1419 | C1419T | E | 3.2e-06 | 3.9e-06 | 2.2e-06 | 3.100E-06 | 8.544E-07 | 2.200E-06 | 3.900E-06 |
| 28378 | 1419 | C1419W | E | 1.5e-06 | 2e-06 | 2.2e-06 | 1.900E-06 | 3.606E-07 | 1.500E-06 | 2.200E-06 |
| 28379 | 1419 | C1419Y | E | 2.3e-06 | 1.4e-06 | 2.1e-06 | 1.933E-06 | 4.726E-07 | 1.400E-06 | 2.300E-06 |
| 28380 | 1419 | C1419V | E | 2.3e-06 | 1.8e-06 | 2.7e-06 | 2.267E-06 | 4.509E-07 | 1.800E-06 | 2.700E-06 |
| 28381 | 1420 | L1420A | E | 1.5e-06 | 2.3e-06 | 3.9e-06 | 2.567E-06 | 1.222E-06 | 1.500E-06 | 3.900E-06 |
| 28382 | 1420 | L1420R | E | 2.4e-06 | 2.5e-06 | 3.2e-06 | 2.700E-06 | 4.359E-07 | 2.400E-06 | 3.200E-06 |
| 28383 | 1420 | L1420N | E | 1.1e-06 | 2.2e-06 | 3.4e-06 | 2.233E-06 | 1.150E-06 | 1.100E-06 | 3.400E-06 |
| 28384 | 1420 | L1420D | E | 2.2e-06 | 2.4e-06 | 1.8e-06 | 2.133E-06 | 3.055E-07 | 1.800E-06 | 2.400E-06 |
| 28385 | 1420 | L1420C | E | 3.1e-06 | 2.3e-06 | 1.6e-06 | 2.333E-06 | 7.506E-07 | 1.600E-06 | 3.100E-06 |
| 28386 | 1420 | L1420E | E | 3.6e-06 | 2.5e-06 | 3.1e-06 | 3.067E-06 | 5.508E-07 | 2.500E-06 | 3.600E-06 |
| 28387 | 1420 | L1420Q | E | 2.3e-06 | 3.1e-06 | 2.7e-06 | 2.700E-06 | 4.000E-07 | 2.300E-06 | 3.100E-06 |
| 28388 | 1420 | L1420G | E | 1.5e-06 | 2e-06 | 1.8e-06 | 1.767E-06 | 2.517E-07 | 1.500E-06 | 2.000E-06 |
| 28389 | 1420 | L1420H | E | 3.3e-06 | 3.5e-06 | 1.7e-06 | 2.833E-06 | 9.866E-07 | 1.700E-06 | 3.500E-06 |

| | | | | | | | | | | |
|---|---|---|---|---|---|---|---|---|---|---|
| 28390 | 1420 | L1420I | E | 1.4e-06 | 3.6e-06 | 3.4e-06 | 2.800E-06 | 1.217E-06 | 1.400E-06 | 3.600E-06 |
| 28391 | 1420 | L1420L | E | 2.5e-06 | 2.7e-06 | 3e-06 | 2.733E-06 | 2.517E-07 | 2.500E-06 | 3.000E-06 |
| 28392 | 1420 | L1420K | E | 1.9e-06 | 2.9e-06 | 3.5e-06 | 2.767E-06 | 8.083E-07 | 1.900E-06 | 3.500E-06 |
| 28393 | 1420 | L1420M | E | 2.8e-06 | 2.2e-06 | 2.3e-06 | 2.433E-06 | 3.215E-07 | 2.200E-06 | 2.800E-06 |
| 28394 | 1420 | L1420F | E | 1.6e-06 | 2.3e-06 | 2.9e-06 | 2.267E-06 | 6.506E-07 | 1.600E-06 | 2.900E-06 |
| 28395 | 1420 | L1420P | E | 2.9e-06 | 2.6e-06 | 2.7e-06 | 2.733E-06 | 1.528E-07 | 2.600E-06 | 2.900E-06 |
| 28396 | 1420 | L1420S | E | 3e-06 | 2.2e-06 | 3e-06 | 2.733E-06 | 4.619E-07 | 2.200E-06 | 3.000E-06 |
| 28397 | 1420 | L1420T | E | 3.1e-06 | 1.6e-06 | 2.1e-06 | 2.267E-06 | 7.638E-07 | 1.600E-06 | 3.100E-06 |
| 28398 | 1420 | L1420W | E | 2.6e-06 | 2.3e-06 | 3.4e-06 | 2.767E-06 | 5.686E-07 | 2.300E-06 | 3.400E-06 |
| 28399 | 1420 | L1420Y | E | 2.3e-06 | 4.7e-06 | 3.8e-06 | 3.600E-06 | 1.212E-06 | 2.300E-06 | 4.700E-06 |
| 28400 | 1420 | L1420V | E | 4.5e-06 | 2.1e-06 | 2.2e-06 | 2.933E-06 | 1.358E-06 | 2.100E-06 | 4.500E-06 |
| 28401 | 1421 | V1421A | E | 2.1e-06 | 1.5e-06 | 3.1e-06 | 2.233E-06 | 8.083E-07 | 1.500E-06 | 3.100E-06 |
| 28402 | 1421 | V1421R | E | 1.5e-06 | 3.6e-06 | 2.5e-06 | 2.533E-06 | 1.050E-06 | 1.500E-06 | 3.600E-06 |
| 28403 | 1421 | V1421N | E | 2e-06 | 2.6e-06 | 2.4e-06 | 2.333E-06 | 3.055E-07 | 2.000E-06 | 2.600E-06 |
| 28404 | 1421 | V1421D | E | 2.9e-06 | 3e-06 | 3e-06 | 2.967E-06 | 5.774E-08 | 2.900E-06 | 3.000E-06 |
| 28405 | 1421 | V1421C | E | 2.5e-06 | 2.5e-06 | 2e-06 | 2.333E-06 | 2.887E-07 | 2.000E-06 | 2.500E-06 |
| 28406 | 1421 | V1421E | E | 1.8e-06 | 2.7e-06 | 2.8e-06 | 2.433E-06 | 5.508E-07 | 1.800E-06 | 2.800E-06 |
| 28407 | 1421 | V1421Q | E | 3.1e-06 | 2e-06 | 2.9e-06 | 2.667E-06 | 5.859E-07 | 2.000E-06 | 3.100E-06 |
| 28408 | 1421 | V1421G | E | 1.9e-06 | 2.2e-06 | 3e-06 | 2.367E-06 | 5.686E-07 | 1.900E-06 | 3.000E-06 |
| 28409 | 1421 | V1421H | E | 4e-06 | 1.5e-06 | 3.4e-06 | 2.967E-06 | 1.305E-06 | 1.500E-06 | 4.000E-06 |
| 28410 | 1421 | V1421I | E | 3.4e-06 | 2.7e-06 | 1.6e-06 | 2.567E-06 | 9.074E-07 | 1.600E-06 | 3.400E-06 |
| 28411 | 1421 | V1421L | E | 2.4e-06 | 1.9e-06 | 3.2e-06 | 2.500E-06 | 6.557E-07 | 1.900E-06 | 3.200E-06 |
| 28412 | 1421 | V1421K | E | 3.1e-06 | 2.2e-06 | 1.7e-06 | 2.333E-06 | 7.095E-07 | 1.700E-06 | 3.100E-06 |
| 28413 | 1421 | V1421M | E | 2.9e-06 | 2.9e-06 | 3e-06 | 2.933E-06 | 5.774E-08 | 2.900E-06 | 3.000E-06 |
| 28414 | 1421 | V1421F | E | 4.4e-06 | 1.8e-06 | 2.1e-06 | 2.767E-06 | 1.422E-06 | 1.800E-06 | 4.400E-06 |
| 28415 | 1421 | V1421P | E | 3.2e-06 | 2e-06 | 2e-06 | 2.400E-06 | 6.928E-07 | 2.000E-06 | 3.200E-06 |
| 28416 | 1421 | V1421S | E | 1.5e-06 | 1.6e-06 | 2.3e-06 | 1.800E-06 | 4.359E-07 | 1.500E-06 | 2.300E-06 |
| 28417 | 1421 | V1421T | E | 2.4e-06 | 3.5e-06 | 2.3e-06 | 2.733E-06 | 6.658E-07 | 2.300E-06 | 3.500E-06 |
| 28418 | 1421 | V1421W | E | 1.6e-06 | 2.3e-06 | 1.1e-06 | 1.667E-06 | 6.028E-07 | 1.100E-06 | 2.300E-06 |
| 28419 | 1421 | V1421Y | E | 2.4e-06 | 1.3e-06 | 1.6e-06 | 1.767E-06 | 5.686E-07 | 1.300E-06 | 2.400E-06 |
| 28420 | 1421 | V1421V | E | 1.9e-06 | 3.4e-06 | 2.5e-06 | 2.600E-06 | 7.550E-07 | 1.900E-06 | 3.400E-06 |
| 28421 | 1422 | K1422A | E | 2e-06 | 3.6e-06 | 1.9e-06 | 2.500E-06 | 9.539E-07 | 1.900E-06 | 3.600E-06 |
| 28422 | 1422 | K1422R | E | 1.6e-06 | 2.2e-06 | 2.1e-06 | 1.967E-06 | 3.215E-07 | 1.600E-06 | 2.200E-06 |
| 28423 | 1422 | K1422N | E | 1.6e-06 | 1.1e-06 | 2.7e-06 | 1.800E-06 | 8.185E-07 | 1.100E-06 | 2.700E-06 |
| 28424 | 1422 | K1422D | E | 2.6e-06 | 2.3e-06 | 3.5e-06 | 2.800E-06 | 6.245E-07 | 2.300E-06 | 3.500E-06 |
| 28425 | 1422 | K1422C | E | 2e-06 | 3.6e-06 | 1.6e-06 | 2.400E-06 | 1.058E-06 | 1.600E-06 | 3.600E-06 |
| 28426 | 1422 | K1422E | E | 2.1e-06 | 3.6e-06 | 1.9e-06 | 2.533E-06 | 9.292E-07 | 1.900E-06 | 3.600E-06 |
| 28427 | 1422 | K1422Q | E | 2e-06 | 2.1e-06 | 3.6e-06 | 2.567E-06 | 8.963E-07 | 2.000E-06 | 3.600E-06 |
| 28428 | 1422 | K1422G | E | 2.2e-06 | 1.9e-06 | 2.3e-06 | 2.133E-06 | 2.082E-07 | 1.900E-06 | 2.300E-06 |
| 28429 | 1422 | K1422H | E | 2.2e-06 | 4.5e-06 | 3.3e-06 | 3.333E-06 | 1.150E-06 | 2.200E-06 | 4.500E-06 |
| 28430 | 1422 | K1422I | E | 2e-06 | 3e-06 | 2e-06 | 2.333E-06 | 5.774E-07 | 2.000E-06 | 3.000E-06 |
| 28431 | 1422 | K1422L | E | 2.5e-06 | 1.7e-06 | 3e-06 | 2.400E-06 | 6.557E-07 | 1.700E-06 | 3.000E-06 |
| 28432 | 1422 | K1422K | E | 2.1e-06 | 1.7e-06 | 2e-06 | 1.933E-06 | 2.082E-07 | 1.700E-06 | 2.100E-06 |
| 28433 | 1422 | K1422M | E | 3e-06 | 2e-06 | 2.9e-06 | 2.633E-06 | 5.508E-07 | 2.000E-06 | 3.000E-06 |
| 28434 | 1422 | K1422F | E | 2.5e-06 | 1.8e-06 | 1.1e-06 | 1.800E-06 | 7.000E-07 | 1.100E-06 | 2.500E-06 |
| 28435 | 1422 | K1422P | E | 1.5e-06 | 1.6e-06 | 2.2e-06 | 1.767E-06 | 3.786E-07 | 1.500E-06 | 2.200E-06 |
| 28436 | 1422 | K1422S | E | 2.2e-06 | 1.9e-06 | 2.5e-06 | 2.200E-06 | 3.000E-07 | 1.900E-06 | 2.500E-06 |
| 28437 | 1422 | K1422T | E | 3.7e-06 | 2.8e-06 | 2.6e-06 | 3.033E-06 | 5.859E-07 | 2.600E-06 | 3.700E-06 |
| 28438 | 1422 | K1422W | E | 2.6e-06 | 3.3e-06 | 2.5e-06 | 2.800E-06 | 4.359E-07 | 2.500E-06 | 3.300E-06 |
| 28439 | 1422 | K1422Y | E | 2e-06 | 1.8e-06 | 3.1e-06 | 2.300E-06 | 7.000E-07 | 1.800E-06 | 3.100E-06 |
| 28440 | 1422 | K1422V | E | 4.3e-06 | 2.1e-06 | 1.7e-06 | 2.700E-06 | 1.400E-06 | 1.700E-06 | 4.300E-06 |
| 28441 | 1423 | D1423A | E | 2e-06 | 2.3e-06 | 3.1e-06 | 2.467E-06 | 5.686E-07 | 2.000E-06 | 3.100E-06 |
| 28442 | 1423 | D1423R | E | 2.6e-06 | 3.1e-06 | 2.9e-06 | 2.867E-06 | 2.517E-07 | 2.600E-06 | 3.100E-06 |
| 28443 | 1423 | D1423N | E | 2.4e-06 | 2e-06 | 2.4e-06 | 2.267E-06 | 2.309E-07 | 2.000E-06 | 2.400E-06 |
| 28444 | 1423 | D1423D | E | 1.8e-06 | 2.7e-06 | 2.7e-06 | 2.400E-06 | 5.196E-07 | 1.800E-06 | 2.700E-06 |
| 28445 | 1423 | D1423C | E | 1.3e-06 | 1.3e-06 | 1.7e-06 | 1.433E-06 | 2.309E-07 | 1.300E-06 | 1.700E-06 |
| 28446 | 1423 | D1423E | E | 1.6e-06 | 1.8e-06 | 1.7e-06 | 1.700E-06 | 1.000E-07 | 1.600E-06 | 1.800E-06 |
| 28447 | 1423 | D1423Q | E | 9.3e-07 | 2.5e-06 | 1.9e-06 | 1.777E-06 | 7.922E-07 | 9.300E-07 | 2.500E-06 |
| 28448 | 1423 | D1423G | E | 2.5e-06 | 2.1e-06 | 2.6e-06 | 2.400E-06 | 2.646E-07 | 2.100E-06 | 2.600E-06 |
| 28449 | 1423 | D1423H | E | 2.8e-06 | 3e-06 | 2.5e-06 | 2.767E-06 | 2.517E-07 | 2.500E-06 | 3.000E-06 |
| 28450 | 1423 | D1423I | E | 3.1e-06 | 2e-06 | 2.6e-06 | 2.567E-06 | 5.508E-07 | 2.000E-06 | 3.100E-06 |
| 28451 | 1423 | D1423L | E | 1.7e-06 | 3.5e-06 | 2.3e-06 | 2.500E-06 | 9.165E-07 | 1.700E-06 | 3.500E-06 |
| 28452 | 1423 | D1423K | E | 2.8e-06 | 3.1e-06 | 1.3e-06 | 2.400E-06 | 9.644E-07 | 1.300E-06 | 3.100E-06 |
| 28453 | 1423 | D1423M | E | 3.1e-06 | 5.8e-06 | 3.1e-06 | 4.000E-06 | 1.559E-06 | 3.100E-06 | 5.800E-06 |
| 28454 | 1423 | D1423F | E | 2.1e-06 | 1.8e-06 | 2e-06 | 1.967E-06 | 1.528E-07 | 1.800E-06 | 2.100E-06 |
| 28455 | 1423 | D1423P | E | 1.8e-06 | 3.1e-06 | 1.2e-06 | 2.033E-06 | 9.713E-07 | 1.200E-06 | 3.100E-06 |
| 28456 | 1423 | D1423S | E | 2.7e-06 | 1.2e-06 | 2.3e-06 | 2.067E-06 | 7.767E-07 | 1.200E-06 | 2.700E-06 |
| 28457 | 1423 | D1423T | E | 2.2e-06 | 4.7e-06 | 4.7e-06 | 3.867E-06 | 1.443E-06 | 2.200E-06 | 4.700E-06 |
| 28458 | 1423 | D1423W | E | 4e-06 | 1.6e-06 | 9e-07 | 2.167E-06 | 1.626E-06 | 9.000E-07 | 4.000E-06 |
| 28459 | 1423 | D1423Y | E | 3e-06 | 1.7e-06 | 2e-06 | 2.233E-06 | 6.807E-07 | 1.700E-06 | 3.000E-06 |
| 28460 | 1423 | D1423V | E | 3.4e-06 | 2.9e-06 | 2.8e-06 | 3.033E-06 | 3.215E-07 | 2.800E-06 | 3.400E-06 |
| 28461 | 1424 | Y1424A | E | 1.9e-06 | 2.4e-06 | 1.7e-06 | 2.000E-06 | 3.606E-07 | 1.700E-06 | 2.400E-06 |
| 28462 | 1424 | Y1424R | E | 2.5e-06 | 1.4e-06 | 2.5e-06 | 2.133E-06 | 6.351E-07 | 1.400E-06 | 2.500E-06 |
| 28463 | 1424 | Y1424N | E | 1.6e-06 | 4e-06 | 3.2e-06 | 2.933E-06 | 1.222E-06 | 1.600E-06 | 4.000E-06 |
| 28464 | 1424 | Y1424D | E | 2.4e-06 | 2.1e-06 | 1.7e-06 | 2.067E-06 | 3.512E-07 | 1.700E-06 | 2.400E-06 |
| 28465 | 1424 | Y1424C | E | 1.5e-06 | 1.5e-06 | 3.1e-06 | 2.033E-06 | 9.238E-07 | 1.500E-06 | 3.100E-06 |
| 28466 | 1424 | Y1424E | E | 1.4e-06 | 2.1e-06 | 2e-06 | 1.833E-06 | 3.786E-07 | 1.400E-06 | 2.100E-06 |
| 28467 | 1424 | Y1424Q | E | 4e-06 | 2.5e-06 | 1.1e-06 | 2.533E-06 | 1.450E-06 | 1.100E-06 | 4.000E-06 |
| 28468 | 1424 | Y1424G | E | 1.9e-06 | 2.4e-06 | 2.6e-06 | 2.300E-06 | 3.606E-07 | 1.900E-06 | 2.600E-06 |
| 28469 | 1424 | Y1424H | E | 2.9e-06 | 2.7e-06 | 2.6e-06 | 2.733E-06 | 1.528E-07 | 2.600E-06 | 2.900E-06 |
| 28470 | 1424 | Y1424I | E | 1.9e-06 | 2.6e-06 | 1.6e-06 | 2.033E-06 | 5.132E-07 | 1.600E-06 | 2.600E-06 |
| 28471 | 1424 | Y1424L | E | 1.9e-06 | 3.1e-06 | 2.5e-06 | 2.500E-06 | 6.000E-07 | 1.900E-06 | 3.100E-06 |
| 28472 | 1424 | Y1424K | E | 2.1e-06 | 4.1e-06 | 2.2e-06 | 2.800E-06 | 1.127E-06 | 2.100E-06 | 4.100E-06 |
| 28473 | 1424 | Y1424M | E | 2.5e-06 | 2.3e-06 | 2.5e-06 | 2.433E-06 | 1.155E-07 | 2.300E-06 | 2.500E-06 |
| 28474 | 1424 | Y1424F | E | 3e-06 | 2.8e-06 | 1.1e-06 | 2.300E-06 | 1.044E-06 | 1.100E-06 | 3.000E-06 |

| | | | | | | | | | | |
|---|---|---|---|---|---|---|---|---|---|---|
| 28475 | 1424 | Y1424P | E | 2.2e-06 | 1.3e-06 | 2.1e-06 | 1.867E-06 | 4.933E-07 | 1.300E-06 | 2.200E-06 |
| 28476 | 1424 | Y1424S | E | 2.7e-06 | 1.5e-06 | 1.9e-06 | 2.033E-06 | 6.110E-07 | 1.500E-06 | 2.700E-06 |
| 28477 | 1424 | Y1424T | E | 4.9e-06 | 2.3e-06 | 1.9e-06 | 3.033E-06 | 1.629E-06 | 1.900E-06 | 4.900E-06 |
| 28478 | 1424 | Y1424W | E | 2.6e-06 | 1.1e-06 | 2.8e-06 | 2.167E-06 | 9.292E-07 | 1.100E-06 | 2.800E-06 |
| 28479 | 1424 | Y1424Y | E | 1.8e-06 | 1.8e-06 | 4.1e-06 | 2.567E-06 | 1.328E-06 | 1.800E-06 | 4.100E-06 |
| 28480 | 1424 | Y1424V | E | 2.8e-06 | 1.7e-06 | 3.5e-06 | 2.667E-06 | 9.074E-07 | 1.700E-06 | 3.500E-06 |
| 28481 | 1425 | F1425A | E | 1.8e-06 | 2.3e-06 | 1.8e-06 | 1.967E-06 | 2.887E-07 | 1.800E-06 | 2.300E-06 |
| 28482 | 1425 | F1425R | E | 9.6e-07 | 1.3e-06 | 3.1e-06 | 1.787E-06 | 1.150E-06 | 9.600E-07 | 3.100E-06 |
| 28483 | 1425 | F1425N | E | 1.4e-06 | 3.6e-06 | 2.7e-06 | 2.567E-06 | 1.106E-06 | 1.400E-06 | 3.600E-06 |
| 28484 | 1425 | F1425D | E | 1.5e-06 | 1.5e-06 | 2.6e-06 | 1.867E-06 | 6.351E-07 | 1.500E-06 | 2.600E-06 |
| 28485 | 1425 | F1425C | E | 8.5e-07 | 4.3e-06 | 2.8e-06 | 2.650E-06 | 1.730E-06 | 8.500E-07 | 4.300E-06 |
| 28486 | 1425 | F1425E | E | 2.9e-06 | 2.2e-06 | 4.4e-06 | 3.167E-06 | 1.124E-06 | 2.200E-06 | 4.400E-06 |
| 28487 | 1425 | F1425Q | E | 4.5e-06 | 3.6e-06 | 1.7e-06 | 3.267E-06 | 1.429E-06 | 1.700E-06 | 4.500E-06 |
| 28488 | 1425 | F1425G | E | 2.5e-06 | 1.9e-06 | 2e-06 | 2.133E-06 | 3.215E-07 | 1.900E-06 | 2.500E-06 |
| 28489 | 1425 | F1425H | E | 1.8e-06 | 3e-06 | 2.1e-06 | 2.300E-06 | 6.245E-07 | 1.800E-06 | 3.000E-06 |
| 28490 | 1425 | F1425I | E | 1.7e-06 | 1.4e-06 | 1.4e-06 | 1.500E-06 | 1.732E-07 | 1.400E-06 | 1.700E-06 |
| 28491 | 1425 | F1425L | E | 2.6e-06 | 2.1e-06 | 2.6e-06 | 2.433E-06 | 2.887E-07 | 2.100E-06 | 2.600E-06 |
| 28492 | 1425 | F1425K | E | 2.8e-06 | 3e-06 | 2.5e-06 | 2.767E-06 | 2.517E-07 | 2.500E-06 | 3.000E-06 |
| 28493 | 1425 | F1425M | E | 2.1e-06 | 3.2e-06 | 2.5e-06 | 2.600E-06 | 5.568E-07 | 2.100E-06 | 3.200E-06 |
| 28494 | 1425 | F1425F | E | 2.1e-06 | 2.3e-06 | 3.1e-06 | 2.500E-06 | 5.292E-07 | 2.100E-06 | 3.100E-06 |
| 28495 | 1425 | F1425P | E | 3.2e-06 | 2e-06 | 2.4e-06 | 2.533E-06 | 6.110E-07 | 2.000E-06 | 3.200E-06 |
| 28496 | 1425 | F1425S | E | 1.7e-06 | 2.7e-06 | 2.3e-06 | 2.233E-06 | 5.033E-07 | 1.700E-06 | 2.700E-06 |
| 28497 | 1425 | F1425T | E | 1.3e-06 | 1.6e-06 | 3.4e-06 | 2.100E-06 | 1.136E-06 | 1.300E-06 | 3.400E-06 |
| 28498 | 1425 | F1425W | E | 1.6e-06 | 3.3e-06 | 2.4e-06 | 2.433E-06 | 8.505E-07 | 1.600E-06 | 3.300E-06 |
| 28499 | 1425 | F1425Y | E | 2.8e-06 | 1.9e-06 | 2.5e-06 | 2.400E-06 | 4.583E-07 | 1.900E-06 | 2.800E-06 |
| 28500 | 1425 | F1425V | E | 3.9e-06 | 2.7e-06 | 1.4e-06 | 2.667E-06 | 1.250E-06 | 1.400E-06 | 3.900E-06 |
| 28501 | 1426 | P1426A | E | 4.5e-06 | 3.8e-06 | 2e-06 | 3.433E-06 | 1.290E-06 | 2.000E-06 | 4.500E-06 |
| 28502 | 1426 | P1426R | E | 3.5e-06 | 1.9e-06 | 1.9e-06 | 2.433E-06 | 9.238E-07 | 1.900E-06 | 3.500E-06 |
| 28503 | 1426 | P1426N | E | 2.4e-06 | 2.7e-06 | 1.9e-06 | 2.333E-06 | 4.041E-07 | 1.900E-06 | 2.700E-06 |
| 28504 | 1426 | P1426D | E | 2.7e-06 | 2.2e-06 | 1.5e-06 | 2.133E-06 | 6.028E-07 | 1.500E-06 | 2.700E-06 |
| 28505 | 1426 | P1426C | E | 2.2e-06 | 2.9e-06 | 1.2e-06 | 2.100E-06 | 8.544E-07 | 1.200E-06 | 2.900E-06 |
| 28506 | 1426 | P1426E | E | 2.8e-06 | 2.8e-06 | 3.4e-06 | 3.000E-06 | 3.464E-07 | 2.800E-06 | 3.400E-06 |
| 28507 | 1426 | P1426Q | E | 2.7e-06 | 2.8e-06 | 3.2e-06 | 2.900E-06 | 2.646E-07 | 2.700E-06 | 3.200E-06 |
| 28508 | 1426 | P1426G | E | 8.5e-07 | 2.7e-06 | 1.6e-06 | 1.717E-06 | 9.305E-07 | 8.500E-07 | 2.700E-06 |
| 28509 | 1426 | P1426H | E | 1.1e-06 | 2.5e-06 | 3.1e-06 | 2.233E-06 | 1.026E-06 | 1.100E-06 | 3.100E-06 |
| 28510 | 1426 | P1426I | E | 2.6e-06 | 1.6e-06 | 3.1e-06 | 2.433E-06 | 7.638E-07 | 1.600E-06 | 3.100E-06 |
| 28511 | 1426 | P1426L | E | 2.3e-06 | 1.4e-06 | 2.2e-06 | 1.967E-06 | 4.933E-07 | 1.400E-06 | 2.300E-06 |
| 28512 | 1426 | P1426K | E | 2.5e-06 | 4e-06 | 3.3e-06 | 3.267E-06 | 7.506E-07 | 2.500E-06 | 4.000E-06 |
| 28513 | 1426 | P1426M | E | 3.1e-06 | 1.4e-06 | 1.2e-06 | 1.900E-06 | 1.044E-06 | 1.200E-06 | 3.100E-06 |
| 28514 | 1426 | P1426F | E | 2.1e-06 | 2.3e-06 | 2.3e-06 | 2.233E-06 | 1.155E-07 | 2.100E-06 | 2.300E-06 |
| 28515 | 1426 | P1426P | E | 2.4e-06 | 2.9e-06 | 2.2e-06 | 2.500E-06 | 3.606E-07 | 2.200E-06 | 2.900E-06 |
| 28516 | 1426 | P1426S | E | 2.2e-06 | 1.6e-06 | 1.8e-06 | 1.867E-06 | 3.055E-07 | 1.600E-06 | 2.200E-06 |
| 28517 | 1426 | P1426T | E | 3.5e-06 | 1.3e-06 | 3.1e-06 | 2.633E-06 | 1.172E-06 | 1.300E-06 | 3.500E-06 |
| 28518 | 1426 | P1426W | E | 2.4e-06 | 2.5e-06 | 2.7e-06 | 2.533E-06 | 1.528E-07 | 2.400E-06 | 2.700E-06 |
| 28519 | 1426 | P1426Y | E | 1.9e-06 | 1.3e-06 | 2.7e-06 | 1.967E-06 | 7.024E-07 | 1.300E-06 | 2.700E-06 |
| 28520 | 1426 | P1426V | E | 1.6e-06 | 1.5e-06 | 3e-06 | 2.033E-06 | 8.386E-07 | 1.500E-06 | 3.000E-06 |
| 28521 | 1427 | E1427A | E | 1.3e-06 | 1.6e-06 | 2e-06 | 1.633E-06 | 3.512E-07 | 1.300E-06 | 2.000E-06 |
| 28522 | 1427 | E1427R | E | 1.8e-06 | 3e-06 | 4.3e-06 | 3.033E-06 | 1.250E-06 | 1.800E-06 | 4.300E-06 |
| 28523 | 1427 | E1427N | E | 2.3e-06 | 1.9e-06 | 3.1e-06 | 2.433E-06 | 6.110E-07 | 1.900E-06 | 3.100E-06 |
| 28524 | 1427 | E1427D | E | 1.8e-06 | 3.1e-06 | 3.9e-06 | 2.933E-06 | 1.060E-06 | 1.800E-06 | 3.900E-06 |
| 28525 | 1427 | E1427C | E | 3.1e-06 | 4.7e-06 | 4.7e-06 | 4.167E-06 | 9.238E-07 | 3.100E-06 | 4.700E-06 |
| 28526 | 1427 | E1427E | E | 1.9e-06 | 2.5e-06 | 3e-06 | 2.467E-06 | 5.508E-07 | 1.900E-06 | 3.000E-06 |
| 28527 | 1427 | E1427Q | E | 1.7e-06 | 2.1e-06 | 3.9e-06 | 2.567E-06 | 1.172E-06 | 1.700E-06 | 3.900E-06 |
| 28528 | 1427 | E1427G | E | 2.6e-06 | 3e-06 | 2.6e-06 | 2.733E-06 | 2.309E-07 | 2.600E-06 | 3.000E-06 |
| 28529 | 1427 | E1427H | E | 2.6e-06 | 2.1e-06 | 2e-06 | 2.233E-06 | 3.215E-07 | 2.000E-06 | 2.600E-06 |
| 28530 | 1427 | E1427I | E | 3.1e-06 | 2e-06 | 3.1e-06 | 2.733E-06 | 6.351E-07 | 2.000E-06 | 3.100E-06 |
| 28531 | 1427 | E1427L | E | 1.5e-06 | 2.3e-06 | 1.4e-06 | 1.733E-06 | 4.933E-07 | 1.400E-06 | 2.300E-06 |
| 28532 | 1427 | E1427K | E | 2.3e-06 | 2.2e-06 | 1.9e-06 | 2.133E-06 | 2.082E-07 | 1.900E-06 | 2.300E-06 |
| 28533 | 1427 | E1427M | E | 1.3e-06 | 2.3e-06 | 1.7e-06 | 1.767E-06 | 5.033E-07 | 1.300E-06 | 2.300E-06 |
| 28534 | 1427 | E1427F | E | 2.1e-06 | 2.1e-06 | 1.9e-06 | 2.033E-06 | 1.155E-07 | 1.900E-06 | 2.100E-06 |
| 28535 | 1427 | E1427P | E | 3.4e-06 | 1.6e-06 | 1.5e-06 | 2.167E-06 | 1.069E-06 | 1.500E-06 | 3.400E-06 |
| 28536 | 1427 | E1427S | E | 2.3e-06 | 2e-06 | 2.5e-06 | 2.267E-06 | 2.517E-07 | 2.000E-06 | 2.500E-06 |
| 28537 | 1427 | E1427T | E | 1.6e-06 | 2e-06 | 3e-06 | 2.200E-06 | 7.211E-07 | 1.600E-06 | 3.000E-06 |
| 28538 | 1427 | E1427W | E | 1.8e-06 | 2.8e-06 | 1.9e-06 | 2.167E-06 | 5.508E-07 | 1.800E-06 | 2.800E-06 |
| 28539 | 1427 | E1427Y | E | 1.6e-06 | 1.7e-06 | 2.1e-06 | 1.800E-06 | 2.646E-07 | 1.600E-06 | 2.100E-06 |
| 28540 | 1427 | E1427V | E | 2e-06 | 3.2e-06 | 4.1e-06 | 3.100E-06 | 1.054E-06 | 2.000E-06 | 4.100E-06 |
| 28541 | 1428 | P1428A | E | 1.6e-06 | 8e-07 | 3.3e-06 | 1.900E-06 | 1.277E-06 | 8.000E-07 | 3.300E-06 |
| 28542 | 1428 | P1428R | E | 2.4e-06 | 1.5e-06 | 4.8e-06 | 2.900E-06 | 1.706E-06 | 1.500E-06 | 4.800E-06 |
| 28543 | 1428 | P1428N | E | 2.5e-06 | 2.8e-06 | 1.8e-06 | 2.367E-06 | 5.132E-07 | 1.800E-06 | 2.800E-06 |
| 28544 | 1428 | P1428D | E | 3.1e-06 | 1.5e-06 | 2.5e-06 | 2.367E-06 | 8.083E-07 | 1.500E-06 | 3.100E-06 |
| 28545 | 1428 | P1428C | E | 2.7e-06 | 1.4e-06 | 2.1e-06 | 2.067E-06 | 6.506E-07 | 1.400E-06 | 2.700E-06 |
| 28546 | 1428 | P1428E | E | 2.1e-06 | 1.4e-06 | 1.9e-06 | 1.800E-06 | 3.606E-07 | 1.400E-06 | 2.100E-06 |
| 28547 | 1428 | P1428Q | E | 3.1e-06 | 2e-06 | 3.5e-06 | 2.867E-06 | 7.767E-07 | 2.000E-06 | 3.500E-06 |
| 28548 | 1428 | P1428G | E | 2.6e-06 | 1.4e-06 | 3.1e-06 | 2.367E-06 | 8.737E-07 | 1.400E-06 | 3.100E-06 |
| 28549 | 1428 | P1428H | E | 2.7e-06 | 2.6e-06 | 1.7e-06 | 2.333E-06 | 5.508E-07 | 1.700E-06 | 2.700E-06 |
| 28550 | 1428 | P1428I | E | 2.1e-06 | 2.5e-06 | 2.3e-06 | 2.300E-06 | 2.000E-07 | 2.100E-06 | 2.500E-06 |
| 28551 | 1428 | P1428L | E | 1.9e-06 | 4.8e-06 | 3e-06 | 3.233E-06 | 1.464E-06 | 1.900E-06 | 4.800E-06 |
| 28552 | 1428 | P1428K | E | 2.9e-06 | 3.1e-06 | 2.7e-06 | 2.900E-06 | 2.000E-07 | 2.700E-06 | 3.100E-06 |
| 28553 | 1428 | P1428M | E | 3.3e-06 | 2.8e-06 | 2.2e-06 | 2.767E-06 | 5.508E-07 | 2.200E-06 | 3.300E-06 |
| 28554 | 1428 | P1428F | E | 3.1e-06 | 2.6e-06 | 3e-06 | 2.900E-06 | 2.646E-07 | 2.600E-06 | 3.100E-06 |
| 28555 | 1428 | P1428P | E | 2.3e-06 | 2.4e-06 | 2.5e-06 | 2.400E-06 | 1.000E-07 | 2.300E-06 | 2.500E-06 |
| 28556 | 1428 | P1428S | E | 2.9e-06 | 2.1e-06 | 1.2e-06 | 2.067E-06 | 8.505E-07 | 1.200E-06 | 2.900E-06 |
| 28557 | 1428 | P1428T | E | 2.9e-06 | 2.1e-06 | 1.6e-06 | 2.200E-06 | 6.557E-07 | 1.600E-06 | 2.900E-06 |
| 28558 | 1428 | P1428W | E | 2.7e-06 | 3e-06 | 2e-06 | 2.567E-06 | 5.132E-07 | 2.000E-06 | 3.000E-06 |
| 28559 | 1428 | P1428Y | E | 2.1e-06 | 2.1e-06 | 3.1e-06 | 2.433E-06 | 5.774E-07 | 2.100E-06 | 3.100E-06 |

| | | | | | | | | | | |
|---|---|---|---|---|---|---|---|---|---|---|
| 28560 | 1428 | P1428V | E | 3.2e-06 | 2.6e-06 | 2.7e-06 | 2.833E-06 | 3.215E-07 | 2.600E-06 | 3.200E-06 |
| 28561 | 1429 | V1429A | E | 2.3e-06 | 2.7e-06 | 8.7e-07 | 1.957E-06 | 9.621E-07 | 8.700E-07 | 2.700E-06 |
| 28562 | 1429 | V1429R | E | 2.6e-06 | 1.8e-06 | 3e-06 | 2.467E-06 | 6.110E-07 | 1.800E-06 | 3.000E-06 |
| 28563 | 1429 | V1429N | E | 2.3e-06 | 3.1e-06 | 2.2e-06 | 2.533E-06 | 4.933E-07 | 2.200E-06 | 3.100E-06 |
| 28564 | 1429 | V1429D | E | 3.5e-06 | 3.2e-06 | 3.4e-06 | 3.367E-06 | 1.528E-07 | 3.200E-06 | 3.500E-06 |
| 28565 | 1429 | V1429C | E | 1.8e-06 | 1.5e-06 | 1.4e-06 | 1.567E-06 | 2.082E-07 | 1.400E-06 | 1.800E-06 |
| 28566 | 1429 | V1429E | E | 1.2e-06 | 2.6e-06 | 2.1e-06 | 1.967E-06 | 7.095E-07 | 1.200E-06 | 2.600E-06 |
| 28567 | 1429 | V1429Q | E | 1.5e-06 | 2.8e-06 | 2.5e-06 | 2.267E-06 | 6.807E-07 | 1.500E-06 | 2.800E-06 |
| 28568 | 1429 | V1429G | E | 2.4e-06 | 2.8e-06 | 8.7e-07 | 2.023E-06 | 1.019E-06 | 8.700E-07 | 2.800E-06 |
| 28569 | 1429 | V1429H | E | 1.7e-06 | 3.6e-06 | 2.4e-06 | 2.567E-06 | 9.609E-07 | 1.700E-06 | 3.600E-06 |
| 28570 | 1429 | V1429I | E | 1.6e-06 | 1.5e-06 | 3.8e-06 | 2.300E-06 | 1.300E-06 | 1.500E-06 | 3.800E-06 |
| 28571 | 1429 | V1429L | E | 2.4e-06 | 2.1e-06 | 2.6e-06 | 2.367E-06 | 2.517E-07 | 2.100E-06 | 2.600E-06 |
| 28572 | 1429 | V1429K | E | 3e-06 | 1.4e-06 | 1.9e-06 | 2.100E-06 | 8.185E-07 | 1.400E-06 | 3.000E-06 |
| 28573 | 1429 | V1429M | E | 2.7e-06 | 1.8e-06 | 2.5e-06 | 2.333E-06 | 4.726E-07 | 1.800E-06 | 2.700E-06 |
| 28574 | 1429 | V1429F | E | 1.7e-06 | 2.2e-06 | 3.2e-06 | 2.367E-06 | 7.638E-07 | 1.700E-06 | 3.200E-06 |
| 28575 | 1429 | V1429P | E | 1.2e-06 | 2e-06 | 9.2e-07 | 1.373E-06 | 5.605E-07 | 9.200E-07 | 2.000E-06 |
| 28576 | 1429 | V1429S | E | 3.4e-06 | 1.9e-06 | 1.7e-06 | 2.333E-06 | 9.292E-07 | 1.700E-06 | 3.400E-06 |
| 28577 | 1429 | V1429T | E | 2.2e-06 | 3.2e-06 | 1.7e-06 | 2.367E-06 | 7.638E-07 | 1.700E-06 | 3.200E-06 |
| 28578 | 1429 | V1429W | E | 1.5e-06 | 2.1e-06 | 2.2e-06 | 1.933E-06 | 3.786E-07 | 1.500E-06 | 2.200E-06 |
| 28579 | 1429 | V1429Y | E | 3.6e-06 | 2e-06 | 2e-06 | 2.533E-06 | 9.238E-07 | 2.000E-06 | 3.600E-06 |
| 28580 | 1429 | V1429V | E | 1.8e-06 | 1.5e-06 | 4.2e-06 | 2.500E-06 | 1.480E-06 | 1.500E-06 | 4.200E-06 |
| 28581 | 1430 | T1430A | E | 2.5e-06 | 2.3e-06 | 2.8e-06 | 2.533E-06 | 2.517E-07 | 2.300E-06 | 2.800E-06 |
| 28582 | 1430 | T1430R | E | 2e-06 | 2.3e-06 | 3.8e-06 | 2.700E-06 | 9.644E-07 | 2.000E-06 | 3.800E-06 |
| 28583 | 1430 | T1430N | E | 3.1e-06 | 2.6e-06 | 2.1e-06 | 2.600E-06 | 5.000E-07 | 2.100E-06 | 3.100E-06 |
| 28584 | 1430 | T1430D | E | 2.2e-06 | 2.8e-06 | 1.6e-06 | 2.200E-06 | 6.000E-07 | 1.600E-06 | 2.800E-06 |
| 28585 | 1430 | T1430C | E | 2.9e-06 | 3.1e-06 | 2.7e-06 | 2.900E-06 | 2.000E-07 | 2.700E-06 | 3.100E-06 |
| 28586 | 1430 | T1430E | E | 1.5e-06 | 2.3e-06 | 2.2e-06 | 2.000E-06 | 4.359E-07 | 1.500E-06 | 2.300E-06 |
| 28587 | 1430 | T1430Q | E | 1.5e-06 | 3.6e-06 | 3.5e-06 | 2.867E-06 | 1.185E-06 | 1.500E-06 | 3.600E-06 |
| 28588 | 1430 | T1430G | E | 4.8e-06 | 2.2e-06 | 2e-06 | 3.000E-06 | 1.562E-06 | 2.000E-06 | 4.800E-06 |
| 28589 | 1430 | T1430H | E | 2.7e-06 | 2.8e-06 | 1.6e-06 | 2.367E-06 | 6.658E-07 | 1.600E-06 | 2.800E-06 |
| 28590 | 1430 | T1430I | E | 1.7e-06 | 2.4e-06 | 1.8e-06 | 1.967E-06 | 3.786E-07 | 1.700E-06 | 2.400E-06 |
| 28591 | 1430 | T1430L | E | 2.8e-06 | 2.3e-06 | 1.5e-06 | 2.200E-06 | 6.557E-07 | 1.500E-06 | 2.800E-06 |
| 28592 | 1430 | T1430K | E | 2.1e-06 | 3.1e-06 | 2.9e-06 | 2.700E-06 | 5.292E-07 | 2.100E-06 | 3.100E-06 |
| 28593 | 1430 | T1430M | E | 2.1e-06 | 3.2e-06 | 2.4e-06 | 2.567E-06 | 5.686E-07 | 2.100E-06 | 3.200E-06 |
| 28594 | 1430 | T1430F | E | 1.8e-06 | 3.9e-06 | 3.3e-06 | 3.000E-06 | 1.082E-06 | 1.800E-06 | 3.900E-06 |
| 28595 | 1430 | T1430P | E | 2.7e-06 | 1.2e-06 | 2.8e-06 | 2.233E-06 | 8.963E-07 | 1.200E-06 | 2.800E-06 |
| 28596 | 1430 | T1430S | E | 1.3e-06 | 2.4e-06 | 1.7e-06 | 1.800E-06 | 5.568E-07 | 1.300E-06 | 2.400E-06 |
| 28597 | 1430 | T1430T | E | 2.3e-06 | 2.8e-06 | 1.7e-06 | 2.267E-06 | 5.508E-07 | 1.700E-06 | 2.800E-06 |
| 28598 | 1430 | T1430W | E | 4.9e-06 | 1.4e-06 | 2.3e-06 | 2.867E-06 | 1.818E-06 | 1.400E-06 | 4.900E-06 |
| 28599 | 1430 | T1430Y | E | 2e-06 | 1.3e-06 | 2.1e-06 | 1.800E-06 | 4.359E-07 | 1.300E-06 | 2.100E-06 |
| 28600 | 1430 | T1430V | E | 2.3e-06 | 1.4e-06 | 1.7e-06 | 1.800E-06 | 4.583E-07 | 1.400E-06 | 2.300E-06 |
| 28601 | 1431 | V1431A | E | 2.7e-06 | 2.7e-06 | 2.2e-06 | 2.533E-06 | 2.887E-07 | 2.200E-06 | 2.700E-06 |
| 28602 | 1431 | V1431R | E | 3.4e-06 | 1.7e-06 | 2.4e-06 | 2.500E-06 | 8.544E-07 | 1.700E-06 | 3.400E-06 |
| 28603 | 1431 | V1431N | E | 2e-06 | 1.3e-06 | 3.2e-06 | 2.167E-06 | 9.609E-07 | 1.300E-06 | 3.200E-06 |
| 28604 | 1431 | V1431D | E | 2e-06 | 2.9e-06 | 2.2e-06 | 2.367E-06 | 4.726E-07 | 2.000E-06 | 2.900E-06 |
| 28605 | 1431 | V1431C | E | 4e-06 | 2.2e-06 | 3.5e-06 | 3.233E-06 | 9.292E-07 | 2.200E-06 | 4.000E-06 |
| 28606 | 1431 | V1431E | E | 1.6e-06 | 1.9e-06 | 1.9e-06 | 1.800E-06 | 1.732E-07 | 1.600E-06 | 1.900E-06 |
| 28607 | 1431 | V1431Q | E | 1.9e-06 | 2.3e-06 | 3.7e-06 | 2.633E-06 | 9.452E-07 | 1.900E-06 | 3.700E-06 |
| 28608 | 1431 | V1431G | E | 2.6e-06 | 2.3e-06 | 2.5e-06 | 2.467E-06 | 1.528E-07 | 2.300E-06 | 2.600E-06 |
| 28609 | 1431 | V1431H | E | 3.5e-06 | 3.1e-06 | 1.3e-06 | 2.633E-06 | 1.172E-06 | 1.300E-06 | 3.500E-06 |
| 28610 | 1431 | V1431I | E | 1.9e-06 | 2.8e-06 | 1.3e-06 | 2.000E-06 | 7.550E-07 | 1.300E-06 | 2.800E-06 |
| 28611 | 1431 | V1431L | E | 2e-06 | 1.7e-06 | 2.1e-06 | 1.933E-06 | 2.082E-07 | 1.700E-06 | 2.100E-06 |
| 28612 | 1431 | V1431K | E | 2.6e-06 | 3.4e-06 | 3.2e-06 | 3.067E-06 | 4.163E-07 | 2.600E-06 | 3.400E-06 |
| 28613 | 1431 | V1431M | E | 2.3e-06 | 1.3e-06 | 2.1e-06 | 1.900E-06 | 5.292E-07 | 1.300E-06 | 2.300E-06 |
| 28614 | 1431 | V1431F | E | 2.1e-06 | 2.2e-06 | 1.7e-06 | 2.000E-06 | 2.646E-07 | 1.700E-06 | 2.200E-06 |
| 28615 | 1431 | V1431P | E | 2.1e-06 | 3.6e-06 | 3e-06 | 2.900E-06 | 7.550E-07 | 2.100E-06 | 3.600E-06 |
| 28616 | 1431 | V1431S | E | 2.8e-06 | 1.5e-06 | 2.5e-06 | 2.267E-06 | 6.807E-07 | 1.500E-06 | 2.800E-06 |
| 28617 | 1431 | V1431T | E | 4.4e-06 | 2.4e-06 | 3.4e-06 | 3.400E-06 | 1.000E-06 | 2.400E-06 | 4.400E-06 |
| 28618 | 1431 | V1431W | E | 1.9e-06 | 1.9e-06 | 3.6e-06 | 2.467E-06 | 9.815E-07 | 1.900E-06 | 3.600E-06 |
| 28619 | 1431 | V1431Y | E | 2.2e-06 | 3.1e-06 | 2.7e-06 | 2.667E-06 | 4.509E-07 | 2.200E-06 | 3.100E-06 |
| 28620 | 1431 | V1431V | E | 3.1e-06 | 3e-06 | 2.1e-06 | 2.733E-06 | 5.508E-07 | 2.100E-06 | 3.100E-06 |
| 28621 | 1432 | S1432A | E | 1.9e-06 | 2e-06 | 2.1e-06 | 2.000E-06 | 1.000E-07 | 1.900E-06 | 2.100E-06 |
| 28622 | 1432 | S1432R | E | 2e-06 | 2.2e-06 | 1.3e-06 | 1.833E-06 | 4.726E-07 | 1.300E-06 | 2.200E-06 |
| 28623 | 1432 | S1432N | E | 1.1e-06 | 2.7e-06 | 3.9e-06 | 2.567E-06 | 1.405E-06 | 1.100E-06 | 3.900E-06 |
| 28624 | 1432 | S1432D | E | 1.4e-06 | 1.4e-06 | 1.8e-06 | 1.533E-06 | 2.309E-07 | 1.400E-06 | 1.800E-06 |
| 28625 | 1432 | S1432C | E | 1.5e-06 | 2.8e-06 | 9.9e-07 | 1.763E-06 | 9.333E-07 | 9.900E-07 | 2.800E-06 |
| 28626 | 1432 | S1432E | E | 2.1e-06 | 1.3e-06 | 3.5e-06 | 2.300E-06 | 1.114E-06 | 1.300E-06 | 3.500E-06 |
| 28627 | 1432 | S1432Q | E | 2.8e-06 | 2.4e-06 | 1.6e-06 | 2.267E-06 | 6.110E-07 | 1.600E-06 | 2.800E-06 |
| 28628 | 1432 | S1432G | E | 2e-06 | 1.2e-06 | 1.8e-06 | 1.667E-06 | 4.163E-07 | 1.200E-06 | 2.000E-06 |
| 28629 | 1432 | S1432H | E | 2.4e-06 | 4.1e-06 | 1.6e-06 | 2.700E-06 | 1.277E-06 | 1.600E-06 | 4.100E-06 |
| 28630 | 1432 | S1432I | E | 1.8e-06 | 2.8e-06 | 1.3e-06 | 1.967E-06 | 7.638E-07 | 1.300E-06 | 2.800E-06 |
| 28631 | 1432 | S1432L | E | 2.2e-06 | 1.7e-06 | 3.4e-06 | 2.433E-06 | 8.737E-07 | 1.700E-06 | 3.400E-06 |
| 28632 | 1432 | S1432K | E | 3.2e-06 | 2.9e-06 | 3.1e-06 | 3.067E-06 | 1.528E-07 | 2.900E-06 | 3.200E-06 |
| 28633 | 1432 | S1432M | E | 2e-06 | 2.2e-06 | 3.2e-06 | 2.467E-06 | 6.429E-07 | 2.000E-06 | 3.200E-06 |
| 28634 | 1432 | S1432F | E | 3.6e-06 | 2.6e-06 | 2.4e-06 | 2.867E-06 | 6.429E-07 | 2.400E-06 | 3.600E-06 |
| 28635 | 1432 | S1432P | E | 1.7e-06 | 2e-06 | 3.9e-06 | 2.533E-06 | 1.193E-06 | 1.700E-06 | 3.900E-06 |
| 28636 | 1432 | S1432S | E | 2.5e-06 | 1.5e-06 | 4.9e-06 | 2.967E-06 | 1.747E-06 | 1.500E-06 | 4.900E-06 |
| 28637 | 1432 | S1432T | E | 1.7e-06 | 2.8e-06 | 1.9e-06 | 2.133E-06 | 5.859E-07 | 1.700E-06 | 2.800E-06 |
| 28638 | 1432 | S1432W | E | 1.9e-06 | 2.4e-06 | 1.6e-06 | 1.967E-06 | 4.041E-07 | 1.600E-06 | 2.400E-06 |
| 28639 | 1432 | S1432Y | E | 2.7e-06 | 3.2e-06 | 3.8e-06 | 3.233E-06 | 5.508E-07 | 2.700E-06 | 3.800E-06 |
| 28640 | 1432 | S1432V | E | 1.6e-06 | 2e-06 | 3.1e-06 | 2.233E-06 | 7.767E-07 | 1.600E-06 | 3.100E-06 |
| 28641 | 1433 | W1433A | E | 1.8e-06 | 2.5e-06 | 2.4e-06 | 2.233E-06 | 3.786E-07 | 1.800E-06 | 2.500E-06 |
| 28642 | 1433 | W1433R | E | 2.5e-06 | 2.4e-06 | 1.7e-06 | 2.200E-06 | 4.359E-07 | 1.700E-06 | 2.500E-06 |
| 28643 | 1433 | W1433N | E | 1.9e-06 | 1.9e-06 | 2.8e-06 | 2.200E-06 | 5.196E-07 | 1.900E-06 | 2.800E-06 |
| 28644 | 1433 | W1433D | E | 1.3e-06 | 2.5e-06 | 2.4e-06 | 2.067E-06 | 6.658E-07 | 1.300E-06 | 2.500E-06 |

| | | | | | | | | | | |
|---|---|---|---|---|---|---|---|---|---|---|
| 28645 | 1433 | W1433C | E | 2.3e-06 | 2.5e-06 | 3.5e-06 | 2.767E-06 | 6.429E-07 | 2.300E-06 | 3.500E-06 |
| 28646 | 1433 | W1433E | E | 4.3e-06 | 2e-06 | 1.7e-06 | 2.667E-06 | 1.422E-06 | 1.700E-06 | 4.300E-06 |
| 28647 | 1433 | W1433Q | E | 2.5e-06 | 2.6e-06 | 2.3e-06 | 2.467E-06 | 1.528E-07 | 2.300E-06 | 2.600E-06 |
| 28648 | 1433 | W1433G | E | 1.4e-06 | 2.3e-06 | 2e-06 | 1.900E-06 | 4.583E-07 | 1.400E-06 | 2.300E-06 |
| 28649 | 1433 | W1433H | E | 1.8e-06 | 2.2e-06 | 1.8e-06 | 1.933E-06 | 2.309E-07 | 1.800E-06 | 2.200E-06 |
| 28650 | 1433 | W1433I | E | 2.7e-06 | 2.7e-06 | 2.4e-06 | 2.600E-06 | 1.732E-07 | 2.400E-06 | 2.700E-06 |
| 28651 | 1433 | W1433L | E | 1.9e-06 | 4.9e-06 | 1.4e-06 | 2.733E-06 | 1.893E-06 | 1.400E-06 | 4.900E-06 |
| 28652 | 1433 | W1433K | E | 5.3e-06 | 1.4e-06 | 1.5e-06 | 2.733E-06 | 2.223E-06 | 1.400E-06 | 5.300E-06 |
| 28653 | 1433 | W1433M | E | 2.5e-06 | 4.3e-06 | 3.2e-06 | 3.333E-06 | 9.074E-07 | 2.500E-06 | 4.300E-06 |
| 28654 | 1433 | W1433F | E | 3.2e-06 | 3.9e-06 | 2.5e-06 | 3.200E-06 | 7.000E-07 | 2.500E-06 | 3.900E-06 |
| 28655 | 1433 | W1433P | E | 2.9e-06 | 2e-06 | 2.9e-06 | 2.600E-06 | 5.196E-07 | 2.000E-06 | 2.900E-06 |
| 28656 | 1433 | W1433S | E | 2e-06 | 4.6e-06 | 1.4e-06 | 2.667E-06 | 1.701E-06 | 1.400E-06 | 4.600E-06 |
| 28657 | 1433 | W1433T | E | 2.6e-06 | 2.4e-06 | 3.2e-06 | 2.733E-06 | 4.163E-07 | 2.400E-06 | 3.200E-06 |
| 28658 | 1433 | W1433W | E | 4.1e-06 | 2.4e-06 | 3.7e-06 | 3.400E-06 | 8.888E-07 | 2.400E-06 | 4.100E-06 |
| 28659 | 1433 | W1433Y | E | 2.4e-06 | 3.2e-06 | 1.9e-06 | 2.500E-06 | 6.557E-07 | 1.900E-06 | 3.200E-06 |
| 28660 | 1433 | W1433V | E | 2.9e-06 | 2.6e-06 | 2.2e-06 | 2.567E-06 | 3.512E-07 | 2.200E-06 | 2.900E-06 |
| 28661 | 1434 | N1434A | E | 3.4e-06 | 1.3e-06 | 2e-06 | 2.233E-06 | 1.069E-06 | 1.300E-06 | 3.400E-06 |
| 28662 | 1434 | N1434R | E | 2.4e-06 | 2.2e-06 | 1e-06 | 1.867E-06 | 7.572E-07 | 1.000E-06 | 2.400E-06 |
| 28663 | 1434 | N1434N | E | 3.2e-06 | 3.6e-06 | 1.7e-06 | 2.833E-06 | 1.002E-06 | 1.700E-06 | 3.600E-06 |
| 28664 | 1434 | N1434D | E | 2.1e-06 | 1.5e-06 | 2.5e-06 | 2.033E-06 | 5.033E-07 | 1.500E-06 | 2.500E-06 |
| 28665 | 1434 | N1434C | E | 2.3e-06 | 2.4e-06 | 1.6e-06 | 2.100E-06 | 4.359E-07 | 1.600E-06 | 2.400E-06 |
| 28666 | 1434 | N1434E | E | 1e-06 | 3.1e-06 | 1.9e-06 | 2.000E-06 | 1.054E-06 | 1.000E-06 | 3.100E-06 |
| 28667 | 1434 | N1434Q | E | 1.9e-06 | 1.3e-06 | 2.6e-06 | 1.933E-06 | 6.506E-07 | 1.300E-06 | 2.600E-06 |
| 28668 | 1434 | N1434G | E | 2.1e-06 | 1.9e-06 | 1.8e-06 | 1.933E-06 | 1.528E-07 | 1.800E-06 | 2.100E-06 |
| 28669 | 1434 | N1434H | E | 2.8e-06 | 2.9e-06 | 2e-06 | 2.567E-06 | 4.933E-07 | 2.000E-06 | 2.900E-06 |
| 28670 | 1434 | N1434I | E | 2.5e-06 | 3.4e-06 | 1.4e-06 | 2.433E-06 | 1.002E-06 | 1.400E-06 | 3.400E-06 |
| 28671 | 1434 | N1434L | E | 3.7e-06 | 1.8e-06 | 2.5e-06 | 2.667E-06 | 9.609E-07 | 1.800E-06 | 3.700E-06 |
| 28672 | 1434 | N1434K | E | 2.8e-06 | 2.8e-06 | 1.7e-06 | 2.433E-06 | 6.351E-07 | 1.700E-06 | 2.800E-06 |
| 28673 | 1434 | N1434M | E | 2.6e-06 | 1.2e-06 | 2.7e-06 | 2.167E-06 | 8.386E-07 | 1.200E-06 | 2.700E-06 |
| 28674 | 1434 | N1434F | E | 2.2e-06 | 3.8e-06 | 2.8e-06 | 2.933E-06 | 8.083E-07 | 2.200E-06 | 3.800E-06 |
| 28675 | 1434 | N1434P | E | 2.7e-06 | 4.1e-06 | 4.1e-06 | 3.633E-06 | 8.083E-07 | 2.700E-06 | 4.100E-06 |
| 28676 | 1434 | N1434S | E | 2.1e-06 | 3.2e-06 | 4.6e-06 | 3.300E-06 | 1.253E-06 | 2.100E-06 | 4.600E-06 |
| 28677 | 1434 | N1434T | E | 3.3e-06 | 2.1e-06 | 3.9e-06 | 3.100E-06 | 9.165E-07 | 2.100E-06 | 3.900E-06 |
| 28678 | 1434 | N1434W | E | 2.4e-06 | 3.2e-06 | 2.1e-06 | 2.567E-06 | 5.686E-07 | 2.100E-06 | 3.200E-06 |
| 28679 | 1434 | N1434Y | E | 1.9e-06 | 2.4e-06 | 3.8e-06 | 2.700E-06 | 9.849E-07 | 1.900E-06 | 3.800E-06 |
| 28680 | 1434 | N1434V | E | 2.7e-06 | 1.4e-06 | 2.1e-06 | 2.067E-06 | 6.506E-07 | 1.400E-06 | 2.700E-06 |
| 28681 | 1435 | S1435A | E | 2.7e-06 | 2.5e-06 | 1.7e-06 | 2.300E-06 | 5.292E-07 | 1.700E-06 | 2.700E-06 |
| 28682 | 1435 | S1435R | E | 1.8e-06 | 2.3e-06 | 1.8e-06 | 1.967E-06 | 2.887E-07 | 1.800E-06 | 2.300E-06 |
| 28683 | 1435 | S1435N | E | 2.5e-06 | 1.5e-06 | 2.4e-06 | 2.133E-06 | 5.508E-07 | 1.500E-06 | 2.500E-06 |
| 28684 | 1435 | S1435D | E | 3.6e-06 | 1.9e-06 | 3.6e-06 | 3.033E-06 | 9.815E-07 | 1.900E-06 | 3.600E-06 |
| 28685 | 1435 | S1435C | E | 2.1e-06 | 2.8e-06 | 3.8e-06 | 2.900E-06 | 8.544E-07 | 2.100E-06 | 3.800E-06 |
| 28686 | 1435 | S1435E | E | 2.9e-06 | 2.4e-06 | 1.8e-06 | 2.367E-06 | 5.508E-07 | 1.800E-06 | 2.900E-06 |
| 28687 | 1435 | S1435Q | E | 2.1e-06 | 2.6e-06 | 1.5e-06 | 2.067E-06 | 5.508E-07 | 1.500E-06 | 2.600E-06 |
| 28688 | 1435 | S1435G | E | 3.1e-06 | 1.7e-06 | 3e-06 | 2.600E-06 | 7.810E-07 | 1.700E-06 | 3.100E-06 |
| 28689 | 1435 | S1435H | E | 3.2e-06 | 2.3e-06 | 2.5e-06 | 2.667E-06 | 4.726E-07 | 2.300E-06 | 3.200E-06 |
| 28690 | 1435 | S1435I | E | 1.8e-06 | 1.4e-06 | 2e-06 | 1.733E-06 | 3.055E-07 | 1.400E-06 | 2.000E-06 |
| 28691 | 1435 | S1435L | E | 2e-06 | 2.3e-06 | 3.8e-06 | 2.700E-06 | 9.644E-07 | 2.000E-06 | 3.800E-06 |
| 28692 | 1435 | S1435K | E | 2.7e-06 | 1.7e-06 | 1.6e-06 | 2.000E-06 | 6.083E-07 | 1.600E-06 | 2.700E-06 |
| 28693 | 1435 | S1435M | E | 2.7e-06 | 4.2e-06 | 4.1e-06 | 3.667E-06 | 8.386E-07 | 2.700E-06 | 4.200E-06 |
| 28694 | 1435 | S1435F | E | 2.6e-06 | 1.6e-06 | 2.3e-06 | 2.167E-06 | 5.132E-07 | 1.600E-06 | 2.600E-06 |
| 28695 | 1435 | S1435P | E | 2.6e-06 | 2.1e-06 | 3.1e-06 | 2.600E-06 | 5.000E-07 | 2.100E-06 | 3.100E-06 |
| 28696 | 1435 | S1435S | E | 2.2e-06 | 4.3e-06 | 1.6e-06 | 2.700E-06 | 1.418E-06 | 1.600E-06 | 4.300E-06 |
| 28697 | 1435 | S1435T | E | 2.9e-06 | 2.1e-06 | 4.8e-06 | 3.267E-06 | 1.387E-06 | 2.100E-06 | 4.800E-06 |
| 28698 | 1435 | S1435W | E | 2.2e-06 | 1.8e-06 | 1.8e-06 | 1.933E-06 | 2.309E-07 | 1.800E-06 | 2.200E-06 |
| 28699 | 1435 | S1435Y | E | 3.2e-06 | 1.1e-06 | 1.5e-06 | 1.933E-06 | 1.115E-06 | 1.100E-06 | 3.200E-06 |
| 28700 | 1435 | S1435V | E | 1.8e-06 | 2.4e-06 | 3.2e-06 | 2.467E-06 | 7.024E-07 | 1.800E-06 | 3.200E-06 |
| 28701 | 1436 | G1436A | E | 2.4e-06 | 2.4e-06 | 2.5e-06 | 2.433E-06 | 5.774E-08 | 2.400E-06 | 2.500E-06 |
| 28702 | 1436 | G1436R | E | 2.5e-06 | 3.6e-06 | 1.9e-06 | 2.667E-06 | 8.622E-07 | 1.900E-06 | 3.600E-06 |
| 28703 | 1436 | G1436N | E | 2.9e-06 | 2.3e-06 | 3.4e-06 | 2.867E-06 | 5.508E-07 | 2.300E-06 | 3.400E-06 |
| 28704 | 1436 | G1436D | E | 2.3e-06 | 2.9e-06 | 1.3e-06 | 2.167E-06 | 8.083E-07 | 1.300E-06 | 2.900E-06 |
| 28705 | 1436 | G1436C | E | 2.3e-06 | 3.1e-06 | 2.7e-06 | 2.700E-06 | 4.000E-07 | 2.300E-06 | 3.100E-06 |
| 28706 | 1436 | G1436E | E | 2.6e-06 | 3.6e-06 | 3.3e-06 | 3.167E-06 | 5.132E-07 | 2.600E-06 | 3.600E-06 |
| 28707 | 1436 | G1436Q | E | 3.1e-06 | 2.1e-06 | 4.2e-06 | 3.133E-06 | 1.050E-06 | 2.100E-06 | 4.200E-06 |
| 28708 | 1436 | G1436G | E | 2.9e-06 | 2.5e-06 | 1.8e-06 | 2.400E-06 | 5.568E-07 | 1.800E-06 | 2.900E-06 |
| 28709 | 1436 | G1436H | E | 1.6e-06 | 3.6e-06 | 1.8e-06 | 2.333E-06 | 1.102E-06 | 1.600E-06 | 3.600E-06 |
| 28710 | 1436 | G1436I | E | 3.2e-06 | 2.2e-06 | 1.3e-06 | 2.233E-06 | 9.504E-07 | 1.300E-06 | 3.200E-06 |
| 28711 | 1436 | G1436L | E | 2.7e-06 | 2.6e-06 | 1.7e-06 | 2.333E-06 | 5.508E-07 | 1.700E-06 | 2.700E-06 |
| 28712 | 1436 | G1436K | E | 1.1e-06 | 1.5e-06 | 1.8e-06 | 1.467E-06 | 3.512E-07 | 1.100E-06 | 1.800E-06 |
| 28713 | 1436 | G1436M | E | 2.9e-06 | 2.6e-06 | 1.8e-06 | 2.433E-06 | 5.686E-07 | 1.800E-06 | 2.900E-06 |
| 28714 | 1436 | G1436F | E | 1.8e-06 | 2.9e-06 | 1.9e-06 | 2.200E-06 | 6.083E-07 | 1.800E-06 | 2.900E-06 |
| 28715 | 1436 | G1436P | E | 2.3e-06 | 2.3e-06 | 1.3e-06 | 1.967E-06 | 5.774E-07 | 1.300E-06 | 2.300E-06 |
| 28716 | 1436 | G1436S | E | 3e-06 | 3.9e-06 | 2.2e-06 | 3.033E-06 | 8.505E-07 | 2.200E-06 | 3.900E-06 |
| 28717 | 1436 | G1436T | E | 3.7e-06 | 2.7e-06 | 2.8e-06 | 3.067E-06 | 5.508E-07 | 2.700E-06 | 3.700E-06 |
| 28718 | 1436 | G1436W | E | 2e-06 | 2.4e-06 | 1.5e-06 | 1.967E-06 | 4.509E-07 | 1.500E-06 | 2.400E-06 |
| 28719 | 1436 | G1436Y | E | 2.2e-06 | 2e-06 | 1.6e-06 | 1.933E-06 | 3.055E-07 | 1.600E-06 | 2.200E-06 |
| 28720 | 1436 | G1436V | E | 3.3e-06 | 2.2e-06 | 3.6e-06 | 3.033E-06 | 7.371E-07 | 2.200E-06 | 3.600E-06 |
| 28721 | 1437 | A1437A | E | 3.4e-06 | 3.2e-06 | 1.8e-06 | 2.800E-06 | 8.718E-07 | 1.800E-06 | 3.400E-06 |
| 28722 | 1437 | A1437R | E | 2.5e-06 | 2.2e-06 | 2e-06 | 2.233E-06 | 2.517E-07 | 2.000E-06 | 2.500E-06 |
| 28723 | 1437 | A1437N | E | 1.6e-06 | 1.9e-06 | 2.7e-06 | 2.067E-06 | 5.686E-07 | 1.600E-06 | 2.700E-06 |
| 28724 | 1437 | A1437D | E | 2.1e-06 | 1.8e-06 | 2e-06 | 1.967E-06 | 1.528E-07 | 1.800E-06 | 2.100E-06 |
| 28725 | 1437 | A1437C | E | 2.3e-06 | 2.5e-06 | 2.6e-06 | 2.467E-06 | 1.528E-07 | 2.300E-06 | 2.600E-06 |
| 28726 | 1437 | A1437E | E | 2e-06 | 1.8e-06 | 1.7e-06 | 1.833E-06 | 1.528E-07 | 1.700E-06 | 2.000E-06 |
| 28727 | 1437 | A1437Q | E | 2.4e-06 | 2.6e-06 | 2.1e-06 | 2.367E-06 | 2.517E-07 | 2.100E-06 | 2.600E-06 |
| 28728 | 1437 | A1437G | E | 2.9e-06 | 3.4e-06 | 2.8e-06 | 3.033E-06 | 3.215E-07 | 2.800E-06 | 3.400E-06 |
| 28729 | 1437 | A1437H | E | 2.2e-06 | 3.5e-06 | 2.2e-06 | 2.633E-06 | 7.506E-07 | 2.200E-06 | 3.500E-06 |

| | | | | | | | | | | |
|---|---|---|---|---|---|---|---|---|---|---|
| 28730 | 1437 | A1437I | E | 1.8e-06 | 3.1e-06 | 3.2e-06 | 2.700E-06 | 7.810E-07 | 1.800E-06 | 3.200E-06 |
| 28731 | 1437 | A1437L | E | 1.9e-06 | 3.4e-06 | 2.5e-06 | 2.600E-06 | 7.550E-07 | 1.900E-06 | 3.400E-06 |
| 28732 | 1437 | A1437K | E | 1.8e-06 | 2.6e-06 | 2e-06 | 2.133E-06 | 4.163E-07 | 1.800E-06 | 2.600E-06 |
| 28733 | 1437 | A1437M | E | 1.3e-06 | 2.3e-06 | 2.3e-06 | 1.967E-06 | 5.774E-07 | 1.300E-06 | 2.300E-06 |
| 28734 | 1437 | A1437F | E | 2.5e-06 | 1.7e-06 | 1.8e-06 | 2.000E-06 | 4.359E-07 | 1.700E-06 | 2.500E-06 |
| 28735 | 1437 | A1437P | E | 1.4e-06 | 3.3e-06 | 2.6e-06 | 2.433E-06 | 9.609E-07 | 1.400E-06 | 3.300E-06 |
| 28736 | 1437 | A1437S | E | 9.1e-07 | 3.6e-06 | 2.2e-06 | 2.237E-06 | 1.345E-06 | 9.100E-07 | 3.600E-06 |
| 28737 | 1437 | A1437T | E | 2.2e-06 | 3.4e-06 | 3.1e-06 | 2.900E-06 | 6.245E-07 | 2.200E-06 | 3.400E-06 |
| 28738 | 1437 | A1437W | E | 1e-06 | 2.7e-06 | 2.3e-06 | 2.000E-06 | 8.888E-07 | 1.000E-06 | 2.700E-06 |
| 28739 | 1437 | A1437Y | E | 1.7e-06 | 2.2e-06 | 3.8e-06 | 2.567E-06 | 1.097E-06 | 1.700E-06 | 3.800E-06 |
| 28740 | 1437 | A1437V | E | 2.8e-06 | 2.7e-06 | 1.5e-06 | 2.333E-06 | 7.234E-07 | 1.500E-06 | 2.800E-06 |
| 28741 | 1438 | L1438A | E | 2.3e-06 | 1.1e-06 | 2.3e-06 | 1.900E-06 | 6.928E-07 | 1.100E-06 | 2.300E-06 |
| 28742 | 1438 | L1438R | E | 1.4e-06 | 1.4e-06 | 2.2e-06 | 1.667E-06 | 4.619E-07 | 1.400E-06 | 2.200E-06 |
| 28743 | 1438 | L1438N | E | 2.5e-06 | 3.5e-06 | 2.1e-06 | 2.700E-06 | 7.211E-07 | 2.100E-06 | 3.500E-06 |
| 28744 | 1438 | L1438D | E | 1.9e-06 | 3.3e-06 | 2.1e-06 | 2.433E-06 | 7.572E-07 | 1.900E-06 | 3.300E-06 |
| 28745 | 1438 | L1438C | E | 2.8e-06 | 2.7e-06 | 4.1e-06 | 3.200E-06 | 7.810E-07 | 2.700E-06 | 4.100E-06 |
| 28746 | 1438 | L1438E | E | 2.8e-06 | 1.6e-06 | 2.1e-06 | 2.167E-06 | 6.028E-07 | 1.600E-06 | 2.800E-06 |
| 28747 | 1438 | L1438Q | E | 2.6e-06 | 2.5e-06 | 1.4e-06 | 2.167E-06 | 6.658E-07 | 1.400E-06 | 2.600E-06 |
| 28748 | 1438 | L1438G | E | 1.9e-06 | 1.9e-06 | 3.5e-06 | 2.433E-06 | 9.238E-07 | 1.900E-06 | 3.500E-06 |
| 28749 | 1438 | L1438H | E | 2.5e-06 | 2.6e-06 | 2.4e-06 | 2.500E-06 | 1.000E-07 | 2.400E-06 | 2.600E-06 |
| 28750 | 1438 | L1438I | E | 2.5e-06 | 3.6e-06 | 3.4e-06 | 3.167E-06 | 5.859E-07 | 2.500E-06 | 3.600E-06 |
| 28751 | 1438 | L1438L | E | 2.1e-06 | 2.6e-06 | 2.1e-06 | 2.267E-06 | 2.887E-07 | 2.100E-06 | 2.600E-06 |
| 28752 | 1438 | L1438K | E | 2.5e-06 | 5.6e-06 | 2.2e-06 | 3.433E-06 | 1.882E-06 | 2.200E-06 | 5.600E-06 |
| 28753 | 1438 | L1438M | E | 1.9e-06 | 2.7e-06 | 2.4e-06 | 2.333E-06 | 4.041E-07 | 1.900E-06 | 2.700E-06 |
| 28754 | 1438 | L1438F | E | 3.1e-06 | 2.5e-06 | 1.9e-06 | 2.500E-06 | 6.000E-07 | 1.900E-06 | 3.100E-06 |
| 28755 | 1438 | L1438P | E | 1.6e-06 | 1.9e-06 | 1.8e-06 | 1.767E-06 | 1.528E-07 | 1.600E-06 | 1.900E-06 |
| 28756 | 1438 | L1438S | E | 3e-06 | 3.4e-06 | 3.5e-06 | 3.300E-06 | 2.646E-07 | 3.000E-06 | 3.500E-06 |
| 28757 | 1438 | L1438T | E | 2.7e-06 | 4.3e-06 | 2.7e-06 | 3.233E-06 | 9.238E-07 | 2.700E-06 | 4.300E-06 |
| 28758 | 1438 | L1438W | E | 2.7e-06 | 2.9e-06 | 2.4e-06 | 2.667E-06 | 2.517E-07 | 2.400E-06 | 2.900E-06 |
| 28759 | 1438 | L1438Y | E | 3.4e-06 | 2.9e-06 | 9.6e-07 | 2.420E-06 | 1.289E-06 | 9.600E-07 | 3.400E-06 |
| 28760 | 1438 | L1438V | E | 1.4e-06 | 3.1e-06 | 4.3e-06 | 2.933E-06 | 1.457E-06 | 1.400E-06 | 4.300E-06 |
| 28761 | 1439 | T1439A | E | 2.5e-06 | 2.3e-06 | 2.5e-06 | 2.433E-06 | 1.155E-07 | 2.300E-06 | 2.500E-06 |
| 28762 | 1439 | T1439R | E | 4.1e-06 | 1.9e-06 | 2.3e-06 | 2.767E-06 | 1.172E-06 | 1.900E-06 | 4.100E-06 |
| 28763 | 1439 | T1439N | E | 2e-06 | 2.3e-06 | 2.9e-06 | 2.400E-06 | 4.583E-07 | 2.000E-06 | 2.900E-06 |
| 28764 | 1439 | T1439D | E | 1.6e-06 | 2.1e-06 | 3.9e-06 | 2.533E-06 | 1.210E-06 | 1.600E-06 | 3.900E-06 |
| 28765 | 1439 | T1439C | E | 2.9e-06 | 3.9e-06 | 2.7e-06 | 3.167E-06 | 6.429E-07 | 2.700E-06 | 3.900E-06 |
| 28766 | 1439 | T1439E | E | 2.6e-06 | 2e-06 | 2.2e-06 | 2.267E-06 | 3.055E-07 | 2.000E-06 | 2.600E-06 |
| 28767 | 1439 | T1439Q | E | 1.8e-06 | 1.1e-06 | 1.7e-06 | 1.533E-06 | 3.786E-07 | 1.100E-06 | 1.800E-06 |
| 28768 | 1439 | T1439G | E | 2e-06 | 2.2e-06 | 3.8e-06 | 2.667E-06 | 9.866E-07 | 2.000E-06 | 3.800E-06 |
| 28769 | 1439 | T1439H | E | 3.3e-06 | 1.6e-06 | 2.1e-06 | 2.333E-06 | 8.737E-07 | 1.600E-06 | 3.300E-06 |
| 28770 | 1439 | T1439I | E | 2.4e-06 | 2.6e-06 | 2.5e-06 | 2.500E-06 | 1.000E-07 | 2.400E-06 | 2.600E-06 |
| 28771 | 1439 | T1439L | E | 6.4e-07 | 2.9e-06 | 2.8e-06 | 2.113E-06 | 1.277E-06 | 6.400E-07 | 2.900E-06 |
| 28772 | 1439 | T1439K | E | 1.6e-06 | 1.3e-06 | 3.1e-06 | 2.000E-06 | 9.644E-07 | 1.300E-06 | 3.100E-06 |
| 28773 | 1439 | T1439M | E | 2.8e-06 | 1.3e-06 | 1.8e-06 | 1.967E-06 | 7.638E-07 | 1.300E-06 | 2.800E-06 |
| 28774 | 1439 | T1439F | E | 1.4e-06 | 2.6e-06 | 3.1e-06 | 2.367E-06 | 8.737E-07 | 1.400E-06 | 3.100E-06 |
| 28775 | 1439 | T1439P | E | 3.7e-06 | 2.2e-06 | 2.9e-06 | 2.933E-06 | 7.506E-07 | 2.200E-06 | 3.700E-06 |
| 28776 | 1439 | T1439S | E | 2.7e-06 | 1.6e-06 | 3.1e-06 | 2.467E-06 | 7.767E-07 | 1.600E-06 | 3.100E-06 |
| 28777 | 1439 | T1439T | E | 3.6e-06 | 1.4e-06 | 1.8e-06 | 2.267E-06 | 1.172E-06 | 1.400E-06 | 3.600E-06 |
| 28778 | 1439 | T1439W | E | 2.3e-06 | 1.8e-06 | 2.2e-06 | 2.100E-06 | 2.646E-07 | 1.800E-06 | 2.300E-06 |
| 28779 | 1439 | T1439Y | E | 2.1e-06 | 2.2e-06 | 1.2e-06 | 1.833E-06 | 5.508E-07 | 1.200E-06 | 2.200E-06 |
| 28780 | 1439 | T1439V | E | 3e-06 | 1.8e-06 | 2.5e-06 | 2.433E-06 | 6.028E-07 | 1.800E-06 | 3.000E-06 |
| 28781 | 1440 | S1440A | E | 1.6e-06 | 2.3e-06 | 3.3e-06 | 2.400E-06 | 8.544E-07 | 1.600E-06 | 3.300E-06 |
| 28782 | 1440 | S1440R | E | 1.8e-06 | 2.4e-06 | 2.1e-06 | 2.100E-06 | 3.000E-07 | 1.800E-06 | 2.400E-06 |
| 28783 | 1440 | S1440N | E | 3.5e-06 | 4e-06 | 2.3e-06 | 3.267E-06 | 8.737E-07 | 2.300E-06 | 4.000E-06 |
| 28784 | 1440 | S1440D | E | 1.9e-06 | 3e-06 | 3.9e-06 | 2.933E-06 | 1.002E-06 | 1.900E-06 | 3.900E-06 |
| 28785 | 1440 | S1440C | E | 2.7e-06 | 3.1e-06 | 3e-06 | 2.933E-06 | 2.082E-07 | 2.700E-06 | 3.100E-06 |
| 28786 | 1440 | S1440E | E | 3.7e-06 | 2.8e-06 | 2.3e-06 | 2.933E-06 | 7.095E-07 | 2.300E-06 | 3.700E-06 |
| 28787 | 1440 | S1440Q | E | 2.3e-06 | 4.3e-06 | 2e-06 | 2.867E-06 | 1.250E-06 | 2.000E-06 | 4.300E-06 |
| 28788 | 1440 | S1440G | E | 3.4e-06 | 2.2e-06 | 4.8e-06 | 3.467E-06 | 1.301E-06 | 2.200E-06 | 4.800E-06 |
| 28789 | 1440 | S1440H | E | 2.3e-06 | 3.4e-06 | 2.7e-06 | 2.800E-06 | 5.568E-07 | 2.300E-06 | 3.400E-06 |
| 28790 | 1440 | S1440I | E | 3.7e-06 | 2.6e-06 | 2.1e-06 | 2.800E-06 | 8.185E-07 | 2.100E-06 | 3.700E-06 |
| 28791 | 1440 | S1440L | E | 3.6e-06 | 3.1e-06 | 3.1e-06 | 3.267E-06 | 2.887E-07 | 3.100E-06 | 3.600E-06 |
| 28792 | 1440 | S1440K | E | 2.6e-06 | 3.4e-06 | 2.9e-06 | 2.967E-06 | 4.041E-07 | 2.600E-06 | 3.400E-06 |
| 28793 | 1440 | S1440M | E | 2.6e-06 | 2.8e-06 | 3e-06 | 2.800E-06 | 2.000E-07 | 2.600E-06 | 3.000E-06 |
| 28794 | 1440 | S1440F | E | 1.7e-06 | 2.4e-06 | 2.7e-06 | 2.267E-06 | 5.132E-07 | 1.700E-06 | 2.700E-06 |
| 28795 | 1440 | S1440P | E | 1.6e-06 | 1.7e-06 | 2.2e-06 | 1.833E-06 | 3.215E-07 | 1.600E-06 | 2.200E-06 |
| 28796 | 1440 | S1440S | E | 4.8e-06 | 2.2e-06 | 2.7e-06 | 3.233E-06 | 1.380E-06 | 2.200E-06 | 4.800E-06 |
| 28797 | 1440 | S1440T | E | 3.3e-06 | 1.8e-06 | 3e-06 | 2.700E-06 | 7.937E-07 | 1.800E-06 | 3.300E-06 |
| 28798 | 1440 | S1440W | E | 1.4e-06 | 3e-06 | 2.3e-06 | 2.233E-06 | 8.021E-07 | 1.400E-06 | 3.000E-06 |
| 28799 | 1440 | S1440Y | E | 2e-06 | 3.6e-06 | 2.6e-06 | 2.733E-06 | 8.083E-07 | 2.000E-06 | 3.600E-06 |
| 28800 | 1440 | S1440V | E | 3.3e-06 | 2.8e-06 | 1.7e-06 | 2.600E-06 | 8.185E-07 | 1.700E-06 | 3.300E-06 |
| 28801 | 1441 | G1441A | E | 1.7e-06 | 2.2e-06 | 2.1e-06 | 2.000E-06 | 2.646E-07 | 1.700E-06 | 2.200E-06 |
| 28802 | 1441 | G1441R | E | 1.5e-06 | 3e-06 | 1.9e-06 | 2.133E-06 | 7.767E-07 | 1.500E-06 | 3.000E-06 |
| 28803 | 1441 | G1441N | E | 2.2e-06 | 1.4e-06 | 1.8e-06 | 1.800E-06 | 4.000E-07 | 1.400E-06 | 2.200E-06 |
| 28804 | 1441 | G1441D | E | 1.7e-06 | 2.5e-06 | 3e-06 | 2.400E-06 | 6.557E-07 | 1.700E-06 | 3.000E-06 |
| 28805 | 1441 | G1441C | E | 2.1e-06 | 2.6e-06 | 3.2e-06 | 2.633E-06 | 5.508E-07 | 2.100E-06 | 3.200E-06 |
| 28806 | 1441 | G1441E | E | 3.3e-06 | 2.7e-06 | 2.2e-06 | 2.733E-06 | 5.508E-07 | 2.200E-06 | 3.300E-06 |
| 28807 | 1441 | G1441Q | E | 1.9e-06 | 3.4e-06 | 2.8e-06 | 2.700E-06 | 7.550E-07 | 1.900E-06 | 3.400E-06 |
| 28808 | 1441 | G1441G | E | 1.2e-06 | 1.6e-06 | 4.3e-06 | 2.367E-06 | 1.686E-06 | 1.200E-06 | 4.300E-06 |
| 28809 | 1441 | G1441H | E | 1.6e-06 | 3.1e-06 | 2.4e-06 | 2.367E-06 | 7.506E-07 | 1.600E-06 | 3.100E-06 |
| 28810 | 1441 | G1441I | E | 3.8e-06 | 3.1e-06 | 2.9e-06 | 3.267E-06 | 4.726E-07 | 2.900E-06 | 3.800E-06 |
| 28811 | 1441 | G1441L | E | 2.2e-06 | 1.8e-06 | 2.6e-06 | 2.200E-06 | 4.000E-07 | 1.800E-06 | 2.600E-06 |
| 28812 | 1441 | G1441K | E | 3.9e-06 | 2.1e-06 | 2.2e-06 | 2.733E-06 | 1.012E-06 | 2.100E-06 | 3.900E-06 |
| 28813 | 1441 | G1441M | E | 3.5e-06 | 4.6e-06 | 4.1e-06 | 4.067E-06 | 5.508E-07 | 3.500E-06 | 4.600E-06 |
| 28814 | 1441 | G1441F | E | 1.1e-06 | 1.7e-06 | 1.5e-06 | 1.433E-06 | 3.055E-07 | 1.100E-06 | 1.700E-06 |

| | | | | | | | | | | |
|---|---|---|---|---|---|---|---|---|---|---|
| 28815 | 1441 | G1441P | E | 2.7e-06 | 1.9e-06 | 1e-06 | 1.867E-06 | 8.505E-07 | 1.000E-06 | 2.700E-06 |
| 28816 | 1441 | G1441S | E | 1.6e-06 | 2.3e-06 | 3.6e-06 | 2.500E-06 | 1.015E-06 | 1.600E-06 | 3.600E-06 |
| 28817 | 1441 | G1441T | E | 1.2e-06 | 4.2e-06 | 1.8e-06 | 2.400E-06 | 1.587E-06 | 1.200E-06 | 4.200E-06 |
| 28818 | 1441 | G1441W | E | 2.5e-06 | 2.4e-06 | 2e-06 | 2.300E-06 | 2.646E-07 | 2.000E-06 | 2.500E-06 |
| 28819 | 1441 | G1441Y | E | 2.4e-06 | 1.7e-06 | 1.8e-06 | 1.967E-06 | 3.786E-07 | 1.700E-06 | 2.400E-06 |
| 28820 | 1441 | G1441V | E | 1.5e-06 | 2.3e-06 | 2.3e-06 | 2.033E-06 | 4.619E-07 | 1.500E-06 | 2.300E-06 |
| 28821 | 1442 | V1442A | E | 3e-06 | 2.7e-06 | 2e-06 | 2.567E-06 | 5.132E-07 | 2.000E-06 | 3.000E-06 |
| 28822 | 1442 | V1442R | E | 2.5e-06 | 2.3e-06 | 2.6e-06 | 2.467E-06 | 1.528E-07 | 2.300E-06 | 2.600E-06 |
| 28823 | 1442 | V1442N | E | 3e-06 | 2.6e-06 | 2.8e-06 | 2.800E-06 | 2.000E-07 | 2.600E-06 | 3.000E-06 |
| 28824 | 1442 | V1442D | E | 1.5e-06 | 2.9e-06 | 2.6e-06 | 2.333E-06 | 7.371E-07 | 1.500E-06 | 2.900E-06 |
| 28825 | 1442 | V1442C | E | 2.4e-06 | 2.9e-06 | 1.5e-06 | 2.267E-06 | 7.095E-07 | 1.500E-06 | 2.900E-06 |
| 28826 | 1442 | V1442E | E | 3.3e-06 | 1.6e-06 | 1.7e-06 | 2.200E-06 | 9.539E-07 | 1.600E-06 | 3.300E-06 |
| 28827 | 1442 | V1442Q | E | 1.4e-06 | 2.2e-06 | 3.7e-06 | 2.433E-06 | 1.168E-06 | 1.400E-06 | 3.700E-06 |
| 28828 | 1442 | V1442G | E | 2.4e-06 | 3.5e-06 | 1.1e-06 | 2.333E-06 | 1.201E-06 | 1.100E-06 | 3.500E-06 |
| 28829 | 1442 | V1442H | E | 1.6e-06 | 2.4e-06 | 2.3e-06 | 2.100E-06 | 4.359E-07 | 1.600E-06 | 2.400E-06 |
| 28830 | 1442 | V1442I | E | 2.6e-06 | 2.1e-06 | 4.4e-06 | 3.033E-06 | 1.210E-06 | 2.100E-06 | 4.400E-06 |
| 28831 | 1442 | V1442L | E | 1.8e-06 | 1.3e-06 | 3.6e-06 | 2.233E-06 | 1.210E-06 | 1.300E-06 | 3.600E-06 |
| 28832 | 1442 | V1442K | E | 3.1e-06 | 2.3e-06 | 3.2e-06 | 2.867E-06 | 4.933E-07 | 2.300E-06 | 3.200E-06 |
| 28833 | 1442 | V1442M | E | 1.1e-06 | 2.4e-06 | 3e-06 | 2.167E-06 | 9.713E-07 | 1.100E-06 | 3.000E-06 |
| 28834 | 1442 | V1442F | E | 2.5e-06 | 2.8e-06 | 2.5e-06 | 2.600E-06 | 1.732E-07 | 2.500E-06 | 2.800E-06 |
| 28835 | 1442 | V1442P | E | 1.5e-06 | 2.9e-06 | 2.5e-06 | 2.300E-06 | 7.211E-07 | 1.500E-06 | 2.900E-06 |
| 28836 | 1442 | V1442S | E | 2e-06 | 2.2e-06 | 2e-06 | 2.067E-06 | 1.155E-07 | 2.000E-06 | 2.200E-06 |
| 28837 | 1442 | V1442T | E | 2.8e-06 | 3e-06 | 2.9e-06 | 2.900E-06 | 1.000E-07 | 2.800E-06 | 3.000E-06 |
| 28838 | 1442 | V1442W | E | 1.7e-06 | 1.7e-06 | 2.6e-06 | 2.000E-06 | 5.196E-07 | 1.700E-06 | 2.600E-06 |
| 28839 | 1442 | V1442Y | E | 2e-06 | 2.7e-06 | 2.8e-06 | 2.500E-06 | 4.359E-07 | 2.000E-06 | 2.800E-06 |
| 28840 | 1442 | V1442V | E | 2.1e-06 | 2.7e-06 | 2.6e-06 | 2.467E-06 | 3.215E-07 | 2.100E-06 | 2.700E-06 |
| 28841 | 1443 | H1443A | E | 2.1e-06 | 1.2e-06 | 3.5e-06 | 2.267E-06 | 1.159E-06 | 1.200E-06 | 3.500E-06 |
| 28842 | 1443 | H1443R | E | 3.1e-06 | 2.6e-06 | 1.3e-06 | 2.333E-06 | 9.292E-07 | 1.300E-06 | 3.100E-06 |
| 28843 | 1443 | H1443N | E | 2.6e-06 | 1.6e-06 | 1.9e-06 | 2.033E-06 | 5.132E-07 | 1.600E-06 | 2.600E-06 |
| 28844 | 1443 | H1443D | E | 2.6e-06 | 1.5e-06 | 2.7e-06 | 2.267E-06 | 6.658E-07 | 1.500E-06 | 2.700E-06 |
| 28845 | 1443 | H1443C | E | 2.6e-06 | 1.9e-06 | 4.5e-06 | 3.000E-06 | 1.345E-06 | 1.900E-06 | 4.500E-06 |
| 28846 | 1443 | H1443E | E | 5.2e-06 | 2.5e-06 | 2.6e-06 | 3.433E-06 | 1.531E-06 | 2.500E-06 | 5.200E-06 |
| 28847 | 1443 | H1443Q | E | 2.7e-06 | 1.9e-06 | 2.7e-06 | 2.433E-06 | 4.619E-07 | 1.900E-06 | 2.700E-06 |
| 28848 | 1443 | H1443G | E | 3.3e-06 | 2e-06 | 2e-06 | 2.433E-06 | 7.506E-07 | 2.000E-06 | 3.300E-06 |
| 28849 | 1443 | H1443H | E | 2.2e-06 | 1.7e-06 | 2.6e-06 | 2.167E-06 | 4.509E-07 | 1.700E-06 | 2.600E-06 |
| 28850 | 1443 | H1443I | E | 2.8e-06 | 2e-06 | 2.2e-06 | 2.333E-06 | 4.163E-07 | 2.000E-06 | 2.800E-06 |
| 28851 | 1443 | H1443L | E | 1.6e-06 | 2e-06 | 2.4e-06 | 2.000E-06 | 4.000E-07 | 1.600E-06 | 2.400E-06 |
| 28852 | 1443 | H1443K | E | 3.3e-06 | 1.2e-06 | 4.1e-06 | 2.867E-06 | 1.498E-06 | 1.200E-06 | 4.100E-06 |
| 28853 | 1443 | H1443M | E | 3.3e-06 | 3.3e-06 | 2.4e-06 | 3.000E-06 | 5.196E-07 | 2.400E-06 | 3.300E-06 |
| 28854 | 1443 | H1443F | E | 3.2e-06 | 2.5e-06 | 3.3e-06 | 3.000E-06 | 4.359E-07 | 2.500E-06 | 3.300E-06 |
| 28855 | 1443 | H1443P | E | 3.2e-06 | 2.4e-06 | 2.9e-06 | 2.833E-06 | 4.041E-07 | 2.400E-06 | 3.200E-06 |
| 28856 | 1443 | H1443S | E | 2.6e-06 | 1.8e-06 | 1.8e-06 | 2.067E-06 | 4.619E-07 | 1.800E-06 | 2.600E-06 |
| 28857 | 1443 | H1443T | E | 4e-06 | 1.3e-06 | 1.9e-06 | 2.400E-06 | 1.418E-06 | 1.300E-06 | 4.000E-06 |
| 28858 | 1443 | H1443W | E | 2.4e-06 | 3.4e-06 | 3.5e-06 | 3.100E-06 | 6.083E-07 | 2.400E-06 | 3.500E-06 |
| 28859 | 1443 | H1443Y | E | 2.6e-06 | 2.8e-06 | 2e-06 | 2.467E-06 | 4.163E-07 | 2.000E-06 | 2.800E-06 |
| 28860 | 1443 | H1443V | E | 3.1e-06 | 1.9e-06 | 2e-06 | 2.333E-06 | 6.658E-07 | 1.900E-06 | 3.100E-06 |
| 28861 | 1444 | T1444A | E | 2.7e-06 | 2.5e-06 | 1.8e-06 | 2.333E-06 | 4.726E-07 | 1.800E-06 | 2.700E-06 |
| 28862 | 1444 | T1444R | E | 3.4e-06 | 3.8e-06 | 2.5e-06 | 3.233E-06 | 6.658E-07 | 2.500E-06 | 3.800E-06 |
| 28863 | 1444 | T1444N | E | 1.3e-06 | 1.6e-06 | 2.2e-06 | 1.700E-06 | 4.583E-07 | 1.300E-06 | 2.200E-06 |
| 28864 | 1444 | T1444D | E | 2.5e-06 | 2.1e-06 | 2.1e-06 | 2.233E-06 | 2.309E-07 | 2.100E-06 | 2.500E-06 |
| 28865 | 1444 | T1444C | E | 2.5e-06 | 2.9e-06 | 2.5e-06 | 2.633E-06 | 2.309E-07 | 2.500E-06 | 2.900E-06 |
| 28866 | 1444 | T1444E | E | 3e-06 | 2.9e-06 | 2.8e-06 | 2.900E-06 | 1.000E-07 | 2.800E-06 | 3.000E-06 |
| 28867 | 1444 | T1444Q | E | 1e-06 | 1.4e-06 | 3.1e-06 | 1.833E-06 | 1.115E-06 | 1.000E-06 | 3.100E-06 |
| 28868 | 1444 | T1444G | E | 4e-06 | 1.6e-06 | 3.7e-06 | 3.100E-06 | 1.308E-06 | 1.600E-06 | 4.000E-06 |
| 28869 | 1444 | T1444H | E | 2.5e-06 | 2.6e-06 | 2.6e-06 | 2.567E-06 | 5.774E-08 | 2.500E-06 | 2.600E-06 |
| 28870 | 1444 | T1444I | E | 4.3e-06 | 1.7e-06 | 1.4e-06 | 2.467E-06 | 1.595E-06 | 1.400E-06 | 4.300E-06 |
| 28871 | 1444 | T1444L | E | 2.7e-06 | 2.4e-06 | 2.7e-06 | 2.600E-06 | 1.732E-07 | 2.400E-06 | 2.700E-06 |
| 28872 | 1444 | T1444K | E | 1.6e-06 | 1.4e-06 | 3.3e-06 | 2.100E-06 | 1.044E-06 | 1.400E-06 | 3.300E-06 |
| 28873 | 1444 | T1444M | E | 3.2e-06 | 2.7e-06 | 2.6e-06 | 2.833E-06 | 3.215E-07 | 2.600E-06 | 3.200E-06 |
| 28874 | 1444 | T1444F | E | 2.8e-06 | 2.2e-06 | 2.7e-06 | 2.567E-06 | 3.215E-07 | 2.200E-06 | 2.800E-06 |
| 28875 | 1444 | T1444P | E | 4.2e-06 | 2e-06 | 2.6e-06 | 2.933E-06 | 1.137E-06 | 2.000E-06 | 4.200E-06 |
| 28876 | 1444 | T1444S | E | 2.4e-06 | 2.5e-06 | 2.3e-06 | 2.400E-06 | 1.000E-07 | 2.300E-06 | 2.500E-06 |
| 28877 | 1444 | T1444T | E | 2.2e-06 | 2.4e-06 | 2.7e-06 | 2.433E-06 | 2.517E-07 | 2.200E-06 | 2.700E-06 |
| 28878 | 1444 | T1444W | E | 3.1e-06 | 2.8e-06 | 2.8e-06 | 2.900E-06 | 1.732E-07 | 2.800E-06 | 3.100E-06 |
| 28879 | 1444 | T1444Y | E | 2.7e-06 | 2.5e-06 | 3e-06 | 2.733E-06 | 2.517E-07 | 2.500E-06 | 3.000E-06 |
| 28880 | 1444 | T1444V | E | 2.6e-06 | 1.9e-06 | 1.5e-06 | 2.000E-06 | 5.568E-07 | 1.500E-06 | 2.600E-06 |
| 28881 | 1445 | F1445A | E | 2e-06 | 3.3e-06 | 2.9e-06 | 2.733E-06 | 6.658E-07 | 2.000E-06 | 3.300E-06 |
| 28882 | 1445 | F1445R | E | 3.2e-06 | 2.1e-06 | 4.1e-06 | 3.133E-06 | 1.002E-06 | 2.100E-06 | 4.100E-06 |
| 28883 | 1445 | F1445N | E | 1.5e-06 | 1.9e-06 | 2.7e-06 | 2.033E-06 | 6.110E-07 | 1.500E-06 | 2.700E-06 |
| 28884 | 1445 | F1445D | E | 2.9e-06 | 2.3e-06 | 3.3e-06 | 2.833E-06 | 5.033E-07 | 2.300E-06 | 3.300E-06 |
| 28885 | 1445 | F1445C | E | 3.4e-06 | 3.1e-06 | 2.4e-06 | 2.967E-06 | 5.132E-07 | 2.400E-06 | 3.400E-06 |
| 28886 | 1445 | F1445E | E | 1.5e-06 | 1.6e-06 | 3e-06 | 2.033E-06 | 8.386E-07 | 1.500E-06 | 3.000E-06 |
| 28887 | 1445 | F1445Q | E | 1.9e-06 | 2.2e-06 | 2.2e-06 | 2.100E-06 | 1.732E-07 | 1.900E-06 | 2.200E-06 |
| 28888 | 1445 | F1445G | E | 1.8e-06 | 2.6e-06 | 3e-06 | 2.467E-06 | 6.110E-07 | 1.800E-06 | 3.000E-06 |
| 28889 | 1445 | F1445H | E | 3.1e-06 | 2.1e-06 | 4.7e-06 | 3.300E-06 | 1.311E-06 | 2.100E-06 | 4.700E-06 |
| 28890 | 1445 | F1445I | E | 7.1e-07 | 1.8e-06 | 2.5e-06 | 1.670E-06 | 9.021E-07 | 7.100E-07 | 2.500E-06 |
| 28891 | 1445 | F1445L | E | 2.4e-06 | 2.9e-06 | 1.7e-06 | 2.333E-06 | 6.028E-07 | 1.700E-06 | 2.900E-06 |
| 28892 | 1445 | F1445K | E | 3.1e-06 | 3.5e-06 | 2.9e-06 | 3.167E-06 | 3.055E-07 | 2.900E-06 | 3.500E-06 |
| 28893 | 1445 | F1445M | E | 1.6e-06 | 3.1e-06 | 4.8e-06 | 3.167E-06 | 1.601E-06 | 1.600E-06 | 4.800E-06 |
| 28894 | 1445 | F1445F | E | 1.9e-06 | 1.5e-06 | 1.4e-06 | 1.600E-06 | 2.646E-07 | 1.400E-06 | 1.900E-06 |
| 28895 | 1445 | F1445P | E | 1.9e-06 | 3e-06 | 2.7e-06 | 2.533E-06 | 5.686E-07 | 1.900E-06 | 3.000E-06 |
| 28896 | 1445 | F1445S | E | 2.5e-06 | 2.3e-06 | 1.5e-06 | 2.100E-06 | 5.292E-07 | 1.500E-06 | 2.500E-06 |
| 28897 | 1445 | F1445T | E | 2.3e-06 | 1.4e-06 | 2.3e-06 | 2.000E-06 | 5.196E-07 | 1.400E-06 | 2.300E-06 |
| 28898 | 1445 | F1445W | E | 1.4e-06 | 2.8e-06 | 2.5e-06 | 2.233E-06 | 7.371E-07 | 1.400E-06 | 2.800E-06 |
| 28899 | 1445 | F1445Y | E | 4.9e-06 | 3.5e-06 | 2.3e-06 | 3.567E-06 | 1.301E-06 | 2.300E-06 | 4.900E-06 |

| | | | | | | | | | | |
|---|---|---|---|---|---|---|---|---|---|---|
| 28900 | 1445 | F1445V | E | 2.3e-06 | 2.9e-06 | 3e-06 | 2.733E-06 | 3.786E-07 | 2.300E-06 | 3.000E-06 |
| 28901 | 1446 | P1446A | E | 2.4e-06 | 1.7e-06 | 2.9e-06 | 2.333E-06 | 6.028E-07 | 1.700E-06 | 2.900E-06 |
| 28902 | 1446 | P1446R | E | 3.1e-06 | 2.1e-06 | 2.8e-06 | 2.667E-06 | 5.132E-07 | 2.100E-06 | 3.100E-06 |
| 28903 | 1446 | P1446N | E | 1.8e-06 | 1.9e-06 | 1.2e-06 | 1.633E-06 | 3.786E-07 | 1.200E-06 | 1.900E-06 |
| 28904 | 1446 | P1446D | E | 3.2e-06 | 3.1e-06 | 2.2e-06 | 2.833E-06 | 5.508E-07 | 2.200E-06 | 3.200E-06 |
| 28905 | 1446 | P1446C | E | 2.2e-06 | 1.7e-06 | 2.1e-06 | 2.000E-06 | 2.646E-07 | 1.700E-06 | 2.200E-06 |
| 28906 | 1446 | P1446E | E | 1.3e-06 | 2.7e-06 | 1.8e-06 | 1.933E-06 | 7.095E-07 | 1.300E-06 | 2.700E-06 |
| 28907 | 1446 | P1446Q | E | 1.4e-06 | 2e-06 | 1.6e-06 | 1.667E-06 | 3.055E-07 | 1.400E-06 | 2.000E-06 |
| 28908 | 1446 | P1446G | E | 2.5e-06 | 2.6e-06 | 3.2e-06 | 2.767E-06 | 3.786E-07 | 2.500E-06 | 3.200E-06 |
| 28909 | 1446 | P1446H | E | 1.5e-06 | 2.5e-06 | 2.3e-06 | 2.100E-06 | 5.292E-07 | 1.500E-06 | 2.500E-06 |
| 28910 | 1446 | P1446I | E | 1.3e-06 | 2.5e-06 | 3.3e-06 | 2.367E-06 | 1.007E-06 | 1.300E-06 | 3.300E-06 |
| 28911 | 1446 | P1446L | E | 1.4e-06 | 3e-06 | 2.2e-06 | 2.200E-06 | 8.000E-07 | 1.400E-06 | 3.000E-06 |
| 28912 | 1446 | P1446K | E | 1.3e-06 | 1.8e-06 | 3.2e-06 | 2.100E-06 | 9.849E-07 | 1.300E-06 | 3.200E-06 |
| 28913 | 1446 | P1446M | E | 3.3e-06 | 1.6e-06 | 1.8e-06 | 2.233E-06 | 9.292E-07 | 1.600E-06 | 3.300E-06 |
| 28914 | 1446 | P1446F | E | 4e-06 | 1.9e-06 | 3.1e-06 | 3.000E-06 | 1.054E-06 | 1.900E-06 | 4.000E-06 |
| 28915 | 1446 | P1446P | E | 2.3e-06 | 3e-06 | 3.2e-06 | 2.833E-06 | 4.726E-07 | 2.300E-06 | 3.200E-06 |
| 28916 | 1446 | P1446S | E | 2.7e-06 | 2.3e-06 | 1.9e-06 | 2.300E-06 | 4.000E-07 | 1.900E-06 | 2.700E-06 |
| 28917 | 1446 | P1446T | E | 2e-06 | 2.5e-06 | 3.1e-06 | 2.533E-06 | 5.508E-07 | 2.000E-06 | 3.100E-06 |
| 28918 | 1446 | P1446W | E | 1.8e-06 | 3.7e-06 | 1.2e-06 | 2.233E-06 | 1.305E-06 | 1.200E-06 | 3.700E-06 |
| 28919 | 1446 | P1446Y | E | 5.2e-06 | 1.8e-06 | 1.9e-06 | 2.967E-06 | 1.935E-06 | 1.800E-06 | 5.200E-06 |
| 28920 | 1446 | P1446V | E | 3.1e-06 | 2.7e-06 | 2.1e-06 | 2.633E-06 | 5.033E-07 | 2.100E-06 | 3.100E-06 |
| 28921 | 1447 | A1447A | E | 3.7e-06 | 3.7e-06 | 2.5e-06 | 3.300E-06 | 6.928E-07 | 2.500E-06 | 3.700E-06 |
| 28922 | 1447 | A1447R | E | 1.6e-06 | 2.5e-06 | 2.7e-06 | 2.267E-06 | 5.859E-07 | 1.600E-06 | 2.700E-06 |
| 28923 | 1447 | A1447N | E | 1.6e-06 | 2e-06 | 2e-06 | 1.867E-06 | 2.309E-07 | 1.600E-06 | 2.000E-06 |
| 28924 | 1447 | A1447D | E | 3.2e-06 | 1.4e-06 | 4.4e-06 | 3.000E-06 | 1.510E-06 | 1.400E-06 | 4.400E-06 |
| 28925 | 1447 | A1447C | E | 2.8e-06 | 3.2e-06 | 2.4e-06 | 2.800E-06 | 4.000E-07 | 2.400E-06 | 3.200E-06 |
| 28926 | 1447 | A1447E | E | 2.9e-06 | 2.4e-06 | 2.2e-06 | 2.500E-06 | 3.606E-07 | 2.200E-06 | 2.900E-06 |
| 28927 | 1447 | A1447Q | E | 2.4e-06 | 4.7e-06 | 2.3e-06 | 3.133E-06 | 1.358E-06 | 2.300E-06 | 4.700E-06 |
| 28928 | 1447 | A1447G | E | 3.3e-06 | 2e-06 | 2.1e-06 | 2.467E-06 | 7.234E-07 | 2.000E-06 | 3.300E-06 |
| 28929 | 1447 | A1447H | E | 1.9e-06 | 2.9e-06 | 2.6e-06 | 2.467E-06 | 5.132E-07 | 1.900E-06 | 2.900E-06 |
| 28930 | 1447 | A1447I | E | 2.3e-06 | 2.2e-06 | 2.6e-06 | 2.367E-06 | 2.082E-07 | 2.200E-06 | 2.600E-06 |
| 28931 | 1447 | A1447L | E | 3.9e-06 | 2.5e-06 | 4.4e-06 | 3.600E-06 | 9.849E-07 | 2.500E-06 | 4.400E-06 |
| 28932 | 1447 | A1447K | E | 2.7e-06 | 2.5e-06 | 2.5e-06 | 2.567E-06 | 1.155E-07 | 2.500E-06 | 2.700E-06 |
| 28933 | 1447 | A1447M | E | 2.7e-06 | 2.3e-06 | 2.5e-06 | 2.500E-06 | 2.000E-07 | 2.300E-06 | 2.700E-06 |
| 28934 | 1447 | A1447F | E | 1.4e-06 | 2.3e-06 | 2e-06 | 1.900E-06 | 4.583E-07 | 1.400E-06 | 2.300E-06 |
| 28935 | 1447 | A1447P | E | 2.3e-06 | 2.7e-06 | 2.2e-06 | 2.400E-06 | 2.646E-07 | 2.200E-06 | 2.700E-06 |
| 28936 | 1447 | A1447S | E | 2e-06 | 2.3e-06 | 2.6e-06 | 2.300E-06 | 3.000E-07 | 2.000E-06 | 2.600E-06 |
| 28937 | 1447 | A1447T | E | 3.6e-06 | 3.1e-06 | 1.8e-06 | 2.833E-06 | 9.292E-07 | 1.800E-06 | 3.600E-06 |
| 28938 | 1447 | A1447W | E | 2.4e-06 | 3.2e-06 | 3.9e-06 | 3.167E-06 | 7.506E-07 | 2.400E-06 | 3.900E-06 |
| 28939 | 1447 | A1447Y | E | 3.4e-06 | 3.4e-06 | 2.1e-06 | 2.967E-06 | 7.506E-07 | 2.100E-06 | 3.400E-06 |
| 28940 | 1447 | A1447V | E | 2.4e-06 | 2.7e-06 | 2.7e-06 | 2.600E-06 | 1.732E-07 | 2.400E-06 | 2.700E-06 |
| 28941 | 1448 | V1448A | E | 2.4e-06 | 2.2e-06 | 2.2e-06 | 2.267E-06 | 1.155E-07 | 2.200E-06 | 2.400E-06 |
| 28942 | 1448 | V1448R | E | 2.8e-06 | 2.8e-06 | 2.3e-06 | 2.633E-06 | 2.887E-07 | 2.300E-06 | 2.800E-06 |
| 28943 | 1448 | V1448N | E | 1.8e-06 | 2e-06 | 2.2e-06 | 2.000E-06 | 2.000E-07 | 1.800E-06 | 2.200E-06 |
| 28944 | 1448 | V1448D | E | 1.9e-06 | 5.1e-06 | 1.9e-06 | 2.967E-06 | 1.848E-06 | 1.900E-06 | 5.100E-06 |
| 28945 | 1448 | V1448C | E | 2.6e-06 | 2.5e-06 | 4.2e-06 | 3.100E-06 | 9.539E-07 | 2.500E-06 | 4.200E-06 |
| 28946 | 1448 | V1448E | E | 1.9e-06 | 1.9e-06 | 3.4e-06 | 2.400E-06 | 8.660E-07 | 1.900E-06 | 3.400E-06 |
| 28947 | 1448 | V1448Q | E | 1.9e-06 | 2.8e-06 | 3.1e-06 | 2.600E-06 | 6.245E-07 | 1.900E-06 | 3.100E-06 |
| 28948 | 1448 | V1448G | E | 1.5e-06 | 2.9e-06 | 2e-06 | 2.133E-06 | 7.095E-07 | 1.500E-06 | 2.900E-06 |
| 28949 | 1448 | V1448H | E | 1.9e-06 | 2e-06 | 2e-06 | 1.967E-06 | 5.774E-08 | 1.900E-06 | 2.000E-06 |
| 28950 | 1448 | V1448I | E | 2.6e-06 | 3.9e-06 | 2.3e-06 | 2.933E-06 | 8.505E-07 | 2.300E-06 | 3.900E-06 |
| 28951 | 1448 | V1448L | E | 1.1e-06 | 2.2e-06 | 1.9e-06 | 1.733E-06 | 5.686E-07 | 1.100E-06 | 2.200E-06 |
| 28952 | 1448 | V1448K | E | 2.4e-06 | 4.3e-06 | 1.2e-06 | 2.633E-06 | 1.563E-06 | 1.200E-06 | 4.300E-06 |
| 28953 | 1448 | V1448M | E | 1.8e-06 | 3.5e-06 | 1.9e-06 | 2.400E-06 | 9.539E-07 | 1.800E-06 | 3.500E-06 |
| 28954 | 1448 | V1448F | E | 3.6e-06 | 1.3e-06 | 2.2e-06 | 2.367E-06 | 1.159E-06 | 1.300E-06 | 3.600E-06 |
| 28955 | 1448 | V1448P | E | 1.4e-06 | 3.2e-06 | 2e-06 | 2.200E-06 | 9.165E-07 | 1.400E-06 | 3.200E-06 |
| 28956 | 1448 | V1448S | E | 1.5e-06 | 3.1e-06 | 4.9e-06 | 3.167E-06 | 1.701E-06 | 1.500E-06 | 4.900E-06 |
| 28957 | 1448 | V1448T | E | 2.9e-06 | 2.4e-06 | 1.8e-06 | 2.367E-06 | 5.508E-07 | 1.800E-06 | 2.900E-06 |
| 28958 | 1448 | V1448W | E | 2.5e-06 | 2e-06 | 2.6e-06 | 2.367E-06 | 3.215E-07 | 2.000E-06 | 2.600E-06 |
| 28959 | 1448 | V1448Y | E | 2.5e-06 | 2.4e-06 | 1.7e-06 | 2.200E-06 | 4.359E-07 | 1.700E-06 | 2.500E-06 |
| 28960 | 1448 | V1448V | E | 1.4e-06 | 2.7e-06 | 1.4e-06 | 1.833E-06 | 7.506E-07 | 1.400E-06 | 2.700E-06 |
| 28961 | 1449 | L1449A | E | 1.8e-06 | 2.9e-06 | 2.9e-06 | 2.533E-06 | 6.351E-07 | 1.800E-06 | 2.900E-06 |
| 28962 | 1449 | L1449R | E | 2.7e-06 | 1.8e-06 | 2.8e-06 | 2.433E-06 | 5.508E-07 | 1.800E-06 | 2.800E-06 |
| 28963 | 1449 | L1449N | E | 5.6e-06 | 1.8e-06 | 2.1e-06 | 3.167E-06 | 2.113E-06 | 1.800E-06 | 5.600E-06 |
| 28964 | 1449 | L1449D | E | 2.1e-06 | 3.4e-06 | 2.1e-06 | 2.533E-06 | 7.506E-07 | 2.100E-06 | 3.400E-06 |
| 28965 | 1449 | L1449C | E | 2.5e-06 | 1.7e-06 | 1.8e-06 | 2.000E-06 | 4.359E-07 | 1.700E-06 | 2.500E-06 |
| 28966 | 1449 | L1449E | E | 1.9e-06 | 2.6e-06 | 5.3e-06 | 3.267E-06 | 1.795E-06 | 1.900E-06 | 5.300E-06 |
| 28967 | 1449 | L1449Q | E | 2.5e-06 | 2.1e-06 | 3.2e-06 | 2.600E-06 | 5.568E-07 | 2.100E-06 | 3.200E-06 |
| 28968 | 1449 | L1449G | E | 3.3e-06 | 1.9e-06 | 3e-06 | 2.733E-06 | 7.371E-07 | 1.900E-06 | 3.300E-06 |
| 28969 | 1449 | L1449H | E | 3e-06 | 2.4e-06 | 3e-06 | 2.800E-06 | 3.464E-07 | 2.400E-06 | 3.000E-06 |
| 28970 | 1449 | L1449I | E | 2.5e-06 | 3e-06 | 2.1e-06 | 2.533E-06 | 4.509E-07 | 2.100E-06 | 3.000E-06 |
| 28971 | 1449 | L1449L | E | 2e-06 | 2.3e-06 | 2.6e-06 | 2.300E-06 | 3.000E-07 | 2.000E-06 | 2.600E-06 |
| 28972 | 1449 | L1449K | E | 3.9e-06 | 2.1e-06 | 2.3e-06 | 2.767E-06 | 9.866E-07 | 2.100E-06 | 3.900E-06 |
| 28973 | 1449 | L1449M | E | 4.9e-06 | 2.2e-06 | 2.7e-06 | 3.267E-06 | 1.436E-06 | 2.200E-06 | 4.900E-06 |
| 28974 | 1449 | L1449F | E | 2.8e-06 | 1.5e-06 | 2.6e-06 | 2.300E-06 | 7.000E-07 | 1.500E-06 | 2.800E-06 |
| 28975 | 1449 | L1449P | E | 1.4e-06 | 1.6e-06 | 2.1e-06 | 1.700E-06 | 3.606E-07 | 1.400E-06 | 2.100E-06 |
| 28976 | 1449 | L1449S | E | 2.6e-06 | 3.1e-06 | 2.1e-06 | 2.600E-06 | 5.000E-07 | 2.100E-06 | 3.100E-06 |
| 28977 | 1449 | L1449T | E | 2.3e-06 | 2.9e-06 | 1.6e-06 | 2.267E-06 | 6.506E-07 | 1.600E-06 | 2.900E-06 |
| 28978 | 1449 | L1449W | E | 1.6e-06 | 2.6e-06 | 2.9e-06 | 2.367E-06 | 6.807E-07 | 1.600E-06 | 2.900E-06 |
| 28979 | 1449 | L1449Y | E | 1.9e-06 | 2.2e-06 | 3.1e-06 | 2.400E-06 | 6.245E-07 | 1.900E-06 | 3.100E-06 |
| 28980 | 1449 | L1449V | E | 3.5e-06 | 2e-06 | 2e-06 | 2.500E-06 | 8.660E-07 | 2.000E-06 | 3.500E-06 |
| 28981 | 1450 | Q1450A | E | 4.3e-06 | 2.2e-06 | 2.3e-06 | 2.933E-06 | 1.185E-06 | 2.200E-06 | 4.300E-06 |
| 28982 | 1450 | Q1450R | E | 1.8e-06 | 3.6e-06 | 2.1e-06 | 2.500E-06 | 9.644E-07 | 1.800E-06 | 3.600E-06 |
| 28983 | 1450 | Q1450N | E | 1.6e-06 | 2.4e-06 | 3.9e-06 | 2.633E-06 | 1.168E-06 | 1.600E-06 | 3.900E-06 |
| 28984 | 1450 | Q1450D | E | 2e-06 | 2.7e-06 | 2e-06 | 2.233E-06 | 4.041E-07 | 2.000E-06 | 2.700E-06 |

| | | | | | | | | | | |
|---|---|---|---|---|---|---|---|---|---|---|
| 28985 | 1450 | Q1450C | E | 2.5e-06 | 1.7e-06 | 2.3e-06 | 2.167E-06 | 4.163E-07 | 1.700E-06 | 2.500E-06 |
| 28986 | 1450 | Q1450E | E | 2e-06 | 1.2e-06 | 3.3e-06 | 2.167E-06 | 1.060E-06 | 1.200E-06 | 3.300E-06 |
| 28987 | 1450 | Q1450Q | E | 3.5e-06 | 3.2e-06 | 2.5e-06 | 3.067E-06 | 5.132E-07 | 2.500E-06 | 3.500E-06 |
| 28988 | 1450 | Q1450G | E | 2.4e-06 | 2.7e-06 | 3.8e-06 | 2.967E-06 | 7.371E-07 | 2.400E-06 | 3.800E-06 |
| 28989 | 1450 | Q1450H | E | 1.5e-06 | 2.6e-06 | 1.7e-06 | 1.933E-06 | 5.859E-07 | 1.500E-06 | 2.600E-06 |
| 28990 | 1450 | Q1450I | E | 2e-06 | 3.3e-06 | 2.6e-06 | 2.633E-06 | 6.506E-07 | 2.000E-06 | 3.300E-06 |
| 28991 | 1450 | Q1450L | E | 3.2e-06 | 3.1e-06 | 2.4e-06 | 2.900E-06 | 4.359E-07 | 2.400E-06 | 3.200E-06 |
| 28992 | 1450 | Q1450K | E | 2.2e-06 | 2.2e-06 | 3.2e-06 | 2.533E-06 | 5.774E-07 | 2.200E-06 | 3.200E-06 |
| 28993 | 1450 | Q1450M | E | 3.6e-06 | 2e-06 | 2.7e-06 | 2.767E-06 | 8.021E-07 | 2.000E-06 | 3.600E-06 |
| 28994 | 1450 | Q1450F | E | 2.5e-06 | 2.2e-06 | 2.6e-06 | 2.433E-06 | 2.082E-07 | 2.200E-06 | 2.600E-06 |
| 28995 | 1450 | Q1450P | E | 4.2e-06 | 2.5e-06 | 1.9e-06 | 2.867E-06 | 1.193E-06 | 1.900E-06 | 4.200E-06 |
| 28996 | 1450 | Q1450S | E | 2.1e-06 | 2e-06 | 1.9e-06 | 2.000E-06 | 1.000E-07 | 1.900E-06 | 2.100E-06 |
| 28997 | 1450 | Q1450T | E | 2.4e-06 | 1.4e-06 | 2.3e-06 | 2.033E-06 | 5.508E-07 | 1.400E-06 | 2.400E-06 |
| 28998 | 1450 | Q1450W | E | 2.1e-06 | 1.4e-06 | 3.4e-06 | 2.300E-06 | 1.015E-06 | 1.400E-06 | 3.400E-06 |
| 28999 | 1450 | Q1450Y | E | 2.6e-06 | 2.1e-06 | 3.8e-06 | 2.833E-06 | 8.737E-07 | 2.100E-06 | 3.800E-06 |
| 29000 | 1450 | Q1450V | E | 2.8e-06 | 2.2e-06 | 3.7e-06 | 2.900E-06 | 7.550E-07 | 2.200E-06 | 3.700E-06 |
| 29001 | 1451 | S1451A | E | 2.7e-06 | 4.7e-06 | 1.5e-06 | 2.967E-06 | 1.617E-06 | 1.500E-06 | 4.700E-06 |
| 29002 | 1451 | S1451R | E | 2.1e-06 | 2.3e-06 | 3.2e-06 | 2.533E-06 | 5.859E-07 | 2.100E-06 | 3.200E-06 |
| 29003 | 1451 | S1451N | E | 3.8e-06 | 2.3e-06 | 1.5e-06 | 2.533E-06 | 1.168E-06 | 1.500E-06 | 3.800E-06 |
| 29004 | 1451 | S1451D | E | 3.6e-06 | 2.2e-06 | 2.1e-06 | 2.633E-06 | 8.386E-07 | 2.100E-06 | 3.600E-06 |
| 29005 | 1451 | S1451C | E | 2.7e-06 | 2.3e-06 | 3.1e-06 | 2.700E-06 | 4.000E-07 | 2.300E-06 | 3.100E-06 |
| 29006 | 1451 | S1451E | E | 1.7e-06 | 3.7e-06 | 3e-06 | 2.800E-06 | 1.015E-06 | 1.700E-06 | 3.700E-06 |
| 29007 | 1451 | S1451Q | E | 2.7e-06 | 2.4e-06 | 2.4e-06 | 2.500E-06 | 1.732E-07 | 2.400E-06 | 2.700E-06 |
| 29008 | 1451 | S1451G | E | 2.4e-06 | 3.2e-06 | 5.1e-06 | 3.567E-06 | 1.387E-06 | 2.400E-06 | 5.100E-06 |
| 29009 | 1451 | S1451H | E | 2.8e-06 | 2.2e-06 | 2.2e-06 | 2.400E-06 | 3.464E-07 | 2.200E-06 | 2.800E-06 |
| 29010 | 1451 | S1451I | E | 3.7e-06 | 2.7e-06 | 3.2e-06 | 3.200E-06 | 5.000E-07 | 2.700E-06 | 3.700E-06 |
| 29011 | 1451 | S1451L | E | 1.8e-06 | 2.6e-06 | 1.7e-06 | 2.033E-06 | 4.933E-07 | 1.700E-06 | 2.600E-06 |
| 29012 | 1451 | S1451K | E | 1.9e-06 | 1.5e-06 | 1.3e-06 | 1.567E-06 | 3.055E-07 | 1.300E-06 | 1.900E-06 |
| 29013 | 1451 | S1451M | E | 2.2e-06 | 1.7e-06 | 2.1e-06 | 2.000E-06 | 2.646E-07 | 1.700E-06 | 2.200E-06 |
| 29014 | 1451 | S1451F | E | 2.2e-06 | 3.3e-06 | 2.6e-06 | 2.700E-06 | 5.568E-07 | 2.200E-06 | 3.300E-06 |
| 29015 | 1451 | S1451P | E | 2e-06 | 3.3e-06 | 1.4e-06 | 2.233E-06 | 9.713E-07 | 1.400E-06 | 3.300E-06 |
| 29016 | 1451 | S1451S | E | 4.8e-06 | 2.3e-06 | 1.9e-06 | 3.000E-06 | 1.572E-06 | 1.900E-06 | 4.800E-06 |
| 29017 | 1451 | S1451T | E | 2.6e-06 | 4.3e-06 | 1.4e-06 | 2.767E-06 | 1.457E-06 | 1.400E-06 | 4.300E-06 |
| 29018 | 1451 | S1451W | E | 3.5e-06 | 1.7e-06 | 2.4e-06 | 2.533E-06 | 9.074E-07 | 1.700E-06 | 3.500E-06 |
| 29019 | 1451 | S1451Y | E | 1.3e-06 | 1.8e-06 | 4e-06 | 2.367E-06 | 1.436E-06 | 1.300E-06 | 4.000E-06 |
| 29020 | 1451 | S1451V | E | 2.9e-06 | 2.2e-06 | 3.4e-06 | 2.833E-06 | 6.028E-07 | 2.200E-06 | 3.400E-06 |
| 29021 | 1452 | S1452A | E | 3e-06 | 2.5e-06 | 2.5e-06 | 2.667E-06 | 2.887E-07 | 2.500E-06 | 3.000E-06 |
| 29022 | 1452 | S1452R | E | 2.7e-06 | 3.4e-06 | 2.5e-06 | 2.867E-06 | 4.726E-07 | 2.500E-06 | 3.400E-06 |
| 29023 | 1452 | S1452N | E | 3.1e-06 | 2.6e-06 | 3.8e-06 | 3.167E-06 | 6.028E-07 | 2.600E-06 | 3.800E-06 |
| 29024 | 1452 | S1452D | E | 2.7e-06 | 3.8e-06 | 1.5e-06 | 2.667E-06 | 1.150E-06 | 1.500E-06 | 3.800E-06 |
| 29025 | 1452 | S1452C | E | 2.9e-06 | 2.3e-06 | 2.2e-06 | 2.467E-06 | 3.786E-07 | 2.200E-06 | 2.900E-06 |
| 29026 | 1452 | S1452E | E | 1.6e-06 | 2.4e-06 | 2.4e-06 | 2.133E-06 | 4.619E-07 | 1.600E-06 | 2.400E-06 |
| 29027 | 1452 | S1452Q | E | 2e-06 | 1.5e-06 | 2.1e-06 | 1.867E-06 | 3.215E-07 | 1.500E-06 | 2.100E-06 |
| 29028 | 1452 | S1452G | E | 2.7e-06 | 3.6e-06 | 2.2e-06 | 2.833E-06 | 7.095E-07 | 2.200E-06 | 3.600E-06 |
| 29029 | 1452 | S1452H | E | 1.5e-06 | 2.4e-06 | 3.2e-06 | 2.367E-06 | 8.505E-07 | 1.500E-06 | 3.200E-06 |
| 29030 | 1452 | S1452I | E | 1.8e-06 | 2.2e-06 | 2.4e-06 | 2.133E-06 | 3.055E-07 | 1.800E-06 | 2.400E-06 |
| 29031 | 1452 | S1452L | E | 2.5e-06 | 2.5e-06 | 2.2e-06 | 2.400E-06 | 1.732E-07 | 2.200E-06 | 2.500E-06 |
| 29032 | 1452 | S1452K | E | 3.1e-06 | 1.9e-06 | 2.3e-06 | 2.433E-06 | 6.110E-07 | 1.900E-06 | 3.100E-06 |
| 29033 | 1452 | S1452M | E | 1.7e-06 | 1.3e-06 | 2.5e-06 | 1.833E-06 | 6.110E-07 | 1.300E-06 | 2.500E-06 |
| 29034 | 1452 | S1452F | E | 2.8e-06 | 2.9e-06 | 2.5e-06 | 2.733E-06 | 2.082E-07 | 2.500E-06 | 2.900E-06 |
| 29035 | 1452 | S1452P | E | 1.9e-06 | 2.4e-06 | 3.7e-06 | 2.667E-06 | 9.292E-07 | 1.900E-06 | 3.700E-06 |
| 29036 | 1452 | S1452S | E | 1.3e-06 | 1.8e-06 | 1.9e-06 | 1.667E-06 | 3.215E-07 | 1.300E-06 | 1.900E-06 |
| 29037 | 1452 | S1452T | E | 2.4e-06 | 5.5e-06 | 1.7e-06 | 3.200E-06 | 2.022E-06 | 1.700E-06 | 5.500E-06 |
| 29038 | 1452 | S1452W | E | 2.7e-06 | 1.7e-06 | 2.2e-06 | 2.200E-06 | 5.000E-07 | 1.700E-06 | 2.700E-06 |
| 29039 | 1452 | S1452Y | E | 2.9e-06 | 2.5e-06 | 2.3e-06 | 2.567E-06 | 3.055E-07 | 2.300E-06 | 2.900E-06 |
| 29040 | 1452 | S1452V | E | 1.9e-06 | 1.8e-06 | 3.9e-06 | 2.533E-06 | 1.185E-06 | 1.800E-06 | 3.900E-06 |
| 29041 | 1453 | G1453A | E | 2.1e-06 | 3.7e-06 | 2.5e-06 | 2.767E-06 | 8.327E-07 | 2.100E-06 | 3.700E-06 |
| 29042 | 1453 | G1453R | E | 2.6e-06 | 2.6e-06 | 1.7e-06 | 2.300E-06 | 5.196E-07 | 1.700E-06 | 2.600E-06 |
| 29043 | 1453 | G1453N | E | 2.5e-06 | 1.8e-06 | 1.1e-06 | 1.800E-06 | 7.000E-07 | 1.100E-06 | 2.500E-06 |
| 29044 | 1453 | G1453D | E | 3e-06 | 1.5e-06 | 1.5e-06 | 2.000E-06 | 8.660E-07 | 1.500E-06 | 3.000E-06 |
| 29045 | 1453 | G1453C | E | 2.8e-06 | 3.3e-06 | 1.6e-06 | 2.567E-06 | 8.737E-07 | 1.600E-06 | 3.300E-06 |
| 29046 | 1453 | G1453E | E | 2e-06 | 2.4e-06 | 2.2e-06 | 2.200E-06 | 2.000E-07 | 2.000E-06 | 2.400E-06 |
| 29047 | 1453 | G1453Q | E | 2.6e-06 | 3.6e-06 | 2.8e-06 | 3.000E-06 | 5.292E-07 | 2.600E-06 | 3.600E-06 |
| 29048 | 1453 | G1453G | E | 2.8e-06 | 1.8e-06 | 2.9e-06 | 2.500E-06 | 6.083E-07 | 1.800E-06 | 2.900E-06 |
| 29049 | 1453 | G1453H | E | 4.1e-06 | 2.9e-06 | 1.5e-06 | 2.833E-06 | 1.301E-06 | 1.500E-06 | 4.100E-06 |
| 29050 | 1453 | G1453I | E | 2.3e-06 | 1.6e-06 | 3.1e-06 | 2.333E-06 | 7.506E-07 | 1.600E-06 | 3.100E-06 |
| 29051 | 1453 | G1453L | E | 3.2e-06 | 2.4e-06 | 4.5e-06 | 3.367E-06 | 1.060E-06 | 2.400E-06 | 4.500E-06 |
| 29052 | 1453 | G1453K | E | 2.8e-06 | 3.1e-06 | 2.3e-06 | 2.733E-06 | 4.041E-07 | 2.300E-06 | 3.100E-06 |
| 29053 | 1453 | G1453M | E | 3e-06 | 4.8e-06 | 1.3e-06 | 3.033E-06 | 1.750E-06 | 1.300E-06 | 4.800E-06 |
| 29054 | 1453 | G1453F | E | 1.7e-06 | 2.9e-06 | 3e-06 | 2.533E-06 | 7.234E-07 | 1.700E-06 | 3.000E-06 |
| 29055 | 1453 | G1453P | E | 2.1e-06 | 2.4e-06 | 2.3e-06 | 2.267E-06 | 1.528E-07 | 2.100E-06 | 2.400E-06 |
| 29056 | 1453 | G1453S | E | 1.7e-06 | 1.9e-06 | 3.4e-06 | 2.333E-06 | 9.292E-07 | 1.700E-06 | 3.400E-06 |
| 29057 | 1453 | G1453T | E | 1.9e-06 | 2.4e-06 | 2.3e-06 | 2.200E-06 | 2.646E-07 | 1.900E-06 | 2.400E-06 |
| 29058 | 1453 | G1453W | E | 2.9e-06 | 4.4e-06 | 2.4e-06 | 3.233E-06 | 1.041E-06 | 2.400E-06 | 4.400E-06 |
| 29059 | 1453 | G1453Y | E | 1.5e-06 | 2.2e-06 | 2.7e-06 | 2.133E-06 | 6.028E-07 | 1.500E-06 | 2.700E-06 |
| 29060 | 1453 | G1453V | E | 2.5e-06 | 1.9e-06 | 1.6e-06 | 2.000E-06 | 4.583E-07 | 1.600E-06 | 2.500E-06 |
| 29061 | 1454 | L1454A | E | 2.6e-06 | 2.1e-06 | 4.6e-06 | 3.100E-06 | 1.323E-06 | 2.100E-06 | 4.600E-06 |
| 29062 | 1454 | L1454R | E | 2e-06 | 1.7e-06 | 2.1e-06 | 1.933E-06 | 2.082E-07 | 1.700E-06 | 2.100E-06 |
| 29063 | 1454 | L1454N | E | 3.7e-06 | 2.1e-06 | 2.9e-06 | 2.900E-06 | 8.000E-07 | 2.100E-06 | 3.700E-06 |
| 29064 | 1454 | L1454D | E | 1.6e-06 | 2.6e-06 | 3.1e-06 | 2.433E-06 | 7.638E-07 | 1.600E-06 | 3.100E-06 |
| 29065 | 1454 | L1454C | E | 3.2e-06 | 2.4e-06 | 3.8e-06 | 3.133E-06 | 7.024E-07 | 2.400E-06 | 3.800E-06 |
| 29066 | 1454 | L1454E | E | 2.5e-06 | 2.9e-06 | 1.3e-06 | 2.233E-06 | 8.327E-07 | 1.300E-06 | 2.900E-06 |
| 29067 | 1454 | L1454Q | E | 3e-06 | 1.8e-06 | 2.3e-06 | 2.367E-06 | 6.028E-07 | 1.800E-06 | 3.000E-06 |
| 29068 | 1454 | L1454G | E | 2.3e-06 | 2.1e-06 | 3.2e-06 | 2.533E-06 | 5.859E-07 | 2.100E-06 | 3.200E-06 |
| 29069 | 1454 | L1454H | E | 2.2e-06 | 2.3e-06 | 1.8e-06 | 2.100E-06 | 2.646E-07 | 1.800E-06 | 2.300E-06 |

| | | | | | | | | | | |
|---|---|---|---|---|---|---|---|---|---|---|
| 29070 | 1454 | L1454I | E | 2.4e-06 | 2.1e-06 | 1.1e-06 | 1.867E-06 | 6.807E-07 | 1.100E-06 | 2.400E-06 |
| 29071 | 1454 | L1454L | E | 1.4e-06 | 1.6e-06 | 4.1e-06 | 2.367E-06 | 1.504E-06 | 1.400E-06 | 4.100E-06 |
| 29072 | 1454 | L1454K | E | 1.7e-06 | 2.2e-06 | 2.1e-06 | 2.000E-06 | 2.646E-07 | 1.700E-06 | 2.200E-06 |
| 29073 | 1454 | L1454M | E | 2.5e-06 | 1.3e-06 | 5.8e-06 | 3.200E-06 | 2.330E-06 | 1.300E-06 | 5.800E-06 |
| 29074 | 1454 | L1454F | E | 2.5e-06 | 1.7e-06 | 3.4e-06 | 2.533E-06 | 8.505E-07 | 1.700E-06 | 3.400E-06 |
| 29075 | 1454 | L1454P | E | 2.6e-06 | 2e-06 | 2.4e-06 | 2.333E-06 | 3.055E-07 | 2.000E-06 | 2.600E-06 |
| 29076 | 1454 | L1454S | E | 2.1e-06 | 2.9e-06 | 2.4e-06 | 2.467E-06 | 4.041E-07 | 2.100E-06 | 2.900E-06 |
| 29077 | 1454 | L1454T | E | 2.3e-06 | 1.5e-06 | 1.8e-06 | 1.867E-06 | 4.041E-07 | 1.500E-06 | 2.300E-06 |
| 29078 | 1454 | L1454W | E | 3.3e-06 | 1.9e-06 | 1.6e-06 | 2.267E-06 | 9.074E-07 | 1.600E-06 | 3.300E-06 |
| 29079 | 1454 | L1454Y | E | 2.2e-06 | 2.2e-06 | 3e-06 | 2.467E-06 | 4.619E-07 | 2.200E-06 | 3.000E-06 |
| 29080 | 1454 | L1454V | E | 2.1e-06 | 2.6e-06 | 4.4e-06 | 3.033E-06 | 1.210E-06 | 2.100E-06 | 4.400E-06 |
| 29081 | 1455 | Y1455A | E | 1.3e-06 | 2.5e-06 | 2.2e-06 | 2.000E-06 | 6.245E-07 | 1.300E-06 | 2.500E-06 |
| 29082 | 1455 | Y1455R | E | 2.1e-06 | 4.9e-06 | 2.2e-06 | 3.067E-06 | 1.589E-06 | 2.100E-06 | 4.900E-06 |
| 29083 | 1455 | Y1455N | E | 2.3e-06 | 3.2e-06 | 1.1e-06 | 2.200E-06 | 1.054E-06 | 1.100E-06 | 3.200E-06 |
| 29084 | 1455 | Y1455D | E | 2.4e-06 | 2.6e-06 | 3.4e-06 | 2.800E-06 | 5.292E-07 | 2.400E-06 | 3.400E-06 |
| 29085 | 1455 | Y1455C | E | 1.1e-06 | 2.1e-06 | 2e-06 | 1.733E-06 | 5.508E-07 | 1.100E-06 | 2.100E-06 |
| 29086 | 1455 | Y1455E | E | 1.4e-06 | 2e-06 | 2.9e-06 | 2.100E-06 | 7.550E-07 | 1.400E-06 | 2.900E-06 |
| 29087 | 1455 | Y1455Q | E | 3.4e-06 | 1.7e-06 | 3.1e-06 | 2.733E-06 | 9.074E-07 | 1.700E-06 | 3.400E-06 |
| 29088 | 1455 | Y1455G | E | 2.5e-06 | 1.4e-06 | 2.4e-06 | 2.100E-06 | 6.083E-07 | 1.400E-06 | 2.500E-06 |
| 29089 | 1455 | Y1455H | E | 1.8e-06 | 2.3e-06 | 3.4e-06 | 2.500E-06 | 8.185E-07 | 1.800E-06 | 3.400E-06 |
| 29090 | 1455 | Y1455I | E | 2.3e-06 | 2.2e-06 | 3.6e-06 | 2.700E-06 | 7.810E-07 | 2.200E-06 | 3.600E-06 |
| 29091 | 1455 | Y1455L | E | 2.8e-06 | 2.6e-06 | 2e-06 | 2.467E-06 | 4.163E-07 | 2.000E-06 | 2.800E-06 |
| 29092 | 1455 | Y1455K | E | 2.2e-06 | 3e-06 | 1.5e-06 | 2.233E-06 | 7.506E-07 | 1.500E-06 | 3.000E-06 |
| 29093 | 1455 | Y1455M | E | 2.3e-06 | 2.5e-06 | 2.3e-06 | 2.367E-06 | 1.155E-07 | 2.300E-06 | 2.500E-06 |
| 29094 | 1455 | Y1455F | E | 1.2e-06 | 3.2e-06 | 3.1e-06 | 2.500E-06 | 1.127E-06 | 1.200E-06 | 3.200E-06 |
| 29095 | 1455 | Y1455P | E | 1.4e-06 | 2e-06 | 1.1e-06 | 1.500E-06 | 4.583E-07 | 1.100E-06 | 2.000E-06 |
| 29096 | 1455 | Y1455S | E | 2.7e-06 | 4.4e-06 | 1.9e-06 | 3.000E-06 | 1.277E-06 | 1.900E-06 | 4.400E-06 |
| 29097 | 1455 | Y1455T | E | 3.4e-06 | 1.9e-06 | 2.8e-06 | 2.700E-06 | 7.550E-07 | 1.900E-06 | 3.400E-06 |
| 29098 | 1455 | Y1455W | E | 3e-06 | 2.5e-06 | 3e-06 | 2.833E-06 | 2.887E-07 | 2.500E-06 | 3.000E-06 |
| 29099 | 1455 | Y1455Y | E | 1.7e-06 | 3.5e-06 | 2.7e-06 | 2.633E-06 | 9.018E-07 | 1.700E-06 | 3.500E-06 |
| 29100 | 1455 | Y1455V | E | 4e-06 | 3.3e-06 | 1.3e-06 | 2.867E-06 | 1.401E-06 | 1.300E-06 | 4.000E-06 |
| 29101 | 1456 | S1456A | E | 1.8e-06 | 3.2e-06 | 3e-06 | 2.667E-06 | 7.572E-07 | 1.800E-06 | 3.200E-06 |
| 29102 | 1456 | S1456R | E | 2.3e-06 | 2e-06 | 2.1e-06 | 2.133E-06 | 1.528E-07 | 2.000E-06 | 2.300E-06 |
| 29103 | 1456 | S1456N | E | 1.9e-06 | 2.2e-06 | 1.8e-06 | 1.967E-06 | 2.082E-07 | 1.800E-06 | 2.200E-06 |
| 29104 | 1456 | S1456D | E | 8.7e-07 | 1.8e-06 | 2.4e-06 | 1.690E-06 | 7.709E-07 | 8.700E-07 | 2.400E-06 |
| 29105 | 1456 | S1456C | E | 2.6e-06 | 2.7e-06 | 2.3e-06 | 2.533E-06 | 2.082E-07 | 2.300E-06 | 2.700E-06 |
| 29106 | 1456 | S1456E | E | 2.3e-06 | 3.6e-06 | 2e-06 | 2.633E-06 | 8.505E-07 | 2.000E-06 | 3.600E-06 |
| 29107 | 1456 | S1456Q | E | 3.7e-06 | 2.7e-06 | 2e-06 | 2.800E-06 | 8.544E-07 | 2.000E-06 | 3.700E-06 |
| 29108 | 1456 | S1456G | E | 1.9e-06 | 2.5e-06 | 2.2e-06 | 2.200E-06 | 3.000E-07 | 1.900E-06 | 2.500E-06 |
| 29109 | 1456 | S1456H | E | 1.4e-06 | 3.9e-06 | 4.7e-06 | 3.333E-06 | 1.721E-06 | 1.400E-06 | 4.700E-06 |
| 29110 | 1456 | S1456I | E | 1.5e-06 | 2.7e-06 | 2.6e-06 | 2.267E-06 | 6.658E-07 | 1.500E-06 | 2.700E-06 |
| 29111 | 1456 | S1456L | E | 3.3e-06 | 4.6e-06 | 2e-06 | 3.300E-06 | 1.300E-06 | 2.000E-06 | 4.600E-06 |
| 29112 | 1456 | S1456K | E | 3.3e-06 | 1.5e-06 | 3.6e-06 | 2.800E-06 | 1.136E-06 | 1.500E-06 | 3.600E-06 |
| 29113 | 1456 | S1456M | E | 3.3e-06 | 1.5e-06 | 1.9e-06 | 2.233E-06 | 9.452E-07 | 1.500E-06 | 3.300E-06 |
| 29114 | 1456 | S1456F | E | 2.1e-06 | 1.2e-06 | 1.7e-06 | 1.667E-06 | 4.509E-07 | 1.200E-06 | 2.100E-06 |
| 29115 | 1456 | S1456P | E | 2.1e-06 | 2.5e-06 | 2.8e-06 | 2.467E-06 | 3.512E-07 | 2.100E-06 | 2.800E-06 |
| 29116 | 1456 | S1456S | E | 4.3e-06 | 2.4e-06 | 2.1e-06 | 2.933E-06 | 1.193E-06 | 2.100E-06 | 4.300E-06 |
| 29117 | 1456 | S1456T | E | 4.8e-06 | 3.1e-06 | 2e-06 | 3.300E-06 | 1.411E-06 | 2.000E-06 | 4.800E-06 |
| 29118 | 1456 | S1456W | E | 3.4e-06 | 2.1e-06 | 2.7e-06 | 2.733E-06 | 6.506E-07 | 2.100E-06 | 3.400E-06 |
| 29119 | 1456 | S1456Y | E | 2.4e-06 | 2.1e-06 | 1.7e-06 | 2.067E-06 | 3.512E-07 | 1.700E-06 | 2.400E-06 |
| 29120 | 1456 | S1456V | E | 3.4e-06 | 3e-06 | 2.8e-06 | 3.067E-06 | 3.055E-07 | 2.800E-06 | 3.400E-06 |
| 29121 | 1457 | L1457A | E | 2.3e-06 | 3.6e-06 | 5.7e-06 | 3.867E-06 | 1.716E-06 | 2.300E-06 | 5.700E-06 |
| 29122 | 1457 | L1457R | E | 1.4e-06 | 3.5e-06 | 2.3e-06 | 2.400E-06 | 1.054E-06 | 1.400E-06 | 3.500E-06 |
| 29123 | 1457 | L1457N | E | 2.3e-06 | 2.8e-06 | 2.5e-06 | 2.533E-06 | 2.517E-07 | 2.300E-06 | 2.800E-06 |
| 29124 | 1457 | L1457D | E | 1.5e-06 | 1.8e-06 | 2.3e-06 | 1.867E-06 | 4.041E-07 | 1.500E-06 | 2.300E-06 |
| 29125 | 1457 | L1457C | E | 2.1e-06 | 2.6e-06 | 2.9e-06 | 2.533E-06 | 4.041E-07 | 2.100E-06 | 2.900E-06 |
| 29126 | 1457 | L1457E | E | 1.8e-06 | 2.5e-06 | 1.9e-06 | 2.067E-06 | 3.786E-07 | 1.800E-06 | 2.500E-06 |
| 29127 | 1457 | L1457Q | E | 2.5e-06 | 2.2e-06 | 3.4e-06 | 2.700E-06 | 6.245E-07 | 2.200E-06 | 3.400E-06 |
| 29128 | 1457 | L1457G | E | 2.6e-06 | 2.1e-06 | 2.7e-06 | 2.467E-06 | 3.215E-07 | 2.100E-06 | 2.700E-06 |
| 29129 | 1457 | L1457H | E | 1.2e-06 | 2.3e-06 | 2.4e-06 | 1.967E-06 | 6.658E-07 | 1.200E-06 | 2.400E-06 |
| 29130 | 1457 | L1457I | E | 2.4e-06 | 1.9e-06 | 1.8e-06 | 2.033E-06 | 3.215E-07 | 1.800E-06 | 2.400E-06 |
| 29131 | 1457 | L1457L | E | 3.5e-06 | 2.3e-06 | 4.2e-06 | 3.333E-06 | 9.609E-07 | 2.300E-06 | 4.200E-06 |
| 29132 | 1457 | L1457K | E | 3.2e-06 | 2.4e-06 | 2.6e-06 | 2.733E-06 | 4.163E-07 | 2.400E-06 | 3.200E-06 |
| 29133 | 1457 | L1457M | E | 3.2e-06 | 1.4e-06 | 3.5e-06 | 2.700E-06 | 1.136E-06 | 1.400E-06 | 3.500E-06 |
| 29134 | 1457 | L1457F | E | 2.7e-06 | 2.5e-06 | 2.9e-06 | 2.700E-06 | 2.000E-07 | 2.500E-06 | 2.900E-06 |
| 29135 | 1457 | L1457P | E | 2.2e-06 | 1.8e-06 | 2.1e-06 | 2.033E-06 | 2.082E-07 | 1.800E-06 | 2.200E-06 |
| 29136 | 1457 | L1457S | E | 2.6e-06 | 2.3e-06 | 4.4e-06 | 3.100E-06 | 1.136E-06 | 2.300E-06 | 4.400E-06 |
| 29137 | 1457 | L1457T | E | 2.5e-06 | 2.2e-06 | 3.1e-06 | 2.600E-06 | 4.583E-07 | 2.200E-06 | 3.100E-06 |
| 29138 | 1457 | L1457W | E | 2.9e-06 | 1.8e-06 | 4.6e-06 | 3.100E-06 | 1.411E-06 | 1.800E-06 | 4.600E-06 |
| 29139 | 1457 | L1457Y | E | 1.7e-06 | 1.7e-06 | 1.9e-06 | 1.767E-06 | 1.155E-07 | 1.700E-06 | 1.900E-06 |
| 29140 | 1457 | L1457V | E | 2.6e-06 | 2.6e-06 | 4.2e-06 | 3.133E-06 | 9.238E-07 | 2.600E-06 | 4.200E-06 |
| 29141 | 1458 | S1458A | E | 2.1e-06 | 2.7e-06 | 1.6e-06 | 2.133E-06 | 5.508E-07 | 1.600E-06 | 2.700E-06 |
| 29142 | 1458 | S1458R | E | 1.6e-06 | 2.7e-06 | 3.6e-06 | 2.633E-06 | 1.002E-06 | 1.600E-06 | 3.600E-06 |
| 29143 | 1458 | S1458N | E | 3.2e-06 | 1.8e-06 | 2.3e-06 | 2.433E-06 | 7.095E-07 | 1.800E-06 | 3.200E-06 |
| 29144 | 1458 | S1458D | E | 2.3e-06 | 2.7e-06 | 1.2e-06 | 2.067E-06 | 7.767E-07 | 1.200E-06 | 2.700E-06 |
| 29145 | 1458 | S1458C | E | 8.4e-07 | 2.7e-06 | 1.7e-06 | 1.747E-06 | 9.309E-07 | 8.400E-07 | 2.700E-06 |
| 29146 | 1458 | S1458E | E | 2.5e-06 | 3.4e-06 | 2.6e-06 | 2.833E-06 | 4.933E-07 | 2.500E-06 | 3.400E-06 |
| 29147 | 1458 | S1458Q | E | 1e-06 | 3.3e-06 | 1.9e-06 | 2.067E-06 | 1.159E-06 | 1.000E-06 | 3.300E-06 |
| 29148 | 1458 | S1458G | E | 3.4e-06 | 2.7e-06 | 4e-06 | 3.367E-06 | 6.506E-07 | 2.700E-06 | 4.000E-06 |
| 29149 | 1458 | S1458H | E | 2.3e-06 | 2.5e-06 | 2.4e-06 | 2.400E-06 | 1.000E-07 | 2.300E-06 | 2.500E-06 |
| 29150 | 1458 | S1458I | E | 2.6e-06 | 2.5e-06 | 2.7e-06 | 2.600E-06 | 1.000E-07 | 2.500E-06 | 2.700E-06 |
| 29151 | 1458 | S1458L | E | 4e-06 | 3.2e-06 | 4.6e-06 | 3.933E-06 | 7.024E-07 | 3.200E-06 | 4.600E-06 |
| 29152 | 1458 | S1458K | E | 3.3e-06 | 3.2e-06 | 3.5e-06 | 3.333E-06 | 1.528E-07 | 3.200E-06 | 3.500E-06 |
| 29153 | 1458 | S1458M | E | 2.6e-06 | 2.3e-06 | 2.9e-06 | 2.600E-06 | 3.000E-07 | 2.300E-06 | 2.900E-06 |
| 29154 | 1458 | S1458F | E | 2.8e-06 | 3.5e-06 | 4.6e-06 | 3.633E-06 | 9.074E-07 | 2.800E-06 | 4.600E-06 |

| | | | | | | | | | | |
|---|---|---|---|---|---|---|---|---|---|---|
| 29155 | 1458 | S1458P | E | 2.7e-06 | 2.8e-06 | 2e-06 | 2.500E-06 | 4.359E-07 | 2.000E-06 | 2.800E-06 |
| 29156 | 1458 | S1458S | E | 4e-06 | 5.5e-06 | 2.5e-06 | 4.000E-06 | 1.500E-06 | 2.500E-06 | 5.500E-06 |
| 29157 | 1458 | S1458T | E | 1.7e-06 | 4.9e-06 | 1.6e-06 | 2.733E-06 | 1.877E-06 | 1.600E-06 | 4.900E-06 |
| 29158 | 1458 | S1458W | E | 1.2e-06 | 1.5e-06 | 3.4e-06 | 2.033E-06 | 1.193E-06 | 1.200E-06 | 3.400E-06 |
| 29159 | 1458 | S1458Y | E | 3.6e-06 | 2.1e-06 | 3.6e-06 | 3.100E-06 | 8.660E-07 | 2.100E-06 | 3.600E-06 |
| 29160 | 1458 | S1458V | E | 2.1e-06 | 2.1e-06 | 2.3e-06 | 2.167E-06 | 1.155E-07 | 2.100E-06 | 2.300E-06 |
| 29161 | 1459 | S1459A | E | 2.5e-06 | 2e-06 | 1.9e-06 | 2.133E-06 | 3.215E-07 | 1.900E-06 | 2.500E-06 |
| 29162 | 1459 | S1459R | E | 2.6e-06 | 2.5e-06 | 5e-06 | 3.367E-06 | 1.415E-06 | 2.500E-06 | 5.000E-06 |
| 29163 | 1459 | S1459N | E | 1.7e-06 | 2.4e-06 | 2.2e-06 | 2.100E-06 | 3.606E-07 | 1.700E-06 | 2.400E-06 |
| 29164 | 1459 | S1459D | E | 1e-06 | 2e-06 | 2.8e-06 | 1.933E-06 | 9.018E-07 | 1.000E-06 | 2.800E-06 |
| 29165 | 1459 | S1459C | E | 3.4e-06 | 2.4e-06 | 2.3e-06 | 2.700E-06 | 6.083E-07 | 2.300E-06 | 3.400E-06 |
| 29166 | 1459 | S1459E | E | 2.5e-06 | 2.3e-06 | 2e-06 | 2.267E-06 | 2.517E-07 | 2.000E-06 | 2.500E-06 |
| 29167 | 1459 | S1459Q | E | 2.6e-06 | 1.7e-06 | 3.3e-06 | 2.533E-06 | 8.021E-07 | 1.700E-06 | 3.300E-06 |
| 29168 | 1459 | S1459G | E | 1.7e-06 | 2.4e-06 | 1.8e-06 | 1.967E-06 | 3.786E-07 | 1.700E-06 | 2.400E-06 |
| 29169 | 1459 | S1459H | E | 2.2e-06 | 3.7e-06 | 5.1e-06 | 3.667E-06 | 1.450E-06 | 2.200E-06 | 5.100E-06 |
| 29170 | 1459 | S1459I | E | 3e-06 | 3e-06 | 1.8e-06 | 2.600E-06 | 6.928E-07 | 1.800E-06 | 3.000E-06 |
| 29171 | 1459 | S1459L | E | 4.1e-06 | 2.6e-06 | 1.5e-06 | 2.733E-06 | 1.305E-06 | 1.500E-06 | 4.100E-06 |
| 29172 | 1459 | S1459K | E | 2.5e-06 | 2.3e-06 | 1.2e-06 | 2.000E-06 | 7.000E-07 | 1.200E-06 | 2.500E-06 |
| 29173 | 1459 | S1459M | E | 2.7e-06 | 2.8e-06 | 2.2e-06 | 2.567E-06 | 3.215E-07 | 2.200E-06 | 2.800E-06 |
| 29174 | 1459 | S1459F | E | 2e-06 | 1.8e-06 | 6.3e-06 | 3.367E-06 | 2.542E-06 | 1.800E-06 | 6.300E-06 |
| 29175 | 1459 | S1459P | E | 1.8e-06 | 1.1e-06 | 1.5e-06 | 1.467E-06 | 3.512E-07 | 1.100E-06 | 1.800E-06 |
| 29176 | 1459 | S1459S | E | 3.8e-06 | 2.7e-06 | 1.7e-06 | 2.733E-06 | 1.050E-06 | 1.700E-06 | 3.800E-06 |
| 29177 | 1459 | S1459T | E | 2.4e-06 | 2.7e-06 | 2.5e-06 | 2.533E-06 | 1.528E-07 | 2.400E-06 | 2.700E-06 |
| 29178 | 1459 | S1459W | E | 9.5e-07 | 3e-06 | 1.1e-06 | 1.683E-06 | 1.143E-06 | 9.500E-07 | 3.000E-06 |
| 29179 | 1459 | S1459Y | E | 3e-06 | 2.2e-06 | 1.6e-06 | 2.267E-06 | 7.024E-07 | 1.600E-06 | 3.000E-06 |
| 29180 | 1459 | S1459V | E | 2.6e-06 | 2.5e-06 | 3.2e-06 | 2.767E-06 | 3.786E-07 | 2.500E-06 | 3.200E-06 |
| 29181 | 1460 | V1460A | E | 3.8e-06 | 1.5e-06 | 2.8e-06 | 2.700E-06 | 1.153E-06 | 1.500E-06 | 3.800E-06 |
| 29182 | 1460 | V1460R | E | 3.7e-06 | 1.7e-06 | 2.4e-06 | 2.600E-06 | 1.015E-06 | 1.700E-06 | 3.700E-06 |
| 29183 | 1460 | V1460N | E | 2.2e-06 | 2.4e-06 | 2.3e-06 | 2.300E-06 | 1.000E-07 | 2.200E-06 | 2.400E-06 |
| 29184 | 1460 | V1460D | E | 3.3e-06 | 2.7e-06 | 3.2e-06 | 3.067E-06 | 3.215E-07 | 2.700E-06 | 3.300E-06 |
| 29185 | 1460 | V1460C | E | 2.3e-06 | 2.6e-06 | 2e-06 | 2.300E-06 | 3.000E-07 | 2.000E-06 | 2.600E-06 |
| 29186 | 1460 | V1460E | E | 1.8e-06 | 2.2e-06 | 1.1e-06 | 1.700E-06 | 5.568E-07 | 1.100E-06 | 2.200E-06 |
| 29187 | 1460 | V1460Q | E | 3e-06 | 4e-06 | 1.9e-06 | 2.967E-06 | 1.050E-06 | 1.900E-06 | 4.000E-06 |
| 29188 | 1460 | V1460G | E | 1.2e-06 | 2.6e-06 | 2.8e-06 | 2.200E-06 | 8.718E-07 | 1.200E-06 | 2.800E-06 |
| 29189 | 1460 | V1460H | E | 1.2e-06 | 2.5e-06 | 3e-06 | 2.233E-06 | 9.292E-07 | 1.200E-06 | 3.000E-06 |
| 29190 | 1460 | V1460I | E | 3e-06 | 2.4e-06 | 3.1e-06 | 2.833E-06 | 3.786E-07 | 2.400E-06 | 3.100E-06 |
| 29191 | 1460 | V1460L | E | 1.9e-06 | 2.6e-06 | 1.5e-06 | 2.000E-06 | 5.568E-07 | 1.500E-06 | 2.600E-06 |
| 29192 | 1460 | V1460K | E | 2e-06 | 2.7e-06 | 2.5e-06 | 2.400E-06 | 3.606E-07 | 2.000E-06 | 2.700E-06 |
| 29193 | 1460 | V1460M | E | 1.9e-06 | 2.1e-06 | 2.3e-06 | 2.100E-06 | 2.000E-07 | 1.900E-06 | 2.300E-06 |
| 29194 | 1460 | V1460F | E | 3.3e-06 | 1.8e-06 | 1.8e-06 | 2.300E-06 | 8.660E-07 | 1.800E-06 | 3.300E-06 |
| 29195 | 1460 | V1460P | E | 2.3e-06 | 2.9e-06 | 1.5e-06 | 2.233E-06 | 7.024E-07 | 1.500E-06 | 2.900E-06 |
| 29196 | 1460 | V1460S | E | 3.4e-06 | 1.9e-06 | 2.7e-06 | 2.667E-06 | 7.506E-07 | 1.900E-06 | 3.400E-06 |
| 29197 | 1460 | V1460T | E | 2.6e-06 | 8.5e-07 | 1.6e-06 | 1.683E-06 | 8.780E-07 | 8.500E-07 | 2.600E-06 |
| 29198 | 1460 | V1460W | E | 2.4e-06 | 2.3e-06 | 4.2e-06 | 2.967E-06 | 1.069E-06 | 2.300E-06 | 4.200E-06 |
| 29199 | 1460 | V1460Y | E | 2.1e-06 | 6.2e-06 | 1.7e-06 | 3.333E-06 | 2.491E-06 | 1.700E-06 | 6.200E-06 |
| 29200 | 1460 | V1460V | E | 3.2e-06 | 2e-06 | 2.1e-06 | 2.433E-06 | 6.658E-07 | 2.000E-06 | 3.200E-06 |
| 29201 | 1461 | V1461A | E | 2e-06 | 2.6e-06 | 2.4e-06 | 2.333E-06 | 3.055E-07 | 2.000E-06 | 2.600E-06 |
| 29202 | 1461 | V1461R | E | 2.2e-06 | 2.9e-06 | 1.3e-06 | 2.133E-06 | 8.021E-07 | 1.300E-06 | 2.900E-06 |
| 29203 | 1461 | V1461N | E | 1.5e-06 | 5e-06 | 1.9e-06 | 2.800E-06 | 1.916E-06 | 1.500E-06 | 5.000E-06 |
| 29204 | 1461 | V1461D | E | 2.2e-06 | 3.3e-06 | 1.6e-06 | 2.367E-06 | 8.622E-07 | 1.600E-06 | 3.300E-06 |
| 29205 | 1461 | V1461C | E | 2.4e-06 | 2.7e-06 | 3e-06 | 2.700E-06 | 3.000E-07 | 2.400E-06 | 3.000E-06 |
| 29206 | 1461 | V1461E | E | 3.3e-06 | 2.5e-06 | 2.1e-06 | 2.633E-06 | 6.110E-07 | 2.100E-06 | 3.300E-06 |
| 29207 | 1461 | V1461Q | E | 3.8e-06 | 2.2e-06 | 3.4e-06 | 3.133E-06 | 8.327E-07 | 2.200E-06 | 3.800E-06 |
| 29208 | 1461 | V1461G | E | 3e-06 | 2.1e-06 | 1.7e-06 | 2.267E-06 | 6.658E-07 | 1.700E-06 | 3.000E-06 |
| 29209 | 1461 | V1461H | E | 3.9e-06 | 2e-06 | 2.6e-06 | 2.833E-06 | 9.713E-07 | 2.000E-06 | 3.900E-06 |
| 29210 | 1461 | V1461I | E | 1.2e-06 | 1.4e-06 | 4.3e-06 | 2.300E-06 | 1.735E-06 | 1.200E-06 | 4.300E-06 |
| 29211 | 1461 | V1461L | E | 2.4e-06 | 1.6e-06 | 2.7e-06 | 2.233E-06 | 5.686E-07 | 1.600E-06 | 2.700E-06 |
| 29212 | 1461 | V1461K | E | 2.2e-06 | 2.3e-06 | 2.6e-06 | 2.367E-06 | 2.082E-07 | 2.200E-06 | 2.600E-06 |
| 29213 | 1461 | V1461M | E | 2.3e-06 | 2.1e-06 | 3.2e-06 | 2.533E-06 | 5.859E-07 | 2.100E-06 | 3.200E-06 |
| 29214 | 1461 | V1461F | E | 1.8e-06 | 2.4e-06 | 2e-06 | 2.067E-06 | 3.055E-07 | 1.800E-06 | 2.400E-06 |
| 29215 | 1461 | V1461P | E | 3.2e-06 | 2e-06 | 1.7e-06 | 2.300E-06 | 7.937E-07 | 1.700E-06 | 3.200E-06 |
| 29216 | 1461 | V1461S | E | 1.3e-06 | 2e-06 | 3.8e-06 | 2.367E-06 | 1.290E-06 | 1.300E-06 | 3.800E-06 |
| 29217 | 1461 | V1461T | E | 2.1e-06 | 1.7e-06 | 2.5e-06 | 2.100E-06 | 4.000E-07 | 1.700E-06 | 2.500E-06 |
| 29218 | 1461 | V1461W | E | 2.1e-06 | 2.8e-06 | 2.2e-06 | 2.367E-06 | 3.786E-07 | 2.100E-06 | 2.800E-06 |
| 29219 | 1461 | V1461Y | E | 2.3e-06 | 2.6e-06 | 2e-06 | 2.300E-06 | 3.000E-07 | 2.000E-06 | 2.600E-06 |
| 29220 | 1461 | V1461V | E | 2.6e-06 | 1.9e-06 | 3.2e-06 | 2.567E-06 | 6.506E-07 | 1.900E-06 | 3.200E-06 |
| 29221 | 1462 | T1462A | E | 5.4e-06 | 2.9e-06 | 3.1e-06 | 3.800E-06 | 1.389E-06 | 2.900E-06 | 5.400E-06 |
| 29222 | 1462 | T1462R | E | 3.1e-06 | 2.2e-06 | 2.7e-06 | 2.667E-06 | 4.509E-07 | 2.200E-06 | 3.100E-06 |
| 29223 | 1462 | T1462N | E | 6.8e-06 | 3.1e-06 | 1.4e-06 | 3.767E-06 | 2.761E-06 | 1.400E-06 | 6.800E-06 |
| 29224 | 1462 | T1462D | E | 3.1e-06 | 1.9e-06 | 2.5e-06 | 2.500E-06 | 6.000E-07 | 1.900E-06 | 3.100E-06 |
| 29225 | 1462 | T1462C | E | 1.3e-06 | 2.2e-06 | 9.9e-07 | 1.497E-06 | 6.285E-07 | 9.900E-07 | 2.200E-06 |
| 29226 | 1462 | T1462E | E | 3.5e-06 | 2.7e-06 | 4e-06 | 3.400E-06 | 6.557E-07 | 2.700E-06 | 4.000E-06 |
| 29227 | 1462 | T1462Q | E | 2.6e-06 | 1.4e-06 | 2.7e-06 | 2.233E-06 | 7.234E-07 | 1.400E-06 | 2.700E-06 |
| 29228 | 1462 | T1462G | E | 2e-06 | 3.5e-06 | 4.2e-06 | 3.233E-06 | 1.124E-06 | 2.000E-06 | 4.200E-06 |
| 29229 | 1462 | T1462H | E | 2.3e-06 | 2.5e-06 | 2.3e-06 | 2.367E-06 | 1.155E-07 | 2.300E-06 | 2.500E-06 |
| 29230 | 1462 | T1462I | E | 1.6e-06 | 1.9e-06 | 2.8e-06 | 2.100E-06 | 6.245E-07 | 1.600E-06 | 2.800E-06 |
| 29231 | 1462 | T1462L | E | 4e-06 | 4.1e-06 | 2e-06 | 3.367E-06 | 1.185E-06 | 2.000E-06 | 4.100E-06 |
| 29232 | 1462 | T1462K | E | 1.3e-06 | 3.8e-06 | 2.3e-06 | 2.467E-06 | 1.258E-06 | 1.300E-06 | 3.800E-06 |
| 29233 | 1462 | T1462M | E | 2e-06 | 2.6e-06 | 2.8e-06 | 2.467E-06 | 4.163E-07 | 2.000E-06 | 2.800E-06 |
| 29234 | 1462 | T1462F | E | 3.3e-06 | 3.4e-06 | 2.5e-06 | 3.067E-06 | 4.933E-07 | 2.500E-06 | 3.400E-06 |
| 29235 | 1462 | T1462P | E | 2.8e-06 | 1.3e-06 | 1.3e-06 | 1.800E-06 | 8.660E-07 | 1.300E-06 | 2.800E-06 |
| 29236 | 1462 | T1462S | E | 3.5e-06 | 2.2e-06 | 2.3e-06 | 2.667E-06 | 7.234E-07 | 2.200E-06 | 3.500E-06 |
| 29237 | 1462 | T1462T | E | 7.6e-07 | 1.6e-06 | 1.9e-06 | 1.420E-06 | 5.909E-07 | 7.600E-07 | 1.900E-06 |
| 29238 | 1462 | T1462W | E | 1.3e-06 | 1e-06 | 3.1e-06 | 1.800E-06 | 1.136E-06 | 1.000E-06 | 3.100E-06 |
| 29239 | 1462 | T1462Y | E | 1.6e-06 | 1.4e-06 | 2.3e-06 | 1.767E-06 | 4.726E-07 | 1.400E-06 | 2.300E-06 |

| | | | | | | | | | | |
|---|---|---|---|---|---|---|---|---|---|---|
| 29240 | 1462 | T1462V | E | 1.8e-06 | 1.6e-06 | 2.2e-06 | 1.867E-06 | 3.055E-07 | 1.600E-06 | 2.200E-06 |
| 29241 | 1463 | V1463A | E | 2e-06 | 2.3e-06 | 2.1e-06 | 2.133E-06 | 1.528E-07 | 2.000E-06 | 2.300E-06 |
| 29242 | 1463 | V1463R | E | 1.5e-06 | 3.2e-06 | 2.5e-06 | 2.400E-06 | 8.544E-07 | 1.500E-06 | 3.200E-06 |
| 29243 | 1463 | V1463N | E | 2.5e-06 | 2e-06 | 1.7e-06 | 2.067E-06 | 4.041E-07 | 1.700E-06 | 2.500E-06 |
| 29244 | 1463 | V1463D | E | 1.3e-06 | 2.6e-06 | 2.4e-06 | 2.100E-06 | 7.000E-07 | 1.300E-06 | 2.600E-06 |
| 29245 | 1463 | V1463C | E | 3.4e-06 | 2.1e-06 | 2.2e-06 | 2.567E-06 | 7.234E-07 | 2.100E-06 | 3.400E-06 |
| 29246 | 1463 | V1463E | E | 2.5e-06 | 1.8e-06 | 2e-06 | 2.100E-06 | 3.606E-07 | 1.800E-06 | 2.500E-06 |
| 29247 | 1463 | V1463Q | E | 2.6e-06 | 2.3e-06 | 2.8e-06 | 2.567E-06 | 2.517E-07 | 2.300E-06 | 2.800E-06 |
| 29248 | 1463 | V1463G | E | 2.4e-06 | 7.8e-07 | 1.9e-06 | 1.693E-06 | 8.295E-07 | 7.800E-07 | 2.400E-06 |
| 29249 | 1463 | V1463H | E | 2.1e-06 | 2.3e-06 | 2.9e-06 | 2.433E-06 | 4.163E-07 | 2.100E-06 | 2.900E-06 |
| 29250 | 1463 | V1463I | E | 2e-06 | 1.9e-06 | 1.2e-06 | 1.700E-06 | 4.359E-07 | 1.200E-06 | 2.000E-06 |
| 29251 | 1463 | V1463L | E | 3e-06 | 3.6e-06 | 2.4e-06 | 3.000E-06 | 6.000E-07 | 2.400E-06 | 3.600E-06 |
| 29252 | 1463 | V1463K | E | 2e-06 | 2.5e-06 | 1.9e-06 | 2.133E-06 | 3.215E-07 | 1.900E-06 | 2.500E-06 |
| 29253 | 1463 | V1463M | E | 1.3e-06 | 3.2e-06 | 2.6e-06 | 2.367E-06 | 9.713E-07 | 1.300E-06 | 3.200E-06 |
| 29254 | 1463 | V1463F | E | 1.2e-06 | 3e-06 | 3.5e-06 | 2.567E-06 | 1.210E-06 | 1.200E-06 | 3.500E-06 |
| 29255 | 1463 | V1463P | E | 4.2e-06 | 5.2e-06 | 1.3e-06 | 3.567E-06 | 2.026E-06 | 1.300E-06 | 5.200E-06 |
| 29256 | 1463 | V1463S | E | 1.3e-06 | 3.5e-06 | 2.1e-06 | 2.300E-06 | 1.114E-06 | 1.300E-06 | 3.500E-06 |
| 29257 | 1463 | V1463T | E | 1.9e-06 | 3.9e-06 | 2.6e-06 | 2.800E-06 | 1.015E-06 | 1.900E-06 | 3.900E-06 |
| 29258 | 1463 | V1463W | E | 1.5e-06 | 1.8e-06 | 3.6e-06 | 2.300E-06 | 1.136E-06 | 1.500E-06 | 3.600E-06 |
| 29259 | 1463 | V1463Y | E | 2.2e-06 | 1.1e-06 | 2.1e-06 | 1.800E-06 | 6.083E-07 | 1.100E-06 | 2.200E-06 |
| 29260 | 1463 | V1463V | E | 2.4e-06 | 2.9e-06 | 3e-06 | 2.767E-06 | 3.215E-07 | 2.400E-06 | 3.000E-06 |
| 29261 | 1464 | P1464A | E | 2.4e-06 | 1.9e-06 | 1.8e-06 | 2.033E-06 | 3.215E-07 | 1.800E-06 | 2.400E-06 |
| 29262 | 1464 | P1464R | E | 2.9e-06 | 1.7e-06 | 1.3e-06 | 1.967E-06 | 8.327E-07 | 1.300E-06 | 2.900E-06 |
| 29263 | 1464 | P1464N | E | 2e-06 | 2.9e-06 | 3.3e-06 | 2.733E-06 | 6.658E-07 | 2.000E-06 | 3.300E-06 |
| 29264 | 1464 | P1464D | E | 1.6e-06 | 3.7e-06 | 2e-06 | 2.433E-06 | 1.115E-06 | 1.600E-06 | 3.700E-06 |
| 29265 | 1464 | P1464C | E | 2.2e-06 | 1.9e-06 | 1.7e-06 | 1.933E-06 | 2.517E-07 | 1.700E-06 | 2.200E-06 |
| 29266 | 1464 | P1464E | E | 2.1e-06 | 1.9e-06 | 2.4e-06 | 2.133E-06 | 2.517E-07 | 1.900E-06 | 2.400E-06 |
| 29267 | 1464 | P1464Q | E | 2.1e-06 | 4e-06 | 1.4e-06 | 2.500E-06 | 1.345E-06 | 1.400E-06 | 4.000E-06 |
| 29268 | 1464 | P1464G | E | 2e-06 | 1.9e-06 | 1.7e-06 | 1.867E-06 | 1.528E-07 | 1.700E-06 | 2.000E-06 |
| 29269 | 1464 | P1464H | E | 3e-06 | 2.9e-06 | 2.4e-06 | 2.767E-06 | 3.215E-07 | 2.400E-06 | 3.000E-06 |
| 29270 | 1464 | P1464I | E | 1.8e-06 | 3.5e-06 | 1.4e-06 | 2.233E-06 | 1.115E-06 | 1.400E-06 | 3.500E-06 |
| 29271 | 1464 | P1464L | E | 2.3e-06 | 2.7e-06 | 2.1e-06 | 2.367E-06 | 3.055E-07 | 2.100E-06 | 2.700E-06 |
| 29272 | 1464 | P1464K | E | 2.9e-06 | 3.5e-06 | 1.1e-06 | 2.500E-06 | 1.249E-06 | 1.100E-06 | 3.500E-06 |
| 29273 | 1464 | P1464M | E | 2e-06 | 3.9e-06 | 3.2e-06 | 3.033E-06 | 9.609E-07 | 2.000E-06 | 3.900E-06 |
| 29274 | 1464 | P1464F | E | 2.4e-06 | 2.8e-06 | 2.8e-06 | 2.667E-06 | 2.309E-07 | 2.400E-06 | 2.800E-06 |
| 29275 | 1464 | P1464P | E | 2.4e-06 | 3e-06 | 3.5e-06 | 2.967E-06 | 5.508E-07 | 2.400E-06 | 3.500E-06 |
| 29276 | 1464 | P1464S | E | 2.2e-06 | 8.8e-07 | 2.8e-06 | 1.960E-06 | 9.822E-07 | 8.800E-07 | 2.800E-06 |
| 29277 | 1464 | P1464T | E | 2.3e-06 | 2.4e-06 | 2.2e-06 | 2.300E-06 | 1.000E-07 | 2.200E-06 | 2.400E-06 |
| 29278 | 1464 | P1464W | E | 2.1e-06 | 1.8e-06 | 1.4e-06 | 1.767E-06 | 3.512E-07 | 1.400E-06 | 2.100E-06 |
| 29279 | 1464 | P1464Y | E | 3.3e-06 | 9.5e-07 | 2.4e-06 | 2.217E-06 | 1.186E-06 | 9.500E-07 | 3.300E-06 |
| 29280 | 1464 | P1464V | E | 3e-06 | 2.3e-06 | 2.7e-06 | 2.667E-06 | 3.512E-07 | 2.300E-06 | 3.000E-06 |
| 29281 | 1465 | S1465A | E | 1.7e-06 | 1.9e-06 | 2.1e-06 | 1.900E-06 | 2.000E-07 | 1.700E-06 | 2.100E-06 |
| 29282 | 1465 | S1465R | E | 3.1e-06 | 2e-06 | 4.5e-06 | 3.200E-06 | 1.253E-06 | 2.000E-06 | 4.500E-06 |
| 29283 | 1465 | S1465N | E | 3.3e-06 | 2.8e-06 | 3.1e-06 | 3.067E-06 | 2.517E-07 | 2.800E-06 | 3.300E-06 |
| 29284 | 1465 | S1465D | E | 2.4e-06 | 1.9e-06 | 1.3e-06 | 1.867E-06 | 5.508E-07 | 1.300E-06 | 2.400E-06 |
| 29285 | 1465 | S1465C | E | 2.6e-06 | 2.2e-06 | 2.2e-06 | 2.333E-06 | 2.309E-07 | 2.200E-06 | 2.600E-06 |
| 29286 | 1465 | S1465E | E | 1.6e-06 | 1.9e-06 | 2e-06 | 1.833E-06 | 2.082E-07 | 1.600E-06 | 2.000E-06 |
| 29287 | 1465 | S1465Q | E | 2.7e-06 | 2.6e-06 | 2.8e-06 | 2.700E-06 | 1.000E-07 | 2.600E-06 | 2.800E-06 |
| 29288 | 1465 | S1465G | E | 1.7e-06 | 2.4e-06 | 2.2e-06 | 2.100E-06 | 3.606E-07 | 1.700E-06 | 2.400E-06 |
| 29289 | 1465 | S1465H | E | 3e-06 | 2.1e-06 | 3.1e-06 | 2.733E-06 | 5.508E-07 | 2.100E-06 | 3.100E-06 |
| 29290 | 1465 | S1465I | E | 3.4e-06 | 1.8e-06 | 2.7e-06 | 2.633E-06 | 8.021E-07 | 1.800E-06 | 3.400E-06 |
| 29291 | 1465 | S1465L | E | 3e-06 | 3.1e-06 | 2.5e-06 | 2.867E-06 | 3.215E-07 | 2.500E-06 | 3.100E-06 |
| 29292 | 1465 | S1465K | E | 2.3e-06 | 2.4e-06 | 2.7e-06 | 2.467E-06 | 2.082E-07 | 2.300E-06 | 2.700E-06 |
| 29293 | 1465 | S1465M | E | 3.1e-06 | 2e-06 | 2.7e-06 | 2.600E-06 | 5.568E-07 | 2.000E-06 | 3.100E-06 |
| 29294 | 1465 | S1465F | E | 5.1e-06 | 1.4e-06 | 2.8e-06 | 3.100E-06 | 1.868E-06 | 1.400E-06 | 5.100E-06 |
| 29295 | 1465 | S1465P | E | 3.6e-06 | 2.3e-06 | 2.2e-06 | 2.700E-06 | 7.810E-07 | 2.200E-06 | 3.600E-06 |
| 29296 | 1465 | S1465S | E | 2.5e-06 | 2.6e-06 | 3.1e-06 | 2.733E-06 | 3.215E-07 | 2.500E-06 | 3.100E-06 |
| 29297 | 1465 | S1465T | E | 3.6e-06 | 3.7e-06 | 3.3e-06 | 3.533E-06 | 2.082E-07 | 3.300E-06 | 3.700E-06 |
| 29298 | 1465 | S1465W | E | 2.5e-06 | 2.7e-06 | 1.8e-06 | 2.333E-06 | 4.726E-07 | 1.800E-06 | 2.700E-06 |
| 29299 | 1465 | S1465Y | E | 1.7e-06 | 1.9e-06 | 2.2e-06 | 1.933E-06 | 2.517E-07 | 1.700E-06 | 2.200E-06 |
| 29300 | 1465 | S1465V | E | 2.1e-06 | 4.3e-06 | 2.1e-06 | 2.833E-06 | 1.270E-06 | 2.100E-06 | 4.300E-06 |
| 29301 | 1466 | S1466A | E | 2.6e-06 | 2.1e-06 | 1.9e-06 | 2.200E-06 | 3.606E-07 | 1.900E-06 | 2.600E-06 |
| 29302 | 1466 | S1466R | E | 1.9e-06 | 3.1e-06 | 2e-06 | 2.333E-06 | 6.658E-07 | 1.900E-06 | 3.100E-06 |
| 29303 | 1466 | S1466N | E | 2.6e-06 | 2.9e-06 | 3.3e-06 | 2.933E-06 | 3.512E-07 | 2.600E-06 | 3.300E-06 |
| 29304 | 1466 | S1466D | E | 1.7e-06 | 2.8e-06 | 1.7e-06 | 2.067E-06 | 6.351E-07 | 1.700E-06 | 2.800E-06 |
| 29305 | 1466 | S1466C | E | 2.9e-06 | 3.1e-06 | 1.4e-06 | 2.467E-06 | 9.292E-07 | 1.400E-06 | 3.100E-06 |
| 29306 | 1466 | S1466E | E | 2.1e-06 | 2.1e-06 | 1.3e-06 | 1.833E-06 | 4.619E-07 | 1.300E-06 | 2.100E-06 |
| 29307 | 1466 | S1466Q | E | 1.8e-06 | 2.1e-06 | 1.3e-06 | 1.733E-06 | 4.041E-07 | 1.300E-06 | 2.100E-06 |
| 29308 | 1466 | S1466G | E | 1.8e-06 | 3.5e-06 | 2.4e-06 | 2.567E-06 | 8.622E-07 | 1.800E-06 | 3.500E-06 |
| 29309 | 1466 | S1466H | E | 3.3e-06 | 1.7e-06 | 2.5e-06 | 2.500E-06 | 8.000E-07 | 1.700E-06 | 3.300E-06 |
| 29310 | 1466 | S1466I | E | 1.7e-06 | 2.8e-06 | 2.4e-06 | 2.300E-06 | 5.568E-07 | 1.700E-06 | 2.800E-06 |
| 29311 | 1466 | S1466L | E | 5.3e-06 | 2e-06 | 2.4e-06 | 3.233E-06 | 1.801E-06 | 2.000E-06 | 5.300E-06 |
| 29312 | 1466 | S1466K | E | 2.6e-06 | 4e-06 | 1.2e-06 | 2.600E-06 | 1.400E-06 | 1.200E-06 | 4.000E-06 |
| 29313 | 1466 | S1466M | E | 2.3e-06 | 1.6e-06 | 4.4e-06 | 2.767E-06 | 1.457E-06 | 1.600E-06 | 4.400E-06 |
| 29314 | 1466 | S1466F | E | 3e-06 | 1.9e-06 | 2.5e-06 | 2.467E-06 | 5.508E-07 | 1.900E-06 | 3.000E-06 |
| 29315 | 1466 | S1466P | E | 1.7e-06 | 2.4e-06 | 2.1e-06 | 2.067E-06 | 3.512E-07 | 1.700E-06 | 2.400E-06 |
| 29316 | 1466 | S1466S | E | 2.8e-06 | 2.3e-06 | 2.7e-06 | 2.600E-06 | 2.646E-07 | 2.300E-06 | 2.800E-06 |
| 29317 | 1466 | S1466T | E | 2.7e-06 | 2e-06 | 3.1e-06 | 2.600E-06 | 5.568E-07 | 2.000E-06 | 3.100E-06 |
| 29318 | 1466 | S1466W | E | 2.1e-06 | 3.5e-06 | 2.8e-06 | 2.800E-06 | 7.000E-07 | 2.100E-06 | 3.500E-06 |
| 29319 | 1466 | S1466Y | E | 2e-06 | 2e-06 | 2.4e-06 | 2.133E-06 | 2.309E-07 | 2.000E-06 | 2.400E-06 |
| 29320 | 1466 | S1466V | E | 1.4e-06 | 2e-06 | 2.4e-06 | 1.933E-06 | 5.033E-07 | 1.400E-06 | 2.400E-06 |
| 29321 | 1467 | S1467A | E | 4.2e-06 | 1.2e-06 | 4.2e-06 | 3.200E-06 | 1.732E-06 | 1.200E-06 | 4.200E-06 |
| 29322 | 1467 | S1467R | E | 1.1e-06 | 1.2e-06 | 2.7e-06 | 1.667E-06 | 8.963E-07 | 1.100E-06 | 2.700E-06 |
| 29323 | 1467 | S1467N | E | 2.6e-06 | 2.1e-06 | 3e-06 | 2.567E-06 | 4.509E-07 | 2.100E-06 | 3.000E-06 |
| 29324 | 1467 | S1467D | E | 3.4e-06 | 1.8e-06 | 1.8e-06 | 2.333E-06 | 9.238E-07 | 1.800E-06 | 3.400E-06 |

| | | | | | | | | | | |
|---|---|---|---|---|---|---|---|---|---|---|
| 29325 | 1467 | S1467C | E | 2.3e-06 | 4.5e-06 | 2.1e-06 | 2.967E-06 | 1.332E-06 | 2.100E-06 | 4.500E-06 |
| 29326 | 1467 | S1467E | E | 2e-06 | 2.2e-06 | 2.1e-06 | 2.100E-06 | 1.000E-07 | 2.000E-06 | 2.200E-06 |
| 29327 | 1467 | S1467Q | E | 2.9e-06 | 3.7e-06 | 1.9e-06 | 2.833E-06 | 9.018E-07 | 1.900E-06 | 3.700E-06 |
| 29328 | 1467 | S1467G | E | 4.3e-06 | 2.7e-06 | 2.7e-06 | 3.233E-06 | 9.238E-07 | 2.700E-06 | 4.300E-06 |
| 29329 | 1467 | S1467H | E | 2.6e-06 | 2.2e-06 | 1.4e-06 | 2.067E-06 | 6.110E-07 | 1.400E-06 | 2.600E-06 |
| 29330 | 1467 | S1467I | E | 2.8e-06 | 1.8e-06 | 1.5e-06 | 2.033E-06 | 6.807E-07 | 1.500E-06 | 2.800E-06 |
| 29331 | 1467 | S1467L | E | 1.9e-06 | 2.8e-06 | 2.2e-06 | 2.300E-06 | 4.583E-07 | 1.900E-06 | 2.800E-06 |
| 29332 | 1467 | S1467K | E | 1.4e-06 | 3.1e-06 | 2.2e-06 | 2.233E-06 | 8.505E-07 | 1.400E-06 | 3.100E-06 |
| 29333 | 1467 | S1467M | E | 3e-06 | 2.8e-06 | 3.4e-06 | 3.067E-06 | 3.055E-07 | 2.800E-06 | 3.400E-06 |
| 29334 | 1467 | S1467F | E | 1.9e-06 | 1.9e-06 | 3.8e-06 | 2.533E-06 | 1.097E-06 | 1.900E-06 | 3.800E-06 |
| 29335 | 1467 | S1467P | E | 5e-06 | 3.4e-06 | 2.3e-06 | 3.567E-06 | 1.358E-06 | 2.300E-06 | 5.000E-06 |
| 29336 | 1467 | S1467S | E | 2.4e-06 | 3.7e-06 | 2.1e-06 | 2.733E-06 | 8.505E-07 | 2.100E-06 | 3.700E-06 |
| 29337 | 1467 | S1467T | E | 2.9e-06 | 3.7e-06 | 1.6e-06 | 2.733E-06 | 1.060E-06 | 1.600E-06 | 3.700E-06 |
| 29338 | 1467 | S1467W | E | 2.2e-06 | 4e-06 | 2.3e-06 | 2.833E-06 | 1.012E-06 | 2.200E-06 | 4.000E-06 |
| 29339 | 1467 | S1467Y | E | 4.6e-06 | 1.5e-06 | 2.4e-06 | 2.833E-06 | 1.595E-06 | 1.500E-06 | 4.600E-06 |
| 29340 | 1467 | S1467V | E | 2e-06 | 2.8e-06 | 2.8e-06 | 2.533E-06 | 4.619E-07 | 2.000E-06 | 2.800E-06 |
| 29341 | 1468 | L1468A | E | 2.9e-06 | 2.9e-06 | 2.1e-06 | 2.633E-06 | 4.619E-07 | 2.100E-06 | 2.900E-06 |
| 29342 | 1468 | L1468R | E | 3.9e-06 | 2.3e-06 | 2.9e-06 | 3.033E-06 | 8.083E-07 | 2.300E-06 | 3.900E-06 |
| 29343 | 1468 | L1468N | E | 2.6e-06 | 2.3e-06 | 2.5e-06 | 2.467E-06 | 1.528E-07 | 2.300E-06 | 2.600E-06 |
| 29344 | 1468 | L1468D | E | 3.1e-06 | 2.7e-06 | 4.9e-06 | 3.567E-06 | 1.172E-06 | 2.700E-06 | 4.900E-06 |
| 29345 | 1468 | L1468C | E | 1.7e-06 | 2.7e-06 | 1.7e-06 | 2.033E-06 | 5.774E-07 | 1.700E-06 | 2.700E-06 |
| 29346 | 1468 | L1468E | E | 3.3e-06 | 2.1e-06 | 2.9e-06 | 2.767E-06 | 6.110E-07 | 2.100E-06 | 3.300E-06 |
| 29347 | 1468 | L1468Q | E | 1.9e-06 | 5e-06 | 1.6e-06 | 2.833E-06 | 1.882E-06 | 1.600E-06 | 5.000E-06 |
| 29348 | 1468 | L1468G | E | 2.4e-06 | 2.1e-06 | 3.1e-06 | 2.533E-06 | 5.132E-07 | 2.100E-06 | 3.100E-06 |
| 29349 | 1468 | L1468H | E | 2e-06 | 2e-06 | 2.4e-06 | 2.133E-06 | 2.309E-07 | 2.000E-06 | 2.400E-06 |
| 29350 | 1468 | L1468I | E | 4.5e-06 | 2.1e-06 | 1.7e-06 | 2.767E-06 | 1.514E-06 | 1.700E-06 | 4.500E-06 |
| 29351 | 1468 | L1468L | E | 3.2e-06 | 3e-06 | 3.2e-06 | 3.133E-06 | 1.155E-07 | 3.000E-06 | 3.200E-06 |
| 29352 | 1468 | L1468K | E | 4.5e-06 | 2.9e-06 | 1.9e-06 | 3.100E-06 | 1.311E-06 | 1.900E-06 | 4.500E-06 |
| 29353 | 1468 | L1468M | E | 3.1e-06 | 2.8e-06 | 2.6e-06 | 2.833E-06 | 2.517E-07 | 2.600E-06 | 3.100E-06 |
| 29354 | 1468 | L1468F | E | 3.5e-06 | 2.1e-06 | 2.5e-06 | 2.700E-06 | 7.211E-07 | 2.100E-06 | 3.500E-06 |
| 29355 | 1468 | L1468P | E | 2e-06 | 3.7e-06 | 2.7e-06 | 2.800E-06 | 8.544E-07 | 2.000E-06 | 3.700E-06 |
| 29356 | 1468 | L1468S | E | 2.1e-06 | 2.3e-06 | 2.3e-06 | 2.233E-06 | 1.155E-07 | 2.100E-06 | 2.300E-06 |
| 29357 | 1468 | L1468T | E | 2.7e-06 | 2.8e-06 | 1.6e-06 | 2.367E-06 | 6.658E-07 | 1.600E-06 | 2.800E-06 |
| 29358 | 1468 | L1468W | E | 3.9e-06 | 3.6e-06 | 2.5e-06 | 3.333E-06 | 7.371E-07 | 2.500E-06 | 3.900E-06 |
| 29359 | 1468 | L1468Y | E | 3.8e-06 | 3e-06 | 2.7e-06 | 3.167E-06 | 5.686E-07 | 2.700E-06 | 3.800E-06 |
| 29360 | 1468 | L1468V | E | 2.1e-06 | 1.4e-06 | 2.7e-06 | 2.067E-06 | 6.506E-07 | 1.400E-06 | 2.700E-06 |
| 29361 | 1469 | G1469A | E | 1.7e-06 | 2.3e-06 | 4.9e-06 | 2.967E-06 | 1.701E-06 | 1.700E-06 | 4.900E-06 |
| 29362 | 1469 | G1469R | E | 1.6e-06 | 2.8e-06 | 2.2e-06 | 2.200E-06 | 6.000E-07 | 1.600E-06 | 2.800E-06 |
| 29363 | 1469 | G1469N | E | 1.7e-06 | 1.6e-06 | 1.3e-06 | 1.533E-06 | 2.082E-07 | 1.300E-06 | 1.700E-06 |
| 29364 | 1469 | G1469D | E | 2.6e-06 | 3e-06 | 1.2e-06 | 2.267E-06 | 9.452E-07 | 1.200E-06 | 3.000E-06 |
| 29365 | 1469 | G1469C | E | 2.9e-06 | 2e-06 | 2.7e-06 | 2.533E-06 | 4.726E-07 | 2.000E-06 | 2.900E-06 |
| 29366 | 1469 | G1469E | E | 3.5e-06 | 3.5e-06 | 2.1e-06 | 3.033E-06 | 8.083E-07 | 2.100E-06 | 3.500E-06 |
| 29367 | 1469 | G1469Q | E | 2e-06 | 1.9e-06 | 2.5e-06 | 2.133E-06 | 3.215E-07 | 1.900E-06 | 2.500E-06 |
| 29368 | 1469 | G1469G | E | 2.2e-06 | 2.1e-06 | 2.7e-06 | 2.333E-06 | 3.215E-07 | 2.100E-06 | 2.700E-06 |
| 29369 | 1469 | G1469H | E | 2.7e-06 | 2.4e-06 | 1.7e-06 | 2.267E-06 | 5.132E-07 | 1.700E-06 | 2.700E-06 |
| 29370 | 1469 | G1469I | E | 2.5e-06 | 2.6e-06 | 1.9e-06 | 2.333E-06 | 3.786E-07 | 1.900E-06 | 2.600E-06 |
| 29371 | 1469 | G1469L | E | 2.9e-06 | 1.6e-06 | 2.6e-06 | 2.367E-06 | 6.807E-07 | 1.600E-06 | 2.900E-06 |
| 29372 | 1469 | G1469K | E | 1.3e-06 | 1.8e-06 | 1.4e-06 | 1.500E-06 | 2.646E-07 | 1.300E-06 | 1.800E-06 |
| 29373 | 1469 | G1469M | E | 2.2e-06 | 2.2e-06 | 2.1e-06 | 2.167E-06 | 5.774E-08 | 2.100E-06 | 2.200E-06 |
| 29374 | 1469 | G1469F | E | 4.3e-06 | 3.1e-06 | 2.5e-06 | 3.300E-06 | 9.165E-07 | 2.500E-06 | 4.300E-06 |
| 29375 | 1469 | G1469P | E | 1.6e-06 | 1.9e-06 | 4.4e-06 | 2.633E-06 | 1.537E-06 | 1.600E-06 | 4.400E-06 |
| 29376 | 1469 | G1469S | E | 4e-06 | 2.9e-06 | 2e-06 | 2.967E-06 | 1.002E-06 | 2.000E-06 | 4.000E-06 |
| 29377 | 1469 | G1469T | E | 2.9e-06 | 1.6e-06 | 2.3e-06 | 2.267E-06 | 6.506E-07 | 1.600E-06 | 2.900E-06 |
| 29378 | 1469 | G1469W | E | 3.2e-06 | 2.2e-06 | 2.8e-06 | 2.733E-06 | 5.033E-07 | 2.200E-06 | 3.200E-06 |
| 29379 | 1469 | G1469Y | E | 1.8e-06 | 3.1e-06 | 1.3e-06 | 2.067E-06 | 9.292E-07 | 1.300E-06 | 3.100E-06 |
| 29380 | 1469 | G1469V | E | 2.9e-06 | 1.4e-06 | 2.1e-06 | 2.133E-06 | 7.506E-07 | 1.400E-06 | 2.900E-06 |
| 29381 | 1470 | T1470A | E | 2.3e-06 | 2.8e-06 | 2.7e-06 | 2.600E-06 | 2.646E-07 | 2.300E-06 | 2.800E-06 |
| 29382 | 1470 | T1470R | E | 2.5e-06 | 2.9e-06 | 2.5e-06 | 2.633E-06 | 2.309E-07 | 2.500E-06 | 2.900E-06 |
| 29383 | 1470 | T1470N | E | 2.4e-06 | 1.2e-06 | 1.4e-06 | 1.667E-06 | 6.429E-07 | 1.200E-06 | 2.400E-06 |
| 29384 | 1470 | T1470D | E | 1.6e-06 | 2.5e-06 | 2.5e-06 | 2.200E-06 | 5.196E-07 | 1.600E-06 | 2.500E-06 |
| 29385 | 1470 | T1470C | E | 2.1e-06 | 5e-06 | 3.2e-06 | 3.433E-06 | 1.464E-06 | 2.100E-06 | 5.000E-06 |
| 29386 | 1470 | T1470E | E | 2.1e-06 | 2.2e-06 | 1.4e-06 | 1.900E-06 | 4.359E-07 | 1.400E-06 | 2.200E-06 |
| 29387 | 1470 | T1470Q | E | 2.9e-06 | 3.4e-06 | 3.3e-06 | 3.200E-06 | 2.646E-07 | 2.900E-06 | 3.400E-06 |
| 29388 | 1470 | T1470G | E | 1.3e-06 | 2.6e-06 | 1.3e-06 | 1.733E-06 | 7.506E-07 | 1.300E-06 | 2.600E-06 |
| 29389 | 1470 | T1470H | E | 3.2e-06 | 1.9e-06 | 3e-06 | 2.700E-06 | 7.000E-07 | 1.900E-06 | 3.200E-06 |
| 29390 | 1470 | T1470I | E | 2.1e-06 | 2.1e-06 | 1.5e-06 | 1.900E-06 | 3.464E-07 | 1.500E-06 | 2.100E-06 |
| 29391 | 1470 | T1470L | E | 2.9e-06 | 2.3e-06 | 2.1e-06 | 2.433E-06 | 4.163E-07 | 2.100E-06 | 2.900E-06 |
| 29392 | 1470 | T1470K | E | 1.9e-06 | 2.2e-06 | 4.1e-06 | 2.733E-06 | 1.193E-06 | 1.900E-06 | 4.100E-06 |
| 29393 | 1470 | T1470M | E | 2.8e-06 | 2.5e-06 | 2e-06 | 2.433E-06 | 4.041E-07 | 2.000E-06 | 2.800E-06 |
| 29394 | 1470 | T1470F | E | 4.8e-06 | 2.4e-06 | 2.1e-06 | 3.100E-06 | 1.480E-06 | 2.100E-06 | 4.800E-06 |
| 29395 | 1470 | T1470P | E | 3.6e-06 | 2.7e-06 | 1.6e-06 | 2.633E-06 | 1.002E-06 | 1.600E-06 | 3.600E-06 |
| 29396 | 1470 | T1470S | E | 1.9e-06 | 9.9e-07 | 2.3e-06 | 1.730E-06 | 6.713E-07 | 9.900E-07 | 2.300E-06 |
| 29397 | 1470 | T1470T | E | 1.6e-06 | 1.9e-06 | 2.5e-06 | 2.000E-06 | 4.583E-07 | 1.600E-06 | 2.500E-06 |
| 29398 | 1470 | T1470W | E | 1.4e-06 | 2.1e-06 | 3e-06 | 2.167E-06 | 8.021E-07 | 1.400E-06 | 3.000E-06 |
| 29399 | 1470 | T1470Y | E | 2.6e-06 | 2.8e-06 | 2.5e-06 | 2.633E-06 | 1.528E-07 | 2.500E-06 | 2.800E-06 |
| 29400 | 1470 | T1470V | E | 1.3e-06 | 2.6e-06 | 2e-06 | 1.967E-06 | 6.506E-07 | 1.300E-06 | 2.600E-06 |
| 29401 | 1471 | Q1471A | E | 1.7e-06 | 1.9e-06 | 1.8e-06 | 1.800E-06 | 1.000E-07 | 1.700E-06 | 1.900E-06 |
| 29402 | 1471 | Q1471R | E | 3.3e-06 | 2.5e-06 | 3.1e-06 | 2.967E-06 | 4.163E-07 | 2.500E-06 | 3.300E-06 |
| 29403 | 1471 | Q1471N | E | 2.1e-06 | 4.5e-06 | 3.2e-06 | 3.267E-06 | 1.201E-06 | 2.100E-06 | 4.500E-06 |
| 29404 | 1471 | Q1471D | E | 2.4e-06 | 3.3e-06 | 1.7e-06 | 2.467E-06 | 8.021E-07 | 1.700E-06 | 3.300E-06 |
| 29405 | 1471 | Q1471C | E | 4.8e-06 | 2.2e-06 | 2.7e-06 | 3.233E-06 | 1.380E-06 | 2.200E-06 | 4.800E-06 |
| 29406 | 1471 | Q1471E | E | 3.2e-06 | 3.1e-06 | 2.4e-06 | 2.900E-06 | 4.359E-07 | 2.400E-06 | 3.200E-06 |
| 29407 | 1471 | Q1471Q | E | 2.6e-06 | 2.3e-06 | 1.6e-06 | 2.167E-06 | 5.132E-07 | 1.600E-06 | 2.600E-06 |
| 29408 | 1471 | Q1471G | E | 3e-06 | 2.2e-06 | 2.6e-06 | 2.600E-06 | 4.000E-07 | 2.200E-06 | 3.000E-06 |
| 29409 | 1471 | Q1471H | E | 3.2e-06 | 2.1e-06 | 2.3e-06 | 2.533E-06 | 5.859E-07 | 2.100E-06 | 3.200E-06 |

| | | | | | | | | | | |
|---|---|---|---|---|---|---|---|---|---|---|
| 29410 | 1471 | Q1471I | E | 4.6e-06 | 2.3e-06 | 2.2e-06 | 3.033E-06 | 1.358E-06 | 2.200E-06 | 4.600E-06 |
| 29411 | 1471 | Q1471L | E | 2.3e-06 | 2.5e-06 | 2e-06 | 2.267E-06 | 2.517E-07 | 2.000E-06 | 2.500E-06 |
| 29412 | 1471 | Q1471K | E | 2.8e-06 | 3.6e-06 | 1.9e-06 | 2.767E-06 | 8.505E-07 | 1.900E-06 | 3.600E-06 |
| 29413 | 1471 | Q1471M | E | 4e-06 | 2.4e-06 | 7.6e-07 | 2.387E-06 | 1.620E-06 | 7.600E-07 | 4.000E-06 |
| 29414 | 1471 | Q1471F | E | 1.8e-06 | 2.5e-06 | 3.7e-06 | 2.667E-06 | 9.609E-07 | 1.800E-06 | 3.700E-06 |
| 29415 | 1471 | Q1471P | E | 3.8e-06 | 2.9e-06 | 1.9e-06 | 2.867E-06 | 9.504E-07 | 1.900E-06 | 3.800E-06 |
| 29416 | 1471 | Q1471S | E | 3.8e-06 | 2.6e-06 | 1.4e-06 | 2.600E-06 | 1.200E-06 | 1.400E-06 | 3.800E-06 |
| 29417 | 1471 | Q1471T | E | 3.4e-06 | 3.4e-06 | 2.7e-06 | 3.167E-06 | 4.041E-07 | 2.700E-06 | 3.400E-06 |
| 29418 | 1471 | Q1471W | E | 2e-06 | 3.1e-06 | 1.5e-06 | 2.200E-06 | 8.185E-07 | 1.500E-06 | 3.100E-06 |
| 29419 | 1471 | Q1471Y | E | 3e-06 | 1.3e-06 | 2.1e-06 | 2.133E-06 | 8.505E-07 | 1.300E-06 | 3.000E-06 |
| 29420 | 1471 | Q1471V | E | 1.6e-06 | 2.2e-06 | 5.2e-06 | 3.000E-06 | 1.929E-06 | 1.600E-06 | 5.200E-06 |
| 29421 | 1472 | T1472A | E | 2.7e-06 | 2.5e-06 | 2.7e-06 | 2.633E-06 | 1.155E-07 | 2.500E-06 | 2.700E-06 |
| 29422 | 1472 | T1472R | E | 2.7e-06 | 2.7e-06 | 4.9e-06 | 3.433E-06 | 1.270E-06 | 2.700E-06 | 4.900E-06 |
| 29423 | 1472 | T1472N | E | 1.8e-06 | 2.6e-06 | 9.3e-07 | 1.777E-06 | 8.352E-07 | 9.300E-07 | 2.600E-06 |
| 29424 | 1472 | T1472D | E | 2.2e-06 | 2.1e-06 | 1.5e-06 | 1.933E-06 | 3.786E-07 | 1.500E-06 | 2.200E-06 |
| 29425 | 1472 | T1472C | E | 1.5e-06 | 1.8e-06 | 2.6e-06 | 1.967E-06 | 5.686E-07 | 1.500E-06 | 2.600E-06 |
| 29426 | 1472 | T1472E | E | 3.8e-06 | 2.5e-06 | 3.9e-06 | 3.400E-06 | 7.810E-07 | 2.500E-06 | 3.900E-06 |
| 29427 | 1472 | T1472Q | E | 2e-06 | 2.1e-06 | 4.3e-06 | 2.800E-06 | 1.300E-06 | 2.000E-06 | 4.300E-06 |
| 29428 | 1472 | T1472G | E | 3.1e-06 | 3.4e-06 | 3.1e-06 | 3.200E-06 | 1.732E-07 | 3.100E-06 | 3.400E-06 |
| 29429 | 1472 | T1472H | E | 2.8e-06 | 2.6e-06 | 2.3e-06 | 2.567E-06 | 2.517E-07 | 2.300E-06 | 2.800E-06 |
| 29430 | 1472 | T1472I | E | 2.3e-06 | 2.6e-06 | 4.9e-06 | 3.267E-06 | 1.422E-06 | 2.300E-06 | 4.900E-06 |
| 29431 | 1472 | T1472L | E | 1.7e-06 | 2.6e-06 | 2.3e-06 | 2.200E-06 | 4.583E-07 | 1.700E-06 | 2.600E-06 |
| 29432 | 1472 | T1472K | E | 2.2e-06 | 1.7e-06 | 3.3e-06 | 2.400E-06 | 8.185E-07 | 1.700E-06 | 3.300E-06 |
| 29433 | 1472 | T1472M | E | 3.6e-06 | 3.3e-06 | 2.5e-06 | 3.133E-06 | 5.686E-07 | 2.500E-06 | 3.600E-06 |
| 29434 | 1472 | T1472F | E | 2.3e-06 | 2.5e-06 | 1.3e-06 | 2.033E-06 | 6.429E-07 | 1.300E-06 | 2.500E-06 |
| 29435 | 1472 | T1472P | E | 4.3e-06 | 2.4e-06 | 1.2e-06 | 2.633E-06 | 1.563E-06 | 1.200E-06 | 4.300E-06 |
| 29436 | 1472 | T1472S | E | 2.4e-06 | 2.9e-06 | 2.7e-06 | 2.667E-06 | 2.517E-07 | 2.400E-06 | 2.900E-06 |
| 29437 | 1472 | T1472T | E | 3.2e-06 | 1.4e-06 | 2.7e-06 | 2.433E-06 | 9.292E-07 | 1.400E-06 | 3.200E-06 |
| 29438 | 1472 | T1472W | E | 3e-06 | 1.7e-06 | 1.1e-06 | 1.933E-06 | 9.713E-07 | 1.100E-06 | 3.000E-06 |
| 29439 | 1472 | T1472Y | E | 2.9e-06 | 2.4e-06 | 2.1e-06 | 2.467E-06 | 4.041E-07 | 2.100E-06 | 2.900E-06 |
| 29440 | 1472 | T1472V | E | 3.5e-06 | 2.3e-06 | 2.2e-06 | 2.667E-06 | 7.234E-07 | 2.200E-06 | 3.500E-06 |
| 29441 | 1473 | Y1473A | E | 2.6e-06 | 3e-06 | 3.3e-06 | 2.967E-06 | 3.512E-07 | 2.600E-06 | 3.300E-06 |
| 29442 | 1473 | Y1473R | E | 3.6e-06 | 1.5e-06 | 2.1e-06 | 2.400E-06 | 1.082E-06 | 1.500E-06 | 3.600E-06 |
| 29443 | 1473 | Y1473N | E | 3e-06 | 1.5e-06 | 2.6e-06 | 2.367E-06 | 7.767E-07 | 1.500E-06 | 3.000E-06 |
| 29444 | 1473 | Y1473D | E | 2.5e-06 | 3.1e-06 | 1.5e-06 | 2.367E-06 | 8.083E-07 | 1.500E-06 | 3.100E-06 |
| 29445 | 1473 | Y1473C | E | 3e-06 | 1.9e-06 | 2.8e-06 | 2.567E-06 | 5.859E-07 | 1.900E-06 | 3.000E-06 |
| 29446 | 1473 | Y1473E | E | 1.9e-06 | 2.9e-06 | 3.3e-06 | 2.700E-06 | 7.211E-07 | 1.900E-06 | 3.300E-06 |
| 29447 | 1473 | Y1473Q | E | 2.2e-06 | 2.6e-06 | 2.4e-06 | 2.400E-06 | 2.000E-07 | 2.200E-06 | 2.600E-06 |
| 29448 | 1473 | Y1473G | E | 2.3e-06 | 2.3e-06 | 2.2e-06 | 2.267E-06 | 5.774E-08 | 2.200E-06 | 2.300E-06 |
| 29449 | 1473 | Y1473H | E | 3.8e-06 | 2.1e-06 | 2.7e-06 | 2.867E-06 | 8.622E-07 | 2.100E-06 | 3.800E-06 |
| 29450 | 1473 | Y1473I | E | 2.1e-06 | 3.4e-06 | 3.5e-06 | 3.000E-06 | 7.810E-07 | 2.100E-06 | 3.500E-06 |
| 29451 | 1473 | Y1473L | E | 2.5e-06 | 1.2e-06 | 3.3e-06 | 2.333E-06 | 1.060E-06 | 1.200E-06 | 3.300E-06 |
| 29452 | 1473 | Y1473K | E | 2.4e-06 | 2.4e-06 | 1.4e-06 | 2.067E-06 | 5.774E-07 | 1.400E-06 | 2.400E-06 |
| 29453 | 1473 | Y1473M | E | 3.3e-06 | 2.8e-06 | 4.1e-06 | 3.400E-06 | 6.557E-07 | 2.800E-06 | 4.100E-06 |
| 29454 | 1473 | Y1473F | E | 1.9e-06 | 2.4e-06 | 1.7e-06 | 2.000E-06 | 3.606E-07 | 1.700E-06 | 2.400E-06 |
| 29455 | 1473 | Y1473P | E | 2.4e-06 | 2e-06 | 2.5e-06 | 2.300E-06 | 2.646E-07 | 2.000E-06 | 2.500E-06 |
| 29456 | 1473 | Y1473S | E | 2.3e-06 | 1.6e-06 | 2.3e-06 | 2.067E-06 | 4.041E-07 | 1.600E-06 | 2.300E-06 |
| 29457 | 1473 | Y1473T | E | 2e-06 | 4e-06 | 2.1e-06 | 2.700E-06 | 1.127E-06 | 2.000E-06 | 4.000E-06 |
| 29458 | 1473 | Y1473W | E | 1.9e-06 | 2.7e-06 | 1.8e-06 | 2.133E-06 | 4.933E-07 | 1.800E-06 | 2.700E-06 |
| 29459 | 1473 | Y1473Y | E | 3.1e-06 | 3e-06 | 2e-06 | 2.700E-06 | 6.083E-07 | 2.000E-06 | 3.100E-06 |
| 29460 | 1473 | Y1473V | E | 1.5e-06 | 3.9e-06 | 2.6e-06 | 2.667E-06 | 1.201E-06 | 1.500E-06 | 3.900E-06 |
| 29461 | 1474 | I1474A | E | 2.6e-06 | 2.5e-06 | 2e-06 | 2.367E-06 | 3.215E-07 | 2.000E-06 | 2.600E-06 |
| 29462 | 1474 | I1474R | E | 1.5e-06 | 1.9e-06 | 1.7e-06 | 1.700E-06 | 2.000E-07 | 1.500E-06 | 1.900E-06 |
| 29463 | 1474 | I1474N | E | 2.2e-06 | 3.1e-06 | 3.7e-06 | 3.000E-06 | 7.550E-07 | 2.200E-06 | 3.700E-06 |
| 29464 | 1474 | I1474D | E | 2.1e-06 | 1.9e-06 | 1.7e-06 | 1.900E-06 | 2.000E-07 | 1.700E-06 | 2.100E-06 |
| 29465 | 1474 | I1474C | E | 1.5e-06 | 2.2e-06 | 1.6e-06 | 1.767E-06 | 3.786E-07 | 1.500E-06 | 2.200E-06 |
| 29466 | 1474 | I1474E | E | 1.5e-06 | 3.3e-06 | 2e-06 | 2.267E-06 | 9.292E-07 | 1.500E-06 | 3.300E-06 |
| 29467 | 1474 | I1474Q | E | 1.9e-06 | 1.8e-06 | 3e-06 | 2.233E-06 | 6.658E-07 | 1.800E-06 | 3.000E-06 |
| 29468 | 1474 | I1474G | E | 1.6e-06 | 2.9e-06 | 2.2e-06 | 2.233E-06 | 6.506E-07 | 1.600E-06 | 2.900E-06 |
| 29469 | 1474 | I1474H | E | 3.7e-06 | 1.8e-06 | 2.8e-06 | 2.767E-06 | 9.504E-07 | 1.800E-06 | 3.700E-06 |
| 29470 | 1474 | I1474I | E | 2.5e-06 | 2.6e-06 | 1.6e-06 | 2.233E-06 | 5.508E-07 | 1.600E-06 | 2.600E-06 |
| 29471 | 1474 | I1474L | E | 2.2e-06 | 2.8e-06 | 2e-06 | 2.333E-06 | 4.163E-07 | 2.000E-06 | 2.800E-06 |
| 29472 | 1474 | I1474K | E | 1.4e-06 | 3.3e-06 | 2.3e-06 | 2.333E-06 | 9.504E-07 | 1.400E-06 | 3.300E-06 |
| 29473 | 1474 | I1474M | E | 1.9e-06 | 2.2e-06 | 3.9e-06 | 2.667E-06 | 1.079E-06 | 1.900E-06 | 3.900E-06 |
| 29474 | 1474 | I1474F | E | 1.5e-06 | 1.7e-06 | 1.2e-06 | 1.467E-06 | 2.517E-07 | 1.200E-06 | 1.700E-06 |
| 29475 | 1474 | I1474P | E | 2.5e-06 | 2.3e-06 | 2.5e-06 | 2.433E-06 | 1.155E-07 | 2.300E-06 | 2.500E-06 |
| 29476 | 1474 | I1474S | E | 2.7e-06 | 2.3e-06 | 3.6e-06 | 2.867E-06 | 6.658E-07 | 2.300E-06 | 3.600E-06 |
| 29477 | 1474 | I1474T | E | 2.8e-06 | 2.9e-06 | 2.3e-06 | 2.667E-06 | 3.215E-07 | 2.300E-06 | 2.900E-06 |
| 29478 | 1474 | I1474W | E | 2.3e-06 | 3.2e-06 | 2.4e-06 | 2.633E-06 | 4.933E-07 | 2.300E-06 | 3.200E-06 |
| 29479 | 1474 | I1474Y | E | 2.2e-06 | 1.7e-06 | 4.8e-06 | 2.900E-06 | 1.664E-06 | 1.700E-06 | 4.800E-06 |
| 29480 | 1474 | I1474V | E | 2.8e-06 | 3.7e-06 | 1.5e-06 | 2.667E-06 | 1.106E-06 | 1.500E-06 | 3.700E-06 |
| 29481 | 1475 | C1475A | E | 1.9e-06 | 2.6e-06 | 1.8e-06 | 2.100E-06 | 4.359E-07 | 1.800E-06 | 2.600E-06 |
| 29482 | 1475 | C1475R | E | 2.1e-06 | 2.9e-06 | 2.7e-06 | 2.567E-06 | 4.163E-07 | 2.100E-06 | 2.900E-06 |
| 29483 | 1475 | C1475N | E | 1.8e-06 | 2.5e-06 | 1.5e-06 | 1.933E-06 | 5.132E-07 | 1.500E-06 | 2.500E-06 |
| 29484 | 1475 | C1475D | E | 2.3e-06 | 2.5e-06 | 1.6e-06 | 2.133E-06 | 4.726E-07 | 1.600E-06 | 2.500E-06 |
| 29485 | 1475 | C1475C | E | 2.2e-06 | 2.3e-06 | 2.3e-06 | 2.267E-06 | 5.774E-08 | 2.200E-06 | 2.300E-06 |
| 29486 | 1475 | C1475E | E | 3.4e-06 | 2.1e-06 | 2.9e-06 | 2.800E-06 | 6.557E-07 | 2.100E-06 | 3.400E-06 |
| 29487 | 1475 | C1475Q | E | 2.5e-06 | 4.1e-06 | 9.7e-07 | 2.523E-06 | 1.565E-06 | 9.700E-07 | 4.100E-06 |
| 29488 | 1475 | C1475G | E | 2.6e-06 | 2.6e-06 | 1.5e-06 | 2.233E-06 | 6.351E-07 | 1.500E-06 | 2.600E-06 |
| 29489 | 1475 | C1475H | E | 1.8e-06 | 8.2e-07 | 2.4e-06 | 1.673E-06 | 7.976E-07 | 8.200E-07 | 2.400E-06 |
| 29490 | 1475 | C1475I | E | 3e-06 | 2.5e-06 | 3.7e-06 | 3.067E-06 | 6.028E-07 | 2.500E-06 | 3.700E-06 |
| 29491 | 1475 | C1475L | E | 1.5e-06 | 3.3e-06 | 2.6e-06 | 2.467E-06 | 9.074E-07 | 1.500E-06 | 3.300E-06 |
| 29492 | 1475 | C1475K | E | 2.4e-06 | 2e-06 | 1.7e-06 | 2.033E-06 | 3.512E-07 | 1.700E-06 | 2.400E-06 |
| 29493 | 1475 | C1475M | E | 2e-06 | 3.5e-06 | 5.1e-06 | 3.533E-06 | 1.550E-06 | 2.000E-06 | 5.100E-06 |
| 29494 | 1475 | C1475F | E | 1.5e-06 | 1.7e-06 | 1.4e-06 | 1.533E-06 | 1.528E-07 | 1.400E-06 | 1.700E-06 |

| | | | | | | | | | | |
|---|---|---|---|---|---|---|---|---|---|---|
| 29495 | 1475 | C1475P | E | 2.6e-06 | 2.1e-06 | 2.2e-06 | 2.300E-06 | 2.646E-07 | 2.100E-06 | 2.600E-06 |
| 29496 | 1475 | C1475S | E | 1.4e-06 | 2.2e-06 | 3.2e-06 | 2.267E-06 | 9.018E-07 | 1.400E-06 | 3.200E-06 |
| 29497 | 1475 | C1475T | E | 3.4e-06 | 2.6e-06 | 2.7e-06 | 2.900E-06 | 4.359E-07 | 2.600E-06 | 3.400E-06 |
| 29498 | 1475 | C1475W | E | 1.9e-06 | 2.7e-06 | 2.2e-06 | 2.267E-06 | 4.041E-07 | 1.900E-06 | 2.700E-06 |
| 29499 | 1475 | C1475Y | E | 1.9e-06 | 2e-06 | 2.7e-06 | 2.200E-06 | 4.359E-07 | 1.900E-06 | 2.700E-06 |
| 29500 | 1475 | C1475V | E | 2.7e-06 | 2.7e-06 | 4.4e-06 | 3.267E-06 | 9.815E-07 | 2.700E-06 | 4.400E-06 |
| 29501 | 1476 | N1476A | E | 3.4e-06 | 4.1e-06 | 3.2e-06 | 3.567E-06 | 4.726E-07 | 3.200E-06 | 4.100E-06 |
| 29502 | 1476 | N1476R | E | 3e-06 | 3.1e-06 | 2.4e-06 | 2.833E-06 | 3.786E-07 | 2.400E-06 | 3.100E-06 |
| 29503 | 1476 | N1476N | E | 2.9e-06 | 2.4e-06 | 2.4e-06 | 2.567E-06 | 2.887E-07 | 2.400E-06 | 2.900E-06 |
| 29504 | 1476 | N1476D | E | 1.5e-06 | 2.3e-06 | 1.8e-06 | 1.867E-06 | 4.041E-07 | 1.500E-06 | 2.300E-06 |
| 29505 | 1476 | N1476C | E | 4.8e-06 | 3.1e-06 | 3.4e-06 | 3.767E-06 | 9.074E-07 | 3.100E-06 | 4.800E-06 |
| 29506 | 1476 | N1476E | E | 1.5e-06 | 3.4e-06 | 1.9e-06 | 2.267E-06 | 1.002E-06 | 1.500E-06 | 3.400E-06 |
| 29507 | 1476 | N1476Q | E | 2.6e-06 | 2e-06 | 1.6e-06 | 2.067E-06 | 5.033E-07 | 1.600E-06 | 2.600E-06 |
| 29508 | 1476 | N1476G | E | 3e-06 | 3.1e-06 | 2e-06 | 2.700E-06 | 6.083E-07 | 2.000E-06 | 3.100E-06 |
| 29509 | 1476 | N1476H | E | 4.8e-06 | 1.7e-06 | 3.2e-06 | 3.233E-06 | 1.550E-06 | 1.700E-06 | 4.800E-06 |
| 29510 | 1476 | N1476I | E | 3e-06 | 1.4e-06 | 2.4e-06 | 2.267E-06 | 8.083E-07 | 1.400E-06 | 3.000E-06 |
| 29511 | 1476 | N1476L | E | 2.2e-06 | 3.2e-06 | 2.8e-06 | 2.733E-06 | 5.033E-07 | 2.200E-06 | 3.200E-06 |
| 29512 | 1476 | N1476K | E | 3.3e-06 | 2.2e-06 | 2.2e-06 | 2.567E-06 | 6.351E-07 | 2.200E-06 | 3.300E-06 |
| 29513 | 1476 | N1476M | E | 2.2e-06 | 2.4e-06 | 3.6e-06 | 2.733E-06 | 7.572E-07 | 2.200E-06 | 3.600E-06 |
| 29514 | 1476 | N1476F | E | 3.4e-06 | 2.8e-06 | 3.4e-06 | 3.200E-06 | 3.464E-07 | 2.800E-06 | 3.400E-06 |
| 29515 | 1476 | N1476P | E | 2.8e-06 | 3.3e-06 | 1.3e-06 | 2.467E-06 | 1.041E-06 | 1.300E-06 | 3.300E-06 |
| 29516 | 1476 | N1476S | E | 2.6e-06 | 2.5e-06 | 2e-06 | 2.367E-06 | 3.215E-07 | 2.000E-06 | 2.600E-06 |
| 29517 | 1476 | N1476T | E | 3.7e-06 | 2e-06 | 3e-06 | 2.900E-06 | 8.544E-07 | 2.000E-06 | 3.700E-06 |
| 29518 | 1476 | N1476W | E | 1.6e-06 | 3.8e-06 | 2.1e-06 | 2.500E-06 | 1.153E-06 | 1.600E-06 | 3.800E-06 |
| 29519 | 1476 | N1476Y | E | 6.5e-06 | 1.9e-06 | 2.2e-06 | 3.533E-06 | 2.574E-06 | 1.900E-06 | 6.500E-06 |
| 29520 | 1476 | N1476V | E | 1.5e-06 | 3.3e-06 | 1.4e-06 | 2.067E-06 | 1.069E-06 | 1.400E-06 | 3.300E-06 |
| 29521 | 1477 | V1477A | E | 2.3e-06 | 4.5e-06 | 2.3e-06 | 3.033E-06 | 1.270E-06 | 2.300E-06 | 4.500E-06 |
| 29522 | 1477 | V1477R | E | 2.5e-06 | 2.2e-06 | 3.2e-06 | 2.633E-06 | 5.132E-07 | 2.200E-06 | 3.200E-06 |
| 29523 | 1477 | V1477N | E | 2.3e-06 | 2.2e-06 | 1.9e-06 | 2.133E-06 | 2.082E-07 | 1.900E-06 | 2.300E-06 |
| 29524 | 1477 | V1477D | E | 3.8e-06 | 2.2e-06 | 1.9e-06 | 2.633E-06 | 1.021E-06 | 1.900E-06 | 3.800E-06 |
| 29525 | 1477 | V1477C | E | 2.4e-06 | 2.8e-06 | 1.6e-06 | 2.267E-06 | 6.110E-07 | 1.600E-06 | 2.800E-06 |
| 29526 | 1477 | V1477E | E | 1.6e-06 | 1.4e-06 | 2.1e-06 | 1.700E-06 | 3.606E-07 | 1.400E-06 | 2.100E-06 |
| 29527 | 1477 | V1477Q | E | 1.3e-06 | 2.5e-06 | 3.3e-06 | 2.367E-06 | 1.007E-06 | 1.300E-06 | 3.300E-06 |
| 29528 | 1477 | V1477G | E | 1.9e-06 | 1.5e-06 | 3.4e-06 | 2.267E-06 | 1.002E-06 | 1.500E-06 | 3.400E-06 |
| 29529 | 1477 | V1477H | E | 1.4e-06 | 2.9e-06 | 2e-06 | 2.100E-06 | 7.550E-07 | 1.400E-06 | 2.900E-06 |
| 29530 | 1477 | V1477I | E | 3.5e-06 | 4e-06 | 2.3e-06 | 3.267E-06 | 8.737E-07 | 2.300E-06 | 4.000E-06 |
| 29531 | 1477 | V1477L | E | 1.1e-06 | 3.2e-06 | 2.3e-06 | 2.200E-06 | 1.054E-06 | 1.100E-06 | 3.200E-06 |
| 29532 | 1477 | V1477K | E | 2.6e-06 | 2e-06 | 1.7e-06 | 2.100E-06 | 4.583E-07 | 1.700E-06 | 2.600E-06 |
| 29533 | 1477 | V1477M | E | 1.7e-06 | 2.8e-06 | 3.6e-06 | 2.700E-06 | 9.539E-07 | 1.700E-06 | 3.600E-06 |
| 29534 | 1477 | V1477F | E | 1.8e-06 | 2.1e-06 | 2.7e-06 | 2.200E-06 | 4.583E-07 | 1.800E-06 | 2.700E-06 |
| 29535 | 1477 | V1477P | E | 3.5e-06 | 3e-06 | 1.7e-06 | 2.733E-06 | 9.292E-07 | 1.700E-06 | 3.500E-06 |
| 29536 | 1477 | V1477S | E | 6.5e-06 | 2.9e-06 | 2.5e-06 | 3.967E-06 | 2.203E-06 | 2.500E-06 | 6.500E-06 |
| 29537 | 1477 | V1477T | E | 2e-06 | 2.4e-06 | 3.9e-06 | 2.767E-06 | 1.002E-06 | 2.000E-06 | 3.900E-06 |
| 29538 | 1477 | V1477W | E | 3.2e-06 | 3e-06 | 3e-06 | 3.067E-06 | 1.155E-07 | 3.000E-06 | 3.200E-06 |
| 29539 | 1477 | V1477Y | E | 2.1e-06 | 2.3e-06 | 2.8e-06 | 2.400E-06 | 3.606E-07 | 2.100E-06 | 2.800E-06 |
| 29540 | 1477 | V1477V | E | 2.4e-06 | 1.2e-06 | 2.4e-06 | 2.000E-06 | 6.928E-07 | 1.200E-06 | 2.400E-06 |
| 29541 | 1478 | N1478A | E | 2.5e-06 | 4.1e-06 | 5.5e-06 | 4.033E-06 | 1.501E-06 | 2.500E-06 | 5.500E-06 |
| 29542 | 1478 | N1478R | E | 1.8e-06 | 2e-06 | 1.9e-06 | 1.900E-06 | 1.000E-07 | 1.800E-06 | 2.000E-06 |
| 29543 | 1478 | N1478N | E | 2.5e-06 | 1.6e-06 | 2.9e-06 | 2.333E-06 | 6.658E-07 | 1.600E-06 | 2.900E-06 |
| 29544 | 1478 | N1478D | E | 1.9e-06 | 1.5e-06 | 1.8e-06 | 1.733E-06 | 2.082E-07 | 1.500E-06 | 1.900E-06 |
| 29545 | 1478 | N1478C | E | 3.3e-06 | 2.6e-06 | 4.4e-06 | 3.433E-06 | 9.074E-07 | 2.600E-06 | 4.400E-06 |
| 29546 | 1478 | N1478E | E | 2e-06 | 2.3e-06 | 1.8e-06 | 2.033E-06 | 2.517E-07 | 1.800E-06 | 2.300E-06 |
| 29547 | 1478 | N1478Q | E | 1.6e-06 | 2.2e-06 | 2.5e-06 | 2.100E-06 | 4.583E-07 | 1.600E-06 | 2.500E-06 |
| 29548 | 1478 | N1478G | E | 2e-06 | 1.6e-06 | 1.9e-06 | 1.833E-06 | 2.082E-07 | 1.600E-06 | 2.000E-06 |
| 29549 | 1478 | N1478H | E | 1.5e-06 | 2.6e-06 | 1.8e-06 | 1.967E-06 | 5.686E-07 | 1.500E-06 | 2.600E-06 |
| 29550 | 1478 | N1478I | E | 2.2e-06 | 1.3e-06 | 2.5e-06 | 2.000E-06 | 6.245E-07 | 1.300E-06 | 2.500E-06 |
| 29551 | 1478 | N1478L | E | 2.8e-06 | 2.8e-06 | 3.5e-06 | 3.033E-06 | 4.041E-07 | 2.800E-06 | 3.500E-06 |
| 29552 | 1478 | N1478K | E | 2e-06 | 2.7e-06 | 2.5e-06 | 2.400E-06 | 3.606E-07 | 2.000E-06 | 2.700E-06 |
| 29553 | 1478 | N1478M | E | 2.2e-06 | 3e-06 | 3.8e-06 | 3.000E-06 | 8.000E-07 | 2.200E-06 | 3.800E-06 |
| 29554 | 1478 | N1478F | E | 3.7e-06 | 1e-06 | 3.1e-06 | 2.600E-06 | 1.418E-06 | 1.000E-06 | 3.700E-06 |
| 29555 | 1478 | N1478P | E | 1.9e-06 | 3.2e-06 | 1.9e-06 | 2.333E-06 | 7.506E-07 | 1.900E-06 | 3.200E-06 |
| 29556 | 1478 | N1478S | E | 3.7e-06 | 2e-06 | 3.1e-06 | 2.933E-06 | 8.622E-07 | 2.000E-06 | 3.700E-06 |
| 29557 | 1478 | N1478T | E | 3.5e-06 | 1.9e-06 | 1.8e-06 | 2.400E-06 | 9.539E-07 | 1.800E-06 | 3.500E-06 |
| 29558 | 1478 | N1478W | E | 3.4e-06 | 3.1e-06 | 2e-06 | 2.833E-06 | 7.371E-07 | 2.000E-06 | 3.400E-06 |
| 29559 | 1478 | N1478Y | E | 4.4e-06 | 3.5e-06 | 1.6e-06 | 3.167E-06 | 1.429E-06 | 1.600E-06 | 4.400E-06 |
| 29560 | 1478 | N1478V | E | 3.9e-06 | 4e-06 | 2.4e-06 | 3.433E-06 | 8.963E-07 | 2.400E-06 | 4.000E-06 |
| 29561 | 1479 | H1479A | E | 2.1e-06 | 1.9e-06 | 1.3e-06 | 1.767E-06 | 4.163E-07 | 1.300E-06 | 2.100E-06 |
| 29562 | 1479 | H1479R | E | 3e-06 | 4e-06 | 2.7e-06 | 3.233E-06 | 6.807E-07 | 2.700E-06 | 4.000E-06 |
| 29563 | 1479 | H1479N | E | 2.6e-06 | 2.8e-06 | 2.4e-06 | 2.600E-06 | 2.000E-07 | 2.400E-06 | 2.800E-06 |
| 29564 | 1479 | H1479D | E | 2.3e-06 | 2e-06 | 2.6e-06 | 2.300E-06 | 3.000E-07 | 2.000E-06 | 2.600E-06 |
| 29565 | 1479 | H1479C | E | 1.6e-06 | 1.8e-06 | 2.2e-06 | 1.867E-06 | 3.055E-07 | 1.600E-06 | 2.200E-06 |
| 29566 | 1479 | H1479E | E | 1.6e-06 | 2.3e-06 | 1.6e-06 | 1.833E-06 | 4.041E-07 | 1.600E-06 | 2.300E-06 |
| 29567 | 1479 | H1479Q | E | 2.5e-06 | 4e-06 | 2.2e-06 | 2.900E-06 | 9.644E-07 | 2.200E-06 | 4.000E-06 |
| 29568 | 1479 | H1479G | E | 3.3e-06 | 2.5e-06 | 2.7e-06 | 2.833E-06 | 4.163E-07 | 2.500E-06 | 3.300E-06 |
| 29569 | 1479 | H1479H | E | 1.7e-06 | 1.5e-06 | 2.6e-06 | 1.933E-06 | 5.859E-07 | 1.500E-06 | 2.600E-06 |
| 29570 | 1479 | H1479I | E | 1.6e-06 | 2.3e-06 | 2.1e-06 | 2.000E-06 | 3.606E-07 | 1.600E-06 | 2.300E-06 |
| 29571 | 1479 | H1479L | E | 2.9e-06 | 2.1e-06 | 2.6e-06 | 2.533E-06 | 4.041E-07 | 2.100E-06 | 2.900E-06 |
| 29572 | 1479 | H1479K | E | 2.8e-06 | 8.1e-07 | 2.9e-06 | 2.170E-06 | 1.179E-06 | 8.100E-07 | 2.900E-06 |
| 29573 | 1479 | H1479M | E | 3.2e-06 | 1.9e-06 | 1.5e-06 | 2.200E-06 | 8.888E-07 | 1.500E-06 | 3.200E-06 |
| 29574 | 1479 | H1479F | E | 2e-06 | 3e-06 | 1.2e-06 | 2.067E-06 | 9.018E-07 | 1.200E-06 | 3.000E-06 |
| 29575 | 1479 | H1479P | E | 2.9e-06 | 2.4e-06 | 1.3e-06 | 2.200E-06 | 8.185E-07 | 1.300E-06 | 2.900E-06 |
| 29576 | 1479 | H1479S | E | 2.4e-06 | 2.3e-06 | 2.3e-06 | 2.333E-06 | 5.774E-08 | 2.300E-06 | 2.400E-06 |
| 29577 | 1479 | H1479T | E | 2.4e-06 | 2.2e-06 | 2.4e-06 | 2.333E-06 | 1.155E-07 | 2.200E-06 | 2.400E-06 |
| 29578 | 1479 | H1479W | E | 2.8e-06 | 2.2e-06 | 2.8e-06 | 2.600E-06 | 3.464E-07 | 2.200E-06 | 2.800E-06 |
| 29579 | 1479 | H1479Y | E | 2.7e-06 | 1.9e-06 | 2.1e-06 | 2.233E-06 | 4.163E-07 | 1.900E-06 | 2.700E-06 |

| | | | | | | | | | | |
|---|---|---|---|---|---|---|---|---|---|---|
| 29580 | 1479 | H1479V | E | 3.7e-06 | 2.6e-06 | 2.7e-06 | 3.000E-06 | 6.083E-07 | 2.600E-06 | 3.700E-06 |
| 29581 | 1480 | K1480A | E | 3.1e-06 | 1.6e-06 | 4.7e-06 | 3.133E-06 | 1.550E-06 | 1.600E-06 | 4.700E-06 |
| 29582 | 1480 | K1480R | E | 2.1e-06 | 2.2e-06 | 3.1e-06 | 2.467E-06 | 5.508E-07 | 2.100E-06 | 3.100E-06 |
| 29583 | 1480 | K1480N | E | 2.8e-06 | 2.1e-06 | 2.2e-06 | 2.367E-06 | 3.786E-07 | 2.100E-06 | 2.800E-06 |
| 29584 | 1480 | K1480D | E | 1.8e-06 | 2.9e-06 | 3.4e-06 | 2.700E-06 | 8.185E-07 | 1.800E-06 | 3.400E-06 |
| 29585 | 1480 | K1480C | E | 2.4e-06 | 2.1e-06 | 2.1e-06 | 2.200E-06 | 1.732E-07 | 2.100E-06 | 2.400E-06 |
| 29586 | 1480 | K1480E | E | 2.1e-06 | 2.6e-06 | 3.3e-06 | 2.667E-06 | 6.028E-07 | 2.100E-06 | 3.300E-06 |
| 29587 | 1480 | K1480Q | E | 2.5e-06 | 1.4e-06 | 2.8e-06 | 2.233E-06 | 7.371E-07 | 1.400E-06 | 2.800E-06 |
| 29588 | 1480 | K1480G | E | 3e-06 | 2.3e-06 | 2.5e-06 | 2.600E-06 | 3.606E-07 | 2.300E-06 | 3.000E-06 |
| 29589 | 1480 | K1480H | E | 2.6e-06 | 2.8e-06 | 1.5e-06 | 2.300E-06 | 7.000E-07 | 1.500E-06 | 2.800E-06 |
| 29590 | 1480 | K1480I | E | 2.3e-06 | 3.6e-06 | 2.7e-06 | 2.867E-06 | 6.658E-07 | 2.300E-06 | 3.600E-06 |
| 29591 | 1480 | K1480L | E | 3.6e-06 | 4.8e-06 | 1.3e-06 | 3.233E-06 | 1.779E-06 | 1.300E-06 | 4.800E-06 |
| 29592 | 1480 | K1480K | E | 2.8e-06 | 2.4e-06 | 2.7e-06 | 2.633E-06 | 2.082E-07 | 2.400E-06 | 2.800E-06 |
| 29593 | 1480 | K1480M | E | 3.1e-06 | 1.9e-06 | 1.3e-06 | 2.100E-06 | 9.165E-07 | 1.300E-06 | 3.100E-06 |
| 29594 | 1480 | K1480F | E | 1.7e-06 | 2.7e-06 | 1.2e-06 | 1.867E-06 | 7.638E-07 | 1.200E-06 | 2.700E-06 |
| 29595 | 1480 | K1480P | E | 2.1e-06 | 2.4e-06 | 1.2e-06 | 1.900E-06 | 6.245E-07 | 1.200E-06 | 2.400E-06 |
| 29596 | 1480 | K1480S | E | 2.4e-06 | 4e-06 | 3.1e-06 | 3.167E-06 | 8.021E-07 | 2.400E-06 | 4.000E-06 |
| 29597 | 1480 | K1480T | E | 1.4e-06 | 2.1e-06 | 2.3e-06 | 1.933E-06 | 4.726E-07 | 1.400E-06 | 2.300E-06 |
| 29598 | 1480 | K1480W | E | 2.4e-06 | 2.6e-06 | 1.5e-06 | 2.167E-06 | 5.859E-07 | 1.500E-06 | 2.600E-06 |
| 29599 | 1480 | K1480Y | E | 3.5e-06 | 1.4e-06 | 2.5e-06 | 2.467E-06 | 1.050E-06 | 1.400E-06 | 3.500E-06 |
| 29600 | 1480 | K1480V | E | 2.4e-06 | 1.6e-06 | 2.4e-06 | 2.133E-06 | 4.619E-07 | 1.600E-06 | 2.400E-06 |
| 29601 | 1481 | P1481A | E | 2.1e-06 | 1.4e-06 | 1.7e-06 | 1.733E-06 | 3.512E-07 | 1.400E-06 | 2.100E-06 |
| 29602 | 1481 | P1481R | E | 4.6e-06 | 2.1e-06 | 1.8e-06 | 2.833E-06 | 1.537E-06 | 1.800E-06 | 4.600E-06 |
| 29603 | 1481 | P1481N | E | 2.1e-06 | 1.3e-06 | 2.2e-06 | 1.867E-06 | 4.933E-07 | 1.300E-06 | 2.200E-06 |
| 29604 | 1481 | P1481D | E | 2.4e-06 | 1.9e-06 | 3.2e-06 | 2.500E-06 | 6.557E-07 | 1.900E-06 | 3.200E-06 |
| 29605 | 1481 | P1481C | E | 1.4e-06 | 2.5e-06 | 1.9e-06 | 1.933E-06 | 5.508E-07 | 1.400E-06 | 2.500E-06 |
| 29606 | 1481 | P1481E | E | 2.1e-06 | 4.3e-06 | 3e-06 | 3.133E-06 | 1.106E-06 | 2.100E-06 | 4.300E-06 |
| 29607 | 1481 | P1481Q | E | 2.6e-06 | 2.6e-06 | 3e-06 | 2.733E-06 | 2.309E-07 | 2.600E-06 | 3.000E-06 |
| 29608 | 1481 | P1481G | E | 2.9e-06 | 2.8e-06 | 3.6e-06 | 3.100E-06 | 4.359E-07 | 2.800E-06 | 3.600E-06 |
| 29609 | 1481 | P1481H | E | 2.3e-06 | 2e-06 | 1.2e-06 | 1.833E-06 | 5.686E-07 | 1.200E-06 | 2.300E-06 |
| 29610 | 1481 | P1481I | E | 2.2e-06 | 2.2e-06 | 1.6e-06 | 2.000E-06 | 3.464E-07 | 1.600E-06 | 2.200E-06 |
| 29611 | 1481 | P1481L | E | 3e-06 | 3.3e-06 | 2.4e-06 | 2.900E-06 | 4.583E-07 | 2.400E-06 | 3.300E-06 |
| 29612 | 1481 | P1481K | E | 2.6e-06 | 2.7e-06 | 2.3e-06 | 2.533E-06 | 2.082E-07 | 2.300E-06 | 2.700E-06 |
| 29613 | 1481 | P1481M | E | 2.6e-06 | 2.8e-06 | 2.1e-06 | 2.500E-06 | 3.606E-07 | 2.100E-06 | 2.800E-06 |
| 29614 | 1481 | P1481F | E | 3.1e-06 | 9.3e-07 | 1.6e-06 | 1.877E-06 | 1.111E-06 | 9.300E-07 | 3.100E-06 |
| 29615 | 1481 | P1481P | E | 3.4e-06 | 1.6e-06 | 3.3e-06 | 2.767E-06 | 1.012E-06 | 1.600E-06 | 3.400E-06 |
| 29616 | 1481 | P1481S | E | 2.1e-06 | 2.7e-06 | 1.7e-06 | 2.167E-06 | 5.033E-07 | 1.700E-06 | 2.700E-06 |
| 29617 | 1481 | P1481T | E | 2.8e-06 | 1.6e-06 | 4.4e-06 | 2.933E-06 | 1.405E-06 | 1.600E-06 | 4.400E-06 |
| 29618 | 1481 | P1481W | E | 1.6e-06 | 4.2e-06 | 1.6e-06 | 2.467E-06 | 1.501E-06 | 1.600E-06 | 4.200E-06 |
| 29619 | 1481 | P1481Y | E | 3.3e-06 | 1.7e-06 | 2.8e-06 | 2.600E-06 | 8.185E-07 | 1.700E-06 | 3.300E-06 |
| 29620 | 1481 | P1481V | E | 3.1e-06 | 2.5e-06 | 2.6e-06 | 2.733E-06 | 3.215E-07 | 2.500E-06 | 3.100E-06 |
| 29621 | 1482 | S1482A | E | 2.9e-06 | 1.2e-06 | 2.3e-06 | 2.133E-06 | 8.622E-07 | 1.200E-06 | 2.900E-06 |
| 29622 | 1482 | S1482R | E | 2.1e-06 | 4.2e-06 | 2.2e-06 | 2.833E-06 | 1.185E-06 | 2.100E-06 | 4.200E-06 |
| 29623 | 1482 | S1482N | E | 2.2e-06 | 3.8e-06 | 3.3e-06 | 3.100E-06 | 8.185E-07 | 2.200E-06 | 3.800E-06 |
| 29624 | 1482 | S1482D | E | 3e-06 | 1.8e-06 | 2.7e-06 | 2.500E-06 | 6.245E-07 | 1.800E-06 | 3.000E-06 |
| 29625 | 1482 | S1482C | E | 2.1e-06 | 3.1e-06 | 2.3e-06 | 2.500E-06 | 5.292E-07 | 2.100E-06 | 3.100E-06 |
| 29626 | 1482 | S1482E | E | 4.6e-06 | 2.6e-06 | 2.7e-06 | 3.300E-06 | 1.127E-06 | 2.600E-06 | 4.600E-06 |
| 29627 | 1482 | S1482Q | E | 2.3e-06 | 1.2e-06 | 1.6e-06 | 1.700E-06 | 5.568E-07 | 1.200E-06 | 2.300E-06 |
| 29628 | 1482 | S1482G | E | 2.6e-06 | 2.5e-06 | 1.5e-06 | 2.200E-06 | 6.083E-07 | 1.500E-06 | 2.600E-06 |
| 29629 | 1482 | S1482H | E | 2.6e-06 | 2.6e-06 | 1.7e-06 | 2.300E-06 | 5.196E-07 | 1.700E-06 | 2.600E-06 |
| 29630 | 1482 | S1482I | E | 2.7e-06 | 3.6e-06 | 3.4e-06 | 3.233E-06 | 4.726E-07 | 2.700E-06 | 3.600E-06 |
| 29631 | 1482 | S1482L | E | 2.2e-06 | 1.7e-06 | 2.5e-06 | 2.133E-06 | 4.041E-07 | 1.700E-06 | 2.500E-06 |
| 29632 | 1482 | S1482K | E | 2.4e-06 | 3.4e-06 | 2.2e-06 | 2.667E-06 | 6.429E-07 | 2.200E-06 | 3.400E-06 |
| 29633 | 1482 | S1482M | E | 5.8e-06 | 2.6e-06 | 1.7e-06 | 3.367E-06 | 2.155E-06 | 1.700E-06 | 5.800E-06 |
| 29634 | 1482 | S1482F | E | 2.3e-06 | 4.4e-06 | 2.4e-06 | 3.033E-06 | 1.185E-06 | 2.300E-06 | 4.400E-06 |
| 29635 | 1482 | S1482P | E | 2.2e-06 | 2.3e-06 | 2.3e-06 | 2.267E-06 | 5.774E-08 | 2.200E-06 | 2.300E-06 |
| 29636 | 1482 | S1482S | E | 2e-06 | 3e-06 | 3e-06 | 2.667E-06 | 5.774E-07 | 2.000E-06 | 3.000E-06 |
| 29637 | 1482 | S1482T | E | 2.1e-06 | 3.6e-06 | 2.6e-06 | 2.767E-06 | 7.638E-07 | 2.100E-06 | 3.600E-06 |
| 29638 | 1482 | S1482W | E | 4.5e-06 | 1.7e-06 | 2.4e-06 | 2.867E-06 | 1.457E-06 | 1.700E-06 | 4.500E-06 |
| 29639 | 1482 | S1482Y | E | 1.5e-06 | 3.2e-06 | 2.2e-06 | 2.300E-06 | 8.544E-07 | 1.500E-06 | 3.200E-06 |
| 29640 | 1482 | S1482V | E | 1.6e-06 | 3e-06 | 2.3e-06 | 2.300E-06 | 7.000E-07 | 1.600E-06 | 3.000E-06 |
| 29641 | 1483 | N1483A | E | 1.6e-06 | 2.6e-06 | 1.6e-06 | 1.933E-06 | 5.774E-07 | 1.600E-06 | 2.600E-06 |
| 29642 | 1483 | N1483R | E | 1.6e-06 | 3.3e-06 | 2.1e-06 | 2.333E-06 | 8.737E-07 | 1.600E-06 | 3.300E-06 |
| 29643 | 1483 | N1483N | E | 5.1e-06 | 2.1e-06 | 2.3e-06 | 3.167E-06 | 1.677E-06 | 2.100E-06 | 5.100E-06 |
| 29644 | 1483 | N1483D | E | 1.1e-06 | 2.9e-06 | 2.2e-06 | 2.067E-06 | 9.074E-07 | 1.100E-06 | 2.900E-06 |
| 29645 | 1483 | N1483C | E | 2.7e-06 | 2.2e-06 | 2.5e-06 | 2.467E-06 | 2.517E-07 | 2.200E-06 | 2.700E-06 |
| 29646 | 1483 | N1483E | E | 2.3e-06 | 3.4e-06 | 3.3e-06 | 3.000E-06 | 6.083E-07 | 2.300E-06 | 3.400E-06 |
| 29647 | 1483 | N1483Q | E | 3e-06 | 2e-06 | 3.2e-06 | 2.733E-06 | 6.429E-07 | 2.000E-06 | 3.200E-06 |
| 29648 | 1483 | N1483G | E | 3.2e-06 | 3.8e-06 | 5.4e-06 | 4.133E-06 | 1.137E-06 | 3.200E-06 | 5.400E-06 |
| 29649 | 1483 | N1483H | E | 2.2e-06 | 2.2e-06 | 2e-06 | 2.133E-06 | 1.155E-07 | 2.000E-06 | 2.200E-06 |
| 29650 | 1483 | N1483I | E | 2.1e-06 | 2e-06 | 1.9e-06 | 2.000E-06 | 1.000E-07 | 1.900E-06 | 2.100E-06 |
| 29651 | 1483 | N1483L | E | 2.7e-06 | 1.8e-06 | 2.2e-06 | 2.233E-06 | 4.509E-07 | 1.800E-06 | 2.700E-06 |
| 29652 | 1483 | N1483K | E | 2.2e-06 | 2e-06 | 2.9e-06 | 2.367E-06 | 4.726E-07 | 2.000E-06 | 2.900E-06 |
| 29653 | 1483 | N1483M | E | 2.6e-06 | 2.3e-06 | 1.7e-06 | 2.200E-06 | 4.583E-07 | 1.700E-06 | 2.600E-06 |
| 29654 | 1483 | N1483F | E | 1.9e-06 | 2.8e-06 | 3e-06 | 2.567E-06 | 5.859E-07 | 1.900E-06 | 3.000E-06 |
| 29655 | 1483 | N1483P | E | 3e-06 | 3.6e-06 | 1.8e-06 | 2.800E-06 | 9.165E-07 | 1.800E-06 | 3.600E-06 |
| 29656 | 1483 | N1483S | E | 1.4e-06 | 1.9e-06 | 1.3e-06 | 1.533E-06 | 3.215E-07 | 1.300E-06 | 1.900E-06 |
| 29657 | 1483 | N1483T | E | 2.8e-06 | 2.4e-06 | 1.3e-06 | 2.167E-06 | 7.767E-07 | 1.300E-06 | 2.800E-06 |
| 29658 | 1483 | N1483W | E | 2.1e-06 | 1.7e-06 | 5.3e-06 | 3.033E-06 | 1.973E-06 | 1.700E-06 | 5.300E-06 |
| 29659 | 1483 | N1483Y | E | 2.7e-06 | 2.5e-06 | 2.9e-06 | 2.700E-06 | 2.000E-07 | 2.500E-06 | 2.900E-06 |
| 29660 | 1483 | N1483V | E | 2.6e-06 | 1.7e-06 | 1.3e-06 | 1.867E-06 | 6.658E-07 | 1.300E-06 | 2.600E-06 |
| 29661 | 1484 | T1484A | E | 2.5e-06 | 2.3e-06 | 2.3e-06 | 2.367E-06 | 1.155E-07 | 2.300E-06 | 2.500E-06 |
| 29662 | 1484 | T1484R | E | 4.5e-06 | 2.1e-06 | 1.5e-06 | 2.700E-06 | 1.587E-06 | 1.500E-06 | 4.500E-06 |
| 29663 | 1484 | T1484N | E | 3.5e-06 | 1.8e-06 | 1.8e-06 | 2.367E-06 | 9.815E-07 | 1.800E-06 | 3.500E-06 |
| 29664 | 1484 | T1484D | E | 2.4e-06 | 2.8e-06 | 2.9e-06 | 2.700E-06 | 2.646E-07 | 2.400E-06 | 2.900E-06 |

| | | | | | | | | | | |
|---|---|---|---|---|---|---|---|---|---|---|
| 29665 | 1484 | T1484C | E | 1.7e-06 | 2.7e-06 | 2.5e-06 | 2.300E-06 | 5.292E-07 | 1.700E-06 | 2.700E-06 |
| 29666 | 1484 | T1484E | E | 2.4e-06 | 2.2e-06 | 2.1e-06 | 2.233E-06 | 1.528E-07 | 2.100E-06 | 2.400E-06 |
| 29667 | 1484 | T1484Q | E | 2.7e-06 | 2.6e-06 | 1.9e-06 | 2.400E-06 | 4.359E-07 | 1.900E-06 | 2.700E-06 |
| 29668 | 1484 | T1484G | E | 2.9e-06 | 1.9e-06 | 3.2e-06 | 2.667E-06 | 6.807E-07 | 1.900E-06 | 3.200E-06 |
| 29669 | 1484 | T1484H | E | 3.8e-06 | 4.7e-06 | 1.8e-06 | 3.433E-06 | 1.484E-06 | 1.800E-06 | 4.700E-06 |
| 29670 | 1484 | T1484I | E | 3.1e-06 | 2.4e-06 | 1.9e-06 | 2.467E-06 | 6.028E-07 | 1.900E-06 | 3.100E-06 |
| 29671 | 1484 | T1484L | E | 3.6e-06 | 1.9e-06 | 1.4e-06 | 2.300E-06 | 1.153E-06 | 1.400E-06 | 3.600E-06 |
| 29672 | 1484 | T1484K | E | 1.9e-06 | 1.3e-06 | 2.7e-06 | 1.967E-06 | 7.024E-07 | 1.300E-06 | 2.700E-06 |
| 29673 | 1484 | T1484M | E | 2.1e-06 | 3.4e-06 | 2.7e-06 | 2.733E-06 | 6.506E-07 | 2.100E-06 | 3.400E-06 |
| 29674 | 1484 | T1484F | E | 3.3e-06 | 2.3e-06 | 3.7e-06 | 3.100E-06 | 7.211E-07 | 2.300E-06 | 3.700E-06 |
| 29675 | 1484 | T1484P | E | 3.9e-06 | 2e-06 | 4.7e-06 | 3.533E-06 | 1.387E-06 | 2.000E-06 | 4.700E-06 |
| 29676 | 1484 | T1484S | E | 1.1e-06 | 3.2e-06 | 4.4e-06 | 2.900E-06 | 1.670E-06 | 1.100E-06 | 4.400E-06 |
| 29677 | 1484 | T1484T | E | 2.6e-06 | 2.3e-06 | 2.6e-06 | 2.500E-06 | 1.732E-07 | 2.300E-06 | 2.600E-06 |
| 29678 | 1484 | T1484W | E | 2.6e-06 | 3e-06 | 3.1e-06 | 2.900E-06 | 2.646E-07 | 2.600E-06 | 3.100E-06 |
| 29679 | 1484 | T1484Y | E | 3e-06 | 2.5e-06 | 3.9e-06 | 3.133E-06 | 7.095E-07 | 2.500E-06 | 3.900E-06 |
| 29680 | 1484 | T1484V | E | 2.1e-06 | 3.4e-06 | 1.6e-06 | 2.367E-06 | 9.292E-07 | 1.600E-06 | 3.400E-06 |
| 29681 | 1485 | K1485A | E | 2.6e-06 | 1.7e-06 | 2.6e-06 | 2.300E-06 | 5.196E-07 | 1.700E-06 | 2.600E-06 |
| 29682 | 1485 | K1485R | E | 2e-06 | 3.8e-06 | 2.5e-06 | 2.767E-06 | 9.292E-07 | 2.000E-06 | 3.800E-06 |
| 29683 | 1485 | K1485N | E | 3.6e-06 | 2.2e-06 | 1.3e-06 | 2.367E-06 | 1.159E-06 | 1.300E-06 | 3.600E-06 |
| 29684 | 1485 | K1485D | E | 4.8e-06 | 1.3e-06 | 2.7e-06 | 2.933E-06 | 1.762E-06 | 1.300E-06 | 4.800E-06 |
| 29685 | 1485 | K1485C | E | 1.7e-06 | 2.3e-06 | 3.3e-06 | 2.433E-06 | 8.083E-07 | 1.700E-06 | 3.300E-06 |
| 29686 | 1485 | K1485E | E | 2e-06 | 2.7e-06 | 2.2e-06 | 2.300E-06 | 3.606E-07 | 2.000E-06 | 2.700E-06 |
| 29687 | 1485 | K1485Q | E | 3.5e-06 | 3.2e-06 | 2.9e-06 | 3.200E-06 | 3.000E-07 | 2.900E-06 | 3.500E-06 |
| 29688 | 1485 | K1485G | E | 1.8e-06 | 1.4e-06 | 2.1e-06 | 1.767E-06 | 3.512E-07 | 1.400E-06 | 2.100E-06 |
| 29689 | 1485 | K1485H | E | 1.6e-06 | 2.5e-06 | 1.9e-06 | 2.000E-06 | 4.583E-07 | 1.600E-06 | 2.500E-06 |
| 29690 | 1485 | K1485I | E | 3.9e-06 | 2.7e-06 | 2.9e-06 | 3.167E-06 | 6.429E-07 | 2.700E-06 | 3.900E-06 |
| 29691 | 1485 | K1485L | E | 3.2e-06 | 1.9e-06 | 2.8e-06 | 2.633E-06 | 6.658E-07 | 1.900E-06 | 3.200E-06 |
| 29692 | 1485 | K1485K | E | 4.5e-06 | 1.9e-06 | 2.5e-06 | 2.967E-06 | 1.361E-06 | 1.900E-06 | 4.500E-06 |
| 29693 | 1485 | K1485M | E | 3.9e-06 | 1.2e-06 | 1.9e-06 | 2.333E-06 | 1.401E-06 | 1.200E-06 | 3.900E-06 |
| 29694 | 1485 | K1485F | E | 3e-06 | 3.2e-06 | 2.5e-06 | 2.900E-06 | 3.606E-07 | 2.500E-06 | 3.200E-06 |
| 29695 | 1485 | K1485P | E | 2e-06 | 3.1e-06 | 9.8e-07 | 2.027E-06 | 1.060E-06 | 9.800E-07 | 3.100E-06 |
| 29696 | 1485 | K1485S | E | 1.6e-06 | 2.5e-06 | 3.6e-06 | 2.567E-06 | 1.002E-06 | 1.600E-06 | 3.600E-06 |
| 29697 | 1485 | K1485T | E | 3.9e-06 | 2.5e-06 | 2.9e-06 | 3.100E-06 | 7.211E-07 | 2.500E-06 | 3.900E-06 |
| 29698 | 1485 | K1485W | E | 3.1e-06 | 2.1e-06 | 1.4e-06 | 2.200E-06 | 8.544E-07 | 1.400E-06 | 3.100E-06 |
| 29699 | 1485 | K1485Y | E | 1.8e-06 | 1.4e-06 | 2.8e-06 | 2.000E-06 | 7.211E-07 | 1.400E-06 | 2.800E-06 |
| 29700 | 1485 | K1485V | E | 2.4e-06 | 3.1e-06 | 4.5e-06 | 3.333E-06 | 1.069E-06 | 2.400E-06 | 4.500E-06 |
| 29701 | 1486 | V1486A | E | 1.5e-06 | 1.5e-06 | 3.4e-06 | 2.133E-06 | 1.097E-06 | 1.500E-06 | 3.400E-06 |
| 29702 | 1486 | V1486R | E | 1.6e-06 | 2.4e-06 | 2.1e-06 | 2.033E-06 | 4.041E-07 | 1.600E-06 | 2.400E-06 |
| 29703 | 1486 | V1486N | E | 1.8e-06 | 2.2e-06 | 2.1e-06 | 2.033E-06 | 2.082E-07 | 1.800E-06 | 2.200E-06 |
| 29704 | 1486 | V1486D | E | 2.2e-06 | 2.7e-06 | 5.3e-06 | 3.400E-06 | 1.664E-06 | 2.200E-06 | 5.300E-06 |
| 29705 | 1486 | V1486C | E | 4e-06 | 1.4e-06 | 1.6e-06 | 2.333E-06 | 1.447E-06 | 1.400E-06 | 4.000E-06 |
| 29706 | 1486 | V1486E | E | 2.3e-06 | 2e-06 | 1.9e-06 | 2.067E-06 | 2.082E-07 | 1.900E-06 | 2.300E-06 |
| 29707 | 1486 | V1486Q | E | 2.2e-06 | 2.4e-06 | 1.8e-06 | 2.133E-06 | 3.055E-07 | 1.800E-06 | 2.400E-06 |
| 29708 | 1486 | V1486G | E | 2.5e-06 | 1.9e-06 | 2.2e-06 | 2.200E-06 | 3.000E-07 | 1.900E-06 | 2.500E-06 |
| 29709 | 1486 | V1486H | E | 2.5e-06 | 2.9e-06 | 1.3e-06 | 2.233E-06 | 8.327E-07 | 1.300E-06 | 2.900E-06 |
| 29710 | 1486 | V1486I | E | 2.1e-06 | 2.4e-06 | 3.3e-06 | 2.600E-06 | 6.245E-07 | 2.100E-06 | 3.300E-06 |
| 29711 | 1486 | V1486L | E | 2.1e-06 | 1.9e-06 | 2.9e-06 | 2.300E-06 | 5.292E-07 | 1.900E-06 | 2.900E-06 |
| 29712 | 1486 | V1486K | E | 2.6e-06 | 1.8e-06 | 3.1e-06 | 2.500E-06 | 6.557E-07 | 1.800E-06 | 3.100E-06 |
| 29713 | 1486 | V1486M | E | 2.6e-06 | 1.8e-06 | 2.1e-06 | 2.167E-06 | 4.041E-07 | 1.800E-06 | 2.600E-06 |
| 29714 | 1486 | V1486F | E | 3.4e-06 | 1.6e-06 | 3.6e-06 | 2.867E-06 | 1.102E-06 | 1.600E-06 | 3.600E-06 |
| 29715 | 1486 | V1486P | E | 2.2e-06 | 4.9e-06 | 3.1e-06 | 3.400E-06 | 1.375E-06 | 2.200E-06 | 4.900E-06 |
| 29716 | 1486 | V1486S | E | 2.5e-06 | 2.2e-06 | 2.4e-06 | 2.367E-06 | 1.528E-07 | 2.200E-06 | 2.500E-06 |
| 29717 | 1486 | V1486T | E | 2.2e-06 | 3e-06 | 2.6e-06 | 2.600E-06 | 4.000E-07 | 2.200E-06 | 3.000E-06 |
| 29718 | 1486 | V1486W | E | 2.5e-06 | 2.1e-06 | 2.7e-06 | 2.433E-06 | 3.055E-07 | 2.100E-06 | 2.700E-06 |
| 29719 | 1486 | V1486Y | E | 3.5e-06 | 3.2e-06 | 2.4e-06 | 3.033E-06 | 5.686E-07 | 2.400E-06 | 3.500E-06 |
| 29720 | 1486 | V1486V | E | 2.5e-06 | 2.7e-06 | 1.3e-06 | 2.167E-06 | 7.572E-07 | 1.300E-06 | 2.700E-06 |
| 29721 | 1487 | D1487A | E | 2.5e-06 | 2.7e-06 | 1.6e-06 | 2.267E-06 | 5.859E-07 | 1.600E-06 | 2.700E-06 |
| 29722 | 1487 | D1487R | E | 3.5e-06 | 2.4e-06 | 1.1e-06 | 2.333E-06 | 1.201E-06 | 1.100E-06 | 3.500E-06 |
| 29723 | 1487 | D1487N | E | 1.9e-06 | 2.2e-06 | 2.8e-06 | 2.300E-06 | 4.583E-07 | 1.900E-06 | 2.800E-06 |
| 29724 | 1487 | D1487D | E | 4.5e-06 | 2.2e-06 | 2.3e-06 | 3.000E-06 | 1.300E-06 | 2.200E-06 | 4.500E-06 |
| 29725 | 1487 | D1487C | E | 2.3e-06 | 3.6e-06 | 2.5e-06 | 2.800E-06 | 7.000E-07 | 2.300E-06 | 3.600E-06 |
| 29726 | 1487 | D1487E | E | 2.4e-06 | 3.8e-06 | 2.4e-06 | 2.867E-06 | 8.083E-07 | 2.400E-06 | 3.800E-06 |
| 29727 | 1487 | D1487Q | E | 3.2e-06 | 2.4e-06 | 1.5e-06 | 2.367E-06 | 8.505E-07 | 1.500E-06 | 3.200E-06 |
| 29728 | 1487 | D1487G | E | 2.1e-06 | 2.3e-06 | 1.7e-06 | 2.033E-06 | 3.055E-07 | 1.700E-06 | 2.300E-06 |
| 29729 | 1487 | D1487H | E | 2.1e-06 | 2.5e-06 | 4.1e-06 | 2.900E-06 | 1.058E-06 | 2.100E-06 | 4.100E-06 |
| 29730 | 1487 | D1487I | E | 2.2e-06 | 3.4e-06 | 2.1e-06 | 2.567E-06 | 7.234E-07 | 2.100E-06 | 3.400E-06 |
| 29731 | 1487 | D1487L | E | 1.7e-06 | 2.3e-06 | 2.7e-06 | 2.233E-06 | 5.033E-07 | 1.700E-06 | 2.700E-06 |
| 29732 | 1487 | D1487K | E | 2.2e-06 | 2.1e-06 | 2.6e-06 | 2.300E-06 | 2.646E-07 | 2.100E-06 | 2.600E-06 |
| 29733 | 1487 | D1487M | E | 2.6e-06 | 2.7e-06 | 1.9e-06 | 2.400E-06 | 4.359E-07 | 1.900E-06 | 2.700E-06 |
| 29734 | 1487 | D1487F | E | 1.5e-06 | 4.8e-06 | 2.4e-06 | 2.900E-06 | 1.706E-06 | 1.500E-06 | 4.800E-06 |
| 29735 | 1487 | D1487P | E | 2.5e-06 | 3.1e-06 | 3.2e-06 | 2.933E-06 | 3.786E-07 | 2.500E-06 | 3.200E-06 |
| 29736 | 1487 | D1487S | E | 1.5e-06 | 3.5e-06 | 3.2e-06 | 2.733E-06 | 1.079E-06 | 1.500E-06 | 3.500E-06 |
| 29737 | 1487 | D1487T | E | 1.6e-06 | 2.2e-06 | 1.9e-06 | 1.900E-06 | 3.000E-07 | 1.600E-06 | 2.200E-06 |
| 29738 | 1487 | D1487W | E | 3.5e-06 | 1.7e-06 | 1.6e-06 | 2.267E-06 | 1.069E-06 | 1.600E-06 | 3.500E-06 |
| 29739 | 1487 | D1487Y | E | 4.3e-06 | 2.9e-06 | 2.3e-06 | 3.167E-06 | 1.026E-06 | 2.300E-06 | 4.300E-06 |
| 29740 | 1487 | D1487V | E | 1.9e-06 | 2.5e-06 | 2.1e-06 | 2.167E-06 | 3.055E-07 | 1.900E-06 | 2.500E-06 |
| 29741 | 1488 | K1488A | E | 1.9e-06 | 2.6e-06 | 1.8e-06 | 2.100E-06 | 4.359E-07 | 1.800E-06 | 2.600E-06 |
| 29742 | 1488 | K1488R | E | 9.2e-07 | 4.7e-06 | 1.6e-06 | 2.407E-06 | 2.015E-06 | 9.200E-07 | 4.700E-06 |
| 29743 | 1488 | K1488N | E | 2.9e-06 | 2.3e-06 | 2.1e-06 | 2.433E-06 | 4.163E-07 | 2.100E-06 | 2.900E-06 |
| 29744 | 1488 | K1488D | E | 2.8e-06 | 3e-06 | 2.3e-06 | 2.700E-06 | 3.606E-07 | 2.300E-06 | 3.000E-06 |
| 29745 | 1488 | K1488C | E | 3.4e-06 | 1.9e-06 | 2.1e-06 | 2.467E-06 | 8.145E-07 | 1.900E-06 | 3.400E-06 |
| 29746 | 1488 | K1488E | E | 2.5e-06 | 2.5e-06 | 2.3e-06 | 2.433E-06 | 1.155E-07 | 2.300E-06 | 2.500E-06 |
| 29747 | 1488 | K1488Q | E | 2.5e-06 | 2.6e-06 | 1.1e-06 | 2.067E-06 | 8.386E-07 | 1.100E-06 | 2.600E-06 |
| 29748 | 1488 | K1488G | E | 1.8e-06 | 3.5e-06 | 1.7e-06 | 2.333E-06 | 1.012E-06 | 1.700E-06 | 3.500E-06 |
| 29749 | 1488 | K1488H | E | 1.2e-06 | 2.1e-06 | 2.1e-06 | 1.800E-06 | 5.196E-07 | 1.200E-06 | 2.100E-06 |

| | | | | | | | | | | |
|---|---|---|---|---|---|---|---|---|---|---|
| 29750 | 1488 | K1488I | E | 1e-06 | 3.3e-06 | 4.5e-06 | 2.933E-06 | 1.779E-06 | 1.000E-06 | 4.500E-06 |
| 29751 | 1488 | K1488L | E | 2.7e-06 | 5.6e-06 | 1.1e-06 | 3.133E-06 | 2.281E-06 | 1.100E-06 | 5.600E-06 |
| 29752 | 1488 | K1488K | E | 3.2e-06 | 1.6e-06 | 2e-06 | 2.267E-06 | 8.327E-07 | 1.600E-06 | 3.200E-06 |
| 29753 | 1488 | K1488M | E | 1.6e-06 | 2.5e-06 | 2e-06 | 2.033E-06 | 4.509E-07 | 1.600E-06 | 2.500E-06 |
| 29754 | 1488 | K1488F | E | 1.1e-06 | 3e-06 | 1.7e-06 | 1.933E-06 | 9.713E-07 | 1.100E-06 | 3.000E-06 |
| 29755 | 1488 | K1488P | E | 2.6e-06 | 2.2e-06 | 2.6e-06 | 2.467E-06 | 2.309E-07 | 2.200E-06 | 2.600E-06 |
| 29756 | 1488 | K1488S | E | 4.2e-06 | 3e-06 | 1.6e-06 | 2.933E-06 | 1.301E-06 | 1.600E-06 | 4.200E-06 |
| 29757 | 1488 | K1488T | E | 2.1e-06 | 1.5e-06 | 1.3e-06 | 1.633E-06 | 4.163E-07 | 1.300E-06 | 2.100E-06 |
| 29758 | 1488 | K1488W | E | 1.8e-06 | 3.1e-06 | 3.1e-06 | 2.667E-06 | 7.506E-07 | 1.800E-06 | 3.100E-06 |
| 29759 | 1488 | K1488Y | E | 1.9e-06 | 3.2e-06 | 2.3e-06 | 2.467E-06 | 6.658E-07 | 1.900E-06 | 3.200E-06 |
| 29760 | 1488 | K1488V | E | 2.2e-06 | 1.2e-06 | 1.9e-06 | 1.767E-06 | 5.132E-07 | 1.200E-06 | 2.200E-06 |
| 29761 | 1489 | K1489A | E | 2.4e-06 | 1.9e-06 | 2.4e-06 | 2.233E-06 | 2.887E-07 | 1.900E-06 | 2.400E-06 |
| 29762 | 1489 | K1489R | E | 2.4e-06 | 2.4e-06 | 2.6e-06 | 2.467E-06 | 1.155E-07 | 2.400E-06 | 2.600E-06 |
| 29763 | 1489 | K1489N | E | 1.7e-06 | 2.3e-06 | 1.4e-06 | 1.800E-06 | 4.583E-07 | 1.400E-06 | 2.300E-06 |
| 29764 | 1489 | K1489D | E | 1.8e-06 | 1.8e-06 | 2e-06 | 1.867E-06 | 1.155E-07 | 1.800E-06 | 2.000E-06 |
| 29765 | 1489 | K1489C | E | 1.7e-06 | 2.2e-06 | 2.3e-06 | 2.067E-06 | 3.215E-07 | 1.700E-06 | 2.300E-06 |
| 29766 | 1489 | K1489E | E | 3.1e-06 | 2.5e-06 | 2.1e-06 | 2.567E-06 | 5.033E-07 | 2.100E-06 | 3.100E-06 |
| 29767 | 1489 | K1489Q | E | 2.3e-06 | 1.7e-06 | 2.9e-06 | 2.300E-06 | 6.000E-07 | 1.700E-06 | 2.900E-06 |
| 29768 | 1489 | K1489G | E | 3.2e-06 | 4.4e-06 | 1.9e-06 | 3.167E-06 | 1.250E-06 | 1.900E-06 | 4.400E-06 |
| 29769 | 1489 | K1489H | E | 2.2e-06 | 2e-06 | 2.3e-06 | 2.167E-06 | 1.528E-07 | 2.000E-06 | 2.300E-06 |
| 29770 | 1489 | K1489I | E | 3.6e-06 | 3.2e-06 | 4.7e-06 | 3.833E-06 | 7.767E-07 | 3.200E-06 | 4.700E-06 |
| 29771 | 1489 | K1489L | E | 1.9e-06 | 3.2e-06 | 1.8e-06 | 2.300E-06 | 7.810E-07 | 1.800E-06 | 3.200E-06 |
| 29772 | 1489 | K1489K | E | 2.9e-06 | 2.8e-06 | 1.8e-06 | 2.500E-06 | 6.083E-07 | 1.800E-06 | 2.900E-06 |
| 29773 | 1489 | K1489M | E | 1.8e-06 | 1.4e-06 | 2.8e-06 | 2.000E-06 | 7.211E-07 | 1.400E-06 | 2.800E-06 |
| 29774 | 1489 | K1489F | E | 1.7e-06 | 1.9e-06 | 4.6e-06 | 2.733E-06 | 1.620E-06 | 1.700E-06 | 4.600E-06 |
| 29775 | 1489 | K1489P | E | 2.1e-06 | 2.8e-06 | 2e-06 | 2.300E-06 | 4.359E-07 | 2.000E-06 | 2.800E-06 |
| 29776 | 1489 | K1489S | E | 2.4e-06 | 1.8e-06 | 3e-06 | 2.400E-06 | 6.000E-07 | 1.800E-06 | 3.000E-06 |
| 29777 | 1489 | K1489T | E | 3.9e-06 | 2.1e-06 | 1.6e-06 | 2.533E-06 | 1.210E-06 | 1.600E-06 | 3.900E-06 |
| 29778 | 1489 | K1489W | E | 1.8e-06 | 4.3e-06 | 3.7e-06 | 3.267E-06 | 1.305E-06 | 1.800E-06 | 4.300E-06 |
| 29779 | 1489 | K1489Y | E | 2.4e-06 | 2.4e-06 | 2.4e-06 | 2.400E-06 | 0.000E+00 | 2.400E-06 | 2.400E-06 |
| 29780 | 1489 | K1489V | E | 2.5e-06 | 1.8e-06 | 2.3e-06 | 2.200E-06 | 3.606E-07 | 1.800E-06 | 2.500E-06 |
| 29781 | 1490 | V1490A | E | 4.4e-06 | 1.9e-06 | 1.7e-06 | 2.667E-06 | 1.504E-06 | 1.700E-06 | 4.400E-06 |
| 29782 | 1490 | V1490R | E | 2e-06 | 3e-06 | 1.9e-06 | 2.300E-06 | 6.083E-07 | 1.900E-06 | 3.000E-06 |
| 29783 | 1490 | V1490N | E | 1.8e-06 | 1.8e-06 | 2e-06 | 1.867E-06 | 1.155E-07 | 1.800E-06 | 2.000E-06 |
| 29784 | 1490 | V1490D | E | 2.2e-06 | 1.9e-06 | 2.7e-06 | 2.267E-06 | 4.041E-07 | 1.900E-06 | 2.700E-06 |
| 29785 | 1490 | V1490C | E | 1.8e-06 | 2.2e-06 | 3.3e-06 | 2.433E-06 | 7.767E-07 | 1.800E-06 | 3.300E-06 |
| 29786 | 1490 | V1490E | E | 2.8e-06 | 1.5e-06 | 2.9e-06 | 2.400E-06 | 7.810E-07 | 1.500E-06 | 2.900E-06 |
| 29787 | 1490 | V1490Q | E | 2.5e-06 | 3.1e-06 | 2.5e-06 | 2.700E-06 | 3.464E-07 | 2.500E-06 | 3.100E-06 |
| 29788 | 1490 | V1490G | E | 2.6e-06 | 2.4e-06 | 2.8e-06 | 2.600E-06 | 2.000E-07 | 2.400E-06 | 2.800E-06 |
| 29789 | 1490 | V1490H | E | 1.7e-06 | 4.8e-06 | 2.7e-06 | 3.067E-06 | 1.582E-06 | 1.700E-06 | 4.800E-06 |
| 29790 | 1490 | V1490I | E | 1.8e-06 | 4.2e-06 | 1.9e-06 | 2.633E-06 | 1.358E-06 | 1.800E-06 | 4.200E-06 |
| 29791 | 1490 | V1490L | E | 2.8e-06 | 2.9e-06 | 4.3e-06 | 3.333E-06 | 8.386E-07 | 2.800E-06 | 4.300E-06 |
| 29792 | 1490 | V1490K | E | 2.5e-06 | 1.8e-06 | 1.9e-06 | 2.067E-06 | 3.786E-07 | 1.800E-06 | 2.500E-06 |
| 29793 | 1490 | V1490M | E | 1.3e-06 | 2.1e-06 | 1.4e-06 | 1.600E-06 | 4.359E-07 | 1.300E-06 | 2.100E-06 |
| 29794 | 1490 | V1490F | E | 3e-06 | 2.7e-06 | 1.6e-06 | 2.433E-06 | 7.371E-07 | 1.600E-06 | 3.000E-06 |
| 29795 | 1490 | V1490P | E | 1.4e-06 | 2.8e-06 | 2.5e-06 | 2.233E-06 | 7.371E-07 | 1.400E-06 | 2.800E-06 |
| 29796 | 1490 | V1490S | E | 1.1e-06 | 2.4e-06 | 2.8e-06 | 2.100E-06 | 8.888E-07 | 1.100E-06 | 2.800E-06 |
| 29797 | 1490 | V1490T | E | 1.3e-06 | 2e-06 | 1.9e-06 | 1.733E-06 | 3.786E-07 | 1.300E-06 | 2.000E-06 |
| 29798 | 1490 | V1490W | E | 3.1e-06 | 2.2e-06 | 2.7e-06 | 2.667E-06 | 4.509E-07 | 2.200E-06 | 3.100E-06 |
| 29799 | 1490 | V1490Y | E | 1.8e-06 | 3.1e-06 | 5e-06 | 3.300E-06 | 1.609E-06 | 1.800E-06 | 5.000E-06 |
| 29800 | 1490 | V1490V | E | 2.1e-06 | 3.1e-06 | 2.9e-06 | 2.700E-06 | 5.292E-07 | 2.100E-06 | 3.100E-06 |
| 29801 | 1491 | E1491A | E | 1.7e-06 | 2.8e-06 | 2e-06 | 2.167E-06 | 5.686E-07 | 1.700E-06 | 2.800E-06 |
| 29802 | 1491 | E1491R | E | 1.9e-06 | 3.1e-06 | 2.1e-06 | 2.367E-06 | 6.429E-07 | 1.900E-06 | 3.100E-06 |
| 29803 | 1491 | E1491N | E | 1.7e-06 | 1.7e-06 | 2.2e-06 | 1.867E-06 | 2.887E-07 | 1.700E-06 | 2.200E-06 |
| 29804 | 1491 | E1491D | E | 2.4e-06 | 3.2e-06 | 1.5e-06 | 2.367E-06 | 8.505E-07 | 1.500E-06 | 3.200E-06 |
| 29805 | 1491 | E1491C | E | 1.9e-06 | 3.1e-06 | 2.2e-06 | 2.400E-06 | 6.245E-07 | 1.900E-06 | 3.100E-06 |
| 29806 | 1491 | E1491E | E | 2.7e-06 | 2.9e-06 | 2.1e-06 | 2.567E-06 | 4.163E-07 | 2.100E-06 | 2.900E-06 |
| 29807 | 1491 | E1491Q | E | 4.3e-06 | 3.3e-06 | 2.8e-06 | 3.467E-06 | 7.638E-07 | 2.800E-06 | 4.300E-06 |
| 29808 | 1491 | E1491G | E | 3.1e-06 | 2.1e-06 | 3.9e-06 | 3.033E-06 | 9.018E-07 | 2.100E-06 | 3.900E-06 |
| 29809 | 1491 | E1491H | E | 2e-06 | 2.2e-06 | 2.6e-06 | 2.267E-06 | 3.055E-07 | 2.000E-06 | 2.600E-06 |
| 29810 | 1491 | E1491I | E | 2.1e-06 | 2.2e-06 | 2.5e-06 | 2.267E-06 | 2.082E-07 | 2.100E-06 | 2.500E-06 |
| 29811 | 1491 | E1491L | E | 1.7e-06 | 2.6e-06 | 2e-06 | 2.100E-06 | 4.583E-07 | 1.700E-06 | 2.600E-06 |
| 29812 | 1491 | E1491K | E | 1.5e-06 | 2.3e-06 | 3.6e-06 | 2.467E-06 | 1.060E-06 | 1.500E-06 | 3.600E-06 |
| 29813 | 1491 | E1491M | E | 1.9e-06 | 1.6e-06 | 2.1e-06 | 1.867E-06 | 2.517E-07 | 1.600E-06 | 2.100E-06 |
| 29814 | 1491 | E1491F | E | 3e-06 | 2.3e-06 | 1.6e-06 | 2.300E-06 | 7.000E-07 | 1.600E-06 | 3.000E-06 |
| 29815 | 1491 | E1491P | E | 2.5e-06 | 1.3e-06 | 2.9e-06 | 2.233E-06 | 8.327E-07 | 1.300E-06 | 2.900E-06 |
| 29816 | 1491 | E1491S | E | 3.3e-06 | 3.2e-06 | 1.2e-06 | 2.567E-06 | 1.185E-06 | 1.200E-06 | 3.300E-06 |
| 29817 | 1491 | E1491T | E | 2.9e-06 | 1.9e-06 | 2.9e-06 | 2.567E-06 | 5.774E-07 | 1.900E-06 | 2.900E-06 |
| 29818 | 1491 | E1491W | E | 2.7e-06 | 2.5e-06 | 2.7e-06 | 2.633E-06 | 1.155E-07 | 2.500E-06 | 2.700E-06 |
| 29819 | 1491 | E1491Y | E | 2e-06 | 1.8e-06 | 2.2e-06 | 2.000E-06 | 2.000E-07 | 1.800E-06 | 2.200E-06 |
| 29820 | 1491 | E1491V | E | 3.1e-06 | 1.9e-06 | 3.5e-06 | 2.833E-06 | 8.327E-07 | 1.900E-06 | 3.500E-06 |
| 29821 | 1492 | P1492A | E | 2.7e-06 | 2.6e-06 | 1.7e-06 | 2.333E-06 | 5.508E-07 | 1.700E-06 | 2.700E-06 |
| 29822 | 1492 | P1492R | E | 3.3e-06 | 3.4e-06 | 2.8e-06 | 3.167E-06 | 3.215E-07 | 2.800E-06 | 3.400E-06 |
| 29823 | 1492 | P1492N | E | 1.7e-06 | 3.7e-06 | 2.7e-06 | 2.700E-06 | 1.000E-06 | 1.700E-06 | 3.700E-06 |
| 29824 | 1492 | P1492D | E | 1.7e-06 | 3.3e-06 | 2.1e-06 | 2.367E-06 | 8.327E-07 | 1.700E-06 | 3.300E-06 |
| 29825 | 1492 | P1492C | E | 2e-06 | 2.1e-06 | 1.6e-06 | 1.900E-06 | 2.646E-07 | 1.600E-06 | 2.100E-06 |
| 29826 | 1492 | P1492E | E | 3.1e-06 | 2.7e-06 | 2.1e-06 | 2.633E-06 | 5.033E-07 | 2.100E-06 | 3.100E-06 |
| 29827 | 1492 | P1492Q | E | 4.3e-06 | 2.4e-06 | 3e-06 | 3.233E-06 | 9.713E-07 | 2.400E-06 | 4.300E-06 |
| 29828 | 1492 | P1492G | E | 1.8e-06 | 2.9e-06 | 1.2e-06 | 1.967E-06 | 8.622E-07 | 1.200E-06 | 2.900E-06 |
| 29829 | 1492 | P1492H | E | 2.2e-06 | 1.4e-06 | 1.3e-06 | 1.633E-06 | 4.933E-07 | 1.300E-06 | 2.200E-06 |
| 29830 | 1492 | P1492I | E | 2.1e-06 | 2.2e-06 | 1.5e-06 | 1.933E-06 | 3.786E-07 | 1.500E-06 | 2.200E-06 |
| 29831 | 1492 | P1492L | E | 2.6e-06 | 4.3e-06 | 2.6e-06 | 3.167E-06 | 9.815E-07 | 2.600E-06 | 4.300E-06 |
| 29832 | 1492 | P1492K | E | 2.1e-06 | 1.6e-06 | 1.4e-06 | 1.700E-06 | 3.606E-07 | 1.400E-06 | 2.100E-06 |
| 29833 | 1492 | P1492M | E | 2.1e-06 | 1.8e-06 | 2.3e-06 | 2.067E-06 | 2.517E-07 | 1.800E-06 | 2.300E-06 |
| 29834 | 1492 | P1492F | E | 2.8e-06 | 2e-06 | 3.3e-06 | 2.700E-06 | 6.557E-07 | 2.000E-06 | 3.300E-06 |

| | | | | | | | | | | |
|---|---|---|---|---|---|---|---|---|---|---|
| 29835 | 1492 | P1492P | E | 2.2e-06 | 3.6e-06 | 3.5e-06 | 3.100E-06 | 7.810E-07 | 2.200E-06 | 3.600E-06 |
| 29836 | 1492 | P1492S | E | 1.5e-06 | 1.5e-06 | 1.6e-06 | 1.533E-06 | 5.774E-08 | 1.500E-06 | 1.600E-06 |
| 29837 | 1492 | P1492T | E | 2.6e-06 | 2.2e-06 | 2.4e-06 | 2.400E-06 | 2.000E-07 | 2.200E-06 | 2.600E-06 |
| 29838 | 1492 | P1492W | E | 2.3e-06 | 2.1e-06 | 5.3e-06 | 3.233E-06 | 1.793E-06 | 2.100E-06 | 5.300E-06 |
| 29839 | 1492 | P1492Y | E | 2.7e-06 | 1.6e-06 | 2.3e-06 | 2.200E-06 | 5.568E-07 | 1.600E-06 | 2.700E-06 |
| 29840 | 1492 | P1492V | E | 4.8e-06 | 1.6e-06 | 3e-06 | 3.133E-06 | 1.604E-06 | 1.600E-06 | 4.800E-06 |

Table 6: A scalable Her2-Trastuzumab-Pertuzumab binding affinity landscape. In this table, **ResID** represents the amino acid residue number of PDB entry 6OGE, while **Muta** represents the site-specific missense mutation introduced into the 1492 amino acid residues in PDB entry 6OGE, **Min**, **Max**, **Mean** and **Std** represent the minimum, the maximum, the average and the standard deviation of the $K_d$ values calculated using Prodigy, respectively.

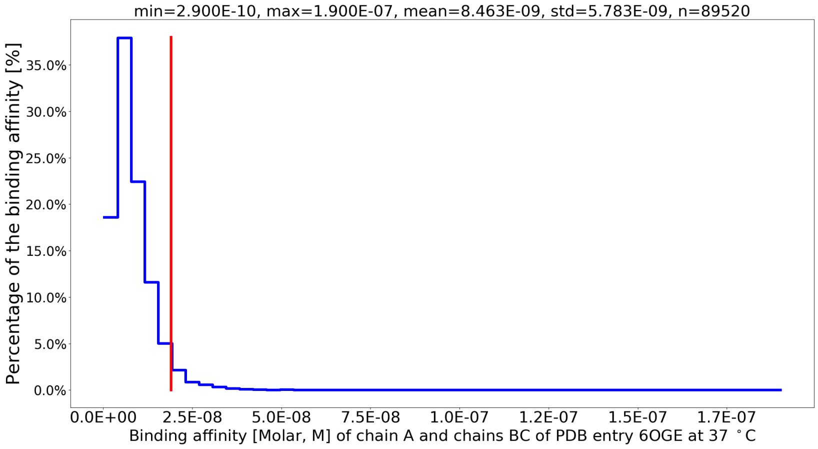


Figure 5: A histogram of the distribution pattern of the Her2-Trastuzumab-Pertuzumab binding affinities between chain A and chain BC (listed in Table 1) of PDB entry 6OGE (with one site-specific missense mutation). In this figure, the vertical red line represents the $K_d$ between chain A (Her2) and chain BC (Pertuzumab FAB, Table 1) of PDB entry 6OGE, i.e., native complex structure of Her2-Trastuzumab-Pertuzumab.

| No. | ResID | Muta | ChainID | $K_{d1}$ (M) | $K_{d2}$ (M) | $K_{d3}$ (M) | Mean (M) | Std (M) | Min (M) | Max (M) |
|---|---|---|---|---|---|---|---|---|---|---|
| 1 | 1 | T1A | A | 1.6e-08 | 1.6e-08 | 1.6e-08 | 1.600E-08 | 0.000E+00 | 1.600E-08 | 1.600E-08 |
| 2 | 1 | T1R | A | 1.2e-08 | 1.2e-08 | 1.2e-08 | 1.200E-08 | 0.000E+00 | 1.200E-08 | 1.200E-08 |
| 3 | 1 | T1N | A | 9e-09 | 9e-09 | 9e-09 | 9.000E-09 | 0.000E+00 | 9.000E-09 | 9.000E-09 |
| 4 | 1 | T1D | A | 4e-09 | 4e-09 | 4e-09 | 4.000E-09 | 0.000E+00 | 4.000E-09 | 4.000E-09 |
| 5 | 1 | T1C | A | 3.8e-09 | 3.8e-09 | 3.8e-09 | 3.800E-09 | 0.000E+00 | 3.800E-09 | 3.800E-09 |
| 6 | 1 | T1E | A | 1.7e-08 | 1.7e-08 | 1.7e-08 | 1.700E-08 | 0.000E+00 | 1.700E-08 | 1.700E-08 |
| 7 | 1 | T1Q | A | 1.1e-08 | 1.1e-08 | 1.1e-08 | 1.100E-08 | 0.000E+00 | 1.100E-08 | 1.100E-08 |
| 8 | 1 | T1G | A | 1.5e-08 | 1.5e-08 | 1.5e-08 | 1.500E-08 | 0.000E+00 | 1.500E-08 | 1.500E-08 |
| 9 | 1 | T1H | A | 9.4e-09 | 9.4e-09 | 9.4e-09 | 9.400E-09 | 0.000E+00 | 9.400E-09 | 9.400E-09 |
| 10 | 1 | T1I | A | 1.6e-08 | 1.6e-08 | 1.6e-08 | 1.600E-08 | 0.000E+00 | 1.600E-08 | 1.600E-08 |
| 11 | 1 | T1L | A | 8.6e-09 | 8.6e-09 | 8.6e-09 | 8.600E-09 | 0.000E+00 | 8.600E-09 | 8.600E-09 |
| 12 | 1 | T1K | A | 1.3e-08 | 1.3e-08 | 1.3e-08 | 1.300E-08 | 2.026E-24 | 1.300E-08 | 1.300E-08 |
| 13 | 1 | T1M | A | 3.1e-09 | 3.1e-09 | 3.1e-09 | 3.100E-09 | 0.000E+00 | 3.100E-09 | 3.100E-09 |
| 14 | 1 | T1F | A | 3.3e-09 | 3.3e-09 | 3.3e-09 | 3.300E-09 | 0.000E+00 | 3.300E-09 | 3.300E-09 |
| 15 | 1 | T1P | A | 2.3e-08 | 2.3e-08 | 2.3e-08 | 2.300E-08 | 4.052E-24 | 2.300E-08 | 2.300E-08 |
| 16 | 1 | T1S | A | 3.1e-09 | 3.1e-09 | 3.1e-09 | 3.100E-09 | 0.000E+00 | 3.100E-09 | 3.100E-09 |
| 17 | 1 | T1T | A | 1.1e-08 | 1.1e-08 | 1.1e-08 | 1.100E-08 | 0.000E+00 | 1.100E-08 | 1.100E-08 |
| 18 | 1 | T1W | A | 9.2e-09 | 9.2e-09 | 9.2e-09 | 9.200E-09 | 0.000E+00 | 9.200E-09 | 9.200E-09 |
| 19 | 1 | T1Y | A | 2.2e-08 | 2.2e-08 | 2.2e-08 | 2.200E-08 | 0.000E+00 | 2.200E-08 | 2.200E-08 |
| 20 | 1 | T1V | A | 1.3e-08 | 1.3e-08 | 1.3e-08 | 1.300E-08 | 2.026E-24 | 1.300E-08 | 1.300E-08 |
| 21 | 2 | Q2A | A | 8.3e-09 | 8.3e-09 | 8.3e-09 | 8.300E-09 | 0.000E+00 | 8.300E-09 | 8.300E-09 |
| 22 | 2 | Q2R | A | 3.1e-08 | 3.1e-08 | 3.1e-08 | 3.100E-08 | 0.000E+00 | 3.100E-08 | 3.100E-08 |
| 23 | 2 | Q2N | A | 7.1e-10 | 7.1e-10 | 7.1e-10 | 7.100E-10 | 1.266E-25 | 7.100E-10 | 7.100E-10 |
| 24 | 2 | Q2D | A | 4e-09 | 4e-09 | 4e-09 | 4.000E-09 | 0.000E+00 | 4.000E-09 | 4.000E-09 |
| 25 | 2 | Q2C | A | 1.1e-08 | 4.5e-09 | 4.5e-09 | 6.667E-09 | 3.753E-09 | 4.500E-09 | 1.100E-08 |
| 26 | 2 | Q2E | A | 5.7e-09 | 5.4e-09 | 8.9e-09 | 6.667E-09 | 1.940E-09 | 5.400E-09 | 8.900E-09 |
| 27 | 2 | Q2Q | A | 2.9e-09 | 9.9e-09 | 1.6e-09 | 4.800E-09 | 4.464E-09 | 1.600E-09 | 9.900E-09 |
| 28 | 2 | Q2G | A | 2.2e-08 | 6e-09 | 1.3e-08 | 1.367E-08 | 8.021E-09 | 6.000E-09 | 2.200E-08 |
| 29 | 2 | Q2H | A | 2.3e-08 | 7.9e-09 | 7.9e-09 | 1.293E-08 | 8.718E-09 | 7.900E-09 | 2.300E-08 |
| 30 | 2 | Q2I | A | 5.1e-09 | 2e-09 | 5.1e-09 | 4.067E-09 | 1.790E-09 | 2.000E-09 | 5.100E-09 |
| 31 | 2 | Q2L | A | 2.4e-09 | 8.5e-09 | 2.4e-09 | 4.433E-09 | 3.522E-09 | 2.400E-09 | 8.500E-09 |
| 32 | 2 | Q2K | A | 6.6e-09 | 6.6e-09 | 1.4e-08 | 9.067E-09 | 4.272E-09 | 6.600E-09 | 1.400E-08 |
| 33 | 2 | Q2M | A | 2.5e-08 | 2.5e-08 | 1e-08 | 2.000E-08 | 8.660E-09 | 1.000E-08 | 2.500E-08 |
| 34 | 2 | Q2F | A | 3.8e-09 | 7e-09 | 3.8e-09 | 4.867E-09 | 1.848E-09 | 3.800E-09 | 7.000E-09 |
| 35 | 2 | Q2P | A | 1.3e-08 | 1.7e-08 | 1.3e-08 | 1.433E-08 | 2.309E-09 | 1.300E-08 | 1.700E-08 |
| 36 | 2 | Q2S | A | 1.3e-08 | 1.3e-08 | 1.3e-08 | 1.300E-08 | 2.026E-24 | 1.300E-08 | 1.300E-08 |
| 37 | 2 | Q2T | A | 8.9e-09 | 9.4e-09 | 8.7e-09 | 9.000E-09 | 3.606E-10 | 8.700E-09 | 9.400E-09 |
| 38 | 2 | Q2W | A | 8.3e-09 | 3.6e-09 | 1.2e-08 | 7.967E-09 | 4.210E-09 | 3.600E-09 | 1.200E-08 |
| 39 | 2 | Q2Y | A | 4.4e-09 | 4.2e-09 | 4e-09 | 4.200E-09 | 2.000E-10 | 4.000E-09 | 4.400E-09 |
| 40 | 2 | Q2V | A | 6.2e-09 | 1.9e-08 | 1.9e-08 | 1.473E-08 | 7.390E-09 | 6.200E-09 | 1.900E-08 |
| 41 | 3 | V3A | A | 1.9e-08 | 1.9e-08 | 4.4e-09 | 1.413E-08 | 8.429E-09 | 4.400E-09 | 1.900E-08 |
| 42 | 3 | V3R | A | 2.1e-09 | 6.6e-09 | 6.6e-09 | 5.100E-09 | 2.598E-09 | 2.100E-09 | 6.600E-09 |
| 43 | 3 | V3N | A | 6.6e-09 | 6.6e-09 | 6.6e-09 | 6.600E-09 | 0.000E+00 | 6.600E-09 | 6.600E-09 |
| 44 | 3 | V3D | A | 5.6e-09 | 5.6e-09 | 5.6e-09 | 5.600E-09 | 0.000E+00 | 5.600E-09 | 5.600E-09 |
| 45 | 3 | V3C | A | 1.6e-08 | 1.6e-08 | 6.8e-09 | 1.293E-08 | 5.312E-09 | 6.800E-09 | 1.600E-08 |
| 46 | 3 | V3E | A | 1.1e-08 | 1.1e-08 | 5.3e-09 | 9.100E-09 | 3.291E-09 | 5.300E-09 | 1.100E-08 |
| 47 | 3 | V3Q | A | 8e-09 | 8e-09 | 1.1e-08 | 9.000E-09 | 1.732E-09 | 8.000E-09 | 1.100E-08 |
| 48 | 3 | V3G | A | 5.5e-09 | 5.5e-09 | 5.5e-09 | 5.500E-09 | 0.000E+00 | 5.500E-09 | 5.500E-09 |
| 49 | 3 | V3H | A | 1e-08 | 8.7e-09 | 1.4e-08 | 1.090E-08 | 2.762E-09 | 8.700E-09 | 1.400E-08 |
| 50 | 3 | V3I | A | 1.6e-08 | 4e-09 | 1.6e-08 | 1.200E-08 | 6.928E-09 | 4.000E-09 | 1.600E-08 |
| 51 | 3 | V3L | A | 9.9e-09 | 6.8e-09 | 6.8e-09 | 7.833E-09 | 1.790E-09 | 6.800E-09 | 9.900E-09 |
| 52 | 3 | V3K | A | 5.1e-09 | 5.9e-09 | 3.9e-09 | 4.967E-09 | 1.007E-09 | 3.900E-09 | 5.900E-09 |
| 53 | 3 | V3M | A | 2.2e-08 | 4.3e-09 | 4.1e-09 | 1.013E-08 | 1.028E-08 | 4.100E-09 | 2.200E-08 |
| 54 | 3 | V3F | A | 5e-09 | 6.4e-09 | 1.1e-08 | 7.467E-09 | 3.139E-09 | 5.000E-09 | 1.100E-08 |
| 55 | 3 | V3P | A | 6.4e-09 | 3.7e-09 | 3.7e-09 | 4.600E-09 | 1.559E-09 | 3.700E-09 | 6.400E-09 |
| 56 | 3 | V3S | A | 1.9e-09 | 1.9e-09 | 1.6e-08 | 6.600E-09 | 8.141E-09 | 1.900E-09 | 1.600E-08 |
| 57 | 3 | V3T | A | 1.4e-08 | 7.9e-09 | 4.3e-09 | 8.733E-09 | 4.903E-09 | 4.300E-09 | 1.400E-08 |
| 58 | 3 | V3W | A | 1.1e-08 | 1.1e-08 | 4.8e-09 | 8.933E-09 | 3.580E-09 | 4.800E-09 | 1.100E-08 |
| 59 | 3 | V3Y | A | 5e-09 | 1.4e-08 | 3.2e-09 | 7.400E-09 | 5.786E-09 | 3.200E-09 | 1.400E-08 |
| 60 | 3 | V3V | A | 2.3e-09 | 2.9e-09 | 1.2e-08 | 5.733E-09 | 5.435E-09 | 2.300E-09 | 1.200E-08 |
| 61 | 4 | C4A | A | 3.7e-09 | 1.3e-08 | 2.4e-09 | 6.367E-09 | 5.781E-09 | 2.400E-09 | 1.300E-08 |
| 62 | 4 | C4R | A | 1.3e-09 | 7.9e-09 | 6.5e-09 | 5.233E-09 | 3.478E-09 | 1.300E-09 | 7.900E-09 |
| 63 | 4 | C4N | A | 8.8e-09 | 5.4e-09 | 4e-09 | 6.067E-09 | 2.468E-09 | 4.000E-09 | 8.800E-09 |
| 64 | 4 | C4D | A | 1.6e-09 | 2.2e-08 | 8.4e-09 | 1.067E-08 | 1.039E-08 | 1.600E-09 | 2.200E-08 |
| 65 | 4 | C4C | A | 2.4e-08 | 2.4e-08 | 2.4e-08 | 2.400E-08 | 0.000E+00 | 2.400E-08 | 2.400E-08 |
| 66 | 4 | C4E | A | 9.7e-09 | 9.1e-09 | 6.1e-09 | 8.300E-09 | 1.929E-09 | 6.100E-09 | 9.700E-09 |
| 67 | 4 | C4Q | A | 6.5e-09 | 1.7e-08 | 2.5e-08 | 1.617E-08 | 9.278E-09 | 6.500E-09 | 2.500E-08 |
| 68 | 4 | C4G | A | 1.4e-08 | 1.4e-08 | 9.2e-09 | 1.240E-08 | 2.771E-09 | 9.200E-09 | 1.400E-08 |
| 69 | 4 | C4H | A | 8.3e-09 | 2.2e-09 | 9e-09 | 6.500E-09 | 3.740E-09 | 2.200E-09 | 9.000E-09 |
| 70 | 4 | C4I | A | 5.3e-09 | 5.3e-09 | 4.6e-09 | 5.067E-09 | 4.041E-10 | 4.600E-09 | 5.300E-09 |
| 71 | 4 | C4L | A | 7.3e-09 | 7e-09 | 2.8e-09 | 5.700E-09 | 2.516E-09 | 2.800E-09 | 7.300E-09 |
| 72 | 4 | C4K | A | 1.9e-08 | 1.9e-08 | 5.4e-09 | 1.447E-08 | 7.852E-09 | 5.400E-09 | 1.900E-08 |
| 73 | 4 | C4M | A | 1.7e-08 | 4.2e-09 | 6.2e-09 | 9.133E-09 | 6.886E-09 | 4.200E-09 | 1.700E-08 |
| 74 | 4 | C4F | A | 6.7e-09 | 3.9e-09 | 2.9e-09 | 4.500E-09 | 1.970E-09 | 2.900E-09 | 6.700E-09 |
| 75 | 4 | C4P | A | 6.4e-09 | 1.1e-08 | 2.1e-08 | 1.280E-08 | 7.465E-09 | 6.400E-09 | 2.100E-08 |
| 76 | 4 | C4S | A | 1.1e-08 | 6.4e-09 | 1.4e-08 | 1.047E-08 | 3.828E-09 | 6.400E-09 | 1.400E-08 |
| 77 | 4 | C4T | A | 1.4e-08 | 9.3e-09 | 1.1e-08 | 1.143E-08 | 2.380E-09 | 9.300E-09 | 1.400E-08 |
| 78 | 4 | C4W | A | 1.3e-08 | 1.5e-08 | 6.2e-09 | 1.140E-08 | 4.613E-09 | 6.200E-09 | 1.500E-08 |
| 79 | 4 | C4Y | A | 8.1e-09 | 5.6e-09 | 7.4e-09 | 7.033E-09 | 1.290E-09 | 5.600E-09 | 8.100E-09 |
| 80 | 4 | C4V | A | 2.8e-09 | 8.2e-09 | 6.2e-09 | 5.733E-09 | 2.730E-09 | 2.800E-09 | 8.200E-09 |
| 81 | 5 | T5A | A | 1e-08 | 1.5e-08 | 8e-09 | 1.100E-08 | 3.606E-09 | 8.000E-09 | 1.500E-08 |
| 82 | 5 | T5R | A | 8.3e-09 | 3.3e-09 | 6.2e-09 | 5.933E-09 | 2.511E-09 | 3.300E-09 | 8.300E-09 |
| 83 | 5 | T5N | A | 4.3e-09 | 3.8e-09 | 1.3e-08 | 7.033E-09 | 5.173E-09 | 3.800E-09 | 1.300E-08 |
| 84 | 5 | T5D | A | 4.4e-09 | 2.7e-09 | 2.7e-09 | 3.267E-09 | 9.815E-10 | 2.700E-09 | 4.400E-09 |

| | | | | | | | | | | |
|---|---|---|---|---|---|---|---|---|---|---|
| 85 | 5 | T5C | A | 3.5e-09 | 3.3e-09 | 1.9e-09 | 2.900E-09 | 8.718E-10 | 1.900E-09 | 3.500E-09 |
| 86 | 5 | T5E | A | 4.2e-09 | 1.5e-08 | 3.1e-09 | 7.433E-09 | 6.576E-09 | 3.100E-09 | 1.500E-08 |
| 87 | 5 | T5Q | A | 6e-09 | 4.4e-09 | 6.8e-09 | 5.733E-09 | 1.222E-09 | 4.400E-09 | 6.800E-09 |
| 88 | 5 | T5G | A | 6.2e-09 | 7.1e-09 | 1.4e-08 | 9.100E-09 | 4.267E-09 | 6.200E-09 | 1.400E-08 |
| 89 | 5 | T5H | A | 2.5e-09 | 1.1e-08 | 5.9e-09 | 6.467E-09 | 4.278E-09 | 2.500E-09 | 1.100E-08 |
| 90 | 5 | T5I | A | 1.8e-08 | 3e-09 | 2.1e-08 | 1.400E-08 | 9.644E-09 | 3.000E-09 | 2.100E-08 |
| 91 | 5 | T5L | A | 3.1e-09 | 3.1e-09 | 9e-09 | 5.067E-09 | 3.406E-09 | 3.100E-09 | 9.000E-09 |
| 92 | 5 | T5K | A | 1e-08 | 3.6e-09 | 6.4e-09 | 6.667E-09 | 3.208E-09 | 3.600E-09 | 1.000E-08 |
| 93 | 5 | T5M | A | 3.5e-09 | 4.4e-09 | 9.5e-09 | 5.800E-09 | 3.236E-09 | 3.500E-09 | 9.500E-09 |
| 94 | 5 | T5F | A | 1.1e-08 | 3.7e-09 | 3.7e-09 | 6.133E-09 | 4.215E-09 | 3.700E-09 | 1.100E-08 |
| 95 | 5 | T5P | A | 1.7e-08 | 6.6e-09 | 4.9e-09 | 9.500E-09 | 6.551E-09 | 4.900E-09 | 1.700E-08 |
| 96 | 5 | T5S | A | 1.2e-08 | 3e-09 | 7.6e-09 | 7.533E-09 | 4.500E-09 | 3.000E-09 | 1.200E-08 |
| 97 | 5 | T5T | A | 1.5e-08 | 1.1e-08 | 1.3e-08 | 1.300E-08 | 2.000E-09 | 1.100E-08 | 1.500E-08 |
| 98 | 5 | T5W | A | 3.9e-09 | 1.2e-08 | 2.4e-08 | 1.330E-08 | 1.011E-08 | 3.900E-09 | 2.400E-08 |
| 99 | 5 | T5Y | A | 4.6e-09 | 7.7e-09 | 4.3e-09 | 5.533E-09 | 1.882E-09 | 4.300E-09 | 7.700E-09 |
| 100 | 5 | T5V | A | 8.1e-09 | 6.3e-09 | 3.7e-09 | 6.033E-09 | 2.212E-09 | 3.700E-09 | 8.100E-09 |
| 101 | 6 | G6A | A | 1.2e-08 | 4.5e-09 | 5.8e-09 | 7.433E-09 | 4.008E-09 | 4.500E-09 | 1.200E-08 |
| 102 | 6 | G6R | A | 1.4e-08 | 6.8e-09 | 3e-09 | 7.933E-09 | 5.587E-09 | 3.000E-09 | 1.400E-08 |
| 103 | 6 | G6N | A | 5.8e-09 | 8.2e-09 | 3.1e-09 | 5.700E-09 | 2.551E-09 | 3.100E-09 | 8.200E-09 |
| 104 | 6 | G6D | A | 1.5e-09 | 1.2e-08 | 2.9e-09 | 5.467E-09 | 5.701E-09 | 1.500E-09 | 1.200E-08 |
| 105 | 6 | G6C | A | 3.6e-09 | 8.1e-09 | 8.9e-09 | 6.867E-09 | 2.857E-09 | 3.600E-09 | 8.900E-09 |
| 106 | 6 | G6E | A | 3.9e-09 | 7.5e-09 | 7.3e-09 | 6.233E-09 | 2.023E-09 | 3.900E-09 | 7.500E-09 |
| 107 | 6 | G6Q | A | 7.3e-09 | 1.4e-08 | 3.8e-09 | 8.367E-09 | 5.183E-09 | 3.800E-09 | 1.400E-08 |
| 108 | 6 | G6G | A | 5.3e-09 | 9.9e-09 | 7.5e-09 | 7.567E-09 | 2.301E-09 | 5.300E-09 | 9.900E-09 |
| 109 | 6 | G6H | A | 8.4e-09 | 4.9e-09 | 6e-09 | 6.433E-09 | 1.790E-09 | 4.900E-09 | 8.400E-09 |
| 110 | 6 | G6I | A | 1.4e-08 | 5.4e-09 | 8.6e-09 | 9.333E-09 | 4.347E-09 | 5.400E-09 | 1.400E-08 |
| 111 | 6 | G6L | A | 7.2e-09 | 5.8e-09 | 1.3e-08 | 8.667E-09 | 3.818E-09 | 5.800E-09 | 1.300E-08 |
| 112 | 6 | G6K | A | 4.8e-09 | 1.4e-08 | 1.9e-08 | 1.260E-08 | 7.203E-09 | 4.800E-09 | 1.900E-08 |
| 113 | 6 | G6M | A | 1.3e-08 | 1.8e-08 | 1.5e-08 | 1.533E-08 | 2.517E-09 | 1.300E-08 | 1.800E-08 |
| 114 | 6 | G6F | A | 6.5e-09 | 2e-09 | 4.1e-09 | 4.200E-09 | 2.252E-09 | 2.000E-09 | 6.500E-09 |
| 115 | 6 | G6P | A | 1e-08 | 5.4e-09 | 3.2e-09 | 6.200E-09 | 3.470E-09 | 3.200E-09 | 1.000E-08 |
| 116 | 6 | G6S | A | 5.7e-09 | 1.6e-08 | 1.1e-08 | 1.090E-08 | 5.151E-09 | 5.700E-09 | 1.600E-08 |
| 117 | 6 | G6T | A | 1.2e-08 | 5.2e-09 | 1.6e-08 | 1.107E-08 | 5.460E-09 | 5.200E-09 | 1.600E-08 |
| 118 | 6 | G6W | A | 5.9e-09 | 8.7e-09 | 5.6e-09 | 6.733E-09 | 1.710E-09 | 5.600E-09 | 8.700E-09 |
| 119 | 6 | G6Y | A | 3.7e-09 | 9.7e-09 | 9e-09 | 7.467E-09 | 3.281E-09 | 3.700E-09 | 9.700E-09 |
| 120 | 6 | G6V | A | 1.1e-08 | 1.1e-08 | 4.5e-09 | 8.833E-09 | 3.753E-09 | 4.500E-09 | 1.100E-08 |
| 121 | 7 | T7A | A | 8.9e-09 | 2.3e-09 | 3.1e-09 | 4.767E-09 | 3.602E-09 | 2.300E-09 | 8.900E-09 |
| 122 | 7 | T7R | A | 5.5e-09 | 1.1e-08 | 9.9e-09 | 8.800E-09 | 2.910E-09 | 5.500E-09 | 1.100E-08 |
| 123 | 7 | T7N | A | 5.3e-09 | 1.7e-08 | 1e-08 | 1.077E-08 | 5.888E-09 | 5.300E-09 | 1.700E-08 |
| 124 | 7 | T7D | A | 2.3e-08 | 8.2e-09 | 5e-09 | 1.207E-08 | 9.603E-09 | 5.000E-09 | 2.300E-08 |
| 125 | 7 | T7C | A | 7.4e-09 | 1.3e-08 | 1.7e-08 | 1.247E-08 | 4.822E-09 | 7.400E-09 | 1.700E-08 |
| 126 | 7 | T7E | A | 5.7e-09 | 5.4e-09 | 8.2e-09 | 6.433E-09 | 1.537E-09 | 5.400E-09 | 8.200E-09 |
| 127 | 7 | T7Q | A | 6.1e-09 | 1.2e-08 | 7.5e-09 | 8.533E-09 | 3.083E-09 | 6.100E-09 | 1.200E-08 |
| 128 | 7 | T7G | A | 3.6e-09 | 1.7e-08 | 1.4e-08 | 1.153E-08 | 7.032E-09 | 3.600E-09 | 1.700E-08 |
| 129 | 7 | T7H | A | 4.7e-09 | 6.6e-09 | 6.6e-09 | 5.967E-09 | 1.097E-09 | 4.700E-09 | 6.600E-09 |
| 130 | 7 | T7I | A | 7.2e-09 | 6.9e-09 | 1.1e-08 | 8.367E-09 | 2.285E-09 | 6.900E-09 | 1.100E-08 |
| 131 | 7 | T7L | A | 8.1e-09 | 3e-09 | 2.9e-09 | 4.667E-09 | 2.974E-09 | 2.900E-09 | 8.100E-09 |
| 132 | 7 | T7K | A | 6.2e-09 | 8.9e-09 | 7.9e-09 | 7.667E-09 | 1.365E-09 | 6.200E-09 | 8.900E-09 |
| 133 | 7 | T7M | A | 9.4e-09 | 6.3e-09 | 6.2e-09 | 7.300E-09 | 1.819E-09 | 6.200E-09 | 9.400E-09 |
| 134 | 7 | T7F | A | 2.1e-09 | 3.2e-09 | 1.6e-08 | 7.100E-09 | 7.727E-09 | 2.100E-09 | 1.600E-08 |
| 135 | 7 | T7P | A | 1.1e-08 | 9e-09 | 7.4e-09 | 9.133E-09 | 1.804E-09 | 7.400E-09 | 1.100E-08 |
| 136 | 7 | T7S | A | 6.2e-09 | 2.2e-09 | 1.3e-08 | 7.133E-09 | 5.460E-09 | 2.200E-09 | 1.300E-08 |
| 137 | 7 | T7T | A | 8.5e-09 | 3.6e-08 | 1e-08 | 1.817E-08 | 1.546E-08 | 8.500E-09 | 3.600E-08 |
| 138 | 7 | T7W | A | 8.7e-09 | 1.5e-08 | 5.2e-09 | 9.633E-09 | 4.966E-09 | 5.200E-09 | 1.500E-08 |
| 139 | 7 | T7Y | A | 5.6e-09 | 1.1e-08 | 6.1e-09 | 7.567E-09 | 2.984E-09 | 5.600E-09 | 1.100E-08 |
| 140 | 7 | T7V | A | 7.5e-09 | 1.6e-08 | 4.1e-09 | 9.200E-09 | 6.129E-09 | 4.100E-09 | 1.600E-08 |
| 141 | 8 | D8A | A | 4.8e-09 | 8.3e-09 | 6.8e-09 | 6.633E-09 | 1.756E-09 | 4.800E-09 | 8.300E-09 |
| 142 | 8 | D8R | A | 9.7e-09 | 6.9e-09 | 6.9e-09 | 7.833E-09 | 1.617E-09 | 6.900E-09 | 9.700E-09 |
| 143 | 8 | D8N | A | 6.3e-09 | 1.9e-09 | 5.9e-09 | 4.700E-09 | 2.433E-09 | 1.900E-09 | 6.300E-09 |
| 144 | 8 | D8D | A | 8.6e-09 | 6.8e-09 | 1.5e-08 | 1.013E-08 | 4.310E-09 | 6.800E-09 | 1.500E-08 |
| 145 | 8 | D8C | A | 3.3e-09 | 9.1e-09 | 4.6e-09 | 5.667E-09 | 3.044E-09 | 3.300E-09 | 9.100E-09 |
| 146 | 8 | D8E | A | 1.4e-08 | 6.4e-09 | 1.9e-08 | 1.313E-08 | 6.345E-09 | 6.400E-09 | 1.900E-08 |
| 147 | 8 | D8Q | A | 5.7e-09 | 1.9e-08 | 4.5e-09 | 9.733E-09 | 8.048E-09 | 4.500E-09 | 1.900E-08 |
| 148 | 8 | D8G | A | 3.7e-09 | 9.2e-09 | 5.2e-09 | 6.033E-09 | 2.843E-09 | 3.700E-09 | 9.200E-09 |
| 149 | 8 | D8H | A | 7.9e-09 | 5.4e-09 | 6.3e-09 | 6.533E-09 | 1.266E-09 | 5.400E-09 | 7.900E-09 |
| 150 | 8 | D8I | A | 9.2e-09 | 1.1e-08 | 5.4e-09 | 8.533E-09 | 2.859E-09 | 5.400E-09 | 1.100E-08 |
| 151 | 8 | D8L | A | 4.1e-09 | 5e-09 | 1.5e-09 | 3.533E-09 | 1.818E-09 | 1.500E-09 | 5.000E-09 |
| 152 | 8 | D8K | A | 7.7e-09 | 3.6e-09 | 4e-09 | 5.100E-09 | 2.261E-09 | 3.600E-09 | 7.700E-09 |
| 153 | 8 | D8M | A | 2.9e-09 | 7e-09 | 8.3e-09 | 6.067E-09 | 2.818E-09 | 2.900E-09 | 8.300E-09 |
| 154 | 8 | D8F | A | 1.2e-08 | 8.1e-09 | 9.9e-09 | 1.000E-08 | 1.952E-09 | 8.100E-09 | 1.200E-08 |
| 155 | 8 | D8P | A | 1.1e-08 | 1.5e-08 | 6.4e-09 | 1.080E-08 | 4.303E-09 | 6.400E-09 | 1.500E-08 |
| 156 | 8 | D8S | A | 9e-09 | 3.9e-09 | 7.3e-09 | 6.733E-09 | 2.597E-09 | 3.900E-09 | 9.000E-09 |
| 157 | 8 | D8T | A | 3.6e-09 | 6.4e-09 | 1.2e-08 | 7.333E-09 | 4.277E-09 | 3.600E-09 | 1.200E-08 |
| 158 | 8 | D8W | A | 1.5e-08 | 7.5e-09 | 3.5e-09 | 8.667E-09 | 5.838E-09 | 3.500E-09 | 1.500E-08 |
| 159 | 8 | D8Y | A | 2.3e-08 | 2.8e-09 | 9.7e-09 | 1.183E-08 | 1.027E-08 | 2.800E-09 | 2.300E-08 |
| 160 | 8 | D8V | A | 4e-09 | 2.9e-09 | 6.2e-09 | 4.367E-09 | 1.680E-09 | 2.900E-09 | 6.200E-09 |
| 161 | 9 | M9A | A | 6.3e-09 | 7.1e-09 | 5.6e-09 | 6.333E-09 | 7.506E-10 | 5.600E-09 | 7.100E-09 |
| 162 | 9 | M9R | A | 9.5e-09 | 9.5e-09 | 1.7e-09 | 6.900E-09 | 4.503E-09 | 1.700E-09 | 9.500E-09 |
| 163 | 9 | M9N | A | 5.5e-09 | 6.2e-09 | 6.4e-09 | 6.033E-09 | 4.726E-10 | 5.500E-09 | 6.400E-09 |
| 164 | 9 | M9D | A | 1.2e-08 | 2.9e-09 | 1.1e-08 | 8.633E-09 | 4.990E-09 | 2.900E-09 | 1.200E-08 |
| 165 | 9 | M9C | A | 6.7e-09 | 3.7e-09 | 4e-09 | 4.800E-09 | 1.652E-09 | 3.700E-09 | 6.700E-09 |
| 166 | 9 | M9E | A | 8.1e-09 | 7.1e-09 | 7.4e-09 | 7.533E-09 | 5.132E-10 | 7.100E-09 | 8.100E-09 |
| 167 | 9 | M9Q | A | 4.6e-09 | 2.5e-09 | 1.5e-08 | 7.367E-09 | 6.694E-09 | 2.500E-09 | 1.500E-08 |
| 168 | 9 | M9G | A | 9e-09 | 7.7e-09 | 2.3e-09 | 6.333E-09 | 3.553E-09 | 2.300E-09 | 9.000E-09 |
| 169 | 9 | M9H | A | 1e-08 | 6.1e-09 | 8.5e-09 | 8.200E-09 | 1.967E-09 | 6.100E-09 | 1.000E-08 |

| | | | | | | | | | | |
|---|---|---|---|---|---|---|---|---|---|---|
| 170 | 9 | M9I | A | 1.2e-08 | 5.1e-09 | 6.4e-09 | 7.833E-09 | 3.667E-09 | 5.100E-09 | 1.200E-08 |
| 171 | 9 | M9L | A | 5.9e-09 | 2.8e-09 | 7.4e-09 | 5.367E-09 | 2.346E-09 | 2.800E-09 | 7.400E-09 |
| 172 | 9 | M9K | A | 1.3e-08 | 1.3e-08 | 1.7e-09 | 9.233E-09 | 6.524E-09 | 1.700E-09 | 1.300E-08 |
| 173 | 9 | M9M | A | 1.9e-08 | 7.9e-09 | 1e-08 | 1.230E-08 | 5.897E-09 | 7.900E-09 | 1.900E-08 |
| 174 | 9 | M9F | A | 7.6e-09 | 8.8e-09 | 6.8e-09 | 7.733E-09 | 1.007E-09 | 6.800E-09 | 8.800E-09 |
| 175 | 9 | M9P | A | 2.8e-09 | 6.2e-09 | 5.9e-09 | 4.967E-09 | 1.882E-09 | 2.800E-09 | 6.200E-09 |
| 176 | 9 | M9S | A | 5.8e-09 | 4.4e-09 | 4.4e-09 | 4.867E-09 | 8.083E-10 | 4.400E-09 | 5.800E-09 |
| 177 | 9 | M9T | A | 1.9e-08 | 1.1e-09 | 6.2e-09 | 8.767E-09 | 9.222E-09 | 1.100E-09 | 1.900E-08 |
| 178 | 9 | M9W | A | 1e-08 | 1.8e-08 | 9.9e-09 | 1.263E-08 | 4.648E-09 | 9.900E-09 | 1.800E-08 |
| 179 | 9 | M9Y | A | 7e-09 | 9.9e-09 | 1.4e-08 | 1.030E-08 | 3.517E-09 | 7.000E-09 | 1.400E-08 |
| 180 | 9 | M9V | A | 3.5e-09 | 1.1e-08 | 6.2e-09 | 6.900E-09 | 3.799E-09 | 3.500E-09 | 1.100E-08 |
| 181 | 10 | K10A | A | 2.2e-09 | 1.5e-08 | 1.1e-08 | 9.400E-09 | 6.548E-09 | 2.200E-09 | 1.500E-08 |
| 182 | 10 | K10R | A | 2.4e-09 | 1.7e-08 | 1e-08 | 9.800E-09 | 7.302E-09 | 2.400E-09 | 1.700E-08 |
| 183 | 10 | K10N | A | 3.5e-09 | 8.8e-09 | 1e-08 | 7.433E-09 | 3.459E-09 | 3.500E-09 | 1.000E-08 |
| 184 | 10 | K10D | A | 2.4e-09 | 1.1e-08 | 4.4e-09 | 5.933E-09 | 4.500E-09 | 2.400E-09 | 1.100E-08 |
| 185 | 10 | K10C | A | 5.4e-09 | 4.4e-09 | 4.8e-09 | 4.867E-09 | 5.033E-10 | 4.400E-09 | 5.400E-09 |
| 186 | 10 | K10E | A | 4.8e-09 | 5.9e-09 | 3e-09 | 4.567E-09 | 1.464E-09 | 3.000E-09 | 5.900E-09 |
| 187 | 10 | K10Q | A | 1.1e-08 | 1.1e-08 | 6.5e-09 | 9.500E-09 | 2.598E-09 | 6.500E-09 | 1.100E-08 |
| 188 | 10 | K10G | A | 6.8e-09 | 4e-09 | 6.8e-09 | 5.867E-09 | 1.617E-09 | 4.000E-09 | 6.800E-09 |
| 189 | 10 | K10H | A | 6e-09 | 4.7e-09 | 2e-08 | 1.023E-08 | 8.483E-09 | 4.700E-09 | 2.000E-08 |
| 190 | 10 | K10I | A | 2.6e-09 | 2.4e-09 | 5.4e-09 | 3.467E-09 | 1.677E-09 | 2.400E-09 | 5.400E-09 |
| 191 | 10 | K10L | A | 2e-09 | 5.7e-09 | 6.3e-09 | 4.667E-09 | 2.329E-09 | 2.000E-09 | 6.300E-09 |
| 192 | 10 | K10K | A | 1.3e-08 | 7.8e-09 | 3e-09 | 7.933E-09 | 5.001E-09 | 3.000E-09 | 1.300E-08 |
| 193 | 10 | K10M | A | 3.5e-09 | 7.5e-09 | 8.2e-09 | 6.400E-09 | 2.536E-09 | 3.500E-09 | 8.200E-09 |
| 194 | 10 | K10F | A | 3.8e-09 | 2e-09 | 5.9e-09 | 3.900E-09 | 1.952E-09 | 2.000E-09 | 5.900E-09 |
| 195 | 10 | K10P | A | 1.9e-08 | 4.9e-09 | 6.4e-09 | 1.010E-08 | 7.744E-09 | 4.900E-09 | 1.900E-08 |
| 196 | 10 | K10S | A | 1.2e-08 | 1.4e-08 | 3.4e-09 | 9.800E-09 | 5.632E-09 | 3.400E-09 | 1.400E-08 |
| 197 | 10 | K10T | A | 7.6e-09 | 9.7e-09 | 7.9e-09 | 8.400E-09 | 1.136E-09 | 7.600E-09 | 9.700E-09 |
| 198 | 10 | K10W | A | 9.1e-09 | 3.7e-09 | 4.9e-09 | 5.900E-09 | 2.835E-09 | 3.700E-09 | 9.100E-09 |
| 199 | 10 | K10Y | A | 4.4e-09 | 3.7e-09 | 6.5e-09 | 4.867E-09 | 1.457E-09 | 3.700E-09 | 6.500E-09 |
| 200 | 10 | K10V | A | 7.4e-09 | 2.9e-09 | 4.9e-09 | 5.067E-09 | 2.255E-09 | 2.900E-09 | 7.400E-09 |
| 201 | 11 | L11A | A | 1.5e-08 | 1.9e-09 | 4.2e-09 | 7.033E-09 | 6.995E-09 | 1.900E-09 | 1.500E-08 |
| 202 | 11 | L11R | A | 3.2e-09 | 2.3e-08 | 2.2e-08 | 1.607E-08 | 1.115E-08 | 3.200E-09 | 2.300E-08 |
| 203 | 11 | L11N | A | 5.7e-09 | 6e-09 | 4.2e-09 | 5.300E-09 | 9.644E-10 | 4.200E-09 | 6.000E-09 |
| 204 | 11 | L11D | A | 3e-09 | 3.4e-09 | 3.3e-09 | 3.233E-09 | 2.082E-10 | 3.000E-09 | 3.400E-09 |
| 205 | 11 | L11C | A | 3.1e-09 | 2.3e-09 | 1.3e-08 | 6.133E-09 | 5.960E-09 | 2.300E-09 | 1.300E-08 |
| 206 | 11 | L11E | A | 1.2e-08 | 1.7e-08 | 9.3e-09 | 1.277E-08 | 3.907E-09 | 9.300E-09 | 1.700E-08 |
| 207 | 11 | L11Q | A | 4.4e-09 | 7.9e-09 | 7.2e-09 | 6.500E-09 | 1.852E-09 | 4.400E-09 | 7.900E-09 |
| 208 | 11 | L11G | A | 1.2e-08 | 1.2e-08 | 8e-09 | 1.067E-08 | 2.309E-09 | 8.000E-09 | 1.200E-08 |
| 209 | 11 | L11H | A | 5e-09 | 3e-09 | 3e-09 | 3.667E-09 | 1.155E-09 | 3.000E-09 | 5.000E-09 |
| 210 | 11 | L11I | A | 6.8e-09 | 6.6e-09 | 3e-09 | 5.467E-09 | 2.139E-09 | 3.000E-09 | 6.800E-09 |
| 211 | 11 | L11L | A | 3.3e-09 | 1.5e-08 | 3.1e-09 | 7.133E-09 | 6.813E-09 | 3.100E-09 | 1.500E-08 |
| 212 | 11 | L11K | A | 3.4e-09 | 9.6e-09 | 1.3e-08 | 8.667E-09 | 4.868E-09 | 3.400E-09 | 1.300E-08 |
| 213 | 11 | L11M | A | 1.7e-08 | 4.1e-09 | 4.5e-09 | 8.533E-09 | 7.335E-09 | 4.100E-09 | 1.700E-08 |
| 214 | 11 | L11F | A | 9.2e-09 | 2.7e-09 | 4.6e-09 | 5.500E-09 | 3.342E-09 | 2.700E-09 | 9.200E-09 |
| 215 | 11 | L11P | A | 5.4e-09 | 1.2e-08 | 8.4e-09 | 8.600E-09 | 3.305E-09 | 5.400E-09 | 1.200E-08 |
| 216 | 11 | L11S | A | 4.7e-09 | 1.1e-08 | 1.3e-08 | 9.567E-09 | 4.332E-09 | 4.700E-09 | 1.300E-08 |
| 217 | 11 | L11T | A | 1.3e-08 | 4.1e-09 | 8.8e-09 | 8.633E-09 | 4.452E-09 | 4.100E-09 | 1.300E-08 |
| 218 | 11 | L11W | A | 4.4e-09 | 1.6e-08 | 6.2e-09 | 8.867E-09 | 6.243E-09 | 4.400E-09 | 1.600E-08 |
| 219 | 11 | L11Y | A | 3.4e-09 | 1.4e-08 | 8.6e-09 | 8.667E-09 | 5.300E-09 | 3.400E-09 | 1.400E-08 |
| 220 | 11 | L11V | A | 2.6e-09 | 4.5e-09 | 6.7e-09 | 4.600E-09 | 2.052E-09 | 2.600E-09 | 6.700E-09 |
| 221 | 12 | R12A | A | 2.8e-09 | 1.7e-09 | 9.5e-09 | 4.667E-09 | 4.222E-09 | 1.700E-09 | 9.500E-09 |
| 222 | 12 | R12R | A | 3.6e-09 | 6.2e-09 | 3.5e-09 | 4.433E-09 | 1.531E-09 | 3.500E-09 | 6.200E-09 |
| 223 | 12 | R12N | A | 1.7e-08 | 4.1e-09 | 2e-08 | 1.370E-08 | 8.448E-09 | 4.100E-09 | 2.000E-08 |
| 224 | 12 | R12D | A | 7.1e-09 | 4.8e-09 | 6.1e-09 | 6.000E-09 | 1.153E-09 | 4.800E-09 | 7.100E-09 |
| 225 | 12 | R12C | A | 7.3e-09 | 1.5e-08 | 5e-09 | 9.100E-09 | 5.237E-09 | 5.000E-09 | 1.500E-08 |
| 226 | 12 | R12E | A | 4.9e-09 | 1.1e-08 | 3.9e-09 | 6.600E-09 | 3.843E-09 | 3.900E-09 | 1.100E-08 |
| 227 | 12 | R12Q | A | 8.9e-09 | 1.6e-08 | 1e-08 | 1.163E-08 | 3.821E-09 | 8.900E-09 | 1.600E-08 |
| 228 | 12 | R12G | A | 5.6e-09 | 6.7e-09 | 2.2e-09 | 4.833E-09 | 2.346E-09 | 2.200E-09 | 6.700E-09 |
| 229 | 12 | R12H | A | 3e-09 | 1.5e-08 | 7.8e-09 | 8.600E-09 | 6.040E-09 | 3.000E-09 | 1.500E-08 |
| 230 | 12 | R12I | A | 5.4e-09 | 3.7e-09 | 8.2e-09 | 5.767E-09 | 2.272E-09 | 3.700E-09 | 8.200E-09 |
| 231 | 12 | R12L | A | 2e-09 | 2e-08 | 8.4e-09 | 1.013E-08 | 9.124E-09 | 2.000E-09 | 2.000E-08 |
| 232 | 12 | R12K | A | 7.9e-09 | 2.4e-08 | 1.1e-08 | 1.430E-08 | 8.542E-09 | 7.900E-09 | 2.400E-08 |
| 233 | 12 | R12M | A | 4.9e-09 | 2.3e-09 | 3.9e-09 | 3.700E-09 | 1.311E-09 | 2.300E-09 | 4.900E-09 |
| 234 | 12 | R12F | A | 2.2e-09 | 6.4e-09 | 3.8e-09 | 4.133E-09 | 2.120E-09 | 2.200E-09 | 6.400E-09 |
| 235 | 12 | R12P | A | 1.2e-08 | 5.1e-09 | 1.3e-08 | 1.003E-08 | 4.302E-09 | 5.100E-09 | 1.300E-08 |
| 236 | 12 | R12S | A | 7.1e-09 | 4.2e-09 | 9.9e-09 | 7.067E-09 | 2.850E-09 | 4.200E-09 | 9.900E-09 |
| 237 | 12 | R12T | A | 9.4e-09 | 1.1e-08 | 1.3e-08 | 1.113E-08 | 1.804E-09 | 9.400E-09 | 1.300E-08 |
| 238 | 12 | R12W | A | 1.2e-08 | 8.6e-09 | 6.6e-09 | 9.067E-09 | 2.730E-09 | 6.600E-09 | 1.200E-08 |
| 239 | 12 | R12Y | A | 1e-08 | 2.4e-09 | 1.8e-08 | 1.013E-08 | 7.801E-09 | 2.400E-09 | 1.800E-08 |
| 240 | 12 | R12V | A | 2e-09 | 9.4e-09 | 5e-09 | 5.467E-09 | 3.722E-09 | 2.000E-09 | 9.400E-09 |
| 241 | 13 | L13A | A | 6e-09 | 3.6e-09 | 1.1e-08 | 6.867E-09 | 3.775E-09 | 3.600E-09 | 1.100E-08 |
| 242 | 13 | L13R | A | 2.2e-08 | 3.8e-09 | 4.4e-09 | 1.007E-08 | 1.034E-08 | 3.800E-09 | 2.200E-08 |
| 243 | 13 | L13N | A | 6.3e-09 | 1e-08 | 4e-09 | 6.767E-09 | 3.027E-09 | 4.000E-09 | 1.000E-08 |
| 244 | 13 | L13D | A | 5.2e-09 | 1.3e-08 | 1.4e-08 | 1.073E-08 | 4.818E-09 | 5.200E-09 | 1.400E-08 |
| 245 | 13 | L13C | A | 7e-09 | 3.7e-09 | 5e-09 | 5.233E-09 | 1.662E-09 | 3.700E-09 | 7.000E-09 |
| 246 | 13 | L13E | A | 5.5e-09 | 4.2e-09 | 5.8e-09 | 5.167E-09 | 8.505E-10 | 4.200E-09 | 5.800E-09 |
| 247 | 13 | L13Q | A | 9.5e-09 | 7.8e-09 | 2.3e-08 | 1.343E-08 | 8.328E-09 | 7.800E-09 | 2.300E-08 |
| 248 | 13 | L13G | A | 3.8e-09 | 4.2e-09 | 5e-09 | 4.333E-09 | 6.110E-10 | 3.800E-09 | 5.000E-09 |
| 249 | 13 | L13H | A | 9.5e-09 | 6.7e-09 | 1.1e-08 | 9.067E-09 | 2.183E-09 | 6.700E-09 | 1.100E-08 |
| 250 | 13 | L13I | A | 1.2e-08 | 6.6e-09 | 1.1e-08 | 9.867E-09 | 2.873E-09 | 6.600E-09 | 1.200E-08 |
| 251 | 13 | L13L | A | 8.9e-09 | 9.2e-09 | 8.7e-09 | 8.933E-09 | 2.517E-10 | 8.700E-09 | 9.200E-09 |
| 252 | 13 | L13K | A | 1.7e-08 | 4.4e-09 | 8.8e-09 | 1.007E-08 | 6.395E-09 | 4.400E-09 | 1.700E-08 |
| 253 | 13 | L13M | A | 6.4e-09 | 3.6e-09 | 1.3e-08 | 7.667E-09 | 4.826E-09 | 3.600E-09 | 1.300E-08 |
| 254 | 13 | L13F | A | 3.6e-09 | 1.1e-08 | 1.5e-08 | 9.867E-09 | 5.784E-09 | 3.600E-09 | 1.500E-08 |

| | | | | | | | | | | |
|---|---|---|---|---|---|---|---|---|---|---|
| 255 | 13 | L13P | A | 3.9e-09 | 8.7e-09 | 8e-09 | 6.867E-09 | 2.593E-09 | 3.900E-09 | 8.700E-09 |
| 256 | 13 | L13S | A | 9.3e-09 | 9.8e-09 | 7.5e-09 | 8.867E-09 | 1.210E-09 | 7.500E-09 | 9.800E-09 |
| 257 | 13 | L13T | A | 8.2e-09 | 2.3e-08 | 4e-09 | 1.173E-08 | 9.981E-09 | 4.000E-09 | 2.300E-08 |
| 258 | 13 | L13W | A | 1.2e-08 | 5.1e-09 | 6.9e-09 | 8.000E-09 | 3.579E-09 | 5.100E-09 | 1.200E-08 |
| 259 | 13 | L13Y | A | 1e-08 | 6.4e-09 | 9.8e-09 | 8.733E-09 | 2.023E-09 | 6.400E-09 | 1.000E-08 |
| 260 | 13 | L13V | A | 3.3e-09 | 1.6e-08 | 2.5e-09 | 7.267E-09 | 7.574E-09 | 2.500E-09 | 1.600E-08 |
| 261 | 14 | P14A | A | 1.7e-08 | 4.7e-09 | 6.5e-09 | 9.400E-09 | 6.643E-09 | 4.700E-09 | 1.700E-08 |
| 262 | 14 | P14R | A | 7.2e-09 | 9e-09 | 4.2e-09 | 6.800E-09 | 2.425E-09 | 4.200E-09 | 9.000E-09 |
| 263 | 14 | P14N | A | 1.4e-08 | 1.3e-08 | 2e-08 | 1.567E-08 | 3.786E-09 | 1.300E-08 | 2.000E-08 |
| 264 | 14 | P14D | A | 3e-09 | 2.8e-09 | 1.2e-08 | 5.933E-09 | 5.255E-09 | 2.800E-09 | 1.200E-08 |
| 265 | 14 | P14C | A | 4.7e-09 | 9.5e-09 | 3.5e-09 | 5.900E-09 | 3.175E-09 | 3.500E-09 | 9.500E-09 |
| 266 | 14 | P14E | A | 4.3e-09 | 7.3e-09 | 6.7e-09 | 6.100E-09 | 1.587E-09 | 4.300E-09 | 7.300E-09 |
| 267 | 14 | P14Q | A | 1.9e-09 | 7e-09 | 3.6e-09 | 4.167E-09 | 2.597E-09 | 1.900E-09 | 7.000E-09 |
| 268 | 14 | P14G | A | 6.5e-09 | 3.5e-09 | 1.1e-08 | 7.000E-09 | 3.775E-09 | 3.500E-09 | 1.100E-08 |
| 269 | 14 | P14H | A | 5.3e-09 | 2.9e-08 | 1.1e-08 | 1.510E-08 | 1.237E-08 | 5.300E-09 | 2.900E-08 |
| 270 | 14 | P14I | A | 1.2e-08 | 1.3e-08 | 8.9e-09 | 1.130E-08 | 2.138E-09 | 8.900E-09 | 1.300E-08 |
| 271 | 14 | P14L | A | 4.9e-09 | 8.2e-09 | 1.3e-08 | 8.700E-09 | 4.073E-09 | 4.900E-09 | 1.300E-08 |
| 272 | 14 | P14K | A | 9.8e-10 | 1.5e-08 | 6.6e-09 | 7.527E-09 | 7.056E-09 | 9.800E-10 | 1.500E-08 |
| 273 | 14 | P14M | A | 1.5e-08 | 7.8e-09 | 3.2e-09 | 8.667E-09 | 5.948E-09 | 3.200E-09 | 1.500E-08 |
| 274 | 14 | P14F | A | 4e-09 | 3.5e-09 | 1e-08 | 5.833E-09 | 3.617E-09 | 3.500E-09 | 1.000E-08 |
| 275 | 14 | P14P | A | 1.1e-08 | 1e-08 | 1e-08 | 1.033E-08 | 5.774E-10 | 1.000E-08 | 1.100E-08 |
| 276 | 14 | P14S | A | 3.4e-09 | 5.6e-09 | 3.5e-09 | 4.167E-09 | 1.242E-09 | 3.400E-09 | 5.600E-09 |
| 277 | 14 | P14T | A | 5.3e-10 | 6.2e-09 | 4.7e-09 | 3.810E-09 | 2.938E-09 | 5.300E-10 | 6.200E-09 |
| 278 | 14 | P14W | A | 7.1e-09 | 2.6e-09 | 8.4e-09 | 6.033E-09 | 3.044E-09 | 2.600E-09 | 8.400E-09 |
| 279 | 14 | P14Y | A | 6e-09 | 5.5e-09 | 8.3e-09 | 6.600E-09 | 1.493E-09 | 5.500E-09 | 8.300E-09 |
| 280 | 14 | P14V | A | 5.8e-09 | 1.3e-08 | 1.3e-08 | 1.060E-08 | 4.157E-09 | 5.800E-09 | 1.300E-08 |
| 281 | 15 | A15A | A | 5.2e-09 | 1e-08 | 9.3e-09 | 8.167E-09 | 2.593E-09 | 5.200E-09 | 1.000E-08 |
| 282 | 15 | A15R | A | 2.8e-09 | 1.1e-08 | 1e-08 | 7.933E-09 | 4.474E-09 | 2.800E-09 | 1.100E-08 |
| 283 | 15 | A15N | A | 2.3e-08 | 6.9e-09 | 5.5e-09 | 1.180E-08 | 9.725E-09 | 5.500E-09 | 2.300E-08 |
| 284 | 15 | A15D | A | 6.9e-09 | 3.6e-09 | 6.6e-09 | 5.700E-09 | 1.825E-09 | 3.600E-09 | 6.900E-09 |
| 285 | 15 | A15C | A | 4e-09 | 1.3e-08 | 8.4e-09 | 8.467E-09 | 4.500E-09 | 4.000E-09 | 1.300E-08 |
| 286 | 15 | A15E | A | 7.7e-09 | 4.4e-09 | 2.7e-08 | 1.303E-08 | 1.221E-08 | 4.400E-09 | 2.700E-08 |
| 287 | 15 | A15Q | A | 9.1e-08 | 7.4e-09 | 1.7e-08 | 3.847E-08 | 4.575E-08 | 7.400E-09 | 9.100E-08 |
| 288 | 15 | A15G | A | 9.3e-09 | 3e-09 | 1.1e-08 | 7.767E-09 | 4.215E-09 | 3.000E-09 | 1.100E-08 |
| 289 | 15 | A15H | A | 7.5e-09 | 8.2e-09 | 6.4e-09 | 7.367E-09 | 9.074E-10 | 6.400E-09 | 8.200E-09 |
| 290 | 15 | A15I | A | 3.9e-09 | 3.7e-09 | 7.7e-09 | 5.100E-09 | 2.254E-09 | 3.700E-09 | 7.700E-09 |
| 291 | 15 | A15L | A | 1.5e-08 | 2e-09 | 2.3e-09 | 6.433E-09 | 7.420E-09 | 2.000E-09 | 1.500E-08 |
| 292 | 15 | A15K | A | 4.7e-09 | 7.1e-09 | 3.4e-08 | 1.527E-08 | 1.627E-08 | 4.700E-09 | 3.400E-08 |
| 293 | 15 | A15M | A | 5.6e-09 | 3.8e-09 | 1.2e-08 | 7.133E-09 | 4.310E-09 | 3.800E-09 | 1.200E-08 |
| 294 | 15 | A15F | A | 4.3e-09 | 3e-09 | 1.2e-08 | 6.433E-09 | 4.864E-09 | 3.000E-09 | 1.200E-08 |
| 295 | 15 | A15P | A | 2.7e-08 | 6e-09 | 1.1e-08 | 1.467E-08 | 1.097E-08 | 6.000E-09 | 2.700E-08 |
| 296 | 15 | A15S | A | 8.4e-09 | 4e-09 | 7e-09 | 6.467E-09 | 2.248E-09 | 4.000E-09 | 8.400E-09 |
| 297 | 15 | A15T | A | 1.5e-08 | 7.4e-09 | 4.9e-09 | 9.100E-09 | 5.260E-09 | 4.900E-09 | 1.500E-08 |
| 298 | 15 | A15W | A | 9.8e-09 | 2.9e-09 | 5.5e-09 | 6.067E-09 | 3.485E-09 | 2.900E-09 | 9.800E-09 |
| 299 | 15 | A15Y | A | 1.4e-08 | 1.9e-09 | 8.5e-09 | 8.133E-09 | 6.058E-09 | 1.900E-09 | 1.400E-08 |
| 300 | 15 | A15V | A | 6.7e-09 | 1.9e-08 | 7.9e-09 | 1.120E-08 | 6.782E-09 | 6.700E-09 | 1.900E-08 |
| 301 | 16 | S16A | A | 4e-09 | 1.7e-08 | 1.7e-08 | 1.267E-08 | 7.506E-09 | 4.000E-09 | 1.700E-08 |
| 302 | 16 | S16R | A | 8.1e-09 | 8.2e-09 | 7.8e-09 | 8.033E-09 | 2.082E-10 | 7.800E-09 | 8.200E-09 |
| 303 | 16 | S16N | A | 1.2e-08 | 2.6e-08 | 1.8e-08 | 1.867E-08 | 7.024E-09 | 1.200E-08 | 2.600E-08 |
| 304 | 16 | S16D | A | 1.1e-08 | 2.3e-09 | 9e-09 | 7.433E-09 | 4.557E-09 | 2.300E-09 | 1.100E-08 |
| 305 | 16 | S16C | A | 8.6e-09 | 1.3e-08 | 8.4e-09 | 1.000E-08 | 2.600E-09 | 8.400E-09 | 1.300E-08 |
| 306 | 16 | S16E | A | 4.1e-09 | 3.1e-09 | 8e-09 | 5.067E-09 | 2.589E-09 | 3.100E-09 | 8.000E-09 |
| 307 | 16 | S16Q | A | 1.5e-08 | 1.4e-08 | 1.3e-08 | 1.400E-08 | 1.000E-09 | 1.300E-08 | 1.500E-08 |
| 308 | 16 | S16G | A | 8.3e-09 | 1.5e-08 | 1e-08 | 1.110E-08 | 3.483E-09 | 8.300E-09 | 1.500E-08 |
| 309 | 16 | S16H | A | 4.9e-09 | 2.9e-09 | 5e-09 | 4.267E-09 | 1.185E-09 | 2.900E-09 | 5.000E-09 |
| 310 | 16 | S16I | A | 4.5e-09 | 1.2e-08 | 4e-09 | 6.833E-09 | 4.481E-09 | 4.000E-09 | 1.200E-08 |
| 311 | 16 | S16L | A | 6.1e-09 | 3.1e-09 | 5.2e-09 | 4.800E-09 | 1.539E-09 | 3.100E-09 | 6.100E-09 |
| 312 | 16 | S16K | A | 3.2e-09 | 1.2e-08 | 6.8e-09 | 7.333E-09 | 4.424E-09 | 3.200E-09 | 1.200E-08 |
| 313 | 16 | S16M | A | 6.8e-08 | 4.1e-09 | 1.2e-08 | 2.803E-08 | 3.484E-08 | 4.100E-09 | 6.800E-08 |
| 314 | 16 | S16F | A | 1.9e-08 | 1.6e-08 | 4.9e-09 | 1.330E-08 | 7.428E-09 | 4.900E-09 | 1.900E-08 |
| 315 | 16 | S16P | A | 7.2e-09 | 1.2e-08 | 4e-09 | 7.733E-09 | 4.027E-09 | 4.000E-09 | 1.200E-08 |
| 316 | 16 | S16S | A | 2.7e-08 | 5.5e-09 | 1.1e-08 | 1.450E-08 | 1.117E-08 | 5.500E-09 | 2.700E-08 |
| 317 | 16 | S16T | A | 5.7e-09 | 8.7e-09 | 1e-08 | 8.133E-09 | 2.205E-09 | 5.700E-09 | 1.000E-08 |
| 318 | 16 | S16W | A | 7.8e-09 | 5e-09 | 5.9e-09 | 6.233E-09 | 1.429E-09 | 5.000E-09 | 7.800E-09 |
| 319 | 16 | S16Y | A | 1.5e-08 | 6.4e-09 | 5.9e-09 | 9.100E-09 | 5.116E-09 | 5.900E-09 | 1.500E-08 |
| 320 | 16 | S16V | A | 5.6e-09 | 2.4e-08 | 1.7e-09 | 1.043E-08 | 1.191E-08 | 1.700E-09 | 2.400E-08 |
| 321 | 17 | P17A | A | 2.2e-09 | 8.9e-09 | 1.4e-08 | 8.367E-09 | 5.918E-09 | 2.200E-09 | 1.400E-08 |
| 322 | 17 | P17R | A | 6.5e-09 | 9e-09 | 1.1e-08 | 8.833E-09 | 2.255E-09 | 6.500E-09 | 1.100E-08 |
| 323 | 17 | P17N | A | 4e-09 | 1.1e-08 | 2.5e-09 | 5.833E-09 | 4.537E-09 | 2.500E-09 | 1.100E-08 |
| 324 | 17 | P17D | A | 6.2e-09 | 1e-08 | 4.3e-09 | 6.833E-09 | 2.902E-09 | 4.300E-09 | 1.000E-08 |
| 325 | 17 | P17C | A | 9.7e-09 | 4.1e-09 | 4e-09 | 5.933E-09 | 3.262E-09 | 4.000E-09 | 9.700E-09 |
| 326 | 17 | P17E | A | 9.6e-09 | 1.2e-08 | 6e-09 | 9.200E-09 | 3.020E-09 | 6.000E-09 | 1.200E-08 |
| 327 | 17 | P17Q | A | 4.1e-09 | 8.9e-09 | 2.3e-08 | 1.200E-08 | 9.824E-09 | 4.100E-09 | 2.300E-08 |
| 328 | 17 | P17G | A | 1e-08 | 5.7e-09 | 5.1e-09 | 6.933E-09 | 2.673E-09 | 5.100E-09 | 1.000E-08 |
| 329 | 17 | P17H | A | 9.9e-09 | 2.6e-08 | 3.7e-09 | 1.320E-08 | 1.151E-08 | 3.700E-09 | 2.600E-08 |
| 330 | 17 | P17I | A | 6.1e-09 | 4.5e-09 | 2.8e-09 | 4.467E-09 | 1.650E-09 | 2.800E-09 | 6.100E-09 |
| 331 | 17 | P17L | A | 2.2e-09 | 2.7e-09 | 4.5e-09 | 3.133E-09 | 1.210E-09 | 2.200E-09 | 4.500E-09 |
| 332 | 17 | P17K | A | 1.8e-08 | 1.1e-08 | 5.7e-09 | 1.157E-08 | 6.170E-09 | 5.700E-09 | 1.800E-08 |
| 333 | 17 | P17M | A | 2.2e-09 | 6.8e-09 | 7.4e-09 | 5.467E-09 | 2.845E-09 | 2.200E-09 | 7.400E-09 |
| 334 | 17 | P17F | A | 9e-09 | 4.1e-09 | 1e-08 | 7.700E-09 | 3.158E-09 | 4.100E-09 | 1.000E-08 |
| 335 | 17 | P17P | A | 4.1e-08 | 3.2e-09 | 5.5e-09 | 1.657E-08 | 2.119E-08 | 3.200E-09 | 4.100E-08 |
| 336 | 17 | P17S | A | 3.1e-09 | 5.9e-09 | 3.6e-09 | 4.200E-09 | 1.493E-09 | 3.100E-09 | 5.900E-09 |
| 337 | 17 | P17T | A | 3.2e-09 | 1.8e-08 | 3.1e-09 | 8.100E-09 | 8.574E-09 | 3.100E-09 | 1.800E-08 |
| 338 | 17 | P17W | A | 2.9e-09 | 3.4e-09 | 8.7e-10 | 2.390E-09 | 1.340E-09 | 8.700E-10 | 3.400E-09 |
| 339 | 17 | P17Y | A | 7.7e-09 | 9.6e-09 | 1.8e-08 | 1.177E-08 | 5.481E-09 | 7.700E-09 | 1.800E-08 |

| | | | | | | | | | | |
|---|---|---|---|---|---|---|---|---|---|---|
| 340 | 17 | P17V | A | 9e-09 | 7.9e-09 | 3.3e-09 | 6.733E-09 | 3.024E-09 | 3.300E-09 | 9.000E-09 |
| 341 | 18 | E18A | A | 7.7e-09 | 1.2e-08 | 8e-09 | 9.233E-09 | 2.401E-09 | 7.700E-09 | 1.200E-08 |
| 342 | 18 | E18R | A | 6.1e-09 | 8.8e-09 | 8.9e-09 | 7.933E-09 | 1.589E-09 | 6.100E-09 | 8.900E-09 |
| 343 | 18 | E18N | A | 9e-09 | 8.6e-09 | 3.9e-09 | 7.167E-09 | 2.836E-09 | 3.900E-09 | 9.000E-09 |
| 344 | 18 | E18D | A | 2.4e-09 | 8.9e-09 | 5.9e-09 | 5.733E-09 | 3.253E-09 | 2.400E-09 | 8.900E-09 |
| 345 | 18 | E18C | A | 4.8e-09 | 1.1e-08 | 1.1e-08 | 8.933E-09 | 3.580E-09 | 4.800E-09 | 1.100E-08 |
| 346 | 18 | E18E | A | 5.6e-09 | 5.1e-09 | 2.8e-09 | 4.500E-09 | 1.493E-09 | 2.800E-09 | 5.600E-09 |
| 347 | 18 | E18Q | A | 1.9e-09 | 7.6e-09 | 7.2e-09 | 5.567E-09 | 3.182E-09 | 1.900E-09 | 7.600E-09 |
| 348 | 18 | E18G | A | 2.1e-08 | 1.6e-08 | 6e-09 | 1.433E-08 | 7.638E-09 | 6.000E-09 | 2.100E-08 |
| 349 | 18 | E18H | A | 9.4e-09 | 6.1e-09 | 1.1e-08 | 8.833E-09 | 2.499E-09 | 6.100E-09 | 1.100E-08 |
| 350 | 18 | E18I | A | 3.1e-09 | 9.4e-09 | 1.7e-08 | 9.833E-09 | 6.960E-09 | 3.100E-09 | 1.700E-08 |
| 351 | 18 | E18L | A | 1.1e-08 | 3.2e-09 | 7.2e-09 | 7.133E-09 | 3.900E-09 | 3.200E-09 | 1.100E-08 |
| 352 | 18 | E18K | A | 4e-09 | 1.3e-09 | 1.2e-08 | 5.767E-09 | 5.564E-09 | 1.300E-09 | 1.200E-08 |
| 353 | 18 | E18M | A | 1.5e-08 | 4.4e-09 | 1.8e-08 | 1.247E-08 | 7.145E-09 | 4.400E-09 | 1.800E-08 |
| 354 | 18 | E18F | A | 4.5e-09 | 6.7e-09 | 4.1e-09 | 5.100E-09 | 1.400E-09 | 4.100E-09 | 6.700E-09 |
| 355 | 18 | E18P | A | 7.7e-09 | 8.3e-09 | 1.3e-08 | 9.667E-09 | 2.902E-09 | 7.700E-09 | 1.300E-08 |
| 356 | 18 | E18S | A | 4.3e-09 | 4.1e-09 | 8.2e-09 | 5.533E-09 | 2.312E-09 | 4.100E-09 | 8.200E-09 |
| 357 | 18 | E18T | A | 2.8e-09 | 9.2e-09 | 6e-09 | 6.000E-09 | 3.200E-09 | 2.800E-09 | 9.200E-09 |
| 358 | 18 | E18W | A | 2.5e-08 | 1.1e-08 | 8e-09 | 1.467E-08 | 9.074E-09 | 8.000E-09 | 2.500E-08 |
| 359 | 18 | E18Y | A | 3.3e-09 | 5e-09 | 1.4e-08 | 7.433E-09 | 5.750E-09 | 3.300E-09 | 1.400E-08 |
| 360 | 18 | E18V | A | 3.2e-09 | 6e-09 | 7.1e-09 | 5.433E-09 | 2.011E-09 | 3.200E-09 | 7.100E-09 |
| 361 | 19 | T19A | A | 4.5e-09 | 4.7e-09 | 4.3e-08 | 1.740E-08 | 2.217E-08 | 4.500E-09 | 4.300E-08 |
| 362 | 19 | T19R | A | 1.7e-08 | 3.6e-09 | 6.3e-09 | 8.967E-09 | 7.087E-09 | 3.600E-09 | 1.700E-08 |
| 363 | 19 | T19N | A | 7.1e-09 | 5.6e-09 | 5.9e-09 | 6.200E-09 | 7.937E-10 | 5.600E-09 | 7.100E-09 |
| 364 | 19 | T19D | A | 5e-09 | 1.2e-08 | 6.6e-09 | 7.867E-09 | 3.668E-09 | 5.000E-09 | 1.200E-08 |
| 365 | 19 | T19C | A | 1.6e-08 | 1.6e-08 | 1.4e-08 | 1.533E-08 | 1.155E-09 | 1.400E-08 | 1.600E-08 |
| 366 | 19 | T19E | A | 6e-09 | 1.2e-08 | 1.1e-08 | 9.667E-09 | 3.215E-09 | 6.000E-09 | 1.200E-08 |
| 367 | 19 | T19Q | A | 8.4e-09 | 7.4e-09 | 1.1e-08 | 8.933E-09 | 1.858E-09 | 7.400E-09 | 1.100E-08 |
| 368 | 19 | T19G | A | 3.3e-09 | 1e-08 | 6.2e-09 | 6.500E-09 | 3.360E-09 | 3.300E-09 | 1.000E-08 |
| 369 | 19 | T19H | A | 8.9e-09 | 8.1e-09 | 5.9e-09 | 7.633E-09 | 1.553E-09 | 5.900E-09 | 8.900E-09 |
| 370 | 19 | T19I | A | 3.6e-09 | 1.1e-08 | 8.9e-09 | 7.833E-09 | 3.814E-09 | 3.600E-09 | 1.100E-08 |
| 371 | 19 | T19L | A | 1.1e-08 | 3.7e-08 | 8.1e-09 | 1.870E-08 | 1.591E-08 | 8.100E-09 | 3.700E-08 |
| 372 | 19 | T19K | A | 7.2e-09 | 2e-08 | 9.5e-09 | 1.223E-08 | 6.824E-09 | 7.200E-09 | 2.000E-08 |
| 373 | 19 | T19M | A | 6.8e-09 | 5.1e-09 | 3.5e-09 | 5.133E-09 | 1.650E-09 | 3.500E-09 | 6.800E-09 |
| 374 | 19 | T19F | A | 2.3e-09 | 7e-09 | 1e-08 | 6.433E-09 | 3.881E-09 | 2.300E-09 | 1.000E-08 |
| 375 | 19 | T19P | A | 9.5e-09 | 4e-09 | 1.1e-08 | 8.167E-09 | 3.686E-09 | 4.000E-09 | 1.100E-08 |
| 376 | 19 | T19S | A | 2.1e-09 | 2.6e-09 | 5.8e-09 | 3.500E-09 | 2.007E-09 | 2.100E-09 | 5.800E-09 |
| 377 | 19 | T19T | A | 9.8e-09 | 8.2e-09 | 6.8e-09 | 8.267E-09 | 1.501E-09 | 6.800E-09 | 9.800E-09 |
| 378 | 19 | T19W | A | 5.8e-09 | 2.3e-08 | 8.9e-09 | 1.257E-08 | 9.168E-09 | 5.800E-09 | 2.300E-08 |
| 379 | 19 | T19Y | A | 6.4e-09 | 3.9e-09 | 4.6e-09 | 4.967E-09 | 1.290E-09 | 3.900E-09 | 6.400E-09 |
| 380 | 19 | T19V | A | 7.5e-09 | 3.6e-09 | 3.3e-09 | 4.800E-09 | 2.343E-09 | 3.300E-09 | 7.500E-09 |
| 381 | 20 | H20A | A | 9.7e-09 | 1.1e-08 | 1.6e-08 | 1.223E-08 | 3.326E-09 | 9.700E-09 | 1.600E-08 |
| 382 | 20 | H20R | A | 6.1e-09 | 1.3e-08 | 1.4e-08 | 1.103E-08 | 4.302E-09 | 6.100E-09 | 1.400E-08 |
| 383 | 20 | H20N | A | 7.4e-09 | 4.6e-09 | 7.1e-09 | 6.367E-09 | 1.537E-09 | 4.600E-09 | 7.400E-09 |
| 384 | 20 | H20D | A | 1.1e-08 | 4e-09 | 5.4e-09 | 6.800E-09 | 3.704E-09 | 4.000E-09 | 1.100E-08 |
| 385 | 20 | H20C | A | 5.8e-09 | 9.2e-09 | 6.6e-09 | 7.200E-09 | 1.778E-09 | 5.800E-09 | 9.200E-09 |
| 386 | 20 | H20E | A | 7.9e-09 | 8.2e-09 | 1.9e-09 | 6.000E-09 | 3.554E-09 | 1.900E-09 | 8.200E-09 |
| 387 | 20 | H20Q | A | 7.1e-09 | 6.8e-09 | 1.2e-08 | 8.633E-09 | 2.919E-09 | 6.800E-09 | 1.200E-08 |
| 388 | 20 | H20G | A | 1.5e-08 | 5.6e-09 | 4.2e-09 | 8.267E-09 | 5.873E-09 | 4.200E-09 | 1.500E-08 |
| 389 | 20 | H20H | A | 6.4e-09 | 1.2e-08 | 5e-09 | 7.800E-09 | 3.704E-09 | 5.000E-09 | 1.200E-08 |
| 390 | 20 | H20I | A | 1.7e-08 | 5.3e-09 | 1.5e-08 | 1.243E-08 | 6.258E-09 | 5.300E-09 | 1.700E-08 |
| 391 | 20 | H20L | A | 7.4e-09 | 2.3e-08 | 7.6e-09 | 1.267E-08 | 8.949E-09 | 7.400E-09 | 2.300E-08 |
| 392 | 20 | H20K | A | 5.6e-09 | 2.3e-09 | 5.7e-09 | 4.533E-09 | 1.935E-09 | 2.300E-09 | 5.700E-09 |
| 393 | 20 | H20M | A | 9.5e-09 | 8.1e-09 | 6.9e-09 | 8.167E-09 | 1.301E-09 | 6.900E-09 | 9.500E-09 |
| 394 | 20 | H20F | A | 4.1e-09 | 6.5e-09 | 9.4e-09 | 6.667E-09 | 2.654E-09 | 4.100E-09 | 9.400E-09 |
| 395 | 20 | H20P | A | 6.4e-09 | 5.7e-09 | 1e-08 | 7.367E-09 | 2.307E-09 | 5.700E-09 | 1.000E-08 |
| 396 | 20 | H20S | A | 6.1e-09 | 8.3e-09 | 9.1e-09 | 7.833E-09 | 1.553E-09 | 6.100E-09 | 9.100E-09 |
| 397 | 20 | H20T | A | 1e-08 | 6.9e-09 | 2.7e-09 | 6.533E-09 | 3.664E-09 | 2.700E-09 | 1.000E-08 |
| 398 | 20 | H20W | A | 5e-09 | 8e-09 | 1.5e-08 | 9.333E-09 | 5.132E-09 | 5.000E-09 | 1.500E-08 |
| 399 | 20 | H20Y | A | 8.5e-09 | 5.2e-09 | 1.7e-08 | 1.023E-08 | 6.088E-09 | 5.200E-09 | 1.700E-08 |
| 400 | 20 | H20V | A | 6.6e-09 | 6.1e-09 | 7.7e-09 | 6.800E-09 | 8.185E-10 | 6.100E-09 | 7.700E-09 |
| 401 | 21 | L21A | A | 8.4e-09 | 1.6e-08 | 9.2e-09 | 1.120E-08 | 4.176E-09 | 8.400E-09 | 1.600E-08 |
| 402 | 21 | L21R | A | 6.4e-09 | 1.1e-08 | 7.4e-09 | 8.267E-09 | 2.419E-09 | 6.400E-09 | 1.100E-08 |
| 403 | 21 | L21N | A | 3.4e-09 | 4.3e-09 | 1.1e-08 | 6.233E-09 | 4.153E-09 | 3.400E-09 | 1.100E-08 |
| 404 | 21 | L21D | A | 1e-08 | 1.1e-08 | 3.4e-09 | 8.133E-09 | 4.130E-09 | 3.400E-09 | 1.100E-08 |
| 405 | 21 | L21C | A | 9.9e-09 | 3.8e-09 | 1.5e-08 | 9.567E-09 | 5.607E-09 | 3.800E-09 | 1.500E-08 |
| 406 | 21 | L21E | A | 1e-08 | 4.2e-09 | 2e-09 | 5.400E-09 | 4.133E-09 | 2.000E-09 | 1.000E-08 |
| 407 | 21 | L21Q | A | 5.2e-09 | 4e-09 | 1.2e-08 | 7.067E-09 | 4.314E-09 | 4.000E-09 | 1.200E-08 |
| 408 | 21 | L21G | A | 7e-09 | 4.8e-09 | 6.2e-09 | 6.000E-09 | 1.114E-09 | 4.800E-09 | 7.000E-09 |
| 409 | 21 | L21H | A | 1.4e-08 | 6.1e-09 | 1.2e-08 | 1.070E-08 | 4.107E-09 | 6.100E-09 | 1.400E-08 |
| 410 | 21 | L21I | A | 3.7e-09 | 4.8e-09 | 1.5e-08 | 7.833E-09 | 6.231E-09 | 3.700E-09 | 1.500E-08 |
| 411 | 21 | L21L | A | 3.9e-09 | 6.5e-09 | 3.5e-09 | 4.633E-09 | 1.629E-09 | 3.500E-09 | 6.500E-09 |
| 412 | 21 | L21K | A | 1.4e-08 | 6.8e-09 | 3.7e-09 | 8.167E-09 | 5.284E-09 | 3.700E-09 | 1.400E-08 |
| 413 | 21 | L21M | A | 7.4e-09 | 8.4e-09 | 9.7e-09 | 8.500E-09 | 1.153E-09 | 7.400E-09 | 9.700E-09 |
| 414 | 21 | L21F | A | 3.6e-08 | 1.2e-08 | 7.7e-09 | 1.857E-08 | 1.525E-08 | 7.700E-09 | 3.600E-08 |
| 415 | 21 | L21P | A | 8.6e-09 | 6.7e-09 | 6.4e-09 | 7.233E-09 | 1.193E-09 | 6.400E-09 | 8.600E-09 |
| 416 | 21 | L21S | A | 3.8e-09 | 7.4e-09 | 5.9e-09 | 5.700E-09 | 1.808E-09 | 3.800E-09 | 7.400E-09 |
| 417 | 21 | L21T | A | 2.7e-09 | 1e-08 | 1.2e-08 | 8.233E-09 | 4.895E-09 | 2.700E-09 | 1.200E-08 |
| 418 | 21 | L21W | A | 6.7e-09 | 1.2e-08 | 1.6e-08 | 1.157E-08 | 4.665E-09 | 6.700E-09 | 1.600E-08 |
| 419 | 21 | L21Y | A | 9.8e-09 | 1.5e-08 | 5.3e-09 | 1.003E-08 | 4.854E-09 | 5.300E-09 | 1.500E-08 |
| 420 | 21 | L21V | A | 4.2e-09 | 5.2e-09 | 4.5e-09 | 4.633E-09 | 5.132E-10 | 4.200E-09 | 5.200E-09 |
| 421 | 22 | D22A | A | 9.6e-09 | 6.3e-09 | 1.4e-08 | 9.967E-09 | 3.863E-09 | 6.300E-09 | 1.400E-08 |
| 422 | 22 | D22R | A | 1.5e-08 | 2e-08 | 4.8e-09 | 1.327E-08 | 7.747E-09 | 4.800E-09 | 2.000E-08 |
| 423 | 22 | D22N | A | 7.1e-09 | 1.1e-08 | 1.3e-08 | 1.037E-08 | 3.001E-09 | 7.100E-09 | 1.300E-08 |
| 424 | 22 | D22D | A | 1e-08 | 2.5e-08 | 3.6e-09 | 1.287E-08 | 1.098E-08 | 3.600E-09 | 2.500E-08 |

| | | | | | | | | | | |
|---|---|---|---|---|---|---|---|---|---|---|
| 425 | 22 | D22C | A | 7.3e-09 | 7e-09 | 9.8e-09 | 8.033E-09 | 1.537E-09 | 7.000E-09 | 9.800E-09 |
| 426 | 22 | D22E | A | 1.3e-08 | 6.9e-09 | 4.3e-09 | 8.067E-09 | 4.466E-09 | 4.300E-09 | 1.300E-08 |
| 427 | 22 | D22Q | A | 1.4e-08 | 7.1e-09 | 1.5e-08 | 1.203E-08 | 4.302E-09 | 7.100E-09 | 1.500E-08 |
| 428 | 22 | D22G | A | 1.6e-09 | 5.4e-09 | 5.7e-09 | 4.233E-09 | 2.285E-09 | 1.600E-09 | 5.700E-09 |
| 429 | 22 | D22H | A | 4.7e-09 | 4.8e-09 | 4.2e-09 | 4.567E-09 | 3.215E-10 | 4.200E-09 | 4.800E-09 |
| 430 | 22 | D22I | A | 5.3e-09 | 1.2e-08 | 2.1e-08 | 1.277E-08 | 7.878E-09 | 5.300E-09 | 2.100E-08 |
| 431 | 22 | D22L | A | 7e-09 | 1.7e-08 | 4.8e-09 | 9.600E-09 | 6.502E-09 | 4.800E-09 | 1.700E-08 |
| 432 | 22 | D22K | A | 9.1e-09 | 4.7e-09 | 9.6e-09 | 7.800E-09 | 2.696E-09 | 4.700E-09 | 9.600E-09 |
| 433 | 22 | D22M | A | 3.1e-09 | 1.6e-08 | 4.7e-09 | 7.933E-09 | 7.032E-09 | 3.100E-09 | 1.600E-08 |
| 434 | 22 | D22F | A | 4.8e-09 | 4.6e-09 | 4.9e-09 | 4.767E-09 | 1.528E-10 | 4.600E-09 | 4.900E-09 |
| 435 | 22 | D22P | A | 6.6e-09 | 7.5e-09 | 6.1e-09 | 6.733E-09 | 7.095E-10 | 6.100E-09 | 7.500E-09 |
| 436 | 22 | D22S | A | 4.7e-09 | 7.5e-09 | 6.7e-09 | 6.300E-09 | 1.442E-09 | 4.700E-09 | 7.500E-09 |
| 437 | 22 | D22T | A | 6.3e-09 | 6.8e-09 | 3.7e-09 | 5.600E-09 | 1.664E-09 | 3.700E-09 | 6.800E-09 |
| 438 | 22 | D22W | A | 2.9e-08 | 1e-08 | 8.2e-09 | 1.573E-08 | 1.152E-08 | 8.200E-09 | 2.900E-08 |
| 439 | 22 | D22Y | A | 2.1e-08 | 9.7e-09 | 1.7e-08 | 1.590E-08 | 5.730E-09 | 9.700E-09 | 2.100E-08 |
| 440 | 22 | D22V | A | 1.5e-08 | 5.5e-09 | 3.4e-09 | 7.967E-09 | 6.181E-09 | 3.400E-09 | 1.500E-08 |
| 441 | 23 | M23A | A | 7.6e-09 | 8.7e-09 | 1e-08 | 8.767E-09 | 1.201E-09 | 7.600E-09 | 1.000E-08 |
| 442 | 23 | M23R | A | 1.3e-08 | 8.3e-09 | 1.8e-08 | 1.310E-08 | 4.851E-09 | 8.300E-09 | 1.800E-08 |
| 443 | 23 | M23N | A | 1.2e-08 | 1.6e-08 | 4.1e-09 | 1.070E-08 | 6.056E-09 | 4.100E-09 | 1.600E-08 |
| 444 | 23 | M23D | A | 4.4e-09 | 3.2e-09 | 5.3e-09 | 4.300E-09 | 1.054E-09 | 3.200E-09 | 5.300E-09 |
| 445 | 23 | M23C | A | 7.2e-09 | 5.6e-09 | 1e-08 | 7.600E-09 | 2.227E-09 | 5.600E-09 | 1.000E-08 |
| 446 | 23 | M23E | A | 4e-09 | 6.9e-09 | 1.7e-08 | 9.300E-09 | 6.824E-09 | 4.000E-09 | 1.700E-08 |
| 447 | 23 | M23Q | A | 1e-08 | 1.4e-08 | 9.4e-09 | 1.113E-08 | 2.501E-09 | 9.400E-09 | 1.400E-08 |
| 448 | 23 | M23G | A | 1.2e-08 | 7.4e-09 | 6.5e-09 | 8.633E-09 | 2.950E-09 | 6.500E-09 | 1.200E-08 |
| 449 | 23 | M23H | A | 1.6e-08 | 1.1e-08 | 2.1e-08 | 1.600E-08 | 5.000E-09 | 1.100E-08 | 2.100E-08 |
| 450 | 23 | M23I | A | 5.9e-09 | 2.7e-08 | 5.1e-09 | 1.267E-08 | 1.242E-08 | 5.100E-09 | 2.700E-08 |
| 451 | 23 | M23L | A | 2.4e-08 | 8.6e-09 | 2.6e-09 | 1.173E-08 | 1.104E-08 | 2.600E-09 | 2.400E-08 |
| 452 | 23 | M23K | A | 8.5e-09 | 5.2e-09 | 6.3e-09 | 6.667E-09 | 1.680E-09 | 5.200E-09 | 8.500E-09 |
| 453 | 23 | M23M | A | 1.1e-08 | 9.2e-09 | 1.3e-08 | 1.107E-08 | 1.901E-09 | 9.200E-09 | 1.300E-08 |
| 454 | 23 | M23F | A | 7.7e-09 | 5.2e-09 | 3.9e-09 | 5.600E-09 | 1.931E-09 | 3.900E-09 | 7.700E-09 |
| 455 | 23 | M23P | A | 6.5e-09 | 2.4e-08 | 6.8e-09 | 1.243E-08 | 1.002E-08 | 6.500E-09 | 2.400E-08 |
| 456 | 23 | M23S | A | 7.7e-09 | 6.5e-09 | 1.5e-08 | 9.733E-09 | 4.600E-09 | 6.500E-09 | 1.500E-08 |
| 457 | 23 | M23T | A | 5.1e-09 | 1.2e-08 | 1.1e-08 | 9.367E-09 | 3.729E-09 | 5.100E-09 | 1.200E-08 |
| 458 | 23 | M23W | A | 9.4e-09 | 4.3e-09 | 1.4e-08 | 9.233E-09 | 4.852E-09 | 4.300E-09 | 1.400E-08 |
| 459 | 23 | M23Y | A | 8.5e-09 | 3.6e-09 | 1.4e-08 | 8.700E-09 | 5.203E-09 | 3.600E-09 | 1.400E-08 |
| 460 | 23 | M23V | A | 4.3e-09 | 2.5e-08 | 1.5e-08 | 1.477E-08 | 1.035E-08 | 4.300E-09 | 2.500E-08 |
| 461 | 24 | L24A | A | 1.5e-08 | 1.4e-09 | 2.6e-08 | 1.413E-08 | 1.232E-08 | 1.400E-09 | 2.600E-08 |
| 462 | 24 | L24R | A | 9.9e-09 | 4.5e-09 | 8.3e-09 | 7.567E-09 | 2.774E-09 | 4.500E-09 | 9.900E-09 |
| 463 | 24 | L24N | A | 1e-08 | 7.7e-09 | 1.5e-08 | 1.090E-08 | 3.732E-09 | 7.700E-09 | 1.500E-08 |
| 464 | 24 | L24D | A | 8.1e-09 | 8.8e-09 | 1.3e-08 | 9.967E-09 | 2.650E-09 | 8.100E-09 | 1.300E-08 |
| 465 | 24 | L24C | A | 8.9e-09 | 9.1e-09 | 1.8e-08 | 1.200E-08 | 5.197E-09 | 8.900E-09 | 1.800E-08 |
| 466 | 24 | L24E | A | 5.9e-09 | 6.7e-09 | 1.3e-08 | 8.533E-09 | 3.889E-09 | 5.900E-09 | 1.300E-08 |
| 467 | 24 | L24Q | A | 8.9e-09 | 7.7e-09 | 5.8e-09 | 7.467E-09 | 1.563E-09 | 5.800E-09 | 8.900E-09 |
| 468 | 24 | L24G | A | 6.8e-09 | 2.6e-09 | 5e-09 | 4.800E-09 | 2.107E-09 | 2.600E-09 | 6.800E-09 |
| 469 | 24 | L24H | A | 2.3e-09 | 9.5e-09 | 1.1e-08 | 7.600E-09 | 4.651E-09 | 2.300E-09 | 1.100E-08 |
| 470 | 24 | L24I | A | 7.1e-09 | 8.7e-09 | 5.2e-09 | 7.000E-09 | 1.752E-09 | 5.200E-09 | 8.700E-09 |
| 471 | 24 | L24L | A | 1e-08 | 6.3e-09 | 5.1e-09 | 7.133E-09 | 2.554E-09 | 5.100E-09 | 1.000E-08 |
| 472 | 24 | L24K | A | 5.2e-09 | 5.7e-09 | 4.2e-09 | 5.033E-09 | 7.638E-10 | 4.200E-09 | 5.700E-09 |
| 473 | 24 | L24M | A | 1e-08 | 4.4e-09 | 6.3e-09 | 6.900E-09 | 2.848E-09 | 4.400E-09 | 1.000E-08 |
| 474 | 24 | L24F | A | 5.6e-09 | 8.9e-09 | 9.2e-09 | 7.900E-09 | 1.997E-09 | 5.600E-09 | 9.200E-09 |
| 475 | 24 | L24P | A | 6e-09 | 9.4e-09 | 1.6e-08 | 1.047E-08 | 5.085E-09 | 6.000E-09 | 1.600E-08 |
| 476 | 24 | L24S | A | 7.4e-09 | 2.9e-09 | 3.4e-09 | 4.567E-09 | 2.466E-09 | 2.900E-09 | 7.400E-09 |
| 477 | 24 | L24T | A | 1.7e-08 | 1.4e-08 | 3.3e-09 | 1.143E-08 | 7.202E-09 | 3.300E-09 | 1.700E-08 |
| 478 | 24 | L24W | A | 9.1e-09 | 5e-09 | 4.9e-09 | 6.333E-09 | 2.397E-09 | 4.900E-09 | 9.100E-09 |
| 479 | 24 | L24Y | A | 1.1e-08 | 1.9e-09 | 5.3e-09 | 6.067E-09 | 4.598E-09 | 1.900E-09 | 1.100E-08 |
| 480 | 24 | L24V | A | 8.6e-09 | 5.7e-09 | 5.4e-09 | 6.567E-09 | 1.767E-09 | 5.400E-09 | 8.600E-09 |
| 481 | 25 | R25A | A | 4.6e-09 | 1.2e-08 | 7.5e-09 | 8.033E-09 | 3.729E-09 | 4.600E-09 | 1.200E-08 |
| 482 | 25 | R25R | A | 1.2e-09 | 9.6e-09 | 7.1e-09 | 5.967E-09 | 4.313E-09 | 1.200E-09 | 9.600E-09 |
| 483 | 25 | R25N | A | 1.1e-08 | 3.3e-09 | 4.5e-09 | 6.267E-09 | 4.143E-09 | 3.300E-09 | 1.100E-08 |
| 484 | 25 | R25D | A | 6.7e-09 | 2.5e-08 | 5.9e-09 | 1.253E-08 | 1.080E-08 | 5.900E-09 | 2.500E-08 |
| 485 | 25 | R25C | A | 1e-08 | 1.4e-08 | 1e-08 | 1.133E-08 | 2.309E-09 | 1.000E-08 | 1.400E-08 |
| 486 | 25 | R25E | A | 7.6e-09 | 9.6e-09 | 6.7e-09 | 7.967E-09 | 1.484E-09 | 6.700E-09 | 9.600E-09 |
| 487 | 25 | R25Q | A | 4.1e-09 | 8.4e-09 | 5.5e-09 | 6.000E-09 | 2.193E-09 | 4.100E-09 | 8.400E-09 |
| 488 | 25 | R25G | A | 3.3e-09 | 1.3e-08 | 1.8e-08 | 1.143E-08 | 7.474E-09 | 3.300E-09 | 1.800E-08 |
| 489 | 25 | R25H | A | 4.9e-09 | 8.2e-09 | 4.8e-09 | 5.967E-09 | 1.935E-09 | 4.800E-09 | 8.200E-09 |
| 490 | 25 | R25I | A | 8.4e-09 | 3.9e-09 | 1.1e-08 | 7.767E-09 | 3.592E-09 | 3.900E-09 | 1.100E-08 |
| 491 | 25 | R25L | A | 7.1e-09 | 9.5e-09 | 1.1e-08 | 9.200E-09 | 1.967E-09 | 7.100E-09 | 1.100E-08 |
| 492 | 25 | R25K | A | 1.4e-08 | 1.8e-08 | 5.7e-09 | 1.257E-08 | 6.274E-09 | 5.700E-09 | 1.800E-08 |
| 493 | 25 | R25M | A | 6.7e-09 | 3.8e-09 | 6e-09 | 5.500E-09 | 1.513E-09 | 3.800E-09 | 6.700E-09 |
| 494 | 25 | R25F | A | 2.3e-09 | 8.6e-09 | 9.2e-10 | 3.940E-09 | 4.094E-09 | 9.200E-10 | 8.600E-09 |
| 495 | 25 | R25P | A | 5.6e-09 | 6e-09 | 1.6e-09 | 4.400E-09 | 2.433E-09 | 1.600E-09 | 6.000E-09 |
| 496 | 25 | R25S | A | 1.7e-08 | 1.1e-08 | 6.8e-09 | 1.160E-08 | 5.126E-09 | 6.800E-09 | 1.700E-08 |
| 497 | 25 | R25T | A | 1.5e-08 | 8.7e-09 | 8.7e-09 | 1.080E-08 | 3.637E-09 | 8.700E-09 | 1.500E-08 |
| 498 | 25 | R25W | A | 1.3e-08 | 1.5e-09 | 4.5e-09 | 6.333E-09 | 5.965E-09 | 1.500E-09 | 1.300E-08 |
| 499 | 25 | R25Y | A | 1.9e-09 | 1.2e-08 | 8.2e-09 | 7.367E-09 | 5.101E-09 | 1.900E-09 | 1.200E-08 |
| 500 | 25 | R25V | A | 2.4e-09 | 1.6e-09 | 5.6e-09 | 3.200E-09 | 2.117E-09 | 1.600E-09 | 5.600E-09 |
| 501 | 26 | H26A | A | 1.3e-08 | 5.2e-09 | 8e-09 | 8.733E-09 | 3.951E-09 | 5.200E-09 | 1.300E-08 |
| 502 | 26 | H26R | A | 5.9e-09 | 3.9e-09 | 9.1e-09 | 6.300E-09 | 2.623E-09 | 3.900E-09 | 9.100E-09 |
| 503 | 26 | H26N | A | 1.1e-08 | 4.9e-09 | 6.4e-09 | 7.433E-09 | 3.179E-09 | 4.900E-09 | 1.100E-08 |
| 504 | 26 | H26D | A | 2.2e-09 | 1.1e-08 | 1.2e-08 | 8.400E-09 | 5.393E-09 | 2.200E-09 | 1.200E-08 |
| 505 | 26 | H26C | A | 2.2e-08 | 2.6e-09 | 1.4e-08 | 1.287E-08 | 9.750E-09 | 2.600E-09 | 2.200E-08 |
| 506 | 26 | H26E | A | 5.3e-09 | 7.8e-09 | 6.9e-09 | 6.667E-09 | 1.266E-09 | 5.300E-09 | 7.800E-09 |
| 507 | 26 | H26Q | A | 1e-08 | 9.1e-09 | 1.2e-08 | 1.037E-08 | 1.484E-09 | 9.100E-09 | 1.200E-08 |
| 508 | 26 | H26G | A | 3.9e-09 | 1.2e-08 | 9.1e-09 | 8.333E-09 | 4.104E-09 | 3.900E-09 | 1.200E-08 |
| 509 | 26 | H26H | A | 1.6e-08 | 1.3e-08 | 2.8e-08 | 1.900E-08 | 7.937E-09 | 1.300E-08 | 2.800E-08 |

| | | | | | | | | | | |
|---|---|---|---|---|---|---|---|---|---|---|
| 510 | 26 | H26I | A | 9.4e-09 | 1.6e-08 | 5.8e-09 | 1.040E-08 | 5.173E-09 | 5.800E-09 | 1.600E-08 |
| 511 | 26 | H26L | A | 8.1e-09 | 1.2e-08 | 4.5e-09 | 8.200E-09 | 3.751E-09 | 4.500E-09 | 1.200E-08 |
| 512 | 26 | H26K | A | 3.7e-08 | 6.9e-09 | 4.2e-09 | 1.603E-08 | 1.821E-08 | 4.200E-09 | 3.700E-08 |
| 513 | 26 | H26M | A | 7.9e-09 | 7.1e-09 | 9.3e-09 | 8.100E-09 | 1.114E-09 | 7.100E-09 | 9.300E-09 |
| 514 | 26 | H26F | A | 7.5e-09 | 3e-09 | 4.2e-09 | 4.900E-09 | 2.330E-09 | 3.000E-09 | 7.500E-09 |
| 515 | 26 | H26P | A | 8.5e-09 | 1.7e-08 | 1.2e-08 | 1.250E-08 | 4.272E-09 | 8.500E-09 | 1.700E-08 |
| 516 | 26 | H26S | A | 8.9e-09 | 1.1e-08 | 2e-09 | 7.300E-09 | 4.709E-09 | 2.000E-09 | 1.100E-08 |
| 517 | 26 | H26T | A | 6.6e-09 | 3.6e-09 | 6.1e-09 | 5.433E-09 | 1.607E-09 | 3.600E-09 | 6.600E-09 |
| 518 | 26 | H26W | A | 6.8e-09 | 1.6e-08 | 8.6e-09 | 1.047E-08 | 4.876E-09 | 6.800E-09 | 1.600E-08 |
| 519 | 26 | H26Y | A | 7.8e-09 | 1.1e-08 | 2.3e-09 | 7.033E-09 | 4.400E-09 | 2.300E-09 | 1.100E-08 |
| 520 | 26 | H26V | A | 8.8e-09 | 7.3e-09 | 2e-09 | 6.033E-09 | 3.573E-09 | 2.000E-09 | 8.800E-09 |
| 521 | 27 | L27A | A | 6.9e-09 | 1.4e-08 | 7.8e-09 | 9.567E-09 | 3.866E-09 | 6.900E-09 | 1.400E-08 |
| 522 | 27 | L27R | A | 6.3e-09 | 2.7e-09 | 7.7e-09 | 5.567E-09 | 2.579E-09 | 2.700E-09 | 7.700E-09 |
| 523 | 27 | L27N | A | 2.1e-08 | 4.8e-09 | 4.1e-09 | 9.967E-09 | 9.562E-09 | 4.100E-09 | 2.100E-08 |
| 524 | 27 | L27D | A | 5.4e-09 | 4.6e-09 | 9.2e-09 | 6.400E-09 | 2.458E-09 | 4.600E-09 | 9.200E-09 |
| 525 | 27 | L27C | A | 6e-09 | 6.2e-09 | 5.8e-09 | 6.000E-09 | 2.000E-10 | 5.800E-09 | 6.200E-09 |
| 526 | 27 | L27E | A | 8.3e-09 | 8.8e-09 | 1.9e-08 | 1.203E-08 | 6.038E-09 | 8.300E-09 | 1.900E-08 |
| 527 | 27 | L27Q | A | 2e-08 | 6e-09 | 5e-09 | 1.033E-08 | 8.386E-09 | 5.000E-09 | 2.000E-08 |
| 528 | 27 | L27G | A | 5.9e-09 | 5.3e-09 | 1.3e-08 | 8.067E-09 | 4.283E-09 | 5.300E-09 | 1.300E-08 |
| 529 | 27 | L27H | A | 1.1e-08 | 2.1e-08 | 1.4e-08 | 1.533E-08 | 5.132E-09 | 1.100E-08 | 2.100E-08 |
| 530 | 27 | L27I | A | 8.1e-09 | 7.7e-09 | 3.4e-09 | 6.400E-09 | 2.606E-09 | 3.400E-09 | 8.100E-09 |
| 531 | 27 | L27L | A | 2.3e-08 | 9.7e-09 | 3.1e-09 | 1.193E-08 | 1.014E-08 | 3.100E-09 | 2.300E-08 |
| 532 | 27 | L27K | A | 1.9e-09 | 1.1e-08 | 3e-09 | 5.300E-09 | 4.967E-09 | 1.900E-09 | 1.100E-08 |
| 533 | 27 | L27M | A | 3.2e-09 | 2.5e-09 | 4.3e-09 | 3.333E-09 | 9.074E-10 | 2.500E-09 | 4.300E-09 |
| 534 | 27 | L27F | A | 3.8e-09 | 6.1e-09 | 2.5e-09 | 4.133E-09 | 1.823E-09 | 2.500E-09 | 6.100E-09 |
| 535 | 27 | L27P | A | 5.1e-09 | 2.3e-09 | 1.1e-08 | 6.133E-09 | 4.441E-09 | 2.300E-09 | 1.100E-08 |
| 536 | 27 | L27S | A | 1.5e-08 | 5.9e-09 | 8.9e-09 | 9.933E-09 | 4.637E-09 | 5.900E-09 | 1.500E-08 |
| 537 | 27 | L27T | A | 9.7e-09 | 2e-09 | 1.8e-08 | 9.900E-09 | 8.002E-09 | 2.000E-09 | 1.800E-08 |
| 538 | 27 | L27W | A | 4.2e-09 | 7.7e-09 | 1.7e-08 | 9.633E-09 | 6.615E-09 | 4.200E-09 | 1.700E-08 |
| 539 | 27 | L27Y | A | 4.4e-09 | 6.1e-09 | 1e-08 | 6.833E-09 | 2.871E-09 | 4.400E-09 | 1.000E-08 |
| 540 | 27 | L27V | A | 5.7e-09 | 3.6e-09 | 6.1e-09 | 5.133E-09 | 1.343E-09 | 3.600E-09 | 6.100E-09 |
| 541 | 28 | Y28A | A | 2.2e-08 | 1.5e-08 | 3.4e-09 | 1.347E-08 | 9.394E-09 | 3.400E-09 | 2.200E-08 |
| 542 | 28 | Y28R | A | 2.5e-09 | 2.5e-09 | 8.5e-09 | 4.500E-09 | 3.464E-09 | 2.500E-09 | 8.500E-09 |
| 543 | 28 | Y28N | A | 2e-09 | 4.5e-09 | 9.3e-09 | 5.267E-09 | 3.710E-09 | 2.000E-09 | 9.300E-09 |
| 544 | 28 | Y28D | A | 2.2e-08 | 2.4e-09 | 2.9e-09 | 9.100E-09 | 1.117E-08 | 2.400E-09 | 2.200E-08 |
| 545 | 28 | Y28C | A | 3.4e-09 | 7.6e-09 | 5.4e-09 | 5.467E-09 | 2.101E-09 | 3.400E-09 | 7.600E-09 |
| 546 | 28 | Y28E | A | 1.9e-08 | 1.3e-08 | 1.1e-08 | 1.433E-08 | 4.163E-09 | 1.100E-08 | 1.900E-08 |
| 547 | 28 | Y28Q | A | 3.3e-09 | 1.1e-08 | 5.5e-09 | 6.600E-09 | 3.966E-09 | 3.300E-09 | 1.100E-08 |
| 548 | 28 | Y28G | A | 3.2e-09 | 1.2e-08 | 1.7e-08 | 1.073E-08 | 6.987E-09 | 3.200E-09 | 1.700E-08 |
| 549 | 28 | Y28H | A | 1e-08 | 3.9e-09 | 1.5e-08 | 9.633E-09 | 5.559E-09 | 3.900E-09 | 1.500E-08 |
| 550 | 28 | Y28I | A | 2.2e-08 | 1.3e-08 | 5.9e-09 | 1.363E-08 | 8.069E-09 | 5.900E-09 | 2.200E-08 |
| 551 | 28 | Y28L | A | 4.1e-09 | 4.7e-09 | 3e-09 | 3.933E-09 | 8.622E-10 | 3.000E-09 | 4.700E-09 |
| 552 | 28 | Y28K | A | 1.4e-08 | 3.8e-09 | 5.9e-09 | 7.900E-09 | 5.386E-09 | 3.800E-09 | 1.400E-08 |
| 553 | 28 | Y28M | A | 2.7e-09 | 7.8e-09 | 2.8e-09 | 4.433E-09 | 2.916E-09 | 2.700E-09 | 7.800E-09 |
| 554 | 28 | Y28F | A | 5.1e-09 | 7.5e-09 | 2.7e-09 | 5.100E-09 | 2.400E-09 | 2.700E-09 | 7.500E-09 |
| 555 | 28 | Y28P | A | 6.9e-09 | 9.8e-09 | 1.1e-08 | 9.233E-09 | 2.108E-09 | 6.900E-09 | 1.100E-08 |
| 556 | 28 | Y28S | A | 4.4e-09 | 7.2e-09 | 2.3e-09 | 4.633E-09 | 2.458E-09 | 2.300E-09 | 7.200E-09 |
| 557 | 28 | Y28T | A | 6.9e-09 | 1.6e-08 | 4.6e-09 | 9.167E-09 | 6.029E-09 | 4.600E-09 | 1.600E-08 |
| 558 | 28 | Y28W | A | 4.8e-09 | 6.6e-09 | 1.3e-08 | 8.133E-09 | 4.310E-09 | 4.800E-09 | 1.300E-08 |
| 559 | 28 | Y28Y | A | 8.4e-09 | 5.8e-09 | 1.2e-08 | 8.733E-09 | 3.113E-09 | 5.800E-09 | 1.200E-08 |
| 560 | 28 | Y28V | A | 4.6e-09 | 3.3e-09 | 2.5e-09 | 3.467E-09 | 1.060E-09 | 2.500E-09 | 4.600E-09 |
| 561 | 29 | Q29A | A | 1.1e-08 | 3.5e-09 | 7.9e-09 | 7.467E-09 | 3.769E-09 | 3.500E-09 | 1.100E-08 |
| 562 | 29 | Q29R | A | 2e-09 | 5.8e-09 | 1.1e-08 | 6.267E-09 | 4.518E-09 | 2.000E-09 | 1.100E-08 |
| 563 | 29 | Q29N | A | 1.4e-08 | 8.7e-09 | 4.4e-09 | 9.033E-09 | 4.809E-09 | 4.400E-09 | 1.400E-08 |
| 564 | 29 | Q29D | A | 1e-08 | 1.4e-08 | 1.3e-08 | 1.233E-08 | 2.082E-09 | 1.000E-08 | 1.400E-08 |
| 565 | 29 | Q29C | A | 2.1e-09 | 8.3e-09 | 3.3e-09 | 4.567E-09 | 3.288E-09 | 2.100E-09 | 8.300E-09 |
| 566 | 29 | Q29E | A | 1.5e-08 | 1.3e-08 | 2e-09 | 1.000E-08 | 7.000E-09 | 2.000E-09 | 1.500E-08 |
| 567 | 29 | Q29Q | A | 6.9e-09 | 6.2e-09 | 6.4e-09 | 6.500E-09 | 3.606E-10 | 6.200E-09 | 6.900E-09 |
| 568 | 29 | Q29G | A | 1.6e-08 | 4.9e-09 | 1.1e-08 | 1.063E-08 | 5.559E-09 | 4.900E-09 | 1.600E-08 |
| 569 | 29 | Q29H | A | 8.5e-09 | 1.2e-08 | 5.6e-09 | 8.700E-09 | 3.205E-09 | 5.600E-09 | 1.200E-08 |
| 570 | 29 | Q29I | A | 3e-09 | 8.8e-09 | 5.2e-09 | 5.667E-09 | 2.928E-09 | 3.000E-09 | 8.800E-09 |
| 571 | 29 | Q29L | A | 4.3e-09 | 7.5e-09 | 5.2e-09 | 5.667E-09 | 1.650E-09 | 4.300E-09 | 7.500E-09 |
| 572 | 29 | Q29K | A | 2.8e-09 | 6.9e-09 | 4e-09 | 4.567E-09 | 2.108E-09 | 2.800E-09 | 6.900E-09 |
| 573 | 29 | Q29M | A | 1.2e-08 | 1.3e-08 | 1.1e-08 | 1.200E-08 | 1.000E-09 | 1.100E-08 | 1.300E-08 |
| 574 | 29 | Q29F | A | 6.7e-09 | 2.1e-09 | 7.9e-09 | 5.567E-09 | 3.062E-09 | 2.100E-09 | 7.900E-09 |
| 575 | 29 | Q29P | A | 3e-09 | 9.1e-09 | 2.5e-08 | 1.237E-08 | 1.136E-08 | 3.000E-09 | 2.500E-08 |
| 576 | 29 | Q29S | A | 1.4e-08 | 6.4e-09 | 3.6e-09 | 8.000E-09 | 5.381E-09 | 3.600E-09 | 1.400E-08 |
| 577 | 29 | Q29T | A | 9.5e-09 | 6.5e-09 | 3.5e-09 | 6.500E-09 | 3.000E-09 | 3.500E-09 | 9.500E-09 |
| 578 | 29 | Q29W | A | 6.6e-09 | 1.4e-09 | 3.7e-09 | 3.900E-09 | 2.606E-09 | 1.400E-09 | 6.600E-09 |
| 579 | 29 | Q29Y | A | 1.4e-08 | 2e-09 | 6.6e-09 | 7.533E-09 | 6.054E-09 | 2.000E-09 | 1.400E-08 |
| 580 | 29 | Q29V | A | 7.5e-09 | 9.1e-09 | 6.2e-09 | 7.600E-09 | 1.453E-09 | 6.200E-09 | 9.100E-09 |
| 581 | 30 | G30A | A | 3.5e-09 | 5.9e-09 | 6.1e-09 | 5.167E-09 | 1.447E-09 | 3.500E-09 | 6.100E-09 |
| 582 | 30 | G30R | A | 6.3e-09 | 6e-09 | 4.4e-09 | 5.567E-09 | 1.021E-09 | 4.400E-09 | 6.300E-09 |
| 583 | 30 | G30N | A | 8.1e-09 | 3.1e-09 | 9.2e-09 | 6.800E-09 | 3.251E-09 | 3.100E-09 | 9.200E-09 |
| 584 | 30 | G30D | A | 1.6e-08 | 1.4e-08 | 3.2e-09 | 1.107E-08 | 6.886E-09 | 3.200E-09 | 1.600E-08 |
| 585 | 30 | G30C | A | 2.8e-08 | 6.8e-09 | 3.3e-09 | 1.270E-08 | 1.337E-08 | 3.300E-09 | 2.800E-08 |
| 586 | 30 | G30E | A | 4.8e-09 | 1.1e-08 | 3.3e-09 | 6.367E-09 | 4.082E-09 | 3.300E-09 | 1.100E-08 |
| 587 | 30 | G30Q | A | 6e-09 | 1.7e-09 | 3.6e-09 | 3.767E-09 | 2.155E-09 | 1.700E-09 | 6.000E-09 |
| 588 | 30 | G30G | A | 8.8e-09 | 4.8e-09 | 1.5e-08 | 9.533E-09 | 5.139E-09 | 4.800E-09 | 1.500E-08 |
| 589 | 30 | G30H | A | 1.1e-08 | 8.4e-10 | 4.2e-09 | 5.347E-09 | 5.176E-09 | 8.400E-10 | 1.100E-08 |
| 590 | 30 | G30I | A | 2.2e-09 | 1.2e-08 | 1.8e-08 | 1.073E-08 | 7.976E-09 | 2.200E-09 | 1.800E-08 |
| 591 | 30 | G30L | A | 3.3e-09 | 8.7e-09 | 5.2e-09 | 5.733E-09 | 2.739E-09 | 3.300E-09 | 8.700E-09 |
| 592 | 30 | G30K | A | 8.4e-09 | 9.5e-09 | 3.4e-09 | 7.100E-09 | 3.251E-09 | 3.400E-09 | 9.500E-09 |
| 593 | 30 | G30M | A | 6.9e-09 | 1.1e-08 | 1e-08 | 9.300E-09 | 2.138E-09 | 6.900E-09 | 1.100E-08 |
| 594 | 30 | G30F | A | 1.5e-08 | 9.4e-09 | 1.8e-08 | 1.413E-08 | 4.365E-09 | 9.400E-09 | 1.800E-08 |

| | | | | | | | | | | |
|---|---|---|---|---|---|---|---|---|---|---|
| 595 | 30 | G30P | A | 1.1e-08 | 6.9e-09 | 7.7e-09 | 8.533E-09 | 2.173E-09 | 6.900E-09 | 1.100E-08 |
| 596 | 30 | G30S | A | 3e-08 | 6.9e-09 | 6.8e-09 | 1.457E-08 | 1.337E-08 | 6.800E-09 | 3.000E-08 |
| 597 | 30 | G30T | A | 8.5e-09 | 6.8e-09 | 4.7e-09 | 6.667E-09 | 1.904E-09 | 4.700E-09 | 8.500E-09 |
| 598 | 30 | G30W | A | 3.9e-09 | 1.1e-08 | 4.4e-09 | 6.433E-09 | 3.963E-09 | 3.900E-09 | 1.100E-08 |
| 599 | 30 | G30Y | A | 7.6e-09 | 1.2e-08 | 1.8e-09 | 7.133E-09 | 5.116E-09 | 1.800E-09 | 1.200E-08 |
| 600 | 30 | G30V | A | 8.4e-09 | 3.1e-09 | 7.8e-09 | 6.433E-09 | 2.902E-09 | 3.100E-09 | 8.400E-09 |
| 601 | 31 | C31A | A | 1e-08 | 9.7e-09 | 1.1e-08 | 1.023E-08 | 6.807E-10 | 9.700E-09 | 1.100E-08 |
| 602 | 31 | C31R | A | 4.1e-08 | 1e-08 | 3e-09 | 1.800E-08 | 2.022E-08 | 3.000E-09 | 4.100E-08 |
| 603 | 31 | C31N | A | 1.6e-08 | 1.5e-08 | 1.2e-08 | 1.433E-08 | 2.082E-09 | 1.200E-08 | 1.600E-08 |
| 604 | 31 | C31D | A | 1.5e-08 | 1.1e-08 | 1.5e-08 | 1.367E-08 | 2.309E-09 | 1.100E-08 | 1.500E-08 |
| 605 | 31 | C31C | A | 7.9e-09 | 1.5e-08 | 1.2e-08 | 1.163E-08 | 3.564E-09 | 7.900E-09 | 1.500E-08 |
| 606 | 31 | C31E | A | 1.2e-08 | 7.3e-09 | 3.6e-09 | 7.633E-09 | 4.210E-09 | 3.600E-09 | 1.200E-08 |
| 607 | 31 | C31Q | A | 3.2e-09 | 6.2e-09 | 8.7e-09 | 6.033E-09 | 2.754E-09 | 3.200E-09 | 8.700E-09 |
| 608 | 31 | C31G | A | 1.5e-08 | 2.3e-08 | 3.2e-09 | 1.373E-08 | 9.961E-09 | 3.200E-09 | 2.300E-08 |
| 609 | 31 | C31H | A | 2.9e-09 | 5.8e-09 | 1.6e-08 | 8.233E-09 | 6.881E-09 | 2.900E-09 | 1.600E-08 |
| 610 | 31 | C31I | A | 1.1e-08 | 1.4e-08 | 4.3e-09 | 9.767E-09 | 4.966E-09 | 4.300E-09 | 1.400E-08 |
| 611 | 31 | C31L | A | 8.3e-09 | 1.8e-08 | 3.4e-09 | 9.900E-09 | 7.430E-09 | 3.400E-09 | 1.800E-08 |
| 612 | 31 | C31K | A | 8.3e-09 | 2e-08 | 9.1e-09 | 1.247E-08 | 6.536E-09 | 8.300E-09 | 2.000E-08 |
| 613 | 31 | C31M | A | 5.8e-09 | 5.9e-09 | 1.2e-08 | 7.900E-09 | 3.551E-09 | 5.800E-09 | 1.200E-08 |
| 614 | 31 | C31F | A | 1.1e-08 | 1.2e-08 | 1.1e-08 | 1.133E-08 | 5.774E-10 | 1.100E-08 | 1.200E-08 |
| 615 | 31 | C31P | A | 8.2e-09 | 6.7e-09 | 6.8e-09 | 7.233E-09 | 8.386E-10 | 6.700E-09 | 8.200E-09 |
| 616 | 31 | C31S | A | 1.7e-08 | 5.5e-09 | 5.3e-09 | 9.267E-09 | 6.698E-09 | 5.300E-09 | 1.700E-08 |
| 617 | 31 | C31T | A | 3.8e-09 | 2.7e-09 | 1.4e-08 | 6.833E-09 | 6.231E-09 | 2.700E-09 | 1.400E-08 |
| 618 | 31 | C31W | A | 1.1e-08 | 1.3e-08 | 4.6e-09 | 9.533E-09 | 4.388E-09 | 4.600E-09 | 1.300E-08 |
| 619 | 31 | C31Y | A | 1.4e-08 | 1.2e-08 | 1.2e-08 | 1.267E-08 | 1.155E-09 | 1.200E-08 | 1.400E-08 |
| 620 | 31 | C31V | A | 6.4e-09 | 6.9e-09 | 4.6e-09 | 5.967E-09 | 1.210E-09 | 4.600E-09 | 6.900E-09 |
| 621 | 32 | Q32A | A | 2.5e-08 | 6.3e-09 | 2.6e-08 | 1.910E-08 | 1.110E-08 | 6.300E-09 | 2.600E-08 |
| 622 | 32 | Q32R | A | 1.4e-08 | 1.1e-08 | 6.8e-09 | 1.060E-08 | 3.617E-09 | 6.800E-09 | 1.400E-08 |
| 623 | 32 | Q32N | A | 8.2e-09 | 1e-08 | 4.7e-09 | 7.633E-09 | 2.695E-09 | 4.700E-09 | 1.000E-08 |
| 624 | 32 | Q32D | A | 3.6e-08 | 4.1e-09 | 2.4e-08 | 2.137E-08 | 1.611E-08 | 4.100E-09 | 3.600E-08 |
| 625 | 32 | Q32C | A | 1.1e-08 | 8.8e-09 | 6.2e-09 | 8.667E-09 | 2.403E-09 | 6.200E-09 | 1.100E-08 |
| 626 | 32 | Q32E | A | 3.1e-09 | 1e-08 | 8.5e-09 | 7.200E-09 | 3.629E-09 | 3.100E-09 | 1.000E-08 |
| 627 | 32 | Q32Q | A | 5.1e-09 | 9.4e-09 | 1.6e-08 | 1.017E-08 | 5.490E-09 | 5.100E-09 | 1.600E-08 |
| 628 | 32 | Q32G | A | 1.9e-08 | 7e-09 | 7.8e-09 | 1.127E-08 | 6.709E-09 | 7.000E-09 | 1.900E-08 |
| 629 | 32 | Q32H | A | 7.6e-09 | 9.3e-09 | 6.8e-09 | 7.900E-09 | 1.277E-09 | 6.800E-09 | 9.300E-09 |
| 630 | 32 | Q32I | A | 4e-09 | 5.9e-09 | 1.1e-08 | 6.967E-09 | 3.620E-09 | 4.000E-09 | 1.100E-08 |
| 631 | 32 | Q32L | A | 7.6e-09 | 5.4e-09 | 1.1e-08 | 8.000E-09 | 2.821E-09 | 5.400E-09 | 1.100E-08 |
| 632 | 32 | Q32K | A | 2.1e-08 | 1.9e-08 | 6.2e-09 | 1.540E-08 | 8.030E-09 | 6.200E-09 | 2.100E-08 |
| 633 | 32 | Q32M | A | 9.5e-09 | 1.9e-08 | 7.6e-09 | 1.203E-08 | 6.108E-09 | 7.600E-09 | 1.900E-08 |
| 634 | 32 | Q32F | A | 3.6e-09 | 4.3e-09 | 3.7e-09 | 3.867E-09 | 3.786E-10 | 3.600E-09 | 4.300E-09 |
| 635 | 32 | Q32P | A | 6.1e-09 | 4.1e-09 | 6.7e-09 | 5.633E-09 | 1.361E-09 | 4.100E-09 | 6.700E-09 |
| 636 | 32 | Q32S | A | 4.4e-09 | 4.9e-09 | 6e-09 | 5.100E-09 | 8.185E-10 | 4.400E-09 | 6.000E-09 |
| 637 | 32 | Q32T | A | 4.1e-09 | 5.8e-09 | 9.9e-10 | 3.630E-09 | 2.439E-09 | 9.900E-10 | 5.800E-09 |
| 638 | 32 | Q32W | A | 8e-09 | 5.3e-09 | 7.9e-09 | 7.067E-09 | 1.531E-09 | 5.300E-09 | 8.000E-09 |
| 639 | 32 | Q32Y | A | 5.4e-09 | 4.8e-09 | 5.9e-09 | 5.367E-09 | 5.508E-10 | 4.800E-09 | 5.900E-09 |
| 640 | 32 | Q32V | A | 8e-09 | 4.7e-09 | 9.4e-09 | 7.367E-09 | 2.413E-09 | 4.700E-09 | 9.400E-09 |
| 641 | 33 | V33A | A | 3.9e-09 | 3e-09 | 4e-09 | 3.633E-09 | 5.508E-10 | 3.000E-09 | 4.000E-09 |
| 642 | 33 | V33R | A | 3.9e-09 | 1.8e-08 | 7.7e-09 | 9.867E-09 | 7.295E-09 | 3.900E-09 | 1.800E-08 |
| 643 | 33 | V33N | A | 4.8e-09 | 1.2e-08 | 7e-09 | 7.933E-09 | 3.690E-09 | 4.800E-09 | 1.200E-08 |
| 644 | 33 | V33D | A | 1.2e-08 | 6.6e-09 | 8.9e-09 | 9.167E-09 | 2.710E-09 | 6.600E-09 | 1.200E-08 |
| 645 | 33 | V33C | A | 2.9e-09 | 8e-09 | 6.2e-09 | 5.700E-09 | 2.587E-09 | 2.900E-09 | 8.000E-09 |
| 646 | 33 | V33E | A | 6.4e-09 | 3.3e-09 | 6.2e-09 | 5.300E-09 | 1.735E-09 | 3.300E-09 | 6.400E-09 |
| 647 | 33 | V33Q | A | 1e-08 | 3.7e-09 | 8.4e-09 | 7.367E-09 | 3.275E-09 | 3.700E-09 | 1.000E-08 |
| 648 | 33 | V33G | A | 1.3e-08 | 3e-09 | 1.2e-08 | 9.333E-09 | 5.508E-09 | 3.000E-09 | 1.300E-08 |
| 649 | 33 | V33H | A | 9.7e-09 | 3.4e-09 | 4.9e-09 | 6.000E-09 | 3.291E-09 | 3.400E-09 | 9.700E-09 |
| 650 | 33 | V33I | A | 4.7e-09 | 3.5e-09 | 1.2e-08 | 6.733E-09 | 4.600E-09 | 3.500E-09 | 1.200E-08 |
| 651 | 33 | V33L | A | 4.8e-09 | 7.1e-09 | 1.4e-08 | 8.633E-09 | 4.788E-09 | 4.800E-09 | 1.400E-08 |
| 652 | 33 | V33K | A | 1.4e-08 | 4.6e-09 | 6.4e-09 | 8.333E-09 | 4.989E-09 | 4.600E-09 | 1.400E-08 |
| 653 | 33 | V33M | A | 8.4e-09 | 4.6e-09 | 2.9e-09 | 5.300E-09 | 2.816E-09 | 2.900E-09 | 8.400E-09 |
| 654 | 33 | V33F | A | 4e-09 | 5.1e-09 | 1.5e-08 | 8.033E-09 | 6.058E-09 | 4.000E-09 | 1.500E-08 |
| 655 | 33 | V33P | A | 2.1e-08 | 5.2e-09 | 3.4e-09 | 9.867E-09 | 9.684E-09 | 3.400E-09 | 2.100E-08 |
| 656 | 33 | V33S | A | 1e-08 | 3e-09 | 3.9e-09 | 5.633E-09 | 3.808E-09 | 3.000E-09 | 1.000E-08 |
| 657 | 33 | V33T | A | 3.4e-09 | 7.4e-09 | 9.4e-09 | 6.733E-09 | 3.055E-09 | 3.400E-09 | 9.400E-09 |
| 658 | 33 | V33W | A | 5.6e-09 | 3.7e-09 | 7.2e-09 | 5.500E-09 | 1.752E-09 | 3.700E-09 | 7.200E-09 |
| 659 | 33 | V33Y | A | 5.6e-09 | 7.9e-09 | 4.8e-09 | 6.100E-09 | 1.609E-09 | 4.800E-09 | 7.900E-09 |
| 660 | 33 | V33V | A | 1.2e-08 | 7e-09 | 4.1e-09 | 7.700E-09 | 3.996E-09 | 4.100E-09 | 1.200E-08 |
| 661 | 34 | V34A | A | 9.7e-09 | 3.7e-09 | 6.7e-09 | 6.700E-09 | 3.000E-09 | 3.700E-09 | 9.700E-09 |
| 662 | 34 | V34R | A | 5.8e-09 | 5.5e-09 | 2.1e-08 | 1.077E-08 | 8.864E-09 | 5.500E-09 | 2.100E-08 |
| 663 | 34 | V34N | A | 3.6e-09 | 1.6e-08 | 3.9e-09 | 7.833E-09 | 7.074E-09 | 3.600E-09 | 1.600E-08 |
| 664 | 34 | V34D | A | 9.3e-09 | 9.6e-09 | 9.2e-09 | 9.367E-09 | 2.082E-10 | 9.200E-09 | 9.600E-09 |
| 665 | 34 | V34C | A | 4.1e-09 | 4.6e-09 | 5.1e-09 | 4.600E-09 | 5.000E-10 | 4.100E-09 | 5.100E-09 |
| 666 | 34 | V34E | A | 4.8e-09 | 8.9e-09 | 5.3e-09 | 6.333E-09 | 2.237E-09 | 4.800E-09 | 8.900E-09 |
| 667 | 34 | V34Q | A | 6.3e-09 | 4.2e-09 | 8e-09 | 6.167E-09 | 1.904E-09 | 4.200E-09 | 8.000E-09 |
| 668 | 34 | V34G | A | 5.3e-09 | 7.3e-09 | 1.3e-08 | 8.533E-09 | 3.995E-09 | 5.300E-09 | 1.300E-08 |
| 669 | 34 | V34H | A | 1.1e-08 | 7e-09 | 3.6e-09 | 7.200E-09 | 3.704E-09 | 3.600E-09 | 1.100E-08 |
| 670 | 34 | V34I | A | 7e-09 | 1e-08 | 5.5e-09 | 7.500E-09 | 2.291E-09 | 5.500E-09 | 1.000E-08 |
| 671 | 34 | V34L | A | 7.2e-09 | 5.7e-09 | 5.9e-09 | 6.267E-09 | 8.145E-10 | 5.700E-09 | 7.200E-09 |
| 672 | 34 | V34K | A | 2.9e-09 | 9.1e-09 | 6.9e-09 | 6.300E-09 | 3.143E-09 | 2.900E-09 | 9.100E-09 |
| 673 | 34 | V34M | A | 1e-08 | 1.7e-08 | 7.2e-09 | 1.140E-08 | 5.048E-09 | 7.200E-09 | 1.700E-08 |
| 674 | 34 | V34F | A | 4.4e-09 | 2.4e-09 | 7.4e-09 | 4.733E-09 | 2.517E-09 | 2.400E-09 | 7.400E-09 |
| 675 | 34 | V34P | A | 1.8e-08 | 7.5e-09 | 6e-09 | 1.050E-08 | 6.538E-09 | 6.000E-09 | 1.800E-08 |
| 676 | 34 | V34S | A | 2.1e-09 | 7.1e-09 | 3.3e-09 | 4.167E-09 | 2.610E-09 | 2.100E-09 | 7.100E-09 |
| 677 | 34 | V34T | A | 3.6e-09 | 1.1e-08 | 1.9e-08 | 1.120E-08 | 7.702E-09 | 3.600E-09 | 1.900E-08 |
| 678 | 34 | V34W | A | 8.1e-09 | 5.9e-09 | 1.1e-08 | 8.333E-09 | 2.558E-09 | 5.900E-09 | 1.100E-08 |
| 679 | 34 | V34Y | A | 7.6e-09 | 2e-08 | 8.6e-09 | 1.207E-08 | 6.889E-09 | 7.600E-09 | 2.000E-08 |

| | | | | | | | | | | |
|---|---|---|---|---|---|---|---|---|---|---|
| 680 | 34 | V34V | A | 9.5e-09 | 5.8e-09 | 5e-09 | 6.767E-09 | 2.401E-09 | 5.000E-09 | 9.500E-09 |
| 681 | 35 | Q35A | A | 9.5e-09 | 1.3e-08 | 1.3e-08 | 1.183E-08 | 2.021E-09 | 9.500E-09 | 1.300E-08 |
| 682 | 35 | Q35R | A | 7.3e-09 | 8.1e-09 | 1e-08 | 8.467E-09 | 1.387E-09 | 7.300E-09 | 1.000E-08 |
| 683 | 35 | Q35N | A | 3.4e-09 | 1e-08 | 2.6e-09 | 5.333E-09 | 4.061E-09 | 2.600E-09 | 1.000E-08 |
| 684 | 35 | Q35D | A | 8.8e-09 | 4e-09 | 2.9e-09 | 5.233E-09 | 3.137E-09 | 2.900E-09 | 8.800E-09 |
| 685 | 35 | Q35C | A | 9.1e-09 | 1.4e-08 | 8.3e-09 | 1.047E-08 | 3.086E-09 | 8.300E-09 | 1.400E-08 |
| 686 | 35 | Q35E | A | 8.3e-09 | 1.7e-08 | 1.1e-08 | 1.210E-08 | 4.453E-09 | 8.300E-09 | 1.700E-08 |
| 687 | 35 | Q35Q | A | 8.3e-09 | 6.5e-09 | 2e-09 | 5.600E-09 | 3.245E-09 | 2.000E-09 | 8.300E-09 |
| 688 | 35 | Q35G | A | 6e-09 | 3.5e-09 | 1.1e-08 | 6.833E-09 | 3.819E-09 | 3.500E-09 | 1.100E-08 |
| 689 | 35 | Q35H | A | 7.1e-09 | 9.4e-09 | 1.2e-08 | 9.500E-09 | 2.452E-09 | 7.100E-09 | 1.200E-08 |
| 690 | 35 | Q35I | A | 9.1e-09 | 1.1e-08 | 7.2e-09 | 9.100E-09 | 1.900E-09 | 7.200E-09 | 1.100E-08 |
| 691 | 35 | Q35L | A | 1.1e-08 | 1.3e-08 | 5.5e-09 | 9.833E-09 | 3.884E-09 | 5.500E-09 | 1.300E-08 |
| 692 | 35 | Q35K | A | 5.5e-09 | 2.9e-09 | 7.4e-09 | 5.267E-09 | 2.259E-09 | 2.900E-09 | 7.400E-09 |
| 693 | 35 | Q35M | A | 1.6e-08 | 1e-08 | 5.8e-09 | 1.060E-08 | 5.126E-09 | 5.800E-09 | 1.600E-08 |
| 694 | 35 | Q35F | A | 1.8e-09 | 4.6e-09 | 9.9e-09 | 5.433E-09 | 4.114E-09 | 1.800E-09 | 9.900E-09 |
| 695 | 35 | Q35P | A | 3.6e-09 | 6.8e-09 | 4.5e-09 | 4.967E-09 | 1.650E-09 | 3.600E-09 | 6.800E-09 |
| 696 | 35 | Q35S | A | 1.9e-08 | 5.2e-09 | 4e-09 | 9.400E-09 | 8.335E-09 | 4.000E-09 | 1.900E-08 |
| 697 | 35 | Q35T | A | 1e-08 | 3.5e-09 | 1.4e-08 | 9.167E-09 | 5.299E-09 | 3.500E-09 | 1.400E-08 |
| 698 | 35 | Q35W | A | 6.5e-09 | 1.5e-08 | 5e-09 | 8.833E-09 | 5.393E-09 | 5.000E-09 | 1.500E-08 |
| 699 | 35 | Q35Y | A | 1.2e-08 | 1.2e-08 | 7.2e-09 | 1.040E-08 | 2.771E-09 | 7.200E-09 | 1.200E-08 |
| 700 | 35 | Q35V | A | 7.9e-09 | 1.5e-08 | 5.6e-09 | 9.500E-09 | 4.900E-09 | 5.600E-09 | 1.500E-08 |
| 701 | 36 | G36A | A | 5.4e-09 | 5.3e-09 | 2.3e-09 | 4.333E-09 | 1.762E-09 | 2.300E-09 | 5.400E-09 |
| 702 | 36 | G36R | A | 7.9e-09 | 1.8e-09 | 9.5e-09 | 6.400E-09 | 4.063E-09 | 1.800E-09 | 9.500E-09 |
| 703 | 36 | G36N | A | 6.3e-09 | 1.3e-08 | 2e-09 | 7.100E-09 | 5.543E-09 | 2.000E-09 | 1.300E-08 |
| 704 | 36 | G36D | A | 6.2e-09 | 5.2e-09 | 9.8e-09 | 7.067E-09 | 2.419E-09 | 5.200E-09 | 9.800E-09 |
| 705 | 36 | G36C | A | 5.8e-09 | 1.8e-08 | 4.7e-09 | 9.500E-09 | 7.382E-09 | 4.700E-09 | 1.800E-08 |
| 706 | 36 | G36E | A | 7.2e-09 | 8.1e-09 | 6.9e-09 | 7.400E-09 | 6.245E-10 | 6.900E-09 | 8.100E-09 |
| 707 | 36 | G36Q | A | 4.7e-09 | 2.1e-09 | 4.4e-09 | 3.733E-09 | 1.422E-09 | 2.100E-09 | 4.700E-09 |
| 708 | 36 | G36G | A | 9.2e-09 | 1.3e-08 | 3e-09 | 8.400E-09 | 5.048E-09 | 3.000E-09 | 1.300E-08 |
| 709 | 36 | G36H | A | 8.9e-09 | 3.9e-09 | 1.1e-08 | 7.933E-09 | 3.647E-09 | 3.900E-09 | 1.100E-08 |
| 710 | 36 | G36I | A | 2.6e-08 | 8.5e-09 | 3.6e-09 | 1.270E-08 | 1.178E-08 | 3.600E-09 | 2.600E-08 |
| 711 | 36 | G36L | A | 2.1e-08 | 9.5e-09 | 9.2e-09 | 1.323E-08 | 6.728E-09 | 9.200E-09 | 2.100E-08 |
| 712 | 36 | G36K | A | 2.6e-08 | 5.2e-09 | 5.9e-09 | 1.237E-08 | 1.181E-08 | 5.200E-09 | 2.600E-08 |
| 713 | 36 | G36M | A | 2.6e-09 | 1.1e-08 | 6.6e-09 | 6.733E-09 | 4.202E-09 | 2.600E-09 | 1.100E-08 |
| 714 | 36 | G36F | A | 7e-09 | 6.1e-09 | 4.4e-09 | 5.833E-09 | 1.320E-09 | 4.400E-09 | 7.000E-09 |
| 715 | 36 | G36P | A | 2.6e-08 | 8.7e-09 | 5.2e-09 | 1.330E-08 | 1.114E-08 | 5.200E-09 | 2.600E-08 |
| 716 | 36 | G36S | A | 4.8e-09 | 9.3e-09 | 7.1e-09 | 7.067E-09 | 2.250E-09 | 4.800E-09 | 9.300E-09 |
| 717 | 36 | G36T | A | 5.6e-09 | 3e-08 | 7.7e-09 | 1.443E-08 | 1.352E-08 | 5.600E-09 | 3.000E-08 |
| 718 | 36 | G36W | A | 6.4e-09 | 4.8e-09 | 7.1e-09 | 6.100E-09 | 1.179E-09 | 4.800E-09 | 7.100E-09 |
| 719 | 36 | G36Y | A | 3.8e-09 | 2.9e-09 | 1.1e-08 | 5.900E-09 | 4.440E-09 | 2.900E-09 | 1.100E-08 |
| 720 | 36 | G36V | A | 7.1e-09 | 4.6e-09 | 1.2e-08 | 7.900E-09 | 3.764E-09 | 4.600E-09 | 1.200E-08 |
| 721 | 37 | N37A | A | 1e-08 | 1e-08 | 3.5e-09 | 7.833E-09 | 3.753E-09 | 3.500E-09 | 1.000E-08 |
| 722 | 37 | N37R | A | 5.7e-09 | 9.5e-09 | 1e-08 | 8.400E-09 | 2.352E-09 | 5.700E-09 | 1.000E-08 |
| 723 | 37 | N37N | A | 8.4e-09 | 2.4e-09 | 1.1e-08 | 7.267E-09 | 4.411E-09 | 2.400E-09 | 1.100E-08 |
| 724 | 37 | N37D | A | 1e-08 | 9.7e-09 | 2.8e-09 | 7.500E-09 | 4.073E-09 | 2.800E-09 | 1.000E-08 |
| 725 | 37 | N37C | A | 7.3e-09 | 1.3e-08 | 6.5e-09 | 8.933E-09 | 3.544E-09 | 6.500E-09 | 1.300E-08 |
| 726 | 37 | N37E | A | 5.2e-09 | 7.2e-09 | 1e-08 | 7.467E-09 | 2.411E-09 | 5.200E-09 | 1.000E-08 |
| 727 | 37 | N37Q | A | 9.1e-09 | 8.5e-09 | 1.6e-08 | 1.120E-08 | 4.168E-09 | 8.500E-09 | 1.600E-08 |
| 728 | 37 | N37G | A | 2.7e-09 | 5.4e-09 | 6.2e-09 | 4.767E-09 | 1.834E-09 | 2.700E-09 | 6.200E-09 |
| 729 | 37 | N37H | A | 4.2e-09 | 9.5e-09 | 2.4e-08 | 1.257E-08 | 1.025E-08 | 4.200E-09 | 2.400E-08 |
| 730 | 37 | N37I | A | 4.6e-09 | 3e-09 | 3.7e-09 | 3.767E-09 | 8.021E-10 | 3.000E-09 | 4.600E-09 |
| 731 | 37 | N37L | A | 7.5e-09 | 3.4e-09 | 7.3e-09 | 6.067E-09 | 2.312E-09 | 3.400E-09 | 7.500E-09 |
| 732 | 37 | N37K | A | 5.9e-09 | 8e-09 | 4.1e-09 | 6.000E-09 | 1.952E-09 | 4.100E-09 | 8.000E-09 |
| 733 | 37 | N37M | A | 6e-09 | 3.7e-09 | 8.9e-09 | 6.200E-09 | 2.606E-09 | 3.700E-09 | 8.900E-09 |
| 734 | 37 | N37F | A | 4.9e-09 | 2.3e-09 | 1.3e-08 | 6.733E-09 | 5.581E-09 | 2.300E-09 | 1.300E-08 |
| 735 | 37 | N37P | A | 3.8e-09 | 1.4e-08 | 6.3e-09 | 8.033E-09 | 5.316E-09 | 3.800E-09 | 1.400E-08 |
| 736 | 37 | N37S | A | 4.1e-09 | 4.7e-09 | 2.6e-09 | 3.800E-09 | 1.082E-09 | 2.600E-09 | 4.700E-09 |
| 737 | 37 | N37T | A | 8.5e-09 | 8.2e-09 | 6.1e-09 | 7.600E-09 | 1.308E-09 | 6.100E-09 | 8.500E-09 |
| 738 | 37 | N37W | A | 7.4e-09 | 2.1e-09 | 2.8e-09 | 4.100E-09 | 2.879E-09 | 2.100E-09 | 7.400E-09 |
| 739 | 37 | N37Y | A | 3.6e-09 | 5.6e-09 | 8e-09 | 5.733E-09 | 2.203E-09 | 3.600E-09 | 8.000E-09 |
| 740 | 37 | N37V | A | 7.7e-09 | 9e-09 | 8.5e-09 | 8.400E-09 | 6.557E-10 | 7.700E-09 | 9.000E-09 |
| 741 | 38 | L38A | A | 5.4e-09 | 4.7e-09 | 1.1e-08 | 7.033E-09 | 3.453E-09 | 4.700E-09 | 1.100E-08 |
| 742 | 38 | L38R | A | 1.2e-08 | 6.7e-09 | 3e-09 | 7.233E-09 | 4.524E-09 | 3.000E-09 | 1.200E-08 |
| 743 | 38 | L38N | A | 7.1e-09 | 9.1e-09 | 7.9e-09 | 8.033E-09 | 1.007E-09 | 7.100E-09 | 9.100E-09 |
| 744 | 38 | L38D | A | 5.1e-09 | 1.7e-08 | 1e-08 | 1.070E-08 | 5.981E-09 | 5.100E-09 | 1.700E-08 |
| 745 | 38 | L38C | A | 7.8e-09 | 1.1e-08 | 6.2e-09 | 8.333E-09 | 2.444E-09 | 6.200E-09 | 1.100E-08 |
| 746 | 38 | L38E | A | 5.2e-09 | 2.5e-09 | 1.4e-08 | 7.233E-09 | 6.014E-09 | 2.500E-09 | 1.400E-08 |
| 747 | 38 | L38Q | A | 1.3e-08 | 7.5e-09 | 1.2e-08 | 1.083E-08 | 2.930E-09 | 7.500E-09 | 1.300E-08 |
| 748 | 38 | L38G | A | 5.4e-09 | 3.3e-09 | 2e-09 | 3.567E-09 | 1.716E-09 | 2.000E-09 | 5.400E-09 |
| 749 | 38 | L38H | A | 1.6e-08 | 9.2e-09 | 3.2e-09 | 9.467E-09 | 6.404E-09 | 3.200E-09 | 1.600E-08 |
| 750 | 38 | L38I | A | 8.1e-09 | 5.6e-09 | 5.9e-09 | 6.533E-09 | 1.365E-09 | 5.600E-09 | 8.100E-09 |
| 751 | 38 | L38L | A | 6.3e-09 | 1.2e-08 | 1.5e-08 | 1.110E-08 | 4.419E-09 | 6.300E-09 | 1.500E-08 |
| 752 | 38 | L38K | A | 2.1e-09 | 2.9e-08 | 7.1e-09 | 1.273E-08 | 1.431E-08 | 2.100E-09 | 2.900E-08 |
| 753 | 38 | L38M | A | 6.8e-09 | 1.1e-08 | 9.7e-09 | 9.167E-09 | 2.150E-09 | 6.800E-09 | 1.100E-08 |
| 754 | 38 | L38F | A | 4.4e-09 | 2.5e-09 | 4.5e-09 | 3.800E-09 | 1.127E-09 | 2.500E-09 | 4.500E-09 |
| 755 | 38 | L38P | A | 1.6e-08 | 8.3e-09 | 2.3e-09 | 8.867E-09 | 6.868E-09 | 2.300E-09 | 1.600E-08 |
| 756 | 38 | L38S | A | 1.2e-08 | 2.5e-09 | 2e-08 | 1.150E-08 | 8.761E-09 | 2.500E-09 | 2.000E-08 |
| 757 | 38 | L38T | A | 1.2e-08 | 8e-09 | 6e-09 | 8.667E-09 | 3.055E-09 | 6.000E-09 | 1.200E-08 |
| 758 | 38 | L38W | A | 1.8e-08 | 6.3e-09 | 4.5e-09 | 9.600E-09 | 7.330E-09 | 4.500E-09 | 1.800E-08 |
| 759 | 38 | L38Y | A | 6.4e-09 | 1.4e-08 | 1.3e-08 | 1.113E-08 | 4.130E-09 | 6.400E-09 | 1.400E-08 |
| 760 | 38 | L38V | A | 1.1e-08 | 1.1e-08 | 4.1e-09 | 8.700E-09 | 3.984E-09 | 4.100E-09 | 1.100E-08 |
| 761 | 39 | E39A | A | 5.8e-09 | 4.8e-09 | 5.2e-09 | 5.267E-09 | 5.033E-10 | 4.800E-09 | 5.800E-09 |
| 762 | 39 | E39R | A | 2.7e-09 | 1.1e-08 | 9.6e-09 | 7.767E-09 | 4.443E-09 | 2.700E-09 | 1.100E-08 |
| 763 | 39 | E39N | A | 3.4e-09 | 1e-08 | 9.6e-09 | 7.667E-09 | 3.700E-09 | 3.400E-09 | 1.000E-08 |
| 764 | 39 | E39D | A | 1.7e-08 | 5.5e-09 | 9.4e-09 | 1.063E-08 | 5.848E-09 | 5.500E-09 | 1.700E-08 |

| | | | | | | | | | | |
|---|---|---|---|---|---|---|---|---|---|---|
| 765 | 39 | E39C | A | 2.2e-09 | 9.8e-09 | 6.3e-09 | 6.100E-09 | 3.804E-09 | 2.200E-09 | 9.800E-09 |
| 766 | 39 | E39E | A | 4.7e-09 | 1.9e-09 | 5.7e-09 | 4.100E-09 | 1.970E-09 | 1.900E-09 | 5.700E-09 |
| 767 | 39 | E39Q | A | 2.1e-09 | 6.6e-09 | 1e-08 | 6.233E-09 | 3.963E-09 | 2.100E-09 | 1.000E-08 |
| 768 | 39 | E39G | A | 1.3e-08 | 6.5e-09 | 6.7e-09 | 8.733E-09 | 3.696E-09 | 6.500E-09 | 1.300E-08 |
| 769 | 39 | E39H | A | 2.7e-08 | 1.4e-08 | 9.1e-09 | 1.670E-08 | 9.250E-09 | 9.100E-09 | 2.700E-08 |
| 770 | 39 | E39I | A | 9.3e-09 | 3.5e-09 | 3.3e-09 | 5.367E-09 | 3.408E-09 | 3.300E-09 | 9.300E-09 |
| 771 | 39 | E39L | A | 8.3e-09 | 5e-09 | 4.5e-09 | 5.933E-09 | 2.065E-09 | 4.500E-09 | 8.300E-09 |
| 772 | 39 | E39K | A | 1.8e-08 | 5.3e-09 | 4.5e-09 | 9.267E-09 | 7.574E-09 | 4.500E-09 | 1.800E-08 |
| 773 | 39 | E39M | A | 3.7e-09 | 5.5e-09 | 9.8e-09 | 6.333E-09 | 3.134E-09 | 3.700E-09 | 9.800E-09 |
| 774 | 39 | E39F | A | 1.4e-08 | 1.4e-08 | 4.3e-09 | 1.077E-08 | 5.600E-09 | 4.300E-09 | 1.400E-08 |
| 775 | 39 | E39P | A | 7.3e-09 | 5.6e-09 | 3e-09 | 5.300E-09 | 2.166E-09 | 3.000E-09 | 7.300E-09 |
| 776 | 39 | E39S | A | 5.7e-09 | 1.1e-08 | 8.5e-09 | 8.400E-09 | 2.651E-09 | 5.700E-09 | 1.100E-08 |
| 777 | 39 | E39T | A | 3.5e-09 | 5.5e-09 | 4.2e-09 | 4.400E-09 | 1.015E-09 | 3.500E-09 | 5.500E-09 |
| 778 | 39 | E39W | A | 1.4e-08 | 3.3e-09 | 3.3e-09 | 6.867E-09 | 6.178E-09 | 3.300E-09 | 1.400E-08 |
| 779 | 39 | E39Y | A | 1.5e-08 | 3.1e-09 | 1.4e-08 | 1.070E-08 | 6.601E-09 | 3.100E-09 | 1.500E-08 |
| 780 | 39 | E39V | A | 8.1e-09 | 3.5e-09 | 2.4e-09 | 4.667E-09 | 3.024E-09 | 2.400E-09 | 8.100E-09 |
| 781 | 40 | L40A | A | 3.3e-09 | 2.5e-09 | 6.1e-09 | 3.967E-09 | 1.890E-09 | 2.500E-09 | 6.100E-09 |
| 782 | 40 | L40R | A | 1.3e-08 | 1.9e-09 | 3.5e-09 | 6.133E-09 | 6.000E-09 | 1.900E-09 | 1.300E-08 |
| 783 | 40 | L40N | A | 1.2e-09 | 5.3e-09 | 1.2e-08 | 6.167E-09 | 5.452E-09 | 1.200E-09 | 1.200E-08 |
| 784 | 40 | L40D | A | 8.3e-09 | 1.2e-08 | 1.1e-08 | 1.043E-08 | 1.914E-09 | 8.300E-09 | 1.200E-08 |
| 785 | 40 | L40C | A | 4.3e-09 | 6.1e-09 | 9.4e-09 | 6.600E-09 | 2.587E-09 | 4.300E-09 | 9.400E-09 |
| 786 | 40 | L40E | A | 5.9e-09 | 6.4e-09 | 4e-09 | 5.433E-09 | 1.266E-09 | 4.000E-09 | 6.400E-09 |
| 787 | 40 | L40Q | A | 4e-09 | 7e-09 | 7.8e-09 | 6.267E-09 | 2.003E-09 | 4.000E-09 | 7.800E-09 |
| 788 | 40 | L40G | A | 6.5e-09 | 5.4e-09 | 5.2e-09 | 5.700E-09 | 7.000E-10 | 5.200E-09 | 6.500E-09 |
| 789 | 40 | L40H | A | 4e-09 | 8.2e-09 | 9.6e-09 | 7.267E-09 | 2.914E-09 | 4.000E-09 | 9.600E-09 |
| 790 | 40 | L40I | A | 2.8e-09 | 9e-09 | 8.3e-09 | 6.700E-09 | 3.396E-09 | 2.800E-09 | 9.000E-09 |
| 791 | 40 | L40L | A | 4.6e-09 | 8e-09 | 1.4e-08 | 8.867E-09 | 4.760E-09 | 4.600E-09 | 1.400E-08 |
| 792 | 40 | L40K | A | 3.3e-09 | 1.7e-08 | 5.5e-09 | 8.600E-09 | 7.357E-09 | 3.300E-09 | 1.700E-08 |
| 793 | 40 | L40M | A | 3.3e-09 | 2e-09 | 2.8e-09 | 2.700E-09 | 6.557E-10 | 2.000E-09 | 3.300E-09 |
| 794 | 40 | L40F | A | 7.8e-09 | 6e-09 | 5e-09 | 6.267E-09 | 1.419E-09 | 5.000E-09 | 7.800E-09 |
| 795 | 40 | L40P | A | 6.8e-09 | 7.5e-09 | 1.7e-09 | 5.333E-09 | 3.166E-09 | 1.700E-09 | 7.500E-09 |
| 796 | 40 | L40S | A | 9e-09 | 7.1e-09 | 4.4e-09 | 6.833E-09 | 2.312E-09 | 4.400E-09 | 9.000E-09 |
| 797 | 40 | L40T | A | 9.9e-09 | 1.5e-08 | 5.4e-09 | 1.010E-08 | 4.803E-09 | 5.400E-09 | 1.500E-08 |
| 798 | 40 | L40W | A | 2.2e-08 | 1.7e-08 | 1.4e-08 | 1.767E-08 | 4.041E-09 | 1.400E-08 | 2.200E-08 |
| 799 | 40 | L40Y | A | 1.2e-08 | 9.3e-09 | 8.4e-09 | 9.900E-09 | 1.873E-09 | 8.400E-09 | 1.200E-08 |
| 800 | 40 | L40V | A | 7.6e-09 | 1e-08 | 1.1e-08 | 9.533E-09 | 1.747E-09 | 7.600E-09 | 1.100E-08 |
| 801 | 41 | T41A | A | 5.3e-09 | 3.4e-09 | 9e-09 | 5.900E-09 | 2.848E-09 | 3.400E-09 | 9.000E-09 |
| 802 | 41 | T41R | A | 7.4e-09 | 1.1e-08 | 2.8e-08 | 1.547E-08 | 1.100E-08 | 7.400E-09 | 2.800E-08 |
| 803 | 41 | T41N | A | 5.7e-09 | 1.6e-09 | 4.3e-09 | 3.867E-09 | 2.084E-09 | 1.600E-09 | 5.700E-09 |
| 804 | 41 | T41D | A | 6.7e-09 | 7.1e-09 | 7.6e-09 | 7.133E-09 | 4.509E-10 | 6.700E-09 | 7.600E-09 |
| 805 | 41 | T41C | A | 3.6e-09 | 5.9e-09 | 8.8e-09 | 6.100E-09 | 2.606E-09 | 3.600E-09 | 8.800E-09 |
| 806 | 41 | T41E | A | 5.9e-09 | 5.9e-09 | 9.9e-09 | 7.233E-09 | 2.309E-09 | 5.900E-09 | 9.900E-09 |
| 807 | 41 | T41Q | A | 2e-08 | 1.1e-08 | 1.5e-08 | 1.533E-08 | 4.509E-09 | 1.100E-08 | 2.000E-08 |
| 808 | 41 | T41G | A | 6.3e-09 | 1.5e-08 | 2e-08 | 1.377E-08 | 6.933E-09 | 6.300E-09 | 2.000E-08 |
| 809 | 41 | T41H | A | 9.8e-09 | 1.3e-08 | 9.6e-09 | 1.080E-08 | 1.908E-09 | 9.600E-09 | 1.300E-08 |
| 810 | 41 | T41I | A | 3.6e-09 | 5.9e-09 | 6.4e-09 | 5.300E-09 | 1.493E-09 | 3.600E-09 | 6.400E-09 |
| 811 | 41 | T41L | A | 5.8e-09 | 1.7e-08 | 6.9e-09 | 9.900E-09 | 6.173E-09 | 5.800E-09 | 1.700E-08 |
| 812 | 41 | T41K | A | 9.3e-09 | 6.3e-09 | 1.2e-08 | 9.200E-09 | 2.851E-09 | 6.300E-09 | 1.200E-08 |
| 813 | 41 | T41M | A | 7.4e-09 | 1.7e-08 | 3.3e-09 | 9.233E-09 | 7.032E-09 | 3.300E-09 | 1.700E-08 |
| 814 | 41 | T41F | A | 8e-09 | 1.8e-08 | 2.7e-09 | 9.567E-09 | 7.769E-09 | 2.700E-09 | 1.800E-08 |
| 815 | 41 | T41P | A | 1.2e-08 | 1.2e-08 | 6.1e-09 | 1.003E-08 | 3.406E-09 | 6.100E-09 | 1.200E-08 |
| 816 | 41 | T41S | A | 4.7e-09 | 5.6e-09 | 3.5e-09 | 4.600E-09 | 1.054E-09 | 3.500E-09 | 5.600E-09 |
| 817 | 41 | T41T | A | 6.2e-09 | 1.6e-08 | 8.3e-09 | 1.017E-08 | 5.160E-09 | 6.200E-09 | 1.600E-08 |
| 818 | 41 | T41W | A | 1.4e-08 | 1.1e-08 | 5.7e-09 | 1.023E-08 | 4.203E-09 | 5.700E-09 | 1.400E-08 |
| 819 | 41 | T41Y | A | 9.4e-09 | 4.9e-09 | 1.2e-08 | 8.767E-09 | 3.592E-09 | 4.900E-09 | 1.200E-08 |
| 820 | 41 | T41V | A | 5.2e-09 | 7e-09 | 2.8e-09 | 5.000E-09 | 2.107E-09 | 2.800E-09 | 7.000E-09 |
| 821 | 42 | Y42A | A | 4.6e-09 | 8e-09 | 1.4e-08 | 8.867E-09 | 4.760E-09 | 4.600E-09 | 1.400E-08 |
| 822 | 42 | Y42R | A | 1e-08 | 3.2e-08 | 7.8e-09 | 1.660E-08 | 1.338E-08 | 7.800E-09 | 3.200E-08 |
| 823 | 42 | Y42N | A | 3.3e-09 | 1.4e-08 | 5.3e-09 | 7.533E-09 | 5.689E-09 | 3.300E-09 | 1.400E-08 |
| 824 | 42 | Y42D | A | 8.9e-09 | 5.2e-09 | 7.6e-09 | 7.233E-09 | 1.877E-09 | 5.200E-09 | 8.900E-09 |
| 825 | 42 | Y42C | A | 8e-09 | 2.3e-09 | 8.4e-09 | 6.233E-09 | 3.412E-09 | 2.300E-09 | 8.400E-09 |
| 826 | 42 | Y42E | A | 9.9e-09 | 1.1e-08 | 1.1e-08 | 1.063E-08 | 6.351E-10 | 9.900E-09 | 1.100E-08 |
| 827 | 42 | Y42Q | A | 6.1e-09 | 3.2e-09 | 4.2e-09 | 4.500E-09 | 1.473E-09 | 3.200E-09 | 6.100E-09 |
| 828 | 42 | Y42G | A | 1.6e-08 | 2.6e-08 | 1.2e-08 | 1.800E-08 | 7.211E-09 | 1.200E-08 | 2.600E-08 |
| 829 | 42 | Y42H | A | 4.9e-09 | 8.6e-09 | 4.8e-09 | 6.100E-09 | 2.166E-09 | 4.800E-09 | 8.600E-09 |
| 830 | 42 | Y42I | A | 1.4e-08 | 9.6e-09 | 3.6e-09 | 9.067E-09 | 5.220E-09 | 3.600E-09 | 1.400E-08 |
| 831 | 42 | Y42L | A | 1.9e-08 | 7.3e-09 | 1e-08 | 1.210E-08 | 6.126E-09 | 7.300E-09 | 1.900E-08 |
| 832 | 42 | Y42K | A | 2e-08 | 2e-09 | 6.3e-09 | 9.433E-09 | 9.400E-09 | 2.000E-09 | 2.000E-08 |
| 833 | 42 | Y42M | A | 3.3e-08 | 1.7e-08 | 1.6e-08 | 2.200E-08 | 9.539E-09 | 1.600E-08 | 3.300E-08 |
| 834 | 42 | Y42F | A | 1.9e-09 | 3.5e-09 | 6.1e-09 | 3.833E-09 | 2.120E-09 | 1.900E-09 | 6.100E-09 |
| 835 | 42 | Y42P | A | 1.2e-08 | 9.7e-09 | 3.7e-09 | 8.467E-09 | 4.285E-09 | 3.700E-09 | 1.200E-08 |
| 836 | 42 | Y42S | A | 7.6e-09 | 7.2e-09 | 2e-08 | 1.160E-08 | 7.277E-09 | 7.200E-09 | 2.000E-08 |
| 837 | 42 | Y42T | A | 8.2e-09 | 7.4e-09 | 8.2e-09 | 7.933E-09 | 4.619E-10 | 7.400E-09 | 8.200E-09 |
| 838 | 42 | Y42W | A | 1.1e-08 | 2.9e-09 | 1.4e-08 | 9.300E-09 | 5.742E-09 | 2.900E-09 | 1.400E-08 |
| 839 | 42 | Y42Y | A | 3.8e-09 | 1.7e-08 | 1.3e-08 | 1.127E-08 | 6.769E-09 | 3.800E-09 | 1.700E-08 |
| 840 | 42 | Y42V | A | 7e-09 | 2.2e-08 | 1.1e-08 | 1.333E-08 | 7.767E-09 | 7.000E-09 | 2.200E-08 |
| 841 | 43 | L43A | A | 9.6e-09 | 1e-08 | 1.5e-08 | 1.153E-08 | 3.009E-09 | 9.600E-09 | 1.500E-08 |
| 842 | 43 | L43R | A | 5.4e-09 | 4.5e-09 | 1.2e-08 | 7.300E-09 | 4.095E-09 | 4.500E-09 | 1.200E-08 |
| 843 | 43 | L43N | A | 5.7e-09 | 9.9e-09 | 4.6e-09 | 6.733E-09 | 2.797E-09 | 4.600E-09 | 9.900E-09 |
| 844 | 43 | L43D | A | 1e-08 | 7.8e-09 | 6.1e-09 | 7.967E-09 | 1.955E-09 | 6.100E-09 | 1.000E-08 |
| 845 | 43 | L43C | A | 2.3e-09 | 3.9e-09 | 4.5e-09 | 3.567E-09 | 1.137E-09 | 2.300E-09 | 4.500E-09 |
| 846 | 43 | L43E | A | 4.9e-09 | 8.3e-09 | 3.5e-09 | 5.567E-09 | 2.468E-09 | 3.500E-09 | 8.300E-09 |
| 847 | 43 | L43Q | A | 1.5e-08 | 4.8e-09 | 5e-09 | 8.267E-09 | 5.832E-09 | 4.800E-09 | 1.500E-08 |
| 848 | 43 | L43G | A | 6.2e-09 | 7.3e-09 | 3.7e-09 | 5.733E-09 | 1.845E-09 | 3.700E-09 | 7.300E-09 |
| 849 | 43 | L43H | A | 7.4e-09 | 1.6e-08 | 1.1e-08 | 1.147E-08 | 4.319E-09 | 7.400E-09 | 1.600E-08 |

| | | | | | | | | | | |
|---|---|---|---|---|---|---|---|---|---|---|
| 850 | 43 | L43I | A | 1e-08 | 3.9e-09 | 8.3e-09 | 7.400E-09 | 3.148E-09 | 3.900E-09 | 1.000E-08 |
| 851 | 43 | L43L | A | 7.9e-09 | 3.5e-09 | 5.8e-09 | 5.733E-09 | 2.201E-09 | 3.500E-09 | 7.900E-09 |
| 852 | 43 | L43K | A | 1.2e-08 | 4.2e-09 | 6.3e-09 | 7.500E-09 | 4.036E-09 | 4.200E-09 | 1.200E-08 |
| 853 | 43 | L43M | A | 1.4e-08 | 1.1e-08 | 2.4e-09 | 9.133E-09 | 6.021E-09 | 2.400E-09 | 1.400E-08 |
| 854 | 43 | L43F | A | 8.4e-09 | 1e-08 | 5.8e-09 | 8.067E-09 | 2.120E-09 | 5.800E-09 | 1.000E-08 |
| 855 | 43 | L43P | A | 9.7e-09 | 7e-09 | 1.2e-08 | 9.567E-09 | 2.503E-09 | 7.000E-09 | 1.200E-08 |
| 856 | 43 | L43S | A | 1.4e-08 | 7.3e-09 | 6.9e-09 | 9.400E-09 | 3.989E-09 | 6.900E-09 | 1.400E-08 |
| 857 | 43 | L43T | A | 1e-08 | 9.2e-09 | 5.5e-09 | 8.233E-09 | 2.401E-09 | 5.500E-09 | 1.000E-08 |
| 858 | 43 | L43W | A | 2.1e-08 | 2.1e-09 | 4.5e-09 | 9.200E-09 | 1.029E-08 | 2.100E-09 | 2.100E-08 |
| 859 | 43 | L43Y | A | 1.1e-08 | 2e-09 | 8.2e-09 | 7.067E-09 | 4.606E-09 | 2.000E-09 | 1.100E-08 |
| 860 | 43 | L43V | A | 6.7e-09 | 6.4e-09 | 2.4e-08 | 1.237E-08 | 1.008E-08 | 6.400E-09 | 2.400E-08 |
| 861 | 44 | P44A | A | 6.9e-09 | 2.9e-09 | 5.5e-09 | 5.100E-09 | 2.030E-09 | 2.900E-09 | 6.900E-09 |
| 862 | 44 | P44R | A | 7.1e-09 | 7.8e-09 | 9.1e-09 | 8.000E-09 | 1.015E-09 | 7.100E-09 | 9.100E-09 |
| 863 | 44 | P44N | A | 7.3e-09 | 5.1e-09 | 5e-09 | 5.800E-09 | 1.300E-09 | 5.000E-09 | 7.300E-09 |
| 864 | 44 | P44D | A | 5.2e-09 | 1.1e-08 | 6.7e-09 | 7.633E-09 | 3.011E-09 | 5.200E-09 | 1.100E-08 |
| 865 | 44 | P44C | A | 1e-08 | 8.6e-09 | 7.3e-09 | 8.633E-09 | 1.350E-09 | 7.300E-09 | 1.000E-08 |
| 866 | 44 | P44E | A | 1.2e-08 | 8.6e-09 | 5.6e-09 | 8.733E-09 | 3.202E-09 | 5.600E-09 | 1.200E-08 |
| 867 | 44 | P44Q | A | 7.3e-09 | 4.1e-09 | 1.1e-08 | 7.467E-09 | 3.453E-09 | 4.100E-09 | 1.100E-08 |
| 868 | 44 | P44G | A | 1.3e-08 | 1.7e-08 | 4.4e-09 | 1.147E-08 | 6.438E-09 | 4.400E-09 | 1.700E-08 |
| 869 | 44 | P44H | A | 1e-08 | 6.9e-09 | 7.1e-09 | 8.000E-09 | 1.735E-09 | 6.900E-09 | 1.000E-08 |
| 870 | 44 | P44I | A | 6.6e-09 | 6e-09 | 1.5e-08 | 9.200E-09 | 5.032E-09 | 6.000E-09 | 1.500E-08 |
| 871 | 44 | P44L | A | 2e-09 | 9e-09 | 9.2e-09 | 6.733E-09 | 4.100E-09 | 2.000E-09 | 9.200E-09 |
| 872 | 44 | P44K | A | 9.3e-09 | 8.1e-09 | 3.2e-09 | 6.867E-09 | 3.232E-09 | 3.200E-09 | 9.300E-09 |
| 873 | 44 | P44M | A | 4.8e-09 | 1.6e-08 | 6.8e-09 | 9.200E-09 | 5.973E-09 | 4.800E-09 | 1.600E-08 |
| 874 | 44 | P44F | A | 5.6e-09 | 9.3e-09 | 7.8e-09 | 7.567E-09 | 1.861E-09 | 5.600E-09 | 9.300E-09 |
| 875 | 44 | P44P | A | 1.1e-08 | 4.7e-09 | 5e-09 | 6.900E-09 | 3.554E-09 | 4.700E-09 | 1.100E-08 |
| 876 | 44 | P44S | A | 7e-09 | 1.2e-08 | 1.3e-08 | 1.067E-08 | 3.215E-09 | 7.000E-09 | 1.300E-08 |
| 877 | 44 | P44T | A | 1.3e-08 | 8.9e-09 | 4.9e-09 | 8.933E-09 | 4.050E-09 | 4.900E-09 | 1.300E-08 |
| 878 | 44 | P44W | A | 3.9e-09 | 5.9e-09 | 8.4e-09 | 6.067E-09 | 2.255E-09 | 3.900E-09 | 8.400E-09 |
| 879 | 44 | P44Y | A | 2.8e-09 | 3.4e-09 | 1.7e-08 | 7.733E-09 | 8.031E-09 | 2.800E-09 | 1.700E-08 |
| 880 | 44 | P44V | A | 6.7e-09 | 6.4e-09 | 5.6e-09 | 6.233E-09 | 5.686E-10 | 5.600E-09 | 6.700E-09 |
| 881 | 45 | T45A | A | 5.3e-09 | 9.6e-09 | 1.8e-08 | 1.097E-08 | 6.459E-09 | 5.300E-09 | 1.800E-08 |
| 882 | 45 | T45R | A | 1.1e-08 | 1.6e-08 | 1.2e-08 | 1.300E-08 | 2.646E-09 | 1.100E-08 | 1.600E-08 |
| 883 | 45 | T45N | A | 6.4e-09 | 3.7e-09 | 5.3e-09 | 5.133E-09 | 1.358E-09 | 3.700E-09 | 6.400E-09 |
| 884 | 45 | T45D | A | 4.5e-09 | 9.9e-09 | 1.2e-08 | 8.800E-09 | 3.869E-09 | 4.500E-09 | 1.200E-08 |
| 885 | 45 | T45C | A | 8.2e-10 | 7.9e-09 | 8.3e-09 | 5.673E-09 | 4.208E-09 | 8.200E-10 | 8.300E-09 |
| 886 | 45 | T45E | A | 5.5e-09 | 1.5e-08 | 7e-09 | 9.167E-09 | 5.107E-09 | 5.500E-09 | 1.500E-08 |
| 887 | 45 | T45Q | A | 9.8e-09 | 8.7e-09 | 7.2e-09 | 8.567E-09 | 1.305E-09 | 7.200E-09 | 9.800E-09 |
| 888 | 45 | T45G | A | 6.2e-09 | 9.7e-09 | 9.3e-09 | 8.400E-09 | 1.916E-09 | 6.200E-09 | 9.700E-09 |
| 889 | 45 | T45H | A | 7.7e-09 | 1.5e-08 | 4.4e-09 | 9.033E-09 | 5.424E-09 | 4.400E-09 | 1.500E-08 |
| 890 | 45 | T45I | A | 9.8e-09 | 1e-08 | 2.7e-09 | 7.500E-09 | 4.158E-09 | 2.700E-09 | 1.000E-08 |
| 891 | 45 | T45L | A | 5.9e-09 | 9.9e-09 | 2.5e-08 | 1.360E-08 | 1.007E-08 | 5.900E-09 | 2.500E-08 |
| 892 | 45 | T45K | A | 5.5e-09 | 1.5e-08 | 7.5e-09 | 9.333E-09 | 5.008E-09 | 5.500E-09 | 1.500E-08 |
| 893 | 45 | T45M | A | 8.8e-09 | 6.6e-09 | 5e-09 | 6.800E-09 | 1.908E-09 | 5.000E-09 | 8.800E-09 |
| 894 | 45 | T45F | A | 5.7e-09 | 4.5e-09 | 7e-09 | 5.733E-09 | 1.250E-09 | 4.500E-09 | 7.000E-09 |
| 895 | 45 | T45P | A | 8.7e-09 | 1.2e-08 | 8.2e-09 | 9.633E-09 | 2.065E-09 | 8.200E-09 | 1.200E-08 |
| 896 | 45 | T45S | A | 1e-08 | 1.3e-08 | 2.7e-09 | 8.567E-09 | 5.297E-09 | 2.700E-09 | 1.300E-08 |
| 897 | 45 | T45T | A | 8.6e-09 | 1e-08 | 2.6e-09 | 7.067E-09 | 3.931E-09 | 2.600E-09 | 1.000E-08 |
| 898 | 45 | T45W | A | 1.5e-08 | 2.3e-09 | 5.5e-09 | 7.600E-09 | 6.605E-09 | 2.300E-09 | 1.500E-08 |
| 899 | 45 | T45Y | A | 6.8e-09 | 9.6e-09 | 5.9e-09 | 7.433E-09 | 1.930E-09 | 5.900E-09 | 9.600E-09 |
| 900 | 45 | T45V | A | 6.4e-09 | 3.1e-08 | 4.8e-09 | 1.407E-08 | 1.469E-08 | 4.800E-09 | 3.100E-08 |
| 901 | 46 | N46A | A | 7.1e-09 | 9.8e-09 | 1.2e-08 | 9.633E-09 | 2.454E-09 | 7.100E-09 | 1.200E-08 |
| 902 | 46 | N46R | A | 5.4e-09 | 1.4e-08 | 3.8e-09 | 7.733E-09 | 5.486E-09 | 3.800E-09 | 1.400E-08 |
| 903 | 46 | N46N | A | 4.2e-09 | 6.7e-09 | 1.2e-08 | 7.633E-09 | 3.983E-09 | 4.200E-09 | 1.200E-08 |
| 904 | 46 | N46D | A | 2.4e-09 | 4.4e-09 | 9.9e-09 | 5.567E-09 | 3.884E-09 | 2.400E-09 | 9.900E-09 |
| 905 | 46 | N46C | A | 1e-08 | 9.3e-09 | 1.7e-08 | 1.210E-08 | 4.258E-09 | 9.300E-09 | 1.700E-08 |
| 906 | 46 | N46E | A | 5.3e-09 | 8.9e-09 | 8.4e-09 | 7.533E-09 | 1.950E-09 | 5.300E-09 | 8.900E-09 |
| 907 | 46 | N46Q | A | 8.5e-09 | 8.6e-09 | 5.7e-09 | 7.600E-09 | 1.646E-09 | 5.700E-09 | 8.600E-09 |
| 908 | 46 | N46G | A | 6.5e-09 | 7.3e-09 | 2.2e-09 | 5.333E-09 | 2.743E-09 | 2.200E-09 | 7.300E-09 |
| 909 | 46 | N46H | A | 6.3e-09 | 2.5e-09 | 1.3e-08 | 7.267E-09 | 5.316E-09 | 2.500E-09 | 1.300E-08 |
| 910 | 46 | N46I | A | 7.6e-09 | 8.9e-09 | 4.5e-09 | 7.000E-09 | 2.261E-09 | 4.500E-09 | 8.900E-09 |
| 911 | 46 | N46L | A | 1.9e-09 | 8.6e-09 | 6.1e-09 | 5.533E-09 | 3.386E-09 | 1.900E-09 | 8.600E-09 |
| 912 | 46 | N46K | A | 5.9e-09 | 9.6e-09 | 4.5e-09 | 6.667E-09 | 2.635E-09 | 4.500E-09 | 9.600E-09 |
| 913 | 46 | N46M | A | 7.1e-09 | 8.3e-09 | 1.2e-08 | 9.133E-09 | 2.554E-09 | 7.100E-09 | 1.200E-08 |
| 914 | 46 | N46F | A | 9.5e-09 | 6.5e-09 | 4.3e-09 | 6.767E-09 | 2.610E-09 | 4.300E-09 | 9.500E-09 |
| 915 | 46 | N46P | A | 9.9e-09 | 5.4e-09 | 5.5e-09 | 6.933E-09 | 2.570E-09 | 5.400E-09 | 9.900E-09 |
| 916 | 46 | N46S | A | 1.1e-08 | 1.4e-08 | 5.6e-09 | 1.020E-08 | 4.257E-09 | 5.600E-09 | 1.400E-08 |
| 917 | 46 | N46T | A | 9e-09 | 4.4e-09 | 5.5e-09 | 6.300E-09 | 2.402E-09 | 4.400E-09 | 9.000E-09 |
| 918 | 46 | N46W | A | 8.3e-09 | 4.8e-09 | 3.9e-09 | 5.667E-09 | 2.325E-09 | 3.900E-09 | 8.300E-09 |
| 919 | 46 | N46Y | A | 7.2e-09 | 1.5e-08 | 1.5e-08 | 1.240E-08 | 4.503E-09 | 7.200E-09 | 1.500E-08 |
| 920 | 46 | N46V | A | 1.5e-08 | 1.5e-08 | 5.8e-09 | 1.193E-08 | 5.312E-09 | 5.800E-09 | 1.500E-08 |
| 921 | 47 | A47A | A | 2.4e-09 | 4.9e-09 | 1.6e-08 | 7.767E-09 | 7.239E-09 | 2.400E-09 | 1.600E-08 |
| 922 | 47 | A47R | A | 1.3e-08 | 5.5e-09 | 5e-09 | 7.833E-09 | 4.481E-09 | 5.000E-09 | 1.300E-08 |
| 923 | 47 | A47N | A | 7.5e-09 | 7.6e-09 | 8.3e-09 | 7.800E-09 | 4.359E-10 | 7.500E-09 | 8.300E-09 |
| 924 | 47 | A47D | A | 2.7e-09 | 6.4e-09 | 9.8e-09 | 6.300E-09 | 3.551E-09 | 2.700E-09 | 9.800E-09 |
| 925 | 47 | A47C | A | 8.1e-09 | 1e-08 | 1.3e-08 | 1.037E-08 | 2.470E-09 | 8.100E-09 | 1.300E-08 |
| 926 | 47 | A47E | A | 2.4e-08 | 1.7e-08 | 5.8e-09 | 1.560E-08 | 9.180E-09 | 5.800E-09 | 2.400E-08 |
| 927 | 47 | A47Q | A | 7.3e-09 | 3.1e-09 | 7.7e-09 | 6.033E-09 | 2.548E-09 | 3.100E-09 | 7.700E-09 |
| 928 | 47 | A47G | A | 5.2e-09 | 1.2e-08 | 3.5e-09 | 6.900E-09 | 4.498E-09 | 3.500E-09 | 1.200E-08 |
| 929 | 47 | A47H | A | 6.4e-09 | 8.5e-09 | 6.9e-09 | 7.267E-09 | 1.097E-09 | 6.400E-09 | 8.500E-09 |
| 930 | 47 | A47I | A | 1.8e-08 | 3.4e-09 | 1.5e-08 | 1.213E-08 | 7.711E-09 | 3.400E-09 | 1.800E-08 |
| 931 | 47 | A47L | A | 8.1e-09 | 5.3e-09 | 6.6e-09 | 6.667E-09 | 1.401E-09 | 5.300E-09 | 8.100E-09 |
| 932 | 47 | A47K | A | 1.7e-09 | 4.2e-09 | 9e-09 | 4.967E-09 | 3.710E-09 | 1.700E-09 | 9.000E-09 |
| 933 | 47 | A47M | A | 1.6e-08 | 1e-08 | 1.3e-08 | 1.300E-08 | 3.000E-09 | 1.000E-08 | 1.600E-08 |
| 934 | 47 | A47F | A | 2.1e-09 | 5.1e-09 | 6.6e-09 | 4.600E-09 | 2.291E-09 | 2.100E-09 | 6.600E-09 |

| | | | | | | | | | | |
|---|---|---|---|---|---|---|---|---|---|---|
| 935 | 47 | A47P | A | 1.2e-08 | 1.2e-08 | 1.6e-08 | 1.333E-08 | 2.309E-09 | 1.200E-08 | 1.600E-08 |
| 936 | 47 | A47S | A | 4.9e-09 | 1.4e-08 | 3.6e-09 | 7.500E-09 | 5.667E-09 | 3.600E-09 | 1.400E-08 |
| 937 | 47 | A47T | A | 1.7e-09 | 5.6e-09 | 1.7e-08 | 8.100E-09 | 7.950E-09 | 1.700E-09 | 1.700E-08 |
| 938 | 47 | A47W | A | 8.5e-09 | 1.9e-08 | 7e-09 | 1.150E-08 | 6.538E-09 | 7.000E-09 | 1.900E-08 |
| 939 | 47 | A47Y | A | 3.3e-09 | 1.1e-08 | 1.9e-08 | 1.110E-08 | 7.850E-09 | 3.300E-09 | 1.900E-08 |
| 940 | 47 | A47V | A | 7.5e-09 | 4.1e-09 | 1.2e-08 | 7.867E-09 | 3.963E-09 | 4.100E-09 | 1.200E-08 |
| 941 | 48 | S48A | A | 5.1e-09 | 1.2e-08 | 1e-08 | 9.033E-09 | 3.550E-09 | 5.100E-09 | 1.200E-08 |
| 942 | 48 | S48R | A | 3e-09 | 1.2e-08 | 3.4e-09 | 6.133E-09 | 5.085E-09 | 3.000E-09 | 1.200E-08 |
| 943 | 48 | S48N | A | 4.9e-09 | 9.9e-09 | 1.5e-08 | 9.933E-09 | 5.050E-09 | 4.900E-09 | 1.500E-08 |
| 944 | 48 | S48D | A | 3.7e-09 | 9.9e-09 | 1e-08 | 7.867E-09 | 3.609E-09 | 3.700E-09 | 1.000E-08 |
| 945 | 48 | S48C | A | 2e-08 | 7.1e-09 | 9.3e-09 | 1.213E-08 | 6.901E-09 | 7.100E-09 | 2.000E-08 |
| 946 | 48 | S48E | A | 1.1e-08 | 9.5e-09 | 1e-08 | 1.017E-08 | 7.638E-10 | 9.500E-09 | 1.100E-08 |
| 947 | 48 | S48Q | A | 2.9e-09 | 1.3e-08 | 1.1e-08 | 8.967E-09 | 5.348E-09 | 2.900E-09 | 1.300E-08 |
| 948 | 48 | S48G | A | 3.9e-08 | 1e-08 | 6.3e-09 | 1.843E-08 | 1.791E-08 | 6.300E-09 | 3.900E-08 |
| 949 | 48 | S48H | A | 6.1e-09 | 1.6e-08 | 2e-09 | 8.033E-09 | 7.197E-09 | 2.000E-09 | 1.600E-08 |
| 950 | 48 | S48I | A | 8e-09 | 5.3e-09 | 1.5e-08 | 9.433E-09 | 5.006E-09 | 5.300E-09 | 1.500E-08 |
| 951 | 48 | S48L | A | 6.9e-09 | 5.3e-09 | 1.6e-08 | 9.400E-09 | 5.771E-09 | 5.300E-09 | 1.600E-08 |
| 952 | 48 | S48K | A | 7.8e-09 | 2.8e-09 | 1.1e-08 | 7.200E-09 | 4.133E-09 | 2.800E-09 | 1.100E-08 |
| 953 | 48 | S48M | A | 2.6e-08 | 5.9e-09 | 6.2e-09 | 1.270E-08 | 1.152E-08 | 5.900E-09 | 2.600E-08 |
| 954 | 48 | S48F | A | 3e-09 | 6e-09 | 1.4e-08 | 7.667E-09 | 5.686E-09 | 3.000E-09 | 1.400E-08 |
| 955 | 48 | S48P | A | 2.5e-09 | 3.4e-09 | 3.1e-09 | 3.000E-09 | 4.583E-10 | 2.500E-09 | 3.400E-09 |
| 956 | 48 | S48S | A | 1.9e-09 | 7.9e-09 | 1e-08 | 6.600E-09 | 4.204E-09 | 1.900E-09 | 1.000E-08 |
| 957 | 48 | S48T | A | 4.6e-09 | 7.9e-09 | 5.2e-09 | 5.900E-09 | 1.758E-09 | 4.600E-09 | 7.900E-09 |
| 958 | 48 | S48W | A | 1.1e-08 | 2.6e-09 | 1.4e-08 | 9.200E-09 | 5.909E-09 | 2.600E-09 | 1.400E-08 |
| 959 | 48 | S48Y | A | 8.8e-09 | 5.7e-09 | 3.1e-09 | 5.867E-09 | 2.854E-09 | 3.100E-09 | 8.800E-09 |
| 960 | 48 | S48V | A | 1.7e-08 | 4.9e-09 | 9.7e-09 | 1.053E-08 | 6.093E-09 | 4.900E-09 | 1.700E-08 |
| 961 | 49 | L49A | A | 9.8e-09 | 2e-09 | 4.2e-09 | 5.333E-09 | 4.022E-09 | 2.000E-09 | 9.800E-09 |
| 962 | 49 | L49R | A | 6.5e-09 | 5.9e-09 | 8.3e-09 | 6.900E-09 | 1.249E-09 | 5.900E-09 | 8.300E-09 |
| 963 | 49 | L49N | A | 1.7e-08 | 7.4e-09 | 8.2e-09 | 1.087E-08 | 5.327E-09 | 7.400E-09 | 1.700E-08 |
| 964 | 49 | L49D | A | 1.2e-09 | 5.5e-09 | 1.2e-08 | 6.233E-09 | 5.437E-09 | 1.200E-09 | 1.200E-08 |
| 965 | 49 | L49C | A | 3.1e-09 | 7.8e-09 | 2e-09 | 4.300E-09 | 3.081E-09 | 2.000E-09 | 7.800E-09 |
| 966 | 49 | L49E | A | 3.7e-09 | 8.6e-09 | 8.9e-09 | 7.067E-09 | 2.919E-09 | 3.700E-09 | 8.900E-09 |
| 967 | 49 | L49Q | A | 1.4e-09 | 8.4e-09 | 5.2e-09 | 5.000E-09 | 3.504E-09 | 1.400E-09 | 8.400E-09 |
| 968 | 49 | L49G | A | 4.2e-09 | 5.8e-09 | 1.5e-08 | 8.333E-09 | 5.829E-09 | 4.200E-09 | 1.500E-08 |
| 969 | 49 | L49H | A | 5.7e-09 | 3.5e-09 | 5e-09 | 4.733E-09 | 1.124E-09 | 3.500E-09 | 5.700E-09 |
| 970 | 49 | L49I | A | 8.2e-09 | 7.6e-09 | 6e-09 | 7.267E-09 | 1.137E-09 | 6.000E-09 | 8.200E-09 |
| 971 | 49 | L49L | A | 2.1e-08 | 5.4e-09 | 5.7e-09 | 1.070E-08 | 8.921E-09 | 5.400E-09 | 2.100E-08 |
| 972 | 49 | L49K | A | 1.8e-08 | 6.7e-09 | 8.3e-09 | 1.100E-08 | 6.115E-09 | 6.700E-09 | 1.800E-08 |
| 973 | 49 | L49M | A | 5.1e-09 | 1.3e-08 | 4.8e-09 | 7.633E-09 | 4.650E-09 | 4.800E-09 | 1.300E-08 |
| 974 | 49 | L49F | A | 6.6e-09 | 3e-09 | 5.2e-09 | 4.933E-09 | 1.815E-09 | 3.000E-09 | 6.600E-09 |
| 975 | 49 | L49P | A | 3.6e-09 | 6.5e-09 | 7.2e-09 | 5.767E-09 | 1.909E-09 | 3.600E-09 | 7.200E-09 |
| 976 | 49 | L49S | A | 7.3e-09 | 1.7e-08 | 1.1e-08 | 1.177E-08 | 4.895E-09 | 7.300E-09 | 1.700E-08 |
| 977 | 49 | L49T | A | 1.2e-08 | 8e-09 | 9.2e-09 | 9.733E-09 | 2.053E-09 | 8.000E-09 | 1.200E-08 |
| 978 | 49 | L49W | A | 9.8e-09 | 7.3e-09 | 6e-09 | 7.700E-09 | 1.931E-09 | 6.000E-09 | 9.800E-09 |
| 979 | 49 | L49Y | A | 1e-08 | 1.5e-08 | 3e-08 | 1.833E-08 | 1.041E-08 | 1.000E-08 | 3.000E-08 |
| 980 | 49 | L49V | A | 6.4e-09 | 9e-09 | 1e-08 | 8.467E-09 | 1.858E-09 | 6.400E-09 | 1.000E-08 |
| 981 | 50 | S50A | A | 8.8e-09 | 4.8e-09 | 3e-09 | 5.533E-09 | 2.969E-09 | 3.000E-09 | 8.800E-09 |
| 982 | 50 | S50R | A | 5e-09 | 5.1e-09 | 1.2e-08 | 7.367E-09 | 4.013E-09 | 5.000E-09 | 1.200E-08 |
| 983 | 50 | S50N | A | 3e-09 | 5.2e-09 | 1.5e-08 | 7.733E-09 | 6.389E-09 | 3.000E-09 | 1.500E-08 |
| 984 | 50 | S50D | A | 2.4e-09 | 9.4e-09 | 3.4e-09 | 5.067E-09 | 3.786E-09 | 2.400E-09 | 9.400E-09 |
| 985 | 50 | S50C | A | 3e-09 | 4.6e-09 | 1.8e-09 | 3.133E-09 | 1.405E-09 | 1.800E-09 | 4.600E-09 |
| 986 | 50 | S50E | A | 3.6e-09 | 2.9e-09 | 2.4e-08 | 1.017E-08 | 1.199E-08 | 2.900E-09 | 2.400E-08 |
| 987 | 50 | S50Q | A | 2.3e-09 | 2.7e-09 | 2e-09 | 2.333E-09 | 3.512E-10 | 2.000E-09 | 2.700E-09 |
| 988 | 50 | S50G | A | 2e-09 | 2.8e-08 | 7.8e-09 | 1.260E-08 | 1.365E-08 | 2.000E-09 | 2.800E-08 |
| 989 | 50 | S50H | A | 4.2e-09 | 1.8e-09 | 8.9e-09 | 4.967E-09 | 3.612E-09 | 1.800E-09 | 8.900E-09 |
| 990 | 50 | S50I | A | 6.8e-09 | 1.4e-08 | 5.5e-08 | 2.527E-08 | 2.600E-08 | 6.800E-09 | 5.500E-08 |
| 991 | 50 | S50L | A | 3.6e-09 | 5e-09 | 8.5e-09 | 5.700E-09 | 2.524E-09 | 3.600E-09 | 8.500E-09 |
| 992 | 50 | S50K | A | 1.6e-08 | 8.7e-09 | 4.4e-09 | 9.700E-09 | 5.864E-09 | 4.400E-09 | 1.600E-08 |
| 993 | 50 | S50M | A | 4.6e-09 | 5.9e-09 | 1.6e-08 | 8.833E-09 | 6.240E-09 | 4.600E-09 | 1.600E-08 |
| 994 | 50 | S50F | A | 5.4e-09 | 4.3e-09 | 8.5e-09 | 6.067E-09 | 2.178E-09 | 4.300E-09 | 8.500E-09 |
| 995 | 50 | S50P | A | 1.3e-08 | 1.9e-08 | 1.1e-08 | 1.433E-08 | 4.163E-09 | 1.100E-08 | 1.900E-08 |
| 996 | 50 | S50S | A | 4e-09 | 1.6e-09 | 2.9e-08 | 1.153E-08 | 1.517E-08 | 1.600E-09 | 2.900E-08 |
| 997 | 50 | S50T | A | 3.2e-09 | 1e-08 | 9.6e-09 | 7.600E-09 | 3.816E-09 | 3.200E-09 | 1.000E-08 |
| 998 | 50 | S50W | A | 1.2e-08 | 5.9e-09 | 4.8e-09 | 7.567E-09 | 3.879E-09 | 4.800E-09 | 1.200E-08 |
| 999 | 50 | S50Y | A | 2.3e-08 | 2.7e-09 | 9.4e-09 | 1.170E-08 | 1.034E-08 | 2.700E-09 | 2.300E-08 |
| 1000 | 50 | S50V | A | 1.5e-08 | 5.9e-09 | 4.9e-09 | 8.600E-09 | 5.565E-09 | 4.900E-09 | 1.500E-08 |
| 1001 | 51 | F51A | A | 9.8e-09 | 4.8e-09 | 3.5e-09 | 6.033E-09 | 3.326E-09 | 3.500E-09 | 9.800E-09 |
| 1002 | 51 | F51R | A | 1.4e-08 | 1.1e-08 | 5.9e-09 | 1.030E-08 | 4.095E-09 | 5.900E-09 | 1.400E-08 |
| 1003 | 51 | F51N | A | 5.4e-09 | 3.6e-09 | 5.2e-09 | 4.733E-09 | 9.866E-10 | 3.600E-09 | 5.400E-09 |
| 1004 | 51 | F51D | A | 1.2e-08 | 4e-09 | 2.4e-08 | 1.333E-08 | 1.007E-08 | 4.000E-09 | 2.400E-08 |
| 1005 | 51 | F51C | A | 9.7e-09 | 1.4e-08 | 5e-09 | 9.567E-09 | 4.501E-09 | 5.000E-09 | 1.400E-08 |
| 1006 | 51 | F51E | A | 8.5e-09 | 5.8e-09 | 4.5e-09 | 6.267E-09 | 2.040E-09 | 4.500E-09 | 8.500E-09 |
| 1007 | 51 | F51Q | A | 5.8e-09 | 6.3e-09 | 2.3e-08 | 1.170E-08 | 9.789E-09 | 5.800E-09 | 2.300E-08 |
| 1008 | 51 | F51G | A | 1.1e-08 | 7.3e-09 | 4.7e-09 | 7.667E-09 | 3.166E-09 | 4.700E-09 | 1.100E-08 |
| 1009 | 51 | F51H | A | 1.8e-09 | 2.1e-09 | 1e-08 | 4.633E-09 | 4.650E-09 | 1.800E-09 | 1.000E-08 |
| 1010 | 51 | F51I | A | 4.7e-09 | 1.1e-08 | 5.9e-09 | 7.200E-09 | 3.345E-09 | 4.700E-09 | 1.100E-08 |
| 1011 | 51 | F51L | A | 7.8e-09 | 2.8e-09 | 2.2e-08 | 1.087E-08 | 9.961E-09 | 2.800E-09 | 2.200E-08 |
| 1012 | 51 | F51K | A | 7.2e-09 | 2e-08 | 6.6e-09 | 1.127E-08 | 7.569E-09 | 6.600E-09 | 2.000E-08 |
| 1013 | 51 | F51M | A | 1.1e-08 | 6.3e-09 | 4.7e-08 | 2.143E-08 | 2.227E-08 | 6.300E-09 | 4.700E-08 |
| 1014 | 51 | F51F | A | 2e-08 | 3e-09 | 2.6e-09 | 8.533E-09 | 9.932E-09 | 2.600E-09 | 2.000E-08 |
| 1015 | 51 | F51P | A | 4.6e-09 | 1e-08 | 9.5e-09 | 8.033E-09 | 2.984E-09 | 4.600E-09 | 1.000E-08 |
| 1016 | 51 | F51S | A | 2.9e-09 | 6.5e-09 | 9.5e-09 | 6.300E-09 | 3.305E-09 | 2.900E-09 | 9.500E-09 |
| 1017 | 51 | F51T | A | 1e-08 | 6e-09 | 1.5e-09 | 5.833E-09 | 4.252E-09 | 1.500E-09 | 1.000E-08 |
| 1018 | 51 | F51W | A | 9.5e-09 | 8.2e-09 | 1.5e-08 | 1.090E-08 | 3.610E-09 | 8.200E-09 | 1.500E-08 |
| 1019 | 51 | F51Y | A | 7.5e-09 | 1e-08 | 5.4e-09 | 7.633E-09 | 2.303E-09 | 5.400E-09 | 1.000E-08 |

| | | | | | | | | | | |
|---|---|---|---|---|---|---|---|---|---|---|
| 1020 | 51 | F51V | A | 6.1e-09 | 7.6e-09 | 2.7e-09 | 5.467E-09 | 2.511E-09 | 2.700E-09 | 7.600E-09 |
| 1021 | 52 | L52A | A | 7.1e-09 | 1.2e-08 | 1.2e-08 | 1.037E-08 | 2.829E-09 | 7.100E-09 | 1.200E-08 |
| 1022 | 52 | L52R | A | 1e-08 | 3.3e-08 | 5.2e-09 | 1.607E-08 | 1.486E-08 | 5.200E-09 | 3.300E-08 |
| 1023 | 52 | L52N | A | 4.4e-09 | 1.1e-08 | 3.6e-09 | 6.333E-09 | 4.061E-09 | 3.600E-09 | 1.100E-08 |
| 1024 | 52 | L52D | A | 1.3e-08 | 6e-09 | 1.3e-08 | 1.067E-08 | 4.041E-09 | 6.000E-09 | 1.300E-08 |
| 1025 | 52 | L52C | A | 9.9e-09 | 5.1e-09 | 1.4e-08 | 9.667E-09 | 4.455E-09 | 5.100E-09 | 1.400E-08 |
| 1026 | 52 | L52E | A | 1.7e-09 | 4.1e-09 | 3.9e-09 | 3.233E-09 | 1.332E-09 | 1.700E-09 | 4.100E-09 |
| 1027 | 52 | L52Q | A | 4.2e-09 | 8.7e-09 | 8.2e-09 | 7.033E-09 | 2.466E-09 | 4.200E-09 | 8.700E-09 |
| 1028 | 52 | L52G | A | 8.8e-09 | 1.4e-08 | 4.8e-09 | 9.200E-09 | 4.613E-09 | 4.800E-09 | 1.400E-08 |
| 1029 | 52 | L52H | A | 3.4e-08 | 1.2e-08 | 7.1e-09 | 1.770E-08 | 1.433E-08 | 7.100E-09 | 3.400E-08 |
| 1030 | 52 | L52I | A | 6.8e-09 | 1e-08 | 5.9e-09 | 7.567E-09 | 2.155E-09 | 5.900E-09 | 1.000E-08 |
| 1031 | 52 | L52L | A | 7e-09 | 6.9e-09 | 5.2e-09 | 6.367E-09 | 1.012E-09 | 5.200E-09 | 7.000E-09 |
| 1032 | 52 | L52K | A | 5.4e-09 | 7.9e-09 | 5e-09 | 6.100E-09 | 1.572E-09 | 5.000E-09 | 7.900E-09 |
| 1033 | 52 | L52M | A | 4.1e-09 | 2.4e-08 | 5.3e-09 | 1.113E-08 | 1.116E-08 | 4.100E-09 | 2.400E-08 |
| 1034 | 52 | L52F | A | 7.4e-09 | 9.3e-09 | 5.1e-09 | 7.267E-09 | 2.103E-09 | 5.100E-09 | 9.300E-09 |
| 1035 | 52 | L52P | A | 4.9e-09 | 4.8e-09 | 5.7e-09 | 5.133E-09 | 4.933E-10 | 4.800E-09 | 5.700E-09 |
| 1036 | 52 | L52S | A | 1.7e-09 | 8.1e-09 | 5.4e-09 | 5.067E-09 | 3.213E-09 | 1.700E-09 | 8.100E-09 |
| 1037 | 52 | L52T | A | 3.6e-09 | 1.3e-08 | 3.8e-09 | 6.800E-09 | 5.370E-09 | 3.600E-09 | 1.300E-08 |
| 1038 | 52 | L52W | A | 8.8e-09 | 1.3e-08 | 1.3e-08 | 1.160E-08 | 2.425E-09 | 8.800E-09 | 1.300E-08 |
| 1039 | 52 | L52Y | A | 7.2e-09 | 1.1e-08 | 2.2e-09 | 6.800E-09 | 4.414E-09 | 2.200E-09 | 1.100E-08 |
| 1040 | 52 | L52V | A | 2.8e-09 | 4.1e-09 | 9.1e-09 | 5.333E-09 | 3.326E-09 | 2.800E-09 | 9.100E-09 |
| 1041 | 53 | Q53A | A | 6.7e-09 | 7.6e-09 | 6e-09 | 6.767E-09 | 8.021E-10 | 6.000E-09 | 7.600E-09 |
| 1042 | 53 | Q53R | A | 1.4e-08 | 1.1e-08 | 2.3e-09 | 9.100E-09 | 6.077E-09 | 2.300E-09 | 1.400E-08 |
| 1043 | 53 | Q53N | A | 5.2e-09 | 8.5e-09 | 4.6e-09 | 6.100E-09 | 2.100E-09 | 4.600E-09 | 8.500E-09 |
| 1044 | 53 | Q53D | A | 3.9e-09 | 1e-08 | 6.7e-09 | 6.867E-09 | 3.053E-09 | 3.900E-09 | 1.000E-08 |
| 1045 | 53 | Q53C | A | 4.3e-09 | 9.4e-09 | 1.6e-08 | 9.900E-09 | 5.866E-09 | 4.300E-09 | 1.600E-08 |
| 1046 | 53 | Q53E | A | 2e-08 | 2.9e-09 | 1.9e-09 | 8.267E-09 | 1.017E-08 | 1.900E-09 | 2.000E-08 |
| 1047 | 53 | Q53Q | A | 2.3e-09 | 4.9e-09 | 2.6e-08 | 1.107E-08 | 1.300E-08 | 2.300E-09 | 2.600E-08 |
| 1048 | 53 | Q53G | A | 1.2e-08 | 1.3e-08 | 7.5e-09 | 1.083E-08 | 2.930E-09 | 7.500E-09 | 1.300E-08 |
| 1049 | 53 | Q53H | A | 5.2e-09 | 5.4e-09 | 1.1e-08 | 7.200E-09 | 3.292E-09 | 5.200E-09 | 1.100E-08 |
| 1050 | 53 | Q53I | A | 6.5e-09 | 1.5e-09 | 1.1e-08 | 6.333E-09 | 4.752E-09 | 1.500E-09 | 1.100E-08 |
| 1051 | 53 | Q53L | A | 1e-08 | 8.1e-09 | 1.1e-08 | 9.700E-09 | 1.473E-09 | 8.100E-09 | 1.100E-08 |
| 1052 | 53 | Q53K | A | 1.6e-09 | 1.3e-08 | 4.4e-09 | 6.333E-09 | 5.941E-09 | 1.600E-09 | 1.300E-08 |
| 1053 | 53 | Q53M | A | 1.3e-08 | 2.7e-09 | 1.2e-08 | 9.233E-09 | 5.680E-09 | 2.700E-09 | 1.300E-08 |
| 1054 | 53 | Q53F | A | 5.2e-09 | 9.4e-09 | 7.5e-09 | 7.367E-09 | 2.103E-09 | 5.200E-09 | 9.400E-09 |
| 1055 | 53 | Q53P | A | 4.3e-09 | 1e-08 | 3.8e-09 | 6.033E-09 | 3.444E-09 | 3.800E-09 | 1.000E-08 |
| 1056 | 53 | Q53S | A | 7.8e-09 | 1.6e-09 | 6.2e-09 | 5.200E-09 | 3.219E-09 | 1.600E-09 | 7.800E-09 |
| 1057 | 53 | Q53T | A | 3.6e-08 | 8.5e-09 | 3.9e-09 | 1.613E-08 | 1.736E-08 | 3.900E-09 | 3.600E-08 |
| 1058 | 53 | Q53W | A | 1.2e-08 | 3e-09 | 1.6e-08 | 1.033E-08 | 6.658E-09 | 3.000E-09 | 1.600E-08 |
| 1059 | 53 | Q53Y | A | 1.9e-08 | 1.5e-08 | 6.6e-09 | 1.353E-08 | 6.329E-09 | 6.600E-09 | 1.900E-08 |
| 1060 | 53 | Q53V | A | 4.3e-09 | 4.9e-09 | 7.7e-09 | 5.633E-09 | 1.815E-09 | 4.300E-09 | 7.700E-09 |
| 1061 | 54 | D54A | A | 2.9e-09 | 4.2e-09 | 1.7e-08 | 8.033E-09 | 7.793E-09 | 2.900E-09 | 1.700E-08 |
| 1062 | 54 | D54R | A | 1.4e-08 | 1.3e-08 | 7.2e-09 | 1.140E-08 | 3.672E-09 | 7.200E-09 | 1.400E-08 |
| 1063 | 54 | D54N | A | 1.9e-08 | 2.9e-08 | 6.1e-09 | 1.803E-08 | 1.148E-08 | 6.100E-09 | 2.900E-08 |
| 1064 | 54 | D54D | A | 4.9e-09 | 4.4e-09 | 1.7e-08 | 8.767E-09 | 7.135E-09 | 4.400E-09 | 1.700E-08 |
| 1065 | 54 | D54C | A | 1.2e-08 | 1.2e-08 | 4.7e-09 | 9.567E-09 | 4.215E-09 | 4.700E-09 | 1.200E-08 |
| 1066 | 54 | D54E | A | 5.8e-09 | 1.1e-08 | 1.1e-08 | 9.267E-09 | 3.002E-09 | 5.800E-09 | 1.100E-08 |
| 1067 | 54 | D54Q | A | 1.5e-08 | 5.1e-09 | 5e-10 | 6.867E-09 | 7.410E-09 | 5.000E-10 | 1.500E-08 |
| 1068 | 54 | D54G | A | 1.1e-08 | 3.8e-09 | 8.7e-09 | 7.833E-09 | 3.677E-09 | 3.800E-09 | 1.100E-08 |
| 1069 | 54 | D54H | A | 4.4e-09 | 1.8e-09 | 9e-09 | 5.067E-09 | 3.646E-09 | 1.800E-09 | 9.000E-09 |
| 1070 | 54 | D54I | A | 2.7e-09 | 1.3e-08 | 1.5e-08 | 1.023E-08 | 6.600E-09 | 2.700E-09 | 1.500E-08 |
| 1071 | 54 | D54L | A | 8.7e-09 | 6.7e-09 | 3.3e-09 | 6.233E-09 | 2.730E-09 | 3.300E-09 | 8.700E-09 |
| 1072 | 54 | D54K | A | 1.9e-09 | 4.6e-09 | 3.7e-09 | 3.400E-09 | 1.375E-09 | 1.900E-09 | 4.600E-09 |
| 1073 | 54 | D54M | A | 9.7e-09 | 4e-09 | 9.8e-09 | 7.833E-09 | 3.320E-09 | 4.000E-09 | 9.800E-09 |
| 1074 | 54 | D54F | A | 4.5e-09 | 3.5e-08 | 1.1e-08 | 1.683E-08 | 1.606E-08 | 4.500E-09 | 3.500E-08 |
| 1075 | 54 | D54P | A | 2.1e-09 | 7.6e-09 | 1.1e-08 | 6.900E-09 | 4.491E-09 | 2.100E-09 | 1.100E-08 |
| 1076 | 54 | D54S | A | 1.5e-08 | 3.9e-09 | 1.8e-08 | 1.230E-08 | 7.428E-09 | 3.900E-09 | 1.800E-08 |
| 1077 | 54 | D54T | A | 3.6e-09 | 1.3e-08 | 8.4e-09 | 8.333E-09 | 4.700E-09 | 3.600E-09 | 1.300E-08 |
| 1078 | 54 | D54W | A | 6.1e-09 | 7.8e-09 | 7.2e-09 | 7.033E-09 | 8.622E-10 | 6.100E-09 | 7.800E-09 |
| 1079 | 54 | D54Y | A | 1.1e-08 | 8.1e-09 | 5.4e-09 | 8.167E-09 | 2.801E-09 | 5.400E-09 | 1.100E-08 |
| 1080 | 54 | D54V | A | 7.2e-09 | 8.6e-09 | 1.6e-08 | 1.060E-08 | 4.729E-09 | 7.200E-09 | 1.600E-08 |
| 1081 | 55 | I55A | A | 4.4e-09 | 9.2e-09 | 1.3e-08 | 8.867E-09 | 4.310E-09 | 4.400E-09 | 1.300E-08 |
| 1082 | 55 | I55R | A | 1.1e-08 | 4.6e-09 | 4.2e-09 | 6.600E-09 | 3.816E-09 | 4.200E-09 | 1.100E-08 |
| 1083 | 55 | I55N | A | 1.1e-08 | 1.1e-08 | 1.7e-08 | 1.300E-08 | 3.464E-09 | 1.100E-08 | 1.700E-08 |
| 1084 | 55 | I55D | A | 5.8e-09 | 5.2e-09 | 5.4e-09 | 5.467E-09 | 3.055E-10 | 5.200E-09 | 5.800E-09 |
| 1085 | 55 | I55C | A | 6.1e-09 | 5.5e-09 | 4.4e-09 | 5.333E-09 | 8.622E-10 | 4.400E-09 | 6.100E-09 |
| 1086 | 55 | I55E | A | 4.8e-09 | 1.2e-08 | 2.9e-08 | 1.527E-08 | 1.243E-08 | 4.800E-09 | 2.900E-08 |
| 1087 | 55 | I55Q | A | 8e-09 | 5.8e-09 | 1.5e-08 | 9.600E-09 | 4.804E-09 | 5.800E-09 | 1.500E-08 |
| 1088 | 55 | I55G | A | 1.4e-08 | 1.7e-09 | 1.2e-08 | 9.233E-09 | 6.600E-09 | 1.700E-09 | 1.400E-08 |
| 1089 | 55 | I55H | A | 6.7e-09 | 6.5e-09 | 1.2e-08 | 8.400E-09 | 3.119E-09 | 6.500E-09 | 1.200E-08 |
| 1090 | 55 | I55I | A | 9.1e-09 | 1.5e-08 | 7.6e-09 | 1.057E-08 | 3.912E-09 | 7.600E-09 | 1.500E-08 |
| 1091 | 55 | I55L | A | 4.3e-09 | 2.7e-09 | 4.2e-09 | 3.733E-09 | 8.963E-10 | 2.700E-09 | 4.300E-09 |
| 1092 | 55 | I55K | A | 6.2e-09 | 7.9e-09 | 1e-08 | 8.033E-09 | 1.904E-09 | 6.200E-09 | 1.000E-08 |
| 1093 | 55 | I55M | A | 8.9e-09 | 8.4e-09 | 4.4e-09 | 7.233E-09 | 2.466E-09 | 4.400E-09 | 8.900E-09 |
| 1094 | 55 | I55F | A | 5.6e-09 | 8.5e-09 | 4.8e-09 | 6.300E-09 | 1.947E-09 | 4.800E-09 | 8.500E-09 |
| 1095 | 55 | I55P | A | 1.4e-08 | 1.3e-08 | 1.4e-08 | 1.367E-08 | 5.774E-10 | 1.300E-08 | 1.400E-08 |
| 1096 | 55 | I55S | A | 1.5e-08 | 1e-08 | 7.7e-09 | 1.090E-08 | 3.732E-09 | 7.700E-09 | 1.500E-08 |
| 1097 | 55 | I55T | A | 6.6e-09 | 3.9e-09 | 1.7e-09 | 4.067E-09 | 2.454E-09 | 1.700E-09 | 6.600E-09 |
| 1098 | 55 | I55W | A | 6.2e-09 | 3.7e-09 | 6e-09 | 5.300E-09 | 1.389E-09 | 3.700E-09 | 6.200E-09 |
| 1099 | 55 | I55Y | A | 4.1e-09 | 3.7e-09 | 1.1e-08 | 6.267E-09 | 4.104E-09 | 3.700E-09 | 1.100E-08 |
| 1100 | 55 | I55V | A | 3.1e-09 | 7.9e-09 | 4.4e-09 | 5.133E-09 | 2.483E-09 | 3.100E-09 | 7.900E-09 |
| 1101 | 56 | Q56A | A | 8.1e-09 | 3.4e-09 | 5.2e-09 | 5.567E-09 | 2.371E-09 | 3.400E-09 | 8.100E-09 |
| 1102 | 56 | Q56R | A | 1.7e-08 | 1.2e-08 | 6.3e-09 | 1.177E-08 | 5.354E-09 | 6.300E-09 | 1.700E-08 |
| 1103 | 56 | Q56N | A | 1.4e-08 | 5.3e-09 | 3.8e-09 | 7.700E-09 | 5.507E-09 | 3.800E-09 | 1.400E-08 |
| 1104 | 56 | Q56D | A | 1.6e-08 | 4.5e-09 | 2.2e-08 | 1.417E-08 | 8.893E-09 | 4.500E-09 | 2.200E-08 |

| | | | | | | | | | | |
|---|---|---|---|---|---|---|---|---|---|---|
| 1105 | 56 | Q56C | A | 7.4e-09 | 5.5e-09 | 6.9e-09 | 6.600E-09 | 9.849E-10 | 5.500E-09 | 7.400E-09 |
| 1106 | 56 | Q56E | A | 3.5e-09 | 6.3e-09 | 9.7e-09 | 6.500E-09 | 3.105E-09 | 3.500E-09 | 9.700E-09 |
| 1107 | 56 | Q56Q | A | 6.4e-09 | 1.2e-08 | 5.2e-09 | 7.867E-09 | 3.630E-09 | 5.200E-09 | 1.200E-08 |
| 1108 | 56 | Q56G | A | 6.1e-09 | 7.5e-09 | 1.6e-08 | 9.867E-09 | 5.358E-09 | 6.100E-09 | 1.600E-08 |
| 1109 | 56 | Q56H | A | 6.7e-09 | 4.6e-09 | 1.4e-08 | 8.433E-09 | 4.934E-09 | 4.600E-09 | 1.400E-08 |
| 1110 | 56 | Q56I | A | 7.8e-09 | 5.9e-09 | 3.8e-09 | 5.833E-09 | 2.001E-09 | 3.800E-09 | 7.800E-09 |
| 1111 | 56 | Q56L | A | 1.2e-09 | 7.2e-09 | 2.4e-08 | 1.080E-08 | 1.182E-08 | 1.200E-09 | 2.400E-08 |
| 1112 | 56 | Q56K | A | 9e-09 | 1.4e-08 | 1.2e-08 | 1.167E-08 | 2.517E-09 | 9.000E-09 | 1.400E-08 |
| 1113 | 56 | Q56M | A | 9.7e-09 | 1.8e-08 | 6.4e-09 | 1.137E-08 | 5.977E-09 | 6.400E-09 | 1.800E-08 |
| 1114 | 56 | Q56F | A | 6.8e-09 | 4.4e-09 | 6.1e-09 | 5.767E-09 | 1.234E-09 | 4.400E-09 | 6.800E-09 |
| 1115 | 56 | Q56P | A | 3.5e-09 | 4.4e-09 | 6.5e-09 | 4.800E-09 | 1.539E-09 | 3.500E-09 | 6.500E-09 |
| 1116 | 56 | Q56S | A | 7.9e-09 | 3.6e-09 | 4.5e-09 | 5.333E-09 | 2.268E-09 | 3.600E-09 | 7.900E-09 |
| 1117 | 56 | Q56T | A | 8.3e-09 | 2.2e-08 | 2.7e-08 | 1.910E-08 | 9.681E-09 | 8.300E-09 | 2.700E-08 |
| 1118 | 56 | Q56W | A | 5.9e-09 | 1.1e-08 | 1.2e-08 | 9.633E-09 | 3.272E-09 | 5.900E-09 | 1.200E-08 |
| 1119 | 56 | Q56Y | A | 4.3e-09 | 1e-08 | 7.5e-09 | 7.267E-09 | 2.857E-09 | 4.300E-09 | 1.000E-08 |
| 1120 | 56 | Q56V | A | 3.9e-09 | 3.5e-09 | 5.8e-09 | 4.400E-09 | 1.229E-09 | 3.500E-09 | 5.800E-09 |
| 1121 | 57 | E57A | A | 1.3e-08 | 4.6e-09 | 4.9e-09 | 7.500E-09 | 4.766E-09 | 4.600E-09 | 1.300E-08 |
| 1122 | 57 | E57R | A | 8.9e-09 | 9.3e-09 | 1.8e-08 | 1.207E-08 | 5.142E-09 | 8.900E-09 | 1.800E-08 |
| 1123 | 57 | E57N | A | 3.7e-09 | 4.5e-09 | 7.2e-09 | 5.133E-09 | 1.834E-09 | 3.700E-09 | 7.200E-09 |
| 1124 | 57 | E57D | A | 5.4e-09 | 3.1e-09 | 1.2e-08 | 6.833E-09 | 4.620E-09 | 3.100E-09 | 1.200E-08 |
| 1125 | 57 | E57C | A | 8.9e-09 | 6.1e-09 | 2.1e-08 | 1.200E-08 | 7.919E-09 | 6.100E-09 | 2.100E-08 |
| 1126 | 57 | E57E | A | 7e-09 | 1.1e-08 | 1e-08 | 9.333E-09 | 2.082E-09 | 7.000E-09 | 1.100E-08 |
| 1127 | 57 | E57Q | A | 1.3e-08 | 9.1e-09 | 6e-09 | 9.367E-09 | 3.508E-09 | 6.000E-09 | 1.300E-08 |
| 1128 | 57 | E57G | A | 1.9e-09 | 5.7e-09 | 1.4e-08 | 7.200E-09 | 6.188E-09 | 1.900E-09 | 1.400E-08 |
| 1129 | 57 | E57H | A | 6.3e-09 | 1.2e-08 | 5.8e-09 | 8.033E-09 | 3.444E-09 | 5.800E-09 | 1.200E-08 |
| 1130 | 57 | E57I | A | 5.1e-09 | 7.2e-09 | 4.5e-09 | 5.600E-09 | 1.418E-09 | 4.500E-09 | 7.200E-09 |
| 1131 | 57 | E57L | A | 6.9e-09 | 1.2e-08 | 4e-09 | 7.633E-09 | 4.050E-09 | 4.000E-09 | 1.200E-08 |
| 1132 | 57 | E57K | A | 2.3e-08 | 5e-09 | 1.2e-08 | 1.333E-08 | 9.074E-09 | 5.000E-09 | 2.300E-08 |
| 1133 | 57 | E57M | A | 1.2e-08 | 4.7e-09 | 1.1e-08 | 9.233E-09 | 3.958E-09 | 4.700E-09 | 1.200E-08 |
| 1134 | 57 | E57F | A | 5e-09 | 4.5e-09 | 1.8e-08 | 9.167E-09 | 7.654E-09 | 4.500E-09 | 1.800E-08 |
| 1135 | 57 | E57P | A | 7e-09 | 6.2e-09 | 1e-08 | 7.733E-09 | 2.003E-09 | 6.200E-09 | 1.000E-08 |
| 1136 | 57 | E57S | A | 4.2e-09 | 1.4e-08 | 4.4e-09 | 7.533E-09 | 5.601E-09 | 4.200E-09 | 1.400E-08 |
| 1137 | 57 | E57T | A | 8.8e-09 | 2.7e-09 | 1.5e-08 | 8.833E-09 | 6.150E-09 | 2.700E-09 | 1.500E-08 |
| 1138 | 57 | E57W | A | 2.7e-09 | 4.8e-09 | 3.3e-09 | 3.600E-09 | 1.082E-09 | 2.700E-09 | 4.800E-09 |
| 1139 | 57 | E57Y | A | 3.3e-09 | 6.5e-09 | 5.3e-09 | 5.033E-09 | 1.617E-09 | 3.300E-09 | 6.500E-09 |
| 1140 | 57 | E57V | A | 5.8e-09 | 5.7e-09 | 6.1e-09 | 5.867E-09 | 2.082E-10 | 5.700E-09 | 6.100E-09 |
| 1141 | 58 | V58A | A | 6e-09 | 5e-09 | 1.7e-08 | 9.333E-09 | 6.658E-09 | 5.000E-09 | 1.700E-08 |
| 1142 | 58 | V58R | A | 9.8e-09 | 5.5e-09 | 6.1e-09 | 7.133E-09 | 2.329E-09 | 5.500E-09 | 9.800E-09 |
| 1143 | 58 | V58N | A | 6.5e-09 | 9.3e-09 | 8.9e-09 | 8.233E-09 | 1.514E-09 | 6.500E-09 | 9.300E-09 |
| 1144 | 58 | V58D | A | 3.1e-09 | 1.4e-09 | 6.8e-09 | 3.767E-09 | 2.761E-09 | 1.400E-09 | 6.800E-09 |
| 1145 | 58 | V58C | A | 1.7e-08 | 5.4e-09 | 9.6e-09 | 1.067E-08 | 5.873E-09 | 5.400E-09 | 1.700E-08 |
| 1146 | 58 | V58E | A | 1.7e-08 | 5.2e-09 | 5e-09 | 9.067E-09 | 6.871E-09 | 5.000E-09 | 1.700E-08 |
| 1147 | 58 | V58Q | A | 6.2e-09 | 1.1e-08 | 1.4e-08 | 1.040E-08 | 3.934E-09 | 6.200E-09 | 1.400E-08 |
| 1148 | 58 | V58G | A | 1.9e-08 | 2.6e-08 | 4.5e-09 | 1.650E-08 | 1.097E-08 | 4.500E-09 | 2.600E-08 |
| 1149 | 58 | V58H | A | 4.3e-09 | 5.7e-09 | 5.5e-09 | 5.167E-09 | 7.572E-10 | 4.300E-09 | 5.700E-09 |
| 1150 | 58 | V58I | A | 1.8e-08 | 8.3e-09 | 2.7e-09 | 9.667E-09 | 7.741E-09 | 2.700E-09 | 1.800E-08 |
| 1151 | 58 | V58L | A | 8.7e-09 | 7.4e-09 | 1.3e-08 | 9.700E-09 | 2.931E-09 | 7.400E-09 | 1.300E-08 |
| 1152 | 58 | V58K | A | 4e-09 | 1.2e-08 | 8e-09 | 8.000E-09 | 4.000E-09 | 4.000E-09 | 1.200E-08 |
| 1153 | 58 | V58M | A | 2.6e-09 | 8.1e-09 | 1.4e-09 | 4.033E-09 | 3.573E-09 | 1.400E-09 | 8.100E-09 |
| 1154 | 58 | V58F | A | 7.8e-09 | 6.1e-09 | 1.7e-09 | 5.200E-09 | 3.148E-09 | 1.700E-09 | 7.800E-09 |
| 1155 | 58 | V58P | A | 1.1e-08 | 3e-09 | 4.5e-09 | 6.167E-09 | 4.252E-09 | 3.000E-09 | 1.100E-08 |
| 1156 | 58 | V58S | A | 6.4e-09 | 1.6e-08 | 1.3e-08 | 1.180E-08 | 4.911E-09 | 6.400E-09 | 1.600E-08 |
| 1157 | 58 | V58T | A | 8.6e-09 | 8.3e-09 | 2.8e-09 | 6.567E-09 | 3.265E-09 | 2.800E-09 | 8.600E-09 |
| 1158 | 58 | V58W | A | 4.4e-09 | 1.4e-08 | 1.4e-08 | 1.080E-08 | 5.543E-09 | 4.400E-09 | 1.400E-08 |
| 1159 | 58 | V58Y | A | 6.9e-09 | 4e-09 | 4.1e-09 | 5.000E-09 | 1.646E-09 | 4.000E-09 | 6.900E-09 |
| 1160 | 58 | V58V | A | 1.4e-08 | 8.9e-09 | 8.7e-09 | 1.053E-08 | 3.004E-09 | 8.700E-09 | 1.400E-08 |
| 1161 | 59 | Q59A | A | 1.1e-08 | 2.7e-09 | 6.2e-09 | 6.633E-09 | 4.167E-09 | 2.700E-09 | 1.100E-08 |
| 1162 | 59 | Q59R | A | 5e-09 | 9.3e-09 | 8.2e-09 | 7.500E-09 | 2.234E-09 | 5.000E-09 | 9.300E-09 |
| 1163 | 59 | Q59N | A | 8e-09 | 3.7e-09 | 1.1e-08 | 7.567E-09 | 3.669E-09 | 3.700E-09 | 1.100E-08 |
| 1164 | 59 | Q59D | A | 3.8e-09 | 2.4e-09 | 9.8e-09 | 5.333E-09 | 3.931E-09 | 2.400E-09 | 9.800E-09 |
| 1165 | 59 | Q59C | A | 5e-09 | 1.5e-08 | 5.3e-09 | 8.433E-09 | 5.689E-09 | 5.000E-09 | 1.500E-08 |
| 1166 | 59 | Q59E | A | 9.2e-09 | 2.5e-09 | 8.2e-09 | 6.633E-09 | 3.614E-09 | 2.500E-09 | 9.200E-09 |
| 1167 | 59 | Q59Q | A | 1.1e-08 | 3.1e-09 | 5.6e-09 | 6.567E-09 | 4.038E-09 | 3.100E-09 | 1.100E-08 |
| 1168 | 59 | Q59G | A | 6.1e-09 | 1.6e-08 | 8e-09 | 1.003E-08 | 5.254E-09 | 6.100E-09 | 1.600E-08 |
| 1169 | 59 | Q59H | A | 1.8e-08 | 1.8e-08 | 1.1e-08 | 1.567E-08 | 4.041E-09 | 1.100E-08 | 1.800E-08 |
| 1170 | 59 | Q59I | A | 2.5e-09 | 2.2e-08 | 1.1e-08 | 1.183E-08 | 9.777E-09 | 2.500E-09 | 2.200E-08 |
| 1171 | 59 | Q59L | A | 4.5e-09 | 5.2e-09 | 7.5e-09 | 5.733E-09 | 1.570E-09 | 4.500E-09 | 7.500E-09 |
| 1172 | 59 | Q59K | A | 5.1e-09 | 1.1e-08 | 2.3e-09 | 6.133E-09 | 4.441E-09 | 2.300E-09 | 1.100E-08 |
| 1173 | 59 | Q59M | A | 7.2e-09 | 9.5e-09 | 9.8e-09 | 8.833E-09 | 1.422E-09 | 7.200E-09 | 9.800E-09 |
| 1174 | 59 | Q59F | A | 8.4e-09 | 7.8e-09 | 6.3e-09 | 7.500E-09 | 1.082E-09 | 6.300E-09 | 8.400E-09 |
| 1175 | 59 | Q59P | A | 7.1e-09 | 1.1e-08 | 4.9e-09 | 7.667E-09 | 3.089E-09 | 4.900E-09 | 1.100E-08 |
| 1176 | 59 | Q59S | A | 1.7e-09 | 9.9e-09 | 3.5e-09 | 5.033E-09 | 4.310E-09 | 1.700E-09 | 9.900E-09 |
| 1177 | 59 | Q59T | A | 6.8e-09 | 1e-08 | 7.9e-09 | 8.233E-09 | 1.626E-09 | 6.800E-09 | 1.000E-08 |
| 1178 | 59 | Q59W | A | 7.7e-09 | 1.1e-09 | 4.7e-09 | 4.500E-09 | 3.305E-09 | 1.100E-09 | 7.700E-09 |
| 1179 | 59 | Q59Y | A | 1.2e-08 | 3.4e-09 | 8e-09 | 7.800E-09 | 4.303E-09 | 3.400E-09 | 1.200E-08 |
| 1180 | 59 | Q59V | A | 7.2e-09 | 6.4e-09 | 3.1e-08 | 1.487E-08 | 1.398E-08 | 6.400E-09 | 3.100E-08 |
| 1181 | 60 | G60A | A | 1.4e-09 | 9.7e-09 | 3.8e-09 | 4.967E-09 | 4.271E-09 | 1.400E-09 | 9.700E-09 |
| 1182 | 60 | G60R | A | 1.2e-08 | 5.3e-09 | 1.6e-08 | 1.110E-08 | 5.406E-09 | 5.300E-09 | 1.600E-08 |
| 1183 | 60 | G60N | A | 9e-09 | 8.1e-09 | 3.1e-09 | 6.733E-09 | 3.179E-09 | 3.100E-09 | 9.000E-09 |
| 1184 | 60 | G60D | A | 4.3e-09 | 3.1e-09 | 2.1e-08 | 9.467E-09 | 1.001E-08 | 3.100E-09 | 2.100E-08 |
| 1185 | 60 | G60C | A | 2.2e-09 | 7.5e-09 | 8.8e-09 | 6.167E-09 | 3.496E-09 | 2.200E-09 | 8.800E-09 |
| 1186 | 60 | G60E | A | 4.8e-09 | 1.6e-08 | 4.7e-09 | 8.500E-09 | 6.495E-09 | 4.700E-09 | 1.600E-08 |
| 1187 | 60 | G60Q | A | 1.9e-09 | 7.2e-09 | 8.2e-09 | 5.767E-09 | 3.386E-09 | 1.900E-09 | 8.200E-09 |
| 1188 | 60 | G60G | A | 9e-09 | 1.4e-08 | 7.9e-09 | 1.030E-08 | 3.251E-09 | 7.900E-09 | 1.400E-08 |
| 1189 | 60 | G60H | A | 5.1e-09 | 2.2e-08 | 6.6e-09 | 1.123E-08 | 9.354E-09 | 5.100E-09 | 2.200E-08 |

| | | | | | | | | | | |
|---|---|---|---|---|---|---|---|---|---|---|
| 1190 | 60 | G60I | A | 1.1e-08 | 1e-08 | 6.7e-09 | 9.233E-09 | 2.250E-09 | 6.700E-09 | 1.100E-08 |
| 1191 | 60 | G60L | A | 5.3e-09 | 4.1e-08 | 3.1e-09 | 1.647E-08 | 2.127E-08 | 3.100E-09 | 4.100E-08 |
| 1192 | 60 | G60K | A | 4e-09 | 2.6e-08 | 2.6e-09 | 1.087E-08 | 1.312E-08 | 2.600E-09 | 2.600E-08 |
| 1193 | 60 | G60M | A | 5.2e-09 | 4.6e-09 | 6.2e-09 | 5.333E-09 | 8.083E-10 | 4.600E-09 | 6.200E-09 |
| 1194 | 60 | G60F | A | 1.9e-09 | 7.3e-09 | 3.8e-09 | 4.333E-09 | 2.739E-09 | 1.900E-09 | 7.300E-09 |
| 1195 | 60 | G60P | A | 8.5e-09 | 1.8e-08 | 3.7e-09 | 1.007E-08 | 7.278E-09 | 3.700E-09 | 1.800E-08 |
| 1196 | 60 | G60S | A | 3.8e-09 | 8.6e-09 | 1.5e-08 | 9.133E-09 | 5.619E-09 | 3.800E-09 | 1.500E-08 |
| 1197 | 60 | G60T | A | 7.4e-09 | 1.2e-08 | 5.1e-09 | 8.167E-09 | 3.513E-09 | 5.100E-09 | 1.200E-08 |
| 1198 | 60 | G60W | A | 8.3e-09 | 1.9e-08 | 6.9e-09 | 1.140E-08 | 6.619E-09 | 6.900E-09 | 1.900E-08 |
| 1199 | 60 | G60Y | A | 1.8e-09 | 3.2e-09 | 8.8e-09 | 4.600E-09 | 3.704E-09 | 1.800E-09 | 8.800E-09 |
| 1200 | 60 | G60V | A | 6.8e-09 | 1.3e-08 | 1.1e-08 | 1.027E-08 | 3.164E-09 | 6.800E-09 | 1.300E-08 |
| 1201 | 61 | Y61A | A | 7.7e-09 | 6.2e-09 | 2.7e-09 | 5.533E-09 | 2.566E-09 | 2.700E-09 | 7.700E-09 |
| 1202 | 61 | Y61R | A | 5.7e-09 | 1.3e-08 | 4.5e-09 | 7.733E-09 | 4.600E-09 | 4.500E-09 | 1.300E-08 |
| 1203 | 61 | Y61N | A | 4.5e-09 | 6.5e-09 | 1.7e-08 | 9.333E-09 | 6.714E-09 | 4.500E-09 | 1.700E-08 |
| 1204 | 61 | Y61D | A | 2e-08 | 4.5e-09 | 8.3e-09 | 1.093E-08 | 8.079E-09 | 4.500E-09 | 2.000E-08 |
| 1205 | 61 | Y61C | A | 3.6e-09 | 1.3e-08 | 6.7e-09 | 7.767E-09 | 4.790E-09 | 3.600E-09 | 1.300E-08 |
| 1206 | 61 | Y61E | A | 7.7e-09 | 5.2e-09 | 2.7e-09 | 5.200E-09 | 2.500E-09 | 2.700E-09 | 7.700E-09 |
| 1207 | 61 | Y61Q | A | 8.8e-09 | 1.3e-08 | 1e-08 | 1.060E-08 | 2.163E-09 | 8.800E-09 | 1.300E-08 |
| 1208 | 61 | Y61G | A | 5.4e-09 | 5.8e-09 | 7.2e-09 | 6.133E-09 | 9.452E-10 | 5.400E-09 | 7.200E-09 |
| 1209 | 61 | Y61H | A | 1.6e-08 | 1.3e-08 | 1.1e-08 | 1.333E-08 | 2.517E-09 | 1.100E-08 | 1.600E-08 |
| 1210 | 61 | Y61I | A | 8e-09 | 1e-08 | 1.1e-08 | 9.667E-09 | 1.528E-09 | 8.000E-09 | 1.100E-08 |
| 1211 | 61 | Y61L | A | 1.3e-08 | 1.1e-08 | 4.9e-09 | 9.633E-09 | 4.219E-09 | 4.900E-09 | 1.300E-08 |
| 1212 | 61 | Y61K | A | 6.1e-09 | 1.1e-08 | 1.1e-08 | 9.367E-09 | 2.829E-09 | 6.100E-09 | 1.100E-08 |
| 1213 | 61 | Y61M | A | 4.7e-09 | 1.1e-08 | 4.3e-09 | 6.667E-09 | 3.758E-09 | 4.300E-09 | 1.100E-08 |
| 1214 | 61 | Y61F | A | 1.1e-08 | 1.6e-08 | 4.8e-09 | 1.060E-08 | 5.611E-09 | 4.800E-09 | 1.600E-08 |
| 1215 | 61 | Y61P | A | 8.9e-09 | 7.1e-09 | 6.8e-09 | 7.600E-09 | 1.136E-09 | 6.800E-09 | 8.900E-09 |
| 1216 | 61 | Y61S | A | 5.2e-09 | 6.7e-09 | 1.3e-08 | 8.300E-09 | 4.139E-09 | 5.200E-09 | 1.300E-08 |
| 1217 | 61 | Y61T | A | 9.3e-09 | 4e-09 | 1.1e-08 | 8.100E-09 | 3.651E-09 | 4.000E-09 | 1.100E-08 |
| 1218 | 61 | Y61W | A | 4.5e-09 | 4e-09 | 4e-09 | 4.167E-09 | 2.887E-10 | 4.000E-09 | 4.500E-09 |
| 1219 | 61 | Y61Y | A | 1.3e-08 | 1.5e-08 | 4.6e-09 | 1.087E-08 | 5.518E-09 | 4.600E-09 | 1.500E-08 |
| 1220 | 61 | Y61V | A | 1e-08 | 2.4e-09 | 1.3e-08 | 8.467E-09 | 5.464E-09 | 2.400E-09 | 1.300E-08 |
| 1221 | 62 | V62A | A | 8.8e-09 | 4.9e-09 | 7.3e-09 | 7.000E-09 | 1.967E-09 | 4.900E-09 | 8.800E-09 |
| 1222 | 62 | V62R | A | 3.9e-09 | 2.2e-08 | 1.1e-08 | 1.230E-08 | 9.120E-09 | 3.900E-09 | 2.200E-08 |
| 1223 | 62 | V62N | A | 8.8e-09 | 3.7e-08 | 6.2e-09 | 1.733E-08 | 1.708E-08 | 6.200E-09 | 3.700E-08 |
| 1224 | 62 | V62D | A | 4e-09 | 1.1e-08 | 6.7e-09 | 7.233E-09 | 3.530E-09 | 4.000E-09 | 1.100E-08 |
| 1225 | 62 | V62C | A | 6.3e-09 | 6.8e-09 | 4.6e-09 | 5.900E-09 | 1.153E-09 | 4.600E-09 | 6.800E-09 |
| 1226 | 62 | V62E | A | 4e-09 | 2.7e-09 | 5e-09 | 3.900E-09 | 1.153E-09 | 2.700E-09 | 5.000E-09 |
| 1227 | 62 | V62Q | A | 7.1e-09 | 5.3e-09 | 1e-08 | 7.467E-09 | 2.371E-09 | 5.300E-09 | 1.000E-08 |
| 1228 | 62 | V62G | A | 5.4e-09 | 9.3e-09 | 5.5e-09 | 6.733E-09 | 2.223E-09 | 5.400E-09 | 9.300E-09 |
| 1229 | 62 | V62H | A | 6.8e-09 | 4.2e-09 | 4.4e-09 | 5.133E-09 | 1.447E-09 | 4.200E-09 | 6.800E-09 |
| 1230 | 62 | V62I | A | 5.1e-09 | 4.1e-09 | 8e-09 | 5.733E-09 | 2.026E-09 | 4.100E-09 | 8.000E-09 |
| 1231 | 62 | V62L | A | 9.9e-09 | 4.9e-09 | 6.8e-09 | 7.200E-09 | 2.524E-09 | 4.900E-09 | 9.900E-09 |
| 1232 | 62 | V62K | A | 3.8e-09 | 1.1e-08 | 3.2e-09 | 6.000E-09 | 4.341E-09 | 3.200E-09 | 1.100E-08 |
| 1233 | 62 | V62M | A | 4.7e-09 | 1.5e-08 | 4.7e-09 | 8.133E-09 | 5.947E-09 | 4.700E-09 | 1.500E-08 |
| 1234 | 62 | V62F | A | 1e-08 | 2.5e-08 | 4.7e-09 | 1.323E-08 | 1.053E-08 | 4.700E-09 | 2.500E-08 |
| 1235 | 62 | V62P | A | 8.9e-09 | 8.5e-09 | 8.8e-09 | 8.733E-09 | 2.082E-10 | 8.500E-09 | 8.900E-09 |
| 1236 | 62 | V62S | A | 6.8e-09 | 6e-09 | 1.6e-08 | 9.600E-09 | 5.557E-09 | 6.000E-09 | 1.600E-08 |
| 1237 | 62 | V62T | A | 5.8e-09 | 1.2e-08 | 7.5e-09 | 8.433E-09 | 3.204E-09 | 5.800E-09 | 1.200E-08 |
| 1238 | 62 | V62W | A | 1.9e-09 | 3.5e-09 | 4.5e-09 | 3.300E-09 | 1.311E-09 | 1.900E-09 | 4.500E-09 |
| 1239 | 62 | V62Y | A | 6.6e-09 | 2e-09 | 7.9e-09 | 5.500E-09 | 3.100E-09 | 2.000E-09 | 7.900E-09 |
| 1240 | 62 | V62V | A | 9.9e-09 | 3.6e-09 | 1.4e-08 | 9.167E-09 | 5.239E-09 | 3.600E-09 | 1.400E-08 |
| 1241 | 63 | L63A | A | 3.3e-09 | 2.7e-09 | 1e-08 | 5.333E-09 | 4.053E-09 | 2.700E-09 | 1.000E-08 |
| 1242 | 63 | L63R | A | 3e-08 | 9.1e-09 | 6.8e-09 | 1.530E-08 | 1.278E-08 | 6.800E-09 | 3.000E-08 |
| 1243 | 63 | L63N | A | 1e-08 | 8.7e-09 | 6.4e-09 | 8.367E-09 | 1.823E-09 | 6.400E-09 | 1.000E-08 |
| 1244 | 63 | L63D | A | 1.1e-08 | 1.1e-08 | 1.3e-08 | 1.167E-08 | 1.155E-09 | 1.100E-08 | 1.300E-08 |
| 1245 | 63 | L63C | A | 6e-09 | 1.1e-08 | 3.9e-09 | 6.967E-09 | 3.647E-09 | 3.900E-09 | 1.100E-08 |
| 1246 | 63 | L63E | A | 1.2e-08 | 3.4e-09 | 4.9e-09 | 6.767E-09 | 4.594E-09 | 3.400E-09 | 1.200E-08 |
| 1247 | 63 | L63Q | A | 1.8e-08 | 9.3e-09 | 4.2e-09 | 1.050E-08 | 6.978E-09 | 4.200E-09 | 1.800E-08 |
| 1248 | 63 | L63G | A | 1.9e-09 | 2.8e-09 | 9e-09 | 4.567E-09 | 3.866E-09 | 1.900E-09 | 9.000E-09 |
| 1249 | 63 | L63H | A | 1.2e-08 | 1.2e-08 | 6.9e-09 | 1.030E-08 | 2.944E-09 | 6.900E-09 | 1.200E-08 |
| 1250 | 63 | L63I | A | 4.6e-09 | 7.9e-09 | 8.1e-09 | 6.867E-09 | 1.966E-09 | 4.600E-09 | 8.100E-09 |
| 1251 | 63 | L63L | A | 1.3e-08 | 1e-08 | 1.2e-08 | 1.167E-08 | 1.528E-09 | 1.000E-08 | 1.300E-08 |
| 1252 | 63 | L63K | A | 5.9e-09 | 5.9e-09 | 1e-08 | 7.267E-09 | 2.367E-09 | 5.900E-09 | 1.000E-08 |
| 1253 | 63 | L63M | A | 9.2e-09 | 1.5e-08 | 8.2e-09 | 1.080E-08 | 3.672E-09 | 8.200E-09 | 1.500E-08 |
| 1254 | 63 | L63F | A | 1.3e-08 | 1.3e-08 | 4.7e-09 | 1.023E-08 | 4.792E-09 | 4.700E-09 | 1.300E-08 |
| 1255 | 63 | L63P | A | 7.1e-09 | 1.4e-08 | 7.9e-09 | 9.667E-09 | 3.774E-09 | 7.100E-09 | 1.400E-08 |
| 1256 | 63 | L63S | A | 6e-09 | 1.6e-08 | 4.5e-09 | 8.833E-09 | 6.252E-09 | 4.500E-09 | 1.600E-08 |
| 1257 | 63 | L63T | A | 1.1e-08 | 4.5e-09 | 1.2e-08 | 9.167E-09 | 4.072E-09 | 4.500E-09 | 1.200E-08 |
| 1258 | 63 | L63W | A | 1.3e-08 | 6.4e-09 | 8.6e-09 | 9.333E-09 | 3.361E-09 | 6.400E-09 | 1.300E-08 |
| 1259 | 63 | L63Y | A | 9.3e-09 | 8e-09 | 3.7e-09 | 7.000E-09 | 2.931E-09 | 3.700E-09 | 9.300E-09 |
| 1260 | 63 | L63V | A | 8.3e-09 | 5.1e-09 | 4e-09 | 5.800E-09 | 2.234E-09 | 4.000E-09 | 8.300E-09 |
| 1261 | 64 | I64A | A | 1.4e-08 | 4.5e-09 | 9.4e-09 | 9.300E-09 | 4.751E-09 | 4.500E-09 | 1.400E-08 |
| 1262 | 64 | I64R | A | 4e-09 | 1e-08 | 8.1e-09 | 7.367E-09 | 3.066E-09 | 4.000E-09 | 1.000E-08 |
| 1263 | 64 | I64N | A | 7.5e-09 | 8.8e-09 | 6.4e-09 | 7.567E-09 | 1.201E-09 | 6.400E-09 | 8.800E-09 |
| 1264 | 64 | I64D | A | 1.3e-08 | 5.6e-09 | 1.7e-08 | 1.187E-08 | 5.784E-09 | 5.600E-09 | 1.700E-08 |
| 1265 | 64 | I64C | A | 1.1e-08 | 1.5e-08 | 8.1e-09 | 1.137E-08 | 3.465E-09 | 8.100E-09 | 1.500E-08 |
| 1266 | 64 | I64E | A | 9.8e-09 | 1e-08 | 7.6e-09 | 9.133E-09 | 1.332E-09 | 7.600E-09 | 1.000E-08 |
| 1267 | 64 | I64Q | A | 8.1e-09 | 1e-08 | 1.8e-08 | 1.203E-08 | 5.254E-09 | 8.100E-09 | 1.800E-08 |
| 1268 | 64 | I64G | A | 5.7e-09 | 7.4e-09 | 4.8e-09 | 5.967E-09 | 1.320E-09 | 4.800E-09 | 7.400E-09 |
| 1269 | 64 | I64H | A | 6.1e-09 | 1.3e-08 | 1.7e-08 | 1.203E-08 | 5.514E-09 | 6.100E-09 | 1.700E-08 |
| 1270 | 64 | I64I | A | 1.9e-08 | 9.9e-09 | 1.1e-08 | 1.330E-08 | 4.967E-09 | 9.900E-09 | 1.900E-08 |
| 1271 | 64 | I64L | A | 7e-09 | 1.9e-09 | 6.5e-09 | 5.133E-09 | 2.811E-09 | 1.900E-09 | 7.000E-09 |
| 1272 | 64 | I64K | A | 1.1e-08 | 1.2e-08 | 5.4e-09 | 9.467E-09 | 3.557E-09 | 5.400E-09 | 1.200E-08 |
| 1273 | 64 | I64M | A | 6.6e-09 | 4.6e-09 | 9.1e-09 | 6.767E-09 | 2.255E-09 | 4.600E-09 | 9.100E-09 |
| 1274 | 64 | I64F | A | 5.5e-09 | 5.8e-09 | 4.8e-09 | 5.367E-09 | 5.132E-10 | 4.800E-09 | 5.800E-09 |

| | | | | | | | | | | |
|---|---|---|---|---|---|---|---|---|---|---|
| 1275 | 64 | I64P | A | 2.9e-09 | 5.9e-09 | 1.9e-08 | 9.267E-09 | 8.562E-09 | 2.900E-09 | 1.900E-08 |
| 1276 | 64 | I64S | A | 2.4e-09 | 1.5e-08 | 9.1e-09 | 8.833E-09 | 6.304E-09 | 2.400E-09 | 1.500E-08 |
| 1277 | 64 | I64T | A | 3.9e-09 | 8.5e-09 | 7.3e-09 | 6.567E-09 | 2.386E-09 | 3.900E-09 | 8.500E-09 |
| 1278 | 64 | I64W | A | 6.2e-09 | 8.6e-09 | 4.9e-09 | 6.567E-09 | 1.877E-09 | 4.900E-09 | 8.600E-09 |
| 1279 | 64 | I64Y | A | 8e-09 | 2.6e-08 | 1.1e-08 | 1.500E-08 | 9.644E-09 | 8.000E-09 | 2.600E-08 |
| 1280 | 64 | I64V | A | 6e-09 | 3.3e-09 | 7.4e-09 | 5.567E-09 | 2.084E-09 | 3.300E-09 | 7.400E-09 |
| 1281 | 65 | A65A | A | 1.6e-08 | 5.6e-09 | 1.1e-08 | 1.087E-08 | 5.201E-09 | 5.600E-09 | 1.600E-08 |
| 1282 | 65 | A65R | A | 1.3e-08 | 9.3e-09 | 1.4e-08 | 1.210E-08 | 2.476E-09 | 9.300E-09 | 1.400E-08 |
| 1283 | 65 | A65N | A | 9.1e-09 | 2.9e-08 | 5.5e-09 | 1.453E-08 | 1.266E-08 | 5.500E-09 | 2.900E-08 |
| 1284 | 65 | A65D | A | 3.8e-09 | 1.4e-08 | 5.9e-09 | 7.900E-09 | 5.386E-09 | 3.800E-09 | 1.400E-08 |
| 1285 | 65 | A65C | A | 4.1e-09 | 3.2e-09 | 5.1e-09 | 4.133E-09 | 9.504E-10 | 3.200E-09 | 5.100E-09 |
| 1286 | 65 | A65E | A | 3.2e-09 | 1.4e-08 | 1.1e-08 | 9.400E-09 | 5.575E-09 | 3.200E-09 | 1.400E-08 |
| 1287 | 65 | A65Q | A | 4.5e-09 | 1.3e-08 | 6.1e-09 | 7.867E-09 | 4.517E-09 | 4.500E-09 | 1.300E-08 |
| 1288 | 65 | A65G | A | 1.2e-08 | 1.5e-08 | 4.8e-09 | 1.060E-08 | 5.242E-09 | 4.800E-09 | 1.500E-08 |
| 1289 | 65 | A65H | A | 1.7e-08 | 6.6e-09 | 3.9e-09 | 9.167E-09 | 6.917E-09 | 3.900E-09 | 1.700E-08 |
| 1290 | 65 | A65I | A | 7e-09 | 6.2e-09 | 1.1e-08 | 8.067E-09 | 2.572E-09 | 6.200E-09 | 1.100E-08 |
| 1291 | 65 | A65L | A | 9.3e-09 | 3.2e-09 | 4.9e-09 | 5.800E-09 | 3.148E-09 | 3.200E-09 | 9.300E-09 |
| 1292 | 65 | A65K | A | 8e-09 | 2.1e-09 | 7.2e-09 | 5.767E-09 | 3.201E-09 | 2.100E-09 | 8.000E-09 |
| 1293 | 65 | A65M | A | 8.3e-09 | 2.6e-09 | 6.8e-09 | 5.900E-09 | 2.955E-09 | 2.600E-09 | 8.300E-09 |
| 1294 | 65 | A65F | A | 1.7e-08 | 4.1e-09 | 4.1e-09 | 8.400E-09 | 7.448E-09 | 4.100E-09 | 1.700E-08 |
| 1295 | 65 | A65P | A | 1.1e-08 | 9.5e-09 | 3.6e-09 | 8.033E-09 | 3.912E-09 | 3.600E-09 | 1.100E-08 |
| 1296 | 65 | A65S | A | 1.6e-08 | 1.7e-08 | 1.1e-08 | 1.467E-08 | 3.215E-09 | 1.100E-08 | 1.700E-08 |
| 1297 | 65 | A65T | A | 2e-09 | 8.1e-09 | 1.6e-08 | 8.700E-09 | 7.019E-09 | 2.000E-09 | 1.600E-08 |
| 1298 | 65 | A65W | A | 5.2e-09 | 8.9e-09 | 3.1e-08 | 1.503E-08 | 1.395E-08 | 5.200E-09 | 3.100E-08 |
| 1299 | 65 | A65Y | A | 1.6e-08 | 4.9e-09 | 1.5e-08 | 1.197E-08 | 6.140E-09 | 4.900E-09 | 1.600E-08 |
| 1300 | 65 | A65V | A | 1.2e-08 | 1e-08 | 8.2e-09 | 1.007E-08 | 1.901E-09 | 8.200E-09 | 1.200E-08 |
| 1301 | 66 | H66A | A | 5.2e-09 | 5.3e-09 | 2.4e-09 | 4.300E-09 | 1.646E-09 | 2.400E-09 | 5.300E-09 |
| 1302 | 66 | H66R | A | 3.8e-09 | 8e-09 | 1.1e-08 | 7.600E-09 | 3.617E-09 | 3.800E-09 | 1.100E-08 |
| 1303 | 66 | H66N | A | 7.4e-09 | 6.6e-09 | 1.4e-09 | 5.133E-09 | 3.258E-09 | 1.400E-09 | 7.400E-09 |
| 1304 | 66 | H66D | A | 6.1e-09 | 7.3e-09 | 8.4e-09 | 7.267E-09 | 1.150E-09 | 6.100E-09 | 8.400E-09 |
| 1305 | 66 | H66C | A | 4.4e-09 | 1.1e-08 | 6.7e-09 | 7.367E-09 | 3.350E-09 | 4.400E-09 | 1.100E-08 |
| 1306 | 66 | H66E | A | 9.9e-09 | 1.8e-08 | 1e-08 | 1.263E-08 | 4.648E-09 | 9.900E-09 | 1.800E-08 |
| 1307 | 66 | H66Q | A | 3.3e-09 | 1.7e-08 | 1.2e-08 | 1.077E-08 | 6.933E-09 | 3.300E-09 | 1.700E-08 |
| 1308 | 66 | H66G | A | 6.8e-09 | 5e-09 | 6.5e-09 | 6.100E-09 | 9.644E-10 | 5.000E-09 | 6.800E-09 |
| 1309 | 66 | H66H | A | 5.9e-09 | 2.1e-08 | 7.7e-09 | 1.153E-08 | 8.248E-09 | 5.900E-09 | 2.100E-08 |
| 1310 | 66 | H66I | A | 5.6e-09 | 5.1e-09 | 5.2e-09 | 5.300E-09 | 2.646E-10 | 5.100E-09 | 5.600E-09 |
| 1311 | 66 | H66L | A | 9.7e-09 | 6.9e-09 | 7.7e-09 | 8.100E-09 | 1.442E-09 | 6.900E-09 | 9.700E-09 |
| 1312 | 66 | H66K | A | 9.5e-09 | 8.8e-09 | 1.8e-08 | 1.210E-08 | 5.122E-09 | 8.800E-09 | 1.800E-08 |
| 1313 | 66 | H66M | A | 6.4e-09 | 4.4e-09 | 7.8e-09 | 6.200E-09 | 1.709E-09 | 4.400E-09 | 7.800E-09 |
| 1314 | 66 | H66F | A | 2.4e-09 | 5.4e-09 | 7e-09 | 4.933E-09 | 2.335E-09 | 2.400E-09 | 7.000E-09 |
| 1315 | 66 | H66P | A | 8.7e-09 | 1.9e-08 | 6e-09 | 1.123E-08 | 6.860E-09 | 6.000E-09 | 1.900E-08 |
| 1316 | 66 | H66S | A | 5.1e-09 | 9.9e-09 | 5.3e-09 | 6.767E-09 | 2.715E-09 | 5.100E-09 | 9.900E-09 |
| 1317 | 66 | H66T | A | 4.6e-09 | 1.7e-08 | 2.7e-09 | 8.100E-09 | 7.766E-09 | 2.700E-09 | 1.700E-08 |
| 1318 | 66 | H66W | A | 4.9e-09 | 1.1e-08 | 1.3e-08 | 9.633E-09 | 4.219E-09 | 4.900E-09 | 1.300E-08 |
| 1319 | 66 | H66Y | A | 1.8e-09 | 8.2e-09 | 3.4e-09 | 4.467E-09 | 3.331E-09 | 1.800E-09 | 8.200E-09 |
| 1320 | 66 | H66V | A | 9.2e-09 | 8.5e-09 | 1.3e-08 | 1.023E-08 | 2.421E-09 | 8.500E-09 | 1.300E-08 |
| 1321 | 67 | N67A | A | 8.3e-09 | 1.1e-08 | 2.4e-08 | 1.443E-08 | 8.394E-09 | 8.300E-09 | 2.400E-08 |
| 1322 | 67 | N67R | A | 8.2e-09 | 5e-09 | 6.5e-09 | 6.567E-09 | 1.601E-09 | 5.000E-09 | 8.200E-09 |
| 1323 | 67 | N67N | A | 3.3e-09 | 4.6e-09 | 9.9e-09 | 5.933E-09 | 3.496E-09 | 3.300E-09 | 9.900E-09 |
| 1324 | 67 | N67D | A | 3.2e-09 | 1.1e-08 | 8.5e-09 | 7.567E-09 | 3.983E-09 | 3.200E-09 | 1.100E-08 |
| 1325 | 67 | N67C | A | 8.6e-09 | 1.3e-08 | 7.8e-09 | 9.800E-09 | 2.800E-09 | 7.800E-09 | 1.300E-08 |
| 1326 | 67 | N67E | A | 6.9e-09 | 8.3e-09 | 1.3e-08 | 9.400E-09 | 3.195E-09 | 6.900E-09 | 1.300E-08 |
| 1327 | 67 | N67Q | A | 1.3e-08 | 5.9e-09 | 5.2e-09 | 8.033E-09 | 4.315E-09 | 5.200E-09 | 1.300E-08 |
| 1328 | 67 | N67G | A | 4.3e-09 | 4.1e-09 | 8.1e-09 | 5.500E-09 | 2.254E-09 | 4.100E-09 | 8.100E-09 |
| 1329 | 67 | N67H | A | 2.5e-09 | 8.7e-09 | 4.2e-09 | 5.133E-09 | 3.204E-09 | 2.500E-09 | 8.700E-09 |
| 1330 | 67 | N67I | A | 5.1e-09 | 1.7e-08 | 2e-09 | 8.033E-09 | 7.919E-09 | 2.000E-09 | 1.700E-08 |
| 1331 | 67 | N67L | A | 9.8e-09 | 2.9e-09 | 1.1e-08 | 7.900E-09 | 4.371E-09 | 2.900E-09 | 1.100E-08 |
| 1332 | 67 | N67K | A | 7.7e-09 | 2.1e-09 | 6.2e-09 | 5.333E-09 | 2.899E-09 | 2.100E-09 | 7.700E-09 |
| 1333 | 67 | N67M | A | 4.9e-09 | 5.4e-09 | 1.2e-08 | 7.433E-09 | 3.963E-09 | 4.900E-09 | 1.200E-08 |
| 1334 | 67 | N67F | A | 5.9e-09 | 1.2e-08 | 4.9e-09 | 7.600E-09 | 3.843E-09 | 4.900E-09 | 1.200E-08 |
| 1335 | 67 | N67P | A | 2.4e-09 | 5.3e-09 | 1.9e-08 | 8.900E-09 | 8.866E-09 | 2.400E-09 | 1.900E-08 |
| 1336 | 67 | N67S | A | 7.8e-09 | 4.6e-09 | 9e-09 | 7.133E-09 | 2.274E-09 | 4.600E-09 | 9.000E-09 |
| 1337 | 67 | N67T | A | 4e-09 | 1e-08 | 7.4e-09 | 7.133E-09 | 3.009E-09 | 4.000E-09 | 1.000E-08 |
| 1338 | 67 | N67W | A | 3.8e-09 | 2.1e-08 | 8.6e-09 | 1.113E-08 | 8.875E-09 | 3.800E-09 | 2.100E-08 |
| 1339 | 67 | N67Y | A | 2.3e-08 | 8.2e-09 | 2.3e-09 | 1.117E-08 | 1.066E-08 | 2.300E-09 | 2.300E-08 |
| 1340 | 67 | N67V | A | 5.3e-09 | 3.5e-09 | 2e-08 | 9.600E-09 | 9.052E-09 | 3.500E-09 | 2.000E-08 |
| 1341 | 68 | Q68A | A | 1.1e-08 | 2.8e-08 | 1.4e-08 | 1.767E-08 | 9.074E-09 | 1.100E-08 | 2.800E-08 |
| 1342 | 68 | Q68R | A | 6.4e-09 | 2.4e-09 | 7.5e-09 | 5.433E-09 | 2.684E-09 | 2.400E-09 | 7.500E-09 |
| 1343 | 68 | Q68N | A | 5.6e-09 | 2.6e-09 | 4.2e-09 | 4.133E-09 | 1.501E-09 | 2.600E-09 | 5.600E-09 |
| 1344 | 68 | Q68D | A | 4.7e-09 | 1e-08 | 3.2e-08 | 1.557E-08 | 1.448E-08 | 4.700E-09 | 3.200E-08 |
| 1345 | 68 | Q68C | A | 7.1e-09 | 2.2e-09 | 1e-08 | 6.433E-09 | 3.943E-09 | 2.200E-09 | 1.000E-08 |
| 1346 | 68 | Q68E | A | 6.4e-09 | 6.4e-09 | 1.1e-08 | 7.933E-09 | 2.656E-09 | 6.400E-09 | 1.100E-08 |
| 1347 | 68 | Q68Q | A | 6.9e-09 | 3.3e-09 | 3.6e-09 | 4.600E-09 | 1.997E-09 | 3.300E-09 | 6.900E-09 |
| 1348 | 68 | Q68G | A | 7.8e-09 | 1.1e-08 | 2.3e-09 | 7.033E-09 | 4.400E-09 | 2.300E-09 | 1.100E-08 |
| 1349 | 68 | Q68H | A | 5.7e-09 | 1e-08 | 1.4e-08 | 9.900E-09 | 4.151E-09 | 5.700E-09 | 1.400E-08 |
| 1350 | 68 | Q68I | A | 1.5e-08 | 3.6e-09 | 4.7e-09 | 7.767E-09 | 6.288E-09 | 3.600E-09 | 1.500E-08 |
| 1351 | 68 | Q68L | A | 1.4e-08 | 5e-09 | 4.3e-09 | 7.767E-09 | 5.410E-09 | 4.300E-09 | 1.400E-08 |
| 1352 | 68 | Q68K | A | 2.3e-08 | 1.4e-08 | 1.4e-08 | 1.700E-08 | 5.196E-09 | 1.400E-08 | 2.300E-08 |
| 1353 | 68 | Q68M | A | 6.3e-09 | 1.2e-08 | 4.7e-09 | 7.667E-09 | 3.837E-09 | 4.700E-09 | 1.200E-08 |
| 1354 | 68 | Q68F | A | 1e-08 | 1.4e-08 | 9.5e-09 | 1.117E-08 | 2.466E-09 | 9.500E-09 | 1.400E-08 |
| 1355 | 68 | Q68P | A | 1.3e-08 | 7.9e-09 | 1.3e-08 | 1.130E-08 | 2.944E-09 | 7.900E-09 | 1.300E-08 |
| 1356 | 68 | Q68S | A | 1.7e-08 | 1.1e-08 | 1.6e-08 | 1.467E-08 | 3.215E-09 | 1.100E-08 | 1.700E-08 |
| 1357 | 68 | Q68T | A | 6.2e-09 | 2.9e-09 | 5.5e-09 | 4.867E-09 | 1.739E-09 | 2.900E-09 | 6.200E-09 |
| 1358 | 68 | Q68W | A | 4.9e-09 | 1.2e-09 | 1.5e-08 | 7.033E-09 | 7.143E-09 | 1.200E-09 | 1.500E-08 |
| 1359 | 68 | Q68Y | A | 6.8e-08 | 3.4e-09 | 3.5e-09 | 2.497E-08 | 3.727E-08 | 3.400E-09 | 6.800E-08 |

| | | | | | | | | | | |
|---|---|---|---|---|---|---|---|---|---|---|
| 1360 | 68 | Q68V | A | 5.4e-09 | 1.1e-08 | 9.8e-09 | 8.733E-09 | 2.948E-09 | 5.400E-09 | 1.100E-08 |
| 1361 | 69 | V69A | A | 9.6e-09 | 1.6e-08 | 8.6e-09 | 1.140E-08 | 4.015E-09 | 8.600E-09 | 1.600E-08 |
| 1362 | 69 | V69R | A | 9e-09 | 4.4e-09 | 2.1e-09 | 5.167E-09 | 3.513E-09 | 2.100E-09 | 9.000E-09 |
| 1363 | 69 | V69N | A | 8.4e-09 | 8.6e-09 | 1.2e-08 | 9.667E-09 | 2.023E-09 | 8.400E-09 | 1.200E-08 |
| 1364 | 69 | V69D | A | 3.9e-09 | 6.6e-09 | 1.2e-08 | 7.500E-09 | 4.124E-09 | 3.900E-09 | 1.200E-08 |
| 1365 | 69 | V69C | A | 1.2e-08 | 7e-09 | 4.1e-09 | 7.700E-09 | 3.996E-09 | 4.100E-09 | 1.200E-08 |
| 1366 | 69 | V69E | A | 4.9e-09 | 4.6e-09 | 1.5e-08 | 8.167E-09 | 5.920E-09 | 4.600E-09 | 1.500E-08 |
| 1367 | 69 | V69Q | A | 1.2e-08 | 2.8e-09 | 1.1e-08 | 8.600E-09 | 5.048E-09 | 2.800E-09 | 1.200E-08 |
| 1368 | 69 | V69G | A | 5.5e-09 | 2.2e-08 | 1.4e-08 | 1.383E-08 | 8.251E-09 | 5.500E-09 | 2.200E-08 |
| 1369 | 69 | V69H | A | 7.2e-09 | 1.8e-08 | 2.8e-08 | 1.773E-08 | 1.040E-08 | 7.200E-09 | 2.800E-08 |
| 1370 | 69 | V69I | A | 1.1e-08 | 1.7e-08 | 1.3e-08 | 1.367E-08 | 3.055E-09 | 1.100E-08 | 1.700E-08 |
| 1371 | 69 | V69L | A | 3.8e-09 | 1e-08 | 7.1e-09 | 6.967E-09 | 3.102E-09 | 3.800E-09 | 1.000E-08 |
| 1372 | 69 | V69K | A | 3.9e-09 | 6.6e-09 | 4.3e-09 | 4.933E-09 | 1.457E-09 | 3.900E-09 | 6.600E-09 |
| 1373 | 69 | V69M | A | 1.5e-08 | 8.9e-09 | 4.5e-09 | 9.467E-09 | 5.273E-09 | 4.500E-09 | 1.500E-08 |
| 1374 | 69 | V69F | A | 2.7e-08 | 1.5e-08 | 1.2e-08 | 1.800E-08 | 7.937E-09 | 1.200E-08 | 2.700E-08 |
| 1375 | 69 | V69P | A | 3.7e-09 | 4.3e-09 | 7.3e-09 | 5.100E-09 | 1.929E-09 | 3.700E-09 | 7.300E-09 |
| 1376 | 69 | V69S | A | 7e-09 | 1.8e-08 | 1.4e-08 | 1.300E-08 | 5.568E-09 | 7.000E-09 | 1.800E-08 |
| 1377 | 69 | V69T | A | 8.4e-09 | 2e-08 | 2.9e-09 | 1.043E-08 | 8.729E-09 | 2.900E-09 | 2.000E-08 |
| 1378 | 69 | V69W | A | 4.2e-09 | 6.2e-09 | 7.8e-09 | 6.067E-09 | 1.804E-09 | 4.200E-09 | 7.800E-09 |
| 1379 | 69 | V69Y | A | 5.6e-09 | 3.4e-09 | 2.1e-08 | 1.000E-08 | 9.590E-09 | 3.400E-09 | 2.100E-08 |
| 1380 | 69 | V69V | A | 1.1e-08 | 1.3e-08 | 1.5e-08 | 1.300E-08 | 2.000E-09 | 1.100E-08 | 1.500E-08 |
| 1381 | 70 | R70A | A | 1.6e-09 | 3.9e-09 | 1.7e-09 | 2.400E-09 | 1.300E-09 | 1.600E-09 | 3.900E-09 |
| 1382 | 70 | R70R | A | 1.2e-08 | 6.4e-09 | 2.7e-09 | 7.033E-09 | 4.682E-09 | 2.700E-09 | 1.200E-08 |
| 1383 | 70 | R70N | A | 8e-09 | 1.1e-08 | 2.6e-08 | 1.500E-08 | 9.644E-09 | 8.000E-09 | 2.600E-08 |
| 1384 | 70 | R70D | A | 3e-09 | 9e-09 | 4e-09 | 5.333E-09 | 3.215E-09 | 3.000E-09 | 9.000E-09 |
| 1385 | 70 | R70C | A | 1.1e-08 | 9.2e-09 | 1.8e-08 | 1.273E-08 | 4.649E-09 | 9.200E-09 | 1.800E-08 |
| 1386 | 70 | R70E | A | 5.5e-09 | 1.1e-08 | 2.7e-08 | 1.450E-08 | 1.117E-08 | 5.500E-09 | 2.700E-08 |
| 1387 | 70 | R70Q | A | 2.4e-08 | 3.9e-08 | 5e-09 | 2.267E-08 | 1.704E-08 | 5.000E-09 | 3.900E-08 |
| 1388 | 70 | R70G | A | 9.1e-09 | 7.9e-09 | 6.7e-09 | 7.900E-09 | 1.200E-09 | 6.700E-09 | 9.100E-09 |
| 1389 | 70 | R70H | A | 1.1e-08 | 1.7e-08 | 1.6e-08 | 1.467E-08 | 3.215E-09 | 1.100E-08 | 1.700E-08 |
| 1390 | 70 | R70I | A | 7.2e-09 | 6.9e-09 | 4.5e-09 | 6.200E-09 | 1.480E-09 | 4.500E-09 | 7.200E-09 |
| 1391 | 70 | R70L | A | 6.9e-09 | 8.3e-09 | 1.4e-08 | 9.733E-09 | 3.761E-09 | 6.900E-09 | 1.400E-08 |
| 1392 | 70 | R70K | A | 3.1e-08 | 4.2e-09 | 1.6e-08 | 1.707E-08 | 1.343E-08 | 4.200E-09 | 3.100E-08 |
| 1393 | 70 | R70M | A | 1.5e-09 | 4.4e-09 | 5.8e-09 | 3.900E-09 | 2.193E-09 | 1.500E-09 | 5.800E-09 |
| 1394 | 70 | R70F | A | 2.9e-09 | 1.8e-08 | 8.6e-09 | 9.833E-09 | 7.625E-09 | 2.900E-09 | 1.800E-08 |
| 1395 | 70 | R70P | A | 8.6e-09 | 8.6e-09 | 3.5e-09 | 6.900E-09 | 2.944E-09 | 3.500E-09 | 8.600E-09 |
| 1396 | 70 | R70S | A | 6e-09 | 3.2e-09 | 5.4e-09 | 4.867E-09 | 1.474E-09 | 3.200E-09 | 6.000E-09 |
| 1397 | 70 | R70T | A | 8.7e-09 | 1.2e-08 | 2.4e-09 | 7.700E-09 | 4.877E-09 | 2.400E-09 | 1.200E-08 |
| 1398 | 70 | R70W | A | 1.7e-08 | 5.7e-09 | 5.2e-09 | 9.300E-09 | 6.673E-09 | 5.200E-09 | 1.700E-08 |
| 1399 | 70 | R70Y | A | 1.1e-08 | 7.5e-09 | 1e-08 | 9.500E-09 | 1.803E-09 | 7.500E-09 | 1.100E-08 |
| 1400 | 70 | R70V | A | 1e-08 | 6.8e-09 | 2.2e-08 | 1.293E-08 | 8.013E-09 | 6.800E-09 | 2.200E-08 |
| 1401 | 71 | Q71A | A | 8.8e-09 | 8.7e-09 | 5.7e-09 | 7.733E-09 | 1.762E-09 | 5.700E-09 | 8.800E-09 |
| 1402 | 71 | Q71R | A | 4.4e-09 | 1.5e-08 | 1.7e-08 | 1.213E-08 | 6.772E-09 | 4.400E-09 | 1.700E-08 |
| 1403 | 71 | Q71N | A | 5.9e-09 | 1.1e-08 | 1.3e-08 | 9.967E-09 | 3.661E-09 | 5.900E-09 | 1.300E-08 |
| 1404 | 71 | Q71D | A | 3.7e-09 | 1.3e-09 | 3e-08 | 1.167E-08 | 1.592E-08 | 1.300E-09 | 3.000E-08 |
| 1405 | 71 | Q71C | A | 1e-08 | 2.6e-09 | 1.2e-08 | 8.200E-09 | 4.952E-09 | 2.600E-09 | 1.200E-08 |
| 1406 | 71 | Q71E | A | 8.1e-09 | 3.6e-09 | 1.2e-08 | 7.900E-09 | 4.204E-09 | 3.600E-09 | 1.200E-08 |
| 1407 | 71 | Q71Q | A | 5.7e-09 | 5.1e-09 | 7.3e-09 | 6.033E-09 | 1.137E-09 | 5.100E-09 | 7.300E-09 |
| 1408 | 71 | Q71G | A | 6.3e-09 | 1.7e-08 | 5.9e-09 | 9.733E-09 | 6.296E-09 | 5.900E-09 | 1.700E-08 |
| 1409 | 71 | Q71H | A | 3.8e-09 | 5.6e-09 | 5e-09 | 4.800E-09 | 9.165E-10 | 3.800E-09 | 5.600E-09 |
| 1410 | 71 | Q71I | A | 3.7e-09 | 7.2e-09 | 9.4e-09 | 6.767E-09 | 2.875E-09 | 3.700E-09 | 9.400E-09 |
| 1411 | 71 | Q71L | A | 6.2e-09 | 5e-09 | 1.6e-08 | 9.067E-09 | 6.034E-09 | 5.000E-09 | 1.600E-08 |
| 1412 | 71 | Q71K | A | 9.6e-09 | 6.8e-09 | 7.3e-09 | 7.900E-09 | 1.493E-09 | 6.800E-09 | 9.600E-09 |
| 1413 | 71 | Q71M | A | 3.4e-09 | 7.7e-09 | 3.1e-09 | 4.733E-09 | 2.574E-09 | 3.100E-09 | 7.700E-09 |
| 1414 | 71 | Q71F | A | 8.5e-09 | 7.8e-09 | 2.7e-09 | 6.333E-09 | 3.166E-09 | 2.700E-09 | 8.500E-09 |
| 1415 | 71 | Q71P | A | 6.5e-09 | 1.1e-08 | 1.6e-08 | 1.117E-08 | 4.752E-09 | 6.500E-09 | 1.600E-08 |
| 1416 | 71 | Q71S | A | 1e-08 | 3.1e-08 | 1.6e-08 | 1.900E-08 | 1.082E-08 | 1.000E-08 | 3.100E-08 |
| 1417 | 71 | Q71T | A | 4.5e-09 | 8.2e-09 | 3.1e-09 | 5.267E-09 | 2.635E-09 | 3.100E-09 | 8.200E-09 |
| 1418 | 71 | Q71W | A | 6e-09 | 5.2e-09 | 8e-09 | 6.400E-09 | 1.442E-09 | 5.200E-09 | 8.000E-09 |
| 1419 | 71 | Q71Y | A | 1e-08 | 7.9e-09 | 7e-09 | 8.300E-09 | 1.539E-09 | 7.000E-09 | 1.000E-08 |
| 1420 | 71 | Q71V | A | 2.1e-08 | 2.5e-09 | 1.3e-08 | 1.217E-08 | 9.278E-09 | 2.500E-09 | 2.100E-08 |
| 1421 | 72 | V72A | A | 6.6e-09 | 3.8e-08 | 1.2e-08 | 1.887E-08 | 1.679E-08 | 6.600E-09 | 3.800E-08 |
| 1422 | 72 | V72R | A | 5.1e-09 | 2.5e-09 | 4e-09 | 3.867E-09 | 1.305E-09 | 2.500E-09 | 5.100E-09 |
| 1423 | 72 | V72N | A | 4.3e-09 | 4.4e-09 | 1e-08 | 6.233E-09 | 3.262E-09 | 4.300E-09 | 1.000E-08 |
| 1424 | 72 | V72D | A | 1.3e-08 | 1.4e-08 | 3.5e-09 | 1.017E-08 | 5.795E-09 | 3.500E-09 | 1.400E-08 |
| 1425 | 72 | V72C | A | 3e-09 | 4.4e-09 | 3.8e-09 | 3.733E-09 | 7.024E-10 | 3.000E-09 | 4.400E-09 |
| 1426 | 72 | V72E | A | 2.9e-08 | 2.8e-09 | 1.8e-08 | 1.660E-08 | 1.316E-08 | 2.800E-09 | 2.900E-08 |
| 1427 | 72 | V72Q | A | 1e-08 | 2.2e-08 | 1e-08 | 1.400E-08 | 6.928E-09 | 1.000E-08 | 2.200E-08 |
| 1428 | 72 | V72G | A | 1.5e-08 | 1.6e-08 | 3.1e-08 | 2.067E-08 | 8.963E-09 | 1.500E-08 | 3.100E-08 |
| 1429 | 72 | V72H | A | 1.7e-08 | 7.6e-09 | 1.2e-08 | 1.220E-08 | 4.703E-09 | 7.600E-09 | 1.700E-08 |
| 1430 | 72 | V72I | A | 1.1e-08 | 5.7e-09 | 8.6e-09 | 8.433E-09 | 2.654E-09 | 5.700E-09 | 1.100E-08 |
| 1431 | 72 | V72L | A | 4.9e-09 | 4.2e-09 | 1e-08 | 6.367E-09 | 3.166E-09 | 4.200E-09 | 1.000E-08 |
| 1432 | 72 | V72K | A | 6.3e-09 | 2.9e-08 | 4.1e-09 | 1.313E-08 | 1.378E-08 | 4.100E-09 | 2.900E-08 |
| 1433 | 72 | V72M | A | 3.1e-09 | 8.6e-09 | 1e-08 | 7.233E-09 | 3.647E-09 | 3.100E-09 | 1.000E-08 |
| 1434 | 72 | V72F | A | 2.3e-09 | 7.7e-09 | 4.4e-09 | 4.800E-09 | 2.722E-09 | 2.300E-09 | 7.700E-09 |
| 1435 | 72 | V72P | A | 6.8e-09 | 1.4e-08 | 3.5e-09 | 8.100E-09 | 5.369E-09 | 3.500E-09 | 1.400E-08 |
| 1436 | 72 | V72S | A | 2.3e-08 | 4.8e-09 | 9.9e-09 | 1.257E-08 | 9.388E-09 | 4.800E-09 | 2.300E-08 |
| 1437 | 72 | V72T | A | 2.8e-08 | 5.4e-09 | 8.6e-09 | 1.400E-08 | 1.223E-08 | 5.400E-09 | 2.800E-08 |
| 1438 | 72 | V72W | A | 1e-08 | 4.4e-09 | 5.4e-09 | 6.600E-09 | 2.987E-09 | 4.400E-09 | 1.000E-08 |
| 1439 | 72 | V72Y | A | 8.9e-09 | 5.8e-09 | 7.5e-09 | 7.400E-09 | 1.552E-09 | 5.800E-09 | 8.900E-09 |
| 1440 | 72 | V72V | A | 2.2e-09 | 6.8e-09 | 9e-09 | 6.000E-09 | 3.470E-09 | 2.200E-09 | 9.000E-09 |
| 1441 | 73 | P73A | A | 1.1e-08 | 1.5e-08 | 2.7e-09 | 9.567E-09 | 6.274E-09 | 2.700E-09 | 1.500E-08 |
| 1442 | 73 | P73R | A | 2e-09 | 9.9e-09 | 2.6e-09 | 4.833E-09 | 4.398E-09 | 2.000E-09 | 9.900E-09 |
| 1443 | 73 | P73N | A | 7.3e-09 | 6.9e-09 | 6.2e-09 | 6.800E-09 | 5.568E-10 | 6.200E-09 | 7.300E-09 |
| 1444 | 73 | P73D | A | 8.2e-09 | 4.3e-09 | 5.2e-09 | 5.900E-09 | 2.042E-09 | 4.300E-09 | 8.200E-09 |

| | | | | | | | | | | |
|---|---|---|---|---|---|---|---|---|---|---|
| 1445 | 73 | P73C | A | 2e-08 | 1.5e-08 | 1.2e-08 | 1.567E-08 | 4.041E-09 | 1.200E-08 | 2.000E-08 |
| 1446 | 73 | P73E | A | 3.8e-09 | 5.7e-09 | 1.3e-08 | 7.500E-09 | 4.857E-09 | 3.800E-09 | 1.300E-08 |
| 1447 | 73 | P73Q | A | 4e-09 | 1.3e-09 | 2.6e-09 | 2.633E-09 | 1.350E-09 | 1.300E-09 | 4.000E-09 |
| 1448 | 73 | P73G | A | 1.8e-09 | 6.9e-09 | 1.1e-09 | 3.267E-09 | 3.166E-09 | 1.100E-09 | 6.900E-09 |
| 1449 | 73 | P73H | A | 6e-09 | 4.2e-09 | 4.3e-09 | 4.833E-09 | 1.012E-09 | 4.200E-09 | 6.000E-09 |
| 1450 | 73 | P73I | A | 4.6e-09 | 1.5e-08 | 1.1e-08 | 1.020E-08 | 5.246E-09 | 4.600E-09 | 1.500E-08 |
| 1451 | 73 | P73L | A | 2e-09 | 1.9e-09 | 7.3e-09 | 3.733E-09 | 3.089E-09 | 1.900E-09 | 7.300E-09 |
| 1452 | 73 | P73K | A | 8.6e-09 | 9.1e-09 | 1.7e-08 | 1.157E-08 | 4.712E-09 | 8.600E-09 | 1.700E-08 |
| 1453 | 73 | P73M | A | 7.7e-09 | 5e-09 | 4.2e-09 | 5.633E-09 | 1.834E-09 | 4.200E-09 | 7.700E-09 |
| 1454 | 73 | P73F | A | 2.2e-09 | 6.9e-09 | 6.1e-09 | 5.067E-09 | 2.515E-09 | 2.200E-09 | 6.900E-09 |
| 1455 | 73 | P73P | A | 1.5e-08 | 1.1e-08 | 5.4e-09 | 1.047E-08 | 4.822E-09 | 5.400E-09 | 1.500E-08 |
| 1456 | 73 | P73S | A | 4.5e-09 | 2.8e-09 | 3.5e-09 | 3.600E-09 | 8.544E-10 | 2.800E-09 | 4.500E-09 |
| 1457 | 73 | P73T | A | 6.7e-09 | 7.5e-09 | 6.6e-09 | 6.933E-09 | 4.933E-10 | 6.600E-09 | 7.500E-09 |
| 1458 | 73 | P73W | A | 4.5e-09 | 7.3e-09 | 9e-09 | 6.933E-09 | 2.272E-09 | 4.500E-09 | 9.000E-09 |
| 1459 | 73 | P73Y | A | 2.7e-09 | 1.2e-09 | 1e-08 | 4.633E-09 | 4.708E-09 | 1.200E-09 | 1.000E-08 |
| 1460 | 73 | P73V | A | 9.1e-09 | 3.2e-09 | 7.9e-09 | 6.733E-09 | 3.118E-09 | 3.200E-09 | 9.100E-09 |
| 1461 | 74 | L74A | A | 5.1e-09 | 1.2e-08 | 5.6e-09 | 7.567E-09 | 3.848E-09 | 5.100E-09 | 1.200E-08 |
| 1462 | 74 | L74R | A | 3.3e-09 | 5.7e-09 | 5.5e-09 | 4.833E-09 | 1.332E-09 | 3.300E-09 | 5.700E-09 |
| 1463 | 74 | L74N | A | 3.3e-09 | 9.2e-09 | 4.4e-09 | 5.633E-09 | 3.137E-09 | 3.300E-09 | 9.200E-09 |
| 1464 | 74 | L74D | A | 4.2e-09 | 2e-09 | 4.7e-09 | 3.633E-09 | 1.436E-09 | 2.000E-09 | 4.700E-09 |
| 1465 | 74 | L74C | A | 4.9e-09 | 5.7e-09 | 1.8e-08 | 9.533E-09 | 7.343E-09 | 4.900E-09 | 1.800E-08 |
| 1466 | 74 | L74E | A | 7.4e-09 | 1e-08 | 3.3e-09 | 6.900E-09 | 3.378E-09 | 3.300E-09 | 1.000E-08 |
| 1467 | 74 | L74Q | A | 1e-08 | 1e-08 | 7.8e-09 | 9.267E-09 | 1.270E-09 | 7.800E-09 | 1.000E-08 |
| 1468 | 74 | L74G | A | 5.7e-09 | 1.1e-08 | 1.6e-08 | 1.090E-08 | 5.151E-09 | 5.700E-09 | 1.600E-08 |
| 1469 | 74 | L74H | A | 2.4e-09 | 2.2e-09 | 5.3e-09 | 3.300E-09 | 1.735E-09 | 2.200E-09 | 5.300E-09 |
| 1470 | 74 | L74I | A | 3.3e-09 | 5.3e-09 | 1.2e-08 | 6.867E-09 | 4.557E-09 | 3.300E-09 | 1.200E-08 |
| 1471 | 74 | L74L | A | 2.2e-09 | 1.8e-09 | 7.9e-09 | 3.967E-09 | 3.412E-09 | 1.800E-09 | 7.900E-09 |
| 1472 | 74 | L74K | A | 9.7e-09 | 4.2e-09 | 9.6e-09 | 7.833E-09 | 3.147E-09 | 4.200E-09 | 9.700E-09 |
| 1473 | 74 | L74M | A | 8.4e-09 | 5.7e-09 | 1.7e-09 | 5.267E-09 | 3.371E-09 | 1.700E-09 | 8.400E-09 |
| 1474 | 74 | L74F | A | 4.4e-09 | 5e-09 | 1.2e-08 | 7.133E-09 | 4.225E-09 | 4.400E-09 | 1.200E-08 |
| 1475 | 74 | L74P | A | 3.8e-09 | 5.3e-09 | 5.8e-09 | 4.967E-09 | 1.041E-09 | 3.800E-09 | 5.800E-09 |
| 1476 | 74 | L74S | A | 6.2e-09 | 3e-09 | 9.9e-09 | 6.367E-09 | 3.453E-09 | 3.000E-09 | 9.900E-09 |
| 1477 | 74 | L74T | A | 2.1e-09 | 1.1e-08 | 9.3e-09 | 7.467E-09 | 4.725E-09 | 2.100E-09 | 1.100E-08 |
| 1478 | 74 | L74W | A | 5.7e-09 | 7.7e-09 | 3.1e-09 | 5.500E-09 | 2.307E-09 | 3.100E-09 | 7.700E-09 |
| 1479 | 74 | L74Y | A | 6.2e-09 | 1.4e-08 | 4.3e-09 | 8.167E-09 | 5.140E-09 | 4.300E-09 | 1.400E-08 |
| 1480 | 74 | L74V | A | 7.8e-09 | 9.3e-09 | 1e-08 | 9.033E-09 | 1.124E-09 | 7.800E-09 | 1.000E-08 |
| 1481 | 75 | Q75A | A | 5.6e-09 | 1.2e-08 | 9.4e-09 | 9.000E-09 | 3.219E-09 | 5.600E-09 | 1.200E-08 |
| 1482 | 75 | Q75R | A | 6e-09 | 6e-09 | 8.5e-09 | 6.833E-09 | 1.443E-09 | 6.000E-09 | 8.500E-09 |
| 1483 | 75 | Q75N | A | 1.7e-08 | 2.9e-08 | 4.7e-09 | 1.690E-08 | 1.215E-08 | 4.700E-09 | 2.900E-08 |
| 1484 | 75 | Q75D | A | 4.1e-09 | 1.5e-08 | 3.6e-09 | 7.567E-09 | 6.442E-09 | 3.600E-09 | 1.500E-08 |
| 1485 | 75 | Q75C | A | 1.2e-08 | 2.3e-08 | 1.8e-09 | 1.227E-08 | 1.060E-08 | 1.800E-09 | 2.300E-08 |
| 1486 | 75 | Q75E | A | 7.1e-09 | 4.9e-09 | 2.8e-09 | 4.933E-09 | 2.150E-09 | 2.800E-09 | 7.100E-09 |
| 1487 | 75 | Q75Q | A | 3.2e-09 | 9.4e-09 | 3.1e-09 | 5.233E-09 | 3.609E-09 | 3.100E-09 | 9.400E-09 |
| 1488 | 75 | Q75G | A | 8.7e-09 | 1e-08 | 1e-08 | 9.567E-09 | 7.506E-10 | 8.700E-09 | 1.000E-08 |
| 1489 | 75 | Q75H | A | 2e-09 | 2e-08 | 1e-08 | 1.067E-08 | 9.018E-09 | 2.000E-09 | 2.000E-08 |
| 1490 | 75 | Q75I | A | 6.6e-09 | 5.7e-09 | 8.4e-09 | 6.900E-09 | 1.375E-09 | 5.700E-09 | 8.400E-09 |
| 1491 | 75 | Q75L | A | 8.3e-09 | 3.1e-09 | 1.9e-08 | 1.013E-08 | 8.107E-09 | 3.100E-09 | 1.900E-08 |
| 1492 | 75 | Q75K | A | 1.8e-08 | 1.1e-08 | 8.2e-09 | 1.240E-08 | 5.048E-09 | 8.200E-09 | 1.800E-08 |
| 1493 | 75 | Q75M | A | 6.3e-09 | 3.1e-09 | 2.5e-08 | 1.147E-08 | 1.183E-08 | 3.100E-09 | 2.500E-08 |
| 1494 | 75 | Q75F | A | 3.5e-09 | 1.1e-08 | 7.2e-09 | 7.233E-09 | 3.750E-09 | 3.500E-09 | 1.100E-08 |
| 1495 | 75 | Q75P | A | 4e-09 | 8.4e-09 | 5.4e-09 | 5.933E-09 | 2.248E-09 | 4.000E-09 | 8.400E-09 |
| 1496 | 75 | Q75S | A | 2.3e-09 | 7.3e-09 | 8.9e-09 | 6.167E-09 | 3.443E-09 | 2.300E-09 | 8.900E-09 |
| 1497 | 75 | Q75T | A | 5.2e-09 | 9.1e-09 | 9.2e-09 | 7.833E-09 | 2.281E-09 | 5.200E-09 | 9.200E-09 |
| 1498 | 75 | Q75W | A | 1.6e-09 | 4.5e-09 | 1e-08 | 5.367E-09 | 4.267E-09 | 1.600E-09 | 1.000E-08 |
| 1499 | 75 | Q75Y | A | 4.1e-09 | 1e-08 | 1.2e-08 | 8.700E-09 | 4.107E-09 | 4.100E-09 | 1.200E-08 |
| 1500 | 75 | Q75V | A | 1.8e-08 | 1.1e-08 | 6.7e-09 | 1.190E-08 | 5.704E-09 | 6.700E-09 | 1.800E-08 |
| 1501 | 76 | R76A | A | 1.6e-08 | 5.3e-09 | 3.8e-09 | 8.367E-09 | 6.653E-09 | 3.800E-09 | 1.600E-08 |
| 1502 | 76 | R76R | A | 6e-09 | 4.4e-09 | 4.3e-09 | 4.900E-09 | 9.539E-10 | 4.300E-09 | 6.000E-09 |
| 1503 | 76 | R76N | A | 5.9e-09 | 1.2e-08 | 1.4e-08 | 1.063E-08 | 4.219E-09 | 5.900E-09 | 1.400E-08 |
| 1504 | 76 | R76D | A | 2.1e-09 | 8.4e-09 | 1.3e-08 | 7.833E-09 | 5.472E-09 | 2.100E-09 | 1.300E-08 |
| 1505 | 76 | R76C | A | 6.6e-09 | 3.8e-09 | 7.2e-09 | 5.867E-09 | 1.815E-09 | 3.800E-09 | 7.200E-09 |
| 1506 | 76 | R76E | A | 5.1e-09 | 7.2e-09 | 9.7e-09 | 7.333E-09 | 2.303E-09 | 5.100E-09 | 9.700E-09 |
| 1507 | 76 | R76Q | A | 4.7e-09 | 4.7e-09 | 1.1e-08 | 6.800E-09 | 3.637E-09 | 4.700E-09 | 1.100E-08 |
| 1508 | 76 | R76G | A | 1.2e-08 | 1.1e-08 | 4e-09 | 9.000E-09 | 4.359E-09 | 4.000E-09 | 1.200E-08 |
| 1509 | 76 | R76H | A | 6.1e-09 | 6.5e-09 | 1.2e-09 | 4.600E-09 | 2.951E-09 | 1.200E-09 | 6.500E-09 |
| 1510 | 76 | R76I | A | 7.4e-09 | 1.7e-09 | 7.1e-09 | 5.400E-09 | 3.208E-09 | 1.700E-09 | 7.400E-09 |
| 1511 | 76 | R76L | A | 9.1e-09 | 9.7e-09 | 3.5e-09 | 7.433E-09 | 3.420E-09 | 3.500E-09 | 9.700E-09 |
| 1512 | 76 | R76K | A | 8.6e-09 | 9.6e-09 | 5.3e-09 | 7.833E-09 | 2.250E-09 | 5.300E-09 | 9.600E-09 |
| 1513 | 76 | R76M | A | 5.8e-09 | 3.5e-09 | 1.3e-08 | 7.433E-09 | 4.956E-09 | 3.500E-09 | 1.300E-08 |
| 1514 | 76 | R76F | A | 7.4e-09 | 8.7e-09 | 1.2e-08 | 9.367E-09 | 2.371E-09 | 7.400E-09 | 1.200E-08 |
| 1515 | 76 | R76P | A | 1.5e-08 | 1.4e-08 | 1.5e-08 | 1.467E-08 | 5.774E-10 | 1.400E-08 | 1.500E-08 |
| 1516 | 76 | R76S | A | 5.4e-09 | 6.1e-09 | 1.4e-08 | 8.500E-09 | 4.776E-09 | 5.400E-09 | 1.400E-08 |
| 1517 | 76 | R76T | A | 3.4e-09 | 1.3e-08 | 2e-09 | 6.133E-09 | 5.988E-09 | 2.000E-09 | 1.300E-08 |
| 1518 | 76 | R76W | A | 8e-09 | 4.9e-09 | 3.6e-09 | 5.500E-09 | 2.261E-09 | 3.600E-09 | 8.000E-09 |
| 1519 | 76 | R76Y | A | 1.6e-08 | 1.2e-08 | 5.7e-09 | 1.123E-08 | 5.193E-09 | 5.700E-09 | 1.600E-08 |
| 1520 | 76 | R76V | A | 5.3e-09 | 4.8e-09 | 1.3e-08 | 7.700E-09 | 4.597E-09 | 4.800E-09 | 1.300E-08 |
| 1521 | 77 | L77A | A | 1.4e-08 | 7.6e-09 | 8.5e-09 | 1.003E-08 | 3.465E-09 | 7.600E-09 | 1.400E-08 |
| 1522 | 77 | L77R | A | 1.3e-08 | 7.9e-09 | 1.3e-08 | 1.130E-08 | 2.944E-09 | 7.900E-09 | 1.300E-08 |
| 1523 | 77 | L77N | A | 1e-08 | 4e-09 | 1.2e-08 | 8.667E-09 | 4.163E-09 | 4.000E-09 | 1.200E-08 |
| 1524 | 77 | L77D | A | 4.1e-09 | 2.2e-09 | 1e-08 | 5.433E-09 | 4.067E-09 | 2.200E-09 | 1.000E-08 |
| 1525 | 77 | L77C | A | 7.5e-09 | 9.4e-09 | 6.1e-09 | 7.667E-09 | 1.656E-09 | 6.100E-09 | 9.400E-09 |
| 1526 | 77 | L77E | A | 5.7e-09 | 6.9e-09 | 4.3e-09 | 5.633E-09 | 1.301E-09 | 4.300E-09 | 6.900E-09 |
| 1527 | 77 | L77Q | A | 4e-09 | 3.8e-09 | 1.3e-08 | 6.933E-09 | 5.255E-09 | 3.800E-09 | 1.300E-08 |
| 1528 | 77 | L77G | A | 6.2e-09 | 5.8e-09 | 1.4e-08 | 8.667E-09 | 4.623E-09 | 5.800E-09 | 1.400E-08 |
| 1529 | 77 | L77H | A | 4.3e-09 | 1.3e-08 | 2.5e-09 | 6.600E-09 | 5.615E-09 | 2.500E-09 | 1.300E-08 |

| | | | | | | | | | | |
|---|---|---|---|---|---|---|---|---|---|---|
| 1530 | 77 | L77I | A | 5.4e-09 | 7.2e-09 | 3.7e-08 | 1.653E-08 | 1.775E-08 | 5.400E-09 | 3.700E-08 |
| 1531 | 77 | L77L | A | 6.4e-09 | 7.5e-09 | 3.6e-09 | 5.833E-09 | 2.011E-09 | 3.600E-09 | 7.500E-09 |
| 1532 | 77 | L77K | A | 1.7e-09 | 3.8e-09 | 2.1e-08 | 8.833E-09 | 1.059E-08 | 1.700E-09 | 2.100E-08 |
| 1533 | 77 | L77M | A | 1e-08 | 2.3e-09 | 3.4e-09 | 5.233E-09 | 4.165E-09 | 2.300E-09 | 1.000E-08 |
| 1534 | 77 | L77F | A | 1e-08 | 6.6e-09 | 9.2e-09 | 8.600E-09 | 1.778E-09 | 6.600E-09 | 1.000E-08 |
| 1535 | 77 | L77P | A | 1.1e-08 | 6.7e-09 | 1.4e-08 | 1.057E-08 | 3.669E-09 | 6.700E-09 | 1.400E-08 |
| 1536 | 77 | L77S | A | 3.2e-09 | 5.9e-09 | 4.7e-09 | 4.600E-09 | 1.353E-09 | 3.200E-09 | 5.900E-09 |
| 1537 | 77 | L77T | A | 1.8e-08 | 4.9e-09 | 9.5e-09 | 1.080E-08 | 6.646E-09 | 4.900E-09 | 1.800E-08 |
| 1538 | 77 | L77W | A | 6.1e-09 | 2.3e-09 | 5.7e-09 | 4.700E-09 | 2.088E-09 | 2.300E-09 | 6.100E-09 |
| 1539 | 77 | L77Y | A | 3.5e-09 | 1.2e-08 | 1.1e-08 | 8.833E-09 | 4.646E-09 | 3.500E-09 | 1.200E-08 |
| 1540 | 77 | L77V | A | 9.1e-09 | 8.2e-09 | 9.4e-09 | 8.900E-09 | 6.245E-10 | 8.200E-09 | 9.400E-09 |
| 1541 | 78 | R78A | A | 1.3e-08 | 4.8e-09 | 6.2e-09 | 8.000E-09 | 4.386E-09 | 4.800E-09 | 1.300E-08 |
| 1542 | 78 | R78R | A | 7.7e-09 | 5.8e-09 | 5.5e-09 | 6.333E-09 | 1.193E-09 | 5.500E-09 | 7.700E-09 |
| 1543 | 78 | R78N | A | 1.3e-08 | 6.2e-09 | 2.2e-09 | 7.133E-09 | 5.460E-09 | 2.200E-09 | 1.300E-08 |
| 1544 | 78 | R78D | A | 1.6e-08 | 1.2e-09 | 8.4e-09 | 8.533E-09 | 7.401E-09 | 1.200E-09 | 1.600E-08 |
| 1545 | 78 | R78C | A | 1.3e-08 | 2.8e-09 | 4.8e-09 | 6.867E-09 | 5.405E-09 | 2.800E-09 | 1.300E-08 |
| 1546 | 78 | R78E | A | 1.4e-08 | 9.4e-09 | 1.3e-08 | 1.213E-08 | 2.419E-09 | 9.400E-09 | 1.400E-08 |
| 1547 | 78 | R78Q | A | 9.1e-09 | 1.1e-08 | 4.4e-09 | 8.167E-09 | 3.398E-09 | 4.400E-09 | 1.100E-08 |
| 1548 | 78 | R78G | A | 9.2e-09 | 6.9e-09 | 9.1e-09 | 8.400E-09 | 1.300E-09 | 6.900E-09 | 9.200E-09 |
| 1549 | 78 | R78H | A | 4.4e-09 | 6.2e-09 | 1.1e-08 | 7.200E-09 | 3.412E-09 | 4.400E-09 | 1.100E-08 |
| 1550 | 78 | R78I | A | 3.3e-09 | 7e-09 | 4.2e-09 | 4.833E-09 | 1.930E-09 | 3.300E-09 | 7.000E-09 |
| 1551 | 78 | R78L | A | 1.2e-08 | 3.9e-09 | 8.1e-09 | 8.000E-09 | 4.051E-09 | 3.900E-09 | 1.200E-08 |
| 1552 | 78 | R78K | A | 6.6e-09 | 5.5e-09 | 2.9e-09 | 5.000E-09 | 1.900E-09 | 2.900E-09 | 6.600E-09 |
| 1553 | 78 | R78M | A | 5.2e-09 | 4.2e-09 | 5.9e-09 | 5.100E-09 | 8.544E-10 | 4.200E-09 | 5.900E-09 |
| 1554 | 78 | R78F | A | 2.4e-09 | 5.9e-09 | 1.7e-08 | 8.433E-09 | 7.623E-09 | 2.400E-09 | 1.700E-08 |
| 1555 | 78 | R78P | A | 5.2e-09 | 3.2e-08 | 5.5e-09 | 1.423E-08 | 1.539E-08 | 5.200E-09 | 3.200E-08 |
| 1556 | 78 | R78S | A | 6.9e-09 | 6.3e-09 | 1.4e-08 | 9.067E-09 | 4.283E-09 | 6.300E-09 | 1.400E-08 |
| 1557 | 78 | R78T | A | 5.3e-09 | 1.5e-08 | 5e-09 | 8.433E-09 | 5.689E-09 | 5.000E-09 | 1.500E-08 |
| 1558 | 78 | R78W | A | 2.7e-08 | 3.7e-09 | 3.1e-09 | 1.127E-08 | 1.363E-08 | 3.100E-09 | 2.700E-08 |
| 1559 | 78 | R78Y | A | 3.7e-09 | 6.3e-09 | 1e-08 | 6.667E-09 | 3.166E-09 | 3.700E-09 | 1.000E-08 |
| 1560 | 78 | R78V | A | 3e-09 | 6.3e-09 | 9e-09 | 6.100E-09 | 3.005E-09 | 3.000E-09 | 9.000E-09 |
| 1561 | 79 | I79A | A | 1.3e-08 | 7.4e-09 | 3.5e-09 | 7.967E-09 | 4.775E-09 | 3.500E-09 | 1.300E-08 |
| 1562 | 79 | I79R | A | 8.7e-09 | 4.5e-09 | 2.8e-08 | 1.373E-08 | 1.253E-08 | 4.500E-09 | 2.800E-08 |
| 1563 | 79 | I79N | A | 2.5e-08 | 6.4e-09 | 8.2e-09 | 1.320E-08 | 1.026E-08 | 6.400E-09 | 2.500E-08 |
| 1564 | 79 | I79D | A | 1.6e-08 | 5.7e-09 | 7.4e-09 | 9.700E-09 | 5.522E-09 | 5.700E-09 | 1.600E-08 |
| 1565 | 79 | I79C | A | 6.2e-09 | 1.4e-08 | 7.5e-09 | 9.233E-09 | 4.179E-09 | 6.200E-09 | 1.400E-08 |
| 1566 | 79 | I79E | A | 1.2e-08 | 7.5e-09 | 1.1e-08 | 1.017E-08 | 2.363E-09 | 7.500E-09 | 1.200E-08 |
| 1567 | 79 | I79Q | A | 9.7e-09 | 8.1e-09 | 7.5e-09 | 8.433E-09 | 1.137E-09 | 7.500E-09 | 9.700E-09 |
| 1568 | 79 | I79G | A | 4.5e-09 | 5.7e-09 | 1.5e-08 | 8.400E-09 | 5.747E-09 | 4.500E-09 | 1.500E-08 |
| 1569 | 79 | I79H | A | 2.1e-08 | 8.2e-09 | 6.7e-09 | 1.197E-08 | 7.859E-09 | 6.700E-09 | 2.100E-08 |
| 1570 | 79 | I79I | A | 4.1e-09 | 4e-09 | 1e-08 | 6.033E-09 | 3.436E-09 | 4.000E-09 | 1.000E-08 |
| 1571 | 79 | I79L | A | 8.1e-09 | 9.6e-09 | 6.9e-09 | 8.200E-09 | 1.353E-09 | 6.900E-09 | 9.600E-09 |
| 1572 | 79 | I79K | A | 5.7e-09 | 5.4e-09 | 3.1e-09 | 4.733E-09 | 1.422E-09 | 3.100E-09 | 5.700E-09 |
| 1573 | 79 | I79M | A | 1.3e-08 | 5e-09 | 1.1e-08 | 9.667E-09 | 4.163E-09 | 5.000E-09 | 1.300E-08 |
| 1574 | 79 | I79F | A | 1.2e-08 | 5.5e-09 | 4.4e-09 | 7.300E-09 | 4.107E-09 | 4.400E-09 | 1.200E-08 |
| 1575 | 79 | I79P | A | 1.5e-08 | 3.4e-09 | 2.6e-08 | 1.480E-08 | 1.130E-08 | 3.400E-09 | 2.600E-08 |
| 1576 | 79 | I79S | A | 9e-09 | 1.4e-08 | 1.1e-08 | 1.133E-08 | 2.517E-09 | 9.000E-09 | 1.400E-08 |
| 1577 | 79 | I79T | A | 6.8e-09 | 1.8e-08 | 2e-08 | 1.493E-08 | 7.114E-09 | 6.800E-09 | 2.000E-08 |
| 1578 | 79 | I79W | A | 2.6e-08 | 7.9e-09 | 3.8e-09 | 1.257E-08 | 1.181E-08 | 3.800E-09 | 2.600E-08 |
| 1579 | 79 | I79Y | A | 7.1e-09 | 5e-09 | 3.5e-09 | 5.200E-09 | 1.808E-09 | 3.500E-09 | 7.100E-09 |
| 1580 | 79 | I79V | A | 1.1e-08 | 9.1e-09 | 1.1e-08 | 1.037E-08 | 1.097E-09 | 9.100E-09 | 1.100E-08 |
| 1581 | 80 | V80A | A | 8.5e-09 | 4.4e-09 | 5.5e-09 | 6.133E-09 | 2.122E-09 | 4.400E-09 | 8.500E-09 |
| 1582 | 80 | V80R | A | 9.4e-09 | 5.7e-09 | 9.4e-09 | 8.167E-09 | 2.136E-09 | 5.700E-09 | 9.400E-09 |
| 1583 | 80 | V80N | A | 4.9e-09 | 1.4e-08 | 5.1e-09 | 8.000E-09 | 5.197E-09 | 4.900E-09 | 1.400E-08 |
| 1584 | 80 | V80D | A | 1.1e-08 | 1.6e-08 | 5.4e-09 | 1.080E-08 | 5.303E-09 | 5.400E-09 | 1.600E-08 |
| 1585 | 80 | V80C | A | 2.4e-09 | 3.3e-09 | 2.4e-09 | 2.700E-09 | 5.196E-10 | 2.400E-09 | 3.300E-09 |
| 1586 | 80 | V80E | A | 3.9e-09 | 7.6e-09 | 9.7e-09 | 7.067E-09 | 2.937E-09 | 3.900E-09 | 9.700E-09 |
| 1587 | 80 | V80Q | A | 8.6e-09 | 1.1e-08 | 8.9e-09 | 9.500E-09 | 1.308E-09 | 8.600E-09 | 1.100E-08 |
| 1588 | 80 | V80G | A | 5.8e-09 | 1.2e-09 | 8.3e-09 | 5.100E-09 | 3.601E-09 | 1.200E-09 | 8.300E-09 |
| 1589 | 80 | V80H | A | 8e-09 | 3.1e-09 | 5.9e-09 | 5.667E-09 | 2.458E-09 | 3.100E-09 | 8.000E-09 |
| 1590 | 80 | V80I | A | 8e-09 | 1e-08 | 3.5e-09 | 7.167E-09 | 3.329E-09 | 3.500E-09 | 1.000E-08 |
| 1591 | 80 | V80L | A | 4.7e-09 | 3.1e-09 | 5.3e-09 | 4.367E-09 | 1.137E-09 | 3.100E-09 | 5.300E-09 |
| 1592 | 80 | V80K | A | 6e-09 | 1.2e-08 | 2.4e-09 | 6.800E-09 | 4.850E-09 | 2.400E-09 | 1.200E-08 |
| 1593 | 80 | V80M | A | 1.2e-08 | 1.2e-08 | 9.2e-09 | 1.107E-08 | 1.617E-09 | 9.200E-09 | 1.200E-08 |
| 1594 | 80 | V80F | A | 5.7e-09 | 3.9e-09 | 8.7e-09 | 6.100E-09 | 2.425E-09 | 3.900E-09 | 8.700E-09 |
| 1595 | 80 | V80P | A | 1.9e-09 | 9e-09 | 7.1e-09 | 6.000E-09 | 3.676E-09 | 1.900E-09 | 9.000E-09 |
| 1596 | 80 | V80S | A | 6.2e-09 | 7e-09 | 4.2e-09 | 5.800E-09 | 1.442E-09 | 4.200E-09 | 7.000E-09 |
| 1597 | 80 | V80T | A | 1.3e-08 | 1.5e-08 | 3.3e-09 | 1.043E-08 | 6.258E-09 | 3.300E-09 | 1.500E-08 |
| 1598 | 80 | V80W | A | 4.2e-09 | 6.9e-09 | 1.2e-08 | 7.700E-09 | 3.961E-09 | 4.200E-09 | 1.200E-08 |
| 1599 | 80 | V80Y | A | 7.5e-09 | 8.6e-09 | 1.5e-09 | 5.867E-09 | 3.821E-09 | 1.500E-09 | 8.600E-09 |
| 1600 | 80 | V80V | A | 1e-08 | 9.4e-09 | 5.1e-09 | 8.167E-09 | 2.673E-09 | 5.100E-09 | 1.000E-08 |
| 1601 | 81 | R81A | A | 7.5e-09 | 6.9e-09 | 6.5e-09 | 6.967E-09 | 5.033E-10 | 6.500E-09 | 7.500E-09 |
| 1602 | 81 | R81R | A | 2.2e-09 | 9.3e-09 | 3.2e-09 | 4.900E-09 | 3.843E-09 | 2.200E-09 | 9.300E-09 |
| 1603 | 81 | R81N | A | 1.2e-08 | 4.3e-09 | 6.9e-09 | 7.733E-09 | 3.917E-09 | 4.300E-09 | 1.200E-08 |
| 1604 | 81 | R81D | A | 1.7e-09 | 1.3e-08 | 9.4e-09 | 8.033E-09 | 5.773E-09 | 1.700E-09 | 1.300E-08 |
| 1605 | 81 | R81C | A | 4.4e-09 | 4.5e-09 | 4.1e-09 | 4.333E-09 | 2.082E-10 | 4.100E-09 | 4.500E-09 |
| 1606 | 81 | R81E | A | 2.3e-09 | 2.3e-08 | 1e-08 | 1.177E-08 | 1.046E-08 | 2.300E-09 | 2.300E-08 |
| 1607 | 81 | R81Q | A | 1.1e-08 | 1.2e-08 | 5e-09 | 9.333E-09 | 3.786E-09 | 5.000E-09 | 1.200E-08 |
| 1608 | 81 | R81G | A | 4.8e-09 | 7.8e-09 | 6.8e-09 | 6.467E-09 | 1.528E-09 | 4.800E-09 | 7.800E-09 |
| 1609 | 81 | R81H | A | 1.6e-08 | 7.1e-09 | 4.7e-09 | 9.267E-09 | 5.953E-09 | 4.700E-09 | 1.600E-08 |
| 1610 | 81 | R81I | A | 3.4e-09 | 7e-09 | 1.3e-08 | 7.800E-09 | 4.850E-09 | 3.400E-09 | 1.300E-08 |
| 1611 | 81 | R81L | A | 2.4e-09 | 4.5e-09 | 2.6e-09 | 3.167E-09 | 1.159E-09 | 2.400E-09 | 4.500E-09 |
| 1612 | 81 | R81K | A | 8.4e-09 | 2.3e-09 | 7.9e-09 | 6.200E-09 | 3.387E-09 | 2.300E-09 | 8.400E-09 |
| 1613 | 81 | R81M | A | 4.6e-09 | 1.9e-09 | 8.6e-09 | 5.033E-09 | 3.371E-09 | 1.900E-09 | 8.600E-09 |
| 1614 | 81 | R81F | A | 8.8e-09 | 1.1e-08 | 2.9e-09 | 7.567E-09 | 4.188E-09 | 2.900E-09 | 1.100E-08 |

| | | | | | | | | | | |
|---|---|---|---|---|---|---|---|---|---|---|
| 1615 | 81 | R81P | A | 5.2e-09 | 1.1e-08 | 6.5e-09 | 7.567E-09 | 3.044E-09 | 5.200E-09 | 1.100E-08 |
| 1616 | 81 | R81S | A | 9.3e-09 | 3.8e-09 | 1e-08 | 7.700E-09 | 3.396E-09 | 3.800E-09 | 1.000E-08 |
| 1617 | 81 | R81T | A | 1.5e-08 | 3.7e-09 | 1.2e-08 | 1.023E-08 | 5.853E-09 | 3.700E-09 | 1.500E-08 |
| 1618 | 81 | R81W | A | 1.2e-08 | 1e-08 | 7.1e-09 | 9.700E-09 | 2.464E-09 | 7.100E-09 | 1.200E-08 |
| 1619 | 81 | R81Y | A | 6e-09 | 3.9e-09 | 3.6e-09 | 4.500E-09 | 1.308E-09 | 3.600E-09 | 6.000E-09 |
| 1620 | 81 | R81V | A | 9.9e-09 | 6.3e-09 | 2.9e-08 | 1.507E-08 | 1.220E-08 | 6.300E-09 | 2.900E-08 |
| 1621 | 82 | G82A | A | 3.3e-09 | 6.7e-09 | 1.2e-08 | 7.333E-09 | 4.384E-09 | 3.300E-09 | 1.200E-08 |
| 1622 | 82 | G82R | A | 9.6e-09 | 2.4e-08 | 5.2e-09 | 1.293E-08 | 9.833E-09 | 5.200E-09 | 2.400E-08 |
| 1623 | 82 | G82N | A | 8.5e-09 | 6.9e-09 | 1.2e-08 | 9.133E-09 | 2.608E-09 | 6.900E-09 | 1.200E-08 |
| 1624 | 82 | G82D | A | 9.8e-09 | 1.3e-08 | 5.3e-09 | 9.367E-09 | 3.868E-09 | 5.300E-09 | 1.300E-08 |
| 1625 | 82 | G82C | A | 8.9e-09 | 6.2e-09 | 1.5e-08 | 1.003E-08 | 4.508E-09 | 6.200E-09 | 1.500E-08 |
| 1626 | 82 | G82E | A | 9.2e-09 | 9.3e-09 | 7.4e-09 | 8.633E-09 | 1.069E-09 | 7.400E-09 | 9.300E-09 |
| 1627 | 82 | G82Q | A | 6.4e-09 | 1e-08 | 1.2e-08 | 9.467E-09 | 2.838E-09 | 6.400E-09 | 1.200E-08 |
| 1628 | 82 | G82G | A | 2.6e-08 | 7.3e-09 | 1.7e-08 | 1.677E-08 | 9.352E-09 | 7.300E-09 | 2.600E-08 |
| 1629 | 82 | G82H | A | 2.4e-09 | 6.9e-09 | 8.4e-09 | 5.900E-09 | 3.122E-09 | 2.400E-09 | 8.400E-09 |
| 1630 | 82 | G82I | A | 4.6e-09 | 3e-09 | 1.4e-08 | 7.200E-09 | 5.943E-09 | 3.000E-09 | 1.400E-08 |
| 1631 | 82 | G82L | A | 5.2e-09 | 1.2e-08 | 1.1e-08 | 9.400E-09 | 3.672E-09 | 5.200E-09 | 1.200E-08 |
| 1632 | 82 | G82K | A | 8.4e-10 | 5.7e-09 | 5.2e-09 | 3.913E-09 | 2.673E-09 | 8.400E-10 | 5.700E-09 |
| 1633 | 82 | G82M | A | 1.1e-09 | 4.1e-09 | 7.6e-09 | 4.267E-09 | 3.253E-09 | 1.100E-09 | 7.600E-09 |
| 1634 | 82 | G82F | A | 7.9e-09 | 7.8e-09 | 8.7e-09 | 8.133E-09 | 4.933E-10 | 7.800E-09 | 8.700E-09 |
| 1635 | 82 | G82P | A | 1.2e-08 | 6e-08 | 4.7e-09 | 2.557E-08 | 3.004E-08 | 4.700E-09 | 6.000E-08 |
| 1636 | 82 | G82S | A | 1.9e-08 | 1.3e-08 | 7.4e-09 | 1.313E-08 | 5.801E-09 | 7.400E-09 | 1.900E-08 |
| 1637 | 82 | G82T | A | 1.9e-09 | 6.4e-09 | 3.4e-09 | 3.900E-09 | 2.291E-09 | 1.900E-09 | 6.400E-09 |
| 1638 | 82 | G82W | A | 9.2e-09 | 3.9e-09 | 1.1e-08 | 8.033E-09 | 3.691E-09 | 3.900E-09 | 1.100E-08 |
| 1639 | 82 | G82Y | A | 6.6e-09 | 1.1e-08 | 1e-08 | 9.200E-09 | 2.307E-09 | 6.600E-09 | 1.100E-08 |
| 1640 | 82 | G82V | A | 1.1e-08 | 1.3e-08 | 7.9e-09 | 1.063E-08 | 2.570E-09 | 7.900E-09 | 1.300E-08 |
| 1641 | 83 | T83A | A | 5.6e-09 | 8.9e-09 | 7.3e-09 | 7.267E-09 | 1.650E-09 | 5.600E-09 | 8.900E-09 |
| 1642 | 83 | T83R | A | 4.3e-09 | 5.5e-09 | 4.9e-09 | 4.900E-09 | 6.000E-10 | 4.300E-09 | 5.500E-09 |
| 1643 | 83 | T83N | A | 1.1e-08 | 1.1e-08 | 1.5e-08 | 1.233E-08 | 2.309E-09 | 1.100E-08 | 1.500E-08 |
| 1644 | 83 | T83D | A | 6.4e-09 | 7.8e-09 | 1e-08 | 8.067E-09 | 1.815E-09 | 6.400E-09 | 1.000E-08 |
| 1645 | 83 | T83C | A | 6.9e-09 | 1.5e-08 | 8.9e-09 | 1.027E-08 | 4.219E-09 | 6.900E-09 | 1.500E-08 |
| 1646 | 83 | T83E | A | 1.3e-08 | 9.4e-09 | 4.5e-09 | 8.967E-09 | 4.267E-09 | 4.500E-09 | 1.300E-08 |
| 1647 | 83 | T83Q | A | 1.1e-08 | 3.4e-09 | 9.7e-09 | 8.033E-09 | 4.065E-09 | 3.400E-09 | 1.100E-08 |
| 1648 | 83 | T83G | A | 5.2e-09 | 4e-09 | 8.7e-09 | 5.967E-09 | 2.442E-09 | 4.000E-09 | 8.700E-09 |
| 1649 | 83 | T83H | A | 9.8e-09 | 2.6e-08 | 9.3e-09 | 1.503E-08 | 9.501E-09 | 9.300E-09 | 2.600E-08 |
| 1650 | 83 | T83I | A | 1.1e-08 | 4.8e-09 | 4.6e-09 | 6.800E-09 | 3.639E-09 | 4.600E-09 | 1.100E-08 |
| 1651 | 83 | T83L | A | 1.5e-08 | 4.1e-09 | 3e-09 | 7.367E-09 | 6.634E-09 | 3.000E-09 | 1.500E-08 |
| 1652 | 83 | T83K | A | 5.3e-09 | 9.5e-09 | 4.1e-09 | 6.300E-09 | 2.835E-09 | 4.100E-09 | 9.500E-09 |
| 1653 | 83 | T83M | A | 1e-08 | 2.3e-09 | 4.2e-09 | 5.500E-09 | 4.011E-09 | 2.300E-09 | 1.000E-08 |
| 1654 | 83 | T83F | A | 4.5e-09 | 8.9e-09 | 6.1e-09 | 6.500E-09 | 2.227E-09 | 4.500E-09 | 8.900E-09 |
| 1655 | 83 | T83P | A | 2.5e-08 | 1.4e-08 | 7.1e-09 | 1.537E-08 | 9.028E-09 | 7.100E-09 | 2.500E-08 |
| 1656 | 83 | T83S | A | 2.1e-08 | 3.7e-09 | 1e-08 | 1.157E-08 | 8.756E-09 | 3.700E-09 | 2.100E-08 |
| 1657 | 83 | T83T | A | 1.5e-08 | 1e-08 | 1.3e-08 | 1.267E-08 | 2.517E-09 | 1.000E-08 | 1.500E-08 |
| 1658 | 83 | T83W | A | 4.2e-09 | 6.5e-09 | 5.7e-09 | 5.467E-09 | 1.168E-09 | 4.200E-09 | 6.500E-09 |
| 1659 | 83 | T83Y | A | 5.9e-09 | 1e-08 | 4.5e-09 | 6.800E-09 | 2.858E-09 | 4.500E-09 | 1.000E-08 |
| 1660 | 83 | T83V | A | 9.6e-09 | 1.4e-08 | 6.8e-09 | 1.013E-08 | 3.630E-09 | 6.800E-09 | 1.400E-08 |
| 1661 | 84 | Q84A | A | 3e-09 | 7.1e-09 | 6e-09 | 5.367E-09 | 2.122E-09 | 3.000E-09 | 7.100E-09 |
| 1662 | 84 | Q84R | A | 7.2e-09 | 6.9e-09 | 1.1e-08 | 8.367E-09 | 2.285E-09 | 6.900E-09 | 1.100E-08 |
| 1663 | 84 | Q84N | A | 4.7e-09 | 8.7e-09 | 1.9e-09 | 5.100E-09 | 3.418E-09 | 1.900E-09 | 8.700E-09 |
| 1664 | 84 | Q84D | A | 8.7e-09 | 5.9e-09 | 7.4e-09 | 7.333E-09 | 1.401E-09 | 5.900E-09 | 8.700E-09 |
| 1665 | 84 | Q84C | A | 1.5e-08 | 1.1e-08 | 2.4e-09 | 9.467E-09 | 6.438E-09 | 2.400E-09 | 1.500E-08 |
| 1666 | 84 | Q84E | A | 1.3e-08 | 5.6e-09 | 1.1e-08 | 9.867E-09 | 3.828E-09 | 5.600E-09 | 1.300E-08 |
| 1667 | 84 | Q84Q | A | 8.8e-09 | 1.2e-08 | 8.1e-09 | 9.633E-09 | 2.079E-09 | 8.100E-09 | 1.200E-08 |
| 1668 | 84 | Q84G | A | 5.2e-09 | 9.7e-09 | 4.4e-09 | 6.433E-09 | 2.857E-09 | 4.400E-09 | 9.700E-09 |
| 1669 | 84 | Q84H | A | 6.5e-09 | 9e-09 | 2.6e-08 | 1.383E-08 | 1.061E-08 | 6.500E-09 | 2.600E-08 |
| 1670 | 84 | Q84I | A | 8.2e-09 | 1.3e-08 | 7.3e-09 | 9.500E-09 | 3.064E-09 | 7.300E-09 | 1.300E-08 |
| 1671 | 84 | Q84L | A | 1.5e-08 | 9e-09 | 2.7e-09 | 8.900E-09 | 6.151E-09 | 2.700E-09 | 1.500E-08 |
| 1672 | 84 | Q84K | A | 8.3e-09 | 8.2e-09 | 9.6e-09 | 8.700E-09 | 7.810E-10 | 8.200E-09 | 9.600E-09 |
| 1673 | 84 | Q84M | A | 5.8e-09 | 7.1e-09 | 5.8e-09 | 6.233E-09 | 7.506E-10 | 5.800E-09 | 7.100E-09 |
| 1674 | 84 | Q84F | A | 1.3e-08 | 4.1e-09 | 5.1e-09 | 7.400E-09 | 4.875E-09 | 4.100E-09 | 1.300E-08 |
| 1675 | 84 | Q84P | A | 5e-09 | 2.6e-09 | 2.6e-09 | 3.400E-09 | 1.386E-09 | 2.600E-09 | 5.000E-09 |
| 1676 | 84 | Q84S | A | 4.5e-09 | 6.7e-09 | 3.7e-09 | 4.967E-09 | 1.553E-09 | 3.700E-09 | 6.700E-09 |
| 1677 | 84 | Q84T | A | 1.8e-08 | 1.9e-09 | 9.2e-09 | 9.700E-09 | 8.062E-09 | 1.900E-09 | 1.800E-08 |
| 1678 | 84 | Q84W | A | 1.1e-08 | 4.2e-08 | 8.2e-09 | 2.040E-08 | 1.876E-08 | 8.200E-09 | 4.200E-08 |
| 1679 | 84 | Q84Y | A | 1.4e-08 | 1.4e-08 | 2.5e-09 | 1.017E-08 | 6.640E-09 | 2.500E-09 | 1.400E-08 |
| 1680 | 84 | Q84V | A | 2.5e-08 | 1.1e-08 | 4.7e-09 | 1.357E-08 | 1.039E-08 | 4.700E-09 | 2.500E-08 |
| 1681 | 85 | L85A | A | 3.3e-09 | 3.3e-09 | 4.2e-09 | 3.600E-09 | 5.196E-10 | 3.300E-09 | 4.200E-09 |
| 1682 | 85 | L85R | A | 1.8e-08 | 1.2e-08 | 3.5e-09 | 1.117E-08 | 7.286E-09 | 3.500E-09 | 1.800E-08 |
| 1683 | 85 | L85N | A | 4.3e-09 | 4.8e-09 | 7.6e-09 | 5.567E-09 | 1.779E-09 | 4.300E-09 | 7.600E-09 |
| 1684 | 85 | L85D | A | 3.5e-09 | 1.5e-08 | 9.4e-09 | 9.300E-09 | 5.751E-09 | 3.500E-09 | 1.500E-08 |
| 1685 | 85 | L85C | A | 4.1e-09 | 1.3e-08 | 2.1e-08 | 1.270E-08 | 8.454E-09 | 4.100E-09 | 2.100E-08 |
| 1686 | 85 | L85E | A | 3.4e-08 | 7.4e-09 | 6.2e-09 | 1.587E-08 | 1.572E-08 | 6.200E-09 | 3.400E-08 |
| 1687 | 85 | L85Q | A | 5.9e-09 | 8.9e-09 | 1.7e-09 | 5.500E-09 | 3.617E-09 | 1.700E-09 | 8.900E-09 |
| 1688 | 85 | L85G | A | 6e-09 | 1.1e-08 | 1.5e-08 | 1.067E-08 | 4.509E-09 | 6.000E-09 | 1.500E-08 |
| 1689 | 85 | L85H | A | 8.8e-09 | 7.8e-09 | 1e-08 | 8.867E-09 | 1.102E-09 | 7.800E-09 | 1.000E-08 |
| 1690 | 85 | L85I | A | 7.3e-09 | 7.7e-09 | 6.8e-09 | 7.267E-09 | 4.509E-10 | 6.800E-09 | 7.700E-09 |
| 1691 | 85 | L85L | A | 1.3e-08 | 9.3e-09 | 4e-09 | 8.767E-09 | 4.524E-09 | 4.000E-09 | 1.300E-08 |
| 1692 | 85 | L85K | A | 7.8e-09 | 1.2e-08 | 2.8e-09 | 7.533E-09 | 4.606E-09 | 2.800E-09 | 1.200E-08 |
| 1693 | 85 | L85M | A | 6.1e-09 | 8.4e-09 | 3.6e-09 | 6.033E-09 | 2.401E-09 | 3.600E-09 | 8.400E-09 |
| 1694 | 85 | L85F | A | 1e-08 | 1.2e-08 | 6.7e-09 | 9.567E-09 | 2.676E-09 | 6.700E-09 | 1.200E-08 |
| 1695 | 85 | L85P | A | 1.3e-08 | 6.5e-09 | 5.4e-09 | 8.300E-09 | 4.107E-09 | 5.400E-09 | 1.300E-08 |
| 1696 | 85 | L85S | A | 2.4e-09 | 6.9e-09 | 5.7e-09 | 5.000E-09 | 2.330E-09 | 2.400E-09 | 6.900E-09 |
| 1697 | 85 | L85T | A | 9.1e-09 | 5e-09 | 5.2e-09 | 6.433E-09 | 2.312E-09 | 5.000E-09 | 9.100E-09 |
| 1698 | 85 | L85W | A | 6.1e-09 | 1.6e-08 | 6e-09 | 9.367E-09 | 5.745E-09 | 6.000E-09 | 1.600E-08 |
| 1699 | 85 | L85Y | A | 7.7e-09 | 6.4e-09 | 4.9e-09 | 6.333E-09 | 1.401E-09 | 4.900E-09 | 7.700E-09 |

| | | | | | | | | | | |
|---|---|---|---|---|---|---|---|---|---|---|
| 1700 | 85 | L85V | A | 1.3e-08 | 1e-08 | 7.6e-09 | 1.020E-08 | 2.706E-09 | 7.600E-09 | 1.300E-08 |
| 1701 | 86 | F86A | A | 3.5e-09 | 6.1e-09 | 1.4e-08 | 7.867E-09 | 5.468E-09 | 3.500E-09 | 1.400E-08 |
| 1702 | 86 | F86R | A | 2.5e-08 | 1.1e-08 | 2.5e-08 | 2.033E-08 | 8.083E-09 | 1.100E-08 | 2.500E-08 |
| 1703 | 86 | F86N | A | 7.8e-09 | 6.9e-09 | 1.3e-08 | 9.233E-09 | 3.293E-09 | 6.900E-09 | 1.300E-08 |
| 1704 | 86 | F86D | A | 4.5e-09 | 2.3e-09 | 5e-09 | 3.933E-09 | 1.436E-09 | 2.300E-09 | 5.000E-09 |
| 1705 | 86 | F86C | A | 6.2e-09 | 3.5e-09 | 9.5e-09 | 6.400E-09 | 3.005E-09 | 3.500E-09 | 9.500E-09 |
| 1706 | 86 | F86E | A | 8.8e-09 | 6.8e-09 | 1.5e-08 | 1.020E-08 | 4.276E-09 | 6.800E-09 | 1.500E-08 |
| 1707 | 86 | F86Q | A | 2.9e-09 | 2.6e-09 | 1.2e-08 | 5.833E-09 | 5.343E-09 | 2.600E-09 | 1.200E-08 |
| 1708 | 86 | F86G | A | 2e-08 | 1.8e-08 | 1.3e-08 | 1.700E-08 | 3.606E-09 | 1.300E-08 | 2.000E-08 |
| 1709 | 86 | F86H | A | 1.2e-08 | 5.8e-09 | 2.1e-09 | 6.633E-09 | 5.002E-09 | 2.100E-09 | 1.200E-08 |
| 1710 | 86 | F86I | A | 8.6e-09 | 1.1e-08 | 4e-09 | 7.867E-09 | 3.557E-09 | 4.000E-09 | 1.100E-08 |
| 1711 | 86 | F86L | A | 7.6e-09 | 6.8e-09 | 1e-08 | 8.133E-09 | 1.665E-09 | 6.800E-09 | 1.000E-08 |
| 1712 | 86 | F86K | A | 5.6e-09 | 9.5e-09 | 4e-09 | 6.367E-09 | 2.829E-09 | 4.000E-09 | 9.500E-09 |
| 1713 | 86 | F86M | A | 5.3e-09 | 1.3e-08 | 2.6e-09 | 6.967E-09 | 5.397E-09 | 2.600E-09 | 1.300E-08 |
| 1714 | 86 | F86F | A | 4.4e-09 | 8e-09 | 6e-09 | 6.133E-09 | 1.804E-09 | 4.400E-09 | 8.000E-09 |
| 1715 | 86 | F86P | A | 7.9e-09 | 5.3e-09 | 9.5e-09 | 7.567E-09 | 2.120E-09 | 5.300E-09 | 9.500E-09 |
| 1716 | 86 | F86S | A | 4.8e-09 | 7.2e-09 | 6.1e-09 | 6.033E-09 | 1.201E-09 | 4.800E-09 | 7.200E-09 |
| 1717 | 86 | F86T | A | 1.7e-08 | 4.2e-09 | 4.6e-09 | 8.600E-09 | 7.277E-09 | 4.200E-09 | 1.700E-08 |
| 1718 | 86 | F86W | A | 4.6e-09 | 9.8e-09 | 4.5e-09 | 6.300E-09 | 3.032E-09 | 4.500E-09 | 9.800E-09 |
| 1719 | 86 | F86Y | A | 3.2e-09 | 7.2e-09 | 4e-09 | 4.800E-09 | 2.117E-09 | 3.200E-09 | 7.200E-09 |
| 1720 | 86 | F86V | A | 5.2e-09 | 1.4e-08 | 6.6e-09 | 8.600E-09 | 4.729E-09 | 5.200E-09 | 1.400E-08 |
| 1721 | 87 | E87A | A | 8.7e-09 | 7.2e-09 | 1.7e-08 | 1.097E-08 | 5.279E-09 | 7.200E-09 | 1.700E-08 |
| 1722 | 87 | E87R | A | 1.3e-08 | 5.3e-09 | 9.6e-09 | 9.300E-09 | 3.859E-09 | 5.300E-09 | 1.300E-08 |
| 1723 | 87 | E87N | A | 8.7e-09 | 4.9e-09 | 1.9e-08 | 1.087E-08 | 7.295E-09 | 4.900E-09 | 1.900E-08 |
| 1724 | 87 | E87D | A | 6.5e-09 | 6.5e-09 | 6.7e-09 | 6.567E-09 | 1.155E-10 | 6.500E-09 | 6.700E-09 |
| 1725 | 87 | E87C | A | 1.2e-08 | 1.7e-08 | 6.2e-09 | 1.173E-08 | 5.405E-09 | 6.200E-09 | 1.700E-08 |
| 1726 | 87 | E87E | A | 9.9e-09 | 7.5e-09 | 1.4e-08 | 1.047E-08 | 3.287E-09 | 7.500E-09 | 1.400E-08 |
| 1727 | 87 | E87Q | A | 4e-09 | 2.7e-09 | 1.5e-08 | 7.233E-09 | 6.757E-09 | 2.700E-09 | 1.500E-08 |
| 1728 | 87 | E87G | A | 7.3e-09 | 6.2e-09 | 8.3e-09 | 7.267E-09 | 1.050E-09 | 6.200E-09 | 8.300E-09 |
| 1729 | 87 | E87H | A | 3.9e-09 | 5.6e-09 | 1.2e-08 | 7.167E-09 | 4.271E-09 | 3.900E-09 | 1.200E-08 |
| 1730 | 87 | E87I | A | 8.5e-09 | 1.4e-08 | 5.6e-09 | 9.367E-09 | 4.267E-09 | 5.600E-09 | 1.400E-08 |
| 1731 | 87 | E87L | A | 5.1e-09 | 8.6e-09 | 1.7e-08 | 1.023E-08 | 6.116E-09 | 5.100E-09 | 1.700E-08 |
| 1732 | 87 | E87K | A | 1.3e-08 | 6.7e-09 | 1.6e-08 | 1.190E-08 | 4.747E-09 | 6.700E-09 | 1.600E-08 |
| 1733 | 87 | E87M | A | 6.8e-09 | 4.5e-09 | 5.6e-09 | 5.633E-09 | 1.150E-09 | 4.500E-09 | 6.800E-09 |
| 1734 | 87 | E87F | A | 1.1e-08 | 1.1e-08 | 1.2e-08 | 1.133E-08 | 5.774E-10 | 1.100E-08 | 1.200E-08 |
| 1735 | 87 | E87P | A | 4.3e-09 | 7.6e-09 | 6.5e-09 | 6.133E-09 | 1.680E-09 | 4.300E-09 | 7.600E-09 |
| 1736 | 87 | E87S | A | 3.5e-09 | 7.2e-09 | 3.9e-09 | 4.867E-09 | 2.031E-09 | 3.500E-09 | 7.200E-09 |
| 1737 | 87 | E87T | A | 8e-09 | 3.6e-09 | 7.3e-09 | 6.300E-09 | 2.364E-09 | 3.600E-09 | 8.000E-09 |
| 1738 | 87 | E87W | A | 3.7e-09 | 3.6e-09 | 1e-08 | 5.767E-09 | 3.667E-09 | 3.600E-09 | 1.000E-08 |
| 1739 | 87 | E87Y | A | 8.3e-09 | 1e-08 | 1.6e-08 | 1.143E-08 | 4.045E-09 | 8.300E-09 | 1.600E-08 |
| 1740 | 87 | E87V | A | 3.6e-09 | 7.4e-09 | 1.4e-08 | 8.333E-09 | 5.262E-09 | 3.600E-09 | 1.400E-08 |
| 1741 | 88 | D88A | A | 1.2e-08 | 1.2e-08 | 6.1e-09 | 1.003E-08 | 3.406E-09 | 6.100E-09 | 1.200E-08 |
| 1742 | 88 | D88R | A | 1.2e-08 | 6.6e-09 | 3e-09 | 7.200E-09 | 4.530E-09 | 3.000E-09 | 1.200E-08 |
| 1743 | 88 | D88N | A | 6.5e-09 | 2.9e-09 | 4.3e-09 | 4.567E-09 | 1.815E-09 | 2.900E-09 | 6.500E-09 |
| 1744 | 88 | D88D | A | 7.5e-09 | 9e-09 | 2e-09 | 6.167E-09 | 3.686E-09 | 2.000E-09 | 9.000E-09 |
| 1745 | 88 | D88C | A | 6.7e-09 | 4.7e-09 | 1.7e-08 | 9.467E-09 | 6.600E-09 | 4.700E-09 | 1.700E-08 |
| 1746 | 88 | D88E | A | 6.9e-09 | 7.8e-09 | 6.5e-09 | 7.067E-09 | 6.658E-10 | 6.500E-09 | 7.800E-09 |
| 1747 | 88 | D88Q | A | 3.8e-09 | 1.1e-08 | 4.2e-09 | 6.333E-09 | 4.046E-09 | 3.800E-09 | 1.100E-08 |
| 1748 | 88 | D88G | A | 1.7e-08 | 6.5e-09 | 9.1e-09 | 1.087E-08 | 5.468E-09 | 6.500E-09 | 1.700E-08 |
| 1749 | 88 | D88H | A | 7.6e-09 | 5.4e-09 | 6.4e-09 | 6.467E-09 | 1.102E-09 | 5.400E-09 | 7.600E-09 |
| 1750 | 88 | D88I | A | 1.5e-08 | 4.3e-09 | 3.9e-09 | 7.733E-09 | 6.296E-09 | 3.900E-09 | 1.500E-08 |
| 1751 | 88 | D88L | A | 4e-09 | 7.7e-09 | 1.5e-08 | 8.900E-09 | 5.597E-09 | 4.000E-09 | 1.500E-08 |
| 1752 | 88 | D88K | A | 8.7e-09 | 9.9e-09 | 3.1e-09 | 7.233E-09 | 3.630E-09 | 3.100E-09 | 9.900E-09 |
| 1753 | 88 | D88M | A | 3.4e-09 | 6.4e-09 | 6.5e-09 | 5.433E-09 | 1.762E-09 | 3.400E-09 | 6.500E-09 |
| 1754 | 88 | D88F | A | 1.2e-08 | 4.6e-09 | 9.6e-09 | 8.733E-09 | 3.775E-09 | 4.600E-09 | 1.200E-08 |
| 1755 | 88 | D88P | A | 2.3e-08 | 6.5e-09 | 8.2e-09 | 1.257E-08 | 9.075E-09 | 6.500E-09 | 2.300E-08 |
| 1756 | 88 | D88S | A | 5.2e-09 | 1.3e-09 | 9.8e-09 | 5.433E-09 | 4.255E-09 | 1.300E-09 | 9.800E-09 |
| 1757 | 88 | D88T | A | 3.6e-09 | 1.8e-08 | 8.6e-09 | 1.007E-08 | 7.311E-09 | 3.600E-09 | 1.800E-08 |
| 1758 | 88 | D88W | A | 2.7e-09 | 4.1e-09 | 8.5e-09 | 5.100E-09 | 3.027E-09 | 2.700E-09 | 8.500E-09 |
| 1759 | 88 | D88Y | A | 6.2e-09 | 9e-09 | 6.7e-09 | 7.300E-09 | 1.493E-09 | 6.200E-09 | 9.000E-09 |
| 1760 | 88 | D88V | A | 3.5e-09 | 3.1e-09 | 4.8e-08 | 1.820E-08 | 2.581E-08 | 3.100E-09 | 4.800E-08 |
| 1761 | 89 | N89A | A | 6.1e-09 | 3.1e-09 | 2.8e-09 | 4.000E-09 | 1.825E-09 | 2.800E-09 | 6.100E-09 |
| 1762 | 89 | N89R | A | 1e-08 | 1.8e-08 | 7.1e-09 | 1.170E-08 | 5.645E-09 | 7.100E-09 | 1.800E-08 |
| 1763 | 89 | N89N | A | 4.1e-09 | 2.1e-08 | 1e-08 | 1.170E-08 | 8.577E-09 | 4.100E-09 | 2.100E-08 |
| 1764 | 89 | N89D | A | 4.6e-09 | 4e-09 | 5.6e-09 | 4.733E-09 | 8.083E-10 | 4.000E-09 | 5.600E-09 |
| 1765 | 89 | N89C | A | 4.9e-09 | 8.4e-09 | 1.5e-08 | 9.433E-09 | 5.129E-09 | 4.900E-09 | 1.500E-08 |
| 1766 | 89 | N89E | A | 2.8e-09 | 1.3e-08 | 3.6e-09 | 6.467E-09 | 5.672E-09 | 2.800E-09 | 1.300E-08 |
| 1767 | 89 | N89Q | A | 1.5e-08 | 2.1e-08 | 4.1e-09 | 1.337E-08 | 8.568E-09 | 4.100E-09 | 2.100E-08 |
| 1768 | 89 | N89G | A | 5.3e-09 | 1.2e-08 | 5.3e-09 | 7.533E-09 | 3.868E-09 | 5.300E-09 | 1.200E-08 |
| 1769 | 89 | N89H | A | 3.7e-09 | 1.1e-08 | 3.3e-08 | 1.590E-08 | 1.525E-08 | 3.700E-09 | 3.300E-08 |
| 1770 | 89 | N89I | A | 4.6e-09 | 1.7e-08 | 1.6e-08 | 1.253E-08 | 6.889E-09 | 4.600E-09 | 1.700E-08 |
| 1771 | 89 | N89L | A | 1.5e-08 | 8.6e-09 | 7.1e-09 | 1.023E-08 | 4.196E-09 | 7.100E-09 | 1.500E-08 |
| 1772 | 89 | N89K | A | 1.4e-08 | 3.2e-09 | 8e-09 | 8.400E-09 | 5.411E-09 | 3.200E-09 | 1.400E-08 |
| 1773 | 89 | N89M | A | 2.4e-09 | 1.1e-08 | 5.6e-09 | 6.333E-09 | 4.347E-09 | 2.400E-09 | 1.100E-08 |
| 1774 | 89 | N89F | A | 8.4e-09 | 4.2e-09 | 2.6e-08 | 1.287E-08 | 1.157E-08 | 4.200E-09 | 2.600E-08 |
| 1775 | 89 | N89P | A | 2.5e-08 | 6.9e-09 | 6.4e-09 | 1.277E-08 | 1.060E-08 | 6.400E-09 | 2.500E-08 |
| 1776 | 89 | N89S | A | 1.2e-08 | 6.9e-09 | 3.8e-09 | 7.567E-09 | 4.140E-09 | 3.800E-09 | 1.200E-08 |
| 1777 | 89 | N89T | A | 4.8e-09 | 4.3e-09 | 4.8e-09 | 4.633E-09 | 2.887E-10 | 4.300E-09 | 4.800E-09 |
| 1778 | 89 | N89W | A | 5.9e-09 | 7.5e-09 | 6.9e-09 | 6.767E-09 | 8.083E-10 | 5.900E-09 | 7.500E-09 |
| 1779 | 89 | N89Y | A | 1.2e-08 | 1.2e-08 | 1.7e-08 | 1.367E-08 | 2.887E-09 | 1.200E-08 | 1.700E-08 |
| 1780 | 89 | N89V | A | 8.3e-09 | 2e-08 | 9.3e-09 | 1.253E-08 | 6.486E-09 | 8.300E-09 | 2.000E-08 |
| 1781 | 90 | Y90A | A | 1.1e-08 | 6.7e-09 | 6.4e-09 | 8.033E-09 | 2.574E-09 | 6.400E-09 | 1.100E-08 |
| 1782 | 90 | Y90R | A | 1.3e-08 | 1.2e-08 | 3.9e-09 | 9.633E-09 | 4.990E-09 | 3.900E-09 | 1.300E-08 |
| 1783 | 90 | Y90N | A | 1.2e-08 | 9e-09 | 7.6e-09 | 9.533E-09 | 2.248E-09 | 7.600E-09 | 1.200E-08 |
| 1784 | 90 | Y90D | A | 1.7e-08 | 2.3e-08 | 1.1e-08 | 1.700E-08 | 6.000E-09 | 1.100E-08 | 2.300E-08 |

| | | | | | | | | | | |
|---|---|---|---|---|---|---|---|---|---|---|
| 1785 | 90 | Y90C | A | 7.1e-09 | 5.4e-09 | 6.9e-09 | 6.467E-09 | 9.292E-10 | 5.400E-09 | 7.100E-09 |
| 1786 | 90 | Y90E | A | 1.2e-08 | 3e-09 | 9.5e-09 | 8.167E-09 | 4.646E-09 | 3.000E-09 | 1.200E-08 |
| 1787 | 90 | Y90Q | A | 7.3e-09 | 1.4e-08 | 2.1e-08 | 1.410E-08 | 6.851E-09 | 7.300E-09 | 2.100E-08 |
| 1788 | 90 | Y90G | A | 9.5e-09 | 5.1e-09 | 2.6e-09 | 5.733E-09 | 3.493E-09 | 2.600E-09 | 9.500E-09 |
| 1789 | 90 | Y90H | A | 7.4e-09 | 1.2e-08 | 5.6e-09 | 8.333E-09 | 3.301E-09 | 5.600E-09 | 1.200E-08 |
| 1790 | 90 | Y90I | A | 2.2e-08 | 5.1e-09 | 8.5e-09 | 1.187E-08 | 8.939E-09 | 5.100E-09 | 2.200E-08 |
| 1791 | 90 | Y90L | A | 6.4e-09 | 1.5e-08 | 7.3e-09 | 9.567E-09 | 4.727E-09 | 6.400E-09 | 1.500E-08 |
| 1792 | 90 | Y90K | A | 1.2e-08 | 1.1e-08 | 4e-09 | 9.000E-09 | 4.359E-09 | 4.000E-09 | 1.200E-08 |
| 1793 | 90 | Y90M | A | 2.3e-09 | 3.9e-09 | 5.9e-09 | 4.033E-09 | 1.804E-09 | 2.300E-09 | 5.900E-09 |
| 1794 | 90 | Y90F | A | 5.6e-09 | 3.4e-09 | 2.8e-09 | 3.933E-09 | 1.474E-09 | 2.800E-09 | 5.600E-09 |
| 1795 | 90 | Y90P | A | 4.5e-09 | 3.6e-09 | 7e-09 | 5.033E-09 | 1.762E-09 | 3.600E-09 | 7.000E-09 |
| 1796 | 90 | Y90S | A | 3.3e-09 | 2.6e-09 | 4.2e-09 | 3.367E-09 | 8.021E-10 | 2.600E-09 | 4.200E-09 |
| 1797 | 90 | Y90T | A | 2.8e-08 | 6.5e-09 | 5.1e-09 | 1.320E-08 | 1.284E-08 | 5.100E-09 | 2.800E-08 |
| 1798 | 90 | Y90W | A | 1.2e-08 | 4.7e-09 | 2.7e-09 | 6.467E-09 | 4.895E-09 | 2.700E-09 | 1.200E-08 |
| 1799 | 90 | Y90Y | A | 6.9e-09 | 4.4e-09 | 1.3e-08 | 8.100E-09 | 4.424E-09 | 4.400E-09 | 1.300E-08 |
| 1800 | 90 | Y90V | A | 3e-09 | 1.6e-08 | 8.3e-09 | 9.100E-09 | 6.537E-09 | 3.000E-09 | 1.600E-08 |
| 1801 | 91 | A91A | A | 1.8e-08 | 1.1e-08 | 1.3e-08 | 1.400E-08 | 3.606E-09 | 1.100E-08 | 1.800E-08 |
| 1802 | 91 | A91R | A | 6.6e-09 | 1.4e-08 | 2.7e-09 | 7.767E-09 | 5.740E-09 | 2.700E-09 | 1.400E-08 |
| 1803 | 91 | A91N | A | 1.3e-08 | 1.1e-08 | 1.2e-08 | 1.200E-08 | 1.000E-09 | 1.100E-08 | 1.300E-08 |
| 1804 | 91 | A91D | A | 1.1e-08 | 5.8e-09 | 5.8e-09 | 7.533E-09 | 3.002E-09 | 5.800E-09 | 1.100E-08 |
| 1805 | 91 | A91C | A | 1.9e-08 | 1.2e-08 | 3.6e-09 | 1.153E-08 | 7.711E-09 | 3.600E-09 | 1.900E-08 |
| 1806 | 91 | A91E | A | 9e-09 | 1.4e-08 | 5.7e-09 | 9.567E-09 | 4.179E-09 | 5.700E-09 | 1.400E-08 |
| 1807 | 91 | A91Q | A | 9.9e-09 | 1.1e-09 | 9.4e-09 | 6.800E-09 | 4.943E-09 | 1.100E-09 | 9.900E-09 |
| 1808 | 91 | A91G | A | 2e-08 | 5.6e-09 | 7.4e-09 | 1.100E-08 | 7.846E-09 | 5.600E-09 | 2.000E-08 |
| 1809 | 91 | A91H | A | 3.4e-08 | 7.8e-09 | 1.3e-08 | 1.827E-08 | 1.387E-08 | 7.800E-09 | 3.400E-08 |
| 1810 | 91 | A91I | A | 4.9e-09 | 9.9e-09 | 8.2e-09 | 7.667E-09 | 2.542E-09 | 4.900E-09 | 9.900E-09 |
| 1811 | 91 | A91L | A | 7.5e-09 | 1.9e-08 | 8.9e-09 | 1.180E-08 | 6.275E-09 | 7.500E-09 | 1.900E-08 |
| 1812 | 91 | A91K | A | 5.2e-09 | 1.7e-08 | 1.1e-08 | 1.107E-08 | 5.900E-09 | 5.200E-09 | 1.700E-08 |
| 1813 | 91 | A91M | A | 2.9e-09 | 1.2e-08 | 4.3e-09 | 6.400E-09 | 4.900E-09 | 2.900E-09 | 1.200E-08 |
| 1814 | 91 | A91F | A | 2.6e-09 | 6e-09 | 4.7e-09 | 4.433E-09 | 1.716E-09 | 2.600E-09 | 6.000E-09 |
| 1815 | 91 | A91P | A | 1.4e-08 | 5.1e-09 | 5.2e-09 | 8.100E-09 | 5.110E-09 | 5.100E-09 | 1.400E-08 |
| 1816 | 91 | A91S | A | 8.2e-09 | 1e-08 | 1.2e-08 | 1.007E-08 | 1.901E-09 | 8.200E-09 | 1.200E-08 |
| 1817 | 91 | A91T | A | 3.6e-09 | 9.5e-09 | 3.8e-09 | 5.633E-09 | 3.350E-09 | 3.600E-09 | 9.500E-09 |
| 1818 | 91 | A91W | A | 6.2e-09 | 6.8e-09 | 1.7e-08 | 1.000E-08 | 6.070E-09 | 6.200E-09 | 1.700E-08 |
| 1819 | 91 | A91Y | A | 9.1e-09 | 9.4e-09 | 1.2e-08 | 1.017E-08 | 1.595E-09 | 9.100E-09 | 1.200E-08 |
| 1820 | 91 | A91V | A | 1.9e-08 | 3.7e-09 | 5.4e-09 | 9.367E-09 | 8.386E-09 | 3.700E-09 | 1.900E-08 |
| 1821 | 92 | L92A | A | 7.9e-09 | 3.7e-09 | 7e-09 | 6.200E-09 | 2.211E-09 | 3.700E-09 | 7.900E-09 |
| 1822 | 92 | L92R | A | 4.9e-09 | 9.3e-09 | 6.8e-09 | 7.000E-09 | 2.207E-09 | 4.900E-09 | 9.300E-09 |
| 1823 | 92 | L92N | A | 7.8e-09 | 2.4e-09 | 1.4e-09 | 3.867E-09 | 3.443E-09 | 1.400E-09 | 7.800E-09 |
| 1824 | 92 | L92D | A | 5.8e-09 | 4.5e-08 | 7.7e-09 | 1.950E-08 | 2.210E-08 | 5.800E-09 | 4.500E-08 |
| 1825 | 92 | L92C | A | 3.2e-09 | 6.5e-09 | 6.5e-09 | 5.400E-09 | 1.905E-09 | 3.200E-09 | 6.500E-09 |
| 1826 | 92 | L92E | A | 8.3e-09 | 3.2e-09 | 1.5e-08 | 8.833E-09 | 5.918E-09 | 3.200E-09 | 1.500E-08 |
| 1827 | 92 | L92Q | A | 1.2e-08 | 1.1e-08 | 3.4e-09 | 8.800E-09 | 4.703E-09 | 3.400E-09 | 1.200E-08 |
| 1828 | 92 | L92G | A | 1.3e-08 | 1.2e-08 | 4.9e-09 | 9.967E-09 | 4.416E-09 | 4.900E-09 | 1.300E-08 |
| 1829 | 92 | L92H | A | 5.9e-09 | 7.5e-09 | 1e-08 | 7.800E-09 | 2.066E-09 | 5.900E-09 | 1.000E-08 |
| 1830 | 92 | L92I | A | 4.5e-09 | 5e-09 | 7.4e-09 | 5.633E-09 | 1.550E-09 | 4.500E-09 | 7.400E-09 |
| 1831 | 92 | L92L | A | 5.4e-09 | 5.2e-09 | 8.8e-09 | 6.467E-09 | 2.023E-09 | 5.200E-09 | 8.800E-09 |
| 1832 | 92 | L92K | A | 6.3e-09 | 6e-09 | 1.5e-08 | 9.100E-09 | 5.112E-09 | 6.000E-09 | 1.500E-08 |
| 1833 | 92 | L92M | A | 1e-08 | 8.9e-09 | 9.2e-09 | 9.367E-09 | 5.686E-10 | 8.900E-09 | 1.000E-08 |
| 1834 | 92 | L92F | A | 8.6e-09 | 3.6e-09 | 5.6e-09 | 5.933E-09 | 2.517E-09 | 3.600E-09 | 8.600E-09 |
| 1835 | 92 | L92P | A | 1.5e-09 | 7.7e-09 | 1.1e-08 | 6.733E-09 | 4.823E-09 | 1.500E-09 | 1.100E-08 |
| 1836 | 92 | L92S | A | 2.7e-09 | 1.5e-08 | 5.7e-09 | 7.800E-09 | 6.413E-09 | 2.700E-09 | 1.500E-08 |
| 1837 | 92 | L92T | A | 5.9e-09 | 1.1e-08 | 4.1e-09 | 7.000E-09 | 3.579E-09 | 4.100E-09 | 1.100E-08 |
| 1838 | 92 | L92W | A | 5.2e-09 | 3.1e-09 | 8.4e-09 | 5.567E-09 | 2.669E-09 | 3.100E-09 | 8.400E-09 |
| 1839 | 92 | L92Y | A | 7.3e-09 | 3.7e-09 | 6.3e-09 | 5.767E-09 | 1.858E-09 | 3.700E-09 | 7.300E-09 |
| 1840 | 92 | L92V | A | 4.4e-09 | 1.1e-08 | 9.8e-10 | 5.460E-09 | 5.093E-09 | 9.800E-10 | 1.100E-08 |
| 1841 | 93 | A93A | A | 7.5e-09 | 4.1e-09 | 1.8e-08 | 9.867E-09 | 7.246E-09 | 4.100E-09 | 1.800E-08 |
| 1842 | 93 | A93R | A | 2.3e-09 | 5.2e-09 | 1.8e-08 | 8.500E-09 | 8.354E-09 | 2.300E-09 | 1.800E-08 |
| 1843 | 93 | A93N | A | 5e-09 | 1.1e-08 | 5.1e-09 | 7.033E-09 | 3.436E-09 | 5.000E-09 | 1.100E-08 |
| 1844 | 93 | A93D | A | 3.9e-09 | 3.6e-08 | 6.2e-09 | 1.537E-08 | 1.791E-08 | 3.900E-09 | 3.600E-08 |
| 1845 | 93 | A93C | A | 6.5e-09 | 6.8e-09 | 7.3e-09 | 6.867E-09 | 4.041E-10 | 6.500E-09 | 7.300E-09 |
| 1846 | 93 | A93E | A | 1.7e-08 | 6.3e-09 | 1.1e-08 | 1.143E-08 | 5.363E-09 | 6.300E-09 | 1.700E-08 |
| 1847 | 93 | A93Q | A | 7.6e-09 | 7.2e-09 | 6.2e-09 | 7.000E-09 | 7.211E-10 | 6.200E-09 | 7.600E-09 |
| 1848 | 93 | A93G | A | 1.2e-08 | 7.9e-09 | 1.4e-08 | 1.130E-08 | 3.110E-09 | 7.900E-09 | 1.400E-08 |
| 1849 | 93 | A93H | A | 7.5e-09 | 1.3e-08 | 4.7e-09 | 8.400E-09 | 4.223E-09 | 4.700E-09 | 1.300E-08 |
| 1850 | 93 | A93I | A | 6.1e-09 | 5.4e-09 | 3.4e-09 | 4.967E-09 | 1.401E-09 | 3.400E-09 | 6.100E-09 |
| 1851 | 93 | A93L | A | 3.8e-09 | 4.2e-09 | 1.1e-08 | 6.333E-09 | 4.046E-09 | 3.800E-09 | 1.100E-08 |
| 1852 | 93 | A93K | A | 9.4e-09 | 1.9e-09 | 7e-09 | 6.100E-09 | 3.830E-09 | 1.900E-09 | 9.400E-09 |
| 1853 | 93 | A93M | A | 4.6e-09 | 6.2e-09 | 1.7e-08 | 9.267E-09 | 6.745E-09 | 4.600E-09 | 1.700E-08 |
| 1854 | 93 | A93F | A | 7.7e-09 | 7.1e-09 | 5.9e-09 | 6.900E-09 | 9.165E-10 | 5.900E-09 | 7.700E-09 |
| 1855 | 93 | A93P | A | 2.7e-09 | 2.6e-09 | 1.1e-08 | 5.433E-09 | 4.821E-09 | 2.600E-09 | 1.100E-08 |
| 1856 | 93 | A93S | A | 2.2e-09 | 1.6e-08 | 2.5e-09 | 6.900E-09 | 7.882E-09 | 2.200E-09 | 1.600E-08 |
| 1857 | 93 | A93T | A | 5.4e-09 | 4.4e-09 | 8.2e-09 | 6.000E-09 | 1.970E-09 | 4.400E-09 | 8.200E-09 |
| 1858 | 93 | A93W | A | 1.7e-08 | 4.9e-09 | 1.3e-08 | 1.163E-08 | 6.165E-09 | 4.900E-09 | 1.700E-08 |
| 1859 | 93 | A93Y | A | 6e-09 | 4.8e-09 | 5.6e-09 | 5.467E-09 | 6.110E-10 | 4.800E-09 | 6.000E-09 |
| 1860 | 93 | A93V | A | 9.9e-09 | 6.2e-09 | 8.7e-09 | 8.267E-09 | 1.888E-09 | 6.200E-09 | 9.900E-09 |
| 1861 | 94 | V94A | A | 1e-08 | 6.3e-09 | 1.1e-08 | 9.100E-09 | 2.476E-09 | 6.300E-09 | 1.100E-08 |
| 1862 | 94 | V94R | A | 4.8e-09 | 4.2e-09 | 2.1e-08 | 1.000E-08 | 9.531E-09 | 4.200E-09 | 2.100E-08 |
| 1863 | 94 | V94N | A | 6.9e-09 | 1.1e-08 | 1.1e-08 | 9.633E-09 | 2.367E-09 | 6.900E-09 | 1.100E-08 |
| 1864 | 94 | V94D | A | 7.8e-09 | 6.5e-09 | 3.8e-09 | 6.033E-09 | 2.040E-09 | 3.800E-09 | 7.800E-09 |
| 1865 | 94 | V94C | A | 7.5e-09 | 1e-08 | 4.9e-09 | 7.467E-09 | 2.550E-09 | 4.900E-09 | 1.000E-08 |
| 1866 | 94 | V94E | A | 3.7e-09 | 7.3e-09 | 1.1e-08 | 7.333E-09 | 3.650E-09 | 3.700E-09 | 1.100E-08 |
| 1867 | 94 | V94Q | A | 3.3e-09 | 4.1e-09 | 6e-09 | 4.467E-09 | 1.387E-09 | 3.300E-09 | 6.000E-09 |
| 1868 | 94 | V94G | A | 8.5e-09 | 2.3e-09 | 8.4e-09 | 6.400E-09 | 3.551E-09 | 2.300E-09 | 8.500E-09 |
| 1869 | 94 | V94H | A | 5.9e-09 | 5.5e-09 | 8.4e-09 | 6.600E-09 | 1.572E-09 | 5.500E-09 | 8.400E-09 |

| | | | | | | | | | | |
|---|---|---|---|---|---|---|---|---|---|---|
| 1870 | 94 | V94I | A | 1.2e-08 | 2.3e-09 | 8.4e-09 | 7.567E-09 | 4.903E-09 | 2.300E-09 | 1.200E-08 |
| 1871 | 94 | V94L | A | 2e-09 | 1e-08 | 5e-09 | 5.667E-09 | 4.041E-09 | 2.000E-09 | 1.000E-08 |
| 1872 | 94 | V94K | A | 5.8e-09 | 9.9e-09 | 4.4e-09 | 6.700E-09 | 2.858E-09 | 4.400E-09 | 9.900E-09 |
| 1873 | 94 | V94M | A | 3.5e-09 | 5.6e-09 | 1.5e-08 | 8.033E-09 | 6.124E-09 | 3.500E-09 | 1.500E-08 |
| 1874 | 94 | V94F | A | 1e-08 | 6.4e-09 | 2.6e-08 | 1.413E-08 | 1.043E-08 | 6.400E-09 | 2.600E-08 |
| 1875 | 94 | V94P | A | 5.5e-09 | 2.5e-09 | 1e-08 | 6.000E-09 | 3.775E-09 | 2.500E-09 | 1.000E-08 |
| 1876 | 94 | V94S | A | 7.1e-09 | 7.7e-09 | 6.9e-09 | 7.233E-09 | 4.163E-10 | 6.900E-09 | 7.700E-09 |
| 1877 | 94 | V94T | A | 3.8e-09 | 2.6e-09 | 6e-09 | 4.133E-09 | 1.724E-09 | 2.600E-09 | 6.000E-09 |
| 1878 | 94 | V94W | A | 3.4e-09 | 7.3e-09 | 4.7e-09 | 5.133E-09 | 1.986E-09 | 3.400E-09 | 7.300E-09 |
| 1879 | 94 | V94Y | A | 1.4e-08 | 9.1e-09 | 1.1e-08 | 1.137E-08 | 2.470E-09 | 9.100E-09 | 1.400E-08 |
| 1880 | 94 | V94V | A | 9.8e-09 | 4.1e-09 | 5.1e-09 | 6.333E-09 | 3.044E-09 | 4.100E-09 | 9.800E-09 |
| 1881 | 95 | L95A | A | 4e-09 | 4.2e-09 | 1.1e-09 | 3.100E-09 | 1.735E-09 | 1.100E-09 | 4.200E-09 |
| 1882 | 95 | L95R | A | 4.9e-09 | 7.2e-09 | 3.7e-09 | 5.267E-09 | 1.779E-09 | 3.700E-09 | 7.200E-09 |
| 1883 | 95 | L95N | A | 6.6e-09 | 8.9e-09 | 8.7e-09 | 8.067E-09 | 1.274E-09 | 6.600E-09 | 8.900E-09 |
| 1884 | 95 | L95D | A | 7.8e-09 | 2.3e-09 | 7.4e-09 | 5.833E-09 | 3.066E-09 | 2.300E-09 | 7.800E-09 |
| 1885 | 95 | L95C | A | 3.1e-09 | 1.6e-08 | 8e-09 | 9.033E-09 | 6.512E-09 | 3.100E-09 | 1.600E-08 |
| 1886 | 95 | L95E | A | 5.3e-09 | 4.1e-09 | 5.7e-09 | 5.033E-09 | 8.327E-10 | 4.100E-09 | 5.700E-09 |
| 1887 | 95 | L95Q | A | 6.2e-09 | 2.7e-09 | 2.4e-09 | 3.767E-09 | 2.113E-09 | 2.400E-09 | 6.200E-09 |
| 1888 | 95 | L95G | A | 2e-08 | 6.9e-09 | 1e-08 | 1.230E-08 | 6.846E-09 | 6.900E-09 | 2.000E-08 |
| 1889 | 95 | L95H | A | 6.2e-09 | 9.5e-09 | 2.4e-09 | 6.033E-09 | 3.553E-09 | 2.400E-09 | 9.500E-09 |
| 1890 | 95 | L95I | A | 4.3e-09 | 2.7e-09 | 1e-08 | 5.667E-09 | 3.837E-09 | 2.700E-09 | 1.000E-08 |
| 1891 | 95 | L95L | A | 9.4e-09 | 4.2e-09 | 7.2e-09 | 6.933E-09 | 2.610E-09 | 4.200E-09 | 9.400E-09 |
| 1892 | 95 | L95K | A | 2.6e-09 | 2.2e-08 | 6.3e-09 | 1.030E-08 | 1.030E-08 | 2.600E-09 | 2.200E-08 |
| 1893 | 95 | L95M | A | 2.3e-08 | 1.7e-08 | 1e-08 | 1.667E-08 | 6.506E-09 | 1.000E-08 | 2.300E-08 |
| 1894 | 95 | L95F | A | 4.4e-09 | 1.2e-08 | 7.8e-09 | 8.067E-09 | 3.807E-09 | 4.400E-09 | 1.200E-08 |
| 1895 | 95 | L95P | A | 1.4e-08 | 6.6e-09 | 1e-08 | 1.020E-08 | 3.704E-09 | 6.600E-09 | 1.400E-08 |
| 1896 | 95 | L95S | A | 6.3e-09 | 1.7e-08 | 6.6e-09 | 9.967E-09 | 6.093E-09 | 6.300E-09 | 1.700E-08 |
| 1897 | 95 | L95T | A | 2.7e-09 | 5.7e-09 | 5.4e-09 | 4.600E-09 | 1.652E-09 | 2.700E-09 | 5.700E-09 |
| 1898 | 95 | L95W | A | 2.6e-09 | 7.3e-09 | 1.4e-08 | 7.967E-09 | 5.729E-09 | 2.600E-09 | 1.400E-08 |
| 1899 | 95 | L95Y | A | 1.2e-08 | 1.4e-08 | 3.8e-09 | 9.933E-09 | 5.405E-09 | 3.800E-09 | 1.400E-08 |
| 1900 | 95 | L95V | A | 1.3e-08 | 1.6e-08 | 3.1e-09 | 1.070E-08 | 6.751E-09 | 3.100E-09 | 1.600E-08 |
| 1901 | 96 | D96A | A | 1.3e-08 | 6.5e-09 | 9e-09 | 9.500E-09 | 3.279E-09 | 6.500E-09 | 1.300E-08 |
| 1902 | 96 | D96R | A | 9.4e-09 | 3.3e-09 | 5.8e-09 | 6.167E-09 | 3.066E-09 | 3.300E-09 | 9.400E-09 |
| 1903 | 96 | D96N | A | 7.8e-09 | 3.4e-09 | 5.6e-09 | 5.600E-09 | 2.200E-09 | 3.400E-09 | 7.800E-09 |
| 1904 | 96 | D96D | A | 6.1e-09 | 1e-08 | 3e-09 | 6.367E-09 | 3.508E-09 | 3.000E-09 | 1.000E-08 |
| 1905 | 96 | D96C | A | 6.6e-09 | 1e-08 | 8.8e-09 | 8.467E-09 | 1.724E-09 | 6.600E-09 | 1.000E-08 |
| 1906 | 96 | D96E | A | 8.3e-09 | 7.2e-09 | 9.4e-09 | 8.300E-09 | 1.100E-09 | 7.200E-09 | 9.400E-09 |
| 1907 | 96 | D96Q | A | 6e-09 | 1e-08 | 6.6e-09 | 7.533E-09 | 2.157E-09 | 6.000E-09 | 1.000E-08 |
| 1908 | 96 | D96G | A | 3.8e-09 | 1.3e-08 | 4.7e-09 | 7.167E-09 | 5.072E-09 | 3.800E-09 | 1.300E-08 |
| 1909 | 96 | D96H | A | 3e-09 | 8.8e-09 | 4.2e-09 | 5.333E-09 | 3.062E-09 | 3.000E-09 | 8.800E-09 |
| 1910 | 96 | D96I | A | 2.7e-09 | 4.7e-09 | 1.5e-08 | 7.467E-09 | 6.600E-09 | 2.700E-09 | 1.500E-08 |
| 1911 | 96 | D96L | A | 1e-08 | 1e-08 | 4.7e-09 | 8.233E-09 | 3.060E-09 | 4.700E-09 | 1.000E-08 |
| 1912 | 96 | D96K | A | 9.5e-09 | 1.6e-08 | 2e-09 | 9.167E-09 | 7.006E-09 | 2.000E-09 | 1.600E-08 |
| 1913 | 96 | D96M | A | 1.8e-09 | 9.4e-09 | 6.6e-09 | 5.933E-09 | 3.844E-09 | 1.800E-09 | 9.400E-09 |
| 1914 | 96 | D96F | A | 1.2e-08 | 2e-08 | 9.6e-09 | 1.387E-08 | 5.445E-09 | 9.600E-09 | 2.000E-08 |
| 1915 | 96 | D96P | A | 5e-09 | 1.6e-08 | 7.3e-09 | 9.433E-09 | 5.802E-09 | 5.000E-09 | 1.600E-08 |
| 1916 | 96 | D96S | A | 4.1e-09 | 1.2e-09 | 8.9e-09 | 4.733E-09 | 3.889E-09 | 1.200E-09 | 8.900E-09 |
| 1917 | 96 | D96T | A | 2.7e-09 | 1.6e-08 | 5.4e-09 | 8.033E-09 | 7.030E-09 | 2.700E-09 | 1.600E-08 |
| 1918 | 96 | D96W | A | 7.6e-09 | 2.4e-09 | 1.2e-08 | 7.333E-09 | 4.806E-09 | 2.400E-09 | 1.200E-08 |
| 1919 | 96 | D96Y | A | 5.8e-09 | 7.8e-09 | 6.7e-09 | 6.767E-09 | 1.002E-09 | 5.800E-09 | 7.800E-09 |
| 1920 | 96 | D96V | A | 9.3e-09 | 6.2e-09 | 6.5e-09 | 7.333E-09 | 1.710E-09 | 6.200E-09 | 9.300E-09 |
| 1921 | 97 | N97A | A | 4.6e-08 | 2.2e-09 | 9.4e-09 | 1.920E-08 | 2.349E-08 | 2.200E-09 | 4.600E-08 |
| 1922 | 97 | N97R | A | 1.4e-08 | 3.9e-09 | 4.1e-09 | 7.333E-09 | 5.774E-09 | 3.900E-09 | 1.400E-08 |
| 1923 | 97 | N97N | A | 1.1e-08 | 6e-09 | 5.3e-09 | 7.433E-09 | 3.109E-09 | 5.300E-09 | 1.100E-08 |
| 1924 | 97 | N97D | A | 5.1e-09 | 7.2e-09 | 2.4e-08 | 1.210E-08 | 1.036E-08 | 5.100E-09 | 2.400E-08 |
| 1925 | 97 | N97C | A | 2.2e-08 | 1.2e-08 | 3.4e-09 | 1.247E-08 | 9.309E-09 | 3.400E-09 | 2.200E-08 |
| 1926 | 97 | N97E | A | 9.7e-09 | 3.7e-09 | 1.1e-08 | 8.133E-09 | 3.894E-09 | 3.700E-09 | 1.100E-08 |
| 1927 | 97 | N97Q | A | 5.5e-09 | 7.2e-09 | 4.5e-09 | 5.733E-09 | 1.365E-09 | 4.500E-09 | 7.200E-09 |
| 1928 | 97 | N97G | A | 1.7e-08 | 1.1e-08 | 4.3e-09 | 1.077E-08 | 6.353E-09 | 4.300E-09 | 1.700E-08 |
| 1929 | 97 | N97H | A | 4.4e-09 | 1.5e-08 | 1.3e-08 | 1.080E-08 | 5.632E-09 | 4.400E-09 | 1.500E-08 |
| 1930 | 97 | N97I | A | 2.1e-08 | 2.2e-08 | 2.3e-09 | 1.510E-08 | 1.110E-08 | 2.300E-09 | 2.200E-08 |
| 1931 | 97 | N97L | A | 1e-08 | 8.8e-09 | 6.5e-09 | 8.433E-09 | 1.779E-09 | 6.500E-09 | 1.000E-08 |
| 1932 | 97 | N97K | A | 8.6e-09 | 6.9e-09 | 4.1e-09 | 6.533E-09 | 2.272E-09 | 4.100E-09 | 8.600E-09 |
| 1933 | 97 | N97M | A | 1.2e-08 | 5.8e-09 | 3.2e-09 | 7.000E-09 | 4.521E-09 | 3.200E-09 | 1.200E-08 |
| 1934 | 97 | N97F | A | 1e-08 | 3.2e-09 | 1.9e-08 | 1.073E-08 | 7.925E-09 | 3.200E-09 | 1.900E-08 |
| 1935 | 97 | N97P | A | 3e-09 | 6.1e-09 | 1.1e-08 | 6.700E-09 | 4.034E-09 | 3.000E-09 | 1.100E-08 |
| 1936 | 97 | N97S | A | 7e-09 | 6.6e-09 | 7.4e-09 | 7.000E-09 | 4.000E-10 | 6.600E-09 | 7.400E-09 |
| 1937 | 97 | N97T | A | 4.5e-09 | 2.8e-09 | 5.6e-09 | 4.300E-09 | 1.411E-09 | 2.800E-09 | 5.600E-09 |
| 1938 | 97 | N97W | A | 8.9e-09 | 5.8e-09 | 8.1e-09 | 7.600E-09 | 1.609E-09 | 5.800E-09 | 8.900E-09 |
| 1939 | 97 | N97Y | A | 8.8e-09 | 7.8e-09 | 1.1e-08 | 9.200E-09 | 1.637E-09 | 7.800E-09 | 1.100E-08 |
| 1940 | 97 | N97V | A | 3.7e-09 | 1.1e-08 | 8.5e-09 | 7.733E-09 | 3.710E-09 | 3.700E-09 | 1.100E-08 |
| 1941 | 98 | G98A | A | 1.5e-08 | 6.9e-09 | 3.7e-09 | 8.533E-09 | 5.824E-09 | 3.700E-09 | 1.500E-08 |
| 1942 | 98 | G98R | A | 8.5e-09 | 9.1e-09 | 6.1e-09 | 7.900E-09 | 1.587E-09 | 6.100E-09 | 9.100E-09 |
| 1943 | 98 | G98N | A | 2.9e-09 | 6.6e-09 | 8.1e-09 | 5.867E-09 | 2.676E-09 | 2.900E-09 | 8.100E-09 |
| 1944 | 98 | G98D | A | 4.2e-09 | 1.6e-09 | 5.6e-09 | 3.800E-09 | 2.030E-09 | 1.600E-09 | 5.600E-09 |
| 1945 | 98 | G98C | A | 3e-09 | 2.2e-09 | 3.4e-09 | 2.867E-09 | 6.110E-10 | 2.200E-09 | 3.400E-09 |
| 1946 | 98 | G98E | A | 7.8e-09 | 1.3e-08 | 5.2e-09 | 8.667E-09 | 3.972E-09 | 5.200E-09 | 1.300E-08 |
| 1947 | 98 | G98Q | A | 1.8e-09 | 1.3e-08 | 1.1e-08 | 8.600E-09 | 5.973E-09 | 1.800E-09 | 1.300E-08 |
| 1948 | 98 | G98G | A | 1.1e-08 | 1.9e-08 | 3.9e-08 | 2.300E-08 | 1.442E-08 | 1.100E-08 | 3.900E-08 |
| 1949 | 98 | G98H | A | 2.8e-09 | 2.8e-09 | 1.2e-08 | 5.867E-09 | 5.312E-09 | 2.800E-09 | 1.200E-08 |
| 1950 | 98 | G98I | A | 1.3e-08 | 8e-09 | 1.7e-08 | 1.267E-08 | 4.509E-09 | 8.000E-09 | 1.700E-08 |
| 1951 | 98 | G98L | A | 4.2e-09 | 2.6e-09 | 5.9e-09 | 4.233E-09 | 1.650E-09 | 2.600E-09 | 5.900E-09 |
| 1952 | 98 | G98K | A | 7e-09 | 1.6e-08 | 7.6e-09 | 1.020E-08 | 5.032E-09 | 7.000E-09 | 1.600E-08 |
| 1953 | 98 | G98M | A | 1.6e-08 | 3.5e-09 | 3.7e-09 | 7.733E-09 | 7.160E-09 | 3.500E-09 | 1.600E-08 |
| 1954 | 98 | G98F | A | 2e-08 | 5.3e-09 | 1.5e-08 | 1.343E-08 | 7.474E-09 | 5.300E-09 | 2.000E-08 |

| | | | | | | | | | | |
|---|---|---|---|---|---|---|---|---|---|---|
| 1955 | 98 | G98P | A | 4e-09 | 1.8e-08 | 3.3e-09 | 8.433E-09 | 8.292E-09 | 3.300E-09 | 1.800E-08 |
| 1956 | 98 | G98S | A | 5.9e-09 | 3.9e-09 | 9.8e-09 | 6.533E-09 | 3.001E-09 | 3.900E-09 | 9.800E-09 |
| 1957 | 98 | G98T | A | 6.1e-09 | 2.8e-09 | 7.7e-09 | 5.533E-09 | 2.499E-09 | 2.800E-09 | 7.700E-09 |
| 1958 | 98 | G98W | A | 1e-08 | 1.9e-08 | 3.6e-09 | 1.087E-08 | 7.736E-09 | 3.600E-09 | 1.900E-08 |
| 1959 | 98 | G98Y | A | 4.8e-09 | 2.1e-09 | 1.1e-08 | 5.967E-09 | 4.563E-09 | 2.100E-09 | 1.100E-08 |
| 1960 | 98 | G98V | A | 6.2e-09 | 5.6e-09 | 2.2e-09 | 4.667E-09 | 2.157E-09 | 2.200E-09 | 6.200E-09 |
| 1961 | 99 | D99A | A | 4.6e-09 | 3.9e-09 | 1.2e-08 | 6.833E-09 | 4.488E-09 | 3.900E-09 | 1.200E-08 |
| 1962 | 99 | D99R | A | 3.5e-08 | 8.4e-09 | 7.4e-09 | 1.693E-08 | 1.565E-08 | 7.400E-09 | 3.500E-08 |
| 1963 | 99 | D99N | A | 5e-09 | 2.2e-08 | 1.8e-09 | 9.600E-09 | 1.086E-08 | 1.800E-09 | 2.200E-08 |
| 1964 | 99 | D99D | A | 2.8e-09 | 4.9e-09 | 6e-09 | 4.567E-09 | 1.626E-09 | 2.800E-09 | 6.000E-09 |
| 1965 | 99 | D99C | A | 6.6e-09 | 5.2e-09 | 9.7e-09 | 7.167E-09 | 2.303E-09 | 5.200E-09 | 9.700E-09 |
| 1966 | 99 | D99E | A | 9.3e-09 | 1.6e-08 | 8.1e-09 | 1.113E-08 | 4.257E-09 | 8.100E-09 | 1.600E-08 |
| 1967 | 99 | D99Q | A | 5.2e-09 | 2e-08 | 1.3e-08 | 1.273E-08 | 7.404E-09 | 5.200E-09 | 2.000E-08 |
| 1968 | 99 | D99G | A | 2.1e-08 | 1.8e-08 | 8.8e-09 | 1.593E-08 | 6.357E-09 | 8.800E-09 | 2.100E-08 |
| 1969 | 99 | D99H | A | 5.2e-09 | 2e-09 | 1.3e-08 | 6.733E-09 | 5.658E-09 | 2.000E-09 | 1.300E-08 |
| 1970 | 99 | D99I | A | 3.2e-09 | 2.5e-08 | 7.2e-09 | 1.180E-08 | 1.161E-08 | 3.200E-09 | 2.500E-08 |
| 1971 | 99 | D99L | A | 3.9e-09 | 2.7e-09 | 7.4e-09 | 4.667E-09 | 2.442E-09 | 2.700E-09 | 7.400E-09 |
| 1972 | 99 | D99K | A | 9.6e-09 | 5.7e-09 | 9.5e-09 | 8.267E-09 | 2.223E-09 | 5.700E-09 | 9.600E-09 |
| 1973 | 99 | D99M | A | 1e-08 | 7.4e-09 | 9e-09 | 8.800E-09 | 1.311E-09 | 7.400E-09 | 1.000E-08 |
| 1974 | 99 | D99F | A | 7e-09 | 1.7e-08 | 8.3e-09 | 1.077E-08 | 5.437E-09 | 7.000E-09 | 1.700E-08 |
| 1975 | 99 | D99P | A | 1.5e-08 | 1.1e-08 | 1.1e-08 | 1.233E-08 | 2.309E-09 | 1.100E-08 | 1.500E-08 |
| 1976 | 99 | D99S | A | 3e-09 | 2e-08 | 1.8e-09 | 8.267E-09 | 1.018E-08 | 1.800E-09 | 2.000E-08 |
| 1977 | 99 | D99T | A | 4.5e-09 | 2.2e-08 | 7.1e-09 | 1.120E-08 | 9.443E-09 | 4.500E-09 | 2.200E-08 |
| 1978 | 99 | D99W | A | 4.2e-09 | 5.4e-09 | 4.8e-09 | 4.800E-09 | 6.000E-10 | 4.200E-09 | 5.400E-09 |
| 1979 | 99 | D99Y | A | 1.6e-08 | 1.7e-08 | 8.5e-09 | 1.383E-08 | 4.646E-09 | 8.500E-09 | 1.700E-08 |
| 1980 | 99 | D99V | A | 8.5e-09 | 1.1e-08 | 6e-09 | 8.500E-09 | 2.500E-09 | 6.000E-09 | 1.100E-08 |
| 1981 | 100 | P100A | A | 7.5e-09 | 7.7e-09 | 1.2e-08 | 9.067E-09 | 2.542E-09 | 7.500E-09 | 1.200E-08 |
| 1982 | 100 | P100R | A | 6.2e-09 | 8.2e-09 | 5.5e-09 | 6.633E-09 | 1.401E-09 | 5.500E-09 | 8.200E-09 |
| 1983 | 100 | P100N | A | 1.3e-08 | 5.2e-09 | 1e-08 | 9.400E-09 | 3.934E-09 | 5.200E-09 | 1.300E-08 |
| 1984 | 100 | P100D | A | 5.2e-09 | 9.8e-09 | 3.2e-09 | 6.067E-09 | 3.384E-09 | 3.200E-09 | 9.800E-09 |
| 1985 | 100 | P100C | A | 1.1e-08 | 8e-09 | 5.9e-09 | 8.300E-09 | 2.563E-09 | 5.900E-09 | 1.100E-08 |
| 1986 | 100 | P100E | A | 7.5e-09 | 9.3e-09 | 2.5e-09 | 6.433E-09 | 3.523E-09 | 2.500E-09 | 9.300E-09 |
| 1987 | 100 | P100Q | A | 7.1e-09 | 7e-09 | 8.3e-09 | 7.467E-09 | 7.234E-10 | 7.000E-09 | 8.300E-09 |
| 1988 | 100 | P100G | A | 1.1e-08 | 1.8e-08 | 1.2e-08 | 1.367E-08 | 3.786E-09 | 1.100E-08 | 1.800E-08 |
| 1989 | 100 | P100H | A | 6e-09 | 1.7e-08 | 5e-09 | 9.333E-09 | 6.658E-09 | 5.000E-09 | 1.700E-08 |
| 1990 | 100 | P100I | A | 9.2e-09 | 8.9e-09 | 4.7e-09 | 7.600E-09 | 2.516E-09 | 4.700E-09 | 9.200E-09 |
| 1991 | 100 | P100L | A | 7.2e-09 | 1.8e-08 | 1.6e-08 | 1.373E-08 | 5.746E-09 | 7.200E-09 | 1.800E-08 |
| 1992 | 100 | P100K | A | 5.5e-09 | 8.5e-09 | 4.3e-09 | 6.100E-09 | 2.163E-09 | 4.300E-09 | 8.500E-09 |
| 1993 | 100 | P100M | A | 6e-09 | 9.8e-09 | 7.5e-09 | 7.767E-09 | 1.914E-09 | 6.000E-09 | 9.800E-09 |
| 1994 | 100 | P100F | A | 6.1e-09 | 7.7e-09 | 3.8e-09 | 5.867E-09 | 1.960E-09 | 3.800E-09 | 7.700E-09 |
| 1995 | 100 | P100P | A | 1e-08 | 3e-09 | 2e-09 | 5.000E-09 | 4.359E-09 | 2.000E-09 | 1.000E-08 |
| 1996 | 100 | P100S | A | 8.2e-09 | 5.6e-09 | 1e-08 | 7.933E-09 | 2.212E-09 | 5.600E-09 | 1.000E-08 |
| 1997 | 100 | P100T | A | 9.5e-09 | 4.4e-09 | 1.6e-08 | 9.967E-09 | 5.814E-09 | 4.400E-09 | 1.600E-08 |
| 1998 | 100 | P100W | A | 2.1e-08 | 2e-08 | 7.4e-09 | 1.613E-08 | 7.580E-09 | 7.400E-09 | 2.100E-08 |
| 1999 | 100 | P100Y | A | 6e-09 | 1.1e-08 | 4.1e-09 | 7.033E-09 | 3.564E-09 | 4.100E-09 | 1.100E-08 |
| 2000 | 100 | P100V | A | 9e-09 | 6.9e-09 | 9e-09 | 8.300E-09 | 1.212E-09 | 6.900E-09 | 9.000E-09 |
| 2001 | 101 | L101A | A | 5.5e-09 | 2.1e-09 | 5.7e-09 | 4.433E-09 | 2.023E-09 | 2.100E-09 | 5.700E-09 |
| 2002 | 101 | L101R | A | 9.5e-09 | 1.1e-08 | 5.7e-09 | 8.733E-09 | 2.732E-09 | 5.700E-09 | 1.100E-08 |
| 2003 | 101 | L101N | A | 8.9e-09 | 1.8e-08 | 1.5e-08 | 1.397E-08 | 4.637E-09 | 8.900E-09 | 1.800E-08 |
| 2004 | 101 | L101D | A | 6e-09 | 2.9e-09 | 2.9e-09 | 3.933E-09 | 1.790E-09 | 2.900E-09 | 6.000E-09 |
| 2005 | 101 | L101C | A | 1.2e-08 | 1.1e-08 | 6.4e-09 | 9.800E-09 | 2.987E-09 | 6.400E-09 | 1.200E-08 |
| 2006 | 101 | L101E | A | 3.9e-09 | 5.2e-09 | 9.7e-09 | 6.267E-09 | 3.044E-09 | 3.900E-09 | 9.700E-09 |
| 2007 | 101 | L101Q | A | 5.6e-09 | 6.8e-09 | 8.1e-09 | 6.833E-09 | 1.250E-09 | 5.600E-09 | 8.100E-09 |
| 2008 | 101 | L101G | A | 6.3e-09 | 1.5e-09 | 7e-09 | 4.933E-09 | 2.994E-09 | 1.500E-09 | 7.000E-09 |
| 2009 | 101 | L101H | A | 5.1e-09 | 6.9e-09 | 4.2e-09 | 5.400E-09 | 1.375E-09 | 4.200E-09 | 6.900E-09 |
| 2010 | 101 | L101I | A | 5.4e-09 | 8.9e-09 | 1.5e-08 | 9.767E-09 | 4.858E-09 | 5.400E-09 | 1.500E-08 |
| 2011 | 101 | L101L | A | 1.5e-08 | 3.4e-08 | 4.9e-09 | 1.797E-08 | 1.478E-08 | 4.900E-09 | 3.400E-08 |
| 2012 | 101 | L101K | A | 1.5e-08 | 1.6e-08 | 3.2e-09 | 1.140E-08 | 7.119E-09 | 3.200E-09 | 1.600E-08 |
| 2013 | 101 | L101M | A | 7.1e-09 | 6.5e-09 | 1.4e-08 | 9.200E-09 | 4.168E-09 | 6.500E-09 | 1.400E-08 |
| 2014 | 101 | L101F | A | 2.2e-09 | 9.8e-09 | 1.6e-08 | 9.333E-09 | 6.912E-09 | 2.200E-09 | 1.600E-08 |
| 2015 | 101 | L101P | A | 3e-09 | 1.7e-08 | 1.2e-08 | 1.067E-08 | 7.095E-09 | 3.000E-09 | 1.700E-08 |
| 2016 | 101 | L101S | A | 4.6e-09 | 8.4e-09 | 1.5e-08 | 9.333E-09 | 5.262E-09 | 4.600E-09 | 1.500E-08 |
| 2017 | 101 | L101T | A | 8.2e-09 | 1.2e-08 | 1.3e-08 | 1.107E-08 | 2.532E-09 | 8.200E-09 | 1.300E-08 |
| 2018 | 101 | L101W | A | 1.3e-08 | 3.8e-09 | 6.2e-09 | 7.667E-09 | 4.772E-09 | 3.800E-09 | 1.300E-08 |
| 2019 | 101 | L101Y | A | 1.1e-08 | 1.3e-08 | 6.4e-09 | 1.013E-08 | 3.384E-09 | 6.400E-09 | 1.300E-08 |
| 2020 | 101 | L101V | A | 8.3e-09 | 3.5e-09 | 2.6e-09 | 4.800E-09 | 3.064E-09 | 2.600E-09 | 8.300E-09 |
| 2021 | 102 | N102A | A | 1.2e-08 | 5.5e-09 | 9.2e-09 | 8.900E-09 | 3.260E-09 | 5.500E-09 | 1.200E-08 |
| 2022 | 102 | N102R | A | 4.7e-09 | 4.5e-09 | 8.2e-09 | 5.800E-09 | 2.081E-09 | 4.500E-09 | 8.200E-09 |
| 2023 | 102 | N102N | A | 6.2e-09 | 5.3e-09 | 6.2e-09 | 5.900E-09 | 5.196E-10 | 5.300E-09 | 6.200E-09 |
| 2024 | 102 | N102D | A | 9.1e-09 | 6.6e-09 | 8.7e-09 | 8.133E-09 | 1.343E-09 | 6.600E-09 | 9.100E-09 |
| 2025 | 102 | N102C | A | 1.1e-08 | 1.4e-08 | 2.4e-08 | 1.633E-08 | 6.807E-09 | 1.100E-08 | 2.400E-08 |
| 2026 | 102 | N102E | A | 4.1e-09 | 1.5e-08 | 1.8e-09 | 6.967E-09 | 7.051E-09 | 1.800E-09 | 1.500E-08 |
| 2027 | 102 | N102Q | A | 1.8e-08 | 8.1e-09 | 1.6e-08 | 1.403E-08 | 5.235E-09 | 8.100E-09 | 1.800E-08 |
| 2028 | 102 | N102G | A | 1.7e-08 | 8.5e-09 | 6.1e-09 | 1.053E-08 | 5.727E-09 | 6.100E-09 | 1.700E-08 |
| 2029 | 102 | N102H | A | 3.8e-09 | 9.2e-09 | 2.9e-09 | 5.300E-09 | 3.407E-09 | 2.900E-09 | 9.200E-09 |
| 2030 | 102 | N102I | A | 3.7e-09 | 1e-08 | 2.5e-09 | 5.400E-09 | 4.029E-09 | 2.500E-09 | 1.000E-08 |
| 2031 | 102 | N102L | A | 1.4e-09 | 1.7e-08 | 9.1e-09 | 9.167E-09 | 7.800E-09 | 1.400E-09 | 1.700E-08 |
| 2032 | 102 | N102K | A | 5.3e-09 | 5.7e-09 | 6.5e-09 | 5.833E-09 | 6.110E-10 | 5.300E-09 | 6.500E-09 |
| 2033 | 102 | N102M | A | 2.2e-08 | 1.3e-08 | 7.2e-09 | 1.407E-08 | 7.457E-09 | 7.200E-09 | 2.200E-08 |
| 2034 | 102 | N102F | A | 6.5e-09 | 1.6e-08 | 2.6e-09 | 8.367E-09 | 6.892E-09 | 2.600E-09 | 1.600E-08 |
| 2035 | 102 | N102P | A | 1.2e-08 | 9e-09 | 2.2e-08 | 1.433E-08 | 6.807E-09 | 9.000E-09 | 2.200E-08 |
| 2036 | 102 | N102S | A | 1e-08 | 5.1e-09 | 1.6e-08 | 1.037E-08 | 5.459E-09 | 5.100E-09 | 1.600E-08 |
| 2037 | 102 | N102T | A | 5e-09 | 8.9e-09 | 3.9e-08 | 1.763E-08 | 1.861E-08 | 5.000E-09 | 3.900E-08 |
| 2038 | 102 | N102W | A | 6.3e-09 | 4.5e-09 | 1.3e-08 | 7.933E-09 | 4.479E-09 | 4.500E-09 | 1.300E-08 |
| 2039 | 102 | N102Y | A | 5.9e-09 | 4.1e-09 | 1.9e-08 | 9.667E-09 | 8.133E-09 | 4.100E-09 | 1.900E-08 |

| | | | | | | | | | | |
|---|---|---|---|---|---|---|---|---|---|---|
| 2040 | 102 | N102V | A | 9e-09 | 1.4e-08 | 4.5e-09 | 9.167E-09 | 4.752E-09 | 4.500E-09 | 1.400E-08 |
| 2041 | 103 | N103A | A | 2e-09 | 5.9e-09 | 1.4e-08 | 7.300E-09 | 6.121E-09 | 2.000E-09 | 1.400E-08 |
| 2042 | 103 | N103R | A | 1.1e-08 | 5e-09 | 1.7e-08 | 1.100E-08 | 6.000E-09 | 5.000E-09 | 1.700E-08 |
| 2043 | 103 | N103N | A | 7.1e-09 | 1.3e-08 | 4e-09 | 8.033E-09 | 4.572E-09 | 4.000E-09 | 1.300E-08 |
| 2044 | 103 | N103D | A | 1.4e-08 | 2.9e-09 | 9.8e-09 | 8.900E-09 | 5.604E-09 | 2.900E-09 | 1.400E-08 |
| 2045 | 103 | N103C | A | 4.9e-09 | 9.8e-09 | 4.8e-09 | 6.500E-09 | 2.858E-09 | 4.800E-09 | 9.800E-09 |
| 2046 | 103 | N103E | A | 1.3e-08 | 2e-09 | 6e-09 | 7.000E-09 | 5.568E-09 | 2.000E-09 | 1.300E-08 |
| 2047 | 103 | N103Q | A | 1.4e-08 | 4.5e-09 | 5.5e-09 | 8.000E-09 | 5.220E-09 | 4.500E-09 | 1.400E-08 |
| 2048 | 103 | N103G | A | 1.4e-08 | 4.3e-09 | 1.7e-08 | 1.177E-08 | 6.638E-09 | 4.300E-09 | 1.700E-08 |
| 2049 | 103 | N103H | A | 7.7e-09 | 7.1e-09 | 1.9e-08 | 1.127E-08 | 6.704E-09 | 7.100E-09 | 1.900E-08 |
| 2050 | 103 | N103I | A | 4.6e-09 | 6.8e-09 | 4.7e-09 | 5.367E-09 | 1.242E-09 | 4.600E-09 | 6.800E-09 |
| 2051 | 103 | N103L | A | 7.9e-09 | 6.2e-09 | 8.2e-09 | 7.433E-09 | 1.079E-09 | 6.200E-09 | 8.200E-09 |
| 2052 | 103 | N103K | A | 6.7e-09 | 6.4e-09 | 6.2e-09 | 6.433E-09 | 2.517E-10 | 6.200E-09 | 6.700E-09 |
| 2053 | 103 | N103M | A | 6e-09 | 5.2e-09 | 3.7e-09 | 4.967E-09 | 1.168E-09 | 3.700E-09 | 6.000E-09 |
| 2054 | 103 | N103F | A | 5.6e-09 | 6e-09 | 8.9e-09 | 6.833E-09 | 1.801E-09 | 5.600E-09 | 8.900E-09 |
| 2055 | 103 | N103P | A | 1.7e-08 | 6e-09 | 7.1e-09 | 1.003E-08 | 6.058E-09 | 6.000E-09 | 1.700E-08 |
| 2056 | 103 | N103S | A | 4.7e-09 | 4.7e-09 | 5.8e-09 | 5.067E-09 | 6.351E-10 | 4.700E-09 | 5.800E-09 |
| 2057 | 103 | N103T | A | 3e-09 | 6.8e-09 | 2.8e-08 | 1.260E-08 | 1.347E-08 | 3.000E-09 | 2.800E-08 |
| 2058 | 103 | N103W | A | 1.3e-08 | 1.7e-08 | 7.9e-09 | 1.263E-08 | 4.561E-09 | 7.900E-09 | 1.700E-08 |
| 2059 | 103 | N103Y | A | 2.5e-09 | 2.1e-08 | 5.5e-09 | 9.667E-09 | 9.929E-09 | 2.500E-09 | 2.100E-08 |
| 2060 | 103 | N103V | A | 3.4e-09 | 7.1e-09 | 3.4e-09 | 4.633E-09 | 2.136E-09 | 3.400E-09 | 7.100E-09 |
| 2061 | 104 | T104A | A | 1.2e-08 | 3.7e-09 | 1.2e-08 | 9.233E-09 | 4.792E-09 | 3.700E-09 | 1.200E-08 |
| 2062 | 104 | T104R | A | 1e-08 | 6e-09 | 3.3e-09 | 6.433E-09 | 3.371E-09 | 3.300E-09 | 1.000E-08 |
| 2063 | 104 | T104N | A | 5.7e-09 | 8.9e-09 | 1.1e-08 | 8.533E-09 | 2.669E-09 | 5.700E-09 | 1.100E-08 |
| 2064 | 104 | T104D | A | 1.4e-08 | 5.9e-09 | 7.6e-09 | 9.167E-09 | 4.271E-09 | 5.900E-09 | 1.400E-08 |
| 2065 | 104 | T104C | A | 6.7e-09 | 6.8e-09 | 4.1e-09 | 5.867E-09 | 1.531E-09 | 4.100E-09 | 6.800E-09 |
| 2066 | 104 | T104E | A | 1.2e-08 | 1.1e-08 | 5.1e-09 | 9.367E-09 | 3.729E-09 | 5.100E-09 | 1.200E-08 |
| 2067 | 104 | T104Q | A | 9.6e-09 | 1.5e-08 | 1.7e-09 | 8.767E-09 | 6.689E-09 | 1.700E-09 | 1.500E-08 |
| 2068 | 104 | T104G | A | 6.9e-09 | 1.6e-08 | 1.1e-08 | 1.130E-08 | 4.557E-09 | 6.900E-09 | 1.600E-08 |
| 2069 | 104 | T104H | A | 4.1e-09 | 1.9e-08 | 8.8e-09 | 1.063E-08 | 7.617E-09 | 4.100E-09 | 1.900E-08 |
| 2070 | 104 | T104I | A | 4.7e-09 | 3.9e-09 | 5.7e-09 | 4.767E-09 | 9.018E-10 | 3.900E-09 | 5.700E-09 |
| 2071 | 104 | T104L | A | 1.7e-09 | 9.6e-09 | 5.5e-09 | 5.600E-09 | 3.951E-09 | 1.700E-09 | 9.600E-09 |
| 2072 | 104 | T104K | A | 9.5e-09 | 1.7e-08 | 6.1e-09 | 1.087E-08 | 5.577E-09 | 6.100E-09 | 1.700E-08 |
| 2073 | 104 | T104M | A | 6.9e-09 | 1.7e-08 | 7.6e-09 | 1.050E-08 | 5.640E-09 | 6.900E-09 | 1.700E-08 |
| 2074 | 104 | T104F | A | 9.4e-09 | 4.1e-09 | 6.5e-09 | 6.667E-09 | 2.654E-09 | 4.100E-09 | 9.400E-09 |
| 2075 | 104 | T104P | A | 7.1e-09 | 2.4e-08 | 9.7e-09 | 1.360E-08 | 9.100E-09 | 7.100E-09 | 2.400E-08 |
| 2076 | 104 | T104S | A | 1.4e-08 | 4.7e-09 | 1.6e-08 | 1.157E-08 | 6.030E-09 | 4.700E-09 | 1.600E-08 |
| 2077 | 104 | T104T | A | 3.9e-09 | 1.1e-08 | 5.9e-09 | 6.933E-09 | 3.661E-09 | 3.900E-09 | 1.100E-08 |
| 2078 | 104 | T104W | A | 1.5e-08 | 1e-08 | 6.1e-09 | 1.037E-08 | 4.461E-09 | 6.100E-09 | 1.500E-08 |
| 2079 | 104 | T104Y | A | 4.9e-09 | 2.6e-09 | 3.6e-09 | 3.700E-09 | 1.153E-09 | 2.600E-09 | 4.900E-09 |
| 2080 | 104 | T104V | A | 4e-09 | 4.1e-09 | 5.2e-08 | 2.003E-08 | 2.768E-08 | 4.000E-09 | 5.200E-08 |
| 2081 | 105 | T105A | A | 1.1e-08 | 8.1e-09 | 5.5e-09 | 8.200E-09 | 2.751E-09 | 5.500E-09 | 1.100E-08 |
| 2082 | 105 | T105R | A | 3.8e-09 | 6.6e-09 | 4.7e-09 | 5.033E-09 | 1.429E-09 | 3.800E-09 | 6.600E-09 |
| 2083 | 105 | T105N | A | 6.8e-09 | 4.9e-09 | 6.5e-09 | 6.067E-09 | 1.021E-09 | 4.900E-09 | 6.800E-09 |
| 2084 | 105 | T105D | A | 1.6e-08 | 7.7e-09 | 2.9e-09 | 8.867E-09 | 6.627E-09 | 2.900E-09 | 1.600E-08 |
| 2085 | 105 | T105C | A | 1.3e-08 | 9.8e-09 | 6.1e-09 | 9.633E-09 | 3.453E-09 | 6.100E-09 | 1.300E-08 |
| 2086 | 105 | T105E | A | 9.6e-09 | 2e-08 | 9.2e-09 | 1.293E-08 | 6.123E-09 | 9.200E-09 | 2.000E-08 |
| 2087 | 105 | T105Q | A | 1.5e-08 | 1.6e-09 | 3.6e-09 | 6.733E-09 | 7.229E-09 | 1.600E-09 | 1.500E-08 |
| 2088 | 105 | T105G | A | 3.3e-08 | 1.2e-08 | 1.7e-09 | 1.557E-08 | 1.595E-08 | 1.700E-09 | 3.300E-08 |
| 2089 | 105 | T105H | A | 3.4e-09 | 6.4e-09 | 1.4e-08 | 7.933E-09 | 5.464E-09 | 3.400E-09 | 1.400E-08 |
| 2090 | 105 | T105I | A | 7.2e-09 | 1.2e-08 | 4.1e-09 | 7.767E-09 | 3.980E-09 | 4.100E-09 | 1.200E-08 |
| 2091 | 105 | T105L | A | 7.8e-09 | 6.1e-09 | 4.1e-09 | 6.000E-09 | 1.852E-09 | 4.100E-09 | 7.800E-09 |
| 2092 | 105 | T105K | A | 1e-08 | 6.3e-09 | 5.7e-09 | 7.333E-09 | 2.329E-09 | 5.700E-09 | 1.000E-08 |
| 2093 | 105 | T105M | A | 1.4e-08 | 3.2e-09 | 9.5e-09 | 8.900E-09 | 5.425E-09 | 3.200E-09 | 1.400E-08 |
| 2094 | 105 | T105F | A | 2e-09 | 7.3e-09 | 1.4e-09 | 3.567E-09 | 3.247E-09 | 1.400E-09 | 7.300E-09 |
| 2095 | 105 | T105P | A | 8.9e-09 | 4e-09 | 3.7e-09 | 5.533E-09 | 2.919E-09 | 3.700E-09 | 8.900E-09 |
| 2096 | 105 | T105S | A | 1.9e-09 | 5.7e-09 | 2.8e-09 | 3.467E-09 | 1.986E-09 | 1.900E-09 | 5.700E-09 |
| 2097 | 105 | T105T | A | 2.1e-09 | 8.6e-09 | 1e-08 | 6.900E-09 | 4.215E-09 | 2.100E-09 | 1.000E-08 |
| 2098 | 105 | T105W | A | 5.5e-09 | 9.2e-09 | 8.7e-09 | 7.800E-09 | 2.007E-09 | 5.500E-09 | 9.200E-09 |
| 2099 | 105 | T105Y | A | 2e-08 | 3.4e-09 | 6.5e-09 | 9.967E-09 | 8.826E-09 | 3.400E-09 | 2.000E-08 |
| 2100 | 105 | T105V | A | 7.4e-09 | 8.9e-09 | 8.7e-09 | 8.333E-09 | 8.145E-10 | 7.400E-09 | 8.900E-09 |
| 2101 | 106 | P106A | A | 3.3e-09 | 9.2e-09 | 7.5e-09 | 6.667E-09 | 3.037E-09 | 3.300E-09 | 9.200E-09 |
| 2102 | 106 | P106R | A | 1.2e-08 | 3.4e-09 | 5.7e-09 | 7.033E-09 | 4.452E-09 | 3.400E-09 | 1.200E-08 |
| 2103 | 106 | P106N | A | 9.9e-09 | 6.3e-09 | 6.3e-09 | 7.500E-09 | 2.078E-09 | 6.300E-09 | 9.900E-09 |
| 2104 | 106 | P106D | A | 4.4e-09 | 9.1e-09 | 4.3e-09 | 5.933E-09 | 2.743E-09 | 4.300E-09 | 9.100E-09 |
| 2105 | 106 | P106C | A | 6e-09 | 2.6e-09 | 5.7e-09 | 4.767E-09 | 1.882E-09 | 2.600E-09 | 6.000E-09 |
| 2106 | 106 | P106E | A | 9.8e-09 | 1.1e-08 | 4.9e-09 | 8.567E-09 | 3.232E-09 | 4.900E-09 | 1.100E-08 |
| 2107 | 106 | P106Q | A | 2.7e-09 | 1.6e-08 | 1.8e-09 | 6.833E-09 | 7.951E-09 | 1.800E-09 | 1.600E-08 |
| 2108 | 106 | P106G | A | 4.5e-09 | 2.2e-08 | 1.3e-08 | 1.317E-08 | 8.751E-09 | 4.500E-09 | 2.200E-08 |
| 2109 | 106 | P106H | A | 7.5e-09 | 7.7e-09 | 1.1e-08 | 8.733E-09 | 1.966E-09 | 7.500E-09 | 1.100E-08 |
| 2110 | 106 | P106I | A | 1.2e-08 | 1.4e-08 | 9.9e-09 | 1.197E-08 | 2.050E-09 | 9.900E-09 | 1.400E-08 |
| 2111 | 106 | P106L | A | 3.3e-09 | 7.7e-09 | 1e-08 | 7.000E-09 | 3.404E-09 | 3.300E-09 | 1.000E-08 |
| 2112 | 106 | P106K | A | 4.5e-09 | 6.7e-09 | 2.7e-09 | 4.633E-09 | 2.003E-09 | 2.700E-09 | 6.700E-09 |
| 2113 | 106 | P106M | A | 1.7e-08 | 3.3e-09 | 7e-09 | 9.100E-09 | 7.087E-09 | 3.300E-09 | 1.700E-08 |
| 2114 | 106 | P106F | A | 1.9e-08 | 1.5e-08 | 1.4e-08 | 1.600E-08 | 2.646E-09 | 1.400E-08 | 1.900E-08 |
| 2115 | 106 | P106P | A | 6e-09 | 2.6e-09 | 3.4e-09 | 4.000E-09 | 1.778E-09 | 2.600E-09 | 6.000E-09 |
| 2116 | 106 | P106S | A | 3.9e-09 | 8.3e-09 | 7.5e-09 | 6.567E-09 | 2.344E-09 | 3.900E-09 | 8.300E-09 |
| 2117 | 106 | P106T | A | 1.1e-08 | 9.4e-09 | 4e-09 | 8.133E-09 | 3.668E-09 | 4.000E-09 | 1.100E-08 |
| 2118 | 106 | P106W | A | 3.1e-08 | 1.2e-08 | 4.9e-09 | 1.597E-08 | 1.349E-08 | 4.900E-09 | 3.100E-08 |
| 2119 | 106 | P106Y | A | 1.9e-08 | 1.1e-08 | 6.1e-09 | 1.203E-08 | 6.512E-09 | 6.100E-09 | 1.900E-08 |
| 2120 | 106 | P106V | A | 4.2e-09 | 2.3e-09 | 1.6e-09 | 2.700E-09 | 1.345E-09 | 1.600E-09 | 4.200E-09 |
| 2121 | 107 | V107A | A | 1e-08 | 1.4e-08 | 9.2e-09 | 1.107E-08 | 2.572E-09 | 9.200E-09 | 1.400E-08 |
| 2122 | 107 | V107R | A | 5.2e-09 | 7.6e-09 | 1.4e-08 | 8.933E-09 | 4.549E-09 | 5.200E-09 | 1.400E-08 |
| 2123 | 107 | V107N | A | 5.6e-09 | 7.3e-09 | 3.3e-09 | 5.400E-09 | 2.007E-09 | 3.300E-09 | 7.300E-09 |
| 2124 | 107 | V107D | A | 1.4e-08 | 7.5e-09 | 3.9e-09 | 8.467E-09 | 5.119E-09 | 3.900E-09 | 1.400E-08 |

| | | | | | | | | | | |
|---|---|---|---|---|---|---|---|---|---|---|
| 2125 | 107 | V107C | A | 3.4e-09 | 3.4e-09 | 2.1e-09 | 2.967E-09 | 7.506E-10 | 2.100E-09 | 3.400E-09 |
| 2126 | 107 | V107E | A | 1.7e-08 | 8.8e-09 | 1.1e-08 | 1.227E-08 | 4.244E-09 | 8.800E-09 | 1.700E-08 |
| 2127 | 107 | V107Q | A | 5.3e-09 | 4.1e-09 | 1.5e-08 | 8.133E-09 | 5.977E-09 | 4.100E-09 | 1.500E-08 |
| 2128 | 107 | V107G | A | 1.7e-09 | 8e-09 | 3.6e-09 | 4.433E-09 | 3.232E-09 | 1.700E-09 | 8.000E-09 |
| 2129 | 107 | V107H | A | 4.2e-09 | 5.4e-09 | 1.4e-08 | 7.867E-09 | 5.345E-09 | 4.200E-09 | 1.400E-08 |
| 2130 | 107 | V107I | A | 7.5e-09 | 6.8e-09 | 1.5e-08 | 9.767E-09 | 4.546E-09 | 6.800E-09 | 1.500E-08 |
| 2131 | 107 | V107L | A | 1.8e-08 | 6.8e-09 | 8.4e-09 | 1.107E-08 | 6.058E-09 | 6.800E-09 | 1.800E-08 |
| 2132 | 107 | V107K | A | 3e-08 | 2.3e-08 | 1.2e-08 | 2.167E-08 | 9.074E-09 | 1.200E-08 | 3.000E-08 |
| 2133 | 107 | V107M | A | 2.6e-09 | 4.1e-09 | 1.4e-08 | 6.900E-09 | 6.194E-09 | 2.600E-09 | 1.400E-08 |
| 2134 | 107 | V107F | A | 4.6e-09 | 8.7e-09 | 7.4e-09 | 6.900E-09 | 2.095E-09 | 4.600E-09 | 8.700E-09 |
| 2135 | 107 | V107P | A | 1e-08 | 8.9e-09 | 2.3e-08 | 1.397E-08 | 7.842E-09 | 8.900E-09 | 2.300E-08 |
| 2136 | 107 | V107S | A | 7.8e-09 | 8.8e-09 | 1.7e-08 | 1.120E-08 | 5.048E-09 | 7.800E-09 | 1.700E-08 |
| 2137 | 107 | V107T | A | 5.6e-09 | 1.7e-08 | 5.3e-09 | 9.300E-09 | 6.670E-09 | 5.300E-09 | 1.700E-08 |
| 2138 | 107 | V107W | A | 5.2e-09 | 1.2e-08 | 3e-09 | 6.733E-09 | 4.692E-09 | 3.000E-09 | 1.200E-08 |
| 2139 | 107 | V107Y | A | 8e-09 | 4.2e-09 | 1.2e-08 | 8.067E-09 | 3.900E-09 | 4.200E-09 | 1.200E-08 |
| 2140 | 107 | V107V | A | 1.4e-08 | 6.3e-09 | 4e-09 | 8.100E-09 | 5.237E-09 | 4.000E-09 | 1.400E-08 |
| 2141 | 108 | T108A | A | 2.4e-09 | 1.5e-08 | 1.2e-08 | 9.800E-09 | 6.582E-09 | 2.400E-09 | 1.500E-08 |
| 2142 | 108 | T108R | A | 4.8e-09 | 1.3e-08 | 1.5e-08 | 1.093E-08 | 5.405E-09 | 4.800E-09 | 1.500E-08 |
| 2143 | 108 | T108N | A | 1.1e-08 | 1.2e-08 | 1.2e-08 | 1.167E-08 | 5.774E-10 | 1.100E-08 | 1.200E-08 |
| 2144 | 108 | T108D | A | 8.7e-09 | 8.1e-09 | 9.9e-09 | 8.900E-09 | 9.165E-10 | 8.100E-09 | 9.900E-09 |
| 2145 | 108 | T108C | A | 9.8e-09 | 4.8e-09 | 5.4e-09 | 6.667E-09 | 2.730E-09 | 4.800E-09 | 9.800E-09 |
| 2146 | 108 | T108E | A | 5.9e-09 | 2e-08 | 1.1e-08 | 1.230E-08 | 7.139E-09 | 5.900E-09 | 2.000E-08 |
| 2147 | 108 | T108Q | A | 7.2e-09 | 2.2e-09 | 2.1e-08 | 1.013E-08 | 9.737E-09 | 2.200E-09 | 2.100E-08 |
| 2148 | 108 | T108G | A | 1.2e-08 | 7.6e-09 | 1.6e-08 | 1.187E-08 | 4.202E-09 | 7.600E-09 | 1.600E-08 |
| 2149 | 108 | T108H | A | 2.5e-09 | 1.1e-08 | 8.2e-09 | 7.233E-09 | 4.332E-09 | 2.500E-09 | 1.100E-08 |
| 2150 | 108 | T108I | A | 1.2e-08 | 1.2e-08 | 6.5e-09 | 1.017E-08 | 3.175E-09 | 6.500E-09 | 1.200E-08 |
| 2151 | 108 | T108L | A | 1.4e-09 | 5.7e-09 | 2.6e-09 | 3.233E-09 | 2.219E-09 | 1.400E-09 | 5.700E-09 |
| 2152 | 108 | T108K | A | 2.5e-09 | 5.5e-09 | 5.8e-09 | 4.600E-09 | 1.825E-09 | 2.500E-09 | 5.800E-09 |
| 2153 | 108 | T108M | A | 8.8e-09 | 1.1e-08 | 7.8e-09 | 9.200E-09 | 1.637E-09 | 7.800E-09 | 1.100E-08 |
| 2154 | 108 | T108F | A | 8.7e-09 | 1.7e-08 | 7.3e-09 | 1.100E-08 | 5.243E-09 | 7.300E-09 | 1.700E-08 |
| 2155 | 108 | T108P | A | 9.2e-09 | 3.7e-09 | 1e-08 | 7.633E-09 | 3.430E-09 | 3.700E-09 | 1.000E-08 |
| 2156 | 108 | T108S | A | 7.4e-09 | 7e-09 | 9.3e-09 | 7.900E-09 | 1.229E-09 | 7.000E-09 | 9.300E-09 |
| 2157 | 108 | T108T | A | 6.6e-09 | 8.1e-09 | 1.3e-08 | 9.233E-09 | 3.347E-09 | 6.600E-09 | 1.300E-08 |
| 2158 | 108 | T108W | A | 1.1e-08 | 1.3e-08 | 8.3e-09 | 1.077E-08 | 2.359E-09 | 8.300E-09 | 1.300E-08 |
| 2159 | 108 | T108Y | A | 2.4e-09 | 3.3e-09 | 8e-09 | 4.567E-09 | 3.007E-09 | 2.400E-09 | 8.000E-09 |
| 2160 | 108 | T108V | A | 3.1e-09 | 7e-09 | 6e-09 | 5.367E-09 | 2.026E-09 | 3.100E-09 | 7.000E-09 |
| 2161 | 109 | G109A | A | 3.7e-09 | 4.1e-09 | 1e-08 | 5.933E-09 | 3.528E-09 | 3.700E-09 | 1.000E-08 |
| 2162 | 109 | G109R | A | 6.6e-09 | 8e-09 | 3e-09 | 5.867E-09 | 2.579E-09 | 3.000E-09 | 8.000E-09 |
| 2163 | 109 | G109N | A | 4.6e-09 | 9.4e-09 | 2.5e-09 | 5.500E-09 | 3.537E-09 | 2.500E-09 | 9.400E-09 |
| 2164 | 109 | G109D | A | 3e-09 | 1.2e-08 | 4.8e-09 | 6.600E-09 | 4.762E-09 | 3.000E-09 | 1.200E-08 |
| 2165 | 109 | G109C | A | 1.1e-08 | 1.3e-08 | 2.5e-08 | 1.633E-08 | 7.572E-09 | 1.100E-08 | 2.500E-08 |
| 2166 | 109 | G109E | A | 6.4e-09 | 9.9e-09 | 4.3e-09 | 6.867E-09 | 2.829E-09 | 4.300E-09 | 9.900E-09 |
| 2167 | 109 | G109Q | A | 9.1e-09 | 1.6e-09 | 7.6e-09 | 6.100E-09 | 3.969E-09 | 1.600E-09 | 9.100E-09 |
| 2168 | 109 | G109G | A | 5.2e-09 | 4.3e-09 | 6.6e-09 | 5.367E-09 | 1.159E-09 | 4.300E-09 | 6.600E-09 |
| 2169 | 109 | G109H | A | 1.2e-08 | 3.5e-09 | 2.1e-09 | 5.867E-09 | 5.358E-09 | 2.100E-09 | 1.200E-08 |
| 2170 | 109 | G109I | A | 7.1e-09 | 3.4e-09 | 6.9e-09 | 5.800E-09 | 2.081E-09 | 3.400E-09 | 7.100E-09 |
| 2171 | 109 | G109L | A | 3e-09 | 2.7e-09 | 1.1e-08 | 5.567E-09 | 4.708E-09 | 2.700E-09 | 1.100E-08 |
| 2172 | 109 | G109K | A | 1.3e-08 | 2.8e-09 | 1.3e-09 | 5.700E-09 | 6.366E-09 | 1.300E-09 | 1.300E-08 |
| 2173 | 109 | G109M | A | 1.9e-08 | 5.1e-09 | 8.1e-09 | 1.073E-08 | 7.315E-09 | 5.100E-09 | 1.900E-08 |
| 2174 | 109 | G109F | A | 8.8e-09 | 9.3e-09 | 1.1e-08 | 9.700E-09 | 1.153E-09 | 8.800E-09 | 1.100E-08 |
| 2175 | 109 | G109P | A | 3.6e-08 | 4.1e-09 | 7.8e-09 | 1.597E-08 | 1.745E-08 | 4.100E-09 | 3.600E-08 |
| 2176 | 109 | G109S | A | 1.2e-08 | 1e-08 | 4.1e-09 | 8.700E-09 | 4.107E-09 | 4.100E-09 | 1.200E-08 |
| 2177 | 109 | G109T | A | 1e-08 | 2.8e-09 | 2.7e-09 | 5.167E-09 | 4.186E-09 | 2.700E-09 | 1.000E-08 |
| 2178 | 109 | G109W | A | 6.4e-09 | 2.1e-08 | 9.7e-09 | 1.237E-08 | 7.657E-09 | 6.400E-09 | 2.100E-08 |
| 2179 | 109 | G109Y | A | 7.7e-09 | 4.8e-09 | 7.2e-09 | 6.567E-09 | 1.550E-09 | 4.800E-09 | 7.700E-09 |
| 2180 | 109 | G109V | A | 5.3e-09 | 4.6e-09 | 1.1e-08 | 6.967E-09 | 3.510E-09 | 4.600E-09 | 1.100E-08 |
| 2181 | 110 | A110A | A | 9.9e-09 | 3.7e-09 | 2.4e-09 | 5.333E-09 | 4.008E-09 | 2.400E-09 | 9.900E-09 |
| 2182 | 110 | A110R | A | 5.7e-09 | 1.2e-08 | 1.2e-08 | 9.900E-09 | 3.637E-09 | 5.700E-09 | 1.200E-08 |
| 2183 | 110 | A110N | A | 3.5e-09 | 5.1e-09 | 3.3e-09 | 3.967E-09 | 9.866E-10 | 3.300E-09 | 5.100E-09 |
| 2184 | 110 | A110D | A | 1.3e-08 | 2.5e-08 | 9.2e-09 | 1.573E-08 | 8.247E-09 | 9.200E-09 | 2.500E-08 |
| 2185 | 110 | A110C | A | 3.1e-09 | 9.6e-09 | 1e-08 | 7.567E-09 | 3.873E-09 | 3.100E-09 | 1.000E-08 |
| 2186 | 110 | A110E | A | 9.6e-09 | 6.8e-09 | 4.5e-09 | 6.967E-09 | 2.554E-09 | 4.500E-09 | 9.600E-09 |
| 2187 | 110 | A110Q | A | 1.7e-09 | 7.7e-09 | 4.1e-09 | 4.500E-09 | 3.020E-09 | 1.700E-09 | 7.700E-09 |
| 2188 | 110 | A110G | A | 7.7e-09 | 4.3e-09 | 8.9e-09 | 6.967E-09 | 2.386E-09 | 4.300E-09 | 8.900E-09 |
| 2189 | 110 | A110H | A | 9e-09 | 6.7e-09 | 8.6e-09 | 8.100E-09 | 1.229E-09 | 6.700E-09 | 9.000E-09 |
| 2190 | 110 | A110I | A | 1.4e-08 | 5.7e-09 | 6.8e-09 | 8.833E-09 | 4.508E-09 | 5.700E-09 | 1.400E-08 |
| 2191 | 110 | A110L | A | 1.7e-08 | 2.3e-09 | 1.2e-08 | 1.043E-08 | 7.474E-09 | 2.300E-09 | 1.700E-08 |
| 2192 | 110 | A110K | A | 1.8e-09 | 8.9e-09 | 1.2e-09 | 3.967E-09 | 4.283E-09 | 1.200E-09 | 8.900E-09 |
| 2193 | 110 | A110M | A | 9.2e-09 | 5.3e-09 | 4.1e-09 | 6.200E-09 | 2.666E-09 | 4.100E-09 | 9.200E-09 |
| 2194 | 110 | A110F | A | 8.9e-09 | 1.4e-08 | 1.3e-08 | 1.197E-08 | 2.702E-09 | 8.900E-09 | 1.400E-08 |
| 2195 | 110 | A110P | A | 8.5e-09 | 9.6e-09 | 4.4e-09 | 7.500E-09 | 2.740E-09 | 4.400E-09 | 9.600E-09 |
| 2196 | 110 | A110S | A | 1.3e-08 | 1.4e-08 | 3.5e-09 | 1.017E-08 | 5.795E-09 | 3.500E-09 | 1.400E-08 |
| 2197 | 110 | A110T | A | 6.9e-09 | 2.2e-09 | 7.3e-09 | 5.467E-09 | 2.836E-09 | 2.200E-09 | 7.300E-09 |
| 2198 | 110 | A110W | A | 3.9e-09 | 2.1e-09 | 1.4e-08 | 6.667E-09 | 6.414E-09 | 2.100E-09 | 1.400E-08 |
| 2199 | 110 | A110Y | A | 5e-09 | 1.5e-09 | 1e-08 | 5.500E-09 | 4.272E-09 | 1.500E-09 | 1.000E-08 |
| 2200 | 110 | A110V | A | 1e-08 | 1.5e-08 | 1.8e-08 | 1.433E-08 | 4.041E-09 | 1.000E-08 | 1.800E-08 |
| 2201 | 111 | S111A | A | 2.5e-09 | 4.4e-09 | 1.3e-08 | 6.633E-09 | 5.595E-09 | 2.500E-09 | 1.300E-08 |
| 2202 | 111 | S111R | A | 9.9e-09 | 1.7e-08 | 4.1e-08 | 2.263E-08 | 1.630E-08 | 9.900E-09 | 4.100E-08 |
| 2203 | 111 | S111N | A | 7.1e-09 | 1.7e-08 | 2.1e-09 | 8.733E-09 | 7.583E-09 | 2.100E-09 | 1.700E-08 |
| 2204 | 111 | S111D | A | 6e-09 | 1.3e-08 | 9.2e-09 | 9.400E-09 | 3.504E-09 | 6.000E-09 | 1.300E-08 |
| 2205 | 111 | S111C | A | 1.2e-08 | 5.3e-09 | 1.9e-08 | 1.210E-08 | 6.851E-09 | 5.300E-09 | 1.900E-08 |
| 2206 | 111 | S111E | A | 8.1e-09 | 1e-08 | 1.7e-08 | 1.170E-08 | 4.687E-09 | 8.100E-09 | 1.700E-08 |
| 2207 | 111 | S111Q | A | 3.4e-09 | 2.5e-09 | 4e-09 | 3.300E-09 | 7.550E-10 | 2.500E-09 | 4.000E-09 |
| 2208 | 111 | S111G | A | 9.5e-09 | 1.2e-08 | 2.2e-08 | 1.450E-08 | 6.614E-09 | 9.500E-09 | 2.200E-08 |
| 2209 | 111 | S111H | A | 6.2e-09 | 8.5e-09 | 9e-09 | 7.900E-09 | 1.493E-09 | 6.200E-09 | 9.000E-09 |

| | | | | | | | | | | |
|---|---|---|---|---|---|---|---|---|---|---|
| 2210 | 111 | S111I | A | 4.9e-09 | 9.6e-09 | 1.4e-08 | 9.500E-09 | 4.551E-09 | 4.900E-09 | 1.400E-08 |
| 2211 | 111 | S111L | A | 8.3e-09 | 7.4e-09 | 3.9e-09 | 6.533E-09 | 2.325E-09 | 3.900E-09 | 8.300E-09 |
| 2212 | 111 | S111K | A | 5.1e-09 | 6.5e-09 | 4.6e-09 | 5.400E-09 | 9.849E-10 | 4.600E-09 | 6.500E-09 |
| 2213 | 111 | S111M | A | 1.7e-09 | 2e-09 | 9.8e-09 | 4.500E-09 | 4.592E-09 | 1.700E-09 | 9.800E-09 |
| 2214 | 111 | S111F | A | 4.9e-09 | 5.1e-09 | 4.6e-09 | 4.867E-09 | 2.517E-10 | 4.600E-09 | 5.100E-09 |
| 2215 | 111 | S111P | A | 9e-09 | 5.7e-09 | 7.7e-09 | 7.467E-09 | 1.662E-09 | 5.700E-09 | 9.000E-09 |
| 2216 | 111 | S111S | A | 5.7e-09 | 6.3e-09 | 8.2e-09 | 6.733E-09 | 1.305E-09 | 5.700E-09 | 8.200E-09 |
| 2217 | 111 | S111T | A | 7.7e-09 | 7.5e-09 | 4.8e-09 | 6.667E-09 | 1.620E-09 | 4.800E-09 | 7.700E-09 |
| 2218 | 111 | S111W | A | 6.9e-09 | 3.1e-09 | 1.2e-08 | 7.333E-09 | 4.466E-09 | 3.100E-09 | 1.200E-08 |
| 2219 | 111 | S111Y | A | 1.2e-08 | 5e-09 | 5.5e-09 | 7.500E-09 | 3.905E-09 | 5.000E-09 | 1.200E-08 |
| 2220 | 111 | S111V | A | 7.6e-09 | 6.8e-09 | 1e-08 | 8.133E-09 | 1.665E-09 | 6.800E-09 | 1.000E-08 |
| 2221 | 112 | P112A | A | 1e-08 | 7.6e-09 | 5.5e-09 | 7.700E-09 | 2.252E-09 | 5.500E-09 | 1.000E-08 |
| 2222 | 112 | P112R | A | 8.5e-09 | 1.4e-09 | 4.3e-09 | 4.733E-09 | 3.570E-09 | 1.400E-09 | 8.500E-09 |
| 2223 | 112 | P112N | A | 5.4e-09 | 3.7e-09 | 4e-09 | 4.367E-09 | 9.074E-10 | 3.700E-09 | 5.400E-09 |
| 2224 | 112 | P112D | A | 1.1e-08 | 9.7e-09 | 1.6e-08 | 1.223E-08 | 3.326E-09 | 9.700E-09 | 1.600E-08 |
| 2225 | 112 | P112C | A | 9.4e-09 | 4.6e-09 | 1e-08 | 8.000E-09 | 2.960E-09 | 4.600E-09 | 1.000E-08 |
| 2226 | 112 | P112E | A | 1.3e-08 | 5e-09 | 2.6e-09 | 6.867E-09 | 5.445E-09 | 2.600E-09 | 1.300E-08 |
| 2227 | 112 | P112Q | A | 5.2e-09 | 4.1e-09 | 6.6e-09 | 5.300E-09 | 1.253E-09 | 4.100E-09 | 6.600E-09 |
| 2228 | 112 | P112G | A | 5.6e-09 | 1e-08 | 1.2e-08 | 9.200E-09 | 3.274E-09 | 5.600E-09 | 1.200E-08 |
| 2229 | 112 | P112H | A | 5.8e-09 | 1.6e-08 | 3e-08 | 1.727E-08 | 1.215E-08 | 5.800E-09 | 3.000E-08 |
| 2230 | 112 | P112I | A | 1.4e-08 | 4.9e-09 | 5.1e-09 | 8.000E-09 | 5.197E-09 | 4.900E-09 | 1.400E-08 |
| 2231 | 112 | P112L | A | 9.9e-09 | 1.2e-08 | 7.7e-09 | 9.867E-09 | 2.150E-09 | 7.700E-09 | 1.200E-08 |
| 2232 | 112 | P112K | A | 8.7e-09 | 7.3e-09 | 1.3e-08 | 9.667E-09 | 2.970E-09 | 7.300E-09 | 1.300E-08 |
| 2233 | 112 | P112M | A | 7.8e-09 | 7.8e-09 | 2.6e-08 | 1.387E-08 | 1.051E-08 | 7.800E-09 | 2.600E-08 |
| 2234 | 112 | P112F | A | 9.1e-09 | 5.1e-09 | 4e-09 | 6.067E-09 | 2.684E-09 | 4.000E-09 | 9.100E-09 |
| 2235 | 112 | P112P | A | 9.4e-09 | 2.1e-08 | 8.9e-09 | 1.310E-08 | 6.846E-09 | 8.900E-09 | 2.100E-08 |
| 2236 | 112 | P112S | A | 6.1e-09 | 1.4e-08 | 5e-09 | 8.367E-09 | 4.910E-09 | 5.000E-09 | 1.400E-08 |
| 2237 | 112 | P112T | A | 1.1e-08 | 7.2e-09 | 9.1e-09 | 9.100E-09 | 1.900E-09 | 7.200E-09 | 1.100E-08 |
| 2238 | 112 | P112W | A | 9.9e-09 | 4.1e-09 | 6.8e-09 | 6.933E-09 | 2.902E-09 | 4.100E-09 | 9.900E-09 |
| 2239 | 112 | P112Y | A | 1.6e-08 | 5.4e-09 | 2.6e-09 | 8.000E-09 | 7.068E-09 | 2.600E-09 | 1.600E-08 |
| 2240 | 112 | P112V | A | 4e-09 | 8e-09 | 5e-09 | 5.667E-09 | 2.082E-09 | 4.000E-09 | 8.000E-09 |
| 2241 | 113 | G113A | A | 9.1e-09 | 7.4e-09 | 5.5e-09 | 7.333E-09 | 1.801E-09 | 5.500E-09 | 9.100E-09 |
| 2242 | 113 | G113R | A | 6.2e-09 | 2.9e-09 | 5.4e-09 | 4.833E-09 | 1.721E-09 | 2.900E-09 | 6.200E-09 |
| 2243 | 113 | G113N | A | 6.1e-09 | 3.4e-09 | 5.8e-09 | 5.100E-09 | 1.480E-09 | 3.400E-09 | 6.100E-09 |
| 2244 | 113 | G113D | A | 4.5e-09 | 4e-09 | 9.5e-09 | 6.000E-09 | 3.041E-09 | 4.000E-09 | 9.500E-09 |
| 2245 | 113 | G113C | A | 8e-09 | 5.8e-09 | 1.7e-08 | 1.027E-08 | 5.934E-09 | 5.800E-09 | 1.700E-08 |
| 2246 | 113 | G113E | A | 5.1e-09 | 3.1e-09 | 7.8e-09 | 5.333E-09 | 2.359E-09 | 3.100E-09 | 7.800E-09 |
| 2247 | 113 | G113Q | A | 1.5e-08 | 1.6e-08 | 1.3e-08 | 1.467E-08 | 1.528E-09 | 1.300E-08 | 1.600E-08 |
| 2248 | 113 | G113G | A | 7e-09 | 3.9e-09 | 1.6e-08 | 8.967E-09 | 6.285E-09 | 3.900E-09 | 1.600E-08 |
| 2249 | 113 | G113H | A | 6.9e-09 | 7.1e-09 | 8.1e-09 | 7.367E-09 | 6.429E-10 | 6.900E-09 | 8.100E-09 |
| 2250 | 113 | G113I | A | 3.9e-09 | 5.1e-09 | 5.5e-09 | 4.833E-09 | 8.327E-10 | 3.900E-09 | 5.500E-09 |
| 2251 | 113 | G113L | A | 4.3e-09 | 5.5e-09 | 4.9e-09 | 4.900E-09 | 6.000E-10 | 4.300E-09 | 5.500E-09 |
| 2252 | 113 | G113K | A | 3.4e-09 | 5.8e-09 | 5.1e-09 | 4.767E-09 | 1.234E-09 | 3.400E-09 | 5.800E-09 |
| 2253 | 113 | G113M | A | 7.4e-09 | 1.2e-08 | 9.7e-09 | 9.700E-09 | 2.300E-09 | 7.400E-09 | 1.200E-08 |
| 2254 | 113 | G113F | A | 1.3e-08 | 1.7e-08 | 1.3e-08 | 1.433E-08 | 2.309E-09 | 1.300E-08 | 1.700E-08 |
| 2255 | 113 | G113P | A | 4.4e-09 | 1.7e-09 | 3.5e-09 | 3.200E-09 | 1.375E-09 | 1.700E-09 | 4.400E-09 |
| 2256 | 113 | G113S | A | 3.5e-09 | 1.2e-08 | 6.3e-09 | 7.267E-09 | 4.332E-09 | 3.500E-09 | 1.200E-08 |
| 2257 | 113 | G113T | A | 4.7e-09 | 3e-09 | 8.6e-09 | 5.433E-09 | 2.871E-09 | 3.000E-09 | 8.600E-09 |
| 2258 | 113 | G113W | A | 1.2e-08 | 3.4e-09 | 3.9e-09 | 6.433E-09 | 4.827E-09 | 3.400E-09 | 1.200E-08 |
| 2259 | 113 | G113Y | A | 1.4e-08 | 9.5e-09 | 3.4e-09 | 8.967E-09 | 5.320E-09 | 3.400E-09 | 1.400E-08 |
| 2260 | 113 | G113V | A | 4.5e-09 | 1.1e-08 | 4.6e-09 | 6.700E-09 | 3.724E-09 | 4.500E-09 | 1.100E-08 |
| 2261 | 114 | G114A | A | 6.2e-09 | 6.9e-09 | 6.2e-09 | 6.433E-09 | 4.041E-10 | 6.200E-09 | 6.900E-09 |
| 2262 | 114 | G114R | A | 7.4e-09 | 9e-09 | 2.6e-08 | 1.413E-08 | 1.031E-08 | 7.400E-09 | 2.600E-08 |
| 2263 | 114 | G114N | A | 1.1e-08 | 4.9e-09 | 2e-08 | 1.197E-08 | 7.596E-09 | 4.900E-09 | 2.000E-08 |
| 2264 | 114 | G114D | A | 8e-09 | 3.8e-09 | 1.4e-08 | 8.600E-09 | 5.126E-09 | 3.800E-09 | 1.400E-08 |
| 2265 | 114 | G114C | A | 6.8e-09 | 4.7e-09 | 1.1e-08 | 7.500E-09 | 3.208E-09 | 4.700E-09 | 1.100E-08 |
| 2266 | 114 | G114E | A | 1.3e-08 | 3.7e-09 | 5.6e-09 | 7.433E-09 | 4.914E-09 | 3.700E-09 | 1.300E-08 |
| 2267 | 114 | G114Q | A | 8.7e-09 | 1e-08 | 3.4e-09 | 7.367E-09 | 3.496E-09 | 3.400E-09 | 1.000E-08 |
| 2268 | 114 | G114G | A | 8.5e-09 | 3.1e-09 | 9.2e-09 | 6.933E-09 | 3.338E-09 | 3.100E-09 | 9.200E-09 |
| 2269 | 114 | G114H | A | 4.3e-09 | 7.1e-09 | 4.2e-09 | 5.200E-09 | 1.646E-09 | 4.200E-09 | 7.100E-09 |
| 2270 | 114 | G114I | A | 1e-08 | 9.1e-09 | 7.5e-09 | 8.867E-09 | 1.266E-09 | 7.500E-09 | 1.000E-08 |
| 2271 | 114 | G114L | A | 4.7e-09 | 1.8e-08 | 6.9e-09 | 9.867E-09 | 7.129E-09 | 4.700E-09 | 1.800E-08 |
| 2272 | 114 | G114K | A | 2.3e-09 | 8.2e-09 | 5e-09 | 5.167E-09 | 2.954E-09 | 2.300E-09 | 8.200E-09 |
| 2273 | 114 | G114M | A | 3.4e-09 | 1.1e-08 | 8.3e-09 | 7.567E-09 | 3.853E-09 | 3.400E-09 | 1.100E-08 |
| 2274 | 114 | G114F | A | 1.2e-08 | 4.8e-09 | 1.2e-08 | 9.600E-09 | 4.157E-09 | 4.800E-09 | 1.200E-08 |
| 2275 | 114 | G114P | A | 8.5e-09 | 9.1e-09 | 1.4e-08 | 1.053E-08 | 3.017E-09 | 8.500E-09 | 1.400E-08 |
| 2276 | 114 | G114S | A | 6.3e-09 | 8.7e-09 | 2.8e-09 | 5.933E-09 | 2.967E-09 | 2.800E-09 | 8.700E-09 |
| 2277 | 114 | G114T | A | 3.7e-08 | 2.6e-09 | 4.2e-09 | 1.460E-08 | 1.942E-08 | 2.600E-09 | 3.700E-08 |
| 2278 | 114 | G114W | A | 1.8e-09 | 1.3e-09 | 1.1e-08 | 4.700E-09 | 5.462E-09 | 1.300E-09 | 1.100E-08 |
| 2279 | 114 | G114Y | A | 8.9e-09 | 5.2e-09 | 8.9e-09 | 7.667E-09 | 2.136E-09 | 5.200E-09 | 8.900E-09 |
| 2280 | 114 | G114V | A | 2.3e-09 | 9.9e-09 | 7e-09 | 6.400E-09 | 3.835E-09 | 2.300E-09 | 9.900E-09 |
| 2281 | 115 | L115A | A | 1.4e-08 | 7.4e-09 | 8.2e-09 | 9.867E-09 | 3.602E-09 | 7.400E-09 | 1.400E-08 |
| 2282 | 115 | L115R | A | 1.1e-08 | 3.1e-09 | 1e-08 | 8.033E-09 | 4.302E-09 | 3.100E-09 | 1.100E-08 |
| 2283 | 115 | L115N | A | 1e-08 | 1.4e-08 | 7.2e-09 | 1.040E-08 | 3.418E-09 | 7.200E-09 | 1.400E-08 |
| 2284 | 115 | L115D | A | 1.7e-08 | 1.9e-08 | 7.6e-09 | 1.453E-08 | 6.087E-09 | 7.600E-09 | 1.900E-08 |
| 2285 | 115 | L115C | A | 5e-09 | 1.4e-08 | 6.9e-09 | 8.633E-09 | 4.744E-09 | 5.000E-09 | 1.400E-08 |
| 2286 | 115 | L115E | A | 8.5e-09 | 9.8e-09 | 9.6e-09 | 9.300E-09 | 7.000E-10 | 8.500E-09 | 9.800E-09 |
| 2287 | 115 | L115Q | A | 1.2e-08 | 5.7e-09 | 6.1e-09 | 7.933E-09 | 3.528E-09 | 5.700E-09 | 1.200E-08 |
| 2288 | 115 | L115G | A | 8.3e-09 | 5.2e-09 | 6.5e-09 | 6.667E-09 | 1.557E-09 | 5.200E-09 | 8.300E-09 |
| 2289 | 115 | L115H | A | 7.7e-09 | 5.9e-09 | 2.5e-09 | 5.367E-09 | 2.641E-09 | 2.500E-09 | 7.700E-09 |
| 2290 | 115 | L115I | A | 6.4e-09 | 1e-08 | 4.4e-09 | 6.933E-09 | 2.838E-09 | 4.400E-09 | 1.000E-08 |
| 2291 | 115 | L115L | A | 7.8e-09 | 6.3e-09 | 2.4e-09 | 5.500E-09 | 2.787E-09 | 2.400E-09 | 7.800E-09 |
| 2292 | 115 | L115K | A | 8.1e-09 | 4.9e-09 | 2.1e-09 | 5.033E-09 | 3.002E-09 | 2.100E-09 | 8.100E-09 |
| 2293 | 115 | L115M | A | 2.9e-09 | 8.6e-09 | 1.5e-08 | 8.833E-09 | 6.053E-09 | 2.900E-09 | 1.500E-08 |
| 2294 | 115 | L115F | A | 4.9e-09 | 6.1e-09 | 4.4e-09 | 5.133E-09 | 8.737E-10 | 4.400E-09 | 6.100E-09 |

| | | | | | | | | | | |
|---|---|---|---|---|---|---|---|---|---|---|
| 2295 | 115 | L115P | A | 4.5e-09 | 7.6e-09 | 5.4e-09 | 5.833E-09 | 1.595E-09 | 4.500E-09 | 7.600E-09 |
| 2296 | 115 | L115S | A | 1.5e-08 | 1.6e-08 | 1.2e-08 | 1.433E-08 | 2.082E-09 | 1.200E-08 | 1.600E-08 |
| 2297 | 115 | L115T | A | 3.3e-09 | 9.7e-09 | 2.8e-08 | 1.367E-08 | 1.282E-08 | 3.300E-09 | 2.800E-08 |
| 2298 | 115 | L115W | A | 4.4e-09 | 4e-09 | 6.5e-09 | 4.967E-09 | 1.343E-09 | 4.000E-09 | 6.500E-09 |
| 2299 | 115 | L115Y | A | 6.6e-09 | 4.6e-09 | 1.1e-08 | 7.400E-09 | 3.274E-09 | 4.600E-09 | 1.100E-08 |
| 2300 | 115 | L115V | A | 5.7e-09 | 1.1e-08 | 4.3e-09 | 7.000E-09 | 3.534E-09 | 4.300E-09 | 1.100E-08 |
| 2301 | 116 | R116A | A | 6e-09 | 4.5e-09 | 1.1e-08 | 7.167E-09 | 3.403E-09 | 4.500E-09 | 1.100E-08 |
| 2302 | 116 | R116R | A | 5.1e-09 | 1.5e-08 | 8.2e-09 | 9.433E-09 | 5.064E-09 | 5.100E-09 | 1.500E-08 |
| 2303 | 116 | R116N | A | 9.7e-09 | 6.6e-09 | 9.6e-09 | 8.633E-09 | 1.762E-09 | 6.600E-09 | 9.700E-09 |
| 2304 | 116 | R116D | A | 1.2e-08 | 3.6e-09 | 6.5e-09 | 7.367E-09 | 4.267E-09 | 3.600E-09 | 1.200E-08 |
| 2305 | 116 | R116C | A | 1.4e-08 | 6.6e-09 | 9.5e-09 | 1.003E-08 | 3.729E-09 | 6.600E-09 | 1.400E-08 |
| 2306 | 116 | R116E | A | 6e-09 | 1.4e-08 | 6.4e-09 | 8.800E-09 | 4.508E-09 | 6.000E-09 | 1.400E-08 |
| 2307 | 116 | R116Q | A | 9.7e-09 | 1.3e-08 | 1.4e-08 | 1.223E-08 | 2.250E-09 | 9.700E-09 | 1.400E-08 |
| 2308 | 116 | R116G | A | 3.1e-09 | 9.8e-10 | 6e-09 | 3.360E-09 | 2.520E-09 | 9.800E-10 | 6.000E-09 |
| 2309 | 116 | R116H | A | 4.2e-09 | 1.2e-08 | 9.2e-09 | 8.467E-09 | 3.951E-09 | 4.200E-09 | 1.200E-08 |
| 2310 | 116 | R116I | A | 3.6e-09 | 2.5e-08 | 5.5e-09 | 1.137E-08 | 1.184E-08 | 3.600E-09 | 2.500E-08 |
| 2311 | 116 | R116L | A | 1.4e-08 | 7.1e-09 | 6.2e-09 | 9.100E-09 | 4.267E-09 | 6.200E-09 | 1.400E-08 |
| 2312 | 116 | R116K | A | 2.3e-09 | 8.2e-09 | 2.2e-08 | 1.083E-08 | 1.011E-08 | 2.300E-09 | 2.200E-08 |
| 2313 | 116 | R116M | A | 2.4e-09 | 4.4e-09 | 2.5e-08 | 1.060E-08 | 1.251E-08 | 2.400E-09 | 2.500E-08 |
| 2314 | 116 | R116F | A | 1.9e-09 | 3.3e-09 | 8.7e-09 | 4.633E-09 | 3.591E-09 | 1.900E-09 | 8.700E-09 |
| 2315 | 116 | R116P | A | 4e-09 | 6.2e-09 | 8.2e-09 | 6.133E-09 | 2.101E-09 | 4.000E-09 | 8.200E-09 |
| 2316 | 116 | R116S | A | 1.6e-08 | 1.1e-08 | 8.5e-09 | 1.183E-08 | 3.819E-09 | 8.500E-09 | 1.600E-08 |
| 2317 | 116 | R116T | A | 3.9e-09 | 5.7e-09 | 1.3e-08 | 7.533E-09 | 4.819E-09 | 3.900E-09 | 1.300E-08 |
| 2318 | 116 | R116W | A | 2.5e-08 | 3.5e-09 | 6.2e-09 | 1.157E-08 | 1.171E-08 | 3.500E-09 | 2.500E-08 |
| 2319 | 116 | R116Y | A | 5.5e-09 | 1.2e-09 | 8.6e-09 | 5.100E-09 | 3.716E-09 | 1.200E-09 | 8.600E-09 |
| 2320 | 116 | R116V | A | 8.5e-09 | 3.8e-09 | 1.4e-08 | 8.767E-09 | 5.105E-09 | 3.800E-09 | 1.400E-08 |
| 2321 | 117 | E117A | A | 1.1e-08 | 1.1e-08 | 6.7e-09 | 9.567E-09 | 2.483E-09 | 6.700E-09 | 1.100E-08 |
| 2322 | 117 | E117R | A | 1e-08 | 7.9e-09 | 1e-08 | 9.300E-09 | 1.212E-09 | 7.900E-09 | 1.000E-08 |
| 2323 | 117 | E117N | A | 1.3e-08 | 1.2e-08 | 1.2e-08 | 1.233E-08 | 5.774E-10 | 1.200E-08 | 1.300E-08 |
| 2324 | 117 | E117D | A | 7.6e-09 | 2.1e-08 | 2e-08 | 1.620E-08 | 7.465E-09 | 7.600E-09 | 2.100E-08 |
| 2325 | 117 | E117C | A | 9.6e-09 | 2.5e-09 | 1.9e-08 | 1.037E-08 | 8.277E-09 | 2.500E-09 | 1.900E-08 |
| 2326 | 117 | E117E | A | 1.6e-08 | 1.9e-08 | 1.1e-08 | 1.533E-08 | 4.041E-09 | 1.100E-08 | 1.900E-08 |
| 2327 | 117 | E117Q | A | 2e-08 | 5.4e-09 | 1.4e-08 | 1.313E-08 | 7.338E-09 | 5.400E-09 | 2.000E-08 |
| 2328 | 117 | E117G | A | 2.4e-09 | 1.1e-08 | 7.6e-09 | 7.000E-09 | 4.331E-09 | 2.400E-09 | 1.100E-08 |
| 2329 | 117 | E117H | A | 9.9e-09 | 8.4e-09 | 8.6e-09 | 8.967E-09 | 8.145E-10 | 8.400E-09 | 9.900E-09 |
| 2330 | 117 | E117I | A | 9.9e-09 | 9.2e-09 | 1e-08 | 9.700E-09 | 4.359E-10 | 9.200E-09 | 1.000E-08 |
| 2331 | 117 | E117L | A | 3.4e-09 | 7.2e-09 | 3.7e-09 | 4.767E-09 | 2.113E-09 | 3.400E-09 | 7.200E-09 |
| 2332 | 117 | E117K | A | 2.6e-09 | 7e-09 | 2.7e-09 | 4.100E-09 | 2.512E-09 | 2.600E-09 | 7.000E-09 |
| 2333 | 117 | E117M | A | 5.9e-09 | 6.7e-09 | 1.8e-08 | 1.020E-08 | 6.767E-09 | 5.900E-09 | 1.800E-08 |
| 2334 | 117 | E117F | A | 7.5e-09 | 1.9e-08 | 8.9e-09 | 1.180E-08 | 6.275E-09 | 7.500E-09 | 1.900E-08 |
| 2335 | 117 | E117P | A | 3.6e-09 | 4.7e-09 | 1.3e-08 | 7.100E-09 | 5.139E-09 | 3.600E-09 | 1.300E-08 |
| 2336 | 117 | E117S | A | 8.4e-09 | 1.1e-08 | 1.3e-08 | 1.080E-08 | 2.307E-09 | 8.400E-09 | 1.300E-08 |
| 2337 | 117 | E117T | A | 9.5e-09 | 1.4e-08 | 3e-09 | 8.833E-09 | 5.530E-09 | 3.000E-09 | 1.400E-08 |
| 2338 | 117 | E117W | A | 7.4e-09 | 9.4e-09 | 1.3e-08 | 9.933E-09 | 2.838E-09 | 7.400E-09 | 1.300E-08 |
| 2339 | 117 | E117Y | A | 4.4e-09 | 3.9e-09 | 4.1e-09 | 4.133E-09 | 2.517E-10 | 3.900E-09 | 4.400E-09 |
| 2340 | 117 | E117V | A | 6e-09 | 5.6e-09 | 8.9e-09 | 6.833E-09 | 1.801E-09 | 5.600E-09 | 8.900E-09 |
| 2341 | 118 | L118A | A | 1.7e-08 | 2.3e-08 | 2.4e-09 | 1.413E-08 | 1.059E-08 | 2.400E-09 | 2.300E-08 |
| 2342 | 118 | L118R | A | 3.8e-09 | 8.8e-09 | 2.4e-08 | 1.220E-08 | 1.052E-08 | 3.800E-09 | 2.400E-08 |
| 2343 | 118 | L118N | A | 2.5e-09 | 1.7e-08 | 2.6e-09 | 7.367E-09 | 8.343E-09 | 2.500E-09 | 1.700E-08 |
| 2344 | 118 | L118D | A | 8.4e-09 | 1.2e-08 | 9.8e-09 | 1.007E-08 | 1.815E-09 | 8.400E-09 | 1.200E-08 |
| 2345 | 118 | L118C | A | 7.8e-09 | 8.1e-09 | 1.1e-08 | 8.967E-09 | 1.767E-09 | 7.800E-09 | 1.100E-08 |
| 2346 | 118 | L118E | A | 1.1e-08 | 1.1e-08 | 8.6e-09 | 1.020E-08 | 1.386E-09 | 8.600E-09 | 1.100E-08 |
| 2347 | 118 | L118Q | A | 6.8e-09 | 7.8e-09 | 2.8e-09 | 5.800E-09 | 2.646E-09 | 2.800E-09 | 7.800E-09 |
| 2348 | 118 | L118G | A | 4.9e-09 | 2.9e-09 | 6.4e-09 | 4.733E-09 | 1.756E-09 | 2.900E-09 | 6.400E-09 |
| 2349 | 118 | L118H | A | 6.3e-09 | 1.2e-09 | 1.1e-08 | 6.167E-09 | 4.901E-09 | 1.200E-09 | 1.100E-08 |
| 2350 | 118 | L118I | A | 3.6e-09 | 5.9e-09 | 1.7e-09 | 3.733E-09 | 2.103E-09 | 1.700E-09 | 5.900E-09 |
| 2351 | 118 | L118L | A | 5.4e-09 | 2e-08 | 5.6e-09 | 1.033E-08 | 8.372E-09 | 5.400E-09 | 2.000E-08 |
| 2352 | 118 | L118K | A | 6.2e-09 | 4e-09 | 6.2e-09 | 5.467E-09 | 1.270E-09 | 4.000E-09 | 6.200E-09 |
| 2353 | 118 | L118M | A | 1.3e-08 | 1.1e-08 | 1.4e-09 | 8.467E-09 | 6.201E-09 | 1.400E-09 | 1.300E-08 |
| 2354 | 118 | L118F | A | 1.8e-08 | 3.5e-09 | 6.9e-09 | 9.467E-09 | 7.583E-09 | 3.500E-09 | 1.800E-08 |
| 2355 | 118 | L118P | A | 2.1e-08 | 1.2e-08 | 4.8e-09 | 1.260E-08 | 8.117E-09 | 4.800E-09 | 2.100E-08 |
| 2356 | 118 | L118S | A | 2.2e-08 | 3.7e-09 | 1.8e-08 | 1.457E-08 | 9.621E-09 | 3.700E-09 | 2.200E-08 |
| 2357 | 118 | L118T | A | 1.8e-08 | 6.6e-09 | 6.2e-09 | 1.027E-08 | 6.700E-09 | 6.200E-09 | 1.800E-08 |
| 2358 | 118 | L118W | A | 3e-09 | 1.1e-08 | 6.8e-09 | 6.933E-09 | 4.002E-09 | 3.000E-09 | 1.100E-08 |
| 2359 | 118 | L118Y | A | 5.9e-09 | 9.8e-09 | 3.4e-09 | 6.367E-09 | 3.225E-09 | 3.400E-09 | 9.800E-09 |
| 2360 | 118 | L118V | A | 5.7e-09 | 3.4e-09 | 1.3e-08 | 7.367E-09 | 5.012E-09 | 3.400E-09 | 1.300E-08 |
| 2361 | 119 | Q119A | A | 7.8e-09 | 7.7e-09 | 2.4e-09 | 5.967E-09 | 3.089E-09 | 2.400E-09 | 7.800E-09 |
| 2362 | 119 | Q119R | A | 5.5e-09 | 4.3e-09 | 1.5e-08 | 8.267E-09 | 5.862E-09 | 4.300E-09 | 1.500E-08 |
| 2363 | 119 | Q119N | A | 9.1e-09 | 9.9e-09 | 8.5e-09 | 9.167E-09 | 7.024E-10 | 8.500E-09 | 9.900E-09 |
| 2364 | 119 | Q119D | A | 3.2e-09 | 9.1e-09 | 6.1e-09 | 6.133E-09 | 2.950E-09 | 3.200E-09 | 9.100E-09 |
| 2365 | 119 | Q119C | A | 6.7e-09 | 7.4e-09 | 6.5e-09 | 6.867E-09 | 4.726E-10 | 6.500E-09 | 7.400E-09 |
| 2366 | 119 | Q119E | A | 4.1e-09 | 4.8e-09 | 5.8e-09 | 4.900E-09 | 8.544E-10 | 4.100E-09 | 5.800E-09 |
| 2367 | 119 | Q119Q | A | 5e-09 | 7e-09 | 1.2e-08 | 8.000E-09 | 3.606E-09 | 5.000E-09 | 1.200E-08 |
| 2368 | 119 | Q119G | A | 6.7e-09 | 1.4e-08 | 1.2e-08 | 1.090E-08 | 3.772E-09 | 6.700E-09 | 1.400E-08 |
| 2369 | 119 | Q119H | A | 1.4e-09 | 1.3e-08 | 9.4e-09 | 7.933E-09 | 5.937E-09 | 1.400E-09 | 1.300E-08 |
| 2370 | 119 | Q119I | A | 9.3e-09 | 6e-09 | 3.8e-09 | 6.367E-09 | 2.768E-09 | 3.800E-09 | 9.300E-09 |
| 2371 | 119 | Q119L | A | 5.2e-09 | 3.8e-09 | 1.1e-08 | 6.667E-09 | 3.818E-09 | 3.800E-09 | 1.100E-08 |
| 2372 | 119 | Q119K | A | 4.1e-09 | 3.6e-09 | 7.2e-09 | 4.967E-09 | 1.950E-09 | 3.600E-09 | 7.200E-09 |
| 2373 | 119 | Q119M | A | 1.1e-08 | 6.4e-09 | 3.9e-09 | 7.100E-09 | 3.601E-09 | 3.900E-09 | 1.100E-08 |
| 2374 | 119 | Q119F | A | 1.5e-09 | 4.1e-09 | 9.4e-09 | 5.000E-09 | 4.026E-09 | 1.500E-09 | 9.400E-09 |
| 2375 | 119 | Q119P | A | 4.9e-09 | 5.3e-09 | 2.2e-09 | 4.133E-09 | 1.686E-09 | 2.200E-09 | 5.300E-09 |
| 2376 | 119 | Q119S | A | 3e-09 | 7.4e-09 | 4.3e-09 | 4.900E-09 | 2.261E-09 | 3.000E-09 | 7.400E-09 |
| 2377 | 119 | Q119T | A | 4.2e-09 | 1.2e-08 | 3e-09 | 6.400E-09 | 4.887E-09 | 3.000E-09 | 1.200E-08 |
| 2378 | 119 | Q119W | A | 5.5e-09 | 1.5e-08 | 3e-09 | 7.833E-09 | 6.331E-09 | 3.000E-09 | 1.500E-08 |
| 2379 | 119 | Q119Y | A | 7.1e-09 | 1.5e-08 | 1.5e-08 | 1.237E-08 | 4.561E-09 | 7.100E-09 | 1.500E-08 |

| | | | | | | | | | | |
|---|---|---|---|---|---|---|---|---|---|---|
| 2380 | 119 | Q119V | A | 9.7e-09 | 2.2e-08 | 5.5e-09 | 1.240E-08 | 8.575E-09 | 5.500E-09 | 2.200E-08 |
| 2381 | 120 | L120A | A | 5e-09 | 5.7e-09 | 6.9e-09 | 5.867E-09 | 9.609E-10 | 5.000E-09 | 6.900E-09 |
| 2382 | 120 | L120R | A | 4.8e-09 | 9.6e-09 | 4.7e-09 | 6.367E-09 | 2.801E-09 | 4.700E-09 | 9.600E-09 |
| 2383 | 120 | L120N | A | 5e-09 | 1.7e-08 | 4.9e-09 | 8.967E-09 | 6.957E-09 | 4.900E-09 | 1.700E-08 |
| 2384 | 120 | L120D | A | 1.2e-08 | 4.5e-09 | 1.5e-08 | 1.050E-08 | 5.408E-09 | 4.500E-09 | 1.500E-08 |
| 2385 | 120 | L120C | A | 2.1e-08 | 4.1e-09 | 7.8e-09 | 1.097E-08 | 8.884E-09 | 4.100E-09 | 2.100E-08 |
| 2386 | 120 | L120E | A | 7.5e-09 | 1.2e-08 | 7.2e-09 | 8.900E-09 | 2.689E-09 | 7.200E-09 | 1.200E-08 |
| 2387 | 120 | L120Q | A | 1e-08 | 1.3e-08 | 6.2e-09 | 9.733E-09 | 3.408E-09 | 6.200E-09 | 1.300E-08 |
| 2388 | 120 | L120G | A | 8.6e-09 | 5.9e-09 | 6.4e-09 | 6.967E-09 | 1.436E-09 | 5.900E-09 | 8.600E-09 |
| 2389 | 120 | L120H | A | 1e-08 | 4.2e-09 | 6.9e-09 | 7.033E-09 | 2.902E-09 | 4.200E-09 | 1.000E-08 |
| 2390 | 120 | L120I | A | 1.3e-08 | 7.6e-09 | 6.6e-09 | 9.067E-09 | 3.443E-09 | 6.600E-09 | 1.300E-08 |
| 2391 | 120 | L120L | A | 1.2e-08 | 9.5e-09 | 7.4e-09 | 9.633E-09 | 2.303E-09 | 7.400E-09 | 1.200E-08 |
| 2392 | 120 | L120K | A | 1.4e-08 | 7.8e-09 | 9.7e-09 | 1.050E-08 | 3.176E-09 | 7.800E-09 | 1.400E-08 |
| 2393 | 120 | L120M | A | 3e-09 | 1.1e-08 | 5.1e-09 | 6.367E-09 | 4.148E-09 | 3.000E-09 | 1.100E-08 |
| 2394 | 120 | L120F | A | 3.1e-09 | 7.1e-09 | 1.4e-08 | 8.067E-09 | 5.514E-09 | 3.100E-09 | 1.400E-08 |
| 2395 | 120 | L120P | A | 1.7e-08 | 8.9e-09 | 5e-09 | 1.030E-08 | 6.121E-09 | 5.000E-09 | 1.700E-08 |
| 2396 | 120 | L120S | A | 6.8e-09 | 3.4e-09 | 9.9e-09 | 6.700E-09 | 3.251E-09 | 3.400E-09 | 9.900E-09 |
| 2397 | 120 | L120T | A | 1.2e-08 | 8.5e-09 | 6.1e-09 | 8.867E-09 | 2.967E-09 | 6.100E-09 | 1.200E-08 |
| 2398 | 120 | L120W | A | 5.6e-09 | 6.7e-09 | 8.1e-09 | 6.800E-09 | 1.253E-09 | 5.600E-09 | 8.100E-09 |
| 2399 | 120 | L120Y | A | 8e-09 | 3.4e-09 | 1.9e-09 | 4.433E-09 | 3.179E-09 | 1.900E-09 | 8.000E-09 |
| 2400 | 120 | L120V | A | 3.9e-09 | 9.6e-09 | 8.4e-09 | 7.300E-09 | 3.005E-09 | 3.900E-09 | 9.600E-09 |
| 2401 | 121 | R121A | A | 8.1e-09 | 8.1e-09 | 1.1e-08 | 9.067E-09 | 1.674E-09 | 8.100E-09 | 1.100E-08 |
| 2402 | 121 | R121R | A | 3.1e-09 | 1.5e-08 | 7.9e-09 | 8.667E-09 | 5.987E-09 | 3.100E-09 | 1.500E-08 |
| 2403 | 121 | R121N | A | 3.8e-09 | 1.2e-08 | 2.1e-08 | 1.227E-08 | 8.603E-09 | 3.800E-09 | 2.100E-08 |
| 2404 | 121 | R121D | A | 1.2e-08 | 3.9e-09 | 7.3e-09 | 7.733E-09 | 4.067E-09 | 3.900E-09 | 1.200E-08 |
| 2405 | 121 | R121C | A | 1.7e-08 | 4.1e-09 | 5.6e-09 | 8.900E-09 | 7.055E-09 | 4.100E-09 | 1.700E-08 |
| 2406 | 121 | R121E | A | 4.2e-09 | 8.3e-09 | 8.1e-09 | 6.867E-09 | 2.312E-09 | 4.200E-09 | 8.300E-09 |
| 2407 | 121 | R121Q | A | 3.2e-09 | 1.1e-08 | 1.5e-08 | 9.733E-09 | 6.001E-09 | 3.200E-09 | 1.500E-08 |
| 2408 | 121 | R121G | A | 2e-09 | 2.7e-09 | 5.6e-09 | 3.433E-09 | 1.909E-09 | 2.000E-09 | 5.600E-09 |
| 2409 | 121 | R121H | A | 6e-09 | 3.6e-09 | 9.9e-09 | 6.500E-09 | 3.180E-09 | 3.600E-09 | 9.900E-09 |
| 2410 | 121 | R121I | A | 5.2e-09 | 7.8e-09 | 9.1e-09 | 7.367E-09 | 1.986E-09 | 5.200E-09 | 9.100E-09 |
| 2411 | 121 | R121L | A | 9.5e-09 | 4.2e-09 | 3.4e-09 | 5.700E-09 | 3.315E-09 | 3.400E-09 | 9.500E-09 |
| 2412 | 121 | R121K | A | 6e-09 | 6.9e-09 | 1.1e-08 | 7.967E-09 | 2.665E-09 | 6.000E-09 | 1.100E-08 |
| 2413 | 121 | R121M | A | 8.6e-09 | 9.7e-09 | 6.6e-09 | 8.300E-09 | 1.572E-09 | 6.600E-09 | 9.700E-09 |
| 2414 | 121 | R121F | A | 6.5e-09 | 5.5e-09 | 4e-09 | 5.333E-09 | 1.258E-09 | 4.000E-09 | 6.500E-09 |
| 2415 | 121 | R121P | A | 2.9e-09 | 1.8e-08 | 8.4e-09 | 9.767E-09 | 7.642E-09 | 2.900E-09 | 1.800E-08 |
| 2416 | 121 | R121S | A | 4.5e-09 | 2e-09 | 7.4e-09 | 4.633E-09 | 2.702E-09 | 2.000E-09 | 7.400E-09 |
| 2417 | 121 | R121T | A | 5e-09 | 1.9e-09 | 4.5e-09 | 3.800E-09 | 1.664E-09 | 1.900E-09 | 5.000E-09 |
| 2418 | 121 | R121W | A | 1.5e-08 | 4.9e-09 | 3.8e-09 | 7.900E-09 | 6.173E-09 | 3.800E-09 | 1.500E-08 |
| 2419 | 121 | R121Y | A | 3.4e-09 | 1.1e-08 | 5.5e-09 | 6.633E-09 | 3.925E-09 | 3.400E-09 | 1.100E-08 |
| 2420 | 121 | R121V | A | 3.5e-09 | 8.8e-09 | 1.1e-08 | 7.767E-09 | 3.855E-09 | 3.500E-09 | 1.100E-08 |
| 2421 | 122 | S122A | A | 7e-09 | 3.7e-09 | 1.7e-08 | 9.233E-09 | 6.926E-09 | 3.700E-09 | 1.700E-08 |
| 2422 | 122 | S122R | A | 3.7e-09 | 6.5e-09 | 6.8e-09 | 5.667E-09 | 1.710E-09 | 3.700E-09 | 6.800E-09 |
| 2423 | 122 | S122N | A | 4.8e-09 | 5e-09 | 5.2e-09 | 5.000E-09 | 2.000E-10 | 4.800E-09 | 5.200E-09 |
| 2424 | 122 | S122D | A | 3.5e-09 | 6.1e-09 | 1.3e-09 | 3.633E-09 | 2.403E-09 | 1.300E-09 | 6.100E-09 |
| 2425 | 122 | S122C | A | 8.7e-09 | 8.9e-09 | 5.9e-09 | 7.833E-09 | 1.677E-09 | 5.900E-09 | 8.900E-09 |
| 2426 | 122 | S122E | A | 7.3e-09 | 1.2e-08 | 6.2e-09 | 8.500E-09 | 3.081E-09 | 6.200E-09 | 1.200E-08 |
| 2427 | 122 | S122Q | A | 4.1e-09 | 6.9e-09 | 1.1e-08 | 7.333E-09 | 3.470E-09 | 4.100E-09 | 1.100E-08 |
| 2428 | 122 | S122G | A | 3e-09 | 5e-09 | 6.4e-09 | 4.800E-09 | 1.709E-09 | 3.000E-09 | 6.400E-09 |
| 2429 | 122 | S122H | A | 2.2e-08 | 5e-09 | 2.4e-08 | 1.700E-08 | 1.044E-08 | 5.000E-09 | 2.400E-08 |
| 2430 | 122 | S122I | A | 1.1e-08 | 1.7e-08 | 4.9e-09 | 1.097E-08 | 6.050E-09 | 4.900E-09 | 1.700E-08 |
| 2431 | 122 | S122L | A | 5.6e-09 | 8.3e-09 | 1.2e-08 | 8.633E-09 | 3.213E-09 | 5.600E-09 | 1.200E-08 |
| 2432 | 122 | S122K | A | 5.7e-09 | 1.4e-08 | 4.9e-09 | 8.200E-09 | 5.039E-09 | 4.900E-09 | 1.400E-08 |
| 2433 | 122 | S122M | A | 3e-09 | 5.1e-09 | 5.6e-09 | 4.567E-09 | 1.380E-09 | 3.000E-09 | 5.600E-09 |
| 2434 | 122 | S122F | A | 1.2e-08 | 2.8e-08 | 9.1e-09 | 1.637E-08 | 1.018E-08 | 9.100E-09 | 2.800E-08 |
| 2435 | 122 | S122P | A | 3.6e-09 | 4.9e-09 | 3.8e-09 | 4.100E-09 | 7.000E-10 | 3.600E-09 | 4.900E-09 |
| 2436 | 122 | S122S | A | 1e-08 | 6.9e-09 | 9.2e-09 | 8.700E-09 | 1.609E-09 | 6.900E-09 | 1.000E-08 |
| 2437 | 122 | S122T | A | 7.6e-09 | 5.9e-09 | 8e-09 | 7.167E-09 | 1.115E-09 | 5.900E-09 | 8.000E-09 |
| 2438 | 122 | S122W | A | 1e-08 | 7.6e-09 | 1.6e-08 | 1.120E-08 | 4.327E-09 | 7.600E-09 | 1.600E-08 |
| 2439 | 122 | S122Y | A | 7.1e-09 | 6.7e-09 | 4.2e-09 | 6.000E-09 | 1.572E-09 | 4.200E-09 | 7.100E-09 |
| 2440 | 122 | S122V | A | 7.7e-09 | 5.8e-09 | 9.7e-09 | 7.733E-09 | 1.950E-09 | 5.800E-09 | 9.700E-09 |
| 2441 | 123 | L123A | A | 3.8e-09 | 1.2e-08 | 8.8e-09 | 8.200E-09 | 4.133E-09 | 3.800E-09 | 1.200E-08 |
| 2442 | 123 | L123R | A | 2.2e-08 | 1.1e-08 | 9.1e-09 | 1.403E-08 | 6.964E-09 | 9.100E-09 | 2.200E-08 |
| 2443 | 123 | L123N | A | 4.7e-09 | 1.5e-08 | 7.1e-09 | 8.933E-09 | 5.389E-09 | 4.700E-09 | 1.500E-08 |
| 2444 | 123 | L123D | A | 3.8e-09 | 9.3e-09 | 1.3e-08 | 8.700E-09 | 4.629E-09 | 3.800E-09 | 1.300E-08 |
| 2445 | 123 | L123C | A | 6.8e-09 | 4.4e-09 | 5.3e-09 | 5.500E-09 | 1.212E-09 | 4.400E-09 | 6.800E-09 |
| 2446 | 123 | L123E | A | 6.8e-09 | 5.4e-09 | 2.7e-09 | 4.967E-09 | 2.084E-09 | 2.700E-09 | 6.800E-09 |
| 2447 | 123 | L123Q | A | 5.2e-09 | 6.8e-09 | 1.5e-08 | 9.000E-09 | 5.257E-09 | 5.200E-09 | 1.500E-08 |
| 2448 | 123 | L123G | A | 7.1e-09 | 1.1e-08 | 6.2e-09 | 8.100E-09 | 2.551E-09 | 6.200E-09 | 1.100E-08 |
| 2449 | 123 | L123H | A | 4.3e-09 | 1.1e-08 | 7e-09 | 7.433E-09 | 3.371E-09 | 4.300E-09 | 1.100E-08 |
| 2450 | 123 | L123I | A | 7.8e-09 | 5.8e-09 | 9.7e-09 | 7.767E-09 | 1.950E-09 | 5.800E-09 | 9.700E-09 |
| 2451 | 123 | L123L | A | 7.3e-09 | 9.5e-09 | 6.8e-09 | 7.867E-09 | 1.436E-09 | 6.800E-09 | 9.500E-09 |
| 2452 | 123 | L123K | A | 1.7e-09 | 8.3e-09 | 4.8e-09 | 4.933E-09 | 3.302E-09 | 1.700E-09 | 8.300E-09 |
| 2453 | 123 | L123M | A | 6.1e-09 | 8.1e-09 | 6.7e-09 | 6.967E-09 | 1.026E-09 | 6.100E-09 | 8.100E-09 |
| 2454 | 123 | L123F | A | 5.8e-09 | 1.3e-08 | 4e-09 | 7.600E-09 | 4.762E-09 | 4.000E-09 | 1.300E-08 |
| 2455 | 123 | L123P | A | 2e-09 | 5.9e-09 | 2.4e-09 | 3.433E-09 | 2.146E-09 | 2.000E-09 | 5.900E-09 |
| 2456 | 123 | L123S | A | 4.2e-09 | 1.6e-08 | 7.5e-09 | 9.233E-09 | 6.088E-09 | 4.200E-09 | 1.600E-08 |
| 2457 | 123 | L123T | A | 5.8e-09 | 2.8e-09 | 7.8e-09 | 5.467E-09 | 2.517E-09 | 2.800E-09 | 7.800E-09 |
| 2458 | 123 | L123W | A | 1.1e-08 | 3.7e-09 | 3.7e-09 | 6.133E-09 | 4.215E-09 | 3.700E-09 | 1.100E-08 |
| 2459 | 123 | L123Y | A | 1.1e-08 | 1.4e-08 | 7.8e-09 | 1.093E-08 | 3.101E-09 | 7.800E-09 | 1.400E-08 |
| 2460 | 123 | L123V | A | 2.5e-09 | 1e-08 | 3.5e-09 | 5.333E-09 | 4.072E-09 | 2.500E-09 | 1.000E-08 |
| 2461 | 124 | T124A | A | 3.6e-09 | 6e-09 | 6.2e-09 | 5.267E-09 | 1.447E-09 | 3.600E-09 | 6.200E-09 |
| 2462 | 124 | T124R | A | 5.4e-09 | 4.1e-09 | 8.5e-09 | 6.000E-09 | 2.261E-09 | 4.100E-09 | 8.500E-09 |
| 2463 | 124 | T124N | A | 3.6e-09 | 4.8e-09 | 9.2e-09 | 5.867E-09 | 2.948E-09 | 3.600E-09 | 9.200E-09 |
| 2464 | 124 | T124D | A | 7.5e-09 | 8.5e-09 | 6.3e-09 | 7.433E-09 | 1.102E-09 | 6.300E-09 | 8.500E-09 |

| | | | | | | | | | | |
|---|---|---|---|---|---|---|---|---|---|---|
| 2465 | 124 | T124C | A | 1e-08 | 6.9e-09 | 8e-09 | 8.300E-09 | 1.572E-09 | 6.900E-09 | 1.000E-08 |
| 2466 | 124 | T124E | A | 3.1e-09 | 7.6e-09 | 7.9e-09 | 6.200E-09 | 2.689E-09 | 3.100E-09 | 7.900E-09 |
| 2467 | 124 | T124Q | A | 7e-09 | 8.5e-09 | 5.4e-09 | 6.967E-09 | 1.550E-09 | 5.400E-09 | 8.500E-09 |
| 2468 | 124 | T124G | A | 6.1e-09 | 2.9e-09 | 1e-08 | 6.333E-09 | 3.556E-09 | 2.900E-09 | 1.000E-08 |
| 2469 | 124 | T124H | A | 7.8e-09 | 2.1e-09 | 9.2e-09 | 6.367E-09 | 3.761E-09 | 2.100E-09 | 9.200E-09 |
| 2470 | 124 | T124I | A | 5.6e-09 | 1.7e-08 | 6.6e-09 | 9.733E-09 | 6.313E-09 | 5.600E-09 | 1.700E-08 |
| 2471 | 124 | T124L | A | 1.8e-09 | 5.4e-09 | 3.8e-09 | 3.667E-09 | 1.804E-09 | 1.800E-09 | 5.400E-09 |
| 2472 | 124 | T124K | A | 4.6e-09 | 8e-09 | 4e-08 | 1.753E-08 | 1.953E-08 | 4.600E-09 | 4.000E-08 |
| 2473 | 124 | T124M | A | 1.2e-08 | 1.3e-08 | 3.9e-09 | 9.633E-09 | 4.990E-09 | 3.900E-09 | 1.300E-08 |
| 2474 | 124 | T124F | A | 8.1e-09 | 9.6e-09 | 1.7e-08 | 1.157E-08 | 4.765E-09 | 8.100E-09 | 1.700E-08 |
| 2475 | 124 | T124P | A | 1.3e-08 | 3e-09 | 4.6e-09 | 6.867E-09 | 5.372E-09 | 3.000E-09 | 1.300E-08 |
| 2476 | 124 | T124S | A | 4.2e-09 | 4.1e-09 | 1.7e-09 | 3.333E-09 | 1.415E-09 | 1.700E-09 | 4.200E-09 |
| 2477 | 124 | T124T | A | 4.2e-09 | 3.9e-09 | 4.5e-09 | 4.200E-09 | 3.000E-10 | 3.900E-09 | 4.500E-09 |
| 2478 | 124 | T124W | A | 3.6e-09 | 2.7e-09 | 3e-09 | 3.100E-09 | 4.583E-10 | 2.700E-09 | 3.600E-09 |
| 2479 | 124 | T124Y | A | 5.1e-09 | 8.9e-09 | 2.8e-09 | 5.600E-09 | 3.081E-09 | 2.800E-09 | 8.900E-09 |
| 2480 | 124 | T124V | A | 2.3e-08 | 2.6e-08 | 3.3e-09 | 1.743E-08 | 1.233E-08 | 3.300E-09 | 2.600E-08 |
| 2481 | 125 | E125A | A | 7.3e-09 | 1.6e-08 | 1.1e-08 | 1.143E-08 | 4.366E-09 | 7.300E-09 | 1.600E-08 |
| 2482 | 125 | E125R | A | 6.9e-09 | 5.1e-09 | 2e-08 | 1.067E-08 | 8.133E-09 | 5.100E-09 | 2.000E-08 |
| 2483 | 125 | E125N | A | 7e-09 | 7.7e-09 | 2.4e-08 | 1.290E-08 | 9.619E-09 | 7.000E-09 | 2.400E-08 |
| 2484 | 125 | E125D | A | 3.4e-09 | 9.2e-09 | 1.1e-08 | 7.867E-09 | 3.972E-09 | 3.400E-09 | 1.100E-08 |
| 2485 | 125 | E125C | A | 4.9e-09 | 8.8e-09 | 9.3e-09 | 7.667E-09 | 2.409E-09 | 4.900E-09 | 9.300E-09 |
| 2486 | 125 | E125E | A | 7.5e-09 | 1.8e-08 | 6.8e-09 | 1.077E-08 | 6.274E-09 | 6.800E-09 | 1.800E-08 |
| 2487 | 125 | E125Q | A | 1.4e-08 | 5e-09 | 5.7e-09 | 8.233E-09 | 5.006E-09 | 5.000E-09 | 1.400E-08 |
| 2488 | 125 | E125G | A | 1.5e-08 | 8.5e-09 | 8.3e-09 | 1.060E-08 | 3.812E-09 | 8.300E-09 | 1.500E-08 |
| 2489 | 125 | E125H | A | 3.7e-09 | 9.5e-09 | 8.3e-09 | 7.167E-09 | 3.062E-09 | 3.700E-09 | 9.500E-09 |
| 2490 | 125 | E125I | A | 3.8e-09 | 4.5e-09 | 9e-09 | 5.767E-09 | 2.822E-09 | 3.800E-09 | 9.000E-09 |
| 2491 | 125 | E125L | A | 6.8e-09 | 1.2e-08 | 7.4e-09 | 8.733E-09 | 2.845E-09 | 6.800E-09 | 1.200E-08 |
| 2492 | 125 | E125K | A | 8.8e-09 | 3.2e-08 | 1.4e-08 | 1.827E-08 | 1.217E-08 | 8.800E-09 | 3.200E-08 |
| 2493 | 125 | E125M | A | 1.9e-09 | 8.4e-09 | 4.8e-09 | 5.033E-09 | 3.256E-09 | 1.900E-09 | 8.400E-09 |
| 2494 | 125 | E125F | A | 1.6e-08 | 8.1e-09 | 1.7e-08 | 1.370E-08 | 4.875E-09 | 8.100E-09 | 1.700E-08 |
| 2495 | 125 | E125P | A | 4.4e-09 | 6.3e-09 | 7.6e-09 | 6.100E-09 | 1.609E-09 | 4.400E-09 | 7.600E-09 |
| 2496 | 125 | E125S | A | 7.3e-09 | 7.5e-09 | 9.8e-09 | 8.200E-09 | 1.389E-09 | 7.300E-09 | 9.800E-09 |
| 2497 | 125 | E125T | A | 2.7e-09 | 3.8e-09 | 8.8e-09 | 5.100E-09 | 3.251E-09 | 2.700E-09 | 8.800E-09 |
| 2498 | 125 | E125W | A | 6.3e-09 | 4e-09 | 4.8e-09 | 5.033E-09 | 1.168E-09 | 4.000E-09 | 6.300E-09 |
| 2499 | 125 | E125Y | A | 1.1e-08 | 1.9e-09 | 2.8e-09 | 5.233E-09 | 5.014E-09 | 1.900E-09 | 1.100E-08 |
| 2500 | 125 | E125V | A | 2e-08 | 1.4e-08 | 4.2e-09 | 1.273E-08 | 7.976E-09 | 4.200E-09 | 2.000E-08 |
| 2501 | 126 | I126A | A | 7.7e-09 | 1.1e-08 | 4.5e-09 | 7.733E-09 | 3.250E-09 | 4.500E-09 | 1.100E-08 |
| 2502 | 126 | I126R | A | 1.2e-08 | 5.8e-09 | 5.6e-09 | 7.800E-09 | 3.639E-09 | 5.600E-09 | 1.200E-08 |
| 2503 | 126 | I126N | A | 5.2e-09 | 1.6e-08 | 1.5e-08 | 1.207E-08 | 5.968E-09 | 5.200E-09 | 1.600E-08 |
| 2504 | 126 | I126D | A | 1.7e-08 | 1.1e-08 | 9.7e-09 | 1.257E-08 | 3.894E-09 | 9.700E-09 | 1.700E-08 |
| 2505 | 126 | I126C | A | 3.1e-09 | 5.3e-09 | 1.5e-08 | 7.800E-09 | 6.332E-09 | 3.100E-09 | 1.500E-08 |
| 2506 | 126 | I126E | A | 4.1e-09 | 7.5e-09 | 5.4e-09 | 5.667E-09 | 1.716E-09 | 4.100E-09 | 7.500E-09 |
| 2507 | 126 | I126Q | A | 5.6e-09 | 4.6e-09 | 3.9e-09 | 4.700E-09 | 8.544E-10 | 3.900E-09 | 5.600E-09 |
| 2508 | 126 | I126G | A | 3.8e-09 | 7.5e-09 | 1.1e-08 | 7.433E-09 | 3.600E-09 | 3.800E-09 | 1.100E-08 |
| 2509 | 126 | I126H | A | 1.5e-08 | 6e-09 | 6.7e-09 | 9.233E-09 | 5.006E-09 | 6.000E-09 | 1.500E-08 |
| 2510 | 126 | I126I | A | 5.1e-09 | 4.2e-09 | 3e-09 | 4.100E-09 | 1.054E-09 | 3.000E-09 | 5.100E-09 |
| 2511 | 126 | I126L | A | 2.2e-08 | 8.5e-09 | 6.7e-09 | 1.240E-08 | 8.362E-09 | 6.700E-09 | 2.200E-08 |
| 2512 | 126 | I126K | A | 8.5e-09 | 5.5e-09 | 9e-09 | 7.667E-09 | 1.893E-09 | 5.500E-09 | 9.000E-09 |
| 2513 | 126 | I126M | A | 6.1e-09 | 4.4e-09 | 2.2e-08 | 1.083E-08 | 9.708E-09 | 4.400E-09 | 2.200E-08 |
| 2514 | 126 | I126F | A | 2.1e-09 | 1.5e-08 | 5.5e-09 | 7.533E-09 | 6.686E-09 | 2.100E-09 | 1.500E-08 |
| 2515 | 126 | I126P | A | 1.1e-08 | 6.2e-09 | 1e-08 | 9.067E-09 | 2.532E-09 | 6.200E-09 | 1.100E-08 |
| 2516 | 126 | I126S | A | 4.6e-09 | 4.3e-09 | 1.6e-08 | 8.300E-09 | 6.670E-09 | 4.300E-09 | 1.600E-08 |
| 2517 | 126 | I126T | A | 9.6e-09 | 4.1e-09 | 2.6e-08 | 1.323E-08 | 1.139E-08 | 4.100E-09 | 2.600E-08 |
| 2518 | 126 | I126W | A | 2.8e-09 | 6.3e-09 | 5.7e-09 | 4.933E-09 | 1.872E-09 | 2.800E-09 | 6.300E-09 |
| 2519 | 126 | I126Y | A | 4.7e-09 | 1e-08 | 9.2e-09 | 7.967E-09 | 2.857E-09 | 4.700E-09 | 1.000E-08 |
| 2520 | 126 | I126V | A | 1.7e-08 | 3.3e-09 | 4.3e-09 | 8.200E-09 | 7.637E-09 | 3.300E-09 | 1.700E-08 |
| 2521 | 127 | L127A | A | 4.5e-09 | 7.6e-09 | 1e-08 | 7.367E-09 | 2.757E-09 | 4.500E-09 | 1.000E-08 |
| 2522 | 127 | L127R | A | 3.8e-09 | 2.6e-09 | 1.1e-08 | 5.800E-09 | 4.543E-09 | 2.600E-09 | 1.100E-08 |
| 2523 | 127 | L127N | A | 3.6e-09 | 1.1e-08 | 6.8e-09 | 7.133E-09 | 3.711E-09 | 3.600E-09 | 1.100E-08 |
| 2524 | 127 | L127D | A | 1.4e-08 | 7.1e-09 | 7.1e-09 | 9.400E-09 | 3.984E-09 | 7.100E-09 | 1.400E-08 |
| 2525 | 127 | L127C | A | 1e-08 | 1.5e-08 | 7.2e-09 | 1.073E-08 | 3.951E-09 | 7.200E-09 | 1.500E-08 |
| 2526 | 127 | L127E | A | 1.1e-08 | 3.6e-09 | 5.5e-09 | 6.700E-09 | 3.843E-09 | 3.600E-09 | 1.100E-08 |
| 2527 | 127 | L127Q | A | 6e-09 | 2.2e-08 | 5e-09 | 1.100E-08 | 9.539E-09 | 5.000E-09 | 2.200E-08 |
| 2528 | 127 | L127G | A | 1.2e-08 | 8.9e-09 | 8.7e-09 | 9.867E-09 | 1.850E-09 | 8.700E-09 | 1.200E-08 |
| 2529 | 127 | L127H | A | 7e-09 | 1.6e-08 | 6.6e-09 | 9.867E-09 | 5.315E-09 | 6.600E-09 | 1.600E-08 |
| 2530 | 127 | L127I | A | 7.6e-09 | 1.1e-08 | 1.3e-08 | 1.053E-08 | 2.730E-09 | 7.600E-09 | 1.300E-08 |
| 2531 | 127 | L127L | A | 1.2e-08 | 1.9e-08 | 5e-09 | 1.200E-08 | 7.000E-09 | 5.000E-09 | 1.900E-08 |
| 2532 | 127 | L127K | A | 1.3e-08 | 5.2e-09 | 5.6e-09 | 7.933E-09 | 4.392E-09 | 5.200E-09 | 1.300E-08 |
| 2533 | 127 | L127M | A | 1.3e-08 | 2.3e-09 | 1.5e-08 | 1.010E-08 | 6.829E-09 | 2.300E-09 | 1.500E-08 |
| 2534 | 127 | L127F | A | 4.7e-09 | 1.5e-08 | 1.4e-08 | 1.123E-08 | 5.680E-09 | 4.700E-09 | 1.500E-08 |
| 2535 | 127 | L127P | A | 2.9e-09 | 3.9e-09 | 6.4e-09 | 4.400E-09 | 1.803E-09 | 2.900E-09 | 6.400E-09 |
| 2536 | 127 | L127S | A | 2.6e-08 | 1e-08 | 4e-09 | 1.333E-08 | 1.137E-08 | 4.000E-09 | 2.600E-08 |
| 2537 | 127 | L127T | A | 1.4e-08 | 4.3e-09 | 5.8e-09 | 8.033E-09 | 5.221E-09 | 4.300E-09 | 1.400E-08 |
| 2538 | 127 | L127W | A | 1.4e-09 | 5.3e-09 | 1.3e-08 | 6.567E-09 | 5.903E-09 | 1.400E-09 | 1.300E-08 |
| 2539 | 127 | L127Y | A | 1.4e-08 | 3.2e-09 | 6.8e-09 | 8.000E-09 | 5.499E-09 | 3.200E-09 | 1.400E-08 |
| 2540 | 127 | L127V | A | 5.3e-09 | 3.3e-09 | 1.2e-08 | 6.867E-09 | 4.557E-09 | 3.300E-09 | 1.200E-08 |
| 2541 | 128 | K128A | A | 6.4e-09 | 1.6e-08 | 1.5e-08 | 1.247E-08 | 5.278E-09 | 6.400E-09 | 1.600E-08 |
| 2542 | 128 | K128R | A | 1.3e-08 | 5.1e-09 | 6.1e-09 | 8.067E-09 | 4.302E-09 | 5.100E-09 | 1.300E-08 |
| 2543 | 128 | K128N | A | 3.4e-09 | 7.9e-09 | 8.6e-09 | 6.633E-09 | 2.822E-09 | 3.400E-09 | 8.600E-09 |
| 2544 | 128 | K128D | A | 6.9e-09 | 8.5e-09 | 3.6e-09 | 6.333E-09 | 2.499E-09 | 3.600E-09 | 8.500E-09 |
| 2545 | 128 | K128C | A | 4.6e-09 | 1.7e-08 | 1.2e-08 | 1.120E-08 | 6.239E-09 | 4.600E-09 | 1.700E-08 |
| 2546 | 128 | K128E | A | 6.9e-09 | 6.2e-09 | 1.3e-08 | 8.700E-09 | 3.740E-09 | 6.200E-09 | 1.300E-08 |
| 2547 | 128 | K128Q | A | 5e-08 | 8.6e-09 | 2e-08 | 2.620E-08 | 2.139E-08 | 8.600E-09 | 5.000E-08 |
| 2548 | 128 | K128G | A | 9.4e-09 | 1e-08 | 3.7e-09 | 7.700E-09 | 3.477E-09 | 3.700E-09 | 1.000E-08 |
| 2549 | 128 | K128H | A | 1.2e-08 | 4.9e-09 | 9.2e-09 | 8.700E-09 | 3.576E-09 | 4.900E-09 | 1.200E-08 |

| | | | | | | | | | | |
|---|---|---|---|---|---|---|---|---|---|---|
| 2550 | 128 | K128I | A | 5.4e-09 | 6.7e-09 | 5.4e-09 | 5.833E-09 | 7.506E-10 | 5.400E-09 | 6.700E-09 |
| 2551 | 128 | K128L | A | 1.2e-08 | 8.9e-09 | 4e-09 | 8.300E-09 | 4.034E-09 | 4.000E-09 | 1.200E-08 |
| 2552 | 128 | K128K | A | 4.4e-09 | 9.5e-09 | 5.9e-09 | 6.600E-09 | 2.621E-09 | 4.400E-09 | 9.500E-09 |
| 2553 | 128 | K128M | A | 1.3e-08 | 3.3e-09 | 4.1e-09 | 6.800E-09 | 5.384E-09 | 3.300E-09 | 1.300E-08 |
| 2554 | 128 | K128F | A | 3.2e-09 | 5e-09 | 1.2e-08 | 6.733E-09 | 4.649E-09 | 3.200E-09 | 1.200E-08 |
| 2555 | 128 | K128P | A | 1.1e-08 | 1.4e-08 | 6.1e-09 | 1.037E-08 | 3.988E-09 | 6.100E-09 | 1.400E-08 |
| 2556 | 128 | K128S | A | 7.3e-09 | 8.3e-09 | 7.2e-09 | 7.600E-09 | 6.083E-10 | 7.200E-09 | 8.300E-09 |
| 2557 | 128 | K128T | A | 1e-08 | 7.1e-09 | 5.6e-09 | 7.567E-09 | 2.237E-09 | 5.600E-09 | 1.000E-08 |
| 2558 | 128 | K128W | A | 1.6e-08 | 1.2e-08 | 2.4e-09 | 1.013E-08 | 6.990E-09 | 2.400E-09 | 1.600E-08 |
| 2559 | 128 | K128Y | A | 9.5e-09 | 1.8e-09 | 3.4e-09 | 4.900E-09 | 4.063E-09 | 1.800E-09 | 9.500E-09 |
| 2560 | 128 | K128V | A | 5e-09 | 3.1e-09 | 2.6e-08 | 1.137E-08 | 1.271E-08 | 3.100E-09 | 2.600E-08 |
| 2561 | 129 | G129A | A | 7.1e-09 | 5.3e-09 | 1.6e-08 | 9.467E-09 | 5.729E-09 | 5.300E-09 | 1.600E-08 |
| 2562 | 129 | G129R | A | 1.9e-08 | 6.4e-09 | 4.4e-09 | 9.933E-09 | 7.915E-09 | 4.400E-09 | 1.900E-08 |
| 2563 | 129 | G129N | A | 2e-09 | 5.1e-09 | 2.5e-09 | 3.200E-09 | 1.664E-09 | 2.000E-09 | 5.100E-09 |
| 2564 | 129 | G129D | A | 7.2e-09 | 2.6e-08 | 9e-09 | 1.407E-08 | 1.037E-08 | 7.200E-09 | 2.600E-08 |
| 2565 | 129 | G129C | A | 9.2e-09 | 1.7e-08 | 1e-08 | 1.207E-08 | 4.291E-09 | 9.200E-09 | 1.700E-08 |
| 2566 | 129 | G129E | A | 4.5e-09 | 1.3e-08 | 7.2e-09 | 8.233E-09 | 4.343E-09 | 4.500E-09 | 1.300E-08 |
| 2567 | 129 | G129Q | A | 1.1e-08 | 4.8e-09 | 2.5e-09 | 6.100E-09 | 4.397E-09 | 2.500E-09 | 1.100E-08 |
| 2568 | 129 | G129G | A | 1.9e-09 | 8.4e-09 | 9.7e-09 | 6.667E-09 | 4.179E-09 | 1.900E-09 | 9.700E-09 |
| 2569 | 129 | G129H | A | 5.1e-09 | 2.6e-09 | 4.7e-09 | 4.133E-09 | 1.343E-09 | 2.600E-09 | 5.100E-09 |
| 2570 | 129 | G129I | A | 8.2e-09 | 9.8e-09 | 2.8e-09 | 6.933E-09 | 3.668E-09 | 2.800E-09 | 9.800E-09 |
| 2571 | 129 | G129L | A | 3.2e-09 | 6e-09 | 4.4e-09 | 4.533E-09 | 1.405E-09 | 3.200E-09 | 6.000E-09 |
| 2572 | 129 | G129K | A | 3.5e-08 | 2.8e-09 | 2.7e-09 | 1.350E-08 | 1.862E-08 | 2.700E-09 | 3.500E-08 |
| 2573 | 129 | G129M | A | 3.2e-09 | 2.7e-09 | 8.7e-09 | 4.867E-09 | 3.329E-09 | 2.700E-09 | 8.700E-09 |
| 2574 | 129 | G129F | A | 9.7e-09 | 1.5e-08 | 1.1e-08 | 1.190E-08 | 2.762E-09 | 9.700E-09 | 1.500E-08 |
| 2575 | 129 | G129P | A | 2.3e-08 | 2e-08 | 6.1e-09 | 1.637E-08 | 9.017E-09 | 6.100E-09 | 2.300E-08 |
| 2576 | 129 | G129S | A | 3.9e-09 | 5.1e-08 | 6.2e-09 | 2.037E-08 | 2.655E-08 | 3.900E-09 | 5.100E-08 |
| 2577 | 129 | G129T | A | 2.2e-08 | 1.4e-08 | 5.7e-09 | 1.390E-08 | 8.150E-09 | 5.700E-09 | 2.200E-08 |
| 2578 | 129 | G129W | A | 4.7e-09 | 1.2e-08 | 6.2e-09 | 7.633E-09 | 3.855E-09 | 4.700E-09 | 1.200E-08 |
| 2579 | 129 | G129Y | A | 1.1e-08 | 7e-09 | 1.2e-08 | 1.000E-08 | 2.646E-09 | 7.000E-09 | 1.200E-08 |
| 2580 | 129 | G129V | A | 1.1e-08 | 3.9e-09 | 1.3e-08 | 9.300E-09 | 4.782E-09 | 3.900E-09 | 1.300E-08 |
| 2581 | 130 | G130A | A | 1.6e-08 | 2.5e-08 | 4.8e-09 | 1.527E-08 | 1.012E-08 | 4.800E-09 | 2.500E-08 |
| 2582 | 130 | G130R | A | 7.3e-09 | 2.9e-09 | 3.1e-08 | 1.373E-08 | 1.511E-08 | 2.900E-09 | 3.100E-08 |
| 2583 | 130 | G130N | A | 1.2e-08 | 8.9e-09 | 9.6e-09 | 1.017E-08 | 1.626E-09 | 8.900E-09 | 1.200E-08 |
| 2584 | 130 | G130D | A | 9.4e-09 | 9.9e-09 | 7.1e-09 | 8.800E-09 | 1.493E-09 | 7.100E-09 | 9.900E-09 |
| 2585 | 130 | G130C | A | 3.7e-09 | 9.9e-09 | 6.4e-09 | 6.667E-09 | 3.109E-09 | 3.700E-09 | 9.900E-09 |
| 2586 | 130 | G130E | A | 1.9e-08 | 3.8e-09 | 7.3e-09 | 1.003E-08 | 7.960E-09 | 3.800E-09 | 1.900E-08 |
| 2587 | 130 | G130Q | A | 9.4e-09 | 1.6e-08 | 4.1e-09 | 9.833E-09 | 5.962E-09 | 4.100E-09 | 1.600E-08 |
| 2588 | 130 | G130G | A | 1.4e-08 | 1.8e-08 | 1.9e-09 | 1.130E-08 | 8.383E-09 | 1.900E-09 | 1.800E-08 |
| 2589 | 130 | G130H | A | 1.5e-08 | 6.6e-09 | 9e-09 | 1.020E-08 | 4.327E-09 | 6.600E-09 | 1.500E-08 |
| 2590 | 130 | G130I | A | 9.3e-09 | 1.7e-08 | 9e-09 | 1.177E-08 | 4.535E-09 | 9.000E-09 | 1.700E-08 |
| 2591 | 130 | G130L | A | 1.1e-08 | 5.3e-09 | 5.7e-09 | 7.333E-09 | 3.182E-09 | 5.300E-09 | 1.100E-08 |
| 2592 | 130 | G130K | A | 4.3e-09 | 3.5e-09 | 1.7e-08 | 8.267E-09 | 7.574E-09 | 3.500E-09 | 1.700E-08 |
| 2593 | 130 | G130M | A | 6.4e-09 | 6.7e-09 | 7.6e-09 | 6.900E-09 | 6.245E-10 | 6.400E-09 | 7.600E-09 |
| 2594 | 130 | G130F | A | 1.3e-08 | 2.3e-09 | 8.8e-09 | 8.033E-09 | 5.391E-09 | 2.300E-09 | 1.300E-08 |
| 2595 | 130 | G130P | A | 1.1e-08 | 1e-08 | 9e-09 | 1.000E-08 | 1.000E-09 | 9.000E-09 | 1.100E-08 |
| 2596 | 130 | G130S | A | 1.1e-08 | 4.5e-09 | 5.4e-09 | 6.967E-09 | 3.522E-09 | 4.500E-09 | 1.100E-08 |
| 2597 | 130 | G130T | A | 8.5e-09 | 1.4e-08 | 1.2e-08 | 1.150E-08 | 2.784E-09 | 8.500E-09 | 1.400E-08 |
| 2598 | 130 | G130W | A | 1.3e-08 | 6.2e-09 | 5.7e-09 | 8.300E-09 | 4.078E-09 | 5.700E-09 | 1.300E-08 |
| 2599 | 130 | G130Y | A | 4.1e-09 | 1e-08 | 1.7e-09 | 5.267E-09 | 4.271E-09 | 1.700E-09 | 1.000E-08 |
| 2600 | 130 | G130V | A | 1.9e-09 | 4.9e-09 | 1e-08 | 5.600E-09 | 4.095E-09 | 1.900E-09 | 1.000E-08 |
| 2601 | 131 | V131A | A | 3.2e-09 | 4.4e-09 | 1.1e-08 | 6.200E-09 | 4.200E-09 | 3.200E-09 | 1.100E-08 |
| 2602 | 131 | V131R | A | 9.3e-09 | 6.1e-09 | 5.7e-09 | 7.033E-09 | 1.973E-09 | 5.700E-09 | 9.300E-09 |
| 2603 | 131 | V131N | A | 9.1e-09 | 9.7e-09 | 9.3e-09 | 9.367E-09 | 3.055E-10 | 9.100E-09 | 9.700E-09 |
| 2604 | 131 | V131D | A | 7.5e-09 | 1.8e-09 | 8.4e-09 | 5.900E-09 | 3.579E-09 | 1.800E-09 | 8.400E-09 |
| 2605 | 131 | V131C | A | 9.3e-09 | 7.5e-09 | 1.3e-08 | 9.933E-09 | 2.804E-09 | 7.500E-09 | 1.300E-08 |
| 2606 | 131 | V131E | A | 4e-09 | 6.6e-09 | 7e-09 | 5.867E-09 | 1.629E-09 | 4.000E-09 | 7.000E-09 |
| 2607 | 131 | V131Q | A | 6.2e-09 | 2.4e-08 | 1e-08 | 1.340E-08 | 9.374E-09 | 6.200E-09 | 2.400E-08 |
| 2608 | 131 | V131G | A | 1.4e-08 | 8.5e-09 | 8.9e-09 | 1.047E-08 | 3.066E-09 | 8.500E-09 | 1.400E-08 |
| 2609 | 131 | V131H | A | 2.5e-09 | 4e-09 | 2.7e-09 | 3.067E-09 | 8.145E-10 | 2.500E-09 | 4.000E-09 |
| 2610 | 131 | V131I | A | 8e-09 | 1.7e-09 | 4.7e-09 | 4.800E-09 | 3.151E-09 | 1.700E-09 | 8.000E-09 |
| 2611 | 131 | V131L | A | 1.1e-08 | 3.2e-08 | 1.2e-08 | 1.833E-08 | 1.185E-08 | 1.100E-08 | 3.200E-08 |
| 2612 | 131 | V131K | A | 1.1e-08 | 1e-08 | 7.3e-09 | 9.433E-09 | 1.914E-09 | 7.300E-09 | 1.100E-08 |
| 2613 | 131 | V131M | A | 5.8e-09 | 6.9e-09 | 9.1e-09 | 7.267E-09 | 1.680E-09 | 5.800E-09 | 9.100E-09 |
| 2614 | 131 | V131F | A | 1.2e-08 | 4e-09 | 1.6e-08 | 1.067E-08 | 6.110E-09 | 4.000E-09 | 1.600E-08 |
| 2615 | 131 | V131P | A | 5.9e-09 | 6.7e-09 | 7.1e-09 | 6.567E-09 | 6.110E-10 | 5.900E-09 | 7.100E-09 |
| 2616 | 131 | V131S | A | 6.9e-09 | 1e-08 | 4.6e-09 | 7.167E-09 | 2.710E-09 | 4.600E-09 | 1.000E-08 |
| 2617 | 131 | V131T | A | 3.6e-09 | 1e-08 | 3.9e-09 | 5.833E-09 | 3.612E-09 | 3.600E-09 | 1.000E-08 |
| 2618 | 131 | V131W | A | 1.6e-08 | 8.4e-09 | 1.5e-08 | 1.313E-08 | 4.130E-09 | 8.400E-09 | 1.600E-08 |
| 2619 | 131 | V131Y | A | 7.2e-09 | 9e-09 | 5.7e-09 | 7.300E-09 | 1.652E-09 | 5.700E-09 | 9.000E-09 |
| 2620 | 131 | V131V | A | 5.4e-09 | 9.3e-09 | 1.1e-08 | 8.567E-09 | 2.871E-09 | 5.400E-09 | 1.100E-08 |
| 2621 | 132 | L132A | A | 3.3e-09 | 3e-09 | 1.1e-08 | 5.767E-09 | 4.535E-09 | 3.000E-09 | 1.100E-08 |
| 2622 | 132 | L132R | A | 7e-09 | 6e-09 | 5.4e-09 | 6.133E-09 | 8.083E-10 | 5.400E-09 | 7.000E-09 |
| 2623 | 132 | L132N | A | 2.4e-09 | 5.8e-09 | 2.4e-08 | 1.073E-08 | 1.161E-08 | 2.400E-09 | 2.400E-08 |
| 2624 | 132 | L132D | A | 4e-09 | 5.8e-09 | 2.9e-09 | 4.233E-09 | 1.464E-09 | 2.900E-09 | 5.800E-09 |
| 2625 | 132 | L132C | A | 4.8e-09 | 8.1e-09 | 9.5e-09 | 7.467E-09 | 2.413E-09 | 4.800E-09 | 9.500E-09 |
| 2626 | 132 | L132E | A | 1.8e-08 | 7.9e-09 | 8e-09 | 1.130E-08 | 5.803E-09 | 7.900E-09 | 1.800E-08 |
| 2627 | 132 | L132Q | A | 3.5e-09 | 1.9e-08 | 1e-08 | 1.083E-08 | 7.784E-09 | 3.500E-09 | 1.900E-08 |
| 2628 | 132 | L132G | A | 5.4e-09 | 2.2e-09 | 5e-09 | 4.200E-09 | 1.744E-09 | 2.200E-09 | 5.400E-09 |
| 2629 | 132 | L132H | A | 9.7e-09 | 2.8e-09 | 7.8e-09 | 6.767E-09 | 3.564E-09 | 2.800E-09 | 9.700E-09 |
| 2630 | 132 | L132I | A | 1.6e-09 | 9.9e-09 | 8.3e-09 | 6.600E-09 | 4.403E-09 | 1.600E-09 | 9.900E-09 |
| 2631 | 132 | L132L | A | 6.3e-09 | 1.4e-08 | 5.2e-09 | 8.500E-09 | 4.795E-09 | 5.200E-09 | 1.400E-08 |
| 2632 | 132 | L132K | A | 1.3e-08 | 1.4e-08 | 1.5e-08 | 1.400E-08 | 1.000E-09 | 1.300E-08 | 1.500E-08 |
| 2633 | 132 | L132M | A | 6.6e-09 | 3.2e-09 | 5e-09 | 4.933E-09 | 1.701E-09 | 3.200E-09 | 6.600E-09 |
| 2634 | 132 | L132F | A | 2.2e-09 | 2.9e-09 | 7.3e-09 | 4.133E-09 | 2.765E-09 | 2.200E-09 | 7.300E-09 |

| | | | | | | | | | | |
|---|---|---|---|---|---|---|---|---|---|---|
| 2635 | 132 | L132P | A | 1e-08 | 2.2e-09 | 8.9e-09 | 7.033E-09 | 4.222E-09 | 2.200E-09 | 1.000E-08 |
| 2636 | 132 | L132S | A | 1.8e-08 | 3.3e-09 | 2.7e-09 | 8.000E-09 | 8.665E-09 | 2.700E-09 | 1.800E-08 |
| 2637 | 132 | L132T | A | 1e-08 | 1.2e-08 | 3.6e-09 | 8.533E-09 | 4.388E-09 | 3.600E-09 | 1.200E-08 |
| 2638 | 132 | L132W | A | 9.5e-09 | 5.7e-09 | 6.3e-09 | 7.167E-09 | 2.043E-09 | 5.700E-09 | 9.500E-09 |
| 2639 | 132 | L132Y | A | 5.2e-09 | 3.3e-09 | 5.7e-09 | 4.733E-09 | 1.266E-09 | 3.300E-09 | 5.700E-09 |
| 2640 | 132 | L132V | A | 1.3e-08 | 4.6e-09 | 1.7e-08 | 1.153E-08 | 6.329E-09 | 4.600E-09 | 1.700E-08 |
| 2641 | 133 | I133A | A | 7.5e-09 | 1.9e-09 | 1e-08 | 6.467E-09 | 4.148E-09 | 1.900E-09 | 1.000E-08 |
| 2642 | 133 | I133R | A | 4.6e-09 | 1.5e-09 | 8.1e-09 | 4.733E-09 | 3.302E-09 | 1.500E-09 | 8.100E-09 |
| 2643 | 133 | I133N | A | 5.6e-09 | 6.9e-09 | 7e-09 | 6.500E-09 | 7.810E-10 | 5.600E-09 | 7.000E-09 |
| 2644 | 133 | I133D | A | 4.5e-09 | 1.5e-08 | 4.1e-09 | 7.867E-09 | 6.181E-09 | 4.100E-09 | 1.500E-08 |
| 2645 | 133 | I133C | A | 5.5e-09 | 4e-09 | 1.9e-09 | 3.800E-09 | 1.808E-09 | 1.900E-09 | 5.500E-09 |
| 2646 | 133 | I133E | A | 1.3e-08 | 6.5e-09 | 5.3e-09 | 8.267E-09 | 4.143E-09 | 5.300E-09 | 1.300E-08 |
| 2647 | 133 | I133Q | A | 1.2e-08 | 1.7e-08 | 1.2e-09 | 1.007E-08 | 8.075E-09 | 1.200E-09 | 1.700E-08 |
| 2648 | 133 | I133G | A | 1.3e-08 | 6.4e-09 | 3e-09 | 7.467E-09 | 5.085E-09 | 3.000E-09 | 1.300E-08 |
| 2649 | 133 | I133H | A | 2.6e-09 | 5.5e-09 | 8.1e-09 | 5.400E-09 | 2.751E-09 | 2.600E-09 | 8.100E-09 |
| 2650 | 133 | I133I | A | 4.1e-09 | 5.2e-09 | 4.1e-09 | 4.467E-09 | 6.351E-10 | 4.100E-09 | 5.200E-09 |
| 2651 | 133 | I133L | A | 3.3e-09 | 5.5e-09 | 1.9e-08 | 9.267E-09 | 8.501E-09 | 3.300E-09 | 1.900E-08 |
| 2652 | 133 | I133K | A | 7e-09 | 5.1e-09 | 1.2e-08 | 8.033E-09 | 3.564E-09 | 5.100E-09 | 1.200E-08 |
| 2653 | 133 | I133M | A | 1.3e-08 | 5.2e-09 | 4.6e-09 | 7.600E-09 | 4.686E-09 | 4.600E-09 | 1.300E-08 |
| 2654 | 133 | I133F | A | 1.2e-08 | 3.8e-09 | 9.3e-09 | 8.367E-09 | 4.179E-09 | 3.800E-09 | 1.200E-08 |
| 2655 | 133 | I133P | A | 2.3e-08 | 5.8e-09 | 5.6e-09 | 1.147E-08 | 9.989E-09 | 5.600E-09 | 2.300E-08 |
| 2656 | 133 | I133S | A | 2.1e-08 | 5.9e-09 | 1.3e-08 | 1.330E-08 | 7.554E-09 | 5.900E-09 | 2.100E-08 |
| 2657 | 133 | I133T | A | 5.6e-09 | 1.1e-08 | 4.1e-09 | 6.900E-09 | 3.629E-09 | 4.100E-09 | 1.100E-08 |
| 2658 | 133 | I133W | A | 2.4e-08 | 9.1e-09 | 5.6e-09 | 1.290E-08 | 9.771E-09 | 5.600E-09 | 2.400E-08 |
| 2659 | 133 | I133Y | A | 1.6e-09 | 7.1e-09 | 5.9e-09 | 4.867E-09 | 2.892E-09 | 1.600E-09 | 7.100E-09 |
| 2660 | 133 | I133V | A | 7.8e-09 | 7.8e-09 | 1.1e-08 | 8.867E-09 | 1.848E-09 | 7.800E-09 | 1.100E-08 |
| 2661 | 134 | Q134A | A | 5.7e-09 | 8.1e-09 | 1.6e-08 | 9.933E-09 | 5.389E-09 | 5.700E-09 | 1.600E-08 |
| 2662 | 134 | Q134R | A | 5e-09 | 9e-09 | 1.9e-08 | 1.100E-08 | 7.211E-09 | 5.000E-09 | 1.900E-08 |
| 2663 | 134 | Q134N | A | 3.9e-09 | 3.5e-09 | 8e-09 | 5.133E-09 | 2.491E-09 | 3.500E-09 | 8.000E-09 |
| 2664 | 134 | Q134D | A | 5.3e-09 | 5.7e-09 | 5e-09 | 5.333E-09 | 3.512E-10 | 5.000E-09 | 5.700E-09 |
| 2665 | 134 | Q134C | A | 8.5e-09 | 4.4e-09 | 1.1e-08 | 7.967E-09 | 3.332E-09 | 4.400E-09 | 1.100E-08 |
| 2666 | 134 | Q134E | A | 2e-09 | 1.1e-08 | 8.4e-09 | 7.133E-09 | 4.632E-09 | 2.000E-09 | 1.100E-08 |
| 2667 | 134 | Q134Q | A | 2.7e-08 | 2.3e-09 | 1.2e-08 | 1.377E-08 | 1.244E-08 | 2.300E-09 | 2.700E-08 |
| 2668 | 134 | Q134G | A | 9.8e-09 | 9e-09 | 1.2e-08 | 1.027E-08 | 1.553E-09 | 9.000E-09 | 1.200E-08 |
| 2669 | 134 | Q134H | A | 1.2e-09 | 5.1e-09 | 2.8e-09 | 3.033E-09 | 1.960E-09 | 1.200E-09 | 5.100E-09 |
| 2670 | 134 | Q134I | A | 1.1e-08 | 3.5e-09 | 5.5e-09 | 6.667E-09 | 3.884E-09 | 3.500E-09 | 1.100E-08 |
| 2671 | 134 | Q134L | A | 8.6e-09 | 1.8e-08 | 3.4e-09 | 1.000E-08 | 7.400E-09 | 3.400E-09 | 1.800E-08 |
| 2672 | 134 | Q134K | A | 4.6e-09 | 6.4e-09 | 5.1e-09 | 5.367E-09 | 9.292E-10 | 4.600E-09 | 6.400E-09 |
| 2673 | 134 | Q134M | A | 1.6e-08 | 1e-08 | 8.3e-09 | 1.143E-08 | 4.045E-09 | 8.300E-09 | 1.600E-08 |
| 2674 | 134 | Q134F | A | 2.1e-08 | 1.5e-08 | 8.9e-09 | 1.497E-08 | 6.050E-09 | 8.900E-09 | 2.100E-08 |
| 2675 | 134 | Q134P | A | 7.5e-09 | 3.8e-09 | 3.2e-09 | 4.833E-09 | 2.329E-09 | 3.200E-09 | 7.500E-09 |
| 2676 | 134 | Q134S | A | 1.3e-08 | 3.9e-09 | 3.9e-09 | 6.933E-09 | 5.254E-09 | 3.900E-09 | 1.300E-08 |
| 2677 | 134 | Q134T | A | 7.6e-09 | 1.8e-08 | 3.4e-09 | 9.667E-09 | 7.516E-09 | 3.400E-09 | 1.800E-08 |
| 2678 | 134 | Q134W | A | 3.4e-09 | 1.8e-08 | 3.2e-09 | 8.200E-09 | 8.488E-09 | 3.200E-09 | 1.800E-08 |
| 2679 | 134 | Q134Y | A | 1.3e-08 | 2.2e-09 | 8.7e-09 | 7.967E-09 | 5.437E-09 | 2.200E-09 | 1.300E-08 |
| 2680 | 134 | Q134V | A | 1.4e-08 | 5.4e-09 | 6.4e-09 | 8.600E-09 | 4.703E-09 | 5.400E-09 | 1.400E-08 |
| 2681 | 135 | R135A | A | 6.2e-09 | 1.2e-08 | 9.9e-09 | 9.367E-09 | 2.937E-09 | 6.200E-09 | 1.200E-08 |
| 2682 | 135 | R135R | A | 1.9e-08 | 5.1e-09 | 1e-08 | 1.137E-08 | 7.050E-09 | 5.100E-09 | 1.900E-08 |
| 2683 | 135 | R135N | A | 5.6e-09 | 3.3e-09 | 4.1e-09 | 4.333E-09 | 1.168E-09 | 3.300E-09 | 5.600E-09 |
| 2684 | 135 | R135D | A | 2.2e-08 | 1e-08 | 5.3e-09 | 1.243E-08 | 8.612E-09 | 5.300E-09 | 2.200E-08 |
| 2685 | 135 | R135C | A | 8.4e-09 | 1.4e-08 | 5.6e-09 | 9.333E-09 | 4.277E-09 | 5.600E-09 | 1.400E-08 |
| 2686 | 135 | R135E | A | 1.4e-08 | 5.2e-09 | 1.1e-08 | 1.007E-08 | 4.474E-09 | 5.200E-09 | 1.400E-08 |
| 2687 | 135 | R135Q | A | 4.4e-09 | 3.8e-09 | 3.1e-09 | 3.767E-09 | 6.506E-10 | 3.100E-09 | 4.400E-09 |
| 2688 | 135 | R135G | A | 1.8e-08 | 2.2e-08 | 8.6e-09 | 1.620E-08 | 6.879E-09 | 8.600E-09 | 2.200E-08 |
| 2689 | 135 | R135H | A | 1.4e-08 | 2.1e-09 | 6.2e-09 | 7.433E-09 | 6.045E-09 | 2.100E-09 | 1.400E-08 |
| 2690 | 135 | R135I | A | 6.4e-09 | 6.1e-09 | 7.1e-09 | 6.533E-09 | 5.132E-10 | 6.100E-09 | 7.100E-09 |
| 2691 | 135 | R135L | A | 1.3e-08 | 1.3e-08 | 9.3e-09 | 1.177E-08 | 2.136E-09 | 9.300E-09 | 1.300E-08 |
| 2692 | 135 | R135K | A | 1.3e-08 | 1.4e-08 | 1.6e-09 | 9.533E-09 | 6.889E-09 | 1.600E-09 | 1.400E-08 |
| 2693 | 135 | R135M | A | 5.2e-09 | 8.6e-09 | 6.5e-09 | 6.767E-09 | 1.716E-09 | 5.200E-09 | 8.600E-09 |
| 2694 | 135 | R135F | A | 5.4e-09 | 1.1e-08 | 6.7e-09 | 7.700E-09 | 2.931E-09 | 5.400E-09 | 1.100E-08 |
| 2695 | 135 | R135P | A | 6.1e-09 | 2.2e-09 | 2.4e-09 | 3.567E-09 | 2.196E-09 | 2.200E-09 | 6.100E-09 |
| 2696 | 135 | R135S | A | 1.9e-08 | 4.4e-09 | 2.4e-09 | 8.600E-09 | 9.062E-09 | 2.400E-09 | 1.900E-08 |
| 2697 | 135 | R135T | A | 9.3e-09 | 1.6e-09 | 3.2e-09 | 4.700E-09 | 4.063E-09 | 1.600E-09 | 9.300E-09 |
| 2698 | 135 | R135W | A | 2.1e-08 | 3.7e-09 | 1.2e-08 | 1.223E-08 | 8.652E-09 | 3.700E-09 | 2.100E-08 |
| 2699 | 135 | R135Y | A | 6.4e-09 | 8.3e-09 | 8.9e-09 | 7.867E-09 | 1.305E-09 | 6.400E-09 | 8.900E-09 |
| 2700 | 135 | R135V | A | 8e-09 | 7.4e-09 | 5.8e-09 | 7.067E-09 | 1.137E-09 | 5.800E-09 | 8.000E-09 |
| 2701 | 136 | N136A | A | 1e-08 | 8.6e-09 | 3.6e-09 | 7.400E-09 | 3.365E-09 | 3.600E-09 | 1.000E-08 |
| 2702 | 136 | N136R | A | 1.1e-08 | 1.2e-08 | 6.2e-09 | 9.733E-09 | 3.101E-09 | 6.200E-09 | 1.200E-08 |
| 2703 | 136 | N136N | A | 3.1e-08 | 1.1e-08 | 4.1e-09 | 1.537E-08 | 1.397E-08 | 4.100E-09 | 3.100E-08 |
| 2704 | 136 | N136D | A | 5.3e-09 | 4e-08 | 6.7e-09 | 1.733E-08 | 1.964E-08 | 5.300E-09 | 4.000E-08 |
| 2705 | 136 | N136C | A | 9.3e-09 | 1.1e-08 | 4e-09 | 8.100E-09 | 3.651E-09 | 4.000E-09 | 1.100E-08 |
| 2706 | 136 | N136E | A | 7.2e-09 | 6.3e-09 | 4.2e-09 | 5.900E-09 | 1.539E-09 | 4.200E-09 | 7.200E-09 |
| 2707 | 136 | N136Q | A | 8e-09 | 5.5e-09 | 5.3e-09 | 6.267E-09 | 1.504E-09 | 5.300E-09 | 8.000E-09 |
| 2708 | 136 | N136G | A | 8.1e-09 | 4.5e-09 | 4.1e-09 | 5.567E-09 | 2.203E-09 | 4.100E-09 | 8.100E-09 |
| 2709 | 136 | N136H | A | 1.2e-08 | 4.4e-09 | 2.8e-09 | 6.400E-09 | 4.915E-09 | 2.800E-09 | 1.200E-08 |
| 2710 | 136 | N136I | A | 2.5e-09 | 4.8e-09 | 3.4e-08 | 1.377E-08 | 1.756E-08 | 2.500E-09 | 3.400E-08 |
| 2711 | 136 | N136L | A | 8.3e-09 | 8.4e-09 | 3.2e-09 | 6.633E-09 | 2.974E-09 | 3.200E-09 | 8.400E-09 |
| 2712 | 136 | N136K | A | 1.4e-08 | 3.7e-09 | 1.1e-08 | 9.567E-09 | 5.297E-09 | 3.700E-09 | 1.400E-08 |
| 2713 | 136 | N136M | A | 9.6e-09 | 7.1e-09 | 6.5e-09 | 7.733E-09 | 1.644E-09 | 6.500E-09 | 9.600E-09 |
| 2714 | 136 | N136F | A | 1.3e-08 | 5.2e-09 | 2.6e-09 | 6.933E-09 | 5.412E-09 | 2.600E-09 | 1.300E-08 |
| 2715 | 136 | N136P | A | 5.5e-09 | 2.6e-09 | 7e-09 | 5.033E-09 | 2.237E-09 | 2.600E-09 | 7.000E-09 |
| 2716 | 136 | N136S | A | 6.4e-09 | 1.6e-08 | 1.2e-08 | 1.147E-08 | 4.822E-09 | 6.400E-09 | 1.600E-08 |
| 2717 | 136 | N136T | A | 4.3e-09 | 6.9e-09 | 2.8e-09 | 4.667E-09 | 2.074E-09 | 2.800E-09 | 6.900E-09 |
| 2718 | 136 | N136W | A | 2.5e-09 | 8e-09 | 1e-08 | 6.833E-09 | 3.884E-09 | 2.500E-09 | 1.000E-08 |
| 2719 | 136 | N136Y | A | 1e-08 | 6.7e-09 | 1.1e-08 | 9.233E-09 | 2.250E-09 | 6.700E-09 | 1.100E-08 |

| | | | | | | | | | | |
|---|---|---|---|---|---|---|---|---|---|---|
| 2720 | 136 | N136V | A | 5.6e-09 | 4.4e-09 | 8.5e-09 | 6.167E-09 | 2.108E-09 | 4.400E-09 | 8.500E-09 |
| 2721 | 137 | P137A | A | 1.8e-08 | 6e-09 | 1.9e-08 | 1.433E-08 | 7.234E-09 | 6.000E-09 | 1.900E-08 |
| 2722 | 137 | P137R | A | 1.6e-09 | 6.7e-09 | 5.8e-09 | 4.700E-09 | 2.722E-09 | 1.600E-09 | 6.700E-09 |
| 2723 | 137 | P137N | A | 3.1e-08 | 9.8e-09 | 1.7e-08 | 1.927E-08 | 1.078E-08 | 9.800E-09 | 3.100E-08 |
| 2724 | 137 | P137D | A | 3.1e-08 | 1.1e-08 | 6e-09 | 1.600E-08 | 1.323E-08 | 6.000E-09 | 3.100E-08 |
| 2725 | 137 | P137C | A | 6.6e-09 | 2.7e-09 | 8.9e-09 | 6.067E-09 | 3.134E-09 | 2.700E-09 | 8.900E-09 |
| 2726 | 137 | P137E | A | 5.3e-09 | 8e-09 | 4.3e-09 | 5.867E-09 | 1.914E-09 | 4.300E-09 | 8.000E-09 |
| 2727 | 137 | P137Q | A | 1.1e-08 | 8.1e-09 | 1.7e-08 | 1.203E-08 | 4.539E-09 | 8.100E-09 | 1.700E-08 |
| 2728 | 137 | P137G | A | 1.1e-08 | 3.9e-09 | 1.2e-08 | 8.967E-09 | 4.416E-09 | 3.900E-09 | 1.200E-08 |
| 2729 | 137 | P137H | A | 5.6e-09 | 3.6e-09 | 7.4e-09 | 5.533E-09 | 1.901E-09 | 3.600E-09 | 7.400E-09 |
| 2730 | 137 | P137I | A | 7e-09 | 8.9e-09 | 1e-08 | 8.633E-09 | 1.518E-09 | 7.000E-09 | 1.000E-08 |
| 2731 | 137 | P137L | A | 1.4e-08 | 4.8e-09 | 7.5e-09 | 8.767E-09 | 4.729E-09 | 4.800E-09 | 1.400E-08 |
| 2732 | 137 | P137K | A | 6.2e-09 | 8.4e-09 | 5.9e-09 | 6.833E-09 | 1.365E-09 | 5.900E-09 | 8.400E-09 |
| 2733 | 137 | P137M | A | 8.3e-09 | 1.4e-08 | 2.5e-08 | 1.577E-08 | 8.489E-09 | 8.300E-09 | 2.500E-08 |
| 2734 | 137 | P137F | A | 1.7e-08 | 1.3e-08 | 8.4e-09 | 1.280E-08 | 4.303E-09 | 8.400E-09 | 1.700E-08 |
| 2735 | 137 | P137P | A | 6.6e-09 | 1.1e-08 | 2.7e-09 | 6.767E-09 | 4.153E-09 | 2.700E-09 | 1.100E-08 |
| 2736 | 137 | P137S | A | 6.3e-09 | 3.4e-08 | 1.3e-08 | 1.777E-08 | 1.445E-08 | 6.300E-09 | 3.400E-08 |
| 2737 | 137 | P137T | A | 7e-09 | 2.8e-09 | 1.2e-08 | 7.267E-09 | 4.606E-09 | 2.800E-09 | 1.200E-08 |
| 2738 | 137 | P137W | A | 5e-09 | 2.3e-08 | 4.5e-09 | 1.083E-08 | 1.054E-08 | 4.500E-09 | 2.300E-08 |
| 2739 | 137 | P137Y | A | 7.1e-09 | 6.6e-09 | 4.9e-09 | 6.200E-09 | 1.153E-09 | 4.900E-09 | 7.100E-09 |
| 2740 | 137 | P137V | A | 9.1e-09 | 1.7e-08 | 1.2e-08 | 1.270E-08 | 3.996E-09 | 9.100E-09 | 1.700E-08 |
| 2741 | 138 | Q138A | A | 9.9e-09 | 9.6e-09 | 5.1e-09 | 8.200E-09 | 2.689E-09 | 5.100E-09 | 9.900E-09 |
| 2742 | 138 | Q138R | A | 6.6e-09 | 1.5e-09 | 5.1e-09 | 4.400E-09 | 2.621E-09 | 1.500E-09 | 6.600E-09 |
| 2743 | 138 | Q138N | A | 3e-09 | 6.5e-09 | 3e-09 | 4.167E-09 | 2.021E-09 | 3.000E-09 | 6.500E-09 |
| 2744 | 138 | Q138D | A | 4.1e-09 | 9.7e-09 | 7.1e-09 | 6.967E-09 | 2.802E-09 | 4.100E-09 | 9.700E-09 |
| 2745 | 138 | Q138C | A | 9.8e-09 | 5.9e-09 | 3.4e-09 | 6.367E-09 | 3.225E-09 | 3.400E-09 | 9.800E-09 |
| 2746 | 138 | Q138E | A | 3.6e-09 | 8.9e-09 | 1.3e-08 | 8.500E-09 | 4.713E-09 | 3.600E-09 | 1.300E-08 |
| 2747 | 138 | Q138Q | A | 7.1e-09 | 5.5e-09 | 8.5e-09 | 7.033E-09 | 1.501E-09 | 5.500E-09 | 8.500E-09 |
| 2748 | 138 | Q138G | A | 3.9e-09 | 9.3e-09 | 3.6e-09 | 5.600E-09 | 3.208E-09 | 3.600E-09 | 9.300E-09 |
| 2749 | 138 | Q138H | A | 1.8e-08 | 8.1e-09 | 1.9e-09 | 9.333E-09 | 8.121E-09 | 1.900E-09 | 1.800E-08 |
| 2750 | 138 | Q138I | A | 8.3e-09 | 7.5e-09 | 1.6e-08 | 1.060E-08 | 4.694E-09 | 7.500E-09 | 1.600E-08 |
| 2751 | 138 | Q138L | A | 6.1e-09 | 7e-09 | 2.1e-09 | 5.067E-09 | 2.608E-09 | 2.100E-09 | 7.000E-09 |
| 2752 | 138 | Q138K | A | 2.7e-09 | 4.1e-09 | 2.4e-09 | 3.067E-09 | 9.074E-10 | 2.400E-09 | 4.100E-09 |
| 2753 | 138 | Q138M | A | 1.3e-09 | 4.6e-09 | 2.7e-09 | 2.867E-09 | 1.656E-09 | 1.300E-09 | 4.600E-09 |
| 2754 | 138 | Q138F | A | 9.5e-09 | 6.4e-09 | 1e-08 | 8.633E-09 | 1.950E-09 | 6.400E-09 | 1.000E-08 |
| 2755 | 138 | Q138P | A | 3.4e-09 | 7.5e-09 | 4.9e-09 | 5.267E-09 | 2.074E-09 | 3.400E-09 | 7.500E-09 |
| 2756 | 138 | Q138S | A | 1.1e-08 | 7.4e-09 | 1.9e-08 | 1.247E-08 | 5.937E-09 | 7.400E-09 | 1.900E-08 |
| 2757 | 138 | Q138T | A | 5.4e-09 | 7.1e-09 | 4.2e-09 | 5.567E-09 | 1.457E-09 | 4.200E-09 | 7.100E-09 |
| 2758 | 138 | Q138W | A | 3.2e-09 | 3.7e-09 | 6e-09 | 4.300E-09 | 1.493E-09 | 3.200E-09 | 6.000E-09 |
| 2759 | 138 | Q138Y | A | 9.8e-09 | 2.2e-09 | 5.5e-09 | 5.833E-09 | 3.811E-09 | 2.200E-09 | 9.800E-09 |
| 2760 | 138 | Q138V | A | 3.6e-09 | 4.8e-09 | 6.7e-09 | 5.033E-09 | 1.563E-09 | 3.600E-09 | 6.700E-09 |
| 2761 | 139 | L139A | A | 4.2e-09 | 1e-08 | 9.9e-09 | 8.033E-09 | 3.320E-09 | 4.200E-09 | 1.000E-08 |
| 2762 | 139 | L139R | A | 7.8e-09 | 5e-09 | 6.3e-09 | 6.367E-09 | 1.401E-09 | 5.000E-09 | 7.800E-09 |
| 2763 | 139 | L139N | A | 1.7e-08 | 2.9e-09 | 2.5e-08 | 1.497E-08 | 1.119E-08 | 2.900E-09 | 2.500E-08 |
| 2764 | 139 | L139D | A | 2.6e-09 | 2e-09 | 6e-09 | 3.533E-09 | 2.157E-09 | 2.000E-09 | 6.000E-09 |
| 2765 | 139 | L139C | A | 8.8e-09 | 8.1e-09 | 4.5e-09 | 7.133E-09 | 2.307E-09 | 4.500E-09 | 8.800E-09 |
| 2766 | 139 | L139E | A | 7.3e-09 | 4.4e-09 | 6.1e-09 | 5.933E-09 | 1.457E-09 | 4.400E-09 | 7.300E-09 |
| 2767 | 139 | L139Q | A | 7.8e-09 | 7.4e-09 | 3.2e-08 | 1.573E-08 | 1.409E-08 | 7.400E-09 | 3.200E-08 |
| 2768 | 139 | L139G | A | 4.6e-09 | 5.2e-09 | 9.1e-09 | 6.300E-09 | 2.443E-09 | 4.600E-09 | 9.100E-09 |
| 2769 | 139 | L139H | A | 3.4e-09 | 1.5e-08 | 7.3e-09 | 8.567E-09 | 5.903E-09 | 3.400E-09 | 1.500E-08 |
| 2770 | 139 | L139I | A | 8.7e-09 | 1.2e-08 | 4e-09 | 8.233E-09 | 4.020E-09 | 4.000E-09 | 1.200E-08 |
| 2771 | 139 | L139L | A | 4.2e-09 | 4.6e-09 | 4.3e-09 | 4.367E-09 | 2.082E-10 | 4.200E-09 | 4.600E-09 |
| 2772 | 139 | L139K | A | 3.5e-09 | 1.2e-08 | 3.6e-09 | 6.367E-09 | 4.879E-09 | 3.500E-09 | 1.200E-08 |
| 2773 | 139 | L139M | A | 1e-08 | 4.5e-09 | 6.5e-09 | 7.000E-09 | 2.784E-09 | 4.500E-09 | 1.000E-08 |
| 2774 | 139 | L139F | A | 6.1e-09 | 1.3e-08 | 1.5e-08 | 1.137E-08 | 4.669E-09 | 6.100E-09 | 1.500E-08 |
| 2775 | 139 | L139P | A | 8.7e-09 | 4.9e-09 | 6.5e-09 | 6.700E-09 | 1.908E-09 | 4.900E-09 | 8.700E-09 |
| 2776 | 139 | L139S | A | 8.7e-09 | 1.2e-08 | 8.1e-09 | 9.600E-09 | 2.100E-09 | 8.100E-09 | 1.200E-08 |
| 2777 | 139 | L139T | A | 9.2e-09 | 1.7e-08 | 6.5e-09 | 1.090E-08 | 5.453E-09 | 6.500E-09 | 1.700E-08 |
| 2778 | 139 | L139W | A | 7.7e-09 | 6.1e-09 | 9.6e-09 | 7.800E-09 | 1.752E-09 | 6.100E-09 | 9.600E-09 |
| 2779 | 139 | L139Y | A | 9.6e-09 | 3.1e-09 | 5e-09 | 5.900E-09 | 3.342E-09 | 3.100E-09 | 9.600E-09 |
| 2780 | 139 | L139V | A | 5.1e-09 | 3.6e-09 | 1.9e-09 | 3.533E-09 | 1.601E-09 | 1.900E-09 | 5.100E-09 |
| 2781 | 140 | C140A | A | 6.6e-09 | 3.1e-09 | 6.4e-09 | 5.367E-09 | 1.966E-09 | 3.100E-09 | 6.600E-09 |
| 2782 | 140 | C140R | A | 4.7e-09 | 5.1e-09 | 2.2e-09 | 4.000E-09 | 1.572E-09 | 2.200E-09 | 5.100E-09 |
| 2783 | 140 | C140N | A | 1.5e-08 | 1.3e-08 | 2.2e-08 | 1.667E-08 | 4.726E-09 | 1.300E-08 | 2.200E-08 |
| 2784 | 140 | C140D | A | 7.5e-09 | 2.1e-09 | 5.9e-09 | 5.167E-09 | 2.774E-09 | 2.100E-09 | 7.500E-09 |
| 2785 | 140 | C140C | A | 7.4e-09 | 1.5e-08 | 3.4e-09 | 8.600E-09 | 5.892E-09 | 3.400E-09 | 1.500E-08 |
| 2786 | 140 | C140E | A | 8.9e-09 | 1.9e-08 | 7.8e-09 | 1.190E-08 | 6.173E-09 | 7.800E-09 | 1.900E-08 |
| 2787 | 140 | C140Q | A | 6.2e-09 | 8.4e-09 | 8.2e-09 | 7.600E-09 | 1.217E-09 | 6.200E-09 | 8.400E-09 |
| 2788 | 140 | C140G | A | 6.1e-09 | 5.8e-09 | 7e-09 | 6.300E-09 | 6.245E-10 | 5.800E-09 | 7.000E-09 |
| 2789 | 140 | C140H | A | 1.1e-08 | 9.6e-09 | 1.3e-08 | 1.120E-08 | 1.709E-09 | 9.600E-09 | 1.300E-08 |
| 2790 | 140 | C140I | A | 5.8e-09 | 2.8e-09 | 7.9e-09 | 5.500E-09 | 2.563E-09 | 2.800E-09 | 7.900E-09 |
| 2791 | 140 | C140L | A | 1e-08 | 4e-09 | 3.3e-08 | 1.567E-08 | 1.531E-08 | 4.000E-09 | 3.300E-08 |
| 2792 | 140 | C140K | A | 5.9e-09 | 1.1e-08 | 1.9e-08 | 1.197E-08 | 6.603E-09 | 5.900E-09 | 1.900E-08 |
| 2793 | 140 | C140M | A | 1.2e-08 | 2.2e-09 | 9.7e-09 | 7.967E-09 | 5.125E-09 | 2.200E-09 | 1.200E-08 |
| 2794 | 140 | C140F | A | 1.6e-08 | 8.7e-09 | 2.2e-09 | 8.967E-09 | 6.904E-09 | 2.200E-09 | 1.600E-08 |
| 2795 | 140 | C140P | A | 9.8e-09 | 5.1e-09 | 5.2e-09 | 6.700E-09 | 2.685E-09 | 5.100E-09 | 9.800E-09 |
| 2796 | 140 | C140S | A | 1.3e-08 | 5e-09 | 5.6e-09 | 7.867E-09 | 4.456E-09 | 5.000E-09 | 1.300E-08 |
| 2797 | 140 | C140T | A | 2.7e-09 | 1.4e-08 | 9.6e-09 | 8.767E-09 | 5.696E-09 | 2.700E-09 | 1.400E-08 |
| 2798 | 140 | C140W | A | 6e-09 | 2.8e-09 | 3.6e-09 | 4.133E-09 | 1.665E-09 | 2.800E-09 | 6.000E-09 |
| 2799 | 140 | C140Y | A | 9.5e-09 | 8.2e-09 | 8.4e-09 | 8.700E-09 | 7.000E-10 | 8.200E-09 | 9.500E-09 |
| 2800 | 140 | C140V | A | 3.9e-09 | 5.5e-09 | 9.5e-09 | 6.300E-09 | 2.884E-09 | 3.900E-09 | 9.500E-09 |
| 2801 | 141 | Y141A | A | 3.3e-09 | 2e-08 | 9.3e-09 | 1.087E-08 | 8.460E-09 | 3.300E-09 | 2.000E-08 |
| 2802 | 141 | Y141R | A | 7e-09 | 1.3e-08 | 2.1e-08 | 1.367E-08 | 7.024E-09 | 7.000E-09 | 2.100E-08 |
| 2803 | 141 | Y141N | A | 5.2e-09 | 7.1e-09 | 6.4e-09 | 6.233E-09 | 9.609E-10 | 5.200E-09 | 7.100E-09 |
| 2804 | 141 | Y141D | A | 6.9e-09 | 7.3e-09 | 3.9e-09 | 6.033E-09 | 1.858E-09 | 3.900E-09 | 7.300E-09 |

| | | | | | | | | | | |
|---|---|---|---|---|---|---|---|---|---|---|
| 2805 | 141 | Y141C | A | 9.8e-09 | 5.1e-09 | 1.2e-08 | 8.967E-09 | 3.525E-09 | 5.100E-09 | 1.200E-08 |
| 2806 | 141 | Y141E | A | 6e-09 | 4e-09 | 2.5e-08 | 1.167E-08 | 1.159E-08 | 4.000E-09 | 2.500E-08 |
| 2807 | 141 | Y141Q | A | 6.3e-09 | 5.8e-09 | 4.1e-09 | 5.400E-09 | 1.153E-09 | 4.100E-09 | 6.300E-09 |
| 2808 | 141 | Y141G | A | 5.2e-09 | 9.9e-09 | 3.9e-09 | 6.333E-09 | 3.156E-09 | 3.900E-09 | 9.900E-09 |
| 2809 | 141 | Y141H | A | 9.5e-09 | 7e-09 | 2.9e-09 | 6.467E-09 | 3.332E-09 | 2.900E-09 | 9.500E-09 |
| 2810 | 141 | Y141I | A | 1.9e-09 | 9.3e-09 | 7.8e-09 | 6.333E-09 | 3.912E-09 | 1.900E-09 | 9.300E-09 |
| 2811 | 141 | Y141L | A | 1.4e-08 | 1.1e-08 | 5.9e-09 | 1.030E-08 | 4.095E-09 | 5.900E-09 | 1.400E-08 |
| 2812 | 141 | Y141K | A | 7.2e-09 | 5.2e-09 | 1.3e-08 | 8.467E-09 | 4.051E-09 | 5.200E-09 | 1.300E-08 |
| 2813 | 141 | Y141M | A | 9.1e-09 | 4e-09 | 8.2e-09 | 7.100E-09 | 2.722E-09 | 4.000E-09 | 9.100E-09 |
| 2814 | 141 | Y141F | A | 1.3e-08 | 1.9e-08 | 6.7e-09 | 1.290E-08 | 6.151E-09 | 6.700E-09 | 1.900E-08 |
| 2815 | 141 | Y141P | A | 6.5e-09 | 2.5e-09 | 6.7e-09 | 5.233E-09 | 2.369E-09 | 2.500E-09 | 6.700E-09 |
| 2816 | 141 | Y141S | A | 1.3e-08 | 9.4e-09 | 1.4e-08 | 1.213E-08 | 2.419E-09 | 9.400E-09 | 1.400E-08 |
| 2817 | 141 | Y141T | A | 3.2e-09 | 1.7e-08 | 1.4e-08 | 1.140E-08 | 7.258E-09 | 3.200E-09 | 1.700E-08 |
| 2818 | 141 | Y141W | A | 7e-09 | 9.5e-09 | 4.4e-09 | 6.967E-09 | 2.550E-09 | 4.400E-09 | 9.500E-09 |
| 2819 | 141 | Y141Y | A | 5.2e-09 | 1.2e-08 | 9.4e-09 | 8.867E-09 | 3.431E-09 | 5.200E-09 | 1.200E-08 |
| 2820 | 141 | Y141V | A | 1.1e-08 | 1e-08 | 7.6e-09 | 9.533E-09 | 1.747E-09 | 7.600E-09 | 1.100E-08 |
| 2821 | 142 | Q142A | A | 8.6e-09 | 1.2e-08 | 1.1e-08 | 1.053E-08 | 1.747E-09 | 8.600E-09 | 1.200E-08 |
| 2822 | 142 | Q142R | A | 8.5e-09 | 6.4e-09 | 1.3e-08 | 9.300E-09 | 3.372E-09 | 6.400E-09 | 1.300E-08 |
| 2823 | 142 | Q142N | A | 4.8e-09 | 1.1e-08 | 9.9e-09 | 8.567E-09 | 3.308E-09 | 4.800E-09 | 1.100E-08 |
| 2824 | 142 | Q142D | A | 5.1e-09 | 1.3e-08 | 1.5e-08 | 1.103E-08 | 5.235E-09 | 5.100E-09 | 1.500E-08 |
| 2825 | 142 | Q142C | A | 8.9e-09 | 7.1e-09 | 2.2e-08 | 1.267E-08 | 8.133E-09 | 7.100E-09 | 2.200E-08 |
| 2826 | 142 | Q142E | A | 5.9e-09 | 1.1e-08 | 3e-09 | 6.633E-09 | 4.050E-09 | 3.000E-09 | 1.100E-08 |
| 2827 | 142 | Q142Q | A | 6e-09 | 7.9e-09 | 5.2e-09 | 6.367E-09 | 1.387E-09 | 5.200E-09 | 7.900E-09 |
| 2828 | 142 | Q142G | A | 4.5e-09 | 8.8e-09 | 7.3e-09 | 6.867E-09 | 2.183E-09 | 4.500E-09 | 8.800E-09 |
| 2829 | 142 | Q142H | A | 6.8e-09 | 1.1e-08 | 1.9e-09 | 6.567E-09 | 4.554E-09 | 1.900E-09 | 1.100E-08 |
| 2830 | 142 | Q142I | A | 8.2e-09 | 3.3e-09 | 2e-08 | 1.050E-08 | 8.584E-09 | 3.300E-09 | 2.000E-08 |
| 2831 | 142 | Q142L | A | 2e-08 | 2.8e-09 | 4.2e-09 | 9.000E-09 | 9.552E-09 | 2.800E-09 | 2.000E-08 |
| 2832 | 142 | Q142K | A | 4.3e-09 | 4.8e-09 | 5.1e-09 | 4.733E-09 | 4.041E-10 | 4.300E-09 | 5.100E-09 |
| 2833 | 142 | Q142M | A | 1.6e-08 | 7.2e-09 | 1.5e-08 | 1.273E-08 | 4.818E-09 | 7.200E-09 | 1.600E-08 |
| 2834 | 142 | Q142F | A | 9.6e-09 | 4.8e-09 | 3.9e-09 | 6.100E-09 | 3.064E-09 | 3.900E-09 | 9.600E-09 |
| 2835 | 142 | Q142P | A | 8.6e-09 | 7.8e-09 | 1.7e-09 | 6.033E-09 | 3.774E-09 | 1.700E-09 | 8.600E-09 |
| 2836 | 142 | Q142S | A | 7.4e-09 | 1.8e-09 | 1.4e-09 | 3.533E-09 | 3.355E-09 | 1.400E-09 | 7.400E-09 |
| 2837 | 142 | Q142T | A | 1e-08 | 1.3e-08 | 1.1e-08 | 1.133E-08 | 1.528E-09 | 1.000E-08 | 1.300E-08 |
| 2838 | 142 | Q142W | A | 7.7e-09 | 5.4e-09 | 8.3e-09 | 7.133E-09 | 1.531E-09 | 5.400E-09 | 8.300E-09 |
| 2839 | 142 | Q142Y | A | 7.5e-09 | 6.3e-09 | 7.4e-09 | 7.067E-09 | 6.658E-10 | 6.300E-09 | 7.500E-09 |
| 2840 | 142 | Q142V | A | 7.2e-09 | 5.6e-09 | 4e-09 | 5.600E-09 | 1.600E-09 | 4.000E-09 | 7.200E-09 |
| 2841 | 143 | D143A | A | 6.8e-09 | 4.8e-09 | 6.7e-09 | 6.100E-09 | 1.127E-09 | 4.800E-09 | 6.800E-09 |
| 2842 | 143 | D143R | A | 4.9e-09 | 2.8e-09 | 1e-08 | 5.900E-09 | 3.703E-09 | 2.800E-09 | 1.000E-08 |
| 2843 | 143 | D143N | A | 8.9e-09 | 5.5e-09 | 9.8e-09 | 8.067E-09 | 2.268E-09 | 5.500E-09 | 9.800E-09 |
| 2844 | 143 | D143D | A | 1.3e-08 | 6.2e-09 | 9.7e-09 | 9.633E-09 | 3.400E-09 | 6.200E-09 | 1.300E-08 |
| 2845 | 143 | D143C | A | 1.1e-08 | 5e-09 | 1.6e-08 | 1.067E-08 | 5.508E-09 | 5.000E-09 | 1.600E-08 |
| 2846 | 143 | D143E | A | 3.3e-09 | 3.4e-09 | 7.4e-09 | 4.700E-09 | 2.339E-09 | 3.300E-09 | 7.400E-09 |
| 2847 | 143 | D143Q | A | 4.7e-09 | 6.1e-09 | 4.4e-09 | 5.067E-09 | 9.074E-10 | 4.400E-09 | 6.100E-09 |
| 2848 | 143 | D143G | A | 1.1e-08 | 3.6e-09 | 1.3e-08 | 9.200E-09 | 4.952E-09 | 3.600E-09 | 1.300E-08 |
| 2849 | 143 | D143H | A | 1.4e-09 | 1.3e-08 | 7.2e-09 | 7.200E-09 | 5.800E-09 | 1.400E-09 | 1.300E-08 |
| 2850 | 143 | D143I | A | 6.6e-09 | 1.9e-08 | 7.2e-09 | 1.093E-08 | 6.992E-09 | 6.600E-09 | 1.900E-08 |
| 2851 | 143 | D143L | A | 5.9e-09 | 5.5e-09 | 2.1e-08 | 1.080E-08 | 8.836E-09 | 5.500E-09 | 2.100E-08 |
| 2852 | 143 | D143K | A | 1e-08 | 1e-08 | 5.9e-09 | 8.633E-09 | 2.367E-09 | 5.900E-09 | 1.000E-08 |
| 2853 | 143 | D143M | A | 4.9e-09 | 1.1e-08 | 6.9e-09 | 7.600E-09 | 3.110E-09 | 4.900E-09 | 1.100E-08 |
| 2854 | 143 | D143F | A | 5.1e-09 | 7.4e-09 | 7.6e-09 | 6.700E-09 | 1.389E-09 | 5.100E-09 | 7.600E-09 |
| 2855 | 143 | D143P | A | 8.7e-09 | 7.4e-09 | 1.3e-08 | 9.700E-09 | 2.931E-09 | 7.400E-09 | 1.300E-08 |
| 2856 | 143 | D143S | A | 1.8e-08 | 6.1e-09 | 7.6e-09 | 1.057E-08 | 6.481E-09 | 6.100E-09 | 1.800E-08 |
| 2857 | 143 | D143T | A | 1.9e-09 | 7.5e-09 | 5.3e-09 | 4.900E-09 | 2.821E-09 | 1.900E-09 | 7.500E-09 |
| 2858 | 143 | D143W | A | 3.3e-09 | 3.3e-09 | 4.4e-09 | 3.667E-09 | 6.351E-10 | 3.300E-09 | 4.400E-09 |
| 2859 | 143 | D143Y | A | 6.3e-09 | 1.1e-08 | 1.1e-08 | 9.433E-09 | 2.714E-09 | 6.300E-09 | 1.100E-08 |
| 2860 | 143 | D143V | A | 4.6e-09 | 7.7e-09 | 6.8e-09 | 6.367E-09 | 1.595E-09 | 4.600E-09 | 7.700E-09 |
| 2861 | 144 | T144A | A | 1.1e-08 | 2.1e-09 | 1.2e-08 | 8.367E-09 | 5.450E-09 | 2.100E-09 | 1.200E-08 |
| 2862 | 144 | T144R | A | 8.1e-09 | 4.5e-09 | 1.1e-08 | 7.867E-09 | 3.256E-09 | 4.500E-09 | 1.100E-08 |
| 2863 | 144 | T144N | A | 1.1e-08 | 5.3e-09 | 1.1e-08 | 9.100E-09 | 3.291E-09 | 5.300E-09 | 1.100E-08 |
| 2864 | 144 | T144D | A | 5.4e-09 | 8.8e-09 | 9e-09 | 7.733E-09 | 2.023E-09 | 5.400E-09 | 9.000E-09 |
| 2865 | 144 | T144C | A | 3.8e-09 | 3.8e-09 | 1.4e-08 | 7.200E-09 | 5.889E-09 | 3.800E-09 | 1.400E-08 |
| 2866 | 144 | T144E | A | 4.6e-09 | 4.7e-09 | 1.7e-08 | 8.767E-09 | 7.130E-09 | 4.600E-09 | 1.700E-08 |
| 2867 | 144 | T144Q | A | 1.7e-08 | 6.5e-09 | 3.9e-09 | 9.133E-09 | 6.936E-09 | 3.900E-09 | 1.700E-08 |
| 2868 | 144 | T144G | A | 5e-09 | 8.4e-09 | 5.6e-09 | 6.333E-09 | 1.815E-09 | 5.000E-09 | 8.400E-09 |
| 2869 | 144 | T144H | A | 8e-09 | 1.6e-08 | 8.7e-09 | 1.090E-08 | 4.431E-09 | 8.000E-09 | 1.600E-08 |
| 2870 | 144 | T144I | A | 5.8e-09 | 3.9e-09 | 5.8e-09 | 5.167E-09 | 1.097E-09 | 3.900E-09 | 5.800E-09 |
| 2871 | 144 | T144L | A | 8.6e-09 | 1.5e-08 | 1e-08 | 1.120E-08 | 3.365E-09 | 8.600E-09 | 1.500E-08 |
| 2872 | 144 | T144K | A | 1.1e-08 | 3.5e-09 | 2.3e-09 | 5.600E-09 | 4.715E-09 | 2.300E-09 | 1.100E-08 |
| 2873 | 144 | T144M | A | 1.1e-08 | 1e-08 | 3e-09 | 8.000E-09 | 4.359E-09 | 3.000E-09 | 1.100E-08 |
| 2874 | 144 | T144F | A | 5.7e-09 | 1.5e-09 | 2.2e-09 | 3.133E-09 | 2.250E-09 | 1.500E-09 | 5.700E-09 |
| 2875 | 144 | T144P | A | 4.4e-09 | 8.7e-09 | 1.4e-08 | 9.033E-09 | 4.809E-09 | 4.400E-09 | 1.400E-08 |
| 2876 | 144 | T144S | A | 9.8e-09 | 7.7e-09 | 4.9e-09 | 7.467E-09 | 2.458E-09 | 4.900E-09 | 9.800E-09 |
| 2877 | 144 | T144T | A | 5.2e-09 | 3.7e-09 | 7.1e-09 | 5.333E-09 | 1.704E-09 | 3.700E-09 | 7.100E-09 |
| 2878 | 144 | T144W | A | 5.1e-09 | 1e-08 | 1.7e-08 | 1.070E-08 | 5.981E-09 | 5.100E-09 | 1.700E-08 |
| 2879 | 144 | T144Y | A | 2.3e-09 | 1.5e-08 | 7.8e-09 | 8.367E-09 | 6.369E-09 | 2.300E-09 | 1.500E-08 |
| 2880 | 144 | T144V | A | 4.7e-09 | 5.8e-09 | 8.4e-09 | 6.300E-09 | 1.900E-09 | 4.700E-09 | 8.400E-09 |
| 2881 | 145 | I145A | A | 6.1e-09 | 1.1e-08 | 6e-09 | 7.700E-09 | 2.858E-09 | 6.000E-09 | 1.100E-08 |
| 2882 | 145 | I145R | A | 2.8e-09 | 5.9e-09 | 5.8e-09 | 4.833E-09 | 1.762E-09 | 2.800E-09 | 5.900E-09 |
| 2883 | 145 | I145N | A | 7.6e-09 | 1.6e-08 | 2.5e-09 | 8.700E-09 | 6.817E-09 | 2.500E-09 | 1.600E-08 |
| 2884 | 145 | I145D | A | 8.7e-09 | 3e-09 | 1.8e-08 | 9.900E-09 | 7.572E-09 | 3.000E-09 | 1.800E-08 |
| 2885 | 145 | I145C | A | 3.6e-08 | 1e-08 | 1.1e-08 | 1.900E-08 | 1.473E-08 | 1.000E-08 | 3.600E-08 |
| 2886 | 145 | I145E | A | 6.5e-09 | 1.2e-08 | 6.1e-09 | 8.200E-09 | 3.297E-09 | 6.100E-09 | 1.200E-08 |
| 2887 | 145 | I145Q | A | 1.1e-08 | 1.6e-09 | 7.8e-09 | 6.800E-09 | 4.779E-09 | 1.600E-09 | 1.100E-08 |
| 2888 | 145 | I145G | A | 1.4e-08 | 3.8e-09 | 1.4e-08 | 1.060E-08 | 5.889E-09 | 3.800E-09 | 1.400E-08 |
| 2889 | 145 | I145H | A | 6.6e-09 | 3.5e-09 | 3.8e-09 | 4.633E-09 | 1.710E-09 | 3.500E-09 | 6.600E-09 |

| | | | | | | | | | | |
|---|---|---|---|---|---|---|---|---|---|---|
| 2890 | 145 | I145I | A | 3.9e-09 | 1.6e-08 | 7.4e-09 | 9.100E-09 | 6.227E-09 | 3.900E-09 | 1.600E-08 |
| 2891 | 145 | I145L | A | 1.5e-08 | 8.9e-09 | 1.5e-08 | 1.297E-08 | 3.522E-09 | 8.900E-09 | 1.500E-08 |
| 2892 | 145 | I145K | A | 1.4e-08 | 1.2e-08 | 6.9e-09 | 1.097E-08 | 3.661E-09 | 6.900E-09 | 1.400E-08 |
| 2893 | 145 | I145M | A | 7.6e-09 | 9.7e-09 | 5.4e-09 | 7.567E-09 | 2.150E-09 | 5.400E-09 | 9.700E-09 |
| 2894 | 145 | I145F | A | 6.4e-09 | 1.3e-08 | 1.3e-08 | 1.080E-08 | 3.811E-09 | 6.400E-09 | 1.300E-08 |
| 2895 | 145 | I145P | A | 4.1e-09 | 2.1e-09 | 8.6e-09 | 4.933E-09 | 3.329E-09 | 2.100E-09 | 8.600E-09 |
| 2896 | 145 | I145S | A | 7.3e-09 | 1.7e-08 | 1.2e-08 | 1.210E-08 | 4.851E-09 | 7.300E-09 | 1.700E-08 |
| 2897 | 145 | I145T | A | 5.3e-09 | 3e-09 | 8.8e-09 | 5.700E-09 | 2.921E-09 | 3.000E-09 | 8.800E-09 |
| 2898 | 145 | I145W | A | 5.3e-09 | 6.7e-09 | 1.2e-08 | 8.000E-09 | 3.534E-09 | 5.300E-09 | 1.200E-08 |
| 2899 | 145 | I145Y | A | 1.1e-08 | 8.2e-09 | 9.2e-09 | 9.467E-09 | 1.419E-09 | 8.200E-09 | 1.100E-08 |
| 2900 | 145 | I145V | A | 7.4e-09 | 7.4e-09 | 6.8e-09 | 7.200E-09 | 3.464E-10 | 6.800E-09 | 7.400E-09 |
| 2901 | 146 | L146A | A | 5.6e-09 | 3.6e-08 | 4.3e-09 | 1.530E-08 | 1.794E-08 | 4.300E-09 | 3.600E-08 |
| 2902 | 146 | L146R | A | 3.5e-09 | 6.4e-09 | 1.7e-08 | 8.967E-09 | 7.107E-09 | 3.500E-09 | 1.700E-08 |
| 2903 | 146 | L146N | A | 8.3e-09 | 3.7e-09 | 1.4e-08 | 8.667E-09 | 5.160E-09 | 3.700E-09 | 1.400E-08 |
| 2904 | 146 | L146D | A | 5.9e-09 | 9.5e-09 | 8.6e-09 | 8.000E-09 | 1.873E-09 | 5.900E-09 | 9.500E-09 |
| 2905 | 146 | L146C | A | 3.1e-09 | 1e-08 | 2.4e-09 | 5.167E-09 | 4.200E-09 | 2.400E-09 | 1.000E-08 |
| 2906 | 146 | L146E | A | 2.2e-09 | 1.4e-08 | 7.9e-09 | 8.033E-09 | 5.901E-09 | 2.200E-09 | 1.400E-08 |
| 2907 | 146 | L146Q | A | 3e-09 | 5.1e-09 | 1.6e-08 | 8.033E-09 | 6.979E-09 | 3.000E-09 | 1.600E-08 |
| 2908 | 146 | L146G | A | 4.8e-09 | 7.8e-09 | 1e-08 | 7.533E-09 | 2.610E-09 | 4.800E-09 | 1.000E-08 |
| 2909 | 146 | L146H | A | 1e-08 | 6.9e-09 | 7.9e-09 | 8.267E-09 | 1.582E-09 | 6.900E-09 | 1.000E-08 |
| 2910 | 146 | L146I | A | 1e-08 | 1.4e-08 | 2.4e-09 | 8.800E-09 | 5.892E-09 | 2.400E-09 | 1.400E-08 |
| 2911 | 146 | L146L | A | 7.3e-09 | 4e-09 | 1.1e-08 | 7.433E-09 | 3.502E-09 | 4.000E-09 | 1.100E-08 |
| 2912 | 146 | L146K | A | 7.7e-09 | 1.6e-08 | 4.4e-09 | 9.367E-09 | 5.977E-09 | 4.400E-09 | 1.600E-08 |
| 2913 | 146 | L146M | A | 1.2e-08 | 1.5e-08 | 7e-09 | 1.133E-08 | 4.041E-09 | 7.000E-09 | 1.500E-08 |
| 2914 | 146 | L146F | A | 1e-08 | 1.1e-08 | 1.6e-08 | 1.233E-08 | 3.215E-09 | 1.000E-08 | 1.600E-08 |
| 2915 | 146 | L146P | A | 4.4e-09 | 3.5e-08 | 2.2e-08 | 2.047E-08 | 1.536E-08 | 4.400E-09 | 3.500E-08 |
| 2916 | 146 | L146S | A | 1.3e-08 | 4.9e-09 | 5.9e-09 | 7.933E-09 | 4.416E-09 | 4.900E-09 | 1.300E-08 |
| 2917 | 146 | L146T | A | 5.4e-09 | 1.1e-08 | 2.8e-09 | 6.400E-09 | 4.190E-09 | 2.800E-09 | 1.100E-08 |
| 2918 | 146 | L146W | A | 1.1e-08 | 1.7e-08 | 1.5e-08 | 1.433E-08 | 3.055E-09 | 1.100E-08 | 1.700E-08 |
| 2919 | 146 | L146Y | A | 1.1e-08 | 5.3e-09 | 5.1e-09 | 7.133E-09 | 3.350E-09 | 5.100E-09 | 1.100E-08 |
| 2920 | 146 | L146V | A | 1.6e-08 | 4.4e-09 | 1.5e-08 | 1.180E-08 | 6.428E-09 | 4.400E-09 | 1.600E-08 |
| 2921 | 147 | W147A | A | 8e-09 | 3.2e-09 | 1.3e-08 | 8.067E-09 | 4.900E-09 | 3.200E-09 | 1.300E-08 |
| 2922 | 147 | W147R | A | 8.9e-09 | 1.7e-08 | 5.7e-09 | 1.053E-08 | 5.824E-09 | 5.700E-09 | 1.700E-08 |
| 2923 | 147 | W147N | A | 8.3e-09 | 4.7e-09 | 1.2e-08 | 8.333E-09 | 3.650E-09 | 4.700E-09 | 1.200E-08 |
| 2924 | 147 | W147D | A | 1.1e-09 | 9.8e-09 | 9.7e-09 | 6.867E-09 | 4.994E-09 | 1.100E-09 | 9.800E-09 |
| 2925 | 147 | W147C | A | 1e-08 | 1.8e-08 | 2.3e-08 | 1.700E-08 | 6.557E-09 | 1.000E-08 | 2.300E-08 |
| 2926 | 147 | W147E | A | 7.3e-09 | 8.6e-09 | 8e-09 | 7.967E-09 | 6.506E-10 | 7.300E-09 | 8.600E-09 |
| 2927 | 147 | W147Q | A | 8.9e-09 | 6e-09 | 3.9e-09 | 6.267E-09 | 2.511E-09 | 3.900E-09 | 8.900E-09 |
| 2928 | 147 | W147G | A | 3.7e-09 | 1.5e-08 | 2.8e-09 | 7.167E-09 | 6.799E-09 | 2.800E-09 | 1.500E-08 |
| 2929 | 147 | W147H | A | 7.5e-09 | 1.3e-08 | 7e-09 | 9.167E-09 | 3.329E-09 | 7.000E-09 | 1.300E-08 |
| 2930 | 147 | W147I | A | 6.4e-09 | 1.1e-08 | 9.5e-09 | 8.967E-09 | 2.346E-09 | 6.400E-09 | 1.100E-08 |
| 2931 | 147 | W147L | A | 1.3e-08 | 5.7e-09 | 5.2e-09 | 7.967E-09 | 4.366E-09 | 5.200E-09 | 1.300E-08 |
| 2932 | 147 | W147K | A | 7.8e-09 | 6.2e-09 | 4e-09 | 6.000E-09 | 1.908E-09 | 4.000E-09 | 7.800E-09 |
| 2933 | 147 | W147M | A | 2.1e-08 | 9.8e-09 | 2.2e-09 | 1.100E-08 | 9.457E-09 | 2.200E-09 | 2.100E-08 |
| 2934 | 147 | W147F | A | 1e-08 | 5.1e-09 | 6.4e-09 | 7.167E-09 | 2.538E-09 | 5.100E-09 | 1.000E-08 |
| 2935 | 147 | W147P | A | 6.9e-09 | 7.3e-09 | 2.3e-09 | 5.500E-09 | 2.778E-09 | 2.300E-09 | 7.300E-09 |
| 2936 | 147 | W147S | A | 3.1e-08 | 4.7e-09 | 7.1e-09 | 1.427E-08 | 1.454E-08 | 4.700E-09 | 3.100E-08 |
| 2937 | 147 | W147T | A | 6.7e-09 | 4.2e-09 | 5.2e-09 | 5.367E-09 | 1.258E-09 | 4.200E-09 | 6.700E-09 |
| 2938 | 147 | W147W | A | 5e-09 | 8e-09 | 9.4e-09 | 7.467E-09 | 2.248E-09 | 5.000E-09 | 9.400E-09 |
| 2939 | 147 | W147Y | A | 1.9e-08 | 5.4e-09 | 5.5e-09 | 9.967E-09 | 7.823E-09 | 5.400E-09 | 1.900E-08 |
| 2940 | 147 | W147V | A | 3.3e-09 | 4.9e-09 | 6.7e-09 | 4.967E-09 | 1.701E-09 | 3.300E-09 | 6.700E-09 |
| 2941 | 148 | K148A | A | 7.2e-09 | 7.2e-09 | 1.9e-08 | 1.113E-08 | 6.813E-09 | 7.200E-09 | 1.900E-08 |
| 2942 | 148 | K148R | A | 6.1e-09 | 1.2e-08 | 6.6e-09 | 8.233E-09 | 3.272E-09 | 6.100E-09 | 1.200E-08 |
| 2943 | 148 | K148N | A | 1.6e-08 | 4e-09 | 4.9e-09 | 8.300E-09 | 6.684E-09 | 4.000E-09 | 1.600E-08 |
| 2944 | 148 | K148D | A | 6.7e-09 | 4.4e-09 | 3e-09 | 4.700E-09 | 1.868E-09 | 3.000E-09 | 6.700E-09 |
| 2945 | 148 | K148C | A | 1e-08 | 1.8e-08 | 6.9e-09 | 1.163E-08 | 5.727E-09 | 6.900E-09 | 1.800E-08 |
| 2946 | 148 | K148E | A | 3.6e-09 | 1.1e-08 | 1.4e-08 | 9.533E-09 | 5.353E-09 | 3.600E-09 | 1.400E-08 |
| 2947 | 148 | K148Q | A | 5.8e-09 | 1e-08 | 6.1e-09 | 7.300E-09 | 2.343E-09 | 5.800E-09 | 1.000E-08 |
| 2948 | 148 | K148G | A | 1.9e-09 | 1e-08 | 1.3e-08 | 8.300E-09 | 5.742E-09 | 1.900E-09 | 1.300E-08 |
| 2949 | 148 | K148H | A | 7.4e-09 | 1.7e-08 | 1.1e-08 | 1.180E-08 | 4.850E-09 | 7.400E-09 | 1.700E-08 |
| 2950 | 148 | K148I | A | 8.9e-09 | 3.5e-09 | 4e-09 | 5.467E-09 | 2.984E-09 | 3.500E-09 | 8.900E-09 |
| 2951 | 148 | K148L | A | 2.5e-09 | 1.2e-08 | 3.5e-09 | 6.000E-09 | 5.220E-09 | 2.500E-09 | 1.200E-08 |
| 2952 | 148 | K148K | A | 7.2e-09 | 5e-09 | 3.3e-09 | 5.167E-09 | 1.955E-09 | 3.300E-09 | 7.200E-09 |
| 2953 | 148 | K148M | A | 2.1e-08 | 7.3e-09 | 8e-09 | 1.210E-08 | 7.716E-09 | 7.300E-09 | 2.100E-08 |
| 2954 | 148 | K148F | A | 1.3e-08 | 2.6e-08 | 1.8e-09 | 1.360E-08 | 1.211E-08 | 1.800E-09 | 2.600E-08 |
| 2955 | 148 | K148P | A | 5e-09 | 5.5e-09 | 4.2e-09 | 4.900E-09 | 6.557E-10 | 4.200E-09 | 5.500E-09 |
| 2956 | 148 | K148S | A | 1.1e-08 | 8.2e-09 | 4.9e-09 | 8.033E-09 | 3.053E-09 | 4.900E-09 | 1.100E-08 |
| 2957 | 148 | K148T | A | 5.6e-09 | 6.6e-09 | 3.1e-09 | 5.100E-09 | 1.803E-09 | 3.100E-09 | 6.600E-09 |
| 2958 | 148 | K148W | A | 5.1e-09 | 1.3e-08 | 1e-08 | 9.367E-09 | 3.988E-09 | 5.100E-09 | 1.300E-08 |
| 2959 | 148 | K148Y | A | 2.5e-09 | 4.4e-09 | 2.4e-09 | 3.100E-09 | 1.127E-09 | 2.400E-09 | 4.400E-09 |
| 2960 | 148 | K148V | A | 7.9e-09 | 1e-08 | 2.7e-08 | 1.497E-08 | 1.047E-08 | 7.900E-09 | 2.700E-08 |
| 2961 | 149 | D149A | A | 2.1e-08 | 4.8e-09 | 6.3e-09 | 1.070E-08 | 8.952E-09 | 4.800E-09 | 2.100E-08 |
| 2962 | 149 | D149R | A | 3.3e-09 | 1.8e-09 | 6.4e-09 | 3.833E-09 | 2.346E-09 | 1.800E-09 | 6.400E-09 |
| 2963 | 149 | D149N | A | 3.5e-09 | 4.1e-09 | 1.3e-08 | 6.867E-09 | 5.320E-09 | 3.500E-09 | 1.300E-08 |
| 2964 | 149 | D149D | A | 3.9e-09 | 7.3e-10 | 1.2e-08 | 5.543E-09 | 5.812E-09 | 7.300E-10 | 1.200E-08 |
| 2965 | 149 | D149C | A | 5.4e-09 | 7.7e-09 | 4.8e-09 | 5.967E-09 | 1.531E-09 | 4.800E-09 | 7.700E-09 |
| 2966 | 149 | D149E | A | 8.2e-09 | 7e-09 | 8.7e-09 | 7.967E-09 | 8.737E-10 | 7.000E-09 | 8.700E-09 |
| 2967 | 149 | D149Q | A | 3.8e-09 | 7.1e-09 | 1.2e-08 | 7.633E-09 | 4.126E-09 | 3.800E-09 | 1.200E-08 |
| 2968 | 149 | D149G | A | 6.5e-09 | 5.4e-09 | 1.6e-08 | 9.300E-09 | 5.828E-09 | 5.400E-09 | 1.600E-08 |
| 2969 | 149 | D149H | A | 5.2e-09 | 7.1e-09 | 1.7e-08 | 9.767E-09 | 6.336E-09 | 5.200E-09 | 1.700E-08 |
| 2970 | 149 | D149I | A | 8.7e-09 | 9.2e-09 | 2e-08 | 1.263E-08 | 6.385E-09 | 8.700E-09 | 2.000E-08 |
| 2971 | 149 | D149L | A | 5e-09 | 1.3e-08 | 1.2e-08 | 1.000E-08 | 4.359E-09 | 5.000E-09 | 1.300E-08 |
| 2972 | 149 | D149K | A | 4e-09 | 4.4e-09 | 9.1e-09 | 5.833E-09 | 2.836E-09 | 4.000E-09 | 9.100E-09 |
| 2973 | 149 | D149M | A | 3.3e-09 | 1.2e-09 | 1.3e-08 | 5.833E-09 | 6.295E-09 | 1.200E-09 | 1.300E-08 |
| 2974 | 149 | D149F | A | 1.3e-08 | 9.3e-09 | 2.9e-09 | 8.400E-09 | 5.110E-09 | 2.900E-09 | 1.300E-08 |

| | | | | | | | | | | |
|---|---|---|---|---|---|---|---|---|---|---|
| 2975 | 149 | D149P | A | 4.1e-09 | 1.2e-08 | 5.8e-09 | 7.300E-09 | 4.158E-09 | 4.100E-09 | 1.200E-08 |
| 2976 | 149 | D149S | A | 1.6e-08 | 2.3e-09 | 8.4e-09 | 8.900E-09 | 6.864E-09 | 2.300E-09 | 1.600E-08 |
| 2977 | 149 | D149T | A | 5e-09 | 1.6e-08 | 4e-09 | 8.333E-09 | 6.658E-09 | 4.000E-09 | 1.600E-08 |
| 2978 | 149 | D149W | A | 3.6e-09 | 5.7e-09 | 1.1e-08 | 6.767E-09 | 3.814E-09 | 3.600E-09 | 1.100E-08 |
| 2979 | 149 | D149Y | A | 5e-09 | 9.9e-09 | 1.3e-08 | 9.300E-09 | 4.034E-09 | 5.000E-09 | 1.300E-08 |
| 2980 | 149 | D149V | A | 1.1e-08 | 3e-09 | 7.6e-09 | 7.200E-09 | 4.015E-09 | 3.000E-09 | 1.100E-08 |
| 2981 | 150 | I150A | A | 4.6e-09 | 7.5e-09 | 2.5e-09 | 4.867E-09 | 2.511E-09 | 2.500E-09 | 7.500E-09 |
| 2982 | 150 | I150R | A | 8.5e-09 | 4.5e-09 | 2e-08 | 1.100E-08 | 8.047E-09 | 4.500E-09 | 2.000E-08 |
| 2983 | 150 | I150N | A | 1.5e-08 | 7e-09 | 1.2e-08 | 1.133E-08 | 4.041E-09 | 7.000E-09 | 1.500E-08 |
| 2984 | 150 | I150D | A | 1.6e-08 | 4.7e-09 | 4e-09 | 8.233E-09 | 6.735E-09 | 4.000E-09 | 1.600E-08 |
| 2985 | 150 | I150C | A | 2.9e-09 | 1.4e-08 | 1.7e-08 | 1.130E-08 | 7.428E-09 | 2.900E-09 | 1.700E-08 |
| 2986 | 150 | I150E | A | 4.7e-09 | 4.7e-09 | 6.3e-09 | 5.233E-09 | 9.238E-10 | 4.700E-09 | 6.300E-09 |
| 2987 | 150 | I150Q | A | 6.4e-09 | 9.2e-09 | 3.4e-09 | 6.333E-09 | 2.901E-09 | 3.400E-09 | 9.200E-09 |
| 2988 | 150 | I150G | A | 1.2e-08 | 1e-08 | 3e-08 | 1.733E-08 | 1.102E-08 | 1.000E-08 | 3.000E-08 |
| 2989 | 150 | I150H | A | 2.2e-08 | 3.8e-09 | 4.3e-09 | 1.003E-08 | 1.037E-08 | 3.800E-09 | 2.200E-08 |
| 2990 | 150 | I150I | A | 6.1e-09 | 6.4e-09 | 1.7e-09 | 4.733E-09 | 2.631E-09 | 1.700E-09 | 6.400E-09 |
| 2991 | 150 | I150L | A | 6.4e-09 | 3.1e-09 | 1.1e-08 | 6.833E-09 | 3.968E-09 | 3.100E-09 | 1.100E-08 |
| 2992 | 150 | I150K | A | 4.2e-09 | 2.7e-09 | 2.2e-08 | 9.633E-09 | 1.074E-08 | 2.700E-09 | 2.200E-08 |
| 2993 | 150 | I150M | A | 5.2e-09 | 1.1e-08 | 1.3e-08 | 9.733E-09 | 4.051E-09 | 5.200E-09 | 1.300E-08 |
| 2994 | 150 | I150F | A | 8.8e-09 | 3.2e-09 | 1.6e-08 | 9.333E-09 | 6.417E-09 | 3.200E-09 | 1.600E-08 |
| 2995 | 150 | I150P | A | 8.4e-09 | 6.1e-09 | 1e-08 | 8.167E-09 | 1.960E-09 | 6.100E-09 | 1.000E-08 |
| 2996 | 150 | I150S | A | 6.3e-09 | 8.3e-09 | 6.9e-09 | 7.167E-09 | 1.026E-09 | 6.300E-09 | 8.300E-09 |
| 2997 | 150 | I150T | A | 1e-08 | 8.7e-09 | 6.7e-09 | 8.467E-09 | 1.662E-09 | 6.700E-09 | 1.000E-08 |
| 2998 | 150 | I150W | A | 5.6e-09 | 3.9e-09 | 5.5e-09 | 5.000E-09 | 9.539E-10 | 3.900E-09 | 5.600E-09 |
| 2999 | 150 | I150Y | A | 4e-09 | 6.1e-09 | 5.9e-09 | 5.333E-09 | 1.159E-09 | 4.000E-09 | 6.100E-09 |
| 3000 | 150 | I150V | A | 1.2e-08 | 8.1e-09 | 1.2e-08 | 1.070E-08 | 2.252E-09 | 8.100E-09 | 1.200E-08 |
| 3001 | 151 | F151A | A | 7e-09 | 4.5e-09 | 4.5e-09 | 5.333E-09 | 1.443E-09 | 4.500E-09 | 7.000E-09 |
| 3002 | 151 | F151R | A | 4.4e-09 | 8.6e-09 | 1.7e-09 | 4.900E-09 | 3.477E-09 | 1.700E-09 | 8.600E-09 |
| 3003 | 151 | F151N | A | 7.4e-09 | 3.5e-09 | 5.9e-09 | 5.600E-09 | 1.967E-09 | 3.500E-09 | 7.400E-09 |
| 3004 | 151 | F151D | A | 4.5e-09 | 1.3e-08 | 7.2e-09 | 8.233E-09 | 4.343E-09 | 4.500E-09 | 1.300E-08 |
| 3005 | 151 | F151C | A | 1.6e-08 | 1.2e-08 | 1.8e-08 | 1.533E-08 | 3.055E-09 | 1.200E-08 | 1.800E-08 |
| 3006 | 151 | F151E | A | 1.2e-08 | 7.3e-09 | 4.2e-09 | 7.833E-09 | 3.927E-09 | 4.200E-09 | 1.200E-08 |
| 3007 | 151 | F151Q | A | 5.3e-09 | 1.9e-09 | 4.5e-09 | 3.900E-09 | 1.778E-09 | 1.900E-09 | 5.300E-09 |
| 3008 | 151 | F151G | A | 6.7e-09 | 3.3e-09 | 9.1e-09 | 6.367E-09 | 2.914E-09 | 3.300E-09 | 9.100E-09 |
| 3009 | 151 | F151H | A | 3.1e-08 | 7.3e-09 | 6.4e-09 | 1.490E-08 | 1.395E-08 | 6.400E-09 | 3.100E-08 |
| 3010 | 151 | F151I | A | 5.9e-09 | 9e-09 | 1.5e-08 | 9.967E-09 | 4.626E-09 | 5.900E-09 | 1.500E-08 |
| 3011 | 151 | F151L | A | 6.8e-09 | 4.9e-09 | 8.3e-09 | 6.667E-09 | 1.704E-09 | 4.900E-09 | 8.300E-09 |
| 3012 | 151 | F151K | A | 5e-09 | 1.6e-08 | 3.8e-09 | 8.267E-09 | 6.724E-09 | 3.800E-09 | 1.600E-08 |
| 3013 | 151 | F151M | A | 1.2e-08 | 1.3e-08 | 4.9e-09 | 9.967E-09 | 4.416E-09 | 4.900E-09 | 1.300E-08 |
| 3014 | 151 | F151F | A | 3.5e-09 | 5e-09 | 9.1e-09 | 5.867E-09 | 2.899E-09 | 3.500E-09 | 9.100E-09 |
| 3015 | 151 | F151P | A | 2.3e-08 | 7.1e-09 | 3.6e-09 | 1.123E-08 | 1.034E-08 | 3.600E-09 | 2.300E-08 |
| 3016 | 151 | F151S | A | 7.3e-09 | 9e-09 | 6e-09 | 7.433E-09 | 1.504E-09 | 6.000E-09 | 9.000E-09 |
| 3017 | 151 | F151T | A | 3.7e-09 | 6.6e-09 | 7.2e-09 | 5.833E-09 | 1.872E-09 | 3.700E-09 | 7.200E-09 |
| 3018 | 151 | F151W | A | 5.3e-09 | 1.5e-08 | 2.1e-08 | 1.377E-08 | 7.922E-09 | 5.300E-09 | 2.100E-08 |
| 3019 | 151 | F151Y | A | 1.9e-08 | 5.5e-09 | 8.1e-09 | 1.087E-08 | 7.163E-09 | 5.500E-09 | 1.900E-08 |
| 3020 | 151 | F151V | A | 3.6e-09 | 1.3e-08 | 4.3e-09 | 6.967E-09 | 5.237E-09 | 3.600E-09 | 1.300E-08 |
| 3021 | 152 | H152A | A | 1.6e-08 | 1e-08 | 1.2e-08 | 1.267E-08 | 3.055E-09 | 1.000E-08 | 1.600E-08 |
| 3022 | 152 | H152R | A | 3.8e-09 | 7.5e-09 | 3.5e-09 | 4.933E-09 | 2.228E-09 | 3.500E-09 | 7.500E-09 |
| 3023 | 152 | H152N | A | 5.2e-09 | 8.8e-09 | 1.2e-08 | 8.667E-09 | 3.402E-09 | 5.200E-09 | 1.200E-08 |
| 3024 | 152 | H152D | A | 2.9e-09 | 2.8e-09 | 3.4e-09 | 3.033E-09 | 3.215E-10 | 2.800E-09 | 3.400E-09 |
| 3025 | 152 | H152C | A | 3.4e-09 | 1.5e-08 | 1.1e-08 | 9.800E-09 | 5.892E-09 | 3.400E-09 | 1.500E-08 |
| 3026 | 152 | H152E | A | 2.6e-09 | 4.5e-09 | 1.2e-08 | 6.367E-09 | 4.970E-09 | 2.600E-09 | 1.200E-08 |
| 3027 | 152 | H152Q | A | 2.8e-09 | 7.1e-09 | 1e-08 | 6.633E-09 | 3.623E-09 | 2.800E-09 | 1.000E-08 |
| 3028 | 152 | H152G | A | 6.2e-09 | 1.1e-08 | 5.4e-09 | 7.533E-09 | 3.029E-09 | 5.400E-09 | 1.100E-08 |
| 3029 | 152 | H152H | A | 5.6e-09 | 1.9e-08 | 1.1e-09 | 8.567E-09 | 9.311E-09 | 1.100E-09 | 1.900E-08 |
| 3030 | 152 | H152I | A | 4.7e-09 | 9.3e-09 | 1.2e-08 | 8.667E-09 | 3.691E-09 | 4.700E-09 | 1.200E-08 |
| 3031 | 152 | H152L | A | 7.8e-09 | 6.6e-09 | 8.2e-09 | 7.533E-09 | 8.327E-10 | 6.600E-09 | 8.200E-09 |
| 3032 | 152 | H152K | A | 2.3e-08 | 5.4e-09 | 2.3e-08 | 1.713E-08 | 1.016E-08 | 5.400E-09 | 2.300E-08 |
| 3033 | 152 | H152M | A | 2.7e-09 | 6.2e-09 | 1.5e-08 | 7.967E-09 | 6.337E-09 | 2.700E-09 | 1.500E-08 |
| 3034 | 152 | H152F | A | 1.3e-08 | 9.6e-09 | 1.3e-08 | 1.187E-08 | 1.963E-09 | 9.600E-09 | 1.300E-08 |
| 3035 | 152 | H152P | A | 6.3e-09 | 1.2e-08 | 4.9e-09 | 7.733E-09 | 3.761E-09 | 4.900E-09 | 1.200E-08 |
| 3036 | 152 | H152S | A | 4.6e-09 | 1.1e-08 | 8.6e-09 | 8.067E-09 | 3.233E-09 | 4.600E-09 | 1.100E-08 |
| 3037 | 152 | H152T | A | 1.1e-08 | 2.3e-09 | 6.4e-09 | 6.567E-09 | 4.352E-09 | 2.300E-09 | 1.100E-08 |
| 3038 | 152 | H152W | A | 1e-08 | 4.8e-09 | 9.1e-09 | 7.967E-09 | 2.779E-09 | 4.800E-09 | 1.000E-08 |
| 3039 | 152 | H152Y | A | 7.2e-09 | 2.6e-08 | 2.3e-08 | 1.873E-08 | 1.010E-08 | 7.200E-09 | 2.600E-08 |
| 3040 | 152 | H152V | A | 4.2e-09 | 4.5e-09 | 2.9e-09 | 3.867E-09 | 8.505E-10 | 2.900E-09 | 4.500E-09 |
| 3041 | 153 | K153A | A | 3.8e-09 | 3e-09 | 1.1e-08 | 5.933E-09 | 4.406E-09 | 3.000E-09 | 1.100E-08 |
| 3042 | 153 | K153R | A | 5.8e-09 | 1.4e-09 | 1.1e-08 | 6.067E-09 | 4.806E-09 | 1.400E-09 | 1.100E-08 |
| 3043 | 153 | K153N | A | 2e-09 | 5.6e-09 | 2.8e-09 | 3.467E-09 | 1.890E-09 | 2.000E-09 | 5.600E-09 |
| 3044 | 153 | K153D | A | 6e-09 | 9.5e-09 | 3.9e-09 | 6.467E-09 | 2.829E-09 | 3.900E-09 | 9.500E-09 |
| 3045 | 153 | K153C | A | 5.8e-09 | 5.2e-09 | 1.6e-08 | 9.000E-09 | 6.070E-09 | 5.200E-09 | 1.600E-08 |
| 3046 | 153 | K153E | A | 4e-09 | 1.6e-09 | 7.6e-09 | 4.400E-09 | 3.020E-09 | 1.600E-09 | 7.600E-09 |
| 3047 | 153 | K153Q | A | 2.6e-09 | 9.8e-09 | 5.1e-09 | 5.833E-09 | 3.656E-09 | 2.600E-09 | 9.800E-09 |
| 3048 | 153 | K153G | A | 1.5e-08 | 9.5e-09 | 1.2e-08 | 1.217E-08 | 2.754E-09 | 9.500E-09 | 1.500E-08 |
| 3049 | 153 | K153H | A | 4.5e-09 | 1.4e-08 | 5e-09 | 7.833E-09 | 5.346E-09 | 4.500E-09 | 1.400E-08 |
| 3050 | 153 | K153I | A | 1e-08 | 1.6e-08 | 9.6e-09 | 1.187E-08 | 3.585E-09 | 9.600E-09 | 1.600E-08 |
| 3051 | 153 | K153L | A | 3.5e-09 | 1.3e-08 | 9e-09 | 8.500E-09 | 4.770E-09 | 3.500E-09 | 1.300E-08 |
| 3052 | 153 | K153K | A | 1.6e-08 | 1.1e-08 | 9.5e-09 | 1.217E-08 | 3.403E-09 | 9.500E-09 | 1.600E-08 |
| 3053 | 153 | K153M | A | 1.1e-08 | 1e-08 | 1.4e-08 | 1.167E-08 | 2.082E-09 | 1.000E-08 | 1.400E-08 |
| 3054 | 153 | K153F | A | 7.5e-09 | 3.5e-08 | 1e-08 | 1.750E-08 | 1.521E-08 | 7.500E-09 | 3.500E-08 |
| 3055 | 153 | K153P | A | 3.5e-09 | 2.5e-09 | 4.5e-09 | 3.500E-09 | 1.000E-09 | 2.500E-09 | 4.500E-09 |
| 3056 | 153 | K153S | A | 3.5e-09 | 2.5e-08 | 6.6e-09 | 1.170E-08 | 1.162E-08 | 3.500E-09 | 2.500E-08 |
| 3057 | 153 | K153T | A | 5.7e-09 | 4e-09 | 1.5e-08 | 8.233E-09 | 5.921E-09 | 4.000E-09 | 1.500E-08 |
| 3058 | 153 | K153W | A | 6.8e-09 | 4.7e-09 | 2.6e-09 | 4.700E-09 | 2.100E-09 | 2.600E-09 | 6.800E-09 |
| 3059 | 153 | K153Y | A | 3.6e-09 | 9.1e-09 | 2.9e-09 | 5.200E-09 | 3.396E-09 | 2.900E-09 | 9.100E-09 |

| | | | | | | | | | | |
|---|---|---|---|---|---|---|---|---|---|---|
| 3060 | 153 | K153V | A | 2.8e-09 | 8.5e-09 | 6.9e-09 | 6.067E-09 | 2.940E-09 | 2.800E-09 | 8.500E-09 |
| 3061 | 154 | N154A | A | 1.5e-08 | 5.2e-09 | 2.9e-09 | 7.700E-09 | 6.426E-09 | 2.900E-09 | 1.500E-08 |
| 3062 | 154 | N154R | A | 2.5e-09 | 5.8e-09 | 5.4e-09 | 4.567E-09 | 1.801E-09 | 2.500E-09 | 5.800E-09 |
| 3063 | 154 | N154N | A | 5.2e-09 | 4.7e-09 | 9.5e-09 | 6.467E-09 | 2.639E-09 | 4.700E-09 | 9.500E-09 |
| 3064 | 154 | N154D | A | 8.7e-09 | 5.1e-09 | 2.3e-08 | 1.227E-08 | 9.468E-09 | 5.100E-09 | 2.300E-08 |
| 3065 | 154 | N154C | A | 7.6e-09 | 2e-08 | 7.6e-09 | 1.173E-08 | 7.159E-09 | 7.600E-09 | 2.000E-08 |
| 3066 | 154 | N154E | A | 1.6e-08 | 4.1e-09 | 7.3e-09 | 9.133E-09 | 6.158E-09 | 4.100E-09 | 1.600E-08 |
| 3067 | 154 | N154Q | A | 5.6e-09 | 6e-09 | 8.7e-09 | 6.767E-09 | 1.686E-09 | 5.600E-09 | 8.700E-09 |
| 3068 | 154 | N154G | A | 8.9e-09 | 6.4e-09 | 1.2e-08 | 9.100E-09 | 2.805E-09 | 6.400E-09 | 1.200E-08 |
| 3069 | 154 | N154H | A | 4.3e-09 | 8.8e-09 | 9.1e-09 | 7.400E-09 | 2.689E-09 | 4.300E-09 | 9.100E-09 |
| 3070 | 154 | N154I | A | 8.5e-09 | 1.3e-09 | 2.4e-09 | 4.067E-09 | 3.879E-09 | 1.300E-09 | 8.500E-09 |
| 3071 | 154 | N154L | A | 3.4e-09 | 5.2e-09 | 7.1e-09 | 5.233E-09 | 1.850E-09 | 3.400E-09 | 7.100E-09 |
| 3072 | 154 | N154K | A | 8.1e-09 | 1.1e-08 | 2.4e-09 | 7.167E-09 | 4.375E-09 | 2.400E-09 | 1.100E-08 |
| 3073 | 154 | N154M | A | 3.4e-09 | 4.3e-09 | 7.1e-09 | 4.933E-09 | 1.930E-09 | 3.400E-09 | 7.100E-09 |
| 3074 | 154 | N154F | A | 4.1e-09 | 2.1e-08 | 8.2e-09 | 1.110E-08 | 8.815E-09 | 4.100E-09 | 2.100E-08 |
| 3075 | 154 | N154P | A | 1.3e-08 | 1.4e-08 | 5.2e-09 | 1.073E-08 | 4.818E-09 | 5.200E-09 | 1.400E-08 |
| 3076 | 154 | N154S | A | 6.2e-09 | 2.9e-09 | 4.2e-09 | 4.433E-09 | 1.662E-09 | 2.900E-09 | 6.200E-09 |
| 3077 | 154 | N154T | A | 1.3e-08 | 4.1e-09 | 1.4e-08 | 1.037E-08 | 5.450E-09 | 4.100E-09 | 1.400E-08 |
| 3078 | 154 | N154W | A | 5.7e-09 | 8.6e-09 | 3.3e-09 | 5.867E-09 | 2.654E-09 | 3.300E-09 | 8.600E-09 |
| 3079 | 154 | N154Y | A | 4e-09 | 9.1e-09 | 4.5e-09 | 5.867E-09 | 2.811E-09 | 4.000E-09 | 9.100E-09 |
| 3080 | 154 | N154V | A | 2.1e-09 | 9.2e-09 | 2e-09 | 4.433E-09 | 4.128E-09 | 2.000E-09 | 9.200E-09 |
| 3081 | 155 | N155A | A | 4.7e-09 | 8.5e-09 | 5.5e-09 | 6.233E-09 | 2.003E-09 | 4.700E-09 | 8.500E-09 |
| 3082 | 155 | N155R | A | 9.2e-09 | 6.7e-09 | 8.9e-09 | 8.267E-09 | 1.365E-09 | 6.700E-09 | 9.200E-09 |
| 3083 | 155 | N155N | A | 2.8e-09 | 6.3e-09 | 6.3e-09 | 5.133E-09 | 2.021E-09 | 2.800E-09 | 6.300E-09 |
| 3084 | 155 | N155D | A | 1.9e-08 | 7.1e-09 | 5.8e-09 | 1.063E-08 | 7.275E-09 | 5.800E-09 | 1.900E-08 |
| 3085 | 155 | N155C | A | 1.1e-08 | 3.3e-09 | 1.8e-08 | 1.077E-08 | 7.353E-09 | 3.300E-09 | 1.800E-08 |
| 3086 | 155 | N155E | A | 7.9e-09 | 5.6e-09 | 9.9e-09 | 7.800E-09 | 2.152E-09 | 5.600E-09 | 9.900E-09 |
| 3087 | 155 | N155Q | A | 3e-09 | 7e-09 | 2.3e-08 | 1.100E-08 | 1.058E-08 | 3.000E-09 | 2.300E-08 |
| 3088 | 155 | N155G | A | 2.8e-09 | 2.3e-08 | 2.5e-09 | 9.433E-09 | 1.175E-08 | 2.500E-09 | 2.300E-08 |
| 3089 | 155 | N155H | A | 1.6e-08 | 5.2e-09 | 7.6e-09 | 9.600E-09 | 5.671E-09 | 5.200E-09 | 1.600E-08 |
| 3090 | 155 | N155I | A | 2.8e-09 | 2.6e-09 | 3e-09 | 2.800E-09 | 2.000E-10 | 2.600E-09 | 3.000E-09 |
| 3091 | 155 | N155L | A | 6.6e-09 | 1.4e-09 | 2.9e-09 | 3.633E-09 | 2.676E-09 | 1.400E-09 | 6.600E-09 |
| 3092 | 155 | N155K | A | 5.4e-09 | 5.1e-09 | 1.4e-09 | 3.967E-09 | 2.228E-09 | 1.400E-09 | 5.400E-09 |
| 3093 | 155 | N155M | A | 1.3e-08 | 9.7e-09 | 3.3e-09 | 8.667E-09 | 4.932E-09 | 3.300E-09 | 1.300E-08 |
| 3094 | 155 | N155F | A | 4.1e-09 | 2.4e-09 | 3e-09 | 3.167E-09 | 8.622E-10 | 2.400E-09 | 4.100E-09 |
| 3095 | 155 | N155P | A | 7.1e-09 | 9.3e-09 | 1e-08 | 8.800E-09 | 1.513E-09 | 7.100E-09 | 1.000E-08 |
| 3096 | 155 | N155S | A | 1e-08 | 6.8e-09 | 4.9e-09 | 7.233E-09 | 2.577E-09 | 4.900E-09 | 1.000E-08 |
| 3097 | 155 | N155T | A | 3.7e-09 | 1.2e-08 | 6.3e-09 | 7.333E-09 | 4.245E-09 | 3.700E-09 | 1.200E-08 |
| 3098 | 155 | N155W | A | 1.1e-08 | 8.7e-09 | 2.9e-09 | 7.533E-09 | 4.174E-09 | 2.900E-09 | 1.100E-08 |
| 3099 | 155 | N155Y | A | 1.5e-08 | 7.5e-09 | 4.8e-09 | 9.100E-09 | 5.285E-09 | 4.800E-09 | 1.500E-08 |
| 3100 | 155 | N155V | A | 8.8e-09 | 7.7e-09 | 2.3e-09 | 6.267E-09 | 3.479E-09 | 2.300E-09 | 8.800E-09 |
| 3101 | 156 | Q156A | A | 3.4e-09 | 4e-09 | 4.1e-09 | 3.833E-09 | 3.786E-10 | 3.400E-09 | 4.100E-09 |
| 3102 | 156 | Q156R | A | 8.5e-09 | 1.3e-08 | 3.2e-09 | 8.233E-09 | 4.905E-09 | 3.200E-09 | 1.300E-08 |
| 3103 | 156 | Q156N | A | 5.2e-09 | 3.1e-09 | 1.2e-08 | 6.767E-09 | 4.652E-09 | 3.100E-09 | 1.200E-08 |
| 3104 | 156 | Q156D | A | 1.5e-08 | 1.2e-08 | 6.7e-09 | 1.123E-08 | 4.203E-09 | 6.700E-09 | 1.500E-08 |
| 3105 | 156 | Q156C | A | 3e-09 | 5.6e-09 | 1.1e-08 | 6.533E-09 | 4.081E-09 | 3.000E-09 | 1.100E-08 |
| 3106 | 156 | Q156E | A | 3.1e-09 | 5.7e-09 | 5e-09 | 4.600E-09 | 1.345E-09 | 3.100E-09 | 5.700E-09 |
| 3107 | 156 | Q156Q | A | 4.6e-09 | 2.3e-08 | 1.3e-08 | 1.353E-08 | 9.212E-09 | 4.600E-09 | 2.300E-08 |
| 3108 | 156 | Q156G | A | 8.5e-09 | 5e-09 | 6.3e-09 | 6.600E-09 | 1.769E-09 | 5.000E-09 | 8.500E-09 |
| 3109 | 156 | Q156H | A | 7e-09 | 8.9e-09 | 3.7e-09 | 6.533E-09 | 2.631E-09 | 3.700E-09 | 8.900E-09 |
| 3110 | 156 | Q156I | A | 6.1e-09 | 8.2e-09 | 6.3e-09 | 6.867E-09 | 1.159E-09 | 6.100E-09 | 8.200E-09 |
| 3111 | 156 | Q156L | A | 4.9e-09 | 1.2e-08 | 9.5e-09 | 8.800E-09 | 3.601E-09 | 4.900E-09 | 1.200E-08 |
| 3112 | 156 | Q156K | A | 9.3e-09 | 4e-09 | 1.7e-09 | 5.000E-09 | 3.897E-09 | 1.700E-09 | 9.300E-09 |
| 3113 | 156 | Q156M | A | 4.6e-09 | 4.1e-09 | 5.3e-09 | 4.667E-09 | 6.028E-10 | 4.100E-09 | 5.300E-09 |
| 3114 | 156 | Q156F | A | 8.3e-09 | 1.8e-08 | 7.6e-09 | 1.130E-08 | 5.813E-09 | 7.600E-09 | 1.800E-08 |
| 3115 | 156 | Q156P | A | 8.6e-09 | 7.9e-09 | 8.7e-09 | 8.400E-09 | 4.359E-10 | 7.900E-09 | 8.700E-09 |
| 3116 | 156 | Q156S | A | 1.6e-08 | 1.3e-08 | 1.2e-08 | 1.367E-08 | 2.082E-09 | 1.200E-08 | 1.600E-08 |
| 3117 | 156 | Q156T | A | 3.1e-09 | 8.1e-09 | 7.8e-09 | 6.333E-09 | 2.804E-09 | 3.100E-09 | 8.100E-09 |
| 3118 | 156 | Q156W | A | 2.7e-09 | 1.3e-08 | 8.1e-09 | 7.933E-09 | 5.152E-09 | 2.700E-09 | 1.300E-08 |
| 3119 | 156 | Q156Y | A | 6.4e-09 | 5.9e-09 | 7.3e-09 | 6.533E-09 | 7.095E-10 | 5.900E-09 | 7.300E-09 |
| 3120 | 156 | Q156V | A | 2e-09 | 8.4e-09 | 2.9e-09 | 4.433E-09 | 3.465E-09 | 2.000E-09 | 8.400E-09 |
| 3121 | 157 | L157A | A | 7.7e-09 | 1.2e-08 | 7.1e-09 | 8.933E-09 | 2.673E-09 | 7.100E-09 | 1.200E-08 |
| 3122 | 157 | L157R | A | 5.9e-09 | 9.7e-09 | 7.9e-09 | 7.833E-09 | 1.901E-09 | 5.900E-09 | 9.700E-09 |
| 3123 | 157 | L157N | A | 4.8e-09 | 4.5e-09 | 5.5e-09 | 4.933E-09 | 5.132E-10 | 4.500E-09 | 5.500E-09 |
| 3124 | 157 | L157D | A | 6.1e-09 | 3.5e-09 | 1.3e-08 | 7.533E-09 | 4.910E-09 | 3.500E-09 | 1.300E-08 |
| 3125 | 157 | L157C | A | 5.6e-09 | 9.1e-09 | 5.3e-09 | 6.667E-09 | 2.113E-09 | 5.300E-09 | 9.100E-09 |
| 3126 | 157 | L157E | A | 2.7e-09 | 4.4e-09 | 2.5e-09 | 3.200E-09 | 1.044E-09 | 2.500E-09 | 4.400E-09 |
| 3127 | 157 | L157Q | A | 2.9e-09 | 7.2e-09 | 4.5e-09 | 4.867E-09 | 2.173E-09 | 2.900E-09 | 7.200E-09 |
| 3128 | 157 | L157G | A | 7.7e-09 | 7.2e-09 | 9.5e-09 | 8.133E-09 | 1.210E-09 | 7.200E-09 | 9.500E-09 |
| 3129 | 157 | L157H | A | 9.9e-09 | 3.6e-09 | 9.8e-09 | 7.767E-09 | 3.609E-09 | 3.600E-09 | 9.900E-09 |
| 3130 | 157 | L157I | A | 1.9e-08 | 8.5e-09 | 4.4e-09 | 1.063E-08 | 7.530E-09 | 4.400E-09 | 1.900E-08 |
| 3131 | 157 | L157L | A | 8.3e-09 | 7.4e-09 | 3.1e-09 | 6.267E-09 | 2.779E-09 | 3.100E-09 | 8.300E-09 |
| 3132 | 157 | L157K | A | 1.7e-08 | 2.7e-09 | 1e-08 | 9.900E-09 | 7.151E-09 | 2.700E-09 | 1.700E-08 |
| 3133 | 157 | L157M | A | 2.2e-09 | 4.3e-09 | 1.4e-08 | 6.833E-09 | 6.295E-09 | 2.200E-09 | 1.400E-08 |
| 3134 | 157 | L157F | A | 1.1e-08 | 1e-08 | 1.1e-09 | 7.367E-09 | 5.450E-09 | 1.100E-09 | 1.100E-08 |
| 3135 | 157 | L157P | A | 1e-08 | 5.4e-09 | 3.1e-09 | 6.167E-09 | 3.513E-09 | 3.100E-09 | 1.000E-08 |
| 3136 | 157 | L157S | A | 1.4e-08 | 1.6e-08 | 7.3e-09 | 1.243E-08 | 4.557E-09 | 7.300E-09 | 1.600E-08 |
| 3137 | 157 | L157T | A | 7.8e-09 | 3.8e-09 | 5.4e-09 | 5.667E-09 | 2.013E-09 | 3.800E-09 | 7.800E-09 |
| 3138 | 157 | L157W | A | 7.2e-09 | 5.8e-09 | 1.3e-08 | 8.667E-09 | 3.818E-09 | 5.800E-09 | 1.300E-08 |
| 3139 | 157 | L157Y | A | 1.9e-08 | 2.2e-08 | 3.6e-09 | 1.487E-08 | 9.872E-09 | 3.600E-09 | 2.200E-08 |
| 3140 | 157 | L157V | A | 5e-09 | 7.8e-09 | 2.4e-09 | 5.067E-09 | 2.701E-09 | 2.400E-09 | 7.800E-09 |
| 3141 | 158 | A158A | A | 9.7e-09 | 7.4e-09 | 9.3e-09 | 8.800E-09 | 1.229E-09 | 7.400E-09 | 9.700E-09 |
| 3142 | 158 | A158R | A | 1.2e-08 | 9.5e-09 | 1.4e-08 | 1.183E-08 | 2.255E-09 | 9.500E-09 | 1.400E-08 |
| 3143 | 158 | A158N | A | 3.3e-09 | 2.6e-09 | 1.1e-08 | 5.633E-09 | 4.661E-09 | 2.600E-09 | 1.100E-08 |
| 3144 | 158 | A158D | A | 4.7e-09 | 7.4e-09 | 4.7e-09 | 5.600E-09 | 1.559E-09 | 4.700E-09 | 7.400E-09 |

| | | | | | | | | | | |
|---|---|---|---|---|---|---|---|---|---|---|
| 3145 | 158 | A158C | A | 8.5e-09 | 1.6e-09 | 1.5e-08 | 8.367E-09 | 6.701E-09 | 1.600E-09 | 1.500E-08 |
| 3146 | 158 | A158E | A | 1.4e-08 | 5.7e-09 | 1.5e-08 | 1.157E-08 | 5.105E-09 | 5.700E-09 | 1.500E-08 |
| 3147 | 158 | A158Q | A | 2.9e-09 | 5e-09 | 7.2e-09 | 5.033E-09 | 2.150E-09 | 2.900E-09 | 7.200E-09 |
| 3148 | 158 | A158G | A | 2.6e-09 | 9.8e-09 | 1e-08 | 7.467E-09 | 4.216E-09 | 2.600E-09 | 1.000E-08 |
| 3149 | 158 | A158H | A | 1.5e-09 | 1.4e-08 | 8.6e-09 | 8.033E-09 | 6.269E-09 | 1.500E-09 | 1.400E-08 |
| 3150 | 158 | A158I | A | 1.7e-08 | 6.4e-09 | 1.6e-08 | 1.313E-08 | 5.853E-09 | 6.400E-09 | 1.700E-08 |
| 3151 | 158 | A158L | A | 5.5e-09 | 2.6e-09 | 8.1e-09 | 5.400E-09 | 2.751E-09 | 2.600E-09 | 8.100E-09 |
| 3152 | 158 | A158K | A | 4.5e-09 | 7.7e-09 | 3.4e-09 | 5.200E-09 | 2.234E-09 | 3.400E-09 | 7.700E-09 |
| 3153 | 158 | A158M | A | 3.6e-09 | 5.1e-09 | 1.8e-08 | 8.900E-09 | 7.916E-09 | 3.600E-09 | 1.800E-08 |
| 3154 | 158 | A158F | A | 7.1e-09 | 1.3e-08 | 7.8e-09 | 9.300E-09 | 3.223E-09 | 7.100E-09 | 1.300E-08 |
| 3155 | 158 | A158P | A | 9.5e-09 | 8.5e-09 | 8e-09 | 8.667E-09 | 7.638E-10 | 8.000E-09 | 9.500E-09 |
| 3156 | 158 | A158S | A | 9.5e-10 | 1.3e-08 | 6.3e-09 | 6.750E-09 | 6.038E-09 | 9.500E-10 | 1.300E-08 |
| 3157 | 158 | A158T | A | 8.5e-09 | 1.3e-08 | 5.3e-09 | 8.933E-09 | 3.868E-09 | 5.300E-09 | 1.300E-08 |
| 3158 | 158 | A158W | A | 2.5e-09 | 6.9e-09 | 1.7e-08 | 8.800E-09 | 7.434E-09 | 2.500E-09 | 1.700E-08 |
| 3159 | 158 | A158Y | A | 6.9e-09 | 2.7e-09 | 7.3e-09 | 5.633E-09 | 2.548E-09 | 2.700E-09 | 7.300E-09 |
| 3160 | 158 | A158V | A | 4.8e-09 | 8.1e-09 | 1.3e-08 | 8.633E-09 | 4.126E-09 | 4.800E-09 | 1.300E-08 |
| 3161 | 159 | L159A | A | 4.9e-09 | 3.6e-09 | 9.6e-09 | 6.033E-09 | 3.156E-09 | 3.600E-09 | 9.600E-09 |
| 3162 | 159 | L159R | A | 1.2e-08 | 1.3e-08 | 1.1e-08 | 1.200E-08 | 1.000E-09 | 1.100E-08 | 1.300E-08 |
| 3163 | 159 | L159N | A | 2.9e-09 | 6.6e-09 | 7.8e-09 | 5.767E-09 | 2.554E-09 | 2.900E-09 | 7.800E-09 |
| 3164 | 159 | L159D | A | 7.6e-09 | 1.5e-08 | 6.1e-09 | 9.567E-09 | 4.765E-09 | 6.100E-09 | 1.500E-08 |
| 3165 | 159 | L159C | A | 1.2e-08 | 6.8e-09 | 7.4e-09 | 8.733E-09 | 2.845E-09 | 6.800E-09 | 1.200E-08 |
| 3166 | 159 | L159E | A | 7.6e-09 | 1.6e-08 | 1.4e-08 | 1.253E-08 | 4.388E-09 | 7.600E-09 | 1.600E-08 |
| 3167 | 159 | L159Q | A | 1.4e-08 | 8.5e-09 | 1.2e-08 | 1.150E-08 | 2.784E-09 | 8.500E-09 | 1.400E-08 |
| 3168 | 159 | L159G | A | 3e-09 | 1.1e-08 | 6.5e-09 | 6.833E-09 | 4.010E-09 | 3.000E-09 | 1.100E-08 |
| 3169 | 159 | L159H | A | 1.2e-09 | 5.7e-09 | 4.8e-09 | 3.900E-09 | 2.381E-09 | 1.200E-09 | 5.700E-09 |
| 3170 | 159 | L159I | A | 2.5e-09 | 7.8e-09 | 3.9e-09 | 4.733E-09 | 2.747E-09 | 2.500E-09 | 7.800E-09 |
| 3171 | 159 | L159L | A | 1.3e-08 | 6.4e-09 | 2e-08 | 1.313E-08 | 6.801E-09 | 6.400E-09 | 2.000E-08 |
| 3172 | 159 | L159K | A | 6.5e-09 | 7.3e-09 | 7.8e-09 | 7.200E-09 | 6.557E-10 | 6.500E-09 | 7.800E-09 |
| 3173 | 159 | L159M | A | 3.8e-09 | 8.8e-09 | 4.4e-08 | 1.887E-08 | 2.191E-08 | 3.800E-09 | 4.400E-08 |
| 3174 | 159 | L159F | A | 3.4e-09 | 9.6e-09 | 3.5e-09 | 5.500E-09 | 3.551E-09 | 3.400E-09 | 9.600E-09 |
| 3175 | 159 | L159P | A | 4.6e-09 | 1.1e-08 | 3.9e-09 | 6.500E-09 | 3.913E-09 | 3.900E-09 | 1.100E-08 |
| 3176 | 159 | L159S | A | 4.1e-09 | 1.3e-08 | 1e-08 | 9.033E-09 | 4.528E-09 | 4.100E-09 | 1.300E-08 |
| 3177 | 159 | L159T | A | 1.3e-08 | 4.3e-09 | 1.3e-08 | 1.010E-08 | 5.023E-09 | 4.300E-09 | 1.300E-08 |
| 3178 | 159 | L159W | A | 7.6e-09 | 1.2e-08 | 1.5e-08 | 1.153E-08 | 3.722E-09 | 7.600E-09 | 1.500E-08 |
| 3179 | 159 | L159Y | A | 1.8e-08 | 2.8e-09 | 7.2e-09 | 9.333E-09 | 7.821E-09 | 2.800E-09 | 1.800E-08 |
| 3180 | 159 | L159V | A | 1.3e-08 | 3.6e-09 | 1.1e-08 | 9.200E-09 | 4.952E-09 | 3.600E-09 | 1.300E-08 |
| 3181 | 160 | T160A | A | 6.3e-09 | 3.9e-09 | 9.7e-09 | 6.633E-09 | 2.914E-09 | 3.900E-09 | 9.700E-09 |
| 3182 | 160 | T160R | A | 4.7e-09 | 6e-08 | 8.5e-09 | 2.440E-08 | 3.089E-08 | 4.700E-09 | 6.000E-08 |
| 3183 | 160 | T160N | A | 7.7e-09 | 9.7e-09 | 4e-09 | 7.133E-09 | 2.892E-09 | 4.000E-09 | 9.700E-09 |
| 3184 | 160 | T160D | A | 4.1e-09 | 4.8e-09 | 2.7e-08 | 1.197E-08 | 1.302E-08 | 4.100E-09 | 2.700E-08 |
| 3185 | 160 | T160C | A | 1.3e-08 | 5.8e-09 | 3.2e-09 | 7.333E-09 | 5.077E-09 | 3.200E-09 | 1.300E-08 |
| 3186 | 160 | T160E | A | 8.5e-09 | 9e-09 | 1.5e-08 | 1.083E-08 | 3.617E-09 | 8.500E-09 | 1.500E-08 |
| 3187 | 160 | T160Q | A | 2.2e-09 | 8.3e-09 | 6.3e-09 | 5.600E-09 | 3.110E-09 | 2.200E-09 | 8.300E-09 |
| 3188 | 160 | T160G | A | 2e-08 | 5.9e-09 | 8e-09 | 1.130E-08 | 7.607E-09 | 5.900E-09 | 2.000E-08 |
| 3189 | 160 | T160H | A | 1e-08 | 2.7e-09 | 6.3e-09 | 6.333E-09 | 3.650E-09 | 2.700E-09 | 1.000E-08 |
| 3190 | 160 | T160I | A | 2.6e-09 | 5.7e-09 | 1.8e-09 | 3.367E-09 | 2.060E-09 | 1.800E-09 | 5.700E-09 |
| 3191 | 160 | T160L | A | 8.2e-09 | 7.8e-09 | 9.3e-09 | 8.433E-09 | 7.767E-10 | 7.800E-09 | 9.300E-09 |
| 3192 | 160 | T160K | A | 3.4e-09 | 2.8e-09 | 1.3e-08 | 6.400E-09 | 5.724E-09 | 2.800E-09 | 1.300E-08 |
| 3193 | 160 | T160M | A | 6e-09 | 4.8e-09 | 5.6e-09 | 5.467E-09 | 6.110E-10 | 4.800E-09 | 6.000E-09 |
| 3194 | 160 | T160F | A | 2.6e-09 | 1.1e-08 | 1.5e-09 | 5.033E-09 | 5.196E-09 | 1.500E-09 | 1.100E-08 |
| 3195 | 160 | T160P | A | 6.3e-09 | 1.1e-09 | 8.3e-09 | 5.233E-09 | 3.717E-09 | 1.100E-09 | 8.300E-09 |
| 3196 | 160 | T160S | A | 3.7e-09 | 5.6e-09 | 4.5e-09 | 4.600E-09 | 9.539E-10 | 3.700E-09 | 5.600E-09 |
| 3197 | 160 | T160T | A | 1.1e-08 | 3.7e-09 | 4.5e-09 | 6.400E-09 | 4.004E-09 | 3.700E-09 | 1.100E-08 |
| 3198 | 160 | T160W | A | 1.6e-08 | 1.1e-08 | 5.5e-09 | 1.083E-08 | 5.252E-09 | 5.500E-09 | 1.600E-08 |
| 3199 | 160 | T160Y | A | 9.8e-09 | 5.5e-09 | 1.1e-08 | 8.767E-09 | 2.892E-09 | 5.500E-09 | 1.100E-08 |
| 3200 | 160 | T160V | A | 3.2e-09 | 6.3e-09 | 6.2e-09 | 5.233E-09 | 1.762E-09 | 3.200E-09 | 6.300E-09 |
| 3201 | 161 | L161A | A | 7.3e-09 | 9.8e-09 | 5e-09 | 7.367E-09 | 2.401E-09 | 5.000E-09 | 9.800E-09 |
| 3202 | 161 | L161R | A | 5.7e-09 | 8.8e-09 | 8.7e-09 | 7.733E-09 | 1.762E-09 | 5.700E-09 | 8.800E-09 |
| 3203 | 161 | L161N | A | 1e-08 | 2.6e-09 | 2.7e-09 | 5.100E-09 | 4.244E-09 | 2.600E-09 | 1.000E-08 |
| 3204 | 161 | L161D | A | 3e-09 | 7.5e-09 | 7.4e-09 | 5.967E-09 | 2.570E-09 | 3.000E-09 | 7.500E-09 |
| 3205 | 161 | L161C | A | 6.4e-09 | 6.3e-09 | 6.8e-09 | 6.500E-09 | 2.646E-10 | 6.300E-09 | 6.800E-09 |
| 3206 | 161 | L161E | A | 5.7e-09 | 1.3e-08 | 4.7e-09 | 7.800E-09 | 4.531E-09 | 4.700E-09 | 1.300E-08 |
| 3207 | 161 | L161Q | A | 6.5e-09 | 6.6e-09 | 1.6e-09 | 4.900E-09 | 2.858E-09 | 1.600E-09 | 6.600E-09 |
| 3208 | 161 | L161G | A | 2e-09 | 7.7e-09 | 5.4e-09 | 5.033E-09 | 2.868E-09 | 2.000E-09 | 7.700E-09 |
| 3209 | 161 | L161H | A | 2.2e-08 | 7e-09 | 1.2e-09 | 1.007E-08 | 1.073E-08 | 1.200E-09 | 2.200E-08 |
| 3210 | 161 | L161I | A | 4.9e-09 | 7.3e-09 | 8.4e-09 | 6.867E-09 | 1.790E-09 | 4.900E-09 | 8.400E-09 |
| 3211 | 161 | L161L | A | 8.8e-09 | 7.8e-09 | 5.3e-09 | 7.300E-09 | 1.803E-09 | 5.300E-09 | 8.800E-09 |
| 3212 | 161 | L161K | A | 1.2e-08 | 8.6e-09 | 6.2e-09 | 8.933E-09 | 2.914E-09 | 6.200E-09 | 1.200E-08 |
| 3213 | 161 | L161M | A | 3e-09 | 5e-09 | 8.3e-09 | 5.433E-09 | 2.676E-09 | 3.000E-09 | 8.300E-09 |
| 3214 | 161 | L161F | A | 6.1e-09 | 1.3e-08 | 6.1e-09 | 8.400E-09 | 3.984E-09 | 6.100E-09 | 1.300E-08 |
| 3215 | 161 | L161P | A | 2.5e-09 | 6.7e-09 | 6.7e-09 | 5.300E-09 | 2.425E-09 | 2.500E-09 | 6.700E-09 |
| 3216 | 161 | L161S | A | 5.3e-09 | 5.9e-09 | 1.4e-08 | 8.400E-09 | 4.859E-09 | 5.300E-09 | 1.400E-08 |
| 3217 | 161 | L161T | A | 8.7e-09 | 3.3e-09 | 8.1e-09 | 6.700E-09 | 2.960E-09 | 3.300E-09 | 8.700E-09 |
| 3218 | 161 | L161W | A | 6.9e-09 | 1.1e-08 | 1e-08 | 9.300E-09 | 2.138E-09 | 6.900E-09 | 1.100E-08 |
| 3219 | 161 | L161Y | A | 1.1e-08 | 3.3e-09 | 1.5e-08 | 9.767E-09 | 5.947E-09 | 3.300E-09 | 1.500E-08 |
| 3220 | 161 | L161V | A | 1.5e-08 | 1.2e-08 | 5.8e-09 | 1.093E-08 | 4.692E-09 | 5.800E-09 | 1.500E-08 |
| 3221 | 162 | I162A | A | 1.1e-08 | 8.6e-09 | 5.8e-09 | 8.467E-09 | 2.603E-09 | 5.800E-09 | 1.100E-08 |
| 3222 | 162 | I162R | A | 7.6e-09 | 9.1e-09 | 3.3e-09 | 6.667E-09 | 3.011E-09 | 3.300E-09 | 9.100E-09 |
| 3223 | 162 | I162N | A | 4.4e-09 | 6.9e-09 | 3.2e-09 | 4.833E-09 | 1.888E-09 | 3.200E-09 | 6.900E-09 |
| 3224 | 162 | I162D | A | 6e-09 | 9.7e-09 | 7.6e-09 | 7.767E-09 | 1.856E-09 | 6.000E-09 | 9.700E-09 |
| 3225 | 162 | I162C | A | 4.2e-09 | 4.2e-08 | 3.5e-09 | 1.657E-08 | 2.203E-08 | 3.500E-09 | 4.200E-08 |
| 3226 | 162 | I162E | A | 7.4e-09 | 1.3e-08 | 2.2e-09 | 7.533E-09 | 5.401E-09 | 2.200E-09 | 1.300E-08 |
| 3227 | 162 | I162Q | A | 5.5e-09 | 1.8e-08 | 3.6e-09 | 9.033E-09 | 7.823E-09 | 3.600E-09 | 1.800E-08 |
| 3228 | 162 | I162G | A | 2.9e-09 | 7e-09 | 4.6e-09 | 4.833E-09 | 2.060E-09 | 2.900E-09 | 7.000E-09 |
| 3229 | 162 | I162H | A | 4.3e-09 | 4.3e-09 | 3.6e-09 | 4.067E-09 | 4.041E-10 | 3.600E-09 | 4.300E-09 |

| | | | | | | | | | | |
|---|---|---|---|---|---|---|---|---|---|---|
| 3230 | 162 | I162I | A | 7.8e-09 | 5.1e-09 | 4.2e-09 | 5.700E-09 | 1.873E-09 | 4.200E-09 | 7.800E-09 |
| 3231 | 162 | I162L | A | 2e-09 | 4.4e-09 | 2e-09 | 2.800E-09 | 1.386E-09 | 2.000E-09 | 4.400E-09 |
| 3232 | 162 | I162K | A | 1.8e-08 | 1.8e-08 | 4e-09 | 1.333E-08 | 8.083E-09 | 4.000E-09 | 1.800E-08 |
| 3233 | 162 | I162M | A | 1.7e-08 | 4.9e-09 | 6.8e-09 | 9.567E-09 | 6.507E-09 | 4.900E-09 | 1.700E-08 |
| 3234 | 162 | I162F | A | 8.4e-09 | 5.9e-09 | 1.2e-08 | 8.767E-09 | 3.066E-09 | 5.900E-09 | 1.200E-08 |
| 3235 | 162 | I162P | A | 5.4e-09 | 2.6e-09 | 4.7e-09 | 4.233E-09 | 1.457E-09 | 2.600E-09 | 5.400E-09 |
| 3236 | 162 | I162S | A | 6e-09 | 7.3e-09 | 3.7e-09 | 5.667E-09 | 1.823E-09 | 3.700E-09 | 7.300E-09 |
| 3237 | 162 | I162T | A | 1e-08 | 1.5e-08 | 1.4e-08 | 1.300E-08 | 2.646E-09 | 1.000E-08 | 1.500E-08 |
| 3238 | 162 | I162W | A | 2.7e-09 | 7.4e-09 | 3.8e-09 | 4.633E-09 | 2.458E-09 | 2.700E-09 | 7.400E-09 |
| 3239 | 162 | I162Y | A | 2e-08 | 1e-08 | 1.1e-08 | 1.367E-08 | 5.508E-09 | 1.000E-08 | 2.000E-08 |
| 3240 | 162 | I162V | A | 3.4e-09 | 1e-08 | 4.3e-09 | 5.900E-09 | 3.579E-09 | 3.400E-09 | 1.000E-08 |
| 3241 | 163 | D163A | A | 1.8e-09 | 1e-08 | 1e-08 | 7.267E-09 | 4.734E-09 | 1.800E-09 | 1.000E-08 |
| 3242 | 163 | D163R | A | 8.3e-09 | 2e-08 | 3.9e-09 | 1.073E-08 | 8.321E-09 | 3.900E-09 | 2.000E-08 |
| 3243 | 163 | D163N | A | 4.4e-09 | 5e-09 | 2.8e-08 | 1.247E-08 | 1.346E-08 | 4.400E-09 | 2.800E-08 |
| 3244 | 163 | D163D | A | 9.1e-09 | 7.6e-09 | 2.3e-08 | 1.323E-08 | 8.491E-09 | 7.600E-09 | 2.300E-08 |
| 3245 | 163 | D163C | A | 7e-09 | 3.1e-09 | 8.3e-09 | 6.133E-09 | 2.706E-09 | 3.100E-09 | 8.300E-09 |
| 3246 | 163 | D163E | A | 1.2e-08 | 6.8e-09 | 1.6e-08 | 1.160E-08 | 4.613E-09 | 6.800E-09 | 1.600E-08 |
| 3247 | 163 | D163Q | A | 1.3e-08 | 4.6e-09 | 3.5e-09 | 7.033E-09 | 5.196E-09 | 3.500E-09 | 1.300E-08 |
| 3248 | 163 | D163G | A | 1.8e-08 | 4.1e-09 | 1.8e-08 | 1.337E-08 | 8.025E-09 | 4.100E-09 | 1.800E-08 |
| 3249 | 163 | D163H | A | 4.4e-09 | 4.1e-09 | 5.6e-09 | 4.700E-09 | 7.937E-10 | 4.100E-09 | 5.600E-09 |
| 3250 | 163 | D163I | A | 2.7e-09 | 1e-08 | 1.7e-08 | 9.900E-09 | 7.151E-09 | 2.700E-09 | 1.700E-08 |
| 3251 | 163 | D163L | A | 7.5e-09 | 9.8e-09 | 8.5e-09 | 8.600E-09 | 1.153E-09 | 7.500E-09 | 9.800E-09 |
| 3252 | 163 | D163K | A | 1.2e-08 | 3.1e-09 | 1.4e-08 | 9.700E-09 | 5.803E-09 | 3.100E-09 | 1.400E-08 |
| 3253 | 163 | D163M | A | 8.8e-09 | 1.1e-08 | 6.1e-09 | 8.633E-09 | 2.454E-09 | 6.100E-09 | 1.100E-08 |
| 3254 | 163 | D163F | A | 2.2e-08 | 7.2e-09 | 4.9e-09 | 1.137E-08 | 9.280E-09 | 4.900E-09 | 2.200E-08 |
| 3255 | 163 | D163P | A | 4.3e-09 | 8.5e-09 | 1.5e-08 | 9.267E-09 | 5.391E-09 | 4.300E-09 | 1.500E-08 |
| 3256 | 163 | D163S | A | 3.3e-09 | 3.3e-09 | 9.8e-09 | 5.467E-09 | 3.753E-09 | 3.300E-09 | 9.800E-09 |
| 3257 | 163 | D163T | A | 1.5e-08 | 6.3e-09 | 3.3e-09 | 8.200E-09 | 6.077E-09 | 3.300E-09 | 1.500E-08 |
| 3258 | 163 | D163W | A | 4e-09 | 2.4e-09 | 8.8e-09 | 5.067E-09 | 3.331E-09 | 2.400E-09 | 8.800E-09 |
| 3259 | 163 | D163Y | A | 4.3e-09 | 5.8e-09 | 5.9e-09 | 5.333E-09 | 8.963E-10 | 4.300E-09 | 5.900E-09 |
| 3260 | 163 | D163V | A | 1.5e-08 | 1.9e-09 | 6.1e-09 | 7.667E-09 | 6.689E-09 | 1.900E-09 | 1.500E-08 |
| 3261 | 164 | T164A | A | 3.1e-09 | 4.6e-09 | 4e-09 | 3.900E-09 | 7.550E-10 | 3.100E-09 | 4.600E-09 |
| 3262 | 164 | T164R | A | 4.8e-09 | 7.4e-09 | 8.7e-09 | 6.967E-09 | 1.986E-09 | 4.800E-09 | 8.700E-09 |
| 3263 | 164 | T164N | A | 7.8e-09 | 1.7e-09 | 6.8e-09 | 5.433E-09 | 3.272E-09 | 1.700E-09 | 7.800E-09 |
| 3264 | 164 | T164D | A | 9.6e-09 | 2e-08 | 1.5e-09 | 1.037E-08 | 9.274E-09 | 1.500E-09 | 2.000E-08 |
| 3265 | 164 | T164C | A | 5.8e-09 | 5.8e-09 | 4.9e-09 | 5.500E-09 | 5.196E-10 | 4.900E-09 | 5.800E-09 |
| 3266 | 164 | T164E | A | 8e-09 | 3.5e-09 | 2.3e-09 | 4.600E-09 | 3.005E-09 | 2.300E-09 | 8.000E-09 |
| 3267 | 164 | T164Q | A | 9.7e-09 | 9.7e-09 | 4.6e-09 | 8.000E-09 | 2.944E-09 | 4.600E-09 | 9.700E-09 |
| 3268 | 164 | T164G | A | 6.6e-09 | 1.1e-08 | 1e-08 | 9.200E-09 | 2.307E-09 | 6.600E-09 | 1.100E-08 |
| 3269 | 164 | T164H | A | 8.6e-09 | 5e-09 | 6.7e-09 | 6.767E-09 | 1.801E-09 | 5.000E-09 | 8.600E-09 |
| 3270 | 164 | T164I | A | 7.4e-09 | 9.3e-09 | 8.7e-09 | 8.467E-09 | 9.713E-10 | 7.400E-09 | 9.300E-09 |
| 3271 | 164 | T164L | A | 3.7e-09 | 5.8e-09 | 8.1e-09 | 5.867E-09 | 2.201E-09 | 3.700E-09 | 8.100E-09 |
| 3272 | 164 | T164K | A | 7e-09 | 4.7e-09 | 1.1e-08 | 7.567E-09 | 3.188E-09 | 4.700E-09 | 1.100E-08 |
| 3273 | 164 | T164M | A | 7.1e-09 | 2.1e-08 | 3.6e-09 | 1.057E-08 | 9.203E-09 | 3.600E-09 | 2.100E-08 |
| 3274 | 164 | T164F | A | 4.5e-09 | 5.4e-09 | 1.8e-08 | 9.300E-09 | 7.548E-09 | 4.500E-09 | 1.800E-08 |
| 3275 | 164 | T164P | A | 3.8e-09 | 2.9e-09 | 1.6e-09 | 2.767E-09 | 1.106E-09 | 1.600E-09 | 3.800E-09 |
| 3276 | 164 | T164S | A | 2.9e-09 | 4e-09 | 4e-09 | 3.633E-09 | 6.351E-10 | 2.900E-09 | 4.000E-09 |
| 3277 | 164 | T164T | A | 5.3e-09 | 1.4e-08 | 7.1e-09 | 8.800E-09 | 4.592E-09 | 5.300E-09 | 1.400E-08 |
| 3278 | 164 | T164W | A | 9.6e-09 | 2.2e-08 | 1.1e-08 | 1.420E-08 | 6.791E-09 | 9.600E-09 | 2.200E-08 |
| 3279 | 164 | T164Y | A | 5.7e-09 | 8.3e-09 | 4.9e-09 | 6.300E-09 | 1.778E-09 | 4.900E-09 | 8.300E-09 |
| 3280 | 164 | T164V | A | 5.7e-09 | 1.6e-08 | 1e-08 | 1.057E-08 | 5.173E-09 | 5.700E-09 | 1.600E-08 |
| 3281 | 165 | N165A | A | 4.9e-09 | 6.8e-09 | 2.4e-09 | 4.700E-09 | 2.207E-09 | 2.400E-09 | 6.800E-09 |
| 3282 | 165 | N165R | A | 6.4e-09 | 1.8e-08 | 1.3e-08 | 1.247E-08 | 5.818E-09 | 6.400E-09 | 1.800E-08 |
| 3283 | 165 | N165N | A | 4.9e-09 | 6.6e-09 | 5.2e-09 | 5.567E-09 | 9.074E-10 | 4.900E-09 | 6.600E-09 |
| 3284 | 165 | N165D | A | 6.8e-09 | 5.6e-09 | 4.6e-09 | 5.667E-09 | 1.102E-09 | 4.600E-09 | 6.800E-09 |
| 3285 | 165 | N165C | A | 1.4e-08 | 5.6e-09 | 9.5e-09 | 9.700E-09 | 4.204E-09 | 5.600E-09 | 1.400E-08 |
| 3286 | 165 | N165E | A | 1.3e-08 | 3.4e-09 | 4.8e-09 | 7.067E-09 | 5.186E-09 | 3.400E-09 | 1.300E-08 |
| 3287 | 165 | N165Q | A | 9.3e-09 | 1.2e-08 | 7e-09 | 9.433E-09 | 2.503E-09 | 7.000E-09 | 1.200E-08 |
| 3288 | 165 | N165G | A | 2.4e-09 | 4.7e-09 | 6.3e-09 | 4.467E-09 | 1.960E-09 | 2.400E-09 | 6.300E-09 |
| 3289 | 165 | N165H | A | 3.9e-09 | 2.3e-08 | 1.6e-08 | 1.430E-08 | 9.663E-09 | 3.900E-09 | 2.300E-08 |
| 3290 | 165 | N165I | A | 7.2e-09 | 7.6e-09 | 2.3e-08 | 1.260E-08 | 9.009E-09 | 7.200E-09 | 2.300E-08 |
| 3291 | 165 | N165L | A | 1.2e-08 | 3.5e-09 | 2.1e-09 | 5.867E-09 | 5.358E-09 | 2.100E-09 | 1.200E-08 |
| 3292 | 165 | N165K | A | 9.8e-09 | 7.1e-09 | 3.7e-09 | 6.867E-09 | 3.057E-09 | 3.700E-09 | 9.800E-09 |
| 3293 | 165 | N165M | A | 6.3e-09 | 8.8e-09 | 5.7e-09 | 6.933E-09 | 1.644E-09 | 5.700E-09 | 8.800E-09 |
| 3294 | 165 | N165F | A | 5.3e-09 | 1.2e-08 | 3e-09 | 6.767E-09 | 4.676E-09 | 3.000E-09 | 1.200E-08 |
| 3295 | 165 | N165P | A | 1.3e-08 | 1.4e-08 | 1.9e-08 | 1.533E-08 | 3.215E-09 | 1.300E-08 | 1.900E-08 |
| 3296 | 165 | N165S | A | 2.2e-08 | 1.6e-09 | 4.2e-09 | 9.267E-09 | 1.110E-08 | 1.600E-09 | 2.200E-08 |
| 3297 | 165 | N165T | A | 6.4e-09 | 1.8e-09 | 1.2e-08 | 6.733E-09 | 5.108E-09 | 1.800E-09 | 1.200E-08 |
| 3298 | 165 | N165W | A | 9e-09 | 7.5e-09 | 4.1e-09 | 6.867E-09 | 2.511E-09 | 4.100E-09 | 9.000E-09 |
| 3299 | 165 | N165Y | A | 1.2e-08 | 2.6e-09 | 6e-09 | 6.867E-09 | 4.760E-09 | 2.600E-09 | 1.200E-08 |
| 3300 | 165 | N165V | A | 8.5e-09 | 1.9e-09 | 3.1e-08 | 1.380E-08 | 1.526E-08 | 1.900E-09 | 3.100E-08 |
| 3301 | 166 | R166A | A | 5.5e-09 | 1e-08 | 4.8e-09 | 6.767E-09 | 2.822E-09 | 4.800E-09 | 1.000E-08 |
| 3302 | 166 | R166R | A | 2.9e-09 | 7.1e-09 | 1.8e-09 | 3.933E-09 | 2.797E-09 | 1.800E-09 | 7.100E-09 |
| 3303 | 166 | R166N | A | 7e-09 | 9.4e-09 | 5.1e-09 | 7.167E-09 | 2.155E-09 | 5.100E-09 | 9.400E-09 |
| 3304 | 166 | R166D | A | 1.8e-08 | 1.8e-08 | 6e-09 | 1.400E-08 | 6.928E-09 | 6.000E-09 | 1.800E-08 |
| 3305 | 166 | R166C | A | 6.4e-09 | 5.3e-09 | 2.1e-09 | 4.600E-09 | 2.234E-09 | 2.100E-09 | 6.400E-09 |
| 3306 | 166 | R166E | A | 4.5e-09 | 1e-08 | 4.2e-09 | 6.233E-09 | 3.265E-09 | 4.200E-09 | 1.000E-08 |
| 3307 | 166 | R166Q | A | 1.2e-08 | 4.1e-09 | 1.2e-08 | 9.367E-09 | 4.561E-09 | 4.100E-09 | 1.200E-08 |
| 3308 | 166 | R166G | A | 7.8e-09 | 8.3e-09 | 1.3e-08 | 9.700E-09 | 2.869E-09 | 7.800E-09 | 1.300E-08 |
| 3309 | 166 | R166H | A | 8.1e-09 | 5.7e-09 | 1.4e-09 | 5.067E-09 | 3.395E-09 | 1.400E-09 | 8.100E-09 |
| 3310 | 166 | R166I | A | 1e-08 | 6.5e-09 | 3.5e-09 | 6.667E-09 | 3.253E-09 | 3.500E-09 | 1.000E-08 |
| 3311 | 166 | R166L | A | 1.4e-09 | 9.8e-09 | 4e-09 | 5.067E-09 | 4.300E-09 | 1.400E-09 | 9.800E-09 |
| 3312 | 166 | R166K | A | 7.6e-09 | 6.8e-09 | 4.4e-09 | 6.267E-09 | 1.665E-09 | 4.400E-09 | 7.600E-09 |
| 3313 | 166 | R166M | A | 6.6e-09 | 2.2e-09 | 1.6e-08 | 8.267E-09 | 7.049E-09 | 2.200E-09 | 1.600E-08 |
| 3314 | 166 | R166F | A | 5.8e-09 | 5.7e-09 | 5e-09 | 5.500E-09 | 4.359E-10 | 5.000E-09 | 5.800E-09 |

| | | | | | | | | | | |
|---|---|---|---|---|---|---|---|---|---|---|
| 3315 | 166 | R166P | A | 6e-09 | 5.2e-09 | 1.3e-08 | 8.067E-09 | 4.291E-09 | 5.200E-09 | 1.300E-08 |
| 3316 | 166 | R166S | A | 5.2e-09 | 4.5e-09 | 5.1e-09 | 4.933E-09 | 3.786E-10 | 4.500E-09 | 5.200E-09 |
| 3317 | 166 | R166T | A | 4.6e-09 | 1e-08 | 1.4e-08 | 9.533E-09 | 4.717E-09 | 4.600E-09 | 1.400E-08 |
| 3318 | 166 | R166W | A | 6.4e-09 | 2.5e-09 | 3.1e-09 | 4.000E-09 | 2.100E-09 | 2.500E-09 | 6.400E-09 |
| 3319 | 166 | R166Y | A | 7.1e-09 | 9e-09 | 4.1e-09 | 6.733E-09 | 2.470E-09 | 4.100E-09 | 9.000E-09 |
| 3320 | 166 | R166V | A | 2.1e-08 | 4.8e-09 | 5.7e-09 | 1.050E-08 | 9.104E-09 | 4.800E-09 | 2.100E-08 |
| 3321 | 167 | S167A | A | 4.1e-09 | 2.7e-09 | 2.5e-09 | 3.100E-09 | 8.718E-10 | 2.500E-09 | 4.100E-09 |
| 3322 | 167 | S167R | A | 5.3e-09 | 3.3e-09 | 4.4e-09 | 4.333E-09 | 1.002E-09 | 3.300E-09 | 5.300E-09 |
| 3323 | 167 | S167N | A | 5.6e-09 | 1.1e-08 | 1.3e-08 | 9.867E-09 | 3.828E-09 | 5.600E-09 | 1.300E-08 |
| 3324 | 167 | S167D | A | 8.8e-09 | 4e-09 | 1.1e-08 | 7.933E-09 | 3.580E-09 | 4.000E-09 | 1.100E-08 |
| 3325 | 167 | S167C | A | 2.3e-08 | 7.3e-09 | 9.1e-09 | 1.313E-08 | 8.592E-09 | 7.300E-09 | 2.300E-08 |
| 3326 | 167 | S167E | A | 5.1e-09 | 5.5e-09 | 6.6e-09 | 5.733E-09 | 7.767E-10 | 5.100E-09 | 6.600E-09 |
| 3327 | 167 | S167Q | A | 1.5e-08 | 5.8e-09 | 3.5e-09 | 8.100E-09 | 6.085E-09 | 3.500E-09 | 1.500E-08 |
| 3328 | 167 | S167G | A | 6.3e-09 | 1.8e-08 | 6.8e-09 | 1.037E-08 | 6.615E-09 | 6.300E-09 | 1.800E-08 |
| 3329 | 167 | S167H | A | 1e-08 | 3.9e-09 | 2.4e-09 | 5.433E-09 | 4.025E-09 | 2.400E-09 | 1.000E-08 |
| 3330 | 167 | S167I | A | 1.5e-09 | 3.1e-08 | 2.8e-08 | 2.017E-08 | 1.624E-08 | 1.500E-09 | 3.100E-08 |
| 3331 | 167 | S167L | A | 8.9e-09 | 5.7e-09 | 8.4e-09 | 7.667E-09 | 1.721E-09 | 5.700E-09 | 8.900E-09 |
| 3332 | 167 | S167K | A | 3.1e-09 | 8.7e-09 | 1.8e-08 | 9.933E-09 | 7.526E-09 | 3.100E-09 | 1.800E-08 |
| 3333 | 167 | S167M | A | 1.1e-08 | 6.1e-09 | 3.1e-09 | 6.733E-09 | 3.988E-09 | 3.100E-09 | 1.100E-08 |
| 3334 | 167 | S167F | A | 4.6e-09 | 9.4e-09 | 5.1e-09 | 6.367E-09 | 2.639E-09 | 4.600E-09 | 9.400E-09 |
| 3335 | 167 | S167P | A | 8.3e-09 | 6e-09 | 1.8e-08 | 1.077E-08 | 6.369E-09 | 6.000E-09 | 1.800E-08 |
| 3336 | 167 | S167S | A | 6.5e-09 | 7.2e-09 | 9.6e-09 | 7.767E-09 | 1.626E-09 | 6.500E-09 | 9.600E-09 |
| 3337 | 167 | S167T | A | 2e-09 | 1e-08 | 1.2e-08 | 8.000E-09 | 5.292E-09 | 2.000E-09 | 1.200E-08 |
| 3338 | 167 | S167W | A | 1.1e-08 | 6.6e-09 | 8.4e-09 | 8.667E-09 | 2.212E-09 | 6.600E-09 | 1.100E-08 |
| 3339 | 167 | S167Y | A | 1.2e-09 | 8.1e-09 | 7.1e-09 | 5.467E-09 | 3.729E-09 | 1.200E-09 | 8.100E-09 |
| 3340 | 167 | S167V | A | 7.9e-09 | 3.1e-09 | 7e-09 | 6.000E-09 | 2.551E-09 | 3.100E-09 | 7.900E-09 |
| 3341 | 168 | R168A | A | 6.9e-09 | 5.5e-09 | 3.7e-09 | 5.367E-09 | 1.604E-09 | 3.700E-09 | 6.900E-09 |
| 3342 | 168 | R168R | A | 4.3e-09 | 7.9e-09 | 1.6e-08 | 9.400E-09 | 5.992E-09 | 4.300E-09 | 1.600E-08 |
| 3343 | 168 | R168N | A | 1.8e-08 | 8.7e-09 | 6.7e-09 | 1.113E-08 | 6.030E-09 | 6.700E-09 | 1.800E-08 |
| 3344 | 168 | R168D | A | 1.1e-08 | 5.3e-09 | 9.9e-09 | 8.733E-09 | 3.024E-09 | 5.300E-09 | 1.100E-08 |
| 3345 | 168 | R168C | A | 3e-09 | 1.1e-08 | 9.1e-09 | 7.700E-09 | 4.180E-09 | 3.000E-09 | 1.100E-08 |
| 3346 | 168 | R168E | A | 6.8e-09 | 1e-08 | 6.3e-09 | 7.700E-09 | 2.007E-09 | 6.300E-09 | 1.000E-08 |
| 3347 | 168 | R168Q | A | 9.1e-09 | 7.7e-09 | 8.6e-09 | 8.467E-09 | 7.095E-10 | 7.700E-09 | 9.100E-09 |
| 3348 | 168 | R168G | A | 7e-09 | 7.6e-09 | 9.7e-09 | 8.100E-09 | 1.418E-09 | 7.000E-09 | 9.700E-09 |
| 3349 | 168 | R168H | A | 4.1e-09 | 4.1e-09 | 1.7e-08 | 8.400E-09 | 7.448E-09 | 4.100E-09 | 1.700E-08 |
| 3350 | 168 | R168I | A | 1.1e-08 | 7.2e-09 | 1.1e-08 | 9.733E-09 | 2.194E-09 | 7.200E-09 | 1.100E-08 |
| 3351 | 168 | R168L | A | 1.4e-08 | 8.5e-09 | 3.5e-09 | 8.667E-09 | 5.252E-09 | 3.500E-09 | 1.400E-08 |
| 3352 | 168 | R168K | A | 9e-09 | 8.3e-09 | 2.9e-09 | 6.733E-09 | 3.338E-09 | 2.900E-09 | 9.000E-09 |
| 3353 | 168 | R168M | A | 3e-09 | 5.8e-09 | 2.8e-09 | 3.867E-09 | 1.677E-09 | 2.800E-09 | 5.800E-09 |
| 3354 | 168 | R168F | A | 1.8e-08 | 5.1e-09 | 4.8e-09 | 9.300E-09 | 7.536E-09 | 4.800E-09 | 1.800E-08 |
| 3355 | 168 | R168P | A | 1.2e-08 | 1e-08 | 1.3e-09 | 7.767E-09 | 5.689E-09 | 1.300E-09 | 1.200E-08 |
| 3356 | 168 | R168S | A | 9.9e-09 | 9.7e-09 | 3.8e-09 | 7.800E-09 | 3.466E-09 | 3.800E-09 | 9.900E-09 |
| 3357 | 168 | R168T | A | 2.3e-09 | 1.5e-08 | 6.2e-09 | 7.833E-09 | 6.506E-09 | 2.300E-09 | 1.500E-08 |
| 3358 | 168 | R168W | A | 7e-09 | 5.1e-09 | 6.6e-09 | 6.233E-09 | 1.002E-09 | 5.100E-09 | 7.000E-09 |
| 3359 | 168 | R168Y | A | 6.8e-09 | 6.8e-09 | 5.5e-09 | 6.367E-09 | 7.506E-10 | 5.500E-09 | 6.800E-09 |
| 3360 | 168 | R168V | A | 3.4e-09 | 1.2e-08 | 1e-08 | 8.467E-09 | 4.500E-09 | 3.400E-09 | 1.200E-08 |
| 3361 | 169 | A169A | A | 6e-09 | 8.4e-09 | 3.8e-09 | 6.067E-09 | 2.301E-09 | 3.800E-09 | 8.400E-09 |
| 3362 | 169 | A169R | A | 4.4e-09 | 1.1e-08 | 2.8e-09 | 6.067E-09 | 4.347E-09 | 2.800E-09 | 1.100E-08 |
| 3363 | 169 | A169N | A | 1.4e-08 | 4.5e-09 | 2.1e-08 | 1.317E-08 | 8.282E-09 | 4.500E-09 | 2.100E-08 |
| 3364 | 169 | A169D | A | 2.2e-09 | 1.6e-08 | 5.4e-09 | 7.867E-09 | 7.223E-09 | 2.200E-09 | 1.600E-08 |
| 3365 | 169 | A169C | A | 1.2e-08 | 6e-09 | 4.2e-09 | 7.400E-09 | 4.084E-09 | 4.200E-09 | 1.200E-08 |
| 3366 | 169 | A169E | A | 7.6e-09 | 1.7e-08 | 3.9e-09 | 9.500E-09 | 6.754E-09 | 3.900E-09 | 1.700E-08 |
| 3367 | 169 | A169Q | A | 3.9e-09 | 4.9e-09 | 1.5e-08 | 7.933E-09 | 6.140E-09 | 3.900E-09 | 1.500E-08 |
| 3368 | 169 | A169G | A | 7e-09 | 1.6e-09 | 2.4e-09 | 3.667E-09 | 2.914E-09 | 1.600E-09 | 7.000E-09 |
| 3369 | 169 | A169H | A | 3.5e-09 | 5.4e-09 | 6.8e-09 | 5.233E-09 | 1.656E-09 | 3.500E-09 | 6.800E-09 |
| 3370 | 169 | A169I | A | 4.9e-09 | 1.9e-09 | 7.3e-09 | 4.700E-09 | 2.706E-09 | 1.900E-09 | 7.300E-09 |
| 3371 | 169 | A169L | A | 8.9e-09 | 1.6e-08 | 5.1e-09 | 1.000E-08 | 5.533E-09 | 5.100E-09 | 1.600E-08 |
| 3372 | 169 | A169K | A | 2.3e-08 | 1.4e-08 | 2e-08 | 1.900E-08 | 4.583E-09 | 1.400E-08 | 2.300E-08 |
| 3373 | 169 | A169M | A | 1.1e-08 | 1.1e-08 | 1.7e-08 | 1.300E-08 | 3.464E-09 | 1.100E-08 | 1.700E-08 |
| 3374 | 169 | A169F | A | 5.1e-09 | 6e-09 | 9.1e-09 | 6.733E-09 | 2.098E-09 | 5.100E-09 | 9.100E-09 |
| 3375 | 169 | A169P | A | 3.9e-09 | 3e-09 | 4.3e-09 | 3.733E-09 | 6.658E-10 | 3.000E-09 | 4.300E-09 |
| 3376 | 169 | A169S | A | 6.3e-09 | 3.6e-09 | 1.9e-08 | 9.633E-09 | 8.223E-09 | 3.600E-09 | 1.900E-08 |
| 3377 | 169 | A169T | A | 5.2e-09 | 7.6e-09 | 2.8e-09 | 5.200E-09 | 2.400E-09 | 2.800E-09 | 7.600E-09 |
| 3378 | 169 | A169W | A | 1.6e-08 | 1.5e-08 | 1.3e-08 | 1.467E-08 | 1.528E-09 | 1.300E-08 | 1.600E-08 |
| 3379 | 169 | A169Y | A | 1.6e-08 | 2.3e-09 | 1.3e-08 | 1.043E-08 | 7.202E-09 | 2.300E-09 | 1.600E-08 |
| 3380 | 169 | A169V | A | 3.3e-09 | 1.6e-08 | 6.3e-09 | 8.533E-09 | 6.638E-09 | 3.300E-09 | 1.600E-08 |
| 3381 | 170 | C170A | A | 4.5e-09 | 1.1e-08 | 7e-09 | 7.500E-09 | 3.279E-09 | 4.500E-09 | 1.100E-08 |
| 3382 | 170 | C170R | A | 1.9e-08 | 1.8e-09 | 3.9e-09 | 8.233E-09 | 9.383E-09 | 1.800E-09 | 1.900E-08 |
| 3383 | 170 | C170N | A | 1.1e-08 | 1.7e-08 | 1.7e-09 | 9.900E-09 | 7.709E-09 | 1.700E-09 | 1.700E-08 |
| 3384 | 170 | C170D | A | 1.4e-08 | 1.4e-08 | 1.5e-08 | 1.433E-08 | 5.774E-10 | 1.400E-08 | 1.500E-08 |
| 3385 | 170 | C170C | A | 2.2e-08 | 4.4e-09 | 9.1e-09 | 1.183E-08 | 9.113E-09 | 4.400E-09 | 2.200E-08 |
| 3386 | 170 | C170E | A | 4e-09 | 3.4e-09 | 1.2e-08 | 6.467E-09 | 4.801E-09 | 3.400E-09 | 1.200E-08 |
| 3387 | 170 | C170Q | A | 4.6e-09 | 8.7e-09 | 1.4e-08 | 9.100E-09 | 4.713E-09 | 4.600E-09 | 1.400E-08 |
| 3388 | 170 | C170G | A | 2.6e-09 | 5e-09 | 2.2e-09 | 3.267E-09 | 1.514E-09 | 2.200E-09 | 5.000E-09 |
| 3389 | 170 | C170H | A | 5.6e-09 | 5e-09 | 1.5e-09 | 4.033E-09 | 2.214E-09 | 1.500E-09 | 5.600E-09 |
| 3390 | 170 | C170I | A | 1.4e-08 | 7.7e-09 | 2.8e-09 | 8.167E-09 | 5.615E-09 | 2.800E-09 | 1.400E-08 |
| 3391 | 170 | C170L | A | 2.4e-09 | 1.8e-09 | 7.1e-09 | 3.767E-09 | 2.902E-09 | 1.800E-09 | 7.100E-09 |
| 3392 | 170 | C170K | A | 6.2e-09 | 5.6e-09 | 2.5e-08 | 1.227E-08 | 1.103E-08 | 5.600E-09 | 2.500E-08 |
| 3393 | 170 | C170M | A | 4.1e-09 | 6.2e-09 | 7.9e-09 | 6.067E-09 | 1.904E-09 | 4.100E-09 | 7.900E-09 |
| 3394 | 170 | C170F | A | 4.9e-09 | 8.5e-09 | 4.6e-09 | 6.000E-09 | 2.170E-09 | 4.600E-09 | 8.500E-09 |
| 3395 | 170 | C170P | A | 1.1e-08 | 1.9e-08 | 4.4e-09 | 1.147E-08 | 7.311E-09 | 4.400E-09 | 1.900E-08 |
| 3396 | 170 | C170S | A | 4.8e-09 | 5.2e-09 | 3.4e-09 | 4.467E-09 | 9.452E-10 | 3.400E-09 | 5.200E-09 |
| 3397 | 170 | C170T | A | 4.4e-09 | 7e-09 | 6.8e-09 | 6.067E-09 | 1.447E-09 | 4.400E-09 | 7.000E-09 |
| 3398 | 170 | C170W | A | 1.3e-08 | 1.1e-08 | 6.6e-09 | 1.020E-08 | 3.274E-09 | 6.600E-09 | 1.300E-08 |
| 3399 | 170 | C170Y | A | 1.2e-08 | 6.7e-09 | 2.9e-09 | 7.200E-09 | 4.571E-09 | 2.900E-09 | 1.200E-08 |

| | | | | | | | | | | |
|---|---|---|---|---|---|---|---|---|---|---|
| 3400 | 170 | C170V | A | 1.3e-08 | 1.1e-08 | 4.8e-09 | 9.600E-09 | 4.276E-09 | 4.800E-09 | 1.300E-08 |
| 3401 | 171 | H171A | A | 3.2e-09 | 7.9e-09 | 9.2e-09 | 6.767E-09 | 3.156E-09 | 3.200E-09 | 9.200E-09 |
| 3402 | 171 | H171R | A | 7.4e-09 | 4.2e-09 | 7.6e-09 | 6.400E-09 | 1.908E-09 | 4.200E-09 | 7.600E-09 |
| 3403 | 171 | H171N | A | 6.9e-09 | 3.5e-09 | 6.7e-09 | 5.700E-09 | 1.908E-09 | 3.500E-09 | 6.900E-09 |
| 3404 | 171 | H171D | A | 2.1e-09 | 4e-09 | 3.5e-09 | 3.200E-09 | 9.849E-10 | 2.100E-09 | 4.000E-09 |
| 3405 | 171 | H171C | A | 5.9e-09 | 1.2e-08 | 5e-09 | 7.633E-09 | 3.808E-09 | 5.000E-09 | 1.200E-08 |
| 3406 | 171 | H171E | A | 9.1e-09 | 1.1e-08 | 5.9e-09 | 8.667E-09 | 2.577E-09 | 5.900E-09 | 1.100E-08 |
| 3407 | 171 | H171Q | A | 4.4e-09 | 9.7e-09 | 1e-08 | 8.033E-09 | 3.150E-09 | 4.400E-09 | 1.000E-08 |
| 3408 | 171 | H171G | A | 5.3e-09 | 2.5e-09 | 1.9e-09 | 3.233E-09 | 1.815E-09 | 1.900E-09 | 5.300E-09 |
| 3409 | 171 | H171H | A | 6.4e-09 | 7.8e-09 | 2e-08 | 1.140E-08 | 7.481E-09 | 6.400E-09 | 2.000E-08 |
| 3410 | 171 | H171I | A | 1.3e-08 | 8.4e-09 | 6.4e-09 | 9.267E-09 | 3.384E-09 | 6.400E-09 | 1.300E-08 |
| 3411 | 171 | H171L | A | 8.4e-09 | 1.1e-08 | 1e-08 | 9.800E-09 | 1.311E-09 | 8.400E-09 | 1.100E-08 |
| 3412 | 171 | H171K | A | 5e-09 | 7.1e-09 | 1.1e-08 | 7.700E-09 | 3.045E-09 | 5.000E-09 | 1.100E-08 |
| 3413 | 171 | H171M | A | 5.9e-09 | 4.3e-09 | 6.3e-09 | 5.500E-09 | 1.058E-09 | 4.300E-09 | 6.300E-09 |
| 3414 | 171 | H171F | A | 6.8e-09 | 4.1e-09 | 7.2e-09 | 6.033E-09 | 1.686E-09 | 4.100E-09 | 7.200E-09 |
| 3415 | 171 | H171P | A | 9.8e-09 | 5.6e-09 | 4.9e-09 | 6.767E-09 | 2.650E-09 | 4.900E-09 | 9.800E-09 |
| 3416 | 171 | H171S | A | 3e-09 | 1.1e-08 | 4e-09 | 6.000E-09 | 4.359E-09 | 3.000E-09 | 1.100E-08 |
| 3417 | 171 | H171T | A | 7.5e-09 | 2.2e-08 | 6.1e-09 | 1.187E-08 | 8.804E-09 | 6.100E-09 | 2.200E-08 |
| 3418 | 171 | H171W | A | 6.9e-09 | 1.3e-08 | 4.2e-09 | 8.033E-09 | 4.508E-09 | 4.200E-09 | 1.300E-08 |
| 3419 | 171 | H171Y | A | 4e-09 | 8.5e-09 | 9.4e-09 | 7.300E-09 | 2.893E-09 | 4.000E-09 | 9.400E-09 |
| 3420 | 171 | H171V | A | 1.4e-08 | 1.8e-09 | 5.3e-09 | 7.033E-09 | 6.282E-09 | 1.800E-09 | 1.400E-08 |
| 3421 | 172 | P172A | A | 3.5e-09 | 1.7e-08 | 6.6e-09 | 9.033E-09 | 7.071E-09 | 3.500E-09 | 1.700E-08 |
| 3422 | 172 | P172R | A | 3.4e-09 | 1.7e-08 | 2.2e-09 | 7.533E-09 | 8.220E-09 | 2.200E-09 | 1.700E-08 |
| 3423 | 172 | P172N | A | 9.3e-09 | 6.1e-09 | 1.2e-08 | 9.133E-09 | 2.954E-09 | 6.100E-09 | 1.200E-08 |
| 3424 | 172 | P172D | A | 4e-09 | 1.1e-08 | 1.1e-08 | 8.667E-09 | 4.041E-09 | 4.000E-09 | 1.100E-08 |
| 3425 | 172 | P172C | A | 7.6e-09 | 4.7e-09 | 4e-09 | 5.433E-09 | 1.909E-09 | 4.000E-09 | 7.600E-09 |
| 3426 | 172 | P172E | A | 1.7e-09 | 1.1e-08 | 2.7e-09 | 5.133E-09 | 5.105E-09 | 1.700E-09 | 1.100E-08 |
| 3427 | 172 | P172Q | A | 2.5e-09 | 3.5e-09 | 6.3e-09 | 4.100E-09 | 1.970E-09 | 2.500E-09 | 6.300E-09 |
| 3428 | 172 | P172G | A | 1.4e-08 | 1e-08 | 2.6e-08 | 1.667E-08 | 8.327E-09 | 1.000E-08 | 2.600E-08 |
| 3429 | 172 | P172H | A | 6.3e-09 | 8.5e-09 | 1.1e-08 | 8.600E-09 | 2.352E-09 | 6.300E-09 | 1.100E-08 |
| 3430 | 172 | P172I | A | 5e-09 | 2.5e-09 | 7e-09 | 4.833E-09 | 2.255E-09 | 2.500E-09 | 7.000E-09 |
| 3431 | 172 | P172L | A | 8.2e-09 | 1.4e-08 | 5.4e-09 | 9.200E-09 | 4.386E-09 | 5.400E-09 | 1.400E-08 |
| 3432 | 172 | P172K | A | 1.5e-08 | 1.2e-08 | 5.3e-09 | 1.077E-08 | 4.966E-09 | 5.300E-09 | 1.500E-08 |
| 3433 | 172 | P172M | A | 9.3e-09 | 1.5e-08 | 8e-09 | 1.077E-08 | 3.723E-09 | 8.000E-09 | 1.500E-08 |
| 3434 | 172 | P172F | A | 5.6e-09 | 3.5e-09 | 4e-09 | 4.367E-09 | 1.097E-09 | 3.500E-09 | 5.600E-09 |
| 3435 | 172 | P172P | A | 5.9e-09 | 1.1e-08 | 6.2e-09 | 7.700E-09 | 2.862E-09 | 5.900E-09 | 1.100E-08 |
| 3436 | 172 | P172S | A | 1.4e-08 | 6.6e-09 | 4.3e-09 | 8.300E-09 | 5.069E-09 | 4.300E-09 | 1.400E-08 |
| 3437 | 172 | P172T | A | 5.1e-09 | 1.7e-09 | 4.2e-09 | 3.667E-09 | 1.762E-09 | 1.700E-09 | 5.100E-09 |
| 3438 | 172 | P172W | A | 4.9e-09 | 5.5e-09 | 6.2e-09 | 5.533E-09 | 6.506E-10 | 4.900E-09 | 6.200E-09 |
| 3439 | 172 | P172Y | A | 6.1e-09 | 8.9e-09 | 4.2e-09 | 6.400E-09 | 2.364E-09 | 4.200E-09 | 8.900E-09 |
| 3440 | 172 | P172V | A | 4.4e-09 | 3.6e-09 | 2.8e-09 | 3.600E-09 | 8.000E-10 | 2.800E-09 | 4.400E-09 |
| 3441 | 173 | C173A | A | 2.3e-09 | 2e-09 | 1.3e-08 | 5.767E-09 | 6.266E-09 | 2.000E-09 | 1.300E-08 |
| 3442 | 173 | C173R | A | 8.7e-09 | 4.4e-09 | 2.3e-09 | 5.133E-09 | 3.262E-09 | 2.300E-09 | 8.700E-09 |
| 3443 | 173 | C173N | A | 1e-08 | 4.6e-09 | 3.4e-09 | 6.000E-09 | 3.516E-09 | 3.400E-09 | 1.000E-08 |
| 3444 | 173 | C173D | A | 1.1e-08 | 3.4e-08 | 5.7e-09 | 1.690E-08 | 1.504E-08 | 5.700E-09 | 3.400E-08 |
| 3445 | 173 | C173C | A | 7.6e-09 | 1.3e-08 | 5.7e-09 | 8.767E-09 | 3.787E-09 | 5.700E-09 | 1.300E-08 |
| 3446 | 173 | C173E | A | 9.4e-09 | 3.6e-09 | 6.5e-09 | 6.500E-09 | 2.900E-09 | 3.600E-09 | 9.400E-09 |
| 3447 | 173 | C173Q | A | 1.9e-09 | 1e-08 | 4.2e-09 | 5.367E-09 | 4.174E-09 | 1.900E-09 | 1.000E-08 |
| 3448 | 173 | C173G | A | 6.5e-09 | 7.6e-09 | 9.3e-09 | 7.800E-09 | 1.411E-09 | 6.500E-09 | 9.300E-09 |
| 3449 | 173 | C173H | A | 7.5e-09 | 5.2e-09 | 7.7e-09 | 6.800E-09 | 1.389E-09 | 5.200E-09 | 7.700E-09 |
| 3450 | 173 | C173I | A | 7.1e-09 | 1.6e-08 | 3.3e-08 | 1.870E-08 | 1.316E-08 | 7.100E-09 | 3.300E-08 |
| 3451 | 173 | C173L | A | 5.1e-09 | 2.4e-09 | 9.6e-09 | 5.700E-09 | 3.637E-09 | 2.400E-09 | 9.600E-09 |
| 3452 | 173 | C173K | A | 8.3e-09 | 1e-08 | 3.5e-08 | 1.777E-08 | 1.495E-08 | 8.300E-09 | 3.500E-08 |
| 3453 | 173 | C173M | A | 4.7e-09 | 1e-08 | 5.8e-09 | 6.833E-09 | 2.797E-09 | 4.700E-09 | 1.000E-08 |
| 3454 | 173 | C173F | A | 3.6e-09 | 5.5e-09 | 2.1e-09 | 3.733E-09 | 1.704E-09 | 2.100E-09 | 5.500E-09 |
| 3455 | 173 | C173P | A | 1.4e-08 | 9.1e-09 | 6.2e-09 | 9.767E-09 | 3.943E-09 | 6.200E-09 | 1.400E-08 |
| 3456 | 173 | C173S | A | 1.5e-09 | 2.1e-08 | 1.6e-08 | 1.283E-08 | 1.013E-08 | 1.500E-09 | 2.100E-08 |
| 3457 | 173 | C173T | A | 2.9e-08 | 5.4e-09 | 1.7e-09 | 1.203E-08 | 1.481E-08 | 1.700E-09 | 2.900E-08 |
| 3458 | 173 | C173W | A | 1.9e-08 | 7.7e-09 | 1.5e-08 | 1.390E-08 | 5.730E-09 | 7.700E-09 | 1.900E-08 |
| 3459 | 173 | C173Y | A | 6.9e-09 | 2.6e-08 | 3.9e-09 | 1.227E-08 | 1.199E-08 | 3.900E-09 | 2.600E-08 |
| 3460 | 173 | C173V | A | 4.4e-09 | 1.5e-08 | 3.9e-09 | 7.767E-09 | 6.269E-09 | 3.900E-09 | 1.500E-08 |
| 3461 | 174 | S174A | A | 1e-08 | 1.3e-08 | 5.1e-09 | 9.367E-09 | 3.988E-09 | 5.100E-09 | 1.300E-08 |
| 3462 | 174 | S174R | A | 1.8e-09 | 4.7e-09 | 3.4e-08 | 1.350E-08 | 1.781E-08 | 1.800E-09 | 3.400E-08 |
| 3463 | 174 | S174N | A | 1.1e-08 | 1.5e-08 | 2.1e-09 | 9.367E-09 | 6.603E-09 | 2.100E-09 | 1.500E-08 |
| 3464 | 174 | S174D | A | 3.3e-09 | 7.9e-09 | 1.2e-08 | 7.733E-09 | 4.352E-09 | 3.300E-09 | 1.200E-08 |
| 3465 | 174 | S174C | A | 1.5e-08 | 8.4e-09 | 5e-09 | 9.467E-09 | 5.085E-09 | 5.000E-09 | 1.500E-08 |
| 3466 | 174 | S174E | A | 8.7e-09 | 1e-08 | 9.7e-09 | 9.467E-09 | 6.807E-10 | 8.700E-09 | 1.000E-08 |
| 3467 | 174 | S174Q | A | 1.4e-08 | 7.2e-09 | 3.7e-09 | 8.300E-09 | 5.237E-09 | 3.700E-09 | 1.400E-08 |
| 3468 | 174 | S174G | A | 6.6e-09 | 6.9e-09 | 7e-09 | 6.833E-09 | 2.082E-10 | 6.600E-09 | 7.000E-09 |
| 3469 | 174 | S174H | A | 2.7e-09 | 1e-08 | 4.2e-09 | 5.633E-09 | 3.855E-09 | 2.700E-09 | 1.000E-08 |
| 3470 | 174 | S174I | A | 8e-09 | 5.2e-09 | 4.3e-09 | 5.833E-09 | 1.930E-09 | 4.300E-09 | 8.000E-09 |
| 3471 | 174 | S174L | A | 5.9e-09 | 1.2e-08 | 4.8e-09 | 7.567E-09 | 3.879E-09 | 4.800E-09 | 1.200E-08 |
| 3472 | 174 | S174K | A | 2e-08 | 9.8e-09 | 7e-09 | 1.227E-08 | 6.842E-09 | 7.000E-09 | 2.000E-08 |
| 3473 | 174 | S174M | A | 1.2e-08 | 1.2e-08 | 8.6e-09 | 1.087E-08 | 1.963E-09 | 8.600E-09 | 1.200E-08 |
| 3474 | 174 | S174F | A | 9.6e-09 | 9.1e-09 | 1.1e-08 | 9.900E-09 | 9.849E-10 | 9.100E-09 | 1.100E-08 |
| 3475 | 174 | S174P | A | 1.2e-08 | 4.2e-09 | 1.4e-08 | 1.007E-08 | 5.178E-09 | 4.200E-09 | 1.400E-08 |
| 3476 | 174 | S174S | A | 5.3e-09 | 9.2e-09 | 3e-09 | 5.833E-09 | 3.134E-09 | 3.000E-09 | 9.200E-09 |
| 3477 | 174 | S174T | A | 3.6e-09 | 5e-09 | 1.4e-08 | 7.533E-09 | 5.644E-09 | 3.600E-09 | 1.400E-08 |
| 3478 | 174 | S174W | A | 1.2e-08 | 8.9e-09 | 2.7e-08 | 1.597E-08 | 9.680E-09 | 8.900E-09 | 2.700E-08 |
| 3479 | 174 | S174Y | A | 5.1e-09 | 5.3e-09 | 7.7e-09 | 6.033E-09 | 1.447E-09 | 5.100E-09 | 7.700E-09 |
| 3480 | 174 | S174V | A | 7.6e-09 | 1.5e-08 | 1.1e-08 | 1.120E-08 | 3.704E-09 | 7.600E-09 | 1.500E-08 |
| 3481 | 175 | P175A | A | 6.4e-09 | 6.6e-09 | 8.9e-09 | 7.300E-09 | 1.389E-09 | 6.400E-09 | 8.900E-09 |
| 3482 | 175 | P175R | A | 1.1e-08 | 5.6e-09 | 1e-08 | 8.867E-09 | 2.873E-09 | 5.600E-09 | 1.100E-08 |
| 3483 | 175 | P175N | A | 4e-09 | 8.4e-09 | 1.3e-08 | 8.467E-09 | 4.500E-09 | 4.000E-09 | 1.300E-08 |
| 3484 | 175 | P175D | A | 1.1e-08 | 6.4e-09 | 3.9e-09 | 7.100E-09 | 3.601E-09 | 3.900E-09 | 1.100E-08 |

| | | | | | | | | | | |
|---|---|---|---|---|---|---|---|---|---|---|
| 3485 | 175 | P175C | A | 7.3e-09 | 1.3e-08 | 3.1e-09 | 7.800E-09 | 4.969E-09 | 3.100E-09 | 1.300E-08 |
| 3486 | 175 | P175E | A | 1.5e-09 | 8.1e-09 | 9.5e-09 | 6.367E-09 | 4.272E-09 | 1.500E-09 | 9.500E-09 |
| 3487 | 175 | P175Q | A | 6.1e-09 | 7.5e-09 | 1.5e-08 | 9.533E-09 | 4.786E-09 | 6.100E-09 | 1.500E-08 |
| 3488 | 175 | P175G | A | 6.6e-09 | 1.3e-08 | 5.5e-09 | 8.367E-09 | 4.050E-09 | 5.500E-09 | 1.300E-08 |
| 3489 | 175 | P175H | A | 1.5e-08 | 5.4e-09 | 8.4e-09 | 9.600E-09 | 4.911E-09 | 5.400E-09 | 1.500E-08 |
| 3490 | 175 | P175I | A | 4.9e-09 | 6.1e-09 | 3.2e-09 | 4.733E-09 | 1.457E-09 | 3.200E-09 | 6.100E-09 |
| 3491 | 175 | P175L | A | 3.4e-09 | 1e-08 | 1.7e-08 | 1.013E-08 | 6.801E-09 | 3.400E-09 | 1.700E-08 |
| 3492 | 175 | P175K | A | 9e-09 | 8.1e-09 | 7.7e-09 | 8.267E-09 | 6.658E-10 | 7.700E-09 | 9.000E-09 |
| 3493 | 175 | P175M | A | 7.4e-09 | 6.9e-09 | 5.6e-09 | 6.633E-09 | 9.292E-10 | 5.600E-09 | 7.400E-09 |
| 3494 | 175 | P175F | A | 2.8e-08 | 5.4e-09 | 1.3e-08 | 1.547E-08 | 1.150E-08 | 5.400E-09 | 2.800E-08 |
| 3495 | 175 | P175P | A | 1.4e-08 | 2.5e-08 | 1.5e-08 | 1.800E-08 | 6.083E-09 | 1.400E-08 | 2.500E-08 |
| 3496 | 175 | P175S | A | 1.6e-08 | 6.7e-09 | 1.1e-09 | 7.933E-09 | 7.526E-09 | 1.100E-09 | 1.600E-08 |
| 3497 | 175 | P175T | A | 6.5e-09 | 4.2e-09 | 6.9e-09 | 5.867E-09 | 1.457E-09 | 4.200E-09 | 6.900E-09 |
| 3498 | 175 | P175W | A | 4.4e-09 | 5.2e-09 | 2.8e-09 | 4.133E-09 | 1.222E-09 | 2.800E-09 | 5.200E-09 |
| 3499 | 175 | P175Y | A | 5e-09 | 1.6e-09 | 6e-09 | 4.200E-09 | 2.307E-09 | 1.600E-09 | 6.000E-09 |
| 3500 | 175 | P175V | A | 3.4e-09 | 4e-08 | 3.7e-09 | 1.570E-08 | 2.104E-08 | 3.400E-09 | 4.000E-08 |
| 3501 | 176 | M176A | A | 3.7e-09 | 1e-08 | 6.4e-09 | 6.700E-09 | 3.161E-09 | 3.700E-09 | 1.000E-08 |
| 3502 | 176 | M176R | A | 3.8e-09 | 1.4e-08 | 7.2e-09 | 8.333E-09 | 5.194E-09 | 3.800E-09 | 1.400E-08 |
| 3503 | 176 | M176N | A | 9.7e-09 | 4.6e-09 | 3.9e-09 | 6.067E-09 | 3.166E-09 | 3.900E-09 | 9.700E-09 |
| 3504 | 176 | M176D | A | 1.9e-08 | 6.8e-09 | 3.2e-09 | 9.667E-09 | 8.281E-09 | 3.200E-09 | 1.900E-08 |
| 3505 | 176 | M176C | A | 4.4e-09 | 5.1e-09 | 3.7e-09 | 4.400E-09 | 7.000E-10 | 3.700E-09 | 5.100E-09 |
| 3506 | 176 | M176E | A | 4.8e-09 | 1.3e-08 | 8.9e-09 | 8.900E-09 | 4.100E-09 | 4.800E-09 | 1.300E-08 |
| 3507 | 176 | M176Q | A | 7.7e-09 | 7.7e-09 | 3.7e-09 | 6.367E-09 | 2.309E-09 | 3.700E-09 | 7.700E-09 |
| 3508 | 176 | M176G | A | 7.8e-09 | 8.5e-09 | 1.4e-08 | 1.010E-08 | 3.396E-09 | 7.800E-09 | 1.400E-08 |
| 3509 | 176 | M176H | A | 2.6e-09 | 1.4e-09 | 8.4e-09 | 4.133E-09 | 3.743E-09 | 1.400E-09 | 8.400E-09 |
| 3510 | 176 | M176I | A | 2.8e-09 | 3.1e-09 | 8.9e-09 | 4.933E-09 | 3.439E-09 | 2.800E-09 | 8.900E-09 |
| 3511 | 176 | M176L | A | 9.4e-09 | 3.6e-09 | 5e-09 | 6.000E-09 | 3.027E-09 | 3.600E-09 | 9.400E-09 |
| 3512 | 176 | M176K | A | 6.2e-09 | 9.8e-09 | 8.8e-09 | 8.267E-09 | 1.858E-09 | 6.200E-09 | 9.800E-09 |
| 3513 | 176 | M176M | A | 8.1e-09 | 3.4e-09 | 3.3e-09 | 4.933E-09 | 2.743E-09 | 3.300E-09 | 8.100E-09 |
| 3514 | 176 | M176F | A | 1.2e-08 | 9.9e-09 | 4.3e-09 | 8.733E-09 | 3.980E-09 | 4.300E-09 | 1.200E-08 |
| 3515 | 176 | M176P | A | 6.1e-09 | 3.2e-09 | 1.8e-08 | 9.100E-09 | 7.843E-09 | 3.200E-09 | 1.800E-08 |
| 3516 | 176 | M176S | A | 7.6e-09 | 1.1e-08 | 1.2e-09 | 6.600E-09 | 4.976E-09 | 1.200E-09 | 1.100E-08 |
| 3517 | 176 | M176T | A | 5.2e-09 | 1.4e-08 | 1.1e-08 | 1.007E-08 | 4.474E-09 | 5.200E-09 | 1.400E-08 |
| 3518 | 176 | M176W | A | 2.6e-09 | 1.4e-08 | 1.9e-09 | 6.167E-09 | 6.793E-09 | 1.900E-09 | 1.400E-08 |
| 3519 | 176 | M176Y | A | 4.8e-09 | 4.3e-09 | 3.1e-09 | 4.067E-09 | 8.737E-10 | 3.100E-09 | 4.800E-09 |
| 3520 | 176 | M176V | A | 6.5e-09 | 5.9e-09 | 1e-08 | 7.467E-09 | 2.214E-09 | 5.900E-09 | 1.000E-08 |
| 3521 | 177 | C177A | A | 2.7e-09 | 4.2e-09 | 5e-09 | 3.967E-09 | 1.168E-09 | 2.700E-09 | 5.000E-09 |
| 3522 | 177 | C177R | A | 3.1e-08 | 3.3e-09 | 4.2e-09 | 1.283E-08 | 1.574E-08 | 3.300E-09 | 3.100E-08 |
| 3523 | 177 | C177N | A | 6.6e-09 | 5.4e-09 | 7.5e-09 | 6.500E-09 | 1.054E-09 | 5.400E-09 | 7.500E-09 |
| 3524 | 177 | C177D | A | 9.8e-09 | 7.2e-09 | 1.2e-08 | 9.667E-09 | 2.403E-09 | 7.200E-09 | 1.200E-08 |
| 3525 | 177 | C177C | A | 5.6e-09 | 1e-08 | 5.2e-09 | 6.933E-09 | 2.663E-09 | 5.200E-09 | 1.000E-08 |
| 3526 | 177 | C177E | A | 6.3e-09 | 5.7e-09 | 2.8e-08 | 1.333E-08 | 1.271E-08 | 5.700E-09 | 2.800E-08 |
| 3527 | 177 | C177Q | A | 1.5e-08 | 4.4e-09 | 1.6e-09 | 7.000E-09 | 7.068E-09 | 1.600E-09 | 1.500E-08 |
| 3528 | 177 | C177G | A | 2.1e-09 | 6e-09 | 1.7e-09 | 3.267E-09 | 2.376E-09 | 1.700E-09 | 6.000E-09 |
| 3529 | 177 | C177H | A | 1.1e-08 | 8e-09 | 1.8e-08 | 1.233E-08 | 5.132E-09 | 8.000E-09 | 1.800E-08 |
| 3530 | 177 | C177I | A | 6.5e-09 | 1.4e-08 | 4.8e-09 | 8.433E-09 | 4.895E-09 | 4.800E-09 | 1.400E-08 |
| 3531 | 177 | C177L | A | 4.3e-09 | 9.5e-09 | 5.5e-09 | 6.433E-09 | 2.723E-09 | 4.300E-09 | 9.500E-09 |
| 3532 | 177 | C177K | A | 7.1e-09 | 5.1e-09 | 7.3e-09 | 6.500E-09 | 1.217E-09 | 5.100E-09 | 7.300E-09 |
| 3533 | 177 | C177M | A | 7.1e-09 | 5.4e-09 | 4e-09 | 5.500E-09 | 1.552E-09 | 4.000E-09 | 7.100E-09 |
| 3534 | 177 | C177F | A | 7.1e-09 | 2.7e-09 | 1.3e-08 | 7.600E-09 | 5.168E-09 | 2.700E-09 | 1.300E-08 |
| 3535 | 177 | C177P | A | 1.4e-08 | 1.3e-08 | 7.1e-09 | 1.137E-08 | 3.729E-09 | 7.100E-09 | 1.400E-08 |
| 3536 | 177 | C177S | A | 8.5e-09 | 6.4e-09 | 6.6e-09 | 7.167E-09 | 1.159E-09 | 6.400E-09 | 8.500E-09 |
| 3537 | 177 | C177T | A | 1.1e-08 | 1e-08 | 3.9e-09 | 8.300E-09 | 3.843E-09 | 3.900E-09 | 1.100E-08 |
| 3538 | 177 | C177W | A | 7.7e-09 | 7.2e-09 | 2.9e-09 | 5.933E-09 | 2.639E-09 | 2.900E-09 | 7.700E-09 |
| 3539 | 177 | C177Y | A | 5.9e-09 | 9.6e-09 | 7.1e-09 | 7.533E-09 | 1.888E-09 | 5.900E-09 | 9.600E-09 |
| 3540 | 177 | C177V | A | 4.3e-09 | 4.1e-09 | 7.2e-09 | 5.200E-09 | 1.735E-09 | 4.100E-09 | 7.200E-09 |
| 3541 | 178 | K178A | A | 1.1e-08 | 3.8e-09 | 5.5e-09 | 6.767E-09 | 3.763E-09 | 3.800E-09 | 1.100E-08 |
| 3542 | 178 | K178R | A | 1e-08 | 2.1e-09 | 8.6e-09 | 6.900E-09 | 4.215E-09 | 2.100E-09 | 1.000E-08 |
| 3543 | 178 | K178N | A | 1.1e-08 | 8.6e-09 | 6.3e-09 | 8.633E-09 | 2.350E-09 | 6.300E-09 | 1.100E-08 |
| 3544 | 178 | K178D | A | 6.1e-09 | 1.1e-08 | 7.3e-09 | 8.133E-09 | 2.554E-09 | 6.100E-09 | 1.100E-08 |
| 3545 | 178 | K178C | A | 7.3e-09 | 7.1e-09 | 7.6e-09 | 7.333E-09 | 2.517E-10 | 7.100E-09 | 7.600E-09 |
| 3546 | 178 | K178E | A | 3.1e-09 | 8.2e-09 | 8.9e-09 | 6.733E-09 | 3.166E-09 | 3.100E-09 | 8.900E-09 |
| 3547 | 178 | K178Q | A | 4.4e-09 | 5e-09 | 3.5e-09 | 4.300E-09 | 7.550E-10 | 3.500E-09 | 5.000E-09 |
| 3548 | 178 | K178G | A | 3.9e-09 | 4.5e-09 | 4.9e-09 | 4.433E-09 | 5.033E-10 | 3.900E-09 | 4.900E-09 |
| 3549 | 178 | K178H | A | 8.6e-10 | 2.9e-09 | 1.6e-08 | 6.587E-09 | 8.216E-09 | 8.600E-10 | 1.600E-08 |
| 3550 | 178 | K178I | A | 1.2e-08 | 3.1e-09 | 1.6e-08 | 1.037E-08 | 6.603E-09 | 3.100E-09 | 1.600E-08 |
| 3551 | 178 | K178L | A | 9.2e-09 | 6.6e-09 | 2.1e-09 | 5.967E-09 | 3.592E-09 | 2.100E-09 | 9.200E-09 |
| 3552 | 178 | K178K | A | 7.6e-09 | 8.8e-09 | 2.4e-09 | 6.267E-09 | 3.402E-09 | 2.400E-09 | 8.800E-09 |
| 3553 | 178 | K178M | A | 3.3e-09 | 4.9e-09 | 3.4e-09 | 3.867E-09 | 8.963E-10 | 3.300E-09 | 4.900E-09 |
| 3554 | 178 | K178F | A | 4.4e-09 | 1.5e-08 | 4.9e-09 | 8.100E-09 | 5.981E-09 | 4.400E-09 | 1.500E-08 |
| 3555 | 178 | K178P | A | 1.6e-08 | 7.2e-09 | 1.4e-09 | 8.200E-09 | 7.351E-09 | 1.400E-09 | 1.600E-08 |
| 3556 | 178 | K178S | A | 2.9e-09 | 4.4e-09 | 2.7e-09 | 3.333E-09 | 9.292E-10 | 2.700E-09 | 4.400E-09 |
| 3557 | 178 | K178T | A | 2.3e-09 | 4.5e-09 | 2.3e-09 | 3.033E-09 | 1.270E-09 | 2.300E-09 | 4.500E-09 |
| 3558 | 178 | K178W | A | 1.6e-08 | 7.2e-09 | 1e-08 | 1.107E-08 | 4.496E-09 | 7.200E-09 | 1.600E-08 |
| 3559 | 178 | K178Y | A | 6.1e-09 | 7.9e-09 | 9e-09 | 7.667E-09 | 1.464E-09 | 6.100E-09 | 9.000E-09 |
| 3560 | 178 | K178V | A | 1.3e-08 | 9.5e-09 | 5.7e-09 | 9.400E-09 | 3.651E-09 | 5.700E-09 | 1.300E-08 |
| 3561 | 179 | G179A | A | 7.4e-09 | 1.5e-08 | 9.5e-09 | 1.063E-08 | 3.925E-09 | 7.400E-09 | 1.500E-08 |
| 3562 | 179 | G179R | A | 3.8e-09 | 7.1e-09 | 7.5e-09 | 6.133E-09 | 2.031E-09 | 3.800E-09 | 7.500E-09 |
| 3563 | 179 | G179N | A | 8.2e-09 | 7.3e-09 | 1.1e-08 | 8.833E-09 | 1.930E-09 | 7.300E-09 | 1.100E-08 |
| 3564 | 179 | G179D | A | 3.9e-09 | 2e-09 | 3.8e-09 | 3.233E-09 | 1.069E-09 | 2.000E-09 | 3.900E-09 |
| 3565 | 179 | G179C | A | 6.2e-09 | 5.3e-09 | 1.3e-08 | 8.167E-09 | 4.210E-09 | 5.300E-09 | 1.300E-08 |
| 3566 | 179 | G179E | A | 1.6e-09 | 2.6e-09 | 8.8e-09 | 4.333E-09 | 3.900E-09 | 1.600E-09 | 8.800E-09 |
| 3567 | 179 | G179Q | A | 2.5e-08 | 1.1e-08 | 1e-08 | 1.533E-08 | 8.386E-09 | 1.000E-08 | 2.500E-08 |
| 3568 | 179 | G179G | A | 1e-08 | 1.2e-08 | 1.2e-08 | 1.133E-08 | 1.155E-09 | 1.000E-08 | 1.200E-08 |
| 3569 | 179 | G179H | A | 2.1e-08 | 7.5e-09 | 1.8e-08 | 1.550E-08 | 7.089E-09 | 7.500E-09 | 2.100E-08 |

| | | | | | | | | | | |
|---|---|---|---|---|---|---|---|---|---|---|
| 3570 | 179 | G179I | A | 6.5e-09 | 9.2e-09 | 2.5e-09 | 6.067E-09 | 3.371E-09 | 2.500E-09 | 9.200E-09 |
| 3571 | 179 | G179L | A | 1.4e-08 | 7.9e-09 | 6.3e-09 | 9.400E-09 | 4.063E-09 | 6.300E-09 | 1.400E-08 |
| 3572 | 179 | G179K | A | 1.6e-08 | 7.4e-09 | 1.7e-08 | 1.347E-08 | 5.278E-09 | 7.400E-09 | 1.700E-08 |
| 3573 | 179 | G179M | A | 4.1e-09 | 5.4e-09 | 5.8e-09 | 5.100E-09 | 8.888E-10 | 4.100E-09 | 5.800E-09 |
| 3574 | 179 | G179F | A | 4.3e-09 | 1.2e-08 | 8.3e-09 | 8.200E-09 | 3.851E-09 | 4.300E-09 | 1.200E-08 |
| 3575 | 179 | G179P | A | 1.8e-09 | 4.9e-09 | 3.9e-09 | 3.533E-09 | 1.582E-09 | 1.800E-09 | 4.900E-09 |
| 3576 | 179 | G179S | A | 8.4e-09 | 1.5e-08 | 2.7e-09 | 8.700E-09 | 6.155E-09 | 2.700E-09 | 1.500E-08 |
| 3577 | 179 | G179T | A | 4.9e-09 | 7e-09 | 3.4e-09 | 5.100E-09 | 1.808E-09 | 3.400E-09 | 7.000E-09 |
| 3578 | 179 | G179W | A | 3.5e-09 | 9.5e-09 | 1.7e-08 | 1.000E-08 | 6.764E-09 | 3.500E-09 | 1.700E-08 |
| 3579 | 179 | G179Y | A | 2.6e-09 | 7.6e-09 | 8.8e-09 | 6.333E-09 | 3.288E-09 | 2.600E-09 | 8.800E-09 |
| 3580 | 179 | G179V | A | 4.9e-09 | 9.1e-09 | 4.9e-09 | 6.300E-09 | 2.425E-09 | 4.900E-09 | 9.100E-09 |
| 3581 | 180 | S180A | A | 9.4e-09 | 6.5e-09 | 2.5e-09 | 6.133E-09 | 3.465E-09 | 2.500E-09 | 9.400E-09 |
| 3582 | 180 | S180R | A | 1.1e-08 | 9.2e-09 | 1.9e-08 | 1.307E-08 | 5.217E-09 | 9.200E-09 | 1.900E-08 |
| 3583 | 180 | S180N | A | 6.8e-09 | 6.2e-09 | 6.6e-09 | 6.533E-09 | 3.055E-10 | 6.200E-09 | 6.800E-09 |
| 3584 | 180 | S180D | A | 6.8e-09 | 6.6e-09 | 2.2e-08 | 1.180E-08 | 8.834E-09 | 6.600E-09 | 2.200E-08 |
| 3585 | 180 | S180C | A | 5.3e-09 | 5.5e-09 | 4.7e-09 | 5.167E-09 | 4.163E-10 | 4.700E-09 | 5.500E-09 |
| 3586 | 180 | S180E | A | 4.2e-09 | 9.5e-09 | 5.6e-09 | 6.433E-09 | 2.747E-09 | 4.200E-09 | 9.500E-09 |
| 3587 | 180 | S180Q | A | 3.6e-09 | 6.8e-09 | 9.8e-09 | 6.733E-09 | 3.101E-09 | 3.600E-09 | 9.800E-09 |
| 3588 | 180 | S180G | A | 1.2e-08 | 1.7e-08 | 1.2e-08 | 1.367E-08 | 2.887E-09 | 1.200E-08 | 1.700E-08 |
| 3589 | 180 | S180H | A | 2.3e-08 | 7e-09 | 8.6e-09 | 1.287E-08 | 8.812E-09 | 7.000E-09 | 2.300E-08 |
| 3590 | 180 | S180I | A | 7.1e-09 | 3.8e-09 | 9.6e-09 | 6.833E-09 | 2.909E-09 | 3.800E-09 | 9.600E-09 |
| 3591 | 180 | S180L | A | 1.2e-08 | 6.9e-09 | 2.3e-09 | 7.067E-09 | 4.852E-09 | 2.300E-09 | 1.200E-08 |
| 3592 | 180 | S180K | A | 6.8e-09 | 1.1e-08 | 9.7e-09 | 9.167E-09 | 2.150E-09 | 6.800E-09 | 1.100E-08 |
| 3593 | 180 | S180M | A | 5.7e-09 | 7e-09 | 7.9e-09 | 6.867E-09 | 1.106E-09 | 5.700E-09 | 7.900E-09 |
| 3594 | 180 | S180F | A | 6.7e-09 | 8.6e-09 | 2e-09 | 5.767E-09 | 3.398E-09 | 2.000E-09 | 8.600E-09 |
| 3595 | 180 | S180P | A | 4e-09 | 6.6e-09 | 4.7e-09 | 5.100E-09 | 1.345E-09 | 4.000E-09 | 6.600E-09 |
| 3596 | 180 | S180S | A | 7.9e-09 | 8.7e-09 | 9.1e-09 | 8.567E-09 | 6.110E-10 | 7.900E-09 | 9.100E-09 |
| 3597 | 180 | S180T | A | 3.3e-09 | 2.7e-09 | 2.4e-09 | 2.800E-09 | 4.583E-10 | 2.400E-09 | 3.300E-09 |
| 3598 | 180 | S180W | A | 9e-09 | 6.2e-09 | 1.1e-08 | 8.733E-09 | 2.411E-09 | 6.200E-09 | 1.100E-08 |
| 3599 | 180 | S180Y | A | 4.2e-09 | 7.1e-09 | 3.7e-09 | 5.000E-09 | 1.836E-09 | 3.700E-09 | 7.100E-09 |
| 3600 | 180 | S180V | A | 3.7e-08 | 1.3e-08 | 7.7e-09 | 1.923E-08 | 1.561E-08 | 7.700E-09 | 3.700E-08 |
| 3601 | 181 | R181A | A | 2e-08 | 1.5e-08 | 3.5e-09 | 1.283E-08 | 8.461E-09 | 3.500E-09 | 2.000E-08 |
| 3602 | 181 | R181R | A | 3e-09 | 1.1e-08 | 8e-09 | 7.333E-09 | 4.041E-09 | 3.000E-09 | 1.100E-08 |
| 3603 | 181 | R181N | A | 9.2e-09 | 1.4e-08 | 1e-08 | 1.107E-08 | 2.572E-09 | 9.200E-09 | 1.400E-08 |
| 3604 | 181 | R181D | A | 1.5e-08 | 4.3e-09 | 6.5e-09 | 8.600E-09 | 5.651E-09 | 4.300E-09 | 1.500E-08 |
| 3605 | 181 | R181C | A | 4.7e-09 | 7.2e-09 | 9.4e-09 | 7.100E-09 | 2.352E-09 | 4.700E-09 | 9.400E-09 |
| 3606 | 181 | R181E | A | 6.4e-09 | 2.9e-09 | 1.1e-08 | 6.767E-09 | 4.062E-09 | 2.900E-09 | 1.100E-08 |
| 3607 | 181 | R181Q | A | 3.5e-09 | 8.2e-09 | 1.2e-08 | 7.900E-09 | 4.258E-09 | 3.500E-09 | 1.200E-08 |
| 3608 | 181 | R181G | A | 5.6e-09 | 1.3e-08 | 6.6e-09 | 8.400E-09 | 4.015E-09 | 5.600E-09 | 1.300E-08 |
| 3609 | 181 | R181H | A | 1.3e-08 | 8.5e-09 | 7e-09 | 9.500E-09 | 3.122E-09 | 7.000E-09 | 1.300E-08 |
| 3610 | 181 | R181I | A | 5e-09 | 1.5e-08 | 3.3e-09 | 7.767E-09 | 6.322E-09 | 3.300E-09 | 1.500E-08 |
| 3611 | 181 | R181L | A | 8e-09 | 7.3e-09 | 5.8e-09 | 7.033E-09 | 1.124E-09 | 5.800E-09 | 8.000E-09 |
| 3612 | 181 | R181K | A | 1.5e-08 | 5.6e-09 | 1.7e-08 | 1.253E-08 | 6.087E-09 | 5.600E-09 | 1.700E-08 |
| 3613 | 181 | R181M | A | 4.1e-09 | 1.7e-08 | 1.4e-08 | 1.170E-08 | 6.751E-09 | 4.100E-09 | 1.700E-08 |
| 3614 | 181 | R181F | A | 1.1e-08 | 7.8e-09 | 8.4e-09 | 9.067E-09 | 1.701E-09 | 7.800E-09 | 1.100E-08 |
| 3615 | 181 | R181P | A | 1.2e-08 | 7.3e-09 | 2.9e-09 | 7.400E-09 | 4.551E-09 | 2.900E-09 | 1.200E-08 |
| 3616 | 181 | R181S | A | 3.8e-09 | 5.1e-09 | 6.5e-09 | 5.133E-09 | 1.350E-09 | 3.800E-09 | 6.500E-09 |
| 3617 | 181 | R181T | A | 4.5e-09 | 7e-09 | 7.4e-09 | 6.300E-09 | 1.572E-09 | 4.500E-09 | 7.400E-09 |
| 3618 | 181 | R181W | A | 1.3e-09 | 8.7e-09 | 4.9e-09 | 4.967E-09 | 3.700E-09 | 1.300E-09 | 8.700E-09 |
| 3619 | 181 | R181Y | A | 1.2e-08 | 4.2e-09 | 1.1e-08 | 9.067E-09 | 4.244E-09 | 4.200E-09 | 1.200E-08 |
| 3620 | 181 | R181V | A | 2.7e-08 | 2.4e-09 | 6.3e-09 | 1.190E-08 | 1.322E-08 | 2.400E-09 | 2.700E-08 |
| 3621 | 182 | C182A | A | 1.8e-08 | 7.1e-09 | 4.2e-09 | 9.767E-09 | 7.276E-09 | 4.200E-09 | 1.800E-08 |
| 3622 | 182 | C182R | A | 4.7e-09 | 1.6e-08 | 7.7e-09 | 9.467E-09 | 5.853E-09 | 4.700E-09 | 1.600E-08 |
| 3623 | 182 | C182N | A | 9.1e-09 | 8.3e-09 | 7.4e-09 | 8.267E-09 | 8.505E-10 | 7.400E-09 | 9.100E-09 |
| 3624 | 182 | C182D | A | 6.4e-09 | 5.3e-09 | 7.8e-09 | 6.500E-09 | 1.253E-09 | 5.300E-09 | 7.800E-09 |
| 3625 | 182 | C182C | A | 4.5e-09 | 7.9e-09 | 1.1e-08 | 7.800E-09 | 3.251E-09 | 4.500E-09 | 1.100E-08 |
| 3626 | 182 | C182E | A | 9.2e-09 | 1.1e-08 | 1.8e-09 | 7.333E-09 | 4.876E-09 | 1.800E-09 | 1.100E-08 |
| 3627 | 182 | C182Q | A | 1e-08 | 7.2e-09 | 5.1e-09 | 7.433E-09 | 2.458E-09 | 5.100E-09 | 1.000E-08 |
| 3628 | 182 | C182G | A | 1e-09 | 3.1e-09 | 1e-08 | 4.700E-09 | 4.709E-09 | 1.000E-09 | 1.000E-08 |
| 3629 | 182 | C182H | A | 9.9e-09 | 7.3e-09 | 3e-09 | 6.733E-09 | 3.485E-09 | 3.000E-09 | 9.900E-09 |
| 3630 | 182 | C182I | A | 4.6e-09 | 9.7e-09 | 7.5e-09 | 7.267E-09 | 2.558E-09 | 4.600E-09 | 9.700E-09 |
| 3631 | 182 | C182L | A | 4.2e-09 | 3e-09 | 6e-09 | 4.400E-09 | 1.510E-09 | 3.000E-09 | 6.000E-09 |
| 3632 | 182 | C182K | A | 1.8e-09 | 6.5e-09 | 5e-09 | 4.433E-09 | 2.401E-09 | 1.800E-09 | 6.500E-09 |
| 3633 | 182 | C182M | A | 1.2e-08 | 8.7e-09 | 6.1e-09 | 8.933E-09 | 2.957E-09 | 6.100E-09 | 1.200E-08 |
| 3634 | 182 | C182F | A | 6.9e-09 | 2.5e-09 | 1.1e-08 | 6.800E-09 | 4.251E-09 | 2.500E-09 | 1.100E-08 |
| 3635 | 182 | C182P | A | 6.8e-09 | 7.9e-09 | 6.7e-09 | 7.133E-09 | 6.658E-10 | 6.700E-09 | 7.900E-09 |
| 3636 | 182 | C182S | A | 1.2e-08 | 6.8e-09 | 6.1e-09 | 8.300E-09 | 3.223E-09 | 6.100E-09 | 1.200E-08 |
| 3637 | 182 | C182T | A | 6.8e-09 | 3.7e-09 | 1.7e-08 | 9.167E-09 | 6.959E-09 | 3.700E-09 | 1.700E-08 |
| 3638 | 182 | C182W | A | 8.2e-09 | 6.4e-09 | 5.8e-09 | 6.800E-09 | 1.249E-09 | 5.800E-09 | 8.200E-09 |
| 3639 | 182 | C182Y | A | 5.6e-09 | 2.5e-09 | 8.8e-09 | 5.633E-09 | 3.150E-09 | 2.500E-09 | 8.800E-09 |
| 3640 | 182 | C182V | A | 9.7e-09 | 1.3e-08 | 1.3e-08 | 1.190E-08 | 1.905E-09 | 9.700E-09 | 1.300E-08 |
| 3641 | 183 | W183A | A | 1.8e-08 | 5.5e-09 | 8.4e-09 | 1.063E-08 | 6.542E-09 | 5.500E-09 | 1.800E-08 |
| 3642 | 183 | W183R | A | 5e-09 | 3.9e-09 | 1.1e-08 | 6.633E-09 | 3.821E-09 | 3.900E-09 | 1.100E-08 |
| 3643 | 183 | W183N | A | 1.5e-08 | 2.9e-09 | 2.2e-09 | 6.700E-09 | 7.197E-09 | 2.200E-09 | 1.500E-08 |
| 3644 | 183 | W183D | A | 1.8e-08 | 1.1e-08 | 4.3e-09 | 1.110E-08 | 6.851E-09 | 4.300E-09 | 1.800E-08 |
| 3645 | 183 | W183C | A | 1e-08 | 8.7e-09 | 9e-09 | 9.233E-09 | 6.807E-10 | 8.700E-09 | 1.000E-08 |
| 3646 | 183 | W183E | A | 4.9e-09 | 2.1e-09 | 6.2e-09 | 4.400E-09 | 2.095E-09 | 2.100E-09 | 6.200E-09 |
| 3647 | 183 | W183Q | A | 8.1e-09 | 1.6e-08 | 7e-09 | 1.037E-08 | 4.910E-09 | 7.000E-09 | 1.600E-08 |
| 3648 | 183 | W183G | A | 6.6e-09 | 1.6e-08 | 1.4e-08 | 1.220E-08 | 4.952E-09 | 6.600E-09 | 1.600E-08 |
| 3649 | 183 | W183H | A | 7.8e-09 | 7.1e-09 | 4.8e-09 | 6.567E-09 | 1.570E-09 | 4.800E-09 | 7.800E-09 |
| 3650 | 183 | W183I | A | 1.8e-08 | 2.1e-08 | 4.9e-09 | 1.463E-08 | 8.562E-09 | 4.900E-09 | 2.100E-08 |
| 3651 | 183 | W183L | A | 4e-08 | 2.1e-09 | 1.2e-08 | 1.803E-08 | 1.966E-08 | 2.100E-09 | 4.000E-08 |
| 3652 | 183 | W183K | A | 6.2e-09 | 8.1e-09 | 5.8e-09 | 6.700E-09 | 1.229E-09 | 5.800E-09 | 8.100E-09 |
| 3653 | 183 | W183M | A | 9.7e-09 | 2.4e-08 | 1.7e-08 | 1.690E-08 | 7.151E-09 | 9.700E-09 | 2.400E-08 |
| 3654 | 183 | W183F | A | 1.2e-08 | 4e-09 | 7.5e-09 | 7.833E-09 | 4.010E-09 | 4.000E-09 | 1.200E-08 |

| | | | | | | | | | | |
|---|---|---|---|---|---|---|---|---|---|---|
| 3655 | 183 | W183P | A | 1.2e-08 | 3.7e-09 | 1.8e-08 | 1.123E-08 | 7.181E-09 | 3.700E-09 | 1.800E-08 |
| 3656 | 183 | W183S | A | 6.5e-09 | 4.3e-09 | 1.1e-08 | 7.267E-09 | 3.415E-09 | 4.300E-09 | 1.100E-08 |
| 3657 | 183 | W183T | A | 2.7e-08 | 2.2e-08 | 7.4e-09 | 1.880E-08 | 1.018E-08 | 7.400E-09 | 2.700E-08 |
| 3658 | 183 | W183W | A | 6.5e-09 | 7.5e-09 | 1.2e-08 | 8.667E-09 | 2.930E-09 | 6.500E-09 | 1.200E-08 |
| 3659 | 183 | W183Y | A | 1.2e-08 | 7.4e-09 | 8.2e-09 | 9.200E-09 | 2.458E-09 | 7.400E-09 | 1.200E-08 |
| 3660 | 183 | W183V | A | 2.7e-09 | 1.1e-08 | 1.2e-08 | 8.567E-09 | 5.105E-09 | 2.700E-09 | 1.200E-08 |
| 3661 | 184 | G184A | A | 7.9e-09 | 1.4e-09 | 2.3e-08 | 1.077E-08 | 1.108E-08 | 1.400E-09 | 2.300E-08 |
| 3662 | 184 | G184R | A | 1.7e-08 | 5.1e-09 | 1.7e-08 | 1.303E-08 | 6.870E-09 | 5.100E-09 | 1.700E-08 |
| 3663 | 184 | G184N | A | 4.5e-09 | 4.1e-09 | 9.3e-09 | 5.967E-09 | 2.894E-09 | 4.100E-09 | 9.300E-09 |
| 3664 | 184 | G184D | A | 1.8e-08 | 1.3e-08 | 3.5e-08 | 2.200E-08 | 1.153E-08 | 1.300E-08 | 3.500E-08 |
| 3665 | 184 | G184C | A | 4.8e-09 | 9.7e-09 | 3.5e-09 | 6.000E-09 | 3.270E-09 | 3.500E-09 | 9.700E-09 |
| 3666 | 184 | G184E | A | 6.1e-09 | 1.2e-08 | 1.2e-08 | 1.003E-08 | 3.406E-09 | 6.100E-09 | 1.200E-08 |
| 3667 | 184 | G184Q | A | 4.4e-09 | 3.6e-09 | 6.5e-09 | 4.833E-09 | 1.498E-09 | 3.600E-09 | 6.500E-09 |
| 3668 | 184 | G184G | A | 8.8e-09 | 6.4e-09 | 2.6e-09 | 5.933E-09 | 3.126E-09 | 2.600E-09 | 8.800E-09 |
| 3669 | 184 | G184H | A | 1e-08 | 7.9e-09 | 7.1e-09 | 8.333E-09 | 1.498E-09 | 7.100E-09 | 1.000E-08 |
| 3670 | 184 | G184I | A | 7.8e-09 | 1.8e-08 | 4.9e-09 | 1.023E-08 | 6.881E-09 | 4.900E-09 | 1.800E-08 |
| 3671 | 184 | G184L | A | 6.8e-09 | 4.6e-09 | 6.2e-09 | 5.867E-09 | 1.137E-09 | 4.600E-09 | 6.800E-09 |
| 3672 | 184 | G184K | A | 1.1e-08 | 2.3e-09 | 3.3e-09 | 5.533E-09 | 4.761E-09 | 2.300E-09 | 1.100E-08 |
| 3673 | 184 | G184M | A | 2.2e-08 | 6.2e-09 | 1.2e-08 | 1.340E-08 | 7.992E-09 | 6.200E-09 | 2.200E-08 |
| 3674 | 184 | G184F | A | 5.5e-09 | 6.3e-09 | 9.4e-09 | 7.067E-09 | 2.060E-09 | 5.500E-09 | 9.400E-09 |
| 3675 | 184 | G184P | A | 5.7e-09 | 1.1e-08 | 6e-09 | 7.567E-09 | 2.977E-09 | 5.700E-09 | 1.100E-08 |
| 3676 | 184 | G184S | A | 9e-09 | 8.8e-09 | 3.2e-08 | 1.660E-08 | 1.334E-08 | 8.800E-09 | 3.200E-08 |
| 3677 | 184 | G184T | A | 1e-08 | 8.1e-09 | 6.1e-09 | 8.067E-09 | 1.950E-09 | 6.100E-09 | 1.000E-08 |
| 3678 | 184 | G184W | A | 4.2e-09 | 6.3e-09 | 4.7e-09 | 5.067E-09 | 1.097E-09 | 4.200E-09 | 6.300E-09 |
| 3679 | 184 | G184Y | A | 5.1e-09 | 3e-09 | 1.7e-08 | 8.367E-09 | 7.550E-09 | 3.000E-09 | 1.700E-08 |
| 3680 | 184 | G184V | A | 1.1e-08 | 1.2e-08 | 7.4e-09 | 1.013E-08 | 2.419E-09 | 7.400E-09 | 1.200E-08 |
| 3681 | 185 | E185A | A | 7.8e-09 | 6.6e-09 | 3.6e-09 | 6.000E-09 | 2.163E-09 | 3.600E-09 | 7.800E-09 |
| 3682 | 185 | E185R | A | 1.2e-08 | 4.8e-09 | 9.7e-09 | 8.833E-09 | 3.677E-09 | 4.800E-09 | 1.200E-08 |
| 3683 | 185 | E185N | A | 2.7e-09 | 1.5e-08 | 7e-09 | 8.233E-09 | 6.242E-09 | 2.700E-09 | 1.500E-08 |
| 3684 | 185 | E185D | A | 5.4e-09 | 7.4e-09 | 4.3e-09 | 5.700E-09 | 1.572E-09 | 4.300E-09 | 7.400E-09 |
| 3685 | 185 | E185C | A | 2.6e-09 | 9.7e-09 | 3.7e-09 | 5.333E-09 | 3.821E-09 | 2.600E-09 | 9.700E-09 |
| 3686 | 185 | E185E | A | 1.2e-08 | 2.9e-09 | 5.5e-09 | 6.800E-09 | 4.687E-09 | 2.900E-09 | 1.200E-08 |
| 3687 | 185 | E185Q | A | 1e-08 | 2.2e-08 | 4.1e-09 | 1.203E-08 | 9.122E-09 | 4.100E-09 | 2.200E-08 |
| 3688 | 185 | E185G | A | 5.2e-09 | 1.3e-08 | 3.8e-09 | 7.333E-09 | 4.957E-09 | 3.800E-09 | 1.300E-08 |
| 3689 | 185 | E185H | A | 7.4e-09 | 3.9e-09 | 6.1e-09 | 5.800E-09 | 1.769E-09 | 3.900E-09 | 7.400E-09 |
| 3690 | 185 | E185I | A | 6.6e-09 | 6.5e-09 | 9.7e-09 | 7.600E-09 | 1.819E-09 | 6.500E-09 | 9.700E-09 |
| 3691 | 185 | E185L | A | 2.1e-08 | 4e-09 | 1.3e-08 | 1.267E-08 | 8.505E-09 | 4.000E-09 | 2.100E-08 |
| 3692 | 185 | E185K | A | 7.5e-09 | 2.8e-09 | 2.3e-08 | 1.110E-08 | 1.057E-08 | 2.800E-09 | 2.300E-08 |
| 3693 | 185 | E185M | A | 3.6e-09 | 1.2e-08 | 6.3e-09 | 7.300E-09 | 4.288E-09 | 3.600E-09 | 1.200E-08 |
| 3694 | 185 | E185F | A | 5.9e-09 | 1.4e-08 | 5.4e-09 | 8.433E-09 | 4.827E-09 | 5.400E-09 | 1.400E-08 |
| 3695 | 185 | E185P | A | 1.1e-08 | 1.2e-08 | 1e-08 | 1.100E-08 | 1.000E-09 | 1.000E-08 | 1.200E-08 |
| 3696 | 185 | E185S | A | 3.5e-09 | 6.1e-09 | 9.2e-10 | 3.507E-09 | 2.590E-09 | 9.200E-10 | 6.100E-09 |
| 3697 | 185 | E185T | A | 4.6e-09 | 4.2e-09 | 2.7e-09 | 3.833E-09 | 1.002E-09 | 2.700E-09 | 4.600E-09 |
| 3698 | 185 | E185W | A | 7.2e-09 | 7.6e-09 | 8e-09 | 7.600E-09 | 4.000E-10 | 7.200E-09 | 8.000E-09 |
| 3699 | 185 | E185Y | A | 6.2e-09 | 6.1e-09 | 1.1e-08 | 7.767E-09 | 2.801E-09 | 6.100E-09 | 1.100E-08 |
| 3700 | 185 | E185V | A | 1.2e-08 | 1.3e-08 | 7.6e-09 | 1.087E-08 | 2.873E-09 | 7.600E-09 | 1.300E-08 |
| 3701 | 186 | S186A | A | 4.4e-09 | 4.8e-09 | 6.2e-09 | 5.133E-09 | 9.452E-10 | 4.400E-09 | 6.200E-09 |
| 3702 | 186 | S186R | A | 1.9e-09 | 1e-08 | 6e-09 | 5.967E-09 | 4.050E-09 | 1.900E-09 | 1.000E-08 |
| 3703 | 186 | S186N | A | 4.1e-09 | 1.5e-08 | 6.3e-09 | 8.467E-09 | 5.764E-09 | 4.100E-09 | 1.500E-08 |
| 3704 | 186 | S186D | A | 1.2e-08 | 1.2e-08 | 1.3e-08 | 1.233E-08 | 5.774E-10 | 1.200E-08 | 1.300E-08 |
| 3705 | 186 | S186C | A | 2.6e-09 | 1.6e-08 | 8.5e-09 | 9.033E-09 | 6.716E-09 | 2.600E-09 | 1.600E-08 |
| 3706 | 186 | S186E | A | 1.1e-08 | 1.2e-08 | 1.8e-08 | 1.367E-08 | 3.786E-09 | 1.100E-08 | 1.800E-08 |
| 3707 | 186 | S186Q | A | 1.8e-09 | 9e-09 | 2.4e-09 | 4.400E-09 | 3.995E-09 | 1.800E-09 | 9.000E-09 |
| 3708 | 186 | S186G | A | 2.5e-08 | 5e-09 | 1.8e-08 | 1.600E-08 | 1.015E-08 | 5.000E-09 | 2.500E-08 |
| 3709 | 186 | S186H | A | 3.4e-09 | 6.3e-09 | 5.4e-09 | 5.033E-09 | 1.484E-09 | 3.400E-09 | 6.300E-09 |
| 3710 | 186 | S186I | A | 1.4e-08 | 9.1e-09 | 1.4e-08 | 1.237E-08 | 2.829E-09 | 9.100E-09 | 1.400E-08 |
| 3711 | 186 | S186L | A | 1.1e-08 | 2.7e-09 | 7.6e-09 | 7.100E-09 | 4.173E-09 | 2.700E-09 | 1.100E-08 |
| 3712 | 186 | S186K | A | 6.5e-09 | 1e-08 | 1.5e-08 | 1.050E-08 | 4.272E-09 | 6.500E-09 | 1.500E-08 |
| 3713 | 186 | S186M | A | 9e-09 | 6.5e-09 | 2.6e-09 | 6.033E-09 | 3.225E-09 | 2.600E-09 | 9.000E-09 |
| 3714 | 186 | S186F | A | 8.3e-09 | 2.2e-08 | 9e-09 | 1.310E-08 | 7.716E-09 | 8.300E-09 | 2.200E-08 |
| 3715 | 186 | S186P | A | 1.2e-08 | 4.1e-09 | 5e-09 | 7.033E-09 | 4.325E-09 | 4.100E-09 | 1.200E-08 |
| 3716 | 186 | S186S | A | 1.1e-09 | 9.5e-09 | 1.5e-08 | 8.533E-09 | 7.000E-09 | 1.100E-09 | 1.500E-08 |
| 3717 | 186 | S186T | A | 3.3e-09 | 6.2e-09 | 6.1e-09 | 5.200E-09 | 1.646E-09 | 3.300E-09 | 6.200E-09 |
| 3718 | 186 | S186W | A | 2.9e-08 | 2.8e-09 | 4.1e-09 | 1.197E-08 | 1.477E-08 | 2.800E-09 | 2.900E-08 |
| 3719 | 186 | S186Y | A | 1.2e-08 | 1.6e-09 | 1.6e-08 | 9.867E-09 | 7.433E-09 | 1.600E-09 | 1.600E-08 |
| 3720 | 186 | S186V | A | 9e-09 | 9.2e-09 | 1.6e-08 | 1.140E-08 | 3.985E-09 | 9.000E-09 | 1.600E-08 |
| 3721 | 187 | S187A | A | 5.3e-09 | 1.2e-08 | 3.3e-09 | 6.867E-09 | 4.557E-09 | 3.300E-09 | 1.200E-08 |
| 3722 | 187 | S187R | A | 6.6e-09 | 2.4e-09 | 4.1e-09 | 4.367E-09 | 2.113E-09 | 2.400E-09 | 6.600E-09 |
| 3723 | 187 | S187N | A | 7.7e-09 | 1.1e-08 | 1.1e-08 | 9.900E-09 | 1.905E-09 | 7.700E-09 | 1.100E-08 |
| 3724 | 187 | S187D | A | 1.2e-08 | 8.4e-09 | 2.7e-09 | 7.700E-09 | 4.689E-09 | 2.700E-09 | 1.200E-08 |
| 3725 | 187 | S187C | A | 1.1e-08 | 1.3e-08 | 3.6e-09 | 9.200E-09 | 4.952E-09 | 3.600E-09 | 1.300E-08 |
| 3726 | 187 | S187E | A | 3.5e-09 | 7.8e-09 | 1.9e-09 | 4.400E-09 | 3.051E-09 | 1.900E-09 | 7.800E-09 |
| 3727 | 187 | S187Q | A | 2.1e-08 | 2.1e-08 | 4e-09 | 1.533E-08 | 9.815E-09 | 4.000E-09 | 2.100E-08 |
| 3728 | 187 | S187G | A | 3.4e-08 | 5.6e-09 | 1.8e-08 | 1.920E-08 | 1.424E-08 | 5.600E-09 | 3.400E-08 |
| 3729 | 187 | S187H | A | 6.2e-09 | 1.9e-08 | 6.8e-09 | 1.067E-08 | 7.223E-09 | 6.200E-09 | 1.900E-08 |
| 3730 | 187 | S187I | A | 7.2e-09 | 1.4e-08 | 8.8e-09 | 1.000E-08 | 3.555E-09 | 7.200E-09 | 1.400E-08 |
| 3731 | 187 | S187L | A | 3e-09 | 6.3e-09 | 1.5e-08 | 8.100E-09 | 6.199E-09 | 3.000E-09 | 1.500E-08 |
| 3732 | 187 | S187K | A | 1.4e-08 | 8.5e-09 | 7.3e-09 | 9.933E-09 | 3.573E-09 | 7.300E-09 | 1.400E-08 |
| 3733 | 187 | S187M | A | 1.5e-08 | 3.3e-09 | 7.4e-09 | 8.567E-09 | 5.937E-09 | 3.300E-09 | 1.500E-08 |
| 3734 | 187 | S187F | A | 8e-09 | 7.3e-09 | 1e-08 | 8.433E-09 | 1.401E-09 | 7.300E-09 | 1.000E-08 |
| 3735 | 187 | S187P | A | 8.9e-09 | 5.6e-09 | 3.7e-09 | 6.067E-09 | 2.631E-09 | 3.700E-09 | 8.900E-09 |
| 3736 | 187 | S187S | A | 2.2e-09 | 7.1e-09 | 7.8e-09 | 5.700E-09 | 3.051E-09 | 2.200E-09 | 7.800E-09 |
| 3737 | 187 | S187T | A | 1.8e-08 | 1.1e-08 | 1.1e-08 | 1.333E-08 | 4.041E-09 | 1.100E-08 | 1.800E-08 |
| 3738 | 187 | S187W | A | 2.2e-08 | 7.6e-09 | 4.2e-09 | 1.127E-08 | 9.450E-09 | 4.200E-09 | 2.200E-08 |
| 3739 | 187 | S187Y | A | 4.8e-09 | 2.3e-09 | 1.2e-08 | 6.367E-09 | 5.036E-09 | 2.300E-09 | 1.200E-08 |

| | | | | | | | | | | |
|---|---|---|---|---|---|---|---|---|---|---|
| 3740 | 187 | S187V | A | 7.2e-09 | 8.8e-09 | 1.5e-08 | 1.033E-08 | 4.120E-09 | 7.200E-09 | 1.500E-08 |
| 3741 | 188 | E188A | A | 4.3e-09 | 4.6e-09 | 2.9e-08 | 1.263E-08 | 1.417E-08 | 4.300E-09 | 2.900E-08 |
| 3742 | 188 | E188R | A | 7e-09 | 7.5e-09 | 1.5e-08 | 9.833E-09 | 4.481E-09 | 7.000E-09 | 1.500E-08 |
| 3743 | 188 | E188N | A | 1.4e-08 | 2.8e-09 | 1.1e-08 | 9.267E-09 | 5.798E-09 | 2.800E-09 | 1.400E-08 |
| 3744 | 188 | E188D | A | 5.7e-09 | 2.2e-08 | 1.3e-08 | 1.357E-08 | 8.165E-09 | 5.700E-09 | 2.200E-08 |
| 3745 | 188 | E188C | A | 7.6e-09 | 1.1e-08 | 7.2e-09 | 8.600E-09 | 2.088E-09 | 7.200E-09 | 1.100E-08 |
| 3746 | 188 | E188E | A | 1.4e-08 | 4.5e-09 | 4.1e-09 | 7.533E-09 | 5.604E-09 | 4.100E-09 | 1.400E-08 |
| 3747 | 188 | E188Q | A | 7e-09 | 8.7e-09 | 1.8e-08 | 1.123E-08 | 5.921E-09 | 7.000E-09 | 1.800E-08 |
| 3748 | 188 | E188G | A | 8.8e-09 | 7.7e-09 | 3.4e-08 | 1.683E-08 | 1.488E-08 | 7.700E-09 | 3.400E-08 |
| 3749 | 188 | E188H | A | 7.3e-09 | 8.7e-09 | 1.2e-08 | 9.333E-09 | 2.413E-09 | 7.300E-09 | 1.200E-08 |
| 3750 | 188 | E188I | A | 1.6e-08 | 7.7e-09 | 1.1e-08 | 1.157E-08 | 4.179E-09 | 7.700E-09 | 1.600E-08 |
| 3751 | 188 | E188L | A | 1.1e-08 | 7.6e-09 | 3.7e-09 | 7.433E-09 | 3.653E-09 | 3.700E-09 | 1.100E-08 |
| 3752 | 188 | E188K | A | 6.3e-09 | 1.4e-08 | 1.7e-08 | 1.243E-08 | 5.519E-09 | 6.300E-09 | 1.700E-08 |
| 3753 | 188 | E188M | A | 1e-08 | 5.5e-09 | 1e-08 | 8.500E-09 | 2.598E-09 | 5.500E-09 | 1.000E-08 |
| 3754 | 188 | E188F | A | 6.8e-09 | 2e-08 | 1.3e-08 | 1.327E-08 | 6.604E-09 | 6.800E-09 | 2.000E-08 |
| 3755 | 188 | E188P | A | 3.5e-09 | 1.8e-08 | 4.2e-09 | 8.567E-09 | 8.177E-09 | 3.500E-09 | 1.800E-08 |
| 3756 | 188 | E188S | A | 3e-08 | 6.5e-09 | 1e-08 | 1.550E-08 | 1.268E-08 | 6.500E-09 | 3.000E-08 |
| 3757 | 188 | E188T | A | 3.7e-08 | 7.4e-09 | 6.2e-09 | 1.687E-08 | 1.745E-08 | 6.200E-09 | 3.700E-08 |
| 3758 | 188 | E188W | A | 6.1e-09 | 4.2e-09 | 7.1e-09 | 5.800E-09 | 1.473E-09 | 4.200E-09 | 7.100E-09 |
| 3759 | 188 | E188Y | A | 8.2e-09 | 7.2e-09 | 8.3e-09 | 7.900E-09 | 6.083E-10 | 7.200E-09 | 8.300E-09 |
| 3760 | 188 | E188V | A | 1.2e-08 | 3.9e-09 | 5.2e-09 | 7.033E-09 | 4.350E-09 | 3.900E-09 | 1.200E-08 |
| 3761 | 189 | D189A | A | 1.4e-08 | 2.8e-09 | 1.1e-08 | 9.267E-09 | 5.798E-09 | 2.800E-09 | 1.400E-08 |
| 3762 | 189 | D189R | A | 1e-08 | 4e-09 | 9.3e-09 | 7.767E-09 | 3.281E-09 | 4.000E-09 | 1.000E-08 |
| 3763 | 189 | D189N | A | 6.7e-09 | 1.2e-08 | 4.9e-09 | 7.867E-09 | 3.691E-09 | 4.900E-09 | 1.200E-08 |
| 3764 | 189 | D189D | A | 6.7e-09 | 4.4e-09 | 8e-09 | 6.367E-09 | 1.823E-09 | 4.400E-09 | 8.000E-09 |
| 3765 | 189 | D189C | A | 6.6e-09 | 7.6e-09 | 1.2e-08 | 8.733E-09 | 2.873E-09 | 6.600E-09 | 1.200E-08 |
| 3766 | 189 | D189E | A | 3.9e-09 | 7.8e-09 | 1.6e-08 | 9.233E-09 | 6.176E-09 | 3.900E-09 | 1.600E-08 |
| 3767 | 189 | D189Q | A | 1.3e-08 | 7.8e-09 | 5.2e-09 | 8.667E-09 | 3.972E-09 | 5.200E-09 | 1.300E-08 |
| 3768 | 189 | D189G | A | 1e-08 | 2.2e-08 | 2.3e-09 | 1.143E-08 | 9.928E-09 | 2.300E-09 | 2.200E-08 |
| 3769 | 189 | D189H | A | 1.9e-08 | 2.6e-08 | 1.1e-08 | 1.867E-08 | 7.506E-09 | 1.100E-08 | 2.600E-08 |
| 3770 | 189 | D189I | A | 1.1e-08 | 1.1e-08 | 6.5e-09 | 9.500E-09 | 2.598E-09 | 6.500E-09 | 1.100E-08 |
| 3771 | 189 | D189L | A | 5.4e-09 | 1.3e-08 | 7.9e-09 | 8.767E-09 | 3.873E-09 | 5.400E-09 | 1.300E-08 |
| 3772 | 189 | D189K | A | 2.3e-09 | 3.3e-09 | 6.2e-09 | 3.933E-09 | 2.026E-09 | 2.300E-09 | 6.200E-09 |
| 3773 | 189 | D189M | A | 1.9e-08 | 7.2e-09 | 5.1e-09 | 1.043E-08 | 7.493E-09 | 5.100E-09 | 1.900E-08 |
| 3774 | 189 | D189F | A | 7.7e-09 | 7.9e-09 | 6.2e-09 | 7.267E-09 | 9.292E-10 | 6.200E-09 | 7.900E-09 |
| 3775 | 189 | D189P | A | 4.4e-09 | 1.4e-09 | 1e-08 | 5.267E-09 | 4.365E-09 | 1.400E-09 | 1.000E-08 |
| 3776 | 189 | D189S | A | 6.4e-09 | 8.7e-09 | 1.2e-08 | 9.033E-09 | 2.815E-09 | 6.400E-09 | 1.200E-08 |
| 3777 | 189 | D189T | A | 3.6e-08 | 1.2e-08 | 9.7e-09 | 1.923E-08 | 1.457E-08 | 9.700E-09 | 3.600E-08 |
| 3778 | 189 | D189W | A | 1.5e-08 | 6.6e-09 | 1.5e-08 | 1.220E-08 | 4.850E-09 | 6.600E-09 | 1.500E-08 |
| 3779 | 189 | D189Y | A | 1.1e-08 | 8.7e-09 | 6.9e-09 | 8.867E-09 | 2.055E-09 | 6.900E-09 | 1.100E-08 |
| 3780 | 189 | D189V | A | 2.1e-08 | 1e-08 | 1.2e-08 | 1.433E-08 | 5.859E-09 | 1.000E-08 | 2.100E-08 |
| 3781 | 190 | C190A | A | 1.4e-08 | 6.3e-09 | 5.3e-09 | 8.533E-09 | 4.761E-09 | 5.300E-09 | 1.400E-08 |
| 3782 | 190 | C190R | A | 5.5e-09 | 2.5e-09 | 7.2e-09 | 5.067E-09 | 2.380E-09 | 2.500E-09 | 7.200E-09 |
| 3783 | 190 | C190N | A | 4.7e-09 | 8.4e-09 | 2.2e-09 | 5.100E-09 | 3.119E-09 | 2.200E-09 | 8.400E-09 |
| 3784 | 190 | C190D | A | 6.4e-09 | 5.7e-09 | 4.6e-09 | 5.567E-09 | 9.074E-10 | 4.600E-09 | 6.400E-09 |
| 3785 | 190 | C190C | A | 6e-09 | 4.5e-09 | 7.5e-09 | 6.000E-09 | 1.500E-09 | 4.500E-09 | 7.500E-09 |
| 3786 | 190 | C190E | A | 4.7e-09 | 4.3e-09 | 7.3e-09 | 5.433E-09 | 1.629E-09 | 4.300E-09 | 7.300E-09 |
| 3787 | 190 | C190Q | A | 3.2e-09 | 2e-08 | 1.1e-08 | 1.140E-08 | 8.407E-09 | 3.200E-09 | 2.000E-08 |
| 3788 | 190 | C190G | A | 1.1e-08 | 2.9e-09 | 9.9e-09 | 7.933E-09 | 4.394E-09 | 2.900E-09 | 1.100E-08 |
| 3789 | 190 | C190H | A | 9.2e-09 | 3.8e-09 | 4.4e-09 | 5.800E-09 | 2.960E-09 | 3.800E-09 | 9.200E-09 |
| 3790 | 190 | C190I | A | 5.5e-09 | 6.3e-09 | 5.3e-09 | 5.700E-09 | 5.292E-10 | 5.300E-09 | 6.300E-09 |
| 3791 | 190 | C190L | A | 1.5e-08 | 5.7e-09 | 4.1e-09 | 8.267E-09 | 5.886E-09 | 4.100E-09 | 1.500E-08 |
| 3792 | 190 | C190K | A | 1.2e-08 | 3.2e-09 | 2.4e-08 | 1.307E-08 | 1.044E-08 | 3.200E-09 | 2.400E-08 |
| 3793 | 190 | C190M | A | 5.5e-09 | 3e-09 | 3.9e-09 | 4.133E-09 | 1.266E-09 | 3.000E-09 | 5.500E-09 |
| 3794 | 190 | C190F | A | 2.1e-09 | 2.6e-09 | 3.7e-09 | 2.800E-09 | 8.185E-10 | 2.100E-09 | 3.700E-09 |
| 3795 | 190 | C190P | A | 3.9e-09 | 6.7e-09 | 1.2e-08 | 7.533E-09 | 4.114E-09 | 3.900E-09 | 1.200E-08 |
| 3796 | 190 | C190S | A | 4.9e-09 | 8.4e-09 | 9e-09 | 7.433E-09 | 2.214E-09 | 4.900E-09 | 9.000E-09 |
| 3797 | 190 | C190T | A | 3.3e-09 | 5.7e-09 | 5.1e-09 | 4.700E-09 | 1.249E-09 | 3.300E-09 | 5.700E-09 |
| 3798 | 190 | C190W | A | 6.8e-09 | 2.6e-09 | 8.9e-09 | 6.100E-09 | 3.208E-09 | 2.600E-09 | 8.900E-09 |
| 3799 | 190 | C190Y | A | 9.2e-09 | 6.5e-09 | 8.3e-09 | 8.000E-09 | 1.375E-09 | 6.500E-09 | 9.200E-09 |
| 3800 | 190 | C190V | A | 6.4e-09 | 9.2e-09 | 4.1e-09 | 6.567E-09 | 2.554E-09 | 4.100E-09 | 9.200E-09 |
| 3801 | 191 | Q191A | A | 6.7e-09 | 9.9e-09 | 1.7e-08 | 1.120E-08 | 5.272E-09 | 6.700E-09 | 1.700E-08 |
| 3802 | 191 | Q191R | A | 9.8e-09 | 5.3e-09 | 1.6e-08 | 1.037E-08 | 5.372E-09 | 5.300E-09 | 1.600E-08 |
| 3803 | 191 | Q191N | A | 1.1e-08 | 5.6e-09 | 6.4e-09 | 7.667E-09 | 2.914E-09 | 5.600E-09 | 1.100E-08 |
| 3804 | 191 | Q191D | A | 5.2e-09 | 4.8e-09 | 4.9e-09 | 4.967E-09 | 2.082E-10 | 4.800E-09 | 5.200E-09 |
| 3805 | 191 | Q191C | A | 3.6e-09 | 4.2e-09 | 3.3e-09 | 3.700E-09 | 4.583E-10 | 3.300E-09 | 4.200E-09 |
| 3806 | 191 | Q191E | A | 4.8e-09 | 8.9e-09 | 6.1e-09 | 6.600E-09 | 2.095E-09 | 4.800E-09 | 8.900E-09 |
| 3807 | 191 | Q191Q | A | 1.1e-08 | 4.2e-09 | 9.6e-09 | 8.267E-09 | 3.591E-09 | 4.200E-09 | 1.100E-08 |
| 3808 | 191 | Q191G | A | 9.6e-09 | 1.2e-08 | 3.6e-09 | 8.400E-09 | 4.327E-09 | 3.600E-09 | 1.200E-08 |
| 3809 | 191 | Q191H | A | 5.2e-09 | 4.9e-09 | 9e-09 | 6.367E-09 | 2.285E-09 | 4.900E-09 | 9.000E-09 |
| 3810 | 191 | Q191I | A | 3.4e-09 | 3.2e-09 | 7.8e-09 | 4.800E-09 | 2.600E-09 | 3.200E-09 | 7.800E-09 |
| 3811 | 191 | Q191L | A | 6.4e-09 | 1.6e-08 | 4.2e-09 | 8.867E-09 | 6.275E-09 | 4.200E-09 | 1.600E-08 |
| 3812 | 191 | Q191K | A | 6.9e-09 | 2.8e-09 | 6.4e-09 | 5.367E-09 | 2.237E-09 | 2.800E-09 | 6.900E-09 |
| 3813 | 191 | Q191M | A | 5.4e-09 | 1.1e-08 | 1.8e-08 | 1.147E-08 | 6.313E-09 | 5.400E-09 | 1.800E-08 |
| 3814 | 191 | Q191F | A | 2.3e-09 | 7.5e-09 | 1.2e-08 | 7.267E-09 | 4.854E-09 | 2.300E-09 | 1.200E-08 |
| 3815 | 191 | Q191P | A | 6.3e-09 | 1.1e-08 | 4e-09 | 7.100E-09 | 3.568E-09 | 4.000E-09 | 1.100E-08 |
| 3816 | 191 | Q191S | A | 3e-08 | 5.3e-09 | 7.5e-09 | 1.427E-08 | 1.367E-08 | 5.300E-09 | 3.000E-08 |
| 3817 | 191 | Q191T | A | 6.2e-09 | 1e-08 | 4.2e-09 | 6.800E-09 | 2.946E-09 | 4.200E-09 | 1.000E-08 |
| 3818 | 191 | Q191W | A | 2e-09 | 1.8e-08 | 6e-09 | 8.667E-09 | 8.327E-09 | 2.000E-09 | 1.800E-08 |
| 3819 | 191 | Q191Y | A | 6.1e-09 | 1.2e-08 | 6.4e-09 | 8.167E-09 | 3.323E-09 | 6.100E-09 | 1.200E-08 |
| 3820 | 191 | Q191V | A | 2.1e-09 | 2.5e-08 | 6.9e-09 | 1.133E-08 | 1.208E-08 | 2.100E-09 | 2.500E-08 |
| 3821 | 192 | S192A | A | 4.7e-09 | 3.4e-09 | 5.8e-09 | 4.633E-09 | 1.201E-09 | 3.400E-09 | 5.800E-09 |
| 3822 | 192 | S192R | A | 1.8e-09 | 3.1e-09 | 4.4e-09 | 3.100E-09 | 1.300E-09 | 1.800E-09 | 4.400E-09 |
| 3823 | 192 | S192N | A | 6.2e-09 | 1.1e-08 | 1.1e-08 | 9.400E-09 | 2.771E-09 | 6.200E-09 | 1.100E-08 |
| 3824 | 192 | S192D | A | 1.3e-08 | 4.2e-09 | 1.2e-09 | 6.133E-09 | 6.133E-09 | 1.200E-09 | 1.300E-08 |

| | | | | | | | | | | |
|---|---|---|---|---|---|---|---|---|---|---|
| 3825 | 192 | S192C | A | 8e-09 | 3.8e-09 | 1e-08 | 7.267E-09 | 3.164E-09 | 3.800E-09 | 1.000E-08 |
| 3826 | 192 | S192E | A | 1.2e-08 | 4.5e-09 | 8.4e-09 | 8.300E-09 | 3.751E-09 | 4.500E-09 | 1.200E-08 |
| 3827 | 192 | S192Q | A | 3.4e-09 | 1e-08 | 3.8e-09 | 5.733E-09 | 3.700E-09 | 3.400E-09 | 1.000E-08 |
| 3828 | 192 | S192G | A | 5.7e-09 | 8.2e-09 | 1.4e-08 | 9.300E-09 | 4.258E-09 | 5.700E-09 | 1.400E-08 |
| 3829 | 192 | S192H | A | 1.1e-08 | 4.8e-09 | 9.7e-09 | 8.500E-09 | 3.270E-09 | 4.800E-09 | 1.100E-08 |
| 3830 | 192 | S192I | A | 6e-09 | 1.1e-08 | 8.4e-09 | 8.467E-09 | 2.501E-09 | 6.000E-09 | 1.100E-08 |
| 3831 | 192 | S192L | A | 1.1e-08 | 7.5e-09 | 9.9e-09 | 9.467E-09 | 1.790E-09 | 7.500E-09 | 1.100E-08 |
| 3832 | 192 | S192K | A | 5e-09 | 5.7e-09 | 6.8e-09 | 5.833E-09 | 9.074E-10 | 5.000E-09 | 6.800E-09 |
| 3833 | 192 | S192M | A | 1.2e-08 | 3.7e-09 | 5.9e-09 | 7.200E-09 | 4.300E-09 | 3.700E-09 | 1.200E-08 |
| 3834 | 192 | S192F | A | 8.9e-09 | 9.5e-09 | 5.3e-09 | 7.900E-09 | 2.272E-09 | 5.300E-09 | 9.500E-09 |
| 3835 | 192 | S192P | A | 8.8e-09 | 1e-08 | 6.9e-09 | 8.567E-09 | 1.563E-09 | 6.900E-09 | 1.000E-08 |
| 3836 | 192 | S192S | A | 2.8e-08 | 1.4e-09 | 6.5e-09 | 1.197E-08 | 1.412E-08 | 1.400E-09 | 2.800E-08 |
| 3837 | 192 | S192T | A | 5.7e-09 | 9.9e-09 | 6.5e-09 | 7.367E-09 | 2.230E-09 | 5.700E-09 | 9.900E-09 |
| 3838 | 192 | S192W | A | 2.6e-08 | 7e-09 | 1.2e-08 | 1.500E-08 | 9.849E-09 | 7.000E-09 | 2.600E-08 |
| 3839 | 192 | S192Y | A | 3.3e-09 | 1.4e-09 | 6.3e-09 | 3.667E-09 | 2.470E-09 | 1.400E-09 | 6.300E-09 |
| 3840 | 192 | S192V | A | 4.5e-09 | 1e-08 | 7e-09 | 7.167E-09 | 2.754E-09 | 4.500E-09 | 1.000E-08 |
| 3841 | 193 | L193A | A | 5.8e-09 | 1.2e-08 | 1.3e-08 | 1.027E-08 | 3.900E-09 | 5.800E-09 | 1.300E-08 |
| 3842 | 193 | L193R | A | 4.8e-09 | 4.2e-09 | 1.8e-09 | 3.600E-09 | 1.587E-09 | 1.800E-09 | 4.800E-09 |
| 3843 | 193 | L193N | A | 6.3e-09 | 5.4e-09 | 5.6e-09 | 5.767E-09 | 4.726E-10 | 5.400E-09 | 6.300E-09 |
| 3844 | 193 | L193D | A | 8.3e-09 | 3.3e-09 | 3.9e-09 | 5.167E-09 | 2.730E-09 | 3.300E-09 | 8.300E-09 |
| 3845 | 193 | L193C | A | 7e-09 | 6.3e-09 | 3.8e-09 | 5.700E-09 | 1.682E-09 | 3.800E-09 | 7.000E-09 |
| 3846 | 193 | L193E | A | 3e-09 | 7.3e-09 | 1.2e-08 | 7.433E-09 | 4.501E-09 | 3.000E-09 | 1.200E-08 |
| 3847 | 193 | L193Q | A | 5e-09 | 6.5e-09 | 9e-09 | 6.833E-09 | 2.021E-09 | 5.000E-09 | 9.000E-09 |
| 3848 | 193 | L193G | A | 5.4e-09 | 1.4e-08 | 2.5e-09 | 7.300E-09 | 5.981E-09 | 2.500E-09 | 1.400E-08 |
| 3849 | 193 | L193H | A | 6e-09 | 1.8e-08 | 3.4e-09 | 9.133E-09 | 7.788E-09 | 3.400E-09 | 1.800E-08 |
| 3850 | 193 | L193I | A | 1.3e-08 | 1e-08 | 1.5e-08 | 1.267E-08 | 2.517E-09 | 1.000E-08 | 1.500E-08 |
| 3851 | 193 | L193L | A | 5e-09 | 1.4e-08 | 1.1e-08 | 1.000E-08 | 4.583E-09 | 5.000E-09 | 1.400E-08 |
| 3852 | 193 | L193K | A | 1.2e-08 | 1.2e-08 | 1.3e-08 | 1.233E-08 | 5.774E-10 | 1.200E-08 | 1.300E-08 |
| 3853 | 193 | L193M | A | 3.8e-09 | 1.5e-08 | 3.6e-09 | 7.467E-09 | 6.525E-09 | 3.600E-09 | 1.500E-08 |
| 3854 | 193 | L193F | A | 1.3e-08 | 2.7e-09 | 7.1e-09 | 7.600E-09 | 5.168E-09 | 2.700E-09 | 1.300E-08 |
| 3855 | 193 | L193P | A | 6.9e-09 | 2.8e-09 | 2.3e-09 | 4.000E-09 | 2.524E-09 | 2.300E-09 | 6.900E-09 |
| 3856 | 193 | L193S | A | 4.6e-09 | 4.9e-09 | 8.8e-09 | 6.100E-09 | 2.343E-09 | 4.600E-09 | 8.800E-09 |
| 3857 | 193 | L193T | A | 5.3e-09 | 5.6e-09 | 4.8e-09 | 5.233E-09 | 4.041E-10 | 4.800E-09 | 5.600E-09 |
| 3858 | 193 | L193W | A | 7.9e-09 | 1.7e-08 | 8.2e-09 | 1.103E-08 | 5.169E-09 | 7.900E-09 | 1.700E-08 |
| 3859 | 193 | L193Y | A | 3.8e-09 | 7.7e-09 | 8e-09 | 6.500E-09 | 2.343E-09 | 3.800E-09 | 8.000E-09 |
| 3860 | 193 | L193V | A | 3.9e-09 | 7.1e-09 | 2.2e-09 | 4.400E-09 | 2.488E-09 | 2.200E-09 | 7.100E-09 |
| 3861 | 194 | T194A | A | 2e-09 | 7.5e-09 | 2e-09 | 3.833E-09 | 3.175E-09 | 2.000E-09 | 7.500E-09 |
| 3862 | 194 | T194R | A | 1.9e-08 | 2.7e-09 | 6.7e-09 | 9.467E-09 | 8.495E-09 | 2.700E-09 | 1.900E-08 |
| 3863 | 194 | T194N | A | 8.7e-09 | 6.6e-09 | 9.6e-09 | 8.300E-09 | 1.539E-09 | 6.600E-09 | 9.600E-09 |
| 3864 | 194 | T194D | A | 7.3e-09 | 1e-09 | 8.3e-09 | 5.533E-09 | 3.958E-09 | 1.000E-09 | 8.300E-09 |
| 3865 | 194 | T194C | A | 3.9e-09 | 2.8e-09 | 3.9e-09 | 3.533E-09 | 6.351E-10 | 2.800E-09 | 3.900E-09 |
| 3866 | 194 | T194E | A | 2.5e-08 | 4.7e-09 | 6.1e-09 | 1.193E-08 | 1.134E-08 | 4.700E-09 | 2.500E-08 |
| 3867 | 194 | T194Q | A | 1.2e-08 | 1.1e-08 | 2.9e-09 | 8.633E-09 | 4.990E-09 | 2.900E-09 | 1.200E-08 |
| 3868 | 194 | T194G | A | 4.9e-09 | 9e-09 | 5.2e-09 | 6.367E-09 | 2.285E-09 | 4.900E-09 | 9.000E-09 |
| 3869 | 194 | T194H | A | 7e-09 | 2.2e-08 | 7.8e-09 | 1.227E-08 | 8.439E-09 | 7.000E-09 | 2.200E-08 |
| 3870 | 194 | T194I | A | 7.3e-09 | 6.2e-09 | 1e-08 | 7.833E-09 | 1.955E-09 | 6.200E-09 | 1.000E-08 |
| 3871 | 194 | T194L | A | 4e-09 | 6.1e-09 | 1.2e-08 | 7.367E-09 | 4.148E-09 | 4.000E-09 | 1.200E-08 |
| 3872 | 194 | T194K | A | 1.8e-08 | 3.1e-09 | 1.2e-08 | 1.103E-08 | 7.497E-09 | 3.100E-09 | 1.800E-08 |
| 3873 | 194 | T194M | A | 7.6e-09 | 8.4e-09 | 4.4e-09 | 6.800E-09 | 2.117E-09 | 4.400E-09 | 8.400E-09 |
| 3874 | 194 | T194F | A | 7.4e-09 | 7.3e-09 | 8.2e-09 | 7.633E-09 | 4.933E-10 | 7.300E-09 | 8.200E-09 |
| 3875 | 194 | T194P | A | 1.3e-08 | 3.8e-08 | 7.1e-09 | 1.937E-08 | 1.640E-08 | 7.100E-09 | 3.800E-08 |
| 3876 | 194 | T194S | A | 7.4e-09 | 6.6e-09 | 1.6e-08 | 1.000E-08 | 5.212E-09 | 6.600E-09 | 1.600E-08 |
| 3877 | 194 | T194T | A | 5.9e-09 | 7e-09 | 5.5e-09 | 6.133E-09 | 7.767E-10 | 5.500E-09 | 7.000E-09 |
| 3878 | 194 | T194W | A | 6.4e-09 | 9.2e-09 | 1.1e-08 | 8.867E-09 | 2.318E-09 | 6.400E-09 | 1.100E-08 |
| 3879 | 194 | T194Y | A | 3.5e-09 | 7.4e-09 | 2.5e-09 | 4.467E-09 | 2.589E-09 | 2.500E-09 | 7.400E-09 |
| 3880 | 194 | T194V | A | 2.3e-09 | 7.5e-09 | 5.4e-09 | 5.067E-09 | 2.616E-09 | 2.300E-09 | 7.500E-09 |
| 3881 | 195 | R195A | A | 4.2e-09 | 1.8e-08 | 9.1e-09 | 1.043E-08 | 6.996E-09 | 4.200E-09 | 1.800E-08 |
| 3882 | 195 | R195R | A | 1.7e-08 | 5.6e-09 | 3.3e-09 | 8.633E-09 | 7.336E-09 | 3.300E-09 | 1.700E-08 |
| 3883 | 195 | R195N | A | 2.9e-09 | 1.6e-09 | 5.7e-09 | 3.400E-09 | 2.095E-09 | 1.600E-09 | 5.700E-09 |
| 3884 | 195 | R195D | A | 1.5e-08 | 1.4e-08 | 4.4e-09 | 1.113E-08 | 5.853E-09 | 4.400E-09 | 1.500E-08 |
| 3885 | 195 | R195C | A | 9.6e-09 | 2e-08 | 9.6e-09 | 1.307E-08 | 6.004E-09 | 9.600E-09 | 2.000E-08 |
| 3886 | 195 | R195E | A | 8.6e-09 | 1.1e-08 | 2.5e-08 | 1.487E-08 | 8.857E-09 | 8.600E-09 | 2.500E-08 |
| 3887 | 195 | R195Q | A | 7.7e-09 | 5.7e-09 | 7.8e-09 | 7.067E-09 | 1.185E-09 | 5.700E-09 | 7.800E-09 |
| 3888 | 195 | R195G | A | 3.2e-09 | 4.2e-09 | 1.2e-08 | 6.467E-09 | 4.818E-09 | 3.200E-09 | 1.200E-08 |
| 3889 | 195 | R195H | A | 9.2e-09 | 5.6e-09 | 4e-09 | 6.267E-09 | 2.663E-09 | 4.000E-09 | 9.200E-09 |
| 3890 | 195 | R195I | A | 1.1e-08 | 6.5e-09 | 5.9e-09 | 7.800E-09 | 2.787E-09 | 5.900E-09 | 1.100E-08 |
| 3891 | 195 | R195L | A | 6.2e-09 | 1.8e-09 | 4.4e-09 | 4.133E-09 | 2.212E-09 | 1.800E-09 | 6.200E-09 |
| 3892 | 195 | R195K | A | 8.6e-09 | 3.9e-09 | 6.8e-09 | 6.433E-09 | 2.371E-09 | 3.900E-09 | 8.600E-09 |
| 3893 | 195 | R195M | A | 3e-09 | 7.5e-09 | 3.5e-09 | 4.667E-09 | 2.466E-09 | 3.000E-09 | 7.500E-09 |
| 3894 | 195 | R195F | A | 9.6e-09 | 6.7e-09 | 8.6e-09 | 8.300E-09 | 1.473E-09 | 6.700E-09 | 9.600E-09 |
| 3895 | 195 | R195P | A | 6.3e-09 | 8.4e-09 | 2e-08 | 1.157E-08 | 7.379E-09 | 6.300E-09 | 2.000E-08 |
| 3896 | 195 | R195S | A | 7.9e-09 | 4.7e-09 | 6.8e-09 | 6.467E-09 | 1.626E-09 | 4.700E-09 | 7.900E-09 |
| 3897 | 195 | R195T | A | 1e-08 | 4.4e-09 | 6.3e-09 | 6.900E-09 | 2.848E-09 | 4.400E-09 | 1.000E-08 |
| 3898 | 195 | R195W | A | 1.4e-08 | 5e-09 | 1.1e-08 | 1.000E-08 | 4.583E-09 | 5.000E-09 | 1.400E-08 |
| 3899 | 195 | R195Y | A | 5.9e-09 | 1.1e-08 | 6.2e-09 | 7.700E-09 | 2.862E-09 | 5.900E-09 | 1.100E-08 |
| 3900 | 195 | R195V | A | 7.6e-09 | 4.4e-09 | 7.5e-09 | 6.500E-09 | 1.819E-09 | 4.400E-09 | 7.600E-09 |
| 3901 | 196 | T196A | A | 1.1e-08 | 1.1e-08 | 5.6e-09 | 9.200E-09 | 3.118E-09 | 5.600E-09 | 1.100E-08 |
| 3902 | 196 | T196R | A | 9.5e-09 | 1.6e-08 | 1.5e-09 | 9.000E-09 | 7.263E-09 | 1.500E-09 | 1.600E-08 |
| 3903 | 196 | T196N | A | 6e-09 | 6.1e-09 | 3.5e-09 | 5.200E-09 | 1.473E-09 | 3.500E-09 | 6.100E-09 |
| 3904 | 196 | T196D | A | 1.6e-08 | 1.1e-08 | 1.1e-08 | 1.267E-08 | 2.887E-09 | 1.100E-08 | 1.600E-08 |
| 3905 | 196 | T196C | A | 6.3e-09 | 4.6e-09 | 3.5e-09 | 4.800E-09 | 1.411E-09 | 3.500E-09 | 6.300E-09 |
| 3906 | 196 | T196E | A | 5.5e-09 | 7.5e-09 | 3.7e-09 | 5.567E-09 | 1.901E-09 | 3.700E-09 | 7.500E-09 |
| 3907 | 196 | T196Q | A | 1.1e-08 | 5.4e-09 | 2e-09 | 6.133E-09 | 4.545E-09 | 2.000E-09 | 1.100E-08 |
| 3908 | 196 | T196G | A | 5.8e-09 | 3.9e-09 | 6.5e-09 | 5.400E-09 | 1.345E-09 | 3.900E-09 | 6.500E-09 |
| 3909 | 196 | T196H | A | 2.5e-08 | 1.8e-08 | 2e-08 | 2.100E-08 | 3.606E-09 | 1.800E-08 | 2.500E-08 |

| | | | | | | | | | | |
|---|---|---|---|---|---|---|---|---|---|---|
| 3910 | 196 | T196I | A | 7.1e-09 | 2.1e-08 | 4.8e-09 | 1.097E-08 | 8.765E-09 | 4.800E-09 | 2.100E-08 |
| 3911 | 196 | T196L | A | 4.5e-09 | 4.5e-09 | 1.1e-08 | 6.667E-09 | 3.753E-09 | 4.500E-09 | 1.100E-08 |
| 3912 | 196 | T196K | A | 5.1e-09 | 1e-09 | 3.8e-09 | 3.300E-09 | 2.095E-09 | 1.000E-09 | 5.100E-09 |
| 3913 | 196 | T196M | A | 4.6e-09 | 3.6e-09 | 7.3e-09 | 5.167E-09 | 1.914E-09 | 3.600E-09 | 7.300E-09 |
| 3914 | 196 | T196F | A | 4.3e-09 | 2.2e-08 | 2.3e-09 | 9.533E-09 | 1.084E-08 | 2.300E-09 | 2.200E-08 |
| 3915 | 196 | T196P | A | 5e-09 | 7.5e-09 | 9.5e-09 | 7.333E-09 | 2.255E-09 | 5.000E-09 | 9.500E-09 |
| 3916 | 196 | T196S | A | 8.4e-09 | 1e-08 | 4.1e-09 | 7.500E-09 | 3.051E-09 | 4.100E-09 | 1.000E-08 |
| 3917 | 196 | T196T | A | 7.5e-09 | 5.9e-09 | 7.3e-09 | 6.900E-09 | 8.718E-10 | 5.900E-09 | 7.500E-09 |
| 3918 | 196 | T196W | A | 7.2e-09 | 4.7e-09 | 4.2e-09 | 5.367E-09 | 1.607E-09 | 4.200E-09 | 7.200E-09 |
| 3919 | 196 | T196Y | A | 1.3e-08 | 4.5e-09 | 2.2e-09 | 6.567E-09 | 5.689E-09 | 2.200E-09 | 1.300E-08 |
| 3920 | 196 | T196V | A | 1.1e-08 | 2.4e-09 | 1.9e-09 | 5.100E-09 | 5.116E-09 | 1.900E-09 | 1.100E-08 |
| 3921 | 197 | V197A | A | 9.1e-09 | 7.1e-09 | 6.5e-09 | 7.567E-09 | 1.361E-09 | 6.500E-09 | 9.100E-09 |
| 3922 | 197 | V197R | A | 3.3e-09 | 8.6e-09 | 6.6e-09 | 6.167E-09 | 2.676E-09 | 3.300E-09 | 8.600E-09 |
| 3923 | 197 | V197N | A | 5.1e-09 | 4.9e-09 | 1.5e-08 | 8.333E-09 | 5.774E-09 | 4.900E-09 | 1.500E-08 |
| 3924 | 197 | V197D | A | 7.4e-09 | 4.2e-09 | 6.6e-09 | 6.067E-09 | 1.665E-09 | 4.200E-09 | 7.400E-09 |
| 3925 | 197 | V197C | A | 2.8e-09 | 5.5e-09 | 4.6e-09 | 4.300E-09 | 1.375E-09 | 2.800E-09 | 5.500E-09 |
| 3926 | 197 | V197E | A | 7.7e-09 | 9.9e-09 | 7.3e-09 | 8.300E-09 | 1.400E-09 | 7.300E-09 | 9.900E-09 |
| 3927 | 197 | V197Q | A | 7.4e-09 | 3.7e-09 | 4.5e-09 | 5.200E-09 | 1.947E-09 | 3.700E-09 | 7.400E-09 |
| 3928 | 197 | V197G | A | 8.3e-09 | 2.3e-08 | 2.6e-09 | 1.130E-08 | 1.053E-08 | 2.600E-09 | 2.300E-08 |
| 3929 | 197 | V197H | A | 6.6e-09 | 1.4e-08 | 7.3e-09 | 9.300E-09 | 4.085E-09 | 6.600E-09 | 1.400E-08 |
| 3930 | 197 | V197I | A | 2.9e-09 | 2.9e-09 | 9.8e-10 | 2.260E-09 | 1.109E-09 | 9.800E-10 | 2.900E-09 |
| 3931 | 197 | V197L | A | 7.6e-09 | 5.2e-09 | 1.2e-08 | 8.267E-09 | 3.449E-09 | 5.200E-09 | 1.200E-08 |
| 3932 | 197 | V197K | A | 2.9e-09 | 5.1e-09 | 8.6e-09 | 5.533E-09 | 2.875E-09 | 2.900E-09 | 8.600E-09 |
| 3933 | 197 | V197M | A | 1.1e-08 | 1.2e-08 | 3.1e-09 | 8.700E-09 | 4.875E-09 | 3.100E-09 | 1.200E-08 |
| 3934 | 197 | V197F | A | 1.9e-09 | 7.1e-09 | 3.9e-09 | 4.300E-09 | 2.623E-09 | 1.900E-09 | 7.100E-09 |
| 3935 | 197 | V197P | A | 1.1e-08 | 1.2e-08 | 3.5e-09 | 8.833E-09 | 4.646E-09 | 3.500E-09 | 1.200E-08 |
| 3936 | 197 | V197S | A | 6.3e-09 | 4.2e-09 | 5.1e-09 | 5.200E-09 | 1.054E-09 | 4.200E-09 | 6.300E-09 |
| 3937 | 197 | V197T | A | 2.5e-09 | 7e-09 | 3.2e-09 | 4.233E-09 | 2.421E-09 | 2.500E-09 | 7.000E-09 |
| 3938 | 197 | V197W | A | 6.3e-09 | 6.7e-09 | 8e-09 | 7.000E-09 | 8.888E-10 | 6.300E-09 | 8.000E-09 |
| 3939 | 197 | V197Y | A | 8.5e-09 | 5.1e-09 | 1.8e-08 | 1.053E-08 | 6.686E-09 | 5.100E-09 | 1.800E-08 |
| 3940 | 197 | V197V | A | 6.2e-09 | 5.7e-09 | 6.8e-09 | 6.233E-09 | 5.508E-10 | 5.700E-09 | 6.800E-09 |
| 3941 | 198 | C198A | A | 1.9e-08 | 1.3e-08 | 2.9e-09 | 1.163E-08 | 8.137E-09 | 2.900E-09 | 1.900E-08 |
| 3942 | 198 | C198R | A | 6.4e-09 | 1.9e-08 | 5.9e-09 | 1.043E-08 | 7.423E-09 | 5.900E-09 | 1.900E-08 |
| 3943 | 198 | C198N | A | 5e-09 | 1.1e-08 | 4.7e-09 | 6.900E-09 | 3.554E-09 | 4.700E-09 | 1.100E-08 |
| 3944 | 198 | C198D | A | 1.8e-08 | 7.6e-09 | 1.4e-08 | 1.320E-08 | 5.246E-09 | 7.600E-09 | 1.800E-08 |
| 3945 | 198 | C198C | A | 3.5e-09 | 5.5e-09 | 1.5e-08 | 8.000E-09 | 6.144E-09 | 3.500E-09 | 1.500E-08 |
| 3946 | 198 | C198E | A | 6.9e-09 | 1.3e-08 | 9.9e-09 | 9.933E-09 | 3.050E-09 | 6.900E-09 | 1.300E-08 |
| 3947 | 198 | C198Q | A | 8.8e-09 | 1.1e-08 | 7.7e-09 | 9.167E-09 | 1.680E-09 | 7.700E-09 | 1.100E-08 |
| 3948 | 198 | C198G | A | 4.7e-09 | 4.3e-09 | 6.6e-09 | 5.200E-09 | 1.229E-09 | 4.300E-09 | 6.600E-09 |
| 3949 | 198 | C198H | A | 5.2e-09 | 7.4e-09 | 7.5e-09 | 6.700E-09 | 1.300E-09 | 5.200E-09 | 7.500E-09 |
| 3950 | 198 | C198I | A | 5e-09 | 9.2e-09 | 9.2e-09 | 7.800E-09 | 2.425E-09 | 5.000E-09 | 9.200E-09 |
| 3951 | 198 | C198L | A | 8.4e-09 | 1.1e-08 | 1.9e-08 | 1.280E-08 | 5.524E-09 | 8.400E-09 | 1.900E-08 |
| 3952 | 198 | C198K | A | 4.1e-09 | 1.5e-08 | 1.9e-08 | 1.270E-08 | 7.712E-09 | 4.100E-09 | 1.900E-08 |
| 3953 | 198 | C198M | A | 7.5e-09 | 6.7e-09 | 4e-09 | 6.067E-09 | 1.834E-09 | 4.000E-09 | 7.500E-09 |
| 3954 | 198 | C198F | A | 9.5e-09 | 8e-09 | 3.6e-09 | 7.033E-09 | 3.066E-09 | 3.600E-09 | 9.500E-09 |
| 3955 | 198 | C198P | A | 3.9e-09 | 2.1e-08 | 1.3e-08 | 1.263E-08 | 8.556E-09 | 3.900E-09 | 2.100E-08 |
| 3956 | 198 | C198S | A | 8.5e-09 | 6.8e-09 | 1.6e-09 | 5.633E-09 | 3.595E-09 | 1.600E-09 | 8.500E-09 |
| 3957 | 198 | C198T | A | 1.1e-08 | 1.6e-08 | 7.7e-09 | 1.157E-08 | 4.179E-09 | 7.700E-09 | 1.600E-08 |
| 3958 | 198 | C198W | A | 1.3e-09 | 7.8e-09 | 7.1e-09 | 5.400E-09 | 3.568E-09 | 1.300E-09 | 7.800E-09 |
| 3959 | 198 | C198Y | A | 8.3e-09 | 3.7e-09 | 2.3e-09 | 4.767E-09 | 3.139E-09 | 2.300E-09 | 8.300E-09 |
| 3960 | 198 | C198V | A | 7.2e-09 | 6.1e-09 | 4.3e-09 | 5.867E-09 | 1.464E-09 | 4.300E-09 | 7.200E-09 |
| 3961 | 199 | A199A | A | 3.7e-09 | 2.8e-08 | 3.3e-09 | 1.167E-08 | 1.415E-08 | 3.300E-09 | 2.800E-08 |
| 3962 | 199 | A199R | A | 1.4e-08 | 5.8e-09 | 3.6e-09 | 7.800E-09 | 5.481E-09 | 3.600E-09 | 1.400E-08 |
| 3963 | 199 | A199N | A | 4.6e-09 | 6.7e-09 | 3.1e-09 | 4.800E-09 | 1.808E-09 | 3.100E-09 | 6.700E-09 |
| 3964 | 199 | A199D | A | 1.5e-08 | 3.8e-09 | 2.1e-09 | 6.967E-09 | 7.009E-09 | 2.100E-09 | 1.500E-08 |
| 3965 | 199 | A199C | A | 1.1e-08 | 8.6e-09 | 9.6e-09 | 9.733E-09 | 1.206E-09 | 8.600E-09 | 1.100E-08 |
| 3966 | 199 | A199E | A | 9.2e-09 | 6.2e-09 | 1.4e-08 | 9.800E-09 | 3.934E-09 | 6.200E-09 | 1.400E-08 |
| 3967 | 199 | A199Q | A | 1.1e-08 | 5.7e-09 | 6.3e-09 | 7.667E-09 | 2.902E-09 | 5.700E-09 | 1.100E-08 |
| 3968 | 199 | A199G | A | 3.6e-08 | 4e-09 | 4.8e-09 | 1.493E-08 | 1.825E-08 | 4.000E-09 | 3.600E-08 |
| 3969 | 199 | A199H | A | 1.1e-08 | 2.1e-09 | 8.7e-09 | 7.267E-09 | 4.620E-09 | 2.100E-09 | 1.100E-08 |
| 3970 | 199 | A199I | A | 7.7e-09 | 1.4e-09 | 2.1e-08 | 1.003E-08 | 1.001E-08 | 1.400E-09 | 2.100E-08 |
| 3971 | 199 | A199L | A | 7.5e-09 | 1.3e-08 | 2.3e-09 | 7.600E-09 | 5.351E-09 | 2.300E-09 | 1.300E-08 |
| 3972 | 199 | A199K | A | 5.4e-09 | 4.5e-09 | 7.9e-09 | 5.933E-09 | 1.762E-09 | 4.500E-09 | 7.900E-09 |
| 3973 | 199 | A199M | A | 1.2e-08 | 4e-09 | 1.1e-08 | 9.000E-09 | 4.359E-09 | 4.000E-09 | 1.200E-08 |
| 3974 | 199 | A199F | A | 1.4e-08 | 4.4e-09 | 1.3e-08 | 1.047E-08 | 5.278E-09 | 4.400E-09 | 1.400E-08 |
| 3975 | 199 | A199P | A | 4.1e-09 | 9.3e-09 | 5.3e-09 | 6.233E-09 | 2.723E-09 | 4.100E-09 | 9.300E-09 |
| 3976 | 199 | A199S | A | 7e-09 | 1e-08 | 3.5e-09 | 6.833E-09 | 3.253E-09 | 3.500E-09 | 1.000E-08 |
| 3977 | 199 | A199T | A | 3.1e-09 | 9.6e-09 | 2.5e-08 | 1.257E-08 | 1.125E-08 | 3.100E-09 | 2.500E-08 |
| 3978 | 199 | A199W | A | 1.5e-08 | 3e-09 | 3.6e-09 | 7.200E-09 | 6.762E-09 | 3.000E-09 | 1.500E-08 |
| 3979 | 199 | A199Y | A | 9.9e-09 | 4.4e-08 | 1.2e-08 | 2.197E-08 | 1.911E-08 | 9.900E-09 | 4.400E-08 |
| 3980 | 199 | A199V | A | 8.5e-09 | 2.2e-09 | 2.1e-08 | 1.057E-08 | 9.569E-09 | 2.200E-09 | 2.100E-08 |
| 3981 | 200 | G200A | A | 4.9e-09 | 1.2e-08 | 1.9e-08 | 1.197E-08 | 7.050E-09 | 4.900E-09 | 1.900E-08 |
| 3982 | 200 | G200R | A | 5.6e-09 | 4.4e-09 | 2.2e-08 | 1.067E-08 | 9.833E-09 | 4.400E-09 | 2.200E-08 |
| 3983 | 200 | G200N | A | 7.2e-09 | 2.5e-08 | 3.5e-09 | 1.190E-08 | 1.149E-08 | 3.500E-09 | 2.500E-08 |
| 3984 | 200 | G200D | A | 4e-09 | 1e-08 | 1.1e-08 | 8.333E-09 | 3.786E-09 | 4.000E-09 | 1.100E-08 |
| 3985 | 200 | G200C | A | 6.5e-09 | 9.8e-09 | 4.7e-09 | 7.000E-09 | 2.587E-09 | 4.700E-09 | 9.800E-09 |
| 3986 | 200 | G200E | A | 3.1e-09 | 1.1e-08 | 1.9e-08 | 1.103E-08 | 7.950E-09 | 3.100E-09 | 1.900E-08 |
| 3987 | 200 | G200Q | A | 5.7e-09 | 6e-09 | 8e-09 | 6.567E-09 | 1.250E-09 | 5.700E-09 | 8.000E-09 |
| 3988 | 200 | G200G | A | 4.2e-09 | 5.8e-09 | 2.9e-09 | 4.300E-09 | 1.453E-09 | 2.900E-09 | 5.800E-09 |
| 3989 | 200 | G200H | A | 8e-09 | 3.1e-09 | 1.7e-08 | 9.367E-09 | 7.050E-09 | 3.100E-09 | 1.700E-08 |
| 3990 | 200 | G200I | A | 1.1e-08 | 3.1e-09 | 4.2e-09 | 6.100E-09 | 4.279E-09 | 3.100E-09 | 1.100E-08 |
| 3991 | 200 | G200L | A | 4.6e-09 | 3e-09 | 2.3e-08 | 1.020E-08 | 1.111E-08 | 3.000E-09 | 2.300E-08 |
| 3992 | 200 | G200K | A | 4.3e-09 | 1.2e-08 | 2.8e-09 | 6.367E-09 | 4.936E-09 | 2.800E-09 | 1.200E-08 |
| 3993 | 200 | G200M | A | 4.8e-09 | 3.6e-09 | 5.4e-09 | 4.600E-09 | 9.165E-10 | 3.600E-09 | 5.400E-09 |
| 3994 | 200 | G200F | A | 2.8e-09 | 3.8e-09 | 6.1e-09 | 4.233E-09 | 1.692E-09 | 2.800E-09 | 6.100E-09 |

| | | | | | | | | | | |
|---|---|---|---|---|---|---|---|---|---|---|
| 3995 | 200 | G200P | A | 5.4e-09 | 9.2e-09 | 7e-09 | 7.200E-09 | 1.908E-09 | 5.400E-09 | 9.200E-09 |
| 3996 | 200 | G200S | A | 5.6e-09 | 1.4e-08 | 4.5e-09 | 8.033E-09 | 5.196E-09 | 4.500E-09 | 1.400E-08 |
| 3997 | 200 | G200T | A | 5.1e-09 | 8.3e-09 | 1e-08 | 7.800E-09 | 2.488E-09 | 5.100E-09 | 1.000E-08 |
| 3998 | 200 | G200W | A | 8.5e-09 | 1.2e-08 | 5.6e-09 | 8.700E-09 | 3.205E-09 | 5.600E-09 | 1.200E-08 |
| 3999 | 200 | G200Y | A | 2.2e-08 | 6.4e-09 | 8.8e-09 | 1.240E-08 | 8.400E-09 | 6.400E-09 | 2.200E-08 |
| 4000 | 200 | G200V | A | 8e-09 | 1e-08 | 1.2e-08 | 1.000E-08 | 2.000E-09 | 8.000E-09 | 1.200E-08 |
| 4001 | 201 | G201A | A | 1.4e-08 | 7.2e-09 | 5.5e-09 | 8.900E-09 | 4.498E-09 | 5.500E-09 | 1.400E-08 |
| 4002 | 201 | G201R | A | 6.4e-09 | 6.8e-09 | 6.7e-09 | 6.633E-09 | 2.082E-10 | 6.400E-09 | 6.800E-09 |
| 4003 | 201 | G201N | A | 4.3e-09 | 2.8e-09 | 1e-08 | 5.700E-09 | 3.799E-09 | 2.800E-09 | 1.000E-08 |
| 4004 | 201 | G201D | A | 1.1e-08 | 6.5e-09 | 4e-09 | 7.167E-09 | 3.547E-09 | 4.000E-09 | 1.100E-08 |
| 4005 | 201 | G201C | A | 1.2e-08 | 1.5e-08 | 1.2e-08 | 1.300E-08 | 1.732E-09 | 1.200E-08 | 1.500E-08 |
| 4006 | 201 | G201E | A | 1.1e-08 | 4.6e-09 | 1.6e-08 | 1.053E-08 | 5.714E-09 | 4.600E-09 | 1.600E-08 |
| 4007 | 201 | G201Q | A | 5.6e-09 | 9.4e-09 | 1.4e-08 | 9.667E-09 | 4.206E-09 | 5.600E-09 | 1.400E-08 |
| 4008 | 201 | G201G | A | 5.8e-09 | 2.9e-08 | 6.3e-09 | 1.370E-08 | 1.325E-08 | 5.800E-09 | 2.900E-08 |
| 4009 | 201 | G201H | A | 1.2e-08 | 9.3e-09 | 8.6e-09 | 9.967E-09 | 1.795E-09 | 8.600E-09 | 1.200E-08 |
| 4010 | 201 | G201I | A | 6.9e-09 | 3.8e-08 | 4.9e-09 | 1.660E-08 | 1.856E-08 | 4.900E-09 | 3.800E-08 |
| 4011 | 201 | G201L | A | 4.3e-09 | 1.1e-08 | 9e-09 | 8.100E-09 | 3.439E-09 | 4.300E-09 | 1.100E-08 |
| 4012 | 201 | G201K | A | 2.1e-08 | 7.2e-09 | 5.3e-09 | 1.117E-08 | 8.569E-09 | 5.300E-09 | 2.100E-08 |
| 4013 | 201 | G201M | A | 2.2e-09 | 2.5e-09 | 1e-08 | 4.900E-09 | 4.419E-09 | 2.200E-09 | 1.000E-08 |
| 4014 | 201 | G201F | A | 1.2e-08 | 1.8e-08 | 9.4e-09 | 1.313E-08 | 4.411E-09 | 9.400E-09 | 1.800E-08 |
| 4015 | 201 | G201P | A | 9.6e-09 | 4.6e-09 | 5.1e-09 | 6.433E-09 | 2.754E-09 | 4.600E-09 | 9.600E-09 |
| 4016 | 201 | G201S | A | 8.7e-09 | 2.8e-08 | 4.1e-09 | 1.360E-08 | 1.268E-08 | 4.100E-09 | 2.800E-08 |
| 4017 | 201 | G201T | A | 4.5e-09 | 1.6e-08 | 8.8e-09 | 9.767E-09 | 5.811E-09 | 4.500E-09 | 1.600E-08 |
| 4018 | 201 | G201W | A | 5.4e-09 | 1.4e-08 | 4.1e-09 | 7.833E-09 | 5.380E-09 | 4.100E-09 | 1.400E-08 |
| 4019 | 201 | G201Y | A | 4.4e-09 | 3.2e-09 | 1.3e-08 | 6.867E-09 | 5.345E-09 | 3.200E-09 | 1.300E-08 |
| 4020 | 201 | G201V | A | 7.2e-09 | 5.3e-09 | 5.6e-09 | 6.033E-09 | 1.021E-09 | 5.300E-09 | 7.200E-09 |
| 4021 | 202 | C202A | A | 1e-08 | 9.1e-09 | 1.1e-08 | 1.003E-08 | 9.504E-10 | 9.100E-09 | 1.100E-08 |
| 4022 | 202 | C202R | A | 6.9e-09 | 7.7e-09 | 6.6e-09 | 7.067E-09 | 5.686E-10 | 6.600E-09 | 7.700E-09 |
| 4023 | 202 | C202N | A | 9e-09 | 2e-08 | 1e-08 | 1.300E-08 | 6.083E-09 | 9.000E-09 | 2.000E-08 |
| 4024 | 202 | C202D | A | 8.7e-09 | 9.8e-09 | 1.8e-08 | 1.217E-08 | 5.082E-09 | 8.700E-09 | 1.800E-08 |
| 4025 | 202 | C202C | A | 9.5e-09 | 5.1e-09 | 7.4e-09 | 7.333E-09 | 2.201E-09 | 5.100E-09 | 9.500E-09 |
| 4026 | 202 | C202E | A | 5.7e-09 | 5.6e-09 | 1.7e-08 | 9.433E-09 | 6.553E-09 | 5.600E-09 | 1.700E-08 |
| 4027 | 202 | C202Q | A | 2.6e-08 | 1.7e-08 | 2.2e-09 | 1.507E-08 | 1.202E-08 | 2.200E-09 | 2.600E-08 |
| 4028 | 202 | C202G | A | 9e-09 | 2.6e-09 | 1.3e-08 | 8.200E-09 | 5.246E-09 | 2.600E-09 | 1.300E-08 |
| 4029 | 202 | C202H | A | 5.1e-09 | 6.2e-09 | 9.5e-09 | 6.933E-09 | 2.290E-09 | 5.100E-09 | 9.500E-09 |
| 4030 | 202 | C202I | A | 7.1e-09 | 5.4e-09 | 1.1e-08 | 7.833E-09 | 2.871E-09 | 5.400E-09 | 1.100E-08 |
| 4031 | 202 | C202L | A | 7.4e-09 | 6.3e-09 | 3.3e-09 | 5.667E-09 | 2.122E-09 | 3.300E-09 | 7.400E-09 |
| 4032 | 202 | C202K | A | 8.1e-09 | 1.2e-08 | 2.2e-09 | 7.433E-09 | 4.934E-09 | 2.200E-09 | 1.200E-08 |
| 4033 | 202 | C202M | A | 1.1e-08 | 2.5e-09 | 6.2e-09 | 6.567E-09 | 4.262E-09 | 2.500E-09 | 1.100E-08 |
| 4034 | 202 | C202F | A | 3e-09 | 5.4e-09 | 5.7e-09 | 4.700E-09 | 1.480E-09 | 3.000E-09 | 5.700E-09 |
| 4035 | 202 | C202P | A | 3.2e-08 | 1.5e-08 | 5e-09 | 1.733E-08 | 1.365E-08 | 5.000E-09 | 3.200E-08 |
| 4036 | 202 | C202S | A | 2e-08 | 2.6e-08 | 1.7e-08 | 2.100E-08 | 4.583E-09 | 1.700E-08 | 2.600E-08 |
| 4037 | 202 | C202T | A | 1.4e-08 | 4.6e-09 | 2.6e-09 | 7.067E-09 | 6.087E-09 | 2.600E-09 | 1.400E-08 |
| 4038 | 202 | C202W | A | 7.1e-09 | 3.8e-09 | 6.1e-09 | 5.667E-09 | 1.692E-09 | 3.800E-09 | 7.100E-09 |
| 4039 | 202 | C202Y | A | 9.5e-09 | 1.5e-08 | 6.4e-09 | 1.030E-08 | 4.355E-09 | 6.400E-09 | 1.500E-08 |
| 4040 | 202 | C202V | A | 1.5e-08 | 1.4e-09 | 5.5e-09 | 7.300E-09 | 6.976E-09 | 1.400E-09 | 1.500E-08 |
| 4041 | 203 | A203A | A | 7.3e-09 | 1.5e-08 | 7.2e-09 | 9.833E-09 | 4.475E-09 | 7.200E-09 | 1.500E-08 |
| 4042 | 203 | A203R | A | 5.8e-09 | 1.4e-08 | 3.2e-09 | 7.667E-09 | 5.637E-09 | 3.200E-09 | 1.400E-08 |
| 4043 | 203 | A203N | A | 6e-09 | 1.1e-08 | 1.2e-08 | 9.667E-09 | 3.215E-09 | 6.000E-09 | 1.200E-08 |
| 4044 | 203 | A203D | A | 2.5e-08 | 6.4e-09 | 6.6e-09 | 1.267E-08 | 1.068E-08 | 6.400E-09 | 2.500E-08 |
| 4045 | 203 | A203C | A | 5.3e-09 | 1.9e-08 | 1.3e-08 | 1.243E-08 | 6.868E-09 | 5.300E-09 | 1.900E-08 |
| 4046 | 203 | A203E | A | 5e-09 | 6.8e-09 | 5.6e-09 | 5.800E-09 | 9.165E-10 | 5.000E-09 | 6.800E-09 |
| 4047 | 203 | A203Q | A | 2.8e-08 | 1.4e-08 | 6.1e-09 | 1.603E-08 | 1.109E-08 | 6.100E-09 | 2.800E-08 |
| 4048 | 203 | A203G | A | 1.1e-08 | 6e-09 | 3.4e-09 | 6.800E-09 | 3.863E-09 | 3.400E-09 | 1.100E-08 |
| 4049 | 203 | A203H | A | 6.2e-09 | 1.8e-08 | 5.8e-09 | 1.000E-08 | 6.931E-09 | 5.800E-09 | 1.800E-08 |
| 4050 | 203 | A203I | A | 4.7e-09 | 7.4e-09 | 4.9e-09 | 5.667E-09 | 1.504E-09 | 4.700E-09 | 7.400E-09 |
| 4051 | 203 | A203L | A | 2.6e-08 | 1.2e-08 | 8.6e-09 | 1.553E-08 | 9.222E-09 | 8.600E-09 | 2.600E-08 |
| 4052 | 203 | A203K | A | 6.2e-09 | 7.9e-09 | 3.3e-08 | 1.570E-08 | 1.501E-08 | 6.200E-09 | 3.300E-08 |
| 4053 | 203 | A203M | A | 1.2e-08 | 1.1e-08 | 1.7e-08 | 1.333E-08 | 3.215E-09 | 1.100E-08 | 1.700E-08 |
| 4054 | 203 | A203F | A | 4.3e-09 | 5.3e-09 | 6.5e-09 | 5.367E-09 | 1.102E-09 | 4.300E-09 | 6.500E-09 |
| 4055 | 203 | A203P | A | 1.7e-08 | 4e-09 | 2.3e-08 | 1.467E-08 | 9.713E-09 | 4.000E-09 | 2.300E-08 |
| 4056 | 203 | A203S | A | 3.8e-09 | 4e-09 | 3.2e-09 | 3.667E-09 | 4.163E-10 | 3.200E-09 | 4.000E-09 |
| 4057 | 203 | A203T | A | 3.5e-09 | 1.4e-08 | 3.8e-09 | 7.100E-09 | 5.977E-09 | 3.500E-09 | 1.400E-08 |
| 4058 | 203 | A203W | A | 9.2e-09 | 9.6e-09 | 3.6e-09 | 7.467E-09 | 3.355E-09 | 3.600E-09 | 9.600E-09 |
| 4059 | 203 | A203Y | A | 6.7e-09 | 8.3e-09 | 2.3e-09 | 5.767E-09 | 3.107E-09 | 2.300E-09 | 8.300E-09 |
| 4060 | 203 | A203V | A | 9.1e-09 | 1.1e-08 | 2.7e-09 | 7.600E-09 | 4.349E-09 | 2.700E-09 | 1.100E-08 |
| 4061 | 204 | R204A | A | 5e-09 | 2.4e-08 | 9.8e-09 | 1.293E-08 | 9.880E-09 | 5.000E-09 | 2.400E-08 |
| 4062 | 204 | R204R | A | 6.4e-09 | 1e-08 | 6.6e-09 | 7.667E-09 | 2.023E-09 | 6.400E-09 | 1.000E-08 |
| 4063 | 204 | R204N | A | 1e-08 | 6.2e-09 | 2.6e-09 | 6.267E-09 | 3.700E-09 | 2.600E-09 | 1.000E-08 |
| 4064 | 204 | R204D | A | 6.2e-09 | 1.4e-08 | 3.8e-09 | 8.000E-09 | 5.333E-09 | 3.800E-09 | 1.400E-08 |
| 4065 | 204 | R204C | A | 2.5e-09 | 4.1e-09 | 1.4e-08 | 6.867E-09 | 6.229E-09 | 2.500E-09 | 1.400E-08 |
| 4066 | 204 | R204E | A | 1.3e-08 | 1.4e-09 | 5.7e-09 | 6.700E-09 | 5.864E-09 | 1.400E-09 | 1.300E-08 |
| 4067 | 204 | R204Q | A | 1.4e-08 | 5.6e-09 | 6.1e-09 | 8.567E-09 | 4.712E-09 | 5.600E-09 | 1.400E-08 |
| 4068 | 204 | R204G | A | 7.2e-09 | 1.2e-08 | 6.2e-09 | 8.467E-09 | 3.101E-09 | 6.200E-09 | 1.200E-08 |
| 4069 | 204 | R204H | A | 1.1e-08 | 3.7e-09 | 2.2e-08 | 1.223E-08 | 9.212E-09 | 3.700E-09 | 2.200E-08 |
| 4070 | 204 | R204I | A | 6.1e-09 | 6.5e-09 | 5.3e-09 | 5.967E-09 | 6.110E-10 | 5.300E-09 | 6.500E-09 |
| 4071 | 204 | R204L | A | 1.7e-08 | 7.1e-09 | 2.8e-09 | 8.967E-09 | 7.282E-09 | 2.800E-09 | 1.700E-08 |
| 4072 | 204 | R204K | A | 8.2e-09 | 7.1e-09 | 1e-08 | 8.433E-09 | 1.464E-09 | 7.100E-09 | 1.000E-08 |
| 4073 | 204 | R204M | A | 1.6e-08 | 5.9e-09 | 5.3e-09 | 9.067E-09 | 6.012E-09 | 5.300E-09 | 1.600E-08 |
| 4074 | 204 | R204F | A | 4.5e-09 | 1e-08 | 1.4e-08 | 9.500E-09 | 4.770E-09 | 4.500E-09 | 1.400E-08 |
| 4075 | 204 | R204P | A | 5.1e-09 | 1.2e-08 | 7.6e-09 | 8.233E-09 | 3.493E-09 | 5.100E-09 | 1.200E-08 |
| 4076 | 204 | R204S | A | 8.5e-09 | 1.2e-08 | 5.7e-09 | 8.733E-09 | 3.156E-09 | 5.700E-09 | 1.200E-08 |
| 4077 | 204 | R204T | A | 8e-09 | 4.2e-09 | 8e-09 | 6.733E-09 | 2.194E-09 | 4.200E-09 | 8.000E-09 |
| 4078 | 204 | R204W | A | 7.4e-09 | 5e-09 | 1.8e-08 | 1.013E-08 | 6.918E-09 | 5.000E-09 | 1.800E-08 |
| 4079 | 204 | R204Y | A | 7.8e-09 | 1.3e-08 | 4.9e-09 | 8.567E-09 | 4.104E-09 | 4.900E-09 | 1.300E-08 |

| | | | | | | | | | | |
|---|---|---|---|---|---|---|---|---|---|---|
| 4080 | 204 | R204V | A | 1.9e-08 | 2.5e-08 | 1.1e-08 | 1.833E-08 | 7.024E-09 | 1.100E-08 | 2.500E-08 |
| 4081 | 205 | C205A | A | 9.1e-09 | 4.9e-09 | 4.2e-09 | 6.067E-09 | 2.650E-09 | 4.200E-09 | 9.100E-09 |
| 4082 | 205 | C205R | A | 6.6e-09 | 1.3e-08 | 4.6e-09 | 8.067E-09 | 4.388E-09 | 4.600E-09 | 1.300E-08 |
| 4083 | 205 | C205N | A | 5.6e-09 | 5.2e-09 | 4.1e-09 | 4.967E-09 | 7.767E-10 | 4.100E-09 | 5.600E-09 |
| 4084 | 205 | C205D | A | 2.2e-09 | 1.8e-08 | 9.9e-09 | 1.003E-08 | 7.901E-09 | 2.200E-09 | 1.800E-08 |
| 4085 | 205 | C205C | A | 1.7e-08 | 8.6e-09 | 4.5e-09 | 1.003E-08 | 6.372E-09 | 4.500E-09 | 1.700E-08 |
| 4086 | 205 | C205E | A | 8e-09 | 2.3e-08 | 6.2e-09 | 1.240E-08 | 9.224E-09 | 6.200E-09 | 2.300E-08 |
| 4087 | 205 | C205Q | A | 6.8e-09 | 1.3e-08 | 1.5e-08 | 1.160E-08 | 4.276E-09 | 6.800E-09 | 1.500E-08 |
| 4088 | 205 | C205G | A | 8.4e-09 | 6.1e-09 | 1.2e-08 | 8.833E-09 | 2.974E-09 | 6.100E-09 | 1.200E-08 |
| 4089 | 205 | C205H | A | 1.4e-08 | 7.4e-09 | 8.7e-09 | 1.003E-08 | 3.496E-09 | 7.400E-09 | 1.400E-08 |
| 4090 | 205 | C205I | A | 3.6e-09 | 1.6e-08 | 4.3e-09 | 7.967E-09 | 6.966E-09 | 3.600E-09 | 1.600E-08 |
| 4091 | 205 | C205L | A | 7.4e-09 | 3.5e-09 | 6.3e-09 | 5.733E-09 | 2.011E-09 | 3.500E-09 | 7.400E-09 |
| 4092 | 205 | C205K | A | 1.7e-08 | 4.3e-09 | 4.4e-09 | 8.567E-09 | 7.304E-09 | 4.300E-09 | 1.700E-08 |
| 4093 | 205 | C205M | A | 2.4e-08 | 8.9e-09 | 4.1e-09 | 1.233E-08 | 1.038E-08 | 4.100E-09 | 2.400E-08 |
| 4094 | 205 | C205F | A | 5.3e-09 | 5.9e-09 | 6.7e-09 | 5.967E-09 | 7.024E-10 | 5.300E-09 | 6.700E-09 |
| 4095 | 205 | C205P | A | 9e-09 | 7.7e-09 | 6.9e-09 | 7.867E-09 | 1.060E-09 | 6.900E-09 | 9.000E-09 |
| 4096 | 205 | C205S | A | 5.5e-09 | 5.8e-09 | 1.7e-08 | 9.433E-09 | 6.555E-09 | 5.500E-09 | 1.700E-08 |
| 4097 | 205 | C205T | A | 9.7e-09 | 8.2e-09 | 6.1e-09 | 8.000E-09 | 1.808E-09 | 6.100E-09 | 9.700E-09 |
| 4098 | 205 | C205W | A | 6.4e-09 | 5.2e-09 | 8.3e-09 | 6.633E-09 | 1.563E-09 | 5.200E-09 | 8.300E-09 |
| 4099 | 205 | C205Y | A | 2.9e-09 | 6e-09 | 1.5e-08 | 7.967E-09 | 6.285E-09 | 2.900E-09 | 1.500E-08 |
| 4100 | 205 | C205V | A | 1.3e-08 | 1.2e-08 | 4.1e-09 | 9.700E-09 | 4.875E-09 | 4.100E-09 | 1.300E-08 |
| 4101 | 206 | K206A | A | 5.1e-09 | 2.7e-09 | 9.2e-09 | 5.667E-09 | 3.287E-09 | 2.700E-09 | 9.200E-09 |
| 4102 | 206 | K206R | A | 2.6e-09 | 3.5e-09 | 6.3e-09 | 4.133E-09 | 1.930E-09 | 2.600E-09 | 6.300E-09 |
| 4103 | 206 | K206N | A | 1.1e-08 | 4.3e-09 | 6.4e-09 | 7.233E-09 | 3.427E-09 | 4.300E-09 | 1.100E-08 |
| 4104 | 206 | K206D | A | 7.6e-09 | 5.6e-09 | 8.5e-09 | 7.233E-09 | 1.484E-09 | 5.600E-09 | 8.500E-09 |
| 4105 | 206 | K206C | A | 8.3e-09 | 3.5e-09 | 8.1e-09 | 6.633E-09 | 2.715E-09 | 3.500E-09 | 8.300E-09 |
| 4106 | 206 | K206E | A | 1e-08 | 1.4e-08 | 3.5e-08 | 1.967E-08 | 1.343E-08 | 1.000E-08 | 3.500E-08 |
| 4107 | 206 | K206Q | A | 6.2e-09 | 7.7e-09 | 4.8e-09 | 6.233E-09 | 1.450E-09 | 4.800E-09 | 7.700E-09 |
| 4108 | 206 | K206G | A | 1.2e-08 | 1.4e-08 | 1e-08 | 1.200E-08 | 2.000E-09 | 1.000E-08 | 1.400E-08 |
| 4109 | 206 | K206H | A | 1.9e-09 | 8e-09 | 9.1e-09 | 6.333E-09 | 3.879E-09 | 1.900E-09 | 9.100E-09 |
| 4110 | 206 | K206I | A | 1.2e-08 | 9.8e-09 | 7.4e-09 | 9.733E-09 | 2.301E-09 | 7.400E-09 | 1.200E-08 |
| 4111 | 206 | K206L | A | 1.9e-08 | 7.7e-09 | 7.5e-09 | 1.140E-08 | 6.583E-09 | 7.500E-09 | 1.900E-08 |
| 4112 | 206 | K206K | A | 9.1e-09 | 1e-08 | 3.2e-09 | 7.433E-09 | 3.694E-09 | 3.200E-09 | 1.000E-08 |
| 4113 | 206 | K206M | A | 4.6e-09 | 6.2e-09 | 3.1e-08 | 1.393E-08 | 1.480E-08 | 4.600E-09 | 3.100E-08 |
| 4114 | 206 | K206F | A | 3.6e-09 | 1.4e-08 | 1.1e-08 | 9.533E-09 | 5.353E-09 | 3.600E-09 | 1.400E-08 |
| 4115 | 206 | K206P | A | 6.9e-09 | 1.4e-08 | 1.3e-08 | 1.130E-08 | 3.843E-09 | 6.900E-09 | 1.400E-08 |
| 4116 | 206 | K206S | A | 3.6e-09 | 1.8e-09 | 9.6e-09 | 5.000E-09 | 4.084E-09 | 1.800E-09 | 9.600E-09 |
| 4117 | 206 | K206T | A | 1.4e-08 | 6.6e-09 | 1.1e-08 | 1.053E-08 | 3.722E-09 | 6.600E-09 | 1.400E-08 |
| 4118 | 206 | K206W | A | 1.9e-09 | 5.7e-09 | 5.8e-09 | 4.467E-09 | 2.223E-09 | 1.900E-09 | 5.800E-09 |
| 4119 | 206 | K206Y | A | 9.2e-09 | 1.2e-08 | 5.5e-09 | 8.900E-09 | 3.260E-09 | 5.500E-09 | 1.200E-08 |
| 4120 | 206 | K206V | A | 9.6e-09 | 2.3e-09 | 3.2e-09 | 5.033E-09 | 3.980E-09 | 2.300E-09 | 9.600E-09 |
| 4121 | 207 | G207A | A | 1.4e-08 | 2.5e-09 | 4e-09 | 6.833E-09 | 6.252E-09 | 2.500E-09 | 1.400E-08 |
| 4122 | 207 | G207R | A | 7.8e-09 | 2.8e-09 | 6.1e-09 | 5.567E-09 | 2.542E-09 | 2.800E-09 | 7.800E-09 |
| 4123 | 207 | G207N | A | 9e-09 | 3.9e-09 | 6.7e-09 | 6.533E-09 | 2.554E-09 | 3.900E-09 | 9.000E-09 |
| 4124 | 207 | G207D | A | 1.7e-08 | 5.3e-09 | 8e-09 | 1.010E-08 | 6.126E-09 | 5.300E-09 | 1.700E-08 |
| 4125 | 207 | G207C | A | 3.3e-09 | 7.9e-09 | 7.5e-09 | 6.233E-09 | 2.548E-09 | 3.300E-09 | 7.900E-09 |
| 4126 | 207 | G207E | A | 6e-09 | 3.8e-09 | 7.1e-09 | 5.633E-09 | 1.680E-09 | 3.800E-09 | 7.100E-09 |
| 4127 | 207 | G207Q | A | 4.4e-09 | 9.8e-09 | 7.7e-09 | 7.300E-09 | 2.722E-09 | 4.400E-09 | 9.800E-09 |
| 4128 | 207 | G207G | A | 2.7e-09 | 4e-09 | 1.2e-08 | 6.233E-09 | 5.036E-09 | 2.700E-09 | 1.200E-08 |
| 4129 | 207 | G207H | A | 1e-08 | 5.5e-09 | 7.4e-09 | 7.633E-09 | 2.259E-09 | 5.500E-09 | 1.000E-08 |
| 4130 | 207 | G207I | A | 4.3e-09 | 1.2e-08 | 7.6e-09 | 7.967E-09 | 3.863E-09 | 4.300E-09 | 1.200E-08 |
| 4131 | 207 | G207L | A | 1.6e-08 | 6.7e-09 | 2.5e-09 | 8.400E-09 | 6.909E-09 | 2.500E-09 | 1.600E-08 |
| 4132 | 207 | G207K | A | 5.4e-09 | 5.9e-09 | 1.3e-08 | 8.100E-09 | 4.251E-09 | 5.400E-09 | 1.300E-08 |
| 4133 | 207 | G207M | A | 4.5e-09 | 5.2e-09 | 7.8e-09 | 5.833E-09 | 1.739E-09 | 4.500E-09 | 7.800E-09 |
| 4134 | 207 | G207F | A | 1.3e-09 | 6.2e-09 | 1.4e-08 | 7.167E-09 | 6.405E-09 | 1.300E-09 | 1.400E-08 |
| 4135 | 207 | G207P | A | 6.5e-09 | 5.3e-09 | 7.4e-09 | 6.400E-09 | 1.054E-09 | 5.300E-09 | 7.400E-09 |
| 4136 | 207 | G207S | A | 9.1e-09 | 3.9e-09 | 5.6e-09 | 6.200E-09 | 2.651E-09 | 3.900E-09 | 9.100E-09 |
| 4137 | 207 | G207T | A | 1.3e-08 | 8.8e-09 | 6e-09 | 9.267E-09 | 3.523E-09 | 6.000E-09 | 1.300E-08 |
| 4138 | 207 | G207W | A | 7.3e-09 | 5.4e-09 | 3.3e-09 | 5.333E-09 | 2.001E-09 | 3.300E-09 | 7.300E-09 |
| 4139 | 207 | G207Y | A | 5.3e-09 | 1.1e-08 | 8.7e-09 | 8.333E-09 | 2.868E-09 | 5.300E-09 | 1.100E-08 |
| 4140 | 207 | G207V | A | 2.4e-09 | 1.2e-08 | 1.1e-08 | 8.467E-09 | 5.278E-09 | 2.400E-09 | 1.200E-08 |
| 4141 | 208 | P208A | A | 1.9e-08 | 1.1e-08 | 4e-09 | 1.133E-08 | 7.506E-09 | 4.000E-09 | 1.900E-08 |
| 4142 | 208 | P208R | A | 5.6e-09 | 1.3e-08 | 4e-09 | 7.533E-09 | 4.801E-09 | 4.000E-09 | 1.300E-08 |
| 4143 | 208 | P208N | A | 4e-09 | 5.3e-09 | 2.1e-09 | 3.800E-09 | 1.609E-09 | 2.100E-09 | 5.300E-09 |
| 4144 | 208 | P208D | A | 2.8e-08 | 1.1e-08 | 7.2e-09 | 1.540E-08 | 1.108E-08 | 7.200E-09 | 2.800E-08 |
| 4145 | 208 | P208C | A | 1.6e-08 | 1.2e-08 | 4.1e-09 | 1.070E-08 | 6.056E-09 | 4.100E-09 | 1.600E-08 |
| 4146 | 208 | P208E | A | 4.9e-09 | 6e-09 | 3.6e-09 | 4.833E-09 | 1.201E-09 | 3.600E-09 | 6.000E-09 |
| 4147 | 208 | P208Q | A | 2.5e-08 | 1e-08 | 7.9e-09 | 1.430E-08 | 9.326E-09 | 7.900E-09 | 2.500E-08 |
| 4148 | 208 | P208G | A | 9e-09 | 4.9e-09 | 1.9e-08 | 1.097E-08 | 7.253E-09 | 4.900E-09 | 1.900E-08 |
| 4149 | 208 | P208H | A | 6.3e-09 | 1.2e-08 | 3.9e-09 | 7.400E-09 | 4.161E-09 | 3.900E-09 | 1.200E-08 |
| 4150 | 208 | P208I | A | 7.1e-09 | 5.2e-09 | 1.4e-08 | 8.767E-09 | 4.631E-09 | 5.200E-09 | 1.400E-08 |
| 4151 | 208 | P208L | A | 6.6e-09 | 5.5e-09 | 1.5e-08 | 9.033E-09 | 5.196E-09 | 5.500E-09 | 1.500E-08 |
| 4152 | 208 | P208K | A | 3.6e-09 | 9e-09 | 4.8e-09 | 5.800E-09 | 2.835E-09 | 3.600E-09 | 9.000E-09 |
| 4153 | 208 | P208M | A | 5.7e-09 | 3.4e-09 | 3.7e-09 | 4.267E-09 | 1.250E-09 | 3.400E-09 | 5.700E-09 |
| 4154 | 208 | P208F | A | 6.3e-09 | 1.6e-08 | 1.3e-08 | 1.177E-08 | 4.966E-09 | 6.300E-09 | 1.600E-08 |
| 4155 | 208 | P208P | A | 7.4e-09 | 2.7e-09 | 5.2e-09 | 5.100E-09 | 2.352E-09 | 2.700E-09 | 7.400E-09 |
| 4156 | 208 | P208S | A | 3.7e-09 | 7.7e-09 | 5.8e-09 | 5.733E-09 | 2.001E-09 | 3.700E-09 | 7.700E-09 |
| 4157 | 208 | P208T | A | 4.5e-09 | 5.5e-09 | 1.1e-08 | 7.000E-09 | 3.500E-09 | 4.500E-09 | 1.100E-08 |
| 4158 | 208 | P208W | A | 7.7e-09 | 9.4e-09 | 1e-08 | 9.033E-09 | 1.193E-09 | 7.700E-09 | 1.000E-08 |
| 4159 | 208 | P208Y | A | 1.2e-08 | 4.7e-09 | 1.2e-08 | 9.567E-09 | 4.215E-09 | 4.700E-09 | 1.200E-08 |
| 4160 | 208 | P208V | A | 7.7e-09 | 9.3e-09 | 5e-09 | 7.333E-09 | 2.173E-09 | 5.000E-09 | 9.300E-09 |
| 4161 | 209 | L209A | A | 1.1e-08 | 2e-08 | 5.2e-09 | 1.207E-08 | 7.457E-09 | 5.200E-09 | 2.000E-08 |
| 4162 | 209 | L209R | A | 6.5e-10 | 8.8e-09 | 1.5e-08 | 8.150E-09 | 7.197E-09 | 6.500E-10 | 1.500E-08 |
| 4163 | 209 | L209N | A | 2.2e-08 | 9.2e-09 | 1.6e-08 | 1.573E-08 | 6.404E-09 | 9.200E-09 | 2.200E-08 |
| 4164 | 209 | L209D | A | 4.1e-09 | 3.3e-09 | 2e-09 | 3.133E-09 | 1.060E-09 | 2.000E-09 | 4.100E-09 |

| | | | | | | | | | | |
|---|---|---|---|---|---|---|---|---|---|---|
| 4165 | 209 | L209C | A | 5.4e-09 | 4.4e-09 | 2.5e-08 | 1.160E-08 | 1.162E-08 | 4.400E-09 | 2.500E-08 |
| 4166 | 209 | L209E | A | 5.4e-09 | 1.5e-08 | 6.8e-09 | 9.067E-09 | 5.186E-09 | 5.400E-09 | 1.500E-08 |
| 4167 | 209 | L209Q | A | 5e-09 | 4.2e-09 | 9.1e-09 | 6.100E-09 | 2.629E-09 | 4.200E-09 | 9.100E-09 |
| 4168 | 209 | L209G | A | 2.5e-09 | 9.7e-09 | 9e-09 | 7.067E-09 | 3.970E-09 | 2.500E-09 | 9.700E-09 |
| 4169 | 209 | L209H | A | 3.8e-09 | 8.7e-09 | 7.4e-09 | 6.633E-09 | 2.538E-09 | 3.800E-09 | 8.700E-09 |
| 4170 | 209 | L209I | A | 6.1e-09 | 3.6e-09 | 8.3e-09 | 6.000E-09 | 2.352E-09 | 3.600E-09 | 8.300E-09 |
| 4171 | 209 | L209L | A | 7.7e-09 | 4.8e-09 | 4.2e-09 | 5.567E-09 | 1.872E-09 | 4.200E-09 | 7.700E-09 |
| 4172 | 209 | L209K | A | 1.1e-08 | 7.3e-09 | 3.8e-09 | 7.367E-09 | 3.600E-09 | 3.800E-09 | 1.100E-08 |
| 4173 | 209 | L209M | A | 4.8e-09 | 2.7e-08 | 5.7e-09 | 1.250E-08 | 1.257E-08 | 4.800E-09 | 2.700E-08 |
| 4174 | 209 | L209F | A | 7.2e-09 | 3.2e-08 | 3.2e-09 | 1.413E-08 | 1.560E-08 | 3.200E-09 | 3.200E-08 |
| 4175 | 209 | L209P | A | 7.3e-09 | 6e-09 | 1.6e-08 | 9.767E-09 | 5.437E-09 | 6.000E-09 | 1.600E-08 |
| 4176 | 209 | L209S | A | 9.1e-09 | 1.7e-08 | 5.9e-09 | 1.067E-08 | 5.713E-09 | 5.900E-09 | 1.700E-08 |
| 4177 | 209 | L209T | A | 9.3e-09 | 1.3e-08 | 5.5e-09 | 9.267E-09 | 3.750E-09 | 5.500E-09 | 1.300E-08 |
| 4178 | 209 | L209W | A | 1.2e-08 | 6.1e-09 | 2.1e-08 | 1.303E-08 | 7.504E-09 | 6.100E-09 | 2.100E-08 |
| 4179 | 209 | L209Y | A | 5.4e-09 | 4.9e-09 | 4.8e-09 | 5.033E-09 | 3.215E-10 | 4.800E-09 | 5.400E-09 |
| 4180 | 209 | L209V | A | 3.4e-09 | 1.5e-08 | 5e-09 | 7.800E-09 | 6.286E-09 | 3.400E-09 | 1.500E-08 |
| 4181 | 210 | P210A | A | 4.3e-09 | 8.2e-09 | 4.6e-09 | 5.700E-09 | 2.170E-09 | 4.300E-09 | 8.200E-09 |
| 4182 | 210 | P210R | A | 4.7e-09 | 4.1e-09 | 6.5e-09 | 5.100E-09 | 1.249E-09 | 4.100E-09 | 6.500E-09 |
| 4183 | 210 | P210N | A | 3.7e-09 | 1.1e-08 | 2.1e-09 | 5.600E-09 | 4.744E-09 | 2.100E-09 | 1.100E-08 |
| 4184 | 210 | P210D | A | 6.4e-09 | 1.9e-08 | 8e-09 | 1.113E-08 | 6.860E-09 | 6.400E-09 | 1.900E-08 |
| 4185 | 210 | P210C | A | 1.4e-08 | 6.4e-09 | 1.3e-08 | 1.113E-08 | 4.130E-09 | 6.400E-09 | 1.400E-08 |
| 4186 | 210 | P210E | A | 1e-08 | 1.1e-08 | 7.1e-09 | 9.367E-09 | 2.026E-09 | 7.100E-09 | 1.100E-08 |
| 4187 | 210 | P210Q | A | 8.4e-09 | 3.9e-09 | 3.2e-08 | 1.477E-08 | 1.509E-08 | 3.900E-09 | 3.200E-08 |
| 4188 | 210 | P210G | A | 1.4e-08 | 4.6e-09 | 4.9e-09 | 7.833E-09 | 5.343E-09 | 4.600E-09 | 1.400E-08 |
| 4189 | 210 | P210H | A | 6.9e-09 | 2.2e-09 | 3.8e-09 | 4.300E-09 | 2.390E-09 | 2.200E-09 | 6.900E-09 |
| 4190 | 210 | P210I | A | 3.3e-09 | 7.3e-09 | 1.9e-08 | 9.867E-09 | 8.159E-09 | 3.300E-09 | 1.900E-08 |
| 4191 | 210 | P210L | A | 6.6e-09 | 9.5e-09 | 7.9e-09 | 8.000E-09 | 1.453E-09 | 6.600E-09 | 9.500E-09 |
| 4192 | 210 | P210K | A | 7.9e-09 | 5.3e-09 | 8.1e-09 | 7.100E-09 | 1.562E-09 | 5.300E-09 | 8.100E-09 |
| 4193 | 210 | P210M | A | 9.5e-10 | 4.6e-09 | 3.3e-09 | 2.950E-09 | 1.850E-09 | 9.500E-10 | 4.600E-09 |
| 4194 | 210 | P210F | A | 6.9e-09 | 7.2e-09 | 3.9e-09 | 6.000E-09 | 1.825E-09 | 3.900E-09 | 7.200E-09 |
| 4195 | 210 | P210P | A | 1.5e-08 | 2.6e-09 | 5.1e-09 | 7.567E-09 | 6.558E-09 | 2.600E-09 | 1.500E-08 |
| 4196 | 210 | P210S | A | 8.2e-09 | 3.2e-09 | 4.8e-09 | 5.400E-09 | 2.553E-09 | 3.200E-09 | 8.200E-09 |
| 4197 | 210 | P210T | A | 1.4e-08 | 1.1e-09 | 2.4e-08 | 1.303E-08 | 1.148E-08 | 1.100E-09 | 2.400E-08 |
| 4198 | 210 | P210W | A | 4e-09 | 1.1e-08 | 1e-08 | 8.333E-09 | 3.786E-09 | 4.000E-09 | 1.100E-08 |
| 4199 | 210 | P210Y | A | 2.4e-09 | 3.7e-09 | 7.7e-09 | 4.600E-09 | 2.762E-09 | 2.400E-09 | 7.700E-09 |
| 4200 | 210 | P210V | A | 5e-09 | 1.6e-08 | 1.4e-08 | 1.167E-08 | 5.859E-09 | 5.000E-09 | 1.600E-08 |
| 4201 | 211 | T211A | A | 1.3e-08 | 2.5e-09 | 8.2e-09 | 7.900E-09 | 5.256E-09 | 2.500E-09 | 1.300E-08 |
| 4202 | 211 | T211R | A | 1.6e-08 | 3.2e-09 | 7.3e-09 | 8.833E-09 | 6.536E-09 | 3.200E-09 | 1.600E-08 |
| 4203 | 211 | T211N | A | 9.6e-09 | 6.9e-09 | 3.9e-09 | 6.800E-09 | 2.851E-09 | 3.900E-09 | 9.600E-09 |
| 4204 | 211 | T211D | A | 2.6e-09 | 8.1e-09 | 4.7e-09 | 5.133E-09 | 2.775E-09 | 2.600E-09 | 8.100E-09 |
| 4205 | 211 | T211C | A | 4.4e-09 | 9.8e-09 | 1.5e-08 | 9.733E-09 | 5.300E-09 | 4.400E-09 | 1.500E-08 |
| 4206 | 211 | T211E | A | 3e-09 | 6.1e-09 | 1e-08 | 6.367E-09 | 3.508E-09 | 3.000E-09 | 1.000E-08 |
| 4207 | 211 | T211Q | A | 1.1e-08 | 6.6e-09 | 5.6e-09 | 7.733E-09 | 2.873E-09 | 5.600E-09 | 1.100E-08 |
| 4208 | 211 | T211G | A | 9.4e-09 | 1.7e-08 | 6.6e-09 | 1.100E-08 | 5.381E-09 | 6.600E-09 | 1.700E-08 |
| 4209 | 211 | T211H | A | 1.1e-08 | 9e-09 | 2.5e-09 | 7.500E-09 | 4.444E-09 | 2.500E-09 | 1.100E-08 |
| 4210 | 211 | T211I | A | 1.7e-08 | 4e-09 | 6.9e-09 | 9.300E-09 | 6.824E-09 | 4.000E-09 | 1.700E-08 |
| 4211 | 211 | T211L | A | 7.8e-09 | 8.2e-09 | 6.5e-09 | 7.500E-09 | 8.888E-10 | 6.500E-09 | 8.200E-09 |
| 4212 | 211 | T211K | A | 2.8e-09 | 5.1e-09 | 9.8e-09 | 5.900E-09 | 3.568E-09 | 2.800E-09 | 9.800E-09 |
| 4213 | 211 | T211M | A | 4.6e-09 | 5.3e-09 | 4.6e-09 | 4.833E-09 | 4.041E-10 | 4.600E-09 | 5.300E-09 |
| 4214 | 211 | T211F | A | 1.7e-08 | 1.1e-08 | 1e-08 | 1.267E-08 | 3.786E-09 | 1.000E-08 | 1.700E-08 |
| 4215 | 211 | T211P | A | 7.1e-09 | 8.9e-09 | 4.9e-09 | 6.967E-09 | 2.003E-09 | 4.900E-09 | 8.900E-09 |
| 4216 | 211 | T211S | A | 5.8e-09 | 1.2e-08 | 3.5e-09 | 7.100E-09 | 4.397E-09 | 3.500E-09 | 1.200E-08 |
| 4217 | 211 | T211T | A | 4.3e-09 | 2.3e-08 | 1e-08 | 1.243E-08 | 9.585E-09 | 4.300E-09 | 2.300E-08 |
| 4218 | 211 | T211W | A | 1.6e-08 | 1.3e-08 | 6.6e-09 | 1.187E-08 | 4.801E-09 | 6.600E-09 | 1.600E-08 |
| 4219 | 211 | T211Y | A | 1.2e-08 | 2.7e-09 | 7.9e-09 | 7.533E-09 | 4.661E-09 | 2.700E-09 | 1.200E-08 |
| 4220 | 211 | T211V | A | 6.4e-09 | 5.9e-09 | 9e-09 | 7.100E-09 | 1.664E-09 | 5.900E-09 | 9.000E-09 |
| 4221 | 212 | D212A | A | 1.3e-08 | 1e-08 | 6.6e-09 | 9.867E-09 | 3.202E-09 | 6.600E-09 | 1.300E-08 |
| 4222 | 212 | D212R | A | 7.1e-09 | 8e-09 | 7.2e-09 | 7.433E-09 | 4.933E-10 | 7.100E-09 | 8.000E-09 |
| 4223 | 212 | D212N | A | 1.1e-08 | 3.5e-09 | 5.2e-09 | 6.567E-09 | 3.932E-09 | 3.500E-09 | 1.100E-08 |
| 4224 | 212 | D212D | A | 8.7e-09 | 3.5e-08 | 5.1e-09 | 1.627E-08 | 1.632E-08 | 5.100E-09 | 3.500E-08 |
| 4225 | 212 | D212C | A | 7.8e-09 | 4.4e-09 | 7e-09 | 6.400E-09 | 1.778E-09 | 4.400E-09 | 7.800E-09 |
| 4226 | 212 | D212E | A | 6.3e-09 | 6.1e-09 | 4.5e-09 | 5.633E-09 | 9.866E-10 | 4.500E-09 | 6.300E-09 |
| 4227 | 212 | D212Q | A | 8.4e-09 | 8e-09 | 4.9e-09 | 7.100E-09 | 1.916E-09 | 4.900E-09 | 8.400E-09 |
| 4228 | 212 | D212G | A | 3.7e-09 | 6.6e-09 | 4.9e-09 | 5.067E-09 | 1.457E-09 | 3.700E-09 | 6.600E-09 |
| 4229 | 212 | D212H | A | 6.6e-09 | 5.5e-09 | 6.7e-09 | 6.267E-09 | 6.658E-10 | 5.500E-09 | 6.700E-09 |
| 4230 | 212 | D212I | A | 7.6e-09 | 1e-08 | 5.8e-09 | 7.800E-09 | 2.107E-09 | 5.800E-09 | 1.000E-08 |
| 4231 | 212 | D212L | A | 3.7e-09 | 6.5e-09 | 1.4e-08 | 8.067E-09 | 5.326E-09 | 3.700E-09 | 1.400E-08 |
| 4232 | 212 | D212K | A | 1.3e-08 | 1.3e-08 | 2e-08 | 1.533E-08 | 4.041E-09 | 1.300E-08 | 2.000E-08 |
| 4233 | 212 | D212M | A | 4e-09 | 1.5e-08 | 1.5e-08 | 1.133E-08 | 6.351E-09 | 4.000E-09 | 1.500E-08 |
| 4234 | 212 | D212F | A | 4.7e-09 | 4.6e-09 | 1.3e-08 | 7.433E-09 | 4.821E-09 | 4.600E-09 | 1.300E-08 |
| 4235 | 212 | D212P | A | 7.1e-09 | 3.5e-09 | 2e-08 | 1.020E-08 | 8.676E-09 | 3.500E-09 | 2.000E-08 |
| 4236 | 212 | D212S | A | 8.2e-09 | 9.2e-09 | 4.5e-09 | 7.300E-09 | 2.476E-09 | 4.500E-09 | 9.200E-09 |
| 4237 | 212 | D212T | A | 5.6e-09 | 5.6e-09 | 6e-09 | 5.733E-09 | 2.309E-10 | 5.600E-09 | 6.000E-09 |
| 4238 | 212 | D212W | A | 1.3e-08 | 1.1e-08 | 8e-09 | 1.067E-08 | 2.517E-09 | 8.000E-09 | 1.300E-08 |
| 4239 | 212 | D212Y | A | 2.7e-08 | 1.5e-08 | 5e-09 | 1.567E-08 | 1.102E-08 | 5.000E-09 | 2.700E-08 |
| 4240 | 212 | D212V | A | 4.8e-09 | 2.8e-09 | 5.1e-09 | 4.233E-09 | 1.250E-09 | 2.800E-09 | 5.100E-09 |
| 4241 | 213 | C213A | A | 5.9e-09 | 6.4e-09 | 5e-09 | 5.767E-09 | 7.095E-10 | 5.000E-09 | 6.400E-09 |
| 4242 | 213 | C213R | A | 5.2e-09 | 4.9e-09 | 6.5e-09 | 5.533E-09 | 8.505E-10 | 4.900E-09 | 6.500E-09 |
| 4243 | 213 | C213N | A | 1.2e-08 | 5.7e-09 | 2.5e-08 | 1.423E-08 | 9.842E-09 | 5.700E-09 | 2.500E-08 |
| 4244 | 213 | C213D | A | 9.1e-09 | 4.6e-09 | 4.2e-09 | 5.967E-09 | 2.721E-09 | 4.200E-09 | 9.100E-09 |
| 4245 | 213 | C213C | A | 6e-09 | 9e-09 | 4.9e-09 | 6.633E-09 | 2.122E-09 | 4.900E-09 | 9.000E-09 |
| 4246 | 213 | C213E | A | 3.9e-09 | 1.6e-08 | 1.4e-08 | 1.130E-08 | 6.486E-09 | 3.900E-09 | 1.600E-08 |
| 4247 | 213 | C213Q | A | 3.6e-08 | 4e-09 | 1.5e-08 | 1.833E-08 | 1.626E-08 | 4.000E-09 | 3.600E-08 |
| 4248 | 213 | C213G | A | 3e-09 | 5.2e-09 | 8.4e-09 | 5.533E-09 | 2.715E-09 | 3.000E-09 | 8.400E-09 |
| 4249 | 213 | C213H | A | 7.4e-09 | 4.7e-09 | 8.4e-09 | 6.833E-09 | 1.914E-09 | 4.700E-09 | 8.400E-09 |

| | | | | | | | | | | |
|---|---|---|---|---|---|---|---|---|---|---|
| 4250 | 213 | C213I | A | 7.5e-09 | 5.3e-09 | 1.5e-08 | 9.267E-09 | 5.086E-09 | 5.300E-09 | 1.500E-08 |
| 4251 | 213 | C213L | A | 3.7e-09 | 5.8e-09 | 3.5e-09 | 4.333E-09 | 1.274E-09 | 3.500E-09 | 5.800E-09 |
| 4252 | 213 | C213K | A | 4e-09 | 6e-09 | 1.1e-08 | 7.000E-09 | 3.606E-09 | 4.000E-09 | 1.100E-08 |
| 4253 | 213 | C213M | A | 1.2e-08 | 1.7e-08 | 6.7e-09 | 1.190E-08 | 5.151E-09 | 6.700E-09 | 1.700E-08 |
| 4254 | 213 | C213F | A | 9.3e-09 | 7.9e-09 | 6.1e-09 | 7.767E-09 | 1.604E-09 | 6.100E-09 | 9.300E-09 |
| 4255 | 213 | C213P | A | 4.8e-09 | 3.4e-09 | 9.8e-09 | 6.000E-09 | 3.365E-09 | 3.400E-09 | 9.800E-09 |
| 4256 | 213 | C213S | A | 1.2e-08 | 1.1e-08 | 4.3e-09 | 9.100E-09 | 4.187E-09 | 4.300E-09 | 1.200E-08 |
| 4257 | 213 | C213T | A | 6e-09 | 2.2e-08 | 4.8e-09 | 1.093E-08 | 9.603E-09 | 4.800E-09 | 2.200E-08 |
| 4258 | 213 | C213W | A | 1.5e-08 | 6.8e-09 | 1.3e-08 | 1.160E-08 | 4.276E-09 | 6.800E-09 | 1.500E-08 |
| 4259 | 213 | C213Y | A | 5.2e-09 | 1e-08 | 9.5e-09 | 8.233E-09 | 2.639E-09 | 5.200E-09 | 1.000E-08 |
| 4260 | 213 | C213V | A | 8.7e-09 | 6e-09 | 2.6e-09 | 5.767E-09 | 3.057E-09 | 2.600E-09 | 8.700E-09 |
| 4261 | 214 | C214A | A | 3.1e-09 | 7.7e-09 | 8.2e-09 | 6.333E-09 | 2.811E-09 | 3.100E-09 | 8.200E-09 |
| 4262 | 214 | C214R | A | 7.5e-09 | 4.3e-09 | 3.8e-09 | 5.200E-09 | 2.007E-09 | 3.800E-09 | 7.500E-09 |
| 4263 | 214 | C214N | A | 1.7e-08 | 4.7e-09 | 8.6e-09 | 1.010E-08 | 6.286E-09 | 4.700E-09 | 1.700E-08 |
| 4264 | 214 | C214D | A | 3.7e-09 | 7.9e-09 | 7.3e-09 | 6.300E-09 | 2.272E-09 | 3.700E-09 | 7.900E-09 |
| 4265 | 214 | C214C | A | 1.1e-08 | 8.3e-09 | 1.1e-08 | 1.010E-08 | 1.559E-09 | 8.300E-09 | 1.100E-08 |
| 4266 | 214 | C214E | A | 3.3e-09 | 1.2e-08 | 5.8e-09 | 7.033E-09 | 4.479E-09 | 3.300E-09 | 1.200E-08 |
| 4267 | 214 | C214Q | A | 1.1e-08 | 1.1e-08 | 4.6e-09 | 8.867E-09 | 3.695E-09 | 4.600E-09 | 1.100E-08 |
| 4268 | 214 | C214G | A | 5.1e-09 | 3e-09 | 4.9e-09 | 4.333E-09 | 1.159E-09 | 3.000E-09 | 5.100E-09 |
| 4269 | 214 | C214H | A | 8.5e-09 | 5.8e-09 | 1.3e-08 | 9.100E-09 | 3.637E-09 | 5.800E-09 | 1.300E-08 |
| 4270 | 214 | C214I | A | 1e-08 | 2.9e-09 | 9.9e-09 | 7.600E-09 | 4.071E-09 | 2.900E-09 | 1.000E-08 |
| 4271 | 214 | C214L | A | 5.1e-09 | 3.2e-09 | 4.1e-09 | 4.133E-09 | 9.504E-10 | 3.200E-09 | 5.100E-09 |
| 4272 | 214 | C214K | A | 6.1e-09 | 1.9e-08 | 1.3e-08 | 1.270E-08 | 6.455E-09 | 6.100E-09 | 1.900E-08 |
| 4273 | 214 | C214M | A | 7.1e-09 | 1.8e-08 | 2e-08 | 1.503E-08 | 6.943E-09 | 7.100E-09 | 2.000E-08 |
| 4274 | 214 | C214F | A | 9.2e-09 | 1.7e-08 | 3.8e-09 | 1.000E-08 | 6.636E-09 | 3.800E-09 | 1.700E-08 |
| 4275 | 214 | C214P | A | 4.4e-09 | 1.1e-08 | 1e-08 | 8.467E-09 | 3.557E-09 | 4.400E-09 | 1.100E-08 |
| 4276 | 214 | C214S | A | 1.9e-08 | 1.4e-08 | 2.9e-08 | 2.067E-08 | 7.638E-09 | 1.400E-08 | 2.900E-08 |
| 4277 | 214 | C214T | A | 5.8e-09 | 8e-09 | 1.1e-08 | 8.267E-09 | 2.610E-09 | 5.800E-09 | 1.100E-08 |
| 4278 | 214 | C214W | A | 5.7e-09 | 2.4e-09 | 9.8e-09 | 5.967E-09 | 3.707E-09 | 2.400E-09 | 9.800E-09 |
| 4279 | 214 | C214Y | A | 5.6e-09 | 1.9e-08 | 2.3e-09 | 8.967E-09 | 8.844E-09 | 2.300E-09 | 1.900E-08 |
| 4280 | 214 | C214V | A | 3e-09 | 1.2e-08 | 8.2e-09 | 7.733E-09 | 4.518E-09 | 3.000E-09 | 1.200E-08 |
| 4281 | 215 | H215A | A | 9.4e-09 | 4.4e-09 | 1.3e-08 | 8.933E-09 | 4.319E-09 | 4.400E-09 | 1.300E-08 |
| 4282 | 215 | H215R | A | 4.7e-09 | 3.5e-09 | 4.4e-09 | 4.200E-09 | 6.245E-10 | 3.500E-09 | 4.700E-09 |
| 4283 | 215 | H215N | A | 4e-09 | 1.1e-08 | 3.9e-09 | 6.300E-09 | 4.071E-09 | 3.900E-09 | 1.100E-08 |
| 4284 | 215 | H215D | A | 6.9e-09 | 2e-09 | 5.8e-09 | 4.900E-09 | 2.571E-09 | 2.000E-09 | 6.900E-09 |
| 4285 | 215 | H215C | A | 6.9e-09 | 1.1e-08 | 1.8e-08 | 1.197E-08 | 5.613E-09 | 6.900E-09 | 1.800E-08 |
| 4286 | 215 | H215E | A | 1.9e-08 | 5.6e-09 | 1.3e-08 | 1.253E-08 | 6.712E-09 | 5.600E-09 | 1.900E-08 |
| 4287 | 215 | H215Q | A | 2.4e-09 | 4.6e-09 | 4.6e-09 | 3.867E-09 | 1.270E-09 | 2.400E-09 | 4.600E-09 |
| 4288 | 215 | H215G | A | 2e-08 | 8.8e-09 | 1.1e-08 | 1.327E-08 | 5.934E-09 | 8.800E-09 | 2.000E-08 |
| 4289 | 215 | H215H | A | 2.6e-08 | 3.4e-09 | 6.1e-09 | 1.183E-08 | 1.234E-08 | 3.400E-09 | 2.600E-08 |
| 4290 | 215 | H215I | A | 8.8e-09 | 2.1e-08 | 1.1e-08 | 1.360E-08 | 6.502E-09 | 8.800E-09 | 2.100E-08 |
| 4291 | 215 | H215L | A | 1.1e-08 | 9.7e-09 | 7.4e-09 | 9.367E-09 | 1.823E-09 | 7.400E-09 | 1.100E-08 |
| 4292 | 215 | H215K | A | 2.2e-08 | 7.1e-09 | 2.5e-09 | 1.053E-08 | 1.019E-08 | 2.500E-09 | 2.200E-08 |
| 4293 | 215 | H215M | A | 8.8e-09 | 3.1e-09 | 3e-09 | 4.967E-09 | 3.320E-09 | 3.000E-09 | 8.800E-09 |
| 4294 | 215 | H215F | A | 2.2e-09 | 1.3e-08 | 9.1e-09 | 8.100E-09 | 5.469E-09 | 2.200E-09 | 1.300E-08 |
| 4295 | 215 | H215P | A | 8.7e-09 | 2.1e-08 | 1e-08 | 1.323E-08 | 6.757E-09 | 8.700E-09 | 2.100E-08 |
| 4296 | 215 | H215S | A | 7.2e-09 | 7.6e-09 | 1.2e-08 | 8.933E-09 | 2.663E-09 | 7.200E-09 | 1.200E-08 |
| 4297 | 215 | H215T | A | 3.7e-09 | 8e-09 | 7.4e-09 | 6.367E-09 | 2.329E-09 | 3.700E-09 | 8.000E-09 |
| 4298 | 215 | H215W | A | 3.2e-09 | 1e-08 | 1e-08 | 7.733E-09 | 3.926E-09 | 3.200E-09 | 1.000E-08 |
| 4299 | 215 | H215Y | A | 1.5e-08 | 2.1e-09 | 3.1e-09 | 6.733E-09 | 7.177E-09 | 2.100E-09 | 1.500E-08 |
| 4300 | 215 | H215V | A | 1e-08 | 5.1e-09 | 7.6e-09 | 7.567E-09 | 2.450E-09 | 5.100E-09 | 1.000E-08 |
| 4301 | 216 | E216A | A | 4.3e-09 | 9.5e-09 | 3.1e-09 | 5.633E-09 | 3.402E-09 | 3.100E-09 | 9.500E-09 |
| 4302 | 216 | E216R | A | 8.1e-09 | 1e-08 | 2.1e-08 | 1.303E-08 | 6.964E-09 | 8.100E-09 | 2.100E-08 |
| 4303 | 216 | E216N | A | 3.3e-09 | 3.8e-09 | 2.6e-09 | 3.233E-09 | 6.028E-10 | 2.600E-09 | 3.800E-09 |
| 4304 | 216 | E216D | A | 2.6e-09 | 2.2e-08 | 4.6e-09 | 9.733E-09 | 1.067E-08 | 2.600E-09 | 2.200E-08 |
| 4305 | 216 | E216C | A | 7.1e-09 | 4.8e-09 | 2.1e-08 | 1.097E-08 | 8.765E-09 | 4.800E-09 | 2.100E-08 |
| 4306 | 216 | E216E | A | 6.2e-09 | 6.9e-09 | 6.9e-09 | 6.667E-09 | 4.041E-10 | 6.200E-09 | 6.900E-09 |
| 4307 | 216 | E216Q | A | 7.7e-09 | 2.2e-08 | 8.1e-09 | 1.260E-08 | 8.143E-09 | 7.700E-09 | 2.200E-08 |
| 4308 | 216 | E216G | A | 1.7e-08 | 1e-08 | 4.6e-09 | 1.053E-08 | 6.217E-09 | 4.600E-09 | 1.700E-08 |
| 4309 | 216 | E216H | A | 5.6e-09 | 8.3e-09 | 1.4e-08 | 9.300E-09 | 4.288E-09 | 5.600E-09 | 1.400E-08 |
| 4310 | 216 | E216I | A | 9.4e-09 | 2.2e-09 | 4.7e-09 | 5.433E-09 | 3.656E-09 | 2.200E-09 | 9.400E-09 |
| 4311 | 216 | E216L | A | 3.5e-09 | 1.7e-08 | 1.1e-08 | 1.050E-08 | 6.764E-09 | 3.500E-09 | 1.700E-08 |
| 4312 | 216 | E216K | A | 3e-09 | 1.2e-08 | 9.1e-09 | 8.033E-09 | 4.594E-09 | 3.000E-09 | 1.200E-08 |
| 4313 | 216 | E216M | A | 5.2e-09 | 3.6e-09 | 1.1e-08 | 6.600E-09 | 3.894E-09 | 3.600E-09 | 1.100E-08 |
| 4314 | 216 | E216F | A | 4.5e-09 | 3.4e-09 | 3.6e-09 | 3.833E-09 | 5.859E-10 | 3.400E-09 | 4.500E-09 |
| 4315 | 216 | E216P | A | 1.4e-08 | 3.3e-09 | 1.8e-08 | 1.177E-08 | 7.600E-09 | 3.300E-09 | 1.800E-08 |
| 4316 | 216 | E216S | A | 3.1e-08 | 4.6e-09 | 1.8e-08 | 1.787E-08 | 1.320E-08 | 4.600E-09 | 3.100E-08 |
| 4317 | 216 | E216T | A | 1e-08 | 6.1e-09 | 1.4e-08 | 1.003E-08 | 3.950E-09 | 6.100E-09 | 1.400E-08 |
| 4318 | 216 | E216W | A | 4.8e-09 | 5.9e-09 | 7.3e-09 | 6.000E-09 | 1.253E-09 | 4.800E-09 | 7.300E-09 |
| 4319 | 216 | E216Y | A | 1.2e-08 | 3.9e-09 | 3.1e-09 | 6.333E-09 | 4.924E-09 | 3.100E-09 | 1.200E-08 |
| 4320 | 216 | E216V | A | 1.3e-08 | 8.4e-09 | 1.1e-08 | 1.080E-08 | 2.307E-09 | 8.400E-09 | 1.300E-08 |
| 4321 | 217 | Q217A | A | 3.9e-09 | 1e-08 | 9.1e-09 | 7.667E-09 | 3.293E-09 | 3.900E-09 | 1.000E-08 |
| 4322 | 217 | Q217R | A | 1.2e-09 | 1.1e-08 | 1e-08 | 7.400E-09 | 5.393E-09 | 1.200E-09 | 1.100E-08 |
| 4323 | 217 | Q217N | A | 2.8e-09 | 1.1e-08 | 2.4e-08 | 1.260E-08 | 1.069E-08 | 2.800E-09 | 2.400E-08 |
| 4324 | 217 | Q217D | A | 1.1e-08 | 5.5e-09 | 8.1e-09 | 8.200E-09 | 2.751E-09 | 5.500E-09 | 1.100E-08 |
| 4325 | 217 | Q217C | A | 5.7e-09 | 9.2e-09 | 4.8e-09 | 6.567E-09 | 2.325E-09 | 4.800E-09 | 9.200E-09 |
| 4326 | 217 | Q217E | A | 5.7e-09 | 6.6e-09 | 5.9e-09 | 6.067E-09 | 4.726E-10 | 5.700E-09 | 6.600E-09 |
| 4327 | 217 | Q217Q | A | 8e-09 | 5.2e-09 | 3.5e-09 | 5.567E-09 | 2.272E-09 | 3.500E-09 | 8.000E-09 |
| 4328 | 217 | Q217G | A | 8.2e-09 | 7.9e-09 | 9.7e-09 | 8.600E-09 | 9.644E-10 | 7.900E-09 | 9.700E-09 |
| 4329 | 217 | Q217H | A | 2.4e-08 | 4e-09 | 1.1e-09 | 9.700E-09 | 1.247E-08 | 1.100E-09 | 2.400E-08 |
| 4330 | 217 | Q217I | A | 4.9e-09 | 8.3e-09 | 5.1e-09 | 6.100E-09 | 1.908E-09 | 4.900E-09 | 8.300E-09 |
| 4331 | 217 | Q217L | A | 1.1e-08 | 9.6e-09 | 4.8e-09 | 8.467E-09 | 3.252E-09 | 4.800E-09 | 1.100E-08 |
| 4332 | 217 | Q217K | A | 4.4e-09 | 5e-09 | 8.4e-09 | 5.933E-09 | 2.157E-09 | 4.400E-09 | 8.400E-09 |
| 4333 | 217 | Q217M | A | 7e-09 | 1.6e-08 | 7.4e-09 | 1.013E-08 | 5.085E-09 | 7.000E-09 | 1.600E-08 |
| 4334 | 217 | Q217F | A | 6.1e-09 | 1.1e-08 | 6.6e-09 | 7.900E-09 | 2.696E-09 | 6.100E-09 | 1.100E-08 |

| | | | | | | | | | | |
|---|---|---|---|---|---|---|---|---|---|---|
| 4335 | 217 | Q217P | A | 4.4e-09 | 6.8e-09 | 4.2e-09 | 5.133E-09 | 1.447E-09 | 4.200E-09 | 6.800E-09 |
| 4336 | 217 | Q217S | A | 1.1e-08 | 9.6e-09 | 1.5e-08 | 1.187E-08 | 2.802E-09 | 9.600E-09 | 1.500E-08 |
| 4337 | 217 | Q217T | A | 3.3e-09 | 5.5e-09 | 8.4e-09 | 5.733E-09 | 2.558E-09 | 3.300E-09 | 8.400E-09 |
| 4338 | 217 | Q217W | A | 4.2e-09 | 9e-09 | 1.6e-08 | 9.733E-09 | 5.934E-09 | 4.200E-09 | 1.600E-08 |
| 4339 | 217 | Q217Y | A | 6.4e-09 | 6.4e-09 | 1.1e-08 | 7.933E-09 | 2.656E-09 | 6.400E-09 | 1.100E-08 |
| 4340 | 217 | Q217V | A | 2e-09 | 8.3e-09 | 4.8e-09 | 5.033E-09 | 3.156E-09 | 2.000E-09 | 8.300E-09 |
| 4341 | 218 | C218A | A | 8.2e-09 | 3.4e-08 | 1.1e-08 | 1.773E-08 | 1.416E-08 | 8.200E-09 | 3.400E-08 |
| 4342 | 218 | C218R | A | 9.5e-09 | 3.7e-09 | 1.2e-08 | 8.400E-09 | 4.258E-09 | 3.700E-09 | 1.200E-08 |
| 4343 | 218 | C218N | A | 1.9e-08 | 6.8e-09 | 5.1e-09 | 1.030E-08 | 7.582E-09 | 5.100E-09 | 1.900E-08 |
| 4344 | 218 | C218D | A | 1.6e-08 | 2.4e-09 | 9.5e-09 | 9.300E-09 | 6.802E-09 | 2.400E-09 | 1.600E-08 |
| 4345 | 218 | C218C | A | 5.8e-09 | 5.9e-09 | 9.4e-09 | 7.033E-09 | 2.050E-09 | 5.800E-09 | 9.400E-09 |
| 4346 | 218 | C218E | A | 3.4e-09 | 8.6e-09 | 5.6e-09 | 5.867E-09 | 2.610E-09 | 3.400E-09 | 8.600E-09 |
| 4347 | 218 | C218Q | A | 1.1e-08 | 5e-09 | 7.3e-09 | 7.767E-09 | 3.027E-09 | 5.000E-09 | 1.100E-08 |
| 4348 | 218 | C218G | A | 3.4e-09 | 1.5e-08 | 9.9e-09 | 9.433E-09 | 5.814E-09 | 3.400E-09 | 1.500E-08 |
| 4349 | 218 | C218H | A | 6.2e-09 | 3.1e-08 | 1.6e-08 | 1.773E-08 | 1.249E-08 | 6.200E-09 | 3.100E-08 |
| 4350 | 218 | C218I | A | 7.5e-09 | 2.1e-09 | 5.5e-09 | 5.033E-09 | 2.730E-09 | 2.100E-09 | 7.500E-09 |
| 4351 | 218 | C218L | A | 1e-08 | 2e-08 | 1.4e-08 | 1.467E-08 | 5.033E-09 | 1.000E-08 | 2.000E-08 |
| 4352 | 218 | C218K | A | 1.3e-08 | 1.2e-08 | 1.1e-08 | 1.200E-08 | 1.000E-09 | 1.100E-08 | 1.300E-08 |
| 4353 | 218 | C218M | A | 3.2e-09 | 6.8e-09 | 1.4e-08 | 8.000E-09 | 5.499E-09 | 3.200E-09 | 1.400E-08 |
| 4354 | 218 | C218F | A | 6.4e-09 | 7.6e-09 | 1.2e-08 | 8.667E-09 | 2.948E-09 | 6.400E-09 | 1.200E-08 |
| 4355 | 218 | C218P | A | 8.2e-09 | 1.9e-09 | 7.5e-09 | 5.867E-09 | 3.453E-09 | 1.900E-09 | 8.200E-09 |
| 4356 | 218 | C218S | A | 5.4e-09 | 5.5e-09 | 1.7e-08 | 9.300E-09 | 6.669E-09 | 5.400E-09 | 1.700E-08 |
| 4357 | 218 | C218T | A | 8.3e-09 | 1.2e-08 | 1.4e-08 | 1.143E-08 | 2.892E-09 | 8.300E-09 | 1.400E-08 |
| 4358 | 218 | C218W | A | 7.2e-09 | 8.6e-09 | 3.7e-09 | 6.500E-09 | 2.524E-09 | 3.700E-09 | 8.600E-09 |
| 4359 | 218 | C218Y | A | 1.4e-08 | 9.1e-09 | 1.3e-08 | 1.203E-08 | 2.589E-09 | 9.100E-09 | 1.400E-08 |
| 4360 | 218 | C218V | A | 3.1e-09 | 1.6e-09 | 4.9e-09 | 3.200E-09 | 1.652E-09 | 1.600E-09 | 4.900E-09 |
| 4361 | 219 | A219A | A | 2.4e-09 | 4.8e-09 | 4.8e-09 | 4.000E-09 | 1.386E-09 | 2.400E-09 | 4.800E-09 |
| 4362 | 219 | A219R | A | 1.8e-08 | 2.2e-09 | 3.9e-09 | 8.033E-09 | 8.673E-09 | 2.200E-09 | 1.800E-08 |
| 4363 | 219 | A219N | A | 3.3e-09 | 1e-08 | 2.8e-09 | 5.367E-09 | 4.020E-09 | 2.800E-09 | 1.000E-08 |
| 4364 | 219 | A219D | A | 8.4e-09 | 7.3e-09 | 4.7e-09 | 6.800E-09 | 1.900E-09 | 4.700E-09 | 8.400E-09 |
| 4365 | 219 | A219C | A | 1.3e-08 | 1.5e-08 | 2.3e-08 | 1.700E-08 | 5.292E-09 | 1.300E-08 | 2.300E-08 |
| 4366 | 219 | A219E | A | 1.2e-08 | 4.6e-09 | 6.1e-09 | 7.567E-09 | 3.912E-09 | 4.600E-09 | 1.200E-08 |
| 4367 | 219 | A219Q | A | 7.5e-09 | 1e-08 | 7.7e-09 | 8.400E-09 | 1.389E-09 | 7.500E-09 | 1.000E-08 |
| 4368 | 219 | A219G | A | 1.2e-08 | 1e-08 | 9.7e-09 | 1.057E-08 | 1.250E-09 | 9.700E-09 | 1.200E-08 |
| 4369 | 219 | A219H | A | 2.3e-09 | 1.1e-08 | 4.2e-09 | 5.833E-09 | 4.574E-09 | 2.300E-09 | 1.100E-08 |
| 4370 | 219 | A219I | A | 1.3e-08 | 3.7e-09 | 6.6e-09 | 7.767E-09 | 4.759E-09 | 3.700E-09 | 1.300E-08 |
| 4371 | 219 | A219L | A | 7.4e-09 | 1e-08 | 1.6e-09 | 6.333E-09 | 4.300E-09 | 1.600E-09 | 1.000E-08 |
| 4372 | 219 | A219K | A | 6.7e-09 | 3.7e-09 | 3.7e-09 | 4.700E-09 | 1.732E-09 | 3.700E-09 | 6.700E-09 |
| 4373 | 219 | A219M | A | 1.7e-08 | 4.3e-09 | 6.5e-09 | 9.267E-09 | 6.787E-09 | 4.300E-09 | 1.700E-08 |
| 4374 | 219 | A219F | A | 7.7e-09 | 6.1e-09 | 3.8e-09 | 5.867E-09 | 1.960E-09 | 3.800E-09 | 7.700E-09 |
| 4375 | 219 | A219P | A | 1.4e-08 | 6.8e-09 | 1.3e-08 | 1.127E-08 | 3.900E-09 | 6.800E-09 | 1.400E-08 |
| 4376 | 219 | A219S | A | 8.3e-09 | 7.2e-09 | 1.4e-08 | 9.833E-09 | 3.650E-09 | 7.200E-09 | 1.400E-08 |
| 4377 | 219 | A219T | A | 1.4e-08 | 2e-08 | 7.5e-09 | 1.383E-08 | 6.252E-09 | 7.500E-09 | 2.000E-08 |
| 4378 | 219 | A219W | A | 2.3e-08 | 1e-08 | 1e-08 | 1.433E-08 | 7.506E-09 | 1.000E-08 | 2.300E-08 |
| 4379 | 219 | A219Y | A | 1.8e-09 | 9.1e-09 | 2e-08 | 1.030E-08 | 9.159E-09 | 1.800E-09 | 2.000E-08 |
| 4380 | 219 | A219V | A | 4.5e-08 | 3.6e-09 | 2.9e-09 | 1.717E-08 | 2.411E-08 | 2.900E-09 | 4.500E-08 |
| 4381 | 220 | A220A | A | 4.4e-09 | 1.1e-09 | 8.7e-09 | 4.733E-09 | 3.811E-09 | 1.100E-09 | 8.700E-09 |
| 4382 | 220 | A220R | A | 8.3e-09 | 3.2e-09 | 6.4e-09 | 5.967E-09 | 2.577E-09 | 3.200E-09 | 8.300E-09 |
| 4383 | 220 | A220N | A | 3.2e-09 | 3.3e-09 | 1.2e-08 | 6.167E-09 | 5.052E-09 | 3.200E-09 | 1.200E-08 |
| 4384 | 220 | A220D | A | 1.2e-08 | 5.1e-09 | 4.6e-09 | 7.233E-09 | 4.136E-09 | 4.600E-09 | 1.200E-08 |
| 4385 | 220 | A220C | A | 1.1e-08 | 1.2e-08 | 9.1e-09 | 1.070E-08 | 1.473E-09 | 9.100E-09 | 1.200E-08 |
| 4386 | 220 | A220E | A | 1.1e-08 | 4.4e-09 | 1.7e-08 | 1.080E-08 | 6.302E-09 | 4.400E-09 | 1.700E-08 |
| 4387 | 220 | A220Q | A | 7.2e-09 | 5.8e-09 | 8.6e-09 | 7.200E-09 | 1.400E-09 | 5.800E-09 | 8.600E-09 |
| 4388 | 220 | A220G | A | 2.2e-09 | 3.9e-09 | 7.8e-09 | 4.633E-09 | 2.871E-09 | 2.200E-09 | 7.800E-09 |
| 4389 | 220 | A220H | A | 5.3e-09 | 5.6e-09 | 9.6e-09 | 6.833E-09 | 2.401E-09 | 5.300E-09 | 9.600E-09 |
| 4390 | 220 | A220I | A | 2.6e-09 | 6.9e-09 | 6.4e-09 | 5.300E-09 | 2.352E-09 | 2.600E-09 | 6.900E-09 |
| 4391 | 220 | A220L | A | 2.6e-09 | 7.7e-09 | 4.2e-09 | 4.833E-09 | 2.608E-09 | 2.600E-09 | 7.700E-09 |
| 4392 | 220 | A220K | A | 7e-10 | 1.3e-08 | 3.9e-09 | 5.867E-09 | 6.381E-09 | 7.000E-10 | 1.300E-08 |
| 4393 | 220 | A220M | A | 8.3e-09 | 3.8e-09 | 1.5e-08 | 9.033E-09 | 5.636E-09 | 3.800E-09 | 1.500E-08 |
| 4394 | 220 | A220F | A | 2.6e-09 | 6.5e-09 | 3.9e-09 | 4.333E-09 | 1.986E-09 | 2.600E-09 | 6.500E-09 |
| 4395 | 220 | A220P | A | 8e-09 | 5.6e-09 | 3.8e-09 | 5.800E-09 | 2.107E-09 | 3.800E-09 | 8.000E-09 |
| 4396 | 220 | A220S | A | 5.6e-09 | 2.5e-09 | 7.6e-09 | 5.233E-09 | 2.570E-09 | 2.500E-09 | 7.600E-09 |
| 4397 | 220 | A220T | A | 4e-09 | 6.4e-09 | 2.5e-09 | 4.300E-09 | 1.967E-09 | 2.500E-09 | 6.400E-09 |
| 4398 | 220 | A220W | A | 3.1e-09 | 1.2e-08 | 3.8e-09 | 6.300E-09 | 4.949E-09 | 3.100E-09 | 1.200E-08 |
| 4399 | 220 | A220Y | A | 1.1e-08 | 7.4e-09 | 4e-09 | 7.467E-09 | 3.500E-09 | 4.000E-09 | 1.100E-08 |
| 4400 | 220 | A220V | A | 4e-09 | 1.4e-08 | 3.6e-09 | 7.200E-09 | 5.892E-09 | 3.600E-09 | 1.400E-08 |
| 4401 | 221 | G221A | A | 1.3e-08 | 7.8e-09 | 9.4e-09 | 1.007E-08 | 2.663E-09 | 7.800E-09 | 1.300E-08 |
| 4402 | 221 | G221R | A | 1.1e-08 | 3e-09 | 2.7e-09 | 5.567E-09 | 4.708E-09 | 2.700E-09 | 1.100E-08 |
| 4403 | 221 | G221N | A | 1e-08 | 7.3e-09 | 3.2e-09 | 6.833E-09 | 3.424E-09 | 3.200E-09 | 1.000E-08 |
| 4404 | 221 | G221D | A | 9.6e-09 | 9e-09 | 1.4e-08 | 1.087E-08 | 2.730E-09 | 9.000E-09 | 1.400E-08 |
| 4405 | 221 | G221C | A | 5.8e-09 | 8.3e-09 | 7.6e-09 | 7.233E-09 | 1.290E-09 | 5.800E-09 | 8.300E-09 |
| 4406 | 221 | G221E | A | 1.1e-08 | 4.3e-09 | 4.2e-09 | 6.500E-09 | 3.897E-09 | 4.200E-09 | 1.100E-08 |
| 4407 | 221 | G221Q | A | 3.9e-09 | 1.8e-09 | 6.3e-09 | 4.000E-09 | 2.252E-09 | 1.800E-09 | 6.300E-09 |
| 4408 | 221 | G221G | A | 6.6e-09 | 7.9e-09 | 4.3e-09 | 6.267E-09 | 1.823E-09 | 4.300E-09 | 7.900E-09 |
| 4409 | 221 | G221H | A | 6.6e-09 | 1.4e-08 | 1.2e-08 | 1.087E-08 | 3.828E-09 | 6.600E-09 | 1.400E-08 |
| 4410 | 221 | G221I | A | 2.3e-08 | 1.1e-08 | 1.1e-08 | 1.500E-08 | 6.928E-09 | 1.100E-08 | 2.300E-08 |
| 4411 | 221 | G221L | A | 4.8e-09 | 1e-08 | 8.6e-09 | 7.800E-09 | 2.691E-09 | 4.800E-09 | 1.000E-08 |
| 4412 | 221 | G221K | A | 9.8e-09 | 4.2e-09 | 6.1e-09 | 6.700E-09 | 2.848E-09 | 4.200E-09 | 9.800E-09 |
| 4413 | 221 | G221M | A | 8.1e-09 | 2e-08 | 2e-08 | 1.603E-08 | 6.870E-09 | 8.100E-09 | 2.000E-08 |
| 4414 | 221 | G221F | A | 5.2e-09 | 3.6e-09 | 6.3e-09 | 5.033E-09 | 1.358E-09 | 3.600E-09 | 6.300E-09 |
| 4415 | 221 | G221P | A | 1.2e-08 | 2e-08 | 6.3e-09 | 1.277E-08 | 6.882E-09 | 6.300E-09 | 2.000E-08 |
| 4416 | 221 | G221S | A | 3.2e-09 | 9.9e-09 | 1.1e-08 | 8.033E-09 | 4.222E-09 | 3.200E-09 | 1.100E-08 |
| 4417 | 221 | G221T | A | 4.4e-09 | 8.3e-09 | 7.6e-09 | 6.767E-09 | 2.079E-09 | 4.400E-09 | 8.300E-09 |
| 4418 | 221 | G221W | A | 1e-08 | 2.9e-09 | 1e-08 | 7.633E-09 | 4.099E-09 | 2.900E-09 | 1.000E-08 |
| 4419 | 221 | G221Y | A | 2.3e-09 | 1.3e-08 | 7.3e-09 | 7.533E-09 | 5.354E-09 | 2.300E-09 | 1.300E-08 |

| | | | | | | | | | | |
|---|---|---|---|---|---|---|---|---|---|---|
| 4420 | 221 | G221V | A | 4.6e-09 | 2.9e-09 | 1.2e-08 | 6.500E-09 | 4.838E-09 | 2.900E-09 | 1.200E-08 |
| 4421 | 222 | C222A | A | 1.5e-08 | 2e-08 | 4.9e-09 | 1.330E-08 | 7.692E-09 | 4.900E-09 | 2.000E-08 |
| 4422 | 222 | C222R | A | 2.9e-09 | 3.1e-09 | 6.9e-09 | 4.300E-09 | 2.254E-09 | 2.900E-09 | 6.900E-09 |
| 4423 | 222 | C222N | A | 2.9e-08 | 7.6e-09 | 6.1e-09 | 1.423E-08 | 1.281E-08 | 6.100E-09 | 2.900E-08 |
| 4424 | 222 | C222D | A | 2.3e-09 | 1.2e-08 | 2.5e-08 | 1.310E-08 | 1.139E-08 | 2.300E-09 | 2.500E-08 |
| 4425 | 222 | C222C | A | 2.1e-08 | 3.9e-09 | 3.8e-09 | 9.567E-09 | 9.902E-09 | 3.800E-09 | 2.100E-08 |
| 4426 | 222 | C222E | A | 2.1e-09 | 5.3e-09 | 4.6e-09 | 4.000E-09 | 1.682E-09 | 2.100E-09 | 5.300E-09 |
| 4427 | 222 | C222Q | A | 6.8e-09 | 9.4e-09 | 3.9e-09 | 6.700E-09 | 2.751E-09 | 3.900E-09 | 9.400E-09 |
| 4428 | 222 | C222G | A | 9e-09 | 3.5e-09 | 6.5e-09 | 6.333E-09 | 2.754E-09 | 3.500E-09 | 9.000E-09 |
| 4429 | 222 | C222H | A | 1.1e-08 | 2.8e-09 | 4.1e-09 | 5.967E-09 | 4.407E-09 | 2.800E-09 | 1.100E-08 |
| 4430 | 222 | C222I | A | 5.2e-09 | 7e-09 | 6.4e-09 | 6.200E-09 | 9.165E-10 | 5.200E-09 | 7.000E-09 |
| 4431 | 222 | C222L | A | 6.5e-09 | 1.4e-08 | 6.3e-09 | 8.933E-09 | 4.389E-09 | 6.300E-09 | 1.400E-08 |
| 4432 | 222 | C222K | A | 2.6e-09 | 2e-08 | 1.5e-08 | 1.253E-08 | 8.958E-09 | 2.600E-09 | 2.000E-08 |
| 4433 | 222 | C222M | A | 3.6e-09 | 1e-08 | 1.2e-08 | 8.533E-09 | 4.388E-09 | 3.600E-09 | 1.200E-08 |
| 4434 | 222 | C222F | A | 8.1e-09 | 6.4e-09 | 1.4e-08 | 9.500E-09 | 3.989E-09 | 6.400E-09 | 1.400E-08 |
| 4435 | 222 | C222P | A | 2e-08 | 5.4e-09 | 6.7e-09 | 1.070E-08 | 8.080E-09 | 5.400E-09 | 2.000E-08 |
| 4436 | 222 | C222S | A | 1.1e-08 | 1e-08 | 3e-09 | 8.000E-09 | 4.359E-09 | 3.000E-09 | 1.100E-08 |
| 4437 | 222 | C222T | A | 4.9e-09 | 4.5e-09 | 8.6e-09 | 6.000E-09 | 2.261E-09 | 4.500E-09 | 8.600E-09 |
| 4438 | 222 | C222W | A | 1.1e-08 | 1.7e-08 | 3.4e-09 | 1.047E-08 | 6.816E-09 | 3.400E-09 | 1.700E-08 |
| 4439 | 222 | C222Y | A | 4.4e-09 | 4.4e-09 | 7.4e-09 | 5.400E-09 | 1.732E-09 | 4.400E-09 | 7.400E-09 |
| 4440 | 222 | C222V | A | 5.2e-09 | 4.9e-09 | 1.2e-08 | 7.367E-09 | 4.015E-09 | 4.900E-09 | 1.200E-08 |
| 4441 | 223 | T223A | A | 9.5e-09 | 8.7e-09 | 2e-08 | 1.273E-08 | 6.306E-09 | 8.700E-09 | 2.000E-08 |
| 4442 | 223 | T223R | A | 1.3e-08 | 2e-08 | 3.6e-09 | 1.220E-08 | 8.229E-09 | 3.600E-09 | 2.000E-08 |
| 4443 | 223 | T223N | A | 4.2e-09 | 4.1e-09 | 2.6e-09 | 3.633E-09 | 8.963E-10 | 2.600E-09 | 4.200E-09 |
| 4444 | 223 | T223D | A | 2.4e-09 | 9.6e-09 | 6.9e-09 | 6.300E-09 | 3.637E-09 | 2.400E-09 | 9.600E-09 |
| 4445 | 223 | T223C | A | 7e-09 | 3.5e-09 | 4.8e-09 | 5.100E-09 | 1.769E-09 | 3.500E-09 | 7.000E-09 |
| 4446 | 223 | T223E | A | 8.8e-09 | 5.7e-09 | 5.2e-09 | 6.567E-09 | 1.950E-09 | 5.200E-09 | 8.800E-09 |
| 4447 | 223 | T223Q | A | 4e-09 | 1.7e-09 | 6.3e-09 | 4.000E-09 | 2.300E-09 | 1.700E-09 | 6.300E-09 |
| 4448 | 223 | T223G | A | 7.6e-09 | 2.9e-08 | 7.4e-09 | 1.467E-08 | 1.241E-08 | 7.400E-09 | 2.900E-08 |
| 4449 | 223 | T223H | A | 1.5e-09 | 6.8e-09 | 7.6e-09 | 5.300E-09 | 3.315E-09 | 1.500E-09 | 7.600E-09 |
| 4450 | 223 | T223I | A | 8.7e-09 | 1e-08 | 5.5e-09 | 8.067E-09 | 2.316E-09 | 5.500E-09 | 1.000E-08 |
| 4451 | 223 | T223L | A | 9.3e-09 | 2.7e-09 | 2.4e-09 | 4.800E-09 | 3.900E-09 | 2.400E-09 | 9.300E-09 |
| 4452 | 223 | T223K | A | 7.1e-09 | 4.4e-09 | 1.1e-08 | 7.500E-09 | 3.318E-09 | 4.400E-09 | 1.100E-08 |
| 4453 | 223 | T223M | A | 8.4e-09 | 8.3e-09 | 1.5e-08 | 1.057E-08 | 3.840E-09 | 8.300E-09 | 1.500E-08 |
| 4454 | 223 | T223F | A | 4.8e-09 | 2.6e-09 | 7.2e-09 | 4.867E-09 | 2.301E-09 | 2.600E-09 | 7.200E-09 |
| 4455 | 223 | T223P | A | 3.4e-09 | 6.2e-09 | 3.4e-09 | 4.333E-09 | 1.617E-09 | 3.400E-09 | 6.200E-09 |
| 4456 | 223 | T223S | A | 3.5e-09 | 6.9e-09 | 4.9e-09 | 5.100E-09 | 1.709E-09 | 3.500E-09 | 6.900E-09 |
| 4457 | 223 | T223T | A | 1.5e-08 | 5.2e-09 | 7.1e-09 | 9.100E-09 | 5.197E-09 | 5.200E-09 | 1.500E-08 |
| 4458 | 223 | T223W | A | 8.5e-09 | 6.1e-09 | 2e-08 | 1.153E-08 | 7.430E-09 | 6.100E-09 | 2.000E-08 |
| 4459 | 223 | T223Y | A | 2.6e-09 | 6.4e-09 | 3.2e-09 | 4.067E-09 | 2.043E-09 | 2.600E-09 | 6.400E-09 |
| 4460 | 223 | T223V | A | 9.8e-09 | 2.3e-09 | 6e-09 | 6.033E-09 | 3.750E-09 | 2.300E-09 | 9.800E-09 |
| 4461 | 224 | G224A | A | 3.5e-09 | 5.2e-09 | 6e-09 | 4.900E-09 | 1.277E-09 | 3.500E-09 | 6.000E-09 |
| 4462 | 224 | G224R | A | 7.1e-09 | 2.4e-09 | 2.1e-08 | 1.017E-08 | 9.672E-09 | 2.400E-09 | 2.100E-08 |
| 4463 | 224 | G224N | A | 6.6e-09 | 4.8e-09 | 1e-08 | 7.133E-09 | 2.641E-09 | 4.800E-09 | 1.000E-08 |
| 4464 | 224 | G224D | A | 8.3e-09 | 3.5e-09 | 3.4e-09 | 5.067E-09 | 2.801E-09 | 3.400E-09 | 8.300E-09 |
| 4465 | 224 | G224C | A | 8.7e-09 | 4.9e-09 | 5.3e-09 | 6.300E-09 | 2.088E-09 | 4.900E-09 | 8.700E-09 |
| 4466 | 224 | G224E | A | 7.9e-09 | 9.1e-09 | 6.8e-09 | 7.933E-09 | 1.150E-09 | 6.800E-09 | 9.100E-09 |
| 4467 | 224 | G224Q | A | 3.8e-09 | 9.2e-09 | 1.4e-08 | 9.000E-09 | 5.103E-09 | 3.800E-09 | 1.400E-08 |
| 4468 | 224 | G224G | A | 1.9e-09 | 3.4e-09 | 7.7e-09 | 4.333E-09 | 3.011E-09 | 1.900E-09 | 7.700E-09 |
| 4469 | 224 | G224H | A | 4.5e-09 | 1.9e-08 | 8.7e-09 | 1.073E-08 | 7.461E-09 | 4.500E-09 | 1.900E-08 |
| 4470 | 224 | G224I | A | 4.1e-09 | 9.5e-09 | 4.9e-09 | 6.167E-09 | 2.914E-09 | 4.100E-09 | 9.500E-09 |
| 4471 | 224 | G224L | A | 4.6e-09 | 7.3e-09 | 2.3e-09 | 4.733E-09 | 2.503E-09 | 2.300E-09 | 7.300E-09 |
| 4472 | 224 | G224K | A | 5.1e-09 | 6.1e-09 | 2.6e-09 | 4.600E-09 | 1.803E-09 | 2.600E-09 | 6.100E-09 |
| 4473 | 224 | G224M | A | 8.7e-09 | 1.2e-08 | 9.6e-09 | 1.010E-08 | 1.706E-09 | 8.700E-09 | 1.200E-08 |
| 4474 | 224 | G224F | A | 3.3e-09 | 1.4e-08 | 5.4e-09 | 7.567E-09 | 5.670E-09 | 3.300E-09 | 1.400E-08 |
| 4475 | 224 | G224P | A | 7.6e-09 | 2e-09 | 1e-08 | 6.533E-09 | 4.105E-09 | 2.000E-09 | 1.000E-08 |
| 4476 | 224 | G224S | A | 4.2e-09 | 2.3e-09 | 1e-08 | 5.500E-09 | 4.011E-09 | 2.300E-09 | 1.000E-08 |
| 4477 | 224 | G224T | A | 1.9e-08 | 9.3e-09 | 1.1e-08 | 1.310E-08 | 5.180E-09 | 9.300E-09 | 1.900E-08 |
| 4478 | 224 | G224W | A | 1.3e-09 | 3.1e-09 | 3.7e-09 | 2.700E-09 | 1.249E-09 | 1.300E-09 | 3.700E-09 |
| 4479 | 224 | G224Y | A | 3.4e-09 | 2.7e-08 | 8e-09 | 1.280E-08 | 1.251E-08 | 3.400E-09 | 2.700E-08 |
| 4480 | 224 | G224V | A | 5.6e-09 | 3.8e-09 | 6.5e-09 | 5.300E-09 | 1.375E-09 | 3.800E-09 | 6.500E-09 |
| 4481 | 225 | P225A | A | 4.4e-09 | 8e-09 | 8.6e-09 | 7.000E-09 | 2.272E-09 | 4.400E-09 | 8.600E-09 |
| 4482 | 225 | P225R | A | 3.7e-09 | 6.4e-09 | 5.5e-09 | 5.200E-09 | 1.375E-09 | 3.700E-09 | 6.400E-09 |
| 4483 | 225 | P225N | A | 1.3e-08 | 9.6e-09 | 9e-09 | 1.053E-08 | 2.157E-09 | 9.000E-09 | 1.300E-08 |
| 4484 | 225 | P225D | A | 2e-08 | 3.9e-09 | 6e-09 | 9.967E-09 | 8.752E-09 | 3.900E-09 | 2.000E-08 |
| 4485 | 225 | P225C | A | 8.5e-09 | 6.8e-09 | 8.2e-09 | 7.833E-09 | 9.074E-10 | 6.800E-09 | 8.500E-09 |
| 4486 | 225 | P225E | A | 1.3e-08 | 2.5e-09 | 9.4e-09 | 8.300E-09 | 5.336E-09 | 2.500E-09 | 1.300E-08 |
| 4487 | 225 | P225Q | A | 7.2e-09 | 2.2e-09 | 3.6e-09 | 4.333E-09 | 2.579E-09 | 2.200E-09 | 7.200E-09 |
| 4488 | 225 | P225G | A | 2.2e-08 | 5.8e-09 | 3.5e-09 | 1.043E-08 | 1.008E-08 | 3.500E-09 | 2.200E-08 |
| 4489 | 225 | P225H | A | 8.6e-09 | 4.3e-09 | 1.9e-09 | 4.933E-09 | 3.395E-09 | 1.900E-09 | 8.600E-09 |
| 4490 | 225 | P225I | A | 2.5e-09 | 8.1e-09 | 5.1e-09 | 5.233E-09 | 2.802E-09 | 2.500E-09 | 8.100E-09 |
| 4491 | 225 | P225L | A | 1.3e-09 | 6.9e-09 | 1.3e-08 | 7.067E-09 | 5.852E-09 | 1.300E-09 | 1.300E-08 |
| 4492 | 225 | P225K | A | 8.4e-09 | 3.8e-09 | 6.8e-09 | 6.333E-09 | 2.335E-09 | 3.800E-09 | 8.400E-09 |
| 4493 | 225 | P225M | A | 1.7e-09 | 3.6e-09 | 1.5e-08 | 6.767E-09 | 7.193E-09 | 1.700E-09 | 1.500E-08 |
| 4494 | 225 | P225F | A | 9.3e-09 | 5.1e-09 | 6.1e-09 | 6.833E-09 | 2.194E-09 | 5.100E-09 | 9.300E-09 |
| 4495 | 225 | P225P | A | 1.5e-08 | 1.1e-08 | 1.5e-08 | 1.367E-08 | 2.309E-09 | 1.100E-08 | 1.500E-08 |
| 4496 | 225 | P225S | A | 4.7e-09 | 4.4e-09 | 8.7e-09 | 5.933E-09 | 2.401E-09 | 4.400E-09 | 8.700E-09 |
| 4497 | 225 | P225T | A | 8.2e-09 | 3.6e-09 | 5.3e-09 | 5.700E-09 | 2.326E-09 | 3.600E-09 | 8.200E-09 |
| 4498 | 225 | P225W | A | 8.6e-09 | 5e-09 | 4.9e-09 | 6.167E-09 | 2.108E-09 | 4.900E-09 | 8.600E-09 |
| 4499 | 225 | P225Y | A | 3.9e-09 | 1.1e-08 | 4.3e-09 | 6.400E-09 | 3.989E-09 | 3.900E-09 | 1.100E-08 |
| 4500 | 225 | P225V | A | 3.2e-09 | 7e-09 | 1.4e-08 | 8.067E-09 | 5.478E-09 | 3.200E-09 | 1.400E-08 |
| 4501 | 226 | K226A | A | 4.5e-09 | 6.1e-09 | 4.3e-09 | 4.967E-09 | 9.866E-10 | 4.300E-09 | 6.100E-09 |
| 4502 | 226 | K226R | A | 1e-08 | 9.1e-09 | 4.5e-09 | 7.867E-09 | 2.950E-09 | 4.500E-09 | 1.000E-08 |
| 4503 | 226 | K226N | A | 8.9e-09 | 1.1e-08 | 2.5e-09 | 7.467E-09 | 4.428E-09 | 2.500E-09 | 1.100E-08 |
| 4504 | 226 | K226D | A | 1.4e-08 | 6.3e-09 | 4.9e-09 | 8.400E-09 | 4.900E-09 | 4.900E-09 | 1.400E-08 |

| | | | | | | | | | | |
|---|---|---|---|---|---|---|---|---|---|---|
| 4505 | 226 | K226C | A | 8.2e-09 | 2.3e-08 | 8.2e-09 | 1.313E-08 | 8.545E-09 | 8.200E-09 | 2.300E-08 |
| 4506 | 226 | K226E | A | 5.4e-09 | 6.8e-09 | 5.1e-09 | 5.767E-09 | 9.074E-10 | 5.100E-09 | 6.800E-09 |
| 4507 | 226 | K226Q | A | 1.2e-08 | 2.7e-09 | 5.6e-09 | 6.767E-09 | 4.759E-09 | 2.700E-09 | 1.200E-08 |
| 4508 | 226 | K226G | A | 2.9e-09 | 5.9e-09 | 4.9e-09 | 4.567E-09 | 1.528E-09 | 2.900E-09 | 5.900E-09 |
| 4509 | 226 | K226H | A | 1.2e-08 | 6.5e-09 | 1.2e-08 | 1.017E-08 | 3.175E-09 | 6.500E-09 | 1.200E-08 |
| 4510 | 226 | K226I | A | 4.4e-09 | 4.9e-09 | 1.4e-08 | 7.767E-09 | 5.404E-09 | 4.400E-09 | 1.400E-08 |
| 4511 | 226 | K226L | A | 2.2e-08 | 2.1e-08 | 2.8e-09 | 1.527E-08 | 1.081E-08 | 2.800E-09 | 2.200E-08 |
| 4512 | 226 | K226K | A | 1.2e-08 | 8.1e-09 | 1.5e-08 | 1.170E-08 | 3.460E-09 | 8.100E-09 | 1.500E-08 |
| 4513 | 226 | K226M | A | 6.7e-09 | 3.2e-09 | 1.4e-09 | 3.767E-09 | 2.695E-09 | 1.400E-09 | 6.700E-09 |
| 4514 | 226 | K226F | A | 4.1e-09 | 1.9e-08 | 5.9e-09 | 9.667E-09 | 8.133E-09 | 4.100E-09 | 1.900E-08 |
| 4515 | 226 | K226P | A | 1.8e-08 | 6.5e-09 | 6.5e-09 | 1.033E-08 | 6.640E-09 | 6.500E-09 | 1.800E-08 |
| 4516 | 226 | K226S | A | 2.8e-09 | 4.5e-09 | 2.1e-08 | 9.433E-09 | 1.005E-08 | 2.800E-09 | 2.100E-08 |
| 4517 | 226 | K226T | A | 4.7e-09 | 1.7e-08 | 6.2e-09 | 9.300E-09 | 6.710E-09 | 4.700E-09 | 1.700E-08 |
| 4518 | 226 | K226W | A | 1e-08 | 7.6e-09 | 5.8e-09 | 7.800E-09 | 2.107E-09 | 5.800E-09 | 1.000E-08 |
| 4519 | 226 | K226Y | A | 3.7e-09 | 1.5e-08 | 2.9e-09 | 7.200E-09 | 6.767E-09 | 2.900E-09 | 1.500E-08 |
| 4520 | 226 | K226V | A | 1.3e-08 | 5.7e-09 | 7e-09 | 8.567E-09 | 3.894E-09 | 5.700E-09 | 1.300E-08 |
| 4521 | 227 | H227A | A | 9.9e-09 | 7.3e-09 | 8e-09 | 8.400E-09 | 1.345E-09 | 7.300E-09 | 9.900E-09 |
| 4522 | 227 | H227R | A | 8.2e-09 | 8.6e-09 | 5.7e-09 | 7.500E-09 | 1.572E-09 | 5.700E-09 | 8.600E-09 |
| 4523 | 227 | H227N | A | 6.4e-09 | 2e-09 | 4.7e-09 | 4.367E-09 | 2.219E-09 | 2.000E-09 | 6.400E-09 |
| 4524 | 227 | H227D | A | 6.2e-09 | 8.4e-09 | 8.8e-09 | 7.800E-09 | 1.400E-09 | 6.200E-09 | 8.800E-09 |
| 4525 | 227 | H227C | A | 3.9e-09 | 7.5e-09 | 9e-09 | 6.800E-09 | 2.621E-09 | 3.900E-09 | 9.000E-09 |
| 4526 | 227 | H227E | A | 9.7e-09 | 1.3e-08 | 8.4e-09 | 1.037E-08 | 2.371E-09 | 8.400E-09 | 1.300E-08 |
| 4527 | 227 | H227Q | A | 1e-08 | 1e-08 | 1.5e-09 | 7.167E-09 | 4.907E-09 | 1.500E-09 | 1.000E-08 |
| 4528 | 227 | H227G | A | 6.3e-09 | 5.1e-09 | 1.5e-08 | 8.800E-09 | 5.403E-09 | 5.100E-09 | 1.500E-08 |
| 4529 | 227 | H227H | A | 4e-09 | 4.8e-09 | 1.6e-08 | 8.267E-09 | 6.709E-09 | 4.000E-09 | 1.600E-08 |
| 4530 | 227 | H227I | A | 7.3e-09 | 2.5e-09 | 8.8e-09 | 6.200E-09 | 3.291E-09 | 2.500E-09 | 8.800E-09 |
| 4531 | 227 | H227L | A | 1.6e-08 | 1.1e-08 | 2.6e-09 | 9.867E-09 | 6.772E-09 | 2.600E-09 | 1.600E-08 |
| 4532 | 227 | H227K | A | 6.8e-09 | 1e-08 | 5.1e-09 | 7.300E-09 | 2.488E-09 | 5.100E-09 | 1.000E-08 |
| 4533 | 227 | H227M | A | 6.8e-09 | 1.3e-08 | 6.6e-09 | 8.800E-09 | 3.639E-09 | 6.600E-09 | 1.300E-08 |
| 4534 | 227 | H227F | A | 9.3e-09 | 1.1e-08 | 5.1e-09 | 8.467E-09 | 3.037E-09 | 5.100E-09 | 1.100E-08 |
| 4535 | 227 | H227P | A | 7.3e-09 | 1.3e-08 | 2.8e-08 | 1.610E-08 | 1.069E-08 | 7.300E-09 | 2.800E-08 |
| 4536 | 227 | H227S | A | 1.4e-08 | 9.2e-09 | 2.4e-09 | 8.533E-09 | 5.829E-09 | 2.400E-09 | 1.400E-08 |
| 4537 | 227 | H227T | A | 5.4e-09 | 8.4e-09 | 9.6e-09 | 7.800E-09 | 2.163E-09 | 5.400E-09 | 9.600E-09 |
| 4538 | 227 | H227W | A | 8.3e-09 | 7.3e-09 | 4.4e-09 | 6.667E-09 | 2.026E-09 | 4.400E-09 | 8.300E-09 |
| 4539 | 227 | H227Y | A | 7e-09 | 9.5e-09 | 1.5e-08 | 1.050E-08 | 4.093E-09 | 7.000E-09 | 1.500E-08 |
| 4540 | 227 | H227V | A | 4.5e-09 | 1.1e-08 | 4.2e-09 | 6.567E-09 | 3.842E-09 | 4.200E-09 | 1.100E-08 |
| 4541 | 228 | S228A | A | 5.2e-09 | 1.7e-08 | 1.7e-08 | 1.307E-08 | 6.813E-09 | 5.200E-09 | 1.700E-08 |
| 4542 | 228 | S228R | A | 1.5e-08 | 6.4e-09 | 7e-09 | 9.467E-09 | 4.801E-09 | 6.400E-09 | 1.500E-08 |
| 4543 | 228 | S228N | A | 4.3e-09 | 4.6e-09 | 9.3e-09 | 6.067E-09 | 2.804E-09 | 4.300E-09 | 9.300E-09 |
| 4544 | 228 | S228D | A | 1.3e-08 | 2.9e-09 | 1.2e-08 | 9.300E-09 | 5.565E-09 | 2.900E-09 | 1.300E-08 |
| 4545 | 228 | S228C | A | 3.4e-09 | 5.5e-09 | 1.5e-08 | 7.967E-09 | 6.181E-09 | 3.400E-09 | 1.500E-08 |
| 4546 | 228 | S228E | A | 2.8e-09 | 5.4e-09 | 1.2e-08 | 6.733E-09 | 4.743E-09 | 2.800E-09 | 1.200E-08 |
| 4547 | 228 | S228Q | A | 4.5e-09 | 4.2e-09 | 1.2e-08 | 6.900E-09 | 4.419E-09 | 4.200E-09 | 1.200E-08 |
| 4548 | 228 | S228G | A | 8e-09 | 2.5e-09 | 3.9e-09 | 4.800E-09 | 2.858E-09 | 2.500E-09 | 8.000E-09 |
| 4549 | 228 | S228H | A | 9.3e-09 | 1.7e-08 | 1.5e-08 | 1.377E-08 | 3.995E-09 | 9.300E-09 | 1.700E-08 |
| 4550 | 228 | S228I | A | 8.4e-09 | 6.3e-09 | 4.8e-09 | 6.500E-09 | 1.808E-09 | 4.800E-09 | 8.400E-09 |
| 4551 | 228 | S228L | A | 7e-09 | 1.1e-08 | 2.5e-09 | 6.833E-09 | 4.252E-09 | 2.500E-09 | 1.100E-08 |
| 4552 | 228 | S228K | A | 7.5e-09 | 7.4e-09 | 1.2e-08 | 8.967E-09 | 2.627E-09 | 7.400E-09 | 1.200E-08 |
| 4553 | 228 | S228M | A | 4.7e-09 | 8.3e-09 | 6.1e-09 | 6.367E-09 | 1.815E-09 | 4.700E-09 | 8.300E-09 |
| 4554 | 228 | S228F | A | 5.5e-09 | 1.1e-08 | 7.6e-09 | 8.033E-09 | 2.775E-09 | 5.500E-09 | 1.100E-08 |
| 4555 | 228 | S228P | A | 2e-09 | 1.6e-08 | 3e-09 | 7.000E-09 | 7.810E-09 | 2.000E-09 | 1.600E-08 |
| 4556 | 228 | S228S | A | 9.7e-09 | 7e-09 | 6e-09 | 7.567E-09 | 1.914E-09 | 6.000E-09 | 9.700E-09 |
| 4557 | 228 | S228T | A | 1.7e-08 | 1.6e-08 | 3e-09 | 1.200E-08 | 7.810E-09 | 3.000E-09 | 1.700E-08 |
| 4558 | 228 | S228W | A | 1.3e-08 | 1.4e-08 | 1.5e-08 | 1.400E-08 | 1.000E-09 | 1.300E-08 | 1.500E-08 |
| 4559 | 228 | S228Y | A | 2.5e-08 | 6.2e-09 | 9e-09 | 1.340E-08 | 1.014E-08 | 6.200E-09 | 2.500E-08 |
| 4560 | 228 | S228V | A | 3.3e-09 | 1.4e-08 | 9.1e-09 | 8.800E-09 | 5.356E-09 | 3.300E-09 | 1.400E-08 |
| 4561 | 229 | D229A | A | 8e-09 | 7.5e-09 | 5.8e-09 | 7.100E-09 | 1.153E-09 | 5.800E-09 | 8.000E-09 |
| 4562 | 229 | D229R | A | 7e-09 | 6.3e-09 | 1.3e-09 | 4.867E-09 | 3.109E-09 | 1.300E-09 | 7.000E-09 |
| 4563 | 229 | D229N | A | 4.6e-09 | 4e-09 | 4.9e-09 | 4.500E-09 | 4.583E-10 | 4.000E-09 | 4.900E-09 |
| 4564 | 229 | D229D | A | 1e-08 | 8.9e-09 | 8.2e-09 | 9.033E-09 | 9.074E-10 | 8.200E-09 | 1.000E-08 |
| 4565 | 229 | D229C | A | 7.4e-09 | 2.7e-08 | 4e-09 | 1.280E-08 | 1.241E-08 | 4.000E-09 | 2.700E-08 |
| 4566 | 229 | D229E | A | 1.1e-08 | 9.2e-09 | 2.5e-09 | 7.567E-09 | 4.479E-09 | 2.500E-09 | 1.100E-08 |
| 4567 | 229 | D229Q | A | 7.8e-09 | 3.3e-09 | 9.2e-09 | 6.767E-09 | 3.083E-09 | 3.300E-09 | 9.200E-09 |
| 4568 | 229 | D229G | A | 3.1e-09 | 1e-08 | 1.6e-08 | 9.700E-09 | 6.455E-09 | 3.100E-09 | 1.600E-08 |
| 4569 | 229 | D229H | A | 8.4e-09 | 3.4e-09 | 4.6e-09 | 5.467E-09 | 2.610E-09 | 3.400E-09 | 8.400E-09 |
| 4570 | 229 | D229I | A | 9.4e-09 | 5.9e-09 | 1.6e-08 | 1.043E-08 | 5.129E-09 | 5.900E-09 | 1.600E-08 |
| 4571 | 229 | D229L | A | 1.5e-08 | 1.6e-08 | 1.3e-08 | 1.467E-08 | 1.528E-09 | 1.300E-08 | 1.600E-08 |
| 4572 | 229 | D229K | A | 4.9e-09 | 4.6e-09 | 1.1e-08 | 6.833E-09 | 3.612E-09 | 4.600E-09 | 1.100E-08 |
| 4573 | 229 | D229M | A | 1.3e-08 | 2.5e-08 | 8.4e-09 | 1.547E-08 | 8.570E-09 | 8.400E-09 | 2.500E-08 |
| 4574 | 229 | D229F | A | 3.4e-09 | 3.7e-09 | 9.1e-09 | 5.400E-09 | 3.208E-09 | 3.400E-09 | 9.100E-09 |
| 4575 | 229 | D229P | A | 1.8e-08 | 1.1e-08 | 4.8e-09 | 1.127E-08 | 6.604E-09 | 4.800E-09 | 1.800E-08 |
| 4576 | 229 | D229S | A | 3.2e-08 | 3.6e-09 | 4.7e-09 | 1.343E-08 | 1.609E-08 | 3.600E-09 | 3.200E-08 |
| 4577 | 229 | D229T | A | 1.2e-09 | 6.2e-09 | 8.8e-09 | 5.400E-09 | 3.863E-09 | 1.200E-09 | 8.800E-09 |
| 4578 | 229 | D229W | A | 9.8e-09 | 1.1e-08 | 1.7e-09 | 7.500E-09 | 5.059E-09 | 1.700E-09 | 1.100E-08 |
| 4579 | 229 | D229Y | A | 1.1e-08 | 1.8e-09 | 4.7e-09 | 5.833E-09 | 4.704E-09 | 1.800E-09 | 1.100E-08 |
| 4580 | 229 | D229V | A | 5.9e-09 | 6.3e-09 | 1.3e-08 | 8.400E-09 | 3.989E-09 | 5.900E-09 | 1.300E-08 |
| 4581 | 230 | C230A | A | 5.9e-09 | 1e-08 | 9.6e-09 | 8.500E-09 | 2.261E-09 | 5.900E-09 | 1.000E-08 |
| 4582 | 230 | C230R | A | 7.6e-09 | 6.2e-09 | 4.1e-09 | 5.967E-09 | 1.762E-09 | 4.100E-09 | 7.600E-09 |
| 4583 | 230 | C230N | A | 6.5e-09 | 6.9e-09 | 5.2e-09 | 6.200E-09 | 8.888E-10 | 5.200E-09 | 6.900E-09 |
| 4584 | 230 | C230D | A | 3.7e-09 | 1.2e-08 | 8.8e-09 | 8.167E-09 | 4.186E-09 | 3.700E-09 | 1.200E-08 |
| 4585 | 230 | C230C | A | 8e-09 | 7.4e-09 | 2.3e-08 | 1.280E-08 | 8.839E-09 | 7.400E-09 | 2.300E-08 |
| 4586 | 230 | C230E | A | 4.7e-09 | 1.1e-08 | 1.2e-09 | 5.633E-09 | 4.966E-09 | 1.200E-09 | 1.100E-08 |
| 4587 | 230 | C230Q | A | 8.4e-09 | 3.2e-09 | 2e-08 | 1.053E-08 | 8.601E-09 | 3.200E-09 | 2.000E-08 |
| 4588 | 230 | C230G | A | 1.7e-08 | 8.3e-09 | 2.1e-08 | 1.543E-08 | 6.493E-09 | 8.300E-09 | 2.100E-08 |
| 4589 | 230 | C230H | A | 1.3e-08 | 1.7e-09 | 2.9e-09 | 5.867E-09 | 6.207E-09 | 1.700E-09 | 1.300E-08 |

| | | | | | | | | | | |
|---|---|---|---|---|---|---|---|---|---|---|
| 4590 | 230 | C230I | A | 3.8e-09 | 9e-09 | 6.3e-09 | 6.367E-09 | 2.601E-09 | 3.800E-09 | 9.000E-09 |
| 4591 | 230 | C230L | A | 2.9e-09 | 4.3e-09 | 9.2e-09 | 5.467E-09 | 3.308E-09 | 2.900E-09 | 9.200E-09 |
| 4592 | 230 | C230K | A | 8.6e-09 | 1.4e-08 | 1.4e-08 | 1.220E-08 | 3.118E-09 | 8.600E-09 | 1.400E-08 |
| 4593 | 230 | C230M | A | 9.3e-09 | 8.2e-09 | 4.3e-09 | 7.267E-09 | 2.627E-09 | 4.300E-09 | 9.300E-09 |
| 4594 | 230 | C230F | A | 6.3e-09 | 1.7e-08 | 9.4e-09 | 1.090E-08 | 5.505E-09 | 6.300E-09 | 1.700E-08 |
| 4595 | 230 | C230P | A | 6.4e-09 | 2e-08 | 3.1e-09 | 9.833E-09 | 8.958E-09 | 3.100E-09 | 2.000E-08 |
| 4596 | 230 | C230S | A | 4.8e-09 | 7.9e-09 | 7.1e-09 | 6.600E-09 | 1.609E-09 | 4.800E-09 | 7.900E-09 |
| 4597 | 230 | C230T | A | 3.7e-09 | 4.1e-09 | 1.4e-08 | 7.267E-09 | 5.835E-09 | 3.700E-09 | 1.400E-08 |
| 4598 | 230 | C230W | A | 1.1e-08 | 9.4e-09 | 1.1e-08 | 1.047E-08 | 9.238E-10 | 9.400E-09 | 1.100E-08 |
| 4599 | 230 | C230Y | A | 1.4e-08 | 1.3e-08 | 1.1e-08 | 1.267E-08 | 1.528E-09 | 1.100E-08 | 1.400E-08 |
| 4600 | 230 | C230V | A | 1.2e-08 | 4.4e-09 | 2.7e-08 | 1.447E-08 | 1.150E-08 | 4.400E-09 | 2.700E-08 |
| 4601 | 231 | L231A | A | 6.8e-09 | 1e-08 | 1.8e-09 | 6.200E-09 | 4.133E-09 | 1.800E-09 | 1.000E-08 |
| 4602 | 231 | L231R | A | 4.2e-09 | 1.1e-08 | 1.7e-08 | 1.073E-08 | 6.404E-09 | 4.200E-09 | 1.700E-08 |
| 4603 | 231 | L231N | A | 1.8e-08 | 1.2e-08 | 2.6e-09 | 1.087E-08 | 7.762E-09 | 2.600E-09 | 1.800E-08 |
| 4604 | 231 | L231D | A | 6.1e-09 | 1.8e-08 | 1.3e-08 | 1.237E-08 | 5.975E-09 | 6.100E-09 | 1.800E-08 |
| 4605 | 231 | L231C | A | 8.6e-09 | 3.6e-09 | 8.8e-09 | 7.000E-09 | 2.946E-09 | 3.600E-09 | 8.800E-09 |
| 4606 | 231 | L231E | A | 4.4e-09 | 2.3e-09 | 4.1e-09 | 3.600E-09 | 1.136E-09 | 2.300E-09 | 4.400E-09 |
| 4607 | 231 | L231Q | A | 5e-09 | 2.1e-09 | 6e-09 | 4.367E-09 | 2.026E-09 | 2.100E-09 | 6.000E-09 |
| 4608 | 231 | L231G | A | 6e-09 | 3.5e-09 | 2e-08 | 9.833E-09 | 8.893E-09 | 3.500E-09 | 2.000E-08 |
| 4609 | 231 | L231H | A | 5.9e-09 | 9.5e-09 | 1.3e-08 | 9.467E-09 | 3.550E-09 | 5.900E-09 | 1.300E-08 |
| 4610 | 231 | L231I | A | 9.6e-09 | 1.8e-08 | 5.3e-09 | 1.097E-08 | 6.459E-09 | 5.300E-09 | 1.800E-08 |
| 4611 | 231 | L231L | A | 2.9e-09 | 7.5e-09 | 7.8e-09 | 6.067E-09 | 2.747E-09 | 2.900E-09 | 7.800E-09 |
| 4612 | 231 | L231K | A | 4.6e-09 | 1.3e-08 | 8.5e-09 | 8.700E-09 | 4.204E-09 | 4.600E-09 | 1.300E-08 |
| 4613 | 231 | L231M | A | 6.4e-09 | 3.8e-09 | 5e-09 | 5.067E-09 | 1.301E-09 | 3.800E-09 | 6.400E-09 |
| 4614 | 231 | L231F | A | 7.9e-09 | 1.2e-08 | 1.5e-08 | 1.163E-08 | 3.564E-09 | 7.900E-09 | 1.500E-08 |
| 4615 | 231 | L231P | A | 4.6e-09 | 4e-09 | 1e-08 | 6.200E-09 | 3.305E-09 | 4.000E-09 | 1.000E-08 |
| 4616 | 231 | L231S | A | 5.5e-09 | 1e-08 | 7e-09 | 7.500E-09 | 2.291E-09 | 5.500E-09 | 1.000E-08 |
| 4617 | 231 | L231T | A | 4.4e-09 | 1.5e-08 | 5.7e-09 | 8.367E-09 | 5.781E-09 | 4.400E-09 | 1.500E-08 |
| 4618 | 231 | L231W | A | 6.6e-09 | 5e-09 | 5.9e-09 | 5.833E-09 | 8.021E-10 | 5.000E-09 | 6.600E-09 |
| 4619 | 231 | L231Y | A | 5.9e-09 | 1.1e-08 | 7.8e-09 | 8.233E-09 | 2.577E-09 | 5.900E-09 | 1.100E-08 |
| 4620 | 231 | L231V | A | 7e-09 | 2.3e-08 | 9.3e-09 | 1.310E-08 | 8.650E-09 | 7.000E-09 | 2.300E-08 |
| 4621 | 232 | A232A | A | 1.8e-08 | 5e-09 | 9.7e-09 | 1.090E-08 | 6.583E-09 | 5.000E-09 | 1.800E-08 |
| 4622 | 232 | A232R | A | 1e-08 | 6e-09 | 3e-09 | 6.333E-09 | 3.512E-09 | 3.000E-09 | 1.000E-08 |
| 4623 | 232 | A232N | A | 6.6e-09 | 5e-09 | 3.6e-09 | 5.067E-09 | 1.501E-09 | 3.600E-09 | 6.600E-09 |
| 4624 | 232 | A232D | A | 9.6e-09 | 6.5e-09 | 7.8e-09 | 7.967E-09 | 1.557E-09 | 6.500E-09 | 9.600E-09 |
| 4625 | 232 | A232C | A | 1.7e-09 | 3.5e-09 | 5.7e-09 | 3.633E-09 | 2.003E-09 | 1.700E-09 | 5.700E-09 |
| 4626 | 232 | A232E | A | 1.8e-09 | 3.6e-08 | 6.1e-09 | 1.463E-08 | 1.863E-08 | 1.800E-09 | 3.600E-08 |
| 4627 | 232 | A232Q | A | 4.2e-09 | 1.2e-08 | 3.7e-09 | 6.633E-09 | 4.654E-09 | 3.700E-09 | 1.200E-08 |
| 4628 | 232 | A232G | A | 4.7e-09 | 1.4e-08 | 1.6e-08 | 1.157E-08 | 6.030E-09 | 4.700E-09 | 1.600E-08 |
| 4629 | 232 | A232H | A | 3.9e-09 | 4.4e-09 | 5.8e-09 | 4.700E-09 | 9.849E-10 | 3.900E-09 | 5.800E-09 |
| 4630 | 232 | A232I | A | 6.7e-09 | 2.6e-09 | 9.1e-09 | 6.133E-09 | 3.287E-09 | 2.600E-09 | 9.100E-09 |
| 4631 | 232 | A232L | A | 1.2e-08 | 5.7e-09 | 9.7e-09 | 9.133E-09 | 3.188E-09 | 5.700E-09 | 1.200E-08 |
| 4632 | 232 | A232K | A | 1.4e-08 | 4.1e-09 | 5.5e-09 | 7.867E-09 | 5.358E-09 | 4.100E-09 | 1.400E-08 |
| 4633 | 232 | A232M | A | 4.2e-09 | 8.2e-09 | 2e-08 | 1.080E-08 | 8.215E-09 | 4.200E-09 | 2.000E-08 |
| 4634 | 232 | A232F | A | 1.2e-08 | 1.8e-08 | 1.1e-08 | 1.367E-08 | 3.786E-09 | 1.100E-08 | 1.800E-08 |
| 4635 | 232 | A232P | A | 9.5e-09 | 1.5e-08 | 6.1e-09 | 1.020E-08 | 4.491E-09 | 6.100E-09 | 1.500E-08 |
| 4636 | 232 | A232S | A | 7.5e-09 | 7e-09 | 1.6e-08 | 1.017E-08 | 5.058E-09 | 7.000E-09 | 1.600E-08 |
| 4637 | 232 | A232T | A | 6.4e-09 | 6.9e-09 | 8.5e-09 | 7.267E-09 | 1.097E-09 | 6.400E-09 | 8.500E-09 |
| 4638 | 232 | A232W | A | 8.5e-09 | 5.1e-09 | 7.1e-09 | 6.900E-09 | 1.709E-09 | 5.100E-09 | 8.500E-09 |
| 4639 | 232 | A232Y | A | 1.8e-08 | 5.9e-09 | 1.4e-08 | 1.263E-08 | 6.165E-09 | 5.900E-09 | 1.800E-08 |
| 4640 | 232 | A232V | A | 1.1e-08 | 7.1e-09 | 5.6e-09 | 7.900E-09 | 2.787E-09 | 5.600E-09 | 1.100E-08 |
| 4641 | 233 | C233A | A | 5.8e-09 | 3.7e-09 | 3.5e-09 | 4.333E-09 | 1.274E-09 | 3.500E-09 | 5.800E-09 |
| 4642 | 233 | C233R | A | 1.5e-09 | 6.9e-09 | 8e-09 | 5.467E-09 | 3.479E-09 | 1.500E-09 | 8.000E-09 |
| 4643 | 233 | C233N | A | 8.4e-09 | 4.7e-09 | 6.7e-09 | 6.600E-09 | 1.852E-09 | 4.700E-09 | 8.400E-09 |
| 4644 | 233 | C233D | A | 3.2e-09 | 6.7e-09 | 6.2e-09 | 5.367E-09 | 1.893E-09 | 3.200E-09 | 6.700E-09 |
| 4645 | 233 | C233C | A | 1.5e-08 | 7.2e-09 | 6.2e-09 | 9.467E-09 | 4.818E-09 | 6.200E-09 | 1.500E-08 |
| 4646 | 233 | C233E | A | 5.5e-09 | 6.2e-09 | 1.2e-08 | 7.900E-09 | 3.568E-09 | 5.500E-09 | 1.200E-08 |
| 4647 | 233 | C233Q | A | 5.9e-09 | 1.4e-09 | 8.9e-09 | 5.400E-09 | 3.775E-09 | 1.400E-09 | 8.900E-09 |
| 4648 | 233 | C233G | A | 1e-08 | 1.3e-08 | 4.9e-09 | 9.300E-09 | 4.095E-09 | 4.900E-09 | 1.300E-08 |
| 4649 | 233 | C233H | A | 5.7e-09 | 1.1e-08 | 5.9e-09 | 7.533E-09 | 3.004E-09 | 5.700E-09 | 1.100E-08 |
| 4650 | 233 | C233I | A | 1.8e-09 | 1.2e-08 | 3.9e-09 | 5.900E-09 | 5.386E-09 | 1.800E-09 | 1.200E-08 |
| 4651 | 233 | C233L | A | 3.5e-09 | 2.1e-08 | 3.8e-09 | 9.433E-09 | 1.002E-08 | 3.500E-09 | 2.100E-08 |
| 4652 | 233 | C233K | A | 5e-09 | 6.5e-09 | 7.2e-09 | 6.233E-09 | 1.124E-09 | 5.000E-09 | 7.200E-09 |
| 4653 | 233 | C233M | A | 7.9e-09 | 6.7e-09 | 3.7e-09 | 6.100E-09 | 2.163E-09 | 3.700E-09 | 7.900E-09 |
| 4654 | 233 | C233F | A | 3e-08 | 9.9e-09 | 5.4e-09 | 1.510E-08 | 1.310E-08 | 5.400E-09 | 3.000E-08 |
| 4655 | 233 | C233P | A | 3.8e-09 | 1.1e-08 | 1.1e-08 | 8.600E-09 | 4.157E-09 | 3.800E-09 | 1.100E-08 |
| 4656 | 233 | C233S | A | 2.7e-09 | 1.8e-08 | 1.5e-08 | 1.190E-08 | 8.107E-09 | 2.700E-09 | 1.800E-08 |
| 4657 | 233 | C233T | A | 7.8e-09 | 9.8e-09 | 7.2e-09 | 8.267E-09 | 1.361E-09 | 7.200E-09 | 9.800E-09 |
| 4658 | 233 | C233W | A | 2.7e-09 | 1.1e-08 | 1.9e-08 | 1.090E-08 | 8.150E-09 | 2.700E-09 | 1.900E-08 |
| 4659 | 233 | C233Y | A | 5.5e-09 | 9.4e-09 | 5.8e-09 | 6.900E-09 | 2.170E-09 | 5.500E-09 | 9.400E-09 |
| 4660 | 233 | C233V | A | 3.2e-09 | 2.2e-08 | 7e-09 | 1.073E-08 | 9.940E-09 | 3.200E-09 | 2.200E-08 |
| 4661 | 234 | L234A | A | 1.8e-08 | 8e-09 | 9.8e-09 | 1.193E-08 | 5.330E-09 | 8.000E-09 | 1.800E-08 |
| 4662 | 234 | L234R | A | 1.8e-08 | 2.1e-08 | 8.8e-09 | 1.593E-08 | 6.357E-09 | 8.800E-09 | 2.100E-08 |
| 4663 | 234 | L234N | A | 1.9e-09 | 7.1e-09 | 5.3e-09 | 4.767E-09 | 2.641E-09 | 1.900E-09 | 7.100E-09 |
| 4664 | 234 | L234D | A | 4.1e-09 | 5.7e-09 | 4.9e-09 | 4.900E-09 | 8.000E-10 | 4.100E-09 | 5.700E-09 |
| 4665 | 234 | L234C | A | 1.2e-08 | 2.3e-08 | 2.1e-09 | 1.237E-08 | 1.045E-08 | 2.100E-09 | 2.300E-08 |
| 4666 | 234 | L234E | A | 8.7e-09 | 9.2e-09 | 1.3e-08 | 1.030E-08 | 2.352E-09 | 8.700E-09 | 1.300E-08 |
| 4667 | 234 | L234Q | A | 1.3e-08 | 1.1e-08 | 8.8e-09 | 1.093E-08 | 2.101E-09 | 8.800E-09 | 1.300E-08 |
| 4668 | 234 | L234G | A | 3.6e-09 | 5.6e-09 | 3.8e-09 | 4.333E-09 | 1.102E-09 | 3.600E-09 | 5.600E-09 |
| 4669 | 234 | L234H | A | 7.5e-09 | 9.7e-09 | 4.5e-09 | 7.233E-09 | 2.610E-09 | 4.500E-09 | 9.700E-09 |
| 4670 | 234 | L234I | A | 7e-09 | 2.4e-09 | 7.9e-09 | 5.767E-09 | 2.950E-09 | 2.400E-09 | 7.900E-09 |
| 4671 | 234 | L234L | A | 7e-09 | 3.3e-09 | 2.5e-08 | 1.177E-08 | 1.161E-08 | 3.300E-09 | 2.500E-08 |
| 4672 | 234 | L234K | A | 1e-08 | 1.3e-08 | 1.1e-08 | 1.133E-08 | 1.528E-09 | 1.000E-08 | 1.300E-08 |
| 4673 | 234 | L234M | A | 7.5e-09 | 9.4e-09 | 3.3e-09 | 6.733E-09 | 3.121E-09 | 3.300E-09 | 9.400E-09 |
| 4674 | 234 | L234F | A | 4.3e-09 | 2.8e-09 | 7.5e-09 | 4.867E-09 | 2.401E-09 | 2.800E-09 | 7.500E-09 |

| | | | | | | | | | | |
|---|---|---|---|---|---|---|---|---|---|---|
| 4675 | 234 | L234P | A | 4.6e-09 | 5.4e-09 | 4.1e-09 | 4.700E-09 | 6.557E-10 | 4.100E-09 | 5.400E-09 |
| 4676 | 234 | L234S | A | 2.3e-09 | 7.2e-09 | 3.1e-09 | 4.200E-09 | 2.629E-09 | 2.300E-09 | 7.200E-09 |
| 4677 | 234 | L234T | A | 4.9e-09 | 3.7e-09 | 9.9e-09 | 6.167E-09 | 3.288E-09 | 3.700E-09 | 9.900E-09 |
| 4678 | 234 | L234W | A | 9.2e-09 | 9e-09 | 6.1e-09 | 8.100E-09 | 1.735E-09 | 6.100E-09 | 9.200E-09 |
| 4679 | 234 | L234Y | A | 1.4e-08 | 3.1e-09 | 6.4e-09 | 7.833E-09 | 5.590E-09 | 3.100E-09 | 1.400E-08 |
| 4680 | 234 | L234V | A | 5.2e-09 | 8.7e-09 | 5.5e-09 | 6.467E-09 | 1.940E-09 | 5.200E-09 | 8.700E-09 |
| 4681 | 235 | H235A | A | 8.2e-09 | 6.8e-09 | 4.8e-09 | 6.600E-09 | 1.709E-09 | 4.800E-09 | 8.200E-09 |
| 4682 | 235 | H235R | A | 3.1e-09 | 5.7e-09 | 4.1e-09 | 4.300E-09 | 1.311E-09 | 3.100E-09 | 5.700E-09 |
| 4683 | 235 | H235N | A | 9.5e-09 | 1.4e-08 | 5.2e-09 | 9.567E-09 | 4.400E-09 | 5.200E-09 | 1.400E-08 |
| 4684 | 235 | H235D | A | 5.8e-09 | 3.8e-09 | 1.7e-09 | 3.767E-09 | 2.050E-09 | 1.700E-09 | 5.800E-09 |
| 4685 | 235 | H235C | A | 6.6e-09 | 3.3e-09 | 1.1e-08 | 6.967E-09 | 3.863E-09 | 3.300E-09 | 1.100E-08 |
| 4686 | 235 | H235E | A | 3.8e-09 | 8.2e-09 | 1.2e-08 | 8.000E-09 | 4.104E-09 | 3.800E-09 | 1.200E-08 |
| 4687 | 235 | H235Q | A | 3.9e-09 | 4.8e-09 | 1.9e-08 | 9.233E-09 | 8.470E-09 | 3.900E-09 | 1.900E-08 |
| 4688 | 235 | H235G | A | 3e-09 | 1e-08 | 1.4e-08 | 9.000E-09 | 5.568E-09 | 3.000E-09 | 1.400E-08 |
| 4689 | 235 | H235H | A | 1.8e-08 | 8.6e-09 | 6.4e-09 | 1.100E-08 | 6.161E-09 | 6.400E-09 | 1.800E-08 |
| 4690 | 235 | H235I | A | 8.8e-09 | 2e-09 | 5.7e-09 | 5.500E-09 | 3.404E-09 | 2.000E-09 | 8.800E-09 |
| 4691 | 235 | H235L | A | 3.7e-09 | 1.1e-08 | 6.4e-09 | 7.033E-09 | 3.691E-09 | 3.700E-09 | 1.100E-08 |
| 4692 | 235 | H235K | A | 1.1e-08 | 6.9e-09 | 4.2e-09 | 7.367E-09 | 3.424E-09 | 4.200E-09 | 1.100E-08 |
| 4693 | 235 | H235M | A | 2.4e-09 | 5.3e-09 | 4.2e-09 | 3.967E-09 | 1.464E-09 | 2.400E-09 | 5.300E-09 |
| 4694 | 235 | H235F | A | 2.1e-09 | 1.5e-08 | 4.7e-09 | 7.267E-09 | 6.822E-09 | 2.100E-09 | 1.500E-08 |
| 4695 | 235 | H235P | A | 1.1e-08 | 7.5e-09 | 3.6e-09 | 7.367E-09 | 3.702E-09 | 3.600E-09 | 1.100E-08 |
| 4696 | 235 | H235S | A | 6.7e-09 | 3.7e-09 | 6.1e-09 | 5.500E-09 | 1.587E-09 | 3.700E-09 | 6.700E-09 |
| 4697 | 235 | H235T | A | 3.4e-09 | 1.8e-08 | 9.7e-09 | 1.037E-08 | 7.323E-09 | 3.400E-09 | 1.800E-08 |
| 4698 | 235 | H235W | A | 2.6e-09 | 2.9e-09 | 2e-09 | 2.500E-09 | 4.583E-10 | 2.000E-09 | 2.900E-09 |
| 4699 | 235 | H235Y | A | 7e-09 | 1e-08 | 9.7e-09 | 8.900E-09 | 1.652E-09 | 7.000E-09 | 1.000E-08 |
| 4700 | 235 | H235V | A | 7.6e-09 | 3.8e-09 | 1.1e-08 | 7.467E-09 | 3.602E-09 | 3.800E-09 | 1.100E-08 |
| 4701 | 236 | F236A | A | 9.1e-09 | 1.1e-08 | 1e-08 | 1.003E-08 | 9.504E-10 | 9.100E-09 | 1.100E-08 |
| 4702 | 236 | F236R | A | 7.3e-09 | 5.8e-09 | 9.9e-09 | 7.667E-09 | 2.074E-09 | 5.800E-09 | 9.900E-09 |
| 4703 | 236 | F236N | A | 6.1e-09 | 1.1e-08 | 2.1e-08 | 1.270E-08 | 7.594E-09 | 6.100E-09 | 2.100E-08 |
| 4704 | 236 | F236D | A | 2.5e-08 | 1.9e-08 | 1.4e-08 | 1.933E-08 | 5.508E-09 | 1.400E-08 | 2.500E-08 |
| 4705 | 236 | F236C | A | 2e-09 | 8.9e-09 | 2.4e-08 | 1.163E-08 | 1.125E-08 | 2.000E-09 | 2.400E-08 |
| 4706 | 236 | F236E | A | 1.4e-08 | 2.6e-09 | 1.3e-08 | 9.867E-09 | 6.313E-09 | 2.600E-09 | 1.400E-08 |
| 4707 | 236 | F236Q | A | 3.9e-08 | 7.6e-09 | 1.7e-08 | 2.120E-08 | 1.612E-08 | 7.600E-09 | 3.900E-08 |
| 4708 | 236 | F236G | A | 1.3e-08 | 5.9e-09 | 9.2e-09 | 9.367E-09 | 3.553E-09 | 5.900E-09 | 1.300E-08 |
| 4709 | 236 | F236H | A | 6.8e-09 | 3.9e-09 | 2.8e-08 | 1.290E-08 | 1.316E-08 | 3.900E-09 | 2.800E-08 |
| 4710 | 236 | F236I | A | 6.8e-09 | 1.1e-08 | 1.1e-08 | 9.600E-09 | 2.425E-09 | 6.800E-09 | 1.100E-08 |
| 4711 | 236 | F236L | A | 6.6e-09 | 6.4e-09 | 6.8e-09 | 6.600E-09 | 2.000E-10 | 6.400E-09 | 6.800E-09 |
| 4712 | 236 | F236K | A | 5.1e-09 | 5e-09 | 5.8e-09 | 5.300E-09 | 4.359E-10 | 5.000E-09 | 5.800E-09 |
| 4713 | 236 | F236M | A | 7.7e-09 | 3.7e-09 | 4e-09 | 5.133E-09 | 2.228E-09 | 3.700E-09 | 7.700E-09 |
| 4714 | 236 | F236F | A | 5.4e-09 | 8.1e-09 | 2.2e-09 | 5.233E-09 | 2.954E-09 | 2.200E-09 | 8.100E-09 |
| 4715 | 236 | F236P | A | 2.3e-08 | 2.3e-08 | 1.7e-08 | 2.100E-08 | 3.464E-09 | 1.700E-08 | 2.300E-08 |
| 4716 | 236 | F236S | A | 1.5e-08 | 1.1e-08 | 1.9e-08 | 1.500E-08 | 4.000E-09 | 1.100E-08 | 1.900E-08 |
| 4717 | 236 | F236T | A | 4.6e-09 | 6.9e-09 | 1e-08 | 7.167E-09 | 2.710E-09 | 4.600E-09 | 1.000E-08 |
| 4718 | 236 | F236W | A | 8.6e-09 | 2.7e-09 | 8.3e-09 | 6.533E-09 | 3.323E-09 | 2.700E-09 | 8.600E-09 |
| 4719 | 236 | F236Y | A | 5e-09 | 3.6e-09 | 8.6e-09 | 5.733E-09 | 2.579E-09 | 3.600E-09 | 8.600E-09 |
| 4720 | 236 | F236V | A | 7.7e-09 | 8.5e-09 | 9.3e-09 | 8.500E-09 | 8.000E-10 | 7.700E-09 | 9.300E-09 |
| 4721 | 237 | N237A | A | 1e-08 | 9.5e-09 | 1.5e-08 | 1.150E-08 | 3.041E-09 | 9.500E-09 | 1.500E-08 |
| 4722 | 237 | N237R | A | 4.5e-09 | 2.2e-08 | 8.8e-09 | 1.177E-08 | 9.119E-09 | 4.500E-09 | 2.200E-08 |
| 4723 | 237 | N237N | A | 1e-08 | 7e-09 | 4.3e-09 | 7.100E-09 | 2.851E-09 | 4.300E-09 | 1.000E-08 |
| 4724 | 237 | N237D | A | 3.5e-09 | 1.7e-08 | 2.8e-09 | 7.767E-09 | 8.004E-09 | 2.800E-09 | 1.700E-08 |
| 4725 | 237 | N237C | A | 8.8e-09 | 1.2e-08 | 6.6e-09 | 9.133E-09 | 2.715E-09 | 6.600E-09 | 1.200E-08 |
| 4726 | 237 | N237E | A | 6.1e-09 | 5.5e-09 | 4.4e-09 | 5.333E-09 | 8.622E-10 | 4.400E-09 | 6.100E-09 |
| 4727 | 237 | N237Q | A | 7.4e-09 | 1.6e-09 | 3.5e-09 | 4.167E-09 | 2.957E-09 | 1.600E-09 | 7.400E-09 |
| 4728 | 237 | N237G | A | 1.7e-08 | 4.2e-09 | 3.7e-09 | 8.300E-09 | 7.539E-09 | 3.700E-09 | 1.700E-08 |
| 4729 | 237 | N237H | A | 3.6e-09 | 5.8e-09 | 1.1e-08 | 6.800E-09 | 3.800E-09 | 3.600E-09 | 1.100E-08 |
| 4730 | 237 | N237I | A | 5.3e-09 | 1.1e-08 | 1e-08 | 8.767E-09 | 3.044E-09 | 5.300E-09 | 1.100E-08 |
| 4731 | 237 | N237L | A | 7.4e-09 | 1.8e-08 | 9.5e-10 | 8.783E-09 | 8.609E-09 | 9.500E-10 | 1.800E-08 |
| 4732 | 237 | N237K | A | 9.5e-09 | 6.2e-09 | 9.7e-09 | 8.467E-09 | 1.966E-09 | 6.200E-09 | 9.700E-09 |
| 4733 | 237 | N237M | A | 9.4e-09 | 6.2e-09 | 4.4e-09 | 6.667E-09 | 2.532E-09 | 4.400E-09 | 9.400E-09 |
| 4734 | 237 | N237F | A | 8.8e-09 | 1e-08 | 9.4e-09 | 9.400E-09 | 6.000E-10 | 8.800E-09 | 1.000E-08 |
| 4735 | 237 | N237P | A | 5.9e-09 | 5.7e-09 | 5e-09 | 5.533E-09 | 4.726E-10 | 5.000E-09 | 5.900E-09 |
| 4736 | 237 | N237S | A | 6.3e-09 | 3.3e-09 | 5.7e-09 | 5.100E-09 | 1.587E-09 | 3.300E-09 | 6.300E-09 |
| 4737 | 237 | N237T | A | 7.2e-09 | 9.1e-09 | 9.5e-09 | 8.600E-09 | 1.229E-09 | 7.200E-09 | 9.500E-09 |
| 4738 | 237 | N237W | A | 7.9e-09 | 4.7e-09 | 1.2e-08 | 8.200E-09 | 3.659E-09 | 4.700E-09 | 1.200E-08 |
| 4739 | 237 | N237Y | A | 7.3e-09 | 1e-08 | 1.4e-08 | 1.043E-08 | 3.371E-09 | 7.300E-09 | 1.400E-08 |
| 4740 | 237 | N237V | A | 3.6e-09 | 2.9e-09 | 9.3e-09 | 5.267E-09 | 3.510E-09 | 2.900E-09 | 9.300E-09 |
| 4741 | 238 | H238A | A | 1.5e-08 | 1.1e-08 | 1.5e-08 | 1.367E-08 | 2.309E-09 | 1.100E-08 | 1.500E-08 |
| 4742 | 238 | H238R | A | 1e-08 | 3.7e-09 | 7.5e-09 | 7.067E-09 | 3.172E-09 | 3.700E-09 | 1.000E-08 |
| 4743 | 238 | H238N | A | 9.5e-09 | 1.3e-08 | 7e-09 | 9.833E-09 | 3.014E-09 | 7.000E-09 | 1.300E-08 |
| 4744 | 238 | H238D | A | 3.8e-09 | 7.9e-09 | 7.1e-09 | 6.267E-09 | 2.173E-09 | 3.800E-09 | 7.900E-09 |
| 4745 | 238 | H238C | A | 3.4e-09 | 2.9e-09 | 2.9e-09 | 3.067E-09 | 2.887E-10 | 2.900E-09 | 3.400E-09 |
| 4746 | 238 | H238E | A | 1.1e-08 | 3.3e-09 | 8.9e-09 | 7.733E-09 | 3.980E-09 | 3.300E-09 | 1.100E-08 |
| 4747 | 238 | H238Q | A | 3.9e-09 | 8.6e-09 | 1.6e-08 | 9.500E-09 | 6.100E-09 | 3.900E-09 | 1.600E-08 |
| 4748 | 238 | H238G | A | 4.3e-09 | 1.6e-08 | 6.8e-09 | 9.033E-09 | 6.161E-09 | 4.300E-09 | 1.600E-08 |
| 4749 | 238 | H238H | A | 7.1e-09 | 8e-09 | 9.6e-09 | 8.233E-09 | 1.266E-09 | 7.100E-09 | 9.600E-09 |
| 4750 | 238 | H238I | A | 1.3e-08 | 1.2e-08 | 3.2e-09 | 9.400E-09 | 5.393E-09 | 3.200E-09 | 1.300E-08 |
| 4751 | 238 | H238L | A | 6.7e-09 | 8.1e-09 | 1.2e-08 | 8.933E-09 | 2.747E-09 | 6.700E-09 | 1.200E-08 |
| 4752 | 238 | H238K | A | 8e-09 | 4.4e-09 | 5.3e-09 | 5.900E-09 | 1.873E-09 | 4.400E-09 | 8.000E-09 |
| 4753 | 238 | H238M | A | 5.6e-09 | 7.8e-09 | 7.3e-09 | 6.900E-09 | 1.153E-09 | 5.600E-09 | 7.800E-09 |
| 4754 | 238 | H238F | A | 9e-09 | 2.8e-09 | 7.4e-09 | 6.400E-09 | 3.219E-09 | 2.800E-09 | 9.000E-09 |
| 4755 | 238 | H238P | A | 5.8e-09 | 1.1e-08 | 1.1e-08 | 9.267E-09 | 3.002E-09 | 5.800E-09 | 1.100E-08 |
| 4756 | 238 | H238S | A | 2.4e-08 | 1.3e-08 | 1.5e-08 | 1.733E-08 | 5.859E-09 | 1.300E-08 | 2.400E-08 |
| 4757 | 238 | H238T | A | 5e-09 | 1.1e-09 | 8.7e-09 | 4.933E-09 | 3.800E-09 | 1.100E-09 | 8.700E-09 |
| 4758 | 238 | H238W | A | 7.2e-09 | 5.6e-09 | 1.7e-09 | 4.833E-09 | 2.829E-09 | 1.700E-09 | 7.200E-09 |
| 4759 | 238 | H238Y | A | 8.8e-09 | 1.2e-08 | 1.6e-08 | 1.227E-08 | 3.607E-09 | 8.800E-09 | 1.600E-08 |

| | | | | | | | | | | |
|---|---|---|---|---|---|---|---|---|---|---|
| 4760 | 238 | H238V | A | 1.7e-09 | 7.5e-09 | 5.6e-09 | 4.933E-09 | 2.957E-09 | 1.700E-09 | 7.500E-09 |
| 4761 | 239 | S239A | A | 1.1e-08 | 1.9e-08 | 6.9e-09 | 1.230E-08 | 6.154E-09 | 6.900E-09 | 1.900E-08 |
| 4762 | 239 | S239R | A | 1.4e-08 | 6.7e-09 | 2.1e-08 | 1.390E-08 | 7.151E-09 | 6.700E-09 | 2.100E-08 |
| 4763 | 239 | S239N | A | 5.5e-09 | 1.3e-08 | 5.6e-09 | 8.033E-09 | 4.302E-09 | 5.500E-09 | 1.300E-08 |
| 4764 | 239 | S239D | A | 9e-09 | 1e-08 | 1.5e-08 | 1.133E-08 | 3.215E-09 | 9.000E-09 | 1.500E-08 |
| 4765 | 239 | S239C | A | 1.1e-08 | 6e-09 | 4.7e-09 | 7.233E-09 | 3.326E-09 | 4.700E-09 | 1.100E-08 |
| 4766 | 239 | S239E | A | 8.7e-09 | 2.4e-08 | 5.3e-09 | 1.267E-08 | 9.961E-09 | 5.300E-09 | 2.400E-08 |
| 4767 | 239 | S239Q | A | 1.1e-08 | 6.7e-09 | 3.8e-09 | 7.167E-09 | 3.623E-09 | 3.800E-09 | 1.100E-08 |
| 4768 | 239 | S239G | A | 5e-09 | 3.7e-09 | 1e-08 | 6.233E-09 | 3.326E-09 | 3.700E-09 | 1.000E-08 |
| 4769 | 239 | S239H | A | 2.1e-09 | 2.4e-09 | 7.3e-09 | 3.933E-09 | 2.919E-09 | 2.100E-09 | 7.300E-09 |
| 4770 | 239 | S239I | A | 6e-09 | 7.4e-09 | 6.2e-09 | 6.533E-09 | 7.572E-10 | 6.000E-09 | 7.400E-09 |
| 4771 | 239 | S239L | A | 2.3e-09 | 5.8e-09 | 8.6e-09 | 5.567E-09 | 3.156E-09 | 2.300E-09 | 8.600E-09 |
| 4772 | 239 | S239K | A | 6.5e-09 | 6.4e-09 | 8.2e-09 | 7.033E-09 | 1.012E-09 | 6.400E-09 | 8.200E-09 |
| 4773 | 239 | S239M | A | 1.6e-08 | 1.1e-08 | 7.3e-09 | 1.143E-08 | 4.366E-09 | 7.300E-09 | 1.600E-08 |
| 4774 | 239 | S239F | A | 9e-09 | 3e-09 | 5.7e-09 | 5.900E-09 | 3.005E-09 | 3.000E-09 | 9.000E-09 |
| 4775 | 239 | S239P | A | 8.4e-09 | 1.3e-08 | 9.7e-09 | 1.037E-08 | 2.371E-09 | 8.400E-09 | 1.300E-08 |
| 4776 | 239 | S239S | A | 1.2e-08 | 1.2e-08 | 5.2e-09 | 9.733E-09 | 3.926E-09 | 5.200E-09 | 1.200E-08 |
| 4777 | 239 | S239T | A | 2.2e-09 | 1.8e-08 | 3.3e-09 | 7.833E-09 | 8.822E-09 | 2.200E-09 | 1.800E-08 |
| 4778 | 239 | S239W | A | 4.8e-09 | 2.1e-08 | 1.6e-08 | 1.393E-08 | 8.295E-09 | 4.800E-09 | 2.100E-08 |
| 4779 | 239 | S239Y | A | 2.7e-08 | 3.4e-09 | 6.3e-09 | 1.223E-08 | 1.287E-08 | 3.400E-09 | 2.700E-08 |
| 4780 | 239 | S239V | A | 2.4e-09 | 2.1e-09 | 8.7e-09 | 4.400E-09 | 3.727E-09 | 2.100E-09 | 8.700E-09 |
| 4781 | 240 | G240A | A | 4.4e-09 | 5.6e-09 | 1.1e-08 | 7.000E-09 | 3.516E-09 | 4.400E-09 | 1.100E-08 |
| 4782 | 240 | G240R | A | 4.1e-09 | 1.7e-08 | 1.7e-09 | 7.600E-09 | 8.229E-09 | 1.700E-09 | 1.700E-08 |
| 4783 | 240 | G240N | A | 7e-09 | 3.9e-09 | 3e-09 | 4.633E-09 | 2.098E-09 | 3.000E-09 | 7.000E-09 |
| 4784 | 240 | G240D | A | 4.4e-09 | 4.7e-09 | 1.2e-08 | 7.033E-09 | 4.304E-09 | 4.400E-09 | 1.200E-08 |
| 4785 | 240 | G240C | A | 5.5e-09 | 1.1e-08 | 2.8e-09 | 6.433E-09 | 4.179E-09 | 2.800E-09 | 1.100E-08 |
| 4786 | 240 | G240E | A | 2.3e-09 | 1.1e-08 | 4.7e-09 | 6.000E-09 | 4.493E-09 | 2.300E-09 | 1.100E-08 |
| 4787 | 240 | G240Q | A | 3e-09 | 1.9e-08 | 2.4e-09 | 8.133E-09 | 9.416E-09 | 2.400E-09 | 1.900E-08 |
| 4788 | 240 | G240G | A | 1.1e-08 | 6.3e-09 | 9.6e-09 | 8.967E-09 | 2.413E-09 | 6.300E-09 | 1.100E-08 |
| 4789 | 240 | G240H | A | 9e-09 | 2.9e-08 | 1.1e-08 | 1.633E-08 | 1.102E-08 | 9.000E-09 | 2.900E-08 |
| 4790 | 240 | G240I | A | 7.4e-09 | 6.5e-09 | 1.5e-08 | 9.633E-09 | 4.669E-09 | 6.500E-09 | 1.500E-08 |
| 4791 | 240 | G240L | A | 9.4e-09 | 5.8e-09 | 1.3e-08 | 9.400E-09 | 3.600E-09 | 5.800E-09 | 1.300E-08 |
| 4792 | 240 | G240K | A | 1e-08 | 1.2e-08 | 6.1e-09 | 9.367E-09 | 3.001E-09 | 6.100E-09 | 1.200E-08 |
| 4793 | 240 | G240M | A | 9.3e-09 | 7.3e-09 | 2.2e-09 | 6.267E-09 | 3.661E-09 | 2.200E-09 | 9.300E-09 |
| 4794 | 240 | G240F | A | 8.2e-09 | 3.8e-09 | 2.2e-09 | 4.733E-09 | 3.107E-09 | 2.200E-09 | 8.200E-09 |
| 4795 | 240 | G240P | A | 9.7e-09 | 3.1e-09 | 9.1e-09 | 7.300E-09 | 3.650E-09 | 3.100E-09 | 9.700E-09 |
| 4796 | 240 | G240S | A | 8.2e-09 | 1.4e-09 | 1.1e-08 | 6.867E-09 | 4.937E-09 | 1.400E-09 | 1.100E-08 |
| 4797 | 240 | G240T | A | 8.1e-09 | 6e-09 | 9.4e-09 | 7.833E-09 | 1.716E-09 | 6.000E-09 | 9.400E-09 |
| 4798 | 240 | G240W | A | 4e-09 | 2.4e-08 | 3.2e-09 | 1.040E-08 | 1.178E-08 | 3.200E-09 | 2.400E-08 |
| 4799 | 240 | G240Y | A | 5.8e-09 | 6e-09 | 3.5e-09 | 5.100E-09 | 1.389E-09 | 3.500E-09 | 6.000E-09 |
| 4800 | 240 | G240V | A | 1e-08 | 1.1e-08 | 4.3e-09 | 8.433E-09 | 3.614E-09 | 4.300E-09 | 1.100E-08 |
| 4801 | 241 | I241A | A | 6.4e-09 | 4.5e-09 | 3.7e-09 | 4.867E-09 | 1.387E-09 | 3.700E-09 | 6.400E-09 |
| 4802 | 241 | I241R | A | 4.6e-09 | 8.2e-09 | 9.8e-09 | 7.533E-09 | 2.663E-09 | 4.600E-09 | 9.800E-09 |
| 4803 | 241 | I241N | A | 1.7e-08 | 1.3e-08 | 1.3e-08 | 1.433E-08 | 2.309E-09 | 1.300E-08 | 1.700E-08 |
| 4804 | 241 | I241D | A | 7.2e-09 | 8.4e-09 | 5e-09 | 6.867E-09 | 1.724E-09 | 5.000E-09 | 8.400E-09 |
| 4805 | 241 | I241C | A | 5.7e-09 | 3.3e-09 | 3e-08 | 1.300E-08 | 1.477E-08 | 3.300E-09 | 3.000E-08 |
| 4806 | 241 | I241E | A | 1.8e-08 | 5.8e-09 | 8.6e-10 | 8.220E-09 | 8.823E-09 | 8.600E-10 | 1.800E-08 |
| 4807 | 241 | I241Q | A | 2.6e-09 | 1.1e-08 | 3.1e-09 | 5.567E-09 | 4.712E-09 | 2.600E-09 | 1.100E-08 |
| 4808 | 241 | I241G | A | 8.2e-09 | 1.1e-08 | 1.7e-09 | 6.967E-09 | 4.771E-09 | 1.700E-09 | 1.100E-08 |
| 4809 | 241 | I241H | A | 5.9e-09 | 7.1e-09 | 3.3e-09 | 5.433E-09 | 1.943E-09 | 3.300E-09 | 7.100E-09 |
| 4810 | 241 | I241I | A | 9.6e-09 | 5.4e-09 | 1.1e-08 | 8.667E-09 | 2.914E-09 | 5.400E-09 | 1.100E-08 |
| 4811 | 241 | I241L | A | 1.1e-08 | 2.2e-09 | 4e-09 | 5.733E-09 | 4.649E-09 | 2.200E-09 | 1.100E-08 |
| 4812 | 241 | I241K | A | 4.3e-09 | 7.5e-09 | 6.3e-09 | 6.033E-09 | 1.617E-09 | 4.300E-09 | 7.500E-09 |
| 4813 | 241 | I241M | A | 7.3e-09 | 1.4e-08 | 2.5e-09 | 7.933E-09 | 5.776E-09 | 2.500E-09 | 1.400E-08 |
| 4814 | 241 | I241F | A | 1.9e-08 | 3.3e-09 | 7.1e-09 | 9.800E-09 | 8.191E-09 | 3.300E-09 | 1.900E-08 |
| 4815 | 241 | I241P | A | 3e-09 | 1.1e-08 | 1.2e-08 | 8.667E-09 | 4.933E-09 | 3.000E-09 | 1.200E-08 |
| 4816 | 241 | I241S | A | 4.7e-09 | 1.1e-08 | 2.7e-08 | 1.423E-08 | 1.150E-08 | 4.700E-09 | 2.700E-08 |
| 4817 | 241 | I241T | A | 4.9e-09 | 4.4e-09 | 1.9e-08 | 9.433E-09 | 8.289E-09 | 4.400E-09 | 1.900E-08 |
| 4818 | 241 | I241W | A | 3.8e-09 | 3.2e-09 | 1.2e-08 | 6.333E-09 | 4.917E-09 | 3.200E-09 | 1.200E-08 |
| 4819 | 241 | I241Y | A | 5.6e-09 | 7.6e-09 | 1e-08 | 7.733E-09 | 2.203E-09 | 5.600E-09 | 1.000E-08 |
| 4820 | 241 | I241V | A | 8.9e-09 | 1.1e-08 | 3.5e-09 | 7.800E-09 | 3.869E-09 | 3.500E-09 | 1.100E-08 |
| 4821 | 242 | C242A | A | 1.1e-08 | 4.2e-09 | 6.3e-09 | 7.167E-09 | 3.482E-09 | 4.200E-09 | 1.100E-08 |
| 4822 | 242 | C242R | A | 8.4e-09 | 1.5e-08 | 8.1e-09 | 1.050E-08 | 3.900E-09 | 8.100E-09 | 1.500E-08 |
| 4823 | 242 | C242N | A | 3.6e-09 | 8.1e-09 | 4.1e-09 | 5.267E-09 | 2.466E-09 | 3.600E-09 | 8.100E-09 |
| 4824 | 242 | C242D | A | 9.7e-09 | 5.9e-09 | 6.1e-09 | 7.233E-09 | 2.139E-09 | 5.900E-09 | 9.700E-09 |
| 4825 | 242 | C242C | A | 3.7e-09 | 1.1e-08 | 1.1e-08 | 8.567E-09 | 4.215E-09 | 3.700E-09 | 1.100E-08 |
| 4826 | 242 | C242E | A | 1.9e-08 | 7.3e-09 | 6.6e-09 | 1.097E-08 | 6.966E-09 | 6.600E-09 | 1.900E-08 |
| 4827 | 242 | C242Q | A | 1.8e-09 | 2.4e-09 | 9.2e-09 | 4.467E-09 | 4.110E-09 | 1.800E-09 | 9.200E-09 |
| 4828 | 242 | C242G | A | 1.2e-08 | 1.6e-08 | 1.1e-08 | 1.300E-08 | 2.646E-09 | 1.100E-08 | 1.600E-08 |
| 4829 | 242 | C242H | A | 7.6e-09 | 1e-08 | 1.2e-08 | 9.867E-09 | 2.203E-09 | 7.600E-09 | 1.200E-08 |
| 4830 | 242 | C242I | A | 5.4e-09 | 1.5e-08 | 4.7e-09 | 8.367E-09 | 5.755E-09 | 4.700E-09 | 1.500E-08 |
| 4831 | 242 | C242L | A | 5.4e-09 | 4.5e-09 | 7.2e-09 | 5.700E-09 | 1.375E-09 | 4.500E-09 | 7.200E-09 |
| 4832 | 242 | C242K | A | 4.7e-09 | 6.4e-09 | 1.3e-08 | 8.033E-09 | 4.384E-09 | 4.700E-09 | 1.300E-08 |
| 4833 | 242 | C242M | A | 5.9e-09 | 1.8e-08 | 6.6e-09 | 1.017E-08 | 6.793E-09 | 5.900E-09 | 1.800E-08 |
| 4834 | 242 | C242F | A | 7.6e-09 | 8.5e-09 | 7.3e-09 | 7.800E-09 | 6.245E-10 | 7.300E-09 | 8.500E-09 |
| 4835 | 242 | C242P | A | 5.4e-09 | 9.4e-09 | 6.3e-09 | 7.033E-09 | 2.098E-09 | 5.400E-09 | 9.400E-09 |
| 4836 | 242 | C242S | A | 5e-09 | 9.1e-09 | 5.7e-09 | 6.600E-09 | 2.193E-09 | 5.000E-09 | 9.100E-09 |
| 4837 | 242 | C242T | A | 6.8e-09 | 4.1e-09 | 3e-09 | 4.633E-09 | 1.955E-09 | 3.000E-09 | 6.800E-09 |
| 4838 | 242 | C242W | A | 1.4e-08 | 5.6e-09 | 7.8e-09 | 9.133E-09 | 4.356E-09 | 5.600E-09 | 1.400E-08 |
| 4839 | 242 | C242Y | A | 1.4e-08 | 1.6e-08 | 5e-09 | 1.167E-08 | 5.859E-09 | 5.000E-09 | 1.600E-08 |
| 4840 | 242 | C242V | A | 1e-08 | 1.2e-08 | 4.8e-09 | 8.933E-09 | 3.717E-09 | 4.800E-09 | 1.200E-08 |
| 4841 | 243 | E243A | A | 4.1e-09 | 4.7e-09 | 3.4e-09 | 4.067E-09 | 6.506E-10 | 3.400E-09 | 4.700E-09 |
| 4842 | 243 | E243R | A | 4e-09 | 8.6e-09 | 1.3e-08 | 8.533E-09 | 4.500E-09 | 4.000E-09 | 1.300E-08 |
| 4843 | 243 | E243N | A | 7.5e-09 | 5.1e-09 | 4e-09 | 5.533E-09 | 1.790E-09 | 4.000E-09 | 7.500E-09 |
| 4844 | 243 | E243D | A | 4.6e-08 | 7.6e-09 | 5.1e-09 | 1.957E-08 | 2.293E-08 | 5.100E-09 | 4.600E-08 |

| | | | | | | | | | | |
|---|---|---|---|---|---|---|---|---|---|---|
| 4845 | 243 | E243C | A | 1.1e-08 | 7.6e-09 | 2.5e-09 | 7.033E-09 | 4.278E-09 | 2.500E-09 | 1.100E-08 |
| 4846 | 243 | E243E | A | 6.6e-09 | 2e-08 | 8.6e-09 | 1.173E-08 | 7.229E-09 | 6.600E-09 | 2.000E-08 |
| 4847 | 243 | E243Q | A | 1.3e-08 | 9.8e-09 | 5.4e-09 | 9.400E-09 | 3.816E-09 | 5.400E-09 | 1.300E-08 |
| 4848 | 243 | E243G | A | 5e-09 | 3.7e-09 | 9.9e-09 | 6.200E-09 | 3.270E-09 | 3.700E-09 | 9.900E-09 |
| 4849 | 243 | E243H | A | 3.5e-09 | 5.4e-09 | 1.2e-08 | 6.967E-09 | 4.461E-09 | 3.500E-09 | 1.200E-08 |
| 4850 | 243 | E243I | A | 5.4e-09 | 3.3e-09 | 7.6e-09 | 5.433E-09 | 2.150E-09 | 3.300E-09 | 7.600E-09 |
| 4851 | 243 | E243L | A | 7.7e-09 | 6.1e-09 | 6.6e-09 | 6.800E-09 | 8.185E-10 | 6.100E-09 | 7.700E-09 |
| 4852 | 243 | E243K | A | 9.2e-09 | 9.8e-09 | 5e-09 | 8.000E-09 | 2.615E-09 | 5.000E-09 | 9.800E-09 |
| 4853 | 243 | E243M | A | 1.3e-08 | 1.4e-08 | 8.9e-09 | 1.197E-08 | 2.702E-09 | 8.900E-09 | 1.400E-08 |
| 4854 | 243 | E243F | A | 1.6e-08 | 7.2e-09 | 5.1e-09 | 9.433E-09 | 5.783E-09 | 5.100E-09 | 1.600E-08 |
| 4855 | 243 | E243P | A | 7.5e-09 | 8.2e-09 | 1.3e-08 | 9.567E-09 | 2.994E-09 | 7.500E-09 | 1.300E-08 |
| 4856 | 243 | E243S | A | 1e-08 | 3.5e-09 | 5.1e-09 | 6.200E-09 | 3.387E-09 | 3.500E-09 | 1.000E-08 |
| 4857 | 243 | E243T | A | 1.5e-08 | 5.4e-09 | 8.9e-09 | 9.767E-09 | 4.858E-09 | 5.400E-09 | 1.500E-08 |
| 4858 | 243 | E243W | A | 3.6e-09 | 1.1e-08 | 1.5e-08 | 9.867E-09 | 5.784E-09 | 3.600E-09 | 1.500E-08 |
| 4859 | 243 | E243Y | A | 1.6e-08 | 7.8e-09 | 6.2e-09 | 1.000E-08 | 5.257E-09 | 6.200E-09 | 1.600E-08 |
| 4860 | 243 | E243V | A | 8.5e-09 | 7.3e-09 | 7e-09 | 7.600E-09 | 7.937E-10 | 7.000E-09 | 8.500E-09 |
| 4861 | 244 | L244A | A | 4.6e-09 | 3e-09 | 6.6e-09 | 4.733E-09 | 1.804E-09 | 3.000E-09 | 6.600E-09 |
| 4862 | 244 | L244R | A | 5.7e-09 | 2.6e-08 | 1.9e-08 | 1.690E-08 | 1.031E-08 | 5.700E-09 | 2.600E-08 |
| 4863 | 244 | L244N | A | 1.9e-08 | 1.8e-08 | 2.9e-08 | 2.200E-08 | 6.083E-09 | 1.800E-08 | 2.900E-08 |
| 4864 | 244 | L244D | A | 4.2e-09 | 2.7e-09 | 5.3e-09 | 4.067E-09 | 1.305E-09 | 2.700E-09 | 5.300E-09 |
| 4865 | 244 | L244C | A | 8.9e-09 | 3.1e-09 | 4.6e-09 | 5.533E-09 | 3.011E-09 | 3.100E-09 | 8.900E-09 |
| 4866 | 244 | L244E | A | 1.6e-08 | 1e-08 | 6e-09 | 1.067E-08 | 5.033E-09 | 6.000E-09 | 1.600E-08 |
| 4867 | 244 | L244Q | A | 1.8e-08 | 4.8e-09 | 2.9e-08 | 1.727E-08 | 1.212E-08 | 4.800E-09 | 2.900E-08 |
| 4868 | 244 | L244G | A | 1.1e-08 | 8.9e-09 | 7.2e-09 | 9.033E-09 | 1.904E-09 | 7.200E-09 | 1.100E-08 |
| 4869 | 244 | L244H | A | 1.1e-08 | 9.8e-09 | 1.2e-08 | 1.093E-08 | 1.102E-09 | 9.800E-09 | 1.200E-08 |
| 4870 | 244 | L244I | A | 3e-09 | 1.7e-08 | 1.3e-09 | 7.100E-09 | 8.616E-09 | 1.300E-09 | 1.700E-08 |
| 4871 | 244 | L244L | A | 7.2e-09 | 9.5e-09 | 1.6e-08 | 1.090E-08 | 4.564E-09 | 7.200E-09 | 1.600E-08 |
| 4872 | 244 | L244K | A | 1.8e-08 | 7e-09 | 1.6e-08 | 1.367E-08 | 5.859E-09 | 7.000E-09 | 1.800E-08 |
| 4873 | 244 | L244M | A | 1.3e-08 | 4.4e-09 | 2.1e-09 | 6.500E-09 | 5.745E-09 | 2.100E-09 | 1.300E-08 |
| 4874 | 244 | L244F | A | 5.6e-09 | 3.4e-09 | 1.4e-08 | 7.667E-09 | 5.594E-09 | 3.400E-09 | 1.400E-08 |
| 4875 | 244 | L244P | A | 9.2e-09 | 4.8e-09 | 1.4e-08 | 9.333E-09 | 4.601E-09 | 4.800E-09 | 1.400E-08 |
| 4876 | 244 | L244S | A | 2.4e-08 | 1.7e-08 | 3.3e-08 | 2.467E-08 | 8.021E-09 | 1.700E-08 | 3.300E-08 |
| 4877 | 244 | L244T | A | 2e-08 | 1.3e-08 | 3.1e-08 | 2.133E-08 | 9.074E-09 | 1.300E-08 | 3.100E-08 |
| 4878 | 244 | L244W | A | 5.5e-09 | 1.7e-08 | 2e-09 | 8.167E-09 | 7.848E-09 | 2.000E-09 | 1.700E-08 |
| 4879 | 244 | L244Y | A | 1e-08 | 4e-09 | 8.5e-09 | 7.500E-09 | 3.122E-09 | 4.000E-09 | 1.000E-08 |
| 4880 | 244 | L244V | A | 8.9e-09 | 2.3e-09 | 1.6e-08 | 9.067E-09 | 6.852E-09 | 2.300E-09 | 1.600E-08 |
| 4881 | 245 | H245A | A | 1.3e-08 | 7.1e-09 | 2.9e-09 | 7.667E-09 | 5.074E-09 | 2.900E-09 | 1.300E-08 |
| 4882 | 245 | H245R | A | 1.3e-08 | 6.6e-09 | 1.4e-08 | 1.120E-08 | 4.015E-09 | 6.600E-09 | 1.400E-08 |
| 4883 | 245 | H245N | A | 2.3e-08 | 8.1e-09 | 7.9e-09 | 1.300E-08 | 8.661E-09 | 7.900E-09 | 2.300E-08 |
| 4884 | 245 | H245D | A | 6.2e-09 | 1.4e-08 | 1.7e-08 | 1.240E-08 | 5.575E-09 | 6.200E-09 | 1.700E-08 |
| 4885 | 245 | H245C | A | 7.8e-09 | 7.5e-09 | 1.4e-08 | 9.767E-09 | 3.669E-09 | 7.500E-09 | 1.400E-08 |
| 4886 | 245 | H245E | A | 1.2e-08 | 1.1e-08 | 8.8e-09 | 1.060E-08 | 1.637E-09 | 8.800E-09 | 1.200E-08 |
| 4887 | 245 | H245Q | A | 1.7e-08 | 1.4e-08 | 8.2e-09 | 1.307E-08 | 4.474E-09 | 8.200E-09 | 1.700E-08 |
| 4888 | 245 | H245G | A | 6.2e-09 | 2.2e-09 | 2.9e-09 | 3.767E-09 | 2.136E-09 | 2.200E-09 | 6.200E-09 |
| 4889 | 245 | H245H | A | 5.3e-09 | 1.4e-08 | 1.7e-08 | 1.210E-08 | 6.077E-09 | 5.300E-09 | 1.700E-08 |
| 4890 | 245 | H245I | A | 5e-09 | 7.8e-09 | 3.6e-09 | 5.467E-09 | 2.139E-09 | 3.600E-09 | 7.800E-09 |
| 4891 | 245 | H245L | A | 1.2e-09 | 6.5e-09 | 5.6e-09 | 4.433E-09 | 2.836E-09 | 1.200E-09 | 6.500E-09 |
| 4892 | 245 | H245K | A | 2.6e-09 | 8e-09 | 9.8e-09 | 6.800E-09 | 3.747E-09 | 2.600E-09 | 9.800E-09 |
| 4893 | 245 | H245M | A | 6.6e-09 | 4.1e-09 | 1.8e-09 | 4.167E-09 | 2.401E-09 | 1.800E-09 | 6.600E-09 |
| 4894 | 245 | H245F | A | 3.4e-09 | 1.3e-09 | 4.8e-09 | 3.167E-09 | 1.762E-09 | 1.300E-09 | 4.800E-09 |
| 4895 | 245 | H245P | A | 9.8e-09 | 1.1e-08 | 4.4e-09 | 8.400E-09 | 3.516E-09 | 4.400E-09 | 1.100E-08 |
| 4896 | 245 | H245S | A | 8.4e-09 | 9.2e-09 | 3.7e-08 | 1.820E-08 | 1.629E-08 | 8.400E-09 | 3.700E-08 |
| 4897 | 245 | H245T | A | 9.5e-09 | 2.2e-08 | 1.3e-08 | 1.483E-08 | 6.449E-09 | 9.500E-09 | 2.200E-08 |
| 4898 | 245 | H245W | A | 3.4e-09 | 2e-09 | 8.8e-09 | 4.733E-09 | 3.591E-09 | 2.000E-09 | 8.800E-09 |
| 4899 | 245 | H245Y | A | 2.2e-09 | 2e-09 | 2.7e-09 | 2.300E-09 | 3.606E-10 | 2.000E-09 | 2.700E-09 |
| 4900 | 245 | H245V | A | 7.7e-09 | 1e-08 | 1.5e-08 | 1.090E-08 | 3.732E-09 | 7.700E-09 | 1.500E-08 |
| 4901 | 246 | C246A | A | 7.9e-09 | 2.2e-09 | 2.3e-08 | 1.103E-08 | 1.075E-08 | 2.200E-09 | 2.300E-08 |
| 4902 | 246 | C246R | A | 1.2e-08 | 1.7e-08 | 9.4e-09 | 1.280E-08 | 3.863E-09 | 9.400E-09 | 1.700E-08 |
| 4903 | 246 | C246N | A | 2e-08 | 2.4e-08 | 4.8e-08 | 3.067E-08 | 1.514E-08 | 2.000E-08 | 4.800E-08 |
| 4904 | 246 | C246D | A | 9.2e-09 | 3.2e-08 | 2.3e-08 | 2.140E-08 | 1.148E-08 | 9.200E-09 | 3.200E-08 |
| 4905 | 246 | C246C | A | 7e-09 | 1.4e-08 | 6.6e-09 | 9.200E-09 | 4.162E-09 | 6.600E-09 | 1.400E-08 |
| 4906 | 246 | C246E | A | 5e-09 | 5e-08 | 2.1e-09 | 1.903E-08 | 2.686E-08 | 2.100E-09 | 5.000E-08 |
| 4907 | 246 | C246Q | A | 9.9e-09 | 2.8e-08 | 6.2e-08 | 3.330E-08 | 2.645E-08 | 9.900E-09 | 6.200E-08 |
| 4908 | 246 | C246G | A | 1.6e-08 | 1.1e-08 | 6.2e-09 | 1.107E-08 | 4.900E-09 | 6.200E-09 | 1.600E-08 |
| 4909 | 246 | C246H | A | 1.8e-08 | 2.6e-08 | 7.2e-08 | 3.867E-08 | 2.914E-08 | 1.800E-08 | 7.200E-08 |
| 4910 | 246 | C246I | A | 1.1e-08 | 3e-09 | 1.2e-08 | 8.667E-09 | 4.933E-09 | 3.000E-09 | 1.200E-08 |
| 4911 | 246 | C246L | A | 1.7e-09 | 3.4e-09 | 5.1e-09 | 3.400E-09 | 1.700E-09 | 1.700E-09 | 5.100E-09 |
| 4912 | 246 | C246K | A | 1.1e-08 | 5e-08 | 3e-08 | 3.033E-08 | 1.950E-08 | 1.100E-08 | 5.000E-08 |
| 4913 | 246 | C246M | A | 1.6e-08 | 1.5e-08 | 2.1e-08 | 1.733E-08 | 3.215E-09 | 1.500E-08 | 2.100E-08 |
| 4914 | 246 | C246F | A | 5.1e-09 | 2.5e-09 | 9.6e-09 | 5.733E-09 | 3.592E-09 | 2.500E-09 | 9.600E-09 |
| 4915 | 246 | C246P | A | 2.1e-08 | 1.2e-08 | 1.9e-08 | 1.733E-08 | 4.726E-09 | 1.200E-08 | 2.100E-08 |
| 4916 | 246 | C246S | A | 5.9e-08 | 8.1e-08 | 2.3e-08 | 5.433E-08 | 2.928E-08 | 2.300E-08 | 8.100E-08 |
| 4917 | 246 | C246T | A | 4.2e-08 | 4.1e-08 | 2.2e-08 | 3.500E-08 | 1.127E-08 | 2.200E-08 | 4.200E-08 |
| 4918 | 246 | C246W | A | 7.7e-09 | 8.9e-09 | 2.1e-08 | 1.253E-08 | 7.357E-09 | 7.700E-09 | 2.100E-08 |
| 4919 | 246 | C246Y | A | 6e-09 | 3.5e-08 | 5e-09 | 1.533E-08 | 1.704E-08 | 5.000E-09 | 3.500E-08 |
| 4920 | 246 | C246V | A | 1.9e-08 | 2.1e-08 | 2.3e-09 | 1.410E-08 | 1.027E-08 | 2.300E-09 | 2.100E-08 |
| 4921 | 247 | P247A | A | 6.9e-09 | 7.4e-09 | 9.7e-09 | 8.000E-09 | 1.493E-09 | 6.900E-09 | 9.700E-09 |
| 4922 | 247 | P247R | A | 1.3e-08 | 1.2e-08 | 2.4e-08 | 1.633E-08 | 6.658E-09 | 1.200E-08 | 2.400E-08 |
| 4923 | 247 | P247N | A | 2e-08 | 2.8e-08 | 1.8e-08 | 2.200E-08 | 5.292E-09 | 1.800E-08 | 2.800E-08 |
| 4924 | 247 | P247D | A | 6.8e-09 | 1.8e-08 | 7e-09 | 1.060E-08 | 6.409E-09 | 6.800E-09 | 1.800E-08 |
| 4925 | 247 | P247C | A | 1e-08 | 1.1e-08 | 1.7e-08 | 1.267E-08 | 3.786E-09 | 1.000E-08 | 1.700E-08 |
| 4926 | 247 | P247E | A | 1.1e-08 | 3.5e-09 | 6.4e-09 | 6.967E-09 | 3.782E-09 | 3.500E-09 | 1.100E-08 |
| 4927 | 247 | P247Q | A | 1.9e-08 | 9.2e-09 | 2.3e-08 | 1.707E-08 | 7.100E-09 | 9.200E-09 | 2.300E-08 |
| 4928 | 247 | P247G | A | 7.9e-09 | 6.1e-09 | 6.4e-09 | 6.800E-09 | 9.644E-10 | 6.100E-09 | 7.900E-09 |
| 4929 | 247 | P247H | A | 1.7e-08 | 3.1e-08 | 1.3e-08 | 2.033E-08 | 9.452E-09 | 1.300E-08 | 3.100E-08 |

| | | | | | | | | | | |
|---|---|---|---|---|---|---|---|---|---|---|
| 4930 | 247 | P247I | A | 1.8e-08 | 1.4e-08 | 3.5e-09 | 1.183E-08 | 7.489E-09 | 3.500E-09 | 1.800E-08 |
| 4931 | 247 | P247L | A | 2.4e-09 | 4.8e-09 | 7e-09 | 4.733E-09 | 2.301E-09 | 2.400E-09 | 7.000E-09 |
| 4932 | 247 | P247K | A | 1.2e-08 | 1.7e-08 | 1.8e-08 | 1.567E-08 | 3.215E-09 | 1.200E-08 | 1.800E-08 |
| 4933 | 247 | P247M | A | 6.1e-09 | 5e-09 | 5.4e-09 | 5.500E-09 | 5.568E-10 | 5.000E-09 | 6.100E-09 |
| 4934 | 247 | P247F | A | 1.1e-08 | 1.9e-08 | 6.6e-09 | 1.220E-08 | 6.286E-09 | 6.600E-09 | 1.900E-08 |
| 4935 | 247 | P247P | A | 7.6e-09 | 5e-09 | 5.8e-09 | 6.133E-09 | 1.332E-09 | 5.000E-09 | 7.600E-09 |
| 4936 | 247 | P247S | A | 1.8e-08 | 1.1e-08 | 1e-08 | 1.300E-08 | 4.359E-09 | 1.000E-08 | 1.800E-08 |
| 4937 | 247 | P247T | A | 1.9e-08 | 1.7e-08 | 1e-08 | 1.533E-08 | 4.726E-09 | 1.000E-08 | 1.900E-08 |
| 4938 | 247 | P247W | A | 2.3e-09 | 1.9e-09 | 1.3e-08 | 5.733E-09 | 6.296E-09 | 1.900E-09 | 1.300E-08 |
| 4939 | 247 | P247Y | A | 5e-09 | 1.2e-08 | 3e-09 | 6.667E-09 | 4.726E-09 | 3.000E-09 | 1.200E-08 |
| 4940 | 247 | P247V | A | 4.2e-09 | 5.6e-09 | 8.6e-09 | 6.133E-09 | 2.248E-09 | 4.200E-09 | 8.600E-09 |
| 4941 | 248 | A248A | A | 7.7e-09 | 8e-09 | 8.7e-09 | 8.133E-09 | 5.132E-10 | 7.700E-09 | 8.700E-09 |
| 4942 | 248 | A248R | A | 6.8e-09 | 5.8e-09 | 3.9e-09 | 5.500E-09 | 1.473E-09 | 3.900E-09 | 6.800E-09 |
| 4943 | 248 | A248N | A | 4.1e-08 | 2.4e-08 | 2e-08 | 2.833E-08 | 1.115E-08 | 2.000E-08 | 4.100E-08 |
| 4944 | 248 | A248D | A | 1.6e-08 | 1.1e-08 | 2.5e-08 | 1.733E-08 | 7.095E-09 | 1.100E-08 | 2.500E-08 |
| 4945 | 248 | A248C | A | 6.1e-09 | 7.4e-09 | 6.8e-09 | 6.767E-09 | 6.506E-10 | 6.100E-09 | 7.400E-09 |
| 4946 | 248 | A248E | A | 6.7e-09 | 1.3e-08 | 1e-08 | 9.900E-09 | 3.151E-09 | 6.700E-09 | 1.300E-08 |
| 4947 | 248 | A248Q | A | 2.2e-08 | 3.7e-08 | 1e-08 | 2.300E-08 | 1.353E-08 | 1.000E-08 | 3.700E-08 |
| 4948 | 248 | A248G | A | 1.4e-08 | 1.3e-08 | 2.2e-08 | 1.633E-08 | 4.933E-09 | 1.300E-08 | 2.200E-08 |
| 4949 | 248 | A248H | A | 7.5e-09 | 5.7e-09 | 3.5e-09 | 5.567E-09 | 2.003E-09 | 3.500E-09 | 7.500E-09 |
| 4950 | 248 | A248I | A | 6.4e-09 | 1.2e-08 | 1e-08 | 9.467E-09 | 2.838E-09 | 6.400E-09 | 1.200E-08 |
| 4951 | 248 | A248L | A | 4.1e-09 | 3e-09 | 1.9e-09 | 3.000E-09 | 1.100E-09 | 1.900E-09 | 4.100E-09 |
| 4952 | 248 | A248K | A | 1.4e-08 | 2.9e-08 | 9.6e-09 | 1.753E-08 | 1.017E-08 | 9.600E-09 | 2.900E-08 |
| 4953 | 248 | A248M | A | 1.1e-08 | 9.2e-09 | 2.2e-08 | 1.407E-08 | 6.929E-09 | 9.200E-09 | 2.200E-08 |
| 4954 | 248 | A248F | A | 1.4e-08 | 1.2e-08 | 2.4e-09 | 9.467E-09 | 6.201E-09 | 2.400E-09 | 1.400E-08 |
| 4955 | 248 | A248P | A | 9.3e-09 | 8.8e-09 | 5.3e-09 | 7.800E-09 | 2.179E-09 | 5.300E-09 | 9.300E-09 |
| 4956 | 248 | A248S | A | 1.7e-08 | 1.6e-08 | 1.2e-08 | 1.500E-08 | 2.646E-09 | 1.200E-08 | 1.700E-08 |
| 4957 | 248 | A248T | A | 4.3e-09 | 1.2e-08 | 1.7e-08 | 1.110E-08 | 6.398E-09 | 4.300E-09 | 1.700E-08 |
| 4958 | 248 | A248W | A | 1.7e-08 | 1.5e-08 | 8.1e-09 | 1.337E-08 | 4.669E-09 | 8.100E-09 | 1.700E-08 |
| 4959 | 248 | A248Y | A | 3.2e-09 | 8e-09 | 1.3e-08 | 8.067E-09 | 4.900E-09 | 3.200E-09 | 1.300E-08 |
| 4960 | 248 | A248V | A | 3.6e-09 | 1.8e-08 | 5.9e-09 | 9.167E-09 | 7.736E-09 | 3.600E-09 | 1.800E-08 |
| 4961 | 249 | L249A | A | 1.6e-08 | 5.4e-09 | 7e-09 | 9.467E-09 | 5.714E-09 | 5.400E-09 | 1.600E-08 |
| 4962 | 249 | L249R | A | 2.9e-08 | 1.5e-08 | 1.5e-08 | 1.967E-08 | 8.083E-09 | 1.500E-08 | 2.900E-08 |
| 4963 | 249 | L249N | A | 2e-08 | 3.2e-08 | 3.5e-09 | 1.850E-08 | 1.431E-08 | 3.500E-09 | 3.200E-08 |
| 4964 | 249 | L249D | A | 1.7e-08 | 4.4e-09 | 2.8e-09 | 8.067E-09 | 7.778E-09 | 2.800E-09 | 1.700E-08 |
| 4965 | 249 | L249C | A | 4.7e-09 | 6.8e-09 | 4.1e-09 | 5.200E-09 | 1.418E-09 | 4.100E-09 | 6.800E-09 |
| 4966 | 249 | L249E | A | 1.6e-08 | 1.1e-08 | 1.7e-08 | 1.467E-08 | 3.215E-09 | 1.100E-08 | 1.700E-08 |
| 4967 | 249 | L249Q | A | 3e-08 | 5.4e-08 | 1.7e-08 | 3.367E-08 | 1.877E-08 | 1.700E-08 | 5.400E-08 |
| 4968 | 249 | L249G | A | 1.8e-08 | 1.3e-08 | 5.6e-09 | 1.220E-08 | 6.239E-09 | 5.600E-09 | 1.800E-08 |
| 4969 | 249 | L249H | A | 2.3e-08 | 1.3e-08 | 9.5e-09 | 1.517E-08 | 7.006E-09 | 9.500E-09 | 2.300E-08 |
| 4970 | 249 | L249I | A | 1.2e-08 | 3.9e-09 | 1.3e-08 | 9.633E-09 | 4.990E-09 | 3.900E-09 | 1.300E-08 |
| 4971 | 249 | L249L | A | 3.1e-09 | 5.1e-09 | 9e-09 | 5.733E-09 | 3.001E-09 | 3.100E-09 | 9.000E-09 |
| 4972 | 249 | L249K | A | 2.7e-08 | 1.7e-08 | 5.6e-09 | 1.653E-08 | 1.071E-08 | 5.600E-09 | 2.700E-08 |
| 4973 | 249 | L249M | A | 1.3e-08 | 6.6e-09 | 4.5e-09 | 8.033E-09 | 4.428E-09 | 4.500E-09 | 1.300E-08 |
| 4974 | 249 | L249F | A | 4.5e-09 | 3.2e-08 | 9.6e-09 | 1.537E-08 | 1.463E-08 | 4.500E-09 | 3.200E-08 |
| 4975 | 249 | L249P | A | 2.4e-09 | 3e-09 | 4.8e-09 | 3.400E-09 | 1.249E-09 | 2.400E-09 | 4.800E-09 |
| 4976 | 249 | L249S | A | 1.2e-08 | 2.7e-08 | 5.3e-08 | 3.067E-08 | 2.074E-08 | 1.200E-08 | 5.300E-08 |
| 4977 | 249 | L249T | A | 1.1e-08 | 1.6e-08 | 2.3e-08 | 1.667E-08 | 6.028E-09 | 1.100E-08 | 2.300E-08 |
| 4978 | 249 | L249W | A | 1.3e-08 | 1.8e-08 | 3.8e-09 | 1.160E-08 | 7.203E-09 | 3.800E-09 | 1.800E-08 |
| 4979 | 249 | L249Y | A | 1.4e-09 | 1.1e-08 | 4.6e-09 | 5.667E-09 | 4.888E-09 | 1.400E-09 | 1.100E-08 |
| 4980 | 249 | L249V | A | 1.5e-08 | 4.8e-09 | 7.8e-09 | 9.200E-09 | 5.242E-09 | 4.800E-09 | 1.500E-08 |
| 4981 | 250 | V250A | A | 7.2e-09 | 6.9e-09 | 3.6e-09 | 5.900E-09 | 1.997E-09 | 3.600E-09 | 7.200E-09 |
| 4982 | 250 | V250R | A | 5.5e-09 | 9.9e-09 | 5.6e-09 | 7.000E-09 | 2.512E-09 | 5.500E-09 | 9.900E-09 |
| 4983 | 250 | V250N | A | 1.4e-08 | 5.9e-09 | 6e-09 | 8.633E-09 | 4.648E-09 | 5.900E-09 | 1.400E-08 |
| 4984 | 250 | V250D | A | 1.6e-08 | 7.1e-09 | 2.8e-09 | 8.633E-09 | 6.732E-09 | 2.800E-09 | 1.600E-08 |
| 4985 | 250 | V250C | A | 5.6e-09 | 4.7e-09 | 6.3e-09 | 5.533E-09 | 8.021E-10 | 4.700E-09 | 6.300E-09 |
| 4986 | 250 | V250E | A | 5.1e-08 | 9.3e-09 | 1.9e-09 | 2.073E-08 | 2.647E-08 | 1.900E-09 | 5.100E-08 |
| 4987 | 250 | V250Q | A | 1.5e-08 | 1.3e-08 | 1.2e-08 | 1.333E-08 | 1.528E-09 | 1.200E-08 | 1.500E-08 |
| 4988 | 250 | V250G | A | 7.6e-09 | 5.8e-09 | 8.7e-09 | 7.367E-09 | 1.464E-09 | 5.800E-09 | 8.700E-09 |
| 4989 | 250 | V250H | A | 9.3e-09 | 9.8e-09 | 3.3e-09 | 7.467E-09 | 3.617E-09 | 3.300E-09 | 9.800E-09 |
| 4990 | 250 | V250I | A | 1.1e-08 | 5.4e-09 | 1.7e-08 | 1.113E-08 | 5.801E-09 | 5.400E-09 | 1.700E-08 |
| 4991 | 250 | V250L | A | 4.6e-09 | 2.4e-09 | 1.6e-09 | 2.867E-09 | 1.553E-09 | 1.600E-09 | 4.600E-09 |
| 4992 | 250 | V250K | A | 8.9e-09 | 7.2e-09 | 3.7e-09 | 6.600E-09 | 2.651E-09 | 3.700E-09 | 8.900E-09 |
| 4993 | 250 | V250M | A | 8.5e-09 | 1e-08 | 6.9e-09 | 8.467E-09 | 1.550E-09 | 6.900E-09 | 1.000E-08 |
| 4994 | 250 | V250F | A | 2.4e-08 | 1.1e-08 | 3.1e-09 | 1.270E-08 | 1.055E-08 | 3.100E-09 | 2.400E-08 |
| 4995 | 250 | V250P | A | 1.4e-08 | 3.9e-09 | 6.3e-09 | 8.067E-09 | 5.277E-09 | 3.900E-09 | 1.400E-08 |
| 4996 | 250 | V250S | A | 6.7e-09 | 1.2e-08 | 3.6e-09 | 7.433E-09 | 4.248E-09 | 3.600E-09 | 1.200E-08 |
| 4997 | 250 | V250T | A | 2e-08 | 5.3e-09 | 6e-09 | 1.043E-08 | 8.292E-09 | 5.300E-09 | 2.000E-08 |
| 4998 | 250 | V250W | A | 5.4e-09 | 6.3e-09 | 1e-08 | 7.233E-09 | 2.438E-09 | 5.400E-09 | 1.000E-08 |
| 4999 | 250 | V250Y | A | 2.5e-09 | 4.5e-09 | 1.3e-09 | 2.767E-09 | 1.617E-09 | 1.300E-09 | 4.500E-09 |
| 5000 | 250 | V250V | A | 8.1e-09 | 4.7e-09 | 1.1e-08 | 7.933E-09 | 3.153E-09 | 4.700E-09 | 1.100E-08 |
| 5001 | 251 | T251A | A | 6.5e-09 | 4.8e-09 | 1e-08 | 7.100E-09 | 2.651E-09 | 4.800E-09 | 1.000E-08 |
| 5002 | 251 | T251R | A | 5.3e-09 | 2.3e-08 | 1.6e-08 | 1.477E-08 | 8.914E-09 | 5.300E-09 | 2.300E-08 |
| 5003 | 251 | T251N | A | 1.7e-09 | 8.1e-09 | 7.7e-09 | 5.833E-09 | 3.585E-09 | 1.700E-09 | 8.100E-09 |
| 5004 | 251 | T251D | A | 4.8e-09 | 3.4e-09 | 5.6e-09 | 4.600E-09 | 1.114E-09 | 3.400E-09 | 5.600E-09 |
| 5005 | 251 | T251C | A | 1.5e-08 | 2.8e-09 | 8.2e-09 | 8.667E-09 | 6.113E-09 | 2.800E-09 | 1.500E-08 |
| 5006 | 251 | T251E | A | 5.5e-09 | 3.7e-09 | 7.1e-09 | 5.433E-09 | 1.701E-09 | 3.700E-09 | 7.100E-09 |
| 5007 | 251 | T251Q | A | 9e-09 | 7.4e-09 | 1.2e-08 | 9.467E-09 | 2.335E-09 | 7.400E-09 | 1.200E-08 |
| 5008 | 251 | T251G | A | 5.9e-09 | 1e-08 | 6.2e-09 | 7.367E-09 | 2.285E-09 | 5.900E-09 | 1.000E-08 |
| 5009 | 251 | T251H | A | 1.5e-09 | 3.4e-09 | 5.6e-09 | 3.500E-09 | 2.052E-09 | 1.500E-09 | 5.600E-09 |
| 5010 | 251 | T251I | A | 1.4e-08 | 3.2e-08 | 7.3e-09 | 1.777E-08 | 1.277E-08 | 7.300E-09 | 3.200E-08 |
| 5011 | 251 | T251L | A | 1.2e-08 | 1.7e-08 | 1.5e-08 | 1.467E-08 | 2.517E-09 | 1.200E-08 | 1.700E-08 |
| 5012 | 251 | T251K | A | 8.1e-09 | 2.3e-08 | 7.4e-09 | 1.283E-08 | 8.812E-09 | 7.400E-09 | 2.300E-08 |
| 5013 | 251 | T251M | A | 7e-09 | 1.5e-08 | 1.8e-08 | 1.333E-08 | 5.686E-09 | 7.000E-09 | 1.800E-08 |
| 5014 | 251 | T251F | A | 8.1e-09 | 3e-09 | 6.8e-09 | 5.967E-09 | 2.650E-09 | 3.000E-09 | 8.100E-09 |

| | | | | | | | | | | |
|---|---|---|---|---|---|---|---|---|---|---|
| 5015 | 251 | T251P | A | 1.2e-08 | 5.2e-09 | 5.9e-09 | 7.700E-09 | 3.740E-09 | 5.200E-09 | 1.200E-08 |
| 5016 | 251 | T251S | A | 1e-08 | 5.1e-09 | 1.1e-08 | 8.700E-09 | 3.158E-09 | 5.100E-09 | 1.100E-08 |
| 5017 | 251 | T251T | A | 2.9e-09 | 1e-08 | 3.1e-09 | 5.333E-09 | 4.043E-09 | 2.900E-09 | 1.000E-08 |
| 5018 | 251 | T251W | A | 1.4e-09 | 2.2e-08 | 3.5e-09 | 8.967E-09 | 1.134E-08 | 1.400E-09 | 2.200E-08 |
| 5019 | 251 | T251Y | A | 2.3e-09 | 5.6e-09 | 3.1e-10 | 2.737E-09 | 2.672E-09 | 3.100E-10 | 5.600E-09 |
| 5020 | 251 | T251V | A | 6.6e-09 | 3.1e-09 | 6.3e-09 | 5.333E-09 | 1.940E-09 | 3.100E-09 | 6.600E-09 |
| 5021 | 252 | Y252A | A | 1.4e-08 | 1.1e-08 | 2.1e-08 | 1.533E-08 | 5.132E-09 | 1.100E-08 | 2.100E-08 |
| 5022 | 252 | Y252R | A | 4.7e-09 | 3.3e-09 | 4.2e-09 | 4.067E-09 | 7.095E-10 | 3.300E-09 | 4.700E-09 |
| 5023 | 252 | Y252N | A | 4.5e-09 | 2.9e-09 | 3.5e-09 | 3.633E-09 | 8.083E-10 | 2.900E-09 | 4.500E-09 |
| 5024 | 252 | Y252D | A | 1.2e-08 | 3.3e-09 | 5.8e-09 | 7.033E-09 | 4.479E-09 | 3.300E-09 | 1.200E-08 |
| 5025 | 252 | Y252C | A | 1.8e-08 | 2.6e-09 | 9.7e-10 | 7.190E-09 | 9.397E-09 | 9.700E-10 | 1.800E-08 |
| 5026 | 252 | Y252E | A | 5e-09 | 5.1e-09 | 6.2e-09 | 5.433E-09 | 6.658E-10 | 5.000E-09 | 6.200E-09 |
| 5027 | 252 | Y252Q | A | 5.6e-09 | 2.1e-09 | 2e-09 | 3.233E-09 | 2.050E-09 | 2.000E-09 | 5.600E-09 |
| 5028 | 252 | Y252G | A | 4.2e-09 | 4e-09 | 5.4e-09 | 4.533E-09 | 7.572E-10 | 4.000E-09 | 5.400E-09 |
| 5029 | 252 | Y252H | A | 1.1e-08 | 4.2e-09 | 9e-09 | 8.067E-09 | 3.495E-09 | 4.200E-09 | 1.100E-08 |
| 5030 | 252 | Y252I | A | 2.7e-09 | 7.5e-09 | 4.5e-09 | 4.900E-09 | 2.425E-09 | 2.700E-09 | 7.500E-09 |
| 5031 | 252 | Y252L | A | 9.3e-09 | 1.1e-08 | 4.4e-09 | 8.233E-09 | 3.427E-09 | 4.400E-09 | 1.100E-08 |
| 5032 | 252 | Y252K | A | 5.5e-09 | 4.3e-09 | 4.8e-09 | 4.867E-09 | 6.028E-10 | 4.300E-09 | 5.500E-09 |
| 5033 | 252 | Y252M | A | 3.1e-09 | 2.3e-09 | 6.8e-09 | 4.067E-09 | 2.401E-09 | 2.300E-09 | 6.800E-09 |
| 5034 | 252 | Y252F | A | 3.5e-09 | 1.5e-08 | 9.1e-09 | 9.200E-09 | 5.751E-09 | 3.500E-09 | 1.500E-08 |
| 5035 | 252 | Y252P | A | 5.7e-09 | 5.1e-09 | 1.6e-08 | 8.933E-09 | 6.127E-09 | 5.100E-09 | 1.600E-08 |
| 5036 | 252 | Y252S | A | 6.2e-09 | 8.3e-09 | 6.7e-09 | 7.067E-09 | 1.097E-09 | 6.200E-09 | 8.300E-09 |
| 5037 | 252 | Y252T | A | 1.8e-09 | 3.9e-09 | 2.8e-09 | 2.833E-09 | 1.050E-09 | 1.800E-09 | 3.900E-09 |
| 5038 | 252 | Y252W | A | 7.1e-09 | 2.5e-09 | 1.9e-08 | 9.533E-09 | 8.515E-09 | 2.500E-09 | 1.900E-08 |
| 5039 | 252 | Y252Y | A | 1.1e-08 | 1.1e-08 | 1.2e-08 | 1.133E-08 | 5.774E-10 | 1.100E-08 | 1.200E-08 |
| 5040 | 252 | Y252V | A | 5.2e-09 | 4.7e-09 | 7.9e-09 | 5.933E-09 | 1.721E-09 | 4.700E-09 | 7.900E-09 |
| 5041 | 253 | N253A | A | 4.6e-09 | 6.3e-09 | 5.1e-09 | 5.333E-09 | 8.737E-10 | 4.600E-09 | 6.300E-09 |
| 5042 | 253 | N253R | A | 1e-08 | 6.2e-09 | 1.7e-08 | 1.107E-08 | 5.478E-09 | 6.200E-09 | 1.700E-08 |
| 5043 | 253 | N253N | A | 6.8e-09 | 9.1e-09 | 7.9e-09 | 7.933E-09 | 1.150E-09 | 6.800E-09 | 9.100E-09 |
| 5044 | 253 | N253D | A | 5.8e-09 | 1e-08 | 1.2e-08 | 9.267E-09 | 3.164E-09 | 5.800E-09 | 1.200E-08 |
| 5045 | 253 | N253C | A | 1.2e-09 | 1.3e-08 | 6.3e-09 | 6.833E-09 | 5.918E-09 | 1.200E-09 | 1.300E-08 |
| 5046 | 253 | N253E | A | 2.4e-08 | 4.8e-09 | 1.2e-08 | 1.360E-08 | 9.699E-09 | 4.800E-09 | 2.400E-08 |
| 5047 | 253 | N253Q | A | 5.7e-09 | 3e-08 | 8.9e-09 | 1.487E-08 | 1.320E-08 | 5.700E-09 | 3.000E-08 |
| 5048 | 253 | N253G | A | 2.1e-08 | 8.8e-09 | 3e-09 | 1.093E-08 | 9.188E-09 | 3.000E-09 | 2.100E-08 |
| 5049 | 253 | N253H | A | 1.3e-08 | 2.2e-08 | 4.8e-09 | 1.327E-08 | 8.603E-09 | 4.800E-09 | 2.200E-08 |
| 5050 | 253 | N253I | A | 7.8e-09 | 1.4e-08 | 8e-09 | 9.933E-09 | 3.523E-09 | 7.800E-09 | 1.400E-08 |
| 5051 | 253 | N253L | A | 3.4e-09 | 1.5e-08 | 1.1e-08 | 9.800E-09 | 5.892E-09 | 3.400E-09 | 1.500E-08 |
| 5052 | 253 | N253K | A | 9.4e-09 | 5.7e-09 | 1.2e-08 | 9.033E-09 | 3.166E-09 | 5.700E-09 | 1.200E-08 |
| 5053 | 253 | N253M | A | 8.2e-09 | 6.5e-09 | 3.2e-08 | 1.557E-08 | 1.426E-08 | 6.500E-09 | 3.200E-08 |
| 5054 | 253 | N253F | A | 5.1e-09 | 7.7e-09 | 1.2e-08 | 8.267E-09 | 3.485E-09 | 5.100E-09 | 1.200E-08 |
| 5055 | 253 | N253P | A | 2.6e-08 | 4.9e-09 | 4e-09 | 1.163E-08 | 1.245E-08 | 4.000E-09 | 2.600E-08 |
| 5056 | 253 | N253S | A | 3.2e-09 | 1.5e-08 | 1.1e-08 | 9.733E-09 | 6.001E-09 | 3.200E-09 | 1.500E-08 |
| 5057 | 253 | N253T | A | 3.1e-09 | 3.2e-09 | 2.6e-08 | 1.077E-08 | 1.319E-08 | 3.100E-09 | 2.600E-08 |
| 5058 | 253 | N253W | A | 6.3e-09 | 5.6e-09 | 6.9e-09 | 6.267E-09 | 6.506E-10 | 5.600E-09 | 6.900E-09 |
| 5059 | 253 | N253Y | A | 1.5e-08 | 5.4e-09 | 5.6e-09 | 8.667E-09 | 5.486E-09 | 5.400E-09 | 1.500E-08 |
| 5060 | 253 | N253V | A | 1.3e-08 | 1.1e-08 | 4.8e-09 | 9.600E-09 | 4.276E-09 | 4.800E-09 | 1.300E-08 |
| 5061 | 254 | T254A | A | 3.2e-09 | 2.7e-09 | 2e-09 | 2.633E-09 | 6.028E-10 | 2.000E-09 | 3.200E-09 |
| 5062 | 254 | T254R | A | 1.6e-08 | 2.3e-09 | 2.6e-09 | 6.967E-09 | 7.825E-09 | 2.300E-09 | 1.600E-08 |
| 5063 | 254 | T254N | A | 9.8e-09 | 8.5e-09 | 5e-09 | 7.767E-09 | 2.483E-09 | 5.000E-09 | 9.800E-09 |
| 5064 | 254 | T254D | A | 6.9e-09 | 3.8e-09 | 7.6e-09 | 6.100E-09 | 2.022E-09 | 3.800E-09 | 7.600E-09 |
| 5065 | 254 | T254C | A | 2.4e-09 | 3e-09 | 6.1e-09 | 3.833E-09 | 1.986E-09 | 2.400E-09 | 6.100E-09 |
| 5066 | 254 | T254E | A | 9.9e-09 | 1.1e-08 | 1.5e-09 | 7.467E-09 | 5.196E-09 | 1.500E-09 | 1.100E-08 |
| 5067 | 254 | T254Q | A | 6.5e-09 | 3.3e-09 | 7e-09 | 5.600E-09 | 2.007E-09 | 3.300E-09 | 7.000E-09 |
| 5068 | 254 | T254G | A | 3.5e-09 | 1.8e-09 | 5.6e-09 | 3.633E-09 | 1.904E-09 | 1.800E-09 | 5.600E-09 |
| 5069 | 254 | T254H | A | 2.6e-09 | 3.6e-09 | 1.8e-09 | 2.667E-09 | 9.018E-10 | 1.800E-09 | 3.600E-09 |
| 5070 | 254 | T254I | A | 3.6e-09 | 2e-09 | 6e-09 | 3.867E-09 | 2.013E-09 | 2.000E-09 | 6.000E-09 |
| 5071 | 254 | T254L | A | 2.6e-09 | 1e-09 | 2.6e-09 | 2.067E-09 | 9.238E-10 | 1.000E-09 | 2.600E-09 |
| 5072 | 254 | T254K | A | 2.3e-09 | 4.1e-09 | 3.4e-09 | 3.267E-09 | 9.074E-10 | 2.300E-09 | 4.100E-09 |
| 5073 | 254 | T254M | A | 2.1e-09 | 4.1e-09 | 6.6e-09 | 4.267E-09 | 2.255E-09 | 2.100E-09 | 6.600E-09 |
| 5074 | 254 | T254F | A | 1.1e-09 | 3.9e-09 | 8e-09 | 4.333E-09 | 3.470E-09 | 1.100E-09 | 8.000E-09 |
| 5075 | 254 | T254P | A | 2.6e-09 | 4.7e-09 | 4.5e-09 | 3.933E-09 | 1.159E-09 | 2.600E-09 | 4.700E-09 |
| 5076 | 254 | T254S | A | 6.6e-09 | 7.8e-09 | 2e-08 | 1.147E-08 | 7.414E-09 | 6.600E-09 | 2.000E-08 |
| 5077 | 254 | T254T | A | 1.1e-08 | 3.3e-09 | 9.8e-09 | 8.033E-09 | 4.143E-09 | 3.300E-09 | 1.100E-08 |
| 5078 | 254 | T254W | A | 3.4e-09 | 3.6e-09 | 2.5e-09 | 3.167E-09 | 5.859E-10 | 2.500E-09 | 3.600E-09 |
| 5079 | 254 | T254Y | A | 3.6e-09 | 4.8e-09 | 7e-09 | 5.133E-09 | 1.724E-09 | 3.600E-09 | 7.000E-09 |
| 5080 | 254 | T254V | A | 2.4e-09 | 1.2e-08 | 4.3e-09 | 6.233E-09 | 5.084E-09 | 2.400E-09 | 1.200E-08 |
| 5081 | 255 | D255A | A | 1.2e-08 | 3.9e-09 | 3.1e-09 | 6.333E-09 | 4.924E-09 | 3.100E-09 | 1.200E-08 |
| 5082 | 255 | D255R | A | 8.5e-09 | 5.2e-09 | 5.9e-09 | 6.533E-09 | 1.739E-09 | 5.200E-09 | 8.500E-09 |
| 5083 | 255 | D255N | A | 5e-08 | 6.4e-09 | 4.6e-09 | 2.033E-08 | 2.571E-08 | 4.600E-09 | 5.000E-08 |
| 5084 | 255 | D255D | A | 4.4e-09 | 9.5e-09 | 2.5e-09 | 5.467E-09 | 3.620E-09 | 2.500E-09 | 9.500E-09 |
| 5085 | 255 | D255C | A | 4.2e-09 | 1.6e-08 | 1e-08 | 1.007E-08 | 5.900E-09 | 4.200E-09 | 1.600E-08 |
| 5086 | 255 | D255E | A | 9.8e-09 | 9.5e-09 | 7.6e-09 | 8.967E-09 | 1.193E-09 | 7.600E-09 | 9.800E-09 |
| 5087 | 255 | D255Q | A | 6.6e-09 | 8.9e-09 | 1.4e-08 | 9.833E-09 | 3.787E-09 | 6.600E-09 | 1.400E-08 |
| 5088 | 255 | D255G | A | 1.8e-09 | 1.1e-08 | 4.7e-09 | 5.833E-09 | 4.704E-09 | 1.800E-09 | 1.100E-08 |
| 5089 | 255 | D255H | A | 2.1e-09 | 1.3e-08 | 6.7e-09 | 7.267E-09 | 5.472E-09 | 2.100E-09 | 1.300E-08 |
| 5090 | 255 | D255I | A | 8.2e-09 | 4.5e-09 | 2.6e-09 | 5.100E-09 | 2.848E-09 | 2.600E-09 | 8.200E-09 |
| 5091 | 255 | D255L | A | 9.9e-09 | 2.4e-09 | 1.5e-08 | 9.100E-09 | 6.338E-09 | 2.400E-09 | 1.500E-08 |
| 5092 | 255 | D255K | A | 3.9e-09 | 5.7e-09 | 8.5e-09 | 6.033E-09 | 2.318E-09 | 3.900E-09 | 8.500E-09 |
| 5093 | 255 | D255M | A | 6.8e-09 | 7e-09 | 1.1e-08 | 8.267E-09 | 2.369E-09 | 6.800E-09 | 1.100E-08 |
| 5094 | 255 | D255F | A | 6e-09 | 1.2e-08 | 3.3e-09 | 7.100E-09 | 4.453E-09 | 3.300E-09 | 1.200E-08 |
| 5095 | 255 | D255P | A | 6.1e-09 | 1.6e-08 | 3.1e-09 | 8.400E-09 | 6.751E-09 | 3.100E-09 | 1.600E-08 |
| 5096 | 255 | D255S | A | 1.2e-08 | 6.3e-09 | 1.7e-08 | 1.177E-08 | 5.354E-09 | 6.300E-09 | 1.700E-08 |
| 5097 | 255 | D255T | A | 1.8e-08 | 8.4e-09 | 4.3e-09 | 1.023E-08 | 7.032E-09 | 4.300E-09 | 1.800E-08 |
| 5098 | 255 | D255W | A | 1.7e-08 | 9.7e-09 | 6.2e-09 | 1.097E-08 | 5.510E-09 | 6.200E-09 | 1.700E-08 |
| 5099 | 255 | D255Y | A | 7.4e-09 | 2.4e-09 | 4.4e-09 | 4.733E-09 | 2.517E-09 | 2.400E-09 | 7.400E-09 |

| | | | | | | | | | | |
|---|---|---|---|---|---|---|---|---|---|---|
| 5100 | 255 | D255V | A | 4e-09 | 1.4e-08 | 3.4e-09 | 7.133E-09 | 5.954E-09 | 3.400E-09 | 1.400E-08 |
| 5101 | 256 | T256A | A | 7.7e-09 | 1.6e-08 | 8.7e-09 | 1.080E-08 | 4.531E-09 | 7.700E-09 | 1.600E-08 |
| 5102 | 256 | T256R | A | 7.5e-09 | 7.6e-09 | 9.2e-09 | 8.100E-09 | 9.539E-10 | 7.500E-09 | 9.200E-09 |
| 5103 | 256 | T256N | A | 1.2e-08 | 6.6e-09 | 1.3e-08 | 1.053E-08 | 3.443E-09 | 6.600E-09 | 1.300E-08 |
| 5104 | 256 | T256D | A | 2.5e-09 | 1.7e-08 | 1.1e-08 | 1.017E-08 | 7.286E-09 | 2.500E-09 | 1.700E-08 |
| 5105 | 256 | T256C | A | 3.6e-09 | 6.3e-09 | 1.8e-08 | 9.300E-09 | 7.654E-09 | 3.600E-09 | 1.800E-08 |
| 5106 | 256 | T256E | A | 3.5e-09 | 7e-09 | 9.4e-09 | 6.633E-09 | 2.967E-09 | 3.500E-09 | 9.400E-09 |
| 5107 | 256 | T256Q | A | 9.8e-09 | 1.9e-08 | 1.1e-08 | 1.327E-08 | 5.001E-09 | 9.800E-09 | 1.900E-08 |
| 5108 | 256 | T256G | A | 6e-09 | 5e-09 | 4.7e-09 | 5.233E-09 | 6.807E-10 | 4.700E-09 | 6.000E-09 |
| 5109 | 256 | T256H | A | 6e-09 | 4e-09 | 1.4e-08 | 8.000E-09 | 5.292E-09 | 4.000E-09 | 1.400E-08 |
| 5110 | 256 | T256I | A | 4.2e-09 | 4.4e-09 | 4e-09 | 4.200E-09 | 2.000E-10 | 4.000E-09 | 4.400E-09 |
| 5111 | 256 | T256L | A | 4.8e-09 | 8.7e-09 | 9.1e-09 | 7.533E-09 | 2.376E-09 | 4.800E-09 | 9.100E-09 |
| 5112 | 256 | T256K | A | 8.2e-09 | 1.2e-08 | 7e-09 | 9.067E-09 | 2.610E-09 | 7.000E-09 | 1.200E-08 |
| 5113 | 256 | T256M | A | 6.9e-09 | 2.7e-09 | 5.2e-09 | 4.933E-09 | 2.113E-09 | 2.700E-09 | 6.900E-09 |
| 5114 | 256 | T256F | A | 9e-09 | 2e-09 | 4.1e-09 | 5.033E-09 | 3.592E-09 | 2.000E-09 | 9.000E-09 |
| 5115 | 256 | T256P | A | 2.9e-09 | 4.3e-09 | 5.6e-09 | 4.267E-09 | 1.350E-09 | 2.900E-09 | 5.600E-09 |
| 5116 | 256 | T256S | A | 6.4e-09 | 8.8e-09 | 3.8e-09 | 6.333E-09 | 2.501E-09 | 3.800E-09 | 8.800E-09 |
| 5117 | 256 | T256T | A | 2.2e-09 | 5.1e-09 | 6.5e-09 | 4.600E-09 | 2.193E-09 | 2.200E-09 | 6.500E-09 |
| 5118 | 256 | T256W | A | 1.2e-08 | 4.7e-09 | 3.1e-08 | 1.590E-08 | 1.358E-08 | 4.700E-09 | 3.100E-08 |
| 5119 | 256 | T256Y | A | 4.3e-09 | 5.8e-09 | 3.7e-09 | 4.600E-09 | 1.082E-09 | 3.700E-09 | 5.800E-09 |
| 5120 | 256 | T256V | A | 2.4e-09 | 5.5e-09 | 9.5e-09 | 5.800E-09 | 3.559E-09 | 2.400E-09 | 9.500E-09 |
| 5121 | 257 | F257A | A | 2e-08 | 9.1e-09 | 1.4e-08 | 1.437E-08 | 5.459E-09 | 9.100E-09 | 2.000E-08 |
| 5122 | 257 | F257R | A | 1.4e-08 | 1.6e-08 | 9.7e-09 | 1.323E-08 | 3.219E-09 | 9.700E-09 | 1.600E-08 |
| 5123 | 257 | F257N | A | 8.9e-09 | 5.2e-09 | 3.7e-09 | 5.933E-09 | 2.676E-09 | 3.700E-09 | 8.900E-09 |
| 5124 | 257 | F257D | A | 1.3e-08 | 2.7e-09 | 1.6e-08 | 1.057E-08 | 6.976E-09 | 2.700E-09 | 1.600E-08 |
| 5125 | 257 | F257C | A | 1.7e-08 | 8.2e-09 | 1.3e-08 | 1.273E-08 | 4.406E-09 | 8.200E-09 | 1.700E-08 |
| 5126 | 257 | F257E | A | 3.1e-09 | 9.1e-09 | 6e-09 | 6.067E-09 | 3.001E-09 | 3.100E-09 | 9.100E-09 |
| 5127 | 257 | F257Q | A | 3e-08 | 1.1e-08 | 4.9e-09 | 1.530E-08 | 1.309E-08 | 4.900E-09 | 3.000E-08 |
| 5128 | 257 | F257G | A | 1.5e-08 | 6.2e-09 | 1.3e-08 | 1.140E-08 | 4.613E-09 | 6.200E-09 | 1.500E-08 |
| 5129 | 257 | F257H | A | 3.2e-09 | 1.8e-08 | 6.9e-09 | 9.367E-09 | 7.702E-09 | 3.200E-09 | 1.800E-08 |
| 5130 | 257 | F257I | A | 1.1e-08 | 2e-08 | 1.9e-08 | 1.667E-08 | 4.933E-09 | 1.100E-08 | 2.000E-08 |
| 5131 | 257 | F257L | A | 2.8e-08 | 1.1e-08 | 1.1e-08 | 1.667E-08 | 9.815E-09 | 1.100E-08 | 2.800E-08 |
| 5132 | 257 | F257K | A | 1e-08 | 1.6e-08 | 7.8e-09 | 1.127E-08 | 4.244E-09 | 7.800E-09 | 1.600E-08 |
| 5133 | 257 | F257M | A | 2.2e-08 | 3.5e-09 | 5.9e-09 | 1.047E-08 | 1.006E-08 | 3.500E-09 | 2.200E-08 |
| 5134 | 257 | F257F | A | 7.8e-09 | 8.7e-09 | 9.1e-09 | 8.533E-09 | 6.658E-10 | 7.800E-09 | 9.100E-09 |
| 5135 | 257 | F257P | A | 6.2e-09 | 1e-08 | 7.3e-09 | 7.833E-09 | 1.955E-09 | 6.200E-09 | 1.000E-08 |
| 5136 | 257 | F257S | A | 5.2e-09 | 7.3e-09 | 8e-09 | 6.833E-09 | 1.457E-09 | 5.200E-09 | 8.000E-09 |
| 5137 | 257 | F257T | A | 1.1e-08 | 9.8e-09 | 7.1e-09 | 9.300E-09 | 1.997E-09 | 7.100E-09 | 1.100E-08 |
| 5138 | 257 | F257W | A | 4.1e-09 | 5.4e-09 | 5.7e-09 | 5.067E-09 | 8.505E-10 | 4.100E-09 | 5.700E-09 |
| 5139 | 257 | F257Y | A | 4.5e-09 | 1.1e-08 | 1.5e-08 | 1.017E-08 | 5.299E-09 | 4.500E-09 | 1.500E-08 |
| 5140 | 257 | F257V | A | 7.5e-09 | 1e-08 | 5.5e-09 | 7.667E-09 | 2.255E-09 | 5.500E-09 | 1.000E-08 |
| 5141 | 258 | E258A | A | 1.4e-08 | 7.8e-09 | 7.3e-09 | 9.700E-09 | 3.732E-09 | 7.300E-09 | 1.400E-08 |
| 5142 | 258 | E258R | A | 6.3e-09 | 1e-08 | 4.8e-09 | 7.033E-09 | 2.676E-09 | 4.800E-09 | 1.000E-08 |
| 5143 | 258 | E258N | A | 3.7e-09 | 7.7e-09 | 2.5e-09 | 4.633E-09 | 2.723E-09 | 2.500E-09 | 7.700E-09 |
| 5144 | 258 | E258D | A | 1.1e-08 | 8.8e-09 | 5.1e-09 | 8.300E-09 | 2.982E-09 | 5.100E-09 | 1.100E-08 |
| 5145 | 258 | E258C | A | 8.7e-09 | 7e-09 | 9.9e-09 | 8.533E-09 | 1.457E-09 | 7.000E-09 | 9.900E-09 |
| 5146 | 258 | E258E | A | 1e-08 | 5.2e-09 | 8e-09 | 7.733E-09 | 2.411E-09 | 5.200E-09 | 1.000E-08 |
| 5147 | 258 | E258Q | A | 2.2e-08 | 6.6e-09 | 7.7e-09 | 1.210E-08 | 8.591E-09 | 6.600E-09 | 2.200E-08 |
| 5148 | 258 | E258G | A | 6.9e-09 | 4e-09 | 4.1e-09 | 5.000E-09 | 1.646E-09 | 4.000E-09 | 6.900E-09 |
| 5149 | 258 | E258H | A | 1e-08 | 4e-09 | 5.4e-09 | 6.467E-09 | 3.139E-09 | 4.000E-09 | 1.000E-08 |
| 5150 | 258 | E258I | A | 7.5e-09 | 4.7e-09 | 8.8e-09 | 7.000E-09 | 2.095E-09 | 4.700E-09 | 8.800E-09 |
| 5151 | 258 | E258L | A | 9.1e-09 | 1.3e-09 | 1.2e-08 | 7.467E-09 | 5.534E-09 | 1.300E-09 | 1.200E-08 |
| 5152 | 258 | E258K | A | 1.4e-08 | 3.7e-09 | 1.1e-08 | 9.567E-09 | 5.297E-09 | 3.700E-09 | 1.400E-08 |
| 5153 | 258 | E258M | A | 1.4e-08 | 1.3e-08 | 1.5e-08 | 1.400E-08 | 1.000E-09 | 1.300E-08 | 1.500E-08 |
| 5154 | 258 | E258F | A | 5.6e-09 | 1.2e-08 | 5.6e-09 | 7.733E-09 | 3.695E-09 | 5.600E-09 | 1.200E-08 |
| 5155 | 258 | E258P | A | 4.6e-09 | 6.1e-09 | 2.9e-08 | 1.323E-08 | 1.367E-08 | 4.600E-09 | 2.900E-08 |
| 5156 | 258 | E258S | A | 9.4e-09 | 3.7e-09 | 9.6e-09 | 7.567E-09 | 3.350E-09 | 3.700E-09 | 9.600E-09 |
| 5157 | 258 | E258T | A | 3.2e-09 | 1.9e-08 | 7.3e-09 | 9.833E-09 | 8.199E-09 | 3.200E-09 | 1.900E-08 |
| 5158 | 258 | E258W | A | 3.7e-09 | 7.6e-09 | 1.5e-09 | 4.267E-09 | 3.089E-09 | 1.500E-09 | 7.600E-09 |
| 5159 | 258 | E258Y | A | 4.7e-09 | 5.5e-09 | 3.3e-09 | 4.500E-09 | 1.114E-09 | 3.300E-09 | 5.500E-09 |
| 5160 | 258 | E258V | A | 2.6e-09 | 7.2e-09 | 3.9e-09 | 4.567E-09 | 2.371E-09 | 2.600E-09 | 7.200E-09 |
| 5161 | 259 | S259A | A | 1.8e-08 | 5.7e-09 | 7e-09 | 1.023E-08 | 6.757E-09 | 5.700E-09 | 1.800E-08 |
| 5162 | 259 | S259R | A | 2e-08 | 1.1e-08 | 2.4e-09 | 1.113E-08 | 8.801E-09 | 2.400E-09 | 2.000E-08 |
| 5163 | 259 | S259N | A | 1.3e-08 | 1.2e-08 | 4.4e-09 | 9.800E-09 | 4.703E-09 | 4.400E-09 | 1.300E-08 |
| 5164 | 259 | S259D | A | 8e-09 | 7.5e-09 | 7.7e-09 | 7.733E-09 | 2.517E-10 | 7.500E-09 | 8.000E-09 |
| 5165 | 259 | S259C | A | 9.8e-09 | 9.3e-09 | 1.6e-08 | 1.170E-08 | 3.732E-09 | 9.300E-09 | 1.600E-08 |
| 5166 | 259 | S259E | A | 2.6e-08 | 7.7e-09 | 6.8e-09 | 1.350E-08 | 1.083E-08 | 6.800E-09 | 2.600E-08 |
| 5167 | 259 | S259Q | A | 2.5e-09 | 3.1e-09 | 9.4e-09 | 5.000E-09 | 3.822E-09 | 2.500E-09 | 9.400E-09 |
| 5168 | 259 | S259G | A | 6.5e-09 | 1.1e-08 | 7.4e-09 | 8.300E-09 | 2.381E-09 | 6.500E-09 | 1.100E-08 |
| 5169 | 259 | S259H | A | 4.4e-09 | 1.6e-08 | 6.4e-09 | 8.933E-09 | 6.201E-09 | 4.400E-09 | 1.600E-08 |
| 5170 | 259 | S259I | A | 7.6e-09 | 4.1e-09 | 1e-08 | 7.233E-09 | 2.967E-09 | 4.100E-09 | 1.000E-08 |
| 5171 | 259 | S259L | A | 2.8e-09 | 7.2e-09 | 4.9e-09 | 4.967E-09 | 2.201E-09 | 2.800E-09 | 7.200E-09 |
| 5172 | 259 | S259K | A | 3.6e-09 | 2.5e-09 | 1.2e-08 | 6.033E-09 | 5.196E-09 | 2.500E-09 | 1.200E-08 |
| 5173 | 259 | S259M | A | 7.2e-09 | 5.8e-09 | 5.1e-09 | 6.033E-09 | 1.069E-09 | 5.100E-09 | 7.200E-09 |
| 5174 | 259 | S259F | A | 7e-09 | 2.3e-08 | 1.7e-09 | 1.057E-08 | 1.109E-08 | 1.700E-09 | 2.300E-08 |
| 5175 | 259 | S259P | A | 6.9e-09 | 7.2e-09 | 3.8e-09 | 5.967E-09 | 1.882E-09 | 3.800E-09 | 7.200E-09 |
| 5176 | 259 | S259S | A | 5e-09 | 5.3e-09 | 1.2e-08 | 7.433E-09 | 3.958E-09 | 5.000E-09 | 1.200E-08 |
| 5177 | 259 | S259T | A | 4.9e-09 | 1e-08 | 5.2e-09 | 6.700E-09 | 2.862E-09 | 4.900E-09 | 1.000E-08 |
| 5178 | 259 | S259W | A | 2.6e-09 | 6.3e-09 | 5.7e-09 | 4.867E-09 | 1.986E-09 | 2.600E-09 | 6.300E-09 |
| 5179 | 259 | S259Y | A | 1e-08 | 6.4e-09 | 3.2e-08 | 1.613E-08 | 1.386E-08 | 6.400E-09 | 3.200E-08 |
| 5180 | 259 | S259V | A | 7e-09 | 1.7e-08 | 7.8e-09 | 1.060E-08 | 5.557E-09 | 7.000E-09 | 1.700E-08 |
| 5181 | 260 | M260A | A | 5e-09 | 3.2e-09 | 9.9e-09 | 6.033E-09 | 3.467E-09 | 3.200E-09 | 9.900E-09 |
| 5182 | 260 | M260R | A | 1.6e-08 | 3.4e-09 | 5.1e-09 | 8.167E-09 | 6.837E-09 | 3.400E-09 | 1.600E-08 |
| 5183 | 260 | M260N | A | 9.4e-09 | 3.6e-09 | 4.4e-09 | 5.800E-09 | 3.143E-09 | 3.600E-09 | 9.400E-09 |
| 5184 | 260 | M260D | A | 7.2e-09 | 3e-09 | 1.1e-08 | 7.067E-09 | 4.002E-09 | 3.000E-09 | 1.100E-08 |

| | | | | | | | | | | |
|---|---|---|---|---|---|---|---|---|---|---|
| 5185 | 260 | M260C | A | 1.7e-08 | 4e-09 | 2.7e-09 | 7.900E-09 | 7.908E-09 | 2.700E-09 | 1.700E-08 |
| 5186 | 260 | M260E | A | 1.6e-08 | 6.5e-09 | 7e-09 | 9.833E-09 | 5.346E-09 | 6.500E-09 | 1.600E-08 |
| 5187 | 260 | M260Q | A | 4.2e-09 | 2.6e-09 | 8.8e-09 | 5.200E-09 | 3.219E-09 | 2.600E-09 | 8.800E-09 |
| 5188 | 260 | M260G | A | 9.6e-09 | 2.7e-09 | 1.4e-08 | 8.767E-09 | 5.696E-09 | 2.700E-09 | 1.400E-08 |
| 5189 | 260 | M260H | A | 5.4e-09 | 4.9e-09 | 6.5e-09 | 5.600E-09 | 8.185E-10 | 4.900E-09 | 6.500E-09 |
| 5190 | 260 | M260I | A | 7.2e-09 | 1.8e-09 | 8.1e-09 | 5.700E-09 | 3.407E-09 | 1.800E-09 | 8.100E-09 |
| 5191 | 260 | M260L | A | 1.1e-08 | 7.2e-09 | 4.5e-09 | 7.567E-09 | 3.265E-09 | 4.500E-09 | 1.100E-08 |
| 5192 | 260 | M260K | A | 7.3e-09 | 8.8e-09 | 2.9e-09 | 6.333E-09 | 3.066E-09 | 2.900E-09 | 8.800E-09 |
| 5193 | 260 | M260M | A | 2.2e-09 | 3.9e-09 | 6.7e-09 | 4.267E-09 | 2.272E-09 | 2.200E-09 | 6.700E-09 |
| 5194 | 260 | M260F | A | 5.3e-09 | 8.2e-09 | 3.1e-08 | 1.483E-08 | 1.408E-08 | 5.300E-09 | 3.100E-08 |
| 5195 | 260 | M260P | A | 4.2e-09 | 4.2e-09 | 6.8e-09 | 5.067E-09 | 1.501E-09 | 4.200E-09 | 6.800E-09 |
| 5196 | 260 | M260S | A | 3.7e-09 | 2.2e-08 | 6.7e-09 | 1.080E-08 | 9.815E-09 | 3.700E-09 | 2.200E-08 |
| 5197 | 260 | M260T | A | 5.4e-09 | 4.7e-09 | 4.7e-09 | 4.933E-09 | 4.041E-10 | 4.700E-09 | 5.400E-09 |
| 5198 | 260 | M260W | A | 3.4e-09 | 7.5e-09 | 4.1e-09 | 5.000E-09 | 2.193E-09 | 3.400E-09 | 7.500E-09 |
| 5199 | 260 | M260Y | A | 3.4e-09 | 1.3e-08 | 1.2e-08 | 9.467E-09 | 5.278E-09 | 3.400E-09 | 1.300E-08 |
| 5200 | 260 | M260V | A | 1e-08 | 4.9e-09 | 3.1e-08 | 1.530E-08 | 1.383E-08 | 4.900E-09 | 3.100E-08 |
| 5201 | 261 | P261A | A | 3.1e-09 | 3.3e-09 | 3.1e-09 | 3.167E-09 | 1.155E-10 | 3.100E-09 | 3.300E-09 |
| 5202 | 261 | P261R | A | 6.6e-09 | 1.1e-08 | 6.3e-09 | 7.967E-09 | 2.631E-09 | 6.300E-09 | 1.100E-08 |
| 5203 | 261 | P261N | A | 3.4e-09 | 2.2e-08 | 3.9e-09 | 9.767E-09 | 1.060E-08 | 3.400E-09 | 2.200E-08 |
| 5204 | 261 | P261D | A | 5.9e-09 | 1e-08 | 6.4e-09 | 7.433E-09 | 2.237E-09 | 5.900E-09 | 1.000E-08 |
| 5205 | 261 | P261C | A | 3.5e-09 | 2.6e-09 | 6.7e-09 | 4.267E-09 | 2.155E-09 | 2.600E-09 | 6.700E-09 |
| 5206 | 261 | P261E | A | 3.9e-09 | 4.6e-09 | 3.4e-09 | 3.967E-09 | 6.028E-10 | 3.400E-09 | 4.600E-09 |
| 5207 | 261 | P261Q | A | 2.3e-09 | 7.3e-09 | 5.3e-09 | 4.967E-09 | 2.517E-09 | 2.300E-09 | 7.300E-09 |
| 5208 | 261 | P261G | A | 5.6e-09 | 3.1e-09 | 4e-09 | 4.233E-09 | 1.266E-09 | 3.100E-09 | 5.600E-09 |
| 5209 | 261 | P261H | A | 8.6e-09 | 8.6e-09 | 7.6e-09 | 8.267E-09 | 5.774E-10 | 7.600E-09 | 8.600E-09 |
| 5210 | 261 | P261I | A | 6.1e-09 | 1.2e-08 | 1.5e-08 | 1.103E-08 | 4.528E-09 | 6.100E-09 | 1.500E-08 |
| 5211 | 261 | P261L | A | 2e-09 | 5.8e-09 | 3.2e-09 | 3.667E-09 | 1.943E-09 | 2.000E-09 | 5.800E-09 |
| 5212 | 261 | P261K | A | 6e-09 | 1.2e-08 | 1e-08 | 9.333E-09 | 3.055E-09 | 6.000E-09 | 1.200E-08 |
| 5213 | 261 | P261M | A | 5.7e-09 | 6.2e-09 | 4.2e-09 | 5.367E-09 | 1.041E-09 | 4.200E-09 | 6.200E-09 |
| 5214 | 261 | P261F | A | 1.1e-08 | 2.4e-08 | 3.5e-09 | 1.283E-08 | 1.037E-08 | 3.500E-09 | 2.400E-08 |
| 5215 | 261 | P261P | A | 2.1e-09 | 8e-09 | 6.1e-09 | 5.400E-09 | 3.012E-09 | 2.100E-09 | 8.000E-09 |
| 5216 | 261 | P261S | A | 7.5e-09 | 1.1e-08 | 5.5e-09 | 8.000E-09 | 2.784E-09 | 5.500E-09 | 1.100E-08 |
| 5217 | 261 | P261T | A | 9.1e-09 | 3.8e-09 | 1.1e-08 | 7.967E-09 | 3.731E-09 | 3.800E-09 | 1.100E-08 |
| 5218 | 261 | P261W | A | 5.8e-09 | 8.7e-09 | 4.4e-09 | 6.300E-09 | 2.193E-09 | 4.400E-09 | 8.700E-09 |
| 5219 | 261 | P261Y | A | 9.6e-09 | 4.8e-09 | 4.2e-09 | 6.200E-09 | 2.960E-09 | 4.200E-09 | 9.600E-09 |
| 5220 | 261 | P261V | A | 4.9e-09 | 1.4e-08 | 9.9e-09 | 9.600E-09 | 4.557E-09 | 4.900E-09 | 1.400E-08 |
| 5221 | 262 | N262A | A | 2.3e-09 | 4.4e-09 | 6.3e-09 | 4.333E-09 | 2.001E-09 | 2.300E-09 | 6.300E-09 |
| 5222 | 262 | N262R | A | 2.9e-09 | 1.1e-08 | 7.8e-09 | 7.233E-09 | 4.080E-09 | 2.900E-09 | 1.100E-08 |
| 5223 | 262 | N262N | A | 1.1e-08 | 5.1e-09 | 6.1e-09 | 7.400E-09 | 3.158E-09 | 5.100E-09 | 1.100E-08 |
| 5224 | 262 | N262D | A | 2.2e-08 | 8.4e-09 | 1.5e-08 | 1.513E-08 | 6.801E-09 | 8.400E-09 | 2.200E-08 |
| 5225 | 262 | N262C | A | 2.2e-09 | 6.3e-09 | 1.4e-08 | 7.500E-09 | 5.991E-09 | 2.200E-09 | 1.400E-08 |
| 5226 | 262 | N262E | A | 5.2e-09 | 3.8e-09 | 2.8e-09 | 3.933E-09 | 1.206E-09 | 2.800E-09 | 5.200E-09 |
| 5227 | 262 | N262Q | A | 2e-08 | 2.5e-09 | 6.2e-09 | 9.567E-09 | 9.223E-09 | 2.500E-09 | 2.000E-08 |
| 5228 | 262 | N262G | A | 7.3e-09 | 1e-08 | 9.8e-09 | 9.033E-09 | 1.504E-09 | 7.300E-09 | 1.000E-08 |
| 5229 | 262 | N262H | A | 6.1e-09 | 2.8e-09 | 8.1e-09 | 5.667E-09 | 2.676E-09 | 2.800E-09 | 8.100E-09 |
| 5230 | 262 | N262I | A | 1.2e-08 | 4.2e-09 | 1.8e-08 | 1.140E-08 | 6.920E-09 | 4.200E-09 | 1.800E-08 |
| 5231 | 262 | N262L | A | 1.1e-08 | 3.4e-09 | 2.8e-09 | 5.733E-09 | 4.571E-09 | 2.800E-09 | 1.100E-08 |
| 5232 | 262 | N262K | A | 3.9e-09 | 1.1e-08 | 1.1e-08 | 8.633E-09 | 4.099E-09 | 3.900E-09 | 1.100E-08 |
| 5233 | 262 | N262M | A | 2.7e-09 | 6.6e-09 | 1.4e-08 | 7.767E-09 | 5.740E-09 | 2.700E-09 | 1.400E-08 |
| 5234 | 262 | N262F | A | 1.3e-08 | 2.8e-09 | 3.8e-09 | 6.533E-09 | 5.623E-09 | 2.800E-09 | 1.300E-08 |
| 5235 | 262 | N262P | A | 3.7e-08 | 1e-08 | 1.2e-08 | 1.967E-08 | 1.504E-08 | 1.000E-08 | 3.700E-08 |
| 5236 | 262 | N262S | A | 1.6e-08 | 1.1e-08 | 6.6e-09 | 1.120E-08 | 4.703E-09 | 6.600E-09 | 1.600E-08 |
| 5237 | 262 | N262T | A | 4.4e-09 | 3.4e-09 | 1.1e-08 | 6.267E-09 | 4.130E-09 | 3.400E-09 | 1.100E-08 |
| 5238 | 262 | N262W | A | 5.9e-09 | 1.3e-08 | 3.9e-09 | 7.600E-09 | 4.782E-09 | 3.900E-09 | 1.300E-08 |
| 5239 | 262 | N262Y | A | 5.4e-09 | 1.1e-08 | 7.9e-09 | 8.100E-09 | 2.805E-09 | 5.400E-09 | 1.100E-08 |
| 5240 | 262 | N262V | A | 1.5e-08 | 1.3e-08 | 1.4e-08 | 1.400E-08 | 1.000E-09 | 1.300E-08 | 1.500E-08 |
| 5241 | 263 | P263A | A | 5.6e-09 | 1.9e-08 | 7.7e-09 | 1.077E-08 | 7.207E-09 | 5.600E-09 | 1.900E-08 |
| 5242 | 263 | P263R | A | 1.8e-08 | 4.9e-09 | 2.8e-09 | 8.567E-09 | 8.237E-09 | 2.800E-09 | 1.800E-08 |
| 5243 | 263 | P263N | A | 8.1e-09 | 3.2e-09 | 5.2e-09 | 5.500E-09 | 2.464E-09 | 3.200E-09 | 8.100E-09 |
| 5244 | 263 | P263D | A | 9.3e-09 | 1.9e-08 | 6.3e-09 | 1.153E-08 | 6.638E-09 | 6.300E-09 | 1.900E-08 |
| 5245 | 263 | P263C | A | 5.4e-09 | 1.3e-08 | 1.4e-08 | 1.080E-08 | 4.703E-09 | 5.400E-09 | 1.400E-08 |
| 5246 | 263 | P263E | A | 2.3e-09 | 1e-08 | 1.5e-08 | 9.100E-09 | 6.398E-09 | 2.300E-09 | 1.500E-08 |
| 5247 | 263 | P263Q | A | 2.4e-08 | 3.3e-09 | 1.1e-08 | 1.277E-08 | 1.046E-08 | 3.300E-09 | 2.400E-08 |
| 5248 | 263 | P263G | A | 7.6e-09 | 8.4e-09 | 8.7e-09 | 8.233E-09 | 5.686E-10 | 7.600E-09 | 8.700E-09 |
| 5249 | 263 | P263H | A | 4.2e-09 | 1.2e-08 | 5.4e-09 | 7.200E-09 | 4.200E-09 | 4.200E-09 | 1.200E-08 |
| 5250 | 263 | P263I | A | 3.6e-09 | 7.3e-09 | 1.3e-09 | 4.067E-09 | 3.027E-09 | 1.300E-09 | 7.300E-09 |
| 5251 | 263 | P263L | A | 6.8e-09 | 4.2e-09 | 1.2e-09 | 4.067E-09 | 2.802E-09 | 1.200E-09 | 6.800E-09 |
| 5252 | 263 | P263K | A | 4.2e-09 | 3.4e-09 | 1.3e-08 | 6.867E-09 | 5.327E-09 | 3.400E-09 | 1.300E-08 |
| 5253 | 263 | P263M | A | 4e-09 | 1.3e-08 | 2.4e-08 | 1.367E-08 | 1.002E-08 | 4.000E-09 | 2.400E-08 |
| 5254 | 263 | P263F | A | 7.7e-09 | 1.5e-08 | 6.9e-09 | 9.867E-09 | 4.464E-09 | 6.900E-09 | 1.500E-08 |
| 5255 | 263 | P263P | A | 8.5e-09 | 5e-09 | 1.1e-08 | 8.167E-09 | 3.014E-09 | 5.000E-09 | 1.100E-08 |
| 5256 | 263 | P263S | A | 8.5e-09 | 4.2e-09 | 5e-09 | 5.900E-09 | 2.287E-09 | 4.200E-09 | 8.500E-09 |
| 5257 | 263 | P263T | A | 5.8e-09 | 1.1e-08 | 3.3e-09 | 6.700E-09 | 3.928E-09 | 3.300E-09 | 1.100E-08 |
| 5258 | 263 | P263W | A | 1.9e-08 | 8.4e-09 | 1.5e-08 | 1.413E-08 | 5.353E-09 | 8.400E-09 | 1.900E-08 |
| 5259 | 263 | P263Y | A | 3e-08 | 1.1e-08 | 1.3e-08 | 1.800E-08 | 1.044E-08 | 1.100E-08 | 3.000E-08 |
| 5260 | 263 | P263V | A | 1.3e-08 | 5e-09 | 9.7e-09 | 9.233E-09 | 4.020E-09 | 5.000E-09 | 1.300E-08 |
| 5261 | 264 | E264A | A | 1.3e-08 | 3.4e-09 | 1.8e-08 | 1.147E-08 | 7.420E-09 | 3.400E-09 | 1.800E-08 |
| 5262 | 264 | E264R | A | 7.9e-09 | 5.7e-09 | 7.4e-09 | 7.000E-09 | 1.153E-09 | 5.700E-09 | 7.900E-09 |
| 5263 | 264 | E264N | A | 2.7e-09 | 2.6e-09 | 6.9e-09 | 4.067E-09 | 2.454E-09 | 2.600E-09 | 6.900E-09 |
| 5264 | 264 | E264D | A | 7.8e-09 | 2.2e-09 | 1.6e-08 | 8.667E-09 | 6.941E-09 | 2.200E-09 | 1.600E-08 |
| 5265 | 264 | E264C | A | 1.6e-08 | 9.4e-09 | 3e-09 | 9.467E-09 | 6.500E-09 | 3.000E-09 | 1.600E-08 |
| 5266 | 264 | E264E | A | 6.8e-09 | 8.3e-09 | 1.1e-08 | 8.700E-09 | 2.128E-09 | 6.800E-09 | 1.100E-08 |
| 5267 | 264 | E264Q | A | 9.5e-09 | 1.2e-08 | 1.1e-08 | 1.083E-08 | 1.258E-09 | 9.500E-09 | 1.200E-08 |
| 5268 | 264 | E264G | A | 2.5e-09 | 9.2e-09 | 1.2e-08 | 7.900E-09 | 4.882E-09 | 2.500E-09 | 1.200E-08 |
| 5269 | 264 | E264H | A | 4.6e-09 | 1.2e-08 | 1.5e-08 | 1.053E-08 | 5.353E-09 | 4.600E-09 | 1.500E-08 |

| | | | | | | | | | | |
|---|---|---|---|---|---|---|---|---|---|---|
| 5270 | 264 | E264I | A | 8.2e-09 | 8.9e-09 | 3.9e-09 | 7.000E-09 | 2.707E-09 | 3.900E-09 | 8.900E-09 |
| 5271 | 264 | E264L | A | 1.7e-08 | 4.5e-09 | 1.5e-08 | 1.217E-08 | 6.714E-09 | 4.500E-09 | 1.700E-08 |
| 5272 | 264 | E264K | A | 6.9e-09 | 1.5e-08 | 5e-09 | 8.967E-09 | 5.311E-09 | 5.000E-09 | 1.500E-08 |
| 5273 | 264 | E264M | A | 1.1e-08 | 2.5e-09 | 1.4e-08 | 9.167E-09 | 5.965E-09 | 2.500E-09 | 1.400E-08 |
| 5274 | 264 | E264F | A | 3.6e-09 | 1e-08 | 1.9e-08 | 1.087E-08 | 7.736E-09 | 3.600E-09 | 1.900E-08 |
| 5275 | 264 | E264P | A | 2.6e-09 | 1.1e-08 | 7.8e-09 | 7.133E-09 | 4.239E-09 | 2.600E-09 | 1.100E-08 |
| 5276 | 264 | E264S | A | 5.7e-09 | 2.6e-08 | 7.7e-09 | 1.313E-08 | 1.119E-08 | 5.700E-09 | 2.600E-08 |
| 5277 | 264 | E264T | A | 1.2e-08 | 9.1e-09 | 8.6e-09 | 9.900E-09 | 1.836E-09 | 8.600E-09 | 1.200E-08 |
| 5278 | 264 | E264W | A | 9e-09 | 7.5e-09 | 7e-09 | 7.833E-09 | 1.041E-09 | 7.000E-09 | 9.000E-09 |
| 5279 | 264 | E264Y | A | 9.6e-09 | 5.9e-09 | 8.1e-09 | 7.867E-09 | 1.861E-09 | 5.900E-09 | 9.600E-09 |
| 5280 | 264 | E264V | A | 5.6e-09 | 1.1e-08 | 2e-08 | 1.220E-08 | 7.275E-09 | 5.600E-09 | 2.000E-08 |
| 5281 | 265 | G265A | A | 3e-09 | 9.5e-09 | 9.7e-09 | 7.400E-09 | 3.812E-09 | 3.000E-09 | 9.700E-09 |
| 5282 | 265 | G265R | A | 5.9e-09 | 1.6e-08 | 9.8e-09 | 1.057E-08 | 5.093E-09 | 5.900E-09 | 1.600E-08 |
| 5283 | 265 | G265N | A | 9.6e-09 | 3.3e-09 | 5e-09 | 5.967E-09 | 3.259E-09 | 3.300E-09 | 9.600E-09 |
| 5284 | 265 | G265D | A | 1.6e-08 | 1.7e-08 | 8.5e-09 | 1.383E-08 | 4.646E-09 | 8.500E-09 | 1.700E-08 |
| 5285 | 265 | G265C | A | 1.2e-08 | 5.7e-09 | 1.3e-09 | 6.333E-09 | 5.378E-09 | 1.300E-09 | 1.200E-08 |
| 5286 | 265 | G265E | A | 3.8e-09 | 1.1e-08 | 4.1e-09 | 6.300E-09 | 4.073E-09 | 3.800E-09 | 1.100E-08 |
| 5287 | 265 | G265Q | A | 6.7e-09 | 1.4e-08 | 5.9e-09 | 8.867E-09 | 4.464E-09 | 5.900E-09 | 1.400E-08 |
| 5288 | 265 | G265G | A | 8.5e-09 | 2.7e-09 | 4.9e-09 | 5.367E-09 | 2.928E-09 | 2.700E-09 | 8.500E-09 |
| 5289 | 265 | G265H | A | 6.7e-09 | 5e-09 | 8.1e-09 | 6.600E-09 | 1.552E-09 | 5.000E-09 | 8.100E-09 |
| 5290 | 265 | G265I | A | 3.2e-09 | 5e-09 | 4.3e-09 | 4.167E-09 | 9.074E-10 | 3.200E-09 | 5.000E-09 |
| 5291 | 265 | G265L | A | 2.9e-09 | 7.3e-09 | 5.6e-09 | 5.267E-09 | 2.219E-09 | 2.900E-09 | 7.300E-09 |
| 5292 | 265 | G265K | A | 5.2e-09 | 7.6e-09 | 1.2e-08 | 8.267E-09 | 3.449E-09 | 5.200E-09 | 1.200E-08 |
| 5293 | 265 | G265M | A | 1.1e-08 | 6.3e-09 | 1.1e-08 | 9.433E-09 | 2.714E-09 | 6.300E-09 | 1.100E-08 |
| 5294 | 265 | G265F | A | 6.2e-09 | 7.5e-09 | 1.3e-09 | 5.000E-09 | 3.270E-09 | 1.300E-09 | 7.500E-09 |
| 5295 | 265 | G265P | A | 3.5e-09 | 4e-09 | 5.6e-09 | 4.367E-09 | 1.097E-09 | 3.500E-09 | 5.600E-09 |
| 5296 | 265 | G265S | A | 1.9e-08 | 1.3e-08 | 1.3e-08 | 1.500E-08 | 3.464E-09 | 1.300E-08 | 1.900E-08 |
| 5297 | 265 | G265T | A | 5.9e-09 | 9.3e-09 | 1.1e-08 | 8.733E-09 | 2.597E-09 | 5.900E-09 | 1.100E-08 |
| 5298 | 265 | G265W | A | 1.3e-09 | 5.2e-09 | 1.8e-09 | 2.767E-09 | 2.122E-09 | 1.300E-09 | 5.200E-09 |
| 5299 | 265 | G265Y | A | 3.5e-09 | 6.3e-09 | 1.3e-08 | 7.600E-09 | 4.882E-09 | 3.500E-09 | 1.300E-08 |
| 5300 | 265 | G265V | A | 5.9e-09 | 3.2e-09 | 1.1e-08 | 6.700E-09 | 3.961E-09 | 3.200E-09 | 1.100E-08 |
| 5301 | 266 | R266A | A | 6.7e-09 | 8.5e-09 | 9.9e-09 | 8.367E-09 | 1.604E-09 | 6.700E-09 | 9.900E-09 |
| 5302 | 266 | R266R | A | 7.4e-09 | 4.5e-09 | 7e-09 | 6.300E-09 | 1.572E-09 | 4.500E-09 | 7.400E-09 |
| 5303 | 266 | R266N | A | 1e-08 | 2.1e-09 | 6.4e-09 | 6.167E-09 | 3.955E-09 | 2.100E-09 | 1.000E-08 |
| 5304 | 266 | R266D | A | 1.3e-08 | 4.6e-08 | 3e-08 | 2.967E-08 | 1.650E-08 | 1.300E-08 | 4.600E-08 |
| 5305 | 266 | R266C | A | 2.8e-09 | 5.9e-09 | 7.9e-09 | 5.533E-09 | 2.570E-09 | 2.800E-09 | 7.900E-09 |
| 5306 | 266 | R266E | A | 2.6e-09 | 7.9e-09 | 1.7e-08 | 9.167E-09 | 7.283E-09 | 2.600E-09 | 1.700E-08 |
| 5307 | 266 | R266Q | A | 1.4e-08 | 2e-08 | 1.6e-08 | 1.667E-08 | 3.055E-09 | 1.400E-08 | 2.000E-08 |
| 5308 | 266 | R266G | A | 7.2e-09 | 3.7e-09 | 4.1e-09 | 5.000E-09 | 1.916E-09 | 3.700E-09 | 7.200E-09 |
| 5309 | 266 | R266H | A | 8.8e-09 | 2.8e-09 | 3.1e-09 | 4.900E-09 | 3.381E-09 | 2.800E-09 | 8.800E-09 |
| 5310 | 266 | R266I | A | 2.8e-09 | 6.9e-09 | 3e-09 | 4.233E-09 | 2.312E-09 | 2.800E-09 | 6.900E-09 |
| 5311 | 266 | R266L | A | 7.1e-09 | 4.7e-09 | 4.2e-09 | 5.333E-09 | 1.550E-09 | 4.200E-09 | 7.100E-09 |
| 5312 | 266 | R266K | A | 7.6e-09 | 5.3e-09 | 3.2e-09 | 5.367E-09 | 2.201E-09 | 3.200E-09 | 7.600E-09 |
| 5313 | 266 | R266M | A | 8e-09 | 1.4e-09 | 5.5e-09 | 4.967E-09 | 3.332E-09 | 1.400E-09 | 8.000E-09 |
| 5314 | 266 | R266F | A | 9.9e-09 | 4.3e-09 | 7.4e-09 | 7.200E-09 | 2.805E-09 | 4.300E-09 | 9.900E-09 |
| 5315 | 266 | R266P | A | 8.8e-09 | 6.4e-09 | 3.9e-09 | 6.367E-09 | 2.450E-09 | 3.900E-09 | 8.800E-09 |
| 5316 | 266 | R266S | A | 1.2e-08 | 7.2e-09 | 4.6e-09 | 7.933E-09 | 3.754E-09 | 4.600E-09 | 1.200E-08 |
| 5317 | 266 | R266T | A | 1.7e-08 | 4.9e-09 | 1.4e-08 | 1.197E-08 | 6.301E-09 | 4.900E-09 | 1.700E-08 |
| 5318 | 266 | R266W | A | 8e-09 | 9.5e-09 | 2.5e-09 | 6.667E-09 | 3.686E-09 | 2.500E-09 | 9.500E-09 |
| 5319 | 266 | R266Y | A | 5e-09 | 1.7e-09 | 3.5e-09 | 3.400E-09 | 1.652E-09 | 1.700E-09 | 5.000E-09 |
| 5320 | 266 | R266V | A | 6e-09 | 8.3e-09 | 6.8e-09 | 7.033E-09 | 1.168E-09 | 6.000E-09 | 8.300E-09 |
| 5321 | 267 | Y267A | A | 3.7e-09 | 1e-08 | 5.6e-09 | 6.433E-09 | 3.232E-09 | 3.700E-09 | 1.000E-08 |
| 5322 | 267 | Y267R | A | 3.6e-09 | 2.5e-08 | 3.5e-09 | 1.070E-08 | 1.238E-08 | 3.500E-09 | 2.500E-08 |
| 5323 | 267 | Y267N | A | 7.8e-09 | 6.7e-09 | 8.1e-09 | 7.533E-09 | 7.371E-10 | 6.700E-09 | 8.100E-09 |
| 5324 | 267 | Y267D | A | 1.9e-08 | 1.5e-08 | 1.4e-08 | 1.600E-08 | 2.646E-09 | 1.400E-08 | 1.900E-08 |
| 5325 | 267 | Y267C | A | 1.8e-08 | 1.1e-08 | 7.3e-09 | 1.210E-08 | 5.434E-09 | 7.300E-09 | 1.800E-08 |
| 5326 | 267 | Y267E | A | 3.7e-09 | 8.8e-09 | 9.2e-09 | 7.233E-09 | 3.066E-09 | 3.700E-09 | 9.200E-09 |
| 5327 | 267 | Y267Q | A | 1.9e-08 | 6.7e-09 | 2e-08 | 1.523E-08 | 7.407E-09 | 6.700E-09 | 2.000E-08 |
| 5328 | 267 | Y267G | A | 6e-09 | 1.4e-08 | 1.2e-08 | 1.067E-08 | 4.163E-09 | 6.000E-09 | 1.400E-08 |
| 5329 | 267 | Y267H | A | 1.5e-08 | 3e-09 | 7.5e-09 | 8.500E-09 | 6.062E-09 | 3.000E-09 | 1.500E-08 |
| 5330 | 267 | Y267I | A | 5.3e-09 | 4.1e-09 | 1.7e-08 | 8.800E-09 | 7.127E-09 | 4.100E-09 | 1.700E-08 |
| 5331 | 267 | Y267L | A | 4.7e-09 | 1.4e-08 | 7e-09 | 8.567E-09 | 4.844E-09 | 4.700E-09 | 1.400E-08 |
| 5332 | 267 | Y267K | A | 9.7e-09 | 3.9e-09 | 7.2e-09 | 6.933E-09 | 2.909E-09 | 3.900E-09 | 9.700E-09 |
| 5333 | 267 | Y267M | A | 4.2e-09 | 2.9e-08 | 6.4e-09 | 1.320E-08 | 1.373E-08 | 4.200E-09 | 2.900E-08 |
| 5334 | 267 | Y267F | A | 6.9e-09 | 5.6e-09 | 9.6e-09 | 7.367E-09 | 2.040E-09 | 5.600E-09 | 9.600E-09 |
| 5335 | 267 | Y267P | A | 8.3e-09 | 8.8e-09 | 1.1e-08 | 9.367E-09 | 1.436E-09 | 8.300E-09 | 1.100E-08 |
| 5336 | 267 | Y267S | A | 1.1e-08 | 6.3e-09 | 8.8e-09 | 8.700E-09 | 2.352E-09 | 6.300E-09 | 1.100E-08 |
| 5337 | 267 | Y267T | A | 9.7e-09 | 1.8e-08 | 3.2e-08 | 1.990E-08 | 1.127E-08 | 9.700E-09 | 3.200E-08 |
| 5338 | 267 | Y267W | A | 2.5e-09 | 8.2e-09 | 2.3e-09 | 4.333E-09 | 3.350E-09 | 2.300E-09 | 8.200E-09 |
| 5339 | 267 | Y267Y | A | 3e-09 | 9.4e-09 | 1.8e-09 | 4.733E-09 | 4.086E-09 | 1.800E-09 | 9.400E-09 |
| 5340 | 267 | Y267V | A | 1.1e-08 | 1.2e-08 | 3.4e-09 | 8.800E-09 | 4.703E-09 | 3.400E-09 | 1.200E-08 |
| 5341 | 268 | T268A | A | 1.4e-09 | 1.4e-09 | 1.9e-09 | 1.567E-09 | 2.887E-10 | 1.400E-09 | 1.900E-09 |
| 5342 | 268 | T268R | A | 3.5e-09 | 2.4e-09 | 4.1e-09 | 3.333E-09 | 8.622E-10 | 2.400E-09 | 4.100E-09 |
| 5343 | 268 | T268N | A | 1.1e-08 | 1.4e-08 | 8e-09 | 1.100E-08 | 3.000E-09 | 8.000E-09 | 1.400E-08 |
| 5344 | 268 | T268D | A | 2.9e-09 | 7.2e-09 | 3.4e-09 | 4.500E-09 | 2.352E-09 | 2.900E-09 | 7.200E-09 |
| 5345 | 268 | T268C | A | 9.8e-10 | 1.2e-09 | 1.7e-09 | 1.293E-09 | 3.690E-10 | 9.800E-10 | 1.700E-09 |
| 5346 | 268 | T268E | A | 3.3e-09 | 2.4e-09 | 4.3e-09 | 3.333E-09 | 9.504E-10 | 2.400E-09 | 4.300E-09 |
| 5347 | 268 | T268Q | A | 1.5e-09 | 5.8e-09 | 1.2e-08 | 6.433E-09 | 5.279E-09 | 1.500E-09 | 1.200E-08 |
| 5348 | 268 | T268G | A | 1.8e-09 | 2.8e-09 | 3.1e-09 | 2.567E-09 | 6.807E-10 | 1.800E-09 | 3.100E-09 |
| 5349 | 268 | T268H | A | 6.4e-09 | 3.4e-09 | 2.6e-09 | 4.133E-09 | 2.003E-09 | 2.600E-09 | 6.400E-09 |
| 5350 | 268 | T268I | A | 1.4e-09 | 3.8e-09 | 1.8e-09 | 2.333E-09 | 1.286E-09 | 1.400E-09 | 3.800E-09 |
| 5351 | 268 | T268L | A | 2.6e-09 | 2e-09 | 1.2e-09 | 1.933E-09 | 7.024E-10 | 1.200E-09 | 2.600E-09 |
| 5352 | 268 | T268K | A | 7.6e-09 | 6.8e-09 | 5.7e-10 | 4.990E-09 | 3.849E-09 | 5.700E-10 | 7.600E-09 |
| 5353 | 268 | T268M | A | 5.1e-09 | 4.1e-09 | 2.2e-09 | 3.800E-09 | 1.473E-09 | 2.200E-09 | 5.100E-09 |
| 5354 | 268 | T268F | A | 1.6e-09 | 2.1e-09 | 1.1e-09 | 1.600E-09 | 5.000E-10 | 1.100E-09 | 2.100E-09 |

| | | | | | | | | | | |
|---|---|---|---|---|---|---|---|---|---|---|
| 5355 | 268 | T268P | A | 6.9e-10 | 8.5e-10 | 5e-09 | 2.180E-09 | 2.444E-09 | 6.900E-10 | 5.000E-09 |
| 5356 | 268 | T268S | A | 1.4e-09 | 1.7e-09 | 2.9e-09 | 2.000E-09 | 7.937E-10 | 1.400E-09 | 2.900E-09 |
| 5357 | 268 | T268T | A | 8.9e-09 | 4.2e-09 | 4.4e-09 | 5.833E-09 | 2.658E-09 | 4.200E-09 | 8.900E-09 |
| 5358 | 268 | T268W | A | 1.6e-09 | 1.9e-09 | 1.8e-09 | 1.767E-09 | 1.528E-10 | 1.600E-09 | 1.900E-09 |
| 5359 | 268 | T268Y | A | 9.4e-10 | 5.4e-09 | 7.7e-10 | 2.370E-09 | 2.625E-09 | 7.700E-10 | 5.400E-09 |
| 5360 | 268 | T268V | A | 7e-10 | 1.6e-09 | 1.4e-09 | 1.233E-09 | 4.726E-10 | 7.000E-10 | 1.600E-09 |
| 5361 | 269 | F269A | A | 1.5e-08 | 5.3e-09 | 2.3e-09 | 7.533E-09 | 6.638E-09 | 2.300E-09 | 1.500E-08 |
| 5362 | 269 | F269R | A | 1.7e-08 | 1.1e-08 | 6.3e-09 | 1.143E-08 | 5.363E-09 | 6.300E-09 | 1.700E-08 |
| 5363 | 269 | F269N | A | 3e-09 | 6.9e-09 | 6.8e-09 | 5.567E-09 | 2.223E-09 | 3.000E-09 | 6.900E-09 |
| 5364 | 269 | F269D | A | 1.2e-08 | 8.6e-09 | 1.1e-08 | 1.053E-08 | 1.747E-09 | 8.600E-09 | 1.200E-08 |
| 5365 | 269 | F269C | A | 1.3e-08 | 1.4e-08 | 1.3e-08 | 1.333E-08 | 5.774E-10 | 1.300E-08 | 1.400E-08 |
| 5366 | 269 | F269E | A | 4.4e-09 | 1.6e-08 | 9.7e-09 | 1.003E-08 | 5.807E-09 | 4.400E-09 | 1.600E-08 |
| 5367 | 269 | F269Q | A | 1.5e-08 | 1.7e-08 | 5e-09 | 1.233E-08 | 6.429E-09 | 5.000E-09 | 1.700E-08 |
| 5368 | 269 | F269G | A | 1.3e-09 | 1e-08 | 3e-09 | 4.767E-09 | 4.611E-09 | 1.300E-09 | 1.000E-08 |
| 5369 | 269 | F269H | A | 3.6e-09 | 5.7e-09 | 1.3e-08 | 7.433E-09 | 4.934E-09 | 3.600E-09 | 1.300E-08 |
| 5370 | 269 | F269I | A | 2.3e-08 | 1.1e-08 | 6.7e-09 | 1.357E-08 | 8.448E-09 | 6.700E-09 | 2.300E-08 |
| 5371 | 269 | F269L | A | 1.6e-08 | 1.2e-08 | 3.6e-09 | 1.053E-08 | 6.329E-09 | 3.600E-09 | 1.600E-08 |
| 5372 | 269 | F269K | A | 1.7e-08 | 6.6e-09 | 6e-09 | 9.867E-09 | 6.185E-09 | 6.000E-09 | 1.700E-08 |
| 5373 | 269 | F269M | A | 4.3e-09 | 7.8e-09 | 9.3e-09 | 7.133E-09 | 2.566E-09 | 4.300E-09 | 9.300E-09 |
| 5374 | 269 | F269F | A | 5.2e-09 | 2.8e-09 | 1.5e-08 | 7.667E-09 | 6.463E-09 | 2.800E-09 | 1.500E-08 |
| 5375 | 269 | F269P | A | 6.3e-09 | 6.8e-09 | 1.2e-08 | 8.367E-09 | 3.156E-09 | 6.300E-09 | 1.200E-08 |
| 5376 | 269 | F269S | A | 4.7e-09 | 5.1e-09 | 2.7e-09 | 4.167E-09 | 1.286E-09 | 2.700E-09 | 5.100E-09 |
| 5377 | 269 | F269T | A | 4.8e-09 | 1.7e-08 | 5.2e-09 | 9.000E-09 | 6.931E-09 | 4.800E-09 | 1.700E-08 |
| 5378 | 269 | F269W | A | 7.2e-09 | 1.1e-08 | 4.8e-09 | 7.667E-09 | 3.126E-09 | 4.800E-09 | 1.100E-08 |
| 5379 | 269 | F269Y | A | 4.9e-09 | 1.2e-08 | 8.6e-09 | 8.500E-09 | 3.551E-09 | 4.900E-09 | 1.200E-08 |
| 5380 | 269 | F269V | A | 2.5e-09 | 9.5e-09 | 7.2e-09 | 6.400E-09 | 3.568E-09 | 2.500E-09 | 9.500E-09 |
| 5381 | 270 | G270A | A | 7.2e-09 | 6.3e-09 | 4.6e-09 | 6.033E-09 | 1.320E-09 | 4.600E-09 | 7.200E-09 |
| 5382 | 270 | G270R | A | 1.1e-08 | 1.3e-08 | 9.8e-09 | 1.127E-08 | 1.617E-09 | 9.800E-09 | 1.300E-08 |
| 5383 | 270 | G270N | A | 1e-08 | 1.8e-08 | 1.2e-08 | 1.333E-08 | 4.163E-09 | 1.000E-08 | 1.800E-08 |
| 5384 | 270 | G270D | A | 1.4e-08 | 1e-08 | 2.5e-08 | 1.633E-08 | 7.767E-09 | 1.000E-08 | 2.500E-08 |
| 5385 | 270 | G270C | A | 1.6e-08 | 1.5e-08 | 8.6e-09 | 1.320E-08 | 4.015E-09 | 8.600E-09 | 1.600E-08 |
| 5386 | 270 | G270E | A | 9.2e-09 | 7.3e-09 | 3.3e-09 | 6.600E-09 | 3.012E-09 | 3.300E-09 | 9.200E-09 |
| 5387 | 270 | G270Q | A | 9.1e-09 | 8.4e-09 | 2.7e-09 | 6.733E-09 | 3.510E-09 | 2.700E-09 | 9.100E-09 |
| 5388 | 270 | G270G | A | 1.2e-08 | 9.1e-09 | 1.3e-08 | 1.137E-08 | 2.026E-09 | 9.100E-09 | 1.300E-08 |
| 5389 | 270 | G270H | A | 4.6e-09 | 6.9e-09 | 6.9e-09 | 6.133E-09 | 1.328E-09 | 4.600E-09 | 6.900E-09 |
| 5390 | 270 | G270I | A | 2e-08 | 3.9e-09 | 2e-08 | 1.463E-08 | 9.295E-09 | 3.900E-09 | 2.000E-08 |
| 5391 | 270 | G270L | A | 6.9e-09 | 6.6e-10 | 6.9e-09 | 4.820E-09 | 3.603E-09 | 6.600E-10 | 6.900E-09 |
| 5392 | 270 | G270K | A | 2.4e-09 | 4.5e-09 | 4.2e-09 | 3.700E-09 | 1.136E-09 | 2.400E-09 | 4.500E-09 |
| 5393 | 270 | G270M | A | 9.3e-09 | 1.8e-09 | 1.4e-08 | 8.367E-09 | 6.153E-09 | 1.800E-09 | 1.400E-08 |
| 5394 | 270 | G270F | A | 2.2e-08 | 6.5e-09 | 3.5e-09 | 1.067E-08 | 9.929E-09 | 3.500E-09 | 2.200E-08 |
| 5395 | 270 | G270P | A | 6.1e-09 | 3.4e-09 | 5.2e-09 | 4.900E-09 | 1.375E-09 | 3.400E-09 | 6.100E-09 |
| 5396 | 270 | G270S | A | 9.7e-09 | 1.2e-08 | 8.9e-09 | 1.020E-08 | 1.609E-09 | 8.900E-09 | 1.200E-08 |
| 5397 | 270 | G270T | A | 1.4e-08 | 1.2e-08 | 1.1e-08 | 1.233E-08 | 1.528E-09 | 1.100E-08 | 1.400E-08 |
| 5398 | 270 | G270W | A | 2.9e-09 | 7.3e-09 | 8.9e-09 | 6.367E-09 | 3.107E-09 | 2.900E-09 | 8.900E-09 |
| 5399 | 270 | G270Y | A | 1.1e-08 | 5.2e-09 | 3e-09 | 6.400E-09 | 4.133E-09 | 3.000E-09 | 1.100E-08 |
| 5400 | 270 | G270V | A | 3.4e-09 | 7.1e-09 | 4.4e-09 | 4.967E-09 | 1.914E-09 | 3.400E-09 | 7.100E-09 |
| 5401 | 271 | A271A | A | 1.3e-08 | 1e-08 | 5.1e-09 | 9.367E-09 | 3.988E-09 | 5.100E-09 | 1.300E-08 |
| 5402 | 271 | A271R | A | 1.3e-08 | 4.2e-09 | 3.8e-09 | 7.000E-09 | 5.200E-09 | 3.800E-09 | 1.300E-08 |
| 5403 | 271 | A271N | A | 5.1e-09 | 3.1e-09 | 3.7e-09 | 3.967E-09 | 1.026E-09 | 3.100E-09 | 5.100E-09 |
| 5404 | 271 | A271D | A | 1.4e-08 | 6.5e-09 | 5.3e-09 | 8.600E-09 | 4.715E-09 | 5.300E-09 | 1.400E-08 |
| 5405 | 271 | A271C | A | 5.9e-09 | 9.3e-09 | 6.8e-09 | 7.333E-09 | 1.762E-09 | 5.900E-09 | 9.300E-09 |
| 5406 | 271 | A271E | A | 5.8e-09 | 5.4e-09 | 1.2e-08 | 7.733E-09 | 3.700E-09 | 5.400E-09 | 1.200E-08 |
| 5407 | 271 | A271Q | A | 1.9e-09 | 5.9e-09 | 2.3e-09 | 3.367E-09 | 2.203E-09 | 1.900E-09 | 5.900E-09 |
| 5408 | 271 | A271G | A | 1.1e-08 | 5.1e-09 | 6.2e-09 | 7.433E-09 | 3.137E-09 | 5.100E-09 | 1.100E-08 |
| 5409 | 271 | A271H | A | 1.3e-08 | 5.3e-09 | 4.6e-09 | 7.633E-09 | 4.661E-09 | 4.600E-09 | 1.300E-08 |
| 5410 | 271 | A271I | A | 8.9e-09 | 2.2e-09 | 5.5e-09 | 5.533E-09 | 3.350E-09 | 2.200E-09 | 8.900E-09 |
| 5411 | 271 | A271L | A | 1.3e-09 | 5.3e-09 | 7.1e-09 | 4.567E-09 | 2.969E-09 | 1.300E-09 | 7.100E-09 |
| 5412 | 271 | A271K | A | 1.6e-08 | 6.5e-09 | 8.3e-09 | 1.027E-08 | 5.046E-09 | 6.500E-09 | 1.600E-08 |
| 5413 | 271 | A271M | A | 2.9e-09 | 7.3e-09 | 1.1e-08 | 7.067E-09 | 4.055E-09 | 2.900E-09 | 1.100E-08 |
| 5414 | 271 | A271F | A | 1e-08 | 7.9e-09 | 5.9e-09 | 7.933E-09 | 2.050E-09 | 5.900E-09 | 1.000E-08 |
| 5415 | 271 | A271P | A | 1.7e-08 | 6e-09 | 1.1e-08 | 1.133E-08 | 5.508E-09 | 6.000E-09 | 1.700E-08 |
| 5416 | 271 | A271S | A | 1e-08 | 5.4e-09 | 8.2e-09 | 7.867E-09 | 2.318E-09 | 5.400E-09 | 1.000E-08 |
| 5417 | 271 | A271T | A | 2.6e-09 | 9.7e-09 | 6.8e-09 | 6.367E-09 | 3.570E-09 | 2.600E-09 | 9.700E-09 |
| 5418 | 271 | A271W | A | 9.8e-09 | 4.2e-09 | 4e-09 | 6.000E-09 | 3.292E-09 | 4.000E-09 | 9.800E-09 |
| 5419 | 271 | A271Y | A | 1.7e-08 | 6.7e-09 | 1.7e-08 | 1.357E-08 | 5.947E-09 | 6.700E-09 | 1.700E-08 |
| 5420 | 271 | A271V | A | 6.9e-09 | 8.8e-09 | 3.3e-09 | 6.333E-09 | 2.793E-09 | 3.300E-09 | 8.800E-09 |
| 5421 | 272 | S272A | A | 1.4e-08 | 7.7e-09 | 3e-09 | 8.233E-09 | 5.519E-09 | 3.000E-09 | 1.400E-08 |
| 5422 | 272 | S272R | A | 3.8e-09 | 7.6e-09 | 4.6e-09 | 5.333E-09 | 2.003E-09 | 3.800E-09 | 7.600E-09 |
| 5423 | 272 | S272N | A | 6.6e-09 | 8e-09 | 2.6e-08 | 1.353E-08 | 1.082E-08 | 6.600E-09 | 2.600E-08 |
| 5424 | 272 | S272D | A | 6.5e-09 | 1.4e-08 | 7.6e-09 | 9.367E-09 | 4.050E-09 | 6.500E-09 | 1.400E-08 |
| 5425 | 272 | S272C | A | 1.1e-08 | 4.5e-09 | 2.3e-09 | 5.933E-09 | 4.524E-09 | 2.300E-09 | 1.100E-08 |
| 5426 | 272 | S272E | A | 3.1e-09 | 4.1e-09 | 7.8e-09 | 5.000E-09 | 2.476E-09 | 3.100E-09 | 7.800E-09 |
| 5427 | 272 | S272Q | A | 5.5e-09 | 1.2e-08 | 9.1e-09 | 8.867E-09 | 3.256E-09 | 5.500E-09 | 1.200E-08 |
| 5428 | 272 | S272G | A | 6.1e-09 | 6.8e-09 | 4.9e-09 | 5.933E-09 | 9.609E-10 | 4.900E-09 | 6.800E-09 |
| 5429 | 272 | S272H | A | 9.6e-09 | 7.4e-09 | 5.4e-09 | 7.467E-09 | 2.101E-09 | 5.400E-09 | 9.600E-09 |
| 5430 | 272 | S272I | A | 1.2e-08 | 2.6e-08 | 4.3e-09 | 1.410E-08 | 1.100E-08 | 4.300E-09 | 2.600E-08 |
| 5431 | 272 | S272L | A | 1.1e-08 | 2.2e-09 | 8e-09 | 7.067E-09 | 4.474E-09 | 2.200E-09 | 1.100E-08 |
| 5432 | 272 | S272K | A | 9.3e-09 | 1.3e-09 | 1.7e-08 | 9.200E-09 | 7.850E-09 | 1.300E-09 | 1.700E-08 |
| 5433 | 272 | S272M | A | 2.4e-08 | 4.8e-09 | 8e-09 | 1.227E-08 | 1.029E-08 | 4.800E-09 | 2.400E-08 |
| 5434 | 272 | S272F | A | 6e-09 | 1.4e-08 | 6e-09 | 8.667E-09 | 4.619E-09 | 6.000E-09 | 1.400E-08 |
| 5435 | 272 | S272P | A | 1.6e-09 | 1.3e-08 | 1e-08 | 8.200E-09 | 5.909E-09 | 1.600E-09 | 1.300E-08 |
| 5436 | 272 | S272S | A | 1.2e-08 | 8.7e-09 | 6e-09 | 8.900E-09 | 3.005E-09 | 6.000E-09 | 1.200E-08 |
| 5437 | 272 | S272T | A | 1.7e-08 | 4.6e-09 | 7.3e-09 | 9.633E-09 | 6.521E-09 | 4.600E-09 | 1.700E-08 |
| 5438 | 272 | S272W | A | 5.2e-09 | 4.8e-09 | 9.5e-09 | 6.500E-09 | 2.606E-09 | 4.800E-09 | 9.500E-09 |
| 5439 | 272 | S272Y | A | 7.3e-09 | 1.2e-08 | 3.5e-09 | 7.600E-09 | 4.258E-09 | 3.500E-09 | 1.200E-08 |

| | | | | | | | | | | |
|---|---|---|---|---|---|---|---|---|---|---|
| 5440 | 272 | S272V | A | 7e-09 | 4.3e-09 | 4.6e-09 | 5.300E-09 | 1.480E-09 | 4.300E-09 | 7.000E-09 |
| 5441 | 273 | C273A | A | 9.4e-09 | 3.3e-09 | 1.3e-08 | 8.567E-09 | 4.903E-09 | 3.300E-09 | 1.300E-08 |
| 5442 | 273 | C273R | A | 1.3e-08 | 1.8e-08 | 1.7e-08 | 1.600E-08 | 2.646E-09 | 1.300E-08 | 1.800E-08 |
| 5443 | 273 | C273N | A | 7.9e-09 | 1e-08 | 2.2e-08 | 1.330E-08 | 7.607E-09 | 7.900E-09 | 2.200E-08 |
| 5444 | 273 | C273D | A | 3e-09 | 1.3e-08 | 2.3e-08 | 1.300E-08 | 1.000E-08 | 3.000E-09 | 2.300E-08 |
| 5445 | 273 | C273C | A | 6.8e-09 | 1.3e-08 | 1.1e-08 | 1.027E-08 | 3.164E-09 | 6.800E-09 | 1.300E-08 |
| 5446 | 273 | C273E | A | 2.5e-08 | 1.8e-08 | 1e-08 | 1.767E-08 | 7.506E-09 | 1.000E-08 | 2.500E-08 |
| 5447 | 273 | C273Q | A | 2.4e-08 | 9e-09 | 9.5e-09 | 1.417E-08 | 8.520E-09 | 9.000E-09 | 2.400E-08 |
| 5448 | 273 | C273G | A | 1.8e-08 | 8.1e-09 | 3.7e-09 | 9.933E-09 | 7.324E-09 | 3.700E-09 | 1.800E-08 |
| 5449 | 273 | C273H | A | 3.4e-09 | 8.3e-09 | 1.8e-08 | 9.900E-09 | 7.430E-09 | 3.400E-09 | 1.800E-08 |
| 5450 | 273 | C273I | A | 9.2e-09 | 5.5e-09 | 4.2e-09 | 6.300E-09 | 2.594E-09 | 4.200E-09 | 9.200E-09 |
| 5451 | 273 | C273L | A | 5.2e-09 | 2.8e-08 | 5.7e-09 | 1.297E-08 | 1.302E-08 | 5.200E-09 | 2.800E-08 |
| 5452 | 273 | C273K | A | 3.7e-09 | 5.1e-09 | 8.1e-09 | 5.633E-09 | 2.248E-09 | 3.700E-09 | 8.100E-09 |
| 5453 | 273 | C273M | A | 4.5e-09 | 4.1e-09 | 7.5e-09 | 5.367E-09 | 1.858E-09 | 4.100E-09 | 7.500E-09 |
| 5454 | 273 | C273F | A | 8.9e-09 | 1e-08 | 5.9e-09 | 8.267E-09 | 2.122E-09 | 5.900E-09 | 1.000E-08 |
| 5455 | 273 | C273P | A | 5.6e-09 | 9.1e-09 | 1.3e-08 | 9.233E-09 | 3.702E-09 | 5.600E-09 | 1.300E-08 |
| 5456 | 273 | C273S | A | 8e-09 | 7e-09 | 1.2e-08 | 9.000E-09 | 2.646E-09 | 7.000E-09 | 1.200E-08 |
| 5457 | 273 | C273T | A | 6.7e-09 | 7.1e-09 | 3.4e-09 | 5.733E-09 | 2.031E-09 | 3.400E-09 | 7.100E-09 |
| 5458 | 273 | C273W | A | 6.3e-09 | 7.5e-09 | 9.9e-09 | 7.900E-09 | 1.833E-09 | 6.300E-09 | 9.900E-09 |
| 5459 | 273 | C273Y | A | 6.3e-09 | 8.5e-09 | 4.9e-09 | 6.567E-09 | 1.815E-09 | 4.900E-09 | 8.500E-09 |
| 5460 | 273 | C273V | A | 4.1e-09 | 1e-08 | 1.5e-08 | 9.700E-09 | 5.456E-09 | 4.100E-09 | 1.500E-08 |
| 5461 | 274 | V274A | A | 1.3e-08 | 1e-08 | 4.9e-09 | 9.300E-09 | 4.095E-09 | 4.900E-09 | 1.300E-08 |
| 5462 | 274 | V274R | A | 4.2e-09 | 8.6e-09 | 1.7e-08 | 9.933E-09 | 6.503E-09 | 4.200E-09 | 1.700E-08 |
| 5463 | 274 | V274N | A | 1.3e-08 | 5e-09 | 6.2e-09 | 8.067E-09 | 4.314E-09 | 5.000E-09 | 1.300E-08 |
| 5464 | 274 | V274D | A | 5.9e-09 | 4.6e-09 | 1.2e-08 | 7.500E-09 | 3.951E-09 | 4.600E-09 | 1.200E-08 |
| 5465 | 274 | V274C | A | 5.7e-09 | 1.6e-08 | 6.2e-09 | 9.300E-09 | 5.808E-09 | 5.700E-09 | 1.600E-08 |
| 5466 | 274 | V274E | A | 6.3e-09 | 1.1e-08 | 6.4e-09 | 7.900E-09 | 2.685E-09 | 6.300E-09 | 1.100E-08 |
| 5467 | 274 | V274Q | A | 2.4e-08 | 5.7e-09 | 8.7e-09 | 1.280E-08 | 9.815E-09 | 5.700E-09 | 2.400E-08 |
| 5468 | 274 | V274G | A | 5.8e-09 | 1.1e-08 | 3e-09 | 6.600E-09 | 4.060E-09 | 3.000E-09 | 1.100E-08 |
| 5469 | 274 | V274H | A | 3.7e-09 | 6.3e-09 | 2.9e-09 | 4.300E-09 | 1.778E-09 | 2.900E-09 | 6.300E-09 |
| 5470 | 274 | V274I | A | 1.8e-08 | 1.5e-09 | 8.6e-09 | 9.367E-09 | 8.277E-09 | 1.500E-09 | 1.800E-08 |
| 5471 | 274 | V274L | A | 7.6e-09 | 4.7e-09 | 3.6e-09 | 5.300E-09 | 2.066E-09 | 3.600E-09 | 7.600E-09 |
| 5472 | 274 | V274K | A | 9.1e-09 | 5.4e-09 | 2.3e-08 | 1.250E-08 | 9.280E-09 | 5.400E-09 | 2.300E-08 |
| 5473 | 274 | V274M | A | 1.2e-08 | 1.3e-08 | 1.7e-09 | 8.900E-09 | 6.255E-09 | 1.700E-09 | 1.300E-08 |
| 5474 | 274 | V274F | A | 4.7e-09 | 5.5e-09 | 6.9e-09 | 5.700E-09 | 1.114E-09 | 4.700E-09 | 6.900E-09 |
| 5475 | 274 | V274P | A | 2.5e-08 | 1.1e-08 | 7.2e-09 | 1.440E-08 | 9.374E-09 | 7.200E-09 | 2.500E-08 |
| 5476 | 274 | V274S | A | 1.5e-08 | 8.3e-09 | 1.6e-08 | 1.310E-08 | 4.187E-09 | 8.300E-09 | 1.600E-08 |
| 5477 | 274 | V274T | A | 1.4e-08 | 1e-08 | 5.3e-09 | 9.767E-09 | 4.355E-09 | 5.300E-09 | 1.400E-08 |
| 5478 | 274 | V274W | A | 1.4e-08 | 5.2e-09 | 3.9e-09 | 7.700E-09 | 5.495E-09 | 3.900E-09 | 1.400E-08 |
| 5479 | 274 | V274Y | A | 5.5e-09 | 5.2e-09 | 8.2e-09 | 6.300E-09 | 1.652E-09 | 5.200E-09 | 8.200E-09 |
| 5480 | 274 | V274V | A | 1.5e-08 | 5.2e-09 | 1e-08 | 1.007E-08 | 4.900E-09 | 5.200E-09 | 1.500E-08 |
| 5481 | 275 | T275A | A | 3e-08 | 7.5e-09 | 7.1e-09 | 1.487E-08 | 1.311E-08 | 7.100E-09 | 3.000E-08 |
| 5482 | 275 | T275R | A | 7.2e-09 | 1.2e-08 | 1.9e-08 | 1.273E-08 | 5.934E-09 | 7.200E-09 | 1.900E-08 |
| 5483 | 275 | T275N | A | 5.8e-09 | 4.2e-09 | 6e-09 | 5.333E-09 | 9.866E-10 | 4.200E-09 | 6.000E-09 |
| 5484 | 275 | T275D | A | 2.4e-09 | 4.4e-09 | 4.4e-09 | 3.733E-09 | 1.155E-09 | 2.400E-09 | 4.400E-09 |
| 5485 | 275 | T275C | A | 8.6e-09 | 6.5e-09 | 6.1e-09 | 7.067E-09 | 1.343E-09 | 6.100E-09 | 8.600E-09 |
| 5486 | 275 | T275E | A | 1.7e-08 | 6.1e-09 | 1.5e-08 | 1.270E-08 | 5.803E-09 | 6.100E-09 | 1.700E-08 |
| 5487 | 275 | T275Q | A | 3.1e-09 | 6.1e-09 | 3e-09 | 4.067E-09 | 1.762E-09 | 3.000E-09 | 6.100E-09 |
| 5488 | 275 | T275G | A | 1.2e-08 | 4.7e-09 | 1.1e-08 | 9.233E-09 | 3.958E-09 | 4.700E-09 | 1.200E-08 |
| 5489 | 275 | T275H | A | 9.7e-09 | 4.4e-09 | 1.1e-08 | 8.367E-09 | 3.496E-09 | 4.400E-09 | 1.100E-08 |
| 5490 | 275 | T275I | A | 1.9e-08 | 3.9e-09 | 8.5e-09 | 1.047E-08 | 7.740E-09 | 3.900E-09 | 1.900E-08 |
| 5491 | 275 | T275L | A | 8.2e-09 | 5.2e-09 | 7.6e-09 | 7.000E-09 | 1.587E-09 | 5.200E-09 | 8.200E-09 |
| 5492 | 275 | T275K | A | 2.4e-09 | 4e-09 | 1.1e-08 | 5.800E-09 | 4.574E-09 | 2.400E-09 | 1.100E-08 |
| 5493 | 275 | T275M | A | 5.1e-09 | 1.6e-08 | 6.4e-09 | 9.167E-09 | 5.953E-09 | 5.100E-09 | 1.600E-08 |
| 5494 | 275 | T275F | A | 6.6e-09 | 5.1e-09 | 5.3e-09 | 5.667E-09 | 8.145E-10 | 5.100E-09 | 6.600E-09 |
| 5495 | 275 | T275P | A | 3.1e-09 | 2.7e-09 | 2.2e-08 | 9.267E-09 | 1.103E-08 | 2.700E-09 | 2.200E-08 |
| 5496 | 275 | T275S | A | 7.4e-09 | 7.3e-09 | 5.2e-09 | 6.633E-09 | 1.242E-09 | 5.200E-09 | 7.400E-09 |
| 5497 | 275 | T275T | A | 4.8e-09 | 7e-09 | 9.8e-09 | 7.200E-09 | 2.506E-09 | 4.800E-09 | 9.800E-09 |
| 5498 | 275 | T275W | A | 2.5e-09 | 3.4e-09 | 6.2e-09 | 4.033E-09 | 1.930E-09 | 2.500E-09 | 6.200E-09 |
| 5499 | 275 | T275Y | A | 9.5e-09 | 6.2e-09 | 2.7e-09 | 6.133E-09 | 3.400E-09 | 2.700E-09 | 9.500E-09 |
| 5500 | 275 | T275V | A | 9.1e-09 | 7.9e-09 | 8.5e-09 | 8.500E-09 | 6.000E-10 | 7.900E-09 | 9.100E-09 |
| 5501 | 276 | A276A | A | 5.6e-09 | 1.4e-08 | 7.5e-09 | 9.033E-09 | 4.405E-09 | 5.600E-09 | 1.400E-08 |
| 5502 | 276 | A276R | A | 1.4e-09 | 5.5e-09 | 7.5e-09 | 4.800E-09 | 3.110E-09 | 1.400E-09 | 7.500E-09 |
| 5503 | 276 | A276N | A | 1.7e-08 | 1.1e-08 | 3.1e-09 | 1.037E-08 | 6.972E-09 | 3.100E-09 | 1.700E-08 |
| 5504 | 276 | A276D | A | 8.6e-09 | 9.2e-09 | 1.1e-08 | 9.600E-09 | 1.249E-09 | 8.600E-09 | 1.100E-08 |
| 5505 | 276 | A276C | A | 3e-08 | 1.2e-09 | 2.7e-09 | 1.130E-08 | 1.621E-08 | 1.200E-09 | 3.000E-08 |
| 5506 | 276 | A276E | A | 9e-09 | 1e-08 | 1.1e-08 | 1.000E-08 | 1.000E-09 | 9.000E-09 | 1.100E-08 |
| 5507 | 276 | A276Q | A | 1.2e-08 | 7e-09 | 1.1e-08 | 1.000E-08 | 2.646E-09 | 7.000E-09 | 1.200E-08 |
| 5508 | 276 | A276G | A | 5.7e-09 | 4.7e-09 | 1.7e-08 | 9.133E-09 | 6.831E-09 | 4.700E-09 | 1.700E-08 |
| 5509 | 276 | A276H | A | 1.2e-08 | 3.5e-08 | 1.2e-08 | 1.967E-08 | 1.328E-08 | 1.200E-08 | 3.500E-08 |
| 5510 | 276 | A276I | A | 7.3e-09 | 4.2e-09 | 9.8e-09 | 7.100E-09 | 2.805E-09 | 4.200E-09 | 9.800E-09 |
| 5511 | 276 | A276L | A | 6.1e-09 | 5.2e-09 | 6.3e-09 | 5.867E-09 | 5.859E-10 | 5.200E-09 | 6.300E-09 |
| 5512 | 276 | A276K | A | 2.2e-08 | 6.5e-09 | 3.8e-08 | 2.217E-08 | 1.575E-08 | 6.500E-09 | 3.800E-08 |
| 5513 | 276 | A276M | A | 5.5e-09 | 6.1e-09 | 4.2e-09 | 5.267E-09 | 9.713E-10 | 4.200E-09 | 6.100E-09 |
| 5514 | 276 | A276F | A | 2.4e-09 | 9.9e-09 | 7.2e-09 | 6.500E-09 | 3.799E-09 | 2.400E-09 | 9.900E-09 |
| 5515 | 276 | A276P | A | 9.5e-09 | 5.6e-09 | 8e-09 | 7.700E-09 | 1.967E-09 | 5.600E-09 | 9.500E-09 |
| 5516 | 276 | A276S | A | 3.4e-09 | 2.7e-09 | 1.4e-08 | 6.700E-09 | 6.332E-09 | 2.700E-09 | 1.400E-08 |
| 5517 | 276 | A276T | A | 7.6e-09 | 1.1e-08 | 7.8e-09 | 8.800E-09 | 1.908E-09 | 7.600E-09 | 1.100E-08 |
| 5518 | 276 | A276W | A | 8.6e-09 | 5.4e-09 | 4.9e-09 | 6.300E-09 | 2.007E-09 | 4.900E-09 | 8.600E-09 |
| 5519 | 276 | A276Y | A | 6.6e-09 | 4.5e-09 | 1.5e-08 | 8.700E-09 | 5.556E-09 | 4.500E-09 | 1.500E-08 |
| 5520 | 276 | A276V | A | 2.9e-09 | 6.5e-09 | 4e-09 | 4.467E-09 | 1.845E-09 | 2.900E-09 | 6.500E-09 |
| 5521 | 277 | C277A | A | 3.1e-09 | 9.5e-09 | 2.8e-09 | 5.133E-09 | 3.785E-09 | 2.800E-09 | 9.500E-09 |
| 5522 | 277 | C277R | A | 9.8e-09 | 1.9e-08 | 2.9e-08 | 1.927E-08 | 9.603E-09 | 9.800E-09 | 2.900E-08 |
| 5523 | 277 | C277N | A | 1.2e-08 | 7.1e-09 | 5.2e-09 | 8.100E-09 | 3.509E-09 | 5.200E-09 | 1.200E-08 |
| 5524 | 277 | C277D | A | 6.9e-09 | 4.9e-09 | 9.3e-09 | 7.033E-09 | 2.203E-09 | 4.900E-09 | 9.300E-09 |

| | | | | | | | | | | |
|---|---|---|---|---|---|---|---|---|---|---|
| 5525 | 277 | C277C | A | 5.3e-09 | 5.6e-09 | 2.1e-09 | 4.333E-09 | 1.940E-09 | 2.100E-09 | 5.600E-09 |
| 5526 | 277 | C277E | A | 5.8e-09 | 6e-09 | 7e-09 | 6.267E-09 | 6.429E-10 | 5.800E-09 | 7.000E-09 |
| 5527 | 277 | C277Q | A | 2.6e-09 | 5.1e-09 | 1.1e-08 | 6.233E-09 | 4.313E-09 | 2.600E-09 | 1.100E-08 |
| 5528 | 277 | C277G | A | 1.3e-08 | 9.4e-09 | 1e-08 | 1.080E-08 | 1.929E-09 | 9.400E-09 | 1.300E-08 |
| 5529 | 277 | C277H | A | 1.2e-08 | 3.1e-09 | 4.3e-09 | 6.467E-09 | 4.829E-09 | 3.100E-09 | 1.200E-08 |
| 5530 | 277 | C277I | A | 2.4e-09 | 5e-09 | 3.3e-09 | 3.567E-09 | 1.320E-09 | 2.400E-09 | 5.000E-09 |
| 5531 | 277 | C277L | A | 1e-08 | 1.5e-08 | 4.9e-09 | 9.967E-09 | 5.050E-09 | 4.900E-09 | 1.500E-08 |
| 5532 | 277 | C277K | A | 9.4e-09 | 5.7e-09 | 7.9e-09 | 7.667E-09 | 1.861E-09 | 5.700E-09 | 9.400E-09 |
| 5533 | 277 | C277M | A | 5e-09 | 3e-09 | 3.8e-09 | 3.933E-09 | 1.007E-09 | 3.000E-09 | 5.000E-09 |
| 5534 | 277 | C277F | A | 2.9e-08 | 1.1e-08 | 1.7e-08 | 1.900E-08 | 9.165E-09 | 1.100E-08 | 2.900E-08 |
| 5535 | 277 | C277P | A | 9.2e-09 | 2.5e-09 | 1.2e-08 | 7.900E-09 | 4.882E-09 | 2.500E-09 | 1.200E-08 |
| 5536 | 277 | C277S | A | 7e-09 | 8.1e-09 | 1e-08 | 8.367E-09 | 1.518E-09 | 7.000E-09 | 1.000E-08 |
| 5537 | 277 | C277T | A | 4.5e-09 | 1.6e-08 | 1.7e-08 | 1.250E-08 | 6.946E-09 | 4.500E-09 | 1.700E-08 |
| 5538 | 277 | C277W | A | 7.2e-09 | 7.2e-09 | 3.7e-09 | 6.033E-09 | 2.021E-09 | 3.700E-09 | 7.200E-09 |
| 5539 | 277 | C277Y | A | 5.9e-09 | 5.7e-09 | 4.6e-09 | 5.400E-09 | 7.000E-10 | 4.600E-09 | 5.900E-09 |
| 5540 | 277 | C277V | A | 7.4e-09 | 1.1e-09 | 8.3e-09 | 5.600E-09 | 3.923E-09 | 1.100E-09 | 8.300E-09 |
| 5541 | 278 | P278A | A | 4.6e-09 | 1.6e-09 | 3.7e-09 | 3.300E-09 | 1.539E-09 | 1.600E-09 | 4.600E-09 |
| 5542 | 278 | P278R | A | 7.8e-09 | 5.8e-09 | 6e-09 | 6.533E-09 | 1.102E-09 | 5.800E-09 | 7.800E-09 |
| 5543 | 278 | P278N | A | 9.7e-09 | 8.9e-09 | 9.7e-09 | 9.433E-09 | 4.619E-10 | 8.900E-09 | 9.700E-09 |
| 5544 | 278 | P278D | A | 6.5e-09 | 9.2e-09 | 7.7e-09 | 7.800E-09 | 1.353E-09 | 6.500E-09 | 9.200E-09 |
| 5545 | 278 | P278C | A | 8.9e-09 | 4.3e-09 | 6.9e-09 | 6.700E-09 | 2.307E-09 | 4.300E-09 | 8.900E-09 |
| 5546 | 278 | P278E | A | 1.3e-08 | 1.3e-08 | 1.8e-08 | 1.467E-08 | 2.887E-09 | 1.300E-08 | 1.800E-08 |
| 5547 | 278 | P278Q | A | 1.2e-08 | 1.9e-08 | 3.8e-09 | 1.160E-08 | 7.608E-09 | 3.800E-09 | 1.900E-08 |
| 5548 | 278 | P278G | A | 6.3e-09 | 4.3e-09 | 8.7e-09 | 6.433E-09 | 2.203E-09 | 4.300E-09 | 8.700E-09 |
| 5549 | 278 | P278H | A | 8.6e-09 | 3.3e-09 | 2.3e-08 | 1.163E-08 | 1.019E-08 | 3.300E-09 | 2.300E-08 |
| 5550 | 278 | P278I | A | 7.4e-09 | 4.1e-09 | 2.1e-09 | 4.533E-09 | 2.676E-09 | 2.100E-09 | 7.400E-09 |
| 5551 | 278 | P278L | A | 4.8e-09 | 9.6e-09 | 2.7e-09 | 5.700E-09 | 3.537E-09 | 2.700E-09 | 9.600E-09 |
| 5552 | 278 | P278K | A | 4.9e-09 | 8e-09 | 7.2e-09 | 6.700E-09 | 1.609E-09 | 4.900E-09 | 8.000E-09 |
| 5553 | 278 | P278M | A | 1.1e-08 | 4.8e-09 | 4.4e-09 | 6.733E-09 | 3.700E-09 | 4.400E-09 | 1.100E-08 |
| 5554 | 278 | P278F | A | 3.2e-09 | 5e-09 | 5.3e-09 | 4.500E-09 | 1.136E-09 | 3.200E-09 | 5.300E-09 |
| 5555 | 278 | P278P | A | 5.6e-09 | 8.6e-09 | 3.5e-09 | 5.900E-09 | 2.563E-09 | 3.500E-09 | 8.600E-09 |
| 5556 | 278 | P278S | A | 9.4e-09 | 2.8e-09 | 3.2e-09 | 5.133E-09 | 3.700E-09 | 2.800E-09 | 9.400E-09 |
| 5557 | 278 | P278T | A | 2.3e-09 | 1.1e-09 | 4.6e-09 | 2.667E-09 | 1.779E-09 | 1.100E-09 | 4.600E-09 |
| 5558 | 278 | P278W | A | 3.2e-09 | 9.4e-09 | 5.9e-09 | 6.167E-09 | 3.109E-09 | 3.200E-09 | 9.400E-09 |
| 5559 | 278 | P278Y | A | 2.8e-09 | 1e-08 | 1.2e-08 | 8.267E-09 | 4.839E-09 | 2.800E-09 | 1.200E-08 |
| 5560 | 278 | P278V | A | 2.4e-09 | 4.2e-09 | 4.4e-09 | 3.667E-09 | 1.102E-09 | 2.400E-09 | 4.400E-09 |
| 5561 | 279 | Y279A | A | 4.7e-09 | 1.2e-08 | 4.5e-09 | 7.067E-09 | 4.274E-09 | 4.500E-09 | 1.200E-08 |
| 5562 | 279 | Y279R | A | 1.6e-08 | 1e-08 | 4.2e-09 | 1.007E-08 | 5.900E-09 | 4.200E-09 | 1.600E-08 |
| 5563 | 279 | Y279N | A | 6.2e-09 | 7.6e-09 | 6.1e-09 | 6.633E-09 | 8.386E-10 | 6.100E-09 | 7.600E-09 |
| 5564 | 279 | Y279D | A | 7.4e-09 | 1.7e-08 | 3e-09 | 9.133E-09 | 7.159E-09 | 3.000E-09 | 1.700E-08 |
| 5565 | 279 | Y279C | A | 5e-09 | 1.7e-08 | 1.6e-08 | 1.267E-08 | 6.658E-09 | 5.000E-09 | 1.700E-08 |
| 5566 | 279 | Y279E | A | 1.7e-08 | 1e-08 | 6.6e-09 | 1.120E-08 | 5.303E-09 | 6.600E-09 | 1.700E-08 |
| 5567 | 279 | Y279Q | A | 4.7e-09 | 4.1e-09 | 2.6e-09 | 3.800E-09 | 1.082E-09 | 2.600E-09 | 4.700E-09 |
| 5568 | 279 | Y279G | A | 1.1e-08 | 1e-08 | 1.2e-08 | 1.100E-08 | 1.000E-09 | 1.000E-08 | 1.200E-08 |
| 5569 | 279 | Y279H | A | 4.2e-09 | 1.1e-08 | 5.2e-09 | 6.800E-09 | 3.672E-09 | 4.200E-09 | 1.100E-08 |
| 5570 | 279 | Y279I | A | 8e-09 | 2e-08 | 3.5e-08 | 2.100E-08 | 1.353E-08 | 8.000E-09 | 3.500E-08 |
| 5571 | 279 | Y279L | A | 4.7e-09 | 3.4e-09 | 1.6e-08 | 8.033E-09 | 6.930E-09 | 3.400E-09 | 1.600E-08 |
| 5572 | 279 | Y279K | A | 1.1e-08 | 1.2e-08 | 6.2e-09 | 9.733E-09 | 3.101E-09 | 6.200E-09 | 1.200E-08 |
| 5573 | 279 | Y279M | A | 1.4e-08 | 4.2e-09 | 2.9e-08 | 1.573E-08 | 1.249E-08 | 4.200E-09 | 2.900E-08 |
| 5574 | 279 | Y279F | A | 8.3e-09 | 8.2e-09 | 3.6e-09 | 6.700E-09 | 2.685E-09 | 3.600E-09 | 8.300E-09 |
| 5575 | 279 | Y279P | A | 7e-09 | 9.3e-09 | 1.1e-08 | 9.100E-09 | 2.007E-09 | 7.000E-09 | 1.100E-08 |
| 5576 | 279 | Y279S | A | 1.5e-08 | 1.5e-08 | 8.7e-09 | 1.290E-08 | 3.637E-09 | 8.700E-09 | 1.500E-08 |
| 5577 | 279 | Y279T | A | 4.3e-09 | 1.3e-08 | 1.2e-08 | 9.767E-09 | 4.761E-09 | 4.300E-09 | 1.300E-08 |
| 5578 | 279 | Y279W | A | 7.3e-09 | 5.8e-09 | 8e-09 | 7.033E-09 | 1.124E-09 | 5.800E-09 | 8.000E-09 |
| 5579 | 279 | Y279Y | A | 2.1e-09 | 7.7e-09 | 9.8e-09 | 6.533E-09 | 3.980E-09 | 2.100E-09 | 9.800E-09 |
| 5580 | 279 | Y279V | A | 1.5e-08 | 8.3e-09 | 5e-09 | 9.433E-09 | 5.095E-09 | 5.000E-09 | 1.500E-08 |
| 5581 | 280 | N280A | A | 1.2e-08 | 4.3e-09 | 4.4e-09 | 6.900E-09 | 4.417E-09 | 4.300E-09 | 1.200E-08 |
| 5582 | 280 | N280R | A | 1.2e-08 | 5.1e-09 | 3.9e-09 | 7.000E-09 | 4.371E-09 | 3.900E-09 | 1.200E-08 |
| 5583 | 280 | N280N | A | 5.5e-09 | 1.6e-08 | 6.4e-09 | 9.300E-09 | 5.820E-09 | 5.500E-09 | 1.600E-08 |
| 5584 | 280 | N280D | A | 4e-09 | 3.7e-09 | 1.9e-08 | 8.900E-09 | 8.748E-09 | 3.700E-09 | 1.900E-08 |
| 5585 | 280 | N280C | A | 1.3e-08 | 1.4e-08 | 2.8e-09 | 9.933E-09 | 6.198E-09 | 2.800E-09 | 1.400E-08 |
| 5586 | 280 | N280E | A | 3.3e-09 | 2.8e-09 | 2.4e-08 | 1.003E-08 | 1.210E-08 | 2.800E-09 | 2.400E-08 |
| 5587 | 280 | N280Q | A | 1.2e-08 | 4.7e-09 | 6.4e-09 | 7.700E-09 | 3.820E-09 | 4.700E-09 | 1.200E-08 |
| 5588 | 280 | N280G | A | 1.1e-08 | 2.1e-08 | 8.5e-09 | 1.350E-08 | 6.614E-09 | 8.500E-09 | 2.100E-08 |
| 5589 | 280 | N280H | A | 5.6e-09 | 7.8e-09 | 7.1e-09 | 6.833E-09 | 1.124E-09 | 5.600E-09 | 7.800E-09 |
| 5590 | 280 | N280I | A | 6e-09 | 2.9e-09 | 1.2e-08 | 6.967E-09 | 4.626E-09 | 2.900E-09 | 1.200E-08 |
| 5591 | 280 | N280L | A | 2.1e-08 | 5.9e-09 | 2.1e-08 | 1.597E-08 | 8.718E-09 | 5.900E-09 | 2.100E-08 |
| 5592 | 280 | N280K | A | 1.1e-08 | 5.8e-09 | 3.2e-09 | 6.667E-09 | 3.972E-09 | 3.200E-09 | 1.100E-08 |
| 5593 | 280 | N280M | A | 8.7e-09 | 8.6e-09 | 5.7e-09 | 7.667E-09 | 1.704E-09 | 5.700E-09 | 8.700E-09 |
| 5594 | 280 | N280F | A | 7.2e-09 | 2.1e-08 | 5.8e-09 | 1.133E-08 | 8.401E-09 | 5.800E-09 | 2.100E-08 |
| 5595 | 280 | N280P | A | 5.7e-09 | 8.7e-09 | 2.6e-09 | 5.667E-09 | 3.050E-09 | 2.600E-09 | 8.700E-09 |
| 5596 | 280 | N280S | A | 3.1e-09 | 1.5e-08 | 2e-08 | 1.270E-08 | 8.682E-09 | 3.100E-09 | 2.000E-08 |
| 5597 | 280 | N280T | A | 5.7e-09 | 1.4e-08 | 4.5e-09 | 8.067E-09 | 5.173E-09 | 4.500E-09 | 1.400E-08 |
| 5598 | 280 | N280W | A | 3.4e-09 | 5.8e-09 | 9.8e-09 | 6.333E-09 | 3.233E-09 | 3.400E-09 | 9.800E-09 |
| 5599 | 280 | N280Y | A | 1.1e-08 | 1.6e-08 | 5.4e-09 | 1.080E-08 | 5.303E-09 | 5.400E-09 | 1.600E-08 |
| 5600 | 280 | N280V | A | 6.5e-09 | 3.1e-09 | 7.7e-09 | 5.767E-09 | 2.386E-09 | 3.100E-09 | 7.700E-09 |
| 5601 | 281 | Y281A | A | 2e-09 | 3e-09 | 6.7e-09 | 3.900E-09 | 2.476E-09 | 2.000E-09 | 6.700E-09 |
| 5602 | 281 | Y281R | A | 5.8e-09 | 9.5e-09 | 5.4e-09 | 6.900E-09 | 2.261E-09 | 5.400E-09 | 9.500E-09 |
| 5603 | 281 | Y281N | A | 9.3e-09 | 5e-09 | 3.3e-09 | 5.867E-09 | 3.092E-09 | 3.300E-09 | 9.300E-09 |
| 5604 | 281 | Y281D | A | 2.2e-09 | 7.8e-09 | 2.3e-09 | 4.100E-09 | 3.205E-09 | 2.200E-09 | 7.800E-09 |
| 5605 | 281 | Y281C | A | 6.3e-09 | 7.2e-09 | 1.8e-09 | 5.100E-09 | 2.893E-09 | 1.800E-09 | 7.200E-09 |
| 5606 | 281 | Y281E | A | 2.6e-09 | 6.4e-09 | 1.4e-08 | 7.667E-09 | 5.805E-09 | 2.600E-09 | 1.400E-08 |
| 5607 | 281 | Y281Q | A | 4e-09 | 9.9e-09 | 3.4e-09 | 5.767E-09 | 3.592E-09 | 3.400E-09 | 9.900E-09 |
| 5608 | 281 | Y281G | A | 9.1e-09 | 2.6e-09 | 1.9e-08 | 1.023E-08 | 8.259E-09 | 2.600E-09 | 1.900E-08 |
| 5609 | 281 | Y281H | A | 6.1e-09 | 7.4e-09 | 8.1e-09 | 7.200E-09 | 1.015E-09 | 6.100E-09 | 8.100E-09 |

| | | | | | | | | | | |
|---|---|---|---|---|---|---|---|---|---|---|
| 5610 | 281 | Y281I | A | 4.8e-09 | 6.1e-09 | 1.2e-08 | 7.633E-09 | 3.837E-09 | 4.800E-09 | 1.200E-08 |
| 5611 | 281 | Y281L | A | 1.6e-08 | 5.3e-09 | 6.7e-09 | 9.333E-09 | 5.816E-09 | 5.300E-09 | 1.600E-08 |
| 5612 | 281 | Y281K | A | 1e-08 | 4.8e-09 | 9.1e-09 | 7.967E-09 | 2.779E-09 | 4.800E-09 | 1.000E-08 |
| 5613 | 281 | Y281M | A | 4.5e-09 | 7e-09 | 4.4e-09 | 5.300E-09 | 1.473E-09 | 4.400E-09 | 7.000E-09 |
| 5614 | 281 | Y281F | A | 1.1e-08 | 3e-09 | 4.6e-09 | 6.200E-09 | 4.233E-09 | 3.000E-09 | 1.100E-08 |
| 5615 | 281 | Y281P | A | 8.3e-09 | 5.4e-09 | 7.9e-09 | 7.200E-09 | 1.572E-09 | 5.400E-09 | 8.300E-09 |
| 5616 | 281 | Y281S | A | 1.5e-08 | 3.5e-09 | 9e-09 | 9.167E-09 | 5.752E-09 | 3.500E-09 | 1.500E-08 |
| 5617 | 281 | Y281T | A | 5.9e-09 | 4.2e-09 | 8e-09 | 6.033E-09 | 1.904E-09 | 4.200E-09 | 8.000E-09 |
| 5618 | 281 | Y281W | A | 5.2e-09 | 1.8e-08 | 5.3e-09 | 9.500E-09 | 7.361E-09 | 5.200E-09 | 1.800E-08 |
| 5619 | 281 | Y281Y | A | 4.7e-09 | 3.2e-09 | 6.7e-09 | 4.867E-09 | 1.756E-09 | 3.200E-09 | 6.700E-09 |
| 5620 | 281 | Y281V | A | 8.1e-09 | 6.4e-09 | 5.2e-09 | 6.567E-09 | 1.457E-09 | 5.200E-09 | 8.100E-09 |
| 5621 | 282 | L282A | A | 1.4e-08 | 7.6e-09 | 2.2e-09 | 7.933E-09 | 5.907E-09 | 2.200E-09 | 1.400E-08 |
| 5622 | 282 | L282R | A | 5.6e-09 | 9.5e-09 | 1.1e-08 | 8.700E-09 | 2.787E-09 | 5.600E-09 | 1.100E-08 |
| 5623 | 282 | L282N | A | 1.2e-08 | 8.9e-09 | 3.2e-09 | 8.033E-09 | 4.464E-09 | 3.200E-09 | 1.200E-08 |
| 5624 | 282 | L282D | A | 7.8e-09 | 1.2e-08 | 2.4e-08 | 1.460E-08 | 8.407E-09 | 7.800E-09 | 2.400E-08 |
| 5625 | 282 | L282C | A | 4.2e-09 | 1.2e-08 | 7.2e-09 | 7.800E-09 | 3.934E-09 | 4.200E-09 | 1.200E-08 |
| 5626 | 282 | L282E | A | 7.8e-09 | 7.7e-09 | 7e-09 | 7.500E-09 | 4.359E-10 | 7.000E-09 | 7.800E-09 |
| 5627 | 282 | L282Q | A | 1.3e-09 | 6.4e-09 | 7e-09 | 4.900E-09 | 3.132E-09 | 1.300E-09 | 7.000E-09 |
| 5628 | 282 | L282G | A | 6.5e-09 | 5.3e-09 | 6.4e-09 | 6.067E-09 | 6.658E-10 | 5.300E-09 | 6.500E-09 |
| 5629 | 282 | L282H | A | 2.3e-08 | 5e-09 | 7.6e-09 | 1.187E-08 | 9.729E-09 | 5.000E-09 | 2.300E-08 |
| 5630 | 282 | L282I | A | 1e-08 | 1.5e-08 | 1.5e-08 | 1.333E-08 | 2.887E-09 | 1.000E-08 | 1.500E-08 |
| 5631 | 282 | L282L | A | 4.8e-09 | 1.4e-08 | 1e-08 | 9.600E-09 | 4.613E-09 | 4.800E-09 | 1.400E-08 |
| 5632 | 282 | L282K | A | 8.3e-09 | 2e-08 | 4.3e-09 | 1.087E-08 | 8.159E-09 | 4.300E-09 | 2.000E-08 |
| 5633 | 282 | L282M | A | 9.8e-09 | 4e-09 | 9.6e-10 | 4.920E-09 | 4.491E-09 | 9.600E-10 | 9.800E-09 |
| 5634 | 282 | L282F | A | 7.8e-09 | 1.1e-08 | 6.5e-09 | 8.433E-09 | 2.316E-09 | 6.500E-09 | 1.100E-08 |
| 5635 | 282 | L282P | A | 1.2e-08 | 6.5e-09 | 2.3e-08 | 1.383E-08 | 8.401E-09 | 6.500E-09 | 2.300E-08 |
| 5636 | 282 | L282S | A | 9.3e-09 | 1e-08 | 1.1e-08 | 1.010E-08 | 8.544E-10 | 9.300E-09 | 1.100E-08 |
| 5637 | 282 | L282T | A | 6.5e-09 | 7.8e-09 | 5.1e-09 | 6.467E-09 | 1.350E-09 | 5.100E-09 | 7.800E-09 |
| 5638 | 282 | L282W | A | 1.3e-08 | 5.1e-09 | 1.5e-08 | 1.103E-08 | 5.235E-09 | 5.100E-09 | 1.500E-08 |
| 5639 | 282 | L282Y | A | 3.6e-09 | 8.3e-09 | 7e-09 | 6.300E-09 | 2.427E-09 | 3.600E-09 | 8.300E-09 |
| 5640 | 282 | L282V | A | 8.9e-09 | 6.3e-09 | 9.3e-09 | 8.167E-09 | 1.629E-09 | 6.300E-09 | 9.300E-09 |
| 5641 | 283 | S283A | A | 1.3e-08 | 2.6e-09 | 5e-09 | 6.867E-09 | 5.445E-09 | 2.600E-09 | 1.300E-08 |
| 5642 | 283 | S283R | A | 1.3e-08 | 9.4e-09 | 4.2e-08 | 2.147E-08 | 1.787E-08 | 9.400E-09 | 4.200E-08 |
| 5643 | 283 | S283N | A | 8e-09 | 4.8e-09 | 4.5e-09 | 5.767E-09 | 1.940E-09 | 4.500E-09 | 8.000E-09 |
| 5644 | 283 | S283D | A | 7.4e-09 | 3.8e-08 | 7.5e-09 | 1.763E-08 | 1.764E-08 | 7.400E-09 | 3.800E-08 |
| 5645 | 283 | S283C | A | 6.3e-09 | 3.8e-09 | 1.9e-08 | 9.700E-09 | 8.150E-09 | 3.800E-09 | 1.900E-08 |
| 5646 | 283 | S283E | A | 1.1e-08 | 4.1e-09 | 1.1e-08 | 8.700E-09 | 3.984E-09 | 4.100E-09 | 1.100E-08 |
| 5647 | 283 | S283Q | A | 1.1e-08 | 5.5e-09 | 3e-09 | 6.500E-09 | 4.093E-09 | 3.000E-09 | 1.100E-08 |
| 5648 | 283 | S283G | A | 4.6e-09 | 3.8e-09 | 5.7e-09 | 4.700E-09 | 9.539E-10 | 3.800E-09 | 5.700E-09 |
| 5649 | 283 | S283H | A | 8.3e-09 | 3.3e-09 | 6.5e-09 | 6.033E-09 | 2.532E-09 | 3.300E-09 | 8.300E-09 |
| 5650 | 283 | S283I | A | 3.8e-09 | 2e-08 | 1.6e-08 | 1.327E-08 | 8.439E-09 | 3.800E-09 | 2.000E-08 |
| 5651 | 283 | S283L | A | 7.6e-09 | 7.4e-09 | 2.4e-09 | 5.800E-09 | 2.946E-09 | 2.400E-09 | 7.600E-09 |
| 5652 | 283 | S283K | A | 5.1e-09 | 8.6e-09 | 4.9e-09 | 6.200E-09 | 2.081E-09 | 4.900E-09 | 8.600E-09 |
| 5653 | 283 | S283M | A | 1.1e-08 | 2.7e-09 | 6.5e-09 | 6.733E-09 | 4.155E-09 | 2.700E-09 | 1.100E-08 |
| 5654 | 283 | S283F | A | 3.2e-09 | 6.8e-09 | 7.9e-09 | 5.967E-09 | 2.458E-09 | 3.200E-09 | 7.900E-09 |
| 5655 | 283 | S283P | A | 1.5e-08 | 4.4e-09 | 1.6e-08 | 1.180E-08 | 6.428E-09 | 4.400E-09 | 1.600E-08 |
| 5656 | 283 | S283S | A | 2.7e-09 | 3e-09 | 1.9e-08 | 8.233E-09 | 9.325E-09 | 2.700E-09 | 1.900E-08 |
| 5657 | 283 | S283T | A | 1.9e-08 | 3.6e-09 | 2.3e-09 | 8.300E-09 | 9.289E-09 | 2.300E-09 | 1.900E-08 |
| 5658 | 283 | S283W | A | 2.3e-08 | 7.8e-09 | 2.7e-08 | 1.927E-08 | 1.013E-08 | 7.800E-09 | 2.700E-08 |
| 5659 | 283 | S283Y | A | 7.4e-09 | 7.8e-09 | 4.4e-09 | 6.533E-09 | 1.858E-09 | 4.400E-09 | 7.800E-09 |
| 5660 | 283 | S283V | A | 2.3e-09 | 2.2e-08 | 8.3e-09 | 1.087E-08 | 1.010E-08 | 2.300E-09 | 2.200E-08 |
| 5661 | 284 | T284A | A | 1.1e-08 | 1.6e-08 | 4.7e-09 | 1.057E-08 | 5.662E-09 | 4.700E-09 | 1.600E-08 |
| 5662 | 284 | T284R | A | 4.1e-09 | 4.2e-09 | 1.4e-08 | 7.433E-09 | 5.687E-09 | 4.100E-09 | 1.400E-08 |
| 5663 | 284 | T284N | A | 1.2e-08 | 6.1e-09 | 1.7e-08 | 1.170E-08 | 5.456E-09 | 6.100E-09 | 1.700E-08 |
| 5664 | 284 | T284D | A | 3.4e-09 | 3.5e-09 | 2.8e-09 | 3.233E-09 | 3.786E-10 | 2.800E-09 | 3.500E-09 |
| 5665 | 284 | T284C | A | 3.5e-09 | 7.5e-09 | 5.1e-09 | 5.367E-09 | 2.013E-09 | 3.500E-09 | 7.500E-09 |
| 5666 | 284 | T284E | A | 9.9e-09 | 5.4e-09 | 8.8e-09 | 8.033E-09 | 2.346E-09 | 5.400E-09 | 9.900E-09 |
| 5667 | 284 | T284Q | A | 5.7e-09 | 7e-09 | 6.7e-09 | 6.467E-09 | 6.807E-10 | 5.700E-09 | 7.000E-09 |
| 5668 | 284 | T284G | A | 5.8e-09 | 5.3e-09 | 1.5e-08 | 8.700E-09 | 5.462E-09 | 5.300E-09 | 1.500E-08 |
| 5669 | 284 | T284H | A | 7.5e-09 | 3.7e-09 | 9.4e-09 | 6.867E-09 | 2.902E-09 | 3.700E-09 | 9.400E-09 |
| 5670 | 284 | T284I | A | 1.1e-08 | 1e-08 | 3.1e-09 | 8.033E-09 | 4.302E-09 | 3.100E-09 | 1.100E-08 |
| 5671 | 284 | T284L | A | 7.4e-09 | 5.5e-09 | 4.1e-09 | 5.667E-09 | 1.656E-09 | 4.100E-09 | 7.400E-09 |
| 5672 | 284 | T284K | A | 1.4e-08 | 2.9e-09 | 9.5e-09 | 8.800E-09 | 5.583E-09 | 2.900E-09 | 1.400E-08 |
| 5673 | 284 | T284M | A | 4.2e-09 | 1.3e-09 | 5e-09 | 3.500E-09 | 1.947E-09 | 1.300E-09 | 5.000E-09 |
| 5674 | 284 | T284F | A | 2.1e-09 | 2.8e-09 | 7.2e-09 | 4.033E-09 | 2.765E-09 | 2.100E-09 | 7.200E-09 |
| 5675 | 284 | T284P | A | 6.5e-09 | 9.2e-09 | 1.6e-08 | 1.057E-08 | 4.895E-09 | 6.500E-09 | 1.600E-08 |
| 5676 | 284 | T284S | A | 1.9e-09 | 4.3e-09 | 2.4e-08 | 1.007E-08 | 1.213E-08 | 1.900E-09 | 2.400E-08 |
| 5677 | 284 | T284T | A | 1.8e-08 | 1.3e-08 | 7.3e-09 | 1.277E-08 | 5.354E-09 | 7.300E-09 | 1.800E-08 |
| 5678 | 284 | T284W | A | 5.4e-09 | 4.1e-09 | 3.9e-09 | 4.467E-09 | 8.145E-10 | 3.900E-09 | 5.400E-09 |
| 5679 | 284 | T284Y | A | 2.8e-09 | 5.9e-09 | 7.8e-09 | 5.500E-09 | 2.524E-09 | 2.800E-09 | 7.800E-09 |
| 5680 | 284 | T284V | A | 3.6e-09 | 7.1e-09 | 8.3e-09 | 6.333E-09 | 2.442E-09 | 3.600E-09 | 8.300E-09 |
| 5681 | 285 | D285A | A | 5.9e-09 | 1.9e-08 | 1.2e-08 | 1.230E-08 | 6.555E-09 | 5.900E-09 | 1.900E-08 |
| 5682 | 285 | D285R | A | 4.1e-09 | 4.2e-09 | 7.9e-09 | 5.400E-09 | 2.166E-09 | 4.100E-09 | 7.900E-09 |
| 5683 | 285 | D285N | A | 8.6e-09 | 6.3e-09 | 6.3e-09 | 7.067E-09 | 1.328E-09 | 6.300E-09 | 8.600E-09 |
| 5684 | 285 | D285D | A | 7.8e-09 | 1.7e-08 | 7.9e-09 | 1.090E-08 | 5.283E-09 | 7.800E-09 | 1.700E-08 |
| 5685 | 285 | D285C | A | 7.2e-09 | 9e-09 | 2.9e-09 | 6.367E-09 | 3.134E-09 | 2.900E-09 | 9.000E-09 |
| 5686 | 285 | D285E | A | 2.4e-08 | 3.1e-09 | 8.2e-09 | 1.177E-08 | 1.090E-08 | 3.100E-09 | 2.400E-08 |
| 5687 | 285 | D285Q | A | 2.8e-09 | 4.2e-09 | 1.3e-08 | 6.667E-09 | 5.529E-09 | 2.800E-09 | 1.300E-08 |
| 5688 | 285 | D285G | A | 5.3e-09 | 1.2e-08 | 9.8e-09 | 9.033E-09 | 3.415E-09 | 5.300E-09 | 1.200E-08 |
| 5689 | 285 | D285H | A | 5.4e-09 | 1.2e-08 | 6.3e-09 | 7.900E-09 | 3.579E-09 | 5.400E-09 | 1.200E-08 |
| 5690 | 285 | D285I | A | 2e-08 | 5.8e-08 | 1.2e-08 | 3.000E-08 | 2.458E-08 | 1.200E-08 | 5.800E-08 |
| 5691 | 285 | D285L | A | 1.2e-08 | 3.9e-09 | 1.1e-09 | 5.667E-09 | 5.661E-09 | 1.100E-09 | 1.200E-08 |
| 5692 | 285 | D285K | A | 1.3e-09 | 1e-08 | 2.6e-09 | 4.633E-09 | 4.693E-09 | 1.300E-09 | 1.000E-08 |
| 5693 | 285 | D285M | A | 2.3e-09 | 7.3e-09 | 1.8e-08 | 9.200E-09 | 8.021E-09 | 2.300E-09 | 1.800E-08 |
| 5694 | 285 | D285F | A | 2.1e-08 | 6.2e-09 | 5e-09 | 1.073E-08 | 8.911E-09 | 5.000E-09 | 2.100E-08 |

| | | | | | | | | | | |
|---|---|---|---|---|---|---|---|---|---|---|
| 5695 | 285 | D285P | A | 1.9e-09 | 1.4e-08 | 1.2e-08 | 9.300E-09 | 6.486E-09 | 1.900E-09 | 1.400E-08 |
| 5696 | 285 | D285S | A | 2.5e-09 | 1.5e-08 | 7.6e-09 | 8.367E-09 | 6.285E-09 | 2.500E-09 | 1.500E-08 |
| 5697 | 285 | D285T | A | 9.5e-09 | 4.5e-09 | 2.4e-09 | 5.467E-09 | 3.647E-09 | 2.400E-09 | 9.500E-09 |
| 5698 | 285 | D285W | A | 1.5e-08 | 4.8e-09 | 7.9e-09 | 9.233E-09 | 5.229E-09 | 4.800E-09 | 1.500E-08 |
| 5699 | 285 | D285Y | A | 1.5e-08 | 9.6e-09 | 1.1e-08 | 1.187E-08 | 2.802E-09 | 9.600E-09 | 1.500E-08 |
| 5700 | 285 | D285V | A | 4.3e-09 | 4.3e-09 | 4.4e-09 | 4.333E-09 | 5.774E-11 | 4.300E-09 | 4.400E-09 |
| 5701 | 286 | V286A | A | 2e-08 | 8.3e-09 | 4.2e-09 | 1.083E-08 | 8.199E-09 | 4.200E-09 | 2.000E-08 |
| 5702 | 286 | V286R | A | 2e-08 | 3.5e-08 | 6.5e-09 | 2.050E-08 | 1.426E-08 | 6.500E-09 | 3.500E-08 |
| 5703 | 286 | V286N | A | 1e-07 | 1.2e-07 | 1.3e-07 | 1.167E-07 | 1.528E-08 | 1.000E-07 | 1.300E-07 |
| 5704 | 286 | V286D | A | 2.8e-08 | 8.9e-09 | 9.4e-09 | 1.543E-08 | 1.089E-08 | 8.900E-09 | 2.800E-08 |
| 5705 | 286 | V286C | A | 2.3e-09 | 1.1e-08 | 3.1e-09 | 5.467E-09 | 4.809E-09 | 2.300E-09 | 1.100E-08 |
| 5706 | 286 | V286E | A | 2.7e-08 | 1.7e-08 | 2.3e-08 | 2.233E-08 | 5.033E-09 | 1.700E-08 | 2.700E-08 |
| 5707 | 286 | V286Q | A | 5.9e-08 | 8.7e-08 | 6.4e-08 | 7.000E-08 | 1.493E-08 | 5.900E-08 | 8.700E-08 |
| 5708 | 286 | V286G | A | 1.1e-08 | 2.2e-08 | 2.1e-08 | 1.800E-08 | 6.083E-09 | 1.100E-08 | 2.200E-08 |
| 5709 | 286 | V286H | A | 1.2e-08 | 1.2e-08 | 2.7e-08 | 1.700E-08 | 8.660E-09 | 1.200E-08 | 2.700E-08 |
| 5710 | 286 | V286I | A | 5.6e-09 | 3.4e-09 | 2.8e-08 | 1.233E-08 | 1.361E-08 | 3.400E-09 | 2.800E-08 |
| 5711 | 286 | V286L | A | 3.2e-09 | 4.3e-09 | 1.5e-08 | 7.500E-09 | 6.518E-09 | 3.200E-09 | 1.500E-08 |
| 5712 | 286 | V286K | A | 2.9e-08 | 1.7e-08 | 2e-08 | 2.200E-08 | 6.245E-09 | 1.700E-08 | 2.900E-08 |
| 5713 | 286 | V286M | A | 5.6e-09 | 4.2e-09 | 9.1e-09 | 6.300E-09 | 2.524E-09 | 4.200E-09 | 9.100E-09 |
| 5714 | 286 | V286F | A | 8.1e-09 | 2.3e-09 | 1.4e-08 | 8.133E-09 | 5.850E-09 | 2.300E-09 | 1.400E-08 |
| 5715 | 286 | V286P | A | 5.6e-09 | 1.9e-09 | 3.2e-09 | 3.567E-09 | 1.877E-09 | 1.900E-09 | 5.600E-09 |
| 5716 | 286 | V286S | A | 5.5e-08 | 1.1e-07 | 3e-08 | 6.500E-08 | 4.093E-08 | 3.000E-08 | 1.100E-07 |
| 5717 | 286 | V286T | A | 9.8e-08 | 5.6e-08 | 1.9e-07 | 1.147E-07 | 6.854E-08 | 5.600E-08 | 1.900E-07 |
| 5718 | 286 | V286W | A | 2.5e-09 | 1.4e-09 | 8.4e-09 | 4.100E-09 | 3.764E-09 | 1.400E-09 | 8.400E-09 |
| 5719 | 286 | V286Y | A | 7.3e-09 | 4.7e-09 | 2.9e-09 | 4.967E-09 | 2.212E-09 | 2.900E-09 | 7.300E-09 |
| 5720 | 286 | V286V | A | 9.6e-09 | 3.3e-09 | 7e-09 | 6.633E-09 | 3.166E-09 | 3.300E-09 | 9.600E-09 |
| 5721 | 287 | G287A | A | 1.9e-08 | 3.7e-09 | 1.2e-08 | 1.157E-08 | 7.659E-09 | 3.700E-09 | 1.900E-08 |
| 5722 | 287 | G287R | A | 6e-09 | 8.9e-09 | 1.9e-08 | 1.130E-08 | 6.824E-09 | 6.000E-09 | 1.900E-08 |
| 5723 | 287 | G287N | A | 6.8e-09 | 1.4e-08 | 1.3e-08 | 1.127E-08 | 3.900E-09 | 6.800E-09 | 1.400E-08 |
| 5724 | 287 | G287D | A | 1.8e-08 | 9.9e-09 | 3.7e-08 | 2.163E-08 | 1.391E-08 | 9.900E-09 | 3.700E-08 |
| 5725 | 287 | G287C | A | 3.1e-09 | 3.9e-09 | 4.6e-09 | 3.867E-09 | 7.506E-10 | 3.100E-09 | 4.600E-09 |
| 5726 | 287 | G287E | A | 3.1e-08 | 1.8e-08 | 2.5e-08 | 2.467E-08 | 6.506E-09 | 1.800E-08 | 3.100E-08 |
| 5727 | 287 | G287Q | A | 4.8e-08 | 1.2e-08 | 3.5e-08 | 3.167E-08 | 1.823E-08 | 1.200E-08 | 4.800E-08 |
| 5728 | 287 | G287G | A | 3.2e-08 | 3.5e-09 | 2.4e-09 | 1.263E-08 | 1.678E-08 | 2.400E-09 | 3.200E-08 |
| 5729 | 287 | G287H | A | 5.8e-09 | 2e-08 | 1.8e-08 | 1.460E-08 | 7.686E-09 | 5.800E-09 | 2.000E-08 |
| 5730 | 287 | G287I | A | 6.3e-09 | 2.7e-09 | 5.7e-09 | 4.900E-09 | 1.929E-09 | 2.700E-09 | 6.300E-09 |
| 5731 | 287 | G287L | A | 9.8e-09 | 3.6e-09 | 8.4e-09 | 7.267E-09 | 3.252E-09 | 3.600E-09 | 9.800E-09 |
| 5732 | 287 | G287K | A | 5.4e-09 | 1.8e-08 | 3.6e-09 | 9.000E-09 | 7.846E-09 | 3.600E-09 | 1.800E-08 |
| 5733 | 287 | G287M | A | 5.9e-09 | 9e-09 | 9.4e-09 | 8.100E-09 | 1.916E-09 | 5.900E-09 | 9.400E-09 |
| 5734 | 287 | G287F | A | 6.5e-09 | 6.7e-09 | 8.7e-09 | 7.300E-09 | 1.217E-09 | 6.500E-09 | 8.700E-09 |
| 5735 | 287 | G287P | A | 2e-09 | 2.8e-09 | 1.3e-09 | 2.033E-09 | 7.506E-10 | 1.300E-09 | 2.800E-09 |
| 5736 | 287 | G287S | A | 5.8e-09 | 9.2e-09 | 3.3e-08 | 1.600E-08 | 1.482E-08 | 5.800E-09 | 3.300E-08 |
| 5737 | 287 | G287T | A | 2e-08 | 1e-08 | 1e-07 | 4.333E-08 | 4.933E-08 | 1.000E-08 | 1.000E-07 |
| 5738 | 287 | G287W | A | 5.1e-09 | 4.9e-09 | 8.7e-09 | 6.233E-09 | 2.139E-09 | 4.900E-09 | 8.700E-09 |
| 5739 | 287 | G287Y | A | 2.3e-09 | 8.9e-09 | 9.4e-09 | 6.867E-09 | 3.963E-09 | 2.300E-09 | 9.400E-09 |
| 5740 | 287 | G287V | A | 1.1e-08 | 1.1e-08 | 8.4e-09 | 1.013E-08 | 1.501E-09 | 8.400E-09 | 1.100E-08 |
| 5741 | 288 | S288A | A | 1.1e-09 | 8.4e-10 | 1e-09 | 9.800E-10 | 1.311E-10 | 8.400E-10 | 1.100E-09 |
| 5742 | 288 | S288R | A | 9.4e-09 | 4.1e-09 | 4.2e-09 | 5.900E-09 | 3.032E-09 | 4.100E-09 | 9.400E-09 |
| 5743 | 288 | S288N | A | 3.2e-08 | 2.8e-09 | 9.5e-09 | 1.477E-08 | 1.530E-08 | 2.800E-09 | 3.200E-08 |
| 5744 | 288 | S288D | A | 6.5e-09 | 4.8e-09 | 2e-09 | 4.433E-09 | 2.272E-09 | 2.000E-09 | 6.500E-09 |
| 5745 | 288 | S288C | A | 1.5e-09 | 1.2e-09 | 1.5e-09 | 1.400E-09 | 1.732E-10 | 1.200E-09 | 1.500E-09 |
| 5746 | 288 | S288E | A | 1.6e-09 | 3.9e-09 | 6e-09 | 3.833E-09 | 2.201E-09 | 1.600E-09 | 6.000E-09 |
| 5747 | 288 | S288Q | A | 7e-09 | 7.6e-09 | 1e-08 | 8.200E-09 | 1.587E-09 | 7.000E-09 | 1.000E-08 |
| 5748 | 288 | S288G | A | 2.8e-09 | 1.4e-09 | 3.9e-09 | 2.700E-09 | 1.253E-09 | 1.400E-09 | 3.900E-09 |
| 5749 | 288 | S288H | A | 6.7e-09 | 2.1e-09 | 2.1e-09 | 3.633E-09 | 2.656E-09 | 2.100E-09 | 6.700E-09 |
| 5750 | 288 | S288I | A | 1.9e-09 | 5.6e-09 | 2.9e-09 | 3.467E-09 | 1.914E-09 | 1.900E-09 | 5.600E-09 |
| 5751 | 288 | S288L | A | 2.3e-09 | 1.8e-09 | 5.9e-09 | 3.333E-09 | 2.237E-09 | 1.800E-09 | 5.900E-09 |
| 5752 | 288 | S288K | A | 3.4e-09 | 5e-09 | 3.2e-10 | 2.907E-09 | 2.379E-09 | 3.200E-10 | 5.000E-09 |
| 5753 | 288 | S288M | A | 1.4e-09 | 5.5e-09 | 1.2e-09 | 2.700E-09 | 2.427E-09 | 1.200E-09 | 5.500E-09 |
| 5754 | 288 | S288F | A | 3.8e-09 | 7.5e-10 | 1.3e-09 | 1.950E-09 | 1.626E-09 | 7.500E-10 | 3.800E-09 |
| 5755 | 288 | S288P | A | 1.8e-09 | 1.6e-09 | 3.3e-09 | 2.233E-09 | 9.292E-10 | 1.600E-09 | 3.300E-09 |
| 5756 | 288 | S288S | A | 6.8e-09 | 3e-09 | 1.2e-08 | 7.267E-09 | 4.518E-09 | 3.000E-09 | 1.200E-08 |
| 5757 | 288 | S288T | A | 4.4e-09 | 5e-09 | 1.2e-08 | 7.133E-09 | 4.225E-09 | 4.400E-09 | 1.200E-08 |
| 5758 | 288 | S288W | A | 2e-09 | 1.4e-09 | 1.6e-09 | 1.667E-09 | 3.055E-10 | 1.400E-09 | 2.000E-09 |
| 5759 | 288 | S288Y | A | 1.3e-09 | 6.2e-10 | 3.2e-09 | 1.707E-09 | 1.337E-09 | 6.200E-10 | 3.200E-09 |
| 5760 | 288 | S288V | A | 5.6e-10 | 6.7e-10 | 1.6e-09 | 9.433E-10 | 5.713E-10 | 5.600E-10 | 1.600E-09 |
| 5761 | 289 | C289A | A | 8.8e-09 | 2.2e-08 | 7.1e-09 | 1.263E-08 | 8.156E-09 | 7.100E-09 | 2.200E-08 |
| 5762 | 289 | C289R | A | 7.6e-09 | 9.9e-09 | 9.7e-09 | 9.067E-09 | 1.274E-09 | 7.600E-09 | 9.900E-09 |
| 5763 | 289 | C289N | A | 2.3e-08 | 1.6e-08 | 1.2e-08 | 1.700E-08 | 5.568E-09 | 1.200E-08 | 2.300E-08 |
| 5764 | 289 | C289D | A | 4.5e-09 | 1.4e-08 | 9.9e-09 | 9.467E-09 | 4.765E-09 | 4.500E-09 | 1.400E-08 |
| 5765 | 289 | C289C | A | 1.6e-08 | 6.4e-09 | 1.5e-08 | 1.247E-08 | 5.278E-09 | 6.400E-09 | 1.600E-08 |
| 5766 | 289 | C289E | A | 1.2e-08 | 5e-09 | 6.5e-09 | 7.833E-09 | 3.686E-09 | 5.000E-09 | 1.200E-08 |
| 5767 | 289 | C289Q | A | 1.8e-09 | 1.1e-08 | 1.9e-08 | 1.060E-08 | 8.607E-09 | 1.800E-09 | 1.900E-08 |
| 5768 | 289 | C289G | A | 4.5e-09 | 9.8e-09 | 1e-08 | 8.100E-09 | 3.119E-09 | 4.500E-09 | 1.000E-08 |
| 5769 | 289 | C289H | A | 8.3e-09 | 2.4e-08 | 5.8e-09 | 1.270E-08 | 9.866E-09 | 5.800E-09 | 2.400E-08 |
| 5770 | 289 | C289I | A | 8.4e-09 | 4.1e-09 | 1.2e-08 | 8.167E-09 | 3.955E-09 | 4.100E-09 | 1.200E-08 |
| 5771 | 289 | C289L | A | 9.3e-09 | 3.5e-09 | 4.4e-09 | 5.733E-09 | 3.121E-09 | 3.500E-09 | 9.300E-09 |
| 5772 | 289 | C289K | A | 6.5e-09 | 6.5e-09 | 9.5e-09 | 7.500E-09 | 1.732E-09 | 6.500E-09 | 9.500E-09 |
| 5773 | 289 | C289M | A | 7.5e-09 | 5.4e-09 | 7.3e-09 | 6.733E-09 | 1.159E-09 | 5.400E-09 | 7.500E-09 |
| 5774 | 289 | C289F | A | 4.3e-09 | 4.8e-09 | 6.4e-09 | 5.167E-09 | 1.097E-09 | 4.300E-09 | 6.400E-09 |
| 5775 | 289 | C289P | A | 1.6e-08 | 8.3e-09 | 5.5e-09 | 9.933E-09 | 5.437E-09 | 5.500E-09 | 1.600E-08 |
| 5776 | 289 | C289S | A | 4.1e-09 | 4.9e-09 | 4e-09 | 4.333E-09 | 4.933E-10 | 4.000E-09 | 4.900E-09 |
| 5777 | 289 | C289T | A | 3.4e-09 | 5e-09 | 8e-09 | 5.467E-09 | 2.335E-09 | 3.400E-09 | 8.000E-09 |
| 5778 | 289 | C289W | A | 9.4e-09 | 3.8e-09 | 2.5e-09 | 5.233E-09 | 3.667E-09 | 2.500E-09 | 9.400E-09 |
| 5779 | 289 | C289Y | A | 8.4e-09 | 1.6e-08 | 4.5e-09 | 9.633E-09 | 5.848E-09 | 4.500E-09 | 1.600E-08 |

| | | | | | | | | | | |
|---|---|---|---|---|---|---|---|---|---|---|
| 5780 | 289 | C289V | A | 1.6e-08 | 1.7e-08 | 2.5e-09 | 1.183E-08 | 8.098E-09 | 2.500E-09 | 1.700E-08 |
| 5781 | 290 | T290A | A | 1e-09 | 2.5e-09 | 9.6e-10 | 1.487E-09 | 8.778E-10 | 9.600E-10 | 2.500E-09 |
| 5782 | 290 | T290R | A | 2.9e-09 | 2.2e-09 | 1.5e-09 | 2.200E-09 | 7.000E-10 | 1.500E-09 | 2.900E-09 |
| 5783 | 290 | T290N | A | 4.3e-09 | 2e-09 | 1.4e-08 | 6.767E-09 | 6.369E-09 | 2.000E-09 | 1.400E-08 |
| 5784 | 290 | T290D | A | 3.1e-09 | 2.1e-09 | 3.3e-09 | 2.833E-09 | 6.429E-10 | 2.100E-09 | 3.300E-09 |
| 5785 | 290 | T290C | A | 3.2e-09 | 1.5e-09 | 3.5e-10 | 1.683E-09 | 1.434E-09 | 3.500E-10 | 3.200E-09 |
| 5786 | 290 | T290E | A | 4.9e-09 | 3.5e-09 | 8.2e-09 | 5.533E-09 | 2.413E-09 | 3.500E-09 | 8.200E-09 |
| 5787 | 290 | T290Q | A | 5.9e-09 | 1e-08 | 1.7e-08 | 1.097E-08 | 5.613E-09 | 5.900E-09 | 1.700E-08 |
| 5788 | 290 | T290G | A | 1.5e-09 | 2.1e-09 | 1.5e-09 | 1.700E-09 | 3.464E-10 | 1.500E-09 | 2.100E-09 |
| 5789 | 290 | T290H | A | 6.6e-10 | 1.4e-09 | 1.1e-08 | 4.353E-09 | 5.768E-09 | 6.600E-10 | 1.100E-08 |
| 5790 | 290 | T290I | A | 8e-10 | 9.6e-10 | 3.5e-10 | 7.033E-10 | 3.163E-10 | 3.500E-10 | 9.600E-10 |
| 5791 | 290 | T290L | A | 2.2e-09 | 2.4e-09 | 1.6e-09 | 2.067E-09 | 4.163E-10 | 1.600E-09 | 2.400E-09 |
| 5792 | 290 | T290K | A | 2.3e-09 | 2.6e-09 | 6.5e-09 | 3.800E-09 | 2.343E-09 | 2.300E-09 | 6.500E-09 |
| 5793 | 290 | T290M | A | 1.9e-09 | 2.3e-09 | 2.2e-09 | 2.133E-09 | 2.082E-10 | 1.900E-09 | 2.300E-09 |
| 5794 | 290 | T290F | A | 7.5e-10 | 3.6e-09 | 2.5e-09 | 2.283E-09 | 1.437E-09 | 7.500E-10 | 3.600E-09 |
| 5795 | 290 | T290P | A | 1.8e-09 | 1.9e-09 | 3.6e-09 | 2.433E-09 | 1.012E-09 | 1.800E-09 | 3.600E-09 |
| 5796 | 290 | T290S | A | 4.2e-09 | 6.5e-09 | 5e-09 | 5.233E-09 | 1.168E-09 | 4.200E-09 | 6.500E-09 |
| 5797 | 290 | T290T | A | 7.2e-09 | 7.7e-09 | 1.3e-08 | 9.300E-09 | 3.214E-09 | 7.200E-09 | 1.300E-08 |
| 5798 | 290 | T290W | A | 3e-09 | 1.8e-09 | 1.3e-09 | 2.033E-09 | 8.737E-10 | 1.300E-09 | 3.000E-09 |
| 5799 | 290 | T290Y | A | 3.9e-09 | 1.5e-09 | 1e-09 | 2.133E-09 | 1.550E-09 | 1.000E-09 | 3.900E-09 |
| 5800 | 290 | T290V | A | 8.4e-09 | 3.7e-09 | 1.6e-09 | 4.567E-09 | 3.482E-09 | 1.600E-09 | 8.400E-09 |
| 5801 | 291 | L291A | A | 5.5e-09 | 9.4e-09 | 1.6e-09 | 5.500E-09 | 3.900E-09 | 1.600E-09 | 9.400E-09 |
| 5802 | 291 | L291R | A | 1.5e-08 | 4.5e-09 | 7.1e-09 | 8.867E-09 | 5.468E-09 | 4.500E-09 | 1.500E-08 |
| 5803 | 291 | L291N | A | 1.8e-08 | 4.3e-09 | 1.4e-08 | 1.210E-08 | 7.045E-09 | 4.300E-09 | 1.800E-08 |
| 5804 | 291 | L291D | A | 1.3e-09 | 1.4e-08 | 4.3e-09 | 6.533E-09 | 6.638E-09 | 1.300E-09 | 1.400E-08 |
| 5805 | 291 | L291C | A | 4.4e-09 | 9e-09 | 2e-08 | 1.113E-08 | 8.016E-09 | 4.400E-09 | 2.000E-08 |
| 5806 | 291 | L291E | A | 1.2e-08 | 4.7e-09 | 5.7e-09 | 7.467E-09 | 3.958E-09 | 4.700E-09 | 1.200E-08 |
| 5807 | 291 | L291Q | A | 7.7e-09 | 6.5e-09 | 4.4e-09 | 6.200E-09 | 1.670E-09 | 4.400E-09 | 7.700E-09 |
| 5808 | 291 | L291G | A | 1.2e-08 | 9.6e-09 | 1.7e-08 | 1.287E-08 | 3.775E-09 | 9.600E-09 | 1.700E-08 |
| 5809 | 291 | L291H | A | 1.4e-08 | 8.6e-09 | 7.3e-09 | 9.967E-09 | 3.553E-09 | 7.300E-09 | 1.400E-08 |
| 5810 | 291 | L291I | A | 8e-09 | 1e-08 | 2.4e-09 | 6.800E-09 | 3.940E-09 | 2.400E-09 | 1.000E-08 |
| 5811 | 291 | L291L | A | 1e-08 | 7.9e-09 | 5.5e-09 | 7.800E-09 | 2.252E-09 | 5.500E-09 | 1.000E-08 |
| 5812 | 291 | L291K | A | 4.3e-09 | 4.8e-09 | 1.2e-08 | 7.033E-09 | 4.309E-09 | 4.300E-09 | 1.200E-08 |
| 5813 | 291 | L291M | A | 4.9e-08 | 2.6e-08 | 1.2e-08 | 2.900E-08 | 1.868E-08 | 1.200E-08 | 4.900E-08 |
| 5814 | 291 | L291F | A | 9.2e-09 | 3.5e-09 | 7.1e-09 | 6.600E-09 | 2.883E-09 | 3.500E-09 | 9.200E-09 |
| 5815 | 291 | L291P | A | 3.6e-09 | 1e-08 | 9.5e-09 | 7.700E-09 | 3.559E-09 | 3.600E-09 | 1.000E-08 |
| 5816 | 291 | L291S | A | 4.7e-09 | 1.3e-08 | 2.9e-09 | 6.867E-09 | 5.387E-09 | 2.900E-09 | 1.300E-08 |
| 5817 | 291 | L291T | A | 4.3e-09 | 3.6e-09 | 4.5e-09 | 4.133E-09 | 4.726E-10 | 3.600E-09 | 4.500E-09 |
| 5818 | 291 | L291W | A | 2.4e-08 | 1.4e-09 | 1.3e-08 | 1.280E-08 | 1.130E-08 | 1.400E-09 | 2.400E-08 |
| 5819 | 291 | L291Y | A | 5.7e-09 | 5.6e-09 | 8.7e-09 | 6.667E-09 | 1.762E-09 | 5.600E-09 | 8.700E-09 |
| 5820 | 291 | L291V | A | 5.6e-09 | 4.8e-09 | 3.8e-09 | 4.733E-09 | 9.018E-10 | 3.800E-09 | 5.600E-09 |
| 5821 | 292 | V292A | A | 6.2e-09 | 3.3e-09 | 4.2e-09 | 4.567E-09 | 1.484E-09 | 3.300E-09 | 6.200E-09 |
| 5822 | 292 | V292R | A | 5.8e-09 | 1.6e-08 | 1.2e-08 | 1.127E-08 | 5.139E-09 | 5.800E-09 | 1.600E-08 |
| 5823 | 292 | V292N | A | 1.4e-08 | 1.1e-08 | 1.2e-08 | 1.233E-08 | 1.528E-09 | 1.100E-08 | 1.400E-08 |
| 5824 | 292 | V292D | A | 8.4e-09 | 1.8e-08 | 8.7e-09 | 1.170E-08 | 5.458E-09 | 8.400E-09 | 1.800E-08 |
| 5825 | 292 | V292C | A | 1.1e-08 | 1.1e-08 | 4.6e-09 | 8.867E-09 | 3.695E-09 | 4.600E-09 | 1.100E-08 |
| 5826 | 292 | V292E | A | 6.5e-09 | 2.9e-09 | 4.8e-09 | 4.733E-09 | 1.801E-09 | 2.900E-09 | 6.500E-09 |
| 5827 | 292 | V292Q | A | 4.8e-09 | 9.8e-09 | 2.7e-09 | 5.767E-09 | 3.647E-09 | 2.700E-09 | 9.800E-09 |
| 5828 | 292 | V292G | A | 1.2e-08 | 4.8e-09 | 1.5e-08 | 1.060E-08 | 5.242E-09 | 4.800E-09 | 1.500E-08 |
| 5829 | 292 | V292H | A | 1.2e-08 | 8.2e-09 | 8e-09 | 9.400E-09 | 2.254E-09 | 8.000E-09 | 1.200E-08 |
| 5830 | 292 | V292I | A | 1.3e-08 | 7.4e-09 | 5.9e-09 | 8.767E-09 | 3.742E-09 | 5.900E-09 | 1.300E-08 |
| 5831 | 292 | V292L | A | 4.3e-09 | 5.9e-09 | 7e-09 | 5.733E-09 | 1.358E-09 | 4.300E-09 | 7.000E-09 |
| 5832 | 292 | V292K | A | 1.6e-08 | 6e-09 | 9.8e-10 | 7.660E-09 | 7.646E-09 | 9.800E-10 | 1.600E-08 |
| 5833 | 292 | V292M | A | 3.8e-09 | 8.3e-09 | 7.6e-09 | 6.567E-09 | 2.421E-09 | 3.800E-09 | 8.300E-09 |
| 5834 | 292 | V292F | A | 1e-08 | 5.3e-09 | 1e-08 | 8.433E-09 | 2.714E-09 | 5.300E-09 | 1.000E-08 |
| 5835 | 292 | V292P | A | 8e-09 | 2.4e-08 | 5e-09 | 1.233E-08 | 1.021E-08 | 5.000E-09 | 2.400E-08 |
| 5836 | 292 | V292S | A | 5.4e-09 | 1.1e-08 | 1.4e-08 | 1.013E-08 | 4.365E-09 | 5.400E-09 | 1.400E-08 |
| 5837 | 292 | V292T | A | 1.7e-08 | 1e-08 | 4.8e-08 | 2.500E-08 | 2.022E-08 | 1.000E-08 | 4.800E-08 |
| 5838 | 292 | V292W | A | 7.5e-09 | 8.4e-09 | 1.5e-08 | 1.030E-08 | 4.095E-09 | 7.500E-09 | 1.500E-08 |
| 5839 | 292 | V292Y | A | 1.2e-08 | 1.1e-08 | 6.7e-09 | 9.900E-09 | 2.816E-09 | 6.700E-09 | 1.200E-08 |
| 5840 | 292 | V292V | A | 8.1e-09 | 3e-09 | 1.3e-08 | 8.033E-09 | 5.000E-09 | 3.000E-09 | 1.300E-08 |
| 5841 | 293 | C293A | A | 3.3e-09 | 2.8e-09 | 7.7e-09 | 4.600E-09 | 2.696E-09 | 2.800E-09 | 7.700E-09 |
| 5842 | 293 | C293R | A | 4.4e-09 | 5.6e-09 | 5.2e-09 | 5.067E-09 | 6.110E-10 | 4.400E-09 | 5.600E-09 |
| 5843 | 293 | C293N | A | 7.7e-09 | 1.6e-08 | 5.6e-09 | 9.767E-09 | 5.499E-09 | 5.600E-09 | 1.600E-08 |
| 5844 | 293 | C293D | A | 5e-09 | 2.2e-08 | 1.5e-08 | 1.400E-08 | 8.544E-09 | 5.000E-09 | 2.200E-08 |
| 5845 | 293 | C293C | A | 1.1e-08 | 2e-08 | 1.3e-08 | 1.467E-08 | 4.726E-09 | 1.100E-08 | 2.000E-08 |
| 5846 | 293 | C293E | A | 9.5e-09 | 4.3e-09 | 7.5e-09 | 7.100E-09 | 2.623E-09 | 4.300E-09 | 9.500E-09 |
| 5847 | 293 | C293Q | A | 6.6e-09 | 6.3e-09 | 7.2e-09 | 6.700E-09 | 4.583E-10 | 6.300E-09 | 7.200E-09 |
| 5848 | 293 | C293G | A | 6.3e-09 | 4.3e-09 | 7.8e-09 | 6.133E-09 | 1.756E-09 | 4.300E-09 | 7.800E-09 |
| 5849 | 293 | C293H | A | 5.8e-09 | 3.6e-09 | 6.1e-09 | 5.167E-09 | 1.365E-09 | 3.600E-09 | 6.100E-09 |
| 5850 | 293 | C293I | A | 5.6e-09 | 5e-09 | 2.1e-09 | 4.233E-09 | 1.872E-09 | 2.100E-09 | 5.600E-09 |
| 5851 | 293 | C293L | A | 4.6e-09 | 1.4e-08 | 2.1e-08 | 1.320E-08 | 8.229E-09 | 4.600E-09 | 2.100E-08 |
| 5852 | 293 | C293K | A | 6.1e-09 | 5.7e-09 | 6e-09 | 5.933E-09 | 2.082E-10 | 5.700E-09 | 6.100E-09 |
| 5853 | 293 | C293M | A | 3.9e-09 | 5.3e-09 | 1.7e-08 | 8.733E-09 | 7.193E-09 | 3.900E-09 | 1.700E-08 |
| 5854 | 293 | C293F | A | 1.4e-08 | 4e-09 | 6.8e-09 | 8.267E-09 | 5.159E-09 | 4.000E-09 | 1.400E-08 |
| 5855 | 293 | C293P | A | 1.2e-08 | 6.8e-09 | 4.4e-09 | 7.733E-09 | 3.885E-09 | 4.400E-09 | 1.200E-08 |
| 5856 | 293 | C293S | A | 1.4e-09 | 2e-08 | 8.1e-09 | 9.833E-09 | 9.420E-09 | 1.400E-09 | 2.000E-08 |
| 5857 | 293 | C293T | A | 1.1e-08 | 1e-08 | 7.8e-09 | 9.600E-09 | 1.637E-09 | 7.800E-09 | 1.100E-08 |
| 5858 | 293 | C293W | A | 9.7e-09 | 1.3e-08 | 1.2e-08 | 1.157E-08 | 1.692E-09 | 9.700E-09 | 1.300E-08 |
| 5859 | 293 | C293Y | A | 4.1e-09 | 5.5e-09 | 7.6e-09 | 5.733E-09 | 1.762E-09 | 4.100E-09 | 7.600E-09 |
| 5860 | 293 | C293V | A | 8e-09 | 3.8e-08 | 7.4e-09 | 1.780E-08 | 1.750E-08 | 7.400E-09 | 3.800E-08 |
| 5861 | 294 | P294A | A | 1.1e-08 | 1.2e-08 | 1.6e-08 | 1.300E-08 | 2.646E-09 | 1.100E-08 | 1.600E-08 |
| 5862 | 294 | P294R | A | 4.4e-09 | 8.2e-09 | 5.6e-09 | 6.067E-09 | 1.943E-09 | 4.400E-09 | 8.200E-09 |
| 5863 | 294 | P294N | A | 3.6e-09 | 1.2e-08 | 3.7e-09 | 6.433E-09 | 4.821E-09 | 3.600E-09 | 1.200E-08 |
| 5864 | 294 | P294D | A | 1.3e-08 | 2.5e-08 | 6.2e-09 | 1.473E-08 | 9.519E-09 | 6.200E-09 | 2.500E-08 |

| | | | | | | | | | | |
|---|---|---|---|---|---|---|---|---|---|---|
| 5865 | 294 | P294C | A | 7.9e-09 | 9.3e-09 | 2.4e-08 | 1.373E-08 | 8.919E-09 | 7.900E-09 | 2.400E-08 |
| 5866 | 294 | P294E | A | 2.6e-09 | 3.1e-09 | 4.9e-09 | 3.533E-09 | 1.210E-09 | 2.600E-09 | 4.900E-09 |
| 5867 | 294 | P294Q | A | 7.6e-09 | 1.2e-08 | 3.7e-09 | 7.767E-09 | 4.153E-09 | 3.700E-09 | 1.200E-08 |
| 5868 | 294 | P294G | A | 4.8e-09 | 9e-09 | 7.7e-09 | 7.167E-09 | 2.150E-09 | 4.800E-09 | 9.000E-09 |
| 5869 | 294 | P294H | A | 3.2e-09 | 1.9e-09 | 1.9e-09 | 2.333E-09 | 7.506E-10 | 1.900E-09 | 3.200E-09 |
| 5870 | 294 | P294I | A | 6.5e-09 | 2e-08 | 7.6e-09 | 1.137E-08 | 7.497E-09 | 6.500E-09 | 2.000E-08 |
| 5871 | 294 | P294L | A | 3.6e-09 | 5.1e-09 | 4.7e-09 | 4.467E-09 | 7.767E-10 | 3.600E-09 | 5.100E-09 |
| 5872 | 294 | P294K | A | 1.2e-08 | 1e-08 | 7.3e-09 | 9.767E-09 | 2.359E-09 | 7.300E-09 | 1.200E-08 |
| 5873 | 294 | P294M | A | 3.7e-09 | 8e-09 | 2.5e-09 | 4.733E-09 | 2.892E-09 | 2.500E-09 | 8.000E-09 |
| 5874 | 294 | P294F | A | 1.3e-08 | 7.4e-09 | 5.1e-09 | 8.500E-09 | 4.063E-09 | 5.100E-09 | 1.300E-08 |
| 5875 | 294 | P294P | A | 1e-08 | 7.2e-09 | 6e-09 | 7.733E-09 | 2.053E-09 | 6.000E-09 | 1.000E-08 |
| 5876 | 294 | P294S | A | 1.6e-09 | 5.8e-09 | 5.5e-09 | 4.300E-09 | 2.343E-09 | 1.600E-09 | 5.800E-09 |
| 5877 | 294 | P294T | A | 6.2e-09 | 2.4e-09 | 5.6e-09 | 4.733E-09 | 2.043E-09 | 2.400E-09 | 6.200E-09 |
| 5878 | 294 | P294W | A | 4.5e-09 | 3.4e-09 | 1.7e-09 | 3.200E-09 | 1.411E-09 | 1.700E-09 | 4.500E-09 |
| 5879 | 294 | P294Y | A | 6.5e-09 | 8.7e-09 | 1.7e-08 | 1.073E-08 | 5.537E-09 | 6.500E-09 | 1.700E-08 |
| 5880 | 294 | P294V | A | 8.5e-09 | 3.6e-08 | 1e-08 | 1.817E-08 | 1.546E-08 | 8.500E-09 | 3.600E-08 |
| 5881 | 295 | L295A | A | 9.4e-09 | 8.1e-09 | 1.1e-08 | 9.500E-09 | 1.453E-09 | 8.100E-09 | 1.100E-08 |
| 5882 | 295 | L295R | A | 6.2e-09 | 1e-08 | 2.4e-09 | 6.200E-09 | 3.800E-09 | 2.400E-09 | 1.000E-08 |
| 5883 | 295 | L295N | A | 8.9e-10 | 2.3e-09 | 1.2e-09 | 1.463E-09 | 7.410E-10 | 8.900E-10 | 2.300E-09 |
| 5884 | 295 | L295D | A | 6.6e-09 | 2.7e-09 | 3.5e-09 | 4.267E-09 | 2.060E-09 | 2.700E-09 | 6.600E-09 |
| 5885 | 295 | L295C | A | 8.9e-09 | 9.1e-09 | 6.7e-09 | 8.233E-09 | 1.332E-09 | 6.700E-09 | 9.100E-09 |
| 5886 | 295 | L295E | A | 1.6e-09 | 2.4e-09 | 3.5e-09 | 2.500E-09 | 9.539E-10 | 1.600E-09 | 3.500E-09 |
| 5887 | 295 | L295Q | A | 1.3e-09 | 2.4e-09 | 2e-09 | 1.900E-09 | 5.568E-10 | 1.300E-09 | 2.400E-09 |
| 5888 | 295 | L295G | A | 4.5e-09 | 1.3e-08 | 2.2e-08 | 1.317E-08 | 8.751E-09 | 4.500E-09 | 2.200E-08 |
| 5889 | 295 | L295H | A | 4.4e-09 | 1.6e-08 | 8e-09 | 9.467E-09 | 5.937E-09 | 4.400E-09 | 1.600E-08 |
| 5890 | 295 | L295I | A | 3.5e-09 | 9.1e-09 | 5.5e-09 | 6.033E-09 | 2.838E-09 | 3.500E-09 | 9.100E-09 |
| 5891 | 295 | L295L | A | 6.3e-09 | 7.8e-09 | 1.9e-08 | 1.103E-08 | 6.940E-09 | 6.300E-09 | 1.900E-08 |
| 5892 | 295 | L295K | A | 4.6e-09 | 1.4e-09 | 1.2e-08 | 6.000E-09 | 5.437E-09 | 1.400E-09 | 1.200E-08 |
| 5893 | 295 | L295M | A | 1.3e-08 | 6.2e-09 | 3.4e-09 | 7.533E-09 | 4.937E-09 | 3.400E-09 | 1.300E-08 |
| 5894 | 295 | L295F | A | 1.1e-08 | 3.3e-09 | 2.5e-09 | 5.600E-09 | 4.694E-09 | 2.500E-09 | 1.100E-08 |
| 5895 | 295 | L295P | A | 9.1e-09 | 5.1e-09 | 4.2e-09 | 6.133E-09 | 2.608E-09 | 4.200E-09 | 9.100E-09 |
| 5896 | 295 | L295S | A | 4.1e-09 | 1.7e-09 | 3.2e-09 | 3.000E-09 | 1.212E-09 | 1.700E-09 | 4.100E-09 |
| 5897 | 295 | L295T | A | 2.3e-09 | 3.3e-09 | 1.7e-09 | 2.433E-09 | 8.083E-10 | 1.700E-09 | 3.300E-09 |
| 5898 | 295 | L295W | A | 3e-09 | 1.3e-08 | 2.5e-08 | 1.367E-08 | 1.102E-08 | 3.000E-09 | 2.500E-08 |
| 5899 | 295 | L295Y | A | 1e-08 | 5.1e-09 | 1e-08 | 8.367E-09 | 2.829E-09 | 5.100E-09 | 1.000E-08 |
| 5900 | 295 | L295V | A | 1.1e-08 | 6.8e-09 | 5.2e-09 | 7.667E-09 | 2.996E-09 | 5.200E-09 | 1.100E-08 |
| 5901 | 296 | H296A | A | 3e-08 | 2.9e-08 | 1.1e-08 | 2.333E-08 | 1.069E-08 | 1.100E-08 | 3.000E-08 |
| 5902 | 296 | H296R | A | 1.6e-08 | 7.4e-09 | 8.6e-09 | 1.067E-08 | 4.658E-09 | 7.400E-09 | 1.600E-08 |
| 5903 | 296 | H296N | A | 7.3e-10 | 1.4e-09 | 7e-10 | 9.433E-10 | 3.958E-10 | 7.000E-10 | 1.400E-09 |
| 5904 | 296 | H296D | A | 6.9e-09 | 8.5e-09 | 9.1e-09 | 8.167E-09 | 1.137E-09 | 6.900E-09 | 9.100E-09 |
| 5905 | 296 | H296C | A | 1.7e-08 | 3.8e-08 | 1.7e-08 | 2.400E-08 | 1.212E-08 | 1.700E-08 | 3.800E-08 |
| 5906 | 296 | H296E | A | 4.2e-09 | 4e-09 | 2.1e-09 | 3.433E-09 | 1.159E-09 | 2.100E-09 | 4.200E-09 |
| 5907 | 296 | H296Q | A | 3.1e-09 | 9.3e-10 | 3.7e-09 | 2.577E-09 | 1.457E-09 | 9.300E-10 | 3.700E-09 |
| 5908 | 296 | H296G | A | 5.1e-09 | 1.2e-08 | 1.1e-08 | 9.367E-09 | 3.729E-09 | 5.100E-09 | 1.200E-08 |
| 5909 | 296 | H296H | A | 1.2e-08 | 3.5e-09 | 1.2e-08 | 9.167E-09 | 4.907E-09 | 3.500E-09 | 1.200E-08 |
| 5910 | 296 | H296I | A | 1.6e-08 | 6.1e-09 | 2.3e-08 | 1.503E-08 | 8.491E-09 | 6.100E-09 | 2.300E-08 |
| 5911 | 296 | H296L | A | 1.4e-08 | 3.6e-08 | 1.2e-08 | 2.067E-08 | 1.332E-08 | 1.200E-08 | 3.600E-08 |
| 5912 | 296 | H296K | A | 7.2e-09 | 4.5e-09 | 7.5e-09 | 6.400E-09 | 1.652E-09 | 4.500E-09 | 7.500E-09 |
| 5913 | 296 | H296M | A | 2.7e-08 | 1.2e-08 | 4e-08 | 2.633E-08 | 1.401E-08 | 1.200E-08 | 4.000E-08 |
| 5914 | 296 | H296F | A | 2e-08 | 1.8e-08 | 1.6e-08 | 1.800E-08 | 2.000E-09 | 1.600E-08 | 2.000E-08 |
| 5915 | 296 | H296P | A | 2.8e-08 | 1.2e-08 | 9e-08 | 4.333E-08 | 4.120E-08 | 1.200E-08 | 9.000E-08 |
| 5916 | 296 | H296S | A | 5.1e-09 | 3.5e-09 | 3.2e-09 | 3.933E-09 | 1.021E-09 | 3.200E-09 | 5.100E-09 |
| 5917 | 296 | H296T | A | 5.6e-10 | 4.3e-09 | 3.9e-09 | 2.920E-09 | 2.054E-09 | 5.600E-10 | 4.300E-09 |
| 5918 | 296 | H296W | A | 9.8e-09 | 1.8e-08 | 2.5e-08 | 1.760E-08 | 7.608E-09 | 9.800E-09 | 2.500E-08 |
| 5919 | 296 | H296Y | A | 1.4e-08 | 9.8e-09 | 1.7e-08 | 1.360E-08 | 3.617E-09 | 9.800E-09 | 1.700E-08 |
| 5920 | 296 | H296V | A | 1.3e-08 | 1e-08 | 5.1e-08 | 2.467E-08 | 2.285E-08 | 1.000E-08 | 5.100E-08 |
| 5921 | 297 | N297A | A | 1e-08 | 4.6e-08 | 2.6e-08 | 2.733E-08 | 1.804E-08 | 1.000E-08 | 4.600E-08 |
| 5922 | 297 | N297R | A | 5.3e-08 | 7.4e-09 | 6.2e-09 | 2.220E-08 | 2.668E-08 | 6.200E-09 | 5.300E-08 |
| 5923 | 297 | N297N | A | 3.6e-09 | 2.8e-09 | 7.3e-09 | 4.567E-09 | 2.401E-09 | 2.800E-09 | 7.300E-09 |
| 5924 | 297 | N297D | A | 5.1e-09 | 9.9e-09 | 1.3e-08 | 9.333E-09 | 3.980E-09 | 5.100E-09 | 1.300E-08 |
| 5925 | 297 | N297C | A | 1.5e-08 | 5.8e-08 | 3.1e-08 | 3.467E-08 | 2.173E-08 | 1.500E-08 | 5.800E-08 |
| 5926 | 297 | N297E | A | 1.1e-08 | 1.7e-08 | 5.9e-09 | 1.130E-08 | 5.556E-09 | 5.900E-09 | 1.700E-08 |
| 5927 | 297 | N297Q | A | 3.8e-09 | 6.5e-09 | 1.3e-08 | 7.767E-09 | 4.729E-09 | 3.800E-09 | 1.300E-08 |
| 5928 | 297 | N297G | A | 1.6e-08 | 5.8e-08 | 3.1e-08 | 3.500E-08 | 2.128E-08 | 1.600E-08 | 5.800E-08 |
| 5929 | 297 | N297H | A | 2.4e-08 | 2.9e-08 | 1.3e-08 | 2.200E-08 | 8.185E-09 | 1.300E-08 | 2.900E-08 |
| 5930 | 297 | N297I | A | 2.1e-08 | 4.3e-08 | 2.4e-08 | 2.933E-08 | 1.193E-08 | 2.100E-08 | 4.300E-08 |
| 5931 | 297 | N297L | A | 2.2e-08 | 1.5e-08 | 2e-08 | 1.900E-08 | 3.606E-09 | 1.500E-08 | 2.200E-08 |
| 5932 | 297 | N297K | A | 1.6e-08 | 2.2e-08 | 1.1e-08 | 1.633E-08 | 5.508E-09 | 1.100E-08 | 2.200E-08 |
| 5933 | 297 | N297M | A | 1.6e-08 | 1.2e-08 | 2.4e-08 | 1.733E-08 | 6.110E-09 | 1.200E-08 | 2.400E-08 |
| 5934 | 297 | N297F | A | 1e-08 | 1.1e-08 | 4.6e-08 | 2.233E-08 | 2.050E-08 | 1.000E-08 | 4.600E-08 |
| 5935 | 297 | N297P | A | 1.4e-08 | 6.7e-08 | 1.6e-08 | 3.233E-08 | 3.004E-08 | 1.400E-08 | 6.700E-08 |
| 5936 | 297 | N297S | A | 7.4e-09 | 1.8e-08 | 1.5e-08 | 1.347E-08 | 5.464E-09 | 7.400E-09 | 1.800E-08 |
| 5937 | 297 | N297T | A | 1.6e-08 | 3.3e-08 | 6.8e-09 | 1.860E-08 | 1.329E-08 | 6.800E-09 | 3.300E-08 |
| 5938 | 297 | N297W | A | 6.9e-09 | 1.8e-08 | 1.8e-08 | 1.430E-08 | 6.409E-09 | 6.900E-09 | 1.800E-08 |
| 5939 | 297 | N297Y | A | 1.2e-08 | 1.3e-08 | 8.9e-09 | 1.130E-08 | 2.138E-09 | 8.900E-09 | 1.300E-08 |
| 5940 | 297 | N297V | A | 1.2e-08 | 1.9e-08 | 3.2e-08 | 2.100E-08 | 1.015E-08 | 1.200E-08 | 3.200E-08 |
| 5941 | 298 | Q298A | A | 5.8e-09 | 3e-09 | 1.2e-08 | 6.933E-09 | 4.606E-09 | 3.000E-09 | 1.200E-08 |
| 5942 | 298 | Q298R | A | 7.6e-09 | 8e-09 | 3e-09 | 6.200E-09 | 2.778E-09 | 3.000E-09 | 8.000E-09 |
| 5943 | 298 | Q298N | A | 1.1e-08 | 3e-09 | 5.7e-09 | 6.567E-09 | 4.070E-09 | 3.000E-09 | 1.100E-08 |
| 5944 | 298 | Q298D | A | 1.6e-08 | 3.9e-08 | 1.1e-08 | 2.200E-08 | 1.493E-08 | 1.100E-08 | 3.900E-08 |
| 5945 | 298 | Q298C | A | 4.3e-09 | 7.7e-09 | 3.6e-09 | 5.200E-09 | 2.193E-09 | 3.600E-09 | 7.700E-09 |
| 5946 | 298 | Q298E | A | 7.8e-09 | 1.2e-08 | 1.1e-08 | 1.027E-08 | 2.194E-09 | 7.800E-09 | 1.200E-08 |
| 5947 | 298 | Q298Q | A | 1.1e-08 | 2.5e-09 | 5.5e-09 | 6.333E-09 | 4.311E-09 | 2.500E-09 | 1.100E-08 |
| 5948 | 298 | Q298G | A | 6.1e-09 | 7.5e-09 | 6.4e-09 | 6.667E-09 | 7.371E-10 | 6.100E-09 | 7.500E-09 |
| 5949 | 298 | Q298H | A | 2.4e-08 | 1.1e-08 | 1e-08 | 1.500E-08 | 7.810E-09 | 1.000E-08 | 2.400E-08 |

| | | | | | | | | | | |
|---|---|---|---|---|---|---|---|---|---|---|
| 5950 | 298 | Q298I | A | 6.3e-09 | 2e-08 | 2.1e-08 | 1.577E-08 | 8.214E-09 | 6.300E-09 | 2.100E-08 |
| 5951 | 298 | Q298L | A | 2e-08 | 8.2e-09 | 8.5e-09 | 1.223E-08 | 6.728E-09 | 8.200E-09 | 2.000E-08 |
| 5952 | 298 | Q298K | A | 7e-09 | 3.4e-08 | 9.9e-09 | 1.697E-08 | 1.482E-08 | 7.000E-09 | 3.400E-08 |
| 5953 | 298 | Q298M | A | 1.1e-08 | 5e-09 | 2.1e-08 | 1.233E-08 | 8.083E-09 | 5.000E-09 | 2.100E-08 |
| 5954 | 298 | Q298F | A | 3.6e-08 | 1.4e-08 | 6.8e-09 | 1.893E-08 | 1.521E-08 | 6.800E-09 | 3.600E-08 |
| 5955 | 298 | Q298P | A | 7.5e-09 | 4.7e-09 | 2e-08 | 1.073E-08 | 8.146E-09 | 4.700E-09 | 2.000E-08 |
| 5956 | 298 | Q298S | A | 1.2e-08 | 3.1e-09 | 2.5e-09 | 5.867E-09 | 5.320E-09 | 2.500E-09 | 1.200E-08 |
| 5957 | 298 | Q298T | A | 1.5e-09 | 2e-08 | 3.1e-09 | 8.200E-09 | 1.025E-08 | 1.500E-09 | 2.000E-08 |
| 5958 | 298 | Q298W | A | 8.7e-09 | 1e-08 | 9.2e-09 | 9.300E-09 | 6.557E-10 | 8.700E-09 | 1.000E-08 |
| 5959 | 298 | Q298Y | A | 5.1e-09 | 8e-09 | 2.2e-09 | 5.100E-09 | 2.900E-09 | 2.200E-09 | 8.000E-09 |
| 5960 | 298 | Q298V | A | 5.3e-09 | 1e-08 | 8.8e-09 | 8.033E-09 | 2.442E-09 | 5.300E-09 | 1.000E-08 |
| 5961 | 299 | E299A | A | 1.5e-09 | 5.2e-09 | 2.7e-09 | 3.133E-09 | 1.888E-09 | 1.500E-09 | 5.200E-09 |
| 5962 | 299 | E299R | A | 1.1e-08 | 1.3e-08 | 6e-09 | 1.000E-08 | 3.606E-09 | 6.000E-09 | 1.300E-08 |
| 5963 | 299 | E299N | A | 1.4e-08 | 9.7e-09 | 2.8e-09 | 8.833E-09 | 5.650E-09 | 2.800E-09 | 1.400E-08 |
| 5964 | 299 | E299D | A | 1.4e-08 | 5.4e-09 | 1.6e-08 | 1.180E-08 | 5.632E-09 | 5.400E-09 | 1.600E-08 |
| 5965 | 299 | E299C | A | 6.2e-09 | 4.9e-09 | 1.6e-08 | 9.033E-09 | 6.068E-09 | 4.900E-09 | 1.600E-08 |
| 5966 | 299 | E299E | A | 1.4e-08 | 4.8e-08 | 1.3e-08 | 2.500E-08 | 1.992E-08 | 1.300E-08 | 4.800E-08 |
| 5967 | 299 | E299Q | A | 5.3e-09 | 7e-09 | 1.6e-08 | 9.433E-09 | 5.750E-09 | 5.300E-09 | 1.600E-08 |
| 5968 | 299 | E299G | A | 2.5e-09 | 2.9e-09 | 6e-09 | 3.800E-09 | 1.916E-09 | 2.500E-09 | 6.000E-09 |
| 5969 | 299 | E299H | A | 5e-09 | 7.9e-09 | 6.4e-09 | 6.433E-09 | 1.450E-09 | 5.000E-09 | 7.900E-09 |
| 5970 | 299 | E299I | A | 3.5e-09 | 5.4e-09 | 4.2e-09 | 4.367E-09 | 9.609E-10 | 3.500E-09 | 5.400E-09 |
| 5971 | 299 | E299L | A | 3.9e-09 | 4.8e-09 | 5.5e-09 | 4.733E-09 | 8.021E-10 | 3.900E-09 | 5.500E-09 |
| 5972 | 299 | E299K | A | 5.2e-09 | 3.1e-09 | 4.6e-09 | 4.300E-09 | 1.082E-09 | 3.100E-09 | 5.200E-09 |
| 5973 | 299 | E299M | A | 8e-09 | 2e-08 | 4.1e-09 | 1.070E-08 | 8.287E-09 | 4.100E-09 | 2.000E-08 |
| 5974 | 299 | E299F | A | 8.3e-09 | 1.3e-09 | 6.4e-09 | 5.333E-09 | 3.620E-09 | 1.300E-09 | 8.300E-09 |
| 5975 | 299 | E299P | A | 2.4e-09 | 1.9e-08 | 6.5e-09 | 9.300E-09 | 8.647E-09 | 2.400E-09 | 1.900E-08 |
| 5976 | 299 | E299S | A | 5.8e-09 | 8.4e-09 | 6.5e-09 | 6.900E-09 | 1.345E-09 | 5.800E-09 | 8.400E-09 |
| 5977 | 299 | E299T | A | 7.9e-10 | 7.5e-09 | 6.4e-09 | 4.897E-09 | 3.599E-09 | 7.900E-10 | 7.500E-09 |
| 5978 | 299 | E299W | A | 1.3e-08 | 6.1e-09 | 4.6e-09 | 7.900E-09 | 4.480E-09 | 4.600E-09 | 1.300E-08 |
| 5979 | 299 | E299Y | A | 2.6e-09 | 7.1e-09 | 1.7e-08 | 8.900E-09 | 7.367E-09 | 2.600E-09 | 1.700E-08 |
| 5980 | 299 | E299V | A | 3.7e-09 | 8.1e-09 | 1.1e-08 | 7.600E-09 | 3.676E-09 | 3.700E-09 | 1.100E-08 |
| 5981 | 300 | V300A | A | 3.3e-09 | 6.2e-09 | 2.6e-09 | 4.033E-09 | 1.909E-09 | 2.600E-09 | 6.200E-09 |
| 5982 | 300 | V300R | A | 1.6e-08 | 1.9e-08 | 8.5e-09 | 1.450E-08 | 5.408E-09 | 8.500E-09 | 1.900E-08 |
| 5983 | 300 | V300N | A | 1.2e-08 | 3e-09 | 2.8e-08 | 1.433E-08 | 1.266E-08 | 3.000E-09 | 2.800E-08 |
| 5984 | 300 | V300D | A | 2e-08 | 1.2e-08 | 9e-09 | 1.367E-08 | 5.686E-09 | 9.000E-09 | 2.000E-08 |
| 5985 | 300 | V300C | A | 8.5e-09 | 5e-09 | 6.7e-09 | 6.733E-09 | 1.750E-09 | 5.000E-09 | 8.500E-09 |
| 5986 | 300 | V300E | A | 1e-08 | 3.4e-09 | 9.4e-09 | 7.600E-09 | 3.650E-09 | 3.400E-09 | 1.000E-08 |
| 5987 | 300 | V300Q | A | 1.3e-08 | 5.4e-09 | 1.2e-08 | 1.013E-08 | 4.130E-09 | 5.400E-09 | 1.300E-08 |
| 5988 | 300 | V300G | A | 5.9e-09 | 5.7e-09 | 2.5e-08 | 1.220E-08 | 1.109E-08 | 5.700E-09 | 2.500E-08 |
| 5989 | 300 | V300H | A | 7.6e-09 | 9.5e-09 | 3.8e-09 | 6.967E-09 | 2.902E-09 | 3.800E-09 | 9.500E-09 |
| 5990 | 300 | V300I | A | 6.2e-09 | 7.9e-09 | 8.2e-09 | 7.433E-09 | 1.079E-09 | 6.200E-09 | 8.200E-09 |
| 5991 | 300 | V300L | A | 2.9e-09 | 9.8e-09 | 7.1e-09 | 6.600E-09 | 3.477E-09 | 2.900E-09 | 9.800E-09 |
| 5992 | 300 | V300K | A | 4e-09 | 3.6e-09 | 2.7e-09 | 3.433E-09 | 6.658E-10 | 2.700E-09 | 4.000E-09 |
| 5993 | 300 | V300M | A | 1.1e-08 | 7.3e-09 | 2.6e-09 | 6.967E-09 | 4.210E-09 | 2.600E-09 | 1.100E-08 |
| 5994 | 300 | V300F | A | 7.6e-09 | 3.7e-09 | 1.9e-08 | 1.010E-08 | 7.950E-09 | 3.700E-09 | 1.900E-08 |
| 5995 | 300 | V300P | A | 1.6e-08 | 8.7e-09 | 5.8e-09 | 1.017E-08 | 5.256E-09 | 5.800E-09 | 1.600E-08 |
| 5996 | 300 | V300S | A | 5.3e-09 | 8.2e-09 | 8.4e-09 | 7.300E-09 | 1.735E-09 | 5.300E-09 | 8.400E-09 |
| 5997 | 300 | V300T | A | 5.1e-09 | 5.9e-09 | 3.5e-09 | 4.833E-09 | 1.222E-09 | 3.500E-09 | 5.900E-09 |
| 5998 | 300 | V300W | A | 8.6e-09 | 8.6e-09 | 2.8e-09 | 6.667E-09 | 3.349E-09 | 2.800E-09 | 8.600E-09 |
| 5999 | 300 | V300Y | A | 5.3e-09 | 8.5e-09 | 9.4e-09 | 7.733E-09 | 2.155E-09 | 5.300E-09 | 9.400E-09 |
| 6000 | 300 | V300V | A | 5.5e-09 | 1.1e-08 | 9.4e-09 | 8.633E-09 | 2.829E-09 | 5.500E-09 | 1.100E-08 |
| 6001 | 301 | T301A | A | 1e-08 | 1.5e-08 | 7.3e-09 | 1.077E-08 | 3.907E-09 | 7.300E-09 | 1.500E-08 |
| 6002 | 301 | T301R | A | 5.6e-09 | 1.4e-08 | 4.4e-09 | 8.000E-09 | 5.231E-09 | 4.400E-09 | 1.400E-08 |
| 6003 | 301 | T301N | A | 1.5e-08 | 6.1e-09 | 7.5e-09 | 9.533E-09 | 4.786E-09 | 6.100E-09 | 1.500E-08 |
| 6004 | 301 | T301D | A | 7.1e-09 | 3.2e-09 | 1.3e-08 | 7.767E-09 | 4.934E-09 | 3.200E-09 | 1.300E-08 |
| 6005 | 301 | T301C | A | 1.3e-08 | 4.9e-09 | 2.3e-08 | 1.363E-08 | 9.067E-09 | 4.900E-09 | 2.300E-08 |
| 6006 | 301 | T301E | A | 8.4e-09 | 4.9e-09 | 1.4e-08 | 9.100E-09 | 4.590E-09 | 4.900E-09 | 1.400E-08 |
| 6007 | 301 | T301Q | A | 8.6e-09 | 1.6e-08 | 4.6e-09 | 9.733E-09 | 5.784E-09 | 4.600E-09 | 1.600E-08 |
| 6008 | 301 | T301G | A | 9.9e-09 | 9e-09 | 6.2e-09 | 8.367E-09 | 1.930E-09 | 6.200E-09 | 9.900E-09 |
| 6009 | 301 | T301H | A | 3.6e-08 | 8.7e-09 | 9.1e-09 | 1.793E-08 | 1.565E-08 | 8.700E-09 | 3.600E-08 |
| 6010 | 301 | T301I | A | 1.1e-08 | 3.5e-09 | 2.7e-08 | 1.383E-08 | 1.200E-08 | 3.500E-09 | 2.700E-08 |
| 6011 | 301 | T301L | A | 1.3e-08 | 1.3e-08 | 1.7e-08 | 1.433E-08 | 2.309E-09 | 1.300E-08 | 1.700E-08 |
| 6012 | 301 | T301K | A | 7.1e-09 | 1.3e-08 | 9e-09 | 9.700E-09 | 3.012E-09 | 7.100E-09 | 1.300E-08 |
| 6013 | 301 | T301M | A | 1.8e-09 | 5.5e-09 | 5e-09 | 4.100E-09 | 2.007E-09 | 1.800E-09 | 5.500E-09 |
| 6014 | 301 | T301F | A | 2.1e-08 | 8.7e-09 | 8.6e-09 | 1.277E-08 | 7.130E-09 | 8.600E-09 | 2.100E-08 |
| 6015 | 301 | T301P | A | 4.1e-09 | 2.8e-09 | 5.8e-09 | 4.233E-09 | 1.504E-09 | 2.800E-09 | 5.800E-09 |
| 6016 | 301 | T301S | A | 1.7e-08 | 9.9e-09 | 4.8e-09 | 1.057E-08 | 6.127E-09 | 4.800E-09 | 1.700E-08 |
| 6017 | 301 | T301T | A | 9.7e-09 | 5.7e-09 | 1.1e-08 | 8.800E-09 | 2.762E-09 | 5.700E-09 | 1.100E-08 |
| 6018 | 301 | T301W | A | 5e-09 | 8.7e-09 | 9.8e-09 | 7.833E-09 | 2.515E-09 | 5.000E-09 | 9.800E-09 |
| 6019 | 301 | T301Y | A | 7.2e-09 | 4.2e-09 | 1.1e-08 | 7.467E-09 | 3.408E-09 | 4.200E-09 | 1.100E-08 |
| 6020 | 301 | T301V | A | 6.5e-09 | 4.4e-09 | 4.6e-09 | 5.167E-09 | 1.159E-09 | 4.400E-09 | 6.500E-09 |
| 6021 | 302 | A302A | A | 1.7e-08 | 1e-08 | 9.1e-09 | 1.203E-08 | 4.325E-09 | 9.100E-09 | 1.700E-08 |
| 6022 | 302 | A302R | A | 4.2e-09 | 5.5e-09 | 8.8e-09 | 6.167E-09 | 2.371E-09 | 4.200E-09 | 8.800E-09 |
| 6023 | 302 | A302N | A | 7.4e-09 | 3.1e-09 | 9.2e-09 | 6.567E-09 | 3.134E-09 | 3.100E-09 | 9.200E-09 |
| 6024 | 302 | A302D | A | 2e-08 | 1.3e-08 | 1.2e-08 | 1.500E-08 | 4.359E-09 | 1.200E-08 | 2.000E-08 |
| 6025 | 302 | A302C | A | 2.2e-08 | 1.1e-08 | 7e-09 | 1.333E-08 | 7.767E-09 | 7.000E-09 | 2.200E-08 |
| 6026 | 302 | A302E | A | 6.6e-09 | 1e-08 | 5.2e-09 | 7.267E-09 | 2.468E-09 | 5.200E-09 | 1.000E-08 |
| 6027 | 302 | A302Q | A | 2.7e-08 | 3.1e-09 | 4.9e-09 | 1.167E-08 | 1.331E-08 | 3.100E-09 | 2.700E-08 |
| 6028 | 302 | A302G | A | 2.2e-08 | 8.6e-09 | 9.1e-09 | 1.323E-08 | 7.596E-09 | 8.600E-09 | 2.200E-08 |
| 6029 | 302 | A302H | A | 5.6e-09 | 7.9e-09 | 8.9e-09 | 7.467E-09 | 1.692E-09 | 5.600E-09 | 8.900E-09 |
| 6030 | 302 | A302I | A | 7.8e-09 | 2.3e-09 | 7.4e-09 | 5.833E-09 | 3.066E-09 | 2.300E-09 | 7.800E-09 |
| 6031 | 302 | A302L | A | 1.2e-08 | 2.2e-09 | 3.3e-09 | 5.833E-09 | 5.369E-09 | 2.200E-09 | 1.200E-08 |
| 6032 | 302 | A302K | A | 1.8e-08 | 6.2e-09 | 1.3e-08 | 1.240E-08 | 5.923E-09 | 6.200E-09 | 1.800E-08 |
| 6033 | 302 | A302M | A | 4.5e-09 | 1.3e-08 | 1.6e-08 | 1.117E-08 | 5.965E-09 | 4.500E-09 | 1.600E-08 |
| 6034 | 302 | A302F | A | 1.3e-08 | 4.3e-09 | 1.2e-08 | 9.767E-09 | 4.761E-09 | 4.300E-09 | 1.300E-08 |

| | | | | | | | | | | |
|---|---|---|---|---|---|---|---|---|---|---|
| 6035 | 302 | A302P | A | 1.7e-08 | 1.3e-08 | 8e-09 | 1.267E-08 | 4.509E-09 | 8.000E-09 | 1.700E-08 |
| 6036 | 302 | A302S | A | 9.3e-09 | 6e-09 | 3.4e-09 | 6.233E-09 | 2.957E-09 | 3.400E-09 | 9.300E-09 |
| 6037 | 302 | A302T | A | 9.6e-09 | 1.3e-08 | 1e-08 | 1.087E-08 | 1.858E-09 | 9.600E-09 | 1.300E-08 |
| 6038 | 302 | A302W | A | 4.7e-09 | 7.5e-09 | 2e-09 | 4.733E-09 | 2.750E-09 | 2.000E-09 | 7.500E-09 |
| 6039 | 302 | A302Y | A | 2.5e-09 | 3.1e-09 | 4.8e-09 | 3.467E-09 | 1.193E-09 | 2.500E-09 | 4.800E-09 |
| 6040 | 302 | A302V | A | 1.3e-08 | 1.4e-08 | 5e-09 | 1.067E-08 | 4.933E-09 | 5.000E-09 | 1.400E-08 |
| 6041 | 303 | E303A | A | 4.7e-09 | 4.2e-09 | 7e-09 | 5.300E-09 | 1.493E-09 | 4.200E-09 | 7.000E-09 |
| 6042 | 303 | E303R | A | 6.6e-09 | 3.5e-09 | 2e-09 | 4.033E-09 | 2.346E-09 | 2.000E-09 | 6.600E-09 |
| 6043 | 303 | E303N | A | 6.8e-09 | 2.6e-09 | 1.1e-08 | 6.800E-09 | 4.200E-09 | 2.600E-09 | 1.100E-08 |
| 6044 | 303 | E303D | A | 9.6e-09 | 3.9e-09 | 7.8e-09 | 7.100E-09 | 2.914E-09 | 3.900E-09 | 9.600E-09 |
| 6045 | 303 | E303C | A | 1.3e-08 | 9.2e-09 | 6.7e-09 | 9.633E-09 | 3.172E-09 | 6.700E-09 | 1.300E-08 |
| 6046 | 303 | E303E | A | 4.4e-08 | 2.6e-09 | 2.1e-09 | 1.623E-08 | 2.405E-08 | 2.100E-09 | 4.400E-08 |
| 6047 | 303 | E303Q | A | 4.8e-09 | 9.9e-09 | 9e-09 | 7.900E-09 | 2.722E-09 | 4.800E-09 | 9.900E-09 |
| 6048 | 303 | E303G | A | 3.1e-09 | 9.6e-09 | 2.3e-08 | 1.190E-08 | 1.015E-08 | 3.100E-09 | 2.300E-08 |
| 6049 | 303 | E303H | A | 3.6e-09 | 6.3e-09 | 1.3e-08 | 7.633E-09 | 4.840E-09 | 3.600E-09 | 1.300E-08 |
| 6050 | 303 | E303I | A | 3.7e-09 | 7.6e-09 | 8.5e-09 | 6.600E-09 | 2.551E-09 | 3.700E-09 | 8.500E-09 |
| 6051 | 303 | E303L | A | 3.9e-09 | 5.3e-09 | 5.9e-09 | 5.033E-09 | 1.026E-09 | 3.900E-09 | 5.900E-09 |
| 6052 | 303 | E303K | A | 6.1e-09 | 2.1e-08 | 6.5e-09 | 1.120E-08 | 8.489E-09 | 6.100E-09 | 2.100E-08 |
| 6053 | 303 | E303M | A | 3.3e-09 | 8.8e-09 | 9.6e-09 | 7.233E-09 | 3.430E-09 | 3.300E-09 | 9.600E-09 |
| 6054 | 303 | E303F | A | 2.4e-09 | 1.5e-09 | 5e-09 | 2.967E-09 | 1.818E-09 | 1.500E-09 | 5.000E-09 |
| 6055 | 303 | E303P | A | 7.7e-09 | 2.5e-09 | 5.2e-09 | 5.133E-09 | 2.601E-09 | 2.500E-09 | 7.700E-09 |
| 6056 | 303 | E303S | A | 1.8e-08 | 9.6e-09 | 2.6e-09 | 1.007E-08 | 7.711E-09 | 2.600E-09 | 1.800E-08 |
| 6057 | 303 | E303T | A | 1.1e-08 | 3.4e-09 | 1.5e-08 | 9.800E-09 | 5.892E-09 | 3.400E-09 | 1.500E-08 |
| 6058 | 303 | E303W | A | 3.2e-09 | 6.5e-09 | 4e-09 | 4.567E-09 | 1.721E-09 | 3.200E-09 | 6.500E-09 |
| 6059 | 303 | E303Y | A | 2.7e-09 | 1.7e-09 | 1.4e-08 | 6.133E-09 | 6.831E-09 | 1.700E-09 | 1.400E-08 |
| 6060 | 303 | E303V | A | 5.6e-09 | 3e-08 | 1.8e-08 | 1.787E-08 | 1.220E-08 | 5.600E-09 | 3.000E-08 |
| 6061 | 304 | D304A | A | 1.6e-08 | 3.7e-09 | 2.8e-08 | 1.590E-08 | 1.215E-08 | 3.700E-09 | 2.800E-08 |
| 6062 | 304 | D304R | A | 4.3e-09 | 1.2e-08 | 3.6e-09 | 6.633E-09 | 4.661E-09 | 3.600E-09 | 1.200E-08 |
| 6063 | 304 | D304N | A | 9.6e-09 | 1.4e-08 | 4.3e-09 | 9.300E-09 | 4.857E-09 | 4.300E-09 | 1.400E-08 |
| 6064 | 304 | D304D | A | 1.7e-08 | 1.1e-08 | 5.6e-09 | 1.120E-08 | 5.703E-09 | 5.600E-09 | 1.700E-08 |
| 6065 | 304 | D304C | A | 4.7e-09 | 2.6e-08 | 7.2e-09 | 1.263E-08 | 1.164E-08 | 4.700E-09 | 2.600E-08 |
| 6066 | 304 | D304E | A | 2.1e-08 | 7.8e-09 | 8.5e-09 | 1.243E-08 | 7.427E-09 | 7.800E-09 | 2.100E-08 |
| 6067 | 304 | D304Q | A | 2.2e-09 | 2.6e-09 | 2e-09 | 2.267E-09 | 3.055E-10 | 2.000E-09 | 2.600E-09 |
| 6068 | 304 | D304G | A | 1.2e-08 | 7.5e-09 | 4.9e-09 | 8.133E-09 | 3.592E-09 | 4.900E-09 | 1.200E-08 |
| 6069 | 304 | D304H | A | 1.1e-08 | 5.1e-09 | 1.6e-08 | 1.070E-08 | 5.456E-09 | 5.100E-09 | 1.600E-08 |
| 6070 | 304 | D304I | A | 2.6e-09 | 1e-08 | 4.4e-09 | 5.667E-09 | 3.859E-09 | 2.600E-09 | 1.000E-08 |
| 6071 | 304 | D304L | A | 3e-09 | 8.4e-09 | 3.8e-09 | 5.067E-09 | 2.914E-09 | 3.000E-09 | 8.400E-09 |
| 6072 | 304 | D304K | A | 3.5e-09 | 6.6e-09 | 4.8e-09 | 4.967E-09 | 1.557E-09 | 3.500E-09 | 6.600E-09 |
| 6073 | 304 | D304M | A | 6e-09 | 1.7e-08 | 3.5e-09 | 8.833E-09 | 7.182E-09 | 3.500E-09 | 1.700E-08 |
| 6074 | 304 | D304F | A | 8e-09 | 8.5e-09 | 2.1e-08 | 1.250E-08 | 7.365E-09 | 8.000E-09 | 2.100E-08 |
| 6075 | 304 | D304P | A | 3.1e-09 | 8.6e-09 | 1e-08 | 7.233E-09 | 3.647E-09 | 3.100E-09 | 1.000E-08 |
| 6076 | 304 | D304S | A | 8.5e-09 | 5e-09 | 4.2e-09 | 5.900E-09 | 2.287E-09 | 4.200E-09 | 8.500E-09 |
| 6077 | 304 | D304T | A | 1e-08 | 6e-09 | 5.1e-09 | 7.033E-09 | 2.608E-09 | 5.100E-09 | 1.000E-08 |
| 6078 | 304 | D304W | A | 1.1e-08 | 1.8e-08 | 8.1e-09 | 1.237E-08 | 5.090E-09 | 8.100E-09 | 1.800E-08 |
| 6079 | 304 | D304Y | A | 1e-08 | 6.8e-09 | 8.2e-09 | 8.333E-09 | 1.604E-09 | 6.800E-09 | 1.000E-08 |
| 6080 | 304 | D304V | A | 5.1e-09 | 3.1e-09 | 8.5e-09 | 5.567E-09 | 2.730E-09 | 3.100E-09 | 8.500E-09 |
| 6081 | 305 | G305A | A | 5.4e-09 | 1e-08 | 1.7e-09 | 5.700E-09 | 4.158E-09 | 1.700E-09 | 1.000E-08 |
| 6082 | 305 | G305R | A | 1.3e-08 | 7.1e-09 | 4.8e-09 | 8.300E-09 | 4.230E-09 | 4.800E-09 | 1.300E-08 |
| 6083 | 305 | G305N | A | 6.4e-09 | 7e-09 | 8.7e-09 | 7.367E-09 | 1.193E-09 | 6.400E-09 | 8.700E-09 |
| 6084 | 305 | G305D | A | 1.8e-08 | 5.7e-09 | 1.1e-08 | 1.157E-08 | 6.170E-09 | 5.700E-09 | 1.800E-08 |
| 6085 | 305 | G305C | A | 1.1e-08 | 1.1e-08 | 2.5e-09 | 8.167E-09 | 4.907E-09 | 2.500E-09 | 1.100E-08 |
| 6086 | 305 | G305E | A | 2.6e-09 | 4e-09 | 8.7e-09 | 5.100E-09 | 3.195E-09 | 2.600E-09 | 8.700E-09 |
| 6087 | 305 | G305Q | A | 2.2e-08 | 2.3e-08 | 1.2e-08 | 1.900E-08 | 6.083E-09 | 1.200E-08 | 2.300E-08 |
| 6088 | 305 | G305G | A | 2e-09 | 7.7e-09 | 2.6e-08 | 1.190E-08 | 1.254E-08 | 2.000E-09 | 2.600E-08 |
| 6089 | 305 | G305H | A | 4.4e-09 | 1.3e-08 | 2.5e-08 | 1.413E-08 | 1.035E-08 | 4.400E-09 | 2.500E-08 |
| 6090 | 305 | G305I | A | 7.2e-09 | 7.2e-09 | 2.1e-09 | 5.500E-09 | 2.944E-09 | 2.100E-09 | 7.200E-09 |
| 6091 | 305 | G305L | A | 1.7e-08 | 6e-09 | 2.1e-08 | 1.467E-08 | 7.767E-09 | 6.000E-09 | 2.100E-08 |
| 6092 | 305 | G305K | A | 1.7e-08 | 6.4e-09 | 2.5e-08 | 1.613E-08 | 9.330E-09 | 6.400E-09 | 2.500E-08 |
| 6093 | 305 | G305M | A | 8.3e-09 | 3.8e-09 | 1.1e-08 | 7.700E-09 | 3.637E-09 | 3.800E-09 | 1.100E-08 |
| 6094 | 305 | G305F | A | 7.8e-09 | 1.6e-08 | 5.7e-09 | 9.833E-09 | 5.443E-09 | 5.700E-09 | 1.600E-08 |
| 6095 | 305 | G305P | A | 5.5e-09 | 9.2e-09 | 5e-09 | 6.567E-09 | 2.294E-09 | 5.000E-09 | 9.200E-09 |
| 6096 | 305 | G305S | A | 1.2e-08 | 3.5e-09 | 1.8e-08 | 1.117E-08 | 7.286E-09 | 3.500E-09 | 1.800E-08 |
| 6097 | 305 | G305T | A | 1.7e-08 | 1.5e-08 | 3e-09 | 1.167E-08 | 7.572E-09 | 3.000E-09 | 1.700E-08 |
| 6098 | 305 | G305W | A | 1.6e-08 | 3.1e-09 | 8.3e-09 | 9.133E-09 | 6.490E-09 | 3.100E-09 | 1.600E-08 |
| 6099 | 305 | G305Y | A | 3.7e-09 | 1e-08 | 8.7e-09 | 7.467E-09 | 3.326E-09 | 3.700E-09 | 1.000E-08 |
| 6100 | 305 | G305V | A | 2e-09 | 6.6e-09 | 7.7e-09 | 5.433E-09 | 3.024E-09 | 2.000E-09 | 7.700E-09 |
| 6101 | 306 | T306A | A | 4.6e-09 | 5.2e-09 | 6.7e-09 | 5.500E-09 | 1.082E-09 | 4.600E-09 | 6.700E-09 |
| 6102 | 306 | T306R | A | 1.8e-08 | 9.1e-09 | 7.2e-09 | 1.143E-08 | 5.766E-09 | 7.200E-09 | 1.800E-08 |
| 6103 | 306 | T306N | A | 1.2e-08 | 1.2e-08 | 3.3e-09 | 9.100E-09 | 5.023E-09 | 3.300E-09 | 1.200E-08 |
| 6104 | 306 | T306D | A | 4.9e-09 | 3.6e-09 | 9.6e-09 | 6.033E-09 | 3.156E-09 | 3.600E-09 | 9.600E-09 |
| 6105 | 306 | T306C | A | 5.7e-09 | 6e-09 | 1.2e-08 | 7.900E-09 | 3.554E-09 | 5.700E-09 | 1.200E-08 |
| 6106 | 306 | T306E | A | 9.4e-09 | 5.1e-09 | 7.6e-09 | 7.367E-09 | 2.159E-09 | 5.100E-09 | 9.400E-09 |
| 6107 | 306 | T306Q | A | 1.2e-08 | 3.4e-09 | 1e-08 | 8.467E-09 | 4.500E-09 | 3.400E-09 | 1.200E-08 |
| 6108 | 306 | T306G | A | 6.9e-09 | 2.6e-09 | 5.8e-09 | 5.100E-09 | 2.234E-09 | 2.600E-09 | 6.900E-09 |
| 6109 | 306 | T306H | A | 4.4e-09 | 1.1e-08 | 6.1e-09 | 7.167E-09 | 3.427E-09 | 4.400E-09 | 1.100E-08 |
| 6110 | 306 | T306I | A | 8.7e-09 | 3.9e-09 | 7.8e-09 | 6.800E-09 | 2.551E-09 | 3.900E-09 | 8.700E-09 |
| 6111 | 306 | T306L | A | 5.2e-09 | 2.4e-09 | 2e-09 | 3.200E-09 | 1.744E-09 | 2.000E-09 | 5.200E-09 |
| 6112 | 306 | T306K | A | 1.3e-08 | 7.9e-09 | 4.3e-09 | 8.400E-09 | 4.371E-09 | 4.300E-09 | 1.300E-08 |
| 6113 | 306 | T306M | A | 8.5e-09 | 5.8e-09 | 8.7e-09 | 7.667E-09 | 1.620E-09 | 5.800E-09 | 8.700E-09 |
| 6114 | 306 | T306F | A | 6.1e-09 | 3.9e-09 | 6.2e-09 | 5.400E-09 | 1.300E-09 | 3.900E-09 | 6.200E-09 |
| 6115 | 306 | T306P | A | 8.7e-09 | 7.5e-09 | 8.5e-09 | 8.233E-09 | 6.429E-10 | 7.500E-09 | 8.700E-09 |
| 6116 | 306 | T306S | A | 5e-09 | 7.2e-09 | 1.9e-08 | 1.040E-08 | 7.529E-09 | 5.000E-09 | 1.900E-08 |
| 6117 | 306 | T306T | A | 4.2e-09 | 1.7e-08 | 8.2e-09 | 9.800E-09 | 6.548E-09 | 4.200E-09 | 1.700E-08 |
| 6118 | 306 | T306W | A | 1.4e-08 | 7.6e-09 | 3.4e-09 | 8.333E-09 | 5.338E-09 | 3.400E-09 | 1.400E-08 |
| 6119 | 306 | T306Y | A | 7.4e-09 | 4.1e-09 | 9.6e-09 | 7.033E-09 | 2.768E-09 | 4.100E-09 | 9.600E-09 |

| | | | | | | | | | | |
|---|---|---|---|---|---|---|---|---|---|---|
| 6120 | 306 | T306V | A | 1.4e-08 | 1.2e-08 | 3e-09 | 9.667E-09 | 5.859E-09 | 3.000E-09 | 1.400E-08 |
| 6121 | 307 | Q307A | A | 2.9e-09 | 3.8e-09 | 7e-09 | 4.567E-09 | 2.155E-09 | 2.900E-09 | 7.000E-09 |
| 6122 | 307 | Q307R | A | 4.9e-09 | 6.5e-09 | 7.7e-09 | 6.367E-09 | 1.405E-09 | 4.900E-09 | 7.700E-09 |
| 6123 | 307 | Q307N | A | 3.7e-09 | 5.1e-09 | 1.8e-08 | 8.933E-09 | 7.883E-09 | 3.700E-09 | 1.800E-08 |
| 6124 | 307 | Q307D | A | 1.3e-08 | 2.7e-08 | 3.8e-09 | 1.460E-08 | 1.168E-08 | 3.800E-09 | 2.700E-08 |
| 6125 | 307 | Q307C | A | 7.6e-09 | 2.8e-09 | 6.8e-09 | 5.733E-09 | 2.572E-09 | 2.800E-09 | 7.600E-09 |
| 6126 | 307 | Q307E | A | 5.8e-09 | 6.8e-09 | 7.5e-09 | 6.700E-09 | 8.544E-10 | 5.800E-09 | 7.500E-09 |
| 6127 | 307 | Q307Q | A | 5.1e-09 | 9.5e-09 | 8.8e-09 | 7.800E-09 | 2.364E-09 | 5.100E-09 | 9.500E-09 |
| 6128 | 307 | Q307G | A | 4.9e-09 | 8.5e-09 | 1.8e-08 | 1.047E-08 | 6.768E-09 | 4.900E-09 | 1.800E-08 |
| 6129 | 307 | Q307H | A | 3.3e-09 | 5.5e-09 | 2e-08 | 9.600E-09 | 9.074E-09 | 3.300E-09 | 2.000E-08 |
| 6130 | 307 | Q307I | A | 9.4e-09 | 6.5e-09 | 7.5e-09 | 7.800E-09 | 1.473E-09 | 6.500E-09 | 9.400E-09 |
| 6131 | 307 | Q307L | A | 5e-09 | 4.9e-09 | 1.3e-08 | 7.633E-09 | 4.648E-09 | 4.900E-09 | 1.300E-08 |
| 6132 | 307 | Q307K | A | 7.9e-09 | 1e-08 | 4.7e-09 | 7.533E-09 | 2.669E-09 | 4.700E-09 | 1.000E-08 |
| 6133 | 307 | Q307M | A | 2.8e-09 | 6.8e-09 | 9.9e-09 | 6.500E-09 | 3.559E-09 | 2.800E-09 | 9.900E-09 |
| 6134 | 307 | Q307F | A | 4.3e-09 | 8.8e-09 | 5.8e-09 | 6.300E-09 | 2.291E-09 | 4.300E-09 | 8.800E-09 |
| 6135 | 307 | Q307P | A | 5.8e-09 | 8.9e-09 | 1e-09 | 5.233E-09 | 3.980E-09 | 1.000E-09 | 8.900E-09 |
| 6136 | 307 | Q307S | A | 6.4e-09 | 9e-09 | 1.9e-08 | 1.147E-08 | 6.652E-09 | 6.400E-09 | 1.900E-08 |
| 6137 | 307 | Q307T | A | 5.4e-09 | 1.4e-08 | 7.4e-09 | 8.933E-09 | 4.500E-09 | 5.400E-09 | 1.400E-08 |
| 6138 | 307 | Q307W | A | 1.9e-09 | 6e-09 | 3.5e-09 | 3.800E-09 | 2.066E-09 | 1.900E-09 | 6.000E-09 |
| 6139 | 307 | Q307Y | A | 4.1e-09 | 1.3e-08 | 1.7e-08 | 1.137E-08 | 6.603E-09 | 4.100E-09 | 1.700E-08 |
| 6140 | 307 | Q307V | A | 2.6e-09 | 7.7e-09 | 6.3e-09 | 5.533E-09 | 2.635E-09 | 2.600E-09 | 7.700E-09 |
| 6141 | 308 | R308A | A | 1.5e-08 | 5.4e-09 | 9.2e-09 | 9.867E-09 | 4.835E-09 | 5.400E-09 | 1.500E-08 |
| 6142 | 308 | R308R | A | 1.5e-08 | 4.2e-09 | 3.9e-09 | 7.700E-09 | 6.324E-09 | 3.900E-09 | 1.500E-08 |
| 6143 | 308 | R308N | A | 1.1e-08 | 7.3e-09 | 7.8e-09 | 8.700E-09 | 2.007E-09 | 7.300E-09 | 1.100E-08 |
| 6144 | 308 | R308D | A | 1.2e-08 | 6.2e-09 | 2.4e-09 | 6.867E-09 | 4.835E-09 | 2.400E-09 | 1.200E-08 |
| 6145 | 308 | R308C | A | 1.3e-08 | 2.4e-08 | 4.1e-09 | 1.370E-08 | 9.968E-09 | 4.100E-09 | 2.400E-08 |
| 6146 | 308 | R308E | A | 7.2e-09 | 3e-09 | 1.7e-08 | 9.067E-09 | 7.184E-09 | 3.000E-09 | 1.700E-08 |
| 6147 | 308 | R308Q | A | 7.4e-09 | 7e-09 | 2.9e-09 | 5.767E-09 | 2.491E-09 | 2.900E-09 | 7.400E-09 |
| 6148 | 308 | R308G | A | 6.4e-09 | 1.4e-08 | 3.1e-08 | 1.713E-08 | 1.260E-08 | 6.400E-09 | 3.100E-08 |
| 6149 | 308 | R308H | A | 5e-09 | 7.5e-09 | 1.2e-08 | 8.167E-09 | 3.547E-09 | 5.000E-09 | 1.200E-08 |
| 6150 | 308 | R308I | A | 1.5e-08 | 8.6e-09 | 1.4e-08 | 1.253E-08 | 3.443E-09 | 8.600E-09 | 1.500E-08 |
| 6151 | 308 | R308L | A | 4.4e-09 | 3.5e-09 | 8.2e-09 | 5.367E-09 | 2.495E-09 | 3.500E-09 | 8.200E-09 |
| 6152 | 308 | R308K | A | 3.5e-09 | 8e-09 | 5.1e-09 | 5.533E-09 | 2.281E-09 | 3.500E-09 | 8.000E-09 |
| 6153 | 308 | R308M | A | 4.2e-09 | 8.5e-09 | 5.6e-09 | 6.100E-09 | 2.193E-09 | 4.200E-09 | 8.500E-09 |
| 6154 | 308 | R308F | A | 1e-08 | 1.4e-08 | 4.5e-09 | 9.500E-09 | 4.770E-09 | 4.500E-09 | 1.400E-08 |
| 6155 | 308 | R308P | A | 7e-09 | 3.2e-09 | 3.2e-08 | 1.407E-08 | 1.565E-08 | 3.200E-09 | 3.200E-08 |
| 6156 | 308 | R308S | A | 3e-09 | 9.5e-09 | 5.3e-09 | 5.933E-09 | 3.296E-09 | 3.000E-09 | 9.500E-09 |
| 6157 | 308 | R308T | A | 3.1e-09 | 4.3e-09 | 5.9e-09 | 4.433E-09 | 1.405E-09 | 3.100E-09 | 5.900E-09 |
| 6158 | 308 | R308W | A | 8.4e-09 | 5.6e-09 | 1.9e-08 | 1.100E-08 | 7.068E-09 | 5.600E-09 | 1.900E-08 |
| 6159 | 308 | R308Y | A | 8.1e-09 | 8.2e-09 | 1.8e-08 | 1.143E-08 | 5.687E-09 | 8.100E-09 | 1.800E-08 |
| 6160 | 308 | R308V | A | 6.7e-09 | 1.2e-08 | 2.1e-09 | 6.933E-09 | 4.954E-09 | 2.100E-09 | 1.200E-08 |
| 6161 | 309 | C309A | A | 8.9e-09 | 1.2e-08 | 6.8e-09 | 9.233E-09 | 2.616E-09 | 6.800E-09 | 1.200E-08 |
| 6162 | 309 | C309R | A | 6.8e-09 | 9.6e-09 | 4.9e-09 | 7.100E-09 | 2.364E-09 | 4.900E-09 | 9.600E-09 |
| 6163 | 309 | C309N | A | 1.3e-08 | 6.7e-09 | 7.1e-09 | 8.933E-09 | 3.528E-09 | 6.700E-09 | 1.300E-08 |
| 6164 | 309 | C309D | A | 8.4e-09 | 2.2e-08 | 1.7e-08 | 1.580E-08 | 6.879E-09 | 8.400E-09 | 2.200E-08 |
| 6165 | 309 | C309C | A | 1e-08 | 4.1e-09 | 2.7e-09 | 5.600E-09 | 3.874E-09 | 2.700E-09 | 1.000E-08 |
| 6166 | 309 | C309E | A | 1.2e-08 | 4.8e-09 | 1.5e-09 | 6.100E-09 | 5.369E-09 | 1.500E-09 | 1.200E-08 |
| 6167 | 309 | C309Q | A | 1e-08 | 1.3e-08 | 4.5e-09 | 9.167E-09 | 4.311E-09 | 4.500E-09 | 1.300E-08 |
| 6168 | 309 | C309G | A | 2.5e-09 | 5.2e-09 | 7.2e-09 | 4.967E-09 | 2.359E-09 | 2.500E-09 | 7.200E-09 |
| 6169 | 309 | C309H | A | 1.3e-08 | 1.3e-08 | 5.8e-09 | 1.060E-08 | 4.157E-09 | 5.800E-09 | 1.300E-08 |
| 6170 | 309 | C309I | A | 4.9e-09 | 6.8e-09 | 4.2e-09 | 5.300E-09 | 1.345E-09 | 4.200E-09 | 6.800E-09 |
| 6171 | 309 | C309L | A | 6.8e-09 | 3.3e-09 | 1.3e-08 | 7.700E-09 | 4.912E-09 | 3.300E-09 | 1.300E-08 |
| 6172 | 309 | C309K | A | 5e-09 | 1.6e-09 | 4.9e-09 | 3.833E-09 | 1.935E-09 | 1.600E-09 | 5.000E-09 |
| 6173 | 309 | C309M | A | 1.7e-08 | 3e-09 | 4.5e-09 | 8.167E-09 | 7.687E-09 | 3.000E-09 | 1.700E-08 |
| 6174 | 309 | C309F | A | 7.2e-09 | 8.6e-09 | 3.7e-09 | 6.500E-09 | 2.524E-09 | 3.700E-09 | 8.600E-09 |
| 6175 | 309 | C309P | A | 8.5e-09 | 1.6e-08 | 4.1e-09 | 9.533E-09 | 6.017E-09 | 4.100E-09 | 1.600E-08 |
| 6176 | 309 | C309S | A | 2.2e-08 | 1.3e-08 | 3.5e-09 | 1.283E-08 | 9.251E-09 | 3.500E-09 | 2.200E-08 |
| 6177 | 309 | C309T | A | 3.6e-09 | 6.7e-09 | 2.4e-09 | 4.233E-09 | 2.219E-09 | 2.400E-09 | 6.700E-09 |
| 6178 | 309 | C309W | A | 4.9e-09 | 3.2e-09 | 1.4e-08 | 7.367E-09 | 5.807E-09 | 3.200E-09 | 1.400E-08 |
| 6179 | 309 | C309Y | A | 4e-09 | 3.6e-08 | 6.1e-09 | 1.537E-08 | 1.790E-08 | 4.000E-09 | 3.600E-08 |
| 6180 | 309 | C309V | A | 1.6e-08 | 1e-08 | 3.4e-09 | 9.800E-09 | 6.302E-09 | 3.400E-09 | 1.600E-08 |
| 6181 | 310 | E310A | A | 4.8e-09 | 8.4e-09 | 8e-09 | 7.067E-09 | 1.973E-09 | 4.800E-09 | 8.400E-09 |
| 6182 | 310 | E310R | A | 6e-09 | 8e-09 | 1.2e-08 | 8.667E-09 | 3.055E-09 | 6.000E-09 | 1.200E-08 |
| 6183 | 310 | E310N | A | 9.1e-09 | 7.4e-09 | 3.2e-09 | 6.567E-09 | 3.037E-09 | 3.200E-09 | 9.100E-09 |
| 6184 | 310 | E310D | A | 2.8e-09 | 1.3e-08 | 6.6e-09 | 7.467E-09 | 5.155E-09 | 2.800E-09 | 1.300E-08 |
| 6185 | 310 | E310C | A | 1.6e-08 | 4.3e-09 | 5.1e-09 | 8.467E-09 | 6.536E-09 | 4.300E-09 | 1.600E-08 |
| 6186 | 310 | E310E | A | 1.2e-08 | 2.7e-09 | 9.9e-09 | 8.200E-09 | 4.877E-09 | 2.700E-09 | 1.200E-08 |
| 6187 | 310 | E310Q | A | 1.2e-08 | 1.2e-08 | 3.6e-09 | 9.200E-09 | 4.850E-09 | 3.600E-09 | 1.200E-08 |
| 6188 | 310 | E310G | A | 5.4e-09 | 8.6e-09 | 7.3e-09 | 7.100E-09 | 1.609E-09 | 5.400E-09 | 8.600E-09 |
| 6189 | 310 | E310H | A | 5.8e-09 | 1e-08 | 8.1e-09 | 7.967E-09 | 2.103E-09 | 5.800E-09 | 1.000E-08 |
| 6190 | 310 | E310I | A | 2.1e-08 | 1.1e-09 | 9.2e-09 | 1.043E-08 | 1.001E-08 | 1.100E-09 | 2.100E-08 |
| 6191 | 310 | E310L | A | 8.9e-09 | 3.9e-09 | 7e-09 | 6.600E-09 | 2.524E-09 | 3.900E-09 | 8.900E-09 |
| 6192 | 310 | E310K | A | 4.8e-09 | 4.7e-09 | 7.3e-09 | 5.600E-09 | 1.473E-09 | 4.700E-09 | 7.300E-09 |
| 6193 | 310 | E310M | A | 1.1e-08 | 4.2e-09 | 8.9e-09 | 8.033E-09 | 3.482E-09 | 4.200E-09 | 1.100E-08 |
| 6194 | 310 | E310F | A | 7.8e-09 | 3.2e-09 | 5.6e-09 | 5.533E-09 | 2.301E-09 | 3.200E-09 | 7.800E-09 |
| 6195 | 310 | E310P | A | 5.5e-09 | 5.4e-09 | 1.7e-08 | 9.300E-09 | 6.669E-09 | 5.400E-09 | 1.700E-08 |
| 6196 | 310 | E310S | A | 1.2e-08 | 1.7e-08 | 1.4e-08 | 1.433E-08 | 2.517E-09 | 1.200E-08 | 1.700E-08 |
| 6197 | 310 | E310T | A | 5.6e-09 | 1e-08 | 1.2e-08 | 9.200E-09 | 3.274E-09 | 5.600E-09 | 1.200E-08 |
| 6198 | 310 | E310W | A | 8e-09 | 6.6e-09 | 8e-09 | 7.533E-09 | 8.083E-10 | 6.600E-09 | 8.000E-09 |
| 6199 | 310 | E310Y | A | 8.1e-09 | 2.8e-09 | 8.1e-09 | 6.333E-09 | 3.060E-09 | 2.800E-09 | 8.100E-09 |
| 6200 | 310 | E310V | A | 5.6e-09 | 1.2e-08 | 1.8e-08 | 1.187E-08 | 6.201E-09 | 5.600E-09 | 1.800E-08 |
| 6201 | 311 | K311A | A | 1.4e-08 | 1.4e-08 | 7.1e-09 | 1.170E-08 | 3.984E-09 | 7.100E-09 | 1.400E-08 |
| 6202 | 311 | K311R | A | 6.5e-09 | 2.9e-09 | 5e-09 | 4.800E-09 | 1.808E-09 | 2.900E-09 | 6.500E-09 |
| 6203 | 311 | K311N | A | 4.8e-09 | 3.5e-09 | 7.9e-09 | 5.400E-09 | 2.261E-09 | 3.500E-09 | 7.900E-09 |
| 6204 | 311 | K311D | A | 1.5e-08 | 6.8e-09 | 4.2e-09 | 8.667E-09 | 5.637E-09 | 4.200E-09 | 1.500E-08 |

| | | | | | | | | | | |
|---|---|---|---|---|---|---|---|---|---|---|
| 6205 | 311 | K311C | A | 4.6e-09 | 2e-08 | 2e-08 | 1.487E-08 | 8.891E-09 | 4.600E-09 | 2.000E-08 |
| 6206 | 311 | K311E | A | 4.2e-09 | 6.6e-09 | 1.4e-08 | 8.267E-09 | 5.108E-09 | 4.200E-09 | 1.400E-08 |
| 6207 | 311 | K311Q | A | 2.1e-08 | 6.8e-09 | 2.8e-09 | 1.020E-08 | 9.565E-09 | 2.800E-09 | 2.100E-08 |
| 6208 | 311 | K311G | A | 7.3e-09 | 2.3e-08 | 2.9e-09 | 1.107E-08 | 1.057E-08 | 2.900E-09 | 2.300E-08 |
| 6209 | 311 | K311H | A | 8.5e-09 | 6.6e-09 | 6.7e-09 | 7.267E-09 | 1.069E-09 | 6.600E-09 | 8.500E-09 |
| 6210 | 311 | K311I | A | 8.3e-09 | 1.5e-08 | 4.8e-09 | 9.367E-09 | 5.183E-09 | 4.800E-09 | 1.500E-08 |
| 6211 | 311 | K311L | A | 3.1e-08 | 3.9e-09 | 1.1e-08 | 1.530E-08 | 1.405E-08 | 3.900E-09 | 3.100E-08 |
| 6212 | 311 | K311K | A | 3e-09 | 9.5e-09 | 1.1e-08 | 7.833E-09 | 4.252E-09 | 3.000E-09 | 1.100E-08 |
| 6213 | 311 | K311M | A | 2.7e-09 | 3.8e-09 | 1.3e-09 | 2.600E-09 | 1.253E-09 | 1.300E-09 | 3.800E-09 |
| 6214 | 311 | K311F | A | 4.2e-08 | 5.1e-09 | 1.3e-08 | 2.003E-08 | 1.943E-08 | 5.100E-09 | 4.200E-08 |
| 6215 | 311 | K311P | A | 8.8e-09 | 5.5e-09 | 9.1e-09 | 7.800E-09 | 1.997E-09 | 5.500E-09 | 9.100E-09 |
| 6216 | 311 | K311S | A | 4.1e-09 | 3.7e-09 | 1.3e-08 | 6.933E-09 | 5.258E-09 | 3.700E-09 | 1.300E-08 |
| 6217 | 311 | K311T | A | 3.7e-09 | 5.8e-09 | 1.5e-08 | 8.167E-09 | 6.010E-09 | 3.700E-09 | 1.500E-08 |
| 6218 | 311 | K311W | A | 1.1e-08 | 3.8e-09 | 2.9e-09 | 5.900E-09 | 4.440E-09 | 2.900E-09 | 1.100E-08 |
| 6219 | 311 | K311Y | A | 2.3e-08 | 6e-09 | 4.4e-09 | 1.113E-08 | 1.031E-08 | 4.400E-09 | 2.300E-08 |
| 6220 | 311 | K311V | A | 3.2e-09 | 7.4e-09 | 3.3e-09 | 4.633E-09 | 2.397E-09 | 3.200E-09 | 7.400E-09 |
| 6221 | 312 | C312A | A | 4.3e-09 | 4.8e-09 | 4.2e-09 | 4.433E-09 | 3.215E-10 | 4.200E-09 | 4.800E-09 |
| 6222 | 312 | C312R | A | 1.1e-08 | 9.9e-09 | 2.8e-09 | 7.900E-09 | 4.451E-09 | 2.800E-09 | 1.100E-08 |
| 6223 | 312 | C312N | A | 8.6e-09 | 7.9e-09 | 3.6e-09 | 6.700E-09 | 2.707E-09 | 3.600E-09 | 8.600E-09 |
| 6224 | 312 | C312D | A | 3.7e-09 | 8.9e-09 | 1.1e-08 | 7.867E-09 | 3.758E-09 | 3.700E-09 | 1.100E-08 |
| 6225 | 312 | C312C | A | 8.7e-09 | 5.9e-09 | 1.7e-08 | 1.053E-08 | 5.773E-09 | 5.900E-09 | 1.700E-08 |
| 6226 | 312 | C312E | A | 9.7e-09 | 4.9e-09 | 3e-09 | 5.867E-09 | 3.453E-09 | 3.000E-09 | 9.700E-09 |
| 6227 | 312 | C312Q | A | 3.6e-09 | 5.1e-09 | 3.2e-09 | 3.967E-09 | 1.002E-09 | 3.200E-09 | 5.100E-09 |
| 6228 | 312 | C312G | A | 2e-09 | 1.8e-08 | 6.7e-09 | 8.900E-09 | 8.224E-09 | 2.000E-09 | 1.800E-08 |
| 6229 | 312 | C312H | A | 2.7e-08 | 6.7e-09 | 1.2e-08 | 1.523E-08 | 1.053E-08 | 6.700E-09 | 2.700E-08 |
| 6230 | 312 | C312I | A | 5.3e-09 | 2.9e-08 | 4.6e-09 | 1.297E-08 | 1.389E-08 | 4.600E-09 | 2.900E-08 |
| 6231 | 312 | C312L | A | 9.6e-09 | 5.6e-09 | 1.1e-08 | 8.733E-09 | 2.802E-09 | 5.600E-09 | 1.100E-08 |
| 6232 | 312 | C312K | A | 1.5e-08 | 2.9e-09 | 4.7e-09 | 7.533E-09 | 6.529E-09 | 2.900E-09 | 1.500E-08 |
| 6233 | 312 | C312M | A | 5.5e-09 | 1.2e-08 | 3e-09 | 6.833E-09 | 4.646E-09 | 3.000E-09 | 1.200E-08 |
| 6234 | 312 | C312F | A | 1.9e-09 | 9.1e-09 | 9.4e-09 | 6.800E-09 | 4.246E-09 | 1.900E-09 | 9.400E-09 |
| 6235 | 312 | C312P | A | 7.7e-09 | 6.4e-09 | 2e-09 | 5.367E-09 | 2.987E-09 | 2.000E-09 | 7.700E-09 |
| 6236 | 312 | C312S | A | 5.6e-09 | 1.2e-08 | 1e-08 | 9.200E-09 | 3.274E-09 | 5.600E-09 | 1.200E-08 |
| 6237 | 312 | C312T | A | 1e-08 | 1.6e-08 | 4.8e-09 | 1.027E-08 | 5.605E-09 | 4.800E-09 | 1.600E-08 |
| 6238 | 312 | C312W | A | 1.5e-08 | 9.2e-09 | 8.9e-09 | 1.103E-08 | 3.439E-09 | 8.900E-09 | 1.500E-08 |
| 6239 | 312 | C312Y | A | 8.6e-09 | 5.7e-09 | 8.1e-09 | 7.467E-09 | 1.550E-09 | 5.700E-09 | 8.600E-09 |
| 6240 | 312 | C312V | A | 5.7e-09 | 1.1e-08 | 9.7e-09 | 8.800E-09 | 2.762E-09 | 5.700E-09 | 1.100E-08 |
| 6241 | 313 | S313A | A | 5.4e-09 | 9.8e-09 | 4.4e-08 | 1.973E-08 | 2.113E-08 | 5.400E-09 | 4.400E-08 |
| 6242 | 313 | S313R | A | 4.2e-09 | 6.3e-09 | 1.2e-08 | 7.500E-09 | 4.036E-09 | 4.200E-09 | 1.200E-08 |
| 6243 | 313 | S313N | A | 3.9e-09 | 9.7e-09 | 5.5e-09 | 6.367E-09 | 2.996E-09 | 3.900E-09 | 9.700E-09 |
| 6244 | 313 | S313D | A | 2.4e-08 | 1.8e-08 | 4.9e-09 | 1.563E-08 | 9.767E-09 | 4.900E-09 | 2.400E-08 |
| 6245 | 313 | S313C | A | 8.1e-09 | 2.4e-09 | 2.3e-09 | 4.267E-09 | 3.320E-09 | 2.300E-09 | 8.100E-09 |
| 6246 | 313 | S313E | A | 5.8e-09 | 3.6e-09 | 5.9e-09 | 5.100E-09 | 1.300E-09 | 3.600E-09 | 5.900E-09 |
| 6247 | 313 | S313Q | A | 2.5e-09 | 1.1e-08 | 4.5e-09 | 6.000E-09 | 4.444E-09 | 2.500E-09 | 1.100E-08 |
| 6248 | 313 | S313G | A | 1.7e-09 | 2.4e-09 | 1.3e-08 | 5.700E-09 | 6.332E-09 | 1.700E-09 | 1.300E-08 |
| 6249 | 313 | S313H | A | 1.8e-09 | 6.4e-09 | 3.9e-09 | 4.033E-09 | 2.303E-09 | 1.800E-09 | 6.400E-09 |
| 6250 | 313 | S313I | A | 5.9e-09 | 8.3e-10 | 9.4e-09 | 5.377E-09 | 4.309E-09 | 8.300E-10 | 9.400E-09 |
| 6251 | 313 | S313L | A | 2e-08 | 1.8e-08 | 6.2e-09 | 1.473E-08 | 7.457E-09 | 6.200E-09 | 2.000E-08 |
| 6252 | 313 | S313K | A | 4.1e-09 | 1.1e-08 | 7.2e-09 | 7.433E-09 | 3.456E-09 | 4.100E-09 | 1.100E-08 |
| 6253 | 313 | S313M | A | 8e-09 | 2.1e-08 | 1.6e-08 | 1.500E-08 | 6.557E-09 | 8.000E-09 | 2.100E-08 |
| 6254 | 313 | S313F | A | 8e-09 | 5.7e-09 | 9.2e-09 | 7.633E-09 | 1.779E-09 | 5.700E-09 | 9.200E-09 |
| 6255 | 313 | S313P | A | 3.2e-08 | 6.8e-09 | 3.7e-09 | 1.417E-08 | 1.552E-08 | 3.700E-09 | 3.200E-08 |
| 6256 | 313 | S313S | A | 5.7e-09 | 6.9e-09 | 4.7e-09 | 5.767E-09 | 1.102E-09 | 4.700E-09 | 6.900E-09 |
| 6257 | 313 | S313T | A | 7.2e-09 | 2.1e-08 | 1.1e-08 | 1.307E-08 | 7.128E-09 | 7.200E-09 | 2.100E-08 |
| 6258 | 313 | S313W | A | 7.5e-09 | 5.8e-09 | 8.7e-09 | 7.333E-09 | 1.457E-09 | 5.800E-09 | 8.700E-09 |
| 6259 | 313 | S313Y | A | 5.2e-09 | 4.7e-09 | 1.4e-08 | 7.967E-09 | 5.231E-09 | 4.700E-09 | 1.400E-08 |
| 6260 | 313 | S313V | A | 2.9e-09 | 8.6e-09 | 1.1e-08 | 7.500E-09 | 4.161E-09 | 2.900E-09 | 1.100E-08 |
| 6261 | 314 | K314A | A | 6.7e-09 | 2e-08 | 9e-09 | 1.190E-08 | 7.108E-09 | 6.700E-09 | 2.000E-08 |
| 6262 | 314 | K314R | A | 3.4e-08 | 7.1e-09 | 7.7e-09 | 1.627E-08 | 1.536E-08 | 7.100E-09 | 3.400E-08 |
| 6263 | 314 | K314N | A | 2e-09 | 1.3e-08 | 8.2e-09 | 7.733E-09 | 5.515E-09 | 2.000E-09 | 1.300E-08 |
| 6264 | 314 | K314D | A | 1.1e-08 | 1.5e-08 | 1e-08 | 1.200E-08 | 2.646E-09 | 1.000E-08 | 1.500E-08 |
| 6265 | 314 | K314C | A | 1.6e-08 | 1.5e-08 | 2.5e-08 | 1.867E-08 | 5.508E-09 | 1.500E-08 | 2.500E-08 |
| 6266 | 314 | K314E | A | 1.2e-08 | 2.2e-08 | 1.2e-08 | 1.533E-08 | 5.774E-09 | 1.200E-08 | 2.200E-08 |
| 6267 | 314 | K314Q | A | 1.5e-08 | 7.3e-09 | 7.9e-09 | 1.007E-08 | 4.283E-09 | 7.300E-09 | 1.500E-08 |
| 6268 | 314 | K314G | A | 8.6e-09 | 2.5e-09 | 1.4e-08 | 8.367E-09 | 5.754E-09 | 2.500E-09 | 1.400E-08 |
| 6269 | 314 | K314H | A | 6.6e-09 | 9.3e-09 | 8.1e-09 | 8.000E-09 | 1.353E-09 | 6.600E-09 | 9.300E-09 |
| 6270 | 314 | K314I | A | 1.7e-09 | 5.6e-09 | 4.8e-09 | 4.033E-09 | 2.060E-09 | 1.700E-09 | 5.600E-09 |
| 6271 | 314 | K314L | A | 9.2e-09 | 6e-09 | 9.1e-09 | 8.100E-09 | 1.819E-09 | 6.000E-09 | 9.200E-09 |
| 6272 | 314 | K314K | A | 1e-08 | 1e-08 | 6.1e-09 | 8.700E-09 | 2.252E-09 | 6.100E-09 | 1.000E-08 |
| 6273 | 314 | K314M | A | 1.6e-08 | 8.5e-09 | 1.9e-08 | 1.450E-08 | 5.408E-09 | 8.500E-09 | 1.900E-08 |
| 6274 | 314 | K314F | A | 6.5e-09 | 7.9e-09 | 1.4e-08 | 9.467E-09 | 3.988E-09 | 6.500E-09 | 1.400E-08 |
| 6275 | 314 | K314P | A | 3.2e-09 | 1.6e-08 | 2.8e-08 | 1.573E-08 | 1.240E-08 | 3.200E-09 | 2.800E-08 |
| 6276 | 314 | K314S | A | 4e-09 | 2e-09 | 1e-08 | 5.333E-09 | 4.163E-09 | 2.000E-09 | 1.000E-08 |
| 6277 | 314 | K314T | A | 6e-09 | 8.8e-09 | 1.7e-09 | 5.500E-09 | 3.576E-09 | 1.700E-09 | 8.800E-09 |
| 6278 | 314 | K314W | A | 4e-09 | 1.2e-08 | 2e-08 | 1.200E-08 | 8.000E-09 | 4.000E-09 | 2.000E-08 |
| 6279 | 314 | K314Y | A | 1.6e-08 | 1.5e-08 | 1.3e-08 | 1.467E-08 | 1.528E-09 | 1.300E-08 | 1.600E-08 |
| 6280 | 314 | K314V | A | 1.1e-08 | 6.2e-09 | 1.7e-09 | 6.300E-09 | 4.651E-09 | 1.700E-09 | 1.100E-08 |
| 6281 | 315 | P315A | A | 6.2e-09 | 1.1e-08 | 1.1e-08 | 9.400E-09 | 2.771E-09 | 6.200E-09 | 1.100E-08 |
| 6282 | 315 | P315R | A | 3.9e-09 | 2.8e-09 | 3.6e-09 | 3.433E-09 | 5.686E-10 | 2.800E-09 | 3.900E-09 |
| 6283 | 315 | P315N | A | 3.3e-09 | 5e-09 | 2.6e-09 | 3.633E-09 | 1.234E-09 | 2.600E-09 | 5.000E-09 |
| 6284 | 315 | P315D | A | 6.9e-09 | 7.6e-09 | 8.4e-09 | 7.633E-09 | 7.506E-10 | 6.900E-09 | 8.400E-09 |
| 6285 | 315 | P315C | A | 3e-09 | 9.2e-09 | 8e-09 | 6.733E-09 | 3.288E-09 | 3.000E-09 | 9.200E-09 |
| 6286 | 315 | P315E | A | 4.6e-09 | 6.7e-09 | 2.2e-09 | 4.500E-09 | 2.252E-09 | 2.200E-09 | 6.700E-09 |
| 6287 | 315 | P315Q | A | 4.1e-09 | 1.4e-09 | 5.3e-09 | 3.600E-09 | 1.997E-09 | 1.400E-09 | 5.300E-09 |
| 6288 | 315 | P315G | A | 4.8e-09 | 9.5e-09 | 6e-09 | 6.767E-09 | 2.442E-09 | 4.800E-09 | 9.500E-09 |
| 6289 | 315 | P315H | A | 2.6e-09 | 3.7e-09 | 1.8e-09 | 2.700E-09 | 9.539E-10 | 1.800E-09 | 3.700E-09 |

| | | | | | | | | | | |
|---|---|---|---|---|---|---|---|---|---|---|
| 6290 | 315 | P315I | A | 2.7e-08 | 7.2e-09 | 5.6e-09 | 1.327E-08 | 1.192E-08 | 5.600E-09 | 2.700E-08 |
| 6291 | 315 | P315L | A | 6.2e-09 | 4.9e-09 | 6.1e-09 | 5.733E-09 | 7.234E-10 | 4.900E-09 | 6.200E-09 |
| 6292 | 315 | P315K | A | 5.1e-09 | 4e-09 | 2.7e-09 | 3.933E-09 | 1.201E-09 | 2.700E-09 | 5.100E-09 |
| 6293 | 315 | P315M | A | 1e-08 | 1e-08 | 4.3e-09 | 8.100E-09 | 3.291E-09 | 4.300E-09 | 1.000E-08 |
| 6294 | 315 | P315F | A | 4.6e-09 | 6.8e-09 | 2.5e-09 | 4.633E-09 | 2.150E-09 | 2.500E-09 | 6.800E-09 |
| 6295 | 315 | P315P | A | 8.7e-09 | 4.2e-09 | 6.1e-09 | 6.333E-09 | 2.259E-09 | 4.200E-09 | 8.700E-09 |
| 6296 | 315 | P315S | A | 4.4e-09 | 3.4e-09 | 3.5e-09 | 3.767E-09 | 5.508E-10 | 3.400E-09 | 4.400E-09 |
| 6297 | 315 | P315T | A | 3.3e-09 | 2.4e-09 | 2.5e-09 | 2.733E-09 | 4.933E-10 | 2.400E-09 | 3.300E-09 |
| 6298 | 315 | P315W | A | 6.2e-09 | 4.4e-09 | 4e-09 | 4.867E-09 | 1.172E-09 | 4.000E-09 | 6.200E-09 |
| 6299 | 315 | P315Y | A | 9e-09 | 8.7e-09 | 7.1e-09 | 8.267E-09 | 1.021E-09 | 7.100E-09 | 9.000E-09 |
| 6300 | 315 | P315V | A | 1.3e-08 | 3.9e-09 | 9.3e-09 | 8.733E-09 | 4.576E-09 | 3.900E-09 | 1.300E-08 |
| 6301 | 316 | C316A | A | 7.7e-09 | 9.9e-09 | 1.1e-08 | 9.533E-09 | 1.680E-09 | 7.700E-09 | 1.100E-08 |
| 6302 | 316 | C316R | A | 1.2e-08 | 1e-08 | 9.2e-09 | 1.040E-08 | 1.442E-09 | 9.200E-09 | 1.200E-08 |
| 6303 | 316 | C316N | A | 3.6e-08 | 8.4e-09 | 3e-09 | 1.580E-08 | 1.770E-08 | 3.000E-09 | 3.600E-08 |
| 6304 | 316 | C316D | A | 7.8e-09 | 4.9e-09 | 1.3e-08 | 8.567E-09 | 4.104E-09 | 4.900E-09 | 1.300E-08 |
| 6305 | 316 | C316C | A | 1.6e-08 | 1.4e-08 | 4.4e-09 | 1.147E-08 | 6.201E-09 | 4.400E-09 | 1.600E-08 |
| 6306 | 316 | C316E | A | 1.6e-08 | 5.4e-09 | 5.7e-09 | 9.033E-09 | 6.035E-09 | 5.400E-09 | 1.600E-08 |
| 6307 | 316 | C316Q | A | 6e-09 | 3.8e-09 | 1.2e-08 | 7.267E-09 | 4.244E-09 | 3.800E-09 | 1.200E-08 |
| 6308 | 316 | C316G | A | 3.2e-09 | 5.4e-09 | 6.8e-09 | 5.133E-09 | 1.815E-09 | 3.200E-09 | 6.800E-09 |
| 6309 | 316 | C316H | A | 3.7e-09 | 1.1e-08 | 5.8e-09 | 6.833E-09 | 3.758E-09 | 3.700E-09 | 1.100E-08 |
| 6310 | 316 | C316I | A | 3.1e-08 | 1.7e-08 | 4.8e-09 | 1.760E-08 | 1.311E-08 | 4.800E-09 | 3.100E-08 |
| 6311 | 316 | C316L | A | 1.3e-09 | 2.6e-09 | 7.8e-09 | 3.900E-09 | 3.439E-09 | 1.300E-09 | 7.800E-09 |
| 6312 | 316 | C316K | A | 3.1e-09 | 2.1e-08 | 1.6e-08 | 1.337E-08 | 9.236E-09 | 3.100E-09 | 2.100E-08 |
| 6313 | 316 | C316M | A | 8.9e-09 | 1.2e-08 | 4.9e-09 | 8.600E-09 | 3.559E-09 | 4.900E-09 | 1.200E-08 |
| 6314 | 316 | C316F | A | 3.6e-09 | 6.7e-09 | 1.7e-08 | 9.100E-09 | 7.015E-09 | 3.600E-09 | 1.700E-08 |
| 6315 | 316 | C316P | A | 8.8e-09 | 5.1e-09 | 2.1e-08 | 1.163E-08 | 8.320E-09 | 5.100E-09 | 2.100E-08 |
| 6316 | 316 | C316S | A | 9.8e-09 | 5.4e-09 | 9.6e-09 | 8.267E-09 | 2.485E-09 | 5.400E-09 | 9.800E-09 |
| 6317 | 316 | C316T | A | 9.9e-09 | 3e-09 | 5.6e-09 | 6.167E-09 | 3.485E-09 | 3.000E-09 | 9.900E-09 |
| 6318 | 316 | C316W | A | 3.7e-09 | 1.6e-08 | 1.2e-08 | 1.057E-08 | 6.274E-09 | 3.700E-09 | 1.600E-08 |
| 6319 | 316 | C316Y | A | 9.8e-09 | 7.4e-09 | 1.4e-08 | 1.040E-08 | 3.341E-09 | 7.400E-09 | 1.400E-08 |
| 6320 | 316 | C316V | A | 3.6e-09 | 6.4e-09 | 6.6e-09 | 5.533E-09 | 1.677E-09 | 3.600E-09 | 6.600E-09 |
| 6321 | 317 | A317A | A | 3.9e-09 | 4.6e-09 | 1.5e-08 | 7.833E-09 | 6.216E-09 | 3.900E-09 | 1.500E-08 |
| 6322 | 317 | A317R | A | 8e-09 | 2.1e-08 | 4.3e-09 | 1.110E-08 | 8.771E-09 | 4.300E-09 | 2.100E-08 |
| 6323 | 317 | A317N | A | 5.3e-09 | 9.7e-09 | 1.9e-09 | 5.633E-09 | 3.911E-09 | 1.900E-09 | 9.700E-09 |
| 6324 | 317 | A317D | A | 5.3e-09 | 4.3e-09 | 5.3e-09 | 4.967E-09 | 5.774E-10 | 4.300E-09 | 5.300E-09 |
| 6325 | 317 | A317C | A | 9.6e-09 | 9.6e-09 | 1.3e-08 | 1.073E-08 | 1.963E-09 | 9.600E-09 | 1.300E-08 |
| 6326 | 317 | A317E | A | 1.7e-09 | 9.1e-09 | 6.6e-09 | 5.800E-09 | 3.764E-09 | 1.700E-09 | 9.100E-09 |
| 6327 | 317 | A317Q | A | 6.5e-09 | 3.7e-09 | 1.1e-08 | 7.067E-09 | 3.683E-09 | 3.700E-09 | 1.100E-08 |
| 6328 | 317 | A317G | A | 7.6e-09 | 9.4e-09 | 7.9e-09 | 8.300E-09 | 9.644E-10 | 7.600E-09 | 9.400E-09 |
| 6329 | 317 | A317H | A | 4.4e-09 | 3.9e-09 | 3.6e-09 | 3.967E-09 | 4.041E-10 | 3.600E-09 | 4.400E-09 |
| 6330 | 317 | A317I | A | 6.7e-09 | 6e-09 | 1e-08 | 7.567E-09 | 2.136E-09 | 6.000E-09 | 1.000E-08 |
| 6331 | 317 | A317L | A | 1.1e-08 | 7e-09 | 1.8e-08 | 1.200E-08 | 5.568E-09 | 7.000E-09 | 1.800E-08 |
| 6332 | 317 | A317K | A | 5.1e-09 | 1.3e-08 | 8.4e-09 | 8.833E-09 | 3.968E-09 | 5.100E-09 | 1.300E-08 |
| 6333 | 317 | A317M | A | 6.2e-09 | 5.4e-09 | 2e-09 | 4.533E-09 | 2.230E-09 | 2.000E-09 | 6.200E-09 |
| 6334 | 317 | A317F | A | 2.1e-09 | 9.2e-09 | 9.9e-09 | 7.067E-09 | 4.315E-09 | 2.100E-09 | 9.900E-09 |
| 6335 | 317 | A317P | A | 9.1e-09 | 1.2e-08 | 2.3e-09 | 7.800E-09 | 4.979E-09 | 2.300E-09 | 1.200E-08 |
| 6336 | 317 | A317S | A | 4.5e-09 | 1.7e-09 | 3.3e-09 | 3.167E-09 | 1.405E-09 | 1.700E-09 | 4.500E-09 |
| 6337 | 317 | A317T | A | 1.2e-08 | 5.2e-09 | 4.8e-09 | 7.333E-09 | 4.046E-09 | 4.800E-09 | 1.200E-08 |
| 6338 | 317 | A317W | A | 1.2e-08 | 7.4e-09 | 1.7e-08 | 1.213E-08 | 4.801E-09 | 7.400E-09 | 1.700E-08 |
| 6339 | 317 | A317Y | A | 2.1e-08 | 9.9e-09 | 1.2e-08 | 1.430E-08 | 5.897E-09 | 9.900E-09 | 2.100E-08 |
| 6340 | 317 | A317V | A | 7.2e-09 | 1.2e-08 | 3.2e-09 | 7.467E-09 | 4.406E-09 | 3.200E-09 | 1.200E-08 |
| 6341 | 318 | R318A | A | 1.1e-08 | 4.7e-09 | 1.9e-08 | 1.157E-08 | 7.167E-09 | 4.700E-09 | 1.900E-08 |
| 6342 | 318 | R318R | A | 1.2e-08 | 8.6e-09 | 9.2e-09 | 9.933E-09 | 1.815E-09 | 8.600E-09 | 1.200E-08 |
| 6343 | 318 | R318N | A | 9.7e-09 | 1.4e-08 | 6.4e-09 | 1.003E-08 | 3.811E-09 | 6.400E-09 | 1.400E-08 |
| 6344 | 318 | R318D | A | 1.2e-08 | 3.1e-08 | 1.2e-08 | 1.833E-08 | 1.097E-08 | 1.200E-08 | 3.100E-08 |
| 6345 | 318 | R318C | A | 9.7e-09 | 9.8e-09 | 7.6e-09 | 9.033E-09 | 1.242E-09 | 7.600E-09 | 9.800E-09 |
| 6346 | 318 | R318E | A | 1.4e-08 | 1.3e-08 | 1.2e-08 | 1.300E-08 | 1.000E-09 | 1.200E-08 | 1.400E-08 |
| 6347 | 318 | R318Q | A | 2.3e-08 | 9.9e-09 | 1.2e-08 | 1.497E-08 | 7.036E-09 | 9.900E-09 | 2.300E-08 |
| 6348 | 318 | R318G | A | 1.4e-08 | 2.4e-08 | 2.6e-09 | 1.353E-08 | 1.071E-08 | 2.600E-09 | 2.400E-08 |
| 6349 | 318 | R318H | A | 8.2e-09 | 1.7e-08 | 1.1e-08 | 1.207E-08 | 4.496E-09 | 8.200E-09 | 1.700E-08 |
| 6350 | 318 | R318I | A | 6.6e-09 | 4.3e-09 | 7.4e-09 | 6.100E-09 | 1.609E-09 | 4.300E-09 | 7.400E-09 |
| 6351 | 318 | R318L | A | 9.8e-09 | 6.5e-09 | 2.3e-09 | 6.200E-09 | 3.759E-09 | 2.300E-09 | 9.800E-09 |
| 6352 | 318 | R318K | A | 1.2e-08 | 1.1e-08 | 1.4e-08 | 1.233E-08 | 1.528E-09 | 1.100E-08 | 1.400E-08 |
| 6353 | 318 | R318M | A | 4.5e-09 | 1.9e-08 | 3e-09 | 8.833E-09 | 8.836E-09 | 3.000E-09 | 1.900E-08 |
| 6354 | 318 | R318F | A | 1.3e-08 | 1.6e-08 | 6.9e-09 | 1.197E-08 | 4.637E-09 | 6.900E-09 | 1.600E-08 |
| 6355 | 318 | R318P | A | 9.4e-09 | 1.5e-08 | 6.4e-09 | 1.027E-08 | 4.365E-09 | 6.400E-09 | 1.500E-08 |
| 6356 | 318 | R318S | A | 3.3e-09 | 1.6e-08 | 6.5e-09 | 8.600E-09 | 6.605E-09 | 3.300E-09 | 1.600E-08 |
| 6357 | 318 | R318T | A | 1.8e-09 | 8.6e-09 | 2e-08 | 1.013E-08 | 9.196E-09 | 1.800E-09 | 2.000E-08 |
| 6358 | 318 | R318W | A | 1.8e-08 | 5.2e-09 | 1.4e-08 | 1.240E-08 | 6.548E-09 | 5.200E-09 | 1.800E-08 |
| 6359 | 318 | R318Y | A | 2.3e-08 | 2.5e-08 | 4e-09 | 1.733E-08 | 1.159E-08 | 4.000E-09 | 2.500E-08 |
| 6360 | 318 | R318V | A | 8e-09 | 7.2e-09 | 4.4e-09 | 6.533E-09 | 1.890E-09 | 4.400E-09 | 8.000E-09 |
| 6361 | 319 | V319A | A | 2.5e-09 | 4.7e-09 | 1.7e-09 | 2.967E-09 | 1.553E-09 | 1.700E-09 | 4.700E-09 |
| 6362 | 319 | V319R | A | 5.7e-09 | 5.4e-09 | 9.3e-09 | 6.800E-09 | 2.170E-09 | 5.400E-09 | 9.300E-09 |
| 6363 | 319 | V319N | A | 3.7e-09 | 9.6e-09 | 9e-09 | 7.433E-09 | 3.247E-09 | 3.700E-09 | 9.600E-09 |
| 6364 | 319 | V319D | A | 8.6e-09 | 4.4e-09 | 5.6e-09 | 6.200E-09 | 2.163E-09 | 4.400E-09 | 8.600E-09 |
| 6365 | 319 | V319C | A | 1.2e-08 | 4.2e-09 | 3.6e-09 | 6.600E-09 | 4.686E-09 | 3.600E-09 | 1.200E-08 |
| 6366 | 319 | V319E | A | 5.8e-09 | 3.6e-09 | 5e-09 | 4.800E-09 | 1.114E-09 | 3.600E-09 | 5.800E-09 |
| 6367 | 319 | V319Q | A | 6.1e-09 | 8e-09 | 7.8e-09 | 7.300E-09 | 1.044E-09 | 6.100E-09 | 8.000E-09 |
| 6368 | 319 | V319G | A | 8.5e-09 | 9.7e-09 | 3e-08 | 1.607E-08 | 1.208E-08 | 8.500E-09 | 3.000E-08 |
| 6369 | 319 | V319H | A | 2.5e-09 | 1.3e-08 | 1.7e-08 | 1.083E-08 | 7.489E-09 | 2.500E-09 | 1.700E-08 |
| 6370 | 319 | V319I | A | 1.4e-08 | 5.4e-09 | 4.9e-09 | 8.100E-09 | 5.116E-09 | 4.900E-09 | 1.400E-08 |
| 6371 | 319 | V319L | A | 6.7e-09 | 1.2e-08 | 1.1e-08 | 9.900E-09 | 2.816E-09 | 6.700E-09 | 1.200E-08 |
| 6372 | 319 | V319K | A | 4.4e-09 | 5.8e-09 | 1.2e-08 | 7.400E-09 | 4.045E-09 | 4.400E-09 | 1.200E-08 |
| 6373 | 319 | V319M | A | 3.2e-09 | 3.5e-09 | 8.1e-09 | 4.933E-09 | 2.747E-09 | 3.200E-09 | 8.100E-09 |
| 6374 | 319 | V319F | A | 7.1e-09 | 6.7e-09 | 9.8e-09 | 7.867E-09 | 1.686E-09 | 6.700E-09 | 9.800E-09 |

| | | | | | | | | | | |
|---|---|---|---|---|---|---|---|---|---|---|
| 6375 | 319 | V319P | A | 1.2e-08 | 4.8e-09 | 4.8e-09 | 7.200E-09 | 4.157E-09 | 4.800E-09 | 1.200E-08 |
| 6376 | 319 | V319S | A | 8.9e-09 | 8.9e-09 | 8.8e-09 | 8.867E-09 | 5.774E-11 | 8.800E-09 | 8.900E-09 |
| 6377 | 319 | V319T | A | 1.6e-08 | 1.4e-08 | 4.6e-09 | 1.153E-08 | 6.087E-09 | 4.600E-09 | 1.600E-08 |
| 6378 | 319 | V319W | A | 4.8e-09 | 7.4e-09 | 7.5e-09 | 6.567E-09 | 1.531E-09 | 4.800E-09 | 7.500E-09 |
| 6379 | 319 | V319Y | A | 2.6e-09 | 1.2e-08 | 4.2e-09 | 6.267E-09 | 5.029E-09 | 2.600E-09 | 1.200E-08 |
| 6380 | 319 | V319V | A | 7e-09 | 7.4e-09 | 2.9e-09 | 5.767E-09 | 2.491E-09 | 2.900E-09 | 7.400E-09 |
| 6381 | 320 | C320A | A | 5.7e-09 | 4.4e-09 | 3.6e-09 | 4.567E-09 | 1.060E-09 | 3.600E-09 | 5.700E-09 |
| 6382 | 320 | C320R | A | 4e-09 | 8.5e-09 | 1e-08 | 7.500E-09 | 3.122E-09 | 4.000E-09 | 1.000E-08 |
| 6383 | 320 | C320N | A | 3.3e-09 | 4.5e-09 | 9.6e-09 | 5.800E-09 | 3.345E-09 | 3.300E-09 | 9.600E-09 |
| 6384 | 320 | C320D | A | 1.5e-08 | 7.8e-09 | 5.1e-09 | 9.300E-09 | 5.118E-09 | 5.100E-09 | 1.500E-08 |
| 6385 | 320 | C320C | A | 1.8e-08 | 8.3e-09 | 1.2e-08 | 1.277E-08 | 4.895E-09 | 8.300E-09 | 1.800E-08 |
| 6386 | 320 | C320E | A | 4.6e-09 | 5.7e-09 | 1.8e-08 | 9.433E-09 | 7.439E-09 | 4.600E-09 | 1.800E-08 |
| 6387 | 320 | C320Q | A | 7.5e-09 | 1.4e-08 | 1e-08 | 1.050E-08 | 3.279E-09 | 7.500E-09 | 1.400E-08 |
| 6388 | 320 | C320G | A | 1.3e-08 | 1.1e-09 | 3.6e-09 | 5.900E-09 | 6.275E-09 | 1.100E-09 | 1.300E-08 |
| 6389 | 320 | C320H | A | 4.2e-09 | 2.3e-08 | 1.1e-09 | 9.433E-09 | 1.185E-08 | 1.100E-09 | 2.300E-08 |
| 6390 | 320 | C320I | A | 3.6e-09 | 8.2e-09 | 7.4e-09 | 6.400E-09 | 2.458E-09 | 3.600E-09 | 8.200E-09 |
| 6391 | 320 | C320L | A | 6.6e-09 | 5.8e-09 | 1e-08 | 7.467E-09 | 2.230E-09 | 5.800E-09 | 1.000E-08 |
| 6392 | 320 | C320K | A | 5.4e-09 | 8.2e-09 | 2.3e-09 | 5.300E-09 | 2.951E-09 | 2.300E-09 | 8.200E-09 |
| 6393 | 320 | C320M | A | 1.1e-08 | 6.2e-09 | 1e-08 | 9.067E-09 | 2.532E-09 | 6.200E-09 | 1.100E-08 |
| 6394 | 320 | C320F | A | 2.8e-09 | 6e-09 | 7e-09 | 5.267E-09 | 2.194E-09 | 2.800E-09 | 7.000E-09 |
| 6395 | 320 | C320P | A | 1.5e-08 | 1.5e-08 | 7e-09 | 1.233E-08 | 4.619E-09 | 7.000E-09 | 1.500E-08 |
| 6396 | 320 | C320S | A | 2.6e-09 | 1.2e-08 | 8.8e-09 | 7.800E-09 | 4.779E-09 | 2.600E-09 | 1.200E-08 |
| 6397 | 320 | C320T | A | 1e-08 | 7.7e-09 | 1.2e-08 | 9.900E-09 | 2.152E-09 | 7.700E-09 | 1.200E-08 |
| 6398 | 320 | C320W | A | 6.4e-09 | 1.1e-08 | 3e-09 | 6.800E-09 | 4.015E-09 | 3.000E-09 | 1.100E-08 |
| 6399 | 320 | C320Y | A | 7.9e-09 | 4.4e-09 | 8.4e-09 | 6.900E-09 | 2.179E-09 | 4.400E-09 | 8.400E-09 |
| 6400 | 320 | C320V | A | 1.1e-08 | 5.8e-09 | 1.1e-08 | 9.267E-09 | 3.002E-09 | 5.800E-09 | 1.100E-08 |
| 6401 | 321 | Y321A | A | 6.9e-09 | 7.9e-09 | 2.5e-09 | 5.767E-09 | 2.873E-09 | 2.500E-09 | 7.900E-09 |
| 6402 | 321 | Y321R | A | 3.2e-09 | 2.7e-09 | 6.3e-09 | 4.067E-09 | 1.950E-09 | 2.700E-09 | 6.300E-09 |
| 6403 | 321 | Y321N | A | 4.3e-09 | 4.6e-09 | 3e-09 | 3.967E-09 | 8.505E-10 | 3.000E-09 | 4.600E-09 |
| 6404 | 321 | Y321D | A | 2.2e-08 | 3.7e-09 | 4.7e-09 | 1.013E-08 | 1.029E-08 | 3.700E-09 | 2.200E-08 |
| 6405 | 321 | Y321C | A | 1.8e-08 | 6.8e-09 | 7.9e-09 | 1.090E-08 | 6.173E-09 | 6.800E-09 | 1.800E-08 |
| 6406 | 321 | Y321E | A | 3.1e-09 | 2.2e-09 | 7.5e-09 | 4.267E-09 | 2.836E-09 | 2.200E-09 | 7.500E-09 |
| 6407 | 321 | Y321Q | A | 1.7e-09 | 4.4e-09 | 2.6e-09 | 2.900E-09 | 1.375E-09 | 1.700E-09 | 4.400E-09 |
| 6408 | 321 | Y321G | A | 8.1e-09 | 1.6e-08 | 1.4e-08 | 1.270E-08 | 4.107E-09 | 8.100E-09 | 1.600E-08 |
| 6409 | 321 | Y321H | A | 2.8e-08 | 2.4e-09 | 3.3e-09 | 1.123E-08 | 1.453E-08 | 2.400E-09 | 2.800E-08 |
| 6410 | 321 | Y321I | A | 2.1e-09 | 1.8e-08 | 6.5e-09 | 8.867E-09 | 8.210E-09 | 2.100E-09 | 1.800E-08 |
| 6411 | 321 | Y321L | A | 1.1e-08 | 5.2e-09 | 5.4e-09 | 7.200E-09 | 3.292E-09 | 5.200E-09 | 1.100E-08 |
| 6412 | 321 | Y321K | A | 9e-09 | 2.7e-09 | 3.2e-09 | 4.967E-09 | 3.502E-09 | 2.700E-09 | 9.000E-09 |
| 6413 | 321 | Y321M | A | 2.8e-09 | 1.2e-08 | 3.6e-09 | 6.133E-09 | 5.096E-09 | 2.800E-09 | 1.200E-08 |
| 6414 | 321 | Y321F | A | 1.1e-08 | 1e-08 | 4e-09 | 8.333E-09 | 3.786E-09 | 4.000E-09 | 1.100E-08 |
| 6415 | 321 | Y321P | A | 1.2e-08 | 1.4e-08 | 1.1e-08 | 1.233E-08 | 1.528E-09 | 1.100E-08 | 1.400E-08 |
| 6416 | 321 | Y321S | A | 1.3e-08 | 1.1e-08 | 1.3e-08 | 1.233E-08 | 1.155E-09 | 1.100E-08 | 1.300E-08 |
| 6417 | 321 | Y321T | A | 7.5e-09 | 4e-09 | 3.2e-09 | 4.900E-09 | 2.287E-09 | 3.200E-09 | 7.500E-09 |
| 6418 | 321 | Y321W | A | 9.5e-09 | 1.5e-08 | 2.5e-09 | 9.000E-09 | 6.265E-09 | 2.500E-09 | 1.500E-08 |
| 6419 | 321 | Y321Y | A | 1.9e-09 | 3.2e-09 | 9e-09 | 4.700E-09 | 3.780E-09 | 1.900E-09 | 9.000E-09 |
| 6420 | 321 | Y321V | A | 8.3e-09 | 1.1e-08 | 1.3e-08 | 1.077E-08 | 2.359E-09 | 8.300E-09 | 1.300E-08 |
| 6421 | 322 | G322A | A | 3.8e-09 | 4.3e-09 | 5.9e-09 | 4.667E-09 | 1.097E-09 | 3.800E-09 | 5.900E-09 |
| 6422 | 322 | G322R | A | 2.8e-09 | 4.3e-09 | 2.1e-09 | 3.067E-09 | 1.124E-09 | 2.100E-09 | 4.300E-09 |
| 6423 | 322 | G322N | A | 4.6e-09 | 2e-08 | 2.4e-08 | 1.620E-08 | 1.024E-08 | 4.600E-09 | 2.400E-08 |
| 6424 | 322 | G322D | A | 6.1e-09 | 2.1e-09 | 6e-09 | 4.733E-09 | 2.281E-09 | 2.100E-09 | 6.100E-09 |
| 6425 | 322 | G322C | A | 1.2e-08 | 1.1e-08 | 1.2e-08 | 1.167E-08 | 5.774E-10 | 1.100E-08 | 1.200E-08 |
| 6426 | 322 | G322E | A | 6.4e-09 | 7.7e-09 | 1.3e-08 | 9.033E-09 | 3.496E-09 | 6.400E-09 | 1.300E-08 |
| 6427 | 322 | G322Q | A | 1.4e-08 | 4.9e-09 | 6e-09 | 8.300E-09 | 4.967E-09 | 4.900E-09 | 1.400E-08 |
| 6428 | 322 | G322G | A | 5.6e-09 | 3.6e-09 | 6.9e-09 | 5.367E-09 | 1.662E-09 | 3.600E-09 | 6.900E-09 |
| 6429 | 322 | G322H | A | 1e-08 | 1.5e-08 | 1e-08 | 1.167E-08 | 2.887E-09 | 1.000E-08 | 1.500E-08 |
| 6430 | 322 | G322I | A | 8.5e-09 | 7.6e-09 | 1.2e-08 | 9.367E-09 | 2.325E-09 | 7.600E-09 | 1.200E-08 |
| 6431 | 322 | G322L | A | 5.1e-09 | 1.5e-08 | 4.5e-09 | 8.200E-09 | 5.897E-09 | 4.500E-09 | 1.500E-08 |
| 6432 | 322 | G322K | A | 9.4e-09 | 8e-09 | 1.4e-08 | 1.047E-08 | 3.139E-09 | 8.000E-09 | 1.400E-08 |
| 6433 | 322 | G322M | A | 9.7e-09 | 3.2e-09 | 1.4e-08 | 8.967E-09 | 5.437E-09 | 3.200E-09 | 1.400E-08 |
| 6434 | 322 | G322F | A | 9.2e-09 | 1e-08 | 6.8e-09 | 8.667E-09 | 1.665E-09 | 6.800E-09 | 1.000E-08 |
| 6435 | 322 | G322P | A | 1.1e-08 | 5.4e-09 | 1.3e-08 | 9.800E-09 | 3.940E-09 | 5.400E-09 | 1.300E-08 |
| 6436 | 322 | G322S | A | 6.5e-09 | 7.5e-09 | 1.4e-08 | 9.333E-09 | 4.072E-09 | 6.500E-09 | 1.400E-08 |
| 6437 | 322 | G322T | A | 6.8e-09 | 7.8e-09 | 7.5e-09 | 7.367E-09 | 5.132E-10 | 6.800E-09 | 7.800E-09 |
| 6438 | 322 | G322W | A | 8e-10 | 7.2e-09 | 6.5e-09 | 4.833E-09 | 3.510E-09 | 8.000E-10 | 7.200E-09 |
| 6439 | 322 | G322Y | A | 5.5e-09 | 2.4e-09 | 7.1e-09 | 5.000E-09 | 2.390E-09 | 2.400E-09 | 7.100E-09 |
| 6440 | 322 | G322V | A | 8.9e-09 | 8e-09 | 8.4e-09 | 8.433E-09 | 4.509E-10 | 8.000E-09 | 8.900E-09 |
| 6441 | 323 | L323A | A | 2.7e-09 | 8.5e-09 | 1.4e-08 | 8.400E-09 | 5.651E-09 | 2.700E-09 | 1.400E-08 |
| 6442 | 323 | L323R | A | 8.2e-09 | 7.7e-09 | 7.2e-09 | 7.700E-09 | 5.000E-10 | 7.200E-09 | 8.200E-09 |
| 6443 | 323 | L323N | A | 6.3e-09 | 1e-08 | 3.6e-09 | 6.633E-09 | 3.213E-09 | 3.600E-09 | 1.000E-08 |
| 6444 | 323 | L323D | A | 8.1e-09 | 8.1e-09 | 3.2e-09 | 6.467E-09 | 2.829E-09 | 3.200E-09 | 8.100E-09 |
| 6445 | 323 | L323C | A | 7.1e-09 | 9.1e-09 | 1e-08 | 8.733E-09 | 1.484E-09 | 7.100E-09 | 1.000E-08 |
| 6446 | 323 | L323E | A | 5.2e-09 | 1.1e-08 | 3.4e-09 | 6.533E-09 | 3.972E-09 | 3.400E-09 | 1.100E-08 |
| 6447 | 323 | L323Q | A | 7.3e-09 | 1e-08 | 6e-09 | 7.767E-09 | 2.040E-09 | 6.000E-09 | 1.000E-08 |
| 6448 | 323 | L323G | A | 2.1e-09 | 3.3e-09 | 5.7e-09 | 3.700E-09 | 1.833E-09 | 2.100E-09 | 5.700E-09 |
| 6449 | 323 | L323H | A | 8.3e-09 | 6.7e-09 | 7.4e-09 | 7.467E-09 | 8.021E-10 | 6.700E-09 | 8.300E-09 |
| 6450 | 323 | L323I | A | 1.5e-08 | 6.9e-09 | 6.6e-09 | 9.500E-09 | 4.766E-09 | 6.600E-09 | 1.500E-08 |
| 6451 | 323 | L323L | A | 5.8e-09 | 6e-09 | 2e-09 | 4.600E-09 | 2.254E-09 | 2.000E-09 | 6.000E-09 |
| 6452 | 323 | L323K | A | 5.1e-09 | 1.2e-08 | 1.5e-08 | 1.070E-08 | 5.076E-09 | 5.100E-09 | 1.500E-08 |
| 6453 | 323 | L323M | A | 1.2e-08 | 3.4e-09 | 4.3e-09 | 6.567E-09 | 4.727E-09 | 3.400E-09 | 1.200E-08 |
| 6454 | 323 | L323F | A | 5.7e-09 | 9.4e-09 | 9.3e-09 | 8.133E-09 | 2.108E-09 | 5.700E-09 | 9.400E-09 |
| 6455 | 323 | L323P | A | 6.7e-09 | 1e-08 | 8.3e-09 | 8.333E-09 | 1.650E-09 | 6.700E-09 | 1.000E-08 |
| 6456 | 323 | L323S | A | 1.4e-08 | 8.5e-09 | 3e-08 | 1.750E-08 | 1.117E-08 | 8.500E-09 | 3.000E-08 |
| 6457 | 323 | L323T | A | 1e-08 | 1.6e-08 | 2.4e-09 | 9.467E-09 | 6.816E-09 | 2.400E-09 | 1.600E-08 |
| 6458 | 323 | L323W | A | 2.6e-08 | 4.9e-09 | 2.3e-09 | 1.107E-08 | 1.300E-08 | 2.300E-09 | 2.600E-08 |
| 6459 | 323 | L323Y | A | 6.5e-09 | 7.2e-09 | 7.5e-09 | 7.067E-09 | 5.132E-10 | 6.500E-09 | 7.500E-09 |

| | | | | | | | | | | |
|---|---|---|---|---|---|---|---|---|---|---|
| 6460 | 323 | L323V | A | 6.7e-09 | 3.7e-09 | 1.3e-08 | 7.800E-09 | 4.747E-09 | 3.700E-09 | 1.300E-08 |
| 6461 | 324 | G324A | A | 2.9e-09 | 5.1e-09 | 3.6e-09 | 3.867E-09 | 1.124E-09 | 2.900E-09 | 5.100E-09 |
| 6462 | 324 | G324R | A | 7.5e-09 | 7.2e-09 | 3.5e-09 | 6.067E-09 | 2.228E-09 | 3.500E-09 | 7.500E-09 |
| 6463 | 324 | G324N | A | 7.7e-09 | 4e-09 | 5.6e-09 | 5.767E-09 | 1.856E-09 | 4.000E-09 | 7.700E-09 |
| 6464 | 324 | G324D | A | 3.1e-09 | 2.3e-08 | 8.5e-09 | 1.153E-08 | 1.029E-08 | 3.100E-09 | 2.300E-08 |
| 6465 | 324 | G324C | A | 1.6e-09 | 1.5e-08 | 7.7e-09 | 8.100E-09 | 6.709E-09 | 1.600E-09 | 1.500E-08 |
| 6466 | 324 | G324E | A | 1.7e-08 | 3.2e-09 | 1.1e-08 | 1.040E-08 | 6.920E-09 | 3.200E-09 | 1.700E-08 |
| 6467 | 324 | G324Q | A | 1.7e-08 | 6.9e-09 | 6.3e-09 | 1.007E-08 | 6.012E-09 | 6.300E-09 | 1.700E-08 |
| 6468 | 324 | G324G | A | 5.8e-09 | 1.9e-09 | 5.6e-09 | 4.433E-09 | 2.196E-09 | 1.900E-09 | 5.800E-09 |
| 6469 | 324 | G324H | A | 6e-09 | 2.8e-09 | 1.5e-08 | 7.933E-09 | 6.326E-09 | 2.800E-09 | 1.500E-08 |
| 6470 | 324 | G324I | A | 8.6e-09 | 1.6e-08 | 3e-09 | 9.200E-09 | 6.521E-09 | 3.000E-09 | 1.600E-08 |
| 6471 | 324 | G324L | A | 9e-09 | 7.4e-09 | 6.3e-09 | 7.567E-09 | 1.358E-09 | 6.300E-09 | 9.000E-09 |
| 6472 | 324 | G324K | A | 4.4e-09 | 9e-09 | 1.2e-08 | 8.467E-09 | 3.828E-09 | 4.400E-09 | 1.200E-08 |
| 6473 | 324 | G324M | A | 1.4e-08 | 9.1e-09 | 6.8e-09 | 9.967E-09 | 3.677E-09 | 6.800E-09 | 1.400E-08 |
| 6474 | 324 | G324F | A | 6.3e-09 | 4.2e-09 | 4.5e-09 | 5.000E-09 | 1.136E-09 | 4.200E-09 | 6.300E-09 |
| 6475 | 324 | G324P | A | 7e-09 | 9.7e-09 | 1.2e-08 | 9.567E-09 | 2.503E-09 | 7.000E-09 | 1.200E-08 |
| 6476 | 324 | G324S | A | 7.6e-09 | 9.7e-09 | 9e-09 | 8.767E-09 | 1.069E-09 | 7.600E-09 | 9.700E-09 |
| 6477 | 324 | G324T | A | 6.2e-09 | 1.7e-09 | 3.1e-09 | 3.667E-09 | 2.303E-09 | 1.700E-09 | 6.200E-09 |
| 6478 | 324 | G324W | A | 8e-09 | 1.8e-08 | 8e-09 | 1.133E-08 | 5.774E-09 | 8.000E-09 | 1.800E-08 |
| 6479 | 324 | G324Y | A | 3.6e-09 | 1e-08 | 1.3e-08 | 8.867E-09 | 4.801E-09 | 3.600E-09 | 1.300E-08 |
| 6480 | 324 | G324V | A | 9.3e-09 | 3.6e-09 | 5.9e-09 | 6.267E-09 | 2.868E-09 | 3.600E-09 | 9.300E-09 |
| 6481 | 325 | M325A | A | 1.8e-08 | 4.6e-09 | 5.8e-09 | 9.467E-09 | 7.414E-09 | 4.600E-09 | 1.800E-08 |
| 6482 | 325 | M325R | A | 3.1e-08 | 5.8e-09 | 1.7e-08 | 1.793E-08 | 1.263E-08 | 5.800E-09 | 3.100E-08 |
| 6483 | 325 | M325N | A | 8e-09 | 1.3e-08 | 3.8e-09 | 8.267E-09 | 4.606E-09 | 3.800E-09 | 1.300E-08 |
| 6484 | 325 | M325D | A | 1.4e-08 | 6.6e-09 | 3e-09 | 7.867E-09 | 5.608E-09 | 3.000E-09 | 1.400E-08 |
| 6485 | 325 | M325C | A | 3.4e-09 | 9.5e-09 | 5.9e-09 | 6.267E-09 | 3.066E-09 | 3.400E-09 | 9.500E-09 |
| 6486 | 325 | M325E | A | 1.2e-08 | 5.3e-09 | 7.7e-09 | 8.333E-09 | 3.395E-09 | 5.300E-09 | 1.200E-08 |
| 6487 | 325 | M325Q | A | 6.3e-09 | 4.8e-09 | 3.2e-09 | 4.767E-09 | 1.550E-09 | 3.200E-09 | 6.300E-09 |
| 6488 | 325 | M325G | A | 2.7e-09 | 7.1e-09 | 9.2e-09 | 6.333E-09 | 3.317E-09 | 2.700E-09 | 9.200E-09 |
| 6489 | 325 | M325H | A | 1.4e-09 | 6.6e-09 | 3e-09 | 3.667E-09 | 2.663E-09 | 1.400E-09 | 6.600E-09 |
| 6490 | 325 | M325I | A | 9.8e-09 | 7.2e-09 | 5.4e-09 | 7.467E-09 | 2.212E-09 | 5.400E-09 | 9.800E-09 |
| 6491 | 325 | M325L | A | 9.4e-09 | 5e-09 | 9.3e-09 | 7.900E-09 | 2.512E-09 | 5.000E-09 | 9.400E-09 |
| 6492 | 325 | M325K | A | 6.7e-09 | 2.1e-08 | 2.2e-09 | 9.967E-09 | 9.816E-09 | 2.200E-09 | 2.100E-08 |
| 6493 | 325 | M325M | A | 5.5e-09 | 7.7e-09 | 1.3e-08 | 8.733E-09 | 3.855E-09 | 5.500E-09 | 1.300E-08 |
| 6494 | 325 | M325F | A | 7.1e-09 | 8e-09 | 7.2e-09 | 7.433E-09 | 4.933E-10 | 7.100E-09 | 8.000E-09 |
| 6495 | 325 | M325P | A | 1.7e-09 | 4.5e-09 | 6.7e-09 | 4.300E-09 | 2.506E-09 | 1.700E-09 | 6.700E-09 |
| 6496 | 325 | M325S | A | 4.8e-09 | 8.9e-09 | 1.1e-08 | 8.233E-09 | 3.153E-09 | 4.800E-09 | 1.100E-08 |
| 6497 | 325 | M325T | A | 3.4e-09 | 1.3e-08 | 9.7e-09 | 8.700E-09 | 4.877E-09 | 3.400E-09 | 1.300E-08 |
| 6498 | 325 | M325W | A | 1.2e-08 | 3e-08 | 1.7e-08 | 1.967E-08 | 9.292E-09 | 1.200E-08 | 3.000E-08 |
| 6499 | 325 | M325Y | A | 2.9e-09 | 3.5e-09 | 1.2e-08 | 6.133E-09 | 5.090E-09 | 2.900E-09 | 1.200E-08 |
| 6500 | 325 | M325V | A | 6.8e-09 | 2.2e-08 | 1.2e-08 | 1.360E-08 | 7.725E-09 | 6.800E-09 | 2.200E-08 |
| 6501 | 326 | E326A | A | 3.5e-09 | 5.8e-09 | 5.6e-09 | 4.967E-09 | 1.274E-09 | 3.500E-09 | 5.800E-09 |
| 6502 | 326 | E326R | A | 1.6e-08 | 8.1e-09 | 5.9e-09 | 1.000E-08 | 5.311E-09 | 5.900E-09 | 1.600E-08 |
| 6503 | 326 | E326N | A | 1.1e-08 | 1e-08 | 1.4e-08 | 1.167E-08 | 2.082E-09 | 1.000E-08 | 1.400E-08 |
| 6504 | 326 | E326D | A | 2.5e-09 | 1.9e-08 | 1.1e-08 | 1.083E-08 | 8.251E-09 | 2.500E-09 | 1.900E-08 |
| 6505 | 326 | E326C | A | 7.1e-09 | 6.8e-09 | 1e-08 | 7.967E-09 | 1.767E-09 | 6.800E-09 | 1.000E-08 |
| 6506 | 326 | E326E | A | 1.1e-08 | 4.2e-09 | 1.5e-08 | 1.007E-08 | 5.460E-09 | 4.200E-09 | 1.500E-08 |
| 6507 | 326 | E326Q | A | 9.4e-09 | 4.7e-09 | 1.3e-08 | 9.033E-09 | 4.162E-09 | 4.700E-09 | 1.300E-08 |
| 6508 | 326 | E326G | A | 1.5e-09 | 6.6e-09 | 7.1e-09 | 5.067E-09 | 3.099E-09 | 1.500E-09 | 7.100E-09 |
| 6509 | 326 | E326H | A | 5.7e-09 | 1.7e-08 | 4.3e-09 | 9.000E-09 | 6.963E-09 | 4.300E-09 | 1.700E-08 |
| 6510 | 326 | E326I | A | 4.9e-09 | 2.6e-08 | 1e-08 | 1.363E-08 | 1.101E-08 | 4.900E-09 | 2.600E-08 |
| 6511 | 326 | E326L | A | 1e-08 | 2.9e-09 | 1.1e-08 | 7.967E-09 | 4.416E-09 | 2.900E-09 | 1.100E-08 |
| 6512 | 326 | E326K | A | 7.7e-09 | 1.8e-08 | 3.5e-09 | 9.733E-09 | 7.461E-09 | 3.500E-09 | 1.800E-08 |
| 6513 | 326 | E326M | A | 2.1e-09 | 1.8e-08 | 2.6e-09 | 7.567E-09 | 9.039E-09 | 2.100E-09 | 1.800E-08 |
| 6514 | 326 | E326F | A | 6.8e-09 | 2.6e-09 | 4.3e-09 | 4.567E-09 | 2.113E-09 | 2.600E-09 | 6.800E-09 |
| 6515 | 326 | E326P | A | 1.4e-08 | 9.1e-09 | 6.4e-09 | 9.833E-09 | 3.853E-09 | 6.400E-09 | 1.400E-08 |
| 6516 | 326 | E326S | A | 6.1e-09 | 1.2e-08 | 4.9e-09 | 7.667E-09 | 3.800E-09 | 4.900E-09 | 1.200E-08 |
| 6517 | 326 | E326T | A | 1.4e-08 | 5.1e-09 | 9.4e-09 | 9.500E-09 | 4.451E-09 | 5.100E-09 | 1.400E-08 |
| 6518 | 326 | E326W | A | 1e-08 | 8.5e-09 | 2.9e-09 | 7.133E-09 | 3.742E-09 | 2.900E-09 | 1.000E-08 |
| 6519 | 326 | E326Y | A | 3.4e-09 | 1e-08 | 8.7e-09 | 7.367E-09 | 3.496E-09 | 3.400E-09 | 1.000E-08 |
| 6520 | 326 | E326V | A | 4.6e-09 | 1.1e-08 | 3.1e-09 | 6.233E-09 | 4.196E-09 | 3.100E-09 | 1.100E-08 |
| 6521 | 327 | H327A | A | 7.7e-09 | 8.9e-09 | 1.2e-08 | 9.533E-09 | 2.219E-09 | 7.700E-09 | 1.200E-08 |
| 6522 | 327 | H327R | A | 1.2e-08 | 1.9e-08 | 3.9e-09 | 1.163E-08 | 7.557E-09 | 3.900E-09 | 1.900E-08 |
| 6523 | 327 | H327N | A | 6.6e-09 | 5.3e-09 | 1.1e-08 | 7.633E-09 | 2.987E-09 | 5.300E-09 | 1.100E-08 |
| 6524 | 327 | H327D | A | 2.9e-09 | 2.6e-09 | 1.2e-08 | 5.833E-09 | 5.343E-09 | 2.600E-09 | 1.200E-08 |
| 6525 | 327 | H327C | A | 6.1e-09 | 5.1e-09 | 4.9e-09 | 5.367E-09 | 6.429E-10 | 4.900E-09 | 6.100E-09 |
| 6526 | 327 | H327E | A | 4.1e-09 | 2.2e-09 | 2.8e-09 | 3.033E-09 | 9.713E-10 | 2.200E-09 | 4.100E-09 |
| 6527 | 327 | H327Q | A | 4.3e-09 | 3.6e-09 | 1.1e-08 | 6.300E-09 | 4.085E-09 | 3.600E-09 | 1.100E-08 |
| 6528 | 327 | H327G | A | 7.9e-09 | 5.9e-09 | 2.7e-09 | 5.500E-09 | 2.623E-09 | 2.700E-09 | 7.900E-09 |
| 6529 | 327 | H327H | A | 7.3e-09 | 4.3e-09 | 1.1e-08 | 7.533E-09 | 3.356E-09 | 4.300E-09 | 1.100E-08 |
| 6530 | 327 | H327I | A | 3.9e-09 | 1e-08 | 3.9e-09 | 5.933E-09 | 3.522E-09 | 3.900E-09 | 1.000E-08 |
| 6531 | 327 | H327L | A | 5.5e-09 | 5.4e-09 | 1.8e-08 | 9.633E-09 | 7.246E-09 | 5.400E-09 | 1.800E-08 |
| 6532 | 327 | H327K | A | 1.7e-08 | 3e-08 | 2.6e-09 | 1.653E-08 | 1.371E-08 | 2.600E-09 | 3.000E-08 |
| 6533 | 327 | H327M | A | 7.5e-09 | 7.8e-09 | 6.6e-09 | 7.300E-09 | 6.245E-10 | 6.600E-09 | 7.800E-09 |
| 6534 | 327 | H327F | A | 1.2e-08 | 1.4e-08 | 1.4e-08 | 1.333E-08 | 1.155E-09 | 1.200E-08 | 1.400E-08 |
| 6535 | 327 | H327P | A | 2.7e-09 | 1.2e-08 | 2e-08 | 1.157E-08 | 8.658E-09 | 2.700E-09 | 2.000E-08 |
| 6536 | 327 | H327S | A | 4.7e-09 | 6.3e-09 | 5.4e-09 | 5.467E-09 | 8.021E-10 | 4.700E-09 | 6.300E-09 |
| 6537 | 327 | H327T | A | 1.6e-08 | 1.4e-08 | 6.8e-09 | 1.227E-08 | 4.839E-09 | 6.800E-09 | 1.600E-08 |
| 6538 | 327 | H327W | A | 6.7e-09 | 3.4e-09 | 3.7e-08 | 1.570E-08 | 1.852E-08 | 3.400E-09 | 3.700E-08 |
| 6539 | 327 | H327Y | A | 6.3e-09 | 5e-09 | 4.7e-09 | 5.333E-09 | 8.505E-10 | 4.700E-09 | 6.300E-09 |
| 6540 | 327 | H327V | A | 5.2e-09 | 5.5e-09 | 1e-08 | 6.900E-09 | 2.689E-09 | 5.200E-09 | 1.000E-08 |
| 6541 | 328 | L328A | A | 2e-08 | 9.7e-09 | 5.2e-09 | 1.163E-08 | 7.587E-09 | 5.200E-09 | 2.000E-08 |
| 6542 | 328 | L328R | A | 9.8e-09 | 1.1e-08 | 2.8e-09 | 7.867E-09 | 4.429E-09 | 2.800E-09 | 1.100E-08 |
| 6543 | 328 | L328N | A | 1.9e-08 | 8.8e-09 | 1.9e-08 | 1.560E-08 | 5.889E-09 | 8.800E-09 | 1.900E-08 |
| 6544 | 328 | L328D | A | 2.6e-09 | 1.1e-08 | 3.9e-09 | 5.833E-09 | 4.521E-09 | 2.600E-09 | 1.100E-08 |

| | | | | | | | | | | |
|---|---|---|---|---|---|---|---|---|---|---|
| 6545 | 328 | L328C | A | 5.7e-09 | 9.6e-09 | 6.6e-09 | 7.300E-09 | 2.042E-09 | 5.700E-09 | 9.600E-09 |
| 6546 | 328 | L328E | A | 4e-09 | 8.4e-09 | 8.6e-09 | 7.000E-09 | 2.600E-09 | 4.000E-09 | 8.600E-09 |
| 6547 | 328 | L328Q | A | 5.4e-09 | 2e-09 | 2.7e-09 | 3.367E-09 | 1.795E-09 | 2.000E-09 | 5.400E-09 |
| 6548 | 328 | L328G | A | 5.1e-09 | 4e-08 | 9.9e-09 | 1.833E-08 | 1.892E-08 | 5.100E-09 | 4.000E-08 |
| 6549 | 328 | L328H | A | 6.4e-09 | 1.3e-08 | 2.7e-09 | 7.367E-09 | 5.218E-09 | 2.700E-09 | 1.300E-08 |
| 6550 | 328 | L328I | A | 3.5e-09 | 7.7e-09 | 2.6e-08 | 1.240E-08 | 1.196E-08 | 3.500E-09 | 2.600E-08 |
| 6551 | 328 | L328L | A | 1.3e-08 | 1.5e-08 | 6.3e-09 | 1.143E-08 | 4.557E-09 | 6.300E-09 | 1.500E-08 |
| 6552 | 328 | L328K | A | 1.3e-08 | 5.4e-09 | 3.5e-09 | 7.300E-09 | 5.027E-09 | 3.500E-09 | 1.300E-08 |
| 6553 | 328 | L328M | A | 5.8e-09 | 3.6e-08 | 9.9e-09 | 1.723E-08 | 1.638E-08 | 5.800E-09 | 3.600E-08 |
| 6554 | 328 | L328F | A | 6.4e-09 | 7.3e-09 | 6.5e-09 | 6.733E-09 | 4.933E-10 | 6.400E-09 | 7.300E-09 |
| 6555 | 328 | L328P | A | 4.4e-09 | 7.6e-09 | 5.7e-09 | 5.900E-09 | 1.609E-09 | 4.400E-09 | 7.600E-09 |
| 6556 | 328 | L328S | A | 3.9e-09 | 1e-08 | 9.4e-09 | 7.767E-09 | 3.362E-09 | 3.900E-09 | 1.000E-08 |
| 6557 | 328 | L328T | A | 1.3e-08 | 8.6e-09 | 4.4e-09 | 8.667E-09 | 4.300E-09 | 4.400E-09 | 1.300E-08 |
| 6558 | 328 | L328W | A | 1.1e-08 | 1.1e-08 | 8.6e-09 | 1.020E-08 | 1.386E-09 | 8.600E-09 | 1.100E-08 |
| 6559 | 328 | L328Y | A | 5.2e-09 | 2.6e-09 | 1.7e-08 | 8.267E-09 | 7.674E-09 | 2.600E-09 | 1.700E-08 |
| 6560 | 328 | L328V | A | 6.9e-09 | 1.1e-08 | 6.2e-09 | 8.033E-09 | 2.593E-09 | 6.200E-09 | 1.100E-08 |
| 6561 | 329 | R329A | A | 8.1e-09 | 7.8e-09 | 8.2e-09 | 8.033E-09 | 2.082E-10 | 7.800E-09 | 8.200E-09 |
| 6562 | 329 | R329R | A | 1e-08 | 5.1e-09 | 1e-08 | 8.367E-09 | 2.829E-09 | 5.100E-09 | 1.000E-08 |
| 6563 | 329 | R329N | A | 8.1e-09 | 1.5e-08 | 3.7e-09 | 8.933E-09 | 5.696E-09 | 3.700E-09 | 1.500E-08 |
| 6564 | 329 | R329D | A | 8e-09 | 9.8e-09 | 3e-09 | 6.933E-09 | 3.523E-09 | 3.000E-09 | 9.800E-09 |
| 6565 | 329 | R329C | A | 5.3e-09 | 3e-09 | 3.3e-09 | 3.867E-09 | 1.250E-09 | 3.000E-09 | 5.300E-09 |
| 6566 | 329 | R329E | A | 5.2e-09 | 5.8e-09 | 1.2e-08 | 7.667E-09 | 3.765E-09 | 5.200E-09 | 1.200E-08 |
| 6567 | 329 | R329Q | A | 7.4e-09 | 5.5e-09 | 1.3e-08 | 8.633E-09 | 3.899E-09 | 5.500E-09 | 1.300E-08 |
| 6568 | 329 | R329G | A | 4.2e-09 | 1.8e-08 | 6.2e-09 | 9.467E-09 | 7.457E-09 | 4.200E-09 | 1.800E-08 |
| 6569 | 329 | R329H | A | 1.1e-08 | 6e-09 | 1.1e-08 | 9.333E-09 | 2.887E-09 | 6.000E-09 | 1.100E-08 |
| 6570 | 329 | R329I | A | 5.6e-09 | 8e-09 | 2e-08 | 1.120E-08 | 7.715E-09 | 5.600E-09 | 2.000E-08 |
| 6571 | 329 | R329L | A | 1.2e-08 | 9.5e-09 | 3.9e-09 | 8.467E-09 | 4.148E-09 | 3.900E-09 | 1.200E-08 |
| 6572 | 329 | R329K | A | 2.1e-09 | 1.4e-08 | 5.7e-09 | 7.267E-09 | 6.103E-09 | 2.100E-09 | 1.400E-08 |
| 6573 | 329 | R329M | A | 5.3e-09 | 8.8e-09 | 7.1e-09 | 7.067E-09 | 1.750E-09 | 5.300E-09 | 8.800E-09 |
| 6574 | 329 | R329F | A | 2.4e-09 | 7.8e-09 | 3.4e-09 | 4.533E-09 | 2.873E-09 | 2.400E-09 | 7.800E-09 |
| 6575 | 329 | R329P | A | 5.2e-09 | 5e-09 | 4e-09 | 4.733E-09 | 6.429E-10 | 4.000E-09 | 5.200E-09 |
| 6576 | 329 | R329S | A | 6.7e-09 | 3.7e-09 | 9.5e-09 | 6.633E-09 | 2.901E-09 | 3.700E-09 | 9.500E-09 |
| 6577 | 329 | R329T | A | 1.3e-08 | 6e-09 | 1.4e-08 | 1.100E-08 | 4.359E-09 | 6.000E-09 | 1.400E-08 |
| 6578 | 329 | R329W | A | 6.9e-09 | 6.2e-09 | 3.3e-09 | 5.467E-09 | 1.909E-09 | 3.300E-09 | 6.900E-09 |
| 6579 | 329 | R329Y | A | 2e-08 | 5e-09 | 8.4e-09 | 1.113E-08 | 7.865E-09 | 5.000E-09 | 2.000E-08 |
| 6580 | 329 | R329V | A | 6.2e-09 | 3.5e-09 | 2.8e-09 | 4.167E-09 | 1.795E-09 | 2.800E-09 | 6.200E-09 |
| 6581 | 330 | E330A | A | 1.1e-08 | 1e-08 | 5e-09 | 8.667E-09 | 3.215E-09 | 5.000E-09 | 1.100E-08 |
| 6582 | 330 | E330R | A | 4.9e-09 | 4.2e-09 | 3.9e-09 | 4.333E-09 | 5.132E-10 | 3.900E-09 | 4.900E-09 |
| 6583 | 330 | E330N | A | 7.6e-09 | 2.7e-09 | 1.1e-08 | 7.100E-09 | 4.173E-09 | 2.700E-09 | 1.100E-08 |
| 6584 | 330 | E330D | A | 4.4e-09 | 3.5e-09 | 7e-09 | 4.967E-09 | 1.818E-09 | 3.500E-09 | 7.000E-09 |
| 6585 | 330 | E330C | A | 7.2e-09 | 3.1e-09 | 7.4e-09 | 5.900E-09 | 2.427E-09 | 3.100E-09 | 7.400E-09 |
| 6586 | 330 | E330E | A | 3.6e-09 | 1.5e-08 | 1.6e-09 | 6.733E-09 | 7.229E-09 | 1.600E-09 | 1.500E-08 |
| 6587 | 330 | E330Q | A | 6e-09 | 5.9e-09 | 1.6e-08 | 9.300E-09 | 5.803E-09 | 5.900E-09 | 1.600E-08 |
| 6588 | 330 | E330G | A | 7.5e-09 | 8.5e-09 | 8.7e-09 | 8.233E-09 | 6.429E-10 | 7.500E-09 | 8.700E-09 |
| 6589 | 330 | E330H | A | 3.4e-09 | 2.2e-09 | 3.9e-09 | 3.167E-09 | 8.737E-10 | 2.200E-09 | 3.900E-09 |
| 6590 | 330 | E330I | A | 8.1e-09 | 7.8e-09 | 4.7e-09 | 6.867E-09 | 1.882E-09 | 4.700E-09 | 8.100E-09 |
| 6591 | 330 | E330L | A | 1.2e-08 | 4.1e-09 | 7e-09 | 7.700E-09 | 3.996E-09 | 4.100E-09 | 1.200E-08 |
| 6592 | 330 | E330K | A | 7.7e-09 | 6e-09 | 7.4e-09 | 7.033E-09 | 9.074E-10 | 6.000E-09 | 7.700E-09 |
| 6593 | 330 | E330M | A | 9.7e-09 | 1.8e-08 | 2.8e-09 | 1.017E-08 | 7.611E-09 | 2.800E-09 | 1.800E-08 |
| 6594 | 330 | E330F | A | 6.8e-09 | 5.3e-09 | 7e-09 | 6.367E-09 | 9.292E-10 | 5.300E-09 | 7.000E-09 |
| 6595 | 330 | E330P | A | 9.9e-09 | 2.5e-08 | 9e-09 | 1.463E-08 | 8.989E-09 | 9.000E-09 | 2.500E-08 |
| 6596 | 330 | E330S | A | 8.7e-09 | 1.5e-08 | 4.9e-09 | 9.533E-09 | 5.101E-09 | 4.900E-09 | 1.500E-08 |
| 6597 | 330 | E330T | A | 1.1e-08 | 3.7e-09 | 2e-08 | 1.157E-08 | 8.165E-09 | 3.700E-09 | 2.000E-08 |
| 6598 | 330 | E330W | A | 2.7e-09 | 4.2e-09 | 1.2e-08 | 6.300E-09 | 4.993E-09 | 2.700E-09 | 1.200E-08 |
| 6599 | 330 | E330Y | A | 6e-09 | 1.8e-08 | 1.1e-08 | 1.167E-08 | 6.028E-09 | 6.000E-09 | 1.800E-08 |
| 6600 | 330 | E330V | A | 3.4e-09 | 1.2e-08 | 9.4e-09 | 8.267E-09 | 4.411E-09 | 3.400E-09 | 1.200E-08 |
| 6601 | 331 | V331A | A | 8.2e-09 | 7.4e-09 | 1.3e-08 | 9.533E-09 | 3.029E-09 | 7.400E-09 | 1.300E-08 |
| 6602 | 331 | V331R | A | 5.7e-09 | 1.2e-08 | 3.3e-09 | 7.000E-09 | 4.493E-09 | 3.300E-09 | 1.200E-08 |
| 6603 | 331 | V331N | A | 1.3e-08 | 3.2e-09 | 8.2e-09 | 8.133E-09 | 4.900E-09 | 3.200E-09 | 1.300E-08 |
| 6604 | 331 | V331D | A | 2.1e-08 | 1.7e-08 | 5.8e-09 | 1.460E-08 | 7.879E-09 | 5.800E-09 | 2.100E-08 |
| 6605 | 331 | V331C | A | 1.2e-08 | 7.8e-09 | 1.5e-08 | 1.160E-08 | 3.617E-09 | 7.800E-09 | 1.500E-08 |
| 6606 | 331 | V331E | A | 2.9e-09 | 7e-09 | 5.9e-09 | 5.267E-09 | 2.122E-09 | 2.900E-09 | 7.000E-09 |
| 6607 | 331 | V331Q | A | 7.1e-09 | 4.4e-09 | 1e-08 | 7.167E-09 | 2.801E-09 | 4.400E-09 | 1.000E-08 |
| 6608 | 331 | V331G | A | 6.3e-09 | 5.1e-09 | 1.7e-08 | 9.467E-09 | 6.552E-09 | 5.100E-09 | 1.700E-08 |
| 6609 | 331 | V331H | A | 7.2e-09 | 3.9e-09 | 5.8e-09 | 5.633E-09 | 1.656E-09 | 3.900E-09 | 7.200E-09 |
| 6610 | 331 | V331I | A | 8.4e-09 | 1.3e-08 | 2.5e-09 | 7.967E-09 | 5.263E-09 | 2.500E-09 | 1.300E-08 |
| 6611 | 331 | V331L | A | 5.9e-09 | 6.1e-09 | 7.8e-09 | 6.600E-09 | 1.044E-09 | 5.900E-09 | 7.800E-09 |
| 6612 | 331 | V331K | A | 1.2e-08 | 3.3e-09 | 5.9e-09 | 7.067E-09 | 4.466E-09 | 3.300E-09 | 1.200E-08 |
| 6613 | 331 | V331M | A | 7e-09 | 4.6e-09 | 1.2e-08 | 7.867E-09 | 3.775E-09 | 4.600E-09 | 1.200E-08 |
| 6614 | 331 | V331F | A | 5.3e-09 | 7.2e-09 | 1.5e-08 | 9.167E-09 | 5.140E-09 | 5.300E-09 | 1.500E-08 |
| 6615 | 331 | V331P | A | 7.8e-09 | 5e-09 | 9.8e-09 | 7.533E-09 | 2.411E-09 | 5.000E-09 | 9.800E-09 |
| 6616 | 331 | V331S | A | 5.5e-09 | 5.1e-09 | 5.5e-09 | 5.367E-09 | 2.309E-10 | 5.100E-09 | 5.500E-09 |
| 6617 | 331 | V331T | A | 3.7e-09 | 7.1e-09 | 6.9e-09 | 5.900E-09 | 1.908E-09 | 3.700E-09 | 7.100E-09 |
| 6618 | 331 | V331W | A | 4.2e-09 | 1e-08 | 5e-09 | 6.400E-09 | 3.143E-09 | 4.200E-09 | 1.000E-08 |
| 6619 | 331 | V331Y | A | 8.5e-09 | 5e-09 | 3.9e-09 | 5.800E-09 | 2.402E-09 | 3.900E-09 | 8.500E-09 |
| 6620 | 331 | V331V | A | 7.8e-09 | 3.7e-09 | 4.9e-09 | 5.467E-09 | 2.108E-09 | 3.700E-09 | 7.800E-09 |
| 6621 | 332 | R332A | A | 9.7e-09 | 5.1e-09 | 7.7e-09 | 7.500E-09 | 2.307E-09 | 5.100E-09 | 9.700E-09 |
| 6622 | 332 | R332R | A | 2.9e-09 | 5.8e-09 | 2.7e-08 | 1.190E-08 | 1.316E-08 | 2.900E-09 | 2.700E-08 |
| 6623 | 332 | R332N | A | 1.1e-08 | 7.4e-09 | 9.9e-09 | 9.433E-09 | 1.845E-09 | 7.400E-09 | 1.100E-08 |
| 6624 | 332 | R332D | A | 1.8e-08 | 3.9e-09 | 3.1e-08 | 1.763E-08 | 1.355E-08 | 3.900E-09 | 3.100E-08 |
| 6625 | 332 | R332C | A | 2.8e-09 | 3e-09 | 4.1e-09 | 3.300E-09 | 7.000E-10 | 2.800E-09 | 4.100E-09 |
| 6626 | 332 | R332E | A | 8.4e-09 | 2e-08 | 9e-09 | 1.247E-08 | 6.531E-09 | 8.400E-09 | 2.000E-08 |
| 6627 | 332 | R332Q | A | 3.6e-09 | 1.1e-08 | 1.6e-08 | 1.020E-08 | 6.239E-09 | 3.600E-09 | 1.600E-08 |
| 6628 | 332 | R332G | A | 1.3e-08 | 7.6e-09 | 4.5e-09 | 8.367E-09 | 4.302E-09 | 4.500E-09 | 1.300E-08 |
| 6629 | 332 | R332H | A | 6.4e-09 | 5.6e-09 | 7e-09 | 6.333E-09 | 7.024E-10 | 5.600E-09 | 7.000E-09 |

| | | | | | | | | | | |
|---|---|---|---|---|---|---|---|---|---|---|
| 6630 | 332 | R332I | A | 8.5e-09 | 1.3e-08 | 5.7e-09 | 9.067E-09 | 3.683E-09 | 5.700E-09 | 1.300E-08 |
| 6631 | 332 | R332L | A | 7.9e-09 | 1.2e-08 | 7.9e-09 | 9.267E-09 | 2.367E-09 | 7.900E-09 | 1.200E-08 |
| 6632 | 332 | R332K | A | 5.2e-09 | 1.8e-09 | 6.1e-09 | 4.367E-09 | 2.268E-09 | 1.800E-09 | 6.100E-09 |
| 6633 | 332 | R332M | A | 2.1e-09 | 1.3e-08 | 2.5e-08 | 1.337E-08 | 1.145E-08 | 2.100E-09 | 2.500E-08 |
| 6634 | 332 | R332F | A | 7.3e-09 | 1.5e-08 | 4.6e-09 | 8.967E-09 | 5.397E-09 | 4.600E-09 | 1.500E-08 |
| 6635 | 332 | R332P | A | 9.5e-09 | 7.2e-09 | 1.1e-08 | 9.233E-09 | 1.914E-09 | 7.200E-09 | 1.100E-08 |
| 6636 | 332 | R332S | A | 6.3e-09 | 4.7e-09 | 4.9e-09 | 5.300E-09 | 8.718E-10 | 4.700E-09 | 6.300E-09 |
| 6637 | 332 | R332T | A | 1.6e-08 | 4.4e-09 | 7.4e-09 | 9.267E-09 | 6.021E-09 | 4.400E-09 | 1.600E-08 |
| 6638 | 332 | R332W | A | 1.8e-09 | 5.1e-09 | 6.5e-09 | 4.467E-09 | 2.413E-09 | 1.800E-09 | 6.500E-09 |
| 6639 | 332 | R332Y | A | 1e-08 | 6.3e-09 | 2.4e-08 | 1.343E-08 | 9.336E-09 | 6.300E-09 | 2.400E-08 |
| 6640 | 332 | R332V | A | 1.2e-08 | 6e-09 | 1.4e-08 | 1.067E-08 | 4.163E-09 | 6.000E-09 | 1.400E-08 |
| 6641 | 333 | A333A | A | 8.8e-09 | 7.2e-09 | 7.3e-09 | 7.767E-09 | 8.963E-10 | 7.200E-09 | 8.800E-09 |
| 6642 | 333 | A333R | A | 4.9e-09 | 5.6e-09 | 7.7e-09 | 6.067E-09 | 1.457E-09 | 4.900E-09 | 7.700E-09 |
| 6643 | 333 | A333N | A | 3.7e-09 | 1e-09 | 3.5e-09 | 2.733E-09 | 1.504E-09 | 1.000E-09 | 3.700E-09 |
| 6644 | 333 | A333D | A | 8.4e-09 | 3.2e-09 | 1.1e-08 | 7.533E-09 | 3.972E-09 | 3.200E-09 | 1.100E-08 |
| 6645 | 333 | A333C | A | 8.3e-09 | 2.2e-09 | 5.4e-09 | 5.300E-09 | 3.051E-09 | 2.200E-09 | 8.300E-09 |
| 6646 | 333 | A333E | A | 3.3e-09 | 2.4e-08 | 2.4e-08 | 1.710E-08 | 1.195E-08 | 3.300E-09 | 2.400E-08 |
| 6647 | 333 | A333Q | A | 6.8e-09 | 6.6e-09 | 3.3e-09 | 5.567E-09 | 1.966E-09 | 3.300E-09 | 6.800E-09 |
| 6648 | 333 | A333G | A | 6.4e-09 | 1.8e-09 | 1.3e-08 | 7.067E-09 | 5.630E-09 | 1.800E-09 | 1.300E-08 |
| 6649 | 333 | A333H | A | 2.3e-08 | 1.3e-08 | 5.1e-09 | 1.370E-08 | 8.971E-09 | 5.100E-09 | 2.300E-08 |
| 6650 | 333 | A333I | A | 4.4e-09 | 5.8e-09 | 1.5e-08 | 8.400E-09 | 5.758E-09 | 4.400E-09 | 1.500E-08 |
| 6651 | 333 | A333L | A | 9e-09 | 1.7e-08 | 5.3e-09 | 1.043E-08 | 5.980E-09 | 5.300E-09 | 1.700E-08 |
| 6652 | 333 | A333K | A | 5.4e-09 | 9.9e-09 | 8.9e-09 | 8.067E-09 | 2.363E-09 | 5.400E-09 | 9.900E-09 |
| 6653 | 333 | A333M | A | 6e-09 | 1.2e-08 | 6.1e-09 | 8.033E-09 | 3.436E-09 | 6.000E-09 | 1.200E-08 |
| 6654 | 333 | A333F | A | 1.4e-08 | 6e-09 | 1.5e-09 | 7.167E-09 | 6.331E-09 | 1.500E-09 | 1.400E-08 |
| 6655 | 333 | A333P | A | 2.7e-09 | 8.1e-09 | 1.1e-08 | 7.267E-09 | 4.212E-09 | 2.700E-09 | 1.100E-08 |
| 6656 | 333 | A333S | A | 5.2e-09 | 7.5e-09 | 8.7e-09 | 7.133E-09 | 1.779E-09 | 5.200E-09 | 8.700E-09 |
| 6657 | 333 | A333T | A | 1.1e-08 | 7.8e-09 | 4.1e-09 | 7.633E-09 | 3.453E-09 | 4.100E-09 | 1.100E-08 |
| 6658 | 333 | A333W | A | 7.4e-09 | 1.3e-08 | 1.4e-08 | 1.147E-08 | 3.557E-09 | 7.400E-09 | 1.400E-08 |
| 6659 | 333 | A333Y | A | 8.9e-09 | 8e-09 | 1.3e-08 | 9.967E-09 | 2.665E-09 | 8.000E-09 | 1.300E-08 |
| 6660 | 333 | A333V | A | 9.4e-09 | 7.7e-09 | 1.2e-08 | 9.700E-09 | 2.166E-09 | 7.700E-09 | 1.200E-08 |
| 6661 | 334 | V334A | A | 1.2e-08 | 6.1e-09 | 1.5e-08 | 1.103E-08 | 4.528E-09 | 6.100E-09 | 1.500E-08 |
| 6662 | 334 | V334R | A | 1.6e-09 | 5.6e-09 | 7e-09 | 4.733E-09 | 2.802E-09 | 1.600E-09 | 7.000E-09 |
| 6663 | 334 | V334N | A | 2.5e-09 | 4.8e-09 | 7e-09 | 4.767E-09 | 2.250E-09 | 2.500E-09 | 7.000E-09 |
| 6664 | 334 | V334D | A | 1.1e-08 | 4.3e-09 | 7.9e-09 | 7.733E-09 | 3.353E-09 | 4.300E-09 | 1.100E-08 |
| 6665 | 334 | V334C | A | 3.3e-09 | 7.7e-09 | 8e-09 | 6.333E-09 | 2.631E-09 | 3.300E-09 | 8.000E-09 |
| 6666 | 334 | V334E | A | 1.5e-08 | 4.7e-09 | 7.6e-09 | 9.100E-09 | 5.311E-09 | 4.700E-09 | 1.500E-08 |
| 6667 | 334 | V334Q | A | 1.2e-08 | 2.9e-09 | 5.4e-09 | 6.767E-09 | 4.701E-09 | 2.900E-09 | 1.200E-08 |
| 6668 | 334 | V334G | A | 5.1e-09 | 1.5e-08 | 6.3e-09 | 8.800E-09 | 5.403E-09 | 5.100E-09 | 1.500E-08 |
| 6669 | 334 | V334H | A | 4.9e-09 | 2e-09 | 1.9e-08 | 8.633E-09 | 9.094E-09 | 2.000E-09 | 1.900E-08 |
| 6670 | 334 | V334I | A | 6.5e-09 | 6.2e-09 | 7.6e-09 | 6.767E-09 | 7.371E-10 | 6.200E-09 | 7.600E-09 |
| 6671 | 334 | V334L | A | 7.2e-09 | 6.6e-09 | 1.2e-08 | 8.600E-09 | 2.960E-09 | 6.600E-09 | 1.200E-08 |
| 6672 | 334 | V334K | A | 7.9e-09 | 1.3e-08 | 6.7e-09 | 9.200E-09 | 3.345E-09 | 6.700E-09 | 1.300E-08 |
| 6673 | 334 | V334M | A | 6.2e-09 | 3.4e-09 | 4.4e-09 | 4.667E-09 | 1.419E-09 | 3.400E-09 | 6.200E-09 |
| 6674 | 334 | V334F | A | 5e-09 | 8.2e-09 | 4.2e-09 | 5.800E-09 | 2.117E-09 | 4.200E-09 | 8.200E-09 |
| 6675 | 334 | V334P | A | 3.7e-08 | 4.9e-09 | 1e-08 | 1.730E-08 | 1.725E-08 | 4.900E-09 | 3.700E-08 |
| 6676 | 334 | V334S | A | 7.4e-09 | 2.7e-09 | 1.4e-08 | 8.033E-09 | 5.677E-09 | 2.700E-09 | 1.400E-08 |
| 6677 | 334 | V334T | A | 2.5e-09 | 4e-09 | 1.1e-08 | 5.833E-09 | 4.537E-09 | 2.500E-09 | 1.100E-08 |
| 6678 | 334 | V334W | A | 5.2e-09 | 8.4e-09 | 1.1e-08 | 8.200E-09 | 2.905E-09 | 5.200E-09 | 1.100E-08 |
| 6679 | 334 | V334Y | A | 8.6e-09 | 1e-08 | 3.1e-09 | 7.233E-09 | 3.647E-09 | 3.100E-09 | 1.000E-08 |
| 6680 | 334 | V334V | A | 7.1e-09 | 5.1e-09 | 6.8e-09 | 6.333E-09 | 1.079E-09 | 5.100E-09 | 7.100E-09 |
| 6681 | 335 | T335A | A | 6.9e-09 | 3.4e-09 | 9e-09 | 6.433E-09 | 2.829E-09 | 3.400E-09 | 9.000E-09 |
| 6682 | 335 | T335R | A | 5.7e-09 | 4.4e-09 | 1.1e-08 | 7.033E-09 | 3.496E-09 | 4.400E-09 | 1.100E-08 |
| 6683 | 335 | T335N | A | 4.3e-09 | 1.1e-08 | 1.4e-08 | 9.767E-09 | 4.966E-09 | 4.300E-09 | 1.400E-08 |
| 6684 | 335 | T335D | A | 1.4e-08 | 7.4e-09 | 4.7e-09 | 8.700E-09 | 4.784E-09 | 4.700E-09 | 1.400E-08 |
| 6685 | 335 | T335C | A | 1e-08 | 1.2e-08 | 3.5e-09 | 8.500E-09 | 4.444E-09 | 3.500E-09 | 1.200E-08 |
| 6686 | 335 | T335E | A | 2.3e-09 | 3.7e-09 | 4.9e-09 | 3.633E-09 | 1.301E-09 | 2.300E-09 | 4.900E-09 |
| 6687 | 335 | T335Q | A | 8.2e-09 | 8.4e-09 | 8.9e-09 | 8.500E-09 | 3.606E-10 | 8.200E-09 | 8.900E-09 |
| 6688 | 335 | T335G | A | 2.3e-08 | 1e-08 | 5.1e-09 | 1.270E-08 | 9.250E-09 | 5.100E-09 | 2.300E-08 |
| 6689 | 335 | T335H | A | 1.2e-08 | 1.2e-08 | 1.9e-08 | 1.433E-08 | 4.041E-09 | 1.200E-08 | 1.900E-08 |
| 6690 | 335 | T335I | A | 4.3e-09 | 1.5e-08 | 1.2e-08 | 1.043E-08 | 5.519E-09 | 4.300E-09 | 1.500E-08 |
| 6691 | 335 | T335L | A | 9.4e-09 | 1e-08 | 1.4e-08 | 1.113E-08 | 2.501E-09 | 9.400E-09 | 1.400E-08 |
| 6692 | 335 | T335K | A | 7.1e-09 | 1e-08 | 1.7e-08 | 1.137E-08 | 5.090E-09 | 7.100E-09 | 1.700E-08 |
| 6693 | 335 | T335M | A | 4.1e-09 | 1.5e-08 | 1e-08 | 9.700E-09 | 5.456E-09 | 4.100E-09 | 1.500E-08 |
| 6694 | 335 | T335F | A | 1.1e-08 | 1.2e-08 | 1.1e-08 | 1.133E-08 | 5.774E-10 | 1.100E-08 | 1.200E-08 |
| 6695 | 335 | T335P | A | 4.7e-09 | 1.5e-08 | 1.3e-08 | 1.090E-08 | 5.462E-09 | 4.700E-09 | 1.500E-08 |
| 6696 | 335 | T335S | A | 3e-09 | 9.4e-09 | 8.6e-09 | 7.000E-09 | 3.487E-09 | 3.000E-09 | 9.400E-09 |
| 6697 | 335 | T335T | A | 3.2e-09 | 5.8e-09 | 7.1e-09 | 5.367E-09 | 1.986E-09 | 3.200E-09 | 7.100E-09 |
| 6698 | 335 | T335W | A | 4.7e-09 | 8.6e-09 | 7.8e-09 | 7.033E-09 | 2.060E-09 | 4.700E-09 | 8.600E-09 |
| 6699 | 335 | T335Y | A | 9.8e-09 | 4.1e-09 | 3.3e-09 | 5.733E-09 | 3.544E-09 | 3.300E-09 | 9.800E-09 |
| 6700 | 335 | T335V | A | 9.4e-09 | 4.6e-09 | 4.9e-09 | 6.300E-09 | 2.689E-09 | 4.600E-09 | 9.400E-09 |
| 6701 | 336 | S336A | A | 5e-09 | 3.4e-09 | 4.2e-09 | 4.200E-09 | 8.000E-10 | 3.400E-09 | 5.000E-09 |
| 6702 | 336 | S336R | A | 1.1e-08 | 1.8e-08 | 1.5e-08 | 1.467E-08 | 3.512E-09 | 1.100E-08 | 1.800E-08 |
| 6703 | 336 | S336N | A | 6.3e-09 | 2.5e-09 | 1.2e-08 | 6.933E-09 | 4.782E-09 | 2.500E-09 | 1.200E-08 |
| 6704 | 336 | S336D | A | 3.9e-09 | 4.9e-09 | 3.5e-08 | 1.460E-08 | 1.767E-08 | 3.900E-09 | 3.500E-08 |
| 6705 | 336 | S336C | A | 1e-08 | 9e-09 | 4.6e-08 | 2.167E-08 | 2.108E-08 | 9.000E-09 | 4.600E-08 |
| 6706 | 336 | S336E | A | 9.2e-09 | 4e-09 | 7.1e-09 | 6.767E-09 | 2.616E-09 | 4.000E-09 | 9.200E-09 |
| 6707 | 336 | S336Q | A | 7.6e-09 | 9.1e-09 | 1.3e-08 | 9.900E-09 | 2.787E-09 | 7.600E-09 | 1.300E-08 |
| 6708 | 336 | S336G | A | 3.8e-09 | 1.6e-08 | 2.8e-08 | 1.593E-08 | 1.210E-08 | 3.800E-09 | 2.800E-08 |
| 6709 | 336 | S336H | A | 1e-08 | 8.1e-09 | 1.5e-09 | 6.533E-09 | 4.461E-09 | 1.500E-09 | 1.000E-08 |
| 6710 | 336 | S336I | A | 6.1e-09 | 9.3e-09 | 7.1e-09 | 7.500E-09 | 1.637E-09 | 6.100E-09 | 9.300E-09 |
| 6711 | 336 | S336L | A | 4.9e-09 | 3.7e-09 | 9.6e-09 | 6.067E-09 | 3.118E-09 | 3.700E-09 | 9.600E-09 |
| 6712 | 336 | S336K | A | 1.3e-08 | 1.7e-08 | 8.8e-09 | 1.293E-08 | 4.100E-09 | 8.800E-09 | 1.700E-08 |
| 6713 | 336 | S336M | A | 4.2e-09 | 4.9e-09 | 2.1e-09 | 3.733E-09 | 1.457E-09 | 2.100E-09 | 4.900E-09 |
| 6714 | 336 | S336F | A | 8.9e-09 | 3.2e-09 | 4.7e-09 | 5.600E-09 | 2.955E-09 | 3.200E-09 | 8.900E-09 |

| | | | | | | | | | | |
|---|---|---|---|---|---|---|---|---|---|---|
| 6715 | 336 | S336P | A | 5.4e-09 | 1.2e-08 | 8e-09 | 8.467E-09 | 3.325E-09 | 5.400E-09 | 1.200E-08 |
| 6716 | 336 | S336S | A | 9.1e-09 | 4.2e-09 | 6.6e-09 | 6.633E-09 | 2.450E-09 | 4.200E-09 | 9.100E-09 |
| 6717 | 336 | S336T | A | 6.8e-09 | 5.6e-09 | 1.5e-08 | 9.133E-09 | 5.116E-09 | 5.600E-09 | 1.500E-08 |
| 6718 | 336 | S336W | A | 2.3e-08 | 9e-09 | 2.8e-09 | 1.160E-08 | 1.035E-08 | 2.800E-09 | 2.300E-08 |
| 6719 | 336 | S336Y | A | 1.2e-08 | 3.5e-09 | 8.5e-09 | 8.000E-09 | 4.272E-09 | 3.500E-09 | 1.200E-08 |
| 6720 | 336 | S336V | A | 8.5e-09 | 1e-08 | 4.5e-09 | 7.667E-09 | 2.843E-09 | 4.500E-09 | 1.000E-08 |
| 6721 | 337 | A337A | A | 5.6e-09 | 1.2e-08 | 1.4e-09 | 6.333E-09 | 5.338E-09 | 1.400E-09 | 1.200E-08 |
| 6722 | 337 | A337R | A | 7e-09 | 8.8e-09 | 1.7e-08 | 1.093E-08 | 5.330E-09 | 7.000E-09 | 1.700E-08 |
| 6723 | 337 | A337N | A | 5.1e-09 | 4.7e-09 | 8.1e-09 | 5.967E-09 | 1.858E-09 | 4.700E-09 | 8.100E-09 |
| 6724 | 337 | A337D | A | 6e-09 | 6.4e-09 | 1.5e-08 | 9.133E-09 | 5.085E-09 | 6.000E-09 | 1.500E-08 |
| 6725 | 337 | A337C | A | 7.4e-09 | 3.3e-09 | 2.6e-09 | 4.433E-09 | 2.593E-09 | 2.600E-09 | 7.400E-09 |
| 6726 | 337 | A337E | A | 6.7e-09 | 1.6e-08 | 1.4e-08 | 1.223E-08 | 4.895E-09 | 6.700E-09 | 1.600E-08 |
| 6727 | 337 | A337Q | A | 1.5e-08 | 5.2e-09 | 5.3e-09 | 8.500E-09 | 5.629E-09 | 5.200E-09 | 1.500E-08 |
| 6728 | 337 | A337G | A | 4.5e-09 | 1.5e-08 | 3.1e-09 | 7.533E-09 | 6.504E-09 | 3.100E-09 | 1.500E-08 |
| 6729 | 337 | A337H | A | 1.2e-08 | 2.9e-09 | 1.4e-08 | 9.633E-09 | 5.916E-09 | 2.900E-09 | 1.400E-08 |
| 6730 | 337 | A337I | A | 5.5e-09 | 9e-09 | 3.1e-09 | 5.867E-09 | 2.967E-09 | 3.100E-09 | 9.000E-09 |
| 6731 | 337 | A337L | A | 6.2e-09 | 9e-09 | 6.8e-09 | 7.333E-09 | 1.474E-09 | 6.200E-09 | 9.000E-09 |
| 6732 | 337 | A337K | A | 4e-09 | 3.1e-09 | 1e-08 | 5.700E-09 | 3.751E-09 | 3.100E-09 | 1.000E-08 |
| 6733 | 337 | A337M | A | 6.4e-09 | 8.6e-09 | 1.3e-08 | 9.333E-09 | 3.361E-09 | 6.400E-09 | 1.300E-08 |
| 6734 | 337 | A337F | A | 4.8e-09 | 3.8e-09 | 6.7e-09 | 5.100E-09 | 1.473E-09 | 3.800E-09 | 6.700E-09 |
| 6735 | 337 | A337P | A | 8.1e-09 | 3.9e-09 | 2.9e-09 | 4.967E-09 | 2.759E-09 | 2.900E-09 | 8.100E-09 |
| 6736 | 337 | A337S | A | 3.7e-09 | 3.4e-09 | 1e-08 | 5.700E-09 | 3.727E-09 | 3.400E-09 | 1.000E-08 |
| 6737 | 337 | A337T | A | 9e-09 | 1.4e-08 | 4.8e-09 | 9.267E-09 | 4.606E-09 | 4.800E-09 | 1.400E-08 |
| 6738 | 337 | A337W | A | 2.7e-08 | 2.5e-09 | 6.5e-09 | 1.200E-08 | 1.314E-08 | 2.500E-09 | 2.700E-08 |
| 6739 | 337 | A337Y | A | 5.3e-09 | 2.9e-09 | 9.4e-09 | 5.867E-09 | 3.287E-09 | 2.900E-09 | 9.400E-09 |
| 6740 | 337 | A337V | A | 1.5e-08 | 3.3e-09 | 4.7e-09 | 7.667E-09 | 6.389E-09 | 3.300E-09 | 1.500E-08 |
| 6741 | 338 | N338A | A | 8.9e-09 | 2.3e-09 | 7.6e-09 | 6.267E-09 | 3.496E-09 | 2.300E-09 | 8.900E-09 |
| 6742 | 338 | N338R | A | 8.8e-09 | 3.4e-09 | 4.1e-09 | 5.433E-09 | 2.937E-09 | 3.400E-09 | 8.800E-09 |
| 6743 | 338 | N338N | A | 1.3e-08 | 1.2e-08 | 8.8e-09 | 1.127E-08 | 2.194E-09 | 8.800E-09 | 1.300E-08 |
| 6744 | 338 | N338D | A | 1.2e-08 | 6.2e-09 | 2.4e-09 | 6.867E-09 | 4.835E-09 | 2.400E-09 | 1.200E-08 |
| 6745 | 338 | N338C | A | 1.2e-08 | 1.2e-08 | 4.6e-09 | 9.533E-09 | 4.272E-09 | 4.600E-09 | 1.200E-08 |
| 6746 | 338 | N338E | A | 6.4e-09 | 2e-08 | 3.7e-09 | 1.003E-08 | 8.736E-09 | 3.700E-09 | 2.000E-08 |
| 6747 | 338 | N338Q | A | 6e-09 | 3.1e-09 | 1.2e-08 | 7.033E-09 | 4.539E-09 | 3.100E-09 | 1.200E-08 |
| 6748 | 338 | N338G | A | 3.1e-09 | 2.5e-09 | 1.4e-08 | 6.533E-09 | 6.473E-09 | 2.500E-09 | 1.400E-08 |
| 6749 | 338 | N338H | A | 1.1e-08 | 2.1e-09 | 1.5e-08 | 9.367E-09 | 6.603E-09 | 2.100E-09 | 1.500E-08 |
| 6750 | 338 | N338I | A | 1.2e-08 | 1.1e-08 | 4.7e-09 | 9.233E-09 | 3.958E-09 | 4.700E-09 | 1.200E-08 |
| 6751 | 338 | N338L | A | 5e-09 | 5.4e-09 | 2.4e-08 | 1.147E-08 | 1.086E-08 | 5.000E-09 | 2.400E-08 |
| 6752 | 338 | N338K | A | 1e-08 | 5.3e-09 | 2e-09 | 5.767E-09 | 4.020E-09 | 2.000E-09 | 1.000E-08 |
| 6753 | 338 | N338M | A | 1.3e-08 | 1.1e-09 | 3e-09 | 5.700E-09 | 6.393E-09 | 1.100E-09 | 1.300E-08 |
| 6754 | 338 | N338F | A | 7.2e-09 | 8.1e-09 | 5.4e-09 | 6.900E-09 | 1.375E-09 | 5.400E-09 | 8.100E-09 |
| 6755 | 338 | N338P | A | 3.6e-09 | 3.2e-09 | 1.4e-08 | 6.933E-09 | 6.123E-09 | 3.200E-09 | 1.400E-08 |
| 6756 | 338 | N338S | A | 1.3e-08 | 2.3e-09 | 3.7e-09 | 6.333E-09 | 5.816E-09 | 2.300E-09 | 1.300E-08 |
| 6757 | 338 | N338T | A | 1.3e-08 | 1.1e-08 | 1.3e-08 | 1.233E-08 | 1.155E-09 | 1.100E-08 | 1.300E-08 |
| 6758 | 338 | N338W | A | 1.2e-08 | 5.1e-09 | 9e-09 | 8.700E-09 | 3.460E-09 | 5.100E-09 | 1.200E-08 |
| 6759 | 338 | N338Y | A | 1.1e-08 | 1.3e-08 | 5.2e-09 | 9.733E-09 | 4.051E-09 | 5.200E-09 | 1.300E-08 |
| 6760 | 338 | N338V | A | 1.7e-08 | 5.9e-09 | 8.8e-09 | 1.057E-08 | 5.757E-09 | 5.900E-09 | 1.700E-08 |
| 6761 | 339 | I339A | A | 9.4e-09 | 3.9e-09 | 1.2e-08 | 8.433E-09 | 4.136E-09 | 3.900E-09 | 1.200E-08 |
| 6762 | 339 | I339R | A | 1.5e-08 | 8.2e-09 | 1.1e-08 | 1.140E-08 | 3.418E-09 | 8.200E-09 | 1.500E-08 |
| 6763 | 339 | I339N | A | 4.6e-09 | 7.1e-09 | 5.4e-09 | 5.700E-09 | 1.277E-09 | 4.600E-09 | 7.100E-09 |
| 6764 | 339 | I339D | A | 1e-08 | 5e-09 | 5.8e-09 | 6.933E-09 | 2.686E-09 | 5.000E-09 | 1.000E-08 |
| 6765 | 339 | I339C | A | 7.6e-09 | 7.9e-09 | 9.2e-09 | 8.233E-09 | 8.505E-10 | 7.600E-09 | 9.200E-09 |
| 6766 | 339 | I339E | A | 5.4e-09 | 4e-09 | 7.9e-09 | 5.767E-09 | 1.976E-09 | 4.000E-09 | 7.900E-09 |
| 6767 | 339 | I339Q | A | 1.1e-08 | 1.3e-08 | 5.5e-09 | 9.833E-09 | 3.884E-09 | 5.500E-09 | 1.300E-08 |
| 6768 | 339 | I339G | A | 1.9e-08 | 4.8e-09 | 1.8e-08 | 1.393E-08 | 7.925E-09 | 4.800E-09 | 1.900E-08 |
| 6769 | 339 | I339H | A | 1.6e-08 | 7.7e-09 | 3e-09 | 8.900E-09 | 6.583E-09 | 3.000E-09 | 1.600E-08 |
| 6770 | 339 | I339I | A | 5.9e-09 | 6.1e-09 | 9e-09 | 7.000E-09 | 1.735E-09 | 5.900E-09 | 9.000E-09 |
| 6771 | 339 | I339L | A | 7.9e-09 | 3.8e-09 | 8.8e-09 | 6.833E-09 | 2.665E-09 | 3.800E-09 | 8.800E-09 |
| 6772 | 339 | I339K | A | 8.3e-09 | 1.5e-08 | 2.5e-09 | 8.600E-09 | 6.255E-09 | 2.500E-09 | 1.500E-08 |
| 6773 | 339 | I339M | A | 5.7e-09 | 8.5e-09 | 4.5e-09 | 6.233E-09 | 2.053E-09 | 4.500E-09 | 8.500E-09 |
| 6774 | 339 | I339F | A | 9.4e-09 | 1.9e-09 | 8.9e-09 | 6.733E-09 | 4.193E-09 | 1.900E-09 | 9.400E-09 |
| 6775 | 339 | I339P | A | 1.5e-08 | 1.6e-08 | 1.4e-09 | 1.080E-08 | 8.156E-09 | 1.400E-09 | 1.600E-08 |
| 6776 | 339 | I339S | A | 8.6e-09 | 3e-09 | 2.1e-09 | 4.567E-09 | 3.522E-09 | 2.100E-09 | 8.600E-09 |
| 6777 | 339 | I339T | A | 9.2e-09 | 3.7e-09 | 5.8e-09 | 6.233E-09 | 2.775E-09 | 3.700E-09 | 9.200E-09 |
| 6778 | 339 | I339W | A | 1.7e-09 | 3.6e-09 | 2.4e-08 | 9.767E-09 | 1.236E-08 | 1.700E-09 | 2.400E-08 |
| 6779 | 339 | I339Y | A | 2.8e-09 | 9.4e-09 | 5.3e-09 | 5.833E-09 | 3.332E-09 | 2.800E-09 | 9.400E-09 |
| 6780 | 339 | I339V | A | 3.9e-09 | 5.9e-09 | 4.8e-09 | 4.867E-09 | 1.002E-09 | 3.900E-09 | 5.900E-09 |
| 6781 | 340 | Q340A | A | 1.1e-08 | 1.8e-09 | 7.8e-09 | 6.867E-09 | 4.670E-09 | 1.800E-09 | 1.100E-08 |
| 6782 | 340 | Q340R | A | 7.2e-09 | 6.7e-09 | 8e-09 | 7.300E-09 | 6.557E-10 | 6.700E-09 | 8.000E-09 |
| 6783 | 340 | Q340N | A | 4.3e-08 | 4.7e-09 | 9.8e-09 | 1.917E-08 | 2.080E-08 | 4.700E-09 | 4.300E-08 |
| 6784 | 340 | Q340D | A | 2.5e-09 | 3.5e-09 | 6.9e-09 | 4.300E-09 | 2.307E-09 | 2.500E-09 | 6.900E-09 |
| 6785 | 340 | Q340C | A | 8.3e-09 | 6.5e-09 | 2e-08 | 1.160E-08 | 7.330E-09 | 6.500E-09 | 2.000E-08 |
| 6786 | 340 | Q340E | A | 1.2e-08 | 7.1e-09 | 2.8e-09 | 7.300E-09 | 4.603E-09 | 2.800E-09 | 1.200E-08 |
| 6787 | 340 | Q340Q | A | 5.8e-09 | 5.7e-09 | 3.5e-09 | 5.000E-09 | 1.300E-09 | 3.500E-09 | 5.800E-09 |
| 6788 | 340 | Q340G | A | 4.4e-09 | 1.6e-08 | 8.3e-08 | 3.447E-08 | 4.243E-08 | 4.400E-09 | 8.300E-08 |
| 6789 | 340 | Q340H | A | 8.4e-09 | 1.6e-08 | 7e-09 | 1.047E-08 | 4.843E-09 | 7.000E-09 | 1.600E-08 |
| 6790 | 340 | Q340I | A | 7.8e-09 | 1e-08 | 9.3e-09 | 9.033E-09 | 1.124E-09 | 7.800E-09 | 1.000E-08 |
| 6791 | 340 | Q340L | A | 8.2e-09 | 4.5e-09 | 4.7e-09 | 5.800E-09 | 2.081E-09 | 4.500E-09 | 8.200E-09 |
| 6792 | 340 | Q340K | A | 1.5e-08 | 3.1e-09 | 2.1e-09 | 6.733E-09 | 7.177E-09 | 2.100E-09 | 1.500E-08 |
| 6793 | 340 | Q340M | A | 1.2e-08 | 9.8e-09 | 2.4e-09 | 8.067E-09 | 5.029E-09 | 2.400E-09 | 1.200E-08 |
| 6794 | 340 | Q340F | A | 4.6e-09 | 1.8e-08 | 6.8e-09 | 9.800E-09 | 7.186E-09 | 4.600E-09 | 1.800E-08 |
| 6795 | 340 | Q340P | A | 2.5e-09 | 1.2e-08 | 1.4e-09 | 5.300E-09 | 5.828E-09 | 1.400E-09 | 1.200E-08 |
| 6796 | 340 | Q340S | A | 1.5e-08 | 1.1e-08 | 9.7e-09 | 1.190E-08 | 2.762E-09 | 9.700E-09 | 1.500E-08 |
| 6797 | 340 | Q340T | A | 2.2e-09 | 1.4e-08 | 1.2e-08 | 9.400E-09 | 6.315E-09 | 2.200E-09 | 1.400E-08 |
| 6798 | 340 | Q340W | A | 6.5e-09 | 1e-08 | 1e-08 | 8.833E-09 | 2.021E-09 | 6.500E-09 | 1.000E-08 |
| 6799 | 340 | Q340Y | A | 3.5e-09 | 3.3e-09 | 1.8e-08 | 8.267E-09 | 8.430E-09 | 3.300E-09 | 1.800E-08 |

| | | | | | | | | | | |
|---|---|---|---|---|---|---|---|---|---|---|
| 6800 | 340 | Q340V | A | 2.4e-09 | 1.7e-09 | 8.3e-09 | 4.133E-09 | 3.625E-09 | 1.700E-09 | 8.300E-09 |
| 6801 | 341 | E341A | A | 3.5e-09 | 1.1e-08 | 7.9e-09 | 7.467E-09 | 3.769E-09 | 3.500E-09 | 1.100E-08 |
| 6802 | 341 | E341R | A | 8.9e-09 | 6.4e-09 | 1.7e-08 | 1.077E-08 | 5.541E-09 | 6.400E-09 | 1.700E-08 |
| 6803 | 341 | E341N | A | 7.5e-09 | 3.7e-09 | 8.9e-09 | 6.700E-09 | 2.691E-09 | 3.700E-09 | 8.900E-09 |
| 6804 | 341 | E341D | A | 5.6e-09 | 5.3e-09 | 2.3e-09 | 4.400E-09 | 1.825E-09 | 2.300E-09 | 5.600E-09 |
| 6805 | 341 | E341C | A | 3.9e-09 | 6e-09 | 7.8e-09 | 5.900E-09 | 1.952E-09 | 3.900E-09 | 7.800E-09 |
| 6806 | 341 | E341E | A | 7.4e-09 | 1.5e-08 | 1.5e-08 | 1.247E-08 | 4.388E-09 | 7.400E-09 | 1.500E-08 |
| 6807 | 341 | E341Q | A | 3.7e-09 | 5.3e-09 | 4.7e-09 | 4.567E-09 | 8.083E-10 | 3.700E-09 | 5.300E-09 |
| 6808 | 341 | E341G | A | 1.2e-08 | 1.4e-09 | 7.8e-09 | 7.067E-09 | 5.338E-09 | 1.400E-09 | 1.200E-08 |
| 6809 | 341 | E341H | A | 5.2e-09 | 3.2e-09 | 6.8e-09 | 5.067E-09 | 1.804E-09 | 3.200E-09 | 6.800E-09 |
| 6810 | 341 | E341I | A | 3e-09 | 7.2e-09 | 1.7e-08 | 9.067E-09 | 7.184E-09 | 3.000E-09 | 1.700E-08 |
| 6811 | 341 | E341L | A | 7.3e-09 | 4.3e-09 | 1.1e-08 | 7.533E-09 | 3.356E-09 | 4.300E-09 | 1.100E-08 |
| 6812 | 341 | E341K | A | 2.6e-08 | 7.2e-09 | 1.3e-08 | 1.540E-08 | 9.627E-09 | 7.200E-09 | 2.600E-08 |
| 6813 | 341 | E341M | A | 5.6e-09 | 8.8e-09 | 9.7e-09 | 8.033E-09 | 2.155E-09 | 5.600E-09 | 9.700E-09 |
| 6814 | 341 | E341F | A | 2e-09 | 1.5e-08 | 4.4e-09 | 7.133E-09 | 6.918E-09 | 2.000E-09 | 1.500E-08 |
| 6815 | 341 | E341P | A | 3.5e-09 | 1.1e-08 | 2.7e-09 | 5.733E-09 | 4.579E-09 | 2.700E-09 | 1.100E-08 |
| 6816 | 341 | E341S | A | 5.1e-09 | 1.7e-08 | 2.3e-08 | 1.503E-08 | 9.111E-09 | 5.100E-09 | 2.300E-08 |
| 6817 | 341 | E341T | A | 7.1e-09 | 9.3e-09 | 4.8e-09 | 7.067E-09 | 2.250E-09 | 4.800E-09 | 9.300E-09 |
| 6818 | 341 | E341W | A | 1.1e-09 | 6e-09 | 6.9e-09 | 4.667E-09 | 3.121E-09 | 1.100E-09 | 6.900E-09 |
| 6819 | 341 | E341Y | A | 4.5e-09 | 4.6e-09 | 1.1e-08 | 6.700E-09 | 3.724E-09 | 4.500E-09 | 1.100E-08 |
| 6820 | 341 | E341V | A | 8.8e-09 | 1.5e-08 | 2.6e-09 | 8.800E-09 | 6.200E-09 | 2.600E-09 | 1.500E-08 |
| 6821 | 342 | F342A | A | 1.1e-08 | 8.9e-09 | 6.5e-09 | 8.800E-09 | 2.252E-09 | 6.500E-09 | 1.100E-08 |
| 6822 | 342 | F342R | A | 1.5e-08 | 1.2e-09 | 5.1e-09 | 7.100E-09 | 7.114E-09 | 1.200E-09 | 1.500E-08 |
| 6823 | 342 | F342N | A | 7.1e-09 | 7.9e-09 | 4.2e-09 | 6.400E-09 | 1.947E-09 | 4.200E-09 | 7.900E-09 |
| 6824 | 342 | F342D | A | 1.3e-08 | 1.2e-08 | 6.8e-09 | 1.060E-08 | 3.329E-09 | 6.800E-09 | 1.300E-08 |
| 6825 | 342 | F342C | A | 3.2e-08 | 2.2e-09 | 1.4e-08 | 1.607E-08 | 1.501E-08 | 2.200E-09 | 3.200E-08 |
| 6826 | 342 | F342E | A | 4.3e-09 | 5.3e-09 | 5.4e-09 | 5.000E-09 | 6.083E-10 | 4.300E-09 | 5.400E-09 |
| 6827 | 342 | F342Q | A | 2.2e-09 | 4.4e-09 | 2.3e-09 | 2.967E-09 | 1.242E-09 | 2.200E-09 | 4.400E-09 |
| 6828 | 342 | F342G | A | 1.9e-08 | 3.2e-08 | 6e-09 | 1.900E-08 | 1.300E-08 | 6.000E-09 | 3.200E-08 |
| 6829 | 342 | F342H | A | 3.5e-09 | 2.5e-09 | 4.4e-09 | 3.467E-09 | 9.504E-10 | 2.500E-09 | 4.400E-09 |
| 6830 | 342 | F342I | A | 8.9e-09 | 6.1e-09 | 1.2e-08 | 9.000E-09 | 2.951E-09 | 6.100E-09 | 1.200E-08 |
| 6831 | 342 | F342L | A | 9.2e-09 | 1.1e-08 | 7.5e-09 | 9.233E-09 | 1.750E-09 | 7.500E-09 | 1.100E-08 |
| 6832 | 342 | F342K | A | 1.2e-08 | 1.2e-08 | 2e-09 | 8.667E-09 | 5.774E-09 | 2.000E-09 | 1.200E-08 |
| 6833 | 342 | F342M | A | 6.7e-09 | 1.6e-08 | 1.4e-08 | 1.223E-08 | 4.895E-09 | 6.700E-09 | 1.600E-08 |
| 6834 | 342 | F342F | A | 8e-09 | 1.1e-08 | 1.2e-09 | 6.733E-09 | 5.021E-09 | 1.200E-09 | 1.100E-08 |
| 6835 | 342 | F342P | A | 9.4e-09 | 3e-09 | 7.4e-09 | 6.600E-09 | 3.274E-09 | 3.000E-09 | 9.400E-09 |
| 6836 | 342 | F342S | A | 1e-08 | 1.8e-08 | 2.2e-08 | 1.667E-08 | 6.110E-09 | 1.000E-08 | 2.200E-08 |
| 6837 | 342 | F342T | A | 4.5e-09 | 8.9e-09 | 3.9e-09 | 5.767E-09 | 2.730E-09 | 3.900E-09 | 8.900E-09 |
| 6838 | 342 | F342W | A | 9.6e-09 | 3e-09 | 1.1e-08 | 7.867E-09 | 4.272E-09 | 3.000E-09 | 1.100E-08 |
| 6839 | 342 | F342Y | A | 1.1e-08 | 1.4e-08 | 5.5e-09 | 1.017E-08 | 4.311E-09 | 5.500E-09 | 1.400E-08 |
| 6840 | 342 | F342V | A | 1.2e-08 | 1.3e-08 | 4.5e-09 | 9.833E-09 | 4.646E-09 | 4.500E-09 | 1.300E-08 |
| 6841 | 343 | A343A | A | 4e-09 | 2.7e-09 | 6.4e-09 | 4.367E-09 | 1.877E-09 | 2.700E-09 | 6.400E-09 |
| 6842 | 343 | A343R | A | 2e-09 | 7.7e-09 | 6.3e-09 | 5.333E-09 | 2.970E-09 | 2.000E-09 | 7.700E-09 |
| 6843 | 343 | A343N | A | 3.4e-09 | 9.2e-10 | 2e-08 | 8.107E-09 | 1.037E-08 | 9.200E-10 | 2.000E-08 |
| 6844 | 343 | A343D | A | 5.4e-09 | 3e-09 | 5e-09 | 4.467E-09 | 1.286E-09 | 3.000E-09 | 5.400E-09 |
| 6845 | 343 | A343C | A | 9.5e-09 | 3.6e-09 | 4.6e-09 | 5.900E-09 | 3.158E-09 | 3.600E-09 | 9.500E-09 |
| 6846 | 343 | A343E | A | 1.1e-08 | 1.5e-08 | 8e-09 | 1.133E-08 | 3.512E-09 | 8.000E-09 | 1.500E-08 |
| 6847 | 343 | A343Q | A | 7.7e-09 | 3.8e-09 | 5.6e-09 | 5.700E-09 | 1.952E-09 | 3.800E-09 | 7.700E-09 |
| 6848 | 343 | A343G | A | 5.2e-09 | 9.9e-09 | 8.8e-09 | 7.967E-09 | 2.458E-09 | 5.200E-09 | 9.900E-09 |
| 6849 | 343 | A343H | A | 6.8e-09 | 6.7e-09 | 5.7e-09 | 6.400E-09 | 6.083E-10 | 5.700E-09 | 6.800E-09 |
| 6850 | 343 | A343I | A | 2.9e-09 | 2.8e-08 | 2.3e-08 | 1.797E-08 | 1.329E-08 | 2.900E-09 | 2.800E-08 |
| 6851 | 343 | A343L | A | 3.1e-09 | 1.2e-08 | 7.7e-09 | 7.600E-09 | 4.451E-09 | 3.100E-09 | 1.200E-08 |
| 6852 | 343 | A343K | A | 6.8e-09 | 6.6e-09 | 4.4e-09 | 5.933E-09 | 1.332E-09 | 4.400E-09 | 6.800E-09 |
| 6853 | 343 | A343M | A | 5.5e-09 | 3.5e-09 | 3.5e-09 | 4.167E-09 | 1.155E-09 | 3.500E-09 | 5.500E-09 |
| 6854 | 343 | A343F | A | 4.5e-09 | 7.3e-09 | 2.2e-09 | 4.667E-09 | 2.554E-09 | 2.200E-09 | 7.300E-09 |
| 6855 | 343 | A343P | A | 5e-09 | 1.4e-08 | 4.1e-09 | 7.700E-09 | 5.474E-09 | 4.100E-09 | 1.400E-08 |
| 6856 | 343 | A343S | A | 1.1e-08 | 1e-08 | 6e-09 | 9.000E-09 | 2.646E-09 | 6.000E-09 | 1.100E-08 |
| 6857 | 343 | A343T | A | 1.3e-08 | 5.3e-09 | 5e-09 | 7.767E-09 | 4.535E-09 | 5.000E-09 | 1.300E-08 |
| 6858 | 343 | A343W | A | 1.8e-08 | 8.7e-09 | 4.5e-09 | 1.040E-08 | 6.909E-09 | 4.500E-09 | 1.800E-08 |
| 6859 | 343 | A343Y | A | 1.3e-08 | 3.5e-09 | 6.8e-09 | 7.767E-09 | 4.823E-09 | 3.500E-09 | 1.300E-08 |
| 6860 | 343 | A343V | A | 4.6e-09 | 3.6e-09 | 1.3e-08 | 7.067E-09 | 5.163E-09 | 3.600E-09 | 1.300E-08 |
| 6861 | 344 | G344A | A | 6e-08 | 8.2e-09 | 1.2e-08 | 2.673E-08 | 2.887E-08 | 8.200E-09 | 6.000E-08 |
| 6862 | 344 | G344R | A | 7.7e-09 | 1.8e-08 | 1.5e-08 | 1.357E-08 | 5.297E-09 | 7.700E-09 | 1.800E-08 |
| 6863 | 344 | G344N | A | 6.9e-09 | 8.3e-09 | 1.1e-08 | 8.733E-09 | 2.084E-09 | 6.900E-09 | 1.100E-08 |
| 6864 | 344 | G344D | A | 2.1e-08 | 7e-09 | 2.2e-09 | 1.007E-08 | 9.768E-09 | 2.200E-09 | 2.100E-08 |
| 6865 | 344 | G344C | A | 1.1e-08 | 6.1e-09 | 7.4e-09 | 8.167E-09 | 2.538E-09 | 6.100E-09 | 1.100E-08 |
| 6866 | 344 | G344E | A | 4.9e-09 | 5.2e-09 | 3.4e-09 | 4.500E-09 | 9.644E-10 | 3.400E-09 | 5.200E-09 |
| 6867 | 344 | G344Q | A | 3.4e-09 | 1e-08 | 7.2e-09 | 6.867E-09 | 3.313E-09 | 3.400E-09 | 1.000E-08 |
| 6868 | 344 | G344G | A | 8.8e-09 | 1.3e-08 | 9.1e-09 | 1.030E-08 | 2.343E-09 | 8.800E-09 | 1.300E-08 |
| 6869 | 344 | G344H | A | 3.4e-09 | 4.8e-09 | 6.2e-09 | 4.800E-09 | 1.400E-09 | 3.400E-09 | 6.200E-09 |
| 6870 | 344 | G344I | A | 8.9e-09 | 9e-09 | 2e-09 | 6.633E-09 | 4.013E-09 | 2.000E-09 | 9.000E-09 |
| 6871 | 344 | G344L | A | 1e-08 | 4.9e-09 | 7.6e-09 | 7.500E-09 | 2.551E-09 | 4.900E-09 | 1.000E-08 |
| 6872 | 344 | G344K | A | 4e-09 | 1.5e-09 | 7e-09 | 4.167E-09 | 2.754E-09 | 1.500E-09 | 7.000E-09 |
| 6873 | 344 | G344M | A | 1.1e-08 | 2.3e-09 | 4.8e-09 | 6.033E-09 | 4.479E-09 | 2.300E-09 | 1.100E-08 |
| 6874 | 344 | G344F | A | 2.8e-09 | 2e-08 | 6.3e-09 | 9.700E-09 | 9.090E-09 | 2.800E-09 | 2.000E-08 |
| 6875 | 344 | G344P | A | 2.1e-08 | 4.7e-09 | 4.7e-09 | 1.013E-08 | 9.411E-09 | 4.700E-09 | 2.100E-08 |
| 6876 | 344 | G344S | A | 9.9e-09 | 9.2e-09 | 6.4e-09 | 8.500E-09 | 1.852E-09 | 6.400E-09 | 9.900E-09 |
| 6877 | 344 | G344T | A | 6.7e-09 | 5.4e-09 | 5.4e-09 | 5.833E-09 | 7.506E-10 | 5.400E-09 | 6.700E-09 |
| 6878 | 344 | G344W | A | 1.1e-08 | 4.8e-09 | 1.3e-08 | 9.600E-09 | 4.276E-09 | 4.800E-09 | 1.300E-08 |
| 6879 | 344 | G344Y | A | 6.7e-09 | 5.5e-09 | 1.3e-09 | 4.500E-09 | 2.835E-09 | 1.300E-09 | 6.700E-09 |
| 6880 | 344 | G344V | A | 8.7e-09 | 3.6e-09 | 5.7e-09 | 6.000E-09 | 2.563E-09 | 3.600E-09 | 8.700E-09 |
| 6881 | 345 | C345A | A | 4.4e-09 | 4.1e-09 | 9.8e-09 | 6.100E-09 | 3.208E-09 | 4.100E-09 | 9.800E-09 |
| 6882 | 345 | C345R | A | 3.5e-09 | 7.9e-09 | 6.8e-09 | 6.067E-09 | 2.290E-09 | 3.500E-09 | 7.900E-09 |
| 6883 | 345 | C345N | A | 1.2e-08 | 3e-08 | 4e-09 | 1.533E-08 | 1.332E-08 | 4.000E-09 | 3.000E-08 |
| 6884 | 345 | C345D | A | 7e-09 | 1.5e-08 | 2.5e-09 | 8.167E-09 | 6.331E-09 | 2.500E-09 | 1.500E-08 |

| | | | | | | | | | | |
|---|---|---|---|---|---|---|---|---|---|---|
| 6885 | 345 | C345C | A | 7.5e-09 | 2.2e-09 | 4.2e-09 | 4.633E-09 | 2.676E-09 | 2.200E-09 | 7.500E-09 |
| 6886 | 345 | C345E | A | 3.6e-09 | 8e-09 | 3.7e-09 | 5.100E-09 | 2.512E-09 | 3.600E-09 | 8.000E-09 |
| 6887 | 345 | C345Q | A | 1.6e-08 | 5.4e-09 | 2.4e-09 | 7.933E-09 | 7.145E-09 | 2.400E-09 | 1.600E-08 |
| 6888 | 345 | C345G | A | 1.3e-08 | 4.6e-09 | 1.2e-09 | 6.267E-09 | 6.074E-09 | 1.200E-09 | 1.300E-08 |
| 6889 | 345 | C345H | A | 5.1e-09 | 6.9e-09 | 8.1e-09 | 6.700E-09 | 1.510E-09 | 5.100E-09 | 8.100E-09 |
| 6890 | 345 | C345I | A | 6.9e-09 | 1.6e-08 | 3.3e-09 | 8.733E-09 | 6.545E-09 | 3.300E-09 | 1.600E-08 |
| 6891 | 345 | C345L | A | 1.3e-08 | 3e-09 | 8.6e-09 | 8.200E-09 | 5.012E-09 | 3.000E-09 | 1.300E-08 |
| 6892 | 345 | C345K | A | 7.3e-09 | 6.7e-09 | 3.8e-09 | 5.933E-09 | 1.872E-09 | 3.800E-09 | 7.300E-09 |
| 6893 | 345 | C345M | A | 8.7e-09 | 2.6e-09 | 5.7e-09 | 5.667E-09 | 3.050E-09 | 2.600E-09 | 8.700E-09 |
| 6894 | 345 | C345F | A | 6.7e-09 | 1.9e-08 | 7e-09 | 1.090E-08 | 7.016E-09 | 6.700E-09 | 1.900E-08 |
| 6895 | 345 | C345P | A | 3.5e-09 | 6.7e-09 | 5.4e-09 | 5.200E-09 | 1.609E-09 | 3.500E-09 | 6.700E-09 |
| 6896 | 345 | C345S | A | 2.1e-09 | 4.2e-09 | 1.6e-08 | 7.433E-09 | 7.493E-09 | 2.100E-09 | 1.600E-08 |
| 6897 | 345 | C345T | A | 5.9e-09 | 6.1e-09 | 8.4e-09 | 6.800E-09 | 1.389E-09 | 5.900E-09 | 8.400E-09 |
| 6898 | 345 | C345W | A | 8e-09 | 5.2e-09 | 3e-09 | 5.400E-09 | 2.506E-09 | 3.000E-09 | 8.000E-09 |
| 6899 | 345 | C345Y | A | 5.8e-09 | 1.6e-08 | 2.1e-09 | 7.967E-09 | 7.199E-09 | 2.100E-09 | 1.600E-08 |
| 6900 | 345 | C345V | A | 2e-08 | 3.6e-09 | 7.5e-09 | 1.037E-08 | 8.568E-09 | 3.600E-09 | 2.000E-08 |
| 6901 | 346 | K346A | A | 5.2e-09 | 8.9e-09 | 1.1e-08 | 8.367E-09 | 2.937E-09 | 5.200E-09 | 1.100E-08 |
| 6902 | 346 | K346R | A | 1.2e-08 | 5.5e-09 | 5.4e-09 | 7.633E-09 | 3.782E-09 | 5.400E-09 | 1.200E-08 |
| 6903 | 346 | K346N | A | 7.8e-09 | 8.4e-09 | 2.1e-08 | 1.240E-08 | 7.454E-09 | 7.800E-09 | 2.100E-08 |
| 6904 | 346 | K346D | A | 4e-09 | 1.1e-08 | 1.9e-08 | 1.133E-08 | 7.506E-09 | 4.000E-09 | 1.900E-08 |
| 6905 | 346 | K346C | A | 3.3e-09 | 4e-09 | 2.4e-09 | 3.233E-09 | 8.021E-10 | 2.400E-09 | 4.000E-09 |
| 6906 | 346 | K346E | A | 6.6e-09 | 1e-08 | 3e-09 | 6.533E-09 | 3.500E-09 | 3.000E-09 | 1.000E-08 |
| 6907 | 346 | K346Q | A | 1.1e-08 | 7.8e-09 | 5.7e-09 | 8.167E-09 | 2.669E-09 | 5.700E-09 | 1.100E-08 |
| 6908 | 346 | K346G | A | 1.4e-08 | 6.4e-09 | 1.2e-08 | 1.080E-08 | 3.940E-09 | 6.400E-09 | 1.400E-08 |
| 6909 | 346 | K346H | A | 4.5e-09 | 1.3e-08 | 2e-08 | 1.250E-08 | 7.762E-09 | 4.500E-09 | 2.000E-08 |
| 6910 | 346 | K346I | A | 5.1e-09 | 1.9e-09 | 4.1e-09 | 3.700E-09 | 1.637E-09 | 1.900E-09 | 5.100E-09 |
| 6911 | 346 | K346L | A | 8.5e-09 | 1.4e-08 | 2.6e-09 | 8.367E-09 | 5.701E-09 | 2.600E-09 | 1.400E-08 |
| 6912 | 346 | K346K | A | 3.2e-09 | 1e-08 | 7.2e-09 | 6.800E-09 | 3.418E-09 | 3.200E-09 | 1.000E-08 |
| 6913 | 346 | K346M | A | 4.8e-09 | 1e-08 | 9.7e-09 | 8.167E-09 | 2.919E-09 | 4.800E-09 | 1.000E-08 |
| 6914 | 346 | K346F | A | 8.7e-09 | 4e-09 | 7.8e-09 | 6.833E-09 | 2.495E-09 | 4.000E-09 | 8.700E-09 |
| 6915 | 346 | K346P | A | 1.3e-08 | 7.5e-09 | 3e-09 | 7.833E-09 | 5.008E-09 | 3.000E-09 | 1.300E-08 |
| 6916 | 346 | K346S | A | 4.6e-09 | 7.1e-09 | 7.1e-09 | 6.267E-09 | 1.443E-09 | 4.600E-09 | 7.100E-09 |
| 6917 | 346 | K346T | A | 1.1e-08 | 7.1e-09 | 2.8e-09 | 6.967E-09 | 4.102E-09 | 2.800E-09 | 1.100E-08 |
| 6918 | 346 | K346W | A | 1.5e-08 | 5.3e-09 | 5.2e-09 | 8.500E-09 | 5.629E-09 | 5.200E-09 | 1.500E-08 |
| 6919 | 346 | K346Y | A | 1.1e-08 | 4.9e-09 | 1.6e-08 | 1.063E-08 | 5.559E-09 | 4.900E-09 | 1.600E-08 |
| 6920 | 346 | K346V | A | 4.4e-09 | 5.2e-09 | 1.8e-08 | 9.200E-09 | 7.632E-09 | 4.400E-09 | 1.800E-08 |
| 6921 | 347 | K347A | A | 1.8e-08 | 5.2e-09 | 2.5e-08 | 1.607E-08 | 1.004E-08 | 5.200E-09 | 2.500E-08 |
| 6922 | 347 | K347R | A | 4.3e-09 | 3.6e-09 | 9.7e-09 | 5.867E-09 | 3.338E-09 | 3.600E-09 | 9.700E-09 |
| 6923 | 347 | K347N | A | 1.3e-08 | 1.1e-08 | 3.5e-09 | 9.167E-09 | 5.008E-09 | 3.500E-09 | 1.300E-08 |
| 6924 | 347 | K347D | A | 1.7e-08 | 7.8e-09 | 1.7e-08 | 1.393E-08 | 5.312E-09 | 7.800E-09 | 1.700E-08 |
| 6925 | 347 | K347C | A | 6.3e-09 | 7.9e-09 | 4.1e-09 | 6.100E-09 | 1.908E-09 | 4.100E-09 | 7.900E-09 |
| 6926 | 347 | K347E | A | 1.4e-08 | 1.7e-08 | 7.5e-09 | 1.283E-08 | 4.856E-09 | 7.500E-09 | 1.700E-08 |
| 6927 | 347 | K347Q | A | 1.7e-08 | 7.6e-09 | 3.6e-09 | 9.400E-09 | 6.879E-09 | 3.600E-09 | 1.700E-08 |
| 6928 | 347 | K347G | A | 1.4e-08 | 1.6e-08 | 6.9e-09 | 1.230E-08 | 4.782E-09 | 6.900E-09 | 1.600E-08 |
| 6929 | 347 | K347H | A | 2.6e-09 | 4.4e-09 | 4.4e-09 | 3.800E-09 | 1.039E-09 | 2.600E-09 | 4.400E-09 |
| 6930 | 347 | K347I | A | 5.8e-09 | 5.2e-09 | 6.5e-09 | 5.833E-09 | 6.506E-10 | 5.200E-09 | 6.500E-09 |
| 6931 | 347 | K347L | A | 1e-08 | 5.5e-09 | 6.1e-09 | 7.200E-09 | 2.443E-09 | 5.500E-09 | 1.000E-08 |
| 6932 | 347 | K347K | A | 8.4e-09 | 5.4e-09 | 6.6e-09 | 6.800E-09 | 1.510E-09 | 5.400E-09 | 8.400E-09 |
| 6933 | 347 | K347M | A | 4.7e-09 | 6e-09 | 9e-09 | 6.567E-09 | 2.205E-09 | 4.700E-09 | 9.000E-09 |
| 6934 | 347 | K347F | A | 8.7e-09 | 4.8e-09 | 7.9e-09 | 7.133E-09 | 2.060E-09 | 4.800E-09 | 8.700E-09 |
| 6935 | 347 | K347P | A | 7.6e-09 | 1.3e-08 | 4.3e-09 | 8.300E-09 | 4.392E-09 | 4.300E-09 | 1.300E-08 |
| 6936 | 347 | K347S | A | 4.9e-09 | 6.2e-09 | 8.9e-09 | 6.667E-09 | 2.040E-09 | 4.900E-09 | 8.900E-09 |
| 6937 | 347 | K347T | A | 1.1e-08 | 5.8e-09 | 7.3e-09 | 8.033E-09 | 2.676E-09 | 5.800E-09 | 1.100E-08 |
| 6938 | 347 | K347W | A | 6.7e-09 | 4.3e-09 | 7.7e-09 | 6.233E-09 | 1.747E-09 | 4.300E-09 | 7.700E-09 |
| 6939 | 347 | K347Y | A | 1.5e-08 | 6.9e-09 | 6.6e-09 | 9.500E-09 | 4.766E-09 | 6.600E-09 | 1.500E-08 |
| 6940 | 347 | K347V | A | 5.5e-09 | 6.6e-09 | 8.8e-09 | 6.967E-09 | 1.680E-09 | 5.500E-09 | 8.800E-09 |
| 6941 | 348 | I348A | A | 5.5e-09 | 1.5e-08 | 8.3e-09 | 9.600E-09 | 4.882E-09 | 5.500E-09 | 1.500E-08 |
| 6942 | 348 | I348R | A | 5.2e-09 | 4.6e-09 | 9.8e-09 | 6.533E-09 | 2.845E-09 | 4.600E-09 | 9.800E-09 |
| 6943 | 348 | I348N | A | 9.7e-09 | 5.8e-09 | 2.9e-09 | 6.133E-09 | 3.412E-09 | 2.900E-09 | 9.700E-09 |
| 6944 | 348 | I348D | A | 9e-09 | 9.9e-09 | 1.5e-08 | 1.130E-08 | 3.236E-09 | 9.000E-09 | 1.500E-08 |
| 6945 | 348 | I348C | A | 9.1e-09 | 1.2e-08 | 2e-08 | 1.370E-08 | 5.645E-09 | 9.100E-09 | 2.000E-08 |
| 6946 | 348 | I348E | A | 4.3e-09 | 9.7e-09 | 5.7e-09 | 6.567E-09 | 2.802E-09 | 4.300E-09 | 9.700E-09 |
| 6947 | 348 | I348Q | A | 3.3e-09 | 9.7e-09 | 8e-09 | 7.000E-09 | 3.315E-09 | 3.300E-09 | 9.700E-09 |
| 6948 | 348 | I348G | A | 1.7e-09 | 5.8e-09 | 1.2e-08 | 6.500E-09 | 5.186E-09 | 1.700E-09 | 1.200E-08 |
| 6949 | 348 | I348H | A | 4.7e-09 | 1.7e-08 | 9.5e-09 | 1.040E-08 | 6.199E-09 | 4.700E-09 | 1.700E-08 |
| 6950 | 348 | I348I | A | 2e-09 | 1.8e-08 | 1e-08 | 1.000E-08 | 8.000E-09 | 2.000E-09 | 1.800E-08 |
| 6951 | 348 | I348L | A | 6.1e-09 | 2.7e-09 | 7.1e-09 | 5.300E-09 | 2.307E-09 | 2.700E-09 | 7.100E-09 |
| 6952 | 348 | I348K | A | 3.3e-09 | 6.4e-09 | 4.6e-09 | 4.767E-09 | 1.557E-09 | 3.300E-09 | 6.400E-09 |
| 6953 | 348 | I348M | A | 3.2e-08 | 1.9e-08 | 4e-09 | 1.833E-08 | 1.401E-08 | 4.000E-09 | 3.200E-08 |
| 6954 | 348 | I348F | A | 5.3e-09 | 1e-08 | 8.5e-09 | 7.933E-09 | 2.401E-09 | 5.300E-09 | 1.000E-08 |
| 6955 | 348 | I348P | A | 4.1e-09 | 3.8e-09 | 6.2e-09 | 4.700E-09 | 1.308E-09 | 3.800E-09 | 6.200E-09 |
| 6956 | 348 | I348S | A | 3.4e-09 | 3.4e-09 | 1.2e-08 | 6.267E-09 | 4.965E-09 | 3.400E-09 | 1.200E-08 |
| 6957 | 348 | I348T | A | 7.2e-09 | 4.6e-09 | 1.2e-08 | 7.933E-09 | 3.754E-09 | 4.600E-09 | 1.200E-08 |
| 6958 | 348 | I348W | A | 3.2e-09 | 1.6e-08 | 1.3e-08 | 1.073E-08 | 6.694E-09 | 3.200E-09 | 1.600E-08 |
| 6959 | 348 | I348Y | A | 9.6e-09 | 7.4e-09 | 8.3e-09 | 8.433E-09 | 1.106E-09 | 7.400E-09 | 9.600E-09 |
| 6960 | 348 | I348V | A | 6.5e-09 | 1.2e-08 | 8.3e-09 | 8.933E-09 | 2.804E-09 | 6.500E-09 | 1.200E-08 |
| 6961 | 349 | F349A | A | 5.7e-09 | 2.7e-09 | 5.6e-09 | 4.667E-09 | 1.704E-09 | 2.700E-09 | 5.700E-09 |
| 6962 | 349 | F349R | A | 5.5e-09 | 1e-08 | 2.4e-09 | 5.967E-09 | 3.821E-09 | 2.400E-09 | 1.000E-08 |
| 6963 | 349 | F349N | A | 4.8e-09 | 2.6e-09 | 1.3e-08 | 6.800E-09 | 5.481E-09 | 2.600E-09 | 1.300E-08 |
| 6964 | 349 | F349D | A | 5e-09 | 1e-08 | 1.8e-08 | 1.100E-08 | 6.557E-09 | 5.000E-09 | 1.800E-08 |
| 6965 | 349 | F349C | A | 1.1e-08 | 7.1e-09 | 1.5e-08 | 1.103E-08 | 3.950E-09 | 7.100E-09 | 1.500E-08 |
| 6966 | 349 | F349E | A | 4.1e-09 | 9.1e-09 | 1.6e-09 | 4.933E-09 | 3.819E-09 | 1.600E-09 | 9.100E-09 |
| 6967 | 349 | F349Q | A | 1.3e-08 | 4.4e-09 | 4.7e-09 | 7.367E-09 | 4.881E-09 | 4.400E-09 | 1.300E-08 |
| 6968 | 349 | F349G | A | 6.9e-09 | 2.7e-09 | 5.7e-09 | 5.100E-09 | 2.163E-09 | 2.700E-09 | 6.900E-09 |
| 6969 | 349 | F349H | A | 3.9e-09 | 8.9e-09 | 4.3e-09 | 5.700E-09 | 2.778E-09 | 3.900E-09 | 8.900E-09 |

| | | | | | | | | | | |
|---|---|---|---|---|---|---|---|---|---|---|
| 6970 | 349 | F349I | A | 1.1e-08 | 6.5e-09 | 7.3e-09 | 8.267E-09 | 2.401E-09 | 6.500E-09 | 1.100E-08 |
| 6971 | 349 | F349L | A | 1.3e-08 | 4.1e-09 | 1.3e-08 | 1.003E-08 | 5.138E-09 | 4.100E-09 | 1.300E-08 |
| 6972 | 349 | F349K | A | 4.3e-09 | 7.4e-09 | 5.7e-09 | 5.800E-09 | 1.552E-09 | 4.300E-09 | 7.400E-09 |
| 6973 | 349 | F349M | A | 1.2e-08 | 3.1e-09 | 3.6e-09 | 6.233E-09 | 5.000E-09 | 3.100E-09 | 1.200E-08 |
| 6974 | 349 | F349F | A | 5.5e-09 | 6.5e-09 | 1.1e-08 | 7.667E-09 | 2.930E-09 | 5.500E-09 | 1.100E-08 |
| 6975 | 349 | F349P | A | 6.7e-09 | 3.9e-09 | 7.6e-09 | 6.067E-09 | 1.930E-09 | 3.900E-09 | 7.600E-09 |
| 6976 | 349 | F349S | A | 2e-09 | 2.2e-08 | 7.8e-09 | 1.060E-08 | 1.029E-08 | 2.000E-09 | 2.200E-08 |
| 6977 | 349 | F349T | A | 9.1e-09 | 4.5e-09 | 7e-09 | 6.867E-09 | 2.303E-09 | 4.500E-09 | 9.100E-09 |
| 6978 | 349 | F349W | A | 4.8e-09 | 4.9e-09 | 5.6e-09 | 5.100E-09 | 4.359E-10 | 4.800E-09 | 5.600E-09 |
| 6979 | 349 | F349Y | A | 9.9e-09 | 9.3e-09 | 1.6e-08 | 1.173E-08 | 3.707E-09 | 9.300E-09 | 1.600E-08 |
| 6980 | 349 | F349V | A | 3.3e-09 | 9.8e-09 | 9.2e-09 | 7.433E-09 | 3.592E-09 | 3.300E-09 | 9.800E-09 |
| 6981 | 350 | G350A | A | 7.6e-09 | 1.2e-08 | 6.7e-09 | 8.767E-09 | 2.836E-09 | 6.700E-09 | 1.200E-08 |
| 6982 | 350 | G350R | A | 3.5e-09 | 2.8e-09 | 1.2e-08 | 6.100E-09 | 5.122E-09 | 2.800E-09 | 1.200E-08 |
| 6983 | 350 | G350N | A | 1.5e-08 | 8.6e-09 | 1.8e-08 | 1.387E-08 | 4.801E-09 | 8.600E-09 | 1.800E-08 |
| 6984 | 350 | G350D | A | 9.5e-09 | 8.2e-09 | 7.1e-10 | 6.137E-09 | 4.744E-09 | 7.100E-10 | 9.500E-09 |
| 6985 | 350 | G350C | A | 5.6e-09 | 1.1e-08 | 3.4e-09 | 6.667E-09 | 3.911E-09 | 3.400E-09 | 1.100E-08 |
| 6986 | 350 | G350E | A | 4.5e-09 | 5.6e-09 | 1.2e-08 | 7.367E-09 | 4.050E-09 | 4.500E-09 | 1.200E-08 |
| 6987 | 350 | G350Q | A | 1.7e-08 | 7.9e-09 | 1.2e-08 | 1.230E-08 | 4.557E-09 | 7.900E-09 | 1.700E-08 |
| 6988 | 350 | G350G | A | 5.3e-09 | 4.8e-09 | 1.1e-08 | 7.033E-09 | 3.444E-09 | 4.800E-09 | 1.100E-08 |
| 6989 | 350 | G350H | A | 3.8e-09 | 1.1e-08 | 9.2e-09 | 8.000E-09 | 3.747E-09 | 3.800E-09 | 1.100E-08 |
| 6990 | 350 | G350I | A | 8.6e-09 | 4.7e-09 | 7.7e-09 | 7.000E-09 | 2.042E-09 | 4.700E-09 | 8.600E-09 |
| 6991 | 350 | G350L | A | 3.3e-09 | 1.6e-08 | 6.6e-09 | 8.633E-09 | 6.590E-09 | 3.300E-09 | 1.600E-08 |
| 6992 | 350 | G350K | A | 6.8e-09 | 1.9e-09 | 3.4e-09 | 4.033E-09 | 2.511E-09 | 1.900E-09 | 6.800E-09 |
| 6993 | 350 | G350M | A | 1.4e-08 | 6.1e-09 | 7.6e-09 | 9.233E-09 | 4.196E-09 | 6.100E-09 | 1.400E-08 |
| 6994 | 350 | G350F | A | 5.7e-09 | 5.3e-09 | 8e-09 | 6.333E-09 | 1.457E-09 | 5.300E-09 | 8.000E-09 |
| 6995 | 350 | G350P | A | 8.2e-09 | 2.9e-09 | 1.2e-08 | 7.700E-09 | 4.571E-09 | 2.900E-09 | 1.200E-08 |
| 6996 | 350 | G350S | A | 1.2e-08 | 1.5e-08 | 3.9e-09 | 1.030E-08 | 5.742E-09 | 3.900E-09 | 1.500E-08 |
| 6997 | 350 | G350T | A | 1.2e-08 | 1.5e-08 | 7.4e-09 | 1.147E-08 | 3.828E-09 | 7.400E-09 | 1.500E-08 |
| 6998 | 350 | G350W | A | 6.5e-09 | 5.3e-09 | 2.5e-08 | 1.227E-08 | 1.104E-08 | 5.300E-09 | 2.500E-08 |
| 6999 | 350 | G350Y | A | 4.9e-09 | 2.4e-08 | 8.7e-09 | 1.253E-08 | 1.011E-08 | 4.900E-09 | 2.400E-08 |
| 7000 | 350 | G350V | A | 1.8e-08 | 1.7e-08 | 5.8e-09 | 1.360E-08 | 6.773E-09 | 5.800E-09 | 1.800E-08 |
| 7001 | 351 | S351A | A | 2.2e-09 | 2e-08 | 8.7e-09 | 1.030E-08 | 9.007E-09 | 2.200E-09 | 2.000E-08 |
| 7002 | 351 | S351R | A | 1.2e-08 | 3.2e-09 | 1.3e-08 | 9.400E-09 | 5.393E-09 | 3.200E-09 | 1.300E-08 |
| 7003 | 351 | S351N | A | 5.9e-09 | 2.3e-09 | 7.7e-09 | 5.300E-09 | 2.750E-09 | 2.300E-09 | 7.700E-09 |
| 7004 | 351 | S351D | A | 1.3e-08 | 1.1e-08 | 9.1e-09 | 1.103E-08 | 1.950E-09 | 9.100E-09 | 1.300E-08 |
| 7005 | 351 | S351C | A | 1.6e-08 | 6.6e-09 | 4.1e-09 | 8.900E-09 | 6.275E-09 | 4.100E-09 | 1.600E-08 |
| 7006 | 351 | S351E | A | 7.2e-09 | 7.4e-09 | 1.1e-08 | 8.533E-09 | 2.139E-09 | 7.200E-09 | 1.100E-08 |
| 7007 | 351 | S351Q | A | 4.8e-09 | 8.3e-09 | 9e-09 | 7.367E-09 | 2.250E-09 | 4.800E-09 | 9.000E-09 |
| 7008 | 351 | S351G | A | 8.8e-09 | 1.8e-08 | 9.5e-09 | 1.210E-08 | 5.122E-09 | 8.800E-09 | 1.800E-08 |
| 7009 | 351 | S351H | A | 9.4e-09 | 3.8e-09 | 1.9e-08 | 1.073E-08 | 7.687E-09 | 3.800E-09 | 1.900E-08 |
| 7010 | 351 | S351I | A | 4e-09 | 4.1e-09 | 4.1e-09 | 4.067E-09 | 5.774E-11 | 4.000E-09 | 4.100E-09 |
| 7011 | 351 | S351L | A | 6.4e-09 | 1.7e-08 | 5.7e-09 | 9.700E-09 | 6.332E-09 | 5.700E-09 | 1.700E-08 |
| 7012 | 351 | S351K | A | 2.6e-08 | 1.2e-08 | 7.2e-09 | 1.507E-08 | 9.768E-09 | 7.200E-09 | 2.600E-08 |
| 7013 | 351 | S351M | A | 1.6e-08 | 9.8e-09 | 4.5e-09 | 1.010E-08 | 5.756E-09 | 4.500E-09 | 1.600E-08 |
| 7014 | 351 | S351F | A | 1.1e-08 | 1.2e-08 | 1.7e-08 | 1.333E-08 | 3.215E-09 | 1.100E-08 | 1.700E-08 |
| 7015 | 351 | S351P | A | 6e-09 | 2e-08 | 1.4e-08 | 1.333E-08 | 7.024E-09 | 6.000E-09 | 2.000E-08 |
| 7016 | 351 | S351S | A | 7.2e-09 | 1.2e-08 | 4.9e-09 | 8.033E-09 | 3.623E-09 | 4.900E-09 | 1.200E-08 |
| 7017 | 351 | S351T | A | 1.1e-08 | 6.5e-09 | 3.7e-09 | 7.067E-09 | 3.683E-09 | 3.700E-09 | 1.100E-08 |
| 7018 | 351 | S351W | A | 6.7e-09 | 3.1e-08 | 2.1e-08 | 1.957E-08 | 1.221E-08 | 6.700E-09 | 3.100E-08 |
| 7019 | 351 | S351Y | A | 1.8e-09 | 2.7e-09 | 6e-09 | 3.500E-09 | 2.211E-09 | 1.800E-09 | 6.000E-09 |
| 7020 | 351 | S351V | A | 5.5e-09 | 3.5e-09 | 1.2e-08 | 7.000E-09 | 4.444E-09 | 3.500E-09 | 1.200E-08 |
| 7021 | 352 | L352A | A | 3.4e-09 | 4.1e-09 | 9e-09 | 5.500E-09 | 3.051E-09 | 3.400E-09 | 9.000E-09 |
| 7022 | 352 | L352R | A | 2.5e-09 | 7.9e-09 | 4.4e-09 | 4.933E-09 | 2.739E-09 | 2.500E-09 | 7.900E-09 |
| 7023 | 352 | L352N | A | 1.7e-08 | 6.2e-09 | 1.2e-08 | 1.173E-08 | 5.405E-09 | 6.200E-09 | 1.700E-08 |
| 7024 | 352 | L352D | A | 1.4e-08 | 1.4e-08 | 4.5e-09 | 1.083E-08 | 5.485E-09 | 4.500E-09 | 1.400E-08 |
| 7025 | 352 | L352C | A | 1.2e-08 | 9.1e-09 | 7e-09 | 9.367E-09 | 2.511E-09 | 7.000E-09 | 1.200E-08 |
| 7026 | 352 | L352E | A | 2.7e-09 | 1.3e-08 | 8e-09 | 7.900E-09 | 5.151E-09 | 2.700E-09 | 1.300E-08 |
| 7027 | 352 | L352Q | A | 7.3e-09 | 1.4e-08 | 4.3e-09 | 8.533E-09 | 4.966E-09 | 4.300E-09 | 1.400E-08 |
| 7028 | 352 | L352G | A | 3.5e-09 | 1.7e-08 | 7e-09 | 9.167E-09 | 7.006E-09 | 3.500E-09 | 1.700E-08 |
| 7029 | 352 | L352H | A | 6.1e-09 | 4.3e-09 | 5.9e-09 | 5.433E-09 | 9.866E-10 | 4.300E-09 | 6.100E-09 |
| 7030 | 352 | L352I | A | 3.8e-09 | 4.9e-09 | 8.2e-09 | 5.633E-09 | 2.290E-09 | 3.800E-09 | 8.200E-09 |
| 7031 | 352 | L352L | A | 1e-08 | 4.4e-09 | 1.1e-08 | 8.467E-09 | 3.557E-09 | 4.400E-09 | 1.100E-08 |
| 7032 | 352 | L352K | A | 6.1e-09 | 1.9e-09 | 9e-09 | 5.667E-09 | 3.570E-09 | 1.900E-09 | 9.000E-09 |
| 7033 | 352 | L352M | A | 9.4e-09 | 9.9e-09 | 2.2e-09 | 7.167E-09 | 4.309E-09 | 2.200E-09 | 9.900E-09 |
| 7034 | 352 | L352F | A | 3.5e-09 | 3.1e-09 | 4.4e-09 | 3.667E-09 | 6.658E-10 | 3.100E-09 | 4.400E-09 |
| 7035 | 352 | L352P | A | 4.1e-09 | 1e-08 | 3.3e-09 | 5.800E-09 | 3.659E-09 | 3.300E-09 | 1.000E-08 |
| 7036 | 352 | L352S | A | 6.1e-09 | 7.8e-09 | 6.6e-09 | 6.833E-09 | 8.737E-10 | 6.100E-09 | 7.800E-09 |
| 7037 | 352 | L352T | A | 7.6e-09 | 1.3e-08 | 1.8e-08 | 1.287E-08 | 5.201E-09 | 7.600E-09 | 1.800E-08 |
| 7038 | 352 | L352W | A | 3.1e-09 | 5.7e-09 | 4.4e-09 | 4.400E-09 | 1.300E-09 | 3.100E-09 | 5.700E-09 |
| 7039 | 352 | L352Y | A | 6.2e-09 | 5.2e-09 | 5.2e-09 | 5.533E-09 | 5.774E-10 | 5.200E-09 | 6.200E-09 |
| 7040 | 352 | L352V | A | 4.1e-09 | 1.5e-08 | 7.8e-09 | 8.967E-09 | 5.543E-09 | 4.100E-09 | 1.500E-08 |
| 7041 | 353 | A353A | A | 1.4e-08 | 7.4e-09 | 8.5e-09 | 9.967E-09 | 3.536E-09 | 7.400E-09 | 1.400E-08 |
| 7042 | 353 | A353R | A | 7.6e-09 | 7.5e-09 | 2.7e-09 | 5.933E-09 | 2.801E-09 | 2.700E-09 | 7.600E-09 |
| 7043 | 353 | A353N | A | 4.9e-09 | 8.1e-09 | 1.5e-08 | 9.333E-09 | 5.162E-09 | 4.900E-09 | 1.500E-08 |
| 7044 | 353 | A353D | A | 7.2e-09 | 3.7e-09 | 6.1e-09 | 5.667E-09 | 1.790E-09 | 3.700E-09 | 7.200E-09 |
| 7045 | 353 | A353C | A | 1.4e-09 | 1.4e-08 | 3.7e-09 | 6.367E-09 | 6.710E-09 | 1.400E-09 | 1.400E-08 |
| 7046 | 353 | A353E | A | 9.4e-09 | 5.4e-09 | 7.5e-09 | 7.433E-09 | 2.001E-09 | 5.400E-09 | 9.400E-09 |
| 7047 | 353 | A353Q | A | 3.9e-09 | 7e-09 | 2.1e-09 | 4.333E-09 | 2.479E-09 | 2.100E-09 | 7.000E-09 |
| 7048 | 353 | A353G | A | 5.1e-09 | 1.9e-08 | 9.6e-09 | 1.123E-08 | 7.092E-09 | 5.100E-09 | 1.900E-08 |
| 7049 | 353 | A353H | A | 3.8e-09 | 4.5e-09 | 9.7e-09 | 6.000E-09 | 3.223E-09 | 3.800E-09 | 9.700E-09 |
| 7050 | 353 | A353I | A | 9e-09 | 7.1e-09 | 1.2e-08 | 9.367E-09 | 2.470E-09 | 7.100E-09 | 1.200E-08 |
| 7051 | 353 | A353L | A | 1.5e-08 | 2e-08 | 8.3e-09 | 1.443E-08 | 5.871E-09 | 8.300E-09 | 2.000E-08 |
| 7052 | 353 | A353K | A | 1.6e-08 | 1.5e-08 | 6.9e-09 | 1.263E-08 | 4.990E-09 | 6.900E-09 | 1.600E-08 |
| 7053 | 353 | A353M | A | 1.2e-08 | 2.4e-08 | 7.3e-09 | 1.443E-08 | 8.612E-09 | 7.300E-09 | 2.400E-08 |
| 7054 | 353 | A353F | A | 1.2e-08 | 2.5e-09 | 7.2e-09 | 7.233E-09 | 4.750E-09 | 2.500E-09 | 1.200E-08 |

| | | | | | | | | | | |
|---|---|---|---|---|---|---|---|---|---|---|
| 7055 | 353 | A353P | A | 1.1e-08 | 1.1e-08 | 1.1e-08 | 1.100E-08 | 0.000E+00 | 1.100E-08 | 1.100E-08 |
| 7056 | 353 | A353S | A | 3.5e-09 | 1e-08 | 5.6e-09 | 6.367E-09 | 3.317E-09 | 3.500E-09 | 1.000E-08 |
| 7057 | 353 | A353T | A | 1e-08 | 4.5e-09 | 2.3e-08 | 1.250E-08 | 9.500E-09 | 4.500E-09 | 2.300E-08 |
| 7058 | 353 | A353W | A | 1.4e-09 | 4.7e-09 | 2.2e-09 | 2.767E-09 | 1.721E-09 | 1.400E-09 | 4.700E-09 |
| 7059 | 353 | A353Y | A | 6.7e-09 | 1.1e-08 | 6.5e-09 | 8.067E-09 | 2.542E-09 | 6.500E-09 | 1.100E-08 |
| 7060 | 353 | A353V | A | 1.2e-08 | 2.1e-08 | 7.5e-09 | 1.350E-08 | 6.874E-09 | 7.500E-09 | 2.100E-08 |
| 7061 | 354 | F354A | A | 1.7e-08 | 6.5e-09 | 7.1e-09 | 1.020E-08 | 5.897E-09 | 6.500E-09 | 1.700E-08 |
| 7062 | 354 | F354R | A | 5e-09 | 1.1e-08 | 6.1e-09 | 7.367E-09 | 3.194E-09 | 5.000E-09 | 1.100E-08 |
| 7063 | 354 | F354N | A | 1.1e-08 | 8e-09 | 4e-09 | 7.667E-09 | 3.512E-09 | 4.000E-09 | 1.100E-08 |
| 7064 | 354 | F354D | A | 3.1e-08 | 1e-08 | 1.7e-08 | 1.933E-08 | 1.069E-08 | 1.000E-08 | 3.100E-08 |
| 7065 | 354 | F354C | A | 2.1e-08 | 8.9e-09 | 4.2e-09 | 1.137E-08 | 8.667E-09 | 4.200E-09 | 2.100E-08 |
| 7066 | 354 | F354E | A | 2.2e-08 | 2e-08 | 7.7e-09 | 1.657E-08 | 7.744E-09 | 7.700E-09 | 2.200E-08 |
| 7067 | 354 | F354Q | A | 2.4e-09 | 5.3e-09 | 1.2e-08 | 6.567E-09 | 4.924E-09 | 2.400E-09 | 1.200E-08 |
| 7068 | 354 | F354G | A | 3.9e-09 | 3.4e-09 | 8e-09 | 5.100E-09 | 2.524E-09 | 3.400E-09 | 8.000E-09 |
| 7069 | 354 | F354H | A | 8e-09 | 6.2e-09 | 8.2e-09 | 7.467E-09 | 1.102E-09 | 6.200E-09 | 8.200E-09 |
| 7070 | 354 | F354I | A | 8.8e-09 | 3.5e-09 | 4.6e-09 | 5.633E-09 | 2.797E-09 | 3.500E-09 | 8.800E-09 |
| 7071 | 354 | F354L | A | 9.1e-09 | 6.7e-09 | 4.9e-09 | 6.900E-09 | 2.107E-09 | 4.900E-09 | 9.100E-09 |
| 7072 | 354 | F354K | A | 1e-08 | 7.2e-09 | 4.3e-09 | 7.167E-09 | 2.850E-09 | 4.300E-09 | 1.000E-08 |
| 7073 | 354 | F354M | A | 1.3e-08 | 1.2e-08 | 5.5e-09 | 1.017E-08 | 4.072E-09 | 5.500E-09 | 1.300E-08 |
| 7074 | 354 | F354F | A | 9.4e-09 | 4.5e-09 | 1.3e-08 | 8.967E-09 | 4.267E-09 | 4.500E-09 | 1.300E-08 |
| 7075 | 354 | F354P | A | 3e-08 | 1.3e-08 | 8.3e-09 | 1.710E-08 | 1.142E-08 | 8.300E-09 | 3.000E-08 |
| 7076 | 354 | F354S | A | 3.2e-09 | 5.2e-09 | 4.1e-09 | 4.167E-09 | 1.002E-09 | 3.200E-09 | 5.200E-09 |
| 7077 | 354 | F354T | A | 5e-09 | 1.7e-08 | 1.2e-08 | 1.133E-08 | 6.028E-09 | 5.000E-09 | 1.700E-08 |
| 7078 | 354 | F354W | A | 2e-09 | 8.5e-09 | 2.4e-09 | 4.300E-09 | 3.643E-09 | 2.000E-09 | 8.500E-09 |
| 7079 | 354 | F354Y | A | 2.2e-09 | 1e-08 | 3.9e-09 | 5.367E-09 | 4.102E-09 | 2.200E-09 | 1.000E-08 |
| 7080 | 354 | F354V | A | 6.9e-09 | 1.1e-08 | 7e-09 | 8.300E-09 | 2.339E-09 | 6.900E-09 | 1.100E-08 |
| 7081 | 355 | L355A | A | 8.6e-09 | 7.3e-09 | 5.1e-09 | 7.000E-09 | 1.769E-09 | 5.100E-09 | 8.600E-09 |
| 7082 | 355 | L355R | A | 3.6e-09 | 7.2e-09 | 9.9e-09 | 6.900E-09 | 3.161E-09 | 3.600E-09 | 9.900E-09 |
| 7083 | 355 | L355N | A | 1.4e-08 | 3.9e-09 | 5e-09 | 7.633E-09 | 5.541E-09 | 3.900E-09 | 1.400E-08 |
| 7084 | 355 | L355D | A | 1.2e-08 | 3.7e-09 | 8.4e-09 | 8.033E-09 | 4.162E-09 | 3.700E-09 | 1.200E-08 |
| 7085 | 355 | L355C | A | 1.3e-08 | 9.1e-09 | 2.4e-09 | 8.167E-09 | 5.361E-09 | 2.400E-09 | 1.300E-08 |
| 7086 | 355 | L355E | A | 3.6e-09 | 6.1e-09 | 6.2e-09 | 5.300E-09 | 1.473E-09 | 3.600E-09 | 6.200E-09 |
| 7087 | 355 | L355Q | A | 3.9e-09 | 1.3e-08 | 9.2e-09 | 8.700E-09 | 4.571E-09 | 3.900E-09 | 1.300E-08 |
| 7088 | 355 | L355G | A | 1.5e-08 | 3e-08 | 9.6e-09 | 1.820E-08 | 1.057E-08 | 9.600E-09 | 3.000E-08 |
| 7089 | 355 | L355H | A | 6.7e-09 | 1.2e-08 | 6.4e-09 | 8.367E-09 | 3.150E-09 | 6.400E-09 | 1.200E-08 |
| 7090 | 355 | L355I | A | 1.9e-08 | 8.7e-09 | 4.5e-09 | 1.073E-08 | 7.461E-09 | 4.500E-09 | 1.900E-08 |
| 7091 | 355 | L355L | A | 8e-09 | 1.4e-08 | 5.7e-09 | 9.233E-09 | 4.285E-09 | 5.700E-09 | 1.400E-08 |
| 7092 | 355 | L355K | A | 1e-08 | 3.4e-09 | 9.4e-09 | 7.600E-09 | 3.650E-09 | 3.400E-09 | 1.000E-08 |
| 7093 | 355 | L355M | A | 7.8e-09 | 9.1e-09 | 5.1e-09 | 7.333E-09 | 2.040E-09 | 5.100E-09 | 9.100E-09 |
| 7094 | 355 | L355F | A | 8.1e-09 | 1.4e-08 | 1.7e-09 | 7.933E-09 | 6.152E-09 | 1.700E-09 | 1.400E-08 |
| 7095 | 355 | L355P | A | 1.3e-08 | 1.4e-08 | 5e-09 | 1.067E-08 | 4.933E-09 | 5.000E-09 | 1.400E-08 |
| 7096 | 355 | L355S | A | 6.9e-09 | 1.1e-08 | 1.2e-08 | 9.967E-09 | 2.702E-09 | 6.900E-09 | 1.200E-08 |
| 7097 | 355 | L355T | A | 9.5e-09 | 1e-08 | 1.8e-08 | 1.250E-08 | 4.770E-09 | 9.500E-09 | 1.800E-08 |
| 7098 | 355 | L355W | A | 4.4e-09 | 7.7e-09 | 3.5e-09 | 5.200E-09 | 2.211E-09 | 3.500E-09 | 7.700E-09 |
| 7099 | 355 | L355Y | A | 7.4e-09 | 3.7e-09 | 9.9e-09 | 7.000E-09 | 3.119E-09 | 3.700E-09 | 9.900E-09 |
| 7100 | 355 | L355V | A | 7.3e-09 | 1e-08 | 4.2e-09 | 7.167E-09 | 2.902E-09 | 4.200E-09 | 1.000E-08 |
| 7101 | 356 | P356A | A | 6.2e-09 | 3.2e-09 | 4.1e-09 | 4.500E-09 | 1.539E-09 | 3.200E-09 | 6.200E-09 |
| 7102 | 356 | P356R | A | 6.2e-09 | 6.5e-09 | 3.9e-09 | 5.533E-09 | 1.422E-09 | 3.900E-09 | 6.500E-09 |
| 7103 | 356 | P356N | A | 2.9e-09 | 6.3e-09 | 1.1e-08 | 6.733E-09 | 4.067E-09 | 2.900E-09 | 1.100E-08 |
| 7104 | 356 | P356D | A | 7.7e-09 | 2e-08 | 9.3e-09 | 1.233E-08 | 6.688E-09 | 7.700E-09 | 2.000E-08 |
| 7105 | 356 | P356C | A | 5.1e-09 | 6e-09 | 9.6e-09 | 6.900E-09 | 2.381E-09 | 5.100E-09 | 9.600E-09 |
| 7106 | 356 | P356E | A | 4.8e-09 | 1.8e-08 | 4.3e-09 | 9.033E-09 | 7.769E-09 | 4.300E-09 | 1.800E-08 |
| 7107 | 356 | P356Q | A | 6.8e-09 | 1.3e-08 | 6.6e-09 | 8.800E-09 | 3.639E-09 | 6.600E-09 | 1.300E-08 |
| 7108 | 356 | P356G | A | 1.2e-08 | 4.8e-09 | 7.7e-09 | 8.167E-09 | 3.623E-09 | 4.800E-09 | 1.200E-08 |
| 7109 | 356 | P356H | A | 9e-09 | 8.8e-09 | 6.3e-09 | 8.033E-09 | 1.504E-09 | 6.300E-09 | 9.000E-09 |
| 7110 | 356 | P356I | A | 1.5e-08 | 1.4e-08 | 1.6e-08 | 1.500E-08 | 1.000E-09 | 1.400E-08 | 1.600E-08 |
| 7111 | 356 | P356L | A | 7.7e-09 | 3.7e-09 | 9e-09 | 6.800E-09 | 2.762E-09 | 3.700E-09 | 9.000E-09 |
| 7112 | 356 | P356K | A | 6.1e-09 | 1.3e-08 | 7.3e-09 | 8.800E-09 | 3.686E-09 | 6.100E-09 | 1.300E-08 |
| 7113 | 356 | P356M | A | 9.2e-09 | 1.1e-08 | 3.5e-09 | 7.900E-09 | 3.915E-09 | 3.500E-09 | 1.100E-08 |
| 7114 | 356 | P356F | A | 5.6e-08 | 1.5e-08 | 6.2e-09 | 2.573E-08 | 2.658E-08 | 6.200E-09 | 5.600E-08 |
| 7115 | 356 | P356P | A | 9.4e-09 | 6.8e-09 | 5.1e-09 | 7.100E-09 | 2.166E-09 | 5.100E-09 | 9.400E-09 |
| 7116 | 356 | P356S | A | 1.6e-08 | 1.4e-08 | 8.1e-09 | 1.270E-08 | 4.107E-09 | 8.100E-09 | 1.600E-08 |
| 7117 | 356 | P356T | A | 3.6e-09 | 1.6e-08 | 7.5e-09 | 9.033E-09 | 6.341E-09 | 3.600E-09 | 1.600E-08 |
| 7118 | 356 | P356W | A | 4.9e-09 | 9.4e-09 | 8.7e-09 | 7.667E-09 | 2.421E-09 | 4.900E-09 | 9.400E-09 |
| 7119 | 356 | P356Y | A | 7.2e-09 | 1.5e-08 | 6.3e-09 | 9.500E-09 | 4.784E-09 | 6.300E-09 | 1.500E-08 |
| 7120 | 356 | P356V | A | 4.9e-09 | 7.5e-09 | 1.2e-08 | 8.133E-09 | 3.592E-09 | 4.900E-09 | 1.200E-08 |
| 7121 | 357 | E357A | A | 2.4e-09 | 5.7e-09 | 8.2e-09 | 5.433E-09 | 2.909E-09 | 2.400E-09 | 8.200E-09 |
| 7122 | 357 | E357R | A | 7.7e-09 | 3.8e-09 | 4.6e-09 | 5.367E-09 | 2.060E-09 | 3.800E-09 | 7.700E-09 |
| 7123 | 357 | E357N | A | 6.5e-09 | 1.3e-08 | 5.9e-09 | 8.467E-09 | 3.937E-09 | 5.900E-09 | 1.300E-08 |
| 7124 | 357 | E357D | A | 1.6e-08 | 3e-09 | 4e-09 | 7.667E-09 | 7.234E-09 | 3.000E-09 | 1.600E-08 |
| 7125 | 357 | E357C | A | 3.5e-09 | 5.7e-09 | 5.7e-09 | 4.967E-09 | 1.270E-09 | 3.500E-09 | 5.700E-09 |
| 7126 | 357 | E357E | A | 9.7e-09 | 1.9e-08 | 5.3e-09 | 1.133E-08 | 6.995E-09 | 5.300E-09 | 1.900E-08 |
| 7127 | 357 | E357Q | A | 6.7e-09 | 9.9e-09 | 6.9e-09 | 7.833E-09 | 1.793E-09 | 6.700E-09 | 9.900E-09 |
| 7128 | 357 | E357G | A | 1.8e-08 | 2.7e-08 | 5.8e-09 | 1.693E-08 | 1.064E-08 | 5.800E-09 | 2.700E-08 |
| 7129 | 357 | E357H | A | 1.3e-08 | 9e-09 | 9.3e-09 | 1.043E-08 | 2.228E-09 | 9.000E-09 | 1.300E-08 |
| 7130 | 357 | E357I | A | 9.6e-09 | 1.2e-08 | 2.7e-08 | 1.620E-08 | 9.430E-09 | 9.600E-09 | 2.700E-08 |
| 7131 | 357 | E357L | A | 1.5e-08 | 1.4e-08 | 1.8e-08 | 1.567E-08 | 2.082E-09 | 1.400E-08 | 1.800E-08 |
| 7132 | 357 | E357K | A | 3.6e-09 | 3.7e-09 | 1e-08 | 5.767E-09 | 3.667E-09 | 3.600E-09 | 1.000E-08 |
| 7133 | 357 | E357M | A | 1e-08 | 6.8e-09 | 8.9e-09 | 8.567E-09 | 1.626E-09 | 6.800E-09 | 1.000E-08 |
| 7134 | 357 | E357F | A | 6.9e-09 | 2.7e-09 | 1.1e-08 | 6.867E-09 | 4.150E-09 | 2.700E-09 | 1.100E-08 |
| 7135 | 357 | E357P | A | 8.5e-09 | 1.1e-08 | 5.2e-09 | 8.233E-09 | 2.909E-09 | 5.200E-09 | 1.100E-08 |
| 7136 | 357 | E357S | A | 2.9e-08 | 1e-08 | 8.9e-09 | 1.597E-08 | 1.130E-08 | 8.900E-09 | 2.900E-08 |
| 7137 | 357 | E357T | A | 1.9e-08 | 5.6e-09 | 6.6e-09 | 1.040E-08 | 7.465E-09 | 5.600E-09 | 1.900E-08 |
| 7138 | 357 | E357W | A | 1.7e-08 | 1.6e-09 | 1.5e-08 | 1.120E-08 | 8.374E-09 | 1.600E-09 | 1.700E-08 |
| 7139 | 357 | E357Y | A | 1.2e-08 | 1.6e-08 | 6.5e-09 | 1.150E-08 | 4.770E-09 | 6.500E-09 | 1.600E-08 |

| | | | | | | | | | | |
|---|---|---|---|---|---|---|---|---|---|---|
| 7140 | 357 | E357V | A | 7.6e-09 | 4.1e-09 | 4.5e-09 | 5.400E-09 | 1.916E-09 | 4.100E-09 | 7.600E-09 |
| 7141 | 358 | S358A | A | 6.8e-09 | 2.8e-09 | 2.6e-08 | 1.187E-08 | 1.240E-08 | 2.800E-09 | 2.600E-08 |
| 7142 | 358 | S358R | A | 3.7e-09 | 1.5e-08 | 4.3e-09 | 7.667E-09 | 6.358E-09 | 3.700E-09 | 1.500E-08 |
| 7143 | 358 | S358N | A | 2.8e-09 | 7.2e-09 | 3.4e-09 | 4.467E-09 | 2.386E-09 | 2.800E-09 | 7.200E-09 |
| 7144 | 358 | S358D | A | 1.1e-08 | 6.8e-09 | 7.3e-09 | 8.367E-09 | 2.294E-09 | 6.800E-09 | 1.100E-08 |
| 7145 | 358 | S358C | A | 6.8e-09 | 2.3e-09 | 6.5e-09 | 5.200E-09 | 2.516E-09 | 2.300E-09 | 6.800E-09 |
| 7146 | 358 | S358E | A | 1.1e-08 | 2.7e-09 | 5.8e-09 | 6.500E-09 | 4.194E-09 | 2.700E-09 | 1.100E-08 |
| 7147 | 358 | S358Q | A | 9.4e-09 | 3.2e-09 | 1.1e-08 | 7.867E-09 | 4.120E-09 | 3.200E-09 | 1.100E-08 |
| 7148 | 358 | S358G | A | 4.4e-09 | 3.3e-09 | 2.4e-09 | 3.367E-09 | 1.002E-09 | 2.400E-09 | 4.400E-09 |
| 7149 | 358 | S358H | A | 9.3e-09 | 1.2e-08 | 1.3e-08 | 1.143E-08 | 1.914E-09 | 9.300E-09 | 1.300E-08 |
| 7150 | 358 | S358I | A | 9.3e-09 | 1.8e-08 | 1.1e-08 | 1.277E-08 | 4.611E-09 | 9.300E-09 | 1.800E-08 |
| 7151 | 358 | S358L | A | 1e-08 | 5.7e-09 | 6.3e-09 | 7.333E-09 | 2.329E-09 | 5.700E-09 | 1.000E-08 |
| 7152 | 358 | S358K | A | 2.4e-08 | 3.1e-09 | 4.5e-09 | 1.053E-08 | 1.168E-08 | 3.100E-09 | 2.400E-08 |
| 7153 | 358 | S358M | A | 6.2e-09 | 2.4e-08 | 2.8e-09 | 1.100E-08 | 1.139E-08 | 2.800E-09 | 2.400E-08 |
| 7154 | 358 | S358F | A | 7.5e-09 | 5.3e-09 | 2.6e-09 | 5.133E-09 | 2.454E-09 | 2.600E-09 | 7.500E-09 |
| 7155 | 358 | S358P | A | 7e-09 | 4.4e-09 | 6e-09 | 5.800E-09 | 1.311E-09 | 4.400E-09 | 7.000E-09 |
| 7156 | 358 | S358S | A | 3.9e-09 | 1.9e-08 | 5e-09 | 9.300E-09 | 8.418E-09 | 3.900E-09 | 1.900E-08 |
| 7157 | 358 | S358T | A | 5.1e-09 | 3.9e-08 | 6.4e-09 | 1.683E-08 | 1.921E-08 | 5.100E-09 | 3.900E-08 |
| 7158 | 358 | S358W | A | 8.1e-09 | 1e-08 | 1.1e-08 | 9.700E-09 | 1.473E-09 | 8.100E-09 | 1.100E-08 |
| 7159 | 358 | S358Y | A | 5.1e-09 | 7.1e-09 | 6.3e-09 | 6.167E-09 | 1.007E-09 | 5.100E-09 | 7.100E-09 |
| 7160 | 358 | S358V | A | 7.1e-09 | 3.7e-09 | 1e-08 | 6.933E-09 | 3.153E-09 | 3.700E-09 | 1.000E-08 |
| 7161 | 359 | F359A | A | 4.9e-09 | 7.9e-09 | 1.8e-08 | 1.027E-08 | 6.863E-09 | 4.900E-09 | 1.800E-08 |
| 7162 | 359 | F359R | A | 7.6e-09 | 3.7e-09 | 4e-09 | 5.100E-09 | 2.170E-09 | 3.700E-09 | 7.600E-09 |
| 7163 | 359 | F359N | A | 4.8e-09 | 8.6e-09 | 4.6e-09 | 6.000E-09 | 2.254E-09 | 4.600E-09 | 8.600E-09 |
| 7164 | 359 | F359D | A | 6.3e-09 | 2.7e-09 | 4.4e-09 | 4.467E-09 | 1.801E-09 | 2.700E-09 | 6.300E-09 |
| 7165 | 359 | F359C | A | 2.2e-08 | 3.9e-09 | 1.5e-08 | 1.363E-08 | 9.127E-09 | 3.900E-09 | 2.200E-08 |
| 7166 | 359 | F359E | A | 8.2e-09 | 4.7e-09 | 3.5e-09 | 5.467E-09 | 2.442E-09 | 3.500E-09 | 8.200E-09 |
| 7167 | 359 | F359Q | A | 4.1e-09 | 5e-09 | 1e-08 | 6.367E-09 | 3.179E-09 | 4.100E-09 | 1.000E-08 |
| 7168 | 359 | F359G | A | 1.7e-08 | 5.1e-09 | 6.4e-09 | 9.500E-09 | 6.528E-09 | 5.100E-09 | 1.700E-08 |
| 7169 | 359 | F359H | A | 2.3e-09 | 2.1e-09 | 2.3e-08 | 9.133E-09 | 1.201E-08 | 2.100E-09 | 2.300E-08 |
| 7170 | 359 | F359I | A | 5.6e-09 | 7.5e-09 | 1.1e-08 | 8.033E-09 | 2.739E-09 | 5.600E-09 | 1.100E-08 |
| 7171 | 359 | F359L | A | 6.1e-09 | 7.1e-09 | 5.8e-09 | 6.333E-09 | 6.807E-10 | 5.800E-09 | 7.100E-09 |
| 7172 | 359 | F359K | A | 7.4e-09 | 1.3e-08 | 1.4e-08 | 1.147E-08 | 3.557E-09 | 7.400E-09 | 1.400E-08 |
| 7173 | 359 | F359M | A | 7.4e-09 | 5.1e-09 | 8.9e-09 | 7.133E-09 | 1.914E-09 | 5.100E-09 | 8.900E-09 |
| 7174 | 359 | F359F | A | 1.1e-08 | 1.4e-09 | 2e-09 | 4.800E-09 | 5.378E-09 | 1.400E-09 | 1.100E-08 |
| 7175 | 359 | F359P | A | 6.4e-09 | 9.5e-09 | 1e-08 | 8.633E-09 | 1.950E-09 | 6.400E-09 | 1.000E-08 |
| 7176 | 359 | F359S | A | 4.1e-09 | 1.3e-08 | 2.6e-09 | 6.567E-09 | 5.622E-09 | 2.600E-09 | 1.300E-08 |
| 7177 | 359 | F359T | A | 1.6e-08 | 3.6e-09 | 5.7e-09 | 8.433E-09 | 6.637E-09 | 3.600E-09 | 1.600E-08 |
| 7178 | 359 | F359W | A | 1.5e-08 | 8.6e-09 | 6.6e-09 | 1.007E-08 | 4.388E-09 | 6.600E-09 | 1.500E-08 |
| 7179 | 359 | F359Y | A | 9.6e-09 | 8.4e-09 | 5.4e-09 | 7.800E-09 | 2.163E-09 | 5.400E-09 | 9.600E-09 |
| 7180 | 359 | F359V | A | 4.5e-09 | 2.2e-08 | 4.5e-09 | 1.033E-08 | 1.010E-08 | 4.500E-09 | 2.200E-08 |
| 7181 | 360 | D360A | A | 6.1e-09 | 7.8e-09 | 3.2e-09 | 5.700E-09 | 2.326E-09 | 3.200E-09 | 7.800E-09 |
| 7182 | 360 | D360R | A | 1e-08 | 1e-08 | 1.9e-08 | 1.300E-08 | 5.196E-09 | 1.000E-08 | 1.900E-08 |
| 7183 | 360 | D360N | A | 3.5e-09 | 5e-09 | 8.2e-09 | 5.567E-09 | 2.401E-09 | 3.500E-09 | 8.200E-09 |
| 7184 | 360 | D360D | A | 1.2e-08 | 6.3e-09 | 1.9e-08 | 1.243E-08 | 6.361E-09 | 6.300E-09 | 1.900E-08 |
| 7185 | 360 | D360C | A | 9.5e-09 | 2e-08 | 4.3e-09 | 1.127E-08 | 7.998E-09 | 4.300E-09 | 2.000E-08 |
| 7186 | 360 | D360E | A | 5.9e-09 | 3.2e-09 | 7.9e-09 | 5.667E-09 | 2.359E-09 | 3.200E-09 | 7.900E-09 |
| 7187 | 360 | D360Q | A | 4.4e-09 | 1.1e-08 | 4.2e-09 | 6.533E-09 | 3.870E-09 | 4.200E-09 | 1.100E-08 |
| 7188 | 360 | D360G | A | 8.3e-09 | 7.2e-09 | 2.4e-09 | 5.967E-09 | 3.137E-09 | 2.400E-09 | 8.300E-09 |
| 7189 | 360 | D360H | A | 1.2e-08 | 7.2e-09 | 7.9e-09 | 9.033E-09 | 2.593E-09 | 7.200E-09 | 1.200E-08 |
| 7190 | 360 | D360I | A | 1.2e-08 | 1.6e-08 | 4.2e-09 | 1.073E-08 | 6.001E-09 | 4.200E-09 | 1.600E-08 |
| 7191 | 360 | D360L | A | 4e-09 | 6.2e-09 | 7.6e-09 | 5.933E-09 | 1.815E-09 | 4.000E-09 | 7.600E-09 |
| 7192 | 360 | D360K | A | 3.8e-09 | 1.7e-08 | 2.7e-09 | 7.833E-09 | 7.958E-09 | 2.700E-09 | 1.700E-08 |
| 7193 | 360 | D360M | A | 8e-09 | 1.4e-08 | 4.8e-09 | 8.933E-09 | 4.670E-09 | 4.800E-09 | 1.400E-08 |
| 7194 | 360 | D360F | A | 4.3e-09 | 5e-09 | 7e-09 | 5.433E-09 | 1.401E-09 | 4.300E-09 | 7.000E-09 |
| 7195 | 360 | D360P | A | 6.9e-09 | 2e-08 | 6e-09 | 1.097E-08 | 7.836E-09 | 6.000E-09 | 2.000E-08 |
| 7196 | 360 | D360S | A | 8.6e-09 | 9.8e-09 | 6.3e-09 | 8.233E-09 | 1.779E-09 | 6.300E-09 | 9.800E-09 |
| 7197 | 360 | D360T | A | 2.6e-09 | 2e-08 | 4.2e-09 | 8.933E-09 | 9.617E-09 | 2.600E-09 | 2.000E-08 |
| 7198 | 360 | D360W | A | 1.2e-08 | 6e-09 | 9.5e-09 | 9.167E-09 | 3.014E-09 | 6.000E-09 | 1.200E-08 |
| 7199 | 360 | D360Y | A | 4.3e-09 | 2.1e-09 | 2.2e-09 | 2.867E-09 | 1.242E-09 | 2.100E-09 | 4.300E-09 |
| 7200 | 360 | D360V | A | 3.3e-09 | 5.7e-09 | 8.1e-09 | 5.700E-09 | 2.400E-09 | 3.300E-09 | 8.100E-09 |
| 7201 | 361 | G361A | A | 1.3e-08 | 5.6e-09 | 9e-09 | 9.200E-09 | 3.704E-09 | 5.600E-09 | 1.300E-08 |
| 7202 | 361 | G361R | A | 5e-09 | 9e-09 | 4.2e-09 | 6.067E-09 | 2.572E-09 | 4.200E-09 | 9.000E-09 |
| 7203 | 361 | G361N | A | 3.8e-09 | 8.8e-09 | 4.6e-09 | 5.733E-09 | 2.686E-09 | 3.800E-09 | 8.800E-09 |
| 7204 | 361 | G361D | A | 2.4e-09 | 6e-09 | 7.2e-09 | 5.200E-09 | 2.498E-09 | 2.400E-09 | 7.200E-09 |
| 7205 | 361 | G361C | A | 1.2e-08 | 1.9e-09 | 1.2e-08 | 8.633E-09 | 5.831E-09 | 1.900E-09 | 1.200E-08 |
| 7206 | 361 | G361E | A | 8e-09 | 8.4e-09 | 1.6e-08 | 1.080E-08 | 4.508E-09 | 8.000E-09 | 1.600E-08 |
| 7207 | 361 | G361Q | A | 1.8e-09 | 8.5e-09 | 1.4e-08 | 8.100E-09 | 6.110E-09 | 1.800E-09 | 1.400E-08 |
| 7208 | 361 | G361G | A | 1.3e-08 | 9.7e-09 | 2.8e-09 | 8.500E-09 | 5.205E-09 | 2.800E-09 | 1.300E-08 |
| 7209 | 361 | G361H | A | 1.4e-08 | 6.8e-09 | 1e-08 | 1.027E-08 | 3.607E-09 | 6.800E-09 | 1.400E-08 |
| 7210 | 361 | G361I | A | 1.2e-08 | 2.7e-09 | 2.9e-09 | 5.867E-09 | 5.313E-09 | 2.700E-09 | 1.200E-08 |
| 7211 | 361 | G361L | A | 8.7e-09 | 7.2e-09 | 7.1e-09 | 7.667E-09 | 8.963E-10 | 7.100E-09 | 8.700E-09 |
| 7212 | 361 | G361K | A | 6.2e-09 | 7.9e-09 | 5.2e-09 | 6.433E-09 | 1.365E-09 | 5.200E-09 | 7.900E-09 |
| 7213 | 361 | G361M | A | 1e-08 | 1.2e-08 | 3.3e-09 | 8.433E-09 | 4.557E-09 | 3.300E-09 | 1.200E-08 |
| 7214 | 361 | G361F | A | 5e-09 | 2.1e-09 | 3.7e-09 | 3.600E-09 | 1.453E-09 | 2.100E-09 | 5.000E-09 |
| 7215 | 361 | G361P | A | 1.3e-08 | 7.5e-09 | 1.3e-08 | 1.117E-08 | 3.175E-09 | 7.500E-09 | 1.300E-08 |
| 7216 | 361 | G361S | A | 2.4e-08 | 7.3e-09 | 8.6e-09 | 1.330E-08 | 9.289E-09 | 7.300E-09 | 2.400E-08 |
| 7217 | 361 | G361T | A | 1.5e-08 | 6.7e-09 | 4e-09 | 8.567E-09 | 5.733E-09 | 4.000E-09 | 1.500E-08 |
| 7218 | 361 | G361W | A | 7.3e-09 | 5.2e-09 | 8.9e-09 | 7.133E-09 | 1.856E-09 | 5.200E-09 | 8.900E-09 |
| 7219 | 361 | G361Y | A | 6.5e-09 | 1.1e-08 | 1.2e-08 | 9.833E-09 | 2.930E-09 | 6.500E-09 | 1.200E-08 |
| 7220 | 361 | G361V | A | 1e-08 | 1.2e-08 | 3.3e-09 | 8.433E-09 | 4.557E-09 | 3.300E-09 | 1.200E-08 |
| 7221 | 362 | D362A | A | 9.1e-09 | 1.3e-08 | 4.3e-09 | 8.800E-09 | 4.358E-09 | 4.300E-09 | 1.300E-08 |
| 7222 | 362 | D362R | A | 7.4e-09 | 1.2e-08 | 5.8e-09 | 8.400E-09 | 3.219E-09 | 5.800E-09 | 1.200E-08 |
| 7223 | 362 | D362N | A | 5.7e-09 | 8.5e-09 | 6.4e-09 | 6.867E-09 | 1.457E-09 | 5.700E-09 | 8.500E-09 |
| 7224 | 362 | D362D | A | 7.5e-09 | 3e-09 | 1.5e-08 | 8.500E-09 | 6.062E-09 | 3.000E-09 | 1.500E-08 |

| | | | | | | | | | | |
|---|---|---|---|---|---|---|---|---|---|---|
| 7225 | 362 | D362C | A | 1.9e-08 | 4.2e-09 | 8.3e-09 | 1.050E-08 | 7.641E-09 | 4.200E-09 | 1.900E-08 |
| 7226 | 362 | D362E | A | 1e-08 | 1.8e-08 | 4e-08 | 2.267E-08 | 1.553E-08 | 1.000E-08 | 4.000E-08 |
| 7227 | 362 | D362Q | A | 6.3e-09 | 6.5e-09 | 2.7e-08 | 1.327E-08 | 1.189E-08 | 6.300E-09 | 2.700E-08 |
| 7228 | 362 | D362G | A | 3.2e-09 | 1.4e-08 | 2.5e-08 | 1.407E-08 | 1.090E-08 | 3.200E-09 | 2.500E-08 |
| 7229 | 362 | D362H | A | 1e-08 | 2.3e-08 | 4.5e-09 | 1.250E-08 | 9.500E-09 | 4.500E-09 | 2.300E-08 |
| 7230 | 362 | D362I | A | 4e-09 | 3.4e-09 | 8.7e-09 | 5.367E-09 | 2.902E-09 | 3.400E-09 | 8.700E-09 |
| 7231 | 362 | D362L | A | 3.6e-09 | 7e-09 | 1.4e-08 | 8.200E-09 | 5.303E-09 | 3.600E-09 | 1.400E-08 |
| 7232 | 362 | D362K | A | 7.9e-09 | 1e-08 | 1e-08 | 9.300E-09 | 1.212E-09 | 7.900E-09 | 1.000E-08 |
| 7233 | 362 | D362M | A | 7.6e-09 | 6.9e-09 | 7.5e-09 | 7.333E-09 | 3.786E-10 | 6.900E-09 | 7.600E-09 |
| 7234 | 362 | D362F | A | 6.1e-09 | 1.2e-08 | 6.6e-09 | 8.233E-09 | 3.272E-09 | 6.100E-09 | 1.200E-08 |
| 7235 | 362 | D362P | A | 6.7e-09 | 1.9e-08 | 7.9e-09 | 1.120E-08 | 6.782E-09 | 6.700E-09 | 1.900E-08 |
| 7236 | 362 | D362S | A | 9.4e-09 | 6e-09 | 1.8e-08 | 1.113E-08 | 6.185E-09 | 6.000E-09 | 1.800E-08 |
| 7237 | 362 | D362T | A | 3.1e-08 | 1.9e-08 | 9.1e-09 | 1.970E-08 | 1.097E-08 | 9.100E-09 | 3.100E-08 |
| 7238 | 362 | D362W | A | 4.4e-09 | 5.6e-09 | 1.5e-08 | 8.333E-09 | 5.805E-09 | 4.400E-09 | 1.500E-08 |
| 7239 | 362 | D362Y | A | 6.8e-09 | 4.1e-09 | 1.7e-08 | 9.300E-09 | 6.804E-09 | 4.100E-09 | 1.700E-08 |
| 7240 | 362 | D362V | A | 4.4e-09 | 1.1e-08 | 8.6e-09 | 8.000E-09 | 3.341E-09 | 4.400E-09 | 1.100E-08 |
| 7241 | 363 | P363A | A | 4.8e-09 | 9.5e-09 | 1.6e-09 | 5.300E-09 | 3.974E-09 | 1.600E-09 | 9.500E-09 |
| 7242 | 363 | P363R | A | 5.8e-09 | 6.1e-09 | 1.5e-09 | 4.467E-09 | 2.574E-09 | 1.500E-09 | 6.100E-09 |
| 7243 | 363 | P363N | A | 3.7e-09 | 8.7e-09 | 1.3e-08 | 8.467E-09 | 4.654E-09 | 3.700E-09 | 1.300E-08 |
| 7244 | 363 | P363D | A | 1e-08 | 1.4e-08 | 1e-08 | 1.133E-08 | 2.309E-09 | 1.000E-08 | 1.400E-08 |
| 7245 | 363 | P363C | A | 1.5e-08 | 9.9e-09 | 5.8e-09 | 1.023E-08 | 4.609E-09 | 5.800E-09 | 1.500E-08 |
| 7246 | 363 | P363E | A | 1e-08 | 1.1e-08 | 4.1e-09 | 8.367E-09 | 3.729E-09 | 4.100E-09 | 1.100E-08 |
| 7247 | 363 | P363Q | A | 5.8e-09 | 8.7e-09 | 5.8e-09 | 6.767E-09 | 1.674E-09 | 5.800E-09 | 8.700E-09 |
| 7248 | 363 | P363G | A | 9.6e-09 | 5.2e-09 | 1.2e-08 | 8.933E-09 | 3.449E-09 | 5.200E-09 | 1.200E-08 |
| 7249 | 363 | P363H | A | 1.3e-08 | 6.7e-09 | 2.6e-09 | 7.433E-09 | 5.239E-09 | 2.600E-09 | 1.300E-08 |
| 7250 | 363 | P363I | A | 5.6e-09 | 5.8e-09 | 4.9e-09 | 5.433E-09 | 4.726E-10 | 4.900E-09 | 5.800E-09 |
| 7251 | 363 | P363L | A | 6.7e-09 | 1.3e-08 | 3.2e-08 | 1.723E-08 | 1.317E-08 | 6.700E-09 | 3.200E-08 |
| 7252 | 363 | P363K | A | 5.6e-09 | 4.3e-09 | 8.2e-09 | 6.033E-09 | 1.986E-09 | 4.300E-09 | 8.200E-09 |
| 7253 | 363 | P363M | A | 1.1e-08 | 9.9e-09 | 3e-09 | 7.967E-09 | 4.336E-09 | 3.000E-09 | 1.100E-08 |
| 7254 | 363 | P363F | A | 7.2e-09 | 4.4e-09 | 4.4e-09 | 5.333E-09 | 1.617E-09 | 4.400E-09 | 7.200E-09 |
| 7255 | 363 | P363P | A | 1.1e-08 | 9.9e-09 | 6.9e-09 | 9.267E-09 | 2.122E-09 | 6.900E-09 | 1.100E-08 |
| 7256 | 363 | P363S | A | 1.6e-08 | 1.4e-09 | 6e-09 | 7.800E-09 | 7.465E-09 | 1.400E-09 | 1.600E-08 |
| 7257 | 363 | P363T | A | 5.6e-09 | 7.9e-09 | 7.2e-09 | 6.900E-09 | 1.179E-09 | 5.600E-09 | 7.900E-09 |
| 7258 | 363 | P363W | A | 6.2e-09 | 7.1e-09 | 9.9e-09 | 7.733E-09 | 1.930E-09 | 6.200E-09 | 9.900E-09 |
| 7259 | 363 | P363Y | A | 8.3e-09 | 1.6e-08 | 2.2e-09 | 8.833E-09 | 6.915E-09 | 2.200E-09 | 1.600E-08 |
| 7260 | 363 | P363V | A | 3.9e-09 | 8.8e-09 | 6.5e-09 | 6.400E-09 | 2.452E-09 | 3.900E-09 | 8.800E-09 |
| 7261 | 364 | A364A | A | 1.2e-08 | 6.3e-09 | 3.7e-09 | 7.333E-09 | 4.245E-09 | 3.700E-09 | 1.200E-08 |
| 7262 | 364 | A364R | A | 7.5e-09 | 8.5e-09 | 4e-09 | 6.667E-09 | 2.363E-09 | 4.000E-09 | 8.500E-09 |
| 7263 | 364 | A364N | A | 4.2e-09 | 1.9e-08 | 2e-08 | 1.440E-08 | 8.848E-09 | 4.200E-09 | 2.000E-08 |
| 7264 | 364 | A364D | A | 1.2e-08 | 4.5e-09 | 3.5e-09 | 6.667E-09 | 4.646E-09 | 3.500E-09 | 1.200E-08 |
| 7265 | 364 | A364C | A | 8.1e-09 | 1.1e-08 | 1.6e-09 | 6.900E-09 | 4.814E-09 | 1.600E-09 | 1.100E-08 |
| 7266 | 364 | A364E | A | 1.7e-08 | 9.6e-09 | 4.1e-09 | 1.023E-08 | 6.473E-09 | 4.100E-09 | 1.700E-08 |
| 7267 | 364 | A364Q | A | 1.7e-08 | 6.1e-09 | 2.4e-08 | 1.570E-08 | 9.021E-09 | 6.100E-09 | 2.400E-08 |
| 7268 | 364 | A364G | A | 7.5e-09 | 9e-09 | 6.8e-09 | 7.767E-09 | 1.124E-09 | 6.800E-09 | 9.000E-09 |
| 7269 | 364 | A364H | A | 1e-08 | 3e-09 | 9.5e-09 | 7.500E-09 | 3.905E-09 | 3.000E-09 | 1.000E-08 |
| 7270 | 364 | A364I | A | 8.1e-09 | 1e-08 | 1.1e-08 | 9.700E-09 | 1.473E-09 | 8.100E-09 | 1.100E-08 |
| 7271 | 364 | A364L | A | 5.1e-09 | 7.5e-09 | 4.6e-09 | 5.733E-09 | 1.550E-09 | 4.600E-09 | 7.500E-09 |
| 7272 | 364 | A364K | A | 5.4e-09 | 4.2e-09 | 6.5e-09 | 5.367E-09 | 1.150E-09 | 4.200E-09 | 6.500E-09 |
| 7273 | 364 | A364M | A | 2.9e-09 | 4.1e-09 | 8.4e-09 | 5.133E-09 | 2.892E-09 | 2.900E-09 | 8.400E-09 |
| 7274 | 364 | A364F | A | 4.1e-09 | 6.2e-09 | 1.2e-08 | 7.433E-09 | 4.092E-09 | 4.100E-09 | 1.200E-08 |
| 7275 | 364 | A364P | A | 9e-09 | 7.5e-09 | 1e-08 | 8.833E-09 | 1.258E-09 | 7.500E-09 | 1.000E-08 |
| 7276 | 364 | A364S | A | 1.3e-09 | 2.3e-09 | 5.9e-09 | 3.167E-09 | 2.419E-09 | 1.300E-09 | 5.900E-09 |
| 7277 | 364 | A364T | A | 1.1e-08 | 9.9e-09 | 1.8e-08 | 1.297E-08 | 4.394E-09 | 9.900E-09 | 1.800E-08 |
| 7278 | 364 | A364W | A | 4.3e-09 | 1.3e-08 | 3.3e-09 | 6.867E-09 | 5.335E-09 | 3.300E-09 | 1.300E-08 |
| 7279 | 364 | A364Y | A | 6.1e-09 | 1.1e-08 | 5.2e-09 | 7.433E-09 | 3.121E-09 | 5.200E-09 | 1.100E-08 |
| 7280 | 364 | A364V | A | 7.9e-09 | 5.4e-09 | 6e-09 | 6.433E-09 | 1.305E-09 | 5.400E-09 | 7.900E-09 |
| 7281 | 365 | S365A | A | 2.8e-09 | 5e-09 | 6.3e-09 | 4.700E-09 | 1.769E-09 | 2.800E-09 | 6.300E-09 |
| 7282 | 365 | S365R | A | 2.6e-09 | 5.5e-09 | 4.7e-09 | 4.267E-09 | 1.498E-09 | 2.600E-09 | 5.500E-09 |
| 7283 | 365 | S365N | A | 9.8e-09 | 6.9e-09 | 7.8e-09 | 8.167E-09 | 1.484E-09 | 6.900E-09 | 9.800E-09 |
| 7284 | 365 | S365D | A | 4.5e-09 | 1.4e-08 | 1.6e-08 | 1.150E-08 | 6.144E-09 | 4.500E-09 | 1.600E-08 |
| 7285 | 365 | S365C | A | 8.5e-09 | 1e-09 | 4.9e-09 | 4.800E-09 | 3.751E-09 | 1.000E-09 | 8.500E-09 |
| 7286 | 365 | S365E | A | 4.9e-09 | 9e-09 | 9.1e-09 | 7.667E-09 | 2.397E-09 | 4.900E-09 | 9.100E-09 |
| 7287 | 365 | S365Q | A | 3.6e-09 | 3.7e-09 | 4.7e-09 | 4.000E-09 | 6.083E-10 | 3.600E-09 | 4.700E-09 |
| 7288 | 365 | S365G | A | 9.6e-09 | 2.3e-08 | 6.3e-09 | 1.297E-08 | 8.844E-09 | 6.300E-09 | 2.300E-08 |
| 7289 | 365 | S365H | A | 5.5e-09 | 3.3e-09 | 6.3e-09 | 5.033E-09 | 1.553E-09 | 3.300E-09 | 6.300E-09 |
| 7290 | 365 | S365I | A | 3.3e-09 | 1.3e-08 | 7.6e-09 | 7.967E-09 | 4.860E-09 | 3.300E-09 | 1.300E-08 |
| 7291 | 365 | S365L | A | 9.5e-09 | 2e-09 | 4.1e-09 | 5.200E-09 | 3.869E-09 | 2.000E-09 | 9.500E-09 |
| 7292 | 365 | S365K | A | 1.1e-08 | 1.3e-08 | 5.1e-09 | 9.700E-09 | 4.107E-09 | 5.100E-09 | 1.300E-08 |
| 7293 | 365 | S365M | A | 4.4e-09 | 3.3e-08 | 2.8e-09 | 1.340E-08 | 1.699E-08 | 2.800E-09 | 3.300E-08 |
| 7294 | 365 | S365F | A | 9.7e-09 | 1.3e-08 | 2e-08 | 1.423E-08 | 5.260E-09 | 9.700E-09 | 2.000E-08 |
| 7295 | 365 | S365P | A | 6.3e-09 | 8.9e-09 | 1.1e-08 | 8.733E-09 | 2.354E-09 | 6.300E-09 | 1.100E-08 |
| 7296 | 365 | S365S | A | 9.9e-09 | 8.3e-09 | 5.1e-09 | 7.767E-09 | 2.444E-09 | 5.100E-09 | 9.900E-09 |
| 7297 | 365 | S365T | A | 5.7e-09 | 9.6e-09 | 1.3e-08 | 9.433E-09 | 3.653E-09 | 5.700E-09 | 1.300E-08 |
| 7298 | 365 | S365W | A | 5.9e-09 | 1.1e-08 | 2.2e-08 | 1.297E-08 | 8.228E-09 | 5.900E-09 | 2.200E-08 |
| 7299 | 365 | S365Y | A | 8.4e-09 | 3.4e-09 | 4.6e-09 | 5.467E-09 | 2.610E-09 | 3.400E-09 | 8.400E-09 |
| 7300 | 365 | S365V | A | 6.4e-09 | 6.4e-09 | 5.2e-09 | 6.000E-09 | 6.928E-10 | 5.200E-09 | 6.400E-09 |
| 7301 | 366 | N366A | A | 1.2e-08 | 4.1e-09 | 6e-09 | 7.367E-09 | 4.124E-09 | 4.100E-09 | 1.200E-08 |
| 7302 | 366 | N366R | A | 1.1e-08 | 1.6e-08 | 3.3e-09 | 1.010E-08 | 6.398E-09 | 3.300E-09 | 1.600E-08 |
| 7303 | 366 | N366N | A | 5.1e-09 | 8.2e-09 | 5.1e-09 | 6.133E-09 | 1.790E-09 | 5.100E-09 | 8.200E-09 |
| 7304 | 366 | N366D | A | 9.4e-09 | 4.4e-09 | 3.2e-09 | 5.667E-09 | 3.288E-09 | 3.200E-09 | 9.400E-09 |
| 7305 | 366 | N366C | A | 5e-09 | 1.3e-08 | 4.6e-09 | 7.533E-09 | 4.738E-09 | 4.600E-09 | 1.300E-08 |
| 7306 | 366 | N366E | A | 5.1e-09 | 9.1e-09 | 1e-08 | 8.067E-09 | 2.608E-09 | 5.100E-09 | 1.000E-08 |
| 7307 | 366 | N366Q | A | 9.4e-09 | 1.4e-09 | 6.6e-09 | 5.800E-09 | 4.060E-09 | 1.400E-09 | 9.400E-09 |
| 7308 | 366 | N366G | A | 5.5e-08 | 2e-08 | 4.9e-09 | 2.663E-08 | 2.570E-08 | 4.900E-09 | 5.500E-08 |
| 7309 | 366 | N366H | A | 8.5e-09 | 1.1e-08 | 4.8e-09 | 8.100E-09 | 3.119E-09 | 4.800E-09 | 1.100E-08 |

| | | | | | | | | | | |
|---|---|---|---|---|---|---|---|---|---|---|
| 7310 | 366 | N366I | A | 5.1e-09 | 8.4e-09 | 2.1e-09 | 5.200E-09 | 3.151E-09 | 2.100E-09 | 8.400E-09 |
| 7311 | 366 | N366L | A | 6.1e-09 | 4e-09 | 1.2e-08 | 7.367E-09 | 4.148E-09 | 4.000E-09 | 1.200E-08 |
| 7312 | 366 | N366K | A | 8.8e-09 | 3.4e-09 | 5.9e-09 | 6.033E-09 | 2.702E-09 | 3.400E-09 | 8.800E-09 |
| 7313 | 366 | N366M | A | 2.6e-09 | 1.2e-08 | 2.8e-09 | 5.800E-09 | 5.370E-09 | 2.600E-09 | 1.200E-08 |
| 7314 | 366 | N366F | A | 3.6e-09 | 6.2e-09 | 4.6e-09 | 4.800E-09 | 1.311E-09 | 3.600E-09 | 6.200E-09 |
| 7315 | 366 | N366P | A | 3e-09 | 1.4e-08 | 9e-09 | 8.667E-09 | 5.508E-09 | 3.000E-09 | 1.400E-08 |
| 7316 | 366 | N366S | A | 2.7e-09 | 1.9e-08 | 1e-08 | 1.057E-08 | 8.165E-09 | 2.700E-09 | 1.900E-08 |
| 7317 | 366 | N366T | A | 4.9e-09 | 2.4e-09 | 3.8e-09 | 3.700E-09 | 1.253E-09 | 2.400E-09 | 4.900E-09 |
| 7318 | 366 | N366W | A | 4.4e-09 | 2.2e-09 | 2.1e-09 | 2.900E-09 | 1.300E-09 | 2.100E-09 | 4.400E-09 |
| 7319 | 366 | N366Y | A | 3.2e-09 | 1.2e-08 | 5.2e-09 | 6.800E-09 | 4.613E-09 | 3.200E-09 | 1.200E-08 |
| 7320 | 366 | N366V | A | 1.3e-08 | 3.2e-09 | 8.5e-09 | 8.233E-09 | 4.905E-09 | 3.200E-09 | 1.300E-08 |
| 7321 | 367 | T367A | A | 7.2e-09 | 6.9e-09 | 8.4e-09 | 7.500E-09 | 7.937E-10 | 6.900E-09 | 8.400E-09 |
| 7322 | 367 | T367R | A | 4.8e-09 | 9e-09 | 8.8e-09 | 7.533E-09 | 2.369E-09 | 4.800E-09 | 9.000E-09 |
| 7323 | 367 | T367N | A | 1.6e-09 | 3.8e-09 | 9.6e-09 | 5.000E-09 | 4.133E-09 | 1.600E-09 | 9.600E-09 |
| 7324 | 367 | T367D | A | 6.9e-09 | 7.7e-09 | 1.2e-08 | 8.867E-09 | 2.743E-09 | 6.900E-09 | 1.200E-08 |
| 7325 | 367 | T367C | A | 6.4e-09 | 2.4e-09 | 5.1e-09 | 4.633E-09 | 2.040E-09 | 2.400E-09 | 6.400E-09 |
| 7326 | 367 | T367E | A | 4.6e-09 | 1.5e-08 | 5.3e-09 | 8.300E-09 | 5.813E-09 | 4.600E-09 | 1.500E-08 |
| 7327 | 367 | T367Q | A | 7.4e-09 | 6.4e-09 | 9.1e-09 | 7.633E-09 | 1.365E-09 | 6.400E-09 | 9.100E-09 |
| 7328 | 367 | T367G | A | 2.1e-08 | 1.4e-08 | 1.7e-08 | 1.733E-08 | 3.512E-09 | 1.400E-08 | 2.100E-08 |
| 7329 | 367 | T367H | A | 5.3e-09 | 4.3e-09 | 8.7e-09 | 6.100E-09 | 2.307E-09 | 4.300E-09 | 8.700E-09 |
| 7330 | 367 | T367I | A | 4.4e-09 | 3.4e-09 | 5e-09 | 4.267E-09 | 8.083E-10 | 3.400E-09 | 5.000E-09 |
| 7331 | 367 | T367L | A | 8.7e-09 | 4.3e-09 | 6.3e-09 | 6.433E-09 | 2.203E-09 | 4.300E-09 | 8.700E-09 |
| 7332 | 367 | T367K | A | 1.1e-08 | 1e-08 | 8.6e-09 | 9.867E-09 | 1.206E-09 | 8.600E-09 | 1.100E-08 |
| 7333 | 367 | T367M | A | 1.7e-08 | 2.4e-08 | 2.2e-08 | 2.100E-08 | 3.606E-09 | 1.700E-08 | 2.400E-08 |
| 7334 | 367 | T367F | A | 4.5e-09 | 9e-09 | 5.1e-09 | 6.200E-09 | 2.443E-09 | 4.500E-09 | 9.000E-09 |
| 7335 | 367 | T367P | A | 1.4e-08 | 6.1e-09 | 4.2e-09 | 8.100E-09 | 5.197E-09 | 4.200E-09 | 1.400E-08 |
| 7336 | 367 | T367S | A | 2.8e-09 | 8.2e-09 | 3.6e-09 | 4.867E-09 | 2.914E-09 | 2.800E-09 | 8.200E-09 |
| 7337 | 367 | T367T | A | 8.4e-09 | 9.5e-09 | 2.5e-09 | 6.800E-09 | 3.764E-09 | 2.500E-09 | 9.500E-09 |
| 7338 | 367 | T367W | A | 7.7e-09 | 1.5e-08 | 4.1e-09 | 8.933E-09 | 5.554E-09 | 4.100E-09 | 1.500E-08 |
| 7339 | 367 | T367Y | A | 6.2e-09 | 8.2e-09 | 2.6e-09 | 5.667E-09 | 2.838E-09 | 2.600E-09 | 8.200E-09 |
| 7340 | 367 | T367V | A | 3.2e-09 | 3.8e-09 | 1.5e-08 | 7.333E-09 | 6.646E-09 | 3.200E-09 | 1.500E-08 |
| 7341 | 368 | A368A | A | 7.7e-09 | 1.5e-08 | 5.5e-09 | 9.400E-09 | 4.973E-09 | 5.500E-09 | 1.500E-08 |
| 7342 | 368 | A368R | A | 8.2e-09 | 3.9e-09 | 2e-08 | 1.070E-08 | 8.336E-09 | 3.900E-09 | 2.000E-08 |
| 7343 | 368 | A368N | A | 2.8e-09 | 4e-09 | 4.6e-09 | 3.800E-09 | 9.165E-10 | 2.800E-09 | 4.600E-09 |
| 7344 | 368 | A368D | A | 2.7e-09 | 8.4e-09 | 2.3e-09 | 4.467E-09 | 3.412E-09 | 2.300E-09 | 8.400E-09 |
| 7345 | 368 | A368C | A | 2.7e-09 | 7.9e-09 | 1.2e-08 | 7.533E-09 | 4.661E-09 | 2.700E-09 | 1.200E-08 |
| 7346 | 368 | A368E | A | 1.2e-08 | 4.6e-09 | 1.3e-08 | 9.867E-09 | 4.588E-09 | 4.600E-09 | 1.300E-08 |
| 7347 | 368 | A368Q | A | 7.8e-09 | 5.9e-09 | 6.2e-09 | 6.633E-09 | 1.021E-09 | 5.900E-09 | 7.800E-09 |
| 7348 | 368 | A368G | A | 7.5e-09 | 2.1e-08 | 1.3e-08 | 1.383E-08 | 6.788E-09 | 7.500E-09 | 2.100E-08 |
| 7349 | 368 | A368H | A | 3.8e-09 | 5.1e-09 | 5.5e-09 | 4.800E-09 | 8.888E-10 | 3.800E-09 | 5.500E-09 |
| 7350 | 368 | A368I | A | 3.8e-09 | 2.2e-09 | 7.3e-09 | 4.433E-09 | 2.608E-09 | 2.200E-09 | 7.300E-09 |
| 7351 | 368 | A368L | A | 1.2e-08 | 4.8e-09 | 2.8e-09 | 6.533E-09 | 4.839E-09 | 2.800E-09 | 1.200E-08 |
| 7352 | 368 | A368K | A | 4.7e-09 | 7.2e-09 | 2.1e-08 | 1.097E-08 | 8.779E-09 | 4.700E-09 | 2.100E-08 |
| 7353 | 368 | A368M | A | 1.1e-08 | 1.1e-08 | 9.4e-09 | 1.047E-08 | 9.238E-10 | 9.400E-09 | 1.100E-08 |
| 7354 | 368 | A368F | A | 1e-08 | 1.3e-08 | 1.7e-09 | 8.233E-09 | 5.853E-09 | 1.700E-09 | 1.300E-08 |
| 7355 | 368 | A368P | A | 6.1e-09 | 5.2e-09 | 6.4e-09 | 5.900E-09 | 6.245E-10 | 5.200E-09 | 6.400E-09 |
| 7356 | 368 | A368S | A | 2.8e-09 | 5.1e-09 | 5.4e-09 | 4.433E-09 | 1.422E-09 | 2.800E-09 | 5.400E-09 |
| 7357 | 368 | A368T | A | 4.3e-09 | 6.5e-09 | 4.4e-09 | 5.067E-09 | 1.242E-09 | 4.300E-09 | 6.500E-09 |
| 7358 | 368 | A368W | A | 3e-09 | 2.1e-08 | 1e-08 | 1.133E-08 | 9.074E-09 | 3.000E-09 | 2.100E-08 |
| 7359 | 368 | A368Y | A | 1.9e-09 | 1.9e-09 | 4.9e-09 | 2.900E-09 | 1.732E-09 | 1.900E-09 | 4.900E-09 |
| 7360 | 368 | A368V | A | 1.4e-08 | 5.3e-09 | 1e-08 | 9.767E-09 | 4.355E-09 | 5.300E-09 | 1.400E-08 |
| 7361 | 369 | P369A | A | 5.9e-09 | 1.1e-08 | 1.9e-09 | 6.267E-09 | 4.561E-09 | 1.900E-09 | 1.100E-08 |
| 7362 | 369 | P369R | A | 6.4e-09 | 9.2e-09 | 8.1e-09 | 7.900E-09 | 1.411E-09 | 6.400E-09 | 9.200E-09 |
| 7363 | 369 | P369N | A | 7.2e-09 | 6.5e-09 | 8.9e-09 | 7.533E-09 | 1.234E-09 | 6.500E-09 | 8.900E-09 |
| 7364 | 369 | P369D | A | 2.7e-08 | 6.5e-09 | 9.8e-09 | 1.443E-08 | 1.101E-08 | 6.500E-09 | 2.700E-08 |
| 7365 | 369 | P369C | A | 5.3e-09 | 1.6e-08 | 4.7e-09 | 8.667E-09 | 6.358E-09 | 4.700E-09 | 1.600E-08 |
| 7366 | 369 | P369E | A | 9.7e-09 | 1.7e-08 | 1.9e-08 | 1.523E-08 | 4.895E-09 | 9.700E-09 | 1.900E-08 |
| 7367 | 369 | P369Q | A | 6.7e-09 | 3.3e-09 | 7.2e-09 | 5.733E-09 | 2.122E-09 | 3.300E-09 | 7.200E-09 |
| 7368 | 369 | P369G | A | 9.8e-09 | 9.9e-09 | 1.2e-08 | 1.057E-08 | 1.242E-09 | 9.800E-09 | 1.200E-08 |
| 7369 | 369 | P369H | A | 5e-09 | 9.7e-09 | 3.6e-09 | 6.100E-09 | 3.195E-09 | 3.600E-09 | 9.700E-09 |
| 7370 | 369 | P369I | A | 1.7e-08 | 1.2e-08 | 1e-08 | 1.300E-08 | 3.606E-09 | 1.000E-08 | 1.700E-08 |
| 7371 | 369 | P369L | A | 7.7e-09 | 7.3e-09 | 9e-09 | 8.000E-09 | 8.888E-10 | 7.300E-09 | 9.000E-09 |
| 7372 | 369 | P369K | A | 1e-08 | 5.9e-09 | 4.1e-09 | 6.667E-09 | 3.024E-09 | 4.100E-09 | 1.000E-08 |
| 7373 | 369 | P369M | A | 1.6e-08 | 5.8e-09 | 5.9e-09 | 9.233E-09 | 5.860E-09 | 5.800E-09 | 1.600E-08 |
| 7374 | 369 | P369F | A | 7.8e-09 | 1.5e-08 | 1.1e-08 | 1.127E-08 | 3.607E-09 | 7.800E-09 | 1.500E-08 |
| 7375 | 369 | P369P | A | 4.9e-09 | 9.8e-09 | 8.1e-09 | 7.600E-09 | 2.488E-09 | 4.900E-09 | 9.800E-09 |
| 7376 | 369 | P369S | A | 3.1e-09 | 1e-08 | 5.1e-09 | 6.067E-09 | 3.550E-09 | 3.100E-09 | 1.000E-08 |
| 7377 | 369 | P369T | A | 1e-08 | 8.9e-09 | 1.6e-08 | 1.163E-08 | 3.821E-09 | 8.900E-09 | 1.600E-08 |
| 7378 | 369 | P369W | A | 9.4e-09 | 2.5e-09 | 9.7e-09 | 7.200E-09 | 4.073E-09 | 2.500E-09 | 9.700E-09 |
| 7379 | 369 | P369Y | A | 7.8e-09 | 3.4e-09 | 8.5e-09 | 6.567E-09 | 2.765E-09 | 3.400E-09 | 8.500E-09 |
| 7380 | 369 | P369V | A | 2.1e-08 | 9.4e-09 | 9.2e-09 | 1.320E-08 | 6.756E-09 | 9.200E-09 | 2.100E-08 |
| 7381 | 370 | L370A | A | 6.2e-09 | 5.2e-09 | 1.2e-08 | 7.800E-09 | 3.672E-09 | 5.200E-09 | 1.200E-08 |
| 7382 | 370 | L370R | A | 8.9e-09 | 1e-08 | 1.2e-08 | 1.030E-08 | 1.572E-09 | 8.900E-09 | 1.200E-08 |
| 7383 | 370 | L370N | A | 8.5e-09 | 3.9e-09 | 7.6e-09 | 6.667E-09 | 2.438E-09 | 3.900E-09 | 8.500E-09 |
| 7384 | 370 | L370D | A | 2.6e-09 | 5.1e-09 | 6.7e-09 | 4.800E-09 | 2.066E-09 | 2.600E-09 | 6.700E-09 |
| 7385 | 370 | L370C | A | 1.3e-08 | 7.5e-09 | 1.1e-08 | 1.050E-08 | 2.784E-09 | 7.500E-09 | 1.300E-08 |
| 7386 | 370 | L370E | A | 1.2e-08 | 4.3e-09 | 5.3e-09 | 7.200E-09 | 4.187E-09 | 4.300E-09 | 1.200E-08 |
| 7387 | 370 | L370Q | A | 6.8e-09 | 3.5e-09 | 5.5e-09 | 5.267E-09 | 1.662E-09 | 3.500E-09 | 6.800E-09 |
| 7388 | 370 | L370G | A | 8.1e-09 | 7.2e-09 | 7e-09 | 7.433E-09 | 5.859E-10 | 7.000E-09 | 8.100E-09 |
| 7389 | 370 | L370H | A | 3.4e-09 | 1.1e-08 | 7.6e-09 | 7.333E-09 | 3.807E-09 | 3.400E-09 | 1.100E-08 |
| 7390 | 370 | L370I | A | 4.9e-09 | 5.1e-09 | 7e-09 | 5.667E-09 | 1.159E-09 | 4.900E-09 | 7.000E-09 |
| 7391 | 370 | L370L | A | 4.5e-08 | 8.3e-09 | 7.2e-09 | 2.017E-08 | 2.151E-08 | 7.200E-09 | 4.500E-08 |
| 7392 | 370 | L370K | A | 4.5e-09 | 5.3e-09 | 3.2e-09 | 4.333E-09 | 1.060E-09 | 3.200E-09 | 5.300E-09 |
| 7393 | 370 | L370M | A | 9.9e-09 | 6.9e-09 | 3.5e-09 | 6.767E-09 | 3.202E-09 | 3.500E-09 | 9.900E-09 |
| 7394 | 370 | L370F | A | 6.1e-09 | 8.7e-09 | 4.3e-09 | 6.367E-09 | 2.212E-09 | 4.300E-09 | 8.700E-09 |

| | | | | | | | | | | |
|---|---|---|---|---|---|---|---|---|---|---|
| 7395 | 370 | L370P | A | 2.2e-08 | 1.2e-08 | 3e-09 | 1.233E-08 | 9.504E-09 | 3.000E-09 | 2.200E-08 |
| 7396 | 370 | L370S | A | 6.2e-09 | 4.1e-09 | 5.2e-09 | 5.167E-09 | 1.050E-09 | 4.100E-09 | 6.200E-09 |
| 7397 | 370 | L370T | A | 5.3e-09 | 1.6e-08 | 3.9e-09 | 8.400E-09 | 6.619E-09 | 3.900E-09 | 1.600E-08 |
| 7398 | 370 | L370W | A | 5.3e-09 | 4.2e-09 | 4.1e-09 | 4.533E-09 | 6.658E-10 | 4.100E-09 | 5.300E-09 |
| 7399 | 370 | L370Y | A | 1.7e-08 | 7e-09 | 2.8e-09 | 8.933E-09 | 7.295E-09 | 2.800E-09 | 1.700E-08 |
| 7400 | 370 | L370V | A | 8.5e-09 | 1.1e-08 | 1.7e-08 | 1.217E-08 | 4.368E-09 | 8.500E-09 | 1.700E-08 |
| 7401 | 371 | Q371A | A | 2.4e-08 | 1.1e-08 | 2.2e-09 | 1.240E-08 | 1.097E-08 | 2.200E-09 | 2.400E-08 |
| 7402 | 371 | Q371R | A | 6.7e-09 | 7.8e-09 | 1.4e-08 | 9.500E-09 | 3.936E-09 | 6.700E-09 | 1.400E-08 |
| 7403 | 371 | Q371N | A | 6.9e-09 | 8e-09 | 7.5e-09 | 7.467E-09 | 5.508E-10 | 6.900E-09 | 8.000E-09 |
| 7404 | 371 | Q371D | A | 6.6e-09 | 4.9e-09 | 5.7e-09 | 5.733E-09 | 8.505E-10 | 4.900E-09 | 6.600E-09 |
| 7405 | 371 | Q371C | A | 8.4e-09 | 1e-09 | 2.1e-08 | 1.013E-08 | 1.011E-08 | 1.000E-09 | 2.100E-08 |
| 7406 | 371 | Q371E | A | 5.9e-09 | 9.4e-09 | 1.3e-08 | 9.433E-09 | 3.550E-09 | 5.900E-09 | 1.300E-08 |
| 7407 | 371 | Q371Q | A | 1.1e-08 | 1.1e-08 | 3.3e-08 | 1.833E-08 | 1.270E-08 | 1.100E-08 | 3.300E-08 |
| 7408 | 371 | Q371G | A | 7.6e-09 | 2.4e-09 | 6.2e-09 | 5.400E-09 | 2.691E-09 | 2.400E-09 | 7.600E-09 |
| 7409 | 371 | Q371H | A | 9.9e-09 | 2.4e-08 | 2.2e-09 | 1.203E-08 | 1.106E-08 | 2.200E-09 | 2.400E-08 |
| 7410 | 371 | Q371I | A | 6e-09 | 1.7e-08 | 1.5e-08 | 1.267E-08 | 5.859E-09 | 6.000E-09 | 1.700E-08 |
| 7411 | 371 | Q371L | A | 4.3e-09 | 9.1e-09 | 6.3e-09 | 6.567E-09 | 2.411E-09 | 4.300E-09 | 9.100E-09 |
| 7412 | 371 | Q371K | A | 8.8e-09 | 1.2e-08 | 9.2e-09 | 1.000E-08 | 1.744E-09 | 8.800E-09 | 1.200E-08 |
| 7413 | 371 | Q371M | A | 9.8e-09 | 6.2e-09 | 4.3e-09 | 6.767E-09 | 2.793E-09 | 4.300E-09 | 9.800E-09 |
| 7414 | 371 | Q371F | A | 3.8e-09 | 1.4e-08 | 3.5e-09 | 7.100E-09 | 5.977E-09 | 3.500E-09 | 1.400E-08 |
| 7415 | 371 | Q371P | A | 6.2e-09 | 3.2e-09 | 1.4e-08 | 7.800E-09 | 5.575E-09 | 3.200E-09 | 1.400E-08 |
| 7416 | 371 | Q371S | A | 8.8e-09 | 2.1e-08 | 9.7e-09 | 1.317E-08 | 6.799E-09 | 8.800E-09 | 2.100E-08 |
| 7417 | 371 | Q371T | A | 3.2e-09 | 6.7e-09 | 4.7e-09 | 4.867E-09 | 1.756E-09 | 3.200E-09 | 6.700E-09 |
| 7418 | 371 | Q371W | A | 5.2e-09 | 8.7e-09 | 1.1e-08 | 8.300E-09 | 2.921E-09 | 5.200E-09 | 1.100E-08 |
| 7419 | 371 | Q371Y | A | 7.3e-09 | 5.3e-09 | 7.7e-09 | 6.767E-09 | 1.286E-09 | 5.300E-09 | 7.700E-09 |
| 7420 | 371 | Q371V | A | 1.7e-09 | 6.1e-09 | 1.1e-08 | 6.267E-09 | 4.652E-09 | 1.700E-09 | 1.100E-08 |
| 7421 | 372 | P372A | A | 2e-08 | 1.6e-08 | 6.1e-09 | 1.403E-08 | 7.156E-09 | 6.100E-09 | 2.000E-08 |
| 7422 | 372 | P372R | A | 3.3e-09 | 1.6e-08 | 8.3e-09 | 9.200E-09 | 6.398E-09 | 3.300E-09 | 1.600E-08 |
| 7423 | 372 | P372N | A | 4.4e-09 | 3.1e-09 | 6.3e-09 | 4.600E-09 | 1.609E-09 | 3.100E-09 | 6.300E-09 |
| 7424 | 372 | P372D | A | 5.7e-09 | 2.6e-09 | 6.6e-09 | 4.967E-09 | 2.098E-09 | 2.600E-09 | 6.600E-09 |
| 7425 | 372 | P372C | A | 1.1e-08 | 9.7e-09 | 6.6e-09 | 9.100E-09 | 2.261E-09 | 6.600E-09 | 1.100E-08 |
| 7426 | 372 | P372E | A | 6.4e-09 | 7.3e-09 | 1.2e-08 | 8.567E-09 | 3.007E-09 | 6.400E-09 | 1.200E-08 |
| 7427 | 372 | P372Q | A | 6.2e-09 | 7.1e-09 | 1.1e-08 | 8.100E-09 | 2.551E-09 | 6.200E-09 | 1.100E-08 |
| 7428 | 372 | P372G | A | 3.6e-09 | 1.4e-08 | 6e-09 | 7.867E-09 | 5.445E-09 | 3.600E-09 | 1.400E-08 |
| 7429 | 372 | P372H | A | 1.9e-09 | 1.3e-08 | 1e-08 | 8.300E-09 | 5.742E-09 | 1.900E-09 | 1.300E-08 |
| 7430 | 372 | P372I | A | 3e-09 | 6.6e-09 | 6.5e-09 | 5.367E-09 | 2.050E-09 | 3.000E-09 | 6.600E-09 |
| 7431 | 372 | P372L | A | 4.7e-09 | 1.7e-08 | 1.1e-08 | 1.090E-08 | 6.151E-09 | 4.700E-09 | 1.700E-08 |
| 7432 | 372 | P372K | A | 1.1e-08 | 4.8e-09 | 1.5e-08 | 1.027E-08 | 5.139E-09 | 4.800E-09 | 1.500E-08 |
| 7433 | 372 | P372M | A | 2.4e-09 | 1.2e-08 | 3.9e-09 | 6.100E-09 | 5.164E-09 | 2.400E-09 | 1.200E-08 |
| 7434 | 372 | P372F | A | 7.5e-09 | 8.7e-09 | 4.6e-09 | 6.933E-09 | 2.108E-09 | 4.600E-09 | 8.700E-09 |
| 7435 | 372 | P372P | A | 1.4e-08 | 1.5e-08 | 3.7e-09 | 1.090E-08 | 6.255E-09 | 3.700E-09 | 1.500E-08 |
| 7436 | 372 | P372S | A | 3.6e-09 | 1.4e-08 | 1.2e-08 | 9.867E-09 | 5.518E-09 | 3.600E-09 | 1.400E-08 |
| 7437 | 372 | P372T | A | 3.2e-09 | 2.1e-09 | 6.8e-09 | 4.033E-09 | 2.458E-09 | 2.100E-09 | 6.800E-09 |
| 7438 | 372 | P372W | A | 4.6e-09 | 4.4e-09 | 1e-08 | 6.333E-09 | 3.177E-09 | 4.400E-09 | 1.000E-08 |
| 7439 | 372 | P372Y | A | 2.7e-08 | 1.1e-08 | 4.1e-09 | 1.403E-08 | 1.175E-08 | 4.100E-09 | 2.700E-08 |
| 7440 | 372 | P372V | A | 3.2e-09 | 2.2e-08 | 1.2e-08 | 1.240E-08 | 9.406E-09 | 3.200E-09 | 2.200E-08 |
| 7441 | 373 | E373A | A | 1.5e-08 | 9.1e-09 | 7.6e-09 | 1.057E-08 | 3.912E-09 | 7.600E-09 | 1.500E-08 |
| 7442 | 373 | E373R | A | 4.2e-09 | 4.2e-09 | 7.2e-09 | 5.200E-09 | 1.732E-09 | 4.200E-09 | 7.200E-09 |
| 7443 | 373 | E373N | A | 1.4e-08 | 2e-08 | 4.9e-09 | 1.297E-08 | 7.603E-09 | 4.900E-09 | 2.000E-08 |
| 7444 | 373 | E373D | A | 8.2e-09 | 2.5e-09 | 1.5e-08 | 8.567E-09 | 6.258E-09 | 2.500E-09 | 1.500E-08 |
| 7445 | 373 | E373C | A | 1.6e-08 | 9e-09 | 6.2e-09 | 1.040E-08 | 5.048E-09 | 6.200E-09 | 1.600E-08 |
| 7446 | 373 | E373E | A | 1.2e-08 | 1e-08 | 7.5e-09 | 9.833E-09 | 2.255E-09 | 7.500E-09 | 1.200E-08 |
| 7447 | 373 | E373Q | A | 1.2e-08 | 4.8e-09 | 1.1e-08 | 9.267E-09 | 3.900E-09 | 4.800E-09 | 1.200E-08 |
| 7448 | 373 | E373G | A | 1.5e-08 | 5.1e-09 | 1e-08 | 1.003E-08 | 4.950E-09 | 5.100E-09 | 1.500E-08 |
| 7449 | 373 | E373H | A | 2.7e-09 | 1.4e-08 | 2e-08 | 1.223E-08 | 8.784E-09 | 2.700E-09 | 2.000E-08 |
| 7450 | 373 | E373I | A | 4.5e-09 | 3.1e-08 | 4e-09 | 1.317E-08 | 1.545E-08 | 4.000E-09 | 3.100E-08 |
| 7451 | 373 | E373L | A | 1.2e-08 | 5.4e-09 | 3.9e-09 | 7.100E-09 | 4.309E-09 | 3.900E-09 | 1.200E-08 |
| 7452 | 373 | E373K | A | 8.3e-09 | 7.7e-09 | 2.4e-08 | 1.333E-08 | 9.242E-09 | 7.700E-09 | 2.400E-08 |
| 7453 | 373 | E373M | A | 3.1e-09 | 9e-09 | 4.4e-09 | 5.500E-09 | 3.100E-09 | 3.100E-09 | 9.000E-09 |
| 7454 | 373 | E373F | A | 2.1e-08 | 5.7e-09 | 3.4e-09 | 1.003E-08 | 9.567E-09 | 3.400E-09 | 2.100E-08 |
| 7455 | 373 | E373P | A | 2.4e-09 | 1.1e-08 | 5.2e-09 | 6.200E-09 | 4.386E-09 | 2.400E-09 | 1.100E-08 |
| 7456 | 373 | E373S | A | 8.9e-09 | 7.9e-09 | 8.6e-09 | 8.467E-09 | 5.132E-10 | 7.900E-09 | 8.900E-09 |
| 7457 | 373 | E373T | A | 5.8e-09 | 1.7e-08 | 3.2e-09 | 8.667E-09 | 7.333E-09 | 3.200E-09 | 1.700E-08 |
| 7458 | 373 | E373W | A | 9.7e-09 | 5.5e-09 | 8.1e-09 | 7.767E-09 | 2.120E-09 | 5.500E-09 | 9.700E-09 |
| 7459 | 373 | E373Y | A | 1.3e-08 | 2.9e-09 | 6.5e-09 | 7.467E-09 | 5.119E-09 | 2.900E-09 | 1.300E-08 |
| 7460 | 373 | E373V | A | 8.2e-09 | 2.8e-09 | 1.4e-08 | 8.333E-09 | 5.601E-09 | 2.800E-09 | 1.400E-08 |
| 7461 | 374 | Q374A | A | 5.5e-09 | 8.2e-09 | 1.3e-08 | 8.900E-09 | 3.799E-09 | 5.500E-09 | 1.300E-08 |
| 7462 | 374 | Q374R | A | 6e-09 | 9.9e-09 | 2.9e-09 | 6.267E-09 | 3.508E-09 | 2.900E-09 | 9.900E-09 |
| 7463 | 374 | Q374N | A | 4.4e-09 | 6.9e-09 | 4.8e-09 | 5.367E-09 | 1.343E-09 | 4.400E-09 | 6.900E-09 |
| 7464 | 374 | Q374D | A | 1.3e-08 | 7.9e-09 | 8e-09 | 9.633E-09 | 2.916E-09 | 7.900E-09 | 1.300E-08 |
| 7465 | 374 | Q374C | A | 1.4e-08 | 3.5e-09 | 2.3e-08 | 1.350E-08 | 9.760E-09 | 3.500E-09 | 2.300E-08 |
| 7466 | 374 | Q374E | A | 8.4e-09 | 6.7e-09 | 1e-08 | 8.367E-09 | 1.650E-09 | 6.700E-09 | 1.000E-08 |
| 7467 | 374 | Q374Q | A | 8.8e-09 | 6e-09 | 7e-09 | 7.267E-09 | 1.419E-09 | 6.000E-09 | 8.800E-09 |
| 7468 | 374 | Q374G | A | 1.4e-08 | 7.5e-09 | 5.1e-09 | 8.867E-09 | 4.605E-09 | 5.100E-09 | 1.400E-08 |
| 7469 | 374 | Q374H | A | 1.3e-08 | 8e-09 | 9.8e-09 | 1.027E-08 | 2.532E-09 | 8.000E-09 | 1.300E-08 |
| 7470 | 374 | Q374I | A | 1.1e-08 | 1.3e-08 | 9.3e-09 | 1.110E-08 | 1.852E-09 | 9.300E-09 | 1.300E-08 |
| 7471 | 374 | Q374L | A | 8.5e-09 | 1.6e-09 | 6.1e-09 | 5.400E-09 | 3.503E-09 | 1.600E-09 | 8.500E-09 |
| 7472 | 374 | Q374K | A | 4.7e-09 | 3e-09 | 4e-09 | 3.900E-09 | 8.544E-10 | 3.000E-09 | 4.700E-09 |
| 7473 | 374 | Q374M | A | 8.9e-09 | 9.1e-09 | 9e-09 | 9.000E-09 | 1.000E-10 | 8.900E-09 | 9.100E-09 |
| 7474 | 374 | Q374F | A | 7.5e-09 | 4.1e-09 | 1.6e-08 | 9.200E-09 | 6.129E-09 | 4.100E-09 | 1.600E-08 |
| 7475 | 374 | Q374P | A | 5.3e-09 | 1.3e-08 | 1.1e-08 | 9.767E-09 | 3.995E-09 | 5.300E-09 | 1.300E-08 |
| 7476 | 374 | Q374S | A | 1e-08 | 1e-08 | 1.3e-08 | 1.100E-08 | 1.732E-09 | 1.000E-08 | 1.300E-08 |
| 7477 | 374 | Q374T | A | 6.5e-09 | 5e-09 | 5.1e-09 | 5.533E-09 | 8.386E-10 | 5.000E-09 | 6.500E-09 |
| 7478 | 374 | Q374W | A | 1.1e-08 | 3e-09 | 2.8e-09 | 5.600E-09 | 4.678E-09 | 2.800E-09 | 1.100E-08 |
| 7479 | 374 | Q374Y | A | 8.5e-09 | 4e-09 | 9.3e-09 | 7.267E-09 | 2.857E-09 | 4.000E-09 | 9.300E-09 |

| | | | | | | | | | | |
|---|---|---|---|---|---|---|---|---|---|---|
| 7480 | 374 | Q374V | A | 7e-09 | 1.4e-08 | 4.7e-09 | 8.567E-09 | 4.844E-09 | 4.700E-09 | 1.400E-08 |
| 7481 | 375 | L375A | A | 4.6e-09 | 6.8e-09 | 5.3e-09 | 5.567E-09 | 1.124E-09 | 4.600E-09 | 6.800E-09 |
| 7482 | 375 | L375R | A | 5e-09 | 9.6e-09 | 3.6e-09 | 6.067E-09 | 3.139E-09 | 3.600E-09 | 9.600E-09 |
| 7483 | 375 | L375N | A | 6.3e-09 | 7.7e-09 | 8.3e-09 | 7.433E-09 | 1.026E-09 | 6.300E-09 | 8.300E-09 |
| 7484 | 375 | L375D | A | 2.7e-09 | 4.9e-09 | 6.7e-09 | 4.767E-09 | 2.003E-09 | 2.700E-09 | 6.700E-09 |
| 7485 | 375 | L375C | A | 5.7e-09 | 1.4e-08 | 1.8e-08 | 1.257E-08 | 6.274E-09 | 5.700E-09 | 1.800E-08 |
| 7486 | 375 | L375E | A | 4.2e-09 | 6.2e-09 | 1.4e-08 | 8.133E-09 | 5.178E-09 | 4.200E-09 | 1.400E-08 |
| 7487 | 375 | L375Q | A | 1.4e-08 | 4.3e-09 | 6.2e-09 | 8.167E-09 | 5.140E-09 | 4.300E-09 | 1.400E-08 |
| 7488 | 375 | L375G | A | 2.3e-09 | 1.3e-08 | 1e-08 | 8.433E-09 | 5.519E-09 | 2.300E-09 | 1.300E-08 |
| 7489 | 375 | L375H | A | 5.8e-09 | 7.6e-09 | 2.3e-08 | 1.213E-08 | 9.454E-09 | 5.800E-09 | 2.300E-08 |
| 7490 | 375 | L375I | A | 5.7e-09 | 1.2e-08 | 1e-08 | 9.233E-09 | 3.219E-09 | 5.700E-09 | 1.200E-08 |
| 7491 | 375 | L375L | A | 6.9e-09 | 1.3e-08 | 7.1e-09 | 9.000E-09 | 3.466E-09 | 6.900E-09 | 1.300E-08 |
| 7492 | 375 | L375K | A | 4.3e-09 | 1.5e-08 | 7.9e-09 | 9.067E-09 | 5.445E-09 | 4.300E-09 | 1.500E-08 |
| 7493 | 375 | L375M | A | 2.4e-09 | 6.4e-09 | 9.7e-09 | 6.167E-09 | 3.656E-09 | 2.400E-09 | 9.700E-09 |
| 7494 | 375 | L375F | A | 1e-08 | 4e-09 | 6.9e-09 | 6.967E-09 | 3.001E-09 | 4.000E-09 | 1.000E-08 |
| 7495 | 375 | L375P | A | 1.3e-08 | 5.2e-09 | 1.1e-08 | 9.733E-09 | 4.051E-09 | 5.200E-09 | 1.300E-08 |
| 7496 | 375 | L375S | A | 7.9e-09 | 7.8e-09 | 3.1e-08 | 1.557E-08 | 1.337E-08 | 7.800E-09 | 3.100E-08 |
| 7497 | 375 | L375T | A | 9.9e-09 | 8e-09 | 5.3e-09 | 7.733E-09 | 2.312E-09 | 5.300E-09 | 9.900E-09 |
| 7498 | 375 | L375W | A | 3.4e-09 | 1.2e-08 | 6.5e-09 | 7.300E-09 | 4.355E-09 | 3.400E-09 | 1.200E-08 |
| 7499 | 375 | L375Y | A | 3.2e-09 | 6.6e-09 | 5.5e-09 | 5.100E-09 | 1.735E-09 | 3.200E-09 | 6.600E-09 |
| 7500 | 375 | L375V | A | 7.9e-09 | 1e-08 | 8.9e-09 | 8.933E-09 | 1.050E-09 | 7.900E-09 | 1.000E-08 |
| 7501 | 376 | Q376A | A | 2.2e-09 | 9.4e-09 | 4.3e-09 | 5.300E-09 | 3.703E-09 | 2.200E-09 | 9.400E-09 |
| 7502 | 376 | Q376R | A | 1e-08 | 1.1e-08 | 1.2e-08 | 1.100E-08 | 1.000E-09 | 1.000E-08 | 1.200E-08 |
| 7503 | 376 | Q376N | A | 1.7e-08 | 1e-08 | 3.8e-09 | 1.027E-08 | 6.604E-09 | 3.800E-09 | 1.700E-08 |
| 7504 | 376 | Q376D | A | 6e-09 | 8.3e-09 | 1.1e-08 | 8.433E-09 | 2.503E-09 | 6.000E-09 | 1.100E-08 |
| 7505 | 376 | Q376C | A | 9e-09 | 5.4e-09 | 1.1e-08 | 8.467E-09 | 2.838E-09 | 5.400E-09 | 1.100E-08 |
| 7506 | 376 | Q376E | A | 5.7e-09 | 3.2e-09 | 3.8e-09 | 4.233E-09 | 1.305E-09 | 3.200E-09 | 5.700E-09 |
| 7507 | 376 | Q376Q | A | 7.1e-09 | 2.2e-09 | 7.9e-09 | 5.733E-09 | 3.086E-09 | 2.200E-09 | 7.900E-09 |
| 7508 | 376 | Q376G | A | 1.6e-08 | 8.7e-09 | 4.1e-09 | 9.600E-09 | 6.001E-09 | 4.100E-09 | 1.600E-08 |
| 7509 | 376 | Q376H | A | 1.4e-08 | 4.1e-09 | 7.1e-09 | 8.400E-09 | 5.076E-09 | 4.100E-09 | 1.400E-08 |
| 7510 | 376 | Q376I | A | 3.8e-08 | 9.8e-09 | 5.8e-09 | 1.787E-08 | 1.755E-08 | 5.800E-09 | 3.800E-08 |
| 7511 | 376 | Q376L | A | 5e-09 | 1.8e-08 | 1.7e-08 | 1.333E-08 | 7.234E-09 | 5.000E-09 | 1.800E-08 |
| 7512 | 376 | Q376K | A | 8.7e-09 | 8.3e-09 | 6.4e-09 | 7.800E-09 | 1.229E-09 | 6.400E-09 | 8.700E-09 |
| 7513 | 376 | Q376M | A | 1.2e-08 | 9e-09 | 3.8e-09 | 8.267E-09 | 4.149E-09 | 3.800E-09 | 1.200E-08 |
| 7514 | 376 | Q376F | A | 5.5e-09 | 7.7e-09 | 4.8e-09 | 6.000E-09 | 1.513E-09 | 4.800E-09 | 7.700E-09 |
| 7515 | 376 | Q376P | A | 7.2e-09 | 7.2e-09 | 7e-09 | 7.133E-09 | 1.155E-10 | 7.000E-09 | 7.200E-09 |
| 7516 | 376 | Q376S | A | 6.5e-09 | 1.2e-08 | 5.9e-09 | 8.133E-09 | 3.362E-09 | 5.900E-09 | 1.200E-08 |
| 7517 | 376 | Q376T | A | 9.1e-09 | 4.2e-09 | 2.8e-09 | 5.367E-09 | 3.308E-09 | 2.800E-09 | 9.100E-09 |
| 7518 | 376 | Q376W | A | 7e-09 | 2.8e-09 | 5.2e-09 | 5.000E-09 | 2.107E-09 | 2.800E-09 | 7.000E-09 |
| 7519 | 376 | Q376Y | A | 1.1e-08 | 1.8e-09 | 2.6e-09 | 5.133E-09 | 5.096E-09 | 1.800E-09 | 1.100E-08 |
| 7520 | 376 | Q376V | A | 1.5e-08 | 9.6e-09 | 3.1e-08 | 1.853E-08 | 1.113E-08 | 9.600E-09 | 3.100E-08 |
| 7521 | 377 | V377A | A | 3.1e-09 | 8.5e-09 | 6.6e-09 | 6.067E-09 | 2.739E-09 | 3.100E-09 | 8.500E-09 |
| 7522 | 377 | V377R | A | 3.3e-09 | 6.8e-09 | 5.4e-09 | 5.167E-09 | 1.762E-09 | 3.300E-09 | 6.800E-09 |
| 7523 | 377 | V377N | A | 6.2e-09 | 4.4e-09 | 4.2e-09 | 4.933E-09 | 1.102E-09 | 4.200E-09 | 6.200E-09 |
| 7524 | 377 | V377D | A | 4e-09 | 1.4e-08 | 1.2e-08 | 1.000E-08 | 5.292E-09 | 4.000E-09 | 1.400E-08 |
| 7525 | 377 | V377C | A | 1.3e-08 | 5.1e-09 | 5.9e-09 | 8.000E-09 | 4.349E-09 | 5.100E-09 | 1.300E-08 |
| 7526 | 377 | V377E | A | 2.1e-08 | 5.8e-09 | 1.1e-08 | 1.260E-08 | 7.725E-09 | 5.800E-09 | 2.100E-08 |
| 7527 | 377 | V377Q | A | 3.6e-09 | 2.8e-09 | 8.4e-09 | 4.933E-09 | 3.029E-09 | 2.800E-09 | 8.400E-09 |
| 7528 | 377 | V377G | A | 1.2e-08 | 4.7e-09 | 1.7e-09 | 6.133E-09 | 5.297E-09 | 1.700E-09 | 1.200E-08 |
| 7529 | 377 | V377H | A | 8.5e-09 | 4.2e-09 | 1.3e-08 | 8.567E-09 | 4.400E-09 | 4.200E-09 | 1.300E-08 |
| 7530 | 377 | V377I | A | 4.8e-09 | 1.3e-08 | 3.9e-09 | 7.233E-09 | 5.014E-09 | 3.900E-09 | 1.300E-08 |
| 7531 | 377 | V377L | A | 8.8e-09 | 1.2e-08 | 3.3e-09 | 8.033E-09 | 4.400E-09 | 3.300E-09 | 1.200E-08 |
| 7532 | 377 | V377K | A | 2.7e-08 | 4.9e-09 | 1.3e-08 | 1.497E-08 | 1.118E-08 | 4.900E-09 | 2.700E-08 |
| 7533 | 377 | V377M | A | 2.6e-09 | 5.4e-09 | 9.7e-09 | 5.900E-09 | 3.576E-09 | 2.600E-09 | 9.700E-09 |
| 7534 | 377 | V377F | A | 4.3e-09 | 2.3e-09 | 2.7e-09 | 3.100E-09 | 1.058E-09 | 2.300E-09 | 4.300E-09 |
| 7535 | 377 | V377P | A | 8.5e-09 | 1.5e-08 | 6e-09 | 9.833E-09 | 4.646E-09 | 6.000E-09 | 1.500E-08 |
| 7536 | 377 | V377S | A | 9.2e-09 | 5.4e-09 | 1.7e-08 | 1.053E-08 | 5.914E-09 | 5.400E-09 | 1.700E-08 |
| 7537 | 377 | V377T | A | 5.6e-09 | 2.5e-09 | 6.6e-09 | 4.900E-09 | 2.138E-09 | 2.500E-09 | 6.600E-09 |
| 7538 | 377 | V377W | A | 8.1e-09 | 1.3e-09 | 1.2e-08 | 7.133E-09 | 5.415E-09 | 1.300E-09 | 1.200E-08 |
| 7539 | 377 | V377Y | A | 9.6e-09 | 1e-08 | 5.2e-09 | 8.267E-09 | 2.663E-09 | 5.200E-09 | 1.000E-08 |
| 7540 | 377 | V377V | A | 1.5e-08 | 1.2e-08 | 8.2e-09 | 1.173E-08 | 3.408E-09 | 8.200E-09 | 1.500E-08 |
| 7541 | 378 | F378A | A | 1.3e-08 | 1.7e-08 | 3.4e-09 | 1.113E-08 | 6.990E-09 | 3.400E-09 | 1.700E-08 |
| 7542 | 378 | F378R | A | 4.7e-09 | 4.2e-09 | 2.1e-09 | 3.667E-09 | 1.380E-09 | 2.100E-09 | 4.700E-09 |
| 7543 | 378 | F378N | A | 1.6e-08 | 2.2e-09 | 8.5e-09 | 8.900E-09 | 6.909E-09 | 2.200E-09 | 1.600E-08 |
| 7544 | 378 | F378D | A | 7.9e-09 | 1.8e-08 | 1.1e-08 | 1.230E-08 | 5.174E-09 | 7.900E-09 | 1.800E-08 |
| 7545 | 378 | F378C | A | 3.3e-09 | 4.7e-09 | 1e-08 | 6.000E-09 | 3.534E-09 | 3.300E-09 | 1.000E-08 |
| 7546 | 378 | F378E | A | 3.1e-09 | 9.7e-09 | 6.3e-09 | 6.367E-09 | 3.301E-09 | 3.100E-09 | 9.700E-09 |
| 7547 | 378 | F378Q | A | 5.4e-09 | 7.2e-09 | 9.7e-09 | 7.433E-09 | 2.159E-09 | 5.400E-09 | 9.700E-09 |
| 7548 | 378 | F378G | A | 1.5e-08 | 6.4e-09 | 1.3e-08 | 1.147E-08 | 4.500E-09 | 6.400E-09 | 1.500E-08 |
| 7549 | 378 | F378H | A | 9.2e-09 | 8.3e-09 | 8.5e-09 | 8.667E-09 | 4.726E-10 | 8.300E-09 | 9.200E-09 |
| 7550 | 378 | F378I | A | 3.3e-09 | 3.1e-09 | 2e-08 | 8.800E-09 | 9.700E-09 | 3.100E-09 | 2.000E-08 |
| 7551 | 378 | F378L | A | 2e-08 | 6.1e-09 | 9.4e-09 | 1.183E-08 | 7.262E-09 | 6.100E-09 | 2.000E-08 |
| 7552 | 378 | F378K | A | 2.1e-08 | 7.2e-09 | 8e-09 | 1.207E-08 | 7.747E-09 | 7.200E-09 | 2.100E-08 |
| 7553 | 378 | F378M | A | 4.3e-09 | 3.7e-09 | 1.4e-08 | 7.333E-09 | 5.781E-09 | 3.700E-09 | 1.400E-08 |
| 7554 | 378 | F378F | A | 1.8e-09 | 8.3e-09 | 7.5e-09 | 5.867E-09 | 3.544E-09 | 1.800E-09 | 8.300E-09 |
| 7555 | 378 | F378P | A | 3.7e-09 | 8.7e-09 | 6.9e-09 | 6.433E-09 | 2.532E-09 | 3.700E-09 | 8.700E-09 |
| 7556 | 378 | F378S | A | 6.9e-09 | 5.8e-09 | 1.5e-08 | 9.233E-09 | 5.024E-09 | 5.800E-09 | 1.500E-08 |
| 7557 | 378 | F378T | A | 6e-09 | 1.2e-08 | 7.6e-09 | 8.533E-09 | 3.107E-09 | 6.000E-09 | 1.200E-08 |
| 7558 | 378 | F378W | A | 3e-09 | 5.9e-09 | 3e-09 | 3.967E-09 | 1.674E-09 | 3.000E-09 | 5.900E-09 |
| 7559 | 378 | F378Y | A | 1.3e-09 | 7.7e-09 | 1.3e-08 | 7.333E-09 | 5.859E-09 | 1.300E-09 | 1.300E-08 |
| 7560 | 378 | F378V | A | 1.7e-08 | 9.3e-09 | 6.7e-09 | 1.100E-08 | 5.356E-09 | 6.700E-09 | 1.700E-08 |
| 7561 | 379 | E379A | A | 5e-09 | 1.2e-08 | 2.7e-09 | 6.567E-09 | 4.844E-09 | 2.700E-09 | 1.200E-08 |
| 7562 | 379 | E379R | A | 5e-09 | 5.1e-09 | 2.9e-09 | 4.333E-09 | 1.242E-09 | 2.900E-09 | 5.100E-09 |
| 7563 | 379 | E379N | A | 1.1e-09 | 5.9e-09 | 3.4e-08 | 1.367E-08 | 1.777E-08 | 1.100E-09 | 3.400E-08 |
| 7564 | 379 | E379D | A | 6.4e-09 | 1.1e-08 | 1.8e-08 | 1.180E-08 | 5.841E-09 | 6.400E-09 | 1.800E-08 |

| | | | | | | | | | | |
|---|---|---|---|---|---|---|---|---|---|---|
| 7565 | 379 | E379C | A | 5.2e-09 | 6.7e-09 | 1.9e-08 | 1.030E-08 | 7.572E-09 | 5.200E-09 | 1.900E-08 |
| 7566 | 379 | E379E | A | 1.3e-08 | 2.6e-09 | 6.2e-09 | 7.267E-09 | 5.281E-09 | 2.600E-09 | 1.300E-08 |
| 7567 | 379 | E379Q | A | 4.5e-09 | 1.2e-08 | 1.1e-08 | 9.167E-09 | 4.072E-09 | 4.500E-09 | 1.200E-08 |
| 7568 | 379 | E379G | A | 5e-09 | 5.4e-09 | 9.5e-09 | 6.633E-09 | 2.491E-09 | 5.000E-09 | 9.500E-09 |
| 7569 | 379 | E379H | A | 1.4e-08 | 5.5e-09 | 1.2e-08 | 1.050E-08 | 4.444E-09 | 5.500E-09 | 1.400E-08 |
| 7570 | 379 | E379I | A | 6e-09 | 6.8e-09 | 1.4e-08 | 8.933E-09 | 4.406E-09 | 6.000E-09 | 1.400E-08 |
| 7571 | 379 | E379L | A | 8.6e-09 | 5.2e-09 | 4.4e-09 | 6.067E-09 | 2.230E-09 | 4.400E-09 | 8.600E-09 |
| 7572 | 379 | E379K | A | 6e-09 | 1e-08 | 5e-09 | 7.000E-09 | 2.646E-09 | 5.000E-09 | 1.000E-08 |
| 7573 | 379 | E379M | A | 8.8e-09 | 6.7e-09 | 9.1e-09 | 8.200E-09 | 1.308E-09 | 6.700E-09 | 9.100E-09 |
| 7574 | 379 | E379F | A | 3.8e-09 | 6.1e-09 | 8.6e-09 | 6.167E-09 | 2.401E-09 | 3.800E-09 | 8.600E-09 |
| 7575 | 379 | E379P | A | 5.7e-09 | 6.3e-09 | 1.5e-08 | 9.000E-09 | 5.205E-09 | 5.700E-09 | 1.500E-08 |
| 7576 | 379 | E379S | A | 8.3e-09 | 1.2e-08 | 6.3e-09 | 8.867E-09 | 2.892E-09 | 6.300E-09 | 1.200E-08 |
| 7577 | 379 | E379T | A | 7.9e-09 | 1.1e-08 | 9.4e-09 | 9.433E-09 | 1.550E-09 | 7.900E-09 | 1.100E-08 |
| 7578 | 379 | E379W | A | 7.6e-09 | 2.9e-08 | 6.6e-09 | 1.440E-08 | 1.265E-08 | 6.600E-09 | 2.900E-08 |
| 7579 | 379 | E379Y | A | 3.6e-08 | 8.2e-09 | 5.6e-09 | 1.660E-08 | 1.685E-08 | 5.600E-09 | 3.600E-08 |
| 7580 | 379 | E379V | A | 8.4e-09 | 1.1e-08 | 1.3e-08 | 1.080E-08 | 2.307E-09 | 8.400E-09 | 1.300E-08 |
| 7581 | 380 | T380A | A | 4.3e-08 | 1.2e-08 | 1.2e-08 | 2.233E-08 | 1.790E-08 | 1.200E-08 | 4.300E-08 |
| 7582 | 380 | T380R | A | 7.3e-09 | 6.2e-09 | 2.2e-08 | 1.183E-08 | 8.822E-09 | 6.200E-09 | 2.200E-08 |
| 7583 | 380 | T380N | A | 6.3e-09 | 7.8e-09 | 8.4e-09 | 7.500E-09 | 1.082E-09 | 6.300E-09 | 8.400E-09 |
| 7584 | 380 | T380D | A | 3.7e-09 | 1.8e-08 | 6.1e-09 | 9.267E-09 | 7.658E-09 | 3.700E-09 | 1.800E-08 |
| 7585 | 380 | T380C | A | 7.5e-09 | 3.1e-09 | 9.6e-09 | 6.733E-09 | 3.317E-09 | 3.100E-09 | 9.600E-09 |
| 7586 | 380 | T380E | A | 3.6e-09 | 6.5e-09 | 4.1e-09 | 4.733E-09 | 1.550E-09 | 3.600E-09 | 6.500E-09 |
| 7587 | 380 | T380Q | A | 1.1e-08 | 8.2e-09 | 5.2e-09 | 8.133E-09 | 2.901E-09 | 5.200E-09 | 1.100E-08 |
| 7588 | 380 | T380G | A | 1.9e-09 | 1e-08 | 1.7e-08 | 9.633E-09 | 7.557E-09 | 1.900E-09 | 1.700E-08 |
| 7589 | 380 | T380H | A | 4.7e-09 | 1.5e-09 | 3.1e-09 | 3.100E-09 | 1.600E-09 | 1.500E-09 | 4.700E-09 |
| 7590 | 380 | T380I | A | 1.2e-08 | 8.8e-09 | 3.4e-09 | 8.067E-09 | 4.347E-09 | 3.400E-09 | 1.200E-08 |
| 7591 | 380 | T380L | A | 6.8e-09 | 9.6e-09 | 6.9e-09 | 7.767E-09 | 1.589E-09 | 6.800E-09 | 9.600E-09 |
| 7592 | 380 | T380K | A | 6.3e-09 | 6.9e-09 | 1e-09 | 4.733E-09 | 3.247E-09 | 1.000E-09 | 6.900E-09 |
| 7593 | 380 | T380M | A | 3.5e-09 | 8.9e-09 | 7e-09 | 6.467E-09 | 2.739E-09 | 3.500E-09 | 8.900E-09 |
| 7594 | 380 | T380F | A | 7.2e-09 | 1.2e-08 | 2.3e-08 | 1.407E-08 | 8.100E-09 | 7.200E-09 | 2.300E-08 |
| 7595 | 380 | T380P | A | 7.6e-09 | 7.9e-09 | 3.2e-09 | 6.233E-09 | 2.631E-09 | 3.200E-09 | 7.900E-09 |
| 7596 | 380 | T380S | A | 5.6e-09 | 9.7e-09 | 5.4e-09 | 6.900E-09 | 2.427E-09 | 5.400E-09 | 9.700E-09 |
| 7597 | 380 | T380T | A | 8.7e-09 | 1.1e-08 | 9.3e-09 | 9.667E-09 | 1.193E-09 | 8.700E-09 | 1.100E-08 |
| 7598 | 380 | T380W | A | 3.8e-09 | 2.1e-09 | 6.9e-09 | 4.267E-09 | 2.434E-09 | 2.100E-09 | 6.900E-09 |
| 7599 | 380 | T380Y | A | 9.9e-09 | 1.2e-08 | 1.1e-08 | 1.097E-08 | 1.050E-09 | 9.900E-09 | 1.200E-08 |
| 7600 | 380 | T380V | A | 1e-08 | 5.1e-09 | 6.4e-09 | 7.167E-09 | 2.538E-09 | 5.100E-09 | 1.000E-08 |
| 7601 | 381 | L381A | A | 7e-09 | 9.4e-09 | 6.1e-09 | 7.500E-09 | 1.706E-09 | 6.100E-09 | 9.400E-09 |
| 7602 | 381 | L381R | A | 2.9e-09 | 1.1e-08 | 3.9e-09 | 5.933E-09 | 4.416E-09 | 2.900E-09 | 1.100E-08 |
| 7603 | 381 | L381N | A | 6.6e-09 | 4.1e-09 | 6.1e-09 | 5.600E-09 | 1.323E-09 | 4.100E-09 | 6.600E-09 |
| 7604 | 381 | L381D | A | 6.3e-09 | 3.1e-09 | 6.6e-09 | 5.333E-09 | 1.940E-09 | 3.100E-09 | 6.600E-09 |
| 7605 | 381 | L381C | A | 2.6e-09 | 3.8e-09 | 1.5e-08 | 7.133E-09 | 6.839E-09 | 2.600E-09 | 1.500E-08 |
| 7606 | 381 | L381E | A | 6.6e-09 | 6e-09 | 4.8e-09 | 5.800E-09 | 9.165E-10 | 4.800E-09 | 6.600E-09 |
| 7607 | 381 | L381Q | A | 2.5e-09 | 6.5e-09 | 1.4e-08 | 7.667E-09 | 5.838E-09 | 2.500E-09 | 1.400E-08 |
| 7608 | 381 | L381G | A | 6.1e-09 | 7.5e-09 | 7e-09 | 6.867E-09 | 7.095E-10 | 6.100E-09 | 7.500E-09 |
| 7609 | 381 | L381H | A | 5e-09 | 2.1e-08 | 8.2e-09 | 1.140E-08 | 8.466E-09 | 5.000E-09 | 2.100E-08 |
| 7610 | 381 | L381I | A | 1.4e-08 | 7.6e-09 | 6.5e-09 | 9.367E-09 | 4.050E-09 | 6.500E-09 | 1.400E-08 |
| 7611 | 381 | L381L | A | 3.1e-09 | 1.3e-08 | 1e-08 | 8.700E-09 | 5.076E-09 | 3.100E-09 | 1.300E-08 |
| 7612 | 381 | L381K | A | 4.6e-09 | 4.9e-09 | 1.5e-08 | 8.167E-09 | 5.920E-09 | 4.600E-09 | 1.500E-08 |
| 7613 | 381 | L381M | A | 5.9e-09 | 1.6e-08 | 1.5e-08 | 1.230E-08 | 5.565E-09 | 5.900E-09 | 1.600E-08 |
| 7614 | 381 | L381F | A | 3.8e-09 | 9.5e-09 | 1.4e-08 | 9.100E-09 | 5.112E-09 | 3.800E-09 | 1.400E-08 |
| 7615 | 381 | L381P | A | 8.6e-09 | 1.1e-08 | 3.7e-09 | 7.767E-09 | 3.721E-09 | 3.700E-09 | 1.100E-08 |
| 7616 | 381 | L381S | A | 2.3e-08 | 9e-09 | 6.4e-09 | 1.280E-08 | 8.929E-09 | 6.400E-09 | 2.300E-08 |
| 7617 | 381 | L381T | A | 9.5e-09 | 5.1e-09 | 1.2e-08 | 8.867E-09 | 3.493E-09 | 5.100E-09 | 1.200E-08 |
| 7618 | 381 | L381W | A | 9.3e-09 | 6.2e-09 | 7.8e-09 | 7.767E-09 | 1.550E-09 | 6.200E-09 | 9.300E-09 |
| 7619 | 381 | L381Y | A | 1.1e-08 | 1.4e-08 | 1.8e-09 | 8.933E-09 | 6.357E-09 | 1.800E-09 | 1.400E-08 |
| 7620 | 381 | L381V | A | 1.9e-09 | 4.2e-09 | 9e-09 | 5.033E-09 | 3.623E-09 | 1.900E-09 | 9.000E-09 |
| 7621 | 382 | E382A | A | 1e-08 | 6e-09 | 3e-09 | 6.333E-09 | 3.512E-09 | 3.000E-09 | 1.000E-08 |
| 7622 | 382 | E382R | A | 4.7e-09 | 5.6e-09 | 5.1e-09 | 5.133E-09 | 4.509E-10 | 4.700E-09 | 5.600E-09 |
| 7623 | 382 | E382N | A | 2.2e-09 | 5.5e-09 | 4.7e-09 | 4.133E-09 | 1.721E-09 | 2.200E-09 | 5.500E-09 |
| 7624 | 382 | E382D | A | 1.6e-09 | 4.9e-09 | 4.9e-09 | 3.800E-09 | 1.905E-09 | 1.600E-09 | 4.900E-09 |
| 7625 | 382 | E382C | A | 3.8e-09 | 2.8e-09 | 8.2e-09 | 4.933E-09 | 2.873E-09 | 2.800E-09 | 8.200E-09 |
| 7626 | 382 | E382E | A | 9.2e-09 | 6.6e-09 | 2.3e-08 | 1.293E-08 | 8.814E-09 | 6.600E-09 | 2.300E-08 |
| 7627 | 382 | E382Q | A | 6.1e-09 | 2.8e-09 | 1.1e-08 | 6.633E-09 | 4.126E-09 | 2.800E-09 | 1.100E-08 |
| 7628 | 382 | E382G | A | 4.9e-09 | 7.6e-09 | 1.5e-08 | 9.167E-09 | 5.229E-09 | 4.900E-09 | 1.500E-08 |
| 7629 | 382 | E382H | A | 3.7e-09 | 5.3e-09 | 4.6e-09 | 4.533E-09 | 8.021E-10 | 3.700E-09 | 5.300E-09 |
| 7630 | 382 | E382I | A | 5.1e-09 | 6.1e-09 | 7.3e-09 | 6.167E-09 | 1.102E-09 | 5.100E-09 | 7.300E-09 |
| 7631 | 382 | E382L | A | 1e-08 | 6.3e-09 | 2.2e-09 | 6.167E-09 | 3.902E-09 | 2.200E-09 | 1.000E-08 |
| 7632 | 382 | E382K | A | 9.9e-09 | 2.9e-09 | 5.1e-09 | 5.967E-09 | 3.580E-09 | 2.900E-09 | 9.900E-09 |
| 7633 | 382 | E382M | A | 4.5e-09 | 2e-08 | 8.7e-09 | 1.107E-08 | 8.016E-09 | 4.500E-09 | 2.000E-08 |
| 7634 | 382 | E382F | A | 2.8e-08 | 9.7e-09 | 1.3e-08 | 1.690E-08 | 9.753E-09 | 9.700E-09 | 2.800E-08 |
| 7635 | 382 | E382P | A | 9.4e-09 | 8.6e-09 | 7.7e-09 | 8.567E-09 | 8.505E-10 | 7.700E-09 | 9.400E-09 |
| 7636 | 382 | E382S | A | 5.8e-09 | 1.6e-08 | 9.2e-09 | 1.033E-08 | 5.194E-09 | 5.800E-09 | 1.600E-08 |
| 7637 | 382 | E382T | A | 8.1e-09 | 4e-09 | 8.2e-09 | 6.767E-09 | 2.397E-09 | 4.000E-09 | 8.200E-09 |
| 7638 | 382 | E382W | A | 9.4e-09 | 2e-08 | 7.9e-09 | 1.243E-08 | 6.596E-09 | 7.900E-09 | 2.000E-08 |
| 7639 | 382 | E382Y | A | 1.8e-08 | 1.2e-08 | 1.8e-08 | 1.600E-08 | 3.464E-09 | 1.200E-08 | 1.800E-08 |
| 7640 | 382 | E382V | A | 7.3e-09 | 1e-08 | 5.9e-09 | 7.733E-09 | 2.084E-09 | 5.900E-09 | 1.000E-08 |
| 7641 | 383 | E383A | A | 5.2e-09 | 6.4e-09 | 1.1e-08 | 7.533E-09 | 3.062E-09 | 5.200E-09 | 1.100E-08 |
| 7642 | 383 | E383R | A | 1.1e-08 | 1.5e-08 | 3.6e-09 | 9.867E-09 | 5.784E-09 | 3.600E-09 | 1.500E-08 |
| 7643 | 383 | E383N | A | 6.1e-09 | 6.7e-09 | 3e-09 | 5.267E-09 | 1.986E-09 | 3.000E-09 | 6.700E-09 |
| 7644 | 383 | E383D | A | 3.6e-09 | 1.1e-08 | 1.4e-08 | 9.533E-09 | 5.353E-09 | 3.600E-09 | 1.400E-08 |
| 7645 | 383 | E383C | A | 1.2e-08 | 1.1e-08 | 9e-09 | 1.067E-08 | 1.528E-09 | 9.000E-09 | 1.200E-08 |
| 7646 | 383 | E383E | A | 4.9e-09 | 5.6e-09 | 8e-09 | 6.167E-09 | 1.626E-09 | 4.900E-09 | 8.000E-09 |
| 7647 | 383 | E383Q | A | 1.7e-08 | 6.5e-09 | 8.3e-09 | 1.060E-08 | 5.615E-09 | 6.500E-09 | 1.700E-08 |
| 7648 | 383 | E383G | A | 4.2e-08 | 5.7e-09 | 1e-08 | 1.923E-08 | 1.983E-08 | 5.700E-09 | 4.200E-08 |
| 7649 | 383 | E383H | A | 1.3e-08 | 6.7e-09 | 8.5e-09 | 9.400E-09 | 3.245E-09 | 6.700E-09 | 1.300E-08 |

| | | | | | | | | | | |
|---|---|---|---|---|---|---|---|---|---|---|
| 7650 | 383 | E383I | A | 3e-09 | 4.7e-09 | 6.5e-09 | 4.733E-09 | 1.750E-09 | 3.000E-09 | 6.500E-09 |
| 7651 | 383 | E383L | A | 7e-09 | 6.6e-09 | 2.2e-08 | 1.187E-08 | 8.778E-09 | 6.600E-09 | 2.200E-08 |
| 7652 | 383 | E383K | A | 3.9e-09 | 7.4e-09 | 7.4e-09 | 6.233E-09 | 2.021E-09 | 3.900E-09 | 7.400E-09 |
| 7653 | 383 | E383M | A | 9.3e-09 | 3.4e-08 | 8.5e-09 | 1.727E-08 | 1.450E-08 | 8.500E-09 | 3.400E-08 |
| 7654 | 383 | E383F | A | 5.1e-09 | 8.5e-09 | 1.3e-08 | 8.867E-09 | 3.963E-09 | 5.100E-09 | 1.300E-08 |
| 7655 | 383 | E383P | A | 3e-09 | 1.1e-08 | 4.7e-09 | 6.233E-09 | 4.215E-09 | 3.000E-09 | 1.100E-08 |
| 7656 | 383 | E383S | A | 1e-08 | 3.6e-09 | 1.3e-09 | 4.967E-09 | 4.508E-09 | 1.300E-09 | 1.000E-08 |
| 7657 | 383 | E383T | A | 1.1e-08 | 2.7e-09 | 5.9e-09 | 6.533E-09 | 4.186E-09 | 2.700E-09 | 1.100E-08 |
| 7658 | 383 | E383W | A | 3.8e-09 | 2.4e-09 | 4.3e-09 | 3.500E-09 | 9.849E-10 | 2.400E-09 | 4.300E-09 |
| 7659 | 383 | E383Y | A | 4.5e-09 | 6.4e-09 | 6.7e-09 | 5.867E-09 | 1.193E-09 | 4.500E-09 | 6.700E-09 |
| 7660 | 383 | E383V | A | 8.5e-09 | 1.1e-08 | 1.1e-08 | 1.017E-08 | 1.443E-09 | 8.500E-09 | 1.100E-08 |
| 7661 | 384 | I384A | A | 1.2e-08 | 6.7e-09 | 1.4e-08 | 1.090E-08 | 3.772E-09 | 6.700E-09 | 1.400E-08 |
| 7662 | 384 | I384R | A | 6.9e-09 | 1.4e-08 | 3.1e-09 | 8.000E-09 | 5.533E-09 | 3.100E-09 | 1.400E-08 |
| 7663 | 384 | I384N | A | 2.6e-09 | 1.2e-08 | 3.1e-09 | 5.900E-09 | 5.289E-09 | 2.600E-09 | 1.200E-08 |
| 7664 | 384 | I384D | A | 6.9e-09 | 4.9e-09 | 7.6e-09 | 6.467E-09 | 1.401E-09 | 4.900E-09 | 7.600E-09 |
| 7665 | 384 | I384C | A | 1.1e-08 | 1.2e-08 | 1.2e-08 | 1.167E-08 | 5.774E-10 | 1.100E-08 | 1.200E-08 |
| 7666 | 384 | I384E | A | 3.5e-09 | 8.3e-09 | 2.3e-08 | 1.160E-08 | 1.016E-08 | 3.500E-09 | 2.300E-08 |
| 7667 | 384 | I384Q | A | 4.8e-09 | 1.5e-08 | 4.4e-09 | 8.067E-09 | 6.008E-09 | 4.400E-09 | 1.500E-08 |
| 7668 | 384 | I384G | A | 5e-09 | 6.3e-09 | 8.9e-09 | 6.733E-09 | 1.986E-09 | 5.000E-09 | 8.900E-09 |
| 7669 | 384 | I384H | A | 4.7e-09 | 7.8e-09 | 7.4e-09 | 6.633E-09 | 1.686E-09 | 4.700E-09 | 7.800E-09 |
| 7670 | 384 | I384I | A | 6.7e-09 | 9.1e-09 | 1.1e-08 | 8.933E-09 | 2.155E-09 | 6.700E-09 | 1.100E-08 |
| 7671 | 384 | I384L | A | 6.5e-09 | 7.3e-09 | 9.2e-09 | 7.667E-09 | 1.387E-09 | 6.500E-09 | 9.200E-09 |
| 7672 | 384 | I384K | A | 6.6e-09 | 5.7e-09 | 6.4e-09 | 6.233E-09 | 4.726E-10 | 5.700E-09 | 6.600E-09 |
| 7673 | 384 | I384M | A | 4.4e-09 | 5.3e-09 | 5.5e-09 | 5.067E-09 | 5.859E-10 | 4.400E-09 | 5.500E-09 |
| 7674 | 384 | I384F | A | 3.2e-09 | 7.8e-09 | 1.5e-08 | 8.667E-09 | 5.948E-09 | 3.200E-09 | 1.500E-08 |
| 7675 | 384 | I384P | A | 2e-08 | 7e-09 | 5.1e-09 | 1.070E-08 | 8.110E-09 | 5.100E-09 | 2.000E-08 |
| 7676 | 384 | I384S | A | 4.9e-09 | 5.1e-09 | 8.4e-09 | 6.133E-09 | 1.966E-09 | 4.900E-09 | 8.400E-09 |
| 7677 | 384 | I384T | A | 5.2e-09 | 8.2e-09 | 1e-08 | 7.800E-09 | 2.425E-09 | 5.200E-09 | 1.000E-08 |
| 7678 | 384 | I384W | A | 8e-09 | 3.3e-09 | 2.7e-09 | 4.667E-09 | 2.902E-09 | 2.700E-09 | 8.000E-09 |
| 7679 | 384 | I384Y | A | 5.7e-09 | 7.4e-09 | 3e-09 | 5.367E-09 | 2.219E-09 | 3.000E-09 | 7.400E-09 |
| 7680 | 384 | I384V | A | 7.7e-09 | 1.1e-08 | 6.8e-09 | 8.500E-09 | 2.211E-09 | 6.800E-09 | 1.100E-08 |
| 7681 | 385 | T385A | A | 3e-09 | 9.1e-09 | 9.6e-09 | 7.233E-09 | 3.675E-09 | 3.000E-09 | 9.600E-09 |
| 7682 | 385 | T385R | A | 4.9e-09 | 7.7e-09 | 7.3e-09 | 6.633E-09 | 1.514E-09 | 4.900E-09 | 7.700E-09 |
| 7683 | 385 | T385N | A | 1.2e-08 | 6.4e-09 | 2.8e-09 | 7.067E-09 | 4.636E-09 | 2.800E-09 | 1.200E-08 |
| 7684 | 385 | T385D | A | 5.9e-09 | 9.2e-09 | 5.3e-09 | 6.800E-09 | 2.100E-09 | 5.300E-09 | 9.200E-09 |
| 7685 | 385 | T385C | A | 1.3e-08 | 6e-09 | 1e-08 | 9.667E-09 | 3.512E-09 | 6.000E-09 | 1.300E-08 |
| 7686 | 385 | T385E | A | 4.3e-09 | 8.3e-09 | 5.8e-09 | 6.133E-09 | 2.021E-09 | 4.300E-09 | 8.300E-09 |
| 7687 | 385 | T385Q | A | 7.2e-09 | 5e-09 | 8.8e-09 | 7.000E-09 | 1.908E-09 | 5.000E-09 | 8.800E-09 |
| 7688 | 385 | T385G | A | 8.9e-09 | 1.1e-08 | 3.5e-09 | 7.800E-09 | 3.869E-09 | 3.500E-09 | 1.100E-08 |
| 7689 | 385 | T385H | A | 1.4e-09 | 5.9e-09 | 3.2e-09 | 3.500E-09 | 2.265E-09 | 1.400E-09 | 5.900E-09 |
| 7690 | 385 | T385I | A | 3.1e-09 | 1.7e-08 | 8e-09 | 9.367E-09 | 7.050E-09 | 3.100E-09 | 1.700E-08 |
| 7691 | 385 | T385L | A | 1.7e-08 | 3.8e-09 | 3.6e-09 | 8.133E-09 | 7.679E-09 | 3.600E-09 | 1.700E-08 |
| 7692 | 385 | T385K | A | 4.6e-09 | 2.9e-09 | 8.8e-09 | 5.433E-09 | 3.037E-09 | 2.900E-09 | 8.800E-09 |
| 7693 | 385 | T385M | A | 7.2e-09 | 3.3e-09 | 1.3e-08 | 7.833E-09 | 4.881E-09 | 3.300E-09 | 1.300E-08 |
| 7694 | 385 | T385F | A | 2e-09 | 1.2e-08 | 1.6e-09 | 5.200E-09 | 5.892E-09 | 1.600E-09 | 1.200E-08 |
| 7695 | 385 | T385P | A | 1.5e-08 | 1.5e-08 | 2.7e-09 | 1.090E-08 | 7.101E-09 | 2.700E-09 | 1.500E-08 |
| 7696 | 385 | T385S | A | 7e-09 | 7.2e-09 | 1e-08 | 8.067E-09 | 1.677E-09 | 7.000E-09 | 1.000E-08 |
| 7697 | 385 | T385T | A | 1.3e-08 | 3.3e-09 | 4.6e-09 | 6.967E-09 | 5.265E-09 | 3.300E-09 | 1.300E-08 |
| 7698 | 385 | T385W | A | 4.5e-09 | 1e-08 | 1.3e-08 | 9.167E-09 | 4.311E-09 | 4.500E-09 | 1.300E-08 |
| 7699 | 385 | T385Y | A | 9.4e-09 | 1.6e-09 | 6.1e-09 | 5.700E-09 | 3.915E-09 | 1.600E-09 | 9.400E-09 |
| 7700 | 385 | T385V | A | 3.4e-09 | 1.4e-08 | 2.9e-09 | 6.767E-09 | 6.269E-09 | 2.900E-09 | 1.400E-08 |
| 7701 | 386 | G386A | A | 1.9e-08 | 1.3e-08 | 1.9e-08 | 1.700E-08 | 3.464E-09 | 1.300E-08 | 1.900E-08 |
| 7702 | 386 | G386R | A | 4.8e-09 | 7.9e-09 | 1.4e-08 | 8.900E-09 | 4.681E-09 | 4.800E-09 | 1.400E-08 |
| 7703 | 386 | G386N | A | 1.7e-08 | 4.6e-09 | 3.1e-09 | 8.233E-09 | 7.629E-09 | 3.100E-09 | 1.700E-08 |
| 7704 | 386 | G386D | A | 7e-09 | 9.3e-09 | 6.3e-09 | 7.533E-09 | 1.570E-09 | 6.300E-09 | 9.300E-09 |
| 7705 | 386 | G386C | A | 1.2e-08 | 1.5e-08 | 9.3e-09 | 1.210E-08 | 2.851E-09 | 9.300E-09 | 1.500E-08 |
| 7706 | 386 | G386E | A | 7.5e-09 | 1.7e-08 | 7.2e-09 | 1.057E-08 | 5.573E-09 | 7.200E-09 | 1.700E-08 |
| 7707 | 386 | G386Q | A | 7.2e-08 | 6.8e-09 | 4.3e-09 | 2.770E-08 | 3.839E-08 | 4.300E-09 | 7.200E-08 |
| 7708 | 386 | G386G | A | 1.6e-08 | 3.8e-09 | 8e-09 | 9.267E-09 | 6.198E-09 | 3.800E-09 | 1.600E-08 |
| 7709 | 386 | G386H | A | 6.6e-09 | 6.3e-09 | 1e-08 | 7.633E-09 | 2.055E-09 | 6.300E-09 | 1.000E-08 |
| 7710 | 386 | G386I | A | 1.1e-08 | 7.9e-09 | 2.5e-09 | 7.133E-09 | 4.302E-09 | 2.500E-09 | 1.100E-08 |
| 7711 | 386 | G386L | A | 1.5e-08 | 1e-08 | 5.6e-09 | 1.020E-08 | 4.703E-09 | 5.600E-09 | 1.500E-08 |
| 7712 | 386 | G386K | A | 4.6e-09 | 1.2e-08 | 2.3e-08 | 1.320E-08 | 9.259E-09 | 4.600E-09 | 2.300E-08 |
| 7713 | 386 | G386M | A | 1.1e-09 | 7.7e-09 | 4.4e-09 | 4.400E-09 | 3.300E-09 | 1.100E-09 | 7.700E-09 |
| 7714 | 386 | G386F | A | 3.8e-09 | 1.5e-08 | 6.2e-09 | 8.333E-09 | 5.897E-09 | 3.800E-09 | 1.500E-08 |
| 7715 | 386 | G386P | A | 1.2e-08 | 1.4e-08 | 6.5e-09 | 1.083E-08 | 3.884E-09 | 6.500E-09 | 1.400E-08 |
| 7716 | 386 | G386S | A | 1.8e-08 | 3.6e-09 | 8.8e-09 | 1.013E-08 | 7.292E-09 | 3.600E-09 | 1.800E-08 |
| 7717 | 386 | G386T | A | 2.3e-09 | 1.4e-08 | 4.2e-09 | 6.833E-09 | 6.279E-09 | 2.300E-09 | 1.400E-08 |
| 7718 | 386 | G386W | A | 1e-08 | 9.2e-09 | 1.5e-08 | 1.140E-08 | 3.143E-09 | 9.200E-09 | 1.500E-08 |
| 7719 | 386 | G386Y | A | 7.9e-09 | 3.9e-09 | 2.2e-09 | 4.667E-09 | 2.926E-09 | 2.200E-09 | 7.900E-09 |
| 7720 | 386 | G386V | A | 1.7e-08 | 1e-08 | 1.6e-08 | 1.433E-08 | 3.786E-09 | 1.000E-08 | 1.700E-08 |
| 7721 | 387 | Y387A | A | 6.3e-09 | 1.9e-08 | 6.3e-09 | 1.053E-08 | 7.332E-09 | 6.300E-09 | 1.900E-08 |
| 7722 | 387 | Y387R | A | 1.1e-08 | 8.8e-09 | 6.1e-09 | 8.633E-09 | 2.454E-09 | 6.100E-09 | 1.100E-08 |
| 7723 | 387 | Y387N | A | 2e-08 | 7.4e-09 | 1.5e-09 | 9.633E-09 | 9.450E-09 | 1.500E-09 | 2.000E-08 |
| 7724 | 387 | Y387D | A | 5e-09 | 6.4e-09 | 5.8e-09 | 5.733E-09 | 7.024E-10 | 5.000E-09 | 6.400E-09 |
| 7725 | 387 | Y387C | A | 7.5e-09 | 2.2e-09 | 8.3e-09 | 6.000E-09 | 3.315E-09 | 2.200E-09 | 8.300E-09 |
| 7726 | 387 | Y387E | A | 3.9e-08 | 1.5e-08 | 1.1e-08 | 2.167E-08 | 1.514E-08 | 1.100E-08 | 3.900E-08 |
| 7727 | 387 | Y387Q | A | 6.6e-09 | 1.2e-08 | 1.1e-08 | 9.867E-09 | 2.873E-09 | 6.600E-09 | 1.200E-08 |
| 7728 | 387 | Y387G | A | 2.5e-09 | 7.5e-09 | 7e-09 | 5.667E-09 | 2.754E-09 | 2.500E-09 | 7.500E-09 |
| 7729 | 387 | Y387H | A | 4.4e-09 | 1e-08 | 4.8e-09 | 6.400E-09 | 3.124E-09 | 4.400E-09 | 1.000E-08 |
| 7730 | 387 | Y387I | A | 2e-09 | 1.5e-08 | 6.9e-09 | 7.967E-09 | 6.565E-09 | 2.000E-09 | 1.500E-08 |
| 7731 | 387 | Y387L | A | 6.7e-09 | 8.4e-09 | 6.1e-09 | 7.067E-09 | 1.193E-09 | 6.100E-09 | 8.400E-09 |
| 7732 | 387 | Y387K | A | 6e-09 | 2.1e-08 | 1.4e-08 | 1.367E-08 | 7.506E-09 | 6.000E-09 | 2.100E-08 |
| 7733 | 387 | Y387M | A | 2.9e-09 | 7.6e-09 | 1.6e-09 | 4.033E-09 | 3.156E-09 | 1.600E-09 | 7.600E-09 |
| 7734 | 387 | Y387F | A | 3e-09 | 3.2e-09 | 1.4e-08 | 6.733E-09 | 6.294E-09 | 3.000E-09 | 1.400E-08 |

| | | | | | | | | | | |
|---|---|---|---|---|---|---|---|---|---|---|
| 7735 | 387 | Y387P | A | 4.7e-09 | 1.1e-08 | 5.5e-09 | 7.067E-09 | 3.430E-09 | 4.700E-09 | 1.100E-08 |
| 7736 | 387 | Y387S | A | 1.6e-09 | 1.8e-08 | 1.3e-08 | 1.087E-08 | 8.406E-09 | 1.600E-09 | 1.800E-08 |
| 7737 | 387 | Y387T | A | 7.3e-09 | 1.1e-08 | 7.5e-09 | 8.600E-09 | 2.081E-09 | 7.300E-09 | 1.100E-08 |
| 7738 | 387 | Y387W | A | 8e-09 | 9.5e-09 | 7.6e-09 | 8.367E-09 | 1.002E-09 | 7.600E-09 | 9.500E-09 |
| 7739 | 387 | Y387Y | A | 8e-09 | 1.2e-08 | 9.4e-09 | 9.800E-09 | 2.030E-09 | 8.000E-09 | 1.200E-08 |
| 7740 | 387 | Y387V | A | 6e-09 | 6.5e-09 | 1.1e-08 | 7.833E-09 | 2.754E-09 | 6.000E-09 | 1.100E-08 |
| 7741 | 388 | L388A | A | 2.5e-08 | 1.3e-08 | 1.1e-08 | 1.633E-08 | 7.572E-09 | 1.100E-08 | 2.500E-08 |
| 7742 | 388 | L388R | A | 5.4e-09 | 4.9e-09 | 1.9e-09 | 4.067E-09 | 1.893E-09 | 1.900E-09 | 5.400E-09 |
| 7743 | 388 | L388N | A | 4.9e-09 | 3.5e-09 | 5.1e-09 | 4.500E-09 | 8.718E-10 | 3.500E-09 | 5.100E-09 |
| 7744 | 388 | L388D | A | 7.4e-09 | 5.3e-09 | 4.4e-09 | 5.700E-09 | 1.539E-09 | 4.400E-09 | 7.400E-09 |
| 7745 | 388 | L388C | A | 5.5e-09 | 6.9e-09 | 1.3e-09 | 4.567E-09 | 2.914E-09 | 1.300E-09 | 6.900E-09 |
| 7746 | 388 | L388E | A | 5.7e-09 | 7e-09 | 1.3e-08 | 8.567E-09 | 3.894E-09 | 5.700E-09 | 1.300E-08 |
| 7747 | 388 | L388Q | A | 7.1e-09 | 2.7e-09 | 4.5e-09 | 4.767E-09 | 2.212E-09 | 2.700E-09 | 7.100E-09 |
| 7748 | 388 | L388G | A | 1.1e-08 | 8.9e-09 | 8.2e-09 | 9.367E-09 | 1.457E-09 | 8.200E-09 | 1.100E-08 |
| 7749 | 388 | L388H | A | 1.1e-08 | 8.1e-09 | 3e-09 | 7.367E-09 | 4.050E-09 | 3.000E-09 | 1.100E-08 |
| 7750 | 388 | L388I | A | 3.6e-08 | 7e-09 | 4.1e-09 | 1.570E-08 | 1.764E-08 | 4.100E-09 | 3.600E-08 |
| 7751 | 388 | L388L | A | 1.2e-08 | 3.8e-09 | 9.1e-10 | 5.570E-09 | 5.753E-09 | 9.100E-10 | 1.200E-08 |
| 7752 | 388 | L388K | A | 6.2e-09 | 7.3e-09 | 2.4e-09 | 5.300E-09 | 2.571E-09 | 2.400E-09 | 7.300E-09 |
| 7753 | 388 | L388M | A | 6.3e-09 | 7.4e-09 | 6e-08 | 2.457E-08 | 3.069E-08 | 6.300E-09 | 6.000E-08 |
| 7754 | 388 | L388F | A | 6.3e-09 | 9.3e-09 | 1.3e-08 | 9.533E-09 | 3.356E-09 | 6.300E-09 | 1.300E-08 |
| 7755 | 388 | L388P | A | 1.8e-08 | 4.1e-09 | 6.3e-09 | 9.467E-09 | 7.472E-09 | 4.100E-09 | 1.800E-08 |
| 7756 | 388 | L388S | A | 1.3e-08 | 5.8e-09 | 3.5e-09 | 7.433E-09 | 4.956E-09 | 3.500E-09 | 1.300E-08 |
| 7757 | 388 | L388T | A | 4.7e-09 | 2.3e-09 | 1.3e-08 | 6.667E-09 | 5.615E-09 | 2.300E-09 | 1.300E-08 |
| 7758 | 388 | L388W | A | 5.1e-09 | 1.1e-08 | 4.6e-09 | 6.900E-09 | 3.559E-09 | 4.600E-09 | 1.100E-08 |
| 7759 | 388 | L388Y | A | 5e-09 | 4.7e-09 | 5.4e-09 | 5.033E-09 | 3.512E-10 | 4.700E-09 | 5.400E-09 |
| 7760 | 388 | L388V | A | 6.8e-09 | 8.6e-09 | 1.8e-09 | 5.733E-09 | 3.523E-09 | 1.800E-09 | 8.600E-09 |
| 7761 | 389 | Y389A | A | 2.9e-09 | 3.7e-09 | 4.2e-09 | 3.600E-09 | 6.557E-10 | 2.900E-09 | 4.200E-09 |
| 7762 | 389 | Y389R | A | 1.1e-08 | 7e-09 | 1.7e-08 | 1.167E-08 | 5.033E-09 | 7.000E-09 | 1.700E-08 |
| 7763 | 389 | Y389N | A | 7e-09 | 1.6e-08 | 1.3e-08 | 1.200E-08 | 4.583E-09 | 7.000E-09 | 1.600E-08 |
| 7764 | 389 | Y389D | A | 5.1e-09 | 8.8e-09 | 9e-09 | 7.633E-09 | 2.196E-09 | 5.100E-09 | 9.000E-09 |
| 7765 | 389 | Y389C | A | 1.2e-08 | 3.5e-09 | 4.6e-09 | 6.700E-09 | 4.623E-09 | 3.500E-09 | 1.200E-08 |
| 7766 | 389 | Y389E | A | 2.6e-09 | 4.6e-09 | 8.1e-09 | 5.100E-09 | 2.784E-09 | 2.600E-09 | 8.100E-09 |
| 7767 | 389 | Y389Q | A | 1.1e-08 | 3.9e-09 | 6.3e-09 | 7.067E-09 | 3.612E-09 | 3.900E-09 | 1.100E-08 |
| 7768 | 389 | Y389G | A | 1.4e-08 | 9.1e-09 | 7.9e-09 | 1.033E-08 | 3.232E-09 | 7.900E-09 | 1.400E-08 |
| 7769 | 389 | Y389H | A | 4.5e-09 | 4.2e-09 | 1.2e-08 | 6.900E-09 | 4.419E-09 | 4.200E-09 | 1.200E-08 |
| 7770 | 389 | Y389I | A | 1.8e-08 | 2.2e-08 | 1e-08 | 1.667E-08 | 6.110E-09 | 1.000E-08 | 2.200E-08 |
| 7771 | 389 | Y389L | A | 1.8e-08 | 1e-08 | 6.2e-09 | 1.140E-08 | 6.023E-09 | 6.200E-09 | 1.800E-08 |
| 7772 | 389 | Y389K | A | 4.6e-09 | 8e-09 | 9.1e-09 | 7.233E-09 | 2.346E-09 | 4.600E-09 | 9.100E-09 |
| 7773 | 389 | Y389M | A | 9.2e-09 | 5.2e-09 | 8.9e-09 | 7.767E-09 | 2.228E-09 | 5.200E-09 | 9.200E-09 |
| 7774 | 389 | Y389F | A | 1.2e-08 | 5.9e-09 | 4e-09 | 7.300E-09 | 4.180E-09 | 4.000E-09 | 1.200E-08 |
| 7775 | 389 | Y389P | A | 1.9e-08 | 1.2e-08 | 2.2e-08 | 1.767E-08 | 5.132E-09 | 1.200E-08 | 2.200E-08 |
| 7776 | 389 | Y389S | A | 8.8e-09 | 7.9e-09 | 1.6e-08 | 1.090E-08 | 4.440E-09 | 7.900E-09 | 1.600E-08 |
| 7777 | 389 | Y389T | A | 3.6e-09 | 1.3e-08 | 4.2e-09 | 6.933E-09 | 5.262E-09 | 3.600E-09 | 1.300E-08 |
| 7778 | 389 | Y389W | A | 6.8e-09 | 2.9e-09 | 4.3e-09 | 4.667E-09 | 1.976E-09 | 2.900E-09 | 6.800E-09 |
| 7779 | 389 | Y389Y | A | 8.3e-09 | 1e-08 | 2.6e-09 | 6.967E-09 | 3.876E-09 | 2.600E-09 | 1.000E-08 |
| 7780 | 389 | Y389V | A | 1e-08 | 2.6e-09 | 2.4e-09 | 5.000E-09 | 4.331E-09 | 2.400E-09 | 1.000E-08 |
| 7781 | 390 | I390A | A | 3.1e-09 | 5.2e-09 | 7.3e-09 | 5.200E-09 | 2.100E-09 | 3.100E-09 | 7.300E-09 |
| 7782 | 390 | I390R | A | 3.6e-09 | 8.7e-09 | 1.1e-08 | 7.767E-09 | 3.787E-09 | 3.600E-09 | 1.100E-08 |
| 7783 | 390 | I390N | A | 2.1e-09 | 5.8e-09 | 2.2e-09 | 3.367E-09 | 2.108E-09 | 2.100E-09 | 5.800E-09 |
| 7784 | 390 | I390D | A | 1.5e-08 | 3.3e-09 | 2.5e-09 | 6.933E-09 | 6.997E-09 | 2.500E-09 | 1.500E-08 |
| 7785 | 390 | I390C | A | 2.8e-09 | 7e-09 | 1e-08 | 6.600E-09 | 3.617E-09 | 2.800E-09 | 1.000E-08 |
| 7786 | 390 | I390E | A | 1.4e-08 | 6.9e-09 | 5.3e-09 | 8.733E-09 | 4.631E-09 | 5.300E-09 | 1.400E-08 |
| 7787 | 390 | I390Q | A | 1.1e-08 | 1.4e-08 | 1.1e-08 | 1.200E-08 | 1.732E-09 | 1.100E-08 | 1.400E-08 |
| 7788 | 390 | I390G | A | 1.1e-09 | 8.2e-09 | 7.3e-09 | 5.533E-09 | 3.866E-09 | 1.100E-09 | 8.200E-09 |
| 7789 | 390 | I390H | A | 2.4e-09 | 1.2e-08 | 1.9e-09 | 5.433E-09 | 5.692E-09 | 1.900E-09 | 1.200E-08 |
| 7790 | 390 | I390I | A | 7.9e-09 | 4.4e-09 | 1.6e-08 | 9.433E-09 | 5.950E-09 | 4.400E-09 | 1.600E-08 |
| 7791 | 390 | I390L | A | 7.9e-09 | 2.2e-09 | 6.6e-09 | 5.567E-09 | 2.987E-09 | 2.200E-09 | 7.900E-09 |
| 7792 | 390 | I390K | A | 8.5e-09 | 8.8e-09 | 5.7e-09 | 7.667E-09 | 1.710E-09 | 5.700E-09 | 8.800E-09 |
| 7793 | 390 | I390M | A | 3e-09 | 7e-09 | 3.7e-09 | 4.567E-09 | 2.136E-09 | 3.000E-09 | 7.000E-09 |
| 7794 | 390 | I390F | A | 6.2e-09 | 9.5e-09 | 1.1e-08 | 8.900E-09 | 2.456E-09 | 6.200E-09 | 1.100E-08 |
| 7795 | 390 | I390P | A | 1.4e-08 | 3.6e-09 | 4.4e-09 | 7.333E-09 | 5.787E-09 | 3.600E-09 | 1.400E-08 |
| 7796 | 390 | I390S | A | 1.6e-08 | 1.8e-09 | 4e-09 | 7.267E-09 | 7.643E-09 | 1.800E-09 | 1.600E-08 |
| 7797 | 390 | I390T | A | 5.1e-09 | 4.1e-09 | 1.1e-08 | 6.733E-09 | 3.729E-09 | 4.100E-09 | 1.100E-08 |
| 7798 | 390 | I390W | A | 7.3e-09 | 6.2e-09 | 2.9e-08 | 1.417E-08 | 1.286E-08 | 6.200E-09 | 2.900E-08 |
| 7799 | 390 | I390Y | A | 3.8e-09 | 1.1e-08 | 1.3e-08 | 9.267E-09 | 4.839E-09 | 3.800E-09 | 1.300E-08 |
| 7800 | 390 | I390V | A | 6.6e-09 | 1.4e-08 | 7.1e-09 | 9.233E-09 | 4.136E-09 | 6.600E-09 | 1.400E-08 |
| 7801 | 391 | S391A | A | 5.6e-09 | 1.2e-08 | 5.6e-09 | 7.733E-09 | 3.695E-09 | 5.600E-09 | 1.200E-08 |
| 7802 | 391 | S391R | A | 1.5e-08 | 3.9e-09 | 1e-08 | 9.633E-09 | 5.559E-09 | 3.900E-09 | 1.500E-08 |
| 7803 | 391 | S391N | A | 1.5e-08 | 2.7e-09 | 1.2e-08 | 9.900E-09 | 6.413E-09 | 2.700E-09 | 1.500E-08 |
| 7804 | 391 | S391D | A | 1.5e-08 | 5e-09 | 1e-08 | 1.000E-08 | 5.000E-09 | 5.000E-09 | 1.500E-08 |
| 7805 | 391 | S391C | A | 6.2e-09 | 1.8e-08 | 8e-09 | 1.073E-08 | 6.357E-09 | 6.200E-09 | 1.800E-08 |
| 7806 | 391 | S391E | A | 6e-09 | 1.8e-08 | 5.6e-09 | 9.867E-09 | 7.047E-09 | 5.600E-09 | 1.800E-08 |
| 7807 | 391 | S391Q | A | 8.7e-09 | 1.8e-09 | 4.6e-09 | 5.033E-09 | 3.470E-09 | 1.800E-09 | 8.700E-09 |
| 7808 | 391 | S391G | A | 1.3e-08 | 1.3e-08 | 2.7e-08 | 1.767E-08 | 8.083E-09 | 1.300E-08 | 2.700E-08 |
| 7809 | 391 | S391H | A | 4.9e-09 | 4.5e-09 | 9.7e-09 | 6.367E-09 | 2.894E-09 | 4.500E-09 | 9.700E-09 |
| 7810 | 391 | S391I | A | 9.2e-09 | 1e-08 | 4e-09 | 7.733E-09 | 3.258E-09 | 4.000E-09 | 1.000E-08 |
| 7811 | 391 | S391L | A | 6.9e-09 | 6.1e-09 | 1.8e-08 | 1.033E-08 | 6.652E-09 | 6.100E-09 | 1.800E-08 |
| 7812 | 391 | S391K | A | 5.9e-09 | 3.1e-08 | 1e-08 | 1.563E-08 | 1.346E-08 | 5.900E-09 | 3.100E-08 |
| 7813 | 391 | S391M | A | 5.5e-09 | 6.6e-09 | 3e-09 | 5.033E-09 | 1.845E-09 | 3.000E-09 | 6.600E-09 |
| 7814 | 391 | S391F | A | 2.2e-08 | 8.8e-09 | 4.8e-09 | 1.187E-08 | 9.001E-09 | 4.800E-09 | 2.200E-08 |
| 7815 | 391 | S391P | A | 9e-09 | 1.3e-08 | 1.3e-08 | 1.167E-08 | 2.309E-09 | 9.000E-09 | 1.300E-08 |
| 7816 | 391 | S391S | A | 5.7e-09 | 3.1e-09 | 1.4e-08 | 7.600E-09 | 5.693E-09 | 3.100E-09 | 1.400E-08 |
| 7817 | 391 | S391T | A | 5.3e-09 | 2.1e-09 | 2.7e-09 | 3.367E-09 | 1.701E-09 | 2.100E-09 | 5.300E-09 |
| 7818 | 391 | S391W | A | 6.7e-09 | 1.2e-08 | 1.9e-09 | 6.867E-09 | 5.052E-09 | 1.900E-09 | 1.200E-08 |
| 7819 | 391 | S391Y | A | 4.7e-09 | 1.1e-08 | 1.4e-08 | 9.900E-09 | 4.747E-09 | 4.700E-09 | 1.400E-08 |

| | | | | | | | | | | |
|---|---|---|---|---|---|---|---|---|---|---|
| 7820 | 391 | S391V | A | 7.2e-09 | 4.2e-09 | 2.5e-09 | 4.633E-09 | 2.380E-09 | 2.500E-09 | 7.200E-09 |
| 7821 | 392 | A392A | A | 8.9e-09 | 5.1e-09 | 3.4e-09 | 5.800E-09 | 2.816E-09 | 3.400E-09 | 8.900E-09 |
| 7822 | 392 | A392R | A | 1.2e-08 | 6.6e-09 | 2.4e-09 | 7.000E-09 | 4.812E-09 | 2.400E-09 | 1.200E-08 |
| 7823 | 392 | A392N | A | 7.5e-09 | 6.3e-09 | 6e-09 | 6.600E-09 | 7.937E-10 | 6.000E-09 | 7.500E-09 |
| 7824 | 392 | A392D | A | 4.5e-09 | 5.4e-09 | 5e-09 | 4.967E-09 | 4.509E-10 | 4.500E-09 | 5.400E-09 |
| 7825 | 392 | A392C | A | 2.1e-09 | 2.4e-08 | 2.2e-09 | 9.433E-09 | 1.262E-08 | 2.100E-09 | 2.400E-08 |
| 7826 | 392 | A392E | A | 5.8e-09 | 5.7e-09 | 2e-08 | 1.050E-08 | 8.227E-09 | 5.700E-09 | 2.000E-08 |
| 7827 | 392 | A392Q | A | 8.8e-09 | 8.4e-09 | 4.3e-09 | 7.167E-09 | 2.491E-09 | 4.300E-09 | 8.800E-09 |
| 7828 | 392 | A392G | A | 3.2e-09 | 6e-09 | 4.9e-09 | 4.700E-09 | 1.411E-09 | 3.200E-09 | 6.000E-09 |
| 7829 | 392 | A392H | A | 1.5e-08 | 6.6e-09 | 3.6e-09 | 8.400E-09 | 5.909E-09 | 3.600E-09 | 1.500E-08 |
| 7830 | 392 | A392I | A | 4.3e-09 | 5.5e-09 | 6.9e-09 | 5.567E-09 | 1.301E-09 | 4.300E-09 | 6.900E-09 |
| 7831 | 392 | A392L | A | 8e-09 | 4.5e-09 | 6.2e-09 | 6.233E-09 | 1.750E-09 | 4.500E-09 | 8.000E-09 |
| 7832 | 392 | A392K | A | 2.2e-08 | 5.6e-09 | 1.1e-08 | 1.287E-08 | 8.358E-09 | 5.600E-09 | 2.200E-08 |
| 7833 | 392 | A392M | A | 8.9e-09 | 4.7e-09 | 4.1e-09 | 5.900E-09 | 2.615E-09 | 4.100E-09 | 8.900E-09 |
| 7834 | 392 | A392F | A | 8.8e-09 | 8.3e-09 | 3.5e-09 | 6.867E-09 | 2.926E-09 | 3.500E-09 | 8.800E-09 |
| 7835 | 392 | A392P | A | 4.7e-09 | 4.8e-09 | 4.3e-09 | 4.600E-09 | 2.646E-10 | 4.300E-09 | 4.800E-09 |
| 7836 | 392 | A392S | A | 5.6e-09 | 3.1e-09 | 9.5e-10 | 3.217E-09 | 2.327E-09 | 9.500E-10 | 5.600E-09 |
| 7837 | 392 | A392T | A | 1.7e-09 | 1.3e-08 | 3.9e-09 | 6.200E-09 | 5.991E-09 | 1.700E-09 | 1.300E-08 |
| 7838 | 392 | A392W | A | 1.1e-08 | 2.4e-09 | 1.5e-08 | 9.467E-09 | 6.438E-09 | 2.400E-09 | 1.500E-08 |
| 7839 | 392 | A392Y | A | 2.3e-09 | 1.2e-08 | 1.1e-08 | 8.433E-09 | 5.335E-09 | 2.300E-09 | 1.200E-08 |
| 7840 | 392 | A392V | A | 3.2e-09 | 1.6e-08 | 1.5e-09 | 6.900E-09 | 7.927E-09 | 1.500E-09 | 1.600E-08 |
| 7841 | 393 | W393A | A | 1.1e-08 | 2.4e-08 | 4.8e-09 | 1.327E-08 | 9.799E-09 | 4.800E-09 | 2.400E-08 |
| 7842 | 393 | W393R | A | 1.4e-08 | 1e-08 | 1.1e-08 | 1.167E-08 | 2.082E-09 | 1.000E-08 | 1.400E-08 |
| 7843 | 393 | W393N | A | 4.6e-09 | 1.1e-08 | 6.7e-09 | 7.433E-09 | 3.262E-09 | 4.600E-09 | 1.100E-08 |
| 7844 | 393 | W393D | A | 5.9e-09 | 9.9e-09 | 8.3e-09 | 8.033E-09 | 2.013E-09 | 5.900E-09 | 9.900E-09 |
| 7845 | 393 | W393C | A | 8.4e-09 | 1.1e-08 | 2e-08 | 1.313E-08 | 6.087E-09 | 8.400E-09 | 2.000E-08 |
| 7846 | 393 | W393E | A | 5.1e-09 | 2.7e-09 | 5.5e-09 | 4.433E-09 | 1.514E-09 | 2.700E-09 | 5.500E-09 |
| 7847 | 393 | W393Q | A | 2.9e-09 | 7.4e-09 | 2.4e-08 | 1.143E-08 | 1.111E-08 | 2.900E-09 | 2.400E-08 |
| 7848 | 393 | W393G | A | 1.1e-08 | 1.1e-08 | 2.9e-09 | 8.300E-09 | 4.677E-09 | 2.900E-09 | 1.100E-08 |
| 7849 | 393 | W393H | A | 8.4e-09 | 1.7e-08 | 3.7e-09 | 9.700E-09 | 6.745E-09 | 3.700E-09 | 1.700E-08 |
| 7850 | 393 | W393I | A | 5.6e-09 | 8.3e-09 | 5.9e-09 | 6.600E-09 | 1.480E-09 | 5.600E-09 | 8.300E-09 |
| 7851 | 393 | W393L | A | 1.8e-09 | 5e-09 | 2.2e-08 | 9.600E-09 | 1.086E-08 | 1.800E-09 | 2.200E-08 |
| 7852 | 393 | W393K | A | 6.3e-09 | 6.7e-09 | 4.7e-09 | 5.900E-09 | 1.058E-09 | 4.700E-09 | 6.700E-09 |
| 7853 | 393 | W393M | A | 6.1e-09 | 2.2e-08 | 1.1e-08 | 1.303E-08 | 8.143E-09 | 6.100E-09 | 2.200E-08 |
| 7854 | 393 | W393F | A | 8.5e-09 | 1.5e-08 | 5.5e-09 | 9.667E-09 | 4.856E-09 | 5.500E-09 | 1.500E-08 |
| 7855 | 393 | W393P | A | 9.2e-09 | 1e-08 | 1e-08 | 9.733E-09 | 4.619E-10 | 9.200E-09 | 1.000E-08 |
| 7856 | 393 | W393S | A | 7.1e-09 | 6.5e-09 | 1.4e-08 | 9.200E-09 | 4.168E-09 | 6.500E-09 | 1.400E-08 |
| 7857 | 393 | W393T | A | 5.3e-09 | 3.6e-09 | 8.6e-09 | 5.833E-09 | 2.542E-09 | 3.600E-09 | 8.600E-09 |
| 7858 | 393 | W393W | A | 3.7e-09 | 3.3e-08 | 6.3e-09 | 1.433E-08 | 1.622E-08 | 3.700E-09 | 3.300E-08 |
| 7859 | 393 | W393Y | A | 1.6e-09 | 5e-09 | 7.8e-09 | 4.800E-09 | 3.105E-09 | 1.600E-09 | 7.800E-09 |
| 7860 | 393 | W393V | A | 1.1e-08 | 1.5e-08 | 1.5e-08 | 1.367E-08 | 2.309E-09 | 1.100E-08 | 1.500E-08 |
| 7861 | 394 | P394A | A | 2.7e-08 | 6.9e-09 | 3.7e-09 | 1.253E-08 | 1.263E-08 | 3.700E-09 | 2.700E-08 |
| 7862 | 394 | P394R | A | 1.7e-08 | 2.4e-09 | 7.6e-09 | 9.000E-09 | 7.400E-09 | 2.400E-09 | 1.700E-08 |
| 7863 | 394 | P394N | A | 5.7e-09 | 1.2e-08 | 9.2e-09 | 8.967E-09 | 3.156E-09 | 5.700E-09 | 1.200E-08 |
| 7864 | 394 | P394D | A | 9.3e-09 | 7.4e-09 | 8.3e-09 | 8.333E-09 | 9.504E-10 | 7.400E-09 | 9.300E-09 |
| 7865 | 394 | P394C | A | 6e-09 | 6.4e-09 | 4.9e-09 | 5.767E-09 | 7.767E-10 | 4.900E-09 | 6.400E-09 |
| 7866 | 394 | P394E | A | 2.5e-09 | 1e-08 | 7.1e-09 | 6.533E-09 | 3.782E-09 | 2.500E-09 | 1.000E-08 |
| 7867 | 394 | P394Q | A | 7.3e-09 | 3e-09 | 2.4e-09 | 4.233E-09 | 2.673E-09 | 2.400E-09 | 7.300E-09 |
| 7868 | 394 | P394G | A | 9.4e-09 | 5.9e-09 | 6.8e-09 | 7.367E-09 | 1.818E-09 | 5.900E-09 | 9.400E-09 |
| 7869 | 394 | P394H | A | 5e-09 | 7.4e-09 | 1.2e-08 | 8.133E-09 | 3.557E-09 | 5.000E-09 | 1.200E-08 |
| 7870 | 394 | P394I | A | 5.8e-09 | 8.2e-09 | 2.1e-08 | 1.167E-08 | 8.171E-09 | 5.800E-09 | 2.100E-08 |
| 7871 | 394 | P394L | A | 2.2e-08 | 1.1e-08 | 1e-08 | 1.433E-08 | 6.658E-09 | 1.000E-08 | 2.200E-08 |
| 7872 | 394 | P394K | A | 5e-09 | 7.5e-09 | 6.2e-09 | 6.233E-09 | 1.250E-09 | 5.000E-09 | 7.500E-09 |
| 7873 | 394 | P394M | A | 7.3e-09 | 2.4e-09 | 7.2e-09 | 5.633E-09 | 2.801E-09 | 2.400E-09 | 7.300E-09 |
| 7874 | 394 | P394F | A | 4.7e-09 | 1.1e-08 | 5.9e-09 | 7.200E-09 | 3.345E-09 | 4.700E-09 | 1.100E-08 |
| 7875 | 394 | P394P | A | 8.6e-09 | 4.7e-09 | 3.6e-09 | 5.633E-09 | 2.627E-09 | 3.600E-09 | 8.600E-09 |
| 7876 | 394 | P394S | A | 1.2e-08 | 1.2e-08 | 5.3e-09 | 9.767E-09 | 3.868E-09 | 5.300E-09 | 1.200E-08 |
| 7877 | 394 | P394T | A | 8.7e-09 | 6.2e-09 | 7.2e-09 | 7.367E-09 | 1.258E-09 | 6.200E-09 | 8.700E-09 |
| 7878 | 394 | P394W | A | 9.7e-09 | 1.1e-08 | 2.8e-09 | 7.833E-09 | 4.407E-09 | 2.800E-09 | 1.100E-08 |
| 7879 | 394 | P394Y | A | 9e-09 | 6.4e-09 | 4e-09 | 6.467E-09 | 2.501E-09 | 4.000E-09 | 9.000E-09 |
| 7880 | 394 | P394V | A | 4.3e-09 | 5.5e-09 | 1.2e-08 | 7.267E-09 | 4.143E-09 | 4.300E-09 | 1.200E-08 |
| 7881 | 395 | D395A | A | 9.2e-09 | 6.9e-09 | 4.5e-09 | 6.867E-09 | 2.350E-09 | 4.500E-09 | 9.200E-09 |
| 7882 | 395 | D395R | A | 7.2e-09 | 3.4e-09 | 9.6e-09 | 6.733E-09 | 3.126E-09 | 3.400E-09 | 9.600E-09 |
| 7883 | 395 | D395N | A | 7.4e-09 | 4.9e-09 | 3e-09 | 5.100E-09 | 2.207E-09 | 3.000E-09 | 7.400E-09 |
| 7884 | 395 | D395D | A | 1.2e-08 | 7.8e-09 | 7.1e-09 | 8.967E-09 | 2.650E-09 | 7.100E-09 | 1.200E-08 |
| 7885 | 395 | D395C | A | 2.6e-09 | 3.9e-09 | 4.8e-09 | 3.767E-09 | 1.106E-09 | 2.600E-09 | 4.800E-09 |
| 7886 | 395 | D395E | A | 4.6e-09 | 3e-09 | 5e-09 | 4.200E-09 | 1.058E-09 | 3.000E-09 | 5.000E-09 |
| 7887 | 395 | D395Q | A | 2.5e-08 | 5.4e-09 | 2.5e-09 | 1.097E-08 | 1.224E-08 | 2.500E-09 | 2.500E-08 |
| 7888 | 395 | D395G | A | 7.8e-09 | 8.1e-09 | 8.7e-09 | 8.200E-09 | 4.583E-10 | 7.800E-09 | 8.700E-09 |
| 7889 | 395 | D395H | A | 8.9e-09 | 1.1e-08 | 7.9e-09 | 9.267E-09 | 1.582E-09 | 7.900E-09 | 1.100E-08 |
| 7890 | 395 | D395I | A | 3.1e-09 | 1.1e-08 | 8.2e-09 | 7.433E-09 | 4.005E-09 | 3.100E-09 | 1.100E-08 |
| 7891 | 395 | D395L | A | 2.7e-09 | 1.4e-08 | 1.6e-08 | 1.090E-08 | 7.171E-09 | 2.700E-09 | 1.600E-08 |
| 7892 | 395 | D395K | A | 1.1e-08 | 7.1e-09 | 5.9e-09 | 8.000E-09 | 2.666E-09 | 5.900E-09 | 1.100E-08 |
| 7893 | 395 | D395M | A | 7.3e-09 | 1.3e-08 | 1.5e-08 | 1.177E-08 | 3.995E-09 | 7.300E-09 | 1.500E-08 |
| 7894 | 395 | D395F | A | 6.1e-09 | 1.1e-08 | 4.4e-09 | 7.167E-09 | 3.427E-09 | 4.400E-09 | 1.100E-08 |
| 7895 | 395 | D395P | A | 1e-08 | 6.4e-09 | 9.9e-09 | 8.767E-09 | 2.050E-09 | 6.400E-09 | 1.000E-08 |
| 7896 | 395 | D395S | A | 2.1e-08 | 7.3e-09 | 4.7e-09 | 1.100E-08 | 8.757E-09 | 4.700E-09 | 2.100E-08 |
| 7897 | 395 | D395T | A | 7.5e-09 | 5.5e-09 | 1.4e-08 | 9.000E-09 | 4.444E-09 | 5.500E-09 | 1.400E-08 |
| 7898 | 395 | D395W | A | 3.3e-09 | 8e-09 | 7.3e-09 | 6.200E-09 | 2.536E-09 | 3.300E-09 | 8.000E-09 |
| 7899 | 395 | D395Y | A | 6.6e-09 | 3e-09 | 4.6e-09 | 4.733E-09 | 1.804E-09 | 3.000E-09 | 6.600E-09 |
| 7900 | 395 | D395V | A | 1.2e-08 | 9.2e-09 | 1.4e-08 | 1.173E-08 | 2.411E-09 | 9.200E-09 | 1.400E-08 |
| 7901 | 396 | S396A | A | 6.6e-09 | 4.3e-09 | 9.3e-09 | 6.733E-09 | 2.503E-09 | 4.300E-09 | 9.300E-09 |
| 7902 | 396 | S396R | A | 6.3e-09 | 1.1e-08 | 6.2e-09 | 7.833E-09 | 2.743E-09 | 6.200E-09 | 1.100E-08 |
| 7903 | 396 | S396N | A | 1.2e-08 | 2.3e-08 | 9e-09 | 1.467E-08 | 7.371E-09 | 9.000E-09 | 2.300E-08 |
| 7904 | 396 | S396D | A | 1.6e-08 | 1.8e-08 | 1.1e-08 | 1.500E-08 | 3.606E-09 | 1.100E-08 | 1.800E-08 |

| | | | | | | | | | | |
|---|---|---|---|---|---|---|---|---|---|---|
| 7905 | 396 | S396C | A | 4.1e-09 | 6.3e-09 | 5.9e-09 | 5.433E-09 | 1.172E-09 | 4.100E-09 | 6.300E-09 |
| 7906 | 396 | S396E | A | 3.6e-09 | 1.6e-08 | 5.3e-09 | 8.300E-09 | 6.722E-09 | 3.600E-09 | 1.600E-08 |
| 7907 | 396 | S396Q | A | 7.6e-09 | 4.7e-09 | 2.3e-09 | 4.867E-09 | 2.654E-09 | 2.300E-09 | 7.600E-09 |
| 7908 | 396 | S396G | A | 7.2e-09 | 4.4e-09 | 7.5e-09 | 6.367E-09 | 1.710E-09 | 4.400E-09 | 7.500E-09 |
| 7909 | 396 | S396H | A | 1.1e-08 | 4.8e-09 | 1.4e-08 | 9.933E-09 | 4.692E-09 | 4.800E-09 | 1.400E-08 |
| 7910 | 396 | S396I | A | 1.3e-08 | 5.9e-09 | 6.9e-09 | 8.600E-09 | 3.843E-09 | 5.900E-09 | 1.300E-08 |
| 7911 | 396 | S396L | A | 2.4e-09 | 4.3e-09 | 1.6e-08 | 7.567E-09 | 7.365E-09 | 2.400E-09 | 1.600E-08 |
| 7912 | 396 | S396K | A | 5.3e-09 | 9.7e-09 | 5.7e-09 | 6.900E-09 | 2.433E-09 | 5.300E-09 | 9.700E-09 |
| 7913 | 396 | S396M | A | 3.9e-09 | 2.6e-08 | 1.9e-08 | 1.630E-08 | 1.129E-08 | 3.900E-09 | 2.600E-08 |
| 7914 | 396 | S396F | A | 4e-09 | 6e-09 | 8e-09 | 6.000E-09 | 2.000E-09 | 4.000E-09 | 8.000E-09 |
| 7915 | 396 | S396P | A | 8.7e-09 | 5.2e-09 | 1e-08 | 7.967E-09 | 2.483E-09 | 5.200E-09 | 1.000E-08 |
| 7916 | 396 | S396S | A | 4.2e-09 | 4.9e-09 | 3.6e-09 | 4.233E-09 | 6.506E-10 | 3.600E-09 | 4.900E-09 |
| 7917 | 396 | S396T | A | 3.5e-09 | 5.7e-09 | 1.5e-08 | 8.067E-09 | 6.104E-09 | 3.500E-09 | 1.500E-08 |
| 7918 | 396 | S396W | A | 6.1e-09 | 1.9e-08 | 7.8e-09 | 1.097E-08 | 7.009E-09 | 6.100E-09 | 1.900E-08 |
| 7919 | 396 | S396Y | A | 3.5e-09 | 2e-08 | 2.6e-09 | 8.700E-09 | 9.796E-09 | 2.600E-09 | 2.000E-08 |
| 7920 | 396 | S396V | A | 4.3e-09 | 8.6e-09 | 8e-09 | 6.967E-09 | 2.329E-09 | 4.300E-09 | 8.600E-09 |
| 7921 | 397 | L397A | A | 7.2e-09 | 9.8e-09 | 3.8e-09 | 6.933E-09 | 3.009E-09 | 3.800E-09 | 9.800E-09 |
| 7922 | 397 | L397R | A | 1.7e-08 | 1.5e-08 | 1.8e-08 | 1.667E-08 | 1.528E-09 | 1.500E-08 | 1.800E-08 |
| 7923 | 397 | L397N | A | 4.3e-09 | 4.2e-09 | 4e-09 | 4.167E-09 | 1.528E-10 | 4.000E-09 | 4.300E-09 |
| 7924 | 397 | L397D | A | 7e-09 | 9e-09 | 8.3e-09 | 8.100E-09 | 1.015E-09 | 7.000E-09 | 9.000E-09 |
| 7925 | 397 | L397C | A | 1.7e-08 | 2.8e-09 | 6.3e-09 | 8.700E-09 | 7.398E-09 | 2.800E-09 | 1.700E-08 |
| 7926 | 397 | L397E | A | 6.2e-09 | 9.1e-09 | 4.2e-09 | 6.500E-09 | 2.464E-09 | 4.200E-09 | 9.100E-09 |
| 7927 | 397 | L397Q | A | 3.1e-09 | 7.2e-09 | 3.8e-09 | 4.700E-09 | 2.193E-09 | 3.100E-09 | 7.200E-09 |
| 7928 | 397 | L397G | A | 3.1e-09 | 5e-09 | 1.1e-08 | 6.367E-09 | 4.124E-09 | 3.100E-09 | 1.100E-08 |
| 7929 | 397 | L397H | A | 5.7e-09 | 5.4e-09 | 3.5e-09 | 4.867E-09 | 1.193E-09 | 3.500E-09 | 5.700E-09 |
| 7930 | 397 | L397I | A | 8.9e-09 | 1e-08 | 5.8e-09 | 8.233E-09 | 2.178E-09 | 5.800E-09 | 1.000E-08 |
| 7931 | 397 | L397L | A | 5.1e-09 | 6.9e-09 | 1.7e-08 | 9.667E-09 | 6.414E-09 | 5.100E-09 | 1.700E-08 |
| 7932 | 397 | L397K | A | 4.6e-09 | 3e-09 | 5e-09 | 4.200E-09 | 1.058E-09 | 3.000E-09 | 5.000E-09 |
| 7933 | 397 | L397M | A | 3e-09 | 1.3e-08 | 9.7e-09 | 8.567E-09 | 5.095E-09 | 3.000E-09 | 1.300E-08 |
| 7934 | 397 | L397F | A | 2.2e-08 | 1.7e-08 | 2.6e-08 | 2.167E-08 | 4.509E-09 | 1.700E-08 | 2.600E-08 |
| 7935 | 397 | L397P | A | 9.1e-09 | 5.9e-09 | 1.3e-08 | 9.333E-09 | 3.556E-09 | 5.900E-09 | 1.300E-08 |
| 7936 | 397 | L397S | A | 9e-09 | 3.9e-09 | 3.3e-09 | 5.400E-09 | 3.132E-09 | 3.300E-09 | 9.000E-09 |
| 7937 | 397 | L397T | A | 8.1e-09 | 1e-08 | 5.5e-09 | 7.867E-09 | 2.259E-09 | 5.500E-09 | 1.000E-08 |
| 7938 | 397 | L397W | A | 3.2e-09 | 5.9e-09 | 6e-09 | 5.033E-09 | 1.589E-09 | 3.200E-09 | 6.000E-09 |
| 7939 | 397 | L397Y | A | 1.5e-08 | 1.4e-08 | 1.2e-08 | 1.367E-08 | 1.528E-09 | 1.200E-08 | 1.500E-08 |
| 7940 | 397 | L397V | A | 9.9e-09 | 5.5e-09 | 1.3e-08 | 9.467E-09 | 3.769E-09 | 5.500E-09 | 1.300E-08 |
| 7941 | 398 | P398A | A | 8.5e-09 | 1.2e-08 | 1e-08 | 1.017E-08 | 1.756E-09 | 8.500E-09 | 1.200E-08 |
| 7942 | 398 | P398R | A | 6.2e-09 | 6.9e-09 | 4.7e-09 | 5.933E-09 | 1.124E-09 | 4.700E-09 | 6.900E-09 |
| 7943 | 398 | P398N | A | 9e-09 | 6.5e-09 | 4.4e-09 | 6.633E-09 | 2.303E-09 | 4.400E-09 | 9.000E-09 |
| 7944 | 398 | P398D | A | 1e-08 | 3.1e-08 | 7.2e-09 | 1.607E-08 | 1.301E-08 | 7.200E-09 | 3.100E-08 |
| 7945 | 398 | P398C | A | 1e-08 | 1.4e-08 | 5.7e-09 | 9.900E-09 | 4.151E-09 | 5.700E-09 | 1.400E-08 |
| 7946 | 398 | P398E | A | 1.4e-09 | 6.1e-09 | 5.4e-09 | 4.300E-09 | 2.536E-09 | 1.400E-09 | 6.100E-09 |
| 7947 | 398 | P398Q | A | 1.9e-09 | 4.8e-09 | 4.6e-09 | 3.767E-09 | 1.620E-09 | 1.900E-09 | 4.800E-09 |
| 7948 | 398 | P398G | A | 6.1e-09 | 1.3e-08 | 5.2e-09 | 8.100E-09 | 4.267E-09 | 5.200E-09 | 1.300E-08 |
| 7949 | 398 | P398H | A | 1.2e-08 | 5.1e-09 | 5.1e-09 | 7.400E-09 | 3.984E-09 | 5.100E-09 | 1.200E-08 |
| 7950 | 398 | P398I | A | 1.1e-08 | 3.6e-09 | 4.4e-09 | 6.333E-09 | 4.061E-09 | 3.600E-09 | 1.100E-08 |
| 7951 | 398 | P398L | A | 2.6e-09 | 4.4e-09 | 5.7e-09 | 4.233E-09 | 1.557E-09 | 2.600E-09 | 5.700E-09 |
| 7952 | 398 | P398K | A | 5.1e-09 | 1.4e-08 | 8.6e-09 | 9.233E-09 | 4.484E-09 | 5.100E-09 | 1.400E-08 |
| 7953 | 398 | P398M | A | 5.4e-09 | 9e-09 | 6.7e-09 | 7.033E-09 | 1.823E-09 | 5.400E-09 | 9.000E-09 |
| 7954 | 398 | P398F | A | 3.2e-08 | 6.8e-09 | 1.6e-08 | 1.827E-08 | 1.275E-08 | 6.800E-09 | 3.200E-08 |
| 7955 | 398 | P398P | A | 2.8e-09 | 4.8e-09 | 5.8e-09 | 4.467E-09 | 1.528E-09 | 2.800E-09 | 5.800E-09 |
| 7956 | 398 | P398S | A | 4.4e-09 | 1.4e-08 | 9.8e-09 | 9.400E-09 | 4.812E-09 | 4.400E-09 | 1.400E-08 |
| 7957 | 398 | P398T | A | 1.7e-08 | 6e-09 | 3.4e-09 | 8.800E-09 | 7.219E-09 | 3.400E-09 | 1.700E-08 |
| 7958 | 398 | P398W | A | 6.4e-09 | 4e-09 | 5.7e-09 | 5.367E-09 | 1.234E-09 | 4.000E-09 | 6.400E-09 |
| 7959 | 398 | P398Y | A | 2.5e-09 | 6.4e-09 | 1.1e-08 | 6.633E-09 | 4.255E-09 | 2.500E-09 | 1.100E-08 |
| 7960 | 398 | P398V | A | 1.3e-08 | 8.2e-09 | 3.4e-09 | 8.200E-09 | 4.800E-09 | 3.400E-09 | 1.300E-08 |
| 7961 | 399 | D399A | A | 4.7e-09 | 1.3e-08 | 1e-08 | 9.233E-09 | 4.203E-09 | 4.700E-09 | 1.300E-08 |
| 7962 | 399 | D399R | A | 7.7e-09 | 2.1e-08 | 1.1e-08 | 1.323E-08 | 6.926E-09 | 7.700E-09 | 2.100E-08 |
| 7963 | 399 | D399N | A | 6.2e-09 | 3.7e-09 | 1.1e-08 | 6.967E-09 | 3.710E-09 | 3.700E-09 | 1.100E-08 |
| 7964 | 399 | D399D | A | 2e-08 | 7.7e-09 | 1.2e-08 | 1.323E-08 | 6.242E-09 | 7.700E-09 | 2.000E-08 |
| 7965 | 399 | D399C | A | 4.9e-09 | 6.2e-09 | 8.4e-09 | 6.500E-09 | 1.769E-09 | 4.900E-09 | 8.400E-09 |
| 7966 | 399 | D399E | A | 1.1e-08 | 1e-08 | 1.6e-08 | 1.233E-08 | 3.215E-09 | 1.000E-08 | 1.600E-08 |
| 7967 | 399 | D399Q | A | 1.1e-08 | 1.7e-08 | 3.1e-09 | 1.037E-08 | 6.972E-09 | 3.100E-09 | 1.700E-08 |
| 7968 | 399 | D399G | A | 1.4e-08 | 3.2e-09 | 5.3e-09 | 7.500E-09 | 5.726E-09 | 3.200E-09 | 1.400E-08 |
| 7969 | 399 | D399H | A | 5.6e-09 | 5.1e-09 | 3.5e-09 | 4.733E-09 | 1.097E-09 | 3.500E-09 | 5.600E-09 |
| 7970 | 399 | D399I | A | 1.3e-08 | 3.4e-09 | 1e-08 | 8.800E-09 | 4.911E-09 | 3.400E-09 | 1.300E-08 |
| 7971 | 399 | D399L | A | 4.5e-09 | 6.1e-09 | 1.2e-09 | 3.933E-09 | 2.499E-09 | 1.200E-09 | 6.100E-09 |
| 7972 | 399 | D399K | A | 3.2e-09 | 1.2e-08 | 4.2e-09 | 6.467E-09 | 4.818E-09 | 3.200E-09 | 1.200E-08 |
| 7973 | 399 | D399M | A | 7.2e-09 | 5.8e-09 | 1e-08 | 7.667E-09 | 2.139E-09 | 5.800E-09 | 1.000E-08 |
| 7974 | 399 | D399F | A | 7.2e-09 | 1e-08 | 5.8e-09 | 7.667E-09 | 2.139E-09 | 5.800E-09 | 1.000E-08 |
| 7975 | 399 | D399P | A | 1.1e-08 | 8.1e-09 | 1.3e-08 | 1.070E-08 | 2.464E-09 | 8.100E-09 | 1.300E-08 |
| 7976 | 399 | D399S | A | 1.2e-08 | 4.9e-09 | 1.2e-08 | 9.633E-09 | 4.099E-09 | 4.900E-09 | 1.200E-08 |
| 7977 | 399 | D399T | A | 1.3e-08 | 8.6e-09 | 8.4e-09 | 1.000E-08 | 2.600E-09 | 8.400E-09 | 1.300E-08 |
| 7978 | 399 | D399W | A | 1.9e-08 | 6.4e-09 | 5e-09 | 1.013E-08 | 7.711E-09 | 5.000E-09 | 1.900E-08 |
| 7979 | 399 | D399Y | A | 7.2e-09 | 5.6e-09 | 6.3e-09 | 6.367E-09 | 8.021E-10 | 5.600E-09 | 7.200E-09 |
| 7980 | 399 | D399V | A | 4.2e-09 | 6.4e-09 | 3.4e-09 | 4.667E-09 | 1.553E-09 | 3.400E-09 | 6.400E-09 |
| 7981 | 400 | L400A | A | 5.7e-09 | 1.2e-08 | 1.2e-08 | 9.900E-09 | 3.637E-09 | 5.700E-09 | 1.200E-08 |
| 7982 | 400 | L400R | A | 2.6e-09 | 3.8e-09 | 5e-09 | 3.800E-09 | 1.200E-09 | 2.600E-09 | 5.000E-09 |
| 7983 | 400 | L400N | A | 7.3e-09 | 3.8e-09 | 1.1e-08 | 7.367E-09 | 3.600E-09 | 3.800E-09 | 1.100E-08 |
| 7984 | 400 | L400D | A | 1e-08 | 8.4e-09 | 8.4e-09 | 8.933E-09 | 9.238E-10 | 8.400E-09 | 1.000E-08 |
| 7985 | 400 | L400C | A | 6.4e-09 | 3.6e-09 | 1.5e-08 | 8.333E-09 | 5.941E-09 | 3.600E-09 | 1.500E-08 |
| 7986 | 400 | L400E | A | 4.7e-09 | 4.8e-09 | 5.6e-09 | 5.033E-09 | 4.933E-10 | 4.700E-09 | 5.600E-09 |
| 7987 | 400 | L400Q | A | 4.6e-09 | 5.8e-09 | 5.5e-09 | 5.300E-09 | 6.245E-10 | 4.600E-09 | 5.800E-09 |
| 7988 | 400 | L400G | A | 1e-08 | 1.2e-08 | 6.7e-09 | 9.567E-09 | 2.676E-09 | 6.700E-09 | 1.200E-08 |
| 7989 | 400 | L400H | A | 7.6e-09 | 8.3e-09 | 8.6e-09 | 8.167E-09 | 5.132E-10 | 7.600E-09 | 8.600E-09 |

| | | | | | | | | | | |
|---|---|---|---|---|---|---|---|---|---|---|
| 7990 | 400 | L400I | A | 1.4e-08 | 4.4e-08 | 1.3e-08 | 2.367E-08 | 1.762E-08 | 1.300E-08 | 4.400E-08 |
| 7991 | 400 | L400L | A | 7.4e-09 | 1.6e-08 | 5.4e-09 | 9.600E-09 | 5.632E-09 | 5.400E-09 | 1.600E-08 |
| 7992 | 400 | L400K | A | 4.3e-09 | 4.8e-09 | 1.4e-08 | 7.700E-09 | 5.462E-09 | 4.300E-09 | 1.400E-08 |
| 7993 | 400 | L400M | A | 8.8e-09 | 4.8e-09 | 1.2e-08 | 8.533E-09 | 3.607E-09 | 4.800E-09 | 1.200E-08 |
| 7994 | 400 | L400F | A | 1e-08 | 1e-08 | 2.1e-08 | 1.367E-08 | 6.351E-09 | 1.000E-08 | 2.100E-08 |
| 7995 | 400 | L400P | A | 1e-08 | 3.6e-09 | 6.8e-09 | 6.800E-09 | 3.200E-09 | 3.600E-09 | 1.000E-08 |
| 7996 | 400 | L400S | A | 4.9e-09 | 5.6e-09 | 7.6e-09 | 6.033E-09 | 1.401E-09 | 4.900E-09 | 7.600E-09 |
| 7997 | 400 | L400T | A | 9e-09 | 9.5e-09 | 6.6e-09 | 8.367E-09 | 1.550E-09 | 6.600E-09 | 9.500E-09 |
| 7998 | 400 | L400W | A | 6.6e-09 | 6.7e-09 | 1.5e-09 | 4.933E-09 | 2.974E-09 | 1.500E-09 | 6.700E-09 |
| 7999 | 400 | L400Y | A | 5.4e-09 | 7e-09 | 1.4e-08 | 8.800E-09 | 4.574E-09 | 5.400E-09 | 1.400E-08 |
| 8000 | 400 | L400V | A | 1.6e-09 | 1.5e-08 | 1.5e-08 | 1.053E-08 | 7.736E-09 | 1.600E-09 | 1.500E-08 |
| 8001 | 401 | S401A | A | 6.3e-09 | 1.4e-08 | 6.2e-09 | 8.833E-09 | 4.475E-09 | 6.200E-09 | 1.400E-08 |
| 8002 | 401 | S401R | A | 5.3e-09 | 3.5e-09 | 7.9e-09 | 5.567E-09 | 2.212E-09 | 3.500E-09 | 7.900E-09 |
| 8003 | 401 | S401N | A | 3.2e-09 | 4.8e-09 | 8.8e-09 | 5.600E-09 | 2.884E-09 | 3.200E-09 | 8.800E-09 |
| 8004 | 401 | S401D | A | 5.4e-09 | 8.4e-09 | 9.1e-09 | 7.633E-09 | 1.966E-09 | 5.400E-09 | 9.100E-09 |
| 8005 | 401 | S401C | A | 1.5e-08 | 3.8e-09 | 1.3e-08 | 1.060E-08 | 5.973E-09 | 3.800E-09 | 1.500E-08 |
| 8006 | 401 | S401E | A | 7.4e-09 | 9e-09 | 3.7e-09 | 6.700E-09 | 2.718E-09 | 3.700E-09 | 9.000E-09 |
| 8007 | 401 | S401Q | A | 5.5e-09 | 1.1e-08 | 6.8e-09 | 7.767E-09 | 2.875E-09 | 5.500E-09 | 1.100E-08 |
| 8008 | 401 | S401G | A | 7.6e-09 | 5.2e-09 | 1.5e-08 | 9.267E-09 | 5.108E-09 | 5.200E-09 | 1.500E-08 |
| 8009 | 401 | S401H | A | 5.9e-09 | 3.6e-09 | 8.1e-09 | 5.867E-09 | 2.250E-09 | 3.600E-09 | 8.100E-09 |
| 8010 | 401 | S401I | A | 1e-08 | 1.3e-08 | 2.6e-08 | 1.633E-08 | 8.505E-09 | 1.000E-08 | 2.600E-08 |
| 8011 | 401 | S401L | A | 7.5e-09 | 9.4e-09 | 8.9e-09 | 8.600E-09 | 9.849E-10 | 7.500E-09 | 9.400E-09 |
| 8012 | 401 | S401K | A | 1.1e-08 | 5.3e-09 | 9.6e-09 | 8.633E-09 | 2.970E-09 | 5.300E-09 | 1.100E-08 |
| 8013 | 401 | S401M | A | 2.8e-09 | 3.5e-09 | 8.9e-09 | 5.067E-09 | 3.338E-09 | 2.800E-09 | 8.900E-09 |
| 8014 | 401 | S401F | A | 1.9e-08 | 1.3e-08 | 6.4e-09 | 1.280E-08 | 6.302E-09 | 6.400E-09 | 1.900E-08 |
| 8015 | 401 | S401P | A | 1.1e-08 | 2.4e-09 | 9.2e-09 | 7.533E-09 | 4.536E-09 | 2.400E-09 | 1.100E-08 |
| 8016 | 401 | S401S | A | 5e-09 | 1.4e-08 | 1e-08 | 9.667E-09 | 4.509E-09 | 5.000E-09 | 1.400E-08 |
| 8017 | 401 | S401T | A | 4.2e-09 | 1.1e-08 | 6e-09 | 7.067E-09 | 3.523E-09 | 4.200E-09 | 1.100E-08 |
| 8018 | 401 | S401W | A | 4.7e-09 | 1.8e-08 | 5.9e-09 | 9.533E-09 | 7.357E-09 | 4.700E-09 | 1.800E-08 |
| 8019 | 401 | S401Y | A | 5e-09 | 2.7e-09 | 1.4e-08 | 7.233E-09 | 5.972E-09 | 2.700E-09 | 1.400E-08 |
| 8020 | 401 | S401V | A | 1e-08 | 4.1e-09 | 1.1e-08 | 8.367E-09 | 3.729E-09 | 4.100E-09 | 1.100E-08 |
| 8021 | 402 | V402A | A | 7.2e-09 | 1.7e-08 | 4.7e-09 | 9.633E-09 | 6.501E-09 | 4.700E-09 | 1.700E-08 |
| 8022 | 402 | V402R | A | 9.6e-09 | 1.3e-08 | 3.8e-09 | 8.800E-09 | 4.652E-09 | 3.800E-09 | 1.300E-08 |
| 8023 | 402 | V402N | A | 1e-08 | 6.9e-09 | 1.4e-08 | 1.030E-08 | 3.559E-09 | 6.900E-09 | 1.400E-08 |
| 8024 | 402 | V402D | A | 7.7e-09 | 7.7e-09 | 5e-09 | 6.800E-09 | 1.559E-09 | 5.000E-09 | 7.700E-09 |
| 8025 | 402 | V402C | A | 1.8e-08 | 2e-08 | 9.9e-09 | 1.597E-08 | 5.348E-09 | 9.900E-09 | 2.000E-08 |
| 8026 | 402 | V402E | A | 9.1e-09 | 8.2e-09 | 1.5e-08 | 1.077E-08 | 3.694E-09 | 8.200E-09 | 1.500E-08 |
| 8027 | 402 | V402Q | A | 1.7e-08 | 9.5e-09 | 1e-08 | 1.217E-08 | 4.193E-09 | 9.500E-09 | 1.700E-08 |
| 8028 | 402 | V402G | A | 1.6e-08 | 5.8e-09 | 3.7e-09 | 8.500E-09 | 6.580E-09 | 3.700E-09 | 1.600E-08 |
| 8029 | 402 | V402H | A | 5.3e-09 | 9.6e-09 | 4.7e-09 | 6.533E-09 | 2.673E-09 | 4.700E-09 | 9.600E-09 |
| 8030 | 402 | V402I | A | 2e-08 | 1.8e-08 | 5.1e-09 | 1.437E-08 | 8.087E-09 | 5.100E-09 | 2.000E-08 |
| 8031 | 402 | V402L | A | 8.4e-09 | 1.8e-08 | 4e-09 | 1.013E-08 | 7.159E-09 | 4.000E-09 | 1.800E-08 |
| 8032 | 402 | V402K | A | 1e-08 | 2.2e-08 | 7.3e-09 | 1.310E-08 | 7.825E-09 | 7.300E-09 | 2.200E-08 |
| 8033 | 402 | V402M | A | 1.1e-08 | 3.5e-09 | 4.1e-09 | 6.200E-09 | 4.168E-09 | 3.500E-09 | 1.100E-08 |
| 8034 | 402 | V402F | A | 2.5e-09 | 9.1e-09 | 8.4e-09 | 6.667E-09 | 3.625E-09 | 2.500E-09 | 9.100E-09 |
| 8035 | 402 | V402P | A | 7.9e-09 | 6.4e-09 | 9.9e-09 | 8.067E-09 | 1.756E-09 | 6.400E-09 | 9.900E-09 |
| 8036 | 402 | V402S | A | 3.1e-09 | 1.5e-08 | 3.1e-09 | 7.067E-09 | 6.870E-09 | 3.100E-09 | 1.500E-08 |
| 8037 | 402 | V402T | A | 8e-09 | 1.3e-08 | 4.9e-09 | 8.633E-09 | 4.087E-09 | 4.900E-09 | 1.300E-08 |
| 8038 | 402 | V402W | A | 4.3e-09 | 6.2e-09 | 5.5e-09 | 5.333E-09 | 9.609E-10 | 4.300E-09 | 6.200E-09 |
| 8039 | 402 | V402Y | A | 6.8e-09 | 5.3e-09 | 3.5e-09 | 5.200E-09 | 1.652E-09 | 3.500E-09 | 6.800E-09 |
| 8040 | 402 | V402V | A | 1.8e-08 | 4.2e-09 | 5.5e-09 | 9.233E-09 | 7.620E-09 | 4.200E-09 | 1.800E-08 |
| 8041 | 403 | F403A | A | 1.8e-08 | 3.4e-09 | 6.4e-09 | 9.267E-09 | 7.711E-09 | 3.400E-09 | 1.800E-08 |
| 8042 | 403 | F403R | A | 8.4e-09 | 3.7e-09 | 5.2e-09 | 5.767E-09 | 2.401E-09 | 3.700E-09 | 8.400E-09 |
| 8043 | 403 | F403N | A | 9.4e-09 | 5.7e-09 | 1e-08 | 8.367E-09 | 2.329E-09 | 5.700E-09 | 1.000E-08 |
| 8044 | 403 | F403D | A | 4e-09 | 1.8e-08 | 1.8e-08 | 1.333E-08 | 8.083E-09 | 4.000E-09 | 1.800E-08 |
| 8045 | 403 | F403C | A | 1e-08 | 4.8e-09 | 1.1e-08 | 8.600E-09 | 3.329E-09 | 4.800E-09 | 1.100E-08 |
| 8046 | 403 | F403E | A | 1.2e-08 | 5.2e-09 | 2.5e-09 | 6.567E-09 | 4.895E-09 | 2.500E-09 | 1.200E-08 |
| 8047 | 403 | F403Q | A | 2.4e-08 | 6.3e-09 | 3.5e-08 | 2.177E-08 | 1.448E-08 | 6.300E-09 | 3.500E-08 |
| 8048 | 403 | F403G | A | 7.7e-09 | 1.2e-08 | 2.6e-09 | 7.433E-09 | 4.706E-09 | 2.600E-09 | 1.200E-08 |
| 8049 | 403 | F403H | A | 7.6e-09 | 5.6e-09 | 4.9e-09 | 6.033E-09 | 1.401E-09 | 4.900E-09 | 7.600E-09 |
| 8050 | 403 | F403I | A | 2.1e-08 | 1.7e-08 | 6.8e-09 | 1.493E-08 | 7.322E-09 | 6.800E-09 | 2.100E-08 |
| 8051 | 403 | F403L | A | 6.7e-09 | 1.7e-08 | 9e-09 | 1.090E-08 | 5.406E-09 | 6.700E-09 | 1.700E-08 |
| 8052 | 403 | F403K | A | 1.2e-08 | 4e-09 | 5.3e-09 | 7.100E-09 | 4.293E-09 | 4.000E-09 | 1.200E-08 |
| 8053 | 403 | F403M | A | 5.9e-09 | 5.6e-09 | 8.1e-09 | 6.533E-09 | 1.365E-09 | 5.600E-09 | 8.100E-09 |
| 8054 | 403 | F403F | A | 3.1e-08 | 1e-08 | 5.4e-09 | 1.547E-08 | 1.365E-08 | 5.400E-09 | 3.100E-08 |
| 8055 | 403 | F403P | A | 2.5e-09 | 6.7e-09 | 1.6e-08 | 8.400E-09 | 6.909E-09 | 2.500E-09 | 1.600E-08 |
| 8056 | 403 | F403S | A | 7.2e-09 | 3e-09 | 4.2e-09 | 4.800E-09 | 2.163E-09 | 3.000E-09 | 7.200E-09 |
| 8057 | 403 | F403T | A | 2.3e-08 | 1.4e-08 | 4.2e-09 | 1.373E-08 | 9.403E-09 | 4.200E-09 | 2.300E-08 |
| 8058 | 403 | F403W | A | 3.4e-09 | 6.3e-09 | 7.8e-09 | 5.833E-09 | 2.237E-09 | 3.400E-09 | 7.800E-09 |
| 8059 | 403 | F403Y | A | 8.6e-09 | 6.1e-09 | 8.9e-09 | 7.867E-09 | 1.537E-09 | 6.100E-09 | 8.900E-09 |
| 8060 | 403 | F403V | A | 4.1e-09 | 4.7e-09 | 8.8e-09 | 5.867E-09 | 2.558E-09 | 4.100E-09 | 8.800E-09 |
| 8061 | 404 | Q404A | A | 3.6e-09 | 7.9e-09 | 4.6e-09 | 5.367E-09 | 2.250E-09 | 3.600E-09 | 7.900E-09 |
| 8062 | 404 | Q404R | A | 7.7e-09 | 5.4e-09 | 8.4e-09 | 7.167E-09 | 1.570E-09 | 5.400E-09 | 8.400E-09 |
| 8063 | 404 | Q404N | A | 1.1e-08 | 7.7e-09 | 3.1e-09 | 7.267E-09 | 3.968E-09 | 3.100E-09 | 1.100E-08 |
| 8064 | 404 | Q404D | A | 6.8e-09 | 7e-09 | 1.2e-08 | 8.600E-09 | 2.946E-09 | 6.800E-09 | 1.200E-08 |
| 8065 | 404 | Q404C | A | 4.9e-09 | 1.6e-08 | 9e-09 | 9.967E-09 | 5.613E-09 | 4.900E-09 | 1.600E-08 |
| 8066 | 404 | Q404E | A | 1.5e-08 | 4.7e-09 | 7.3e-09 | 9.000E-09 | 5.356E-09 | 4.700E-09 | 1.500E-08 |
| 8067 | 404 | Q404Q | A | 3.3e-09 | 4.3e-09 | 5.3e-09 | 4.300E-09 | 1.000E-09 | 3.300E-09 | 5.300E-09 |
| 8068 | 404 | Q404G | A | 1.1e-08 | 4e-09 | 5.9e-09 | 6.967E-09 | 3.620E-09 | 4.000E-09 | 1.100E-08 |
| 8069 | 404 | Q404H | A | 8.9e-09 | 1.2e-08 | 2.9e-09 | 7.933E-09 | 4.626E-09 | 2.900E-09 | 1.200E-08 |
| 8070 | 404 | Q404I | A | 4.7e-09 | 1.4e-08 | 1.1e-08 | 9.900E-09 | 4.747E-09 | 4.700E-09 | 1.400E-08 |
| 8071 | 404 | Q404L | A | 6.2e-09 | 1.2e-08 | 3.8e-09 | 7.333E-09 | 4.216E-09 | 3.800E-09 | 1.200E-08 |
| 8072 | 404 | Q404K | A | 7.8e-09 | 1.5e-09 | 5.2e-09 | 4.833E-09 | 3.166E-09 | 1.500E-09 | 7.800E-09 |
| 8073 | 404 | Q404M | A | 8.2e-09 | 2e-08 | 5.6e-09 | 1.127E-08 | 7.674E-09 | 5.600E-09 | 2.000E-08 |
| 8074 | 404 | Q404F | A | 1.8e-08 | 2.6e-08 | 1e-08 | 1.800E-08 | 8.000E-09 | 1.000E-08 | 2.600E-08 |

| | | | | | | | | | | |
|---|---|---|---|---|---|---|---|---|---|---|
| 8075 | 404 | Q404P | A | 2e-08 | 6.3e-09 | 4.9e-09 | 1.040E-08 | 8.343E-09 | 4.900E-09 | 2.000E-08 |
| 8076 | 404 | Q404S | A | 7.6e-09 | 4.2e-09 | 1.9e-08 | 1.027E-08 | 7.752E-09 | 4.200E-09 | 1.900E-08 |
| 8077 | 404 | Q404T | A | 9.5e-09 | 8.4e-09 | 6.4e-09 | 8.100E-09 | 1.572E-09 | 6.400E-09 | 9.500E-09 |
| 8078 | 404 | Q404W | A | 1.9e-08 | 6.9e-09 | 5e-09 | 1.030E-08 | 7.594E-09 | 5.000E-09 | 1.900E-08 |
| 8079 | 404 | Q404Y | A | 1.1e-08 | 9.5e-09 | 8.4e-09 | 9.633E-09 | 1.305E-09 | 8.400E-09 | 1.100E-08 |
| 8080 | 404 | Q404V | A | 8e-09 | 7.6e-09 | 1.5e-08 | 1.020E-08 | 4.162E-09 | 7.600E-09 | 1.500E-08 |
| 8081 | 405 | N405A | A | 5e-09 | 4.4e-09 | 3.3e-09 | 4.233E-09 | 8.622E-10 | 3.300E-09 | 5.000E-09 |
| 8082 | 405 | N405R | A | 1.2e-08 | 1.1e-08 | 1.4e-08 | 1.233E-08 | 1.528E-09 | 1.100E-08 | 1.400E-08 |
| 8083 | 405 | N405N | A | 4e-09 | 3.6e-09 | 9.6e-09 | 5.733E-09 | 3.355E-09 | 3.600E-09 | 9.600E-09 |
| 8084 | 405 | N405D | A | 6e-09 | 1.2e-08 | 3.1e-08 | 1.633E-08 | 1.305E-08 | 6.000E-09 | 3.100E-08 |
| 8085 | 405 | N405C | A | 1.5e-08 | 3.2e-09 | 5.5e-09 | 7.900E-09 | 6.255E-09 | 3.200E-09 | 1.500E-08 |
| 8086 | 405 | N405E | A | 7.1e-09 | 8.9e-09 | 2.8e-09 | 6.267E-09 | 3.134E-09 | 2.800E-09 | 8.900E-09 |
| 8087 | 405 | N405Q | A | 1.2e-09 | 6.8e-09 | 3.9e-09 | 3.967E-09 | 2.801E-09 | 1.200E-09 | 6.800E-09 |
| 8088 | 405 | N405G | A | 1.2e-08 | 9.7e-09 | 4.5e-09 | 8.733E-09 | 3.842E-09 | 4.500E-09 | 1.200E-08 |
| 8089 | 405 | N405H | A | 6.7e-09 | 5.1e-09 | 5.5e-09 | 5.767E-09 | 8.327E-10 | 5.100E-09 | 6.700E-09 |
| 8090 | 405 | N405I | A | 1.2e-08 | 4.2e-09 | 8.7e-09 | 8.300E-09 | 3.915E-09 | 4.200E-09 | 1.200E-08 |
| 8091 | 405 | N405L | A | 1.6e-08 | 7.6e-09 | 7.2e-09 | 1.027E-08 | 4.969E-09 | 7.200E-09 | 1.600E-08 |
| 8092 | 405 | N405K | A | 8.5e-09 | 1.5e-08 | 2e-08 | 1.450E-08 | 5.766E-09 | 8.500E-09 | 2.000E-08 |
| 8093 | 405 | N405M | A | 3.1e-09 | 2.9e-08 | 6.9e-09 | 1.300E-08 | 1.399E-08 | 3.100E-09 | 2.900E-08 |
| 8094 | 405 | N405F | A | 6.8e-09 | 1.3e-08 | 2.1e-08 | 1.360E-08 | 7.119E-09 | 6.800E-09 | 2.100E-08 |
| 8095 | 405 | N405P | A | 6.8e-09 | 5.5e-09 | 1.1e-08 | 7.767E-09 | 2.875E-09 | 5.500E-09 | 1.100E-08 |
| 8096 | 405 | N405S | A | 1.1e-08 | 2.1e-08 | 5.3e-09 | 1.243E-08 | 7.948E-09 | 5.300E-09 | 2.100E-08 |
| 8097 | 405 | N405T | A | 1.3e-08 | 1.1e-08 | 1.4e-08 | 1.267E-08 | 1.528E-09 | 1.100E-08 | 1.400E-08 |
| 8098 | 405 | N405W | A | 1.7e-09 | 1.7e-08 | 8e-09 | 8.900E-09 | 7.690E-09 | 1.700E-09 | 1.700E-08 |
| 8099 | 405 | N405Y | A | 1.7e-08 | 3.4e-09 | 8.2e-09 | 9.533E-09 | 6.897E-09 | 3.400E-09 | 1.700E-08 |
| 8100 | 405 | N405V | A | 1.2e-08 | 7.6e-09 | 1.5e-08 | 1.153E-08 | 3.722E-09 | 7.600E-09 | 1.500E-08 |
| 8101 | 406 | L406A | A | 5.1e-09 | 1.3e-08 | 2.9e-08 | 1.570E-08 | 1.218E-08 | 5.100E-09 | 2.900E-08 |
| 8102 | 406 | L406R | A | 6.1e-09 | 1e-08 | 1e-08 | 8.700E-09 | 2.252E-09 | 6.100E-09 | 1.000E-08 |
| 8103 | 406 | L406N | A | 2.6e-09 | 1.9e-08 | 5.4e-09 | 9.000E-09 | 8.773E-09 | 2.600E-09 | 1.900E-08 |
| 8104 | 406 | L406D | A | 1.1e-08 | 1.1e-08 | 3.7e-09 | 8.567E-09 | 4.215E-09 | 3.700E-09 | 1.100E-08 |
| 8105 | 406 | L406C | A | 4.6e-09 | 4.1e-09 | 8.7e-09 | 5.800E-09 | 2.524E-09 | 4.100E-09 | 8.700E-09 |
| 8106 | 406 | L406E | A | 8.9e-09 | 3.9e-09 | 1.2e-08 | 8.267E-09 | 4.087E-09 | 3.900E-09 | 1.200E-08 |
| 8107 | 406 | L406Q | A | 5.2e-09 | 1.1e-08 | 4.2e-09 | 6.800E-09 | 3.672E-09 | 4.200E-09 | 1.100E-08 |
| 8108 | 406 | L406G | A | 1.6e-08 | 8.7e-09 | 1.2e-08 | 1.223E-08 | 3.656E-09 | 8.700E-09 | 1.600E-08 |
| 8109 | 406 | L406H | A | 6.5e-09 | 6.5e-09 | 1.1e-08 | 8.000E-09 | 2.598E-09 | 6.500E-09 | 1.100E-08 |
| 8110 | 406 | L406I | A | 8.8e-09 | 4.4e-09 | 4.4e-09 | 5.867E-09 | 2.540E-09 | 4.400E-09 | 8.800E-09 |
| 8111 | 406 | L406L | A | 6.1e-09 | 6e-09 | 1.9e-08 | 1.037E-08 | 7.477E-09 | 6.000E-09 | 1.900E-08 |
| 8112 | 406 | L406K | A | 6.2e-09 | 3.1e-09 | 1.1e-08 | 6.767E-09 | 3.980E-09 | 3.100E-09 | 1.100E-08 |
| 8113 | 406 | L406M | A | 1e-08 | 1.7e-08 | 4.7e-09 | 1.057E-08 | 6.170E-09 | 4.700E-09 | 1.700E-08 |
| 8114 | 406 | L406F | A | 1.5e-08 | 1e-08 | 1.1e-08 | 1.200E-08 | 2.646E-09 | 1.000E-08 | 1.500E-08 |
| 8115 | 406 | L406P | A | 7.6e-09 | 7.7e-09 | 1.4e-08 | 9.767E-09 | 3.667E-09 | 7.600E-09 | 1.400E-08 |
| 8116 | 406 | L406S | A | 5.3e-09 | 1.4e-08 | 5e-09 | 8.100E-09 | 5.112E-09 | 5.000E-09 | 1.400E-08 |
| 8117 | 406 | L406T | A | 7.2e-09 | 4.3e-09 | 4e-09 | 5.167E-09 | 1.767E-09 | 4.000E-09 | 7.200E-09 |
| 8118 | 406 | L406W | A | 1.2e-08 | 6e-09 | 7.4e-09 | 8.467E-09 | 3.139E-09 | 6.000E-09 | 1.200E-08 |
| 8119 | 406 | L406Y | A | 2.4e-08 | 1.2e-08 | 2.1e-08 | 1.900E-08 | 6.245E-09 | 1.200E-08 | 2.400E-08 |
| 8120 | 406 | L406V | A | 1.3e-08 | 1.3e-08 | 9.4e-09 | 1.180E-08 | 2.078E-09 | 9.400E-09 | 1.300E-08 |
| 8121 | 407 | Q407A | A | 7.1e-09 | 1.7e-09 | 8e-09 | 5.600E-09 | 3.407E-09 | 1.700E-09 | 8.000E-09 |
| 8122 | 407 | Q407R | A | 4.3e-09 | 5.5e-09 | 7.8e-09 | 5.867E-09 | 1.779E-09 | 4.300E-09 | 7.800E-09 |
| 8123 | 407 | Q407N | A | 6.1e-09 | 7e-09 | 8.9e-09 | 7.333E-09 | 1.429E-09 | 6.100E-09 | 8.900E-09 |
| 8124 | 407 | Q407D | A | 3.4e-09 | 1.3e-08 | 4.8e-09 | 7.067E-09 | 5.186E-09 | 3.400E-09 | 1.300E-08 |
| 8125 | 407 | Q407C | A | 8e-09 | 8.5e-09 | 1.2e-08 | 9.500E-09 | 2.179E-09 | 8.000E-09 | 1.200E-08 |
| 8126 | 407 | Q407E | A | 7.2e-09 | 5.8e-09 | 9e-09 | 7.333E-09 | 1.604E-09 | 5.800E-09 | 9.000E-09 |
| 8127 | 407 | Q407Q | A | 9.5e-09 | 6.1e-09 | 4.4e-09 | 6.667E-09 | 2.597E-09 | 4.400E-09 | 9.500E-09 |
| 8128 | 407 | Q407G | A | 2.4e-09 | 2.4e-09 | 4.2e-09 | 3.000E-09 | 1.039E-09 | 2.400E-09 | 4.200E-09 |
| 8129 | 407 | Q407H | A | 1.2e-08 | 1.2e-08 | 3.1e-09 | 9.033E-09 | 5.138E-09 | 3.100E-09 | 1.200E-08 |
| 8130 | 407 | Q407I | A | 5.1e-09 | 1.2e-08 | 6.1e-09 | 7.733E-09 | 3.729E-09 | 5.100E-09 | 1.200E-08 |
| 8131 | 407 | Q407L | A | 4.7e-09 | 4.8e-09 | 1.1e-08 | 6.833E-09 | 3.609E-09 | 4.700E-09 | 1.100E-08 |
| 8132 | 407 | Q407K | A | 8.3e-09 | 4.3e-09 | 1.3e-08 | 8.533E-09 | 4.355E-09 | 4.300E-09 | 1.300E-08 |
| 8133 | 407 | Q407M | A | 2.2e-08 | 1.6e-08 | 2.3e-08 | 2.033E-08 | 3.786E-09 | 1.600E-08 | 2.300E-08 |
| 8134 | 407 | Q407F | A | 2.2e-09 | 7.1e-09 | 3.4e-09 | 4.233E-09 | 2.554E-09 | 2.200E-09 | 7.100E-09 |
| 8135 | 407 | Q407P | A | 9.8e-09 | 1e-08 | 6e-09 | 8.600E-09 | 2.254E-09 | 6.000E-09 | 1.000E-08 |
| 8136 | 407 | Q407S | A | 1.2e-08 | 1.4e-08 | 7.3e-09 | 1.110E-08 | 3.439E-09 | 7.300E-09 | 1.400E-08 |
| 8137 | 407 | Q407T | A | 6e-09 | 1.5e-08 | 3.3e-09 | 8.100E-09 | 6.126E-09 | 3.300E-09 | 1.500E-08 |
| 8138 | 407 | Q407W | A | 5.8e-09 | 1.4e-08 | 1.2e-08 | 1.060E-08 | 4.276E-09 | 5.800E-09 | 1.400E-08 |
| 8139 | 407 | Q407Y | A | 5.9e-09 | 6.2e-09 | 3.6e-09 | 5.233E-09 | 1.422E-09 | 3.600E-09 | 6.200E-09 |
| 8140 | 407 | Q407V | A | 5.1e-09 | 7.4e-09 | 8e-09 | 6.833E-09 | 1.531E-09 | 5.100E-09 | 8.000E-09 |
| 8141 | 408 | V408A | A | 7.3e-09 | 1.1e-08 | 7.4e-09 | 8.567E-09 | 2.108E-09 | 7.300E-09 | 1.100E-08 |
| 8142 | 408 | V408R | A | 5e-09 | 2.8e-09 | 5.6e-09 | 4.467E-09 | 1.474E-09 | 2.800E-09 | 5.600E-09 |
| 8143 | 408 | V408N | A | 7e-09 | 8.7e-09 | 4.8e-09 | 6.833E-09 | 1.955E-09 | 4.800E-09 | 8.700E-09 |
| 8144 | 408 | V408D | A | 1.4e-09 | 1.7e-08 | 1.2e-08 | 1.013E-08 | 7.966E-09 | 1.400E-09 | 1.700E-08 |
| 8145 | 408 | V408C | A | 8e-09 | 5.2e-09 | 4.5e-09 | 5.900E-09 | 1.852E-09 | 4.500E-09 | 8.000E-09 |
| 8146 | 408 | V408E | A | 9.6e-09 | 5.4e-09 | 1e-08 | 8.333E-09 | 2.548E-09 | 5.400E-09 | 1.000E-08 |
| 8147 | 408 | V408Q | A | 4.6e-09 | 8.4e-09 | 4.9e-09 | 5.967E-09 | 2.113E-09 | 4.600E-09 | 8.400E-09 |
| 8148 | 408 | V408G | A | 9e-09 | 9.2e-09 | 6e-09 | 8.067E-09 | 1.793E-09 | 6.000E-09 | 9.200E-09 |
| 8149 | 408 | V408H | A | 3.2e-08 | 1.1e-08 | 6.1e-09 | 1.637E-08 | 1.376E-08 | 6.100E-09 | 3.200E-08 |
| 8150 | 408 | V408I | A | 6.4e-09 | 2.9e-09 | 6.3e-09 | 5.200E-09 | 1.992E-09 | 2.900E-09 | 6.400E-09 |
| 8151 | 408 | V408L | A | 3.8e-09 | 8.9e-09 | 4.1e-09 | 5.600E-09 | 2.862E-09 | 3.800E-09 | 8.900E-09 |
| 8152 | 408 | V408K | A | 2.4e-09 | 3.8e-09 | 3e-08 | 1.207E-08 | 1.555E-08 | 2.400E-09 | 3.000E-08 |
| 8153 | 408 | V408M | A | 3.2e-09 | 2.7e-09 | 4e-09 | 3.300E-09 | 6.557E-10 | 2.700E-09 | 4.000E-09 |
| 8154 | 408 | V408F | A | 7e-09 | 6.1e-09 | 5.1e-09 | 6.067E-09 | 9.504E-10 | 5.100E-09 | 7.000E-09 |
| 8155 | 408 | V408P | A | 5.5e-09 | 1.9e-08 | 4.3e-09 | 9.600E-09 | 8.163E-09 | 4.300E-09 | 1.900E-08 |
| 8156 | 408 | V408S | A | 2.1e-09 | 7.3e-09 | 2.4e-09 | 3.933E-09 | 2.919E-09 | 2.100E-09 | 7.300E-09 |
| 8157 | 408 | V408T | A | 6.6e-09 | 8.3e-09 | 6.1e-09 | 7.000E-09 | 1.153E-09 | 6.100E-09 | 8.300E-09 |
| 8158 | 408 | V408W | A | 4.1e-09 | 1.5e-08 | 7.3e-09 | 8.800E-09 | 5.603E-09 | 4.100E-09 | 1.500E-08 |
| 8159 | 408 | V408Y | A | 4.7e-09 | 9e-09 | 2.9e-09 | 5.533E-09 | 3.134E-09 | 2.900E-09 | 9.000E-09 |

| | | | | | | | | | | |
|---|---|---|---|---|---|---|---|---|---|---|
| 8160 | 408 | V408V | A | 1.1e-08 | 1.7e-08 | 5.9e-09 | 1.130E-08 | 5.556E-09 | 5.900E-09 | 1.700E-08 |
| 8161 | 409 | I409A | A | 9.2e-09 | 7.3e-09 | 1.9e-08 | 1.183E-08 | 6.279E-09 | 7.300E-09 | 1.900E-08 |
| 8162 | 409 | I409R | A | 5.6e-09 | 6.1e-09 | 1.5e-08 | 8.900E-09 | 5.289E-09 | 5.600E-09 | 1.500E-08 |
| 8163 | 409 | I409N | A | 2.4e-09 | 1.5e-08 | 2.5e-09 | 6.633E-09 | 7.246E-09 | 2.400E-09 | 1.500E-08 |
| 8164 | 409 | I409D | A | 5.2e-09 | 8.5e-09 | 2.1e-08 | 1.157E-08 | 8.334E-09 | 5.200E-09 | 2.100E-08 |
| 8165 | 409 | I409C | A | 1.8e-08 | 1.3e-08 | 8.9e-09 | 1.330E-08 | 4.557E-09 | 8.900E-09 | 1.800E-08 |
| 8166 | 409 | I409E | A | 9.5e-09 | 4.7e-09 | 5.2e-09 | 6.467E-09 | 2.639E-09 | 4.700E-09 | 9.500E-09 |
| 8167 | 409 | I409Q | A | 8.3e-09 | 6.2e-09 | 8.8e-09 | 7.767E-09 | 1.380E-09 | 6.200E-09 | 8.800E-09 |
| 8168 | 409 | I409G | A | 4.8e-09 | 7.1e-09 | 1e-08 | 7.300E-09 | 2.606E-09 | 4.800E-09 | 1.000E-08 |
| 8169 | 409 | I409H | A | 1.1e-08 | 1.3e-08 | 2.5e-08 | 1.633E-08 | 7.572E-09 | 1.100E-08 | 2.500E-08 |
| 8170 | 409 | I409I | A | 1.2e-08 | 9.3e-09 | 9.7e-09 | 1.033E-08 | 1.457E-09 | 9.300E-09 | 1.200E-08 |
| 8171 | 409 | I409L | A | 7.2e-09 | 9.4e-09 | 1e-08 | 8.867E-09 | 1.474E-09 | 7.200E-09 | 1.000E-08 |
| 8172 | 409 | I409K | A | 2.9e-09 | 1.3e-08 | 8e-09 | 7.967E-09 | 5.050E-09 | 2.900E-09 | 1.300E-08 |
| 8173 | 409 | I409M | A | 2.4e-09 | 1e-08 | 5.4e-09 | 5.933E-09 | 3.828E-09 | 2.400E-09 | 1.000E-08 |
| 8174 | 409 | I409F | A | 5.7e-09 | 7.4e-09 | 4.6e-09 | 5.900E-09 | 1.411E-09 | 4.600E-09 | 7.400E-09 |
| 8175 | 409 | I409P | A | 8e-09 | 2.7e-08 | 5.9e-09 | 1.363E-08 | 1.162E-08 | 5.900E-09 | 2.700E-08 |
| 8176 | 409 | I409S | A | 8.9e-09 | 2.2e-09 | 7.1e-09 | 6.067E-09 | 3.467E-09 | 2.200E-09 | 8.900E-09 |
| 8177 | 409 | I409T | A | 1.2e-08 | 9.5e-09 | 3.1e-08 | 1.750E-08 | 1.176E-08 | 9.500E-09 | 3.100E-08 |
| 8178 | 409 | I409W | A | 4.1e-09 | 6.1e-09 | 3.7e-09 | 4.633E-09 | 1.286E-09 | 3.700E-09 | 6.100E-09 |
| 8179 | 409 | I409Y | A | 1.2e-08 | 4.2e-09 | 5.9e-09 | 7.367E-09 | 4.102E-09 | 4.200E-09 | 1.200E-08 |
| 8180 | 409 | I409V | A | 4.2e-09 | 7.7e-09 | 4e-09 | 5.300E-09 | 2.081E-09 | 4.000E-09 | 7.700E-09 |
| 8181 | 410 | R410A | A | 6.4e-09 | 9.6e-09 | 7.4e-09 | 7.800E-09 | 1.637E-09 | 6.400E-09 | 9.600E-09 |
| 8182 | 410 | R410R | A | 3.9e-09 | 6.3e-09 | 2.2e-09 | 4.133E-09 | 2.060E-09 | 2.200E-09 | 6.300E-09 |
| 8183 | 410 | R410N | A | 9.7e-09 | 1.6e-08 | 8.2e-09 | 1.130E-08 | 4.139E-09 | 8.200E-09 | 1.600E-08 |
| 8184 | 410 | R410D | A | 7.9e-09 | 3.2e-09 | 9e-09 | 6.700E-09 | 3.081E-09 | 3.200E-09 | 9.000E-09 |
| 8185 | 410 | R410C | A | 6.5e-09 | 1.4e-08 | 4.9e-09 | 8.467E-09 | 4.858E-09 | 4.900E-09 | 1.400E-08 |
| 8186 | 410 | R410E | A | 1.1e-08 | 5.2e-09 | 9.2e-09 | 8.467E-09 | 2.969E-09 | 5.200E-09 | 1.100E-08 |
| 8187 | 410 | R410Q | A | 2.7e-09 | 2.4e-09 | 4.2e-09 | 3.100E-09 | 9.644E-10 | 2.400E-09 | 4.200E-09 |
| 8188 | 410 | R410G | A | 3.8e-09 | 7.2e-09 | 1.4e-08 | 8.333E-09 | 5.194E-09 | 3.800E-09 | 1.400E-08 |
| 8189 | 410 | R410H | A | 8.4e-09 | 1.6e-08 | 3.2e-09 | 9.200E-09 | 6.437E-09 | 3.200E-09 | 1.600E-08 |
| 8190 | 410 | R410I | A | 9.1e-09 | 4.4e-09 | 5.6e-09 | 6.367E-09 | 2.442E-09 | 4.400E-09 | 9.100E-09 |
| 8191 | 410 | R410L | A | 2.6e-09 | 4.2e-09 | 2.9e-09 | 3.233E-09 | 8.505E-10 | 2.600E-09 | 4.200E-09 |
| 8192 | 410 | R410K | A | 1.1e-08 | 2.8e-09 | 1.3e-08 | 8.933E-09 | 5.405E-09 | 2.800E-09 | 1.300E-08 |
| 8193 | 410 | R410M | A | 1.1e-08 | 7.2e-09 | 1e-08 | 9.400E-09 | 1.970E-09 | 7.200E-09 | 1.100E-08 |
| 8194 | 410 | R410F | A | 7.5e-09 | 4.9e-09 | 8.3e-09 | 6.900E-09 | 1.778E-09 | 4.900E-09 | 8.300E-09 |
| 8195 | 410 | R410P | A | 2.6e-09 | 4.4e-09 | 1e-08 | 5.667E-09 | 3.859E-09 | 2.600E-09 | 1.000E-08 |
| 8196 | 410 | R410S | A | 1.7e-08 | 9.8e-09 | 3.5e-09 | 1.010E-08 | 6.755E-09 | 3.500E-09 | 1.700E-08 |
| 8197 | 410 | R410T | A | 3.8e-08 | 4.7e-09 | 2.2e-08 | 2.157E-08 | 1.665E-08 | 4.700E-09 | 3.800E-08 |
| 8198 | 410 | R410W | A | 5.2e-09 | 2.1e-08 | 8.6e-09 | 1.160E-08 | 8.316E-09 | 5.200E-09 | 2.100E-08 |
| 8199 | 410 | R410Y | A | 8.1e-09 | 6e-09 | 1.1e-08 | 8.367E-09 | 2.511E-09 | 6.000E-09 | 1.100E-08 |
| 8200 | 410 | R410V | A | 2.2e-09 | 5.8e-09 | 9.4e-09 | 5.800E-09 | 3.600E-09 | 2.200E-09 | 9.400E-09 |
| 8201 | 411 | G411A | A | 1.5e-08 | 7.7e-09 | 1.7e-09 | 8.133E-09 | 6.661E-09 | 1.700E-09 | 1.500E-08 |
| 8202 | 411 | G411R | A | 3.2e-09 | 7.3e-09 | 2.7e-09 | 4.400E-09 | 2.524E-09 | 2.700E-09 | 7.300E-09 |
| 8203 | 411 | G411N | A | 2.8e-09 | 5.3e-09 | 3.5e-09 | 3.867E-09 | 1.290E-09 | 2.800E-09 | 5.300E-09 |
| 8204 | 411 | G411D | A | 5.3e-09 | 8.2e-09 | 4.5e-09 | 6.000E-09 | 1.947E-09 | 4.500E-09 | 8.200E-09 |
| 8205 | 411 | G411C | A | 5.7e-09 | 4.4e-09 | 1.3e-08 | 7.700E-09 | 4.636E-09 | 4.400E-09 | 1.300E-08 |
| 8206 | 411 | G411E | A | 8.8e-09 | 5.8e-09 | 8.2e-09 | 7.600E-09 | 1.587E-09 | 5.800E-09 | 8.800E-09 |
| 8207 | 411 | G411Q | A | 9.3e-09 | 7.9e-09 | 8.6e-09 | 8.600E-09 | 7.000E-10 | 7.900E-09 | 9.300E-09 |
| 8208 | 411 | G411G | A | 7.3e-09 | 3.7e-09 | 2.9e-09 | 4.633E-09 | 2.344E-09 | 2.900E-09 | 7.300E-09 |
| 8209 | 411 | G411H | A | 7.5e-09 | 1.5e-09 | 6.4e-09 | 5.133E-09 | 3.194E-09 | 1.500E-09 | 7.500E-09 |
| 8210 | 411 | G411I | A | 4.4e-09 | 5.1e-09 | 1e-08 | 6.500E-09 | 3.051E-09 | 4.400E-09 | 1.000E-08 |
| 8211 | 411 | G411L | A | 3.8e-09 | 7.3e-09 | 5.3e-09 | 5.467E-09 | 1.756E-09 | 3.800E-09 | 7.300E-09 |
| 8212 | 411 | G411K | A | 2.1e-08 | 4.6e-09 | 8.5e-09 | 1.137E-08 | 8.568E-09 | 4.600E-09 | 2.100E-08 |
| 8213 | 411 | G411M | A | 4.3e-09 | 1.8e-08 | 1.3e-08 | 1.177E-08 | 6.933E-09 | 4.300E-09 | 1.800E-08 |
| 8214 | 411 | G411F | A | 8.6e-10 | 8.2e-09 | 1.2e-08 | 7.020E-09 | 5.663E-09 | 8.600E-10 | 1.200E-08 |
| 8215 | 411 | G411P | A | 4.5e-09 | 1e-08 | 3.5e-09 | 6.000E-09 | 3.500E-09 | 3.500E-09 | 1.000E-08 |
| 8216 | 411 | G411S | A | 7.3e-09 | 8.7e-09 | 3e-09 | 6.333E-09 | 2.970E-09 | 3.000E-09 | 8.700E-09 |
| 8217 | 411 | G411T | A | 1e-08 | 1.3e-08 | 9.2e-09 | 1.073E-08 | 2.003E-09 | 9.200E-09 | 1.300E-08 |
| 8218 | 411 | G411W | A | 6.2e-09 | 8e-09 | 1.7e-08 | 1.040E-08 | 5.786E-09 | 6.200E-09 | 1.700E-08 |
| 8219 | 411 | G411Y | A | 2.7e-09 | 1.2e-08 | 1.3e-08 | 9.233E-09 | 5.680E-09 | 2.700E-09 | 1.300E-08 |
| 8220 | 411 | G411V | A | 6e-09 | 9.9e-09 | 1.5e-09 | 5.800E-09 | 4.204E-09 | 1.500E-09 | 9.900E-09 |
| 8221 | 412 | R412A | A | 8.4e-09 | 2.6e-08 | 5.2e-09 | 1.320E-08 | 1.120E-08 | 5.200E-09 | 2.600E-08 |
| 8222 | 412 | R412R | A | 3.6e-09 | 3.8e-09 | 2.5e-09 | 3.300E-09 | 7.000E-10 | 2.500E-09 | 3.800E-09 |
| 8223 | 412 | R412N | A | 1.2e-08 | 9.9e-09 | 5.2e-09 | 9.033E-09 | 3.482E-09 | 5.200E-09 | 1.200E-08 |
| 8224 | 412 | R412D | A | 1.3e-08 | 8.1e-09 | 4.3e-09 | 8.467E-09 | 4.362E-09 | 4.300E-09 | 1.300E-08 |
| 8225 | 412 | R412C | A | 1.9e-08 | 1.3e-08 | 2.6e-09 | 1.153E-08 | 8.298E-09 | 2.600E-09 | 1.900E-08 |
| 8226 | 412 | R412E | A | 1.1e-08 | 1.2e-08 | 1.7e-08 | 1.333E-08 | 3.215E-09 | 1.100E-08 | 1.700E-08 |
| 8227 | 412 | R412Q | A | 2.1e-09 | 8.2e-09 | 6.3e-09 | 5.533E-09 | 3.121E-09 | 2.100E-09 | 8.200E-09 |
| 8228 | 412 | R412G | A | 1.2e-08 | 6e-09 | 1.5e-08 | 1.100E-08 | 4.583E-09 | 6.000E-09 | 1.500E-08 |
| 8229 | 412 | R412H | A | 9.6e-09 | 5e-09 | 5.5e-09 | 6.700E-09 | 2.524E-09 | 5.000E-09 | 9.600E-09 |
| 8230 | 412 | R412I | A | 1.2e-08 | 2e-09 | 9.9e-09 | 7.967E-09 | 5.273E-09 | 2.000E-09 | 1.200E-08 |
| 8231 | 412 | R412L | A | 3.9e-09 | 1.5e-08 | 5.1e-09 | 8.000E-09 | 6.092E-09 | 3.900E-09 | 1.500E-08 |
| 8232 | 412 | R412K | A | 5.8e-09 | 6.6e-09 | 3e-09 | 5.133E-09 | 1.890E-09 | 3.000E-09 | 6.600E-09 |
| 8233 | 412 | R412M | A | 8.4e-09 | 2e-09 | 2.8e-09 | 4.400E-09 | 3.487E-09 | 2.000E-09 | 8.400E-09 |
| 8234 | 412 | R412F | A | 8e-09 | 1.4e-09 | 1.6e-09 | 3.667E-09 | 3.754E-09 | 1.400E-09 | 8.000E-09 |
| 8235 | 412 | R412P | A | 4.7e-09 | 1.7e-08 | 1.3e-08 | 1.157E-08 | 6.274E-09 | 4.700E-09 | 1.700E-08 |
| 8236 | 412 | R412S | A | 1.2e-08 | 7.9e-09 | 8.6e-09 | 9.500E-09 | 2.193E-09 | 7.900E-09 | 1.200E-08 |
| 8237 | 412 | R412T | A | 1.5e-08 | 1.8e-08 | 5.3e-09 | 1.277E-08 | 6.638E-09 | 5.300E-09 | 1.800E-08 |
| 8238 | 412 | R412W | A | 6.6e-09 | 1.3e-08 | 1.2e-08 | 1.053E-08 | 3.443E-09 | 6.600E-09 | 1.300E-08 |
| 8239 | 412 | R412Y | A | 5.4e-09 | 4.7e-09 | 1e-08 | 6.700E-09 | 2.879E-09 | 4.700E-09 | 1.000E-08 |
| 8240 | 412 | R412V | A | 5.2e-09 | 4e-09 | 5.4e-09 | 4.867E-09 | 7.572E-10 | 4.000E-09 | 5.400E-09 |
| 8241 | 413 | I413A | A | 1.2e-08 | 1.8e-08 | 5.5e-09 | 1.183E-08 | 6.252E-09 | 5.500E-09 | 1.800E-08 |
| 8242 | 413 | I413R | A | 1.1e-08 | 1.5e-08 | 1e-08 | 1.200E-08 | 2.646E-09 | 1.000E-08 | 1.500E-08 |
| 8243 | 413 | I413N | A | 1.5e-08 | 1.2e-08 | 6.8e-09 | 1.127E-08 | 4.149E-09 | 6.800E-09 | 1.500E-08 |
| 8244 | 413 | I413D | A | 1.3e-08 | 7e-09 | 4e-09 | 8.000E-09 | 4.583E-09 | 4.000E-09 | 1.300E-08 |

| | | | | | | | | | | |
|---|---|---|---|---|---|---|---|---|---|---|
| 8245 | 413 | I413C | A | 6.7e-09 | 1.1e-08 | 2.1e-09 | 6.600E-09 | 4.451E-09 | 2.100E-09 | 1.100E-08 |
| 8246 | 413 | I413E | A | 3e-09 | 3.6e-09 | 2.2e-08 | 9.533E-09 | 1.080E-08 | 3.000E-09 | 2.200E-08 |
| 8247 | 413 | I413Q | A | 1.1e-08 | 1.4e-08 | 1.1e-08 | 1.200E-08 | 1.732E-09 | 1.100E-08 | 1.400E-08 |
| 8248 | 413 | I413G | A | 1.2e-08 | 4.9e-09 | 1.1e-08 | 9.300E-09 | 3.843E-09 | 4.900E-09 | 1.200E-08 |
| 8249 | 413 | I413H | A | 4e-09 | 1.3e-08 | 1.4e-08 | 1.033E-08 | 5.508E-09 | 4.000E-09 | 1.400E-08 |
| 8250 | 413 | I413I | A | 5.9e-09 | 1.2e-09 | 1.3e-08 | 6.700E-09 | 5.941E-09 | 1.200E-09 | 1.300E-08 |
| 8251 | 413 | I413L | A | 1.2e-08 | 1.7e-09 | 1.2e-08 | 8.567E-09 | 5.947E-09 | 1.700E-09 | 1.200E-08 |
| 8252 | 413 | I413K | A | 5.6e-09 | 1.9e-08 | 4e-09 | 9.533E-09 | 8.237E-09 | 4.000E-09 | 1.900E-08 |
| 8253 | 413 | I413M | A | 2.8e-09 | 2.2e-08 | 6e-09 | 1.027E-08 | 1.029E-08 | 2.800E-09 | 2.200E-08 |
| 8254 | 413 | I413F | A | 1.2e-09 | 8.7e-09 | 9.9e-09 | 6.600E-09 | 4.715E-09 | 1.200E-09 | 9.900E-09 |
| 8255 | 413 | I413P | A | 8.5e-09 | 4.2e-09 | 7e-09 | 6.567E-09 | 2.183E-09 | 4.200E-09 | 8.500E-09 |
| 8256 | 413 | I413S | A | 3.6e-09 | 6.9e-09 | 4.2e-09 | 4.900E-09 | 1.758E-09 | 3.600E-09 | 6.900E-09 |
| 8257 | 413 | I413T | A | 4e-09 | 9.8e-09 | 6.8e-09 | 6.867E-09 | 2.901E-09 | 4.000E-09 | 9.800E-09 |
| 8258 | 413 | I413W | A | 3.5e-08 | 9.3e-09 | 1.2e-08 | 1.877E-08 | 1.412E-08 | 9.300E-09 | 3.500E-08 |
| 8259 | 413 | I413Y | A | 2e-09 | 1.6e-08 | 3.6e-09 | 7.200E-09 | 7.663E-09 | 2.000E-09 | 1.600E-08 |
| 8260 | 413 | I413V | A | 3.5e-09 | 1e-08 | 1.1e-08 | 8.167E-09 | 4.072E-09 | 3.500E-09 | 1.100E-08 |
| 8261 | 414 | L414A | A | 1.2e-08 | 4.8e-09 | 3.6e-09 | 6.800E-09 | 4.543E-09 | 3.600E-09 | 1.200E-08 |
| 8262 | 414 | L414R | A | 3e-09 | 2.4e-08 | 1.4e-08 | 1.367E-08 | 1.050E-08 | 3.000E-09 | 2.400E-08 |
| 8263 | 414 | L414N | A | 6e-09 | 3e-09 | 5.3e-09 | 4.767E-09 | 1.570E-09 | 3.000E-09 | 6.000E-09 |
| 8264 | 414 | L414D | A | 6.9e-09 | 8.8e-09 | 1.3e-08 | 9.567E-09 | 3.121E-09 | 6.900E-09 | 1.300E-08 |
| 8265 | 414 | L414C | A | 1.3e-08 | 1.5e-08 | 6.5e-09 | 1.150E-08 | 4.444E-09 | 6.500E-09 | 1.500E-08 |
| 8266 | 414 | L414E | A | 5.4e-09 | 5.1e-09 | 5e-09 | 5.167E-09 | 2.082E-10 | 5.000E-09 | 5.400E-09 |
| 8267 | 414 | L414Q | A | 7.7e-09 | 4.8e-09 | 4.1e-09 | 5.533E-09 | 1.909E-09 | 4.100E-09 | 7.700E-09 |
| 8268 | 414 | L414G | A | 8.1e-09 | 6.1e-09 | 9.1e-09 | 7.767E-09 | 1.528E-09 | 6.100E-09 | 9.100E-09 |
| 8269 | 414 | L414H | A | 6.2e-09 | 1.2e-08 | 3e-09 | 7.067E-09 | 4.562E-09 | 3.000E-09 | 1.200E-08 |
| 8270 | 414 | L414I | A | 3.8e-09 | 5.7e-09 | 2.5e-09 | 4.000E-09 | 1.609E-09 | 2.500E-09 | 5.700E-09 |
| 8271 | 414 | L414L | A | 7e-09 | 8.6e-09 | 9.7e-09 | 8.433E-09 | 1.358E-09 | 7.000E-09 | 9.700E-09 |
| 8272 | 414 | L414K | A | 3.1e-09 | 4.2e-09 | 1.5e-09 | 2.933E-09 | 1.358E-09 | 1.500E-09 | 4.200E-09 |
| 8273 | 414 | L414M | A | 6.3e-09 | 8.7e-09 | 1e-09 | 5.333E-09 | 3.940E-09 | 1.000E-09 | 8.700E-09 |
| 8274 | 414 | L414F | A | 3.6e-09 | 9.2e-09 | 1e-08 | 7.600E-09 | 3.487E-09 | 3.600E-09 | 1.000E-08 |
| 8275 | 414 | L414P | A | 9.5e-09 | 4.7e-09 | 7.7e-09 | 7.300E-09 | 2.425E-09 | 4.700E-09 | 9.500E-09 |
| 8276 | 414 | L414S | A | 5.1e-09 | 7.6e-09 | 1e-08 | 7.567E-09 | 2.450E-09 | 5.100E-09 | 1.000E-08 |
| 8277 | 414 | L414T | A | 4e-09 | 1.8e-09 | 1.7e-09 | 2.500E-09 | 1.300E-09 | 1.700E-09 | 4.000E-09 |
| 8278 | 414 | L414W | A | 1.4e-08 | 5.8e-09 | 3.9e-09 | 7.900E-09 | 5.367E-09 | 3.900E-09 | 1.400E-08 |
| 8279 | 414 | L414Y | A | 3.9e-09 | 1.2e-08 | 6.8e-09 | 7.567E-09 | 4.104E-09 | 3.900E-09 | 1.200E-08 |
| 8280 | 414 | L414V | A | 9.9e-09 | 1.8e-08 | 6.5e-09 | 1.147E-08 | 5.908E-09 | 6.500E-09 | 1.800E-08 |
| 8281 | 415 | H415A | A | 3.7e-09 | 8.1e-09 | 5.2e-09 | 5.667E-09 | 2.237E-09 | 3.700E-09 | 8.100E-09 |
| 8282 | 415 | H415R | A | 1.8e-08 | 1.1e-08 | 2.6e-09 | 1.053E-08 | 7.711E-09 | 2.600E-09 | 1.800E-08 |
| 8283 | 415 | H415N | A | 8.6e-09 | 9e-09 | 1.4e-08 | 1.053E-08 | 3.009E-09 | 8.600E-09 | 1.400E-08 |
| 8284 | 415 | H415D | A | 1.3e-08 | 5.4e-09 | 5.6e-09 | 8.000E-09 | 4.331E-09 | 5.400E-09 | 1.300E-08 |
| 8285 | 415 | H415C | A | 2.7e-08 | 5.6e-09 | 9e-09 | 1.387E-08 | 1.150E-08 | 5.600E-09 | 2.700E-08 |
| 8286 | 415 | H415E | A | 2.8e-09 | 7e-09 | 3.5e-09 | 4.433E-09 | 2.250E-09 | 2.800E-09 | 7.000E-09 |
| 8287 | 415 | H415Q | A | 1.2e-09 | 6.5e-09 | 7.4e-09 | 5.033E-09 | 3.350E-09 | 1.200E-09 | 7.400E-09 |
| 8288 | 415 | H415G | A | 4.5e-09 | 6.6e-09 | 6.7e-09 | 5.933E-09 | 1.242E-09 | 4.500E-09 | 6.700E-09 |
| 8289 | 415 | H415H | A | 5.7e-09 | 1.4e-08 | 7.2e-09 | 8.967E-09 | 4.423E-09 | 5.700E-09 | 1.400E-08 |
| 8290 | 415 | H415I | A | 1e-08 | 6e-09 | 9.6e-09 | 8.533E-09 | 2.203E-09 | 6.000E-09 | 1.000E-08 |
| 8291 | 415 | H415L | A | 5.5e-09 | 1.2e-08 | 8.6e-09 | 8.700E-09 | 3.251E-09 | 5.500E-09 | 1.200E-08 |
| 8292 | 415 | H415K | A | 3.3e-09 | 4.9e-09 | 2.4e-09 | 3.533E-09 | 1.266E-09 | 2.400E-09 | 4.900E-09 |
| 8293 | 415 | H415M | A | 1.5e-08 | 8e-09 | 7.7e-09 | 1.023E-08 | 4.131E-09 | 7.700E-09 | 1.500E-08 |
| 8294 | 415 | H415F | A | 5.7e-09 | 8.2e-09 | 1.1e-08 | 8.300E-09 | 2.651E-09 | 5.700E-09 | 1.100E-08 |
| 8295 | 415 | H415P | A | 2.9e-08 | 4.9e-09 | 1.3e-08 | 1.563E-08 | 1.226E-08 | 4.900E-09 | 2.900E-08 |
| 8296 | 415 | H415S | A | 7.9e-09 | 2.9e-09 | 3.5e-09 | 4.767E-09 | 2.730E-09 | 2.900E-09 | 7.900E-09 |
| 8297 | 415 | H415T | A | 6e-09 | 2.7e-09 | 4.5e-09 | 4.400E-09 | 1.652E-09 | 2.700E-09 | 6.000E-09 |
| 8298 | 415 | H415W | A | 1.3e-08 | 6.5e-09 | 7.1e-09 | 8.867E-09 | 3.592E-09 | 6.500E-09 | 1.300E-08 |
| 8299 | 415 | H415Y | A | 4.3e-09 | 4.9e-09 | 3.4e-09 | 4.200E-09 | 7.550E-10 | 3.400E-09 | 4.900E-09 |
| 8300 | 415 | H415V | A | 3.5e-09 | 3.8e-09 | 1.9e-08 | 8.767E-09 | 8.864E-09 | 3.500E-09 | 1.900E-08 |
| 8301 | 416 | N416A | A | 3.8e-09 | 5.9e-09 | 3.3e-09 | 4.333E-09 | 1.380E-09 | 3.300E-09 | 5.900E-09 |
| 8302 | 416 | N416R | A | 1.3e-08 | 5.5e-09 | 1.4e-08 | 1.083E-08 | 4.646E-09 | 5.500E-09 | 1.400E-08 |
| 8303 | 416 | N416N | A | 5.2e-09 | 1.3e-08 | 8.2e-09 | 8.800E-09 | 3.934E-09 | 5.200E-09 | 1.300E-08 |
| 8304 | 416 | N416D | A | 8.8e-09 | 5.3e-09 | 1.4e-08 | 9.367E-09 | 4.378E-09 | 5.300E-09 | 1.400E-08 |
| 8305 | 416 | N416C | A | 3.7e-09 | 1.3e-08 | 2.6e-09 | 6.433E-09 | 5.713E-09 | 2.600E-09 | 1.300E-08 |
| 8306 | 416 | N416E | A | 5.1e-09 | 4.1e-09 | 5.4e-09 | 4.867E-09 | 6.807E-10 | 4.100E-09 | 5.400E-09 |
| 8307 | 416 | N416Q | A | 2.8e-09 | 1.9e-08 | 1.4e-08 | 1.193E-08 | 8.295E-09 | 2.800E-09 | 1.900E-08 |
| 8308 | 416 | N416G | A | 1.9e-08 | 3.8e-09 | 6.3e-09 | 9.700E-09 | 8.150E-09 | 3.800E-09 | 1.900E-08 |
| 8309 | 416 | N416H | A | 3.5e-09 | 2.8e-09 | 6.2e-09 | 4.167E-09 | 1.795E-09 | 2.800E-09 | 6.200E-09 |
| 8310 | 416 | N416I | A | 6.5e-09 | 5.3e-09 | 1.7e-08 | 9.600E-09 | 6.437E-09 | 5.300E-09 | 1.700E-08 |
| 8311 | 416 | N416L | A | 4.4e-09 | 9.8e-09 | 1.4e-08 | 9.400E-09 | 4.812E-09 | 4.400E-09 | 1.400E-08 |
| 8312 | 416 | N416K | A | 1.1e-08 | 3.3e-08 | 9e-09 | 1.767E-08 | 1.332E-08 | 9.000E-09 | 3.300E-08 |
| 8313 | 416 | N416M | A | 5.3e-09 | 1.4e-08 | 1.3e-08 | 1.077E-08 | 4.761E-09 | 5.300E-09 | 1.400E-08 |
| 8314 | 416 | N416F | A | 5.2e-09 | 9.7e-09 | 7.4e-09 | 7.433E-09 | 2.250E-09 | 5.200E-09 | 9.700E-09 |
| 8315 | 416 | N416P | A | 2.9e-09 | 1e-08 | 5.2e-09 | 6.033E-09 | 3.623E-09 | 2.900E-09 | 1.000E-08 |
| 8316 | 416 | N416S | A | 7.2e-09 | 7e-09 | 5.7e-09 | 6.633E-09 | 8.145E-10 | 5.700E-09 | 7.200E-09 |
| 8317 | 416 | N416T | A | 1.7e-08 | 6.1e-09 | 5.9e-09 | 9.667E-09 | 6.352E-09 | 5.900E-09 | 1.700E-08 |
| 8318 | 416 | N416W | A | 6.4e-09 | 1.9e-09 | 1.8e-09 | 3.367E-09 | 2.627E-09 | 1.800E-09 | 6.400E-09 |
| 8319 | 416 | N416Y | A | 7.8e-09 | 1.2e-08 | 5.8e-09 | 8.533E-09 | 3.164E-09 | 5.800E-09 | 1.200E-08 |
| 8320 | 416 | N416V | A | 2.5e-09 | 4.7e-09 | 4.7e-09 | 3.967E-09 | 1.270E-09 | 2.500E-09 | 4.700E-09 |
| 8321 | 417 | G417A | A | 1.4e-08 | 1.5e-08 | 9.7e-09 | 1.290E-08 | 2.816E-09 | 9.700E-09 | 1.500E-08 |
| 8322 | 417 | G417R | A | 4.5e-09 | 4.7e-09 | 6.8e-09 | 5.333E-09 | 1.274E-09 | 4.500E-09 | 6.800E-09 |
| 8323 | 417 | G417N | A | 5.7e-09 | 1.2e-08 | 7.4e-09 | 8.367E-09 | 3.259E-09 | 5.700E-09 | 1.200E-08 |
| 8324 | 417 | G417D | A | 7.8e-09 | 1.4e-08 | 1.9e-08 | 1.360E-08 | 5.611E-09 | 7.800E-09 | 1.900E-08 |
| 8325 | 417 | G417C | A | 8.9e-09 | 3.5e-09 | 4.6e-09 | 5.667E-09 | 2.854E-09 | 3.500E-09 | 8.900E-09 |
| 8326 | 417 | G417E | A | 8.8e-09 | 7e-09 | 9.2e-09 | 8.333E-09 | 1.172E-09 | 7.000E-09 | 9.200E-09 |
| 8327 | 417 | G417Q | A | 1.1e-08 | 9e-09 | 4e-09 | 8.000E-09 | 3.606E-09 | 4.000E-09 | 1.100E-08 |
| 8328 | 417 | G417G | A | 5.7e-09 | 2.6e-09 | 7.3e-09 | 5.200E-09 | 2.390E-09 | 2.600E-09 | 7.300E-09 |
| 8329 | 417 | G417H | A | 1.2e-08 | 8.5e-09 | 3.2e-08 | 1.750E-08 | 1.268E-08 | 8.500E-09 | 3.200E-08 |

| | | | | | | | | | | |
|---|---|---|---|---|---|---|---|---|---|---|
| 8330 | 417 | G417I | A | 2.6e-09 | 2.9e-09 | 3.6e-09 | 3.033E-09 | 5.132E-10 | 2.600E-09 | 3.600E-09 |
| 8331 | 417 | G417L | A | 8e-09 | 5.9e-09 | 5.2e-09 | 6.367E-09 | 1.457E-09 | 5.200E-09 | 8.000E-09 |
| 8332 | 417 | G417K | A | 9.8e-09 | 6.3e-09 | 8.6e-09 | 8.233E-09 | 1.779E-09 | 6.300E-09 | 9.800E-09 |
| 8333 | 417 | G417M | A | 9.2e-09 | 1.2e-08 | 2.7e-09 | 7.967E-09 | 4.771E-09 | 2.700E-09 | 1.200E-08 |
| 8334 | 417 | G417F | A | 1.9e-08 | 1.4e-08 | 2.6e-09 | 1.187E-08 | 8.406E-09 | 2.600E-09 | 1.900E-08 |
| 8335 | 417 | G417P | A | 7.2e-09 | 1.4e-08 | 1.1e-08 | 1.073E-08 | 3.408E-09 | 7.200E-09 | 1.400E-08 |
| 8336 | 417 | G417S | A | 6.5e-09 | 1.6e-08 | 2.7e-09 | 8.400E-09 | 6.851E-09 | 2.700E-09 | 1.600E-08 |
| 8337 | 417 | G417T | A | 3.8e-09 | 6.9e-09 | 8.8e-09 | 6.500E-09 | 2.524E-09 | 3.800E-09 | 8.800E-09 |
| 8338 | 417 | G417W | A | 4.1e-09 | 3.1e-09 | 8.5e-09 | 5.233E-09 | 2.873E-09 | 3.100E-09 | 8.500E-09 |
| 8339 | 417 | G417Y | A | 1.2e-08 | 5.7e-09 | 6.4e-09 | 8.033E-09 | 3.453E-09 | 5.700E-09 | 1.200E-08 |
| 8340 | 417 | G417V | A | 1.1e-08 | 5.9e-09 | 3.5e-09 | 6.800E-09 | 3.830E-09 | 3.500E-09 | 1.100E-08 |
| 8341 | 418 | A418A | A | 7.5e-09 | 9.6e-09 | 1.5e-08 | 1.070E-08 | 3.869E-09 | 7.500E-09 | 1.500E-08 |
| 8342 | 418 | A418R | A | 2.7e-09 | 8.3e-09 | 8.1e-09 | 6.367E-09 | 3.177E-09 | 2.700E-09 | 8.300E-09 |
| 8343 | 418 | A418N | A | 5e-09 | 7.7e-09 | 7.8e-09 | 6.833E-09 | 1.589E-09 | 5.000E-09 | 7.800E-09 |
| 8344 | 418 | A418D | A | 6.4e-09 | 5.8e-09 | 3.3e-09 | 5.167E-09 | 1.644E-09 | 3.300E-09 | 6.400E-09 |
| 8345 | 418 | A418C | A | 2.8e-09 | 7.6e-09 | 8.5e-09 | 6.300E-09 | 3.064E-09 | 2.800E-09 | 8.500E-09 |
| 8346 | 418 | A418E | A | 4.5e-09 | 8.7e-09 | 6e-09 | 6.400E-09 | 2.128E-09 | 4.500E-09 | 8.700E-09 |
| 8347 | 418 | A418Q | A | 4.7e-09 | 2.9e-09 | 6e-09 | 4.533E-09 | 1.557E-09 | 2.900E-09 | 6.000E-09 |
| 8348 | 418 | A418G | A | 2.2e-09 | 5.2e-09 | 2.8e-09 | 3.400E-09 | 1.587E-09 | 2.200E-09 | 5.200E-09 |
| 8349 | 418 | A418H | A | 1e-08 | 7.4e-09 | 7.6e-09 | 8.333E-09 | 1.447E-09 | 7.400E-09 | 1.000E-08 |
| 8350 | 418 | A418I | A | 4e-09 | 1e-08 | 3.5e-09 | 5.833E-09 | 3.617E-09 | 3.500E-09 | 1.000E-08 |
| 8351 | 418 | A418L | A | 8.2e-09 | 1.2e-08 | 7.6e-09 | 9.267E-09 | 2.386E-09 | 7.600E-09 | 1.200E-08 |
| 8352 | 418 | A418K | A | 7.6e-09 | 2.1e-08 | 5.2e-09 | 1.127E-08 | 8.514E-09 | 5.200E-09 | 2.100E-08 |
| 8353 | 418 | A418M | A | 7.5e-09 | 6.1e-09 | 1.2e-08 | 8.533E-09 | 3.083E-09 | 6.100E-09 | 1.200E-08 |
| 8354 | 418 | A418F | A | 6.1e-09 | 7e-09 | 9.9e-09 | 7.667E-09 | 1.986E-09 | 6.100E-09 | 9.900E-09 |
| 8355 | 418 | A418P | A | 2.8e-09 | 9.5e-09 | 6.8e-09 | 6.367E-09 | 3.371E-09 | 2.800E-09 | 9.500E-09 |
| 8356 | 418 | A418S | A | 7.4e-09 | 1e-08 | 1.5e-08 | 1.080E-08 | 3.863E-09 | 7.400E-09 | 1.500E-08 |
| 8357 | 418 | A418T | A | 5.3e-09 | 4e-09 | 1.3e-08 | 7.433E-09 | 4.864E-09 | 4.000E-09 | 1.300E-08 |
| 8358 | 418 | A418W | A | 9.3e-09 | 9e-09 | 4.5e-09 | 7.600E-09 | 2.689E-09 | 4.500E-09 | 9.300E-09 |
| 8359 | 418 | A418Y | A | 6e-09 | 8.3e-09 | 9.2e-09 | 7.833E-09 | 1.650E-09 | 6.000E-09 | 9.200E-09 |
| 8360 | 418 | A418V | A | 3.2e-09 | 6.4e-09 | 8.5e-09 | 6.033E-09 | 2.669E-09 | 3.200E-09 | 8.500E-09 |
| 8361 | 419 | Y419A | A | 1.3e-09 | 3.2e-09 | 1.1e-08 | 5.167E-09 | 5.140E-09 | 1.300E-09 | 1.100E-08 |
| 8362 | 419 | Y419R | A | 3.7e-09 | 4.3e-09 | 6.2e-09 | 4.733E-09 | 1.305E-09 | 3.700E-09 | 6.200E-09 |
| 8363 | 419 | Y419N | A | 5.9e-09 | 7.5e-09 | 5.4e-09 | 6.267E-09 | 1.097E-09 | 5.400E-09 | 7.500E-09 |
| 8364 | 419 | Y419D | A | 1.1e-08 | 8.3e-09 | 1.5e-08 | 1.143E-08 | 3.371E-09 | 8.300E-09 | 1.500E-08 |
| 8365 | 419 | Y419C | A | 7.7e-09 | 2.7e-09 | 1.9e-08 | 9.800E-09 | 8.350E-09 | 2.700E-09 | 1.900E-08 |
| 8366 | 419 | Y419E | A | 3e-09 | 1.7e-08 | 1.3e-08 | 1.100E-08 | 7.211E-09 | 3.000E-09 | 1.700E-08 |
| 8367 | 419 | Y419Q | A | 1.1e-08 | 6.8e-09 | 4.6e-09 | 7.467E-09 | 3.252E-09 | 4.600E-09 | 1.100E-08 |
| 8368 | 419 | Y419G | A | 8.2e-09 | 6.4e-09 | 1.4e-08 | 9.533E-09 | 3.972E-09 | 6.400E-09 | 1.400E-08 |
| 8369 | 419 | Y419H | A | 9.2e-09 | 9e-09 | 1.2e-08 | 1.007E-08 | 1.677E-09 | 9.000E-09 | 1.200E-08 |
| 8370 | 419 | Y419I | A | 7.3e-09 | 6.2e-09 | 2.7e-09 | 5.400E-09 | 2.402E-09 | 2.700E-09 | 7.300E-09 |
| 8371 | 419 | Y419L | A | 7.2e-09 | 4.4e-09 | 4.9e-09 | 5.500E-09 | 1.493E-09 | 4.400E-09 | 7.200E-09 |
| 8372 | 419 | Y419K | A | 9e-09 | 8.3e-09 | 1.2e-08 | 9.767E-09 | 1.966E-09 | 8.300E-09 | 1.200E-08 |
| 8373 | 419 | Y419M | A | 1.7e-08 | 6.7e-09 | 6.7e-09 | 1.013E-08 | 5.947E-09 | 6.700E-09 | 1.700E-08 |
| 8374 | 419 | Y419F | A | 9e-09 | 2.6e-09 | 6.1e-09 | 5.900E-09 | 3.205E-09 | 2.600E-09 | 9.000E-09 |
| 8375 | 419 | Y419P | A | 4.9e-09 | 3.2e-09 | 5.6e-09 | 4.567E-09 | 1.234E-09 | 3.200E-09 | 5.600E-09 |
| 8376 | 419 | Y419S | A | 1.3e-08 | 4.8e-09 | 4.6e-09 | 7.467E-09 | 4.793E-09 | 4.600E-09 | 1.300E-08 |
| 8377 | 419 | Y419T | A | 1.3e-08 | 9.7e-09 | 9.6e-09 | 1.077E-08 | 1.935E-09 | 9.600E-09 | 1.300E-08 |
| 8378 | 419 | Y419W | A | 1.5e-08 | 8.6e-09 | 3.3e-09 | 8.967E-09 | 5.859E-09 | 3.300E-09 | 1.500E-08 |
| 8379 | 419 | Y419Y | A | 3.2e-09 | 3.9e-09 | 7.4e-09 | 4.833E-09 | 2.250E-09 | 3.200E-09 | 7.400E-09 |
| 8380 | 419 | Y419V | A | 7.7e-09 | 7e-09 | 5.6e-09 | 6.767E-09 | 1.069E-09 | 5.600E-09 | 7.700E-09 |
| 8381 | 420 | S420A | A | 1.6e-09 | 1.4e-08 | 1.2e-08 | 9.200E-09 | 6.657E-09 | 1.600E-09 | 1.400E-08 |
| 8382 | 420 | S420R | A | 3.7e-08 | 1.5e-08 | 4.3e-09 | 1.877E-08 | 1.667E-08 | 4.300E-09 | 3.700E-08 |
| 8383 | 420 | S420N | A | 4.9e-09 | 7.6e-09 | 6e-09 | 6.167E-09 | 1.358E-09 | 4.900E-09 | 7.600E-09 |
| 8384 | 420 | S420D | A | 1.1e-08 | 7.9e-09 | 1.8e-09 | 6.900E-09 | 4.681E-09 | 1.800E-09 | 1.100E-08 |
| 8385 | 420 | S420C | A | 1.1e-08 | 3e-09 | 5.3e-09 | 6.433E-09 | 4.119E-09 | 3.000E-09 | 1.100E-08 |
| 8386 | 420 | S420E | A | 1.3e-08 | 7.8e-09 | 1.6e-08 | 1.227E-08 | 4.149E-09 | 7.800E-09 | 1.600E-08 |
| 8387 | 420 | S420Q | A | 5.1e-09 | 1e-08 | 5.2e-09 | 6.767E-09 | 2.801E-09 | 5.100E-09 | 1.000E-08 |
| 8388 | 420 | S420G | A | 3.8e-09 | 1.3e-08 | 1.5e-08 | 1.060E-08 | 5.973E-09 | 3.800E-09 | 1.500E-08 |
| 8389 | 420 | S420H | A | 1e-08 | 5.6e-09 | 9.6e-09 | 8.400E-09 | 2.433E-09 | 5.600E-09 | 1.000E-08 |
| 8390 | 420 | S420I | A | 1.3e-08 | 4.8e-09 | 1.5e-09 | 6.433E-09 | 5.921E-09 | 1.500E-09 | 1.300E-08 |
| 8391 | 420 | S420L | A | 1.1e-08 | 5.7e-09 | 9.4e-09 | 8.700E-09 | 2.718E-09 | 5.700E-09 | 1.100E-08 |
| 8392 | 420 | S420K | A | 1e-08 | 9.5e-09 | 2.5e-08 | 1.483E-08 | 8.808E-09 | 9.500E-09 | 2.500E-08 |
| 8393 | 420 | S420M | A | 5e-09 | 1.8e-08 | 3.3e-09 | 8.767E-09 | 8.041E-09 | 3.300E-09 | 1.800E-08 |
| 8394 | 420 | S420F | A | 9.1e-09 | 8.1e-09 | 6e-09 | 7.733E-09 | 1.582E-09 | 6.000E-09 | 9.100E-09 |
| 8395 | 420 | S420P | A | 2.5e-09 | 8.5e-09 | 4.5e-09 | 5.167E-09 | 3.055E-09 | 2.500E-09 | 8.500E-09 |
| 8396 | 420 | S420S | A | 5.2e-09 | 2e-08 | 5.7e-09 | 1.030E-08 | 8.404E-09 | 5.200E-09 | 2.000E-08 |
| 8397 | 420 | S420T | A | 9.5e-09 | 5.9e-09 | 7.7e-09 | 7.700E-09 | 1.800E-09 | 5.900E-09 | 9.500E-09 |
| 8398 | 420 | S420W | A | 1.5e-09 | 1.7e-08 | 3.7e-09 | 7.400E-09 | 8.386E-09 | 1.500E-09 | 1.700E-08 |
| 8399 | 420 | S420Y | A | 2e-08 | 3e-09 | 5.2e-09 | 9.400E-09 | 9.246E-09 | 3.000E-09 | 2.000E-08 |
| 8400 | 420 | S420V | A | 3.5e-09 | 4.6e-09 | 8.5e-09 | 5.533E-09 | 2.627E-09 | 3.500E-09 | 8.500E-09 |
| 8401 | 421 | L421A | A | 4.3e-09 | 6.7e-09 | 2e-09 | 4.333E-09 | 2.350E-09 | 2.000E-09 | 6.700E-09 |
| 8402 | 421 | L421R | A | 4.8e-09 | 1.3e-08 | 7.2e-09 | 8.333E-09 | 4.216E-09 | 4.800E-09 | 1.300E-08 |
| 8403 | 421 | L421N | A | 5.3e-09 | 1.1e-08 | 2.3e-08 | 1.310E-08 | 9.035E-09 | 5.300E-09 | 2.300E-08 |
| 8404 | 421 | L421D | A | 1.3e-08 | 4.4e-09 | 5.9e-09 | 7.767E-09 | 4.594E-09 | 4.400E-09 | 1.300E-08 |
| 8405 | 421 | L421C | A | 1.8e-09 | 4.3e-09 | 3.2e-09 | 3.100E-09 | 1.253E-09 | 1.800E-09 | 4.300E-09 |
| 8406 | 421 | L421E | A | 7.1e-09 | 3.8e-09 | 8.7e-09 | 6.533E-09 | 2.499E-09 | 3.800E-09 | 8.700E-09 |
| 8407 | 421 | L421Q | A | 6.1e-09 | 1.1e-08 | 5.6e-09 | 7.567E-09 | 2.984E-09 | 5.600E-09 | 1.100E-08 |
| 8408 | 421 | L421G | A | 5.1e-09 | 4.6e-09 | 3.4e-09 | 4.367E-09 | 8.737E-10 | 3.400E-09 | 5.100E-09 |
| 8409 | 421 | L421H | A | 3.7e-09 | 1.4e-09 | 7.3e-09 | 4.133E-09 | 2.974E-09 | 1.400E-09 | 7.300E-09 |
| 8410 | 421 | L421I | A | 7.4e-09 | 7.7e-09 | 7.3e-09 | 7.467E-09 | 2.082E-10 | 7.300E-09 | 7.700E-09 |
| 8411 | 421 | L421L | A | 1.5e-08 | 6.9e-09 | 5.8e-09 | 9.233E-09 | 5.024E-09 | 5.800E-09 | 1.500E-08 |
| 8412 | 421 | L421K | A | 5.4e-09 | 5e-09 | 3.6e-09 | 4.667E-09 | 9.452E-10 | 3.600E-09 | 5.400E-09 |
| 8413 | 421 | L421M | A | 5.3e-09 | 4.7e-09 | 1.1e-08 | 7.000E-09 | 3.477E-09 | 4.700E-09 | 1.100E-08 |
| 8414 | 421 | L421F | A | 1.2e-08 | 1.4e-08 | 1.9e-09 | 9.300E-09 | 6.486E-09 | 1.900E-09 | 1.400E-08 |

| | | | | | | | | | | |
|---|---|---|---|---|---|---|---|---|---|---|
| 8415 | 421 | L421P | A | 5.1e-09 | 8.2e-09 | 9e-09 | 7.433E-09 | 2.060E-09 | 5.100E-09 | 9.000E-09 |
| 8416 | 421 | L421S | A | 7.7e-09 | 1.5e-08 | 1.2e-08 | 1.157E-08 | 3.669E-09 | 7.700E-09 | 1.500E-08 |
| 8417 | 421 | L421T | A | 1.1e-08 | 1e-08 | 4.5e-09 | 8.500E-09 | 3.500E-09 | 4.500E-09 | 1.100E-08 |
| 8418 | 421 | L421W | A | 7.6e-09 | 1.3e-08 | 1e-08 | 1.020E-08 | 2.706E-09 | 7.600E-09 | 1.300E-08 |
| 8419 | 421 | L421Y | A | 1.4e-09 | 5.4e-09 | 1.7e-08 | 7.933E-09 | 8.103E-09 | 1.400E-09 | 1.700E-08 |
| 8420 | 421 | L421V | A | 2.9e-09 | 4.2e-09 | 5.7e-09 | 4.267E-09 | 1.401E-09 | 2.900E-09 | 5.700E-09 |
| 8421 | 422 | T422A | A | 8.4e-09 | 6.6e-09 | 4.8e-09 | 6.600E-09 | 1.800E-09 | 4.800E-09 | 8.400E-09 |
| 8422 | 422 | T422R | A | 6e-09 | 5.8e-09 | 2.2e-09 | 4.667E-09 | 2.139E-09 | 2.200E-09 | 6.000E-09 |
| 8423 | 422 | T422N | A | 2e-09 | 5.9e-09 | 1.2e-08 | 6.633E-09 | 5.040E-09 | 2.000E-09 | 1.200E-08 |
| 8424 | 422 | T422D | A | 4.7e-09 | 7.8e-09 | 6e-09 | 6.167E-09 | 1.557E-09 | 4.700E-09 | 7.800E-09 |
| 8425 | 422 | T422C | A | 6.1e-09 | 2.3e-09 | 1.7e-08 | 8.467E-09 | 7.630E-09 | 2.300E-09 | 1.700E-08 |
| 8426 | 422 | T422E | A | 5.9e-09 | 2.7e-09 | 1e-08 | 6.200E-09 | 3.659E-09 | 2.700E-09 | 1.000E-08 |
| 8427 | 422 | T422Q | A | 7.7e-09 | 6.7e-09 | 5.7e-09 | 6.700E-09 | 1.000E-09 | 5.700E-09 | 7.700E-09 |
| 8428 | 422 | T422G | A | 6.8e-09 | 4.5e-09 | 5.3e-09 | 5.533E-09 | 1.168E-09 | 4.500E-09 | 6.800E-09 |
| 8429 | 422 | T422H | A | 1.1e-08 | 1.3e-08 | 5.6e-09 | 9.867E-09 | 3.828E-09 | 5.600E-09 | 1.300E-08 |
| 8430 | 422 | T422I | A | 1.1e-08 | 3.8e-09 | 2.1e-08 | 1.193E-08 | 8.638E-09 | 3.800E-09 | 2.100E-08 |
| 8431 | 422 | T422L | A | 1e-08 | 1.3e-08 | 6.6e-09 | 9.867E-09 | 3.202E-09 | 6.600E-09 | 1.300E-08 |
| 8432 | 422 | T422K | A | 6.6e-09 | 5.8e-09 | 3.3e-09 | 5.233E-09 | 1.721E-09 | 3.300E-09 | 6.600E-09 |
| 8433 | 422 | T422M | A | 3e-09 | 4.2e-09 | 1.4e-08 | 7.067E-09 | 6.034E-09 | 3.000E-09 | 1.400E-08 |
| 8434 | 422 | T422F | A | 8e-09 | 3.8e-09 | 2.3e-09 | 4.700E-09 | 2.955E-09 | 2.300E-09 | 8.000E-09 |
| 8435 | 422 | T422P | A | 2.7e-08 | 1.6e-08 | 4.3e-09 | 1.577E-08 | 1.135E-08 | 4.300E-09 | 2.700E-08 |
| 8436 | 422 | T422S | A | 5.7e-09 | 6.8e-09 | 5.6e-09 | 6.033E-09 | 6.658E-10 | 5.600E-09 | 6.800E-09 |
| 8437 | 422 | T422T | A | 4.2e-09 | 3.5e-09 | 7.3e-09 | 5.000E-09 | 2.022E-09 | 3.500E-09 | 7.300E-09 |
| 8438 | 422 | T422W | A | 2.1e-08 | 5.2e-09 | 9.2e-09 | 1.180E-08 | 8.215E-09 | 5.200E-09 | 2.100E-08 |
| 8439 | 422 | T422Y | A | 9.6e-09 | 4.3e-09 | 6.7e-09 | 6.867E-09 | 2.654E-09 | 4.300E-09 | 9.600E-09 |
| 8440 | 422 | T422V | A | 2.7e-09 | 4.4e-09 | 4.6e-09 | 3.900E-09 | 1.044E-09 | 2.700E-09 | 4.600E-09 |
| 8441 | 423 | L423A | A | 1.2e-08 | 8.4e-09 | 5.3e-09 | 8.567E-09 | 3.353E-09 | 5.300E-09 | 1.200E-08 |
| 8442 | 423 | L423R | A | 3.8e-09 | 2.4e-08 | 5.1e-09 | 1.097E-08 | 1.131E-08 | 3.800E-09 | 2.400E-08 |
| 8443 | 423 | L423N | A | 5.8e-09 | 1.9e-08 | 7e-09 | 1.060E-08 | 7.299E-09 | 5.800E-09 | 1.900E-08 |
| 8444 | 423 | L423D | A | 3.5e-09 | 9.6e-09 | 6.5e-09 | 6.533E-09 | 3.050E-09 | 3.500E-09 | 9.600E-09 |
| 8445 | 423 | L423C | A | 5.7e-09 | 1.2e-08 | 9.4e-09 | 9.033E-09 | 3.166E-09 | 5.700E-09 | 1.200E-08 |
| 8446 | 423 | L423E | A | 6.5e-09 | 1.8e-08 | 5.9e-09 | 1.013E-08 | 6.819E-09 | 5.900E-09 | 1.800E-08 |
| 8447 | 423 | L423Q | A | 7.2e-09 | 3.3e-09 | 6e-09 | 5.500E-09 | 1.997E-09 | 3.300E-09 | 7.200E-09 |
| 8448 | 423 | L423G | A | 1e-08 | 7.5e-09 | 1.5e-08 | 1.083E-08 | 3.819E-09 | 7.500E-09 | 1.500E-08 |
| 8449 | 423 | L423H | A | 5e-09 | 6.9e-09 | 4.1e-09 | 5.333E-09 | 1.429E-09 | 4.100E-09 | 6.900E-09 |
| 8450 | 423 | L423I | A | 9.5e-09 | 4.5e-09 | 1.2e-08 | 8.667E-09 | 3.819E-09 | 4.500E-09 | 1.200E-08 |
| 8451 | 423 | L423L | A | 1.4e-08 | 1.4e-08 | 1e-08 | 1.267E-08 | 2.309E-09 | 1.000E-08 | 1.400E-08 |
| 8452 | 423 | L423K | A | 9.7e-09 | 7.1e-09 | 7.2e-09 | 8.000E-09 | 1.473E-09 | 7.100E-09 | 9.700E-09 |
| 8453 | 423 | L423M | A | 5.9e-09 | 3.1e-09 | 1.4e-08 | 7.667E-09 | 5.661E-09 | 3.100E-09 | 1.400E-08 |
| 8454 | 423 | L423F | A | 1.9e-09 | 7.6e-09 | 8.2e-09 | 5.900E-09 | 3.477E-09 | 1.900E-09 | 8.200E-09 |
| 8455 | 423 | L423P | A | 9.9e-09 | 4.3e-09 | 1.4e-08 | 9.400E-09 | 4.869E-09 | 4.300E-09 | 1.400E-08 |
| 8456 | 423 | L423S | A | 3.8e-09 | 6.6e-09 | 5.2e-09 | 5.200E-09 | 1.400E-09 | 3.800E-09 | 6.600E-09 |
| 8457 | 423 | L423T | A | 8.3e-09 | 3.6e-09 | 6.4e-09 | 6.100E-09 | 2.364E-09 | 3.600E-09 | 8.300E-09 |
| 8458 | 423 | L423W | A | 1.2e-08 | 7.2e-09 | 4.9e-09 | 8.033E-09 | 3.623E-09 | 4.900E-09 | 1.200E-08 |
| 8459 | 423 | L423Y | A | 3.2e-09 | 4.4e-09 | 1e-08 | 5.867E-09 | 3.630E-09 | 3.200E-09 | 1.000E-08 |
| 8460 | 423 | L423V | A | 5.8e-09 | 5.4e-09 | 7.9e-09 | 6.367E-09 | 1.343E-09 | 5.400E-09 | 7.900E-09 |
| 8461 | 424 | Q424A | A | 5.1e-08 | 6.5e-09 | 2.8e-09 | 2.010E-08 | 2.682E-08 | 2.800E-09 | 5.100E-08 |
| 8462 | 424 | Q424R | A | 6.9e-09 | 1e-08 | 2.6e-09 | 6.500E-09 | 3.716E-09 | 2.600E-09 | 1.000E-08 |
| 8463 | 424 | Q424N | A | 5.1e-09 | 7.6e-09 | 1.4e-08 | 8.900E-09 | 4.590E-09 | 5.100E-09 | 1.400E-08 |
| 8464 | 424 | Q424D | A | 6.4e-09 | 8.6e-09 | 3.2e-09 | 6.067E-09 | 2.715E-09 | 3.200E-09 | 8.600E-09 |
| 8465 | 424 | Q424C | A | 3.6e-09 | 1.3e-09 | 9.7e-09 | 4.867E-09 | 4.341E-09 | 1.300E-09 | 9.700E-09 |
| 8466 | 424 | Q424E | A | 4.5e-09 | 6.4e-09 | 1.2e-08 | 7.633E-09 | 3.899E-09 | 4.500E-09 | 1.200E-08 |
| 8467 | 424 | Q424Q | A | 7.2e-09 | 1.1e-08 | 9.7e-09 | 9.300E-09 | 1.931E-09 | 7.200E-09 | 1.100E-08 |
| 8468 | 424 | Q424G | A | 7.1e-09 | 3.2e-09 | 7.3e-09 | 5.867E-09 | 2.312E-09 | 3.200E-09 | 7.300E-09 |
| 8469 | 424 | Q424H | A | 2.4e-09 | 5e-09 | 3.1e-09 | 3.500E-09 | 1.345E-09 | 2.400E-09 | 5.000E-09 |
| 8470 | 424 | Q424I | A | 3.9e-09 | 8.6e-09 | 3.3e-09 | 5.267E-09 | 2.902E-09 | 3.300E-09 | 8.600E-09 |
| 8471 | 424 | Q424L | A | 6.5e-09 | 1.1e-08 | 5.7e-09 | 7.733E-09 | 2.857E-09 | 5.700E-09 | 1.100E-08 |
| 8472 | 424 | Q424K | A | 7.9e-09 | 6.5e-09 | 4.4e-09 | 6.267E-09 | 1.762E-09 | 4.400E-09 | 7.900E-09 |
| 8473 | 424 | Q424M | A | 5.3e-09 | 5.9e-09 | 5.9e-09 | 5.700E-09 | 3.464E-10 | 5.300E-09 | 5.900E-09 |
| 8474 | 424 | Q424F | A | 5.8e-09 | 4.2e-09 | 8.1e-09 | 6.033E-09 | 1.960E-09 | 4.200E-09 | 8.100E-09 |
| 8475 | 424 | Q424P | A | 7.1e-09 | 3e-09 | 3.8e-09 | 4.633E-09 | 2.173E-09 | 3.000E-09 | 7.100E-09 |
| 8476 | 424 | Q424S | A | 1.1e-08 | 3.1e-09 | 3.2e-09 | 5.767E-09 | 4.532E-09 | 3.100E-09 | 1.100E-08 |
| 8477 | 424 | Q424T | A | 4.6e-09 | 4.3e-09 | 6.9e-09 | 5.267E-09 | 1.422E-09 | 4.300E-09 | 6.900E-09 |
| 8478 | 424 | Q424W | A | 1.4e-09 | 5.7e-09 | 8.4e-09 | 5.167E-09 | 3.530E-09 | 1.400E-09 | 8.400E-09 |
| 8479 | 424 | Q424Y | A | 5.3e-09 | 3.8e-09 | 3.9e-09 | 4.333E-09 | 8.386E-10 | 3.800E-09 | 5.300E-09 |
| 8480 | 424 | Q424V | A | 3.6e-09 | 5.6e-09 | 9.4e-09 | 6.200E-09 | 2.946E-09 | 3.600E-09 | 9.400E-09 |
| 8481 | 425 | G425A | A | 1.3e-08 | 5.4e-09 | 3.8e-09 | 7.400E-09 | 4.915E-09 | 3.800E-09 | 1.300E-08 |
| 8482 | 425 | G425R | A | 3.9e-09 | 2.3e-08 | 7.8e-09 | 1.157E-08 | 1.009E-08 | 3.900E-09 | 2.300E-08 |
| 8483 | 425 | G425N | A | 6e-09 | 4.1e-09 | 1.1e-08 | 7.033E-09 | 3.564E-09 | 4.100E-09 | 1.100E-08 |
| 8484 | 425 | G425D | A | 5.7e-09 | 8.5e-09 | 3.7e-09 | 5.967E-09 | 2.411E-09 | 3.700E-09 | 8.500E-09 |
| 8485 | 425 | G425C | A | 6.7e-09 | 1.2e-08 | 5.7e-09 | 8.133E-09 | 3.386E-09 | 5.700E-09 | 1.200E-08 |
| 8486 | 425 | G425E | A | 6.2e-09 | 1.1e-08 | 7.4e-09 | 8.200E-09 | 2.498E-09 | 6.200E-09 | 1.100E-08 |
| 8487 | 425 | G425Q | A | 5.3e-09 | 1.2e-08 | 7.6e-09 | 8.300E-09 | 3.404E-09 | 5.300E-09 | 1.200E-08 |
| 8488 | 425 | G425G | A | 8.8e-09 | 1.7e-08 | 9.3e-09 | 1.170E-08 | 4.597E-09 | 8.800E-09 | 1.700E-08 |
| 8489 | 425 | G425H | A | 1.7e-09 | 1.6e-08 | 7.3e-09 | 8.333E-09 | 7.206E-09 | 1.700E-09 | 1.600E-08 |
| 8490 | 425 | G425I | A | 1.8e-08 | 6.3e-09 | 3e-09 | 9.100E-09 | 7.882E-09 | 3.000E-09 | 1.800E-08 |
| 8491 | 425 | G425L | A | 7.3e-09 | 6.2e-09 | 1.2e-08 | 8.500E-09 | 3.081E-09 | 6.200E-09 | 1.200E-08 |
| 8492 | 425 | G425K | A | 7e-09 | 2.1e-09 | 4.6e-09 | 4.567E-09 | 2.450E-09 | 2.100E-09 | 7.000E-09 |
| 8493 | 425 | G425M | A | 7.1e-09 | 1e-08 | 5.5e-09 | 7.533E-09 | 2.281E-09 | 5.500E-09 | 1.000E-08 |
| 8494 | 425 | G425F | A | 3.7e-09 | 1.4e-09 | 1.6e-08 | 7.033E-09 | 7.850E-09 | 1.400E-09 | 1.600E-08 |
| 8495 | 425 | G425P | A | 9.2e-09 | 3.9e-09 | 2.7e-09 | 5.267E-09 | 3.459E-09 | 2.700E-09 | 9.200E-09 |
| 8496 | 425 | G425S | A | 3.2e-09 | 6.2e-09 | 3e-09 | 4.133E-09 | 1.793E-09 | 3.000E-09 | 6.200E-09 |
| 8497 | 425 | G425T | A | 6.8e-09 | 6e-09 | 6.1e-09 | 6.300E-09 | 4.359E-10 | 6.000E-09 | 6.800E-09 |
| 8498 | 425 | G425W | A | 6e-09 | 1.1e-08 | 5.9e-09 | 7.633E-09 | 2.916E-09 | 5.900E-09 | 1.100E-08 |
| 8499 | 425 | G425Y | A | 1.8e-08 | 7.5e-09 | 9.7e-09 | 1.173E-08 | 5.537E-09 | 7.500E-09 | 1.800E-08 |

| | | | | | | | | | | |
|---|---|---|---|---|---|---|---|---|---|---|
| 8500 | 425 | G425V | A | 1e-08 | 8.8e-09 | 7.8e-09 | 8.867E-09 | 1.102E-09 | 7.800E-09 | 1.000E-08 |
| 8501 | 426 | L426A | A | 6.8e-09 | 3.6e-09 | 1.7e-08 | 9.133E-09 | 6.998E-09 | 3.600E-09 | 1.700E-08 |
| 8502 | 426 | L426R | A | 2.8e-09 | 3.9e-09 | 4.1e-09 | 3.600E-09 | 7.000E-10 | 2.800E-09 | 4.100E-09 |
| 8503 | 426 | L426N | A | 2.3e-09 | 6.8e-09 | 5.9e-09 | 5.000E-09 | 2.381E-09 | 2.300E-09 | 6.800E-09 |
| 8504 | 426 | L426D | A | 8.8e-09 | 1.1e-08 | 8e-09 | 9.267E-09 | 1.553E-09 | 8.000E-09 | 1.100E-08 |
| 8505 | 426 | L426C | A | 7.6e-09 | 1.1e-08 | 8.2e-09 | 8.933E-09 | 1.815E-09 | 7.600E-09 | 1.100E-08 |
| 8506 | 426 | L426E | A | 9.4e-09 | 7.5e-09 | 7.4e-09 | 8.100E-09 | 1.127E-09 | 7.400E-09 | 9.400E-09 |
| 8507 | 426 | L426Q | A | 3.2e-09 | 3.6e-09 | 1.5e-08 | 7.267E-09 | 6.700E-09 | 3.200E-09 | 1.500E-08 |
| 8508 | 426 | L426G | A | 2.5e-09 | 3.1e-09 | 4.8e-09 | 3.467E-09 | 1.193E-09 | 2.500E-09 | 4.800E-09 |
| 8509 | 426 | L426H | A | 5.9e-09 | 1.1e-08 | 3.4e-09 | 6.767E-09 | 3.873E-09 | 3.400E-09 | 1.100E-08 |
| 8510 | 426 | L426I | A | 4e-09 | 2.7e-08 | 6.5e-09 | 1.250E-08 | 1.262E-08 | 4.000E-09 | 2.700E-08 |
| 8511 | 426 | L426L | A | 8.9e-09 | 9.7e-09 | 1.5e-08 | 1.120E-08 | 3.315E-09 | 8.900E-09 | 1.500E-08 |
| 8512 | 426 | L426K | A | 4.4e-09 | 1.4e-08 | 7.3e-09 | 8.567E-09 | 4.924E-09 | 4.400E-09 | 1.400E-08 |
| 8513 | 426 | L426M | A | 5.3e-09 | 6.3e-09 | 3.9e-09 | 5.167E-09 | 1.206E-09 | 3.900E-09 | 6.300E-09 |
| 8514 | 426 | L426F | A | 9.5e-09 | 8.7e-09 | 1.1e-08 | 9.733E-09 | 1.168E-09 | 8.700E-09 | 1.100E-08 |
| 8515 | 426 | L426P | A | 3.8e-09 | 1e-08 | 8.9e-09 | 7.567E-09 | 3.308E-09 | 3.800E-09 | 1.000E-08 |
| 8516 | 426 | L426S | A | 6.7e-09 | 4.4e-09 | 2.7e-09 | 4.600E-09 | 2.007E-09 | 2.700E-09 | 6.700E-09 |
| 8517 | 426 | L426T | A | 5.5e-09 | 7.7e-09 | 2e-08 | 1.107E-08 | 7.814E-09 | 5.500E-09 | 2.000E-08 |
| 8518 | 426 | L426W | A | 2.7e-08 | 2.3e-09 | 4.9e-09 | 1.140E-08 | 1.357E-08 | 2.300E-09 | 2.700E-08 |
| 8519 | 426 | L426Y | A | 1.9e-08 | 6.9e-09 | 7.9e-09 | 1.127E-08 | 6.716E-09 | 6.900E-09 | 1.900E-08 |
| 8520 | 426 | L426V | A | 1.5e-08 | 1.3e-08 | 4.6e-09 | 1.087E-08 | 5.518E-09 | 4.600E-09 | 1.500E-08 |
| 8521 | 427 | G427A | A | 1.2e-08 | 7.6e-09 | 5.2e-09 | 8.267E-09 | 3.449E-09 | 5.200E-09 | 1.200E-08 |
| 8522 | 427 | G427R | A | 5.9e-09 | 1.3e-08 | 1.6e-09 | 6.833E-09 | 5.757E-09 | 1.600E-09 | 1.300E-08 |
| 8523 | 427 | G427N | A | 7.7e-09 | 1.4e-08 | 9.1e-09 | 1.027E-08 | 3.308E-09 | 7.700E-09 | 1.400E-08 |
| 8524 | 427 | G427D | A | 4e-09 | 9.2e-09 | 5.9e-09 | 6.367E-09 | 2.631E-09 | 4.000E-09 | 9.200E-09 |
| 8525 | 427 | G427C | A | 6.4e-09 | 8.5e-09 | 3.3e-09 | 6.067E-09 | 2.616E-09 | 3.300E-09 | 8.500E-09 |
| 8526 | 427 | G427E | A | 6.6e-09 | 7.9e-09 | 3.4e-09 | 5.967E-09 | 2.316E-09 | 3.400E-09 | 7.900E-09 |
| 8527 | 427 | G427Q | A | 9.2e-09 | 1.5e-08 | 4.4e-09 | 9.533E-09 | 5.308E-09 | 4.400E-09 | 1.500E-08 |
| 8528 | 427 | G427G | A | 9.2e-09 | 2.8e-09 | 1.2e-08 | 8.000E-09 | 4.716E-09 | 2.800E-09 | 1.200E-08 |
| 8529 | 427 | G427H | A | 8e-09 | 3.7e-09 | 1.4e-08 | 8.567E-09 | 5.173E-09 | 3.700E-09 | 1.400E-08 |
| 8530 | 427 | G427I | A | 1.5e-08 | 6.9e-09 | 9e-09 | 1.030E-08 | 4.204E-09 | 6.900E-09 | 1.500E-08 |
| 8531 | 427 | G427L | A | 8.4e-09 | 3.7e-09 | 1.5e-08 | 9.033E-09 | 5.677E-09 | 3.700E-09 | 1.500E-08 |
| 8532 | 427 | G427K | A | 6e-09 | 1.3e-08 | 3.3e-08 | 1.733E-08 | 1.401E-08 | 6.000E-09 | 3.300E-08 |
| 8533 | 427 | G427M | A | 9.3e-09 | 2e-08 | 6.2e-09 | 1.183E-08 | 7.240E-09 | 6.200E-09 | 2.000E-08 |
| 8534 | 427 | G427F | A | 5.1e-09 | 2.8e-08 | 6.1e-09 | 1.307E-08 | 1.294E-08 | 5.100E-09 | 2.800E-08 |
| 8535 | 427 | G427P | A | 1.4e-08 | 6.2e-09 | 5.5e-09 | 8.567E-09 | 4.718E-09 | 5.500E-09 | 1.400E-08 |
| 8536 | 427 | G427S | A | 1.7e-08 | 7.7e-09 | 2e-09 | 8.900E-09 | 7.572E-09 | 2.000E-09 | 1.700E-08 |
| 8537 | 427 | G427T | A | 2.4e-09 | 1.4e-08 | 3.3e-09 | 6.567E-09 | 6.453E-09 | 2.400E-09 | 1.400E-08 |
| 8538 | 427 | G427W | A | 1.3e-08 | 3.7e-09 | 2.4e-09 | 6.367E-09 | 5.781E-09 | 2.400E-09 | 1.300E-08 |
| 8539 | 427 | G427Y | A | 1.6e-08 | 7e-09 | 7.5e-09 | 1.017E-08 | 5.058E-09 | 7.000E-09 | 1.600E-08 |
| 8540 | 427 | G427V | A | 8.7e-09 | 1.3e-08 | 9e-09 | 1.023E-08 | 2.401E-09 | 8.700E-09 | 1.300E-08 |
| 8541 | 428 | I428A | A | 1.8e-08 | 8.2e-09 | 5.7e-09 | 1.063E-08 | 6.501E-09 | 5.700E-09 | 1.800E-08 |
| 8542 | 428 | I428R | A | 2.9e-08 | 3.8e-09 | 1e-08 | 1.427E-08 | 1.313E-08 | 3.800E-09 | 2.900E-08 |
| 8543 | 428 | I428N | A | 3.1e-09 | 7.7e-09 | 3.4e-09 | 4.733E-09 | 2.574E-09 | 3.100E-09 | 7.700E-09 |
| 8544 | 428 | I428D | A | 1.2e-08 | 2.7e-09 | 8.5e-09 | 7.733E-09 | 4.697E-09 | 2.700E-09 | 1.200E-08 |
| 8545 | 428 | I428C | A | 1.2e-08 | 5.5e-09 | 6.4e-09 | 7.967E-09 | 3.522E-09 | 5.500E-09 | 1.200E-08 |
| 8546 | 428 | I428E | A | 8.3e-09 | 1.3e-08 | 1.7e-08 | 1.277E-08 | 4.355E-09 | 8.300E-09 | 1.700E-08 |
| 8547 | 428 | I428Q | A | 1.5e-08 | 8.7e-09 | 5.5e-09 | 9.733E-09 | 4.834E-09 | 5.500E-09 | 1.500E-08 |
| 8548 | 428 | I428G | A | 7.4e-09 | 6.4e-09 | 2.1e-08 | 1.160E-08 | 8.156E-09 | 6.400E-09 | 2.100E-08 |
| 8549 | 428 | I428H | A | 1e-08 | 1.3e-08 | 9.4e-09 | 1.080E-08 | 1.929E-09 | 9.400E-09 | 1.300E-08 |
| 8550 | 428 | I428I | A | 4.4e-09 | 8.6e-09 | 1.3e-08 | 8.667E-09 | 4.300E-09 | 4.400E-09 | 1.300E-08 |
| 8551 | 428 | I428L | A | 2.2e-09 | 2e-09 | 3.5e-09 | 2.567E-09 | 8.145E-10 | 2.000E-09 | 3.500E-09 |
| 8552 | 428 | I428K | A | 5.6e-09 | 5.3e-09 | 1.2e-08 | 7.633E-09 | 3.785E-09 | 5.300E-09 | 1.200E-08 |
| 8553 | 428 | I428M | A | 7e-09 | 1.1e-08 | 6e-09 | 8.000E-09 | 2.646E-09 | 6.000E-09 | 1.100E-08 |
| 8554 | 428 | I428F | A | 1.7e-08 | 1.4e-08 | 2.1e-08 | 1.733E-08 | 3.512E-09 | 1.400E-08 | 2.100E-08 |
| 8555 | 428 | I428P | A | 1.2e-08 | 1.2e-08 | 4.1e-09 | 9.367E-09 | 4.561E-09 | 4.100E-09 | 1.200E-08 |
| 8556 | 428 | I428S | A | 2.2e-08 | 3.6e-09 | 4.1e-09 | 9.900E-09 | 1.048E-08 | 3.600E-09 | 2.200E-08 |
| 8557 | 428 | I428T | A | 5.8e-09 | 1e-08 | 6.4e-09 | 7.400E-09 | 2.272E-09 | 5.800E-09 | 1.000E-08 |
| 8558 | 428 | I428W | A | 1.1e-08 | 5.6e-09 | 2.6e-09 | 6.400E-09 | 4.257E-09 | 2.600E-09 | 1.100E-08 |
| 8559 | 428 | I428Y | A | 4.6e-09 | 4.4e-09 | 7.9e-09 | 5.633E-09 | 1.966E-09 | 4.400E-09 | 7.900E-09 |
| 8560 | 428 | I428V | A | 6e-09 | 5.4e-09 | 2.9e-09 | 4.767E-09 | 1.644E-09 | 2.900E-09 | 6.000E-09 |
| 8561 | 429 | S429A | A | 1.2e-09 | 1.4e-08 | 9.7e-09 | 8.300E-09 | 6.514E-09 | 1.200E-09 | 1.400E-08 |
| 8562 | 429 | S429R | A | 3.7e-09 | 7e-09 | 1e-08 | 6.900E-09 | 3.151E-09 | 3.700E-09 | 1.000E-08 |
| 8563 | 429 | S429N | A | 1.1e-08 | 7.6e-09 | 4e-09 | 7.533E-09 | 3.500E-09 | 4.000E-09 | 1.100E-08 |
| 8564 | 429 | S429D | A | 3.7e-09 | 8.2e-09 | 5.3e-09 | 5.733E-09 | 2.281E-09 | 3.700E-09 | 8.200E-09 |
| 8565 | 429 | S429C | A | 4.1e-09 | 7.9e-09 | 4.8e-09 | 5.600E-09 | 2.022E-09 | 4.100E-09 | 7.900E-09 |
| 8566 | 429 | S429E | A | 6.7e-09 | 4.7e-09 | 1.6e-08 | 9.133E-09 | 6.030E-09 | 4.700E-09 | 1.600E-08 |
| 8567 | 429 | S429Q | A | 5.5e-09 | 9e-09 | 1.2e-08 | 8.833E-09 | 3.253E-09 | 5.500E-09 | 1.200E-08 |
| 8568 | 429 | S429G | A | 8.5e-09 | 4.8e-09 | 4.2e-08 | 1.843E-08 | 2.049E-08 | 4.800E-09 | 4.200E-08 |
| 8569 | 429 | S429H | A | 7.5e-09 | 6.5e-09 | 1.1e-08 | 8.333E-09 | 2.363E-09 | 6.500E-09 | 1.100E-08 |
| 8570 | 429 | S429I | A | 8.8e-09 | 1.3e-08 | 1e-08 | 1.060E-08 | 2.163E-09 | 8.800E-09 | 1.300E-08 |
| 8571 | 429 | S429L | A | 1.7e-08 | 4.3e-09 | 1.4e-08 | 1.177E-08 | 6.638E-09 | 4.300E-09 | 1.700E-08 |
| 8572 | 429 | S429K | A | 2.3e-09 | 1.1e-08 | 9.3e-09 | 7.533E-09 | 4.611E-09 | 2.300E-09 | 1.100E-08 |
| 8573 | 429 | S429M | A | 3.9e-09 | 6.8e-09 | 1e-08 | 6.900E-09 | 3.051E-09 | 3.900E-09 | 1.000E-08 |
| 8574 | 429 | S429F | A | 1.1e-08 | 5.5e-09 | 8.3e-09 | 8.267E-09 | 2.750E-09 | 5.500E-09 | 1.100E-08 |
| 8575 | 429 | S429P | A | 4.5e-09 | 1.3e-08 | 2.2e-09 | 6.567E-09 | 5.689E-09 | 2.200E-09 | 1.300E-08 |
| 8576 | 429 | S429S | A | 1.1e-08 | 6.3e-09 | 7e-09 | 8.100E-09 | 2.536E-09 | 6.300E-09 | 1.100E-08 |
| 8577 | 429 | S429T | A | 1.2e-08 | 4.7e-09 | 5.7e-09 | 7.467E-09 | 3.958E-09 | 4.700E-09 | 1.200E-08 |
| 8578 | 429 | S429W | A | 1.4e-08 | 9e-09 | 1.5e-08 | 1.267E-08 | 3.215E-09 | 9.000E-09 | 1.500E-08 |
| 8579 | 429 | S429Y | A | 7.7e-09 | 1.1e-08 | 1.5e-08 | 1.123E-08 | 3.656E-09 | 7.700E-09 | 1.500E-08 |
| 8580 | 429 | S429V | A | 6e-09 | 5.1e-09 | 1.2e-08 | 7.700E-09 | 3.751E-09 | 5.100E-09 | 1.200E-08 |
| 8581 | 430 | W430A | A | 1.3e-08 | 2.7e-08 | 5.8e-09 | 1.527E-08 | 1.078E-08 | 5.800E-09 | 2.700E-08 |
| 8582 | 430 | W430R | A | 1.5e-08 | 5.2e-09 | 6e-09 | 8.733E-09 | 5.442E-09 | 5.200E-09 | 1.500E-08 |
| 8583 | 430 | W430N | A | 1.7e-09 | 3.5e-09 | 6.1e-09 | 3.767E-09 | 2.212E-09 | 1.700E-09 | 6.100E-09 |
| 8584 | 430 | W430D | A | 8.3e-09 | 4.9e-09 | 4.9e-09 | 6.033E-09 | 1.963E-09 | 4.900E-09 | 8.300E-09 |

| | | | | | | | | | | |
|---|---|---|---|---|---|---|---|---|---|---|
| 8585 | 430 | W430C | A | 5.4e-09 | 1.8e-08 | 7e-09 | 1.013E-08 | 6.860E-09 | 5.400E-09 | 1.800E-08 |
| 8586 | 430 | W430E | A | 5.6e-09 | 2e-08 | 6.8e-09 | 1.080E-08 | 7.990E-09 | 5.600E-09 | 2.000E-08 |
| 8587 | 430 | W430Q | A | 1.2e-08 | 7.7e-09 | 3e-09 | 7.567E-09 | 4.501E-09 | 3.000E-09 | 1.200E-08 |
| 8588 | 430 | W430G | A | 2.9e-09 | 7.2e-09 | 6.9e-09 | 5.667E-09 | 2.401E-09 | 2.900E-09 | 7.200E-09 |
| 8589 | 430 | W430H | A | 1.2e-08 | 5.7e-09 | 1.5e-08 | 1.090E-08 | 4.747E-09 | 5.700E-09 | 1.500E-08 |
| 8590 | 430 | W430I | A | 8.7e-09 | 9.9e-09 | 5.1e-09 | 7.900E-09 | 2.498E-09 | 5.100E-09 | 9.900E-09 |
| 8591 | 430 | W430L | A | 1.5e-08 | 1.6e-08 | 1.6e-08 | 1.567E-08 | 5.774E-10 | 1.500E-08 | 1.600E-08 |
| 8592 | 430 | W430K | A | 5.4e-09 | 4.5e-09 | 3.9e-09 | 4.600E-09 | 7.550E-10 | 3.900E-09 | 5.400E-09 |
| 8593 | 430 | W430M | A | 4e-09 | 1.4e-08 | 5.9e-09 | 7.967E-09 | 5.311E-09 | 4.000E-09 | 1.400E-08 |
| 8594 | 430 | W430F | A | 5.5e-09 | 2.3e-09 | 4.6e-09 | 4.133E-09 | 1.650E-09 | 2.300E-09 | 5.500E-09 |
| 8595 | 430 | W430P | A | 1.5e-08 | 3.6e-09 | 1.1e-08 | 9.867E-09 | 5.784E-09 | 3.600E-09 | 1.500E-08 |
| 8596 | 430 | W430S | A | 4.5e-09 | 1.7e-08 | 5.5e-09 | 9.000E-09 | 6.946E-09 | 4.500E-09 | 1.700E-08 |
| 8597 | 430 | W430T | A | 7.4e-09 | 4.7e-09 | 3.4e-08 | 1.537E-08 | 1.619E-08 | 4.700E-09 | 3.400E-08 |
| 8598 | 430 | W430W | A | 2.8e-09 | 2.3e-09 | 3e-09 | 2.700E-09 | 3.606E-10 | 2.300E-09 | 3.000E-09 |
| 8599 | 430 | W430Y | A | 7.1e-09 | 8.5e-09 | 1.3e-08 | 9.533E-09 | 3.083E-09 | 7.100E-09 | 1.300E-08 |
| 8600 | 430 | W430V | A | 5.5e-09 | 1.4e-08 | 9.2e-09 | 9.567E-09 | 4.262E-09 | 5.500E-09 | 1.400E-08 |
| 8601 | 431 | L431A | A | 1.9e-08 | 3.5e-09 | 1.2e-08 | 1.150E-08 | 7.762E-09 | 3.500E-09 | 1.900E-08 |
| 8602 | 431 | L431R | A | 3.3e-09 | 5.8e-09 | 7.9e-09 | 5.667E-09 | 2.303E-09 | 3.300E-09 | 7.900E-09 |
| 8603 | 431 | L431N | A | 3.2e-09 | 1.1e-08 | 3.2e-09 | 5.800E-09 | 4.503E-09 | 3.200E-09 | 1.100E-08 |
| 8604 | 431 | L431D | A | 1.2e-08 | 5.3e-09 | 2.1e-09 | 6.467E-09 | 5.052E-09 | 2.100E-09 | 1.200E-08 |
| 8605 | 431 | L431C | A | 2.3e-09 | 1.5e-08 | 6.6e-09 | 7.967E-09 | 6.459E-09 | 2.300E-09 | 1.500E-08 |
| 8606 | 431 | L431E | A | 1.3e-08 | 7.4e-09 | 4.2e-09 | 8.200E-09 | 4.454E-09 | 4.200E-09 | 1.300E-08 |
| 8607 | 431 | L431Q | A | 4.3e-09 | 1.1e-08 | 1.8e-09 | 5.700E-09 | 4.757E-09 | 1.800E-09 | 1.100E-08 |
| 8608 | 431 | L431G | A | 5e-09 | 9.2e-09 | 1.8e-08 | 1.073E-08 | 6.634E-09 | 5.000E-09 | 1.800E-08 |
| 8609 | 431 | L431H | A | 4.4e-09 | 2.2e-09 | 2.4e-08 | 1.020E-08 | 1.200E-08 | 2.200E-09 | 2.400E-08 |
| 8610 | 431 | L431I | A | 1e-08 | 1.7e-08 | 2.2e-09 | 9.733E-09 | 7.404E-09 | 2.200E-09 | 1.700E-08 |
| 8611 | 431 | L431L | A | 8e-09 | 5.2e-09 | 6.2e-09 | 6.467E-09 | 1.419E-09 | 5.200E-09 | 8.000E-09 |
| 8612 | 431 | L431K | A | 9.1e-09 | 2.9e-09 | 7.1e-09 | 6.367E-09 | 3.164E-09 | 2.900E-09 | 9.100E-09 |
| 8613 | 431 | L431M | A | 4.9e-09 | 5.5e-09 | 7e-09 | 5.800E-09 | 1.082E-09 | 4.900E-09 | 7.000E-09 |
| 8614 | 431 | L431F | A | 2.6e-09 | 1.1e-08 | 6.4e-09 | 6.667E-09 | 4.206E-09 | 2.600E-09 | 1.100E-08 |
| 8615 | 431 | L431P | A | 5.9e-09 | 1.5e-08 | 6.1e-09 | 9.000E-09 | 5.197E-09 | 5.900E-09 | 1.500E-08 |
| 8616 | 431 | L431S | A | 4.3e-09 | 1e-08 | 7.9e-09 | 7.400E-09 | 2.883E-09 | 4.300E-09 | 1.000E-08 |
| 8617 | 431 | L431T | A | 7.7e-09 | 9.3e-09 | 6.9e-09 | 7.967E-09 | 1.222E-09 | 6.900E-09 | 9.300E-09 |
| 8618 | 431 | L431W | A | 4.7e-09 | 7.1e-09 | 1.5e-08 | 8.933E-09 | 5.389E-09 | 4.700E-09 | 1.500E-08 |
| 8619 | 431 | L431Y | A | 1.2e-08 | 4.8e-09 | 8.8e-09 | 8.533E-09 | 3.607E-09 | 4.800E-09 | 1.200E-08 |
| 8620 | 431 | L431V | A | 9.8e-09 | 8.8e-09 | 5.2e-09 | 7.933E-09 | 2.419E-09 | 5.200E-09 | 9.800E-09 |
| 8621 | 432 | G432A | A | 1.4e-08 | 8.6e-09 | 1.2e-08 | 1.153E-08 | 2.730E-09 | 8.600E-09 | 1.400E-08 |
| 8622 | 432 | G432R | A | 4.6e-09 | 4.8e-09 | 3.2e-09 | 4.200E-09 | 8.718E-10 | 3.200E-09 | 4.800E-09 |
| 8623 | 432 | G432N | A | 8e-09 | 4.7e-09 | 1.2e-08 | 8.233E-09 | 3.656E-09 | 4.700E-09 | 1.200E-08 |
| 8624 | 432 | G432D | A | 9.3e-09 | 4.9e-09 | 1.9e-09 | 5.367E-09 | 3.722E-09 | 1.900E-09 | 9.300E-09 |
| 8625 | 432 | G432C | A | 2.6e-09 | 1.1e-08 | 8.6e-09 | 7.400E-09 | 4.327E-09 | 2.600E-09 | 1.100E-08 |
| 8626 | 432 | G432E | A | 6.5e-09 | 7.8e-09 | 4.9e-09 | 6.400E-09 | 1.453E-09 | 4.900E-09 | 7.800E-09 |
| 8627 | 432 | G432Q | A | 1.6e-08 | 1.1e-08 | 1.8e-08 | 1.500E-08 | 3.606E-09 | 1.100E-08 | 1.800E-08 |
| 8628 | 432 | G432G | A | 2.2e-09 | 9.1e-09 | 6.2e-09 | 5.833E-09 | 3.465E-09 | 2.200E-09 | 9.100E-09 |
| 8629 | 432 | G432H | A | 3.9e-09 | 3e-09 | 3.1e-09 | 3.333E-09 | 4.933E-10 | 3.000E-09 | 3.900E-09 |
| 8630 | 432 | G432I | A | 2.4e-09 | 1.1e-08 | 4.6e-09 | 6.000E-09 | 4.468E-09 | 2.400E-09 | 1.100E-08 |
| 8631 | 432 | G432L | A | 1.2e-08 | 1.1e-08 | 7e-09 | 1.000E-08 | 2.646E-09 | 7.000E-09 | 1.200E-08 |
| 8632 | 432 | G432K | A | 1.2e-08 | 4.3e-09 | 1.5e-08 | 1.043E-08 | 5.519E-09 | 4.300E-09 | 1.500E-08 |
| 8633 | 432 | G432M | A | 6.2e-09 | 1e-08 | 6.8e-09 | 7.667E-09 | 2.043E-09 | 6.200E-09 | 1.000E-08 |
| 8634 | 432 | G432F | A | 1.4e-08 | 5.4e-09 | 1.3e-08 | 1.080E-08 | 4.703E-09 | 5.400E-09 | 1.400E-08 |
| 8635 | 432 | G432P | A | 1.1e-08 | 6.8e-09 | 1.5e-08 | 1.093E-08 | 4.100E-09 | 6.800E-09 | 1.500E-08 |
| 8636 | 432 | G432S | A | 1.4e-08 | 1.3e-08 | 8.3e-09 | 1.177E-08 | 3.044E-09 | 8.300E-09 | 1.400E-08 |
| 8637 | 432 | G432T | A | 7e-09 | 8e-09 | 7.5e-09 | 7.500E-09 | 5.000E-10 | 7.000E-09 | 8.000E-09 |
| 8638 | 432 | G432W | A | 6.7e-09 | 4.7e-09 | 1.1e-09 | 4.167E-09 | 2.838E-09 | 1.100E-09 | 6.700E-09 |
| 8639 | 432 | G432Y | A | 5.3e-09 | 7.6e-09 | 1e-08 | 7.633E-09 | 2.350E-09 | 5.300E-09 | 1.000E-08 |
| 8640 | 432 | G432V | A | 1.1e-08 | 5.3e-09 | 5.1e-09 | 7.133E-09 | 3.350E-09 | 5.100E-09 | 1.100E-08 |
| 8641 | 433 | L433A | A | 1.1e-08 | 4.8e-09 | 2.8e-09 | 6.200E-09 | 4.276E-09 | 2.800E-09 | 1.100E-08 |
| 8642 | 433 | L433R | A | 1.7e-08 | 1.7e-08 | 9.1e-10 | 1.164E-08 | 9.290E-09 | 9.100E-10 | 1.700E-08 |
| 8643 | 433 | L433N | A | 1.3e-08 | 8.9e-09 | 3.5e-09 | 8.467E-09 | 4.765E-09 | 3.500E-09 | 1.300E-08 |
| 8644 | 433 | L433D | A | 7.9e-09 | 5.7e-09 | 1.4e-08 | 9.200E-09 | 4.300E-09 | 5.700E-09 | 1.400E-08 |
| 8645 | 433 | L433C | A | 5.4e-09 | 5.1e-09 | 1.2e-08 | 7.500E-09 | 3.900E-09 | 5.100E-09 | 1.200E-08 |
| 8646 | 433 | L433E | A | 7.2e-09 | 6.7e-09 | 1.9e-08 | 1.097E-08 | 6.962E-09 | 6.700E-09 | 1.900E-08 |
| 8647 | 433 | L433Q | A | 5.6e-09 | 4.4e-09 | 6.5e-09 | 5.500E-09 | 1.054E-09 | 4.400E-09 | 6.500E-09 |
| 8648 | 433 | L433G | A | 1.1e-08 | 6.9e-09 | 7.1e-09 | 8.333E-09 | 2.312E-09 | 6.900E-09 | 1.100E-08 |
| 8649 | 433 | L433H | A | 1.5e-08 | 7.4e-09 | 3.1e-09 | 8.500E-09 | 6.026E-09 | 3.100E-09 | 1.500E-08 |
| 8650 | 433 | L433I | A | 7.5e-09 | 8.8e-09 | 2.9e-09 | 6.400E-09 | 3.100E-09 | 2.900E-09 | 8.800E-09 |
| 8651 | 433 | L433L | A | 1.2e-08 | 1.4e-08 | 4.6e-09 | 1.020E-08 | 4.952E-09 | 4.600E-09 | 1.400E-08 |
| 8652 | 433 | L433K | A | 1.4e-09 | 4.3e-09 | 1.1e-08 | 5.567E-09 | 4.924E-09 | 1.400E-09 | 1.100E-08 |
| 8653 | 433 | L433M | A | 1.3e-08 | 2.2e-08 | 6.3e-09 | 1.377E-08 | 7.878E-09 | 6.300E-09 | 2.200E-08 |
| 8654 | 433 | L433F | A | 3.6e-09 | 7.8e-09 | 6.7e-09 | 6.033E-09 | 2.178E-09 | 3.600E-09 | 7.800E-09 |
| 8655 | 433 | L433P | A | 4.3e-09 | 1.4e-08 | 4.3e-09 | 7.533E-09 | 5.600E-09 | 4.300E-09 | 1.400E-08 |
| 8656 | 433 | L433S | A | 4.1e-08 | 5.4e-09 | 2.2e-09 | 1.620E-08 | 2.154E-08 | 2.200E-09 | 4.100E-08 |
| 8657 | 433 | L433T | A | 5.2e-09 | 1.9e-08 | 1.2e-08 | 1.207E-08 | 6.900E-09 | 5.200E-09 | 1.900E-08 |
| 8658 | 433 | L433W | A | 1e-08 | 3.6e-09 | 1.2e-08 | 8.533E-09 | 4.388E-09 | 3.600E-09 | 1.200E-08 |
| 8659 | 433 | L433Y | A | 6.4e-09 | 1.5e-08 | 5.8e-09 | 9.067E-09 | 5.147E-09 | 5.800E-09 | 1.500E-08 |
| 8660 | 433 | L433V | A | 1e-08 | 2.9e-09 | 2e-08 | 1.097E-08 | 8.591E-09 | 2.900E-09 | 2.000E-08 |
| 8661 | 434 | R434A | A | 1.1e-08 | 9.1e-09 | 1.5e-08 | 1.170E-08 | 3.012E-09 | 9.100E-09 | 1.500E-08 |
| 8662 | 434 | R434R | A | 3.3e-09 | 8.4e-09 | 2.1e-09 | 4.600E-09 | 3.345E-09 | 2.100E-09 | 8.400E-09 |
| 8663 | 434 | R434N | A | 1.1e-08 | 5.3e-09 | 1.1e-08 | 9.100E-09 | 3.291E-09 | 5.300E-09 | 1.100E-08 |
| 8664 | 434 | R434D | A | 6.3e-09 | 1.5e-08 | 6.4e-09 | 9.233E-09 | 4.994E-09 | 6.300E-09 | 1.500E-08 |
| 8665 | 434 | R434C | A | 1.1e-08 | 1.1e-08 | 5.3e-09 | 9.100E-09 | 3.291E-09 | 5.300E-09 | 1.100E-08 |
| 8666 | 434 | R434E | A | 9.2e-09 | 8.6e-09 | 7.5e-09 | 8.433E-09 | 8.622E-10 | 7.500E-09 | 9.200E-09 |
| 8667 | 434 | R434Q | A | 2e-09 | 1.6e-08 | 8.2e-09 | 8.733E-09 | 7.015E-09 | 2.000E-09 | 1.600E-08 |
| 8668 | 434 | R434G | A | 1.5e-08 | 5.6e-09 | 5.2e-09 | 8.600E-09 | 5.546E-09 | 5.200E-09 | 1.500E-08 |
| 8669 | 434 | R434H | A | 1.5e-08 | 9.3e-09 | 7.8e-09 | 1.070E-08 | 3.799E-09 | 7.800E-09 | 1.500E-08 |

| | | | | | | | | | | |
|---|---|---|---|---|---|---|---|---|---|---|
| 8670 | 434 | R434I | A | 3e-09 | 6.3e-09 | 2.4e-09 | 3.900E-09 | 2.100E-09 | 2.400E-09 | 6.300E-09 |
| 8671 | 434 | R434L | A | 2.2e-08 | 1.8e-08 | 1e-08 | 1.667E-08 | 6.110E-09 | 1.000E-08 | 2.200E-08 |
| 8672 | 434 | R434K | A | 6.5e-09 | 7.4e-09 | 5.1e-09 | 6.333E-09 | 1.159E-09 | 5.100E-09 | 7.400E-09 |
| 8673 | 434 | R434M | A | 4.2e-09 | 4.8e-09 | 8.4e-09 | 5.800E-09 | 2.272E-09 | 4.200E-09 | 8.400E-09 |
| 8674 | 434 | R434F | A | 3.5e-09 | 6e-09 | 5.7e-09 | 5.067E-09 | 1.365E-09 | 3.500E-09 | 6.000E-09 |
| 8675 | 434 | R434P | A | 1.1e-08 | 4e-09 | 1.9e-08 | 1.133E-08 | 7.506E-09 | 4.000E-09 | 1.900E-08 |
| 8676 | 434 | R434S | A | 1.8e-09 | 5.5e-09 | 3.4e-09 | 3.567E-09 | 1.856E-09 | 1.800E-09 | 5.500E-09 |
| 8677 | 434 | R434T | A | 3.4e-09 | 7.1e-09 | 8.6e-09 | 6.367E-09 | 2.676E-09 | 3.400E-09 | 8.600E-09 |
| 8678 | 434 | R434W | A | 1.2e-08 | 4.6e-09 | 7.3e-09 | 7.967E-09 | 3.745E-09 | 4.600E-09 | 1.200E-08 |
| 8679 | 434 | R434Y | A | 3e-09 | 1.1e-08 | 3.6e-09 | 5.867E-09 | 4.456E-09 | 3.000E-09 | 1.100E-08 |
| 8680 | 434 | R434V | A | 1.1e-08 | 6.4e-09 | 5e-09 | 7.467E-09 | 3.139E-09 | 5.000E-09 | 1.100E-08 |
| 8681 | 435 | S435A | A | 1.3e-08 | 7.7e-09 | 7.9e-09 | 9.533E-09 | 3.004E-09 | 7.700E-09 | 1.300E-08 |
| 8682 | 435 | S435R | A | 3.1e-09 | 9.7e-09 | 1.2e-08 | 8.267E-09 | 4.620E-09 | 3.100E-09 | 1.200E-08 |
| 8683 | 435 | S435N | A | 3.3e-08 | 4.3e-09 | 6.9e-09 | 1.473E-08 | 1.587E-08 | 4.300E-09 | 3.300E-08 |
| 8684 | 435 | S435D | A | 5e-09 | 9.5e-09 | 1.1e-08 | 8.500E-09 | 3.122E-09 | 5.000E-09 | 1.100E-08 |
| 8685 | 435 | S435C | A | 7e-09 | 6.6e-09 | 1.6e-08 | 9.867E-09 | 5.315E-09 | 6.600E-09 | 1.600E-08 |
| 8686 | 435 | S435E | A | 8e-09 | 1.7e-08 | 3.3e-09 | 9.433E-09 | 6.962E-09 | 3.300E-09 | 1.700E-08 |
| 8687 | 435 | S435Q | A | 5.9e-09 | 1.7e-08 | 9.5e-09 | 1.080E-08 | 5.663E-09 | 5.900E-09 | 1.700E-08 |
| 8688 | 435 | S435G | A | 4.8e-09 | 2e-08 | 5.1e-09 | 9.967E-09 | 8.690E-09 | 4.800E-09 | 2.000E-08 |
| 8689 | 435 | S435H | A | 1.1e-08 | 7.4e-09 | 4.9e-09 | 7.767E-09 | 3.066E-09 | 4.900E-09 | 1.100E-08 |
| 8690 | 435 | S435I | A | 7.8e-09 | 3.2e-09 | 5.1e-09 | 5.367E-09 | 2.312E-09 | 3.200E-09 | 7.800E-09 |
| 8691 | 435 | S435L | A | 2.3e-09 | 7.8e-09 | 1.6e-08 | 8.700E-09 | 6.894E-09 | 2.300E-09 | 1.600E-08 |
| 8692 | 435 | S435K | A | 1.2e-08 | 7.5e-09 | 1e-08 | 9.833E-09 | 2.255E-09 | 7.500E-09 | 1.200E-08 |
| 8693 | 435 | S435M | A | 1.8e-08 | 4.2e-09 | 1.1e-08 | 1.107E-08 | 6.900E-09 | 4.200E-09 | 1.800E-08 |
| 8694 | 435 | S435F | A | 3.9e-09 | 7.4e-09 | 5.5e-09 | 5.600E-09 | 1.752E-09 | 3.900E-09 | 7.400E-09 |
| 8695 | 435 | S435P | A | 2e-08 | 3.7e-09 | 8.1e-09 | 1.060E-08 | 8.433E-09 | 3.700E-09 | 2.000E-08 |
| 8696 | 435 | S435S | A | 4.9e-09 | 3e-09 | 1.2e-08 | 6.633E-09 | 4.744E-09 | 3.000E-09 | 1.200E-08 |
| 8697 | 435 | S435T | A | 1e-08 | 8.6e-09 | 4.7e-09 | 7.767E-09 | 2.747E-09 | 4.700E-09 | 1.000E-08 |
| 8698 | 435 | S435W | A | 8.2e-09 | 1.8e-08 | 2.9e-09 | 9.700E-09 | 7.661E-09 | 2.900E-09 | 1.800E-08 |
| 8699 | 435 | S435Y | A | 8.1e-09 | 8.9e-09 | 5.4e-09 | 7.467E-09 | 1.834E-09 | 5.400E-09 | 8.900E-09 |
| 8700 | 435 | S435V | A | 4.5e-09 | 5.5e-09 | 2.5e-08 | 1.167E-08 | 1.156E-08 | 4.500E-09 | 2.500E-08 |
| 8701 | 436 | L436A | A | 3.9e-09 | 9.1e-09 | 5.1e-09 | 6.033E-09 | 2.723E-09 | 3.900E-09 | 9.100E-09 |
| 8702 | 436 | L436R | A | 5e-09 | 2.3e-09 | 7e-09 | 4.767E-09 | 2.359E-09 | 2.300E-09 | 7.000E-09 |
| 8703 | 436 | L436N | A | 1.6e-08 | 7.1e-09 | 4.5e-09 | 9.200E-09 | 6.031E-09 | 4.500E-09 | 1.600E-08 |
| 8704 | 436 | L436D | A | 4e-09 | 1.2e-08 | 7.8e-09 | 7.933E-09 | 4.002E-09 | 4.000E-09 | 1.200E-08 |
| 8705 | 436 | L436C | A | 5.3e-09 | 1.3e-08 | 1.4e-08 | 1.077E-08 | 4.761E-09 | 5.300E-09 | 1.400E-08 |
| 8706 | 436 | L436E | A | 1.3e-08 | 8.2e-09 | 3.8e-09 | 8.333E-09 | 4.601E-09 | 3.800E-09 | 1.300E-08 |
| 8707 | 436 | L436Q | A | 4.4e-09 | 7.6e-09 | 1.6e-08 | 9.333E-09 | 5.991E-09 | 4.400E-09 | 1.600E-08 |
| 8708 | 436 | L436G | A | 6.9e-09 | 6.1e-09 | 1.1e-08 | 8.000E-09 | 2.629E-09 | 6.100E-09 | 1.100E-08 |
| 8709 | 436 | L436H | A | 2.8e-09 | 6.6e-09 | 4.5e-09 | 4.633E-09 | 1.904E-09 | 2.800E-09 | 6.600E-09 |
| 8710 | 436 | L436I | A | 2.1e-09 | 1.1e-08 | 2.6e-08 | 1.303E-08 | 1.208E-08 | 2.100E-09 | 2.600E-08 |
| 8711 | 436 | L436L | A | 7.3e-09 | 1.4e-09 | 4.6e-09 | 4.433E-09 | 2.954E-09 | 1.400E-09 | 7.300E-09 |
| 8712 | 436 | L436K | A | 1.8e-08 | 4.2e-09 | 1.5e-08 | 1.240E-08 | 7.258E-09 | 4.200E-09 | 1.800E-08 |
| 8713 | 436 | L436M | A | 1.2e-08 | 7.3e-09 | 4.1e-09 | 7.800E-09 | 3.974E-09 | 4.100E-09 | 1.200E-08 |
| 8714 | 436 | L436F | A | 3.4e-09 | 5.9e-09 | 6.7e-09 | 5.333E-09 | 1.721E-09 | 3.400E-09 | 6.700E-09 |
| 8715 | 436 | L436P | A | 8.2e-09 | 1.6e-08 | 4.9e-09 | 9.700E-09 | 5.700E-09 | 4.900E-09 | 1.600E-08 |
| 8716 | 436 | L436S | A | 1e-08 | 5.2e-09 | 6.5e-09 | 7.233E-09 | 2.483E-09 | 5.200E-09 | 1.000E-08 |
| 8717 | 436 | L436T | A | 7.2e-09 | 1.8e-08 | 3.8e-08 | 2.107E-08 | 1.563E-08 | 7.200E-09 | 3.800E-08 |
| 8718 | 436 | L436W | A | 1.2e-08 | 4.3e-09 | 6.8e-09 | 7.700E-09 | 3.928E-09 | 4.300E-09 | 1.200E-08 |
| 8719 | 436 | L436Y | A | 9.6e-09 | 9.1e-09 | 7.3e-09 | 8.667E-09 | 1.210E-09 | 7.300E-09 | 9.600E-09 |
| 8720 | 436 | L436V | A | 9.2e-09 | 8.7e-09 | 1.1e-08 | 9.633E-09 | 1.210E-09 | 8.700E-09 | 1.100E-08 |
| 8721 | 437 | R437A | A | 2e-09 | 3e-09 | 5.5e-09 | 3.500E-09 | 1.803E-09 | 2.000E-09 | 5.500E-09 |
| 8722 | 437 | R437R | A | 1.6e-08 | 6.2e-09 | 9e-09 | 1.040E-08 | 5.048E-09 | 6.200E-09 | 1.600E-08 |
| 8723 | 437 | R437N | A | 2.8e-09 | 1.9e-08 | 4.1e-09 | 8.633E-09 | 9.001E-09 | 2.800E-09 | 1.900E-08 |
| 8724 | 437 | R437D | A | 3.1e-09 | 6.8e-09 | 1.1e-08 | 6.967E-09 | 3.953E-09 | 3.100E-09 | 1.100E-08 |
| 8725 | 437 | R437C | A | 2.6e-08 | 5e-09 | 6.6e-09 | 1.253E-08 | 1.169E-08 | 5.000E-09 | 2.600E-08 |
| 8726 | 437 | R437E | A | 1.5e-08 | 2e-08 | 3.1e-08 | 2.200E-08 | 8.185E-09 | 1.500E-08 | 3.100E-08 |
| 8727 | 437 | R437Q | A | 1.7e-09 | 1.8e-08 | 5.5e-09 | 8.400E-09 | 8.528E-09 | 1.700E-09 | 1.800E-08 |
| 8728 | 437 | R437G | A | 9.6e-09 | 8.9e-09 | 1e-08 | 9.500E-09 | 5.568E-10 | 8.900E-09 | 1.000E-08 |
| 8729 | 437 | R437H | A | 1e-08 | 2.3e-09 | 1.1e-08 | 7.767E-09 | 4.761E-09 | 2.300E-09 | 1.100E-08 |
| 8730 | 437 | R437I | A | 7.2e-09 | 3.7e-09 | 8.6e-09 | 6.500E-09 | 2.524E-09 | 3.700E-09 | 8.600E-09 |
| 8731 | 437 | R437L | A | 4.6e-09 | 2.1e-09 | 2.3e-09 | 3.000E-09 | 1.389E-09 | 2.100E-09 | 4.600E-09 |
| 8732 | 437 | R437K | A | 8.3e-09 | 7.5e-09 | 9.7e-09 | 8.500E-09 | 1.114E-09 | 7.500E-09 | 9.700E-09 |
| 8733 | 437 | R437M | A | 6.7e-09 | 2.4e-08 | 2.2e-08 | 1.757E-08 | 9.464E-09 | 6.700E-09 | 2.400E-08 |
| 8734 | 437 | R437F | A | 7.8e-09 | 5.8e-09 | 6.9e-09 | 6.833E-09 | 1.002E-09 | 5.800E-09 | 7.800E-09 |
| 8735 | 437 | R437P | A | 8e-09 | 4.8e-09 | 7.2e-09 | 6.667E-09 | 1.665E-09 | 4.800E-09 | 8.000E-09 |
| 8736 | 437 | R437S | A | 1.2e-08 | 1e-08 | 4e-09 | 8.667E-09 | 4.163E-09 | 4.000E-09 | 1.200E-08 |
| 8737 | 437 | R437T | A | 8.4e-09 | 7.4e-09 | 1.2e-08 | 9.267E-09 | 2.419E-09 | 7.400E-09 | 1.200E-08 |
| 8738 | 437 | R437W | A | 4.1e-09 | 2.5e-08 | 6.2e-09 | 1.177E-08 | 1.151E-08 | 4.100E-09 | 2.500E-08 |
| 8739 | 437 | R437Y | A | 3.3e-09 | 2.9e-09 | 7.5e-09 | 4.567E-09 | 2.548E-09 | 2.900E-09 | 7.500E-09 |
| 8740 | 437 | R437V | A | 5e-09 | 4.4e-09 | 4.2e-09 | 4.533E-09 | 4.163E-10 | 4.200E-09 | 5.000E-09 |
| 8741 | 438 | E438A | A | 4.1e-09 | 1.4e-08 | 1.3e-08 | 1.037E-08 | 5.450E-09 | 4.100E-09 | 1.400E-08 |
| 8742 | 438 | E438R | A | 1.8e-08 | 9.8e-09 | 7.5e-09 | 1.177E-08 | 5.519E-09 | 7.500E-09 | 1.800E-08 |
| 8743 | 438 | E438N | A | 1.9e-09 | 1.2e-08 | 7.1e-09 | 7.000E-09 | 5.051E-09 | 1.900E-09 | 1.200E-08 |
| 8744 | 438 | E438D | A | 2.1e-09 | 8.7e-09 | 1.5e-08 | 8.600E-09 | 6.451E-09 | 2.100E-09 | 1.500E-08 |
| 8745 | 438 | E438C | A | 6.2e-09 | 7.1e-09 | 1.9e-08 | 1.077E-08 | 7.144E-09 | 6.200E-09 | 1.900E-08 |
| 8746 | 438 | E438E | A | 5e-09 | 6.9e-09 | 5.4e-10 | 4.147E-09 | 3.265E-09 | 5.400E-10 | 6.900E-09 |
| 8747 | 438 | E438Q | A | 8.6e-09 | 7.4e-09 | 5.7e-09 | 7.233E-09 | 1.457E-09 | 5.700E-09 | 8.600E-09 |
| 8748 | 438 | E438G | A | 4e-09 | 9.4e-09 | 5.6e-09 | 6.333E-09 | 2.774E-09 | 4.000E-09 | 9.400E-09 |
| 8749 | 438 | E438H | A | 1.3e-09 | 1e-08 | 2.1e-09 | 4.467E-09 | 4.809E-09 | 1.300E-09 | 1.000E-08 |
| 8750 | 438 | E438I | A | 1.7e-08 | 8.4e-09 | 4e-09 | 9.800E-09 | 6.612E-09 | 4.000E-09 | 1.700E-08 |
| 8751 | 438 | E438L | A | 1e-08 | 1.4e-08 | 1.2e-08 | 1.200E-08 | 2.000E-09 | 1.000E-08 | 1.400E-08 |
| 8752 | 438 | E438K | A | 2.2e-09 | 7.5e-09 | 8.6e-09 | 6.100E-09 | 3.422E-09 | 2.200E-09 | 8.600E-09 |
| 8753 | 438 | E438M | A | 6e-09 | 6.4e-09 | 3.1e-09 | 5.167E-09 | 1.801E-09 | 3.100E-09 | 6.400E-09 |
| 8754 | 438 | E438F | A | 4.3e-09 | 4.7e-09 | 2.6e-08 | 1.167E-08 | 1.241E-08 | 4.300E-09 | 2.600E-08 |

| | | | | | | | | | | |
|---|---|---|---|---|---|---|---|---|---|---|
| 8755 | 438 | E438P | A | 6.4e-09 | 1.2e-08 | 7.2e-09 | 8.533E-09 | 3.029E-09 | 6.400E-09 | 1.200E-08 |
| 8756 | 438 | E438S | A | 9.1e-09 | 1.4e-08 | 1.2e-08 | 1.170E-08 | 2.464E-09 | 9.100E-09 | 1.400E-08 |
| 8757 | 438 | E438T | A | 1.4e-08 | 4.6e-09 | 3.9e-09 | 7.500E-09 | 5.640E-09 | 3.900E-09 | 1.400E-08 |
| 8758 | 438 | E438W | A | 9.5e-09 | 7.5e-09 | 1.3e-08 | 1.000E-08 | 2.784E-09 | 7.500E-09 | 1.300E-08 |
| 8759 | 438 | E438Y | A | 7.3e-09 | 2.2e-08 | 5.9e-09 | 1.173E-08 | 8.919E-09 | 5.900E-09 | 2.200E-08 |
| 8760 | 438 | E438V | A | 7.7e-09 | 2.3e-08 | 1.3e-08 | 1.457E-08 | 7.769E-09 | 7.700E-09 | 2.300E-08 |
| 8761 | 439 | L439A | A | 9.3e-09 | 4.8e-09 | 8.4e-09 | 7.500E-09 | 2.381E-09 | 4.800E-09 | 9.300E-09 |
| 8762 | 439 | L439R | A | 8.1e-09 | 1.6e-08 | 3.7e-09 | 9.267E-09 | 6.232E-09 | 3.700E-09 | 1.600E-08 |
| 8763 | 439 | L439N | A | 1.1e-08 | 7.5e-09 | 1.1e-08 | 9.833E-09 | 2.021E-09 | 7.500E-09 | 1.100E-08 |
| 8764 | 439 | L439D | A | 1e-08 | 2.2e-08 | 4.6e-09 | 1.220E-08 | 8.906E-09 | 4.600E-09 | 2.200E-08 |
| 8765 | 439 | L439C | A | 4.3e-09 | 6.7e-09 | 5.7e-09 | 5.567E-09 | 1.206E-09 | 4.300E-09 | 6.700E-09 |
| 8766 | 439 | L439E | A | 1.4e-08 | 7.8e-09 | 5.3e-09 | 9.033E-09 | 4.479E-09 | 5.300E-09 | 1.400E-08 |
| 8767 | 439 | L439Q | A | 3.4e-09 | 8.6e-09 | 8.7e-09 | 6.900E-09 | 3.032E-09 | 3.400E-09 | 8.700E-09 |
| 8768 | 439 | L439G | A | 1.7e-08 | 3e-09 | 7.5e-09 | 9.167E-09 | 7.147E-09 | 3.000E-09 | 1.700E-08 |
| 8769 | 439 | L439H | A | 1.3e-08 | 6e-09 | 4.3e-09 | 7.767E-09 | 4.611E-09 | 4.300E-09 | 1.300E-08 |
| 8770 | 439 | L439I | A | 6.6e-09 | 9.7e-09 | 2.4e-09 | 6.233E-09 | 3.664E-09 | 2.400E-09 | 9.700E-09 |
| 8771 | 439 | L439L | A | 1.3e-08 | 1.7e-08 | 4.3e-09 | 1.143E-08 | 6.493E-09 | 4.300E-09 | 1.700E-08 |
| 8772 | 439 | L439K | A | 4.9e-09 | 4.3e-09 | 6e-09 | 5.067E-09 | 8.622E-10 | 4.300E-09 | 6.000E-09 |
| 8773 | 439 | L439M | A | 2.1e-09 | 5.8e-09 | 9.6e-09 | 5.833E-09 | 3.750E-09 | 2.100E-09 | 9.600E-09 |
| 8774 | 439 | L439F | A | 2.2e-08 | 6.1e-08 | 7.7e-09 | 3.023E-08 | 2.759E-08 | 7.700E-09 | 6.100E-08 |
| 8775 | 439 | L439P | A | 3.3e-09 | 2e-08 | 2.6e-09 | 8.633E-09 | 9.850E-09 | 2.600E-09 | 2.000E-08 |
| 8776 | 439 | L439S | A | 5.7e-09 | 5.5e-09 | 2.2e-08 | 1.107E-08 | 9.469E-09 | 5.500E-09 | 2.200E-08 |
| 8777 | 439 | L439T | A | 1.1e-08 | 5.8e-09 | 7.7e-09 | 8.167E-09 | 2.631E-09 | 5.800E-09 | 1.100E-08 |
| 8778 | 439 | L439W | A | 1.1e-08 | 3.2e-09 | 1.9e-09 | 5.367E-09 | 4.922E-09 | 1.900E-09 | 1.100E-08 |
| 8779 | 439 | L439Y | A | 1e-08 | 2.7e-08 | 8.6e-09 | 1.520E-08 | 1.024E-08 | 8.600E-09 | 2.700E-08 |
| 8780 | 439 | L439V | A | 3.2e-09 | 1.2e-08 | 3.6e-09 | 6.267E-09 | 4.969E-09 | 3.200E-09 | 1.200E-08 |
| 8781 | 440 | G440A | A | 1e-08 | 9.3e-09 | 6e-09 | 8.433E-09 | 2.136E-09 | 6.000E-09 | 1.000E-08 |
| 8782 | 440 | G440R | A | 6.3e-09 | 3.6e-09 | 1.7e-08 | 8.967E-09 | 7.087E-09 | 3.600E-09 | 1.700E-08 |
| 8783 | 440 | G440N | A | 1.5e-08 | 4e-09 | 1.1e-08 | 1.000E-08 | 5.568E-09 | 4.000E-09 | 1.500E-08 |
| 8784 | 440 | G440D | A | 1.9e-08 | 2.6e-08 | 4.5e-09 | 1.650E-08 | 1.097E-08 | 4.500E-09 | 2.600E-08 |
| 8785 | 440 | G440C | A | 2.8e-08 | 5.7e-09 | 3.5e-09 | 1.240E-08 | 1.355E-08 | 3.500E-09 | 2.800E-08 |
| 8786 | 440 | G440E | A | 9.9e-09 | 1.8e-08 | 6.7e-09 | 1.153E-08 | 5.824E-09 | 6.700E-09 | 1.800E-08 |
| 8787 | 440 | G440Q | A | 6.9e-09 | 2.9e-09 | 3.9e-09 | 4.567E-09 | 2.082E-09 | 2.900E-09 | 6.900E-09 |
| 8788 | 440 | G440G | A | 4.7e-09 | 1.3e-08 | 5.3e-09 | 7.667E-09 | 4.629E-09 | 4.700E-09 | 1.300E-08 |
| 8789 | 440 | G440H | A | 1e-08 | 9.5e-09 | 8e-09 | 9.167E-09 | 1.041E-09 | 8.000E-09 | 1.000E-08 |
| 8790 | 440 | G440I | A | 1.5e-08 | 5.4e-09 | 9.8e-09 | 1.007E-08 | 4.806E-09 | 5.400E-09 | 1.500E-08 |
| 8791 | 440 | G440L | A | 1e-08 | 3.2e-09 | 5.7e-09 | 6.300E-09 | 3.439E-09 | 3.200E-09 | 1.000E-08 |
| 8792 | 440 | G440K | A | 1.1e-08 | 4.6e-09 | 5.5e-09 | 7.033E-09 | 3.465E-09 | 4.600E-09 | 1.100E-08 |
| 8793 | 440 | G440M | A | 1.2e-08 | 1.7e-08 | 7.3e-09 | 1.210E-08 | 4.851E-09 | 7.300E-09 | 1.700E-08 |
| 8794 | 440 | G440F | A | 4.4e-09 | 4e-09 | 6.7e-09 | 5.033E-09 | 1.457E-09 | 4.000E-09 | 6.700E-09 |
| 8795 | 440 | G440P | A | 5.1e-09 | 5.7e-09 | 1.2e-08 | 7.600E-09 | 3.822E-09 | 5.100E-09 | 1.200E-08 |
| 8796 | 440 | G440S | A | 6.8e-09 | 3.5e-09 | 5.5e-09 | 5.267E-09 | 1.662E-09 | 3.500E-09 | 6.800E-09 |
| 8797 | 440 | G440T | A | 6.2e-09 | 7.6e-09 | 1.9e-09 | 5.233E-09 | 2.970E-09 | 1.900E-09 | 7.600E-09 |
| 8798 | 440 | G440W | A | 9.8e-09 | 3e-09 | 1.9e-08 | 1.060E-08 | 8.030E-09 | 3.000E-09 | 1.900E-08 |
| 8799 | 440 | G440Y | A | 5e-09 | 8.3e-09 | 8.6e-09 | 7.300E-09 | 1.997E-09 | 5.000E-09 | 8.600E-09 |
| 8800 | 440 | G440V | A | 5.9e-09 | 2.8e-09 | 8.6e-09 | 5.767E-09 | 2.902E-09 | 2.800E-09 | 8.600E-09 |
| 8801 | 441 | S441A | A | 8e-09 | 8.3e-09 | 8.9e-09 | 8.400E-09 | 4.583E-10 | 8.000E-09 | 8.900E-09 |
| 8802 | 441 | S441R | A | 4.9e-09 | 6.3e-09 | 7e-09 | 6.067E-09 | 1.069E-09 | 4.900E-09 | 7.000E-09 |
| 8803 | 441 | S441N | A | 7.1e-09 | 7.3e-09 | 9.4e-09 | 7.933E-09 | 1.274E-09 | 7.100E-09 | 9.400E-09 |
| 8804 | 441 | S441D | A | 7.2e-09 | 1.1e-08 | 1.8e-09 | 6.667E-09 | 4.623E-09 | 1.800E-09 | 1.100E-08 |
| 8805 | 441 | S441C | A | 4.6e-09 | 2.9e-09 | 6.3e-09 | 4.600E-09 | 1.700E-09 | 2.900E-09 | 6.300E-09 |
| 8806 | 441 | S441E | A | 8.6e-09 | 6.6e-09 | 2.5e-09 | 5.900E-09 | 3.110E-09 | 2.500E-09 | 8.600E-09 |
| 8807 | 441 | S441Q | A | 1.8e-09 | 1.5e-08 | 1.8e-09 | 6.200E-09 | 7.621E-09 | 1.800E-09 | 1.500E-08 |
| 8808 | 441 | S441G | A | 4.8e-09 | 1.7e-09 | 1.5e-08 | 7.167E-09 | 6.959E-09 | 1.700E-09 | 1.500E-08 |
| 8809 | 441 | S441H | A | 1.3e-09 | 4.5e-09 | 4.9e-09 | 3.567E-09 | 1.973E-09 | 1.300E-09 | 4.900E-09 |
| 8810 | 441 | S441I | A | 4.3e-09 | 4.2e-09 | 1.7e-08 | 8.500E-09 | 7.361E-09 | 4.200E-09 | 1.700E-08 |
| 8811 | 441 | S441L | A | 5.1e-09 | 8.5e-09 | 1e-08 | 7.867E-09 | 2.511E-09 | 5.100E-09 | 1.000E-08 |
| 8812 | 441 | S441K | A | 5.8e-09 | 6.5e-09 | 4.1e-09 | 5.467E-09 | 1.234E-09 | 4.100E-09 | 6.500E-09 |
| 8813 | 441 | S441M | A | 1.3e-08 | 6.8e-09 | 1e-08 | 9.933E-09 | 3.101E-09 | 6.800E-09 | 1.300E-08 |
| 8814 | 441 | S441F | A | 3.8e-09 | 3.1e-09 | 4.7e-09 | 3.867E-09 | 8.021E-10 | 3.100E-09 | 4.700E-09 |
| 8815 | 441 | S441P | A | 7.2e-09 | 3.3e-09 | 7.6e-09 | 6.033E-09 | 2.376E-09 | 3.300E-09 | 7.600E-09 |
| 8816 | 441 | S441S | A | 8.7e-09 | 1.2e-08 | 1.7e-08 | 1.257E-08 | 4.179E-09 | 8.700E-09 | 1.700E-08 |
| 8817 | 441 | S441T | A | 7.6e-09 | 2.6e-09 | 2.4e-09 | 4.200E-09 | 2.946E-09 | 2.400E-09 | 7.600E-09 |
| 8818 | 441 | S441W | A | 1.3e-08 | 1e-08 | 8.9e-09 | 1.063E-08 | 2.122E-09 | 8.900E-09 | 1.300E-08 |
| 8819 | 441 | S441Y | A | 1.8e-08 | 1.1e-08 | 6.8e-09 | 1.193E-08 | 5.658E-09 | 6.800E-09 | 1.800E-08 |
| 8820 | 441 | S441V | A | 1.3e-08 | 2.9e-09 | 1.7e-08 | 1.097E-08 | 7.267E-09 | 2.900E-09 | 1.700E-08 |
| 8821 | 442 | G442A | A | 4.2e-09 | 4e-09 | 1e-08 | 6.067E-09 | 3.408E-09 | 4.000E-09 | 1.000E-08 |
| 8822 | 442 | G442R | A | 9e-09 | 8.1e-09 | 7e-09 | 8.033E-09 | 1.002E-09 | 7.000E-09 | 9.000E-09 |
| 8823 | 442 | G442N | A | 4.8e-09 | 2.8e-09 | 8.6e-09 | 5.400E-09 | 2.946E-09 | 2.800E-09 | 8.600E-09 |
| 8824 | 442 | G442D | A | 1e-08 | 3.3e-09 | 6e-09 | 6.433E-09 | 3.371E-09 | 3.300E-09 | 1.000E-08 |
| 8825 | 442 | G442C | A | 9e-09 | 5.2e-09 | 3.9e-09 | 6.033E-09 | 2.650E-09 | 3.900E-09 | 9.000E-09 |
| 8826 | 442 | G442E | A | 4.9e-09 | 1.8e-09 | 3.4e-09 | 3.367E-09 | 1.550E-09 | 1.800E-09 | 4.900E-09 |
| 8827 | 442 | G442Q | A | 1.4e-08 | 3.3e-09 | 9.8e-09 | 9.033E-09 | 5.391E-09 | 3.300E-09 | 1.400E-08 |
| 8828 | 442 | G442G | A | 4.3e-09 | 1.6e-08 | 5.7e-09 | 8.667E-09 | 6.389E-09 | 4.300E-09 | 1.600E-08 |
| 8829 | 442 | G442H | A | 4.4e-09 | 8.4e-09 | 3.3e-09 | 5.367E-09 | 2.684E-09 | 3.300E-09 | 8.400E-09 |
| 8830 | 442 | G442I | A | 1.1e-08 | 5.8e-09 | 3.6e-09 | 6.800E-09 | 3.800E-09 | 3.600E-09 | 1.100E-08 |
| 8831 | 442 | G442L | A | 1.5e-08 | 5.7e-09 | 5.5e-09 | 8.733E-09 | 5.428E-09 | 5.500E-09 | 1.500E-08 |
| 8832 | 442 | G442K | A | 7.6e-09 | 2.1e-09 | 3.1e-09 | 4.267E-09 | 2.930E-09 | 2.100E-09 | 7.600E-09 |
| 8833 | 442 | G442M | A | 2.7e-08 | 3.5e-09 | 2.1e-08 | 1.717E-08 | 1.221E-08 | 3.500E-09 | 2.700E-08 |
| 8834 | 442 | G442F | A | 1.2e-08 | 1.1e-08 | 9.6e-09 | 1.087E-08 | 1.206E-09 | 9.600E-09 | 1.200E-08 |
| 8835 | 442 | G442P | A | 9.2e-09 | 1.4e-08 | 6.7e-09 | 9.967E-09 | 3.710E-09 | 6.700E-09 | 1.400E-08 |
| 8836 | 442 | G442S | A | 1.1e-08 | 2.1e-08 | 3.5e-09 | 1.183E-08 | 8.780E-09 | 3.500E-09 | 2.100E-08 |
| 8837 | 442 | G442T | A | 1.5e-08 | 5.6e-09 | 3.5e-09 | 8.033E-09 | 6.124E-09 | 3.500E-09 | 1.500E-08 |
| 8838 | 442 | G442W | A | 1.2e-08 | 5.1e-09 | 1.8e-08 | 1.170E-08 | 6.455E-09 | 5.100E-09 | 1.800E-08 |
| 8839 | 442 | G442Y | A | 1e-08 | 1.3e-08 | 1.6e-08 | 1.300E-08 | 3.000E-09 | 1.000E-08 | 1.600E-08 |

| | | | | | | | | | | |
|---|---|---|---|---|---|---|---|---|---|---|
| 8840 | 442 | G442V | A | 6.8e-09 | 9.2e-09 | 1e-08 | 8.667E-09 | 1.665E-09 | 6.800E-09 | 1.000E-08 |
| 8841 | 443 | L443A | A | 1.1e-08 | 1.2e-08 | 2.6e-08 | 1.633E-08 | 8.386E-09 | 1.100E-08 | 2.600E-08 |
| 8842 | 443 | L443R | A | 9.6e-09 | 2.4e-09 | 1.1e-08 | 7.667E-09 | 4.614E-09 | 2.400E-09 | 1.100E-08 |
| 8843 | 443 | L443N | A | 3.2e-09 | 4.6e-09 | 3.8e-09 | 3.867E-09 | 7.024E-10 | 3.200E-09 | 4.600E-09 |
| 8844 | 443 | L443D | A | 7.2e-09 | 1.7e-09 | 9.2e-09 | 6.033E-09 | 3.884E-09 | 1.700E-09 | 9.200E-09 |
| 8845 | 443 | L443C | A | 1.7e-08 | 7.9e-09 | 8e-09 | 1.097E-08 | 5.225E-09 | 7.900E-09 | 1.700E-08 |
| 8846 | 443 | L443E | A | 4.2e-09 | 1.3e-08 | 1.6e-08 | 1.107E-08 | 6.133E-09 | 4.200E-09 | 1.600E-08 |
| 8847 | 443 | L443Q | A | 2.6e-08 | 9.3e-09 | 6.8e-09 | 1.403E-08 | 1.044E-08 | 6.800E-09 | 2.600E-08 |
| 8848 | 443 | L443G | A | 8.9e-09 | 1.2e-08 | 4e-09 | 8.300E-09 | 4.034E-09 | 4.000E-09 | 1.200E-08 |
| 8849 | 443 | L443H | A | 8.4e-09 | 3.6e-08 | 5.5e-09 | 1.663E-08 | 1.683E-08 | 5.500E-09 | 3.600E-08 |
| 8850 | 443 | L443I | A | 1.5e-08 | 1.1e-08 | 1.2e-08 | 1.267E-08 | 2.082E-09 | 1.100E-08 | 1.500E-08 |
| 8851 | 443 | L443L | A | 3.4e-09 | 2.1e-09 | 6.5e-09 | 4.000E-09 | 2.261E-09 | 2.100E-09 | 6.500E-09 |
| 8852 | 443 | L443K | A | 4.9e-09 | 2e-08 | 1.1e-08 | 1.197E-08 | 7.596E-09 | 4.900E-09 | 2.000E-08 |
| 8853 | 443 | L443M | A | 7.9e-09 | 9.3e-09 | 4.7e-09 | 7.300E-09 | 2.358E-09 | 4.700E-09 | 9.300E-09 |
| 8854 | 443 | L443F | A | 5.4e-09 | 2.5e-09 | 9.4e-09 | 5.767E-09 | 3.465E-09 | 2.500E-09 | 9.400E-09 |
| 8855 | 443 | L443P | A | 5.3e-09 | 9.6e-09 | 1.4e-08 | 9.633E-09 | 4.350E-09 | 5.300E-09 | 1.400E-08 |
| 8856 | 443 | L443S | A | 6.5e-09 | 7.6e-09 | 6.5e-09 | 6.867E-09 | 6.351E-10 | 6.500E-09 | 7.600E-09 |
| 8857 | 443 | L443T | A | 3.2e-09 | 9.2e-09 | 1.2e-08 | 8.133E-09 | 4.496E-09 | 3.200E-09 | 1.200E-08 |
| 8858 | 443 | L443W | A | 1.4e-08 | 8.7e-09 | 1.5e-08 | 1.257E-08 | 3.386E-09 | 8.700E-09 | 1.500E-08 |
| 8859 | 443 | L443Y | A | 6.8e-09 | 1.6e-08 | 3.7e-09 | 8.833E-09 | 6.397E-09 | 3.700E-09 | 1.600E-08 |
| 8860 | 443 | L443V | A | 6.5e-09 | 5.3e-09 | 2.4e-09 | 4.733E-09 | 2.108E-09 | 2.400E-09 | 6.500E-09 |
| 8861 | 444 | A444A | A | 5.7e-09 | 8.1e-09 | 7.2e-09 | 7.000E-09 | 1.212E-09 | 5.700E-09 | 8.100E-09 |
| 8862 | 444 | A444R | A | 8.9e-09 | 6.1e-09 | 4e-09 | 6.333E-09 | 2.458E-09 | 4.000E-09 | 8.900E-09 |
| 8863 | 444 | A444N | A | 1.9e-09 | 6.7e-09 | 2.3e-09 | 3.633E-09 | 2.663E-09 | 1.900E-09 | 6.700E-09 |
| 8864 | 444 | A444D | A | 8.8e-09 | 7.9e-09 | 6.3e-09 | 7.667E-09 | 1.266E-09 | 6.300E-09 | 8.800E-09 |
| 8865 | 444 | A444C | A | 7.8e-09 | 1.2e-08 | 6e-09 | 8.600E-09 | 3.079E-09 | 6.000E-09 | 1.200E-08 |
| 8866 | 444 | A444E | A | 5.3e-09 | 8.5e-09 | 4.3e-09 | 6.033E-09 | 2.194E-09 | 4.300E-09 | 8.500E-09 |
| 8867 | 444 | A444Q | A | 8e-09 | 4.1e-09 | 7e-09 | 6.367E-09 | 2.026E-09 | 4.100E-09 | 8.000E-09 |
| 8868 | 444 | A444G | A | 8e-09 | 3.8e-09 | 4.3e-09 | 5.367E-09 | 2.294E-09 | 3.800E-09 | 8.000E-09 |
| 8869 | 444 | A444H | A | 3.6e-09 | 1.1e-08 | 2.3e-09 | 5.633E-09 | 4.693E-09 | 2.300E-09 | 1.100E-08 |
| 8870 | 444 | A444I | A | 1e-08 | 8.9e-09 | 5.4e-09 | 8.100E-09 | 2.402E-09 | 5.400E-09 | 1.000E-08 |
| 8871 | 444 | A444L | A | 5.6e-09 | 4.9e-09 | 5.1e-09 | 5.200E-09 | 3.606E-10 | 4.900E-09 | 5.600E-09 |
| 8872 | 444 | A444K | A | 7.8e-09 | 4.3e-09 | 5.8e-09 | 5.967E-09 | 1.756E-09 | 4.300E-09 | 7.800E-09 |
| 8873 | 444 | A444M | A | 1.4e-08 | 5.6e-09 | 1.2e-08 | 1.053E-08 | 4.388E-09 | 5.600E-09 | 1.400E-08 |
| 8874 | 444 | A444F | A | 2.1e-09 | 1.5e-08 | 3.1e-09 | 6.733E-09 | 7.177E-09 | 2.100E-09 | 1.500E-08 |
| 8875 | 444 | A444P | A | 1.2e-08 | 8.4e-09 | 3.5e-09 | 7.967E-09 | 4.267E-09 | 3.500E-09 | 1.200E-08 |
| 8876 | 444 | A444S | A | 5.6e-09 | 7.8e-09 | 2.7e-09 | 5.367E-09 | 2.558E-09 | 2.700E-09 | 7.800E-09 |
| 8877 | 444 | A444T | A | 4.1e-09 | 1e-08 | 6.9e-09 | 7.000E-09 | 2.951E-09 | 4.100E-09 | 1.000E-08 |
| 8878 | 444 | A444W | A | 7.2e-09 | 1.2e-08 | 1e-08 | 9.733E-09 | 2.411E-09 | 7.200E-09 | 1.200E-08 |
| 8879 | 444 | A444Y | A | 4.4e-09 | 5.9e-09 | 1.5e-08 | 8.433E-09 | 5.736E-09 | 4.400E-09 | 1.500E-08 |
| 8880 | 444 | A444V | A | 4.1e-09 | 1.4e-08 | 8e-09 | 8.700E-09 | 4.987E-09 | 4.100E-09 | 1.400E-08 |
| 8881 | 445 | L445A | A | 2.1e-08 | 5.1e-09 | 1.6e-08 | 1.403E-08 | 8.130E-09 | 5.100E-09 | 2.100E-08 |
| 8882 | 445 | L445R | A | 2.5e-09 | 9.1e-09 | 6e-09 | 5.867E-09 | 3.302E-09 | 2.500E-09 | 9.100E-09 |
| 8883 | 445 | L445N | A | 9.6e-09 | 3.1e-09 | 5.4e-09 | 6.033E-09 | 3.296E-09 | 3.100E-09 | 9.600E-09 |
| 8884 | 445 | L445D | A | 2.6e-09 | 4.9e-09 | 1.2e-08 | 6.500E-09 | 4.900E-09 | 2.600E-09 | 1.200E-08 |
| 8885 | 445 | L445C | A | 1.2e-08 | 8.9e-09 | 6.5e-09 | 9.133E-09 | 2.757E-09 | 6.500E-09 | 1.200E-08 |
| 8886 | 445 | L445E | A | 7.2e-09 | 1.4e-08 | 3e-09 | 8.067E-09 | 5.551E-09 | 3.000E-09 | 1.400E-08 |
| 8887 | 445 | L445Q | A | 3.6e-09 | 6.1e-09 | 4e-09 | 4.567E-09 | 1.343E-09 | 3.600E-09 | 6.100E-09 |
| 8888 | 445 | L445G | A | 7.3e-09 | 1.3e-09 | 7.5e-09 | 5.367E-09 | 3.523E-09 | 1.300E-09 | 7.500E-09 |
| 8889 | 445 | L445H | A | 5.7e-09 | 8.3e-09 | 9.7e-09 | 7.900E-09 | 2.030E-09 | 5.700E-09 | 9.700E-09 |
| 8890 | 445 | L445I | A | 1.2e-08 | 9.8e-09 | 7.8e-09 | 9.867E-09 | 2.101E-09 | 7.800E-09 | 1.200E-08 |
| 8891 | 445 | L445L | A | 6.5e-09 | 7.9e-09 | 6.3e-09 | 6.900E-09 | 8.718E-10 | 6.300E-09 | 7.900E-09 |
| 8892 | 445 | L445K | A | 2.7e-09 | 6e-09 | 8.7e-09 | 5.800E-09 | 3.005E-09 | 2.700E-09 | 8.700E-09 |
| 8893 | 445 | L445M | A | 1.8e-08 | 7.3e-09 | 4.5e-09 | 9.933E-09 | 7.125E-09 | 4.500E-09 | 1.800E-08 |
| 8894 | 445 | L445F | A | 2.3e-08 | 1.4e-08 | 6.5e-09 | 1.450E-08 | 8.261E-09 | 6.500E-09 | 2.300E-08 |
| 8895 | 445 | L445P | A | 2.1e-09 | 7.2e-09 | 6.7e-09 | 5.333E-09 | 2.811E-09 | 2.100E-09 | 7.200E-09 |
| 8896 | 445 | L445S | A | 1.2e-08 | 8e-09 | 1e-08 | 1.000E-08 | 2.000E-09 | 8.000E-09 | 1.200E-08 |
| 8897 | 445 | L445T | A | 2.8e-09 | 1.7e-08 | 1e-08 | 9.933E-09 | 7.100E-09 | 2.800E-09 | 1.700E-08 |
| 8898 | 445 | L445W | A | 9e-09 | 7.6e-09 | 6.2e-09 | 7.600E-09 | 1.400E-09 | 6.200E-09 | 9.000E-09 |
| 8899 | 445 | L445Y | A | 8.2e-09 | 1.2e-08 | 3.5e-09 | 7.900E-09 | 4.258E-09 | 3.500E-09 | 1.200E-08 |
| 8900 | 445 | L445V | A | 1.4e-08 | 8.1e-09 | 1.2e-08 | 1.137E-08 | 3.001E-09 | 8.100E-09 | 1.400E-08 |
| 8901 | 446 | I446A | A | 4.8e-09 | 4.2e-09 | 1.6e-08 | 8.333E-09 | 6.646E-09 | 4.200E-09 | 1.600E-08 |
| 8902 | 446 | I446R | A | 7.6e-09 | 7e-09 | 9.3e-10 | 5.177E-09 | 3.690E-09 | 9.300E-10 | 7.600E-09 |
| 8903 | 446 | I446N | A | 4.8e-09 | 1.4e-08 | 2.3e-09 | 7.033E-09 | 6.161E-09 | 2.300E-09 | 1.400E-08 |
| 8904 | 446 | I446D | A | 1.8e-08 | 5.8e-09 | 8.9e-09 | 1.090E-08 | 6.341E-09 | 5.800E-09 | 1.800E-08 |
| 8905 | 446 | I446C | A | 1.4e-08 | 6.5e-09 | 2e-08 | 1.350E-08 | 6.764E-09 | 6.500E-09 | 2.000E-08 |
| 8906 | 446 | I446E | A | 1e-08 | 3.7e-09 | 3.1e-09 | 5.600E-09 | 3.822E-09 | 3.100E-09 | 1.000E-08 |
| 8907 | 446 | I446Q | A | 3.3e-09 | 2.8e-09 | 7e-09 | 4.367E-09 | 2.294E-09 | 2.800E-09 | 7.000E-09 |
| 8908 | 446 | I446G | A | 4.1e-09 | 1e-08 | 1.1e-08 | 8.367E-09 | 3.729E-09 | 4.100E-09 | 1.100E-08 |
| 8909 | 446 | I446H | A | 7.9e-09 | 1e-08 | 6.8e-09 | 8.233E-09 | 1.626E-09 | 6.800E-09 | 1.000E-08 |
| 8910 | 446 | I446I | A | 1.1e-08 | 1.1e-08 | 3.5e-09 | 8.500E-09 | 4.330E-09 | 3.500E-09 | 1.100E-08 |
| 8911 | 446 | I446L | A | 6.9e-09 | 8.1e-09 | 1.9e-09 | 5.633E-09 | 3.288E-09 | 1.900E-09 | 8.100E-09 |
| 8912 | 446 | I446K | A | 5.5e-09 | 2e-08 | 1.7e-08 | 1.417E-08 | 7.654E-09 | 5.500E-09 | 2.000E-08 |
| 8913 | 446 | I446M | A | 2.9e-09 | 3.9e-09 | 4.7e-09 | 3.833E-09 | 9.018E-10 | 2.900E-09 | 4.700E-09 |
| 8914 | 446 | I446F | A | 6e-09 | 8.5e-09 | 2.7e-09 | 5.733E-09 | 2.909E-09 | 2.700E-09 | 8.500E-09 |
| 8915 | 446 | I446P | A | 4.9e-09 | 7.2e-09 | 8.5e-09 | 6.867E-09 | 1.823E-09 | 4.900E-09 | 8.500E-09 |
| 8916 | 446 | I446S | A | 1.6e-08 | 4.6e-09 | 1.9e-08 | 1.320E-08 | 7.597E-09 | 4.600E-09 | 1.900E-08 |
| 8917 | 446 | I446T | A | 1e-08 | 5.5e-09 | 7.6e-09 | 7.700E-09 | 2.252E-09 | 5.500E-09 | 1.000E-08 |
| 8918 | 446 | I446W | A | 1.4e-08 | 5.9e-09 | 7.7e-09 | 9.200E-09 | 4.253E-09 | 5.900E-09 | 1.400E-08 |
| 8919 | 446 | I446Y | A | 7.6e-09 | 7.7e-09 | 7.4e-09 | 7.567E-09 | 1.528E-10 | 7.400E-09 | 7.700E-09 |
| 8920 | 446 | I446V | A | 6.4e-09 | 6.2e-09 | 1.2e-08 | 8.200E-09 | 3.292E-09 | 6.200E-09 | 1.200E-08 |
| 8921 | 447 | H447A | A | 7.1e-09 | 6.2e-09 | 6.7e-09 | 6.667E-09 | 4.509E-10 | 6.200E-09 | 7.100E-09 |
| 8922 | 447 | H447R | A | 5.4e-09 | 5.4e-09 | 5.5e-09 | 5.433E-09 | 5.774E-11 | 5.400E-09 | 5.500E-09 |
| 8923 | 447 | H447N | A | 9.6e-09 | 1.3e-08 | 7.5e-09 | 1.003E-08 | 2.775E-09 | 7.500E-09 | 1.300E-08 |
| 8924 | 447 | H447D | A | 3.7e-09 | 2.7e-08 | 1.5e-08 | 1.523E-08 | 1.165E-08 | 3.700E-09 | 2.700E-08 |

| | | | | | | | | | | |
|---|---|---|---|---|---|---|---|---|---|---|
| 8925 | 447 | H447C | A | 6.3e-09 | 5.4e-09 | 8e-09 | 6.567E-09 | 1.320E-09 | 5.400E-09 | 8.000E-09 |
| 8926 | 447 | H447E | A | 4.3e-09 | 1.2e-08 | 8.3e-09 | 8.200E-09 | 3.851E-09 | 4.300E-09 | 1.200E-08 |
| 8927 | 447 | H447Q | A | 3.5e-09 | 4.1e-09 | 5.4e-09 | 4.333E-09 | 9.713E-10 | 3.500E-09 | 5.400E-09 |
| 8928 | 447 | H447G | A | 1e-08 | 7.6e-09 | 7e-09 | 8.200E-09 | 1.587E-09 | 7.000E-09 | 1.000E-08 |
| 8929 | 447 | H447H | A | 7e-09 | 2.4e-08 | 7.1e-09 | 1.270E-08 | 9.786E-09 | 7.000E-09 | 2.400E-08 |
| 8930 | 447 | H447I | A | 2e-09 | 3.7e-09 | 1.2e-08 | 5.900E-09 | 5.351E-09 | 2.000E-09 | 1.200E-08 |
| 8931 | 447 | H447L | A | 2.3e-08 | 4.1e-09 | 5.5e-09 | 1.087E-08 | 1.053E-08 | 4.100E-09 | 2.300E-08 |
| 8932 | 447 | H447K | A | 7.7e-09 | 9.4e-09 | 8e-09 | 8.367E-09 | 9.074E-10 | 7.700E-09 | 9.400E-09 |
| 8933 | 447 | H447M | A | 1.1e-08 | 1.6e-08 | 1.8e-09 | 9.600E-09 | 7.203E-09 | 1.800E-09 | 1.600E-08 |
| 8934 | 447 | H447F | A | 5.2e-09 | 1.1e-08 | 1.1e-08 | 9.067E-09 | 3.349E-09 | 5.200E-09 | 1.100E-08 |
| 8935 | 447 | H447P | A | 7.4e-09 | 5.7e-09 | 1.2e-08 | 8.367E-09 | 3.259E-09 | 5.700E-09 | 1.200E-08 |
| 8936 | 447 | H447S | A | 1.3e-08 | 7.7e-09 | 5.6e-09 | 8.767E-09 | 3.814E-09 | 5.600E-09 | 1.300E-08 |
| 8937 | 447 | H447T | A | 9.4e-09 | 1.2e-08 | 7.4e-09 | 9.600E-09 | 2.307E-09 | 7.400E-09 | 1.200E-08 |
| 8938 | 447 | H447W | A | 6.4e-09 | 7e-09 | 9.9e-09 | 7.767E-09 | 1.872E-09 | 6.400E-09 | 9.900E-09 |
| 8939 | 447 | H447Y | A | 1.1e-08 | 5.8e-09 | 1e-08 | 8.933E-09 | 2.759E-09 | 5.800E-09 | 1.100E-08 |
| 8940 | 447 | H447V | A | 1.3e-08 | 1.2e-08 | 4.7e-09 | 9.900E-09 | 4.531E-09 | 4.700E-09 | 1.300E-08 |
| 8941 | 448 | H448A | A | 2.4e-08 | 8.9e-09 | 2.4e-09 | 1.177E-08 | 1.108E-08 | 2.400E-09 | 2.400E-08 |
| 8942 | 448 | H448R | A | 4.8e-09 | 2.3e-08 | 1.2e-08 | 1.327E-08 | 9.166E-09 | 4.800E-09 | 2.300E-08 |
| 8943 | 448 | H448N | A | 3.7e-09 | 1.2e-08 | 1e-08 | 8.567E-09 | 4.332E-09 | 3.700E-09 | 1.200E-08 |
| 8944 | 448 | H448D | A | 1e-08 | 1.4e-08 | 3.8e-09 | 9.267E-09 | 5.139E-09 | 3.800E-09 | 1.400E-08 |
| 8945 | 448 | H448C | A | 3.6e-09 | 7.1e-09 | 1.1e-08 | 7.233E-09 | 3.702E-09 | 3.600E-09 | 1.100E-08 |
| 8946 | 448 | H448E | A | 8.1e-09 | 2.5e-08 | 8.5e-09 | 1.387E-08 | 9.644E-09 | 8.100E-09 | 2.500E-08 |
| 8947 | 448 | H448Q | A | 5.9e-09 | 2.1e-08 | 1.8e-08 | 1.497E-08 | 7.994E-09 | 5.900E-09 | 2.100E-08 |
| 8948 | 448 | H448G | A | 1.1e-08 | 3.4e-09 | 9.9e-09 | 8.100E-09 | 4.107E-09 | 3.400E-09 | 1.100E-08 |
| 8949 | 448 | H448H | A | 7.4e-09 | 5.6e-09 | 6.6e-09 | 6.533E-09 | 9.018E-10 | 5.600E-09 | 7.400E-09 |
| 8950 | 448 | H448I | A | 9.7e-09 | 7.3e-09 | 3.1e-08 | 1.600E-08 | 1.305E-08 | 7.300E-09 | 3.100E-08 |
| 8951 | 448 | H448L | A | 3.8e-09 | 2.4e-08 | 4e-09 | 1.060E-08 | 1.161E-08 | 3.800E-09 | 2.400E-08 |
| 8952 | 448 | H448K | A | 1.3e-08 | 5.3e-09 | 1.1e-08 | 9.767E-09 | 3.995E-09 | 5.300E-09 | 1.300E-08 |
| 8953 | 448 | H448M | A | 8.1e-09 | 1.3e-08 | 1.2e-08 | 1.103E-08 | 2.589E-09 | 8.100E-09 | 1.300E-08 |
| 8954 | 448 | H448F | A | 6.2e-09 | 7.2e-09 | 1.2e-08 | 8.467E-09 | 3.101E-09 | 6.200E-09 | 1.200E-08 |
| 8955 | 448 | H448P | A | 1.5e-08 | 6.7e-09 | 8.2e-09 | 9.967E-09 | 4.423E-09 | 6.700E-09 | 1.500E-08 |
| 8956 | 448 | H448S | A | 5.5e-09 | 1.5e-08 | 6.7e-09 | 9.067E-09 | 5.173E-09 | 5.500E-09 | 1.500E-08 |
| 8957 | 448 | H448T | A | 8.7e-09 | 7e-09 | 3e-09 | 6.233E-09 | 2.926E-09 | 3.000E-09 | 8.700E-09 |
| 8958 | 448 | H448W | A | 3.7e-09 | 9.4e-09 | 4e-09 | 5.700E-09 | 3.208E-09 | 3.700E-09 | 9.400E-09 |
| 8959 | 448 | H448Y | A | 1e-08 | 6.4e-09 | 6.6e-09 | 7.667E-09 | 2.023E-09 | 6.400E-09 | 1.000E-08 |
| 8960 | 448 | H448V | A | 4.5e-09 | 9.5e-09 | 1.3e-08 | 9.000E-09 | 4.272E-09 | 4.500E-09 | 1.300E-08 |
| 8961 | 449 | N449A | A | 5.8e-09 | 4.4e-09 | 1.1e-08 | 7.067E-09 | 3.478E-09 | 4.400E-09 | 1.100E-08 |
| 8962 | 449 | N449R | A | 2.8e-09 | 2.6e-09 | 6.3e-09 | 3.900E-09 | 2.081E-09 | 2.600E-09 | 6.300E-09 |
| 8963 | 449 | N449N | A | 5.4e-09 | 2.2e-08 | 1.3e-08 | 1.347E-08 | 8.310E-09 | 5.400E-09 | 2.200E-08 |
| 8964 | 449 | N449D | A | 4.2e-09 | 1.7e-08 | 3.5e-09 | 8.233E-09 | 7.600E-09 | 3.500E-09 | 1.700E-08 |
| 8965 | 449 | N449C | A | 7.5e-09 | 3.7e-09 | 3.5e-09 | 4.900E-09 | 2.254E-09 | 3.500E-09 | 7.500E-09 |
| 8966 | 449 | N449E | A | 6.5e-09 | 1.1e-08 | 1.6e-08 | 1.117E-08 | 4.752E-09 | 6.500E-09 | 1.600E-08 |
| 8967 | 449 | N449Q | A | 4.3e-09 | 5.5e-09 | 3.1e-08 | 1.360E-08 | 1.508E-08 | 4.300E-09 | 3.100E-08 |
| 8968 | 449 | N449G | A | 8.1e-09 | 1e-08 | 9.6e-09 | 9.233E-09 | 1.002E-09 | 8.100E-09 | 1.000E-08 |
| 8969 | 449 | N449H | A | 5.2e-09 | 1.8e-08 | 5.7e-09 | 9.633E-09 | 7.250E-09 | 5.200E-09 | 1.800E-08 |
| 8970 | 449 | N449I | A | 1.4e-08 | 1.2e-08 | 6.4e-09 | 1.080E-08 | 3.940E-09 | 6.400E-09 | 1.400E-08 |
| 8971 | 449 | N449L | A | 7.3e-09 | 2e-09 | 1e-08 | 6.433E-09 | 4.070E-09 | 2.000E-09 | 1.000E-08 |
| 8972 | 449 | N449K | A | 1.7e-09 | 4e-09 | 6.8e-09 | 4.167E-09 | 2.554E-09 | 1.700E-09 | 6.800E-09 |
| 8973 | 449 | N449M | A | 2.3e-09 | 3.8e-09 | 3.2e-08 | 1.270E-08 | 1.673E-08 | 2.300E-09 | 3.200E-08 |
| 8974 | 449 | N449F | A | 4.9e-09 | 5.2e-09 | 5.7e-09 | 5.267E-09 | 4.041E-10 | 4.900E-09 | 5.700E-09 |
| 8975 | 449 | N449P | A | 2.9e-09 | 1.4e-08 | 7.6e-09 | 8.167E-09 | 5.572E-09 | 2.900E-09 | 1.400E-08 |
| 8976 | 449 | N449S | A | 4.2e-09 | 2e-08 | 5.8e-09 | 1.000E-08 | 8.697E-09 | 4.200E-09 | 2.000E-08 |
| 8977 | 449 | N449T | A | 1.4e-08 | 9.3e-09 | 8.1e-09 | 1.047E-08 | 3.118E-09 | 8.100E-09 | 1.400E-08 |
| 8978 | 449 | N449W | A | 2.3e-09 | 9.8e-09 | 5.2e-09 | 5.767E-09 | 3.782E-09 | 2.300E-09 | 9.800E-09 |
| 8979 | 449 | N449Y | A | 6.6e-09 | 6.9e-09 | 7.2e-09 | 6.900E-09 | 3.000E-10 | 6.600E-09 | 7.200E-09 |
| 8980 | 449 | N449V | A | 5.7e-09 | 4.8e-09 | 1.9e-08 | 9.833E-09 | 7.951E-09 | 4.800E-09 | 1.900E-08 |
| 8981 | 450 | T450A | A | 8.8e-09 | 9.4e-09 | 1e-08 | 9.400E-09 | 6.000E-10 | 8.800E-09 | 1.000E-08 |
| 8982 | 450 | T450R | A | 7.4e-09 | 8e-09 | 3.7e-09 | 6.367E-09 | 2.329E-09 | 3.700E-09 | 8.000E-09 |
| 8983 | 450 | T450N | A | 6.6e-09 | 4.3e-09 | 5.8e-09 | 5.567E-09 | 1.168E-09 | 4.300E-09 | 6.600E-09 |
| 8984 | 450 | T450D | A | 7.1e-09 | 5.4e-09 | 7e-09 | 6.500E-09 | 9.539E-10 | 5.400E-09 | 7.100E-09 |
| 8985 | 450 | T450C | A | 7.4e-09 | 1e-08 | 5.5e-09 | 7.633E-09 | 2.259E-09 | 5.500E-09 | 1.000E-08 |
| 8986 | 450 | T450E | A | 1.1e-08 | 1.2e-08 | 1.1e-08 | 1.133E-08 | 5.774E-10 | 1.100E-08 | 1.200E-08 |
| 8987 | 450 | T450Q | A | 1.1e-08 | 3e-09 | 1.3e-08 | 9.000E-09 | 5.292E-09 | 3.000E-09 | 1.300E-08 |
| 8988 | 450 | T450G | A | 3.8e-09 | 3.1e-09 | 4.7e-09 | 3.867E-09 | 8.021E-10 | 3.100E-09 | 4.700E-09 |
| 8989 | 450 | T450H | A | 6.5e-09 | 8e-09 | 2.2e-09 | 5.567E-09 | 3.011E-09 | 2.200E-09 | 8.000E-09 |
| 8990 | 450 | T450I | A | 2.6e-09 | 1.2e-08 | 7.9e-09 | 7.500E-09 | 4.713E-09 | 2.600E-09 | 1.200E-08 |
| 8991 | 450 | T450L | A | 1.4e-08 | 4.9e-09 | 4e-09 | 7.633E-09 | 5.532E-09 | 4.000E-09 | 1.400E-08 |
| 8992 | 450 | T450K | A | 2.4e-08 | 5.6e-09 | 7e-09 | 1.220E-08 | 1.024E-08 | 5.600E-09 | 2.400E-08 |
| 8993 | 450 | T450M | A | 3.6e-09 | 1.1e-08 | 1.5e-08 | 9.867E-09 | 5.784E-09 | 3.600E-09 | 1.500E-08 |
| 8994 | 450 | T450F | A | 7.6e-09 | 4.3e-09 | 5.8e-09 | 5.900E-09 | 1.652E-09 | 4.300E-09 | 7.600E-09 |
| 8995 | 450 | T450P | A | 6.2e-09 | 6.1e-09 | 1e-08 | 7.433E-09 | 2.223E-09 | 6.100E-09 | 1.000E-08 |
| 8996 | 450 | T450S | A | 3.7e-09 | 6.8e-09 | 3.9e-09 | 4.800E-09 | 1.735E-09 | 3.700E-09 | 6.800E-09 |
| 8997 | 450 | T450T | A | 2.6e-09 | 7.2e-09 | 1.1e-08 | 6.933E-09 | 4.206E-09 | 2.600E-09 | 1.100E-08 |
| 8998 | 450 | T450W | A | 2e-08 | 7.7e-09 | 1.9e-08 | 1.557E-08 | 6.831E-09 | 7.700E-09 | 2.000E-08 |
| 8999 | 450 | T450Y | A | 7.3e-09 | 1.2e-08 | 1.9e-08 | 1.277E-08 | 5.888E-09 | 7.300E-09 | 1.900E-08 |
| 9000 | 450 | T450V | A | 8e-09 | 7.1e-09 | 1.5e-08 | 1.003E-08 | 4.325E-09 | 7.100E-09 | 1.500E-08 |
| 9001 | 451 | H451A | A | 1.4e-08 | 4.3e-09 | 1.1e-08 | 9.767E-09 | 4.966E-09 | 4.300E-09 | 1.400E-08 |
| 9002 | 451 | H451R | A | 7.5e-09 | 1.4e-08 | 8.9e-09 | 1.013E-08 | 3.421E-09 | 7.500E-09 | 1.400E-08 |
| 9003 | 451 | H451N | A | 5.5e-09 | 1.6e-08 | 7.4e-09 | 9.633E-09 | 5.595E-09 | 5.500E-09 | 1.600E-08 |
| 9004 | 451 | H451D | A | 7.6e-09 | 4.8e-09 | 3.3e-09 | 5.233E-09 | 2.183E-09 | 3.300E-09 | 7.600E-09 |
| 9005 | 451 | H451C | A | 1.3e-08 | 5.1e-09 | 1.5e-08 | 1.103E-08 | 5.235E-09 | 5.100E-09 | 1.500E-08 |
| 9006 | 451 | H451E | A | 3.1e-09 | 6e-09 | 1.2e-08 | 7.033E-09 | 4.539E-09 | 3.100E-09 | 1.200E-08 |
| 9007 | 451 | H451Q | A | 7.1e-09 | 1.2e-08 | 8.3e-09 | 9.133E-09 | 2.554E-09 | 7.100E-09 | 1.200E-08 |
| 9008 | 451 | H451G | A | 2.3e-08 | 8.8e-09 | 9.6e-09 | 1.380E-08 | 7.977E-09 | 8.800E-09 | 2.300E-08 |
| 9009 | 451 | H451H | A | 3.5e-09 | 6.8e-09 | 4.4e-09 | 4.900E-09 | 1.706E-09 | 3.500E-09 | 6.800E-09 |

| | | | | | | | | | | |
|---|---|---|---|---|---|---|---|---|---|---|
| 9010 | 451 | H451I | A | 9.3e-09 | 1.6e-08 | 1.2e-08 | 1.243E-08 | 3.371E-09 | 9.300E-09 | 1.600E-08 |
| 9011 | 451 | H451L | A | 3.5e-09 | 7.8e-09 | 1.6e-08 | 9.100E-09 | 6.351E-09 | 3.500E-09 | 1.600E-08 |
| 9012 | 451 | H451K | A | 1.1e-08 | 8.1e-09 | 3.6e-09 | 7.567E-09 | 3.729E-09 | 3.600E-09 | 1.100E-08 |
| 9013 | 451 | H451M | A | 4.1e-09 | 1.5e-08 | 3.5e-09 | 7.533E-09 | 6.473E-09 | 3.500E-09 | 1.500E-08 |
| 9014 | 451 | H451F | A | 7.9e-09 | 1.6e-08 | 4.2e-09 | 9.367E-09 | 6.035E-09 | 4.200E-09 | 1.600E-08 |
| 9015 | 451 | H451P | A | 9.8e-09 | 1.2e-08 | 1.7e-08 | 1.293E-08 | 3.690E-09 | 9.800E-09 | 1.700E-08 |
| 9016 | 451 | H451S | A | 6.6e-09 | 1.7e-08 | 2.2e-09 | 8.600E-09 | 7.600E-09 | 2.200E-09 | 1.700E-08 |
| 9017 | 451 | H451T | A | 4.2e-09 | 4.2e-09 | 1.4e-08 | 7.467E-09 | 5.658E-09 | 4.200E-09 | 1.400E-08 |
| 9018 | 451 | H451W | A | 4e-09 | 8.9e-09 | 8.5e-09 | 7.133E-09 | 2.721E-09 | 4.000E-09 | 8.900E-09 |
| 9019 | 451 | H451Y | A | 1.5e-08 | 7.6e-09 | 1.5e-08 | 1.253E-08 | 4.272E-09 | 7.600E-09 | 1.500E-08 |
| 9020 | 451 | H451V | A | 1.1e-08 | 1e-08 | 1.1e-08 | 1.067E-08 | 5.774E-10 | 1.000E-08 | 1.100E-08 |
| 9021 | 452 | L452A | A | 6.7e-09 | 9.3e-09 | 7.6e-09 | 7.867E-09 | 1.320E-09 | 6.700E-09 | 9.300E-09 |
| 9022 | 452 | L452R | A | 5.6e-09 | 3e-09 | 9.8e-09 | 6.133E-09 | 3.431E-09 | 3.000E-09 | 9.800E-09 |
| 9023 | 452 | L452N | A | 4.2e-09 | 6.1e-09 | 9.7e-09 | 6.667E-09 | 2.793E-09 | 4.200E-09 | 9.700E-09 |
| 9024 | 452 | L452D | A | 1.7e-08 | 7.8e-09 | 8.9e-09 | 1.123E-08 | 5.024E-09 | 7.800E-09 | 1.700E-08 |
| 9025 | 452 | L452C | A | 2.6e-09 | 3.6e-09 | 6.4e-09 | 4.200E-09 | 1.970E-09 | 2.600E-09 | 6.400E-09 |
| 9026 | 452 | L452E | A | 8.2e-09 | 7.7e-09 | 5e-09 | 6.967E-09 | 1.721E-09 | 5.000E-09 | 8.200E-09 |
| 9027 | 452 | L452Q | A | 6.5e-09 | 3.6e-08 | 9.1e-09 | 1.720E-08 | 1.633E-08 | 6.500E-09 | 3.600E-08 |
| 9028 | 452 | L452G | A | 7.7e-09 | 5.5e-09 | 5.2e-09 | 6.133E-09 | 1.365E-09 | 5.200E-09 | 7.700E-09 |
| 9029 | 452 | L452H | A | 1.2e-08 | 2.4e-08 | 1e-08 | 1.533E-08 | 7.572E-09 | 1.000E-08 | 2.400E-08 |
| 9030 | 452 | L452I | A | 1.5e-09 | 5.9e-09 | 9.1e-09 | 5.500E-09 | 3.816E-09 | 1.500E-09 | 9.100E-09 |
| 9031 | 452 | L452L | A | 1.4e-08 | 4.4e-09 | 3.3e-09 | 7.233E-09 | 5.886E-09 | 3.300E-09 | 1.400E-08 |
| 9032 | 452 | L452K | A | 2.7e-09 | 6.9e-09 | 1.1e-08 | 6.867E-09 | 4.150E-09 | 2.700E-09 | 1.100E-08 |
| 9033 | 452 | L452M | A | 9.2e-09 | 6e-09 | 5.7e-09 | 6.967E-09 | 1.940E-09 | 5.700E-09 | 9.200E-09 |
| 9034 | 452 | L452F | A | 1.3e-08 | 7.5e-09 | 1.2e-09 | 7.233E-09 | 5.905E-09 | 1.200E-09 | 1.300E-08 |
| 9035 | 452 | L452P | A | 7.4e-09 | 2.2e-09 | 1.6e-09 | 3.733E-09 | 3.190E-09 | 1.600E-09 | 7.400E-09 |
| 9036 | 452 | L452S | A | 3.6e-09 | 2.1e-09 | 1.1e-08 | 5.567E-09 | 4.765E-09 | 2.100E-09 | 1.100E-08 |
| 9037 | 452 | L452T | A | 5.6e-09 | 4.3e-09 | 6.7e-09 | 5.533E-09 | 1.201E-09 | 4.300E-09 | 6.700E-09 |
| 9038 | 452 | L452W | A | 2.3e-09 | 7.8e-09 | 8.3e-09 | 6.133E-09 | 3.329E-09 | 2.300E-09 | 8.300E-09 |
| 9039 | 452 | L452Y | A | 1.9e-08 | 1e-08 | 8.3e-09 | 1.243E-08 | 5.750E-09 | 8.300E-09 | 1.900E-08 |
| 9040 | 452 | L452V | A | 7.7e-09 | 3.5e-09 | 8.8e-09 | 6.667E-09 | 2.797E-09 | 3.500E-09 | 8.800E-09 |
| 9041 | 453 | C453A | A | 1.9e-09 | 1.1e-08 | 7.2e-09 | 6.700E-09 | 4.571E-09 | 1.900E-09 | 1.100E-08 |
| 9042 | 453 | C453R | A | 1e-08 | 2e-08 | 2.8e-09 | 1.093E-08 | 8.638E-09 | 2.800E-09 | 2.000E-08 |
| 9043 | 453 | C453N | A | 3.6e-09 | 6.7e-09 | 2.1e-09 | 4.133E-09 | 2.346E-09 | 2.100E-09 | 6.700E-09 |
| 9044 | 453 | C453D | A | 9.1e-09 | 1.1e-08 | 8.1e-09 | 9.400E-09 | 1.473E-09 | 8.100E-09 | 1.100E-08 |
| 9045 | 453 | C453C | A | 3.5e-09 | 6.7e-09 | 1.3e-08 | 7.733E-09 | 4.834E-09 | 3.500E-09 | 1.300E-08 |
| 9046 | 453 | C453E | A | 5.3e-09 | 7.5e-09 | 9.2e-09 | 7.333E-09 | 1.955E-09 | 5.300E-09 | 9.200E-09 |
| 9047 | 453 | C453Q | A | 1.1e-08 | 2.7e-09 | 4.5e-09 | 6.067E-09 | 4.366E-09 | 2.700E-09 | 1.100E-08 |
| 9048 | 453 | C453G | A | 1.8e-08 | 7.5e-09 | 2.9e-09 | 9.467E-09 | 7.740E-09 | 2.900E-09 | 1.800E-08 |
| 9049 | 453 | C453H | A | 1e-08 | 2.5e-09 | 1.5e-08 | 9.167E-09 | 6.292E-09 | 2.500E-09 | 1.500E-08 |
| 9050 | 453 | C453I | A | 6.9e-09 | 2.9e-08 | 4.3e-09 | 1.340E-08 | 1.357E-08 | 4.300E-09 | 2.900E-08 |
| 9051 | 453 | C453L | A | 9.6e-09 | 2.3e-08 | 6e-09 | 1.287E-08 | 8.958E-09 | 6.000E-09 | 2.300E-08 |
| 9052 | 453 | C453K | A | 3.5e-09 | 6.2e-09 | 1.4e-08 | 7.900E-09 | 5.453E-09 | 3.500E-09 | 1.400E-08 |
| 9053 | 453 | C453M | A | 1.8e-08 | 5e-09 | 4.9e-09 | 9.300E-09 | 7.535E-09 | 4.900E-09 | 1.800E-08 |
| 9054 | 453 | C453F | A | 4.4e-09 | 1.7e-08 | 4.1e-09 | 8.500E-09 | 7.363E-09 | 4.100E-09 | 1.700E-08 |
| 9055 | 453 | C453P | A | 5.5e-09 | 6.3e-09 | 7.6e-09 | 6.467E-09 | 1.060E-09 | 5.500E-09 | 7.600E-09 |
| 9056 | 453 | C453S | A | 8.1e-09 | 4.9e-09 | 2.5e-09 | 5.167E-09 | 2.810E-09 | 2.500E-09 | 8.100E-09 |
| 9057 | 453 | C453T | A | 1.8e-08 | 1.2e-08 | 1.6e-08 | 1.533E-08 | 3.055E-09 | 1.200E-08 | 1.800E-08 |
| 9058 | 453 | C453W | A | 5.1e-09 | 1.8e-08 | 6.8e-09 | 9.967E-09 | 7.009E-09 | 5.100E-09 | 1.800E-08 |
| 9059 | 453 | C453Y | A | 4.2e-09 | 9.6e-09 | 6.8e-09 | 6.867E-09 | 2.701E-09 | 4.200E-09 | 9.600E-09 |
| 9060 | 453 | C453V | A | 6.4e-09 | 5.9e-09 | 1.1e-08 | 7.767E-09 | 2.811E-09 | 5.900E-09 | 1.100E-08 |
| 9061 | 454 | F454A | A | 1.2e-08 | 1.4e-08 | 3.1e-09 | 9.700E-09 | 5.803E-09 | 3.100E-09 | 1.400E-08 |
| 9062 | 454 | F454R | A | 1.1e-08 | 2.7e-09 | 4.8e-09 | 6.167E-09 | 4.315E-09 | 2.700E-09 | 1.100E-08 |
| 9063 | 454 | F454N | A | 2.5e-08 | 5.7e-09 | 1.8e-08 | 1.623E-08 | 9.771E-09 | 5.700E-09 | 2.500E-08 |
| 9064 | 454 | F454D | A | 1.8e-09 | 3.6e-09 | 1e-08 | 5.133E-09 | 4.310E-09 | 1.800E-09 | 1.000E-08 |
| 9065 | 454 | F454C | A | 4.3e-09 | 2.2e-08 | 2.9e-09 | 9.733E-09 | 1.065E-08 | 2.900E-09 | 2.200E-08 |
| 9066 | 454 | F454E | A | 3.7e-09 | 3e-09 | 5.6e-09 | 4.100E-09 | 1.345E-09 | 3.000E-09 | 5.600E-09 |
| 9067 | 454 | F454Q | A | 2e-09 | 5.9e-09 | 2.1e-08 | 9.633E-09 | 1.004E-08 | 2.000E-09 | 2.100E-08 |
| 9068 | 454 | F454G | A | 1.3e-08 | 7.5e-09 | 7.8e-09 | 9.433E-09 | 3.092E-09 | 7.500E-09 | 1.300E-08 |
| 9069 | 454 | F454H | A | 9e-09 | 1.1e-08 | 6.1e-09 | 8.700E-09 | 2.464E-09 | 6.100E-09 | 1.100E-08 |
| 9070 | 454 | F454I | A | 2.8e-09 | 6.9e-09 | 8.7e-09 | 6.133E-09 | 3.024E-09 | 2.800E-09 | 8.700E-09 |
| 9071 | 454 | F454L | A | 1.1e-08 | 1.8e-08 | 6.3e-09 | 1.177E-08 | 5.888E-09 | 6.300E-09 | 1.800E-08 |
| 9072 | 454 | F454K | A | 1.6e-08 | 7.2e-09 | 4.4e-09 | 9.200E-09 | 6.053E-09 | 4.400E-09 | 1.600E-08 |
| 9073 | 454 | F454M | A | 1.1e-08 | 2.7e-09 | 8.7e-09 | 7.467E-09 | 4.285E-09 | 2.700E-09 | 1.100E-08 |
| 9074 | 454 | F454F | A | 5.7e-09 | 9.2e-09 | 1.2e-08 | 8.967E-09 | 3.156E-09 | 5.700E-09 | 1.200E-08 |
| 9075 | 454 | F454P | A | 6.7e-09 | 1.9e-08 | 2.3e-09 | 9.333E-09 | 8.656E-09 | 2.300E-09 | 1.900E-08 |
| 9076 | 454 | F454S | A | 8.8e-09 | 7.1e-09 | 3.6e-08 | 1.730E-08 | 1.622E-08 | 7.100E-09 | 3.600E-08 |
| 9077 | 454 | F454T | A | 2e-08 | 6.2e-09 | 6.1e-09 | 1.077E-08 | 7.996E-09 | 6.100E-09 | 2.000E-08 |
| 9078 | 454 | F454W | A | 1.4e-08 | 9.9e-09 | 4.5e-09 | 9.467E-09 | 4.765E-09 | 4.500E-09 | 1.400E-08 |
| 9079 | 454 | F454Y | A | 1e-08 | 3.9e-09 | 4.1e-09 | 6.000E-09 | 3.466E-09 | 3.900E-09 | 1.000E-08 |
| 9080 | 454 | F454V | A | 2.7e-09 | 3.2e-09 | 1.3e-08 | 6.300E-09 | 5.808E-09 | 2.700E-09 | 1.300E-08 |
| 9081 | 455 | V455A | A | 1.4e-08 | 8.4e-09 | 6.3e-09 | 9.567E-09 | 3.980E-09 | 6.300E-09 | 1.400E-08 |
| 9082 | 455 | V455R | A | 7.9e-09 | 8.1e-09 | 8.5e-09 | 8.167E-09 | 3.055E-10 | 7.900E-09 | 8.500E-09 |
| 9083 | 455 | V455N | A | 4.4e-09 | 1.8e-08 | 5.2e-09 | 9.200E-09 | 7.632E-09 | 4.400E-09 | 1.800E-08 |
| 9084 | 455 | V455D | A | 5e-09 | 1.1e-08 | 6.3e-09 | 7.433E-09 | 3.156E-09 | 5.000E-09 | 1.100E-08 |
| 9085 | 455 | V455C | A | 4.2e-09 | 2.9e-09 | 8.5e-09 | 5.200E-09 | 2.931E-09 | 2.900E-09 | 8.500E-09 |
| 9086 | 455 | V455E | A | 6.5e-09 | 6.6e-09 | 3.8e-09 | 5.633E-09 | 1.589E-09 | 3.800E-09 | 6.600E-09 |
| 9087 | 455 | V455Q | A | 1.1e-08 | 9.1e-09 | 4e-09 | 8.033E-09 | 3.620E-09 | 4.000E-09 | 1.100E-08 |
| 9088 | 455 | V455G | A | 7.1e-09 | 4.2e-09 | 1.3e-08 | 8.100E-09 | 4.484E-09 | 4.200E-09 | 1.300E-08 |
| 9089 | 455 | V455H | A | 5.6e-09 | 6.6e-09 | 1.3e-08 | 8.400E-09 | 4.015E-09 | 5.600E-09 | 1.300E-08 |
| 9090 | 455 | V455I | A | 6.5e-09 | 5.8e-09 | 5.8e-09 | 6.033E-09 | 4.041E-10 | 5.800E-09 | 6.500E-09 |
| 9091 | 455 | V455L | A | 1e-08 | 2.1e-08 | 2e-09 | 1.100E-08 | 9.539E-09 | 2.000E-09 | 2.100E-08 |
| 9092 | 455 | V455K | A | 2.1e-08 | 5.6e-09 | 6.8e-09 | 1.113E-08 | 8.566E-09 | 5.600E-09 | 2.100E-08 |
| 9093 | 455 | V455M | A | 5e-09 | 7.7e-09 | 7.1e-09 | 6.600E-09 | 1.418E-09 | 5.000E-09 | 7.700E-09 |
| 9094 | 455 | V455F | A | 3.4e-09 | 6.7e-09 | 5.5e-09 | 5.200E-09 | 1.670E-09 | 3.400E-09 | 6.700E-09 |

| | | | | | | | | | | |
|---|---|---|---|---|---|---|---|---|---|---|
| 9095 | 455 | V455P | A | 7.1e-09 | 2.9e-09 | 1.6e-08 | 8.667E-09 | 6.689E-09 | 2.900E-09 | 1.600E-08 |
| 9096 | 455 | V455S | A | 8.4e-09 | 8.3e-09 | 6.2e-09 | 7.633E-09 | 1.242E-09 | 6.200E-09 | 8.400E-09 |
| 9097 | 455 | V455T | A | 1.2e-08 | 4.7e-09 | 3.6e-09 | 6.767E-09 | 4.565E-09 | 3.600E-09 | 1.200E-08 |
| 9098 | 455 | V455W | A | 3.1e-09 | 9e-09 | 5.1e-09 | 5.733E-09 | 3.001E-09 | 3.100E-09 | 9.000E-09 |
| 9099 | 455 | V455Y | A | 1e-08 | 2e-09 | 1.1e-08 | 7.667E-09 | 4.933E-09 | 2.000E-09 | 1.100E-08 |
| 9100 | 455 | V455V | A | 8.1e-09 | 1.2e-08 | 4.9e-09 | 8.333E-09 | 3.556E-09 | 4.900E-09 | 1.200E-08 |
| 9101 | 456 | H456A | A | 1.3e-08 | 1.6e-08 | 2.5e-08 | 1.800E-08 | 6.245E-09 | 1.300E-08 | 2.500E-08 |
| 9102 | 456 | H456R | A | 2.3e-09 | 1.5e-08 | 3.5e-09 | 6.933E-09 | 7.012E-09 | 2.300E-09 | 1.500E-08 |
| 9103 | 456 | H456N | A | 2.5e-09 | 6.4e-09 | 1.1e-08 | 6.633E-09 | 4.255E-09 | 2.500E-09 | 1.100E-08 |
| 9104 | 456 | H456D | A | 1.6e-08 | 4.8e-09 | 2.2e-08 | 1.427E-08 | 8.730E-09 | 4.800E-09 | 2.200E-08 |
| 9105 | 456 | H456C | A | 2.9e-09 | 1.1e-08 | 1e-08 | 7.967E-09 | 4.416E-09 | 2.900E-09 | 1.100E-08 |
| 9106 | 456 | H456E | A | 9.6e-09 | 7.4e-09 | 5.7e-09 | 7.567E-09 | 1.955E-09 | 5.700E-09 | 9.600E-09 |
| 9107 | 456 | H456Q | A | 4.3e-09 | 1.1e-08 | 4.2e-09 | 6.500E-09 | 3.897E-09 | 4.200E-09 | 1.100E-08 |
| 9108 | 456 | H456G | A | 6.7e-09 | 1.7e-08 | 4.8e-09 | 9.500E-09 | 6.564E-09 | 4.800E-09 | 1.700E-08 |
| 9109 | 456 | H456H | A | 6.3e-09 | 8.5e-09 | 1e-08 | 8.267E-09 | 1.861E-09 | 6.300E-09 | 1.000E-08 |
| 9110 | 456 | H456I | A | 6.7e-09 | 1.1e-08 | 1.6e-09 | 6.433E-09 | 4.706E-09 | 1.600E-09 | 1.100E-08 |
| 9111 | 456 | H456L | A | 2.2e-08 | 1.2e-08 | 2.7e-09 | 1.223E-08 | 9.652E-09 | 2.700E-09 | 2.200E-08 |
| 9112 | 456 | H456K | A | 5.5e-09 | 8e-09 | 6.6e-09 | 6.700E-09 | 1.253E-09 | 5.500E-09 | 8.000E-09 |
| 9113 | 456 | H456M | A | 1.2e-08 | 7.4e-09 | 9.4e-09 | 9.600E-09 | 2.307E-09 | 7.400E-09 | 1.200E-08 |
| 9114 | 456 | H456F | A | 1e-08 | 4.2e-09 | 4.6e-09 | 6.267E-09 | 3.239E-09 | 4.200E-09 | 1.000E-08 |
| 9115 | 456 | H456P | A | 1.1e-08 | 9.1e-09 | 1.8e-08 | 1.270E-08 | 4.687E-09 | 9.100E-09 | 1.800E-08 |
| 9116 | 456 | H456S | A | 8.3e-09 | 1.2e-08 | 2.8e-09 | 7.700E-09 | 4.629E-09 | 2.800E-09 | 1.200E-08 |
| 9117 | 456 | H456T | A | 2e-09 | 3.5e-09 | 3.2e-09 | 2.900E-09 | 7.937E-10 | 2.000E-09 | 3.500E-09 |
| 9118 | 456 | H456W | A | 3.1e-09 | 4.4e-09 | 3e-09 | 3.500E-09 | 7.810E-10 | 3.000E-09 | 4.400E-09 |
| 9119 | 456 | H456Y | A | 9.6e-09 | 5e-09 | 3.8e-09 | 6.133E-09 | 3.062E-09 | 3.800E-09 | 9.600E-09 |
| 9120 | 456 | H456V | A | 4.9e-09 | 1.8e-08 | 1.8e-08 | 1.363E-08 | 7.563E-09 | 4.900E-09 | 1.800E-08 |
| 9121 | 457 | T457A | A | 1.9e-08 | 2.8e-09 | 1.3e-08 | 1.160E-08 | 8.190E-09 | 2.800E-09 | 1.900E-08 |
| 9122 | 457 | T457R | A | 3.1e-09 | 6.2e-09 | 1.1e-08 | 6.767E-09 | 3.980E-09 | 3.100E-09 | 1.100E-08 |
| 9123 | 457 | T457N | A | 6.2e-09 | 9.9e-09 | 1.1e-08 | 9.033E-09 | 2.515E-09 | 6.200E-09 | 1.100E-08 |
| 9124 | 457 | T457D | A | 1.2e-08 | 6.4e-09 | 1.5e-08 | 1.113E-08 | 4.365E-09 | 6.400E-09 | 1.500E-08 |
| 9125 | 457 | T457C | A | 6.9e-09 | 6.9e-09 | 1e-08 | 7.933E-09 | 1.790E-09 | 6.900E-09 | 1.000E-08 |
| 9126 | 457 | T457E | A | 3.8e-09 | 4.9e-09 | 4.6e-09 | 4.433E-09 | 5.686E-10 | 3.800E-09 | 4.900E-09 |
| 9127 | 457 | T457Q | A | 4.1e-09 | 1e-08 | 4.2e-09 | 6.100E-09 | 3.378E-09 | 4.100E-09 | 1.000E-08 |
| 9128 | 457 | T457G | A | 4.4e-09 | 1e-09 | 7.7e-09 | 4.367E-09 | 3.350E-09 | 1.000E-09 | 7.700E-09 |
| 9129 | 457 | T457H | A | 6.3e-09 | 3.2e-08 | 1.1e-08 | 1.643E-08 | 1.368E-08 | 6.300E-09 | 3.200E-08 |
| 9130 | 457 | T457I | A | 1.1e-08 | 1.4e-08 | 9.1e-09 | 1.137E-08 | 2.470E-09 | 9.100E-09 | 1.400E-08 |
| 9131 | 457 | T457L | A | 1e-08 | 2.6e-08 | 5.6e-09 | 1.387E-08 | 1.074E-08 | 5.600E-09 | 2.600E-08 |
| 9132 | 457 | T457K | A | 7.6e-09 | 6.8e-09 | 3.5e-09 | 5.967E-09 | 2.173E-09 | 3.500E-09 | 7.600E-09 |
| 9133 | 457 | T457M | A | 7.1e-09 | 9.1e-09 | 3.6e-09 | 6.600E-09 | 2.784E-09 | 3.600E-09 | 9.100E-09 |
| 9134 | 457 | T457F | A | 1.1e-09 | 2.8e-08 | 7.7e-09 | 1.227E-08 | 1.402E-08 | 1.100E-09 | 2.800E-08 |
| 9135 | 457 | T457P | A | 1.4e-08 | 4.4e-09 | 6.4e-09 | 8.267E-09 | 5.065E-09 | 4.400E-09 | 1.400E-08 |
| 9136 | 457 | T457S | A | 4.1e-09 | 5.6e-09 | 2.3e-09 | 4.000E-09 | 1.652E-09 | 2.300E-09 | 5.600E-09 |
| 9137 | 457 | T457T | A | 5.8e-09 | 3.8e-09 | 7.3e-09 | 5.633E-09 | 1.756E-09 | 3.800E-09 | 7.300E-09 |
| 9138 | 457 | T457W | A | 8e-09 | 8.2e-09 | 3e-09 | 6.400E-09 | 2.946E-09 | 3.000E-09 | 8.200E-09 |
| 9139 | 457 | T457Y | A | 8.5e-09 | 8.2e-09 | 9.2e-09 | 8.633E-09 | 5.132E-10 | 8.200E-09 | 9.200E-09 |
| 9140 | 457 | T457V | A | 9.1e-09 | 4.5e-09 | 7.2e-09 | 6.933E-09 | 2.312E-09 | 4.500E-09 | 9.100E-09 |
| 9141 | 458 | V458A | A | 1.9e-08 | 1.2e-08 | 1.6e-08 | 1.567E-08 | 3.512E-09 | 1.200E-08 | 1.900E-08 |
| 9142 | 458 | V458R | A | 1.7e-08 | 7.8e-09 | 3.9e-09 | 9.567E-09 | 6.726E-09 | 3.900E-09 | 1.700E-08 |
| 9143 | 458 | V458N | A | 4.1e-09 | 2.8e-09 | 9.7e-09 | 5.533E-09 | 3.667E-09 | 2.800E-09 | 9.700E-09 |
| 9144 | 458 | V458D | A | 2.7e-09 | 5e-09 | 1e-08 | 5.900E-09 | 3.732E-09 | 2.700E-09 | 1.000E-08 |
| 9145 | 458 | V458C | A | 1.3e-08 | 1.3e-08 | 4.2e-09 | 1.007E-08 | 5.081E-09 | 4.200E-09 | 1.300E-08 |
| 9146 | 458 | V458E | A | 7.7e-09 | 3.5e-09 | 9.9e-09 | 7.033E-09 | 3.252E-09 | 3.500E-09 | 9.900E-09 |
| 9147 | 458 | V458Q | A | 6.2e-09 | 4.4e-09 | 8.1e-09 | 6.233E-09 | 1.850E-09 | 4.400E-09 | 8.100E-09 |
| 9148 | 458 | V458G | A | 7e-09 | 1.1e-08 | 5e-09 | 7.667E-09 | 3.055E-09 | 5.000E-09 | 1.100E-08 |
| 9149 | 458 | V458H | A | 6.7e-09 | 1.2e-08 | 1.2e-08 | 1.023E-08 | 3.060E-09 | 6.700E-09 | 1.200E-08 |
| 9150 | 458 | V458I | A | 4.8e-09 | 3.1e-09 | 1.3e-08 | 6.967E-09 | 5.294E-09 | 3.100E-09 | 1.300E-08 |
| 9151 | 458 | V458L | A | 7.2e-09 | 1e-08 | 4.7e-09 | 7.300E-09 | 2.651E-09 | 4.700E-09 | 1.000E-08 |
| 9152 | 458 | V458K | A | 1.9e-08 | 8.4e-09 | 6.5e-09 | 1.130E-08 | 6.736E-09 | 6.500E-09 | 1.900E-08 |
| 9153 | 458 | V458M | A | 1.8e-08 | 7.6e-09 | 5.3e-09 | 1.030E-08 | 6.767E-09 | 5.300E-09 | 1.800E-08 |
| 9154 | 458 | V458F | A | 8.8e-09 | 1.5e-08 | 1.4e-08 | 1.260E-08 | 3.329E-09 | 8.800E-09 | 1.500E-08 |
| 9155 | 458 | V458P | A | 9.6e-09 | 9.4e-09 | 6.7e-09 | 8.567E-09 | 1.620E-09 | 6.700E-09 | 9.600E-09 |
| 9156 | 458 | V458S | A | 5.5e-09 | 2.3e-09 | 5e-09 | 4.267E-09 | 1.721E-09 | 2.300E-09 | 5.500E-09 |
| 9157 | 458 | V458T | A | 5.5e-09 | 6e-09 | 1.7e-08 | 9.500E-09 | 6.500E-09 | 5.500E-09 | 1.700E-08 |
| 9158 | 458 | V458W | A | 5.3e-09 | 5.9e-09 | 7.2e-09 | 6.133E-09 | 9.713E-10 | 5.300E-09 | 7.200E-09 |
| 9159 | 458 | V458Y | A | 1.4e-08 | 1.4e-08 | 4.2e-09 | 1.073E-08 | 5.658E-09 | 4.200E-09 | 1.400E-08 |
| 9160 | 458 | V458V | A | 1.6e-08 | 1.3e-08 | 5.8e-09 | 1.160E-08 | 5.242E-09 | 5.800E-09 | 1.600E-08 |
| 9161 | 459 | P459A | A | 1.7e-09 | 2.9e-09 | 1.6e-08 | 6.867E-09 | 7.932E-09 | 1.700E-09 | 1.600E-08 |
| 9162 | 459 | P459R | A | 4.4e-09 | 1.1e-08 | 4.3e-09 | 6.567E-09 | 3.840E-09 | 4.300E-09 | 1.100E-08 |
| 9163 | 459 | P459N | A | 6.4e-09 | 5e-09 | 4.8e-09 | 5.400E-09 | 8.718E-10 | 4.800E-09 | 6.400E-09 |
| 9164 | 459 | P459D | A | 1.2e-08 | 6.6e-09 | 4.7e-09 | 7.767E-09 | 3.787E-09 | 4.700E-09 | 1.200E-08 |
| 9165 | 459 | P459C | A | 6.1e-09 | 5.6e-09 | 5.2e-09 | 5.633E-09 | 4.509E-10 | 5.200E-09 | 6.100E-09 |
| 9166 | 459 | P459E | A | 5.2e-09 | 1.2e-08 | 1.1e-08 | 9.400E-09 | 3.672E-09 | 5.200E-09 | 1.200E-08 |
| 9167 | 459 | P459Q | A | 5.7e-09 | 3.8e-09 | 3.5e-08 | 1.483E-08 | 1.749E-08 | 3.800E-09 | 3.500E-08 |
| 9168 | 459 | P459G | A | 2e-09 | 5.9e-09 | 2.1e-08 | 9.633E-09 | 1.004E-08 | 2.000E-09 | 2.100E-08 |
| 9169 | 459 | P459H | A | 9.4e-09 | 5.4e-09 | 8.2e-09 | 7.667E-09 | 2.053E-09 | 5.400E-09 | 9.400E-09 |
| 9170 | 459 | P459I | A | 9.2e-09 | 4e-09 | 7e-09 | 6.733E-09 | 2.610E-09 | 4.000E-09 | 9.200E-09 |
| 9171 | 459 | P459L | A | 1.8e-08 | 5.6e-09 | 6.9e-09 | 1.017E-08 | 6.815E-09 | 5.600E-09 | 1.800E-08 |
| 9172 | 459 | P459K | A | 3.1e-09 | 5.7e-09 | 4.2e-09 | 4.333E-09 | 1.305E-09 | 3.100E-09 | 5.700E-09 |
| 9173 | 459 | P459M | A | 1.2e-08 | 1.2e-08 | 5.5e-09 | 9.833E-09 | 3.753E-09 | 5.500E-09 | 1.200E-08 |
| 9174 | 459 | P459F | A | 2e-08 | 1e-08 | 5e-09 | 1.167E-08 | 7.638E-09 | 5.000E-09 | 2.000E-08 |
| 9175 | 459 | P459P | A | 7.6e-09 | 5e-09 | 3.6e-09 | 5.400E-09 | 2.030E-09 | 3.600E-09 | 7.600E-09 |
| 9176 | 459 | P459S | A | 6.8e-09 | 7.6e-09 | 4.7e-09 | 6.367E-09 | 1.498E-09 | 4.700E-09 | 7.600E-09 |
| 9177 | 459 | P459T | A | 1.8e-08 | 7.2e-09 | 4.4e-09 | 9.867E-09 | 7.181E-09 | 4.400E-09 | 1.800E-08 |
| 9178 | 459 | P459W | A | 2.8e-09 | 9.7e-09 | 4.4e-09 | 5.633E-09 | 3.612E-09 | 2.800E-09 | 9.700E-09 |
| 9179 | 459 | P459Y | A | 3.8e-09 | 1.4e-08 | 5.8e-09 | 7.867E-09 | 5.405E-09 | 3.800E-09 | 1.400E-08 |

| | | | | | | | | | | |
|---|---|---|---|---|---|---|---|---|---|---|
| 9180 | 459 | P459V | A | 8.2e-09 | 4.1e-09 | 3.3e-09 | 5.200E-09 | 2.629E-09 | 3.300E-09 | 8.200E-09 |
| 9181 | 460 | W460A | A | 3.9e-09 | 1e-08 | 1.3e-08 | 8.967E-09 | 4.637E-09 | 3.900E-09 | 1.300E-08 |
| 9182 | 460 | W460R | A | 9.4e-09 | 5.8e-09 | 4.4e-09 | 6.533E-09 | 2.579E-09 | 4.400E-09 | 9.400E-09 |
| 9183 | 460 | W460N | A | 3.4e-09 | 3.9e-09 | 4.5e-09 | 3.933E-09 | 5.508E-10 | 3.400E-09 | 4.500E-09 |
| 9184 | 460 | W460D | A | 9.2e-09 | 5.7e-09 | 3.7e-09 | 6.200E-09 | 2.784E-09 | 3.700E-09 | 9.200E-09 |
| 9185 | 460 | W460C | A | 1.3e-08 | 1.1e-08 | 9.4e-09 | 1.113E-08 | 1.804E-09 | 9.400E-09 | 1.300E-08 |
| 9186 | 460 | W460E | A | 2.3e-08 | 1.4e-08 | 1.1e-08 | 1.600E-08 | 6.245E-09 | 1.100E-08 | 2.300E-08 |
| 9187 | 460 | W460Q | A | 1e-08 | 5.2e-09 | 4.6e-09 | 6.600E-09 | 2.960E-09 | 4.600E-09 | 1.000E-08 |
| 9188 | 460 | W460G | A | 5.3e-09 | 1.9e-09 | 1.1e-08 | 6.067E-09 | 4.598E-09 | 1.900E-09 | 1.100E-08 |
| 9189 | 460 | W460H | A | 1.4e-08 | 8.5e-09 | 5.7e-09 | 9.400E-09 | 4.223E-09 | 5.700E-09 | 1.400E-08 |
| 9190 | 460 | W460I | A | 4.5e-09 | 4.5e-09 | 4.4e-09 | 4.467E-09 | 5.774E-11 | 4.400E-09 | 4.500E-09 |
| 9191 | 460 | W460L | A | 1.1e-08 | 7.9e-09 | 5.8e-09 | 8.233E-09 | 2.616E-09 | 5.800E-09 | 1.100E-08 |
| 9192 | 460 | W460K | A | 1.5e-08 | 3.3e-09 | 9.5e-09 | 9.267E-09 | 5.853E-09 | 3.300E-09 | 1.500E-08 |
| 9193 | 460 | W460M | A | 7.8e-09 | 3.2e-09 | 7.3e-09 | 6.100E-09 | 2.524E-09 | 3.200E-09 | 7.800E-09 |
| 9194 | 460 | W460F | A | 8.5e-09 | 8.4e-09 | 6.2e-09 | 7.700E-09 | 1.300E-09 | 6.200E-09 | 8.500E-09 |
| 9195 | 460 | W460P | A | 9e-09 | 9.8e-09 | 6.9e-09 | 8.567E-09 | 1.498E-09 | 6.900E-09 | 9.800E-09 |
| 9196 | 460 | W460S | A | 3.7e-09 | 5.5e-09 | 8.7e-09 | 5.967E-09 | 2.532E-09 | 3.700E-09 | 8.700E-09 |
| 9197 | 460 | W460T | A | 6.7e-09 | 6e-09 | 5.1e-09 | 5.933E-09 | 8.021E-10 | 5.100E-09 | 6.700E-09 |
| 9198 | 460 | W460W | A | 4.4e-09 | 5.9e-09 | 3.1e-09 | 4.467E-09 | 1.401E-09 | 3.100E-09 | 5.900E-09 |
| 9199 | 460 | W460Y | A | 1.5e-08 | 3.4e-09 | 3.1e-09 | 7.167E-09 | 6.786E-09 | 3.100E-09 | 1.500E-08 |
| 9200 | 460 | W460V | A | 8.1e-09 | 3.3e-09 | 8.2e-09 | 6.533E-09 | 2.801E-09 | 3.300E-09 | 8.200E-09 |
| 9201 | 461 | D461A | A | 1.9e-09 | 4.2e-09 | 5.7e-09 | 3.933E-09 | 1.914E-09 | 1.900E-09 | 5.700E-09 |
| 9202 | 461 | D461R | A | 5.3e-09 | 6.3e-09 | 6.5e-09 | 6.033E-09 | 6.429E-10 | 5.300E-09 | 6.500E-09 |
| 9203 | 461 | D461N | A | 4.8e-09 | 1.3e-08 | 2.7e-09 | 6.833E-09 | 5.443E-09 | 2.700E-09 | 1.300E-08 |
| 9204 | 461 | D461D | A | 1.3e-08 | 2.3e-09 | 1.5e-08 | 1.010E-08 | 6.829E-09 | 2.300E-09 | 1.500E-08 |
| 9205 | 461 | D461C | A | 5.6e-09 | 7.9e-09 | 1.4e-09 | 4.967E-09 | 3.296E-09 | 1.400E-09 | 7.900E-09 |
| 9206 | 461 | D461E | A | 4.8e-09 | 9.7e-09 | 7.3e-09 | 7.267E-09 | 2.450E-09 | 4.800E-09 | 9.700E-09 |
| 9207 | 461 | D461Q | A | 1.4e-08 | 6.7e-09 | 4.8e-09 | 8.500E-09 | 4.857E-09 | 4.800E-09 | 1.400E-08 |
| 9208 | 461 | D461G | A | 1.4e-08 | 3.7e-09 | 5.1e-09 | 7.600E-09 | 5.587E-09 | 3.700E-09 | 1.400E-08 |
| 9209 | 461 | D461H | A | 1.1e-08 | 6.8e-09 | 4.9e-09 | 7.567E-09 | 3.121E-09 | 4.900E-09 | 1.100E-08 |
| 9210 | 461 | D461I | A | 1.8e-09 | 2.6e-08 | 3.2e-09 | 1.033E-08 | 1.359E-08 | 1.800E-09 | 2.600E-08 |
| 9211 | 461 | D461L | A | 2.5e-09 | 7.4e-09 | 3.4e-09 | 4.433E-09 | 2.608E-09 | 2.500E-09 | 7.400E-09 |
| 9212 | 461 | D461K | A | 5.9e-09 | 4e-09 | 3e-09 | 4.300E-09 | 1.473E-09 | 3.000E-09 | 5.900E-09 |
| 9213 | 461 | D461M | A | 8.3e-09 | 1.5e-08 | 7.7e-09 | 1.033E-08 | 4.053E-09 | 7.700E-09 | 1.500E-08 |
| 9214 | 461 | D461F | A | 1.5e-08 | 1.2e-08 | 5.3e-09 | 1.077E-08 | 4.966E-09 | 5.300E-09 | 1.500E-08 |
| 9215 | 461 | D461P | A | 1.6e-08 | 5.2e-09 | 3.2e-09 | 8.133E-09 | 6.886E-09 | 3.200E-09 | 1.600E-08 |
| 9216 | 461 | D461S | A | 3.6e-09 | 6.1e-09 | 1.5e-08 | 8.233E-09 | 5.992E-09 | 3.600E-09 | 1.500E-08 |
| 9217 | 461 | D461T | A | 9.6e-09 | 8.7e-09 | 3.1e-09 | 7.133E-09 | 3.522E-09 | 3.100E-09 | 9.600E-09 |
| 9218 | 461 | D461W | A | 3e-09 | 6.7e-09 | 2.6e-09 | 4.100E-09 | 2.261E-09 | 2.600E-09 | 6.700E-09 |
| 9219 | 461 | D461Y | A | 6.6e-09 | 6e-09 | 4.6e-09 | 5.733E-09 | 1.026E-09 | 4.600E-09 | 6.600E-09 |
| 9220 | 461 | D461V | A | 4.9e-09 | 4.5e-09 | 5.9e-09 | 5.100E-09 | 7.211E-10 | 4.500E-09 | 5.900E-09 |
| 9221 | 462 | Q462A | A | 2.2e-09 | 4.7e-09 | 8.1e-09 | 5.000E-09 | 2.961E-09 | 2.200E-09 | 8.100E-09 |
| 9222 | 462 | Q462R | A | 3.2e-09 | 3.2e-09 | 5.9e-09 | 4.100E-09 | 1.559E-09 | 3.200E-09 | 5.900E-09 |
| 9223 | 462 | Q462N | A | 1.4e-08 | 2.2e-09 | 2.9e-09 | 6.367E-09 | 6.620E-09 | 2.200E-09 | 1.400E-08 |
| 9224 | 462 | Q462D | A | 2.2e-09 | 1.3e-08 | 3.5e-09 | 6.233E-09 | 5.896E-09 | 2.200E-09 | 1.300E-08 |
| 9225 | 462 | Q462C | A | 1.8e-08 | 1.7e-08 | 2.2e-08 | 1.900E-08 | 2.646E-09 | 1.700E-08 | 2.200E-08 |
| 9226 | 462 | Q462E | A | 3.7e-09 | 1.2e-08 | 9.2e-09 | 8.300E-09 | 4.223E-09 | 3.700E-09 | 1.200E-08 |
| 9227 | 462 | Q462Q | A | 9.1e-09 | 3.2e-09 | 1.4e-08 | 8.767E-09 | 5.408E-09 | 3.200E-09 | 1.400E-08 |
| 9228 | 462 | Q462G | A | 3e-09 | 7.9e-09 | 4e-09 | 4.967E-09 | 2.589E-09 | 3.000E-09 | 7.900E-09 |
| 9229 | 462 | Q462H | A | 5.1e-09 | 1.2e-08 | 3.8e-09 | 6.967E-09 | 4.407E-09 | 3.800E-09 | 1.200E-08 |
| 9230 | 462 | Q462I | A | 4.8e-09 | 1e-08 | 2.5e-09 | 5.767E-09 | 3.842E-09 | 2.500E-09 | 1.000E-08 |
| 9231 | 462 | Q462L | A | 6.1e-09 | 7.9e-09 | 1.5e-08 | 9.667E-09 | 4.706E-09 | 6.100E-09 | 1.500E-08 |
| 9232 | 462 | Q462K | A | 7.7e-09 | 3.5e-09 | 6.1e-09 | 5.767E-09 | 2.120E-09 | 3.500E-09 | 7.700E-09 |
| 9233 | 462 | Q462M | A | 7.5e-09 | 3.3e-09 | 1.2e-08 | 7.600E-09 | 4.351E-09 | 3.300E-09 | 1.200E-08 |
| 9234 | 462 | Q462F | A | 4.1e-09 | 4.4e-09 | 1.7e-08 | 8.500E-09 | 7.363E-09 | 4.100E-09 | 1.700E-08 |
| 9235 | 462 | Q462P | A | 3.6e-09 | 1.6e-09 | 3.2e-09 | 2.800E-09 | 1.058E-09 | 1.600E-09 | 3.600E-09 |
| 9236 | 462 | Q462S | A | 3.6e-09 | 1.1e-08 | 5.1e-09 | 6.567E-09 | 3.912E-09 | 3.600E-09 | 1.100E-08 |
| 9237 | 462 | Q462T | A | 6.1e-09 | 6e-09 | 7.5e-09 | 6.533E-09 | 8.386E-10 | 6.000E-09 | 7.500E-09 |
| 9238 | 462 | Q462W | A | 5.7e-09 | 4.6e-09 | 9e-09 | 6.433E-09 | 2.290E-09 | 4.600E-09 | 9.000E-09 |
| 9239 | 462 | Q462Y | A | 2e-08 | 1.9e-08 | 3.5e-09 | 1.417E-08 | 9.251E-09 | 3.500E-09 | 2.000E-08 |
| 9240 | 462 | Q462V | A | 4.2e-09 | 2.6e-08 | 1.1e-08 | 1.373E-08 | 1.115E-08 | 4.200E-09 | 2.600E-08 |
| 9241 | 463 | L463A | A | 3.9e-09 | 9.1e-09 | 8.5e-09 | 7.167E-09 | 2.845E-09 | 3.900E-09 | 9.100E-09 |
| 9242 | 463 | L463R | A | 2.1e-09 | 1.2e-08 | 1.4e-08 | 9.367E-09 | 6.372E-09 | 2.100E-09 | 1.400E-08 |
| 9243 | 463 | L463N | A | 7.1e-09 | 2.4e-09 | 7.1e-09 | 5.533E-09 | 2.714E-09 | 2.400E-09 | 7.100E-09 |
| 9244 | 463 | L463D | A | 1.9e-09 | 9.1e-09 | 7e-09 | 6.000E-09 | 3.703E-09 | 1.900E-09 | 9.100E-09 |
| 9245 | 463 | L463C | A | 4.3e-08 | 6.5e-09 | 1.2e-08 | 2.050E-08 | 1.968E-08 | 6.500E-09 | 4.300E-08 |
| 9246 | 463 | L463E | A | 7.1e-09 | 1.3e-08 | 3.7e-09 | 7.933E-09 | 4.706E-09 | 3.700E-09 | 1.300E-08 |
| 9247 | 463 | L463Q | A | 7.3e-09 | 1.4e-08 | 1e-08 | 1.043E-08 | 3.371E-09 | 7.300E-09 | 1.400E-08 |
| 9248 | 463 | L463G | A | 4.3e-09 | 8.2e-09 | 1.1e-08 | 7.833E-09 | 3.365E-09 | 4.300E-09 | 1.100E-08 |
| 9249 | 463 | L463H | A | 1.2e-08 | 2.3e-09 | 5.6e-09 | 6.633E-09 | 4.932E-09 | 2.300E-09 | 1.200E-08 |
| 9250 | 463 | L463I | A | 6e-09 | 5.8e-09 | 2.3e-08 | 1.160E-08 | 9.873E-09 | 5.800E-09 | 2.300E-08 |
| 9251 | 463 | L463L | A | 3.7e-09 | 8.8e-09 | 4.8e-09 | 5.767E-09 | 2.684E-09 | 3.700E-09 | 8.800E-09 |
| 9252 | 463 | L463K | A | 3.5e-09 | 1.5e-08 | 1.2e-08 | 1.017E-08 | 5.965E-09 | 3.500E-09 | 1.500E-08 |
| 9253 | 463 | L463M | A | 4.1e-09 | 9.8e-09 | 1.3e-08 | 8.967E-09 | 4.508E-09 | 4.100E-09 | 1.300E-08 |
| 9254 | 463 | L463F | A | 3.3e-09 | 1.4e-08 | 1.9e-08 | 1.210E-08 | 8.021E-09 | 3.300E-09 | 1.900E-08 |
| 9255 | 463 | L463P | A | 5.6e-09 | 3.3e-09 | 5.3e-09 | 4.733E-09 | 1.250E-09 | 3.300E-09 | 5.600E-09 |
| 9256 | 463 | L463S | A | 1.4e-08 | 9.1e-09 | 5.4e-08 | 2.570E-08 | 2.463E-08 | 9.100E-09 | 5.400E-08 |
| 9257 | 463 | L463T | A | 5.8e-09 | 5.1e-09 | 6.6e-09 | 5.833E-09 | 7.506E-10 | 5.100E-09 | 6.600E-09 |
| 9258 | 463 | L463W | A | 1.3e-08 | 5e-09 | 7.6e-09 | 8.533E-09 | 4.081E-09 | 5.000E-09 | 1.300E-08 |
| 9259 | 463 | L463Y | A | 1.1e-08 | 1.4e-08 | 7e-09 | 1.067E-08 | 3.512E-09 | 7.000E-09 | 1.400E-08 |
| 9260 | 463 | L463V | A | 7e-09 | 6.4e-09 | 2.1e-08 | 1.147E-08 | 8.262E-09 | 6.400E-09 | 2.100E-08 |
| 9261 | 464 | F464A | A | 5.8e-09 | 6.8e-09 | 1.1e-08 | 7.867E-09 | 2.759E-09 | 5.800E-09 | 1.100E-08 |
| 9262 | 464 | F464R | A | 1.1e-08 | 6.2e-09 | 5e-09 | 7.400E-09 | 3.175E-09 | 5.000E-09 | 1.100E-08 |
| 9263 | 464 | F464N | A | 2.2e-08 | 1e-08 | 1.9e-08 | 1.700E-08 | 6.245E-09 | 1.000E-08 | 2.200E-08 |
| 9264 | 464 | F464D | A | 8.6e-09 | 8.9e-09 | 6.2e-09 | 7.900E-09 | 1.480E-09 | 6.200E-09 | 8.900E-09 |

| | | | | | | | | | | |
|---|---|---|---|---|---|---|---|---|---|---|
| 9265 | 464 | F464C | A | 4.4e-09 | 8.6e-09 | 1.3e-08 | 8.667E-09 | 4.300E-09 | 4.400E-09 | 1.300E-08 |
| 9266 | 464 | F464E | A | 7e-09 | 3e-09 | 3e-09 | 4.333E-09 | 2.309E-09 | 3.000E-09 | 7.000E-09 |
| 9267 | 464 | F464Q | A | 1.2e-08 | 6.7e-09 | 1.5e-08 | 1.123E-08 | 4.203E-09 | 6.700E-09 | 1.500E-08 |
| 9268 | 464 | F464G | A | 8e-09 | 7.9e-09 | 3.8e-09 | 6.567E-09 | 2.397E-09 | 3.800E-09 | 8.000E-09 |
| 9269 | 464 | F464H | A | 4.5e-09 | 5.7e-09 | 4.5e-09 | 4.900E-09 | 6.928E-10 | 4.500E-09 | 5.700E-09 |
| 9270 | 464 | F464I | A | 4.5e-09 | 1.1e-08 | 1.3e-08 | 9.500E-09 | 4.444E-09 | 4.500E-09 | 1.300E-08 |
| 9271 | 464 | F464L | A | 6.2e-09 | 5.9e-09 | 3.7e-09 | 5.267E-09 | 1.365E-09 | 3.700E-09 | 6.200E-09 |
| 9272 | 464 | F464K | A | 1.4e-08 | 7.4e-09 | 6.2e-09 | 9.200E-09 | 4.200E-09 | 6.200E-09 | 1.400E-08 |
| 9273 | 464 | F464M | A | 5e-09 | 1.1e-08 | 1.4e-08 | 1.000E-08 | 4.583E-09 | 5.000E-09 | 1.400E-08 |
| 9274 | 464 | F464F | A | 4.7e-09 | 3.5e-09 | 7.8e-09 | 5.333E-09 | 2.219E-09 | 3.500E-09 | 7.800E-09 |
| 9275 | 464 | F464P | A | 6.1e-09 | 2.4e-08 | 4.5e-09 | 1.153E-08 | 1.083E-08 | 4.500E-09 | 2.400E-08 |
| 9276 | 464 | F464S | A | 1.5e-08 | 4.2e-09 | 8.6e-09 | 9.267E-09 | 5.431E-09 | 4.200E-09 | 1.500E-08 |
| 9277 | 464 | F464T | A | 5.5e-09 | 6.5e-09 | 1.2e-08 | 8.000E-09 | 3.500E-09 | 5.500E-09 | 1.200E-08 |
| 9278 | 464 | F464W | A | 7.3e-09 | 7.7e-09 | 6.8e-09 | 7.267E-09 | 4.509E-10 | 6.800E-09 | 7.700E-09 |
| 9279 | 464 | F464Y | A | 1e-08 | 6e-09 | 7.3e-09 | 7.767E-09 | 2.040E-09 | 6.000E-09 | 1.000E-08 |
| 9280 | 464 | F464V | A | 1e-08 | 2.8e-09 | 1.3e-08 | 8.600E-09 | 5.242E-09 | 2.800E-09 | 1.300E-08 |
| 9281 | 465 | R465A | A | 1.1e-08 | 4.2e-09 | 2.4e-09 | 5.867E-09 | 4.536E-09 | 2.400E-09 | 1.100E-08 |
| 9282 | 465 | R465R | A | 1.2e-08 | 6.3e-09 | 9.3e-09 | 9.200E-09 | 2.851E-09 | 6.300E-09 | 1.200E-08 |
| 9283 | 465 | R465N | A | 9.1e-09 | 6e-09 | 5.4e-09 | 6.833E-09 | 1.986E-09 | 5.400E-09 | 9.100E-09 |
| 9284 | 465 | R465D | A | 1.3e-08 | 5.7e-09 | 1.6e-08 | 1.157E-08 | 5.297E-09 | 5.700E-09 | 1.600E-08 |
| 9285 | 465 | R465C | A | 9.3e-09 | 3.7e-09 | 6.6e-09 | 6.533E-09 | 2.801E-09 | 3.700E-09 | 9.300E-09 |
| 9286 | 465 | R465E | A | 5.1e-09 | 1.1e-08 | 5.1e-09 | 7.067E-09 | 3.406E-09 | 5.100E-09 | 1.100E-08 |
| 9287 | 465 | R465Q | A | 3.9e-09 | 3e-08 | 7.2e-09 | 1.370E-08 | 1.421E-08 | 3.900E-09 | 3.000E-08 |
| 9288 | 465 | R465G | A | 5.5e-09 | 4.8e-09 | 4.1e-09 | 4.800E-09 | 7.000E-10 | 4.100E-09 | 5.500E-09 |
| 9289 | 465 | R465H | A | 7.5e-09 | 1.3e-08 | 9.2e-09 | 9.900E-09 | 2.816E-09 | 7.500E-09 | 1.300E-08 |
| 9290 | 465 | R465I | A | 1.3e-08 | 8.4e-09 | 1.2e-08 | 1.113E-08 | 2.419E-09 | 8.400E-09 | 1.300E-08 |
| 9291 | 465 | R465L | A | 3.4e-09 | 8.1e-09 | 5.1e-09 | 5.533E-09 | 2.380E-09 | 3.400E-09 | 8.100E-09 |
| 9292 | 465 | R465K | A | 4.5e-09 | 3.6e-09 | 3e-09 | 3.700E-09 | 7.550E-10 | 3.000E-09 | 4.500E-09 |
| 9293 | 465 | R465M | A | 1.5e-08 | 4.9e-09 | 2.6e-09 | 7.500E-09 | 6.596E-09 | 2.600E-09 | 1.500E-08 |
| 9294 | 465 | R465F | A | 5.2e-09 | 1.7e-08 | 6.7e-09 | 9.633E-09 | 6.424E-09 | 5.200E-09 | 1.700E-08 |
| 9295 | 465 | R465P | A | 1e-08 | 2.1e-08 | 3.1e-08 | 2.067E-08 | 1.050E-08 | 1.000E-08 | 3.100E-08 |
| 9296 | 465 | R465S | A | 5.7e-09 | 1e-08 | 2.9e-09 | 6.200E-09 | 3.576E-09 | 2.900E-09 | 1.000E-08 |
| 9297 | 465 | R465T | A | 3.9e-09 | 4.7e-09 | 6.6e-09 | 5.067E-09 | 1.387E-09 | 3.900E-09 | 6.600E-09 |
| 9298 | 465 | R465W | A | 2.3e-09 | 6.7e-09 | 1.2e-08 | 7.000E-09 | 4.857E-09 | 2.300E-09 | 1.200E-08 |
| 9299 | 465 | R465Y | A | 5.2e-09 | 3.7e-09 | 7.7e-09 | 5.533E-09 | 2.021E-09 | 3.700E-09 | 7.700E-09 |
| 9300 | 465 | R465V | A | 1.4e-08 | 8.5e-09 | 9.2e-09 | 1.057E-08 | 2.994E-09 | 8.500E-09 | 1.400E-08 |
| 9301 | 466 | N466A | A | 6.1e-09 | 8e-09 | 7.5e-09 | 7.200E-09 | 9.849E-10 | 6.100E-09 | 8.000E-09 |
| 9302 | 466 | N466R | A | 1.3e-08 | 7.3e-09 | 4e-09 | 8.100E-09 | 4.553E-09 | 4.000E-09 | 1.300E-08 |
| 9303 | 466 | N466N | A | 1.2e-08 | 1.5e-08 | 8.7e-09 | 1.190E-08 | 3.151E-09 | 8.700E-09 | 1.500E-08 |
| 9304 | 466 | N466D | A | 3e-09 | 6.2e-09 | 4.4e-09 | 4.533E-09 | 1.604E-09 | 3.000E-09 | 6.200E-09 |
| 9305 | 466 | N466C | A | 3.2e-09 | 3.3e-09 | 5.2e-09 | 3.900E-09 | 1.127E-09 | 3.200E-09 | 5.200E-09 |
| 9306 | 466 | N466E | A | 5e-09 | 3.8e-09 | 5.9e-09 | 4.900E-09 | 1.054E-09 | 3.800E-09 | 5.900E-09 |
| 9307 | 466 | N466Q | A | 7.9e-09 | 4.9e-09 | 1.4e-08 | 8.933E-09 | 4.637E-09 | 4.900E-09 | 1.400E-08 |
| 9308 | 466 | N466G | A | 7.2e-09 | 3.1e-09 | 9.1e-09 | 6.467E-09 | 3.066E-09 | 3.100E-09 | 9.100E-09 |
| 9309 | 466 | N466H | A | 6.9e-09 | 1.8e-08 | 1.4e-09 | 8.767E-09 | 8.456E-09 | 1.400E-09 | 1.800E-08 |
| 9310 | 466 | N466I | A | 6.6e-09 | 7.1e-09 | 5.7e-09 | 6.467E-09 | 7.095E-10 | 5.700E-09 | 7.100E-09 |
| 9311 | 466 | N466L | A | 1.1e-08 | 6.9e-09 | 5.1e-09 | 7.667E-09 | 3.024E-09 | 5.100E-09 | 1.100E-08 |
| 9312 | 466 | N466K | A | 8.4e-09 | 3.9e-09 | 1e-08 | 7.433E-09 | 3.163E-09 | 3.900E-09 | 1.000E-08 |
| 9313 | 466 | N466M | A | 3.1e-08 | 1.3e-08 | 6.7e-09 | 1.690E-08 | 1.261E-08 | 6.700E-09 | 3.100E-08 |
| 9314 | 466 | N466F | A | 1.6e-08 | 1e-08 | 9.8e-09 | 1.193E-08 | 3.523E-09 | 9.800E-09 | 1.600E-08 |
| 9315 | 466 | N466P | A | 7.4e-09 | 1.5e-09 | 8.7e-09 | 5.867E-09 | 3.837E-09 | 1.500E-09 | 8.700E-09 |
| 9316 | 466 | N466S | A | 3.3e-09 | 4.9e-09 | 4.3e-09 | 4.167E-09 | 8.083E-10 | 3.300E-09 | 4.900E-09 |
| 9317 | 466 | N466T | A | 8.5e-09 | 1.4e-08 | 1.2e-08 | 1.150E-08 | 2.784E-09 | 8.500E-09 | 1.400E-08 |
| 9318 | 466 | N466W | A | 2.9e-09 | 3.2e-09 | 2.9e-09 | 3.000E-09 | 1.732E-10 | 2.900E-09 | 3.200E-09 |
| 9319 | 466 | N466Y | A | 8.4e-09 | 6.8e-09 | 4.5e-09 | 6.567E-09 | 1.960E-09 | 4.500E-09 | 8.400E-09 |
| 9320 | 466 | N466V | A | 2.6e-08 | 5.4e-09 | 3.9e-09 | 1.177E-08 | 1.235E-08 | 3.900E-09 | 2.600E-08 |
| 9321 | 467 | P467A | A | 1.1e-08 | 5.9e-09 | 1.6e-08 | 1.097E-08 | 5.050E-09 | 5.900E-09 | 1.600E-08 |
| 9322 | 467 | P467R | A | 2.9e-09 | 6.1e-09 | 6.6e-09 | 5.200E-09 | 2.007E-09 | 2.900E-09 | 6.600E-09 |
| 9323 | 467 | P467N | A | 1.5e-08 | 9.8e-09 | 4.8e-09 | 9.867E-09 | 5.100E-09 | 4.800E-09 | 1.500E-08 |
| 9324 | 467 | P467D | A | 1.9e-08 | 5.2e-09 | 1.1e-08 | 1.173E-08 | 6.929E-09 | 5.200E-09 | 1.900E-08 |
| 9325 | 467 | P467C | A | 2.2e-08 | 8.7e-09 | 4.5e-09 | 1.173E-08 | 9.136E-09 | 4.500E-09 | 2.200E-08 |
| 9326 | 467 | P467E | A | 6.6e-09 | 4.1e-09 | 7.7e-09 | 6.133E-09 | 1.845E-09 | 4.100E-09 | 7.700E-09 |
| 9327 | 467 | P467Q | A | 2.8e-09 | 6.9e-09 | 6.5e-09 | 5.400E-09 | 2.261E-09 | 2.800E-09 | 6.900E-09 |
| 9328 | 467 | P467G | A | 4e-09 | 4.5e-09 | 1.3e-08 | 7.167E-09 | 5.058E-09 | 4.000E-09 | 1.300E-08 |
| 9329 | 467 | P467H | A | 3.8e-09 | 7.9e-09 | 8.1e-09 | 6.600E-09 | 2.427E-09 | 3.800E-09 | 8.100E-09 |
| 9330 | 467 | P467I | A | 1.2e-08 | 6.1e-09 | 9e-09 | 9.033E-09 | 2.950E-09 | 6.100E-09 | 1.200E-08 |
| 9331 | 467 | P467L | A | 4.1e-09 | 7.1e-09 | 1.1e-08 | 7.400E-09 | 3.460E-09 | 4.100E-09 | 1.100E-08 |
| 9332 | 467 | P467K | A | 1.3e-08 | 1.7e-08 | 5.6e-09 | 1.187E-08 | 5.784E-09 | 5.600E-09 | 1.700E-08 |
| 9333 | 467 | P467M | A | 1.5e-08 | 1.8e-08 | 5.3e-09 | 1.277E-08 | 6.638E-09 | 5.300E-09 | 1.800E-08 |
| 9334 | 467 | P467F | A | 3.2e-09 | 1.1e-08 | 9.3e-09 | 7.833E-09 | 4.102E-09 | 3.200E-09 | 1.100E-08 |
| 9335 | 467 | P467P | A | 1.1e-08 | 1.4e-08 | 3.8e-09 | 9.600E-09 | 5.242E-09 | 3.800E-09 | 1.400E-08 |
| 9336 | 467 | P467S | A | 1.2e-08 | 8.7e-09 | 1.1e-08 | 1.057E-08 | 1.692E-09 | 8.700E-09 | 1.200E-08 |
| 9337 | 467 | P467T | A | 8.5e-09 | 6.4e-09 | 9.4e-09 | 8.100E-09 | 1.539E-09 | 6.400E-09 | 9.400E-09 |
| 9338 | 467 | P467W | A | 9.7e-09 | 1.7e-08 | 5.3e-09 | 1.067E-08 | 5.910E-09 | 5.300E-09 | 1.700E-08 |
| 9339 | 467 | P467Y | A | 8.1e-09 | 2.4e-08 | 7.1e-09 | 1.307E-08 | 9.482E-09 | 7.100E-09 | 2.400E-08 |
| 9340 | 467 | P467V | A | 7.9e-09 | 8.6e-09 | 6.1e-09 | 7.533E-09 | 1.290E-09 | 6.100E-09 | 8.600E-09 |
| 9341 | 468 | H468A | A | 8.2e-09 | 4.3e-09 | 1.1e-08 | 7.833E-09 | 3.365E-09 | 4.300E-09 | 1.100E-08 |
| 9342 | 468 | H468R | A | 1.4e-08 | 7e-09 | 1.1e-08 | 1.067E-08 | 3.512E-09 | 7.000E-09 | 1.400E-08 |
| 9343 | 468 | H468N | A | 5.9e-09 | 1.2e-08 | 1e-08 | 9.300E-09 | 3.110E-09 | 5.900E-09 | 1.200E-08 |
| 9344 | 468 | H468D | A | 4.5e-09 | 1.1e-08 | 5.2e-09 | 6.900E-09 | 3.568E-09 | 4.500E-09 | 1.100E-08 |
| 9345 | 468 | H468C | A | 9.3e-09 | 5.6e-09 | 6.6e-09 | 7.167E-09 | 1.914E-09 | 5.600E-09 | 9.300E-09 |
| 9346 | 468 | H468E | A | 2.1e-09 | 1.6e-08 | 8.5e-09 | 8.867E-09 | 6.957E-09 | 2.100E-09 | 1.600E-08 |
| 9347 | 468 | H468Q | A | 1.1e-09 | 6.3e-09 | 7e-09 | 4.800E-09 | 3.223E-09 | 1.100E-09 | 7.000E-09 |
| 9348 | 468 | H468G | A | 7.8e-09 | 2.9e-09 | 1.3e-08 | 7.900E-09 | 5.051E-09 | 2.900E-09 | 1.300E-08 |
| 9349 | 468 | H468H | A | 3.4e-09 | 6.3e-09 | 2.5e-08 | 1.157E-08 | 1.172E-08 | 3.400E-09 | 2.500E-08 |

| | | | | | | | | | | |
|---|---|---|---|---|---|---|---|---|---|---|
| 9350 | 468 | H468I | A | 2.2e-09 | 8.7e-09 | 8.4e-09 | 6.433E-09 | 3.669E-09 | 2.200E-09 | 8.700E-09 |
| 9351 | 468 | H468L | A | 7.8e-09 | 1.3e-08 | 1.1e-08 | 1.060E-08 | 2.623E-09 | 7.800E-09 | 1.300E-08 |
| 9352 | 468 | H468K | A | 6.8e-09 | 4.5e-09 | 2.5e-09 | 4.600E-09 | 2.152E-09 | 2.500E-09 | 6.800E-09 |
| 9353 | 468 | H468M | A | 3.1e-08 | 5.4e-09 | 1.1e-08 | 1.580E-08 | 1.346E-08 | 5.400E-09 | 3.100E-08 |
| 9354 | 468 | H468F | A | 7e-09 | 4.7e-09 | 5.7e-09 | 5.800E-09 | 1.153E-09 | 4.700E-09 | 7.000E-09 |
| 9355 | 468 | H468P | A | 9.9e-09 | 6e-09 | 2.5e-09 | 6.133E-09 | 3.702E-09 | 2.500E-09 | 9.900E-09 |
| 9356 | 468 | H468S | A | 7.9e-09 | 1.1e-08 | 5.2e-09 | 8.033E-09 | 2.902E-09 | 5.200E-09 | 1.100E-08 |
| 9357 | 468 | H468T | A | 2.1e-08 | 9.1e-09 | 1e-08 | 1.337E-08 | 6.626E-09 | 9.100E-09 | 2.100E-08 |
| 9358 | 468 | H468W | A | 3.9e-09 | 9.9e-09 | 7.7e-09 | 7.167E-09 | 3.035E-09 | 3.900E-09 | 9.900E-09 |
| 9359 | 468 | H468Y | A | 1e-08 | 3e-08 | 7e-09 | 1.567E-08 | 1.250E-08 | 7.000E-09 | 3.000E-08 |
| 9360 | 468 | H468V | A | 4e-09 | 6.7e-09 | 6.1e-09 | 5.600E-09 | 1.418E-09 | 4.000E-09 | 6.700E-09 |
| 9361 | 469 | Q469A | A | 1.2e-08 | 5e-09 | 1.5e-08 | 1.067E-08 | 5.132E-09 | 5.000E-09 | 1.500E-08 |
| 9362 | 469 | Q469R | A | 8.2e-09 | 1e-08 | 8.6e-09 | 8.933E-09 | 9.452E-10 | 8.200E-09 | 1.000E-08 |
| 9363 | 469 | Q469N | A | 9e-09 | 6.7e-09 | 3.4e-09 | 6.367E-09 | 2.815E-09 | 3.400E-09 | 9.000E-09 |
| 9364 | 469 | Q469D | A | 1.4e-08 | 1.8e-09 | 4.1e-09 | 6.633E-09 | 6.483E-09 | 1.800E-09 | 1.400E-08 |
| 9365 | 469 | Q469C | A | 1.2e-08 | 3.3e-09 | 1.1e-08 | 8.767E-09 | 4.761E-09 | 3.300E-09 | 1.200E-08 |
| 9366 | 469 | Q469E | A | 9.3e-09 | 2.9e-09 | 1.2e-08 | 8.067E-09 | 4.674E-09 | 2.900E-09 | 1.200E-08 |
| 9367 | 469 | Q469Q | A | 1.3e-08 | 8.7e-09 | 7.8e-09 | 9.833E-09 | 2.779E-09 | 7.800E-09 | 1.300E-08 |
| 9368 | 469 | Q469G | A | 1.8e-08 | 1.2e-08 | 5.1e-09 | 1.170E-08 | 6.455E-09 | 5.100E-09 | 1.800E-08 |
| 9369 | 469 | Q469H | A | 1.4e-08 | 1.1e-08 | 4.7e-09 | 9.900E-09 | 4.747E-09 | 4.700E-09 | 1.400E-08 |
| 9370 | 469 | Q469I | A | 1.5e-08 | 6.1e-09 | 9.4e-09 | 1.017E-08 | 4.499E-09 | 6.100E-09 | 1.500E-08 |
| 9371 | 469 | Q469L | A | 4.8e-09 | 1.4e-08 | 2.2e-08 | 1.360E-08 | 8.607E-09 | 4.800E-09 | 2.200E-08 |
| 9372 | 469 | Q469K | A | 4.6e-09 | 5.5e-09 | 4.1e-09 | 4.733E-09 | 7.095E-10 | 4.100E-09 | 5.500E-09 |
| 9373 | 469 | Q469M | A | 3e-08 | 7.9e-09 | 4.7e-09 | 1.420E-08 | 1.378E-08 | 4.700E-09 | 3.000E-08 |
| 9374 | 469 | Q469F | A | 8.5e-09 | 1.6e-08 | 8.7e-09 | 1.107E-08 | 4.274E-09 | 8.500E-09 | 1.600E-08 |
| 9375 | 469 | Q469P | A | 3.8e-09 | 2.6e-08 | 1.5e-08 | 1.493E-08 | 1.110E-08 | 3.800E-09 | 2.600E-08 |
| 9376 | 469 | Q469S | A | 1.8e-08 | 1.2e-08 | 6.2e-09 | 1.207E-08 | 5.900E-09 | 6.200E-09 | 1.800E-08 |
| 9377 | 469 | Q469T | A | 2e-08 | 1e-08 | 3.8e-09 | 1.127E-08 | 8.174E-09 | 3.800E-09 | 2.000E-08 |
| 9378 | 469 | Q469W | A | 1.2e-08 | 2.7e-08 | 5.3e-09 | 1.477E-08 | 1.111E-08 | 5.300E-09 | 2.700E-08 |
| 9379 | 469 | Q469Y | A | 2.9e-09 | 1.2e-08 | 1.5e-08 | 9.967E-09 | 6.301E-09 | 2.900E-09 | 1.500E-08 |
| 9380 | 469 | Q469V | A | 1.5e-08 | 2.4e-08 | 3.9e-08 | 2.600E-08 | 1.212E-08 | 1.500E-08 | 3.900E-08 |
| 9381 | 470 | A470A | A | 7.5e-09 | 1.7e-08 | 7.1e-09 | 1.053E-08 | 5.604E-09 | 7.100E-09 | 1.700E-08 |
| 9382 | 470 | A470R | A | 1e-08 | 6.2e-09 | 3.1e-09 | 6.433E-09 | 3.456E-09 | 3.100E-09 | 1.000E-08 |
| 9383 | 470 | A470N | A | 7.5e-09 | 5e-09 | 4.2e-09 | 5.567E-09 | 1.721E-09 | 4.200E-09 | 7.500E-09 |
| 9384 | 470 | A470D | A | 3.6e-09 | 4.5e-09 | 3.4e-09 | 3.833E-09 | 5.859E-10 | 3.400E-09 | 4.500E-09 |
| 9385 | 470 | A470C | A | 7e-09 | 2.3e-08 | 8.9e-09 | 1.297E-08 | 8.741E-09 | 7.000E-09 | 2.300E-08 |
| 9386 | 470 | A470E | A | 2.1e-09 | 6.1e-09 | 4e-09 | 4.067E-09 | 2.001E-09 | 2.100E-09 | 6.100E-09 |
| 9387 | 470 | A470Q | A | 3.9e-09 | 5e-09 | 1.2e-08 | 6.967E-09 | 4.394E-09 | 3.900E-09 | 1.200E-08 |
| 9388 | 470 | A470G | A | 4e-09 | 5.8e-09 | 1.4e-08 | 7.933E-09 | 5.330E-09 | 4.000E-09 | 1.400E-08 |
| 9389 | 470 | A470H | A | 8.8e-09 | 9.2e-09 | 4.2e-09 | 7.400E-09 | 2.778E-09 | 4.200E-09 | 9.200E-09 |
| 9390 | 470 | A470I | A | 8.4e-09 | 1.1e-08 | 5.7e-09 | 8.367E-09 | 2.650E-09 | 5.700E-09 | 1.100E-08 |
| 9391 | 470 | A470L | A | 1.2e-08 | 8.6e-09 | 5.4e-09 | 8.667E-09 | 3.301E-09 | 5.400E-09 | 1.200E-08 |
| 9392 | 470 | A470K | A | 4.5e-09 | 6.3e-09 | 6.3e-09 | 5.700E-09 | 1.039E-09 | 4.500E-09 | 6.300E-09 |
| 9393 | 470 | A470M | A | 1.1e-08 | 2.2e-09 | 1.6e-08 | 9.733E-09 | 6.987E-09 | 2.200E-09 | 1.600E-08 |
| 9394 | 470 | A470F | A | 8.5e-09 | 9.8e-09 | 8.4e-09 | 8.900E-09 | 7.810E-10 | 8.400E-09 | 9.800E-09 |
| 9395 | 470 | A470P | A | 2.5e-09 | 4.1e-09 | 7.9e-09 | 4.833E-09 | 2.774E-09 | 2.500E-09 | 7.900E-09 |
| 9396 | 470 | A470S | A | 2.2e-08 | 3.7e-09 | 1.1e-08 | 1.223E-08 | 9.212E-09 | 3.700E-09 | 2.200E-08 |
| 9397 | 470 | A470T | A | 1.7e-08 | 1.1e-08 | 3.3e-09 | 1.043E-08 | 6.868E-09 | 3.300E-09 | 1.700E-08 |
| 9398 | 470 | A470W | A | 8.6e-09 | 1.2e-08 | 8e-09 | 9.533E-09 | 2.157E-09 | 8.000E-09 | 1.200E-08 |
| 9399 | 470 | A470Y | A | 6.5e-09 | 7.8e-09 | 7.5e-09 | 7.267E-09 | 6.807E-10 | 6.500E-09 | 7.800E-09 |
| 9400 | 470 | A470V | A | 6.9e-09 | 2.4e-08 | 1.3e-08 | 1.463E-08 | 8.666E-09 | 6.900E-09 | 2.400E-08 |
| 9401 | 471 | L471A | A | 6.5e-09 | 4.9e-09 | 1.3e-08 | 8.133E-09 | 4.290E-09 | 4.900E-09 | 1.300E-08 |
| 9402 | 471 | L471R | A | 1.7e-08 | 6.2e-09 | 4.2e-09 | 9.133E-09 | 6.886E-09 | 4.200E-09 | 1.700E-08 |
| 9403 | 471 | L471N | A | 1.2e-08 | 4.2e-09 | 4.6e-09 | 6.933E-09 | 4.392E-09 | 4.200E-09 | 1.200E-08 |
| 9404 | 471 | L471D | A | 4.1e-09 | 6.8e-09 | 3.9e-09 | 4.933E-09 | 1.620E-09 | 3.900E-09 | 6.800E-09 |
| 9405 | 471 | L471C | A | 3.1e-09 | 1.5e-08 | 7.5e-09 | 8.533E-09 | 6.017E-09 | 3.100E-09 | 1.500E-08 |
| 9406 | 471 | L471E | A | 4.6e-09 | 1.7e-08 | 7.2e-09 | 9.600E-09 | 6.539E-09 | 4.600E-09 | 1.700E-08 |
| 9407 | 471 | L471Q | A | 1.7e-09 | 3.9e-09 | 7.4e-09 | 4.333E-09 | 2.875E-09 | 1.700E-09 | 7.400E-09 |
| 9408 | 471 | L471G | A | 1.5e-08 | 8.4e-09 | 9.1e-09 | 1.083E-08 | 3.625E-09 | 8.400E-09 | 1.500E-08 |
| 9409 | 471 | L471H | A | 1.4e-08 | 6.3e-09 | 1.8e-09 | 7.367E-09 | 6.170E-09 | 1.800E-09 | 1.400E-08 |
| 9410 | 471 | L471I | A | 5.2e-09 | 7.3e-09 | 1.3e-08 | 8.500E-09 | 4.036E-09 | 5.200E-09 | 1.300E-08 |
| 9411 | 471 | L471L | A | 1.2e-08 | 4.5e-09 | 1.3e-08 | 9.833E-09 | 4.646E-09 | 4.500E-09 | 1.300E-08 |
| 9412 | 471 | L471K | A | 6.7e-09 | 5.1e-09 | 6e-09 | 5.933E-09 | 8.021E-10 | 5.100E-09 | 6.700E-09 |
| 9413 | 471 | L471M | A | 1.7e-08 | 1.7e-09 | 1.4e-08 | 1.090E-08 | 8.107E-09 | 1.700E-09 | 1.700E-08 |
| 9414 | 471 | L471F | A | 2.4e-09 | 1.2e-08 | 6.8e-09 | 7.067E-09 | 4.806E-09 | 2.400E-09 | 1.200E-08 |
| 9415 | 471 | L471P | A | 1.8e-08 | 3.1e-09 | 5.3e-09 | 8.800E-09 | 8.043E-09 | 3.100E-09 | 1.800E-08 |
| 9416 | 471 | L471S | A | 6.5e-09 | 7.8e-09 | 6.5e-09 | 6.933E-09 | 7.506E-10 | 6.500E-09 | 7.800E-09 |
| 9417 | 471 | L471T | A | 1.1e-08 | 3.2e-09 | 3.4e-09 | 5.867E-09 | 4.447E-09 | 3.200E-09 | 1.100E-08 |
| 9418 | 471 | L471W | A | 1.1e-08 | 6.8e-09 | 6.8e-09 | 8.200E-09 | 2.425E-09 | 6.800E-09 | 1.100E-08 |
| 9419 | 471 | L471Y | A | 1.4e-08 | 1.5e-08 | 5e-09 | 1.133E-08 | 5.508E-09 | 5.000E-09 | 1.500E-08 |
| 9420 | 471 | L471V | A | 5e-09 | 5.6e-09 | 1.6e-08 | 8.867E-09 | 6.185E-09 | 5.000E-09 | 1.600E-08 |
| 9421 | 472 | L472A | A | 9.4e-09 | 5.1e-09 | 4.2e-09 | 6.233E-09 | 2.779E-09 | 4.200E-09 | 9.400E-09 |
| 9422 | 472 | L472R | A | 3.8e-09 | 8.8e-09 | 6.3e-09 | 6.300E-09 | 2.500E-09 | 3.800E-09 | 8.800E-09 |
| 9423 | 472 | L472N | A | 6e-09 | 2.5e-08 | 4.3e-09 | 1.177E-08 | 1.149E-08 | 4.300E-09 | 2.500E-08 |
| 9424 | 472 | L472D | A | 1.1e-08 | 3.3e-09 | 7.7e-09 | 7.333E-09 | 3.863E-09 | 3.300E-09 | 1.100E-08 |
| 9425 | 472 | L472C | A | 9e-09 | 1.7e-08 | 6.7e-09 | 1.090E-08 | 5.406E-09 | 6.700E-09 | 1.700E-08 |
| 9426 | 472 | L472E | A | 5.7e-09 | 5.3e-09 | 6.8e-09 | 5.933E-09 | 7.767E-10 | 5.300E-09 | 6.800E-09 |
| 9427 | 472 | L472Q | A | 1.5e-08 | 1.4e-08 | 1.2e-08 | 1.367E-08 | 1.528E-09 | 1.200E-08 | 1.500E-08 |
| 9428 | 472 | L472G | A | 5.1e-09 | 7.8e-09 | 6.5e-09 | 6.467E-09 | 1.350E-09 | 5.100E-09 | 7.800E-09 |
| 9429 | 472 | L472H | A | 1.7e-09 | 4.8e-09 | 1.2e-08 | 6.167E-09 | 5.284E-09 | 1.700E-09 | 1.200E-08 |
| 9430 | 472 | L472I | A | 7.2e-09 | 7.3e-09 | 5.4e-09 | 6.633E-09 | 1.069E-09 | 5.400E-09 | 7.300E-09 |
| 9431 | 472 | L472L | A | 1e-08 | 8.7e-09 | 9.6e-09 | 9.433E-09 | 6.658E-10 | 8.700E-09 | 1.000E-08 |
| 9432 | 472 | L472K | A | 3.6e-09 | 1.2e-08 | 9.1e-09 | 8.233E-09 | 4.267E-09 | 3.600E-09 | 1.200E-08 |
| 9433 | 472 | L472M | A | 2.4e-09 | 1.3e-08 | 5e-09 | 6.800E-09 | 5.524E-09 | 2.400E-09 | 1.300E-08 |
| 9434 | 472 | L472F | A | 2.8e-09 | 5e-09 | 1.1e-08 | 6.267E-09 | 4.244E-09 | 2.800E-09 | 1.100E-08 |

| | | | | | | | | | | |
|---|---|---|---|---|---|---|---|---|---|---|
| 9435 | 472 | L472P | A | 5.8e-09 | 1.7e-08 | 8.5e-09 | 1.043E-08 | 5.845E-09 | 5.800E-09 | 1.700E-08 |
| 9436 | 472 | L472S | A | 8.1e-09 | 6.1e-09 | 3.2e-08 | 1.540E-08 | 1.441E-08 | 6.100E-09 | 3.200E-08 |
| 9437 | 472 | L472T | A | 4.9e-09 | 1.1e-08 | 9e-09 | 8.300E-09 | 3.110E-09 | 4.900E-09 | 1.100E-08 |
| 9438 | 472 | L472W | A | 3.9e-09 | 1.1e-08 | 3.8e-09 | 6.233E-09 | 4.128E-09 | 3.800E-09 | 1.100E-08 |
| 9439 | 472 | L472Y | A | 2.4e-08 | 3.6e-09 | 2.3e-08 | 1.687E-08 | 1.150E-08 | 3.600E-09 | 2.400E-08 |
| 9440 | 472 | L472V | A | 4.7e-09 | 9.1e-09 | 3.1e-08 | 1.493E-08 | 1.409E-08 | 4.700E-09 | 3.100E-08 |
| 9441 | 473 | H473A | A | 9.3e-09 | 2.5e-09 | 6.2e-09 | 6.000E-09 | 3.404E-09 | 2.500E-09 | 9.300E-09 |
| 9442 | 473 | H473R | A | 7.1e-09 | 3.2e-09 | 5.7e-09 | 5.333E-09 | 1.976E-09 | 3.200E-09 | 7.100E-09 |
| 9443 | 473 | H473N | A | 1.4e-08 | 2.5e-09 | 1.1e-08 | 9.167E-09 | 5.965E-09 | 2.500E-09 | 1.400E-08 |
| 9444 | 473 | H473D | A | 3.1e-09 | 1.8e-08 | 7.3e-09 | 9.467E-09 | 7.683E-09 | 3.100E-09 | 1.800E-08 |
| 9445 | 473 | H473C | A | 5e-09 | 7.8e-09 | 6.9e-09 | 6.567E-09 | 1.429E-09 | 5.000E-09 | 7.800E-09 |
| 9446 | 473 | H473E | A | 5e-09 | 1.5e-09 | 4.4e-09 | 3.633E-09 | 1.872E-09 | 1.500E-09 | 5.000E-09 |
| 9447 | 473 | H473Q | A | 5.8e-09 | 8.8e-09 | 3.4e-09 | 6.000E-09 | 2.706E-09 | 3.400E-09 | 8.800E-09 |
| 9448 | 473 | H473G | A | 4.2e-09 | 5.4e-09 | 7.7e-09 | 5.767E-09 | 1.779E-09 | 4.200E-09 | 7.700E-09 |
| 9449 | 473 | H473H | A | 2e-09 | 6.7e-09 | 6.1e-09 | 4.933E-09 | 2.558E-09 | 2.000E-09 | 6.700E-09 |
| 9450 | 473 | H473I | A | 3.6e-09 | 6.9e-09 | 3.9e-09 | 4.800E-09 | 1.825E-09 | 3.600E-09 | 6.900E-09 |
| 9451 | 473 | H473L | A | 7.7e-09 | 1.9e-09 | 8.4e-09 | 6.000E-09 | 3.568E-09 | 1.900E-09 | 8.400E-09 |
| 9452 | 473 | H473K | A | 6.2e-09 | 3.8e-09 | 1.3e-08 | 7.667E-09 | 4.772E-09 | 3.800E-09 | 1.300E-08 |
| 9453 | 473 | H473M | A | 1.3e-08 | 1.3e-08 | 1.3e-08 | 1.300E-08 | 2.026E-24 | 1.300E-08 | 1.300E-08 |
| 9454 | 473 | H473F | A | 1.4e-08 | 4.1e-09 | 4.7e-09 | 7.600E-09 | 5.551E-09 | 4.100E-09 | 1.400E-08 |
| 9455 | 473 | H473P | A | 5.5e-09 | 2.7e-09 | 2.4e-09 | 3.533E-09 | 1.710E-09 | 2.400E-09 | 5.500E-09 |
| 9456 | 473 | H473S | A | 6.6e-09 | 4.5e-09 | 1.5e-08 | 8.700E-09 | 5.556E-09 | 4.500E-09 | 1.500E-08 |
| 9457 | 473 | H473T | A | 2e-08 | 8.6e-09 | 8.1e-09 | 1.223E-08 | 6.731E-09 | 8.100E-09 | 2.000E-08 |
| 9458 | 473 | H473W | A | 5.7e-09 | 6.6e-09 | 8.2e-09 | 6.833E-09 | 1.266E-09 | 5.700E-09 | 8.200E-09 |
| 9459 | 473 | H473Y | A | 1.3e-08 | 1.1e-08 | 3.8e-09 | 9.267E-09 | 4.839E-09 | 3.800E-09 | 1.300E-08 |
| 9460 | 473 | H473V | A | 2e-08 | 2.8e-09 | 1.3e-08 | 1.193E-08 | 8.649E-09 | 2.800E-09 | 2.000E-08 |
| 9461 | 474 | T474A | A | 1.1e-08 | 5.3e-09 | 3.2e-09 | 6.500E-09 | 4.036E-09 | 3.200E-09 | 1.100E-08 |
| 9462 | 474 | T474R | A | 6.4e-09 | 1e-08 | 6.8e-09 | 7.733E-09 | 1.973E-09 | 6.400E-09 | 1.000E-08 |
| 9463 | 474 | T474N | A | 6.1e-09 | 6.8e-09 | 1.8e-08 | 1.030E-08 | 6.678E-09 | 6.100E-09 | 1.800E-08 |
| 9464 | 474 | T474D | A | 8.6e-09 | 5.5e-09 | 2.5e-09 | 5.533E-09 | 3.050E-09 | 2.500E-09 | 8.600E-09 |
| 9465 | 474 | T474C | A | 1.8e-08 | 1.1e-08 | 5.8e-09 | 1.160E-08 | 6.122E-09 | 5.800E-09 | 1.800E-08 |
| 9466 | 474 | T474E | A | 3.4e-09 | 6.7e-09 | 8.5e-09 | 6.200E-09 | 2.587E-09 | 3.400E-09 | 8.500E-09 |
| 9467 | 474 | T474Q | A | 8.1e-09 | 7.1e-09 | 4.3e-09 | 6.500E-09 | 1.970E-09 | 4.300E-09 | 8.100E-09 |
| 9468 | 474 | T474G | A | 2e-08 | 2.1e-08 | 6.1e-09 | 1.570E-08 | 8.329E-09 | 6.100E-09 | 2.100E-08 |
| 9469 | 474 | T474H | A | 9.6e-09 | 9.4e-09 | 1.3e-08 | 1.067E-08 | 2.023E-09 | 9.400E-09 | 1.300E-08 |
| 9470 | 474 | T474I | A | 5.1e-09 | 5.6e-09 | 1.1e-08 | 7.233E-09 | 3.272E-09 | 5.100E-09 | 1.100E-08 |
| 9471 | 474 | T474L | A | 4.4e-09 | 6e-09 | 5.1e-09 | 5.167E-09 | 8.021E-10 | 4.400E-09 | 6.000E-09 |
| 9472 | 474 | T474K | A | 6.3e-09 | 2.9e-09 | 2.7e-09 | 3.967E-09 | 2.023E-09 | 2.700E-09 | 6.300E-09 |
| 9473 | 474 | T474M | A | 9.6e-09 | 1.2e-08 | 3.9e-09 | 8.500E-09 | 4.161E-09 | 3.900E-09 | 1.200E-08 |
| 9474 | 474 | T474F | A | 8.4e-10 | 6.7e-09 | 5.4e-09 | 4.313E-09 | 3.077E-09 | 8.400E-10 | 6.700E-09 |
| 9475 | 474 | T474P | A | 1.2e-08 | 1.1e-08 | 1.6e-08 | 1.300E-08 | 2.646E-09 | 1.100E-08 | 1.600E-08 |
| 9476 | 474 | T474S | A | 8e-09 | 3.6e-09 | 4.3e-09 | 5.300E-09 | 2.364E-09 | 3.600E-09 | 8.000E-09 |
| 9477 | 474 | T474T | A | 5.5e-09 | 1.1e-08 | 7.2e-09 | 7.900E-09 | 2.816E-09 | 5.500E-09 | 1.100E-08 |
| 9478 | 474 | T474W | A | 2.6e-08 | 1.2e-08 | 5.1e-09 | 1.437E-08 | 1.065E-08 | 5.100E-09 | 2.600E-08 |
| 9479 | 474 | T474Y | A | 3.1e-09 | 4.2e-09 | 3e-09 | 3.433E-09 | 6.658E-10 | 3.000E-09 | 4.200E-09 |
| 9480 | 474 | T474V | A | 8.9e-09 | 1.7e-09 | 8.8e-09 | 6.467E-09 | 4.128E-09 | 1.700E-09 | 8.900E-09 |
| 9481 | 475 | A475A | A | 7.4e-09 | 8.5e-09 | 9.6e-09 | 8.500E-09 | 1.100E-09 | 7.400E-09 | 9.600E-09 |
| 9482 | 475 | A475R | A | 2.9e-09 | 1.9e-08 | 9.8e-09 | 1.057E-08 | 8.077E-09 | 2.900E-09 | 1.900E-08 |
| 9483 | 475 | A475N | A | 1e-08 | 4e-09 | 3.5e-09 | 5.833E-09 | 3.617E-09 | 3.500E-09 | 1.000E-08 |
| 9484 | 475 | A475D | A | 2.1e-08 | 7.8e-09 | 5.3e-09 | 1.137E-08 | 8.436E-09 | 5.300E-09 | 2.100E-08 |
| 9485 | 475 | A475C | A | 6e-09 | 1.2e-08 | 6.2e-09 | 8.067E-09 | 3.408E-09 | 6.000E-09 | 1.200E-08 |
| 9486 | 475 | A475E | A | 2.4e-09 | 3.1e-09 | 3e-09 | 2.833E-09 | 3.786E-10 | 2.400E-09 | 3.100E-09 |
| 9487 | 475 | A475Q | A | 2.4e-09 | 9.2e-09 | 8.1e-09 | 6.567E-09 | 3.650E-09 | 2.400E-09 | 9.200E-09 |
| 9488 | 475 | A475G | A | 1.5e-08 | 4.7e-09 | 9.9e-09 | 9.867E-09 | 5.150E-09 | 4.700E-09 | 1.500E-08 |
| 9489 | 475 | A475H | A | 8.9e-09 | 7.6e-09 | 5.3e-09 | 7.267E-09 | 1.823E-09 | 5.300E-09 | 8.900E-09 |
| 9490 | 475 | A475I | A | 8.8e-09 | 1.3e-08 | 9e-09 | 1.027E-08 | 2.369E-09 | 8.800E-09 | 1.300E-08 |
| 9491 | 475 | A475L | A | 8.2e-09 | 8.4e-09 | 9.5e-09 | 8.700E-09 | 7.000E-10 | 8.200E-09 | 9.500E-09 |
| 9492 | 475 | A475K | A | 1.5e-08 | 1.7e-08 | 4.3e-09 | 1.210E-08 | 6.829E-09 | 4.300E-09 | 1.700E-08 |
| 9493 | 475 | A475M | A | 2.1e-09 | 1.3e-09 | 2.2e-09 | 1.867E-09 | 4.933E-10 | 1.300E-09 | 2.200E-09 |
| 9494 | 475 | A475F | A | 4.7e-09 | 3.5e-09 | 3.8e-09 | 4.000E-09 | 6.245E-10 | 3.500E-09 | 4.700E-09 |
| 9495 | 475 | A475P | A | 5.7e-09 | 7.7e-09 | 3e-08 | 1.447E-08 | 1.349E-08 | 5.700E-09 | 3.000E-08 |
| 9496 | 475 | A475S | A | 1.3e-08 | 7e-09 | 2.1e-09 | 7.367E-09 | 5.459E-09 | 2.100E-09 | 1.300E-08 |
| 9497 | 475 | A475T | A | 8e-09 | 1e-08 | 3.1e-08 | 1.633E-08 | 1.274E-08 | 8.000E-09 | 3.100E-08 |
| 9498 | 475 | A475W | A | 4.8e-09 | 6.4e-09 | 1.6e-08 | 9.067E-09 | 6.058E-09 | 4.800E-09 | 1.600E-08 |
| 9499 | 475 | A475Y | A | 7.6e-09 | 9.1e-09 | 6.3e-09 | 7.667E-09 | 1.401E-09 | 6.300E-09 | 9.100E-09 |
| 9500 | 475 | A475V | A | 4.3e-09 | 6.3e-09 | 1.1e-08 | 7.200E-09 | 3.439E-09 | 4.300E-09 | 1.100E-08 |
| 9501 | 476 | N476A | A | 1.9e-08 | 9.6e-09 | 3.5e-09 | 1.070E-08 | 7.808E-09 | 3.500E-09 | 1.900E-08 |
| 9502 | 476 | N476R | A | 8e-09 | 5.3e-09 | 1.4e-08 | 9.100E-09 | 4.453E-09 | 5.300E-09 | 1.400E-08 |
| 9503 | 476 | N476N | A | 4.4e-09 | 1.2e-08 | 1.4e-08 | 1.013E-08 | 5.065E-09 | 4.400E-09 | 1.400E-08 |
| 9504 | 476 | N476D | A | 1.2e-08 | 3.1e-08 | 5.2e-09 | 1.607E-08 | 1.337E-08 | 5.200E-09 | 3.100E-08 |
| 9505 | 476 | N476C | A | 1.7e-09 | 1.1e-08 | 6e-09 | 6.233E-09 | 4.654E-09 | 1.700E-09 | 1.100E-08 |
| 9506 | 476 | N476E | A | 4.1e-09 | 6.4e-09 | 1.4e-08 | 8.167E-09 | 5.181E-09 | 4.100E-09 | 1.400E-08 |
| 9507 | 476 | N476Q | A | 8.2e-09 | 3.5e-09 | 7.6e-09 | 6.433E-09 | 2.558E-09 | 3.500E-09 | 8.200E-09 |
| 9508 | 476 | N476G | A | 8.7e-09 | 8.9e-09 | 8.3e-09 | 8.633E-09 | 3.055E-10 | 8.300E-09 | 8.900E-09 |
| 9509 | 476 | N476H | A | 4.8e-09 | 5.9e-09 | 4e-09 | 4.900E-09 | 9.539E-10 | 4.000E-09 | 5.900E-09 |
| 9510 | 476 | N476I | A | 2.7e-09 | 6.5e-09 | 8.4e-09 | 5.867E-09 | 2.902E-09 | 2.700E-09 | 8.400E-09 |
| 9511 | 476 | N476L | A | 1.3e-08 | 3.5e-09 | 1.2e-08 | 9.500E-09 | 5.220E-09 | 3.500E-09 | 1.300E-08 |
| 9512 | 476 | N476K | A | 8e-09 | 3.9e-09 | 9.6e-09 | 7.167E-09 | 2.940E-09 | 3.900E-09 | 9.600E-09 |
| 9513 | 476 | N476M | A | 7.5e-09 | 7.5e-09 | 1e-08 | 8.333E-09 | 1.443E-09 | 7.500E-09 | 1.000E-08 |
| 9514 | 476 | N476F | A | 1.1e-08 | 7.8e-09 | 3.3e-09 | 7.367E-09 | 3.868E-09 | 3.300E-09 | 1.100E-08 |
| 9515 | 476 | N476P | A | 8.5e-09 | 5.4e-09 | 7.1e-09 | 7.000E-09 | 1.552E-09 | 5.400E-09 | 8.500E-09 |
| 9516 | 476 | N476S | A | 4.8e-09 | 6.3e-09 | 1.1e-08 | 7.367E-09 | 3.235E-09 | 4.800E-09 | 1.100E-08 |
| 9517 | 476 | N476T | A | 9.4e-09 | 6.1e-09 | 1.2e-08 | 9.167E-09 | 2.957E-09 | 6.100E-09 | 1.200E-08 |
| 9518 | 476 | N476W | A | 7.8e-09 | 4.1e-09 | 9.8e-09 | 7.233E-09 | 2.892E-09 | 4.100E-09 | 9.800E-09 |
| 9519 | 476 | N476Y | A | 9.6e-09 | 1.5e-08 | 3.4e-09 | 9.333E-09 | 5.805E-09 | 3.400E-09 | 1.500E-08 |

| | | | | | | | | | | |
|---|---|---|---|---|---|---|---|---|---|---|
| 9520 | 476 | N476V | A | 5.5e-09 | 5.7e-09 | 4.9e-09 | 5.367E-09 | 4.163E-10 | 4.900E-09 | 5.700E-09 |
| 9521 | 477 | R477A | A | 8.4e-09 | 8.5e-09 | 1.2e-08 | 9.633E-09 | 2.050E-09 | 8.400E-09 | 1.200E-08 |
| 9522 | 477 | R477R | A | 7.7e-09 | 7e-09 | 4.8e-09 | 6.500E-09 | 1.513E-09 | 4.800E-09 | 7.700E-09 |
| 9523 | 477 | R477N | A | 1.2e-08 | 8.5e-09 | 4.9e-09 | 8.467E-09 | 3.550E-09 | 4.900E-09 | 1.200E-08 |
| 9524 | 477 | R477D | A | 5.4e-09 | 4.1e-09 | 8.7e-09 | 6.067E-09 | 2.371E-09 | 4.100E-09 | 8.700E-09 |
| 9525 | 477 | R477C | A | 7.3e-09 | 3.1e-09 | 3.5e-09 | 4.633E-09 | 2.318E-09 | 3.100E-09 | 7.300E-09 |
| 9526 | 477 | R477E | A | 1.8e-08 | 3.6e-09 | 1.3e-08 | 1.153E-08 | 7.311E-09 | 3.600E-09 | 1.800E-08 |
| 9527 | 477 | R477Q | A | 1.1e-08 | 1.2e-08 | 5.5e-09 | 9.500E-09 | 3.500E-09 | 5.500E-09 | 1.200E-08 |
| 9528 | 477 | R477G | A | 4.7e-09 | 8.3e-09 | 7.9e-09 | 6.967E-09 | 1.973E-09 | 4.700E-09 | 8.300E-09 |
| 9529 | 477 | R477H | A | 1.1e-09 | 1.1e-08 | 2.6e-09 | 4.900E-09 | 5.336E-09 | 1.100E-09 | 1.100E-08 |
| 9530 | 477 | R477I | A | 3.1e-09 | 9.6e-09 | 2.1e-09 | 4.933E-09 | 4.072E-09 | 2.100E-09 | 9.600E-09 |
| 9531 | 477 | R477L | A | 1.2e-08 | 1.6e-08 | 6e-09 | 1.133E-08 | 5.033E-09 | 6.000E-09 | 1.600E-08 |
| 9532 | 477 | R477K | A | 7.1e-09 | 2.9e-09 | 1.1e-08 | 7.000E-09 | 4.051E-09 | 2.900E-09 | 1.100E-08 |
| 9533 | 477 | R477M | A | 4.1e-09 | 6e-09 | 5.2e-09 | 5.100E-09 | 9.539E-10 | 4.100E-09 | 6.000E-09 |
| 9534 | 477 | R477F | A | 2.2e-09 | 1.5e-09 | 7.2e-09 | 3.633E-09 | 3.109E-09 | 1.500E-09 | 7.200E-09 |
| 9535 | 477 | R477P | A | 2.1e-08 | 7.6e-09 | 3.6e-09 | 1.073E-08 | 9.113E-09 | 3.600E-09 | 2.100E-08 |
| 9536 | 477 | R477S | A | 1.8e-08 | 4.5e-09 | 4.8e-09 | 9.100E-09 | 7.709E-09 | 4.500E-09 | 1.800E-08 |
| 9537 | 477 | R477T | A | 1.2e-08 | 6e-09 | 1.9e-08 | 1.233E-08 | 6.506E-09 | 6.000E-09 | 1.900E-08 |
| 9538 | 477 | R477W | A | 1e-08 | 1e-08 | 1.3e-08 | 1.100E-08 | 1.732E-09 | 1.000E-08 | 1.300E-08 |
| 9539 | 477 | R477Y | A | 9e-09 | 8.2e-09 | 2.6e-08 | 1.440E-08 | 1.005E-08 | 8.200E-09 | 2.600E-08 |
| 9540 | 477 | R477V | A | 2.4e-08 | 7.3e-09 | 5.1e-09 | 1.213E-08 | 1.034E-08 | 5.100E-09 | 2.400E-08 |
| 9541 | 478 | P478A | A | 1.2e-08 | 1e-08 | 9.6e-09 | 1.053E-08 | 1.286E-09 | 9.600E-09 | 1.200E-08 |
| 9542 | 478 | P478R | A | 8.7e-09 | 3.5e-09 | 7.5e-09 | 6.567E-09 | 2.723E-09 | 3.500E-09 | 8.700E-09 |
| 9543 | 478 | P478N | A | 3.6e-09 | 2.7e-09 | 5.6e-09 | 3.967E-09 | 1.484E-09 | 2.700E-09 | 5.600E-09 |
| 9544 | 478 | P478D | A | 8.5e-09 | 5.4e-09 | 6.6e-09 | 6.833E-09 | 1.563E-09 | 5.400E-09 | 8.500E-09 |
| 9545 | 478 | P478C | A | 2.8e-09 | 2.6e-09 | 6.1e-09 | 3.833E-09 | 1.966E-09 | 2.600E-09 | 6.100E-09 |
| 9546 | 478 | P478E | A | 1.5e-08 | 3.8e-09 | 2.1e-08 | 1.327E-08 | 8.730E-09 | 3.800E-09 | 2.100E-08 |
| 9547 | 478 | P478Q | A | 5.3e-09 | 8.2e-09 | 5.8e-09 | 6.433E-09 | 1.550E-09 | 5.300E-09 | 8.200E-09 |
| 9548 | 478 | P478G | A | 5.6e-09 | 7e-09 | 5.5e-09 | 6.033E-09 | 8.386E-10 | 5.500E-09 | 7.000E-09 |
| 9549 | 478 | P478H | A | 8.7e-09 | 2.6e-09 | 5.6e-09 | 5.633E-09 | 3.050E-09 | 2.600E-09 | 8.700E-09 |
| 9550 | 478 | P478I | A | 4e-09 | 1.9e-08 | 9.2e-09 | 1.073E-08 | 7.617E-09 | 4.000E-09 | 1.900E-08 |
| 9551 | 478 | P478L | A | 5.1e-09 | 9e-09 | 2.4e-08 | 1.270E-08 | 9.978E-09 | 5.100E-09 | 2.400E-08 |
| 9552 | 478 | P478K | A | 1.3e-08 | 2.1e-09 | 1.7e-08 | 1.070E-08 | 7.712E-09 | 2.100E-09 | 1.700E-08 |
| 9553 | 478 | P478M | A | 2.7e-08 | 6.8e-09 | 5.9e-09 | 1.323E-08 | 1.193E-08 | 5.900E-09 | 2.700E-08 |
| 9554 | 478 | P478F | A | 1.3e-08 | 5.5e-09 | 1e-08 | 9.500E-09 | 3.775E-09 | 5.500E-09 | 1.300E-08 |
| 9555 | 478 | P478P | A | 6.3e-09 | 2.3e-09 | 5.6e-09 | 4.733E-09 | 2.136E-09 | 2.300E-09 | 6.300E-09 |
| 9556 | 478 | P478S | A | 1.2e-08 | 8.3e-09 | 8.4e-09 | 9.567E-09 | 2.108E-09 | 8.300E-09 | 1.200E-08 |
| 9557 | 478 | P478T | A | 5.8e-09 | 6.1e-09 | 3e-09 | 4.967E-09 | 1.710E-09 | 3.000E-09 | 6.100E-09 |
| 9558 | 478 | P478W | A | 7.1e-09 | 2.4e-09 | 1.1e-08 | 6.833E-09 | 4.306E-09 | 2.400E-09 | 1.100E-08 |
| 9559 | 478 | P478Y | A | 9.9e-09 | 3.8e-09 | 4.3e-09 | 6.000E-09 | 3.387E-09 | 3.800E-09 | 9.900E-09 |
| 9560 | 478 | P478V | A | 1.1e-08 | 5e-09 | 1.3e-08 | 9.667E-09 | 4.163E-09 | 5.000E-09 | 1.300E-08 |
| 9561 | 479 | E479A | A | 1e-08 | 3.9e-09 | 4.5e-09 | 6.133E-09 | 3.362E-09 | 3.900E-09 | 1.000E-08 |
| 9562 | 479 | E479R | A | 1.1e-08 | 3.9e-09 | 6.7e-09 | 7.200E-09 | 3.576E-09 | 3.900E-09 | 1.100E-08 |
| 9563 | 479 | E479N | A | 1e-09 | 3e-09 | 2e-09 | 2.000E-09 | 1.000E-09 | 1.000E-09 | 3.000E-09 |
| 9564 | 479 | E479D | A | 4.4e-09 | 4.3e-09 | 8.7e-09 | 5.800E-09 | 2.512E-09 | 4.300E-09 | 8.700E-09 |
| 9565 | 479 | E479C | A | 1.4e-08 | 4.4e-09 | 3.4e-09 | 7.267E-09 | 5.853E-09 | 3.400E-09 | 1.400E-08 |
| 9566 | 479 | E479E | A | 6.3e-09 | 9.2e-09 | 5.6e-09 | 7.033E-09 | 1.909E-09 | 5.600E-09 | 9.200E-09 |
| 9567 | 479 | E479Q | A | 8.1e-09 | 2.4e-08 | 1.4e-08 | 1.537E-08 | 8.038E-09 | 8.100E-09 | 2.400E-08 |
| 9568 | 479 | E479G | A | 4.4e-09 | 3.9e-08 | 1e-08 | 1.780E-08 | 1.857E-08 | 4.400E-09 | 3.900E-08 |
| 9569 | 479 | E479H | A | 8.6e-09 | 1.1e-08 | 8.8e-09 | 9.467E-09 | 1.332E-09 | 8.600E-09 | 1.100E-08 |
| 9570 | 479 | E479I | A | 2.5e-09 | 2e-09 | 5e-09 | 3.167E-09 | 1.607E-09 | 2.000E-09 | 5.000E-09 |
| 9571 | 479 | E479L | A | 9.4e-09 | 3.9e-09 | 1.4e-08 | 9.100E-09 | 5.057E-09 | 3.900E-09 | 1.400E-08 |
| 9572 | 479 | E479K | A | 1.2e-08 | 5.3e-09 | 1.6e-08 | 1.110E-08 | 5.406E-09 | 5.300E-09 | 1.600E-08 |
| 9573 | 479 | E479M | A | 7.2e-09 | 8.7e-09 | 7.9e-09 | 7.933E-09 | 7.506E-10 | 7.200E-09 | 8.700E-09 |
| 9574 | 479 | E479F | A | 4.1e-09 | 1e-08 | 8.6e-09 | 7.567E-09 | 3.083E-09 | 4.100E-09 | 1.000E-08 |
| 9575 | 479 | E479P | A | 7.7e-09 | 8.6e-09 | 8.9e-09 | 8.400E-09 | 6.245E-10 | 7.700E-09 | 8.900E-09 |
| 9576 | 479 | E479S | A | 1.1e-08 | 4.8e-09 | 2.9e-09 | 6.233E-09 | 4.236E-09 | 2.900E-09 | 1.100E-08 |
| 9577 | 479 | E479T | A | 3.4e-09 | 1.7e-08 | 1.2e-08 | 1.080E-08 | 6.879E-09 | 3.400E-09 | 1.700E-08 |
| 9578 | 479 | E479W | A | 2.7e-09 | 1.2e-08 | 5.9e-09 | 6.867E-09 | 4.725E-09 | 2.700E-09 | 1.200E-08 |
| 9579 | 479 | E479Y | A | 6.3e-09 | 1e-08 | 1e-08 | 8.767E-09 | 2.136E-09 | 6.300E-09 | 1.000E-08 |
| 9580 | 479 | E479V | A | 1.6e-08 | 8.7e-09 | 7e-09 | 1.057E-08 | 4.782E-09 | 7.000E-09 | 1.600E-08 |
| 9581 | 480 | D480A | A | 4.6e-09 | 3.5e-09 | 5.9e-09 | 4.667E-09 | 1.201E-09 | 3.500E-09 | 5.900E-09 |
| 9582 | 480 | D480R | A | 2.4e-09 | 3.5e-09 | 8.1e-09 | 4.667E-09 | 3.024E-09 | 2.400E-09 | 8.100E-09 |
| 9583 | 480 | D480N | A | 5.7e-09 | 7.4e-09 | 1.1e-08 | 8.033E-09 | 2.706E-09 | 5.700E-09 | 1.100E-08 |
| 9584 | 480 | D480D | A | 7.7e-09 | 1e-08 | 1.2e-08 | 9.900E-09 | 2.152E-09 | 7.700E-09 | 1.200E-08 |
| 9585 | 480 | D480C | A | 1.4e-08 | 3.8e-09 | 4.4e-09 | 7.400E-09 | 5.724E-09 | 3.800E-09 | 1.400E-08 |
| 9586 | 480 | D480E | A | 3.8e-09 | 4.7e-09 | 9e-09 | 5.833E-09 | 2.779E-09 | 3.800E-09 | 9.000E-09 |
| 9587 | 480 | D480Q | A | 1.1e-08 | 8.9e-09 | 3.8e-09 | 7.900E-09 | 3.703E-09 | 3.800E-09 | 1.100E-08 |
| 9588 | 480 | D480G | A | 8.6e-09 | 4.4e-09 | 6.5e-09 | 6.500E-09 | 2.100E-09 | 4.400E-09 | 8.600E-09 |
| 9589 | 480 | D480H | A | 4.8e-09 | 3.5e-09 | 7.6e-09 | 5.300E-09 | 2.095E-09 | 3.500E-09 | 7.600E-09 |
| 9590 | 480 | D480I | A | 5.4e-09 | 1.7e-08 | 1.4e-08 | 1.213E-08 | 6.021E-09 | 5.400E-09 | 1.700E-08 |
| 9591 | 480 | D480L | A | 1.6e-08 | 3.3e-09 | 4.3e-09 | 7.867E-09 | 7.061E-09 | 3.300E-09 | 1.600E-08 |
| 9592 | 480 | D480K | A | 1.4e-08 | 1e-08 | 8.2e-09 | 1.073E-08 | 2.969E-09 | 8.200E-09 | 1.400E-08 |
| 9593 | 480 | D480M | A | 4.2e-09 | 1.9e-08 | 1.2e-08 | 1.173E-08 | 7.404E-09 | 4.200E-09 | 1.900E-08 |
| 9594 | 480 | D480F | A | 8.2e-09 | 4e-09 | 3.7e-09 | 5.300E-09 | 2.516E-09 | 3.700E-09 | 8.200E-09 |
| 9595 | 480 | D480P | A | 1.3e-08 | 6.7e-09 | 3.5e-09 | 7.733E-09 | 4.834E-09 | 3.500E-09 | 1.300E-08 |
| 9596 | 480 | D480S | A | 6.3e-09 | 4.7e-09 | 1.9e-09 | 4.300E-09 | 2.227E-09 | 1.900E-09 | 6.300E-09 |
| 9597 | 480 | D480T | A | 7.1e-09 | 5.8e-09 | 7.1e-09 | 6.667E-09 | 7.506E-10 | 5.800E-09 | 7.100E-09 |
| 9598 | 480 | D480W | A | 8.5e-09 | 4.4e-09 | 5.3e-09 | 6.067E-09 | 2.155E-09 | 4.400E-09 | 8.500E-09 |
| 9599 | 480 | D480Y | A | 8.6e-09 | 1.3e-08 | 1.1e-08 | 1.087E-08 | 2.203E-09 | 8.600E-09 | 1.300E-08 |
| 9600 | 480 | D480V | A | 5.6e-09 | 7.7e-09 | 3.9e-09 | 5.733E-09 | 1.904E-09 | 3.900E-09 | 7.700E-09 |
| 9601 | 481 | E481A | A | 1.1e-08 | 8.3e-09 | 1.8e-08 | 1.243E-08 | 5.006E-09 | 8.300E-09 | 1.800E-08 |
| 9602 | 481 | E481R | A | 5.5e-09 | 7.9e-09 | 4.4e-09 | 5.933E-09 | 1.790E-09 | 4.400E-09 | 7.900E-09 |
| 9603 | 481 | E481N | A | 6e-09 | 7.6e-09 | 1.9e-08 | 1.087E-08 | 7.089E-09 | 6.000E-09 | 1.900E-08 |
| 9604 | 481 | E481D | A | 7.5e-09 | 3.4e-09 | 1.3e-08 | 7.967E-09 | 4.817E-09 | 3.400E-09 | 1.300E-08 |

| | | | | | | | | | | |
|---|---|---|---|---|---|---|---|---|---|---|
| 9605 | 481 | E481C | A | 7.2e-09 | 3.9e-09 | 1.1e-08 | 7.367E-09 | 3.553E-09 | 3.900E-09 | 1.100E-08 |
| 9606 | 481 | E481E | A | 6.9e-09 | 4.1e-09 | 5.7e-09 | 5.567E-09 | 1.405E-09 | 4.100E-09 | 6.900E-09 |
| 9607 | 481 | E481Q | A | 5.2e-09 | 6.7e-09 | 1.1e-08 | 7.633E-09 | 3.011E-09 | 5.200E-09 | 1.100E-08 |
| 9608 | 481 | E481G | A | 3.1e-09 | 2.6e-09 | 3.5e-09 | 3.067E-09 | 4.509E-10 | 2.600E-09 | 3.500E-09 |
| 9609 | 481 | E481H | A | 4e-09 | 4.8e-09 | 4.6e-09 | 4.467E-09 | 4.163E-10 | 4.000E-09 | 4.800E-09 |
| 9610 | 481 | E481I | A | 4e-09 | 4.2e-09 | 3.2e-09 | 3.800E-09 | 5.292E-10 | 3.200E-09 | 4.200E-09 |
| 9611 | 481 | E481L | A | 3.2e-09 | 9.2e-09 | 7.5e-09 | 6.633E-09 | 3.092E-09 | 3.200E-09 | 9.200E-09 |
| 9612 | 481 | E481K | A | 1.1e-08 | 5.9e-09 | 6.2e-09 | 7.700E-09 | 2.862E-09 | 5.900E-09 | 1.100E-08 |
| 9613 | 481 | E481M | A | 3.7e-09 | 6.6e-09 | 1e-08 | 6.767E-09 | 3.153E-09 | 3.700E-09 | 1.000E-08 |
| 9614 | 481 | E481F | A | 5.6e-09 | 5.9e-09 | 2.1e-09 | 4.533E-09 | 2.113E-09 | 2.100E-09 | 5.900E-09 |
| 9615 | 481 | E481P | A | 3.4e-09 | 6.4e-09 | 2.2e-09 | 4.000E-09 | 2.163E-09 | 2.200E-09 | 6.400E-09 |
| 9616 | 481 | E481S | A | 3e-09 | 5.4e-09 | 2.7e-09 | 3.700E-09 | 1.480E-09 | 2.700E-09 | 5.400E-09 |
| 9617 | 481 | E481T | A | 3.6e-09 | 7.9e-09 | 6e-09 | 5.833E-09 | 2.155E-09 | 3.600E-09 | 7.900E-09 |
| 9618 | 481 | E481W | A | 1.6e-08 | 1.1e-08 | 7.3e-09 | 1.143E-08 | 4.366E-09 | 7.300E-09 | 1.600E-08 |
| 9619 | 481 | E481Y | A | 9.9e-09 | 7.9e-09 | 8.2e-09 | 8.667E-09 | 1.079E-09 | 7.900E-09 | 9.900E-09 |
| 9620 | 481 | E481V | A | 5.6e-09 | 5.5e-09 | 1.6e-08 | 9.033E-09 | 6.034E-09 | 5.500E-09 | 1.600E-08 |
| 9621 | 482 | C482A | A | 1e-08 | 7.3e-09 | 7.2e-09 | 8.167E-09 | 1.589E-09 | 7.200E-09 | 1.000E-08 |
| 9622 | 482 | C482R | A | 1.7e-08 | 7.2e-09 | 2.7e-08 | 1.707E-08 | 9.900E-09 | 7.200E-09 | 2.700E-08 |
| 9623 | 482 | C482N | A | 1.2e-08 | 3.3e-09 | 4.2e-09 | 6.500E-09 | 4.784E-09 | 3.300E-09 | 1.200E-08 |
| 9624 | 482 | C482D | A | 7.3e-09 | 6.3e-09 | 3.9e-09 | 5.833E-09 | 1.747E-09 | 3.900E-09 | 7.300E-09 |
| 9625 | 482 | C482C | A | 3e-09 | 9e-09 | 4.6e-09 | 5.533E-09 | 3.107E-09 | 3.000E-09 | 9.000E-09 |
| 9626 | 482 | C482E | A | 9.5e-09 | 5.3e-09 | 1.1e-08 | 8.600E-09 | 2.955E-09 | 5.300E-09 | 1.100E-08 |
| 9627 | 482 | C482Q | A | 4.4e-09 | 3.7e-09 | 5.2e-09 | 4.433E-09 | 7.506E-10 | 3.700E-09 | 5.200E-09 |
| 9628 | 482 | C482G | A | 9.2e-09 | 6e-09 | 5.7e-09 | 6.967E-09 | 1.940E-09 | 5.700E-09 | 9.200E-09 |
| 9629 | 482 | C482H | A | 8.1e-09 | 9.9e-09 | 7.6e-09 | 8.533E-09 | 1.210E-09 | 7.600E-09 | 9.900E-09 |
| 9630 | 482 | C482I | A | 1.2e-08 | 3.5e-09 | 2.4e-08 | 1.317E-08 | 1.030E-08 | 3.500E-09 | 2.400E-08 |
| 9631 | 482 | C482L | A | 6.3e-09 | 3.6e-09 | 8.2e-09 | 6.033E-09 | 2.312E-09 | 3.600E-09 | 8.200E-09 |
| 9632 | 482 | C482K | A | 2e-08 | 8.1e-09 | 1.8e-08 | 1.537E-08 | 6.372E-09 | 8.100E-09 | 2.000E-08 |
| 9633 | 482 | C482M | A | 1.2e-08 | 8.4e-09 | 2.3e-09 | 7.567E-09 | 4.903E-09 | 2.300E-09 | 1.200E-08 |
| 9634 | 482 | C482F | A | 6e-09 | 4.3e-09 | 1.1e-08 | 7.100E-09 | 3.483E-09 | 4.300E-09 | 1.100E-08 |
| 9635 | 482 | C482P | A | 1.8e-08 | 1.3e-08 | 4.2e-09 | 1.173E-08 | 6.987E-09 | 4.200E-09 | 1.800E-08 |
| 9636 | 482 | C482S | A | 8.5e-09 | 3.7e-09 | 5.1e-09 | 5.767E-09 | 2.468E-09 | 3.700E-09 | 8.500E-09 |
| 9637 | 482 | C482T | A | 3.6e-08 | 1.1e-08 | 3.5e-09 | 1.683E-08 | 1.702E-08 | 3.500E-09 | 3.600E-08 |
| 9638 | 482 | C482W | A | 1.1e-08 | 1.4e-08 | 1.3e-08 | 1.267E-08 | 1.528E-09 | 1.100E-08 | 1.400E-08 |
| 9639 | 482 | C482Y | A | 6e-09 | 3.6e-09 | 2.9e-09 | 4.167E-09 | 1.626E-09 | 2.900E-09 | 6.000E-09 |
| 9640 | 482 | C482V | A | 5.1e-09 | 5e-09 | 3.3e-09 | 4.467E-09 | 1.012E-09 | 3.300E-09 | 5.100E-09 |
| 9641 | 483 | V483A | A | 5.4e-09 | 1.2e-08 | 1.3e-08 | 1.013E-08 | 4.130E-09 | 5.400E-09 | 1.300E-08 |
| 9642 | 483 | V483R | A | 2.8e-08 | 6.4e-09 | 3.7e-09 | 1.270E-08 | 1.332E-08 | 3.700E-09 | 2.800E-08 |
| 9643 | 483 | V483N | A | 4.7e-09 | 8.1e-09 | 4e-09 | 5.600E-09 | 2.193E-09 | 4.000E-09 | 8.100E-09 |
| 9644 | 483 | V483D | A | 1e-08 | 1.4e-08 | 6.8e-09 | 1.027E-08 | 3.607E-09 | 6.800E-09 | 1.400E-08 |
| 9645 | 483 | V483C | A | 2.3e-08 | 3.6e-09 | 2.4e-09 | 9.667E-09 | 1.156E-08 | 2.400E-09 | 2.300E-08 |
| 9646 | 483 | V483E | A | 1e-08 | 5.9e-09 | 7.2e-09 | 7.700E-09 | 2.095E-09 | 5.900E-09 | 1.000E-08 |
| 9647 | 483 | V483Q | A | 2.7e-09 | 3.8e-09 | 6.3e-09 | 4.267E-09 | 1.845E-09 | 2.700E-09 | 6.300E-09 |
| 9648 | 483 | V483G | A | 3.6e-09 | 6.2e-09 | 6.5e-09 | 5.433E-09 | 1.595E-09 | 3.600E-09 | 6.500E-09 |
| 9649 | 483 | V483H | A | 7.6e-09 | 9.9e-09 | 3.6e-09 | 7.033E-09 | 3.188E-09 | 3.600E-09 | 9.900E-09 |
| 9650 | 483 | V483I | A | 5.3e-09 | 1.5e-08 | 4.8e-09 | 8.367E-09 | 5.750E-09 | 4.800E-09 | 1.500E-08 |
| 9651 | 483 | V483L | A | 1e-08 | 6.5e-09 | 1.6e-09 | 6.033E-09 | 4.219E-09 | 1.600E-09 | 1.000E-08 |
| 9652 | 483 | V483K | A | 4.5e-09 | 5.7e-09 | 5e-09 | 5.067E-09 | 6.028E-10 | 4.500E-09 | 5.700E-09 |
| 9653 | 483 | V483M | A | 4.7e-09 | 7.5e-09 | 4.5e-09 | 5.567E-09 | 1.677E-09 | 4.500E-09 | 7.500E-09 |
| 9654 | 483 | V483F | A | 1.3e-08 | 2.8e-08 | 7e-09 | 1.600E-08 | 1.082E-08 | 7.000E-09 | 2.800E-08 |
| 9655 | 483 | V483P | A | 8.3e-09 | 6.8e-09 | 1.4e-08 | 9.700E-09 | 3.799E-09 | 6.800E-09 | 1.400E-08 |
| 9656 | 483 | V483S | A | 2.5e-09 | 1.2e-08 | 1.2e-08 | 8.833E-09 | 5.485E-09 | 2.500E-09 | 1.200E-08 |
| 9657 | 483 | V483T | A | 1.3e-08 | 4.5e-09 | 7e-09 | 8.167E-09 | 4.368E-09 | 4.500E-09 | 1.300E-08 |
| 9658 | 483 | V483W | A | 4.2e-09 | 2.5e-09 | 1.8e-08 | 8.233E-09 | 8.501E-09 | 2.500E-09 | 1.800E-08 |
| 9659 | 483 | V483Y | A | 7.5e-09 | 1.1e-08 | 1.8e-08 | 1.217E-08 | 5.346E-09 | 7.500E-09 | 1.800E-08 |
| 9660 | 483 | V483V | A | 1.3e-08 | 5.1e-09 | 8.9e-09 | 9.000E-09 | 3.951E-09 | 5.100E-09 | 1.300E-08 |
| 9661 | 484 | G484A | A | 8.2e-09 | 3.5e-09 | 4.6e-09 | 5.433E-09 | 2.458E-09 | 3.500E-09 | 8.200E-09 |
| 9662 | 484 | G484R | A | 1.5e-08 | 3.4e-09 | 1.7e-08 | 1.180E-08 | 7.343E-09 | 3.400E-09 | 1.700E-08 |
| 9663 | 484 | G484N | A | 8.9e-09 | 7.2e-09 | 1.3e-08 | 9.700E-09 | 2.982E-09 | 7.200E-09 | 1.300E-08 |
| 9664 | 484 | G484D | A | 2.5e-08 | 2.8e-09 | 1.2e-08 | 1.327E-08 | 1.115E-08 | 2.800E-09 | 2.500E-08 |
| 9665 | 484 | G484C | A | 5.9e-09 | 6.5e-09 | 1.7e-08 | 9.800E-09 | 6.243E-09 | 5.900E-09 | 1.700E-08 |
| 9666 | 484 | G484E | A | 7.4e-09 | 5.4e-09 | 3.7e-09 | 5.500E-09 | 1.852E-09 | 3.700E-09 | 7.400E-09 |
| 9667 | 484 | G484Q | A | 3.2e-09 | 4.5e-09 | 1.5e-08 | 7.567E-09 | 6.470E-09 | 3.200E-09 | 1.500E-08 |
| 9668 | 484 | G484G | A | 1.8e-08 | 5.3e-09 | 6.1e-09 | 9.800E-09 | 7.113E-09 | 5.300E-09 | 1.800E-08 |
| 9669 | 484 | G484H | A | 7.8e-09 | 1.2e-08 | 1.8e-09 | 7.200E-09 | 5.126E-09 | 1.800E-09 | 1.200E-08 |
| 9670 | 484 | G484I | A | 2.1e-08 | 8.4e-09 | 1.7e-08 | 1.547E-08 | 6.438E-09 | 8.400E-09 | 2.100E-08 |
| 9671 | 484 | G484L | A | 1.3e-08 | 5e-09 | 9.7e-09 | 9.233E-09 | 4.020E-09 | 5.000E-09 | 1.300E-08 |
| 9672 | 484 | G484K | A | 9.4e-09 | 4e-09 | 1.1e-08 | 8.133E-09 | 3.668E-09 | 4.000E-09 | 1.100E-08 |
| 9673 | 484 | G484M | A | 1.8e-08 | 4.6e-09 | 7.3e-09 | 9.967E-09 | 7.087E-09 | 4.600E-09 | 1.800E-08 |
| 9674 | 484 | G484F | A | 5.3e-09 | 5.5e-09 | 7.2e-09 | 6.000E-09 | 1.044E-09 | 5.300E-09 | 7.200E-09 |
| 9675 | 484 | G484P | A | 1.3e-08 | 2.5e-08 | 8.6e-09 | 1.553E-08 | 8.488E-09 | 8.600E-09 | 2.500E-08 |
| 9676 | 484 | G484S | A | 6e-09 | 1e-08 | 1.2e-08 | 9.333E-09 | 3.055E-09 | 6.000E-09 | 1.200E-08 |
| 9677 | 484 | G484T | A | 6.2e-09 | 3e-09 | 2.6e-09 | 3.933E-09 | 1.973E-09 | 2.600E-09 | 6.200E-09 |
| 9678 | 484 | G484W | A | 3.9e-09 | 9e-09 | 1.1e-08 | 7.967E-09 | 3.661E-09 | 3.900E-09 | 1.100E-08 |
| 9679 | 484 | G484Y | A | 1.3e-08 | 6.4e-09 | 5.8e-09 | 8.400E-09 | 3.995E-09 | 5.800E-09 | 1.300E-08 |
| 9680 | 484 | G484V | A | 6.8e-09 | 5.9e-09 | 1e-08 | 7.567E-09 | 2.155E-09 | 5.900E-09 | 1.000E-08 |
| 9681 | 485 | E485A | A | 1.2e-08 | 6.7e-09 | 2.6e-08 | 1.490E-08 | 9.971E-09 | 6.700E-09 | 2.600E-08 |
| 9682 | 485 | E485R | A | 1.2e-08 | 1.2e-08 | 4.2e-09 | 9.400E-09 | 4.503E-09 | 4.200E-09 | 1.200E-08 |
| 9683 | 485 | E485N | A | 7.6e-09 | 3.3e-09 | 3.4e-09 | 4.767E-09 | 2.454E-09 | 3.300E-09 | 7.600E-09 |
| 9684 | 485 | E485D | A | 7.9e-09 | 1.2e-08 | 1.1e-08 | 1.030E-08 | 2.138E-09 | 7.900E-09 | 1.200E-08 |
| 9685 | 485 | E485C | A | 2.9e-09 | 1.2e-08 | 8.9e-09 | 7.933E-09 | 4.626E-09 | 2.900E-09 | 1.200E-08 |
| 9686 | 485 | E485E | A | 7.7e-09 | 5.4e-09 | 5.8e-09 | 6.300E-09 | 1.229E-09 | 5.400E-09 | 7.700E-09 |
| 9687 | 485 | E485Q | A | 6.8e-09 | 3.3e-09 | 7.7e-09 | 5.933E-09 | 2.325E-09 | 3.300E-09 | 7.700E-09 |
| 9688 | 485 | E485G | A | 1e-08 | 1.3e-08 | 3.6e-09 | 8.867E-09 | 4.801E-09 | 3.600E-09 | 1.300E-08 |
| 9689 | 485 | E485H | A | 3.3e-09 | 4.6e-09 | 2.3e-09 | 3.400E-09 | 1.153E-09 | 2.300E-09 | 4.600E-09 |

| | | | | | | | | | | |
|---|---|---|---|---|---|---|---|---|---|---|
| 9690 | 485 | E485I | A | 1e-08 | 6.6e-09 | 5e-09 | 7.200E-09 | 2.553E-09 | 5.000E-09 | 1.000E-08 |
| 9691 | 485 | E485L | A | 1.1e-08 | 9.8e-09 | 2.9e-09 | 7.900E-09 | 4.371E-09 | 2.900E-09 | 1.100E-08 |
| 9692 | 485 | E485K | A | 1.3e-08 | 5.1e-09 | 7.6e-09 | 8.567E-09 | 4.038E-09 | 5.100E-09 | 1.300E-08 |
| 9693 | 485 | E485M | A | 1.1e-08 | 3.7e-09 | 3.3e-08 | 1.590E-08 | 1.525E-08 | 3.700E-09 | 3.300E-08 |
| 9694 | 485 | E485F | A | 6.2e-09 | 1e-08 | 3.1e-09 | 6.433E-09 | 3.456E-09 | 3.100E-09 | 1.000E-08 |
| 9695 | 485 | E485P | A | 4.4e-09 | 5.6e-09 | 5.3e-09 | 5.100E-09 | 6.245E-10 | 4.400E-09 | 5.600E-09 |
| 9696 | 485 | E485S | A | 1.3e-09 | 9e-09 | 1.1e-08 | 7.100E-09 | 5.122E-09 | 1.300E-09 | 1.100E-08 |
| 9697 | 485 | E485T | A | 3.4e-09 | 1.4e-08 | 9.8e-09 | 9.067E-09 | 5.338E-09 | 3.400E-09 | 1.400E-08 |
| 9698 | 485 | E485W | A | 1.2e-08 | 1.5e-08 | 2.6e-09 | 9.867E-09 | 6.469E-09 | 2.600E-09 | 1.500E-08 |
| 9699 | 485 | E485Y | A | 1.5e-08 | 7.8e-09 | 6.2e-09 | 9.667E-09 | 4.688E-09 | 6.200E-09 | 1.500E-08 |
| 9700 | 485 | E485V | A | 9.9e-09 | 1e-08 | 3.4e-09 | 7.767E-09 | 3.782E-09 | 3.400E-09 | 1.000E-08 |
| 9701 | 486 | G486A | A | 2.7e-09 | 1.3e-08 | 4.2e-09 | 6.633E-09 | 5.564E-09 | 2.700E-09 | 1.300E-08 |
| 9702 | 486 | G486R | A | 5.6e-09 | 2.4e-08 | 1.9e-09 | 1.050E-08 | 1.184E-08 | 1.900E-09 | 2.400E-08 |
| 9703 | 486 | G486N | A | 4.7e-09 | 6.4e-09 | 3.3e-09 | 4.800E-09 | 1.552E-09 | 3.300E-09 | 6.400E-09 |
| 9704 | 486 | G486D | A | 1.5e-08 | 7.7e-09 | 1e-08 | 1.090E-08 | 3.732E-09 | 7.700E-09 | 1.500E-08 |
| 9705 | 486 | G486C | A | 5.6e-09 | 1.1e-08 | 2.1e-08 | 1.253E-08 | 7.814E-09 | 5.600E-09 | 2.100E-08 |
| 9706 | 486 | G486E | A | 7.8e-09 | 3e-09 | 1.9e-08 | 9.933E-09 | 8.211E-09 | 3.000E-09 | 1.900E-08 |
| 9707 | 486 | G486Q | A | 6.9e-09 | 3.4e-09 | 8.7e-09 | 6.333E-09 | 2.695E-09 | 3.400E-09 | 8.700E-09 |
| 9708 | 486 | G486G | A | 4.4e-09 | 1e-08 | 9.8e-09 | 8.067E-09 | 3.177E-09 | 4.400E-09 | 1.000E-08 |
| 9709 | 486 | G486H | A | 1.7e-08 | 1e-08 | 1.5e-08 | 1.400E-08 | 3.606E-09 | 1.000E-08 | 1.700E-08 |
| 9710 | 486 | G486I | A | 3.6e-09 | 1.2e-08 | 3.6e-09 | 6.400E-09 | 4.850E-09 | 3.600E-09 | 1.200E-08 |
| 9711 | 486 | G486L | A | 1.3e-08 | 7e-09 | 6e-09 | 8.667E-09 | 3.786E-09 | 6.000E-09 | 1.300E-08 |
| 9712 | 486 | G486K | A | 9.3e-09 | 8.3e-09 | 5.2e-09 | 7.600E-09 | 2.138E-09 | 5.200E-09 | 9.300E-09 |
| 9713 | 486 | G486M | A | 1.3e-08 | 1.1e-08 | 7.1e-09 | 1.037E-08 | 3.001E-09 | 7.100E-09 | 1.300E-08 |
| 9714 | 486 | G486F | A | 5.9e-09 | 2.3e-09 | 3.5e-09 | 3.900E-09 | 1.833E-09 | 2.300E-09 | 5.900E-09 |
| 9715 | 486 | G486P | A | 8.6e-09 | 7.7e-09 | 3e-09 | 6.433E-09 | 3.007E-09 | 3.000E-09 | 8.600E-09 |
| 9716 | 486 | G486S | A | 4.1e-09 | 1.9e-08 | 5.4e-09 | 9.500E-09 | 8.253E-09 | 4.100E-09 | 1.900E-08 |
| 9717 | 486 | G486T | A | 1.5e-09 | 1.2e-08 | 6.6e-09 | 6.700E-09 | 5.251E-09 | 1.500E-09 | 1.200E-08 |
| 9718 | 486 | G486W | A | 6.8e-09 | 8.5e-09 | 2e-09 | 5.767E-09 | 3.371E-09 | 2.000E-09 | 8.500E-09 |
| 9719 | 486 | G486Y | A | 4e-09 | 4.5e-09 | 4.3e-09 | 4.267E-09 | 2.517E-10 | 4.000E-09 | 4.500E-09 |
| 9720 | 486 | G486V | A | 2e-09 | 4.7e-09 | 1.8e-08 | 8.233E-09 | 8.565E-09 | 2.000E-09 | 1.800E-08 |
| 9721 | 487 | L487A | A | 8.6e-09 | 6.8e-09 | 7e-09 | 7.467E-09 | 9.866E-10 | 6.800E-09 | 8.600E-09 |
| 9722 | 487 | L487R | A | 8.8e-09 | 4.4e-09 | 7.4e-09 | 6.867E-09 | 2.248E-09 | 4.400E-09 | 8.800E-09 |
| 9723 | 487 | L487N | A | 1.3e-08 | 5.3e-09 | 4.2e-09 | 7.500E-09 | 4.795E-09 | 4.200E-09 | 1.300E-08 |
| 9724 | 487 | L487D | A | 1.1e-08 | 1e-08 | 2.5e-09 | 7.833E-09 | 4.646E-09 | 2.500E-09 | 1.100E-08 |
| 9725 | 487 | L487C | A | 3.2e-09 | 2.4e-09 | 5.7e-09 | 3.767E-09 | 1.721E-09 | 2.400E-09 | 5.700E-09 |
| 9726 | 487 | L487E | A | 7.9e-09 | 4.2e-09 | 2.5e-09 | 4.867E-09 | 2.761E-09 | 2.500E-09 | 7.900E-09 |
| 9727 | 487 | L487Q | A | 1.2e-08 | 1.5e-08 | 4.4e-09 | 1.047E-08 | 5.464E-09 | 4.400E-09 | 1.500E-08 |
| 9728 | 487 | L487G | A | 6.1e-09 | 8.1e-09 | 3.1e-09 | 5.767E-09 | 2.517E-09 | 3.100E-09 | 8.100E-09 |
| 9729 | 487 | L487H | A | 2e-09 | 8.5e-09 | 8.7e-09 | 6.400E-09 | 3.812E-09 | 2.000E-09 | 8.700E-09 |
| 9730 | 487 | L487I | A | 5.4e-09 | 1e-08 | 8.3e-09 | 7.900E-09 | 2.326E-09 | 5.400E-09 | 1.000E-08 |
| 9731 | 487 | L487L | A | 8.2e-09 | 1.3e-08 | 3.2e-08 | 1.773E-08 | 1.259E-08 | 8.200E-09 | 3.200E-08 |
| 9732 | 487 | L487K | A | 6.5e-09 | 4.8e-09 | 1.3e-08 | 8.100E-09 | 4.328E-09 | 4.800E-09 | 1.300E-08 |
| 9733 | 487 | L487M | A | 4.7e-09 | 1.3e-08 | 6.5e-09 | 8.067E-09 | 4.366E-09 | 4.700E-09 | 1.300E-08 |
| 9734 | 487 | L487F | A | 1.2e-08 | 6.6e-09 | 4.7e-09 | 7.767E-09 | 3.787E-09 | 4.700E-09 | 1.200E-08 |
| 9735 | 487 | L487P | A | 7e-09 | 1.5e-08 | 6.2e-09 | 9.400E-09 | 4.866E-09 | 6.200E-09 | 1.500E-08 |
| 9736 | 487 | L487S | A | 1.8e-08 | 1.7e-08 | 3.5e-09 | 1.283E-08 | 8.098E-09 | 3.500E-09 | 1.800E-08 |
| 9737 | 487 | L487T | A | 2.5e-09 | 2e-08 | 1.4e-08 | 1.217E-08 | 8.893E-09 | 2.500E-09 | 2.000E-08 |
| 9738 | 487 | L487W | A | 9.2e-09 | 4.6e-09 | 1.5e-08 | 9.600E-09 | 5.212E-09 | 4.600E-09 | 1.500E-08 |
| 9739 | 487 | L487Y | A | 4.3e-09 | 7.5e-09 | 3.3e-09 | 5.033E-09 | 2.194E-09 | 3.300E-09 | 7.500E-09 |
| 9740 | 487 | L487V | A | 6e-09 | 1.3e-08 | 1.1e-08 | 1.000E-08 | 3.606E-09 | 6.000E-09 | 1.300E-08 |
| 9741 | 488 | A488A | A | 3e-09 | 1.4e-08 | 5.4e-09 | 7.467E-09 | 5.784E-09 | 3.000E-09 | 1.400E-08 |
| 9742 | 488 | A488R | A | 2.2e-09 | 6.3e-09 | 8.3e-09 | 5.600E-09 | 3.110E-09 | 2.200E-09 | 8.300E-09 |
| 9743 | 488 | A488N | A | 3.7e-09 | 1e-08 | 6.9e-09 | 6.867E-09 | 3.150E-09 | 3.700E-09 | 1.000E-08 |
| 9744 | 488 | A488D | A | 1.2e-08 | 9.8e-09 | 9.2e-09 | 1.033E-08 | 1.474E-09 | 9.200E-09 | 1.200E-08 |
| 9745 | 488 | A488C | A | 7.7e-09 | 1.1e-08 | 2.6e-08 | 1.490E-08 | 9.753E-09 | 7.700E-09 | 2.600E-08 |
| 9746 | 488 | A488E | A | 5.4e-09 | 1e-08 | 6.6e-09 | 7.333E-09 | 2.386E-09 | 5.400E-09 | 1.000E-08 |
| 9747 | 488 | A488Q | A | 1.3e-08 | 6.8e-09 | 1.3e-08 | 1.093E-08 | 3.580E-09 | 6.800E-09 | 1.300E-08 |
| 9748 | 488 | A488G | A | 1.4e-08 | 7.1e-09 | 1.5e-08 | 1.203E-08 | 4.302E-09 | 7.100E-09 | 1.500E-08 |
| 9749 | 488 | A488H | A | 1.5e-08 | 7.4e-09 | 5.2e-09 | 9.200E-09 | 5.142E-09 | 5.200E-09 | 1.500E-08 |
| 9750 | 488 | A488I | A | 2.4e-09 | 3.3e-09 | 2.8e-09 | 2.833E-09 | 4.509E-10 | 2.400E-09 | 3.300E-09 |
| 9751 | 488 | A488L | A | 2e-08 | 6e-09 | 2.6e-09 | 9.533E-09 | 9.222E-09 | 2.600E-09 | 2.000E-08 |
| 9752 | 488 | A488K | A | 5.9e-09 | 8.2e-09 | 9.2e-09 | 7.767E-09 | 1.692E-09 | 5.900E-09 | 9.200E-09 |
| 9753 | 488 | A488M | A | 3.6e-09 | 1.2e-08 | 1.8e-08 | 1.120E-08 | 7.233E-09 | 3.600E-09 | 1.800E-08 |
| 9754 | 488 | A488F | A | 2.5e-08 | 5e-09 | 6.4e-09 | 1.213E-08 | 1.116E-08 | 5.000E-09 | 2.500E-08 |
| 9755 | 488 | A488P | A | 9.5e-09 | 3.2e-09 | 1.6e-08 | 9.567E-09 | 6.400E-09 | 3.200E-09 | 1.600E-08 |
| 9756 | 488 | A488S | A | 8.8e-09 | 8e-09 | 7.1e-09 | 7.967E-09 | 8.505E-10 | 7.100E-09 | 8.800E-09 |
| 9757 | 488 | A488T | A | 3e-09 | 6.3e-09 | 8.2e-09 | 5.833E-09 | 2.631E-09 | 3.000E-09 | 8.200E-09 |
| 9758 | 488 | A488W | A | 2.5e-08 | 3.7e-09 | 9.9e-09 | 1.287E-08 | 1.096E-08 | 3.700E-09 | 2.500E-08 |
| 9759 | 488 | A488Y | A | 7.7e-09 | 1.1e-08 | 3.1e-09 | 7.267E-09 | 3.968E-09 | 3.100E-09 | 1.100E-08 |
| 9760 | 488 | A488V | A | 1.5e-08 | 3.8e-09 | 9e-09 | 9.267E-09 | 5.605E-09 | 3.800E-09 | 1.500E-08 |
| 9761 | 489 | C489A | A | 1e-08 | 8.1e-09 | 2.9e-09 | 7.000E-09 | 3.676E-09 | 2.900E-09 | 1.000E-08 |
| 9762 | 489 | C489R | A | 6.6e-09 | 7.9e-09 | 1.1e-08 | 8.500E-09 | 2.261E-09 | 6.600E-09 | 1.100E-08 |
| 9763 | 489 | C489N | A | 5.4e-09 | 7.3e-09 | 1.9e-08 | 1.057E-08 | 7.365E-09 | 5.400E-09 | 1.900E-08 |
| 9764 | 489 | C489D | A | 4.1e-09 | 6.1e-09 | 2.4e-08 | 1.140E-08 | 1.096E-08 | 4.100E-09 | 2.400E-08 |
| 9765 | 489 | C489C | A | 5.1e-09 | 2.7e-08 | 5e-09 | 1.237E-08 | 1.267E-08 | 5.000E-09 | 2.700E-08 |
| 9766 | 489 | C489E | A | 5.5e-09 | 5.5e-09 | 7.8e-09 | 6.267E-09 | 1.328E-09 | 5.500E-09 | 7.800E-09 |
| 9767 | 489 | C489Q | A | 1.8e-08 | 4.6e-09 | 7.4e-09 | 1.000E-08 | 7.068E-09 | 4.600E-09 | 1.800E-08 |
| 9768 | 489 | C489G | A | 4e-09 | 1.3e-08 | 5.8e-09 | 7.600E-09 | 4.762E-09 | 4.000E-09 | 1.300E-08 |
| 9769 | 489 | C489H | A | 9.8e-09 | 3.8e-09 | 6.9e-09 | 6.833E-09 | 3.001E-09 | 3.800E-09 | 9.800E-09 |
| 9770 | 489 | C489I | A | 6.3e-09 | 6.9e-09 | 6.8e-09 | 6.667E-09 | 3.215E-10 | 6.300E-09 | 6.900E-09 |
| 9771 | 489 | C489L | A | 5.4e-09 | 8.8e-09 | 3.5e-09 | 5.900E-09 | 2.685E-09 | 3.500E-09 | 8.800E-09 |
| 9772 | 489 | C489K | A | 7.4e-09 | 7.9e-09 | 1.1e-08 | 8.767E-09 | 1.950E-09 | 7.400E-09 | 1.100E-08 |
| 9773 | 489 | C489M | A | 2e-08 | 6.5e-09 | 8.9e-09 | 1.180E-08 | 7.202E-09 | 6.500E-09 | 2.000E-08 |
| 9774 | 489 | C489F | A | 6.3e-09 | 3.9e-09 | 5e-09 | 5.067E-09 | 1.201E-09 | 3.900E-09 | 6.300E-09 |

| | | | | | | | | | | |
|---|---|---|---|---|---|---|---|---|---|---|
| 9775 | 489 | C489P | A | 1.5e-08 | 3.2e-09 | 1.2e-08 | 1.007E-08 | 6.133E-09 | 3.200E-09 | 1.500E-08 |
| 9776 | 489 | C489S | A | 5.1e-09 | 6.2e-09 | 1.3e-08 | 8.100E-09 | 4.279E-09 | 5.100E-09 | 1.300E-08 |
| 9777 | 489 | C489T | A | 1.1e-08 | 1e-08 | 1.3e-08 | 1.133E-08 | 1.528E-09 | 1.000E-08 | 1.300E-08 |
| 9778 | 489 | C489W | A | 4.4e-09 | 8.6e-09 | 1.7e-08 | 1.000E-08 | 6.416E-09 | 4.400E-09 | 1.700E-08 |
| 9779 | 489 | C489Y | A | 1.8e-09 | 1e-08 | 6.2e-09 | 6.000E-09 | 4.104E-09 | 1.800E-09 | 1.000E-08 |
| 9780 | 489 | C489V | A | 6.9e-09 | 4.9e-09 | 8.8e-09 | 6.867E-09 | 1.950E-09 | 4.900E-09 | 8.800E-09 |
| 9781 | 490 | H490A | A | 7.1e-09 | 4.4e-09 | 1.5e-08 | 8.833E-09 | 5.508E-09 | 4.400E-09 | 1.500E-08 |
| 9782 | 490 | H490R | A | 7.6e-09 | 5.2e-09 | 6.9e-09 | 6.567E-09 | 1.234E-09 | 5.200E-09 | 7.600E-09 |
| 9783 | 490 | H490N | A | 6.7e-09 | 4.8e-09 | 8.7e-09 | 6.733E-09 | 1.950E-09 | 4.800E-09 | 8.700E-09 |
| 9784 | 490 | H490D | A | 7.2e-09 | 5.6e-09 | 5.3e-09 | 6.033E-09 | 1.021E-09 | 5.300E-09 | 7.200E-09 |
| 9785 | 490 | H490C | A | 2.8e-08 | 1.3e-08 | 1.3e-08 | 1.800E-08 | 8.660E-09 | 1.300E-08 | 2.800E-08 |
| 9786 | 490 | H490E | A | 2.5e-08 | 6.6e-09 | 2.5e-09 | 1.137E-08 | 1.198E-08 | 2.500E-09 | 2.500E-08 |
| 9787 | 490 | H490Q | A | 2.8e-09 | 7.8e-09 | 8.8e-09 | 6.467E-09 | 3.215E-09 | 2.800E-09 | 8.800E-09 |
| 9788 | 490 | H490G | A | 1e-08 | 2.2e-08 | 1.1e-08 | 1.433E-08 | 6.658E-09 | 1.000E-08 | 2.200E-08 |
| 9789 | 490 | H490H | A | 9e-09 | 9.8e-09 | 6.6e-09 | 8.467E-09 | 1.665E-09 | 6.600E-09 | 9.800E-09 |
| 9790 | 490 | H490I | A | 6e-09 | 9.3e-09 | 2e-09 | 5.767E-09 | 3.656E-09 | 2.000E-09 | 9.300E-09 |
| 9791 | 490 | H490L | A | 5.2e-09 | 1.4e-08 | 1.3e-08 | 1.073E-08 | 4.818E-09 | 5.200E-09 | 1.400E-08 |
| 9792 | 490 | H490K | A | 3.9e-09 | 6.3e-09 | 8.3e-09 | 6.167E-09 | 2.203E-09 | 3.900E-09 | 8.300E-09 |
| 9793 | 490 | H490M | A | 4.7e-09 | 2.8e-09 | 1.3e-08 | 6.833E-09 | 5.424E-09 | 2.800E-09 | 1.300E-08 |
| 9794 | 490 | H490F | A | 2.5e-08 | 7.2e-09 | 3.6e-09 | 1.193E-08 | 1.146E-08 | 3.600E-09 | 2.500E-08 |
| 9795 | 490 | H490P | A | 6.8e-09 | 5.4e-09 | 2.9e-08 | 1.373E-08 | 1.324E-08 | 5.400E-09 | 2.900E-08 |
| 9796 | 490 | H490S | A | 7e-09 | 5.5e-09 | 4e-09 | 5.500E-09 | 1.500E-09 | 4.000E-09 | 7.000E-09 |
| 9797 | 490 | H490T | A | 9.8e-09 | 1.5e-08 | 1.4e-08 | 1.293E-08 | 2.759E-09 | 9.800E-09 | 1.500E-08 |
| 9798 | 490 | H490W | A | 1.7e-08 | 7.5e-09 | 4.3e-09 | 9.600E-09 | 6.605E-09 | 4.300E-09 | 1.700E-08 |
| 9799 | 490 | H490Y | A | 8.5e-09 | 3.5e-09 | 5.8e-09 | 5.933E-09 | 2.503E-09 | 3.500E-09 | 8.500E-09 |
| 9800 | 490 | H490V | A | 6.9e-09 | 8.9e-09 | 5.4e-09 | 7.067E-09 | 1.756E-09 | 5.400E-09 | 8.900E-09 |
| 9801 | 491 | Q491A | A | 1.8e-09 | 5e-09 | 5.9e-09 | 4.233E-09 | 2.155E-09 | 1.800E-09 | 5.900E-09 |
| 9802 | 491 | Q491R | A | 1.9e-08 | 3.6e-08 | 9.5e-09 | 2.150E-08 | 1.343E-08 | 9.500E-09 | 3.600E-08 |
| 9803 | 491 | Q491N | A | 1.8e-08 | 8.2e-09 | 2.9e-09 | 9.700E-09 | 7.661E-09 | 2.900E-09 | 1.800E-08 |
| 9804 | 491 | Q491D | A | 1.1e-08 | 1.1e-08 | 4.1e-09 | 8.700E-09 | 3.984E-09 | 4.100E-09 | 1.100E-08 |
| 9805 | 491 | Q491C | A | 4.2e-09 | 9.5e-09 | 4.2e-09 | 5.967E-09 | 3.060E-09 | 4.200E-09 | 9.500E-09 |
| 9806 | 491 | Q491E | A | 1.2e-08 | 7.3e-09 | 7.1e-09 | 8.800E-09 | 2.773E-09 | 7.100E-09 | 1.200E-08 |
| 9807 | 491 | Q491Q | A | 8.9e-09 | 1.4e-09 | 4.2e-09 | 4.833E-09 | 3.790E-09 | 1.400E-09 | 8.900E-09 |
| 9808 | 491 | Q491G | A | 1.9e-08 | 2.1e-09 | 1.9e-08 | 1.337E-08 | 9.757E-09 | 2.100E-09 | 1.900E-08 |
| 9809 | 491 | Q491H | A | 5.2e-09 | 5.1e-09 | 7.8e-09 | 6.033E-09 | 1.531E-09 | 5.100E-09 | 7.800E-09 |
| 9810 | 491 | Q491I | A | 4.5e-09 | 1.3e-08 | 3.7e-09 | 7.067E-09 | 5.154E-09 | 3.700E-09 | 1.300E-08 |
| 9811 | 491 | Q491L | A | 1.4e-08 | 1.9e-08 | 4.4e-09 | 1.247E-08 | 7.420E-09 | 4.400E-09 | 1.900E-08 |
| 9812 | 491 | Q491K | A | 7.7e-09 | 2.2e-09 | 1.7e-09 | 3.867E-09 | 3.329E-09 | 1.700E-09 | 7.700E-09 |
| 9813 | 491 | Q491M | A | 1e-08 | 5.1e-09 | 1.7e-08 | 1.070E-08 | 5.981E-09 | 5.100E-09 | 1.700E-08 |
| 9814 | 491 | Q491F | A | 5.3e-09 | 6e-09 | 8.1e-09 | 6.467E-09 | 1.457E-09 | 5.300E-09 | 8.100E-09 |
| 9815 | 491 | Q491P | A | 9.7e-09 | 2.4e-09 | 1e-08 | 7.367E-09 | 4.304E-09 | 2.400E-09 | 1.000E-08 |
| 9816 | 491 | Q491S | A | 5.8e-09 | 6.7e-09 | 1.1e-08 | 7.833E-09 | 2.779E-09 | 5.800E-09 | 1.100E-08 |
| 9817 | 491 | Q491T | A | 7.2e-09 | 1.5e-08 | 1.5e-08 | 1.240E-08 | 4.503E-09 | 7.200E-09 | 1.500E-08 |
| 9818 | 491 | Q491W | A | 8.5e-09 | 1.1e-08 | 7e-09 | 8.833E-09 | 2.021E-09 | 7.000E-09 | 1.100E-08 |
| 9819 | 491 | Q491Y | A | 1.1e-08 | 1e-08 | 1.2e-08 | 1.100E-08 | 1.000E-09 | 1.000E-08 | 1.200E-08 |
| 9820 | 491 | Q491V | A | 1.3e-08 | 6.9e-09 | 4.8e-09 | 8.233E-09 | 4.259E-09 | 4.800E-09 | 1.300E-08 |
| 9821 | 492 | L492A | A | 4.2e-09 | 1.2e-08 | 6.6e-09 | 7.600E-09 | 3.995E-09 | 4.200E-09 | 1.200E-08 |
| 9822 | 492 | L492R | A | 2.4e-09 | 3.7e-09 | 1.4e-08 | 6.700E-09 | 6.355E-09 | 2.400E-09 | 1.400E-08 |
| 9823 | 492 | L492N | A | 1.2e-08 | 7.7e-09 | 1.3e-08 | 1.090E-08 | 2.816E-09 | 7.700E-09 | 1.300E-08 |
| 9824 | 492 | L492D | A | 5.9e-09 | 5.9e-09 | 5.9e-09 | 5.900E-09 | 0.000E+00 | 5.900E-09 | 5.900E-09 |
| 9825 | 492 | L492C | A | 4.9e-09 | 5e-09 | 7.1e-09 | 5.667E-09 | 1.242E-09 | 4.900E-09 | 7.100E-09 |
| 9826 | 492 | L492E | A | 2.4e-09 | 1.5e-08 | 7.1e-09 | 8.167E-09 | 6.367E-09 | 2.400E-09 | 1.500E-08 |
| 9827 | 492 | L492Q | A | 2.2e-09 | 8.3e-09 | 4.7e-09 | 5.067E-09 | 3.066E-09 | 2.200E-09 | 8.300E-09 |
| 9828 | 492 | L492G | A | 8.3e-09 | 7.2e-09 | 1.2e-08 | 9.167E-09 | 2.515E-09 | 7.200E-09 | 1.200E-08 |
| 9829 | 492 | L492H | A | 1.2e-08 | 7e-09 | 1.1e-08 | 1.000E-08 | 2.646E-09 | 7.000E-09 | 1.200E-08 |
| 9830 | 492 | L492I | A | 1.3e-08 | 4.7e-09 | 4.3e-09 | 7.333E-09 | 4.912E-09 | 4.300E-09 | 1.300E-08 |
| 9831 | 492 | L492L | A | 1e-08 | 7.7e-09 | 3.2e-09 | 6.967E-09 | 3.459E-09 | 3.200E-09 | 1.000E-08 |
| 9832 | 492 | L492K | A | 4.7e-09 | 8.1e-09 | 1.9e-08 | 1.060E-08 | 7.471E-09 | 4.700E-09 | 1.900E-08 |
| 9833 | 492 | L492M | A | 8.9e-09 | 6.7e-09 | 1.3e-08 | 9.533E-09 | 3.197E-09 | 6.700E-09 | 1.300E-08 |
| 9834 | 492 | L492F | A | 1e-08 | 4.2e-09 | 7.4e-09 | 7.200E-09 | 2.905E-09 | 4.200E-09 | 1.000E-08 |
| 9835 | 492 | L492P | A | 5.4e-09 | 2.6e-09 | 4.1e-09 | 4.033E-09 | 1.401E-09 | 2.600E-09 | 5.400E-09 |
| 9836 | 492 | L492S | A | 2.5e-08 | 3.9e-09 | 3.8e-09 | 1.090E-08 | 1.221E-08 | 3.800E-09 | 2.500E-08 |
| 9837 | 492 | L492T | A | 4.1e-08 | 1.7e-08 | 1.6e-08 | 2.467E-08 | 1.415E-08 | 1.600E-08 | 4.100E-08 |
| 9838 | 492 | L492W | A | 1.7e-08 | 4.6e-09 | 5.9e-09 | 9.167E-09 | 6.815E-09 | 4.600E-09 | 1.700E-08 |
| 9839 | 492 | L492Y | A | 4.6e-09 | 3.9e-09 | 2e-09 | 3.500E-09 | 1.345E-09 | 2.000E-09 | 4.600E-09 |
| 9840 | 492 | L492V | A | 1.2e-08 | 4.8e-09 | 1.1e-08 | 9.267E-09 | 3.900E-09 | 4.800E-09 | 1.200E-08 |
| 9841 | 493 | C493A | A | 1.8e-08 | 9.6e-09 | 4.9e-09 | 1.083E-08 | 6.637E-09 | 4.900E-09 | 1.800E-08 |
| 9842 | 493 | C493R | A | 2.3e-08 | 4.4e-09 | 8.5e-09 | 1.197E-08 | 9.773E-09 | 4.400E-09 | 2.300E-08 |
| 9843 | 493 | C493N | A | 4.6e-09 | 5.9e-09 | 2.4e-09 | 4.300E-09 | 1.769E-09 | 2.400E-09 | 5.900E-09 |
| 9844 | 493 | C493D | A | 3.7e-09 | 5.1e-09 | 2.2e-08 | 1.027E-08 | 1.019E-08 | 3.700E-09 | 2.200E-08 |
| 9845 | 493 | C493C | A | 7.2e-09 | 2.3e-08 | 1.3e-08 | 1.440E-08 | 7.992E-09 | 7.200E-09 | 2.300E-08 |
| 9846 | 493 | C493E | A | 2.5e-09 | 5.8e-09 | 3.5e-09 | 3.933E-09 | 1.692E-09 | 2.500E-09 | 5.800E-09 |
| 9847 | 493 | C493Q | A | 5.3e-09 | 6.4e-09 | 5.8e-09 | 5.833E-09 | 5.508E-10 | 5.300E-09 | 6.400E-09 |
| 9848 | 493 | C493G | A | 9e-09 | 7.3e-09 | 5.8e-09 | 7.367E-09 | 1.601E-09 | 5.800E-09 | 9.000E-09 |
| 9849 | 493 | C493H | A | 2.1e-09 | 7e-09 | 6.2e-09 | 5.100E-09 | 2.629E-09 | 2.100E-09 | 7.000E-09 |
| 9850 | 493 | C493I | A | 1.1e-08 | 7.3e-09 | 1.1e-08 | 9.767E-09 | 2.136E-09 | 7.300E-09 | 1.100E-08 |
| 9851 | 493 | C493L | A | 9.4e-09 | 2.3e-08 | 1.3e-08 | 1.513E-08 | 7.047E-09 | 9.400E-09 | 2.300E-08 |
| 9852 | 493 | C493K | A | 7.9e-09 | 6.4e-09 | 7.3e-09 | 7.200E-09 | 7.550E-10 | 6.400E-09 | 7.900E-09 |
| 9853 | 493 | C493M | A | 7.3e-09 | 6.2e-09 | 1.5e-08 | 9.500E-09 | 4.795E-09 | 6.200E-09 | 1.500E-08 |
| 9854 | 493 | C493F | A | 8.8e-09 | 2.7e-09 | 1e-08 | 7.167E-09 | 3.915E-09 | 2.700E-09 | 1.000E-08 |
| 9855 | 493 | C493P | A | 5.3e-09 | 9.3e-09 | 1e-08 | 8.200E-09 | 2.536E-09 | 5.300E-09 | 1.000E-08 |
| 9856 | 493 | C493S | A | 1.1e-08 | 6.6e-09 | 9e-09 | 8.867E-09 | 2.203E-09 | 6.600E-09 | 1.100E-08 |
| 9857 | 493 | C493T | A | 1.6e-08 | 3.4e-09 | 4.2e-09 | 7.867E-09 | 7.055E-09 | 3.400E-09 | 1.600E-08 |
| 9858 | 493 | C493W | A | 3e-09 | 2.7e-09 | 3.4e-09 | 3.033E-09 | 3.512E-10 | 2.700E-09 | 3.400E-09 |
| 9859 | 493 | C493Y | A | 1.4e-08 | 5.4e-09 | 7.8e-09 | 9.067E-09 | 4.438E-09 | 5.400E-09 | 1.400E-08 |

| | | | | | | | | | | |
|---|---|---|---|---|---|---|---|---|---|---|
| 9860 | 493 | C493V | A | 2.6e-09 | 6.3e-09 | 2.9e-09 | 3.933E-09 | 2.055E-09 | 2.600E-09 | 6.300E-09 |
| 9861 | 494 | A494A | A | 7.8e-09 | 1.2e-08 | 2.1e-08 | 1.360E-08 | 6.744E-09 | 7.800E-09 | 2.100E-08 |
| 9862 | 494 | A494R | A | 3.3e-09 | 3.7e-09 | 5.8e-09 | 4.267E-09 | 1.343E-09 | 3.300E-09 | 5.800E-09 |
| 9863 | 494 | A494N | A | 1.5e-09 | 4.7e-09 | 6.4e-09 | 4.200E-09 | 2.488E-09 | 1.500E-09 | 6.400E-09 |
| 9864 | 494 | A494D | A | 3.6e-09 | 3.3e-09 | 9.6e-09 | 5.500E-09 | 3.554E-09 | 3.300E-09 | 9.600E-09 |
| 9865 | 494 | A494C | A | 7.5e-09 | 9.7e-09 | 4.7e-09 | 7.300E-09 | 2.506E-09 | 4.700E-09 | 9.700E-09 |
| 9866 | 494 | A494E | A | 1.5e-08 | 6.6e-09 | 1.1e-08 | 1.087E-08 | 4.202E-09 | 6.600E-09 | 1.500E-08 |
| 9867 | 494 | A494Q | A | 6.7e-09 | 4.2e-09 | 8.5e-09 | 6.467E-09 | 2.159E-09 | 4.200E-09 | 8.500E-09 |
| 9868 | 494 | A494G | A | 4.9e-09 | 6.8e-09 | 1e-08 | 7.233E-09 | 2.577E-09 | 4.900E-09 | 1.000E-08 |
| 9869 | 494 | A494H | A | 9.2e-09 | 6.4e-09 | 3.7e-09 | 6.433E-09 | 2.750E-09 | 3.700E-09 | 9.200E-09 |
| 9870 | 494 | A494I | A | 7.3e-09 | 2.9e-08 | 1.1e-08 | 1.577E-08 | 1.161E-08 | 7.300E-09 | 2.900E-08 |
| 9871 | 494 | A494L | A | 4.4e-09 | 1.3e-08 | 5.4e-09 | 7.600E-09 | 4.703E-09 | 4.400E-09 | 1.300E-08 |
| 9872 | 494 | A494K | A | 8.6e-09 | 1e-08 | 3.1e-09 | 7.233E-09 | 3.647E-09 | 3.100E-09 | 1.000E-08 |
| 9873 | 494 | A494M | A | 5.3e-09 | 3.9e-09 | 9.2e-09 | 6.133E-09 | 2.747E-09 | 3.900E-09 | 9.200E-09 |
| 9874 | 494 | A494F | A | 1.3e-08 | 8.6e-09 | 1.1e-08 | 1.087E-08 | 2.203E-09 | 8.600E-09 | 1.300E-08 |
| 9875 | 494 | A494P | A | 5.5e-09 | 1.7e-08 | 7.8e-09 | 1.010E-08 | 6.085E-09 | 5.500E-09 | 1.700E-08 |
| 9876 | 494 | A494S | A | 1.7e-08 | 6.6e-09 | 7.8e-09 | 1.047E-08 | 5.690E-09 | 6.600E-09 | 1.700E-08 |
| 9877 | 494 | A494T | A | 1.2e-08 | 1.3e-08 | 8.3e-09 | 1.110E-08 | 2.476E-09 | 8.300E-09 | 1.300E-08 |
| 9878 | 494 | A494W | A | 1.2e-08 | 5.5e-09 | 6.3e-09 | 7.933E-09 | 3.544E-09 | 5.500E-09 | 1.200E-08 |
| 9879 | 494 | A494Y | A | 3.7e-09 | 1.3e-08 | 3.8e-09 | 6.833E-09 | 5.341E-09 | 3.700E-09 | 1.300E-08 |
| 9880 | 494 | A494V | A | 1.5e-08 | 1.4e-08 | 1.1e-08 | 1.333E-08 | 2.082E-09 | 1.100E-08 | 1.500E-08 |
| 9881 | 495 | R495A | A | 1.5e-08 | 3.6e-09 | 7.4e-09 | 8.667E-09 | 5.805E-09 | 3.600E-09 | 1.500E-08 |
| 9882 | 495 | R495R | A | 6.8e-09 | 5.4e-09 | 9.2e-09 | 7.133E-09 | 1.922E-09 | 5.400E-09 | 9.200E-09 |
| 9883 | 495 | R495N | A | 4.4e-09 | 1e-08 | 8.6e-09 | 7.667E-09 | 2.914E-09 | 4.400E-09 | 1.000E-08 |
| 9884 | 495 | R495D | A | 1.5e-08 | 6.1e-09 | 5.7e-09 | 8.933E-09 | 5.258E-09 | 5.700E-09 | 1.500E-08 |
| 9885 | 495 | R495C | A | 2e-08 | 5.6e-09 | 5.1e-09 | 1.023E-08 | 8.462E-09 | 5.100E-09 | 2.000E-08 |
| 9886 | 495 | R495E | A | 2.5e-09 | 1.2e-08 | 2.9e-09 | 5.800E-09 | 5.373E-09 | 2.500E-09 | 1.200E-08 |
| 9887 | 495 | R495Q | A | 5.9e-09 | 1.6e-09 | 2.5e-09 | 3.333E-09 | 2.268E-09 | 1.600E-09 | 5.900E-09 |
| 9888 | 495 | R495G | A | 7.2e-09 | 8.8e-09 | 4.6e-09 | 6.867E-09 | 2.120E-09 | 4.600E-09 | 8.800E-09 |
| 9889 | 495 | R495H | A | 1.1e-08 | 1.2e-08 | 1e-08 | 1.100E-08 | 1.000E-09 | 1.000E-08 | 1.200E-08 |
| 9890 | 495 | R495I | A | 6.8e-09 | 1.9e-08 | 1e-08 | 1.193E-08 | 6.326E-09 | 6.800E-09 | 1.900E-08 |
| 9891 | 495 | R495L | A | 7.5e-09 | 1.5e-08 | 7e-09 | 9.833E-09 | 4.481E-09 | 7.000E-09 | 1.500E-08 |
| 9892 | 495 | R495K | A | 6.6e-09 | 1.3e-08 | 2.3e-09 | 7.300E-09 | 5.384E-09 | 2.300E-09 | 1.300E-08 |
| 9893 | 495 | R495M | A | 8.1e-09 | 5.5e-09 | 5.2e-09 | 6.267E-09 | 1.595E-09 | 5.200E-09 | 8.100E-09 |
| 9894 | 495 | R495F | A | 3.3e-08 | 6.1e-09 | 4.8e-09 | 1.463E-08 | 1.592E-08 | 4.800E-09 | 3.300E-08 |
| 9895 | 495 | R495P | A | 3.2e-09 | 4.5e-09 | 6.2e-09 | 4.633E-09 | 1.504E-09 | 3.200E-09 | 6.200E-09 |
| 9896 | 495 | R495S | A | 1.1e-08 | 7.5e-09 | 1.6e-08 | 1.150E-08 | 4.272E-09 | 7.500E-09 | 1.600E-08 |
| 9897 | 495 | R495T | A | 1.1e-08 | 5.2e-09 | 4.4e-09 | 6.867E-09 | 3.602E-09 | 4.400E-09 | 1.100E-08 |
| 9898 | 495 | R495W | A | 6.2e-09 | 5.6e-09 | 7.8e-09 | 6.533E-09 | 1.137E-09 | 5.600E-09 | 7.800E-09 |
| 9899 | 495 | R495Y | A | 4.1e-09 | 5.7e-09 | 1.7e-08 | 8.933E-09 | 7.032E-09 | 4.100E-09 | 1.700E-08 |
| 9900 | 495 | R495V | A | 1.5e-08 | 1e-08 | 3.3e-08 | 1.933E-08 | 1.210E-08 | 1.000E-08 | 3.300E-08 |
| 9901 | 496 | G496A | A | 5.9e-09 | 1.3e-08 | 1.3e-08 | 1.063E-08 | 4.099E-09 | 5.900E-09 | 1.300E-08 |
| 9902 | 496 | G496R | A | 1.5e-08 | 1.6e-08 | 1e-08 | 1.367E-08 | 3.215E-09 | 1.000E-08 | 1.600E-08 |
| 9903 | 496 | G496N | A | 7.5e-09 | 1e-08 | 7.5e-09 | 8.333E-09 | 1.443E-09 | 7.500E-09 | 1.000E-08 |
| 9904 | 496 | G496D | A | 9.7e-09 | 1.5e-08 | 2.8e-09 | 9.167E-09 | 6.117E-09 | 2.800E-09 | 1.500E-08 |
| 9905 | 496 | G496C | A | 3.4e-09 | 4.9e-09 | 1.4e-08 | 7.433E-09 | 5.736E-09 | 3.400E-09 | 1.400E-08 |
| 9906 | 496 | G496E | A | 5.9e-09 | 8e-09 | 1.3e-08 | 8.967E-09 | 3.647E-09 | 5.900E-09 | 1.300E-08 |
| 9907 | 496 | G496Q | A | 2.3e-08 | 7e-09 | 6.6e-09 | 1.220E-08 | 9.355E-09 | 6.600E-09 | 2.300E-08 |
| 9908 | 496 | G496G | A | 3.5e-09 | 3.6e-09 | 3.7e-09 | 3.600E-09 | 1.000E-10 | 3.500E-09 | 3.700E-09 |
| 9909 | 496 | G496H | A | 1e-08 | 5.9e-09 | 1.5e-08 | 1.030E-08 | 4.557E-09 | 5.900E-09 | 1.500E-08 |
| 9910 | 496 | G496I | A | 1.3e-08 | 4.7e-09 | 3.3e-09 | 7.000E-09 | 5.243E-09 | 3.300E-09 | 1.300E-08 |
| 9911 | 496 | G496L | A | 1.2e-08 | 4.4e-09 | 2.4e-08 | 1.347E-08 | 9.882E-09 | 4.400E-09 | 2.400E-08 |
| 9912 | 496 | G496K | A | 4.1e-09 | 6.8e-09 | 4e-09 | 4.967E-09 | 1.589E-09 | 4.000E-09 | 6.800E-09 |
| 9913 | 496 | G496M | A | 1.3e-08 | 6.3e-09 | 1.3e-08 | 1.077E-08 | 3.868E-09 | 6.300E-09 | 1.300E-08 |
| 9914 | 496 | G496F | A | 5.5e-09 | 6.5e-09 | 1.1e-08 | 7.667E-09 | 2.930E-09 | 5.500E-09 | 1.100E-08 |
| 9915 | 496 | G496P | A | 2.8e-09 | 1.1e-08 | 3.5e-09 | 5.767E-09 | 4.546E-09 | 2.800E-09 | 1.100E-08 |
| 9916 | 496 | G496S | A | 3.1e-08 | 9.3e-09 | 9.1e-09 | 1.647E-08 | 1.259E-08 | 9.100E-09 | 3.100E-08 |
| 9917 | 496 | G496T | A | 8.2e-09 | 1.1e-08 | 4.3e-09 | 7.833E-09 | 3.365E-09 | 4.300E-09 | 1.100E-08 |
| 9918 | 496 | G496W | A | 6.5e-09 | 7.1e-09 | 7e-09 | 6.867E-09 | 3.215E-10 | 6.500E-09 | 7.100E-09 |
| 9919 | 496 | G496Y | A | 3.7e-09 | 6.7e-09 | 3.6e-09 | 4.667E-09 | 1.762E-09 | 3.600E-09 | 6.700E-09 |
| 9920 | 496 | G496V | A | 1.8e-09 | 6.8e-09 | 1.7e-09 | 3.433E-09 | 2.916E-09 | 1.700E-09 | 6.800E-09 |
| 9921 | 497 | H497A | A | 1.9e-08 | 4.5e-09 | 3.9e-09 | 9.133E-09 | 8.550E-09 | 3.900E-09 | 1.900E-08 |
| 9922 | 497 | H497R | A | 6.9e-09 | 1.4e-08 | 1e-08 | 1.030E-08 | 3.559E-09 | 6.900E-09 | 1.400E-08 |
| 9923 | 497 | H497N | A | 6.1e-09 | 2e-09 | 1.1e-09 | 3.067E-09 | 2.665E-09 | 1.100E-09 | 6.100E-09 |
| 9924 | 497 | H497D | A | 2.6e-08 | 1.3e-08 | 1.7e-08 | 1.867E-08 | 6.658E-09 | 1.300E-08 | 2.600E-08 |
| 9925 | 497 | H497C | A | 1.5e-08 | 7.1e-09 | 4.6e-09 | 8.900E-09 | 5.429E-09 | 4.600E-09 | 1.500E-08 |
| 9926 | 497 | H497E | A | 2e-08 | 1.3e-08 | 5.3e-09 | 1.277E-08 | 7.353E-09 | 5.300E-09 | 2.000E-08 |
| 9927 | 497 | H497Q | A | 7.3e-09 | 5e-09 | 5.1e-09 | 5.800E-09 | 1.300E-09 | 5.000E-09 | 7.300E-09 |
| 9928 | 497 | H497G | A | 5.9e-09 | 6.5e-09 | 5e-09 | 5.800E-09 | 7.550E-10 | 5.000E-09 | 6.500E-09 |
| 9929 | 497 | H497H | A | 2.7e-09 | 8.3e-09 | 3.3e-09 | 4.767E-09 | 3.075E-09 | 2.700E-09 | 8.300E-09 |
| 9930 | 497 | H497I | A | 7.7e-09 | 2e-09 | 5.3e-09 | 5.000E-09 | 2.862E-09 | 2.000E-09 | 7.700E-09 |
| 9931 | 497 | H497L | A | 5.5e-09 | 6.4e-09 | 6.3e-09 | 6.067E-09 | 4.933E-10 | 5.500E-09 | 6.400E-09 |
| 9932 | 497 | H497K | A | 1.4e-08 | 5.1e-09 | 3.1e-09 | 7.400E-09 | 5.803E-09 | 3.100E-09 | 1.400E-08 |
| 9933 | 497 | H497M | A | 6.2e-09 | 1.3e-08 | 3.7e-09 | 7.633E-09 | 4.813E-09 | 3.700E-09 | 1.300E-08 |
| 9934 | 497 | H497F | A | 6.4e-09 | 1.2e-08 | 1.1e-08 | 9.800E-09 | 2.987E-09 | 6.400E-09 | 1.200E-08 |
| 9935 | 497 | H497P | A | 5e-09 | 6.6e-09 | 2.3e-08 | 1.153E-08 | 9.963E-09 | 5.000E-09 | 2.300E-08 |
| 9936 | 497 | H497S | A | 3.9e-09 | 6.8e-09 | 6.6e-09 | 5.767E-09 | 1.620E-09 | 3.900E-09 | 6.800E-09 |
| 9937 | 497 | H497T | A | 4.2e-09 | 8.7e-09 | 1.1e-08 | 7.967E-09 | 3.459E-09 | 4.200E-09 | 1.100E-08 |
| 9938 | 497 | H497W | A | 9.6e-09 | 8e-09 | 7.1e-09 | 8.233E-09 | 1.266E-09 | 7.100E-09 | 9.600E-09 |
| 9939 | 497 | H497Y | A | 8e-09 | 8.3e-09 | 8.7e-09 | 8.333E-09 | 3.512E-10 | 8.000E-09 | 8.700E-09 |
| 9940 | 497 | H497V | A | 4e-09 | 6.4e-09 | 1e-08 | 6.800E-09 | 3.020E-09 | 4.000E-09 | 1.000E-08 |
| 9941 | 498 | C498A | A | 4.7e-09 | 1.1e-08 | 4.3e-09 | 6.667E-09 | 3.758E-09 | 4.300E-09 | 1.100E-08 |
| 9942 | 498 | C498R | A | 1e-08 | 4.1e-09 | 1.2e-08 | 8.700E-09 | 4.107E-09 | 4.100E-09 | 1.200E-08 |
| 9943 | 498 | C498N | A | 2.2e-08 | 2.8e-08 | 2.6e-09 | 1.753E-08 | 1.328E-08 | 2.600E-09 | 2.800E-08 |
| 9944 | 498 | C498D | A | 4.9e-09 | 8.2e-09 | 2.9e-09 | 5.333E-09 | 2.676E-09 | 2.900E-09 | 8.200E-09 |

| | | | | | | | | | | |
|---|---|---|---|---|---|---|---|---|---|---|
| 9945 | 498 | C498C | A | 7.6e-09 | 5.4e-09 | 7.5e-09 | 6.833E-09 | 1.242E-09 | 5.400E-09 | 7.600E-09 |
| 9946 | 498 | C498E | A | 1.5e-08 | 7e-09 | 2.5e-08 | 1.567E-08 | 9.018E-09 | 7.000E-09 | 2.500E-08 |
| 9947 | 498 | C498Q | A | 7.1e-09 | 1.3e-08 | 1.9e-08 | 1.303E-08 | 5.950E-09 | 7.100E-09 | 1.900E-08 |
| 9948 | 498 | C498G | A | 5.9e-09 | 1.6e-08 | 2.8e-09 | 8.233E-09 | 6.902E-09 | 2.800E-09 | 1.600E-08 |
| 9949 | 498 | C498H | A | 8.2e-09 | 3.6e-09 | 1.6e-09 | 4.467E-09 | 3.384E-09 | 1.600E-09 | 8.200E-09 |
| 9950 | 498 | C498I | A | 1.3e-08 | 7e-09 | 5.8e-09 | 8.600E-09 | 3.857E-09 | 5.800E-09 | 1.300E-08 |
| 9951 | 498 | C498L | A | 4.3e-09 | 6.8e-09 | 9.3e-09 | 6.800E-09 | 2.500E-09 | 4.300E-09 | 9.300E-09 |
| 9952 | 498 | C498K | A | 7.6e-09 | 2.4e-09 | 2.8e-09 | 4.267E-09 | 2.894E-09 | 2.400E-09 | 7.600E-09 |
| 9953 | 498 | C498M | A | 7.1e-09 | 1.4e-08 | 5.5e-09 | 8.867E-09 | 4.517E-09 | 5.500E-09 | 1.400E-08 |
| 9954 | 498 | C498F | A | 4.4e-09 | 5.4e-09 | 4.5e-09 | 4.767E-09 | 5.508E-10 | 4.400E-09 | 5.400E-09 |
| 9955 | 498 | C498P | A | 2.1e-08 | 1.5e-08 | 5.8e-09 | 1.393E-08 | 7.656E-09 | 5.800E-09 | 2.100E-08 |
| 9956 | 498 | C498S | A | 8.8e-09 | 8.1e-09 | 4.7e-09 | 7.200E-09 | 2.193E-09 | 4.700E-09 | 8.800E-09 |
| 9957 | 498 | C498T | A | 7.8e-09 | 3.9e-09 | 5.2e-09 | 5.633E-09 | 1.986E-09 | 3.900E-09 | 7.800E-09 |
| 9958 | 498 | C498W | A | 1.3e-09 | 1.9e-09 | 1.4e-08 | 5.733E-09 | 7.165E-09 | 1.300E-09 | 1.400E-08 |
| 9959 | 498 | C498Y | A | 2.5e-08 | 1.2e-08 | 1.3e-08 | 1.667E-08 | 7.234E-09 | 1.200E-08 | 2.500E-08 |
| 9960 | 498 | C498V | A | 7.6e-09 | 1.6e-08 | 9.3e-09 | 1.097E-08 | 4.441E-09 | 7.600E-09 | 1.600E-08 |
| 9961 | 499 | W499A | A | 4.4e-09 | 6.4e-09 | 8.8e-09 | 6.533E-09 | 2.203E-09 | 4.400E-09 | 8.800E-09 |
| 9962 | 499 | W499R | A | 3.4e-09 | 5.2e-09 | 5.4e-09 | 4.667E-09 | 1.102E-09 | 3.400E-09 | 5.400E-09 |
| 9963 | 499 | W499N | A | 1.3e-08 | 1.1e-08 | 1.2e-08 | 1.200E-08 | 1.000E-09 | 1.100E-08 | 1.300E-08 |
| 9964 | 499 | W499D | A | 4.3e-08 | 3.1e-09 | 1.1e-08 | 1.903E-08 | 2.113E-08 | 3.100E-09 | 4.300E-08 |
| 9965 | 499 | W499C | A | 1.1e-08 | 3.1e-09 | 5.6e-09 | 6.567E-09 | 4.038E-09 | 3.100E-09 | 1.100E-08 |
| 9966 | 499 | W499E | A | 8.3e-09 | 1.1e-08 | 3.7e-09 | 7.667E-09 | 3.691E-09 | 3.700E-09 | 1.100E-08 |
| 9967 | 499 | W499Q | A | 8.8e-09 | 8e-09 | 4.3e-09 | 7.033E-09 | 2.401E-09 | 4.300E-09 | 8.800E-09 |
| 9968 | 499 | W499G | A | 1e-08 | 3.7e-09 | 2e-08 | 1.123E-08 | 8.220E-09 | 3.700E-09 | 2.000E-08 |
| 9969 | 499 | W499H | A | 1.5e-08 | 1.8e-08 | 3.3e-09 | 1.210E-08 | 7.767E-09 | 3.300E-09 | 1.800E-08 |
| 9970 | 499 | W499I | A | 2.4e-09 | 1.5e-08 | 3.6e-09 | 7.000E-09 | 6.954E-09 | 2.400E-09 | 1.500E-08 |
| 9971 | 499 | W499L | A | 1.8e-08 | 1.1e-08 | 4.6e-09 | 1.120E-08 | 6.702E-09 | 4.600E-09 | 1.800E-08 |
| 9972 | 499 | W499K | A | 2.9e-09 | 1e-08 | 1.5e-08 | 9.300E-09 | 6.080E-09 | 2.900E-09 | 1.500E-08 |
| 9973 | 499 | W499M | A | 7.6e-09 | 7e-09 | 2.2e-09 | 5.600E-09 | 2.960E-09 | 2.200E-09 | 7.600E-09 |
| 9974 | 499 | W499F | A | 3.1e-09 | 8.5e-09 | 6.1e-09 | 5.900E-09 | 2.706E-09 | 3.100E-09 | 8.500E-09 |
| 9975 | 499 | W499P | A | 5.5e-09 | 1.9e-08 | 7.2e-09 | 1.057E-08 | 7.353E-09 | 5.500E-09 | 1.900E-08 |
| 9976 | 499 | W499S | A | 1.6e-08 | 4e-09 | 2.2e-08 | 1.400E-08 | 9.165E-09 | 4.000E-09 | 2.200E-08 |
| 9977 | 499 | W499T | A | 7.4e-09 | 4.3e-09 | 5.1e-09 | 5.600E-09 | 1.609E-09 | 4.300E-09 | 7.400E-09 |
| 9978 | 499 | W499W | A | 3.9e-09 | 8.8e-09 | 1.1e-08 | 7.900E-09 | 3.635E-09 | 3.900E-09 | 1.100E-08 |
| 9979 | 499 | W499Y | A | 1.2e-08 | 4.1e-09 | 5.3e-09 | 7.133E-09 | 4.257E-09 | 4.100E-09 | 1.200E-08 |
| 9980 | 499 | W499V | A | 1.5e-08 | 3.2e-09 | 5.7e-09 | 7.967E-09 | 6.218E-09 | 3.200E-09 | 1.500E-08 |
| 9981 | 500 | G500A | A | 1.4e-08 | 1e-08 | 4.3e-09 | 9.433E-09 | 4.875E-09 | 4.300E-09 | 1.400E-08 |
| 9982 | 500 | G500R | A | 1.2e-08 | 2.3e-09 | 5.5e-09 | 6.600E-09 | 4.943E-09 | 2.300E-09 | 1.200E-08 |
| 9983 | 500 | G500N | A | 6.8e-09 | 4.8e-09 | 4.9e-09 | 5.500E-09 | 1.127E-09 | 4.800E-09 | 6.800E-09 |
| 9984 | 500 | G500D | A | 4.7e-09 | 4.7e-09 | 5.6e-09 | 5.000E-09 | 5.196E-10 | 4.700E-09 | 5.600E-09 |
| 9985 | 500 | G500C | A | 9e-09 | 3.9e-09 | 4.4e-09 | 5.767E-09 | 2.811E-09 | 3.900E-09 | 9.000E-09 |
| 9986 | 500 | G500E | A | 2.2e-09 | 2.2e-09 | 4.1e-09 | 2.833E-09 | 1.097E-09 | 2.200E-09 | 4.100E-09 |
| 9987 | 500 | G500Q | A | 7.3e-09 | 7.3e-09 | 2.7e-09 | 5.767E-09 | 2.656E-09 | 2.700E-09 | 7.300E-09 |
| 9988 | 500 | G500G | A | 1.3e-08 | 7.9e-09 | 9.2e-09 | 1.003E-08 | 2.650E-09 | 7.900E-09 | 1.300E-08 |
| 9989 | 500 | G500H | A | 9.4e-09 | 2.5e-08 | 1e-08 | 1.480E-08 | 8.839E-09 | 9.400E-09 | 2.500E-08 |
| 9990 | 500 | G500I | A | 1.4e-08 | 1.6e-08 | 9.5e-09 | 1.317E-08 | 3.329E-09 | 9.500E-09 | 1.600E-08 |
| 9991 | 500 | G500L | A | 2.1e-09 | 3.5e-09 | 9.5e-09 | 5.033E-09 | 3.931E-09 | 2.100E-09 | 9.500E-09 |
| 9992 | 500 | G500K | A | 1e-08 | 1.1e-08 | 1.5e-08 | 1.200E-08 | 2.646E-09 | 1.000E-08 | 1.500E-08 |
| 9993 | 500 | G500M | A | 1.2e-08 | 3.1e-09 | 6.4e-09 | 7.167E-09 | 4.499E-09 | 3.100E-09 | 1.200E-08 |
| 9994 | 500 | G500F | A | 7.6e-09 | 6.4e-09 | 4.9e-09 | 6.300E-09 | 1.353E-09 | 4.900E-09 | 7.600E-09 |
| 9995 | 500 | G500P | A | 4.4e-09 | 5.8e-09 | 1.3e-08 | 7.733E-09 | 4.614E-09 | 4.400E-09 | 1.300E-08 |
| 9996 | 500 | G500S | A | 8e-09 | 8.5e-09 | 5e-09 | 7.167E-09 | 1.893E-09 | 5.000E-09 | 8.500E-09 |
| 9997 | 500 | G500T | A | 2.5e-09 | 9e-09 | 8e-09 | 6.500E-09 | 3.500E-09 | 2.500E-09 | 9.000E-09 |
| 9998 | 500 | G500W | A | 6.9e-09 | 1.3e-08 | 6.6e-09 | 8.833E-09 | 3.612E-09 | 6.600E-09 | 1.300E-08 |
| 9999 | 500 | G500Y | A | 9.9e-09 | 2.5e-09 | 4.7e-09 | 5.700E-09 | 3.800E-09 | 2.500E-09 | 9.900E-09 |
| 10000 | 500 | G500V | A | 1.1e-08 | 9.9e-09 | 5.4e-09 | 8.767E-09 | 2.967E-09 | 5.400E-09 | 1.100E-08 |
| 10001 | 501 | P501A | A | 4.5e-09 | 2.9e-09 | 2e-09 | 3.133E-09 | 1.266E-09 | 2.000E-09 | 4.500E-09 |
| 10002 | 501 | P501R | A | 9e-09 | 1.5e-08 | 3.9e-09 | 9.300E-09 | 5.556E-09 | 3.900E-09 | 1.500E-08 |
| 10003 | 501 | P501N | A | 3.4e-09 | 7.9e-09 | 1.1e-08 | 7.433E-09 | 3.821E-09 | 3.400E-09 | 1.100E-08 |
| 10004 | 501 | P501D | A | 6.3e-09 | 1.5e-08 | 1.1e-08 | 1.077E-08 | 4.355E-09 | 6.300E-09 | 1.500E-08 |
| 10005 | 501 | P501C | A | 2.8e-09 | 2.9e-09 | 8.2e-09 | 4.633E-09 | 3.089E-09 | 2.800E-09 | 8.200E-09 |
| 10006 | 501 | P501E | A | 3e-08 | 3.8e-09 | 9.3e-09 | 1.437E-08 | 1.382E-08 | 3.800E-09 | 3.000E-08 |
| 10007 | 501 | P501Q | A | 1.5e-08 | 8.7e-09 | 1.1e-08 | 1.157E-08 | 3.188E-09 | 8.700E-09 | 1.500E-08 |
| 10008 | 501 | P501G | A | 6e-09 | 2e-08 | 7.9e-09 | 1.130E-08 | 7.594E-09 | 6.000E-09 | 2.000E-08 |
| 10009 | 501 | P501H | A | 3.2e-09 | 1.5e-08 | 1.6e-08 | 1.140E-08 | 7.119E-09 | 3.200E-09 | 1.600E-08 |
| 10010 | 501 | P501I | A | 4.1e-09 | 6.8e-09 | 7.1e-09 | 6.000E-09 | 1.652E-09 | 4.100E-09 | 7.100E-09 |
| 10011 | 501 | P501L | A | 2.8e-09 | 7.9e-09 | 7.4e-09 | 6.033E-09 | 2.811E-09 | 2.800E-09 | 7.900E-09 |
| 10012 | 501 | P501K | A | 7.4e-09 | 5.2e-09 | 1.7e-09 | 4.767E-09 | 2.875E-09 | 1.700E-09 | 7.400E-09 |
| 10013 | 501 | P501M | A | 1.6e-09 | 2.3e-09 | 7.7e-09 | 3.867E-09 | 3.338E-09 | 1.600E-09 | 7.700E-09 |
| 10014 | 501 | P501F | A | 5.7e-09 | 9.6e-09 | 9.3e-09 | 8.200E-09 | 2.170E-09 | 5.700E-09 | 9.600E-09 |
| 10015 | 501 | P501P | A | 8.4e-09 | 9.7e-09 | 4e-09 | 7.367E-09 | 2.987E-09 | 4.000E-09 | 9.700E-09 |
| 10016 | 501 | P501S | A | 1e-08 | 3.8e-09 | 2.3e-08 | 1.227E-08 | 9.799E-09 | 3.800E-09 | 2.300E-08 |
| 10017 | 501 | P501T | A | 5.6e-09 | 1.1e-08 | 9.2e-09 | 8.600E-09 | 2.750E-09 | 5.600E-09 | 1.100E-08 |
| 10018 | 501 | P501W | A | 8.3e-09 | 4.3e-09 | 1.5e-08 | 9.200E-09 | 5.406E-09 | 4.300E-09 | 1.500E-08 |
| 10019 | 501 | P501Y | A | 5.8e-09 | 1.8e-08 | 8.6e-09 | 1.080E-08 | 6.391E-09 | 5.800E-09 | 1.800E-08 |
| 10020 | 501 | P501V | A | 8.6e-09 | 1e-08 | 8.1e-09 | 8.900E-09 | 9.849E-10 | 8.100E-09 | 1.000E-08 |
| 10021 | 502 | G502A | A | 5.1e-09 | 3.2e-09 | 8.7e-09 | 5.667E-09 | 2.793E-09 | 3.200E-09 | 8.700E-09 |
| 10022 | 502 | G502R | A | 6.3e-09 | 5.6e-09 | 9.4e-09 | 7.100E-09 | 2.022E-09 | 5.600E-09 | 9.400E-09 |
| 10023 | 502 | G502N | A | 1.9e-08 | 7.2e-09 | 2.7e-09 | 9.633E-09 | 8.418E-09 | 2.700E-09 | 1.900E-08 |
| 10024 | 502 | G502D | A | 3.2e-09 | 8.8e-09 | 1.3e-08 | 8.333E-09 | 4.917E-09 | 3.200E-09 | 1.300E-08 |
| 10025 | 502 | G502C | A | 8.2e-09 | 6.4e-09 | 4.3e-09 | 6.300E-09 | 1.952E-09 | 4.300E-09 | 8.200E-09 |
| 10026 | 502 | G502E | A | 6.3e-09 | 7e-09 | 2.3e-09 | 5.200E-09 | 2.536E-09 | 2.300E-09 | 7.000E-09 |
| 10027 | 502 | G502Q | A | 9.4e-09 | 1.4e-08 | 3.3e-09 | 8.900E-09 | 5.367E-09 | 3.300E-09 | 1.400E-08 |
| 10028 | 502 | G502G | A | 3.7e-09 | 2.2e-08 | 8.9e-10 | 8.863E-09 | 1.146E-08 | 8.900E-10 | 2.200E-08 |
| 10029 | 502 | G502H | A | 2.5e-09 | 3.2e-09 | 2.3e-09 | 2.667E-09 | 4.726E-10 | 2.300E-09 | 3.200E-09 |

| | | | | | | | | | | |
|---|---|---|---|---|---|---|---|---|---|---|
| 10030 | 502 | G502I | A | 1.1e-08 | 6.5e-09 | 1.6e-08 | 1.117E-08 | 4.752E-09 | 6.500E-09 | 1.600E-08 |
| 10031 | 502 | G502L | A | 5.5e-09 | 1.5e-08 | 4.1e-09 | 8.200E-09 | 5.930E-09 | 4.100E-09 | 1.500E-08 |
| 10032 | 502 | G502K | A | 2.8e-09 | 7.4e-09 | 8e-09 | 6.067E-09 | 2.845E-09 | 2.800E-09 | 8.000E-09 |
| 10033 | 502 | G502M | A | 1.8e-08 | 8.3e-09 | 1.4e-08 | 1.343E-08 | 4.875E-09 | 8.300E-09 | 1.800E-08 |
| 10034 | 502 | G502F | A | 8.8e-09 | 1.3e-09 | 5.5e-09 | 5.200E-09 | 3.759E-09 | 1.300E-09 | 8.800E-09 |
| 10035 | 502 | G502P | A | 5e-09 | 1.6e-08 | 1.9e-08 | 1.333E-08 | 7.371E-09 | 5.000E-09 | 1.900E-08 |
| 10036 | 502 | G502S | A | 6.4e-09 | 2.4e-09 | 2.2e-09 | 3.667E-09 | 2.369E-09 | 2.200E-09 | 6.400E-09 |
| 10037 | 502 | G502T | A | 1.2e-08 | 7.3e-09 | 2.7e-09 | 7.333E-09 | 4.650E-09 | 2.700E-09 | 1.200E-08 |
| 10038 | 502 | G502W | A | 3.2e-08 | 8.8e-09 | 1.2e-08 | 1.760E-08 | 1.257E-08 | 8.800E-09 | 3.200E-08 |
| 10039 | 502 | G502Y | A | 6.1e-09 | 8.2e-09 | 6.3e-09 | 6.867E-09 | 1.159E-09 | 6.100E-09 | 8.200E-09 |
| 10040 | 502 | G502V | A | 7.3e-09 | 2.5e-09 | 2.7e-09 | 4.167E-09 | 2.715E-09 | 2.500E-09 | 7.300E-09 |
| 10041 | 503 | P503A | A | 1.3e-08 | 6.3e-09 | 9.4e-09 | 9.567E-09 | 3.353E-09 | 6.300E-09 | 1.300E-08 |
| 10042 | 503 | P503R | A | 6.8e-09 | 1e-08 | 1.9e-09 | 6.233E-09 | 4.080E-09 | 1.900E-09 | 1.000E-08 |
| 10043 | 503 | P503N | A | 6e-09 | 6.6e-09 | 1.1e-08 | 7.867E-09 | 2.730E-09 | 6.000E-09 | 1.100E-08 |
| 10044 | 503 | P503D | A | 3.8e-09 | 1.7e-08 | 6.1e-09 | 8.967E-09 | 7.051E-09 | 3.800E-09 | 1.700E-08 |
| 10045 | 503 | P503C | A | 8.2e-09 | 8.8e-09 | 3.1e-09 | 6.700E-09 | 3.132E-09 | 3.100E-09 | 8.800E-09 |
| 10046 | 503 | P503E | A | 1.5e-08 | 2.1e-09 | 1.2e-08 | 9.700E-09 | 6.751E-09 | 2.100E-09 | 1.500E-08 |
| 10047 | 503 | P503Q | A | 6e-09 | 7.4e-09 | 2.4e-09 | 5.267E-09 | 2.579E-09 | 2.400E-09 | 7.400E-09 |
| 10048 | 503 | P503G | A | 6.4e-09 | 7.9e-09 | 5.4e-09 | 6.567E-09 | 1.258E-09 | 5.400E-09 | 7.900E-09 |
| 10049 | 503 | P503H | A | 7.2e-09 | 1.1e-08 | 1.1e-08 | 9.733E-09 | 2.194E-09 | 7.200E-09 | 1.100E-08 |
| 10050 | 503 | P503I | A | 7.2e-09 | 1.5e-08 | 1e-08 | 1.073E-08 | 3.951E-09 | 7.200E-09 | 1.500E-08 |
| 10051 | 503 | P503L | A | 4e-09 | 8.8e-09 | 5.5e-09 | 6.100E-09 | 2.456E-09 | 4.000E-09 | 8.800E-09 |
| 10052 | 503 | P503K | A | 8e-09 | 4.3e-09 | 7.4e-09 | 6.567E-09 | 1.986E-09 | 4.300E-09 | 8.000E-09 |
| 10053 | 503 | P503M | A | 7.3e-09 | 1.4e-08 | 6.3e-09 | 9.200E-09 | 4.187E-09 | 6.300E-09 | 1.400E-08 |
| 10054 | 503 | P503F | A | 7.8e-09 | 7.1e-09 | 1.1e-08 | 8.633E-09 | 2.079E-09 | 7.100E-09 | 1.100E-08 |
| 10055 | 503 | P503P | A | 2.7e-09 | 2.1e-09 | 1e-08 | 4.933E-09 | 4.398E-09 | 2.100E-09 | 1.000E-08 |
| 10056 | 503 | P503S | A | 1.1e-08 | 2.3e-09 | 7.3e-09 | 6.867E-09 | 4.366E-09 | 2.300E-09 | 1.100E-08 |
| 10057 | 503 | P503T | A | 6.3e-09 | 2.6e-08 | 4.4e-09 | 1.223E-08 | 1.196E-08 | 4.400E-09 | 2.600E-08 |
| 10058 | 503 | P503W | A | 3.5e-09 | 4.2e-09 | 1.4e-08 | 7.233E-09 | 5.871E-09 | 3.500E-09 | 1.400E-08 |
| 10059 | 503 | P503Y | A | 4.1e-09 | 1.2e-08 | 5.9e-09 | 7.333E-09 | 4.140E-09 | 4.100E-09 | 1.200E-08 |
| 10060 | 503 | P503V | A | 3.5e-09 | 7.1e-09 | 1e-08 | 6.867E-09 | 3.256E-09 | 3.500E-09 | 1.000E-08 |
| 10061 | 504 | T504A | A | 7.7e-09 | 1.2e-08 | 8e-09 | 9.233E-09 | 2.401E-09 | 7.700E-09 | 1.200E-08 |
| 10062 | 504 | T504R | A | 8.5e-10 | 1.1e-08 | 1.6e-08 | 9.283E-09 | 7.720E-09 | 8.500E-10 | 1.600E-08 |
| 10063 | 504 | T504N | A | 1.3e-08 | 3.1e-09 | 4.8e-09 | 6.967E-09 | 5.294E-09 | 3.100E-09 | 1.300E-08 |
| 10064 | 504 | T504D | A | 7.3e-09 | 1.4e-08 | 7.4e-09 | 9.567E-09 | 3.840E-09 | 7.300E-09 | 1.400E-08 |
| 10065 | 504 | T504C | A | 7.8e-09 | 2.5e-09 | 5.6e-09 | 5.300E-09 | 2.663E-09 | 2.500E-09 | 7.800E-09 |
| 10066 | 504 | T504E | A | 7.2e-09 | 4e-09 | 1.3e-08 | 8.067E-09 | 4.562E-09 | 4.000E-09 | 1.300E-08 |
| 10067 | 504 | T504Q | A | 7.9e-09 | 4.5e-09 | 3.4e-09 | 5.267E-09 | 2.346E-09 | 3.400E-09 | 7.900E-09 |
| 10068 | 504 | T504G | A | 6e-09 | 6.8e-09 | 1e-08 | 7.600E-09 | 2.117E-09 | 6.000E-09 | 1.000E-08 |
| 10069 | 504 | T504H | A | 2.8e-09 | 7.4e-09 | 6.9e-09 | 5.700E-09 | 2.524E-09 | 2.800E-09 | 7.400E-09 |
| 10070 | 504 | T504I | A | 1.8e-08 | 4.3e-09 | 2.4e-09 | 8.233E-09 | 8.511E-09 | 2.400E-09 | 1.800E-08 |
| 10071 | 504 | T504L | A | 6e-09 | 2.2e-08 | 1.3e-08 | 1.367E-08 | 8.021E-09 | 6.000E-09 | 2.200E-08 |
| 10072 | 504 | T504K | A | 2e-08 | 7.4e-09 | 3.4e-09 | 1.027E-08 | 8.663E-09 | 3.400E-09 | 2.000E-08 |
| 10073 | 504 | T504M | A | 4.1e-09 | 5.4e-09 | 1.1e-08 | 6.833E-09 | 3.667E-09 | 4.100E-09 | 1.100E-08 |
| 10074 | 504 | T504F | A | 8.6e-09 | 5e-09 | 5.3e-09 | 6.300E-09 | 1.997E-09 | 5.000E-09 | 8.600E-09 |
| 10075 | 504 | T504P | A | 5.4e-09 | 7e-09 | 1e-08 | 7.467E-09 | 2.335E-09 | 5.400E-09 | 1.000E-08 |
| 10076 | 504 | T504S | A | 9.5e-09 | 3.6e-09 | 1.9e-08 | 1.070E-08 | 7.770E-09 | 3.600E-09 | 1.900E-08 |
| 10077 | 504 | T504T | A | 9e-09 | 6.1e-09 | 1.1e-08 | 8.700E-09 | 2.464E-09 | 6.100E-09 | 1.100E-08 |
| 10078 | 504 | T504W | A | 6.8e-09 | 4.4e-09 | 6.1e-09 | 5.767E-09 | 1.234E-09 | 4.400E-09 | 6.800E-09 |
| 10079 | 504 | T504Y | A | 5.8e-09 | 8.1e-09 | 1.3e-08 | 8.967E-09 | 3.677E-09 | 5.800E-09 | 1.300E-08 |
| 10080 | 504 | T504V | A | 5.8e-09 | 6.8e-09 | 7.8e-09 | 6.800E-09 | 1.000E-09 | 5.800E-09 | 7.800E-09 |
| 10081 | 505 | Q505A | A | 6.8e-09 | 3.4e-09 | 1.6e-08 | 8.733E-09 | 6.519E-09 | 3.400E-09 | 1.600E-08 |
| 10082 | 505 | Q505R | A | 9.1e-09 | 1.1e-08 | 1.5e-08 | 1.170E-08 | 3.012E-09 | 9.100E-09 | 1.500E-08 |
| 10083 | 505 | Q505N | A | 3.7e-09 | 1e-08 | 5.9e-09 | 6.533E-09 | 3.197E-09 | 3.700E-09 | 1.000E-08 |
| 10084 | 505 | Q505D | A | 1.4e-08 | 1.3e-08 | 2.1e-08 | 1.600E-08 | 4.359E-09 | 1.300E-08 | 2.100E-08 |
| 10085 | 505 | Q505C | A | 9.8e-09 | 2.3e-08 | 1.8e-08 | 1.693E-08 | 6.664E-09 | 9.800E-09 | 2.300E-08 |
| 10086 | 505 | Q505E | A | 3.3e-09 | 2.6e-09 | 7.2e-09 | 4.367E-09 | 2.479E-09 | 2.600E-09 | 7.200E-09 |
| 10087 | 505 | Q505Q | A | 1.1e-08 | 1.2e-08 | 1.7e-08 | 1.333E-08 | 3.215E-09 | 1.100E-08 | 1.700E-08 |
| 10088 | 505 | Q505G | A | 6e-09 | 1.3e-09 | 6e-09 | 4.433E-09 | 2.714E-09 | 1.300E-09 | 6.000E-09 |
| 10089 | 505 | Q505H | A | 2.7e-09 | 7.5e-09 | 6.3e-09 | 5.500E-09 | 2.498E-09 | 2.700E-09 | 7.500E-09 |
| 10090 | 505 | Q505I | A | 1.1e-08 | 8.8e-09 | 1.2e-08 | 1.060E-08 | 1.637E-09 | 8.800E-09 | 1.200E-08 |
| 10091 | 505 | Q505L | A | 1.3e-08 | 2.9e-08 | 2.2e-08 | 2.133E-08 | 8.021E-09 | 1.300E-08 | 2.900E-08 |
| 10092 | 505 | Q505K | A | 8.6e-09 | 9.3e-09 | 7.6e-09 | 8.500E-09 | 8.544E-10 | 7.600E-09 | 9.300E-09 |
| 10093 | 505 | Q505M | A | 9.3e-09 | 1.5e-08 | 3.7e-09 | 9.333E-09 | 5.650E-09 | 3.700E-09 | 1.500E-08 |
| 10094 | 505 | Q505F | A | 1e-08 | 9.5e-09 | 9e-09 | 9.500E-09 | 5.000E-10 | 9.000E-09 | 1.000E-08 |
| 10095 | 505 | Q505P | A | 1e-08 | 1e-08 | 9.7e-09 | 9.900E-09 | 1.732E-10 | 9.700E-09 | 1.000E-08 |
| 10096 | 505 | Q505S | A | 4.5e-09 | 1e-08 | 8.4e-09 | 7.633E-09 | 2.829E-09 | 4.500E-09 | 1.000E-08 |
| 10097 | 505 | Q505T | A | 8e-09 | 3e-08 | 2.3e-09 | 1.343E-08 | 1.463E-08 | 2.300E-09 | 3.000E-08 |
| 10098 | 505 | Q505W | A | 1.1e-08 | 9.4e-09 | 1.4e-08 | 1.147E-08 | 2.335E-09 | 9.400E-09 | 1.400E-08 |
| 10099 | 505 | Q505Y | A | 4.2e-09 | 4.3e-09 | 7.8e-09 | 5.433E-09 | 2.050E-09 | 4.200E-09 | 7.800E-09 |
| 10100 | 505 | Q505V | A | 5.3e-09 | 1.2e-08 | 9.3e-09 | 8.867E-09 | 3.371E-09 | 5.300E-09 | 1.200E-08 |
| 10101 | 506 | C506A | A | 1.1e-08 | 1.2e-08 | 1.3e-08 | 1.200E-08 | 1.000E-09 | 1.100E-08 | 1.300E-08 |
| 10102 | 506 | C506R | A | 8.3e-09 | 4.9e-09 | 4.1e-09 | 5.767E-09 | 2.230E-09 | 4.100E-09 | 8.300E-09 |
| 10103 | 506 | C506N | A | 5.5e-09 | 6e-09 | 6.4e-09 | 5.967E-09 | 4.509E-10 | 5.500E-09 | 6.400E-09 |
| 10104 | 506 | C506D | A | 6.9e-09 | 3.3e-09 | 1.7e-08 | 9.067E-09 | 7.102E-09 | 3.300E-09 | 1.700E-08 |
| 10105 | 506 | C506C | A | 9.3e-09 | 2.4e-08 | 9.1e-09 | 1.413E-08 | 8.545E-09 | 9.100E-09 | 2.400E-08 |
| 10106 | 506 | C506E | A | 8.2e-09 | 5.8e-09 | 1.4e-08 | 9.333E-09 | 4.216E-09 | 5.800E-09 | 1.400E-08 |
| 10107 | 506 | C506Q | A | 5.1e-09 | 6.9e-09 | 4.6e-09 | 5.533E-09 | 1.210E-09 | 4.600E-09 | 6.900E-09 |
| 10108 | 506 | C506G | A | 8.8e-09 | 4.2e-09 | 9.6e-09 | 7.533E-09 | 2.914E-09 | 4.200E-09 | 9.600E-09 |
| 10109 | 506 | C506H | A | 7.8e-09 | 1.4e-08 | 5.1e-09 | 8.967E-09 | 4.563E-09 | 5.100E-09 | 1.400E-08 |
| 10110 | 506 | C506I | A | 6.5e-09 | 1.1e-08 | 7.1e-09 | 8.200E-09 | 2.443E-09 | 6.500E-09 | 1.100E-08 |
| 10111 | 506 | C506L | A | 1.3e-08 | 5.4e-09 | 3.8e-09 | 7.400E-09 | 4.915E-09 | 3.800E-09 | 1.300E-08 |
| 10112 | 506 | C506K | A | 6.4e-09 | 6.6e-09 | 3.7e-09 | 5.567E-09 | 1.620E-09 | 3.700E-09 | 6.600E-09 |
| 10113 | 506 | C506M | A | 3.4e-09 | 1.1e-08 | 5.3e-09 | 6.567E-09 | 3.955E-09 | 3.400E-09 | 1.100E-08 |
| 10114 | 506 | C506F | A | 8.1e-09 | 9e-09 | 5e-09 | 7.367E-09 | 2.098E-09 | 5.000E-09 | 9.000E-09 |

| | | | | | | | | | | |
|---|---|---|---|---|---|---|---|---|---|---|
| 10115 | 506 | C506P | A | 6.1e-09 | 4.3e-09 | 5.5e-09 | 5.300E-09 | 9.165E-10 | 4.300E-09 | 6.100E-09 |
| 10116 | 506 | C506S | A | 2.5e-08 | 6.4e-09 | 5.8e-09 | 1.240E-08 | 1.092E-08 | 5.800E-09 | 2.500E-08 |
| 10117 | 506 | C506T | A | 2.6e-09 | 4.4e-08 | 6e-09 | 1.753E-08 | 2.298E-08 | 2.600E-09 | 4.400E-08 |
| 10118 | 506 | C506W | A | 1.2e-08 | 8.2e-09 | 7.1e-09 | 9.100E-09 | 2.571E-09 | 7.100E-09 | 1.200E-08 |
| 10119 | 506 | C506Y | A | 1.9e-08 | 6.9e-09 | 8.2e-09 | 1.137E-08 | 6.643E-09 | 6.900E-09 | 1.900E-08 |
| 10120 | 506 | C506V | A | 8.4e-09 | 6.1e-09 | 6.9e-09 | 7.133E-09 | 1.168E-09 | 6.100E-09 | 8.400E-09 |
| 10121 | 507 | V507A | A | 6.1e-09 | 5.2e-09 | 8.6e-09 | 6.633E-09 | 1.762E-09 | 5.200E-09 | 8.600E-09 |
| 10122 | 507 | V507R | A | 4.9e-09 | 1.5e-08 | 7.7e-09 | 9.200E-09 | 5.214E-09 | 4.900E-09 | 1.500E-08 |
| 10123 | 507 | V507N | A | 4.7e-09 | 1.2e-08 | 7.7e-09 | 8.133E-09 | 3.669E-09 | 4.700E-09 | 1.200E-08 |
| 10124 | 507 | V507D | A | 7.4e-09 | 4.2e-08 | 9.8e-09 | 1.973E-08 | 1.932E-08 | 7.400E-09 | 4.200E-08 |
| 10125 | 507 | V507C | A | 8.1e-09 | 6.5e-09 | 7.7e-09 | 7.433E-09 | 8.327E-10 | 6.500E-09 | 8.100E-09 |
| 10126 | 507 | V507E | A | 5.4e-09 | 1.5e-08 | 9.1e-09 | 9.833E-09 | 4.842E-09 | 5.400E-09 | 1.500E-08 |
| 10127 | 507 | V507Q | A | 9.8e-09 | 4.2e-09 | 6.2e-09 | 6.733E-09 | 2.838E-09 | 4.200E-09 | 9.800E-09 |
| 10128 | 507 | V507G | A | 7.4e-09 | 1.3e-08 | 3.2e-08 | 1.747E-08 | 1.289E-08 | 7.400E-09 | 3.200E-08 |
| 10129 | 507 | V507H | A | 1.2e-08 | 6e-09 | 7.7e-09 | 8.567E-09 | 3.092E-09 | 6.000E-09 | 1.200E-08 |
| 10130 | 507 | V507I | A | 1.3e-08 | 1e-08 | 4.1e-09 | 9.033E-09 | 4.528E-09 | 4.100E-09 | 1.300E-08 |
| 10131 | 507 | V507L | A | 7.1e-09 | 1.2e-08 | 8.2e-09 | 9.100E-09 | 2.571E-09 | 7.100E-09 | 1.200E-08 |
| 10132 | 507 | V507K | A | 7e-09 | 5.7e-09 | 3e-09 | 5.233E-09 | 2.040E-09 | 3.000E-09 | 7.000E-09 |
| 10133 | 507 | V507M | A | 2.3e-09 | 1.1e-09 | 2.8e-09 | 2.067E-09 | 8.737E-10 | 1.100E-09 | 2.800E-09 |
| 10134 | 507 | V507F | A | 4.1e-09 | 6.9e-09 | 1.9e-08 | 1.000E-08 | 7.919E-09 | 4.100E-09 | 1.900E-08 |
| 10135 | 507 | V507P | A | 8.9e-09 | 1.5e-08 | 1.8e-08 | 1.397E-08 | 4.637E-09 | 8.900E-09 | 1.800E-08 |
| 10136 | 507 | V507S | A | 4.4e-09 | 1.4e-08 | 5.3e-09 | 7.900E-09 | 5.302E-09 | 4.400E-09 | 1.400E-08 |
| 10137 | 507 | V507T | A | 5.8e-09 | 6.4e-09 | 3.5e-09 | 5.233E-09 | 1.531E-09 | 3.500E-09 | 6.400E-09 |
| 10138 | 507 | V507W | A | 8.9e-09 | 8.9e-09 | 1.8e-09 | 6.533E-09 | 4.099E-09 | 1.800E-09 | 8.900E-09 |
| 10139 | 507 | V507Y | A | 1.4e-08 | 2e-08 | 4.5e-09 | 1.283E-08 | 7.816E-09 | 4.500E-09 | 2.000E-08 |
| 10140 | 507 | V507V | A | 3.5e-09 | 3.6e-09 | 7.4e-09 | 4.833E-09 | 2.223E-09 | 3.500E-09 | 7.400E-09 |
| 10141 | 508 | N508A | A | 6.4e-09 | 4.7e-09 | 3.1e-09 | 4.733E-09 | 1.650E-09 | 3.100E-09 | 6.400E-09 |
| 10142 | 508 | N508R | A | 1e-08 | 7e-09 | 8.3e-09 | 8.433E-09 | 1.504E-09 | 7.000E-09 | 1.000E-08 |
| 10143 | 508 | N508N | A | 8e-09 | 6.1e-09 | 4.4e-09 | 6.167E-09 | 1.801E-09 | 4.400E-09 | 8.000E-09 |
| 10144 | 508 | N508D | A | 3.7e-09 | 1.1e-08 | 5.2e-09 | 6.633E-09 | 3.855E-09 | 3.700E-09 | 1.100E-08 |
| 10145 | 508 | N508C | A | 2.9e-09 | 4e-09 | 7.1e-09 | 4.667E-09 | 2.178E-09 | 2.900E-09 | 7.100E-09 |
| 10146 | 508 | N508E | A | 3.9e-09 | 1.4e-08 | 8.7e-09 | 8.867E-09 | 5.052E-09 | 3.900E-09 | 1.400E-08 |
| 10147 | 508 | N508Q | A | 7.6e-09 | 5.7e-09 | 8.8e-09 | 7.367E-09 | 1.563E-09 | 5.700E-09 | 8.800E-09 |
| 10148 | 508 | N508G | A | 8.3e-09 | 6.5e-09 | 1.1e-08 | 8.600E-09 | 2.265E-09 | 6.500E-09 | 1.100E-08 |
| 10149 | 508 | N508H | A | 1.9e-09 | 1.3e-08 | 1.6e-08 | 1.030E-08 | 7.428E-09 | 1.900E-09 | 1.600E-08 |
| 10150 | 508 | N508I | A | 1.2e-08 | 3.7e-09 | 9.5e-09 | 8.400E-09 | 4.258E-09 | 3.700E-09 | 1.200E-08 |
| 10151 | 508 | N508L | A | 4.3e-09 | 7.4e-09 | 5.1e-09 | 5.600E-09 | 1.609E-09 | 4.300E-09 | 7.400E-09 |
| 10152 | 508 | N508K | A | 5.2e-09 | 1.2e-08 | 9.2e-09 | 8.800E-09 | 3.418E-09 | 5.200E-09 | 1.200E-08 |
| 10153 | 508 | N508M | A | 6.8e-09 | 5.1e-09 | 5.7e-09 | 5.867E-09 | 8.622E-10 | 5.100E-09 | 6.800E-09 |
| 10154 | 508 | N508F | A | 1.9e-09 | 1.4e-08 | 1.1e-08 | 8.967E-09 | 6.301E-09 | 1.900E-09 | 1.400E-08 |
| 10155 | 508 | N508P | A | 3.7e-09 | 5.9e-09 | 7.1e-09 | 5.567E-09 | 1.724E-09 | 3.700E-09 | 7.100E-09 |
| 10156 | 508 | N508S | A | 7.9e-09 | 4.3e-09 | 3.4e-09 | 5.200E-09 | 2.381E-09 | 3.400E-09 | 7.900E-09 |
| 10157 | 508 | N508T | A | 5.7e-09 | 4.2e-09 | 1.9e-08 | 9.633E-09 | 8.146E-09 | 4.200E-09 | 1.900E-08 |
| 10158 | 508 | N508W | A | 8.2e-09 | 2e-09 | 8.3e-09 | 6.167E-09 | 3.609E-09 | 2.000E-09 | 8.300E-09 |
| 10159 | 508 | N508Y | A | 7.8e-09 | 4.8e-09 | 3.4e-09 | 5.333E-09 | 2.248E-09 | 3.400E-09 | 7.800E-09 |
| 10160 | 508 | N508V | A | 1.5e-08 | 1.1e-08 | 1.1e-08 | 1.233E-08 | 2.309E-09 | 1.100E-08 | 1.500E-08 |
| 10161 | 509 | C509A | A | 1.2e-09 | 6.2e-09 | 1.6e-08 | 7.800E-09 | 7.529E-09 | 1.200E-09 | 1.600E-08 |
| 10162 | 509 | C509R | A | 9e-09 | 1.2e-08 | 9.6e-09 | 1.020E-08 | 1.587E-09 | 9.000E-09 | 1.200E-08 |
| 10163 | 509 | C509N | A | 1e-08 | 2.3e-08 | 9.7e-09 | 1.423E-08 | 7.594E-09 | 9.700E-09 | 2.300E-08 |
| 10164 | 509 | C509D | A | 1.2e-08 | 1.8e-08 | 2.3e-08 | 1.767E-08 | 5.508E-09 | 1.200E-08 | 2.300E-08 |
| 10165 | 509 | C509C | A | 8e-09 | 1.4e-08 | 3.8e-09 | 8.600E-09 | 5.126E-09 | 3.800E-09 | 1.400E-08 |
| 10166 | 509 | C509E | A | 4.5e-09 | 8.7e-09 | 7.7e-09 | 6.967E-09 | 2.194E-09 | 4.500E-09 | 8.700E-09 |
| 10167 | 509 | C509Q | A | 7.8e-09 | 1.1e-08 | 7.5e-09 | 8.767E-09 | 1.940E-09 | 7.500E-09 | 1.100E-08 |
| 10168 | 509 | C509G | A | 5.9e-09 | 3.2e-09 | 3.7e-09 | 4.267E-09 | 1.436E-09 | 3.200E-09 | 5.900E-09 |
| 10169 | 509 | C509H | A | 8.8e-09 | 1.3e-08 | 9.8e-09 | 1.053E-08 | 2.194E-09 | 8.800E-09 | 1.300E-08 |
| 10170 | 509 | C509I | A | 3.6e-09 | 8.4e-09 | 1.1e-08 | 7.667E-09 | 3.754E-09 | 3.600E-09 | 1.100E-08 |
| 10171 | 509 | C509L | A | 8.3e-09 | 1.1e-08 | 6.3e-09 | 8.533E-09 | 2.359E-09 | 6.300E-09 | 1.100E-08 |
| 10172 | 509 | C509K | A | 2.1e-08 | 1.3e-08 | 1.3e-08 | 1.567E-08 | 4.619E-09 | 1.300E-08 | 2.100E-08 |
| 10173 | 509 | C509M | A | 9e-09 | 9.4e-09 | 9.6e-09 | 9.333E-09 | 3.055E-10 | 9.000E-09 | 9.600E-09 |
| 10174 | 509 | C509F | A | 3.5e-09 | 3.6e-09 | 9.3e-09 | 5.467E-09 | 3.320E-09 | 3.500E-09 | 9.300E-09 |
| 10175 | 509 | C509P | A | 4.5e-09 | 7.3e-09 | 9.2e-09 | 7.000E-09 | 2.364E-09 | 4.500E-09 | 9.200E-09 |
| 10176 | 509 | C509S | A | 6.1e-09 | 9.1e-09 | 4.5e-09 | 6.567E-09 | 2.335E-09 | 4.500E-09 | 9.100E-09 |
| 10177 | 509 | C509T | A | 7.6e-09 | 1.4e-08 | 1.5e-08 | 1.220E-08 | 4.015E-09 | 7.600E-09 | 1.500E-08 |
| 10178 | 509 | C509W | A | 1.2e-08 | 1.6e-09 | 1.2e-08 | 8.533E-09 | 6.004E-09 | 1.600E-09 | 1.200E-08 |
| 10179 | 509 | C509Y | A | 1.1e-08 | 7e-09 | 7.9e-09 | 8.633E-09 | 2.098E-09 | 7.000E-09 | 1.100E-08 |
| 10180 | 509 | C509V | A | 6.7e-09 | 2.7e-09 | 1.3e-08 | 7.467E-09 | 5.193E-09 | 2.700E-09 | 1.300E-08 |
| 10181 | 510 | S510A | A | 1.4e-08 | 4.8e-09 | 1.1e-08 | 9.933E-09 | 4.692E-09 | 4.800E-09 | 1.400E-08 |
| 10182 | 510 | S510R | A | 1.3e-08 | 7.6e-09 | 1.1e-08 | 1.053E-08 | 2.730E-09 | 7.600E-09 | 1.300E-08 |
| 10183 | 510 | S510N | A | 2e-08 | 4.6e-09 | 5.6e-09 | 1.007E-08 | 8.617E-09 | 4.600E-09 | 2.000E-08 |
| 10184 | 510 | S510D | A | 5.9e-09 | 1.8e-08 | 1e-08 | 1.130E-08 | 6.154E-09 | 5.900E-09 | 1.800E-08 |
| 10185 | 510 | S510C | A | 4.3e-09 | 9.6e-09 | 8.7e-09 | 7.533E-09 | 2.836E-09 | 4.300E-09 | 9.600E-09 |
| 10186 | 510 | S510E | A | 3.7e-09 | 6e-09 | 2.1e-08 | 1.023E-08 | 9.395E-09 | 3.700E-09 | 2.100E-08 |
| 10187 | 510 | S510Q | A | 2.2e-08 | 2.1e-09 | 1e-08 | 1.137E-08 | 1.002E-08 | 2.100E-09 | 2.200E-08 |
| 10188 | 510 | S510G | A | 6.8e-09 | 4.5e-09 | 7.5e-09 | 6.267E-09 | 1.570E-09 | 4.500E-09 | 7.500E-09 |
| 10189 | 510 | S510H | A | 8.5e-09 | 1.2e-08 | 6.6e-09 | 9.033E-09 | 2.739E-09 | 6.600E-09 | 1.200E-08 |
| 10190 | 510 | S510I | A | 1.2e-08 | 4.4e-09 | 2.8e-09 | 6.400E-09 | 4.915E-09 | 2.800E-09 | 1.200E-08 |
| 10191 | 510 | S510L | A | 4.1e-09 | 2.8e-09 | 3.5e-09 | 3.467E-09 | 6.506E-10 | 2.800E-09 | 4.100E-09 |
| 10192 | 510 | S510K | A | 7.6e-09 | 8.9e-09 | 3.7e-09 | 6.733E-09 | 2.706E-09 | 3.700E-09 | 8.900E-09 |
| 10193 | 510 | S510M | A | 4.3e-09 | 7.7e-09 | 1.9e-08 | 1.033E-08 | 7.696E-09 | 4.300E-09 | 1.900E-08 |
| 10194 | 510 | S510F | A | 1.1e-08 | 3.6e-09 | 8.9e-09 | 7.833E-09 | 3.814E-09 | 3.600E-09 | 1.100E-08 |
| 10195 | 510 | S510P | A | 5.5e-09 | 2.2e-09 | 4.8e-09 | 4.167E-09 | 1.739E-09 | 2.200E-09 | 5.500E-09 |
| 10196 | 510 | S510S | A | 1.3e-08 | 8.3e-09 | 3.2e-09 | 8.167E-09 | 4.901E-09 | 3.200E-09 | 1.300E-08 |
| 10197 | 510 | S510T | A | 6.6e-09 | 1e-08 | 6.8e-09 | 7.800E-09 | 1.908E-09 | 6.600E-09 | 1.000E-08 |
| 10198 | 510 | S510W | A | 5e-09 | 1e-08 | 5.9e-09 | 6.967E-09 | 2.665E-09 | 5.000E-09 | 1.000E-08 |
| 10199 | 510 | S510Y | A | 1e-08 | 2.9e-09 | 7e-09 | 6.633E-09 | 3.564E-09 | 2.900E-09 | 1.000E-08 |

| | | | | | | | | | | |
|---|---|---|---|---|---|---|---|---|---|---|
| 10200 | 510 | S510V | A | 6e-09 | 8.3e-09 | 1e-08 | 8.100E-09 | 2.007E-09 | 6.000E-09 | 1.000E-08 |
| 10201 | 511 | Q511A | A | 7.6e-09 | 1.3e-08 | 4e-09 | 8.200E-09 | 4.530E-09 | 4.000E-09 | 1.300E-08 |
| 10202 | 511 | Q511R | A | 2.2e-08 | 4.4e-09 | 1.6e-08 | 1.413E-08 | 8.947E-09 | 4.400E-09 | 2.200E-08 |
| 10203 | 511 | Q511N | A | 1.8e-08 | 1e-08 | 4.6e-09 | 1.087E-08 | 6.742E-09 | 4.600E-09 | 1.800E-08 |
| 10204 | 511 | Q511D | A | 5.3e-09 | 3.1e-08 | 6.2e-09 | 1.417E-08 | 1.459E-08 | 5.300E-09 | 3.100E-08 |
| 10205 | 511 | Q511C | A | 1.5e-08 | 9.2e-09 | 8.2e-09 | 1.080E-08 | 3.672E-09 | 8.200E-09 | 1.500E-08 |
| 10206 | 511 | Q511E | A | 1e-08 | 5.8e-09 | 7.6e-09 | 7.800E-09 | 2.107E-09 | 5.800E-09 | 1.000E-08 |
| 10207 | 511 | Q511Q | A | 4.9e-09 | 7.3e-09 | 6.1e-09 | 6.100E-09 | 1.200E-09 | 4.900E-09 | 7.300E-09 |
| 10208 | 511 | Q511G | A | 8.6e-09 | 2.4e-09 | 3.5e-09 | 4.833E-09 | 3.308E-09 | 2.400E-09 | 8.600E-09 |
| 10209 | 511 | Q511H | A | 2.2e-09 | 1e-08 | 3.7e-09 | 5.300E-09 | 4.139E-09 | 2.200E-09 | 1.000E-08 |
| 10210 | 511 | Q511I | A | 6.5e-09 | 2.1e-09 | 2.8e-08 | 1.220E-08 | 1.386E-08 | 2.100E-09 | 2.800E-08 |
| 10211 | 511 | Q511L | A | 1.1e-08 | 1.9e-08 | 1.5e-08 | 1.500E-08 | 4.000E-09 | 1.100E-08 | 1.900E-08 |
| 10212 | 511 | Q511K | A | 7.7e-09 | 1.1e-08 | 4.6e-09 | 7.767E-09 | 3.201E-09 | 4.600E-09 | 1.100E-08 |
| 10213 | 511 | Q511M | A | 1.1e-08 | 7.6e-09 | 1.6e-08 | 1.153E-08 | 4.225E-09 | 7.600E-09 | 1.600E-08 |
| 10214 | 511 | Q511F | A | 1.1e-08 | 9.4e-09 | 1.6e-08 | 1.213E-08 | 3.443E-09 | 9.400E-09 | 1.600E-08 |
| 10215 | 511 | Q511P | A | 4.9e-09 | 5.5e-09 | 1.4e-08 | 8.133E-09 | 5.090E-09 | 4.900E-09 | 1.400E-08 |
| 10216 | 511 | Q511S | A | 6.1e-09 | 1.6e-08 | 1.4e-08 | 1.203E-08 | 5.235E-09 | 6.100E-09 | 1.600E-08 |
| 10217 | 511 | Q511T | A | 4.3e-09 | 7.7e-09 | 3.8e-09 | 5.267E-09 | 2.122E-09 | 3.800E-09 | 7.700E-09 |
| 10218 | 511 | Q511W | A | 1.2e-08 | 1.2e-08 | 6.6e-09 | 1.020E-08 | 3.118E-09 | 6.600E-09 | 1.200E-08 |
| 10219 | 511 | Q511Y | A | 9.1e-09 | 1.6e-08 | 5.5e-09 | 1.020E-08 | 5.336E-09 | 5.500E-09 | 1.600E-08 |
| 10220 | 511 | Q511V | A | 6.7e-09 | 1.3e-08 | 8.7e-09 | 9.467E-09 | 3.219E-09 | 6.700E-09 | 1.300E-08 |
| 10221 | 512 | F512A | A | 1.1e-08 | 3.7e-09 | 5.3e-09 | 6.667E-09 | 3.837E-09 | 3.700E-09 | 1.100E-08 |
| 10222 | 512 | F512R | A | 9.5e-09 | 4.1e-09 | 8.4e-09 | 7.333E-09 | 2.854E-09 | 4.100E-09 | 9.500E-09 |
| 10223 | 512 | F512N | A | 4.8e-09 | 6.1e-09 | 1.2e-08 | 7.633E-09 | 3.837E-09 | 4.800E-09 | 1.200E-08 |
| 10224 | 512 | F512D | A | 3.3e-09 | 5.3e-09 | 1.5e-08 | 7.867E-09 | 6.258E-09 | 3.300E-09 | 1.500E-08 |
| 10225 | 512 | F512C | A | 5.6e-09 | 7.2e-09 | 4.3e-09 | 5.700E-09 | 1.453E-09 | 4.300E-09 | 7.200E-09 |
| 10226 | 512 | F512E | A | 2e-08 | 9.3e-09 | 7e-09 | 1.210E-08 | 6.938E-09 | 7.000E-09 | 2.000E-08 |
| 10227 | 512 | F512Q | A | 4e-09 | 2.8e-09 | 7.8e-09 | 4.867E-09 | 2.610E-09 | 2.800E-09 | 7.800E-09 |
| 10228 | 512 | F512G | A | 3.2e-09 | 1.2e-08 | 1.1e-08 | 8.733E-09 | 4.818E-09 | 3.200E-09 | 1.200E-08 |
| 10229 | 512 | F512H | A | 1e-08 | 4.4e-09 | 2e-08 | 1.147E-08 | 7.903E-09 | 4.400E-09 | 2.000E-08 |
| 10230 | 512 | F512I | A | 2e-08 | 4.9e-09 | 4.2e-09 | 9.700E-09 | 8.927E-09 | 4.200E-09 | 2.000E-08 |
| 10231 | 512 | F512L | A | 4.4e-09 | 6.1e-09 | 6.3e-09 | 5.600E-09 | 1.044E-09 | 4.400E-09 | 6.300E-09 |
| 10232 | 512 | F512K | A | 1.3e-08 | 4.7e-09 | 3.1e-09 | 6.933E-09 | 5.314E-09 | 3.100E-09 | 1.300E-08 |
| 10233 | 512 | F512M | A | 2.6e-09 | 7.4e-09 | 1.3e-08 | 7.667E-09 | 5.205E-09 | 2.600E-09 | 1.300E-08 |
| 10234 | 512 | F512F | A | 8e-09 | 5.4e-09 | 3.8e-09 | 5.733E-09 | 2.120E-09 | 3.800E-09 | 8.000E-09 |
| 10235 | 512 | F512P | A | 1.3e-08 | 6.4e-09 | 6.9e-09 | 8.767E-09 | 3.675E-09 | 6.400E-09 | 1.300E-08 |
| 10236 | 512 | F512S | A | 9.6e-09 | 3.8e-09 | 2.1e-09 | 5.167E-09 | 3.932E-09 | 2.100E-09 | 9.600E-09 |
| 10237 | 512 | F512T | A | 3.3e-09 | 2.1e-08 | 1.3e-08 | 1.243E-08 | 8.864E-09 | 3.300E-09 | 2.100E-08 |
| 10238 | 512 | F512W | A | 3.7e-09 | 1.8e-08 | 9.7e-09 | 1.047E-08 | 7.181E-09 | 3.700E-09 | 1.800E-08 |
| 10239 | 512 | F512Y | A | 1.1e-08 | 4.3e-09 | 5.6e-09 | 6.967E-09 | 3.553E-09 | 4.300E-09 | 1.100E-08 |
| 10240 | 512 | F512V | A | 9.5e-09 | 2.8e-09 | 4.2e-09 | 5.500E-09 | 3.534E-09 | 2.800E-09 | 9.500E-09 |
| 10241 | 513 | L513A | A | 4.4e-09 | 7.2e-09 | 1e-08 | 7.200E-09 | 2.800E-09 | 4.400E-09 | 1.000E-08 |
| 10242 | 513 | L513R | A | 8.5e-09 | 6.8e-09 | 6.4e-09 | 7.233E-09 | 1.115E-09 | 6.400E-09 | 8.500E-09 |
| 10243 | 513 | L513N | A | 5.8e-09 | 5.3e-09 | 7e-09 | 6.033E-09 | 8.737E-10 | 5.300E-09 | 7.000E-09 |
| 10244 | 513 | L513D | A | 1.1e-08 | 2.1e-08 | 1.4e-08 | 1.533E-08 | 5.132E-09 | 1.100E-08 | 2.100E-08 |
| 10245 | 513 | L513C | A | 4.2e-08 | 2.5e-09 | 7.1e-09 | 1.720E-08 | 2.160E-08 | 2.500E-09 | 4.200E-08 |
| 10246 | 513 | L513E | A | 1.7e-08 | 3.3e-09 | 6.9e-09 | 9.067E-09 | 7.102E-09 | 3.300E-09 | 1.700E-08 |
| 10247 | 513 | L513Q | A | 9.7e-09 | 7.7e-09 | 1.4e-08 | 1.047E-08 | 3.219E-09 | 7.700E-09 | 1.400E-08 |
| 10248 | 513 | L513G | A | 3.5e-09 | 1.2e-09 | 1.1e-08 | 5.233E-09 | 5.125E-09 | 1.200E-09 | 1.100E-08 |
| 10249 | 513 | L513H | A | 1.4e-08 | 1.1e-08 | 4.8e-09 | 9.933E-09 | 4.692E-09 | 4.800E-09 | 1.400E-08 |
| 10250 | 513 | L513I | A | 7.1e-09 | 1e-08 | 1e-08 | 9.033E-09 | 1.674E-09 | 7.100E-09 | 1.000E-08 |
| 10251 | 513 | L513L | A | 9.5e-09 | 7.3e-09 | 6.9e-09 | 7.900E-09 | 1.400E-09 | 6.900E-09 | 9.500E-09 |
| 10252 | 513 | L513K | A | 7.9e-09 | 1.3e-08 | 6.3e-09 | 9.067E-09 | 3.499E-09 | 6.300E-09 | 1.300E-08 |
| 10253 | 513 | L513M | A | 1.1e-08 | 2.2e-08 | 5.4e-09 | 1.280E-08 | 8.445E-09 | 5.400E-09 | 2.200E-08 |
| 10254 | 513 | L513F | A | 1.1e-08 | 1.2e-08 | 5.6e-09 | 9.533E-09 | 3.443E-09 | 5.600E-09 | 1.200E-08 |
| 10255 | 513 | L513P | A | 5.9e-09 | 6.8e-09 | 3.8e-09 | 5.500E-09 | 1.539E-09 | 3.800E-09 | 6.800E-09 |
| 10256 | 513 | L513S | A | 6.4e-09 | 2.9e-09 | 6.9e-09 | 5.400E-09 | 2.179E-09 | 2.900E-09 | 6.900E-09 |
| 10257 | 513 | L513T | A | 1.8e-08 | 6.6e-09 | 5.8e-09 | 1.013E-08 | 6.824E-09 | 5.800E-09 | 1.800E-08 |
| 10258 | 513 | L513W | A | 4.3e-09 | 1.2e-08 | 8.4e-09 | 8.233E-09 | 3.853E-09 | 4.300E-09 | 1.200E-08 |
| 10259 | 513 | L513Y | A | 1.5e-08 | 9.8e-09 | 9.2e-09 | 1.133E-08 | 3.190E-09 | 9.200E-09 | 1.500E-08 |
| 10260 | 513 | L513V | A | 5.4e-09 | 2.8e-09 | 4.7e-09 | 4.300E-09 | 1.345E-09 | 2.800E-09 | 5.400E-09 |
| 10261 | 514 | R514A | A | 3.5e-09 | 3e-09 | 9.2e-09 | 5.233E-09 | 3.444E-09 | 3.000E-09 | 9.200E-09 |
| 10262 | 514 | R514R | A | 2.2e-09 | 5.8e-09 | 1.4e-08 | 7.333E-09 | 6.048E-09 | 2.200E-09 | 1.400E-08 |
| 10263 | 514 | R514N | A | 7e-09 | 7.5e-09 | 5.9e-09 | 6.800E-09 | 8.185E-10 | 5.900E-09 | 7.500E-09 |
| 10264 | 514 | R514D | A | 6.6e-09 | 5.7e-09 | 4.7e-09 | 5.667E-09 | 9.504E-10 | 4.700E-09 | 6.600E-09 |
| 10265 | 514 | R514C | A | 1.1e-08 | 1.5e-08 | 4.6e-09 | 1.020E-08 | 5.246E-09 | 4.600E-09 | 1.500E-08 |
| 10266 | 514 | R514E | A | 6.8e-09 | 6.5e-09 | 2.6e-09 | 5.300E-09 | 2.343E-09 | 2.600E-09 | 6.800E-09 |
| 10267 | 514 | R514Q | A | 8.7e-09 | 5.2e-09 | 5e-09 | 6.300E-09 | 2.081E-09 | 5.000E-09 | 8.700E-09 |
| 10268 | 514 | R514G | A | 1.4e-09 | 1.1e-08 | 2.6e-08 | 1.280E-08 | 1.240E-08 | 1.400E-09 | 2.600E-08 |
| 10269 | 514 | R514H | A | 7e-09 | 6.4e-09 | 1.6e-09 | 5.000E-09 | 2.960E-09 | 1.600E-09 | 7.000E-09 |
| 10270 | 514 | R514I | A | 3.1e-09 | 2.6e-08 | 5.2e-09 | 1.143E-08 | 1.266E-08 | 3.100E-09 | 2.600E-08 |
| 10271 | 514 | R514L | A | 2e-08 | 1e-08 | 7.1e-09 | 1.237E-08 | 6.768E-09 | 7.100E-09 | 2.000E-08 |
| 10272 | 514 | R514K | A | 3.5e-09 | 7.3e-09 | 8.2e-09 | 6.333E-09 | 2.495E-09 | 3.500E-09 | 8.200E-09 |
| 10273 | 514 | R514M | A | 5.8e-09 | 4.8e-09 | 1.2e-08 | 7.533E-09 | 3.900E-09 | 4.800E-09 | 1.200E-08 |
| 10274 | 514 | R514F | A | 5.7e-09 | 1.5e-08 | 1e-08 | 1.023E-08 | 4.654E-09 | 5.700E-09 | 1.500E-08 |
| 10275 | 514 | R514P | A | 9e-09 | 1e-08 | 9.1e-09 | 9.367E-09 | 5.508E-10 | 9.000E-09 | 1.000E-08 |
| 10276 | 514 | R514S | A | 7.3e-09 | 3.8e-09 | 6.3e-09 | 5.800E-09 | 1.803E-09 | 3.800E-09 | 7.300E-09 |
| 10277 | 514 | R514T | A | 1.6e-08 | 6.3e-09 | 6.4e-09 | 9.567E-09 | 5.572E-09 | 6.300E-09 | 1.600E-08 |
| 10278 | 514 | R514W | A | 7.3e-09 | 8.3e-09 | 1.9e-09 | 5.833E-09 | 3.443E-09 | 1.900E-09 | 8.300E-09 |
| 10279 | 514 | R514Y | A | 1.6e-08 | 8.7e-09 | 6.4e-09 | 1.037E-08 | 5.012E-09 | 6.400E-09 | 1.600E-08 |
| 10280 | 514 | R514V | A | 9.3e-09 | 6.1e-09 | 7.7e-09 | 7.700E-09 | 1.600E-09 | 6.100E-09 | 9.300E-09 |
| 10281 | 515 | G515A | A | 1.3e-08 | 8.8e-09 | 6.2e-09 | 9.333E-09 | 3.431E-09 | 6.200E-09 | 1.300E-08 |
| 10282 | 515 | G515R | A | 1.2e-08 | 4.1e-09 | 1.1e-08 | 9.033E-09 | 4.302E-09 | 4.100E-09 | 1.200E-08 |
| 10283 | 515 | G515N | A | 4.6e-09 | 7.3e-09 | 3.4e-09 | 5.100E-09 | 1.997E-09 | 3.400E-09 | 7.300E-09 |
| 10284 | 515 | G515D | A | 3.2e-09 | 6.5e-09 | 1.7e-08 | 8.900E-09 | 7.206E-09 | 3.200E-09 | 1.700E-08 |

| | | | | | | | | | | |
|---|---|---|---|---|---|---|---|---|---|---|
| 10285 | 515 | G515C | A | 2e-08 | 1e-08 | 4.5e-09 | 1.150E-08 | 7.858E-09 | 4.500E-09 | 2.000E-08 |
| 10286 | 515 | G515E | A | 4.8e-09 | 8.2e-09 | 3.3e-09 | 5.433E-09 | 2.511E-09 | 3.300E-09 | 8.200E-09 |
| 10287 | 515 | G515Q | A | 7.6e-09 | 1.3e-08 | 8.5e-09 | 9.700E-09 | 2.893E-09 | 7.600E-09 | 1.300E-08 |
| 10288 | 515 | G515G | A | 5.2e-09 | 6.4e-09 | 4.8e-09 | 5.467E-09 | 8.327E-10 | 4.800E-09 | 6.400E-09 |
| 10289 | 515 | G515H | A | 9.3e-09 | 4.4e-09 | 9e-09 | 7.567E-09 | 2.747E-09 | 4.400E-09 | 9.300E-09 |
| 10290 | 515 | G515I | A | 5.4e-08 | 9.2e-09 | 1.1e-08 | 2.473E-08 | 2.536E-08 | 9.200E-09 | 5.400E-08 |
| 10291 | 515 | G515L | A | 9.3e-09 | 5.1e-09 | 7.7e-09 | 7.367E-09 | 2.120E-09 | 5.100E-09 | 9.300E-09 |
| 10292 | 515 | G515K | A | 2.1e-08 | 6.9e-09 | 5.2e-09 | 1.103E-08 | 8.673E-09 | 5.200E-09 | 2.100E-08 |
| 10293 | 515 | G515M | A | 1.3e-08 | 1.1e-08 | 1e-08 | 1.133E-08 | 1.528E-09 | 1.000E-08 | 1.300E-08 |
| 10294 | 515 | G515F | A | 3e-09 | 9.6e-09 | 6.9e-09 | 6.500E-09 | 3.318E-09 | 3.000E-09 | 9.600E-09 |
| 10295 | 515 | G515P | A | 7.6e-09 | 1.1e-08 | 7.2e-09 | 8.600E-09 | 2.088E-09 | 7.200E-09 | 1.100E-08 |
| 10296 | 515 | G515S | A | 3.5e-09 | 4.2e-09 | 4.2e-09 | 3.967E-09 | 4.041E-10 | 3.500E-09 | 4.200E-09 |
| 10297 | 515 | G515T | A | 3.9e-09 | 2.1e-08 | 7.2e-09 | 1.070E-08 | 9.071E-09 | 3.900E-09 | 2.100E-08 |
| 10298 | 515 | G515W | A | 1.6e-08 | 8.9e-09 | 5.4e-09 | 1.010E-08 | 5.401E-09 | 5.400E-09 | 1.600E-08 |
| 10299 | 515 | G515Y | A | 7.1e-09 | 2.4e-08 | 7.9e-09 | 1.300E-08 | 9.535E-09 | 7.100E-09 | 2.400E-08 |
| 10300 | 515 | G515V | A | 4.4e-09 | 5.7e-09 | 3e-08 | 1.337E-08 | 1.442E-08 | 4.400E-09 | 3.000E-08 |
| 10301 | 516 | Q516A | A | 1.3e-08 | 1.2e-08 | 9.2e-09 | 1.140E-08 | 1.970E-09 | 9.200E-09 | 1.300E-08 |
| 10302 | 516 | Q516R | A | 9.6e-09 | 4.7e-09 | 3.2e-09 | 5.833E-09 | 3.347E-09 | 3.200E-09 | 9.600E-09 |
| 10303 | 516 | Q516N | A | 9.3e-09 | 1.8e-09 | 1.1e-08 | 7.367E-09 | 4.895E-09 | 1.800E-09 | 1.100E-08 |
| 10304 | 516 | Q516D | A | 9.5e-09 | 1.3e-08 | 1.3e-08 | 1.183E-08 | 2.021E-09 | 9.500E-09 | 1.300E-08 |
| 10305 | 516 | Q516C | A | 2.8e-09 | 2e-09 | 4.4e-09 | 3.067E-09 | 1.222E-09 | 2.000E-09 | 4.400E-09 |
| 10306 | 516 | Q516E | A | 7.9e-09 | 1.5e-08 | 7e-09 | 9.967E-09 | 4.382E-09 | 7.000E-09 | 1.500E-08 |
| 10307 | 516 | Q516Q | A | 9e-09 | 1.6e-08 | 6.2e-10 | 8.540E-09 | 7.700E-09 | 6.200E-10 | 1.600E-08 |
| 10308 | 516 | Q516G | A | 6.1e-09 | 1.9e-08 | 6.6e-09 | 1.057E-08 | 7.308E-09 | 6.100E-09 | 1.900E-08 |
| 10309 | 516 | Q516H | A | 1.3e-08 | 9.4e-09 | 1.1e-08 | 1.113E-08 | 1.804E-09 | 9.400E-09 | 1.300E-08 |
| 10310 | 516 | Q516I | A | 4.3e-09 | 3.8e-09 | 1.3e-08 | 7.033E-09 | 5.173E-09 | 3.800E-09 | 1.300E-08 |
| 10311 | 516 | Q516L | A | 4.4e-09 | 8.6e-09 | 3.3e-09 | 5.433E-09 | 2.797E-09 | 3.300E-09 | 8.600E-09 |
| 10312 | 516 | Q516K | A | 3.1e-09 | 7.5e-09 | 6.2e-09 | 5.600E-09 | 2.261E-09 | 3.100E-09 | 7.500E-09 |
| 10313 | 516 | Q516M | A | 7.5e-09 | 7.9e-09 | 7.4e-09 | 7.600E-09 | 2.646E-10 | 7.400E-09 | 7.900E-09 |
| 10314 | 516 | Q516F | A | 9e-09 | 4e-09 | 6e-09 | 6.333E-09 | 2.517E-09 | 4.000E-09 | 9.000E-09 |
| 10315 | 516 | Q516P | A | 1e-08 | 5.1e-09 | 4.5e-09 | 6.533E-09 | 3.017E-09 | 4.500E-09 | 1.000E-08 |
| 10316 | 516 | Q516S | A | 7.1e-09 | 2.1e-08 | 3.3e-09 | 1.047E-08 | 9.318E-09 | 3.300E-09 | 2.100E-08 |
| 10317 | 516 | Q516T | A | 5.8e-09 | 7.6e-09 | 8.1e-09 | 7.167E-09 | 1.210E-09 | 5.800E-09 | 8.100E-09 |
| 10318 | 516 | Q516W | A | 6.5e-09 | 1.1e-08 | 2.8e-09 | 6.767E-09 | 4.106E-09 | 2.800E-09 | 1.100E-08 |
| 10319 | 516 | Q516Y | A | 6.3e-09 | 5.4e-09 | 7.7e-09 | 6.467E-09 | 1.159E-09 | 5.400E-09 | 7.700E-09 |
| 10320 | 516 | Q516V | A | 9.1e-09 | 9.9e-09 | 4.9e-09 | 7.967E-09 | 2.686E-09 | 4.900E-09 | 9.900E-09 |
| 10321 | 517 | E517A | A | 9.9e-09 | 1.2e-08 | 8.9e-09 | 1.027E-08 | 1.582E-09 | 8.900E-09 | 1.200E-08 |
| 10322 | 517 | E517R | A | 6.2e-09 | 6e-09 | 1.3e-08 | 8.400E-09 | 3.985E-09 | 6.000E-09 | 1.300E-08 |
| 10323 | 517 | E517N | A | 9.6e-09 | 7.3e-09 | 1e-08 | 8.967E-09 | 1.457E-09 | 7.300E-09 | 1.000E-08 |
| 10324 | 517 | E517D | A | 5.9e-09 | 7.5e-09 | 2.4e-09 | 5.267E-09 | 2.608E-09 | 2.400E-09 | 7.500E-09 |
| 10325 | 517 | E517C | A | 2.2e-08 | 8.3e-09 | 4.1e-09 | 1.147E-08 | 9.361E-09 | 4.100E-09 | 2.200E-08 |
| 10326 | 517 | E517E | A | 3.8e-09 | 1.7e-08 | 1.1e-08 | 1.060E-08 | 6.609E-09 | 3.800E-09 | 1.700E-08 |
| 10327 | 517 | E517Q | A | 7.5e-09 | 1e-08 | 8.5e-09 | 8.667E-09 | 1.258E-09 | 7.500E-09 | 1.000E-08 |
| 10328 | 517 | E517G | A | 1.2e-08 | 3.9e-09 | 5.4e-09 | 7.100E-09 | 4.309E-09 | 3.900E-09 | 1.200E-08 |
| 10329 | 517 | E517H | A | 7e-09 | 5.9e-09 | 6.7e-09 | 6.533E-09 | 5.686E-10 | 5.900E-09 | 7.000E-09 |
| 10330 | 517 | E517I | A | 2.9e-08 | 9.1e-09 | 2.2e-09 | 1.343E-08 | 1.392E-08 | 2.200E-09 | 2.900E-08 |
| 10331 | 517 | E517L | A | 6.3e-09 | 8.7e-09 | 4.4e-09 | 6.467E-09 | 2.155E-09 | 4.400E-09 | 8.700E-09 |
| 10332 | 517 | E517K | A | 7.7e-09 | 1.4e-08 | 2.6e-09 | 8.100E-09 | 5.711E-09 | 2.600E-09 | 1.400E-08 |
| 10333 | 517 | E517M | A | 7.3e-09 | 7.4e-09 | 1.6e-08 | 1.023E-08 | 4.994E-09 | 7.300E-09 | 1.600E-08 |
| 10334 | 517 | E517F | A | 1e-08 | 3.4e-09 | 3.1e-09 | 5.500E-09 | 3.900E-09 | 3.100E-09 | 1.000E-08 |
| 10335 | 517 | E517P | A | 4.2e-09 | 6.1e-09 | 3.1e-09 | 4.467E-09 | 1.518E-09 | 3.100E-09 | 6.100E-09 |
| 10336 | 517 | E517S | A | 7.4e-09 | 8.5e-09 | 1.7e-09 | 5.867E-09 | 3.650E-09 | 1.700E-09 | 8.500E-09 |
| 10337 | 517 | E517T | A | 4.8e-09 | 2.8e-09 | 8.7e-09 | 5.433E-09 | 3.001E-09 | 2.800E-09 | 8.700E-09 |
| 10338 | 517 | E517W | A | 6.7e-09 | 3.1e-09 | 5.2e-09 | 5.000E-09 | 1.808E-09 | 3.100E-09 | 6.700E-09 |
| 10339 | 517 | E517Y | A | 4.1e-09 | 8e-09 | 6.4e-09 | 6.167E-09 | 1.960E-09 | 4.100E-09 | 8.000E-09 |
| 10340 | 517 | E517V | A | 1.4e-08 | 4e-09 | 1.8e-08 | 1.200E-08 | 7.211E-09 | 4.000E-09 | 1.800E-08 |
| 10341 | 518 | C518A | A | 2e-08 | 7.5e-09 | 6e-09 | 1.117E-08 | 7.687E-09 | 6.000E-09 | 2.000E-08 |
| 10342 | 518 | C518R | A | 5.9e-09 | 7.9e-09 | 6.9e-09 | 6.900E-09 | 1.000E-09 | 5.900E-09 | 7.900E-09 |
| 10343 | 518 | C518N | A | 6.7e-09 | 1.8e-08 | 2.6e-09 | 9.100E-09 | 7.976E-09 | 2.600E-09 | 1.800E-08 |
| 10344 | 518 | C518D | A | 9.2e-09 | 9.9e-09 | 7.2e-09 | 8.767E-09 | 1.401E-09 | 7.200E-09 | 9.900E-09 |
| 10345 | 518 | C518C | A | 2.4e-08 | 6.7e-09 | 2.8e-08 | 1.957E-08 | 1.132E-08 | 6.700E-09 | 2.800E-08 |
| 10346 | 518 | C518E | A | 7.2e-09 | 5.1e-09 | 5.2e-09 | 5.833E-09 | 1.185E-09 | 5.100E-09 | 7.200E-09 |
| 10347 | 518 | C518Q | A | 1.6e-08 | 7.1e-09 | 1.2e-08 | 1.170E-08 | 4.458E-09 | 7.100E-09 | 1.600E-08 |
| 10348 | 518 | C518G | A | 1.7e-08 | 5.4e-09 | 9.1e-09 | 1.050E-08 | 5.925E-09 | 5.400E-09 | 1.700E-08 |
| 10349 | 518 | C518H | A | 9.4e-09 | 8.9e-09 | 4.7e-09 | 7.667E-09 | 2.581E-09 | 4.700E-09 | 9.400E-09 |
| 10350 | 518 | C518I | A | 1.1e-08 | 1.1e-08 | 6.8e-09 | 9.600E-09 | 2.425E-09 | 6.800E-09 | 1.100E-08 |
| 10351 | 518 | C518L | A | 2.5e-08 | 3.5e-09 | 1.2e-08 | 1.350E-08 | 1.083E-08 | 3.500E-09 | 2.500E-08 |
| 10352 | 518 | C518K | A | 6.4e-09 | 1.5e-08 | 7.3e-09 | 9.567E-09 | 4.727E-09 | 6.400E-09 | 1.500E-08 |
| 10353 | 518 | C518M | A | 1.8e-08 | 1.2e-08 | 1.7e-08 | 1.567E-08 | 3.215E-09 | 1.200E-08 | 1.800E-08 |
| 10354 | 518 | C518F | A | 1.8e-08 | 9e-09 | 3.2e-09 | 1.007E-08 | 7.457E-09 | 3.200E-09 | 1.800E-08 |
| 10355 | 518 | C518P | A | 1.2e-08 | 6.8e-08 | 2.6e-09 | 2.753E-08 | 3.536E-08 | 2.600E-09 | 6.800E-08 |
| 10356 | 518 | C518S | A | 1e-08 | 4.2e-09 | 3e-09 | 5.733E-09 | 3.743E-09 | 3.000E-09 | 1.000E-08 |
| 10357 | 518 | C518T | A | 1e-08 | 4.3e-09 | 5.6e-09 | 6.633E-09 | 2.987E-09 | 4.300E-09 | 1.000E-08 |
| 10358 | 518 | C518W | A | 6.5e-09 | 8.4e-09 | 2.2e-09 | 5.700E-09 | 3.176E-09 | 2.200E-09 | 8.400E-09 |
| 10359 | 518 | C518Y | A | 2.9e-09 | 3.2e-09 | 4.7e-09 | 3.600E-09 | 9.644E-10 | 2.900E-09 | 4.700E-09 |
| 10360 | 518 | C518V | A | 1.1e-08 | 1.3e-08 | 5.5e-09 | 9.833E-09 | 3.884E-09 | 5.500E-09 | 1.300E-08 |
| 10361 | 519 | V519A | A | 5.2e-09 | 1e-08 | 4.3e-09 | 6.500E-09 | 3.064E-09 | 4.300E-09 | 1.000E-08 |
| 10362 | 519 | V519R | A | 5.2e-09 | 6.3e-09 | 7.8e-09 | 6.433E-09 | 1.305E-09 | 5.200E-09 | 7.800E-09 |
| 10363 | 519 | V519N | A | 4.2e-09 | 7.1e-09 | 6.4e-09 | 5.900E-09 | 1.513E-09 | 4.200E-09 | 7.100E-09 |
| 10364 | 519 | V519D | A | 3.7e-09 | 5.8e-09 | 8.7e-09 | 6.067E-09 | 2.511E-09 | 3.700E-09 | 8.700E-09 |
| 10365 | 519 | V519C | A | 6e-09 | 3.7e-09 | 5.1e-09 | 4.933E-09 | 1.159E-09 | 3.700E-09 | 6.000E-09 |
| 10366 | 519 | V519E | A | 7.8e-09 | 3.5e-09 | 3.7e-09 | 5.000E-09 | 2.427E-09 | 3.500E-09 | 7.800E-09 |
| 10367 | 519 | V519Q | A | 3.3e-09 | 3.4e-09 | 1.3e-08 | 6.567E-09 | 5.572E-09 | 3.300E-09 | 1.300E-08 |
| 10368 | 519 | V519G | A | 1.1e-08 | 2.6e-09 | 1.4e-08 | 9.200E-09 | 5.909E-09 | 2.600E-09 | 1.400E-08 |
| 10369 | 519 | V519H | A | 3.9e-09 | 1.9e-08 | 6.6e-09 | 9.833E-09 | 8.053E-09 | 3.900E-09 | 1.900E-08 |

| | | | | | | | | | | |
|---|---|---|---|---|---|---|---|---|---|---|
| 10370 | 519 | V519I | A | 1.2e-08 | 5.2e-09 | 2.2e-09 | 6.467E-09 | 5.021E-09 | 2.200E-09 | 1.200E-08 |
| 10371 | 519 | V519L | A | 8.2e-09 | 6.9e-09 | 9.2e-09 | 8.100E-09 | 1.153E-09 | 6.900E-09 | 9.200E-09 |
| 10372 | 519 | V519K | A | 1.3e-09 | 8.3e-09 | 4.9e-09 | 4.833E-09 | 3.500E-09 | 1.300E-09 | 8.300E-09 |
| 10373 | 519 | V519M | A | 6.6e-09 | 4.2e-09 | 8.2e-09 | 6.333E-09 | 2.013E-09 | 4.200E-09 | 8.200E-09 |
| 10374 | 519 | V519F | A | 7.3e-09 | 4.6e-09 | 2.3e-09 | 4.733E-09 | 2.503E-09 | 2.300E-09 | 7.300E-09 |
| 10375 | 519 | V519P | A | 1.8e-09 | 2.6e-09 | 1.6e-08 | 6.800E-09 | 7.977E-09 | 1.800E-09 | 1.600E-08 |
| 10376 | 519 | V519S | A | 3.3e-09 | 8.8e-09 | 9.6e-09 | 7.233E-09 | 3.430E-09 | 3.300E-09 | 9.600E-09 |
| 10377 | 519 | V519T | A | 1.7e-08 | 8.4e-09 | 6.6e-09 | 1.067E-08 | 5.558E-09 | 6.600E-09 | 1.700E-08 |
| 10378 | 519 | V519W | A | 2.6e-08 | 6e-09 | 1.3e-08 | 1.500E-08 | 1.015E-08 | 6.000E-09 | 2.600E-08 |
| 10379 | 519 | V519Y | A | 3.8e-09 | 6.5e-09 | 4e-09 | 4.767E-09 | 1.504E-09 | 3.800E-09 | 6.500E-09 |
| 10380 | 519 | V519V | A | 1.8e-08 | 9e-09 | 4.3e-09 | 1.043E-08 | 6.962E-09 | 4.300E-09 | 1.800E-08 |
| 10381 | 520 | E520A | A | 7.7e-09 | 5e-09 | 5.6e-09 | 6.100E-09 | 1.418E-09 | 5.000E-09 | 7.700E-09 |
| 10382 | 520 | E520R | A | 1.4e-08 | 5.7e-09 | 7.1e-09 | 8.933E-09 | 4.443E-09 | 5.700E-09 | 1.400E-08 |
| 10383 | 520 | E520N | A | 4.7e-09 | 9.3e-09 | 8.6e-09 | 7.533E-09 | 2.479E-09 | 4.700E-09 | 9.300E-09 |
| 10384 | 520 | E520D | A | 1.9e-09 | 6.7e-09 | 1.2e-08 | 6.867E-09 | 5.052E-09 | 1.900E-09 | 1.200E-08 |
| 10385 | 520 | E520C | A | 1e-08 | 7.4e-09 | 1.6e-08 | 1.113E-08 | 4.411E-09 | 7.400E-09 | 1.600E-08 |
| 10386 | 520 | E520E | A | 4e-09 | 5.3e-09 | 7.4e-09 | 5.567E-09 | 1.716E-09 | 4.000E-09 | 7.400E-09 |
| 10387 | 520 | E520Q | A | 1.3e-08 | 4e-09 | 1.4e-08 | 1.033E-08 | 5.508E-09 | 4.000E-09 | 1.400E-08 |
| 10388 | 520 | E520G | A | 5.8e-09 | 4.4e-09 | 1.2e-08 | 7.400E-09 | 4.045E-09 | 4.400E-09 | 1.200E-08 |
| 10389 | 520 | E520H | A | 5.3e-09 | 5.2e-09 | 8.2e-09 | 6.233E-09 | 1.704E-09 | 5.200E-09 | 8.200E-09 |
| 10390 | 520 | E520I | A | 1e-08 | 2.8e-09 | 1.3e-08 | 8.600E-09 | 5.242E-09 | 2.800E-09 | 1.300E-08 |
| 10391 | 520 | E520L | A | 9.2e-09 | 4.8e-09 | 2.1e-08 | 1.167E-08 | 8.377E-09 | 4.800E-09 | 2.100E-08 |
| 10392 | 520 | E520K | A | 1.8e-08 | 9.3e-09 | 3.7e-09 | 1.033E-08 | 7.206E-09 | 3.700E-09 | 1.800E-08 |
| 10393 | 520 | E520M | A | 1.4e-08 | 8.1e-09 | 5.7e-09 | 9.267E-09 | 4.271E-09 | 5.700E-09 | 1.400E-08 |
| 10394 | 520 | E520F | A | 1.3e-08 | 2.7e-09 | 3.5e-09 | 6.400E-09 | 5.730E-09 | 2.700E-09 | 1.300E-08 |
| 10395 | 520 | E520P | A | 4.9e-09 | 5.4e-09 | 1.5e-08 | 8.433E-09 | 5.692E-09 | 4.900E-09 | 1.500E-08 |
| 10396 | 520 | E520S | A | 2e-09 | 1.1e-08 | 4.2e-09 | 5.733E-09 | 4.692E-09 | 2.000E-09 | 1.100E-08 |
| 10397 | 520 | E520T | A | 5.2e-09 | 3.7e-09 | 8.4e-09 | 5.767E-09 | 2.401E-09 | 3.700E-09 | 8.400E-09 |
| 10398 | 520 | E520W | A | 4.8e-09 | 4.3e-09 | 2.7e-09 | 3.933E-09 | 1.097E-09 | 2.700E-09 | 4.800E-09 |
| 10399 | 520 | E520Y | A | 1.8e-08 | 9.1e-09 | 6.6e-09 | 1.123E-08 | 5.992E-09 | 6.600E-09 | 1.800E-08 |
| 10400 | 520 | E520V | A | 8.8e-09 | 3.9e-09 | 6.6e-09 | 6.433E-09 | 2.454E-09 | 3.900E-09 | 8.800E-09 |
| 10401 | 521 | E521A | A | 1.9e-08 | 2.3e-09 | 6.7e-09 | 9.333E-09 | 8.656E-09 | 2.300E-09 | 1.900E-08 |
| 10402 | 521 | E521R | A | 1.1e-08 | 1.3e-08 | 8.6e-09 | 1.087E-08 | 2.203E-09 | 8.600E-09 | 1.300E-08 |
| 10403 | 521 | E521N | A | 1.1e-08 | 3.1e-09 | 1.2e-08 | 8.700E-09 | 4.875E-09 | 3.100E-09 | 1.200E-08 |
| 10404 | 521 | E521D | A | 7e-09 | 1.3e-08 | 4.8e-09 | 8.267E-09 | 4.244E-09 | 4.800E-09 | 1.300E-08 |
| 10405 | 521 | E521C | A | 9.7e-09 | 1.2e-08 | 1.4e-08 | 1.190E-08 | 2.152E-09 | 9.700E-09 | 1.400E-08 |
| 10406 | 521 | E521E | A | 7.4e-09 | 1.2e-08 | 8.7e-09 | 9.367E-09 | 2.371E-09 | 7.400E-09 | 1.200E-08 |
| 10407 | 521 | E521Q | A | 1.1e-08 | 4.4e-09 | 1.1e-08 | 8.800E-09 | 3.811E-09 | 4.400E-09 | 1.100E-08 |
| 10408 | 521 | E521G | A | 8e-09 | 2.4e-09 | 6.9e-09 | 5.767E-09 | 2.967E-09 | 2.400E-09 | 8.000E-09 |
| 10409 | 521 | E521H | A | 8.2e-09 | 4.8e-09 | 1.2e-08 | 8.333E-09 | 3.602E-09 | 4.800E-09 | 1.200E-08 |
| 10410 | 521 | E521I | A | 8.1e-09 | 1e-08 | 1.9e-09 | 6.667E-09 | 4.236E-09 | 1.900E-09 | 1.000E-08 |
| 10411 | 521 | E521L | A | 1.3e-08 | 3.6e-09 | 8.4e-09 | 8.333E-09 | 4.700E-09 | 3.600E-09 | 1.300E-08 |
| 10412 | 521 | E521K | A | 1.4e-08 | 1.3e-08 | 2e-09 | 9.667E-09 | 6.658E-09 | 2.000E-09 | 1.400E-08 |
| 10413 | 521 | E521M | A | 7.1e-09 | 3.7e-09 | 8.6e-09 | 6.467E-09 | 2.511E-09 | 3.700E-09 | 8.600E-09 |
| 10414 | 521 | E521F | A | 3.4e-09 | 7.5e-09 | 2.3e-08 | 1.130E-08 | 1.034E-08 | 3.400E-09 | 2.300E-08 |
| 10415 | 521 | E521P | A | 6.4e-09 | 1.4e-08 | 3.2e-09 | 7.867E-09 | 5.547E-09 | 3.200E-09 | 1.400E-08 |
| 10416 | 521 | E521S | A | 4.9e-09 | 1e-08 | 5.2e-09 | 6.700E-09 | 2.862E-09 | 4.900E-09 | 1.000E-08 |
| 10417 | 521 | E521T | A | 7.4e-09 | 1.9e-08 | 2.6e-09 | 9.667E-09 | 8.432E-09 | 2.600E-09 | 1.900E-08 |
| 10418 | 521 | E521W | A | 3.2e-09 | 7.8e-09 | 2e-09 | 4.333E-09 | 3.062E-09 | 2.000E-09 | 7.800E-09 |
| 10419 | 521 | E521Y | A | 1.4e-09 | 8.2e-09 | 5.6e-09 | 5.067E-09 | 3.431E-09 | 1.400E-09 | 8.200E-09 |
| 10420 | 521 | E521V | A | 1.2e-08 | 1.4e-08 | 7e-09 | 1.100E-08 | 3.606E-09 | 7.000E-09 | 1.400E-08 |
| 10421 | 522 | C522A | A | 7.7e-09 | 5.5e-09 | 2.6e-09 | 5.267E-09 | 2.558E-09 | 2.600E-09 | 7.700E-09 |
| 10422 | 522 | C522R | A | 3.3e-09 | 1.5e-08 | 8.2e-09 | 8.833E-09 | 5.876E-09 | 3.300E-09 | 1.500E-08 |
| 10423 | 522 | C522N | A | 5.4e-09 | 5.2e-09 | 4.8e-09 | 5.133E-09 | 3.055E-10 | 4.800E-09 | 5.400E-09 |
| 10424 | 522 | C522D | A | 4.4e-09 | 7.6e-09 | 7.8e-09 | 6.600E-09 | 1.908E-09 | 4.400E-09 | 7.800E-09 |
| 10425 | 522 | C522C | A | 7.1e-09 | 6e-09 | 2.3e-08 | 1.203E-08 | 9.513E-09 | 6.000E-09 | 2.300E-08 |
| 10426 | 522 | C522E | A | 7.9e-09 | 7.1e-09 | 4.2e-09 | 6.400E-09 | 1.947E-09 | 4.200E-09 | 7.900E-09 |
| 10427 | 522 | C522Q | A | 7.7e-09 | 6.8e-09 | 8.9e-09 | 7.800E-09 | 1.054E-09 | 6.800E-09 | 8.900E-09 |
| 10428 | 522 | C522G | A | 4.7e-09 | 3.8e-09 | 1.2e-08 | 6.833E-09 | 4.497E-09 | 3.800E-09 | 1.200E-08 |
| 10429 | 522 | C522H | A | 8.1e-09 | 2.5e-08 | 1.2e-08 | 1.503E-08 | 8.849E-09 | 8.100E-09 | 2.500E-08 |
| 10430 | 522 | C522I | A | 7.7e-09 | 5.6e-09 | 1.1e-08 | 8.100E-09 | 2.722E-09 | 5.600E-09 | 1.100E-08 |
| 10431 | 522 | C522L | A | 8.3e-09 | 5.1e-09 | 1e-08 | 7.800E-09 | 2.488E-09 | 5.100E-09 | 1.000E-08 |
| 10432 | 522 | C522K | A | 8.8e-09 | 4.5e-09 | 1.4e-08 | 9.100E-09 | 4.757E-09 | 4.500E-09 | 1.400E-08 |
| 10433 | 522 | C522M | A | 8.8e-09 | 6.3e-09 | 7.1e-09 | 7.400E-09 | 1.277E-09 | 6.300E-09 | 8.800E-09 |
| 10434 | 522 | C522F | A | 4.6e-09 | 5.3e-08 | 5.1e-09 | 2.090E-08 | 2.780E-08 | 4.600E-09 | 5.300E-08 |
| 10435 | 522 | C522P | A | 1.2e-08 | 1.4e-08 | 1.5e-08 | 1.367E-08 | 1.528E-09 | 1.200E-08 | 1.500E-08 |
| 10436 | 522 | C522S | A | 1e-08 | 8.7e-09 | 2.3e-09 | 7.000E-09 | 4.122E-09 | 2.300E-09 | 1.000E-08 |
| 10437 | 522 | C522T | A | 7.8e-09 | 2.8e-09 | 8.2e-09 | 6.267E-09 | 3.009E-09 | 2.800E-09 | 8.200E-09 |
| 10438 | 522 | C522W | A | 2.3e-09 | 6.1e-09 | 4.7e-09 | 4.367E-09 | 1.922E-09 | 2.300E-09 | 6.100E-09 |
| 10439 | 522 | C522Y | A | 1.9e-08 | 4.1e-09 | 6.6e-09 | 9.900E-09 | 7.979E-09 | 4.100E-09 | 1.900E-08 |
| 10440 | 522 | C522V | A | 8.3e-09 | 4e-09 | 6.9e-09 | 6.400E-09 | 2.193E-09 | 4.000E-09 | 8.300E-09 |
| 10441 | 523 | R523A | A | 9.9e-09 | 6.3e-09 | 4.2e-09 | 6.800E-09 | 2.883E-09 | 4.200E-09 | 9.900E-09 |
| 10442 | 523 | R523R | A | 8.4e-09 | 1.6e-08 | 5.7e-09 | 1.003E-08 | 5.341E-09 | 5.700E-09 | 1.600E-08 |
| 10443 | 523 | R523N | A | 4.5e-09 | 7.3e-09 | 7.9e-09 | 6.567E-09 | 1.815E-09 | 4.500E-09 | 7.900E-09 |
| 10444 | 523 | R523D | A | 8.3e-09 | 4.5e-09 | 1.5e-08 | 9.267E-09 | 5.316E-09 | 4.500E-09 | 1.500E-08 |
| 10445 | 523 | R523C | A | 6.5e-09 | 9.6e-09 | 1e-08 | 8.700E-09 | 1.916E-09 | 6.500E-09 | 1.000E-08 |
| 10446 | 523 | R523E | A | 7.5e-09 | 4.8e-09 | 1.1e-08 | 7.767E-09 | 3.109E-09 | 4.800E-09 | 1.100E-08 |
| 10447 | 523 | R523Q | A | 2.4e-09 | 6.5e-09 | 8.6e-09 | 5.833E-09 | 3.153E-09 | 2.400E-09 | 8.600E-09 |
| 10448 | 523 | R523G | A | 1.7e-08 | 6.2e-09 | 1.4e-08 | 1.240E-08 | 5.575E-09 | 6.200E-09 | 1.700E-08 |
| 10449 | 523 | R523H | A | 4.1e-09 | 8.7e-09 | 3.7e-09 | 5.500E-09 | 2.778E-09 | 3.700E-09 | 8.700E-09 |
| 10450 | 523 | R523I | A | 3.9e-09 | 9.5e-09 | 8e-09 | 7.133E-09 | 2.899E-09 | 3.900E-09 | 9.500E-09 |
| 10451 | 523 | R523L | A | 2.6e-09 | 7.5e-09 | 2.8e-09 | 4.300E-09 | 2.773E-09 | 2.600E-09 | 7.500E-09 |
| 10452 | 523 | R523K | A | 1e-08 | 1.5e-08 | 7.1e-09 | 1.070E-08 | 3.996E-09 | 7.100E-09 | 1.500E-08 |
| 10453 | 523 | R523M | A | 2e-08 | 6.4e-09 | 6.3e-09 | 1.090E-08 | 7.881E-09 | 6.300E-09 | 2.000E-08 |
| 10454 | 523 | R523F | A | 1e-08 | 1e-08 | 1.7e-08 | 1.233E-08 | 4.041E-09 | 1.000E-08 | 1.700E-08 |

| | | | | | | | | | | |
|---|---|---|---|---|---|---|---|---|---|---|
| 10455 | 523 | R523P | A | 4.6e-09 | 8.3e-09 | 7.5e-09 | 6.800E-09 | 1.947E-09 | 4.600E-09 | 8.300E-09 |
| 10456 | 523 | R523S | A | 5.7e-09 | 7.7e-09 | 6.9e-09 | 6.767E-09 | 1.007E-09 | 5.700E-09 | 7.700E-09 |
| 10457 | 523 | R523T | A | 8.3e-09 | 5.4e-09 | 1.7e-08 | 1.023E-08 | 6.037E-09 | 5.400E-09 | 1.700E-08 |
| 10458 | 523 | R523W | A | 1.8e-08 | 3.8e-09 | 1.5e-09 | 7.767E-09 | 8.937E-09 | 1.500E-09 | 1.800E-08 |
| 10459 | 523 | R523Y | A | 9.1e-09 | 1.2e-08 | 1.5e-08 | 1.203E-08 | 2.950E-09 | 9.100E-09 | 1.500E-08 |
| 10460 | 523 | R523V | A | 9.3e-09 | 6.9e-09 | 4.8e-09 | 7.000E-09 | 2.252E-09 | 4.800E-09 | 9.300E-09 |
| 10461 | 524 | V524A | A | 1.1e-08 | 1e-08 | 3.2e-09 | 8.067E-09 | 4.244E-09 | 3.200E-09 | 1.100E-08 |
| 10462 | 524 | V524R | A | 3.1e-08 | 8.1e-09 | 2.6e-08 | 2.170E-08 | 1.204E-08 | 8.100E-09 | 3.100E-08 |
| 10463 | 524 | V524N | A | 5.1e-09 | 5.3e-09 | 2.8e-09 | 4.400E-09 | 1.389E-09 | 2.800E-09 | 5.300E-09 |
| 10464 | 524 | V524D | A | 5e-09 | 5.1e-09 | 2.7e-09 | 4.267E-09 | 1.358E-09 | 2.700E-09 | 5.100E-09 |
| 10465 | 524 | V524C | A | 1.5e-08 | 1.2e-08 | 3.1e-09 | 1.003E-08 | 6.189E-09 | 3.100E-09 | 1.500E-08 |
| 10466 | 524 | V524E | A | 2.6e-09 | 4.9e-09 | 1.6e-08 | 7.833E-09 | 7.165E-09 | 2.600E-09 | 1.600E-08 |
| 10467 | 524 | V524Q | A | 5e-09 | 2.9e-08 | 6.6e-09 | 1.353E-08 | 1.342E-08 | 5.000E-09 | 2.900E-08 |
| 10468 | 524 | V524G | A | 1.2e-08 | 5.1e-09 | 7.6e-09 | 8.233E-09 | 3.493E-09 | 5.100E-09 | 1.200E-08 |
| 10469 | 524 | V524H | A | 5.2e-09 | 7.1e-09 | 9.1e-09 | 7.133E-09 | 1.950E-09 | 5.200E-09 | 9.100E-09 |
| 10470 | 524 | V524I | A | 6.4e-09 | 7.3e-09 | 4.5e-09 | 6.067E-09 | 1.429E-09 | 4.500E-09 | 7.300E-09 |
| 10471 | 524 | V524L | A | 7.9e-09 | 1.1e-08 | 1.2e-08 | 1.030E-08 | 2.138E-09 | 7.900E-09 | 1.200E-08 |
| 10472 | 524 | V524K | A | 9.7e-09 | 1e-08 | 4.3e-09 | 8.000E-09 | 3.208E-09 | 4.300E-09 | 1.000E-08 |
| 10473 | 524 | V524M | A | 1e-08 | 1.3e-08 | 5.9e-09 | 9.633E-09 | 3.564E-09 | 5.900E-09 | 1.300E-08 |
| 10474 | 524 | V524F | A | 1.3e-09 | 2.2e-08 | 8.7e-09 | 1.067E-08 | 1.049E-08 | 1.300E-09 | 2.200E-08 |
| 10475 | 524 | V524P | A | 1.5e-08 | 1.2e-08 | 5.9e-09 | 1.097E-08 | 4.637E-09 | 5.900E-09 | 1.500E-08 |
| 10476 | 524 | V524S | A | 5.7e-09 | 2.8e-09 | 1.1e-08 | 6.500E-09 | 4.158E-09 | 2.800E-09 | 1.100E-08 |
| 10477 | 524 | V524T | A | 8.3e-09 | 7.7e-09 | 3.3e-09 | 6.433E-09 | 2.730E-09 | 3.300E-09 | 8.300E-09 |
| 10478 | 524 | V524W | A | 6.4e-09 | 1.1e-08 | 4.6e-09 | 7.333E-09 | 3.301E-09 | 4.600E-09 | 1.100E-08 |
| 10479 | 524 | V524Y | A | 7.7e-09 | 4.6e-09 | 6.7e-09 | 6.333E-09 | 1.582E-09 | 4.600E-09 | 7.700E-09 |
| 10480 | 524 | V524V | A | 4.5e-09 | 2.9e-09 | 1.3e-08 | 6.800E-09 | 5.429E-09 | 2.900E-09 | 1.300E-08 |
| 10481 | 525 | L525A | A | 9.2e-09 | 6.1e-09 | 6.7e-09 | 7.333E-09 | 1.644E-09 | 6.100E-09 | 9.200E-09 |
| 10482 | 525 | L525R | A | 7e-09 | 1.3e-08 | 1.4e-08 | 1.133E-08 | 3.786E-09 | 7.000E-09 | 1.400E-08 |
| 10483 | 525 | L525N | A | 4.1e-09 | 5.6e-09 | 2.7e-09 | 4.133E-09 | 1.450E-09 | 2.700E-09 | 5.600E-09 |
| 10484 | 525 | L525D | A | 8.2e-09 | 1.1e-08 | 5.1e-09 | 8.100E-09 | 2.951E-09 | 5.100E-09 | 1.100E-08 |
| 10485 | 525 | L525C | A | 1.5e-08 | 8e-09 | 2.4e-08 | 1.567E-08 | 8.021E-09 | 8.000E-09 | 2.400E-08 |
| 10486 | 525 | L525E | A | 7.5e-09 | 1.2e-08 | 4e-09 | 7.833E-09 | 4.010E-09 | 4.000E-09 | 1.200E-08 |
| 10487 | 525 | L525Q | A | 7.6e-09 | 6.3e-09 | 3.6e-09 | 5.833E-09 | 2.040E-09 | 3.600E-09 | 7.600E-09 |
| 10488 | 525 | L525G | A | 3.9e-08 | 2.8e-09 | 5.8e-09 | 1.587E-08 | 2.009E-08 | 2.800E-09 | 3.900E-08 |
| 10489 | 525 | L525H | A | 1.2e-08 | 2.1e-08 | 2.2e-08 | 1.833E-08 | 5.508E-09 | 1.200E-08 | 2.200E-08 |
| 10490 | 525 | L525I | A | 8.5e-09 | 5.3e-09 | 5.5e-09 | 6.433E-09 | 1.793E-09 | 5.300E-09 | 8.500E-09 |
| 10491 | 525 | L525L | A | 1e-08 | 5.9e-09 | 4.2e-09 | 6.700E-09 | 2.982E-09 | 4.200E-09 | 1.000E-08 |
| 10492 | 525 | L525K | A | 1.6e-08 | 3.8e-09 | 5e-09 | 8.267E-09 | 6.724E-09 | 3.800E-09 | 1.600E-08 |
| 10493 | 525 | L525M | A | 2.5e-09 | 7.2e-09 | 8e-09 | 5.900E-09 | 2.972E-09 | 2.500E-09 | 8.000E-09 |
| 10494 | 525 | L525F | A | 1.6e-08 | 5.1e-09 | 3e-09 | 8.033E-09 | 6.979E-09 | 3.000E-09 | 1.600E-08 |
| 10495 | 525 | L525P | A | 8.2e-09 | 5.3e-09 | 3.4e-09 | 5.633E-09 | 2.417E-09 | 3.400E-09 | 8.200E-09 |
| 10496 | 525 | L525S | A | 3.3e-09 | 3.6e-09 | 4.1e-09 | 3.667E-09 | 4.041E-10 | 3.300E-09 | 4.100E-09 |
| 10497 | 525 | L525T | A | 2.3e-08 | 7.9e-09 | 1.9e-08 | 1.663E-08 | 7.823E-09 | 7.900E-09 | 2.300E-08 |
| 10498 | 525 | L525W | A | 4.8e-09 | 5.4e-09 | 3.2e-09 | 4.467E-09 | 1.137E-09 | 3.200E-09 | 5.400E-09 |
| 10499 | 525 | L525Y | A | 6.3e-09 | 1.1e-08 | 5.3e-09 | 7.533E-09 | 3.044E-09 | 5.300E-09 | 1.100E-08 |
| 10500 | 525 | L525V | A | 1.7e-08 | 6.1e-09 | 2.4e-09 | 8.500E-09 | 7.590E-09 | 2.400E-09 | 1.700E-08 |
| 10501 | 526 | Q526A | A | 1.3e-08 | 1.4e-08 | 1.1e-08 | 1.267E-08 | 1.528E-09 | 1.100E-08 | 1.400E-08 |
| 10502 | 526 | Q526R | A | 1.7e-08 | 4.3e-09 | 2.7e-08 | 1.610E-08 | 1.138E-08 | 4.300E-09 | 2.700E-08 |
| 10503 | 526 | Q526N | A | 3.2e-09 | 1.4e-08 | 8.7e-10 | 6.023E-09 | 7.006E-09 | 8.700E-10 | 1.400E-08 |
| 10504 | 526 | Q526D | A | 2.4e-09 | 1.5e-08 | 1.3e-08 | 1.013E-08 | 6.772E-09 | 2.400E-09 | 1.500E-08 |
| 10505 | 526 | Q526C | A | 1.2e-08 | 5.9e-09 | 6.2e-09 | 8.033E-09 | 3.439E-09 | 5.900E-09 | 1.200E-08 |
| 10506 | 526 | Q526E | A | 1.8e-08 | 9.3e-09 | 5.4e-09 | 1.090E-08 | 6.451E-09 | 5.400E-09 | 1.800E-08 |
| 10507 | 526 | Q526Q | A | 4.4e-09 | 1.9e-08 | 7.1e-09 | 1.017E-08 | 7.768E-09 | 4.400E-09 | 1.900E-08 |
| 10508 | 526 | Q526G | A | 4.5e-09 | 1.5e-08 | 1.8e-08 | 1.250E-08 | 7.089E-09 | 4.500E-09 | 1.800E-08 |
| 10509 | 526 | Q526H | A | 2.2e-08 | 1.5e-08 | 5.6e-09 | 1.420E-08 | 8.229E-09 | 5.600E-09 | 2.200E-08 |
| 10510 | 526 | Q526I | A | 7.3e-09 | 1.7e-08 | 6.4e-09 | 1.023E-08 | 5.877E-09 | 6.400E-09 | 1.700E-08 |
| 10511 | 526 | Q526L | A | 6.2e-09 | 2.2e-09 | 9.9e-09 | 6.100E-09 | 3.851E-09 | 2.200E-09 | 9.900E-09 |
| 10512 | 526 | Q526K | A | 4.9e-09 | 8.6e-09 | 2.4e-09 | 5.300E-09 | 3.119E-09 | 2.400E-09 | 8.600E-09 |
| 10513 | 526 | Q526M | A | 1.2e-08 | 7.6e-09 | 2.4e-09 | 7.333E-09 | 4.806E-09 | 2.400E-09 | 1.200E-08 |
| 10514 | 526 | Q526F | A | 3.3e-09 | 1.8e-08 | 2.7e-09 | 8.000E-09 | 8.665E-09 | 2.700E-09 | 1.800E-08 |
| 10515 | 526 | Q526P | A | 4.1e-09 | 5e-09 | 4.7e-09 | 4.600E-09 | 4.583E-10 | 4.100E-09 | 5.000E-09 |
| 10516 | 526 | Q526S | A | 1.4e-08 | 8.9e-09 | 6.7e-09 | 9.867E-09 | 3.745E-09 | 6.700E-09 | 1.400E-08 |
| 10517 | 526 | Q526T | A | 2.1e-08 | 7e-09 | 1.1e-08 | 1.300E-08 | 7.211E-09 | 7.000E-09 | 2.100E-08 |
| 10518 | 526 | Q526W | A | 1.2e-08 | 8.1e-09 | 3.9e-09 | 8.000E-09 | 4.051E-09 | 3.900E-09 | 1.200E-08 |
| 10519 | 526 | Q526Y | A | 2.9e-09 | 8.2e-09 | 1.2e-08 | 7.700E-09 | 4.571E-09 | 2.900E-09 | 1.200E-08 |
| 10520 | 526 | Q526V | A | 2.2e-08 | 4.1e-09 | 6.7e-09 | 1.093E-08 | 9.672E-09 | 4.100E-09 | 2.200E-08 |
| 10521 | 527 | G527A | A | 3.9e-09 | 1.9e-08 | 9.9e-09 | 1.093E-08 | 7.603E-09 | 3.900E-09 | 1.900E-08 |
| 10522 | 527 | G527R | A | 6.9e-09 | 4.9e-09 | 4.9e-09 | 5.567E-09 | 1.155E-09 | 4.900E-09 | 6.900E-09 |
| 10523 | 527 | G527N | A | 3.5e-09 | 2e-08 | 8.2e-09 | 1.057E-08 | 8.501E-09 | 3.500E-09 | 2.000E-08 |
| 10524 | 527 | G527D | A | 9.8e-09 | 7.5e-09 | 5.9e-09 | 7.733E-09 | 1.960E-09 | 5.900E-09 | 9.800E-09 |
| 10525 | 527 | G527C | A | 4.2e-09 | 5.7e-09 | 7e-09 | 5.633E-09 | 1.401E-09 | 4.200E-09 | 7.000E-09 |
| 10526 | 527 | G527E | A | 1.1e-08 | 2.5e-09 | 7.2e-09 | 6.900E-09 | 4.258E-09 | 2.500E-09 | 1.100E-08 |
| 10527 | 527 | G527Q | A | 4e-09 | 8.7e-09 | 9.3e-09 | 7.333E-09 | 2.902E-09 | 4.000E-09 | 9.300E-09 |
| 10528 | 527 | G527G | A | 1.6e-08 | 7.3e-09 | 8.9e-09 | 1.073E-08 | 4.631E-09 | 7.300E-09 | 1.600E-08 |
| 10529 | 527 | G527H | A | 1.9e-09 | 6e-09 | 5.8e-09 | 4.567E-09 | 2.312E-09 | 1.900E-09 | 6.000E-09 |
| 10530 | 527 | G527I | A | 1e-08 | 4.1e-09 | 4.2e-09 | 6.100E-09 | 3.378E-09 | 4.100E-09 | 1.000E-08 |
| 10531 | 527 | G527L | A | 8.1e-09 | 4.6e-09 | 1.8e-08 | 1.023E-08 | 6.950E-09 | 4.600E-09 | 1.800E-08 |
| 10532 | 527 | G527K | A | 5.3e-09 | 1.6e-08 | 1.1e-08 | 1.077E-08 | 5.354E-09 | 5.300E-09 | 1.600E-08 |
| 10533 | 527 | G527M | A | 4.9e-09 | 9.1e-09 | 1.6e-08 | 1.000E-08 | 5.604E-09 | 4.900E-09 | 1.600E-08 |
| 10534 | 527 | G527F | A | 1.2e-08 | 9.2e-09 | 6.1e-09 | 9.100E-09 | 2.951E-09 | 6.100E-09 | 1.200E-08 |
| 10535 | 527 | G527P | A | 1e-08 | 4.2e-09 | 8.6e-09 | 7.600E-09 | 3.027E-09 | 4.200E-09 | 1.000E-08 |
| 10536 | 527 | G527S | A | 5.4e-09 | 7.9e-09 | 5.9e-09 | 6.400E-09 | 1.323E-09 | 5.400E-09 | 7.900E-09 |
| 10537 | 527 | G527T | A | 9.7e-09 | 1e-08 | 4.7e-09 | 8.133E-09 | 2.977E-09 | 4.700E-09 | 1.000E-08 |
| 10538 | 527 | G527W | A | 6.8e-09 | 5.2e-09 | 9.6e-09 | 7.200E-09 | 2.227E-09 | 5.200E-09 | 9.600E-09 |
| 10539 | 527 | G527Y | A | 6.8e-09 | 3.4e-09 | 6.2e-09 | 5.467E-09 | 1.815E-09 | 3.400E-09 | 6.800E-09 |

| | | | | | | | | | | |
|---|---|---|---|---|---|---|---|---|---|---|
| 10540 | 527 | G527V | A | 8e-09 | 6.7e-09 | 3.8e-09 | 6.167E-09 | 2.150E-09 | 3.800E-09 | 8.000E-09 |
| 10541 | 528 | L528A | A | 4.7e-09 | 4.2e-09 | 3.5e-09 | 4.133E-09 | 6.028E-10 | 3.500E-09 | 4.700E-09 |
| 10542 | 528 | L528R | A | 5.3e-09 | 4.9e-09 | 3.6e-09 | 4.600E-09 | 8.888E-10 | 3.600E-09 | 5.300E-09 |
| 10543 | 528 | L528N | A | 2.9e-09 | 1.2e-08 | 3.7e-09 | 6.200E-09 | 5.039E-09 | 2.900E-09 | 1.200E-08 |
| 10544 | 528 | L528D | A | 5.2e-09 | 4.9e-09 | 1e-08 | 6.700E-09 | 2.862E-09 | 4.900E-09 | 1.000E-08 |
| 10545 | 528 | L528C | A | 3.2e-08 | 9e-09 | 5.3e-09 | 1.543E-08 | 1.447E-08 | 5.300E-09 | 3.200E-08 |
| 10546 | 528 | L528E | A | 1.5e-08 | 9.1e-09 | 7e-09 | 1.037E-08 | 4.148E-09 | 7.000E-09 | 1.500E-08 |
| 10547 | 528 | L528Q | A | 5.3e-09 | 2.7e-09 | 6.5e-09 | 4.833E-09 | 1.943E-09 | 2.700E-09 | 6.500E-09 |
| 10548 | 528 | L528G | A | 1.6e-08 | 1.5e-08 | 6.9e-09 | 1.263E-08 | 4.990E-09 | 6.900E-09 | 1.600E-08 |
| 10549 | 528 | L528H | A | 8.8e-09 | 7.3e-09 | 3.3e-09 | 6.467E-09 | 2.843E-09 | 3.300E-09 | 8.800E-09 |
| 10550 | 528 | L528I | A | 2e-08 | 5.8e-09 | 8.6e-09 | 1.147E-08 | 7.522E-09 | 5.800E-09 | 2.000E-08 |
| 10551 | 528 | L528L | A | 7.6e-09 | 1.4e-08 | 6.1e-09 | 9.233E-09 | 4.196E-09 | 6.100E-09 | 1.400E-08 |
| 10552 | 528 | L528K | A | 4.6e-09 | 6.9e-09 | 8.3e-09 | 6.600E-09 | 1.868E-09 | 4.600E-09 | 8.300E-09 |
| 10553 | 528 | L528M | A | 9.9e-09 | 7.1e-09 | 6.4e-09 | 7.800E-09 | 1.852E-09 | 6.400E-09 | 9.900E-09 |
| 10554 | 528 | L528F | A | 4.3e-09 | 9.5e-09 | 4.4e-09 | 6.067E-09 | 2.974E-09 | 4.300E-09 | 9.500E-09 |
| 10555 | 528 | L528P | A | 3.7e-09 | 4.6e-09 | 9.3e-09 | 5.867E-09 | 3.007E-09 | 3.700E-09 | 9.300E-09 |
| 10556 | 528 | L528S | A | 5e-09 | 1.4e-08 | 6.9e-09 | 8.633E-09 | 4.744E-09 | 5.000E-09 | 1.400E-08 |
| 10557 | 528 | L528T | A | 1.2e-08 | 1.5e-08 | 2.7e-08 | 1.800E-08 | 7.937E-09 | 1.200E-08 | 2.700E-08 |
| 10558 | 528 | L528W | A | 4.2e-09 | 9.7e-09 | 5e-09 | 6.300E-09 | 2.972E-09 | 4.200E-09 | 9.700E-09 |
| 10559 | 528 | L528Y | A | 8.1e-09 | 7e-09 | 1.5e-08 | 1.003E-08 | 4.336E-09 | 7.000E-09 | 1.500E-08 |
| 10560 | 528 | L528V | A | 1.6e-08 | 8.9e-09 | 1.4e-08 | 1.297E-08 | 3.661E-09 | 8.900E-09 | 1.600E-08 |
| 10561 | 529 | P529A | A | 1.7e-08 | 5.3e-09 | 1.7e-08 | 1.310E-08 | 6.755E-09 | 5.300E-09 | 1.700E-08 |
| 10562 | 529 | P529R | A | 3e-09 | 6.9e-09 | 5.2e-09 | 5.033E-09 | 1.955E-09 | 3.000E-09 | 6.900E-09 |
| 10563 | 529 | P529N | A | 1.3e-08 | 3.4e-09 | 5.5e-09 | 7.300E-09 | 5.047E-09 | 3.400E-09 | 1.300E-08 |
| 10564 | 529 | P529D | A | 6.6e-09 | 5.5e-09 | 1e-08 | 7.367E-09 | 2.346E-09 | 5.500E-09 | 1.000E-08 |
| 10565 | 529 | P529C | A | 5.8e-09 | 4.4e-09 | 7.5e-09 | 5.900E-09 | 1.552E-09 | 4.400E-09 | 7.500E-09 |
| 10566 | 529 | P529E | A | 8.5e-09 | 7.9e-09 | 9.6e-09 | 8.667E-09 | 8.622E-10 | 7.900E-09 | 9.600E-09 |
| 10567 | 529 | P529Q | A | 1.1e-08 | 1.5e-08 | 1e-08 | 1.200E-08 | 2.646E-09 | 1.000E-08 | 1.500E-08 |
| 10568 | 529 | P529G | A | 5.1e-09 | 6.2e-09 | 1.5e-08 | 8.767E-09 | 5.426E-09 | 5.100E-09 | 1.500E-08 |
| 10569 | 529 | P529H | A | 7.2e-09 | 1.1e-08 | 1e-08 | 9.400E-09 | 1.970E-09 | 7.200E-09 | 1.100E-08 |
| 10570 | 529 | P529I | A | 4.8e-09 | 1.2e-08 | 1.1e-08 | 9.267E-09 | 3.900E-09 | 4.800E-09 | 1.200E-08 |
| 10571 | 529 | P529L | A | 6.6e-09 | 3.7e-09 | 2.3e-08 | 1.110E-08 | 1.041E-08 | 3.700E-09 | 2.300E-08 |
| 10572 | 529 | P529K | A | 6.1e-09 | 8.8e-09 | 1.4e-08 | 9.633E-09 | 4.015E-09 | 6.100E-09 | 1.400E-08 |
| 10573 | 529 | P529M | A | 1.1e-08 | 8.1e-09 | 6.7e-09 | 8.600E-09 | 2.193E-09 | 6.700E-09 | 1.100E-08 |
| 10574 | 529 | P529F | A | 3.1e-09 | 9e-09 | 7.3e-09 | 6.467E-09 | 3.037E-09 | 3.100E-09 | 9.000E-09 |
| 10575 | 529 | P529P | A | 8.8e-09 | 6.5e-09 | 2.4e-08 | 1.310E-08 | 9.509E-09 | 6.500E-09 | 2.400E-08 |
| 10576 | 529 | P529S | A | 1.2e-08 | 1.9e-09 | 4.8e-09 | 6.233E-09 | 5.200E-09 | 1.900E-09 | 1.200E-08 |
| 10577 | 529 | P529T | A | 8.6e-09 | 2.9e-09 | 3.9e-09 | 5.133E-09 | 3.044E-09 | 2.900E-09 | 8.600E-09 |
| 10578 | 529 | P529W | A | 1.5e-08 | 7.5e-09 | 3.3e-09 | 8.600E-09 | 5.927E-09 | 3.300E-09 | 1.500E-08 |
| 10579 | 529 | P529Y | A | 1.1e-08 | 1e-08 | 5.7e-09 | 8.900E-09 | 2.816E-09 | 5.700E-09 | 1.100E-08 |
| 10580 | 529 | P529V | A | 4.2e-09 | 8.5e-09 | 5.2e-09 | 5.967E-09 | 2.250E-09 | 4.200E-09 | 8.500E-09 |
| 10581 | 530 | R530A | A | 1.1e-08 | 1.3e-08 | 3.4e-09 | 9.133E-09 | 5.065E-09 | 3.400E-09 | 1.300E-08 |
| 10582 | 530 | R530R | A | 1.6e-08 | 6e-09 | 5.3e-09 | 9.100E-09 | 5.986E-09 | 5.300E-09 | 1.600E-08 |
| 10583 | 530 | R530N | A | 4.6e-09 | 6.6e-09 | 7.8e-09 | 6.333E-09 | 1.617E-09 | 4.600E-09 | 7.800E-09 |
| 10584 | 530 | R530D | A | 4.5e-09 | 5.7e-09 | 1.7e-08 | 9.067E-09 | 6.897E-09 | 4.500E-09 | 1.700E-08 |
| 10585 | 530 | R530C | A | 7.7e-09 | 3.5e-09 | 1.9e-08 | 1.007E-08 | 8.016E-09 | 3.500E-09 | 1.900E-08 |
| 10586 | 530 | R530E | A | 1.3e-08 | 6.7e-09 | 7.4e-09 | 9.033E-09 | 3.453E-09 | 6.700E-09 | 1.300E-08 |
| 10587 | 530 | R530Q | A | 1.9e-09 | 3.6e-09 | 4.5e-09 | 3.333E-09 | 1.320E-09 | 1.900E-09 | 4.500E-09 |
| 10588 | 530 | R530G | A | 3.9e-09 | 8.1e-09 | 5.9e-09 | 5.967E-09 | 2.101E-09 | 3.900E-09 | 8.100E-09 |
| 10589 | 530 | R530H | A | 1.8e-09 | 7.5e-09 | 5.8e-09 | 5.033E-09 | 2.926E-09 | 1.800E-09 | 7.500E-09 |
| 10590 | 530 | R530I | A | 9.6e-09 | 1.7e-08 | 4.6e-09 | 1.040E-08 | 6.239E-09 | 4.600E-09 | 1.700E-08 |
| 10591 | 530 | R530L | A | 1.1e-08 | 5.8e-09 | 6.8e-09 | 7.867E-09 | 2.759E-09 | 5.800E-09 | 1.100E-08 |
| 10592 | 530 | R530K | A | 9.6e-09 | 6.9e-09 | 1.6e-08 | 1.083E-08 | 4.674E-09 | 6.900E-09 | 1.600E-08 |
| 10593 | 530 | R530M | A | 1.3e-08 | 1.8e-09 | 1.8e-09 | 5.533E-09 | 6.466E-09 | 1.800E-09 | 1.300E-08 |
| 10594 | 530 | R530F | A | 2.9e-09 | 1.2e-08 | 8e-09 | 7.633E-09 | 4.561E-09 | 2.900E-09 | 1.200E-08 |
| 10595 | 530 | R530P | A | 7.2e-09 | 6e-09 | 2.2e-08 | 1.173E-08 | 8.911E-09 | 6.000E-09 | 2.200E-08 |
| 10596 | 530 | R530S | A | 2.3e-09 | 6.9e-09 | 4.9e-09 | 4.700E-09 | 2.307E-09 | 2.300E-09 | 6.900E-09 |
| 10597 | 530 | R530T | A | 1.1e-08 | 9.2e-09 | 5.6e-09 | 8.600E-09 | 2.750E-09 | 5.600E-09 | 1.100E-08 |
| 10598 | 530 | R530W | A | 3.9e-09 | 8.6e-09 | 5.9e-09 | 6.133E-09 | 2.359E-09 | 3.900E-09 | 8.600E-09 |
| 10599 | 530 | R530Y | A | 1.3e-08 | 6.7e-09 | 3.5e-09 | 7.733E-09 | 4.834E-09 | 3.500E-09 | 1.300E-08 |
| 10600 | 530 | R530V | A | 4.4e-09 | 9.8e-09 | 1.5e-08 | 9.733E-09 | 5.300E-09 | 4.400E-09 | 1.500E-08 |
| 10601 | 531 | E531A | A | 6.7e-09 | 1.5e-08 | 1.7e-08 | 1.290E-08 | 5.462E-09 | 6.700E-09 | 1.700E-08 |
| 10602 | 531 | E531R | A | 4.9e-09 | 4.7e-09 | 5.6e-09 | 5.067E-09 | 4.726E-10 | 4.700E-09 | 5.600E-09 |
| 10603 | 531 | E531N | A | 5.5e-09 | 1.3e-09 | 1e-08 | 5.600E-09 | 4.351E-09 | 1.300E-09 | 1.000E-08 |
| 10604 | 531 | E531D | A | 6.3e-09 | 7.5e-09 | 7.8e-09 | 7.200E-09 | 7.937E-10 | 6.300E-09 | 7.800E-09 |
| 10605 | 531 | E531C | A | 8.4e-09 | 4.1e-09 | 1e-08 | 7.500E-09 | 3.051E-09 | 4.100E-09 | 1.000E-08 |
| 10606 | 531 | E531E | A | 2.6e-09 | 4.7e-09 | 4.9e-09 | 4.067E-09 | 1.274E-09 | 2.600E-09 | 4.900E-09 |
| 10607 | 531 | E531Q | A | 5.7e-09 | 9.9e-09 | 9.4e-09 | 8.333E-09 | 2.294E-09 | 5.700E-09 | 9.900E-09 |
| 10608 | 531 | E531G | A | 6.7e-09 | 1.9e-08 | 1.7e-08 | 1.423E-08 | 6.600E-09 | 6.700E-09 | 1.900E-08 |
| 10609 | 531 | E531H | A | 1.5e-08 | 9.1e-09 | 1.7e-08 | 1.370E-08 | 4.107E-09 | 9.100E-09 | 1.700E-08 |
| 10610 | 531 | E531I | A | 4.5e-09 | 6.7e-09 | 5.9e-09 | 5.700E-09 | 1.114E-09 | 4.500E-09 | 6.700E-09 |
| 10611 | 531 | E531L | A | 1.2e-08 | 3.3e-09 | 5.3e-09 | 6.867E-09 | 4.557E-09 | 3.300E-09 | 1.200E-08 |
| 10612 | 531 | E531K | A | 6.6e-09 | 6.4e-09 | 8.1e-09 | 7.033E-09 | 9.292E-10 | 6.400E-09 | 8.100E-09 |
| 10613 | 531 | E531M | A | 6.6e-09 | 2.8e-09 | 9.1e-09 | 6.167E-09 | 3.172E-09 | 2.800E-09 | 9.100E-09 |
| 10614 | 531 | E531F | A | 1.9e-08 | 1.2e-08 | 5.5e-09 | 1.217E-08 | 6.752E-09 | 5.500E-09 | 1.900E-08 |
| 10615 | 531 | E531P | A | 8.3e-09 | 4.7e-09 | 5.2e-09 | 6.067E-09 | 1.950E-09 | 4.700E-09 | 8.300E-09 |
| 10616 | 531 | E531S | A | 2.8e-09 | 1.3e-08 | 8e-09 | 7.933E-09 | 5.100E-09 | 2.800E-09 | 1.300E-08 |
| 10617 | 531 | E531T | A | 2.1e-09 | 9.3e-09 | 8.3e-09 | 6.567E-09 | 3.900E-09 | 2.100E-09 | 9.300E-09 |
| 10618 | 531 | E531W | A | 3.2e-09 | 3.7e-09 | 2.2e-09 | 3.033E-09 | 7.638E-10 | 2.200E-09 | 3.700E-09 |
| 10619 | 531 | E531Y | A | 3.8e-09 | 4.4e-09 | 1.1e-08 | 6.400E-09 | 3.995E-09 | 3.800E-09 | 1.100E-08 |
| 10620 | 531 | E531V | A | 2e-08 | 1.1e-08 | 2.7e-09 | 1.123E-08 | 8.652E-09 | 2.700E-09 | 2.000E-08 |
| 10621 | 532 | Y532A | A | 5.3e-09 | 8.2e-09 | 6.1e-09 | 6.533E-09 | 1.498E-09 | 5.300E-09 | 8.200E-09 |
| 10622 | 532 | Y532R | A | 4.2e-09 | 6.1e-09 | 4.8e-09 | 5.033E-09 | 9.713E-10 | 4.200E-09 | 6.100E-09 |
| 10623 | 532 | Y532N | A | 7.6e-09 | 4.4e-09 | 8.4e-09 | 6.800E-09 | 2.117E-09 | 4.400E-09 | 8.400E-09 |
| 10624 | 532 | Y532D | A | 1.2e-08 | 9e-09 | 8.6e-09 | 9.867E-09 | 1.858E-09 | 8.600E-09 | 1.200E-08 |

| | | | | | | | | | | |
|---|---|---|---|---|---|---|---|---|---|---|
| 10625 | 532 | Y532C | A | 8.2e-09 | 6.3e-09 | 5.1e-09 | 6.533E-09 | 1.563E-09 | 5.100E-09 | 8.200E-09 |
| 10626 | 532 | Y532E | A | 3.1e-09 | 1.7e-08 | 6.8e-09 | 8.967E-09 | 7.199E-09 | 3.100E-09 | 1.700E-08 |
| 10627 | 532 | Y532Q | A | 5.3e-09 | 1.3e-08 | 3.4e-09 | 7.233E-09 | 5.084E-09 | 3.400E-09 | 1.300E-08 |
| 10628 | 532 | Y532G | A | 3.8e-09 | 6.8e-09 | 1.5e-08 | 8.533E-09 | 5.798E-09 | 3.800E-09 | 1.500E-08 |
| 10629 | 532 | Y532H | A | 1.2e-08 | 9.8e-09 | 4.7e-09 | 8.833E-09 | 3.745E-09 | 4.700E-09 | 1.200E-08 |
| 10630 | 532 | Y532I | A | 1e-08 | 7.2e-09 | 4.8e-09 | 7.333E-09 | 2.603E-09 | 4.800E-09 | 1.000E-08 |
| 10631 | 532 | Y532L | A | 3.9e-09 | 6.5e-09 | 1.4e-08 | 8.133E-09 | 5.244E-09 | 3.900E-09 | 1.400E-08 |
| 10632 | 532 | Y532K | A | 3.9e-09 | 7.3e-09 | 2.3e-08 | 1.140E-08 | 1.019E-08 | 3.900E-09 | 2.300E-08 |
| 10633 | 532 | Y532M | A | 1.1e-09 | 6.9e-09 | 6.8e-09 | 4.933E-09 | 3.320E-09 | 1.100E-09 | 6.900E-09 |
| 10634 | 532 | Y532F | A | 1.1e-08 | 8.7e-09 | 9.4e-09 | 9.700E-09 | 1.179E-09 | 8.700E-09 | 1.100E-08 |
| 10635 | 532 | Y532P | A | 9.2e-09 | 1.2e-08 | 8.2e-09 | 9.800E-09 | 1.970E-09 | 8.200E-09 | 1.200E-08 |
| 10636 | 532 | Y532S | A | 5e-09 | 3.8e-09 | 1.3e-08 | 7.267E-09 | 5.001E-09 | 3.800E-09 | 1.300E-08 |
| 10637 | 532 | Y532T | A | 7.1e-09 | 5.7e-09 | 1.4e-08 | 8.933E-09 | 4.443E-09 | 5.700E-09 | 1.400E-08 |
| 10638 | 532 | Y532W | A | 2.2e-09 | 2.6e-09 | 1.2e-08 | 5.600E-09 | 5.546E-09 | 2.200E-09 | 1.200E-08 |
| 10639 | 532 | Y532Y | A | 1.4e-08 | 6.9e-09 | 5.2e-09 | 8.700E-09 | 4.668E-09 | 5.200E-09 | 1.400E-08 |
| 10640 | 532 | Y532V | A | 1.2e-08 | 1.1e-08 | 5.5e-09 | 9.500E-09 | 3.500E-09 | 5.500E-09 | 1.200E-08 |
| 10641 | 533 | V533A | A | 1.1e-08 | 6.9e-09 | 6.1e-09 | 8.000E-09 | 2.629E-09 | 6.100E-09 | 1.100E-08 |
| 10642 | 533 | V533R | A | 8.9e-09 | 5.6e-09 | 1.9e-08 | 1.117E-08 | 6.982E-09 | 5.600E-09 | 1.900E-08 |
| 10643 | 533 | V533N | A | 6.5e-09 | 2.9e-09 | 3.7e-09 | 4.367E-09 | 1.890E-09 | 2.900E-09 | 6.500E-09 |
| 10644 | 533 | V533D | A | 4.2e-09 | 3e-09 | 3.6e-09 | 3.600E-09 | 6.000E-10 | 3.000E-09 | 4.200E-09 |
| 10645 | 533 | V533C | A | 2.2e-08 | 4.3e-09 | 7.5e-09 | 1.127E-08 | 9.432E-09 | 4.300E-09 | 2.200E-08 |
| 10646 | 533 | V533E | A | 5e-09 | 5.2e-09 | 1.9e-08 | 9.733E-09 | 8.026E-09 | 5.000E-09 | 1.900E-08 |
| 10647 | 533 | V533Q | A | 1e-08 | 1.6e-09 | 4.7e-09 | 5.433E-09 | 4.248E-09 | 1.600E-09 | 1.000E-08 |
| 10648 | 533 | V533G | A | 3.2e-09 | 3.1e-08 | 2e-08 | 1.807E-08 | 1.400E-08 | 3.200E-09 | 3.100E-08 |
| 10649 | 533 | V533H | A | 8.4e-09 | 8.5e-09 | 1.8e-08 | 1.163E-08 | 5.514E-09 | 8.400E-09 | 1.800E-08 |
| 10650 | 533 | V533I | A | 1.4e-08 | 4.2e-09 | 2.3e-09 | 6.833E-09 | 6.279E-09 | 2.300E-09 | 1.400E-08 |
| 10651 | 533 | V533L | A | 1.4e-08 | 6.4e-09 | 1.1e-08 | 1.047E-08 | 3.828E-09 | 6.400E-09 | 1.400E-08 |
| 10652 | 533 | V533K | A | 4.1e-09 | 5.6e-09 | 1.5e-08 | 8.233E-09 | 5.908E-09 | 4.100E-09 | 1.500E-08 |
| 10653 | 533 | V533M | A | 8.5e-09 | 2.4e-09 | 5.9e-09 | 5.600E-09 | 3.061E-09 | 2.400E-09 | 8.500E-09 |
| 10654 | 533 | V533F | A | 1.5e-08 | 4.4e-09 | 2.8e-09 | 7.400E-09 | 6.630E-09 | 2.800E-09 | 1.500E-08 |
| 10655 | 533 | V533P | A | 8.8e-09 | 9.9e-09 | 6.1e-09 | 8.267E-09 | 1.955E-09 | 6.100E-09 | 9.900E-09 |
| 10656 | 533 | V533S | A | 6.6e-09 | 9.2e-09 | 9.2e-09 | 8.333E-09 | 1.501E-09 | 6.600E-09 | 9.200E-09 |
| 10657 | 533 | V533T | A | 7e-09 | 9.5e-09 | 1.2e-08 | 9.500E-09 | 2.500E-09 | 7.000E-09 | 1.200E-08 |
| 10658 | 533 | V533W | A | 6.8e-09 | 3.5e-09 | 1.8e-08 | 9.433E-09 | 7.600E-09 | 3.500E-09 | 1.800E-08 |
| 10659 | 533 | V533Y | A | 8.4e-09 | 1.4e-08 | 9.6e-09 | 1.067E-08 | 2.948E-09 | 8.400E-09 | 1.400E-08 |
| 10660 | 533 | V533V | A | 1e-08 | 8.9e-09 | 9.4e-09 | 9.433E-09 | 5.508E-10 | 8.900E-09 | 1.000E-08 |
| 10661 | 534 | N534A | A | 3e-08 | 2.4e-09 | 9.5e-09 | 1.397E-08 | 1.433E-08 | 2.400E-09 | 3.000E-08 |
| 10662 | 534 | N534R | A | 3.5e-09 | 1.3e-08 | 9.9e-09 | 8.800E-09 | 4.845E-09 | 3.500E-09 | 1.300E-08 |
| 10663 | 534 | N534N | A | 3.3e-09 | 4e-09 | 4.7e-09 | 4.000E-09 | 7.000E-10 | 3.300E-09 | 4.700E-09 |
| 10664 | 534 | N534D | A | 1.4e-08 | 7.3e-09 | 8.9e-09 | 1.007E-08 | 3.499E-09 | 7.300E-09 | 1.400E-08 |
| 10665 | 534 | N534C | A | 1.5e-09 | 1.9e-09 | 5.3e-09 | 2.900E-09 | 2.088E-09 | 1.500E-09 | 5.300E-09 |
| 10666 | 534 | N534E | A | 1.1e-08 | 1.6e-09 | 5.9e-09 | 6.167E-09 | 4.706E-09 | 1.600E-09 | 1.100E-08 |
| 10667 | 534 | N534Q | A | 5.6e-09 | 4.1e-09 | 6.2e-09 | 5.300E-09 | 1.082E-09 | 4.100E-09 | 6.200E-09 |
| 10668 | 534 | N534G | A | 2.2e-09 | 1.1e-08 | 7.8e-09 | 7.000E-09 | 4.454E-09 | 2.200E-09 | 1.100E-08 |
| 10669 | 534 | N534H | A | 7.5e-09 | 1.2e-08 | 1.2e-08 | 1.050E-08 | 2.598E-09 | 7.500E-09 | 1.200E-08 |
| 10670 | 534 | N534I | A | 6.4e-09 | 7.4e-09 | 3.7e-09 | 5.833E-09 | 1.914E-09 | 3.700E-09 | 7.400E-09 |
| 10671 | 534 | N534L | A | 4.9e-09 | 1.4e-08 | 3.4e-09 | 7.433E-09 | 5.736E-09 | 3.400E-09 | 1.400E-08 |
| 10672 | 534 | N534K | A | 1.3e-08 | 4.8e-09 | 4.1e-09 | 7.300E-09 | 4.949E-09 | 4.100E-09 | 1.300E-08 |
| 10673 | 534 | N534M | A | 4.9e-09 | 1.7e-08 | 9.5e-09 | 1.047E-08 | 6.108E-09 | 4.900E-09 | 1.700E-08 |
| 10674 | 534 | N534F | A | 1.5e-09 | 4.6e-09 | 5.3e-09 | 3.800E-09 | 2.022E-09 | 1.500E-09 | 5.300E-09 |
| 10675 | 534 | N534P | A | 8.4e-09 | 4.3e-09 | 3.7e-09 | 5.467E-09 | 2.558E-09 | 3.700E-09 | 8.400E-09 |
| 10676 | 534 | N534S | A | 1.2e-08 | 9.2e-09 | 5.3e-09 | 8.833E-09 | 3.365E-09 | 5.300E-09 | 1.200E-08 |
| 10677 | 534 | N534T | A | 3.2e-08 | 2.2e-08 | 7.8e-09 | 2.060E-08 | 1.216E-08 | 7.800E-09 | 3.200E-08 |
| 10678 | 534 | N534W | A | 7.9e-09 | 2.2e-08 | 7.3e-09 | 1.240E-08 | 8.319E-09 | 7.300E-09 | 2.200E-08 |
| 10679 | 534 | N534Y | A | 6.8e-09 | 4.5e-09 | 6.8e-09 | 6.033E-09 | 1.328E-09 | 4.500E-09 | 6.800E-09 |
| 10680 | 534 | N534V | A | 2.4e-08 | 8.3e-09 | 1e-08 | 1.410E-08 | 8.616E-09 | 8.300E-09 | 2.400E-08 |
| 10681 | 535 | A535A | A | 1.1e-08 | 4.2e-09 | 5e-09 | 6.733E-09 | 3.717E-09 | 4.200E-09 | 1.100E-08 |
| 10682 | 535 | A535R | A | 5.6e-09 | 8.7e-09 | 6.5e-09 | 6.933E-09 | 1.595E-09 | 5.600E-09 | 8.700E-09 |
| 10683 | 535 | A535N | A | 1.1e-08 | 5.2e-09 | 1.9e-08 | 1.173E-08 | 6.929E-09 | 5.200E-09 | 1.900E-08 |
| 10684 | 535 | A535D | A | 6.8e-09 | 1.8e-09 | 3.2e-08 | 1.353E-08 | 1.619E-08 | 1.800E-09 | 3.200E-08 |
| 10685 | 535 | A535C | A | 2.3e-09 | 1.4e-08 | 8e-09 | 8.100E-09 | 5.851E-09 | 2.300E-09 | 1.400E-08 |
| 10686 | 535 | A535E | A | 7.2e-09 | 1.4e-08 | 1.4e-08 | 1.173E-08 | 3.926E-09 | 7.200E-09 | 1.400E-08 |
| 10687 | 535 | A535Q | A | 1.2e-08 | 2.3e-08 | 5.3e-09 | 1.343E-08 | 8.937E-09 | 5.300E-09 | 2.300E-08 |
| 10688 | 535 | A535G | A | 4.9e-09 | 5.6e-09 | 4.8e-09 | 5.100E-09 | 4.359E-10 | 4.800E-09 | 5.600E-09 |
| 10689 | 535 | A535H | A | 5.7e-09 | 8.7e-09 | 4.1e-09 | 6.167E-09 | 2.335E-09 | 4.100E-09 | 8.700E-09 |
| 10690 | 535 | A535I | A | 4.7e-09 | 8.3e-09 | 7.4e-09 | 6.800E-09 | 1.873E-09 | 4.700E-09 | 8.300E-09 |
| 10691 | 535 | A535L | A | 4e-09 | 1.8e-08 | 1.4e-08 | 1.200E-08 | 7.211E-09 | 4.000E-09 | 1.800E-08 |
| 10692 | 535 | A535K | A | 1.4e-08 | 4.3e-09 | 7.2e-09 | 8.500E-09 | 4.979E-09 | 4.300E-09 | 1.400E-08 |
| 10693 | 535 | A535M | A | 9.2e-09 | 4.6e-09 | 8.2e-09 | 7.333E-09 | 2.419E-09 | 4.600E-09 | 9.200E-09 |
| 10694 | 535 | A535F | A | 9.5e-09 | 6.6e-09 | 9.3e-09 | 8.467E-09 | 1.620E-09 | 6.600E-09 | 9.500E-09 |
| 10695 | 535 | A535P | A | 1.3e-08 | 7.2e-09 | 5.7e-09 | 8.633E-09 | 3.855E-09 | 5.700E-09 | 1.300E-08 |
| 10696 | 535 | A535S | A | 5e-09 | 6e-09 | 3.3e-09 | 4.767E-09 | 1.365E-09 | 3.300E-09 | 6.000E-09 |
| 10697 | 535 | A535T | A | 8.5e-09 | 6.8e-09 | 4.8e-09 | 6.700E-09 | 1.852E-09 | 4.800E-09 | 8.500E-09 |
| 10698 | 535 | A535W | A | 1.2e-08 | 1.5e-08 | 3.3e-09 | 1.010E-08 | 6.077E-09 | 3.300E-09 | 1.500E-08 |
| 10699 | 535 | A535Y | A | 8.1e-09 | 1.6e-08 | 9.3e-09 | 1.113E-08 | 4.257E-09 | 8.100E-09 | 1.600E-08 |
| 10700 | 535 | A535V | A | 9.4e-09 | 7.4e-09 | 9.5e-09 | 8.767E-09 | 1.185E-09 | 7.400E-09 | 9.500E-09 |
| 10701 | 536 | R536A | A | 5e-09 | 7.4e-09 | 1.7e-08 | 9.800E-09 | 6.350E-09 | 5.000E-09 | 1.700E-08 |
| 10702 | 536 | R536R | A | 4.3e-09 | 1.1e-08 | 4.7e-09 | 6.667E-09 | 3.758E-09 | 4.300E-09 | 1.100E-08 |
| 10703 | 536 | R536N | A | 7.6e-09 | 8.5e-09 | 4.8e-09 | 6.967E-09 | 1.930E-09 | 4.800E-09 | 8.500E-09 |
| 10704 | 536 | R536D | A | 1.1e-08 | 7.1e-09 | 1.1e-08 | 9.700E-09 | 2.252E-09 | 7.100E-09 | 1.100E-08 |
| 10705 | 536 | R536C | A | 1.1e-08 | 5.3e-09 | 6.1e-09 | 7.467E-09 | 3.086E-09 | 5.300E-09 | 1.100E-08 |
| 10706 | 536 | R536E | A | 6.4e-09 | 3.7e-09 | 9e-09 | 6.367E-09 | 2.650E-09 | 3.700E-09 | 9.000E-09 |
| 10707 | 536 | R536Q | A | 3.4e-09 | 6.1e-09 | 7.9e-09 | 5.800E-09 | 2.265E-09 | 3.400E-09 | 7.900E-09 |
| 10708 | 536 | R536G | A | 1.7e-08 | 1.2e-08 | 6.1e-09 | 1.170E-08 | 5.456E-09 | 6.100E-09 | 1.700E-08 |
| 10709 | 536 | R536H | A | 6.9e-09 | 1.4e-08 | 8.3e-09 | 9.733E-09 | 3.761E-09 | 6.900E-09 | 1.400E-08 |

| | | | | | | | | | | |
|---|---|---|---|---|---|---|---|---|---|---|
| 10710 | 536 | R536I | A | 3.7e-09 | 7.4e-09 | 1e-08 | 7.033E-09 | 3.166E-09 | 3.700E-09 | 1.000E-08 |
| 10711 | 536 | R536L | A | 6.9e-09 | 5.6e-09 | 4.3e-09 | 5.600E-09 | 1.300E-09 | 4.300E-09 | 6.900E-09 |
| 10712 | 536 | R536K | A | 1.3e-08 | 1.3e-08 | 3.7e-09 | 9.900E-09 | 5.369E-09 | 3.700E-09 | 1.300E-08 |
| 10713 | 536 | R536M | A | 4.9e-09 | 5.9e-09 | 3e-09 | 4.600E-09 | 1.473E-09 | 3.000E-09 | 5.900E-09 |
| 10714 | 536 | R536F | A | 9.7e-09 | 1.7e-08 | 1.6e-09 | 9.433E-09 | 7.703E-09 | 1.600E-09 | 1.700E-08 |
| 10715 | 536 | R536P | A | 5.5e-09 | 2.6e-09 | 1.6e-09 | 3.233E-09 | 2.026E-09 | 1.600E-09 | 5.500E-09 |
| 10716 | 536 | R536S | A | 1.1e-08 | 6.1e-09 | 1.3e-08 | 1.003E-08 | 3.550E-09 | 6.100E-09 | 1.300E-08 |
| 10717 | 536 | R536T | A | 6.1e-09 | 3.5e-09 | 5e-09 | 4.867E-09 | 1.305E-09 | 3.500E-09 | 6.100E-09 |
| 10718 | 536 | R536W | A | 3.6e-09 | 1.4e-08 | 6.8e-09 | 8.133E-09 | 5.327E-09 | 3.600E-09 | 1.400E-08 |
| 10719 | 536 | R536Y | A | 1.3e-08 | 6.3e-09 | 5e-09 | 8.100E-09 | 4.293E-09 | 5.000E-09 | 1.300E-08 |
| 10720 | 536 | R536V | A | 6.3e-09 | 3.7e-09 | 5.2e-09 | 5.067E-09 | 1.305E-09 | 3.700E-09 | 6.300E-09 |
| 10721 | 537 | H537A | A | 5.1e-09 | 1e-08 | 5.1e-09 | 6.733E-09 | 2.829E-09 | 5.100E-09 | 1.000E-08 |
| 10722 | 537 | H537R | A | 4.6e-09 | 2.6e-08 | 3.5e-09 | 1.137E-08 | 1.268E-08 | 3.500E-09 | 2.600E-08 |
| 10723 | 537 | H537N | A | 8.3e-09 | 6.1e-09 | 8.6e-09 | 7.667E-09 | 1.365E-09 | 6.100E-09 | 8.600E-09 |
| 10724 | 537 | H537D | A | 1.3e-08 | 7.1e-09 | 9.2e-09 | 9.767E-09 | 2.991E-09 | 7.100E-09 | 1.300E-08 |
| 10725 | 537 | H537C | A | 7.6e-09 | 3.8e-09 | 4.2e-09 | 5.200E-09 | 2.088E-09 | 3.800E-09 | 7.600E-09 |
| 10726 | 537 | H537E | A | 3.7e-09 | 6.3e-09 | 6.9e-09 | 5.633E-09 | 1.701E-09 | 3.700E-09 | 6.900E-09 |
| 10727 | 537 | H537Q | A | 3.4e-09 | 3e-09 | 8.3e-09 | 4.900E-09 | 2.951E-09 | 3.000E-09 | 8.300E-09 |
| 10728 | 537 | H537G | A | 4.6e-09 | 7.9e-09 | 5.9e-09 | 6.133E-09 | 1.662E-09 | 4.600E-09 | 7.900E-09 |
| 10729 | 537 | H537H | A | 4.3e-09 | 9e-09 | 4.3e-09 | 5.867E-09 | 2.714E-09 | 4.300E-09 | 9.000E-09 |
| 10730 | 537 | H537I | A | 9.2e-09 | 1.4e-08 | 1.6e-08 | 1.307E-08 | 3.495E-09 | 9.200E-09 | 1.600E-08 |
| 10731 | 537 | H537L | A | 5.3e-09 | 1.2e-08 | 4.2e-09 | 7.167E-09 | 4.222E-09 | 4.200E-09 | 1.200E-08 |
| 10732 | 537 | H537K | A | 2.8e-08 | 1.6e-09 | 5.6e-09 | 1.173E-08 | 1.423E-08 | 1.600E-09 | 2.800E-08 |
| 10733 | 537 | H537M | A | 5e-09 | 5.5e-09 | 3.9e-08 | 1.650E-08 | 1.949E-08 | 5.000E-09 | 3.900E-08 |
| 10734 | 537 | H537F | A | 5.8e-09 | 3.8e-09 | 8.9e-09 | 6.167E-09 | 2.570E-09 | 3.800E-09 | 8.900E-09 |
| 10735 | 537 | H537P | A | 2.7e-08 | 8.4e-09 | 2.4e-08 | 1.980E-08 | 9.986E-09 | 8.400E-09 | 2.700E-08 |
| 10736 | 537 | H537S | A | 4.7e-09 | 9.5e-09 | 2.3e-09 | 5.500E-09 | 3.666E-09 | 2.300E-09 | 9.500E-09 |
| 10737 | 537 | H537T | A | 5.4e-09 | 6.9e-09 | 1.7e-08 | 9.767E-09 | 6.309E-09 | 5.400E-09 | 1.700E-08 |
| 10738 | 537 | H537W | A | 1.5e-08 | 2.2e-08 | 7.9e-09 | 1.497E-08 | 7.050E-09 | 7.900E-09 | 2.200E-08 |
| 10739 | 537 | H537Y | A | 3e-09 | 9.1e-09 | 8.2e-09 | 6.767E-09 | 3.293E-09 | 3.000E-09 | 9.100E-09 |
| 10740 | 537 | H537V | A | 1.1e-08 | 1.1e-08 | 2.9e-09 | 8.300E-09 | 4.677E-09 | 2.900E-09 | 1.100E-08 |
| 10741 | 538 | C538A | A | 8.2e-09 | 7.3e-09 | 5.9e-09 | 7.133E-09 | 1.159E-09 | 5.900E-09 | 8.200E-09 |
| 10742 | 538 | C538R | A | 8.5e-09 | 3.3e-09 | 5.1e-09 | 5.633E-09 | 2.641E-09 | 3.300E-09 | 8.500E-09 |
| 10743 | 538 | C538N | A | 5e-09 | 9.5e-09 | 3.4e-09 | 5.967E-09 | 3.163E-09 | 3.400E-09 | 9.500E-09 |
| 10744 | 538 | C538D | A | 5.5e-09 | 3.3e-09 | 3.5e-09 | 4.100E-09 | 1.217E-09 | 3.300E-09 | 5.500E-09 |
| 10745 | 538 | C538C | A | 3.7e-09 | 9.3e-09 | 8.7e-09 | 7.233E-09 | 3.075E-09 | 3.700E-09 | 9.300E-09 |
| 10746 | 538 | C538E | A | 1.2e-08 | 1e-08 | 7.9e-09 | 9.967E-09 | 2.050E-09 | 7.900E-09 | 1.200E-08 |
| 10747 | 538 | C538Q | A | 4.7e-09 | 1.2e-08 | 4.8e-09 | 7.167E-09 | 4.186E-09 | 4.700E-09 | 1.200E-08 |
| 10748 | 538 | C538G | A | 9.3e-09 | 8.1e-09 | 3.2e-09 | 6.867E-09 | 3.232E-09 | 3.200E-09 | 9.300E-09 |
| 10749 | 538 | C538H | A | 8.4e-09 | 1.6e-08 | 9.9e-09 | 1.143E-08 | 4.025E-09 | 8.400E-09 | 1.600E-08 |
| 10750 | 538 | C538I | A | 5.8e-09 | 1e-08 | 6.1e-09 | 7.300E-09 | 2.343E-09 | 5.800E-09 | 1.000E-08 |
| 10751 | 538 | C538L | A | 1.1e-08 | 3.8e-09 | 5.1e-09 | 6.633E-09 | 3.837E-09 | 3.800E-09 | 1.100E-08 |
| 10752 | 538 | C538K | A | 4.3e-09 | 6e-09 | 1.7e-08 | 9.100E-09 | 6.894E-09 | 4.300E-09 | 1.700E-08 |
| 10753 | 538 | C538M | A | 1.3e-08 | 6.1e-09 | 5.8e-09 | 8.300E-09 | 4.073E-09 | 5.800E-09 | 1.300E-08 |
| 10754 | 538 | C538F | A | 1.1e-08 | 1.1e-08 | 5.9e-09 | 9.300E-09 | 2.944E-09 | 5.900E-09 | 1.100E-08 |
| 10755 | 538 | C538P | A | 1.3e-08 | 3.9e-09 | 3.4e-09 | 6.767E-09 | 5.404E-09 | 3.400E-09 | 1.300E-08 |
| 10756 | 538 | C538S | A | 1e-08 | 2.7e-09 | 1.1e-08 | 7.900E-09 | 4.531E-09 | 2.700E-09 | 1.100E-08 |
| 10757 | 538 | C538T | A | 1e-08 | 2.7e-08 | 7.8e-09 | 1.493E-08 | 1.051E-08 | 7.800E-09 | 2.700E-08 |
| 10758 | 538 | C538W | A | 1.2e-08 | 2.5e-08 | 5.5e-09 | 1.417E-08 | 9.929E-09 | 5.500E-09 | 2.500E-08 |
| 10759 | 538 | C538Y | A | 4.3e-09 | 4.5e-09 | 9.7e-09 | 6.167E-09 | 3.062E-09 | 4.300E-09 | 9.700E-09 |
| 10760 | 538 | C538V | A | 4.5e-09 | 5.7e-09 | 4.7e-09 | 4.967E-09 | 6.429E-10 | 4.500E-09 | 5.700E-09 |
| 10761 | 539 | L539A | A | 2e-08 | 1.4e-08 | 2.5e-09 | 1.217E-08 | 8.893E-09 | 2.500E-09 | 2.000E-08 |
| 10762 | 539 | L539R | A | 4.7e-09 | 5.1e-09 | 3.4e-09 | 4.400E-09 | 8.888E-10 | 3.400E-09 | 5.100E-09 |
| 10763 | 539 | L539N | A | 6.3e-09 | 1.4e-08 | 1.5e-08 | 1.177E-08 | 4.761E-09 | 6.300E-09 | 1.500E-08 |
| 10764 | 539 | L539D | A | 2.3e-09 | 1.2e-08 | 4.6e-09 | 6.300E-09 | 5.069E-09 | 2.300E-09 | 1.200E-08 |
| 10765 | 539 | L539C | A | 4.3e-09 | 1.8e-08 | 8.6e-09 | 1.030E-08 | 7.006E-09 | 4.300E-09 | 1.800E-08 |
| 10766 | 539 | L539E | A | 3.8e-09 | 9.6e-09 | 9.9e-09 | 7.767E-09 | 3.439E-09 | 3.800E-09 | 9.900E-09 |
| 10767 | 539 | L539Q | A | 2.7e-09 | 4e-09 | 6.3e-09 | 4.333E-09 | 1.823E-09 | 2.700E-09 | 6.300E-09 |
| 10768 | 539 | L539G | A | 9.4e-09 | 1.2e-08 | 7.5e-09 | 9.633E-09 | 2.259E-09 | 7.500E-09 | 1.200E-08 |
| 10769 | 539 | L539H | A | 2.9e-09 | 8.8e-09 | 8.1e-09 | 6.600E-09 | 3.223E-09 | 2.900E-09 | 8.800E-09 |
| 10770 | 539 | L539I | A | 7.3e-09 | 7.2e-09 | 3.5e-08 | 1.650E-08 | 1.602E-08 | 7.200E-09 | 3.500E-08 |
| 10771 | 539 | L539L | A | 6.5e-09 | 9.2e-09 | 1.3e-08 | 9.567E-09 | 3.265E-09 | 6.500E-09 | 1.300E-08 |
| 10772 | 539 | L539K | A | 9.3e-09 | 9e-09 | 7.9e-09 | 8.733E-09 | 7.371E-10 | 7.900E-09 | 9.300E-09 |
| 10773 | 539 | L539M | A | 1.2e-08 | 2.5e-08 | 9.5e-09 | 1.550E-08 | 8.322E-09 | 9.500E-09 | 2.500E-08 |
| 10774 | 539 | L539F | A | 4.5e-09 | 6.1e-09 | 1.5e-08 | 8.533E-09 | 5.657E-09 | 4.500E-09 | 1.500E-08 |
| 10775 | 539 | L539P | A | 6.7e-09 | 8.1e-09 | 1e-08 | 8.267E-09 | 1.656E-09 | 6.700E-09 | 1.000E-08 |
| 10776 | 539 | L539S | A | 5.6e-09 | 7.3e-09 | 5.6e-09 | 6.167E-09 | 9.815E-10 | 5.600E-09 | 7.300E-09 |
| 10777 | 539 | L539T | A | 1.9e-08 | 5.5e-09 | 1.3e-08 | 1.250E-08 | 6.764E-09 | 5.500E-09 | 1.900E-08 |
| 10778 | 539 | L539W | A | 4.4e-09 | 3.4e-09 | 4.1e-09 | 3.967E-09 | 5.132E-10 | 3.400E-09 | 4.400E-09 |
| 10779 | 539 | L539Y | A | 1e-08 | 1.3e-08 | 5.8e-09 | 9.600E-09 | 3.617E-09 | 5.800E-09 | 1.300E-08 |
| 10780 | 539 | L539V | A | 6e-09 | 1.2e-08 | 4.2e-09 | 7.400E-09 | 4.084E-09 | 4.200E-09 | 1.200E-08 |
| 10781 | 540 | P540A | A | 6e-09 | 5.4e-09 | 6.1e-09 | 5.833E-09 | 3.786E-10 | 5.400E-09 | 6.100E-09 |
| 10782 | 540 | P540R | A | 1.1e-08 | 3.6e-09 | 9e-09 | 7.867E-09 | 3.828E-09 | 3.600E-09 | 1.100E-08 |
| 10783 | 540 | P540N | A | 5.5e-09 | 2.9e-09 | 1.2e-08 | 6.800E-09 | 4.687E-09 | 2.900E-09 | 1.200E-08 |
| 10784 | 540 | P540D | A | 7.5e-09 | 4.3e-09 | 5.5e-09 | 5.767E-09 | 1.617E-09 | 4.300E-09 | 7.500E-09 |
| 10785 | 540 | P540C | A | 2.5e-09 | 6.8e-09 | 1.6e-08 | 8.433E-09 | 6.897E-09 | 2.500E-09 | 1.600E-08 |
| 10786 | 540 | P540E | A | 1.4e-08 | 1e-08 | 1e-08 | 1.133E-08 | 2.309E-09 | 1.000E-08 | 1.400E-08 |
| 10787 | 540 | P540Q | A | 1.9e-08 | 1.2e-08 | 1.8e-08 | 1.633E-08 | 3.786E-09 | 1.200E-08 | 1.900E-08 |
| 10788 | 540 | P540G | A | 6.1e-09 | 8.7e-09 | 3.4e-09 | 6.067E-09 | 2.650E-09 | 3.400E-09 | 8.700E-09 |
| 10789 | 540 | P540H | A | 1.3e-08 | 1e-08 | 6.7e-09 | 9.900E-09 | 3.151E-09 | 6.700E-09 | 1.300E-08 |
| 10790 | 540 | P540I | A | 6.6e-09 | 1.5e-08 | 2.1e-08 | 1.420E-08 | 7.233E-09 | 6.600E-09 | 2.100E-08 |
| 10791 | 540 | P540L | A | 2.6e-09 | 1.3e-08 | 5.2e-09 | 6.933E-09 | 5.412E-09 | 2.600E-09 | 1.300E-08 |
| 10792 | 540 | P540K | A | 1.5e-08 | 8.1e-09 | 1.2e-08 | 1.170E-08 | 3.460E-09 | 8.100E-09 | 1.500E-08 |
| 10793 | 540 | P540M | A | 2.8e-09 | 1.6e-08 | 1.1e-08 | 9.933E-09 | 6.664E-09 | 2.800E-09 | 1.600E-08 |
| 10794 | 540 | P540F | A | 5.6e-09 | 2.1e-08 | 1.6e-08 | 1.420E-08 | 7.856E-09 | 5.600E-09 | 2.100E-08 |

| | | | | | | | | | | |
|---|---|---|---|---|---|---|---|---|---|---|
| 10795 | 540 | P540P | A | 9.3e-09 | 3e-08 | 2.1e-08 | 2.010E-08 | 1.038E-08 | 9.300E-09 | 3.000E-08 |
| 10796 | 540 | P540S | A | 3.5e-09 | 9.3e-09 | 7.8e-09 | 6.867E-09 | 3.011E-09 | 3.500E-09 | 9.300E-09 |
| 10797 | 540 | P540T | A | 4.7e-09 | 6.6e-09 | 1.3e-08 | 8.100E-09 | 4.349E-09 | 4.700E-09 | 1.300E-08 |
| 10798 | 540 | P540W | A | 9.8e-09 | 9.4e-09 | 7.9e-09 | 9.033E-09 | 1.002E-09 | 7.900E-09 | 9.800E-09 |
| 10799 | 540 | P540Y | A | 5.4e-09 | 1.1e-08 | 3.6e-09 | 6.667E-09 | 3.859E-09 | 3.600E-09 | 1.100E-08 |
| 10800 | 540 | P540V | A | 5.6e-09 | 2.4e-09 | 9.8e-09 | 5.933E-09 | 3.711E-09 | 2.400E-09 | 9.800E-09 |
| 10801 | 541 | C541A | A | 1e-08 | 6.1e-09 | 7e-09 | 7.700E-09 | 2.042E-09 | 6.100E-09 | 1.000E-08 |
| 10802 | 541 | C541R | A | 1.8e-08 | 3.5e-09 | 4.5e-09 | 8.667E-09 | 8.098E-09 | 3.500E-09 | 1.800E-08 |
| 10803 | 541 | C541N | A | 3e-09 | 9.2e-09 | 3e-09 | 5.067E-09 | 3.580E-09 | 3.000E-09 | 9.200E-09 |
| 10804 | 541 | C541D | A | 7e-09 | 1.7e-08 | 6.7e-09 | 1.023E-08 | 5.862E-09 | 6.700E-09 | 1.700E-08 |
| 10805 | 541 | C541C | A | 5.6e-09 | 7.5e-09 | 7.7e-09 | 6.933E-09 | 1.159E-09 | 5.600E-09 | 7.700E-09 |
| 10806 | 541 | C541E | A | 1.8e-08 | 5.1e-09 | 2.1e-08 | 1.470E-08 | 8.448E-09 | 5.100E-09 | 2.100E-08 |
| 10807 | 541 | C541Q | A | 7.1e-09 | 6.2e-09 | 5.1e-09 | 6.133E-09 | 1.002E-09 | 5.100E-09 | 7.100E-09 |
| 10808 | 541 | C541G | A | 7.9e-09 | 3.1e-08 | 1.5e-08 | 1.797E-08 | 1.183E-08 | 7.900E-09 | 3.100E-08 |
| 10809 | 541 | C541H | A | 2.5e-09 | 2.9e-08 | 9.5e-09 | 1.367E-08 | 1.373E-08 | 2.500E-09 | 2.900E-08 |
| 10810 | 541 | C541I | A | 7.4e-10 | 7.1e-09 | 3e-09 | 3.613E-09 | 3.224E-09 | 7.400E-10 | 7.100E-09 |
| 10811 | 541 | C541L | A | 7.8e-09 | 1e-08 | 4.8e-09 | 7.533E-09 | 2.610E-09 | 4.800E-09 | 1.000E-08 |
| 10812 | 541 | C541K | A | 6.2e-09 | 8.6e-09 | 6.7e-09 | 7.167E-09 | 1.266E-09 | 6.200E-09 | 8.600E-09 |
| 10813 | 541 | C541M | A | 4.8e-09 | 1e-08 | 5.4e-09 | 6.733E-09 | 2.845E-09 | 4.800E-09 | 1.000E-08 |
| 10814 | 541 | C541F | A | 9.8e-09 | 1.6e-08 | 3.1e-08 | 1.893E-08 | 1.090E-08 | 9.800E-09 | 3.100E-08 |
| 10815 | 541 | C541P | A | 3.4e-09 | 4.2e-09 | 5.7e-09 | 4.433E-09 | 1.168E-09 | 3.400E-09 | 5.700E-09 |
| 10816 | 541 | C541S | A | 7.9e-09 | 7.3e-09 | 7.1e-09 | 7.433E-09 | 4.163E-10 | 7.100E-09 | 7.900E-09 |
| 10817 | 541 | C541T | A | 8.9e-09 | 6.7e-09 | 2.3e-09 | 5.967E-09 | 3.361E-09 | 2.300E-09 | 8.900E-09 |
| 10818 | 541 | C541W | A | 5.2e-09 | 9.4e-09 | 1.3e-08 | 9.200E-09 | 3.904E-09 | 5.200E-09 | 1.300E-08 |
| 10819 | 541 | C541Y | A | 7.7e-09 | 4.4e-09 | 1.4e-08 | 8.700E-09 | 4.877E-09 | 4.400E-09 | 1.400E-08 |
| 10820 | 541 | C541V | A | 5.5e-09 | 8.6e-09 | 9.1e-09 | 7.733E-09 | 1.950E-09 | 5.500E-09 | 9.100E-09 |
| 10821 | 542 | H542A | A | 1e-08 | 9e-09 | 4.1e-09 | 7.700E-09 | 3.158E-09 | 4.100E-09 | 1.000E-08 |
| 10822 | 542 | H542R | A | 1.3e-08 | 8.8e-09 | 2.8e-09 | 8.200E-09 | 5.126E-09 | 2.800E-09 | 1.300E-08 |
| 10823 | 542 | H542N | A | 8.9e-09 | 5.5e-09 | 1e-08 | 8.133E-09 | 2.346E-09 | 5.500E-09 | 1.000E-08 |
| 10824 | 542 | H542D | A | 3.8e-09 | 8.4e-09 | 8.5e-09 | 6.900E-09 | 2.685E-09 | 3.800E-09 | 8.500E-09 |
| 10825 | 542 | H542C | A | 2.4e-09 | 5.3e-09 | 6.1e-09 | 4.600E-09 | 1.947E-09 | 2.400E-09 | 6.100E-09 |
| 10826 | 542 | H542E | A | 5e-09 | 1.5e-08 | 8.9e-09 | 9.633E-09 | 5.040E-09 | 5.000E-09 | 1.500E-08 |
| 10827 | 542 | H542Q | A | 1.1e-08 | 6.5e-09 | 7.9e-09 | 8.467E-09 | 2.303E-09 | 6.500E-09 | 1.100E-08 |
| 10828 | 542 | H542G | A | 5.8e-09 | 1.5e-08 | 4.3e-09 | 8.367E-09 | 5.793E-09 | 4.300E-09 | 1.500E-08 |
| 10829 | 542 | H542H | A | 1e-08 | 5.1e-09 | 6e-09 | 7.033E-09 | 2.608E-09 | 5.100E-09 | 1.000E-08 |
| 10830 | 542 | H542I | A | 8.6e-09 | 1.1e-08 | 4.8e-09 | 8.133E-09 | 3.126E-09 | 4.800E-09 | 1.100E-08 |
| 10831 | 542 | H542L | A | 9e-09 | 4.4e-09 | 5.4e-09 | 6.267E-09 | 2.419E-09 | 4.400E-09 | 9.000E-09 |
| 10832 | 542 | H542K | A | 1e-08 | 7.5e-09 | 8.4e-09 | 8.633E-09 | 1.266E-09 | 7.500E-09 | 1.000E-08 |
| 10833 | 542 | H542M | A | 1.1e-08 | 1.9e-08 | 7e-09 | 1.233E-08 | 6.110E-09 | 7.000E-09 | 1.900E-08 |
| 10834 | 542 | H542F | A | 1.1e-08 | 3.8e-09 | 2e-09 | 5.600E-09 | 4.762E-09 | 2.000E-09 | 1.100E-08 |
| 10835 | 542 | H542P | A | 5.2e-09 | 6.5e-09 | 4.8e-09 | 5.500E-09 | 8.888E-10 | 4.800E-09 | 6.500E-09 |
| 10836 | 542 | H542S | A | 4.6e-09 | 1.5e-08 | 1.4e-08 | 1.120E-08 | 5.738E-09 | 4.600E-09 | 1.500E-08 |
| 10837 | 542 | H542T | A | 1.5e-09 | 1.2e-08 | 5.1e-09 | 6.200E-09 | 5.336E-09 | 1.500E-09 | 1.200E-08 |
| 10838 | 542 | H542W | A | 1.4e-08 | 6.5e-09 | 1.2e-08 | 1.083E-08 | 3.884E-09 | 6.500E-09 | 1.400E-08 |
| 10839 | 542 | H542Y | A | 3.6e-09 | 8.4e-09 | 1e-08 | 7.333E-09 | 3.331E-09 | 3.600E-09 | 1.000E-08 |
| 10840 | 542 | H542V | A | 2.5e-08 | 8e-09 | 7.5e-09 | 1.350E-08 | 9.962E-09 | 7.500E-09 | 2.500E-08 |
| 10841 | 543 | P543A | A | 1.1e-08 | 4.4e-09 | 7.1e-09 | 7.500E-09 | 3.318E-09 | 4.400E-09 | 1.100E-08 |
| 10842 | 543 | P543R | A | 4e-09 | 1.3e-08 | 4.4e-09 | 7.133E-09 | 5.085E-09 | 4.000E-09 | 1.300E-08 |
| 10843 | 543 | P543N | A | 7.1e-09 | 8.2e-09 | 5.6e-09 | 6.967E-09 | 1.305E-09 | 5.600E-09 | 8.200E-09 |
| 10844 | 543 | P543D | A | 5.3e-09 | 5.3e-09 | 5.7e-09 | 5.433E-09 | 2.309E-10 | 5.300E-09 | 5.700E-09 |
| 10845 | 543 | P543C | A | 9.7e-09 | 7.8e-09 | 8.7e-09 | 8.733E-09 | 9.504E-10 | 7.800E-09 | 9.700E-09 |
| 10846 | 543 | P543E | A | 2.6e-09 | 4.3e-09 | 8e-09 | 4.967E-09 | 2.761E-09 | 2.600E-09 | 8.000E-09 |
| 10847 | 543 | P543Q | A | 6.3e-09 | 7.9e-09 | 4.5e-09 | 6.233E-09 | 1.701E-09 | 4.500E-09 | 7.900E-09 |
| 10848 | 543 | P543G | A | 5.1e-09 | 3.2e-08 | 7.6e-09 | 1.490E-08 | 1.486E-08 | 5.100E-09 | 3.200E-08 |
| 10849 | 543 | P543H | A | 1.5e-08 | 5.9e-09 | 8.7e-09 | 9.867E-09 | 4.661E-09 | 5.900E-09 | 1.500E-08 |
| 10850 | 543 | P543I | A | 9.6e-09 | 6.5e-09 | 3e-08 | 1.537E-08 | 1.277E-08 | 6.500E-09 | 3.000E-08 |
| 10851 | 543 | P543L | A | 5.4e-09 | 4.2e-09 | 7.9e-09 | 5.833E-09 | 1.888E-09 | 4.200E-09 | 7.900E-09 |
| 10852 | 543 | P543K | A | 9.9e-09 | 5.5e-09 | 7.2e-09 | 7.533E-09 | 2.219E-09 | 5.500E-09 | 9.900E-09 |
| 10853 | 543 | P543M | A | 6.2e-09 | 2.4e-09 | 4.4e-09 | 4.333E-09 | 1.901E-09 | 2.400E-09 | 6.200E-09 |
| 10854 | 543 | P543F | A | 7.3e-09 | 1.7e-08 | 1.4e-08 | 1.277E-08 | 4.966E-09 | 7.300E-09 | 1.700E-08 |
| 10855 | 543 | P543P | A | 3.1e-09 | 5.5e-09 | 5.4e-09 | 4.667E-09 | 1.358E-09 | 3.100E-09 | 5.500E-09 |
| 10856 | 543 | P543S | A | 6.6e-09 | 3.5e-09 | 4.3e-09 | 4.800E-09 | 1.609E-09 | 3.500E-09 | 6.600E-09 |
| 10857 | 543 | P543T | A | 7.4e-09 | 3.7e-09 | 8.1e-09 | 6.400E-09 | 2.364E-09 | 3.700E-09 | 8.100E-09 |
| 10858 | 543 | P543W | A | 5.1e-09 | 9.6e-09 | 9.3e-09 | 8.000E-09 | 2.516E-09 | 5.100E-09 | 9.600E-09 |
| 10859 | 543 | P543Y | A | 6.7e-09 | 1.4e-08 | 1e-08 | 1.023E-08 | 3.656E-09 | 6.700E-09 | 1.400E-08 |
| 10860 | 543 | P543V | A | 7.3e-09 | 8.1e-09 | 7.6e-09 | 7.667E-09 | 4.041E-10 | 7.300E-09 | 8.100E-09 |
| 10861 | 544 | E544A | A | 2.6e-09 | 1.2e-08 | 2.4e-08 | 1.287E-08 | 1.073E-08 | 2.600E-09 | 2.400E-08 |
| 10862 | 544 | E544R | A | 1.1e-08 | 1.1e-08 | 4e-09 | 8.667E-09 | 4.041E-09 | 4.000E-09 | 1.100E-08 |
| 10863 | 544 | E544N | A | 1.3e-08 | 9.4e-09 | 7.5e-09 | 9.967E-09 | 2.793E-09 | 7.500E-09 | 1.300E-08 |
| 10864 | 544 | E544D | A | 1.2e-08 | 3.3e-09 | 1.6e-08 | 1.043E-08 | 6.493E-09 | 3.300E-09 | 1.600E-08 |
| 10865 | 544 | E544C | A | 1.1e-08 | 1.4e-08 | 1.2e-08 | 1.233E-08 | 1.528E-09 | 1.100E-08 | 1.400E-08 |
| 10866 | 544 | E544E | A | 6.1e-09 | 1.1e-08 | 4.8e-09 | 7.300E-09 | 3.270E-09 | 4.800E-09 | 1.100E-08 |
| 10867 | 544 | E544Q | A | 1.2e-08 | 1e-08 | 9.7e-09 | 1.057E-08 | 1.250E-09 | 9.700E-09 | 1.200E-08 |
| 10868 | 544 | E544G | A | 1e-08 | 1e-08 | 1.2e-08 | 1.067E-08 | 1.155E-09 | 1.000E-08 | 1.200E-08 |
| 10869 | 544 | E544H | A | 3.6e-09 | 4.4e-09 | 7.3e-09 | 5.100E-09 | 1.947E-09 | 3.600E-09 | 7.300E-09 |
| 10870 | 544 | E544I | A | 4.1e-09 | 7.7e-09 | 5.2e-09 | 5.667E-09 | 1.845E-09 | 4.100E-09 | 7.700E-09 |
| 10871 | 544 | E544L | A | 8.1e-09 | 1.5e-08 | 8.5e-09 | 1.053E-08 | 3.873E-09 | 8.100E-09 | 1.500E-08 |
| 10872 | 544 | E544K | A | 7.3e-09 | 5.1e-09 | 1.6e-08 | 9.467E-09 | 5.764E-09 | 5.100E-09 | 1.600E-08 |
| 10873 | 544 | E544M | A | 9.7e-09 | 7.4e-09 | 4.9e-09 | 7.333E-09 | 2.401E-09 | 4.900E-09 | 9.700E-09 |
| 10874 | 544 | E544F | A | 4.4e-09 | 6.6e-09 | 3.6e-08 | 1.567E-08 | 1.764E-08 | 4.400E-09 | 3.600E-08 |
| 10875 | 544 | E544P | A | 8.5e-09 | 5.9e-09 | 4.5e-09 | 6.300E-09 | 2.030E-09 | 4.500E-09 | 8.500E-09 |
| 10876 | 544 | E544S | A | 1.3e-08 | 6.6e-09 | 1.6e-08 | 1.187E-08 | 4.801E-09 | 6.600E-09 | 1.600E-08 |
| 10877 | 544 | E544T | A | 6.8e-09 | 1.2e-08 | 1.3e-08 | 1.060E-08 | 3.329E-09 | 6.800E-09 | 1.300E-08 |
| 10878 | 544 | E544W | A | 1.1e-08 | 1.1e-09 | 2.4e-09 | 4.833E-09 | 5.380E-09 | 1.100E-09 | 1.100E-08 |
| 10879 | 544 | E544Y | A | 1.5e-08 | 7.8e-09 | 1.2e-08 | 1.160E-08 | 3.617E-09 | 7.800E-09 | 1.500E-08 |

| | | | | | | | | | | |
|---|---|---|---|---|---|---|---|---|---|---|
| 10880 | 544 | E544V | A | 9.2e-09 | 3.2e-09 | 9.7e-09 | 7.367E-09 | 3.617E-09 | 3.200E-09 | 9.700E-09 |
| 10881 | 545 | C545A | A | 5.1e-09 | 9.2e-09 | 1.7e-09 | 5.333E-09 | 3.755E-09 | 1.700E-09 | 9.200E-09 |
| 10882 | 545 | C545R | A | 3.7e-08 | 2.2e-08 | 5.9e-09 | 2.163E-08 | 1.555E-08 | 5.900E-09 | 3.700E-08 |
| 10883 | 545 | C545N | A | 1.2e-08 | 1.7e-08 | 8.7e-09 | 1.257E-08 | 4.179E-09 | 8.700E-09 | 1.700E-08 |
| 10884 | 545 | C545D | A | 9.2e-09 | 2.2e-08 | 1.3e-08 | 1.473E-08 | 6.574E-09 | 9.200E-09 | 2.200E-08 |
| 10885 | 545 | C545C | A | 4.9e-09 | 5.6e-09 | 5.4e-09 | 5.300E-09 | 3.606E-10 | 4.900E-09 | 5.600E-09 |
| 10886 | 545 | C545E | A | 9.9e-09 | 1.2e-08 | 4.3e-09 | 8.733E-09 | 3.980E-09 | 4.300E-09 | 1.200E-08 |
| 10887 | 545 | C545Q | A | 3.2e-08 | 2.6e-09 | 5.5e-09 | 1.337E-08 | 1.620E-08 | 2.600E-09 | 3.200E-08 |
| 10888 | 545 | C545G | A | 3.1e-09 | 1.4e-08 | 2.4e-09 | 6.500E-09 | 6.505E-09 | 2.400E-09 | 1.400E-08 |
| 10889 | 545 | C545H | A | 3.7e-09 | 7.1e-09 | 6.5e-09 | 5.767E-09 | 1.815E-09 | 3.700E-09 | 7.100E-09 |
| 10890 | 545 | C545I | A | 8.4e-09 | 6.9e-09 | 7.2e-09 | 7.500E-09 | 7.937E-10 | 6.900E-09 | 8.400E-09 |
| 10891 | 545 | C545L | A | 1.1e-08 | 2.6e-08 | 1.8e-08 | 1.833E-08 | 7.506E-09 | 1.100E-08 | 2.600E-08 |
| 10892 | 545 | C545K | A | 1.4e-08 | 9.9e-09 | 8.2e-09 | 1.070E-08 | 2.982E-09 | 8.200E-09 | 1.400E-08 |
| 10893 | 545 | C545M | A | 7.3e-09 | 3.5e-09 | 7.1e-09 | 5.967E-09 | 2.139E-09 | 3.500E-09 | 7.300E-09 |
| 10894 | 545 | C545F | A | 5.2e-09 | 4.9e-09 | 5.6e-09 | 5.233E-09 | 3.512E-10 | 4.900E-09 | 5.600E-09 |
| 10895 | 545 | C545P | A | 5e-09 | 2.8e-09 | 9.8e-09 | 5.867E-09 | 3.580E-09 | 2.800E-09 | 9.800E-09 |
| 10896 | 545 | C545S | A | 9.6e-09 | 4.1e-09 | 4.7e-09 | 6.133E-09 | 3.017E-09 | 4.100E-09 | 9.600E-09 |
| 10897 | 545 | C545T | A | 2.7e-09 | 5.5e-09 | 1.4e-08 | 7.400E-09 | 5.885E-09 | 2.700E-09 | 1.400E-08 |
| 10898 | 545 | C545W | A | 1.3e-08 | 8.4e-09 | 4.9e-09 | 8.767E-09 | 4.062E-09 | 4.900E-09 | 1.300E-08 |
| 10899 | 545 | C545Y | A | 8.9e-09 | 6.4e-09 | 1e-08 | 8.433E-09 | 1.845E-09 | 6.400E-09 | 1.000E-08 |
| 10900 | 545 | C545V | A | 8.1e-09 | 8.1e-09 | 4.9e-09 | 7.033E-09 | 1.848E-09 | 4.900E-09 | 8.100E-09 |
| 10901 | 546 | Q546A | A | 7.6e-09 | 3e-09 | 2.7e-09 | 4.433E-09 | 2.747E-09 | 2.700E-09 | 7.600E-09 |
| 10902 | 546 | Q546R | A | 7.6e-09 | 3.9e-09 | 6.2e-09 | 5.900E-09 | 1.868E-09 | 3.900E-09 | 7.600E-09 |
| 10903 | 546 | Q546N | A | 7.1e-09 | 6.4e-09 | 1.1e-08 | 8.167E-09 | 2.479E-09 | 6.400E-09 | 1.100E-08 |
| 10904 | 546 | Q546D | A | 5.6e-09 | 7.8e-09 | 6.5e-09 | 6.633E-09 | 1.106E-09 | 5.600E-09 | 7.800E-09 |
| 10905 | 546 | Q546C | A | 1.6e-09 | 1.1e-08 | 4.2e-09 | 5.600E-09 | 4.854E-09 | 1.600E-09 | 1.100E-08 |
| 10906 | 546 | Q546E | A | 1.8e-08 | 1.2e-08 | 2.8e-09 | 1.093E-08 | 7.656E-09 | 2.800E-09 | 1.800E-08 |
| 10907 | 546 | Q546Q | A | 1.2e-08 | 9.1e-09 | 6.6e-09 | 9.233E-09 | 2.702E-09 | 6.600E-09 | 1.200E-08 |
| 10908 | 546 | Q546G | A | 3.8e-09 | 1.1e-08 | 6.3e-09 | 7.033E-09 | 3.656E-09 | 3.800E-09 | 1.100E-08 |
| 10909 | 546 | Q546H | A | 1.9e-08 | 1.5e-08 | 1.7e-08 | 1.700E-08 | 2.000E-09 | 1.500E-08 | 1.900E-08 |
| 10910 | 546 | Q546I | A | 1.1e-08 | 5.4e-09 | 5.8e-09 | 7.400E-09 | 3.124E-09 | 5.400E-09 | 1.100E-08 |
| 10911 | 546 | Q546L | A | 5.3e-09 | 7.8e-09 | 4.8e-09 | 5.967E-09 | 1.607E-09 | 4.800E-09 | 7.800E-09 |
| 10912 | 546 | Q546K | A | 5e-09 | 2.6e-08 | 5.6e-09 | 1.220E-08 | 1.195E-08 | 5.000E-09 | 2.600E-08 |
| 10913 | 546 | Q546M | A | 6.7e-09 | 7.4e-09 | 7.2e-09 | 7.100E-09 | 3.606E-10 | 6.700E-09 | 7.400E-09 |
| 10914 | 546 | Q546F | A | 3.5e-09 | 6e-09 | 3.3e-09 | 4.267E-09 | 1.504E-09 | 3.300E-09 | 6.000E-09 |
| 10915 | 546 | Q546P | A | 3.7e-09 | 6.5e-09 | 3.2e-09 | 4.467E-09 | 1.779E-09 | 3.200E-09 | 6.500E-09 |
| 10916 | 546 | Q546S | A | 1.1e-08 | 1.7e-08 | 8.9e-09 | 1.230E-08 | 4.204E-09 | 8.900E-09 | 1.700E-08 |
| 10917 | 546 | Q546T | A | 5.7e-09 | 5.5e-09 | 9.1e-09 | 6.767E-09 | 2.023E-09 | 5.500E-09 | 9.100E-09 |
| 10918 | 546 | Q546W | A | 3.3e-09 | 4.9e-09 | 1.7e-08 | 8.400E-09 | 7.491E-09 | 3.300E-09 | 1.700E-08 |
| 10919 | 546 | Q546Y | A | 9.7e-09 | 2.5e-09 | 9e-09 | 7.067E-09 | 3.970E-09 | 2.500E-09 | 9.700E-09 |
| 10920 | 546 | Q546V | A | 6.7e-09 | 3.7e-09 | 8e-09 | 6.133E-09 | 2.205E-09 | 3.700E-09 | 8.000E-09 |
| 10921 | 547 | P547A | A | 3.7e-09 | 3.1e-09 | 1.1e-08 | 5.933E-09 | 4.398E-09 | 3.100E-09 | 1.100E-08 |
| 10922 | 547 | P547R | A | 8.8e-09 | 9.5e-09 | 1.6e-08 | 1.143E-08 | 3.970E-09 | 8.800E-09 | 1.600E-08 |
| 10923 | 547 | P547N | A | 3.9e-09 | 1.1e-08 | 9.3e-09 | 8.067E-09 | 3.707E-09 | 3.900E-09 | 1.100E-08 |
| 10924 | 547 | P547D | A | 4.9e-09 | 6.1e-09 | 8.5e-09 | 6.500E-09 | 1.833E-09 | 4.900E-09 | 8.500E-09 |
| 10925 | 547 | P547C | A | 1.1e-09 | 4.3e-09 | 4e-09 | 3.133E-09 | 1.767E-09 | 1.100E-09 | 4.300E-09 |
| 10926 | 547 | P547E | A | 6.8e-09 | 1.6e-09 | 7.3e-09 | 5.233E-09 | 3.156E-09 | 1.600E-09 | 7.300E-09 |
| 10927 | 547 | P547Q | A | 4.1e-09 | 9.4e-09 | 5.8e-09 | 6.433E-09 | 2.706E-09 | 4.100E-09 | 9.400E-09 |
| 10928 | 547 | P547G | A | 2.8e-09 | 1.6e-08 | 1.1e-08 | 9.933E-09 | 6.664E-09 | 2.800E-09 | 1.600E-08 |
| 10929 | 547 | P547H | A | 6.9e-09 | 1.7e-08 | 2.2e-08 | 1.530E-08 | 7.692E-09 | 6.900E-09 | 2.200E-08 |
| 10930 | 547 | P547I | A | 8.1e-09 | 6.5e-09 | 6.7e-09 | 7.100E-09 | 8.718E-10 | 6.500E-09 | 8.100E-09 |
| 10931 | 547 | P547L | A | 1.4e-09 | 1.1e-08 | 6e-09 | 6.133E-09 | 4.801E-09 | 1.400E-09 | 1.100E-08 |
| 10932 | 547 | P547K | A | 3.5e-09 | 6.2e-09 | 2.9e-09 | 4.200E-09 | 1.758E-09 | 2.900E-09 | 6.200E-09 |
| 10933 | 547 | P547M | A | 1.1e-08 | 1.5e-08 | 2.4e-08 | 1.667E-08 | 6.658E-09 | 1.100E-08 | 2.400E-08 |
| 10934 | 547 | P547F | A | 3.7e-09 | 8.4e-09 | 3.2e-08 | 1.470E-08 | 1.517E-08 | 3.700E-09 | 3.200E-08 |
| 10935 | 547 | P547P | A | 4.6e-09 | 7.7e-09 | 7.7e-09 | 6.667E-09 | 1.790E-09 | 4.600E-09 | 7.700E-09 |
| 10936 | 547 | P547S | A | 9.6e-09 | 5.4e-09 | 3.9e-09 | 6.300E-09 | 2.955E-09 | 3.900E-09 | 9.600E-09 |
| 10937 | 547 | P547T | A | 1.7e-08 | 4.7e-09 | 5.2e-09 | 8.967E-09 | 6.962E-09 | 4.700E-09 | 1.700E-08 |
| 10938 | 547 | P547W | A | 3.2e-09 | 1e-08 | 3.3e-09 | 5.500E-09 | 3.897E-09 | 3.200E-09 | 1.000E-08 |
| 10939 | 547 | P547Y | A | 4.5e-09 | 1.1e-08 | 5.3e-09 | 6.933E-09 | 3.544E-09 | 4.500E-09 | 1.100E-08 |
| 10940 | 547 | P547V | A | 2.7e-08 | 5.2e-09 | 1.9e-08 | 1.707E-08 | 1.103E-08 | 5.200E-09 | 2.700E-08 |
| 10941 | 548 | Q548A | A | 5.3e-09 | 5e-09 | 9.1e-09 | 6.467E-09 | 2.285E-09 | 5.000E-09 | 9.100E-09 |
| 10942 | 548 | Q548R | A | 3.3e-09 | 1.2e-08 | 5.5e-09 | 6.933E-09 | 4.524E-09 | 3.300E-09 | 1.200E-08 |
| 10943 | 548 | Q548N | A | 2.6e-09 | 6.1e-09 | 4.9e-09 | 4.533E-09 | 1.779E-09 | 2.600E-09 | 6.100E-09 |
| 10944 | 548 | Q548D | A | 5.3e-09 | 2.9e-09 | 1.9e-08 | 9.067E-09 | 8.686E-09 | 2.900E-09 | 1.900E-08 |
| 10945 | 548 | Q548C | A | 1e-08 | 4.5e-09 | 1.7e-08 | 1.050E-08 | 6.265E-09 | 4.500E-09 | 1.700E-08 |
| 10946 | 548 | Q548E | A | 1.3e-08 | 7.3e-09 | 3e-09 | 7.767E-09 | 5.016E-09 | 3.000E-09 | 1.300E-08 |
| 10947 | 548 | Q548Q | A | 2.1e-09 | 2.1e-09 | 1.4e-08 | 6.067E-09 | 6.870E-09 | 2.100E-09 | 1.400E-08 |
| 10948 | 548 | Q548G | A | 3.8e-09 | 1e-08 | 4.6e-09 | 6.133E-09 | 3.372E-09 | 3.800E-09 | 1.000E-08 |
| 10949 | 548 | Q548H | A | 2.3e-09 | 6.9e-09 | 9.4e-09 | 6.200E-09 | 3.601E-09 | 2.300E-09 | 9.400E-09 |
| 10950 | 548 | Q548I | A | 7.1e-09 | 8.4e-09 | 9.1e-09 | 8.200E-09 | 1.015E-09 | 7.100E-09 | 9.100E-09 |
| 10951 | 548 | Q548L | A | 5.3e-09 | 5.2e-09 | 1.3e-08 | 7.833E-09 | 4.475E-09 | 5.200E-09 | 1.300E-08 |
| 10952 | 548 | Q548K | A | 7.2e-09 | 5.1e-09 | 4.3e-09 | 5.533E-09 | 1.498E-09 | 4.300E-09 | 7.200E-09 |
| 10953 | 548 | Q548M | A | 6.6e-09 | 1e-08 | 4.8e-09 | 7.133E-09 | 2.641E-09 | 4.800E-09 | 1.000E-08 |
| 10954 | 548 | Q548F | A | 5e-09 | 2.3e-09 | 4.9e-09 | 4.067E-09 | 1.531E-09 | 2.300E-09 | 5.000E-09 |
| 10955 | 548 | Q548P | A | 4.4e-09 | 9.2e-09 | 8.6e-09 | 7.400E-09 | 2.615E-09 | 4.400E-09 | 9.200E-09 |
| 10956 | 548 | Q548S | A | 5e-09 | 8.6e-09 | 7.1e-09 | 6.900E-09 | 1.808E-09 | 5.000E-09 | 8.600E-09 |
| 10957 | 548 | Q548T | A | 4.8e-09 | 1.1e-08 | 3.2e-09 | 6.333E-09 | 4.120E-09 | 3.200E-09 | 1.100E-08 |
| 10958 | 548 | Q548W | A | 2.2e-09 | 2.1e-08 | 4.6e-09 | 9.267E-09 | 1.023E-08 | 2.200E-09 | 2.100E-08 |
| 10959 | 548 | Q548Y | A | 6.9e-09 | 3.5e-09 | 8e-09 | 6.133E-09 | 2.346E-09 | 3.500E-09 | 8.000E-09 |
| 10960 | 548 | Q548V | A | 5.8e-09 | 3.4e-09 | 9.5e-09 | 6.233E-09 | 3.073E-09 | 3.400E-09 | 9.500E-09 |
| 10961 | 549 | N549A | A | 7.7e-09 | 1e-08 | 6.6e-09 | 8.100E-09 | 1.735E-09 | 6.600E-09 | 1.000E-08 |
| 10962 | 549 | N549R | A | 1.1e-09 | 6.4e-09 | 1.6e-08 | 7.833E-09 | 7.553E-09 | 1.100E-09 | 1.600E-08 |
| 10963 | 549 | N549N | A | 7.6e-09 | 5.2e-09 | 5.2e-09 | 6.000E-09 | 1.386E-09 | 5.200E-09 | 7.600E-09 |
| 10964 | 549 | N549D | A | 1.2e-08 | 1e-08 | 8.4e-09 | 1.013E-08 | 1.804E-09 | 8.400E-09 | 1.200E-08 |

| | | | | | | | | | | |
|---|---|---|---|---|---|---|---|---|---|---|
| 10965 | 549 | N549C | A | 1.9e-09 | 6.4e-09 | 1.3e-08 | 7.100E-09 | 5.583E-09 | 1.900E-09 | 1.300E-08 |
| 10966 | 549 | N549E | A | 9.4e-09 | 4.4e-09 | 1.5e-08 | 9.600E-09 | 5.303E-09 | 4.400E-09 | 1.500E-08 |
| 10967 | 549 | N549Q | A | 8.3e-09 | 1.5e-08 | 7.7e-09 | 1.033E-08 | 4.053E-09 | 7.700E-09 | 1.500E-08 |
| 10968 | 549 | N549G | A | 9.4e-09 | 7.3e-09 | 8.7e-09 | 8.467E-09 | 1.069E-09 | 7.300E-09 | 9.400E-09 |
| 10969 | 549 | N549H | A | 2.6e-09 | 8.6e-09 | 1e-08 | 7.067E-09 | 3.931E-09 | 2.600E-09 | 1.000E-08 |
| 10970 | 549 | N549I | A | 4.7e-09 | 7.2e-09 | 2.6e-09 | 4.833E-09 | 2.303E-09 | 2.600E-09 | 7.200E-09 |
| 10971 | 549 | N549L | A | 6.7e-09 | 1.2e-08 | 1.1e-08 | 9.900E-09 | 2.816E-09 | 6.700E-09 | 1.200E-08 |
| 10972 | 549 | N549K | A | 7.1e-09 | 2.1e-09 | 7.3e-09 | 5.500E-09 | 2.946E-09 | 2.100E-09 | 7.300E-09 |
| 10973 | 549 | N549M | A | 1.2e-08 | 1.8e-08 | 3.8e-09 | 1.127E-08 | 7.128E-09 | 3.800E-09 | 1.800E-08 |
| 10974 | 549 | N549F | A | 5.5e-09 | 1.9e-08 | 2.4e-09 | 8.967E-09 | 8.826E-09 | 2.400E-09 | 1.900E-08 |
| 10975 | 549 | N549P | A | 4.3e-09 | 5.6e-09 | 1.3e-08 | 7.633E-09 | 4.693E-09 | 4.300E-09 | 1.300E-08 |
| 10976 | 549 | N549S | A | 4e-09 | 2.2e-08 | 1.1e-08 | 1.233E-08 | 9.074E-09 | 4.000E-09 | 2.200E-08 |
| 10977 | 549 | N549T | A | 6.7e-09 | 4.2e-09 | 1e-08 | 6.967E-09 | 2.909E-09 | 4.200E-09 | 1.000E-08 |
| 10978 | 549 | N549W | A | 2.3e-09 | 8.1e-09 | 6e-09 | 5.467E-09 | 2.937E-09 | 2.300E-09 | 8.100E-09 |
| 10979 | 549 | N549Y | A | 1.6e-09 | 4.6e-09 | 7e-09 | 4.400E-09 | 2.706E-09 | 1.600E-09 | 7.000E-09 |
| 10980 | 549 | N549V | A | 1.8e-08 | 5.8e-09 | 8.9e-09 | 1.090E-08 | 6.341E-09 | 5.800E-09 | 1.800E-08 |
| 10981 | 550 | G550A | A | 8.2e-09 | 2.7e-09 | 1.9e-08 | 9.967E-09 | 8.292E-09 | 2.700E-09 | 1.900E-08 |
| 10982 | 550 | G550R | A | 9.3e-09 | 5.2e-09 | 4.9e-09 | 6.467E-09 | 2.458E-09 | 4.900E-09 | 9.300E-09 |
| 10983 | 550 | G550N | A | 7.4e-09 | 1.1e-08 | 5.9e-09 | 8.100E-09 | 2.621E-09 | 5.900E-09 | 1.100E-08 |
| 10984 | 550 | G550D | A | 8.1e-09 | 5.6e-09 | 7.3e-09 | 7.000E-09 | 1.277E-09 | 5.600E-09 | 8.100E-09 |
| 10985 | 550 | G550C | A | 9.1e-09 | 3e-08 | 2.9e-09 | 1.400E-08 | 1.420E-08 | 2.900E-09 | 3.000E-08 |
| 10986 | 550 | G550E | A | 2.1e-09 | 7.8e-09 | 4.1e-09 | 4.667E-09 | 2.892E-09 | 2.100E-09 | 7.800E-09 |
| 10987 | 550 | G550Q | A | 1.7e-08 | 1.4e-09 | 1.2e-08 | 1.013E-08 | 7.966E-09 | 1.400E-09 | 1.700E-08 |
| 10988 | 550 | G550G | A | 6e-09 | 8.3e-09 | 5.4e-09 | 6.567E-09 | 1.531E-09 | 5.400E-09 | 8.300E-09 |
| 10989 | 550 | G550H | A | 9.5e-09 | 3.2e-09 | 1.4e-08 | 8.900E-09 | 5.425E-09 | 3.200E-09 | 1.400E-08 |
| 10990 | 550 | G550I | A | 3.3e-09 | 1.8e-08 | 2.5e-08 | 1.543E-08 | 1.108E-08 | 3.300E-09 | 2.500E-08 |
| 10991 | 550 | G550L | A | 3.8e-09 | 8.5e-09 | 2e-09 | 4.767E-09 | 3.356E-09 | 2.000E-09 | 8.500E-09 |
| 10992 | 550 | G550K | A | 2.7e-08 | 4.4e-09 | 5.6e-09 | 1.233E-08 | 1.272E-08 | 4.400E-09 | 2.700E-08 |
| 10993 | 550 | G550M | A | 1.2e-08 | 1.8e-08 | 2.1e-08 | 1.700E-08 | 4.583E-09 | 1.200E-08 | 2.100E-08 |
| 10994 | 550 | G550F | A | 2.6e-09 | 1.1e-08 | 6e-09 | 6.533E-09 | 4.225E-09 | 2.600E-09 | 1.100E-08 |
| 10995 | 550 | G550P | A | 7e-09 | 7.3e-09 | 7.9e-09 | 7.400E-09 | 4.583E-10 | 7.000E-09 | 7.900E-09 |
| 10996 | 550 | G550S | A | 1.3e-08 | 1.4e-08 | 4.9e-09 | 1.063E-08 | 4.990E-09 | 4.900E-09 | 1.400E-08 |
| 10997 | 550 | G550T | A | 1.4e-08 | 1.1e-08 | 9.8e-09 | 1.160E-08 | 2.163E-09 | 9.800E-09 | 1.400E-08 |
| 10998 | 550 | G550W | A | 1.5e-08 | 1.8e-08 | 5.3e-09 | 1.277E-08 | 6.638E-09 | 5.300E-09 | 1.800E-08 |
| 10999 | 550 | G550Y | A | 1.1e-08 | 2.6e-09 | 8.7e-09 | 7.433E-09 | 4.341E-09 | 2.600E-09 | 1.100E-08 |
| 11000 | 550 | G550V | A | 7.6e-09 | 8.5e-09 | 5.9e-09 | 7.333E-09 | 1.320E-09 | 5.900E-09 | 8.500E-09 |
| 11001 | 551 | S551A | A | 1.6e-08 | 5.9e-09 | 8.5e-09 | 1.013E-08 | 5.244E-09 | 5.900E-09 | 1.600E-08 |
| 11002 | 551 | S551R | A | 1.8e-08 | 1.9e-08 | 4.9e-09 | 1.397E-08 | 7.868E-09 | 4.900E-09 | 1.900E-08 |
| 11003 | 551 | S551N | A | 9.8e-09 | 1.2e-08 | 2.1e-09 | 7.967E-09 | 5.198E-09 | 2.100E-09 | 1.200E-08 |
| 11004 | 551 | S551D | A | 8.7e-09 | 3.1e-09 | 6.6e-09 | 6.133E-09 | 2.829E-09 | 3.100E-09 | 8.700E-09 |
| 11005 | 551 | S551C | A | 1.7e-08 | 1e-08 | 2.6e-09 | 9.867E-09 | 7.201E-09 | 2.600E-09 | 1.700E-08 |
| 11006 | 551 | S551E | A | 4.3e-09 | 9.1e-09 | 2e-08 | 1.113E-08 | 8.045E-09 | 4.300E-09 | 2.000E-08 |
| 11007 | 551 | S551Q | A | 4.1e-09 | 6.3e-09 | 5.4e-09 | 5.267E-09 | 1.106E-09 | 4.100E-09 | 6.300E-09 |
| 11008 | 551 | S551G | A | 1.8e-08 | 9.6e-09 | 4.2e-09 | 1.060E-08 | 6.954E-09 | 4.200E-09 | 1.800E-08 |
| 11009 | 551 | S551H | A | 9.4e-09 | 4e-09 | 4.7e-09 | 6.033E-09 | 2.937E-09 | 4.000E-09 | 9.400E-09 |
| 11010 | 551 | S551I | A | 4.9e-09 | 3.6e-09 | 6.6e-09 | 5.033E-09 | 1.504E-09 | 3.600E-09 | 6.600E-09 |
| 11011 | 551 | S551L | A | 5.2e-09 | 8e-09 | 4.7e-09 | 5.967E-09 | 1.779E-09 | 4.700E-09 | 8.000E-09 |
| 11012 | 551 | S551K | A | 4.4e-09 | 8.8e-09 | 6.9e-09 | 6.700E-09 | 2.207E-09 | 4.400E-09 | 8.800E-09 |
| 11013 | 551 | S551M | A | 4.7e-09 | 1.1e-08 | 5.5e-09 | 7.067E-09 | 3.430E-09 | 4.700E-09 | 1.100E-08 |
| 11014 | 551 | S551F | A | 2.3e-09 | 1.8e-08 | 9.6e-09 | 9.967E-09 | 7.856E-09 | 2.300E-09 | 1.800E-08 |
| 11015 | 551 | S551P | A | 4.8e-09 | 9.9e-09 | 1.3e-08 | 9.233E-09 | 4.140E-09 | 4.800E-09 | 1.300E-08 |
| 11016 | 551 | S551S | A | 4.9e-09 | 8.2e-09 | 4.5e-09 | 5.867E-09 | 2.031E-09 | 4.500E-09 | 8.200E-09 |
| 11017 | 551 | S551T | A | 4e-09 | 4.8e-09 | 1.8e-08 | 8.933E-09 | 7.862E-09 | 4.000E-09 | 1.800E-08 |
| 11018 | 551 | S551W | A | 6.6e-09 | 4.7e-09 | 7.9e-09 | 6.400E-09 | 1.609E-09 | 4.700E-09 | 7.900E-09 |
| 11019 | 551 | S551Y | A | 8.1e-09 | 1.7e-08 | 5.4e-09 | 1.017E-08 | 6.070E-09 | 5.400E-09 | 1.700E-08 |
| 11020 | 551 | S551V | A | 1.2e-08 | 2e-08 | 5e-09 | 1.233E-08 | 7.506E-09 | 5.000E-09 | 2.000E-08 |
| 11021 | 552 | V552A | A | 8e-09 | 7.4e-09 | 1.7e-09 | 5.700E-09 | 3.477E-09 | 1.700E-09 | 8.000E-09 |
| 11022 | 552 | V552R | A | 1.4e-08 | 2.6e-09 | 5.1e-09 | 7.233E-09 | 5.992E-09 | 2.600E-09 | 1.400E-08 |
| 11023 | 552 | V552N | A | 7.3e-09 | 8.3e-09 | 6.1e-09 | 7.233E-09 | 1.102E-09 | 6.100E-09 | 8.300E-09 |
| 11024 | 552 | V552D | A | 9.4e-09 | 1.8e-09 | 1.1e-08 | 7.400E-09 | 4.915E-09 | 1.800E-09 | 1.100E-08 |
| 11025 | 552 | V552C | A | 1.1e-08 | 1.1e-08 | 4.2e-09 | 8.733E-09 | 3.926E-09 | 4.200E-09 | 1.100E-08 |
| 11026 | 552 | V552E | A | 7.2e-09 | 8.9e-09 | 6.3e-09 | 7.467E-09 | 1.320E-09 | 6.300E-09 | 8.900E-09 |
| 11027 | 552 | V552Q | A | 1.2e-08 | 3.8e-09 | 8e-09 | 7.933E-09 | 4.100E-09 | 3.800E-09 | 1.200E-08 |
| 11028 | 552 | V552G | A | 6.1e-09 | 5.7e-09 | 1.7e-08 | 9.600E-09 | 6.412E-09 | 5.700E-09 | 1.700E-08 |
| 11029 | 552 | V552H | A | 6e-09 | 3.7e-09 | 7.1e-09 | 5.600E-09 | 1.735E-09 | 3.700E-09 | 7.100E-09 |
| 11030 | 552 | V552I | A | 1.2e-08 | 3.9e-09 | 8.9e-09 | 8.267E-09 | 4.087E-09 | 3.900E-09 | 1.200E-08 |
| 11031 | 552 | V552L | A | 5e-08 | 4.7e-09 | 9.3e-09 | 2.133E-08 | 2.493E-08 | 4.700E-09 | 5.000E-08 |
| 11032 | 552 | V552K | A | 2.6e-08 | 2.4e-08 | 3.6e-09 | 1.787E-08 | 1.240E-08 | 3.600E-09 | 2.600E-08 |
| 11033 | 552 | V552M | A | 9.5e-09 | 6.9e-09 | 2.3e-09 | 6.233E-09 | 3.646E-09 | 2.300E-09 | 9.500E-09 |
| 11034 | 552 | V552F | A | 7.5e-09 | 1.1e-08 | 7.4e-09 | 8.633E-09 | 2.050E-09 | 7.400E-09 | 1.100E-08 |
| 11035 | 552 | V552P | A | 3.8e-09 | 5.1e-09 | 4e-09 | 4.300E-09 | 7.000E-10 | 3.800E-09 | 5.100E-09 |
| 11036 | 552 | V552S | A | 9.5e-09 | 1.6e-08 | 2.2e-09 | 9.233E-09 | 6.904E-09 | 2.200E-09 | 1.600E-08 |
| 11037 | 552 | V552T | A | 1.2e-08 | 3.5e-09 | 1.1e-08 | 8.833E-09 | 4.646E-09 | 3.500E-09 | 1.200E-08 |
| 11038 | 552 | V552W | A | 8.2e-09 | 7.4e-09 | 7.2e-09 | 7.600E-09 | 5.292E-10 | 7.200E-09 | 8.200E-09 |
| 11039 | 552 | V552Y | A | 6.9e-09 | 7.5e-09 | 3.8e-09 | 6.067E-09 | 1.986E-09 | 3.800E-09 | 7.500E-09 |
| 11040 | 552 | V552V | A | 8e-09 | 1.1e-08 | 1.9e-09 | 6.967E-09 | 4.637E-09 | 1.900E-09 | 1.100E-08 |
| 11041 | 553 | T553A | A | 3.5e-09 | 3.2e-09 | 1.1e-08 | 5.900E-09 | 4.419E-09 | 3.200E-09 | 1.100E-08 |
| 11042 | 553 | T553R | A | 5.6e-09 | 5.4e-09 | 7.1e-09 | 6.033E-09 | 9.292E-10 | 5.400E-09 | 7.100E-09 |
| 11043 | 553 | T553N | A | 3.1e-09 | 9.7e-09 | 4.8e-09 | 5.867E-09 | 3.427E-09 | 3.100E-09 | 9.700E-09 |
| 11044 | 553 | T553D | A | 3.2e-09 | 1.3e-08 | 1.8e-08 | 1.140E-08 | 7.529E-09 | 3.200E-09 | 1.800E-08 |
| 11045 | 553 | T553C | A | 1.3e-08 | 1.5e-08 | 9.9e-09 | 1.263E-08 | 2.570E-09 | 9.900E-09 | 1.500E-08 |
| 11046 | 553 | T553E | A | 8.3e-09 | 7.2e-09 | 1.6e-08 | 1.050E-08 | 4.795E-09 | 7.200E-09 | 1.600E-08 |
| 11047 | 553 | T553Q | A | 2.9e-09 | 9.6e-09 | 7.4e-09 | 6.633E-09 | 3.415E-09 | 2.900E-09 | 9.600E-09 |
| 11048 | 553 | T553G | A | 7.1e-09 | 7.9e-09 | 4.7e-09 | 6.567E-09 | 1.665E-09 | 4.700E-09 | 7.900E-09 |
| 11049 | 553 | T553H | A | 8.9e-09 | 3e-09 | 6.8e-09 | 6.233E-09 | 2.991E-09 | 3.000E-09 | 8.900E-09 |

| | | | | | | | | | | |
|---|---|---|---|---|---|---|---|---|---|---|
| 11050 | 553 | T553I | A | 7.9e-09 | 9.6e-09 | 1.7e-08 | 1.150E-08 | 4.838E-09 | 7.900E-09 | 1.700E-08 |
| 11051 | 553 | T553L | A | 3.8e-09 | 1e-08 | 9.5e-09 | 7.767E-09 | 3.444E-09 | 3.800E-09 | 1.000E-08 |
| 11052 | 553 | T553K | A | 1.1e-08 | 9.2e-09 | 2.8e-09 | 7.667E-09 | 4.310E-09 | 2.800E-09 | 1.100E-08 |
| 11053 | 553 | T553M | A | 8.9e-09 | 1.3e-09 | 9.2e-09 | 6.467E-09 | 4.477E-09 | 1.300E-09 | 9.200E-09 |
| 11054 | 553 | T553F | A | 1.7e-08 | 6.6e-09 | 3e-09 | 8.867E-09 | 7.270E-09 | 3.000E-09 | 1.700E-08 |
| 11055 | 553 | T553P | A | 5.1e-09 | 5e-09 | 1.2e-08 | 7.367E-09 | 4.013E-09 | 5.000E-09 | 1.200E-08 |
| 11056 | 553 | T553S | A | 4.4e-09 | 4.4e-09 | 1.2e-08 | 6.933E-09 | 4.388E-09 | 4.400E-09 | 1.200E-08 |
| 11057 | 553 | T553T | A | 8.5e-09 | 2.1e-09 | 4.2e-09 | 4.933E-09 | 3.262E-09 | 2.100E-09 | 8.500E-09 |
| 11058 | 553 | T553W | A | 4.2e-09 | 7.6e-09 | 7.9e-09 | 6.567E-09 | 2.055E-09 | 4.200E-09 | 7.900E-09 |
| 11059 | 553 | T553Y | A | 4.1e-09 | 4.8e-09 | 4.9e-09 | 4.600E-09 | 4.359E-10 | 4.100E-09 | 4.900E-09 |
| 11060 | 553 | T553V | A | 5.6e-09 | 8.1e-09 | 1.2e-08 | 8.567E-09 | 3.225E-09 | 5.600E-09 | 1.200E-08 |
| 11061 | 554 | C554A | A | 9.7e-09 | 4.9e-09 | 5.8e-09 | 6.800E-09 | 2.551E-09 | 4.900E-09 | 9.700E-09 |
| 11062 | 554 | C554R | A | 1.5e-08 | 7.4e-09 | 1e-08 | 1.080E-08 | 3.863E-09 | 7.400E-09 | 1.500E-08 |
| 11063 | 554 | C554N | A | 1e-08 | 4.7e-09 | 3.3e-09 | 6.000E-09 | 3.534E-09 | 3.300E-09 | 1.000E-08 |
| 11064 | 554 | C554D | A | 6e-09 | 4.5e-09 | 9.6e-09 | 6.700E-09 | 2.621E-09 | 4.500E-09 | 9.600E-09 |
| 11065 | 554 | C554C | A | 2.5e-09 | 9.3e-09 | 4.9e-09 | 5.567E-09 | 3.449E-09 | 2.500E-09 | 9.300E-09 |
| 11066 | 554 | C554E | A | 6e-09 | 5.2e-09 | 7.7e-09 | 6.300E-09 | 1.277E-09 | 5.200E-09 | 7.700E-09 |
| 11067 | 554 | C554Q | A | 8.8e-09 | 1.4e-08 | 8.2e-09 | 1.033E-08 | 3.190E-09 | 8.200E-09 | 1.400E-08 |
| 11068 | 554 | C554G | A | 7.5e-09 | 3.2e-09 | 4.3e-09 | 5.000E-09 | 2.234E-09 | 3.200E-09 | 7.500E-09 |
| 11069 | 554 | C554H | A | 1.8e-08 | 7.4e-09 | 6.1e-09 | 1.050E-08 | 6.528E-09 | 6.100E-09 | 1.800E-08 |
| 11070 | 554 | C554I | A | 2.8e-09 | 1.6e-08 | 1.2e-08 | 1.027E-08 | 6.769E-09 | 2.800E-09 | 1.600E-08 |
| 11071 | 554 | C554L | A | 1.4e-08 | 1.6e-08 | 5.7e-09 | 1.190E-08 | 5.462E-09 | 5.700E-09 | 1.600E-08 |
| 11072 | 554 | C554K | A | 3.7e-09 | 1.1e-08 | 1.4e-08 | 9.567E-09 | 5.297E-09 | 3.700E-09 | 1.400E-08 |
| 11073 | 554 | C554M | A | 3e-09 | 4.7e-09 | 2.3e-08 | 1.023E-08 | 1.109E-08 | 3.000E-09 | 2.300E-08 |
| 11074 | 554 | C554F | A | 5.2e-09 | 5.2e-09 | 5.3e-09 | 5.233E-09 | 5.774E-11 | 5.200E-09 | 5.300E-09 |
| 11075 | 554 | C554P | A | 1.5e-08 | 1.3e-08 | 8.5e-09 | 1.217E-08 | 3.329E-09 | 8.500E-09 | 1.500E-08 |
| 11076 | 554 | C554S | A | 1.1e-08 | 1.2e-08 | 7.4e-09 | 1.013E-08 | 2.419E-09 | 7.400E-09 | 1.200E-08 |
| 11077 | 554 | C554T | A | 5.5e-09 | 6.7e-09 | 7.3e-09 | 6.500E-09 | 9.165E-10 | 5.500E-09 | 7.300E-09 |
| 11078 | 554 | C554W | A | 7.4e-09 | 1.8e-08 | 6.3e-09 | 1.057E-08 | 6.461E-09 | 6.300E-09 | 1.800E-08 |
| 11079 | 554 | C554Y | A | 2.4e-09 | 1.3e-08 | 4.8e-09 | 6.733E-09 | 5.558E-09 | 2.400E-09 | 1.300E-08 |
| 11080 | 554 | C554V | A | 6.5e-09 | 2.9e-09 | 1e-08 | 6.467E-09 | 3.550E-09 | 2.900E-09 | 1.000E-08 |
| 11081 | 555 | F555A | A | 4.8e-09 | 8.9e-09 | 5.6e-09 | 6.433E-09 | 2.173E-09 | 4.800E-09 | 8.900E-09 |
| 11082 | 555 | F555R | A | 1.6e-08 | 4.5e-09 | 1.7e-08 | 1.250E-08 | 6.946E-09 | 4.500E-09 | 1.700E-08 |
| 11083 | 555 | F555N | A | 1.4e-08 | 6.1e-09 | 7.4e-09 | 9.167E-09 | 4.236E-09 | 6.100E-09 | 1.400E-08 |
| 11084 | 555 | F555D | A | 6.1e-09 | 1.3e-08 | 1.4e-08 | 1.103E-08 | 4.302E-09 | 6.100E-09 | 1.400E-08 |
| 11085 | 555 | F555C | A | 2.9e-09 | 4.8e-09 | 8.5e-09 | 5.400E-09 | 2.848E-09 | 2.900E-09 | 8.500E-09 |
| 11086 | 555 | F555E | A | 5.3e-09 | 1.8e-08 | 7.4e-09 | 1.023E-08 | 6.808E-09 | 5.300E-09 | 1.800E-08 |
| 11087 | 555 | F555Q | A | 3.7e-09 | 4.1e-09 | 5.9e-09 | 4.567E-09 | 1.172E-09 | 3.700E-09 | 5.900E-09 |
| 11088 | 555 | F555G | A | 8.2e-09 | 4.1e-09 | 5.8e-09 | 6.033E-09 | 2.060E-09 | 4.100E-09 | 8.200E-09 |
| 11089 | 555 | F555H | A | 9e-09 | 1.4e-08 | 4e-09 | 9.000E-09 | 5.000E-09 | 4.000E-09 | 1.400E-08 |
| 11090 | 555 | F555I | A | 9.3e-09 | 1.4e-08 | 2.2e-08 | 1.510E-08 | 6.421E-09 | 9.300E-09 | 2.200E-08 |
| 11091 | 555 | F555L | A | 8.3e-09 | 5.3e-09 | 2.1e-09 | 5.233E-09 | 3.101E-09 | 2.100E-09 | 8.300E-09 |
| 11092 | 555 | F555K | A | 1.8e-08 | 3.4e-09 | 1.8e-08 | 1.313E-08 | 8.429E-09 | 3.400E-09 | 1.800E-08 |
| 11093 | 555 | F555M | A | 7.5e-09 | 3.4e-09 | 1.2e-08 | 7.633E-09 | 4.302E-09 | 3.400E-09 | 1.200E-08 |
| 11094 | 555 | F555F | A | 3.2e-09 | 8.6e-09 | 1.2e-08 | 7.933E-09 | 4.438E-09 | 3.200E-09 | 1.200E-08 |
| 11095 | 555 | F555P | A | 1.7e-08 | 3.4e-09 | 2.9e-08 | 1.647E-08 | 1.281E-08 | 3.400E-09 | 2.900E-08 |
| 11096 | 555 | F555S | A | 5.6e-09 | 4e-09 | 5.7e-09 | 5.100E-09 | 9.539E-10 | 4.000E-09 | 5.700E-09 |
| 11097 | 555 | F555T | A | 8.5e-09 | 4.6e-09 | 1e-08 | 7.700E-09 | 2.787E-09 | 4.600E-09 | 1.000E-08 |
| 11098 | 555 | F555W | A | 3.1e-09 | 6.5e-09 | 3e-08 | 1.320E-08 | 1.465E-08 | 3.100E-09 | 3.000E-08 |
| 11099 | 555 | F555Y | A | 9.3e-09 | 4e-09 | 6.1e-09 | 6.467E-09 | 2.669E-09 | 4.000E-09 | 9.300E-09 |
| 11100 | 555 | F555V | A | 4.2e-09 | 3.2e-09 | 1.4e-08 | 7.133E-09 | 5.968E-09 | 3.200E-09 | 1.400E-08 |
| 11101 | 556 | G556A | A | 1.2e-08 | 3.6e-09 | 2.4e-09 | 6.000E-09 | 5.231E-09 | 2.400E-09 | 1.200E-08 |
| 11102 | 556 | G556R | A | 2.7e-09 | 9.5e-09 | 1.1e-08 | 7.733E-09 | 4.423E-09 | 2.700E-09 | 1.100E-08 |
| 11103 | 556 | G556N | A | 1.5e-08 | 2.3e-09 | 1.3e-08 | 1.010E-08 | 6.829E-09 | 2.300E-09 | 1.500E-08 |
| 11104 | 556 | G556D | A | 5.3e-09 | 6.3e-09 | 4.3e-08 | 1.820E-08 | 2.148E-08 | 5.300E-09 | 4.300E-08 |
| 11105 | 556 | G556C | A | 3.3e-09 | 9e-09 | 1.1e-08 | 7.767E-09 | 3.995E-09 | 3.300E-09 | 1.100E-08 |
| 11106 | 556 | G556E | A | 7.1e-09 | 6.4e-09 | 5.6e-09 | 6.367E-09 | 7.506E-10 | 5.600E-09 | 7.100E-09 |
| 11107 | 556 | G556Q | A | 6.7e-09 | 2.8e-09 | 6.8e-09 | 5.433E-09 | 2.281E-09 | 2.800E-09 | 6.800E-09 |
| 11108 | 556 | G556G | A | 2e-09 | 3.9e-09 | 9.4e-09 | 5.100E-09 | 3.843E-09 | 2.000E-09 | 9.400E-09 |
| 11109 | 556 | G556H | A | 1.4e-08 | 3.4e-09 | 1e-08 | 9.133E-09 | 5.353E-09 | 3.400E-09 | 1.400E-08 |
| 11110 | 556 | G556I | A | 8.3e-09 | 1.7e-09 | 2.5e-09 | 4.167E-09 | 3.602E-09 | 1.700E-09 | 8.300E-09 |
| 11111 | 556 | G556L | A | 9.6e-09 | 8.3e-09 | 1.4e-08 | 1.063E-08 | 2.987E-09 | 8.300E-09 | 1.400E-08 |
| 11112 | 556 | G556K | A | 3.1e-09 | 7e-09 | 4.9e-09 | 5.000E-09 | 1.952E-09 | 3.100E-09 | 7.000E-09 |
| 11113 | 556 | G556M | A | 1.7e-08 | 8.1e-09 | 1.6e-08 | 1.370E-08 | 4.875E-09 | 8.100E-09 | 1.700E-08 |
| 11114 | 556 | G556F | A | 3.6e-09 | 4.2e-09 | 6.2e-09 | 4.667E-09 | 1.361E-09 | 3.600E-09 | 6.200E-09 |
| 11115 | 556 | G556P | A | 5.4e-09 | 3.9e-09 | 9.2e-09 | 6.167E-09 | 2.732E-09 | 3.900E-09 | 9.200E-09 |
| 11116 | 556 | G556S | A | 3.4e-09 | 4.3e-09 | 8.3e-09 | 5.333E-09 | 2.608E-09 | 3.400E-09 | 8.300E-09 |
| 11117 | 556 | G556T | A | 5.5e-09 | 4.4e-09 | 6.6e-09 | 5.500E-09 | 1.100E-09 | 4.400E-09 | 6.600E-09 |
| 11118 | 556 | G556W | A | 1.1e-08 | 9.2e-09 | 8.1e-09 | 9.433E-09 | 1.464E-09 | 8.100E-09 | 1.100E-08 |
| 11119 | 556 | G556Y | A | 8.2e-09 | 6.7e-09 | 9.7e-09 | 8.200E-09 | 1.500E-09 | 6.700E-09 | 9.700E-09 |
| 11120 | 556 | G556V | A | 8e-09 | 6.2e-09 | 8.3e-09 | 7.500E-09 | 1.136E-09 | 6.200E-09 | 8.300E-09 |
| 11121 | 557 | P557A | A | 4.8e-09 | 4.2e-09 | 8.3e-09 | 5.767E-09 | 2.214E-09 | 4.200E-09 | 8.300E-09 |
| 11122 | 557 | P557R | A | 1.5e-08 | 3.1e-09 | 4.4e-09 | 7.500E-09 | 6.528E-09 | 3.100E-09 | 1.500E-08 |
| 11123 | 557 | P557N | A | 4.1e-09 | 7.8e-09 | 2.9e-08 | 1.363E-08 | 1.344E-08 | 4.100E-09 | 2.900E-08 |
| 11124 | 557 | P557D | A | 2.1e-09 | 7.7e-09 | 5.5e-09 | 5.100E-09 | 2.821E-09 | 2.100E-09 | 7.700E-09 |
| 11125 | 557 | P557C | A | 9.1e-09 | 2.4e-08 | 2.3e-09 | 1.180E-08 | 1.110E-08 | 2.300E-09 | 2.400E-08 |
| 11126 | 557 | P557E | A | 2.1e-09 | 6.7e-09 | 5.5e-09 | 4.767E-09 | 2.386E-09 | 2.100E-09 | 6.700E-09 |
| 11127 | 557 | P557Q | A | 7.4e-09 | 1.1e-08 | 2.3e-08 | 1.380E-08 | 8.168E-09 | 7.400E-09 | 2.300E-08 |
| 11128 | 557 | P557G | A | 1.1e-08 | 1.2e-08 | 2.3e-09 | 8.433E-09 | 5.335E-09 | 2.300E-09 | 1.200E-08 |
| 11129 | 557 | P557H | A | 5e-09 | 9.9e-09 | 7.1e-09 | 7.333E-09 | 2.458E-09 | 5.000E-09 | 9.900E-09 |
| 11130 | 557 | P557I | A | 3.8e-09 | 1.5e-08 | 5.1e-09 | 7.967E-09 | 6.126E-09 | 3.800E-09 | 1.500E-08 |
| 11131 | 557 | P557L | A | 5.4e-09 | 1.8e-09 | 7.3e-09 | 4.833E-09 | 2.793E-09 | 1.800E-09 | 7.300E-09 |
| 11132 | 557 | P557K | A | 6.4e-09 | 1.6e-08 | 6.1e-09 | 9.500E-09 | 5.631E-09 | 6.100E-09 | 1.600E-08 |
| 11133 | 557 | P557M | A | 8.6e-09 | 3.8e-09 | 1e-08 | 7.467E-09 | 3.252E-09 | 3.800E-09 | 1.000E-08 |
| 11134 | 557 | P557F | A | 4.8e-09 | 1.8e-08 | 5.5e-09 | 9.433E-09 | 7.427E-09 | 4.800E-09 | 1.800E-08 |

| | | | | | | | | | | |
|---|---|---|---|---|---|---|---|---|---|---|
| 11135 | 557 | P557P | A | 8.9e-09 | 6e-09 | 5.2e-09 | 6.700E-09 | 1.947E-09 | 5.200E-09 | 8.900E-09 |
| 11136 | 557 | P557S | A | 1.3e-08 | 1.6e-08 | 1.1e-08 | 1.333E-08 | 2.517E-09 | 1.100E-08 | 1.600E-08 |
| 11137 | 557 | P557T | A | 9.2e-09 | 1.3e-08 | 8.1e-09 | 1.010E-08 | 2.571E-09 | 8.100E-09 | 1.300E-08 |
| 11138 | 557 | P557W | A | 4e-09 | 1.4e-08 | 3.7e-09 | 7.233E-09 | 5.862E-09 | 3.700E-09 | 1.400E-08 |
| 11139 | 557 | P557Y | A | 1.1e-08 | 8.2e-09 | 1.1e-08 | 1.007E-08 | 1.617E-09 | 8.200E-09 | 1.100E-08 |
| 11140 | 557 | P557V | A | 2.5e-09 | 5.7e-09 | 5.2e-09 | 4.467E-09 | 1.721E-09 | 2.500E-09 | 5.700E-09 |
| 11141 | 558 | E558A | A | 1.2e-08 | 7.7e-09 | 3.9e-09 | 7.867E-09 | 4.053E-09 | 3.900E-09 | 1.200E-08 |
| 11142 | 558 | E558R | A | 3e-09 | 4.3e-09 | 3.3e-09 | 3.533E-09 | 6.807E-10 | 3.000E-09 | 4.300E-09 |
| 11143 | 558 | E558N | A | 8.9e-09 | 1.5e-09 | 1.1e-08 | 7.133E-09 | 4.990E-09 | 1.500E-09 | 1.100E-08 |
| 11144 | 558 | E558D | A | 8.9e-09 | 2.7e-09 | 6.3e-09 | 5.967E-09 | 3.113E-09 | 2.700E-09 | 8.900E-09 |
| 11145 | 558 | E558C | A | 1.1e-08 | 4.1e-09 | 6e-09 | 7.033E-09 | 3.564E-09 | 4.100E-09 | 1.100E-08 |
| 11146 | 558 | E558E | A | 4.3e-09 | 4.7e-09 | 1.7e-08 | 8.667E-09 | 7.220E-09 | 4.300E-09 | 1.700E-08 |
| 11147 | 558 | E558Q | A | 9.8e-10 | 6.7e-09 | 8.5e-09 | 5.393E-09 | 3.927E-09 | 9.800E-10 | 8.500E-09 |
| 11148 | 558 | E558G | A | 6e-09 | 3.6e-09 | 1.4e-08 | 7.867E-09 | 5.445E-09 | 3.600E-09 | 1.400E-08 |
| 11149 | 558 | E558H | A | 2.6e-09 | 6.9e-09 | 3.5e-09 | 4.333E-09 | 2.268E-09 | 2.600E-09 | 6.900E-09 |
| 11150 | 558 | E558I | A | 5.7e-09 | 4.3e-08 | 3e-09 | 1.723E-08 | 2.236E-08 | 3.000E-09 | 4.300E-08 |
| 11151 | 558 | E558L | A | 1.5e-08 | 8.2e-09 | 8e-09 | 1.040E-08 | 3.985E-09 | 8.000E-09 | 1.500E-08 |
| 11152 | 558 | E558K | A | 5e-09 | 5.4e-09 | 3.1e-09 | 4.500E-09 | 1.229E-09 | 3.100E-09 | 5.400E-09 |
| 11153 | 558 | E558M | A | 6.7e-09 | 5.6e-09 | 3.6e-09 | 5.300E-09 | 1.572E-09 | 3.600E-09 | 6.700E-09 |
| 11154 | 558 | E558F | A | 9e-09 | 6.8e-09 | 5.5e-09 | 7.100E-09 | 1.769E-09 | 5.500E-09 | 9.000E-09 |
| 11155 | 558 | E558P | A | 1e-08 | 2.3e-08 | 4.2e-09 | 1.240E-08 | 9.627E-09 | 4.200E-09 | 2.300E-08 |
| 11156 | 558 | E558S | A | 1.1e-08 | 5.6e-09 | 1.2e-08 | 9.533E-09 | 3.443E-09 | 5.600E-09 | 1.200E-08 |
| 11157 | 558 | E558T | A | 9.6e-09 | 1.6e-08 | 1.2e-08 | 1.253E-08 | 3.233E-09 | 9.600E-09 | 1.600E-08 |
| 11158 | 558 | E558W | A | 3.1e-09 | 9.1e-09 | 1.4e-08 | 8.733E-09 | 5.459E-09 | 3.100E-09 | 1.400E-08 |
| 11159 | 558 | E558Y | A | 5.3e-09 | 1.2e-08 | 1.9e-08 | 1.210E-08 | 6.851E-09 | 5.300E-09 | 1.900E-08 |
| 11160 | 558 | E558V | A | 5.3e-09 | 6.8e-09 | 1.5e-08 | 9.033E-09 | 5.221E-09 | 5.300E-09 | 1.500E-08 |
| 11161 | 559 | A559A | A | 7.4e-09 | 1e-08 | 4.4e-09 | 7.267E-09 | 2.802E-09 | 4.400E-09 | 1.000E-08 |
| 11162 | 559 | A559R | A | 4.9e-09 | 6.8e-09 | 9.8e-09 | 7.167E-09 | 2.470E-09 | 4.900E-09 | 9.800E-09 |
| 11163 | 559 | A559N | A | 5.1e-09 | 3.5e-09 | 1.1e-08 | 6.533E-09 | 3.950E-09 | 3.500E-09 | 1.100E-08 |
| 11164 | 559 | A559D | A | 1.3e-08 | 8.3e-09 | 6.2e-09 | 9.167E-09 | 3.482E-09 | 6.200E-09 | 1.300E-08 |
| 11165 | 559 | A559C | A | 1.6e-08 | 8.6e-09 | 1.2e-09 | 8.600E-09 | 7.400E-09 | 1.200E-09 | 1.600E-08 |
| 11166 | 559 | A559E | A | 8e-09 | 1.1e-08 | 3.7e-09 | 7.567E-09 | 3.669E-09 | 3.700E-09 | 1.100E-08 |
| 11167 | 559 | A559Q | A | 1.3e-09 | 9.3e-09 | 9.7e-09 | 6.767E-09 | 4.738E-09 | 1.300E-09 | 9.700E-09 |
| 11168 | 559 | A559G | A | 5e-09 | 2.5e-09 | 3.7e-09 | 3.733E-09 | 1.250E-09 | 2.500E-09 | 5.000E-09 |
| 11169 | 559 | A559H | A | 1.1e-08 | 1.3e-08 | 1e-08 | 1.133E-08 | 1.528E-09 | 1.000E-08 | 1.300E-08 |
| 11170 | 559 | A559I | A | 9.2e-09 | 7.4e-09 | 1.7e-08 | 1.120E-08 | 5.103E-09 | 7.400E-09 | 1.700E-08 |
| 11171 | 559 | A559L | A | 1.3e-08 | 1.3e-09 | 5.5e-09 | 6.600E-09 | 5.927E-09 | 1.300E-09 | 1.300E-08 |
| 11172 | 559 | A559K | A | 1.9e-08 | 1.2e-08 | 1.9e-08 | 1.667E-08 | 4.041E-09 | 1.200E-08 | 1.900E-08 |
| 11173 | 559 | A559M | A | 7.1e-09 | 1.4e-08 | 3.3e-09 | 8.133E-09 | 5.424E-09 | 3.300E-09 | 1.400E-08 |
| 11174 | 559 | A559F | A | 9.7e-09 | 7e-09 | 7e-09 | 7.900E-09 | 1.559E-09 | 7.000E-09 | 9.700E-09 |
| 11175 | 559 | A559P | A | 1.6e-08 | 1e-08 | 7.8e-09 | 1.127E-08 | 4.244E-09 | 7.800E-09 | 1.600E-08 |
| 11176 | 559 | A559S | A | 1.1e-08 | 5.3e-09 | 2.3e-09 | 6.200E-09 | 4.419E-09 | 2.300E-09 | 1.100E-08 |
| 11177 | 559 | A559T | A | 1.3e-08 | 6.7e-09 | 8.8e-09 | 9.500E-09 | 3.208E-09 | 6.700E-09 | 1.300E-08 |
| 11178 | 559 | A559W | A | 8.3e-09 | 1.4e-09 | 1e-08 | 6.567E-09 | 4.554E-09 | 1.400E-09 | 1.000E-08 |
| 11179 | 559 | A559Y | A | 8.8e-09 | 1.5e-08 | 1.3e-08 | 1.227E-08 | 3.164E-09 | 8.800E-09 | 1.500E-08 |
| 11180 | 559 | A559V | A | 9e-09 | 1e-08 | 1.1e-08 | 1.000E-08 | 1.000E-09 | 9.000E-09 | 1.100E-08 |
| 11181 | 560 | D560A | A | 3.6e-09 | 6.4e-09 | 1.4e-08 | 8.000E-09 | 5.381E-09 | 3.600E-09 | 1.400E-08 |
| 11182 | 560 | D560R | A | 5.1e-09 | 6.7e-09 | 6.2e-09 | 6.000E-09 | 8.185E-10 | 5.100E-09 | 6.700E-09 |
| 11183 | 560 | D560N | A | 7.4e-09 | 9.4e-09 | 4.5e-09 | 7.100E-09 | 2.464E-09 | 4.500E-09 | 9.400E-09 |
| 11184 | 560 | D560D | A | 7.9e-09 | 1.2e-08 | 1.7e-08 | 1.230E-08 | 4.557E-09 | 7.900E-09 | 1.700E-08 |
| 11185 | 560 | D560C | A | 7.7e-09 | 1.1e-08 | 1e-08 | 9.567E-09 | 1.692E-09 | 7.700E-09 | 1.100E-08 |
| 11186 | 560 | D560E | A | 4.3e-09 | 3.5e-09 | 1.1e-08 | 6.267E-09 | 4.119E-09 | 3.500E-09 | 1.100E-08 |
| 11187 | 560 | D560Q | A | 1.2e-08 | 1.1e-08 | 5e-09 | 9.333E-09 | 3.786E-09 | 5.000E-09 | 1.200E-08 |
| 11188 | 560 | D560G | A | 8e-09 | 2.7e-09 | 5.4e-09 | 5.367E-09 | 2.650E-09 | 2.700E-09 | 8.000E-09 |
| 11189 | 560 | D560H | A | 1.9e-08 | 3.7e-09 | 2.4e-09 | 8.367E-09 | 9.232E-09 | 2.400E-09 | 1.900E-08 |
| 11190 | 560 | D560I | A | 1.2e-08 | 9.2e-09 | 3.8e-09 | 8.333E-09 | 4.168E-09 | 3.800E-09 | 1.200E-08 |
| 11191 | 560 | D560L | A | 7.4e-09 | 2.7e-08 | 7.3e-09 | 1.390E-08 | 1.135E-08 | 7.300E-09 | 2.700E-08 |
| 11192 | 560 | D560K | A | 7.3e-09 | 6.3e-09 | 3.4e-09 | 5.667E-09 | 2.026E-09 | 3.400E-09 | 7.300E-09 |
| 11193 | 560 | D560M | A | 1.1e-08 | 4.6e-09 | 1.1e-08 | 8.867E-09 | 3.695E-09 | 4.600E-09 | 1.100E-08 |
| 11194 | 560 | D560F | A | 2e-08 | 1.2e-08 | 6.5e-09 | 1.283E-08 | 6.788E-09 | 6.500E-09 | 2.000E-08 |
| 11195 | 560 | D560P | A | 4.3e-09 | 1.3e-08 | 1.2e-08 | 9.767E-09 | 4.761E-09 | 4.300E-09 | 1.300E-08 |
| 11196 | 560 | D560S | A | 6e-09 | 1.5e-08 | 7.8e-09 | 9.600E-09 | 4.762E-09 | 6.000E-09 | 1.500E-08 |
| 11197 | 560 | D560T | A | 6e-09 | 1.2e-08 | 5.6e-09 | 7.867E-09 | 3.585E-09 | 5.600E-09 | 1.200E-08 |
| 11198 | 560 | D560W | A | 8.4e-09 | 4e-09 | 1.1e-08 | 7.800E-09 | 3.538E-09 | 4.000E-09 | 1.100E-08 |
| 11199 | 560 | D560Y | A | 9.8e-09 | 1.1e-08 | 4.3e-09 | 8.367E-09 | 3.573E-09 | 4.300E-09 | 1.100E-08 |
| 11200 | 560 | D560V | A | 3.5e-09 | 6.1e-09 | 4.1e-09 | 4.567E-09 | 1.361E-09 | 3.500E-09 | 6.100E-09 |
| 11201 | 561 | Q561A | A | 6e-09 | 9.1e-09 | 4.4e-09 | 6.500E-09 | 2.390E-09 | 4.400E-09 | 9.100E-09 |
| 11202 | 561 | Q561R | A | 1.1e-08 | 1.8e-08 | 4.6e-09 | 1.120E-08 | 6.702E-09 | 4.600E-09 | 1.800E-08 |
| 11203 | 561 | Q561N | A | 6.5e-09 | 2.5e-09 | 1.4e-08 | 7.667E-09 | 5.838E-09 | 2.500E-09 | 1.400E-08 |
| 11204 | 561 | Q561D | A | 9.4e-09 | 2.1e-08 | 7.4e-09 | 1.260E-08 | 7.343E-09 | 7.400E-09 | 2.100E-08 |
| 11205 | 561 | Q561C | A | 5.2e-09 | 1.3e-08 | 7e-09 | 8.400E-09 | 4.084E-09 | 5.200E-09 | 1.300E-08 |
| 11206 | 561 | Q561E | A | 4.5e-09 | 7.6e-09 | 4.2e-09 | 5.433E-09 | 1.882E-09 | 4.200E-09 | 7.600E-09 |
| 11207 | 561 | Q561Q | A | 2.1e-09 | 1.5e-08 | 1e-08 | 9.033E-09 | 6.504E-09 | 2.100E-09 | 1.500E-08 |
| 11208 | 561 | Q561G | A | 3.2e-09 | 6.7e-09 | 1.7e-08 | 8.967E-09 | 7.174E-09 | 3.200E-09 | 1.700E-08 |
| 11209 | 561 | Q561H | A | 6.8e-09 | 2.2e-09 | 8.5e-09 | 5.833E-09 | 3.259E-09 | 2.200E-09 | 8.500E-09 |
| 11210 | 561 | Q561I | A | 7.1e-09 | 1.4e-09 | 1e-08 | 6.167E-09 | 4.375E-09 | 1.400E-09 | 1.000E-08 |
| 11211 | 561 | Q561L | A | 1.4e-08 | 1.7e-08 | 9.7e-09 | 1.357E-08 | 3.669E-09 | 9.700E-09 | 1.700E-08 |
| 11212 | 561 | Q561K | A | 9.1e-09 | 1.6e-08 | 4.1e-09 | 9.733E-09 | 5.975E-09 | 4.100E-09 | 1.600E-08 |
| 11213 | 561 | Q561M | A | 5e-09 | 6.2e-09 | 5.2e-09 | 5.467E-09 | 6.429E-10 | 5.000E-09 | 6.200E-09 |
| 11214 | 561 | Q561F | A | 9.8e-09 | 8.7e-09 | 1.1e-08 | 9.833E-09 | 1.150E-09 | 8.700E-09 | 1.100E-08 |
| 11215 | 561 | Q561P | A | 1.1e-08 | 5.9e-09 | 4e-09 | 6.967E-09 | 3.620E-09 | 4.000E-09 | 1.100E-08 |
| 11216 | 561 | Q561S | A | 6.7e-09 | 8.5e-09 | 1.9e-08 | 1.140E-08 | 6.643E-09 | 6.700E-09 | 1.900E-08 |
| 11217 | 561 | Q561T | A | 2.7e-09 | 6.9e-09 | 2.2e-08 | 1.053E-08 | 1.015E-08 | 2.700E-09 | 2.200E-08 |
| 11218 | 561 | Q561W | A | 5.8e-09 | 8.1e-09 | 3.7e-09 | 5.867E-09 | 2.201E-09 | 3.700E-09 | 8.100E-09 |
| 11219 | 561 | Q561Y | A | 1.1e-08 | 1.8e-09 | 4.9e-09 | 5.900E-09 | 4.681E-09 | 1.800E-09 | 1.100E-08 |

| | | | | | | | | | | |
|---|---|---|---|---|---|---|---|---|---|---|
| 11220 | 561 | Q561V | A | 1.5e-08 | 9.4e-09 | 5.2e-09 | 9.867E-09 | 4.917E-09 | 5.200E-09 | 1.500E-08 |
| 11221 | 562 | C562A | A | 3.4e-09 | 3.5e-09 | 9.7e-09 | 5.533E-09 | 3.609E-09 | 3.400E-09 | 9.700E-09 |
| 11222 | 562 | C562R | A | 1e-08 | 2e-08 | 1.4e-08 | 1.467E-08 | 5.033E-09 | 1.000E-08 | 2.000E-08 |
| 11223 | 562 | C562N | A | 3.1e-08 | 1.2e-08 | 9.8e-09 | 1.760E-08 | 1.166E-08 | 9.800E-09 | 3.100E-08 |
| 11224 | 562 | C562D | A | 3e-09 | 7.9e-09 | 3.9e-09 | 4.933E-09 | 2.608E-09 | 3.000E-09 | 7.900E-09 |
| 11225 | 562 | C562C | A | 1.4e-08 | 6.6e-09 | 2.8e-09 | 7.800E-09 | 5.696E-09 | 2.800E-09 | 1.400E-08 |
| 11226 | 562 | C562E | A | 4.8e-09 | 1.8e-08 | 3.6e-09 | 8.800E-09 | 7.990E-09 | 3.600E-09 | 1.800E-08 |
| 11227 | 562 | C562Q | A | 5.8e-09 | 1.6e-08 | 6.5e-09 | 9.433E-09 | 5.698E-09 | 5.800E-09 | 1.600E-08 |
| 11228 | 562 | C562G | A | 1.1e-08 | 4e-08 | 1e-08 | 2.033E-08 | 1.704E-08 | 1.000E-08 | 4.000E-08 |
| 11229 | 562 | C562H | A | 1.6e-08 | 2.4e-08 | 1.1e-08 | 1.700E-08 | 6.557E-09 | 1.100E-08 | 2.400E-08 |
| 11230 | 562 | C562I | A | 8.7e-09 | 1.1e-08 | 8.7e-09 | 9.467E-09 | 1.328E-09 | 8.700E-09 | 1.100E-08 |
| 11231 | 562 | C562L | A | 3e-09 | 3.5e-08 | 1e-08 | 1.600E-08 | 1.682E-08 | 3.000E-09 | 3.500E-08 |
| 11232 | 562 | C562K | A | 1.2e-08 | 6.9e-09 | 2.4e-09 | 7.100E-09 | 4.803E-09 | 2.400E-09 | 1.200E-08 |
| 11233 | 562 | C562M | A | 4.3e-09 | 9.6e-09 | 8.6e-09 | 7.500E-09 | 2.816E-09 | 4.300E-09 | 9.600E-09 |
| 11234 | 562 | C562F | A | 3.8e-09 | 1.3e-08 | 8.8e-09 | 8.533E-09 | 4.606E-09 | 3.800E-09 | 1.300E-08 |
| 11235 | 562 | C562P | A | 9.7e-09 | 9.4e-09 | 1.1e-08 | 1.003E-08 | 8.505E-10 | 9.400E-09 | 1.100E-08 |
| 11236 | 562 | C562S | A | 1.2e-08 | 8.8e-09 | 7.7e-09 | 9.500E-09 | 2.234E-09 | 7.700E-09 | 1.200E-08 |
| 11237 | 562 | C562T | A | 4.9e-09 | 8.8e-09 | 9.3e-09 | 7.667E-09 | 2.409E-09 | 4.900E-09 | 9.300E-09 |
| 11238 | 562 | C562W | A | 2.9e-09 | 7.4e-09 | 4.7e-09 | 5.000E-09 | 2.265E-09 | 2.900E-09 | 7.400E-09 |
| 11239 | 562 | C562Y | A | 1e-08 | 1.2e-08 | 5.4e-09 | 9.133E-09 | 3.384E-09 | 5.400E-09 | 1.200E-08 |
| 11240 | 562 | C562V | A | 1.1e-08 | 2.2e-09 | 6.1e-09 | 6.433E-09 | 4.409E-09 | 2.200E-09 | 1.100E-08 |
| 11241 | 563 | V563A | A | 1.2e-08 | 2.5e-09 | 1.4e-08 | 9.500E-09 | 6.144E-09 | 2.500E-09 | 1.400E-08 |
| 11242 | 563 | V563R | A | 1.1e-08 | 1e-08 | 6.7e-09 | 9.233E-09 | 2.250E-09 | 6.700E-09 | 1.100E-08 |
| 11243 | 563 | V563N | A | 8.7e-09 | 2e-08 | 3.3e-09 | 1.067E-08 | 8.522E-09 | 3.300E-09 | 2.000E-08 |
| 11244 | 563 | V563D | A | 6.3e-09 | 1.2e-08 | 2e-08 | 1.277E-08 | 6.882E-09 | 6.300E-09 | 2.000E-08 |
| 11245 | 563 | V563C | A | 1.1e-08 | 4.1e-09 | 3e-09 | 6.033E-09 | 4.336E-09 | 3.000E-09 | 1.100E-08 |
| 11246 | 563 | V563E | A | 7.5e-09 | 4.9e-09 | 3.6e-09 | 5.333E-09 | 1.986E-09 | 3.600E-09 | 7.500E-09 |
| 11247 | 563 | V563Q | A | 1e-08 | 1.2e-08 | 4.7e-09 | 8.900E-09 | 3.772E-09 | 4.700E-09 | 1.200E-08 |
| 11248 | 563 | V563G | A | 5.3e-09 | 8.8e-09 | 1.2e-08 | 8.700E-09 | 3.351E-09 | 5.300E-09 | 1.200E-08 |
| 11249 | 563 | V563H | A | 2.9e-08 | 1.1e-09 | 5e-09 | 1.170E-08 | 1.511E-08 | 1.100E-09 | 2.900E-08 |
| 11250 | 563 | V563I | A | 8.9e-09 | 4.7e-09 | 3.7e-09 | 5.767E-09 | 2.759E-09 | 3.700E-09 | 8.900E-09 |
| 11251 | 563 | V563L | A | 4.1e-09 | 3e-09 | 1.9e-08 | 8.700E-09 | 8.937E-09 | 3.000E-09 | 1.900E-08 |
| 11252 | 563 | V563K | A | 2.8e-09 | 1.2e-08 | 5.2e-09 | 6.667E-09 | 4.772E-09 | 2.800E-09 | 1.200E-08 |
| 11253 | 563 | V563M | A | 9.1e-09 | 5.8e-09 | 2.3e-09 | 5.733E-09 | 3.400E-09 | 2.300E-09 | 9.100E-09 |
| 11254 | 563 | V563F | A | 9.9e-09 | 1.1e-08 | 4.4e-09 | 8.433E-09 | 3.536E-09 | 4.400E-09 | 1.100E-08 |
| 11255 | 563 | V563P | A | 2.5e-08 | 8.5e-09 | 1.2e-08 | 1.517E-08 | 8.694E-09 | 8.500E-09 | 2.500E-08 |
| 11256 | 563 | V563S | A | 8.8e-09 | 6.7e-09 | 9e-09 | 8.167E-09 | 1.274E-09 | 6.700E-09 | 9.000E-09 |
| 11257 | 563 | V563T | A | 4e-09 | 7.4e-09 | 6.2e-09 | 5.867E-09 | 1.724E-09 | 4.000E-09 | 7.400E-09 |
| 11258 | 563 | V563W | A | 5.1e-09 | 6.5e-09 | 1.3e-08 | 8.200E-09 | 4.215E-09 | 5.100E-09 | 1.300E-08 |
| 11259 | 563 | V563Y | A | 1.1e-08 | 5.7e-08 | 4e-09 | 2.400E-08 | 2.879E-08 | 4.000E-09 | 5.700E-08 |
| 11260 | 563 | V563V | A | 1.7e-08 | 7.9e-09 | 4.1e-08 | 2.197E-08 | 1.710E-08 | 7.900E-09 | 4.100E-08 |
| 11261 | 564 | A564A | A | 2e-09 | 2.6e-09 | 5.4e-09 | 3.333E-09 | 1.815E-09 | 2.000E-09 | 5.400E-09 |
| 11262 | 564 | A564R | A | 1.2e-08 | 1.4e-08 | 1.9e-08 | 1.500E-08 | 3.606E-09 | 1.200E-08 | 1.900E-08 |
| 11263 | 564 | A564N | A | 9.4e-09 | 3.3e-09 | 3.7e-09 | 5.467E-09 | 3.412E-09 | 3.300E-09 | 9.400E-09 |
| 11264 | 564 | A564D | A | 2.9e-09 | 9.1e-09 | 4.8e-09 | 5.600E-09 | 3.176E-09 | 2.900E-09 | 9.100E-09 |
| 11265 | 564 | A564C | A | 2.1e-09 | 1.8e-08 | 9.7e-09 | 9.933E-09 | 7.953E-09 | 2.100E-09 | 1.800E-08 |
| 11266 | 564 | A564E | A | 6.1e-09 | 1.8e-08 | 9e-09 | 1.103E-08 | 6.205E-09 | 6.100E-09 | 1.800E-08 |
| 11267 | 564 | A564Q | A | 1.7e-09 | 6.7e-09 | 2.2e-08 | 1.013E-08 | 1.058E-08 | 1.700E-09 | 2.200E-08 |
| 11268 | 564 | A564G | A | 5.4e-09 | 3.4e-09 | 5.4e-09 | 4.733E-09 | 1.155E-09 | 3.400E-09 | 5.400E-09 |
| 11269 | 564 | A564H | A | 2.6e-08 | 8.4e-09 | 6.1e-09 | 1.350E-08 | 1.089E-08 | 6.100E-09 | 2.600E-08 |
| 11270 | 564 | A564I | A | 8.3e-09 | 1e-08 | 2.2e-08 | 1.343E-08 | 7.467E-09 | 8.300E-09 | 2.200E-08 |
| 11271 | 564 | A564L | A | 2.4e-09 | 2.8e-09 | 8.2e-09 | 4.467E-09 | 3.239E-09 | 2.400E-09 | 8.200E-09 |
| 11272 | 564 | A564K | A | 7.4e-09 | 4.9e-09 | 7.1e-09 | 6.467E-09 | 1.365E-09 | 4.900E-09 | 7.400E-09 |
| 11273 | 564 | A564M | A | 1.7e-08 | 1.2e-08 | 1.5e-09 | 1.017E-08 | 7.911E-09 | 1.500E-09 | 1.700E-08 |
| 11274 | 564 | A564F | A | 4.2e-09 | 8.6e-09 | 1.1e-08 | 7.933E-09 | 3.449E-09 | 4.200E-09 | 1.100E-08 |
| 11275 | 564 | A564P | A | 7.1e-09 | 1.5e-08 | 6.8e-09 | 9.633E-09 | 4.650E-09 | 6.800E-09 | 1.500E-08 |
| 11276 | 564 | A564S | A | 1.1e-08 | 8.1e-09 | 8.7e-09 | 9.267E-09 | 1.531E-09 | 8.100E-09 | 1.100E-08 |
| 11277 | 564 | A564T | A | 3.1e-09 | 3e-09 | 1.3e-08 | 6.367E-09 | 5.745E-09 | 3.000E-09 | 1.300E-08 |
| 11278 | 564 | A564W | A | 8e-09 | 3e-09 | 2e-09 | 4.333E-09 | 3.215E-09 | 2.000E-09 | 8.000E-09 |
| 11279 | 564 | A564Y | A | 5.1e-09 | 8.2e-09 | 9.8e-09 | 7.700E-09 | 2.390E-09 | 5.100E-09 | 9.800E-09 |
| 11280 | 564 | A564V | A | 2.4e-09 | 9.9e-09 | 3.5e-09 | 5.267E-09 | 4.050E-09 | 2.400E-09 | 9.900E-09 |
| 11281 | 565 | C565A | A | 3.8e-09 | 5.8e-09 | 7.5e-09 | 5.700E-09 | 1.852E-09 | 3.800E-09 | 7.500E-09 |
| 11282 | 565 | C565R | A | 1.7e-09 | 8.6e-09 | 7.7e-09 | 6.000E-09 | 3.751E-09 | 1.700E-09 | 8.600E-09 |
| 11283 | 565 | C565N | A | 6.7e-09 | 6.6e-09 | 2.3e-09 | 5.200E-09 | 2.512E-09 | 2.300E-09 | 6.700E-09 |
| 11284 | 565 | C565D | A | 1.4e-08 | 1e-08 | 1.2e-08 | 1.200E-08 | 2.000E-09 | 1.000E-08 | 1.400E-08 |
| 11285 | 565 | C565C | A | 6.1e-09 | 6.5e-09 | 8.2e-09 | 6.933E-09 | 1.115E-09 | 6.100E-09 | 8.200E-09 |
| 11286 | 565 | C565E | A | 2.8e-09 | 7e-09 | 5.5e-09 | 5.100E-09 | 2.128E-09 | 2.800E-09 | 7.000E-09 |
| 11287 | 565 | C565Q | A | 1.3e-08 | 1.1e-08 | 3.1e-09 | 9.033E-09 | 5.235E-09 | 3.100E-09 | 1.300E-08 |
| 11288 | 565 | C565G | A | 9.4e-09 | 1.7e-08 | 4.2e-09 | 1.020E-08 | 6.437E-09 | 4.200E-09 | 1.700E-08 |
| 11289 | 565 | C565H | A | 1.1e-08 | 6.8e-09 | 9e-09 | 8.933E-09 | 2.101E-09 | 6.800E-09 | 1.100E-08 |
| 11290 | 565 | C565I | A | 2e-08 | 1.1e-08 | 5.3e-09 | 1.210E-08 | 7.411E-09 | 5.300E-09 | 2.000E-08 |
| 11291 | 565 | C565L | A | 1.9e-08 | 4.7e-09 | 7.4e-09 | 1.037E-08 | 7.598E-09 | 4.700E-09 | 1.900E-08 |
| 11292 | 565 | C565K | A | 6.8e-09 | 1.2e-08 | 3e-09 | 7.267E-09 | 4.518E-09 | 3.000E-09 | 1.200E-08 |
| 11293 | 565 | C565M | A | 6.6e-09 | 8.5e-09 | 5.5e-09 | 6.867E-09 | 1.518E-09 | 5.500E-09 | 8.500E-09 |
| 11294 | 565 | C565F | A | 6.5e-09 | 3.9e-09 | 5.7e-09 | 5.367E-09 | 1.332E-09 | 3.900E-09 | 6.500E-09 |
| 11295 | 565 | C565P | A | 1.2e-08 | 1.1e-08 | 6.6e-09 | 9.867E-09 | 2.873E-09 | 6.600E-09 | 1.200E-08 |
| 11296 | 565 | C565S | A | 4e-09 | 2.2e-09 | 2.7e-09 | 2.967E-09 | 9.292E-10 | 2.200E-09 | 4.000E-09 |
| 11297 | 565 | C565T | A | 3e-09 | 8.8e-09 | 1.3e-09 | 4.367E-09 | 3.932E-09 | 1.300E-09 | 8.800E-09 |
| 11298 | 565 | C565W | A | 7.5e-09 | 3e-09 | 1.1e-08 | 7.167E-09 | 4.010E-09 | 3.000E-09 | 1.100E-08 |
| 11299 | 565 | C565Y | A | 3.2e-09 | 9.8e-09 | 2.2e-08 | 1.167E-08 | 9.538E-09 | 3.200E-09 | 2.200E-08 |
| 11300 | 565 | C565V | A | 4e-09 | 4.4e-09 | 4.3e-09 | 4.233E-09 | 2.082E-10 | 4.000E-09 | 4.400E-09 |
| 11301 | 566 | A566A | A | 1.1e-08 | 7.1e-09 | 3.8e-09 | 7.300E-09 | 3.604E-09 | 3.800E-09 | 1.100E-08 |
| 11302 | 566 | A566R | A | 4.9e-09 | 7.6e-09 | 1e-08 | 7.500E-09 | 2.551E-09 | 4.900E-09 | 1.000E-08 |
| 11303 | 566 | A566N | A | 7.4e-09 | 5.9e-09 | 5.2e-09 | 6.167E-09 | 1.124E-09 | 5.200E-09 | 7.400E-09 |
| 11304 | 566 | A566D | A | 6.9e-09 | 1.4e-08 | 6.2e-09 | 9.033E-09 | 4.315E-09 | 6.200E-09 | 1.400E-08 |

| | | | | | | | | | | |
|---|---|---|---|---|---|---|---|---|---|---|
| 11305 | 566 | A566C | A | 3.9e-09 | 3.7e-09 | 8.8e-09 | 5.467E-09 | 2.888E-09 | 3.700E-09 | 8.800E-09 |
| 11306 | 566 | A566E | A | 1.8e-08 | 2.4e-09 | 1.2e-08 | 1.080E-08 | 7.869E-09 | 2.400E-09 | 1.800E-08 |
| 11307 | 566 | A566Q | A | 7.2e-09 | 3.7e-09 | 4.3e-09 | 5.067E-09 | 1.872E-09 | 3.700E-09 | 7.200E-09 |
| 11308 | 566 | A566G | A | 3.8e-09 | 1.1e-08 | 7.9e-09 | 7.567E-09 | 3.612E-09 | 3.800E-09 | 1.100E-08 |
| 11309 | 566 | A566H | A | 9.7e-09 | 5.9e-09 | 7.7e-09 | 7.767E-09 | 1.901E-09 | 5.900E-09 | 9.700E-09 |
| 11310 | 566 | A566I | A | 7.6e-09 | 7.8e-09 | 1.6e-08 | 1.047E-08 | 4.793E-09 | 7.600E-09 | 1.600E-08 |
| 11311 | 566 | A566L | A | 2.6e-09 | 9.9e-09 | 4.2e-09 | 5.567E-09 | 3.837E-09 | 2.600E-09 | 9.900E-09 |
| 11312 | 566 | A566K | A | 3.4e-09 | 1.4e-08 | 6e-09 | 7.800E-09 | 5.524E-09 | 3.400E-09 | 1.400E-08 |
| 11313 | 566 | A566M | A | 7.3e-09 | 1.6e-08 | 6.6e-09 | 9.967E-09 | 5.237E-09 | 6.600E-09 | 1.600E-08 |
| 11314 | 566 | A566F | A | 9.8e-09 | 6.8e-09 | 3.1e-09 | 6.567E-09 | 3.356E-09 | 3.100E-09 | 9.800E-09 |
| 11315 | 566 | A566P | A | 2.2e-09 | 5.6e-09 | 4.8e-09 | 4.200E-09 | 1.778E-09 | 2.200E-09 | 5.600E-09 |
| 11316 | 566 | A566S | A | 8.9e-09 | 1.2e-08 | 4.7e-09 | 8.533E-09 | 3.664E-09 | 4.700E-09 | 1.200E-08 |
| 11317 | 566 | A566T | A | 7.9e-09 | 3.1e-09 | 5.2e-09 | 5.400E-09 | 2.406E-09 | 3.100E-09 | 7.900E-09 |
| 11318 | 566 | A566W | A | 5.9e-09 | 3.5e-09 | 7.2e-09 | 5.533E-09 | 1.877E-09 | 3.500E-09 | 7.200E-09 |
| 11319 | 566 | A566Y | A | 5.6e-09 | 1.3e-08 | 4.1e-09 | 7.567E-09 | 4.765E-09 | 4.100E-09 | 1.300E-08 |
| 11320 | 566 | A566V | A | 6e-09 | 5.8e-09 | 2.1e-08 | 1.093E-08 | 8.719E-09 | 5.800E-09 | 2.100E-08 |
| 11321 | 567 | H567A | A | 1e-08 | 5.7e-09 | 6e-09 | 7.233E-09 | 2.401E-09 | 5.700E-09 | 1.000E-08 |
| 11322 | 567 | H567R | A | 3.5e-09 | 6.3e-09 | 1.8e-09 | 3.867E-09 | 2.272E-09 | 1.800E-09 | 6.300E-09 |
| 11323 | 567 | H567N | A | 1.8e-08 | 9.6e-09 | 5.5e-09 | 1.103E-08 | 6.372E-09 | 5.500E-09 | 1.800E-08 |
| 11324 | 567 | H567D | A | 4.3e-09 | 1e-08 | 4.3e-09 | 6.200E-09 | 3.291E-09 | 4.300E-09 | 1.000E-08 |
| 11325 | 567 | H567C | A | 4.9e-09 | 1.4e-08 | 7e-09 | 8.633E-09 | 4.765E-09 | 4.900E-09 | 1.400E-08 |
| 11326 | 567 | H567E | A | 5.2e-09 | 4.8e-09 | 4.4e-09 | 4.800E-09 | 4.000E-10 | 4.400E-09 | 5.200E-09 |
| 11327 | 567 | H567Q | A | 7.1e-09 | 2.3e-08 | 1.8e-08 | 1.603E-08 | 8.130E-09 | 7.100E-09 | 2.300E-08 |
| 11328 | 567 | H567G | A | 8.2e-09 | 7.8e-09 | 5.8e-09 | 7.267E-09 | 1.286E-09 | 5.800E-09 | 8.200E-09 |
| 11329 | 567 | H567H | A | 4.5e-09 | 6e-09 | 6.1e-09 | 5.533E-09 | 8.963E-10 | 4.500E-09 | 6.100E-09 |
| 11330 | 567 | H567I | A | 7.9e-09 | 8.7e-09 | 7.1e-09 | 7.900E-09 | 8.000E-10 | 7.100E-09 | 8.700E-09 |
| 11331 | 567 | H567L | A | 7.5e-09 | 5.3e-09 | 5.7e-09 | 6.167E-09 | 1.172E-09 | 5.300E-09 | 7.500E-09 |
| 11332 | 567 | H567K | A | 1e-08 | 1.4e-08 | 7.1e-09 | 1.037E-08 | 3.465E-09 | 7.100E-09 | 1.400E-08 |
| 11333 | 567 | H567M | A | 2e-08 | 5e-09 | 2.3e-09 | 9.100E-09 | 9.536E-09 | 2.300E-09 | 2.000E-08 |
| 11334 | 567 | H567F | A | 6.7e-09 | 1.3e-08 | 9.3e-09 | 9.667E-09 | 3.166E-09 | 6.700E-09 | 1.300E-08 |
| 11335 | 567 | H567P | A | 6.5e-09 | 5.2e-09 | 6.8e-09 | 6.167E-09 | 8.505E-10 | 5.200E-09 | 6.800E-09 |
| 11336 | 567 | H567S | A | 1e-08 | 1.5e-08 | 1.8e-08 | 1.433E-08 | 4.041E-09 | 1.000E-08 | 1.800E-08 |
| 11337 | 567 | H567T | A | 1.3e-08 | 1.6e-09 | 9.8e-09 | 8.133E-09 | 5.880E-09 | 1.600E-09 | 1.300E-08 |
| 11338 | 567 | H567W | A | 9.9e-09 | 5e-09 | 5.4e-09 | 6.767E-09 | 2.721E-09 | 5.000E-09 | 9.900E-09 |
| 11339 | 567 | H567Y | A | 5.1e-09 | 1e-08 | 6.5e-09 | 7.200E-09 | 2.524E-09 | 5.100E-09 | 1.000E-08 |
| 11340 | 567 | H567V | A | 8.6e-09 | 1.1e-08 | 1.3e-08 | 1.087E-08 | 2.203E-09 | 8.600E-09 | 1.300E-08 |
| 11341 | 568 | Y568A | A | 5.8e-09 | 1.7e-08 | 2.8e-09 | 8.533E-09 | 7.484E-09 | 2.800E-09 | 1.700E-08 |
| 11342 | 568 | Y568R | A | 3.7e-09 | 1.8e-08 | 4.1e-09 | 8.600E-09 | 8.143E-09 | 3.700E-09 | 1.800E-08 |
| 11343 | 568 | Y568N | A | 1e-08 | 1.6e-08 | 8.1e-09 | 1.137E-08 | 4.124E-09 | 8.100E-09 | 1.600E-08 |
| 11344 | 568 | Y568D | A | 2e-08 | 5e-09 | 5.4e-09 | 1.013E-08 | 8.547E-09 | 5.000E-09 | 2.000E-08 |
| 11345 | 568 | Y568C | A | 2.2e-08 | 6.6e-09 | 1.7e-08 | 1.520E-08 | 7.856E-09 | 6.600E-09 | 2.200E-08 |
| 11346 | 568 | Y568E | A | 1.4e-08 | 1.2e-08 | 2.8e-08 | 1.800E-08 | 8.718E-09 | 1.200E-08 | 2.800E-08 |
| 11347 | 568 | Y568Q | A | 4.7e-09 | 1.7e-08 | 6e-09 | 9.233E-09 | 6.757E-09 | 4.700E-09 | 1.700E-08 |
| 11348 | 568 | Y568G | A | 2.8e-09 | 3.1e-09 | 8e-09 | 4.633E-09 | 2.919E-09 | 2.800E-09 | 8.000E-09 |
| 11349 | 568 | Y568H | A | 4.7e-09 | 7.6e-09 | 9.5e-09 | 7.267E-09 | 2.417E-09 | 4.700E-09 | 9.500E-09 |
| 11350 | 568 | Y568I | A | 5.3e-09 | 5.1e-09 | 1.4e-08 | 8.133E-09 | 5.082E-09 | 5.100E-09 | 1.400E-08 |
| 11351 | 568 | Y568L | A | 2.5e-09 | 2.8e-09 | 4.8e-09 | 3.367E-09 | 1.250E-09 | 2.500E-09 | 4.800E-09 |
| 11352 | 568 | Y568K | A | 5.2e-09 | 1e-08 | 1.2e-08 | 9.067E-09 | 3.495E-09 | 5.200E-09 | 1.200E-08 |
| 11353 | 568 | Y568M | A | 7.4e-09 | 8.5e-09 | 1.3e-08 | 9.633E-09 | 2.967E-09 | 7.400E-09 | 1.300E-08 |
| 11354 | 568 | Y568F | A | 3.1e-09 | 5.7e-09 | 1.1e-08 | 6.600E-09 | 4.026E-09 | 3.100E-09 | 1.100E-08 |
| 11355 | 568 | Y568P | A | 1.1e-08 | 1e-09 | 3e-09 | 5.000E-09 | 5.292E-09 | 1.000E-09 | 1.100E-08 |
| 11356 | 568 | Y568S | A | 6.8e-09 | 4e-09 | 3.6e-09 | 4.800E-09 | 1.744E-09 | 3.600E-09 | 6.800E-09 |
| 11357 | 568 | Y568T | A | 4.7e-09 | 1.1e-08 | 1.1e-08 | 8.900E-09 | 3.637E-09 | 4.700E-09 | 1.100E-08 |
| 11358 | 568 | Y568W | A | 6.7e-09 | 1.5e-08 | 6.7e-09 | 9.467E-09 | 4.792E-09 | 6.700E-09 | 1.500E-08 |
| 11359 | 568 | Y568Y | A | 2.3e-08 | 5.5e-09 | 9.1e-09 | 1.253E-08 | 9.241E-09 | 5.500E-09 | 2.300E-08 |
| 11360 | 568 | Y568V | A | 1.2e-08 | 8.1e-09 | 1.1e-08 | 1.037E-08 | 2.026E-09 | 8.100E-09 | 1.200E-08 |
| 11361 | 569 | K569A | A | 1.5e-08 | 7.7e-09 | 3.1e-09 | 8.600E-09 | 6.001E-09 | 3.100E-09 | 1.500E-08 |
| 11362 | 569 | K569R | A | 3.1e-09 | 5.6e-09 | 2.4e-09 | 3.700E-09 | 1.682E-09 | 2.400E-09 | 5.600E-09 |
| 11363 | 569 | K569N | A | 5e-09 | 8.4e-09 | 6e-09 | 6.467E-09 | 1.747E-09 | 5.000E-09 | 8.400E-09 |
| 11364 | 569 | K569D | A | 5.4e-09 | 1.7e-08 | 8.4e-09 | 1.027E-08 | 6.021E-09 | 5.400E-09 | 1.700E-08 |
| 11365 | 569 | K569C | A | 8.9e-09 | 2.9e-09 | 4.9e-09 | 5.567E-09 | 3.055E-09 | 2.900E-09 | 8.900E-09 |
| 11366 | 569 | K569E | A | 7.4e-09 | 2.6e-09 | 4.3e-09 | 4.767E-09 | 2.434E-09 | 2.600E-09 | 7.400E-09 |
| 11367 | 569 | K569Q | A | 8.6e-09 | 7.4e-09 | 3.9e-09 | 6.633E-09 | 2.442E-09 | 3.900E-09 | 8.600E-09 |
| 11368 | 569 | K569G | A | 1.3e-08 | 1.1e-08 | 5.5e-09 | 9.833E-09 | 3.884E-09 | 5.500E-09 | 1.300E-08 |
| 11369 | 569 | K569H | A | 2.8e-09 | 1.1e-08 | 8e-09 | 7.267E-09 | 4.149E-09 | 2.800E-09 | 1.100E-08 |
| 11370 | 569 | K569I | A | 6.2e-09 | 2.9e-09 | 4.6e-09 | 4.567E-09 | 1.650E-09 | 2.900E-09 | 6.200E-09 |
| 11371 | 569 | K569L | A | 5.7e-09 | 1.2e-08 | 1.1e-08 | 9.567E-09 | 3.386E-09 | 5.700E-09 | 1.200E-08 |
| 11372 | 569 | K569K | A | 7e-09 | 5.1e-09 | 1.8e-08 | 1.003E-08 | 6.964E-09 | 5.100E-09 | 1.800E-08 |
| 11373 | 569 | K569M | A | 1.8e-08 | 4.7e-09 | 2.2e-09 | 8.300E-09 | 8.493E-09 | 2.200E-09 | 1.800E-08 |
| 11374 | 569 | K569F | A | 4.6e-09 | 8.5e-09 | 8.2e-09 | 7.100E-09 | 2.170E-09 | 4.600E-09 | 8.500E-09 |
| 11375 | 569 | K569P | A | 3.3e-09 | 7.2e-09 | 1.1e-08 | 7.167E-09 | 3.850E-09 | 3.300E-09 | 1.100E-08 |
| 11376 | 569 | K569S | A | 4.9e-09 | 1.1e-08 | 2.3e-09 | 6.067E-09 | 4.466E-09 | 2.300E-09 | 1.100E-08 |
| 11377 | 569 | K569T | A | 7.5e-09 | 6.1e-09 | 8.3e-09 | 7.300E-09 | 1.114E-09 | 6.100E-09 | 8.300E-09 |
| 11378 | 569 | K569W | A | 1.3e-08 | 9.9e-09 | 6.7e-09 | 9.867E-09 | 3.150E-09 | 6.700E-09 | 1.300E-08 |
| 11379 | 569 | K569Y | A | 9.9e-09 | 2.9e-09 | 4.2e-09 | 5.667E-09 | 3.723E-09 | 2.900E-09 | 9.900E-09 |
| 11380 | 569 | K569V | A | 2.3e-08 | 5.2e-09 | 6.7e-09 | 1.163E-08 | 9.872E-09 | 5.200E-09 | 2.300E-08 |
| 11381 | 570 | D570A | A | 5.7e-09 | 7.5e-09 | 7.9e-09 | 7.033E-09 | 1.172E-09 | 5.700E-09 | 7.900E-09 |
| 11382 | 570 | D570R | A | 1e-08 | 6.4e-09 | 2.1e-08 | 1.247E-08 | 7.606E-09 | 6.400E-09 | 2.100E-08 |
| 11383 | 570 | D570N | A | 1.3e-08 | 1.8e-08 | 5e-09 | 1.200E-08 | 6.557E-09 | 5.000E-09 | 1.800E-08 |
| 11384 | 570 | D570D | A | 1e-08 | 7e-09 | 5.1e-09 | 7.367E-09 | 2.470E-09 | 5.100E-09 | 1.000E-08 |
| 11385 | 570 | D570C | A | 2.1e-09 | 1.2e-08 | 9e-09 | 7.700E-09 | 5.076E-09 | 2.100E-09 | 1.200E-08 |
| 11386 | 570 | D570E | A | 4.9e-09 | 4e-09 | 1.6e-08 | 8.300E-09 | 6.684E-09 | 4.000E-09 | 1.600E-08 |
| 11387 | 570 | D570Q | A | 9.5e-09 | 1.1e-08 | 9.1e-09 | 9.867E-09 | 1.002E-09 | 9.100E-09 | 1.100E-08 |
| 11388 | 570 | D570G | A | 2.6e-08 | 8.8e-09 | 5.5e-08 | 2.993E-08 | 2.335E-08 | 8.800E-09 | 5.500E-08 |
| 11389 | 570 | D570H | A | 6.2e-09 | 7e-09 | 1.2e-08 | 8.400E-09 | 3.143E-09 | 6.200E-09 | 1.200E-08 |

| | | | | | | | | | | |
|---|---|---|---|---|---|---|---|---|---|---|
| 11390 | 570 | D570I | A | 1.2e-08 | 6.2e-09 | 8e-09 | 8.733E-09 | 2.969E-09 | 6.200E-09 | 1.200E-08 |
| 11391 | 570 | D570L | A | 3.3e-09 | 7.2e-09 | 1.2e-08 | 7.500E-09 | 4.358E-09 | 3.300E-09 | 1.200E-08 |
| 11392 | 570 | D570K | A | 1.4e-08 | 7e-09 | 5.6e-09 | 8.867E-09 | 4.500E-09 | 5.600E-09 | 1.400E-08 |
| 11393 | 570 | D570M | A | 1.2e-08 | 8.3e-09 | 9.3e-09 | 9.867E-09 | 1.914E-09 | 8.300E-09 | 1.200E-08 |
| 11394 | 570 | D570F | A | 7.4e-09 | 3.8e-09 | 5.9e-09 | 5.700E-09 | 1.808E-09 | 3.800E-09 | 7.400E-09 |
| 11395 | 570 | D570P | A | 3e-09 | 7.2e-09 | 5.1e-09 | 5.100E-09 | 2.100E-09 | 3.000E-09 | 7.200E-09 |
| 11396 | 570 | D570S | A | 1.1e-08 | 5.3e-09 | 1.3e-08 | 9.767E-09 | 3.995E-09 | 5.300E-09 | 1.300E-08 |
| 11397 | 570 | D570T | A | 8.5e-09 | 5.7e-09 | 6.5e-09 | 6.900E-09 | 1.442E-09 | 5.700E-09 | 8.500E-09 |
| 11398 | 570 | D570W | A | 2.4e-09 | 6.7e-09 | 1.7e-09 | 3.600E-09 | 2.707E-09 | 1.700E-09 | 6.700E-09 |
| 11399 | 570 | D570Y | A | 2.3e-09 | 3.1e-09 | 1.1e-08 | 5.467E-09 | 4.809E-09 | 2.300E-09 | 1.100E-08 |
| 11400 | 570 | D570V | A | 1.1e-08 | 6.6e-09 | 9.6e-09 | 9.067E-09 | 2.248E-09 | 6.600E-09 | 1.100E-08 |
| 11401 | 571 | P571A | A | 3e-09 | 6.4e-09 | 1.5e-08 | 8.133E-09 | 6.185E-09 | 3.000E-09 | 1.500E-08 |
| 11402 | 571 | P571R | A | 8.1e-09 | 1.3e-08 | 1.7e-09 | 7.600E-09 | 5.667E-09 | 1.700E-09 | 1.300E-08 |
| 11403 | 571 | P571N | A | 9.9e-09 | 6.6e-09 | 7.9e-09 | 8.133E-09 | 1.662E-09 | 6.600E-09 | 9.900E-09 |
| 11404 | 571 | P571D | A | 1.4e-08 | 6.2e-09 | 1.6e-08 | 1.207E-08 | 5.178E-09 | 6.200E-09 | 1.600E-08 |
| 11405 | 571 | P571C | A | 7.1e-09 | 5.6e-10 | 3.9e-08 | 1.555E-08 | 2.057E-08 | 5.600E-10 | 3.900E-08 |
| 11406 | 571 | P571E | A | 4.8e-09 | 9.2e-09 | 9.3e-09 | 7.767E-09 | 2.570E-09 | 4.800E-09 | 9.300E-09 |
| 11407 | 571 | P571Q | A | 5.8e-09 | 2.8e-09 | 9.7e-09 | 6.100E-09 | 3.460E-09 | 2.800E-09 | 9.700E-09 |
| 11408 | 571 | P571G | A | 7.5e-09 | 7.7e-09 | 1.8e-08 | 1.107E-08 | 6.005E-09 | 7.500E-09 | 1.800E-08 |
| 11409 | 571 | P571H | A | 4.8e-09 | 2.9e-09 | 5.7e-09 | 4.467E-09 | 1.429E-09 | 2.900E-09 | 5.700E-09 |
| 11410 | 571 | P571I | A | 1.6e-08 | 5.9e-09 | 1.1e-08 | 1.097E-08 | 5.050E-09 | 5.900E-09 | 1.600E-08 |
| 11411 | 571 | P571L | A | 1.4e-08 | 5.6e-09 | 8.2e-09 | 9.267E-09 | 4.300E-09 | 5.600E-09 | 1.400E-08 |
| 11412 | 571 | P571K | A | 4.3e-09 | 1.2e-08 | 1.2e-08 | 9.433E-09 | 4.446E-09 | 4.300E-09 | 1.200E-08 |
| 11413 | 571 | P571M | A | 1.1e-08 | 1.2e-08 | 7.4e-09 | 1.013E-08 | 2.419E-09 | 7.400E-09 | 1.200E-08 |
| 11414 | 571 | P571F | A | 2.4e-08 | 1.3e-09 | 4.6e-09 | 9.967E-09 | 1.226E-08 | 1.300E-09 | 2.400E-08 |
| 11415 | 571 | P571P | A | 3.5e-09 | 1.7e-08 | 7.3e-09 | 9.267E-09 | 6.962E-09 | 3.500E-09 | 1.700E-08 |
| 11416 | 571 | P571S | A | 2.2e-09 | 1.9e-08 | 2.6e-09 | 7.933E-09 | 9.586E-09 | 2.200E-09 | 1.900E-08 |
| 11417 | 571 | P571T | A | 6.1e-09 | 6e-09 | 7.9e-09 | 6.667E-09 | 1.069E-09 | 6.000E-09 | 7.900E-09 |
| 11418 | 571 | P571W | A | 2.9e-09 | 1.9e-08 | 2.1e-08 | 1.430E-08 | 9.923E-09 | 2.900E-09 | 2.100E-08 |
| 11419 | 571 | P571Y | A | 3.6e-09 | 4.1e-09 | 1.1e-08 | 6.233E-09 | 4.136E-09 | 3.600E-09 | 1.100E-08 |
| 11420 | 571 | P571V | A | 2.6e-09 | 4.8e-09 | 7.3e-09 | 4.900E-09 | 2.352E-09 | 2.600E-09 | 7.300E-09 |
| 11421 | 572 | P572A | A | 6e-09 | 1.2e-08 | 5e-09 | 7.667E-09 | 3.786E-09 | 5.000E-09 | 1.200E-08 |
| 11422 | 572 | P572R | A | 8.6e-09 | 7.4e-09 | 4.8e-09 | 6.933E-09 | 1.943E-09 | 4.800E-09 | 8.600E-09 |
| 11423 | 572 | P572N | A | 3.2e-09 | 3e-09 | 9.6e-09 | 5.267E-09 | 3.754E-09 | 3.000E-09 | 9.600E-09 |
| 11424 | 572 | P572D | A | 9.7e-09 | 2.7e-09 | 8.1e-09 | 6.833E-09 | 3.668E-09 | 2.700E-09 | 9.700E-09 |
| 11425 | 572 | P572C | A | 2.5e-09 | 5.8e-09 | 4.8e-09 | 4.367E-09 | 1.692E-09 | 2.500E-09 | 5.800E-09 |
| 11426 | 572 | P572E | A | 6.1e-09 | 7.7e-09 | 9.5e-09 | 7.767E-09 | 1.701E-09 | 6.100E-09 | 9.500E-09 |
| 11427 | 572 | P572Q | A | 4.9e-09 | 2e-08 | 4.6e-09 | 9.833E-09 | 8.806E-09 | 4.600E-09 | 2.000E-08 |
| 11428 | 572 | P572G | A | 2.8e-08 | 1e-08 | 2.9e-08 | 2.233E-08 | 1.069E-08 | 1.000E-08 | 2.900E-08 |
| 11429 | 572 | P572H | A | 4.5e-09 | 1.2e-08 | 1.1e-08 | 9.167E-09 | 4.072E-09 | 4.500E-09 | 1.200E-08 |
| 11430 | 572 | P572I | A | 3e-09 | 3.7e-09 | 5.6e-09 | 4.100E-09 | 1.345E-09 | 3.000E-09 | 5.600E-09 |
| 11431 | 572 | P572L | A | 3.3e-09 | 1e-08 | 6e-09 | 6.433E-09 | 3.371E-09 | 3.300E-09 | 1.000E-08 |
| 11432 | 572 | P572K | A | 1.1e-08 | 1.1e-08 | 9.2e-09 | 1.040E-08 | 1.039E-09 | 9.200E-09 | 1.100E-08 |
| 11433 | 572 | P572M | A | 7.1e-09 | 3.5e-09 | 7.1e-09 | 5.900E-09 | 2.078E-09 | 3.500E-09 | 7.100E-09 |
| 11434 | 572 | P572F | A | 1.6e-08 | 7.6e-09 | 8.1e-09 | 1.057E-08 | 4.712E-09 | 7.600E-09 | 1.600E-08 |
| 11435 | 572 | P572P | A | 7.4e-09 | 4.8e-09 | 7.5e-09 | 6.567E-09 | 1.531E-09 | 4.800E-09 | 7.500E-09 |
| 11436 | 572 | P572S | A | 1.2e-08 | 6.7e-09 | 3.3e-09 | 7.333E-09 | 4.384E-09 | 3.300E-09 | 1.200E-08 |
| 11437 | 572 | P572T | A | 1.1e-08 | 3.1e-09 | 7.5e-09 | 7.200E-09 | 3.959E-09 | 3.100E-09 | 1.100E-08 |
| 11438 | 572 | P572W | A | 7.7e-09 | 2.1e-09 | 6.9e-09 | 5.567E-09 | 3.029E-09 | 2.100E-09 | 7.700E-09 |
| 11439 | 572 | P572Y | A | 1.6e-08 | 1.4e-08 | 7e-09 | 1.233E-08 | 4.726E-09 | 7.000E-09 | 1.600E-08 |
| 11440 | 572 | P572V | A | 1.9e-08 | 1.2e-08 | 1.1e-08 | 1.400E-08 | 4.359E-09 | 1.100E-08 | 1.900E-08 |
| 11441 | 573 | F573A | A | 2.3e-09 | 1.5e-08 | 6.4e-09 | 7.900E-09 | 6.482E-09 | 2.300E-09 | 1.500E-08 |
| 11442 | 573 | F573R | A | 8.7e-09 | 6.8e-09 | 1.7e-08 | 1.083E-08 | 5.424E-09 | 6.800E-09 | 1.700E-08 |
| 11443 | 573 | F573N | A | 4.3e-09 | 1.3e-08 | 1.2e-08 | 9.767E-09 | 4.761E-09 | 4.300E-09 | 1.300E-08 |
| 11444 | 573 | F573D | A | 5e-09 | 1e-08 | 5.1e-09 | 6.700E-09 | 2.858E-09 | 5.000E-09 | 1.000E-08 |
| 11445 | 573 | F573C | A | 2.9e-09 | 4.4e-09 | 7.9e-09 | 5.067E-09 | 2.566E-09 | 2.900E-09 | 7.900E-09 |
| 11446 | 573 | F573E | A | 3.5e-09 | 1e-08 | 6e-09 | 6.500E-09 | 3.279E-09 | 3.500E-09 | 1.000E-08 |
| 11447 | 573 | F573Q | A | 8.8e-09 | 1.1e-08 | 1e-08 | 9.933E-09 | 1.102E-09 | 8.800E-09 | 1.100E-08 |
| 11448 | 573 | F573G | A | 7.7e-09 | 4e-09 | 2.2e-08 | 1.123E-08 | 9.506E-09 | 4.000E-09 | 2.200E-08 |
| 11449 | 573 | F573H | A | 8.4e-09 | 1.3e-08 | 7.2e-09 | 9.533E-09 | 3.062E-09 | 7.200E-09 | 1.300E-08 |
| 11450 | 573 | F573I | A | 5.9e-09 | 8.3e-09 | 3.5e-09 | 5.900E-09 | 2.400E-09 | 3.500E-09 | 8.300E-09 |
| 11451 | 573 | F573L | A | 1.2e-08 | 1.8e-08 | 1.5e-08 | 1.500E-08 | 3.000E-09 | 1.200E-08 | 1.800E-08 |
| 11452 | 573 | F573K | A | 4.6e-09 | 8.8e-09 | 7.4e-09 | 6.933E-09 | 2.139E-09 | 4.600E-09 | 8.800E-09 |
| 11453 | 573 | F573M | A | 2.5e-09 | 3.2e-09 | 1e-08 | 5.233E-09 | 4.143E-09 | 2.500E-09 | 1.000E-08 |
| 11454 | 573 | F573F | A | 1.7e-08 | 4e-09 | 3.7e-09 | 8.233E-09 | 7.594E-09 | 3.700E-09 | 1.700E-08 |
| 11455 | 573 | F573P | A | 1.2e-09 | 1.5e-08 | 1.4e-08 | 1.007E-08 | 7.695E-09 | 1.200E-09 | 1.500E-08 |
| 11456 | 573 | F573S | A | 4.9e-09 | 7.5e-09 | 4.8e-09 | 5.733E-09 | 1.531E-09 | 4.800E-09 | 7.500E-09 |
| 11457 | 573 | F573T | A | 6.2e-09 | 1.2e-08 | 7.3e-09 | 8.500E-09 | 3.081E-09 | 6.200E-09 | 1.200E-08 |
| 11458 | 573 | F573W | A | 1.3e-08 | 3e-09 | 5.7e-09 | 7.233E-09 | 5.173E-09 | 3.000E-09 | 1.300E-08 |
| 11459 | 573 | F573Y | A | 8.7e-09 | 8.6e-09 | 2.4e-09 | 6.567E-09 | 3.609E-09 | 2.400E-09 | 8.700E-09 |
| 11460 | 573 | F573V | A | 7.7e-09 | 1.1e-08 | 7.4e-09 | 8.700E-09 | 1.997E-09 | 7.400E-09 | 1.100E-08 |
| 11461 | 574 | C574A | A | 7.5e-09 | 1.1e-08 | 2.3e-08 | 1.383E-08 | 8.129E-09 | 7.500E-09 | 2.300E-08 |
| 11462 | 574 | C574R | A | 1.5e-08 | 8.6e-09 | 5e-09 | 9.533E-09 | 5.065E-09 | 5.000E-09 | 1.500E-08 |
| 11463 | 574 | C574N | A | 8.4e-09 | 3e-09 | 7.1e-09 | 6.167E-09 | 2.818E-09 | 3.000E-09 | 8.400E-09 |
| 11464 | 574 | C574D | A | 2.4e-09 | 2.2e-09 | 1.5e-08 | 6.533E-09 | 7.333E-09 | 2.200E-09 | 1.500E-08 |
| 11465 | 574 | C574C | A | 5.4e-09 | 1.2e-08 | 3e-09 | 6.800E-09 | 4.660E-09 | 3.000E-09 | 1.200E-08 |
| 11466 | 574 | C574E | A | 4.6e-09 | 6.2e-09 | 1.2e-08 | 7.600E-09 | 3.894E-09 | 4.600E-09 | 1.200E-08 |
| 11467 | 574 | C574Q | A | 1.4e-08 | 6.5e-09 | 7.4e-09 | 9.300E-09 | 4.095E-09 | 6.500E-09 | 1.400E-08 |
| 11468 | 574 | C574G | A | 1e-08 | 2e-09 | 1e-08 | 7.333E-09 | 4.619E-09 | 2.000E-09 | 1.000E-08 |
| 11469 | 574 | C574H | A | 6.8e-09 | 6.4e-09 | 9.5e-09 | 7.567E-09 | 1.686E-09 | 6.400E-09 | 9.500E-09 |
| 11470 | 574 | C574I | A | 8.7e-09 | 6.4e-09 | 2.8e-09 | 5.967E-09 | 2.974E-09 | 2.800E-09 | 8.700E-09 |
| 11471 | 574 | C574L | A | 1.4e-08 | 4.4e-09 | 1.7e-09 | 6.700E-09 | 6.465E-09 | 1.700E-09 | 1.400E-08 |
| 11472 | 574 | C574K | A | 6.9e-09 | 3.7e-09 | 9.4e-09 | 6.667E-09 | 2.857E-09 | 3.700E-09 | 9.400E-09 |
| 11473 | 574 | C574M | A | 1.8e-08 | 1.1e-08 | 6e-09 | 1.167E-08 | 6.028E-09 | 6.000E-09 | 1.800E-08 |
| 11474 | 574 | C574F | A | 1.2e-08 | 2.7e-09 | 5.1e-09 | 6.600E-09 | 4.828E-09 | 2.700E-09 | 1.200E-08 |

| | | | | | | | | | | |
|---|---|---|---|---|---|---|---|---|---|---|
| 11475 | 574 | C574P | A | 1.9e-09 | 6.2e-09 | 1.6e-09 | 3.233E-09 | 2.574E-09 | 1.600E-09 | 6.200E-09 |
| 11476 | 574 | C574S | A | 1.1e-08 | 1.1e-08 | 9.1e-09 | 1.037E-08 | 1.097E-09 | 9.100E-09 | 1.100E-08 |
| 11477 | 574 | C574T | A | 1.4e-08 | 7.7e-09 | 5.6e-09 | 9.100E-09 | 4.371E-09 | 5.600E-09 | 1.400E-08 |
| 11478 | 574 | C574W | A | 3.4e-09 | 1.2e-08 | 1.7e-08 | 1.080E-08 | 6.879E-09 | 3.400E-09 | 1.700E-08 |
| 11479 | 574 | C574Y | A | 1.7e-08 | 4.8e-09 | 1.2e-08 | 1.127E-08 | 6.133E-09 | 4.800E-09 | 1.700E-08 |
| 11480 | 574 | C574V | A | 3.4e-09 | 1.5e-08 | 1.2e-08 | 1.013E-08 | 6.021E-09 | 3.400E-09 | 1.500E-08 |
| 11481 | 575 | V575A | A | 6.5e-09 | 5.3e-09 | 1.6e-08 | 9.267E-09 | 5.862E-09 | 5.300E-09 | 1.600E-08 |
| 11482 | 575 | V575R | A | 4.8e-09 | 8.3e-09 | 6.1e-09 | 6.400E-09 | 1.769E-09 | 4.800E-09 | 8.300E-09 |
| 11483 | 575 | V575N | A | 1.1e-08 | 2e-09 | 4.4e-09 | 5.800E-09 | 4.660E-09 | 2.000E-09 | 1.100E-08 |
| 11484 | 575 | V575D | A | 8.4e-09 | 8e-09 | 2.7e-09 | 6.367E-09 | 3.182E-09 | 2.700E-09 | 8.400E-09 |
| 11485 | 575 | V575C | A | 9.3e-09 | 1.1e-08 | 3.7e-09 | 8.000E-09 | 3.820E-09 | 3.700E-09 | 1.100E-08 |
| 11486 | 575 | V575E | A | 5.8e-09 | 8.2e-09 | 7.8e-09 | 7.267E-09 | 1.286E-09 | 5.800E-09 | 8.200E-09 |
| 11487 | 575 | V575Q | A | 5.1e-09 | 9e-09 | 8.3e-09 | 7.467E-09 | 2.079E-09 | 5.100E-09 | 9.000E-09 |
| 11488 | 575 | V575G | A | 4.3e-09 | 6.4e-09 | 8e-09 | 6.233E-09 | 1.856E-09 | 4.300E-09 | 8.000E-09 |
| 11489 | 575 | V575H | A | 7.4e-09 | 7.1e-09 | 7.3e-09 | 7.267E-09 | 1.528E-10 | 7.100E-09 | 7.400E-09 |
| 11490 | 575 | V575I | A | 5.8e-09 | 5.3e-09 | 2.5e-09 | 4.533E-09 | 1.779E-09 | 2.500E-09 | 5.800E-09 |
| 11491 | 575 | V575L | A | 7.7e-09 | 4.6e-09 | 3.4e-09 | 5.233E-09 | 2.219E-09 | 3.400E-09 | 7.700E-09 |
| 11492 | 575 | V575K | A | 3.6e-09 | 3.5e-09 | 7.2e-09 | 4.767E-09 | 2.108E-09 | 3.500E-09 | 7.200E-09 |
| 11493 | 575 | V575M | A | 2.6e-09 | 9.5e-10 | 6.2e-09 | 3.250E-09 | 2.685E-09 | 9.500E-10 | 6.200E-09 |
| 11494 | 575 | V575F | A | 2.1e-08 | 8.8e-09 | 8.8e-09 | 1.287E-08 | 7.044E-09 | 8.800E-09 | 2.100E-08 |
| 11495 | 575 | V575P | A | 2.9e-09 | 1.2e-08 | 3.7e-09 | 6.200E-09 | 5.039E-09 | 2.900E-09 | 1.200E-08 |
| 11496 | 575 | V575S | A | 7.6e-09 | 6.6e-09 | 3.5e-09 | 5.900E-09 | 2.138E-09 | 3.500E-09 | 7.600E-09 |
| 11497 | 575 | V575T | A | 3.4e-09 | 5.3e-09 | 2e-09 | 3.567E-09 | 1.656E-09 | 2.000E-09 | 5.300E-09 |
| 11498 | 575 | V575W | A | 5.1e-09 | 5.3e-09 | 3.8e-08 | 1.613E-08 | 1.894E-08 | 5.100E-09 | 3.800E-08 |
| 11499 | 575 | V575Y | A | 9.1e-09 | 1.3e-08 | 9.3e-09 | 1.047E-08 | 2.196E-09 | 9.100E-09 | 1.300E-08 |
| 11500 | 575 | V575V | A | 4.2e-09 | 4.5e-09 | 1.2e-08 | 6.900E-09 | 4.419E-09 | 4.200E-09 | 1.200E-08 |
| 11501 | 576 | A576A | A | 7.4e-09 | 1.4e-08 | 5.7e-09 | 9.033E-09 | 4.384E-09 | 5.700E-09 | 1.400E-08 |
| 11502 | 576 | A576R | A | 1.8e-08 | 5.7e-09 | 1.4e-08 | 1.257E-08 | 6.274E-09 | 5.700E-09 | 1.800E-08 |
| 11503 | 576 | A576N | A | 1.9e-08 | 3.6e-09 | 3.9e-09 | 8.833E-09 | 8.806E-09 | 3.600E-09 | 1.900E-08 |
| 11504 | 576 | A576D | A | 1.1e-08 | 5.7e-09 | 6e-09 | 7.567E-09 | 2.977E-09 | 5.700E-09 | 1.100E-08 |
| 11505 | 576 | A576C | A | 1.1e-08 | 7.6e-09 | 2.7e-09 | 7.100E-09 | 4.173E-09 | 2.700E-09 | 1.100E-08 |
| 11506 | 576 | A576E | A | 6e-09 | 5.4e-09 | 7.1e-09 | 6.167E-09 | 8.622E-10 | 5.400E-09 | 7.100E-09 |
| 11507 | 576 | A576Q | A | 4.1e-09 | 2.5e-09 | 1.2e-08 | 6.200E-09 | 5.086E-09 | 2.500E-09 | 1.200E-08 |
| 11508 | 576 | A576G | A | 8.3e-09 | 2.8e-09 | 8.1e-09 | 6.400E-09 | 3.119E-09 | 2.800E-09 | 8.300E-09 |
| 11509 | 576 | A576H | A | 2.6e-09 | 8.4e-09 | 2.9e-09 | 4.633E-09 | 3.265E-09 | 2.600E-09 | 8.400E-09 |
| 11510 | 576 | A576I | A | 1.8e-08 | 7.4e-09 | 6.2e-09 | 1.053E-08 | 6.494E-09 | 6.200E-09 | 1.800E-08 |
| 11511 | 576 | A576L | A | 5.1e-09 | 3.7e-09 | 1.3e-08 | 7.267E-09 | 5.014E-09 | 3.700E-09 | 1.300E-08 |
| 11512 | 576 | A576K | A | 3.4e-09 | 9.2e-09 | 4.4e-09 | 5.667E-09 | 3.101E-09 | 3.400E-09 | 9.200E-09 |
| 11513 | 576 | A576M | A | 2.3e-09 | 7.9e-09 | 1e-08 | 6.733E-09 | 3.980E-09 | 2.300E-09 | 1.000E-08 |
| 11514 | 576 | A576F | A | 8e-09 | 4.1e-09 | 1.6e-08 | 9.367E-09 | 6.067E-09 | 4.100E-09 | 1.600E-08 |
| 11515 | 576 | A576P | A | 6.5e-09 | 1.1e-08 | 3.1e-09 | 6.867E-09 | 3.963E-09 | 3.100E-09 | 1.100E-08 |
| 11516 | 576 | A576S | A | 9.7e-09 | 2.9e-09 | 4.6e-09 | 5.733E-09 | 3.539E-09 | 2.900E-09 | 9.700E-09 |
| 11517 | 576 | A576T | A | 1.1e-08 | 3.8e-09 | 4e-09 | 6.267E-09 | 4.100E-09 | 3.800E-09 | 1.100E-08 |
| 11518 | 576 | A576W | A | 8.5e-09 | 9e-09 | 9.5e-09 | 9.000E-09 | 5.000E-10 | 8.500E-09 | 9.500E-09 |
| 11519 | 576 | A576Y | A | 1.7e-08 | 2.2e-09 | 4.1e-09 | 7.767E-09 | 8.053E-09 | 2.200E-09 | 1.700E-08 |
| 11520 | 576 | A576V | A | 3.4e-09 | 5.1e-09 | 5.8e-09 | 4.767E-09 | 1.234E-09 | 3.400E-09 | 5.800E-09 |
| 11521 | 577 | R577A | A | 1.9e-09 | 2.3e-08 | 6.2e-09 | 1.037E-08 | 1.115E-08 | 1.900E-09 | 2.300E-08 |
| 11522 | 577 | R577R | A | 1e-08 | 2.5e-09 | 9.9e-09 | 7.467E-09 | 4.302E-09 | 2.500E-09 | 1.000E-08 |
| 11523 | 577 | R577N | A | 3.1e-09 | 2.9e-09 | 1.1e-08 | 5.667E-09 | 4.620E-09 | 2.900E-09 | 1.100E-08 |
| 11524 | 577 | R577D | A | 9.9e-09 | 8.1e-09 | 1.1e-08 | 9.667E-09 | 1.464E-09 | 8.100E-09 | 1.100E-08 |
| 11525 | 577 | R577C | A | 1.1e-08 | 1.1e-08 | 7.9e-09 | 9.967E-09 | 1.790E-09 | 7.900E-09 | 1.100E-08 |
| 11526 | 577 | R577E | A | 1e-08 | 1.6e-08 | 6.2e-09 | 1.073E-08 | 4.941E-09 | 6.200E-09 | 1.600E-08 |
| 11527 | 577 | R577Q | A | 1.4e-08 | 3.3e-09 | 3.6e-09 | 6.967E-09 | 6.093E-09 | 3.300E-09 | 1.400E-08 |
| 11528 | 577 | R577G | A | 1.5e-08 | 2.2e-09 | 1.5e-08 | 1.073E-08 | 7.390E-09 | 2.200E-09 | 1.500E-08 |
| 11529 | 577 | R577H | A | 7.5e-09 | 5.6e-09 | 1.4e-08 | 9.033E-09 | 4.405E-09 | 5.600E-09 | 1.400E-08 |
| 11530 | 577 | R577I | A | 2.3e-08 | 1.1e-08 | 1e-08 | 1.467E-08 | 7.234E-09 | 1.000E-08 | 2.300E-08 |
| 11531 | 577 | R577L | A | 3.7e-09 | 6.4e-09 | 3.9e-09 | 4.667E-09 | 1.504E-09 | 3.700E-09 | 6.400E-09 |
| 11532 | 577 | R577K | A | 2.2e-08 | 9.4e-09 | 2.4e-09 | 1.127E-08 | 9.932E-09 | 2.400E-09 | 2.200E-08 |
| 11533 | 577 | R577M | A | 9.9e-09 | 1.6e-09 | 5.2e-09 | 5.567E-09 | 4.162E-09 | 1.600E-09 | 9.900E-09 |
| 11534 | 577 | R577F | A | 9.8e-09 | 1.2e-08 | 9.1e-09 | 1.030E-08 | 1.513E-09 | 9.100E-09 | 1.200E-08 |
| 11535 | 577 | R577P | A | 9.2e-09 | 1.3e-08 | 7.4e-09 | 9.867E-09 | 2.859E-09 | 7.400E-09 | 1.300E-08 |
| 11536 | 577 | R577S | A | 6.7e-09 | 9.4e-09 | 1.8e-08 | 1.137E-08 | 5.901E-09 | 6.700E-09 | 1.800E-08 |
| 11537 | 577 | R577T | A | 6.2e-09 | 7.6e-09 | 7.9e-09 | 7.233E-09 | 9.074E-10 | 6.200E-09 | 7.900E-09 |
| 11538 | 577 | R577W | A | 2.4e-09 | 2.7e-09 | 1.9e-09 | 2.333E-09 | 4.041E-10 | 1.900E-09 | 2.700E-09 |
| 11539 | 577 | R577Y | A | 3.7e-09 | 9.4e-09 | 3.2e-09 | 5.433E-09 | 3.444E-09 | 3.200E-09 | 9.400E-09 |
| 11540 | 577 | R577V | A | 3.8e-09 | 3.5e-09 | 4.3e-09 | 3.867E-09 | 4.041E-10 | 3.500E-09 | 4.300E-09 |
| 11541 | 578 | C578A | A | 9.2e-09 | 7e-09 | 8.6e-09 | 8.267E-09 | 1.137E-09 | 7.000E-09 | 9.200E-09 |
| 11542 | 578 | C578R | A | 3.2e-09 | 2.9e-09 | 7e-09 | 4.367E-09 | 2.285E-09 | 2.900E-09 | 7.000E-09 |
| 11543 | 578 | C578N | A | 5.9e-09 | 3.1e-09 | 3.7e-09 | 4.233E-09 | 1.474E-09 | 3.100E-09 | 5.900E-09 |
| 11544 | 578 | C578D | A | 2e-09 | 3.6e-09 | 2e-08 | 8.533E-09 | 9.963E-09 | 2.000E-09 | 2.000E-08 |
| 11545 | 578 | C578C | A | 4.5e-09 | 1.6e-08 | 1.4e-08 | 1.150E-08 | 6.144E-09 | 4.500E-09 | 1.600E-08 |
| 11546 | 578 | C578E | A | 6.2e-09 | 8.5e-09 | 8.7e-09 | 7.800E-09 | 1.389E-09 | 6.200E-09 | 8.700E-09 |
| 11547 | 578 | C578Q | A | 1.8e-08 | 1.1e-08 | 1e-08 | 1.300E-08 | 4.359E-09 | 1.000E-08 | 1.800E-08 |
| 11548 | 578 | C578G | A | 1.4e-08 | 6.4e-09 | 1e-08 | 1.013E-08 | 3.802E-09 | 6.400E-09 | 1.400E-08 |
| 11549 | 578 | C578H | A | 1.8e-08 | 7.3e-09 | 4.3e-09 | 9.867E-09 | 7.202E-09 | 4.300E-09 | 1.800E-08 |
| 11550 | 578 | C578I | A | 9.2e-09 | 6.2e-09 | 5.3e-09 | 6.900E-09 | 2.042E-09 | 5.300E-09 | 9.200E-09 |
| 11551 | 578 | C578L | A | 4.9e-09 | 6.4e-09 | 8.7e-09 | 6.667E-09 | 1.914E-09 | 4.900E-09 | 8.700E-09 |
| 11552 | 578 | C578K | A | 6.2e-09 | 1.1e-08 | 5.2e-09 | 7.467E-09 | 3.101E-09 | 5.200E-09 | 1.100E-08 |
| 11553 | 578 | C578M | A | 8.1e-09 | 7e-09 | 1.1e-08 | 8.700E-09 | 2.066E-09 | 7.000E-09 | 1.100E-08 |
| 11554 | 578 | C578F | A | 8.2e-09 | 3.4e-09 | 5.1e-09 | 5.567E-09 | 2.434E-09 | 3.400E-09 | 8.200E-09 |
| 11555 | 578 | C578P | A | 1.2e-09 | 1.5e-08 | 2.1e-09 | 6.100E-09 | 7.721E-09 | 1.200E-09 | 1.500E-08 |
| 11556 | 578 | C578S | A | 5.6e-09 | 4.3e-09 | 2.4e-09 | 4.100E-09 | 1.609E-09 | 2.400E-09 | 5.600E-09 |
| 11557 | 578 | C578T | A | 4.5e-09 | 3.3e-09 | 6.1e-09 | 4.633E-09 | 1.405E-09 | 3.300E-09 | 6.100E-09 |
| 11558 | 578 | C578W | A | 1e-08 | 5.6e-09 | 9.4e-09 | 8.333E-09 | 2.386E-09 | 5.600E-09 | 1.000E-08 |
| 11559 | 578 | C578Y | A | 5.8e-09 | 5.9e-09 | 7.8e-09 | 6.500E-09 | 1.127E-09 | 5.800E-09 | 7.800E-09 |

| | | | | | | | | | | |
|---|---|---|---|---|---|---|---|---|---|---|
| 11560 | 578 | C578V | A | 3.7e-09 | 4.1e-09 | 3.9e-09 | 3.900E-09 | 2.000E-10 | 3.700E-09 | 4.100E-09 |
| 11561 | 579 | P579A | A | 2.9e-09 | 1e-08 | 1.8e-08 | 1.030E-08 | 7.554E-09 | 2.900E-09 | 1.800E-08 |
| 11562 | 579 | P579R | A | 1.5e-08 | 4.4e-09 | 8.2e-09 | 9.200E-09 | 5.370E-09 | 4.400E-09 | 1.500E-08 |
| 11563 | 579 | P579N | A | 1.6e-08 | 2.7e-09 | 4.3e-09 | 7.667E-09 | 7.261E-09 | 2.700E-09 | 1.600E-08 |
| 11564 | 579 | P579D | A | 1.1e-08 | 1.8e-08 | 2.4e-09 | 1.047E-08 | 7.814E-09 | 2.400E-09 | 1.800E-08 |
| 11565 | 579 | P579C | A | 5e-09 | 9.9e-09 | 2.9e-09 | 5.933E-09 | 3.592E-09 | 2.900E-09 | 9.900E-09 |
| 11566 | 579 | P579E | A | 5.7e-09 | 6.2e-09 | 6.6e-09 | 6.167E-09 | 4.509E-10 | 5.700E-09 | 6.600E-09 |
| 11567 | 579 | P579Q | A | 1.2e-08 | 4e-09 | 1.4e-08 | 1.000E-08 | 5.292E-09 | 4.000E-09 | 1.400E-08 |
| 11568 | 579 | P579G | A | 1.1e-08 | 1.2e-08 | 1.1e-08 | 1.133E-08 | 5.774E-10 | 1.100E-08 | 1.200E-08 |
| 11569 | 579 | P579H | A | 1e-08 | 2e-09 | 8.4e-09 | 6.800E-09 | 4.233E-09 | 2.000E-09 | 1.000E-08 |
| 11570 | 579 | P579I | A | 4.7e-09 | 1.1e-08 | 3.9e-09 | 6.533E-09 | 3.889E-09 | 3.900E-09 | 1.100E-08 |
| 11571 | 579 | P579L | A | 9.6e-09 | 8.4e-09 | 5.4e-09 | 7.800E-09 | 2.163E-09 | 5.400E-09 | 9.600E-09 |
| 11572 | 579 | P579K | A | 1.9e-08 | 2.6e-08 | 3.8e-09 | 1.627E-08 | 1.135E-08 | 3.800E-09 | 2.600E-08 |
| 11573 | 579 | P579M | A | 1.3e-08 | 4.7e-09 | 1.1e-08 | 9.567E-09 | 4.332E-09 | 4.700E-09 | 1.300E-08 |
| 11574 | 579 | P579F | A | 5e-09 | 7.9e-09 | 1.4e-08 | 8.967E-09 | 4.594E-09 | 5.000E-09 | 1.400E-08 |
| 11575 | 579 | P579P | A | 1.8e-09 | 1.1e-08 | 3.7e-09 | 5.500E-09 | 4.857E-09 | 1.800E-09 | 1.100E-08 |
| 11576 | 579 | P579S | A | 8.9e-09 | 6.6e-09 | 2.8e-09 | 6.100E-09 | 3.081E-09 | 2.800E-09 | 8.900E-09 |
| 11577 | 579 | P579T | A | 1.4e-08 | 8.3e-09 | 4.6e-09 | 8.967E-09 | 4.735E-09 | 4.600E-09 | 1.400E-08 |
| 11578 | 579 | P579W | A | 5e-09 | 1.9e-08 | 3.1e-09 | 9.033E-09 | 8.684E-09 | 3.100E-09 | 1.900E-08 |
| 11579 | 579 | P579Y | A | 6.5e-09 | 6e-09 | 4e-09 | 5.500E-09 | 1.323E-09 | 4.000E-09 | 6.500E-09 |
| 11580 | 579 | P579V | A | 1.3e-08 | 6.5e-09 | 7.4e-09 | 8.967E-09 | 3.522E-09 | 6.500E-09 | 1.300E-08 |
| 11581 | 580 | S580A | A | 8e-09 | 4.5e-09 | 7.6e-09 | 6.700E-09 | 1.916E-09 | 4.500E-09 | 8.000E-09 |
| 11582 | 580 | S580R | A | 1.3e-08 | 2.1e-08 | 6.6e-09 | 1.353E-08 | 7.215E-09 | 6.600E-09 | 2.100E-08 |
| 11583 | 580 | S580N | A | 5.8e-09 | 3.7e-09 | 4.3e-09 | 4.600E-09 | 1.082E-09 | 3.700E-09 | 5.800E-09 |
| 11584 | 580 | S580D | A | 5.9e-09 | 7.5e-09 | 9.4e-09 | 7.600E-09 | 1.752E-09 | 5.900E-09 | 9.400E-09 |
| 11585 | 580 | S580C | A | 2.3e-08 | 4.3e-09 | 6.2e-09 | 1.117E-08 | 1.029E-08 | 4.300E-09 | 2.300E-08 |
| 11586 | 580 | S580E | A | 8.5e-09 | 8.2e-09 | 2.3e-08 | 1.323E-08 | 8.460E-09 | 8.200E-09 | 2.300E-08 |
| 11587 | 580 | S580Q | A | 4.4e-09 | 1e-08 | 8.2e-09 | 7.533E-09 | 2.859E-09 | 4.400E-09 | 1.000E-08 |
| 11588 | 580 | S580G | A | 5.8e-09 | 1.7e-08 | 1.8e-08 | 1.360E-08 | 6.773E-09 | 5.800E-09 | 1.800E-08 |
| 11589 | 580 | S580H | A | 1.9e-09 | 5.9e-09 | 1.7e-08 | 8.267E-09 | 7.823E-09 | 1.900E-09 | 1.700E-08 |
| 11590 | 580 | S580I | A | 2.2e-09 | 7.6e-09 | 2e-08 | 9.933E-09 | 9.127E-09 | 2.200E-09 | 2.000E-08 |
| 11591 | 580 | S580L | A | 7.1e-09 | 4.6e-09 | 7.5e-09 | 6.400E-09 | 1.572E-09 | 4.600E-09 | 7.500E-09 |
| 11592 | 580 | S580K | A | 8.7e-09 | 4e-09 | 2.1e-08 | 1.123E-08 | 8.779E-09 | 4.000E-09 | 2.100E-08 |
| 11593 | 580 | S580M | A | 1.4e-08 | 6.6e-09 | 8.1e-09 | 9.567E-09 | 3.912E-09 | 6.600E-09 | 1.400E-08 |
| 11594 | 580 | S580F | A | 6.5e-09 | 3.1e-09 | 3e-09 | 4.200E-09 | 1.992E-09 | 3.000E-09 | 6.500E-09 |
| 11595 | 580 | S580P | A | 2.2e-09 | 4.2e-09 | 1.2e-08 | 6.133E-09 | 5.178E-09 | 2.200E-09 | 1.200E-08 |
| 11596 | 580 | S580S | A | 4.1e-09 | 9e-09 | 5.4e-09 | 6.167E-09 | 2.538E-09 | 4.100E-09 | 9.000E-09 |
| 11597 | 580 | S580T | A | 8.5e-09 | 7.3e-09 | 2.5e-09 | 6.100E-09 | 3.175E-09 | 2.500E-09 | 8.500E-09 |
| 11598 | 580 | S580W | A | 2.5e-09 | 5.8e-09 | 3.6e-09 | 3.967E-09 | 1.680E-09 | 2.500E-09 | 5.800E-09 |
| 11599 | 580 | S580Y | A | 6.8e-09 | 6.6e-09 | 1.1e-08 | 8.133E-09 | 2.485E-09 | 6.600E-09 | 1.100E-08 |
| 11600 | 580 | S580V | A | 1e-08 | 7.9e-09 | 3.8e-09 | 7.233E-09 | 3.153E-09 | 3.800E-09 | 1.000E-08 |
| 11601 | 581 | G581A | A | 1e-08 | 5.6e-09 | 1.2e-08 | 9.200E-09 | 3.274E-09 | 5.600E-09 | 1.200E-08 |
| 11602 | 581 | G581R | A | 8.6e-09 | 2.1e-08 | 1e-08 | 1.320E-08 | 6.791E-09 | 8.600E-09 | 2.100E-08 |
| 11603 | 581 | G581N | A | 4.2e-09 | 1.1e-08 | 5.9e-09 | 7.033E-09 | 3.539E-09 | 4.200E-09 | 1.100E-08 |
| 11604 | 581 | G581D | A | 5e-09 | 8.3e-09 | 7.5e-09 | 6.933E-09 | 1.721E-09 | 5.000E-09 | 8.300E-09 |
| 11605 | 581 | G581C | A | 9.5e-09 | 6.7e-09 | 1e-08 | 8.733E-09 | 1.779E-09 | 6.700E-09 | 1.000E-08 |
| 11606 | 581 | G581E | A | 8.3e-09 | 1.5e-08 | 7.7e-09 | 1.033E-08 | 4.053E-09 | 7.700E-09 | 1.500E-08 |
| 11607 | 581 | G581Q | A | 2.4e-09 | 2.7e-09 | 9.4e-09 | 4.833E-09 | 3.958E-09 | 2.400E-09 | 9.400E-09 |
| 11608 | 581 | G581G | A | 5.2e-09 | 8.2e-09 | 8.9e-09 | 7.433E-09 | 1.966E-09 | 5.200E-09 | 8.900E-09 |
| 11609 | 581 | G581H | A | 5.6e-09 | 8.3e-09 | 1.6e-08 | 9.967E-09 | 5.397E-09 | 5.600E-09 | 1.600E-08 |
| 11610 | 581 | G581I | A | 4.8e-09 | 2.5e-09 | 3.6e-09 | 3.633E-09 | 1.150E-09 | 2.500E-09 | 4.800E-09 |
| 11611 | 581 | G581L | A | 4.9e-09 | 4.4e-09 | 2.4e-09 | 3.900E-09 | 1.323E-09 | 2.400E-09 | 4.900E-09 |
| 11612 | 581 | G581K | A | 2.9e-09 | 8e-09 | 9.7e-09 | 6.867E-09 | 3.539E-09 | 2.900E-09 | 9.700E-09 |
| 11613 | 581 | G581M | A | 1.9e-09 | 1.1e-08 | 3.9e-09 | 5.600E-09 | 4.782E-09 | 1.900E-09 | 1.100E-08 |
| 11614 | 581 | G581F | A | 1.2e-08 | 8.7e-09 | 1.7e-08 | 1.257E-08 | 4.179E-09 | 8.700E-09 | 1.700E-08 |
| 11615 | 581 | G581P | A | 8.9e-09 | 2.1e-09 | 1.1e-08 | 7.333E-09 | 4.652E-09 | 2.100E-09 | 1.100E-08 |
| 11616 | 581 | G581S | A | 2.1e-08 | 8.8e-09 | 3e-09 | 1.093E-08 | 9.188E-09 | 3.000E-09 | 2.100E-08 |
| 11617 | 581 | G581T | A | 6.9e-09 | 9.6e-09 | 1.2e-08 | 9.500E-09 | 2.551E-09 | 6.900E-09 | 1.200E-08 |
| 11618 | 581 | G581W | A | 3.2e-08 | 4.4e-09 | 7.8e-09 | 1.473E-08 | 1.505E-08 | 4.400E-09 | 3.200E-08 |
| 11619 | 581 | G581Y | A | 5.8e-09 | 1.8e-09 | 3.1e-09 | 3.567E-09 | 2.040E-09 | 1.800E-09 | 5.800E-09 |
| 11620 | 581 | G581V | A | 4.1e-09 | 3.2e-09 | 5.7e-09 | 4.333E-09 | 1.266E-09 | 3.200E-09 | 5.700E-09 |
| 11621 | 582 | V582A | A | 3.4e-09 | 2.9e-09 | 4.5e-09 | 3.600E-09 | 8.185E-10 | 2.900E-09 | 4.500E-09 |
| 11622 | 582 | V582R | A | 8.7e-09 | 8.5e-09 | 6.4e-09 | 7.867E-09 | 1.274E-09 | 6.400E-09 | 8.700E-09 |
| 11623 | 582 | V582N | A | 2.8e-09 | 2.7e-08 | 1.9e-08 | 1.627E-08 | 1.233E-08 | 2.800E-09 | 2.700E-08 |
| 11624 | 582 | V582D | A | 5.1e-09 | 2.4e-09 | 2.8e-09 | 3.433E-09 | 1.457E-09 | 2.400E-09 | 5.100E-09 |
| 11625 | 582 | V582C | A | 6.6e-09 | 5.9e-09 | 2.9e-09 | 5.133E-09 | 1.966E-09 | 2.900E-09 | 6.600E-09 |
| 11626 | 582 | V582E | A | 2.5e-09 | 2.6e-09 | 5.4e-09 | 3.500E-09 | 1.646E-09 | 2.500E-09 | 5.400E-09 |
| 11627 | 582 | V582Q | A | 8.8e-09 | 7.8e-09 | 7.4e-09 | 8.000E-09 | 7.211E-10 | 7.400E-09 | 8.800E-09 |
| 11628 | 582 | V582G | A | 6.7e-09 | 1.3e-08 | 5.9e-09 | 8.533E-09 | 3.889E-09 | 5.900E-09 | 1.300E-08 |
| 11629 | 582 | V582H | A | 3.6e-09 | 4.4e-09 | 5.8e-09 | 4.600E-09 | 1.114E-09 | 3.600E-09 | 5.800E-09 |
| 11630 | 582 | V582I | A | 6.3e-09 | 1.4e-08 | 4.2e-09 | 8.167E-09 | 5.160E-09 | 4.200E-09 | 1.400E-08 |
| 11631 | 582 | V582L | A | 8.5e-09 | 3.5e-09 | 1.2e-08 | 8.000E-09 | 4.272E-09 | 3.500E-09 | 1.200E-08 |
| 11632 | 582 | V582K | A | 6.9e-09 | 4.7e-09 | 1e-08 | 7.200E-09 | 2.663E-09 | 4.700E-09 | 1.000E-08 |
| 11633 | 582 | V582M | A | 1.2e-08 | 4.4e-09 | 2.1e-09 | 6.167E-09 | 5.181E-09 | 2.100E-09 | 1.200E-08 |
| 11634 | 582 | V582F | A | 4.7e-09 | 6.3e-09 | 5.5e-09 | 5.500E-09 | 8.000E-10 | 4.700E-09 | 6.300E-09 |
| 11635 | 582 | V582P | A | 4.2e-09 | 5.3e-09 | 3.7e-09 | 4.400E-09 | 8.185E-10 | 3.700E-09 | 5.300E-09 |
| 11636 | 582 | V582S | A | 1.3e-08 | 1.2e-08 | 1.3e-08 | 1.267E-08 | 5.774E-10 | 1.200E-08 | 1.300E-08 |
| 11637 | 582 | V582T | A | 5.8e-09 | 2.9e-08 | 5.2e-09 | 1.333E-08 | 1.357E-08 | 5.200E-09 | 2.900E-08 |
| 11638 | 582 | V582W | A | 1e-08 | 6.9e-09 | 1.2e-08 | 9.633E-09 | 2.570E-09 | 6.900E-09 | 1.200E-08 |
| 11639 | 582 | V582Y | A | 1.9e-08 | 8.4e-09 | 3.7e-09 | 1.037E-08 | 7.837E-09 | 3.700E-09 | 1.900E-08 |
| 11640 | 582 | V582V | A | 6.9e-09 | 7.8e-09 | 3.8e-09 | 6.167E-09 | 2.098E-09 | 3.800E-09 | 7.800E-09 |
| 11641 | 583 | K583A | A | 5.1e-09 | 6.1e-09 | 3.3e-09 | 4.833E-09 | 1.419E-09 | 3.300E-09 | 6.100E-09 |
| 11642 | 583 | K583R | A | 3.7e-09 | 7.1e-09 | 1.2e-08 | 7.600E-09 | 4.173E-09 | 3.700E-09 | 1.200E-08 |
| 11643 | 583 | K583N | A | 1.4e-08 | 1.5e-08 | 4e-09 | 1.100E-08 | 6.083E-09 | 4.000E-09 | 1.500E-08 |
| 11644 | 583 | K583D | A | 7.7e-09 | 1e-08 | 1.4e-08 | 1.057E-08 | 3.188E-09 | 7.700E-09 | 1.400E-08 |

| | | | | | | | | | | |
|---|---|---|---|---|---|---|---|---|---|---|
| 11645 | 583 | K583C | A | 4.5e-09 | 1.7e-09 | 4.1e-09 | 3.433E-09 | 1.514E-09 | 1.700E-09 | 4.500E-09 |
| 11646 | 583 | K583E | A | 1.6e-08 | 1.3e-08 | 5.9e-09 | 1.163E-08 | 5.187E-09 | 5.900E-09 | 1.600E-08 |
| 11647 | 583 | K583Q | A | 1.9e-09 | 1.4e-08 | 9.4e-09 | 8.433E-09 | 6.108E-09 | 1.900E-09 | 1.400E-08 |
| 11648 | 583 | K583G | A | 2.6e-09 | 1.5e-08 | 7.4e-09 | 8.333E-09 | 6.252E-09 | 2.600E-09 | 1.500E-08 |
| 11649 | 583 | K583H | A | 1.5e-08 | 1.9e-09 | 2.1e-09 | 6.333E-09 | 7.506E-09 | 1.900E-09 | 1.500E-08 |
| 11650 | 583 | K583I | A | 3.3e-09 | 3e-09 | 2.5e-08 | 1.043E-08 | 1.262E-08 | 3.000E-09 | 2.500E-08 |
| 11651 | 583 | K583L | A | 5.9e-09 | 5.3e-09 | 7.6e-09 | 6.267E-09 | 1.193E-09 | 5.300E-09 | 7.600E-09 |
| 11652 | 583 | K583K | A | 4.6e-09 | 3e-09 | 9.6e-09 | 5.733E-09 | 3.443E-09 | 3.000E-09 | 9.600E-09 |
| 11653 | 583 | K583M | A | 7.9e-09 | 2.1e-09 | 1.8e-08 | 9.333E-09 | 8.046E-09 | 2.100E-09 | 1.800E-08 |
| 11654 | 583 | K583F | A | 6.2e-09 | 4.9e-09 | 7.6e-09 | 6.233E-09 | 1.350E-09 | 4.900E-09 | 7.600E-09 |
| 11655 | 583 | K583P | A | 1e-08 | 6.3e-09 | 9e-09 | 8.433E-09 | 1.914E-09 | 6.300E-09 | 1.000E-08 |
| 11656 | 583 | K583S | A | 3.8e-09 | 7.6e-09 | 4.6e-09 | 5.333E-09 | 2.003E-09 | 3.800E-09 | 7.600E-09 |
| 11657 | 583 | K583T | A | 3.8e-09 | 8.5e-09 | 7.8e-09 | 6.700E-09 | 2.536E-09 | 3.800E-09 | 8.500E-09 |
| 11658 | 583 | K583W | A | 2.6e-09 | 8.2e-09 | 5.1e-09 | 5.300E-09 | 2.805E-09 | 2.600E-09 | 8.200E-09 |
| 11659 | 583 | K583Y | A | 5.6e-09 | 1.2e-08 | 1.7e-08 | 1.153E-08 | 5.714E-09 | 5.600E-09 | 1.700E-08 |
| 11660 | 583 | K583V | A | 9.4e-09 | 7.7e-09 | 1.3e-08 | 1.003E-08 | 2.706E-09 | 7.700E-09 | 1.300E-08 |
| 11661 | 584 | P584A | A | 1.2e-08 | 1.4e-08 | 7.6e-09 | 1.120E-08 | 3.274E-09 | 7.600E-09 | 1.400E-08 |
| 11662 | 584 | P584R | A | 2.9e-09 | 1.1e-08 | 2e-08 | 1.130E-08 | 8.554E-09 | 2.900E-09 | 2.000E-08 |
| 11663 | 584 | P584N | A | 6e-09 | 4.1e-09 | 1.1e-08 | 7.033E-09 | 3.564E-09 | 4.100E-09 | 1.100E-08 |
| 11664 | 584 | P584D | A | 1e-08 | 3.7e-09 | 6.9e-09 | 6.867E-09 | 3.150E-09 | 3.700E-09 | 1.000E-08 |
| 11665 | 584 | P584C | A | 2.9e-09 | 7.5e-09 | 1.6e-08 | 8.800E-09 | 6.646E-09 | 2.900E-09 | 1.600E-08 |
| 11666 | 584 | P584E | A | 7.1e-09 | 1e-08 | 3.8e-09 | 6.967E-09 | 3.102E-09 | 3.800E-09 | 1.000E-08 |
| 11667 | 584 | P584Q | A | 1.1e-08 | 1.1e-08 | 1.4e-08 | 1.200E-08 | 1.732E-09 | 1.100E-08 | 1.400E-08 |
| 11668 | 584 | P584G | A | 3.5e-09 | 4.7e-09 | 7.2e-09 | 5.133E-09 | 1.888E-09 | 3.500E-09 | 7.200E-09 |
| 11669 | 584 | P584H | A | 3.3e-08 | 7.3e-09 | 4.6e-09 | 1.497E-08 | 1.568E-08 | 4.600E-09 | 3.300E-08 |
| 11670 | 584 | P584I | A | 9.9e-09 | 6.7e-09 | 8.5e-09 | 8.367E-09 | 1.604E-09 | 6.700E-09 | 9.900E-09 |
| 11671 | 584 | P584L | A | 1.3e-08 | 2e-09 | 6.6e-09 | 7.200E-09 | 5.524E-09 | 2.000E-09 | 1.300E-08 |
| 11672 | 584 | P584K | A | 5.5e-09 | 7.6e-09 | 1.5e-08 | 9.367E-09 | 4.990E-09 | 5.500E-09 | 1.500E-08 |
| 11673 | 584 | P584M | A | 6.6e-09 | 1.4e-08 | 7.8e-09 | 9.467E-09 | 3.972E-09 | 6.600E-09 | 1.400E-08 |
| 11674 | 584 | P584F | A | 8.3e-09 | 6.2e-09 | 7.3e-09 | 7.267E-09 | 1.050E-09 | 6.200E-09 | 8.300E-09 |
| 11675 | 584 | P584P | A | 7.3e-09 | 6e-09 | 2.8e-08 | 1.377E-08 | 1.234E-08 | 6.000E-09 | 2.800E-08 |
| 11676 | 584 | P584S | A | 6.1e-09 | 2.4e-09 | 1.1e-08 | 6.500E-09 | 4.314E-09 | 2.400E-09 | 1.100E-08 |
| 11677 | 584 | P584T | A | 3.5e-09 | 2.6e-09 | 2.8e-09 | 2.967E-09 | 4.726E-10 | 2.600E-09 | 3.500E-09 |
| 11678 | 584 | P584W | A | 5e-09 | 4.5e-09 | 4e-09 | 4.500E-09 | 5.000E-10 | 4.000E-09 | 5.000E-09 |
| 11679 | 584 | P584Y | A | 4e-09 | 1.2e-08 | 8e-09 | 8.000E-09 | 4.000E-09 | 4.000E-09 | 1.200E-08 |
| 11680 | 584 | P584V | A | 3.4e-09 | 9.8e-09 | 8.5e-09 | 7.233E-09 | 3.383E-09 | 3.400E-09 | 9.800E-09 |
| 11681 | 585 | D585A | A | 6.9e-09 | 4.2e-09 | 4.2e-09 | 5.100E-09 | 1.559E-09 | 4.200E-09 | 6.900E-09 |
| 11682 | 585 | D585R | A | 1.2e-08 | 9.9e-09 | 3e-09 | 8.300E-09 | 4.709E-09 | 3.000E-09 | 1.200E-08 |
| 11683 | 585 | D585N | A | 6.3e-09 | 4.3e-09 | 7e-09 | 5.867E-09 | 1.401E-09 | 4.300E-09 | 7.000E-09 |
| 11684 | 585 | D585D | A | 2.5e-09 | 7.8e-09 | 1.4e-08 | 8.100E-09 | 5.756E-09 | 2.500E-09 | 1.400E-08 |
| 11685 | 585 | D585C | A | 1.8e-08 | 1.2e-08 | 1e-08 | 1.333E-08 | 4.163E-09 | 1.000E-08 | 1.800E-08 |
| 11686 | 585 | D585E | A | 7.7e-09 | 4.9e-09 | 6.1e-09 | 6.233E-09 | 1.405E-09 | 4.900E-09 | 7.700E-09 |
| 11687 | 585 | D585Q | A | 2e-08 | 9.2e-09 | 2.8e-09 | 1.067E-08 | 8.693E-09 | 2.800E-09 | 2.000E-08 |
| 11688 | 585 | D585G | A | 1.2e-08 | 1.1e-08 | 5e-09 | 9.333E-09 | 3.786E-09 | 5.000E-09 | 1.200E-08 |
| 11689 | 585 | D585H | A | 3.9e-08 | 5.6e-09 | 1.2e-08 | 1.887E-08 | 1.773E-08 | 5.600E-09 | 3.900E-08 |
| 11690 | 585 | D585I | A | 7.9e-09 | 8.5e-09 | 6.5e-09 | 7.633E-09 | 1.026E-09 | 6.500E-09 | 8.500E-09 |
| 11691 | 585 | D585L | A | 5.3e-09 | 1.4e-09 | 5.4e-09 | 4.033E-09 | 2.281E-09 | 1.400E-09 | 5.400E-09 |
| 11692 | 585 | D585K | A | 1.1e-08 | 5.5e-09 | 1.4e-08 | 1.017E-08 | 4.311E-09 | 5.500E-09 | 1.400E-08 |
| 11693 | 585 | D585M | A | 9.2e-09 | 6.4e-09 | 7.5e-09 | 7.700E-09 | 1.411E-09 | 6.400E-09 | 9.200E-09 |
| 11694 | 585 | D585F | A | 4e-09 | 5.1e-09 | 9.1e-09 | 6.067E-09 | 2.684E-09 | 4.000E-09 | 9.100E-09 |
| 11695 | 585 | D585P | A | 3.9e-09 | 4.5e-09 | 1.3e-08 | 7.133E-09 | 5.090E-09 | 3.900E-09 | 1.300E-08 |
| 11696 | 585 | D585S | A | 3e-09 | 3.6e-09 | 4.6e-09 | 3.733E-09 | 8.083E-10 | 3.000E-09 | 4.600E-09 |
| 11697 | 585 | D585T | A | 1.5e-08 | 7.9e-09 | 8.8e-09 | 1.057E-08 | 3.866E-09 | 7.900E-09 | 1.500E-08 |
| 11698 | 585 | D585W | A | 6.7e-09 | 2.4e-09 | 3.5e-09 | 4.200E-09 | 2.234E-09 | 2.400E-09 | 6.700E-09 |
| 11699 | 585 | D585Y | A | 6.4e-09 | 1e-08 | 3.1e-09 | 6.500E-09 | 3.451E-09 | 3.100E-09 | 1.000E-08 |
| 11700 | 585 | D585V | A | 2.6e-08 | 6.1e-09 | 1.6e-08 | 1.603E-08 | 9.950E-09 | 6.100E-09 | 2.600E-08 |
| 11701 | 586 | L586A | A | 6.6e-09 | 1.6e-09 | 2.2e-09 | 3.467E-09 | 2.730E-09 | 1.600E-09 | 6.600E-09 |
| 11702 | 586 | L586R | A | 6.2e-09 | 9.5e-09 | 8.9e-09 | 8.200E-09 | 1.758E-09 | 6.200E-09 | 9.500E-09 |
| 11703 | 586 | L586N | A | 1.3e-08 | 1.5e-08 | 5.1e-09 | 1.103E-08 | 5.235E-09 | 5.100E-09 | 1.500E-08 |
| 11704 | 586 | L586D | A | 2.1e-08 | 8.5e-09 | 5.7e-09 | 1.173E-08 | 8.146E-09 | 5.700E-09 | 2.100E-08 |
| 11705 | 586 | L586C | A | 3.1e-09 | 6.9e-09 | 1e-08 | 6.667E-09 | 3.456E-09 | 3.100E-09 | 1.000E-08 |
| 11706 | 586 | L586E | A | 7.9e-09 | 9.3e-09 | 1.1e-08 | 9.400E-09 | 1.552E-09 | 7.900E-09 | 1.100E-08 |
| 11707 | 586 | L586Q | A | 9.4e-09 | 5.9e-09 | 1.4e-08 | 9.767E-09 | 4.062E-09 | 5.900E-09 | 1.400E-08 |
| 11708 | 586 | L586G | A | 7.3e-09 | 1.6e-08 | 7.1e-09 | 1.013E-08 | 5.082E-09 | 7.100E-09 | 1.600E-08 |
| 11709 | 586 | L586H | A | 1.4e-08 | 3.6e-09 | 6.2e-09 | 7.933E-09 | 5.412E-09 | 3.600E-09 | 1.400E-08 |
| 11710 | 586 | L586I | A | 7.2e-09 | 4.9e-09 | 6.2e-09 | 6.100E-09 | 1.153E-09 | 4.900E-09 | 7.200E-09 |
| 11711 | 586 | L586L | A | 6.1e-09 | 4.7e-10 | 4.9e-09 | 3.823E-09 | 2.965E-09 | 4.700E-10 | 6.100E-09 |
| 11712 | 586 | L586K | A | 3.9e-09 | 6.6e-09 | 8e-09 | 6.167E-09 | 2.084E-09 | 3.900E-09 | 8.000E-09 |
| 11713 | 586 | L586M | A | 2.4e-08 | 1.1e-08 | 5.5e-09 | 1.350E-08 | 9.500E-09 | 5.500E-09 | 2.400E-08 |
| 11714 | 586 | L586F | A | 3.6e-09 | 9.6e-09 | 1.1e-08 | 8.067E-09 | 3.931E-09 | 3.600E-09 | 1.100E-08 |
| 11715 | 586 | L586P | A | 1.8e-09 | 8.9e-09 | 6.5e-09 | 5.733E-09 | 3.612E-09 | 1.800E-09 | 8.900E-09 |
| 11716 | 586 | L586S | A | 1.7e-08 | 1.6e-08 | 6.8e-09 | 1.327E-08 | 5.623E-09 | 6.800E-09 | 1.700E-08 |
| 11717 | 586 | L586T | A | 1.4e-08 | 1e-08 | 2.6e-08 | 1.667E-08 | 8.327E-09 | 1.000E-08 | 2.600E-08 |
| 11718 | 586 | L586W | A | 8.5e-09 | 5.2e-09 | 3.1e-09 | 5.600E-09 | 2.722E-09 | 3.100E-09 | 8.500E-09 |
| 11719 | 586 | L586Y | A | 1.4e-08 | 4.4e-09 | 3.7e-09 | 7.367E-09 | 5.755E-09 | 3.700E-09 | 1.400E-08 |
| 11720 | 586 | L586V | A | 1.7e-09 | 7.7e-09 | 2.6e-09 | 4.000E-09 | 3.236E-09 | 1.700E-09 | 7.700E-09 |
| 11721 | 587 | S587A | A | 6.1e-09 | 4.1e-09 | 1.1e-08 | 7.067E-09 | 3.550E-09 | 4.100E-09 | 1.100E-08 |
| 11722 | 587 | S587R | A | 8.7e-09 | 1.3e-09 | 1e-08 | 6.667E-09 | 4.693E-09 | 1.300E-09 | 1.000E-08 |
| 11723 | 587 | S587N | A | 1.2e-08 | 1.5e-08 | 6.4e-09 | 1.113E-08 | 4.365E-09 | 6.400E-09 | 1.500E-08 |
| 11724 | 587 | S587D | A | 6.1e-09 | 1.3e-08 | 8.6e-09 | 9.233E-09 | 3.493E-09 | 6.100E-09 | 1.300E-08 |
| 11725 | 587 | S587C | A | 1.2e-09 | 2.9e-09 | 9e-09 | 4.367E-09 | 4.102E-09 | 1.200E-09 | 9.000E-09 |
| 11726 | 587 | S587E | A | 5.9e-09 | 1.2e-08 | 2.7e-08 | 1.497E-08 | 1.086E-08 | 5.900E-09 | 2.700E-08 |
| 11727 | 587 | S587Q | A | 8.8e-09 | 8.3e-09 | 3.8e-09 | 6.967E-09 | 2.754E-09 | 3.800E-09 | 8.800E-09 |
| 11728 | 587 | S587G | A | 6.2e-09 | 5.7e-09 | 2.6e-09 | 4.833E-09 | 1.950E-09 | 2.600E-09 | 6.200E-09 |
| 11729 | 587 | S587H | A | 2.3e-09 | 1.1e-08 | 2e-08 | 1.110E-08 | 8.850E-09 | 2.300E-09 | 2.000E-08 |

| | | | | | | | | | | |
|---|---|---|---|---|---|---|---|---|---|---|
| 11730 | 587 | S587I | A | 1.1e-08 | 5.1e-09 | 5.6e-09 | 7.233E-09 | 3.272E-09 | 5.100E-09 | 1.100E-08 |
| 11731 | 587 | S587L | A | 1.2e-08 | 1.5e-08 | 1.5e-08 | 1.400E-08 | 1.732E-09 | 1.200E-08 | 1.500E-08 |
| 11732 | 587 | S587K | A | 1.6e-08 | 3e-09 | 1.8e-08 | 1.233E-08 | 8.145E-09 | 3.000E-09 | 1.800E-08 |
| 11733 | 587 | S587M | A | 1.5e-08 | 6.1e-09 | 4e-09 | 8.367E-09 | 5.840E-09 | 4.000E-09 | 1.500E-08 |
| 11734 | 587 | S587F | A | 1.6e-08 | 7e-09 | 2.5e-08 | 1.600E-08 | 9.000E-09 | 7.000E-09 | 2.500E-08 |
| 11735 | 587 | S587P | A | 2.8e-09 | 6.6e-09 | 6.4e-09 | 5.267E-09 | 2.139E-09 | 2.800E-09 | 6.600E-09 |
| 11736 | 587 | S587S | A | 1.4e-08 | 5.5e-09 | 1.3e-08 | 1.083E-08 | 4.646E-09 | 5.500E-09 | 1.400E-08 |
| 11737 | 587 | S587T | A | 6.2e-09 | 1.4e-08 | 4.6e-09 | 8.267E-09 | 5.029E-09 | 4.600E-09 | 1.400E-08 |
| 11738 | 587 | S587W | A | 4.9e-09 | 1e-08 | 2.7e-08 | 1.397E-08 | 1.157E-08 | 4.900E-09 | 2.700E-08 |
| 11739 | 587 | S587Y | A | 9.9e-09 | 5.8e-09 | 7.3e-09 | 7.667E-09 | 2.074E-09 | 5.800E-09 | 9.900E-09 |
| 11740 | 587 | S587V | A | 6.9e-09 | 1.6e-09 | 7.9e-09 | 5.467E-09 | 3.386E-09 | 1.600E-09 | 7.900E-09 |
| 11741 | 588 | Y588A | A | 3e-09 | 6.7e-09 | 1.5e-08 | 8.233E-09 | 6.145E-09 | 3.000E-09 | 1.500E-08 |
| 11742 | 588 | Y588R | A | 2e-09 | 6.4e-09 | 1e-08 | 6.133E-09 | 4.007E-09 | 2.000E-09 | 1.000E-08 |
| 11743 | 588 | Y588N | A | 1.6e-08 | 2.2e-08 | 2.8e-09 | 1.360E-08 | 9.822E-09 | 2.800E-09 | 2.200E-08 |
| 11744 | 588 | Y588D | A | 1.6e-08 | 1.4e-08 | 2.2e-08 | 1.733E-08 | 4.163E-09 | 1.400E-08 | 2.200E-08 |
| 11745 | 588 | Y588C | A | 5.8e-09 | 1.1e-08 | 5.9e-09 | 7.567E-09 | 2.974E-09 | 5.800E-09 | 1.100E-08 |
| 11746 | 588 | Y588E | A | 9.8e-09 | 9.3e-09 | 5.2e-09 | 8.100E-09 | 2.524E-09 | 5.200E-09 | 9.800E-09 |
| 11747 | 588 | Y588Q | A | 6.4e-09 | 8.1e-09 | 7.7e-09 | 7.400E-09 | 8.888E-10 | 6.400E-09 | 8.100E-09 |
| 11748 | 588 | Y588G | A | 4.4e-09 | 4.5e-09 | 8.2e-09 | 5.700E-09 | 2.166E-09 | 4.400E-09 | 8.200E-09 |
| 11749 | 588 | Y588H | A | 1.1e-08 | 5e-09 | 1e-08 | 8.667E-09 | 3.215E-09 | 5.000E-09 | 1.100E-08 |
| 11750 | 588 | Y588I | A | 1.1e-08 | 6.8e-09 | 1.8e-09 | 6.533E-09 | 4.606E-09 | 1.800E-09 | 1.100E-08 |
| 11751 | 588 | Y588L | A | 7.6e-09 | 1.5e-08 | 4.7e-09 | 9.100E-09 | 5.311E-09 | 4.700E-09 | 1.500E-08 |
| 11752 | 588 | Y588K | A | 1.4e-08 | 1.1e-08 | 1e-08 | 1.167E-08 | 2.082E-09 | 1.000E-08 | 1.400E-08 |
| 11753 | 588 | Y588M | A | 6.9e-09 | 8.2e-09 | 3.4e-09 | 6.167E-09 | 2.483E-09 | 3.400E-09 | 8.200E-09 |
| 11754 | 588 | Y588F | A | 3.1e-08 | 8.1e-09 | 5.3e-08 | 3.070E-08 | 2.245E-08 | 8.100E-09 | 5.300E-08 |
| 11755 | 588 | Y588P | A | 6e-09 | 1.3e-08 | 1e-08 | 9.667E-09 | 3.512E-09 | 6.000E-09 | 1.300E-08 |
| 11756 | 588 | Y588S | A | 1.8e-08 | 4.4e-09 | 9.5e-09 | 1.063E-08 | 6.870E-09 | 4.400E-09 | 1.800E-08 |
| 11757 | 588 | Y588T | A | 2.8e-09 | 7e-09 | 9.4e-09 | 6.400E-09 | 3.341E-09 | 2.800E-09 | 9.400E-09 |
| 11758 | 588 | Y588W | A | 2.2e-09 | 6.6e-09 | 1.2e-08 | 6.933E-09 | 4.908E-09 | 2.200E-09 | 1.200E-08 |
| 11759 | 588 | Y588Y | A | 7.3e-09 | 7.7e-09 | 5e-09 | 6.667E-09 | 1.457E-09 | 5.000E-09 | 7.700E-09 |
| 11760 | 588 | Y588V | A | 7e-09 | 8.7e-09 | 2e-09 | 5.900E-09 | 3.483E-09 | 2.000E-09 | 8.700E-09 |
| 11761 | 589 | M589A | A | 1.4e-08 | 2.9e-09 | 2.8e-09 | 6.567E-09 | 6.438E-09 | 2.800E-09 | 1.400E-08 |
| 11762 | 589 | M589R | A | 6.2e-09 | 1.9e-09 | 8.7e-10 | 2.990E-09 | 2.827E-09 | 8.700E-10 | 6.200E-09 |
| 11763 | 589 | M589N | A | 3.1e-09 | 5.7e-09 | 7.2e-09 | 5.333E-09 | 2.074E-09 | 3.100E-09 | 7.200E-09 |
| 11764 | 589 | M589D | A | 2.2e-09 | 6.9e-09 | 2.7e-09 | 3.933E-09 | 2.581E-09 | 2.200E-09 | 6.900E-09 |
| 11765 | 589 | M589C | A | 1.2e-08 | 2.1e-08 | 1.1e-08 | 1.467E-08 | 5.508E-09 | 1.100E-08 | 2.100E-08 |
| 11766 | 589 | M589E | A | 3.6e-08 | 3e-09 | 3.4e-09 | 1.413E-08 | 1.894E-08 | 3.000E-09 | 3.600E-08 |
| 11767 | 589 | M589Q | A | 6.7e-09 | 6.9e-09 | 6.9e-09 | 6.833E-09 | 1.155E-10 | 6.700E-09 | 6.900E-09 |
| 11768 | 589 | M589G | A | 1.1e-08 | 1.1e-08 | 5.5e-09 | 9.167E-09 | 3.175E-09 | 5.500E-09 | 1.100E-08 |
| 11769 | 589 | M589H | A | 7.7e-10 | 8.2e-09 | 6.8e-09 | 5.257E-09 | 3.948E-09 | 7.700E-10 | 8.200E-09 |
| 11770 | 589 | M589I | A | 2e-08 | 2.9e-09 | 3.3e-09 | 8.733E-09 | 9.759E-09 | 2.900E-09 | 2.000E-08 |
| 11771 | 589 | M589L | A | 6e-09 | 1.3e-08 | 2e-09 | 7.000E-09 | 5.568E-09 | 2.000E-09 | 1.300E-08 |
| 11772 | 589 | M589K | A | 1.2e-08 | 6.1e-09 | 1.1e-08 | 9.700E-09 | 3.158E-09 | 6.100E-09 | 1.200E-08 |
| 11773 | 589 | M589M | A | 4.7e-09 | 5.8e-09 | 9.5e-09 | 6.667E-09 | 2.515E-09 | 4.700E-09 | 9.500E-09 |
| 11774 | 589 | M589F | A | 1.3e-08 | 5.4e-09 | 2e-09 | 6.800E-09 | 5.632E-09 | 2.000E-09 | 1.300E-08 |
| 11775 | 589 | M589P | A | 1.2e-08 | 5e-09 | 9.4e-09 | 8.800E-09 | 3.538E-09 | 5.000E-09 | 1.200E-08 |
| 11776 | 589 | M589S | A | 3.5e-09 | 3.5e-09 | 3.4e-09 | 3.467E-09 | 5.774E-11 | 3.400E-09 | 3.500E-09 |
| 11777 | 589 | M589T | A | 6.4e-09 | 1.2e-08 | 1.9e-08 | 1.247E-08 | 6.313E-09 | 6.400E-09 | 1.900E-08 |
| 11778 | 589 | M589W | A | 6.3e-09 | 2.1e-08 | 9.8e-09 | 1.237E-08 | 7.679E-09 | 6.300E-09 | 2.100E-08 |
| 11779 | 589 | M589Y | A | 9.8e-09 | 4.2e-09 | 4.9e-09 | 6.300E-09 | 3.051E-09 | 4.200E-09 | 9.800E-09 |
| 11780 | 589 | M589V | A | 1.2e-08 | 4e-09 | 2.2e-08 | 1.267E-08 | 9.018E-09 | 4.000E-09 | 2.200E-08 |
| 11781 | 590 | P590A | A | 2.1e-08 | 9.4e-10 | 9e-09 | 1.031E-08 | 1.009E-08 | 9.400E-10 | 2.100E-08 |
| 11782 | 590 | P590R | A | 1.2e-08 | 6.9e-09 | 1.4e-08 | 1.097E-08 | 3.661E-09 | 6.900E-09 | 1.400E-08 |
| 11783 | 590 | P590N | A | 1.3e-08 | 6.6e-09 | 1.3e-08 | 1.087E-08 | 3.695E-09 | 6.600E-09 | 1.300E-08 |
| 11784 | 590 | P590D | A | 1e-08 | 9.5e-09 | 2e-08 | 1.317E-08 | 5.923E-09 | 9.500E-09 | 2.000E-08 |
| 11785 | 590 | P590C | A | 5.3e-09 | 1e-08 | 4.5e-09 | 6.600E-09 | 2.972E-09 | 4.500E-09 | 1.000E-08 |
| 11786 | 590 | P590E | A | 2.5e-09 | 7.1e-10 | 9.4e-09 | 4.203E-09 | 4.589E-09 | 7.100E-10 | 9.400E-09 |
| 11787 | 590 | P590Q | A | 7.7e-09 | 5.8e-09 | 7.9e-09 | 7.133E-09 | 1.159E-09 | 5.800E-09 | 7.900E-09 |
| 11788 | 590 | P590G | A | 2.9e-09 | 5.2e-09 | 2.4e-09 | 3.500E-09 | 1.493E-09 | 2.400E-09 | 5.200E-09 |
| 11789 | 590 | P590H | A | 4.3e-09 | 2.1e-08 | 6.3e-09 | 1.053E-08 | 9.119E-09 | 4.300E-09 | 2.100E-08 |
| 11790 | 590 | P590I | A | 4.9e-09 | 1.3e-08 | 3.7e-09 | 7.200E-09 | 5.059E-09 | 3.700E-09 | 1.300E-08 |
| 11791 | 590 | P590L | A | 2.3e-08 | 5.5e-09 | 3.6e-09 | 1.070E-08 | 1.069E-08 | 3.600E-09 | 2.300E-08 |
| 11792 | 590 | P590K | A | 8.7e-09 | 7.6e-09 | 1.2e-08 | 9.433E-09 | 2.290E-09 | 7.600E-09 | 1.200E-08 |
| 11793 | 590 | P590M | A | 3.7e-09 | 4.3e-09 | 1.1e-08 | 6.333E-09 | 4.053E-09 | 3.700E-09 | 1.100E-08 |
| 11794 | 590 | P590F | A | 2.9e-09 | 4e-09 | 4.4e-09 | 3.767E-09 | 7.767E-10 | 2.900E-09 | 4.400E-09 |
| 11795 | 590 | P590P | A | 5e-09 | 4.1e-09 | 1.5e-08 | 8.033E-09 | 6.050E-09 | 4.100E-09 | 1.500E-08 |
| 11796 | 590 | P590S | A | 1.7e-08 | 1.3e-08 | 3.8e-09 | 1.127E-08 | 6.769E-09 | 3.800E-09 | 1.700E-08 |
| 11797 | 590 | P590T | A | 8.5e-09 | 3.4e-09 | 2.3e-08 | 1.163E-08 | 1.017E-08 | 3.400E-09 | 2.300E-08 |
| 11798 | 590 | P590W | A | 9.6e-09 | 4.1e-09 | 3.8e-09 | 5.833E-09 | 3.265E-09 | 3.800E-09 | 9.600E-09 |
| 11799 | 590 | P590Y | A | 7.1e-09 | 1.3e-08 | 4.9e-09 | 8.333E-09 | 4.188E-09 | 4.900E-09 | 1.300E-08 |
| 11800 | 590 | P590V | A | 7.2e-09 | 3.8e-09 | 1.3e-08 | 8.000E-09 | 4.652E-09 | 3.800E-09 | 1.300E-08 |
| 11801 | 591 | I591A | A | 6.6e-09 | 2.8e-09 | 2.8e-08 | 1.247E-08 | 1.359E-08 | 2.800E-09 | 2.800E-08 |
| 11802 | 591 | I591R | A | 1.1e-08 | 3.9e-09 | 9.8e-09 | 8.233E-09 | 3.800E-09 | 3.900E-09 | 1.100E-08 |
| 11803 | 591 | I591N | A | 7.2e-09 | 4.9e-09 | 1.2e-08 | 8.033E-09 | 3.623E-09 | 4.900E-09 | 1.200E-08 |
| 11804 | 591 | I591D | A | 5.3e-09 | 7.5e-09 | 8.4e-09 | 7.067E-09 | 1.595E-09 | 5.300E-09 | 8.400E-09 |
| 11805 | 591 | I591C | A | 5.2e-09 | 5.9e-09 | 3.6e-09 | 4.900E-09 | 1.179E-09 | 3.600E-09 | 5.900E-09 |
| 11806 | 591 | I591E | A | 4.4e-09 | 7.6e-09 | 9.4e-09 | 7.133E-09 | 2.532E-09 | 4.400E-09 | 9.400E-09 |
| 11807 | 591 | I591Q | A | 2.3e-09 | 7.4e-09 | 9.4e-09 | 6.367E-09 | 3.661E-09 | 2.300E-09 | 9.400E-09 |
| 11808 | 591 | I591G | A | 7.5e-09 | 1.2e-08 | 8e-09 | 9.167E-09 | 2.466E-09 | 7.500E-09 | 1.200E-08 |
| 11809 | 591 | I591H | A | 3.5e-09 | 1.5e-08 | 2.7e-08 | 1.517E-08 | 1.175E-08 | 3.500E-09 | 2.700E-08 |
| 11810 | 591 | I591I | A | 8.1e-09 | 7.2e-09 | 8.6e-09 | 7.967E-09 | 7.095E-10 | 7.200E-09 | 8.600E-09 |
| 11811 | 591 | I591L | A | 2.9e-08 | 6e-09 | 5e-09 | 1.333E-08 | 1.358E-08 | 5.000E-09 | 2.900E-08 |
| 11812 | 591 | I591K | A | 8.9e-09 | 1.1e-09 | 4.1e-09 | 4.700E-09 | 3.934E-09 | 1.100E-09 | 8.900E-09 |
| 11813 | 591 | I591M | A | 8.9e-09 | 2.5e-08 | 7.2e-09 | 1.370E-08 | 9.823E-09 | 7.200E-09 | 2.500E-08 |
| 11814 | 591 | I591F | A | 1.1e-08 | 4.2e-09 | 4.6e-09 | 6.600E-09 | 3.816E-09 | 4.200E-09 | 1.100E-08 |

| | | | | | | | | | | |
|---|---|---|---|---|---|---|---|---|---|---|
| 11815 | 591 | I591P | A | 1.1e-08 | 4.3e-08 | 1.6e-08 | 2.333E-08 | 1.721E-08 | 1.100E-08 | 4.300E-08 |
| 11816 | 591 | I591S | A | 8.3e-09 | 8.2e-09 | 8.1e-09 | 8.200E-09 | 1.000E-10 | 8.100E-09 | 8.300E-09 |
| 11817 | 591 | I591T | A | 1.4e-08 | 2.7e-09 | 6.7e-09 | 7.800E-09 | 5.730E-09 | 2.700E-09 | 1.400E-08 |
| 11818 | 591 | I591W | A | 5.5e-09 | 8.5e-09 | 4.7e-09 | 6.233E-09 | 2.003E-09 | 4.700E-09 | 8.500E-09 |
| 11819 | 591 | I591Y | A | 5.7e-09 | 5.2e-09 | 7.4e-09 | 6.100E-09 | 1.153E-09 | 5.200E-09 | 7.400E-09 |
| 11820 | 591 | I591V | A | 2e-08 | 4.4e-09 | 9.1e-09 | 1.117E-08 | 8.003E-09 | 4.400E-09 | 2.000E-08 |
| 11821 | 592 | W592A | A | 1.8e-09 | 7.6e-09 | 2.1e-09 | 3.833E-09 | 3.265E-09 | 1.800E-09 | 7.600E-09 |
| 11822 | 592 | W592R | A | 1.7e-08 | 9.7e-09 | 1e-08 | 1.223E-08 | 4.131E-09 | 9.700E-09 | 1.700E-08 |
| 11823 | 592 | W592N | A | 1.6e-09 | 8.9e-09 | 8.4e-09 | 6.300E-09 | 4.078E-09 | 1.600E-09 | 8.900E-09 |
| 11824 | 592 | W592D | A | 1.3e-08 | 1.4e-08 | 4.1e-09 | 1.037E-08 | 5.450E-09 | 4.100E-09 | 1.400E-08 |
| 11825 | 592 | W592C | A | 7.3e-09 | 3.9e-09 | 1.6e-09 | 4.267E-09 | 2.868E-09 | 1.600E-09 | 7.300E-09 |
| 11826 | 592 | W592E | A | 5.3e-09 | 8.2e-09 | 4.9e-09 | 6.133E-09 | 1.801E-09 | 4.900E-09 | 8.200E-09 |
| 11827 | 592 | W592Q | A | 3.4e-09 | 6e-09 | 1.1e-08 | 6.800E-09 | 3.863E-09 | 3.400E-09 | 1.100E-08 |
| 11828 | 592 | W592G | A | 2.9e-09 | 1.4e-08 | 7.9e-09 | 8.267E-09 | 5.559E-09 | 2.900E-09 | 1.400E-08 |
| 11829 | 592 | W592H | A | 2.4e-09 | 3.7e-09 | 7.7e-09 | 4.600E-09 | 2.762E-09 | 2.400E-09 | 7.700E-09 |
| 11830 | 592 | W592I | A | 7.8e-09 | 2.1e-09 | 5.1e-09 | 5.000E-09 | 2.851E-09 | 2.100E-09 | 7.800E-09 |
| 11831 | 592 | W592L | A | 1.3e-08 | 5.4e-09 | 3.5e-09 | 7.300E-09 | 5.027E-09 | 3.500E-09 | 1.300E-08 |
| 11832 | 592 | W592K | A | 8.4e-09 | 1.4e-08 | 4.8e-09 | 9.067E-09 | 4.636E-09 | 4.800E-09 | 1.400E-08 |
| 11833 | 592 | W592M | A | 2.2e-08 | 9.6e-09 | 5.3e-09 | 1.230E-08 | 8.671E-09 | 5.300E-09 | 2.200E-08 |
| 11834 | 592 | W592F | A | 3.4e-09 | 5.8e-09 | 1.7e-08 | 8.733E-09 | 7.259E-09 | 3.400E-09 | 1.700E-08 |
| 11835 | 592 | W592P | A | 3.6e-09 | 1.1e-08 | 8.5e-09 | 7.700E-09 | 3.764E-09 | 3.600E-09 | 1.100E-08 |
| 11836 | 592 | W592S | A | 8.1e-09 | 9.1e-09 | 2.9e-09 | 6.700E-09 | 3.329E-09 | 2.900E-09 | 9.100E-09 |
| 11837 | 592 | W592T | A | 2.9e-09 | 7.9e-09 | 5.5e-09 | 5.433E-09 | 2.501E-09 | 2.900E-09 | 7.900E-09 |
| 11838 | 592 | W592W | A | 1.5e-08 | 3.1e-09 | 1.1e-08 | 9.700E-09 | 6.056E-09 | 3.100E-09 | 1.500E-08 |
| 11839 | 592 | W592Y | A | 2.8e-09 | 5.5e-09 | 1.2e-08 | 6.767E-09 | 4.729E-09 | 2.800E-09 | 1.200E-08 |
| 11840 | 592 | W592V | A | 8.7e-09 | 1.1e-08 | 1.7e-08 | 1.223E-08 | 4.285E-09 | 8.700E-09 | 1.700E-08 |
| 11841 | 593 | K593A | A | 9e-09 | 9.4e-09 | 2.9e-09 | 7.100E-09 | 3.643E-09 | 2.900E-09 | 9.400E-09 |
| 11842 | 593 | K593R | A | 5.3e-09 | 4e-09 | 2e-08 | 9.767E-09 | 8.886E-09 | 4.000E-09 | 2.000E-08 |
| 11843 | 593 | K593N | A | 3.8e-09 | 1.1e-08 | 2.9e-09 | 5.900E-09 | 4.440E-09 | 2.900E-09 | 1.100E-08 |
| 11844 | 593 | K593D | A | 3.7e-09 | 6.3e-09 | 7.6e-09 | 5.867E-09 | 1.986E-09 | 3.700E-09 | 7.600E-09 |
| 11845 | 593 | K593C | A | 7.2e-09 | 4.4e-09 | 6.6e-09 | 6.067E-09 | 1.474E-09 | 4.400E-09 | 7.200E-09 |
| 11846 | 593 | K593E | A | 8.9e-09 | 4.4e-09 | 7.8e-09 | 7.033E-09 | 2.346E-09 | 4.400E-09 | 8.900E-09 |
| 11847 | 593 | K593Q | A | 1.3e-08 | 6.2e-09 | 4.8e-09 | 8.000E-09 | 4.386E-09 | 4.800E-09 | 1.300E-08 |
| 11848 | 593 | K593G | A | 6.8e-09 | 4.6e-09 | 5.5e-09 | 5.633E-09 | 1.106E-09 | 4.600E-09 | 6.800E-09 |
| 11849 | 593 | K593H | A | 5.6e-09 | 3.2e-09 | 6.1e-09 | 4.967E-09 | 1.550E-09 | 3.200E-09 | 6.100E-09 |
| 11850 | 593 | K593I | A | 3.4e-09 | 4.4e-09 | 4.5e-09 | 4.100E-09 | 6.083E-10 | 3.400E-09 | 4.500E-09 |
| 11851 | 593 | K593L | A | 7.9e-09 | 6.5e-09 | 1.4e-08 | 9.467E-09 | 3.988E-09 | 6.500E-09 | 1.400E-08 |
| 11852 | 593 | K593K | A | 6.8e-09 | 6.5e-09 | 2.5e-09 | 5.267E-09 | 2.401E-09 | 2.500E-09 | 6.800E-09 |
| 11853 | 593 | K593M | A | 6.9e-09 | 3e-09 | 1.2e-08 | 7.300E-09 | 4.513E-09 | 3.000E-09 | 1.200E-08 |
| 11854 | 593 | K593F | A | 5.5e-09 | 1e-08 | 7.8e-09 | 7.767E-09 | 2.250E-09 | 5.500E-09 | 1.000E-08 |
| 11855 | 593 | K593P | A | 1.2e-08 | 1e-08 | 3e-09 | 8.333E-09 | 4.726E-09 | 3.000E-09 | 1.200E-08 |
| 11856 | 593 | K593S | A | 1.7e-08 | 4.9e-09 | 2.3e-08 | 1.497E-08 | 9.220E-09 | 4.900E-09 | 2.300E-08 |
| 11857 | 593 | K593T | A | 1.1e-08 | 5.7e-09 | 2.2e-08 | 1.290E-08 | 8.314E-09 | 5.700E-09 | 2.200E-08 |
| 11858 | 593 | K593W | A | 1.1e-08 | 1.2e-08 | 5.6e-09 | 9.533E-09 | 3.443E-09 | 5.600E-09 | 1.200E-08 |
| 11859 | 593 | K593Y | A | 1e-08 | 1.7e-09 | 2.7e-09 | 4.800E-09 | 4.531E-09 | 1.700E-09 | 1.000E-08 |
| 11860 | 593 | K593V | A | 7.5e-09 | 1.3e-08 | 6.6e-09 | 9.033E-09 | 3.465E-09 | 6.600E-09 | 1.300E-08 |
| 11861 | 594 | F594A | A | 8.3e-09 | 6.3e-09 | 3.8e-09 | 6.133E-09 | 2.255E-09 | 3.800E-09 | 8.300E-09 |
| 11862 | 594 | F594R | A | 7.6e-09 | 1.1e-08 | 2e-08 | 1.287E-08 | 6.407E-09 | 7.600E-09 | 2.000E-08 |
| 11863 | 594 | F594N | A | 4.5e-09 | 9.8e-09 | 1e-08 | 8.100E-09 | 3.119E-09 | 4.500E-09 | 1.000E-08 |
| 11864 | 594 | F594D | A | 1.2e-08 | 1.1e-08 | 7.4e-09 | 1.013E-08 | 2.419E-09 | 7.400E-09 | 1.200E-08 |
| 11865 | 594 | F594C | A | 7.7e-09 | 8.7e-09 | 7.9e-09 | 8.100E-09 | 5.292E-10 | 7.700E-09 | 8.700E-09 |
| 11866 | 594 | F594E | A | 3.2e-09 | 1.9e-08 | 1.2e-08 | 1.140E-08 | 7.917E-09 | 3.200E-09 | 1.900E-08 |
| 11867 | 594 | F594Q | A | 1.1e-08 | 4.8e-09 | 2.6e-09 | 6.133E-09 | 4.356E-09 | 2.600E-09 | 1.100E-08 |
| 11868 | 594 | F594G | A | 9.7e-09 | 9.2e-09 | 5.4e-09 | 8.100E-09 | 2.352E-09 | 5.400E-09 | 9.700E-09 |
| 11869 | 594 | F594H | A | 4.1e-09 | 1.3e-08 | 4.7e-09 | 7.267E-09 | 4.974E-09 | 4.100E-09 | 1.300E-08 |
| 11870 | 594 | F594I | A | 6.1e-09 | 5.1e-09 | 7.5e-09 | 6.233E-09 | 1.206E-09 | 5.100E-09 | 7.500E-09 |
| 11871 | 594 | F594L | A | 8.8e-09 | 7.7e-09 | 3e-09 | 6.500E-09 | 3.081E-09 | 3.000E-09 | 8.800E-09 |
| 11872 | 594 | F594K | A | 3.5e-09 | 6.9e-09 | 9.1e-09 | 6.500E-09 | 2.821E-09 | 3.500E-09 | 9.100E-09 |
| 11873 | 594 | F594M | A | 1.1e-08 | 2.6e-08 | 1.7e-08 | 1.800E-08 | 7.550E-09 | 1.100E-08 | 2.600E-08 |
| 11874 | 594 | F594F | A | 3.7e-09 | 1.8e-09 | 6.3e-09 | 3.933E-09 | 2.259E-09 | 1.800E-09 | 6.300E-09 |
| 11875 | 594 | F594P | A | 1.9e-08 | 3.7e-09 | 2.1e-08 | 1.457E-08 | 9.464E-09 | 3.700E-09 | 2.100E-08 |
| 11876 | 594 | F594S | A | 5.5e-09 | 5.4e-09 | 3.4e-09 | 4.767E-09 | 1.185E-09 | 3.400E-09 | 5.500E-09 |
| 11877 | 594 | F594T | A | 1.5e-08 | 7.2e-09 | 4.9e-09 | 9.033E-09 | 5.294E-09 | 4.900E-09 | 1.500E-08 |
| 11878 | 594 | F594W | A | 2.2e-08 | 1.3e-08 | 1.3e-08 | 1.600E-08 | 5.196E-09 | 1.300E-08 | 2.200E-08 |
| 11879 | 594 | F594Y | A | 1.7e-08 | 5.7e-09 | 3.6e-09 | 8.767E-09 | 7.207E-09 | 3.600E-09 | 1.700E-08 |
| 11880 | 594 | F594V | A | 1.1e-08 | 3.8e-09 | 1.4e-08 | 9.600E-09 | 5.242E-09 | 3.800E-09 | 1.400E-08 |
| 11881 | 595 | P595A | A | 3.2e-09 | 6.8e-09 | 2.6e-09 | 4.200E-09 | 2.272E-09 | 2.600E-09 | 6.800E-09 |
| 11882 | 595 | P595R | A | 6.7e-09 | 4.5e-09 | 4.4e-09 | 5.200E-09 | 1.300E-09 | 4.400E-09 | 6.700E-09 |
| 11883 | 595 | P595N | A | 1.3e-08 | 5.2e-09 | 3.1e-09 | 7.100E-09 | 5.216E-09 | 3.100E-09 | 1.300E-08 |
| 11884 | 595 | P595D | A | 1.1e-08 | 2.1e-08 | 7.6e-09 | 1.320E-08 | 6.966E-09 | 7.600E-09 | 2.100E-08 |
| 11885 | 595 | P595C | A | 6.7e-09 | 4.9e-09 | 4.4e-09 | 5.333E-09 | 1.210E-09 | 4.400E-09 | 6.700E-09 |
| 11886 | 595 | P595E | A | 2.3e-08 | 5.7e-09 | 7e-09 | 1.190E-08 | 9.635E-09 | 5.700E-09 | 2.300E-08 |
| 11887 | 595 | P595Q | A | 2.4e-09 | 1.8e-08 | 6.8e-09 | 9.067E-09 | 8.043E-09 | 2.400E-09 | 1.800E-08 |
| 11888 | 595 | P595G | A | 2.4e-09 | 7.1e-09 | 2.9e-09 | 4.133E-09 | 2.581E-09 | 2.400E-09 | 7.100E-09 |
| 11889 | 595 | P595H | A | 4.7e-09 | 1.3e-08 | 6.3e-09 | 8.000E-09 | 4.403E-09 | 4.700E-09 | 1.300E-08 |
| 11890 | 595 | P595I | A | 3.8e-09 | 2.1e-09 | 7e-09 | 4.300E-09 | 2.488E-09 | 2.100E-09 | 7.000E-09 |
| 11891 | 595 | P595L | A | 8.8e-09 | 8e-09 | 6e-09 | 7.600E-09 | 1.442E-09 | 6.000E-09 | 8.800E-09 |
| 11892 | 595 | P595K | A | 2e-09 | 2.6e-09 | 5.7e-09 | 3.433E-09 | 1.986E-09 | 2.000E-09 | 5.700E-09 |
| 11893 | 595 | P595M | A | 9.5e-09 | 3.2e-09 | 2.8e-09 | 5.167E-09 | 3.758E-09 | 2.800E-09 | 9.500E-09 |
| 11894 | 595 | P595F | A | 1e-08 | 1.5e-08 | 4.5e-09 | 9.833E-09 | 5.252E-09 | 4.500E-09 | 1.500E-08 |
| 11895 | 595 | P595P | A | 5.8e-09 | 4.9e-09 | 5.6e-09 | 5.433E-09 | 4.726E-10 | 4.900E-09 | 5.800E-09 |
| 11896 | 595 | P595S | A | 5.5e-09 | 6.4e-09 | 4.7e-09 | 5.533E-09 | 8.505E-10 | 4.700E-09 | 6.400E-09 |
| 11897 | 595 | P595T | A | 2.6e-09 | 9.9e-09 | 6.9e-09 | 6.467E-09 | 3.669E-09 | 2.600E-09 | 9.900E-09 |
| 11898 | 595 | P595W | A | 8.9e-09 | 2.9e-08 | 1.3e-08 | 1.697E-08 | 1.062E-08 | 8.900E-09 | 2.900E-08 |
| 11899 | 595 | P595Y | A | 1.5e-08 | 1.6e-08 | 4.3e-09 | 1.177E-08 | 6.486E-09 | 4.300E-09 | 1.600E-08 |

| | | | | | | | | | | |
|---|---|---|---|---|---|---|---|---|---|---|
| 11900 | 595 | P595V | A | 4.5e-09 | 5.2e-09 | 4e-09 | 4.567E-09 | 6.028E-10 | 4.000E-09 | 5.200E-09 |
| 11901 | 596 | D596A | A | 2e-08 | 1.1e-08 | 3.1e-09 | 1.137E-08 | 8.456E-09 | 3.100E-09 | 2.000E-08 |
| 11902 | 596 | D596R | A | 1.8e-08 | 4.9e-09 | 1.2e-08 | 1.163E-08 | 6.558E-09 | 4.900E-09 | 1.800E-08 |
| 11903 | 596 | D596N | A | 9.5e-10 | 7.1e-09 | 1.1e-08 | 6.350E-09 | 5.067E-09 | 9.500E-10 | 1.100E-08 |
| 11904 | 596 | D596D | A | 4.8e-09 | 5.2e-09 | 5.8e-09 | 5.267E-09 | 5.033E-10 | 4.800E-09 | 5.800E-09 |
| 11905 | 596 | D596C | A | 1.6e-08 | 8.5e-09 | 1e-08 | 1.150E-08 | 3.969E-09 | 8.500E-09 | 1.600E-08 |
| 11906 | 596 | D596E | A | 6.6e-09 | 4.8e-09 | 7.7e-09 | 6.367E-09 | 1.464E-09 | 4.800E-09 | 7.700E-09 |
| 11907 | 596 | D596Q | A | 9.5e-09 | 8.9e-09 | 1.2e-08 | 1.013E-08 | 1.644E-09 | 8.900E-09 | 1.200E-08 |
| 11908 | 596 | D596G | A | 2.8e-09 | 3.4e-09 | 8.5e-09 | 4.900E-09 | 3.132E-09 | 2.800E-09 | 8.500E-09 |
| 11909 | 596 | D596H | A | 1e-08 | 5.1e-09 | 5.1e-09 | 6.733E-09 | 2.829E-09 | 5.100E-09 | 1.000E-08 |
| 11910 | 596 | D596I | A | 5.6e-09 | 4.2e-09 | 1.2e-09 | 3.667E-09 | 2.248E-09 | 1.200E-09 | 5.600E-09 |
| 11911 | 596 | D596L | A | 1.5e-08 | 8.7e-09 | 2.4e-09 | 8.700E-09 | 6.300E-09 | 2.400E-09 | 1.500E-08 |
| 11912 | 596 | D596K | A | 9.9e-09 | 6e-09 | 7.9e-09 | 7.933E-09 | 1.950E-09 | 6.000E-09 | 9.900E-09 |
| 11913 | 596 | D596M | A | 7.6e-09 | 8.3e-09 | 7.4e-09 | 7.767E-09 | 4.726E-10 | 7.400E-09 | 8.300E-09 |
| 11914 | 596 | D596F | A | 2.9e-09 | 5.4e-09 | 7.1e-09 | 5.133E-09 | 2.113E-09 | 2.900E-09 | 7.100E-09 |
| 11915 | 596 | D596P | A | 5e-09 | 6.1e-09 | 1e-08 | 7.033E-09 | 2.627E-09 | 5.000E-09 | 1.000E-08 |
| 11916 | 596 | D596S | A | 7.5e-09 | 4.3e-09 | 1.7e-09 | 4.500E-09 | 2.905E-09 | 1.700E-09 | 7.500E-09 |
| 11917 | 596 | D596T | A | 6.8e-09 | 1.1e-08 | 2.3e-09 | 6.700E-09 | 4.351E-09 | 2.300E-09 | 1.100E-08 |
| 11918 | 596 | D596W | A | 5.2e-09 | 2e-09 | 1e-08 | 5.733E-09 | 4.027E-09 | 2.000E-09 | 1.000E-08 |
| 11919 | 596 | D596Y | A | 6.4e-09 | 4.4e-09 | 3.8e-08 | 1.627E-08 | 1.885E-08 | 4.400E-09 | 3.800E-08 |
| 11920 | 596 | D596V | A | 9.3e-09 | 4.7e-09 | 1.4e-08 | 9.333E-09 | 4.650E-09 | 4.700E-09 | 1.400E-08 |
| 11921 | 597 | E597A | A | 6.4e-09 | 9.6e-09 | 3.1e-09 | 6.367E-09 | 3.250E-09 | 3.100E-09 | 9.600E-09 |
| 11922 | 597 | E597R | A | 1.8e-08 | 1.3e-08 | 4.2e-09 | 1.173E-08 | 6.987E-09 | 4.200E-09 | 1.800E-08 |
| 11923 | 597 | E597N | A | 3.3e-09 | 4.1e-09 | 6.6e-09 | 4.667E-09 | 1.721E-09 | 3.300E-09 | 6.600E-09 |
| 11924 | 597 | E597D | A | 5.2e-09 | 7e-09 | 3e-09 | 5.067E-09 | 2.003E-09 | 3.000E-09 | 7.000E-09 |
| 11925 | 597 | E597C | A | 2e-09 | 4.3e-09 | 1.1e-08 | 5.767E-09 | 4.676E-09 | 2.000E-09 | 1.100E-08 |
| 11926 | 597 | E597E | A | 9.3e-09 | 4.9e-09 | 6.3e-09 | 6.833E-09 | 2.248E-09 | 4.900E-09 | 9.300E-09 |
| 11927 | 597 | E597Q | A | 9.9e-09 | 8.2e-09 | 8.1e-09 | 8.733E-09 | 1.012E-09 | 8.100E-09 | 9.900E-09 |
| 11928 | 597 | E597G | A | 6.8e-09 | 4.6e-09 | 1.8e-08 | 9.800E-09 | 7.186E-09 | 4.600E-09 | 1.800E-08 |
| 11929 | 597 | E597H | A | 5.4e-09 | 1.1e-08 | 7.7e-09 | 8.033E-09 | 2.815E-09 | 5.400E-09 | 1.100E-08 |
| 11930 | 597 | E597I | A | 2.3e-08 | 2.7e-09 | 3.7e-09 | 9.800E-09 | 1.144E-08 | 2.700E-09 | 2.300E-08 |
| 11931 | 597 | E597L | A | 4.2e-09 | 6.7e-09 | 3.8e-09 | 4.900E-09 | 1.572E-09 | 3.800E-09 | 6.700E-09 |
| 11932 | 597 | E597K | A | 7.2e-09 | 3.7e-09 | 5.8e-09 | 5.567E-09 | 1.762E-09 | 3.700E-09 | 7.200E-09 |
| 11933 | 597 | E597M | A | 1.9e-09 | 8.3e-09 | 3.1e-09 | 4.433E-09 | 3.402E-09 | 1.900E-09 | 8.300E-09 |
| 11934 | 597 | E597F | A | 9.9e-09 | 8.2e-09 | 3.1e-09 | 7.067E-09 | 3.539E-09 | 3.100E-09 | 9.900E-09 |
| 11935 | 597 | E597P | A | 7.5e-09 | 8.4e-09 | 6.3e-09 | 7.400E-09 | 1.054E-09 | 6.300E-09 | 8.400E-09 |
| 11936 | 597 | E597S | A | 8.4e-09 | 1.2e-08 | 1e-08 | 1.013E-08 | 1.804E-09 | 8.400E-09 | 1.200E-08 |
| 11937 | 597 | E597T | A | 9.1e-09 | 8.1e-09 | 2.6e-09 | 6.600E-09 | 3.500E-09 | 2.600E-09 | 9.100E-09 |
| 11938 | 597 | E597W | A | 7.8e-09 | 2.5e-08 | 7.2e-09 | 1.333E-08 | 1.011E-08 | 7.200E-09 | 2.500E-08 |
| 11939 | 597 | E597Y | A | 6.4e-09 | 1.7e-09 | 3.9e-09 | 4.000E-09 | 2.352E-09 | 1.700E-09 | 6.400E-09 |
| 11940 | 597 | E597V | A | 1.7e-08 | 2.5e-08 | 7.1e-09 | 1.637E-08 | 8.967E-09 | 7.100E-09 | 2.500E-08 |
| 11941 | 598 | E598A | A | 4.4e-09 | 9.5e-09 | 4.5e-09 | 6.133E-09 | 2.916E-09 | 4.400E-09 | 9.500E-09 |
| 11942 | 598 | E598R | A | 2.9e-08 | 6.4e-09 | 2e-08 | 1.847E-08 | 1.138E-08 | 6.400E-09 | 2.900E-08 |
| 11943 | 598 | E598N | A | 1e-08 | 9.2e-09 | 2.8e-09 | 7.333E-09 | 3.946E-09 | 2.800E-09 | 1.000E-08 |
| 11944 | 598 | E598D | A | 2.2e-09 | 4.3e-09 | 5.7e-09 | 4.067E-09 | 1.762E-09 | 2.200E-09 | 5.700E-09 |
| 11945 | 598 | E598C | A | 6.5e-09 | 2.8e-09 | 8.9e-09 | 6.067E-09 | 3.073E-09 | 2.800E-09 | 8.900E-09 |
| 11946 | 598 | E598E | A | 1.5e-08 | 8.9e-09 | 1.5e-09 | 8.467E-09 | 6.760E-09 | 1.500E-09 | 1.500E-08 |
| 11947 | 598 | E598Q | A | 1.4e-08 | 8e-09 | 1.7e-08 | 1.300E-08 | 4.583E-09 | 8.000E-09 | 1.700E-08 |
| 11948 | 598 | E598G | A | 5.6e-09 | 3e-09 | 1.1e-08 | 6.533E-09 | 4.081E-09 | 3.000E-09 | 1.100E-08 |
| 11949 | 598 | E598H | A | 9.9e-09 | 1.9e-08 | 8.6e-09 | 1.250E-08 | 5.667E-09 | 8.600E-09 | 1.900E-08 |
| 11950 | 598 | E598I | A | 5.4e-09 | 6.2e-09 | 5.4e-09 | 5.667E-09 | 4.619E-10 | 5.400E-09 | 6.200E-09 |
| 11951 | 598 | E598L | A | 8.2e-09 | 5e-09 | 4.2e-09 | 5.800E-09 | 2.117E-09 | 4.200E-09 | 8.200E-09 |
| 11952 | 598 | E598K | A | 1e-08 | 4.4e-09 | 9e-09 | 7.800E-09 | 2.987E-09 | 4.400E-09 | 1.000E-08 |
| 11953 | 598 | E598M | A | 5.3e-09 | 2.4e-09 | 8.9e-09 | 5.533E-09 | 3.256E-09 | 2.400E-09 | 8.900E-09 |
| 11954 | 598 | E598F | A | 5.5e-09 | 6.1e-09 | 8.6e-09 | 6.733E-09 | 1.644E-09 | 5.500E-09 | 8.600E-09 |
| 11955 | 598 | E598P | A | 5.4e-09 | 1.4e-08 | 7.9e-09 | 9.100E-09 | 4.424E-09 | 5.400E-09 | 1.400E-08 |
| 11956 | 598 | E598S | A | 3.1e-09 | 1.3e-08 | 5.6e-09 | 7.233E-09 | 5.148E-09 | 3.100E-09 | 1.300E-08 |
| 11957 | 598 | E598T | A | 6.7e-09 | 9.9e-09 | 7.3e-09 | 7.967E-09 | 1.701E-09 | 6.700E-09 | 9.900E-09 |
| 11958 | 598 | E598W | A | 1e-08 | 1.7e-08 | 2e-08 | 1.567E-08 | 5.132E-09 | 1.000E-08 | 2.000E-08 |
| 11959 | 598 | E598Y | A | 1.2e-08 | 1e-08 | 5.7e-09 | 9.233E-09 | 3.219E-09 | 5.700E-09 | 1.200E-08 |
| 11960 | 598 | E598V | A | 3.4e-08 | 7.9e-09 | 3.6e-09 | 1.517E-08 | 1.645E-08 | 3.600E-09 | 3.400E-08 |
| 11961 | 599 | G599A | A | 1.1e-08 | 7.8e-09 | 4.1e-09 | 7.633E-09 | 3.453E-09 | 4.100E-09 | 1.100E-08 |
| 11962 | 599 | G599R | A | 3.5e-09 | 5e-09 | 1.3e-08 | 7.167E-09 | 5.107E-09 | 3.500E-09 | 1.300E-08 |
| 11963 | 599 | G599N | A | 4.6e-09 | 6.6e-09 | 4.4e-09 | 5.200E-09 | 1.217E-09 | 4.400E-09 | 6.600E-09 |
| 11964 | 599 | G599D | A | 8e-09 | 2.7e-08 | 4.9e-09 | 1.330E-08 | 1.197E-08 | 4.900E-09 | 2.700E-08 |
| 11965 | 599 | G599C | A | 1e-08 | 4e-09 | 3.7e-09 | 5.900E-09 | 3.554E-09 | 3.700E-09 | 1.000E-08 |
| 11966 | 599 | G599E | A | 7.6e-09 | 1.6e-08 | 8.6e-09 | 1.073E-08 | 4.588E-09 | 7.600E-09 | 1.600E-08 |
| 11967 | 599 | G599Q | A | 4e-09 | 7.2e-09 | 1.4e-08 | 8.400E-09 | 5.107E-09 | 4.000E-09 | 1.400E-08 |
| 11968 | 599 | G599G | A | 1.3e-08 | 1.4e-08 | 8e-09 | 1.167E-08 | 3.215E-09 | 8.000E-09 | 1.400E-08 |
| 11969 | 599 | G599H | A | 4.7e-09 | 1e-08 | 6.2e-09 | 6.967E-09 | 2.732E-09 | 4.700E-09 | 1.000E-08 |
| 11970 | 599 | G599I | A | 7.4e-09 | 1.5e-08 | 4.6e-09 | 9.000E-09 | 5.381E-09 | 4.600E-09 | 1.500E-08 |
| 11971 | 599 | G599L | A | 8.2e-09 | 9e-09 | 8.8e-09 | 8.667E-09 | 4.163E-10 | 8.200E-09 | 9.000E-09 |
| 11972 | 599 | G599K | A | 8e-09 | 9.8e-09 | 1.5e-09 | 6.433E-09 | 4.366E-09 | 1.500E-09 | 9.800E-09 |
| 11973 | 599 | G599M | A | 1.4e-08 | 3.8e-09 | 5.9e-09 | 7.900E-09 | 5.386E-09 | 3.800E-09 | 1.400E-08 |
| 11974 | 599 | G599F | A | 5.4e-09 | 5.2e-09 | 5.4e-09 | 5.333E-09 | 1.155E-10 | 5.200E-09 | 5.400E-09 |
| 11975 | 599 | G599P | A | 1.4e-08 | 9.7e-09 | 1.4e-08 | 1.257E-08 | 2.483E-09 | 9.700E-09 | 1.400E-08 |
| 11976 | 599 | G599S | A | 4.6e-09 | 3.5e-09 | 4.1e-09 | 4.067E-09 | 5.508E-10 | 3.500E-09 | 4.600E-09 |
| 11977 | 599 | G599T | A | 7.5e-09 | 7.6e-09 | 1.3e-08 | 9.367E-09 | 3.147E-09 | 7.500E-09 | 1.300E-08 |
| 11978 | 599 | G599W | A | 7.2e-09 | 5.4e-09 | 1.3e-08 | 8.533E-09 | 3.972E-09 | 5.400E-09 | 1.300E-08 |
| 11979 | 599 | G599Y | A | 1.1e-08 | 9e-09 | 4.9e-09 | 8.300E-09 | 3.110E-09 | 4.900E-09 | 1.100E-08 |
| 11980 | 599 | G599V | A | 6.5e-09 | 9.9e-09 | 8.3e-09 | 8.233E-09 | 1.701E-09 | 6.500E-09 | 9.900E-09 |
| 11981 | 600 | A600A | A | 7.7e-09 | 2.7e-08 | 4.4e-09 | 1.303E-08 | 1.221E-08 | 4.400E-09 | 2.700E-08 |
| 11982 | 600 | A600R | A | 6.7e-09 | 8.3e-09 | 6.8e-09 | 7.267E-09 | 8.963E-10 | 6.700E-09 | 8.300E-09 |
| 11983 | 600 | A600N | A | 3.8e-09 | 1.2e-08 | 1.8e-08 | 1.127E-08 | 7.128E-09 | 3.800E-09 | 1.800E-08 |
| 11984 | 600 | A600D | A | 1.9e-08 | 2.2e-09 | 4.9e-09 | 8.700E-09 | 9.022E-09 | 2.200E-09 | 1.900E-08 |

| | | | | | | | | | | |
|---|---|---|---|---|---|---|---|---|---|---|
| 11985 | 600 | A600C | A | 6.3e-09 | 1.1e-08 | 5e-09 | 7.433E-09 | 3.156E-09 | 5.000E-09 | 1.100E-08 |
| 11986 | 600 | A600E | A | 5.1e-09 | 3.7e-08 | 4.5e-09 | 1.553E-08 | 1.859E-08 | 4.500E-09 | 3.700E-08 |
| 11987 | 600 | A600Q | A | 1.3e-08 | 4.3e-09 | 8.4e-09 | 8.567E-09 | 4.352E-09 | 4.300E-09 | 1.300E-08 |
| 11988 | 600 | A600G | A | 1.8e-08 | 8.7e-09 | 8e-09 | 1.157E-08 | 5.582E-09 | 8.000E-09 | 1.800E-08 |
| 11989 | 600 | A600H | A | 5.6e-09 | 7.5e-09 | 2.7e-09 | 5.267E-09 | 2.417E-09 | 2.700E-09 | 7.500E-09 |
| 11990 | 600 | A600I | A | 2.7e-09 | 8.1e-09 | 1.4e-08 | 8.267E-09 | 5.652E-09 | 2.700E-09 | 1.400E-08 |
| 11991 | 600 | A600L | A | 3.3e-09 | 1.7e-08 | 4.7e-09 | 8.333E-09 | 7.538E-09 | 3.300E-09 | 1.700E-08 |
| 11992 | 600 | A600K | A | 1.4e-08 | 2.5e-09 | 3.1e-09 | 6.533E-09 | 6.473E-09 | 2.500E-09 | 1.400E-08 |
| 11993 | 600 | A600M | A | 1e-08 | 6.4e-09 | 1.1e-08 | 9.133E-09 | 2.419E-09 | 6.400E-09 | 1.100E-08 |
| 11994 | 600 | A600F | A | 8.1e-09 | 1.4e-08 | 3.8e-09 | 8.633E-09 | 5.121E-09 | 3.800E-09 | 1.400E-08 |
| 11995 | 600 | A600P | A | 2.8e-09 | 1e-08 | 3.9e-09 | 5.567E-09 | 3.879E-09 | 2.800E-09 | 1.000E-08 |
| 11996 | 600 | A600S | A | 4.6e-09 | 5e-09 | 4.1e-09 | 4.567E-09 | 4.509E-10 | 4.100E-09 | 5.000E-09 |
| 11997 | 600 | A600T | A | 6.4e-09 | 5.9e-09 | 2.5e-08 | 1.243E-08 | 1.089E-08 | 5.900E-09 | 2.500E-08 |
| 11998 | 600 | A600W | A | 4.7e-09 | 5.8e-09 | 1.2e-08 | 7.500E-09 | 3.936E-09 | 4.700E-09 | 1.200E-08 |
| 11999 | 600 | A600Y | A | 2.5e-09 | 1.8e-08 | 5.2e-09 | 8.567E-09 | 8.280E-09 | 2.500E-09 | 1.800E-08 |
| 12000 | 600 | A600V | A | 2.9e-09 | 8.8e-09 | 9.3e-09 | 7.000E-09 | 3.559E-09 | 2.900E-09 | 9.300E-09 |
| 12001 | 601 | C601A | A | 3.5e-09 | 1.5e-08 | 2.1e-09 | 6.867E-09 | 7.078E-09 | 2.100E-09 | 1.500E-08 |
| 12002 | 601 | C601R | A | 7.5e-09 | 2.1e-09 | 2.2e-09 | 3.933E-09 | 3.089E-09 | 2.100E-09 | 7.500E-09 |
| 12003 | 601 | C601N | A | 7e-09 | 2.8e-09 | 3.3e-08 | 1.427E-08 | 1.636E-08 | 2.800E-09 | 3.300E-08 |
| 12004 | 601 | C601D | A | 6e-09 | 2.5e-09 | 6.9e-09 | 5.133E-09 | 2.325E-09 | 2.500E-09 | 6.900E-09 |
| 12005 | 601 | C601C | A | 5.4e-09 | 6.4e-09 | 5.8e-09 | 5.867E-09 | 5.033E-10 | 5.400E-09 | 6.400E-09 |
| 12006 | 601 | C601E | A | 4.3e-09 | 7.3e-09 | 5.6e-09 | 5.733E-09 | 1.504E-09 | 4.300E-09 | 7.300E-09 |
| 12007 | 601 | C601Q | A | 2e-08 | 7.7e-09 | 4.2e-09 | 1.063E-08 | 8.298E-09 | 4.200E-09 | 2.000E-08 |
| 12008 | 601 | C601G | A | 6.9e-09 | 3.6e-09 | 1.2e-08 | 7.500E-09 | 4.232E-09 | 3.600E-09 | 1.200E-08 |
| 12009 | 601 | C601H | A | 4.6e-09 | 2.9e-09 | 7.1e-09 | 4.867E-09 | 2.113E-09 | 2.900E-09 | 7.100E-09 |
| 12010 | 601 | C601I | A | 1.2e-08 | 1.2e-08 | 5.2e-09 | 9.733E-09 | 3.926E-09 | 5.200E-09 | 1.200E-08 |
| 12011 | 601 | C601L | A | 7e-09 | 3.6e-09 | 7.5e-09 | 6.033E-09 | 2.122E-09 | 3.600E-09 | 7.500E-09 |
| 12012 | 601 | C601K | A | 1.1e-08 | 1.7e-08 | 5.3e-09 | 1.110E-08 | 5.851E-09 | 5.300E-09 | 1.700E-08 |
| 12013 | 601 | C601M | A | 1e-08 | 8.9e-09 | 1.3e-08 | 1.063E-08 | 2.122E-09 | 8.900E-09 | 1.300E-08 |
| 12014 | 601 | C601F | A | 4.3e-09 | 1.2e-08 | 1.2e-08 | 9.433E-09 | 4.446E-09 | 4.300E-09 | 1.200E-08 |
| 12015 | 601 | C601P | A | 1.2e-08 | 9.1e-09 | 7.9e-09 | 9.667E-09 | 2.108E-09 | 7.900E-09 | 1.200E-08 |
| 12016 | 601 | C601S | A | 4.2e-09 | 9.8e-09 | 5.5e-09 | 6.500E-09 | 2.931E-09 | 4.200E-09 | 9.800E-09 |
| 12017 | 601 | C601T | A | 1.2e-08 | 7.6e-09 | 1.1e-08 | 1.020E-08 | 2.307E-09 | 7.600E-09 | 1.200E-08 |
| 12018 | 601 | C601W | A | 5.9e-09 | 1.1e-08 | 4.8e-09 | 7.233E-09 | 3.308E-09 | 4.800E-09 | 1.100E-08 |
| 12019 | 601 | C601Y | A | 3.1e-09 | 1.8e-09 | 2.8e-09 | 2.567E-09 | 6.807E-10 | 1.800E-09 | 3.100E-09 |
| 12020 | 601 | C601V | A | 8.7e-09 | 6.2e-09 | 7.2e-09 | 7.367E-09 | 1.258E-09 | 6.200E-09 | 8.700E-09 |
| 12021 | 602 | Q602A | A | 3.3e-08 | 7.6e-09 | 2.4e-09 | 1.433E-08 | 1.637E-08 | 2.400E-09 | 3.300E-08 |
| 12022 | 602 | Q602R | A | 6.6e-09 | 1.5e-08 | 4.1e-09 | 8.567E-09 | 5.710E-09 | 4.100E-09 | 1.500E-08 |
| 12023 | 602 | Q602N | A | 1e-08 | 1.3e-08 | 7e-09 | 1.000E-08 | 3.000E-09 | 7.000E-09 | 1.300E-08 |
| 12024 | 602 | Q602D | A | 5.7e-09 | 5.6e-09 | 1.1e-08 | 7.433E-09 | 3.089E-09 | 5.600E-09 | 1.100E-08 |
| 12025 | 602 | Q602C | A | 3.5e-09 | 3.3e-09 | 1.2e-08 | 6.267E-09 | 4.966E-09 | 3.300E-09 | 1.200E-08 |
| 12026 | 602 | Q602E | A | 3.7e-09 | 3.6e-09 | 7.7e-09 | 5.000E-09 | 2.339E-09 | 3.600E-09 | 7.700E-09 |
| 12027 | 602 | Q602Q | A | 9e-09 | 1.1e-08 | 5.7e-09 | 8.567E-09 | 2.676E-09 | 5.700E-09 | 1.100E-08 |
| 12028 | 602 | Q602G | A | 9.4e-09 | 4.9e-09 | 6.7e-09 | 7.000E-09 | 2.265E-09 | 4.900E-09 | 9.400E-09 |
| 12029 | 602 | Q602H | A | 2.4e-09 | 1.9e-08 | 1.8e-09 | 7.733E-09 | 9.762E-09 | 1.800E-09 | 1.900E-08 |
| 12030 | 602 | Q602I | A | 6.2e-09 | 7.3e-09 | 4.6e-08 | 1.983E-08 | 2.267E-08 | 6.200E-09 | 4.600E-08 |
| 12031 | 602 | Q602L | A | 1.5e-08 | 5.8e-09 | 2.4e-09 | 7.733E-09 | 6.519E-09 | 2.400E-09 | 1.500E-08 |
| 12032 | 602 | Q602K | A | 5.4e-09 | 1.1e-08 | 9.5e-09 | 8.633E-09 | 2.899E-09 | 5.400E-09 | 1.100E-08 |
| 12033 | 602 | Q602M | A | 9.3e-09 | 4e-09 | 9.8e-09 | 7.700E-09 | 3.214E-09 | 4.000E-09 | 9.800E-09 |
| 12034 | 602 | Q602F | A | 2.5e-09 | 1.6e-08 | 2.9e-08 | 1.583E-08 | 1.325E-08 | 2.500E-09 | 2.900E-08 |
| 12035 | 602 | Q602P | A | 1.6e-08 | 7.6e-09 | 5.8e-09 | 9.800E-09 | 5.444E-09 | 5.800E-09 | 1.600E-08 |
| 12036 | 602 | Q602S | A | 1.3e-08 | 6.6e-09 | 1.5e-08 | 1.153E-08 | 4.388E-09 | 6.600E-09 | 1.500E-08 |
| 12037 | 602 | Q602T | A | 4.6e-09 | 1.2e-08 | 4.3e-09 | 6.967E-09 | 4.362E-09 | 4.300E-09 | 1.200E-08 |
| 12038 | 602 | Q602W | A | 1e-08 | 3.5e-09 | 7.1e-09 | 6.867E-09 | 3.256E-09 | 3.500E-09 | 1.000E-08 |
| 12039 | 602 | Q602Y | A | 4.1e-09 | 3.3e-09 | 2.4e-08 | 1.047E-08 | 1.173E-08 | 3.300E-09 | 2.400E-08 |
| 12040 | 602 | Q602V | A | 1.3e-08 | 1.4e-08 | 2e-09 | 9.667E-09 | 6.658E-09 | 2.000E-09 | 1.400E-08 |
| 12041 | 603 | P603A | A | 7.4e-09 | 3.6e-09 | 8.9e-09 | 6.633E-09 | 2.732E-09 | 3.600E-09 | 8.900E-09 |
| 12042 | 603 | P603R | A | 7.7e-09 | 4e-09 | 5.9e-09 | 5.867E-09 | 1.850E-09 | 4.000E-09 | 7.700E-09 |
| 12043 | 603 | P603N | A | 1.2e-08 | 3.5e-09 | 2.4e-08 | 1.317E-08 | 1.030E-08 | 3.500E-09 | 2.400E-08 |
| 12044 | 603 | P603D | A | 1.1e-08 | 6.8e-09 | 5e-09 | 7.600E-09 | 3.079E-09 | 5.000E-09 | 1.100E-08 |
| 12045 | 603 | P603C | A | 2.9e-08 | 6e-09 | 1.3e-08 | 1.600E-08 | 1.179E-08 | 6.000E-09 | 2.900E-08 |
| 12046 | 603 | P603E | A | 6.7e-09 | 2.3e-09 | 6.5e-09 | 5.167E-09 | 2.485E-09 | 2.300E-09 | 6.700E-09 |
| 12047 | 603 | P603Q | A | 9.5e-09 | 4.7e-09 | 4.1e-09 | 6.100E-09 | 2.960E-09 | 4.100E-09 | 9.500E-09 |
| 12048 | 603 | P603G | A | 3.8e-09 | 1.6e-08 | 1.3e-08 | 1.093E-08 | 6.357E-09 | 3.800E-09 | 1.600E-08 |
| 12049 | 603 | P603H | A | 1.4e-09 | 2.9e-09 | 1.1e-08 | 5.100E-09 | 5.164E-09 | 1.400E-09 | 1.100E-08 |
| 12050 | 603 | P603I | A | 2.2e-09 | 5.6e-09 | 7.8e-09 | 5.200E-09 | 2.821E-09 | 2.200E-09 | 7.800E-09 |
| 12051 | 603 | P603L | A | 1.1e-08 | 1.2e-08 | 2.9e-09 | 8.633E-09 | 4.990E-09 | 2.900E-09 | 1.200E-08 |
| 12052 | 603 | P603K | A | 1.1e-08 | 1.3e-08 | 6.1e-09 | 1.003E-08 | 3.550E-09 | 6.100E-09 | 1.300E-08 |
| 12053 | 603 | P603M | A | 1.5e-08 | 1.5e-08 | 1e-08 | 1.333E-08 | 2.887E-09 | 1.000E-08 | 1.500E-08 |
| 12054 | 603 | P603F | A | 5.2e-09 | 7.8e-09 | 4.9e-09 | 5.967E-09 | 1.595E-09 | 4.900E-09 | 7.800E-09 |
| 12055 | 603 | P603P | A | 3.9e-09 | 7.9e-09 | 5e-09 | 5.600E-09 | 2.066E-09 | 3.900E-09 | 7.900E-09 |
| 12056 | 603 | P603S | A | 1.2e-08 | 5.5e-09 | 2.3e-09 | 6.600E-09 | 4.943E-09 | 2.300E-09 | 1.200E-08 |
| 12057 | 603 | P603T | A | 2.4e-08 | 4.9e-09 | 1.4e-08 | 1.430E-08 | 9.554E-09 | 4.900E-09 | 2.400E-08 |
| 12058 | 603 | P603W | A | 7e-09 | 9.4e-09 | 5.3e-09 | 7.233E-09 | 2.060E-09 | 5.300E-09 | 9.400E-09 |
| 12059 | 603 | P603Y | A | 5.3e-09 | 7.6e-09 | 1.1e-08 | 7.967E-09 | 2.868E-09 | 5.300E-09 | 1.100E-08 |
| 12060 | 603 | P603V | A | 1.2e-08 | 6.8e-09 | 1.5e-08 | 1.127E-08 | 4.149E-09 | 6.800E-09 | 1.500E-08 |
| 12061 | 604 | C604A | A | 2.4e-08 | 3.3e-09 | 7.1e-09 | 1.147E-08 | 1.102E-08 | 3.300E-09 | 2.400E-08 |
| 12062 | 604 | C604R | A | 4.8e-09 | 7.5e-09 | 2e-09 | 4.767E-09 | 2.750E-09 | 2.000E-09 | 7.500E-09 |
| 12063 | 604 | C604N | A | 7.4e-09 | 1e-08 | 4.2e-09 | 7.200E-09 | 2.905E-09 | 4.200E-09 | 1.000E-08 |
| 12064 | 604 | C604D | A | 7.8e-09 | 9.9e-09 | 5.3e-09 | 7.667E-09 | 2.303E-09 | 5.300E-09 | 9.900E-09 |
| 12065 | 604 | C604C | A | 4.8e-09 | 1.3e-08 | 9.5e-09 | 9.100E-09 | 4.115E-09 | 4.800E-09 | 1.300E-08 |
| 12066 | 604 | C604E | A | 6.8e-09 | 1e-08 | 3.5e-09 | 6.767E-09 | 3.250E-09 | 3.500E-09 | 1.000E-08 |
| 12067 | 604 | C604Q | A | 2.5e-09 | 1.3e-08 | 1.3e-08 | 9.500E-09 | 6.062E-09 | 2.500E-09 | 1.300E-08 |
| 12068 | 604 | C604G | A | 5.9e-09 | 3.9e-09 | 7.5e-09 | 5.767E-09 | 1.804E-09 | 3.900E-09 | 7.500E-09 |
| 12069 | 604 | C604H | A | 5.5e-09 | 7.8e-09 | 7.8e-09 | 7.033E-09 | 1.328E-09 | 5.500E-09 | 7.800E-09 |

| | | | | | | | | | | |
|---|---|---|---|---|---|---|---|---|---|---|
| 12070 | 604 | C604I | A | 2.8e-09 | 2.2e-08 | 4.4e-09 | 9.733E-09 | 1.065E-08 | 2.800E-09 | 2.200E-08 |
| 12071 | 604 | C604L | A | 5.4e-09 | 2e-08 | 2.3e-09 | 9.233E-09 | 9.452E-09 | 2.300E-09 | 2.000E-08 |
| 12072 | 604 | C604K | A | 6.5e-09 | 3.4e-09 | 5.7e-09 | 5.200E-09 | 1.609E-09 | 3.400E-09 | 6.500E-09 |
| 12073 | 604 | C604M | A | 1.2e-08 | 3.3e-09 | 4.7e-09 | 6.667E-09 | 4.672E-09 | 3.300E-09 | 1.200E-08 |
| 12074 | 604 | C604F | A | 1.3e-08 | 7e-09 | 4.7e-09 | 8.233E-09 | 4.285E-09 | 4.700E-09 | 1.300E-08 |
| 12075 | 604 | C604P | A | 4.9e-09 | 1.5e-08 | 9.4e-09 | 9.767E-09 | 5.060E-09 | 4.900E-09 | 1.500E-08 |
| 12076 | 604 | C604S | A | 3.6e-09 | 7.2e-09 | 7.9e-09 | 6.233E-09 | 2.307E-09 | 3.600E-09 | 7.900E-09 |
| 12077 | 604 | C604T | A | 8.9e-09 | 1.1e-08 | 9.4e-09 | 9.767E-09 | 1.097E-09 | 8.900E-09 | 1.100E-08 |
| 12078 | 604 | C604W | A | 4.4e-09 | 2.3e-08 | 1.1e-08 | 1.280E-08 | 9.430E-09 | 4.400E-09 | 2.300E-08 |
| 12079 | 604 | C604Y | A | 4.2e-09 | 2e-08 | 5.3e-09 | 9.833E-09 | 8.822E-09 | 4.200E-09 | 2.000E-08 |
| 12080 | 604 | C604V | A | 7.6e-09 | 3.8e-09 | 9.5e-09 | 6.967E-09 | 2.902E-09 | 3.800E-09 | 9.500E-09 |
| 12081 | 605 | P605A | A | 6.3e-09 | 6.3e-09 | 6.4e-09 | 6.333E-09 | 5.774E-11 | 6.300E-09 | 6.400E-09 |
| 12082 | 605 | P605R | A | 7.5e-09 | 7.1e-09 | 4.9e-09 | 6.500E-09 | 1.400E-09 | 4.900E-09 | 7.500E-09 |
| 12083 | 605 | P605N | A | 3.9e-09 | 9.5e-09 | 6.1e-09 | 6.500E-09 | 2.821E-09 | 3.900E-09 | 9.500E-09 |
| 12084 | 605 | P605D | A | 1.4e-08 | 2.3e-09 | 6.7e-09 | 7.667E-09 | 5.910E-09 | 2.300E-09 | 1.400E-08 |
| 12085 | 605 | P605C | A | 1.9e-08 | 6.2e-09 | 2.6e-09 | 9.267E-09 | 8.619E-09 | 2.600E-09 | 1.900E-08 |
| 12086 | 605 | P605E | A | 1.7e-09 | 1.2e-08 | 3.2e-08 | 1.523E-08 | 1.541E-08 | 1.700E-09 | 3.200E-08 |
| 12087 | 605 | P605Q | A | 6.2e-09 | 3.2e-09 | 6.8e-09 | 5.400E-09 | 1.929E-09 | 3.200E-09 | 6.800E-09 |
| 12088 | 605 | P605G | A | 6.3e-09 | 3.5e-08 | 2.9e-09 | 1.473E-08 | 1.763E-08 | 2.900E-09 | 3.500E-08 |
| 12089 | 605 | P605H | A | 1.5e-08 | 8.4e-09 | 8.1e-09 | 1.050E-08 | 3.900E-09 | 8.100E-09 | 1.500E-08 |
| 12090 | 605 | P605I | A | 5.2e-09 | 4.7e-09 | 8.7e-09 | 6.200E-09 | 2.179E-09 | 4.700E-09 | 8.700E-09 |
| 12091 | 605 | P605L | A | 3.7e-09 | 5.8e-09 | 4.2e-09 | 4.567E-09 | 1.097E-09 | 3.700E-09 | 5.800E-09 |
| 12092 | 605 | P605K | A | 1.2e-08 | 2.9e-09 | 6.7e-09 | 7.200E-09 | 4.571E-09 | 2.900E-09 | 1.200E-08 |
| 12093 | 605 | P605M | A | 1.2e-08 | 1.3e-08 | 1.5e-08 | 1.333E-08 | 1.528E-09 | 1.200E-08 | 1.500E-08 |
| 12094 | 605 | P605F | A | 7.6e-09 | 4e-09 | 3.3e-09 | 4.967E-09 | 2.307E-09 | 3.300E-09 | 7.600E-09 |
| 12095 | 605 | P605P | A | 1e-08 | 9.1e-09 | 2.6e-08 | 1.503E-08 | 9.508E-09 | 9.100E-09 | 2.600E-08 |
| 12096 | 605 | P605S | A | 2.6e-08 | 8.5e-09 | 5.7e-09 | 1.340E-08 | 1.100E-08 | 5.700E-09 | 2.600E-08 |
| 12097 | 605 | P605T | A | 1.1e-08 | 2.1e-09 | 3.8e-09 | 5.633E-09 | 4.725E-09 | 2.100E-09 | 1.100E-08 |
| 12098 | 605 | P605W | A | 4.1e-09 | 4.5e-09 | 1.1e-08 | 6.533E-09 | 3.873E-09 | 4.100E-09 | 1.100E-08 |
| 12099 | 605 | P605Y | A | 1.9e-09 | 1.9e-09 | 8.8e-09 | 4.200E-09 | 3.984E-09 | 1.900E-09 | 8.800E-09 |
| 12100 | 605 | P605V | A | 5.2e-09 | 1.4e-09 | 1.4e-08 | 6.867E-09 | 6.463E-09 | 1.400E-09 | 1.400E-08 |
| 12101 | 606 | I606A | A | 8.3e-09 | 7.8e-09 | 7.2e-09 | 7.767E-09 | 5.508E-10 | 7.200E-09 | 8.300E-09 |
| 12102 | 606 | I606R | A | 4.3e-09 | 2.5e-09 | 1.9e-08 | 8.600E-09 | 9.052E-09 | 2.500E-09 | 1.900E-08 |
| 12103 | 606 | I606N | A | 6.4e-09 | 2.7e-09 | 3.9e-09 | 4.333E-09 | 1.888E-09 | 2.700E-09 | 6.400E-09 |
| 12104 | 606 | I606D | A | 1.2e-08 | 1.4e-08 | 1.5e-08 | 1.367E-08 | 1.528E-09 | 1.200E-08 | 1.500E-08 |
| 12105 | 606 | I606C | A | 4.3e-09 | 5.2e-09 | 1.8e-08 | 9.167E-09 | 7.663E-09 | 4.300E-09 | 1.800E-08 |
| 12106 | 606 | I606E | A | 3.7e-09 | 6.3e-09 | 7.9e-09 | 5.967E-09 | 2.120E-09 | 3.700E-09 | 7.900E-09 |
| 12107 | 606 | I606Q | A | 8.3e-09 | 6.1e-09 | 4.4e-09 | 6.267E-09 | 1.955E-09 | 4.400E-09 | 8.300E-09 |
| 12108 | 606 | I606G | A | 5.2e-09 | 7.5e-09 | 4.2e-09 | 5.633E-09 | 1.692E-09 | 4.200E-09 | 7.500E-09 |
| 12109 | 606 | I606H | A | 1.4e-08 | 4.9e-09 | 4.3e-09 | 7.733E-09 | 5.435E-09 | 4.300E-09 | 1.400E-08 |
| 12110 | 606 | I606I | A | 1.2e-08 | 7.8e-09 | 3.8e-09 | 7.867E-09 | 4.100E-09 | 3.800E-09 | 1.200E-08 |
| 12111 | 606 | I606L | A | 9.3e-09 | 8e-09 | 8.8e-09 | 8.700E-09 | 6.557E-10 | 8.000E-09 | 9.300E-09 |
| 12112 | 606 | I606K | A | 1e-08 | 2.7e-08 | 9e-09 | 1.533E-08 | 1.012E-08 | 9.000E-09 | 2.700E-08 |
| 12113 | 606 | I606M | A | 2.6e-09 | 3.4e-09 | 1.9e-08 | 8.333E-09 | 9.246E-09 | 2.600E-09 | 1.900E-08 |
| 12114 | 606 | I606F | A | 5.4e-09 | 3.7e-09 | 1.7e-08 | 8.700E-09 | 7.238E-09 | 3.700E-09 | 1.700E-08 |
| 12115 | 606 | I606P | A | 3.9e-09 | 2.1e-09 | 5.8e-09 | 3.933E-09 | 1.850E-09 | 2.100E-09 | 5.800E-09 |
| 12116 | 606 | I606S | A | 4.7e-09 | 1.2e-08 | 7.1e-09 | 7.933E-09 | 3.721E-09 | 4.700E-09 | 1.200E-08 |
| 12117 | 606 | I606T | A | 1.4e-08 | 8.3e-09 | 8.8e-09 | 1.037E-08 | 3.156E-09 | 8.300E-09 | 1.400E-08 |
| 12118 | 606 | I606W | A | 2.2e-09 | 7.4e-09 | 7.1e-09 | 5.567E-09 | 2.919E-09 | 2.200E-09 | 7.400E-09 |
| 12119 | 606 | I606Y | A | 3.2e-09 | 6.1e-09 | 1.2e-09 | 3.500E-09 | 2.464E-09 | 1.200E-09 | 6.100E-09 |
| 12120 | 606 | I606V | A | 4.1e-09 | 9.6e-09 | 4.4e-09 | 6.033E-09 | 3.092E-09 | 4.100E-09 | 9.600E-09 |
| 12121 | 607 | N607A | A | 5.8e-09 | 2.8e-09 | 2.8e-09 | 3.800E-09 | 1.732E-09 | 2.800E-09 | 5.800E-09 |
| 12122 | 607 | N607R | A | 7.7e-09 | 1.4e-08 | 5.1e-09 | 8.933E-09 | 4.576E-09 | 5.100E-09 | 1.400E-08 |
| 12123 | 607 | N607N | A | 8.4e-09 | 5.1e-09 | 6.2e-09 | 6.567E-09 | 1.680E-09 | 5.100E-09 | 8.400E-09 |
| 12124 | 607 | N607D | A | 5e-09 | 3.2e-09 | 1.3e-08 | 7.067E-09 | 5.217E-09 | 3.200E-09 | 1.300E-08 |
| 12125 | 607 | N607C | A | 6.8e-09 | 4.2e-09 | 7.1e-09 | 6.033E-09 | 1.595E-09 | 4.200E-09 | 7.100E-09 |
| 12126 | 607 | N607E | A | 7.6e-09 | 1.2e-08 | 1.1e-08 | 1.020E-08 | 2.307E-09 | 7.600E-09 | 1.200E-08 |
| 12127 | 607 | N607Q | A | 2.8e-09 | 3.6e-09 | 2.2e-08 | 9.467E-09 | 1.086E-08 | 2.800E-09 | 2.200E-08 |
| 12128 | 607 | N607G | A | 1e-08 | 6.8e-09 | 1.4e-08 | 1.027E-08 | 3.607E-09 | 6.800E-09 | 1.400E-08 |
| 12129 | 607 | N607H | A | 1e-08 | 1.4e-08 | 8.4e-09 | 1.080E-08 | 2.884E-09 | 8.400E-09 | 1.400E-08 |
| 12130 | 607 | N607I | A | 6.7e-09 | 5e-09 | 1.6e-08 | 9.233E-09 | 5.921E-09 | 5.000E-09 | 1.600E-08 |
| 12131 | 607 | N607L | A | 1.4e-08 | 7.1e-09 | 1.3e-08 | 1.137E-08 | 3.729E-09 | 7.100E-09 | 1.400E-08 |
| 12132 | 607 | N607K | A | 6.3e-09 | 2e-08 | 5e-09 | 1.043E-08 | 8.310E-09 | 5.000E-09 | 2.000E-08 |
| 12133 | 607 | N607M | A | 4.6e-09 | 5e-09 | 4.2e-09 | 4.600E-09 | 4.000E-10 | 4.200E-09 | 5.000E-09 |
| 12134 | 607 | N607F | A | 2.2e-08 | 6.3e-09 | 1e-08 | 1.277E-08 | 8.208E-09 | 6.300E-09 | 2.200E-08 |
| 12135 | 607 | N607P | A | 1.8e-08 | 6.7e-09 | 2.2e-08 | 1.557E-08 | 7.935E-09 | 6.700E-09 | 2.200E-08 |
| 12136 | 607 | N607S | A | 9.7e-09 | 5e-09 | 1.1e-08 | 8.567E-09 | 3.156E-09 | 5.000E-09 | 1.100E-08 |
| 12137 | 607 | N607T | A | 2.2e-09 | 1.1e-08 | 5.9e-09 | 6.367E-09 | 4.419E-09 | 2.200E-09 | 1.100E-08 |
| 12138 | 607 | N607W | A | 1.1e-09 | 4.2e-09 | 7.4e-09 | 4.233E-09 | 3.150E-09 | 1.100E-09 | 7.400E-09 |
| 12139 | 607 | N607Y | A | 1.1e-08 | 1.1e-08 | 4.1e-09 | 8.700E-09 | 3.984E-09 | 4.100E-09 | 1.100E-08 |
| 12140 | 607 | N607V | A | 1.4e-08 | 5.2e-09 | 1.7e-09 | 6.967E-09 | 6.337E-09 | 1.700E-09 | 1.400E-08 |
| 12141 | 608 | C608A | A | 1.5e-08 | 4.4e-09 | 5e-09 | 8.133E-09 | 5.954E-09 | 4.400E-09 | 1.500E-08 |
| 12142 | 608 | C608R | A | 2.8e-09 | 5.1e-09 | 1.8e-08 | 8.633E-09 | 8.193E-09 | 2.800E-09 | 1.800E-08 |
| 12143 | 608 | C608N | A | 9.5e-09 | 3e-09 | 3.2e-09 | 5.233E-09 | 3.696E-09 | 3.000E-09 | 9.500E-09 |
| 12144 | 608 | C608D | A | 1.1e-08 | 1.8e-08 | 4.1e-09 | 1.103E-08 | 6.950E-09 | 4.100E-09 | 1.800E-08 |
| 12145 | 608 | C608C | A | 9e-09 | 1.5e-09 | 1.3e-08 | 7.833E-09 | 5.838E-09 | 1.500E-09 | 1.300E-08 |
| 12146 | 608 | C608E | A | 2.5e-08 | 1.6e-08 | 8.1e-09 | 1.637E-08 | 8.456E-09 | 8.100E-09 | 2.500E-08 |
| 12147 | 608 | C608Q | A | 9.7e-09 | 6.2e-09 | 1e-08 | 8.633E-09 | 2.113E-09 | 6.200E-09 | 1.000E-08 |
| 12148 | 608 | C608G | A | 6.1e-09 | 1.1e-08 | 3.1e-09 | 6.733E-09 | 3.988E-09 | 3.100E-09 | 1.100E-08 |
| 12149 | 608 | C608H | A | 6.2e-09 | 7.9e-09 | 1.8e-08 | 1.070E-08 | 6.379E-09 | 6.200E-09 | 1.800E-08 |
| 12150 | 608 | C608I | A | 1e-08 | 1.1e-08 | 1.2e-08 | 1.100E-08 | 1.000E-09 | 1.000E-08 | 1.200E-08 |
| 12151 | 608 | C608L | A | 1.2e-08 | 4.2e-09 | 8.2e-09 | 8.133E-09 | 3.900E-09 | 4.200E-09 | 1.200E-08 |
| 12152 | 608 | C608K | A | 1.5e-08 | 1.3e-08 | 1e-08 | 1.267E-08 | 2.517E-09 | 1.000E-08 | 1.500E-08 |
| 12153 | 608 | C608M | A | 2.2e-08 | 5.8e-09 | 5.6e-09 | 1.113E-08 | 9.411E-09 | 5.600E-09 | 2.200E-08 |
| 12154 | 608 | C608F | A | 2.2e-09 | 1.5e-08 | 1.2e-08 | 9.733E-09 | 6.694E-09 | 2.200E-09 | 1.500E-08 |

| | | | | | | | | | | |
|---|---|---|---|---|---|---|---|---|---|---|
| 12155 | 608 | C608P | A | 4.6e-09 | 7e-09 | 5.5e-09 | 5.700E-09 | 1.212E-09 | 4.600E-09 | 7.000E-09 |
| 12156 | 608 | C608S | A | 2.1e-09 | 7.3e-10 | 5.3e-09 | 2.710E-09 | 2.345E-09 | 7.300E-10 | 5.300E-09 |
| 12157 | 608 | C608T | A | 2.6e-08 | 1e-08 | 7.1e-09 | 1.437E-08 | 1.018E-08 | 7.100E-09 | 2.600E-08 |
| 12158 | 608 | C608W | A | 9.4e-09 | 7e-09 | 4.5e-09 | 6.967E-09 | 2.450E-09 | 4.500E-09 | 9.400E-09 |
| 12159 | 608 | C608Y | A | 1.3e-08 | 5.5e-09 | 4.6e-09 | 7.700E-09 | 4.612E-09 | 4.600E-09 | 1.300E-08 |
| 12160 | 608 | C608V | A | 3.7e-08 | 7.6e-09 | 8.8e-09 | 1.780E-08 | 1.664E-08 | 7.600E-09 | 3.700E-08 |
| 12161 | 609 | T609A | A | 7.7e-09 | 2.3e-08 | 1e-08 | 1.357E-08 | 8.250E-09 | 7.700E-09 | 2.300E-08 |
| 12162 | 609 | T609R | A | 7.2e-09 | 5e-09 | 7.7e-09 | 6.633E-09 | 1.436E-09 | 5.000E-09 | 7.700E-09 |
| 12163 | 609 | T609N | A | 5e-09 | 7.3e-09 | 5.3e-09 | 5.867E-09 | 1.250E-09 | 5.000E-09 | 7.300E-09 |
| 12164 | 609 | T609D | A | 6.9e-09 | 5.3e-09 | 3.5e-09 | 5.233E-09 | 1.701E-09 | 3.500E-09 | 6.900E-09 |
| 12165 | 609 | T609C | A | 8.8e-09 | 4.7e-09 | 7.7e-09 | 7.067E-09 | 2.122E-09 | 4.700E-09 | 8.800E-09 |
| 12166 | 609 | T609E | A | 2.3e-08 | 7.7e-09 | 7.1e-09 | 1.260E-08 | 9.012E-09 | 7.100E-09 | 2.300E-08 |
| 12167 | 609 | T609Q | A | 8.8e-09 | 6e-09 | 4.1e-09 | 6.300E-09 | 2.364E-09 | 4.100E-09 | 8.800E-09 |
| 12168 | 609 | T609G | A | 6.2e-09 | 1.2e-08 | 6.4e-09 | 8.200E-09 | 3.292E-09 | 6.200E-09 | 1.200E-08 |
| 12169 | 609 | T609H | A | 8.3e-09 | 3.5e-09 | 2.3e-08 | 1.160E-08 | 1.016E-08 | 3.500E-09 | 2.300E-08 |
| 12170 | 609 | T609I | A | 1.2e-08 | 7.9e-09 | 1.1e-09 | 7.000E-09 | 5.505E-09 | 1.100E-09 | 1.200E-08 |
| 12171 | 609 | T609L | A | 1.5e-08 | 9e-09 | 1e-08 | 1.133E-08 | 3.215E-09 | 9.000E-09 | 1.500E-08 |
| 12172 | 609 | T609K | A | 5.1e-09 | 1.5e-08 | 6.6e-09 | 8.900E-09 | 5.336E-09 | 5.100E-09 | 1.500E-08 |
| 12173 | 609 | T609M | A | 5.3e-09 | 5.9e-09 | 1.4e-08 | 8.400E-09 | 4.859E-09 | 5.300E-09 | 1.400E-08 |
| 12174 | 609 | T609F | A | 1.1e-08 | 2.3e-09 | 3e-09 | 5.433E-09 | 4.834E-09 | 2.300E-09 | 1.100E-08 |
| 12175 | 609 | T609P | A | 7.2e-09 | 5.4e-09 | 6.5e-08 | 2.587E-08 | 3.390E-08 | 5.400E-09 | 6.500E-08 |
| 12176 | 609 | T609S | A | 1.3e-08 | 6.2e-09 | 3.8e-09 | 7.667E-09 | 4.772E-09 | 3.800E-09 | 1.300E-08 |
| 12177 | 609 | T609T | A | 2.1e-08 | 1.7e-08 | 2.8e-08 | 2.200E-08 | 5.568E-09 | 1.700E-08 | 2.800E-08 |
| 12178 | 609 | T609W | A | 9e-09 | 1.8e-08 | 7.5e-09 | 1.150E-08 | 5.679E-09 | 7.500E-09 | 1.800E-08 |
| 12179 | 609 | T609Y | A | 4e-09 | 6.3e-09 | 7.8e-09 | 6.033E-09 | 1.914E-09 | 4.000E-09 | 7.800E-09 |
| 12180 | 609 | T609V | A | 5.9e-09 | 9.7e-09 | 8.5e-09 | 8.033E-09 | 1.943E-09 | 5.900E-09 | 9.700E-09 |
| 12181 | 610 | H610A | A | 4.5e-09 | 4.6e-09 | 1.9e-08 | 9.367E-09 | 8.343E-09 | 4.500E-09 | 1.900E-08 |
| 12182 | 610 | H610R | A | 8.1e-09 | 1.5e-08 | 3.2e-09 | 8.767E-09 | 5.928E-09 | 3.200E-09 | 1.500E-08 |
| 12183 | 610 | H610N | A | 5.3e-09 | 7.6e-09 | 1.6e-08 | 9.633E-09 | 5.632E-09 | 5.300E-09 | 1.600E-08 |
| 12184 | 610 | H610D | A | 8.8e-09 | 3e-09 | 7.4e-09 | 6.400E-09 | 3.027E-09 | 3.000E-09 | 8.800E-09 |
| 12185 | 610 | H610C | A | 3.2e-09 | 9.6e-09 | 4.3e-08 | 1.860E-08 | 2.137E-08 | 3.200E-09 | 4.300E-08 |
| 12186 | 610 | H610E | A | 2e-08 | 8.3e-09 | 1e-08 | 1.277E-08 | 6.322E-09 | 8.300E-09 | 2.000E-08 |
| 12187 | 610 | H610Q | A | 3e-08 | 4.1e-09 | 7.2e-09 | 1.377E-08 | 1.414E-08 | 4.100E-09 | 3.000E-08 |
| 12188 | 610 | H610G | A | 7.8e-09 | 8.2e-09 | 5.2e-09 | 7.067E-09 | 1.629E-09 | 5.200E-09 | 8.200E-09 |
| 12189 | 610 | H610H | A | 3.9e-09 | 3.9e-09 | 3.1e-09 | 3.633E-09 | 4.619E-10 | 3.100E-09 | 3.900E-09 |
| 12190 | 610 | H610I | A | 5.1e-09 | 2e-08 | 4.5e-09 | 9.867E-09 | 8.781E-09 | 4.500E-09 | 2.000E-08 |
| 12191 | 610 | H610L | A | 1.6e-08 | 4.3e-09 | 4.2e-09 | 8.167E-09 | 6.784E-09 | 4.200E-09 | 1.600E-08 |
| 12192 | 610 | H610K | A | 8e-09 | 8.4e-09 | 2e-09 | 6.133E-09 | 3.585E-09 | 2.000E-09 | 8.400E-09 |
| 12193 | 610 | H610M | A | 9.3e-09 | 6.1e-09 | 3.7e-09 | 6.367E-09 | 2.810E-09 | 3.700E-09 | 9.300E-09 |
| 12194 | 610 | H610F | A | 6.7e-09 | 1e-08 | 2.8e-09 | 6.500E-09 | 3.604E-09 | 2.800E-09 | 1.000E-08 |
| 12195 | 610 | H610P | A | 7.7e-09 | 4.9e-09 | 2.6e-08 | 1.287E-08 | 1.146E-08 | 4.900E-09 | 2.600E-08 |
| 12196 | 610 | H610S | A | 9.7e-09 | 3.2e-09 | 7.7e-09 | 6.867E-09 | 3.329E-09 | 3.200E-09 | 9.700E-09 |
| 12197 | 610 | H610T | A | 2.4e-09 | 3.5e-09 | 4.9e-09 | 3.600E-09 | 1.253E-09 | 2.400E-09 | 4.900E-09 |
| 12198 | 610 | H610W | A | 1.1e-08 | 7.6e-09 | 3.6e-09 | 7.400E-09 | 3.704E-09 | 3.600E-09 | 1.100E-08 |
| 12199 | 610 | H610Y | A | 3.4e-09 | 8e-09 | 6.7e-09 | 6.033E-09 | 2.371E-09 | 3.400E-09 | 8.000E-09 |
| 12200 | 610 | H610V | A | 7.2e-09 | 6.6e-09 | 1.3e-08 | 8.933E-09 | 3.535E-09 | 6.600E-09 | 1.300E-08 |
| 12201 | 611 | S611A | A | 5.4e-09 | 5.2e-09 | 1.3e-08 | 7.867E-09 | 4.447E-09 | 5.200E-09 | 1.300E-08 |
| 12202 | 611 | S611R | A | 2.1e-09 | 2.1e-09 | 7.8e-09 | 4.000E-09 | 3.291E-09 | 2.100E-09 | 7.800E-09 |
| 12203 | 611 | S611N | A | 2.1e-09 | 5.1e-09 | 2.4e-09 | 3.200E-09 | 1.652E-09 | 2.100E-09 | 5.100E-09 |
| 12204 | 611 | S611D | A | 4.5e-09 | 7.1e-09 | 9.3e-09 | 6.967E-09 | 2.403E-09 | 4.500E-09 | 9.300E-09 |
| 12205 | 611 | S611C | A | 8.2e-09 | 5.7e-09 | 6.4e-09 | 6.767E-09 | 1.290E-09 | 5.700E-09 | 8.200E-09 |
| 12206 | 611 | S611E | A | 6.8e-09 | 3.8e-09 | 1.7e-08 | 9.200E-09 | 6.920E-09 | 3.800E-09 | 1.700E-08 |
| 12207 | 611 | S611Q | A | 2.4e-09 | 7.9e-09 | 8.1e-09 | 6.133E-09 | 3.235E-09 | 2.400E-09 | 8.100E-09 |
| 12208 | 611 | S611G | A | 5.9e-09 | 3.6e-09 | 6.5e-09 | 5.333E-09 | 1.531E-09 | 3.600E-09 | 6.500E-09 |
| 12209 | 611 | S611H | A | 1.1e-08 | 1.3e-08 | 2.2e-08 | 1.533E-08 | 5.859E-09 | 1.100E-08 | 2.200E-08 |
| 12210 | 611 | S611I | A | 4.2e-09 | 6.8e-09 | 3.5e-09 | 4.833E-09 | 1.739E-09 | 3.500E-09 | 6.800E-09 |
| 12211 | 611 | S611L | A | 8.3e-09 | 4.3e-09 | 9.6e-09 | 7.400E-09 | 2.762E-09 | 4.300E-09 | 9.600E-09 |
| 12212 | 611 | S611K | A | 5e-09 | 7.1e-09 | 6.3e-09 | 6.133E-09 | 1.060E-09 | 5.000E-09 | 7.100E-09 |
| 12213 | 611 | S611M | A | 7.8e-09 | 7.5e-09 | 4.1e-09 | 6.467E-09 | 2.055E-09 | 4.100E-09 | 7.800E-09 |
| 12214 | 611 | S611F | A | 7.2e-09 | 2.7e-09 | 1.6e-08 | 8.633E-09 | 6.765E-09 | 2.700E-09 | 1.600E-08 |
| 12215 | 611 | S611P | A | 1.2e-08 | 4e-09 | 1.6e-08 | 1.067E-08 | 6.110E-09 | 4.000E-09 | 1.600E-08 |
| 12216 | 611 | S611S | A | 1.3e-08 | 2.5e-09 | 7.2e-09 | 7.567E-09 | 5.260E-09 | 2.500E-09 | 1.300E-08 |
| 12217 | 611 | S611T | A | 8.6e-09 | 9.1e-09 | 3.9e-09 | 7.200E-09 | 2.869E-09 | 3.900E-09 | 9.100E-09 |
| 12218 | 611 | S611W | A | 1.2e-08 | 9e-09 | 3.9e-09 | 8.300E-09 | 4.095E-09 | 3.900E-09 | 1.200E-08 |
| 12219 | 611 | S611Y | A | 1.3e-08 | 9.1e-09 | 3.4e-09 | 8.500E-09 | 4.828E-09 | 3.400E-09 | 1.300E-08 |
| 12220 | 611 | S611V | A | 1.2e-08 | 5.8e-09 | 6.5e-09 | 8.100E-09 | 3.396E-09 | 5.800E-09 | 1.200E-08 |
| 12221 | 612 | C612A | A | 1.2e-08 | 5.5e-09 | 7.7e-09 | 8.400E-09 | 3.306E-09 | 5.500E-09 | 1.200E-08 |
| 12222 | 612 | C612R | A | 4.5e-09 | 8.5e-09 | 9.9e-09 | 7.633E-09 | 2.802E-09 | 4.500E-09 | 9.900E-09 |
| 12223 | 612 | C612N | A | 2.7e-09 | 4.5e-09 | 1.5e-08 | 7.400E-09 | 6.643E-09 | 2.700E-09 | 1.500E-08 |
| 12224 | 612 | C612D | A | 9.9e-09 | 7.9e-09 | 9.3e-10 | 6.243E-09 | 4.709E-09 | 9.300E-10 | 9.900E-09 |
| 12225 | 612 | C612C | A | 3.9e-09 | 8.8e-09 | 2.2e-08 | 1.157E-08 | 9.362E-09 | 3.900E-09 | 2.200E-08 |
| 12226 | 612 | C612E | A | 1.5e-08 | 2e-08 | 7e-09 | 1.400E-08 | 6.557E-09 | 7.000E-09 | 2.000E-08 |
| 12227 | 612 | C612Q | A | 1.6e-08 | 1e-08 | 7e-09 | 1.100E-08 | 4.583E-09 | 7.000E-09 | 1.600E-08 |
| 12228 | 612 | C612G | A | 9.4e-09 | 1.7e-08 | 1.8e-09 | 9.400E-09 | 7.600E-09 | 1.800E-09 | 1.700E-08 |
| 12229 | 612 | C612H | A | 6.8e-09 | 9.8e-09 | 1.7e-08 | 1.120E-08 | 5.242E-09 | 6.800E-09 | 1.700E-08 |
| 12230 | 612 | C612I | A | 1.1e-08 | 7.1e-09 | 6.7e-09 | 8.267E-09 | 2.376E-09 | 6.700E-09 | 1.100E-08 |
| 12231 | 612 | C612L | A | 8.3e-09 | 4.1e-09 | 2e-09 | 4.800E-09 | 3.208E-09 | 2.000E-09 | 8.300E-09 |
| 12232 | 612 | C612K | A | 5.9e-09 | 3.7e-09 | 4.2e-09 | 4.600E-09 | 1.153E-09 | 3.700E-09 | 5.900E-09 |
| 12233 | 612 | C612M | A | 9.6e-09 | 5.9e-09 | 1.3e-08 | 9.500E-09 | 3.551E-09 | 5.900E-09 | 1.300E-08 |
| 12234 | 612 | C612F | A | 3.6e-09 | 1.4e-08 | 3.3e-09 | 6.967E-09 | 6.093E-09 | 3.300E-09 | 1.400E-08 |
| 12235 | 612 | C612P | A | 9.6e-09 | 2.9e-09 | 1.3e-08 | 8.500E-09 | 5.139E-09 | 2.900E-09 | 1.300E-08 |
| 12236 | 612 | C612S | A | 6e-09 | 1.5e-08 | 1.4e-08 | 1.167E-08 | 4.933E-09 | 6.000E-09 | 1.500E-08 |
| 12237 | 612 | C612T | A | 1.3e-08 | 1.3e-08 | 1.4e-08 | 1.333E-08 | 5.774E-10 | 1.300E-08 | 1.400E-08 |
| 12238 | 612 | C612W | A | 3.9e-09 | 3e-09 | 1.9e-08 | 8.633E-09 | 8.989E-09 | 3.000E-09 | 1.900E-08 |
| 12239 | 612 | C612Y | A | 2e-08 | 4.8e-09 | 9.6e-09 | 1.147E-08 | 7.770E-09 | 4.800E-09 | 2.000E-08 |

| | | | | | | | | | | |
|---|---|---|---|---|---|---|---|---|---|---|
| 12240 | 612 | C612V | A | 1.1e-08 | 3.1e-09 | 3.3e-09 | 5.800E-09 | 4.504E-09 | 3.100E-09 | 1.100E-08 |
| 12241 | 613 | V613A | A | 5.7e-09 | 2.1e-08 | 3.4e-09 | 1.003E-08 | 9.567E-09 | 3.400E-09 | 2.100E-08 |
| 12242 | 613 | V613R | A | 6.9e-09 | 1.2e-08 | 3.6e-09 | 7.500E-09 | 4.232E-09 | 3.600E-09 | 1.200E-08 |
| 12243 | 613 | V613N | A | 4.2e-09 | 4.3e-09 | 1.2e-08 | 6.833E-09 | 4.475E-09 | 4.200E-09 | 1.200E-08 |
| 12244 | 613 | V613D | A | 2.6e-09 | 6.6e-09 | 7.6e-09 | 5.600E-09 | 2.646E-09 | 2.600E-09 | 7.600E-09 |
| 12245 | 613 | V613C | A | 3.5e-09 | 8.1e-09 | 3.2e-09 | 4.933E-09 | 2.747E-09 | 3.200E-09 | 8.100E-09 |
| 12246 | 613 | V613E | A | 9.8e-09 | 1.5e-08 | 2.4e-09 | 9.067E-09 | 6.332E-09 | 2.400E-09 | 1.500E-08 |
| 12247 | 613 | V613Q | A | 2.8e-09 | 1.1e-08 | 1.1e-08 | 8.267E-09 | 4.734E-09 | 2.800E-09 | 1.100E-08 |
| 12248 | 613 | V613G | A | 3.3e-09 | 8.9e-09 | 1.4e-08 | 8.733E-09 | 5.352E-09 | 3.300E-09 | 1.400E-08 |
| 12249 | 613 | V613H | A | 7.4e-09 | 9.5e-09 | 6.2e-09 | 7.700E-09 | 1.670E-09 | 6.200E-09 | 9.500E-09 |
| 12250 | 613 | V613I | A | 7.7e-09 | 4.2e-09 | 8.8e-09 | 6.900E-09 | 2.402E-09 | 4.200E-09 | 8.800E-09 |
| 12251 | 613 | V613L | A | 9.8e-09 | 9.5e-09 | 1.5e-08 | 1.143E-08 | 3.092E-09 | 9.500E-09 | 1.500E-08 |
| 12252 | 613 | V613K | A | 1.2e-09 | 1.7e-08 | 1.4e-08 | 1.073E-08 | 8.391E-09 | 1.200E-09 | 1.700E-08 |
| 12253 | 613 | V613M | A | 2.4e-09 | 4.1e-09 | 4.3e-09 | 3.600E-09 | 1.044E-09 | 2.400E-09 | 4.300E-09 |
| 12254 | 613 | V613F | A | 8.3e-09 | 8.9e-09 | 1e-08 | 9.067E-09 | 8.622E-10 | 8.300E-09 | 1.000E-08 |
| 12255 | 613 | V613P | A | 4e-09 | 6e-09 | 5.3e-09 | 5.100E-09 | 1.015E-09 | 4.000E-09 | 6.000E-09 |
| 12256 | 613 | V613S | A | 9.2e-09 | 7.4e-09 | 7.4e-09 | 8.000E-09 | 1.039E-09 | 7.400E-09 | 9.200E-09 |
| 12257 | 613 | V613T | A | 1e-08 | 2.4e-09 | 1.2e-08 | 8.133E-09 | 5.065E-09 | 2.400E-09 | 1.200E-08 |
| 12258 | 613 | V613W | A | 1.6e-09 | 6e-09 | 4.8e-09 | 4.133E-09 | 2.274E-09 | 1.600E-09 | 6.000E-09 |
| 12259 | 613 | V613Y | A | 2.8e-09 | 7.6e-09 | 1.7e-09 | 4.033E-09 | 3.137E-09 | 1.700E-09 | 7.600E-09 |
| 12260 | 613 | V613V | A | 1.3e-08 | 6.4e-09 | 5.8e-09 | 8.400E-09 | 3.995E-09 | 5.800E-09 | 1.300E-08 |
| 12261 | 614 | D614A | A | 9.8e-09 | 8.1e-09 | 1.6e-08 | 1.130E-08 | 4.158E-09 | 8.100E-09 | 1.600E-08 |
| 12262 | 614 | D614R | A | 1.1e-08 | 2.4e-09 | 6.6e-09 | 6.667E-09 | 4.300E-09 | 2.400E-09 | 1.100E-08 |
| 12263 | 614 | D614N | A | 6.4e-09 | 5.4e-09 | 6.1e-09 | 5.967E-09 | 5.132E-10 | 5.400E-09 | 6.400E-09 |
| 12264 | 614 | D614D | A | 4e-09 | 2.4e-08 | 5.5e-09 | 1.117E-08 | 1.114E-08 | 4.000E-09 | 2.400E-08 |
| 12265 | 614 | D614C | A | 3.3e-09 | 9.2e-09 | 2.7e-08 | 1.317E-08 | 1.234E-08 | 3.300E-09 | 2.700E-08 |
| 12266 | 614 | D614E | A | 9.2e-09 | 1.1e-08 | 4.6e-09 | 8.267E-09 | 3.301E-09 | 4.600E-09 | 1.100E-08 |
| 12267 | 614 | D614Q | A | 1.3e-08 | 1e-08 | 1.2e-08 | 1.167E-08 | 1.528E-09 | 1.000E-08 | 1.300E-08 |
| 12268 | 614 | D614G | A | 7e-09 | 3.6e-09 | 9.7e-09 | 6.767E-09 | 3.057E-09 | 3.600E-09 | 9.700E-09 |
| 12269 | 614 | D614H | A | 5.5e-09 | 4.3e-09 | 9e-09 | 6.267E-09 | 2.442E-09 | 4.300E-09 | 9.000E-09 |
| 12270 | 614 | D614I | A | 6.4e-09 | 6.2e-09 | 1.1e-08 | 7.867E-09 | 2.715E-09 | 6.200E-09 | 1.100E-08 |
| 12271 | 614 | D614L | A | 1.3e-08 | 7.2e-09 | 6.8e-09 | 9.000E-09 | 3.470E-09 | 6.800E-09 | 1.300E-08 |
| 12272 | 614 | D614K | A | 5.4e-09 | 1.3e-08 | 4.6e-09 | 7.667E-09 | 4.636E-09 | 4.600E-09 | 1.300E-08 |
| 12273 | 614 | D614M | A | 2e-08 | 2e-09 | 6.3e-09 | 9.433E-09 | 9.400E-09 | 2.000E-09 | 2.000E-08 |
| 12274 | 614 | D614F | A | 6.5e-09 | 9.2e-09 | 6.9e-09 | 7.533E-09 | 1.457E-09 | 6.500E-09 | 9.200E-09 |
| 12275 | 614 | D614P | A | 9.1e-09 | 8.3e-09 | 3.4e-09 | 6.933E-09 | 3.086E-09 | 3.400E-09 | 9.100E-09 |
| 12276 | 614 | D614S | A | 1.1e-08 | 3.8e-09 | 2.1e-09 | 5.633E-09 | 4.725E-09 | 2.100E-09 | 1.100E-08 |
| 12277 | 614 | D614T | A | 5.8e-09 | 3.5e-09 | 1.2e-08 | 7.100E-09 | 4.397E-09 | 3.500E-09 | 1.200E-08 |
| 12278 | 614 | D614W | A | 5.9e-09 | 1.4e-08 | 7.5e-09 | 9.133E-09 | 4.290E-09 | 5.900E-09 | 1.400E-08 |
| 12279 | 614 | D614Y | A | 6.6e-09 | 6.1e-09 | 6.1e-09 | 6.267E-09 | 2.887E-10 | 6.100E-09 | 6.600E-09 |
| 12280 | 614 | D614V | A | 1.3e-08 | 5.9e-09 | 1.4e-08 | 1.097E-08 | 4.416E-09 | 5.900E-09 | 1.400E-08 |
| 12281 | 615 | L615A | A | 1.6e-08 | 1.2e-08 | 1.2e-08 | 1.333E-08 | 2.309E-09 | 1.200E-08 | 1.600E-08 |
| 12282 | 615 | L615R | A | 1.8e-09 | 2.1e-09 | 8.2e-09 | 4.033E-09 | 3.612E-09 | 1.800E-09 | 8.200E-09 |
| 12283 | 615 | L615N | A | 2.8e-09 | 5.6e-09 | 3.6e-09 | 4.000E-09 | 1.442E-09 | 2.800E-09 | 5.600E-09 |
| 12284 | 615 | L615D | A | 1.1e-08 | 9.2e-09 | 1.2e-08 | 1.073E-08 | 1.419E-09 | 9.200E-09 | 1.200E-08 |
| 12285 | 615 | L615C | A | 9.4e-09 | 3.8e-09 | 7.5e-09 | 6.900E-09 | 2.848E-09 | 3.800E-09 | 9.400E-09 |
| 12286 | 615 | L615E | A | 3.2e-09 | 3.3e-09 | 8.3e-09 | 4.933E-09 | 2.916E-09 | 3.200E-09 | 8.300E-09 |
| 12287 | 615 | L615Q | A | 1.2e-08 | 1.6e-08 | 8e-09 | 1.200E-08 | 4.000E-09 | 8.000E-09 | 1.600E-08 |
| 12288 | 615 | L615G | A | 1.1e-08 | 5.4e-09 | 1.4e-08 | 1.013E-08 | 4.365E-09 | 5.400E-09 | 1.400E-08 |
| 12289 | 615 | L615H | A | 4.5e-09 | 1.2e-08 | 9.9e-09 | 8.800E-09 | 3.869E-09 | 4.500E-09 | 1.200E-08 |
| 12290 | 615 | L615I | A | 1.3e-08 | 3.2e-09 | 2.9e-09 | 6.367E-09 | 5.747E-09 | 2.900E-09 | 1.300E-08 |
| 12291 | 615 | L615L | A | 3.6e-09 | 8.6e-09 | 6.8e-09 | 6.333E-09 | 2.532E-09 | 3.600E-09 | 8.600E-09 |
| 12292 | 615 | L615K | A | 7.9e-09 | 1.6e-08 | 1.1e-08 | 1.163E-08 | 4.087E-09 | 7.900E-09 | 1.600E-08 |
| 12293 | 615 | L615M | A | 1.2e-08 | 6.1e-09 | 2e-09 | 6.700E-09 | 5.027E-09 | 2.000E-09 | 1.200E-08 |
| 12294 | 615 | L615F | A | 9.8e-09 | 1e-08 | 4.1e-09 | 7.967E-09 | 3.350E-09 | 4.100E-09 | 1.000E-08 |
| 12295 | 615 | L615P | A | 3.3e-09 | 2e-08 | 2.9e-09 | 8.733E-09 | 9.759E-09 | 2.900E-09 | 2.000E-08 |
| 12296 | 615 | L615S | A | 1.7e-08 | 4.8e-09 | 1e-08 | 1.060E-08 | 6.122E-09 | 4.800E-09 | 1.700E-08 |
| 12297 | 615 | L615T | A | 1.5e-08 | 4.8e-09 | 1.3e-08 | 1.093E-08 | 5.405E-09 | 4.800E-09 | 1.500E-08 |
| 12298 | 615 | L615W | A | 6.9e-09 | 5.4e-09 | 4.2e-09 | 5.500E-09 | 1.353E-09 | 4.200E-09 | 6.900E-09 |
| 12299 | 615 | L615Y | A | 5.6e-09 | 7.9e-09 | 1.9e-09 | 5.133E-09 | 3.027E-09 | 1.900E-09 | 7.900E-09 |
| 12300 | 615 | L615V | A | 1.9e-08 | 4.4e-09 | 8.3e-09 | 1.057E-08 | 7.559E-09 | 4.400E-09 | 1.900E-08 |
| 12301 | 616 | D616A | A | 2.1e-08 | 3.7e-09 | 1.7e-08 | 1.390E-08 | 9.057E-09 | 3.700E-09 | 2.100E-08 |
| 12302 | 616 | D616R | A | 1.2e-08 | 2.8e-09 | 1.5e-08 | 9.933E-09 | 6.357E-09 | 2.800E-09 | 1.500E-08 |
| 12303 | 616 | D616N | A | 1.1e-08 | 8.1e-09 | 1.3e-08 | 1.070E-08 | 2.464E-09 | 8.100E-09 | 1.300E-08 |
| 12304 | 616 | D616D | A | 1.7e-09 | 8.1e-09 | 2.1e-09 | 3.967E-09 | 3.585E-09 | 1.700E-09 | 8.100E-09 |
| 12305 | 616 | D616C | A | 9.1e-09 | 1e-08 | 1.3e-09 | 6.800E-09 | 4.784E-09 | 1.300E-09 | 1.000E-08 |
| 12306 | 616 | D616E | A | 8e-09 | 1.6e-08 | 8.2e-09 | 1.073E-08 | 4.562E-09 | 8.000E-09 | 1.600E-08 |
| 12307 | 616 | D616Q | A | 1.1e-08 | 9.5e-09 | 5e-09 | 8.500E-09 | 3.122E-09 | 5.000E-09 | 1.100E-08 |
| 12308 | 616 | D616G | A | 3.6e-09 | 2e-08 | 2.8e-09 | 8.800E-09 | 9.708E-09 | 2.800E-09 | 2.000E-08 |
| 12309 | 616 | D616H | A | 2e-08 | 1e-08 | 9.9e-09 | 1.330E-08 | 5.803E-09 | 9.900E-09 | 2.000E-08 |
| 12310 | 616 | D616I | A | 1.8e-08 | 6.2e-09 | 2e-09 | 8.733E-09 | 8.295E-09 | 2.000E-09 | 1.800E-08 |
| 12311 | 616 | D616L | A | 7.2e-09 | 1.3e-08 | 3.9e-09 | 8.033E-09 | 4.607E-09 | 3.900E-09 | 1.300E-08 |
| 12312 | 616 | D616K | A | 9.9e-09 | 3.5e-09 | 6.3e-09 | 6.567E-09 | 3.208E-09 | 3.500E-09 | 9.900E-09 |
| 12313 | 616 | D616M | A | 2e-08 | 1.1e-08 | 1e-08 | 1.367E-08 | 5.508E-09 | 1.000E-08 | 2.000E-08 |
| 12314 | 616 | D616F | A | 2.3e-09 | 7.7e-09 | 5.9e-09 | 5.300E-09 | 2.750E-09 | 2.300E-09 | 7.700E-09 |
| 12315 | 616 | D616P | A | 1.9e-08 | 3.7e-09 | 1.9e-08 | 1.390E-08 | 8.833E-09 | 3.700E-09 | 1.900E-08 |
| 12316 | 616 | D616S | A | 1.1e-08 | 2.9e-09 | 5.5e-09 | 6.467E-09 | 4.136E-09 | 2.900E-09 | 1.100E-08 |
| 12317 | 616 | D616T | A | 1.2e-09 | 4.1e-09 | 3.6e-09 | 2.967E-09 | 1.550E-09 | 1.200E-09 | 4.100E-09 |
| 12318 | 616 | D616W | A | 6.7e-09 | 1e-08 | 1.2e-08 | 9.567E-09 | 2.676E-09 | 6.700E-09 | 1.200E-08 |
| 12319 | 616 | D616Y | A | 5.5e-09 | 4.5e-09 | 7.8e-09 | 5.933E-09 | 1.692E-09 | 4.500E-09 | 7.800E-09 |
| 12320 | 616 | D616V | A | 1.5e-08 | 6.3e-09 | 5.3e-09 | 8.867E-09 | 5.335E-09 | 5.300E-09 | 1.500E-08 |
| 12321 | 617 | D617A | A | 6.7e-09 | 6.1e-09 | 1.2e-08 | 8.267E-09 | 3.247E-09 | 6.100E-09 | 1.200E-08 |
| 12322 | 617 | D617R | A | 7.3e-09 | 2e-09 | 8.7e-09 | 6.000E-09 | 3.534E-09 | 2.000E-09 | 8.700E-09 |
| 12323 | 617 | D617N | A | 6e-09 | 2.4e-08 | 1.5e-08 | 1.500E-08 | 9.000E-09 | 6.000E-09 | 2.400E-08 |
| 12324 | 617 | D617D | A | 7.1e-09 | 9.1e-09 | 2.7e-09 | 6.300E-09 | 3.274E-09 | 2.700E-09 | 9.100E-09 |

| | | | | | | | | | | |
|---|---|---|---|---|---|---|---|---|---|---|
| 12325 | 617 | D617C | A | 1e-08 | 7.9e-09 | 1.5e-08 | 1.097E-08 | 3.647E-09 | 7.900E-09 | 1.500E-08 |
| 12326 | 617 | D617E | A | 4.4e-09 | 1.6e-08 | 4.6e-09 | 8.333E-09 | 6.640E-09 | 4.400E-09 | 1.600E-08 |
| 12327 | 617 | D617Q | A | 9.2e-09 | 2.1e-08 | 7.3e-09 | 1.250E-08 | 7.422E-09 | 7.300E-09 | 2.100E-08 |
| 12328 | 617 | D617G | A | 9e-09 | 3.6e-09 | 9.5e-09 | 7.367E-09 | 3.272E-09 | 3.600E-09 | 9.500E-09 |
| 12329 | 617 | D617H | A | 8.5e-09 | 1.3e-08 | 5.1e-09 | 8.867E-09 | 3.963E-09 | 5.100E-09 | 1.300E-08 |
| 12330 | 617 | D617I | A | 4.4e-09 | 9.3e-09 | 4.9e-09 | 6.200E-09 | 2.696E-09 | 4.400E-09 | 9.300E-09 |
| 12331 | 617 | D617L | A | 1.6e-08 | 1.7e-08 | 4.2e-09 | 1.240E-08 | 7.119E-09 | 4.200E-09 | 1.700E-08 |
| 12332 | 617 | D617K | A | 9.5e-09 | 1.1e-08 | 8.2e-09 | 9.567E-09 | 1.401E-09 | 8.200E-09 | 1.100E-08 |
| 12333 | 617 | D617M | A | 4.9e-09 | 3.4e-09 | 5.1e-09 | 4.467E-09 | 9.292E-10 | 3.400E-09 | 5.100E-09 |
| 12334 | 617 | D617F | A | 9.8e-09 | 5.7e-09 | 2.8e-09 | 6.100E-09 | 3.517E-09 | 2.800E-09 | 9.800E-09 |
| 12335 | 617 | D617P | A | 7.7e-09 | 9.3e-09 | 4.3e-09 | 7.100E-09 | 2.553E-09 | 4.300E-09 | 9.300E-09 |
| 12336 | 617 | D617S | A | 6.9e-09 | 9.9e-09 | 1.7e-09 | 6.167E-09 | 4.149E-09 | 1.700E-09 | 9.900E-09 |
| 12337 | 617 | D617T | A | 4.9e-09 | 3.6e-09 | 2.2e-08 | 1.017E-08 | 1.027E-08 | 3.600E-09 | 2.200E-08 |
| 12338 | 617 | D617W | A | 9.2e-09 | 1.2e-08 | 9e-09 | 1.007E-08 | 1.677E-09 | 9.000E-09 | 1.200E-08 |
| 12339 | 617 | D617Y | A | 7.6e-09 | 1.5e-08 | 6.1e-09 | 9.567E-09 | 4.765E-09 | 6.100E-09 | 1.500E-08 |
| 12340 | 617 | D617V | A | 7.9e-09 | 3e-09 | 1.4e-08 | 8.300E-09 | 5.511E-09 | 3.000E-09 | 1.400E-08 |
| 12341 | 618 | K618A | A | 7.1e-09 | 3.6e-09 | 7.9e-09 | 6.200E-09 | 2.287E-09 | 3.600E-09 | 7.900E-09 |
| 12342 | 618 | K618R | A | 4.1e-09 | 6.7e-09 | 6.9e-09 | 5.900E-09 | 1.562E-09 | 4.100E-09 | 6.900E-09 |
| 12343 | 618 | K618N | A | 5.4e-09 | 4.5e-09 | 1.4e-08 | 7.967E-09 | 5.244E-09 | 4.500E-09 | 1.400E-08 |
| 12344 | 618 | K618D | A | 1.2e-08 | 1e-08 | 1.6e-08 | 1.267E-08 | 3.055E-09 | 1.000E-08 | 1.600E-08 |
| 12345 | 618 | K618C | A | 5.9e-09 | 1.2e-08 | 9e-09 | 8.967E-09 | 3.050E-09 | 5.900E-09 | 1.200E-08 |
| 12346 | 618 | K618E | A | 1.2e-08 | 1e-08 | 8e-09 | 1.000E-08 | 2.000E-09 | 8.000E-09 | 1.200E-08 |
| 12347 | 618 | K618Q | A | 5.4e-09 | 6.6e-09 | 9.3e-09 | 7.100E-09 | 1.997E-09 | 5.400E-09 | 9.300E-09 |
| 12348 | 618 | K618G | A | 4.2e-09 | 7.4e-09 | 1.3e-08 | 8.200E-09 | 4.454E-09 | 4.200E-09 | 1.300E-08 |
| 12349 | 618 | K618H | A | 5.2e-09 | 1.1e-08 | 1.5e-08 | 1.040E-08 | 4.927E-09 | 5.200E-09 | 1.500E-08 |
| 12350 | 618 | K618I | A | 9e-09 | 2.3e-09 | 4.5e-09 | 5.267E-09 | 3.415E-09 | 2.300E-09 | 9.000E-09 |
| 12351 | 618 | K618L | A | 7.3e-09 | 3.9e-09 | 1e-08 | 7.067E-09 | 3.057E-09 | 3.900E-09 | 1.000E-08 |
| 12352 | 618 | K618K | A | 2.5e-08 | 1.3e-08 | 1.1e-08 | 1.633E-08 | 7.572E-09 | 1.100E-08 | 2.500E-08 |
| 12353 | 618 | K618M | A | 1.7e-08 | 1.6e-08 | 1e-08 | 1.433E-08 | 3.786E-09 | 1.000E-08 | 1.700E-08 |
| 12354 | 618 | K618F | A | 9.7e-09 | 8.9e-09 | 7.9e-09 | 8.833E-09 | 9.018E-10 | 7.900E-09 | 9.700E-09 |
| 12355 | 618 | K618P | A | 3.9e-09 | 5.8e-09 | 8.7e-09 | 6.133E-09 | 2.417E-09 | 3.900E-09 | 8.700E-09 |
| 12356 | 618 | K618S | A | 5.7e-09 | 6.3e-09 | 7.4e-09 | 6.467E-09 | 8.622E-10 | 5.700E-09 | 7.400E-09 |
| 12357 | 618 | K618T | A | 4.2e-09 | 5.4e-09 | 9.5e-09 | 6.367E-09 | 2.779E-09 | 4.200E-09 | 9.500E-09 |
| 12358 | 618 | K618W | A | 5.7e-09 | 8.9e-09 | 1.1e-08 | 8.533E-09 | 2.669E-09 | 5.700E-09 | 1.100E-08 |
| 12359 | 618 | K618Y | A | 9.4e-09 | 5.2e-09 | 5.9e-09 | 6.833E-09 | 2.250E-09 | 5.200E-09 | 9.400E-09 |
| 12360 | 618 | K618V | A | 8.5e-09 | 1.2e-08 | 2.3e-08 | 1.450E-08 | 7.566E-09 | 8.500E-09 | 2.300E-08 |
| 12361 | 619 | G619A | A | 7.4e-09 | 1.1e-08 | 6.1e-09 | 8.167E-09 | 2.538E-09 | 6.100E-09 | 1.100E-08 |
| 12362 | 619 | G619R | A | 7.3e-09 | 4.6e-09 | 4.8e-09 | 5.567E-09 | 1.504E-09 | 4.600E-09 | 7.300E-09 |
| 12363 | 619 | G619N | A | 7e-09 | 4.7e-09 | 3.1e-09 | 4.933E-09 | 1.960E-09 | 3.100E-09 | 7.000E-09 |
| 12364 | 619 | G619D | A | 1.7e-08 | 2.6e-09 | 8.4e-09 | 9.333E-09 | 7.245E-09 | 2.600E-09 | 1.700E-08 |
| 12365 | 619 | G619C | A | 2.2e-09 | 6.3e-09 | 7.9e-09 | 5.467E-09 | 2.940E-09 | 2.200E-09 | 7.900E-09 |
| 12366 | 619 | G619E | A | 1e-08 | 6.2e-09 | 4.8e-09 | 7.000E-09 | 2.691E-09 | 4.800E-09 | 1.000E-08 |
| 12367 | 619 | G619Q | A | 6.7e-09 | 7.5e-09 | 2.8e-09 | 5.667E-09 | 2.515E-09 | 2.800E-09 | 7.500E-09 |
| 12368 | 619 | G619G | A | 3.1e-09 | 1.1e-08 | 3.2e-09 | 5.767E-09 | 4.532E-09 | 3.100E-09 | 1.100E-08 |
| 12369 | 619 | G619H | A | 2e-08 | 1.2e-08 | 3.5e-09 | 1.183E-08 | 8.251E-09 | 3.500E-09 | 2.000E-08 |
| 12370 | 619 | G619I | A | 7.7e-09 | 4.6e-09 | 4e-09 | 5.433E-09 | 1.986E-09 | 4.000E-09 | 7.700E-09 |
| 12371 | 619 | G619L | A | 1.4e-08 | 4.6e-09 | 1.1e-08 | 9.867E-09 | 4.801E-09 | 4.600E-09 | 1.400E-08 |
| 12372 | 619 | G619K | A | 3.1e-08 | 1.1e-08 | 1e-08 | 1.733E-08 | 1.185E-08 | 1.000E-08 | 3.100E-08 |
| 12373 | 619 | G619M | A | 1.1e-08 | 3.3e-08 | 3.8e-09 | 1.593E-08 | 1.521E-08 | 3.800E-09 | 3.300E-08 |
| 12374 | 619 | G619F | A | 8.3e-09 | 8.1e-09 | 2.7e-08 | 1.447E-08 | 1.085E-08 | 8.100E-09 | 2.700E-08 |
| 12375 | 619 | G619P | A | 6.2e-09 | 1.4e-09 | 4.6e-09 | 4.067E-09 | 2.444E-09 | 1.400E-09 | 6.200E-09 |
| 12376 | 619 | G619S | A | 1.2e-08 | 9.9e-09 | 1.9e-09 | 7.933E-09 | 5.329E-09 | 1.900E-09 | 1.200E-08 |
| 12377 | 619 | G619T | A | 1.8e-09 | 2.9e-08 | 2.9e-08 | 1.993E-08 | 1.570E-08 | 1.800E-09 | 2.900E-08 |
| 12378 | 619 | G619W | A | 5e-09 | 7.4e-09 | 5.1e-09 | 5.833E-09 | 1.358E-09 | 5.000E-09 | 7.400E-09 |
| 12379 | 619 | G619Y | A | 5.8e-09 | 3.9e-09 | 1.4e-08 | 7.900E-09 | 5.367E-09 | 3.900E-09 | 1.400E-08 |
| 12380 | 619 | G619V | A | 1.2e-08 | 5e-09 | 5.8e-09 | 7.600E-09 | 3.831E-09 | 5.000E-09 | 1.200E-08 |
| 12381 | 620 | C620A | A | 2.2e-08 | 6.2e-09 | 8.6e-09 | 1.227E-08 | 8.514E-09 | 6.200E-09 | 2.200E-08 |
| 12382 | 620 | C620R | A | 5.5e-09 | 1.6e-08 | 1.7e-08 | 1.283E-08 | 6.371E-09 | 5.500E-09 | 1.700E-08 |
| 12383 | 620 | C620N | A | 1.3e-08 | 2.9e-09 | 8.5e-09 | 8.133E-09 | 5.060E-09 | 2.900E-09 | 1.300E-08 |
| 12384 | 620 | C620D | A | 1.1e-08 | 6.9e-09 | 1.6e-08 | 1.130E-08 | 4.557E-09 | 6.900E-09 | 1.600E-08 |
| 12385 | 620 | C620C | A | 9.2e-09 | 7e-09 | 5.2e-09 | 7.133E-09 | 2.003E-09 | 5.200E-09 | 9.200E-09 |
| 12386 | 620 | C620E | A | 6.7e-09 | 5.7e-09 | 8.4e-09 | 6.933E-09 | 1.365E-09 | 5.700E-09 | 8.400E-09 |
| 12387 | 620 | C620Q | A | 5.1e-09 | 2.2e-09 | 4.4e-09 | 3.900E-09 | 1.513E-09 | 2.200E-09 | 5.100E-09 |
| 12388 | 620 | C620G | A | 7.6e-09 | 2.4e-08 | 1.2e-08 | 1.453E-08 | 8.488E-09 | 7.600E-09 | 2.400E-08 |
| 12389 | 620 | C620H | A | 8.8e-09 | 5.3e-09 | 1.1e-08 | 8.367E-09 | 2.875E-09 | 5.300E-09 | 1.100E-08 |
| 12390 | 620 | C620I | A | 1.1e-08 | 6.2e-09 | 4.2e-09 | 7.133E-09 | 3.495E-09 | 4.200E-09 | 1.100E-08 |
| 12391 | 620 | C620L | A | 5.4e-09 | 8.5e-09 | 4.2e-09 | 6.033E-09 | 2.219E-09 | 4.200E-09 | 8.500E-09 |
| 12392 | 620 | C620K | A | 1.3e-08 | 1.3e-08 | 5.8e-09 | 1.060E-08 | 4.157E-09 | 5.800E-09 | 1.300E-08 |
| 12393 | 620 | C620M | A | 5.4e-09 | 8.6e-09 | 1.1e-08 | 8.333E-09 | 2.810E-09 | 5.400E-09 | 1.100E-08 |
| 12394 | 620 | C620F | A | 9.9e-09 | 5.1e-09 | 5.9e-09 | 6.967E-09 | 2.572E-09 | 5.100E-09 | 9.900E-09 |
| 12395 | 620 | C620P | A | 6.4e-09 | 1.8e-09 | 4.9e-09 | 4.367E-09 | 2.346E-09 | 1.800E-09 | 6.400E-09 |
| 12396 | 620 | C620S | A | 5.2e-09 | 8.6e-09 | 4.5e-09 | 6.100E-09 | 2.193E-09 | 4.500E-09 | 8.600E-09 |
| 12397 | 620 | C620T | A | 3.2e-09 | 4.5e-09 | 6.8e-09 | 4.833E-09 | 1.823E-09 | 3.200E-09 | 6.800E-09 |
| 12398 | 620 | C620W | A | 1.5e-08 | 8.9e-09 | 2.2e-08 | 1.530E-08 | 6.555E-09 | 8.900E-09 | 2.200E-08 |
| 12399 | 620 | C620Y | A | 9e-09 | 8.7e-09 | 1.1e-08 | 9.567E-09 | 1.250E-09 | 8.700E-09 | 1.100E-08 |
| 12400 | 620 | C620V | A | 3.7e-09 | 2e-09 | 6.1e-09 | 3.933E-09 | 2.060E-09 | 2.000E-09 | 6.100E-09 |
| 12401 | 621 | P621A | A | 1.6e-08 | 3e-09 | 1.5e-08 | 1.133E-08 | 7.234E-09 | 3.000E-09 | 1.600E-08 |
| 12402 | 621 | P621R | A | 7.3e-09 | 1.2e-08 | 3.4e-09 | 7.567E-09 | 4.306E-09 | 3.400E-09 | 1.200E-08 |
| 12403 | 621 | P621N | A | 8.3e-09 | 5.1e-09 | 1.2e-08 | 8.467E-09 | 3.453E-09 | 5.100E-09 | 1.200E-08 |
| 12404 | 621 | P621D | A | 5.1e-09 | 2.1e-09 | 9.1e-09 | 5.433E-09 | 3.512E-09 | 2.100E-09 | 9.100E-09 |
| 12405 | 621 | P621C | A | 4.3e-09 | 1.7e-08 | 8e-09 | 9.767E-09 | 6.532E-09 | 4.300E-09 | 1.700E-08 |
| 12406 | 621 | P621E | A | 4.8e-09 | 7.6e-09 | 2.2e-09 | 4.867E-09 | 2.701E-09 | 2.200E-09 | 7.600E-09 |
| 12407 | 621 | P621Q | A | 5.8e-09 | 5.6e-09 | 9.4e-09 | 6.933E-09 | 2.139E-09 | 5.600E-09 | 9.400E-09 |
| 12408 | 621 | P621G | A | 6.2e-09 | 5.3e-09 | 6.1e-09 | 5.867E-09 | 4.933E-10 | 5.300E-09 | 6.200E-09 |
| 12409 | 621 | P621H | A | 7.1e-09 | 7.8e-09 | 5.9e-09 | 6.933E-09 | 9.609E-10 | 5.900E-09 | 7.800E-09 |

| | | | | | | | | | | |
|---|---|---|---|---|---|---|---|---|---|---|
| 12410 | 621 | P621I | A | 4.4e-09 | 3.3e-09 | 4.2e-09 | 3.967E-09 | 5.859E-10 | 3.300E-09 | 4.400E-09 |
| 12411 | 621 | P621L | A | 3.1e-09 | 6.9e-09 | 1.9e-09 | 3.967E-09 | 2.610E-09 | 1.900E-09 | 6.900E-09 |
| 12412 | 621 | P621K | A | 8e-09 | 8.1e-09 | 4.6e-09 | 6.900E-09 | 1.992E-09 | 4.600E-09 | 8.100E-09 |
| 12413 | 621 | P621M | A | 5.1e-09 | 1.3e-08 | 8.3e-09 | 8.800E-09 | 3.974E-09 | 5.100E-09 | 1.300E-08 |
| 12414 | 621 | P621F | A | 6.2e-09 | 6.9e-09 | 2.7e-08 | 1.337E-08 | 1.181E-08 | 6.200E-09 | 2.700E-08 |
| 12415 | 621 | P621P | A | 1.6e-08 | 3.3e-09 | 4.5e-09 | 7.933E-09 | 7.012E-09 | 3.300E-09 | 1.600E-08 |
| 12416 | 621 | P621S | A | 5e-09 | 9.2e-09 | 2.6e-08 | 1.340E-08 | 1.111E-08 | 5.000E-09 | 2.600E-08 |
| 12417 | 621 | P621T | A | 1.7e-09 | 1.2e-08 | 8.9e-09 | 7.533E-09 | 5.284E-09 | 1.700E-09 | 1.200E-08 |
| 12418 | 621 | P621W | A | 9.8e-09 | 2.7e-08 | 4.3e-09 | 1.370E-08 | 1.184E-08 | 4.300E-09 | 2.700E-08 |
| 12419 | 621 | P621Y | A | 3e-09 | 1.1e-08 | 1.4e-08 | 9.333E-09 | 5.686E-09 | 3.000E-09 | 1.400E-08 |
| 12420 | 621 | P621V | A | 6.8e-09 | 8.8e-09 | 8.6e-09 | 8.067E-09 | 1.102E-09 | 6.800E-09 | 8.800E-09 |
| 12421 | 622 | A622A | A | 3e-09 | 3.1e-09 | 1.9e-09 | 2.667E-09 | 6.658E-10 | 1.900E-09 | 3.100E-09 |
| 12422 | 622 | A622R | A | 1.3e-08 | 3.7e-09 | 1.1e-08 | 9.233E-09 | 4.895E-09 | 3.700E-09 | 1.300E-08 |
| 12423 | 622 | A622N | A | 3.2e-09 | 7.1e-09 | 8.4e-09 | 6.233E-09 | 2.706E-09 | 3.200E-09 | 8.400E-09 |
| 12424 | 622 | A622D | A | 6.4e-09 | 5e-09 | 9.1e-09 | 6.833E-09 | 2.084E-09 | 5.000E-09 | 9.100E-09 |
| 12425 | 622 | A622C | A | 4.1e-09 | 5.9e-09 | 8.4e-09 | 6.133E-09 | 2.159E-09 | 4.100E-09 | 8.400E-09 |
| 12426 | 622 | A622E | A | 8e-09 | 6.2e-09 | 1e-08 | 8.067E-09 | 1.901E-09 | 6.200E-09 | 1.000E-08 |
| 12427 | 622 | A622Q | A | 4.6e-09 | 1.4e-08 | 1.1e-08 | 9.867E-09 | 4.801E-09 | 4.600E-09 | 1.400E-08 |
| 12428 | 622 | A622G | A | 5.2e-09 | 6.6e-09 | 2.5e-08 | 1.227E-08 | 1.105E-08 | 5.200E-09 | 2.500E-08 |
| 12429 | 622 | A622H | A | 4e-09 | 7.7e-09 | 4.2e-09 | 5.300E-09 | 2.081E-09 | 4.000E-09 | 7.700E-09 |
| 12430 | 622 | A622I | A | 1.3e-08 | 6.1e-09 | 1.3e-08 | 1.070E-08 | 3.984E-09 | 6.100E-09 | 1.300E-08 |
| 12431 | 622 | A622L | A | 4.7e-09 | 1.8e-08 | 1.7e-09 | 8.133E-09 | 8.675E-09 | 1.700E-09 | 1.800E-08 |
| 12432 | 622 | A622K | A | 8.2e-09 | 1.1e-08 | 8.8e-09 | 9.333E-09 | 1.474E-09 | 8.200E-09 | 1.100E-08 |
| 12433 | 622 | A622M | A | 1.6e-08 | 2.5e-09 | 6.3e-09 | 8.267E-09 | 6.962E-09 | 2.500E-09 | 1.600E-08 |
| 12434 | 622 | A622F | A | 4.2e-09 | 6.9e-09 | 8.7e-09 | 6.600E-09 | 2.265E-09 | 4.200E-09 | 8.700E-09 |
| 12435 | 622 | A622P | A | 2.2e-09 | 7.2e-09 | 5e-09 | 4.800E-09 | 2.506E-09 | 2.200E-09 | 7.200E-09 |
| 12436 | 622 | A622S | A | 1.3e-08 | 3.6e-08 | 5.9e-09 | 1.830E-08 | 1.573E-08 | 5.900E-09 | 3.600E-08 |
| 12437 | 622 | A622T | A | 1e-08 | 2.8e-09 | 1.6e-08 | 9.600E-09 | 6.609E-09 | 2.800E-09 | 1.600E-08 |
| 12438 | 622 | A622W | A | 1.3e-08 | 6.2e-09 | 6.6e-09 | 8.600E-09 | 3.816E-09 | 6.200E-09 | 1.300E-08 |
| 12439 | 622 | A622Y | A | 9.4e-09 | 4.1e-09 | 5.4e-09 | 6.300E-09 | 2.762E-09 | 4.100E-09 | 9.400E-09 |
| 12440 | 622 | A622V | A | 3.5e-09 | 2e-08 | 1.2e-09 | 8.233E-09 | 1.025E-08 | 1.200E-09 | 2.000E-08 |
| 12441 | 623 | D623A | B | 4.4e-09 | 7.8e-09 | 1.2e-08 | 8.067E-09 | 3.807E-09 | 4.400E-09 | 1.200E-08 |
| 12442 | 623 | D623R | B | 7.4e-09 | 1e-08 | 8.6e-09 | 8.667E-09 | 1.301E-09 | 7.400E-09 | 1.000E-08 |
| 12443 | 623 | D623N | B | 1.2e-09 | 6.7e-09 | 4.2e-09 | 4.033E-09 | 2.754E-09 | 1.200E-09 | 6.700E-09 |
| 12444 | 623 | D623D | B | 8.8e-09 | 4.4e-09 | 5e-09 | 6.067E-09 | 2.386E-09 | 4.400E-09 | 8.800E-09 |
| 12445 | 623 | D623C | B | 1.7e-09 | 1e-08 | 1.3e-08 | 8.233E-09 | 5.853E-09 | 1.700E-09 | 1.300E-08 |
| 12446 | 623 | D623E | B | 1.1e-08 | 3.1e-09 | 1e-08 | 8.033E-09 | 4.302E-09 | 3.100E-09 | 1.100E-08 |
| 12447 | 623 | D623Q | B | 1.6e-09 | 3.9e-09 | 3.7e-09 | 3.067E-09 | 1.274E-09 | 1.600E-09 | 3.900E-09 |
| 12448 | 623 | D623G | B | 2.4e-09 | 7.5e-09 | 4.8e-09 | 4.900E-09 | 2.551E-09 | 2.400E-09 | 7.500E-09 |
| 12449 | 623 | D623H | B | 1.1e-08 | 6.7e-09 | 8.2e-09 | 8.633E-09 | 2.183E-09 | 6.700E-09 | 1.100E-08 |
| 12450 | 623 | D623I | B | 1e-08 | 1.2e-08 | 1.7e-08 | 1.300E-08 | 3.606E-09 | 1.000E-08 | 1.700E-08 |
| 12451 | 623 | D623L | B | 4.6e-09 | 3.8e-09 | 1.6e-08 | 8.133E-09 | 6.824E-09 | 3.800E-09 | 1.600E-08 |
| 12452 | 623 | D623K | B | 1.8e-09 | 1.5e-08 | 8.1e-09 | 8.300E-09 | 6.602E-09 | 1.800E-09 | 1.500E-08 |
| 12453 | 623 | D623M | B | 5.1e-09 | 5.3e-09 | 1.6e-08 | 8.800E-09 | 6.236E-09 | 5.100E-09 | 1.600E-08 |
| 12454 | 623 | D623F | B | 6.8e-09 | 2.2e-09 | 8.5e-09 | 5.833E-09 | 3.259E-09 | 2.200E-09 | 8.500E-09 |
| 12455 | 623 | D623P | B | 5.5e-09 | 1.4e-08 | 7.4e-09 | 8.967E-09 | 4.461E-09 | 5.500E-09 | 1.400E-08 |
| 12456 | 623 | D623S | B | 4.7e-09 | 1.4e-08 | 1.7e-08 | 1.190E-08 | 6.413E-09 | 4.700E-09 | 1.700E-08 |
| 12457 | 623 | D623T | B | 2e-08 | 9.9e-09 | 6.6e-09 | 1.217E-08 | 6.982E-09 | 6.600E-09 | 2.000E-08 |
| 12458 | 623 | D623W | B | 1.1e-08 | 7.9e-09 | 7.2e-09 | 8.700E-09 | 2.022E-09 | 7.200E-09 | 1.100E-08 |
| 12459 | 623 | D623Y | B | 1.4e-08 | 4.3e-09 | 6.4e-09 | 8.233E-09 | 5.103E-09 | 4.300E-09 | 1.400E-08 |
| 12460 | 623 | D623V | B | 4e-09 | 5.5e-09 | 1.3e-08 | 7.500E-09 | 4.822E-09 | 4.000E-09 | 1.300E-08 |
| 12461 | 624 | I624A | B | 3.7e-08 | 1.3e-08 | 9e-09 | 1.967E-08 | 1.514E-08 | 9.000E-09 | 3.700E-08 |
| 12462 | 624 | I624R | B | 1.9e-09 | 4.4e-09 | 1.2e-08 | 6.100E-09 | 5.260E-09 | 1.900E-09 | 1.200E-08 |
| 12463 | 624 | I624N | B | 3.6e-09 | 8e-09 | 3.9e-09 | 5.167E-09 | 2.458E-09 | 3.600E-09 | 8.000E-09 |
| 12464 | 624 | I624D | B | 1.3e-08 | 7.7e-09 | 3.3e-09 | 8.000E-09 | 4.857E-09 | 3.300E-09 | 1.300E-08 |
| 12465 | 624 | I624C | B | 1.4e-08 | 2.3e-08 | 6.3e-09 | 1.443E-08 | 8.358E-09 | 6.300E-09 | 2.300E-08 |
| 12466 | 624 | I624E | B | 4.6e-09 | 1.1e-08 | 2e-08 | 1.187E-08 | 7.736E-09 | 4.600E-09 | 2.000E-08 |
| 12467 | 624 | I624Q | B | 1e-08 | 6e-09 | 2.5e-08 | 1.367E-08 | 1.002E-08 | 6.000E-09 | 2.500E-08 |
| 12468 | 624 | I624G | B | 8.9e-09 | 4.3e-09 | 1.4e-08 | 9.067E-09 | 4.852E-09 | 4.300E-09 | 1.400E-08 |
| 12469 | 624 | I624H | B | 8.9e-09 | 4.3e-09 | 8.5e-09 | 7.233E-09 | 2.548E-09 | 4.300E-09 | 8.900E-09 |
| 12470 | 624 | I624I | B | 1.9e-09 | 5e-09 | 7.5e-09 | 4.800E-09 | 2.805E-09 | 1.900E-09 | 7.500E-09 |
| 12471 | 624 | I624L | B | 7.8e-09 | 2.6e-09 | 6.6e-09 | 5.667E-09 | 2.723E-09 | 2.600E-09 | 7.800E-09 |
| 12472 | 624 | I624K | B | 4.1e-09 | 7e-09 | 7e-09 | 6.033E-09 | 1.674E-09 | 4.100E-09 | 7.000E-09 |
| 12473 | 624 | I624M | B | 1.2e-08 | 5.7e-09 | 6e-09 | 7.900E-09 | 3.554E-09 | 5.700E-09 | 1.200E-08 |
| 12474 | 624 | I624F | B | 1e-08 | 7.2e-09 | 9.3e-09 | 8.833E-09 | 1.457E-09 | 7.200E-09 | 1.000E-08 |
| 12475 | 624 | I624P | B | 3.2e-09 | 1.4e-08 | 1.4e-09 | 6.200E-09 | 6.815E-09 | 1.400E-09 | 1.400E-08 |
| 12476 | 624 | I624S | B | 5.9e-09 | 1.1e-08 | 1.3e-08 | 9.967E-09 | 3.661E-09 | 5.900E-09 | 1.300E-08 |
| 12477 | 624 | I624T | B | 5.1e-09 | 3.2e-09 | 4.4e-09 | 4.233E-09 | 9.609E-10 | 3.200E-09 | 5.100E-09 |
| 12478 | 624 | I624W | B | 4.3e-09 | 4e-09 | 1.4e-08 | 7.433E-09 | 5.689E-09 | 4.000E-09 | 1.400E-08 |
| 12479 | 624 | I624Y | B | 9.3e-09 | 1.2e-08 | 8.8e-09 | 1.003E-08 | 1.721E-09 | 8.800E-09 | 1.200E-08 |
| 12480 | 624 | I624V | B | 1.7e-08 | 2.7e-09 | 6.2e-09 | 8.633E-09 | 7.454E-09 | 2.700E-09 | 1.700E-08 |
| 12481 | 625 | Q625A | B | 4.5e-09 | 9.7e-09 | 1.3e-08 | 9.067E-09 | 4.285E-09 | 4.500E-09 | 1.300E-08 |
| 12482 | 625 | Q625R | B | 7.9e-09 | 5.2e-09 | 6.1e-09 | 6.400E-09 | 1.375E-09 | 5.200E-09 | 7.900E-09 |
| 12483 | 625 | Q625N | B | 3.5e-09 | 1.8e-08 | 9.6e-09 | 1.037E-08 | 7.280E-09 | 3.500E-09 | 1.800E-08 |
| 12484 | 625 | Q625D | B | 2.8e-09 | 6.7e-09 | 1.2e-09 | 3.567E-09 | 2.829E-09 | 1.200E-09 | 6.700E-09 |
| 12485 | 625 | Q625C | B | 9.1e-09 | 5.1e-09 | 3.8e-09 | 6.000E-09 | 2.762E-09 | 3.800E-09 | 9.100E-09 |
| 12486 | 625 | Q625E | B | 1.4e-08 | 6.7e-09 | 8.4e-09 | 9.700E-09 | 3.820E-09 | 6.700E-09 | 1.400E-08 |
| 12487 | 625 | Q625Q | B | 5.5e-09 | 1.1e-08 | 7.4e-09 | 7.967E-09 | 2.793E-09 | 5.500E-09 | 1.100E-08 |
| 12488 | 625 | Q625G | B | 9.4e-09 | 4.4e-09 | 2.1e-08 | 1.160E-08 | 8.516E-09 | 4.400E-09 | 2.100E-08 |
| 12489 | 625 | Q625H | B | 7.8e-09 | 3.6e-09 | 1.3e-08 | 8.133E-09 | 4.709E-09 | 3.600E-09 | 1.300E-08 |
| 12490 | 625 | Q625I | B | 8.4e-09 | 5e-09 | 1.5e-08 | 9.467E-09 | 5.085E-09 | 5.000E-09 | 1.500E-08 |
| 12491 | 625 | Q625L | B | 7.5e-09 | 5.7e-09 | 7.9e-09 | 7.033E-09 | 1.172E-09 | 5.700E-09 | 7.900E-09 |
| 12492 | 625 | Q625K | B | 7.6e-09 | 1.4e-08 | 1.9e-08 | 1.353E-08 | 5.714E-09 | 7.600E-09 | 1.900E-08 |
| 12493 | 625 | Q625M | B | 8.4e-09 | 1.4e-08 | 7.1e-09 | 9.833E-09 | 3.667E-09 | 7.100E-09 | 1.400E-08 |
| 12494 | 625 | Q625F | B | 7e-09 | 5.6e-09 | 1.7e-08 | 9.867E-09 | 6.217E-09 | 5.600E-09 | 1.700E-08 |

| | | | | | | | | | | |
|---|---|---|---|---|---|---|---|---|---|---|
| 12495 | 625 | Q625P | B | 1e-08 | 1.3e-08 | 3e-09 | 8.667E-09 | 5.132E-09 | 3.000E-09 | 1.300E-08 |
| 12496 | 625 | Q625S | B | 5.6e-09 | 5.1e-09 | 5.9e-09 | 5.533E-09 | 4.041E-10 | 5.100E-09 | 5.900E-09 |
| 12497 | 625 | Q625T | B | 7.2e-09 | 1e-08 | 3.1e-08 | 1.607E-08 | 1.301E-08 | 7.200E-09 | 3.100E-08 |
| 12498 | 625 | Q625W | B | 8.1e-09 | 1.6e-08 | 7e-09 | 1.037E-08 | 4.910E-09 | 7.000E-09 | 1.600E-08 |
| 12499 | 625 | Q625Y | B | 3.1e-09 | 5.1e-09 | 1.1e-08 | 6.400E-09 | 4.107E-09 | 3.100E-09 | 1.100E-08 |
| 12500 | 625 | Q625V | B | 2.5e-09 | 6.1e-09 | 4.9e-09 | 4.500E-09 | 1.833E-09 | 2.500E-09 | 6.100E-09 |
| 12501 | 626 | M626A | B | 7.2e-09 | 4.1e-09 | 5.6e-09 | 5.633E-09 | 1.550E-09 | 4.100E-09 | 7.200E-09 |
| 12502 | 626 | M626R | B | 3.9e-09 | 1.5e-08 | 9.4e-09 | 9.433E-09 | 5.550E-09 | 3.900E-09 | 1.500E-08 |
| 12503 | 626 | M626N | B | 7.8e-09 | 2.5e-09 | 6e-09 | 5.433E-09 | 2.695E-09 | 2.500E-09 | 7.800E-09 |
| 12504 | 626 | M626D | B | 2e-08 | 6.8e-09 | 1e-08 | 1.227E-08 | 6.886E-09 | 6.800E-09 | 2.000E-08 |
| 12505 | 626 | M626C | B | 1.1e-08 | 1.1e-08 | 3.3e-09 | 8.433E-09 | 4.446E-09 | 3.300E-09 | 1.100E-08 |
| 12506 | 626 | M626E | B | 1.1e-08 | 4.3e-09 | 6.7e-09 | 7.333E-09 | 3.395E-09 | 4.300E-09 | 1.100E-08 |
| 12507 | 626 | M626Q | B | 2.3e-09 | 3.2e-09 | 7.5e-09 | 4.333E-09 | 2.779E-09 | 2.300E-09 | 7.500E-09 |
| 12508 | 626 | M626G | B | 7.9e-09 | 8.8e-09 | 2.9e-09 | 6.533E-09 | 3.179E-09 | 2.900E-09 | 8.800E-09 |
| 12509 | 626 | M626H | B | 3.5e-09 | 4.4e-09 | 7.1e-09 | 5.000E-09 | 1.873E-09 | 3.500E-09 | 7.100E-09 |
| 12510 | 626 | M626I | B | 1.3e-08 | 7.5e-09 | 8.4e-09 | 9.633E-09 | 2.950E-09 | 7.500E-09 | 1.300E-08 |
| 12511 | 626 | M626L | B | 1e-08 | 1.7e-08 | 6.3e-09 | 1.110E-08 | 5.434E-09 | 6.300E-09 | 1.700E-08 |
| 12512 | 626 | M626K | B | 1.9e-08 | 8.8e-09 | 4.5e-09 | 1.077E-08 | 7.447E-09 | 4.500E-09 | 1.900E-08 |
| 12513 | 626 | M626M | B | 7.7e-09 | 1.6e-09 | 3.8e-09 | 4.367E-09 | 3.089E-09 | 1.600E-09 | 7.700E-09 |
| 12514 | 626 | M626F | B | 2.2e-09 | 1.1e-08 | 2.1e-09 | 5.100E-09 | 5.110E-09 | 2.100E-09 | 1.100E-08 |
| 12515 | 626 | M626P | B | 1.3e-08 | 7.4e-09 | 4.1e-09 | 8.167E-09 | 4.499E-09 | 4.100E-09 | 1.300E-08 |
| 12516 | 626 | M626S | B | 6.7e-09 | 2.9e-09 | 5.2e-09 | 4.933E-09 | 1.914E-09 | 2.900E-09 | 6.700E-09 |
| 12517 | 626 | M626T | B | 5.4e-09 | 6.3e-09 | 6.8e-09 | 6.167E-09 | 7.095E-10 | 5.400E-09 | 6.800E-09 |
| 12518 | 626 | M626W | B | 2.6e-09 | 3.1e-09 | 5.9e-09 | 3.867E-09 | 1.779E-09 | 2.600E-09 | 5.900E-09 |
| 12519 | 626 | M626Y | B | 2e-08 | 7.3e-09 | 8.3e-09 | 1.187E-08 | 7.061E-09 | 7.300E-09 | 2.000E-08 |
| 12520 | 626 | M626V | B | 8.5e-09 | 8.1e-09 | 9.1e-09 | 8.567E-09 | 5.033E-10 | 8.100E-09 | 9.100E-09 |
| 12521 | 627 | T627A | B | 4.9e-09 | 3.6e-09 | 5.7e-09 | 4.733E-09 | 1.060E-09 | 3.600E-09 | 5.700E-09 |
| 12522 | 627 | T627R | B | 1e-08 | 4.4e-09 | 1.1e-08 | 8.467E-09 | 3.557E-09 | 4.400E-09 | 1.100E-08 |
| 12523 | 627 | T627N | B | 3.3e-09 | 3.9e-09 | 1.3e-08 | 6.733E-09 | 5.435E-09 | 3.300E-09 | 1.300E-08 |
| 12524 | 627 | T627D | B | 6.6e-09 | 1.3e-08 | 1.3e-08 | 1.087E-08 | 3.695E-09 | 6.600E-09 | 1.300E-08 |
| 12525 | 627 | T627C | B | 3.8e-09 | 1.8e-08 | 4.7e-09 | 8.833E-09 | 7.951E-09 | 3.800E-09 | 1.800E-08 |
| 12526 | 627 | T627E | B | 4.7e-09 | 5.4e-09 | 4.9e-09 | 5.000E-09 | 3.606E-10 | 4.700E-09 | 5.400E-09 |
| 12527 | 627 | T627Q | B | 3.6e-08 | 4.9e-09 | 1.5e-09 | 1.413E-08 | 1.901E-08 | 1.500E-09 | 3.600E-08 |
| 12528 | 627 | T627G | B | 1.1e-08 | 1.3e-08 | 1.6e-08 | 1.333E-08 | 2.517E-09 | 1.100E-08 | 1.600E-08 |
| 12529 | 627 | T627H | B | 1e-08 | 2e-09 | 2.5e-09 | 4.833E-09 | 4.481E-09 | 2.000E-09 | 1.000E-08 |
| 12530 | 627 | T627I | B | 9.4e-09 | 8.1e-09 | 6.6e-09 | 8.033E-09 | 1.401E-09 | 6.600E-09 | 9.400E-09 |
| 12531 | 627 | T627L | B | 2.3e-09 | 3.4e-09 | 7.1e-09 | 4.267E-09 | 2.515E-09 | 2.300E-09 | 7.100E-09 |
| 12532 | 627 | T627K | B | 6.6e-09 | 1.1e-08 | 1e-08 | 9.200E-09 | 2.307E-09 | 6.600E-09 | 1.100E-08 |
| 12533 | 627 | T627M | B | 4.4e-09 | 4.6e-09 | 9.9e-09 | 6.300E-09 | 3.119E-09 | 4.400E-09 | 9.900E-09 |
| 12534 | 627 | T627F | B | 1.1e-08 | 1.3e-08 | 3.1e-09 | 9.033E-09 | 5.235E-09 | 3.100E-09 | 1.300E-08 |
| 12535 | 627 | T627P | B | 3.1e-09 | 6.5e-09 | 3.2e-09 | 4.267E-09 | 1.935E-09 | 3.100E-09 | 6.500E-09 |
| 12536 | 627 | T627S | B | 1.1e-08 | 9.8e-09 | 1.4e-08 | 1.160E-08 | 2.163E-09 | 9.800E-09 | 1.400E-08 |
| 12537 | 627 | T627T | B | 3.8e-09 | 3e-09 | 3.5e-09 | 3.433E-09 | 4.041E-10 | 3.000E-09 | 3.800E-09 |
| 12538 | 627 | T627W | B | 1.1e-08 | 1.3e-08 | 8e-09 | 1.067E-08 | 2.517E-09 | 8.000E-09 | 1.300E-08 |
| 12539 | 627 | T627Y | B | 3.3e-09 | 1.1e-08 | 1.1e-08 | 8.433E-09 | 4.446E-09 | 3.300E-09 | 1.100E-08 |
| 12540 | 627 | T627V | B | 9.9e-09 | 9e-09 | 9.2e-09 | 9.367E-09 | 4.726E-10 | 9.000E-09 | 9.900E-09 |
| 12541 | 628 | Q628A | B | 4.4e-09 | 8.2e-09 | 1.8e-08 | 1.020E-08 | 7.017E-09 | 4.400E-09 | 1.800E-08 |
| 12542 | 628 | Q628R | B | 6.1e-09 | 3e-09 | 5.7e-09 | 4.933E-09 | 1.686E-09 | 3.000E-09 | 6.100E-09 |
| 12543 | 628 | Q628N | B | 7.4e-09 | 1.7e-08 | 8.6e-09 | 1.100E-08 | 5.231E-09 | 7.400E-09 | 1.700E-08 |
| 12544 | 628 | Q628D | B | 7.7e-09 | 1.2e-08 | 7.3e-09 | 9.000E-09 | 2.606E-09 | 7.300E-09 | 1.200E-08 |
| 12545 | 628 | Q628C | B | 9.8e-09 | 1.7e-08 | 4.5e-09 | 1.043E-08 | 6.274E-09 | 4.500E-09 | 1.700E-08 |
| 12546 | 628 | Q628E | B | 9.2e-09 | 5.6e-09 | 1.3e-08 | 9.267E-09 | 3.700E-09 | 5.600E-09 | 1.300E-08 |
| 12547 | 628 | Q628Q | B | 4.5e-09 | 1.3e-09 | 1.2e-08 | 5.933E-09 | 5.492E-09 | 1.300E-09 | 1.200E-08 |
| 12548 | 628 | Q628G | B | 8.3e-09 | 4.1e-09 | 6.3e-09 | 6.233E-09 | 2.101E-09 | 4.100E-09 | 8.300E-09 |
| 12549 | 628 | Q628H | B | 3.7e-09 | 7.4e-09 | 6.3e-09 | 5.800E-09 | 1.900E-09 | 3.700E-09 | 7.400E-09 |
| 12550 | 628 | Q628I | B | 5.9e-09 | 1.3e-08 | 3.7e-09 | 7.533E-09 | 4.860E-09 | 3.700E-09 | 1.300E-08 |
| 12551 | 628 | Q628L | B | 7.9e-09 | 3.9e-09 | 5.4e-09 | 5.733E-09 | 2.021E-09 | 3.900E-09 | 7.900E-09 |
| 12552 | 628 | Q628K | B | 8.4e-09 | 6.1e-09 | 2.2e-08 | 1.217E-08 | 8.593E-09 | 6.100E-09 | 2.200E-08 |
| 12553 | 628 | Q628M | B | 8.2e-09 | 6.9e-09 | 6.3e-09 | 7.133E-09 | 9.713E-10 | 6.300E-09 | 8.200E-09 |
| 12554 | 628 | Q628F | B | 7.6e-09 | 1.2e-08 | 2e-08 | 1.320E-08 | 6.286E-09 | 7.600E-09 | 2.000E-08 |
| 12555 | 628 | Q628P | B | 1e-08 | 1.5e-08 | 7.3e-09 | 1.077E-08 | 3.907E-09 | 7.300E-09 | 1.500E-08 |
| 12556 | 628 | Q628S | B | 2.3e-09 | 6.6e-09 | 5.9e-09 | 4.933E-09 | 2.307E-09 | 2.300E-09 | 6.600E-09 |
| 12557 | 628 | Q628T | B | 1e-08 | 1e-08 | 2.2e-08 | 1.400E-08 | 6.928E-09 | 1.000E-08 | 2.200E-08 |
| 12558 | 628 | Q628W | B | 4.6e-09 | 5.7e-09 | 6e-09 | 5.433E-09 | 7.371E-10 | 4.600E-09 | 6.000E-09 |
| 12559 | 628 | Q628Y | B | 1.2e-08 | 2e-08 | 1.3e-08 | 1.500E-08 | 4.359E-09 | 1.200E-08 | 2.000E-08 |
| 12560 | 628 | Q628V | B | 1.5e-09 | 8.2e-09 | 8.4e-09 | 6.033E-09 | 3.927E-09 | 1.500E-09 | 8.400E-09 |
| 12561 | 629 | S629A | B | 2.5e-09 | 1.1e-08 | 5.4e-09 | 6.300E-09 | 4.321E-09 | 2.500E-09 | 1.100E-08 |
| 12562 | 629 | S629R | B | 2.6e-08 | 8.5e-09 | 4.9e-09 | 1.313E-08 | 1.129E-08 | 4.900E-09 | 2.600E-08 |
| 12563 | 629 | S629N | B | 7.4e-09 | 3e-08 | 1.2e-08 | 1.647E-08 | 1.194E-08 | 7.400E-09 | 3.000E-08 |
| 12564 | 629 | S629D | B | 8.3e-09 | 1.6e-08 | 6.8e-09 | 1.037E-08 | 4.936E-09 | 6.800E-09 | 1.600E-08 |
| 12565 | 629 | S629C | B | 5e-09 | 1.3e-08 | 1.3e-08 | 1.033E-08 | 4.619E-09 | 5.000E-09 | 1.300E-08 |
| 12566 | 629 | S629E | B | 1.8e-08 | 1.1e-08 | 9.5e-09 | 1.283E-08 | 4.537E-09 | 9.500E-09 | 1.800E-08 |
| 12567 | 629 | S629Q | B | 1.6e-08 | 1.4e-08 | 7e-09 | 1.233E-08 | 4.726E-09 | 7.000E-09 | 1.600E-08 |
| 12568 | 629 | S629G | B | 4.5e-09 | 1.8e-08 | 1e-08 | 1.083E-08 | 6.788E-09 | 4.500E-09 | 1.800E-08 |
| 12569 | 629 | S629H | B | 4.5e-09 | 5.1e-09 | 9.6e-09 | 6.400E-09 | 2.787E-09 | 4.500E-09 | 9.600E-09 |
| 12570 | 629 | S629I | B | 5.2e-09 | 9.4e-09 | 3.2e-09 | 5.933E-09 | 3.164E-09 | 3.200E-09 | 9.400E-09 |
| 12571 | 629 | S629L | B | 1.9e-08 | 1e-08 | 6e-09 | 1.167E-08 | 6.658E-09 | 6.000E-09 | 1.900E-08 |
| 12572 | 629 | S629K | B | 1.2e-08 | 7e-09 | 1.3e-08 | 1.067E-08 | 3.215E-09 | 7.000E-09 | 1.300E-08 |
| 12573 | 629 | S629M | B | 9.2e-09 | 2e-09 | 8.9e-09 | 6.700E-09 | 4.073E-09 | 2.000E-09 | 9.200E-09 |
| 12574 | 629 | S629F | B | 3e-09 | 7.1e-09 | 2.6e-09 | 4.233E-09 | 2.491E-09 | 2.600E-09 | 7.100E-09 |
| 12575 | 629 | S629P | B | 1.3e-08 | 8.9e-09 | 5.4e-09 | 9.100E-09 | 3.804E-09 | 5.400E-09 | 1.300E-08 |
| 12576 | 629 | S629S | B | 6.4e-09 | 6.4e-09 | 1e-09 | 4.600E-09 | 3.118E-09 | 1.000E-09 | 6.400E-09 |
| 12577 | 629 | S629T | B | 4.6e-09 | 1.3e-08 | 1.4e-08 | 1.053E-08 | 5.163E-09 | 4.600E-09 | 1.400E-08 |
| 12578 | 629 | S629W | B | 6e-09 | 1.5e-08 | 9.6e-09 | 1.020E-08 | 4.530E-09 | 6.000E-09 | 1.500E-08 |
| 12579 | 629 | S629Y | B | 1.1e-08 | 6.4e-09 | 1.1e-08 | 9.467E-09 | 2.656E-09 | 6.400E-09 | 1.100E-08 |

| | | | | | | | | | | |
|---|---|---|---|---|---|---|---|---|---|---|
| 12580 | 629 | S629V | B | 1.2e-08 | 9.6e-09 | 9.8e-09 | 1.047E-08 | 1.332E-09 | 9.600E-09 | 1.200E-08 |
| 12581 | 630 | P630A | B | 7.2e-09 | 9.4e-09 | 6.4e-09 | 7.667E-09 | 1.553E-09 | 6.400E-09 | 9.400E-09 |
| 12582 | 630 | P630R | B | 3.5e-08 | 7.7e-09 | 1.3e-08 | 1.857E-08 | 1.448E-08 | 7.700E-09 | 3.500E-08 |
| 12583 | 630 | P630N | B | 1.2e-08 | 6.8e-09 | 6.4e-09 | 8.400E-09 | 3.124E-09 | 6.400E-09 | 1.200E-08 |
| 12584 | 630 | P630D | B | 1.2e-08 | 9e-09 | 1e-08 | 1.033E-08 | 1.528E-09 | 9.000E-09 | 1.200E-08 |
| 12585 | 630 | P630C | B | 2.2e-08 | 9.1e-09 | 1.1e-08 | 1.403E-08 | 6.964E-09 | 9.100E-09 | 2.200E-08 |
| 12586 | 630 | P630E | B | 1e-08 | 5.5e-09 | 1.6e-08 | 1.050E-08 | 5.268E-09 | 5.500E-09 | 1.600E-08 |
| 12587 | 630 | P630Q | B | 1e-08 | 3.5e-09 | 1.1e-08 | 8.167E-09 | 4.072E-09 | 3.500E-09 | 1.100E-08 |
| 12588 | 630 | P630G | B | 5.8e-09 | 2e-08 | 4.2e-09 | 1.000E-08 | 8.697E-09 | 4.200E-09 | 2.000E-08 |
| 12589 | 630 | P630H | B | 6.7e-09 | 2.9e-09 | 8.5e-09 | 6.033E-09 | 2.859E-09 | 2.900E-09 | 8.500E-09 |
| 12590 | 630 | P630I | B | 1.5e-08 | 4.2e-09 | 3.2e-08 | 1.707E-08 | 1.401E-08 | 4.200E-09 | 3.200E-08 |
| 12591 | 630 | P630L | B | 4.6e-09 | 5.3e-09 | 1.2e-08 | 7.300E-09 | 4.085E-09 | 4.600E-09 | 1.200E-08 |
| 12592 | 630 | P630K | B | 4.3e-09 | 2.7e-09 | 9.4e-09 | 5.467E-09 | 3.499E-09 | 2.700E-09 | 9.400E-09 |
| 12593 | 630 | P630M | B | 1.8e-09 | 5.9e-09 | 9.4e-09 | 5.700E-09 | 3.804E-09 | 1.800E-09 | 9.400E-09 |
| 12594 | 630 | P630F | B | 6.3e-09 | 2.1e-08 | 4.7e-09 | 1.067E-08 | 8.985E-09 | 4.700E-09 | 2.100E-08 |
| 12595 | 630 | P630P | B | 1.1e-08 | 2.6e-09 | 5.4e-09 | 6.333E-09 | 4.277E-09 | 2.600E-09 | 1.100E-08 |
| 12596 | 630 | P630S | B | 3.5e-09 | 3.3e-09 | 1.2e-08 | 6.267E-09 | 4.966E-09 | 3.300E-09 | 1.200E-08 |
| 12597 | 630 | P630T | B | 8.8e-09 | 5.8e-09 | 8.5e-09 | 7.700E-09 | 1.652E-09 | 5.800E-09 | 8.800E-09 |
| 12598 | 630 | P630W | B | 1e-08 | 9.9e-10 | 4.4e-09 | 5.130E-09 | 4.549E-09 | 9.900E-10 | 1.000E-08 |
| 12599 | 630 | P630Y | B | 6.8e-09 | 1.5e-08 | 1.4e-08 | 1.193E-08 | 4.474E-09 | 6.800E-09 | 1.500E-08 |
| 12600 | 630 | P630V | B | 2.4e-09 | 4.7e-09 | 1.2e-08 | 6.367E-09 | 5.012E-09 | 2.400E-09 | 1.200E-08 |
| 12601 | 631 | S631A | B | 2.3e-09 | 1.2e-08 | 2.5e-08 | 1.310E-08 | 1.139E-08 | 2.300E-09 | 2.500E-08 |
| 12602 | 631 | S631R | B | 5.4e-09 | 1.7e-08 | 1.3e-08 | 1.180E-08 | 5.892E-09 | 5.400E-09 | 1.700E-08 |
| 12603 | 631 | S631N | B | 2.8e-09 | 5.2e-09 | 6.6e-09 | 4.867E-09 | 1.922E-09 | 2.800E-09 | 6.600E-09 |
| 12604 | 631 | S631D | B | 1.9e-08 | 8.7e-09 | 1e-08 | 1.257E-08 | 5.609E-09 | 8.700E-09 | 1.900E-08 |
| 12605 | 631 | S631C | B | 8.8e-09 | 9.8e-09 | 9.6e-09 | 9.400E-09 | 5.292E-10 | 8.800E-09 | 9.800E-09 |
| 12606 | 631 | S631E | B | 7.1e-09 | 9.1e-09 | 7.2e-09 | 7.800E-09 | 1.127E-09 | 7.100E-09 | 9.100E-09 |
| 12607 | 631 | S631Q | B | 1e-08 | 8.3e-09 | 3.2e-09 | 7.167E-09 | 3.539E-09 | 3.200E-09 | 1.000E-08 |
| 12608 | 631 | S631G | B | 1e-08 | 1.4e-08 | 1.3e-08 | 1.233E-08 | 2.082E-09 | 1.000E-08 | 1.400E-08 |
| 12609 | 631 | S631H | B | 1.2e-08 | 1.4e-08 | 1.1e-08 | 1.233E-08 | 1.528E-09 | 1.100E-08 | 1.400E-08 |
| 12610 | 631 | S631I | B | 3.6e-09 | 8e-09 | 3.5e-09 | 5.033E-09 | 2.570E-09 | 3.500E-09 | 8.000E-09 |
| 12611 | 631 | S631L | B | 5.7e-09 | 6.3e-09 | 7.5e-09 | 6.500E-09 | 9.165E-10 | 5.700E-09 | 7.500E-09 |
| 12612 | 631 | S631K | B | 6.2e-09 | 6e-09 | 8.2e-09 | 6.800E-09 | 1.217E-09 | 6.000E-09 | 8.200E-09 |
| 12613 | 631 | S631M | B | 1.2e-08 | 3.5e-09 | 1.4e-08 | 9.833E-09 | 5.575E-09 | 3.500E-09 | 1.400E-08 |
| 12614 | 631 | S631F | B | 1.5e-08 | 3.6e-09 | 6.1e-09 | 8.233E-09 | 5.992E-09 | 3.600E-09 | 1.500E-08 |
| 12615 | 631 | S631P | B | 8.8e-09 | 6.8e-09 | 6.1e-09 | 7.233E-09 | 1.401E-09 | 6.100E-09 | 8.800E-09 |
| 12616 | 631 | S631S | B | 8.3e-09 | 2.9e-09 | 2e-09 | 4.400E-09 | 3.407E-09 | 2.000E-09 | 8.300E-09 |
| 12617 | 631 | S631T | B | 8.1e-09 | 1.3e-08 | 4.1e-09 | 8.400E-09 | 4.458E-09 | 4.100E-09 | 1.300E-08 |
| 12618 | 631 | S631W | B | 8.3e-09 | 4.2e-09 | 8.9e-09 | 7.133E-09 | 2.558E-09 | 4.200E-09 | 8.900E-09 |
| 12619 | 631 | S631Y | B | 1.2e-08 | 2.9e-08 | 7.2e-09 | 1.607E-08 | 1.145E-08 | 7.200E-09 | 2.900E-08 |
| 12620 | 631 | S631V | B | 3.4e-09 | 1e-08 | 2.4e-08 | 1.247E-08 | 1.052E-08 | 3.400E-09 | 2.400E-08 |
| 12621 | 632 | S632A | B | 8e-09 | 1.2e-08 | 4.7e-09 | 8.233E-09 | 3.656E-09 | 4.700E-09 | 1.200E-08 |
| 12622 | 632 | S632R | B | 8.9e-09 | 4.4e-09 | 5.6e-09 | 6.300E-09 | 2.330E-09 | 4.400E-09 | 8.900E-09 |
| 12623 | 632 | S632N | B | 1e-08 | 3.5e-09 | 1.8e-08 | 1.050E-08 | 7.263E-09 | 3.500E-09 | 1.800E-08 |
| 12624 | 632 | S632D | B | 5.8e-09 | 3.9e-09 | 7.1e-09 | 5.600E-09 | 1.609E-09 | 3.900E-09 | 7.100E-09 |
| 12625 | 632 | S632C | B | 8.5e-09 | 8.1e-09 | 9.3e-09 | 8.633E-09 | 6.110E-10 | 8.100E-09 | 9.300E-09 |
| 12626 | 632 | S632E | B | 5.1e-09 | 1.3e-08 | 6.2e-09 | 8.100E-09 | 4.279E-09 | 5.100E-09 | 1.300E-08 |
| 12627 | 632 | S632Q | B | 3.8e-09 | 7.1e-09 | 3.4e-09 | 4.767E-09 | 2.031E-09 | 3.400E-09 | 7.100E-09 |
| 12628 | 632 | S632G | B | 8.9e-09 | 5.8e-09 | 1.2e-08 | 8.900E-09 | 3.100E-09 | 5.800E-09 | 1.200E-08 |
| 12629 | 632 | S632H | B | 5.8e-09 | 9.9e-09 | 5e-09 | 6.900E-09 | 2.629E-09 | 5.000E-09 | 9.900E-09 |
| 12630 | 632 | S632I | B | 6.3e-09 | 2.2e-09 | 3.4e-09 | 3.967E-09 | 2.108E-09 | 2.200E-09 | 6.300E-09 |
| 12631 | 632 | S632L | B | 9.2e-09 | 1.5e-08 | 6.7e-09 | 1.030E-08 | 4.258E-09 | 6.700E-09 | 1.500E-08 |
| 12632 | 632 | S632K | B | 6.4e-09 | 7.9e-09 | 7e-09 | 7.100E-09 | 7.550E-10 | 6.400E-09 | 7.900E-09 |
| 12633 | 632 | S632M | B | 6.3e-09 | 8.8e-09 | 6.6e-09 | 7.233E-09 | 1.365E-09 | 6.300E-09 | 8.800E-09 |
| 12634 | 632 | S632F | B | 1.3e-08 | 5.9e-09 | 7.7e-09 | 8.867E-09 | 3.691E-09 | 5.900E-09 | 1.300E-08 |
| 12635 | 632 | S632P | B | 2.2e-08 | 7.8e-09 | 1.2e-08 | 1.393E-08 | 7.295E-09 | 7.800E-09 | 2.200E-08 |
| 12636 | 632 | S632S | B | 7.6e-09 | 4.1e-09 | 8.8e-09 | 6.833E-09 | 2.442E-09 | 4.100E-09 | 8.800E-09 |
| 12637 | 632 | S632T | B | 8.1e-09 | 6.8e-09 | 8.4e-09 | 7.767E-09 | 8.505E-10 | 6.800E-09 | 8.400E-09 |
| 12638 | 632 | S632W | B | 7.1e-09 | 6.6e-09 | 6e-09 | 6.567E-09 | 5.508E-10 | 6.000E-09 | 7.100E-09 |
| 12639 | 632 | S632Y | B | 6.3e-09 | 3.5e-09 | 6.4e-09 | 5.400E-09 | 1.646E-09 | 3.500E-09 | 6.400E-09 |
| 12640 | 632 | S632V | B | 3.8e-09 | 1.5e-08 | 8.9e-09 | 9.233E-09 | 5.607E-09 | 3.800E-09 | 1.500E-08 |
| 12641 | 633 | L633A | B | 3.9e-09 | 2.1e-09 | 4.5e-09 | 3.500E-09 | 1.249E-09 | 2.100E-09 | 4.500E-09 |
| 12642 | 633 | L633R | B | 1.1e-08 | 2.7e-09 | 6.4e-09 | 6.700E-09 | 4.158E-09 | 2.700E-09 | 1.100E-08 |
| 12643 | 633 | L633N | B | 5.8e-09 | 1e-08 | 7.7e-09 | 7.833E-09 | 2.103E-09 | 5.800E-09 | 1.000E-08 |
| 12644 | 633 | L633D | B | 6.5e-09 | 4e-09 | 5.2e-09 | 5.233E-09 | 1.250E-09 | 4.000E-09 | 6.500E-09 |
| 12645 | 633 | L633C | B | 2.6e-08 | 1e-08 | 2.2e-08 | 1.933E-08 | 8.327E-09 | 1.000E-08 | 2.600E-08 |
| 12646 | 633 | L633E | B | 8.8e-09 | 2.8e-09 | 7.3e-09 | 6.300E-09 | 3.122E-09 | 2.800E-09 | 8.800E-09 |
| 12647 | 633 | L633Q | B | 4e-09 | 6.3e-09 | 7.1e-09 | 5.800E-09 | 1.609E-09 | 4.000E-09 | 7.100E-09 |
| 12648 | 633 | L633G | B | 3.3e-09 | 4.6e-09 | 1.7e-08 | 8.300E-09 | 7.562E-09 | 3.300E-09 | 1.700E-08 |
| 12649 | 633 | L633H | B | 4.5e-09 | 2.2e-09 | 1e-08 | 5.567E-09 | 4.008E-09 | 2.200E-09 | 1.000E-08 |
| 12650 | 633 | L633I | B | 1.6e-08 | 1.7e-08 | 1.2e-08 | 1.500E-08 | 2.646E-09 | 1.200E-08 | 1.700E-08 |
| 12651 | 633 | L633L | B | 9.3e-09 | 1.7e-08 | 5.2e-09 | 1.050E-08 | 5.991E-09 | 5.200E-09 | 1.700E-08 |
| 12652 | 633 | L633K | B | 1.2e-08 | 2.4e-09 | 3.5e-09 | 5.967E-09 | 5.254E-09 | 2.400E-09 | 1.200E-08 |
| 12653 | 633 | L633M | B | 9.7e-09 | 1.1e-08 | 1e-08 | 1.023E-08 | 6.807E-10 | 9.700E-09 | 1.100E-08 |
| 12654 | 633 | L633F | B | 5.2e-09 | 2.8e-09 | 7.4e-09 | 5.133E-09 | 2.301E-09 | 2.800E-09 | 7.400E-09 |
| 12655 | 633 | L633P | B | 1.2e-08 | 6.1e-09 | 7.1e-09 | 8.400E-09 | 3.158E-09 | 6.100E-09 | 1.200E-08 |
| 12656 | 633 | L633S | B | 3.7e-09 | 1e-08 | 4.8e-09 | 6.167E-09 | 3.365E-09 | 3.700E-09 | 1.000E-08 |
| 12657 | 633 | L633T | B | 5.9e-09 | 1.3e-08 | 1.9e-08 | 1.263E-08 | 6.558E-09 | 5.900E-09 | 1.900E-08 |
| 12658 | 633 | L633W | B | 2.8e-09 | 1.9e-08 | 2.4e-09 | 8.067E-09 | 9.471E-09 | 2.400E-09 | 1.900E-08 |
| 12659 | 633 | L633Y | B | 3.6e-09 | 1.6e-08 | 1.1e-08 | 1.020E-08 | 6.239E-09 | 3.600E-09 | 1.600E-08 |
| 12660 | 633 | L633V | B | 6.1e-09 | 4.7e-09 | 4.9e-09 | 5.233E-09 | 7.572E-10 | 4.700E-09 | 6.100E-09 |
| 12661 | 634 | S634A | B | 6.4e-09 | 8.7e-09 | 4e-09 | 6.367E-09 | 2.350E-09 | 4.000E-09 | 8.700E-09 |
| 12662 | 634 | S634R | B | 1.4e-08 | 1.1e-08 | 2.6e-08 | 1.700E-08 | 7.937E-09 | 1.100E-08 | 2.600E-08 |
| 12663 | 634 | S634N | B | 5.5e-09 | 1.2e-08 | 1.8e-08 | 1.183E-08 | 6.252E-09 | 5.500E-09 | 1.800E-08 |
| 12664 | 634 | S634D | B | 6.9e-09 | 2.6e-09 | 4.2e-09 | 4.567E-09 | 2.173E-09 | 2.600E-09 | 6.900E-09 |

| | | | | | | | | | | |
|---|---|---|---|---|---|---|---|---|---|---|
| 12665 | 634 | S634C | B | 5.2e-09 | 2.8e-09 | 7.3e-09 | 5.100E-09 | 2.252E-09 | 2.800E-09 | 7.300E-09 |
| 12666 | 634 | S634E | B | 1.9e-08 | 6.1e-09 | 7.5e-09 | 1.087E-08 | 7.078E-09 | 6.100E-09 | 1.900E-08 |
| 12667 | 634 | S634Q | B | 1.2e-08 | 1.1e-08 | 1.7e-09 | 8.233E-09 | 5.680E-09 | 1.700E-09 | 1.200E-08 |
| 12668 | 634 | S634G | B | 1.1e-08 | 5.2e-09 | 2.2e-08 | 1.273E-08 | 8.533E-09 | 5.200E-09 | 2.200E-08 |
| 12669 | 634 | S634H | B | 5e-09 | 1.1e-08 | 2.2e-09 | 6.067E-09 | 4.496E-09 | 2.200E-09 | 1.100E-08 |
| 12670 | 634 | S634I | B | 7e-09 | 1.4e-08 | 5.2e-09 | 8.733E-09 | 4.649E-09 | 5.200E-09 | 1.400E-08 |
| 12671 | 634 | S634L | B | 1.1e-08 | 2.9e-09 | 7.8e-09 | 7.233E-09 | 4.080E-09 | 2.900E-09 | 1.100E-08 |
| 12672 | 634 | S634K | B | 1.8e-09 | 9.6e-09 | 1.3e-08 | 8.133E-09 | 5.742E-09 | 1.800E-09 | 1.300E-08 |
| 12673 | 634 | S634M | B | 4.8e-09 | 4e-09 | 3.9e-09 | 4.233E-09 | 4.933E-10 | 3.900E-09 | 4.800E-09 |
| 12674 | 634 | S634F | B | 5.3e-09 | 7e-09 | 1.9e-08 | 1.043E-08 | 7.467E-09 | 5.300E-09 | 1.900E-08 |
| 12675 | 634 | S634P | B | 7e-09 | 1.3e-08 | 7.5e-09 | 9.167E-09 | 3.329E-09 | 7.000E-09 | 1.300E-08 |
| 12676 | 634 | S634S | B | 5.5e-09 | 6.3e-09 | 1.4e-08 | 8.600E-09 | 4.694E-09 | 5.500E-09 | 1.400E-08 |
| 12677 | 634 | S634T | B | 1.1e-08 | 7.5e-09 | 3.4e-09 | 7.300E-09 | 3.804E-09 | 3.400E-09 | 1.100E-08 |
| 12678 | 634 | S634W | B | 4.9e-09 | 1.4e-08 | 1e-08 | 9.633E-09 | 4.561E-09 | 4.900E-09 | 1.400E-08 |
| 12679 | 634 | S634Y | B | 1.3e-08 | 1.3e-08 | 3.9e-09 | 9.967E-09 | 5.254E-09 | 3.900E-09 | 1.300E-08 |
| 12680 | 634 | S634V | B | 1.8e-08 | 1.7e-08 | 5.3e-09 | 1.343E-08 | 7.061E-09 | 5.300E-09 | 1.800E-08 |
| 12681 | 635 | A635A | B | 4.7e-09 | 1.4e-08 | 8.6e-09 | 9.100E-09 | 4.670E-09 | 4.700E-09 | 1.400E-08 |
| 12682 | 635 | A635R | B | 2.3e-08 | 2.8e-08 | 1.4e-08 | 2.167E-08 | 7.095E-09 | 1.400E-08 | 2.800E-08 |
| 12683 | 635 | A635N | B | 3.8e-09 | 1.5e-08 | 5.7e-09 | 8.167E-09 | 5.994E-09 | 3.800E-09 | 1.500E-08 |
| 12684 | 635 | A635D | B | 5.9e-09 | 6.1e-09 | 2.6e-09 | 4.867E-09 | 1.966E-09 | 2.600E-09 | 6.100E-09 |
| 12685 | 635 | A635C | B | 4.2e-09 | 2.4e-09 | 6.8e-09 | 4.467E-09 | 2.212E-09 | 2.400E-09 | 6.800E-09 |
| 12686 | 635 | A635E | B | 7.6e-09 | 1.6e-08 | 7.2e-09 | 1.027E-08 | 4.969E-09 | 7.200E-09 | 1.600E-08 |
| 12687 | 635 | A635Q | B | 4.2e-09 | 2e-08 | 1.9e-09 | 8.700E-09 | 9.853E-09 | 1.900E-09 | 2.000E-08 |
| 12688 | 635 | A635G | B | 5.5e-09 | 6.4e-09 | 4.5e-09 | 5.467E-09 | 9.504E-10 | 4.500E-09 | 6.400E-09 |
| 12689 | 635 | A635H | B | 4.7e-09 | 1.5e-08 | 5.6e-09 | 8.433E-09 | 5.705E-09 | 4.700E-09 | 1.500E-08 |
| 12690 | 635 | A635I | B | 9.8e-09 | 1.1e-08 | 5e-09 | 8.600E-09 | 3.175E-09 | 5.000E-09 | 1.100E-08 |
| 12691 | 635 | A635L | B | 1.6e-08 | 3.6e-09 | 3.2e-09 | 7.600E-09 | 7.277E-09 | 3.200E-09 | 1.600E-08 |
| 12692 | 635 | A635K | B | 1.2e-08 | 3.2e-09 | 6.7e-09 | 7.300E-09 | 4.431E-09 | 3.200E-09 | 1.200E-08 |
| 12693 | 635 | A635M | B | 8.4e-09 | 6.7e-09 | 8.8e-09 | 7.967E-09 | 1.115E-09 | 6.700E-09 | 8.800E-09 |
| 12694 | 635 | A635F | B | 9.6e-09 | 1.1e-08 | 7.5e-09 | 9.367E-09 | 1.762E-09 | 7.500E-09 | 1.100E-08 |
| 12695 | 635 | A635P | B | 1.1e-08 | 8.2e-09 | 4.8e-09 | 8.000E-09 | 3.105E-09 | 4.800E-09 | 1.100E-08 |
| 12696 | 635 | A635S | B | 1.8e-09 | 3.2e-09 | 2.2e-09 | 2.400E-09 | 7.211E-10 | 1.800E-09 | 3.200E-09 |
| 12697 | 635 | A635T | B | 8e-09 | 1.4e-08 | 1.1e-08 | 1.100E-08 | 3.000E-09 | 8.000E-09 | 1.400E-08 |
| 12698 | 635 | A635W | B | 1.5e-08 | 8.1e-09 | 3.7e-09 | 8.933E-09 | 5.696E-09 | 3.700E-09 | 1.500E-08 |
| 12699 | 635 | A635Y | B | 1.7e-08 | 5e-09 | 5.2e-09 | 9.067E-09 | 6.871E-09 | 5.000E-09 | 1.700E-08 |
| 12700 | 635 | A635V | B | 6.8e-09 | 4.9e-09 | 4.2e-09 | 5.300E-09 | 1.345E-09 | 4.200E-09 | 6.800E-09 |
| 12701 | 636 | S636A | B | 1.2e-08 | 2e-08 | 9.8e-09 | 1.393E-08 | 5.368E-09 | 9.800E-09 | 2.000E-08 |
| 12702 | 636 | S636R | B | 2.1e-08 | 6.9e-09 | 9.9e-09 | 1.260E-08 | 7.428E-09 | 6.900E-09 | 2.100E-08 |
| 12703 | 636 | S636N | B | 5e-09 | 1.6e-08 | 8.6e-09 | 9.867E-09 | 5.608E-09 | 5.000E-09 | 1.600E-08 |
| 12704 | 636 | S636D | B | 9.5e-09 | 7.7e-09 | 4.3e-09 | 7.167E-09 | 2.641E-09 | 4.300E-09 | 9.500E-09 |
| 12705 | 636 | S636C | B | 4.2e-09 | 6.3e-09 | 9.7e-09 | 6.733E-09 | 2.775E-09 | 4.200E-09 | 9.700E-09 |
| 12706 | 636 | S636E | B | 1.1e-08 | 1e-08 | 7.6e-09 | 9.533E-09 | 1.747E-09 | 7.600E-09 | 1.100E-08 |
| 12707 | 636 | S636Q | B | 3.6e-09 | 1.2e-08 | 4.2e-09 | 6.600E-09 | 4.686E-09 | 3.600E-09 | 1.200E-08 |
| 12708 | 636 | S636G | B | 3.6e-09 | 1.4e-08 | 9.7e-09 | 9.100E-09 | 5.226E-09 | 3.600E-09 | 1.400E-08 |
| 12709 | 636 | S636H | B | 1.7e-08 | 5.9e-09 | 1.8e-08 | 1.363E-08 | 6.716E-09 | 5.900E-09 | 1.800E-08 |
| 12710 | 636 | S636I | B | 1.2e-08 | 8.1e-09 | 1.1e-08 | 1.037E-08 | 2.026E-09 | 8.100E-09 | 1.200E-08 |
| 12711 | 636 | S636L | B | 7.4e-09 | 9.5e-09 | 1.2e-08 | 9.633E-09 | 2.303E-09 | 7.400E-09 | 1.200E-08 |
| 12712 | 636 | S636K | B | 9.8e-09 | 6.3e-09 | 2.4e-09 | 6.167E-09 | 3.702E-09 | 2.400E-09 | 9.800E-09 |
| 12713 | 636 | S636M | B | 1.5e-08 | 1.3e-08 | 1.7e-09 | 9.900E-09 | 7.171E-09 | 1.700E-09 | 1.500E-08 |
| 12714 | 636 | S636F | B | 7.1e-09 | 1.6e-08 | 7.8e-09 | 1.030E-08 | 4.949E-09 | 7.100E-09 | 1.600E-08 |
| 12715 | 636 | S636P | B | 6.2e-09 | 3.8e-09 | 1e-08 | 6.667E-09 | 3.126E-09 | 3.800E-09 | 1.000E-08 |
| 12716 | 636 | S636S | B | 5.7e-09 | 1e-08 | 1.3e-08 | 9.567E-09 | 3.669E-09 | 5.700E-09 | 1.300E-08 |
| 12717 | 636 | S636T | B | 1.5e-08 | 7.4e-09 | 3.2e-09 | 8.533E-09 | 5.981E-09 | 3.200E-09 | 1.500E-08 |
| 12718 | 636 | S636W | B | 6.7e-09 | 3.5e-09 | 6.2e-09 | 5.467E-09 | 1.721E-09 | 3.500E-09 | 6.700E-09 |
| 12719 | 636 | S636Y | B | 8.6e-09 | 3.5e-09 | 2.7e-09 | 4.933E-09 | 3.201E-09 | 2.700E-09 | 8.600E-09 |
| 12720 | 636 | S636V | B | 1.6e-08 | 6.1e-09 | 9.1e-09 | 1.040E-08 | 5.076E-09 | 6.100E-09 | 1.600E-08 |
| 12721 | 637 | V637A | B | 7.8e-09 | 9.2e-09 | 8.6e-09 | 8.533E-09 | 7.024E-10 | 7.800E-09 | 9.200E-09 |
| 12722 | 637 | V637R | B | 7.3e-09 | 7.5e-09 | 1.3e-08 | 9.267E-09 | 3.235E-09 | 7.300E-09 | 1.300E-08 |
| 12723 | 637 | V637N | B | 1e-08 | 6.2e-09 | 1e-08 | 8.733E-09 | 2.194E-09 | 6.200E-09 | 1.000E-08 |
| 12724 | 637 | V637D | B | 1.8e-09 | 3e-09 | 3.9e-09 | 2.900E-09 | 1.054E-09 | 1.800E-09 | 3.900E-09 |
| 12725 | 637 | V637C | B | 9.9e-09 | 1.2e-08 | 9.4e-09 | 1.043E-08 | 1.380E-09 | 9.400E-09 | 1.200E-08 |
| 12726 | 637 | V637E | B | 1.8e-08 | 2.2e-08 | 1.8e-08 | 1.933E-08 | 2.309E-09 | 1.800E-08 | 2.200E-08 |
| 12727 | 637 | V637Q | B | 8e-09 | 2.7e-09 | 1.1e-08 | 7.233E-09 | 4.203E-09 | 2.700E-09 | 1.100E-08 |
| 12728 | 637 | V637G | B | 3.8e-09 | 5.6e-09 | 2.1e-09 | 3.833E-09 | 1.750E-09 | 2.100E-09 | 5.600E-09 |
| 12729 | 637 | V637H | B | 1.3e-08 | 4e-09 | 7.6e-09 | 8.200E-09 | 4.530E-09 | 4.000E-09 | 1.300E-08 |
| 12730 | 637 | V637I | B | 3.5e-09 | 1.2e-08 | 3.7e-09 | 6.400E-09 | 4.851E-09 | 3.500E-09 | 1.200E-08 |
| 12731 | 637 | V637L | B | 1.6e-08 | 7.8e-09 | 2.3e-09 | 8.700E-09 | 6.894E-09 | 2.300E-09 | 1.600E-08 |
| 12732 | 637 | V637K | B | 1.9e-09 | 1.3e-08 | 1.2e-09 | 5.367E-09 | 6.620E-09 | 1.200E-09 | 1.300E-08 |
| 12733 | 637 | V637M | B | 4.6e-09 | 5.4e-09 | 6.2e-09 | 5.400E-09 | 8.000E-10 | 4.600E-09 | 6.200E-09 |
| 12734 | 637 | V637F | B | 5.3e-09 | 1e-08 | 8.6e-09 | 7.967E-09 | 2.413E-09 | 5.300E-09 | 1.000E-08 |
| 12735 | 637 | V637P | B | 6.9e-09 | 6.5e-09 | 8.4e-09 | 7.267E-09 | 1.002E-09 | 6.500E-09 | 8.400E-09 |
| 12736 | 637 | V637S | B | 3e-09 | 9.3e-09 | 7.3e-09 | 6.533E-09 | 3.219E-09 | 3.000E-09 | 9.300E-09 |
| 12737 | 637 | V637T | B | 1.8e-08 | 4.9e-09 | 5.6e-09 | 9.500E-09 | 7.370E-09 | 4.900E-09 | 1.800E-08 |
| 12738 | 637 | V637W | B | 5.1e-09 | 2.7e-09 | 1.5e-08 | 7.600E-09 | 6.520E-09 | 2.700E-09 | 1.500E-08 |
| 12739 | 637 | V637Y | B | 1.6e-08 | 2.9e-09 | 5.8e-09 | 8.233E-09 | 6.881E-09 | 2.900E-09 | 1.600E-08 |
| 12740 | 637 | V637V | B | 6.6e-09 | 1.1e-08 | 2.6e-09 | 6.733E-09 | 4.202E-09 | 2.600E-09 | 1.100E-08 |
| 12741 | 638 | G638A | B | 5.3e-09 | 5.6e-09 | 3.7e-09 | 4.867E-09 | 1.021E-09 | 3.700E-09 | 5.600E-09 |
| 12742 | 638 | G638R | B | 1.6e-08 | 1.2e-08 | 3.3e-09 | 1.043E-08 | 6.493E-09 | 3.300E-09 | 1.600E-08 |
| 12743 | 638 | G638N | B | 1.2e-08 | 7.4e-09 | 2.8e-09 | 7.400E-09 | 4.600E-09 | 2.800E-09 | 1.200E-08 |
| 12744 | 638 | G638D | B | 3.1e-09 | 2e-08 | 1.9e-08 | 1.403E-08 | 9.482E-09 | 3.100E-09 | 2.000E-08 |
| 12745 | 638 | G638C | B | 2.8e-08 | 1.1e-08 | 3.3e-09 | 1.410E-08 | 1.264E-08 | 3.300E-09 | 2.800E-08 |
| 12746 | 638 | G638E | B | 2.7e-09 | 1.4e-08 | 1.3e-08 | 9.900E-09 | 6.255E-09 | 2.700E-09 | 1.400E-08 |
| 12747 | 638 | G638Q | B | 6.6e-09 | 7.2e-09 | 1.6e-08 | 9.933E-09 | 5.262E-09 | 6.600E-09 | 1.600E-08 |
| 12748 | 638 | G638G | B | 7.7e-09 | 3.2e-09 | 7.9e-09 | 6.267E-09 | 2.658E-09 | 3.200E-09 | 7.900E-09 |
| 12749 | 638 | G638H | B | 9.4e-09 | 6.6e-09 | 4.3e-09 | 6.767E-09 | 2.554E-09 | 4.300E-09 | 9.400E-09 |

| | | | | | | | | | | |
|---|---|---|---|---|---|---|---|---|---|---|
| 12750 | 638 | G638I | B | 1e-09 | 3.3e-09 | 7.2e-09 | 3.833E-09 | 3.134E-09 | 1.000E-09 | 7.200E-09 |
| 12751 | 638 | G638L | B | 1.1e-08 | 9.5e-09 | 7.7e-09 | 9.400E-09 | 1.652E-09 | 7.700E-09 | 1.100E-08 |
| 12752 | 638 | G638K | B | 6.7e-09 | 4.4e-09 | 3.6e-09 | 4.900E-09 | 1.609E-09 | 3.600E-09 | 6.700E-09 |
| 12753 | 638 | G638M | B | 4.4e-09 | 1.7e-08 | 5.7e-09 | 9.033E-09 | 6.930E-09 | 4.400E-09 | 1.700E-08 |
| 12754 | 638 | G638F | B | 4.1e-09 | 2.5e-09 | 6.5e-09 | 4.367E-09 | 2.013E-09 | 2.500E-09 | 6.500E-09 |
| 12755 | 638 | G638P | B | 7.9e-09 | 8.7e-09 | 1.2e-08 | 9.533E-09 | 2.173E-09 | 7.900E-09 | 1.200E-08 |
| 12756 | 638 | G638S | B | 5.5e-09 | 4.2e-09 | 1.9e-09 | 3.867E-09 | 1.823E-09 | 1.900E-09 | 5.500E-09 |
| 12757 | 638 | G638T | B | 1.3e-08 | 4.4e-09 | 3.3e-09 | 6.900E-09 | 5.311E-09 | 3.300E-09 | 1.300E-08 |
| 12758 | 638 | G638W | B | 4.8e-09 | 6.1e-09 | 3.2e-09 | 4.700E-09 | 1.453E-09 | 3.200E-09 | 6.100E-09 |
| 12759 | 638 | G638Y | B | 7.7e-09 | 6.5e-09 | 5.7e-09 | 6.633E-09 | 1.007E-09 | 5.700E-09 | 7.700E-09 |
| 12760 | 638 | G638V | B | 3.4e-09 | 1.4e-09 | 4.3e-09 | 3.033E-09 | 1.484E-09 | 1.400E-09 | 4.300E-09 |
| 12761 | 639 | D639A | B | 1e-08 | 5.1e-09 | 6.5e-09 | 7.200E-09 | 2.524E-09 | 5.100E-09 | 1.000E-08 |
| 12762 | 639 | D639R | B | 3.2e-09 | 4e-09 | 7.2e-09 | 4.800E-09 | 2.117E-09 | 3.200E-09 | 7.200E-09 |
| 12763 | 639 | D639N | B | 1.7e-08 | 4.6e-09 | 3.4e-09 | 8.333E-09 | 7.529E-09 | 3.400E-09 | 1.700E-08 |
| 12764 | 639 | D639D | B | 1.1e-08 | 9.1e-09 | 3.8e-09 | 7.967E-09 | 3.731E-09 | 3.800E-09 | 1.100E-08 |
| 12765 | 639 | D639C | B | 3.5e-09 | 3e-09 | 5.3e-09 | 3.933E-09 | 1.210E-09 | 3.000E-09 | 5.300E-09 |
| 12766 | 639 | D639E | B | 2.2e-09 | 2e-09 | 7e-09 | 3.733E-09 | 2.831E-09 | 2.000E-09 | 7.000E-09 |
| 12767 | 639 | D639Q | B | 1.7e-08 | 8e-09 | 1.6e-08 | 1.367E-08 | 4.933E-09 | 8.000E-09 | 1.700E-08 |
| 12768 | 639 | D639G | B | 3.2e-09 | 4.4e-09 | 8.6e-09 | 5.400E-09 | 2.835E-09 | 3.200E-09 | 8.600E-09 |
| 12769 | 639 | D639H | B | 2.5e-08 | 1.7e-08 | 2.5e-09 | 1.483E-08 | 1.141E-08 | 2.500E-09 | 2.500E-08 |
| 12770 | 639 | D639I | B | 2.1e-08 | 5.3e-09 | 1.5e-08 | 1.377E-08 | 7.922E-09 | 5.300E-09 | 2.100E-08 |
| 12771 | 639 | D639L | B | 5.4e-09 | 1.3e-08 | 1.1e-08 | 9.800E-09 | 3.940E-09 | 5.400E-09 | 1.300E-08 |
| 12772 | 639 | D639K | B | 1.5e-09 | 7.5e-09 | 2.5e-09 | 3.833E-09 | 3.215E-09 | 1.500E-09 | 7.500E-09 |
| 12773 | 639 | D639M | B | 3e-09 | 9.8e-09 | 1.1e-08 | 7.933E-09 | 4.314E-09 | 3.000E-09 | 1.100E-08 |
| 12774 | 639 | D639F | B | 2.6e-09 | 1e-08 | 1.6e-08 | 9.533E-09 | 6.712E-09 | 2.600E-09 | 1.600E-08 |
| 12775 | 639 | D639P | B | 1.3e-08 | 7.1e-09 | 7.8e-09 | 9.300E-09 | 3.223E-09 | 7.100E-09 | 1.300E-08 |
| 12776 | 639 | D639S | B | 5.3e-09 | 8.4e-09 | 8e-09 | 7.233E-09 | 1.686E-09 | 5.300E-09 | 8.400E-09 |
| 12777 | 639 | D639T | B | 1.4e-08 | 3.2e-08 | 9.5e-09 | 1.850E-08 | 1.191E-08 | 9.500E-09 | 3.200E-08 |
| 12778 | 639 | D639W | B | 1.2e-08 | 1.9e-08 | 7.3e-09 | 1.277E-08 | 5.888E-09 | 7.300E-09 | 1.900E-08 |
| 12779 | 639 | D639Y | B | 6.1e-09 | 8e-09 | 7.7e-09 | 7.267E-09 | 1.021E-09 | 6.100E-09 | 8.000E-09 |
| 12780 | 639 | D639V | B | 2.4e-09 | 9.9e-09 | 3.8e-09 | 5.367E-09 | 3.988E-09 | 2.400E-09 | 9.900E-09 |
| 12781 | 640 | R640A | B | 7.3e-09 | 6.3e-09 | 1.5e-08 | 9.533E-09 | 4.761E-09 | 6.300E-09 | 1.500E-08 |
| 12782 | 640 | R640R | B | 9.3e-09 | 7.4e-09 | 3.1e-09 | 6.600E-09 | 3.176E-09 | 3.100E-09 | 9.300E-09 |
| 12783 | 640 | R640N | B | 4.7e-09 | 1.2e-08 | 2.1e-08 | 1.257E-08 | 8.165E-09 | 4.700E-09 | 2.100E-08 |
| 12784 | 640 | R640D | B | 2.6e-08 | 1.4e-08 | 1.3e-08 | 1.767E-08 | 7.234E-09 | 1.300E-08 | 2.600E-08 |
| 12785 | 640 | R640C | B | 7.3e-09 | 9.3e-09 | 6.1e-09 | 7.567E-09 | 1.617E-09 | 6.100E-09 | 9.300E-09 |
| 12786 | 640 | R640E | B | 7.2e-09 | 4.7e-09 | 5.2e-09 | 5.700E-09 | 1.323E-09 | 4.700E-09 | 7.200E-09 |
| 12787 | 640 | R640Q | B | 1.1e-08 | 2.9e-09 | 5.9e-09 | 6.600E-09 | 4.095E-09 | 2.900E-09 | 1.100E-08 |
| 12788 | 640 | R640G | B | 9.1e-09 | 5.9e-09 | 5.3e-09 | 6.767E-09 | 2.043E-09 | 5.300E-09 | 9.100E-09 |
| 12789 | 640 | R640H | B | 9.7e-09 | 2.6e-09 | 4.8e-09 | 5.700E-09 | 3.635E-09 | 2.600E-09 | 9.700E-09 |
| 12790 | 640 | R640I | B | 9e-09 | 3.6e-09 | 7.5e-09 | 6.700E-09 | 2.787E-09 | 3.600E-09 | 9.000E-09 |
| 12791 | 640 | R640L | B | 1.3e-08 | 2.1e-08 | 1.8e-08 | 1.733E-08 | 4.041E-09 | 1.300E-08 | 2.100E-08 |
| 12792 | 640 | R640K | B | 9.7e-09 | 1e-08 | 4.9e-09 | 8.200E-09 | 2.862E-09 | 4.900E-09 | 1.000E-08 |
| 12793 | 640 | R640M | B | 4.5e-09 | 1.3e-08 | 4.2e-09 | 7.233E-09 | 4.996E-09 | 4.200E-09 | 1.300E-08 |
| 12794 | 640 | R640F | B | 5.3e-09 | 1.2e-08 | 2.8e-09 | 6.700E-09 | 4.757E-09 | 2.800E-09 | 1.200E-08 |
| 12795 | 640 | R640P | B | 3.2e-09 | 3.2e-09 | 6e-09 | 4.133E-09 | 1.617E-09 | 3.200E-09 | 6.000E-09 |
| 12796 | 640 | R640S | B | 8.1e-09 | 1.2e-08 | 7.5e-09 | 9.200E-09 | 2.443E-09 | 7.500E-09 | 1.200E-08 |
| 12797 | 640 | R640T | B | 1.9e-08 | 5.8e-09 | 9.8e-09 | 1.153E-08 | 6.769E-09 | 5.800E-09 | 1.900E-08 |
| 12798 | 640 | R640W | B | 2.4e-09 | 4.6e-09 | 7e-09 | 4.667E-09 | 2.301E-09 | 2.400E-09 | 7.000E-09 |
| 12799 | 640 | R640Y | B | 6.8e-09 | 7.9e-09 | 3.9e-09 | 6.200E-09 | 2.066E-09 | 3.900E-09 | 7.900E-09 |
| 12800 | 640 | R640V | B | 2e-09 | 7.7e-09 | 1.1e-08 | 6.900E-09 | 4.553E-09 | 2.000E-09 | 1.100E-08 |
| 12801 | 641 | V641A | B | 1.8e-08 | 1.1e-08 | 9.4e-09 | 1.280E-08 | 4.574E-09 | 9.400E-09 | 1.800E-08 |
| 12802 | 641 | V641R | B | 1.1e-08 | 2e-08 | 6.6e-09 | 1.253E-08 | 6.830E-09 | 6.600E-09 | 2.000E-08 |
| 12803 | 641 | V641N | B | 8.1e-09 | 5.2e-08 | 8.3e-09 | 2.280E-08 | 2.529E-08 | 8.100E-09 | 5.200E-08 |
| 12804 | 641 | V641D | B | 4e-09 | 1.5e-08 | 8.4e-09 | 9.133E-09 | 5.537E-09 | 4.000E-09 | 1.500E-08 |
| 12805 | 641 | V641C | B | 6.3e-09 | 4.7e-09 | 6.3e-09 | 5.767E-09 | 9.238E-10 | 4.700E-09 | 6.300E-09 |
| 12806 | 641 | V641E | B | 3.1e-09 | 9.2e-09 | 2e-09 | 4.767E-09 | 3.879E-09 | 2.000E-09 | 9.200E-09 |
| 12807 | 641 | V641Q | B | 5.5e-09 | 6.2e-09 | 3.9e-09 | 5.200E-09 | 1.179E-09 | 3.900E-09 | 6.200E-09 |
| 12808 | 641 | V641G | B | 8.5e-09 | 1.1e-08 | 4e-09 | 7.833E-09 | 3.547E-09 | 4.000E-09 | 1.100E-08 |
| 12809 | 641 | V641H | B | 6.2e-09 | 5.9e-09 | 7.2e-09 | 6.433E-09 | 6.807E-10 | 5.900E-09 | 7.200E-09 |
| 12810 | 641 | V641I | B | 4e-09 | 6e-09 | 6.2e-09 | 5.400E-09 | 1.217E-09 | 4.000E-09 | 6.200E-09 |
| 12811 | 641 | V641L | B | 4.8e-09 | 2.4e-09 | 3.8e-09 | 3.667E-09 | 1.206E-09 | 2.400E-09 | 4.800E-09 |
| 12812 | 641 | V641K | B | 2.4e-09 | 5e-09 | 5.2e-09 | 4.200E-09 | 1.562E-09 | 2.400E-09 | 5.200E-09 |
| 12813 | 641 | V641M | B | 1e-08 | 5.6e-09 | 6.7e-09 | 7.433E-09 | 2.290E-09 | 5.600E-09 | 1.000E-08 |
| 12814 | 641 | V641F | B | 9.1e-09 | 1.5e-08 | 6.3e-09 | 1.013E-08 | 4.441E-09 | 6.300E-09 | 1.500E-08 |
| 12815 | 641 | V641P | B | 4.9e-09 | 1.9e-08 | 2.6e-09 | 8.833E-09 | 8.879E-09 | 2.600E-09 | 1.900E-08 |
| 12816 | 641 | V641S | B | 7.1e-09 | 8.3e-09 | 9e-09 | 8.133E-09 | 9.609E-10 | 7.100E-09 | 9.000E-09 |
| 12817 | 641 | V641T | B | 4e-08 | 8.4e-09 | 2.2e-09 | 1.687E-08 | 2.027E-08 | 2.200E-09 | 4.000E-08 |
| 12818 | 641 | V641W | B | 2.1e-08 | 7.3e-09 | 7.9e-09 | 1.207E-08 | 7.742E-09 | 7.300E-09 | 2.100E-08 |
| 12819 | 641 | V641Y | B | 1.5e-08 | 2.5e-09 | 9e-09 | 8.833E-09 | 6.252E-09 | 2.500E-09 | 1.500E-08 |
| 12820 | 641 | V641V | B | 5.2e-09 | 4.2e-09 | 1.6e-08 | 8.467E-09 | 6.543E-09 | 4.200E-09 | 1.600E-08 |
| 12821 | 642 | T642A | B | 5.9e-09 | 1.5e-08 | 7.5e-09 | 9.467E-09 | 4.858E-09 | 5.900E-09 | 1.500E-08 |
| 12822 | 642 | T642R | B | 5.1e-09 | 4.1e-09 | 3.8e-09 | 4.333E-09 | 6.807E-10 | 3.800E-09 | 5.100E-09 |
| 12823 | 642 | T642N | B | 7.7e-09 | 4.9e-09 | 6.6e-09 | 6.400E-09 | 1.411E-09 | 4.900E-09 | 7.700E-09 |
| 12824 | 642 | T642D | B | 3.4e-09 | 2.7e-09 | 8.7e-09 | 4.933E-09 | 3.281E-09 | 2.700E-09 | 8.700E-09 |
| 12825 | 642 | T642C | B | 8.9e-09 | 1.2e-08 | 5e-09 | 8.633E-09 | 3.508E-09 | 5.000E-09 | 1.200E-08 |
| 12826 | 642 | T642E | B | 5.5e-09 | 1.1e-08 | 7.9e-09 | 8.133E-09 | 2.757E-09 | 5.500E-09 | 1.100E-08 |
| 12827 | 642 | T642Q | B | 5.1e-09 | 8.3e-09 | 4.9e-09 | 6.100E-09 | 1.908E-09 | 4.900E-09 | 8.300E-09 |
| 12828 | 642 | T642G | B | 5.2e-09 | 7.8e-09 | 8.6e-09 | 7.200E-09 | 1.778E-09 | 5.200E-09 | 8.600E-09 |
| 12829 | 642 | T642H | B | 1.7e-08 | 1.2e-08 | 1.2e-08 | 1.367E-08 | 2.887E-09 | 1.200E-08 | 1.700E-08 |
| 12830 | 642 | T642I | B | 1.2e-08 | 3.2e-09 | 1.6e-08 | 1.040E-08 | 6.548E-09 | 3.200E-09 | 1.600E-08 |
| 12831 | 642 | T642L | B | 5.6e-09 | 1e-08 | 5.1e-09 | 6.900E-09 | 2.696E-09 | 5.100E-09 | 1.000E-08 |
| 12832 | 642 | T642K | B | 5.1e-09 | 1.6e-08 | 3.6e-09 | 8.233E-09 | 6.768E-09 | 3.600E-09 | 1.600E-08 |
| 12833 | 642 | T642M | B | 6.8e-09 | 9.1e-09 | 4e-09 | 6.633E-09 | 2.554E-09 | 4.000E-09 | 9.100E-09 |
| 12834 | 642 | T642F | B | 8.4e-10 | 6.5e-09 | 2.4e-09 | 3.247E-09 | 2.923E-09 | 8.400E-10 | 6.500E-09 |

| | | | | | | | | | | |
|---|---|---|---|---|---|---|---|---|---|---|
| 12835 | 642 | T642P | B | 1e-08 | 2.4e-09 | 7.7e-09 | 6.700E-09 | 3.897E-09 | 2.400E-09 | 1.000E-08 |
| 12836 | 642 | T642S | B | 5.1e-09 | 1.5e-08 | 5.8e-09 | 8.633E-09 | 5.525E-09 | 5.100E-09 | 1.500E-08 |
| 12837 | 642 | T642T | B | 1.5e-09 | 1.2e-08 | 3.9e-09 | 5.800E-09 | 5.502E-09 | 1.500E-09 | 1.200E-08 |
| 12838 | 642 | T642W | B | 2e-09 | 7.2e-09 | 3.5e-09 | 4.233E-09 | 2.676E-09 | 2.000E-09 | 7.200E-09 |
| 12839 | 642 | T642Y | B | 5.9e-09 | 5.2e-09 | 5.8e-09 | 5.633E-09 | 3.786E-10 | 5.200E-09 | 5.900E-09 |
| 12840 | 642 | T642V | B | 1.4e-08 | 1e-08 | 8.3e-09 | 1.077E-08 | 2.926E-09 | 8.300E-09 | 1.400E-08 |
| 12841 | 643 | I643A | B | 1.6e-08 | 3.9e-09 | 1.9e-09 | 7.267E-09 | 7.629E-09 | 1.900E-09 | 1.600E-08 |
| 12842 | 643 | I643R | B | 1e-08 | 4.8e-09 | 2e-09 | 5.600E-09 | 4.060E-09 | 2.000E-09 | 1.000E-08 |
| 12843 | 643 | I643N | B | 7.2e-09 | 2.6e-09 | 6.4e-09 | 5.400E-09 | 2.458E-09 | 2.600E-09 | 7.200E-09 |
| 12844 | 643 | I643D | B | 4.2e-09 | 8.9e-09 | 9.7e-09 | 7.600E-09 | 2.972E-09 | 4.200E-09 | 9.700E-09 |
| 12845 | 643 | I643C | B | 1.7e-09 | 7e-10 | 6.9e-09 | 3.100E-09 | 3.329E-09 | 7.000E-10 | 6.900E-09 |
| 12846 | 643 | I643E | B | 1.6e-08 | 4.8e-09 | 2.1e-08 | 1.393E-08 | 8.295E-09 | 4.800E-09 | 2.100E-08 |
| 12847 | 643 | I643Q | B | 4.8e-09 | 1e-08 | 3.5e-09 | 6.100E-09 | 3.439E-09 | 3.500E-09 | 1.000E-08 |
| 12848 | 643 | I643G | B | 6.8e-09 | 6.6e-09 | 1.2e-08 | 8.467E-09 | 3.062E-09 | 6.600E-09 | 1.200E-08 |
| 12849 | 643 | I643H | B | 7e-09 | 6.4e-09 | 2.3e-09 | 5.233E-09 | 2.558E-09 | 2.300E-09 | 7.000E-09 |
| 12850 | 643 | I643I | B | 1.8e-08 | 3.7e-09 | 1e-08 | 1.057E-08 | 7.167E-09 | 3.700E-09 | 1.800E-08 |
| 12851 | 643 | I643L | B | 7e-09 | 1.2e-08 | 3.9e-09 | 7.633E-09 | 4.087E-09 | 3.900E-09 | 1.200E-08 |
| 12852 | 643 | I643K | B | 7.4e-09 | 6.4e-09 | 1.4e-08 | 9.267E-09 | 4.130E-09 | 6.400E-09 | 1.400E-08 |
| 12853 | 643 | I643M | B | 9.2e-09 | 5.2e-09 | 1.6e-08 | 1.013E-08 | 5.460E-09 | 5.200E-09 | 1.600E-08 |
| 12854 | 643 | I643F | B | 1.1e-08 | 5.5e-09 | 7.3e-09 | 7.933E-09 | 2.804E-09 | 5.500E-09 | 1.100E-08 |
| 12855 | 643 | I643P | B | 4.4e-09 | 1e-08 | 5.7e-09 | 6.700E-09 | 2.931E-09 | 4.400E-09 | 1.000E-08 |
| 12856 | 643 | I643S | B | 2.7e-09 | 4.2e-09 | 1.2e-08 | 6.300E-09 | 4.993E-09 | 2.700E-09 | 1.200E-08 |
| 12857 | 643 | I643T | B | 1.2e-08 | 8.4e-09 | 3.1e-09 | 7.833E-09 | 4.477E-09 | 3.100E-09 | 1.200E-08 |
| 12858 | 643 | I643W | B | 9.5e-09 | 3.3e-09 | 1e-08 | 7.600E-09 | 3.732E-09 | 3.300E-09 | 1.000E-08 |
| 12859 | 643 | I643Y | B | 4.4e-09 | 1e-08 | 1.2e-08 | 8.800E-09 | 3.940E-09 | 4.400E-09 | 1.200E-08 |
| 12860 | 643 | I643V | B | 1.6e-08 | 1.1e-08 | 8.5e-09 | 1.183E-08 | 3.819E-09 | 8.500E-09 | 1.600E-08 |
| 12861 | 644 | T644A | B | 3.5e-09 | 8.5e-09 | 1.1e-08 | 7.667E-09 | 3.819E-09 | 3.500E-09 | 1.100E-08 |
| 12862 | 644 | T644R | B | 7.4e-09 | 9.8e-09 | 5e-09 | 7.400E-09 | 2.400E-09 | 5.000E-09 | 9.800E-09 |
| 12863 | 644 | T644N | B | 5e-09 | 7.8e-09 | 7.5e-09 | 6.767E-09 | 1.537E-09 | 5.000E-09 | 7.800E-09 |
| 12864 | 644 | T644D | B | 9e-09 | 1e-08 | 2.2e-09 | 7.067E-09 | 4.244E-09 | 2.200E-09 | 1.000E-08 |
| 12865 | 644 | T644C | B | 4.6e-09 | 4.2e-09 | 1.2e-08 | 6.933E-09 | 4.392E-09 | 4.200E-09 | 1.200E-08 |
| 12866 | 644 | T644E | B | 7.5e-09 | 6.1e-09 | 5.3e-09 | 6.300E-09 | 1.114E-09 | 5.300E-09 | 7.500E-09 |
| 12867 | 644 | T644Q | B | 3.7e-09 | 5.9e-09 | 9e-09 | 6.200E-09 | 2.663E-09 | 3.700E-09 | 9.000E-09 |
| 12868 | 644 | T644G | B | 5.4e-09 | 3.7e-09 | 7.6e-09 | 5.567E-09 | 1.955E-09 | 3.700E-09 | 7.600E-09 |
| 12869 | 644 | T644H | B | 5.7e-09 | 5.3e-09 | 6.4e-09 | 5.800E-09 | 5.568E-10 | 5.300E-09 | 6.400E-09 |
| 12870 | 644 | T644I | B | 7.3e-09 | 8.9e-09 | 1.1e-09 | 5.767E-09 | 4.120E-09 | 1.100E-09 | 8.900E-09 |
| 12871 | 644 | T644L | B | 4.8e-09 | 5.3e-09 | 5.3e-09 | 5.133E-09 | 2.887E-10 | 4.800E-09 | 5.300E-09 |
| 12872 | 644 | T644K | B | 3.6e-09 | 1.6e-08 | 7.4e-09 | 9.000E-09 | 6.353E-09 | 3.600E-09 | 1.600E-08 |
| 12873 | 644 | T644M | B | 6.2e-09 | 6.7e-09 | 2e-08 | 1.097E-08 | 7.827E-09 | 6.200E-09 | 2.000E-08 |
| 12874 | 644 | T644F | B | 9.2e-09 | 1e-08 | 2.8e-08 | 1.573E-08 | 1.063E-08 | 9.200E-09 | 2.800E-08 |
| 12875 | 644 | T644P | B | 1.2e-08 | 6.2e-09 | 5e-09 | 7.733E-09 | 3.743E-09 | 5.000E-09 | 1.200E-08 |
| 12876 | 644 | T644S | B | 5.8e-09 | 7.1e-09 | 4.7e-09 | 5.867E-09 | 1.201E-09 | 4.700E-09 | 7.100E-09 |
| 12877 | 644 | T644T | B | 1.1e-08 | 2.9e-09 | 5.5e-09 | 6.467E-09 | 4.136E-09 | 2.900E-09 | 1.100E-08 |
| 12878 | 644 | T644W | B | 4.4e-09 | 2.8e-09 | 3.5e-09 | 3.567E-09 | 8.021E-10 | 2.800E-09 | 4.400E-09 |
| 12879 | 644 | T644Y | B | 6e-09 | 1.3e-08 | 9.9e-09 | 9.633E-09 | 3.508E-09 | 6.000E-09 | 1.300E-08 |
| 12880 | 644 | T644V | B | 1.4e-08 | 6.8e-09 | 5.1e-09 | 8.633E-09 | 4.725E-09 | 5.100E-09 | 1.400E-08 |
| 12881 | 645 | C645A | B | 9.9e-09 | 4.3e-09 | 1e-08 | 8.067E-09 | 3.262E-09 | 4.300E-09 | 1.000E-08 |
| 12882 | 645 | C645R | B | 3.4e-09 | 8.7e-09 | 1.6e-08 | 9.367E-09 | 6.326E-09 | 3.400E-09 | 1.600E-08 |
| 12883 | 645 | C645N | B | 6.2e-08 | 4e-09 | 5.2e-09 | 2.373E-08 | 3.315E-08 | 4.000E-09 | 6.200E-08 |
| 12884 | 645 | C645D | B | 1.2e-08 | 1e-08 | 1.1e-08 | 1.100E-08 | 1.000E-09 | 1.000E-08 | 1.200E-08 |
| 12885 | 645 | C645C | B | 1.6e-08 | 7.4e-09 | 6.7e-09 | 1.003E-08 | 5.179E-09 | 6.700E-09 | 1.600E-08 |
| 12886 | 645 | C645E | B | 3.6e-09 | 3.2e-09 | 7.8e-09 | 4.867E-09 | 2.548E-09 | 3.200E-09 | 7.800E-09 |
| 12887 | 645 | C645Q | B | 5.1e-09 | 6.6e-09 | 1.1e-08 | 7.567E-09 | 3.066E-09 | 5.100E-09 | 1.100E-08 |
| 12888 | 645 | C645G | B | 1.7e-08 | 7.1e-09 | 1.3e-08 | 1.237E-08 | 4.980E-09 | 7.100E-09 | 1.700E-08 |
| 12889 | 645 | C645H | B | 8.5e-09 | 4.3e-09 | 8.3e-09 | 7.033E-09 | 2.369E-09 | 4.300E-09 | 8.500E-09 |
| 12890 | 645 | C645I | B | 8.3e-09 | 2.9e-09 | 6.7e-09 | 5.967E-09 | 2.774E-09 | 2.900E-09 | 8.300E-09 |
| 12891 | 645 | C645L | B | 4.1e-09 | 1.9e-09 | 8.7e-09 | 4.900E-09 | 3.470E-09 | 1.900E-09 | 8.700E-09 |
| 12892 | 645 | C645K | B | 6.1e-09 | 5.2e-09 | 1.1e-08 | 7.433E-09 | 3.121E-09 | 5.200E-09 | 1.100E-08 |
| 12893 | 645 | C645M | B | 4.8e-09 | 2.8e-09 | 5.2e-09 | 4.267E-09 | 1.286E-09 | 2.800E-09 | 5.200E-09 |
| 12894 | 645 | C645F | B | 6.6e-09 | 6.9e-09 | 1.4e-08 | 9.167E-09 | 4.188E-09 | 6.600E-09 | 1.400E-08 |
| 12895 | 645 | C645P | B | 1.5e-08 | 2.3e-09 | 7.3e-09 | 8.200E-09 | 6.398E-09 | 2.300E-09 | 1.500E-08 |
| 12896 | 645 | C645S | B | 1.1e-08 | 1.1e-08 | 5e-09 | 9.000E-09 | 3.464E-09 | 5.000E-09 | 1.100E-08 |
| 12897 | 645 | C645T | B | 2.1e-09 | 2.1e-08 | 4.3e-09 | 9.133E-09 | 1.034E-08 | 2.100E-09 | 2.100E-08 |
| 12898 | 645 | C645W | B | 2.4e-09 | 1.3e-08 | 2e-08 | 1.180E-08 | 8.861E-09 | 2.400E-09 | 2.000E-08 |
| 12899 | 645 | C645Y | B | 1.4e-08 | 7.1e-09 | 7.1e-09 | 9.400E-09 | 3.984E-09 | 7.100E-09 | 1.400E-08 |
| 12900 | 645 | C645V | B | 1.9e-08 | 8.5e-09 | 1.4e-08 | 1.383E-08 | 5.252E-09 | 8.500E-09 | 1.900E-08 |
| 12901 | 646 | K646A | B | 6.3e-09 | 8.9e-09 | 3.1e-09 | 6.100E-09 | 2.905E-09 | 3.100E-09 | 8.900E-09 |
| 12902 | 646 | K646R | B | 1.2e-08 | 3.1e-09 | 2.2e-08 | 1.237E-08 | 9.455E-09 | 3.100E-09 | 2.200E-08 |
| 12903 | 646 | K646N | B | 1.5e-08 | 1.4e-09 | 2.8e-09 | 6.400E-09 | 7.481E-09 | 1.400E-09 | 1.500E-08 |
| 12904 | 646 | K646D | B | 3.6e-09 | 1.4e-08 | 1.2e-08 | 9.867E-09 | 5.518E-09 | 3.600E-09 | 1.400E-08 |
| 12905 | 646 | K646C | B | 7.9e-09 | 9.7e-09 | 6.8e-09 | 8.133E-09 | 1.464E-09 | 6.800E-09 | 9.700E-09 |
| 12906 | 646 | K646E | B | 4.2e-09 | 8.8e-09 | 2.1e-08 | 1.133E-08 | 8.682E-09 | 4.200E-09 | 2.100E-08 |
| 12907 | 646 | K646Q | B | 2e-09 | 8.9e-09 | 1.2e-08 | 7.633E-09 | 5.119E-09 | 2.000E-09 | 1.200E-08 |
| 12908 | 646 | K646G | B | 1.1e-08 | 1e-08 | 1.3e-08 | 1.133E-08 | 1.528E-09 | 1.000E-08 | 1.300E-08 |
| 12909 | 646 | K646H | B | 5.3e-09 | 8.2e-09 | 1.3e-08 | 8.833E-09 | 3.889E-09 | 5.300E-09 | 1.300E-08 |
| 12910 | 646 | K646I | B | 7.8e-09 | 7.4e-09 | 1.7e-08 | 1.073E-08 | 5.431E-09 | 7.400E-09 | 1.700E-08 |
| 12911 | 646 | K646L | B | 6.4e-09 | 5.4e-09 | 1.1e-08 | 7.600E-09 | 2.987E-09 | 5.400E-09 | 1.100E-08 |
| 12912 | 646 | K646K | B | 2.5e-08 | 1e-08 | 3.8e-09 | 1.293E-08 | 1.090E-08 | 3.800E-09 | 2.500E-08 |
| 12913 | 646 | K646M | B | 3.3e-09 | 4.6e-09 | 4.1e-09 | 4.000E-09 | 6.557E-10 | 3.300E-09 | 4.600E-09 |
| 12914 | 646 | K646F | B | 3.6e-09 | 2.3e-08 | 1.2e-08 | 1.287E-08 | 9.729E-09 | 3.600E-09 | 2.300E-08 |
| 12915 | 646 | K646P | B | 1.5e-08 | 7.9e-09 | 5.2e-09 | 9.367E-09 | 5.062E-09 | 5.200E-09 | 1.500E-08 |
| 12916 | 646 | K646S | B | 2.2e-09 | 4.5e-09 | 5.5e-09 | 4.067E-09 | 1.692E-09 | 2.200E-09 | 5.500E-09 |
| 12917 | 646 | K646T | B | 7.1e-09 | 1.9e-09 | 1.1e-08 | 6.667E-09 | 4.565E-09 | 1.900E-09 | 1.100E-08 |
| 12918 | 646 | K646W | B | 7.2e-09 | 4.8e-09 | 1.1e-08 | 7.667E-09 | 3.126E-09 | 4.800E-09 | 1.100E-08 |
| 12919 | 646 | K646Y | B | 2.7e-09 | 7.8e-09 | 6.2e-09 | 5.567E-09 | 2.608E-09 | 2.700E-09 | 7.800E-09 |

| | | | | | | | | | | |
|---|---|---|---|---|---|---|---|---|---|---|
| 12920 | 646 | K646V | B | 7.2e-09 | 1.2e-08 | 5.1e-09 | 8.100E-09 | 3.537E-09 | 5.100E-09 | 1.200E-08 |
| 12921 | 647 | A647A | B | 5.1e-09 | 8.6e-09 | 1e-08 | 7.900E-09 | 2.524E-09 | 5.100E-09 | 1.000E-08 |
| 12922 | 647 | A647R | B | 6.3e-09 | 1.4e-08 | 8.7e-09 | 9.667E-09 | 3.940E-09 | 6.300E-09 | 1.400E-08 |
| 12923 | 647 | A647N | B | 5.1e-09 | 1e-08 | 5.8e-09 | 6.967E-09 | 2.650E-09 | 5.100E-09 | 1.000E-08 |
| 12924 | 647 | A647D | B | 2.5e-09 | 4.5e-09 | 6.7e-09 | 4.567E-09 | 2.101E-09 | 2.500E-09 | 6.700E-09 |
| 12925 | 647 | A647C | B | 7.1e-09 | 3.8e-09 | 1.5e-08 | 8.633E-09 | 5.755E-09 | 3.800E-09 | 1.500E-08 |
| 12926 | 647 | A647E | B | 9.1e-09 | 4.6e-09 | 6.5e-09 | 6.733E-09 | 2.259E-09 | 4.600E-09 | 9.100E-09 |
| 12927 | 647 | A647Q | B | 4.8e-09 | 4.6e-09 | 1.5e-08 | 8.133E-09 | 5.948E-09 | 4.600E-09 | 1.500E-08 |
| 12928 | 647 | A647G | B | 1.3e-08 | 8e-09 | 2.9e-09 | 7.967E-09 | 5.050E-09 | 2.900E-09 | 1.300E-08 |
| 12929 | 647 | A647H | B | 5.7e-09 | 1.1e-08 | 3.6e-09 | 6.767E-09 | 3.814E-09 | 3.600E-09 | 1.100E-08 |
| 12930 | 647 | A647I | B | 1.7e-08 | 8e-09 | 6.8e-09 | 1.060E-08 | 5.575E-09 | 6.800E-09 | 1.700E-08 |
| 12931 | 647 | A647L | B | 1.8e-08 | 7.1e-09 | 7.2e-09 | 1.077E-08 | 6.264E-09 | 7.100E-09 | 1.800E-08 |
| 12932 | 647 | A647K | B | 1.5e-08 | 6.1e-09 | 5.4e-09 | 8.833E-09 | 5.352E-09 | 5.400E-09 | 1.500E-08 |
| 12933 | 647 | A647M | B | 6.9e-09 | 9.8e-09 | 9e-09 | 8.567E-09 | 1.498E-09 | 6.900E-09 | 9.800E-09 |
| 12934 | 647 | A647F | B | 8.9e-09 | 7.3e-09 | 1.8e-08 | 1.140E-08 | 5.771E-09 | 7.300E-09 | 1.800E-08 |
| 12935 | 647 | A647P | B | 2.1e-08 | 1.5e-08 | 1e-08 | 1.533E-08 | 5.508E-09 | 1.000E-08 | 2.100E-08 |
| 12936 | 647 | A647S | B | 1.6e-08 | 9.9e-09 | 1.4e-08 | 1.330E-08 | 3.110E-09 | 9.900E-09 | 1.600E-08 |
| 12937 | 647 | A647T | B | 6.9e-09 | 7.6e-09 | 1.7e-08 | 1.050E-08 | 5.640E-09 | 6.900E-09 | 1.700E-08 |
| 12938 | 647 | A647W | B | 8.8e-09 | 7e-09 | 6.3e-09 | 7.367E-09 | 1.290E-09 | 6.300E-09 | 8.800E-09 |
| 12939 | 647 | A647Y | B | 3.9e-09 | 6.1e-09 | 1.6e-08 | 8.667E-09 | 6.445E-09 | 3.900E-09 | 1.600E-08 |
| 12940 | 647 | A647V | B | 1.5e-08 | 5.2e-09 | 1.3e-08 | 1.107E-08 | 5.178E-09 | 5.200E-09 | 1.500E-08 |
| 12941 | 648 | S648A | B | 6.5e-09 | 3.7e-09 | 7.8e-09 | 6.000E-09 | 2.095E-09 | 3.700E-09 | 7.800E-09 |
| 12942 | 648 | S648R | B | 3.2e-09 | 3.7e-09 | 3.9e-09 | 3.600E-09 | 3.606E-10 | 3.200E-09 | 3.900E-09 |
| 12943 | 648 | S648N | B | 6.8e-09 | 7.6e-09 | 2.3e-08 | 1.247E-08 | 9.131E-09 | 6.800E-09 | 2.300E-08 |
| 12944 | 648 | S648D | B | 1.4e-08 | 6.3e-09 | 4.9e-09 | 8.400E-09 | 4.900E-09 | 4.900E-09 | 1.400E-08 |
| 12945 | 648 | S648C | B | 1.4e-08 | 1e-08 | 6.6e-09 | 1.020E-08 | 3.704E-09 | 6.600E-09 | 1.400E-08 |
| 12946 | 648 | S648E | B | 1.7e-08 | 7.3e-09 | 1.6e-08 | 1.343E-08 | 5.335E-09 | 7.300E-09 | 1.700E-08 |
| 12947 | 648 | S648Q | B | 1.5e-09 | 7.4e-09 | 5.1e-09 | 4.667E-09 | 2.974E-09 | 1.500E-09 | 7.400E-09 |
| 12948 | 648 | S648G | B | 3.3e-09 | 1.3e-08 | 2.4e-08 | 1.343E-08 | 1.036E-08 | 3.300E-09 | 2.400E-08 |
| 12949 | 648 | S648H | B | 5.7e-09 | 2.4e-09 | 1.5e-08 | 7.700E-09 | 6.534E-09 | 2.400E-09 | 1.500E-08 |
| 12950 | 648 | S648I | B | 3.6e-09 | 1.3e-08 | 1.4e-08 | 1.020E-08 | 5.738E-09 | 3.600E-09 | 1.400E-08 |
| 12951 | 648 | S648L | B | 8.2e-09 | 6e-09 | 2.6e-08 | 1.340E-08 | 1.097E-08 | 6.000E-09 | 2.600E-08 |
| 12952 | 648 | S648K | B | 8.8e-09 | 2.5e-09 | 4.7e-09 | 5.333E-09 | 3.197E-09 | 2.500E-09 | 8.800E-09 |
| 12953 | 648 | S648M | B | 1.3e-09 | 4.2e-09 | 9e-09 | 4.833E-09 | 3.889E-09 | 1.300E-09 | 9.000E-09 |
| 12954 | 648 | S648F | B | 5.6e-09 | 1.3e-08 | 8.7e-09 | 9.100E-09 | 3.716E-09 | 5.600E-09 | 1.300E-08 |
| 12955 | 648 | S648P | B | 1.5e-08 | 1.3e-08 | 5.3e-09 | 1.110E-08 | 5.122E-09 | 5.300E-09 | 1.500E-08 |
| 12956 | 648 | S648S | B | 7.4e-09 | 2.2e-08 | 4.7e-09 | 1.137E-08 | 9.307E-09 | 4.700E-09 | 2.200E-08 |
| 12957 | 648 | S648T | B | 5e-09 | 3.8e-09 | 9.7e-09 | 6.167E-09 | 3.118E-09 | 3.800E-09 | 9.700E-09 |
| 12958 | 648 | S648W | B | 1.2e-08 | 2.8e-09 | 5.6e-09 | 6.800E-09 | 4.716E-09 | 2.800E-09 | 1.200E-08 |
| 12959 | 648 | S648Y | B | 7.6e-09 | 1.2e-08 | 3.7e-09 | 7.767E-09 | 4.153E-09 | 3.700E-09 | 1.200E-08 |
| 12960 | 648 | S648V | B | 2.9e-08 | 5.9e-09 | 1.3e-08 | 1.597E-08 | 1.183E-08 | 5.900E-09 | 2.900E-08 |
| 12961 | 649 | Q649A | B | 7.2e-09 | 9.7e-09 | 6.6e-09 | 7.833E-09 | 1.644E-09 | 6.600E-09 | 9.700E-09 |
| 12962 | 649 | Q649R | B | 9.8e-09 | 4.7e-09 | 4e-09 | 6.167E-09 | 3.166E-09 | 4.000E-09 | 9.800E-09 |
| 12963 | 649 | Q649N | B | 9.7e-09 | 3.5e-09 | 2.9e-09 | 5.367E-09 | 3.765E-09 | 2.900E-09 | 9.700E-09 |
| 12964 | 649 | Q649D | B | 9.4e-09 | 6.1e-09 | 1.7e-08 | 1.083E-08 | 5.590E-09 | 6.100E-09 | 1.700E-08 |
| 12965 | 649 | Q649C | B | 4.5e-09 | 6.3e-09 | 7.3e-09 | 6.033E-09 | 1.419E-09 | 4.500E-09 | 7.300E-09 |
| 12966 | 649 | Q649E | B | 7.4e-09 | 1.7e-08 | 7.2e-09 | 1.053E-08 | 5.601E-09 | 7.200E-09 | 1.700E-08 |
| 12967 | 649 | Q649Q | B | 1.6e-08 | 1.5e-08 | 1.2e-08 | 1.433E-08 | 2.082E-09 | 1.200E-08 | 1.600E-08 |
| 12968 | 649 | Q649G | B | 1.1e-08 | 2.2e-08 | 5.2e-09 | 1.273E-08 | 8.533E-09 | 5.200E-09 | 2.200E-08 |
| 12969 | 649 | Q649H | B | 9.2e-09 | 4.7e-09 | 1.2e-08 | 8.633E-09 | 3.683E-09 | 4.700E-09 | 1.200E-08 |
| 12970 | 649 | Q649I | B | 6.2e-09 | 4.7e-09 | 8.7e-09 | 6.533E-09 | 2.021E-09 | 4.700E-09 | 8.700E-09 |
| 12971 | 649 | Q649L | B | 2.9e-09 | 5.4e-09 | 1.6e-08 | 8.100E-09 | 6.955E-09 | 2.900E-09 | 1.600E-08 |
| 12972 | 649 | Q649K | B | 3.2e-09 | 2.3e-08 | 3e-09 | 9.733E-09 | 1.149E-08 | 3.000E-09 | 2.300E-08 |
| 12973 | 649 | Q649M | B | 5.3e-09 | 9.1e-09 | 3.9e-09 | 6.100E-09 | 2.691E-09 | 3.900E-09 | 9.100E-09 |
| 12974 | 649 | Q649F | B | 6e-09 | 8.9e-09 | 7.5e-09 | 7.467E-09 | 1.450E-09 | 6.000E-09 | 8.900E-09 |
| 12975 | 649 | Q649P | B | 6.1e-09 | 8.7e-09 | 1.1e-08 | 8.600E-09 | 2.452E-09 | 6.100E-09 | 1.100E-08 |
| 12976 | 649 | Q649S | B | 2.1e-08 | 5.1e-09 | 1.3e-08 | 1.303E-08 | 7.950E-09 | 5.100E-09 | 2.100E-08 |
| 12977 | 649 | Q649T | B | 5.4e-09 | 8.5e-09 | 4.8e-09 | 6.233E-09 | 1.986E-09 | 4.800E-09 | 8.500E-09 |
| 12978 | 649 | Q649W | B | 3e-08 | 7.2e-09 | 1.3e-08 | 1.673E-08 | 1.185E-08 | 7.200E-09 | 3.000E-08 |
| 12979 | 649 | Q649Y | B | 1.4e-08 | 4.3e-09 | 9.7e-09 | 9.333E-09 | 4.860E-09 | 4.300E-09 | 1.400E-08 |
| 12980 | 649 | Q649V | B | 7.6e-09 | 4.4e-09 | 5.6e-09 | 5.867E-09 | 1.617E-09 | 4.400E-09 | 7.600E-09 |
| 12981 | 650 | D650A | B | 1.9e-09 | 1.8e-09 | 1.1e-08 | 4.900E-09 | 5.283E-09 | 1.800E-09 | 1.100E-08 |
| 12982 | 650 | D650R | B | 9.8e-09 | 1.9e-08 | 3e-09 | 1.060E-08 | 8.030E-09 | 3.000E-09 | 1.900E-08 |
| 12983 | 650 | D650N | B | 1.9e-08 | 7.1e-09 | 1.1e-08 | 1.237E-08 | 6.067E-09 | 7.100E-09 | 1.900E-08 |
| 12984 | 650 | D650D | B | 9.5e-09 | 8.9e-09 | 4.2e-09 | 7.533E-09 | 2.902E-09 | 4.200E-09 | 9.500E-09 |
| 12985 | 650 | D650C | B | 5e-09 | 1.1e-08 | 1.3e-08 | 9.667E-09 | 4.163E-09 | 5.000E-09 | 1.300E-08 |
| 12986 | 650 | D650E | B | 1.5e-08 | 1.7e-08 | 8.3e-09 | 1.343E-08 | 4.557E-09 | 8.300E-09 | 1.700E-08 |
| 12987 | 650 | D650Q | B | 5.8e-09 | 4.3e-09 | 9.6e-09 | 6.567E-09 | 2.732E-09 | 4.300E-09 | 9.600E-09 |
| 12988 | 650 | D650G | B | 5.2e-09 | 8.5e-09 | 1.3e-09 | 5.000E-09 | 3.604E-09 | 1.300E-09 | 8.500E-09 |
| 12989 | 650 | D650H | B | 6.6e-09 | 1.3e-08 | 1.2e-08 | 1.053E-08 | 3.443E-09 | 6.600E-09 | 1.300E-08 |
| 12990 | 650 | D650I | B | 1.8e-08 | 7.2e-09 | 4.4e-09 | 9.867E-09 | 7.181E-09 | 4.400E-09 | 1.800E-08 |
| 12991 | 650 | D650L | B | 1.1e-08 | 4.8e-09 | 1e-08 | 8.600E-09 | 3.329E-09 | 4.800E-09 | 1.100E-08 |
| 12992 | 650 | D650K | B | 6.6e-09 | 3.7e-09 | 3.6e-09 | 4.633E-09 | 1.704E-09 | 3.600E-09 | 6.600E-09 |
| 12993 | 650 | D650M | B | 1.6e-08 | 4.3e-09 | 1e-08 | 1.010E-08 | 5.851E-09 | 4.300E-09 | 1.600E-08 |
| 12994 | 650 | D650F | B | 4.1e-09 | 1.1e-08 | 2.9e-09 | 6.000E-09 | 4.371E-09 | 2.900E-09 | 1.100E-08 |
| 12995 | 650 | D650P | B | 6.2e-09 | 1.6e-08 | 1.1e-08 | 1.107E-08 | 4.900E-09 | 6.200E-09 | 1.600E-08 |
| 12996 | 650 | D650S | B | 1e-08 | 5.2e-09 | 7.1e-09 | 7.433E-09 | 2.417E-09 | 5.200E-09 | 1.000E-08 |
| 12997 | 650 | D650T | B | 2.5e-09 | 3.6e-09 | 5.7e-09 | 3.933E-09 | 1.626E-09 | 2.500E-09 | 5.700E-09 |
| 12998 | 650 | D650W | B | 2.7e-09 | 4.2e-09 | 5.3e-09 | 4.067E-09 | 1.305E-09 | 2.700E-09 | 5.300E-09 |
| 12999 | 650 | D650Y | B | 9.2e-09 | 3.1e-09 | 7.2e-09 | 6.500E-09 | 3.110E-09 | 3.100E-09 | 9.200E-09 |
| 13000 | 650 | D650V | B | 1.7e-08 | 4.6e-08 | 7.6e-09 | 2.353E-08 | 2.002E-08 | 7.600E-09 | 4.600E-08 |
| 13001 | 651 | V651A | B | 1.6e-08 | 5.6e-09 | 1.3e-08 | 1.153E-08 | 5.353E-09 | 5.600E-09 | 1.600E-08 |
| 13002 | 651 | V651R | B | 7.1e-09 | 4.1e-09 | 3.7e-09 | 4.967E-09 | 1.858E-09 | 3.700E-09 | 7.100E-09 |
| 13003 | 651 | V651N | B | 2.1e-08 | 6.1e-09 | 5.9e-09 | 1.100E-08 | 8.661E-09 | 5.900E-09 | 2.100E-08 |
| 13004 | 651 | V651D | B | 5e-09 | 7.4e-09 | 4.9e-09 | 5.767E-09 | 1.415E-09 | 4.900E-09 | 7.400E-09 |

| | | | | | | | | | | |
|---|---|---|---|---|---|---|---|---|---|---|
| 13005 | 651 | V651C | B | 5e-09 | 4.3e-09 | 2e-08 | 9.767E-09 | 8.869E-09 | 4.300E-09 | 2.000E-08 |
| 13006 | 651 | V651E | B | 8.3e-09 | 1.5e-08 | 4.5e-09 | 9.267E-09 | 5.316E-09 | 4.500E-09 | 1.500E-08 |
| 13007 | 651 | V651Q | B | 1.1e-08 | 1.3e-08 | 1e-08 | 1.133E-08 | 1.528E-09 | 1.000E-08 | 1.300E-08 |
| 13008 | 651 | V651G | B | 5.3e-09 | 8.1e-09 | 9.7e-09 | 7.700E-09 | 2.227E-09 | 5.300E-09 | 9.700E-09 |
| 13009 | 651 | V651H | B | 4e-09 | 3.3e-09 | 1.1e-08 | 6.100E-09 | 4.258E-09 | 3.300E-09 | 1.100E-08 |
| 13010 | 651 | V651I | B | 9.6e-09 | 1.3e-08 | 4e-09 | 8.867E-09 | 4.545E-09 | 4.000E-09 | 1.300E-08 |
| 13011 | 651 | V651L | B | 1.4e-08 | 9e-09 | 4.3e-09 | 9.100E-09 | 4.851E-09 | 4.300E-09 | 1.400E-08 |
| 13012 | 651 | V651K | B | 9.3e-09 | 1.4e-08 | 3.6e-09 | 8.967E-09 | 5.208E-09 | 3.600E-09 | 1.400E-08 |
| 13013 | 651 | V651M | B | 8.3e-09 | 1e-08 | 1.2e-08 | 1.010E-08 | 1.852E-09 | 8.300E-09 | 1.200E-08 |
| 13014 | 651 | V651F | B | 6.7e-09 | 6.3e-09 | 6e-09 | 6.333E-09 | 3.512E-10 | 6.000E-09 | 6.700E-09 |
| 13015 | 651 | V651P | B | 9.9e-09 | 4.3e-09 | 1.2e-08 | 8.733E-09 | 3.980E-09 | 4.300E-09 | 1.200E-08 |
| 13016 | 651 | V651S | B | 2.6e-09 | 4e-09 | 2.3e-08 | 9.867E-09 | 1.140E-08 | 2.600E-09 | 2.300E-08 |
| 13017 | 651 | V651T | B | 3.4e-09 | 1.5e-08 | 1.2e-08 | 1.013E-08 | 6.021E-09 | 3.400E-09 | 1.500E-08 |
| 13018 | 651 | V651W | B | 6.2e-09 | 1e-08 | 9e-09 | 8.400E-09 | 1.970E-09 | 6.200E-09 | 1.000E-08 |
| 13019 | 651 | V651Y | B | 4e-09 | 2.8e-09 | 8.7e-09 | 5.167E-09 | 3.118E-09 | 2.800E-09 | 8.700E-09 |
| 13020 | 651 | V651V | B | 1.2e-08 | 4e-09 | 1.2e-08 | 9.333E-09 | 4.619E-09 | 4.000E-09 | 1.200E-08 |
| 13021 | 652 | S652A | B | 1.2e-08 | 6.1e-09 | 7.5e-09 | 8.533E-09 | 3.083E-09 | 6.100E-09 | 1.200E-08 |
| 13022 | 652 | S652R | B | 6.3e-09 | 6.7e-09 | 2.4e-09 | 5.133E-09 | 2.376E-09 | 2.400E-09 | 6.700E-09 |
| 13023 | 652 | S652N | B | 6.5e-09 | 1e-08 | 4.5e-09 | 7.000E-09 | 2.784E-09 | 4.500E-09 | 1.000E-08 |
| 13024 | 652 | S652D | B | 7.3e-09 | 7e-09 | 5.6e-09 | 6.633E-09 | 9.074E-10 | 5.600E-09 | 7.300E-09 |
| 13025 | 652 | S652C | B | 4.9e-09 | 3.7e-09 | 3.9e-09 | 4.167E-09 | 6.429E-10 | 3.700E-09 | 4.900E-09 |
| 13026 | 652 | S652E | B | 6.8e-09 | 2.7e-09 | 4.5e-09 | 4.667E-09 | 2.055E-09 | 2.700E-09 | 6.800E-09 |
| 13027 | 652 | S652Q | B | 7.9e-09 | 4.7e-09 | 7.7e-09 | 6.767E-09 | 1.793E-09 | 4.700E-09 | 7.900E-09 |
| 13028 | 652 | S652G | B | 1.7e-09 | 9.4e-09 | 1.3e-08 | 8.033E-09 | 5.773E-09 | 1.700E-09 | 1.300E-08 |
| 13029 | 652 | S652H | B | 1.1e-08 | 8.3e-09 | 1.1e-08 | 1.010E-08 | 1.559E-09 | 8.300E-09 | 1.100E-08 |
| 13030 | 652 | S652I | B | 2.9e-09 | 7.5e-09 | 3.7e-09 | 4.700E-09 | 2.458E-09 | 2.900E-09 | 7.500E-09 |
| 13031 | 652 | S652L | B | 1.7e-08 | 9.5e-09 | 6.1e-09 | 1.087E-08 | 5.577E-09 | 6.100E-09 | 1.700E-08 |
| 13032 | 652 | S652K | B | 8.9e-09 | 4.4e-09 | 2.1e-08 | 1.143E-08 | 8.585E-09 | 4.400E-09 | 2.100E-08 |
| 13033 | 652 | S652M | B | 6.7e-09 | 9.7e-09 | 3.2e-09 | 6.533E-09 | 3.253E-09 | 3.200E-09 | 9.700E-09 |
| 13034 | 652 | S652F | B | 1.5e-08 | 1.1e-08 | 6.5e-09 | 1.083E-08 | 4.252E-09 | 6.500E-09 | 1.500E-08 |
| 13035 | 652 | S652P | B | 4.2e-09 | 1.2e-08 | 9.7e-09 | 8.633E-09 | 4.008E-09 | 4.200E-09 | 1.200E-08 |
| 13036 | 652 | S652S | B | 5.6e-09 | 1.9e-08 | 4.3e-09 | 9.633E-09 | 8.138E-09 | 4.300E-09 | 1.900E-08 |
| 13037 | 652 | S652T | B | 1.9e-09 | 5.8e-09 | 7.9e-09 | 5.200E-09 | 3.045E-09 | 1.900E-09 | 7.900E-09 |
| 13038 | 652 | S652W | B | 7.4e-09 | 1.7e-09 | 2.7e-08 | 1.203E-08 | 1.327E-08 | 1.700E-09 | 2.700E-08 |
| 13039 | 652 | S652Y | B | 1e-08 | 1.3e-08 | 1.2e-08 | 1.167E-08 | 1.528E-09 | 1.000E-08 | 1.300E-08 |
| 13040 | 652 | S652V | B | 5.8e-09 | 9.9e-09 | 6e-09 | 7.233E-09 | 2.312E-09 | 5.800E-09 | 9.900E-09 |
| 13041 | 653 | I653A | B | 7.4e-09 | 5.9e-09 | 5.5e-09 | 6.267E-09 | 1.002E-09 | 5.500E-09 | 7.400E-09 |
| 13042 | 653 | I653R | B | 5.8e-09 | 2.8e-08 | 7.6e-09 | 1.380E-08 | 1.233E-08 | 5.800E-09 | 2.800E-08 |
| 13043 | 653 | I653N | B | 6.7e-09 | 1.8e-08 | 7.1e-09 | 1.060E-08 | 6.412E-09 | 6.700E-09 | 1.800E-08 |
| 13044 | 653 | I653D | B | 2.8e-09 | 7.1e-09 | 7.9e-09 | 5.933E-09 | 2.743E-09 | 2.800E-09 | 7.900E-09 |
| 13045 | 653 | I653C | B | 7.9e-09 | 1.2e-08 | 1e-08 | 9.967E-09 | 2.050E-09 | 7.900E-09 | 1.200E-08 |
| 13046 | 653 | I653E | B | 8.7e-09 | 1.7e-08 | 3.1e-09 | 9.600E-09 | 6.994E-09 | 3.100E-09 | 1.700E-08 |
| 13047 | 653 | I653Q | B | 9.2e-09 | 6.5e-09 | 5.4e-09 | 7.033E-09 | 1.955E-09 | 5.400E-09 | 9.200E-09 |
| 13048 | 653 | I653G | B | 7.7e-09 | 4.2e-09 | 1.7e-08 | 9.633E-09 | 6.615E-09 | 4.200E-09 | 1.700E-08 |
| 13049 | 653 | I653H | B | 9.7e-09 | 9.6e-09 | 2.4e-08 | 1.443E-08 | 8.285E-09 | 9.600E-09 | 2.400E-08 |
| 13050 | 653 | I653I | B | 9.3e-09 | 8e-09 | 2.8e-09 | 6.700E-09 | 3.439E-09 | 2.800E-09 | 9.300E-09 |
| 13051 | 653 | I653L | B | 1.1e-08 | 8.1e-09 | 2.2e-09 | 7.100E-09 | 4.484E-09 | 2.200E-09 | 1.100E-08 |
| 13052 | 653 | I653K | B | 8.9e-09 | 2.2e-09 | 6.5e-09 | 5.867E-09 | 3.395E-09 | 2.200E-09 | 8.900E-09 |
| 13053 | 653 | I653M | B | 1e-08 | 7.2e-09 | 2.7e-09 | 6.633E-09 | 3.683E-09 | 2.700E-09 | 1.000E-08 |
| 13054 | 653 | I653F | B | 8.3e-09 | 5.9e-09 | 6.6e-09 | 6.933E-09 | 1.234E-09 | 5.900E-09 | 8.300E-09 |
| 13055 | 653 | I653P | B | 1.1e-08 | 2.5e-09 | 7.3e-09 | 6.933E-09 | 4.262E-09 | 2.500E-09 | 1.100E-08 |
| 13056 | 653 | I653S | B | 5.3e-09 | 6.7e-09 | 3.6e-09 | 5.200E-09 | 1.552E-09 | 3.600E-09 | 6.700E-09 |
| 13057 | 653 | I653T | B | 1.3e-08 | 1.6e-08 | 1.5e-08 | 1.467E-08 | 1.528E-09 | 1.300E-08 | 1.600E-08 |
| 13058 | 653 | I653W | B | 5.1e-09 | 6.5e-09 | 6.8e-09 | 6.133E-09 | 9.074E-10 | 5.100E-09 | 6.800E-09 |
| 13059 | 653 | I653Y | B | 8e-09 | 5e-09 | 3.5e-09 | 5.500E-09 | 2.291E-09 | 3.500E-09 | 8.000E-09 |
| 13060 | 653 | I653V | B | 4.6e-09 | 7.4e-09 | 5.8e-09 | 5.933E-09 | 1.405E-09 | 4.600E-09 | 7.400E-09 |
| 13061 | 654 | G654A | B | 7.6e-09 | 5.9e-09 | 1e-08 | 7.833E-09 | 2.060E-09 | 5.900E-09 | 1.000E-08 |
| 13062 | 654 | G654R | B | 4.8e-09 | 4.7e-09 | 6.5e-09 | 5.333E-09 | 1.012E-09 | 4.700E-09 | 6.500E-09 |
| 13063 | 654 | G654N | B | 4.8e-09 | 3.7e-09 | 5.1e-09 | 4.533E-09 | 7.371E-10 | 3.700E-09 | 5.100E-09 |
| 13064 | 654 | G654D | B | 6.5e-09 | 4.1e-09 | 3.3e-09 | 4.633E-09 | 1.665E-09 | 3.300E-09 | 6.500E-09 |
| 13065 | 654 | G654C | B | 8.3e-09 | 5e-09 | 3.7e-09 | 5.667E-09 | 2.371E-09 | 3.700E-09 | 8.300E-09 |
| 13066 | 654 | G654E | B | 1.3e-08 | 9.1e-09 | 1.2e-08 | 1.137E-08 | 2.026E-09 | 9.100E-09 | 1.300E-08 |
| 13067 | 654 | G654Q | B | 1e-08 | 1.2e-08 | 6.2e-09 | 9.400E-09 | 2.946E-09 | 6.200E-09 | 1.200E-08 |
| 13068 | 654 | G654G | B | 2.6e-09 | 7.3e-09 | 1e-08 | 6.633E-09 | 3.745E-09 | 2.600E-09 | 1.000E-08 |
| 13069 | 654 | G654H | B | 1.1e-08 | 4.3e-09 | 2.6e-08 | 1.377E-08 | 1.111E-08 | 4.300E-09 | 2.600E-08 |
| 13070 | 654 | G654I | B | 1.7e-08 | 1.2e-08 | 7e-09 | 1.200E-08 | 5.000E-09 | 7.000E-09 | 1.700E-08 |
| 13071 | 654 | G654L | B | 6.5e-09 | 7.9e-09 | 6.4e-09 | 6.933E-09 | 8.386E-10 | 6.400E-09 | 7.900E-09 |
| 13072 | 654 | G654K | B | 6.6e-09 | 3.9e-09 | 2.9e-09 | 4.467E-09 | 1.914E-09 | 2.900E-09 | 6.600E-09 |
| 13073 | 654 | G654M | B | 5.6e-09 | 5.3e-09 | 5e-09 | 5.300E-09 | 3.000E-10 | 5.000E-09 | 5.600E-09 |
| 13074 | 654 | G654F | B | 3.2e-09 | 8.1e-09 | 1.3e-08 | 8.100E-09 | 4.900E-09 | 3.200E-09 | 1.300E-08 |
| 13075 | 654 | G654P | B | 1.3e-08 | 3.7e-09 | 4.2e-09 | 6.967E-09 | 5.231E-09 | 3.700E-09 | 1.300E-08 |
| 13076 | 654 | G654S | B | 5e-09 | 5.4e-09 | 2.1e-09 | 4.167E-09 | 1.801E-09 | 2.100E-09 | 5.400E-09 |
| 13077 | 654 | G654T | B | 1.5e-09 | 5e-09 | 3.9e-09 | 3.467E-09 | 1.790E-09 | 1.500E-09 | 5.000E-09 |
| 13078 | 654 | G654W | B | 6.7e-09 | 1.9e-08 | 5.3e-09 | 1.033E-08 | 7.538E-09 | 5.300E-09 | 1.900E-08 |
| 13079 | 654 | G654Y | B | 4.5e-09 | 6.6e-09 | 2.3e-09 | 4.467E-09 | 2.150E-09 | 2.300E-09 | 6.600E-09 |
| 13080 | 654 | G654V | B | 1.1e-08 | 1.3e-08 | 5.1e-09 | 9.700E-09 | 4.107E-09 | 5.100E-09 | 1.300E-08 |
| 13081 | 655 | V655A | B | 1.2e-08 | 1.2e-08 | 1.1e-08 | 1.167E-08 | 5.774E-10 | 1.100E-08 | 1.200E-08 |
| 13082 | 655 | V655R | B | 5e-09 | 3.7e-09 | 3.6e-08 | 1.490E-08 | 1.828E-08 | 3.700E-09 | 3.600E-08 |
| 13083 | 655 | V655N | B | 2.3e-09 | 4.8e-09 | 6.6e-09 | 4.567E-09 | 2.159E-09 | 2.300E-09 | 6.600E-09 |
| 13084 | 655 | V655D | B | 8.1e-09 | 3.9e-09 | 8.2e-09 | 6.733E-09 | 2.454E-09 | 3.900E-09 | 8.200E-09 |
| 13085 | 655 | V655C | B | 2.4e-09 | 9.6e-10 | 2.4e-08 | 9.120E-09 | 1.291E-08 | 9.600E-10 | 2.400E-08 |
| 13086 | 655 | V655E | B | 9.9e-09 | 3.6e-08 | 7.7e-09 | 1.787E-08 | 1.574E-08 | 7.700E-09 | 3.600E-08 |
| 13087 | 655 | V655Q | B | 1.3e-08 | 1.1e-08 | 1.6e-08 | 1.333E-08 | 2.517E-09 | 1.100E-08 | 1.600E-08 |
| 13088 | 655 | V655G | B | 6.8e-09 | 5e-09 | 9.5e-09 | 7.100E-09 | 2.265E-09 | 5.000E-09 | 9.500E-09 |
| 13089 | 655 | V655H | B | 3.9e-09 | 6.4e-09 | 9.4e-09 | 6.567E-09 | 2.754E-09 | 3.900E-09 | 9.400E-09 |

| | | | | | | | | | | |
|---|---|---|---|---|---|---|---|---|---|---|
| 13090 | 655 | V655I | B | 5.5e-09 | 9.1e-09 | 7.2e-09 | 7.267E-09 | 1.801E-09 | 5.500E-09 | 9.100E-09 |
| 13091 | 655 | V655L | B | 1.6e-08 | 6.1e-09 | 2.1e-09 | 8.067E-09 | 7.156E-09 | 2.100E-09 | 1.600E-08 |
| 13092 | 655 | V655K | B | 1.5e-09 | 4.4e-09 | 6.8e-09 | 4.233E-09 | 2.654E-09 | 1.500E-09 | 6.800E-09 |
| 13093 | 655 | V655M | B | 5.4e-09 | 1e-08 | 5.3e-09 | 6.900E-09 | 2.685E-09 | 5.300E-09 | 1.000E-08 |
| 13094 | 655 | V655F | B | 3.3e-09 | 5.6e-09 | 8.3e-09 | 5.733E-09 | 2.503E-09 | 3.300E-09 | 8.300E-09 |
| 13095 | 655 | V655P | B | 9.9e-09 | 7.8e-09 | 8.3e-09 | 8.667E-09 | 1.097E-09 | 7.800E-09 | 9.900E-09 |
| 13096 | 655 | V655S | B | 5.4e-09 | 2.5e-09 | 8e-09 | 5.300E-09 | 2.751E-09 | 2.500E-09 | 8.000E-09 |
| 13097 | 655 | V655T | B | 1.5e-08 | 4.9e-09 | 7.5e-09 | 9.133E-09 | 5.244E-09 | 4.900E-09 | 1.500E-08 |
| 13098 | 655 | V655W | B | 1.5e-08 | 1.8e-08 | 1.7e-08 | 1.667E-08 | 1.528E-09 | 1.500E-08 | 1.800E-08 |
| 13099 | 655 | V655Y | B | 4e-09 | 1.1e-08 | 5.8e-09 | 6.933E-09 | 3.635E-09 | 4.000E-09 | 1.100E-08 |
| 13100 | 655 | V655V | B | 2e-08 | 7.9e-09 | 8.3e-09 | 1.207E-08 | 6.873E-09 | 7.900E-09 | 2.000E-08 |
| 13101 | 656 | A656A | B | 5.3e-09 | 9.7e-09 | 1.7e-08 | 1.067E-08 | 5.910E-09 | 5.300E-09 | 1.700E-08 |
| 13102 | 656 | A656R | B | 4.5e-09 | 1.3e-08 | 9.1e-09 | 8.867E-09 | 4.255E-09 | 4.500E-09 | 1.300E-08 |
| 13103 | 656 | A656N | B | 2.7e-08 | 1.4e-08 | 5.2e-09 | 1.540E-08 | 1.097E-08 | 5.200E-09 | 2.700E-08 |
| 13104 | 656 | A656D | B | 2e-09 | 4.1e-09 | 1.1e-08 | 5.700E-09 | 4.709E-09 | 2.000E-09 | 1.100E-08 |
| 13105 | 656 | A656C | B | 7e-09 | 3.6e-08 | 4.3e-09 | 1.577E-08 | 1.757E-08 | 4.300E-09 | 3.600E-08 |
| 13106 | 656 | A656E | B | 5.4e-08 | 6.2e-09 | 6.8e-09 | 2.233E-08 | 2.743E-08 | 6.200E-09 | 5.400E-08 |
| 13107 | 656 | A656Q | B | 8.9e-09 | 2.9e-08 | 4.8e-09 | 1.423E-08 | 1.295E-08 | 4.800E-09 | 2.900E-08 |
| 13108 | 656 | A656G | B | 2.1e-09 | 7.5e-09 | 1.7e-08 | 8.867E-09 | 7.543E-09 | 2.100E-09 | 1.700E-08 |
| 13109 | 656 | A656H | B | 7.9e-09 | 7.6e-09 | 3.9e-09 | 6.467E-09 | 2.228E-09 | 3.900E-09 | 7.900E-09 |
| 13110 | 656 | A656I | B | 7.3e-09 | 5.5e-09 | 8.5e-09 | 7.100E-09 | 1.510E-09 | 5.500E-09 | 8.500E-09 |
| 13111 | 656 | A656L | B | 1.9e-08 | 9.1e-09 | 8.8e-09 | 1.230E-08 | 5.804E-09 | 8.800E-09 | 1.900E-08 |
| 13112 | 656 | A656K | B | 6.8e-09 | 8.1e-09 | 7.6e-09 | 7.500E-09 | 6.557E-10 | 6.800E-09 | 8.100E-09 |
| 13113 | 656 | A656M | B | 1.3e-08 | 6.1e-09 | 2.3e-09 | 7.133E-09 | 5.424E-09 | 2.300E-09 | 1.300E-08 |
| 13114 | 656 | A656F | B | 2e-08 | 1.8e-08 | 1.7e-08 | 1.833E-08 | 1.528E-09 | 1.700E-08 | 2.000E-08 |
| 13115 | 656 | A656P | B | 5.8e-09 | 9.2e-09 | 1.8e-08 | 1.100E-08 | 6.296E-09 | 5.800E-09 | 1.800E-08 |
| 13116 | 656 | A656S | B | 8.5e-09 | 2.3e-09 | 7.5e-09 | 6.100E-09 | 3.329E-09 | 2.300E-09 | 8.500E-09 |
| 13117 | 656 | A656T | B | 6.9e-09 | 8.1e-09 | 2.1e-09 | 5.700E-09 | 3.175E-09 | 2.100E-09 | 8.100E-09 |
| 13118 | 656 | A656W | B | 5.3e-09 | 8.1e-09 | 3.1e-09 | 5.500E-09 | 2.506E-09 | 3.100E-09 | 8.100E-09 |
| 13119 | 656 | A656Y | B | 4.6e-09 | 2.5e-09 | 4.1e-09 | 3.733E-09 | 1.097E-09 | 2.500E-09 | 4.600E-09 |
| 13120 | 656 | A656V | B | 2.1e-08 | 8.1e-09 | 1.1e-08 | 1.337E-08 | 6.768E-09 | 8.100E-09 | 2.100E-08 |
| 13121 | 657 | W657A | B | 8e-09 | 1.3e-08 | 1.1e-08 | 1.067E-08 | 2.517E-09 | 8.000E-09 | 1.300E-08 |
| 13122 | 657 | W657R | B | 6.2e-09 | 4.1e-09 | 3.7e-09 | 4.667E-09 | 1.343E-09 | 3.700E-09 | 6.200E-09 |
| 13123 | 657 | W657N | B | 5.4e-09 | 1.3e-08 | 1.3e-08 | 1.047E-08 | 4.388E-09 | 5.400E-09 | 1.300E-08 |
| 13124 | 657 | W657D | B | 5.1e-09 | 9.8e-09 | 1.2e-08 | 8.967E-09 | 3.525E-09 | 5.100E-09 | 1.200E-08 |
| 13125 | 657 | W657C | B | 7.6e-09 | 1.8e-08 | 1.3e-08 | 1.287E-08 | 5.201E-09 | 7.600E-09 | 1.800E-08 |
| 13126 | 657 | W657E | B | 1e-08 | 1e-08 | 1.2e-08 | 1.067E-08 | 1.155E-09 | 1.000E-08 | 1.200E-08 |
| 13127 | 657 | W657Q | B | 6.7e-09 | 4.3e-09 | 9.7e-09 | 6.900E-09 | 2.706E-09 | 4.300E-09 | 9.700E-09 |
| 13128 | 657 | W657G | B | 5.3e-09 | 6.5e-09 | 7.7e-09 | 6.500E-09 | 1.200E-09 | 5.300E-09 | 7.700E-09 |
| 13129 | 657 | W657H | B | 1.6e-08 | 5.1e-09 | 4.3e-09 | 8.467E-09 | 6.536E-09 | 4.300E-09 | 1.600E-08 |
| 13130 | 657 | W657I | B | 3.7e-09 | 6.4e-09 | 8.2e-09 | 6.100E-09 | 2.265E-09 | 3.700E-09 | 8.200E-09 |
| 13131 | 657 | W657L | B | 4.6e-09 | 1.3e-08 | 2.1e-08 | 1.287E-08 | 8.201E-09 | 4.600E-09 | 2.100E-08 |
| 13132 | 657 | W657K | B | 6.4e-09 | 2.9e-09 | 4.8e-09 | 4.700E-09 | 1.752E-09 | 2.900E-09 | 6.400E-09 |
| 13133 | 657 | W657M | B | 1.2e-08 | 6.5e-09 | 9.3e-09 | 9.267E-09 | 2.750E-09 | 6.500E-09 | 1.200E-08 |
| 13134 | 657 | W657F | B | 5.3e-09 | 4.6e-09 | 1.4e-08 | 7.967E-09 | 5.237E-09 | 4.600E-09 | 1.400E-08 |
| 13135 | 657 | W657P | B | 1.1e-08 | 1e-08 | 1.4e-08 | 1.167E-08 | 2.082E-09 | 1.000E-08 | 1.400E-08 |
| 13136 | 657 | W657S | B | 2e-08 | 5.9e-09 | 2.1e-09 | 9.333E-09 | 9.431E-09 | 2.100E-09 | 2.000E-08 |
| 13137 | 657 | W657T | B | 2.9e-09 | 1.3e-08 | 7.2e-09 | 7.700E-09 | 5.069E-09 | 2.900E-09 | 1.300E-08 |
| 13138 | 657 | W657W | B | 9.2e-09 | 7.1e-09 | 5.8e-09 | 7.367E-09 | 1.716E-09 | 5.800E-09 | 9.200E-09 |
| 13139 | 657 | W657Y | B | 8.9e-09 | 1.7e-08 | 4.1e-09 | 1.000E-08 | 6.520E-09 | 4.100E-09 | 1.700E-08 |
| 13140 | 657 | W657V | B | 1.5e-08 | 2.2e-08 | 8.8e-09 | 1.527E-08 | 6.604E-09 | 8.800E-09 | 2.200E-08 |
| 13141 | 658 | Y658A | B | 5.6e-09 | 7.1e-09 | 1.8e-09 | 4.833E-09 | 2.732E-09 | 1.800E-09 | 7.100E-09 |
| 13142 | 658 | Y658R | B | 1.4e-08 | 1.6e-08 | 1.1e-08 | 1.367E-08 | 2.517E-09 | 1.100E-08 | 1.600E-08 |
| 13143 | 658 | Y658N | B | 2.5e-09 | 8.7e-09 | 6.1e-09 | 5.767E-09 | 3.113E-09 | 2.500E-09 | 8.700E-09 |
| 13144 | 658 | Y658D | B | 2.2e-08 | 5.9e-09 | 1.1e-08 | 1.297E-08 | 8.228E-09 | 5.900E-09 | 2.200E-08 |
| 13145 | 658 | Y658C | B | 3.4e-09 | 1.8e-08 | 1.7e-08 | 1.280E-08 | 8.156E-09 | 3.400E-09 | 1.800E-08 |
| 13146 | 658 | Y658E | B | 4e-09 | 9.8e-09 | 6.6e-09 | 6.800E-09 | 2.905E-09 | 4.000E-09 | 9.800E-09 |
| 13147 | 658 | Y658Q | B | 3.3e-09 | 6.5e-09 | 4.4e-09 | 4.733E-09 | 1.626E-09 | 3.300E-09 | 6.500E-09 |
| 13148 | 658 | Y658G | B | 1.3e-08 | 2.3e-08 | 8.9e-09 | 1.497E-08 | 7.253E-09 | 8.900E-09 | 2.300E-08 |
| 13149 | 658 | Y658H | B | 1.8e-08 | 6.8e-09 | 2.9e-09 | 9.233E-09 | 7.839E-09 | 2.900E-09 | 1.800E-08 |
| 13150 | 658 | Y658I | B | 1.1e-08 | 7.4e-09 | 7.1e-09 | 8.500E-09 | 2.170E-09 | 7.100E-09 | 1.100E-08 |
| 13151 | 658 | Y658L | B | 5.4e-09 | 1.2e-08 | 1.1e-08 | 9.467E-09 | 3.557E-09 | 5.400E-09 | 1.200E-08 |
| 13152 | 658 | Y658K | B | 8.8e-09 | 1.3e-08 | 6.7e-09 | 9.500E-09 | 3.208E-09 | 6.700E-09 | 1.300E-08 |
| 13153 | 658 | Y658M | B | 9e-09 | 5.2e-09 | 3.2e-08 | 1.540E-08 | 1.450E-08 | 5.200E-09 | 3.200E-08 |
| 13154 | 658 | Y658F | B | 5.6e-09 | 7e-09 | 1.4e-08 | 8.867E-09 | 4.500E-09 | 5.600E-09 | 1.400E-08 |
| 13155 | 658 | Y658P | B | 2.6e-09 | 1.1e-08 | 1.3e-08 | 8.867E-09 | 5.518E-09 | 2.600E-09 | 1.300E-08 |
| 13156 | 658 | Y658S | B | 2.4e-09 | 6.1e-09 | 6.5e-09 | 5.000E-09 | 2.261E-09 | 2.400E-09 | 6.500E-09 |
| 13157 | 658 | Y658T | B | 6.4e-09 | 4.6e-09 | 4.7e-09 | 5.233E-09 | 1.012E-09 | 4.600E-09 | 6.400E-09 |
| 13158 | 658 | Y658W | B | 1.3e-08 | 8.8e-09 | 8.9e-09 | 1.023E-08 | 2.397E-09 | 8.800E-09 | 1.300E-08 |
| 13159 | 658 | Y658Y | B | 4.2e-09 | 1e-08 | 3.7e-09 | 5.967E-09 | 3.502E-09 | 3.700E-09 | 1.000E-08 |
| 13160 | 658 | Y658V | B | 6.2e-09 | 1.3e-08 | 9.6e-09 | 9.600E-09 | 3.400E-09 | 6.200E-09 | 1.300E-08 |
| 13161 | 659 | Q659A | B | 2.9e-09 | 1.2e-08 | 1.6e-09 | 5.500E-09 | 5.667E-09 | 1.600E-09 | 1.200E-08 |
| 13162 | 659 | Q659R | B | 5.1e-09 | 1.2e-08 | 4.6e-09 | 7.233E-09 | 4.136E-09 | 4.600E-09 | 1.200E-08 |
| 13163 | 659 | Q659N | B | 1.4e-08 | 7e-09 | 3.4e-09 | 8.133E-09 | 5.390E-09 | 3.400E-09 | 1.400E-08 |
| 13164 | 659 | Q659D | B | 1e-08 | 1.1e-08 | 1e-08 | 1.033E-08 | 5.774E-10 | 1.000E-08 | 1.100E-08 |
| 13165 | 659 | Q659C | B | 3e-09 | 6.7e-09 | 1.4e-08 | 7.900E-09 | 5.597E-09 | 3.000E-09 | 1.400E-08 |
| 13166 | 659 | Q659E | B | 4e-09 | 6e-09 | 2.1e-08 | 1.033E-08 | 9.292E-09 | 4.000E-09 | 2.100E-08 |
| 13167 | 659 | Q659Q | B | 3.3e-09 | 1.9e-08 | 9.9e-09 | 1.073E-08 | 7.883E-09 | 3.300E-09 | 1.900E-08 |
| 13168 | 659 | Q659G | B | 2.7e-09 | 5.7e-09 | 1.3e-08 | 7.133E-09 | 5.297E-09 | 2.700E-09 | 1.300E-08 |
| 13169 | 659 | Q659H | B | 2.9e-09 | 1.1e-08 | 4.6e-09 | 6.167E-09 | 4.271E-09 | 2.900E-09 | 1.100E-08 |
| 13170 | 659 | Q659I | B | 8.2e-09 | 9.3e-09 | 4.5e-09 | 7.333E-09 | 2.515E-09 | 4.500E-09 | 9.300E-09 |
| 13171 | 659 | Q659L | B | 8.5e-09 | 5.9e-09 | 1.9e-08 | 1.113E-08 | 6.936E-09 | 5.900E-09 | 1.900E-08 |
| 13172 | 659 | Q659K | B | 3.3e-09 | 6.6e-09 | 7.5e-09 | 5.800E-09 | 2.211E-09 | 3.300E-09 | 7.500E-09 |
| 13173 | 659 | Q659M | B | 7.1e-09 | 9.5e-09 | 2.9e-09 | 6.500E-09 | 3.341E-09 | 2.900E-09 | 9.500E-09 |
| 13174 | 659 | Q659F | B | 1.1e-08 | 7.2e-09 | 6.7e-09 | 8.300E-09 | 2.352E-09 | 6.700E-09 | 1.100E-08 |

| | | | | | | | | | | |
|---|---|---|---|---|---|---|---|---|---|---|
| 13175 | 659 | Q659P | B | 1.8e-08 | 4.2e-09 | 1.4e-09 | 7.867E-09 | 8.887E-09 | 1.400E-09 | 1.800E-08 |
| 13176 | 659 | Q659S | B | 1.3e-09 | 3e-09 | 2e-09 | 2.100E-09 | 8.544E-10 | 1.300E-09 | 3.000E-09 |
| 13177 | 659 | Q659T | B | 7e-09 | 2.6e-09 | 2.4e-08 | 1.120E-08 | 1.130E-08 | 2.600E-09 | 2.400E-08 |
| 13178 | 659 | Q659W | B | 9.5e-09 | 4.1e-09 | 1.2e-08 | 8.533E-09 | 4.038E-09 | 4.100E-09 | 1.200E-08 |
| 13179 | 659 | Q659Y | B | 5.3e-09 | 4.8e-09 | 6.8e-09 | 5.633E-09 | 1.041E-09 | 4.800E-09 | 6.800E-09 |
| 13180 | 659 | Q659V | B | 8.1e-09 | 8.2e-09 | 3e-09 | 6.433E-09 | 2.974E-09 | 3.000E-09 | 8.200E-09 |
| 13181 | 660 | Q660A | B | 6.4e-09 | 6.7e-09 | 6e-09 | 6.367E-09 | 3.512E-10 | 6.000E-09 | 6.700E-09 |
| 13182 | 660 | Q660R | B | 1.1e-08 | 1.4e-08 | 3e-09 | 9.333E-09 | 5.686E-09 | 3.000E-09 | 1.400E-08 |
| 13183 | 660 | Q660N | B | 5e-09 | 9.3e-09 | 7.8e-09 | 7.367E-09 | 2.183E-09 | 5.000E-09 | 9.300E-09 |
| 13184 | 660 | Q660D | B | 8.8e-09 | 7.1e-09 | 1.4e-08 | 9.967E-09 | 3.595E-09 | 7.100E-09 | 1.400E-08 |
| 13185 | 660 | Q660C | B | 6.1e-09 | 8.1e-09 | 9.2e-09 | 7.800E-09 | 1.572E-09 | 6.100E-09 | 9.200E-09 |
| 13186 | 660 | Q660E | B | 5.7e-09 | 8.5e-09 | 6.6e-09 | 6.933E-09 | 1.429E-09 | 5.700E-09 | 8.500E-09 |
| 13187 | 660 | Q660Q | B | 8.2e-09 | 8e-09 | 3.3e-09 | 6.500E-09 | 2.773E-09 | 3.300E-09 | 8.200E-09 |
| 13188 | 660 | Q660G | B | 7.6e-09 | 3.1e-09 | 3.8e-09 | 4.833E-09 | 2.421E-09 | 3.100E-09 | 7.600E-09 |
| 13189 | 660 | Q660H | B | 9.4e-09 | 7.2e-09 | 9.7e-09 | 8.767E-09 | 1.365E-09 | 7.200E-09 | 9.700E-09 |
| 13190 | 660 | Q660I | B | 1e-08 | 8.4e-09 | 8.5e-09 | 8.967E-09 | 8.963E-10 | 8.400E-09 | 1.000E-08 |
| 13191 | 660 | Q660L | B | 7.2e-09 | 2.5e-09 | 9.4e-09 | 6.367E-09 | 3.525E-09 | 2.500E-09 | 9.400E-09 |
| 13192 | 660 | Q660K | B | 1.2e-08 | 1e-08 | 8.2e-09 | 1.007E-08 | 1.901E-09 | 8.200E-09 | 1.200E-08 |
| 13193 | 660 | Q660M | B | 1e-08 | 1.8e-09 | 6.4e-09 | 6.067E-09 | 4.110E-09 | 1.800E-09 | 1.000E-08 |
| 13194 | 660 | Q660F | B | 5.9e-09 | 8.2e-09 | 8.7e-09 | 7.600E-09 | 1.493E-09 | 5.900E-09 | 8.700E-09 |
| 13195 | 660 | Q660P | B | 1.3e-08 | 2.6e-08 | 1.1e-08 | 1.667E-08 | 8.145E-09 | 1.100E-08 | 2.600E-08 |
| 13196 | 660 | Q660S | B | 1.7e-08 | 5.6e-09 | 3.7e-09 | 8.767E-09 | 7.193E-09 | 3.700E-09 | 1.700E-08 |
| 13197 | 660 | Q660T | B | 2.8e-09 | 1.4e-08 | 4.7e-09 | 7.167E-09 | 5.994E-09 | 2.800E-09 | 1.400E-08 |
| 13198 | 660 | Q660W | B | 5.4e-09 | 6e-09 | 6.5e-09 | 5.967E-09 | 5.508E-10 | 5.400E-09 | 6.500E-09 |
| 13199 | 660 | Q660Y | B | 1.1e-08 | 5.3e-09 | 9.9e-09 | 8.733E-09 | 3.024E-09 | 5.300E-09 | 1.100E-08 |
| 13200 | 660 | Q660V | B | 3e-09 | 8.5e-09 | 5.7e-09 | 5.733E-09 | 2.750E-09 | 3.000E-09 | 8.500E-09 |
| 13201 | 661 | K661A | B | 1.5e-08 | 1e-08 | 9.6e-09 | 1.153E-08 | 3.009E-09 | 9.600E-09 | 1.500E-08 |
| 13202 | 661 | K661R | B | 4.3e-09 | 1.2e-08 | 8.6e-09 | 8.300E-09 | 3.859E-09 | 4.300E-09 | 1.200E-08 |
| 13203 | 661 | K661N | B | 5.6e-09 | 3.8e-09 | 7e-09 | 5.467E-09 | 1.604E-09 | 3.800E-09 | 7.000E-09 |
| 13204 | 661 | K661D | B | 5e-09 | 5.2e-09 | 8.7e-10 | 3.690E-09 | 2.444E-09 | 8.700E-10 | 5.200E-09 |
| 13205 | 661 | K661C | B | 5.7e-09 | 1e-08 | 5.3e-08 | 2.290E-08 | 2.616E-08 | 5.700E-09 | 5.300E-08 |
| 13206 | 661 | K661E | B | 7.5e-09 | 4.4e-09 | 1.3e-08 | 8.300E-09 | 4.355E-09 | 4.400E-09 | 1.300E-08 |
| 13207 | 661 | K661Q | B | 1.6e-08 | 9.6e-09 | 1.5e-08 | 1.353E-08 | 3.443E-09 | 9.600E-09 | 1.600E-08 |
| 13208 | 661 | K661G | B | 4.9e-09 | 4e-09 | 1.7e-08 | 8.633E-09 | 7.260E-09 | 4.000E-09 | 1.700E-08 |
| 13209 | 661 | K661H | B | 8.8e-09 | 4.8e-09 | 1.3e-08 | 8.867E-09 | 4.100E-09 | 4.800E-09 | 1.300E-08 |
| 13210 | 661 | K661I | B | 9.8e-09 | 2.6e-09 | 1.1e-09 | 4.500E-09 | 4.651E-09 | 1.100E-09 | 9.800E-09 |
| 13211 | 661 | K661L | B | 1.9e-09 | 1.4e-08 | 2.3e-09 | 6.067E-09 | 6.873E-09 | 1.900E-09 | 1.400E-08 |
| 13212 | 661 | K661K | B | 9.7e-10 | 7.5e-09 | 1.5e-08 | 7.823E-09 | 7.021E-09 | 9.700E-10 | 1.500E-08 |
| 13213 | 661 | K661M | B | 4.9e-09 | 4.3e-09 | 1.4e-08 | 7.733E-09 | 5.435E-09 | 4.300E-09 | 1.400E-08 |
| 13214 | 661 | K661F | B | 1.5e-09 | 5.2e-09 | 1.2e-08 | 6.233E-09 | 5.326E-09 | 1.500E-09 | 1.200E-08 |
| 13215 | 661 | K661P | B | 9.2e-09 | 2.8e-08 | 9e-09 | 1.540E-08 | 1.091E-08 | 9.000E-09 | 2.800E-08 |
| 13216 | 661 | K661S | B | 8.5e-09 | 5e-09 | 2e-08 | 1.117E-08 | 7.848E-09 | 5.000E-09 | 2.000E-08 |
| 13217 | 661 | K661T | B | 8.6e-09 | 2.6e-08 | 5.6e-09 | 1.340E-08 | 1.101E-08 | 5.600E-09 | 2.600E-08 |
| 13218 | 661 | K661W | B | 8.2e-09 | 9.6e-09 | 7e-09 | 8.267E-09 | 1.301E-09 | 7.000E-09 | 9.600E-09 |
| 13219 | 661 | K661Y | B | 1.4e-08 | 3.3e-09 | 9.6e-09 | 8.967E-09 | 5.378E-09 | 3.300E-09 | 1.400E-08 |
| 13220 | 661 | K661V | B | 9.7e-09 | 7.3e-09 | 7e-09 | 8.000E-09 | 1.480E-09 | 7.000E-09 | 9.700E-09 |
| 13221 | 662 | P662A | B | 4.9e-09 | 4.2e-09 | 4.8e-09 | 4.633E-09 | 3.786E-10 | 4.200E-09 | 4.900E-09 |
| 13222 | 662 | P662R | B | 9.9e-09 | 4.4e-09 | 9e-09 | 7.767E-09 | 2.950E-09 | 4.400E-09 | 9.900E-09 |
| 13223 | 662 | P662N | B | 1.5e-09 | 4.9e-09 | 2.2e-08 | 9.467E-09 | 1.099E-08 | 1.500E-09 | 2.200E-08 |
| 13224 | 662 | P662D | B | 1.9e-08 | 4.3e-09 | 7e-09 | 1.010E-08 | 7.825E-09 | 4.300E-09 | 1.900E-08 |
| 13225 | 662 | P662C | B | 6.2e-09 | 3.9e-09 | 3.5e-09 | 4.533E-09 | 1.457E-09 | 3.500E-09 | 6.200E-09 |
| 13226 | 662 | P662E | B | 1.7e-08 | 1.1e-08 | 5e-09 | 1.100E-08 | 6.000E-09 | 5.000E-09 | 1.700E-08 |
| 13227 | 662 | P662Q | B | 2.8e-09 | 5.2e-09 | 8.2e-09 | 5.400E-09 | 2.706E-09 | 2.800E-09 | 8.200E-09 |
| 13228 | 662 | P662G | B | 7.4e-09 | 2.7e-09 | 3.5e-09 | 4.533E-09 | 2.515E-09 | 2.700E-09 | 7.400E-09 |
| 13229 | 662 | P662H | B | 1.1e-08 | 5.1e-09 | 9.8e-09 | 8.633E-09 | 3.118E-09 | 5.100E-09 | 1.100E-08 |
| 13230 | 662 | P662I | B | 5.7e-09 | 1.9e-08 | 5.7e-09 | 1.013E-08 | 7.679E-09 | 5.700E-09 | 1.900E-08 |
| 13231 | 662 | P662L | B | 6.3e-09 | 4.6e-09 | 3.6e-09 | 4.833E-09 | 1.365E-09 | 3.600E-09 | 6.300E-09 |
| 13232 | 662 | P662K | B | 4.9e-09 | 2.2e-09 | 7.8e-09 | 4.967E-09 | 2.801E-09 | 2.200E-09 | 7.800E-09 |
| 13233 | 662 | P662M | B | 6.8e-09 | 8.9e-09 | 1.1e-09 | 5.600E-09 | 4.036E-09 | 1.100E-09 | 8.900E-09 |
| 13234 | 662 | P662F | B | 8.8e-09 | 2.5e-09 | 4.2e-09 | 5.167E-09 | 3.259E-09 | 2.500E-09 | 8.800E-09 |
| 13235 | 662 | P662P | B | 5.5e-09 | 5.1e-09 | 7.8e-09 | 6.133E-09 | 1.457E-09 | 5.100E-09 | 7.800E-09 |
| 13236 | 662 | P662S | B | 9.8e-09 | 6.5e-09 | 4.9e-09 | 7.067E-09 | 2.499E-09 | 4.900E-09 | 9.800E-09 |
| 13237 | 662 | P662T | B | 1.1e-08 | 3.4e-09 | 6.1e-09 | 6.833E-09 | 3.853E-09 | 3.400E-09 | 1.100E-08 |
| 13238 | 662 | P662W | B | 2.7e-09 | 1.2e-08 | 2.3e-08 | 1.257E-08 | 1.016E-08 | 2.700E-09 | 2.300E-08 |
| 13239 | 662 | P662Y | B | 4.1e-09 | 6.4e-09 | 5e-09 | 5.167E-09 | 1.159E-09 | 4.100E-09 | 6.400E-09 |
| 13240 | 662 | P662V | B | 2.1e-08 | 5.6e-09 | 9.6e-09 | 1.207E-08 | 7.991E-09 | 5.600E-09 | 2.100E-08 |
| 13241 | 663 | G663A | B | 9.3e-09 | 3e-09 | 1.7e-08 | 9.767E-09 | 7.012E-09 | 3.000E-09 | 1.700E-08 |
| 13242 | 663 | G663R | B | 9.8e-09 | 8.8e-09 | 6.6e-09 | 8.400E-09 | 1.637E-09 | 6.600E-09 | 9.800E-09 |
| 13243 | 663 | G663N | B | 1.6e-08 | 1.6e-09 | 2e-08 | 1.253E-08 | 9.677E-09 | 1.600E-09 | 2.000E-08 |
| 13244 | 663 | G663D | B | 5.5e-09 | 8.4e-09 | 4.4e-09 | 6.100E-09 | 2.066E-09 | 4.400E-09 | 8.400E-09 |
| 13245 | 663 | G663C | B | 3.4e-09 | 1.1e-08 | 1.4e-08 | 9.467E-09 | 5.464E-09 | 3.400E-09 | 1.400E-08 |
| 13246 | 663 | G663E | B | 7.3e-09 | 4e-09 | 6e-09 | 5.767E-09 | 1.662E-09 | 4.000E-09 | 7.300E-09 |
| 13247 | 663 | G663Q | B | 1.7e-08 | 1.3e-08 | 1.9e-08 | 1.633E-08 | 3.055E-09 | 1.300E-08 | 1.900E-08 |
| 13248 | 663 | G663G | B | 3.5e-09 | 5.1e-09 | 5.2e-09 | 4.600E-09 | 9.539E-10 | 3.500E-09 | 5.200E-09 |
| 13249 | 663 | G663H | B | 3.6e-09 | 8.7e-09 | 1e-08 | 7.433E-09 | 3.383E-09 | 3.600E-09 | 1.000E-08 |
| 13250 | 663 | G663I | B | 7.9e-09 | 8.2e-09 | 9.5e-09 | 8.533E-09 | 8.505E-10 | 7.900E-09 | 9.500E-09 |
| 13251 | 663 | G663L | B | 4.6e-09 | 1.2e-08 | 2.3e-09 | 6.300E-09 | 5.069E-09 | 2.300E-09 | 1.200E-08 |
| 13252 | 663 | G663K | B | 5.5e-09 | 4.6e-09 | 7.8e-09 | 5.967E-09 | 1.650E-09 | 4.600E-09 | 7.800E-09 |
| 13253 | 663 | G663M | B | 2.7e-09 | 5.9e-09 | 5.3e-09 | 4.633E-09 | 1.701E-09 | 2.700E-09 | 5.900E-09 |
| 13254 | 663 | G663F | B | 6.8e-09 | 1.1e-08 | 4.2e-09 | 7.333E-09 | 3.431E-09 | 4.200E-09 | 1.100E-08 |
| 13255 | 663 | G663P | B | 1.6e-09 | 4.3e-09 | 5.1e-09 | 3.667E-09 | 1.834E-09 | 1.600E-09 | 5.100E-09 |
| 13256 | 663 | G663S | B | 3.3e-09 | 1.4e-08 | 3.4e-09 | 6.900E-09 | 6.149E-09 | 3.300E-09 | 1.400E-08 |
| 13257 | 663 | G663T | B | 1.5e-08 | 5.3e-09 | 5.9e-09 | 8.733E-09 | 5.435E-09 | 5.300E-09 | 1.500E-08 |
| 13258 | 663 | G663W | B | 7.1e-09 | 7.2e-09 | 9.6e-09 | 7.967E-09 | 1.415E-09 | 7.100E-09 | 9.600E-09 |
| 13259 | 663 | G663Y | B | 3.4e-09 | 5.7e-09 | 9.9e-09 | 6.333E-09 | 3.296E-09 | 3.400E-09 | 9.900E-09 |

| | | | | | | | | | | |
|---|---|---|---|---|---|---|---|---|---|---|
| 13260 | 663 | G663V | B | 8.5e-09 | 1.6e-09 | 1.1e-08 | 7.033E-09 | 4.869E-09 | 1.600E-09 | 1.100E-08 |
| 13261 | 664 | K664A | B | 4.4e-09 | 5.3e-09 | 7.7e-09 | 5.800E-09 | 1.706E-09 | 4.400E-09 | 7.700E-09 |
| 13262 | 664 | K664R | B | 9.5e-09 | 4.7e-09 | 4.7e-09 | 6.300E-09 | 2.771E-09 | 4.700E-09 | 9.500E-09 |
| 13263 | 664 | K664N | B | 3.3e-09 | 1.5e-08 | 8.2e-09 | 8.833E-09 | 5.876E-09 | 3.300E-09 | 1.500E-08 |
| 13264 | 664 | K664D | B | 3.9e-09 | 1.4e-08 | 9.6e-09 | 9.167E-09 | 5.064E-09 | 3.900E-09 | 1.400E-08 |
| 13265 | 664 | K664C | B | 1.5e-08 | 2.2e-09 | 8.1e-09 | 8.433E-09 | 6.407E-09 | 2.200E-09 | 1.500E-08 |
| 13266 | 664 | K664E | B | 4.1e-09 | 1.9e-08 | 1.2e-08 | 1.170E-08 | 7.455E-09 | 4.100E-09 | 1.900E-08 |
| 13267 | 664 | K664Q | B | 8.6e-09 | 5.1e-09 | 1.3e-08 | 8.900E-09 | 3.959E-09 | 5.100E-09 | 1.300E-08 |
| 13268 | 664 | K664G | B | 4.9e-09 | 2.5e-09 | 7.6e-09 | 5.000E-09 | 2.551E-09 | 2.500E-09 | 7.600E-09 |
| 13269 | 664 | K664H | B | 3.5e-09 | 6.5e-09 | 2.9e-09 | 4.300E-09 | 1.929E-09 | 2.900E-09 | 6.500E-09 |
| 13270 | 664 | K664I | B | 1.5e-08 | 9e-09 | 1e-08 | 1.133E-08 | 3.215E-09 | 9.000E-09 | 1.500E-08 |
| 13271 | 664 | K664L | B | 2.8e-09 | 3.1e-09 | 5.2e-09 | 3.700E-09 | 1.308E-09 | 2.800E-09 | 5.200E-09 |
| 13272 | 664 | K664K | B | 1.5e-08 | 2.7e-09 | 1.1e-08 | 9.567E-09 | 6.274E-09 | 2.700E-09 | 1.500E-08 |
| 13273 | 664 | K664M | B | 5.2e-09 | 4e-09 | 7.5e-09 | 5.567E-09 | 1.779E-09 | 4.000E-09 | 7.500E-09 |
| 13274 | 664 | K664F | B | 6e-09 | 9.2e-09 | 5.5e-09 | 6.900E-09 | 2.007E-09 | 5.500E-09 | 9.200E-09 |
| 13275 | 664 | K664P | B | 7.2e-09 | 5.7e-09 | 3.1e-09 | 5.333E-09 | 2.074E-09 | 3.100E-09 | 7.200E-09 |
| 13276 | 664 | K664S | B | 4.8e-09 | 1.1e-08 | 3.7e-09 | 6.500E-09 | 3.936E-09 | 3.700E-09 | 1.100E-08 |
| 13277 | 664 | K664T | B | 4.3e-09 | 6e-09 | 2.5e-08 | 1.177E-08 | 1.149E-08 | 4.300E-09 | 2.500E-08 |
| 13278 | 664 | K664W | B | 3.8e-09 | 8.6e-09 | 8.8e-09 | 7.067E-09 | 2.831E-09 | 3.800E-09 | 8.800E-09 |
| 13279 | 664 | K664Y | B | 1.6e-08 | 5.3e-09 | 6.7e-09 | 9.333E-09 | 5.816E-09 | 5.300E-09 | 1.600E-08 |
| 13280 | 664 | K664V | B | 1.2e-08 | 5.9e-09 | 4e-09 | 7.300E-09 | 4.180E-09 | 4.000E-09 | 1.200E-08 |
| 13281 | 665 | A665A | B | 7.9e-09 | 2e-08 | 2.3e-08 | 1.697E-08 | 7.994E-09 | 7.900E-09 | 2.300E-08 |
| 13282 | 665 | A665R | B | 1.6e-08 | 1.9e-08 | 5.2e-09 | 1.340E-08 | 7.258E-09 | 5.200E-09 | 1.900E-08 |
| 13283 | 665 | A665N | B | 5.3e-09 | 1.1e-08 | 6.8e-09 | 7.700E-09 | 2.955E-09 | 5.300E-09 | 1.100E-08 |
| 13284 | 665 | A665D | B | 1e-08 | 2e-08 | 1.4e-08 | 1.467E-08 | 5.033E-09 | 1.000E-08 | 2.000E-08 |
| 13285 | 665 | A665C | B | 3.5e-09 | 5.2e-09 | 1e-08 | 6.233E-09 | 3.371E-09 | 3.500E-09 | 1.000E-08 |
| 13286 | 665 | A665E | B | 5.7e-09 | 6.2e-09 | 1.7e-08 | 9.633E-09 | 6.385E-09 | 5.700E-09 | 1.700E-08 |
| 13287 | 665 | A665Q | B | 6.2e-09 | 1.6e-08 | 8.3e-09 | 1.017E-08 | 5.160E-09 | 6.200E-09 | 1.600E-08 |
| 13288 | 665 | A665G | B | 1.4e-08 | 9e-09 | 6.8e-09 | 9.933E-09 | 3.690E-09 | 6.800E-09 | 1.400E-08 |
| 13289 | 665 | A665H | B | 4.4e-09 | 3.6e-09 | 8e-09 | 5.333E-09 | 2.344E-09 | 3.600E-09 | 8.000E-09 |
| 13290 | 665 | A665I | B | 3.1e-09 | 4e-09 | 5.5e-09 | 4.200E-09 | 1.212E-09 | 3.100E-09 | 5.500E-09 |
| 13291 | 665 | A665L | B | 7.2e-09 | 1.2e-09 | 3.8e-09 | 4.067E-09 | 3.009E-09 | 1.200E-09 | 7.200E-09 |
| 13292 | 665 | A665K | B | 6.1e-09 | 4.4e-09 | 7.5e-09 | 6.000E-09 | 1.552E-09 | 4.400E-09 | 7.500E-09 |
| 13293 | 665 | A665M | B | 9.2e-09 | 1.8e-08 | 1e-08 | 1.240E-08 | 4.866E-09 | 9.200E-09 | 1.800E-08 |
| 13294 | 665 | A665F | B | 1.4e-08 | 4.4e-09 | 4.2e-09 | 7.533E-09 | 5.601E-09 | 4.200E-09 | 1.400E-08 |
| 13295 | 665 | A665P | B | 2.6e-08 | 8e-09 | 1.7e-08 | 1.700E-08 | 9.000E-09 | 8.000E-09 | 2.600E-08 |
| 13296 | 665 | A665S | B | 2.7e-09 | 1.3e-08 | 1.2e-08 | 9.233E-09 | 5.680E-09 | 2.700E-09 | 1.300E-08 |
| 13297 | 665 | A665T | B | 8.5e-09 | 9.2e-09 | 5.7e-09 | 7.800E-09 | 1.852E-09 | 5.700E-09 | 9.200E-09 |
| 13298 | 665 | A665W | B | 4e-09 | 3.9e-09 | 5.8e-09 | 4.567E-09 | 1.069E-09 | 3.900E-09 | 5.800E-09 |
| 13299 | 665 | A665Y | B | 6.3e-09 | 9.3e-09 | 1.7e-08 | 1.087E-08 | 5.519E-09 | 6.300E-09 | 1.700E-08 |
| 13300 | 665 | A665V | B | 6.7e-09 | 1.2e-08 | 1.5e-08 | 1.123E-08 | 4.203E-09 | 6.700E-09 | 1.500E-08 |
| 13301 | 666 | P666A | B | 3e-09 | 5.4e-09 | 1.3e-08 | 7.133E-09 | 5.220E-09 | 3.000E-09 | 1.300E-08 |
| 13302 | 666 | P666R | B | 1.7e-09 | 1.5e-08 | 5.5e-09 | 7.400E-09 | 6.851E-09 | 1.700E-09 | 1.500E-08 |
| 13303 | 666 | P666N | B | 1.1e-08 | 1.2e-08 | 5.6e-09 | 9.533E-09 | 3.443E-09 | 5.600E-09 | 1.200E-08 |
| 13304 | 666 | P666D | B | 9.3e-09 | 1e-08 | 9.8e-09 | 9.700E-09 | 3.606E-10 | 9.300E-09 | 1.000E-08 |
| 13305 | 666 | P666C | B | 1.3e-08 | 8.9e-09 | 6e-09 | 9.300E-09 | 3.517E-09 | 6.000E-09 | 1.300E-08 |
| 13306 | 666 | P666E | B | 1.4e-08 | 5.8e-09 | 7.8e-09 | 9.200E-09 | 4.276E-09 | 5.800E-09 | 1.400E-08 |
| 13307 | 666 | P666Q | B | 6.6e-09 | 4.6e-09 | 1.1e-08 | 7.400E-09 | 3.274E-09 | 4.600E-09 | 1.100E-08 |
| 13308 | 666 | P666G | B | 7.3e-09 | 5e-09 | 7e-09 | 6.433E-09 | 1.250E-09 | 5.000E-09 | 7.300E-09 |
| 13309 | 666 | P666H | B | 1.1e-08 | 7.8e-09 | 1.6e-09 | 6.800E-09 | 4.779E-09 | 1.600E-09 | 1.100E-08 |
| 13310 | 666 | P666I | B | 1.3e-08 | 2.9e-09 | 4.6e-09 | 6.833E-09 | 5.408E-09 | 2.900E-09 | 1.300E-08 |
| 13311 | 666 | P666L | B | 8.6e-09 | 4.9e-09 | 1.3e-08 | 8.833E-09 | 4.055E-09 | 4.900E-09 | 1.300E-08 |
| 13312 | 666 | P666K | B | 1.1e-08 | 2.4e-09 | 7e-09 | 6.800E-09 | 4.303E-09 | 2.400E-09 | 1.100E-08 |
| 13313 | 666 | P666M | B | 1.4e-08 | 1.3e-08 | 3.1e-09 | 1.003E-08 | 6.025E-09 | 3.100E-09 | 1.400E-08 |
| 13314 | 666 | P666F | B | 2.2e-08 | 1.9e-08 | 1.4e-08 | 1.833E-08 | 4.041E-09 | 1.400E-08 | 2.200E-08 |
| 13315 | 666 | P666P | B | 7.6e-09 | 3.8e-09 | 2.5e-09 | 4.633E-09 | 2.650E-09 | 2.500E-09 | 7.600E-09 |
| 13316 | 666 | P666S | B | 3.8e-09 | 2e-08 | 1.5e-08 | 1.293E-08 | 8.295E-09 | 3.800E-09 | 2.000E-08 |
| 13317 | 666 | P666T | B | 1.4e-08 | 1.7e-08 | 1.1e-08 | 1.400E-08 | 3.000E-09 | 1.100E-08 | 1.700E-08 |
| 13318 | 666 | P666W | B | 6.7e-09 | 5.9e-09 | 1.3e-08 | 8.533E-09 | 3.889E-09 | 5.900E-09 | 1.300E-08 |
| 13319 | 666 | P666Y | B | 5.1e-09 | 1.5e-08 | 1.7e-09 | 7.267E-09 | 6.910E-09 | 1.700E-09 | 1.500E-08 |
| 13320 | 666 | P666V | B | 6.4e-09 | 6.3e-09 | 6.8e-09 | 6.500E-09 | 2.646E-10 | 6.300E-09 | 6.800E-09 |
| 13321 | 667 | K667A | B | 4.5e-09 | 9.6e-09 | 7.4e-09 | 7.167E-09 | 2.558E-09 | 4.500E-09 | 9.600E-09 |
| 13322 | 667 | K667R | B | 7.6e-09 | 4.9e-09 | 5.1e-09 | 5.867E-09 | 1.504E-09 | 4.900E-09 | 7.600E-09 |
| 13323 | 667 | K667N | B | 5.8e-09 | 3e-09 | 8.2e-09 | 5.667E-09 | 2.603E-09 | 3.000E-09 | 8.200E-09 |
| 13324 | 667 | K667D | B | 2.2e-09 | 1.9e-09 | 1.7e-08 | 7.033E-09 | 8.633E-09 | 1.900E-09 | 1.700E-08 |
| 13325 | 667 | K667C | B | 7.6e-09 | 1.3e-08 | 2.2e-09 | 7.600E-09 | 5.400E-09 | 2.200E-09 | 1.300E-08 |
| 13326 | 667 | K667E | B | 3.2e-09 | 9.3e-09 | 1.2e-08 | 8.167E-09 | 4.508E-09 | 3.200E-09 | 1.200E-08 |
| 13327 | 667 | K667Q | B | 1.4e-08 | 1.2e-08 | 1e-08 | 1.200E-08 | 2.000E-09 | 1.000E-08 | 1.400E-08 |
| 13328 | 667 | K667G | B | 4e-09 | 4.7e-09 | 1e-08 | 6.233E-09 | 3.281E-09 | 4.000E-09 | 1.000E-08 |
| 13329 | 667 | K667H | B | 2.3e-09 | 3.7e-09 | 6.4e-09 | 4.133E-09 | 2.084E-09 | 2.300E-09 | 6.400E-09 |
| 13330 | 667 | K667I | B | 3.5e-09 | 1.4e-08 | 6.5e-09 | 8.000E-09 | 5.408E-09 | 3.500E-09 | 1.400E-08 |
| 13331 | 667 | K667L | B | 5.3e-09 | 5.3e-09 | 1.3e-08 | 7.867E-09 | 4.446E-09 | 5.300E-09 | 1.300E-08 |
| 13332 | 667 | K667K | B | 5.3e-09 | 5.6e-09 | 4.9e-09 | 5.267E-09 | 3.512E-10 | 4.900E-09 | 5.600E-09 |
| 13333 | 667 | K667M | B | 4.7e-09 | 1.8e-08 | 4.2e-09 | 8.967E-09 | 7.827E-09 | 4.200E-09 | 1.800E-08 |
| 13334 | 667 | K667F | B | 5e-09 | 1.7e-08 | 3e-09 | 8.333E-09 | 7.572E-09 | 3.000E-09 | 1.700E-08 |
| 13335 | 667 | K667P | B | 7.1e-09 | 8.5e-09 | 1.5e-08 | 1.020E-08 | 4.215E-09 | 7.100E-09 | 1.500E-08 |
| 13336 | 667 | K667S | B | 4.5e-09 | 6.2e-09 | 6.4e-09 | 5.700E-09 | 1.044E-09 | 4.500E-09 | 6.400E-09 |
| 13337 | 667 | K667T | B | 1.4e-08 | 4.2e-09 | 5e-09 | 7.733E-09 | 5.442E-09 | 4.200E-09 | 1.400E-08 |
| 13338 | 667 | K667W | B | 7.3e-09 | 1.3e-08 | 4.3e-09 | 8.200E-09 | 4.419E-09 | 4.300E-09 | 1.300E-08 |
| 13339 | 667 | K667Y | B | 5.7e-09 | 1.5e-08 | 6.5e-09 | 9.067E-09 | 5.154E-09 | 5.700E-09 | 1.500E-08 |
| 13340 | 667 | K667V | B | 6.2e-09 | 6.8e-09 | 5.1e-09 | 6.033E-09 | 8.622E-10 | 5.100E-09 | 6.800E-09 |
| 13341 | 668 | L668A | B | 4.2e-09 | 1.9e-09 | 7.4e-09 | 4.500E-09 | 2.762E-09 | 1.900E-09 | 7.400E-09 |
| 13342 | 668 | L668R | B | 4.3e-09 | 9.5e-09 | 3.4e-09 | 5.733E-09 | 3.293E-09 | 3.400E-09 | 9.500E-09 |
| 13343 | 668 | L668N | B | 4e-09 | 4.8e-09 | 7e-09 | 5.267E-09 | 1.553E-09 | 4.000E-09 | 7.000E-09 |
| 13344 | 668 | L668D | B | 2.8e-09 | 9.8e-09 | 5.4e-09 | 6.000E-09 | 3.538E-09 | 2.800E-09 | 9.800E-09 |

| | | | | | | | | | | |
|---|---|---|---|---|---|---|---|---|---|---|
| 13345 | 668 | L668C | B | 1.5e-08 | 1.2e-08 | 4.5e-09 | 1.050E-08 | 5.408E-09 | 4.500E-09 | 1.500E-08 |
| 13346 | 668 | L668E | B | 1.1e-08 | 6.6e-09 | 5.8e-09 | 7.800E-09 | 2.800E-09 | 5.800E-09 | 1.100E-08 |
| 13347 | 668 | L668Q | B | 1.2e-08 | 8.3e-09 | 6.9e-09 | 9.067E-09 | 2.635E-09 | 6.900E-09 | 1.200E-08 |
| 13348 | 668 | L668G | B | 5.5e-09 | 9.3e-09 | 7.9e-09 | 7.567E-09 | 1.922E-09 | 5.500E-09 | 9.300E-09 |
| 13349 | 668 | L668H | B | 1.2e-08 | 3.4e-08 | 1.7e-08 | 2.100E-08 | 1.153E-08 | 1.200E-08 | 3.400E-08 |
| 13350 | 668 | L668I | B | 6.1e-09 | 5.9e-09 | 1.1e-09 | 4.367E-09 | 2.831E-09 | 1.100E-09 | 6.100E-09 |
| 13351 | 668 | L668L | B | 7.2e-09 | 6.2e-09 | 5.9e-09 | 6.433E-09 | 6.807E-10 | 5.900E-09 | 7.200E-09 |
| 13352 | 668 | L668K | B | 2.5e-09 | 2.1e-08 | 9.1e-09 | 1.087E-08 | 9.376E-09 | 2.500E-09 | 2.100E-08 |
| 13353 | 668 | L668M | B | 2.4e-08 | 1.8e-08 | 2.4e-09 | 1.480E-08 | 1.115E-08 | 2.400E-09 | 2.400E-08 |
| 13354 | 668 | L668F | B | 8.1e-09 | 2.5e-09 | 7.5e-09 | 6.033E-09 | 3.075E-09 | 2.500E-09 | 8.100E-09 |
| 13355 | 668 | L668P | B | 1.6e-08 | 4.4e-09 | 4.1e-09 | 8.167E-09 | 6.786E-09 | 4.100E-09 | 1.600E-08 |
| 13356 | 668 | L668S | B | 9.5e-09 | 4e-09 | 4.9e-09 | 6.133E-09 | 2.950E-09 | 4.000E-09 | 9.500E-09 |
| 13357 | 668 | L668T | B | 1.8e-08 | 2e-08 | 1.5e-08 | 1.767E-08 | 2.517E-09 | 1.500E-08 | 2.000E-08 |
| 13358 | 668 | L668W | B | 4.9e-09 | 3.1e-09 | 2.7e-08 | 1.167E-08 | 1.331E-08 | 3.100E-09 | 2.700E-08 |
| 13359 | 668 | L668Y | B | 1.7e-08 | 2.9e-09 | 6e-09 | 8.633E-09 | 7.410E-09 | 2.900E-09 | 1.700E-08 |
| 13360 | 668 | L668V | B | 6.3e-09 | 1.2e-08 | 1.2e-08 | 1.010E-08 | 3.291E-09 | 6.300E-09 | 1.200E-08 |
| 13361 | 669 | L669A | B | 1.3e-08 | 5.9e-09 | 5.6e-09 | 8.167E-09 | 4.188E-09 | 5.600E-09 | 1.300E-08 |
| 13362 | 669 | L669R | B | 3.2e-09 | 9.7e-09 | 3e-08 | 1.430E-08 | 1.398E-08 | 3.200E-09 | 3.000E-08 |
| 13363 | 669 | L669N | B | 6.9e-10 | 2e-09 | 3e-09 | 1.897E-09 | 1.158E-09 | 6.900E-10 | 3.000E-09 |
| 13364 | 669 | L669D | B | 1.2e-08 | 9.4e-09 | 9.5e-09 | 1.030E-08 | 1.473E-09 | 9.400E-09 | 1.200E-08 |
| 13365 | 669 | L669C | B | 1e-08 | 1.7e-08 | 8.8e-09 | 1.193E-08 | 4.429E-09 | 8.800E-09 | 1.700E-08 |
| 13366 | 669 | L669E | B | 1.7e-08 | 6.1e-09 | 8.9e-09 | 1.067E-08 | 5.661E-09 | 6.100E-09 | 1.700E-08 |
| 13367 | 669 | L669Q | B | 3.9e-09 | 1.7e-08 | 4e-09 | 8.300E-09 | 7.535E-09 | 3.900E-09 | 1.700E-08 |
| 13368 | 669 | L669G | B | 1e-08 | 1e-08 | 5.5e-09 | 8.500E-09 | 2.598E-09 | 5.500E-09 | 1.000E-08 |
| 13369 | 669 | L669H | B | 1.1e-08 | 5.6e-09 | 2e-08 | 1.220E-08 | 7.275E-09 | 5.600E-09 | 2.000E-08 |
| 13370 | 669 | L669I | B | 8.5e-09 | 1.2e-08 | 5.8e-09 | 8.767E-09 | 3.109E-09 | 5.800E-09 | 1.200E-08 |
| 13371 | 669 | L669L | B | 7.5e-09 | 5.7e-09 | 3.1e-08 | 1.473E-08 | 1.412E-08 | 5.700E-09 | 3.100E-08 |
| 13372 | 669 | L669K | B | 2.1e-08 | 5.3e-09 | 4.1e-09 | 1.013E-08 | 9.430E-09 | 4.100E-09 | 2.100E-08 |
| 13373 | 669 | L669M | B | 1.3e-08 | 8.1e-09 | 5.3e-09 | 8.800E-09 | 3.897E-09 | 5.300E-09 | 1.300E-08 |
| 13374 | 669 | L669F | B | 6.3e-09 | 4.9e-09 | 4.1e-09 | 5.100E-09 | 1.114E-09 | 4.100E-09 | 6.300E-09 |
| 13375 | 669 | L669P | B | 1.5e-08 | 4.2e-09 | 2.2e-09 | 7.133E-09 | 6.886E-09 | 2.200E-09 | 1.500E-08 |
| 13376 | 669 | L669S | B | 1.2e-08 | 1.1e-08 | 1.1e-08 | 1.133E-08 | 5.774E-10 | 1.100E-08 | 1.200E-08 |
| 13377 | 669 | L669T | B | 2e-09 | 9.1e-09 | 1.1e-08 | 7.367E-09 | 4.744E-09 | 2.000E-09 | 1.100E-08 |
| 13378 | 669 | L669W | B | 6.5e-09 | 5.2e-09 | 6.4e-09 | 6.033E-09 | 7.234E-10 | 5.200E-09 | 6.500E-09 |
| 13379 | 669 | L669Y | B | 7.7e-09 | 7.6e-09 | 8.1e-09 | 7.800E-09 | 2.646E-10 | 7.600E-09 | 8.100E-09 |
| 13380 | 669 | L669V | B | 2.3e-08 | 7e-09 | 6.7e-09 | 1.223E-08 | 9.325E-09 | 6.700E-09 | 2.300E-08 |
| 13381 | 670 | I670A | B | 4.2e-09 | 3.6e-09 | 1.1e-08 | 6.267E-09 | 4.110E-09 | 3.600E-09 | 1.100E-08 |
| 13382 | 670 | I670R | B | 1.8e-08 | 1.3e-08 | 1.2e-08 | 1.433E-08 | 3.215E-09 | 1.200E-08 | 1.800E-08 |
| 13383 | 670 | I670N | B | 1.7e-09 | 8.6e-09 | 4.1e-09 | 4.800E-09 | 3.503E-09 | 1.700E-09 | 8.600E-09 |
| 13384 | 670 | I670D | B | 3.9e-09 | 1e-08 | 5.3e-09 | 6.400E-09 | 3.195E-09 | 3.900E-09 | 1.000E-08 |
| 13385 | 670 | I670C | B | 2.1e-09 | 7.1e-09 | 6.3e-09 | 5.167E-09 | 2.686E-09 | 2.100E-09 | 7.100E-09 |
| 13386 | 670 | I670E | B | 8.4e-09 | 5.6e-09 | 7e-09 | 7.000E-09 | 1.400E-09 | 5.600E-09 | 8.400E-09 |
| 13387 | 670 | I670Q | B | 9.8e-09 | 1.5e-08 | 1e-08 | 1.160E-08 | 2.946E-09 | 9.800E-09 | 1.500E-08 |
| 13388 | 670 | I670G | B | 9.9e-09 | 7.8e-09 | 6.4e-09 | 8.033E-09 | 1.762E-09 | 6.400E-09 | 9.900E-09 |
| 13389 | 670 | I670H | B | 2.6e-09 | 5.6e-09 | 1.5e-08 | 7.733E-09 | 6.469E-09 | 2.600E-09 | 1.500E-08 |
| 13390 | 670 | I670I | B | 4.4e-09 | 6.2e-09 | 7.1e-09 | 5.900E-09 | 1.375E-09 | 4.400E-09 | 7.100E-09 |
| 13391 | 670 | I670L | B | 6.9e-09 | 3e-09 | 3.5e-09 | 4.467E-09 | 2.122E-09 | 3.000E-09 | 6.900E-09 |
| 13392 | 670 | I670K | B | 2.8e-09 | 4.6e-09 | 3.7e-09 | 3.700E-09 | 9.000E-10 | 2.800E-09 | 4.600E-09 |
| 13393 | 670 | I670M | B | 4e-09 | 7.1e-09 | 5e-09 | 5.367E-09 | 1.582E-09 | 4.000E-09 | 7.100E-09 |
| 13394 | 670 | I670F | B | 9.7e-09 | 7.9e-09 | 8.8e-09 | 8.800E-09 | 9.000E-10 | 7.900E-09 | 9.700E-09 |
| 13395 | 670 | I670P | B | 1.7e-08 | 1.3e-08 | 3.3e-09 | 1.110E-08 | 7.045E-09 | 3.300E-09 | 1.700E-08 |
| 13396 | 670 | I670S | B | 1e-08 | 8.8e-09 | 1e-08 | 9.600E-09 | 6.928E-10 | 8.800E-09 | 1.000E-08 |
| 13397 | 670 | I670T | B | 8.5e-09 | 1.1e-08 | 1.2e-08 | 1.050E-08 | 1.803E-09 | 8.500E-09 | 1.200E-08 |
| 13398 | 670 | I670W | B | 6.2e-09 | 4.1e-09 | 4.8e-09 | 5.033E-09 | 1.069E-09 | 4.100E-09 | 6.200E-09 |
| 13399 | 670 | I670Y | B | 7e-09 | 6.9e-09 | 5.2e-09 | 6.367E-09 | 1.012E-09 | 5.200E-09 | 7.000E-09 |
| 13400 | 670 | I670V | B | 1e-08 | 1.3e-08 | 4.2e-09 | 9.067E-09 | 4.474E-09 | 4.200E-09 | 1.300E-08 |
| 13401 | 671 | Y671A | B | 1.3e-08 | 1.8e-08 | 1.7e-08 | 1.600E-08 | 2.646E-09 | 1.300E-08 | 1.800E-08 |
| 13402 | 671 | Y671R | B | 4.1e-09 | 1.5e-08 | 9.1e-09 | 9.400E-09 | 5.456E-09 | 4.100E-09 | 1.500E-08 |
| 13403 | 671 | Y671N | B | 1.2e-08 | 1.1e-08 | 1.7e-08 | 1.333E-08 | 3.215E-09 | 1.100E-08 | 1.700E-08 |
| 13404 | 671 | Y671D | B | 7.8e-09 | 6.5e-09 | 1.2e-08 | 8.767E-09 | 2.875E-09 | 6.500E-09 | 1.200E-08 |
| 13405 | 671 | Y671C | B | 5.4e-09 | 1.8e-08 | 1.4e-08 | 1.247E-08 | 6.438E-09 | 5.400E-09 | 1.800E-08 |
| 13406 | 671 | Y671E | B | 4.9e-09 | 1e-08 | 1.4e-08 | 9.633E-09 | 4.561E-09 | 4.900E-09 | 1.400E-08 |
| 13407 | 671 | Y671Q | B | 8e-09 | 1.3e-08 | 1.3e-08 | 1.133E-08 | 2.887E-09 | 8.000E-09 | 1.300E-08 |
| 13408 | 671 | Y671G | B | 8.4e-09 | 2.1e-08 | 1.5e-08 | 1.480E-08 | 6.302E-09 | 8.400E-09 | 2.100E-08 |
| 13409 | 671 | Y671H | B | 1.3e-08 | 8.7e-09 | 3.5e-09 | 8.400E-09 | 4.757E-09 | 3.500E-09 | 1.300E-08 |
| 13410 | 671 | Y671I | B | 1.3e-08 | 1.1e-08 | 9.7e-09 | 1.123E-08 | 1.662E-09 | 9.700E-09 | 1.300E-08 |
| 13411 | 671 | Y671L | B | 2e-08 | 1e-08 | 1.2e-08 | 1.400E-08 | 5.292E-09 | 1.000E-08 | 2.000E-08 |
| 13412 | 671 | Y671K | B | 1.9e-08 | 3e-08 | 1.4e-08 | 2.100E-08 | 8.185E-09 | 1.400E-08 | 3.000E-08 |
| 13413 | 671 | Y671M | B | 2.2e-08 | 1.4e-08 | 1.8e-08 | 1.800E-08 | 4.000E-09 | 1.400E-08 | 2.200E-08 |
| 13414 | 671 | Y671F | B | 1.1e-08 | 1.2e-08 | 5.3e-09 | 9.433E-09 | 3.614E-09 | 5.300E-09 | 1.200E-08 |
| 13415 | 671 | Y671P | B | 5.1e-08 | 2.6e-08 | 5.4e-09 | 2.747E-08 | 2.284E-08 | 5.400E-09 | 5.100E-08 |
| 13416 | 671 | Y671S | B | 1.1e-08 | 1.6e-08 | 1.4e-08 | 1.367E-08 | 2.517E-09 | 1.100E-08 | 1.600E-08 |
| 13417 | 671 | Y671T | B | 8.8e-09 | 9.4e-09 | 1.6e-08 | 1.140E-08 | 3.995E-09 | 8.800E-09 | 1.600E-08 |
| 13418 | 671 | Y671W | B | 1.9e-08 | 7.7e-09 | 5.5e-09 | 1.073E-08 | 7.243E-09 | 5.500E-09 | 1.900E-08 |
| 13419 | 671 | Y671Y | B | 5.5e-09 | 2.1e-08 | 4.5e-09 | 1.033E-08 | 9.251E-09 | 4.500E-09 | 2.100E-08 |
| 13420 | 671 | Y671V | B | 6.1e-09 | 3e-08 | 1.7e-08 | 1.770E-08 | 1.197E-08 | 6.100E-09 | 3.000E-08 |
| 13421 | 672 | S672A | B | 2.9e-09 | 1.6e-09 | 6.2e-09 | 3.567E-09 | 2.371E-09 | 1.600E-09 | 6.200E-09 |
| 13422 | 672 | S672R | B | 3.8e-09 | 4.1e-09 | 6e-09 | 4.633E-09 | 1.193E-09 | 3.800E-09 | 6.000E-09 |
| 13423 | 672 | S672N | B | 3.8e-09 | 1.1e-08 | 8.6e-09 | 7.800E-09 | 3.666E-09 | 3.800E-09 | 1.100E-08 |
| 13424 | 672 | S672D | B | 2.2e-09 | 5.4e-09 | 6.8e-09 | 4.800E-09 | 2.358E-09 | 2.200E-09 | 6.800E-09 |
| 13425 | 672 | S672C | B | 6.9e-09 | 1.8e-09 | 3.5e-09 | 4.067E-09 | 2.597E-09 | 1.800E-09 | 6.900E-09 |
| 13426 | 672 | S672E | B | 1.7e-08 | 7.9e-09 | 4.3e-09 | 9.733E-09 | 6.545E-09 | 4.300E-09 | 1.700E-08 |
| 13427 | 672 | S672Q | B | 5.2e-09 | 5e-09 | 7.7e-09 | 5.967E-09 | 1.504E-09 | 5.000E-09 | 7.700E-09 |
| 13428 | 672 | S672G | B | 6.5e-09 | 4.4e-09 | 2e-09 | 4.300E-09 | 2.252E-09 | 2.000E-09 | 6.500E-09 |
| 13429 | 672 | S672H | B | 7.4e-09 | 1.1e-09 | 1.5e-08 | 7.833E-09 | 6.960E-09 | 1.100E-09 | 1.500E-08 |

| | | | | | | | | | | |
|---|---|---|---|---|---|---|---|---|---|---|
| 13430 | 672 | S672I | B | 3.4e-09 | 5.3e-09 | 5.6e-09 | 4.767E-09 | 1.193E-09 | 3.400E-09 | 5.600E-09 |
| 13431 | 672 | S672L | B | 2.5e-09 | 1.4e-09 | 3.4e-09 | 2.433E-09 | 1.002E-09 | 1.400E-09 | 3.400E-09 |
| 13432 | 672 | S672K | B | 4e-09 | 8.8e-09 | 2.3e-09 | 5.033E-09 | 3.371E-09 | 2.300E-09 | 8.800E-09 |
| 13433 | 672 | S672M | B | 3.9e-09 | 2.1e-09 | 2.3e-09 | 2.767E-09 | 9.866E-10 | 2.100E-09 | 3.900E-09 |
| 13434 | 672 | S672F | B | 2.2e-09 | 2.8e-09 | 3.8e-09 | 2.933E-09 | 8.083E-10 | 2.200E-09 | 3.800E-09 |
| 13435 | 672 | S672P | B | 2.7e-09 | 3.2e-09 | 5.1e-09 | 3.667E-09 | 1.266E-09 | 2.700E-09 | 5.100E-09 |
| 13436 | 672 | S672S | B | 9.8e-09 | 7.5e-09 | 1.1e-08 | 9.433E-09 | 1.779E-09 | 7.500E-09 | 1.100E-08 |
| 13437 | 672 | S672T | B | 8.2e-09 | 4.5e-09 | 8.2e-09 | 6.967E-09 | 2.136E-09 | 4.500E-09 | 8.200E-09 |
| 13438 | 672 | S672W | B | 3.5e-09 | 1.6e-09 | 4e-09 | 3.033E-09 | 1.266E-09 | 1.600E-09 | 4.000E-09 |
| 13439 | 672 | S672Y | B | 6.6e-09 | 4.2e-09 | 2.7e-09 | 4.500E-09 | 1.967E-09 | 2.700E-09 | 6.600E-09 |
| 13440 | 672 | S672V | B | 4.4e-09 | 3.8e-09 | 5.5e-09 | 4.567E-09 | 8.622E-10 | 3.800E-09 | 5.500E-09 |
| 13441 | 673 | A673A | B | 5.1e-09 | 1.9e-08 | 6.9e-09 | 1.033E-08 | 7.559E-09 | 5.100E-09 | 1.900E-08 |
| 13442 | 673 | A673R | B | 1.1e-08 | 8.2e-09 | 1.3e-08 | 1.073E-08 | 2.411E-09 | 8.200E-09 | 1.300E-08 |
| 13443 | 673 | A673N | B | 1e-08 | 4.2e-09 | 1.9e-08 | 1.107E-08 | 7.457E-09 | 4.200E-09 | 1.900E-08 |
| 13444 | 673 | A673D | B | 8.8e-09 | 6.6e-09 | 2.5e-08 | 1.347E-08 | 1.005E-08 | 6.600E-09 | 2.500E-08 |
| 13445 | 673 | A673C | B | 9e-09 | 9.6e-09 | 1.1e-08 | 9.867E-09 | 1.026E-09 | 9.000E-09 | 1.100E-08 |
| 13446 | 673 | A673E | B | 8.5e-09 | 3.5e-09 | 2.2e-09 | 4.733E-09 | 3.326E-09 | 2.200E-09 | 8.500E-09 |
| 13447 | 673 | A673Q | B | 6.3e-09 | 7.1e-09 | 7.2e-09 | 6.867E-09 | 4.933E-10 | 6.300E-09 | 7.200E-09 |
| 13448 | 673 | A673G | B | 1.2e-08 | 7.2e-09 | 8.4e-09 | 9.200E-09 | 2.498E-09 | 7.200E-09 | 1.200E-08 |
| 13449 | 673 | A673H | B | 7e-09 | 4e-09 | 6.2e-09 | 5.733E-09 | 1.553E-09 | 4.000E-09 | 7.000E-09 |
| 13450 | 673 | A673I | B | 4.5e-09 | 5e-09 | 1.2e-08 | 7.167E-09 | 4.193E-09 | 4.500E-09 | 1.200E-08 |
| 13451 | 673 | A673L | B | 7.6e-09 | 5.3e-09 | 5.9e-09 | 6.267E-09 | 1.193E-09 | 5.300E-09 | 7.600E-09 |
| 13452 | 673 | A673K | B | 1.2e-08 | 8.5e-09 | 1.2e-08 | 1.083E-08 | 2.021E-09 | 8.500E-09 | 1.200E-08 |
| 13453 | 673 | A673M | B | 5.6e-09 | 6.2e-09 | 1.4e-08 | 8.600E-09 | 4.686E-09 | 5.600E-09 | 1.400E-08 |
| 13454 | 673 | A673F | B | 5.1e-09 | 2.3e-09 | 1.8e-08 | 8.467E-09 | 8.374E-09 | 2.300E-09 | 1.800E-08 |
| 13455 | 673 | A673P | B | 6.6e-09 | 1.6e-08 | 5.7e-09 | 9.433E-09 | 5.705E-09 | 5.700E-09 | 1.600E-08 |
| 13456 | 673 | A673S | B | 1.3e-08 | 9.5e-09 | 8.7e-09 | 1.040E-08 | 2.287E-09 | 8.700E-09 | 1.300E-08 |
| 13457 | 673 | A673T | B | 5.9e-09 | 1.3e-08 | 6.9e-09 | 8.600E-09 | 3.843E-09 | 5.900E-09 | 1.300E-08 |
| 13458 | 673 | A673W | B | 7e-09 | 2.9e-09 | 7.7e-09 | 5.867E-09 | 2.593E-09 | 2.900E-09 | 7.700E-09 |
| 13459 | 673 | A673Y | B | 3e-09 | 9.4e-09 | 4.8e-09 | 5.733E-09 | 3.301E-09 | 3.000E-09 | 9.400E-09 |
| 13460 | 673 | A673V | B | 9.7e-09 | 5.9e-09 | 1.2e-08 | 9.200E-09 | 3.081E-09 | 5.900E-09 | 1.200E-08 |
| 13461 | 674 | S674A | B | 1.5e-08 | 1.9e-08 | 1.7e-09 | 1.190E-08 | 9.057E-09 | 1.700E-09 | 1.900E-08 |
| 13462 | 674 | S674R | B | 7.6e-09 | 2.7e-09 | 3.5e-09 | 4.600E-09 | 2.629E-09 | 2.700E-09 | 7.600E-09 |
| 13463 | 674 | S674N | B | 2.1e-08 | 5.6e-09 | 4e-09 | 1.020E-08 | 9.387E-09 | 4.000E-09 | 2.100E-08 |
| 13464 | 674 | S674D | B | 2.5e-09 | 1.1e-08 | 1e-09 | 4.833E-09 | 5.393E-09 | 1.000E-09 | 1.100E-08 |
| 13465 | 674 | S674C | B | 4.2e-09 | 5.8e-09 | 3.5e-09 | 4.500E-09 | 1.179E-09 | 3.500E-09 | 5.800E-09 |
| 13466 | 674 | S674E | B | 1.4e-08 | 9e-09 | 9.7e-09 | 1.090E-08 | 2.707E-09 | 9.000E-09 | 1.400E-08 |
| 13467 | 674 | S674Q | B | 5.8e-09 | 9.9e-09 | 7.8e-09 | 7.833E-09 | 2.050E-09 | 5.800E-09 | 9.900E-09 |
| 13468 | 674 | S674G | B | 5e-09 | 2.2e-09 | 1.6e-08 | 7.733E-09 | 7.295E-09 | 2.200E-09 | 1.600E-08 |
| 13469 | 674 | S674H | B | 1.7e-08 | 1.5e-08 | 9.2e-09 | 1.373E-08 | 4.051E-09 | 9.200E-09 | 1.700E-08 |
| 13470 | 674 | S674I | B | 3.3e-09 | 9.1e-09 | 5.6e-09 | 6.000E-09 | 2.921E-09 | 3.300E-09 | 9.100E-09 |
| 13471 | 674 | S674L | B | 1.7e-08 | 8.6e-09 | 2.8e-09 | 9.467E-09 | 7.140E-09 | 2.800E-09 | 1.700E-08 |
| 13472 | 674 | S674K | B | 1.3e-09 | 1.8e-09 | 1.1e-08 | 4.700E-09 | 5.462E-09 | 1.300E-09 | 1.100E-08 |
| 13473 | 674 | S674M | B | 4.9e-09 | 4.7e-09 | 3.7e-09 | 4.433E-09 | 6.429E-10 | 3.700E-09 | 4.900E-09 |
| 13474 | 674 | S674F | B | 1.4e-08 | 8.8e-09 | 8.6e-09 | 1.047E-08 | 3.062E-09 | 8.600E-09 | 1.400E-08 |
| 13475 | 674 | S674P | B | 6.6e-09 | 2e-08 | 7.9e-09 | 1.150E-08 | 7.390E-09 | 6.600E-09 | 2.000E-08 |
| 13476 | 674 | S674S | B | 1.2e-08 | 7.7e-09 | 4.3e-09 | 8.000E-09 | 3.859E-09 | 4.300E-09 | 1.200E-08 |
| 13477 | 674 | S674T | B | 4.8e-09 | 3e-09 | 9.6e-09 | 5.800E-09 | 3.412E-09 | 3.000E-09 | 9.600E-09 |
| 13478 | 674 | S674W | B | 1.9e-08 | 1.2e-08 | 4.3e-09 | 1.177E-08 | 7.353E-09 | 4.300E-09 | 1.900E-08 |
| 13479 | 674 | S674Y | B | 3e-09 | 9e-09 | 9.1e-09 | 7.033E-09 | 3.493E-09 | 3.000E-09 | 9.100E-09 |
| 13480 | 674 | S674V | B | 1.1e-08 | 1.5e-08 | 7.7e-09 | 1.123E-08 | 3.656E-09 | 7.700E-09 | 1.500E-08 |
| 13481 | 675 | Y675A | B | 5.3e-09 | 1.9e-08 | 1.2e-08 | 1.210E-08 | 6.851E-09 | 5.300E-09 | 1.900E-08 |
| 13482 | 675 | Y675R | B | 7e-09 | 9.7e-09 | 6.9e-09 | 7.867E-09 | 1.589E-09 | 6.900E-09 | 9.700E-09 |
| 13483 | 675 | Y675N | B | 1.1e-08 | 1.5e-08 | 1.3e-08 | 1.300E-08 | 2.000E-09 | 1.100E-08 | 1.500E-08 |
| 13484 | 675 | Y675D | B | 1.6e-08 | 9.3e-09 | 1.2e-08 | 1.243E-08 | 3.371E-09 | 9.300E-09 | 1.600E-08 |
| 13485 | 675 | Y675C | B | 1.7e-08 | 5.2e-09 | 7.5e-09 | 9.900E-09 | 6.255E-09 | 5.200E-09 | 1.700E-08 |
| 13486 | 675 | Y675E | B | 2.1e-08 | 1.1e-08 | 7.4e-09 | 1.313E-08 | 7.047E-09 | 7.400E-09 | 2.100E-08 |
| 13487 | 675 | Y675Q | B | 2.3e-08 | 1.5e-08 | 1.6e-08 | 1.800E-08 | 4.359E-09 | 1.500E-08 | 2.300E-08 |
| 13488 | 675 | Y675G | B | 8.9e-09 | 3e-09 | 1.7e-08 | 9.633E-09 | 7.029E-09 | 3.000E-09 | 1.700E-08 |
| 13489 | 675 | Y675H | B | 3.8e-09 | 9.1e-09 | 8e-09 | 6.967E-09 | 2.797E-09 | 3.800E-09 | 9.100E-09 |
| 13490 | 675 | Y675I | B | 3.7e-09 | 6.3e-09 | 1.8e-08 | 9.333E-09 | 7.617E-09 | 3.700E-09 | 1.800E-08 |
| 13491 | 675 | Y675L | B | 1.5e-08 | 7.2e-09 | 8.1e-09 | 1.010E-08 | 4.267E-09 | 7.200E-09 | 1.500E-08 |
| 13492 | 675 | Y675K | B | 3e-09 | 9.1e-09 | 5.4e-09 | 5.833E-09 | 3.073E-09 | 3.000E-09 | 9.100E-09 |
| 13493 | 675 | Y675M | B | 4.4e-09 | 1.6e-09 | 1e-08 | 5.333E-09 | 4.277E-09 | 1.600E-09 | 1.000E-08 |
| 13494 | 675 | Y675F | B | 3.1e-09 | 1.1e-08 | 4.4e-09 | 6.167E-09 | 4.236E-09 | 3.100E-09 | 1.100E-08 |
| 13495 | 675 | Y675P | B | 4.8e-09 | 8.2e-09 | 4e-09 | 5.667E-09 | 2.230E-09 | 4.000E-09 | 8.200E-09 |
| 13496 | 675 | Y675S | B | 1.6e-08 | 5e-09 | 2.1e-08 | 1.400E-08 | 8.185E-09 | 5.000E-09 | 2.100E-08 |
| 13497 | 675 | Y675T | B | 1.6e-08 | 3.7e-09 | 1.8e-08 | 1.257E-08 | 7.744E-09 | 3.700E-09 | 1.800E-08 |
| 13498 | 675 | Y675W | B | 1.5e-08 | 6.4e-09 | 1.8e-09 | 7.733E-09 | 6.700E-09 | 1.800E-09 | 1.500E-08 |
| 13499 | 675 | Y675Y | B | 1.3e-08 | 6.6e-09 | 4.3e-09 | 7.967E-09 | 4.508E-09 | 4.300E-09 | 1.300E-08 |
| 13500 | 675 | Y675V | B | 4.8e-09 | 1e-08 | 1.2e-08 | 8.933E-09 | 3.717E-09 | 4.800E-09 | 1.200E-08 |
| 13501 | 676 | R676A | B | 4.2e-09 | 7.3e-09 | 3.5e-08 | 1.550E-08 | 1.696E-08 | 4.200E-09 | 3.500E-08 |
| 13502 | 676 | R676R | B | 6e-09 | 2.8e-09 | 3.9e-09 | 4.233E-09 | 1.626E-09 | 2.800E-09 | 6.000E-09 |
| 13503 | 676 | R676N | B | 1.7e-08 | 4.8e-09 | 6.1e-09 | 9.300E-09 | 6.700E-09 | 4.800E-09 | 1.700E-08 |
| 13504 | 676 | R676D | B | 1.1e-08 | 6.5e-09 | 3.7e-09 | 7.067E-09 | 3.683E-09 | 3.700E-09 | 1.100E-08 |
| 13505 | 676 | R676C | B | 3e-08 | 2.7e-09 | 1.1e-08 | 1.457E-08 | 1.400E-08 | 2.700E-09 | 3.000E-08 |
| 13506 | 676 | R676E | B | 9.1e-09 | 9.6e-09 | 4e-09 | 7.567E-09 | 3.099E-09 | 4.000E-09 | 9.600E-09 |
| 13507 | 676 | R676Q | B | 4.8e-09 | 9.5e-09 | 1.3e-08 | 9.100E-09 | 4.115E-09 | 4.800E-09 | 1.300E-08 |
| 13508 | 676 | R676G | B | 1.9e-08 | 1e-08 | 5.1e-09 | 1.137E-08 | 7.050E-09 | 5.100E-09 | 1.900E-08 |
| 13509 | 676 | R676H | B | 4.2e-09 | 2.7e-08 | 3.4e-09 | 1.153E-08 | 1.340E-08 | 3.400E-09 | 2.700E-08 |
| 13510 | 676 | R676I | B | 7.5e-09 | 8.4e-09 | 4.3e-09 | 6.733E-09 | 2.155E-09 | 4.300E-09 | 8.400E-09 |
| 13511 | 676 | R676L | B | 4.9e-09 | 4.3e-09 | 5.6e-09 | 4.933E-09 | 6.506E-10 | 4.300E-09 | 5.600E-09 |
| 13512 | 676 | R676K | B | 8.1e-09 | 1.3e-08 | 4.5e-09 | 8.533E-09 | 4.267E-09 | 4.500E-09 | 1.300E-08 |
| 13513 | 676 | R676M | B | 7.2e-09 | 1.2e-08 | 4.1e-09 | 7.767E-09 | 3.980E-09 | 4.100E-09 | 1.200E-08 |
| 13514 | 676 | R676F | B | 7.3e-09 | 9.5e-09 | 4.4e-09 | 7.067E-09 | 2.558E-09 | 4.400E-09 | 9.500E-09 |

| | | | | | | | | | | |
|---|---|---|---|---|---|---|---|---|---|---|
| 13515 | 676 | R676P | B | 1.2e-08 | 6.6e-09 | 6.1e-09 | 8.233E-09 | 3.272E-09 | 6.100E-09 | 1.200E-08 |
| 13516 | 676 | R676S | B | 1e-08 | 5.9e-09 | 1e-08 | 8.633E-09 | 2.367E-09 | 5.900E-09 | 1.000E-08 |
| 13517 | 676 | R676T | B | 8e-09 | 2.1e-08 | 4.5e-09 | 1.117E-08 | 8.694E-09 | 4.500E-09 | 2.100E-08 |
| 13518 | 676 | R676W | B | 5.8e-09 | 1.4e-08 | 3.7e-09 | 7.833E-09 | 5.443E-09 | 3.700E-09 | 1.400E-08 |
| 13519 | 676 | R676Y | B | 7.5e-09 | 1.2e-08 | 1.5e-08 | 1.150E-08 | 3.775E-09 | 7.500E-09 | 1.500E-08 |
| 13520 | 676 | R676V | B | 4.6e-09 | 3.9e-09 | 1.1e-08 | 6.500E-09 | 3.913E-09 | 3.900E-09 | 1.100E-08 |
| 13521 | 677 | Y677A | B | 2.1e-08 | 4.3e-09 | 9.6e-09 | 1.163E-08 | 8.534E-09 | 4.300E-09 | 2.100E-08 |
| 13522 | 677 | Y677R | B | 6.3e-09 | 1.8e-08 | 1.6e-08 | 1.343E-08 | 6.258E-09 | 6.300E-09 | 1.800E-08 |
| 13523 | 677 | Y677N | B | 4.4e-09 | 2.6e-08 | 9.1e-09 | 1.317E-08 | 1.136E-08 | 4.400E-09 | 2.600E-08 |
| 13524 | 677 | Y677D | B | 4.9e-09 | 6.1e-09 | 1.2e-08 | 7.667E-09 | 3.800E-09 | 4.900E-09 | 1.200E-08 |
| 13525 | 677 | Y677C | B | 9.6e-09 | 5e-09 | 6.2e-09 | 6.933E-09 | 2.386E-09 | 5.000E-09 | 9.600E-09 |
| 13526 | 677 | Y677E | B | 5.7e-09 | 1.1e-08 | 2.1e-09 | 6.267E-09 | 4.477E-09 | 2.100E-09 | 1.100E-08 |
| 13527 | 677 | Y677Q | B | 1.6e-08 | 4.3e-09 | 1.1e-08 | 1.043E-08 | 5.871E-09 | 4.300E-09 | 1.600E-08 |
| 13528 | 677 | Y677G | B | 1.3e-08 | 1.3e-08 | 1.1e-08 | 1.233E-08 | 1.155E-09 | 1.100E-08 | 1.300E-08 |
| 13529 | 677 | Y677H | B | 1.1e-08 | 9.4e-09 | 4.5e-09 | 8.300E-09 | 3.387E-09 | 4.500E-09 | 1.100E-08 |
| 13530 | 677 | Y677I | B | 7.5e-09 | 2e-09 | 1.2e-08 | 7.167E-09 | 5.008E-09 | 2.000E-09 | 1.200E-08 |
| 13531 | 677 | Y677L | B | 7.7e-09 | 2.2e-08 | 6.9e-09 | 1.220E-08 | 8.496E-09 | 6.900E-09 | 2.200E-08 |
| 13532 | 677 | Y677K | B | 4.5e-09 | 9.8e-09 | 8.9e-09 | 7.733E-09 | 2.836E-09 | 4.500E-09 | 9.800E-09 |
| 13533 | 677 | Y677M | B | 7.8e-09 | 8.1e-09 | 5.7e-09 | 7.200E-09 | 1.308E-09 | 5.700E-09 | 8.100E-09 |
| 13534 | 677 | Y677F | B | 9.4e-09 | 5.4e-09 | 3.4e-09 | 6.067E-09 | 3.055E-09 | 3.400E-09 | 9.400E-09 |
| 13535 | 677 | Y677P | B | 1.7e-08 | 3.3e-09 | 1.4e-08 | 1.143E-08 | 7.202E-09 | 3.300E-09 | 1.700E-08 |
| 13536 | 677 | Y677S | B | 4.8e-09 | 1.2e-08 | 5.8e-09 | 7.533E-09 | 3.900E-09 | 4.800E-09 | 1.200E-08 |
| 13537 | 677 | Y677T | B | 5.6e-09 | 1.7e-08 | 6.2e-09 | 9.600E-09 | 6.416E-09 | 5.600E-09 | 1.700E-08 |
| 13538 | 677 | Y677W | B | 1.3e-08 | 7.7e-09 | 1.4e-08 | 1.157E-08 | 3.386E-09 | 7.700E-09 | 1.400E-08 |
| 13539 | 677 | Y677Y | B | 5.6e-09 | 3.4e-09 | 5.6e-09 | 4.867E-09 | 1.270E-09 | 3.400E-09 | 5.600E-09 |
| 13540 | 677 | Y677V | B | 9.6e-09 | 5e-09 | 5.3e-09 | 6.633E-09 | 2.574E-09 | 5.000E-09 | 9.600E-09 |
| 13541 | 678 | T678A | B | 1.4e-09 | 2.1e-08 | 4.3e-09 | 8.900E-09 | 1.058E-08 | 1.400E-09 | 2.100E-08 |
| 13542 | 678 | T678R | B | 9.4e-09 | 3.7e-09 | 1.7e-08 | 1.003E-08 | 6.673E-09 | 3.700E-09 | 1.700E-08 |
| 13543 | 678 | T678N | B | 5.3e-09 | 5.9e-09 | 1.7e-08 | 9.400E-09 | 6.589E-09 | 5.300E-09 | 1.700E-08 |
| 13544 | 678 | T678D | B | 2.5e-09 | 3.7e-09 | 2e-09 | 2.733E-09 | 8.737E-10 | 2.000E-09 | 3.700E-09 |
| 13545 | 678 | T678C | B | 5e-09 | 1.4e-08 | 1.5e-08 | 1.133E-08 | 5.508E-09 | 5.000E-09 | 1.500E-08 |
| 13546 | 678 | T678E | B | 7.1e-09 | 7.1e-09 | 4e-09 | 6.067E-09 | 1.790E-09 | 4.000E-09 | 7.100E-09 |
| 13547 | 678 | T678Q | B | 2.6e-09 | 4e-09 | 2.7e-09 | 3.100E-09 | 7.810E-10 | 2.600E-09 | 4.000E-09 |
| 13548 | 678 | T678G | B | 7.4e-09 | 6.6e-09 | 1.9e-08 | 1.100E-08 | 6.940E-09 | 6.600E-09 | 1.900E-08 |
| 13549 | 678 | T678H | B | 7.4e-09 | 4.4e-09 | 1.9e-08 | 1.027E-08 | 7.711E-09 | 4.400E-09 | 1.900E-08 |
| 13550 | 678 | T678I | B | 5.5e-09 | 9.1e-09 | 8e-09 | 7.533E-09 | 1.845E-09 | 5.500E-09 | 9.100E-09 |
| 13551 | 678 | T678L | B | 6.3e-09 | 7.7e-09 | 1.6e-08 | 1.000E-08 | 5.243E-09 | 6.300E-09 | 1.600E-08 |
| 13552 | 678 | T678K | B | 6.1e-09 | 9.4e-09 | 5.8e-09 | 7.100E-09 | 1.997E-09 | 5.800E-09 | 9.400E-09 |
| 13553 | 678 | T678M | B | 6.8e-09 | 3.7e-09 | 6.4e-09 | 5.633E-09 | 1.686E-09 | 3.700E-09 | 6.800E-09 |
| 13554 | 678 | T678F | B | 2.8e-09 | 4.4e-08 | 7.1e-09 | 1.797E-08 | 2.265E-08 | 2.800E-09 | 4.400E-08 |
| 13555 | 678 | T678P | B | 3.3e-09 | 7.4e-09 | 1.1e-08 | 7.233E-09 | 3.853E-09 | 3.300E-09 | 1.100E-08 |
| 13556 | 678 | T678S | B | 6.4e-09 | 3.6e-09 | 5.5e-09 | 5.167E-09 | 1.429E-09 | 3.600E-09 | 6.400E-09 |
| 13557 | 678 | T678T | B | 7.2e-09 | 4.5e-09 | 7.7e-09 | 6.467E-09 | 1.721E-09 | 4.500E-09 | 7.700E-09 |
| 13558 | 678 | T678W | B | 6.7e-09 | 5e-09 | 6e-09 | 5.900E-09 | 8.544E-10 | 5.000E-09 | 6.700E-09 |
| 13559 | 678 | T678Y | B | 6e-09 | 7.2e-09 | 1.7e-08 | 1.007E-08 | 6.034E-09 | 6.000E-09 | 1.700E-08 |
| 13560 | 678 | T678V | B | 1.7e-09 | 5.5e-09 | 4.4e-09 | 3.867E-09 | 1.955E-09 | 1.700E-09 | 5.500E-09 |
| 13561 | 679 | G679A | B | 3.1e-09 | 9.8e-09 | 2.9e-09 | 5.267E-09 | 3.927E-09 | 2.900E-09 | 9.800E-09 |
| 13562 | 679 | G679R | B | 5e-09 | 4.2e-09 | 6.1e-09 | 5.100E-09 | 9.539E-10 | 4.200E-09 | 6.100E-09 |
| 13563 | 679 | G679N | B | 9.4e-09 | 3.5e-09 | 4e-09 | 5.633E-09 | 3.272E-09 | 3.500E-09 | 9.400E-09 |
| 13564 | 679 | G679D | B | 8.9e-09 | 4.6e-09 | 8.5e-09 | 7.333E-09 | 2.376E-09 | 4.600E-09 | 8.900E-09 |
| 13565 | 679 | G679C | B | 1.4e-08 | 5.8e-09 | 1.1e-08 | 1.027E-08 | 4.149E-09 | 5.800E-09 | 1.400E-08 |
| 13566 | 679 | G679E | B | 7.4e-09 | 1.9e-08 | 6.8e-09 | 1.107E-08 | 6.877E-09 | 6.800E-09 | 1.900E-08 |
| 13567 | 679 | G679Q | B | 3.4e-09 | 3.6e-09 | 6.6e-09 | 4.533E-09 | 1.793E-09 | 3.400E-09 | 6.600E-09 |
| 13568 | 679 | G679G | B | 5.3e-09 | 6.8e-09 | 8.5e-09 | 6.867E-09 | 1.601E-09 | 5.300E-09 | 8.500E-09 |
| 13569 | 679 | G679H | B | 5.9e-09 | 1.6e-08 | 2.3e-09 | 8.067E-09 | 7.102E-09 | 2.300E-09 | 1.600E-08 |
| 13570 | 679 | G679I | B | 1.2e-08 | 6.9e-09 | 8.6e-09 | 9.167E-09 | 2.597E-09 | 6.900E-09 | 1.200E-08 |
| 13571 | 679 | G679L | B | 6.3e-09 | 4.3e-09 | 3.4e-09 | 4.667E-09 | 1.484E-09 | 3.400E-09 | 6.300E-09 |
| 13572 | 679 | G679K | B | 3.6e-09 | 6.8e-09 | 8.2e-09 | 6.200E-09 | 2.358E-09 | 3.600E-09 | 8.200E-09 |
| 13573 | 679 | G679M | B | 2.8e-09 | 6.9e-09 | 1.5e-08 | 8.233E-09 | 6.208E-09 | 2.800E-09 | 1.500E-08 |
| 13574 | 679 | G679F | B | 1.2e-08 | 7.7e-09 | 5.2e-09 | 8.300E-09 | 3.439E-09 | 5.200E-09 | 1.200E-08 |
| 13575 | 679 | G679P | B | 5.2e-09 | 7.4e-09 | 3.9e-09 | 5.500E-09 | 1.769E-09 | 3.900E-09 | 7.400E-09 |
| 13576 | 679 | G679S | B | 6.6e-09 | 3.6e-09 | 3.7e-09 | 4.633E-09 | 1.704E-09 | 3.600E-09 | 6.600E-09 |
| 13577 | 679 | G679T | B | 1.5e-08 | 8.6e-09 | 3.8e-09 | 9.133E-09 | 5.619E-09 | 3.800E-09 | 1.500E-08 |
| 13578 | 679 | G679W | B | 9.7e-09 | 1.2e-08 | 4.2e-09 | 8.633E-09 | 4.008E-09 | 4.200E-09 | 1.200E-08 |
| 13579 | 679 | G679Y | B | 6.2e-09 | 7.4e-09 | 9e-09 | 7.533E-09 | 1.405E-09 | 6.200E-09 | 9.000E-09 |
| 13580 | 679 | G679V | B | 1.9e-08 | 4.9e-09 | 1.3e-08 | 1.230E-08 | 7.076E-09 | 4.900E-09 | 1.900E-08 |
| 13581 | 680 | V680A | B | 7.9e-09 | 1.1e-08 | 2.9e-09 | 7.267E-09 | 4.087E-09 | 2.900E-09 | 1.100E-08 |
| 13582 | 680 | V680R | B | 4.7e-09 | 9.6e-09 | 7.5e-09 | 7.267E-09 | 2.458E-09 | 4.700E-09 | 9.600E-09 |
| 13583 | 680 | V680N | B | 2.6e-09 | 3.4e-08 | 5.1e-09 | 1.390E-08 | 1.745E-08 | 2.600E-09 | 3.400E-08 |
| 13584 | 680 | V680D | B | 8.3e-09 | 6.1e-09 | 1.1e-08 | 8.467E-09 | 2.454E-09 | 6.100E-09 | 1.100E-08 |
| 13585 | 680 | V680C | B | 9.3e-09 | 1.1e-08 | 1e-08 | 1.010E-08 | 8.544E-10 | 9.300E-09 | 1.100E-08 |
| 13586 | 680 | V680E | B | 1.1e-08 | 7.5e-09 | 5.7e-09 | 8.067E-09 | 2.695E-09 | 5.700E-09 | 1.100E-08 |
| 13587 | 680 | V680Q | B | 7.5e-09 | 1.1e-08 | 2.7e-09 | 7.067E-09 | 4.167E-09 | 2.700E-09 | 1.100E-08 |
| 13588 | 680 | V680G | B | 1.4e-08 | 1.4e-08 | 1.8e-08 | 1.533E-08 | 2.309E-09 | 1.400E-08 | 1.800E-08 |
| 13589 | 680 | V680H | B | 6.1e-09 | 7.1e-09 | 1.4e-09 | 4.867E-09 | 3.044E-09 | 1.400E-09 | 7.100E-09 |
| 13590 | 680 | V680I | B | 9.2e-09 | 3.7e-09 | 5.6e-09 | 6.167E-09 | 2.793E-09 | 3.700E-09 | 9.200E-09 |
| 13591 | 680 | V680L | B | 3.5e-09 | 6.2e-09 | 1.3e-08 | 7.567E-09 | 4.895E-09 | 3.500E-09 | 1.300E-08 |
| 13592 | 680 | V680K | B | 1.7e-08 | 2.3e-08 | 3.3e-09 | 1.443E-08 | 1.010E-08 | 3.300E-09 | 2.300E-08 |
| 13593 | 680 | V680M | B | 1.8e-08 | 6.6e-09 | 4e-09 | 9.533E-09 | 7.447E-09 | 4.000E-09 | 1.800E-08 |
| 13594 | 680 | V680F | B | 1e-08 | 1e-08 | 5.2e-09 | 8.400E-09 | 2.771E-09 | 5.200E-09 | 1.000E-08 |
| 13595 | 680 | V680P | B | 4.7e-09 | 4.8e-09 | 3.8e-09 | 4.433E-09 | 5.508E-10 | 3.800E-09 | 4.800E-09 |
| 13596 | 680 | V680S | B | 9.5e-09 | 2.9e-09 | 1e-08 | 7.467E-09 | 3.963E-09 | 2.900E-09 | 1.000E-08 |
| 13597 | 680 | V680T | B | 1.7e-08 | 9.5e-09 | 4.1e-09 | 1.020E-08 | 6.478E-09 | 4.100E-09 | 1.700E-08 |
| 13598 | 680 | V680W | B | 3.9e-09 | 5.6e-09 | 8.5e-09 | 6.000E-09 | 2.326E-09 | 3.900E-09 | 8.500E-09 |
| 13599 | 680 | V680Y | B | 7.5e-09 | 9.4e-09 | 3.3e-09 | 6.733E-09 | 3.121E-09 | 3.300E-09 | 9.400E-09 |

| | | | | | | | | | | |
|---|---|---|---|---|---|---|---|---|---|---|
| 13600 | 680 | V680V | B | 3e-09 | 1.1e-08 | 7.2e-09 | 7.067E-09 | 4.002E-09 | 3.000E-09 | 1.100E-08 |
| 13601 | 681 | P681A | B | 5.8e-09 | 6.5e-09 | 4.1e-09 | 5.467E-09 | 1.234E-09 | 4.100E-09 | 6.500E-09 |
| 13602 | 681 | P681R | B | 1.5e-08 | 3.9e-09 | 6.3e-09 | 8.400E-09 | 5.840E-09 | 3.900E-09 | 1.500E-08 |
| 13603 | 681 | P681N | B | 5e-09 | 1.2e-08 | 3e-09 | 6.667E-09 | 4.726E-09 | 3.000E-09 | 1.200E-08 |
| 13604 | 681 | P681D | B | 5.6e-09 | 4.6e-09 | 1e-08 | 6.733E-09 | 2.873E-09 | 4.600E-09 | 1.000E-08 |
| 13605 | 681 | P681C | B | 5.9e-09 | 7.4e-09 | 4.9e-09 | 6.067E-09 | 1.258E-09 | 4.900E-09 | 7.400E-09 |
| 13606 | 681 | P681E | B | 8.8e-09 | 2.8e-09 | 7.1e-09 | 6.233E-09 | 3.092E-09 | 2.800E-09 | 8.800E-09 |
| 13607 | 681 | P681Q | B | 6.8e-09 | 8.8e-09 | 7.2e-09 | 7.600E-09 | 1.058E-09 | 6.800E-09 | 8.800E-09 |
| 13608 | 681 | P681G | B | 1e-08 | 3.3e-09 | 4.9e-09 | 6.067E-09 | 3.499E-09 | 3.300E-09 | 1.000E-08 |
| 13609 | 681 | P681H | B | 5e-09 | 7.9e-09 | 4.2e-09 | 5.700E-09 | 1.947E-09 | 4.200E-09 | 7.900E-09 |
| 13610 | 681 | P681I | B | 9.7e-09 | 5.9e-09 | 5.8e-09 | 7.133E-09 | 2.223E-09 | 5.800E-09 | 9.700E-09 |
| 13611 | 681 | P681L | B | 4.3e-09 | 6.9e-09 | 9.6e-09 | 6.933E-09 | 2.650E-09 | 4.300E-09 | 9.600E-09 |
| 13612 | 681 | P681K | B | 5.4e-09 | 5.6e-09 | 6.9e-09 | 5.967E-09 | 8.145E-10 | 5.400E-09 | 6.900E-09 |
| 13613 | 681 | P681M | B | 5e-09 | 4.9e-09 | 4e-09 | 4.633E-09 | 5.508E-10 | 4.000E-09 | 5.000E-09 |
| 13614 | 681 | P681F | B | 6.1e-09 | 1.7e-08 | 6.9e-09 | 1.000E-08 | 6.075E-09 | 6.100E-09 | 1.700E-08 |
| 13615 | 681 | P681P | B | 7e-09 | 1.4e-08 | 2.6e-09 | 7.867E-09 | 5.749E-09 | 2.600E-09 | 1.400E-08 |
| 13616 | 681 | P681S | B | 1.2e-08 | 7.6e-09 | 9.3e-09 | 9.633E-09 | 2.219E-09 | 7.600E-09 | 1.200E-08 |
| 13617 | 681 | P681T | B | 7e-09 | 1.4e-08 | 8.1e-09 | 9.700E-09 | 3.764E-09 | 7.000E-09 | 1.400E-08 |
| 13618 | 681 | P681W | B | 9.2e-09 | 6e-09 | 2.4e-09 | 5.867E-09 | 3.402E-09 | 2.400E-09 | 9.200E-09 |
| 13619 | 681 | P681Y | B | 6.9e-09 | 4.7e-09 | 7.6e-09 | 6.400E-09 | 1.513E-09 | 4.700E-09 | 7.600E-09 |
| 13620 | 681 | P681V | B | 5.5e-09 | 6e-09 | 2.2e-09 | 4.567E-09 | 2.065E-09 | 2.200E-09 | 6.000E-09 |
| 13621 | 682 | S682A | B | 1.1e-08 | 6.8e-09 | 5.2e-09 | 7.667E-09 | 2.996E-09 | 5.200E-09 | 1.100E-08 |
| 13622 | 682 | S682R | B | 8.3e-09 | 1e-08 | 1.7e-08 | 1.177E-08 | 4.611E-09 | 8.300E-09 | 1.700E-08 |
| 13623 | 682 | S682N | B | 7.7e-09 | 6.4e-09 | 1.4e-08 | 9.367E-09 | 4.065E-09 | 6.400E-09 | 1.400E-08 |
| 13624 | 682 | S682D | B | 1.3e-08 | 8.2e-09 | 1.1e-08 | 1.073E-08 | 2.411E-09 | 8.200E-09 | 1.300E-08 |
| 13625 | 682 | S682C | B | 9.6e-09 | 7.1e-09 | 6.3e-09 | 7.667E-09 | 1.721E-09 | 6.300E-09 | 9.600E-09 |
| 13626 | 682 | S682E | B | 1.9e-08 | 5.4e-09 | 3e-09 | 9.133E-09 | 8.629E-09 | 3.000E-09 | 1.900E-08 |
| 13627 | 682 | S682Q | B | 4.8e-09 | 5.1e-09 | 8.1e-09 | 6.000E-09 | 1.825E-09 | 4.800E-09 | 8.100E-09 |
| 13628 | 682 | S682G | B | 4.6e-09 | 8.9e-09 | 1.1e-08 | 8.167E-09 | 3.262E-09 | 4.600E-09 | 1.100E-08 |
| 13629 | 682 | S682H | B | 9.5e-09 | 1.4e-08 | 4.6e-09 | 9.367E-09 | 4.701E-09 | 4.600E-09 | 1.400E-08 |
| 13630 | 682 | S682I | B | 2.9e-08 | 5.9e-09 | 7.8e-09 | 1.423E-08 | 1.282E-08 | 5.900E-09 | 2.900E-08 |
| 13631 | 682 | S682L | B | 5.4e-09 | 7e-09 | 4.3e-09 | 5.567E-09 | 1.358E-09 | 4.300E-09 | 7.000E-09 |
| 13632 | 682 | S682K | B | 9.7e-09 | 8.4e-09 | 4.7e-09 | 7.600E-09 | 2.594E-09 | 4.700E-09 | 9.700E-09 |
| 13633 | 682 | S682M | B | 5.3e-09 | 7.8e-09 | 1.2e-08 | 8.367E-09 | 3.386E-09 | 5.300E-09 | 1.200E-08 |
| 13634 | 682 | S682F | B | 7.7e-09 | 7.4e-09 | 4.6e-09 | 6.567E-09 | 1.710E-09 | 4.600E-09 | 7.700E-09 |
| 13635 | 682 | S682P | B | 6.3e-09 | 7.8e-09 | 2.9e-09 | 5.667E-09 | 2.511E-09 | 2.900E-09 | 7.800E-09 |
| 13636 | 682 | S682S | B | 1.9e-08 | 1.6e-08 | 4.1e-09 | 1.303E-08 | 7.881E-09 | 4.100E-09 | 1.900E-08 |
| 13637 | 682 | S682T | B | 1.3e-08 | 1.7e-08 | 4.9e-09 | 1.163E-08 | 6.165E-09 | 4.900E-09 | 1.700E-08 |
| 13638 | 682 | S682W | B | 9.8e-09 | 1.5e-08 | 9.3e-09 | 1.137E-08 | 3.156E-09 | 9.300E-09 | 1.500E-08 |
| 13639 | 682 | S682Y | B | 1.7e-09 | 4e-09 | 8.7e-09 | 4.800E-09 | 3.568E-09 | 1.700E-09 | 8.700E-09 |
| 13640 | 682 | S682V | B | 3.4e-09 | 1.1e-08 | 4.3e-09 | 6.233E-09 | 4.153E-09 | 3.400E-09 | 1.100E-08 |
| 13641 | 683 | R683A | B | 1.2e-08 | 6.5e-09 | 5.7e-09 | 8.067E-09 | 3.430E-09 | 5.700E-09 | 1.200E-08 |
| 13642 | 683 | R683R | B | 2.2e-08 | 1.6e-08 | 1e-08 | 1.600E-08 | 6.000E-09 | 1.000E-08 | 2.200E-08 |
| 13643 | 683 | R683N | B | 4.3e-09 | 4.2e-09 | 1.9e-08 | 9.167E-09 | 8.516E-09 | 4.200E-09 | 1.900E-08 |
| 13644 | 683 | R683D | B | 4.3e-09 | 1.6e-08 | 5.9e-09 | 8.733E-09 | 6.344E-09 | 4.300E-09 | 1.600E-08 |
| 13645 | 683 | R683C | B | 7.9e-09 | 6.4e-09 | 2e-08 | 1.143E-08 | 7.457E-09 | 6.400E-09 | 2.000E-08 |
| 13646 | 683 | R683E | B | 6.8e-09 | 4e-09 | 6.2e-09 | 5.667E-09 | 1.474E-09 | 4.000E-09 | 6.800E-09 |
| 13647 | 683 | R683Q | B | 5.4e-09 | 2.9e-09 | 3.9e-09 | 4.067E-09 | 1.258E-09 | 2.900E-09 | 5.400E-09 |
| 13648 | 683 | R683G | B | 9.5e-09 | 3.6e-09 | 1.1e-08 | 8.033E-09 | 3.912E-09 | 3.600E-09 | 1.100E-08 |
| 13649 | 683 | R683H | B | 1e-08 | 9e-09 | 3.8e-09 | 7.600E-09 | 3.329E-09 | 3.800E-09 | 1.000E-08 |
| 13650 | 683 | R683I | B | 2.3e-09 | 3.5e-09 | 2.7e-09 | 2.833E-09 | 6.110E-10 | 2.300E-09 | 3.500E-09 |
| 13651 | 683 | R683L | B | 7.7e-09 | 1.5e-08 | 1.1e-08 | 1.123E-08 | 3.656E-09 | 7.700E-09 | 1.500E-08 |
| 13652 | 683 | R683K | B | 2.8e-09 | 8.5e-09 | 4.7e-09 | 5.333E-09 | 2.902E-09 | 2.800E-09 | 8.500E-09 |
| 13653 | 683 | R683M | B | 6.8e-09 | 7.9e-09 | 1.4e-08 | 9.567E-09 | 3.879E-09 | 6.800E-09 | 1.400E-08 |
| 13654 | 683 | R683F | B | 8.8e-09 | 1.5e-08 | 3.3e-09 | 9.033E-09 | 5.853E-09 | 3.300E-09 | 1.500E-08 |
| 13655 | 683 | R683P | B | 8.8e-09 | 9.8e-09 | 1.6e-08 | 1.153E-08 | 3.900E-09 | 8.800E-09 | 1.600E-08 |
| 13656 | 683 | R683S | B | 2.1e-08 | 1e-08 | 7.2e-09 | 1.273E-08 | 7.295E-09 | 7.200E-09 | 2.100E-08 |
| 13657 | 683 | R683T | B | 6.7e-09 | 4.7e-09 | 5.4e-09 | 5.600E-09 | 1.015E-09 | 4.700E-09 | 6.700E-09 |
| 13658 | 683 | R683W | B | 7.6e-09 | 5.4e-09 | 3.6e-09 | 5.533E-09 | 2.003E-09 | 3.600E-09 | 7.600E-09 |
| 13659 | 683 | R683Y | B | 1e-08 | 4.8e-09 | 3.6e-09 | 6.133E-09 | 3.402E-09 | 3.600E-09 | 1.000E-08 |
| 13660 | 683 | R683V | B | 7.3e-09 | 8.5e-09 | 7.2e-09 | 7.667E-09 | 7.234E-10 | 7.200E-09 | 8.500E-09 |
| 13661 | 684 | F684A | B | 1.3e-08 | 1.6e-08 | 7.1e-09 | 1.203E-08 | 4.528E-09 | 7.100E-09 | 1.600E-08 |
| 13662 | 684 | F684R | B | 1e-08 | 2e-09 | 2.5e-09 | 4.833E-09 | 4.481E-09 | 2.000E-09 | 1.000E-08 |
| 13663 | 684 | F684N | B | 1e-08 | 5.6e-09 | 6.1e-09 | 7.233E-09 | 2.409E-09 | 5.600E-09 | 1.000E-08 |
| 13664 | 684 | F684D | B | 1.1e-08 | 2.5e-09 | 5.7e-09 | 6.400E-09 | 4.293E-09 | 2.500E-09 | 1.100E-08 |
| 13665 | 684 | F684C | B | 4.4e-09 | 8.9e-09 | 1.3e-08 | 8.767E-09 | 4.302E-09 | 4.400E-09 | 1.300E-08 |
| 13666 | 684 | F684E | B | 1.2e-08 | 4.5e-09 | 4.6e-09 | 7.033E-09 | 4.302E-09 | 4.500E-09 | 1.200E-08 |
| 13667 | 684 | F684Q | B | 4.7e-09 | 1.1e-08 | 9e-09 | 8.233E-09 | 3.219E-09 | 4.700E-09 | 1.100E-08 |
| 13668 | 684 | F684G | B | 6.5e-09 | 3.9e-09 | 5.2e-09 | 5.200E-09 | 1.300E-09 | 3.900E-09 | 6.500E-09 |
| 13669 | 684 | F684H | B | 5.6e-09 | 5.8e-09 | 4.4e-09 | 5.267E-09 | 7.572E-10 | 4.400E-09 | 5.800E-09 |
| 13670 | 684 | F684I | B | 3.7e-09 | 5.8e-09 | 9e-09 | 6.167E-09 | 2.669E-09 | 3.700E-09 | 9.000E-09 |
| 13671 | 684 | F684L | B | 6.8e-09 | 6e-09 | 6.1e-09 | 6.300E-09 | 4.359E-10 | 6.000E-09 | 6.800E-09 |
| 13672 | 684 | F684K | B | 6.7e-09 | 5.2e-09 | 2.6e-09 | 4.833E-09 | 2.074E-09 | 2.600E-09 | 6.700E-09 |
| 13673 | 684 | F684M | B | 7.7e-09 | 3.2e-09 | 4.6e-09 | 5.167E-09 | 2.303E-09 | 3.200E-09 | 7.700E-09 |
| 13674 | 684 | F684F | B | 1.8e-08 | 7.1e-09 | 1.2e-08 | 1.237E-08 | 5.459E-09 | 7.100E-09 | 1.800E-08 |
| 13675 | 684 | F684P | B | 1.5e-08 | 8.4e-09 | 6.1e-09 | 9.833E-09 | 4.620E-09 | 6.100E-09 | 1.500E-08 |
| 13676 | 684 | F684S | B | 3.7e-09 | 1e-08 | 1.9e-09 | 5.200E-09 | 4.253E-09 | 1.900E-09 | 1.000E-08 |
| 13677 | 684 | F684T | B | 7.3e-09 | 9.6e-09 | 5.9e-09 | 7.600E-09 | 1.868E-09 | 5.900E-09 | 9.600E-09 |
| 13678 | 684 | F684W | B | 2.3e-08 | 5.7e-09 | 1.2e-08 | 1.357E-08 | 8.756E-09 | 5.700E-09 | 2.300E-08 |
| 13679 | 684 | F684Y | B | 3e-08 | 1.1e-08 | 3.2e-09 | 1.473E-08 | 1.378E-08 | 3.200E-09 | 3.000E-08 |
| 13680 | 684 | F684V | B | 4.2e-09 | 7.6e-09 | 3e-09 | 4.933E-09 | 2.386E-09 | 3.000E-09 | 7.600E-09 |
| 13681 | 685 | S685A | B | 7.5e-09 | 1.1e-08 | 8.7e-09 | 9.067E-09 | 1.779E-09 | 7.500E-09 | 1.100E-08 |
| 13682 | 685 | S685R | B | 4.2e-09 | 3.9e-09 | 4.9e-09 | 4.333E-09 | 5.132E-10 | 3.900E-09 | 4.900E-09 |
| 13683 | 685 | S685N | B | 7.4e-09 | 1.6e-08 | 1.3e-08 | 1.213E-08 | 4.365E-09 | 7.400E-09 | 1.600E-08 |
| 13684 | 685 | S685D | B | 6.2e-09 | 2.2e-08 | 4.3e-09 | 1.083E-08 | 9.717E-09 | 4.300E-09 | 2.200E-08 |

| | | | | | | | | | | |
|---|---|---|---|---|---|---|---|---|---|---|
| 13685 | 685 | S685C | B | 5.6e-09 | 5.4e-09 | 5.5e-09 | 5.500E-09 | 1.000E-10 | 5.400E-09 | 5.600E-09 |
| 13686 | 685 | S685E | B | 1.2e-08 | 2.5e-08 | 2e-08 | 1.900E-08 | 6.557E-09 | 1.200E-08 | 2.500E-08 |
| 13687 | 685 | S685Q | B | 3.2e-09 | 4.7e-09 | 5.6e-09 | 4.500E-09 | 1.212E-09 | 3.200E-09 | 5.600E-09 |
| 13688 | 685 | S685G | B | 1.1e-08 | 4.6e-09 | 9.8e-09 | 8.467E-09 | 3.402E-09 | 4.600E-09 | 1.100E-08 |
| 13689 | 685 | S685H | B | 9.4e-09 | 2.7e-09 | 6.5e-09 | 6.200E-09 | 3.360E-09 | 2.700E-09 | 9.400E-09 |
| 13690 | 685 | S685I | B | 1e-08 | 1.7e-08 | 9.6e-09 | 1.220E-08 | 4.162E-09 | 9.600E-09 | 1.700E-08 |
| 13691 | 685 | S685L | B | 7.6e-09 | 1.7e-08 | 9.7e-09 | 1.143E-08 | 4.934E-09 | 7.600E-09 | 1.700E-08 |
| 13692 | 685 | S685K | B | 9.9e-09 | 2.1e-08 | 1e-08 | 1.363E-08 | 6.380E-09 | 9.900E-09 | 2.100E-08 |
| 13693 | 685 | S685M | B | 3.2e-09 | 1.9e-08 | 2.4e-09 | 8.200E-09 | 9.362E-09 | 2.400E-09 | 1.900E-08 |
| 13694 | 685 | S685F | B | 9.4e-09 | 5.2e-09 | 6.3e-09 | 6.967E-09 | 2.178E-09 | 5.200E-09 | 9.400E-09 |
| 13695 | 685 | S685P | B | 2.7e-09 | 4.2e-09 | 1e-08 | 5.633E-09 | 3.855E-09 | 2.700E-09 | 1.000E-08 |
| 13696 | 685 | S685S | B | 6.2e-09 | 7.3e-09 | 7.7e-09 | 7.067E-09 | 7.767E-10 | 6.200E-09 | 7.700E-09 |
| 13697 | 685 | S685T | B | 4.7e-09 | 4.9e-09 | 4.7e-09 | 4.767E-09 | 1.155E-10 | 4.700E-09 | 4.900E-09 |
| 13698 | 685 | S685W | B | 5.3e-09 | 2e-09 | 4.8e-09 | 4.033E-09 | 1.779E-09 | 2.000E-09 | 5.300E-09 |
| 13699 | 685 | S685Y | B | 4.8e-09 | 7.2e-09 | 6.5e-09 | 6.167E-09 | 1.234E-09 | 4.800E-09 | 7.200E-09 |
| 13700 | 685 | S685V | B | 2.3e-08 | 2.1e-09 | 3.1e-09 | 9.400E-09 | 1.179E-08 | 2.100E-09 | 2.300E-08 |
| 13701 | 686 | G686A | B | 1.5e-08 | 1.3e-08 | 1.6e-08 | 1.467E-08 | 1.528E-09 | 1.300E-08 | 1.600E-08 |
| 13702 | 686 | G686R | B | 1.2e-08 | 7.4e-09 | 3.7e-09 | 7.700E-09 | 4.158E-09 | 3.700E-09 | 1.200E-08 |
| 13703 | 686 | G686N | B | 6.6e-09 | 2.1e-08 | 8.8e-09 | 1.213E-08 | 7.757E-09 | 6.600E-09 | 2.100E-08 |
| 13704 | 686 | G686D | B | 1.6e-09 | 8.5e-09 | 6.6e-09 | 5.567E-09 | 3.564E-09 | 1.600E-09 | 8.500E-09 |
| 13705 | 686 | G686C | B | 3.8e-09 | 5.9e-09 | 3.7e-09 | 4.467E-09 | 1.242E-09 | 3.700E-09 | 5.900E-09 |
| 13706 | 686 | G686E | B | 7.2e-09 | 4.7e-09 | 3.3e-08 | 1.497E-08 | 1.567E-08 | 4.700E-09 | 3.300E-08 |
| 13707 | 686 | G686Q | B | 2.6e-09 | 6.3e-09 | 3.1e-09 | 4.000E-09 | 2.007E-09 | 2.600E-09 | 6.300E-09 |
| 13708 | 686 | G686G | B | 3e-09 | 1.2e-08 | 1.6e-08 | 1.033E-08 | 6.658E-09 | 3.000E-09 | 1.600E-08 |
| 13709 | 686 | G686H | B | 4.3e-09 | 7.8e-09 | 4e-09 | 5.367E-09 | 2.113E-09 | 4.000E-09 | 7.800E-09 |
| 13710 | 686 | G686I | B | 5.4e-09 | 2.5e-08 | 2.5e-09 | 1.097E-08 | 1.224E-08 | 2.500E-09 | 2.500E-08 |
| 13711 | 686 | G686L | B | 9.8e-09 | 6.3e-09 | 1.5e-09 | 5.867E-09 | 4.167E-09 | 1.500E-09 | 9.800E-09 |
| 13712 | 686 | G686K | B | 1.3e-08 | 8.2e-09 | 1.2e-08 | 1.107E-08 | 2.532E-09 | 8.200E-09 | 1.300E-08 |
| 13713 | 686 | G686M | B | 5.4e-09 | 1.8e-09 | 2.7e-09 | 3.300E-09 | 1.873E-09 | 1.800E-09 | 5.400E-09 |
| 13714 | 686 | G686F | B | 6e-09 | 7.2e-09 | 6.3e-09 | 6.500E-09 | 6.245E-10 | 6.000E-09 | 7.200E-09 |
| 13715 | 686 | G686P | B | 1.4e-08 | 8.9e-09 | 6.1e-09 | 9.667E-09 | 4.005E-09 | 6.100E-09 | 1.400E-08 |
| 13716 | 686 | G686S | B | 8e-09 | 8e-09 | 1.1e-08 | 9.000E-09 | 1.732E-09 | 8.000E-09 | 1.100E-08 |
| 13717 | 686 | G686T | B | 5.9e-09 | 7.7e-09 | 4.1e-09 | 5.900E-09 | 1.800E-09 | 4.100E-09 | 7.700E-09 |
| 13718 | 686 | G686W | B | 3.4e-08 | 7.3e-09 | 2.6e-09 | 1.463E-08 | 1.694E-08 | 2.600E-09 | 3.400E-08 |
| 13719 | 686 | G686Y | B | 9.9e-09 | 2.6e-09 | 8.4e-09 | 6.967E-09 | 3.855E-09 | 2.600E-09 | 9.900E-09 |
| 13720 | 686 | G686V | B | 3.4e-09 | 9.4e-09 | 3.7e-09 | 5.500E-09 | 3.381E-09 | 3.400E-09 | 9.400E-09 |
| 13721 | 687 | S687A | B | 3.4e-09 | 7.5e-09 | 1.6e-08 | 8.967E-09 | 6.427E-09 | 3.400E-09 | 1.600E-08 |
| 13722 | 687 | S687R | B | 1e-08 | 7e-09 | 3.3e-09 | 6.767E-09 | 3.356E-09 | 3.300E-09 | 1.000E-08 |
| 13723 | 687 | S687N | B | 1.5e-08 | 1.6e-08 | 2.9e-09 | 1.130E-08 | 7.292E-09 | 2.900E-09 | 1.600E-08 |
| 13724 | 687 | S687D | B | 3e-09 | 1.5e-08 | 9.7e-09 | 9.233E-09 | 6.014E-09 | 3.000E-09 | 1.500E-08 |
| 13725 | 687 | S687C | B | 4e-09 | 1e-08 | 4.5e-09 | 6.167E-09 | 3.329E-09 | 4.000E-09 | 1.000E-08 |
| 13726 | 687 | S687E | B | 6.5e-09 | 5e-09 | 3.7e-09 | 5.067E-09 | 1.401E-09 | 3.700E-09 | 6.500E-09 |
| 13727 | 687 | S687Q | B | 3.5e-09 | 8.6e-09 | 3.3e-09 | 5.133E-09 | 3.004E-09 | 3.300E-09 | 8.600E-09 |
| 13728 | 687 | S687G | B | 5.6e-09 | 9.3e-09 | 6.9e-09 | 7.267E-09 | 1.877E-09 | 5.600E-09 | 9.300E-09 |
| 13729 | 687 | S687H | B | 2.4e-08 | 2e-08 | 9.1e-09 | 1.770E-08 | 7.712E-09 | 9.100E-09 | 2.400E-08 |
| 13730 | 687 | S687I | B | 8e-09 | 8.2e-09 | 1.4e-08 | 1.007E-08 | 3.408E-09 | 8.000E-09 | 1.400E-08 |
| 13731 | 687 | S687L | B | 9e-09 | 1.2e-08 | 5.3e-09 | 8.767E-09 | 3.356E-09 | 5.300E-09 | 1.200E-08 |
| 13732 | 687 | S687K | B | 1.2e-08 | 1.4e-08 | 1.7e-08 | 1.433E-08 | 2.517E-09 | 1.200E-08 | 1.700E-08 |
| 13733 | 687 | S687M | B | 1.4e-08 | 3.5e-09 | 3.1e-09 | 6.867E-09 | 6.181E-09 | 3.100E-09 | 1.400E-08 |
| 13734 | 687 | S687F | B | 4.1e-09 | 1.8e-08 | 6.5e-09 | 9.533E-09 | 7.430E-09 | 4.100E-09 | 1.800E-08 |
| 13735 | 687 | S687P | B | 5e-09 | 5.4e-09 | 3.9e-09 | 4.767E-09 | 7.767E-10 | 3.900E-09 | 5.400E-09 |
| 13736 | 687 | S687S | B | 5.2e-08 | 4.8e-09 | 2e-08 | 2.560E-08 | 2.409E-08 | 4.800E-09 | 5.200E-08 |
| 13737 | 687 | S687T | B | 1.4e-08 | 1.1e-08 | 2.2e-09 | 9.067E-09 | 6.133E-09 | 2.200E-09 | 1.400E-08 |
| 13738 | 687 | S687W | B | 1.1e-08 | 2.8e-09 | 3.9e-09 | 5.900E-09 | 4.451E-09 | 2.800E-09 | 1.100E-08 |
| 13739 | 687 | S687Y | B | 1.3e-08 | 2.2e-08 | 6.7e-09 | 1.390E-08 | 7.690E-09 | 6.700E-09 | 2.200E-08 |
| 13740 | 687 | S687V | B | 1.5e-08 | 7.7e-09 | 3e-09 | 8.567E-09 | 6.047E-09 | 3.000E-09 | 1.500E-08 |
| 13741 | 688 | G688A | B | 9.9e-09 | 7.2e-09 | 1.9e-08 | 1.203E-08 | 6.183E-09 | 7.200E-09 | 1.900E-08 |
| 13742 | 688 | G688R | B | 1.7e-09 | 1.6e-09 | 3e-09 | 2.100E-09 | 7.810E-10 | 1.600E-09 | 3.000E-09 |
| 13743 | 688 | G688N | B | 2.6e-09 | 2e-08 | 6.5e-09 | 9.700E-09 | 9.131E-09 | 2.600E-09 | 2.000E-08 |
| 13744 | 688 | G688D | B | 1.2e-08 | 1.4e-08 | 6.6e-09 | 1.087E-08 | 3.828E-09 | 6.600E-09 | 1.400E-08 |
| 13745 | 688 | G688C | B | 8.3e-09 | 1.1e-08 | 1.2e-08 | 1.043E-08 | 1.914E-09 | 8.300E-09 | 1.200E-08 |
| 13746 | 688 | G688E | B | 1.4e-08 | 1e-08 | 2.8e-09 | 8.933E-09 | 5.676E-09 | 2.800E-09 | 1.400E-08 |
| 13747 | 688 | G688Q | B | 1.2e-08 | 6.1e-09 | 4.2e-09 | 7.433E-09 | 4.067E-09 | 4.200E-09 | 1.200E-08 |
| 13748 | 688 | G688G | B | 5e-09 | 3.4e-09 | 7.4e-09 | 5.267E-09 | 2.013E-09 | 3.400E-09 | 7.400E-09 |
| 13749 | 688 | G688H | B | 3.9e-09 | 3.2e-09 | 1.5e-08 | 7.367E-09 | 6.620E-09 | 3.200E-09 | 1.500E-08 |
| 13750 | 688 | G688I | B | 5.1e-09 | 1.5e-08 | 9.4e-09 | 9.833E-09 | 4.964E-09 | 5.100E-09 | 1.500E-08 |
| 13751 | 688 | G688L | B | 4e-09 | 6e-09 | 6.2e-09 | 5.400E-09 | 1.217E-09 | 4.000E-09 | 6.200E-09 |
| 13752 | 688 | G688K | B | 1.9e-08 | 3.1e-09 | 7.1e-09 | 9.733E-09 | 8.271E-09 | 3.100E-09 | 1.900E-08 |
| 13753 | 688 | G688M | B | 5.3e-09 | 5.1e-09 | 1.5e-08 | 8.467E-09 | 5.659E-09 | 5.100E-09 | 1.500E-08 |
| 13754 | 688 | G688F | B | 7.8e-09 | 3.7e-09 | 5.6e-09 | 5.700E-09 | 2.052E-09 | 3.700E-09 | 7.800E-09 |
| 13755 | 688 | G688P | B | 4.5e-09 | 7e-09 | 7.4e-09 | 6.300E-09 | 1.572E-09 | 4.500E-09 | 7.400E-09 |
| 13756 | 688 | G688S | B | 9.4e-09 | 1.2e-08 | 4.9e-09 | 8.767E-09 | 3.592E-09 | 4.900E-09 | 1.200E-08 |
| 13757 | 688 | G688T | B | 1e-08 | 5.3e-09 | 5.7e-09 | 7.000E-09 | 2.606E-09 | 5.300E-09 | 1.000E-08 |
| 13758 | 688 | G688W | B | 1.1e-08 | 6.4e-09 | 3e-08 | 1.580E-08 | 1.251E-08 | 6.400E-09 | 3.000E-08 |
| 13759 | 688 | G688Y | B | 8.2e-09 | 1.9e-08 | 1.3e-08 | 1.340E-08 | 5.411E-09 | 8.200E-09 | 1.900E-08 |
| 13760 | 688 | G688V | B | 3.7e-09 | 8.3e-09 | 7e-09 | 6.333E-09 | 2.371E-09 | 3.700E-09 | 8.300E-09 |
| 13761 | 689 | S689A | B | 3.3e-09 | 7.4e-09 | 4.4e-09 | 5.033E-09 | 2.122E-09 | 3.300E-09 | 7.400E-09 |
| 13762 | 689 | S689R | B | 7.1e-09 | 9.4e-09 | 3.6e-09 | 6.700E-09 | 2.921E-09 | 3.600E-09 | 9.400E-09 |
| 13763 | 689 | S689N | B | 1.1e-08 | 3.6e-09 | 7.2e-09 | 7.267E-09 | 3.700E-09 | 3.600E-09 | 1.100E-08 |
| 13764 | 689 | S689D | B | 2.8e-09 | 3.3e-09 | 4.8e-09 | 3.633E-09 | 1.041E-09 | 2.800E-09 | 4.800E-09 |
| 13765 | 689 | S689C | B | 3.8e-09 | 1.5e-08 | 1e-08 | 9.600E-09 | 5.611E-09 | 3.800E-09 | 1.500E-08 |
| 13766 | 689 | S689E | B | 4.9e-09 | 1.1e-08 | 3.3e-09 | 6.400E-09 | 4.063E-09 | 3.300E-09 | 1.100E-08 |
| 13767 | 689 | S689Q | B | 1.5e-08 | 2.6e-08 | 7.6e-09 | 1.620E-08 | 9.259E-09 | 7.600E-09 | 2.600E-08 |
| 13768 | 689 | S689G | B | 5.8e-09 | 9.7e-09 | 1.7e-08 | 1.083E-08 | 5.685E-09 | 5.800E-09 | 1.700E-08 |
| 13769 | 689 | S689H | B | 1e-08 | 7.8e-09 | 8.3e-09 | 8.700E-09 | 1.153E-09 | 7.800E-09 | 1.000E-08 |

| | | | | | | | | | | |
|---|---|---|---|---|---|---|---|---|---|---|
| 13770 | 689 | S689I | B | 6.5e-09 | 5.6e-09 | 5.9e-09 | 6.000E-09 | 4.583E-10 | 5.600E-09 | 6.500E-09 |
| 13771 | 689 | S689L | B | 3.8e-09 | 5.1e-09 | 3.4e-09 | 4.100E-09 | 8.888E-10 | 3.400E-09 | 5.100E-09 |
| 13772 | 689 | S689K | B | 1.3e-08 | 8.2e-09 | 6e-09 | 9.067E-09 | 3.580E-09 | 6.000E-09 | 1.300E-08 |
| 13773 | 689 | S689M | B | 1.4e-08 | 4.9e-09 | 4.1e-08 | 1.997E-08 | 1.878E-08 | 4.900E-09 | 4.100E-08 |
| 13774 | 689 | S689F | B | 1.4e-08 | 5e-09 | 8.7e-09 | 9.233E-09 | 4.524E-09 | 5.000E-09 | 1.400E-08 |
| 13775 | 689 | S689P | B | 1.6e-09 | 1.3e-08 | 3.4e-09 | 6.000E-09 | 6.129E-09 | 1.600E-09 | 1.300E-08 |
| 13776 | 689 | S689S | B | 6.5e-09 | 2.4e-09 | 7.7e-09 | 5.533E-09 | 2.779E-09 | 2.400E-09 | 7.700E-09 |
| 13777 | 689 | S689T | B | 6e-09 | 7.6e-09 | 7e-09 | 6.867E-09 | 8.083E-10 | 6.000E-09 | 7.600E-09 |
| 13778 | 689 | S689W | B | 7.5e-09 | 6e-09 | 1e-08 | 7.833E-09 | 2.021E-09 | 6.000E-09 | 1.000E-08 |
| 13779 | 689 | S689Y | B | 9.1e-09 | 4.5e-09 | 9.5e-09 | 7.700E-09 | 2.778E-09 | 4.500E-09 | 9.500E-09 |
| 13780 | 689 | S689V | B | 2e-08 | 9.5e-09 | 7.4e-09 | 1.230E-08 | 6.751E-09 | 7.400E-09 | 2.000E-08 |
| 13781 | 690 | G690A | B | 9.6e-09 | 3.7e-09 | 5.1e-09 | 6.133E-09 | 3.083E-09 | 3.700E-09 | 9.600E-09 |
| 13782 | 690 | G690R | B | 1.6e-08 | 1.5e-09 | 1.8e-08 | 1.183E-08 | 9.005E-09 | 1.500E-09 | 1.800E-08 |
| 13783 | 690 | G690N | B | 1.1e-08 | 8.1e-09 | 7.8e-09 | 8.967E-09 | 1.767E-09 | 7.800E-09 | 1.100E-08 |
| 13784 | 690 | G690D | B | 1.6e-08 | 6.8e-09 | 2.8e-09 | 8.533E-09 | 6.769E-09 | 2.800E-09 | 1.600E-08 |
| 13785 | 690 | G690C | B | 1.4e-08 | 1.3e-08 | 4.8e-09 | 1.060E-08 | 5.048E-09 | 4.800E-09 | 1.400E-08 |
| 13786 | 690 | G690E | B | 6.8e-09 | 3.9e-09 | 3.2e-09 | 4.633E-09 | 1.909E-09 | 3.200E-09 | 6.800E-09 |
| 13787 | 690 | G690Q | B | 4.7e-09 | 3.4e-09 | 8e-09 | 5.367E-09 | 2.371E-09 | 3.400E-09 | 8.000E-09 |
| 13788 | 690 | G690G | B | 8.8e-09 | 6.8e-09 | 6.3e-09 | 7.300E-09 | 1.323E-09 | 6.300E-09 | 8.800E-09 |
| 13789 | 690 | G690H | B | 1.9e-08 | 1.2e-08 | 5.8e-09 | 1.227E-08 | 6.604E-09 | 5.800E-09 | 1.900E-08 |
| 13790 | 690 | G690I | B | 8.7e-09 | 4.8e-09 | 3e-08 | 1.450E-08 | 1.356E-08 | 4.800E-09 | 3.000E-08 |
| 13791 | 690 | G690L | B | 5.3e-09 | 5e-09 | 2.9e-09 | 4.400E-09 | 1.308E-09 | 2.900E-09 | 5.300E-09 |
| 13792 | 690 | G690K | B | 5.8e-09 | 6.3e-09 | 1.4e-09 | 4.500E-09 | 2.696E-09 | 1.400E-09 | 6.300E-09 |
| 13793 | 690 | G690M | B | 3.1e-09 | 1.6e-08 | 3.4e-09 | 7.500E-09 | 7.363E-09 | 3.100E-09 | 1.600E-08 |
| 13794 | 690 | G690F | B | 1.8e-08 | 5.6e-09 | 5.7e-09 | 9.767E-09 | 7.130E-09 | 5.600E-09 | 1.800E-08 |
| 13795 | 690 | G690P | B | 1.2e-09 | 9.9e-09 | 4e-09 | 5.033E-09 | 4.441E-09 | 1.200E-09 | 9.900E-09 |
| 13796 | 690 | G690S | B | 1e-08 | 9.7e-09 | 3.1e-08 | 1.690E-08 | 1.221E-08 | 9.700E-09 | 3.100E-08 |
| 13797 | 690 | G690T | B | 3.8e-09 | 5.6e-09 | 8.3e-09 | 5.900E-09 | 2.265E-09 | 3.800E-09 | 8.300E-09 |
| 13798 | 690 | G690W | B | 1.5e-08 | 6.2e-09 | 1.5e-08 | 1.207E-08 | 5.081E-09 | 6.200E-09 | 1.500E-08 |
| 13799 | 690 | G690Y | B | 7.5e-09 | 6.6e-09 | 4.8e-09 | 6.300E-09 | 1.375E-09 | 4.800E-09 | 7.500E-09 |
| 13800 | 690 | G690V | B | 6.4e-09 | 3e-09 | 2.2e-09 | 3.867E-09 | 2.230E-09 | 2.200E-09 | 6.400E-09 |
| 13801 | 691 | T691A | B | 1.1e-08 | 5.3e-09 | 2e-08 | 1.210E-08 | 7.411E-09 | 5.300E-09 | 2.000E-08 |
| 13802 | 691 | T691R | B | 1.7e-09 | 1.3e-08 | 6.3e-09 | 7.000E-09 | 5.682E-09 | 1.700E-09 | 1.300E-08 |
| 13803 | 691 | T691N | B | 3.4e-09 | 1.2e-08 | 1e-08 | 8.467E-09 | 4.500E-09 | 3.400E-09 | 1.200E-08 |
| 13804 | 691 | T691D | B | 2.3e-08 | 1.7e-08 | 9.4e-09 | 1.647E-08 | 6.816E-09 | 9.400E-09 | 2.300E-08 |
| 13805 | 691 | T691C | B | 9.6e-09 | 5.5e-09 | 3.8e-09 | 6.300E-09 | 2.982E-09 | 3.800E-09 | 9.600E-09 |
| 13806 | 691 | T691E | B | 5e-09 | 9.2e-09 | 6.4e-09 | 6.867E-09 | 2.139E-09 | 5.000E-09 | 9.200E-09 |
| 13807 | 691 | T691Q | B | 2.8e-09 | 1.4e-08 | 7.7e-09 | 8.167E-09 | 5.615E-09 | 2.800E-09 | 1.400E-08 |
| 13808 | 691 | T691G | B | 1.1e-08 | 1.3e-08 | 5.2e-09 | 9.733E-09 | 4.051E-09 | 5.200E-09 | 1.300E-08 |
| 13809 | 691 | T691H | B | 8.8e-09 | 6.1e-09 | 1e-08 | 8.300E-09 | 1.997E-09 | 6.100E-09 | 1.000E-08 |
| 13810 | 691 | T691I | B | 2.3e-09 | 7.2e-09 | 4.5e-09 | 4.667E-09 | 2.454E-09 | 2.300E-09 | 7.200E-09 |
| 13811 | 691 | T691L | B | 4e-09 | 6.4e-09 | 1.9e-08 | 9.800E-09 | 8.057E-09 | 4.000E-09 | 1.900E-08 |
| 13812 | 691 | T691K | B | 8.3e-09 | 3.6e-09 | 3.5e-09 | 5.133E-09 | 2.743E-09 | 3.500E-09 | 8.300E-09 |
| 13813 | 691 | T691M | B | 3.5e-09 | 4.2e-09 | 7.9e-09 | 5.200E-09 | 2.364E-09 | 3.500E-09 | 7.900E-09 |
| 13814 | 691 | T691F | B | 1.4e-08 | 1.1e-08 | 9.8e-10 | 8.660E-09 | 6.818E-09 | 9.800E-10 | 1.400E-08 |
| 13815 | 691 | T691P | B | 8.1e-09 | 3.5e-09 | 6.8e-09 | 6.133E-09 | 2.371E-09 | 3.500E-09 | 8.100E-09 |
| 13816 | 691 | T691S | B | 6e-09 | 2.2e-08 | 1.1e-08 | 1.300E-08 | 8.185E-09 | 6.000E-09 | 2.200E-08 |
| 13817 | 691 | T691T | B | 9.7e-09 | 5.5e-09 | 4.6e-09 | 6.600E-09 | 2.722E-09 | 4.600E-09 | 9.700E-09 |
| 13818 | 691 | T691W | B | 9.3e-09 | 6.4e-09 | 2.1e-09 | 5.933E-09 | 3.623E-09 | 2.100E-09 | 9.300E-09 |
| 13819 | 691 | T691Y | B | 4.1e-09 | 6.4e-09 | 7.9e-09 | 6.133E-09 | 1.914E-09 | 4.100E-09 | 7.900E-09 |
| 13820 | 691 | T691V | B | 4.7e-09 | 1.9e-08 | 7.5e-09 | 1.040E-08 | 7.578E-09 | 4.700E-09 | 1.900E-08 |
| 13821 | 692 | D692A | B | 3e-09 | 3.3e-08 | 8.2e-09 | 1.473E-08 | 1.603E-08 | 3.000E-09 | 3.300E-08 |
| 13822 | 692 | D692R | B | 2.5e-08 | 4.3e-09 | 6.2e-09 | 1.183E-08 | 1.144E-08 | 4.300E-09 | 2.500E-08 |
| 13823 | 692 | D692N | B | 7.9e-09 | 9.9e-09 | 3e-09 | 6.933E-09 | 3.550E-09 | 3.000E-09 | 9.900E-09 |
| 13824 | 692 | D692D | B | 3.8e-09 | 1.8e-08 | 4.8e-09 | 8.867E-09 | 7.925E-09 | 3.800E-09 | 1.800E-08 |
| 13825 | 692 | D692C | B | 1.4e-08 | 6.7e-09 | 1.4e-08 | 1.157E-08 | 4.215E-09 | 6.700E-09 | 1.400E-08 |
| 13826 | 692 | D692E | B | 6e-09 | 9.4e-09 | 1.1e-08 | 8.800E-09 | 2.553E-09 | 6.000E-09 | 1.100E-08 |
| 13827 | 692 | D692Q | B | 3.2e-09 | 6.5e-09 | 3.5e-09 | 4.400E-09 | 1.825E-09 | 3.200E-09 | 6.500E-09 |
| 13828 | 692 | D692G | B | 4e-09 | 1.8e-08 | 1.6e-08 | 1.267E-08 | 7.572E-09 | 4.000E-09 | 1.800E-08 |
| 13829 | 692 | D692H | B | 7.2e-09 | 1.1e-08 | 4.7e-09 | 7.633E-09 | 3.172E-09 | 4.700E-09 | 1.100E-08 |
| 13830 | 692 | D692I | B | 8.6e-09 | 1.3e-08 | 6.4e-09 | 9.333E-09 | 3.361E-09 | 6.400E-09 | 1.300E-08 |
| 13831 | 692 | D692L | B | 3.3e-09 | 2.8e-09 | 4.3e-09 | 3.467E-09 | 7.638E-10 | 2.800E-09 | 4.300E-09 |
| 13832 | 692 | D692K | B | 1.3e-08 | 1.6e-08 | 1.3e-08 | 1.400E-08 | 1.732E-09 | 1.300E-08 | 1.600E-08 |
| 13833 | 692 | D692M | B | 6.2e-09 | 8.1e-09 | 6.2e-09 | 6.833E-09 | 1.097E-09 | 6.200E-09 | 8.100E-09 |
| 13834 | 692 | D692F | B | 4.4e-09 | 1.3e-08 | 4.9e-09 | 7.433E-09 | 4.827E-09 | 4.400E-09 | 1.300E-08 |
| 13835 | 692 | D692P | B | 1.5e-09 | 6.4e-09 | 3.1e-09 | 3.667E-09 | 2.499E-09 | 1.500E-09 | 6.400E-09 |
| 13836 | 692 | D692S | B | 2.8e-09 | 2.2e-09 | 5.4e-09 | 3.467E-09 | 1.701E-09 | 2.200E-09 | 5.400E-09 |
| 13837 | 692 | D692T | B | 2.7e-09 | 2.6e-09 | 3.8e-09 | 3.033E-09 | 6.658E-10 | 2.600E-09 | 3.800E-09 |
| 13838 | 692 | D692W | B | 2.1e-09 | 1.3e-08 | 1.3e-08 | 9.367E-09 | 6.293E-09 | 2.100E-09 | 1.300E-08 |
| 13839 | 692 | D692Y | B | 1.8e-08 | 1e-08 | 1.8e-09 | 9.933E-09 | 8.100E-09 | 1.800E-09 | 1.800E-08 |
| 13840 | 692 | D692V | B | 5.9e-09 | 1.9e-08 | 7.9e-09 | 1.093E-08 | 7.057E-09 | 5.900E-09 | 1.900E-08 |
| 13841 | 693 | F693A | B | 6.5e-09 | 2.4e-09 | 7.1e-09 | 5.333E-09 | 2.558E-09 | 2.400E-09 | 7.100E-09 |
| 13842 | 693 | F693R | B | 6.4e-09 | 5.3e-09 | 1.3e-08 | 8.233E-09 | 4.165E-09 | 5.300E-09 | 1.300E-08 |
| 13843 | 693 | F693N | B | 1e-08 | 5.4e-09 | 5.5e-09 | 6.967E-09 | 2.627E-09 | 5.400E-09 | 1.000E-08 |
| 13844 | 693 | F693D | B | 2.7e-09 | 7e-09 | 9.1e-09 | 6.267E-09 | 3.262E-09 | 2.700E-09 | 9.100E-09 |
| 13845 | 693 | F693C | B | 1.5e-08 | 1.2e-08 | 5e-09 | 1.067E-08 | 5.132E-09 | 5.000E-09 | 1.500E-08 |
| 13846 | 693 | F693E | B | 1.1e-08 | 8.5e-09 | 1.8e-08 | 1.250E-08 | 4.924E-09 | 8.500E-09 | 1.800E-08 |
| 13847 | 693 | F693Q | B | 4.4e-09 | 6.1e-09 | 3.9e-09 | 4.800E-09 | 1.153E-09 | 3.900E-09 | 6.100E-09 |
| 13848 | 693 | F693G | B | 3e-09 | 9.9e-09 | 1.6e-08 | 9.633E-09 | 6.504E-09 | 3.000E-09 | 1.600E-08 |
| 13849 | 693 | F693H | B | 5.6e-09 | 8.4e-09 | 8.9e-09 | 7.633E-09 | 1.779E-09 | 5.600E-09 | 8.900E-09 |
| 13850 | 693 | F693I | B | 5.5e-09 | 6.2e-09 | 1.3e-08 | 8.233E-09 | 4.143E-09 | 5.500E-09 | 1.300E-08 |
| 13851 | 693 | F693L | B | 5.3e-09 | 4.2e-09 | 3.9e-09 | 4.467E-09 | 7.371E-10 | 3.900E-09 | 5.300E-09 |
| 13852 | 693 | F693K | B | 4.3e-09 | 7.2e-09 | 9.9e-10 | 4.163E-09 | 3.107E-09 | 9.900E-10 | 7.200E-09 |
| 13853 | 693 | F693M | B | 4.3e-09 | 7.1e-09 | 5.1e-09 | 5.500E-09 | 1.442E-09 | 4.300E-09 | 7.100E-09 |
| 13854 | 693 | F693F | B | 5.8e-09 | 1.5e-08 | 6.1e-09 | 8.967E-09 | 5.227E-09 | 5.800E-09 | 1.500E-08 |

| | | | | | | | | | | |
|---|---|---|---|---|---|---|---|---|---|---|
| 13855 | 693 | F693P | B | 2.2e-09 | 6e-09 | 5.2e-09 | 4.467E-09 | 2.003E-09 | 2.200E-09 | 6.000E-09 |
| 13856 | 693 | F693S | B | 2e-08 | 3.2e-09 | 3.6e-09 | 8.933E-09 | 9.586E-09 | 3.200E-09 | 2.000E-08 |
| 13857 | 693 | F693T | B | 2.4e-09 | 1.4e-08 | 1.3e-08 | 9.800E-09 | 6.428E-09 | 2.400E-09 | 1.400E-08 |
| 13858 | 693 | F693W | B | 1.4e-08 | 1.1e-08 | 1.6e-08 | 1.367E-08 | 2.517E-09 | 1.100E-08 | 1.600E-08 |
| 13859 | 693 | F693Y | B | 8.7e-09 | 8.7e-09 | 1.1e-08 | 9.467E-09 | 1.328E-09 | 8.700E-09 | 1.100E-08 |
| 13860 | 693 | F693V | B | 3.2e-09 | 5e-09 | 3e-09 | 3.733E-09 | 1.102E-09 | 3.000E-09 | 5.000E-09 |
| 13861 | 694 | T694A | B | 1.6e-08 | 3.3e-09 | 1.4e-08 | 1.110E-08 | 6.829E-09 | 3.300E-09 | 1.600E-08 |
| 13862 | 694 | T694R | B | 1.7e-08 | 5.2e-09 | 7e-09 | 9.733E-09 | 6.357E-09 | 5.200E-09 | 1.700E-08 |
| 13863 | 694 | T694N | B | 8.2e-09 | 8.7e-09 | 1.2e-08 | 9.633E-09 | 2.065E-09 | 8.200E-09 | 1.200E-08 |
| 13864 | 694 | T694D | B | 2.3e-09 | 1.2e-08 | 4.6e-09 | 6.300E-09 | 5.069E-09 | 2.300E-09 | 1.200E-08 |
| 13865 | 694 | T694C | B | 6.9e-09 | 5.6e-09 | 2.1e-08 | 1.117E-08 | 8.541E-09 | 5.600E-09 | 2.100E-08 |
| 13866 | 694 | T694E | B | 1.8e-08 | 8.3e-09 | 9.6e-09 | 1.197E-08 | 5.265E-09 | 8.300E-09 | 1.800E-08 |
| 13867 | 694 | T694Q | B | 5.1e-09 | 4.6e-09 | 1.4e-08 | 7.900E-09 | 5.289E-09 | 4.600E-09 | 1.400E-08 |
| 13868 | 694 | T694G | B | 1.6e-08 | 1.8e-08 | 7.6e-09 | 1.387E-08 | 5.518E-09 | 7.600E-09 | 1.800E-08 |
| 13869 | 694 | T694H | B | 5.9e-09 | 6.4e-09 | 5.5e-09 | 5.933E-09 | 4.509E-10 | 5.500E-09 | 6.400E-09 |
| 13870 | 694 | T694I | B | 5.7e-09 | 4.9e-09 | 2.4e-09 | 4.333E-09 | 1.721E-09 | 2.400E-09 | 5.700E-09 |
| 13871 | 694 | T694L | B | 6.2e-09 | 6.1e-09 | 8.1e-09 | 6.800E-09 | 1.127E-09 | 6.100E-09 | 8.100E-09 |
| 13872 | 694 | T694K | B | 5.5e-09 | 1.6e-09 | 1.4e-08 | 7.033E-09 | 6.341E-09 | 1.600E-09 | 1.400E-08 |
| 13873 | 694 | T694M | B | 9e-09 | 9.5e-09 | 1.4e-08 | 1.083E-08 | 2.754E-09 | 9.000E-09 | 1.400E-08 |
| 13874 | 694 | T694F | B | 8.6e-09 | 6.8e-09 | 1e-08 | 8.467E-09 | 1.604E-09 | 6.800E-09 | 1.000E-08 |
| 13875 | 694 | T694P | B | 8.9e-09 | 7.1e-09 | 9.6e-09 | 8.533E-09 | 1.290E-09 | 7.100E-09 | 9.600E-09 |
| 13876 | 694 | T694S | B | 9.9e-09 | 2e-09 | 7.4e-09 | 6.433E-09 | 4.038E-09 | 2.000E-09 | 9.900E-09 |
| 13877 | 694 | T694T | B | 2.7e-09 | 5.4e-09 | 8.4e-09 | 5.500E-09 | 2.851E-09 | 2.700E-09 | 8.400E-09 |
| 13878 | 694 | T694W | B | 1.1e-08 | 2.4e-08 | 2.1e-09 | 1.237E-08 | 1.101E-08 | 2.100E-09 | 2.400E-08 |
| 13879 | 694 | T694Y | B | 1.2e-08 | 1.7e-08 | 8.2e-09 | 1.240E-08 | 4.414E-09 | 8.200E-09 | 1.700E-08 |
| 13880 | 694 | T694V | B | 8.8e-09 | 6.3e-09 | 1.2e-08 | 9.033E-09 | 2.857E-09 | 6.300E-09 | 1.200E-08 |
| 13881 | 695 | L695A | B | 1.2e-08 | 4.9e-09 | 1.5e-08 | 1.063E-08 | 5.187E-09 | 4.900E-09 | 1.500E-08 |
| 13882 | 695 | L695R | B | 5.3e-09 | 3.6e-09 | 1.7e-09 | 3.533E-09 | 1.801E-09 | 1.700E-09 | 5.300E-09 |
| 13883 | 695 | L695N | B | 5e-09 | 5.4e-09 | 5.1e-09 | 5.167E-09 | 2.082E-10 | 5.000E-09 | 5.400E-09 |
| 13884 | 695 | L695D | B | 9.2e-09 | 4.2e-09 | 1.8e-08 | 1.047E-08 | 6.987E-09 | 4.200E-09 | 1.800E-08 |
| 13885 | 695 | L695C | B | 6.8e-09 | 1e-08 | 1e-08 | 8.933E-09 | 1.848E-09 | 6.800E-09 | 1.000E-08 |
| 13886 | 695 | L695E | B | 8e-09 | 3.9e-09 | 4e-08 | 1.730E-08 | 1.977E-08 | 3.900E-09 | 4.000E-08 |
| 13887 | 695 | L695Q | B | 8.7e-09 | 2.8e-09 | 6.6e-09 | 6.033E-09 | 2.991E-09 | 2.800E-09 | 8.700E-09 |
| 13888 | 695 | L695G | B | 1.3e-08 | 2.6e-09 | 1.7e-08 | 1.087E-08 | 7.433E-09 | 2.600E-09 | 1.700E-08 |
| 13889 | 695 | L695H | B | 3.9e-08 | 2.5e-09 | 9.3e-09 | 1.693E-08 | 1.941E-08 | 2.500E-09 | 3.900E-08 |
| 13890 | 695 | L695I | B | 3e-08 | 7.3e-09 | 3.7e-09 | 1.367E-08 | 1.426E-08 | 3.700E-09 | 3.000E-08 |
| 13891 | 695 | L695L | B | 5e-09 | 8e-09 | 1.1e-08 | 8.000E-09 | 3.000E-09 | 5.000E-09 | 1.100E-08 |
| 13892 | 695 | L695K | B | 8e-09 | 6.7e-09 | 1.3e-08 | 9.233E-09 | 3.326E-09 | 6.700E-09 | 1.300E-08 |
| 13893 | 695 | L695M | B | 1e-08 | 7.5e-09 | 8e-09 | 8.500E-09 | 1.323E-09 | 7.500E-09 | 1.000E-08 |
| 13894 | 695 | L695F | B | 5.7e-09 | 4.6e-09 | 1e-08 | 6.767E-09 | 2.854E-09 | 4.600E-09 | 1.000E-08 |
| 13895 | 695 | L695P | B | 1.9e-08 | 7.1e-09 | 3.5e-09 | 9.867E-09 | 8.112E-09 | 3.500E-09 | 1.900E-08 |
| 13896 | 695 | L695S | B | 4.6e-09 | 5.7e-09 | 7.5e-09 | 5.933E-09 | 1.464E-09 | 4.600E-09 | 7.500E-09 |
| 13897 | 695 | L695T | B | 3.1e-09 | 9e-09 | 6.1e-09 | 6.067E-09 | 2.950E-09 | 3.100E-09 | 9.000E-09 |
| 13898 | 695 | L695W | B | 5.9e-09 | 1.3e-08 | 2e-09 | 6.967E-09 | 5.577E-09 | 2.000E-09 | 1.300E-08 |
| 13899 | 695 | L695Y | B | 7.4e-09 | 9.7e-09 | 2.1e-08 | 1.270E-08 | 7.279E-09 | 7.400E-09 | 2.100E-08 |
| 13900 | 695 | L695V | B | 1.3e-08 | 1.5e-08 | 1.1e-08 | 1.300E-08 | 2.000E-09 | 1.100E-08 | 1.500E-08 |
| 13901 | 696 | T696A | B | 5.7e-09 | 1.3e-08 | 2.4e-08 | 1.423E-08 | 9.212E-09 | 5.700E-09 | 2.400E-08 |
| 13902 | 696 | T696R | B | 6.1e-09 | 9e-09 | 3.5e-09 | 6.200E-09 | 2.751E-09 | 3.500E-09 | 9.000E-09 |
| 13903 | 696 | T696N | B | 3.4e-09 | 2.9e-09 | 8.2e-09 | 4.833E-09 | 2.926E-09 | 2.900E-09 | 8.200E-09 |
| 13904 | 696 | T696D | B | 3.8e-09 | 6.4e-09 | 7.7e-09 | 5.967E-09 | 1.986E-09 | 3.800E-09 | 7.700E-09 |
| 13905 | 696 | T696C | B | 6.9e-09 | 3.7e-09 | 5.8e-09 | 5.467E-09 | 1.626E-09 | 3.700E-09 | 6.900E-09 |
| 13906 | 696 | T696E | B | 3.7e-09 | 9.6e-09 | 3.6e-09 | 5.633E-09 | 3.436E-09 | 3.600E-09 | 9.600E-09 |
| 13907 | 696 | T696Q | B | 4e-09 | 9.2e-09 | 9.3e-09 | 7.500E-09 | 3.032E-09 | 4.000E-09 | 9.300E-09 |
| 13908 | 696 | T696G | B | 7.5e-09 | 6.5e-09 | 2.2e-08 | 1.200E-08 | 8.675E-09 | 6.500E-09 | 2.200E-08 |
| 13909 | 696 | T696H | B | 7.6e-09 | 7.4e-09 | 4.1e-09 | 6.367E-09 | 1.966E-09 | 4.100E-09 | 7.600E-09 |
| 13910 | 696 | T696I | B | 5.2e-09 | 7e-09 | 4.9e-09 | 5.700E-09 | 1.136E-09 | 4.900E-09 | 7.000E-09 |
| 13911 | 696 | T696L | B | 1.6e-08 | 9.3e-09 | 3e-09 | 9.433E-09 | 6.501E-09 | 3.000E-09 | 1.600E-08 |
| 13912 | 696 | T696K | B | 1e-08 | 1.2e-08 | 4e-09 | 8.667E-09 | 4.163E-09 | 4.000E-09 | 1.200E-08 |
| 13913 | 696 | T696M | B | 3e-09 | 7.3e-09 | 7.8e-09 | 6.033E-09 | 2.639E-09 | 3.000E-09 | 7.800E-09 |
| 13914 | 696 | T696F | B | 2.1e-08 | 6.7e-09 | 2e-08 | 1.590E-08 | 7.983E-09 | 6.700E-09 | 2.100E-08 |
| 13915 | 696 | T696P | B | 4.5e-09 | 3.9e-09 | 5.9e-09 | 4.767E-09 | 1.026E-09 | 3.900E-09 | 5.900E-09 |
| 13916 | 696 | T696S | B | 1.1e-08 | 8.5e-09 | 9.3e-10 | 6.810E-09 | 5.243E-09 | 9.300E-10 | 1.100E-08 |
| 13917 | 696 | T696T | B | 4.7e-09 | 6.5e-09 | 3.1e-09 | 4.767E-09 | 1.701E-09 | 3.100E-09 | 6.500E-09 |
| 13918 | 696 | T696W | B | 8.9e-09 | 9.7e-09 | 7e-09 | 8.533E-09 | 1.387E-09 | 7.000E-09 | 9.700E-09 |
| 13919 | 696 | T696Y | B | 9.2e-09 | 1.4e-08 | 1.7e-08 | 1.340E-08 | 3.934E-09 | 9.200E-09 | 1.700E-08 |
| 13920 | 696 | T696V | B | 1.2e-08 | 9.2e-09 | 1.4e-08 | 1.173E-08 | 2.411E-09 | 9.200E-09 | 1.400E-08 |
| 13921 | 697 | I697A | B | 1.1e-08 | 4.6e-09 | 9.6e-09 | 8.400E-09 | 3.365E-09 | 4.600E-09 | 1.100E-08 |
| 13922 | 697 | I697R | B | 4.7e-09 | 6e-09 | 1.6e-08 | 8.900E-09 | 6.183E-09 | 4.700E-09 | 1.600E-08 |
| 13923 | 697 | I697N | B | 7.6e-09 | 2.2e-08 | 6e-09 | 1.187E-08 | 8.812E-09 | 6.000E-09 | 2.200E-08 |
| 13924 | 697 | I697D | B | 1.4e-08 | 9.1e-09 | 1.2e-08 | 1.170E-08 | 2.464E-09 | 9.100E-09 | 1.400E-08 |
| 13925 | 697 | I697C | B | 9.6e-09 | 7.8e-09 | 1.1e-08 | 9.467E-09 | 1.604E-09 | 7.800E-09 | 1.100E-08 |
| 13926 | 697 | I697E | B | 5.3e-09 | 1e-08 | 2.1e-09 | 5.800E-09 | 3.974E-09 | 2.100E-09 | 1.000E-08 |
| 13927 | 697 | I697Q | B | 1.3e-08 | 3.2e-08 | 4.6e-09 | 1.653E-08 | 1.404E-08 | 4.600E-09 | 3.200E-08 |
| 13928 | 697 | I697G | B | 4.1e-09 | 1.8e-08 | 5.2e-09 | 9.100E-09 | 7.727E-09 | 4.100E-09 | 1.800E-08 |
| 13929 | 697 | I697H | B | 7.8e-09 | 3.4e-09 | 3.5e-09 | 4.900E-09 | 2.512E-09 | 3.400E-09 | 7.800E-09 |
| 13930 | 697 | I697I | B | 1.2e-08 | 8.1e-09 | 7.4e-09 | 9.167E-09 | 2.479E-09 | 7.400E-09 | 1.200E-08 |
| 13931 | 697 | I697L | B | 5.7e-09 | 8e-09 | 1.8e-08 | 1.057E-08 | 6.539E-09 | 5.700E-09 | 1.800E-08 |
| 13932 | 697 | I697K | B | 2.3e-09 | 3.8e-09 | 6.7e-09 | 4.267E-09 | 2.237E-09 | 2.300E-09 | 6.700E-09 |
| 13933 | 697 | I697M | B | 1.6e-08 | 3.7e-09 | 3.7e-08 | 1.890E-08 | 1.684E-08 | 3.700E-09 | 3.700E-08 |
| 13934 | 697 | I697F | B | 1e-08 | 9.8e-09 | 8.7e-09 | 9.500E-09 | 7.000E-10 | 8.700E-09 | 1.000E-08 |
| 13935 | 697 | I697P | B | 1.5e-08 | 9.3e-09 | 4.2e-09 | 9.500E-09 | 5.403E-09 | 4.200E-09 | 1.500E-08 |
| 13936 | 697 | I697S | B | 7.4e-09 | 1.7e-08 | 7.9e-09 | 1.077E-08 | 5.404E-09 | 7.400E-09 | 1.700E-08 |
| 13937 | 697 | I697T | B | 7.4e-09 | 3.1e-09 | 1.1e-08 | 7.167E-09 | 3.955E-09 | 3.100E-09 | 1.100E-08 |
| 13938 | 697 | I697W | B | 6.4e-09 | 1.2e-08 | 1.2e-08 | 1.013E-08 | 3.233E-09 | 6.400E-09 | 1.200E-08 |
| 13939 | 697 | I697Y | B | 6e-09 | 5e-09 | 1.3e-08 | 8.000E-09 | 4.359E-09 | 5.000E-09 | 1.300E-08 |

| | | | | | | | | | | |
|---|---|---|---|---|---|---|---|---|---|---|
| 13940 | 697 | I697V | B | 4.5e-09 | 4.5e-09 | 1.4e-08 | 7.667E-09 | 5.485E-09 | 4.500E-09 | 1.400E-08 |
| 13941 | 698 | S698A | B | 5.4e-09 | 3.6e-09 | 5.7e-09 | 4.900E-09 | 1.136E-09 | 3.600E-09 | 5.700E-09 |
| 13942 | 698 | S698R | B | 4.3e-08 | 4.8e-09 | 3e-09 | 1.693E-08 | 2.259E-08 | 3.000E-09 | 4.300E-08 |
| 13943 | 698 | S698N | B | 7.7e-09 | 3.4e-09 | 1e-08 | 7.033E-09 | 3.350E-09 | 3.400E-09 | 1.000E-08 |
| 13944 | 698 | S698D | B | 4.5e-09 | 1.9e-08 | 5.9e-09 | 9.800E-09 | 7.998E-09 | 4.500E-09 | 1.900E-08 |
| 13945 | 698 | S698C | B | 8.8e-09 | 2.4e-09 | 2.2e-09 | 4.467E-09 | 3.754E-09 | 2.200E-09 | 8.800E-09 |
| 13946 | 698 | S698E | B | 5.9e-09 | 8.6e-09 | 8.5e-09 | 7.667E-09 | 1.531E-09 | 5.900E-09 | 8.600E-09 |
| 13947 | 698 | S698Q | B | 5e-09 | 4.3e-09 | 5.6e-09 | 4.967E-09 | 6.506E-10 | 4.300E-09 | 5.600E-09 |
| 13948 | 698 | S698G | B | 4.9e-09 | 1.2e-08 | 1.4e-08 | 1.030E-08 | 4.782E-09 | 4.900E-09 | 1.400E-08 |
| 13949 | 698 | S698H | B | 5.3e-09 | 1e-08 | 2.2e-08 | 1.243E-08 | 8.612E-09 | 5.300E-09 | 2.200E-08 |
| 13950 | 698 | S698I | B | 7.7e-09 | 1e-08 | 6.2e-09 | 7.967E-09 | 1.914E-09 | 6.200E-09 | 1.000E-08 |
| 13951 | 698 | S698L | B | 2.9e-09 | 6e-09 | 2.3e-08 | 1.063E-08 | 1.082E-08 | 2.900E-09 | 2.300E-08 |
| 13952 | 698 | S698K | B | 1.6e-09 | 6.8e-09 | 1.3e-09 | 3.233E-09 | 3.092E-09 | 1.300E-09 | 6.800E-09 |
| 13953 | 698 | S698M | B | 1e-08 | 6.5e-09 | 1.5e-08 | 1.050E-08 | 4.272E-09 | 6.500E-09 | 1.500E-08 |
| 13954 | 698 | S698F | B | 3.7e-09 | 1.1e-08 | 7e-09 | 7.233E-09 | 3.656E-09 | 3.700E-09 | 1.100E-08 |
| 13955 | 698 | S698P | B | 1.7e-08 | 2.7e-09 | 7.1e-09 | 8.933E-09 | 7.324E-09 | 2.700E-09 | 1.700E-08 |
| 13956 | 698 | S698S | B | 1.3e-08 | 5.7e-09 | 1.2e-08 | 1.023E-08 | 3.958E-09 | 5.700E-09 | 1.300E-08 |
| 13957 | 698 | S698T | B | 1.7e-08 | 1.3e-08 | 8.5e-09 | 1.283E-08 | 4.252E-09 | 8.500E-09 | 1.700E-08 |
| 13958 | 698 | S698W | B | 8.3e-09 | 1.9e-08 | 1.6e-08 | 1.443E-08 | 5.519E-09 | 8.300E-09 | 1.900E-08 |
| 13959 | 698 | S698Y | B | 5.7e-09 | 2.7e-09 | 8.9e-09 | 5.767E-09 | 3.101E-09 | 2.700E-09 | 8.900E-09 |
| 13960 | 698 | S698V | B | 4.2e-09 | 2e-09 | 3.1e-09 | 3.100E-09 | 1.100E-09 | 2.000E-09 | 4.200E-09 |
| 13961 | 699 | S699A | B | 5.9e-09 | 1.1e-08 | 1.1e-08 | 9.300E-09 | 2.944E-09 | 5.900E-09 | 1.100E-08 |
| 13962 | 699 | S699R | B | 9.3e-09 | 5.1e-09 | 1.3e-08 | 9.133E-09 | 3.953E-09 | 5.100E-09 | 1.300E-08 |
| 13963 | 699 | S699N | B | 5.7e-09 | 5.9e-09 | 5.6e-09 | 5.733E-09 | 1.528E-10 | 5.600E-09 | 5.900E-09 |
| 13964 | 699 | S699D | B | 5.5e-09 | 9.8e-09 | 1.6e-08 | 1.043E-08 | 5.279E-09 | 5.500E-09 | 1.600E-08 |
| 13965 | 699 | S699C | B | 4.4e-09 | 1.1e-08 | 1.1e-08 | 8.800E-09 | 3.811E-09 | 4.400E-09 | 1.100E-08 |
| 13966 | 699 | S699E | B | 1e-08 | 5.8e-09 | 1.4e-08 | 9.933E-09 | 4.100E-09 | 5.800E-09 | 1.400E-08 |
| 13967 | 699 | S699Q | B | 2.4e-09 | 9.1e-09 | 2.6e-09 | 4.700E-09 | 3.812E-09 | 2.400E-09 | 9.100E-09 |
| 13968 | 699 | S699G | B | 1.6e-09 | 8.1e-09 | 4.1e-09 | 4.600E-09 | 3.279E-09 | 1.600E-09 | 8.100E-09 |
| 13969 | 699 | S699H | B | 4.2e-09 | 7.9e-09 | 5.5e-09 | 5.867E-09 | 1.877E-09 | 4.200E-09 | 7.900E-09 |
| 13970 | 699 | S699I | B | 5.6e-09 | 2.2e-09 | 5.9e-09 | 4.567E-09 | 2.055E-09 | 2.200E-09 | 5.900E-09 |
| 13971 | 699 | S699L | B | 1.5e-08 | 1.8e-08 | 5.3e-09 | 1.277E-08 | 6.638E-09 | 5.300E-09 | 1.800E-08 |
| 13972 | 699 | S699K | B | 3.7e-09 | 9e-09 | 7.4e-09 | 6.700E-09 | 2.718E-09 | 3.700E-09 | 9.000E-09 |
| 13973 | 699 | S699M | B | 2.3e-08 | 2.9e-09 | 1e-08 | 1.197E-08 | 1.019E-08 | 2.900E-09 | 2.300E-08 |
| 13974 | 699 | S699F | B | 7.5e-09 | 1e-08 | 1.9e-09 | 6.467E-09 | 4.148E-09 | 1.900E-09 | 1.000E-08 |
| 13975 | 699 | S699P | B | 1.2e-08 | 1.1e-08 | 4.7e-09 | 9.233E-09 | 3.958E-09 | 4.700E-09 | 1.200E-08 |
| 13976 | 699 | S699S | B | 3.6e-09 | 4.2e-09 | 2.3e-09 | 3.367E-09 | 9.713E-10 | 2.300E-09 | 4.200E-09 |
| 13977 | 699 | S699T | B | 3.2e-09 | 7.3e-09 | 6.5e-09 | 5.667E-09 | 2.173E-09 | 3.200E-09 | 7.300E-09 |
| 13978 | 699 | S699W | B | 2.8e-09 | 9.1e-09 | 2.5e-09 | 4.800E-09 | 3.727E-09 | 2.500E-09 | 9.100E-09 |
| 13979 | 699 | S699Y | B | 5.3e-09 | 2.4e-09 | 5.7e-09 | 4.467E-09 | 1.801E-09 | 2.400E-09 | 5.700E-09 |
| 13980 | 699 | S699V | B | 9e-09 | 5.6e-09 | 3.6e-09 | 6.067E-09 | 2.730E-09 | 3.600E-09 | 9.000E-09 |
| 13981 | 700 | L700A | B | 1.1e-08 | 1.5e-08 | 4.5e-09 | 1.017E-08 | 5.299E-09 | 4.500E-09 | 1.500E-08 |
| 13982 | 700 | L700R | B | 1.1e-08 | 1.1e-08 | 4.8e-09 | 8.933E-09 | 3.580E-09 | 4.800E-09 | 1.100E-08 |
| 13983 | 700 | L700N | B | 2e-09 | 1.6e-08 | 1.5e-08 | 1.100E-08 | 7.810E-09 | 2.000E-09 | 1.600E-08 |
| 13984 | 700 | L700D | B | 1.7e-08 | 1.2e-08 | 4.9e-09 | 1.130E-08 | 6.080E-09 | 4.900E-09 | 1.700E-08 |
| 13985 | 700 | L700C | B | 1.3e-08 | 4.3e-09 | 1.8e-09 | 6.367E-09 | 5.879E-09 | 1.800E-09 | 1.300E-08 |
| 13986 | 700 | L700E | B | 9.5e-09 | 1.7e-08 | 2.7e-08 | 1.783E-08 | 8.780E-09 | 9.500E-09 | 2.700E-08 |
| 13987 | 700 | L700Q | B | 4.9e-09 | 1.8e-08 | 1.1e-08 | 1.130E-08 | 6.555E-09 | 4.900E-09 | 1.800E-08 |
| 13988 | 700 | L700G | B | 5.3e-09 | 9.1e-09 | 1.3e-08 | 9.133E-09 | 3.850E-09 | 5.300E-09 | 1.300E-08 |
| 13989 | 700 | L700H | B | 2.6e-09 | 3.8e-09 | 2.7e-09 | 3.033E-09 | 6.658E-10 | 2.600E-09 | 3.800E-09 |
| 13990 | 700 | L700I | B | 3.9e-09 | 7.4e-09 | 1e-08 | 7.100E-09 | 3.061E-09 | 3.900E-09 | 1.000E-08 |
| 13991 | 700 | L700L | B | 3e-09 | 5.6e-09 | 8.4e-09 | 5.667E-09 | 2.701E-09 | 3.000E-09 | 8.400E-09 |
| 13992 | 700 | L700K | B | 8.6e-09 | 3.3e-09 | 1.2e-08 | 7.967E-09 | 4.384E-09 | 3.300E-09 | 1.200E-08 |
| 13993 | 700 | L700M | B | 1.4e-08 | 4.4e-08 | 5.3e-09 | 2.110E-08 | 2.030E-08 | 5.300E-09 | 4.400E-08 |
| 13994 | 700 | L700F | B | 2.5e-09 | 1.5e-08 | 3.6e-09 | 7.033E-09 | 6.921E-09 | 2.500E-09 | 1.500E-08 |
| 13995 | 700 | L700P | B | 9.7e-09 | 8.1e-09 | 7.6e-09 | 8.467E-09 | 1.097E-09 | 7.600E-09 | 9.700E-09 |
| 13996 | 700 | L700S | B | 7.4e-09 | 6.6e-09 | 6.2e-09 | 6.733E-09 | 6.110E-10 | 6.200E-09 | 7.400E-09 |
| 13997 | 700 | L700T | B | 7.6e-09 | 5.2e-09 | 6.6e-09 | 6.467E-09 | 1.206E-09 | 5.200E-09 | 7.600E-09 |
| 13998 | 700 | L700W | B | 1.4e-08 | 9.3e-09 | 3.3e-09 | 8.867E-09 | 5.363E-09 | 3.300E-09 | 1.400E-08 |
| 13999 | 700 | L700Y | B | 4.1e-09 | 7.6e-09 | 7e-09 | 6.233E-09 | 1.872E-09 | 4.100E-09 | 7.600E-09 |
| 14000 | 700 | L700V | B | 1.5e-08 | 1.5e-08 | 1.4e-08 | 1.467E-08 | 5.774E-10 | 1.400E-08 | 1.500E-08 |
| 14001 | 701 | Q701A | B | 3.6e-09 | 5e-09 | 7.6e-09 | 5.400E-09 | 2.030E-09 | 3.600E-09 | 7.600E-09 |
| 14002 | 701 | Q701R | B | 2.4e-09 | 8.3e-09 | 1e-08 | 6.900E-09 | 3.989E-09 | 2.400E-09 | 1.000E-08 |
| 14003 | 701 | Q701N | B | 2.8e-09 | 3.9e-09 | 7.3e-09 | 4.667E-09 | 2.346E-09 | 2.800E-09 | 7.300E-09 |
| 14004 | 701 | Q701D | B | 8.5e-09 | 7.8e-09 | 1.2e-08 | 9.433E-09 | 2.250E-09 | 7.800E-09 | 1.200E-08 |
| 14005 | 701 | Q701C | B | 1.6e-08 | 1.3e-08 | 1.1e-08 | 1.333E-08 | 2.517E-09 | 1.100E-08 | 1.600E-08 |
| 14006 | 701 | Q701E | B | 3.6e-09 | 1.3e-08 | 1.5e-08 | 1.053E-08 | 6.087E-09 | 3.600E-09 | 1.500E-08 |
| 14007 | 701 | Q701Q | B | 8e-09 | 4.8e-09 | 2.4e-08 | 1.227E-08 | 1.029E-08 | 4.800E-09 | 2.400E-08 |
| 14008 | 701 | Q701G | B | 1.2e-08 | 2e-08 | 2.8e-09 | 1.160E-08 | 8.607E-09 | 2.800E-09 | 2.000E-08 |
| 14009 | 701 | Q701H | B | 3e-09 | 3.1e-09 | 8.5e-09 | 4.867E-09 | 3.147E-09 | 3.000E-09 | 8.500E-09 |
| 14010 | 701 | Q701I | B | 2.4e-08 | 6.7e-09 | 5.8e-09 | 1.217E-08 | 1.026E-08 | 5.800E-09 | 2.400E-08 |
| 14011 | 701 | Q701L | B | 2.5e-08 | 4.1e-09 | 4.5e-09 | 1.120E-08 | 1.195E-08 | 4.100E-09 | 2.500E-08 |
| 14012 | 701 | Q701K | B | 5.5e-09 | 1e-08 | 5.8e-09 | 7.100E-09 | 2.516E-09 | 5.500E-09 | 1.000E-08 |
| 14013 | 701 | Q701M | B | 3.6e-09 | 6.4e-09 | 1.2e-08 | 7.333E-09 | 4.277E-09 | 3.600E-09 | 1.200E-08 |
| 14014 | 701 | Q701F | B | 4.5e-09 | 1e-08 | 5.5e-09 | 6.667E-09 | 2.930E-09 | 4.500E-09 | 1.000E-08 |
| 14015 | 701 | Q701P | B | 5.3e-09 | 5.5e-09 | 4.9e-09 | 5.233E-09 | 3.055E-10 | 4.900E-09 | 5.500E-09 |
| 14016 | 701 | Q701S | B | 3.2e-08 | 9.6e-09 | 5.2e-09 | 1.560E-08 | 1.437E-08 | 5.200E-09 | 3.200E-08 |
| 14017 | 701 | Q701T | B | 8.3e-09 | 6.8e-09 | 3.3e-09 | 6.133E-09 | 2.566E-09 | 3.300E-09 | 8.300E-09 |
| 14018 | 701 | Q701W | B | 1.7e-09 | 8.7e-09 | 6.7e-09 | 5.700E-09 | 3.606E-09 | 1.700E-09 | 8.700E-09 |
| 14019 | 701 | Q701Y | B | 8e-09 | 6.5e-09 | 8.8e-09 | 7.767E-09 | 1.168E-09 | 6.500E-09 | 8.800E-09 |
| 14020 | 701 | Q701V | B | 4.3e-09 | 7.8e-09 | 1.3e-08 | 8.367E-09 | 4.378E-09 | 4.300E-09 | 1.300E-08 |
| 14021 | 702 | P702A | B | 8.2e-09 | 4e-09 | 6.8e-09 | 6.333E-09 | 2.139E-09 | 4.000E-09 | 8.200E-09 |
| 14022 | 702 | P702R | B | 3.8e-09 | 4.2e-09 | 4.6e-09 | 4.200E-09 | 4.000E-10 | 3.800E-09 | 4.600E-09 |
| 14023 | 702 | P702N | B | 1.2e-08 | 4.8e-09 | 5e-09 | 7.267E-09 | 4.100E-09 | 4.800E-09 | 1.200E-08 |
| 14024 | 702 | P702D | B | 1.1e-08 | 2.8e-09 | 1e-08 | 7.933E-09 | 4.474E-09 | 2.800E-09 | 1.100E-08 |

| | | | | | | | | | | |
|---|---|---|---|---|---|---|---|---|---|---|
| 14025 | 702 | P702C | B | 4.2e-09 | 5.1e-09 | 7.2e-09 | 5.500E-09 | 1.539E-09 | 4.200E-09 | 7.200E-09 |
| 14026 | 702 | P702E | B | 5e-09 | 1.3e-08 | 6e-09 | 8.000E-09 | 4.359E-09 | 5.000E-09 | 1.300E-08 |
| 14027 | 702 | P702Q | B | 2.2e-08 | 3.1e-09 | 5.1e-09 | 1.007E-08 | 1.038E-08 | 3.100E-09 | 2.200E-08 |
| 14028 | 702 | P702G | B | 2.3e-08 | 1.4e-08 | 6.1e-09 | 1.437E-08 | 8.456E-09 | 6.100E-09 | 2.300E-08 |
| 14029 | 702 | P702H | B | 3.1e-08 | 6e-09 | 7.3e-09 | 1.477E-08 | 1.407E-08 | 6.000E-09 | 3.100E-08 |
| 14030 | 702 | P702I | B | 4.6e-09 | 7e-09 | 7.2e-09 | 6.267E-09 | 1.447E-09 | 4.600E-09 | 7.200E-09 |
| 14031 | 702 | P702L | B | 4.5e-09 | 1.2e-08 | 7.5e-09 | 8.000E-09 | 3.775E-09 | 4.500E-09 | 1.200E-08 |
| 14032 | 702 | P702K | B | 7.9e-09 | 6.4e-09 | 7.2e-09 | 7.167E-09 | 7.506E-10 | 6.400E-09 | 7.900E-09 |
| 14033 | 702 | P702M | B | 4.9e-08 | 9.8e-09 | 3.8e-09 | 2.087E-08 | 2.455E-08 | 3.800E-09 | 4.900E-08 |
| 14034 | 702 | P702F | B | 3.1e-09 | 1e-08 | 2.8e-09 | 5.300E-09 | 4.073E-09 | 2.800E-09 | 1.000E-08 |
| 14035 | 702 | P702P | B | 3.7e-09 | 6.8e-09 | 3.3e-09 | 4.600E-09 | 1.916E-09 | 3.300E-09 | 6.800E-09 |
| 14036 | 702 | P702S | B | 7.5e-09 | 2.8e-09 | 6.5e-09 | 5.600E-09 | 2.476E-09 | 2.800E-09 | 7.500E-09 |
| 14037 | 702 | P702T | B | 6.7e-09 | 4e-09 | 8.2e-09 | 6.300E-09 | 2.128E-09 | 4.000E-09 | 8.200E-09 |
| 14038 | 702 | P702W | B | 6.6e-09 | 4.5e-09 | 7e-09 | 6.033E-09 | 1.343E-09 | 4.500E-09 | 7.000E-09 |
| 14039 | 702 | P702Y | B | 5e-09 | 4.4e-09 | 4e-09 | 4.467E-09 | 5.033E-10 | 4.000E-09 | 5.000E-09 |
| 14040 | 702 | P702V | B | 4.4e-09 | 5.1e-09 | 3.3e-09 | 4.267E-09 | 9.074E-10 | 3.300E-09 | 5.100E-09 |
| 14041 | 703 | E703A | B | 5.9e-09 | 3.4e-09 | 9.4e-09 | 6.233E-09 | 3.014E-09 | 3.400E-09 | 9.400E-09 |
| 14042 | 703 | E703R | B | 4e-09 | 7.6e-09 | 1.4e-08 | 8.533E-09 | 5.065E-09 | 4.000E-09 | 1.400E-08 |
| 14043 | 703 | E703N | B | 4.7e-09 | 5.8e-09 | 5.4e-09 | 5.300E-09 | 5.568E-10 | 4.700E-09 | 5.800E-09 |
| 14044 | 703 | E703D | B | 3.2e-09 | 2.4e-09 | 2.8e-09 | 2.800E-09 | 4.000E-10 | 2.400E-09 | 3.200E-09 |
| 14045 | 703 | E703C | B | 8.4e-09 | 1.6e-08 | 5.5e-09 | 9.967E-09 | 5.422E-09 | 5.500E-09 | 1.600E-08 |
| 14046 | 703 | E703E | B | 6.6e-09 | 6.8e-09 | 1.1e-08 | 8.133E-09 | 2.485E-09 | 6.600E-09 | 1.100E-08 |
| 14047 | 703 | E703Q | B | 5.7e-09 | 1.1e-08 | 6.8e-09 | 7.833E-09 | 2.797E-09 | 5.700E-09 | 1.100E-08 |
| 14048 | 703 | E703G | B | 5.9e-09 | 1.6e-08 | 4.2e-09 | 8.700E-09 | 6.379E-09 | 4.200E-09 | 1.600E-08 |
| 14049 | 703 | E703H | B | 2.6e-09 | 9.6e-09 | 2.2e-08 | 1.140E-08 | 9.824E-09 | 2.600E-09 | 2.200E-08 |
| 14050 | 703 | E703I | B | 6e-09 | 4.6e-09 | 1.6e-08 | 8.867E-09 | 6.217E-09 | 4.600E-09 | 1.600E-08 |
| 14051 | 703 | E703L | B | 1.6e-08 | 1.2e-08 | 2.4e-08 | 1.733E-08 | 6.110E-09 | 1.200E-08 | 2.400E-08 |
| 14052 | 703 | E703K | B | 1.4e-08 | 8.3e-09 | 5.3e-09 | 9.200E-09 | 4.419E-09 | 5.300E-09 | 1.400E-08 |
| 14053 | 703 | E703M | B | 1.8e-08 | 1.6e-08 | 6.2e-09 | 1.340E-08 | 6.315E-09 | 6.200E-09 | 1.800E-08 |
| 14054 | 703 | E703F | B | 7.6e-09 | 5.3e-09 | 9.4e-09 | 7.433E-09 | 2.055E-09 | 5.300E-09 | 9.400E-09 |
| 14055 | 703 | E703P | B | 1.1e-08 | 6.1e-09 | 2.2e-08 | 1.303E-08 | 8.143E-09 | 6.100E-09 | 2.200E-08 |
| 14056 | 703 | E703S | B | 2.1e-09 | 8.8e-09 | 7.6e-09 | 6.167E-09 | 3.573E-09 | 2.100E-09 | 8.800E-09 |
| 14057 | 703 | E703T | B | 1.2e-08 | 6.2e-09 | 5.4e-09 | 7.867E-09 | 3.602E-09 | 5.400E-09 | 1.200E-08 |
| 14058 | 703 | E703W | B | 4.4e-09 | 1.1e-08 | 2e-09 | 5.800E-09 | 4.660E-09 | 2.000E-09 | 1.100E-08 |
| 14059 | 703 | E703Y | B | 9.6e-09 | 6.8e-09 | 8.1e-09 | 8.167E-09 | 1.401E-09 | 6.800E-09 | 9.600E-09 |
| 14060 | 703 | E703V | B | 1.3e-08 | 4e-09 | 5.1e-09 | 7.367E-09 | 4.910E-09 | 4.000E-09 | 1.300E-08 |
| 14061 | 704 | D704A | B | 5e-09 | 1.1e-08 | 4.8e-09 | 6.933E-09 | 3.523E-09 | 4.800E-09 | 1.100E-08 |
| 14062 | 704 | D704R | B | 1.6e-08 | 4.1e-09 | 7.4e-09 | 9.167E-09 | 6.144E-09 | 4.100E-09 | 1.600E-08 |
| 14063 | 704 | D704N | B | 1e-08 | 3.8e-09 | 9.6e-09 | 7.800E-09 | 3.470E-09 | 3.800E-09 | 1.000E-08 |
| 14064 | 704 | D704D | B | 2.5e-09 | 3.4e-09 | 2e-09 | 2.633E-09 | 7.095E-10 | 2.000E-09 | 3.400E-09 |
| 14065 | 704 | D704C | B | 3.6e-09 | 7.5e-09 | 9.8e-09 | 6.967E-09 | 3.134E-09 | 3.600E-09 | 9.800E-09 |
| 14066 | 704 | D704E | B | 1.6e-08 | 9.2e-09 | 3.4e-09 | 9.533E-09 | 6.307E-09 | 3.400E-09 | 1.600E-08 |
| 14067 | 704 | D704Q | B | 5.2e-09 | 6.6e-09 | 5.3e-09 | 5.700E-09 | 7.810E-10 | 5.200E-09 | 6.600E-09 |
| 14068 | 704 | D704G | B | 3.8e-09 | 1.4e-08 | 4.4e-09 | 7.400E-09 | 5.724E-09 | 3.800E-09 | 1.400E-08 |
| 14069 | 704 | D704H | B | 3.5e-09 | 2.1e-08 | 6.7e-09 | 1.040E-08 | 9.318E-09 | 3.500E-09 | 2.100E-08 |
| 14070 | 704 | D704I | B | 1.3e-08 | 2.5e-08 | 6.3e-09 | 1.477E-08 | 9.474E-09 | 6.300E-09 | 2.500E-08 |
| 14071 | 704 | D704L | B | 6.2e-09 | 1.2e-08 | 2.9e-08 | 1.573E-08 | 1.185E-08 | 6.200E-09 | 2.900E-08 |
| 14072 | 704 | D704K | B | 8.4e-09 | 1.4e-08 | 6.9e-09 | 9.767E-09 | 3.742E-09 | 6.900E-09 | 1.400E-08 |
| 14073 | 704 | D704M | B | 5e-09 | 1.3e-08 | 4.1e-08 | 1.967E-08 | 1.890E-08 | 5.000E-09 | 4.100E-08 |
| 14074 | 704 | D704F | B | 7.4e-09 | 2.8e-09 | 4.3e-09 | 4.833E-09 | 2.346E-09 | 2.800E-09 | 7.400E-09 |
| 14075 | 704 | D704P | B | 4.4e-09 | 6.4e-09 | 5.5e-09 | 5.433E-09 | 1.002E-09 | 4.400E-09 | 6.400E-09 |
| 14076 | 704 | D704S | B | 3.2e-08 | 1.2e-08 | 6.7e-09 | 1.690E-08 | 1.334E-08 | 6.700E-09 | 3.200E-08 |
| 14077 | 704 | D704T | B | 4.2e-09 | 6.8e-09 | 9.6e-09 | 6.867E-09 | 2.701E-09 | 4.200E-09 | 9.600E-09 |
| 14078 | 704 | D704W | B | 1.4e-08 | 3.2e-09 | 1e-08 | 9.067E-09 | 5.460E-09 | 3.200E-09 | 1.400E-08 |
| 14079 | 704 | D704Y | B | 6.7e-09 | 1.4e-08 | 7.4e-09 | 9.367E-09 | 4.028E-09 | 6.700E-09 | 1.400E-08 |
| 14080 | 704 | D704V | B | 4.8e-09 | 6.7e-09 | 1.2e-08 | 7.833E-09 | 3.731E-09 | 4.800E-09 | 1.200E-08 |
| 14081 | 705 | F705A | B | 1.3e-08 | 1.9e-08 | 6.7e-09 | 1.290E-08 | 6.151E-09 | 6.700E-09 | 1.900E-08 |
| 14082 | 705 | F705R | B | 5.9e-09 | 1.2e-08 | 3.9e-09 | 7.267E-09 | 4.219E-09 | 3.900E-09 | 1.200E-08 |
| 14083 | 705 | F705N | B | 5.9e-09 | 2.1e-08 | 6e-09 | 1.097E-08 | 8.689E-09 | 5.900E-09 | 2.100E-08 |
| 14084 | 705 | F705D | B | 6.1e-09 | 7.6e-09 | 6.9e-09 | 6.867E-09 | 7.506E-10 | 6.100E-09 | 7.600E-09 |
| 14085 | 705 | F705C | B | 4.4e-09 | 1.3e-08 | 7.7e-09 | 8.367E-09 | 4.339E-09 | 4.400E-09 | 1.300E-08 |
| 14086 | 705 | F705E | B | 7.9e-09 | 6.6e-09 | 1.1e-08 | 8.500E-09 | 2.261E-09 | 6.600E-09 | 1.100E-08 |
| 14087 | 705 | F705Q | B | 9.7e-09 | 6.8e-09 | 1.3e-08 | 9.833E-09 | 3.102E-09 | 6.800E-09 | 1.300E-08 |
| 14088 | 705 | F705G | B | 9.4e-09 | 6.9e-09 | 3.1e-09 | 6.467E-09 | 3.172E-09 | 3.100E-09 | 9.400E-09 |
| 14089 | 705 | F705H | B | 8.5e-09 | 1e-08 | 5e-09 | 7.833E-09 | 2.566E-09 | 5.000E-09 | 1.000E-08 |
| 14090 | 705 | F705I | B | 7.4e-09 | 4.9e-09 | 5e-09 | 5.767E-09 | 1.415E-09 | 4.900E-09 | 7.400E-09 |
| 14091 | 705 | F705L | B | 3.1e-09 | 1e-08 | 6.1e-09 | 6.400E-09 | 3.460E-09 | 3.100E-09 | 1.000E-08 |
| 14092 | 705 | F705K | B | 5.7e-09 | 7e-09 | 4.1e-09 | 5.600E-09 | 1.453E-09 | 4.100E-09 | 7.000E-09 |
| 14093 | 705 | F705M | B | 1.3e-08 | 1.2e-08 | 1.2e-08 | 1.233E-08 | 5.774E-10 | 1.200E-08 | 1.300E-08 |
| 14094 | 705 | F705F | B | 1.2e-08 | 9e-09 | 5.8e-09 | 8.933E-09 | 3.101E-09 | 5.800E-09 | 1.200E-08 |
| 14095 | 705 | F705P | B | 2.3e-08 | 1.4e-08 | 8.6e-09 | 1.520E-08 | 7.275E-09 | 8.600E-09 | 2.300E-08 |
| 14096 | 705 | F705S | B | 1.8e-09 | 1.1e-08 | 1.2e-08 | 8.267E-09 | 5.623E-09 | 1.800E-09 | 1.200E-08 |
| 14097 | 705 | F705T | B | 4.3e-09 | 1.2e-08 | 2.5e-08 | 1.377E-08 | 1.046E-08 | 4.300E-09 | 2.500E-08 |
| 14098 | 705 | F705W | B | 6.6e-09 | 9.8e-09 | 4e-09 | 6.800E-09 | 2.905E-09 | 4.000E-09 | 9.800E-09 |
| 14099 | 705 | F705Y | B | 9.5e-09 | 1.1e-08 | 3.3e-09 | 7.933E-09 | 4.082E-09 | 3.300E-09 | 1.100E-08 |
| 14100 | 705 | F705V | B | 1.3e-08 | 5.2e-09 | 4.7e-09 | 7.633E-09 | 4.654E-09 | 4.700E-09 | 1.300E-08 |
| 14101 | 706 | A706A | B | 5.7e-09 | 1.6e-08 | 4.5e-09 | 8.733E-09 | 6.322E-09 | 4.500E-09 | 1.600E-08 |
| 14102 | 706 | A706R | B | 1.8e-09 | 1.9e-08 | 8.5e-09 | 9.767E-09 | 8.670E-09 | 1.800E-09 | 1.900E-08 |
| 14103 | 706 | A706N | B | 2e-08 | 9e-09 | 3.7e-09 | 1.090E-08 | 8.314E-09 | 3.700E-09 | 2.000E-08 |
| 14104 | 706 | A706D | B | 7.2e-09 | 6.4e-09 | 4.4e-09 | 6.000E-09 | 1.442E-09 | 4.400E-09 | 7.200E-09 |
| 14105 | 706 | A706C | B | 8.9e-09 | 7.7e-09 | 2.4e-09 | 6.333E-09 | 3.459E-09 | 2.400E-09 | 8.900E-09 |
| 14106 | 706 | A706E | B | 4.6e-09 | 5.1e-09 | 1.6e-08 | 8.567E-09 | 6.442E-09 | 4.600E-09 | 1.600E-08 |
| 14107 | 706 | A706Q | B | 5.1e-09 | 8.9e-09 | 1e-08 | 8.000E-09 | 2.571E-09 | 5.100E-09 | 1.000E-08 |
| 14108 | 706 | A706G | B | 4.3e-09 | 5.9e-09 | 1.5e-08 | 8.400E-09 | 5.771E-09 | 4.300E-09 | 1.500E-08 |
| 14109 | 706 | A706H | B | 2.5e-09 | 1.1e-08 | 3.9e-09 | 5.800E-09 | 4.557E-09 | 2.500E-09 | 1.100E-08 |

| | | | | | | | | | | |
|---|---|---|---|---|---|---|---|---|---|---|
| 14110 | 706 | A706I | B | 3.1e-09 | 1.7e-08 | 1.2e-08 | 1.070E-08 | 7.041E-09 | 3.100E-09 | 1.700E-08 |
| 14111 | 706 | A706L | B | 9.8e-09 | 6.7e-09 | 6.2e-09 | 7.567E-09 | 1.950E-09 | 6.200E-09 | 9.800E-09 |
| 14112 | 706 | A706K | B | 4.5e-09 | 8.4e-09 | 1.6e-08 | 9.633E-09 | 5.848E-09 | 4.500E-09 | 1.600E-08 |
| 14113 | 706 | A706M | B | 1.8e-09 | 3.6e-09 | 3.9e-09 | 3.100E-09 | 1.136E-09 | 1.800E-09 | 3.900E-09 |
| 14114 | 706 | A706F | B | 2.8e-09 | 1.1e-08 | 4.4e-09 | 6.067E-09 | 4.347E-09 | 2.800E-09 | 1.100E-08 |
| 14115 | 706 | A706P | B | 7.9e-09 | 1e-08 | 5.6e-09 | 7.833E-09 | 2.201E-09 | 5.600E-09 | 1.000E-08 |
| 14116 | 706 | A706S | B | 3.6e-09 | 1.3e-08 | 1.1e-08 | 9.200E-09 | 4.952E-09 | 3.600E-09 | 1.300E-08 |
| 14117 | 706 | A706T | B | 7.3e-09 | 3.3e-09 | 3.4e-09 | 4.667E-09 | 2.281E-09 | 3.300E-09 | 7.300E-09 |
| 14118 | 706 | A706W | B | 6.9e-09 | 1.6e-08 | 4.2e-09 | 9.033E-09 | 6.183E-09 | 4.200E-09 | 1.600E-08 |
| 14119 | 706 | A706Y | B | 8.6e-09 | 4.4e-09 | 6.1e-09 | 6.367E-09 | 2.113E-09 | 4.400E-09 | 8.600E-09 |
| 14120 | 706 | A706V | B | 1.6e-08 | 6.3e-09 | 1.4e-08 | 1.210E-08 | 5.122E-09 | 6.300E-09 | 1.600E-08 |
| 14121 | 707 | T707A | B | 5.3e-09 | 6.6e-09 | 2.2e-08 | 1.130E-08 | 9.289E-09 | 5.300E-09 | 2.200E-08 |
| 14122 | 707 | T707R | B | 9.1e-10 | 1.4e-08 | 7.3e-09 | 7.403E-09 | 6.546E-09 | 9.100E-10 | 1.400E-08 |
| 14123 | 707 | T707N | B | 5.7e-09 | 5.5e-09 | 8.9e-09 | 6.700E-09 | 1.908E-09 | 5.500E-09 | 8.900E-09 |
| 14124 | 707 | T707D | B | 8.8e-09 | 1.2e-08 | 2.1e-09 | 7.633E-09 | 5.052E-09 | 2.100E-09 | 1.200E-08 |
| 14125 | 707 | T707C | B | 8.5e-09 | 1.2e-08 | 3.8e-09 | 8.100E-09 | 4.115E-09 | 3.800E-09 | 1.200E-08 |
| 14126 | 707 | T707E | B | 7e-09 | 9.6e-09 | 6e-09 | 7.533E-09 | 1.858E-09 | 6.000E-09 | 9.600E-09 |
| 14127 | 707 | T707Q | B | 1e-08 | 5.3e-09 | 3.1e-09 | 6.133E-09 | 3.525E-09 | 3.100E-09 | 1.000E-08 |
| 14128 | 707 | T707G | B | 4.3e-09 | 5.8e-09 | 7.7e-09 | 5.933E-09 | 1.704E-09 | 4.300E-09 | 7.700E-09 |
| 14129 | 707 | T707H | B | 5.4e-09 | 1.1e-08 | 7.1e-09 | 7.833E-09 | 2.871E-09 | 5.400E-09 | 1.100E-08 |
| 14130 | 707 | T707I | B | 5.3e-09 | 9.7e-09 | 1.6e-08 | 1.033E-08 | 5.378E-09 | 5.300E-09 | 1.600E-08 |
| 14131 | 707 | T707L | B | 5.4e-09 | 6.8e-09 | 5.9e-09 | 6.033E-09 | 7.095E-10 | 5.400E-09 | 6.800E-09 |
| 14132 | 707 | T707K | B | 7.7e-09 | 5.8e-09 | 1.6e-08 | 9.833E-09 | 5.424E-09 | 5.800E-09 | 1.600E-08 |
| 14133 | 707 | T707M | B | 7.2e-09 | 2.8e-08 | 2e-08 | 1.840E-08 | 1.049E-08 | 7.200E-09 | 2.800E-08 |
| 14134 | 707 | T707F | B | 3.5e-09 | 8.1e-09 | 1.1e-08 | 7.533E-09 | 3.782E-09 | 3.500E-09 | 1.100E-08 |
| 14135 | 707 | T707P | B | 8.6e-09 | 1.5e-08 | 3.5e-09 | 9.033E-09 | 5.762E-09 | 3.500E-09 | 1.500E-08 |
| 14136 | 707 | T707S | B | 6.1e-09 | 5e-09 | 3.8e-09 | 4.967E-09 | 1.150E-09 | 3.800E-09 | 6.100E-09 |
| 14137 | 707 | T707T | B | 2.9e-09 | 7.9e-09 | 5e-09 | 5.267E-09 | 2.511E-09 | 2.900E-09 | 7.900E-09 |
| 14138 | 707 | T707W | B | 2.5e-09 | 4.9e-09 | 5.2e-09 | 4.200E-09 | 1.480E-09 | 2.500E-09 | 5.200E-09 |
| 14139 | 707 | T707Y | B | 3.9e-09 | 7.5e-09 | 5.1e-09 | 5.500E-09 | 1.833E-09 | 3.900E-09 | 7.500E-09 |
| 14140 | 707 | T707V | B | 1.5e-08 | 3.9e-09 | 5.2e-09 | 8.033E-09 | 6.068E-09 | 3.900E-09 | 1.500E-08 |
| 14141 | 708 | Y708A | B | 6.7e-09 | 1.4e-08 | 4.7e-09 | 8.467E-09 | 4.895E-09 | 4.700E-09 | 1.400E-08 |
| 14142 | 708 | Y708R | B | 5.9e-09 | 1.2e-08 | 1.1e-08 | 9.633E-09 | 3.272E-09 | 5.900E-09 | 1.200E-08 |
| 14143 | 708 | Y708N | B | 1.3e-09 | 5.7e-09 | 3.5e-09 | 3.500E-09 | 2.200E-09 | 1.300E-09 | 5.700E-09 |
| 14144 | 708 | Y708D | B | 2.5e-09 | 6.2e-09 | 6e-09 | 4.900E-09 | 2.081E-09 | 2.500E-09 | 6.200E-09 |
| 14145 | 708 | Y708C | B | 1.3e-08 | 8.9e-09 | 7.3e-09 | 9.733E-09 | 2.940E-09 | 7.300E-09 | 1.300E-08 |
| 14146 | 708 | Y708E | B | 8.3e-09 | 9.6e-09 | 1.2e-08 | 9.967E-09 | 1.877E-09 | 8.300E-09 | 1.200E-08 |
| 14147 | 708 | Y708Q | B | 1.4e-08 | 3.2e-09 | 1.7e-08 | 1.140E-08 | 7.258E-09 | 3.200E-09 | 1.700E-08 |
| 14148 | 708 | Y708G | B | 6.2e-09 | 1.3e-08 | 1.3e-08 | 1.073E-08 | 3.926E-09 | 6.200E-09 | 1.300E-08 |
| 14149 | 708 | Y708H | B | 8.5e-09 | 1e-08 | 9.1e-09 | 9.200E-09 | 7.550E-10 | 8.500E-09 | 1.000E-08 |
| 14150 | 708 | Y708I | B | 8.4e-09 | 5.4e-09 | 5.9e-09 | 6.567E-09 | 1.607E-09 | 5.400E-09 | 8.400E-09 |
| 14151 | 708 | Y708L | B | 9.8e-09 | 2.3e-09 | 7.8e-09 | 6.633E-09 | 3.884E-09 | 2.300E-09 | 9.800E-09 |
| 14152 | 708 | Y708K | B | 1.4e-08 | 1.9e-08 | 2.2e-08 | 1.833E-08 | 4.041E-09 | 1.400E-08 | 2.200E-08 |
| 14153 | 708 | Y708M | B | 4.2e-09 | 1e-08 | 9.3e-09 | 7.833E-09 | 3.166E-09 | 4.200E-09 | 1.000E-08 |
| 14154 | 708 | Y708F | B | 5.4e-09 | 1.3e-08 | 1e-08 | 9.467E-09 | 3.828E-09 | 5.400E-09 | 1.300E-08 |
| 14155 | 708 | Y708P | B | 6.9e-09 | 2.2e-09 | 5.4e-09 | 4.833E-09 | 2.401E-09 | 2.200E-09 | 6.900E-09 |
| 14156 | 708 | Y708S | B | 7.5e-09 | 5.7e-09 | 5.4e-09 | 6.200E-09 | 1.136E-09 | 5.400E-09 | 7.500E-09 |
| 14157 | 708 | Y708T | B | 1.7e-08 | 2.6e-09 | 6.6e-09 | 8.733E-09 | 7.433E-09 | 2.600E-09 | 1.700E-08 |
| 14158 | 708 | Y708W | B | 4.8e-09 | 6.4e-09 | 3.8e-09 | 5.000E-09 | 1.311E-09 | 3.800E-09 | 6.400E-09 |
| 14159 | 708 | Y708Y | B | 6.7e-09 | 3.4e-09 | 3.3e-09 | 4.467E-09 | 1.935E-09 | 3.300E-09 | 6.700E-09 |
| 14160 | 708 | Y708V | B | 7.8e-09 | 3.7e-09 | 4.9e-09 | 5.467E-09 | 2.108E-09 | 3.700E-09 | 7.800E-09 |
| 14161 | 709 | Y709A | B | 6.7e-09 | 2.7e-09 | 6.8e-09 | 5.400E-09 | 2.339E-09 | 2.700E-09 | 6.800E-09 |
| 14162 | 709 | Y709R | B | 7.8e-09 | 5.5e-09 | 8.3e-09 | 7.200E-09 | 1.493E-09 | 5.500E-09 | 8.300E-09 |
| 14163 | 709 | Y709N | B | 8.2e-09 | 1.6e-08 | 7.9e-09 | 1.070E-08 | 4.592E-09 | 7.900E-09 | 1.600E-08 |
| 14164 | 709 | Y709D | B | 2.9e-09 | 1.2e-08 | 2.4e-09 | 5.767E-09 | 5.404E-09 | 2.400E-09 | 1.200E-08 |
| 14165 | 709 | Y709C | B | 5.2e-09 | 7.7e-09 | 1.3e-08 | 8.633E-09 | 3.983E-09 | 5.200E-09 | 1.300E-08 |
| 14166 | 709 | Y709E | B | 1.2e-08 | 5.4e-09 | 7.2e-09 | 8.200E-09 | 3.412E-09 | 5.400E-09 | 1.200E-08 |
| 14167 | 709 | Y709Q | B | 6.1e-09 | 1e-08 | 7.3e-09 | 7.800E-09 | 1.997E-09 | 6.100E-09 | 1.000E-08 |
| 14168 | 709 | Y709G | B | 1.4e-08 | 1.3e-08 | 5.1e-09 | 1.070E-08 | 4.875E-09 | 5.100E-09 | 1.400E-08 |
| 14169 | 709 | Y709H | B | 3.9e-09 | 2.2e-08 | 5.9e-09 | 1.060E-08 | 9.923E-09 | 3.900E-09 | 2.200E-08 |
| 14170 | 709 | Y709I | B | 7.2e-09 | 2.1e-08 | 4e-09 | 1.073E-08 | 9.034E-09 | 4.000E-09 | 2.100E-08 |
| 14171 | 709 | Y709L | B | 6.5e-09 | 7.7e-09 | 1e-08 | 8.067E-09 | 1.779E-09 | 6.500E-09 | 1.000E-08 |
| 14172 | 709 | Y709K | B | 1.2e-08 | 5.1e-09 | 4.2e-09 | 7.100E-09 | 4.267E-09 | 4.200E-09 | 1.200E-08 |
| 14173 | 709 | Y709M | B | 1e-08 | 1.3e-08 | 1.7e-08 | 1.333E-08 | 3.512E-09 | 1.000E-08 | 1.700E-08 |
| 14174 | 709 | Y709F | B | 2.2e-08 | 4.5e-09 | 2.4e-09 | 9.633E-09 | 1.076E-08 | 2.400E-09 | 2.200E-08 |
| 14175 | 709 | Y709P | B | 5.1e-09 | 4.8e-09 | 5.3e-09 | 5.067E-09 | 2.517E-10 | 4.800E-09 | 5.300E-09 |
| 14176 | 709 | Y709S | B | 6.6e-09 | 9.1e-09 | 1.8e-09 | 5.833E-09 | 3.710E-09 | 1.800E-09 | 9.100E-09 |
| 14177 | 709 | Y709T | B | 6.2e-09 | 8.9e-09 | 2.7e-09 | 5.933E-09 | 3.109E-09 | 2.700E-09 | 8.900E-09 |
| 14178 | 709 | Y709W | B | 1.5e-08 | 3e-09 | 6.9e-09 | 8.300E-09 | 6.121E-09 | 3.000E-09 | 1.500E-08 |
| 14179 | 709 | Y709Y | B | 6.9e-09 | 1.1e-08 | 3.3e-09 | 7.067E-09 | 3.853E-09 | 3.300E-09 | 1.100E-08 |
| 14180 | 709 | Y709V | B | 6.9e-09 | 6.3e-09 | 5.7e-09 | 6.300E-09 | 6.000E-10 | 5.700E-09 | 6.900E-09 |
| 14181 | 710 | C710A | B | 4.2e-09 | 7.2e-09 | 1.7e-08 | 9.467E-09 | 6.694E-09 | 4.200E-09 | 1.700E-08 |
| 14182 | 710 | C710R | B | 6.2e-09 | 2.5e-09 | 8.7e-09 | 5.800E-09 | 3.119E-09 | 2.500E-09 | 8.700E-09 |
| 14183 | 710 | C710N | B | 2.7e-08 | 5.1e-09 | 1.3e-08 | 1.503E-08 | 1.109E-08 | 5.100E-09 | 2.700E-08 |
| 14184 | 710 | C710D | B | 2.5e-09 | 2.7e-09 | 5.1e-09 | 3.433E-09 | 1.447E-09 | 2.500E-09 | 5.100E-09 |
| 14185 | 710 | C710C | B | 1.3e-08 | 1.5e-08 | 1.4e-08 | 1.400E-08 | 1.000E-09 | 1.300E-08 | 1.500E-08 |
| 14186 | 710 | C710E | B | 6.2e-09 | 8.4e-09 | 1.2e-08 | 8.867E-09 | 2.928E-09 | 6.200E-09 | 1.200E-08 |
| 14187 | 710 | C710Q | B | 7.1e-09 | 1.9e-09 | 1.7e-08 | 8.667E-09 | 7.671E-09 | 1.900E-09 | 1.700E-08 |
| 14188 | 710 | C710G | B | 1e-08 | 6.8e-09 | 1.3e-09 | 6.033E-09 | 4.400E-09 | 1.300E-09 | 1.000E-08 |
| 14189 | 710 | C710H | B | 6.1e-09 | 3.1e-09 | 1.3e-08 | 7.400E-09 | 5.076E-09 | 3.100E-09 | 1.300E-08 |
| 14190 | 710 | C710I | B | 2.4e-09 | 2.2e-09 | 6.2e-09 | 3.600E-09 | 2.254E-09 | 2.200E-09 | 6.200E-09 |
| 14191 | 710 | C710L | B | 7.5e-09 | 3e-09 | 8.2e-09 | 6.233E-09 | 2.822E-09 | 3.000E-09 | 8.200E-09 |
| 14192 | 710 | C710K | B | 2.7e-08 | 2.1e-09 | 6.9e-09 | 1.200E-08 | 1.321E-08 | 2.100E-09 | 2.700E-08 |
| 14193 | 710 | C710M | B | 9.2e-09 | 6.2e-09 | 1.3e-09 | 5.567E-09 | 3.988E-09 | 1.300E-09 | 9.200E-09 |
| 14194 | 710 | C710F | B | 4.1e-09 | 9.7e-09 | 1.9e-08 | 1.093E-08 | 7.526E-09 | 4.100E-09 | 1.900E-08 |

| | | | | | | | | | | |
|---|---|---|---|---|---|---|---|---|---|---|
| 14195 | 710 | C710P | B | 1.3e-08 | 2.1e-08 | 6e-09 | 1.333E-08 | 7.506E-09 | 6.000E-09 | 2.100E-08 |
| 14196 | 710 | C710S | B | 1e-08 | 6.2e-09 | 6.9e-09 | 7.700E-09 | 2.022E-09 | 6.200E-09 | 1.000E-08 |
| 14197 | 710 | C710T | B | 6.4e-09 | 1.2e-08 | 2.4e-09 | 6.933E-09 | 4.822E-09 | 2.400E-09 | 1.200E-08 |
| 14198 | 710 | C710W | B | 4.2e-09 | 1e-08 | 1.2e-08 | 8.733E-09 | 4.051E-09 | 4.200E-09 | 1.200E-08 |
| 14199 | 710 | C710Y | B | 1.5e-08 | 7e-09 | 2.8e-09 | 8.267E-09 | 6.198E-09 | 2.800E-09 | 1.500E-08 |
| 14200 | 710 | C710V | B | 6.1e-09 | 8e-09 | 6.8e-09 | 6.967E-09 | 9.609E-10 | 6.100E-09 | 8.000E-09 |
| 14201 | 711 | Q711A | B | 3.5e-09 | 4.6e-09 | 2.6e-09 | 3.567E-09 | 1.002E-09 | 2.600E-09 | 4.600E-09 |
| 14202 | 711 | Q711R | B | 1.4e-08 | 5.9e-09 | 1.1e-08 | 1.030E-08 | 4.095E-09 | 5.900E-09 | 1.400E-08 |
| 14203 | 711 | Q711N | B | 5.5e-09 | 4.8e-09 | 8.8e-09 | 6.367E-09 | 2.136E-09 | 4.800E-09 | 8.800E-09 |
| 14204 | 711 | Q711D | B | 7.5e-09 | 5.4e-09 | 4.6e-09 | 5.833E-09 | 1.498E-09 | 4.600E-09 | 7.500E-09 |
| 14205 | 711 | Q711C | B | 1.6e-08 | 2.2e-08 | 3.9e-09 | 1.397E-08 | 9.220E-09 | 3.900E-09 | 2.200E-08 |
| 14206 | 711 | Q711E | B | 5.4e-09 | 4.9e-09 | 2.2e-08 | 1.077E-08 | 9.732E-09 | 4.900E-09 | 2.200E-08 |
| 14207 | 711 | Q711Q | B | 5.8e-09 | 1.8e-08 | 9.4e-09 | 1.107E-08 | 6.268E-09 | 5.800E-09 | 1.800E-08 |
| 14208 | 711 | Q711G | B | 9.2e-09 | 3e-09 | 6e-09 | 6.067E-09 | 3.101E-09 | 3.000E-09 | 9.200E-09 |
| 14209 | 711 | Q711H | B | 5e-09 | 5.8e-09 | 5.7e-09 | 5.500E-09 | 4.359E-10 | 5.000E-09 | 5.800E-09 |
| 14210 | 711 | Q711I | B | 6.9e-09 | 1.9e-09 | 1.8e-09 | 3.533E-09 | 2.916E-09 | 1.800E-09 | 6.900E-09 |
| 14211 | 711 | Q711L | B | 4.9e-09 | 1.1e-08 | 5.3e-09 | 7.067E-09 | 3.412E-09 | 4.900E-09 | 1.100E-08 |
| 14212 | 711 | Q711K | B | 3.9e-09 | 2.3e-09 | 3.4e-09 | 3.200E-09 | 8.185E-10 | 2.300E-09 | 3.900E-09 |
| 14213 | 711 | Q711M | B | 3.5e-09 | 2.6e-09 | 1.4e-08 | 6.700E-09 | 6.338E-09 | 2.600E-09 | 1.400E-08 |
| 14214 | 711 | Q711F | B | 4.1e-09 | 2e-08 | 7.2e-09 | 1.043E-08 | 8.429E-09 | 4.100E-09 | 2.000E-08 |
| 14215 | 711 | Q711P | B | 1.5e-08 | 4.3e-09 | 8.3e-09 | 9.200E-09 | 5.406E-09 | 4.300E-09 | 1.500E-08 |
| 14216 | 711 | Q711S | B | 6.4e-09 | 1.9e-09 | 6.2e-09 | 4.833E-09 | 2.542E-09 | 1.900E-09 | 6.400E-09 |
| 14217 | 711 | Q711T | B | 1.8e-08 | 7.9e-09 | 2.8e-09 | 9.567E-09 | 7.736E-09 | 2.800E-09 | 1.800E-08 |
| 14218 | 711 | Q711W | B | 6e-09 | 7e-09 | 5.5e-09 | 6.167E-09 | 7.638E-10 | 5.500E-09 | 7.000E-09 |
| 14219 | 711 | Q711Y | B | 6.1e-09 | 6.9e-09 | 5.1e-09 | 6.033E-09 | 9.018E-10 | 5.100E-09 | 6.900E-09 |
| 14220 | 711 | Q711V | B | 1.3e-08 | 7.7e-09 | 4.6e-09 | 8.433E-09 | 4.248E-09 | 4.600E-09 | 1.300E-08 |
| 14221 | 712 | Q712A | B | 2.4e-09 | 4.3e-09 | 6.1e-09 | 4.267E-09 | 1.850E-09 | 2.400E-09 | 6.100E-09 |
| 14222 | 712 | Q712R | B | 1e-08 | 5.2e-09 | 5.9e-09 | 7.033E-09 | 2.593E-09 | 5.200E-09 | 1.000E-08 |
| 14223 | 712 | Q712N | B | 7.2e-09 | 8.2e-09 | 1.1e-08 | 8.800E-09 | 1.970E-09 | 7.200E-09 | 1.100E-08 |
| 14224 | 712 | Q712D | B | 7.5e-09 | 7.7e-09 | 2.9e-08 | 1.473E-08 | 1.236E-08 | 7.500E-09 | 2.900E-08 |
| 14225 | 712 | Q712C | B | 2.7e-08 | 3.3e-09 | 8.7e-09 | 1.300E-08 | 1.242E-08 | 3.300E-09 | 2.700E-08 |
| 14226 | 712 | Q712E | B | 4.5e-09 | 5.7e-09 | 7.4e-09 | 5.867E-09 | 1.457E-09 | 4.500E-09 | 7.400E-09 |
| 14227 | 712 | Q712Q | B | 1.5e-09 | 6.1e-09 | 1e-08 | 5.867E-09 | 4.255E-09 | 1.500E-09 | 1.000E-08 |
| 14228 | 712 | Q712G | B | 2.1e-08 | 4.8e-09 | 5.5e-09 | 1.043E-08 | 9.158E-09 | 4.800E-09 | 2.100E-08 |
| 14229 | 712 | Q712H | B | 3.3e-09 | 5.2e-09 | 3e-08 | 1.283E-08 | 1.490E-08 | 3.300E-09 | 3.000E-08 |
| 14230 | 712 | Q712I | B | 1.6e-08 | 1.3e-08 | 2.7e-09 | 1.057E-08 | 6.976E-09 | 2.700E-09 | 1.600E-08 |
| 14231 | 712 | Q712L | B | 1e-08 | 4.4e-09 | 2e-08 | 1.147E-08 | 7.903E-09 | 4.400E-09 | 2.000E-08 |
| 14232 | 712 | Q712K | B | 2.8e-09 | 9.3e-09 | 1.5e-08 | 9.033E-09 | 6.104E-09 | 2.800E-09 | 1.500E-08 |
| 14233 | 712 | Q712M | B | 9.2e-09 | 7.9e-09 | 7.2e-09 | 8.100E-09 | 1.015E-09 | 7.200E-09 | 9.200E-09 |
| 14234 | 712 | Q712F | B | 4.5e-09 | 1.5e-08 | 7.8e-09 | 9.100E-09 | 5.369E-09 | 4.500E-09 | 1.500E-08 |
| 14235 | 712 | Q712P | B | 5.6e-09 | 8.2e-09 | 1.8e-09 | 5.200E-09 | 3.219E-09 | 1.800E-09 | 8.200E-09 |
| 14236 | 712 | Q712S | B | 1.3e-08 | 5.2e-09 | 3.8e-09 | 7.333E-09 | 4.957E-09 | 3.800E-09 | 1.300E-08 |
| 14237 | 712 | Q712T | B | 1.2e-08 | 7.8e-09 | 1.7e-08 | 1.227E-08 | 4.606E-09 | 7.800E-09 | 1.700E-08 |
| 14238 | 712 | Q712W | B | 1.5e-08 | 3.9e-09 | 9.5e-09 | 9.467E-09 | 5.550E-09 | 3.900E-09 | 1.500E-08 |
| 14239 | 712 | Q712Y | B | 1.4e-08 | 3.6e-09 | 1.6e-08 | 1.120E-08 | 6.657E-09 | 3.600E-09 | 1.600E-08 |
| 14240 | 712 | Q712V | B | 2.4e-08 | 9.3e-09 | 1.7e-08 | 1.677E-08 | 7.353E-09 | 9.300E-09 | 2.400E-08 |
| 14241 | 713 | Y713A | B | 5.1e-08 | 4.4e-09 | 5.1e-09 | 2.017E-08 | 2.670E-08 | 4.400E-09 | 5.100E-08 |
| 14242 | 713 | Y713R | B | 6.9e-09 | 4.2e-09 | 2.9e-09 | 4.667E-09 | 2.040E-09 | 2.900E-09 | 6.900E-09 |
| 14243 | 713 | Y713N | B | 7.7e-09 | 1e-08 | 1.2e-08 | 9.900E-09 | 2.152E-09 | 7.700E-09 | 1.200E-08 |
| 14244 | 713 | Y713D | B | 1.2e-08 | 7.3e-09 | 9.6e-09 | 9.633E-09 | 2.350E-09 | 7.300E-09 | 1.200E-08 |
| 14245 | 713 | Y713C | B | 2.6e-08 | 5.9e-09 | 5.1e-09 | 1.233E-08 | 1.184E-08 | 5.100E-09 | 2.600E-08 |
| 14246 | 713 | Y713E | B | 9.9e-09 | 9.5e-09 | 7e-09 | 8.800E-09 | 1.572E-09 | 7.000E-09 | 9.900E-09 |
| 14247 | 713 | Y713Q | B | 2.8e-09 | 5.3e-09 | 4.7e-09 | 4.267E-09 | 1.305E-09 | 2.800E-09 | 5.300E-09 |
| 14248 | 713 | Y713G | B | 7.6e-09 | 7.4e-09 | 6e-09 | 7.000E-09 | 8.718E-10 | 6.000E-09 | 7.600E-09 |
| 14249 | 713 | Y713H | B | 1.1e-08 | 4.4e-09 | 1.2e-08 | 9.133E-09 | 4.130E-09 | 4.400E-09 | 1.200E-08 |
| 14250 | 713 | Y713I | B | 1.3e-08 | 7.8e-09 | 4.9e-09 | 8.567E-09 | 4.104E-09 | 4.900E-09 | 1.300E-08 |
| 14251 | 713 | Y713L | B | 1.8e-08 | 4.4e-09 | 2.6e-09 | 8.333E-09 | 8.420E-09 | 2.600E-09 | 1.800E-08 |
| 14252 | 713 | Y713K | B | 7.6e-09 | 1.5e-08 | 9.5e-09 | 1.070E-08 | 3.843E-09 | 7.600E-09 | 1.500E-08 |
| 14253 | 713 | Y713M | B | 1.6e-08 | 8.9e-09 | 5.2e-09 | 1.003E-08 | 5.488E-09 | 5.200E-09 | 1.600E-08 |
| 14254 | 713 | Y713F | B | 5.7e-09 | 7.6e-09 | 1.7e-08 | 1.010E-08 | 6.051E-09 | 5.700E-09 | 1.700E-08 |
| 14255 | 713 | Y713P | B | 9.9e-09 | 8.7e-09 | 1.1e-08 | 9.867E-09 | 1.150E-09 | 8.700E-09 | 1.100E-08 |
| 14256 | 713 | Y713S | B | 8.8e-09 | 2e-09 | 4.2e-09 | 5.000E-09 | 3.470E-09 | 2.000E-09 | 8.800E-09 |
| 14257 | 713 | Y713T | B | 8.5e-09 | 1.1e-08 | 2.1e-08 | 1.350E-08 | 6.614E-09 | 8.500E-09 | 2.100E-08 |
| 14258 | 713 | Y713W | B | 9.1e-09 | 1e-08 | 5.3e-09 | 8.133E-09 | 2.495E-09 | 5.300E-09 | 1.000E-08 |
| 14259 | 713 | Y713Y | B | 3.9e-09 | 7e-09 | 4.7e-09 | 5.200E-09 | 1.609E-09 | 3.900E-09 | 7.000E-09 |
| 14260 | 713 | Y713V | B | 6.4e-09 | 5.5e-09 | 6.2e-09 | 6.033E-09 | 4.726E-10 | 5.500E-09 | 6.400E-09 |
| 14261 | 714 | Y714A | B | 1.8e-08 | 5.7e-09 | 9.5e-09 | 1.107E-08 | 6.298E-09 | 5.700E-09 | 1.800E-08 |
| 14262 | 714 | Y714R | B | 1e-08 | 2.3e-09 | 1.2e-08 | 8.100E-09 | 5.122E-09 | 2.300E-09 | 1.200E-08 |
| 14263 | 714 | Y714N | B | 1.2e-08 | 2.1e-08 | 2.8e-09 | 1.193E-08 | 9.100E-09 | 2.800E-09 | 2.100E-08 |
| 14264 | 714 | Y714D | B | 4.7e-09 | 5e-09 | 1.2e-08 | 7.233E-09 | 4.131E-09 | 4.700E-09 | 1.200E-08 |
| 14265 | 714 | Y714C | B | 1.4e-08 | 1.3e-08 | 4.4e-09 | 1.047E-08 | 5.278E-09 | 4.400E-09 | 1.400E-08 |
| 14266 | 714 | Y714E | B | 1.7e-09 | 2e-09 | 1.3e-08 | 5.567E-09 | 6.439E-09 | 1.700E-09 | 1.300E-08 |
| 14267 | 714 | Y714Q | B | 7.1e-09 | 1.4e-08 | 8.3e-09 | 9.800E-09 | 3.686E-09 | 7.100E-09 | 1.400E-08 |
| 14268 | 714 | Y714G | B | 9.3e-09 | 1.1e-08 | 2.6e-09 | 7.633E-09 | 4.441E-09 | 2.600E-09 | 1.100E-08 |
| 14269 | 714 | Y714H | B | 1.1e-08 | 5.6e-09 | 1e-08 | 8.867E-09 | 2.873E-09 | 5.600E-09 | 1.100E-08 |
| 14270 | 714 | Y714I | B | 2.1e-08 | 6.9e-09 | 1.1e-08 | 1.297E-08 | 7.253E-09 | 6.900E-09 | 2.100E-08 |
| 14271 | 714 | Y714L | B | 3.6e-09 | 6.3e-09 | 7.4e-09 | 5.767E-09 | 1.955E-09 | 3.600E-09 | 7.400E-09 |
| 14272 | 714 | Y714K | B | 1e-08 | 1.8e-08 | 1e-08 | 1.267E-08 | 4.619E-09 | 1.000E-08 | 1.800E-08 |
| 14273 | 714 | Y714M | B | 8.9e-09 | 1e-08 | 5.7e-09 | 8.200E-09 | 2.234E-09 | 5.700E-09 | 1.000E-08 |
| 14274 | 714 | Y714F | B | 1.1e-08 | 8.2e-09 | 2.4e-09 | 7.200E-09 | 4.386E-09 | 2.400E-09 | 1.100E-08 |
| 14275 | 714 | Y714P | B | 1.1e-08 | 5.4e-09 | 1.7e-08 | 1.113E-08 | 5.801E-09 | 5.400E-09 | 1.700E-08 |
| 14276 | 714 | Y714S | B | 2.9e-09 | 5.5e-09 | 6e-09 | 4.800E-09 | 1.664E-09 | 2.900E-09 | 6.000E-09 |
| 14277 | 714 | Y714T | B | 9.7e-09 | 3.1e-09 | 1e-08 | 7.600E-09 | 3.900E-09 | 3.100E-09 | 1.000E-08 |
| 14278 | 714 | Y714W | B | 1.2e-08 | 9e-09 | 5.8e-09 | 8.933E-09 | 3.101E-09 | 5.800E-09 | 1.200E-08 |
| 14279 | 714 | Y714Y | B | 1e-08 | 3.9e-09 | 1.7e-08 | 1.030E-08 | 6.555E-09 | 3.900E-09 | 1.700E-08 |

| | | | | | | | | | | |
|---|---|---|---|---|---|---|---|---|---|---|
| 14280 | 714 | Y714V | B | 8.2e-09 | 4.1e-09 | 4.7e-09 | 5.667E-09 | 2.214E-09 | 4.100E-09 | 8.200E-09 |
| 14281 | 715 | I715A | B | 1.6e-08 | 8.7e-09 | 3.8e-09 | 9.500E-09 | 6.139E-09 | 3.800E-09 | 1.600E-08 |
| 14282 | 715 | I715R | B | 6.9e-09 | 9e-09 | 3.6e-09 | 6.500E-09 | 2.722E-09 | 3.600E-09 | 9.000E-09 |
| 14283 | 715 | I715N | B | 1.3e-08 | 2.6e-09 | 6.2e-09 | 7.267E-09 | 5.281E-09 | 2.600E-09 | 1.300E-08 |
| 14284 | 715 | I715D | B | 2.2e-08 | 4.3e-09 | 2.6e-09 | 9.633E-09 | 1.074E-08 | 2.600E-09 | 2.200E-08 |
| 14285 | 715 | I715C | B | 3.8e-09 | 1.7e-08 | 8.6e-09 | 9.800E-09 | 6.681E-09 | 3.800E-09 | 1.700E-08 |
| 14286 | 715 | I715E | B | 2.7e-09 | 3.4e-09 | 1.4e-08 | 6.700E-09 | 6.332E-09 | 2.700E-09 | 1.400E-08 |
| 14287 | 715 | I715Q | B | 6.7e-09 | 1.2e-08 | 1.1e-08 | 9.900E-09 | 2.816E-09 | 6.700E-09 | 1.200E-08 |
| 14288 | 715 | I715G | B | 1.3e-08 | 4.9e-09 | 7.5e-09 | 8.467E-09 | 4.136E-09 | 4.900E-09 | 1.300E-08 |
| 14289 | 715 | I715H | B | 2e-09 | 7.6e-09 | 1.1e-08 | 6.867E-09 | 4.545E-09 | 2.000E-09 | 1.100E-08 |
| 14290 | 715 | I715I | B | 2.4e-08 | 2e-09 | 1.6e-09 | 9.200E-09 | 1.282E-08 | 1.600E-09 | 2.400E-08 |
| 14291 | 715 | I715L | B | 8.4e-09 | 2.2e-08 | 3.9e-09 | 1.143E-08 | 9.424E-09 | 3.900E-09 | 2.200E-08 |
| 14292 | 715 | I715K | B | 2.6e-09 | 5.6e-09 | 7.9e-09 | 5.367E-09 | 2.658E-09 | 2.600E-09 | 7.900E-09 |
| 14293 | 715 | I715M | B | 1.3e-08 | 1.5e-08 | 1e-08 | 1.267E-08 | 2.517E-09 | 1.000E-08 | 1.500E-08 |
| 14294 | 715 | I715F | B | 2.9e-09 | 1e-08 | 1.3e-08 | 8.633E-09 | 5.187E-09 | 2.900E-09 | 1.300E-08 |
| 14295 | 715 | I715P | B | 1.2e-08 | 4.7e-09 | 1.1e-08 | 9.233E-09 | 3.958E-09 | 4.700E-09 | 1.200E-08 |
| 14296 | 715 | I715S | B | 8.3e-09 | 5.5e-09 | 1.1e-08 | 8.267E-09 | 2.750E-09 | 5.500E-09 | 1.100E-08 |
| 14297 | 715 | I715T | B | 6e-09 | 5.7e-09 | 3.4e-09 | 5.033E-09 | 1.422E-09 | 3.400E-09 | 6.000E-09 |
| 14298 | 715 | I715W | B | 2.9e-09 | 5e-09 | 2.4e-08 | 1.063E-08 | 1.162E-08 | 2.900E-09 | 2.400E-08 |
| 14299 | 715 | I715Y | B | 5.2e-09 | 3.1e-09 | 6.8e-09 | 5.033E-09 | 1.856E-09 | 3.100E-09 | 6.800E-09 |
| 14300 | 715 | I715V | B | 7.2e-09 | 1.3e-08 | 1.2e-08 | 1.073E-08 | 3.101E-09 | 7.200E-09 | 1.300E-08 |
| 14301 | 716 | Y716A | B | 7.9e-09 | 5.6e-09 | 6.7e-09 | 6.733E-09 | 1.150E-09 | 5.600E-09 | 7.900E-09 |
| 14302 | 716 | Y716R | B | 7.6e-09 | 6e-09 | 4.6e-09 | 6.067E-09 | 1.501E-09 | 4.600E-09 | 7.600E-09 |
| 14303 | 716 | Y716N | B | 1.4e-08 | 1.5e-08 | 2.7e-08 | 1.867E-08 | 7.234E-09 | 1.400E-08 | 2.700E-08 |
| 14304 | 716 | Y716D | B | 9.1e-09 | 1.1e-08 | 7.1e-09 | 9.067E-09 | 1.950E-09 | 7.100E-09 | 1.100E-08 |
| 14305 | 716 | Y716C | B | 9.4e-09 | 1.7e-08 | 7.9e-09 | 1.143E-08 | 4.879E-09 | 7.900E-09 | 1.700E-08 |
| 14306 | 716 | Y716E | B | 1.1e-08 | 1.9e-09 | 1.1e-08 | 7.967E-09 | 5.254E-09 | 1.900E-09 | 1.100E-08 |
| 14307 | 716 | Y716Q | B | 1.4e-08 | 9.8e-09 | 1e-08 | 1.127E-08 | 2.369E-09 | 9.800E-09 | 1.400E-08 |
| 14308 | 716 | Y716G | B | 2.1e-08 | 4.8e-09 | 3.1e-08 | 1.893E-08 | 1.322E-08 | 4.800E-09 | 3.100E-08 |
| 14309 | 716 | Y716H | B | 9.8e-09 | 8.6e-09 | 1.2e-08 | 1.013E-08 | 1.724E-09 | 8.600E-09 | 1.200E-08 |
| 14310 | 716 | Y716I | B | 1.4e-08 | 3.6e-09 | 8.8e-09 | 8.800E-09 | 5.200E-09 | 3.600E-09 | 1.400E-08 |
| 14311 | 716 | Y716L | B | 7.1e-09 | 1.7e-08 | 3.7e-08 | 2.037E-08 | 1.523E-08 | 7.100E-09 | 3.700E-08 |
| 14312 | 716 | Y716K | B | 8.1e-09 | 7.3e-09 | 1.4e-08 | 9.800E-09 | 3.659E-09 | 7.300E-09 | 1.400E-08 |
| 14313 | 716 | Y716M | B | 4.2e-09 | 1.1e-08 | 3.5e-09 | 6.233E-09 | 4.143E-09 | 3.500E-09 | 1.100E-08 |
| 14314 | 716 | Y716F | B | 4.4e-09 | 1e-08 | 1.5e-08 | 9.800E-09 | 5.303E-09 | 4.400E-09 | 1.500E-08 |
| 14315 | 716 | Y716P | B | 3.8e-09 | 9e-09 | 1e-08 | 7.600E-09 | 3.329E-09 | 3.800E-09 | 1.000E-08 |
| 14316 | 716 | Y716S | B | 1.5e-08 | 4.8e-09 | 5.3e-09 | 8.367E-09 | 5.750E-09 | 4.800E-09 | 1.500E-08 |
| 14317 | 716 | Y716T | B | 1.8e-08 | 2.4e-08 | 9.4e-09 | 1.713E-08 | 7.338E-09 | 9.400E-09 | 2.400E-08 |
| 14318 | 716 | Y716W | B | 7.8e-09 | 1.8e-09 | 6.2e-09 | 5.267E-09 | 3.107E-09 | 1.800E-09 | 7.800E-09 |
| 14319 | 716 | Y716Y | B | 5.9e-09 | 1.1e-08 | 7.6e-09 | 8.167E-09 | 2.597E-09 | 5.900E-09 | 1.100E-08 |
| 14320 | 716 | Y716V | B | 1e-08 | 1.4e-08 | 5e-09 | 9.667E-09 | 4.509E-09 | 5.000E-09 | 1.400E-08 |
| 14321 | 717 | P717A | B | 1.7e-09 | 1.2e-08 | 6.2e-09 | 6.633E-09 | 5.164E-09 | 1.700E-09 | 1.200E-08 |
| 14322 | 717 | P717R | B | 4.7e-09 | 9.7e-09 | 8.3e-09 | 7.567E-09 | 2.579E-09 | 4.700E-09 | 9.700E-09 |
| 14323 | 717 | P717N | B | 7.4e-09 | 6.2e-09 | 5.2e-09 | 6.267E-09 | 1.102E-09 | 5.200E-09 | 7.400E-09 |
| 14324 | 717 | P717D | B | 3.4e-09 | 1.1e-08 | 6.3e-09 | 6.900E-09 | 3.835E-09 | 3.400E-09 | 1.100E-08 |
| 14325 | 717 | P717C | B | 6.6e-09 | 8.8e-09 | 3.6e-09 | 6.333E-09 | 2.610E-09 | 3.600E-09 | 8.800E-09 |
| 14326 | 717 | P717E | B | 1.4e-08 | 1.3e-08 | 3e-08 | 1.900E-08 | 9.539E-09 | 1.300E-08 | 3.000E-08 |
| 14327 | 717 | P717Q | B | 8.2e-09 | 3.2e-09 | 1.9e-08 | 1.013E-08 | 8.075E-09 | 3.200E-09 | 1.900E-08 |
| 14328 | 717 | P717G | B | 1e-08 | 9.7e-09 | 8.2e-09 | 9.300E-09 | 9.644E-10 | 8.200E-09 | 1.000E-08 |
| 14329 | 717 | P717H | B | 3.8e-09 | 3e-09 | 1e-08 | 5.600E-09 | 3.831E-09 | 3.000E-09 | 1.000E-08 |
| 14330 | 717 | P717I | B | 7.3e-09 | 5.9e-09 | 5.1e-09 | 6.100E-09 | 1.114E-09 | 5.100E-09 | 7.300E-09 |
| 14331 | 717 | P717L | B | 4.8e-09 | 9.4e-09 | 6.8e-09 | 7.000E-09 | 2.307E-09 | 4.800E-09 | 9.400E-09 |
| 14332 | 717 | P717K | B | 2.6e-09 | 3.1e-09 | 2.4e-08 | 9.900E-09 | 1.221E-08 | 2.600E-09 | 2.400E-08 |
| 14333 | 717 | P717M | B | 1.3e-09 | 5.7e-09 | 4.5e-09 | 3.833E-09 | 2.274E-09 | 1.300E-09 | 5.700E-09 |
| 14334 | 717 | P717F | B | 4.2e-09 | 1.7e-08 | 2.8e-09 | 8.000E-09 | 7.826E-09 | 2.800E-09 | 1.700E-08 |
| 14335 | 717 | P717P | B | 4.5e-09 | 4.4e-09 | 3.4e-09 | 4.100E-09 | 6.083E-10 | 3.400E-09 | 4.500E-09 |
| 14336 | 717 | P717S | B | 5e-09 | 1.1e-08 | 3.5e-09 | 6.500E-09 | 3.969E-09 | 3.500E-09 | 1.100E-08 |
| 14337 | 717 | P717T | B | 8.9e-09 | 2.2e-08 | 1.4e-08 | 1.497E-08 | 6.603E-09 | 8.900E-09 | 2.200E-08 |
| 14338 | 717 | P717W | B | 4.3e-09 | 5.8e-09 | 2.1e-08 | 1.037E-08 | 9.239E-09 | 4.300E-09 | 2.100E-08 |
| 14339 | 717 | P717Y | B | 1.5e-08 | 7.5e-09 | 3.3e-09 | 8.600E-09 | 5.927E-09 | 3.300E-09 | 1.500E-08 |
| 14340 | 717 | P717V | B | 3.1e-09 | 1.1e-08 | 6.2e-09 | 6.767E-09 | 3.980E-09 | 3.100E-09 | 1.100E-08 |
| 14341 | 718 | Y718A | B | 4.5e-09 | 6.4e-09 | 5.5e-09 | 5.467E-09 | 9.504E-10 | 4.500E-09 | 6.400E-09 |
| 14342 | 718 | Y718R | B | 1.5e-09 | 3.2e-09 | 8e-09 | 4.233E-09 | 3.371E-09 | 1.500E-09 | 8.000E-09 |
| 14343 | 718 | Y718N | B | 7.3e-09 | 1e-08 | 1.1e-08 | 9.433E-09 | 1.914E-09 | 7.300E-09 | 1.100E-08 |
| 14344 | 718 | Y718D | B | 4e-09 | 6.4e-09 | 7.8e-09 | 6.067E-09 | 1.922E-09 | 4.000E-09 | 7.800E-09 |
| 14345 | 718 | Y718C | B | 6e-09 | 3.3e-09 | 1.2e-08 | 7.100E-09 | 4.453E-09 | 3.300E-09 | 1.200E-08 |
| 14346 | 718 | Y718E | B | 7.9e-09 | 9.6e-09 | 4.1e-09 | 7.200E-09 | 2.816E-09 | 4.100E-09 | 9.600E-09 |
| 14347 | 718 | Y718Q | B | 4.2e-09 | 5.7e-09 | 4.2e-09 | 4.700E-09 | 8.660E-10 | 4.200E-09 | 5.700E-09 |
| 14348 | 718 | Y718G | B | 2e-09 | 8.7e-09 | 9.7e-09 | 6.800E-09 | 4.187E-09 | 2.000E-09 | 9.700E-09 |
| 14349 | 718 | Y718H | B | 4.2e-09 | 1.1e-08 | 1.8e-09 | 5.667E-09 | 4.772E-09 | 1.800E-09 | 1.100E-08 |
| 14350 | 718 | Y718I | B | 6.5e-09 | 4.1e-09 | 3.1e-09 | 4.567E-09 | 1.747E-09 | 3.100E-09 | 6.500E-09 |
| 14351 | 718 | Y718L | B | 1e-08 | 1e-08 | 6.7e-09 | 8.900E-09 | 1.905E-09 | 6.700E-09 | 1.000E-08 |
| 14352 | 718 | Y718K | B | 5.2e-09 | 1e-08 | 3.2e-09 | 6.133E-09 | 3.495E-09 | 3.200E-09 | 1.000E-08 |
| 14353 | 718 | Y718M | B | 2.9e-09 | 4.4e-09 | 4.7e-09 | 4.000E-09 | 9.644E-10 | 2.900E-09 | 4.700E-09 |
| 14354 | 718 | Y718F | B | 7.6e-09 | 4.9e-09 | 5.7e-09 | 6.067E-09 | 1.387E-09 | 4.900E-09 | 7.600E-09 |
| 14355 | 718 | Y718P | B | 4.3e-09 | 3.3e-09 | 7.9e-09 | 5.167E-09 | 2.419E-09 | 3.300E-09 | 7.900E-09 |
| 14356 | 718 | Y718S | B | 4.4e-09 | 3.7e-09 | 1.3e-08 | 7.033E-09 | 5.179E-09 | 3.700E-09 | 1.300E-08 |
| 14357 | 718 | Y718T | B | 6.5e-09 | 3.4e-09 | 1.3e-08 | 7.633E-09 | 4.899E-09 | 3.400E-09 | 1.300E-08 |
| 14358 | 718 | Y718W | B | 7.2e-09 | 1.9e-08 | 2.7e-09 | 9.633E-09 | 8.418E-09 | 2.700E-09 | 1.900E-08 |
| 14359 | 718 | Y718Y | B | 5.8e-09 | 4.8e-09 | 9.6e-09 | 6.733E-09 | 2.532E-09 | 4.800E-09 | 9.600E-09 |
| 14360 | 718 | Y718V | B | 8.5e-09 | 3.6e-09 | 6.8e-09 | 6.300E-09 | 2.488E-09 | 3.600E-09 | 8.500E-09 |
| 14361 | 719 | T719A | B | 6.4e-09 | 3.7e-09 | 7e-09 | 5.700E-09 | 1.758E-09 | 3.700E-09 | 7.000E-09 |
| 14362 | 719 | T719R | B | 6.7e-09 | 9.9e-09 | 3.3e-09 | 6.633E-09 | 3.301E-09 | 3.300E-09 | 9.900E-09 |
| 14363 | 719 | T719N | B | 4e-09 | 4.4e-09 | 5.6e-09 | 4.667E-09 | 8.327E-10 | 4.000E-09 | 5.600E-09 |
| 14364 | 719 | T719D | B | 8.4e-09 | 5e-09 | 1.3e-08 | 8.800E-09 | 4.015E-09 | 5.000E-09 | 1.300E-08 |

| | | | | | | | | | | |
|---|---|---|---|---|---|---|---|---|---|---|
| 14365 | 719 | T719C | B | 6.3e-09 | 2.1e-09 | 5.7e-09 | 4.700E-09 | 2.272E-09 | 2.100E-09 | 6.300E-09 |
| 14366 | 719 | T719E | B | 4.3e-09 | 5e-09 | 6.4e-09 | 5.233E-09 | 1.069E-09 | 4.300E-09 | 6.400E-09 |
| 14367 | 719 | T719Q | B | 3.6e-09 | 3.7e-09 | 8.8e-09 | 5.367E-09 | 2.974E-09 | 3.600E-09 | 8.800E-09 |
| 14368 | 719 | T719G | B | 3.9e-09 | 6.3e-09 | 1.5e-08 | 8.400E-09 | 5.840E-09 | 3.900E-09 | 1.500E-08 |
| 14369 | 719 | T719H | B | 1.4e-08 | 6.7e-09 | 7.4e-09 | 9.367E-09 | 4.028E-09 | 6.700E-09 | 1.400E-08 |
| 14370 | 719 | T719I | B | 3e-09 | 2.9e-09 | 2.4e-08 | 9.967E-09 | 1.215E-08 | 2.900E-09 | 2.400E-08 |
| 14371 | 719 | T719L | B | 3.5e-09 | 2.5e-09 | 7.6e-09 | 4.533E-09 | 2.702E-09 | 2.500E-09 | 7.600E-09 |
| 14372 | 719 | T719K | B | 2e-08 | 1.1e-08 | 7.6e-09 | 1.287E-08 | 6.407E-09 | 7.600E-09 | 2.000E-08 |
| 14373 | 719 | T719M | B | 4.3e-09 | 6.3e-09 | 6.7e-09 | 5.767E-09 | 1.286E-09 | 4.300E-09 | 6.700E-09 |
| 14374 | 719 | T719F | B | 8e-09 | 7.9e-09 | 3.9e-09 | 6.600E-09 | 2.339E-09 | 3.900E-09 | 8.000E-09 |
| 14375 | 719 | T719P | B | 5.5e-09 | 3.9e-09 | 5.4e-09 | 4.933E-09 | 8.963E-10 | 3.900E-09 | 5.500E-09 |
| 14376 | 719 | T719S | B | 4.8e-09 | 8.1e-09 | 3.5e-09 | 5.467E-09 | 2.371E-09 | 3.500E-09 | 8.100E-09 |
| 14377 | 719 | T719T | B | 1.4e-09 | 1.6e-08 | 4.2e-09 | 7.200E-09 | 7.749E-09 | 1.400E-09 | 1.600E-08 |
| 14378 | 719 | T719W | B | 5.1e-09 | 1.7e-08 | 6e-09 | 9.367E-09 | 6.626E-09 | 5.100E-09 | 1.700E-08 |
| 14379 | 719 | T719Y | B | 9e-09 | 9.2e-09 | 3.8e-09 | 7.333E-09 | 3.062E-09 | 3.800E-09 | 9.200E-09 |
| 14380 | 719 | T719V | B | 8.9e-09 | 7.2e-09 | 1.7e-08 | 1.103E-08 | 5.237E-09 | 7.200E-09 | 1.700E-08 |
| 14381 | 720 | F720A | B | 2e-08 | 1.1e-08 | 8.9e-09 | 1.330E-08 | 5.897E-09 | 8.900E-09 | 2.000E-08 |
| 14382 | 720 | F720R | B | 7.4e-09 | 5.1e-09 | 8.9e-09 | 7.133E-09 | 1.914E-09 | 5.100E-09 | 8.900E-09 |
| 14383 | 720 | F720N | B | 5.5e-09 | 6.2e-09 | 1.1e-08 | 7.567E-09 | 2.994E-09 | 5.500E-09 | 1.100E-08 |
| 14384 | 720 | F720D | B | 1e-08 | 8.5e-09 | 1.2e-08 | 1.017E-08 | 1.756E-09 | 8.500E-09 | 1.200E-08 |
| 14385 | 720 | F720C | B | 1.1e-08 | 3.6e-09 | 9.8e-09 | 8.133E-09 | 3.972E-09 | 3.600E-09 | 1.100E-08 |
| 14386 | 720 | F720E | B | 5.3e-09 | 6.1e-09 | 7.8e-09 | 6.400E-09 | 1.277E-09 | 5.300E-09 | 7.800E-09 |
| 14387 | 720 | F720Q | B | 1e-08 | 1.5e-08 | 1.5e-08 | 1.333E-08 | 2.887E-09 | 1.000E-08 | 1.500E-08 |
| 14388 | 720 | F720G | B | 4.4e-09 | 1.3e-08 | 1.3e-08 | 1.013E-08 | 4.965E-09 | 4.400E-09 | 1.300E-08 |
| 14389 | 720 | F720H | B | 5.2e-09 | 1e-08 | 3.8e-09 | 6.333E-09 | 3.252E-09 | 3.800E-09 | 1.000E-08 |
| 14390 | 720 | F720I | B | 5.3e-09 | 8.3e-09 | 5.6e-09 | 6.400E-09 | 1.652E-09 | 5.300E-09 | 8.300E-09 |
| 14391 | 720 | F720L | B | 6.2e-09 | 1.6e-08 | 3.3e-09 | 8.500E-09 | 6.655E-09 | 3.300E-09 | 1.600E-08 |
| 14392 | 720 | F720K | B | 6.1e-09 | 4.2e-09 | 5.4e-09 | 5.233E-09 | 9.609E-10 | 4.200E-09 | 6.100E-09 |
| 14393 | 720 | F720M | B | 1.1e-08 | 1.2e-08 | 2.1e-08 | 1.467E-08 | 5.508E-09 | 1.100E-08 | 2.100E-08 |
| 14394 | 720 | F720F | B | 5.5e-09 | 7.3e-09 | 4.4e-09 | 5.733E-09 | 1.464E-09 | 4.400E-09 | 7.300E-09 |
| 14395 | 720 | F720P | B | 3.4e-09 | 7.6e-09 | 8.2e-09 | 6.400E-09 | 2.615E-09 | 3.400E-09 | 8.200E-09 |
| 14396 | 720 | F720S | B | 7.4e-09 | 1.2e-08 | 5.7e-09 | 8.367E-09 | 3.259E-09 | 5.700E-09 | 1.200E-08 |
| 14397 | 720 | F720T | B | 5.9e-09 | 1e-08 | 4.8e-09 | 6.900E-09 | 2.740E-09 | 4.800E-09 | 1.000E-08 |
| 14398 | 720 | F720W | B | 6.8e-09 | 8.3e-09 | 6.2e-09 | 7.100E-09 | 1.082E-09 | 6.200E-09 | 8.300E-09 |
| 14399 | 720 | F720Y | B | 8.9e-09 | 6e-09 | 9.6e-09 | 8.167E-09 | 1.909E-09 | 6.000E-09 | 9.600E-09 |
| 14400 | 720 | F720V | B | 9.2e-09 | 2.8e-09 | 4.7e-09 | 5.567E-09 | 3.287E-09 | 2.800E-09 | 9.200E-09 |
| 14401 | 721 | G721A | B | 6.7e-09 | 9.1e-09 | 5.2e-09 | 7.000E-09 | 1.967E-09 | 5.200E-09 | 9.100E-09 |
| 14402 | 721 | G721R | B | 7.4e-09 | 4.3e-09 | 3.7e-09 | 5.133E-09 | 1.986E-09 | 3.700E-09 | 7.400E-09 |
| 14403 | 721 | G721N | B | 3e-09 | 5.6e-09 | 3.3e-09 | 3.967E-09 | 1.422E-09 | 3.000E-09 | 5.600E-09 |
| 14404 | 721 | G721D | B | 2.4e-09 | 1.4e-08 | 9e-09 | 8.467E-09 | 5.818E-09 | 2.400E-09 | 1.400E-08 |
| 14405 | 721 | G721C | B | 5.8e-09 | 7e-09 | 9e-09 | 7.267E-09 | 1.617E-09 | 5.800E-09 | 9.000E-09 |
| 14406 | 721 | G721E | B | 3.8e-09 | 8.2e-09 | 1.2e-08 | 8.000E-09 | 4.104E-09 | 3.800E-09 | 1.200E-08 |
| 14407 | 721 | G721Q | B | 9.5e-09 | 7.1e-09 | 9.6e-09 | 8.733E-09 | 1.415E-09 | 7.100E-09 | 9.600E-09 |
| 14408 | 721 | G721G | B | 1.6e-08 | 4.4e-09 | 5e-09 | 8.467E-09 | 6.531E-09 | 4.400E-09 | 1.600E-08 |
| 14409 | 721 | G721H | B | 5.1e-09 | 1.9e-08 | 3.2e-09 | 9.100E-09 | 8.626E-09 | 3.200E-09 | 1.900E-08 |
| 14410 | 721 | G721I | B | 1.5e-08 | 5.8e-09 | 3.2e-09 | 8.000E-09 | 6.200E-09 | 3.200E-09 | 1.500E-08 |
| 14411 | 721 | G721L | B | 1.9e-08 | 4.2e-09 | 1.9e-08 | 1.407E-08 | 8.545E-09 | 4.200E-09 | 1.900E-08 |
| 14412 | 721 | G721K | B | 2.5e-09 | 7.2e-09 | 1.2e-08 | 7.233E-09 | 4.750E-09 | 2.500E-09 | 1.200E-08 |
| 14413 | 721 | G721M | B | 4.7e-09 | 5.4e-09 | 1.8e-09 | 3.967E-09 | 1.909E-09 | 1.800E-09 | 5.400E-09 |
| 14414 | 721 | G721F | B | 2.5e-09 | 6.4e-09 | 1e-08 | 6.300E-09 | 3.751E-09 | 2.500E-09 | 1.000E-08 |
| 14415 | 721 | G721P | B | 6.3e-09 | 5.5e-09 | 5.1e-09 | 5.633E-09 | 6.110E-10 | 5.100E-09 | 6.300E-09 |
| 14416 | 721 | G721S | B | 8.8e-09 | 3.9e-09 | 2.4e-09 | 5.033E-09 | 3.347E-09 | 2.400E-09 | 8.800E-09 |
| 14417 | 721 | G721T | B | 8.8e-09 | 7e-09 | 8.4e-09 | 8.067E-09 | 9.452E-10 | 7.000E-09 | 8.800E-09 |
| 14418 | 721 | G721W | B | 1.6e-08 | 3.4e-09 | 6.8e-09 | 8.733E-09 | 6.519E-09 | 3.400E-09 | 1.600E-08 |
| 14419 | 721 | G721Y | B | 4.9e-09 | 5.7e-09 | 4.5e-09 | 5.033E-09 | 6.110E-10 | 4.500E-09 | 5.700E-09 |
| 14420 | 721 | G721V | B | 8.2e-09 | 3.4e-09 | 9.3e-09 | 6.967E-09 | 3.137E-09 | 3.400E-09 | 9.300E-09 |
| 14421 | 722 | Q722A | B | 1e-08 | 3.9e-09 | 9.2e-09 | 7.700E-09 | 3.315E-09 | 3.900E-09 | 1.000E-08 |
| 14422 | 722 | Q722R | B | 5.6e-09 | 2.1e-09 | 5.3e-09 | 4.333E-09 | 1.940E-09 | 2.100E-09 | 5.600E-09 |
| 14423 | 722 | Q722N | B | 1.2e-08 | 1.4e-08 | 4.5e-09 | 1.017E-08 | 5.008E-09 | 4.500E-09 | 1.400E-08 |
| 14424 | 722 | Q722D | B | 5.4e-09 | 2.3e-08 | 1.6e-08 | 1.480E-08 | 8.861E-09 | 5.400E-09 | 2.300E-08 |
| 14425 | 722 | Q722C | B | 7.9e-09 | 2.4e-09 | 3.4e-09 | 4.567E-09 | 2.930E-09 | 2.400E-09 | 7.900E-09 |
| 14426 | 722 | Q722E | B | 9.2e-09 | 9.3e-09 | 3.4e-09 | 7.300E-09 | 3.378E-09 | 3.400E-09 | 9.300E-09 |
| 14427 | 722 | Q722Q | B | 8.3e-09 | 1.5e-08 | 3.3e-09 | 8.867E-09 | 5.871E-09 | 3.300E-09 | 1.500E-08 |
| 14428 | 722 | Q722G | B | 1.8e-08 | 1.1e-08 | 4.5e-09 | 1.117E-08 | 6.752E-09 | 4.500E-09 | 1.800E-08 |
| 14429 | 722 | Q722H | B | 7.3e-09 | 1.2e-09 | 3.2e-09 | 3.900E-09 | 3.110E-09 | 1.200E-09 | 7.300E-09 |
| 14430 | 722 | Q722I | B | 2.1e-08 | 4.5e-09 | 9.3e-09 | 1.160E-08 | 8.487E-09 | 4.500E-09 | 2.100E-08 |
| 14431 | 722 | Q722L | B | 5.4e-09 | 6.4e-09 | 7.1e-09 | 6.300E-09 | 8.544E-10 | 5.400E-09 | 7.100E-09 |
| 14432 | 722 | Q722K | B | 6.9e-09 | 1.5e-08 | 6.2e-09 | 9.367E-09 | 4.891E-09 | 6.200E-09 | 1.500E-08 |
| 14433 | 722 | Q722M | B | 4e-09 | 8.1e-09 | 5.9e-09 | 6.000E-09 | 2.052E-09 | 4.000E-09 | 8.100E-09 |
| 14434 | 722 | Q722F | B | 1.3e-08 | 1.4e-08 | 4e-09 | 1.033E-08 | 5.508E-09 | 4.000E-09 | 1.400E-08 |
| 14435 | 722 | Q722P | B | 4.3e-09 | 3.4e-09 | 1.6e-09 | 3.100E-09 | 1.375E-09 | 1.600E-09 | 4.300E-09 |
| 14436 | 722 | Q722S | B | 5.8e-09 | 3.6e-09 | 6.9e-09 | 5.433E-09 | 1.680E-09 | 3.600E-09 | 6.900E-09 |
| 14437 | 722 | Q722T | B | 1.9e-08 | 1.3e-08 | 9.5e-09 | 1.383E-08 | 4.805E-09 | 9.500E-09 | 1.900E-08 |
| 14438 | 722 | Q722W | B | 8.4e-09 | 3.8e-09 | 8.1e-09 | 6.767E-09 | 2.574E-09 | 3.800E-09 | 8.400E-09 |
| 14439 | 722 | Q722Y | B | 2.8e-09 | 1.7e-08 | 7.4e-09 | 9.067E-09 | 7.245E-09 | 2.800E-09 | 1.700E-08 |
| 14440 | 722 | Q722V | B | 6.3e-09 | 3.3e-09 | 5.7e-09 | 5.100E-09 | 1.587E-09 | 3.300E-09 | 6.300E-09 |
| 14441 | 723 | G723A | B | 2.7e-09 | 6.3e-09 | 2.2e-09 | 3.733E-09 | 2.237E-09 | 2.200E-09 | 6.300E-09 |
| 14442 | 723 | G723R | B | 2e-09 | 1.7e-08 | 2.8e-09 | 7.267E-09 | 8.439E-09 | 2.000E-09 | 1.700E-08 |
| 14443 | 723 | G723N | B | 6.3e-09 | 5.5e-09 | 9.2e-09 | 7.000E-09 | 1.947E-09 | 5.500E-09 | 9.200E-09 |
| 14444 | 723 | G723D | B | 5.8e-09 | 2.4e-09 | 7.3e-09 | 5.167E-09 | 2.511E-09 | 2.400E-09 | 7.300E-09 |
| 14445 | 723 | G723C | B | 9e-09 | 8e-09 | 1.9e-08 | 1.200E-08 | 6.083E-09 | 8.000E-09 | 1.900E-08 |
| 14446 | 723 | G723E | B | 2.2e-09 | 4.5e-09 | 1.1e-08 | 5.900E-09 | 4.564E-09 | 2.200E-09 | 1.100E-08 |
| 14447 | 723 | G723Q | B | 1.4e-08 | 7.9e-09 | 1.1e-08 | 1.097E-08 | 3.050E-09 | 7.900E-09 | 1.400E-08 |
| 14448 | 723 | G723G | B | 1.5e-09 | 6e-09 | 9.8e-09 | 5.767E-09 | 4.155E-09 | 1.500E-09 | 9.800E-09 |
| 14449 | 723 | G723H | B | 5.7e-09 | 1e-08 | 1.7e-08 | 1.090E-08 | 5.704E-09 | 5.700E-09 | 1.700E-08 |

| | | | | | | | | | | |
|---|---|---|---|---|---|---|---|---|---|---|
| 14450 | 723 | G723I | B | 1.5e-08 | 1.3e-08 | 2.5e-08 | 1.767E-08 | 6.429E-09 | 1.300E-08 | 2.500E-08 |
| 14451 | 723 | G723L | B | 5.7e-09 | 9.9e-09 | 1.4e-08 | 9.867E-09 | 4.150E-09 | 5.700E-09 | 1.400E-08 |
| 14452 | 723 | G723K | B | 5.5e-09 | 1.1e-08 | 1.2e-08 | 9.500E-09 | 3.500E-09 | 5.500E-09 | 1.200E-08 |
| 14453 | 723 | G723M | B | 3.5e-09 | 1.5e-08 | 4.3e-09 | 7.600E-09 | 6.421E-09 | 3.500E-09 | 1.500E-08 |
| 14454 | 723 | G723F | B | 7.9e-09 | 3.8e-09 | 1.5e-08 | 8.900E-09 | 5.667E-09 | 3.800E-09 | 1.500E-08 |
| 14455 | 723 | G723P | B | 4.2e-09 | 4.7e-09 | 5.6e-09 | 4.833E-09 | 7.095E-10 | 4.200E-09 | 5.600E-09 |
| 14456 | 723 | G723S | B | 1.7e-08 | 1.8e-08 | 3.8e-09 | 1.293E-08 | 7.925E-09 | 3.800E-09 | 1.800E-08 |
| 14457 | 723 | G723T | B | 1.4e-08 | 4.9e-09 | 8.7e-09 | 9.200E-09 | 4.571E-09 | 4.900E-09 | 1.400E-08 |
| 14458 | 723 | G723W | B | 1.2e-08 | 6.5e-09 | 5e-09 | 7.833E-09 | 3.686E-09 | 5.000E-09 | 1.200E-08 |
| 14459 | 723 | G723Y | B | 4.8e-09 | 4.9e-09 | 1.3e-08 | 7.567E-09 | 4.706E-09 | 4.800E-09 | 1.300E-08 |
| 14460 | 723 | G723V | B | 1.3e-08 | 2.6e-09 | 9e-09 | 8.200E-09 | 5.246E-09 | 2.600E-09 | 1.300E-08 |
| 14461 | 724 | T724A | B | 7.8e-09 | 9.9e-09 | 4e-09 | 7.233E-09 | 2.991E-09 | 4.000E-09 | 9.900E-09 |
| 14462 | 724 | T724R | B | 1.5e-08 | 5e-09 | 5.5e-09 | 8.500E-09 | 5.635E-09 | 5.000E-09 | 1.500E-08 |
| 14463 | 724 | T724N | B | 4e-09 | 2.9e-09 | 1e-08 | 5.633E-09 | 3.821E-09 | 2.900E-09 | 1.000E-08 |
| 14464 | 724 | T724D | B | 2.9e-09 | 8e-09 | 2.4e-09 | 4.433E-09 | 3.099E-09 | 2.400E-09 | 8.000E-09 |
| 14465 | 724 | T724C | B | 5.1e-09 | 6.5e-09 | 7.9e-09 | 6.500E-09 | 1.400E-09 | 5.100E-09 | 7.900E-09 |
| 14466 | 724 | T724E | B | 6.6e-09 | 7.2e-09 | 9.9e-09 | 7.900E-09 | 1.758E-09 | 6.600E-09 | 9.900E-09 |
| 14467 | 724 | T724Q | B | 1.4e-08 | 7.8e-09 | 4.4e-09 | 8.733E-09 | 4.868E-09 | 4.400E-09 | 1.400E-08 |
| 14468 | 724 | T724G | B | 5.3e-09 | 7.5e-09 | 4.1e-09 | 5.633E-09 | 1.724E-09 | 4.100E-09 | 7.500E-09 |
| 14469 | 724 | T724H | B | 8.8e-09 | 9.4e-09 | 1.1e-08 | 9.733E-09 | 1.137E-09 | 8.800E-09 | 1.100E-08 |
| 14470 | 724 | T724I | B | 6.1e-09 | 1.7e-08 | 4.4e-09 | 9.167E-09 | 6.837E-09 | 4.400E-09 | 1.700E-08 |
| 14471 | 724 | T724L | B | 5.6e-09 | 7.2e-09 | 1.2e-08 | 8.267E-09 | 3.331E-09 | 5.600E-09 | 1.200E-08 |
| 14472 | 724 | T724K | B | 6.4e-10 | 5.8e-09 | 2.1e-08 | 9.147E-09 | 1.058E-08 | 6.400E-10 | 2.100E-08 |
| 14473 | 724 | T724M | B | 5e-09 | 5.4e-09 | 1.6e-09 | 4.000E-09 | 2.088E-09 | 1.600E-09 | 5.400E-09 |
| 14474 | 724 | T724F | B | 1.9e-08 | 9.8e-09 | 7.5e-09 | 1.210E-08 | 6.085E-09 | 7.500E-09 | 1.900E-08 |
| 14475 | 724 | T724P | B | 3.5e-09 | 8e-09 | 7.2e-09 | 6.233E-09 | 2.401E-09 | 3.500E-09 | 8.000E-09 |
| 14476 | 724 | T724S | B | 2.3e-09 | 6.8e-09 | 6.5e-09 | 5.200E-09 | 2.516E-09 | 2.300E-09 | 6.800E-09 |
| 14477 | 724 | T724T | B | 9.4e-09 | 8e-09 | 1e-08 | 9.133E-09 | 1.026E-09 | 8.000E-09 | 1.000E-08 |
| 14478 | 724 | T724W | B | 6.9e-09 | 2e-08 | 9e-09 | 1.197E-08 | 7.036E-09 | 6.900E-09 | 2.000E-08 |
| 14479 | 724 | T724Y | B | 6.4e-09 | 8.1e-09 | 5.7e-09 | 6.733E-09 | 1.234E-09 | 5.700E-09 | 8.100E-09 |
| 14480 | 724 | T724V | B | 1.4e-08 | 1e-08 | 4.4e-09 | 9.467E-09 | 4.822E-09 | 4.400E-09 | 1.400E-08 |
| 14481 | 725 | K725A | B | 8.5e-09 | 3.3e-09 | 1.2e-08 | 7.933E-09 | 4.378E-09 | 3.300E-09 | 1.200E-08 |
| 14482 | 725 | K725R | B | 1.1e-08 | 1.6e-08 | 8.6e-09 | 1.187E-08 | 3.775E-09 | 8.600E-09 | 1.600E-08 |
| 14483 | 725 | K725N | B | 6.4e-09 | 1.2e-08 | 3.1e-09 | 7.167E-09 | 4.499E-09 | 3.100E-09 | 1.200E-08 |
| 14484 | 725 | K725D | B | 2.2e-08 | 4.9e-09 | 5.7e-09 | 1.087E-08 | 9.650E-09 | 4.900E-09 | 2.200E-08 |
| 14485 | 725 | K725C | B | 1.1e-08 | 3.9e-09 | 6e-09 | 6.967E-09 | 3.647E-09 | 3.900E-09 | 1.100E-08 |
| 14486 | 725 | K725E | B | 6.5e-09 | 7e-09 | 8.7e-09 | 7.400E-09 | 1.153E-09 | 6.500E-09 | 8.700E-09 |
| 14487 | 725 | K725Q | B | 1e-08 | 4.7e-09 | 5.7e-09 | 6.800E-09 | 2.816E-09 | 4.700E-09 | 1.000E-08 |
| 14488 | 725 | K725G | B | 2.1e-08 | 1.1e-08 | 5.1e-09 | 1.237E-08 | 8.038E-09 | 5.100E-09 | 2.100E-08 |
| 14489 | 725 | K725H | B | 9.6e-09 | 6.9e-09 | 5.4e-09 | 7.300E-09 | 2.128E-09 | 5.400E-09 | 9.600E-09 |
| 14490 | 725 | K725I | B | 5.1e-09 | 1.4e-08 | 3.2e-09 | 7.433E-09 | 5.766E-09 | 3.200E-09 | 1.400E-08 |
| 14491 | 725 | K725L | B | 5.1e-09 | 2.5e-09 | 1.7e-08 | 8.200E-09 | 7.731E-09 | 2.500E-09 | 1.700E-08 |
| 14492 | 725 | K725K | B | 6.5e-09 | 6.9e-09 | 1.4e-08 | 9.133E-09 | 4.219E-09 | 6.500E-09 | 1.400E-08 |
| 14493 | 725 | K725M | B | 5.7e-09 | 6.6e-09 | 9.3e-09 | 7.200E-09 | 1.873E-09 | 5.700E-09 | 9.300E-09 |
| 14494 | 725 | K725F | B | 1e-08 | 1.3e-08 | 7.4e-09 | 1.013E-08 | 2.802E-09 | 7.400E-09 | 1.300E-08 |
| 14495 | 725 | K725P | B | 1.1e-08 | 1.3e-08 | 1.3e-08 | 1.233E-08 | 1.155E-09 | 1.100E-08 | 1.300E-08 |
| 14496 | 725 | K725S | B | 3.8e-09 | 8e-09 | 6.2e-09 | 6.000E-09 | 2.107E-09 | 3.800E-09 | 8.000E-09 |
| 14497 | 725 | K725T | B | 1.2e-08 | 2.7e-09 | 8.4e-09 | 7.700E-09 | 4.689E-09 | 2.700E-09 | 1.200E-08 |
| 14498 | 725 | K725W | B | 1.6e-08 | 1.8e-09 | 5.6e-09 | 7.800E-09 | 7.351E-09 | 1.800E-09 | 1.600E-08 |
| 14499 | 725 | K725Y | B | 7.6e-09 | 1.2e-08 | 1.3e-08 | 1.087E-08 | 2.873E-09 | 7.600E-09 | 1.300E-08 |
| 14500 | 725 | K725V | B | 9.4e-09 | 8e-09 | 1e-08 | 9.133E-09 | 1.026E-09 | 8.000E-09 | 1.000E-08 |
| 14501 | 726 | V726A | B | 6.8e-09 | 3.8e-09 | 2.5e-09 | 4.367E-09 | 2.205E-09 | 2.500E-09 | 6.800E-09 |
| 14502 | 726 | V726R | B | 9e-09 | 6.1e-09 | 9.4e-09 | 8.167E-09 | 1.801E-09 | 6.100E-09 | 9.400E-09 |
| 14503 | 726 | V726N | B | 6.8e-09 | 7.6e-09 | 7.3e-09 | 7.233E-09 | 4.041E-10 | 6.800E-09 | 7.600E-09 |
| 14504 | 726 | V726D | B | 7.1e-09 | 5.9e-09 | 7.3e-09 | 6.767E-09 | 7.572E-10 | 5.900E-09 | 7.300E-09 |
| 14505 | 726 | V726C | B | 2.7e-09 | 3.2e-09 | 2.6e-09 | 2.833E-09 | 3.215E-10 | 2.600E-09 | 3.200E-09 |
| 14506 | 726 | V726E | B | 5.6e-09 | 3.8e-09 | 1.3e-09 | 3.567E-09 | 2.159E-09 | 1.300E-09 | 5.600E-09 |
| 14507 | 726 | V726Q | B | 2.9e-09 | 3.5e-09 | 6.3e-09 | 4.233E-09 | 1.815E-09 | 2.900E-09 | 6.300E-09 |
| 14508 | 726 | V726G | B | 6.3e-09 | 1.5e-08 | 1.3e-08 | 1.143E-08 | 4.557E-09 | 6.300E-09 | 1.500E-08 |
| 14509 | 726 | V726H | B | 2.3e-09 | 1.3e-08 | 1.9e-08 | 1.143E-08 | 8.460E-09 | 2.300E-09 | 1.900E-08 |
| 14510 | 726 | V726I | B | 6.9e-09 | 7.5e-09 | 8.2e-09 | 7.533E-09 | 6.506E-10 | 6.900E-09 | 8.200E-09 |
| 14511 | 726 | V726L | B | 1.8e-08 | 5e-09 | 4.6e-09 | 9.200E-09 | 7.624E-09 | 4.600E-09 | 1.800E-08 |
| 14512 | 726 | V726K | B | 6.9e-09 | 5.1e-09 | 1e-09 | 4.333E-09 | 3.024E-09 | 1.000E-09 | 6.900E-09 |
| 14513 | 726 | V726M | B | 6.6e-09 | 3.8e-09 | 9.2e-09 | 6.533E-09 | 2.701E-09 | 3.800E-09 | 9.200E-09 |
| 14514 | 726 | V726F | B | 1.5e-08 | 9e-09 | 2.8e-09 | 8.933E-09 | 6.100E-09 | 2.800E-09 | 1.500E-08 |
| 14515 | 726 | V726P | B | 1.4e-08 | 4.7e-09 | 5.4e-09 | 8.033E-09 | 5.179E-09 | 4.700E-09 | 1.400E-08 |
| 14516 | 726 | V726S | B | 1.3e-08 | 3.6e-09 | 9.1e-09 | 8.567E-09 | 4.723E-09 | 3.600E-09 | 1.300E-08 |
| 14517 | 726 | V726T | B | 7.2e-09 | 4.9e-09 | 4.9e-09 | 5.667E-09 | 1.328E-09 | 4.900E-09 | 7.200E-09 |
| 14518 | 726 | V726W | B | 7.3e-09 | 8.8e-09 | 6.3e-09 | 7.467E-09 | 1.258E-09 | 6.300E-09 | 8.800E-09 |
| 14519 | 726 | V726Y | B | 1.3e-08 | 1.5e-08 | 2.8e-09 | 1.027E-08 | 6.543E-09 | 2.800E-09 | 1.500E-08 |
| 14520 | 726 | V726V | B | 3.2e-09 | 4.6e-09 | 5e-09 | 4.267E-09 | 9.452E-10 | 3.200E-09 | 5.000E-09 |
| 14521 | 727 | E727A | B | 1e-08 | 1.5e-08 | 5.8e-09 | 1.027E-08 | 4.606E-09 | 5.800E-09 | 1.500E-08 |
| 14522 | 727 | E727R | B | 1.2e-08 | 7.5e-09 | 1.1e-08 | 1.017E-08 | 2.363E-09 | 7.500E-09 | 1.200E-08 |
| 14523 | 727 | E727N | B | 7.6e-09 | 1.5e-08 | 1.2e-08 | 1.153E-08 | 3.722E-09 | 7.600E-09 | 1.500E-08 |
| 14524 | 727 | E727D | B | 1e-08 | 1.1e-08 | 8.5e-09 | 9.833E-09 | 1.258E-09 | 8.500E-09 | 1.100E-08 |
| 14525 | 727 | E727C | B | 1e-08 | 5e-09 | 1.2e-08 | 9.000E-09 | 3.606E-09 | 5.000E-09 | 1.200E-08 |
| 14526 | 727 | E727E | B | 7.7e-09 | 2.9e-09 | 5.3e-09 | 5.300E-09 | 2.400E-09 | 2.900E-09 | 7.700E-09 |
| 14527 | 727 | E727Q | B | 5.7e-09 | 3.2e-09 | 7.5e-09 | 5.467E-09 | 2.159E-09 | 3.200E-09 | 7.500E-09 |
| 14528 | 727 | E727G | B | 6.5e-09 | 3.4e-09 | 4.7e-09 | 4.867E-09 | 1.557E-09 | 3.400E-09 | 6.500E-09 |
| 14529 | 727 | E727H | B | 1.7e-08 | 1.1e-08 | 7e-09 | 1.167E-08 | 5.033E-09 | 7.000E-09 | 1.700E-08 |
| 14530 | 727 | E727I | B | 6.3e-09 | 2.2e-08 | 3.3e-09 | 1.053E-08 | 1.004E-08 | 3.300E-09 | 2.200E-08 |
| 14531 | 727 | E727L | B | 5e-09 | 8.2e-09 | 6.1e-09 | 6.433E-09 | 1.626E-09 | 5.000E-09 | 8.200E-09 |
| 14532 | 727 | E727K | B | 1.2e-08 | 5.8e-09 | 1.1e-08 | 9.600E-09 | 3.329E-09 | 5.800E-09 | 1.200E-08 |
| 14533 | 727 | E727M | B | 7.1e-09 | 8.3e-09 | 1.3e-08 | 9.467E-09 | 3.118E-09 | 7.100E-09 | 1.300E-08 |
| 14534 | 727 | E727F | B | 2.2e-09 | 7.1e-09 | 6.4e-09 | 5.233E-09 | 2.650E-09 | 2.200E-09 | 7.100E-09 |

| | | | | | | | | | | |
|---|---|---|---|---|---|---|---|---|---|---|
| 14535 | 727 | E727P | B | 3.1e-09 | 9.1e-09 | 8.6e-09 | 6.933E-09 | 3.329E-09 | 3.100E-09 | 9.100E-09 |
| 14536 | 727 | E727S | B | 1.4e-08 | 7.2e-09 | 5.3e-09 | 8.833E-09 | 4.574E-09 | 5.300E-09 | 1.400E-08 |
| 14537 | 727 | E727T | B | 1.1e-08 | 1.5e-08 | 5.8e-09 | 1.060E-08 | 4.613E-09 | 5.800E-09 | 1.500E-08 |
| 14538 | 727 | E727W | B | 4.3e-09 | 7.2e-09 | 2.3e-08 | 1.150E-08 | 1.006E-08 | 4.300E-09 | 2.300E-08 |
| 14539 | 727 | E727Y | B | 2.7e-09 | 8.2e-09 | 8.9e-09 | 6.600E-09 | 3.396E-09 | 2.700E-09 | 8.900E-09 |
| 14540 | 727 | E727V | B | 7.5e-09 | 5e-09 | 2e-09 | 4.833E-09 | 2.754E-09 | 2.000E-09 | 7.500E-09 |
| 14541 | 728 | I728A | B | 1.2e-08 | 3.1e-09 | 7.9e-09 | 7.667E-09 | 4.455E-09 | 3.100E-09 | 1.200E-08 |
| 14542 | 728 | I728R | B | 5.1e-09 | 9.7e-09 | 1.9e-08 | 1.127E-08 | 7.081E-09 | 5.100E-09 | 1.900E-08 |
| 14543 | 728 | I728N | B | 2.5e-09 | 6.4e-09 | 8.1e-09 | 5.667E-09 | 2.871E-09 | 2.500E-09 | 8.100E-09 |
| 14544 | 728 | I728D | B | 5.1e-09 | 1.1e-08 | 7.6e-09 | 7.900E-09 | 2.961E-09 | 5.100E-09 | 1.100E-08 |
| 14545 | 728 | I728C | B | 8.5e-09 | 2.9e-09 | 6.5e-09 | 5.967E-09 | 2.838E-09 | 2.900E-09 | 8.500E-09 |
| 14546 | 728 | I728E | B | 1.4e-08 | 3.7e-09 | 6.7e-09 | 8.133E-09 | 5.297E-09 | 3.700E-09 | 1.400E-08 |
| 14547 | 728 | I728Q | B | 7.1e-09 | 6.1e-09 | 5.5e-09 | 6.233E-09 | 8.083E-10 | 5.500E-09 | 7.100E-09 |
| 14548 | 728 | I728G | B | 8.8e-09 | 4e-09 | 4.7e-09 | 5.833E-09 | 2.593E-09 | 4.000E-09 | 8.800E-09 |
| 14549 | 728 | I728H | B | 2.1e-09 | 3.1e-09 | 2.7e-09 | 2.633E-09 | 5.033E-10 | 2.100E-09 | 3.100E-09 |
| 14550 | 728 | I728I | B | 4.5e-09 | 8.4e-09 | 5.3e-09 | 6.067E-09 | 2.060E-09 | 4.500E-09 | 8.400E-09 |
| 14551 | 728 | I728L | B | 9.1e-09 | 4e-09 | 5.6e-09 | 6.233E-09 | 2.608E-09 | 4.000E-09 | 9.100E-09 |
| 14552 | 728 | I728K | B | 6.7e-09 | 9.8e-09 | 1.2e-08 | 9.500E-09 | 2.663E-09 | 6.700E-09 | 1.200E-08 |
| 14553 | 728 | I728M | B | 4.1e-09 | 3.2e-08 | 8.9e-09 | 1.500E-08 | 1.492E-08 | 4.100E-09 | 3.200E-08 |
| 14554 | 728 | I728F | B | 3e-09 | 4.9e-09 | 1.2e-08 | 6.633E-09 | 4.744E-09 | 3.000E-09 | 1.200E-08 |
| 14555 | 728 | I728P | B | 9.7e-09 | 5.9e-09 | 2.2e-08 | 1.253E-08 | 8.416E-09 | 5.900E-09 | 2.200E-08 |
| 14556 | 728 | I728S | B | 1.3e-08 | 9.2e-09 | 1e-08 | 1.073E-08 | 2.003E-09 | 9.200E-09 | 1.300E-08 |
| 14557 | 728 | I728T | B | 4.2e-09 | 1e-08 | 1.9e-08 | 1.107E-08 | 7.457E-09 | 4.200E-09 | 1.900E-08 |
| 14558 | 728 | I728W | B | 9.7e-09 | 6.8e-09 | 1.4e-08 | 1.017E-08 | 3.623E-09 | 6.800E-09 | 1.400E-08 |
| 14559 | 728 | I728Y | B | 9.9e-09 | 6.4e-09 | 9.2e-09 | 8.500E-09 | 1.852E-09 | 6.400E-09 | 9.900E-09 |
| 14560 | 728 | I728V | B | 3.9e-09 | 5.3e-09 | 1.6e-08 | 8.400E-09 | 6.619E-09 | 3.900E-09 | 1.600E-08 |
| 14561 | 729 | K729A | B | 1e-08 | 6.3e-09 | 2.4e-09 | 6.233E-09 | 3.800E-09 | 2.400E-09 | 1.000E-08 |
| 14562 | 729 | K729R | B | 3.7e-08 | 3e-09 | 8.5e-09 | 1.617E-08 | 1.825E-08 | 3.000E-09 | 3.700E-08 |
| 14563 | 729 | K729N | B | 9.6e-09 | 3e-09 | 1.1e-08 | 7.867E-09 | 4.272E-09 | 3.000E-09 | 1.100E-08 |
| 14564 | 729 | K729D | B | 5.2e-09 | 8.4e-09 | 3.4e-09 | 5.667E-09 | 2.532E-09 | 3.400E-09 | 8.400E-09 |
| 14565 | 729 | K729C | B | 7.5e-09 | 9.9e-09 | 6.7e-09 | 8.033E-09 | 1.665E-09 | 6.700E-09 | 9.900E-09 |
| 14566 | 729 | K729E | B | 7.2e-09 | 5.5e-09 | 3.6e-09 | 5.433E-09 | 1.801E-09 | 3.600E-09 | 7.200E-09 |
| 14567 | 729 | K729Q | B | 6.4e-09 | 4e-09 | 4.2e-09 | 4.867E-09 | 1.332E-09 | 4.000E-09 | 6.400E-09 |
| 14568 | 729 | K729G | B | 9.9e-09 | 1.1e-08 | 6e-09 | 8.967E-09 | 2.627E-09 | 6.000E-09 | 1.100E-08 |
| 14569 | 729 | K729H | B | 6.5e-09 | 1.5e-08 | 5.7e-09 | 9.067E-09 | 5.154E-09 | 5.700E-09 | 1.500E-08 |
| 14570 | 729 | K729I | B | 1.2e-08 | 1e-08 | 7.5e-09 | 9.833E-09 | 2.255E-09 | 7.500E-09 | 1.200E-08 |
| 14571 | 729 | K729L | B | 2.3e-09 | 8.2e-09 | 6.1e-09 | 5.533E-09 | 2.991E-09 | 2.300E-09 | 8.200E-09 |
| 14572 | 729 | K729K | B | 4.6e-09 | 1.4e-09 | 9.5e-09 | 5.167E-09 | 4.080E-09 | 1.400E-09 | 9.500E-09 |
| 14573 | 729 | K729M | B | 6.8e-09 | 4.7e-09 | 2.8e-08 | 1.317E-08 | 1.289E-08 | 4.700E-09 | 2.800E-08 |
| 14574 | 729 | K729F | B | 1.1e-08 | 1e-08 | 1.1e-08 | 1.067E-08 | 5.774E-10 | 1.000E-08 | 1.100E-08 |
| 14575 | 729 | K729P | B | 1.7e-09 | 4.5e-09 | 5e-09 | 3.733E-09 | 1.779E-09 | 1.700E-09 | 5.000E-09 |
| 14576 | 729 | K729S | B | 1.1e-08 | 7.6e-09 | 1.3e-08 | 1.053E-08 | 2.730E-09 | 7.600E-09 | 1.300E-08 |
| 14577 | 729 | K729T | B | 6.7e-09 | 2.4e-08 | 5.2e-09 | 1.197E-08 | 1.045E-08 | 5.200E-09 | 2.400E-08 |
| 14578 | 729 | K729W | B | 8.1e-09 | 6.8e-09 | 8.5e-09 | 7.800E-09 | 8.888E-10 | 6.800E-09 | 8.500E-09 |
| 14579 | 729 | K729Y | B | 5.8e-09 | 4.4e-09 | 5.7e-09 | 5.300E-09 | 7.810E-10 | 4.400E-09 | 5.800E-09 |
| 14580 | 729 | K729V | B | 2.5e-09 | 3.3e-09 | 1e-08 | 5.267E-09 | 4.119E-09 | 2.500E-09 | 1.000E-08 |
| 14581 | 730 | R730A | B | 5.6e-09 | 4.5e-09 | 9.4e-09 | 6.500E-09 | 2.571E-09 | 4.500E-09 | 9.400E-09 |
| 14582 | 730 | R730R | B | 3.7e-09 | 3.7e-09 | 5.1e-09 | 4.167E-09 | 8.083E-10 | 3.700E-09 | 5.100E-09 |
| 14583 | 730 | R730N | B | 4.4e-09 | 2.3e-08 | 4.4e-09 | 1.060E-08 | 1.074E-08 | 4.400E-09 | 2.300E-08 |
| 14584 | 730 | R730D | B | 7.5e-09 | 3.9e-09 | 8.6e-09 | 6.667E-09 | 2.458E-09 | 3.900E-09 | 8.600E-09 |
| 14585 | 730 | R730C | B | 9.5e-09 | 8.4e-09 | 9.8e-09 | 9.233E-09 | 7.371E-10 | 8.400E-09 | 9.800E-09 |
| 14586 | 730 | R730E | B | 7.3e-09 | 4e-09 | 8.5e-09 | 6.600E-09 | 2.330E-09 | 4.000E-09 | 8.500E-09 |
| 14587 | 730 | R730Q | B | 6.9e-09 | 4.8e-09 | 8.7e-09 | 6.800E-09 | 1.952E-09 | 4.800E-09 | 8.700E-09 |
| 14588 | 730 | R730G | B | 4.8e-09 | 8.2e-09 | 1.4e-08 | 9.000E-09 | 4.652E-09 | 4.800E-09 | 1.400E-08 |
| 14589 | 730 | R730H | B | 6.9e-09 | 3.8e-09 | 8.2e-09 | 6.300E-09 | 2.261E-09 | 3.800E-09 | 8.200E-09 |
| 14590 | 730 | R730I | B | 2.6e-09 | 1.3e-08 | 4.1e-09 | 6.567E-09 | 5.622E-09 | 2.600E-09 | 1.300E-08 |
| 14591 | 730 | R730L | B | 1e-08 | 3e-09 | 7.2e-09 | 6.733E-09 | 3.523E-09 | 3.000E-09 | 1.000E-08 |
| 14592 | 730 | R730K | B | 4.2e-09 | 9.6e-09 | 8.3e-09 | 7.367E-09 | 2.818E-09 | 4.200E-09 | 9.600E-09 |
| 14593 | 730 | R730M | B | 2.2e-08 | 7.3e-09 | 1.2e-08 | 1.377E-08 | 7.508E-09 | 7.300E-09 | 2.200E-08 |
| 14594 | 730 | R730F | B | 8.5e-09 | 7e-09 | 4.7e-09 | 6.733E-09 | 1.914E-09 | 4.700E-09 | 8.500E-09 |
| 14595 | 730 | R730P | B | 9.9e-09 | 1.1e-08 | 7.9e-09 | 9.600E-09 | 1.572E-09 | 7.900E-09 | 1.100E-08 |
| 14596 | 730 | R730S | B | 3.9e-09 | 6.2e-09 | 4.5e-09 | 4.867E-09 | 1.193E-09 | 3.900E-09 | 6.200E-09 |
| 14597 | 730 | R730T | B | 1.1e-08 | 3.3e-09 | 4.3e-09 | 6.200E-09 | 4.187E-09 | 3.300E-09 | 1.100E-08 |
| 14598 | 730 | R730W | B | 6.4e-09 | 2.6e-09 | 1.1e-08 | 6.667E-09 | 4.206E-09 | 2.600E-09 | 1.100E-08 |
| 14599 | 730 | R730Y | B | 1.1e-08 | 2.9e-09 | 5e-09 | 6.300E-09 | 4.204E-09 | 2.900E-09 | 1.100E-08 |
| 14600 | 730 | R730V | B | 1.3e-08 | 7e-09 | 2.5e-09 | 7.500E-09 | 5.268E-09 | 2.500E-09 | 1.300E-08 |
| 14601 | 731 | T731A | B | 3.7e-09 | 6.1e-09 | 3.2e-09 | 4.333E-09 | 1.550E-09 | 3.200E-09 | 6.100E-09 |
| 14602 | 731 | T731R | B | 8.5e-09 | 4.2e-09 | 2e-08 | 1.090E-08 | 8.169E-09 | 4.200E-09 | 2.000E-08 |
| 14603 | 731 | T731N | B | 1.4e-09 | 7.3e-09 | 1.9e-08 | 9.233E-09 | 8.958E-09 | 1.400E-09 | 1.900E-08 |
| 14604 | 731 | T731D | B | 4.2e-09 | 1.1e-08 | 2.9e-09 | 6.033E-09 | 4.350E-09 | 2.900E-09 | 1.100E-08 |
| 14605 | 731 | T731C | B | 1.3e-08 | 2.8e-09 | 5.3e-09 | 7.033E-09 | 5.316E-09 | 2.800E-09 | 1.300E-08 |
| 14606 | 731 | T731E | B | 1.1e-08 | 5.3e-09 | 6.7e-09 | 7.667E-09 | 2.970E-09 | 5.300E-09 | 1.100E-08 |
| 14607 | 731 | T731Q | B | 3.6e-08 | 3.2e-08 | 1e-08 | 2.600E-08 | 1.400E-08 | 1.000E-08 | 3.600E-08 |
| 14608 | 731 | T731G | B | 1.1e-08 | 8.4e-09 | 5.5e-09 | 8.300E-09 | 2.751E-09 | 5.500E-09 | 1.100E-08 |
| 14609 | 731 | T731H | B | 2.3e-09 | 3e-08 | 8.1e-09 | 1.347E-08 | 1.461E-08 | 2.300E-09 | 3.000E-08 |
| 14610 | 731 | T731I | B | 1.6e-08 | 1.8e-08 | 8.6e-09 | 1.420E-08 | 4.952E-09 | 8.600E-09 | 1.800E-08 |
| 14611 | 731 | T731L | B | 6.8e-09 | 1.4e-08 | 2.9e-09 | 7.900E-09 | 5.631E-09 | 2.900E-09 | 1.400E-08 |
| 14612 | 731 | T731K | B | 8e-09 | 7.2e-09 | 4.3e-09 | 6.500E-09 | 1.947E-09 | 4.300E-09 | 8.000E-09 |
| 14613 | 731 | T731M | B | 7.7e-09 | 1.1e-08 | 8.4e-09 | 9.033E-09 | 1.739E-09 | 7.700E-09 | 1.100E-08 |
| 14614 | 731 | T731F | B | 6.3e-09 | 9.3e-09 | 1.6e-08 | 1.053E-08 | 4.966E-09 | 6.300E-09 | 1.600E-08 |
| 14615 | 731 | T731P | B | 2.5e-09 | 1e-08 | 1e-08 | 7.500E-09 | 4.330E-09 | 2.500E-09 | 1.000E-08 |
| 14616 | 731 | T731S | B | 2.2e-08 | 1.3e-08 | 2.9e-09 | 1.263E-08 | 9.555E-09 | 2.900E-09 | 2.200E-08 |
| 14617 | 731 | T731T | B | 6e-09 | 1.2e-08 | 8.8e-09 | 8.933E-09 | 3.002E-09 | 6.000E-09 | 1.200E-08 |
| 14618 | 731 | T731W | B | 9.5e-09 | 1.7e-08 | 3.3e-09 | 9.933E-09 | 6.860E-09 | 3.300E-09 | 1.700E-08 |
| 14619 | 731 | T731Y | B | 2.5e-09 | 3.7e-09 | 8.1e-09 | 4.767E-09 | 2.948E-09 | 2.500E-09 | 8.100E-09 |

| | | | | | | | | | | |
|---|---|---|---|---|---|---|---|---|---|---|
| 14620 | 731 | T731V | B | 7.5e-09 | 7.4e-09 | 9e-09 | 7.967E-09 | 8.963E-10 | 7.400E-09 | 9.000E-09 |
| 14621 | 732 | V732A | B | 2.5e-09 | 8.2e-09 | 8.7e-10 | 3.857E-09 | 3.849E-09 | 8.700E-10 | 8.200E-09 |
| 14622 | 732 | V732R | B | 1.1e-08 | 1.2e-08 | 5.3e-09 | 9.433E-09 | 3.614E-09 | 5.300E-09 | 1.200E-08 |
| 14623 | 732 | V732N | B | 3e-09 | 7.8e-09 | 4.4e-09 | 5.067E-09 | 2.468E-09 | 3.000E-09 | 7.800E-09 |
| 14624 | 732 | V732D | B | 3.2e-09 | 6.5e-09 | 1.2e-08 | 7.233E-09 | 4.446E-09 | 3.200E-09 | 1.200E-08 |
| 14625 | 732 | V732C | B | 6.1e-09 | 5.3e-09 | 1.2e-08 | 7.800E-09 | 3.659E-09 | 5.300E-09 | 1.200E-08 |
| 14626 | 732 | V732E | B | 3.4e-08 | 1.2e-08 | 4.5e-09 | 1.683E-08 | 1.533E-08 | 4.500E-09 | 3.400E-08 |
| 14627 | 732 | V732Q | B | 7.4e-09 | 5.6e-09 | 2e-08 | 1.100E-08 | 7.846E-09 | 5.600E-09 | 2.000E-08 |
| 14628 | 732 | V732G | B | 6.8e-09 | 1e-08 | 5.2e-09 | 7.333E-09 | 2.444E-09 | 5.200E-09 | 1.000E-08 |
| 14629 | 732 | V732H | B | 1.8e-08 | 1.4e-08 | 1e-08 | 1.400E-08 | 4.000E-09 | 1.000E-08 | 1.800E-08 |
| 14630 | 732 | V732I | B | 3.1e-09 | 5.3e-09 | 1.1e-08 | 6.467E-09 | 4.077E-09 | 3.100E-09 | 1.100E-08 |
| 14631 | 732 | V732L | B | 9.5e-09 | 2.5e-08 | 4.9e-09 | 1.313E-08 | 1.053E-08 | 4.900E-09 | 2.500E-08 |
| 14632 | 732 | V732K | B | 6.3e-09 | 9.5e-09 | 5.7e-09 | 7.167E-09 | 2.043E-09 | 5.700E-09 | 9.500E-09 |
| 14633 | 732 | V732M | B | 6.8e-09 | 5.3e-09 | 3.1e-09 | 5.067E-09 | 1.861E-09 | 3.100E-09 | 6.800E-09 |
| 14634 | 732 | V732F | B | 8.7e-09 | 5.1e-09 | 1.8e-08 | 1.060E-08 | 6.657E-09 | 5.100E-09 | 1.800E-08 |
| 14635 | 732 | V732P | B | 4.5e-09 | 1.1e-08 | 7.6e-09 | 7.700E-09 | 3.251E-09 | 4.500E-09 | 1.100E-08 |
| 14636 | 732 | V732S | B | 7.1e-09 | 1.3e-08 | 2.4e-09 | 7.500E-09 | 5.311E-09 | 2.400E-09 | 1.300E-08 |
| 14637 | 732 | V732T | B | 2.1e-08 | 5.9e-09 | 7.4e-09 | 1.143E-08 | 8.319E-09 | 5.900E-09 | 2.100E-08 |
| 14638 | 732 | V732W | B | 1.4e-08 | 8.7e-09 | 7.3e-09 | 1.000E-08 | 3.534E-09 | 7.300E-09 | 1.400E-08 |
| 14639 | 732 | V732Y | B | 2.7e-09 | 5.9e-09 | 1.2e-08 | 6.867E-09 | 4.725E-09 | 2.700E-09 | 1.200E-08 |
| 14640 | 732 | V732V | B | 1e-08 | 1.1e-08 | 6.4e-09 | 9.133E-09 | 2.419E-09 | 6.400E-09 | 1.100E-08 |
| 14641 | 733 | A733A | B | 3.2e-09 | 9.6e-09 | 4.2e-09 | 5.667E-09 | 3.443E-09 | 3.200E-09 | 9.600E-09 |
| 14642 | 733 | A733R | B | 9.4e-09 | 1.2e-08 | 9e-09 | 1.013E-08 | 1.629E-09 | 9.000E-09 | 1.200E-08 |
| 14643 | 733 | A733N | B | 1.2e-08 | 2.6e-09 | 5.2e-09 | 6.600E-09 | 4.854E-09 | 2.600E-09 | 1.200E-08 |
| 14644 | 733 | A733D | B | 1.1e-08 | 3.1e-09 | 6e-09 | 6.700E-09 | 3.996E-09 | 3.100E-09 | 1.100E-08 |
| 14645 | 733 | A733C | B | 1.4e-08 | 6.8e-09 | 9.7e-09 | 1.017E-08 | 3.623E-09 | 6.800E-09 | 1.400E-08 |
| 14646 | 733 | A733E | B | 5.9e-08 | 6.6e-09 | 2.1e-08 | 2.887E-08 | 2.707E-08 | 6.600E-09 | 5.900E-08 |
| 14647 | 733 | A733Q | B | 6.1e-09 | 1.4e-08 | 1.4e-08 | 1.137E-08 | 4.561E-09 | 6.100E-09 | 1.400E-08 |
| 14648 | 733 | A733G | B | 1.2e-08 | 9.8e-09 | 1.5e-08 | 1.227E-08 | 2.610E-09 | 9.800E-09 | 1.500E-08 |
| 14649 | 733 | A733H | B | 4.8e-09 | 7.2e-09 | 1.6e-08 | 9.333E-09 | 5.897E-09 | 4.800E-09 | 1.600E-08 |
| 14650 | 733 | A733I | B | 3.2e-09 | 2.5e-09 | 1.5e-08 | 6.900E-09 | 7.024E-09 | 2.500E-09 | 1.500E-08 |
| 14651 | 733 | A733L | B | 1.1e-08 | 1.2e-08 | 3.3e-09 | 8.767E-09 | 4.761E-09 | 3.300E-09 | 1.200E-08 |
| 14652 | 733 | A733K | B | 8.1e-09 | 5.1e-09 | 9.9e-09 | 7.700E-09 | 2.425E-09 | 5.100E-09 | 9.900E-09 |
| 14653 | 733 | A733M | B | 4.4e-09 | 8.3e-09 | 7.9e-09 | 6.867E-09 | 2.146E-09 | 4.400E-09 | 8.300E-09 |
| 14654 | 733 | A733F | B | 7.3e-09 | 7.2e-09 | 7.4e-09 | 7.300E-09 | 1.000E-10 | 7.200E-09 | 7.400E-09 |
| 14655 | 733 | A733P | B | 6.2e-09 | 1.2e-08 | 5.5e-09 | 7.900E-09 | 3.568E-09 | 5.500E-09 | 1.200E-08 |
| 14656 | 733 | A733S | B | 2.9e-09 | 2e-08 | 1.3e-08 | 1.197E-08 | 8.597E-09 | 2.900E-09 | 2.000E-08 |
| 14657 | 733 | A733T | B | 3.8e-09 | 2e-09 | 8.7e-09 | 4.833E-09 | 3.467E-09 | 2.000E-09 | 8.700E-09 |
| 14658 | 733 | A733W | B | 9.8e-09 | 3.9e-09 | 7e-09 | 6.900E-09 | 2.951E-09 | 3.900E-09 | 9.800E-09 |
| 14659 | 733 | A733Y | B | 7.3e-09 | 2.6e-09 | 6.9e-09 | 5.600E-09 | 2.606E-09 | 2.600E-09 | 7.300E-09 |
| 14660 | 733 | A733V | B | 7.7e-09 | 3.5e-09 | 1e-08 | 7.067E-09 | 3.296E-09 | 3.500E-09 | 1.000E-08 |
| 14661 | 734 | A734A | B | 8.4e-09 | 5.2e-09 | 5.7e-09 | 6.433E-09 | 1.721E-09 | 5.200E-09 | 8.400E-09 |
| 14662 | 734 | A734R | B | 8.3e-09 | 9.2e-09 | 7.6e-09 | 8.367E-09 | 8.021E-10 | 7.600E-09 | 9.200E-09 |
| 14663 | 734 | A734N | B | 3.2e-09 | 5.5e-09 | 5.6e-09 | 4.767E-09 | 1.358E-09 | 3.200E-09 | 5.600E-09 |
| 14664 | 734 | A734D | B | 4e-09 | 3.4e-09 | 1.7e-08 | 8.133E-09 | 7.685E-09 | 3.400E-09 | 1.700E-08 |
| 14665 | 734 | A734C | B | 8.8e-09 | 6.3e-09 | 1.6e-08 | 1.037E-08 | 5.036E-09 | 6.300E-09 | 1.600E-08 |
| 14666 | 734 | A734E | B | 2.6e-08 | 4.4e-09 | 9.2e-09 | 1.320E-08 | 1.134E-08 | 4.400E-09 | 2.600E-08 |
| 14667 | 734 | A734Q | B | 7.1e-09 | 1.4e-08 | 1.4e-08 | 1.170E-08 | 3.984E-09 | 7.100E-09 | 1.400E-08 |
| 14668 | 734 | A734G | B | 3.8e-09 | 2.3e-09 | 1.1e-08 | 5.700E-09 | 4.651E-09 | 2.300E-09 | 1.100E-08 |
| 14669 | 734 | A734H | B | 7.1e-09 | 3.9e-09 | 1.8e-09 | 4.267E-09 | 2.669E-09 | 1.800E-09 | 7.100E-09 |
| 14670 | 734 | A734I | B | 7.7e-09 | 4.6e-09 | 3.7e-09 | 5.333E-09 | 2.098E-09 | 3.700E-09 | 7.700E-09 |
| 14671 | 734 | A734L | B | 1.8e-08 | 8.6e-09 | 2.9e-09 | 9.833E-09 | 7.625E-09 | 2.900E-09 | 1.800E-08 |
| 14672 | 734 | A734K | B | 2e-08 | 7.1e-09 | 7.2e-09 | 1.143E-08 | 7.419E-09 | 7.100E-09 | 2.000E-08 |
| 14673 | 734 | A734M | B | 1.1e-08 | 6.5e-09 | 6.4e-09 | 7.967E-09 | 2.627E-09 | 6.400E-09 | 1.100E-08 |
| 14674 | 734 | A734F | B | 1.8e-08 | 5.2e-09 | 3.2e-09 | 8.800E-09 | 8.030E-09 | 3.200E-09 | 1.800E-08 |
| 14675 | 734 | A734P | B | 5.9e-09 | 1.3e-08 | 2.9e-09 | 7.267E-09 | 5.187E-09 | 2.900E-09 | 1.300E-08 |
| 14676 | 734 | A734S | B | 7.6e-09 | 8.3e-09 | 4.9e-09 | 6.933E-09 | 1.795E-09 | 4.900E-09 | 8.300E-09 |
| 14677 | 734 | A734T | B | 2.1e-09 | 2.9e-09 | 4.6e-09 | 3.200E-09 | 1.277E-09 | 2.100E-09 | 4.600E-09 |
| 14678 | 734 | A734W | B | 4.1e-09 | 7.6e-09 | 2.2e-09 | 4.633E-09 | 2.739E-09 | 2.200E-09 | 7.600E-09 |
| 14679 | 734 | A734Y | B | 9.1e-09 | 1.2e-08 | 1.1e-08 | 1.070E-08 | 1.473E-09 | 9.100E-09 | 1.200E-08 |
| 14680 | 734 | A734V | B | 4.3e-09 | 6.9e-09 | 5.7e-09 | 5.633E-09 | 1.301E-09 | 4.300E-09 | 6.900E-09 |
| 14681 | 735 | P735A | B | 3.9e-09 | 1.3e-08 | 7.2e-09 | 8.033E-09 | 4.607E-09 | 3.900E-09 | 1.300E-08 |
| 14682 | 735 | P735R | B | 2.4e-09 | 5.5e-09 | 7.9e-09 | 5.267E-09 | 2.757E-09 | 2.400E-09 | 7.900E-09 |
| 14683 | 735 | P735N | B | 7.1e-09 | 6.2e-09 | 1.5e-08 | 9.433E-09 | 4.842E-09 | 6.200E-09 | 1.500E-08 |
| 14684 | 735 | P735D | B | 1.8e-08 | 9.8e-09 | 1.3e-08 | 1.360E-08 | 4.133E-09 | 9.800E-09 | 1.800E-08 |
| 14685 | 735 | P735C | B | 8.6e-09 | 3e-09 | 5e-09 | 5.533E-09 | 2.838E-09 | 3.000E-09 | 8.600E-09 |
| 14686 | 735 | P735E | B | 7.5e-09 | 5.7e-09 | 4.2e-09 | 5.800E-09 | 1.652E-09 | 4.200E-09 | 7.500E-09 |
| 14687 | 735 | P735Q | B | 8.8e-09 | 1.4e-08 | 1.2e-08 | 1.160E-08 | 2.623E-09 | 8.800E-09 | 1.400E-08 |
| 14688 | 735 | P735G | B | 7.7e-09 | 6.1e-09 | 4e-09 | 5.933E-09 | 1.856E-09 | 4.000E-09 | 7.700E-09 |
| 14689 | 735 | P735H | B | 7.9e-09 | 5.2e-09 | 6.9e-09 | 6.667E-09 | 1.365E-09 | 5.200E-09 | 7.900E-09 |
| 14690 | 735 | P735I | B | 6.3e-09 | 1.5e-08 | 6.9e-09 | 9.400E-09 | 4.859E-09 | 6.300E-09 | 1.500E-08 |
| 14691 | 735 | P735L | B | 2.7e-09 | 5.9e-09 | 7.9e-09 | 5.500E-09 | 2.623E-09 | 2.700E-09 | 7.900E-09 |
| 14692 | 735 | P735K | B | 5.9e-09 | 3.5e-09 | 7.9e-09 | 5.767E-09 | 2.203E-09 | 3.500E-09 | 7.900E-09 |
| 14693 | 735 | P735M | B | 1.6e-08 | 3.2e-09 | 7.7e-09 | 8.967E-09 | 6.493E-09 | 3.200E-09 | 1.600E-08 |
| 14694 | 735 | P735F | B | 1.1e-08 | 1.9e-08 | 3.8e-09 | 1.127E-08 | 7.604E-09 | 3.800E-09 | 1.900E-08 |
| 14695 | 735 | P735P | B | 4.2e-09 | 8e-09 | 8.5e-09 | 6.900E-09 | 2.352E-09 | 4.200E-09 | 8.500E-09 |
| 14696 | 735 | P735S | B | 3.1e-09 | 7.4e-09 | 1e-08 | 6.833E-09 | 3.485E-09 | 3.100E-09 | 1.000E-08 |
| 14697 | 735 | P735T | B | 1.7e-08 | 1.1e-08 | 1.1e-08 | 1.300E-08 | 3.464E-09 | 1.100E-08 | 1.700E-08 |
| 14698 | 735 | P735W | B | 4.7e-09 | 2e-09 | 5.5e-09 | 4.067E-09 | 1.834E-09 | 2.000E-09 | 5.500E-09 |
| 14699 | 735 | P735Y | B | 7e-09 | 8.2e-09 | 6.8e-09 | 7.333E-09 | 7.572E-10 | 6.800E-09 | 8.200E-09 |
| 14700 | 735 | P735V | B | 9.1e-09 | 1.4e-08 | 1.1e-08 | 1.137E-08 | 2.470E-09 | 9.100E-09 | 1.400E-08 |
| 14701 | 736 | S736A | B | 1.7e-09 | 3.8e-09 | 6.3e-09 | 3.933E-09 | 2.303E-09 | 1.700E-09 | 6.300E-09 |
| 14702 | 736 | S736R | B | 7.9e-09 | 1e-08 | 4.2e-09 | 7.367E-09 | 2.937E-09 | 4.200E-09 | 1.000E-08 |
| 14703 | 736 | S736N | B | 5.6e-09 | 6e-09 | 1.1e-08 | 7.533E-09 | 3.009E-09 | 5.600E-09 | 1.100E-08 |
| 14704 | 736 | S736D | B | 4.8e-09 | 4e-09 | 6e-09 | 4.933E-09 | 1.007E-09 | 4.000E-09 | 6.000E-09 |

| | | | | | | | | | | |
|---|---|---|---|---|---|---|---|---|---|---|
| 14705 | 736 | S736C | B | 1.6e-08 | 3.4e-09 | 3.5e-09 | 7.633E-09 | 7.246E-09 | 3.400E-09 | 1.600E-08 |
| 14706 | 736 | S736E | B | 6e-09 | 5.5e-09 | 5.4e-09 | 5.633E-09 | 3.215E-10 | 5.400E-09 | 6.000E-09 |
| 14707 | 736 | S736Q | B | 7e-09 | 7.2e-09 | 3.9e-09 | 6.033E-09 | 1.850E-09 | 3.900E-09 | 7.200E-09 |
| 14708 | 736 | S736G | B | 1.9e-08 | 6.1e-09 | 1.3e-08 | 1.270E-08 | 6.455E-09 | 6.100E-09 | 1.900E-08 |
| 14709 | 736 | S736H | B | 4.8e-09 | 6.2e-09 | 8.3e-09 | 6.433E-09 | 1.762E-09 | 4.800E-09 | 8.300E-09 |
| 14710 | 736 | S736I | B | 4e-09 | 1.3e-08 | 1e-08 | 9.000E-09 | 4.583E-09 | 4.000E-09 | 1.300E-08 |
| 14711 | 736 | S736L | B | 1e-08 | 3.2e-09 | 2.4e-08 | 1.240E-08 | 1.061E-08 | 3.200E-09 | 2.400E-08 |
| 14712 | 736 | S736K | B | 1.1e-08 | 4.5e-09 | 6.9e-09 | 7.467E-09 | 3.287E-09 | 4.500E-09 | 1.100E-08 |
| 14713 | 736 | S736M | B | 1.8e-08 | 6.2e-09 | 4.8e-09 | 9.667E-09 | 7.251E-09 | 4.800E-09 | 1.800E-08 |
| 14714 | 736 | S736F | B | 1.8e-08 | 7.7e-09 | 4.6e-09 | 1.010E-08 | 7.015E-09 | 4.600E-09 | 1.800E-08 |
| 14715 | 736 | S736P | B | 5.6e-09 | 1e-08 | 3.5e-09 | 6.367E-09 | 3.317E-09 | 3.500E-09 | 1.000E-08 |
| 14716 | 736 | S736S | B | 5.4e-09 | 6.3e-09 | 1.1e-08 | 7.567E-09 | 3.007E-09 | 5.400E-09 | 1.100E-08 |
| 14717 | 736 | S736T | B | 6.4e-09 | 6.5e-09 | 9.2e-09 | 7.367E-09 | 1.589E-09 | 6.400E-09 | 9.200E-09 |
| 14718 | 736 | S736W | B | 7.2e-09 | 1.4e-09 | 3.3e-09 | 3.967E-09 | 2.957E-09 | 1.400E-09 | 7.200E-09 |
| 14719 | 736 | S736Y | B | 2.8e-09 | 6.2e-09 | 7.3e-09 | 5.433E-09 | 2.346E-09 | 2.800E-09 | 7.300E-09 |
| 14720 | 736 | S736V | B | 5.8e-09 | 6.8e-09 | 4.3e-09 | 5.633E-09 | 1.258E-09 | 4.300E-09 | 6.800E-09 |
| 14721 | 737 | V737A | B | 1.1e-08 | 2.1e-08 | 6e-09 | 1.267E-08 | 7.638E-09 | 6.000E-09 | 2.100E-08 |
| 14722 | 737 | V737R | B | 1e-08 | 1.9e-09 | 6.3e-09 | 6.067E-09 | 4.055E-09 | 1.900E-09 | 1.000E-08 |
| 14723 | 737 | V737N | B | 9.9e-09 | 1.4e-08 | 8.6e-09 | 1.083E-08 | 2.818E-09 | 8.600E-09 | 1.400E-08 |
| 14724 | 737 | V737D | B | 1.3e-08 | 9.3e-09 | 7.8e-09 | 1.003E-08 | 2.676E-09 | 7.800E-09 | 1.300E-08 |
| 14725 | 737 | V737C | B | 7.7e-09 | 1.3e-08 | 8.5e-09 | 9.733E-09 | 2.857E-09 | 7.700E-09 | 1.300E-08 |
| 14726 | 737 | V737E | B | 5.8e-09 | 1.4e-08 | 1.8e-08 | 1.260E-08 | 6.219E-09 | 5.800E-09 | 1.800E-08 |
| 14727 | 737 | V737Q | B | 5.1e-09 | 6.3e-09 | 8.2e-09 | 6.533E-09 | 1.563E-09 | 5.100E-09 | 8.200E-09 |
| 14728 | 737 | V737G | B | 1.2e-08 | 6.3e-09 | 6e-09 | 8.100E-09 | 3.381E-09 | 6.000E-09 | 1.200E-08 |
| 14729 | 737 | V737H | B | 6.8e-09 | 2.4e-09 | 6.5e-09 | 5.233E-09 | 2.458E-09 | 2.400E-09 | 6.800E-09 |
| 14730 | 737 | V737I | B | 5.4e-09 | 4e-09 | 1.3e-08 | 7.467E-09 | 4.843E-09 | 4.000E-09 | 1.300E-08 |
| 14731 | 737 | V737L | B | 6.2e-09 | 6.8e-09 | 5.7e-09 | 6.233E-09 | 5.508E-10 | 5.700E-09 | 6.800E-09 |
| 14732 | 737 | V737K | B | 1e-08 | 2.9e-09 | 5.6e-09 | 6.167E-09 | 3.584E-09 | 2.900E-09 | 1.000E-08 |
| 14733 | 737 | V737M | B | 1.4e-08 | 9.5e-09 | 8.1e-09 | 1.053E-08 | 3.083E-09 | 8.100E-09 | 1.400E-08 |
| 14734 | 737 | V737F | B | 9.4e-09 | 1.4e-08 | 1.4e-08 | 1.247E-08 | 2.656E-09 | 9.400E-09 | 1.400E-08 |
| 14735 | 737 | V737P | B | 2.4e-08 | 9.3e-09 | 4.8e-09 | 1.270E-08 | 1.004E-08 | 4.800E-09 | 2.400E-08 |
| 14736 | 737 | V737S | B | 6.4e-09 | 4.3e-09 | 7.1e-09 | 5.933E-09 | 1.457E-09 | 4.300E-09 | 7.100E-09 |
| 14737 | 737 | V737T | B | 5.1e-09 | 5e-09 | 7.5e-09 | 5.867E-09 | 1.415E-09 | 5.000E-09 | 7.500E-09 |
| 14738 | 737 | V737W | B | 1e-08 | 8e-09 | 2e-08 | 1.267E-08 | 6.429E-09 | 8.000E-09 | 2.000E-08 |
| 14739 | 737 | V737Y | B | 1.4e-08 | 1.2e-08 | 3.3e-09 | 9.767E-09 | 5.689E-09 | 3.300E-09 | 1.400E-08 |
| 14740 | 737 | V737V | B | 6.8e-09 | 4.5e-09 | 2.5e-09 | 4.600E-09 | 2.152E-09 | 2.500E-09 | 6.800E-09 |
| 14741 | 738 | F738A | B | 5.6e-09 | 9.1e-09 | 1.1e-08 | 8.567E-09 | 2.739E-09 | 5.600E-09 | 1.100E-08 |
| 14742 | 738 | F738R | B | 3.9e-09 | 4.2e-09 | 1.3e-08 | 7.033E-09 | 5.169E-09 | 3.900E-09 | 1.300E-08 |
| 14743 | 738 | F738N | B | 6.7e-09 | 9.8e-09 | 4.7e-09 | 7.067E-09 | 2.570E-09 | 4.700E-09 | 9.800E-09 |
| 14744 | 738 | F738D | B | 3.7e-09 | 1.5e-08 | 4.6e-09 | 7.767E-09 | 6.280E-09 | 3.700E-09 | 1.500E-08 |
| 14745 | 738 | F738C | B | 1.1e-08 | 5.4e-09 | 4.3e-09 | 6.900E-09 | 3.593E-09 | 4.300E-09 | 1.100E-08 |
| 14746 | 738 | F738E | B | 3e-09 | 4.1e-09 | 4.5e-09 | 3.867E-09 | 7.767E-10 | 3.000E-09 | 4.500E-09 |
| 14747 | 738 | F738Q | B | 1.3e-09 | 1.2e-08 | 4.3e-09 | 5.867E-09 | 5.519E-09 | 1.300E-09 | 1.200E-08 |
| 14748 | 738 | F738G | B | 7e-09 | 1.3e-08 | 8.7e-09 | 9.567E-09 | 3.092E-09 | 7.000E-09 | 1.300E-08 |
| 14749 | 738 | F738H | B | 7.2e-09 | 2.7e-09 | 1.7e-08 | 8.967E-09 | 7.312E-09 | 2.700E-09 | 1.700E-08 |
| 14750 | 738 | F738I | B | 8.9e-09 | 1.2e-08 | 4.2e-09 | 8.367E-09 | 3.927E-09 | 4.200E-09 | 1.200E-08 |
| 14751 | 738 | F738L | B | 6e-09 | 3.9e-09 | 4.7e-09 | 4.867E-09 | 1.060E-09 | 3.900E-09 | 6.000E-09 |
| 14752 | 738 | F738K | B | 8.6e-09 | 3.1e-09 | 6.7e-09 | 6.133E-09 | 2.793E-09 | 3.100E-09 | 8.600E-09 |
| 14753 | 738 | F738M | B | 6.5e-09 | 5.3e-09 | 2.2e-08 | 1.127E-08 | 9.315E-09 | 5.300E-09 | 2.200E-08 |
| 14754 | 738 | F738F | B | 4.3e-09 | 2.7e-08 | 1.2e-08 | 1.443E-08 | 1.154E-08 | 4.300E-09 | 2.700E-08 |
| 14755 | 738 | F738P | B | 1.1e-08 | 2.9e-09 | 2.2e-09 | 5.367E-09 | 4.891E-09 | 2.200E-09 | 1.100E-08 |
| 14756 | 738 | F738S | B | 8.8e-09 | 2.8e-09 | 2.5e-09 | 4.700E-09 | 3.554E-09 | 2.500E-09 | 8.800E-09 |
| 14757 | 738 | F738T | B | 1.5e-08 | 7.2e-09 | 3.4e-09 | 8.533E-09 | 5.914E-09 | 3.400E-09 | 1.500E-08 |
| 14758 | 738 | F738W | B | 4.6e-09 | 8.2e-09 | 5.3e-09 | 6.033E-09 | 1.909E-09 | 4.600E-09 | 8.200E-09 |
| 14759 | 738 | F738Y | B | 7.7e-09 | 2.2e-09 | 5e-09 | 4.967E-09 | 2.750E-09 | 2.200E-09 | 7.700E-09 |
| 14760 | 738 | F738V | B | 9.4e-09 | 1.3e-08 | 9.1e-09 | 1.050E-08 | 2.170E-09 | 9.100E-09 | 1.300E-08 |
| 14761 | 739 | I739A | B | 1.2e-08 | 6.3e-09 | 1.3e-08 | 1.043E-08 | 3.614E-09 | 6.300E-09 | 1.300E-08 |
| 14762 | 739 | I739R | B | 7e-09 | 8.3e-09 | 1.2e-09 | 5.500E-09 | 3.780E-09 | 1.200E-09 | 8.300E-09 |
| 14763 | 739 | I739N | B | 1.3e-08 | 1.4e-08 | 8.9e-09 | 1.197E-08 | 2.702E-09 | 8.900E-09 | 1.400E-08 |
| 14764 | 739 | I739D | B | 7e-09 | 1.6e-08 | 2.6e-09 | 8.533E-09 | 6.830E-09 | 2.600E-09 | 1.600E-08 |
| 14765 | 739 | I739C | B | 1.2e-09 | 1.2e-08 | 2e-09 | 5.067E-09 | 6.018E-09 | 1.200E-09 | 1.200E-08 |
| 14766 | 739 | I739E | B | 4.2e-09 | 1.7e-08 | 9e-09 | 1.007E-08 | 6.466E-09 | 4.200E-09 | 1.700E-08 |
| 14767 | 739 | I739Q | B | 5.3e-09 | 6e-09 | 1e-08 | 7.100E-09 | 2.536E-09 | 5.300E-09 | 1.000E-08 |
| 14768 | 739 | I739G | B | 7.7e-09 | 1.6e-09 | 3.6e-09 | 4.300E-09 | 3.110E-09 | 1.600E-09 | 7.700E-09 |
| 14769 | 739 | I739H | B | 3.7e-09 | 9.7e-09 | 8.3e-09 | 7.233E-09 | 3.139E-09 | 3.700E-09 | 9.700E-09 |
| 14770 | 739 | I739I | B | 8.1e-09 | 7.9e-09 | 8e-09 | 8.000E-09 | 1.000E-10 | 7.900E-09 | 8.100E-09 |
| 14771 | 739 | I739L | B | 1.5e-08 | 1.9e-08 | 6.9e-09 | 1.363E-08 | 6.165E-09 | 6.900E-09 | 1.900E-08 |
| 14772 | 739 | I739K | B | 5.7e-09 | 6.5e-09 | 8.6e-09 | 6.933E-09 | 1.498E-09 | 5.700E-09 | 8.600E-09 |
| 14773 | 739 | I739M | B | 6.9e-09 | 3.6e-09 | 1.1e-08 | 7.167E-09 | 3.707E-09 | 3.600E-09 | 1.100E-08 |
| 14774 | 739 | I739F | B | 1.1e-08 | 7.7e-09 | 8.6e-09 | 9.100E-09 | 1.706E-09 | 7.700E-09 | 1.100E-08 |
| 14775 | 739 | I739P | B | 1.3e-08 | 4.4e-09 | 5e-09 | 7.467E-09 | 4.801E-09 | 4.400E-09 | 1.300E-08 |
| 14776 | 739 | I739S | B | 1.8e-08 | 8.5e-10 | 5.3e-09 | 8.050E-09 | 8.900E-09 | 8.500E-10 | 1.800E-08 |
| 14777 | 739 | I739T | B | 3.7e-09 | 6.5e-09 | 5.2e-09 | 5.133E-09 | 1.401E-09 | 3.700E-09 | 6.500E-09 |
| 14778 | 739 | I739W | B | 3.2e-09 | 1.3e-08 | 2.7e-09 | 6.300E-09 | 5.808E-09 | 2.700E-09 | 1.300E-08 |
| 14779 | 739 | I739Y | B | 9.9e-09 | 1.4e-08 | 4e-09 | 9.300E-09 | 5.027E-09 | 4.000E-09 | 1.400E-08 |
| 14780 | 739 | I739V | B | 1.1e-08 | 4.7e-09 | 4.4e-09 | 6.700E-09 | 3.727E-09 | 4.400E-09 | 1.100E-08 |
| 14781 | 740 | F740A | B | 6.3e-09 | 8.6e-09 | 2.3e-08 | 1.263E-08 | 9.051E-09 | 6.300E-09 | 2.300E-08 |
| 14782 | 740 | F740R | B | 6.8e-09 | 4.8e-09 | 3.9e-09 | 5.167E-09 | 1.484E-09 | 3.900E-09 | 6.800E-09 |
| 14783 | 740 | F740N | B | 9.2e-09 | 4.9e-09 | 5.1e-09 | 6.400E-09 | 2.427E-09 | 4.900E-09 | 9.200E-09 |
| 14784 | 740 | F740D | B | 5.9e-09 | 5e-09 | 9.3e-09 | 6.733E-09 | 2.268E-09 | 5.000E-09 | 9.300E-09 |
| 14785 | 740 | F740C | B | 1.1e-08 | 3.1e-09 | 6.7e-09 | 6.933E-09 | 3.955E-09 | 3.100E-09 | 1.100E-08 |
| 14786 | 740 | F740E | B | 3e-09 | 5.1e-09 | 9.6e-09 | 5.900E-09 | 3.372E-09 | 3.000E-09 | 9.600E-09 |
| 14787 | 740 | F740Q | B | 6.3e-09 | 1.6e-08 | 5.3e-09 | 9.200E-09 | 5.910E-09 | 5.300E-09 | 1.600E-08 |
| 14788 | 740 | F740G | B | 7.9e-09 | 1.5e-08 | 8.9e-09 | 1.060E-08 | 3.843E-09 | 7.900E-09 | 1.500E-08 |
| 14789 | 740 | F740H | B | 2.7e-08 | 1.5e-08 | 3.9e-09 | 1.530E-08 | 1.155E-08 | 3.900E-09 | 2.700E-08 |

| | | | | | | | | | | |
|---|---|---|---|---|---|---|---|---|---|---|
| 14790 | 740 | F740I | B | 7.6e-09 | 7.3e-09 | 3.5e-09 | 6.133E-09 | 2.285E-09 | 3.500E-09 | 7.600E-09 |
| 14791 | 740 | F740L | B | 8.9e-09 | 1.2e-08 | 9.7e-09 | 1.020E-08 | 1.609E-09 | 8.900E-09 | 1.200E-08 |
| 14792 | 740 | F740K | B | 3.8e-09 | 6.1e-09 | 1.4e-08 | 7.967E-09 | 5.350E-09 | 3.800E-09 | 1.400E-08 |
| 14793 | 740 | F740M | B | 7.3e-09 | 1.2e-08 | 1e-08 | 9.767E-09 | 2.359E-09 | 7.300E-09 | 1.200E-08 |
| 14794 | 740 | F740F | B | 8.7e-09 | 6.7e-09 | 8.9e-09 | 8.100E-09 | 1.217E-09 | 6.700E-09 | 8.900E-09 |
| 14795 | 740 | F740P | B | 1.5e-08 | 1e-08 | 1.4e-08 | 1.300E-08 | 2.646E-09 | 1.000E-08 | 1.500E-08 |
| 14796 | 740 | F740S | B | 2.5e-09 | 2.3e-08 | 1.1e-08 | 1.217E-08 | 1.030E-08 | 2.500E-09 | 2.300E-08 |
| 14797 | 740 | F740T | B | 3.7e-09 | 1.5e-08 | 1.2e-08 | 1.023E-08 | 5.853E-09 | 3.700E-09 | 1.500E-08 |
| 14798 | 740 | F740W | B | 2.3e-08 | 7.8e-09 | 1.3e-08 | 1.460E-08 | 7.725E-09 | 7.800E-09 | 2.300E-08 |
| 14799 | 740 | F740Y | B | 6.7e-09 | 1.7e-08 | 1.3e-08 | 1.223E-08 | 5.193E-09 | 6.700E-09 | 1.700E-08 |
| 14800 | 740 | F740V | B | 6.3e-09 | 6.9e-09 | 7.1e-09 | 6.767E-09 | 4.163E-10 | 6.300E-09 | 7.100E-09 |
| 14801 | 741 | P741A | B | 1.5e-08 | 1.1e-08 | 1.8e-08 | 1.467E-08 | 3.512E-09 | 1.100E-08 | 1.800E-08 |
| 14802 | 741 | P741R | B | 2.9e-09 | 4.1e-09 | 5.2e-09 | 4.067E-09 | 1.150E-09 | 2.900E-09 | 5.200E-09 |
| 14803 | 741 | P741N | B | 8.9e-09 | 4.5e-09 | 6.4e-09 | 6.600E-09 | 2.207E-09 | 4.500E-09 | 8.900E-09 |
| 14804 | 741 | P741D | B | 1.2e-08 | 1.7e-08 | 9.6e-09 | 1.287E-08 | 3.775E-09 | 9.600E-09 | 1.700E-08 |
| 14805 | 741 | P741C | B | 1.2e-08 | 5.5e-09 | 1e-08 | 9.167E-09 | 3.329E-09 | 5.500E-09 | 1.200E-08 |
| 14806 | 741 | P741E | B | 6.3e-09 | 1.1e-08 | 9e-09 | 8.767E-09 | 2.359E-09 | 6.300E-09 | 1.100E-08 |
| 14807 | 741 | P741Q | B | 4.6e-09 | 2.8e-08 | 8e-09 | 1.353E-08 | 1.264E-08 | 4.600E-09 | 2.800E-08 |
| 14808 | 741 | P741G | B | 1.7e-08 | 4.5e-09 | 1.3e-08 | 1.150E-08 | 6.384E-09 | 4.500E-09 | 1.700E-08 |
| 14809 | 741 | P741H | B | 6.6e-09 | 9.7e-09 | 4.7e-09 | 7.000E-09 | 2.524E-09 | 4.700E-09 | 9.700E-09 |
| 14810 | 741 | P741I | B | 1.5e-09 | 4.1e-09 | 1.4e-08 | 6.533E-09 | 6.596E-09 | 1.500E-09 | 1.400E-08 |
| 14811 | 741 | P741L | B | 2.5e-09 | 4.7e-09 | 1.5e-08 | 7.400E-09 | 6.673E-09 | 2.500E-09 | 1.500E-08 |
| 14812 | 741 | P741K | B | 1.6e-08 | 1.9e-09 | 4.2e-09 | 7.367E-09 | 7.565E-09 | 1.900E-09 | 1.600E-08 |
| 14813 | 741 | P741M | B | 5.7e-09 | 4.1e-09 | 3.2e-09 | 4.333E-09 | 1.266E-09 | 3.200E-09 | 5.700E-09 |
| 14814 | 741 | P741F | B | 5.1e-09 | 3.5e-09 | 7.9e-09 | 5.500E-09 | 2.227E-09 | 3.500E-09 | 7.900E-09 |
| 14815 | 741 | P741P | B | 8.2e-09 | 7.2e-09 | 5.3e-09 | 6.900E-09 | 1.473E-09 | 5.300E-09 | 8.200E-09 |
| 14816 | 741 | P741S | B | 6.1e-09 | 2.2e-09 | 1.2e-08 | 6.767E-09 | 4.934E-09 | 2.200E-09 | 1.200E-08 |
| 14817 | 741 | P741T | B | 5.7e-09 | 5.5e-09 | 4.3e-09 | 5.167E-09 | 7.572E-10 | 4.300E-09 | 5.700E-09 |
| 14818 | 741 | P741W | B | 6e-09 | 1.2e-08 | 3.5e-09 | 7.167E-09 | 4.368E-09 | 3.500E-09 | 1.200E-08 |
| 14819 | 741 | P741Y | B | 8.5e-09 | 4.6e-09 | 5.8e-09 | 6.300E-09 | 1.997E-09 | 4.600E-09 | 8.500E-09 |
| 14820 | 741 | P741V | B | 9.5e-09 | 1.2e-08 | 3.4e-09 | 8.300E-09 | 4.424E-09 | 3.400E-09 | 1.200E-08 |
| 14821 | 742 | P742A | B | 4.9e-09 | 1.4e-08 | 2.7e-09 | 7.200E-09 | 5.991E-09 | 2.700E-09 | 1.400E-08 |
| 14822 | 742 | P742R | B | 1.5e-08 | 1.3e-08 | 5e-09 | 1.100E-08 | 5.292E-09 | 5.000E-09 | 1.500E-08 |
| 14823 | 742 | P742N | B | 2e-08 | 7.2e-09 | 7.1e-09 | 1.143E-08 | 7.419E-09 | 7.100E-09 | 2.000E-08 |
| 14824 | 742 | P742D | B | 1.5e-08 | 1.2e-08 | 5.6e-09 | 1.087E-08 | 4.801E-09 | 5.600E-09 | 1.500E-08 |
| 14825 | 742 | P742C | B | 3.2e-09 | 5.9e-09 | 8.2e-09 | 5.767E-09 | 2.503E-09 | 3.200E-09 | 8.200E-09 |
| 14826 | 742 | P742E | B | 1.5e-08 | 1.2e-08 | 2.1e-08 | 1.600E-08 | 4.583E-09 | 1.200E-08 | 2.100E-08 |
| 14827 | 742 | P742Q | B | 7.3e-09 | 2.9e-09 | 6.6e-09 | 5.600E-09 | 2.364E-09 | 2.900E-09 | 7.300E-09 |
| 14828 | 742 | P742G | B | 9e-09 | 1.1e-08 | 3.5e-09 | 7.833E-09 | 3.884E-09 | 3.500E-09 | 1.100E-08 |
| 14829 | 742 | P742H | B | 2.8e-09 | 5.5e-09 | 8.5e-09 | 5.600E-09 | 2.851E-09 | 2.800E-09 | 8.500E-09 |
| 14830 | 742 | P742I | B | 3.5e-09 | 3.7e-09 | 1.1e-08 | 6.067E-09 | 4.274E-09 | 3.500E-09 | 1.100E-08 |
| 14831 | 742 | P742L | B | 4.8e-09 | 3.4e-09 | 1.1e-08 | 6.400E-09 | 4.045E-09 | 3.400E-09 | 1.100E-08 |
| 14832 | 742 | P742K | B | 7.7e-09 | 6.8e-09 | 4.8e-09 | 6.433E-09 | 1.484E-09 | 4.800E-09 | 7.700E-09 |
| 14833 | 742 | P742M | B | 1.2e-08 | 8e-09 | 1.5e-08 | 1.167E-08 | 3.512E-09 | 8.000E-09 | 1.500E-08 |
| 14834 | 742 | P742F | B | 4e-09 | 3.6e-09 | 8.3e-09 | 5.300E-09 | 2.606E-09 | 3.600E-09 | 8.300E-09 |
| 14835 | 742 | P742P | B | 1.5e-08 | 1.5e-08 | 6.6e-09 | 1.220E-08 | 4.850E-09 | 6.600E-09 | 1.500E-08 |
| 14836 | 742 | P742S | B | 3.9e-09 | 3e-09 | 1.4e-08 | 6.967E-09 | 6.108E-09 | 3.000E-09 | 1.400E-08 |
| 14837 | 742 | P742T | B | 3.2e-09 | 7.2e-09 | 3.8e-09 | 4.733E-09 | 2.157E-09 | 3.200E-09 | 7.200E-09 |
| 14838 | 742 | P742W | B | 4.5e-09 | 1.7e-08 | 8.7e-09 | 1.007E-08 | 6.361E-09 | 4.500E-09 | 1.700E-08 |
| 14839 | 742 | P742Y | B | 1.2e-08 | 4.3e-09 | 8.1e-09 | 8.133E-09 | 3.850E-09 | 4.300E-09 | 1.200E-08 |
| 14840 | 742 | P742V | B | 6.9e-09 | 3.3e-09 | 2.8e-09 | 4.333E-09 | 2.237E-09 | 2.800E-09 | 6.900E-09 |
| 14841 | 743 | S743A | B | 7.5e-09 | 1.4e-08 | 1.1e-08 | 1.083E-08 | 3.253E-09 | 7.500E-09 | 1.400E-08 |
| 14842 | 743 | S743R | B | 4.6e-09 | 7.4e-09 | 1.6e-08 | 9.333E-09 | 5.941E-09 | 4.600E-09 | 1.600E-08 |
| 14843 | 743 | S743N | B | 1.4e-08 | 1.2e-08 | 3.4e-09 | 9.800E-09 | 5.632E-09 | 3.400E-09 | 1.400E-08 |
| 14844 | 743 | S743D | B | 1.9e-08 | 6.1e-09 | 7.3e-09 | 1.080E-08 | 7.127E-09 | 6.100E-09 | 1.900E-08 |
| 14845 | 743 | S743C | B | 3.1e-09 | 1e-08 | 2.8e-09 | 5.300E-09 | 4.073E-09 | 2.800E-09 | 1.000E-08 |
| 14846 | 743 | S743E | B | 5.8e-09 | 8.1e-09 | 1.1e-08 | 8.300E-09 | 2.606E-09 | 5.800E-09 | 1.100E-08 |
| 14847 | 743 | S743Q | B | 6.4e-09 | 4.8e-09 | 9e-09 | 6.733E-09 | 2.120E-09 | 4.800E-09 | 9.000E-09 |
| 14848 | 743 | S743G | B | 7.6e-09 | 1.3e-08 | 1.5e-08 | 1.187E-08 | 3.828E-09 | 7.600E-09 | 1.500E-08 |
| 14849 | 743 | S743H | B | 2.3e-09 | 2.7e-09 | 5.5e-09 | 3.500E-09 | 1.744E-09 | 2.300E-09 | 5.500E-09 |
| 14850 | 743 | S743I | B | 7.4e-09 | 6.5e-09 | 2.2e-08 | 1.197E-08 | 8.701E-09 | 6.500E-09 | 2.200E-08 |
| 14851 | 743 | S743L | B | 1.4e-08 | 5.7e-09 | 4.8e-09 | 8.167E-09 | 5.072E-09 | 4.800E-09 | 1.400E-08 |
| 14852 | 743 | S743K | B | 5.9e-09 | 2.9e-08 | 2.3e-09 | 1.240E-08 | 1.449E-08 | 2.300E-09 | 2.900E-08 |
| 14853 | 743 | S743M | B | 7.7e-09 | 7.6e-09 | 8.7e-09 | 8.000E-09 | 6.083E-10 | 7.600E-09 | 8.700E-09 |
| 14854 | 743 | S743F | B | 4.6e-09 | 7.4e-09 | 4e-09 | 5.333E-09 | 1.815E-09 | 4.000E-09 | 7.400E-09 |
| 14855 | 743 | S743P | B | 4.5e-09 | 5.5e-09 | 8.9e-09 | 6.300E-09 | 2.307E-09 | 4.500E-09 | 8.900E-09 |
| 14856 | 743 | S743S | B | 5.9e-09 | 9.5e-09 | 5.1e-09 | 6.833E-09 | 2.344E-09 | 5.100E-09 | 9.500E-09 |
| 14857 | 743 | S743T | B | 6.7e-09 | 4.3e-09 | 6.7e-09 | 5.900E-09 | 1.386E-09 | 4.300E-09 | 6.700E-09 |
| 14858 | 743 | S743W | B | 1.8e-08 | 5.9e-09 | 6.3e-09 | 1.007E-08 | 6.873E-09 | 5.900E-09 | 1.800E-08 |
| 14859 | 743 | S743Y | B | 7.3e-09 | 1.6e-08 | 1.5e-08 | 1.277E-08 | 4.761E-09 | 7.300E-09 | 1.600E-08 |
| 14860 | 743 | S743V | B | 5.6e-09 | 7.5e-09 | 9.9e-09 | 7.667E-09 | 2.155E-09 | 5.600E-09 | 9.900E-09 |
| 14861 | 744 | D744A | B | 6.9e-09 | 2.5e-08 | 6.4e-09 | 1.277E-08 | 1.060E-08 | 6.400E-09 | 2.500E-08 |
| 14862 | 744 | D744R | B | 8.2e-09 | 8.4e-09 | 2.3e-08 | 1.320E-08 | 8.488E-09 | 8.200E-09 | 2.300E-08 |
| 14863 | 744 | D744N | B | 6.3e-09 | 8.2e-09 | 1e-08 | 8.167E-09 | 1.850E-09 | 6.300E-09 | 1.000E-08 |
| 14864 | 744 | D744D | B | 4.6e-09 | 7.2e-09 | 4.8e-09 | 5.533E-09 | 1.447E-09 | 4.600E-09 | 7.200E-09 |
| 14865 | 744 | D744C | B | 1e-08 | 5.2e-09 | 1.6e-08 | 1.040E-08 | 5.411E-09 | 5.200E-09 | 1.600E-08 |
| 14866 | 744 | D744E | B | 9.5e-09 | 8.6e-09 | 1.3e-08 | 1.037E-08 | 2.325E-09 | 8.600E-09 | 1.300E-08 |
| 14867 | 744 | D744Q | B | 6.2e-09 | 4e-09 | 9e-09 | 6.400E-09 | 2.506E-09 | 4.000E-09 | 9.000E-09 |
| 14868 | 744 | D744G | B | 2.5e-09 | 3.7e-09 | 1.1e-08 | 5.733E-09 | 4.600E-09 | 2.500E-09 | 1.100E-08 |
| 14869 | 744 | D744H | B | 8.6e-09 | 6.8e-09 | 7.6e-09 | 7.667E-09 | 9.018E-10 | 6.800E-09 | 8.600E-09 |
| 14870 | 744 | D744I | B | 1.9e-08 | 1.5e-08 | 5.3e-09 | 1.310E-08 | 7.045E-09 | 5.300E-09 | 1.900E-08 |
| 14871 | 744 | D744L | B | 1.3e-08 | 2.3e-08 | 1.1e-08 | 1.567E-08 | 6.429E-09 | 1.100E-08 | 2.300E-08 |
| 14872 | 744 | D744K | B | 5.6e-09 | 7.1e-09 | 9.2e-09 | 7.300E-09 | 1.808E-09 | 5.600E-09 | 9.200E-09 |
| 14873 | 744 | D744M | B | 3.3e-09 | 4.7e-09 | 6e-09 | 4.667E-09 | 1.350E-09 | 3.300E-09 | 6.000E-09 |
| 14874 | 744 | D744F | B | 3.7e-09 | 5.8e-09 | 5.4e-09 | 4.967E-09 | 1.115E-09 | 3.700E-09 | 5.800E-09 |

| | | | | | | | | | | |
|---|---|---|---|---|---|---|---|---|---|---|
| 14875 | 744 | D744P | B | 1.9e-09 | 4.6e-08 | 1.2e-08 | 1.997E-08 | 2.310E-08 | 1.900E-09 | 4.600E-08 |
| 14876 | 744 | D744S | B | 5.2e-09 | 8.9e-09 | 8.8e-09 | 7.633E-09 | 2.108E-09 | 5.200E-09 | 8.900E-09 |
| 14877 | 744 | D744T | B | 1.7e-09 | 7.7e-09 | 1.9e-08 | 9.467E-09 | 8.784E-09 | 1.700E-09 | 1.900E-08 |
| 14878 | 744 | D744W | B | 6.8e-09 | 5.3e-09 | 3.1e-09 | 5.067E-09 | 1.861E-09 | 3.100E-09 | 6.800E-09 |
| 14879 | 744 | D744Y | B | 8.6e-09 | 3.9e-09 | 1.1e-08 | 7.833E-09 | 3.612E-09 | 3.900E-09 | 1.100E-08 |
| 14880 | 744 | D744V | B | 7.1e-09 | 6.2e-09 | 7.9e-09 | 7.067E-09 | 8.505E-10 | 6.200E-09 | 7.900E-09 |
| 14881 | 745 | E745A | B | 6.7e-09 | 1.5e-09 | 4.8e-09 | 4.333E-09 | 2.631E-09 | 1.500E-09 | 6.700E-09 |
| 14882 | 745 | E745R | B | 3.1e-09 | 1.2e-08 | 2.1e-09 | 5.733E-09 | 5.450E-09 | 2.100E-09 | 1.200E-08 |
| 14883 | 745 | E745N | B | 1e-08 | 7.5e-09 | 2.6e-08 | 1.450E-08 | 1.004E-08 | 7.500E-09 | 2.600E-08 |
| 14884 | 745 | E745D | B | 3.5e-09 | 5.5e-09 | 3e-09 | 4.000E-09 | 1.323E-09 | 3.000E-09 | 5.500E-09 |
| 14885 | 745 | E745C | B | 4.2e-09 | 5.5e-09 | 4.8e-09 | 4.833E-09 | 6.506E-10 | 4.200E-09 | 5.500E-09 |
| 14886 | 745 | E745E | B | 5e-09 | 2.7e-09 | 2.5e-09 | 3.400E-09 | 1.389E-09 | 2.500E-09 | 5.000E-09 |
| 14887 | 745 | E745Q | B | 8.5e-09 | 3.6e-09 | 1.3e-08 | 8.367E-09 | 4.701E-09 | 3.600E-09 | 1.300E-08 |
| 14888 | 745 | E745G | B | 6.3e-09 | 5.2e-09 | 2.7e-09 | 4.733E-09 | 1.845E-09 | 2.700E-09 | 6.300E-09 |
| 14889 | 745 | E745H | B | 8.4e-09 | 8.2e-09 | 1.5e-08 | 1.053E-08 | 3.870E-09 | 8.200E-09 | 1.500E-08 |
| 14890 | 745 | E745I | B | 5.2e-09 | 7.3e-09 | 9.2e-09 | 7.233E-09 | 2.001E-09 | 5.200E-09 | 9.200E-09 |
| 14891 | 745 | E745L | B | 4.6e-09 | 1.9e-08 | 3.7e-09 | 9.100E-09 | 8.585E-09 | 3.700E-09 | 1.900E-08 |
| 14892 | 745 | E745K | B | 5.9e-09 | 6.8e-09 | 3e-09 | 5.233E-09 | 1.986E-09 | 3.000E-09 | 6.800E-09 |
| 14893 | 745 | E745M | B | 2.2e-09 | 5.7e-09 | 7.3e-09 | 5.067E-09 | 2.608E-09 | 2.200E-09 | 7.300E-09 |
| 14894 | 745 | E745F | B | 4.8e-09 | 4.1e-09 | 3.8e-09 | 4.233E-09 | 5.132E-10 | 3.800E-09 | 4.800E-09 |
| 14895 | 745 | E745P | B | 2.5e-09 | 3.2e-09 | 5.1e-09 | 3.600E-09 | 1.345E-09 | 2.500E-09 | 5.100E-09 |
| 14896 | 745 | E745S | B | 1.3e-08 | 1.8e-08 | 4.7e-09 | 1.190E-08 | 6.718E-09 | 4.700E-09 | 1.800E-08 |
| 14897 | 745 | E745T | B | 1.1e-08 | 1.3e-09 | 1.1e-08 | 7.767E-09 | 5.600E-09 | 1.300E-09 | 1.100E-08 |
| 14898 | 745 | E745W | B | 9.1e-09 | 2.1e-09 | 3.5e-08 | 1.540E-08 | 1.733E-08 | 2.100E-09 | 3.500E-08 |
| 14899 | 745 | E745Y | B | 6.2e-09 | 4.4e-09 | 8.5e-09 | 6.367E-09 | 2.055E-09 | 4.400E-09 | 8.500E-09 |
| 14900 | 745 | E745V | B | 1e-08 | 5.2e-09 | 5.8e-09 | 7.000E-09 | 2.615E-09 | 5.200E-09 | 1.000E-08 |
| 14901 | 746 | Q746A | B | 2.6e-08 | 5.2e-09 | 8.3e-09 | 1.317E-08 | 1.122E-08 | 5.200E-09 | 2.600E-08 |
| 14902 | 746 | Q746R | B | 1.7e-08 | 3.6e-09 | 7.6e-09 | 9.400E-09 | 6.879E-09 | 3.600E-09 | 1.700E-08 |
| 14903 | 746 | Q746N | B | 3.3e-09 | 1.4e-08 | 9.4e-09 | 8.900E-09 | 5.367E-09 | 3.300E-09 | 1.400E-08 |
| 14904 | 746 | Q746D | B | 6e-09 | 1.1e-08 | 2.9e-09 | 6.633E-09 | 4.087E-09 | 2.900E-09 | 1.100E-08 |
| 14905 | 746 | Q746C | B | 8.1e-09 | 7.1e-09 | 5.9e-09 | 7.033E-09 | 1.102E-09 | 5.900E-09 | 8.100E-09 |
| 14906 | 746 | Q746E | B | 6.3e-09 | 4.4e-09 | 6.9e-09 | 5.867E-09 | 1.305E-09 | 4.400E-09 | 6.900E-09 |
| 14907 | 746 | Q746Q | B | 1.1e-08 | 3.4e-09 | 5.2e-09 | 6.533E-09 | 3.972E-09 | 3.400E-09 | 1.100E-08 |
| 14908 | 746 | Q746G | B | 1.2e-08 | 2.7e-09 | 1.1e-08 | 8.567E-09 | 5.105E-09 | 2.700E-09 | 1.200E-08 |
| 14909 | 746 | Q746H | B | 5.8e-09 | 2.6e-08 | 2.6e-09 | 1.147E-08 | 1.269E-08 | 2.600E-09 | 2.600E-08 |
| 14910 | 746 | Q746I | B | 8.1e-09 | 4.1e-09 | 1.5e-08 | 9.067E-09 | 5.514E-09 | 4.100E-09 | 1.500E-08 |
| 14911 | 746 | Q746L | B | 1.3e-08 | 7.9e-09 | 3.4e-09 | 8.100E-09 | 4.803E-09 | 3.400E-09 | 1.300E-08 |
| 14912 | 746 | Q746K | B | 5.8e-09 | 2.8e-09 | 1.2e-08 | 6.867E-09 | 4.692E-09 | 2.800E-09 | 1.200E-08 |
| 14913 | 746 | Q746M | B | 1.5e-09 | 1.3e-08 | 5.9e-09 | 6.800E-09 | 5.803E-09 | 1.500E-09 | 1.300E-08 |
| 14914 | 746 | Q746F | B | 4.8e-09 | 4.1e-08 | 8.7e-09 | 1.817E-08 | 1.987E-08 | 4.800E-09 | 4.100E-08 |
| 14915 | 746 | Q746P | B | 5.8e-09 | 2.3e-08 | 5e-09 | 1.127E-08 | 1.017E-08 | 5.000E-09 | 2.300E-08 |
| 14916 | 746 | Q746S | B | 3.6e-09 | 8e-09 | 1e-08 | 7.200E-09 | 3.274E-09 | 3.600E-09 | 1.000E-08 |
| 14917 | 746 | Q746T | B | 1.9e-08 | 9.6e-09 | 5.9e-09 | 1.150E-08 | 6.754E-09 | 5.900E-09 | 1.900E-08 |
| 14918 | 746 | Q746W | B | 2e-08 | 6.4e-09 | 9.1e-09 | 1.183E-08 | 7.200E-09 | 6.400E-09 | 2.000E-08 |
| 14919 | 746 | Q746Y | B | 2.8e-09 | 1.2e-08 | 2.2e-08 | 1.227E-08 | 9.603E-09 | 2.800E-09 | 2.200E-08 |
| 14920 | 746 | Q746V | B | 7.9e-09 | 8.2e-09 | 1.1e-08 | 9.033E-09 | 1.710E-09 | 7.900E-09 | 1.100E-08 |
| 14921 | 747 | L747A | B | 9.9e-09 | 2e-08 | 4.8e-09 | 1.157E-08 | 7.736E-09 | 4.800E-09 | 2.000E-08 |
| 14922 | 747 | L747R | B | 1.3e-08 | 9.5e-09 | 9.8e-09 | 1.077E-08 | 1.940E-09 | 9.500E-09 | 1.300E-08 |
| 14923 | 747 | L747N | B | 9.3e-09 | 1.2e-08 | 8.6e-09 | 9.967E-09 | 1.795E-09 | 8.600E-09 | 1.200E-08 |
| 14924 | 747 | L747D | B | 4.1e-09 | 6.7e-09 | 8.3e-09 | 6.367E-09 | 2.120E-09 | 4.100E-09 | 8.300E-09 |
| 14925 | 747 | L747C | B | 1.4e-08 | 7.9e-09 | 9.5e-09 | 1.047E-08 | 3.163E-09 | 7.900E-09 | 1.400E-08 |
| 14926 | 747 | L747E | B | 2.7e-09 | 5.5e-09 | 1.3e-08 | 7.067E-09 | 5.326E-09 | 2.700E-09 | 1.300E-08 |
| 14927 | 747 | L747Q | B | 5.6e-09 | 1.1e-08 | 1.9e-08 | 1.187E-08 | 6.742E-09 | 5.600E-09 | 1.900E-08 |
| 14928 | 747 | L747G | B | 2.8e-09 | 4.3e-09 | 8.8e-09 | 5.300E-09 | 3.122E-09 | 2.800E-09 | 8.800E-09 |
| 14929 | 747 | L747H | B | 5e-09 | 4.9e-09 | 7.4e-09 | 5.767E-09 | 1.415E-09 | 4.900E-09 | 7.400E-09 |
| 14930 | 747 | L747I | B | 7e-09 | 1.1e-08 | 1.7e-08 | 1.167E-08 | 5.033E-09 | 7.000E-09 | 1.700E-08 |
| 14931 | 747 | L747L | B | 8.7e-09 | 4.3e-09 | 6.6e-09 | 6.533E-09 | 2.201E-09 | 4.300E-09 | 8.700E-09 |
| 14932 | 747 | L747K | B | 3.5e-09 | 3.5e-09 | 2.6e-08 | 1.100E-08 | 1.299E-08 | 3.500E-09 | 2.600E-08 |
| 14933 | 747 | L747M | B | 5.6e-09 | 7.8e-09 | 4.8e-09 | 6.067E-09 | 1.553E-09 | 4.800E-09 | 7.800E-09 |
| 14934 | 747 | L747F | B | 5.9e-09 | 1.4e-08 | 2.7e-08 | 1.563E-08 | 1.064E-08 | 5.900E-09 | 2.700E-08 |
| 14935 | 747 | L747P | B | 3e-09 | 3.4e-09 | 1.7e-08 | 7.800E-09 | 7.970E-09 | 3.000E-09 | 1.700E-08 |
| 14936 | 747 | L747S | B | 1.7e-09 | 4.6e-09 | 1.4e-08 | 6.767E-09 | 6.430E-09 | 1.700E-09 | 1.400E-08 |
| 14937 | 747 | L747T | B | 4.1e-09 | 1.2e-08 | 1.5e-08 | 1.037E-08 | 5.631E-09 | 4.100E-09 | 1.500E-08 |
| 14938 | 747 | L747W | B | 1.2e-08 | 7.1e-09 | 8.1e-09 | 9.067E-09 | 2.589E-09 | 7.100E-09 | 1.200E-08 |
| 14939 | 747 | L747Y | B | 4.2e-09 | 6.1e-09 | 5.3e-09 | 5.200E-09 | 9.539E-10 | 4.200E-09 | 6.100E-09 |
| 14940 | 747 | L747V | B | 1.2e-08 | 1.8e-08 | 3.3e-09 | 1.110E-08 | 7.391E-09 | 3.300E-09 | 1.800E-08 |
| 14941 | 748 | K748A | B | 4.4e-09 | 4.6e-09 | 9.7e-09 | 6.233E-09 | 3.004E-09 | 4.400E-09 | 9.700E-09 |
| 14942 | 748 | K748R | B | 5.1e-09 | 3.8e-09 | 1.4e-08 | 7.633E-09 | 5.552E-09 | 3.800E-09 | 1.400E-08 |
| 14943 | 748 | K748N | B | 1e-08 | 1.5e-08 | 2.8e-09 | 9.267E-09 | 6.133E-09 | 2.800E-09 | 1.500E-08 |
| 14944 | 748 | K748D | B | 1.5e-08 | 1.4e-08 | 5.5e-09 | 1.150E-08 | 5.220E-09 | 5.500E-09 | 1.500E-08 |
| 14945 | 748 | K748C | B | 3.5e-09 | 6.5e-09 | 4.6e-09 | 4.867E-09 | 1.518E-09 | 3.500E-09 | 6.500E-09 |
| 14946 | 748 | K748E | B | 2.8e-09 | 2.2e-09 | 2.8e-09 | 2.600E-09 | 3.464E-10 | 2.200E-09 | 2.800E-09 |
| 14947 | 748 | K748Q | B | 1.1e-08 | 6.4e-09 | 7.9e-09 | 8.433E-09 | 2.346E-09 | 6.400E-09 | 1.100E-08 |
| 14948 | 748 | K748G | B | 7e-09 | 1.2e-08 | 3.1e-09 | 7.367E-09 | 4.461E-09 | 3.100E-09 | 1.200E-08 |
| 14949 | 748 | K748H | B | 4.2e-09 | 5.7e-09 | 5.4e-09 | 5.100E-09 | 7.937E-10 | 4.200E-09 | 5.700E-09 |
| 14950 | 748 | K748I | B | 1.5e-08 | 8e-09 | 6.7e-09 | 9.900E-09 | 4.464E-09 | 6.700E-09 | 1.500E-08 |
| 14951 | 748 | K748L | B | 5.5e-09 | 1.6e-08 | 6e-09 | 9.167E-09 | 5.923E-09 | 5.500E-09 | 1.600E-08 |
| 14952 | 748 | K748K | B | 9.1e-09 | 2.5e-09 | 4e-09 | 5.200E-09 | 3.460E-09 | 2.500E-09 | 9.100E-09 |
| 14953 | 748 | K748M | B | 4.2e-09 | 1.4e-08 | 6.1e-09 | 8.100E-09 | 5.197E-09 | 4.200E-09 | 1.400E-08 |
| 14954 | 748 | K748F | B | 1.3e-08 | 2.3e-08 | 1e-08 | 1.533E-08 | 6.807E-09 | 1.000E-08 | 2.300E-08 |
| 14955 | 748 | K748P | B | 7e-09 | 3.6e-09 | 1.1e-08 | 7.200E-09 | 3.704E-09 | 3.600E-09 | 1.100E-08 |
| 14956 | 748 | K748S | B | 4e-09 | 3.7e-09 | 1e-08 | 5.900E-09 | 3.554E-09 | 3.700E-09 | 1.000E-08 |
| 14957 | 748 | K748T | B | 8e-09 | 1.3e-08 | 7.1e-09 | 9.367E-09 | 3.179E-09 | 7.100E-09 | 1.300E-08 |
| 14958 | 748 | K748W | B | 1.4e-08 | 1.6e-08 | 5.4e-09 | 1.180E-08 | 5.632E-09 | 5.400E-09 | 1.600E-08 |
| 14959 | 748 | K748Y | B | 1.9e-09 | 1.2e-08 | 1.6e-08 | 9.967E-09 | 7.267E-09 | 1.900E-09 | 1.600E-08 |

| | | | | | | | | | | |
|---|---|---|---|---|---|---|---|---|---|---|
| 14960 | 748 | K748V | B | 1.2e-08 | 3.6e-09 | 7.8e-09 | 7.800E-09 | 4.200E-09 | 3.600E-09 | 1.200E-08 |
| 14961 | 749 | S749A | B | 1.3e-09 | 7.3e-09 | 9.2e-09 | 5.933E-09 | 4.124E-09 | 1.300E-09 | 9.200E-09 |
| 14962 | 749 | S749R | B | 1.6e-08 | 1.6e-08 | 7.2e-09 | 1.307E-08 | 5.081E-09 | 7.200E-09 | 1.600E-08 |
| 14963 | 749 | S749N | B | 1.5e-08 | 6.1e-09 | 2e-09 | 7.700E-09 | 6.646E-09 | 2.000E-09 | 1.500E-08 |
| 14964 | 749 | S749D | B | 3.7e-09 | 8.1e-09 | 6.8e-09 | 6.200E-09 | 2.261E-09 | 3.700E-09 | 8.100E-09 |
| 14965 | 749 | S749C | B | 3.7e-09 | 1.4e-08 | 4.3e-09 | 7.333E-09 | 5.781E-09 | 3.700E-09 | 1.400E-08 |
| 14966 | 749 | S749E | B | 1e-08 | 6.4e-09 | 7e-09 | 7.800E-09 | 1.929E-09 | 6.400E-09 | 1.000E-08 |
| 14967 | 749 | S749Q | B | 9.9e-09 | 9.6e-09 | 5.1e-09 | 8.200E-09 | 2.689E-09 | 5.100E-09 | 9.900E-09 |
| 14968 | 749 | S749G | B | 7.6e-09 | 4.1e-09 | 8.8e-09 | 6.833E-09 | 2.442E-09 | 4.100E-09 | 8.800E-09 |
| 14969 | 749 | S749H | B | 4.6e-09 | 9.2e-09 | 1.8e-09 | 5.200E-09 | 3.736E-09 | 1.800E-09 | 9.200E-09 |
| 14970 | 749 | S749I | B | 6.6e-09 | 1.8e-09 | 1.3e-08 | 7.133E-09 | 5.619E-09 | 1.800E-09 | 1.300E-08 |
| 14971 | 749 | S749L | B | 6.1e-09 | 7.8e-09 | 8.9e-09 | 7.600E-09 | 1.411E-09 | 6.100E-09 | 8.900E-09 |
| 14972 | 749 | S749K | B | 8.9e-09 | 7.1e-09 | 1.3e-08 | 9.667E-09 | 3.024E-09 | 7.100E-09 | 1.300E-08 |
| 14973 | 749 | S749M | B | 3.8e-09 | 1.2e-09 | 3.6e-09 | 2.867E-09 | 1.447E-09 | 1.200E-09 | 3.800E-09 |
| 14974 | 749 | S749F | B | 1.9e-09 | 8.8e-10 | 7.8e-09 | 3.527E-09 | 3.736E-09 | 8.800E-10 | 7.800E-09 |
| 14975 | 749 | S749P | B | 4.8e-09 | 4.6e-08 | 5.4e-09 | 1.873E-08 | 2.362E-08 | 4.800E-09 | 4.600E-08 |
| 14976 | 749 | S749S | B | 3.2e-09 | 7.9e-09 | 7.8e-09 | 6.300E-09 | 2.685E-09 | 3.200E-09 | 7.900E-09 |
| 14977 | 749 | S749T | B | 7.9e-09 | 7.4e-09 | 3.1e-09 | 6.133E-09 | 2.639E-09 | 3.100E-09 | 7.900E-09 |
| 14978 | 749 | S749W | B | 1.2e-08 | 1.7e-08 | 9.3e-09 | 1.277E-08 | 3.907E-09 | 9.300E-09 | 1.700E-08 |
| 14979 | 749 | S749Y | B | 4.4e-08 | 8.4e-09 | 2.2e-09 | 1.820E-08 | 2.256E-08 | 2.200E-09 | 4.400E-08 |
| 14980 | 749 | S749V | B | 3.7e-09 | 6.8e-09 | 1.3e-08 | 7.833E-09 | 4.735E-09 | 3.700E-09 | 1.300E-08 |
| 14981 | 750 | G750A | B | 4.3e-09 | 4.9e-09 | 1.5e-08 | 8.067E-09 | 6.012E-09 | 4.300E-09 | 1.500E-08 |
| 14982 | 750 | G750R | B | 3.8e-09 | 4.8e-09 | 2.5e-09 | 3.700E-09 | 1.153E-09 | 2.500E-09 | 4.800E-09 |
| 14983 | 750 | G750N | B | 2.6e-09 | 1.1e-08 | 4.4e-09 | 6.000E-09 | 4.423E-09 | 2.600E-09 | 1.100E-08 |
| 14984 | 750 | G750D | B | 1e-08 | 2e-09 | 5.5e-09 | 5.833E-09 | 4.010E-09 | 2.000E-09 | 1.000E-08 |
| 14985 | 750 | G750C | B | 5.4e-09 | 1e-08 | 5.3e-09 | 6.900E-09 | 2.685E-09 | 5.300E-09 | 1.000E-08 |
| 14986 | 750 | G750E | B | 1.6e-08 | 7.1e-09 | 6.2e-09 | 9.767E-09 | 5.417E-09 | 6.200E-09 | 1.600E-08 |
| 14987 | 750 | G750Q | B | 4.1e-09 | 2.1e-08 | 4.5e-09 | 9.867E-09 | 9.644E-09 | 4.100E-09 | 2.100E-08 |
| 14988 | 750 | G750G | B | 8.5e-09 | 7.4e-09 | 1.9e-08 | 1.163E-08 | 6.403E-09 | 7.400E-09 | 1.900E-08 |
| 14989 | 750 | G750H | B | 2.2e-08 | 4.2e-09 | 1.8e-08 | 1.473E-08 | 9.339E-09 | 4.200E-09 | 2.200E-08 |
| 14990 | 750 | G750I | B | 5.7e-09 | 1.3e-08 | 6.5e-09 | 8.400E-09 | 4.004E-09 | 5.700E-09 | 1.300E-08 |
| 14991 | 750 | G750L | B | 2.3e-08 | 1.1e-08 | 1.6e-08 | 1.667E-08 | 6.028E-09 | 1.100E-08 | 2.300E-08 |
| 14992 | 750 | G750K | B | 4.3e-09 | 1.4e-08 | 1.3e-08 | 1.043E-08 | 5.335E-09 | 4.300E-09 | 1.400E-08 |
| 14993 | 750 | G750M | B | 9.1e-09 | 8.3e-09 | 6.9e-09 | 8.100E-09 | 1.114E-09 | 6.900E-09 | 9.100E-09 |
| 14994 | 750 | G750F | B | 8.9e-09 | 7.1e-09 | 1.5e-08 | 1.033E-08 | 4.140E-09 | 7.100E-09 | 1.500E-08 |
| 14995 | 750 | G750P | B | 3.5e-09 | 1.3e-08 | 2.7e-08 | 1.450E-08 | 1.182E-08 | 3.500E-09 | 2.700E-08 |
| 14996 | 750 | G750S | B | 4.1e-09 | 1e-08 | 4.5e-09 | 6.200E-09 | 3.297E-09 | 4.100E-09 | 1.000E-08 |
| 14997 | 750 | G750T | B | 4.4e-09 | 9.1e-09 | 1.1e-08 | 8.167E-09 | 3.398E-09 | 4.400E-09 | 1.100E-08 |
| 14998 | 750 | G750W | B | 8.5e-09 | 8.9e-09 | 3e-09 | 6.800E-09 | 3.297E-09 | 3.000E-09 | 8.900E-09 |
| 14999 | 750 | G750Y | B | 3.1e-09 | 2.7e-09 | 8.1e-09 | 4.633E-09 | 3.009E-09 | 2.700E-09 | 8.100E-09 |
| 15000 | 750 | G750V | B | 6.1e-09 | 6e-09 | 2.2e-09 | 4.767E-09 | 2.223E-09 | 2.200E-09 | 6.100E-09 |
| 15001 | 751 | T751A | B | 4.8e-09 | 9.9e-09 | 1e-08 | 8.233E-09 | 2.974E-09 | 4.800E-09 | 1.000E-08 |
| 15002 | 751 | T751R | B | 4.7e-09 | 6.3e-09 | 8.5e-09 | 6.500E-09 | 1.908E-09 | 4.700E-09 | 8.500E-09 |
| 15003 | 751 | T751N | B | 8.7e-09 | 3.1e-09 | 8.8e-09 | 6.867E-09 | 3.262E-09 | 3.100E-09 | 8.800E-09 |
| 15004 | 751 | T751D | B | 1.3e-08 | 2.2e-08 | 5.6e-09 | 1.353E-08 | 8.213E-09 | 5.600E-09 | 2.200E-08 |
| 15005 | 751 | T751C | B | 9.5e-09 | 4.1e-09 | 8.7e-09 | 7.433E-09 | 2.914E-09 | 4.100E-09 | 9.500E-09 |
| 15006 | 751 | T751E | B | 8.4e-09 | 2.9e-09 | 4.7e-09 | 5.333E-09 | 2.804E-09 | 2.900E-09 | 8.400E-09 |
| 15007 | 751 | T751Q | B | 4.9e-09 | 8.2e-09 | 1e-08 | 7.700E-09 | 2.587E-09 | 4.900E-09 | 1.000E-08 |
| 15008 | 751 | T751G | B | 9.5e-09 | 3.2e-09 | 6.3e-09 | 6.333E-09 | 3.150E-09 | 3.200E-09 | 9.500E-09 |
| 15009 | 751 | T751H | B | 6.3e-09 | 1.2e-08 | 5.7e-09 | 8.000E-09 | 3.477E-09 | 5.700E-09 | 1.200E-08 |
| 15010 | 751 | T751I | B | 8.2e-09 | 7.3e-09 | 7.5e-09 | 7.667E-09 | 4.726E-10 | 7.300E-09 | 8.200E-09 |
| 15011 | 751 | T751L | B | 1.8e-08 | 3.2e-09 | 4e-09 | 8.400E-09 | 8.323E-09 | 3.200E-09 | 1.800E-08 |
| 15012 | 751 | T751K | B | 2.7e-09 | 5.1e-09 | 9.9e-09 | 5.900E-09 | 3.666E-09 | 2.700E-09 | 9.900E-09 |
| 15013 | 751 | T751M | B | 6.4e-09 | 5.3e-09 | 1.1e-08 | 7.567E-09 | 3.024E-09 | 5.300E-09 | 1.100E-08 |
| 15014 | 751 | T751F | B | 2.8e-09 | 1.2e-08 | 3.7e-09 | 6.167E-09 | 5.072E-09 | 2.800E-09 | 1.200E-08 |
| 15015 | 751 | T751P | B | 7.5e-09 | 1.5e-09 | 1.5e-08 | 8.000E-09 | 6.764E-09 | 1.500E-09 | 1.500E-08 |
| 15016 | 751 | T751S | B | 2e-08 | 1.4e-08 | 1.4e-08 | 1.600E-08 | 3.464E-09 | 1.400E-08 | 2.000E-08 |
| 15017 | 751 | T751T | B | 1.5e-08 | 4e-09 | 2.4e-09 | 7.133E-09 | 6.860E-09 | 2.400E-09 | 1.500E-08 |
| 15018 | 751 | T751W | B | 9.3e-09 | 7.9e-09 | 6.1e-09 | 7.767E-09 | 1.604E-09 | 6.100E-09 | 9.300E-09 |
| 15019 | 751 | T751Y | B | 1e-08 | 3.9e-09 | 4.5e-09 | 6.133E-09 | 3.362E-09 | 3.900E-09 | 1.000E-08 |
| 15020 | 751 | T751V | B | 8.8e-09 | 6.5e-09 | 3.7e-09 | 6.333E-09 | 2.554E-09 | 3.700E-09 | 8.800E-09 |
| 15021 | 752 | A752A | B | 1.5e-08 | 1.5e-08 | 1.6e-08 | 1.533E-08 | 5.774E-10 | 1.500E-08 | 1.600E-08 |
| 15022 | 752 | A752R | B | 9.6e-09 | 9.5e-09 | 9.5e-09 | 9.533E-09 | 5.774E-11 | 9.500E-09 | 9.600E-09 |
| 15023 | 752 | A752N | B | 4.9e-09 | 1.7e-08 | 2.5e-09 | 8.133E-09 | 7.772E-09 | 2.500E-09 | 1.700E-08 |
| 15024 | 752 | A752D | B | 7.5e-09 | 2.2e-09 | 1.7e-08 | 8.900E-09 | 7.499E-09 | 2.200E-09 | 1.700E-08 |
| 15025 | 752 | A752C | B | 1.4e-08 | 1.3e-08 | 2.9e-09 | 9.967E-09 | 6.140E-09 | 2.900E-09 | 1.400E-08 |
| 15026 | 752 | A752E | B | 5.6e-09 | 7.7e-09 | 1.1e-08 | 8.100E-09 | 2.722E-09 | 5.600E-09 | 1.100E-08 |
| 15027 | 752 | A752Q | B | 1e-08 | 7.6e-09 | 9e-09 | 8.867E-09 | 1.206E-09 | 7.600E-09 | 1.000E-08 |
| 15028 | 752 | A752G | B | 6.1e-09 | 1.4e-08 | 5.4e-09 | 8.500E-09 | 4.776E-09 | 5.400E-09 | 1.400E-08 |
| 15029 | 752 | A752H | B | 7.3e-09 | 1.7e-08 | 1.3e-08 | 1.243E-08 | 4.875E-09 | 7.300E-09 | 1.700E-08 |
| 15030 | 752 | A752I | B | 1.6e-08 | 8.5e-09 | 7.2e-09 | 1.057E-08 | 4.750E-09 | 7.200E-09 | 1.600E-08 |
| 15031 | 752 | A752L | B | 2.5e-08 | 2.3e-08 | 1.3e-08 | 2.033E-08 | 6.429E-09 | 1.300E-08 | 2.500E-08 |
| 15032 | 752 | A752K | B | 6.3e-09 | 6.8e-09 | 1.2e-08 | 8.367E-09 | 3.156E-09 | 6.300E-09 | 1.200E-08 |
| 15033 | 752 | A752M | B | 5.8e-09 | 3.6e-09 | 4.1e-09 | 4.500E-09 | 1.153E-09 | 3.600E-09 | 5.800E-09 |
| 15034 | 752 | A752F | B | 9.6e-09 | 4.4e-09 | 1.9e-08 | 1.100E-08 | 7.400E-09 | 4.400E-09 | 1.900E-08 |
| 15035 | 752 | A752P | B | 1.7e-08 | 8.6e-09 | 1e-08 | 1.187E-08 | 4.500E-09 | 8.600E-09 | 1.700E-08 |
| 15036 | 752 | A752S | B | 1.3e-08 | 6e-09 | 6.5e-09 | 8.500E-09 | 3.905E-09 | 6.000E-09 | 1.300E-08 |
| 15037 | 752 | A752T | B | 1e-08 | 8.9e-09 | 5.5e-09 | 8.133E-09 | 2.346E-09 | 5.500E-09 | 1.000E-08 |
| 15038 | 752 | A752W | B | 4.2e-09 | 8.2e-09 | 7.9e-09 | 6.767E-09 | 2.228E-09 | 4.200E-09 | 8.200E-09 |
| 15039 | 752 | A752Y | B | 1.6e-08 | 4.6e-09 | 3.5e-09 | 8.033E-09 | 6.921E-09 | 3.500E-09 | 1.600E-08 |
| 15040 | 752 | A752V | B | 2.5e-09 | 5.4e-09 | 9.1e-09 | 5.667E-09 | 3.308E-09 | 2.500E-09 | 9.100E-09 |
| 15041 | 753 | S753A | B | 1.4e-08 | 4.6e-09 | 1.4e-08 | 1.087E-08 | 5.427E-09 | 4.600E-09 | 1.400E-08 |
| 15042 | 753 | S753R | B | 1.6e-08 | 4.6e-09 | 2.5e-09 | 7.700E-09 | 7.264E-09 | 2.500E-09 | 1.600E-08 |
| 15043 | 753 | S753N | B | 8.5e-09 | 7.1e-09 | 1.5e-08 | 1.020E-08 | 4.215E-09 | 7.100E-09 | 1.500E-08 |
| 15044 | 753 | S753D | B | 4.3e-09 | 1.6e-08 | 1.1e-08 | 1.043E-08 | 5.871E-09 | 4.300E-09 | 1.600E-08 |

| | | | | | | | | | | |
|---|---|---|---|---|---|---|---|---|---|---|
| 15045 | 753 | S753C | B | 2.3e-08 | 5.8e-09 | 3.1e-09 | 1.063E-08 | 1.079E-08 | 3.100E-09 | 2.300E-08 |
| 15046 | 753 | S753E | B | 8.3e-09 | 4.6e-09 | 4.7e-09 | 5.867E-09 | 2.108E-09 | 4.600E-09 | 8.300E-09 |
| 15047 | 753 | S753Q | B | 4.5e-09 | 1.5e-08 | 1.3e-08 | 1.083E-08 | 5.575E-09 | 4.500E-09 | 1.500E-08 |
| 15048 | 753 | S753G | B | 1.1e-08 | 5.8e-09 | 7.4e-09 | 8.067E-09 | 2.663E-09 | 5.800E-09 | 1.100E-08 |
| 15049 | 753 | S753H | B | 7.1e-09 | 5.7e-09 | 1.1e-08 | 7.933E-09 | 2.747E-09 | 5.700E-09 | 1.100E-08 |
| 15050 | 753 | S753I | B | 6.2e-09 | 8e-09 | 3e-08 | 1.473E-08 | 1.325E-08 | 6.200E-09 | 3.000E-08 |
| 15051 | 753 | S753L | B | 2e-08 | 7.2e-09 | 2.4e-09 | 9.867E-09 | 9.098E-09 | 2.400E-09 | 2.000E-08 |
| 15052 | 753 | S753K | B | 9.9e-09 | 3.3e-08 | 5.6e-09 | 1.617E-08 | 1.474E-08 | 5.600E-09 | 3.300E-08 |
| 15053 | 753 | S753M | B | 7.2e-09 | 1.1e-08 | 6.8e-09 | 8.333E-09 | 2.318E-09 | 6.800E-09 | 1.100E-08 |
| 15054 | 753 | S753F | B | 5.2e-09 | 6.7e-09 | 5.2e-09 | 5.700E-09 | 8.660E-10 | 5.200E-09 | 6.700E-09 |
| 15055 | 753 | S753P | B | 1e-08 | 1.4e-08 | 3.5e-09 | 9.167E-09 | 5.299E-09 | 3.500E-09 | 1.400E-08 |
| 15056 | 753 | S753S | B | 3.1e-09 | 1.5e-08 | 4.4e-09 | 7.500E-09 | 6.528E-09 | 3.100E-09 | 1.500E-08 |
| 15057 | 753 | S753T | B | 6.6e-09 | 4.2e-09 | 8.4e-09 | 6.400E-09 | 2.107E-09 | 4.200E-09 | 8.400E-09 |
| 15058 | 753 | S753W | B | 6.4e-09 | 3.7e-09 | 1.9e-08 | 9.700E-09 | 8.166E-09 | 3.700E-09 | 1.900E-08 |
| 15059 | 753 | S753Y | B | 1.4e-08 | 2.1e-08 | 4.4e-09 | 1.313E-08 | 8.334E-09 | 4.400E-09 | 2.100E-08 |
| 15060 | 753 | S753V | B | 7.8e-09 | 3.1e-09 | 4.2e-09 | 5.033E-09 | 2.458E-09 | 3.100E-09 | 7.800E-09 |
| 15061 | 754 | V754A | B | 9.7e-09 | 5.9e-09 | 7.6e-09 | 7.733E-09 | 1.904E-09 | 5.900E-09 | 9.700E-09 |
| 15062 | 754 | V754R | B | 1.4e-08 | 1.1e-08 | 8.4e-09 | 1.113E-08 | 2.802E-09 | 8.400E-09 | 1.400E-08 |
| 15063 | 754 | V754N | B | 1.7e-08 | 1.5e-08 | 7.2e-09 | 1.307E-08 | 5.178E-09 | 7.200E-09 | 1.700E-08 |
| 15064 | 754 | V754D | B | 2.9e-09 | 5e-09 | 4.5e-09 | 4.133E-09 | 1.097E-09 | 2.900E-09 | 5.000E-09 |
| 15065 | 754 | V754C | B | 5.4e-09 | 5.3e-09 | 5e-09 | 5.233E-09 | 2.082E-10 | 5.000E-09 | 5.400E-09 |
| 15066 | 754 | V754E | B | 1.1e-08 | 3.7e-09 | 1.1e-08 | 8.567E-09 | 4.215E-09 | 3.700E-09 | 1.100E-08 |
| 15067 | 754 | V754Q | B | 6.2e-09 | 5.1e-09 | 5.5e-09 | 5.600E-09 | 5.568E-10 | 5.100E-09 | 6.200E-09 |
| 15068 | 754 | V754G | B | 1.9e-09 | 9.6e-09 | 2.9e-09 | 4.800E-09 | 4.187E-09 | 1.900E-09 | 9.600E-09 |
| 15069 | 754 | V754H | B | 8.7e-09 | 5.1e-09 | 6.3e-09 | 6.700E-09 | 1.833E-09 | 5.100E-09 | 8.700E-09 |
| 15070 | 754 | V754I | B | 6.4e-09 | 1.6e-08 | 4.3e-09 | 8.900E-09 | 6.238E-09 | 4.300E-09 | 1.600E-08 |
| 15071 | 754 | V754L | B | 4.1e-09 | 2.5e-08 | 6.7e-09 | 1.193E-08 | 1.139E-08 | 4.100E-09 | 2.500E-08 |
| 15072 | 754 | V754K | B | 6.9e-09 | 3.1e-09 | 3.8e-09 | 4.600E-09 | 2.022E-09 | 3.100E-09 | 6.900E-09 |
| 15073 | 754 | V754M | B | 8e-09 | 3.2e-09 | 1.1e-08 | 7.400E-09 | 3.934E-09 | 3.200E-09 | 1.100E-08 |
| 15074 | 754 | V754F | B | 1.3e-08 | 4.1e-09 | 1.8e-08 | 1.170E-08 | 7.041E-09 | 4.100E-09 | 1.800E-08 |
| 15075 | 754 | V754P | B | 3.6e-09 | 5e-09 | 4.9e-09 | 4.500E-09 | 7.810E-10 | 3.600E-09 | 5.000E-09 |
| 15076 | 754 | V754S | B | 9.7e-09 | 9.1e-09 | 1.9e-09 | 6.900E-09 | 4.341E-09 | 1.900E-09 | 9.700E-09 |
| 15077 | 754 | V754T | B | 7e-09 | 2.2e-08 | 3.3e-09 | 1.077E-08 | 9.903E-09 | 3.300E-09 | 2.200E-08 |
| 15078 | 754 | V754W | B | 9.8e-09 | 5.2e-09 | 1.2e-08 | 9.000E-09 | 3.470E-09 | 5.200E-09 | 1.200E-08 |
| 15079 | 754 | V754Y | B | 1.4e-08 | 2.6e-08 | 5.6e-09 | 1.520E-08 | 1.025E-08 | 5.600E-09 | 2.600E-08 |
| 15080 | 754 | V754V | B | 1.7e-08 | 7e-09 | 5.9e-09 | 9.967E-09 | 6.116E-09 | 5.900E-09 | 1.700E-08 |
| 15081 | 755 | V755A | B | 4e-09 | 4.2e-09 | 6.9e-09 | 5.033E-09 | 1.620E-09 | 4.000E-09 | 6.900E-09 |
| 15082 | 755 | V755R | B | 6.8e-09 | 4.1e-09 | 1.9e-08 | 9.967E-09 | 7.939E-09 | 4.100E-09 | 1.900E-08 |
| 15083 | 755 | V755N | B | 8.7e-09 | 2.8e-08 | 1.4e-08 | 1.690E-08 | 9.971E-09 | 8.700E-09 | 2.800E-08 |
| 15084 | 755 | V755D | B | 2.2e-08 | 1.8e-08 | 3e-09 | 1.433E-08 | 1.002E-08 | 3.000E-09 | 2.200E-08 |
| 15085 | 755 | V755C | B | 2e-09 | 1.2e-08 | 1.2e-08 | 8.667E-09 | 5.774E-09 | 2.000E-09 | 1.200E-08 |
| 15086 | 755 | V755E | B | 3.4e-09 | 6.2e-09 | 1.3e-08 | 7.533E-09 | 4.937E-09 | 3.400E-09 | 1.300E-08 |
| 15087 | 755 | V755Q | B | 4e-09 | 3.5e-09 | 7.3e-09 | 4.933E-09 | 2.065E-09 | 3.500E-09 | 7.300E-09 |
| 15088 | 755 | V755G | B | 1.1e-08 | 1.6e-08 | 1.9e-08 | 1.533E-08 | 4.041E-09 | 1.100E-08 | 1.900E-08 |
| 15089 | 755 | V755H | B | 3e-09 | 2.1e-09 | 4e-09 | 3.033E-09 | 9.504E-10 | 2.100E-09 | 4.000E-09 |
| 15090 | 755 | V755I | B | 7.2e-09 | 1.2e-08 | 6.3e-09 | 8.500E-09 | 3.064E-09 | 6.300E-09 | 1.200E-08 |
| 15091 | 755 | V755L | B | 5.1e-09 | 7.5e-09 | 6.4e-09 | 6.333E-09 | 1.201E-09 | 5.100E-09 | 7.500E-09 |
| 15092 | 755 | V755K | B | 5.2e-09 | 5.1e-09 | 7.9e-09 | 6.067E-09 | 1.589E-09 | 5.100E-09 | 7.900E-09 |
| 15093 | 755 | V755M | B | 1.7e-09 | 3.3e-09 | 1.5e-08 | 6.667E-09 | 7.261E-09 | 1.700E-09 | 1.500E-08 |
| 15094 | 755 | V755F | B | 1.2e-08 | 8.5e-09 | 6e-09 | 8.833E-09 | 3.014E-09 | 6.000E-09 | 1.200E-08 |
| 15095 | 755 | V755P | B | 9.2e-09 | 1.5e-08 | 1.8e-08 | 1.407E-08 | 4.474E-09 | 9.200E-09 | 1.800E-08 |
| 15096 | 755 | V755S | B | 8.1e-09 | 5.8e-09 | 3.8e-09 | 5.900E-09 | 2.152E-09 | 3.800E-09 | 8.100E-09 |
| 15097 | 755 | V755T | B | 3.6e-09 | 1.2e-08 | 6.2e-09 | 7.267E-09 | 4.300E-09 | 3.600E-09 | 1.200E-08 |
| 15098 | 755 | V755W | B | 1.3e-08 | 1.8e-08 | 1.7e-08 | 1.600E-08 | 2.646E-09 | 1.300E-08 | 1.800E-08 |
| 15099 | 755 | V755Y | B | 1.5e-08 | 2e-08 | 3.8e-09 | 1.293E-08 | 8.295E-09 | 3.800E-09 | 2.000E-08 |
| 15100 | 755 | V755V | B | 1e-08 | 1e-08 | 1.1e-08 | 1.033E-08 | 5.774E-10 | 1.000E-08 | 1.100E-08 |
| 15101 | 756 | C756A | B | 7.2e-09 | 1.2e-08 | 1.8e-08 | 1.240E-08 | 5.411E-09 | 7.200E-09 | 1.800E-08 |
| 15102 | 756 | C756R | B | 1.1e-08 | 8.8e-09 | 7.3e-09 | 9.033E-09 | 1.861E-09 | 7.300E-09 | 1.100E-08 |
| 15103 | 756 | C756N | B | 3.5e-09 | 9.5e-09 | 1.1e-08 | 8.000E-09 | 3.969E-09 | 3.500E-09 | 1.100E-08 |
| 15104 | 756 | C756D | B | 4.5e-09 | 6.5e-09 | 3.6e-09 | 4.867E-09 | 1.484E-09 | 3.600E-09 | 6.500E-09 |
| 15105 | 756 | C756C | B | 5.2e-09 | 7.1e-09 | 8.2e-09 | 6.833E-09 | 1.518E-09 | 5.200E-09 | 8.200E-09 |
| 15106 | 756 | C756E | B | 1.1e-08 | 1.7e-08 | 3.9e-09 | 1.063E-08 | 6.558E-09 | 3.900E-09 | 1.700E-08 |
| 15107 | 756 | C756Q | B | 9.3e-09 | 6.1e-09 | 4.8e-09 | 6.733E-09 | 2.316E-09 | 4.800E-09 | 9.300E-09 |
| 15108 | 756 | C756G | B | 7e-09 | 6.8e-09 | 2.6e-08 | 1.327E-08 | 1.103E-08 | 6.800E-09 | 2.600E-08 |
| 15109 | 756 | C756H | B | 6.3e-09 | 2.6e-09 | 6.1e-09 | 5.000E-09 | 2.081E-09 | 2.600E-09 | 6.300E-09 |
| 15110 | 756 | C756I | B | 1.5e-08 | 5.1e-09 | 8.8e-09 | 9.633E-09 | 5.002E-09 | 5.100E-09 | 1.500E-08 |
| 15111 | 756 | C756L | B | 1e-08 | 4.2e-09 | 8.4e-09 | 7.533E-09 | 2.996E-09 | 4.200E-09 | 1.000E-08 |
| 15112 | 756 | C756K | B | 8.5e-09 | 4.8e-09 | 4.7e-09 | 6.000E-09 | 2.166E-09 | 4.700E-09 | 8.500E-09 |
| 15113 | 756 | C756M | B | 3.7e-09 | 1.1e-08 | 9.3e-10 | 5.210E-09 | 5.202E-09 | 9.300E-10 | 1.100E-08 |
| 15114 | 756 | C756F | B | 1.6e-08 | 1.1e-08 | 9.4e-09 | 1.213E-08 | 3.443E-09 | 9.400E-09 | 1.600E-08 |
| 15115 | 756 | C756P | B | 8.4e-09 | 7e-09 | 3.7e-09 | 6.367E-09 | 2.413E-09 | 3.700E-09 | 8.400E-09 |
| 15116 | 756 | C756S | B | 1.3e-08 | 7.3e-09 | 5.1e-09 | 8.467E-09 | 4.077E-09 | 5.100E-09 | 1.300E-08 |
| 15117 | 756 | C756T | B | 1.5e-08 | 8.7e-09 | 7.4e-09 | 1.037E-08 | 4.065E-09 | 7.400E-09 | 1.500E-08 |
| 15118 | 756 | C756W | B | 1.3e-08 | 2.9e-09 | 4.7e-09 | 6.867E-09 | 5.387E-09 | 2.900E-09 | 1.300E-08 |
| 15119 | 756 | C756Y | B | 1.1e-08 | 9.5e-09 | 4.5e-09 | 8.333E-09 | 3.403E-09 | 4.500E-09 | 1.100E-08 |
| 15120 | 756 | C756V | B | 2.8e-09 | 4.3e-09 | 9.2e-09 | 5.433E-09 | 3.347E-09 | 2.800E-09 | 9.200E-09 |
| 15121 | 757 | L757A | B | 1.2e-08 | 1.6e-08 | 3.4e-09 | 1.047E-08 | 6.438E-09 | 3.400E-09 | 1.600E-08 |
| 15122 | 757 | L757R | B | 6.3e-09 | 1.3e-08 | 4.8e-09 | 8.033E-09 | 4.366E-09 | 4.800E-09 | 1.300E-08 |
| 15123 | 757 | L757N | B | 7.2e-09 | 1.1e-08 | 7.2e-09 | 8.467E-09 | 2.194E-09 | 7.200E-09 | 1.100E-08 |
| 15124 | 757 | L757D | B | 1e-08 | 1.4e-08 | 6.2e-09 | 1.007E-08 | 3.900E-09 | 6.200E-09 | 1.400E-08 |
| 15125 | 757 | L757C | B | 3.7e-09 | 1.2e-08 | 2.5e-09 | 6.067E-09 | 5.173E-09 | 2.500E-09 | 1.200E-08 |
| 15126 | 757 | L757E | B | 1.8e-09 | 9.7e-09 | 1.3e-09 | 4.267E-09 | 4.712E-09 | 1.300E-09 | 9.700E-09 |
| 15127 | 757 | L757Q | B | 3.7e-09 | 7.4e-09 | 1.6e-08 | 9.033E-09 | 6.311E-09 | 3.700E-09 | 1.600E-08 |
| 15128 | 757 | L757G | B | 8.3e-09 | 5.4e-09 | 1.7e-08 | 1.023E-08 | 6.037E-09 | 5.400E-09 | 1.700E-08 |
| 15129 | 757 | L757H | B | 9.2e-09 | 4.7e-09 | 1.2e-08 | 8.633E-09 | 3.683E-09 | 4.700E-09 | 1.200E-08 |

| | | | | | | | | | | |
|---|---|---|---|---|---|---|---|---|---|---|
| 15130 | 757 | L757I | B | 6e-09 | 1.4e-08 | 1.3e-08 | 1.100E-08 | 4.359E-09 | 6.000E-09 | 1.400E-08 |
| 15131 | 757 | L757L | B | 9.4e-09 | 8e-09 | 7.1e-09 | 8.167E-09 | 1.159E-09 | 7.100E-09 | 9.400E-09 |
| 15132 | 757 | L757K | B | 6.3e-09 | 6.8e-09 | 4.4e-09 | 5.833E-09 | 1.266E-09 | 4.400E-09 | 6.800E-09 |
| 15133 | 757 | L757M | B | 1.6e-08 | 2.1e-08 | 8.9e-09 | 1.530E-08 | 6.080E-09 | 8.900E-09 | 2.100E-08 |
| 15134 | 757 | L757F | B | 7.7e-09 | 2.7e-09 | 8e-09 | 6.133E-09 | 2.977E-09 | 2.700E-09 | 8.000E-09 |
| 15135 | 757 | L757P | B | 1.8e-08 | 1.7e-09 | 5.1e-09 | 8.267E-09 | 8.599E-09 | 1.700E-09 | 1.800E-08 |
| 15136 | 757 | L757S | B | 6.7e-09 | 2.2e-09 | 3.6e-09 | 4.167E-09 | 2.303E-09 | 2.200E-09 | 6.700E-09 |
| 15137 | 757 | L757T | B | 9.3e-09 | 5.5e-09 | 4.8e-09 | 6.533E-09 | 2.421E-09 | 4.800E-09 | 9.300E-09 |
| 15138 | 757 | L757W | B | 4.3e-09 | 1.4e-08 | 8e-09 | 8.767E-09 | 4.895E-09 | 4.300E-09 | 1.400E-08 |
| 15139 | 757 | L757Y | B | 9.1e-09 | 4.8e-09 | 7.5e-09 | 7.133E-09 | 2.173E-09 | 4.800E-09 | 9.100E-09 |
| 15140 | 757 | L757V | B | 1.5e-08 | 1.2e-08 | 5.4e-09 | 1.080E-08 | 4.911E-09 | 5.400E-09 | 1.500E-08 |
| 15141 | 758 | L758A | B | 1.6e-09 | 2.7e-09 | 5.5e-09 | 3.267E-09 | 2.011E-09 | 1.600E-09 | 5.500E-09 |
| 15142 | 758 | L758R | B | 9e-09 | 9.3e-09 | 8.2e-09 | 8.833E-09 | 5.686E-10 | 8.200E-09 | 9.300E-09 |
| 15143 | 758 | L758N | B | 7.1e-09 | 4.8e-09 | 2.6e-09 | 4.833E-09 | 2.250E-09 | 2.600E-09 | 7.100E-09 |
| 15144 | 758 | L758D | B | 2.5e-08 | 9e-09 | 1.3e-08 | 1.567E-08 | 8.327E-09 | 9.000E-09 | 2.500E-08 |
| 15145 | 758 | L758C | B | 3.1e-09 | 4.9e-09 | 6.9e-09 | 4.967E-09 | 1.901E-09 | 3.100E-09 | 6.900E-09 |
| 15146 | 758 | L758E | B | 4.8e-09 | 5.7e-09 | 4e-09 | 4.833E-09 | 8.505E-10 | 4.000E-09 | 5.700E-09 |
| 15147 | 758 | L758Q | B | 5.9e-09 | 5.1e-09 | 9.6e-09 | 6.867E-09 | 2.401E-09 | 5.100E-09 | 9.600E-09 |
| 15148 | 758 | L758G | B | 1.2e-08 | 4.9e-09 | 6.6e-09 | 7.833E-09 | 3.707E-09 | 4.900E-09 | 1.200E-08 |
| 15149 | 758 | L758H | B | 2.4e-09 | 2.9e-09 | 4.7e-09 | 3.333E-09 | 1.210E-09 | 2.400E-09 | 4.700E-09 |
| 15150 | 758 | L758I | B | 9.1e-09 | 1.2e-08 | 8e-09 | 9.700E-09 | 2.066E-09 | 8.000E-09 | 1.200E-08 |
| 15151 | 758 | L758L | B | 4.3e-09 | 7.7e-09 | 1.2e-08 | 8.000E-09 | 3.859E-09 | 4.300E-09 | 1.200E-08 |
| 15152 | 758 | L758K | B | 6.1e-09 | 9e-09 | 1.4e-08 | 9.700E-09 | 3.996E-09 | 6.100E-09 | 1.400E-08 |
| 15153 | 758 | L758M | B | 2.8e-09 | 1.5e-08 | 6.7e-09 | 8.167E-09 | 6.231E-09 | 2.800E-09 | 1.500E-08 |
| 15154 | 758 | L758F | B | 1e-08 | 7.4e-09 | 1.9e-08 | 1.213E-08 | 6.087E-09 | 7.400E-09 | 1.900E-08 |
| 15155 | 758 | L758P | B | 4.8e-09 | 2e-08 | 4.1e-09 | 9.633E-09 | 8.985E-09 | 4.100E-09 | 2.000E-08 |
| 15156 | 758 | L758S | B | 1.2e-08 | 9.9e-09 | 1.1e-08 | 1.097E-08 | 1.050E-09 | 9.900E-09 | 1.200E-08 |
| 15157 | 758 | L758T | B | 1.8e-09 | 4.5e-09 | 1.4e-08 | 6.767E-09 | 6.408E-09 | 1.800E-09 | 1.400E-08 |
| 15158 | 758 | L758W | B | 8.8e-09 | 3.7e-09 | 8.1e-09 | 6.867E-09 | 2.765E-09 | 3.700E-09 | 8.800E-09 |
| 15159 | 758 | L758Y | B | 1.2e-08 | 3.5e-09 | 8.7e-09 | 8.067E-09 | 4.285E-09 | 3.500E-09 | 1.200E-08 |
| 15160 | 758 | L758V | B | 1.1e-08 | 3.8e-08 | 8.9e-09 | 1.930E-08 | 1.623E-08 | 8.900E-09 | 3.800E-08 |
| 15161 | 759 | N759A | B | 4.8e-09 | 9.1e-09 | 4.9e-09 | 6.267E-09 | 2.454E-09 | 4.800E-09 | 9.100E-09 |
| 15162 | 759 | N759R | B | 6.3e-09 | 2.3e-09 | 1.6e-08 | 8.200E-09 | 7.045E-09 | 2.300E-09 | 1.600E-08 |
| 15163 | 759 | N759N | B | 4e-09 | 6.4e-09 | 9.4e-09 | 6.600E-09 | 2.706E-09 | 4.000E-09 | 9.400E-09 |
| 15164 | 759 | N759D | B | 1.9e-08 | 2.1e-09 | 9.2e-09 | 1.010E-08 | 8.486E-09 | 2.100E-09 | 1.900E-08 |
| 15165 | 759 | N759C | B | 1e-08 | 4.2e-09 | 6.8e-09 | 7.000E-09 | 2.905E-09 | 4.200E-09 | 1.000E-08 |
| 15166 | 759 | N759E | B | 9.2e-09 | 1.6e-08 | 9e-09 | 1.140E-08 | 3.985E-09 | 9.000E-09 | 1.600E-08 |
| 15167 | 759 | N759Q | B | 9.1e-09 | 1.9e-08 | 1.8e-09 | 9.967E-09 | 8.633E-09 | 1.800E-09 | 1.900E-08 |
| 15168 | 759 | N759G | B | 3e-09 | 5.6e-09 | 8.9e-09 | 5.833E-09 | 2.957E-09 | 3.000E-09 | 8.900E-09 |
| 15169 | 759 | N759H | B | 5.5e-09 | 7.6e-09 | 6.8e-09 | 6.633E-09 | 1.060E-09 | 5.500E-09 | 7.600E-09 |
| 15170 | 759 | N759I | B | 7.1e-09 | 9.2e-09 | 2.1e-09 | 6.133E-09 | 3.647E-09 | 2.100E-09 | 9.200E-09 |
| 15171 | 759 | N759L | B | 3e-09 | 1e-08 | 3e-08 | 1.433E-08 | 1.401E-08 | 3.000E-09 | 3.000E-08 |
| 15172 | 759 | N759K | B | 2.4e-08 | 1.3e-08 | 8.5e-09 | 1.517E-08 | 7.974E-09 | 8.500E-09 | 2.400E-08 |
| 15173 | 759 | N759M | B | 5.1e-09 | 1e-08 | 4.4e-09 | 6.500E-09 | 3.051E-09 | 4.400E-09 | 1.000E-08 |
| 15174 | 759 | N759F | B | 8.7e-09 | 8.4e-09 | 5.6e-09 | 7.567E-09 | 1.710E-09 | 5.600E-09 | 8.700E-09 |
| 15175 | 759 | N759P | B | 6.1e-09 | 5e-09 | 9.5e-09 | 6.867E-09 | 2.346E-09 | 5.000E-09 | 9.500E-09 |
| 15176 | 759 | N759S | B | 8.8e-09 | 3.2e-09 | 6.2e-09 | 6.067E-09 | 2.802E-09 | 3.200E-09 | 8.800E-09 |
| 15177 | 759 | N759T | B | 6.5e-09 | 9e-09 | 8.4e-09 | 7.967E-09 | 1.305E-09 | 6.500E-09 | 9.000E-09 |
| 15178 | 759 | N759W | B | 5.1e-09 | 1.5e-08 | 4.4e-09 | 8.167E-09 | 5.928E-09 | 4.400E-09 | 1.500E-08 |
| 15179 | 759 | N759Y | B | 1.1e-08 | 8.1e-09 | 1.1e-09 | 6.733E-09 | 5.090E-09 | 1.100E-09 | 1.100E-08 |
| 15180 | 759 | N759V | B | 5.6e-09 | 1.1e-08 | 5.8e-09 | 7.467E-09 | 3.062E-09 | 5.600E-09 | 1.100E-08 |
| 15181 | 760 | N760A | B | 5.9e-09 | 9.1e-09 | 1.6e-08 | 1.033E-08 | 5.162E-09 | 5.900E-09 | 1.600E-08 |
| 15182 | 760 | N760R | B | 6e-09 | 8.7e-09 | 4.9e-09 | 6.533E-09 | 1.955E-09 | 4.900E-09 | 8.700E-09 |
| 15183 | 760 | N760N | B | 4.7e-09 | 3.9e-09 | 3.5e-08 | 1.453E-08 | 1.773E-08 | 3.900E-09 | 3.500E-08 |
| 15184 | 760 | N760D | B | 6e-09 | 1.1e-08 | 1.2e-08 | 9.667E-09 | 3.215E-09 | 6.000E-09 | 1.200E-08 |
| 15185 | 760 | N760C | B | 7.4e-09 | 4.4e-09 | 5.8e-09 | 5.867E-09 | 1.501E-09 | 4.400E-09 | 7.400E-09 |
| 15186 | 760 | N760E | B | 5.8e-09 | 5.5e-09 | 3.6e-09 | 4.967E-09 | 1.193E-09 | 3.600E-09 | 5.800E-09 |
| 15187 | 760 | N760Q | B | 5.6e-09 | 9.2e-09 | 4.3e-09 | 6.367E-09 | 2.538E-09 | 4.300E-09 | 9.200E-09 |
| 15188 | 760 | N760G | B | 1.5e-09 | 7.8e-09 | 7.7e-09 | 5.667E-09 | 3.609E-09 | 1.500E-09 | 7.800E-09 |
| 15189 | 760 | N760H | B | 1.8e-09 | 7e-09 | 1.3e-08 | 7.267E-09 | 5.605E-09 | 1.800E-09 | 1.300E-08 |
| 15190 | 760 | N760I | B | 7.1e-09 | 6e-09 | 8.6e-09 | 7.233E-09 | 1.305E-09 | 6.000E-09 | 8.600E-09 |
| 15191 | 760 | N760L | B | 3.8e-09 | 1.3e-08 | 9.2e-09 | 8.667E-09 | 4.623E-09 | 3.800E-09 | 1.300E-08 |
| 15192 | 760 | N760K | B | 6e-09 | 7.1e-09 | 4e-09 | 5.700E-09 | 1.572E-09 | 4.000E-09 | 7.100E-09 |
| 15193 | 760 | N760M | B | 8.8e-09 | 1.7e-08 | 9.6e-09 | 1.180E-08 | 4.521E-09 | 8.800E-09 | 1.700E-08 |
| 15194 | 760 | N760F | B | 4.1e-09 | 1.4e-08 | 1.1e-08 | 9.700E-09 | 5.076E-09 | 4.100E-09 | 1.400E-08 |
| 15195 | 760 | N760P | B | 1.6e-08 | 9.5e-09 | 2.2e-09 | 9.233E-09 | 6.904E-09 | 2.200E-09 | 1.600E-08 |
| 15196 | 760 | N760S | B | 1.7e-08 | 3.4e-09 | 3.1e-09 | 7.833E-09 | 7.940E-09 | 3.100E-09 | 1.700E-08 |
| 15197 | 760 | N760T | B | 5.9e-09 | 6.6e-09 | 5.9e-09 | 6.133E-09 | 4.041E-10 | 5.900E-09 | 6.600E-09 |
| 15198 | 760 | N760W | B | 1.4e-08 | 4.8e-09 | 6e-09 | 8.267E-09 | 5.001E-09 | 4.800E-09 | 1.400E-08 |
| 15199 | 760 | N760Y | B | 4.2e-09 | 4.1e-09 | 2e-08 | 9.433E-09 | 9.151E-09 | 4.100E-09 | 2.000E-08 |
| 15200 | 760 | N760V | B | 1.1e-08 | 1e-08 | 6.1e-09 | 9.033E-09 | 2.589E-09 | 6.100E-09 | 1.100E-08 |
| 15201 | 761 | F761A | B | 8.1e-09 | 5.1e-09 | 1.3e-08 | 8.733E-09 | 3.988E-09 | 5.100E-09 | 1.300E-08 |
| 15202 | 761 | F761R | B | 4.9e-09 | 3e-09 | 8.6e-09 | 5.500E-09 | 2.848E-09 | 3.000E-09 | 8.600E-09 |
| 15203 | 761 | F761N | B | 5.9e-09 | 9.6e-09 | 2.7e-09 | 6.067E-09 | 3.453E-09 | 2.700E-09 | 9.600E-09 |
| 15204 | 761 | F761D | B | 8.5e-09 | 7.7e-09 | 8.7e-09 | 8.300E-09 | 5.292E-10 | 7.700E-09 | 8.700E-09 |
| 15205 | 761 | F761C | B | 1.8e-09 | 3.4e-09 | 5.8e-09 | 3.667E-09 | 2.013E-09 | 1.800E-09 | 5.800E-09 |
| 15206 | 761 | F761E | B | 7.1e-09 | 4e-09 | 6.7e-09 | 5.933E-09 | 1.686E-09 | 4.000E-09 | 7.100E-09 |
| 15207 | 761 | F761Q | B | 4e-09 | 8.8e-09 | 5.4e-09 | 6.067E-09 | 2.468E-09 | 4.000E-09 | 8.800E-09 |
| 15208 | 761 | F761G | B | 2.2e-08 | 4.2e-09 | 7.9e-09 | 1.137E-08 | 9.393E-09 | 4.200E-09 | 2.200E-08 |
| 15209 | 761 | F761H | B | 4.5e-09 | 6.6e-09 | 1.5e-08 | 8.700E-09 | 5.556E-09 | 4.500E-09 | 1.500E-08 |
| 15210 | 761 | F761I | B | 1.6e-08 | 6.3e-09 | 2.3e-08 | 1.510E-08 | 8.386E-09 | 6.300E-09 | 2.300E-08 |
| 15211 | 761 | F761L | B | 6.7e-09 | 6.9e-09 | 6.7e-09 | 6.767E-09 | 1.155E-10 | 6.700E-09 | 6.900E-09 |
| 15212 | 761 | F761K | B | 1.1e-08 | 3.3e-09 | 7.6e-09 | 7.300E-09 | 3.859E-09 | 3.300E-09 | 1.100E-08 |
| 15213 | 761 | F761M | B | 4e-09 | 1.1e-08 | 1.2e-08 | 9.000E-09 | 4.359E-09 | 4.000E-09 | 1.200E-08 |
| 15214 | 761 | F761F | B | 4.8e-09 | 4.3e-09 | 4.5e-09 | 4.533E-09 | 2.517E-10 | 4.300E-09 | 4.800E-09 |

| | | | | | | | | | | |
|---|---|---|---|---|---|---|---|---|---|---|
| 15215 | 761 | F761P | B | 5.3e-09 | 1.9e-09 | 5e-09 | 4.067E-09 | 1.882E-09 | 1.900E-09 | 5.300E-09 |
| 15216 | 761 | F761S | B | 1e-08 | 6.8e-09 | 8.4e-09 | 8.400E-09 | 1.600E-09 | 6.800E-09 | 1.000E-08 |
| 15217 | 761 | F761T | B | 5.5e-09 | 1.8e-09 | 8e-09 | 5.100E-09 | 3.119E-09 | 1.800E-09 | 8.000E-09 |
| 15218 | 761 | F761W | B | 2.2e-08 | 1e-08 | 5.9e-09 | 1.263E-08 | 8.367E-09 | 5.900E-09 | 2.200E-08 |
| 15219 | 761 | F761Y | B | 3.9e-09 | 6.3e-09 | 7.3e-09 | 5.833E-09 | 1.747E-09 | 3.900E-09 | 7.300E-09 |
| 15220 | 761 | F761V | B | 3.5e-09 | 9.1e-09 | 5.4e-09 | 6.000E-09 | 2.848E-09 | 3.500E-09 | 9.100E-09 |
| 15221 | 762 | Y762A | B | 6.3e-09 | 1.1e-08 | 1.2e-08 | 9.767E-09 | 3.044E-09 | 6.300E-09 | 1.200E-08 |
| 15222 | 762 | Y762R | B | 1e-08 | 5e-09 | 7.2e-09 | 7.400E-09 | 2.506E-09 | 5.000E-09 | 1.000E-08 |
| 15223 | 762 | Y762N | B | 4.1e-09 | 1.5e-08 | 2.4e-08 | 1.437E-08 | 9.965E-09 | 4.100E-09 | 2.400E-08 |
| 15224 | 762 | Y762D | B | 6e-09 | 5.2e-09 | 1e-08 | 7.067E-09 | 2.572E-09 | 5.200E-09 | 1.000E-08 |
| 15225 | 762 | Y762C | B | 7.3e-09 | 1.2e-09 | 6.3e-09 | 4.933E-09 | 3.272E-09 | 1.200E-09 | 7.300E-09 |
| 15226 | 762 | Y762E | B | 1.1e-08 | 9.2e-09 | 6.5e-09 | 8.900E-09 | 2.265E-09 | 6.500E-09 | 1.100E-08 |
| 15227 | 762 | Y762Q | B | 1.2e-09 | 3.4e-09 | 1.4e-08 | 6.200E-09 | 6.844E-09 | 1.200E-09 | 1.400E-08 |
| 15228 | 762 | Y762G | B | 4e-09 | 4.1e-09 | 8.1e-09 | 5.400E-09 | 2.339E-09 | 4.000E-09 | 8.100E-09 |
| 15229 | 762 | Y762H | B | 7.4e-09 | 1.7e-08 | 3.4e-09 | 9.267E-09 | 6.990E-09 | 3.400E-09 | 1.700E-08 |
| 15230 | 762 | Y762I | B | 2.4e-08 | 1.6e-08 | 8.8e-09 | 1.627E-08 | 7.604E-09 | 8.800E-09 | 2.400E-08 |
| 15231 | 762 | Y762L | B | 4.4e-09 | 9.2e-09 | 5.6e-08 | 2.320E-08 | 2.851E-08 | 4.400E-09 | 5.600E-08 |
| 15232 | 762 | Y762K | B | 1.3e-08 | 4.8e-09 | 1.1e-08 | 9.600E-09 | 4.276E-09 | 4.800E-09 | 1.300E-08 |
| 15233 | 762 | Y762M | B | 1e-08 | 3.3e-09 | 1.3e-08 | 8.767E-09 | 4.966E-09 | 3.300E-09 | 1.300E-08 |
| 15234 | 762 | Y762F | B | 9.5e-09 | 1.6e-08 | 3.7e-09 | 9.733E-09 | 6.153E-09 | 3.700E-09 | 1.600E-08 |
| 15235 | 762 | Y762P | B | 5.2e-09 | 5e-09 | 7.5e-09 | 5.900E-09 | 1.389E-09 | 5.000E-09 | 7.500E-09 |
| 15236 | 762 | Y762S | B | 8.6e-09 | 1e-08 | 5.9e-09 | 8.167E-09 | 2.084E-09 | 5.900E-09 | 1.000E-08 |
| 15237 | 762 | Y762T | B | 8.9e-09 | 5.5e-09 | 3.7e-09 | 6.033E-09 | 2.641E-09 | 3.700E-09 | 8.900E-09 |
| 15238 | 762 | Y762W | B | 1e-08 | 1.7e-08 | 1.2e-08 | 1.300E-08 | 3.606E-09 | 1.000E-08 | 1.700E-08 |
| 15239 | 762 | Y762Y | B | 7.3e-09 | 1.1e-08 | 5.2e-09 | 7.833E-09 | 2.937E-09 | 5.200E-09 | 1.100E-08 |
| 15240 | 762 | Y762V | B | 1.2e-08 | 6.3e-09 | 1.6e-08 | 1.143E-08 | 4.875E-09 | 6.300E-09 | 1.600E-08 |
| 15241 | 763 | P763A | B | 9.5e-09 | 1.1e-08 | 8.1e-09 | 9.533E-09 | 1.450E-09 | 8.100E-09 | 1.100E-08 |
| 15242 | 763 | P763R | B | 7.6e-09 | 5.4e-09 | 1.3e-08 | 8.667E-09 | 3.911E-09 | 5.400E-09 | 1.300E-08 |
| 15243 | 763 | P763N | B | 1.3e-08 | 7.5e-09 | 3.3e-09 | 7.933E-09 | 4.864E-09 | 3.300E-09 | 1.300E-08 |
| 15244 | 763 | P763D | B | 2.6e-09 | 1.3e-08 | 8.6e-09 | 8.067E-09 | 5.220E-09 | 2.600E-09 | 1.300E-08 |
| 15245 | 763 | P763C | B | 5.1e-09 | 6.9e-09 | 1.8e-08 | 1.000E-08 | 6.986E-09 | 5.100E-09 | 1.800E-08 |
| 15246 | 763 | P763E | B | 7.1e-09 | 7.5e-09 | 1.3e-08 | 9.200E-09 | 3.297E-09 | 7.100E-09 | 1.300E-08 |
| 15247 | 763 | P763Q | B | 4.5e-09 | 6e-09 | 7.2e-09 | 5.900E-09 | 1.353E-09 | 4.500E-09 | 7.200E-09 |
| 15248 | 763 | P763G | B | 4.8e-09 | 3.8e-09 | 1.2e-08 | 6.867E-09 | 4.474E-09 | 3.800E-09 | 1.200E-08 |
| 15249 | 763 | P763H | B | 4.5e-09 | 7.2e-09 | 7.7e-09 | 6.467E-09 | 1.721E-09 | 4.500E-09 | 7.700E-09 |
| 15250 | 763 | P763I | B | 2.1e-08 | 2.7e-09 | 1.3e-08 | 1.223E-08 | 9.174E-09 | 2.700E-09 | 2.100E-08 |
| 15251 | 763 | P763L | B | 3.5e-09 | 1.7e-08 | 1.8e-08 | 1.283E-08 | 8.098E-09 | 3.500E-09 | 1.800E-08 |
| 15252 | 763 | P763K | B | 1.4e-08 | 1.1e-08 | 2.3e-09 | 9.100E-09 | 6.077E-09 | 2.300E-09 | 1.400E-08 |
| 15253 | 763 | P763M | B | 7e-09 | 5.7e-09 | 1.2e-08 | 8.233E-09 | 3.326E-09 | 5.700E-09 | 1.200E-08 |
| 15254 | 763 | P763F | B | 1e-08 | 4.7e-09 | 4.5e-09 | 6.400E-09 | 3.119E-09 | 4.500E-09 | 1.000E-08 |
| 15255 | 763 | P763P | B | 7.4e-09 | 2.5e-08 | 3.9e-09 | 1.210E-08 | 1.131E-08 | 3.900E-09 | 2.500E-08 |
| 15256 | 763 | P763S | B | 7.9e-09 | 9.6e-09 | 1.1e-08 | 9.500E-09 | 1.552E-09 | 7.900E-09 | 1.100E-08 |
| 15257 | 763 | P763T | B | 1.1e-08 | 9.9e-09 | 2.1e-08 | 1.397E-08 | 6.116E-09 | 9.900E-09 | 2.100E-08 |
| 15258 | 763 | P763W | B | 5.8e-09 | 4.1e-09 | 6.8e-09 | 5.567E-09 | 1.365E-09 | 4.100E-09 | 6.800E-09 |
| 15259 | 763 | P763Y | B | 5.1e-09 | 7.7e-09 | 3e-09 | 5.267E-09 | 2.354E-09 | 3.000E-09 | 7.700E-09 |
| 15260 | 763 | P763V | B | 9.2e-09 | 1.9e-08 | 2.1e-09 | 1.010E-08 | 8.486E-09 | 2.100E-09 | 1.900E-08 |
| 15261 | 764 | R764A | B | 1.6e-08 | 7e-09 | 4.3e-09 | 9.100E-09 | 6.126E-09 | 4.300E-09 | 1.600E-08 |
| 15262 | 764 | R764R | B | 5.7e-09 | 9e-09 | 5.8e-09 | 6.833E-09 | 1.877E-09 | 5.700E-09 | 9.000E-09 |
| 15263 | 764 | R764N | B | 3.2e-09 | 1.2e-08 | 3.4e-08 | 1.640E-08 | 1.586E-08 | 3.200E-09 | 3.400E-08 |
| 15264 | 764 | R764D | B | 9.9e-09 | 6.7e-09 | 5.5e-09 | 7.367E-09 | 2.274E-09 | 5.500E-09 | 9.900E-09 |
| 15265 | 764 | R764C | B | 5.5e-09 | 1.4e-08 | 6.6e-09 | 8.700E-09 | 4.623E-09 | 5.500E-09 | 1.400E-08 |
| 15266 | 764 | R764E | B | 1.4e-08 | 3.6e-09 | 1.2e-08 | 9.867E-09 | 5.518E-09 | 3.600E-09 | 1.400E-08 |
| 15267 | 764 | R764Q | B | 7.8e-09 | 8e-09 | 6.6e-09 | 7.467E-09 | 7.572E-10 | 6.600E-09 | 8.000E-09 |
| 15268 | 764 | R764G | B | 8.8e-10 | 5.1e-09 | 8e-09 | 4.660E-09 | 3.580E-09 | 8.800E-10 | 8.000E-09 |
| 15269 | 764 | R764H | B | 3.1e-09 | 1.3e-08 | 6.5e-09 | 7.533E-09 | 5.030E-09 | 3.100E-09 | 1.300E-08 |
| 15270 | 764 | R764I | B | 6.2e-09 | 1.1e-08 | 9.7e-09 | 8.967E-09 | 2.483E-09 | 6.200E-09 | 1.100E-08 |
| 15271 | 764 | R764L | B | 1.4e-08 | 3e-09 | 8.6e-09 | 8.533E-09 | 5.500E-09 | 3.000E-09 | 1.400E-08 |
| 15272 | 764 | R764K | B | 5.7e-09 | 4.6e-09 | 4.9e-09 | 5.067E-09 | 5.686E-10 | 4.600E-09 | 5.700E-09 |
| 15273 | 764 | R764M | B | 2.8e-09 | 6.3e-09 | 9.3e-09 | 6.133E-09 | 3.253E-09 | 2.800E-09 | 9.300E-09 |
| 15274 | 764 | R764F | B | 6.6e-09 | 7.1e-09 | 2.2e-08 | 1.190E-08 | 8.750E-09 | 6.600E-09 | 2.200E-08 |
| 15275 | 764 | R764P | B | 1.8e-08 | 7.4e-09 | 1.4e-08 | 1.313E-08 | 5.353E-09 | 7.400E-09 | 1.800E-08 |
| 15276 | 764 | R764S | B | 4.5e-09 | 3.6e-09 | 6.4e-09 | 4.833E-09 | 1.429E-09 | 3.600E-09 | 6.400E-09 |
| 15277 | 764 | R764T | B | 1.4e-08 | 4.1e-09 | 7.2e-09 | 8.433E-09 | 5.064E-09 | 4.100E-09 | 1.400E-08 |
| 15278 | 764 | R764W | B | 2e-09 | 1.6e-08 | 1.5e-08 | 1.100E-08 | 7.810E-09 | 2.000E-09 | 1.600E-08 |
| 15279 | 764 | R764Y | B | 2.3e-08 | 1.3e-08 | 2.6e-09 | 1.287E-08 | 1.020E-08 | 2.600E-09 | 2.300E-08 |
| 15280 | 764 | R764V | B | 6.1e-09 | 2.6e-09 | 1.1e-08 | 6.567E-09 | 4.219E-09 | 2.600E-09 | 1.100E-08 |
| 15281 | 765 | E765A | B | 9.6e-09 | 1.3e-08 | 1.5e-09 | 8.033E-09 | 5.908E-09 | 1.500E-09 | 1.300E-08 |
| 15282 | 765 | E765R | B | 1.4e-08 | 4e-09 | 3.6e-09 | 7.200E-09 | 5.892E-09 | 3.600E-09 | 1.400E-08 |
| 15283 | 765 | E765N | B | 5.6e-09 | 8.9e-09 | 3.1e-09 | 5.867E-09 | 2.909E-09 | 3.100E-09 | 8.900E-09 |
| 15284 | 765 | E765D | B | 1.5e-09 | 1e-08 | 6.7e-09 | 6.067E-09 | 4.285E-09 | 1.500E-09 | 1.000E-08 |
| 15285 | 765 | E765C | B | 3.5e-09 | 4.4e-09 | 3.4e-09 | 3.767E-09 | 5.508E-10 | 3.400E-09 | 4.400E-09 |
| 15286 | 765 | E765E | B | 2.5e-09 | 6.4e-09 | 1.1e-08 | 6.633E-09 | 4.255E-09 | 2.500E-09 | 1.100E-08 |
| 15287 | 765 | E765Q | B | 6.1e-09 | 6.4e-09 | 1e-08 | 7.500E-09 | 2.170E-09 | 6.100E-09 | 1.000E-08 |
| 15288 | 765 | E765G | B | 9.2e-09 | 3.2e-09 | 8e-09 | 6.800E-09 | 3.175E-09 | 3.200E-09 | 9.200E-09 |
| 15289 | 765 | E765H | B | 8.2e-09 | 8.9e-09 | 9.7e-09 | 8.933E-09 | 7.506E-10 | 8.200E-09 | 9.700E-09 |
| 15290 | 765 | E765I | B | 3.8e-09 | 6.4e-09 | 9.3e-09 | 6.500E-09 | 2.751E-09 | 3.800E-09 | 9.300E-09 |
| 15291 | 765 | E765L | B | 3.5e-09 | 1.2e-08 | 3.3e-09 | 6.267E-09 | 4.966E-09 | 3.300E-09 | 1.200E-08 |
| 15292 | 765 | E765K | B | 9.6e-09 | 7.4e-09 | 6.9e-09 | 7.967E-09 | 1.436E-09 | 6.900E-09 | 9.600E-09 |
| 15293 | 765 | E765M | B | 4e-09 | 2e-09 | 5.3e-09 | 3.767E-09 | 1.662E-09 | 2.000E-09 | 5.300E-09 |
| 15294 | 765 | E765F | B | 1.3e-08 | 3.8e-09 | 8.7e-09 | 8.500E-09 | 4.603E-09 | 3.800E-09 | 1.300E-08 |
| 15295 | 765 | E765P | B | 7.4e-09 | 5.2e-09 | 4.6e-09 | 5.733E-09 | 1.474E-09 | 4.600E-09 | 7.400E-09 |
| 15296 | 765 | E765S | B | 3.8e-09 | 1.6e-08 | 4.5e-09 | 8.100E-09 | 6.851E-09 | 3.800E-09 | 1.600E-08 |
| 15297 | 765 | E765T | B | 6.7e-09 | 7.9e-09 | 5.1e-09 | 6.567E-09 | 1.405E-09 | 5.100E-09 | 7.900E-09 |
| 15298 | 765 | E765W | B | 5.2e-09 | 4.5e-09 | 6e-09 | 5.233E-09 | 7.506E-10 | 4.500E-09 | 6.000E-09 |
| 15299 | 765 | E765Y | B | 8e-09 | 1.5e-08 | 5e-09 | 9.333E-09 | 5.132E-09 | 5.000E-09 | 1.500E-08 |

| | | | | | | | | | | |
|---|---|---|---|---|---|---|---|---|---|---|
| 15300 | 765 | E765V | B | 8.3e-09 | 5.6e-09 | 6.2e-09 | 6.700E-09 | 1.418E-09 | 5.600E-09 | 8.300E-09 |
| 15301 | 766 | A766A | B | 7.1e-09 | 7.6e-09 | 1.1e-08 | 8.567E-09 | 2.122E-09 | 7.100E-09 | 1.100E-08 |
| 15302 | 766 | A766R | B | 4.8e-09 | 1.1e-08 | 6.1e-09 | 7.300E-09 | 3.270E-09 | 4.800E-09 | 1.100E-08 |
| 15303 | 766 | A766N | B | 1.1e-08 | 1.7e-08 | 1.4e-08 | 1.400E-08 | 3.000E-09 | 1.100E-08 | 1.700E-08 |
| 15304 | 766 | A766D | B | 5.8e-09 | 1.9e-08 | 6.8e-09 | 1.053E-08 | 7.349E-09 | 5.800E-09 | 1.900E-08 |
| 15305 | 766 | A766C | B | 4.7e-09 | 7e-09 | 1.2e-08 | 7.900E-09 | 3.732E-09 | 4.700E-09 | 1.200E-08 |
| 15306 | 766 | A766E | B | 6.1e-09 | 1.4e-08 | 1.1e-08 | 1.037E-08 | 3.988E-09 | 6.100E-09 | 1.400E-08 |
| 15307 | 766 | A766Q | B | 1.6e-08 | 9.7e-09 | 8.4e-09 | 1.137E-08 | 4.065E-09 | 8.400E-09 | 1.600E-08 |
| 15308 | 766 | A766G | B | 4e-08 | 6.8e-09 | 7.7e-09 | 1.817E-08 | 1.891E-08 | 6.800E-09 | 4.000E-08 |
| 15309 | 766 | A766H | B | 1.5e-08 | 1.2e-08 | 2.9e-09 | 9.967E-09 | 6.301E-09 | 2.900E-09 | 1.500E-08 |
| 15310 | 766 | A766I | B | 8e-09 | 4.3e-09 | 5.6e-09 | 5.967E-09 | 1.877E-09 | 4.300E-09 | 8.000E-09 |
| 15311 | 766 | A766L | B | 7.8e-09 | 5.7e-09 | 6e-09 | 6.500E-09 | 1.136E-09 | 5.700E-09 | 7.800E-09 |
| 15312 | 766 | A766K | B | 5.4e-09 | 5.1e-09 | 2.5e-09 | 4.333E-09 | 1.595E-09 | 2.500E-09 | 5.400E-09 |
| 15313 | 766 | A766M | B | 3.5e-09 | 1.5e-08 | 4.3e-09 | 7.600E-09 | 6.421E-09 | 3.500E-09 | 1.500E-08 |
| 15314 | 766 | A766F | B | 1.1e-08 | 1.4e-08 | 4.6e-09 | 9.867E-09 | 4.801E-09 | 4.600E-09 | 1.400E-08 |
| 15315 | 766 | A766P | B | 8.3e-09 | 6.9e-09 | 2.5e-08 | 1.340E-08 | 1.007E-08 | 6.900E-09 | 2.500E-08 |
| 15316 | 766 | A766S | B | 4.7e-08 | 5.6e-09 | 2.8e-09 | 1.847E-08 | 2.475E-08 | 2.800E-09 | 4.700E-08 |
| 15317 | 766 | A766T | B | 6.7e-09 | 5.8e-09 | 5.6e-09 | 6.033E-09 | 5.859E-10 | 5.600E-09 | 6.700E-09 |
| 15318 | 766 | A766W | B | 2.6e-09 | 2.5e-08 | 6.7e-09 | 1.143E-08 | 1.193E-08 | 2.600E-09 | 2.500E-08 |
| 15319 | 766 | A766Y | B | 8.4e-09 | 7.5e-09 | 5.5e-09 | 7.133E-09 | 1.484E-09 | 5.500E-09 | 8.400E-09 |
| 15320 | 766 | A766V | B | 1.3e-08 | 4.9e-09 | 3.8e-09 | 7.233E-09 | 5.024E-09 | 3.800E-09 | 1.300E-08 |
| 15321 | 767 | K767A | B | 6.9e-09 | 1.5e-08 | 6.7e-09 | 9.533E-09 | 4.735E-09 | 6.700E-09 | 1.500E-08 |
| 15322 | 767 | K767R | B | 9.8e-09 | 1.5e-08 | 8.4e-09 | 1.107E-08 | 3.478E-09 | 8.400E-09 | 1.500E-08 |
| 15323 | 767 | K767N | B | 2.6e-09 | 5.8e-09 | 1.2e-08 | 6.800E-09 | 4.779E-09 | 2.600E-09 | 1.200E-08 |
| 15324 | 767 | K767D | B | 1.2e-08 | 2.4e-09 | 1.2e-08 | 8.800E-09 | 5.543E-09 | 2.400E-09 | 1.200E-08 |
| 15325 | 767 | K767C | B | 1.6e-08 | 7.3e-09 | 8.1e-09 | 1.047E-08 | 4.809E-09 | 7.300E-09 | 1.600E-08 |
| 15326 | 767 | K767E | B | 4.1e-09 | 1.5e-08 | 1.6e-08 | 1.170E-08 | 6.601E-09 | 4.100E-09 | 1.600E-08 |
| 15327 | 767 | K767Q | B | 2.7e-09 | 3.8e-09 | 4.9e-09 | 3.800E-09 | 1.100E-09 | 2.700E-09 | 4.900E-09 |
| 15328 | 767 | K767G | B | 5.4e-09 | 5.5e-09 | 3.3e-09 | 4.733E-09 | 1.242E-09 | 3.300E-09 | 5.500E-09 |
| 15329 | 767 | K767H | B | 2.4e-09 | 3.5e-09 | 1.7e-08 | 7.633E-09 | 8.130E-09 | 2.400E-09 | 1.700E-08 |
| 15330 | 767 | K767I | B | 4.6e-09 | 1.2e-08 | 5.6e-09 | 7.400E-09 | 4.015E-09 | 4.600E-09 | 1.200E-08 |
| 15331 | 767 | K767L | B | 5e-09 | 9.3e-09 | 6.3e-09 | 6.867E-09 | 2.205E-09 | 5.000E-09 | 9.300E-09 |
| 15332 | 767 | K767K | B | 2.4e-08 | 1.1e-08 | 7.3e-09 | 1.410E-08 | 8.771E-09 | 7.300E-09 | 2.400E-08 |
| 15333 | 767 | K767M | B | 2e-09 | 7.1e-09 | 8.9e-09 | 6.000E-09 | 3.579E-09 | 2.000E-09 | 8.900E-09 |
| 15334 | 767 | K767F | B | 2.5e-09 | 1.3e-08 | 4.5e-09 | 6.667E-09 | 5.575E-09 | 2.500E-09 | 1.300E-08 |
| 15335 | 767 | K767P | B | 8.4e-09 | 8.4e-09 | 5.8e-09 | 7.533E-09 | 1.501E-09 | 5.800E-09 | 8.400E-09 |
| 15336 | 767 | K767S | B | 8e-09 | 2.3e-08 | 1.7e-08 | 1.600E-08 | 7.550E-09 | 8.000E-09 | 2.300E-08 |
| 15337 | 767 | K767T | B | 7.6e-09 | 4.7e-09 | 7.7e-09 | 6.667E-09 | 1.704E-09 | 4.700E-09 | 7.700E-09 |
| 15338 | 767 | K767W | B | 6.1e-09 | 6.1e-09 | 9.5e-09 | 7.233E-09 | 1.963E-09 | 6.100E-09 | 9.500E-09 |
| 15339 | 767 | K767Y | B | 1.4e-08 | 7.4e-09 | 1.4e-08 | 1.180E-08 | 3.811E-09 | 7.400E-09 | 1.400E-08 |
| 15340 | 767 | K767V | B | 1.3e-08 | 1.1e-08 | 4e-09 | 9.333E-09 | 4.726E-09 | 4.000E-09 | 1.300E-08 |
| 15341 | 768 | V768A | B | 2.6e-08 | 6.7e-09 | 7.1e-09 | 1.327E-08 | 1.103E-08 | 6.700E-09 | 2.600E-08 |
| 15342 | 768 | V768R | B | 1.2e-08 | 6.1e-09 | 4.2e-09 | 7.433E-09 | 4.067E-09 | 4.200E-09 | 1.200E-08 |
| 15343 | 768 | V768N | B | 9.5e-09 | 1.2e-08 | 6.6e-09 | 9.367E-09 | 2.702E-09 | 6.600E-09 | 1.200E-08 |
| 15344 | 768 | V768D | B | 9.7e-09 | 5.4e-09 | 1.9e-08 | 1.137E-08 | 6.951E-09 | 5.400E-09 | 1.900E-08 |
| 15345 | 768 | V768C | B | 5.9e-09 | 7.2e-09 | 3.3e-09 | 5.467E-09 | 1.986E-09 | 3.300E-09 | 7.200E-09 |
| 15346 | 768 | V768E | B | 1.1e-08 | 5.7e-09 | 3.2e-09 | 6.633E-09 | 3.983E-09 | 3.200E-09 | 1.100E-08 |
| 15347 | 768 | V768Q | B | 1.5e-08 | 9.6e-09 | 7.6e-09 | 1.073E-08 | 3.828E-09 | 7.600E-09 | 1.500E-08 |
| 15348 | 768 | V768G | B | 2.1e-09 | 2.4e-08 | 1.1e-08 | 1.237E-08 | 1.101E-08 | 2.100E-09 | 2.400E-08 |
| 15349 | 768 | V768H | B | 1.9e-08 | 6.6e-09 | 1.8e-08 | 1.453E-08 | 6.889E-09 | 6.600E-09 | 1.900E-08 |
| 15350 | 768 | V768I | B | 1.8e-08 | 2.4e-08 | 2.3e-08 | 2.167E-08 | 3.215E-09 | 1.800E-08 | 2.400E-08 |
| 15351 | 768 | V768L | B | 8e-09 | 6.7e-09 | 5.9e-09 | 6.867E-09 | 1.060E-09 | 5.900E-09 | 8.000E-09 |
| 15352 | 768 | V768K | B | 3.8e-09 | 1.8e-08 | 6.4e-09 | 9.400E-09 | 7.560E-09 | 3.800E-09 | 1.800E-08 |
| 15353 | 768 | V768M | B | 3.3e-09 | 5.5e-09 | 5.9e-09 | 4.900E-09 | 1.400E-09 | 3.300E-09 | 5.900E-09 |
| 15354 | 768 | V768F | B | 8.4e-09 | 8.7e-09 | 2.8e-09 | 6.633E-09 | 3.323E-09 | 2.800E-09 | 8.700E-09 |
| 15355 | 768 | V768P | B | 3e-09 | 6.6e-09 | 2.1e-09 | 3.900E-09 | 2.381E-09 | 2.100E-09 | 6.600E-09 |
| 15356 | 768 | V768S | B | 5.7e-09 | 8.1e-09 | 8.9e-09 | 7.567E-09 | 1.665E-09 | 5.700E-09 | 8.900E-09 |
| 15357 | 768 | V768T | B | 4.7e-09 | 1.1e-08 | 8.6e-09 | 8.100E-09 | 3.180E-09 | 4.700E-09 | 1.100E-08 |
| 15358 | 768 | V768W | B | 5.6e-09 | 9.6e-09 | 6.9e-09 | 7.367E-09 | 2.040E-09 | 5.600E-09 | 9.600E-09 |
| 15359 | 768 | V768Y | B | 9e-09 | 6.3e-09 | 4.8e-09 | 6.700E-09 | 2.128E-09 | 4.800E-09 | 9.000E-09 |
| 15360 | 768 | V768V | B | 7.6e-09 | 4.3e-09 | 5.5e-09 | 5.800E-09 | 1.670E-09 | 4.300E-09 | 7.600E-09 |
| 15361 | 769 | Q769A | B | 1.5e-08 | 6.4e-09 | 9e-09 | 1.013E-08 | 4.411E-09 | 6.400E-09 | 1.500E-08 |
| 15362 | 769 | Q769R | B | 5.4e-09 | 5.4e-09 | 3.9e-09 | 4.900E-09 | 8.660E-10 | 3.900E-09 | 5.400E-09 |
| 15363 | 769 | Q769N | B | 7.6e-09 | 7.2e-09 | 8.1e-09 | 7.633E-09 | 4.509E-10 | 7.200E-09 | 8.100E-09 |
| 15364 | 769 | Q769D | B | 1.6e-08 | 3.4e-09 | 5.2e-09 | 8.200E-09 | 6.815E-09 | 3.400E-09 | 1.600E-08 |
| 15365 | 769 | Q769C | B | 8.3e-09 | 1.1e-08 | 1.1e-08 | 1.010E-08 | 1.559E-09 | 8.300E-09 | 1.100E-08 |
| 15366 | 769 | Q769E | B | 5.9e-09 | 3.7e-09 | 1.1e-08 | 6.867E-09 | 3.745E-09 | 3.700E-09 | 1.100E-08 |
| 15367 | 769 | Q769Q | B | 9.7e-09 | 8.9e-10 | 1e-08 | 6.863E-09 | 5.175E-09 | 8.900E-10 | 1.000E-08 |
| 15368 | 769 | Q769G | B | 1.1e-08 | 9.4e-09 | 7.7e-09 | 9.367E-09 | 1.650E-09 | 7.700E-09 | 1.100E-08 |
| 15369 | 769 | Q769H | B | 4.1e-09 | 6.7e-09 | 5.1e-09 | 5.300E-09 | 1.311E-09 | 4.100E-09 | 6.700E-09 |
| 15370 | 769 | Q769I | B | 7e-09 | 9.9e-09 | 2.2e-09 | 6.367E-09 | 3.889E-09 | 2.200E-09 | 9.900E-09 |
| 15371 | 769 | Q769L | B | 6.2e-09 | 8.5e-09 | 6.2e-09 | 6.967E-09 | 1.328E-09 | 6.200E-09 | 8.500E-09 |
| 15372 | 769 | Q769K | B | 9.9e-09 | 3.7e-09 | 5.9e-09 | 6.500E-09 | 3.143E-09 | 3.700E-09 | 9.900E-09 |
| 15373 | 769 | Q769M | B | 7.5e-09 | 8.3e-09 | 2.1e-09 | 5.967E-09 | 3.372E-09 | 2.100E-09 | 8.300E-09 |
| 15374 | 769 | Q769F | B | 1.1e-08 | 3.7e-09 | 6.7e-09 | 7.133E-09 | 3.669E-09 | 3.700E-09 | 1.100E-08 |
| 15375 | 769 | Q769P | B | 3.1e-09 | 9.8e-09 | 3.5e-09 | 5.467E-09 | 3.758E-09 | 3.100E-09 | 9.800E-09 |
| 15376 | 769 | Q769S | B | 1.2e-08 | 2.3e-08 | 6.2e-09 | 1.373E-08 | 8.533E-09 | 6.200E-09 | 2.300E-08 |
| 15377 | 769 | Q769T | B | 5.9e-09 | 7.1e-09 | 3.7e-09 | 5.567E-09 | 1.724E-09 | 3.700E-09 | 7.100E-09 |
| 15378 | 769 | Q769W | B | 6.4e-09 | 7.9e-09 | 4.5e-09 | 6.267E-09 | 1.704E-09 | 4.500E-09 | 7.900E-09 |
| 15379 | 769 | Q769Y | B | 5e-09 | 3.9e-09 | 1.6e-08 | 8.300E-09 | 6.691E-09 | 3.900E-09 | 1.600E-08 |
| 15380 | 769 | Q769V | B | 6e-09 | 1.4e-08 | 2.4e-08 | 1.467E-08 | 9.018E-09 | 6.000E-09 | 2.400E-08 |
| 15381 | 770 | W770A | B | 5.9e-09 | 8.3e-09 | 3.4e-09 | 5.867E-09 | 2.450E-09 | 3.400E-09 | 8.300E-09 |
| 15382 | 770 | W770R | B | 1.5e-08 | 8.8e-09 | 1.6e-08 | 1.327E-08 | 3.900E-09 | 8.800E-09 | 1.600E-08 |
| 15383 | 770 | W770N | B | 7.4e-09 | 6.6e-09 | 4.7e-09 | 6.233E-09 | 1.387E-09 | 4.700E-09 | 7.400E-09 |
| 15384 | 770 | W770D | B | 5.6e-09 | 6.7e-09 | 6.9e-09 | 6.400E-09 | 7.000E-10 | 5.600E-09 | 6.900E-09 |

| | | | | | | | | | | |
|---|---|---|---|---|---|---|---|---|---|---|
| 15385 | 770 | W770C | B | 4.3e-09 | 6.7e-09 | 1.2e-08 | 7.667E-09 | 3.940E-09 | 4.300E-09 | 1.200E-08 |
| 15386 | 770 | W770E | B | 6.9e-09 | 9e-09 | 5.1e-09 | 7.000E-09 | 1.952E-09 | 5.100E-09 | 9.000E-09 |
| 15387 | 770 | W770Q | B | 2.1e-08 | 2.5e-09 | 1.8e-08 | 1.383E-08 | 9.929E-09 | 2.500E-09 | 2.100E-08 |
| 15388 | 770 | W770G | B | 3e-09 | 9.2e-09 | 1.7e-08 | 9.733E-09 | 7.015E-09 | 3.000E-09 | 1.700E-08 |
| 15389 | 770 | W770H | B | 3.3e-08 | 9e-09 | 2.1e-08 | 2.100E-08 | 1.200E-08 | 9.000E-09 | 3.300E-08 |
| 15390 | 770 | W770I | B | 4.3e-09 | 1.8e-09 | 1.8e-09 | 2.633E-09 | 1.443E-09 | 1.800E-09 | 4.300E-09 |
| 15391 | 770 | W770L | B | 6.2e-09 | 7.9e-09 | 2.3e-09 | 5.467E-09 | 2.871E-09 | 2.300E-09 | 7.900E-09 |
| 15392 | 770 | W770K | B | 5.9e-09 | 7.2e-09 | 2.2e-08 | 1.170E-08 | 8.944E-09 | 5.900E-09 | 2.200E-08 |
| 15393 | 770 | W770M | B | 9.5e-09 | 5.9e-09 | 1.9e-08 | 1.147E-08 | 6.768E-09 | 5.900E-09 | 1.900E-08 |
| 15394 | 770 | W770F | B | 8.6e-09 | 6.8e-09 | 7.3e-09 | 7.567E-09 | 9.292E-10 | 6.800E-09 | 8.600E-09 |
| 15395 | 770 | W770P | B | 5.2e-09 | 6.1e-09 | 3.9e-09 | 5.067E-09 | 1.106E-09 | 3.900E-09 | 6.100E-09 |
| 15396 | 770 | W770S | B | 9.2e-09 | 5e-09 | 5.1e-09 | 6.433E-09 | 2.397E-09 | 5.000E-09 | 9.200E-09 |
| 15397 | 770 | W770T | B | 5.6e-09 | 3.5e-09 | 3.6e-09 | 4.233E-09 | 1.185E-09 | 3.500E-09 | 5.600E-09 |
| 15398 | 770 | W770W | B | 7e-09 | 8.6e-09 | 2.4e-09 | 6.000E-09 | 3.219E-09 | 2.400E-09 | 8.600E-09 |
| 15399 | 770 | W770Y | B | 9.6e-09 | 9.4e-09 | 5.8e-09 | 8.267E-09 | 2.139E-09 | 5.800E-09 | 9.600E-09 |
| 15400 | 770 | W770V | B | 9.8e-09 | 1.8e-08 | 7e-09 | 1.160E-08 | 5.717E-09 | 7.000E-09 | 1.800E-08 |
| 15401 | 771 | K771A | B | 7.7e-09 | 6.8e-09 | 1.4e-09 | 5.300E-09 | 3.407E-09 | 1.400E-09 | 7.700E-09 |
| 15402 | 771 | K771R | B | 4.6e-09 | 5.7e-09 | 1.6e-08 | 8.767E-09 | 6.288E-09 | 4.600E-09 | 1.600E-08 |
| 15403 | 771 | K771N | B | 5.5e-09 | 3.8e-09 | 1.1e-08 | 6.767E-09 | 3.763E-09 | 3.800E-09 | 1.100E-08 |
| 15404 | 771 | K771D | B | 1.7e-08 | 7.6e-09 | 3.4e-09 | 9.333E-09 | 6.964E-09 | 3.400E-09 | 1.700E-08 |
| 15405 | 771 | K771C | B | 7.2e-09 | 9.9e-09 | 1.2e-08 | 9.700E-09 | 2.406E-09 | 7.200E-09 | 1.200E-08 |
| 15406 | 771 | K771E | B | 3.6e-09 | 4.5e-09 | 1.3e-09 | 3.133E-09 | 1.650E-09 | 1.300E-09 | 4.500E-09 |
| 15407 | 771 | K771Q | B | 1.1e-08 | 3.4e-09 | 1.8e-08 | 1.080E-08 | 7.302E-09 | 3.400E-09 | 1.800E-08 |
| 15408 | 771 | K771G | B | 1.1e-08 | 5.5e-09 | 1.4e-08 | 1.017E-08 | 4.311E-09 | 5.500E-09 | 1.400E-08 |
| 15409 | 771 | K771H | B | 4.6e-09 | 4.8e-09 | 2e-08 | 9.800E-09 | 8.834E-09 | 4.600E-09 | 2.000E-08 |
| 15410 | 771 | K771I | B | 1.1e-08 | 1.4e-08 | 1.6e-08 | 1.367E-08 | 2.517E-09 | 1.100E-08 | 1.600E-08 |
| 15411 | 771 | K771L | B | 4.7e-09 | 5.1e-09 | 7.4e-09 | 5.733E-09 | 1.457E-09 | 4.700E-09 | 7.400E-09 |
| 15412 | 771 | K771K | B | 2.2e-08 | 7.2e-09 | 8.6e-09 | 1.260E-08 | 8.171E-09 | 7.200E-09 | 2.200E-08 |
| 15413 | 771 | K771M | B | 4.6e-09 | 9.8e-09 | 1.5e-08 | 9.800E-09 | 5.200E-09 | 4.600E-09 | 1.500E-08 |
| 15414 | 771 | K771F | B | 8.1e-09 | 2.6e-08 | 1.5e-09 | 1.187E-08 | 1.268E-08 | 1.500E-09 | 2.600E-08 |
| 15415 | 771 | K771P | B | 1.3e-08 | 1.6e-08 | 4.1e-09 | 1.103E-08 | 6.189E-09 | 4.100E-09 | 1.600E-08 |
| 15416 | 771 | K771S | B | 6.3e-09 | 3.4e-09 | 9.6e-09 | 6.433E-09 | 3.102E-09 | 3.400E-09 | 9.600E-09 |
| 15417 | 771 | K771T | B | 4.8e-09 | 8.3e-09 | 1.3e-08 | 8.700E-09 | 4.115E-09 | 4.800E-09 | 1.300E-08 |
| 15418 | 771 | K771W | B | 6.9e-09 | 7.2e-09 | 7.7e-09 | 7.267E-09 | 4.041E-10 | 6.900E-09 | 7.700E-09 |
| 15419 | 771 | K771Y | B | 1.2e-08 | 6.7e-09 | 3.5e-09 | 7.400E-09 | 4.293E-09 | 3.500E-09 | 1.200E-08 |
| 15420 | 771 | K771V | B | 1.8e-08 | 1.1e-08 | 1.6e-08 | 1.500E-08 | 3.606E-09 | 1.100E-08 | 1.800E-08 |
| 15421 | 772 | V772A | B | 5.2e-09 | 7.2e-09 | 1.5e-08 | 9.133E-09 | 5.178E-09 | 5.200E-09 | 1.500E-08 |
| 15422 | 772 | V772R | B | 1.2e-08 | 8.6e-09 | 5e-09 | 8.533E-09 | 3.500E-09 | 5.000E-09 | 1.200E-08 |
| 15423 | 772 | V772N | B | 3.2e-09 | 5.9e-09 | 2.2e-09 | 3.767E-09 | 1.914E-09 | 2.200E-09 | 5.900E-09 |
| 15424 | 772 | V772D | B | 5.1e-09 | 6.3e-09 | 1.3e-09 | 4.233E-09 | 2.610E-09 | 1.300E-09 | 6.300E-09 |
| 15425 | 772 | V772C | B | 1.3e-08 | 3.7e-09 | 3.1e-09 | 6.600E-09 | 5.551E-09 | 3.100E-09 | 1.300E-08 |
| 15426 | 772 | V772E | B | 9.9e-09 | 1e-08 | 4.5e-09 | 8.133E-09 | 3.147E-09 | 4.500E-09 | 1.000E-08 |
| 15427 | 772 | V772Q | B | 9.9e-10 | 4.8e-09 | 1e-08 | 5.263E-09 | 4.523E-09 | 9.900E-10 | 1.000E-08 |
| 15428 | 772 | V772G | B | 7.8e-09 | 1.4e-08 | 9.4e-09 | 1.040E-08 | 3.219E-09 | 7.800E-09 | 1.400E-08 |
| 15429 | 772 | V772H | B | 3.1e-09 | 7.4e-09 | 5.6e-09 | 5.367E-09 | 2.159E-09 | 3.100E-09 | 7.400E-09 |
| 15430 | 772 | V772I | B | 7e-09 | 2e-08 | 5.5e-09 | 1.083E-08 | 7.974E-09 | 5.500E-09 | 2.000E-08 |
| 15431 | 772 | V772L | B | 1.1e-08 | 6.7e-09 | 6.8e-09 | 8.167E-09 | 2.454E-09 | 6.700E-09 | 1.100E-08 |
| 15432 | 772 | V772K | B | 1.4e-08 | 5.1e-09 | 7.3e-09 | 8.800E-09 | 4.636E-09 | 5.100E-09 | 1.400E-08 |
| 15433 | 772 | V772M | B | 7.9e-09 | 9.6e-09 | 2.3e-09 | 6.600E-09 | 3.820E-09 | 2.300E-09 | 9.600E-09 |
| 15434 | 772 | V772F | B | 7.5e-09 | 8.2e-09 | 3.9e-09 | 6.533E-09 | 2.307E-09 | 3.900E-09 | 8.200E-09 |
| 15435 | 772 | V772P | B | 7.4e-09 | 6.3e-09 | 1.2e-08 | 8.567E-09 | 3.024E-09 | 6.300E-09 | 1.200E-08 |
| 15436 | 772 | V772S | B | 1.2e-08 | 1.6e-08 | 3e-09 | 1.033E-08 | 6.658E-09 | 3.000E-09 | 1.600E-08 |
| 15437 | 772 | V772T | B | 5.3e-09 | 9.7e-09 | 6e-09 | 7.000E-09 | 2.364E-09 | 5.300E-09 | 9.700E-09 |
| 15438 | 772 | V772W | B | 1.8e-08 | 2.3e-08 | 9.8e-09 | 1.693E-08 | 6.664E-09 | 9.800E-09 | 2.300E-08 |
| 15439 | 772 | V772Y | B | 9.5e-09 | 1.7e-09 | 4.7e-09 | 5.300E-09 | 3.934E-09 | 1.700E-09 | 9.500E-09 |
| 15440 | 772 | V772V | B | 8e-09 | 9.8e-10 | 6.2e-09 | 5.060E-09 | 3.646E-09 | 9.800E-10 | 8.000E-09 |
| 15441 | 773 | D773A | B | 6.2e-09 | 5.3e-09 | 1.9e-08 | 1.017E-08 | 7.663E-09 | 5.300E-09 | 1.900E-08 |
| 15442 | 773 | D773R | B | 1.4e-09 | 6.4e-09 | 6.6e-09 | 4.800E-09 | 2.946E-09 | 1.400E-09 | 6.600E-09 |
| 15443 | 773 | D773N | B | 7.4e-09 | 2.6e-08 | 7.6e-09 | 1.367E-08 | 1.068E-08 | 7.400E-09 | 2.600E-08 |
| 15444 | 773 | D773D | B | 4.3e-09 | 2.8e-09 | 5.3e-09 | 4.133E-09 | 1.258E-09 | 2.800E-09 | 5.300E-09 |
| 15445 | 773 | D773C | B | 4e-09 | 6.7e-09 | 6.1e-09 | 5.600E-09 | 1.418E-09 | 4.000E-09 | 6.700E-09 |
| 15446 | 773 | D773E | B | 1.4e-08 | 3.8e-09 | 1.3e-08 | 1.027E-08 | 5.623E-09 | 3.800E-09 | 1.400E-08 |
| 15447 | 773 | D773Q | B | 2.4e-08 | 1e-08 | 7.3e-09 | 1.377E-08 | 8.965E-09 | 7.300E-09 | 2.400E-08 |
| 15448 | 773 | D773G | B | 6.3e-09 | 9.3e-09 | 1.1e-08 | 8.867E-09 | 2.380E-09 | 6.300E-09 | 1.100E-08 |
| 15449 | 773 | D773H | B | 6.8e-09 | 1.7e-08 | 6.6e-09 | 1.013E-08 | 5.948E-09 | 6.600E-09 | 1.700E-08 |
| 15450 | 773 | D773I | B | 4.4e-09 | 3.7e-09 | 5.9e-09 | 4.667E-09 | 1.124E-09 | 3.700E-09 | 5.900E-09 |
| 15451 | 773 | D773L | B | 4.2e-09 | 5.5e-09 | 1.8e-09 | 3.833E-09 | 1.877E-09 | 1.800E-09 | 5.500E-09 |
| 15452 | 773 | D773K | B | 1e-08 | 3.6e-09 | 5.4e-09 | 6.333E-09 | 3.301E-09 | 3.600E-09 | 1.000E-08 |
| 15453 | 773 | D773M | B | 1.1e-08 | 5.8e-09 | 7.1e-09 | 7.967E-09 | 2.706E-09 | 5.800E-09 | 1.100E-08 |
| 15454 | 773 | D773F | B | 1.4e-09 | 6.7e-09 | 1.4e-08 | 7.367E-09 | 6.326E-09 | 1.400E-09 | 1.400E-08 |
| 15455 | 773 | D773P | B | 5.4e-09 | 8.6e-09 | 1.7e-08 | 1.033E-08 | 5.991E-09 | 5.400E-09 | 1.700E-08 |
| 15456 | 773 | D773S | B | 1.2e-08 | 1.1e-08 | 1.1e-08 | 1.133E-08 | 5.774E-10 | 1.100E-08 | 1.200E-08 |
| 15457 | 773 | D773T | B | 3.4e-09 | 1.5e-08 | 1e-08 | 9.467E-09 | 5.818E-09 | 3.400E-09 | 1.500E-08 |
| 15458 | 773 | D773W | B | 6.3e-09 | 1.4e-08 | 1e-08 | 1.010E-08 | 3.851E-09 | 6.300E-09 | 1.400E-08 |
| 15459 | 773 | D773Y | B | 6.4e-09 | 3.8e-09 | 5.5e-09 | 5.233E-09 | 1.320E-09 | 3.800E-09 | 6.400E-09 |
| 15460 | 773 | D773V | B | 7.5e-09 | 5e-09 | 4.3e-09 | 5.600E-09 | 1.682E-09 | 4.300E-09 | 7.500E-09 |
| 15461 | 774 | N774A | B | 4.3e-09 | 2.8e-09 | 1e-08 | 5.700E-09 | 3.799E-09 | 2.800E-09 | 1.000E-08 |
| 15462 | 774 | N774R | B | 8.1e-09 | 4.6e-09 | 8.8e-09 | 7.167E-09 | 2.250E-09 | 4.600E-09 | 8.800E-09 |
| 15463 | 774 | N774N | B | 4e-09 | 3.9e-09 | 4.6e-09 | 4.167E-09 | 3.786E-10 | 3.900E-09 | 4.600E-09 |
| 15464 | 774 | N774D | B | 1e-08 | 6.9e-09 | 1.1e-08 | 9.300E-09 | 2.138E-09 | 6.900E-09 | 1.100E-08 |
| 15465 | 774 | N774C | B | 6.9e-09 | 1.7e-09 | 5.4e-09 | 4.667E-09 | 2.676E-09 | 1.700E-09 | 6.900E-09 |
| 15466 | 774 | N774E | B | 7.3e-09 | 4.1e-09 | 1.1e-08 | 7.467E-09 | 3.453E-09 | 4.100E-09 | 1.100E-08 |
| 15467 | 774 | N774Q | B | 1.9e-09 | 8.6e-09 | 1.7e-08 | 9.167E-09 | 7.566E-09 | 1.900E-09 | 1.700E-08 |
| 15468 | 774 | N774G | B | 6.6e-09 | 4.1e-08 | 4.6e-09 | 1.740E-08 | 2.046E-08 | 4.600E-09 | 4.100E-08 |
| 15469 | 774 | N774H | B | 5.1e-09 | 7.3e-09 | 9.3e-09 | 7.233E-09 | 2.101E-09 | 5.100E-09 | 9.300E-09 |

| | | | | | | | | | | |
|---|---|---|---|---|---|---|---|---|---|---|
| 15470 | 774 | N774I | B | 6.3e-09 | 1.3e-08 | 1e-08 | 9.767E-09 | 3.356E-09 | 6.300E-09 | 1.300E-08 |
| 15471 | 774 | N774L | B | 2.2e-09 | 5.7e-09 | 5.5e-09 | 4.467E-09 | 1.966E-09 | 2.200E-09 | 5.700E-09 |
| 15472 | 774 | N774K | B | 6.7e-09 | 1.3e-08 | 8.3e-09 | 9.333E-09 | 3.275E-09 | 6.700E-09 | 1.300E-08 |
| 15473 | 774 | N774M | B | 4.6e-09 | 8.6e-09 | 1.7e-09 | 4.967E-09 | 3.465E-09 | 1.700E-09 | 8.600E-09 |
| 15474 | 774 | N774F | B | 1.1e-08 | 7.8e-09 | 7.5e-09 | 8.767E-09 | 1.940E-09 | 7.500E-09 | 1.100E-08 |
| 15475 | 774 | N774P | B | 8.8e-09 | 1.6e-08 | 7.5e-09 | 1.077E-08 | 4.579E-09 | 7.500E-09 | 1.600E-08 |
| 15476 | 774 | N774S | B | 2.1e-08 | 2.6e-08 | 9.9e-09 | 1.897E-08 | 8.240E-09 | 9.900E-09 | 2.600E-08 |
| 15477 | 774 | N774T | B | 5e-09 | 4e-09 | 1.5e-08 | 8.000E-09 | 6.083E-09 | 4.000E-09 | 1.500E-08 |
| 15478 | 774 | N774W | B | 3.6e-09 | 8.5e-09 | 9.4e-09 | 7.167E-09 | 3.121E-09 | 3.600E-09 | 9.400E-09 |
| 15479 | 774 | N774Y | B | 1e-08 | 1.4e-09 | 4.8e-09 | 5.400E-09 | 4.331E-09 | 1.400E-09 | 1.000E-08 |
| 15480 | 774 | N774V | B | 2.9e-09 | 5.5e-09 | 2.2e-08 | 1.013E-08 | 1.036E-08 | 2.900E-09 | 2.200E-08 |
| 15481 | 775 | A775A | B | 5.6e-09 | 5.2e-09 | 1.2e-08 | 7.600E-09 | 3.816E-09 | 5.200E-09 | 1.200E-08 |
| 15482 | 775 | A775R | B | 7.8e-09 | 6.2e-09 | 7.6e-09 | 7.200E-09 | 8.718E-10 | 6.200E-09 | 7.800E-09 |
| 15483 | 775 | A775N | B | 2.3e-08 | 3.7e-09 | 2.5e-09 | 9.733E-09 | 1.150E-08 | 2.500E-09 | 2.300E-08 |
| 15484 | 775 | A775D | B | 4.9e-09 | 2.7e-09 | 5.2e-09 | 4.267E-09 | 1.365E-09 | 2.700E-09 | 5.200E-09 |
| 15485 | 775 | A775C | B | 6.5e-09 | 6.8e-09 | 3.1e-09 | 5.467E-09 | 2.055E-09 | 3.100E-09 | 6.800E-09 |
| 15486 | 775 | A775E | B | 4e-09 | 6.7e-09 | 4.9e-09 | 5.200E-09 | 1.375E-09 | 4.000E-09 | 6.700E-09 |
| 15487 | 775 | A775Q | B | 1.7e-09 | 2.6e-08 | 2.6e-09 | 1.010E-08 | 1.378E-08 | 1.700E-09 | 2.600E-08 |
| 15488 | 775 | A775G | B | 4.2e-09 | 6.5e-09 | 5.2e-09 | 5.300E-09 | 1.153E-09 | 4.200E-09 | 6.500E-09 |
| 15489 | 775 | A775H | B | 3.6e-09 | 2.9e-09 | 1.4e-08 | 6.833E-09 | 6.216E-09 | 2.900E-09 | 1.400E-08 |
| 15490 | 775 | A775I | B | 9.3e-09 | 1.7e-08 | 3.8e-09 | 1.003E-08 | 6.630E-09 | 3.800E-09 | 1.700E-08 |
| 15491 | 775 | A775L | B | 9.7e-09 | 8.3e-09 | 8.6e-09 | 8.867E-09 | 7.371E-10 | 8.300E-09 | 9.700E-09 |
| 15492 | 775 | A775K | B | 1.3e-08 | 1.2e-08 | 2.3e-08 | 1.600E-08 | 6.083E-09 | 1.200E-08 | 2.300E-08 |
| 15493 | 775 | A775M | B | 8.7e-09 | 3.2e-09 | 1.7e-08 | 9.633E-09 | 6.947E-09 | 3.200E-09 | 1.700E-08 |
| 15494 | 775 | A775F | B | 1e-08 | 4.5e-09 | 5.6e-09 | 6.700E-09 | 2.910E-09 | 4.500E-09 | 1.000E-08 |
| 15495 | 775 | A775P | B | 5.2e-09 | 6.4e-09 | 5.2e-09 | 5.600E-09 | 6.928E-10 | 5.200E-09 | 6.400E-09 |
| 15496 | 775 | A775S | B | 6.1e-09 | 7.3e-09 | 7.4e-09 | 6.933E-09 | 7.234E-10 | 6.100E-09 | 7.400E-09 |
| 15497 | 775 | A775T | B | 1.5e-08 | 3.8e-09 | 6.5e-09 | 8.433E-09 | 5.845E-09 | 3.800E-09 | 1.500E-08 |
| 15498 | 775 | A775W | B | 4.7e-09 | 1.7e-09 | 2.4e-09 | 2.933E-09 | 1.570E-09 | 1.700E-09 | 4.700E-09 |
| 15499 | 775 | A775Y | B | 3.7e-09 | 7e-09 | 2.6e-08 | 1.223E-08 | 1.204E-08 | 3.700E-09 | 2.600E-08 |
| 15500 | 775 | A775V | B | 1.4e-08 | 2.6e-08 | 6.7e-09 | 1.557E-08 | 9.745E-09 | 6.700E-09 | 2.600E-08 |
| 15501 | 776 | L776A | B | 4.8e-09 | 5.6e-09 | 4.2e-09 | 4.867E-09 | 7.024E-10 | 4.200E-09 | 5.600E-09 |
| 15502 | 776 | L776R | B | 4.9e-09 | 5.8e-09 | 3.5e-09 | 4.733E-09 | 1.159E-09 | 3.500E-09 | 5.800E-09 |
| 15503 | 776 | L776N | B | 1.5e-09 | 3.2e-09 | 6.9e-09 | 3.867E-09 | 2.761E-09 | 1.500E-09 | 6.900E-09 |
| 15504 | 776 | L776D | B | 2.4e-09 | 1.6e-08 | 1.1e-08 | 9.800E-09 | 6.879E-09 | 2.400E-09 | 1.600E-08 |
| 15505 | 776 | L776C | B | 8.4e-09 | 2.8e-09 | 1.1e-08 | 7.400E-09 | 4.190E-09 | 2.800E-09 | 1.100E-08 |
| 15506 | 776 | L776E | B | 1.3e-08 | 1.6e-08 | 5.1e-09 | 1.137E-08 | 5.631E-09 | 5.100E-09 | 1.600E-08 |
| 15507 | 776 | L776Q | B | 8.8e-09 | 7.2e-09 | 6.2e-09 | 7.400E-09 | 1.311E-09 | 6.200E-09 | 8.800E-09 |
| 15508 | 776 | L776G | B | 6.1e-09 | 8.2e-09 | 4e-09 | 6.100E-09 | 2.100E-09 | 4.000E-09 | 8.200E-09 |
| 15509 | 776 | L776H | B | 1.3e-08 | 8.3e-09 | 3.7e-09 | 8.333E-09 | 4.650E-09 | 3.700E-09 | 1.300E-08 |
| 15510 | 776 | L776I | B | 2.1e-08 | 3e-09 | 3.8e-09 | 9.267E-09 | 1.017E-08 | 3.000E-09 | 2.100E-08 |
| 15511 | 776 | L776L | B | 1.2e-08 | 4.9e-09 | 8.8e-09 | 8.567E-09 | 3.556E-09 | 4.900E-09 | 1.200E-08 |
| 15512 | 776 | L776K | B | 8.9e-09 | 5e-09 | 4.9e-09 | 6.267E-09 | 2.281E-09 | 4.900E-09 | 8.900E-09 |
| 15513 | 776 | L776M | B | 3e-09 | 4.7e-09 | 1e-08 | 5.900E-09 | 3.651E-09 | 3.000E-09 | 1.000E-08 |
| 15514 | 776 | L776F | B | 5.5e-09 | 1.7e-08 | 3.5e-09 | 8.667E-09 | 7.286E-09 | 3.500E-09 | 1.700E-08 |
| 15515 | 776 | L776P | B | 9.7e-09 | 2.9e-09 | 1.9e-08 | 1.053E-08 | 8.082E-09 | 2.900E-09 | 1.900E-08 |
| 15516 | 776 | L776S | B | 1.1e-08 | 6.3e-09 | 4.2e-09 | 7.167E-09 | 3.482E-09 | 4.200E-09 | 1.100E-08 |
| 15517 | 776 | L776T | B | 6.6e-09 | 4.7e-09 | 3.2e-09 | 4.833E-09 | 1.704E-09 | 3.200E-09 | 6.600E-09 |
| 15518 | 776 | L776W | B | 1.1e-08 | 1.4e-08 | 6.3e-09 | 1.043E-08 | 3.881E-09 | 6.300E-09 | 1.400E-08 |
| 15519 | 776 | L776Y | B | 8.6e-09 | 4.8e-09 | 7.4e-09 | 6.933E-09 | 1.943E-09 | 4.800E-09 | 8.600E-09 |
| 15520 | 776 | L776V | B | 6.4e-09 | 1.8e-08 | 4.2e-09 | 9.533E-09 | 7.414E-09 | 4.200E-09 | 1.800E-08 |
| 15521 | 777 | Q777A | B | 6.9e-09 | 1.1e-08 | 1.2e-08 | 9.967E-09 | 2.702E-09 | 6.900E-09 | 1.200E-08 |
| 15522 | 777 | Q777R | B | 6.5e-09 | 6.4e-09 | 9.5e-09 | 7.467E-09 | 1.762E-09 | 6.400E-09 | 9.500E-09 |
| 15523 | 777 | Q777N | B | 3e-09 | 6.3e-09 | 6.6e-09 | 5.300E-09 | 1.997E-09 | 3.000E-09 | 6.600E-09 |
| 15524 | 777 | Q777D | B | 1.4e-08 | 6.4e-09 | 5.1e-09 | 8.500E-09 | 4.807E-09 | 5.100E-09 | 1.400E-08 |
| 15525 | 777 | Q777C | B | 5.3e-09 | 5.4e-09 | 1.2e-08 | 7.567E-09 | 3.840E-09 | 5.300E-09 | 1.200E-08 |
| 15526 | 777 | Q777E | B | 1.5e-08 | 2e-08 | 1.3e-08 | 1.600E-08 | 3.606E-09 | 1.300E-08 | 2.000E-08 |
| 15527 | 777 | Q777Q | B | 1.2e-08 | 6.3e-09 | 1e-08 | 9.433E-09 | 2.892E-09 | 6.300E-09 | 1.200E-08 |
| 15528 | 777 | Q777G | B | 3.6e-09 | 7.5e-09 | 4.8e-09 | 5.300E-09 | 1.997E-09 | 3.600E-09 | 7.500E-09 |
| 15529 | 777 | Q777H | B | 8.7e-09 | 6.4e-09 | 2.2e-09 | 5.767E-09 | 3.296E-09 | 2.200E-09 | 8.700E-09 |
| 15530 | 777 | Q777I | B | 7.9e-09 | 6.2e-09 | 5.8e-09 | 6.633E-09 | 1.115E-09 | 5.800E-09 | 7.900E-09 |
| 15531 | 777 | Q777L | B | 2.1e-08 | 1e-08 | 7.4e-09 | 1.280E-08 | 7.219E-09 | 7.400E-09 | 2.100E-08 |
| 15532 | 777 | Q777K | B | 1.4e-08 | 1e-08 | 9.2e-09 | 1.107E-08 | 2.572E-09 | 9.200E-09 | 1.400E-08 |
| 15533 | 777 | Q777M | B | 6e-09 | 1.4e-08 | 3.1e-09 | 7.700E-09 | 5.645E-09 | 3.100E-09 | 1.400E-08 |
| 15534 | 777 | Q777F | B | 9.8e-09 | 8.6e-09 | 6e-09 | 8.133E-09 | 1.943E-09 | 6.000E-09 | 9.800E-09 |
| 15535 | 777 | Q777P | B | 8.4e-09 | 1.2e-08 | 2.3e-09 | 7.567E-09 | 4.903E-09 | 2.300E-09 | 1.200E-08 |
| 15536 | 777 | Q777S | B | 1.5e-09 | 1.7e-08 | 2.8e-09 | 7.100E-09 | 8.598E-09 | 1.500E-09 | 1.700E-08 |
| 15537 | 777 | Q777T | B | 9.1e-09 | 3.6e-09 | 4.7e-09 | 5.800E-09 | 2.910E-09 | 3.600E-09 | 9.100E-09 |
| 15538 | 777 | Q777W | B | 4.8e-09 | 3e-09 | 1.3e-09 | 3.033E-09 | 1.750E-09 | 1.300E-09 | 4.800E-09 |
| 15539 | 777 | Q777Y | B | 2.1e-09 | 5.5e-09 | 6.2e-09 | 4.600E-09 | 2.193E-09 | 2.100E-09 | 6.200E-09 |
| 15540 | 777 | Q777V | B | 9.4e-09 | 9.3e-09 | 5.9e-09 | 8.200E-09 | 1.992E-09 | 5.900E-09 | 9.400E-09 |
| 15541 | 778 | S778A | B | 7.1e-09 | 1.3e-08 | 7.4e-09 | 9.167E-09 | 3.323E-09 | 7.100E-09 | 1.300E-08 |
| 15542 | 778 | S778R | B | 1.5e-08 | 6.5e-09 | 6e-09 | 9.167E-09 | 5.058E-09 | 6.000E-09 | 1.500E-08 |
| 15543 | 778 | S778N | B | 1.2e-08 | 4.1e-09 | 6.3e-09 | 7.467E-09 | 4.077E-09 | 4.100E-09 | 1.200E-08 |
| 15544 | 778 | S778D | B | 3.2e-09 | 4e-09 | 6.7e-09 | 4.633E-09 | 1.834E-09 | 3.200E-09 | 6.700E-09 |
| 15545 | 778 | S778C | B | 3.7e-09 | 1e-08 | 1e-08 | 7.900E-09 | 3.637E-09 | 3.700E-09 | 1.000E-08 |
| 15546 | 778 | S778E | B | 6.8e-09 | 1.6e-08 | 1.1e-08 | 1.127E-08 | 4.606E-09 | 6.800E-09 | 1.600E-08 |
| 15547 | 778 | S778Q | B | 6.1e-09 | 1.1e-08 | 9.9e-09 | 9.000E-09 | 2.571E-09 | 6.100E-09 | 1.100E-08 |
| 15548 | 778 | S778G | B | 1.1e-08 | 1.3e-08 | 3.4e-09 | 9.133E-09 | 5.065E-09 | 3.400E-09 | 1.300E-08 |
| 15549 | 778 | S778H | B | 8.4e-09 | 9.9e-09 | 5.8e-09 | 8.033E-09 | 2.074E-09 | 5.800E-09 | 9.900E-09 |
| 15550 | 778 | S778I | B | 2.1e-08 | 1.5e-08 | 2.3e-08 | 1.967E-08 | 4.163E-09 | 1.500E-08 | 2.300E-08 |
| 15551 | 778 | S778L | B | 5.6e-09 | 4.9e-09 | 3.5e-09 | 4.667E-09 | 1.069E-09 | 3.500E-09 | 5.600E-09 |
| 15552 | 778 | S778K | B | 8.8e-09 | 7.8e-09 | 8.6e-09 | 8.400E-09 | 5.292E-10 | 7.800E-09 | 8.800E-09 |
| 15553 | 778 | S778M | B | 5e-09 | 1.4e-08 | 3.8e-09 | 7.600E-09 | 5.575E-09 | 3.800E-09 | 1.400E-08 |
| 15554 | 778 | S778F | B | 7.3e-09 | 1.6e-08 | 7e-09 | 1.010E-08 | 5.112E-09 | 7.000E-09 | 1.600E-08 |

| | | | | | | | | | | |
|---|---|---|---|---|---|---|---|---|---|---|
| 15555 | 778 | S778P | B | 3e-09 | 1.3e-08 | 4.6e-09 | 6.867E-09 | 5.372E-09 | 3.000E-09 | 1.300E-08 |
| 15556 | 778 | S778S | B | 4.1e-09 | 7.2e-09 | 8.7e-09 | 6.667E-09 | 2.346E-09 | 4.100E-09 | 8.700E-09 |
| 15557 | 778 | S778T | B | 1.3e-09 | 5.4e-09 | 4.8e-09 | 3.833E-09 | 2.214E-09 | 1.300E-09 | 5.400E-09 |
| 15558 | 778 | S778W | B | 1.4e-08 | 8.4e-09 | 9e-09 | 1.047E-08 | 3.075E-09 | 8.400E-09 | 1.400E-08 |
| 15559 | 778 | S778Y | B | 5e-09 | 1.2e-08 | 9.6e-09 | 8.867E-09 | 3.557E-09 | 5.000E-09 | 1.200E-08 |
| 15560 | 778 | S778V | B | 7e-09 | 5.3e-09 | 3.9e-09 | 5.400E-09 | 1.552E-09 | 3.900E-09 | 7.000E-09 |
| 15561 | 779 | G779A | B | 5.9e-09 | 1.6e-08 | 7.6e-09 | 9.833E-09 | 5.408E-09 | 5.900E-09 | 1.600E-08 |
| 15562 | 779 | G779R | B | 2.5e-08 | 1.7e-08 | 3.4e-09 | 1.513E-08 | 1.092E-08 | 3.400E-09 | 2.500E-08 |
| 15563 | 779 | G779N | B | 4.4e-09 | 7.6e-09 | 7.6e-09 | 6.533E-09 | 1.848E-09 | 4.400E-09 | 7.600E-09 |
| 15564 | 779 | G779D | B | 1.8e-08 | 3.3e-09 | 5.9e-09 | 9.067E-09 | 7.845E-09 | 3.300E-09 | 1.800E-08 |
| 15565 | 779 | G779C | B | 5.1e-09 | 4.8e-09 | 3.8e-09 | 4.567E-09 | 6.807E-10 | 3.800E-09 | 5.100E-09 |
| 15566 | 779 | G779E | B | 7.2e-09 | 3.3e-09 | 4.2e-09 | 4.900E-09 | 2.042E-09 | 3.300E-09 | 7.200E-09 |
| 15567 | 779 | G779Q | B | 6.2e-09 | 9.7e-09 | 3.3e-09 | 6.400E-09 | 3.205E-09 | 3.300E-09 | 9.700E-09 |
| 15568 | 779 | G779G | B | 1.7e-08 | 7.9e-09 | 1.1e-08 | 1.197E-08 | 4.626E-09 | 7.900E-09 | 1.700E-08 |
| 15569 | 779 | G779H | B | 2.6e-09 | 1.1e-08 | 8.2e-09 | 7.267E-09 | 4.277E-09 | 2.600E-09 | 1.100E-08 |
| 15570 | 779 | G779I | B | 9.2e-09 | 7.8e-09 | 2.5e-09 | 6.500E-09 | 3.534E-09 | 2.500E-09 | 9.200E-09 |
| 15571 | 779 | G779L | B | 1.5e-08 | 4.1e-09 | 6.1e-09 | 8.400E-09 | 5.803E-09 | 4.100E-09 | 1.500E-08 |
| 15572 | 779 | G779K | B | 1.6e-08 | 5e-09 | 4e-09 | 8.333E-09 | 6.658E-09 | 4.000E-09 | 1.600E-08 |
| 15573 | 779 | G779M | B | 1.5e-08 | 2.8e-09 | 9.4e-09 | 9.067E-09 | 6.107E-09 | 2.800E-09 | 1.500E-08 |
| 15574 | 779 | G779F | B | 8.1e-09 | 7e-09 | 2.6e-09 | 5.900E-09 | 2.910E-09 | 2.600E-09 | 8.100E-09 |
| 15575 | 779 | G779P | B | 9.8e-09 | 9.5e-09 | 9e-09 | 9.433E-09 | 4.041E-10 | 9.000E-09 | 9.800E-09 |
| 15576 | 779 | G779S | B | 3.5e-09 | 1.9e-08 | 3.9e-09 | 8.800E-09 | 8.836E-09 | 3.500E-09 | 1.900E-08 |
| 15577 | 779 | G779T | B | 3.6e-09 | 4.7e-09 | 3.7e-09 | 4.000E-09 | 6.083E-10 | 3.600E-09 | 4.700E-09 |
| 15578 | 779 | G779W | B | 8e-09 | 2.4e-09 | 8e-09 | 6.133E-09 | 3.233E-09 | 2.400E-09 | 8.000E-09 |
| 15579 | 779 | G779Y | B | 3e-08 | 4.1e-09 | 6.7e-09 | 1.360E-08 | 1.426E-08 | 4.100E-09 | 3.000E-08 |
| 15580 | 779 | G779V | B | 1.7e-08 | 8.3e-09 | 2.7e-09 | 9.333E-09 | 7.206E-09 | 2.700E-09 | 1.700E-08 |
| 15581 | 780 | N780A | B | 9.1e-09 | 4.7e-09 | 1.1e-08 | 8.267E-09 | 3.232E-09 | 4.700E-09 | 1.100E-08 |
| 15582 | 780 | N780R | B | 4.3e-09 | 9.5e-09 | 6.9e-09 | 6.900E-09 | 2.600E-09 | 4.300E-09 | 9.500E-09 |
| 15583 | 780 | N780N | B | 6e-09 | 4.2e-09 | 1.1e-08 | 7.067E-09 | 3.523E-09 | 4.200E-09 | 1.100E-08 |
| 15584 | 780 | N780D | B | 6.3e-09 | 9.3e-09 | 2.9e-09 | 6.167E-09 | 3.202E-09 | 2.900E-09 | 9.300E-09 |
| 15585 | 780 | N780C | B | 5.3e-09 | 5.5e-09 | 3.3e-09 | 4.700E-09 | 1.217E-09 | 3.300E-09 | 5.500E-09 |
| 15586 | 780 | N780E | B | 1.4e-08 | 1.2e-08 | 6.4e-09 | 1.080E-08 | 3.940E-09 | 6.400E-09 | 1.400E-08 |
| 15587 | 780 | N780Q | B | 9.9e-09 | 5.5e-09 | 3.8e-09 | 6.400E-09 | 3.148E-09 | 3.800E-09 | 9.900E-09 |
| 15588 | 780 | N780G | B | 1.4e-08 | 1.2e-08 | 7.4e-09 | 1.113E-08 | 3.384E-09 | 7.400E-09 | 1.400E-08 |
| 15589 | 780 | N780H | B | 2.8e-08 | 6.9e-09 | 5e-09 | 1.330E-08 | 1.277E-08 | 5.000E-09 | 2.800E-08 |
| 15590 | 780 | N780I | B | 1.1e-08 | 5.5e-09 | 8.2e-09 | 8.233E-09 | 2.750E-09 | 5.500E-09 | 1.100E-08 |
| 15591 | 780 | N780L | B | 8.4e-09 | 2.4e-08 | 3.4e-09 | 1.193E-08 | 1.074E-08 | 3.400E-09 | 2.400E-08 |
| 15592 | 780 | N780K | B | 3e-09 | 6.7e-09 | 4.3e-09 | 4.667E-09 | 1.877E-09 | 3.000E-09 | 6.700E-09 |
| 15593 | 780 | N780M | B | 1.7e-08 | 1.8e-08 | 4.1e-09 | 1.303E-08 | 7.753E-09 | 4.100E-09 | 1.800E-08 |
| 15594 | 780 | N780F | B | 1.1e-08 | 3e-09 | 1.6e-08 | 1.000E-08 | 6.557E-09 | 3.000E-09 | 1.600E-08 |
| 15595 | 780 | N780P | B | 2.7e-09 | 3e-09 | 4.6e-09 | 3.433E-09 | 1.021E-09 | 2.700E-09 | 4.600E-09 |
| 15596 | 780 | N780S | B | 1.2e-08 | 1.7e-08 | 4e-09 | 1.100E-08 | 6.557E-09 | 4.000E-09 | 1.700E-08 |
| 15597 | 780 | N780T | B | 7.1e-09 | 4.9e-09 | 6e-09 | 6.000E-09 | 1.100E-09 | 4.900E-09 | 7.100E-09 |
| 15598 | 780 | N780W | B | 4.2e-09 | 1.1e-08 | 2.1e-08 | 1.207E-08 | 8.451E-09 | 4.200E-09 | 2.100E-08 |
| 15599 | 780 | N780Y | B | 9.4e-09 | 1.1e-08 | 8.3e-09 | 9.567E-09 | 1.358E-09 | 8.300E-09 | 1.100E-08 |
| 15600 | 780 | N780V | B | 4.7e-09 | 1.5e-08 | 2e-08 | 1.323E-08 | 7.801E-09 | 4.700E-09 | 2.000E-08 |
| 15601 | 781 | S781A | B | 5.3e-09 | 1.1e-08 | 3.6e-09 | 6.633E-09 | 3.876E-09 | 3.600E-09 | 1.100E-08 |
| 15602 | 781 | S781R | B | 6.3e-09 | 1.1e-08 | 1.2e-08 | 9.767E-09 | 3.044E-09 | 6.300E-09 | 1.200E-08 |
| 15603 | 781 | S781N | B | 5.7e-09 | 8e-09 | 1.5e-08 | 9.567E-09 | 4.844E-09 | 5.700E-09 | 1.500E-08 |
| 15604 | 781 | S781D | B | 9.1e-09 | 2.7e-09 | 7.5e-09 | 6.433E-09 | 3.331E-09 | 2.700E-09 | 9.100E-09 |
| 15605 | 781 | S781C | B | 8.2e-09 | 1.6e-08 | 4.7e-09 | 9.633E-09 | 5.785E-09 | 4.700E-09 | 1.600E-08 |
| 15606 | 781 | S781E | B | 7.9e-09 | 6.6e-09 | 4e-09 | 6.167E-09 | 1.986E-09 | 4.000E-09 | 7.900E-09 |
| 15607 | 781 | S781Q | B | 9.5e-09 | 1.3e-08 | 5e-09 | 9.167E-09 | 4.010E-09 | 5.000E-09 | 1.300E-08 |
| 15608 | 781 | S781G | B | 1.2e-08 | 9.5e-09 | 2.6e-09 | 8.033E-09 | 4.869E-09 | 2.600E-09 | 1.200E-08 |
| 15609 | 781 | S781H | B | 7.9e-09 | 2.8e-08 | 2.9e-09 | 1.293E-08 | 1.329E-08 | 2.900E-09 | 2.800E-08 |
| 15610 | 781 | S781I | B | 5.3e-09 | 1.8e-08 | 6.9e-09 | 1.007E-08 | 6.917E-09 | 5.300E-09 | 1.800E-08 |
| 15611 | 781 | S781L | B | 8.7e-09 | 1.1e-08 | 9.8e-09 | 9.833E-09 | 1.150E-09 | 8.700E-09 | 1.100E-08 |
| 15612 | 781 | S781K | B | 4.2e-09 | 4e-09 | 1.8e-08 | 8.733E-09 | 8.026E-09 | 4.000E-09 | 1.800E-08 |
| 15613 | 781 | S781M | B | 7.4e-09 | 2e-09 | 1.7e-08 | 8.800E-09 | 7.597E-09 | 2.000E-09 | 1.700E-08 |
| 15614 | 781 | S781F | B | 8.3e-09 | 8.2e-09 | 1.7e-09 | 6.067E-09 | 3.782E-09 | 1.700E-09 | 8.300E-09 |
| 15615 | 781 | S781P | B | 7.2e-09 | 6.2e-09 | 1.2e-08 | 8.467E-09 | 3.101E-09 | 6.200E-09 | 1.200E-08 |
| 15616 | 781 | S781S | B | 3e-09 | 5.6e-09 | 5e-09 | 4.533E-09 | 1.361E-09 | 3.000E-09 | 5.600E-09 |
| 15617 | 781 | S781T | B | 4.9e-09 | 9.1e-09 | 8e-09 | 7.333E-09 | 2.178E-09 | 4.900E-09 | 9.100E-09 |
| 15618 | 781 | S781W | B | 5.4e-09 | 5.2e-08 | 2.2e-09 | 1.987E-08 | 2.787E-08 | 2.200E-09 | 5.200E-08 |
| 15619 | 781 | S781Y | B | 5.5e-09 | 2.7e-08 | 3.7e-09 | 1.207E-08 | 1.296E-08 | 3.700E-09 | 2.700E-08 |
| 15620 | 781 | S781V | B | 4.7e-09 | 7.1e-09 | 8.1e-09 | 6.633E-09 | 1.747E-09 | 4.700E-09 | 8.100E-09 |
| 15621 | 782 | Q782A | B | 1.3e-08 | 1.2e-08 | 3.9e-09 | 9.633E-09 | 4.990E-09 | 3.900E-09 | 1.300E-08 |
| 15622 | 782 | Q782R | B | 7.5e-09 | 8.4e-09 | 4.2e-09 | 6.700E-09 | 2.211E-09 | 4.200E-09 | 8.400E-09 |
| 15623 | 782 | Q782N | B | 4.4e-09 | 3.8e-09 | 8e-09 | 5.400E-09 | 2.272E-09 | 3.800E-09 | 8.000E-09 |
| 15624 | 782 | Q782D | B | 3.5e-09 | 1.8e-08 | 1.7e-09 | 7.733E-09 | 8.937E-09 | 1.700E-09 | 1.800E-08 |
| 15625 | 782 | Q782C | B | 1.6e-08 | 8.8e-09 | 1.3e-08 | 1.260E-08 | 3.617E-09 | 8.800E-09 | 1.600E-08 |
| 15626 | 782 | Q782E | B | 1.6e-08 | 2.1e-08 | 1.4e-08 | 1.700E-08 | 3.606E-09 | 1.400E-08 | 2.100E-08 |
| 15627 | 782 | Q782Q | B | 3.5e-09 | 1.7e-08 | 2.2e-09 | 7.567E-09 | 8.195E-09 | 2.200E-09 | 1.700E-08 |
| 15628 | 782 | Q782G | B | 6.7e-09 | 6.4e-09 | 1e-08 | 7.700E-09 | 1.997E-09 | 6.400E-09 | 1.000E-08 |
| 15629 | 782 | Q782H | B | 1.6e-08 | 3.6e-09 | 2.2e-08 | 1.387E-08 | 9.384E-09 | 3.600E-09 | 2.200E-08 |
| 15630 | 782 | Q782I | B | 9.6e-09 | 9.5e-09 | 1.3e-08 | 1.070E-08 | 1.992E-09 | 9.500E-09 | 1.300E-08 |
| 15631 | 782 | Q782L | B | 8.2e-09 | 4.7e-09 | 5.6e-09 | 6.167E-09 | 1.818E-09 | 4.700E-09 | 8.200E-09 |
| 15632 | 782 | Q782K | B | 1.1e-08 | 5.1e-09 | 2.1e-09 | 6.067E-09 | 4.528E-09 | 2.100E-09 | 1.100E-08 |
| 15633 | 782 | Q782M | B | 1.2e-08 | 3e-09 | 1.4e-08 | 9.667E-09 | 5.859E-09 | 3.000E-09 | 1.400E-08 |
| 15634 | 782 | Q782F | B | 3.7e-09 | 5.5e-09 | 4.8e-09 | 4.667E-09 | 9.074E-10 | 3.700E-09 | 5.500E-09 |
| 15635 | 782 | Q782P | B | 1.4e-08 | 1e-08 | 3.7e-09 | 9.233E-09 | 5.193E-09 | 3.700E-09 | 1.400E-08 |
| 15636 | 782 | Q782S | B | 3.7e-09 | 5.7e-09 | 1.3e-08 | 7.467E-09 | 4.895E-09 | 3.700E-09 | 1.300E-08 |
| 15637 | 782 | Q782T | B | 3.4e-09 | 1.1e-08 | 3.5e-08 | 1.647E-08 | 1.649E-08 | 3.400E-09 | 3.500E-08 |
| 15638 | 782 | Q782W | B | 2.7e-09 | 8.4e-09 | 6.7e-09 | 5.933E-09 | 2.926E-09 | 2.700E-09 | 8.400E-09 |
| 15639 | 782 | Q782Y | B | 3.9e-09 | 3.3e-09 | 6.1e-09 | 4.433E-09 | 1.474E-09 | 3.300E-09 | 6.100E-09 |

| | | | | | | | | | | |
|---|---|---|---|---|---|---|---|---|---|---|
| 15640 | 782 | Q782V | B | 5.1e-09 | 1.1e-08 | 4.5e-09 | 6.867E-09 | 3.592E-09 | 4.500E-09 | 1.100E-08 |
| 15641 | 783 | E783A | B | 1.4e-08 | 1.5e-08 | 1.2e-08 | 1.367E-08 | 1.528E-09 | 1.200E-08 | 1.500E-08 |
| 15642 | 783 | E783R | B | 6.4e-09 | 1.2e-09 | 5.9e-09 | 4.500E-09 | 2.869E-09 | 1.200E-09 | 6.400E-09 |
| 15643 | 783 | E783N | B | 1e-08 | 5.4e-09 | 6.6e-09 | 7.333E-09 | 2.386E-09 | 5.400E-09 | 1.000E-08 |
| 15644 | 783 | E783D | B | 1.1e-08 | 6e-09 | 1.2e-08 | 9.667E-09 | 3.215E-09 | 6.000E-09 | 1.200E-08 |
| 15645 | 783 | E783C | B | 6.4e-09 | 5e-09 | 1.2e-08 | 7.800E-09 | 3.704E-09 | 5.000E-09 | 1.200E-08 |
| 15646 | 783 | E783E | B | 6.5e-09 | 1.1e-08 | 7e-09 | 8.167E-09 | 2.466E-09 | 6.500E-09 | 1.100E-08 |
| 15647 | 783 | E783Q | B | 8.8e-09 | 5.9e-09 | 5.2e-09 | 6.633E-09 | 1.909E-09 | 5.200E-09 | 8.800E-09 |
| 15648 | 783 | E783G | B | 1.3e-08 | 7e-09 | 1.1e-08 | 1.033E-08 | 3.055E-09 | 7.000E-09 | 1.300E-08 |
| 15649 | 783 | E783H | B | 2.7e-09 | 1.8e-08 | 5.3e-09 | 8.667E-09 | 8.187E-09 | 2.700E-09 | 1.800E-08 |
| 15650 | 783 | E783I | B | 7.3e-09 | 3.4e-08 | 2.7e-09 | 1.467E-08 | 1.690E-08 | 2.700E-09 | 3.400E-08 |
| 15651 | 783 | E783L | B | 1.1e-08 | 1.9e-08 | 1.1e-08 | 1.367E-08 | 4.619E-09 | 1.100E-08 | 1.900E-08 |
| 15652 | 783 | E783K | B | 6.1e-09 | 4.3e-09 | 1e-08 | 6.800E-09 | 2.914E-09 | 4.300E-09 | 1.000E-08 |
| 15653 | 783 | E783M | B | 7e-09 | 3e-09 | 1.4e-08 | 8.000E-09 | 5.568E-09 | 3.000E-09 | 1.400E-08 |
| 15654 | 783 | E783F | B | 5.3e-09 | 5.4e-09 | 3.3e-09 | 4.667E-09 | 1.185E-09 | 3.300E-09 | 5.400E-09 |
| 15655 | 783 | E783P | B | 1.3e-08 | 8.8e-09 | 9.8e-09 | 1.053E-08 | 2.194E-09 | 8.800E-09 | 1.300E-08 |
| 15656 | 783 | E783S | B | 2.5e-08 | 5e-09 | 7.2e-09 | 1.240E-08 | 1.097E-08 | 5.000E-09 | 2.500E-08 |
| 15657 | 783 | E783T | B | 5.8e-09 | 1e-08 | 4.5e-09 | 6.767E-09 | 2.875E-09 | 4.500E-09 | 1.000E-08 |
| 15658 | 783 | E783W | B | 1.4e-08 | 1.2e-08 | 5.1e-09 | 1.037E-08 | 4.669E-09 | 5.100E-09 | 1.400E-08 |
| 15659 | 783 | E783Y | B | 4e-09 | 7.1e-09 | 6.1e-09 | 5.733E-09 | 1.582E-09 | 4.000E-09 | 7.100E-09 |
| 15660 | 783 | E783V | B | 4.8e-09 | 7.6e-09 | 6e-09 | 6.133E-09 | 1.405E-09 | 4.800E-09 | 7.600E-09 |
| 15661 | 784 | S784A | B | 1.7e-08 | 7.7e-09 | 1.1e-08 | 1.190E-08 | 4.715E-09 | 7.700E-09 | 1.700E-08 |
| 15662 | 784 | S784R | B | 4.2e-09 | 1.1e-08 | 9.1e-09 | 8.100E-09 | 3.509E-09 | 4.200E-09 | 1.100E-08 |
| 15663 | 784 | S784N | B | 1.3e-08 | 1.3e-08 | 7.9e-09 | 1.130E-08 | 2.944E-09 | 7.900E-09 | 1.300E-08 |
| 15664 | 784 | S784D | B | 8.6e-09 | 4.5e-09 | 4.3e-09 | 5.800E-09 | 2.427E-09 | 4.300E-09 | 8.600E-09 |
| 15665 | 784 | S784C | B | 5.5e-09 | 1.4e-08 | 8.5e-09 | 9.333E-09 | 4.311E-09 | 5.500E-09 | 1.400E-08 |
| 15666 | 784 | S784E | B | 1.6e-08 | 8.2e-09 | 3.7e-09 | 9.300E-09 | 6.223E-09 | 3.700E-09 | 1.600E-08 |
| 15667 | 784 | S784Q | B | 6.6e-09 | 1.4e-08 | 4.2e-09 | 8.267E-09 | 5.108E-09 | 4.200E-09 | 1.400E-08 |
| 15668 | 784 | S784G | B | 3e-09 | 1e-08 | 3.8e-09 | 5.600E-09 | 3.831E-09 | 3.000E-09 | 1.000E-08 |
| 15669 | 784 | S784H | B | 6.8e-09 | 1e-08 | 1.4e-08 | 1.027E-08 | 3.607E-09 | 6.800E-09 | 1.400E-08 |
| 15670 | 784 | S784I | B | 1.7e-08 | 1.1e-08 | 1e-08 | 1.267E-08 | 3.786E-09 | 1.000E-08 | 1.700E-08 |
| 15671 | 784 | S784L | B | 4e-09 | 7.4e-09 | 2.2e-09 | 4.533E-09 | 2.641E-09 | 2.200E-09 | 7.400E-09 |
| 15672 | 784 | S784K | B | 6.8e-09 | 2.2e-08 | 1.2e-08 | 1.360E-08 | 7.725E-09 | 6.800E-09 | 2.200E-08 |
| 15673 | 784 | S784M | B | 8.5e-09 | 6.6e-09 | 9.2e-09 | 8.100E-09 | 1.345E-09 | 6.600E-09 | 9.200E-09 |
| 15674 | 784 | S784F | B | 3.2e-09 | 7.6e-09 | 8.9e-09 | 6.567E-09 | 2.987E-09 | 3.200E-09 | 8.900E-09 |
| 15675 | 784 | S784P | B | 1.2e-08 | 5.9e-09 | 1.7e-08 | 1.163E-08 | 5.559E-09 | 5.900E-09 | 1.700E-08 |
| 15676 | 784 | S784S | B | 1.4e-08 | 5.5e-09 | 5.2e-09 | 8.233E-09 | 4.996E-09 | 5.200E-09 | 1.400E-08 |
| 15677 | 784 | S784T | B | 1.3e-08 | 5.7e-09 | 7.3e-09 | 8.667E-09 | 3.837E-09 | 5.700E-09 | 1.300E-08 |
| 15678 | 784 | S784W | B | 5.8e-09 | 5.2e-09 | 4.1e-09 | 5.033E-09 | 8.622E-10 | 4.100E-09 | 5.800E-09 |
| 15679 | 784 | S784Y | B | 8e-09 | 2.5e-09 | 5.7e-09 | 5.400E-09 | 2.762E-09 | 2.500E-09 | 8.000E-09 |
| 15680 | 784 | S784V | B | 2.8e-09 | 1e-08 | 1.5e-08 | 9.267E-09 | 6.133E-09 | 2.800E-09 | 1.500E-08 |
| 15681 | 785 | V785A | B | 2.3e-09 | 6.6e-09 | 1.2e-08 | 6.967E-09 | 4.860E-09 | 2.300E-09 | 1.200E-08 |
| 15682 | 785 | V785R | B | 7.4e-09 | 4.6e-09 | 5.2e-09 | 5.733E-09 | 1.474E-09 | 4.600E-09 | 7.400E-09 |
| 15683 | 785 | V785N | B | 4.5e-09 | 8.5e-09 | 2.7e-08 | 1.333E-08 | 1.200E-08 | 4.500E-09 | 2.700E-08 |
| 15684 | 785 | V785D | B | 5e-09 | 1e-08 | 1e-08 | 8.333E-09 | 2.887E-09 | 5.000E-09 | 1.000E-08 |
| 15685 | 785 | V785C | B | 7.8e-09 | 1.8e-08 | 4.9e-09 | 1.023E-08 | 6.881E-09 | 4.900E-09 | 1.800E-08 |
| 15686 | 785 | V785E | B | 5.3e-09 | 1.1e-08 | 7.2e-09 | 7.833E-09 | 2.902E-09 | 5.300E-09 | 1.100E-08 |
| 15687 | 785 | V785Q | B | 6.4e-09 | 1.1e-08 | 5.5e-09 | 7.633E-09 | 2.950E-09 | 5.500E-09 | 1.100E-08 |
| 15688 | 785 | V785G | B | 6.9e-09 | 9.3e-09 | 8.5e-09 | 8.233E-09 | 1.222E-09 | 6.900E-09 | 9.300E-09 |
| 15689 | 785 | V785H | B | 5.6e-09 | 6.8e-09 | 9.2e-09 | 7.200E-09 | 1.833E-09 | 5.600E-09 | 9.200E-09 |
| 15690 | 785 | V785I | B | 2.1e-09 | 1.2e-08 | 1.2e-08 | 8.700E-09 | 5.716E-09 | 2.100E-09 | 1.200E-08 |
| 15691 | 785 | V785L | B | 1.2e-08 | 2.9e-09 | 1e-08 | 8.300E-09 | 4.782E-09 | 2.900E-09 | 1.200E-08 |
| 15692 | 785 | V785K | B | 1.5e-08 | 2e-08 | 7.2e-09 | 1.407E-08 | 6.451E-09 | 7.200E-09 | 2.000E-08 |
| 15693 | 785 | V785M | B | 5.8e-09 | 6.5e-09 | 8.8e-09 | 7.033E-09 | 1.570E-09 | 5.800E-09 | 8.800E-09 |
| 15694 | 785 | V785F | B | 7.3e-09 | 6e-09 | 3.2e-09 | 5.500E-09 | 2.095E-09 | 3.200E-09 | 7.300E-09 |
| 15695 | 785 | V785P | B | 6.7e-09 | 4.3e-09 | 6.3e-09 | 5.767E-09 | 1.286E-09 | 4.300E-09 | 6.700E-09 |
| 15696 | 785 | V785S | B | 4.8e-09 | 1.4e-08 | 3.9e-09 | 7.567E-09 | 5.590E-09 | 3.900E-09 | 1.400E-08 |
| 15697 | 785 | V785T | B | 1.6e-08 | 4.4e-09 | 4.7e-09 | 8.367E-09 | 6.612E-09 | 4.400E-09 | 1.600E-08 |
| 15698 | 785 | V785W | B | 3e-09 | 1.3e-08 | 8.4e-09 | 8.133E-09 | 5.005E-09 | 3.000E-09 | 1.300E-08 |
| 15699 | 785 | V785Y | B | 5.2e-09 | 9.4e-09 | 1.3e-08 | 9.200E-09 | 3.904E-09 | 5.200E-09 | 1.300E-08 |
| 15700 | 785 | V785V | B | 3.1e-09 | 2.9e-09 | 4e-09 | 3.333E-09 | 5.859E-10 | 2.900E-09 | 4.000E-09 |
| 15701 | 786 | T786A | B | 6.9e-09 | 4.9e-09 | 1.8e-08 | 9.933E-09 | 7.057E-09 | 4.900E-09 | 1.800E-08 |
| 15702 | 786 | T786R | B | 3.1e-09 | 1.7e-08 | 3.6e-09 | 7.900E-09 | 7.885E-09 | 3.100E-09 | 1.700E-08 |
| 15703 | 786 | T786N | B | 1.5e-09 | 7e-09 | 6.2e-09 | 4.900E-09 | 2.972E-09 | 1.500E-09 | 7.000E-09 |
| 15704 | 786 | T786D | B | 2.3e-08 | 6.9e-09 | 1.2e-09 | 1.037E-08 | 1.131E-08 | 1.200E-09 | 2.300E-08 |
| 15705 | 786 | T786C | B | 1.6e-08 | 3.6e-09 | 1.7e-09 | 7.100E-09 | 7.766E-09 | 1.700E-09 | 1.600E-08 |
| 15706 | 786 | T786E | B | 2.2e-09 | 5.7e-09 | 2.8e-09 | 3.567E-09 | 1.872E-09 | 2.200E-09 | 5.700E-09 |
| 15707 | 786 | T786Q | B | 6.5e-09 | 4.9e-09 | 6.2e-09 | 5.867E-09 | 8.505E-10 | 4.900E-09 | 6.500E-09 |
| 15708 | 786 | T786G | B | 1.9e-08 | 5.2e-09 | 1.1e-08 | 1.173E-08 | 6.929E-09 | 5.200E-09 | 1.900E-08 |
| 15709 | 786 | T786H | B | 1.2e-08 | 4.3e-09 | 3.8e-09 | 6.700E-09 | 4.597E-09 | 3.800E-09 | 1.200E-08 |
| 15710 | 786 | T786I | B | 1e-08 | 1.6e-08 | 9.8e-09 | 1.193E-08 | 3.523E-09 | 9.800E-09 | 1.600E-08 |
| 15711 | 786 | T786L | B | 4.6e-09 | 1.1e-08 | 7e-09 | 7.533E-09 | 3.233E-09 | 4.600E-09 | 1.100E-08 |
| 15712 | 786 | T786K | B | 8.4e-09 | 8.4e-09 | 3.7e-09 | 6.833E-09 | 2.714E-09 | 3.700E-09 | 8.400E-09 |
| 15713 | 786 | T786M | B | 7e-09 | 4.8e-09 | 1.1e-08 | 7.600E-09 | 3.143E-09 | 4.800E-09 | 1.100E-08 |
| 15714 | 786 | T786F | B | 2.7e-09 | 1.1e-08 | 5e-09 | 6.233E-09 | 4.285E-09 | 2.700E-09 | 1.100E-08 |
| 15715 | 786 | T786P | B | 1.2e-08 | 8.2e-09 | 7.5e-09 | 9.233E-09 | 2.421E-09 | 7.500E-09 | 1.200E-08 |
| 15716 | 786 | T786S | B | 1.5e-08 | 1.6e-09 | 2.2e-09 | 6.267E-09 | 7.569E-09 | 1.600E-09 | 1.500E-08 |
| 15717 | 786 | T786T | B | 1.1e-08 | 1.2e-08 | 7.1e-09 | 1.003E-08 | 2.589E-09 | 7.100E-09 | 1.200E-08 |
| 15718 | 786 | T786W | B | 3.3e-09 | 1.3e-08 | 6.9e-09 | 7.733E-09 | 4.903E-09 | 3.300E-09 | 1.300E-08 |
| 15719 | 786 | T786Y | B | 9.6e-09 | 4.8e-09 | 9.6e-09 | 8.000E-09 | 2.771E-09 | 4.800E-09 | 9.600E-09 |
| 15720 | 786 | T786V | B | 4.2e-09 | 8.2e-09 | 2.7e-09 | 5.033E-09 | 2.843E-09 | 2.700E-09 | 8.200E-09 |
| 15721 | 787 | E787A | B | 7.1e-09 | 1.6e-08 | 1.4e-08 | 1.237E-08 | 4.669E-09 | 7.100E-09 | 1.600E-08 |
| 15722 | 787 | E787R | B | 2e-08 | 7.4e-09 | 5.2e-09 | 1.087E-08 | 7.986E-09 | 5.200E-09 | 2.000E-08 |
| 15723 | 787 | E787N | B | 5e-09 | 1.2e-08 | 3.9e-09 | 6.967E-09 | 4.394E-09 | 3.900E-09 | 1.200E-08 |
| 15724 | 787 | E787D | B | 3.2e-09 | 7.7e-09 | 9.7e-09 | 6.867E-09 | 3.329E-09 | 3.200E-09 | 9.700E-09 |

| | | | | | | | | | | |
|---|---|---|---|---|---|---|---|---|---|---|
| 15725 | 787 | E787C | B | 9e-09 | 3.3e-08 | 9.2e-09 | 1.707E-08 | 1.380E-08 | 9.000E-09 | 3.300E-08 |
| 15726 | 787 | E787E | B | 1.8e-09 | 1.2e-08 | 6.4e-09 | 6.733E-09 | 5.108E-09 | 1.800E-09 | 1.200E-08 |
| 15727 | 787 | E787Q | B | 7.1e-09 | 5.9e-09 | 5.8e-09 | 6.267E-09 | 7.234E-10 | 5.800E-09 | 7.100E-09 |
| 15728 | 787 | E787G | B | 6e-09 | 1.6e-08 | 3.5e-09 | 8.500E-09 | 6.614E-09 | 3.500E-09 | 1.600E-08 |
| 15729 | 787 | E787H | B | 4.4e-09 | 2e-09 | 2.1e-09 | 2.833E-09 | 1.358E-09 | 2.000E-09 | 4.400E-09 |
| 15730 | 787 | E787I | B | 4.3e-09 | 4.1e-09 | 8.7e-09 | 5.700E-09 | 2.600E-09 | 4.100E-09 | 8.700E-09 |
| 15731 | 787 | E787L | B | 9.9e-09 | 9e-09 | 6.1e-09 | 8.333E-09 | 1.986E-09 | 6.100E-09 | 9.900E-09 |
| 15732 | 787 | E787K | B | 7.3e-09 | 3e-09 | 8.5e-09 | 6.267E-09 | 2.892E-09 | 3.000E-09 | 8.500E-09 |
| 15733 | 787 | E787M | B | 1e-08 | 6e-09 | 4.7e-09 | 6.900E-09 | 2.762E-09 | 4.700E-09 | 1.000E-08 |
| 15734 | 787 | E787F | B | 1.8e-08 | 1e-08 | 5.3e-09 | 1.110E-08 | 6.421E-09 | 5.300E-09 | 1.800E-08 |
| 15735 | 787 | E787P | B | 3.3e-09 | 1.3e-08 | 1e-08 | 8.767E-09 | 4.966E-09 | 3.300E-09 | 1.300E-08 |
| 15736 | 787 | E787S | B | 4.1e-09 | 4.6e-09 | 4.4e-09 | 4.367E-09 | 2.517E-10 | 4.100E-09 | 4.600E-09 |
| 15737 | 787 | E787T | B | 2.4e-08 | 1e-08 | 2.4e-09 | 1.213E-08 | 1.096E-08 | 2.400E-09 | 2.400E-08 |
| 15738 | 787 | E787W | B | 3.6e-08 | 1.2e-08 | 3.1e-09 | 1.703E-08 | 1.702E-08 | 3.100E-09 | 3.600E-08 |
| 15739 | 787 | E787Y | B | 1e-08 | 2.9e-09 | 2.1e-09 | 5.000E-09 | 4.349E-09 | 2.100E-09 | 1.000E-08 |
| 15740 | 787 | E787V | B | 7.4e-09 | 3.6e-09 | 4.9e-09 | 5.300E-09 | 1.931E-09 | 3.600E-09 | 7.400E-09 |
| 15741 | 788 | Q788A | B | 4.9e-09 | 6.2e-09 | 1.3e-08 | 8.033E-09 | 4.350E-09 | 4.900E-09 | 1.300E-08 |
| 15742 | 788 | Q788R | B | 4.7e-09 | 8.1e-09 | 6e-09 | 6.267E-09 | 1.716E-09 | 4.700E-09 | 8.100E-09 |
| 15743 | 788 | Q788N | B | 8.6e-09 | 1.6e-08 | 8.3e-09 | 1.097E-08 | 4.362E-09 | 8.300E-09 | 1.600E-08 |
| 15744 | 788 | Q788D | B | 7.8e-09 | 6.9e-09 | 4.6e-09 | 6.433E-09 | 1.650E-09 | 4.600E-09 | 7.800E-09 |
| 15745 | 788 | Q788C | B | 1e-08 | 7.2e-09 | 1.5e-09 | 6.233E-09 | 4.332E-09 | 1.500E-09 | 1.000E-08 |
| 15746 | 788 | Q788E | B | 4.9e-09 | 6.2e-09 | 2.2e-08 | 1.103E-08 | 9.520E-09 | 4.900E-09 | 2.200E-08 |
| 15747 | 788 | Q788Q | B | 4e-09 | 6.2e-09 | 5.1e-09 | 5.100E-09 | 1.100E-09 | 4.000E-09 | 6.200E-09 |
| 15748 | 788 | Q788G | B | 5.7e-09 | 6.6e-09 | 1.2e-08 | 8.100E-09 | 3.407E-09 | 5.700E-09 | 1.200E-08 |
| 15749 | 788 | Q788H | B | 5.5e-09 | 1.1e-08 | 1.6e-08 | 1.083E-08 | 5.252E-09 | 5.500E-09 | 1.600E-08 |
| 15750 | 788 | Q788I | B | 1.5e-08 | 1.1e-08 | 5.3e-09 | 1.043E-08 | 4.875E-09 | 5.300E-09 | 1.500E-08 |
| 15751 | 788 | Q788L | B | 1.7e-08 | 2.5e-09 | 9.7e-09 | 9.733E-09 | 7.250E-09 | 2.500E-09 | 1.700E-08 |
| 15752 | 788 | Q788K | B | 6.3e-09 | 5.3e-09 | 4.3e-09 | 5.300E-09 | 1.000E-09 | 4.300E-09 | 6.300E-09 |
| 15753 | 788 | Q788M | B | 5.7e-09 | 7.1e-09 | 5.1e-09 | 5.967E-09 | 1.026E-09 | 5.100E-09 | 7.100E-09 |
| 15754 | 788 | Q788F | B | 1.1e-08 | 9.1e-09 | 7.7e-09 | 9.267E-09 | 1.656E-09 | 7.700E-09 | 1.100E-08 |
| 15755 | 788 | Q788P | B | 1.2e-08 | 3.2e-09 | 6.4e-09 | 7.200E-09 | 4.454E-09 | 3.200E-09 | 1.200E-08 |
| 15756 | 788 | Q788S | B | 7.6e-09 | 8.3e-09 | 2.8e-09 | 6.233E-09 | 2.994E-09 | 2.800E-09 | 8.300E-09 |
| 15757 | 788 | Q788T | B | 1e-08 | 1.3e-08 | 8.7e-09 | 1.057E-08 | 2.205E-09 | 8.700E-09 | 1.300E-08 |
| 15758 | 788 | Q788W | B | 7.2e-09 | 2.7e-09 | 1.2e-08 | 7.300E-09 | 4.651E-09 | 2.700E-09 | 1.200E-08 |
| 15759 | 788 | Q788Y | B | 1.7e-08 | 6.8e-09 | 4.1e-09 | 9.300E-09 | 6.804E-09 | 4.100E-09 | 1.700E-08 |
| 15760 | 788 | Q788V | B | 3.1e-09 | 8.8e-09 | 1.1e-08 | 7.633E-09 | 4.077E-09 | 3.100E-09 | 1.100E-08 |
| 15761 | 789 | D789A | B | 1.3e-08 | 6.9e-09 | 2.7e-09 | 7.533E-09 | 5.179E-09 | 2.700E-09 | 1.300E-08 |
| 15762 | 789 | D789R | B | 3.4e-09 | 1.1e-08 | 1.2e-08 | 8.800E-09 | 4.703E-09 | 3.400E-09 | 1.200E-08 |
| 15763 | 789 | D789N | B | 6.9e-09 | 4.9e-09 | 5.6e-09 | 5.800E-09 | 1.015E-09 | 4.900E-09 | 6.900E-09 |
| 15764 | 789 | D789D | B | 1e-08 | 3.6e-09 | 7e-09 | 6.867E-09 | 3.202E-09 | 3.600E-09 | 1.000E-08 |
| 15765 | 789 | D789C | B | 3.2e-08 | 7.7e-09 | 4.3e-09 | 1.467E-08 | 1.511E-08 | 4.300E-09 | 3.200E-08 |
| 15766 | 789 | D789E | B | 9.1e-09 | 3e-09 | 8.4e-09 | 6.833E-09 | 3.338E-09 | 3.000E-09 | 9.100E-09 |
| 15767 | 789 | D789Q | B | 1.2e-08 | 9.4e-09 | 2e-09 | 7.800E-09 | 5.188E-09 | 2.000E-09 | 1.200E-08 |
| 15768 | 789 | D789G | B | 1.5e-08 | 4e-09 | 1.5e-08 | 1.133E-08 | 6.351E-09 | 4.000E-09 | 1.500E-08 |
| 15769 | 789 | D789H | B | 8.1e-09 | 1.4e-08 | 7.4e-09 | 9.833E-09 | 3.625E-09 | 7.400E-09 | 1.400E-08 |
| 15770 | 789 | D789I | B | 1.7e-08 | 4.7e-09 | 2.2e-08 | 1.457E-08 | 8.903E-09 | 4.700E-09 | 2.200E-08 |
| 15771 | 789 | D789L | B | 4.8e-09 | 5.4e-09 | 2.6e-09 | 4.267E-09 | 1.474E-09 | 2.600E-09 | 5.400E-09 |
| 15772 | 789 | D789K | B | 1.6e-08 | 9.9e-09 | 5.1e-09 | 1.033E-08 | 5.463E-09 | 5.100E-09 | 1.600E-08 |
| 15773 | 789 | D789M | B | 5.6e-09 | 2.1e-09 | 2.4e-09 | 3.367E-09 | 1.940E-09 | 2.100E-09 | 5.600E-09 |
| 15774 | 789 | D789F | B | 1.2e-08 | 5e-09 | 4.6e-09 | 7.200E-09 | 4.162E-09 | 4.600E-09 | 1.200E-08 |
| 15775 | 789 | D789P | B | 1.1e-08 | 1.4e-08 | 4.5e-09 | 9.833E-09 | 4.856E-09 | 4.500E-09 | 1.400E-08 |
| 15776 | 789 | D789S | B | 6.6e-09 | 1.2e-08 | 1.4e-08 | 1.087E-08 | 3.828E-09 | 6.600E-09 | 1.400E-08 |
| 15777 | 789 | D789T | B | 1.8e-09 | 4.5e-09 | 3.3e-09 | 3.200E-09 | 1.353E-09 | 1.800E-09 | 4.500E-09 |
| 15778 | 789 | D789W | B | 4.8e-09 | 3e-09 | 2.8e-09 | 3.533E-09 | 1.102E-09 | 2.800E-09 | 4.800E-09 |
| 15779 | 789 | D789Y | B | 5.7e-09 | 8.9e-09 | 5.7e-09 | 6.767E-09 | 1.848E-09 | 5.700E-09 | 8.900E-09 |
| 15780 | 789 | D789V | B | 2.6e-09 | 3.9e-09 | 7e-09 | 4.500E-09 | 2.261E-09 | 2.600E-09 | 7.000E-09 |
| 15781 | 790 | S790A | B | 5.1e-09 | 9.3e-09 | 8.2e-09 | 7.533E-09 | 2.178E-09 | 5.100E-09 | 9.300E-09 |
| 15782 | 790 | S790R | B | 4.4e-09 | 1.1e-08 | 3.3e-09 | 6.233E-09 | 4.165E-09 | 3.300E-09 | 1.100E-08 |
| 15783 | 790 | S790N | B | 4.4e-09 | 7e-09 | 5.2e-09 | 5.533E-09 | 1.332E-09 | 4.400E-09 | 7.000E-09 |
| 15784 | 790 | S790D | B | 3.3e-09 | 1.7e-08 | 7.4e-09 | 9.233E-09 | 7.032E-09 | 3.300E-09 | 1.700E-08 |
| 15785 | 790 | S790C | B | 7.5e-09 | 3.4e-08 | 3.9e-09 | 1.513E-08 | 1.644E-08 | 3.900E-09 | 3.400E-08 |
| 15786 | 790 | S790E | B | 1.3e-08 | 1.5e-08 | 8.9e-09 | 1.230E-08 | 3.110E-09 | 8.900E-09 | 1.500E-08 |
| 15787 | 790 | S790Q | B | 1.3e-08 | 9.5e-09 | 1.1e-08 | 1.117E-08 | 1.756E-09 | 9.500E-09 | 1.300E-08 |
| 15788 | 790 | S790G | B | 5.9e-09 | 1.3e-08 | 2.8e-09 | 7.233E-09 | 5.229E-09 | 2.800E-09 | 1.300E-08 |
| 15789 | 790 | S790H | B | 7.6e-09 | 8.5e-09 | 8.3e-09 | 8.133E-09 | 4.726E-10 | 7.600E-09 | 8.500E-09 |
| 15790 | 790 | S790I | B | 3.1e-09 | 7.8e-09 | 5.3e-09 | 5.400E-09 | 2.352E-09 | 3.100E-09 | 7.800E-09 |
| 15791 | 790 | S790L | B | 8.8e-09 | 9.3e-09 | 5.6e-09 | 7.900E-09 | 2.007E-09 | 5.600E-09 | 9.300E-09 |
| 15792 | 790 | S790K | B | 9.7e-09 | 1.9e-08 | 4.6e-09 | 1.110E-08 | 7.301E-09 | 4.600E-09 | 1.900E-08 |
| 15793 | 790 | S790M | B | 7.6e-09 | 1.5e-08 | 3.3e-09 | 8.633E-09 | 5.918E-09 | 3.300E-09 | 1.500E-08 |
| 15794 | 790 | S790F | B | 6.7e-09 | 1.5e-08 | 2.9e-09 | 8.200E-09 | 6.188E-09 | 2.900E-09 | 1.500E-08 |
| 15795 | 790 | S790P | B | 1e-08 | 1.2e-08 | 2.4e-09 | 8.133E-09 | 5.065E-09 | 2.400E-09 | 1.200E-08 |
| 15796 | 790 | S790S | B | 6.6e-09 | 9.2e-09 | 1.3e-08 | 9.600E-09 | 3.219E-09 | 6.600E-09 | 1.300E-08 |
| 15797 | 790 | S790T | B | 1.8e-09 | 5.1e-09 | 3.8e-09 | 3.567E-09 | 1.662E-09 | 1.800E-09 | 5.100E-09 |
| 15798 | 790 | S790W | B | 5.1e-09 | 2.9e-09 | 1.1e-08 | 6.333E-09 | 4.188E-09 | 2.900E-09 | 1.100E-08 |
| 15799 | 790 | S790Y | B | 1.7e-08 | 8.4e-09 | 6e-09 | 1.047E-08 | 5.784E-09 | 6.000E-09 | 1.700E-08 |
| 15800 | 790 | S790V | B | 6.4e-09 | 4.4e-09 | 6.9e-09 | 5.900E-09 | 1.323E-09 | 4.400E-09 | 6.900E-09 |
| 15801 | 791 | K791A | B | 8.1e-09 | 4.6e-09 | 1.1e-08 | 7.900E-09 | 3.205E-09 | 4.600E-09 | 1.100E-08 |
| 15802 | 791 | K791R | B | 2.9e-09 | 1.8e-08 | 4.2e-09 | 8.367E-09 | 8.368E-09 | 2.900E-09 | 1.800E-08 |
| 15803 | 791 | K791N | B | 4.4e-09 | 1.7e-08 | 4.5e-09 | 8.633E-09 | 7.246E-09 | 4.400E-09 | 1.700E-08 |
| 15804 | 791 | K791D | B | 1.2e-08 | 9.2e-09 | 1.2e-08 | 1.107E-08 | 1.617E-09 | 9.200E-09 | 1.200E-08 |
| 15805 | 791 | K791C | B | 9e-09 | 1.2e-08 | 1.4e-08 | 1.167E-08 | 2.517E-09 | 9.000E-09 | 1.400E-08 |
| 15806 | 791 | K791E | B | 8.6e-09 | 2.4e-09 | 4.4e-09 | 5.133E-09 | 3.164E-09 | 2.400E-09 | 8.600E-09 |
| 15807 | 791 | K791Q | B | 7.7e-09 | 9.7e-09 | 6e-09 | 7.800E-09 | 1.852E-09 | 6.000E-09 | 9.700E-09 |
| 15808 | 791 | K791G | B | 6.7e-09 | 9.9e-09 | 7.2e-09 | 7.933E-09 | 1.721E-09 | 6.700E-09 | 9.900E-09 |
| 15809 | 791 | K791H | B | 2.3e-08 | 5.7e-09 | 9.5e-09 | 1.273E-08 | 9.092E-09 | 5.700E-09 | 2.300E-08 |

| | | | | | | | | | | |
|---|---|---|---|---|---|---|---|---|---|---|
| 15810 | 791 | K791I | B | 6.8e-09 | 5.4e-09 | 5.6e-09 | 5.933E-09 | 7.572E-10 | 5.400E-09 | 6.800E-09 |
| 15811 | 791 | K791L | B | 2.5e-09 | 5.7e-09 | 8.4e-09 | 5.533E-09 | 2.954E-09 | 2.500E-09 | 8.400E-09 |
| 15812 | 791 | K791K | B | 4.7e-09 | 1.8e-08 | 2.5e-09 | 8.400E-09 | 8.386E-09 | 2.500E-09 | 1.800E-08 |
| 15813 | 791 | K791M | B | 1.6e-08 | 1.3e-09 | 6.9e-09 | 8.067E-09 | 7.419E-09 | 1.300E-09 | 1.600E-08 |
| 15814 | 791 | K791F | B | 3.3e-09 | 7.4e-09 | 9.2e-09 | 6.633E-09 | 3.024E-09 | 3.300E-09 | 9.200E-09 |
| 15815 | 791 | K791P | B | 4e-09 | 2.6e-08 | 3.4e-09 | 1.113E-08 | 1.288E-08 | 3.400E-09 | 2.600E-08 |
| 15816 | 791 | K791S | B | 5.8e-09 | 4.6e-09 | 3.4e-09 | 4.600E-09 | 1.200E-09 | 3.400E-09 | 5.800E-09 |
| 15817 | 791 | K791T | B | 5.6e-09 | 3.1e-09 | 8e-09 | 5.567E-09 | 2.450E-09 | 3.100E-09 | 8.000E-09 |
| 15818 | 791 | K791W | B | 6.8e-09 | 3.7e-09 | 8.6e-09 | 6.367E-09 | 2.479E-09 | 3.700E-09 | 8.600E-09 |
| 15819 | 791 | K791Y | B | 2e-09 | 5.5e-09 | 1.6e-08 | 7.833E-09 | 7.286E-09 | 2.000E-09 | 1.600E-08 |
| 15820 | 791 | K791V | B | 8.3e-09 | 9.6e-09 | 9.4e-09 | 9.100E-09 | 7.000E-10 | 8.300E-09 | 9.600E-09 |
| 15821 | 792 | D792A | B | 8.8e-09 | 5.6e-09 | 8e-09 | 7.467E-09 | 1.665E-09 | 5.600E-09 | 8.800E-09 |
| 15822 | 792 | D792R | B | 6.4e-08 | 8.1e-09 | 7e-09 | 2.637E-08 | 3.260E-08 | 7.000E-09 | 6.400E-08 |
| 15823 | 792 | D792N | B | 8.5e-09 | 2e-08 | 8.6e-09 | 1.237E-08 | 6.611E-09 | 8.500E-09 | 2.000E-08 |
| 15824 | 792 | D792D | B | 1.9e-08 | 2.2e-09 | 1.6e-08 | 1.240E-08 | 8.960E-09 | 2.200E-09 | 1.900E-08 |
| 15825 | 792 | D792C | B | 5.3e-09 | 6.7e-09 | 1.3e-08 | 8.333E-09 | 4.102E-09 | 5.300E-09 | 1.300E-08 |
| 15826 | 792 | D792E | B | 8.1e-09 | 8e-09 | 3.9e-09 | 6.667E-09 | 2.397E-09 | 3.900E-09 | 8.100E-09 |
| 15827 | 792 | D792Q | B | 1.2e-08 | 7e-09 | 6.7e-09 | 8.567E-09 | 2.977E-09 | 6.700E-09 | 1.200E-08 |
| 15828 | 792 | D792G | B | 1.2e-08 | 5.4e-09 | 5.4e-09 | 7.600E-09 | 3.811E-09 | 5.400E-09 | 1.200E-08 |
| 15829 | 792 | D792H | B | 2.6e-09 | 6.4e-09 | 9.1e-09 | 6.033E-09 | 3.265E-09 | 2.600E-09 | 9.100E-09 |
| 15830 | 792 | D792I | B | 1e-08 | 1.4e-08 | 1.2e-08 | 1.200E-08 | 2.000E-09 | 1.000E-08 | 1.400E-08 |
| 15831 | 792 | D792L | B | 2.4e-08 | 6.3e-09 | 6.7e-09 | 1.233E-08 | 1.011E-08 | 6.300E-09 | 2.400E-08 |
| 15832 | 792 | D792K | B | 6.2e-09 | 1.9e-08 | 6e-09 | 1.040E-08 | 7.448E-09 | 6.000E-09 | 1.900E-08 |
| 15833 | 792 | D792M | B | 6.1e-09 | 3.4e-09 | 1.4e-08 | 7.833E-09 | 5.508E-09 | 3.400E-09 | 1.400E-08 |
| 15834 | 792 | D792F | B | 4.7e-09 | 5.7e-09 | 2.3e-09 | 4.233E-09 | 1.747E-09 | 2.300E-09 | 5.700E-09 |
| 15835 | 792 | D792P | B | 7.5e-09 | 3.9e-09 | 7.7e-09 | 6.367E-09 | 2.139E-09 | 3.900E-09 | 7.700E-09 |
| 15836 | 792 | D792S | B | 2.9e-09 | 5.6e-09 | 2.7e-08 | 1.183E-08 | 1.320E-08 | 2.900E-09 | 2.700E-08 |
| 15837 | 792 | D792T | B | 2.2e-08 | 1.9e-08 | 8.2e-09 | 1.640E-08 | 7.258E-09 | 8.200E-09 | 2.200E-08 |
| 15838 | 792 | D792W | B | 6.3e-09 | 8.3e-09 | 8.1e-09 | 7.567E-09 | 1.102E-09 | 6.300E-09 | 8.300E-09 |
| 15839 | 792 | D792Y | B | 9.8e-09 | 2.1e-09 | 7.2e-09 | 6.367E-09 | 3.917E-09 | 2.100E-09 | 9.800E-09 |
| 15840 | 792 | D792V | B | 1.6e-08 | 2.6e-08 | 3e-09 | 1.500E-08 | 1.153E-08 | 3.000E-09 | 2.600E-08 |
| 15841 | 793 | S793A | B | 7.4e-09 | 5.2e-09 | 1.2e-08 | 8.200E-09 | 3.470E-09 | 5.200E-09 | 1.200E-08 |
| 15842 | 793 | S793R | B | 4.8e-09 | 9.4e-09 | 9.6e-09 | 7.933E-09 | 2.715E-09 | 4.800E-09 | 9.600E-09 |
| 15843 | 793 | S793N | B | 3.8e-09 | 2.5e-09 | 3.9e-09 | 3.400E-09 | 7.810E-10 | 2.500E-09 | 3.900E-09 |
| 15844 | 793 | S793D | B | 7.9e-09 | 1.6e-08 | 8.3e-09 | 1.073E-08 | 4.565E-09 | 7.900E-09 | 1.600E-08 |
| 15845 | 793 | S793C | B | 4.3e-09 | 4.2e-09 | 1.1e-08 | 6.500E-09 | 3.897E-09 | 4.200E-09 | 1.100E-08 |
| 15846 | 793 | S793E | B | 9.3e-09 | 8.2e-09 | 4.6e-09 | 7.367E-09 | 2.458E-09 | 4.600E-09 | 9.300E-09 |
| 15847 | 793 | S793Q | B | 5.5e-09 | 1.3e-08 | 6.6e-09 | 8.367E-09 | 4.050E-09 | 5.500E-09 | 1.300E-08 |
| 15848 | 793 | S793G | B | 1.1e-08 | 1.3e-08 | 9.8e-09 | 1.127E-08 | 1.617E-09 | 9.800E-09 | 1.300E-08 |
| 15849 | 793 | S793H | B | 1.4e-08 | 2.1e-09 | 9.9e-09 | 8.667E-09 | 6.045E-09 | 2.100E-09 | 1.400E-08 |
| 15850 | 793 | S793I | B | 1.1e-08 | 6e-09 | 9.5e-09 | 8.833E-09 | 2.566E-09 | 6.000E-09 | 1.100E-08 |
| 15851 | 793 | S793L | B | 5e-09 | 1.4e-08 | 3e-08 | 1.633E-08 | 1.266E-08 | 5.000E-09 | 3.000E-08 |
| 15852 | 793 | S793K | B | 5.5e-09 | 1.6e-08 | 2.4e-08 | 1.517E-08 | 9.278E-09 | 5.500E-09 | 2.400E-08 |
| 15853 | 793 | S793M | B | 9.4e-09 | 1.7e-08 | 9.2e-09 | 1.187E-08 | 4.447E-09 | 9.200E-09 | 1.700E-08 |
| 15854 | 793 | S793F | B | 4.7e-09 | 4.9e-09 | 6.5e-09 | 5.367E-09 | 9.866E-10 | 4.700E-09 | 6.500E-09 |
| 15855 | 793 | S793P | B | 1.1e-08 | 2e-08 | 1.7e-09 | 1.090E-08 | 9.150E-09 | 1.700E-09 | 2.000E-08 |
| 15856 | 793 | S793S | B | 3.6e-08 | 6.6e-09 | 1.8e-08 | 2.020E-08 | 1.482E-08 | 6.600E-09 | 3.600E-08 |
| 15857 | 793 | S793T | B | 1.5e-08 | 2.9e-09 | 4.4e-09 | 7.433E-09 | 6.596E-09 | 2.900E-09 | 1.500E-08 |
| 15858 | 793 | S793W | B | 6.6e-09 | 2.3e-09 | 1.3e-08 | 7.300E-09 | 5.384E-09 | 2.300E-09 | 1.300E-08 |
| 15859 | 793 | S793Y | B | 7.1e-09 | 9.8e-09 | 7.1e-09 | 8.000E-09 | 1.559E-09 | 7.100E-09 | 9.800E-09 |
| 15860 | 793 | S793V | B | 1.7e-08 | 5.6e-09 | 3e-09 | 8.533E-09 | 7.447E-09 | 3.000E-09 | 1.700E-08 |
| 15861 | 794 | T794A | B | 8.8e-09 | 2.4e-09 | 2.4e-08 | 1.173E-08 | 1.109E-08 | 2.400E-09 | 2.400E-08 |
| 15862 | 794 | T794R | B | 3.9e-09 | 5.4e-09 | 2.5e-09 | 3.933E-09 | 1.450E-09 | 2.500E-09 | 5.400E-09 |
| 15863 | 794 | T794N | B | 1.7e-08 | 5.8e-09 | 4.2e-09 | 9.000E-09 | 6.974E-09 | 4.200E-09 | 1.700E-08 |
| 15864 | 794 | T794D | B | 7.6e-09 | 5.9e-09 | 3.6e-09 | 5.700E-09 | 2.007E-09 | 3.600E-09 | 7.600E-09 |
| 15865 | 794 | T794C | B | 8e-09 | 5.2e-09 | 2.9e-09 | 5.367E-09 | 2.554E-09 | 2.900E-09 | 8.000E-09 |
| 15866 | 794 | T794E | B | 8.1e-09 | 7.3e-09 | 1.6e-08 | 1.047E-08 | 4.809E-09 | 7.300E-09 | 1.600E-08 |
| 15867 | 794 | T794Q | B | 4.2e-09 | 2.1e-08 | 3.9e-09 | 9.700E-09 | 9.787E-09 | 3.900E-09 | 2.100E-08 |
| 15868 | 794 | T794G | B | 4.9e-09 | 8.5e-09 | 1.8e-09 | 5.067E-09 | 3.353E-09 | 1.800E-09 | 8.500E-09 |
| 15869 | 794 | T794H | B | 5.3e-09 | 8.2e-09 | 6.6e-09 | 6.700E-09 | 1.453E-09 | 5.300E-09 | 8.200E-09 |
| 15870 | 794 | T794I | B | 6.8e-09 | 5.2e-09 | 1.4e-08 | 8.667E-09 | 4.688E-09 | 5.200E-09 | 1.400E-08 |
| 15871 | 794 | T794L | B | 3.8e-09 | 2.2e-09 | 5.1e-09 | 3.700E-09 | 1.453E-09 | 2.200E-09 | 5.100E-09 |
| 15872 | 794 | T794K | B | 2.7e-09 | 3.5e-09 | 6.5e-09 | 4.233E-09 | 2.003E-09 | 2.700E-09 | 6.500E-09 |
| 15873 | 794 | T794M | B | 1.3e-08 | 2.5e-09 | 8.5e-09 | 8.000E-09 | 5.268E-09 | 2.500E-09 | 1.300E-08 |
| 15874 | 794 | T794F | B | 4.9e-09 | 2.5e-08 | 2e-08 | 1.663E-08 | 1.046E-08 | 4.900E-09 | 2.500E-08 |
| 15875 | 794 | T794P | B | 5.1e-09 | 3.4e-09 | 6.5e-09 | 5.000E-09 | 1.552E-09 | 3.400E-09 | 6.500E-09 |
| 15876 | 794 | T794S | B | 1.2e-08 | 6e-09 | 9.1e-09 | 9.033E-09 | 3.001E-09 | 6.000E-09 | 1.200E-08 |
| 15877 | 794 | T794T | B | 3.4e-08 | 4e-09 | 7.3e-09 | 1.510E-08 | 1.645E-08 | 4.000E-09 | 3.400E-08 |
| 15878 | 794 | T794W | B | 6.2e-09 | 7.5e-09 | 1.7e-08 | 1.023E-08 | 5.896E-09 | 6.200E-09 | 1.700E-08 |
| 15879 | 794 | T794Y | B | 8.1e-09 | 2.2e-09 | 1.6e-08 | 8.767E-09 | 6.924E-09 | 2.200E-09 | 1.600E-08 |
| 15880 | 794 | T794V | B | 2e-08 | 5.8e-09 | 1.1e-08 | 1.227E-08 | 7.184E-09 | 5.800E-09 | 2.000E-08 |
| 15881 | 795 | Y795A | B | 9.5e-09 | 4.6e-09 | 7.1e-09 | 7.067E-09 | 2.450E-09 | 4.600E-09 | 9.500E-09 |
| 15882 | 795 | Y795R | B | 5.6e-09 | 6.6e-09 | 7.9e-09 | 6.700E-09 | 1.153E-09 | 5.600E-09 | 7.900E-09 |
| 15883 | 795 | Y795N | B | 5.8e-09 | 5.5e-09 | 1.2e-08 | 7.767E-09 | 3.669E-09 | 5.500E-09 | 1.200E-08 |
| 15884 | 795 | Y795D | B | 1.2e-08 | 1.3e-08 | 1.1e-08 | 1.200E-08 | 1.000E-09 | 1.100E-08 | 1.300E-08 |
| 15885 | 795 | Y795C | B | 9.2e-09 | 3.9e-09 | 1.1e-08 | 8.033E-09 | 3.691E-09 | 3.900E-09 | 1.100E-08 |
| 15886 | 795 | Y795E | B | 4.3e-09 | 6.1e-09 | 4.9e-09 | 5.100E-09 | 9.165E-10 | 4.300E-09 | 6.100E-09 |
| 15887 | 795 | Y795Q | B | 7.9e-09 | 1.6e-08 | 5.3e-09 | 9.733E-09 | 5.581E-09 | 5.300E-09 | 1.600E-08 |
| 15888 | 795 | Y795G | B | 3.4e-09 | 7.9e-09 | 4.7e-09 | 5.333E-09 | 2.316E-09 | 3.400E-09 | 7.900E-09 |
| 15889 | 795 | Y795H | B | 1.4e-09 | 7.2e-09 | 5.8e-09 | 4.800E-09 | 3.027E-09 | 1.400E-09 | 7.200E-09 |
| 15890 | 795 | Y795I | B | 9.2e-09 | 9.6e-09 | 7.5e-09 | 8.767E-09 | 1.115E-09 | 7.500E-09 | 9.600E-09 |
| 15891 | 795 | Y795L | B | 5.1e-09 | 3.8e-09 | 4.4e-09 | 4.433E-09 | 6.506E-10 | 3.800E-09 | 5.100E-09 |
| 15892 | 795 | Y795K | B | 3.3e-09 | 4.4e-09 | 5.4e-09 | 4.367E-09 | 1.050E-09 | 3.300E-09 | 5.400E-09 |
| 15893 | 795 | Y795M | B | 6.8e-09 | 2.9e-09 | 1.3e-08 | 7.567E-09 | 5.093E-09 | 2.900E-09 | 1.300E-08 |
| 15894 | 795 | Y795F | B | 6.8e-09 | 1.9e-08 | 9e-09 | 1.160E-08 | 6.502E-09 | 6.800E-09 | 1.900E-08 |

| | | | | | | | | | | |
|---|---|---|---|---|---|---|---|---|---|---|
| 15895 | 795 | Y795P | B | 6.7e-09 | 4.9e-09 | 6e-09 | 5.867E-09 | 9.074E-10 | 4.900E-09 | 6.700E-09 |
| 15896 | 795 | Y795S | B | 7.5e-09 | 9.2e-09 | 6e-09 | 7.567E-09 | 1.601E-09 | 6.000E-09 | 9.200E-09 |
| 15897 | 795 | Y795T | B | 3.1e-09 | 2.3e-08 | 1.5e-08 | 1.370E-08 | 1.001E-08 | 3.100E-09 | 2.300E-08 |
| 15898 | 795 | Y795W | B | 6.8e-09 | 1.2e-08 | 1.8e-08 | 1.227E-08 | 5.605E-09 | 6.800E-09 | 1.800E-08 |
| 15899 | 795 | Y795Y | B | 7.6e-09 | 5.5e-09 | 6.6e-09 | 6.567E-09 | 1.050E-09 | 5.500E-09 | 7.600E-09 |
| 15900 | 795 | Y795V | B | 1.2e-08 | 7.1e-09 | 1.1e-08 | 1.003E-08 | 2.589E-09 | 7.100E-09 | 1.200E-08 |
| 15901 | 796 | S796A | B | 7.3e-09 | 4.7e-09 | 6.1e-09 | 6.033E-09 | 1.301E-09 | 4.700E-09 | 7.300E-09 |
| 15902 | 796 | S796R | B | 3.3e-09 | 4.2e-09 | 3.5e-09 | 3.667E-09 | 4.726E-10 | 3.300E-09 | 4.200E-09 |
| 15903 | 796 | S796N | B | 3.3e-09 | 1.2e-08 | 3.1e-09 | 6.133E-09 | 5.082E-09 | 3.100E-09 | 1.200E-08 |
| 15904 | 796 | S796D | B | 4.1e-09 | 4e-09 | 1.3e-08 | 7.033E-09 | 5.168E-09 | 4.000E-09 | 1.300E-08 |
| 15905 | 796 | S796C | B | 1.9e-09 | 3.1e-09 | 1.3e-08 | 6.000E-09 | 6.092E-09 | 1.900E-09 | 1.300E-08 |
| 15906 | 796 | S796E | B | 9.3e-09 | 9.8e-09 | 1.2e-08 | 1.037E-08 | 1.436E-09 | 9.300E-09 | 1.200E-08 |
| 15907 | 796 | S796Q | B | 3.9e-09 | 2.2e-09 | 4.1e-09 | 3.400E-09 | 1.044E-09 | 2.200E-09 | 4.100E-09 |
| 15908 | 796 | S796G | B | 9e-09 | 1.2e-08 | 2.9e-09 | 7.967E-09 | 4.637E-09 | 2.900E-09 | 1.200E-08 |
| 15909 | 796 | S796H | B | 1.9e-08 | 1.4e-08 | 4.1e-09 | 1.237E-08 | 7.583E-09 | 4.100E-09 | 1.900E-08 |
| 15910 | 796 | S796I | B | 3.1e-09 | 6.4e-09 | 2.1e-08 | 1.017E-08 | 9.526E-09 | 3.100E-09 | 2.100E-08 |
| 15911 | 796 | S796L | B | 8.7e-09 | 1.3e-08 | 3e-09 | 8.233E-09 | 5.016E-09 | 3.000E-09 | 1.300E-08 |
| 15912 | 796 | S796K | B | 6.5e-09 | 6.8e-09 | 6.3e-09 | 6.533E-09 | 2.517E-10 | 6.300E-09 | 6.800E-09 |
| 15913 | 796 | S796M | B | 5.9e-09 | 7.2e-09 | 8.2e-09 | 7.100E-09 | 1.153E-09 | 5.900E-09 | 8.200E-09 |
| 15914 | 796 | S796F | B | 9.6e-09 | 5.4e-09 | 5.6e-09 | 6.867E-09 | 2.369E-09 | 5.400E-09 | 9.600E-09 |
| 15915 | 796 | S796P | B | 1e-08 | 3.4e-09 | 9.7e-09 | 7.700E-09 | 3.727E-09 | 3.400E-09 | 1.000E-08 |
| 15916 | 796 | S796S | B | 6.9e-09 | 2.3e-08 | 3.8e-09 | 1.123E-08 | 1.031E-08 | 3.800E-09 | 2.300E-08 |
| 15917 | 796 | S796T | B | 8.6e-09 | 3.6e-09 | 1.2e-08 | 8.067E-09 | 4.225E-09 | 3.600E-09 | 1.200E-08 |
| 15918 | 796 | S796W | B | 4.8e-09 | 8e-09 | 6.5e-09 | 6.433E-09 | 1.601E-09 | 4.800E-09 | 8.000E-09 |
| 15919 | 796 | S796Y | B | 1.1e-08 | 1.6e-08 | 8.8e-09 | 1.193E-08 | 3.690E-09 | 8.800E-09 | 1.600E-08 |
| 15920 | 796 | S796V | B | 1e-08 | 2.7e-09 | 8.5e-09 | 7.067E-09 | 3.855E-09 | 2.700E-09 | 1.000E-08 |
| 15921 | 797 | L797A | B | 9e-09 | 1.3e-08 | 8.3e-09 | 1.010E-08 | 2.536E-09 | 8.300E-09 | 1.300E-08 |
| 15922 | 797 | L797R | B | 7.6e-09 | 1.7e-09 | 3.3e-09 | 4.200E-09 | 3.051E-09 | 1.700E-09 | 7.600E-09 |
| 15923 | 797 | L797N | B | 3.6e-09 | 7.3e-09 | 1e-08 | 6.967E-09 | 3.213E-09 | 3.600E-09 | 1.000E-08 |
| 15924 | 797 | L797D | B | 3.5e-09 | 5.7e-09 | 3.7e-09 | 4.300E-09 | 1.217E-09 | 3.500E-09 | 5.700E-09 |
| 15925 | 797 | L797C | B | 1e-08 | 1.2e-08 | 1.1e-08 | 1.100E-08 | 1.000E-09 | 1.000E-08 | 1.200E-08 |
| 15926 | 797 | L797E | B | 5e-09 | 9.7e-09 | 2.2e-09 | 5.633E-09 | 3.790E-09 | 2.200E-09 | 9.700E-09 |
| 15927 | 797 | L797Q | B | 7.9e-09 | 3.4e-09 | 6.6e-09 | 5.967E-09 | 2.316E-09 | 3.400E-09 | 7.900E-09 |
| 15928 | 797 | L797G | B | 6.9e-09 | 6.9e-09 | 2.6e-08 | 1.327E-08 | 1.103E-08 | 6.900E-09 | 2.600E-08 |
| 15929 | 797 | L797H | B | 1.4e-08 | 5.2e-09 | 7.1e-09 | 8.767E-09 | 4.631E-09 | 5.200E-09 | 1.400E-08 |
| 15930 | 797 | L797I | B | 9.8e-09 | 7.6e-09 | 1e-08 | 9.133E-09 | 1.332E-09 | 7.600E-09 | 1.000E-08 |
| 15931 | 797 | L797L | B | 3.9e-09 | 8.1e-09 | 6.8e-09 | 6.267E-09 | 2.150E-09 | 3.900E-09 | 8.100E-09 |
| 15932 | 797 | L797K | B | 1.1e-08 | 1e-08 | 3.7e-09 | 8.233E-09 | 3.958E-09 | 3.700E-09 | 1.100E-08 |
| 15933 | 797 | L797M | B | 2.3e-09 | 1.3e-08 | 1.1e-08 | 8.767E-09 | 5.689E-09 | 2.300E-09 | 1.300E-08 |
| 15934 | 797 | L797F | B | 6.5e-09 | 8.7e-09 | 9.6e-09 | 8.267E-09 | 1.595E-09 | 6.500E-09 | 9.600E-09 |
| 15935 | 797 | L797P | B | 1.2e-08 | 3.3e-09 | 8.2e-09 | 7.833E-09 | 4.362E-09 | 3.300E-09 | 1.200E-08 |
| 15936 | 797 | L797S | B | 2e-09 | 1.6e-08 | 1.7e-08 | 1.167E-08 | 8.386E-09 | 2.000E-09 | 1.700E-08 |
| 15937 | 797 | L797T | B | 8.1e-09 | 3.8e-09 | 1.2e-08 | 7.967E-09 | 4.102E-09 | 3.800E-09 | 1.200E-08 |
| 15938 | 797 | L797W | B | 1.5e-08 | 4.9e-09 | 7e-09 | 8.967E-09 | 5.329E-09 | 4.900E-09 | 1.500E-08 |
| 15939 | 797 | L797Y | B | 5e-09 | 1.4e-08 | 7.5e-09 | 8.833E-09 | 4.646E-09 | 5.000E-09 | 1.400E-08 |
| 15940 | 797 | L797V | B | 1.1e-08 | 2e-08 | 5.5e-09 | 1.217E-08 | 7.320E-09 | 5.500E-09 | 2.000E-08 |
| 15941 | 798 | S798A | B | 2.8e-09 | 3.9e-09 | 1.2e-08 | 6.233E-09 | 5.024E-09 | 2.800E-09 | 1.200E-08 |
| 15942 | 798 | S798R | B | 2.1e-09 | 8.9e-09 | 8.9e-09 | 6.633E-09 | 3.926E-09 | 2.100E-09 | 8.900E-09 |
| 15943 | 798 | S798N | B | 8.3e-09 | 7.5e-09 | 5.3e-09 | 7.033E-09 | 1.553E-09 | 5.300E-09 | 8.300E-09 |
| 15944 | 798 | S798D | B | 4.5e-09 | 7e-09 | 5.3e-09 | 5.600E-09 | 1.277E-09 | 4.500E-09 | 7.000E-09 |
| 15945 | 798 | S798C | B | 5e-09 | 3.9e-09 | 2.3e-08 | 1.063E-08 | 1.072E-08 | 3.900E-09 | 2.300E-08 |
| 15946 | 798 | S798E | B | 1.5e-08 | 3.3e-09 | 8.4e-09 | 8.900E-09 | 5.866E-09 | 3.300E-09 | 1.500E-08 |
| 15947 | 798 | S798Q | B | 1.1e-08 | 5.5e-09 | 2.2e-08 | 1.283E-08 | 8.401E-09 | 5.500E-09 | 2.200E-08 |
| 15948 | 798 | S798G | B | 3.4e-09 | 7.6e-09 | 4.2e-09 | 5.067E-09 | 2.230E-09 | 3.400E-09 | 7.600E-09 |
| 15949 | 798 | S798H | B | 2.1e-09 | 7.3e-09 | 8.3e-09 | 5.900E-09 | 3.329E-09 | 2.100E-09 | 8.300E-09 |
| 15950 | 798 | S798I | B | 1e-08 | 1.2e-08 | 1.1e-08 | 1.100E-08 | 1.000E-09 | 1.000E-08 | 1.200E-08 |
| 15951 | 798 | S798L | B | 6.3e-09 | 5.4e-09 | 5.5e-09 | 5.733E-09 | 4.933E-10 | 5.400E-09 | 6.300E-09 |
| 15952 | 798 | S798K | B | 2.6e-09 | 7.4e-09 | 5.4e-09 | 5.133E-09 | 2.411E-09 | 2.600E-09 | 7.400E-09 |
| 15953 | 798 | S798M | B | 8.2e-09 | 9.2e-09 | 4e-09 | 7.133E-09 | 2.759E-09 | 4.000E-09 | 9.200E-09 |
| 15954 | 798 | S798F | B | 8.9e-09 | 1.6e-09 | 2.6e-08 | 1.217E-08 | 1.252E-08 | 1.600E-09 | 2.600E-08 |
| 15955 | 798 | S798P | B | 8.5e-09 | 6.7e-09 | 8.9e-09 | 8.033E-09 | 1.172E-09 | 6.700E-09 | 8.900E-09 |
| 15956 | 798 | S798S | B | 1.4e-08 | 1.3e-08 | 1.1e-08 | 1.267E-08 | 1.528E-09 | 1.100E-08 | 1.400E-08 |
| 15957 | 798 | S798T | B | 4.6e-09 | 1.4e-08 | 5.9e-09 | 8.167E-09 | 5.093E-09 | 4.600E-09 | 1.400E-08 |
| 15958 | 798 | S798W | B | 9e-09 | 6.3e-09 | 6.7e-09 | 7.333E-09 | 1.457E-09 | 6.300E-09 | 9.000E-09 |
| 15959 | 798 | S798Y | B | 2.3e-09 | 8e-09 | 1.3e-08 | 7.767E-09 | 5.354E-09 | 2.300E-09 | 1.300E-08 |
| 15960 | 798 | S798V | B | 3.9e-09 | 5.3e-09 | 1.2e-08 | 7.067E-09 | 4.329E-09 | 3.900E-09 | 1.200E-08 |
| 15961 | 799 | S799A | B | 4.9e-09 | 5.8e-09 | 8.4e-09 | 6.367E-09 | 1.818E-09 | 4.900E-09 | 8.400E-09 |
| 15962 | 799 | S799R | B | 1.2e-09 | 4.3e-09 | 8.3e-09 | 4.600E-09 | 3.559E-09 | 1.200E-09 | 8.300E-09 |
| 15963 | 799 | S799N | B | 4.2e-09 | 1.3e-09 | 4.8e-08 | 1.783E-08 | 2.617E-08 | 1.300E-09 | 4.800E-08 |
| 15964 | 799 | S799D | B | 3.8e-09 | 5.6e-09 | 5.8e-09 | 5.067E-09 | 1.102E-09 | 3.800E-09 | 5.800E-09 |
| 15965 | 799 | S799C | B | 9.7e-09 | 9.9e-09 | 2.7e-08 | 1.553E-08 | 9.931E-09 | 9.700E-09 | 2.700E-08 |
| 15966 | 799 | S799E | B | 9.9e-09 | 3.8e-09 | 1.4e-08 | 9.233E-09 | 5.133E-09 | 3.800E-09 | 1.400E-08 |
| 15967 | 799 | S799Q | B | 1.2e-08 | 1.7e-08 | 5.6e-09 | 1.153E-08 | 5.714E-09 | 5.600E-09 | 1.700E-08 |
| 15968 | 799 | S799G | B | 3.9e-09 | 1.1e-08 | 6.3e-09 | 7.067E-09 | 3.612E-09 | 3.900E-09 | 1.100E-08 |
| 15969 | 799 | S799H | B | 5.7e-09 | 3.1e-09 | 4.2e-09 | 4.333E-09 | 1.305E-09 | 3.100E-09 | 5.700E-09 |
| 15970 | 799 | S799I | B | 1e-08 | 5.2e-09 | 8.2e-09 | 7.800E-09 | 2.425E-09 | 5.200E-09 | 1.000E-08 |
| 15971 | 799 | S799L | B | 7.7e-09 | 1.5e-09 | 5.7e-09 | 4.967E-09 | 3.164E-09 | 1.500E-09 | 7.700E-09 |
| 15972 | 799 | S799K | B | 3e-09 | 6.5e-09 | 5e-09 | 4.833E-09 | 1.756E-09 | 3.000E-09 | 6.500E-09 |
| 15973 | 799 | S799M | B | 2.7e-09 | 4.7e-09 | 5.2e-09 | 4.200E-09 | 1.323E-09 | 2.700E-09 | 5.200E-09 |
| 15974 | 799 | S799F | B | 1.4e-08 | 1.9e-08 | 4e-09 | 1.233E-08 | 7.638E-09 | 4.000E-09 | 1.900E-08 |
| 15975 | 799 | S799P | B | 2.5e-08 | 8e-09 | 5.3e-09 | 1.277E-08 | 1.068E-08 | 5.300E-09 | 2.500E-08 |
| 15976 | 799 | S799S | B | 3.5e-09 | 4.5e-09 | 1e-08 | 6.000E-09 | 3.500E-09 | 3.500E-09 | 1.000E-08 |
| 15977 | 799 | S799T | B | 1.1e-08 | 7.3e-09 | 3.1e-09 | 7.133E-09 | 3.953E-09 | 3.100E-09 | 1.100E-08 |
| 15978 | 799 | S799W | B | 4.8e-09 | 3.1e-09 | 6.2e-09 | 4.700E-09 | 1.552E-09 | 3.100E-09 | 6.200E-09 |
| 15979 | 799 | S799Y | B | 4.6e-09 | 5.5e-09 | 1.4e-08 | 8.033E-09 | 5.187E-09 | 4.600E-09 | 1.400E-08 |

| | | | | | | | | | | |
|---|---|---|---|---|---|---|---|---|---|---|
| 15980 | 799 | S799V | B | 4.9e-09 | 2.1e-09 | 8.9e-09 | 5.300E-09 | 3.418E-09 | 2.100E-09 | 8.900E-09 |
| 15981 | 800 | T800A | B | 6.5e-09 | 1.3e-08 | 5.9e-09 | 8.467E-09 | 3.937E-09 | 5.900E-09 | 1.300E-08 |
| 15982 | 800 | T800R | B | 1e-08 | 4.4e-09 | 5.8e-09 | 6.733E-09 | 2.914E-09 | 4.400E-09 | 1.000E-08 |
| 15983 | 800 | T800N | B | 5.9e-09 | 1.5e-08 | 8.8e-09 | 9.900E-09 | 4.649E-09 | 5.900E-09 | 1.500E-08 |
| 15984 | 800 | T800D | B | 3.6e-09 | 3.8e-09 | 7.8e-09 | 5.067E-09 | 2.369E-09 | 3.600E-09 | 7.800E-09 |
| 15985 | 800 | T800C | B | 1.2e-08 | 2e-08 | 4.8e-09 | 1.227E-08 | 7.604E-09 | 4.800E-09 | 2.000E-08 |
| 15986 | 800 | T800E | B | 1.4e-08 | 5.1e-09 | 3e-09 | 7.367E-09 | 5.840E-09 | 3.000E-09 | 1.400E-08 |
| 15987 | 800 | T800Q | B | 9e-09 | 1.1e-08 | 1e-08 | 1.000E-08 | 1.000E-09 | 9.000E-09 | 1.100E-08 |
| 15988 | 800 | T800G | B | 7.5e-09 | 6.6e-09 | 8.3e-09 | 7.467E-09 | 8.505E-10 | 6.600E-09 | 8.300E-09 |
| 15989 | 800 | T800H | B | 1.6e-08 | 7.5e-10 | 4.9e-09 | 7.217E-09 | 7.885E-09 | 7.500E-10 | 1.600E-08 |
| 15990 | 800 | T800I | B | 5.3e-09 | 3.2e-09 | 8.1e-09 | 5.533E-09 | 2.458E-09 | 3.200E-09 | 8.100E-09 |
| 15991 | 800 | T800L | B | 1.4e-08 | 2.6e-09 | 6.5e-09 | 7.700E-09 | 5.794E-09 | 2.600E-09 | 1.400E-08 |
| 15992 | 800 | T800K | B | 2.5e-08 | 1.1e-08 | 5.4e-09 | 1.380E-08 | 1.010E-08 | 5.400E-09 | 2.500E-08 |
| 15993 | 800 | T800M | B | 1.1e-08 | 7.2e-09 | 4.1e-09 | 7.433E-09 | 3.456E-09 | 4.100E-09 | 1.100E-08 |
| 15994 | 800 | T800F | B | 9.9e-09 | 3.3e-09 | 8.7e-09 | 7.300E-09 | 3.516E-09 | 3.300E-09 | 9.900E-09 |
| 15995 | 800 | T800P | B | 1e-08 | 2e-08 | 8.6e-09 | 1.287E-08 | 6.217E-09 | 8.600E-09 | 2.000E-08 |
| 15996 | 800 | T800S | B | 5.4e-09 | 6.1e-09 | 5.6e-09 | 5.700E-09 | 3.606E-10 | 5.400E-09 | 6.100E-09 |
| 15997 | 800 | T800T | B | 1.1e-08 | 5.8e-09 | 1.4e-08 | 1.027E-08 | 4.149E-09 | 5.800E-09 | 1.400E-08 |
| 15998 | 800 | T800W | B | 3.7e-09 | 8.3e-09 | 5.7e-09 | 5.900E-09 | 2.307E-09 | 3.700E-09 | 8.300E-09 |
| 15999 | 800 | T800Y | B | 4.5e-09 | 8.9e-09 | 3.6e-09 | 5.667E-09 | 2.836E-09 | 3.600E-09 | 8.900E-09 |
| 16000 | 800 | T800V | B | 3.7e-09 | 4.6e-09 | 7.8e-09 | 5.367E-09 | 2.155E-09 | 3.700E-09 | 7.800E-09 |
| 16001 | 801 | L801A | B | 1.9e-09 | 4.3e-09 | 2.5e-09 | 2.900E-09 | 1.249E-09 | 1.900E-09 | 4.300E-09 |
| 16002 | 801 | L801R | B | 1.6e-08 | 1.1e-08 | 5.1e-09 | 1.070E-08 | 5.456E-09 | 5.100E-09 | 1.600E-08 |
| 16003 | 801 | L801N | B | 2.6e-09 | 7.7e-09 | 2.8e-09 | 4.367E-09 | 2.888E-09 | 2.600E-09 | 7.700E-09 |
| 16004 | 801 | L801D | B | 1e-08 | 2.7e-09 | 1.6e-08 | 9.567E-09 | 6.661E-09 | 2.700E-09 | 1.600E-08 |
| 16005 | 801 | L801C | B | 3.1e-09 | 7e-09 | 3.6e-09 | 4.567E-09 | 2.122E-09 | 3.100E-09 | 7.000E-09 |
| 16006 | 801 | L801E | B | 8.7e-09 | 2.2e-09 | 4.9e-09 | 5.267E-09 | 3.265E-09 | 2.200E-09 | 8.700E-09 |
| 16007 | 801 | L801Q | B | 1.2e-08 | 2.3e-09 | 6.5e-09 | 6.933E-09 | 4.864E-09 | 2.300E-09 | 1.200E-08 |
| 16008 | 801 | L801G | B | 1.8e-09 | 8.5e-09 | 1.3e-08 | 7.767E-09 | 5.636E-09 | 1.800E-09 | 1.300E-08 |
| 16009 | 801 | L801H | B | 7.4e-09 | 8.1e-09 | 6.7e-09 | 7.400E-09 | 7.000E-10 | 6.700E-09 | 8.100E-09 |
| 16010 | 801 | L801I | B | 5.1e-09 | 8.7e-09 | 1.3e-08 | 8.933E-09 | 3.955E-09 | 5.100E-09 | 1.300E-08 |
| 16011 | 801 | L801L | B | 7.5e-09 | 3.8e-09 | 4.6e-09 | 5.300E-09 | 1.947E-09 | 3.800E-09 | 7.500E-09 |
| 16012 | 801 | L801K | B | 4.2e-09 | 1.4e-08 | 1.3e-08 | 1.040E-08 | 5.393E-09 | 4.200E-09 | 1.400E-08 |
| 16013 | 801 | L801M | B | 2.8e-09 | 6.1e-09 | 8e-09 | 5.633E-09 | 2.631E-09 | 2.800E-09 | 8.000E-09 |
| 16014 | 801 | L801F | B | 3.7e-08 | 2.6e-09 | 9.4e-09 | 1.633E-08 | 1.822E-08 | 2.600E-09 | 3.700E-08 |
| 16015 | 801 | L801P | B | 5.4e-09 | 1e-08 | 1.2e-08 | 9.133E-09 | 3.384E-09 | 5.400E-09 | 1.200E-08 |
| 16016 | 801 | L801S | B | 1.3e-08 | 4e-09 | 1.2e-08 | 9.667E-09 | 4.933E-09 | 4.000E-09 | 1.300E-08 |
| 16017 | 801 | L801T | B | 1.3e-08 | 3.4e-08 | 1.1e-08 | 1.933E-08 | 1.274E-08 | 1.100E-08 | 3.400E-08 |
| 16018 | 801 | L801W | B | 1.4e-08 | 3.1e-09 | 6.9e-09 | 8.000E-09 | 5.533E-09 | 3.100E-09 | 1.400E-08 |
| 16019 | 801 | L801Y | B | 3.6e-09 | 2.9e-09 | 1.8e-08 | 8.167E-09 | 8.523E-09 | 2.900E-09 | 1.800E-08 |
| 16020 | 801 | L801V | B | 4.2e-09 | 1.9e-09 | 6e-09 | 4.033E-09 | 2.055E-09 | 1.900E-09 | 6.000E-09 |
| 16021 | 802 | T802A | B | 6.2e-09 | 8.5e-09 | 9.1e-09 | 7.933E-09 | 1.531E-09 | 6.200E-09 | 9.100E-09 |
| 16022 | 802 | T802R | B | 8.2e-09 | 1.6e-09 | 1.7e-09 | 3.833E-09 | 3.782E-09 | 1.600E-09 | 8.200E-09 |
| 16023 | 802 | T802N | B | 4e-09 | 5.3e-09 | 3.7e-09 | 4.333E-09 | 8.505E-10 | 3.700E-09 | 5.300E-09 |
| 16024 | 802 | T802D | B | 7.4e-09 | 3.6e-09 | 6.3e-09 | 5.767E-09 | 1.955E-09 | 3.600E-09 | 7.400E-09 |
| 16025 | 802 | T802C | B | 5.2e-09 | 9.5e-09 | 1.2e-08 | 8.900E-09 | 3.439E-09 | 5.200E-09 | 1.200E-08 |
| 16026 | 802 | T802E | B | 1.5e-08 | 8.5e-09 | 7.9e-09 | 1.047E-08 | 3.937E-09 | 7.900E-09 | 1.500E-08 |
| 16027 | 802 | T802Q | B | 8.2e-09 | 7e-09 | 5.8e-09 | 7.000E-09 | 1.200E-09 | 5.800E-09 | 8.200E-09 |
| 16028 | 802 | T802G | B | 2.4e-08 | 5.4e-09 | 1.5e-08 | 1.480E-08 | 9.302E-09 | 5.400E-09 | 2.400E-08 |
| 16029 | 802 | T802H | B | 6.4e-09 | 8.1e-09 | 1.9e-08 | 1.117E-08 | 6.837E-09 | 6.400E-09 | 1.900E-08 |
| 16030 | 802 | T802I | B | 8.9e-09 | 4.9e-09 | 8.3e-09 | 7.367E-09 | 2.157E-09 | 4.900E-09 | 8.900E-09 |
| 16031 | 802 | T802L | B | 1.3e-08 | 4.5e-09 | 1.1e-08 | 9.500E-09 | 4.444E-09 | 4.500E-09 | 1.300E-08 |
| 16032 | 802 | T802K | B | 1.1e-08 | 8.5e-09 | 1.2e-08 | 1.050E-08 | 1.803E-09 | 8.500E-09 | 1.200E-08 |
| 16033 | 802 | T802M | B | 4.7e-09 | 6e-09 | 1.4e-09 | 4.033E-09 | 2.371E-09 | 1.400E-09 | 6.000E-09 |
| 16034 | 802 | T802F | B | 4.2e-09 | 6.2e-09 | 2.8e-08 | 1.280E-08 | 1.320E-08 | 4.200E-09 | 2.800E-08 |
| 16035 | 802 | T802P | B | 3.9e-09 | 3.7e-08 | 1e-09 | 1.397E-08 | 2.000E-08 | 1.000E-09 | 3.700E-08 |
| 16036 | 802 | T802S | B | 8.1e-09 | 1.2e-08 | 5e-09 | 8.367E-09 | 3.508E-09 | 5.000E-09 | 1.200E-08 |
| 16037 | 802 | T802T | B | 1.3e-08 | 9.1e-09 | 1.8e-09 | 7.967E-09 | 5.685E-09 | 1.800E-09 | 1.300E-08 |
| 16038 | 802 | T802W | B | 4.4e-09 | 5.8e-09 | 1e-08 | 6.733E-09 | 2.914E-09 | 4.400E-09 | 1.000E-08 |
| 16039 | 802 | T802Y | B | 3.6e-09 | 2.4e-09 | 8e-09 | 4.667E-09 | 2.948E-09 | 2.400E-09 | 8.000E-09 |
| 16040 | 802 | T802V | B | 8.4e-09 | 2.8e-09 | 4.8e-09 | 5.333E-09 | 2.838E-09 | 2.800E-09 | 8.400E-09 |
| 16041 | 803 | L803A | B | 1e-08 | 1.1e-08 | 8.4e-09 | 9.800E-09 | 1.311E-09 | 8.400E-09 | 1.100E-08 |
| 16042 | 803 | L803R | B | 1.1e-08 | 1.7e-08 | 9.2e-09 | 1.240E-08 | 4.084E-09 | 9.200E-09 | 1.700E-08 |
| 16043 | 803 | L803N | B | 5e-09 | 2.6e-09 | 1e-08 | 5.867E-09 | 3.775E-09 | 2.600E-09 | 1.000E-08 |
| 16044 | 803 | L803D | B | 5.7e-09 | 5.3e-09 | 9.2e-09 | 6.733E-09 | 2.146E-09 | 5.300E-09 | 9.200E-09 |
| 16045 | 803 | L803C | B | 1.3e-08 | 1.6e-08 | 8.6e-09 | 1.253E-08 | 3.722E-09 | 8.600E-09 | 1.600E-08 |
| 16046 | 803 | L803E | B | 6.5e-09 | 1.4e-08 | 8e-09 | 9.500E-09 | 3.969E-09 | 6.500E-09 | 1.400E-08 |
| 16047 | 803 | L803Q | B | 5.8e-09 | 5e-09 | 6.9e-09 | 5.900E-09 | 9.539E-10 | 5.000E-09 | 6.900E-09 |
| 16048 | 803 | L803G | B | 2.9e-09 | 6e-09 | 1.1e-08 | 6.633E-09 | 4.087E-09 | 2.900E-09 | 1.100E-08 |
| 16049 | 803 | L803H | B | 3.4e-09 | 1.1e-08 | 4.5e-09 | 6.300E-09 | 4.107E-09 | 3.400E-09 | 1.100E-08 |
| 16050 | 803 | L803I | B | 9.9e-09 | 4e-09 | 5.6e-09 | 6.500E-09 | 3.051E-09 | 4.000E-09 | 9.900E-09 |
| 16051 | 803 | L803L | B | 4.2e-09 | 3.5e-09 | 6.5e-09 | 4.733E-09 | 1.570E-09 | 3.500E-09 | 6.500E-09 |
| 16052 | 803 | L803K | B | 3.2e-09 | 3.2e-09 | 2.9e-09 | 3.100E-09 | 1.732E-10 | 2.900E-09 | 3.200E-09 |
| 16053 | 803 | L803M | B | 1e-08 | 1.5e-08 | 1.5e-08 | 1.333E-08 | 2.887E-09 | 1.000E-08 | 1.500E-08 |
| 16054 | 803 | L803F | B | 5.6e-09 | 6.8e-09 | 2.1e-08 | 1.113E-08 | 8.566E-09 | 5.600E-09 | 2.100E-08 |
| 16055 | 803 | L803P | B | 6.8e-09 | 9.9e-09 | 1.1e-08 | 9.233E-09 | 2.178E-09 | 6.800E-09 | 1.100E-08 |
| 16056 | 803 | L803S | B | 1.2e-08 | 5e-09 | 1.1e-08 | 9.333E-09 | 3.786E-09 | 5.000E-09 | 1.200E-08 |
| 16057 | 803 | L803T | B | 9e-09 | 7.4e-09 | 1.6e-08 | 1.080E-08 | 4.574E-09 | 7.400E-09 | 1.600E-08 |
| 16058 | 803 | L803W | B | 1.4e-09 | 8.9e-09 | 6.4e-09 | 5.567E-09 | 3.819E-09 | 1.400E-09 | 8.900E-09 |
| 16059 | 803 | L803Y | B | 2.7e-09 | 3e-09 | 1.6e-08 | 7.233E-09 | 7.594E-09 | 2.700E-09 | 1.600E-08 |
| 16060 | 803 | L803V | B | 1.1e-08 | 6.1e-09 | 7.2e-09 | 8.100E-09 | 2.571E-09 | 6.100E-09 | 1.100E-08 |
| 16061 | 804 | S804A | B | 1.6e-08 | 8.1e-09 | 5.8e-09 | 9.967E-09 | 5.350E-09 | 5.800E-09 | 1.600E-08 |
| 16062 | 804 | S804R | B | 7e-09 | 1e-08 | 3.3e-09 | 6.767E-09 | 3.356E-09 | 3.300E-09 | 1.000E-08 |
| 16063 | 804 | S804N | B | 4.5e-09 | 6.4e-09 | 1.2e-08 | 7.633E-09 | 3.899E-09 | 4.500E-09 | 1.200E-08 |
| 16064 | 804 | S804D | B | 1.7e-08 | 4.9e-09 | 2.9e-09 | 8.267E-09 | 7.629E-09 | 2.900E-09 | 1.700E-08 |

| | | | | | | | | | | |
|---|---|---|---|---|---|---|---|---|---|---|
| 16065 | 804 | S804C | B | 5.5e-09 | 1.5e-08 | 6.8e-09 | 9.100E-09 | 5.151E-09 | 5.500E-09 | 1.500E-08 |
| 16066 | 804 | S804E | B | 2.6e-08 | 8.8e-09 | 1.1e-08 | 1.527E-08 | 9.360E-09 | 8.800E-09 | 2.600E-08 |
| 16067 | 804 | S804Q | B | 6.4e-09 | 9e-09 | 1.3e-08 | 9.467E-09 | 3.325E-09 | 6.400E-09 | 1.300E-08 |
| 16068 | 804 | S804G | B | 1.8e-08 | 2.8e-09 | 2.4e-08 | 1.493E-08 | 1.093E-08 | 2.800E-09 | 2.400E-08 |
| 16069 | 804 | S804H | B | 2.3e-09 | 5.5e-09 | 4.3e-09 | 4.033E-09 | 1.617E-09 | 2.300E-09 | 5.500E-09 |
| 16070 | 804 | S804I | B | 1.4e-08 | 9.6e-09 | 2.9e-09 | 8.833E-09 | 5.590E-09 | 2.900E-09 | 1.400E-08 |
| 16071 | 804 | S804L | B | 6e-09 | 1.6e-09 | 4.6e-09 | 4.067E-09 | 2.248E-09 | 1.600E-09 | 6.000E-09 |
| 16072 | 804 | S804K | B | 6e-09 | 7.6e-09 | 4.2e-09 | 5.933E-09 | 1.701E-09 | 4.200E-09 | 7.600E-09 |
| 16073 | 804 | S804M | B | 8.4e-09 | 3e-08 | 4.1e-09 | 1.417E-08 | 1.388E-08 | 4.100E-09 | 3.000E-08 |
| 16074 | 804 | S804F | B | 6e-09 | 5.7e-09 | 5.2e-09 | 5.633E-09 | 4.041E-10 | 5.200E-09 | 6.000E-09 |
| 16075 | 804 | S804P | B | 4.4e-09 | 4.4e-09 | 7.1e-09 | 5.300E-09 | 1.559E-09 | 4.400E-09 | 7.100E-09 |
| 16076 | 804 | S804S | B | 1.2e-08 | 1.6e-08 | 4.1e-09 | 1.070E-08 | 6.056E-09 | 4.100E-09 | 1.600E-08 |
| 16077 | 804 | S804T | B | 7.4e-09 | 1.3e-08 | 1.5e-08 | 1.180E-08 | 3.940E-09 | 7.400E-09 | 1.500E-08 |
| 16078 | 804 | S804W | B | 1.4e-08 | 9.9e-09 | 6.4e-09 | 1.010E-08 | 3.804E-09 | 6.400E-09 | 1.400E-08 |
| 16079 | 804 | S804Y | B | 1.2e-08 | 1.1e-08 | 1.1e-08 | 1.133E-08 | 5.774E-10 | 1.100E-08 | 1.200E-08 |
| 16080 | 804 | S804V | B | 1.8e-08 | 4e-09 | 1.1e-08 | 1.100E-08 | 7.000E-09 | 4.000E-09 | 1.800E-08 |
| 16081 | 805 | K805A | B | 1.2e-08 | 1.4e-09 | 5.2e-09 | 6.200E-09 | 5.370E-09 | 1.400E-09 | 1.200E-08 |
| 16082 | 805 | K805R | B | 1e-08 | 1.5e-08 | 6.8e-09 | 1.060E-08 | 4.133E-09 | 6.800E-09 | 1.500E-08 |
| 16083 | 805 | K805N | B | 7.8e-09 | 9.2e-09 | 8.6e-09 | 8.533E-09 | 7.024E-10 | 7.800E-09 | 9.200E-09 |
| 16084 | 805 | K805D | B | 3.4e-09 | 3.8e-09 | 6.7e-09 | 4.633E-09 | 1.801E-09 | 3.400E-09 | 6.700E-09 |
| 16085 | 805 | K805C | B | 7.2e-09 | 4.3e-09 | 5.6e-09 | 5.700E-09 | 1.453E-09 | 4.300E-09 | 7.200E-09 |
| 16086 | 805 | K805E | B | 3.4e-09 | 9.8e-09 | 3e-09 | 5.400E-09 | 3.816E-09 | 3.000E-09 | 9.800E-09 |
| 16087 | 805 | K805Q | B | 3.4e-09 | 4.3e-09 | 6.8e-09 | 4.833E-09 | 1.762E-09 | 3.400E-09 | 6.800E-09 |
| 16088 | 805 | K805G | B | 1.3e-08 | 6.5e-09 | 5.1e-09 | 8.200E-09 | 4.215E-09 | 5.100E-09 | 1.300E-08 |
| 16089 | 805 | K805H | B | 2.2e-09 | 1.2e-08 | 8.6e-09 | 7.600E-09 | 4.976E-09 | 2.200E-09 | 1.200E-08 |
| 16090 | 805 | K805I | B | 8.7e-09 | 2.5e-09 | 1.5e-08 | 8.733E-09 | 6.250E-09 | 2.500E-09 | 1.500E-08 |
| 16091 | 805 | K805L | B | 2.4e-08 | 4e-09 | 7.5e-09 | 1.183E-08 | 1.068E-08 | 4.000E-09 | 2.400E-08 |
| 16092 | 805 | K805K | B | 7e-09 | 6.1e-09 | 1.3e-08 | 8.700E-09 | 3.751E-09 | 6.100E-09 | 1.300E-08 |
| 16093 | 805 | K805M | B | 4.6e-09 | 3e-09 | 1e-08 | 5.867E-09 | 3.668E-09 | 3.000E-09 | 1.000E-08 |
| 16094 | 805 | K805F | B | 5.4e-09 | 5e-09 | 1.6e-08 | 8.800E-09 | 6.239E-09 | 5.000E-09 | 1.600E-08 |
| 16095 | 805 | K805P | B | 6e-09 | 3.5e-09 | 5.8e-09 | 5.100E-09 | 1.389E-09 | 3.500E-09 | 6.000E-09 |
| 16096 | 805 | K805S | B | 5.5e-09 | 3.7e-09 | 4.5e-09 | 4.567E-09 | 9.018E-10 | 3.700E-09 | 5.500E-09 |
| 16097 | 805 | K805T | B | 4.1e-09 | 6.7e-09 | 9.8e-09 | 6.867E-09 | 2.854E-09 | 4.100E-09 | 9.800E-09 |
| 16098 | 805 | K805W | B | 9.1e-09 | 1.4e-09 | 1e-08 | 6.833E-09 | 4.727E-09 | 1.400E-09 | 1.000E-08 |
| 16099 | 805 | K805Y | B | 5.5e-09 | 8.1e-09 | 7.4e-09 | 7.000E-09 | 1.345E-09 | 5.500E-09 | 8.100E-09 |
| 16100 | 805 | K805V | B | 7e-09 | 7.9e-09 | 4.8e-09 | 6.567E-09 | 1.595E-09 | 4.800E-09 | 7.900E-09 |
| 16101 | 806 | A806A | B | 3e-09 | 6.9e-09 | 7.5e-09 | 5.800E-09 | 2.443E-09 | 3.000E-09 | 7.500E-09 |
| 16102 | 806 | A806R | B | 9.8e-09 | 8.1e-09 | 8e-09 | 8.633E-09 | 1.012E-09 | 8.000E-09 | 9.800E-09 |
| 16103 | 806 | A806N | B | 7.7e-09 | 6.8e-09 | 8.4e-09 | 7.633E-09 | 8.021E-10 | 6.800E-09 | 8.400E-09 |
| 16104 | 806 | A806D | B | 1.4e-08 | 8.3e-09 | 2.4e-08 | 1.543E-08 | 7.948E-09 | 8.300E-09 | 2.400E-08 |
| 16105 | 806 | A806C | B | 2.6e-09 | 3.5e-09 | 2.3e-08 | 9.700E-09 | 1.153E-08 | 2.600E-09 | 2.300E-08 |
| 16106 | 806 | A806E | B | 4.7e-09 | 4e-09 | 5e-09 | 4.567E-09 | 5.132E-10 | 4.000E-09 | 5.000E-09 |
| 16107 | 806 | A806Q | B | 6.7e-09 | 7e-09 | 3e-09 | 5.567E-09 | 2.228E-09 | 3.000E-09 | 7.000E-09 |
| 16108 | 806 | A806G | B | 3.9e-09 | 1.8e-08 | 5.6e-09 | 9.167E-09 | 7.697E-09 | 3.900E-09 | 1.800E-08 |
| 16109 | 806 | A806H | B | 1.5e-08 | 7.8e-09 | 4.1e-09 | 8.967E-09 | 5.543E-09 | 4.100E-09 | 1.500E-08 |
| 16110 | 806 | A806I | B | 3.3e-09 | 4.2e-09 | 1.8e-08 | 8.500E-09 | 8.240E-09 | 3.300E-09 | 1.800E-08 |
| 16111 | 806 | A806L | B | 6.3e-09 | 2.2e-09 | 8.3e-09 | 5.600E-09 | 3.110E-09 | 2.200E-09 | 8.300E-09 |
| 16112 | 806 | A806K | B | 7.4e-09 | 2.7e-09 | 5.5e-09 | 5.200E-09 | 2.364E-09 | 2.700E-09 | 7.400E-09 |
| 16113 | 806 | A806M | B | 8.3e-09 | 1.6e-08 | 5.1e-09 | 9.800E-09 | 5.603E-09 | 5.100E-09 | 1.600E-08 |
| 16114 | 806 | A806F | B | 5.1e-09 | 7.1e-09 | 5e-09 | 5.733E-09 | 1.185E-09 | 5.000E-09 | 7.100E-09 |
| 16115 | 806 | A806P | B | 6.4e-09 | 4.9e-09 | 5.3e-09 | 5.533E-09 | 7.767E-10 | 4.900E-09 | 6.400E-09 |
| 16116 | 806 | A806S | B | 9.7e-09 | 5.7e-09 | 1.1e-08 | 8.800E-09 | 2.762E-09 | 5.700E-09 | 1.100E-08 |
| 16117 | 806 | A806T | B | 1.4e-08 | 4.6e-09 | 4.7e-09 | 7.767E-09 | 5.398E-09 | 4.600E-09 | 1.400E-08 |
| 16118 | 806 | A806W | B | 5.8e-09 | 4.9e-09 | 5.8e-09 | 5.500E-09 | 5.196E-10 | 4.900E-09 | 5.800E-09 |
| 16119 | 806 | A806Y | B | 4.7e-09 | 1.2e-08 | 3.4e-09 | 6.700E-09 | 4.636E-09 | 3.400E-09 | 1.200E-08 |
| 16120 | 806 | A806V | B | 3.5e-09 | 7.2e-09 | 5.5e-09 | 5.400E-09 | 1.852E-09 | 3.500E-09 | 7.200E-09 |
| 16121 | 807 | D807A | B | 9.3e-09 | 1.2e-08 | 1.7e-08 | 1.277E-08 | 3.907E-09 | 9.300E-09 | 1.700E-08 |
| 16122 | 807 | D807R | B | 5.6e-09 | 6.4e-09 | 6.8e-09 | 6.267E-09 | 6.110E-10 | 5.600E-09 | 6.800E-09 |
| 16123 | 807 | D807N | B | 8.7e-09 | 2.4e-09 | 2.2e-08 | 1.103E-08 | 1.001E-08 | 2.400E-09 | 2.200E-08 |
| 16124 | 807 | D807D | B | 1.1e-08 | 2e-09 | 1.2e-08 | 8.333E-09 | 5.508E-09 | 2.000E-09 | 1.200E-08 |
| 16125 | 807 | D807C | B | 5.4e-09 | 4.2e-09 | 3.6e-09 | 4.400E-09 | 9.165E-10 | 3.600E-09 | 5.400E-09 |
| 16126 | 807 | D807E | B | 2.7e-08 | 6.6e-09 | 6.2e-09 | 1.327E-08 | 1.190E-08 | 6.200E-09 | 2.700E-08 |
| 16127 | 807 | D807Q | B | 3e-09 | 1.1e-08 | 2.2e-08 | 1.200E-08 | 9.539E-09 | 3.000E-09 | 2.200E-08 |
| 16128 | 807 | D807G | B | 3.4e-09 | 7.7e-09 | 1.2e-08 | 7.700E-09 | 4.300E-09 | 3.400E-09 | 1.200E-08 |
| 16129 | 807 | D807H | B | 3.5e-09 | 3.7e-09 | 1.4e-08 | 7.067E-09 | 6.005E-09 | 3.500E-09 | 1.400E-08 |
| 16130 | 807 | D807I | B | 3.3e-09 | 1e-08 | 1.4e-08 | 9.100E-09 | 5.406E-09 | 3.300E-09 | 1.400E-08 |
| 16131 | 807 | D807L | B | 6.7e-09 | 4.9e-09 | 4.9e-09 | 5.500E-09 | 1.039E-09 | 4.900E-09 | 6.700E-09 |
| 16132 | 807 | D807K | B | 1e-08 | 2.9e-09 | 1e-08 | 7.633E-09 | 4.099E-09 | 2.900E-09 | 1.000E-08 |
| 16133 | 807 | D807M | B | 1.2e-09 | 1.4e-08 | 5.3e-09 | 6.833E-09 | 6.536E-09 | 1.200E-09 | 1.400E-08 |
| 16134 | 807 | D807F | B | 8.7e-09 | 4.8e-09 | 5.9e-09 | 6.467E-09 | 2.011E-09 | 4.800E-09 | 8.700E-09 |
| 16135 | 807 | D807P | B | 3.6e-09 | 2e-08 | 3.2e-09 | 8.933E-09 | 9.586E-09 | 3.200E-09 | 2.000E-08 |
| 16136 | 807 | D807S | B | 5e-09 | 1.1e-08 | 1e-08 | 8.667E-09 | 3.215E-09 | 5.000E-09 | 1.100E-08 |
| 16137 | 807 | D807T | B | 7.5e-09 | 8.5e-09 | 2.4e-08 | 1.333E-08 | 9.251E-09 | 7.500E-09 | 2.400E-08 |
| 16138 | 807 | D807W | B | 1.4e-08 | 6.8e-09 | 5.4e-09 | 8.733E-09 | 4.614E-09 | 5.400E-09 | 1.400E-08 |
| 16139 | 807 | D807Y | B | 5e-09 | 3.4e-09 | 1.2e-08 | 6.800E-09 | 4.574E-09 | 3.400E-09 | 1.200E-08 |
| 16140 | 807 | D807V | B | 6.8e-09 | 5.3e-09 | 7.5e-09 | 6.533E-09 | 1.124E-09 | 5.300E-09 | 7.500E-09 |
| 16141 | 808 | Y808A | B | 1.3e-08 | 1.1e-08 | 1.9e-08 | 1.433E-08 | 4.163E-09 | 1.100E-08 | 1.900E-08 |
| 16142 | 808 | Y808R | B | 1.5e-08 | 1.8e-08 | 1.5e-08 | 1.600E-08 | 1.732E-09 | 1.500E-08 | 1.800E-08 |
| 16143 | 808 | Y808N | B | 5.6e-09 | 8.4e-09 | 1.1e-09 | 5.033E-09 | 3.683E-09 | 1.100E-09 | 8.400E-09 |
| 16144 | 808 | Y808D | B | 1.8e-08 | 8.3e-09 | 4.8e-09 | 1.037E-08 | 6.838E-09 | 4.800E-09 | 1.800E-08 |
| 16145 | 808 | Y808C | B | 2.7e-08 | 6.5e-09 | 2.3e-09 | 1.193E-08 | 1.322E-08 | 2.300E-09 | 2.700E-08 |
| 16146 | 808 | Y808E | B | 1.1e-08 | 1.7e-08 | 9.2e-09 | 1.240E-08 | 4.084E-09 | 9.200E-09 | 1.700E-08 |
| 16147 | 808 | Y808Q | B | 3.9e-09 | 5.7e-09 | 5.6e-09 | 5.067E-09 | 1.012E-09 | 3.900E-09 | 5.700E-09 |
| 16148 | 808 | Y808G | B | 3.7e-09 | 4.1e-09 | 6.3e-09 | 4.700E-09 | 1.400E-09 | 3.700E-09 | 6.300E-09 |
| 16149 | 808 | Y808H | B | 7.4e-09 | 9.3e-09 | 4.3e-09 | 7.000E-09 | 2.524E-09 | 4.300E-09 | 9.300E-09 |

| | | | | | | | | | | |
|---|---|---|---|---|---|---|---|---|---|---|
| 16150 | 808 | Y808I | B | 1.1e-08 | 1.1e-08 | 1.4e-08 | 1.200E-08 | 1.732E-09 | 1.100E-08 | 1.400E-08 |
| 16151 | 808 | Y808L | B | 1.1e-08 | 6.2e-09 | 4.1e-09 | 7.100E-09 | 3.537E-09 | 4.100E-09 | 1.100E-08 |
| 16152 | 808 | Y808K | B | 6.4e-09 | 8.4e-09 | 7.3e-09 | 7.367E-09 | 1.002E-09 | 6.400E-09 | 8.400E-09 |
| 16153 | 808 | Y808M | B | 1.2e-08 | 1.3e-08 | 8.4e-09 | 1.113E-08 | 2.419E-09 | 8.400E-09 | 1.300E-08 |
| 16154 | 808 | Y808F | B | 7.2e-09 | 6e-09 | 1e-08 | 7.733E-09 | 2.053E-09 | 6.000E-09 | 1.000E-08 |
| 16155 | 808 | Y808P | B | 1.3e-08 | 1.4e-08 | 9e-09 | 1.200E-08 | 2.646E-09 | 9.000E-09 | 1.400E-08 |
| 16156 | 808 | Y808S | B | 1.3e-08 | 6.6e-09 | 6.5e-09 | 8.700E-09 | 3.724E-09 | 6.500E-09 | 1.300E-08 |
| 16157 | 808 | Y808T | B | 2.7e-09 | 9.7e-09 | 5.6e-09 | 6.000E-09 | 3.517E-09 | 2.700E-09 | 9.700E-09 |
| 16158 | 808 | Y808W | B | 1.1e-08 | 8.9e-09 | 2.1e-08 | 1.363E-08 | 6.466E-09 | 8.900E-09 | 2.100E-08 |
| 16159 | 808 | Y808Y | B | 1.1e-08 | 4.7e-09 | 5.3e-09 | 7.000E-09 | 3.477E-09 | 4.700E-09 | 1.100E-08 |
| 16160 | 808 | Y808V | B | 5.1e-09 | 2.5e-09 | 7e-09 | 4.867E-09 | 2.259E-09 | 2.500E-09 | 7.000E-09 |
| 16161 | 809 | E809A | B | 5.8e-09 | 1e-08 | 5.2e-09 | 7.000E-09 | 2.615E-09 | 5.200E-09 | 1.000E-08 |
| 16162 | 809 | E809R | B | 3.3e-09 | 8e-09 | 6.4e-09 | 5.900E-09 | 2.390E-09 | 3.300E-09 | 8.000E-09 |
| 16163 | 809 | E809N | B | 1.5e-08 | 5.7e-09 | 4.5e-09 | 8.400E-09 | 5.747E-09 | 4.500E-09 | 1.500E-08 |
| 16164 | 809 | E809D | B | 1.3e-08 | 4.2e-09 | 1.6e-08 | 1.107E-08 | 6.133E-09 | 4.200E-09 | 1.600E-08 |
| 16165 | 809 | E809C | B | 7.9e-09 | 2.2e-09 | 3.2e-09 | 4.433E-09 | 3.044E-09 | 2.200E-09 | 7.900E-09 |
| 16166 | 809 | E809E | B | 9.9e-09 | 2.1e-08 | 5.3e-09 | 1.207E-08 | 8.071E-09 | 5.300E-09 | 2.100E-08 |
| 16167 | 809 | E809Q | B | 8.7e-09 | 2.3e-09 | 8.7e-09 | 6.567E-09 | 3.695E-09 | 2.300E-09 | 8.700E-09 |
| 16168 | 809 | E809G | B | 2.3e-09 | 1.1e-08 | 3.6e-09 | 5.633E-09 | 4.693E-09 | 2.300E-09 | 1.100E-08 |
| 16169 | 809 | E809H | B | 5.7e-09 | 5.5e-09 | 3e-09 | 4.733E-09 | 1.504E-09 | 3.000E-09 | 5.700E-09 |
| 16170 | 809 | E809I | B | 1.1e-08 | 6.9e-09 | 7.4e-09 | 8.433E-09 | 2.237E-09 | 6.900E-09 | 1.100E-08 |
| 16171 | 809 | E809L | B | 6.2e-09 | 3.7e-09 | 5e-09 | 4.967E-09 | 1.250E-09 | 3.700E-09 | 6.200E-09 |
| 16172 | 809 | E809K | B | 2.3e-09 | 4e-09 | 2.8e-09 | 3.033E-09 | 8.737E-10 | 2.300E-09 | 4.000E-09 |
| 16173 | 809 | E809M | B | 2.9e-08 | 1.1e-08 | 5.2e-09 | 1.507E-08 | 1.241E-08 | 5.200E-09 | 2.900E-08 |
| 16174 | 809 | E809F | B | 1.2e-08 | 1.6e-08 | 9.1e-09 | 1.237E-08 | 3.465E-09 | 9.100E-09 | 1.600E-08 |
| 16175 | 809 | E809P | B | 3.8e-09 | 1.5e-08 | 9e-09 | 9.267E-09 | 5.605E-09 | 3.800E-09 | 1.500E-08 |
| 16176 | 809 | E809S | B | 1.8e-08 | 8.2e-09 | 1.3e-08 | 1.307E-08 | 4.900E-09 | 8.200E-09 | 1.800E-08 |
| 16177 | 809 | E809T | B | 6.2e-09 | 4.9e-09 | 2.8e-09 | 4.633E-09 | 1.716E-09 | 2.800E-09 | 6.200E-09 |
| 16178 | 809 | E809W | B | 1e-08 | 1.1e-08 | 5.6e-09 | 8.867E-09 | 2.873E-09 | 5.600E-09 | 1.100E-08 |
| 16179 | 809 | E809Y | B | 1.3e-08 | 1.3e-08 | 8.8e-09 | 1.160E-08 | 2.425E-09 | 8.800E-09 | 1.300E-08 |
| 16180 | 809 | E809V | B | 4.4e-09 | 8e-09 | 6e-09 | 6.133E-09 | 1.804E-09 | 4.400E-09 | 8.000E-09 |
| 16181 | 810 | K810A | B | 1.2e-08 | 8e-09 | 8.1e-09 | 9.367E-09 | 2.281E-09 | 8.000E-09 | 1.200E-08 |
| 16182 | 810 | K810R | B | 6.9e-09 | 1.6e-08 | 4.9e-09 | 9.267E-09 | 5.916E-09 | 4.900E-09 | 1.600E-08 |
| 16183 | 810 | K810N | B | 9.6e-09 | 1.8e-08 | 9.2e-09 | 1.227E-08 | 4.969E-09 | 9.200E-09 | 1.800E-08 |
| 16184 | 810 | K810D | B | 9.3e-09 | 1.4e-08 | 1.2e-08 | 1.177E-08 | 2.359E-09 | 9.300E-09 | 1.400E-08 |
| 16185 | 810 | K810C | B | 6e-09 | 1.3e-08 | 6.9e-09 | 8.633E-09 | 3.808E-09 | 6.000E-09 | 1.300E-08 |
| 16186 | 810 | K810E | B | 2.7e-09 | 6.1e-09 | 3.4e-09 | 4.067E-09 | 1.795E-09 | 2.700E-09 | 6.100E-09 |
| 16187 | 810 | K810Q | B | 4.2e-09 | 4.1e-09 | 8.2e-09 | 5.500E-09 | 2.339E-09 | 4.100E-09 | 8.200E-09 |
| 16188 | 810 | K810G | B | 4.4e-09 | 1.9e-08 | 8.4e-09 | 1.060E-08 | 7.545E-09 | 4.400E-09 | 1.900E-08 |
| 16189 | 810 | K810H | B | 7.4e-09 | 4.5e-08 | 4.2e-09 | 1.887E-08 | 2.269E-08 | 4.200E-09 | 4.500E-08 |
| 16190 | 810 | K810I | B | 2.6e-09 | 1.2e-08 | 6.2e-09 | 6.933E-09 | 4.743E-09 | 2.600E-09 | 1.200E-08 |
| 16191 | 810 | K810L | B | 6.4e-09 | 8.4e-09 | 2.1e-08 | 1.193E-08 | 7.915E-09 | 6.400E-09 | 2.100E-08 |
| 16192 | 810 | K810K | B | 3.5e-09 | 1.4e-08 | 2e-08 | 1.250E-08 | 8.352E-09 | 3.500E-09 | 2.000E-08 |
| 16193 | 810 | K810M | B | 1e-08 | 7.2e-09 | 4.4e-08 | 2.040E-08 | 2.049E-08 | 7.200E-09 | 4.400E-08 |
| 16194 | 810 | K810F | B | 9.7e-09 | 1.9e-08 | 9.9e-09 | 1.287E-08 | 5.313E-09 | 9.700E-09 | 1.900E-08 |
| 16195 | 810 | K810P | B | 3.6e-09 | 1e-08 | 7.5e-09 | 7.033E-09 | 3.225E-09 | 3.600E-09 | 1.000E-08 |
| 16196 | 810 | K810S | B | 6.3e-09 | 8.7e-09 | 4.1e-09 | 6.367E-09 | 2.301E-09 | 4.100E-09 | 8.700E-09 |
| 16197 | 810 | K810T | B | 1.3e-08 | 9e-09 | 1.1e-08 | 1.100E-08 | 2.000E-09 | 9.000E-09 | 1.300E-08 |
| 16198 | 810 | K810W | B | 1.3e-08 | 4.7e-09 | 6.6e-09 | 8.100E-09 | 4.349E-09 | 4.700E-09 | 1.300E-08 |
| 16199 | 810 | K810Y | B | 9e-09 | 1.9e-08 | 2.3e-09 | 1.010E-08 | 8.404E-09 | 2.300E-09 | 1.900E-08 |
| 16200 | 810 | K810V | B | 1.1e-08 | 8.1e-09 | 1.4e-08 | 1.103E-08 | 2.950E-09 | 8.100E-09 | 1.400E-08 |
| 16201 | 811 | H811A | B | 6.8e-09 | 5.9e-09 | 1.1e-08 | 7.900E-09 | 2.722E-09 | 5.900E-09 | 1.100E-08 |
| 16202 | 811 | H811R | B | 4.1e-09 | 7.2e-09 | 6.2e-09 | 5.833E-09 | 1.582E-09 | 4.100E-09 | 7.200E-09 |
| 16203 | 811 | H811N | B | 3.6e-09 | 4e-09 | 7.7e-09 | 5.100E-09 | 2.261E-09 | 3.600E-09 | 7.700E-09 |
| 16204 | 811 | H811D | B | 4.7e-09 | 6.3e-09 | 1.1e-08 | 7.333E-09 | 3.275E-09 | 4.700E-09 | 1.100E-08 |
| 16205 | 811 | H811C | B | 3.7e-09 | 2.6e-09 | 2.5e-09 | 2.933E-09 | 6.658E-10 | 2.500E-09 | 3.700E-09 |
| 16206 | 811 | H811E | B | 7e-09 | 1.9e-08 | 9.5e-09 | 1.183E-08 | 6.331E-09 | 7.000E-09 | 1.900E-08 |
| 16207 | 811 | H811Q | B | 2.7e-09 | 1.4e-08 | 3.2e-09 | 6.633E-09 | 6.385E-09 | 2.700E-09 | 1.400E-08 |
| 16208 | 811 | H811G | B | 5e-09 | 1.9e-08 | 1.5e-08 | 1.300E-08 | 7.211E-09 | 5.000E-09 | 1.900E-08 |
| 16209 | 811 | H811H | B | 4.9e-09 | 5.2e-09 | 8.3e-09 | 6.133E-09 | 1.882E-09 | 4.900E-09 | 8.300E-09 |
| 16210 | 811 | H811I | B | 3.8e-09 | 9.9e-09 | 4.7e-09 | 6.133E-09 | 3.293E-09 | 3.800E-09 | 9.900E-09 |
| 16211 | 811 | H811L | B | 4.4e-09 | 5.5e-09 | 1.1e-08 | 6.967E-09 | 3.536E-09 | 4.400E-09 | 1.100E-08 |
| 16212 | 811 | H811K | B | 1.7e-08 | 1.6e-08 | 1.5e-08 | 1.600E-08 | 1.000E-09 | 1.500E-08 | 1.700E-08 |
| 16213 | 811 | H811M | B | 5.1e-09 | 2.7e-09 | 1.6e-09 | 3.133E-09 | 1.790E-09 | 1.600E-09 | 5.100E-09 |
| 16214 | 811 | H811F | B | 2.1e-08 | 4.6e-09 | 1.9e-08 | 1.487E-08 | 8.947E-09 | 4.600E-09 | 2.100E-08 |
| 16215 | 811 | H811P | B | 1.2e-08 | 3.5e-09 | 7.9e-09 | 7.800E-09 | 4.251E-09 | 3.500E-09 | 1.200E-08 |
| 16216 | 811 | H811S | B | 4.8e-09 | 1.1e-08 | 3.2e-09 | 6.333E-09 | 4.120E-09 | 3.200E-09 | 1.100E-08 |
| 16217 | 811 | H811T | B | 1e-08 | 2e-09 | 7.3e-09 | 6.433E-09 | 4.070E-09 | 2.000E-09 | 1.000E-08 |
| 16218 | 811 | H811W | B | 8.6e-09 | 4.4e-09 | 2.5e-08 | 1.267E-08 | 1.089E-08 | 4.400E-09 | 2.500E-08 |
| 16219 | 811 | H811Y | B | 5.3e-09 | 2.4e-09 | 1e-08 | 5.900E-09 | 3.835E-09 | 2.400E-09 | 1.000E-08 |
| 16220 | 811 | H811V | B | 2.4e-09 | 5.2e-09 | 1.6e-08 | 7.867E-09 | 7.181E-09 | 2.400E-09 | 1.600E-08 |
| 16221 | 812 | K812A | B | 9.6e-09 | 6.3e-09 | 1.3e-08 | 9.633E-09 | 3.350E-09 | 6.300E-09 | 1.300E-08 |
| 16222 | 812 | K812R | B | 1.1e-08 | 1.2e-08 | 5.6e-09 | 9.533E-09 | 3.443E-09 | 5.600E-09 | 1.200E-08 |
| 16223 | 812 | K812N | B | 3e-09 | 2.1e-09 | 1.3e-08 | 6.033E-09 | 6.050E-09 | 2.100E-09 | 1.300E-08 |
| 16224 | 812 | K812D | B | 7e-09 | 1.5e-08 | 9.9e-09 | 1.063E-08 | 4.050E-09 | 7.000E-09 | 1.500E-08 |
| 16225 | 812 | K812C | B | 2.5e-08 | 1.3e-08 | 8.2e-09 | 1.540E-08 | 8.653E-09 | 8.200E-09 | 2.500E-08 |
| 16226 | 812 | K812E | B | 8.7e-09 | 5.4e-09 | 5.5e-09 | 6.533E-09 | 1.877E-09 | 5.400E-09 | 8.700E-09 |
| 16227 | 812 | K812Q | B | 9.8e-09 | 2.9e-09 | 1.1e-08 | 7.900E-09 | 4.371E-09 | 2.900E-09 | 1.100E-08 |
| 16228 | 812 | K812G | B | 3.2e-08 | 8.8e-09 | 5e-08 | 3.027E-08 | 2.065E-08 | 8.800E-09 | 5.000E-08 |
| 16229 | 812 | K812H | B | 2.4e-08 | 1.7e-08 | 1.5e-08 | 1.867E-08 | 4.726E-09 | 1.500E-08 | 2.400E-08 |
| 16230 | 812 | K812I | B | 5.2e-09 | 1.6e-08 | 4.7e-09 | 8.633E-09 | 6.385E-09 | 4.700E-09 | 1.600E-08 |
| 16231 | 812 | K812L | B | 1.1e-08 | 4.1e-09 | 3e-09 | 6.033E-09 | 4.336E-09 | 3.000E-09 | 1.100E-08 |
| 16232 | 812 | K812K | B | 5.8e-09 | 1.3e-08 | 2.4e-08 | 1.427E-08 | 9.166E-09 | 5.800E-09 | 2.400E-08 |
| 16233 | 812 | K812M | B | 8.9e-09 | 5.9e-09 | 5.7e-09 | 6.833E-09 | 1.793E-09 | 5.700E-09 | 8.900E-09 |
| 16234 | 812 | K812F | B | 1.3e-08 | 2.1e-08 | 2.4e-09 | 1.213E-08 | 9.330E-09 | 2.400E-09 | 2.100E-08 |

| | | | | | | | | | | |
|---|---|---|---|---|---|---|---|---|---|---|
| 16235 | 812 | K812P | B | 9.4e-09 | 1.3e-08 | 1.2e-09 | 7.867E-09 | 6.048E-09 | 1.200E-09 | 1.300E-08 |
| 16236 | 812 | K812S | B | 5.1e-09 | 2e-08 | 2.1e-09 | 9.067E-09 | 9.587E-09 | 2.100E-09 | 2.000E-08 |
| 16237 | 812 | K812T | B | 6e-09 | 1.5e-08 | 2e-08 | 1.367E-08 | 7.095E-09 | 6.000E-09 | 2.000E-08 |
| 16238 | 812 | K812W | B | 6.6e-09 | 4.2e-09 | 1.4e-09 | 4.067E-09 | 2.603E-09 | 1.400E-09 | 6.600E-09 |
| 16239 | 812 | K812Y | B | 4.2e-09 | 9.9e-09 | 8.6e-09 | 7.567E-09 | 2.987E-09 | 4.200E-09 | 9.900E-09 |
| 16240 | 812 | K812V | B | 1.3e-08 | 7.2e-09 | 2.6e-09 | 7.600E-09 | 5.212E-09 | 2.600E-09 | 1.300E-08 |
| 16241 | 813 | V813A | B | 5e-09 | 4.4e-09 | 3.9e-09 | 4.433E-09 | 5.508E-10 | 3.900E-09 | 5.000E-09 |
| 16242 | 813 | V813R | B | 1.1e-08 | 5.9e-09 | 1.6e-08 | 1.097E-08 | 5.050E-09 | 5.900E-09 | 1.600E-08 |
| 16243 | 813 | V813N | B | 1.4e-08 | 2.9e-09 | 1.7e-08 | 1.130E-08 | 7.428E-09 | 2.900E-09 | 1.700E-08 |
| 16244 | 813 | V813D | B | 3.7e-09 | 1.4e-08 | 4e-09 | 7.233E-09 | 5.862E-09 | 3.700E-09 | 1.400E-08 |
| 16245 | 813 | V813C | B | 2.4e-08 | 4.7e-09 | 3.9e-09 | 1.087E-08 | 1.138E-08 | 3.900E-09 | 2.400E-08 |
| 16246 | 813 | V813E | B | 4.4e-09 | 4.7e-09 | 5.2e-09 | 4.767E-09 | 4.041E-10 | 4.400E-09 | 5.200E-09 |
| 16247 | 813 | V813Q | B | 1.2e-08 | 2.5e-09 | 9e-09 | 7.833E-09 | 4.856E-09 | 2.500E-09 | 1.200E-08 |
| 16248 | 813 | V813G | B | 5.9e-09 | 1.3e-08 | 5.8e-09 | 8.233E-09 | 4.128E-09 | 5.800E-09 | 1.300E-08 |
| 16249 | 813 | V813H | B | 7.7e-09 | 1e-08 | 3.5e-09 | 7.067E-09 | 3.296E-09 | 3.500E-09 | 1.000E-08 |
| 16250 | 813 | V813I | B | 7.6e-09 | 7.8e-09 | 1e-08 | 8.467E-09 | 1.332E-09 | 7.600E-09 | 1.000E-08 |
| 16251 | 813 | V813L | B | 4.3e-09 | 1.2e-08 | 1.3e-08 | 9.767E-09 | 4.761E-09 | 4.300E-09 | 1.300E-08 |
| 16252 | 813 | V813K | B | 6.6e-09 | 4.7e-09 | 7.2e-09 | 6.167E-09 | 1.305E-09 | 4.700E-09 | 7.200E-09 |
| 16253 | 813 | V813M | B | 7.4e-09 | 1.9e-08 | 2.7e-09 | 9.700E-09 | 8.390E-09 | 2.700E-09 | 1.900E-08 |
| 16254 | 813 | V813F | B | 3e-09 | 5.8e-09 | 1.1e-08 | 6.600E-09 | 4.060E-09 | 3.000E-09 | 1.100E-08 |
| 16255 | 813 | V813P | B | 4.1e-09 | 5.9e-09 | 1.3e-08 | 7.667E-09 | 4.706E-09 | 4.100E-09 | 1.300E-08 |
| 16256 | 813 | V813S | B | 6.5e-09 | 2e-09 | 1e-08 | 6.167E-09 | 4.010E-09 | 2.000E-09 | 1.000E-08 |
| 16257 | 813 | V813T | B | 4.3e-09 | 6.1e-09 | 6.1e-09 | 5.500E-09 | 1.039E-09 | 4.300E-09 | 6.100E-09 |
| 16258 | 813 | V813W | B | 8e-09 | 1.4e-08 | 1.3e-08 | 1.167E-08 | 3.215E-09 | 8.000E-09 | 1.400E-08 |
| 16259 | 813 | V813Y | B | 6.7e-09 | 8.1e-09 | 4.9e-09 | 6.567E-09 | 1.604E-09 | 4.900E-09 | 8.100E-09 |
| 16260 | 813 | V813V | B | 1.1e-08 | 6.4e-09 | 4.9e-09 | 7.433E-09 | 3.179E-09 | 4.900E-09 | 1.100E-08 |
| 16261 | 814 | Y814A | B | 4.4e-09 | 4.1e-09 | 8.2e-09 | 5.567E-09 | 2.285E-09 | 4.100E-09 | 8.200E-09 |
| 16262 | 814 | Y814R | B | 5.6e-09 | 1.3e-08 | 7.5e-09 | 8.700E-09 | 3.843E-09 | 5.600E-09 | 1.300E-08 |
| 16263 | 814 | Y814N | B | 2.2e-09 | 3.1e-09 | 1.3e-08 | 6.100E-09 | 5.992E-09 | 2.200E-09 | 1.300E-08 |
| 16264 | 814 | Y814D | B | 1.2e-08 | 5.3e-09 | 1.6e-08 | 1.110E-08 | 5.406E-09 | 5.300E-09 | 1.600E-08 |
| 16265 | 814 | Y814C | B | 4.5e-09 | 4.6e-09 | 8.7e-09 | 5.933E-09 | 2.397E-09 | 4.500E-09 | 8.700E-09 |
| 16266 | 814 | Y814E | B | 8.4e-09 | 7e-09 | 8.3e-09 | 7.900E-09 | 7.810E-10 | 7.000E-09 | 8.400E-09 |
| 16267 | 814 | Y814Q | B | 8.1e-09 | 3.3e-09 | 1.1e-08 | 7.467E-09 | 3.889E-09 | 3.300E-09 | 1.100E-08 |
| 16268 | 814 | Y814G | B | 1.3e-08 | 5.4e-09 | 4e-09 | 7.467E-09 | 4.843E-09 | 4.000E-09 | 1.300E-08 |
| 16269 | 814 | Y814H | B | 1.4e-08 | 1.8e-08 | 5.3e-09 | 1.243E-08 | 6.493E-09 | 5.300E-09 | 1.800E-08 |
| 16270 | 814 | Y814I | B | 1e-08 | 3.4e-09 | 9.7e-09 | 7.700E-09 | 3.727E-09 | 3.400E-09 | 1.000E-08 |
| 16271 | 814 | Y814L | B | 7.3e-09 | 3.2e-09 | 1.2e-08 | 7.500E-09 | 4.403E-09 | 3.200E-09 | 1.200E-08 |
| 16272 | 814 | Y814K | B | 7e-09 | 9.9e-09 | 4.8e-09 | 7.233E-09 | 2.558E-09 | 4.800E-09 | 9.900E-09 |
| 16273 | 814 | Y814M | B | 3.9e-09 | 1.1e-08 | 4.4e-09 | 6.433E-09 | 3.963E-09 | 3.900E-09 | 1.100E-08 |
| 16274 | 814 | Y814F | B | 9e-09 | 3.2e-09 | 5.1e-09 | 5.767E-09 | 2.957E-09 | 3.200E-09 | 9.000E-09 |
| 16275 | 814 | Y814P | B | 5.8e-09 | 1.3e-08 | 4.7e-09 | 7.833E-09 | 4.508E-09 | 4.700E-09 | 1.300E-08 |
| 16276 | 814 | Y814S | B | 7.2e-09 | 4.4e-09 | 2.6e-09 | 4.733E-09 | 2.318E-09 | 2.600E-09 | 7.200E-09 |
| 16277 | 814 | Y814T | B | 1.1e-08 | 9.2e-09 | 1.1e-08 | 1.040E-08 | 1.039E-09 | 9.200E-09 | 1.100E-08 |
| 16278 | 814 | Y814W | B | 1.5e-08 | 3.6e-09 | 4.4e-09 | 7.667E-09 | 6.363E-09 | 3.600E-09 | 1.500E-08 |
| 16279 | 814 | Y814Y | B | 1.2e-08 | 6.6e-09 | 1.8e-09 | 6.800E-09 | 5.103E-09 | 1.800E-09 | 1.200E-08 |
| 16280 | 814 | Y814V | B | 3.2e-09 | 1.3e-08 | 2.2e-09 | 6.133E-09 | 5.968E-09 | 2.200E-09 | 1.300E-08 |
| 16281 | 815 | A815A | B | 4.4e-09 | 8.5e-09 | 1.5e-08 | 9.300E-09 | 5.345E-09 | 4.400E-09 | 1.500E-08 |
| 16282 | 815 | A815R | B | 1.4e-08 | 5.3e-09 | 8e-09 | 9.100E-09 | 4.453E-09 | 5.300E-09 | 1.400E-08 |
| 16283 | 815 | A815N | B | 5.1e-09 | 6.1e-09 | 1.1e-08 | 7.400E-09 | 3.158E-09 | 5.100E-09 | 1.100E-08 |
| 16284 | 815 | A815D | B | 1e-08 | 8.6e-09 | 2.7e-08 | 1.520E-08 | 1.024E-08 | 8.600E-09 | 2.700E-08 |
| 16285 | 815 | A815C | B | 1.6e-08 | 5.5e-09 | 1.9e-08 | 1.350E-08 | 7.089E-09 | 5.500E-09 | 1.900E-08 |
| 16286 | 815 | A815E | B | 9.3e-09 | 1.7e-09 | 2.8e-09 | 4.600E-09 | 4.107E-09 | 1.700E-09 | 9.300E-09 |
| 16287 | 815 | A815Q | B | 6.1e-09 | 5e-09 | 3.6e-09 | 4.900E-09 | 1.253E-09 | 3.600E-09 | 6.100E-09 |
| 16288 | 815 | A815G | B | 6.8e-09 | 4.6e-09 | 7.9e-09 | 6.433E-09 | 1.680E-09 | 4.600E-09 | 7.900E-09 |
| 16289 | 815 | A815H | B | 2.7e-09 | 6.3e-09 | 8.6e-09 | 5.867E-09 | 2.974E-09 | 2.700E-09 | 8.600E-09 |
| 16290 | 815 | A815I | B | 7e-09 | 9.2e-09 | 5.3e-09 | 7.167E-09 | 1.955E-09 | 5.300E-09 | 9.200E-09 |
| 16291 | 815 | A815L | B | 1.8e-08 | 1e-08 | 1.8e-09 | 9.933E-09 | 8.100E-09 | 1.800E-09 | 1.800E-08 |
| 16292 | 815 | A815K | B | 6.2e-09 | 1.9e-08 | 3.2e-08 | 1.907E-08 | 1.290E-08 | 6.200E-09 | 3.200E-08 |
| 16293 | 815 | A815M | B | 1.1e-08 | 5.3e-09 | 8.4e-09 | 8.233E-09 | 2.854E-09 | 5.300E-09 | 1.100E-08 |
| 16294 | 815 | A815F | B | 1.6e-08 | 1.8e-08 | 9.2e-09 | 1.440E-08 | 4.613E-09 | 9.200E-09 | 1.800E-08 |
| 16295 | 815 | A815P | B | 6e-09 | 1.1e-08 | 4.2e-09 | 7.067E-09 | 3.523E-09 | 4.200E-09 | 1.100E-08 |
| 16296 | 815 | A815S | B | 4e-08 | 4.9e-09 | 5.7e-09 | 1.687E-08 | 2.004E-08 | 4.900E-09 | 4.000E-08 |
| 16297 | 815 | A815T | B | 7.5e-09 | 6.2e-09 | 4.5e-09 | 6.067E-09 | 1.504E-09 | 4.500E-09 | 7.500E-09 |
| 16298 | 815 | A815W | B | 2e-08 | 1e-08 | 2.4e-08 | 1.800E-08 | 7.211E-09 | 1.000E-08 | 2.400E-08 |
| 16299 | 815 | A815Y | B | 4.8e-09 | 1.2e-08 | 1.9e-09 | 6.233E-09 | 5.200E-09 | 1.900E-09 | 1.200E-08 |
| 16300 | 815 | A815V | B | 8e-09 | 5e-09 | 1.6e-08 | 9.667E-09 | 5.686E-09 | 5.000E-09 | 1.600E-08 |
| 16301 | 816 | C816A | B | 2.6e-09 | 8.9e-09 | 2e-08 | 1.050E-08 | 8.810E-09 | 2.600E-09 | 2.000E-08 |
| 16302 | 816 | C816R | B | 9.1e-09 | 9e-09 | 1.7e-08 | 1.170E-08 | 4.590E-09 | 9.000E-09 | 1.700E-08 |
| 16303 | 816 | C816N | B | 4.4e-09 | 1.8e-08 | 6.9e-09 | 9.767E-09 | 7.239E-09 | 4.400E-09 | 1.800E-08 |
| 16304 | 816 | C816D | B | 4.7e-09 | 1.4e-08 | 5.4e-09 | 8.033E-09 | 5.179E-09 | 4.700E-09 | 1.400E-08 |
| 16305 | 816 | C816C | B | 2.9e-08 | 7e-09 | 7.8e-09 | 1.460E-08 | 1.248E-08 | 7.000E-09 | 2.900E-08 |
| 16306 | 816 | C816E | B | 7.6e-09 | 7.9e-09 | 4e-09 | 6.500E-09 | 2.170E-09 | 4.000E-09 | 7.900E-09 |
| 16307 | 816 | C816Q | B | 5.3e-09 | 2.5e-09 | 4.7e-09 | 4.167E-09 | 1.474E-09 | 2.500E-09 | 5.300E-09 |
| 16308 | 816 | C816G | B | 8.2e-09 | 6e-09 | 1.3e-08 | 9.067E-09 | 3.580E-09 | 6.000E-09 | 1.300E-08 |
| 16309 | 816 | C816H | B | 1.9e-08 | 6.6e-09 | 1.9e-09 | 9.167E-09 | 8.834E-09 | 1.900E-09 | 1.900E-08 |
| 16310 | 816 | C816I | B | 9e-09 | 4.4e-09 | 1e-08 | 7.800E-09 | 2.987E-09 | 4.400E-09 | 1.000E-08 |
| 16311 | 816 | C816L | B | 5e-09 | 1.9e-09 | 8e-09 | 4.967E-09 | 3.050E-09 | 1.900E-09 | 8.000E-09 |
| 16312 | 816 | C816K | B | 7.1e-09 | 1.3e-08 | 1.4e-08 | 1.137E-08 | 3.729E-09 | 7.100E-09 | 1.400E-08 |
| 16313 | 816 | C816M | B | 1.8e-08 | 6.2e-09 | 8.3e-09 | 1.083E-08 | 6.295E-09 | 6.200E-09 | 1.800E-08 |
| 16314 | 816 | C816F | B | 3.8e-09 | 6.1e-09 | 1.1e-08 | 6.967E-09 | 3.677E-09 | 3.800E-09 | 1.100E-08 |
| 16315 | 816 | C816P | B | 6.6e-09 | 6.9e-09 | 5.5e-09 | 6.333E-09 | 7.371E-10 | 5.500E-09 | 6.900E-09 |
| 16316 | 816 | C816S | B | 6e-09 | 5.1e-09 | 2e-08 | 1.037E-08 | 8.355E-09 | 5.100E-09 | 2.000E-08 |
| 16317 | 816 | C816T | B | 5.1e-09 | 8.2e-09 | 3.6e-09 | 5.633E-09 | 2.346E-09 | 3.600E-09 | 8.200E-09 |
| 16318 | 816 | C816W | B | 2.7e-09 | 1.2e-08 | 7.1e-09 | 7.267E-09 | 4.652E-09 | 2.700E-09 | 1.200E-08 |
| 16319 | 816 | C816Y | B | 7.9e-09 | 1e-08 | 5e-09 | 7.633E-09 | 2.511E-09 | 5.000E-09 | 1.000E-08 |

| | | | | | | | | | | |
|---|---|---|---|---|---|---|---|---|---|---|
| 16320 | 816 | C816V | B | 6.2e-09 | 2.2e-09 | 4.5e-09 | 4.300E-09 | 2.007E-09 | 2.200E-09 | 6.200E-09 |
| 16321 | 817 | E817A | B | 8.5e-09 | 9.5e-09 | 9.5e-09 | 9.167E-09 | 5.774E-10 | 8.500E-09 | 9.500E-09 |
| 16322 | 817 | E817R | B | 3.3e-09 | 4.3e-09 | 4.7e-09 | 4.100E-09 | 7.211E-10 | 3.300E-09 | 4.700E-09 |
| 16323 | 817 | E817N | B | 7.3e-09 | 6e-09 | 2.1e-08 | 1.143E-08 | 8.310E-09 | 6.000E-09 | 2.100E-08 |
| 16324 | 817 | E817D | B | 2.6e-09 | 1.5e-08 | 3.1e-09 | 6.900E-09 | 7.019E-09 | 2.600E-09 | 1.500E-08 |
| 16325 | 817 | E817C | B | 3.9e-09 | 1.4e-08 | 2e-08 | 1.263E-08 | 8.137E-09 | 3.900E-09 | 2.000E-08 |
| 16326 | 817 | E817E | B | 1.7e-08 | 3.4e-09 | 1.1e-08 | 1.047E-08 | 6.816E-09 | 3.400E-09 | 1.700E-08 |
| 16327 | 817 | E817Q | B | 3.5e-09 | 6.7e-09 | 3.1e-08 | 1.373E-08 | 1.504E-08 | 3.500E-09 | 3.100E-08 |
| 16328 | 817 | E817G | B | 1e-08 | 7.5e-09 | 7.5e-09 | 8.333E-09 | 1.443E-09 | 7.500E-09 | 1.000E-08 |
| 16329 | 817 | E817H | B | 1.1e-08 | 5.4e-09 | 5.8e-09 | 7.400E-09 | 3.124E-09 | 5.400E-09 | 1.100E-08 |
| 16330 | 817 | E817I | B | 9.7e-09 | 5.6e-09 | 3.3e-09 | 6.200E-09 | 3.242E-09 | 3.300E-09 | 9.700E-09 |
| 16331 | 817 | E817L | B | 1e-08 | 2.8e-09 | 6.3e-09 | 6.367E-09 | 3.600E-09 | 2.800E-09 | 1.000E-08 |
| 16332 | 817 | E817K | B | 9.3e-09 | 6.8e-09 | 7.1e-09 | 7.733E-09 | 1.365E-09 | 6.800E-09 | 9.300E-09 |
| 16333 | 817 | E817M | B | 6.3e-09 | 8.4e-09 | 4.4e-09 | 6.367E-09 | 2.001E-09 | 4.400E-09 | 8.400E-09 |
| 16334 | 817 | E817F | B | 1.2e-08 | 2e-09 | 5.8e-09 | 6.600E-09 | 5.048E-09 | 2.000E-09 | 1.200E-08 |
| 16335 | 817 | E817P | B | 3e-09 | 4.5e-09 | 4.4e-09 | 3.967E-09 | 8.386E-10 | 3.000E-09 | 4.500E-09 |
| 16336 | 817 | E817S | B | 1.4e-08 | 5.6e-09 | 9.7e-09 | 9.767E-09 | 4.200E-09 | 5.600E-09 | 1.400E-08 |
| 16337 | 817 | E817T | B | 1.1e-08 | 7.5e-09 | 6.1e-09 | 8.200E-09 | 2.524E-09 | 6.100E-09 | 1.100E-08 |
| 16338 | 817 | E817W | B | 9.4e-09 | 1.9e-08 | 3e-09 | 1.047E-08 | 8.053E-09 | 3.000E-09 | 1.900E-08 |
| 16339 | 817 | E817Y | B | 6.6e-09 | 5.9e-09 | 1.1e-08 | 7.833E-09 | 2.765E-09 | 5.900E-09 | 1.100E-08 |
| 16340 | 817 | E817V | B | 9e-09 | 1.7e-08 | 4.5e-09 | 1.017E-08 | 6.331E-09 | 4.500E-09 | 1.700E-08 |
| 16341 | 818 | V818A | B | 6.4e-09 | 5.8e-09 | 6e-09 | 6.067E-09 | 3.055E-10 | 5.800E-09 | 6.400E-09 |
| 16342 | 818 | V818R | B | 5.8e-09 | 1.2e-08 | 7.4e-09 | 8.400E-09 | 3.219E-09 | 5.800E-09 | 1.200E-08 |
| 16343 | 818 | V818N | B | 7.6e-09 | 4e-09 | 8.9e-09 | 6.833E-09 | 2.538E-09 | 4.000E-09 | 8.900E-09 |
| 16344 | 818 | V818D | B | 2.5e-09 | 5.6e-09 | 1.3e-08 | 7.033E-09 | 5.395E-09 | 2.500E-09 | 1.300E-08 |
| 16345 | 818 | V818C | B | 1.6e-08 | 1.3e-08 | 4.4e-09 | 1.113E-08 | 6.021E-09 | 4.400E-09 | 1.600E-08 |
| 16346 | 818 | V818E | B | 7.3e-09 | 2.7e-09 | 1.2e-08 | 7.333E-09 | 4.650E-09 | 2.700E-09 | 1.200E-08 |
| 16347 | 818 | V818Q | B | 1e-08 | 1.1e-08 | 9e-09 | 1.000E-08 | 1.000E-09 | 9.000E-09 | 1.100E-08 |
| 16348 | 818 | V818G | B | 8.7e-09 | 6.6e-09 | 8.1e-09 | 7.800E-09 | 1.082E-09 | 6.600E-09 | 8.700E-09 |
| 16349 | 818 | V818H | B | 3.8e-09 | 5.4e-09 | 1.8e-08 | 9.067E-09 | 7.778E-09 | 3.800E-09 | 1.800E-08 |
| 16350 | 818 | V818I | B | 4.3e-09 | 1.2e-08 | 4.8e-09 | 7.033E-09 | 4.309E-09 | 4.300E-09 | 1.200E-08 |
| 16351 | 818 | V818L | B | 4.3e-09 | 1.5e-08 | 4.3e-09 | 7.867E-09 | 6.178E-09 | 4.300E-09 | 1.500E-08 |
| 16352 | 818 | V818K | B | 9.8e-09 | 1e-08 | 3.3e-09 | 7.700E-09 | 3.812E-09 | 3.300E-09 | 1.000E-08 |
| 16353 | 818 | V818M | B | 2.4e-08 | 8e-09 | 1.9e-09 | 1.130E-08 | 1.141E-08 | 1.900E-09 | 2.400E-08 |
| 16354 | 818 | V818F | B | 3.5e-09 | 1.1e-08 | 7.2e-09 | 7.233E-09 | 3.750E-09 | 3.500E-09 | 1.100E-08 |
| 16355 | 818 | V818P | B | 2.5e-09 | 5e-09 | 2.3e-08 | 1.017E-08 | 1.118E-08 | 2.500E-09 | 2.300E-08 |
| 16356 | 818 | V818S | B | 7.2e-09 | 2.6e-09 | 1.7e-08 | 8.933E-09 | 7.355E-09 | 2.600E-09 | 1.700E-08 |
| 16357 | 818 | V818T | B | 1.8e-08 | 7.5e-09 | 2e-09 | 9.167E-09 | 8.129E-09 | 2.000E-09 | 1.800E-08 |
| 16358 | 818 | V818W | B | 8.5e-09 | 8.3e-09 | 1.1e-08 | 9.267E-09 | 1.504E-09 | 8.300E-09 | 1.100E-08 |
| 16359 | 818 | V818Y | B | 3.9e-09 | 8.9e-09 | 8.4e-09 | 7.067E-09 | 2.754E-09 | 3.900E-09 | 8.900E-09 |
| 16360 | 818 | V818V | B | 1e-08 | 5.6e-09 | 9.3e-09 | 8.300E-09 | 2.364E-09 | 5.600E-09 | 1.000E-08 |
| 16361 | 819 | T819A | B | 6.6e-09 | 4.8e-09 | 3.2e-09 | 4.867E-09 | 1.701E-09 | 3.200E-09 | 6.600E-09 |
| 16362 | 819 | T819R | B | 8.1e-09 | 7.2e-09 | 7.4e-09 | 7.567E-09 | 4.726E-10 | 7.200E-09 | 8.100E-09 |
| 16363 | 819 | T819N | B | 1e-08 | 5.8e-09 | 1.3e-08 | 9.600E-09 | 3.617E-09 | 5.800E-09 | 1.300E-08 |
| 16364 | 819 | T819D | B | 5.5e-09 | 1.4e-08 | 3.5e-09 | 7.667E-09 | 5.575E-09 | 3.500E-09 | 1.400E-08 |
| 16365 | 819 | T819C | B | 9.2e-09 | 5.3e-09 | 9e-09 | 7.833E-09 | 2.196E-09 | 5.300E-09 | 9.200E-09 |
| 16366 | 819 | T819E | B | 3.5e-09 | 5.4e-09 | 1.6e-08 | 8.300E-09 | 6.736E-09 | 3.500E-09 | 1.600E-08 |
| 16367 | 819 | T819Q | B | 3e-09 | 1.6e-08 | 9.8e-09 | 9.600E-09 | 6.502E-09 | 3.000E-09 | 1.600E-08 |
| 16368 | 819 | T819G | B | 3.1e-09 | 2.9e-09 | 1.1e-08 | 5.667E-09 | 4.620E-09 | 2.900E-09 | 1.100E-08 |
| 16369 | 819 | T819H | B | 7.2e-09 | 8.3e-09 | 2.7e-08 | 1.417E-08 | 1.113E-08 | 7.200E-09 | 2.700E-08 |
| 16370 | 819 | T819I | B | 5.4e-09 | 2.8e-09 | 7e-09 | 5.067E-09 | 2.120E-09 | 2.800E-09 | 7.000E-09 |
| 16371 | 819 | T819L | B | 7.4e-09 | 3.9e-09 | 6.4e-09 | 5.900E-09 | 1.803E-09 | 3.900E-09 | 7.400E-09 |
| 16372 | 819 | T819K | B | 3.7e-09 | 4.6e-09 | 7e-09 | 5.100E-09 | 1.706E-09 | 3.700E-09 | 7.000E-09 |
| 16373 | 819 | T819M | B | 4.5e-09 | 6.4e-09 | 6.1e-09 | 5.667E-09 | 1.021E-09 | 4.500E-09 | 6.400E-09 |
| 16374 | 819 | T819F | B | 1e-08 | 3.4e-09 | 4.4e-09 | 5.933E-09 | 3.557E-09 | 3.400E-09 | 1.000E-08 |
| 16375 | 819 | T819P | B | 2.7e-09 | 7.2e-09 | 1e-08 | 6.633E-09 | 3.683E-09 | 2.700E-09 | 1.000E-08 |
| 16376 | 819 | T819S | B | 8e-09 | 9.2e-09 | 5.2e-09 | 7.467E-09 | 2.053E-09 | 5.200E-09 | 9.200E-09 |
| 16377 | 819 | T819T | B | 6.1e-09 | 2.2e-09 | 6.6e-09 | 4.967E-09 | 2.409E-09 | 2.200E-09 | 6.600E-09 |
| 16378 | 819 | T819W | B | 6e-09 | 3.4e-09 | 7.4e-09 | 5.600E-09 | 2.030E-09 | 3.400E-09 | 7.400E-09 |
| 16379 | 819 | T819Y | B | 6.9e-09 | 4.4e-09 | 7.1e-09 | 6.133E-09 | 1.504E-09 | 4.400E-09 | 7.100E-09 |
| 16380 | 819 | T819V | B | 6.5e-09 | 4.5e-09 | 4.6e-09 | 5.200E-09 | 1.127E-09 | 4.500E-09 | 6.500E-09 |
| 16381 | 820 | H820A | B | 4.3e-09 | 6.8e-09 | 5.4e-09 | 5.500E-09 | 1.253E-09 | 4.300E-09 | 6.800E-09 |
| 16382 | 820 | H820R | B | 8.2e-09 | 9.6e-09 | 9.4e-09 | 9.067E-09 | 7.572E-10 | 8.200E-09 | 9.600E-09 |
| 16383 | 820 | H820N | B | 9.9e-09 | 4.3e-09 | 4.1e-09 | 6.100E-09 | 3.292E-09 | 4.100E-09 | 9.900E-09 |
| 16384 | 820 | H820D | B | 1.7e-08 | 1.3e-08 | 8.9e-09 | 1.297E-08 | 4.050E-09 | 8.900E-09 | 1.700E-08 |
| 16385 | 820 | H820C | B | 4.8e-09 | 1.4e-08 | 7.7e-09 | 8.833E-09 | 4.704E-09 | 4.800E-09 | 1.400E-08 |
| 16386 | 820 | H820E | B | 3.4e-09 | 9.2e-09 | 6.3e-09 | 6.300E-09 | 2.900E-09 | 3.400E-09 | 9.200E-09 |
| 16387 | 820 | H820Q | B | 2.8e-09 | 9.9e-09 | 8.1e-09 | 6.933E-09 | 3.691E-09 | 2.800E-09 | 9.900E-09 |
| 16388 | 820 | H820G | B | 1.7e-08 | 5.7e-09 | 1.1e-08 | 1.123E-08 | 5.654E-09 | 5.700E-09 | 1.700E-08 |
| 16389 | 820 | H820H | B | 2.8e-09 | 7.8e-09 | 2.4e-09 | 4.333E-09 | 3.009E-09 | 2.400E-09 | 7.800E-09 |
| 16390 | 820 | H820I | B | 8.2e-09 | 2.5e-09 | 2.8e-09 | 4.500E-09 | 3.208E-09 | 2.500E-09 | 8.200E-09 |
| 16391 | 820 | H820L | B | 7.9e-09 | 2e-09 | 9e-09 | 6.300E-09 | 3.764E-09 | 2.000E-09 | 9.000E-09 |
| 16392 | 820 | H820K | B | 4.4e-09 | 1.5e-08 | 4.2e-09 | 7.867E-09 | 6.178E-09 | 4.200E-09 | 1.500E-08 |
| 16393 | 820 | H820M | B | 1.4e-08 | 4.8e-09 | 7e-09 | 8.600E-09 | 4.804E-09 | 4.800E-09 | 1.400E-08 |
| 16394 | 820 | H820F | B | 1.1e-08 | 6.2e-09 | 8.9e-09 | 8.700E-09 | 2.406E-09 | 6.200E-09 | 1.100E-08 |
| 16395 | 820 | H820P | B | 1e-08 | 1.7e-09 | 4.6e-09 | 5.433E-09 | 4.212E-09 | 1.700E-09 | 1.000E-08 |
| 16396 | 820 | H820S | B | 3.3e-09 | 3.3e-09 | 7e-09 | 4.533E-09 | 2.136E-09 | 3.300E-09 | 7.000E-09 |
| 16397 | 820 | H820T | B | 3.9e-08 | 3.5e-09 | 5.9e-09 | 1.613E-08 | 1.984E-08 | 3.500E-09 | 3.900E-08 |
| 16398 | 820 | H820W | B | 6.2e-09 | 2.7e-08 | 1.4e-08 | 1.573E-08 | 1.051E-08 | 6.200E-09 | 2.700E-08 |
| 16399 | 820 | H820Y | B | 8.9e-09 | 7.4e-09 | 7.3e-09 | 7.867E-09 | 8.963E-10 | 7.300E-09 | 8.900E-09 |
| 16400 | 820 | H820V | B | 4.3e-09 | 1.2e-08 | 3.6e-09 | 6.633E-09 | 4.661E-09 | 3.600E-09 | 1.200E-08 |
| 16401 | 821 | Q821A | B | 1.2e-08 | 8.9e-09 | 1.1e-08 | 1.063E-08 | 1.582E-09 | 8.900E-09 | 1.200E-08 |
| 16402 | 821 | Q821R | B | 5.8e-09 | 9.6e-09 | 9.9e-09 | 8.433E-09 | 2.285E-09 | 5.800E-09 | 9.900E-09 |
| 16403 | 821 | Q821N | B | 5e-09 | 4.8e-09 | 9.9e-09 | 6.567E-09 | 2.888E-09 | 4.800E-09 | 9.900E-09 |
| 16404 | 821 | Q821D | B | 8.2e-09 | 7.8e-09 | 5e-09 | 7.000E-09 | 1.744E-09 | 5.000E-09 | 8.200E-09 |

| | | | | | | | | | | |
|---|---|---|---|---|---|---|---|---|---|---|
| 16405 | 821 | Q821C | B | 8.1e-09 | 5.2e-09 | 7.3e-09 | 6.867E-09 | 1.498E-09 | 5.200E-09 | 8.100E-09 |
| 16406 | 821 | Q821E | B | 5.5e-09 | 1.2e-08 | 4.9e-09 | 7.467E-09 | 3.937E-09 | 4.900E-09 | 1.200E-08 |
| 16407 | 821 | Q821Q | B | 9.5e-09 | 1.2e-08 | 4.7e-09 | 8.733E-09 | 3.710E-09 | 4.700E-09 | 1.200E-08 |
| 16408 | 821 | Q821G | B | 4.5e-09 | 1.2e-08 | 1.9e-08 | 1.183E-08 | 7.251E-09 | 4.500E-09 | 1.900E-08 |
| 16409 | 821 | Q821H | B | 6e-09 | 1.8e-08 | 1e-08 | 1.133E-08 | 6.110E-09 | 6.000E-09 | 1.800E-08 |
| 16410 | 821 | Q821I | B | 2.3e-08 | 1.2e-08 | 4.5e-09 | 1.317E-08 | 9.305E-09 | 4.500E-09 | 2.300E-08 |
| 16411 | 821 | Q821L | B | 1.6e-08 | 1.2e-08 | 6.4e-09 | 1.147E-08 | 4.822E-09 | 6.400E-09 | 1.600E-08 |
| 16412 | 821 | Q821K | B | 2.4e-09 | 5.6e-09 | 3.3e-09 | 3.767E-09 | 1.650E-09 | 2.400E-09 | 5.600E-09 |
| 16413 | 821 | Q821M | B | 9.8e-09 | 3.3e-09 | 1.2e-08 | 8.367E-09 | 4.524E-09 | 3.300E-09 | 1.200E-08 |
| 16414 | 821 | Q821F | B | 7.2e-10 | 1.5e-08 | 5e-09 | 6.907E-09 | 7.328E-09 | 7.200E-10 | 1.500E-08 |
| 16415 | 821 | Q821P | B | 2.7e-09 | 9e-09 | 7.9e-09 | 6.533E-09 | 3.365E-09 | 2.700E-09 | 9.000E-09 |
| 16416 | 821 | Q821S | B | 1.4e-08 | 9.5e-09 | 4.6e-09 | 9.367E-09 | 4.701E-09 | 4.600E-09 | 1.400E-08 |
| 16417 | 821 | Q821T | B | 9.5e-09 | 8.4e-09 | 8.6e-09 | 8.833E-09 | 5.859E-10 | 8.400E-09 | 9.500E-09 |
| 16418 | 821 | Q821W | B | 3.7e-09 | 8.2e-09 | 1.2e-08 | 7.967E-09 | 4.155E-09 | 3.700E-09 | 1.200E-08 |
| 16419 | 821 | Q821Y | B | 2.1e-09 | 3.9e-09 | 8.4e-09 | 4.800E-09 | 3.245E-09 | 2.100E-09 | 8.400E-09 |
| 16420 | 821 | Q821V | B | 3.3e-09 | 2.8e-08 | 1.3e-08 | 1.477E-08 | 1.244E-08 | 3.300E-09 | 2.800E-08 |
| 16421 | 822 | G822A | B | 1e-08 | 7.8e-09 | 4.6e-09 | 7.467E-09 | 2.715E-09 | 4.600E-09 | 1.000E-08 |
| 16422 | 822 | G822R | B | 5.4e-09 | 3.8e-09 | 1.1e-08 | 6.733E-09 | 3.781E-09 | 3.800E-09 | 1.100E-08 |
| 16423 | 822 | G822N | B | 1e-08 | 1.2e-08 | 3.3e-09 | 8.433E-09 | 4.557E-09 | 3.300E-09 | 1.200E-08 |
| 16424 | 822 | G822D | B | 6e-09 | 4.2e-09 | 3.9e-09 | 4.700E-09 | 1.136E-09 | 3.900E-09 | 6.000E-09 |
| 16425 | 822 | G822C | B | 1.4e-08 | 8.7e-09 | 8.9e-09 | 1.053E-08 | 3.004E-09 | 8.700E-09 | 1.400E-08 |
| 16426 | 822 | G822E | B | 9.9e-09 | 6.9e-09 | 5.3e-09 | 7.367E-09 | 2.335E-09 | 5.300E-09 | 9.900E-09 |
| 16427 | 822 | G822Q | B | 6.3e-09 | 1e-08 | 9.3e-09 | 8.533E-09 | 1.966E-09 | 6.300E-09 | 1.000E-08 |
| 16428 | 822 | G822G | B | 2e-08 | 6.3e-09 | 8.7e-09 | 1.167E-08 | 7.316E-09 | 6.300E-09 | 2.000E-08 |
| 16429 | 822 | G822H | B | 7e-09 | 2.9e-09 | 1.8e-08 | 9.300E-09 | 7.808E-09 | 2.900E-09 | 1.800E-08 |
| 16430 | 822 | G822I | B | 2.6e-09 | 5e-09 | 5.6e-09 | 4.400E-09 | 1.587E-09 | 2.600E-09 | 5.600E-09 |
| 16431 | 822 | G822L | B | 9.8e-09 | 3.6e-09 | 3.4e-09 | 5.600E-09 | 3.639E-09 | 3.400E-09 | 9.800E-09 |
| 16432 | 822 | G822K | B | 5.1e-09 | 7.4e-09 | 4.8e-09 | 5.767E-09 | 1.422E-09 | 4.800E-09 | 7.400E-09 |
| 16433 | 822 | G822M | B | 2.7e-09 | 2.5e-09 | 5.2e-09 | 3.467E-09 | 1.504E-09 | 2.500E-09 | 5.200E-09 |
| 16434 | 822 | G822F | B | 3.7e-09 | 2.8e-09 | 6.1e-09 | 4.200E-09 | 1.706E-09 | 2.800E-09 | 6.100E-09 |
| 16435 | 822 | G822P | B | 2.4e-09 | 1.5e-08 | 4.4e-09 | 7.267E-09 | 6.772E-09 | 2.400E-09 | 1.500E-08 |
| 16436 | 822 | G822S | B | 1e-08 | 8.9e-09 | 6e-09 | 8.300E-09 | 2.066E-09 | 6.000E-09 | 1.000E-08 |
| 16437 | 822 | G822T | B | 1e-08 | 4.1e-09 | 5.5e-09 | 6.533E-09 | 3.083E-09 | 4.100E-09 | 1.000E-08 |
| 16438 | 822 | G822W | B | 4.9e-09 | 1.2e-08 | 2.5e-09 | 6.467E-09 | 4.940E-09 | 2.500E-09 | 1.200E-08 |
| 16439 | 822 | G822Y | B | 1.4e-08 | 6.4e-09 | 3.8e-09 | 8.067E-09 | 5.300E-09 | 3.800E-09 | 1.400E-08 |
| 16440 | 822 | G822V | B | 8e-09 | 7.3e-09 | 7.8e-09 | 7.700E-09 | 3.606E-10 | 7.300E-09 | 8.000E-09 |
| 16441 | 823 | L823A | B | 4.6e-09 | 1.4e-08 | 6.3e-09 | 8.300E-09 | 5.009E-09 | 4.600E-09 | 1.400E-08 |
| 16442 | 823 | L823R | B | 1.1e-08 | 1e-08 | 5.8e-09 | 8.933E-09 | 2.759E-09 | 5.800E-09 | 1.100E-08 |
| 16443 | 823 | L823N | B | 9.4e-09 | 6.4e-09 | 7.8e-09 | 7.867E-09 | 1.501E-09 | 6.400E-09 | 9.400E-09 |
| 16444 | 823 | L823D | B | 5.9e-09 | 3.7e-09 | 7.9e-09 | 5.833E-09 | 2.101E-09 | 3.700E-09 | 7.900E-09 |
| 16445 | 823 | L823C | B | 9.3e-09 | 6e-09 | 8.6e-09 | 7.967E-09 | 1.739E-09 | 6.000E-09 | 9.300E-09 |
| 16446 | 823 | L823E | B | 4.7e-09 | 8.7e-09 | 5.4e-09 | 6.267E-09 | 2.136E-09 | 4.700E-09 | 8.700E-09 |
| 16447 | 823 | L823Q | B | 3.9e-09 | 4.2e-09 | 1.1e-08 | 6.367E-09 | 4.015E-09 | 3.900E-09 | 1.100E-08 |
| 16448 | 823 | L823G | B | 9.1e-10 | 3.6e-09 | 5.1e-09 | 3.203E-09 | 2.123E-09 | 9.100E-10 | 5.100E-09 |
| 16449 | 823 | L823H | B | 5e-09 | 3.6e-09 | 2.5e-09 | 3.700E-09 | 1.253E-09 | 2.500E-09 | 5.000E-09 |
| 16450 | 823 | L823I | B | 6.3e-09 | 1.1e-08 | 2.1e-09 | 6.467E-09 | 4.452E-09 | 2.100E-09 | 1.100E-08 |
| 16451 | 823 | L823L | B | 1.8e-08 | 7e-09 | 5.9e-09 | 1.030E-08 | 6.691E-09 | 5.900E-09 | 1.800E-08 |
| 16452 | 823 | L823K | B | 3.7e-09 | 2.6e-09 | 1e-08 | 5.433E-09 | 3.993E-09 | 2.600E-09 | 1.000E-08 |
| 16453 | 823 | L823M | B | 3.2e-09 | 1.3e-08 | 5.7e-09 | 7.300E-09 | 5.092E-09 | 3.200E-09 | 1.300E-08 |
| 16454 | 823 | L823F | B | 3.4e-09 | 1.1e-08 | 3.8e-09 | 6.067E-09 | 4.277E-09 | 3.400E-09 | 1.100E-08 |
| 16455 | 823 | L823P | B | 1.7e-08 | 6.9e-09 | 7.5e-09 | 1.047E-08 | 5.666E-09 | 6.900E-09 | 1.700E-08 |
| 16456 | 823 | L823S | B | 9.7e-09 | 4e-08 | 2e-08 | 2.323E-08 | 1.541E-08 | 9.700E-09 | 4.000E-08 |
| 16457 | 823 | L823T | B | 7.2e-09 | 8.1e-09 | 9.2e-09 | 8.167E-09 | 1.002E-09 | 7.200E-09 | 9.200E-09 |
| 16458 | 823 | L823W | B | 1.1e-08 | 1.9e-08 | 8.3e-09 | 1.277E-08 | 5.564E-09 | 8.300E-09 | 1.900E-08 |
| 16459 | 823 | L823Y | B | 1.3e-08 | 3.6e-09 | 4.7e-09 | 7.100E-09 | 5.139E-09 | 3.600E-09 | 1.300E-08 |
| 16460 | 823 | L823V | B | 1.5e-08 | 3.7e-09 | 4e-09 | 7.567E-09 | 6.439E-09 | 3.700E-09 | 1.500E-08 |
| 16461 | 824 | S824A | B | 6.8e-09 | 5.4e-09 | 6.6e-09 | 6.267E-09 | 7.572E-10 | 5.400E-09 | 6.800E-09 |
| 16462 | 824 | S824R | B | 9.4e-09 | 4.3e-09 | 8.2e-09 | 7.300E-09 | 2.666E-09 | 4.300E-09 | 9.400E-09 |
| 16463 | 824 | S824N | B | 6.2e-09 | 7.5e-09 | 2.4e-08 | 1.257E-08 | 9.923E-09 | 6.200E-09 | 2.400E-08 |
| 16464 | 824 | S824D | B | 1.1e-08 | 6.2e-09 | 7.1e-09 | 8.100E-09 | 2.551E-09 | 6.200E-09 | 1.100E-08 |
| 16465 | 824 | S824C | B | 2.4e-08 | 5.3e-09 | 1.4e-08 | 1.443E-08 | 9.358E-09 | 5.300E-09 | 2.400E-08 |
| 16466 | 824 | S824E | B | 1.8e-09 | 9.3e-09 | 5.8e-09 | 5.633E-09 | 3.753E-09 | 1.800E-09 | 9.300E-09 |
| 16467 | 824 | S824Q | B | 4.5e-09 | 5.4e-09 | 2.2e-08 | 1.063E-08 | 9.854E-09 | 4.500E-09 | 2.200E-08 |
| 16468 | 824 | S824G | B | 2.3e-09 | 4.5e-09 | 1.3e-08 | 6.600E-09 | 5.651E-09 | 2.300E-09 | 1.300E-08 |
| 16469 | 824 | S824H | B | 9.2e-09 | 4.6e-09 | 7.3e-09 | 7.033E-09 | 2.312E-09 | 4.600E-09 | 9.200E-09 |
| 16470 | 824 | S824I | B | 1e-08 | 1.1e-08 | 7.8e-09 | 9.600E-09 | 1.637E-09 | 7.800E-09 | 1.100E-08 |
| 16471 | 824 | S824L | B | 9.3e-09 | 7.4e-09 | 7e-09 | 7.900E-09 | 1.229E-09 | 7.000E-09 | 9.300E-09 |
| 16472 | 824 | S824K | B | 4.8e-09 | 5.1e-09 | 6.1e-09 | 5.333E-09 | 6.807E-10 | 4.800E-09 | 6.100E-09 |
| 16473 | 824 | S824M | B | 8e-09 | 6.1e-09 | 1.5e-08 | 9.700E-09 | 4.687E-09 | 6.100E-09 | 1.500E-08 |
| 16474 | 824 | S824F | B | 7.7e-09 | 4.3e-09 | 1e-08 | 7.333E-09 | 2.868E-09 | 4.300E-09 | 1.000E-08 |
| 16475 | 824 | S824P | B | 7.5e-09 | 8e-09 | 9.6e-09 | 8.367E-09 | 1.097E-09 | 7.500E-09 | 9.600E-09 |
| 16476 | 824 | S824S | B | 7e-09 | 8e-09 | 7.9e-09 | 7.633E-09 | 5.508E-10 | 7.000E-09 | 8.000E-09 |
| 16477 | 824 | S824T | B | 6.5e-09 | 4.2e-09 | 2.4e-09 | 4.367E-09 | 2.055E-09 | 2.400E-09 | 6.500E-09 |
| 16478 | 824 | S824W | B | 1.5e-08 | 1.4e-08 | 6.6e-09 | 1.187E-08 | 4.588E-09 | 6.600E-09 | 1.500E-08 |
| 16479 | 824 | S824Y | B | 1.1e-08 | 1.3e-09 | 2.8e-09 | 5.033E-09 | 5.221E-09 | 1.300E-09 | 1.100E-08 |
| 16480 | 824 | S824V | B | 3.8e-09 | 1.1e-08 | 1.3e-08 | 9.267E-09 | 4.839E-09 | 3.800E-09 | 1.300E-08 |
| 16481 | 825 | S825A | B | 7.2e-09 | 2.2e-09 | 1.1e-08 | 6.800E-09 | 4.414E-09 | 2.200E-09 | 1.100E-08 |
| 16482 | 825 | S825R | B | 8e-09 | 2.8e-08 | 6.3e-09 | 1.410E-08 | 1.207E-08 | 6.300E-09 | 2.800E-08 |
| 16483 | 825 | S825N | B | 5.7e-09 | 1e-08 | 9.6e-09 | 8.433E-09 | 2.376E-09 | 5.700E-09 | 1.000E-08 |
| 16484 | 825 | S825D | B | 9.9e-09 | 1.9e-09 | 2.1e-08 | 1.093E-08 | 9.592E-09 | 1.900E-09 | 2.100E-08 |
| 16485 | 825 | S825C | B | 1.2e-08 | 6.2e-09 | 4e-09 | 7.400E-09 | 4.133E-09 | 4.000E-09 | 1.200E-08 |
| 16486 | 825 | S825E | B | 6e-09 | 3.7e-09 | 5.5e-09 | 5.067E-09 | 1.210E-09 | 3.700E-09 | 6.000E-09 |
| 16487 | 825 | S825Q | B | 3.6e-09 | 7.9e-09 | 1.9e-08 | 1.017E-08 | 7.946E-09 | 3.600E-09 | 1.900E-08 |
| 16488 | 825 | S825G | B | 6e-09 | 3.7e-09 | 1.8e-08 | 9.233E-09 | 7.679E-09 | 3.700E-09 | 1.800E-08 |
| 16489 | 825 | S825H | B | 4.3e-09 | 1.3e-08 | 9.4e-09 | 8.900E-09 | 4.371E-09 | 4.300E-09 | 1.300E-08 |

| | | | | | | | | | | |
|---|---|---|---|---|---|---|---|---|---|---|
| 16490 | 825 | S825I | B | 1.6e-08 | 3.5e-09 | 4.5e-09 | 8.000E-09 | 6.946E-09 | 3.500E-09 | 1.600E-08 |
| 16491 | 825 | S825L | B | 2.6e-09 | 4.7e-09 | 5.2e-09 | 4.167E-09 | 1.380E-09 | 2.600E-09 | 5.200E-09 |
| 16492 | 825 | S825K | B | 8.7e-09 | 1.1e-08 | 5.8e-09 | 8.500E-09 | 2.606E-09 | 5.800E-09 | 1.100E-08 |
| 16493 | 825 | S825M | B | 8.2e-09 | 1.7e-08 | 1.3e-08 | 1.273E-08 | 4.406E-09 | 8.200E-09 | 1.700E-08 |
| 16494 | 825 | S825F | B | 5.5e-09 | 1.1e-08 | 1.1e-08 | 9.167E-09 | 3.175E-09 | 5.500E-09 | 1.100E-08 |
| 16495 | 825 | S825P | B | 9.1e-09 | 4.9e-09 | 1e-08 | 8.000E-09 | 2.722E-09 | 4.900E-09 | 1.000E-08 |
| 16496 | 825 | S825S | B | 1.8e-09 | 9.7e-10 | 2.6e-08 | 9.590E-09 | 1.422E-08 | 9.700E-10 | 2.600E-08 |
| 16497 | 825 | S825T | B | 5.9e-09 | 6.1e-09 | 9.2e-09 | 7.067E-09 | 1.850E-09 | 5.900E-09 | 9.200E-09 |
| 16498 | 825 | S825W | B | 1.2e-08 | 8.1e-09 | 5.5e-09 | 8.533E-09 | 3.272E-09 | 5.500E-09 | 1.200E-08 |
| 16499 | 825 | S825Y | B | 3e-09 | 1.8e-09 | 4.4e-09 | 3.067E-09 | 1.301E-09 | 1.800E-09 | 4.400E-09 |
| 16500 | 825 | S825V | B | 1.7e-08 | 1.2e-08 | 1.1e-08 | 1.333E-08 | 3.215E-09 | 1.100E-08 | 1.700E-08 |
| 16501 | 826 | P826A | B | 1.5e-08 | 3.7e-08 | 4.6e-09 | 1.887E-08 | 1.654E-08 | 4.600E-09 | 3.700E-08 |
| 16502 | 826 | P826R | B | 3.3e-09 | 9.9e-09 | 2.9e-09 | 5.367E-09 | 3.931E-09 | 2.900E-09 | 9.900E-09 |
| 16503 | 826 | P826N | B | 1.1e-08 | 1.4e-08 | 3.7e-09 | 9.567E-09 | 5.297E-09 | 3.700E-09 | 1.400E-08 |
| 16504 | 826 | P826D | B | 2.3e-08 | 7.4e-09 | 1.5e-08 | 1.513E-08 | 7.801E-09 | 7.400E-09 | 2.300E-08 |
| 16505 | 826 | P826C | B | 6e-09 | 7.9e-09 | 1.2e-08 | 8.633E-09 | 3.066E-09 | 6.000E-09 | 1.200E-08 |
| 16506 | 826 | P826E | B | 1.7e-09 | 9.6e-09 | 1.2e-08 | 7.767E-09 | 5.389E-09 | 1.700E-09 | 1.200E-08 |
| 16507 | 826 | P826Q | B | 3.8e-09 | 1.8e-08 | 6.6e-09 | 9.467E-09 | 7.522E-09 | 3.800E-09 | 1.800E-08 |
| 16508 | 826 | P826G | B | 4.2e-09 | 1.1e-08 | 2.8e-08 | 1.440E-08 | 1.226E-08 | 4.200E-09 | 2.800E-08 |
| 16509 | 826 | P826H | B | 1.6e-08 | 4.5e-09 | 1.1e-08 | 1.050E-08 | 5.766E-09 | 4.500E-09 | 1.600E-08 |
| 16510 | 826 | P826I | B | 2.9e-09 | 1.3e-09 | 8e-09 | 4.067E-09 | 3.499E-09 | 1.300E-09 | 8.000E-09 |
| 16511 | 826 | P826L | B | 2.8e-08 | 1.1e-08 | 3.1e-09 | 1.403E-08 | 1.272E-08 | 3.100E-09 | 2.800E-08 |
| 16512 | 826 | P826K | B | 1.7e-08 | 6.6e-09 | 6.3e-09 | 9.967E-09 | 6.093E-09 | 6.300E-09 | 1.700E-08 |
| 16513 | 826 | P826M | B | 9.2e-09 | 4.8e-09 | 1.1e-08 | 8.333E-09 | 3.190E-09 | 4.800E-09 | 1.100E-08 |
| 16514 | 826 | P826F | B | 4.6e-09 | 1.3e-08 | 5.2e-09 | 7.600E-09 | 4.686E-09 | 4.600E-09 | 1.300E-08 |
| 16515 | 826 | P826P | B | 1.7e-08 | 1.7e-08 | 6.3e-09 | 1.343E-08 | 6.178E-09 | 6.300E-09 | 1.700E-08 |
| 16516 | 826 | P826S | B | 2.5e-09 | 1.8e-08 | 4.6e-09 | 8.367E-09 | 8.409E-09 | 2.500E-09 | 1.800E-08 |
| 16517 | 826 | P826T | B | 8.5e-09 | 1.8e-09 | 8e-09 | 6.100E-09 | 3.732E-09 | 1.800E-09 | 8.500E-09 |
| 16518 | 826 | P826W | B | 4.6e-09 | 3.2e-09 | 8.9e-09 | 5.567E-09 | 2.970E-09 | 3.200E-09 | 8.900E-09 |
| 16519 | 826 | P826Y | B | 1.2e-08 | 2e-08 | 1.1e-08 | 1.433E-08 | 4.933E-09 | 1.100E-08 | 2.000E-08 |
| 16520 | 826 | P826V | B | 4.4e-09 | 1.1e-08 | 2.2e-08 | 1.247E-08 | 8.891E-09 | 4.400E-09 | 2.200E-08 |
| 16521 | 827 | V827A | B | 5.1e-09 | 5.4e-09 | 3.1e-09 | 4.533E-09 | 1.250E-09 | 3.100E-09 | 5.400E-09 |
| 16522 | 827 | V827R | B | 6.9e-09 | 3.3e-09 | 5.4e-09 | 5.200E-09 | 1.808E-09 | 3.300E-09 | 6.900E-09 |
| 16523 | 827 | V827N | B | 1.2e-08 | 1.3e-08 | 7.9e-09 | 1.097E-08 | 2.702E-09 | 7.900E-09 | 1.300E-08 |
| 16524 | 827 | V827D | B | 8.1e-09 | 1.1e-09 | 7.6e-09 | 5.600E-09 | 3.905E-09 | 1.100E-09 | 8.100E-09 |
| 16525 | 827 | V827C | B | 5.3e-09 | 3.2e-09 | 3.4e-09 | 3.967E-09 | 1.159E-09 | 3.200E-09 | 5.300E-09 |
| 16526 | 827 | V827E | B | 5e-09 | 6.4e-09 | 5.5e-09 | 5.633E-09 | 7.095E-10 | 5.000E-09 | 6.400E-09 |
| 16527 | 827 | V827Q | B | 7.2e-09 | 8.8e-09 | 6e-09 | 7.333E-09 | 1.405E-09 | 6.000E-09 | 8.800E-09 |
| 16528 | 827 | V827G | B | 2.2e-08 | 2.6e-08 | 4.2e-09 | 1.740E-08 | 1.161E-08 | 4.200E-09 | 2.600E-08 |
| 16529 | 827 | V827H | B | 2.9e-08 | 5.5e-09 | 8.6e-09 | 1.437E-08 | 1.277E-08 | 5.500E-09 | 2.900E-08 |
| 16530 | 827 | V827I | B | 1.2e-08 | 1e-08 | 2.3e-08 | 1.500E-08 | 7.000E-09 | 1.000E-08 | 2.300E-08 |
| 16531 | 827 | V827L | B | 5.7e-09 | 8.7e-09 | 2.3e-09 | 5.567E-09 | 3.202E-09 | 2.300E-09 | 8.700E-09 |
| 16532 | 827 | V827K | B | 5.5e-09 | 8.6e-09 | 8.9e-09 | 7.667E-09 | 1.882E-09 | 5.500E-09 | 8.900E-09 |
| 16533 | 827 | V827M | B | 2.5e-08 | 6e-09 | 5.2e-09 | 1.207E-08 | 1.121E-08 | 5.200E-09 | 2.500E-08 |
| 16534 | 827 | V827F | B | 1.1e-08 | 8e-09 | 1.8e-08 | 1.233E-08 | 5.132E-09 | 8.000E-09 | 1.800E-08 |
| 16535 | 827 | V827P | B | 1.6e-08 | 2.1e-08 | 9.6e-09 | 1.553E-08 | 5.714E-09 | 9.600E-09 | 2.100E-08 |
| 16536 | 827 | V827S | B | 6.4e-09 | 1.7e-08 | 1.4e-08 | 1.247E-08 | 5.464E-09 | 6.400E-09 | 1.700E-08 |
| 16537 | 827 | V827T | B | 6.2e-09 | 2.6e-09 | 1.7e-08 | 8.600E-09 | 7.494E-09 | 2.600E-09 | 1.700E-08 |
| 16538 | 827 | V827W | B | 6.7e-09 | 7.2e-09 | 8.8e-09 | 7.567E-09 | 1.097E-09 | 6.700E-09 | 8.800E-09 |
| 16539 | 827 | V827Y | B | 1.1e-08 | 6.7e-09 | 1.8e-09 | 6.500E-09 | 4.603E-09 | 1.800E-09 | 1.100E-08 |
| 16540 | 827 | V827V | B | 7.1e-09 | 7.2e-09 | 8e-09 | 7.433E-09 | 4.933E-10 | 7.100E-09 | 8.000E-09 |
| 16541 | 828 | T828A | B | 1e-08 | 2.3e-08 | 2.4e-08 | 1.900E-08 | 7.810E-09 | 1.000E-08 | 2.400E-08 |
| 16542 | 828 | T828R | B | 1e-08 | 4.7e-09 | 4.2e-09 | 6.300E-09 | 3.214E-09 | 4.200E-09 | 1.000E-08 |
| 16543 | 828 | T828N | B | 9.7e-09 | 3.8e-09 | 8.3e-09 | 7.267E-09 | 3.083E-09 | 3.800E-09 | 9.700E-09 |
| 16544 | 828 | T828D | B | 1.9e-08 | 1.9e-08 | 2.6e-09 | 1.353E-08 | 9.469E-09 | 2.600E-09 | 1.900E-08 |
| 16545 | 828 | T828C | B | 7.4e-09 | 9.5e-09 | 4.1e-09 | 7.000E-09 | 2.722E-09 | 4.100E-09 | 9.500E-09 |
| 16546 | 828 | T828E | B | 5.4e-09 | 7.8e-09 | 2.7e-09 | 5.300E-09 | 2.551E-09 | 2.700E-09 | 7.800E-09 |
| 16547 | 828 | T828Q | B | 3.4e-09 | 6.6e-09 | 4.2e-09 | 4.733E-09 | 1.665E-09 | 3.400E-09 | 6.600E-09 |
| 16548 | 828 | T828G | B | 2e-08 | 7.1e-09 | 6.4e-09 | 1.117E-08 | 7.658E-09 | 6.400E-09 | 2.000E-08 |
| 16549 | 828 | T828H | B | 5.5e-09 | 9.7e-09 | 2e-09 | 5.733E-09 | 3.855E-09 | 2.000E-09 | 9.700E-09 |
| 16550 | 828 | T828I | B | 3.2e-08 | 2.4e-09 | 1.8e-08 | 1.747E-08 | 1.481E-08 | 2.400E-09 | 3.200E-08 |
| 16551 | 828 | T828L | B | 2.8e-09 | 7.3e-09 | 1.7e-08 | 9.033E-09 | 7.257E-09 | 2.800E-09 | 1.700E-08 |
| 16552 | 828 | T828K | B | 6.9e-09 | 1e-08 | 3.5e-09 | 6.800E-09 | 3.251E-09 | 3.500E-09 | 1.000E-08 |
| 16553 | 828 | T828M | B | 1.3e-08 | 7.2e-09 | 5.9e-09 | 8.700E-09 | 3.780E-09 | 5.900E-09 | 1.300E-08 |
| 16554 | 828 | T828F | B | 4.2e-09 | 6.3e-09 | 2.5e-09 | 4.333E-09 | 1.904E-09 | 2.500E-09 | 6.300E-09 |
| 16555 | 828 | T828P | B | 2.7e-09 | 1.4e-08 | 9.7e-09 | 8.800E-09 | 5.704E-09 | 2.700E-09 | 1.400E-08 |
| 16556 | 828 | T828S | B | 6.3e-09 | 1e-08 | 1.5e-08 | 1.043E-08 | 4.366E-09 | 6.300E-09 | 1.500E-08 |
| 16557 | 828 | T828T | B | 5.4e-09 | 3.6e-09 | 2.9e-09 | 3.967E-09 | 1.290E-09 | 2.900E-09 | 5.400E-09 |
| 16558 | 828 | T828W | B | 8.1e-09 | 3.5e-09 | 8.3e-09 | 6.633E-09 | 2.715E-09 | 3.500E-09 | 8.300E-09 |
| 16559 | 828 | T828Y | B | 8.5e-09 | 3.4e-09 | 8.5e-09 | 6.800E-09 | 2.944E-09 | 3.400E-09 | 8.500E-09 |
| 16560 | 828 | T828V | B | 7e-09 | 6.2e-09 | 1.5e-09 | 4.900E-09 | 2.972E-09 | 1.500E-09 | 7.000E-09 |
| 16561 | 829 | K829A | B | 2.2e-09 | 5.3e-09 | 2.6e-09 | 3.367E-09 | 1.686E-09 | 2.200E-09 | 5.300E-09 |
| 16562 | 829 | K829R | B | 3.3e-09 | 9.7e-09 | 8.7e-09 | 7.233E-09 | 3.443E-09 | 3.300E-09 | 9.700E-09 |
| 16563 | 829 | K829N | B | 1.2e-08 | 2e-08 | 9.7e-09 | 1.390E-08 | 5.406E-09 | 9.700E-09 | 2.000E-08 |
| 16564 | 829 | K829D | B | 7.7e-09 | 2e-08 | 1.3e-09 | 9.667E-09 | 9.504E-09 | 1.300E-09 | 2.000E-08 |
| 16565 | 829 | K829C | B | 1e-08 | 4.1e-09 | 9.1e-09 | 7.733E-09 | 3.179E-09 | 4.100E-09 | 1.000E-08 |
| 16566 | 829 | K829E | B | 4.5e-09 | 6.5e-09 | 1.3e-08 | 8.000E-09 | 4.444E-09 | 4.500E-09 | 1.300E-08 |
| 16567 | 829 | K829Q | B | 1.3e-08 | 2.2e-09 | 4e-09 | 6.400E-09 | 5.786E-09 | 2.200E-09 | 1.300E-08 |
| 16568 | 829 | K829G | B | 3.9e-09 | 4.7e-09 | 7.4e-09 | 5.333E-09 | 1.834E-09 | 3.900E-09 | 7.400E-09 |
| 16569 | 829 | K829H | B | 8.6e-09 | 5.4e-09 | 4.6e-09 | 6.200E-09 | 2.117E-09 | 4.600E-09 | 8.600E-09 |
| 16570 | 829 | K829I | B | 8.1e-09 | 6e-09 | 4.2e-09 | 6.100E-09 | 1.952E-09 | 4.200E-09 | 8.100E-09 |
| 16571 | 829 | K829L | B | 5.3e-09 | 1.9e-08 | 1.8e-08 | 1.410E-08 | 7.637E-09 | 5.300E-09 | 1.900E-08 |
| 16572 | 829 | K829K | B | 4.8e-09 | 8.3e-09 | 5.5e-09 | 6.200E-09 | 1.852E-09 | 4.800E-09 | 8.300E-09 |
| 16573 | 829 | K829M | B | 3.6e-09 | 3.6e-09 | 2e-09 | 3.067E-09 | 9.238E-10 | 2.000E-09 | 3.600E-09 |
| 16574 | 829 | K829F | B | 1.9e-08 | 3.7e-09 | 3.3e-09 | 8.667E-09 | 8.951E-09 | 3.300E-09 | 1.900E-08 |

| | | | | | | | | | | |
|---|---|---|---|---|---|---|---|---|---|---|
| 16575 | 829 | K829P | B | 1.6e-08 | 3.9e-09 | 1.3e-08 | 1.097E-08 | 6.301E-09 | 3.900E-09 | 1.600E-08 |
| 16576 | 829 | K829S | B | 8.4e-09 | 1.1e-08 | 9.5e-09 | 9.633E-09 | 1.305E-09 | 8.400E-09 | 1.100E-08 |
| 16577 | 829 | K829T | B | 1.3e-08 | 7.1e-09 | 5.2e-09 | 8.433E-09 | 4.067E-09 | 5.200E-09 | 1.300E-08 |
| 16578 | 829 | K829W | B | 8.4e-09 | 5.8e-09 | 6.3e-09 | 6.833E-09 | 1.380E-09 | 5.800E-09 | 8.400E-09 |
| 16579 | 829 | K829Y | B | 1.4e-08 | 7.7e-09 | 8.9e-09 | 1.020E-08 | 3.345E-09 | 7.700E-09 | 1.400E-08 |
| 16580 | 829 | K829V | B | 2.9e-09 | 1.3e-08 | 9.1e-09 | 8.333E-09 | 5.093E-09 | 2.900E-09 | 1.300E-08 |
| 16581 | 830 | S830A | B | 2.5e-08 | 7.5e-09 | 1.2e-08 | 1.483E-08 | 9.088E-09 | 7.500E-09 | 2.500E-08 |
| 16582 | 830 | S830R | B | 1.8e-08 | 9.3e-09 | 4.8e-09 | 1.070E-08 | 6.710E-09 | 4.800E-09 | 1.800E-08 |
| 16583 | 830 | S830N | B | 5.6e-09 | 5.3e-09 | 5.4e-09 | 5.433E-09 | 1.528E-10 | 5.300E-09 | 5.600E-09 |
| 16584 | 830 | S830D | B | 5.1e-09 | 9e-09 | 6.1e-09 | 6.733E-09 | 2.026E-09 | 5.100E-09 | 9.000E-09 |
| 16585 | 830 | S830C | B | 7.9e-09 | 5.7e-09 | 1.3e-08 | 8.867E-09 | 3.745E-09 | 5.700E-09 | 1.300E-08 |
| 16586 | 830 | S830E | B | 4.9e-09 | 4.4e-09 | 2.1e-09 | 3.800E-09 | 1.493E-09 | 2.100E-09 | 4.900E-09 |
| 16587 | 830 | S830Q | B | 4.6e-09 | 3.7e-09 | 1.7e-08 | 8.433E-09 | 7.433E-09 | 3.700E-09 | 1.700E-08 |
| 16588 | 830 | S830G | B | 2.8e-09 | 1.9e-09 | 6.5e-09 | 3.733E-09 | 2.438E-09 | 1.900E-09 | 6.500E-09 |
| 16589 | 830 | S830H | B | 3.6e-09 | 1.6e-08 | 3.2e-09 | 7.600E-09 | 7.277E-09 | 3.200E-09 | 1.600E-08 |
| 16590 | 830 | S830I | B | 5.7e-09 | 9.7e-09 | 6.1e-09 | 7.167E-09 | 2.203E-09 | 5.700E-09 | 9.700E-09 |
| 16591 | 830 | S830L | B | 1.6e-08 | 9.8e-09 | 4.7e-09 | 1.017E-08 | 5.659E-09 | 4.700E-09 | 1.600E-08 |
| 16592 | 830 | S830K | B | 3.6e-09 | 5.2e-09 | 6.5e-09 | 5.100E-09 | 1.453E-09 | 3.600E-09 | 6.500E-09 |
| 16593 | 830 | S830M | B | 2.3e-09 | 4.2e-09 | 8.1e-09 | 4.867E-09 | 2.957E-09 | 2.300E-09 | 8.100E-09 |
| 16594 | 830 | S830F | B | 5.7e-09 | 1.3e-08 | 7.4e-09 | 8.700E-09 | 3.820E-09 | 5.700E-09 | 1.300E-08 |
| 16595 | 830 | S830P | B | 7.7e-09 | 2.8e-09 | 3.9e-09 | 4.800E-09 | 2.571E-09 | 2.800E-09 | 7.700E-09 |
| 16596 | 830 | S830S | B | 1.3e-08 | 4.7e-09 | 9.9e-09 | 9.200E-09 | 4.194E-09 | 4.700E-09 | 1.300E-08 |
| 16597 | 830 | S830T | B | 1.1e-08 | 5.2e-09 | 1e-08 | 8.733E-09 | 3.101E-09 | 5.200E-09 | 1.100E-08 |
| 16598 | 830 | S830W | B | 1.2e-08 | 4.7e-09 | 9.6e-09 | 8.767E-09 | 3.721E-09 | 4.700E-09 | 1.200E-08 |
| 16599 | 830 | S830Y | B | 4.7e-09 | 1.3e-08 | 7.9e-09 | 8.533E-09 | 4.186E-09 | 4.700E-09 | 1.300E-08 |
| 16600 | 830 | S830V | B | 1.8e-08 | 1e-08 | 9e-09 | 1.233E-08 | 4.933E-09 | 9.000E-09 | 1.800E-08 |
| 16601 | 831 | F831A | B | 7.4e-09 | 9.8e-09 | 6.9e-09 | 8.033E-09 | 1.550E-09 | 6.900E-09 | 9.800E-09 |
| 16602 | 831 | F831R | B | 1e-08 | 1e-08 | 4.3e-09 | 8.100E-09 | 3.291E-09 | 4.300E-09 | 1.000E-08 |
| 16603 | 831 | F831N | B | 4.6e-09 | 1e-08 | 3.1e-09 | 5.900E-09 | 3.629E-09 | 3.100E-09 | 1.000E-08 |
| 16604 | 831 | F831D | B | 8e-09 | 1.9e-08 | 4.2e-09 | 1.040E-08 | 7.686E-09 | 4.200E-09 | 1.900E-08 |
| 16605 | 831 | F831C | B | 1.2e-08 | 5.8e-09 | 1.1e-08 | 9.600E-09 | 3.329E-09 | 5.800E-09 | 1.200E-08 |
| 16606 | 831 | F831E | B | 2.8e-09 | 8.5e-09 | 4.1e-09 | 5.133E-09 | 2.987E-09 | 2.800E-09 | 8.500E-09 |
| 16607 | 831 | F831Q | B | 7e-09 | 1.2e-08 | 9.9e-09 | 9.633E-09 | 2.511E-09 | 7.000E-09 | 1.200E-08 |
| 16608 | 831 | F831G | B | 1.6e-08 | 4.8e-09 | 5.7e-09 | 8.833E-09 | 6.223E-09 | 4.800E-09 | 1.600E-08 |
| 16609 | 831 | F831H | B | 4.5e-09 | 9e-09 | 3.5e-09 | 5.667E-09 | 2.930E-09 | 3.500E-09 | 9.000E-09 |
| 16610 | 831 | F831I | B | 2.6e-08 | 1e-08 | 5e-09 | 1.367E-08 | 1.097E-08 | 5.000E-09 | 2.600E-08 |
| 16611 | 831 | F831L | B | 4.1e-09 | 7.1e-09 | 1.3e-08 | 8.067E-09 | 4.528E-09 | 4.100E-09 | 1.300E-08 |
| 16612 | 831 | F831K | B | 3e-09 | 1.2e-08 | 6.9e-09 | 7.300E-09 | 4.513E-09 | 3.000E-09 | 1.200E-08 |
| 16613 | 831 | F831M | B | 6e-09 | 2.3e-08 | 1e-08 | 1.300E-08 | 8.888E-09 | 6.000E-09 | 2.300E-08 |
| 16614 | 831 | F831F | B | 1.1e-08 | 1.6e-08 | 3.9e-09 | 1.030E-08 | 6.080E-09 | 3.900E-09 | 1.600E-08 |
| 16615 | 831 | F831P | B | 3.1e-09 | 1.7e-08 | 1.1e-08 | 1.037E-08 | 6.972E-09 | 3.100E-09 | 1.700E-08 |
| 16616 | 831 | F831S | B | 8.5e-09 | 2.3e-09 | 1.7e-08 | 9.267E-09 | 7.380E-09 | 2.300E-09 | 1.700E-08 |
| 16617 | 831 | F831T | B | 4.4e-09 | 1.3e-08 | 2.4e-09 | 6.600E-09 | 5.632E-09 | 2.400E-09 | 1.300E-08 |
| 16618 | 831 | F831W | B | 1.2e-08 | 7.4e-09 | 7.8e-09 | 9.067E-09 | 2.548E-09 | 7.400E-09 | 1.200E-08 |
| 16619 | 831 | F831Y | B | 1.1e-08 | 9.4e-09 | 2.9e-09 | 7.767E-09 | 4.290E-09 | 2.900E-09 | 1.100E-08 |
| 16620 | 831 | F831V | B | 8.5e-09 | 3.5e-09 | 5.6e-09 | 5.867E-09 | 2.511E-09 | 3.500E-09 | 8.500E-09 |
| 16621 | 832 | N832A | B | 6.7e-09 | 1.4e-08 | 5.1e-09 | 8.600E-09 | 4.744E-09 | 5.100E-09 | 1.400E-08 |
| 16622 | 832 | N832R | B | 6.1e-09 | 1.1e-08 | 6.8e-09 | 7.967E-09 | 2.650E-09 | 6.100E-09 | 1.100E-08 |
| 16623 | 832 | N832N | B | 1.8e-08 | 4.2e-09 | 1.5e-08 | 1.240E-08 | 7.258E-09 | 4.200E-09 | 1.800E-08 |
| 16624 | 832 | N832D | B | 8.3e-09 | 3.8e-09 | 2.7e-08 | 1.303E-08 | 1.230E-08 | 3.800E-09 | 2.700E-08 |
| 16625 | 832 | N832C | B | 8.1e-09 | 2.6e-08 | 2e-08 | 1.803E-08 | 9.111E-09 | 8.100E-09 | 2.600E-08 |
| 16626 | 832 | N832E | B | 1.7e-08 | 2.3e-09 | 1.3e-08 | 1.077E-08 | 7.600E-09 | 2.300E-09 | 1.700E-08 |
| 16627 | 832 | N832Q | B | 2.3e-08 | 6.6e-09 | 5.3e-09 | 1.163E-08 | 9.865E-09 | 5.300E-09 | 2.300E-08 |
| 16628 | 832 | N832G | B | 5.5e-09 | 5.6e-09 | 1.5e-08 | 8.700E-09 | 5.456E-09 | 5.500E-09 | 1.500E-08 |
| 16629 | 832 | N832H | B | 7.9e-09 | 8.7e-09 | 5.3e-09 | 7.300E-09 | 1.778E-09 | 5.300E-09 | 8.700E-09 |
| 16630 | 832 | N832I | B | 7.2e-09 | 4.7e-09 | 8.3e-09 | 6.733E-09 | 1.845E-09 | 4.700E-09 | 8.300E-09 |
| 16631 | 832 | N832L | B | 7.1e-09 | 4.1e-09 | 9.4e-09 | 6.867E-09 | 2.658E-09 | 4.100E-09 | 9.400E-09 |
| 16632 | 832 | N832K | B | 6.8e-09 | 5.4e-09 | 1.6e-08 | 9.400E-09 | 5.758E-09 | 5.400E-09 | 1.600E-08 |
| 16633 | 832 | N832M | B | 7e-09 | 4.1e-09 | 8.3e-09 | 6.467E-09 | 2.150E-09 | 4.100E-09 | 8.300E-09 |
| 16634 | 832 | N832F | B | 7.2e-09 | 1.3e-08 | 2.2e-08 | 1.407E-08 | 7.457E-09 | 7.200E-09 | 2.200E-08 |
| 16635 | 832 | N832P | B | 1e-08 | 8.9e-09 | 9.1e-09 | 9.333E-09 | 5.859E-10 | 8.900E-09 | 1.000E-08 |
| 16636 | 832 | N832S | B | 1.3e-08 | 1.3e-08 | 3.4e-09 | 9.800E-09 | 5.543E-09 | 3.400E-09 | 1.300E-08 |
| 16637 | 832 | N832T | B | 9.4e-09 | 1.4e-08 | 1.6e-08 | 1.313E-08 | 3.384E-09 | 9.400E-09 | 1.600E-08 |
| 16638 | 832 | N832W | B | 2.1e-08 | 2.6e-09 | 3.5e-09 | 9.033E-09 | 1.037E-08 | 2.600E-09 | 2.100E-08 |
| 16639 | 832 | N832Y | B | 1.3e-08 | 1.4e-08 | 1.4e-08 | 1.367E-08 | 5.774E-10 | 1.300E-08 | 1.400E-08 |
| 16640 | 832 | N832V | B | 4.2e-09 | 8.7e-09 | 8.9e-09 | 7.267E-09 | 2.658E-09 | 4.200E-09 | 8.900E-09 |
| 16641 | 833 | R833A | B | 9.3e-09 | 5.8e-09 | 9.1e-09 | 8.067E-09 | 1.966E-09 | 5.800E-09 | 9.300E-09 |
| 16642 | 833 | R833R | B | 1.3e-08 | 1.2e-08 | 5.5e-09 | 1.017E-08 | 4.072E-09 | 5.500E-09 | 1.300E-08 |
| 16643 | 833 | R833N | B | 1.2e-08 | 1.1e-08 | 1.4e-08 | 1.233E-08 | 1.528E-09 | 1.100E-08 | 1.400E-08 |
| 16644 | 833 | R833D | B | 9.3e-09 | 4.4e-09 | 3.5e-09 | 5.733E-09 | 3.121E-09 | 3.500E-09 | 9.300E-09 |
| 16645 | 833 | R833C | B | 5.5e-09 | 4.9e-09 | 4.3e-09 | 4.900E-09 | 6.000E-10 | 4.300E-09 | 5.500E-09 |
| 16646 | 833 | R833E | B | 7.1e-09 | 1.3e-08 | 2.8e-09 | 7.633E-09 | 5.121E-09 | 2.800E-09 | 1.300E-08 |
| 16647 | 833 | R833Q | B | 1.3e-08 | 5.1e-09 | 5.1e-09 | 7.733E-09 | 4.561E-09 | 5.100E-09 | 1.300E-08 |
| 16648 | 833 | R833G | B | 5.2e-09 | 1.3e-08 | 8.5e-09 | 8.900E-09 | 3.915E-09 | 5.200E-09 | 1.300E-08 |
| 16649 | 833 | R833H | B | 1.3e-08 | 3.1e-08 | 1.9e-09 | 1.530E-08 | 1.469E-08 | 1.900E-09 | 3.100E-08 |
| 16650 | 833 | R833I | B | 2.4e-09 | 1.1e-08 | 6.8e-09 | 6.733E-09 | 4.300E-09 | 2.400E-09 | 1.100E-08 |
| 16651 | 833 | R833L | B | 1.4e-08 | 4.2e-09 | 1.3e-08 | 1.040E-08 | 5.393E-09 | 4.200E-09 | 1.400E-08 |
| 16652 | 833 | R833K | B | 1.2e-08 | 4.6e-09 | 9.5e-10 | 5.850E-09 | 5.630E-09 | 9.500E-10 | 1.200E-08 |
| 16653 | 833 | R833M | B | 2.2e-08 | 6.9e-09 | 4.2e-09 | 1.103E-08 | 9.593E-09 | 4.200E-09 | 2.200E-08 |
| 16654 | 833 | R833F | B | 5.3e-09 | 1.1e-08 | 6.9e-09 | 7.733E-09 | 2.940E-09 | 5.300E-09 | 1.100E-08 |
| 16655 | 833 | R833P | B | 6.6e-09 | 7.3e-09 | 3e-09 | 5.633E-09 | 2.307E-09 | 3.000E-09 | 7.300E-09 |
| 16656 | 833 | R833S | B | 6.8e-09 | 5.1e-09 | 5.4e-09 | 5.767E-09 | 9.074E-10 | 5.100E-09 | 6.800E-09 |
| 16657 | 833 | R833T | B | 1.8e-09 | 4.5e-09 | 5.8e-09 | 4.033E-09 | 2.040E-09 | 1.800E-09 | 5.800E-09 |
| 16658 | 833 | R833W | B | 4.6e-09 | 2.2e-09 | 1.7e-08 | 7.933E-09 | 7.943E-09 | 2.200E-09 | 1.700E-08 |
| 16659 | 833 | R833Y | B | 4.4e-09 | 2.4e-09 | 6.4e-09 | 4.400E-09 | 2.000E-09 | 2.400E-09 | 6.400E-09 |

| | | | | | | | | | | |
|---|---|---|---|---|---|---|---|---|---|---|
| 16660 | 833 | R833V | B | 3.8e-09 | 7.9e-09 | 6.2e-09 | 5.967E-09 | 2.060E-09 | 3.800E-09 | 7.900E-09 |
| 16661 | 834 | G834A | B | 4.1e-09 | 1.2e-08 | 5.2e-09 | 7.100E-09 | 4.279E-09 | 4.100E-09 | 1.200E-08 |
| 16662 | 834 | G834R | B | 2.6e-09 | 1.1e-08 | 1.1e-08 | 8.200E-09 | 4.850E-09 | 2.600E-09 | 1.100E-08 |
| 16663 | 834 | G834N | B | 8.4e-09 | 9.3e-09 | 1.5e-08 | 1.090E-08 | 3.579E-09 | 8.400E-09 | 1.500E-08 |
| 16664 | 834 | G834D | B | 1.3e-08 | 8.5e-09 | 3.3e-09 | 8.267E-09 | 4.854E-09 | 3.300E-09 | 1.300E-08 |
| 16665 | 834 | G834C | B | 3.1e-09 | 1.4e-08 | 8.3e-09 | 8.467E-09 | 5.452E-09 | 3.100E-09 | 1.400E-08 |
| 16666 | 834 | G834E | B | 6.6e-09 | 6.5e-09 | 5.3e-09 | 6.133E-09 | 7.234E-10 | 5.300E-09 | 6.600E-09 |
| 16667 | 834 | G834Q | B | 4.3e-09 | 7.1e-09 | 3.2e-09 | 4.867E-09 | 2.011E-09 | 3.200E-09 | 7.100E-09 |
| 16668 | 834 | G834G | B | 5.2e-09 | 2.8e-09 | 1.4e-08 | 7.333E-09 | 5.897E-09 | 2.800E-09 | 1.400E-08 |
| 16669 | 834 | G834H | B | 6e-09 | 5.8e-09 | 8.5e-09 | 6.767E-09 | 1.504E-09 | 5.800E-09 | 8.500E-09 |
| 16670 | 834 | G834I | B | 2e-09 | 1.4e-08 | 7.4e-09 | 7.800E-09 | 6.010E-09 | 2.000E-09 | 1.400E-08 |
| 16671 | 834 | G834L | B | 7.1e-09 | 5.2e-09 | 1e-08 | 7.433E-09 | 2.417E-09 | 5.200E-09 | 1.000E-08 |
| 16672 | 834 | G834K | B | 2.3e-09 | 2e-09 | 7.2e-09 | 3.833E-09 | 2.919E-09 | 2.000E-09 | 7.200E-09 |
| 16673 | 834 | G834M | B | 8.3e-09 | 6.3e-09 | 8.6e-09 | 7.733E-09 | 1.250E-09 | 6.300E-09 | 8.600E-09 |
| 16674 | 834 | G834F | B | 5.8e-09 | 1.3e-08 | 4.3e-09 | 7.700E-09 | 4.651E-09 | 4.300E-09 | 1.300E-08 |
| 16675 | 834 | G834P | B | 9.8e-09 | 1.8e-08 | 7.7e-09 | 1.183E-08 | 5.443E-09 | 7.700E-09 | 1.800E-08 |
| 16676 | 834 | G834S | B | 9.3e-09 | 8.5e-09 | 7.4e-09 | 8.400E-09 | 9.539E-10 | 7.400E-09 | 9.300E-09 |
| 16677 | 834 | G834T | B | 1.9e-08 | 4.3e-09 | 3.3e-09 | 8.867E-09 | 8.790E-09 | 3.300E-09 | 1.900E-08 |
| 16678 | 834 | G834W | B | 7.1e-09 | 1e-08 | 1.3e-08 | 1.003E-08 | 2.950E-09 | 7.100E-09 | 1.300E-08 |
| 16679 | 834 | G834Y | B | 3.1e-09 | 8.5e-09 | 6.7e-09 | 6.100E-09 | 2.750E-09 | 3.100E-09 | 8.500E-09 |
| 16680 | 834 | G834V | B | 1.2e-08 | 8.9e-09 | 4.9e-09 | 8.600E-09 | 3.559E-09 | 4.900E-09 | 1.200E-08 |
| 16681 | 835 | E835A | B | 1.2e-08 | 5.1e-09 | 8.4e-09 | 8.500E-09 | 3.451E-09 | 5.100E-09 | 1.200E-08 |
| 16682 | 835 | E835R | B | 3.9e-08 | 1e-08 | 5.5e-09 | 1.817E-08 | 1.818E-08 | 5.500E-09 | 3.900E-08 |
| 16683 | 835 | E835N | B | 9.4e-09 | 4.9e-09 | 3.3e-09 | 5.867E-09 | 3.163E-09 | 3.300E-09 | 9.400E-09 |
| 16684 | 835 | E835D | B | 5.9e-09 | 6.5e-09 | 1.5e-08 | 9.133E-09 | 5.090E-09 | 5.900E-09 | 1.500E-08 |
| 16685 | 835 | E835C | B | 4.6e-09 | 9.4e-09 | 6.2e-09 | 6.733E-09 | 2.444E-09 | 4.600E-09 | 9.400E-09 |
| 16686 | 835 | E835E | B | 5.2e-09 | 5e-09 | 8.9e-09 | 6.367E-09 | 2.196E-09 | 5.000E-09 | 8.900E-09 |
| 16687 | 835 | E835Q | B | 3e-09 | 1.1e-08 | 7.5e-09 | 7.167E-09 | 4.010E-09 | 3.000E-09 | 1.100E-08 |
| 16688 | 835 | E835G | B | 7.5e-09 | 1.3e-08 | 9.2e-09 | 9.900E-09 | 2.816E-09 | 7.500E-09 | 1.300E-08 |
| 16689 | 835 | E835H | B | 5.4e-09 | 5.3e-09 | 8.4e-09 | 6.367E-09 | 1.762E-09 | 5.300E-09 | 8.400E-09 |
| 16690 | 835 | E835I | B | 2.3e-08 | 9.5e-09 | 4e-09 | 1.217E-08 | 9.777E-09 | 4.000E-09 | 2.300E-08 |
| 16691 | 835 | E835L | B | 6e-09 | 5.9e-09 | 7.6e-09 | 6.500E-09 | 9.539E-10 | 5.900E-09 | 7.600E-09 |
| 16692 | 835 | E835K | B | 1e-08 | 6e-09 | 4.2e-09 | 6.733E-09 | 2.969E-09 | 4.200E-09 | 1.000E-08 |
| 16693 | 835 | E835M | B | 1.9e-08 | 2e-08 | 2.2e-09 | 1.373E-08 | 1.000E-08 | 2.200E-09 | 2.000E-08 |
| 16694 | 835 | E835F | B | 9.9e-09 | 5.4e-09 | 5.4e-09 | 6.900E-09 | 2.598E-09 | 5.400E-09 | 9.900E-09 |
| 16695 | 835 | E835P | B | 7.5e-09 | 1.3e-08 | 1.1e-08 | 1.050E-08 | 2.784E-09 | 7.500E-09 | 1.300E-08 |
| 16696 | 835 | E835S | B | 7.4e-09 | 8.5e-09 | 1.1e-08 | 8.967E-09 | 1.845E-09 | 7.400E-09 | 1.100E-08 |
| 16697 | 835 | E835T | B | 4.3e-09 | 3e-09 | 7.7e-09 | 5.000E-09 | 2.427E-09 | 3.000E-09 | 7.700E-09 |
| 16698 | 835 | E835W | B | 4.9e-09 | 6.6e-09 | 6.8e-09 | 6.100E-09 | 1.044E-09 | 4.900E-09 | 6.800E-09 |
| 16699 | 835 | E835Y | B | 3.3e-09 | 2.4e-09 | 4.1e-09 | 3.267E-09 | 8.505E-10 | 2.400E-09 | 4.100E-09 |
| 16700 | 835 | E835V | B | 2.7e-08 | 1.8e-09 | 9.1e-09 | 1.263E-08 | 1.297E-08 | 1.800E-09 | 2.700E-08 |
| 16701 | 836 | C836A | B | 8.5e-09 | 1e-08 | 6e-09 | 8.167E-09 | 2.021E-09 | 6.000E-09 | 1.000E-08 |
| 16702 | 836 | C836R | B | 6.4e-09 | 3.1e-09 | 1.9e-08 | 9.500E-09 | 8.391E-09 | 3.100E-09 | 1.900E-08 |
| 16703 | 836 | C836N | B | 5.2e-09 | 2.1e-09 | 6.5e-09 | 4.600E-09 | 2.261E-09 | 2.100E-09 | 6.500E-09 |
| 16704 | 836 | C836D | B | 4.7e-09 | 3.9e-09 | 3.1e-09 | 3.900E-09 | 8.000E-10 | 3.100E-09 | 4.700E-09 |
| 16705 | 836 | C836C | B | 8.3e-09 | 3e-09 | 8.8e-09 | 6.700E-09 | 3.214E-09 | 3.000E-09 | 8.800E-09 |
| 16706 | 836 | C836E | B | 1.5e-08 | 4e-09 | 2.6e-09 | 7.200E-09 | 6.791E-09 | 2.600E-09 | 1.500E-08 |
| 16707 | 836 | C836Q | B | 1.4e-08 | 8.8e-09 | 5.9e-09 | 9.567E-09 | 4.104E-09 | 5.900E-09 | 1.400E-08 |
| 16708 | 836 | C836G | B | 6.9e-09 | 8.6e-09 | 3.2e-09 | 6.233E-09 | 2.761E-09 | 3.200E-09 | 8.600E-09 |
| 16709 | 836 | C836H | B | 8.9e-09 | 1.5e-08 | 1.2e-08 | 1.197E-08 | 3.050E-09 | 8.900E-09 | 1.500E-08 |
| 16710 | 836 | C836I | B | 1.1e-08 | 5.2e-09 | 7.9e-09 | 8.033E-09 | 2.902E-09 | 5.200E-09 | 1.100E-08 |
| 16711 | 836 | C836L | B | 4.2e-09 | 1.6e-08 | 9.5e-09 | 9.900E-09 | 5.910E-09 | 4.200E-09 | 1.600E-08 |
| 16712 | 836 | C836K | B | 5e-09 | 6.1e-09 | 5.1e-09 | 5.400E-09 | 6.083E-10 | 5.000E-09 | 6.100E-09 |
| 16713 | 836 | C836M | B | 4.7e-09 | 6.1e-09 | 3.1e-09 | 4.633E-09 | 1.501E-09 | 3.100E-09 | 6.100E-09 |
| 16714 | 836 | C836F | B | 6.5e-09 | 4.3e-09 | 5e-09 | 5.267E-09 | 1.124E-09 | 4.300E-09 | 6.500E-09 |
| 16715 | 836 | C836P | B | 1.4e-08 | 1.1e-08 | 4.3e-09 | 9.767E-09 | 4.966E-09 | 4.300E-09 | 1.400E-08 |
| 16716 | 836 | C836S | B | 1.1e-08 | 8e-09 | 8.5e-09 | 9.167E-09 | 1.607E-09 | 8.000E-09 | 1.100E-08 |
| 16717 | 836 | C836T | B | 4.3e-09 | 7.4e-09 | 4.3e-09 | 5.333E-09 | 1.790E-09 | 4.300E-09 | 7.400E-09 |
| 16718 | 836 | C836W | B | 2.5e-08 | 3.4e-09 | 1.9e-08 | 1.580E-08 | 1.115E-08 | 3.400E-09 | 2.500E-08 |
| 16719 | 836 | C836Y | B | 6.7e-09 | 3.4e-09 | 5.7e-09 | 5.267E-09 | 1.692E-09 | 3.400E-09 | 6.700E-09 |
| 16720 | 836 | C836V | B | 6.5e-09 | 8.6e-09 | 5e-09 | 6.700E-09 | 1.808E-09 | 5.000E-09 | 8.600E-09 |
| 16721 | 837 | E837A | C | 3.5e-09 | 3.8e-09 | 2e-09 | 3.100E-09 | 9.644E-10 | 2.000E-09 | 3.800E-09 |
| 16722 | 837 | E837R | C | 6.2e-09 | 3.4e-09 | 1.3e-08 | 7.533E-09 | 4.937E-09 | 3.400E-09 | 1.300E-08 |
| 16723 | 837 | E837N | C | 1.5e-08 | 3.8e-09 | 5.1e-09 | 7.967E-09 | 6.126E-09 | 3.800E-09 | 1.500E-08 |
| 16724 | 837 | E837D | C | 5.2e-09 | 1.1e-08 | 5.1e-09 | 7.100E-09 | 3.378E-09 | 5.100E-09 | 1.100E-08 |
| 16725 | 837 | E837C | C | 1.3e-08 | 8.2e-09 | 9.1e-09 | 1.010E-08 | 2.551E-09 | 8.200E-09 | 1.300E-08 |
| 16726 | 837 | E837E | C | 5.2e-09 | 5.4e-09 | 7.5e-09 | 6.033E-09 | 1.274E-09 | 5.200E-09 | 7.500E-09 |
| 16727 | 837 | E837Q | C | 4.1e-09 | 1.2e-08 | 8.6e-09 | 8.233E-09 | 3.963E-09 | 4.100E-09 | 1.200E-08 |
| 16728 | 837 | E837G | C | 1e-08 | 1.3e-08 | 5.2e-09 | 9.400E-09 | 3.934E-09 | 5.200E-09 | 1.300E-08 |
| 16729 | 837 | E837H | C | 9.5e-09 | 5.7e-09 | 9.9e-09 | 8.367E-09 | 2.318E-09 | 5.700E-09 | 9.900E-09 |
| 16730 | 837 | E837I | C | 6e-09 | 3.7e-09 | 5.8e-09 | 5.167E-09 | 1.274E-09 | 3.700E-09 | 6.000E-09 |
| 16731 | 837 | E837L | C | 2.3e-08 | 1.9e-08 | 9.5e-09 | 1.717E-08 | 6.934E-09 | 9.500E-09 | 2.300E-08 |
| 16732 | 837 | E837K | C | 1.2e-09 | 7.8e-09 | 4.6e-09 | 4.533E-09 | 3.301E-09 | 1.200E-09 | 7.800E-09 |
| 16733 | 837 | E837M | C | 4.5e-09 | 9.7e-09 | 1.5e-08 | 9.733E-09 | 5.250E-09 | 4.500E-09 | 1.500E-08 |
| 16734 | 837 | E837F | C | 7.7e-09 | 4.8e-09 | 1.4e-08 | 8.833E-09 | 4.704E-09 | 4.800E-09 | 1.400E-08 |
| 16735 | 837 | E837P | C | 5.6e-09 | 7.7e-09 | 8.3e-09 | 7.200E-09 | 1.418E-09 | 5.600E-09 | 8.300E-09 |
| 16736 | 837 | E837S | C | 5.7e-09 | 1.1e-08 | 8.8e-09 | 8.500E-09 | 2.663E-09 | 5.700E-09 | 1.100E-08 |
| 16737 | 837 | E837T | C | 1.1e-08 | 2.8e-09 | 4.7e-09 | 6.167E-09 | 4.292E-09 | 2.800E-09 | 1.100E-08 |
| 16738 | 837 | E837W | C | 3.2e-09 | 4.3e-09 | 7.3e-09 | 4.933E-09 | 2.122E-09 | 3.200E-09 | 7.300E-09 |
| 16739 | 837 | E837Y | C | 6.9e-09 | 8.9e-09 | 6e-09 | 7.267E-09 | 1.484E-09 | 6.000E-09 | 8.900E-09 |
| 16740 | 837 | E837V | C | 2.9e-09 | 3.6e-09 | 1.1e-08 | 5.833E-09 | 4.488E-09 | 2.900E-09 | 1.100E-08 |
| 16741 | 838 | V838A | C | 8.4e-09 | 9.6e-09 | 5.5e-09 | 7.833E-09 | 2.108E-09 | 5.500E-09 | 9.600E-09 |
| 16742 | 838 | V838R | C | 7e-09 | 2.1e-09 | 1.7e-08 | 8.700E-09 | 7.594E-09 | 2.100E-09 | 1.700E-08 |
| 16743 | 838 | V838N | C | 8.8e-09 | 8.5e-09 | 1e-08 | 9.100E-09 | 7.937E-10 | 8.500E-09 | 1.000E-08 |
| 16744 | 838 | V838D | C | 3.7e-09 | 7e-09 | 1.6e-08 | 8.900E-09 | 6.366E-09 | 3.700E-09 | 1.600E-08 |

| | | | | | | | | | | |
|---|---|---|---|---|---|---|---|---|---|---|
| 16745 | 838 | V838C | C | 1.8e-08 | 6e-09 | 8.5e-09 | 1.083E-08 | 6.331E-09 | 6.000E-09 | 1.800E-08 |
| 16746 | 838 | V838E | C | 4.3e-09 | 1.6e-08 | 5.7e-09 | 8.667E-09 | 6.389E-09 | 4.300E-09 | 1.600E-08 |
| 16747 | 838 | V838Q | C | 6.9e-09 | 7e-09 | 7.4e-09 | 7.100E-09 | 2.646E-10 | 6.900E-09 | 7.400E-09 |
| 16748 | 838 | V838G | C | 2.1e-08 | 4.6e-09 | 3.4e-09 | 9.667E-09 | 9.833E-09 | 3.400E-09 | 2.100E-08 |
| 16749 | 838 | V838H | C | 8.8e-09 | 1.2e-08 | 6.7e-09 | 9.167E-09 | 2.669E-09 | 6.700E-09 | 1.200E-08 |
| 16750 | 838 | V838I | C | 3.2e-09 | 3.1e-09 | 4.5e-09 | 3.600E-09 | 7.810E-10 | 3.100E-09 | 4.500E-09 |
| 16751 | 838 | V838L | C | 4.6e-09 | 1.3e-09 | 9.9e-09 | 5.267E-09 | 4.339E-09 | 1.300E-09 | 9.900E-09 |
| 16752 | 838 | V838K | C | 1.3e-08 | 9.4e-09 | 5.5e-09 | 9.300E-09 | 3.751E-09 | 5.500E-09 | 1.300E-08 |
| 16753 | 838 | V838M | C | 5.1e-09 | 1.2e-08 | 1.3e-08 | 1.003E-08 | 4.302E-09 | 5.100E-09 | 1.300E-08 |
| 16754 | 838 | V838F | C | 9.6e-09 | 7.4e-09 | 5.8e-09 | 7.600E-09 | 1.908E-09 | 5.800E-09 | 9.600E-09 |
| 16755 | 838 | V838P | C | 5.6e-09 | 8.8e-09 | 3.4e-09 | 5.933E-09 | 2.715E-09 | 3.400E-09 | 8.800E-09 |
| 16756 | 838 | V838S | C | 5.2e-09 | 5.9e-09 | 8.2e-09 | 6.433E-09 | 1.570E-09 | 5.200E-09 | 8.200E-09 |
| 16757 | 838 | V838T | C | 8.8e-09 | 2.4e-09 | 1.3e-08 | 8.067E-09 | 5.338E-09 | 2.400E-09 | 1.300E-08 |
| 16758 | 838 | V838W | C | 7.3e-09 | 4.1e-09 | 7.9e-09 | 6.433E-09 | 2.043E-09 | 4.100E-09 | 7.900E-09 |
| 16759 | 838 | V838Y | C | 6.3e-09 | 6e-09 | 3.7e-09 | 5.333E-09 | 1.422E-09 | 3.700E-09 | 6.300E-09 |
| 16760 | 838 | V838V | C | 8.7e-09 | 3.5e-09 | 1.4e-08 | 8.733E-09 | 5.250E-09 | 3.500E-09 | 1.400E-08 |
| 16761 | 839 | Q839A | C | 6.7e-09 | 6e-09 | 2.9e-08 | 1.390E-08 | 1.308E-08 | 6.000E-09 | 2.900E-08 |
| 16762 | 839 | Q839R | C | 7.3e-09 | 1e-08 | 2.2e-09 | 6.500E-09 | 3.961E-09 | 2.200E-09 | 1.000E-08 |
| 16763 | 839 | Q839N | C | 1.5e-08 | 1.1e-08 | 4.2e-09 | 1.007E-08 | 5.460E-09 | 4.200E-09 | 1.500E-08 |
| 16764 | 839 | Q839D | C | 1.1e-08 | 6.4e-09 | 5.8e-09 | 7.733E-09 | 2.845E-09 | 5.800E-09 | 1.100E-08 |
| 16765 | 839 | Q839C | C | 6.9e-09 | 2.4e-09 | 4.9e-09 | 4.733E-09 | 2.255E-09 | 2.400E-09 | 6.900E-09 |
| 16766 | 839 | Q839E | C | 3.5e-09 | 1.6e-08 | 2.8e-09 | 7.433E-09 | 7.427E-09 | 2.800E-09 | 1.600E-08 |
| 16767 | 839 | Q839Q | C | 8e-09 | 8.9e-09 | 5.4e-09 | 7.433E-09 | 1.818E-09 | 5.400E-09 | 8.900E-09 |
| 16768 | 839 | Q839G | C | 9.2e-09 | 9.5e-09 | 4.4e-09 | 7.700E-09 | 2.862E-09 | 4.400E-09 | 9.500E-09 |
| 16769 | 839 | Q839H | C | 1.2e-08 | 8.1e-09 | 7.8e-09 | 9.300E-09 | 2.343E-09 | 7.800E-09 | 1.200E-08 |
| 16770 | 839 | Q839I | C | 1e-08 | 1.9e-08 | 1.5e-08 | 1.467E-08 | 4.509E-09 | 1.000E-08 | 1.900E-08 |
| 16771 | 839 | Q839L | C | 3.2e-09 | 1.3e-08 | 1.7e-09 | 5.967E-09 | 6.137E-09 | 1.700E-09 | 1.300E-08 |
| 16772 | 839 | Q839K | C | 1.5e-08 | 7.8e-09 | 1.9e-09 | 8.233E-09 | 6.561E-09 | 1.900E-09 | 1.500E-08 |
| 16773 | 839 | Q839M | C | 3e-09 | 6.1e-09 | 7.1e-09 | 5.400E-09 | 2.138E-09 | 3.000E-09 | 7.100E-09 |
| 16774 | 839 | Q839F | C | 5e-09 | 8.1e-09 | 1.3e-08 | 8.700E-09 | 4.034E-09 | 5.000E-09 | 1.300E-08 |
| 16775 | 839 | Q839P | C | 1.2e-08 | 1.4e-08 | 1.7e-08 | 1.433E-08 | 2.517E-09 | 1.200E-08 | 1.700E-08 |
| 16776 | 839 | Q839S | C | 3.4e-09 | 6.5e-09 | 1.3e-08 | 7.633E-09 | 4.899E-09 | 3.400E-09 | 1.300E-08 |
| 16777 | 839 | Q839T | C | 1.3e-08 | 6.6e-09 | 6.8e-09 | 8.800E-09 | 3.639E-09 | 6.600E-09 | 1.300E-08 |
| 16778 | 839 | Q839W | C | 4.6e-09 | 6e-09 | 2e-09 | 4.200E-09 | 2.030E-09 | 2.000E-09 | 6.000E-09 |
| 16779 | 839 | Q839Y | C | 1.4e-08 | 3.3e-09 | 6.6e-09 | 7.967E-09 | 5.479E-09 | 3.300E-09 | 1.400E-08 |
| 16780 | 839 | Q839V | C | 9.7e-09 | 1.9e-08 | 5.5e-09 | 1.140E-08 | 6.909E-09 | 5.500E-09 | 1.900E-08 |
| 16781 | 840 | L840A | C | 1.4e-08 | 6.3e-09 | 5.6e-09 | 8.633E-09 | 4.661E-09 | 5.600E-09 | 1.400E-08 |
| 16782 | 840 | L840R | C | 9.7e-09 | 1.4e-08 | 9.6e-09 | 1.110E-08 | 2.512E-09 | 9.600E-09 | 1.400E-08 |
| 16783 | 840 | L840N | C | 6e-09 | 9.7e-09 | 3.9e-09 | 6.533E-09 | 2.937E-09 | 3.900E-09 | 9.700E-09 |
| 16784 | 840 | L840D | C | 9.3e-09 | 3.3e-09 | 7.2e-09 | 6.600E-09 | 3.045E-09 | 3.300E-09 | 9.300E-09 |
| 16785 | 840 | L840C | C | 8.4e-09 | 1.4e-08 | 1.4e-08 | 1.213E-08 | 3.233E-09 | 8.400E-09 | 1.400E-08 |
| 16786 | 840 | L840E | C | 8.2e-09 | 1.2e-08 | 6.9e-09 | 9.033E-09 | 2.650E-09 | 6.900E-09 | 1.200E-08 |
| 16787 | 840 | L840Q | C | 1.3e-08 | 6.9e-09 | 1.4e-08 | 1.130E-08 | 3.843E-09 | 6.900E-09 | 1.400E-08 |
| 16788 | 840 | L840G | C | 5.9e-09 | 4e-09 | 3e-09 | 4.300E-09 | 1.473E-09 | 3.000E-09 | 5.900E-09 |
| 16789 | 840 | L840H | C | 8.2e-09 | 3.3e-09 | 8.6e-09 | 6.700E-09 | 2.951E-09 | 3.300E-09 | 8.600E-09 |
| 16790 | 840 | L840I | C | 3.5e-09 | 2.6e-09 | 7e-09 | 4.367E-09 | 2.325E-09 | 2.600E-09 | 7.000E-09 |
| 16791 | 840 | L840L | C | 4.6e-09 | 1e-08 | 1.2e-08 | 8.867E-09 | 3.828E-09 | 4.600E-09 | 1.200E-08 |
| 16792 | 840 | L840K | C | 3.5e-09 | 3.6e-09 | 5e-09 | 4.033E-09 | 8.386E-10 | 3.500E-09 | 5.000E-09 |
| 16793 | 840 | L840M | C | 9.3e-09 | 3.1e-09 | 2.7e-09 | 5.033E-09 | 3.700E-09 | 2.700E-09 | 9.300E-09 |
| 16794 | 840 | L840F | C | 3.8e-09 | 1.3e-08 | 5.3e-09 | 7.367E-09 | 4.936E-09 | 3.800E-09 | 1.300E-08 |
| 16795 | 840 | L840P | C | 4.4e-09 | 6.7e-09 | 7.3e-09 | 6.133E-09 | 1.531E-09 | 4.400E-09 | 7.300E-09 |
| 16796 | 840 | L840S | C | 9.2e-09 | 3.6e-09 | 7e-09 | 6.600E-09 | 2.821E-09 | 3.600E-09 | 9.200E-09 |
| 16797 | 840 | L840T | C | 4.8e-09 | 8.1e-09 | 5.1e-09 | 6.000E-09 | 1.825E-09 | 4.800E-09 | 8.100E-09 |
| 16798 | 840 | L840W | C | 1.4e-08 | 7.6e-09 | 1e-08 | 1.053E-08 | 3.233E-09 | 7.600E-09 | 1.400E-08 |
| 16799 | 840 | L840Y | C | 6e-09 | 1.1e-08 | 9.8e-09 | 8.933E-09 | 2.610E-09 | 6.000E-09 | 1.100E-08 |
| 16800 | 840 | L840V | C | 4.6e-09 | 4.3e-09 | 1e-08 | 6.300E-09 | 3.208E-09 | 4.300E-09 | 1.000E-08 |
| 16801 | 841 | V841A | C | 6.5e-09 | 5.4e-09 | 2.3e-08 | 1.163E-08 | 9.859E-09 | 5.400E-09 | 2.300E-08 |
| 16802 | 841 | V841R | C | 6.4e-09 | 1.8e-08 | 6.3e-09 | 1.023E-08 | 6.726E-09 | 6.300E-09 | 1.800E-08 |
| 16803 | 841 | V841N | C | 6.2e-09 | 3.1e-09 | 7.9e-09 | 5.733E-09 | 2.434E-09 | 3.100E-09 | 7.900E-09 |
| 16804 | 841 | V841D | C | 4.5e-09 | 1e-08 | 4.5e-09 | 6.333E-09 | 3.175E-09 | 4.500E-09 | 1.000E-08 |
| 16805 | 841 | V841C | C | 4.6e-09 | 1.5e-08 | 1.1e-08 | 1.020E-08 | 5.246E-09 | 4.600E-09 | 1.500E-08 |
| 16806 | 841 | V841E | C | 4.3e-09 | 4.7e-09 | 7.2e-09 | 5.400E-09 | 1.572E-09 | 4.300E-09 | 7.200E-09 |
| 16807 | 841 | V841Q | C | 9.3e-09 | 8.3e-09 | 1.1e-08 | 9.533E-09 | 1.365E-09 | 8.300E-09 | 1.100E-08 |
| 16808 | 841 | V841G | C | 2.6e-09 | 2.7e-08 | 7.4e-09 | 1.233E-08 | 1.293E-08 | 2.600E-09 | 2.700E-08 |
| 16809 | 841 | V841H | C | 6.7e-09 | 5e-09 | 8.7e-09 | 6.800E-09 | 1.852E-09 | 5.000E-09 | 8.700E-09 |
| 16810 | 841 | V841I | C | 7.6e-09 | 7.1e-09 | 8.5e-09 | 7.733E-09 | 7.095E-10 | 7.100E-09 | 8.500E-09 |
| 16811 | 841 | V841L | C | 3.7e-09 | 3.7e-09 | 4.5e-09 | 3.967E-09 | 4.619E-10 | 3.700E-09 | 4.500E-09 |
| 16812 | 841 | V841K | C | 5.6e-09 | 1.3e-08 | 4.3e-09 | 7.633E-09 | 4.693E-09 | 4.300E-09 | 1.300E-08 |
| 16813 | 841 | V841M | C | 3.4e-09 | 5.5e-09 | 3.3e-09 | 4.067E-09 | 1.242E-09 | 3.300E-09 | 5.500E-09 |
| 16814 | 841 | V841F | C | 1.8e-08 | 9.1e-09 | 9.4e-09 | 1.217E-08 | 5.054E-09 | 9.100E-09 | 1.800E-08 |
| 16815 | 841 | V841P | C | 5.5e-09 | 8.9e-09 | 7.3e-09 | 7.233E-09 | 1.701E-09 | 5.500E-09 | 8.900E-09 |
| 16816 | 841 | V841S | C | 1e-08 | 3.8e-09 | 1.3e-08 | 8.933E-09 | 4.692E-09 | 3.800E-09 | 1.300E-08 |
| 16817 | 841 | V841T | C | 1.3e-08 | 8.3e-09 | 2e-09 | 7.767E-09 | 5.519E-09 | 2.000E-09 | 1.300E-08 |
| 16818 | 841 | V841W | C | 8.8e-09 | 6.7e-09 | 5.4e-09 | 6.967E-09 | 1.716E-09 | 5.400E-09 | 8.800E-09 |
| 16819 | 841 | V841Y | C | 3.1e-09 | 3.8e-09 | 5.8e-09 | 4.233E-09 | 1.401E-09 | 3.100E-09 | 5.800E-09 |
| 16820 | 841 | V841V | C | 5.3e-09 | 7.8e-09 | 4.1e-09 | 5.733E-09 | 1.888E-09 | 4.100E-09 | 7.800E-09 |
| 16821 | 842 | E842A | C | 6.5e-09 | 6.3e-09 | 4.6e-09 | 5.800E-09 | 1.044E-09 | 4.600E-09 | 6.500E-09 |
| 16822 | 842 | E842R | C | 1.8e-08 | 2.9e-09 | 8.6e-09 | 9.833E-09 | 7.625E-09 | 2.900E-09 | 1.800E-08 |
| 16823 | 842 | E842N | C | 1.7e-08 | 7e-09 | 1.2e-08 | 1.200E-08 | 5.000E-09 | 7.000E-09 | 1.700E-08 |
| 16824 | 842 | E842D | C | 8e-09 | 8.7e-09 | 1.3e-08 | 9.900E-09 | 2.707E-09 | 8.000E-09 | 1.300E-08 |
| 16825 | 842 | E842C | C | 4.9e-09 | 2.8e-09 | 1.1e-08 | 6.233E-09 | 4.259E-09 | 2.800E-09 | 1.100E-08 |
| 16826 | 842 | E842E | C | 1.1e-08 | 2e-09 | 7.7e-09 | 6.900E-09 | 4.553E-09 | 2.000E-09 | 1.100E-08 |
| 16827 | 842 | E842Q | C | 9.4e-09 | 3.1e-09 | 1.6e-08 | 9.500E-09 | 6.451E-09 | 3.100E-09 | 1.600E-08 |
| 16828 | 842 | E842G | C | 7.1e-09 | 1.3e-08 | 1.3e-08 | 1.103E-08 | 3.406E-09 | 7.100E-09 | 1.300E-08 |
| 16829 | 842 | E842H | C | 5.9e-09 | 3.3e-09 | 2.4e-09 | 3.867E-09 | 1.818E-09 | 2.400E-09 | 5.900E-09 |

| | | | | | | | | | | |
|---|---|---|---|---|---|---|---|---|---|---|
| 16830 | 842 | E842I | C | 4.3e-09 | 3.8e-09 | 4.8e-09 | 4.300E-09 | 5.000E-10 | 3.800E-09 | 4.800E-09 |
| 16831 | 842 | E842L | C | 5.1e-09 | 4.1e-08 | 2.5e-08 | 2.370E-08 | 1.799E-08 | 5.100E-09 | 4.100E-08 |
| 16832 | 842 | E842K | C | 9.2e-09 | 7.5e-09 | 1.8e-09 | 6.167E-09 | 3.876E-09 | 1.800E-09 | 9.200E-09 |
| 16833 | 842 | E842M | C | 9e-09 | 4.5e-09 | 5.1e-09 | 6.200E-09 | 2.443E-09 | 4.500E-09 | 9.000E-09 |
| 16834 | 842 | E842F | C | 9e-09 | 1.3e-08 | 1.7e-09 | 7.900E-09 | 5.730E-09 | 1.700E-09 | 1.300E-08 |
| 16835 | 842 | E842P | C | 1.4e-08 | 5.7e-09 | 9.4e-09 | 9.700E-09 | 4.158E-09 | 5.700E-09 | 1.400E-08 |
| 16836 | 842 | E842S | C | 5.3e-09 | 1.2e-08 | 7.6e-09 | 8.300E-09 | 3.404E-09 | 5.300E-09 | 1.200E-08 |
| 16837 | 842 | E842T | C | 4.8e-09 | 6.5e-09 | 5.5e-09 | 5.600E-09 | 8.544E-10 | 4.800E-09 | 6.500E-09 |
| 16838 | 842 | E842W | C | 6.5e-09 | 1.9e-09 | 6.3e-09 | 4.900E-09 | 2.600E-09 | 1.900E-09 | 6.500E-09 |
| 16839 | 842 | E842Y | C | 6e-09 | 6.3e-09 | 4e-09 | 5.433E-09 | 1.250E-09 | 4.000E-09 | 6.300E-09 |
| 16840 | 842 | E842V | C | 2e-08 | 7.3e-09 | 1.8e-08 | 1.510E-08 | 6.829E-09 | 7.300E-09 | 2.000E-08 |
| 16841 | 843 | S843A | C | 3.5e-09 | 2.5e-09 | 2.2e-08 | 9.333E-09 | 1.098E-08 | 2.500E-09 | 2.200E-08 |
| 16842 | 843 | S843R | C | 5.9e-09 | 2.3e-08 | 2.4e-09 | 1.043E-08 | 1.102E-08 | 2.400E-09 | 2.300E-08 |
| 16843 | 843 | S843N | C | 5.8e-09 | 4.6e-09 | 2.4e-08 | 1.147E-08 | 1.087E-08 | 4.600E-09 | 2.400E-08 |
| 16844 | 843 | S843D | C | 4e-09 | 3.8e-09 | 1.7e-08 | 8.267E-09 | 7.564E-09 | 3.800E-09 | 1.700E-08 |
| 16845 | 843 | S843C | C | 4.1e-09 | 8.8e-09 | 4.5e-09 | 5.800E-09 | 2.606E-09 | 4.100E-09 | 8.800E-09 |
| 16846 | 843 | S843E | C | 8.3e-09 | 1.1e-08 | 8.5e-09 | 9.267E-09 | 1.504E-09 | 8.300E-09 | 1.100E-08 |
| 16847 | 843 | S843Q | C | 1.5e-08 | 7.3e-09 | 1.2e-08 | 1.143E-08 | 3.881E-09 | 7.300E-09 | 1.500E-08 |
| 16848 | 843 | S843G | C | 5.6e-09 | 2.3e-09 | 7.2e-09 | 5.033E-09 | 2.499E-09 | 2.300E-09 | 7.200E-09 |
| 16849 | 843 | S843H | C | 2.5e-08 | 8e-09 | 3.3e-09 | 1.210E-08 | 1.142E-08 | 3.300E-09 | 2.500E-08 |
| 16850 | 843 | S843I | C | 5.7e-09 | 3.7e-09 | 8.5e-09 | 5.967E-09 | 2.411E-09 | 3.700E-09 | 8.500E-09 |
| 16851 | 843 | S843L | C | 6.4e-09 | 7.5e-09 | 9.3e-09 | 7.733E-09 | 1.464E-09 | 6.400E-09 | 9.300E-09 |
| 16852 | 843 | S843K | C | 5.9e-09 | 4.3e-09 | 1.8e-08 | 9.400E-09 | 7.491E-09 | 4.300E-09 | 1.800E-08 |
| 16853 | 843 | S843M | C | 2.7e-09 | 7e-09 | 7.4e-09 | 5.700E-09 | 2.606E-09 | 2.700E-09 | 7.400E-09 |
| 16854 | 843 | S843F | C | 7.6e-09 | 2.4e-09 | 4e-09 | 4.667E-09 | 2.663E-09 | 2.400E-09 | 7.600E-09 |
| 16855 | 843 | S843P | C | 5.1e-09 | 5.3e-09 | 5.2e-09 | 5.200E-09 | 1.000E-10 | 5.100E-09 | 5.300E-09 |
| 16856 | 843 | S843S | C | 6.2e-09 | 4.9e-09 | 5.2e-09 | 5.433E-09 | 6.807E-10 | 4.900E-09 | 6.200E-09 |
| 16857 | 843 | S843T | C | 9e-09 | 2.5e-09 | 7.5e-09 | 6.333E-09 | 3.403E-09 | 2.500E-09 | 9.000E-09 |
| 16858 | 843 | S843W | C | 3e-09 | 1.4e-08 | 4e-09 | 7.000E-09 | 6.083E-09 | 3.000E-09 | 1.400E-08 |
| 16859 | 843 | S843Y | C | 8.7e-09 | 3e-09 | 3.6e-09 | 5.100E-09 | 3.132E-09 | 3.000E-09 | 8.700E-09 |
| 16860 | 843 | S843V | C | 1e-08 | 8.5e-09 | 1e-08 | 9.500E-09 | 8.660E-10 | 8.500E-09 | 1.000E-08 |
| 16861 | 844 | G844A | C | 4e-09 | 4.6e-09 | 3.7e-09 | 4.100E-09 | 4.583E-10 | 3.700E-09 | 4.600E-09 |
| 16862 | 844 | G844R | C | 1.1e-08 | 8.3e-09 | 1.8e-08 | 1.243E-08 | 5.006E-09 | 8.300E-09 | 1.800E-08 |
| 16863 | 844 | G844N | C | 9e-09 | 2.5e-09 | 1.6e-08 | 9.167E-09 | 6.752E-09 | 2.500E-09 | 1.600E-08 |
| 16864 | 844 | G844D | C | 5.5e-09 | 4.1e-09 | 4.9e-09 | 4.833E-09 | 7.024E-10 | 4.100E-09 | 5.500E-09 |
| 16865 | 844 | G844C | C | 6e-09 | 7.5e-09 | 1.2e-08 | 8.500E-09 | 3.122E-09 | 6.000E-09 | 1.200E-08 |
| 16866 | 844 | G844E | C | 4.9e-09 | 6.1e-09 | 4.5e-09 | 5.167E-09 | 8.327E-10 | 4.500E-09 | 6.100E-09 |
| 16867 | 844 | G844Q | C | 1e-08 | 7.7e-09 | 1.4e-08 | 1.057E-08 | 3.188E-09 | 7.700E-09 | 1.400E-08 |
| 16868 | 844 | G844G | C | 1.4e-08 | 6.5e-09 | 9.7e-09 | 1.007E-08 | 3.763E-09 | 6.500E-09 | 1.400E-08 |
| 16869 | 844 | G844H | C | 4.2e-09 | 8.9e-09 | 5e-09 | 6.033E-09 | 2.515E-09 | 4.200E-09 | 8.900E-09 |
| 16870 | 844 | G844I | C | 2.6e-09 | 7.1e-09 | 7.4e-09 | 5.700E-09 | 2.689E-09 | 2.600E-09 | 7.400E-09 |
| 16871 | 844 | G844L | C | 2.1e-09 | 4.2e-09 | 5.4e-09 | 3.900E-09 | 1.670E-09 | 2.100E-09 | 5.400E-09 |
| 16872 | 844 | G844K | C | 8e-09 | 1e-08 | 5.2e-09 | 7.733E-09 | 2.411E-09 | 5.200E-09 | 1.000E-08 |
| 16873 | 844 | G844M | C | 6.4e-09 | 8.4e-09 | 6.3e-09 | 7.033E-09 | 1.185E-09 | 6.300E-09 | 8.400E-09 |
| 16874 | 844 | G844F | C | 1.3e-08 | 3.3e-09 | 1.3e-08 | 9.767E-09 | 5.600E-09 | 3.300E-09 | 1.300E-08 |
| 16875 | 844 | G844P | C | 1.2e-08 | 9.7e-09 | 4e-09 | 8.567E-09 | 4.119E-09 | 4.000E-09 | 1.200E-08 |
| 16876 | 844 | G844S | C | 8.3e-09 | 1.1e-08 | 1.6e-08 | 1.177E-08 | 3.907E-09 | 8.300E-09 | 1.600E-08 |
| 16877 | 844 | G844T | C | 7.1e-09 | 4.6e-09 | 3.7e-09 | 5.133E-09 | 1.762E-09 | 3.700E-09 | 7.100E-09 |
| 16878 | 844 | G844W | C | 3.6e-09 | 1.2e-08 | 7.7e-09 | 7.767E-09 | 4.200E-09 | 3.600E-09 | 1.200E-08 |
| 16879 | 844 | G844Y | C | 1.4e-08 | 1.1e-08 | 2e-08 | 1.500E-08 | 4.583E-09 | 1.100E-08 | 2.000E-08 |
| 16880 | 844 | G844V | C | 7.9e-09 | 5.2e-09 | 4.1e-09 | 5.733E-09 | 1.955E-09 | 4.100E-09 | 7.900E-09 |
| 16881 | 845 | G845A | C | 6.6e-09 | 3.6e-09 | 5.9e-09 | 5.367E-09 | 1.570E-09 | 3.600E-09 | 6.600E-09 |
| 16882 | 845 | G845R | C | 2.7e-09 | 1.1e-08 | 1.4e-08 | 9.233E-09 | 5.853E-09 | 2.700E-09 | 1.400E-08 |
| 16883 | 845 | G845N | C | 4.7e-09 | 6.4e-09 | 3.5e-09 | 4.867E-09 | 1.457E-09 | 3.500E-09 | 6.400E-09 |
| 16884 | 845 | G845D | C | 2.9e-09 | 4.9e-09 | 3.3e-09 | 3.700E-09 | 1.058E-09 | 2.900E-09 | 4.900E-09 |
| 16885 | 845 | G845C | C | 1.3e-08 | 2.6e-09 | 2.6e-09 | 6.067E-09 | 6.004E-09 | 2.600E-09 | 1.300E-08 |
| 16886 | 845 | G845E | C | 7.1e-09 | 1.1e-08 | 1.4e-08 | 1.070E-08 | 3.460E-09 | 7.100E-09 | 1.400E-08 |
| 16887 | 845 | G845Q | C | 7.3e-09 | 7e-09 | 1.2e-08 | 8.767E-09 | 2.804E-09 | 7.000E-09 | 1.200E-08 |
| 16888 | 845 | G845G | C | 1.2e-08 | 1.9e-08 | 3.1e-09 | 1.137E-08 | 7.969E-09 | 3.100E-09 | 1.900E-08 |
| 16889 | 845 | G845H | C | 3.7e-09 | 5.1e-09 | 8.3e-09 | 5.700E-09 | 2.358E-09 | 3.700E-09 | 8.300E-09 |
| 16890 | 845 | G845I | C | 4.5e-09 | 8.5e-09 | 9.5e-09 | 7.500E-09 | 2.646E-09 | 4.500E-09 | 9.500E-09 |
| 16891 | 845 | G845L | C | 1.4e-08 | 9e-09 | 1.3e-08 | 1.200E-08 | 2.646E-09 | 9.000E-09 | 1.400E-08 |
| 16892 | 845 | G845K | C | 9e-09 | 8.4e-09 | 5e-09 | 7.467E-09 | 2.157E-09 | 5.000E-09 | 9.000E-09 |
| 16893 | 845 | G845M | C | 8.2e-09 | 6.5e-09 | 1.7e-09 | 5.467E-09 | 3.371E-09 | 1.700E-09 | 8.200E-09 |
| 16894 | 845 | G845F | C | 6.1e-08 | 7.7e-09 | 2.3e-08 | 3.057E-08 | 2.744E-08 | 7.700E-09 | 6.100E-08 |
| 16895 | 845 | G845P | C | 7.1e-09 | 7.8e-09 | 1.5e-08 | 9.967E-09 | 4.373E-09 | 7.100E-09 | 1.500E-08 |
| 16896 | 845 | G845S | C | 5.7e-09 | 1.7e-09 | 1e-08 | 5.800E-09 | 4.151E-09 | 1.700E-09 | 1.000E-08 |
| 16897 | 845 | G845T | C | 2.9e-09 | 5.9e-09 | 1.1e-08 | 6.600E-09 | 4.095E-09 | 2.900E-09 | 1.100E-08 |
| 16898 | 845 | G845W | C | 1.5e-08 | 9.2e-09 | 1.7e-09 | 8.633E-09 | 6.668E-09 | 1.700E-09 | 1.500E-08 |
| 16899 | 845 | G845Y | C | 4.5e-09 | 9.2e-09 | 3.4e-09 | 5.700E-09 | 3.081E-09 | 3.400E-09 | 9.200E-09 |
| 16900 | 845 | G845V | C | 6.1e-09 | 6.2e-09 | 9.6e-09 | 7.300E-09 | 1.992E-09 | 6.100E-09 | 9.600E-09 |
| 16901 | 846 | G846A | C | 2.7e-09 | 5.2e-09 | 1.1e-08 | 6.300E-09 | 4.258E-09 | 2.700E-09 | 1.100E-08 |
| 16902 | 846 | G846R | C | 5.2e-09 | 8e-09 | 7.1e-09 | 6.767E-09 | 1.429E-09 | 5.200E-09 | 8.000E-09 |
| 16903 | 846 | G846N | C | 1.1e-08 | 4.9e-09 | 7.6e-09 | 7.833E-09 | 3.057E-09 | 4.900E-09 | 1.100E-08 |
| 16904 | 846 | G846D | C | 1.3e-08 | 6.8e-09 | 4.8e-09 | 8.200E-09 | 4.276E-09 | 4.800E-09 | 1.300E-08 |
| 16905 | 846 | G846C | C | 2.4e-09 | 5.4e-09 | 4e-09 | 3.933E-09 | 1.501E-09 | 2.400E-09 | 5.400E-09 |
| 16906 | 846 | G846E | C | 6.8e-09 | 9.2e-09 | 8.6e-09 | 8.200E-09 | 1.249E-09 | 6.800E-09 | 9.200E-09 |
| 16907 | 846 | G846Q | C | 4.6e-09 | 1.2e-08 | 1.1e-08 | 9.200E-09 | 4.015E-09 | 4.600E-09 | 1.200E-08 |
| 16908 | 846 | G846G | C | 2e-08 | 5.3e-09 | 4.2e-09 | 9.833E-09 | 8.822E-09 | 4.200E-09 | 2.000E-08 |
| 16909 | 846 | G846H | C | 9.2e-09 | 1.2e-08 | 1.8e-08 | 1.307E-08 | 4.496E-09 | 9.200E-09 | 1.800E-08 |
| 16910 | 846 | G846I | C | 2.3e-09 | 1.1e-08 | 4e-09 | 5.767E-09 | 4.611E-09 | 2.300E-09 | 1.100E-08 |
| 16911 | 846 | G846L | C | 2.8e-09 | 8.2e-09 | 5.6e-09 | 5.533E-09 | 2.701E-09 | 2.800E-09 | 8.200E-09 |
| 16912 | 846 | G846K | C | 9.8e-09 | 1.3e-08 | 6.9e-09 | 9.900E-09 | 3.051E-09 | 6.900E-09 | 1.300E-08 |
| 16913 | 846 | G846M | C | 7e-09 | 4.4e-09 | 7.8e-09 | 6.400E-09 | 1.778E-09 | 4.400E-09 | 7.800E-09 |
| 16914 | 846 | G846F | C | 6.1e-09 | 5.6e-09 | 4.6e-09 | 5.433E-09 | 7.638E-10 | 4.600E-09 | 6.100E-09 |

| | | | | | | | | | | |
|---|---|---|---|---|---|---|---|---|---|---|
| 16915 | 846 | G846P | C | 2.8e-09 | 2.9e-09 | 3.9e-09 | 3.200E-09 | 6.083E-10 | 2.800E-09 | 3.900E-09 |
| 16916 | 846 | G846S | C | 1e-08 | 4.5e-09 | 2.9e-09 | 5.800E-09 | 3.724E-09 | 2.900E-09 | 1.000E-08 |
| 16917 | 846 | G846T | C | 5.4e-09 | 1.4e-08 | 2.9e-09 | 7.433E-09 | 5.823E-09 | 2.900E-09 | 1.400E-08 |
| 16918 | 846 | G846W | C | 1.8e-08 | 7.3e-09 | 1.7e-08 | 1.410E-08 | 5.910E-09 | 7.300E-09 | 1.800E-08 |
| 16919 | 846 | G846Y | C | 5.5e-09 | 1.1e-08 | 6.1e-09 | 7.533E-09 | 3.017E-09 | 5.500E-09 | 1.100E-08 |
| 16920 | 846 | G846V | C | 2.2e-09 | 1.3e-08 | 6.3e-09 | 7.167E-09 | 5.452E-09 | 2.200E-09 | 1.300E-08 |
| 16921 | 847 | L847A | C | 1.3e-08 | 4.4e-09 | 5.7e-09 | 7.700E-09 | 4.636E-09 | 4.400E-09 | 1.300E-08 |
| 16922 | 847 | L847R | C | 4.8e-09 | 4.8e-09 | 6.4e-09 | 5.333E-09 | 9.238E-10 | 4.800E-09 | 6.400E-09 |
| 16923 | 847 | L847N | C | 6.1e-09 | 1.8e-09 | 9.4e-09 | 5.767E-09 | 3.811E-09 | 1.800E-09 | 9.400E-09 |
| 16924 | 847 | L847D | C | 1.2e-08 | 1.2e-08 | 5.8e-09 | 9.933E-09 | 3.580E-09 | 5.800E-09 | 1.200E-08 |
| 16925 | 847 | L847C | C | 7e-08 | 3.5e-09 | 1.2e-08 | 2.850E-08 | 3.619E-08 | 3.500E-09 | 7.000E-08 |
| 16926 | 847 | L847E | C | 1.2e-08 | 1.8e-09 | 9.5e-09 | 7.767E-09 | 5.316E-09 | 1.800E-09 | 1.200E-08 |
| 16927 | 847 | L847Q | C | 1.1e-08 | 7.7e-09 | 1.3e-08 | 1.057E-08 | 2.676E-09 | 7.700E-09 | 1.300E-08 |
| 16928 | 847 | L847G | C | 4e-09 | 1.2e-08 | 4.6e-09 | 6.867E-09 | 4.456E-09 | 4.000E-09 | 1.200E-08 |
| 16929 | 847 | L847H | C | 3.7e-09 | 4e-09 | 3.2e-09 | 3.633E-09 | 4.041E-10 | 3.200E-09 | 4.000E-09 |
| 16930 | 847 | L847I | C | 7.3e-09 | 6.5e-09 | 2.2e-08 | 1.193E-08 | 8.727E-09 | 6.500E-09 | 2.200E-08 |
| 16931 | 847 | L847L | C | 4.5e-09 | 5.9e-09 | 2e-08 | 1.013E-08 | 8.573E-09 | 4.500E-09 | 2.000E-08 |
| 16932 | 847 | L847K | C | 8.6e-09 | 6.4e-09 | 1.3e-08 | 9.333E-09 | 3.361E-09 | 6.400E-09 | 1.300E-08 |
| 16933 | 847 | L847M | C | 4.7e-09 | 5.2e-09 | 3.8e-09 | 4.567E-09 | 7.095E-10 | 3.800E-09 | 5.200E-09 |
| 16934 | 847 | L847F | C | 5.3e-09 | 7.2e-10 | 4e-09 | 3.340E-09 | 2.360E-09 | 7.200E-10 | 5.300E-09 |
| 16935 | 847 | L847P | C | 2e-08 | 5.6e-09 | 1.2e-08 | 1.253E-08 | 7.215E-09 | 5.600E-09 | 2.000E-08 |
| 16936 | 847 | L847S | C | 3e-09 | 7.1e-09 | 9.1e-09 | 6.400E-09 | 3.110E-09 | 3.000E-09 | 9.100E-09 |
| 16937 | 847 | L847T | C | 1.4e-08 | 8.4e-09 | 2.9e-09 | 8.433E-09 | 5.550E-09 | 2.900E-09 | 1.400E-08 |
| 16938 | 847 | L847W | C | 3.1e-09 | 1.1e-08 | 6.7e-09 | 6.933E-09 | 3.955E-09 | 3.100E-09 | 1.100E-08 |
| 16939 | 847 | L847Y | C | 3.4e-09 | 1.5e-08 | 8.7e-09 | 9.033E-09 | 5.807E-09 | 3.400E-09 | 1.500E-08 |
| 16940 | 847 | L847V | C | 2.8e-09 | 1.2e-08 | 5.3e-09 | 6.700E-09 | 4.757E-09 | 2.800E-09 | 1.200E-08 |
| 16941 | 848 | V848A | C | 1.6e-08 | 1.3e-08 | 5.5e-09 | 1.150E-08 | 5.408E-09 | 5.500E-09 | 1.600E-08 |
| 16942 | 848 | V848R | C | 1.4e-09 | 8.5e-09 | 1.5e-08 | 8.300E-09 | 6.802E-09 | 1.400E-09 | 1.500E-08 |
| 16943 | 848 | V848N | C | 1.1e-08 | 1.3e-08 | 7e-09 | 1.033E-08 | 3.055E-09 | 7.000E-09 | 1.300E-08 |
| 16944 | 848 | V848D | C | 1.1e-08 | 1e-08 | 1.4e-09 | 7.467E-09 | 5.278E-09 | 1.400E-09 | 1.100E-08 |
| 16945 | 848 | V848C | C | 1.3e-08 | 1.3e-08 | 1.8e-08 | 1.467E-08 | 2.887E-09 | 1.300E-08 | 1.800E-08 |
| 16946 | 848 | V848E | C | 4.7e-09 | 4.3e-09 | 9.4e-09 | 6.133E-09 | 2.836E-09 | 4.300E-09 | 9.400E-09 |
| 16947 | 848 | V848Q | C | 3.8e-09 | 3.2e-08 | 3.3e-09 | 1.303E-08 | 1.643E-08 | 3.300E-09 | 3.200E-08 |
| 16948 | 848 | V848G | C | 9e-09 | 8.2e-09 | 3.6e-09 | 6.933E-09 | 2.914E-09 | 3.600E-09 | 9.000E-09 |
| 16949 | 848 | V848H | C | 8.3e-09 | 4.1e-09 | 4.6e-09 | 5.667E-09 | 2.294E-09 | 4.100E-09 | 8.300E-09 |
| 16950 | 848 | V848I | C | 9.1e-09 | 1e-08 | 1.1e-08 | 1.003E-08 | 9.504E-10 | 9.100E-09 | 1.100E-08 |
| 16951 | 848 | V848L | C | 3.4e-08 | 1.2e-08 | 1.1e-08 | 1.900E-08 | 1.300E-08 | 1.100E-08 | 3.400E-08 |
| 16952 | 848 | V848K | C | 8.5e-09 | 1e-08 | 9.5e-09 | 9.333E-09 | 7.638E-10 | 8.500E-09 | 1.000E-08 |
| 16953 | 848 | V848M | C | 1.1e-08 | 1.5e-08 | 5.2e-09 | 1.040E-08 | 4.927E-09 | 5.200E-09 | 1.500E-08 |
| 16954 | 848 | V848F | C | 1.9e-09 | 8.2e-09 | 7.4e-09 | 5.833E-09 | 3.430E-09 | 1.900E-09 | 8.200E-09 |
| 16955 | 848 | V848P | C | 6.2e-09 | 6.7e-09 | 3.2e-09 | 5.367E-09 | 1.893E-09 | 3.200E-09 | 6.700E-09 |
| 16956 | 848 | V848S | C | 1.4e-08 | 4.1e-09 | 5.5e-09 | 7.867E-09 | 5.358E-09 | 4.100E-09 | 1.400E-08 |
| 16957 | 848 | V848T | C | 6.3e-09 | 3e-09 | 3.3e-09 | 4.200E-09 | 1.825E-09 | 3.000E-09 | 6.300E-09 |
| 16958 | 848 | V848W | C | 5.1e-09 | 7.4e-09 | 1.5e-08 | 9.167E-09 | 5.181E-09 | 5.100E-09 | 1.500E-08 |
| 16959 | 848 | V848Y | C | 3.7e-09 | 3e-09 | 9.8e-09 | 5.500E-09 | 3.740E-09 | 3.000E-09 | 9.800E-09 |
| 16960 | 848 | V848V | C | 1.9e-08 | 1.5e-09 | 1.1e-08 | 1.050E-08 | 8.761E-09 | 1.500E-09 | 1.900E-08 |
| 16961 | 849 | Q849A | C | 1.9e-08 | 1.6e-09 | 5.2e-09 | 8.600E-09 | 9.185E-09 | 1.600E-09 | 1.900E-08 |
| 16962 | 849 | Q849R | C | 6e-09 | 1.1e-08 | 9.9e-09 | 8.967E-09 | 2.627E-09 | 6.000E-09 | 1.100E-08 |
| 16963 | 849 | Q849N | C | 3.7e-09 | 3.5e-09 | 5.3e-09 | 4.167E-09 | 9.866E-10 | 3.500E-09 | 5.300E-09 |
| 16964 | 849 | Q849D | C | 1.4e-09 | 9.2e-09 | 9.1e-09 | 6.567E-09 | 4.475E-09 | 1.400E-09 | 9.200E-09 |
| 16965 | 849 | Q849C | C | 4.7e-09 | 7.1e-09 | 2.8e-09 | 4.867E-09 | 2.155E-09 | 2.800E-09 | 7.100E-09 |
| 16966 | 849 | Q849E | C | 9.1e-09 | 3.9e-09 | 1.4e-08 | 9.000E-09 | 5.051E-09 | 3.900E-09 | 1.400E-08 |
| 16967 | 849 | Q849Q | C | 5.4e-09 | 5.3e-09 | 4.5e-09 | 5.067E-09 | 4.933E-10 | 4.500E-09 | 5.400E-09 |
| 16968 | 849 | Q849G | C | 1.3e-08 | 6.2e-09 | 7.8e-09 | 9.000E-09 | 3.555E-09 | 6.200E-09 | 1.300E-08 |
| 16969 | 849 | Q849H | C | 1.3e-08 | 9.7e-09 | 1.1e-08 | 1.123E-08 | 1.662E-09 | 9.700E-09 | 1.300E-08 |
| 16970 | 849 | Q849I | C | 6.8e-09 | 4.4e-09 | 5.5e-09 | 5.567E-09 | 1.201E-09 | 4.400E-09 | 6.800E-09 |
| 16971 | 849 | Q849L | C | 1.2e-09 | 3.6e-09 | 6e-09 | 3.600E-09 | 2.400E-09 | 1.200E-09 | 6.000E-09 |
| 16972 | 849 | Q849K | C | 9.4e-09 | 2.1e-08 | 6.5e-09 | 1.230E-08 | 7.673E-09 | 6.500E-09 | 2.100E-08 |
| 16973 | 849 | Q849M | C | 2.8e-09 | 6e-09 | 2.1e-08 | 9.933E-09 | 9.717E-09 | 2.800E-09 | 2.100E-08 |
| 16974 | 849 | Q849F | C | 6e-09 | 5.8e-09 | 5.1e-09 | 5.633E-09 | 4.726E-10 | 5.100E-09 | 6.000E-09 |
| 16975 | 849 | Q849P | C | 4.9e-09 | 5.8e-09 | 2.2e-08 | 1.090E-08 | 9.623E-09 | 4.900E-09 | 2.200E-08 |
| 16976 | 849 | Q849S | C | 5.4e-09 | 7.2e-09 | 8.7e-09 | 7.100E-09 | 1.652E-09 | 5.400E-09 | 8.700E-09 |
| 16977 | 849 | Q849T | C | 7e-09 | 3.2e-09 | 1.4e-08 | 8.067E-09 | 5.478E-09 | 3.200E-09 | 1.400E-08 |
| 16978 | 849 | Q849W | C | 4.2e-09 | 3.1e-09 | 6.7e-09 | 4.667E-09 | 1.845E-09 | 3.100E-09 | 6.700E-09 |
| 16979 | 849 | Q849Y | C | 3.6e-09 | 5.1e-09 | 5.9e-09 | 4.867E-09 | 1.168E-09 | 3.600E-09 | 5.900E-09 |
| 16980 | 849 | Q849V | C | 1.3e-08 | 6.5e-09 | 9.9e-09 | 9.800E-09 | 3.251E-09 | 6.500E-09 | 1.300E-08 |
| 16981 | 850 | P850A | C | 4.1e-09 | 4.6e-09 | 1.8e-08 | 8.900E-09 | 7.885E-09 | 4.100E-09 | 1.800E-08 |
| 16982 | 850 | P850R | C | 4.9e-09 | 6.5e-09 | 7.9e-09 | 6.433E-09 | 1.501E-09 | 4.900E-09 | 7.900E-09 |
| 16983 | 850 | P850N | C | 5.5e-09 | 1.3e-08 | 2.1e-09 | 6.867E-09 | 5.577E-09 | 2.100E-09 | 1.300E-08 |
| 16984 | 850 | P850D | C | 1.4e-09 | 8.6e-09 | 1.3e-08 | 7.667E-09 | 5.856E-09 | 1.400E-09 | 1.300E-08 |
| 16985 | 850 | P850C | C | 2.3e-09 | 8.8e-09 | 1.2e-08 | 7.700E-09 | 4.943E-09 | 2.300E-09 | 1.200E-08 |
| 16986 | 850 | P850E | C | 1.7e-08 | 9e-09 | 7.9e-09 | 1.130E-08 | 4.967E-09 | 7.900E-09 | 1.700E-08 |
| 16987 | 850 | P850Q | C | 7.2e-09 | 6.5e-09 | 8.8e-09 | 7.500E-09 | 1.179E-09 | 6.500E-09 | 8.800E-09 |
| 16988 | 850 | P850G | C | 8.7e-09 | 2.8e-09 | 2.6e-08 | 1.250E-08 | 1.206E-08 | 2.800E-09 | 2.600E-08 |
| 16989 | 850 | P850H | C | 4.3e-09 | 8e-09 | 1.5e-08 | 9.100E-09 | 5.434E-09 | 4.300E-09 | 1.500E-08 |
| 16990 | 850 | P850I | C | 6.7e-09 | 2.7e-08 | 2.6e-09 | 1.210E-08 | 1.307E-08 | 2.600E-09 | 2.700E-08 |
| 16991 | 850 | P850L | C | 4.2e-09 | 3.1e-09 | 4.4e-09 | 3.900E-09 | 7.000E-10 | 3.100E-09 | 4.400E-09 |
| 16992 | 850 | P850K | C | 1.4e-08 | 9e-09 | 1.3e-08 | 1.200E-08 | 2.646E-09 | 9.000E-09 | 1.400E-08 |
| 16993 | 850 | P850M | C | 8.8e-09 | 7.7e-09 | 4.9e-09 | 7.133E-09 | 2.011E-09 | 4.900E-09 | 8.800E-09 |
| 16994 | 850 | P850F | C | 3.9e-09 | 7.1e-09 | 2.6e-09 | 4.533E-09 | 2.316E-09 | 2.600E-09 | 7.100E-09 |
| 16995 | 850 | P850P | C | 6e-09 | 1.9e-08 | 1.3e-08 | 1.267E-08 | 6.506E-09 | 6.000E-09 | 1.900E-08 |
| 16996 | 850 | P850S | C | 2.1e-09 | 1e-08 | 1.3e-08 | 8.367E-09 | 5.631E-09 | 2.100E-09 | 1.300E-08 |
| 16997 | 850 | P850T | C | 1.6e-08 | 6.9e-09 | 6e-09 | 9.633E-09 | 5.532E-09 | 6.000E-09 | 1.600E-08 |
| 16998 | 850 | P850W | C | 4.2e-09 | 4.7e-09 | 1.2e-08 | 6.967E-09 | 4.366E-09 | 4.200E-09 | 1.200E-08 |
| 16999 | 850 | P850Y | C | 2.6e-09 | 2.2e-08 | 1.3e-08 | 1.253E-08 | 9.708E-09 | 2.600E-09 | 2.200E-08 |

| | | | | | | | | | | |
|---|---|---|---|---|---|---|---|---|---|---|
| 17000 | 850 | P850V | C | 1.6e-08 | 1.7e-08 | 1.7e-09 | 1.157E-08 | 8.559E-09 | 1.700E-09 | 1.700E-08 |
| 17001 | 851 | G851A | C | 6.1e-09 | 1.6e-08 | 7.8e-09 | 9.967E-09 | 5.294E-09 | 6.100E-09 | 1.600E-08 |
| 17002 | 851 | G851R | C | 1.5e-08 | 6.5e-09 | 3.4e-09 | 8.300E-09 | 6.006E-09 | 3.400E-09 | 1.500E-08 |
| 17003 | 851 | G851N | C | 4.3e-09 | 7.8e-09 | 6.8e-09 | 6.300E-09 | 1.803E-09 | 4.300E-09 | 7.800E-09 |
| 17004 | 851 | G851D | C | 4e-09 | 1e-08 | 1e-08 | 8.000E-09 | 3.464E-09 | 4.000E-09 | 1.000E-08 |
| 17005 | 851 | G851C | C | 6.3e-09 | 5.7e-09 | 2.4e-09 | 4.800E-09 | 2.100E-09 | 2.400E-09 | 6.300E-09 |
| 17006 | 851 | G851E | C | 1.3e-08 | 1e-08 | 1.3e-08 | 1.200E-08 | 1.732E-09 | 1.000E-08 | 1.300E-08 |
| 17007 | 851 | G851Q | C | 4.4e-09 | 4.5e-09 | 2.7e-08 | 1.197E-08 | 1.302E-08 | 4.400E-09 | 2.700E-08 |
| 17008 | 851 | G851G | C | 1.1e-08 | 6.9e-09 | 1e-08 | 9.300E-09 | 2.138E-09 | 6.900E-09 | 1.100E-08 |
| 17009 | 851 | G851H | C | 9e-09 | 2.8e-09 | 1.4e-08 | 8.600E-09 | 5.611E-09 | 2.800E-09 | 1.400E-08 |
| 17010 | 851 | G851I | C | 6.7e-09 | 8.7e-09 | 5.1e-09 | 6.833E-09 | 1.804E-09 | 5.100E-09 | 8.700E-09 |
| 17011 | 851 | G851L | C | 2.8e-09 | 7.6e-09 | 6.9e-09 | 5.767E-09 | 2.593E-09 | 2.800E-09 | 7.600E-09 |
| 17012 | 851 | G851K | C | 8.9e-09 | 1.1e-08 | 9.8e-09 | 9.900E-09 | 1.054E-09 | 8.900E-09 | 1.100E-08 |
| 17013 | 851 | G851M | C | 4.3e-09 | 3.8e-09 | 9.2e-09 | 5.767E-09 | 2.984E-09 | 3.800E-09 | 9.200E-09 |
| 17014 | 851 | G851F | C | 2.6e-09 | 3.4e-09 | 7.5e-09 | 4.500E-09 | 2.629E-09 | 2.600E-09 | 7.500E-09 |
| 17015 | 851 | G851P | C | 1e-08 | 8.9e-09 | 8.2e-09 | 9.033E-09 | 9.074E-10 | 8.200E-09 | 1.000E-08 |
| 17016 | 851 | G851S | C | 7.4e-09 | 9e-09 | 3.9e-09 | 6.767E-09 | 2.608E-09 | 3.900E-09 | 9.000E-09 |
| 17017 | 851 | G851T | C | 8.6e-09 | 7.6e-09 | 5.8e-09 | 7.333E-09 | 1.419E-09 | 5.800E-09 | 8.600E-09 |
| 17018 | 851 | G851W | C | 1.1e-08 | 1.5e-08 | 2.6e-09 | 9.533E-09 | 6.329E-09 | 2.600E-09 | 1.500E-08 |
| 17019 | 851 | G851Y | C | 1.4e-08 | 7.2e-09 | 1.1e-08 | 1.073E-08 | 3.408E-09 | 7.200E-09 | 1.400E-08 |
| 17020 | 851 | G851V | C | 2.4e-09 | 6.4e-09 | 3.5e-09 | 4.100E-09 | 2.066E-09 | 2.400E-09 | 6.400E-09 |
| 17021 | 852 | G852A | C | 3e-09 | 2.6e-09 | 1.2e-08 | 5.867E-09 | 5.315E-09 | 2.600E-09 | 1.200E-08 |
| 17022 | 852 | G852R | C | 8.8e-09 | 1.7e-08 | 8e-09 | 1.127E-08 | 4.981E-09 | 8.000E-09 | 1.700E-08 |
| 17023 | 852 | G852N | C | 6.8e-09 | 6.5e-09 | 7.8e-09 | 7.033E-09 | 6.807E-10 | 6.500E-09 | 7.800E-09 |
| 17024 | 852 | G852D | C | 3.5e-09 | 3e-09 | 1.1e-08 | 5.833E-09 | 4.481E-09 | 3.000E-09 | 1.100E-08 |
| 17025 | 852 | G852C | C | 1.1e-08 | 1.9e-08 | 1.6e-08 | 1.533E-08 | 4.041E-09 | 1.100E-08 | 1.900E-08 |
| 17026 | 852 | G852E | C | 4.4e-09 | 8.4e-09 | 9.9e-09 | 7.567E-09 | 2.843E-09 | 4.400E-09 | 9.900E-09 |
| 17027 | 852 | G852Q | C | 3.9e-09 | 5.7e-09 | 2.9e-09 | 4.167E-09 | 1.419E-09 | 2.900E-09 | 5.700E-09 |
| 17028 | 852 | G852G | C | 2.8e-09 | 9.7e-09 | 4.5e-09 | 5.667E-09 | 3.595E-09 | 2.800E-09 | 9.700E-09 |
| 17029 | 852 | G852H | C | 2.7e-09 | 1.1e-08 | 1e-08 | 7.900E-09 | 4.531E-09 | 2.700E-09 | 1.100E-08 |
| 17030 | 852 | G852I | C | 4.1e-09 | 9.9e-09 | 7.3e-09 | 7.100E-09 | 2.905E-09 | 4.100E-09 | 9.900E-09 |
| 17031 | 852 | G852L | C | 6.3e-09 | 1.7e-08 | 1.2e-08 | 1.177E-08 | 5.354E-09 | 6.300E-09 | 1.700E-08 |
| 17032 | 852 | G852K | C | 7.8e-09 | 2.9e-09 | 1.1e-08 | 7.233E-09 | 4.080E-09 | 2.900E-09 | 1.100E-08 |
| 17033 | 852 | G852M | C | 5.2e-09 | 5.7e-09 | 8.1e-09 | 6.333E-09 | 1.550E-09 | 5.200E-09 | 8.100E-09 |
| 17034 | 852 | G852F | C | 4.3e-09 | 2e-08 | 4.6e-09 | 9.633E-09 | 8.979E-09 | 4.300E-09 | 2.000E-08 |
| 17035 | 852 | G852P | C | 9.8e-09 | 5.4e-09 | 1.4e-08 | 9.733E-09 | 4.300E-09 | 5.400E-09 | 1.400E-08 |
| 17036 | 852 | G852S | C | 1.8e-08 | 2.8e-09 | 1.2e-08 | 1.093E-08 | 7.656E-09 | 2.800E-09 | 1.800E-08 |
| 17037 | 852 | G852T | C | 2.9e-09 | 1.3e-08 | 1.4e-08 | 9.967E-09 | 6.140E-09 | 2.900E-09 | 1.400E-08 |
| 17038 | 852 | G852W | C | 1.2e-08 | 4.6e-09 | 8e-09 | 8.200E-09 | 3.704E-09 | 4.600E-09 | 1.200E-08 |
| 17039 | 852 | G852Y | C | 3.7e-09 | 5.2e-09 | 8.1e-09 | 5.667E-09 | 2.237E-09 | 3.700E-09 | 8.100E-09 |
| 17040 | 852 | G852V | C | 4.6e-09 | 3.2e-09 | 5.7e-09 | 4.500E-09 | 1.253E-09 | 3.200E-09 | 5.700E-09 |
| 17041 | 853 | S853A | C | 3.3e-09 | 4.6e-09 | 2e-08 | 9.300E-09 | 9.289E-09 | 3.300E-09 | 2.000E-08 |
| 17042 | 853 | S853R | C | 1.2e-08 | 2.9e-09 | 1.1e-08 | 8.633E-09 | 4.990E-09 | 2.900E-09 | 1.200E-08 |
| 17043 | 853 | S853N | C | 1.7e-08 | 7.3e-09 | 1.1e-08 | 1.177E-08 | 4.895E-09 | 7.300E-09 | 1.700E-08 |
| 17044 | 853 | S853D | C | 1.2e-08 | 1.6e-09 | 8.5e-09 | 7.367E-09 | 5.292E-09 | 1.600E-09 | 1.200E-08 |
| 17045 | 853 | S853C | C | 1.1e-08 | 5.2e-09 | 4.2e-09 | 6.800E-09 | 3.672E-09 | 4.200E-09 | 1.100E-08 |
| 17046 | 853 | S853E | C | 6.8e-09 | 4.5e-09 | 5.2e-09 | 5.500E-09 | 1.179E-09 | 4.500E-09 | 6.800E-09 |
| 17047 | 853 | S853Q | C | 1.3e-08 | 2.3e-09 | 4.2e-09 | 6.500E-09 | 5.709E-09 | 2.300E-09 | 1.300E-08 |
| 17048 | 853 | S853G | C | 1.4e-08 | 5.3e-09 | 4.9e-09 | 8.067E-09 | 5.142E-09 | 4.900E-09 | 1.400E-08 |
| 17049 | 853 | S853H | C | 1.1e-08 | 6.3e-09 | 5e-09 | 7.433E-09 | 3.156E-09 | 5.000E-09 | 1.100E-08 |
| 17050 | 853 | S853I | C | 8.7e-09 | 4.6e-09 | 1.4e-08 | 9.100E-09 | 4.713E-09 | 4.600E-09 | 1.400E-08 |
| 17051 | 853 | S853L | C | 5.6e-09 | 4.7e-09 | 8.5e-09 | 6.267E-09 | 1.986E-09 | 4.700E-09 | 8.500E-09 |
| 17052 | 853 | S853K | C | 1.1e-08 | 1.3e-08 | 5.8e-09 | 9.933E-09 | 3.717E-09 | 5.800E-09 | 1.300E-08 |
| 17053 | 853 | S853M | C | 6e-09 | 3.2e-09 | 7.8e-09 | 5.667E-09 | 2.318E-09 | 3.200E-09 | 7.800E-09 |
| 17054 | 853 | S853F | C | 8.1e-09 | 1.6e-08 | 3.6e-09 | 9.233E-09 | 6.277E-09 | 3.600E-09 | 1.600E-08 |
| 17055 | 853 | S853P | C | 4.7e-09 | 1.2e-08 | 7.7e-09 | 8.133E-09 | 3.669E-09 | 4.700E-09 | 1.200E-08 |
| 17056 | 853 | S853S | C | 5.3e-09 | 7.9e-09 | 1e-08 | 7.733E-09 | 2.354E-09 | 5.300E-09 | 1.000E-08 |
| 17057 | 853 | S853T | C | 5.9e-09 | 4e-09 | 3.4e-09 | 4.433E-09 | 1.305E-09 | 3.400E-09 | 5.900E-09 |
| 17058 | 853 | S853W | C | 4.4e-09 | 1.9e-08 | 9.9e-09 | 1.110E-08 | 7.374E-09 | 4.400E-09 | 1.900E-08 |
| 17059 | 853 | S853Y | C | 1.6e-08 | 3.4e-09 | 6.7e-09 | 8.700E-09 | 6.534E-09 | 3.400E-09 | 1.600E-08 |
| 17060 | 853 | S853V | C | 4.8e-09 | 7.6e-09 | 1.6e-09 | 4.667E-09 | 3.002E-09 | 1.600E-09 | 7.600E-09 |
| 17061 | 854 | L854A | C | 6e-09 | 5.2e-09 | 6.7e-09 | 5.967E-09 | 7.506E-10 | 5.200E-09 | 6.700E-09 |
| 17062 | 854 | L854R | C | 2.3e-08 | 7.6e-09 | 6e-09 | 1.220E-08 | 9.387E-09 | 6.000E-09 | 2.300E-08 |
| 17063 | 854 | L854N | C | 1.2e-08 | 4.2e-09 | 4e-09 | 6.733E-09 | 4.562E-09 | 4.000E-09 | 1.200E-08 |
| 17064 | 854 | L854D | C | 4.3e-09 | 1.4e-08 | 4.8e-09 | 7.700E-09 | 5.462E-09 | 4.300E-09 | 1.400E-08 |
| 17065 | 854 | L854C | C | 1.8e-09 | 5.8e-09 | 4.8e-09 | 4.133E-09 | 2.082E-09 | 1.800E-09 | 5.800E-09 |
| 17066 | 854 | L854E | C | 2.7e-08 | 6.5e-09 | 9.5e-09 | 1.433E-08 | 1.107E-08 | 6.500E-09 | 2.700E-08 |
| 17067 | 854 | L854Q | C | 3.7e-09 | 2.9e-09 | 4.3e-09 | 3.633E-09 | 7.024E-10 | 2.900E-09 | 4.300E-09 |
| 17068 | 854 | L854G | C | 2.5e-08 | 4e-08 | 1.2e-08 | 2.567E-08 | 1.401E-08 | 1.200E-08 | 4.000E-08 |
| 17069 | 854 | L854H | C | 3.2e-09 | 5.9e-09 | 1.3e-08 | 7.367E-09 | 5.062E-09 | 3.200E-09 | 1.300E-08 |
| 17070 | 854 | L854I | C | 8e-09 | 4.5e-09 | 4.7e-09 | 5.733E-09 | 1.966E-09 | 4.500E-09 | 8.000E-09 |
| 17071 | 854 | L854L | C | 1.3e-08 | 6.3e-09 | 2.1e-08 | 1.343E-08 | 7.360E-09 | 6.300E-09 | 2.100E-08 |
| 17072 | 854 | L854K | C | 1.2e-08 | 1.5e-08 | 7.2e-09 | 1.140E-08 | 3.934E-09 | 7.200E-09 | 1.500E-08 |
| 17073 | 854 | L854M | C | 2.5e-09 | 8.7e-09 | 2e-08 | 1.040E-08 | 8.873E-09 | 2.500E-09 | 2.000E-08 |
| 17074 | 854 | L854F | C | 3.6e-09 | 2e-08 | 9e-09 | 1.087E-08 | 8.358E-09 | 3.600E-09 | 2.000E-08 |
| 17075 | 854 | L854P | C | 8.6e-09 | 1e-08 | 9.8e-09 | 9.467E-09 | 7.572E-10 | 8.600E-09 | 1.000E-08 |
| 17076 | 854 | L854S | C | 8.1e-09 | 2.2e-09 | 8.9e-09 | 6.400E-09 | 3.659E-09 | 2.200E-09 | 8.900E-09 |
| 17077 | 854 | L854T | C | 1.5e-08 | 6.2e-09 | 4.1e-09 | 8.433E-09 | 5.783E-09 | 4.100E-09 | 1.500E-08 |
| 17078 | 854 | L854W | C | 4.6e-09 | 8.1e-09 | 7.2e-09 | 6.633E-09 | 1.818E-09 | 4.600E-09 | 8.100E-09 |
| 17079 | 854 | L854Y | C | 1.1e-08 | 3.8e-09 | 6.1e-09 | 6.967E-09 | 3.677E-09 | 3.800E-09 | 1.100E-08 |
| 17080 | 854 | L854V | C | 7.2e-09 | 8.6e-09 | 4.5e-09 | 6.767E-09 | 2.084E-09 | 4.500E-09 | 8.600E-09 |
| 17081 | 855 | R855A | C | 1.5e-08 | 2.6e-08 | 5.5e-09 | 1.550E-08 | 1.026E-08 | 5.500E-09 | 2.600E-08 |
| 17082 | 855 | R855R | C | 8.2e-09 | 1.9e-08 | 1.6e-08 | 1.440E-08 | 5.575E-09 | 8.200E-09 | 1.900E-08 |
| 17083 | 855 | R855N | C | 7.4e-09 | 3.4e-09 | 1.1e-08 | 7.267E-09 | 3.802E-09 | 3.400E-09 | 1.100E-08 |
| 17084 | 855 | R855D | C | 5.1e-09 | 9.4e-09 | 3.9e-09 | 6.133E-09 | 2.892E-09 | 3.900E-09 | 9.400E-09 |

| | | | | | | | | | | |
|---|---|---|---|---|---|---|---|---|---|---|
| 17085 | 855 | R855C | C | 5.3e-09 | 5.2e-09 | 4.9e-09 | 5.133E-09 | 2.082E-10 | 4.900E-09 | 5.300E-09 |
| 17086 | 855 | R855E | C | 3.6e-09 | 3.6e-09 | 5.9e-09 | 4.367E-09 | 1.328E-09 | 3.600E-09 | 5.900E-09 |
| 17087 | 855 | R855Q | C | 3.1e-09 | 1.3e-08 | 1.6e-08 | 1.070E-08 | 6.751E-09 | 3.100E-09 | 1.600E-08 |
| 17088 | 855 | R855G | C | 1.1e-08 | 3.2e-09 | 2.9e-09 | 5.700E-09 | 4.592E-09 | 2.900E-09 | 1.100E-08 |
| 17089 | 855 | R855H | C | 4.6e-09 | 9.4e-09 | 3.5e-08 | 1.633E-08 | 1.634E-08 | 4.600E-09 | 3.500E-08 |
| 17090 | 855 | R855I | C | 1.1e-08 | 5.7e-09 | 1.3e-08 | 9.900E-09 | 3.772E-09 | 5.700E-09 | 1.300E-08 |
| 17091 | 855 | R855L | C | 1.6e-09 | 1.6e-08 | 4e-09 | 7.200E-09 | 7.715E-09 | 1.600E-09 | 1.600E-08 |
| 17092 | 855 | R855K | C | 5.1e-09 | 7.1e-09 | 5.5e-09 | 5.900E-09 | 1.058E-09 | 5.100E-09 | 7.100E-09 |
| 17093 | 855 | R855M | C | 1e-08 | 4.5e-09 | 1.4e-08 | 9.500E-09 | 4.770E-09 | 4.500E-09 | 1.400E-08 |
| 17094 | 855 | R855F | C | 9.2e-09 | 2.5e-09 | 8.5e-09 | 6.733E-09 | 3.683E-09 | 2.500E-09 | 9.200E-09 |
| 17095 | 855 | R855P | C | 9.7e-09 | 4.1e-09 | 5.6e-09 | 6.467E-09 | 2.899E-09 | 4.100E-09 | 9.700E-09 |
| 17096 | 855 | R855S | C | 8.2e-09 | 4.5e-09 | 1.6e-08 | 9.567E-09 | 5.871E-09 | 4.500E-09 | 1.600E-08 |
| 17097 | 855 | R855T | C | 1e-08 | 7.6e-09 | 5.7e-09 | 7.767E-09 | 2.155E-09 | 5.700E-09 | 1.000E-08 |
| 17098 | 855 | R855W | C | 8.4e-09 | 5.9e-09 | 1.2e-08 | 8.767E-09 | 3.066E-09 | 5.900E-09 | 1.200E-08 |
| 17099 | 855 | R855Y | C | 3e-09 | 9.3e-09 | 4e-09 | 5.433E-09 | 3.386E-09 | 3.000E-09 | 9.300E-09 |
| 17100 | 855 | R855V | C | 2.2e-08 | 8.4e-09 | 4.3e-09 | 1.157E-08 | 9.265E-09 | 4.300E-09 | 2.200E-08 |
| 17101 | 856 | L856A | C | 4.2e-09 | 6.2e-09 | 4.7e-09 | 5.033E-09 | 1.041E-09 | 4.200E-09 | 6.200E-09 |
| 17102 | 856 | L856R | C | 8.2e-09 | 6e-09 | 5.6e-09 | 6.600E-09 | 1.400E-09 | 5.600E-09 | 8.200E-09 |
| 17103 | 856 | L856N | C | 1.6e-08 | 8.2e-09 | 1e-08 | 1.140E-08 | 4.084E-09 | 8.200E-09 | 1.600E-08 |
| 17104 | 856 | L856D | C | 5.7e-09 | 2.1e-08 | 1.4e-08 | 1.357E-08 | 7.659E-09 | 5.700E-09 | 2.100E-08 |
| 17105 | 856 | L856C | C | 9.4e-09 | 6.3e-09 | 2.6e-09 | 6.100E-09 | 3.404E-09 | 2.600E-09 | 9.400E-09 |
| 17106 | 856 | L856E | C | 8.8e-09 | 4.9e-09 | 5.1e-09 | 6.267E-09 | 2.196E-09 | 4.900E-09 | 8.800E-09 |
| 17107 | 856 | L856Q | C | 9.4e-09 | 2.3e-09 | 1.2e-08 | 7.900E-09 | 5.021E-09 | 2.300E-09 | 1.200E-08 |
| 17108 | 856 | L856G | C | 7.1e-09 | 5.5e-09 | 5.5e-09 | 6.033E-09 | 9.238E-10 | 5.500E-09 | 7.100E-09 |
| 17109 | 856 | L856H | C | 2.3e-08 | 9e-09 | 3.6e-09 | 1.187E-08 | 1.001E-08 | 3.600E-09 | 2.300E-08 |
| 17110 | 856 | L856I | C | 5.3e-09 | 1.7e-08 | 1.3e-08 | 1.177E-08 | 5.947E-09 | 5.300E-09 | 1.700E-08 |
| 17111 | 856 | L856L | C | 4.5e-09 | 1.2e-08 | 6.9e-09 | 7.800E-09 | 3.830E-09 | 4.500E-09 | 1.200E-08 |
| 17112 | 856 | L856K | C | 1e-08 | 2.7e-09 | 8.9e-09 | 7.200E-09 | 3.936E-09 | 2.700E-09 | 1.000E-08 |
| 17113 | 856 | L856M | C | 1.5e-08 | 2.6e-09 | 1.4e-08 | 1.053E-08 | 6.889E-09 | 2.600E-09 | 1.500E-08 |
| 17114 | 856 | L856F | C | 1.1e-08 | 5.8e-09 | 5.2e-09 | 7.333E-09 | 3.190E-09 | 5.200E-09 | 1.100E-08 |
| 17115 | 856 | L856P | C | 5.7e-09 | 5e-09 | 1.4e-08 | 8.233E-09 | 5.006E-09 | 5.000E-09 | 1.400E-08 |
| 17116 | 856 | L856S | C | 4.9e-09 | 1.3e-08 | 2.8e-08 | 1.530E-08 | 1.172E-08 | 4.900E-09 | 2.800E-08 |
| 17117 | 856 | L856T | C | 7.4e-09 | 2e-08 | 7.6e-09 | 1.167E-08 | 7.218E-09 | 7.400E-09 | 2.000E-08 |
| 17118 | 856 | L856W | C | 1.1e-08 | 9.9e-09 | 3e-09 | 7.967E-09 | 4.336E-09 | 3.000E-09 | 1.100E-08 |
| 17119 | 856 | L856Y | C | 1e-08 | 1.5e-08 | 6.6e-09 | 1.053E-08 | 4.225E-09 | 6.600E-09 | 1.500E-08 |
| 17120 | 856 | L856V | C | 2.1e-08 | 3.4e-09 | 3.3e-09 | 9.233E-09 | 1.019E-08 | 3.300E-09 | 2.100E-08 |
| 17121 | 857 | S857A | C | 4.9e-09 | 3.6e-09 | 2.6e-09 | 3.700E-09 | 1.153E-09 | 2.600E-09 | 4.900E-09 |
| 17122 | 857 | S857R | C | 6.1e-09 | 1.8e-08 | 6e-09 | 1.003E-08 | 6.900E-09 | 6.000E-09 | 1.800E-08 |
| 17123 | 857 | S857N | C | 1.8e-09 | 3.6e-09 | 7.1e-09 | 4.167E-09 | 2.695E-09 | 1.800E-09 | 7.100E-09 |
| 17124 | 857 | S857D | C | 1.2e-09 | 1.5e-08 | 6e-09 | 7.400E-09 | 7.006E-09 | 1.200E-09 | 1.500E-08 |
| 17125 | 857 | S857C | C | 6.3e-09 | 5.3e-09 | 6.1e-09 | 5.900E-09 | 5.292E-10 | 5.300E-09 | 6.300E-09 |
| 17126 | 857 | S857E | C | 3.1e-09 | 1.3e-08 | 5.6e-09 | 7.233E-09 | 5.148E-09 | 3.100E-09 | 1.300E-08 |
| 17127 | 857 | S857Q | C | 1.3e-08 | 6.5e-09 | 5.4e-09 | 8.300E-09 | 4.107E-09 | 5.400E-09 | 1.300E-08 |
| 17128 | 857 | S857G | C | 1.2e-08 | 2.3e-09 | 6.8e-09 | 7.033E-09 | 4.854E-09 | 2.300E-09 | 1.200E-08 |
| 17129 | 857 | S857H | C | 1.7e-08 | 7.1e-09 | 8.8e-09 | 1.097E-08 | 5.294E-09 | 7.100E-09 | 1.700E-08 |
| 17130 | 857 | S857I | C | 2.2e-09 | 1.1e-08 | 3.3e-09 | 5.500E-09 | 4.795E-09 | 2.200E-09 | 1.100E-08 |
| 17131 | 857 | S857L | C | 5.6e-09 | 4.3e-09 | 1.3e-08 | 7.633E-09 | 4.693E-09 | 4.300E-09 | 1.300E-08 |
| 17132 | 857 | S857K | C | 1.2e-08 | 7.5e-09 | 1.8e-08 | 1.250E-08 | 5.268E-09 | 7.500E-09 | 1.800E-08 |
| 17133 | 857 | S857M | C | 1e-08 | 1.5e-08 | 1.1e-08 | 1.200E-08 | 2.646E-09 | 1.000E-08 | 1.500E-08 |
| 17134 | 857 | S857F | C | 3.9e-09 | 3.9e-09 | 6.6e-09 | 4.800E-09 | 1.559E-09 | 3.900E-09 | 6.600E-09 |
| 17135 | 857 | S857P | C | 1.6e-08 | 7.7e-09 | 1.2e-08 | 1.190E-08 | 4.151E-09 | 7.700E-09 | 1.600E-08 |
| 17136 | 857 | S857S | C | 1.1e-08 | 3.8e-09 | 1.1e-08 | 8.600E-09 | 4.157E-09 | 3.800E-09 | 1.100E-08 |
| 17137 | 857 | S857T | C | 5.9e-09 | 2.9e-09 | 1.1e-08 | 6.600E-09 | 4.095E-09 | 2.900E-09 | 1.100E-08 |
| 17138 | 857 | S857W | C | 9.1e-09 | 4.4e-09 | 9.7e-09 | 7.733E-09 | 2.902E-09 | 4.400E-09 | 9.700E-09 |
| 17139 | 857 | S857Y | C | 1.1e-08 | 2.8e-09 | 2.8e-09 | 5.533E-09 | 4.734E-09 | 2.800E-09 | 1.100E-08 |
| 17140 | 857 | S857V | C | 4.7e-09 | 1.7e-08 | 8.9e-09 | 1.020E-08 | 6.252E-09 | 4.700E-09 | 1.700E-08 |
| 17141 | 858 | C858A | C | 9.8e-09 | 4.1e-09 | 2.9e-09 | 5.600E-09 | 3.686E-09 | 2.900E-09 | 9.800E-09 |
| 17142 | 858 | C858R | C | 9.8e-09 | 7.9e-09 | 7.2e-09 | 8.300E-09 | 1.345E-09 | 7.200E-09 | 9.800E-09 |
| 17143 | 858 | C858N | C | 4e-09 | 1.7e-09 | 1.3e-08 | 6.233E-09 | 5.972E-09 | 1.700E-09 | 1.300E-08 |
| 17144 | 858 | C858D | C | 4.2e-09 | 8e-09 | 4.8e-09 | 5.667E-09 | 2.043E-09 | 4.200E-09 | 8.000E-09 |
| 17145 | 858 | C858C | C | 7.6e-09 | 9.9e-09 | 5e-09 | 7.500E-09 | 2.452E-09 | 5.000E-09 | 9.900E-09 |
| 17146 | 858 | C858E | C | 3.7e-09 | 1.3e-08 | 8.1e-09 | 8.267E-09 | 4.652E-09 | 3.700E-09 | 1.300E-08 |
| 17147 | 858 | C858Q | C | 1.3e-08 | 4.1e-09 | 1.2e-08 | 9.700E-09 | 4.875E-09 | 4.100E-09 | 1.300E-08 |
| 17148 | 858 | C858G | C | 6.3e-09 | 2.1e-09 | 3.1e-09 | 3.833E-09 | 2.194E-09 | 2.100E-09 | 6.300E-09 |
| 17149 | 858 | C858H | C | 1.7e-08 | 8.7e-09 | 7.2e-09 | 1.097E-08 | 5.279E-09 | 7.200E-09 | 1.700E-08 |
| 17150 | 858 | C858I | C | 1.8e-08 | 7e-09 | 5.1e-09 | 1.003E-08 | 6.964E-09 | 5.100E-09 | 1.800E-08 |
| 17151 | 858 | C858L | C | 2.5e-08 | 5.9e-09 | 1.3e-08 | 1.463E-08 | 9.654E-09 | 5.900E-09 | 2.500E-08 |
| 17152 | 858 | C858K | C | 9.7e-09 | 1.8e-08 | 9.7e-09 | 1.247E-08 | 4.792E-09 | 9.700E-09 | 1.800E-08 |
| 17153 | 858 | C858M | C | 5.3e-09 | 4.2e-09 | 1.9e-08 | 9.500E-09 | 8.246E-09 | 4.200E-09 | 1.900E-08 |
| 17154 | 858 | C858F | C | 1.4e-08 | 7.2e-09 | 5.1e-09 | 8.767E-09 | 4.652E-09 | 5.100E-09 | 1.400E-08 |
| 17155 | 858 | C858P | C | 5.2e-09 | 1.7e-08 | 6.1e-09 | 9.433E-09 | 6.568E-09 | 5.200E-09 | 1.700E-08 |
| 17156 | 858 | C858S | C | 8.1e-09 | 9.2e-09 | 3.6e-09 | 6.967E-09 | 2.967E-09 | 3.600E-09 | 9.200E-09 |
| 17157 | 858 | C858T | C | 6e-09 | 3.6e-09 | 9.9e-09 | 6.500E-09 | 3.180E-09 | 3.600E-09 | 9.900E-09 |
| 17158 | 858 | C858W | C | 4.5e-09 | 1.8e-08 | 3.1e-09 | 8.533E-09 | 8.228E-09 | 3.100E-09 | 1.800E-08 |
| 17159 | 858 | C858Y | C | 5.5e-09 | 3.6e-09 | 3.3e-09 | 4.133E-09 | 1.193E-09 | 3.300E-09 | 5.500E-09 |
| 17160 | 858 | C858V | C | 1.6e-08 | 6.6e-09 | 5.4e-09 | 9.333E-09 | 5.805E-09 | 5.400E-09 | 1.600E-08 |
| 17161 | 859 | A859A | C | 8.6e-09 | 5.8e-09 | 1e-08 | 8.133E-09 | 2.139E-09 | 5.800E-09 | 1.000E-08 |
| 17162 | 859 | A859R | C | 3.3e-09 | 5.7e-09 | 1.4e-08 | 7.667E-09 | 5.615E-09 | 3.300E-09 | 1.400E-08 |
| 17163 | 859 | A859N | C | 3.4e-09 | 4.2e-09 | 5.9e-09 | 4.500E-09 | 1.277E-09 | 3.400E-09 | 5.900E-09 |
| 17164 | 859 | A859D | C | 1.3e-08 | 9.8e-09 | 4.9e-09 | 9.233E-09 | 4.080E-09 | 4.900E-09 | 1.300E-08 |
| 17165 | 859 | A859C | C | 1e-08 | 1.1e-08 | 6.1e-09 | 9.033E-09 | 2.589E-09 | 6.100E-09 | 1.100E-08 |
| 17166 | 859 | A859E | C | 3.5e-09 | 5.2e-08 | 6.6e-09 | 2.070E-08 | 2.715E-08 | 3.500E-09 | 5.200E-08 |
| 17167 | 859 | A859Q | C | 2.7e-09 | 1.5e-09 | 8.8e-09 | 4.333E-09 | 3.915E-09 | 1.500E-09 | 8.800E-09 |
| 17168 | 859 | A859G | C | 1.2e-08 | 1.3e-08 | 2.2e-09 | 9.067E-09 | 5.968E-09 | 2.200E-09 | 1.300E-08 |
| 17169 | 859 | A859H | C | 1.2e-08 | 7.7e-09 | 8.3e-09 | 9.333E-09 | 2.329E-09 | 7.700E-09 | 1.200E-08 |

| | | | | | | | | | | |
|---|---|---|---|---|---|---|---|---|---|---|
| 17170 | 859 | A859I | C | 1.3e-08 | 1e-08 | 8.4e-09 | 1.047E-08 | 2.335E-09 | 8.400E-09 | 1.300E-08 |
| 17171 | 859 | A859L | C | 4.8e-09 | 7e-09 | 3.1e-09 | 4.967E-09 | 1.955E-09 | 3.100E-09 | 7.000E-09 |
| 17172 | 859 | A859K | C | 1.7e-09 | 3.2e-09 | 6.3e-09 | 3.733E-09 | 2.346E-09 | 1.700E-09 | 6.300E-09 |
| 17173 | 859 | A859M | C | 2.4e-09 | 1.1e-09 | 9.5e-09 | 4.333E-09 | 4.521E-09 | 1.100E-09 | 9.500E-09 |
| 17174 | 859 | A859F | C | 1.1e-08 | 4.4e-09 | 5.2e-09 | 6.867E-09 | 3.602E-09 | 4.400E-09 | 1.100E-08 |
| 17175 | 859 | A859P | C | 6.1e-09 | 5.2e-09 | 1.8e-08 | 9.767E-09 | 7.144E-09 | 5.200E-09 | 1.800E-08 |
| 17176 | 859 | A859S | C | 9.9e-09 | 1.4e-08 | 6.2e-09 | 1.003E-08 | 3.902E-09 | 6.200E-09 | 1.400E-08 |
| 17177 | 859 | A859T | C | 1.1e-08 | 3e-09 | 2.4e-09 | 5.467E-09 | 4.801E-09 | 2.400E-09 | 1.100E-08 |
| 17178 | 859 | A859W | C | 9.5e-09 | 6.6e-09 | 2.9e-09 | 6.333E-09 | 3.308E-09 | 2.900E-09 | 9.500E-09 |
| 17179 | 859 | A859Y | C | 1e-08 | 1.3e-08 | 7.2e-09 | 1.007E-08 | 2.901E-09 | 7.200E-09 | 1.300E-08 |
| 17180 | 859 | A859V | C | 9.9e-09 | 3e-09 | 1.2e-08 | 8.300E-09 | 4.709E-09 | 3.000E-09 | 1.200E-08 |
| 17181 | 860 | A860A | C | 7e-09 | 7.9e-09 | 9.6e-09 | 8.167E-09 | 1.320E-09 | 7.000E-09 | 9.600E-09 |
| 17182 | 860 | A860R | C | 1.1e-08 | 6.5e-09 | 1.6e-08 | 1.117E-08 | 4.752E-09 | 6.500E-09 | 1.600E-08 |
| 17183 | 860 | A860N | C | 1.7e-08 | 6.7e-09 | 1e-08 | 1.123E-08 | 5.260E-09 | 6.700E-09 | 1.700E-08 |
| 17184 | 860 | A860D | C | 5.3e-09 | 2.4e-09 | 1e-08 | 5.900E-09 | 3.835E-09 | 2.400E-09 | 1.000E-08 |
| 17185 | 860 | A860C | C | 1.2e-08 | 5.9e-09 | 6.1e-09 | 8.000E-09 | 3.466E-09 | 5.900E-09 | 1.200E-08 |
| 17186 | 860 | A860E | C | 1.9e-09 | 1.9e-08 | 4.9e-09 | 8.600E-09 | 9.131E-09 | 1.900E-09 | 1.900E-08 |
| 17187 | 860 | A860Q | C | 1.1e-08 | 5.3e-09 | 1.6e-08 | 1.077E-08 | 5.354E-09 | 5.300E-09 | 1.600E-08 |
| 17188 | 860 | A860G | C | 5.8e-09 | 1.2e-08 | 7.4e-09 | 8.400E-09 | 3.219E-09 | 5.800E-09 | 1.200E-08 |
| 17189 | 860 | A860H | C | 5.7e-09 | 1.1e-08 | 3.7e-09 | 6.800E-09 | 3.772E-09 | 3.700E-09 | 1.100E-08 |
| 17190 | 860 | A860I | C | 6.4e-09 | 1.2e-08 | 3.8e-09 | 7.400E-09 | 4.190E-09 | 3.800E-09 | 1.200E-08 |
| 17191 | 860 | A860L | C | 5.9e-09 | 1.5e-08 | 6e-09 | 8.967E-09 | 5.225E-09 | 5.900E-09 | 1.500E-08 |
| 17192 | 860 | A860K | C | 1.7e-08 | 1.2e-08 | 1.5e-08 | 1.467E-08 | 2.517E-09 | 1.200E-08 | 1.700E-08 |
| 17193 | 860 | A860M | C | 5.1e-09 | 6.9e-09 | 2.9e-09 | 4.967E-09 | 2.003E-09 | 2.900E-09 | 6.900E-09 |
| 17194 | 860 | A860F | C | 5.9e-09 | 7.3e-09 | 5.7e-09 | 6.300E-09 | 8.718E-10 | 5.700E-09 | 7.300E-09 |
| 17195 | 860 | A860P | C | 6.2e-09 | 4e-09 | 4.3e-09 | 4.833E-09 | 1.193E-09 | 4.000E-09 | 6.200E-09 |
| 17196 | 860 | A860S | C | 6e-09 | 4.5e-09 | 3.2e-09 | 4.567E-09 | 1.401E-09 | 3.200E-09 | 6.000E-09 |
| 17197 | 860 | A860T | C | 5.2e-09 | 3.6e-09 | 7.7e-09 | 5.500E-09 | 2.066E-09 | 3.600E-09 | 7.700E-09 |
| 17198 | 860 | A860W | C | 1.3e-09 | 1.9e-08 | 6.7e-09 | 9.000E-09 | 9.071E-09 | 1.300E-09 | 1.900E-08 |
| 17199 | 860 | A860Y | C | 9.1e-09 | 7e-09 | 9.1e-09 | 8.400E-09 | 1.212E-09 | 7.000E-09 | 9.100E-09 |
| 17200 | 860 | A860V | C | 6.4e-09 | 6.9e-09 | 8e-09 | 7.100E-09 | 8.185E-10 | 6.400E-09 | 8.000E-09 |
| 17201 | 861 | S861A | C | 6.5e-09 | 3.6e-09 | 1.6e-08 | 8.700E-09 | 6.486E-09 | 3.600E-09 | 1.600E-08 |
| 17202 | 861 | S861R | C | 1.1e-08 | 6.7e-09 | 3.2e-09 | 6.967E-09 | 3.907E-09 | 3.200E-09 | 1.100E-08 |
| 17203 | 861 | S861N | C | 6.7e-09 | 1.4e-08 | 4.7e-09 | 8.467E-09 | 4.895E-09 | 4.700E-09 | 1.400E-08 |
| 17204 | 861 | S861D | C | 3.8e-09 | 2e-09 | 6.8e-09 | 4.200E-09 | 2.425E-09 | 2.000E-09 | 6.800E-09 |
| 17205 | 861 | S861C | C | 7.3e-09 | 2.8e-09 | 9.8e-09 | 6.633E-09 | 3.547E-09 | 2.800E-09 | 9.800E-09 |
| 17206 | 861 | S861E | C | 2.5e-09 | 9e-09 | 1.3e-08 | 8.167E-09 | 5.299E-09 | 2.500E-09 | 1.300E-08 |
| 17207 | 861 | S861Q | C | 8.4e-09 | 1.5e-09 | 4.9e-09 | 4.933E-09 | 3.450E-09 | 1.500E-09 | 8.400E-09 |
| 17208 | 861 | S861G | C | 2.5e-08 | 3e-09 | 8e-09 | 1.200E-08 | 1.153E-08 | 3.000E-09 | 2.500E-08 |
| 17209 | 861 | S861H | C | 8.2e-09 | 1.3e-08 | 1.2e-08 | 1.107E-08 | 2.532E-09 | 8.200E-09 | 1.300E-08 |
| 17210 | 861 | S861I | C | 7.8e-09 | 7.2e-09 | 8.9e-09 | 7.967E-09 | 8.622E-10 | 7.200E-09 | 8.900E-09 |
| 17211 | 861 | S861L | C | 5.7e-09 | 1.2e-08 | 5.5e-09 | 7.733E-09 | 3.696E-09 | 5.500E-09 | 1.200E-08 |
| 17212 | 861 | S861K | C | 1.3e-08 | 1.3e-08 | 3.8e-09 | 9.933E-09 | 5.312E-09 | 3.800E-09 | 1.300E-08 |
| 17213 | 861 | S861M | C | 3.5e-09 | 8.5e-09 | 7.1e-09 | 6.367E-09 | 2.579E-09 | 3.500E-09 | 8.500E-09 |
| 17214 | 861 | S861F | C | 7.1e-09 | 9.9e-09 | 6.2e-09 | 7.733E-09 | 1.930E-09 | 6.200E-09 | 9.900E-09 |
| 17215 | 861 | S861P | C | 5.8e-09 | 2.1e-08 | 6.3e-09 | 1.103E-08 | 8.635E-09 | 5.800E-09 | 2.100E-08 |
| 17216 | 861 | S861S | C | 2.8e-08 | 5.6e-09 | 2.9e-09 | 1.217E-08 | 1.378E-08 | 2.900E-09 | 2.800E-08 |
| 17217 | 861 | S861T | C | 7.9e-09 | 2.2e-09 | 3.5e-09 | 4.533E-09 | 2.987E-09 | 2.200E-09 | 7.900E-09 |
| 17218 | 861 | S861W | C | 7.4e-09 | 4.4e-09 | 2.1e-09 | 4.633E-09 | 2.658E-09 | 2.100E-09 | 7.400E-09 |
| 17219 | 861 | S861Y | C | 4.1e-09 | 1.5e-08 | 1.5e-08 | 1.137E-08 | 6.293E-09 | 4.100E-09 | 1.500E-08 |
| 17220 | 861 | S861V | C | 1.6e-08 | 1.6e-08 | 1.5e-08 | 1.567E-08 | 5.774E-10 | 1.500E-08 | 1.600E-08 |
| 17221 | 862 | G862A | C | 8.2e-09 | 7.1e-09 | 4.9e-09 | 6.733E-09 | 1.680E-09 | 4.900E-09 | 8.200E-09 |
| 17222 | 862 | G862R | C | 2.3e-08 | 1.8e-08 | 1.5e-08 | 1.867E-08 | 4.041E-09 | 1.500E-08 | 2.300E-08 |
| 17223 | 862 | G862N | C | 8.6e-09 | 1.1e-08 | 6e-09 | 8.533E-09 | 2.501E-09 | 6.000E-09 | 1.100E-08 |
| 17224 | 862 | G862D | C | 2e-08 | 3.4e-09 | 9e-09 | 1.080E-08 | 8.445E-09 | 3.400E-09 | 2.000E-08 |
| 17225 | 862 | G862C | C | 4.2e-09 | 1.4e-08 | 4e-09 | 7.400E-09 | 5.717E-09 | 4.000E-09 | 1.400E-08 |
| 17226 | 862 | G862E | C | 7.1e-09 | 1e-08 | 3.8e-09 | 6.967E-09 | 3.102E-09 | 3.800E-09 | 1.000E-08 |
| 17227 | 862 | G862Q | C | 1.6e-08 | 1.1e-08 | 7e-09 | 1.133E-08 | 4.509E-09 | 7.000E-09 | 1.600E-08 |
| 17228 | 862 | G862G | C | 7.9e-09 | 4.3e-09 | 1.1e-08 | 7.733E-09 | 3.353E-09 | 4.300E-09 | 1.100E-08 |
| 17229 | 862 | G862H | C | 3.1e-09 | 6.2e-09 | 1e-08 | 6.433E-09 | 3.456E-09 | 3.100E-09 | 1.000E-08 |
| 17230 | 862 | G862I | C | 5.5e-09 | 3.8e-09 | 5.3e-09 | 4.867E-09 | 9.292E-10 | 3.800E-09 | 5.500E-09 |
| 17231 | 862 | G862L | C | 2.7e-09 | 1.5e-08 | 4.3e-09 | 7.333E-09 | 6.688E-09 | 2.700E-09 | 1.500E-08 |
| 17232 | 862 | G862K | C | 2.7e-09 | 3.3e-09 | 5.5e-09 | 3.833E-09 | 1.474E-09 | 2.700E-09 | 5.500E-09 |
| 17233 | 862 | G862M | C | 2.2e-08 | 2.9e-09 | 6.5e-09 | 1.047E-08 | 1.015E-08 | 2.900E-09 | 2.200E-08 |
| 17234 | 862 | G862F | C | 7.8e-09 | 6.6e-09 | 1.2e-08 | 8.800E-09 | 2.835E-09 | 6.600E-09 | 1.200E-08 |
| 17235 | 862 | G862P | C | 1.4e-08 | 5.3e-09 | 5.7e-09 | 8.333E-09 | 4.912E-09 | 5.300E-09 | 1.400E-08 |
| 17236 | 862 | G862S | C | 4.1e-09 | 6.5e-09 | 1.2e-08 | 7.533E-09 | 4.050E-09 | 4.100E-09 | 1.200E-08 |
| 17237 | 862 | G862T | C | 7.8e-09 | 1.5e-08 | 5.9e-09 | 9.567E-09 | 4.800E-09 | 5.900E-09 | 1.500E-08 |
| 17238 | 862 | G862W | C | 1.1e-08 | 3.7e-09 | 6.3e-09 | 7.000E-09 | 3.700E-09 | 3.700E-09 | 1.100E-08 |
| 17239 | 862 | G862Y | C | 1.2e-08 | 5e-09 | 7.4e-09 | 8.133E-09 | 3.557E-09 | 5.000E-09 | 1.200E-08 |
| 17240 | 862 | G862V | C | 2.6e-09 | 5.3e-09 | 7.4e-09 | 5.100E-09 | 2.406E-09 | 2.600E-09 | 7.400E-09 |
| 17241 | 863 | F863A | C | 8.9e-09 | 9.8e-09 | 9.2e-09 | 9.300E-09 | 4.583E-10 | 8.900E-09 | 9.800E-09 |
| 17242 | 863 | F863R | C | 4.8e-09 | 9.1e-09 | 2.8e-09 | 5.567E-09 | 3.219E-09 | 2.800E-09 | 9.100E-09 |
| 17243 | 863 | F863N | C | 1.1e-08 | 9.6e-09 | 1.3e-08 | 1.120E-08 | 1.709E-09 | 9.600E-09 | 1.300E-08 |
| 17244 | 863 | F863D | C | 1e-08 | 5.9e-09 | 1.5e-08 | 1.030E-08 | 4.557E-09 | 5.900E-09 | 1.500E-08 |
| 17245 | 863 | F863C | C | 1.9e-08 | 6e-09 | 7.3e-09 | 1.077E-08 | 7.160E-09 | 6.000E-09 | 1.900E-08 |
| 17246 | 863 | F863E | C | 5.9e-09 | 5.2e-09 | 7.3e-09 | 6.133E-09 | 1.069E-09 | 5.200E-09 | 7.300E-09 |
| 17247 | 863 | F863Q | C | 2.3e-09 | 5.2e-09 | 1.6e-09 | 3.033E-09 | 1.909E-09 | 1.600E-09 | 5.200E-09 |
| 17248 | 863 | F863G | C | 6.8e-09 | 5.2e-09 | 2.7e-09 | 4.900E-09 | 2.066E-09 | 2.700E-09 | 6.800E-09 |
| 17249 | 863 | F863H | C | 1e-08 | 6.9e-09 | 8.3e-09 | 8.400E-09 | 1.552E-09 | 6.900E-09 | 1.000E-08 |
| 17250 | 863 | F863I | C | 6.9e-09 | 7.2e-09 | 6.4e-09 | 6.833E-09 | 4.041E-10 | 6.400E-09 | 7.200E-09 |
| 17251 | 863 | F863L | C | 4.6e-09 | 6.9e-09 | 8.4e-09 | 6.633E-09 | 1.914E-09 | 4.600E-09 | 8.400E-09 |
| 17252 | 863 | F863K | C | 3.3e-09 | 5.1e-09 | 1.2e-08 | 6.800E-09 | 4.592E-09 | 3.300E-09 | 1.200E-08 |
| 17253 | 863 | F863M | C | 5.2e-09 | 5.7e-09 | 5e-09 | 5.300E-09 | 3.606E-10 | 5.000E-09 | 5.700E-09 |
| 17254 | 863 | F863F | C | 4e-09 | 3.1e-09 | 6.9e-09 | 4.667E-09 | 1.986E-09 | 3.100E-09 | 6.900E-09 |

| | | | | | | | | | | |
|---|---|---|---|---|---|---|---|---|---|---|
| 17255 | 863 | F863P | C | 1.1e-08 | 9e-09 | 7.5e-09 | 9.167E-09 | 1.756E-09 | 7.500E-09 | 1.100E-08 |
| 17256 | 863 | F863S | C | 1.7e-08 | 8.5e-09 | 1.3e-09 | 8.933E-09 | 7.859E-09 | 1.300E-09 | 1.700E-08 |
| 17257 | 863 | F863T | C | 3.1e-09 | 3.4e-09 | 4.4e-09 | 3.633E-09 | 6.807E-10 | 3.100E-09 | 4.400E-09 |
| 17258 | 863 | F863W | C | 1.9e-09 | 2.7e-09 | 4.1e-09 | 2.900E-09 | 1.114E-09 | 1.900E-09 | 4.100E-09 |
| 17259 | 863 | F863Y | C | 9e-09 | 6.7e-09 | 6.9e-09 | 7.533E-09 | 1.274E-09 | 6.700E-09 | 9.000E-09 |
| 17260 | 863 | F863V | C | 6.1e-09 | 5.8e-09 | 1.1e-08 | 7.633E-09 | 2.919E-09 | 5.800E-09 | 1.100E-08 |
| 17261 | 864 | T864A | C | 1e-08 | 2.8e-09 | 2.8e-09 | 5.200E-09 | 4.157E-09 | 2.800E-09 | 1.000E-08 |
| 17262 | 864 | T864R | C | 5.3e-09 | 3.1e-09 | 4.8e-09 | 4.400E-09 | 1.153E-09 | 3.100E-09 | 5.300E-09 |
| 17263 | 864 | T864N | C | 1.4e-08 | 2.1e-08 | 1.9e-09 | 1.230E-08 | 9.663E-09 | 1.900E-09 | 2.100E-08 |
| 17264 | 864 | T864D | C | 4e-09 | 6.8e-09 | 7.1e-09 | 5.967E-09 | 1.710E-09 | 4.000E-09 | 7.100E-09 |
| 17265 | 864 | T864C | C | 3.5e-09 | 3e-09 | 5.6e-09 | 4.033E-09 | 1.380E-09 | 3.000E-09 | 5.600E-09 |
| 17266 | 864 | T864E | C | 9.6e-09 | 3.9e-09 | 2.7e-09 | 5.400E-09 | 3.686E-09 | 2.700E-09 | 9.600E-09 |
| 17267 | 864 | T864Q | C | 9.7e-09 | 8.8e-09 | 1e-08 | 9.500E-09 | 6.245E-10 | 8.800E-09 | 1.000E-08 |
| 17268 | 864 | T864G | C | 8.5e-09 | 6.4e-09 | 2.6e-08 | 1.363E-08 | 1.076E-08 | 6.400E-09 | 2.600E-08 |
| 17269 | 864 | T864H | C | 1.3e-08 | 7.1e-09 | 5.4e-09 | 8.500E-09 | 3.989E-09 | 5.400E-09 | 1.300E-08 |
| 17270 | 864 | T864I | C | 6.6e-09 | 1.4e-09 | 3.6e-09 | 3.867E-09 | 2.610E-09 | 1.400E-09 | 6.600E-09 |
| 17271 | 864 | T864L | C | 2.9e-09 | 5.1e-09 | 6.4e-09 | 4.800E-09 | 1.769E-09 | 2.900E-09 | 6.400E-09 |
| 17272 | 864 | T864K | C | 6.8e-09 | 5.9e-09 | 8.4e-09 | 7.033E-09 | 1.266E-09 | 5.900E-09 | 8.400E-09 |
| 17273 | 864 | T864M | C | 3.2e-09 | 7.5e-09 | 2.9e-09 | 4.533E-09 | 2.574E-09 | 2.900E-09 | 7.500E-09 |
| 17274 | 864 | T864F | C | 1.1e-09 | 3.8e-09 | 3.3e-09 | 2.733E-09 | 1.436E-09 | 1.100E-09 | 3.800E-09 |
| 17275 | 864 | T864P | C | 6.1e-09 | 2.7e-08 | 4e-09 | 1.237E-08 | 1.272E-08 | 4.000E-09 | 2.700E-08 |
| 17276 | 864 | T864S | C | 6e-09 | 6.3e-09 | 5.7e-09 | 6.000E-09 | 3.000E-10 | 5.700E-09 | 6.300E-09 |
| 17277 | 864 | T864T | C | 4.6e-09 | 1.5e-08 | 9.4e-09 | 9.667E-09 | 5.205E-09 | 4.600E-09 | 1.500E-08 |
| 17278 | 864 | T864W | C | 6.8e-09 | 8.1e-10 | 6.1e-09 | 4.570E-09 | 3.275E-09 | 8.100E-10 | 6.800E-09 |
| 17279 | 864 | T864Y | C | 4e-09 | 3.3e-09 | 2.4e-09 | 3.233E-09 | 8.021E-10 | 2.400E-09 | 4.000E-09 |
| 17280 | 864 | T864V | C | 9.6e-09 | 4e-09 | 2.1e-09 | 5.233E-09 | 3.899E-09 | 2.100E-09 | 9.600E-09 |
| 17281 | 865 | F865A | C | 3.1e-09 | 2.2e-08 | 4.8e-09 | 9.967E-09 | 1.046E-08 | 3.100E-09 | 2.200E-08 |
| 17282 | 865 | F865R | C | 1.1e-08 | 6.3e-09 | 8.1e-09 | 8.467E-09 | 2.371E-09 | 6.300E-09 | 1.100E-08 |
| 17283 | 865 | F865N | C | 1.3e-08 | 8.3e-09 | 5.5e-09 | 8.933E-09 | 3.790E-09 | 5.500E-09 | 1.300E-08 |
| 17284 | 865 | F865D | C | 3.3e-09 | 9.4e-09 | 1.3e-08 | 8.567E-09 | 4.903E-09 | 3.300E-09 | 1.300E-08 |
| 17285 | 865 | F865C | C | 1.9e-08 | 2.3e-08 | 1.3e-08 | 1.833E-08 | 5.033E-09 | 1.300E-08 | 2.300E-08 |
| 17286 | 865 | F865E | C | 1.1e-08 | 7.3e-09 | 7.2e-09 | 8.500E-09 | 2.166E-09 | 7.200E-09 | 1.100E-08 |
| 17287 | 865 | F865Q | C | 3.5e-09 | 1.3e-08 | 4.5e-09 | 7.000E-09 | 5.220E-09 | 3.500E-09 | 1.300E-08 |
| 17288 | 865 | F865G | C | 8.4e-09 | 3.6e-09 | 1e-08 | 7.333E-09 | 3.331E-09 | 3.600E-09 | 1.000E-08 |
| 17289 | 865 | F865H | C | 3.2e-09 | 8.6e-09 | 8e-09 | 6.600E-09 | 2.960E-09 | 3.200E-09 | 8.600E-09 |
| 17290 | 865 | F865I | C | 5.4e-09 | 7.1e-09 | 9.9e-09 | 7.467E-09 | 2.272E-09 | 5.400E-09 | 9.900E-09 |
| 17291 | 865 | F865L | C | 1.4e-08 | 5.1e-09 | 1.7e-08 | 1.203E-08 | 6.189E-09 | 5.100E-09 | 1.700E-08 |
| 17292 | 865 | F865K | C | 1e-08 | 2.6e-09 | 1.6e-08 | 9.533E-09 | 6.712E-09 | 2.600E-09 | 1.600E-08 |
| 17293 | 865 | F865M | C | 5.8e-09 | 6e-09 | 6e-09 | 5.933E-09 | 1.155E-10 | 5.800E-09 | 6.000E-09 |
| 17294 | 865 | F865F | C | 1.3e-09 | 2.1e-08 | 5.5e-09 | 9.267E-09 | 1.038E-08 | 1.300E-09 | 2.100E-08 |
| 17295 | 865 | F865P | C | 2.1e-08 | 1e-08 | 5.1e-09 | 1.203E-08 | 8.143E-09 | 5.100E-09 | 2.100E-08 |
| 17296 | 865 | F865S | C | 6.2e-09 | 5.5e-09 | 6.1e-09 | 5.933E-09 | 3.786E-10 | 5.500E-09 | 6.200E-09 |
| 17297 | 865 | F865T | C | 6e-09 | 1.2e-08 | 3.4e-09 | 7.133E-09 | 4.411E-09 | 3.400E-09 | 1.200E-08 |
| 17298 | 865 | F865W | C | 6.7e-09 | 7.5e-09 | 7.6e-09 | 7.267E-09 | 4.933E-10 | 6.700E-09 | 7.600E-09 |
| 17299 | 865 | F865Y | C | 1.6e-09 | 1.1e-08 | 2.1e-09 | 4.900E-09 | 5.289E-09 | 1.600E-09 | 1.100E-08 |
| 17300 | 865 | F865V | C | 4.4e-09 | 7.3e-09 | 1e-08 | 7.233E-09 | 2.801E-09 | 4.400E-09 | 1.000E-08 |
| 17301 | 866 | T866A | C | 4.9e-09 | 1.9e-09 | 4.7e-09 | 3.833E-09 | 1.677E-09 | 1.900E-09 | 4.900E-09 |
| 17302 | 866 | T866R | C | 3.6e-09 | 2e-09 | 4.2e-09 | 3.267E-09 | 1.137E-09 | 2.000E-09 | 4.200E-09 |
| 17303 | 866 | T866N | C | 8.9e-09 | 3.6e-09 | 3.6e-09 | 5.367E-09 | 3.060E-09 | 3.600E-09 | 8.900E-09 |
| 17304 | 866 | T866D | C | 1.6e-09 | 3.9e-09 | 5.5e-09 | 3.667E-09 | 1.960E-09 | 1.600E-09 | 5.500E-09 |
| 17305 | 866 | T866C | C | 1.7e-09 | 2.3e-09 | 5.5e-10 | 1.517E-09 | 8.893E-10 | 5.500E-10 | 2.300E-09 |
| 17306 | 866 | T866E | C | 4.8e-10 | 2.1e-09 | 2.6e-09 | 1.727E-09 | 1.108E-09 | 4.800E-10 | 2.600E-09 |
| 17307 | 866 | T866Q | C | 3.5e-09 | 4.4e-09 | 3.3e-09 | 3.733E-09 | 5.859E-10 | 3.300E-09 | 4.400E-09 |
| 17308 | 866 | T866G | C | 2.2e-09 | 3.7e-09 | 1.2e-09 | 2.367E-09 | 1.258E-09 | 1.200E-09 | 3.700E-09 |
| 17309 | 866 | T866H | C | 1.5e-09 | 2.3e-09 | 4.1e-09 | 2.633E-09 | 1.332E-09 | 1.500E-09 | 4.100E-09 |
| 17310 | 866 | T866I | C | 1.1e-09 | 2.7e-09 | 1.4e-09 | 1.733E-09 | 8.505E-10 | 1.100E-09 | 2.700E-09 |
| 17311 | 866 | T866L | C | 2.4e-09 | 1.9e-09 | 2.5e-09 | 2.267E-09 | 3.215E-10 | 1.900E-09 | 2.500E-09 |
| 17312 | 866 | T866K | C | 2.6e-09 | 1.6e-09 | 2.3e-09 | 2.167E-09 | 5.132E-10 | 1.600E-09 | 2.600E-09 |
| 17313 | 866 | T866M | C | 1.3e-09 | 1.8e-09 | 8.5e-10 | 1.317E-09 | 4.752E-10 | 8.500E-10 | 1.800E-09 |
| 17314 | 866 | T866F | C | 2.5e-09 | 9.1e-10 | 1.2e-09 | 1.537E-09 | 8.468E-10 | 9.100E-10 | 2.500E-09 |
| 17315 | 866 | T866P | C | 3.4e-09 | 1.6e-09 | 1.2e-09 | 2.067E-09 | 1.172E-09 | 1.200E-09 | 3.400E-09 |
| 17316 | 866 | T866S | C | 3.8e-09 | 9.5e-09 | 7.9e-09 | 7.067E-09 | 2.940E-09 | 3.800E-09 | 9.500E-09 |
| 17317 | 866 | T866T | C | 1.6e-08 | 2.7e-09 | 2.6e-09 | 7.100E-09 | 7.708E-09 | 2.600E-09 | 1.600E-08 |
| 17318 | 866 | T866W | C | 4.7e-10 | 3.6e-10 | 1.1e-09 | 6.433E-10 | 3.993E-10 | 3.600E-10 | 1.100E-09 |
| 17319 | 866 | T866Y | C | 1.2e-09 | 1e-09 | 2e-09 | 1.400E-09 | 5.292E-10 | 1.000E-09 | 2.000E-09 |
| 17320 | 866 | T866V | C | 1.7e-09 | 1.3e-09 | 5.4e-10 | 1.180E-09 | 5.892E-10 | 5.400E-10 | 1.700E-09 |
| 17321 | 867 | D867A | C | 2.6e-08 | 8.5e-09 | 7.9e-09 | 1.413E-08 | 1.028E-08 | 7.900E-09 | 2.600E-08 |
| 17322 | 867 | D867R | C | 9.5e-09 | 5.1e-09 | 1.4e-08 | 9.533E-09 | 4.450E-09 | 5.100E-09 | 1.400E-08 |
| 17323 | 867 | D867N | C | 6.8e-09 | 1.3e-08 | 3.2e-09 | 7.667E-09 | 4.957E-09 | 3.200E-09 | 1.300E-08 |
| 17324 | 867 | D867D | C | 3.2e-09 | 8.3e-09 | 5.5e-09 | 5.667E-09 | 2.554E-09 | 3.200E-09 | 8.300E-09 |
| 17325 | 867 | D867C | C | 9.7e-09 | 5.6e-09 | 1.4e-08 | 9.767E-09 | 4.200E-09 | 5.600E-09 | 1.400E-08 |
| 17326 | 867 | D867E | C | 2.7e-09 | 8.7e-09 | 3.3e-09 | 4.900E-09 | 3.305E-09 | 2.700E-09 | 8.700E-09 |
| 17327 | 867 | D867Q | C | 1e-09 | 1.1e-08 | 1.7e-09 | 4.567E-09 | 5.582E-09 | 1.000E-09 | 1.100E-08 |
| 17328 | 867 | D867G | C | 2e-08 | 1e-08 | 4.8e-09 | 1.160E-08 | 7.725E-09 | 4.800E-09 | 2.000E-08 |
| 17329 | 867 | D867H | C | 2.6e-09 | 4e-09 | 1.2e-08 | 6.200E-09 | 5.071E-09 | 2.600E-09 | 1.200E-08 |
| 17330 | 867 | D867I | C | 1.3e-08 | 4.1e-09 | 3.6e-09 | 6.900E-09 | 5.289E-09 | 3.600E-09 | 1.300E-08 |
| 17331 | 867 | D867L | C | 1.5e-08 | 4.6e-09 | 7.7e-09 | 9.100E-09 | 5.339E-09 | 4.600E-09 | 1.500E-08 |
| 17332 | 867 | D867K | C | 3.7e-09 | 3.9e-09 | 1.6e-08 | 7.867E-09 | 7.044E-09 | 3.700E-09 | 1.600E-08 |
| 17333 | 867 | D867M | C | 6.1e-09 | 1.6e-08 | 4e-09 | 8.700E-09 | 6.409E-09 | 4.000E-09 | 1.600E-08 |
| 17334 | 867 | D867F | C | 9.8e-09 | 1.2e-08 | 1e-08 | 1.060E-08 | 1.217E-09 | 9.800E-09 | 1.200E-08 |
| 17335 | 867 | D867P | C | 1.6e-08 | 1.2e-09 | 1.9e-09 | 6.367E-09 | 8.350E-09 | 1.200E-09 | 1.600E-08 |
| 17336 | 867 | D867S | C | 1.1e-08 | 2.1e-08 | 4.2e-09 | 1.207E-08 | 8.451E-09 | 4.200E-09 | 2.100E-08 |
| 17337 | 867 | D867T | C | 3.7e-09 | 1.7e-08 | 8.1e-09 | 9.600E-09 | 6.776E-09 | 3.700E-09 | 1.700E-08 |
| 17338 | 867 | D867W | C | 1.2e-08 | 1.4e-08 | 3.3e-09 | 9.767E-09 | 5.689E-09 | 3.300E-09 | 1.400E-08 |
| 17339 | 867 | D867Y | C | 6.4e-09 | 3e-09 | 4.8e-09 | 4.733E-09 | 1.701E-09 | 3.000E-09 | 6.400E-09 |

| | | | | | | | | | | |
|---|---|---|---|---|---|---|---|---|---|---|
| 17340 | 867 | D867V | C | 7.8e-09 | 7.3e-09 | 3.8e-09 | 6.300E-09 | 2.179E-09 | 3.800E-09 | 7.800E-09 |
| 17341 | 868 | Y868A | C | 2.2e-08 | 5.7e-09 | 6.8e-09 | 1.150E-08 | 9.110E-09 | 5.700E-09 | 2.200E-08 |
| 17342 | 868 | Y868R | C | 4.2e-09 | 6.2e-09 | 3.1e-09 | 4.500E-09 | 1.572E-09 | 3.100E-09 | 6.200E-09 |
| 17343 | 868 | Y868N | C | 1e-08 | 5.4e-09 | 4.3e-09 | 6.567E-09 | 3.024E-09 | 4.300E-09 | 1.000E-08 |
| 17344 | 868 | Y868D | C | 8.8e-09 | 1.1e-08 | 1.8e-08 | 1.260E-08 | 4.804E-09 | 8.800E-09 | 1.800E-08 |
| 17345 | 868 | Y868C | C | 1.3e-08 | 1.5e-08 | 1.4e-08 | 1.400E-08 | 1.000E-09 | 1.300E-08 | 1.500E-08 |
| 17346 | 868 | Y868E | C | 9.3e-09 | 6e-09 | 6.4e-09 | 7.233E-09 | 1.801E-09 | 6.000E-09 | 9.300E-09 |
| 17347 | 868 | Y868Q | C | 6.9e-09 | 9.8e-09 | 4.5e-09 | 7.067E-09 | 2.654E-09 | 4.500E-09 | 9.800E-09 |
| 17348 | 868 | Y868G | C | 4.4e-09 | 1.5e-08 | 9.4e-09 | 9.600E-09 | 5.303E-09 | 4.400E-09 | 1.500E-08 |
| 17349 | 868 | Y868H | C | 4.2e-09 | 3.4e-09 | 9.4e-09 | 5.667E-09 | 3.258E-09 | 3.400E-09 | 9.400E-09 |
| 17350 | 868 | Y868I | C | 7.3e-09 | 1.3e-08 | 2.1e-08 | 1.377E-08 | 6.882E-09 | 7.300E-09 | 2.100E-08 |
| 17351 | 868 | Y868L | C | 9.3e-09 | 1.2e-08 | 9.8e-09 | 1.037E-08 | 1.436E-09 | 9.300E-09 | 1.200E-08 |
| 17352 | 868 | Y868K | C | 5.4e-09 | 6.8e-09 | 9.6e-09 | 7.267E-09 | 2.139E-09 | 5.400E-09 | 9.600E-09 |
| 17353 | 868 | Y868M | C | 9.4e-09 | 8.6e-09 | 7.2e-09 | 8.400E-09 | 1.114E-09 | 7.200E-09 | 9.400E-09 |
| 17354 | 868 | Y868F | C | 3.5e-09 | 1.3e-08 | 3.8e-09 | 6.767E-09 | 5.400E-09 | 3.500E-09 | 1.300E-08 |
| 17355 | 868 | Y868P | C | 1.6e-08 | 8.3e-09 | 1.2e-08 | 1.210E-08 | 3.851E-09 | 8.300E-09 | 1.600E-08 |
| 17356 | 868 | Y868S | C | 4.1e-09 | 5.2e-09 | 8.9e-09 | 6.067E-09 | 2.515E-09 | 4.100E-09 | 8.900E-09 |
| 17357 | 868 | Y868T | C | 7.4e-09 | 6e-09 | 8.9e-09 | 7.433E-09 | 1.450E-09 | 6.000E-09 | 8.900E-09 |
| 17358 | 868 | Y868W | C | 3.6e-09 | 6.5e-09 | 7.7e-09 | 5.933E-09 | 2.108E-09 | 3.600E-09 | 7.700E-09 |
| 17359 | 868 | Y868Y | C | 1.4e-08 | 7.5e-09 | 7.7e-09 | 9.733E-09 | 3.696E-09 | 7.500E-09 | 1.400E-08 |
| 17360 | 868 | Y868V | C | 9.2e-09 | 1.5e-08 | 1e-08 | 1.140E-08 | 3.143E-09 | 9.200E-09 | 1.500E-08 |
| 17361 | 869 | T869A | C | 7.2e-09 | 8.2e-09 | 4.8e-09 | 6.733E-09 | 1.747E-09 | 4.800E-09 | 8.200E-09 |
| 17362 | 869 | T869R | C | 3.8e-09 | 1.3e-08 | 6.8e-09 | 7.867E-09 | 4.692E-09 | 3.800E-09 | 1.300E-08 |
| 17363 | 869 | T869N | C | 1.1e-08 | 1.1e-08 | 3.4e-09 | 8.467E-09 | 4.388E-09 | 3.400E-09 | 1.100E-08 |
| 17364 | 869 | T869D | C | 1.6e-08 | 1.9e-08 | 3.6e-09 | 1.287E-08 | 8.164E-09 | 3.600E-09 | 1.900E-08 |
| 17365 | 869 | T869C | C | 1.4e-08 | 1.2e-08 | 8.6e-09 | 1.153E-08 | 2.730E-09 | 8.600E-09 | 1.400E-08 |
| 17366 | 869 | T869E | C | 1.5e-08 | 7.1e-09 | 9.9e-09 | 1.067E-08 | 4.005E-09 | 7.100E-09 | 1.500E-08 |
| 17367 | 869 | T869Q | C | 4.8e-09 | 6.5e-09 | 5.5e-09 | 5.600E-09 | 8.544E-10 | 4.800E-09 | 6.500E-09 |
| 17368 | 869 | T869G | C | 3.9e-09 | 1.5e-09 | 3.7e-09 | 3.033E-09 | 1.332E-09 | 1.500E-09 | 3.900E-09 |
| 17369 | 869 | T869H | C | 9.5e-09 | 1.2e-08 | 9.4e-09 | 1.030E-08 | 1.473E-09 | 9.400E-09 | 1.200E-08 |
| 17370 | 869 | T869I | C | 1.7e-08 | 1.1e-08 | 1.6e-08 | 1.467E-08 | 3.215E-09 | 1.100E-08 | 1.700E-08 |
| 17371 | 869 | T869L | C | 4.3e-09 | 8.1e-09 | 1.3e-08 | 8.467E-09 | 4.362E-09 | 4.300E-09 | 1.300E-08 |
| 17372 | 869 | T869K | C | 7.7e-09 | 1.1e-08 | 1.8e-08 | 1.223E-08 | 5.260E-09 | 7.700E-09 | 1.800E-08 |
| 17373 | 869 | T869M | C | 9.2e-09 | 2e-09 | 2.5e-08 | 1.207E-08 | 1.176E-08 | 2.000E-09 | 2.500E-08 |
| 17374 | 869 | T869F | C | 7.3e-09 | 4.2e-09 | 8.2e-09 | 6.567E-09 | 2.098E-09 | 4.200E-09 | 8.200E-09 |
| 17375 | 869 | T869P | C | 1e-08 | 1.5e-08 | 5.6e-08 | 2.700E-08 | 2.524E-08 | 1.000E-08 | 5.600E-08 |
| 17376 | 869 | T869S | C | 1.3e-08 | 1.2e-08 | 1.4e-08 | 1.300E-08 | 1.000E-09 | 1.200E-08 | 1.400E-08 |
| 17377 | 869 | T869T | C | 2.9e-08 | 5e-09 | 1.1e-08 | 1.500E-08 | 1.249E-08 | 5.000E-09 | 2.900E-08 |
| 17378 | 869 | T869W | C | 5.6e-09 | 5.4e-09 | 1.1e-08 | 7.333E-09 | 3.177E-09 | 5.400E-09 | 1.100E-08 |
| 17379 | 869 | T869Y | C | 1.2e-08 | 5.3e-09 | 4e-09 | 7.100E-09 | 4.293E-09 | 4.000E-09 | 1.200E-08 |
| 17380 | 869 | T869V | C | 1.5e-08 | 1.9e-09 | 4.5e-09 | 7.133E-09 | 6.936E-09 | 1.900E-09 | 1.500E-08 |
| 17381 | 870 | M870A | C | 4.9e-09 | 7.9e-09 | 1e-08 | 7.600E-09 | 2.563E-09 | 4.900E-09 | 1.000E-08 |
| 17382 | 870 | M870R | C | 3.3e-09 | 4.7e-09 | 1.2e-08 | 6.667E-09 | 4.672E-09 | 3.300E-09 | 1.200E-08 |
| 17383 | 870 | M870N | C | 8e-09 | 6.5e-09 | 5.4e-09 | 6.633E-09 | 1.305E-09 | 5.400E-09 | 8.000E-09 |
| 17384 | 870 | M870D | C | 5.4e-09 | 1.1e-08 | 8.9e-09 | 8.433E-09 | 2.829E-09 | 5.400E-09 | 1.100E-08 |
| 17385 | 870 | M870C | C | 5.9e-09 | 8.5e-09 | 8.2e-09 | 7.533E-09 | 1.422E-09 | 5.900E-09 | 8.500E-09 |
| 17386 | 870 | M870E | C | 2.2e-08 | 1.1e-08 | 8.3e-09 | 1.377E-08 | 7.257E-09 | 8.300E-09 | 2.200E-08 |
| 17387 | 870 | M870Q | C | 1.1e-08 | 1.8e-08 | 1.1e-08 | 1.333E-08 | 4.041E-09 | 1.100E-08 | 1.800E-08 |
| 17388 | 870 | M870G | C | 9.9e-09 | 8.9e-09 | 1.1e-08 | 9.933E-09 | 1.050E-09 | 8.900E-09 | 1.100E-08 |
| 17389 | 870 | M870H | C | 1e-08 | 1e-08 | 6.3e-09 | 8.767E-09 | 2.136E-09 | 6.300E-09 | 1.000E-08 |
| 17390 | 870 | M870I | C | 1.1e-08 | 2.2e-09 | 4.2e-09 | 5.800E-09 | 4.613E-09 | 2.200E-09 | 1.100E-08 |
| 17391 | 870 | M870L | C | 7.2e-09 | 2e-09 | 5e-08 | 1.973E-08 | 2.634E-08 | 2.000E-09 | 5.000E-08 |
| 17392 | 870 | M870K | C | 5.9e-09 | 4.8e-09 | 9.1e-09 | 6.600E-09 | 2.234E-09 | 4.800E-09 | 9.100E-09 |
| 17393 | 870 | M870M | C | 3.6e-09 | 9.1e-09 | 1e-08 | 7.567E-09 | 3.465E-09 | 3.600E-09 | 1.000E-08 |
| 17394 | 870 | M870F | C | 5.4e-09 | 4.8e-09 | 7.2e-09 | 5.800E-09 | 1.249E-09 | 4.800E-09 | 7.200E-09 |
| 17395 | 870 | M870P | C | 5.4e-09 | 3.5e-09 | 7.7e-09 | 5.533E-09 | 2.103E-09 | 3.500E-09 | 7.700E-09 |
| 17396 | 870 | M870S | C | 4.3e-09 | 4.8e-09 | 5.3e-09 | 4.800E-09 | 5.000E-10 | 4.300E-09 | 5.300E-09 |
| 17397 | 870 | M870T | C | 1.4e-08 | 1.8e-08 | 1.7e-08 | 1.633E-08 | 2.082E-09 | 1.400E-08 | 1.800E-08 |
| 17398 | 870 | M870W | C | 1.2e-08 | 1.3e-08 | 8.3e-09 | 1.110E-08 | 2.476E-09 | 8.300E-09 | 1.300E-08 |
| 17399 | 870 | M870Y | C | 6.1e-09 | 1.5e-08 | 9.4e-09 | 1.017E-08 | 4.499E-09 | 6.100E-09 | 1.500E-08 |
| 17400 | 870 | M870V | C | 7.7e-09 | 2.9e-09 | 2.4e-09 | 4.333E-09 | 2.926E-09 | 2.400E-09 | 7.700E-09 |
| 17401 | 871 | D871A | C | 1.8e-08 | 4.4e-09 | 7.3e-09 | 9.900E-09 | 7.163E-09 | 4.400E-09 | 1.800E-08 |
| 17402 | 871 | D871R | C | 3.6e-09 | 7.3e-09 | 6.3e-09 | 5.733E-09 | 1.914E-09 | 3.600E-09 | 7.300E-09 |
| 17403 | 871 | D871N | C | 5.1e-09 | 4.5e-09 | 1.5e-08 | 8.200E-09 | 5.897E-09 | 4.500E-09 | 1.500E-08 |
| 17404 | 871 | D871D | C | 4.4e-09 | 1.4e-08 | 1.3e-08 | 1.047E-08 | 5.278E-09 | 4.400E-09 | 1.400E-08 |
| 17405 | 871 | D871C | C | 2.2e-08 | 8.3e-09 | 1.5e-08 | 1.510E-08 | 6.851E-09 | 8.300E-09 | 2.200E-08 |
| 17406 | 871 | D871E | C | 7.8e-09 | 1.3e-08 | 9.9e-09 | 1.023E-08 | 2.616E-09 | 7.800E-09 | 1.300E-08 |
| 17407 | 871 | D871Q | C | 4.4e-09 | 5.4e-09 | 3.6e-09 | 4.467E-09 | 9.018E-10 | 3.600E-09 | 5.400E-09 |
| 17408 | 871 | D871G | C | 6.3e-09 | 7.2e-09 | 2.3e-09 | 5.267E-09 | 2.608E-09 | 2.300E-09 | 7.200E-09 |
| 17409 | 871 | D871H | C | 9.1e-09 | 1.4e-08 | 6.7e-09 | 9.933E-09 | 3.721E-09 | 6.700E-09 | 1.400E-08 |
| 17410 | 871 | D871I | C | 8.6e-09 | 3e-09 | 8.3e-09 | 6.633E-09 | 3.150E-09 | 3.000E-09 | 8.600E-09 |
| 17411 | 871 | D871L | C | 1.1e-08 | 3.8e-09 | 7e-09 | 7.267E-09 | 3.607E-09 | 3.800E-09 | 1.100E-08 |
| 17412 | 871 | D871K | C | 7e-09 | 2.4e-09 | 2.8e-08 | 1.247E-08 | 1.365E-08 | 2.400E-09 | 2.800E-08 |
| 17413 | 871 | D871M | C | 9.6e-09 | 2.1e-08 | 2.4e-08 | 1.820E-08 | 7.597E-09 | 9.600E-09 | 2.400E-08 |
| 17414 | 871 | D871F | C | 8.8e-09 | 6.8e-09 | 2.5e-08 | 1.353E-08 | 9.981E-09 | 6.800E-09 | 2.500E-08 |
| 17415 | 871 | D871P | C | 2.9e-09 | 3e-09 | 4.2e-09 | 3.367E-09 | 7.234E-10 | 2.900E-09 | 4.200E-09 |
| 17416 | 871 | D871S | C | 2e-09 | 4e-09 | 1.3e-08 | 6.333E-09 | 5.859E-09 | 2.000E-09 | 1.300E-08 |
| 17417 | 871 | D871T | C | 1.1e-08 | 2.7e-09 | 1.7e-08 | 1.023E-08 | 7.181E-09 | 2.700E-09 | 1.700E-08 |
| 17418 | 871 | D871W | C | 1.5e-09 | 1.1e-08 | 1e-08 | 7.500E-09 | 5.220E-09 | 1.500E-09 | 1.100E-08 |
| 17419 | 871 | D871Y | C | 1.2e-08 | 1.4e-09 | 9.8e-09 | 7.733E-09 | 5.594E-09 | 1.400E-09 | 1.200E-08 |
| 17420 | 871 | D871V | C | 1e-08 | 8.4e-09 | 4.1e-09 | 7.500E-09 | 3.051E-09 | 4.100E-09 | 1.000E-08 |
| 17421 | 872 | W872A | C | 4.9e-09 | 1.6e-08 | 5.7e-09 | 8.867E-09 | 6.191E-09 | 4.900E-09 | 1.600E-08 |
| 17422 | 872 | W872R | C | 7.1e-09 | 5.1e-09 | 9.5e-09 | 7.233E-09 | 2.203E-09 | 5.100E-09 | 9.500E-09 |
| 17423 | 872 | W872N | C | 4.2e-09 | 3.8e-09 | 5.9e-09 | 4.633E-09 | 1.115E-09 | 3.800E-09 | 5.900E-09 |
| 17424 | 872 | W872D | C | 5.4e-09 | 3.8e-09 | 6.4e-09 | 5.200E-09 | 1.311E-09 | 3.800E-09 | 6.400E-09 |

| | | | | | | | | | | |
|---|---|---|---|---|---|---|---|---|---|---|
| 17425 | 872 | W872C | C | 1.2e-08 | 4.2e-09 | 1.5e-09 | 5.900E-09 | 5.453E-09 | 1.500E-09 | 1.200E-08 |
| 17426 | 872 | W872E | C | 1.1e-08 | 9.9e-09 | 1.2e-08 | 1.097E-08 | 1.050E-09 | 9.900E-09 | 1.200E-08 |
| 17427 | 872 | W872Q | C | 7.1e-09 | 6.8e-09 | 1.1e-08 | 8.300E-09 | 2.343E-09 | 6.800E-09 | 1.100E-08 |
| 17428 | 872 | W872G | C | 7.2e-09 | 4.5e-09 | 2.5e-09 | 4.733E-09 | 2.359E-09 | 2.500E-09 | 7.200E-09 |
| 17429 | 872 | W872H | C | 4.7e-09 | 9e-09 | 5e-09 | 6.233E-09 | 2.401E-09 | 4.700E-09 | 9.000E-09 |
| 17430 | 872 | W872I | C | 2.6e-09 | 1.6e-08 | 5.2e-09 | 7.933E-09 | 7.106E-09 | 2.600E-09 | 1.600E-08 |
| 17431 | 872 | W872L | C | 8.7e-09 | 1.2e-08 | 1.2e-08 | 1.090E-08 | 1.905E-09 | 8.700E-09 | 1.200E-08 |
| 17432 | 872 | W872K | C | 7.9e-09 | 1.7e-08 | 2.1e-09 | 9.000E-09 | 7.511E-09 | 2.100E-09 | 1.700E-08 |
| 17433 | 872 | W872M | C | 1.6e-09 | 1.8e-08 | 1.2e-08 | 1.053E-08 | 8.298E-09 | 1.600E-09 | 1.800E-08 |
| 17434 | 872 | W872F | C | 1.3e-08 | 1.6e-08 | 8.4e-09 | 1.247E-08 | 3.828E-09 | 8.400E-09 | 1.600E-08 |
| 17435 | 872 | W872P | C | 2.2e-08 | 5.6e-09 | 7.9e-09 | 1.183E-08 | 8.879E-09 | 5.600E-09 | 2.200E-08 |
| 17436 | 872 | W872S | C | 6.5e-09 | 8.6e-09 | 5.4e-09 | 6.833E-09 | 1.626E-09 | 5.400E-09 | 8.600E-09 |
| 17437 | 872 | W872T | C | 5.1e-09 | 7.4e-09 | 1.1e-08 | 7.833E-09 | 2.974E-09 | 5.100E-09 | 1.100E-08 |
| 17438 | 872 | W872W | C | 2.7e-08 | 1.6e-08 | 3.8e-09 | 1.560E-08 | 1.161E-08 | 3.800E-09 | 2.700E-08 |
| 17439 | 872 | W872Y | C | 2.5e-08 | 9.6e-09 | 5e-09 | 1.320E-08 | 1.047E-08 | 5.000E-09 | 2.500E-08 |
| 17440 | 872 | W872V | C | 7.6e-09 | 5.7e-09 | 1.1e-08 | 8.100E-09 | 2.685E-09 | 5.700E-09 | 1.100E-08 |
| 17441 | 873 | V873A | C | 9.3e-09 | 5.8e-09 | 5.3e-09 | 6.800E-09 | 2.179E-09 | 5.300E-09 | 9.300E-09 |
| 17442 | 873 | V873R | C | 5.8e-09 | 3.6e-09 | 2e-09 | 3.800E-09 | 1.908E-09 | 2.000E-09 | 5.800E-09 |
| 17443 | 873 | V873N | C | 5.4e-09 | 1.8e-08 | 5.8e-09 | 9.733E-09 | 7.162E-09 | 5.400E-09 | 1.800E-08 |
| 17444 | 873 | V873D | C | 1.8e-08 | 5.5e-09 | 6.5e-09 | 1.000E-08 | 6.946E-09 | 5.500E-09 | 1.800E-08 |
| 17445 | 873 | V873C | C | 3.3e-09 | 1.3e-08 | 6.1e-09 | 7.467E-09 | 4.992E-09 | 3.300E-09 | 1.300E-08 |
| 17446 | 873 | V873E | C | 9.4e-09 | 3.6e-09 | 1.7e-08 | 1.000E-08 | 6.720E-09 | 3.600E-09 | 1.700E-08 |
| 17447 | 873 | V873Q | C | 8.1e-09 | 1.2e-08 | 3.4e-09 | 7.833E-09 | 4.306E-09 | 3.400E-09 | 1.200E-08 |
| 17448 | 873 | V873G | C | 4.4e-09 | 4.7e-09 | 4.4e-09 | 4.500E-09 | 1.732E-10 | 4.400E-09 | 4.700E-09 |
| 17449 | 873 | V873H | C | 4.3e-09 | 8.8e-09 | 4.7e-09 | 5.933E-09 | 2.491E-09 | 4.300E-09 | 8.800E-09 |
| 17450 | 873 | V873I | C | 3e-09 | 1.8e-08 | 6.4e-09 | 9.133E-09 | 7.865E-09 | 3.000E-09 | 1.800E-08 |
| 17451 | 873 | V873L | C | 9e-09 | 2.3e-08 | 1.4e-08 | 1.533E-08 | 7.095E-09 | 9.000E-09 | 2.300E-08 |
| 17452 | 873 | V873K | C | 1.2e-08 | 1.2e-08 | 7.1e-09 | 1.037E-08 | 2.829E-09 | 7.100E-09 | 1.200E-08 |
| 17453 | 873 | V873M | C | 9.9e-09 | 8.1e-09 | 5.4e-09 | 7.800E-09 | 2.265E-09 | 5.400E-09 | 9.900E-09 |
| 17454 | 873 | V873F | C | 1.4e-09 | 2.5e-09 | 5.5e-09 | 3.133E-09 | 2.122E-09 | 1.400E-09 | 5.500E-09 |
| 17455 | 873 | V873P | C | 8.7e-09 | 1.8e-08 | 1.8e-08 | 1.490E-08 | 5.369E-09 | 8.700E-09 | 1.800E-08 |
| 17456 | 873 | V873S | C | 8.4e-09 | 5.2e-09 | 3.6e-09 | 5.733E-09 | 2.444E-09 | 3.600E-09 | 8.400E-09 |
| 17457 | 873 | V873T | C | 8.1e-09 | 4.4e-09 | 3e-09 | 5.167E-09 | 2.635E-09 | 3.000E-09 | 8.100E-09 |
| 17458 | 873 | V873W | C | 9.5e-09 | 1.8e-08 | 7.5e-09 | 1.167E-08 | 5.575E-09 | 7.500E-09 | 1.800E-08 |
| 17459 | 873 | V873Y | C | 9.6e-09 | 9e-09 | 1.3e-08 | 1.053E-08 | 2.157E-09 | 9.000E-09 | 1.300E-08 |
| 17460 | 873 | V873V | C | 1.6e-08 | 5.1e-09 | 4e-09 | 8.367E-09 | 6.634E-09 | 4.000E-09 | 1.600E-08 |
| 17461 | 874 | R874A | C | 2.7e-09 | 6.7e-09 | 1.3e-08 | 7.467E-09 | 5.193E-09 | 2.700E-09 | 1.300E-08 |
| 17462 | 874 | R874R | C | 6.9e-09 | 1e-08 | 8e-09 | 8.300E-09 | 1.572E-09 | 6.900E-09 | 1.000E-08 |
| 17463 | 874 | R874N | C | 1.6e-08 | 2.3e-08 | 8.3e-09 | 1.577E-08 | 7.353E-09 | 8.300E-09 | 2.300E-08 |
| 17464 | 874 | R874D | C | 1.1e-08 | 7.3e-09 | 4.2e-09 | 7.500E-09 | 3.404E-09 | 4.200E-09 | 1.100E-08 |
| 17465 | 874 | R874C | C | 2.2e-09 | 8e-09 | 2.9e-09 | 4.367E-09 | 3.166E-09 | 2.200E-09 | 8.000E-09 |
| 17466 | 874 | R874E | C | 1.2e-08 | 2.7e-08 | 5.4e-09 | 1.480E-08 | 1.107E-08 | 5.400E-09 | 2.700E-08 |
| 17467 | 874 | R874Q | C | 7.7e-09 | 9.5e-09 | 1.4e-08 | 1.040E-08 | 3.245E-09 | 7.700E-09 | 1.400E-08 |
| 17468 | 874 | R874G | C | 5.6e-09 | 5.9e-09 | 1.2e-08 | 7.833E-09 | 3.612E-09 | 5.600E-09 | 1.200E-08 |
| 17469 | 874 | R874H | C | 8.7e-09 | 1e-08 | 6.4e-09 | 8.367E-09 | 1.823E-09 | 6.400E-09 | 1.000E-08 |
| 17470 | 874 | R874I | C | 8.7e-09 | 3.6e-09 | 3.9e-09 | 5.400E-09 | 2.862E-09 | 3.600E-09 | 8.700E-09 |
| 17471 | 874 | R874L | C | 7.2e-09 | 7.7e-09 | 1.2e-08 | 8.967E-09 | 2.639E-09 | 7.200E-09 | 1.200E-08 |
| 17472 | 874 | R874K | C | 8.2e-09 | 6.9e-09 | 6e-09 | 7.033E-09 | 1.106E-09 | 6.000E-09 | 8.200E-09 |
| 17473 | 874 | R874M | C | 6.4e-09 | 1.9e-09 | 9.7e-09 | 6.000E-09 | 3.915E-09 | 1.900E-09 | 9.700E-09 |
| 17474 | 874 | R874F | C | 3.7e-09 | 1.6e-08 | 6.8e-09 | 8.833E-09 | 6.397E-09 | 3.700E-09 | 1.600E-08 |
| 17475 | 874 | R874P | C | 2.4e-08 | 4.4e-09 | 1.2e-08 | 1.347E-08 | 9.882E-09 | 4.400E-09 | 2.400E-08 |
| 17476 | 874 | R874S | C | 9.7e-09 | 3.4e-09 | 6.9e-09 | 6.667E-09 | 3.156E-09 | 3.400E-09 | 9.700E-09 |
| 17477 | 874 | R874T | C | 8.4e-09 | 8.3e-09 | 5.9e-09 | 7.533E-09 | 1.415E-09 | 5.900E-09 | 8.400E-09 |
| 17478 | 874 | R874W | C | 2.4e-09 | 9.7e-09 | 2.8e-09 | 4.967E-09 | 4.104E-09 | 2.400E-09 | 9.700E-09 |
| 17479 | 874 | R874Y | C | 7.6e-09 | 2e-08 | 1.1e-08 | 1.287E-08 | 6.407E-09 | 7.600E-09 | 2.000E-08 |
| 17480 | 874 | R874V | C | 3.2e-09 | 7.5e-09 | 4.6e-09 | 5.100E-09 | 2.193E-09 | 3.200E-09 | 7.500E-09 |
| 17481 | 875 | Q875A | C | 7.3e-09 | 3.8e-09 | 2.5e-08 | 1.203E-08 | 1.137E-08 | 3.800E-09 | 2.500E-08 |
| 17482 | 875 | Q875R | C | 1.3e-08 | 1.2e-08 | 9.4e-09 | 1.147E-08 | 1.858E-09 | 9.400E-09 | 1.300E-08 |
| 17483 | 875 | Q875N | C | 6e-09 | 3e-09 | 8.2e-09 | 5.733E-09 | 2.610E-09 | 3.000E-09 | 8.200E-09 |
| 17484 | 875 | Q875D | C | 1e-08 | 8.3e-10 | 4.8e-09 | 5.210E-09 | 4.599E-09 | 8.300E-10 | 1.000E-08 |
| 17485 | 875 | Q875C | C | 5.6e-09 | 7.9e-09 | 9.6e-09 | 7.700E-09 | 2.007E-09 | 5.600E-09 | 9.600E-09 |
| 17486 | 875 | Q875E | C | 3.6e-08 | 9e-09 | 5.6e-09 | 1.687E-08 | 1.666E-08 | 5.600E-09 | 3.600E-08 |
| 17487 | 875 | Q875Q | C | 1.3e-08 | 4.7e-09 | 7e-09 | 8.233E-09 | 4.285E-09 | 4.700E-09 | 1.300E-08 |
| 17488 | 875 | Q875G | C | 8.2e-09 | 5.6e-09 | 8.9e-09 | 7.567E-09 | 1.739E-09 | 5.600E-09 | 8.900E-09 |
| 17489 | 875 | Q875H | C | 1.2e-08 | 9e-09 | 1.5e-08 | 1.200E-08 | 3.000E-09 | 9.000E-09 | 1.500E-08 |
| 17490 | 875 | Q875I | C | 5.6e-09 | 1.2e-08 | 8.5e-09 | 8.700E-09 | 3.205E-09 | 5.600E-09 | 1.200E-08 |
| 17491 | 875 | Q875L | C | 6e-09 | 9.6e-09 | 4.5e-09 | 6.700E-09 | 2.621E-09 | 4.500E-09 | 9.600E-09 |
| 17492 | 875 | Q875K | C | 8.4e-09 | 7.9e-09 | 5.6e-09 | 7.300E-09 | 1.493E-09 | 5.600E-09 | 8.400E-09 |
| 17493 | 875 | Q875M | C | 4.8e-09 | 7.2e-09 | 9.9e-09 | 7.300E-09 | 2.551E-09 | 4.800E-09 | 9.900E-09 |
| 17494 | 875 | Q875F | C | 8.2e-09 | 1.2e-08 | 1.2e-08 | 1.073E-08 | 2.194E-09 | 8.200E-09 | 1.200E-08 |
| 17495 | 875 | Q875P | C | 5.2e-09 | 3e-09 | 3.2e-09 | 3.800E-09 | 1.217E-09 | 3.000E-09 | 5.200E-09 |
| 17496 | 875 | Q875S | C | 5.8e-09 | 2.2e-09 | 5.9e-08 | 2.233E-08 | 3.181E-08 | 2.200E-09 | 5.900E-08 |
| 17497 | 875 | Q875T | C | 5.8e-09 | 5.1e-09 | 6.6e-09 | 5.833E-09 | 7.506E-10 | 5.100E-09 | 6.600E-09 |
| 17498 | 875 | Q875W | C | 1.1e-08 | 6.3e-09 | 9.2e-09 | 8.833E-09 | 2.371E-09 | 6.300E-09 | 1.100E-08 |
| 17499 | 875 | Q875Y | C | 1e-08 | 3e-09 | 1.6e-08 | 9.667E-09 | 6.506E-09 | 3.000E-09 | 1.600E-08 |
| 17500 | 875 | Q875V | C | 4e-09 | 1.6e-08 | 2.7e-09 | 7.567E-09 | 7.332E-09 | 2.700E-09 | 1.600E-08 |
| 17501 | 876 | A876A | C | 5.1e-09 | 2.4e-09 | 1.4e-08 | 7.167E-09 | 6.070E-09 | 2.400E-09 | 1.400E-08 |
| 17502 | 876 | A876R | C | 8e-09 | 4.9e-09 | 9e-09 | 7.300E-09 | 2.138E-09 | 4.900E-09 | 9.000E-09 |
| 17503 | 876 | A876N | C | 1.4e-09 | 9.5e-09 | 4.5e-09 | 5.133E-09 | 4.087E-09 | 1.400E-09 | 9.500E-09 |
| 17504 | 876 | A876D | C | 2.6e-09 | 1.2e-08 | 2.3e-09 | 5.633E-09 | 5.516E-09 | 2.300E-09 | 1.200E-08 |
| 17505 | 876 | A876C | C | 1e-08 | 5.9e-09 | 4e-09 | 6.633E-09 | 3.066E-09 | 4.000E-09 | 1.000E-08 |
| 17506 | 876 | A876E | C | 4.5e-09 | 3.2e-09 | 2.9e-09 | 3.533E-09 | 8.505E-10 | 2.900E-09 | 4.500E-09 |
| 17507 | 876 | A876Q | C | 4.2e-09 | 6.9e-09 | 1.1e-08 | 7.367E-09 | 3.424E-09 | 4.200E-09 | 1.100E-08 |
| 17508 | 876 | A876G | C | 9.9e-09 | 1.2e-08 | 5.6e-09 | 9.167E-09 | 3.262E-09 | 5.600E-09 | 1.200E-08 |
| 17509 | 876 | A876H | C | 2.6e-08 | 1.2e-08 | 7e-09 | 1.500E-08 | 9.849E-09 | 7.000E-09 | 2.600E-08 |

| | | | | | | | | | | |
|---|---|---|---|---|---|---|---|---|---|---|
| 17510 | 876 | A876I | C | 1e-08 | 6.4e-09 | 5e-09 | 7.133E-09 | 2.579E-09 | 5.000E-09 | 1.000E-08 |
| 17511 | 876 | A876L | C | 2.6e-09 | 5.3e-09 | 6.3e-09 | 4.733E-09 | 1.914E-09 | 2.600E-09 | 6.300E-09 |
| 17512 | 876 | A876K | C | 4.4e-09 | 1.1e-08 | 6.3e-09 | 7.233E-09 | 3.398E-09 | 4.400E-09 | 1.100E-08 |
| 17513 | 876 | A876M | C | 1.2e-08 | 3e-09 | 1.4e-08 | 9.667E-09 | 5.859E-09 | 3.000E-09 | 1.400E-08 |
| 17514 | 876 | A876F | C | 1.1e-08 | 8.1e-09 | 6.1e-09 | 8.400E-09 | 2.464E-09 | 6.100E-09 | 1.100E-08 |
| 17515 | 876 | A876P | C | 9e-09 | 2.9e-09 | 3.6e-09 | 5.167E-09 | 3.338E-09 | 2.900E-09 | 9.000E-09 |
| 17516 | 876 | A876S | C | 5.6e-09 | 1.6e-08 | 6.3e-09 | 9.300E-09 | 5.813E-09 | 5.600E-09 | 1.600E-08 |
| 17517 | 876 | A876T | C | 2.4e-08 | 1.3e-08 | 8.1e-09 | 1.503E-08 | 8.143E-09 | 8.100E-09 | 2.400E-08 |
| 17518 | 876 | A876W | C | 1.7e-09 | 5.5e-09 | 7.2e-09 | 4.800E-09 | 2.816E-09 | 1.700E-09 | 7.200E-09 |
| 17519 | 876 | A876Y | C | 4.8e-09 | 1.2e-08 | 7.1e-09 | 7.967E-09 | 3.677E-09 | 4.800E-09 | 1.200E-08 |
| 17520 | 876 | A876V | C | 1.2e-08 | 1.1e-08 | 1.4e-08 | 1.233E-08 | 1.528E-09 | 1.100E-08 | 1.400E-08 |
| 17521 | 877 | P877A | C | 1.1e-08 | 1.7e-08 | 1.8e-08 | 1.533E-08 | 3.786E-09 | 1.100E-08 | 1.800E-08 |
| 17522 | 877 | P877R | C | 9.4e-09 | 5.2e-09 | 1.2e-08 | 8.867E-09 | 3.431E-09 | 5.200E-09 | 1.200E-08 |
| 17523 | 877 | P877N | C | 1.1e-08 | 2.4e-08 | 5e-09 | 1.333E-08 | 9.713E-09 | 5.000E-09 | 2.400E-08 |
| 17524 | 877 | P877D | C | 3.4e-09 | 7.5e-09 | 7.9e-09 | 6.267E-09 | 2.491E-09 | 3.400E-09 | 7.900E-09 |
| 17525 | 877 | P877C | C | 1.2e-08 | 1.9e-09 | 5.6e-09 | 6.500E-09 | 5.110E-09 | 1.900E-09 | 1.200E-08 |
| 17526 | 877 | P877E | C | 5.4e-09 | 3.8e-09 | 8.3e-09 | 5.833E-09 | 2.281E-09 | 3.800E-09 | 8.300E-09 |
| 17527 | 877 | P877Q | C | 3.5e-09 | 6.3e-09 | 1.2e-08 | 7.267E-09 | 4.332E-09 | 3.500E-09 | 1.200E-08 |
| 17528 | 877 | P877G | C | 1.1e-08 | 1.1e-08 | 9.2e-09 | 1.040E-08 | 1.039E-09 | 9.200E-09 | 1.100E-08 |
| 17529 | 877 | P877H | C | 7.6e-09 | 8.3e-09 | 3e-09 | 6.300E-09 | 2.879E-09 | 3.000E-09 | 8.300E-09 |
| 17530 | 877 | P877I | C | 7.6e-09 | 4e-09 | 2.7e-09 | 4.767E-09 | 2.538E-09 | 2.700E-09 | 7.600E-09 |
| 17531 | 877 | P877L | C | 2.1e-08 | 1.6e-08 | 3.9e-09 | 1.363E-08 | 8.792E-09 | 3.900E-09 | 2.100E-08 |
| 17532 | 877 | P877K | C | 4.2e-09 | 5.5e-09 | 3.9e-09 | 4.533E-09 | 8.505E-10 | 3.900E-09 | 5.500E-09 |
| 17533 | 877 | P877M | C | 9.2e-09 | 1.5e-08 | 1e-08 | 1.140E-08 | 3.143E-09 | 9.200E-09 | 1.500E-08 |
| 17534 | 877 | P877F | C | 4.5e-09 | 9.9e-09 | 8.1e-09 | 7.500E-09 | 2.750E-09 | 4.500E-09 | 9.900E-09 |
| 17535 | 877 | P877P | C | 1.8e-08 | 1.3e-08 | 1e-08 | 1.367E-08 | 4.041E-09 | 1.000E-08 | 1.800E-08 |
| 17536 | 877 | P877S | C | 6.7e-09 | 9.1e-09 | 4.4e-09 | 6.733E-09 | 2.350E-09 | 4.400E-09 | 9.100E-09 |
| 17537 | 877 | P877T | C | 8e-09 | 9.3e-09 | 7.9e-09 | 8.400E-09 | 7.810E-10 | 7.900E-09 | 9.300E-09 |
| 17538 | 877 | P877W | C | 2.9e-09 | 2.5e-09 | 5.9e-09 | 3.767E-09 | 1.858E-09 | 2.500E-09 | 5.900E-09 |
| 17539 | 877 | P877Y | C | 1.3e-08 | 2e-08 | 1.3e-08 | 1.533E-08 | 4.041E-09 | 1.300E-08 | 2.000E-08 |
| 17540 | 877 | P877V | C | 1e-08 | 8.8e-09 | 8.2e-09 | 9.000E-09 | 9.165E-10 | 8.200E-09 | 1.000E-08 |
| 17541 | 878 | G878A | C | 9.3e-09 | 1.1e-08 | 5.9e-09 | 8.733E-09 | 2.597E-09 | 5.900E-09 | 1.100E-08 |
| 17542 | 878 | G878R | C | 7e-09 | 1.1e-08 | 5.3e-09 | 7.767E-09 | 2.926E-09 | 5.300E-09 | 1.100E-08 |
| 17543 | 878 | G878N | C | 1.1e-08 | 4.3e-09 | 6.9e-09 | 7.400E-09 | 3.378E-09 | 4.300E-09 | 1.100E-08 |
| 17544 | 878 | G878D | C | 2.5e-09 | 1.3e-08 | 2.8e-09 | 6.100E-09 | 5.977E-09 | 2.500E-09 | 1.300E-08 |
| 17545 | 878 | G878C | C | 7.1e-09 | 7.5e-09 | 1.4e-08 | 9.533E-09 | 3.873E-09 | 7.100E-09 | 1.400E-08 |
| 17546 | 878 | G878E | C | 4.6e-09 | 1.9e-08 | 1.2e-08 | 1.187E-08 | 7.201E-09 | 4.600E-09 | 1.900E-08 |
| 17547 | 878 | G878Q | C | 1.5e-08 | 1.1e-08 | 7e-09 | 1.100E-08 | 4.000E-09 | 7.000E-09 | 1.500E-08 |
| 17548 | 878 | G878G | C | 8.5e-09 | 8.2e-09 | 6.3e-09 | 7.667E-09 | 1.193E-09 | 6.300E-09 | 8.500E-09 |
| 17549 | 878 | G878H | C | 3.9e-09 | 8.7e-09 | 6.9e-09 | 6.500E-09 | 2.425E-09 | 3.900E-09 | 8.700E-09 |
| 17550 | 878 | G878I | C | 4.4e-09 | 6.7e-09 | 6.1e-09 | 5.733E-09 | 1.193E-09 | 4.400E-09 | 6.700E-09 |
| 17551 | 878 | G878L | C | 6.5e-09 | 1.1e-08 | 1.1e-08 | 9.500E-09 | 2.598E-09 | 6.500E-09 | 1.100E-08 |
| 17552 | 878 | G878K | C | 7.8e-09 | 1.9e-08 | 6e-09 | 1.093E-08 | 7.044E-09 | 6.000E-09 | 1.900E-08 |
| 17553 | 878 | G878M | C | 4e-09 | 7.8e-09 | 1.8e-08 | 9.933E-09 | 7.240E-09 | 4.000E-09 | 1.800E-08 |
| 17554 | 878 | G878F | C | 3.7e-09 | 2.9e-09 | 1.2e-08 | 6.200E-09 | 5.039E-09 | 2.900E-09 | 1.200E-08 |
| 17555 | 878 | G878P | C | 1.1e-08 | 1.6e-08 | 1.7e-08 | 1.467E-08 | 3.215E-09 | 1.100E-08 | 1.700E-08 |
| 17556 | 878 | G878S | C | 4.3e-09 | 2.6e-09 | 6.4e-09 | 4.433E-09 | 1.904E-09 | 2.600E-09 | 6.400E-09 |
| 17557 | 878 | G878T | C | 1.4e-08 | 1.8e-08 | 1.2e-08 | 1.467E-08 | 3.055E-09 | 1.200E-08 | 1.800E-08 |
| 17558 | 878 | G878W | C | 3.9e-09 | 2.5e-08 | 7.2e-09 | 1.203E-08 | 1.135E-08 | 3.900E-09 | 2.500E-08 |
| 17559 | 878 | G878Y | C | 6.5e-09 | 7.8e-09 | 1.2e-08 | 8.767E-09 | 2.875E-09 | 6.500E-09 | 1.200E-08 |
| 17560 | 878 | G878V | C | 1.5e-08 | 3.8e-09 | 8.9e-09 | 9.233E-09 | 5.607E-09 | 3.800E-09 | 1.500E-08 |
| 17561 | 879 | K879A | C | 7.1e-09 | 3.7e-09 | 6.4e-09 | 5.733E-09 | 1.795E-09 | 3.700E-09 | 7.100E-09 |
| 17562 | 879 | K879R | C | 3.4e-09 | 2e-08 | 1e-08 | 1.113E-08 | 8.358E-09 | 3.400E-09 | 2.000E-08 |
| 17563 | 879 | K879N | C | 3.1e-09 | 1.1e-08 | 6e-09 | 6.700E-09 | 3.996E-09 | 3.100E-09 | 1.100E-08 |
| 17564 | 879 | K879D | C | 3.7e-09 | 7.7e-09 | 8.4e-09 | 6.600E-09 | 2.536E-09 | 3.700E-09 | 8.400E-09 |
| 17565 | 879 | K879C | C | 6e-08 | 3.5e-09 | 1.5e-08 | 2.617E-08 | 2.986E-08 | 3.500E-09 | 6.000E-08 |
| 17566 | 879 | K879E | C | 6.7e-09 | 2.7e-09 | 7.3e-09 | 5.567E-09 | 2.501E-09 | 2.700E-09 | 7.300E-09 |
| 17567 | 879 | K879Q | C | 4.1e-09 | 9.2e-09 | 4.2e-09 | 5.833E-09 | 2.916E-09 | 4.100E-09 | 9.200E-09 |
| 17568 | 879 | K879G | C | 2.2e-08 | 6.7e-09 | 5.9e-09 | 1.153E-08 | 9.073E-09 | 5.900E-09 | 2.200E-08 |
| 17569 | 879 | K879H | C | 7.2e-09 | 1.2e-08 | 7.1e-09 | 8.767E-09 | 2.801E-09 | 7.100E-09 | 1.200E-08 |
| 17570 | 879 | K879I | C | 6e-09 | 5.5e-09 | 4.5e-08 | 1.883E-08 | 2.266E-08 | 5.500E-09 | 4.500E-08 |
| 17571 | 879 | K879L | C | 1.2e-09 | 1.1e-08 | 4.2e-09 | 5.467E-09 | 5.021E-09 | 1.200E-09 | 1.100E-08 |
| 17572 | 879 | K879K | C | 1.5e-08 | 4.8e-09 | 7.1e-09 | 8.967E-09 | 5.350E-09 | 4.800E-09 | 1.500E-08 |
| 17573 | 879 | K879M | C | 3.4e-09 | 1.5e-08 | 2.3e-09 | 6.900E-09 | 7.036E-09 | 2.300E-09 | 1.500E-08 |
| 17574 | 879 | K879F | C | 7.6e-09 | 4.9e-09 | 7.7e-09 | 6.733E-09 | 1.589E-09 | 4.900E-09 | 7.700E-09 |
| 17575 | 879 | K879P | C | 5.2e-09 | 2.9e-09 | 3.2e-09 | 3.767E-09 | 1.250E-09 | 2.900E-09 | 5.200E-09 |
| 17576 | 879 | K879S | C | 6.3e-09 | 9.1e-09 | 3.8e-09 | 6.400E-09 | 2.651E-09 | 3.800E-09 | 9.100E-09 |
| 17577 | 879 | K879T | C | 7.9e-09 | 3.2e-09 | 1.1e-08 | 7.367E-09 | 3.927E-09 | 3.200E-09 | 1.100E-08 |
| 17578 | 879 | K879W | C | 8.1e-09 | 7.4e-09 | 7.5e-09 | 7.667E-09 | 3.786E-10 | 7.400E-09 | 8.100E-09 |
| 17579 | 879 | K879Y | C | 5.9e-09 | 1.4e-08 | 1.9e-09 | 7.267E-09 | 6.165E-09 | 1.900E-09 | 1.400E-08 |
| 17580 | 879 | K879V | C | 1.1e-08 | 4.2e-09 | 3.9e-09 | 6.367E-09 | 4.015E-09 | 3.900E-09 | 1.100E-08 |
| 17581 | 880 | G880A | C | 5.9e-09 | 3.2e-09 | 1.3e-08 | 7.367E-09 | 5.062E-09 | 3.200E-09 | 1.300E-08 |
| 17582 | 880 | G880R | C | 5.4e-09 | 5.5e-09 | 5.2e-09 | 5.367E-09 | 1.528E-10 | 5.200E-09 | 5.500E-09 |
| 17583 | 880 | G880N | C | 9.2e-09 | 8.9e-09 | 4.7e-09 | 7.600E-09 | 2.516E-09 | 4.700E-09 | 9.200E-09 |
| 17584 | 880 | G880D | C | 5e-09 | 4.3e-09 | 8e-09 | 5.767E-09 | 1.966E-09 | 4.300E-09 | 8.000E-09 |
| 17585 | 880 | G880C | C | 1.3e-08 | 8.3e-09 | 4.9e-09 | 8.733E-09 | 4.067E-09 | 4.900E-09 | 1.300E-08 |
| 17586 | 880 | G880E | C | 6e-09 | 1.4e-08 | 1.6e-08 | 1.200E-08 | 5.292E-09 | 6.000E-09 | 1.600E-08 |
| 17587 | 880 | G880Q | C | 1.8e-08 | 6.9e-09 | 6.1e-09 | 1.033E-08 | 6.652E-09 | 6.100E-09 | 1.800E-08 |
| 17588 | 880 | G880G | C | 6e-09 | 1.1e-08 | 6.9e-09 | 7.967E-09 | 2.665E-09 | 6.000E-09 | 1.100E-08 |
| 17589 | 880 | G880H | C | 6.1e-09 | 4e-09 | 3.2e-09 | 4.433E-09 | 1.498E-09 | 3.200E-09 | 6.100E-09 |
| 17590 | 880 | G880I | C | 6.9e-09 | 1.2e-08 | 7.8e-09 | 8.900E-09 | 2.722E-09 | 6.900E-09 | 1.200E-08 |
| 17591 | 880 | G880L | C | 7.9e-09 | 2.4e-09 | 1.7e-09 | 4.000E-09 | 3.396E-09 | 1.700E-09 | 7.900E-09 |
| 17592 | 880 | G880K | C | 9e-09 | 3.9e-09 | 1.7e-08 | 9.967E-09 | 6.603E-09 | 3.900E-09 | 1.700E-08 |
| 17593 | 880 | G880M | C | 2e-09 | 4e-09 | 7.1e-09 | 4.367E-09 | 2.570E-09 | 2.000E-09 | 7.100E-09 |
| 17594 | 880 | G880F | C | 1.3e-08 | 6.5e-09 | 9e-09 | 9.500E-09 | 3.279E-09 | 6.500E-09 | 1.300E-08 |

| | | | | | | | | | | |
|---|---|---|---|---|---|---|---|---|---|---|
| 17595 | 880 | G880P | C | 5.5e-09 | 6e-09 | 6.6e-09 | 6.033E-09 | 5.508E-10 | 5.500E-09 | 6.600E-09 |
| 17596 | 880 | G880S | C | 1.1e-08 | 9.6e-09 | 3.4e-09 | 8.000E-09 | 4.045E-09 | 3.400E-09 | 1.100E-08 |
| 17597 | 880 | G880T | C | 1.2e-08 | 4.2e-09 | 1.3e-08 | 9.733E-09 | 4.818E-09 | 4.200E-09 | 1.300E-08 |
| 17598 | 880 | G880W | C | 1.4e-08 | 2.6e-09 | 2.6e-09 | 6.400E-09 | 6.582E-09 | 2.600E-09 | 1.400E-08 |
| 17599 | 880 | G880Y | C | 1e-08 | 4.4e-08 | 6.4e-09 | 2.013E-08 | 2.075E-08 | 6.400E-09 | 4.400E-08 |
| 17600 | 880 | G880V | C | 6.2e-09 | 6.6e-09 | 3.8e-09 | 5.533E-09 | 1.514E-09 | 3.800E-09 | 6.600E-09 |
| 17601 | 881 | L881A | C | 4.9e-09 | 1.8e-08 | 4.5e-09 | 9.133E-09 | 7.681E-09 | 4.500E-09 | 1.800E-08 |
| 17602 | 881 | L881R | C | 6.2e-09 | 4.2e-09 | 1.2e-08 | 7.467E-09 | 4.051E-09 | 4.200E-09 | 1.200E-08 |
| 17603 | 881 | L881N | C | 1e-08 | 4.9e-09 | 1.7e-08 | 1.063E-08 | 6.075E-09 | 4.900E-09 | 1.700E-08 |
| 17604 | 881 | L881D | C | 1.9e-08 | 4.7e-09 | 1.6e-08 | 1.323E-08 | 7.541E-09 | 4.700E-09 | 1.900E-08 |
| 17605 | 881 | L881C | C | 1.3e-08 | 1.5e-08 | 8.2e-09 | 1.207E-08 | 3.495E-09 | 8.200E-09 | 1.500E-08 |
| 17606 | 881 | L881E | C | 7.5e-09 | 7.7e-09 | 5.8e-09 | 7.000E-09 | 1.044E-09 | 5.800E-09 | 7.700E-09 |
| 17607 | 881 | L881Q | C | 1e-08 | 1.8e-08 | 4.9e-09 | 1.097E-08 | 6.603E-09 | 4.900E-09 | 1.800E-08 |
| 17608 | 881 | L881G | C | 7.5e-09 | 6.7e-09 | 1e-08 | 8.067E-09 | 1.721E-09 | 6.700E-09 | 1.000E-08 |
| 17609 | 881 | L881H | C | 3.6e-09 | 1.9e-08 | 1.6e-08 | 1.287E-08 | 8.164E-09 | 3.600E-09 | 1.900E-08 |
| 17610 | 881 | L881I | C | 1.9e-08 | 4.8e-09 | 5.8e-09 | 9.867E-09 | 7.925E-09 | 4.800E-09 | 1.900E-08 |
| 17611 | 881 | L881L | C | 6.3e-09 | 1.3e-08 | 3.4e-09 | 7.567E-09 | 4.924E-09 | 3.400E-09 | 1.300E-08 |
| 17612 | 881 | L881K | C | 2.5e-09 | 5.9e-09 | 7.5e-09 | 5.300E-09 | 2.553E-09 | 2.500E-09 | 7.500E-09 |
| 17613 | 881 | L881M | C | 2.7e-09 | 7.2e-09 | 1.6e-08 | 8.633E-09 | 6.765E-09 | 2.700E-09 | 1.600E-08 |
| 17614 | 881 | L881F | C | 9.8e-09 | 8.1e-09 | 1.2e-08 | 9.967E-09 | 1.955E-09 | 8.100E-09 | 1.200E-08 |
| 17615 | 881 | L881P | C | 1.1e-08 | 1.2e-08 | 4.6e-09 | 9.200E-09 | 4.015E-09 | 4.600E-09 | 1.200E-08 |
| 17616 | 881 | L881S | C | 1.4e-08 | 3.7e-09 | 8.2e-09 | 8.633E-09 | 5.164E-09 | 3.700E-09 | 1.400E-08 |
| 17617 | 881 | L881T | C | 6.2e-09 | 5.9e-09 | 2.1e-08 | 1.103E-08 | 8.633E-09 | 5.900E-09 | 2.100E-08 |
| 17618 | 881 | L881W | C | 2.8e-08 | 4.1e-09 | 7.4e-09 | 1.317E-08 | 1.295E-08 | 4.100E-09 | 2.800E-08 |
| 17619 | 881 | L881Y | C | 1.5e-08 | 1.8e-08 | 6.8e-09 | 1.327E-08 | 5.798E-09 | 6.800E-09 | 1.800E-08 |
| 17620 | 881 | L881V | C | 4.7e-09 | 7.6e-09 | 3.7e-09 | 5.333E-09 | 2.026E-09 | 3.700E-09 | 7.600E-09 |
| 17621 | 882 | E882A | C | 2e-08 | 5.5e-09 | 2.7e-09 | 9.400E-09 | 9.286E-09 | 2.700E-09 | 2.000E-08 |
| 17622 | 882 | E882R | C | 6.2e-09 | 4.1e-08 | 1.2e-08 | 1.973E-08 | 1.864E-08 | 6.200E-09 | 4.100E-08 |
| 17623 | 882 | E882N | C | 1.4e-08 | 5.9e-09 | 8.9e-09 | 9.600E-09 | 4.095E-09 | 5.900E-09 | 1.400E-08 |
| 17624 | 882 | E882D | C | 6e-09 | 1.2e-08 | 7e-09 | 8.333E-09 | 3.215E-09 | 6.000E-09 | 1.200E-08 |
| 17625 | 882 | E882C | C | 8e-09 | 5.1e-09 | 8.6e-09 | 7.233E-09 | 1.872E-09 | 5.100E-09 | 8.600E-09 |
| 17626 | 882 | E882E | C | 1e-08 | 3.8e-09 | 3.1e-09 | 5.633E-09 | 3.798E-09 | 3.100E-09 | 1.000E-08 |
| 17627 | 882 | E882Q | C | 8e-09 | 5.7e-09 | 1e-08 | 7.900E-09 | 2.152E-09 | 5.700E-09 | 1.000E-08 |
| 17628 | 882 | E882G | C | 6.4e-09 | 7.5e-09 | 4.6e-09 | 6.167E-09 | 1.464E-09 | 4.600E-09 | 7.500E-09 |
| 17629 | 882 | E882H | C | 1.2e-08 | 4.4e-09 | 7.1e-09 | 7.833E-09 | 3.853E-09 | 4.400E-09 | 1.200E-08 |
| 17630 | 882 | E882I | C | 1.8e-08 | 7.3e-09 | 1.2e-08 | 1.243E-08 | 5.363E-09 | 7.300E-09 | 1.800E-08 |
| 17631 | 882 | E882L | C | 6.4e-09 | 3.3e-09 | 3.5e-09 | 4.400E-09 | 1.735E-09 | 3.300E-09 | 6.400E-09 |
| 17632 | 882 | E882K | C | 8.2e-09 | 3.1e-09 | 1.1e-08 | 7.433E-09 | 4.005E-09 | 3.100E-09 | 1.100E-08 |
| 17633 | 882 | E882M | C | 2.1e-09 | 1.8e-08 | 1.3e-08 | 1.103E-08 | 8.130E-09 | 2.100E-09 | 1.800E-08 |
| 17634 | 882 | E882F | C | 2.2e-08 | 4.9e-09 | 1.2e-09 | 9.367E-09 | 1.110E-08 | 1.200E-09 | 2.200E-08 |
| 17635 | 882 | E882P | C | 5.4e-09 | 4.3e-09 | 1.1e-08 | 6.900E-09 | 3.593E-09 | 4.300E-09 | 1.100E-08 |
| 17636 | 882 | E882S | C | 4.3e-09 | 6.1e-09 | 8.3e-09 | 6.233E-09 | 2.003E-09 | 4.300E-09 | 8.300E-09 |
| 17637 | 882 | E882T | C | 5.1e-09 | 7.2e-09 | 1.4e-08 | 8.767E-09 | 4.652E-09 | 5.100E-09 | 1.400E-08 |
| 17638 | 882 | E882W | C | 1.5e-08 | 5e-09 | 3.1e-08 | 1.700E-08 | 1.311E-08 | 5.000E-09 | 3.100E-08 |
| 17639 | 882 | E882Y | C | 1e-08 | 1.7e-09 | 1.2e-08 | 7.900E-09 | 5.462E-09 | 1.700E-09 | 1.200E-08 |
| 17640 | 882 | E882V | C | 1e-08 | 5.7e-09 | 5.1e-09 | 6.933E-09 | 2.673E-09 | 5.100E-09 | 1.000E-08 |
| 17641 | 883 | W883A | C | 5.3e-09 | 5.6e-09 | 1.1e-08 | 7.300E-09 | 3.208E-09 | 5.300E-09 | 1.100E-08 |
| 17642 | 883 | W883R | C | 1.5e-09 | 5.2e-09 | 1.2e-08 | 6.233E-09 | 5.326E-09 | 1.500E-09 | 1.200E-08 |
| 17643 | 883 | W883N | C | 3.9e-09 | 7.9e-09 | 4.9e-09 | 5.567E-09 | 2.082E-09 | 3.900E-09 | 7.900E-09 |
| 17644 | 883 | W883D | C | 1.7e-08 | 4.5e-09 | 3.8e-09 | 8.433E-09 | 7.427E-09 | 3.800E-09 | 1.700E-08 |
| 17645 | 883 | W883C | C | 5.4e-09 | 8.7e-09 | 5.7e-09 | 6.600E-09 | 1.825E-09 | 5.400E-09 | 8.700E-09 |
| 17646 | 883 | W883E | C | 7.1e-09 | 3.9e-09 | 7.6e-09 | 6.200E-09 | 2.007E-09 | 3.900E-09 | 7.600E-09 |
| 17647 | 883 | W883Q | C | 8.7e-09 | 1.2e-08 | 6.6e-09 | 9.100E-09 | 2.722E-09 | 6.600E-09 | 1.200E-08 |
| 17648 | 883 | W883G | C | 9.9e-09 | 1.2e-08 | 9e-09 | 1.030E-08 | 1.539E-09 | 9.000E-09 | 1.200E-08 |
| 17649 | 883 | W883H | C | 8.1e-09 | 2.2e-09 | 3.6e-09 | 4.633E-09 | 3.083E-09 | 2.200E-09 | 8.100E-09 |
| 17650 | 883 | W883I | C | 6.3e-09 | 1.2e-08 | 2.1e-09 | 6.800E-09 | 4.969E-09 | 2.100E-09 | 1.200E-08 |
| 17651 | 883 | W883L | C | 2.9e-09 | 1.9e-08 | 1.7e-08 | 1.297E-08 | 8.775E-09 | 2.900E-09 | 1.900E-08 |
| 17652 | 883 | W883K | C | 3.4e-09 | 6.6e-09 | 1.3e-08 | 7.667E-09 | 4.888E-09 | 3.400E-09 | 1.300E-08 |
| 17653 | 883 | W883M | C | 7.1e-09 | 1.1e-08 | 5.4e-09 | 7.833E-09 | 2.871E-09 | 5.400E-09 | 1.100E-08 |
| 17654 | 883 | W883F | C | 2.4e-09 | 9.4e-09 | 1e-08 | 7.267E-09 | 4.225E-09 | 2.400E-09 | 1.000E-08 |
| 17655 | 883 | W883P | C | 6.3e-09 | 2.6e-09 | 1.3e-08 | 7.300E-09 | 5.272E-09 | 2.600E-09 | 1.300E-08 |
| 17656 | 883 | W883S | C | 1.2e-08 | 6.4e-09 | 3.1e-09 | 7.167E-09 | 4.499E-09 | 3.100E-09 | 1.200E-08 |
| 17657 | 883 | W883T | C | 5.8e-09 | 4.9e-09 | 3.6e-09 | 4.767E-09 | 1.106E-09 | 3.600E-09 | 5.800E-09 |
| 17658 | 883 | W883W | C | 1.4e-08 | 7.2e-09 | 7.2e-09 | 9.467E-09 | 3.926E-09 | 7.200E-09 | 1.400E-08 |
| 17659 | 883 | W883Y | C | 1.5e-08 | 3.4e-09 | 6.3e-09 | 8.233E-09 | 6.037E-09 | 3.400E-09 | 1.500E-08 |
| 17660 | 883 | W883V | C | 9.8e-09 | 1.3e-08 | 3.5e-09 | 8.767E-09 | 4.834E-09 | 3.500E-09 | 1.300E-08 |
| 17661 | 884 | V884A | C | 5.7e-09 | 1.3e-08 | 6.4e-09 | 8.367E-09 | 4.028E-09 | 5.700E-09 | 1.300E-08 |
| 17662 | 884 | V884R | C | 9e-09 | 8.9e-09 | 8.3e-09 | 8.733E-09 | 3.786E-10 | 8.300E-09 | 9.000E-09 |
| 17663 | 884 | V884N | C | 1.2e-08 | 4.1e-09 | 5.4e-09 | 7.167E-09 | 4.236E-09 | 4.100E-09 | 1.200E-08 |
| 17664 | 884 | V884D | C | 1.2e-08 | 3.9e-09 | 2.5e-09 | 6.133E-09 | 5.129E-09 | 2.500E-09 | 1.200E-08 |
| 17665 | 884 | V884C | C | 1.5e-08 | 1.2e-08 | 1.8e-08 | 1.500E-08 | 3.000E-09 | 1.200E-08 | 1.800E-08 |
| 17666 | 884 | V884E | C | 1.8e-08 | 4.2e-09 | 4.9e-09 | 9.033E-09 | 7.773E-09 | 4.200E-09 | 1.800E-08 |
| 17667 | 884 | V884Q | C | 6.9e-09 | 1.2e-08 | 8.4e-09 | 9.100E-09 | 2.621E-09 | 6.900E-09 | 1.200E-08 |
| 17668 | 884 | V884G | C | 9.9e-09 | 9.2e-09 | 8.9e-09 | 9.333E-09 | 5.132E-10 | 8.900E-09 | 9.900E-09 |
| 17669 | 884 | V884H | C | 1.7e-08 | 5e-09 | 5.9e-09 | 9.300E-09 | 6.684E-09 | 5.000E-09 | 1.700E-08 |
| 17670 | 884 | V884I | C | 6.4e-09 | 5.9e-09 | 7.5e-09 | 6.600E-09 | 8.185E-10 | 5.900E-09 | 7.500E-09 |
| 17671 | 884 | V884L | C | 8.1e-09 | 6.4e-09 | 1.2e-08 | 8.833E-09 | 2.871E-09 | 6.400E-09 | 1.200E-08 |
| 17672 | 884 | V884K | C | 5.8e-09 | 5.4e-09 | 2.2e-09 | 4.467E-09 | 1.973E-09 | 2.200E-09 | 5.800E-09 |
| 17673 | 884 | V884M | C | 1.3e-08 | 8.1e-09 | 5.7e-09 | 8.933E-09 | 3.721E-09 | 5.700E-09 | 1.300E-08 |
| 17674 | 884 | V884F | C | 8e-09 | 7.1e-09 | 6.3e-09 | 7.133E-09 | 8.505E-10 | 6.300E-09 | 8.000E-09 |
| 17675 | 884 | V884P | C | 1.1e-08 | 7.6e-09 | 1.5e-08 | 1.120E-08 | 3.704E-09 | 7.600E-09 | 1.500E-08 |
| 17676 | 884 | V884S | C | 1.1e-08 | 2.4e-08 | 9.7e-09 | 1.490E-08 | 7.908E-09 | 9.700E-09 | 2.400E-08 |
| 17677 | 884 | V884T | C | 8.2e-09 | 5.7e-09 | 7e-09 | 6.967E-09 | 1.250E-09 | 5.700E-09 | 8.200E-09 |
| 17678 | 884 | V884W | C | 1.5e-08 | 8.7e-09 | 1.7e-09 | 8.467E-09 | 6.653E-09 | 1.700E-09 | 1.500E-08 |
| 17679 | 884 | V884Y | C | 4.5e-09 | 2.2e-09 | 3.3e-09 | 3.333E-09 | 1.150E-09 | 2.200E-09 | 4.500E-09 |

| | | | | | | | | | | |
|---|---|---|---|---|---|---|---|---|---|---|
| 17680 | 884 | V884V | C | 3.2e-09 | 6.6e-09 | 1.2e-08 | 7.267E-09 | 4.438E-09 | 3.200E-09 | 1.200E-08 |
| 17681 | 885 | A885A | C | 1.1e-08 | 5.2e-09 | 1.1e-08 | 9.067E-09 | 3.349E-09 | 5.200E-09 | 1.100E-08 |
| 17682 | 885 | A885R | C | 8.8e-09 | 5.7e-09 | 6.8e-09 | 7.100E-09 | 1.572E-09 | 5.700E-09 | 8.800E-09 |
| 17683 | 885 | A885N | C | 2.1e-09 | 1.3e-08 | 3.4e-09 | 6.167E-09 | 5.953E-09 | 2.100E-09 | 1.300E-08 |
| 17684 | 885 | A885D | C | 4.2e-09 | 7e-09 | 3e-08 | 1.373E-08 | 1.416E-08 | 4.200E-09 | 3.000E-08 |
| 17685 | 885 | A885C | C | 1.1e-08 | 6.9e-09 | 4.7e-09 | 7.533E-09 | 3.197E-09 | 4.700E-09 | 1.100E-08 |
| 17686 | 885 | A885E | C | 1.8e-08 | 3.3e-09 | 2e-09 | 7.767E-09 | 8.886E-09 | 2.000E-09 | 1.800E-08 |
| 17687 | 885 | A885Q | C | 1.4e-08 | 5.8e-09 | 9.8e-09 | 9.867E-09 | 4.100E-09 | 5.800E-09 | 1.400E-08 |
| 17688 | 885 | A885G | C | 7.8e-09 | 3.8e-09 | 3.2e-09 | 4.933E-09 | 2.501E-09 | 3.200E-09 | 7.800E-09 |
| 17689 | 885 | A885H | C | 5.4e-09 | 2.6e-08 | 1.3e-08 | 1.480E-08 | 1.042E-08 | 5.400E-09 | 2.600E-08 |
| 17690 | 885 | A885I | C | 2.2e-09 | 5.8e-09 | 3.8e-09 | 3.933E-09 | 1.804E-09 | 2.200E-09 | 5.800E-09 |
| 17691 | 885 | A885L | C | 4.2e-09 | 7.3e-09 | 9e-09 | 6.833E-09 | 2.434E-09 | 4.200E-09 | 9.000E-09 |
| 17692 | 885 | A885K | C | 6.7e-09 | 4.6e-08 | 6.3e-09 | 1.967E-08 | 2.281E-08 | 6.300E-09 | 4.600E-08 |
| 17693 | 885 | A885M | C | 1.3e-08 | 1e-08 | 4.4e-09 | 9.133E-09 | 4.365E-09 | 4.400E-09 | 1.300E-08 |
| 17694 | 885 | A885F | C | 5.4e-09 | 3.5e-08 | 1.5e-08 | 1.847E-08 | 1.510E-08 | 5.400E-09 | 3.500E-08 |
| 17695 | 885 | A885P | C | 1.4e-08 | 3.9e-09 | 2.6e-09 | 6.833E-09 | 6.240E-09 | 2.600E-09 | 1.400E-08 |
| 17696 | 885 | A885S | C | 7.9e-09 | 9.7e-09 | 1.1e-08 | 9.533E-09 | 1.557E-09 | 7.900E-09 | 1.100E-08 |
| 17697 | 885 | A885T | C | 1e-08 | 1.5e-08 | 2.4e-09 | 9.133E-09 | 6.345E-09 | 2.400E-09 | 1.500E-08 |
| 17698 | 885 | A885W | C | 8.2e-09 | 4.3e-09 | 2.2e-08 | 1.150E-08 | 9.300E-09 | 4.300E-09 | 2.200E-08 |
| 17699 | 885 | A885Y | C | 6.1e-09 | 9.4e-09 | 4.2e-09 | 6.567E-09 | 2.631E-09 | 4.200E-09 | 9.400E-09 |
| 17700 | 885 | A885V | C | 1.8e-08 | 1.3e-08 | 5.5e-09 | 1.217E-08 | 6.292E-09 | 5.500E-09 | 1.800E-08 |
| 17701 | 886 | D886A | C | 3.5e-09 | 6.7e-09 | 5.5e-09 | 5.233E-09 | 1.617E-09 | 3.500E-09 | 6.700E-09 |
| 17702 | 886 | D886R | C | 8.5e-09 | 9.2e-09 | 4.3e-09 | 7.333E-09 | 2.650E-09 | 4.300E-09 | 9.200E-09 |
| 17703 | 886 | D886N | C | 5.4e-09 | 4.4e-09 | 9e-09 | 6.267E-09 | 2.419E-09 | 4.400E-09 | 9.000E-09 |
| 17704 | 886 | D886D | C | 5e-09 | 2.5e-08 | 5.7e-09 | 1.190E-08 | 1.135E-08 | 5.000E-09 | 2.500E-08 |
| 17705 | 886 | D886C | C | 4.5e-09 | 2.6e-09 | 5.2e-09 | 4.100E-09 | 1.345E-09 | 2.600E-09 | 5.200E-09 |
| 17706 | 886 | D886E | C | 9.2e-09 | 1.6e-08 | 4.2e-09 | 9.800E-09 | 5.923E-09 | 4.200E-09 | 1.600E-08 |
| 17707 | 886 | D886Q | C | 2.1e-08 | 8.1e-09 | 6.2e-09 | 1.177E-08 | 8.053E-09 | 6.200E-09 | 2.100E-08 |
| 17708 | 886 | D886G | C | 8.1e-09 | 3.7e-09 | 1.8e-08 | 9.933E-09 | 7.324E-09 | 3.700E-09 | 1.800E-08 |
| 17709 | 886 | D886H | C | 9.5e-09 | 5.2e-09 | 7.4e-09 | 7.367E-09 | 2.150E-09 | 5.200E-09 | 9.500E-09 |
| 17710 | 886 | D886I | C | 4.8e-09 | 4e-09 | 3.9e-09 | 4.233E-09 | 4.933E-10 | 3.900E-09 | 4.800E-09 |
| 17711 | 886 | D886L | C | 2.1e-08 | 3.2e-09 | 8.7e-09 | 1.097E-08 | 9.114E-09 | 3.200E-09 | 2.100E-08 |
| 17712 | 886 | D886K | C | 1.2e-08 | 4.4e-09 | 3.7e-09 | 6.700E-09 | 4.603E-09 | 3.700E-09 | 1.200E-08 |
| 17713 | 886 | D886M | C | 1.3e-08 | 7.2e-09 | 9.8e-09 | 1.000E-08 | 2.905E-09 | 7.200E-09 | 1.300E-08 |
| 17714 | 886 | D886F | C | 9e-09 | 2.8e-08 | 3.2e-09 | 1.340E-08 | 1.297E-08 | 3.200E-09 | 2.800E-08 |
| 17715 | 886 | D886P | C | 9.2e-09 | 6.1e-09 | 4.2e-09 | 6.500E-09 | 2.524E-09 | 4.200E-09 | 9.200E-09 |
| 17716 | 886 | D886S | C | 4.9e-09 | 9.1e-09 | 1.5e-08 | 9.667E-09 | 5.074E-09 | 4.900E-09 | 1.500E-08 |
| 17717 | 886 | D886T | C | 4.3e-09 | 9.5e-09 | 3.8e-09 | 5.867E-09 | 3.156E-09 | 3.800E-09 | 9.500E-09 |
| 17718 | 886 | D886W | C | 9.2e-09 | 4.6e-09 | 9e-09 | 7.600E-09 | 2.600E-09 | 4.600E-09 | 9.200E-09 |
| 17719 | 886 | D886Y | C | 7.5e-09 | 1.7e-08 | 1e-08 | 1.150E-08 | 4.924E-09 | 7.500E-09 | 1.700E-08 |
| 17720 | 886 | D886V | C | 3.2e-09 | 5e-09 | 1e-08 | 6.067E-09 | 3.523E-09 | 3.200E-09 | 1.000E-08 |
| 17721 | 887 | V887A | C | 1.2e-08 | 7.9e-09 | 4.5e-09 | 8.133E-09 | 3.755E-09 | 4.500E-09 | 1.200E-08 |
| 17722 | 887 | V887R | C | 4.6e-09 | 5.6e-09 | 4.9e-09 | 5.033E-09 | 5.132E-10 | 4.600E-09 | 5.600E-09 |
| 17723 | 887 | V887N | C | 1.3e-08 | 1.5e-08 | 4.3e-09 | 1.077E-08 | 5.689E-09 | 4.300E-09 | 1.500E-08 |
| 17724 | 887 | V887D | C | 5.5e-09 | 4.4e-09 | 5.3e-09 | 5.067E-09 | 5.859E-10 | 4.400E-09 | 5.500E-09 |
| 17725 | 887 | V887C | C | 6.3e-09 | 6e-09 | 8.5e-09 | 6.933E-09 | 1.365E-09 | 6.000E-09 | 8.500E-09 |
| 17726 | 887 | V887E | C | 8.8e-09 | 5.2e-09 | 1.4e-08 | 9.333E-09 | 4.424E-09 | 5.200E-09 | 1.400E-08 |
| 17727 | 887 | V887Q | C | 4.8e-09 | 5.2e-09 | 7.1e-09 | 5.700E-09 | 1.229E-09 | 4.800E-09 | 7.100E-09 |
| 17728 | 887 | V887G | C | 4.4e-09 | 5.7e-09 | 1.5e-08 | 8.367E-09 | 5.781E-09 | 4.400E-09 | 1.500E-08 |
| 17729 | 887 | V887H | C | 2.3e-08 | 7.6e-09 | 2.2e-09 | 1.093E-08 | 1.079E-08 | 2.200E-09 | 2.300E-08 |
| 17730 | 887 | V887I | C | 7.9e-09 | 4e-09 | 1.2e-08 | 7.967E-09 | 4.000E-09 | 4.000E-09 | 1.200E-08 |
| 17731 | 887 | V887L | C | 5.8e-09 | 9.9e-09 | 6.6e-09 | 7.433E-09 | 2.173E-09 | 5.800E-09 | 9.900E-09 |
| 17732 | 887 | V887K | C | 9.5e-09 | 1e-08 | 3e-09 | 7.500E-09 | 3.905E-09 | 3.000E-09 | 1.000E-08 |
| 17733 | 887 | V887M | C | 1.7e-08 | 1.3e-08 | 4.6e-09 | 1.153E-08 | 6.329E-09 | 4.600E-09 | 1.700E-08 |
| 17734 | 887 | V887F | C | 5.2e-09 | 3.9e-09 | 3.7e-09 | 4.267E-09 | 8.145E-10 | 3.700E-09 | 5.200E-09 |
| 17735 | 887 | V887P | C | 6.5e-09 | 8.6e-09 | 9.8e-09 | 8.300E-09 | 1.670E-09 | 6.500E-09 | 9.800E-09 |
| 17736 | 887 | V887S | C | 1.1e-08 | 6.3e-09 | 1.6e-08 | 1.110E-08 | 4.851E-09 | 6.300E-09 | 1.600E-08 |
| 17737 | 887 | V887T | C | 3.6e-09 | 5.7e-09 | 5.1e-09 | 4.800E-09 | 1.082E-09 | 3.600E-09 | 5.700E-09 |
| 17738 | 887 | V887W | C | 5.5e-09 | 1.2e-08 | 2.9e-09 | 6.800E-09 | 4.687E-09 | 2.900E-09 | 1.200E-08 |
| 17739 | 887 | V887Y | C | 6.1e-09 | 8e-09 | 2e-09 | 5.367E-09 | 3.066E-09 | 2.000E-09 | 8.000E-09 |
| 17740 | 887 | V887V | C | 2.1e-08 | 2.8e-08 | 1.2e-08 | 2.033E-08 | 8.021E-09 | 1.200E-08 | 2.800E-08 |
| 17741 | 888 | N888A | C | 8.2e-09 | 7.8e-09 | 1.2e-08 | 9.333E-09 | 2.318E-09 | 7.800E-09 | 1.200E-08 |
| 17742 | 888 | N888R | C | 5.6e-09 | 4.8e-09 | 7.4e-09 | 5.933E-09 | 1.332E-09 | 4.800E-09 | 7.400E-09 |
| 17743 | 888 | N888N | C | 9.3e-09 | 4.3e-09 | 3.4e-09 | 5.667E-09 | 3.179E-09 | 3.400E-09 | 9.300E-09 |
| 17744 | 888 | N888D | C | 1e-08 | 3e-09 | 7.3e-09 | 6.767E-09 | 3.530E-09 | 3.000E-09 | 1.000E-08 |
| 17745 | 888 | N888C | C | 9.2e-09 | 1.1e-08 | 2.5e-08 | 1.507E-08 | 8.649E-09 | 9.200E-09 | 2.500E-08 |
| 17746 | 888 | N888E | C | 2.6e-09 | 6.7e-09 | 9.9e-09 | 6.400E-09 | 3.659E-09 | 2.600E-09 | 9.900E-09 |
| 17747 | 888 | N888Q | C | 2.5e-09 | 1.6e-09 | 7.6e-09 | 3.900E-09 | 3.236E-09 | 1.600E-09 | 7.600E-09 |
| 17748 | 888 | N888G | C | 1.6e-08 | 1.2e-08 | 3.5e-09 | 1.050E-08 | 6.384E-09 | 3.500E-09 | 1.600E-08 |
| 17749 | 888 | N888H | C | 3.1e-09 | 7.1e-09 | 3.6e-09 | 4.600E-09 | 2.179E-09 | 3.100E-09 | 7.100E-09 |
| 17750 | 888 | N888I | C | 1.4e-08 | 1.4e-08 | 7.9e-09 | 1.197E-08 | 3.522E-09 | 7.900E-09 | 1.400E-08 |
| 17751 | 888 | N888L | C | 1.2e-08 | 1.2e-08 | 4.2e-09 | 9.400E-09 | 4.503E-09 | 4.200E-09 | 1.200E-08 |
| 17752 | 888 | N888K | C | 1.4e-08 | 2.5e-09 | 3.7e-09 | 6.733E-09 | 6.322E-09 | 2.500E-09 | 1.400E-08 |
| 17753 | 888 | N888M | C | 1.5e-08 | 2.2e-08 | 5.1e-09 | 1.403E-08 | 8.491E-09 | 5.100E-09 | 2.200E-08 |
| 17754 | 888 | N888F | C | 7.1e-09 | 2.9e-08 | 1.3e-08 | 1.637E-08 | 1.133E-08 | 7.100E-09 | 2.900E-08 |
| 17755 | 888 | N888P | C | 6.7e-09 | 7.9e-09 | 1e-08 | 8.200E-09 | 1.670E-09 | 6.700E-09 | 1.000E-08 |
| 17756 | 888 | N888S | C | 1.5e-08 | 6.6e-09 | 1.2e-08 | 1.120E-08 | 4.257E-09 | 6.600E-09 | 1.500E-08 |
| 17757 | 888 | N888T | C | 3.6e-09 | 1.1e-08 | 2.3e-08 | 1.253E-08 | 9.790E-09 | 3.600E-09 | 2.300E-08 |
| 17758 | 888 | N888W | C | 3.8e-09 | 4.2e-08 | 2.1e-08 | 2.227E-08 | 1.913E-08 | 3.800E-09 | 4.200E-08 |
| 17759 | 888 | N888Y | C | 1.1e-08 | 3.5e-08 | 6.1e-09 | 1.737E-08 | 1.547E-08 | 6.100E-09 | 3.500E-08 |
| 17760 | 888 | N888V | C | 4.4e-09 | 4.2e-09 | 1.2e-08 | 6.867E-09 | 4.447E-09 | 4.200E-09 | 1.200E-08 |
| 17761 | 889 | P889A | C | 4.1e-09 | 5.4e-09 | 4.2e-09 | 4.567E-09 | 7.234E-10 | 4.100E-09 | 5.400E-09 |
| 17762 | 889 | P889R | C | 1.3e-08 | 5.4e-09 | 4.4e-09 | 7.600E-09 | 4.703E-09 | 4.400E-09 | 1.300E-08 |
| 17763 | 889 | P889N | C | 6.3e-09 | 1.6e-09 | 1.4e-08 | 7.300E-09 | 6.260E-09 | 1.600E-09 | 1.400E-08 |
| 17764 | 889 | P889D | C | 2.4e-08 | 2.2e-08 | 1.3e-08 | 1.967E-08 | 5.859E-09 | 1.300E-08 | 2.400E-08 |

| | | | | | | | | | | |
|---|---|---|---|---|---|---|---|---|---|---|
| 17765 | 889 | P889C | C | 1.2e-08 | 1e-08 | 1.2e-08 | 1.133E-08 | 1.155E-09 | 1.000E-08 | 1.200E-08 |
| 17766 | 889 | P889E | C | 1.2e-08 | 2.5e-09 | 1.6e-08 | 1.017E-08 | 6.934E-09 | 2.500E-09 | 1.600E-08 |
| 17767 | 889 | P889Q | C | 2.3e-09 | 5.9e-09 | 3.5e-09 | 3.900E-09 | 1.833E-09 | 2.300E-09 | 5.900E-09 |
| 17768 | 889 | P889G | C | 9.3e-09 | 1.8e-08 | 2.9e-08 | 1.877E-08 | 9.872E-09 | 9.300E-09 | 2.900E-08 |
| 17769 | 889 | P889H | C | 8.3e-09 | 7.4e-09 | 4.2e-09 | 6.633E-09 | 2.155E-09 | 4.200E-09 | 8.300E-09 |
| 17770 | 889 | P889I | C | 8.6e-09 | 6.9e-09 | 1.9e-09 | 5.800E-09 | 3.483E-09 | 1.900E-09 | 8.600E-09 |
| 17771 | 889 | P889L | C | 1.2e-08 | 6.4e-09 | 1e-08 | 9.467E-09 | 2.838E-09 | 6.400E-09 | 1.200E-08 |
| 17772 | 889 | P889K | C | 1.1e-08 | 1.2e-08 | 9.8e-09 | 1.093E-08 | 1.102E-09 | 9.800E-09 | 1.200E-08 |
| 17773 | 889 | P889M | C | 1.1e-08 | 4.3e-09 | 5.1e-09 | 6.800E-09 | 3.659E-09 | 4.300E-09 | 1.100E-08 |
| 17774 | 889 | P889F | C | 5.5e-09 | 6e-09 | 7.4e-09 | 6.300E-09 | 9.849E-10 | 5.500E-09 | 7.400E-09 |
| 17775 | 889 | P889P | C | 3.7e-09 | 4.8e-09 | 4.9e-09 | 4.467E-09 | 6.658E-10 | 3.700E-09 | 4.900E-09 |
| 17776 | 889 | P889S | C | 1.4e-08 | 2.9e-08 | 5.7e-09 | 1.623E-08 | 1.181E-08 | 5.700E-09 | 2.900E-08 |
| 17777 | 889 | P889T | C | 5.2e-09 | 5.1e-09 | 1e-08 | 6.767E-09 | 2.801E-09 | 5.100E-09 | 1.000E-08 |
| 17778 | 889 | P889W | C | 4.1e-09 | 2.4e-09 | 4.3e-09 | 3.600E-09 | 1.044E-09 | 2.400E-09 | 4.300E-09 |
| 17779 | 889 | P889Y | C | 1.7e-09 | 5e-09 | 2.5e-09 | 3.067E-09 | 1.721E-09 | 1.700E-09 | 5.000E-09 |
| 17780 | 889 | P889V | C | 8.5e-09 | 2.5e-08 | 3e-09 | 1.217E-08 | 1.145E-08 | 3.000E-09 | 2.500E-08 |
| 17781 | 890 | N890A | C | 3e-08 | 1.7e-08 | 5e-08 | 3.233E-08 | 1.662E-08 | 1.700E-08 | 5.000E-08 |
| 17782 | 890 | N890R | C | 3.7e-09 | 1.5e-08 | 1.1e-08 | 9.900E-09 | 5.730E-09 | 3.700E-09 | 1.500E-08 |
| 17783 | 890 | N890N | C | 2.6e-09 | 5.6e-09 | 5.4e-09 | 4.533E-09 | 1.677E-09 | 2.600E-09 | 5.600E-09 |
| 17784 | 890 | N890D | C | 1.7e-08 | 2.1e-08 | 1.8e-08 | 1.867E-08 | 2.082E-09 | 1.700E-08 | 2.100E-08 |
| 17785 | 890 | N890C | C | 1e-08 | 2.2e-08 | 1.5e-08 | 1.567E-08 | 6.028E-09 | 1.000E-08 | 2.200E-08 |
| 17786 | 890 | N890E | C | 1e-08 | 8.4e-09 | 1.4e-08 | 1.080E-08 | 2.884E-09 | 8.400E-09 | 1.400E-08 |
| 17787 | 890 | N890Q | C | 4.9e-09 | 1.3e-08 | 1.9e-08 | 1.230E-08 | 7.076E-09 | 4.900E-09 | 1.900E-08 |
| 17788 | 890 | N890G | C | 3.2e-08 | 1.2e-08 | 1.8e-08 | 2.067E-08 | 1.026E-08 | 1.200E-08 | 3.200E-08 |
| 17789 | 890 | N890H | C | 6.2e-09 | 1e-08 | 1.2e-08 | 9.400E-09 | 2.946E-09 | 6.200E-09 | 1.200E-08 |
| 17790 | 890 | N890I | C | 4.7e-09 | 1.6e-08 | 1.7e-08 | 1.257E-08 | 6.831E-09 | 4.700E-09 | 1.700E-08 |
| 17791 | 890 | N890L | C | 2.7e-08 | 7.8e-09 | 3.4e-08 | 2.293E-08 | 1.357E-08 | 7.800E-09 | 3.400E-08 |
| 17792 | 890 | N890K | C | 1.2e-08 | 9.1e-09 | 2e-08 | 1.370E-08 | 5.645E-09 | 9.100E-09 | 2.000E-08 |
| 17793 | 890 | N890M | C | 2.9e-08 | 2.4e-08 | 8.8e-08 | 4.700E-08 | 3.559E-08 | 2.400E-08 | 8.800E-08 |
| 17794 | 890 | N890F | C | 1.7e-08 | 6.2e-08 | 3.3e-08 | 3.733E-08 | 2.281E-08 | 1.700E-08 | 6.200E-08 |
| 17795 | 890 | N890P | C | 4.7e-08 | 3.5e-08 | 6e-08 | 4.733E-08 | 1.250E-08 | 3.500E-08 | 6.000E-08 |
| 17796 | 890 | N890S | C | 2.1e-09 | 2.2e-09 | 1.4e-08 | 6.100E-09 | 6.842E-09 | 2.100E-09 | 1.400E-08 |
| 17797 | 890 | N890T | C | 1.8e-08 | 7e-09 | 3.2e-09 | 9.400E-09 | 7.686E-09 | 3.200E-09 | 1.800E-08 |
| 17798 | 890 | N890W | C | 4.3e-08 | 2.3e-08 | 1.6e-08 | 2.733E-08 | 1.401E-08 | 1.600E-08 | 4.300E-08 |
| 17799 | 890 | N890Y | C | 6.3e-08 | 3.1e-08 | 3e-08 | 4.133E-08 | 1.877E-08 | 3.000E-08 | 6.300E-08 |
| 17800 | 890 | N890V | C | 2.2e-08 | 2.3e-08 | 1.4e-08 | 1.967E-08 | 4.933E-09 | 1.400E-08 | 2.300E-08 |
| 17801 | 891 | S891A | C | 1.1e-08 | 2.3e-08 | 6e-09 | 1.333E-08 | 8.737E-09 | 6.000E-09 | 2.300E-08 |
| 17802 | 891 | S891R | C | 1.2e-08 | 1e-08 | 8.6e-09 | 1.020E-08 | 1.709E-09 | 8.600E-09 | 1.200E-08 |
| 17803 | 891 | S891N | C | 6.3e-09 | 2.5e-09 | 3.1e-09 | 3.967E-09 | 2.043E-09 | 2.500E-09 | 6.300E-09 |
| 17804 | 891 | S891D | C | 9.3e-09 | 9.9e-09 | 4.7e-09 | 7.967E-09 | 2.845E-09 | 4.700E-09 | 9.900E-09 |
| 17805 | 891 | S891C | C | 3.1e-08 | 1.1e-08 | 4.5e-09 | 1.550E-08 | 1.381E-08 | 4.500E-09 | 3.100E-08 |
| 17806 | 891 | S891E | C | 5.2e-09 | 1e-08 | 3e-09 | 6.067E-09 | 3.580E-09 | 3.000E-09 | 1.000E-08 |
| 17807 | 891 | S891Q | C | 3.3e-09 | 8e-09 | 8.2e-09 | 6.500E-09 | 2.773E-09 | 3.300E-09 | 8.200E-09 |
| 17808 | 891 | S891G | C | 2.3e-08 | 3.2e-08 | 2e-08 | 2.500E-08 | 6.245E-09 | 2.000E-08 | 3.200E-08 |
| 17809 | 891 | S891H | C | 4.9e-09 | 7e-09 | 2.8e-09 | 4.900E-09 | 2.100E-09 | 2.800E-09 | 7.000E-09 |
| 17810 | 891 | S891I | C | 2e-08 | 8.3e-09 | 3e-09 | 1.043E-08 | 8.698E-09 | 3.000E-09 | 2.000E-08 |
| 17811 | 891 | S891L | C | 1e-08 | 1.5e-08 | 2.3e-08 | 1.600E-08 | 6.557E-09 | 1.000E-08 | 2.300E-08 |
| 17812 | 891 | S891K | C | 1e-08 | 6.3e-09 | 9.5e-09 | 8.600E-09 | 2.007E-09 | 6.300E-09 | 1.000E-08 |
| 17813 | 891 | S891M | C | 7.6e-09 | 8.4e-09 | 1.7e-08 | 1.100E-08 | 5.212E-09 | 7.600E-09 | 1.700E-08 |
| 17814 | 891 | S891F | C | 1.3e-08 | 2.6e-08 | 1.7e-08 | 1.867E-08 | 6.658E-09 | 1.300E-08 | 2.600E-08 |
| 17815 | 891 | S891P | C | 2.5e-08 | 1.2e-08 | 1e-08 | 1.567E-08 | 8.145E-09 | 1.000E-08 | 2.500E-08 |
| 17816 | 891 | S891S | C | 6.2e-09 | 1.3e-08 | 1.2e-08 | 1.040E-08 | 3.672E-09 | 6.200E-09 | 1.300E-08 |
| 17817 | 891 | S891T | C | 1.8e-09 | 3.9e-09 | 1e-08 | 5.233E-09 | 4.259E-09 | 1.800E-09 | 1.000E-08 |
| 17818 | 891 | S891W | C | 8.9e-09 | 9.3e-09 | 1.5e-08 | 1.107E-08 | 3.412E-09 | 8.900E-09 | 1.500E-08 |
| 17819 | 891 | S891Y | C | 7.1e-09 | 1.5e-08 | 1.3e-08 | 1.170E-08 | 4.107E-09 | 7.100E-09 | 1.500E-08 |
| 17820 | 891 | S891V | C | 7.2e-09 | 2e-08 | 1.1e-08 | 1.273E-08 | 6.574E-09 | 7.200E-09 | 2.000E-08 |
| 17821 | 892 | G892A | C | 1.1e-08 | 6e-09 | 3.1e-09 | 6.700E-09 | 3.996E-09 | 3.100E-09 | 1.100E-08 |
| 17822 | 892 | G892R | C | 1.1e-09 | 6.5e-09 | 4.1e-09 | 3.900E-09 | 2.706E-09 | 1.100E-09 | 6.500E-09 |
| 17823 | 892 | G892N | C | 6.9e-09 | 1.7e-09 | 4.2e-09 | 4.267E-09 | 2.601E-09 | 1.700E-09 | 6.900E-09 |
| 17824 | 892 | G892D | C | 3.6e-09 | 3.7e-09 | 2.9e-09 | 3.400E-09 | 4.359E-10 | 2.900E-09 | 3.700E-09 |
| 17825 | 892 | G892C | C | 1.9e-08 | 7.1e-09 | 8.9e-09 | 1.167E-08 | 6.414E-09 | 7.100E-09 | 1.900E-08 |
| 17826 | 892 | G892E | C | 3.3e-09 | 2.3e-09 | 1.1e-08 | 5.533E-09 | 4.761E-09 | 2.300E-09 | 1.100E-08 |
| 17827 | 892 | G892Q | C | 2.3e-08 | 3.3e-09 | 1.3e-08 | 1.310E-08 | 9.850E-09 | 3.300E-09 | 2.300E-08 |
| 17828 | 892 | G892G | C | 6.4e-09 | 4.3e-09 | 4.3e-09 | 5.000E-09 | 1.212E-09 | 4.300E-09 | 6.400E-09 |
| 17829 | 892 | G892H | C | 6.9e-09 | 6.6e-09 | 5.5e-09 | 6.333E-09 | 7.371E-10 | 5.500E-09 | 6.900E-09 |
| 17830 | 892 | G892I | C | 9.7e-09 | 4.9e-09 | 2.7e-09 | 5.767E-09 | 3.580E-09 | 2.700E-09 | 9.700E-09 |
| 17831 | 892 | G892L | C | 1.6e-08 | 4e-09 | 1.1e-08 | 1.033E-08 | 6.028E-09 | 4.000E-09 | 1.600E-08 |
| 17832 | 892 | G892K | C | 8.5e-09 | 6.1e-09 | 2.7e-09 | 5.767E-09 | 2.914E-09 | 2.700E-09 | 8.500E-09 |
| 17833 | 892 | G892M | C | 5.6e-09 | 7.3e-09 | 1.1e-08 | 7.967E-09 | 2.761E-09 | 5.600E-09 | 1.100E-08 |
| 17834 | 892 | G892F | C | 1.1e-08 | 3.6e-09 | 1.3e-09 | 5.300E-09 | 5.069E-09 | 1.300E-09 | 1.100E-08 |
| 17835 | 892 | G892P | C | 1.2e-08 | 2.1e-08 | 1e-08 | 1.433E-08 | 5.859E-09 | 1.000E-08 | 2.100E-08 |
| 17836 | 892 | G892S | C | 1.3e-08 | 9.1e-09 | 3.9e-09 | 8.667E-09 | 4.565E-09 | 3.900E-09 | 1.300E-08 |
| 17837 | 892 | G892T | C | 4.5e-09 | 1.1e-08 | 4.5e-09 | 6.667E-09 | 3.753E-09 | 4.500E-09 | 1.100E-08 |
| 17838 | 892 | G892W | C | 4.3e-09 | 3e-09 | 4.7e-09 | 4.000E-09 | 8.888E-10 | 3.000E-09 | 4.700E-09 |
| 17839 | 892 | G892Y | C | 3.3e-09 | 6e-09 | 1.6e-09 | 3.633E-09 | 2.219E-09 | 1.600E-09 | 6.000E-09 |
| 17840 | 892 | G892V | C | 3.4e-09 | 5.9e-09 | 1.3e-08 | 7.433E-09 | 4.980E-09 | 3.400E-09 | 1.300E-08 |
| 17841 | 893 | G893A | C | 1.4e-08 | 4.1e-09 | 1.1e-08 | 9.700E-09 | 5.076E-09 | 4.100E-09 | 1.400E-08 |
| 17842 | 893 | G893R | C | 8.6e-09 | 6.3e-09 | 3.3e-09 | 6.067E-09 | 2.658E-09 | 3.300E-09 | 8.600E-09 |
| 17843 | 893 | G893N | C | 7.8e-09 | 5.2e-09 | 8.9e-09 | 7.300E-09 | 1.900E-09 | 5.200E-09 | 8.900E-09 |
| 17844 | 893 | G893D | C | 1.3e-08 | 4.3e-09 | 8.1e-09 | 8.467E-09 | 4.362E-09 | 4.300E-09 | 1.300E-08 |
| 17845 | 893 | G893C | C | 5.4e-09 | 6.9e-09 | 4.1e-08 | 1.777E-08 | 2.013E-08 | 5.400E-09 | 4.100E-08 |
| 17846 | 893 | G893E | C | 3.6e-09 | 3.1e-09 | 4.1e-09 | 3.600E-09 | 5.000E-10 | 3.100E-09 | 4.100E-09 |
| 17847 | 893 | G893Q | C | 1.8e-09 | 3.6e-09 | 3.5e-09 | 2.967E-09 | 1.012E-09 | 1.800E-09 | 3.600E-09 |
| 17848 | 893 | G893G | C | 3.7e-09 | 9.2e-09 | 1.1e-08 | 7.967E-09 | 3.803E-09 | 3.700E-09 | 1.100E-08 |
| 17849 | 893 | G893H | C | 3.9e-09 | 3.3e-09 | 6.6e-09 | 4.600E-09 | 1.758E-09 | 3.300E-09 | 6.600E-09 |

| | | | | | | | | | | |
|---|---|---|---|---|---|---|---|---|---|---|
| 17850 | 893 | G893I | C | 4.5e-09 | 6.1e-09 | 6e-09 | 5.533E-09 | 8.963E-10 | 4.500E-09 | 6.100E-09 |
| 17851 | 893 | G893L | C | 2e-09 | 8.8e-09 | 1.4e-08 | 8.267E-09 | 6.018E-09 | 2.000E-09 | 1.400E-08 |
| 17852 | 893 | G893K | C | 1.5e-09 | 1.1e-08 | 2e-09 | 4.833E-09 | 5.346E-09 | 1.500E-09 | 1.100E-08 |
| 17853 | 893 | G893M | C | 1.4e-08 | 2.5e-08 | 5.7e-09 | 1.490E-08 | 9.681E-09 | 5.700E-09 | 2.500E-08 |
| 17854 | 893 | G893F | C | 8.7e-09 | 4e-09 | 1.3e-08 | 8.567E-09 | 4.501E-09 | 4.000E-09 | 1.300E-08 |
| 17855 | 893 | G893P | C | 6e-09 | 2.2e-09 | 5.2e-09 | 4.467E-09 | 2.003E-09 | 2.200E-09 | 6.000E-09 |
| 17856 | 893 | G893S | C | 4.3e-09 | 6.1e-09 | 2.7e-09 | 4.367E-09 | 1.701E-09 | 2.700E-09 | 6.100E-09 |
| 17857 | 893 | G893T | C | 1.9e-08 | 2.9e-09 | 1.9e-09 | 7.933E-09 | 9.597E-09 | 1.900E-09 | 1.900E-08 |
| 17858 | 893 | G893W | C | 8.1e-09 | 1.1e-08 | 1.3e-08 | 1.070E-08 | 2.464E-09 | 8.100E-09 | 1.300E-08 |
| 17859 | 893 | G893Y | C | 4.6e-09 | 5.4e-09 | 1.4e-08 | 8.000E-09 | 5.212E-09 | 4.600E-09 | 1.400E-08 |
| 17860 | 893 | G893V | C | 1.2e-08 | 4.2e-09 | 8.3e-09 | 8.167E-09 | 3.902E-09 | 4.200E-09 | 1.200E-08 |
| 17861 | 894 | S894A | C | 1.1e-08 | 9.9e-10 | 7.8e-09 | 6.597E-09 | 5.112E-09 | 9.900E-10 | 1.100E-08 |
| 17862 | 894 | S894R | C | 9.7e-09 | 4e-09 | 1.7e-08 | 1.023E-08 | 6.516E-09 | 4.000E-09 | 1.700E-08 |
| 17863 | 894 | S894N | C | 1.2e-08 | 9.6e-10 | 2.1e-08 | 1.132E-08 | 1.004E-08 | 9.600E-10 | 2.100E-08 |
| 17864 | 894 | S894D | C | 4.3e-09 | 6.6e-09 | 1.2e-08 | 7.633E-09 | 3.953E-09 | 4.300E-09 | 1.200E-08 |
| 17865 | 894 | S894C | C | 9.4e-09 | 1e-08 | 5e-09 | 8.133E-09 | 2.730E-09 | 5.000E-09 | 1.000E-08 |
| 17866 | 894 | S894E | C | 7.4e-09 | 1.2e-08 | 7e-09 | 8.800E-09 | 2.778E-09 | 7.000E-09 | 1.200E-08 |
| 17867 | 894 | S894Q | C | 7.5e-09 | 1.2e-08 | 5.7e-09 | 8.400E-09 | 3.245E-09 | 5.700E-09 | 1.200E-08 |
| 17868 | 894 | S894G | C | 9.3e-09 | 6e-09 | 1.1e-08 | 8.767E-09 | 2.542E-09 | 6.000E-09 | 1.100E-08 |
| 17869 | 894 | S894H | C | 5.9e-09 | 7.9e-09 | 1.3e-08 | 8.933E-09 | 3.661E-09 | 5.900E-09 | 1.300E-08 |
| 17870 | 894 | S894I | C | 9.1e-09 | 8.7e-09 | 1.2e-08 | 9.933E-09 | 1.801E-09 | 8.700E-09 | 1.200E-08 |
| 17871 | 894 | S894L | C | 2.6e-08 | 4.6e-09 | 1.8e-08 | 1.620E-08 | 1.081E-08 | 4.600E-09 | 2.600E-08 |
| 17872 | 894 | S894K | C | 6.9e-09 | 8.8e-09 | 5.2e-09 | 6.967E-09 | 1.801E-09 | 5.200E-09 | 8.800E-09 |
| 17873 | 894 | S894M | C | 3.5e-08 | 2.4e-08 | 3.4e-08 | 3.100E-08 | 6.083E-09 | 2.400E-08 | 3.500E-08 |
| 17874 | 894 | S894F | C | 1.3e-08 | 2.2e-08 | 2e-09 | 1.233E-08 | 1.002E-08 | 2.000E-09 | 2.200E-08 |
| 17875 | 894 | S894P | C | 3.1e-08 | 2.3e-08 | 1.3e-08 | 2.233E-08 | 9.018E-09 | 1.300E-08 | 3.100E-08 |
| 17876 | 894 | S894S | C | 5.2e-09 | 2.1e-09 | 7.3e-09 | 4.867E-09 | 2.616E-09 | 2.100E-09 | 7.300E-09 |
| 17877 | 894 | S894T | C | 7e-09 | 1e-08 | 1.7e-08 | 1.133E-08 | 5.132E-09 | 7.000E-09 | 1.700E-08 |
| 17878 | 894 | S894W | C | 1.1e-08 | 1.1e-08 | 8e-09 | 1.000E-08 | 1.732E-09 | 8.000E-09 | 1.100E-08 |
| 17879 | 894 | S894Y | C | 5.4e-09 | 1.2e-08 | 1.8e-08 | 1.180E-08 | 6.302E-09 | 5.400E-09 | 1.800E-08 |
| 17880 | 894 | S894V | C | 1.1e-08 | 8.4e-09 | 6.8e-09 | 8.733E-09 | 2.120E-09 | 6.800E-09 | 1.100E-08 |
| 17881 | 895 | I895A | C | 8.7e-10 | 3e-09 | 2.4e-09 | 2.090E-09 | 1.098E-09 | 8.700E-10 | 3.000E-09 |
| 17882 | 895 | I895R | C | 3.6e-09 | 2.6e-09 | 1e-09 | 2.400E-09 | 1.311E-09 | 1.000E-09 | 3.600E-09 |
| 17883 | 895 | I895N | C | 7.2e-09 | 2.2e-09 | 6.3e-09 | 5.233E-09 | 2.665E-09 | 2.200E-09 | 7.200E-09 |
| 17884 | 895 | I895D | C | 4.7e-09 | 6.4e-09 | 9.9e-09 | 7.000E-09 | 2.651E-09 | 4.700E-09 | 9.900E-09 |
| 17885 | 895 | I895C | C | 3.1e-09 | 2.1e-08 | 3.2e-09 | 9.100E-09 | 1.031E-08 | 3.100E-09 | 2.100E-08 |
| 17886 | 895 | I895E | C | 2e-09 | 2.4e-09 | 7.1e-09 | 3.833E-09 | 2.836E-09 | 2.000E-09 | 7.100E-09 |
| 17887 | 895 | I895Q | C | 2.9e-09 | 1.5e-08 | 8.8e-09 | 8.900E-09 | 6.051E-09 | 2.900E-09 | 1.500E-08 |
| 17888 | 895 | I895G | C | 7.5e-09 | 5.6e-09 | 9.4e-09 | 7.500E-09 | 1.900E-09 | 5.600E-09 | 9.400E-09 |
| 17889 | 895 | I895H | C | 4.9e-09 | 5.5e-09 | 8.4e-09 | 6.267E-09 | 1.872E-09 | 4.900E-09 | 8.400E-09 |
| 17890 | 895 | I895I | C | 9.8e-09 | 3.6e-09 | 6.2e-09 | 6.533E-09 | 3.113E-09 | 3.600E-09 | 9.800E-09 |
| 17891 | 895 | I895L | C | 5.2e-09 | 3.2e-09 | 6.1e-09 | 4.833E-09 | 1.484E-09 | 3.200E-09 | 6.100E-09 |
| 17892 | 895 | I895K | C | 5.3e-09 | 4e-09 | 5.1e-09 | 4.800E-09 | 7.000E-10 | 4.000E-09 | 5.300E-09 |
| 17893 | 895 | I895M | C | 8.1e-09 | 1.6e-08 | 1.1e-08 | 1.170E-08 | 3.996E-09 | 8.100E-09 | 1.600E-08 |
| 17894 | 895 | I895F | C | 6.9e-09 | 6.8e-09 | 7.6e-09 | 7.100E-09 | 4.359E-10 | 6.800E-09 | 7.600E-09 |
| 17895 | 895 | I895P | C | 5.5e-09 | 3.1e-09 | 9.8e-09 | 6.133E-09 | 3.395E-09 | 3.100E-09 | 9.800E-09 |
| 17896 | 895 | I895S | C | 5.7e-09 | 1.5e-09 | 2e-09 | 3.067E-09 | 2.294E-09 | 1.500E-09 | 5.700E-09 |
| 17897 | 895 | I895T | C | 5.8e-09 | 3e-09 | 6.8e-09 | 5.200E-09 | 1.970E-09 | 3.000E-09 | 6.800E-09 |
| 17898 | 895 | I895W | C | 2.6e-08 | 6.8e-09 | 5.8e-09 | 1.287E-08 | 1.138E-08 | 5.800E-09 | 2.600E-08 |
| 17899 | 895 | I895Y | C | 2.4e-09 | 4.5e-09 | 1.1e-08 | 5.967E-09 | 4.484E-09 | 2.400E-09 | 1.100E-08 |
| 17900 | 895 | I895V | C | 2e-08 | 1.1e-08 | 2.9e-09 | 1.130E-08 | 8.554E-09 | 2.900E-09 | 2.000E-08 |
| 17901 | 896 | Y896A | C | 1.4e-08 | 1.2e-08 | 1.5e-08 | 1.367E-08 | 1.528E-09 | 1.200E-08 | 1.500E-08 |
| 17902 | 896 | Y896R | C | 4.6e-09 | 6.6e-09 | 3.5e-09 | 4.900E-09 | 1.572E-09 | 3.500E-09 | 6.600E-09 |
| 17903 | 896 | Y896N | C | 3.8e-09 | 7e-09 | 6.9e-09 | 5.900E-09 | 1.819E-09 | 3.800E-09 | 7.000E-09 |
| 17904 | 896 | Y896D | C | 9.5e-09 | 9.3e-09 | 2e-09 | 6.933E-09 | 4.274E-09 | 2.000E-09 | 9.500E-09 |
| 17905 | 896 | Y896C | C | 4.7e-09 | 6.3e-09 | 4.1e-09 | 5.033E-09 | 1.137E-09 | 4.100E-09 | 6.300E-09 |
| 17906 | 896 | Y896E | C | 5e-09 | 1.5e-08 | 6.7e-09 | 8.900E-09 | 5.351E-09 | 5.000E-09 | 1.500E-08 |
| 17907 | 896 | Y896Q | C | 6.4e-09 | 1.9e-09 | 3.9e-09 | 4.067E-09 | 2.255E-09 | 1.900E-09 | 6.400E-09 |
| 17908 | 896 | Y896G | C | 9.2e-09 | 5e-09 | 4.7e-09 | 6.300E-09 | 2.516E-09 | 4.700E-09 | 9.200E-09 |
| 17909 | 896 | Y896H | C | 5.2e-09 | 1.3e-08 | 1.3e-08 | 1.040E-08 | 4.503E-09 | 5.200E-09 | 1.300E-08 |
| 17910 | 896 | Y896I | C | 3.4e-09 | 1.2e-08 | 2.7e-09 | 6.033E-09 | 5.179E-09 | 2.700E-09 | 1.200E-08 |
| 17911 | 896 | Y896L | C | 1.2e-08 | 1e-08 | 4.2e-09 | 8.733E-09 | 4.051E-09 | 4.200E-09 | 1.200E-08 |
| 17912 | 896 | Y896K | C | 8e-09 | 3.2e-09 | 4.5e-09 | 5.233E-09 | 2.483E-09 | 3.200E-09 | 8.000E-09 |
| 17913 | 896 | Y896M | C | 3.9e-09 | 7.1e-09 | 4.7e-09 | 5.233E-09 | 1.665E-09 | 3.900E-09 | 7.100E-09 |
| 17914 | 896 | Y896F | C | 2.4e-08 | 1e-08 | 9.4e-09 | 1.447E-08 | 8.262E-09 | 9.400E-09 | 2.400E-08 |
| 17915 | 896 | Y896P | C | 1.4e-08 | 9.4e-09 | 1.3e-08 | 1.213E-08 | 2.419E-09 | 9.400E-09 | 1.400E-08 |
| 17916 | 896 | Y896S | C | 7.1e-09 | 1.1e-08 | 8.5e-09 | 8.867E-09 | 1.976E-09 | 7.100E-09 | 1.100E-08 |
| 17917 | 896 | Y896T | C | 2.8e-09 | 5.7e-09 | 1.1e-08 | 6.500E-09 | 4.158E-09 | 2.800E-09 | 1.100E-08 |
| 17918 | 896 | Y896W | C | 1.3e-08 | 6.8e-09 | 5.4e-09 | 8.400E-09 | 4.045E-09 | 5.400E-09 | 1.300E-08 |
| 17919 | 896 | Y896Y | C | 2.6e-09 | 7.8e-09 | 2.6e-09 | 4.333E-09 | 3.002E-09 | 2.600E-09 | 7.800E-09 |
| 17920 | 896 | Y896V | C | 1e-08 | 5e-09 | 1.2e-08 | 9.000E-09 | 3.606E-09 | 5.000E-09 | 1.200E-08 |
| 17921 | 897 | N897A | C | 4.3e-09 | 6.9e-09 | 6.7e-09 | 5.967E-09 | 1.447E-09 | 4.300E-09 | 6.900E-09 |
| 17922 | 897 | N897R | C | 5.1e-09 | 8.1e-09 | 1.6e-08 | 9.733E-09 | 5.631E-09 | 5.100E-09 | 1.600E-08 |
| 17923 | 897 | N897N | C | 4.9e-09 | 1.9e-09 | 6.5e-09 | 4.433E-09 | 2.335E-09 | 1.900E-09 | 6.500E-09 |
| 17924 | 897 | N897D | C | 1.9e-08 | 1.7e-08 | 1.4e-08 | 1.667E-08 | 2.517E-09 | 1.400E-08 | 1.900E-08 |
| 17925 | 897 | N897C | C | 4.3e-09 | 1.1e-08 | 7e-09 | 7.433E-09 | 3.371E-09 | 4.300E-09 | 1.100E-08 |
| 17926 | 897 | N897E | C | 7.9e-09 | 8.5e-09 | 3.7e-09 | 6.700E-09 | 2.615E-09 | 3.700E-09 | 8.500E-09 |
| 17927 | 897 | N897Q | C | 5.9e-09 | 1.5e-08 | 4e-09 | 8.300E-09 | 5.880E-09 | 4.000E-09 | 1.500E-08 |
| 17928 | 897 | N897G | C | 8.6e-09 | 5.9e-09 | 4.5e-09 | 6.333E-09 | 2.084E-09 | 4.500E-09 | 8.600E-09 |
| 17929 | 897 | N897H | C | 6.1e-09 | 6.9e-09 | 5.3e-09 | 6.100E-09 | 8.000E-10 | 5.300E-09 | 6.900E-09 |
| 17930 | 897 | N897I | C | 4.4e-09 | 6e-09 | 1.8e-08 | 9.467E-09 | 7.433E-09 | 4.400E-09 | 1.800E-08 |
| 17931 | 897 | N897L | C | 8.3e-09 | 7e-09 | 4.7e-09 | 6.667E-09 | 1.823E-09 | 4.700E-09 | 8.300E-09 |
| 17932 | 897 | N897K | C | 1.4e-08 | 3.8e-09 | 9.4e-09 | 9.067E-09 | 5.108E-09 | 3.800E-09 | 1.400E-08 |
| 17933 | 897 | N897M | C | 1.1e-08 | 9.9e-09 | 5.2e-09 | 8.700E-09 | 3.081E-09 | 5.200E-09 | 1.100E-08 |
| 17934 | 897 | N897F | C | 1.2e-08 | 1.6e-08 | 9.3e-09 | 1.243E-08 | 3.371E-09 | 9.300E-09 | 1.600E-08 |

| | | | | | | | | | | |
|---|---|---|---|---|---|---|---|---|---|---|
| 17935 | 897 | N897P | C | 1.6e-08 | 7.8e-09 | 9.5e-09 | 1.110E-08 | 4.328E-09 | 7.800E-09 | 1.600E-08 |
| 17936 | 897 | N897S | C | 8.8e-09 | 3.6e-09 | 6.4e-09 | 6.267E-09 | 2.603E-09 | 3.600E-09 | 8.800E-09 |
| 17937 | 897 | N897T | C | 4.4e-09 | 1.3e-08 | 7.1e-09 | 8.167E-09 | 4.398E-09 | 4.400E-09 | 1.300E-08 |
| 17938 | 897 | N897W | C | 1e-08 | 5.6e-09 | 2.4e-08 | 1.320E-08 | 9.608E-09 | 5.600E-09 | 2.400E-08 |
| 17939 | 897 | N897Y | C | 8.4e-09 | 1e-08 | 1.4e-08 | 1.080E-08 | 2.884E-09 | 8.400E-09 | 1.400E-08 |
| 17940 | 897 | N897V | C | 1.6e-08 | 2e-08 | 2e-08 | 1.867E-08 | 2.309E-09 | 1.600E-08 | 2.000E-08 |
| 17941 | 898 | Q898A | C | 9.6e-09 | 2.6e-09 | 2.6e-09 | 4.933E-09 | 4.041E-09 | 2.600E-09 | 9.600E-09 |
| 17942 | 898 | Q898R | C | 3.6e-09 | 1.4e-08 | 1.2e-09 | 6.267E-09 | 6.804E-09 | 1.200E-09 | 1.400E-08 |
| 17943 | 898 | Q898N | C | 2.4e-09 | 1.3e-08 | 7.7e-09 | 7.700E-09 | 5.300E-09 | 2.400E-09 | 1.300E-08 |
| 17944 | 898 | Q898D | C | 4.3e-09 | 9.6e-09 | 4.8e-09 | 6.233E-09 | 2.926E-09 | 4.300E-09 | 9.600E-09 |
| 17945 | 898 | Q898C | C | 5e-09 | 5.4e-09 | 4e-09 | 4.800E-09 | 7.211E-10 | 4.000E-09 | 5.400E-09 |
| 17946 | 898 | Q898E | C | 8.3e-09 | 2.1e-09 | 2.8e-09 | 4.400E-09 | 3.396E-09 | 2.100E-09 | 8.300E-09 |
| 17947 | 898 | Q898Q | C | 3.7e-09 | 5.9e-09 | 1.5e-08 | 8.200E-09 | 5.991E-09 | 3.700E-09 | 1.500E-08 |
| 17948 | 898 | Q898G | C | 7.9e-10 | 6.3e-09 | 5.2e-09 | 4.097E-09 | 2.916E-09 | 7.900E-10 | 6.300E-09 |
| 17949 | 898 | Q898H | C | 1.3e-08 | 4.6e-09 | 6.1e-09 | 7.900E-09 | 4.480E-09 | 4.600E-09 | 1.300E-08 |
| 17950 | 898 | Q898I | C | 1.4e-08 | 2.1e-09 | 5.3e-09 | 7.133E-09 | 6.158E-09 | 2.100E-09 | 1.400E-08 |
| 17951 | 898 | Q898L | C | 8.1e-09 | 2.6e-09 | 2.6e-09 | 4.433E-09 | 3.175E-09 | 2.600E-09 | 8.100E-09 |
| 17952 | 898 | Q898K | C | 5.3e-09 | 1.7e-09 | 4.3e-09 | 3.767E-09 | 1.858E-09 | 1.700E-09 | 5.300E-09 |
| 17953 | 898 | Q898M | C | 3.9e-09 | 5.3e-09 | 1.3e-09 | 3.500E-09 | 2.030E-09 | 1.300E-09 | 5.300E-09 |
| 17954 | 898 | Q898F | C | 2.6e-09 | 1.1e-09 | 7.7e-10 | 1.490E-09 | 9.753E-10 | 7.700E-10 | 2.600E-09 |
| 17955 | 898 | Q898P | C | 6e-09 | 2.2e-09 | 3.7e-09 | 3.967E-09 | 1.914E-09 | 2.200E-09 | 6.000E-09 |
| 17956 | 898 | Q898S | C | 5.4e-09 | 1e-08 | 7.7e-09 | 7.700E-09 | 2.300E-09 | 5.400E-09 | 1.000E-08 |
| 17957 | 898 | Q898T | C | 4.9e-09 | 9.1e-09 | 4.2e-09 | 6.067E-09 | 2.650E-09 | 4.200E-09 | 9.100E-09 |
| 17958 | 898 | Q898W | C | 3.1e-09 | 2e-09 | 3.2e-09 | 2.767E-09 | 6.658E-10 | 2.000E-09 | 3.200E-09 |
| 17959 | 898 | Q898Y | C | 3.5e-09 | 3.6e-09 | 3.8e-09 | 3.633E-09 | 1.528E-10 | 3.500E-09 | 3.800E-09 |
| 17960 | 898 | Q898V | C | 2e-09 | 2e-09 | 5.4e-09 | 3.133E-09 | 1.963E-09 | 2.000E-09 | 5.400E-09 |
| 17961 | 899 | R899A | C | 7.4e-09 | 8.5e-09 | 6.9e-09 | 7.600E-09 | 8.185E-10 | 6.900E-09 | 8.500E-09 |
| 17962 | 899 | R899R | C | 3.7e-09 | 6e-09 | 5.6e-09 | 5.100E-09 | 1.229E-09 | 3.700E-09 | 6.000E-09 |
| 17963 | 899 | R899N | C | 2.9e-09 | 1.5e-08 | 8.5e-09 | 8.800E-09 | 6.056E-09 | 2.900E-09 | 1.500E-08 |
| 17964 | 899 | R899D | C | 8.9e-09 | 2.1e-09 | 4.3e-09 | 5.100E-09 | 3.470E-09 | 2.100E-09 | 8.900E-09 |
| 17965 | 899 | R899C | C | 1.9e-08 | 2e-09 | 1.3e-09 | 7.433E-09 | 1.002E-08 | 1.300E-09 | 1.900E-08 |
| 17966 | 899 | R899E | C | 4e-09 | 5.2e-09 | 1.1e-08 | 6.733E-09 | 3.743E-09 | 4.000E-09 | 1.100E-08 |
| 17967 | 899 | R899Q | C | 7.6e-09 | 4.3e-09 | 6.1e-09 | 6.000E-09 | 1.652E-09 | 4.300E-09 | 7.600E-09 |
| 17968 | 899 | R899G | C | 7.4e-09 | 6.5e-09 | 1.3e-08 | 8.967E-09 | 3.522E-09 | 6.500E-09 | 1.300E-08 |
| 17969 | 899 | R899H | C | 6.5e-09 | 1.7e-08 | 6.2e-09 | 9.900E-09 | 6.151E-09 | 6.200E-09 | 1.700E-08 |
| 17970 | 899 | R899I | C | 1e-08 | 2.4e-08 | 7.8e-09 | 1.393E-08 | 8.787E-09 | 7.800E-09 | 2.400E-08 |
| 17971 | 899 | R899L | C | 2e-09 | 1.4e-08 | 8.5e-09 | 8.167E-09 | 6.007E-09 | 2.000E-09 | 1.400E-08 |
| 17972 | 899 | R899K | C | 8.2e-09 | 4.2e-09 | 2.7e-09 | 5.033E-09 | 2.843E-09 | 2.700E-09 | 8.200E-09 |
| 17973 | 899 | R899M | C | 7.4e-09 | 4.2e-09 | 1.7e-09 | 4.433E-09 | 2.857E-09 | 1.700E-09 | 7.400E-09 |
| 17974 | 899 | R899F | C | 1.1e-08 | 4.1e-09 | 1e-09 | 5.367E-09 | 5.119E-09 | 1.000E-09 | 1.100E-08 |
| 17975 | 899 | R899P | C | 1.6e-09 | 9.7e-09 | 1.1e-08 | 7.433E-09 | 5.093E-09 | 1.600E-09 | 1.100E-08 |
| 17976 | 899 | R899S | C | 1.5e-08 | 1.6e-08 | 2e-08 | 1.700E-08 | 2.646E-09 | 1.500E-08 | 2.000E-08 |
| 17977 | 899 | R899T | C | 5.2e-09 | 4.4e-09 | 5.6e-09 | 5.067E-09 | 6.110E-10 | 4.400E-09 | 5.600E-09 |
| 17978 | 899 | R899W | C | 3.4e-09 | 9.4e-09 | 8.6e-09 | 7.133E-09 | 3.258E-09 | 3.400E-09 | 9.400E-09 |
| 17979 | 899 | R899Y | C | 1.1e-08 | 6e-09 | 6.9e-09 | 7.967E-09 | 2.665E-09 | 6.000E-09 | 1.100E-08 |
| 17980 | 899 | R899V | C | 1.2e-08 | 5.8e-09 | 6.2e-09 | 8.000E-09 | 3.470E-09 | 5.800E-09 | 1.200E-08 |
| 17981 | 900 | F900A | C | 8e-09 | 2e-08 | 1.4e-08 | 1.400E-08 | 6.000E-09 | 8.000E-09 | 2.000E-08 |
| 17982 | 900 | F900R | C | 7.9e-09 | 7.9e-09 | 8.8e-09 | 8.200E-09 | 5.196E-10 | 7.900E-09 | 8.800E-09 |
| 17983 | 900 | F900N | C | 1.4e-08 | 1.7e-08 | 1.7e-08 | 1.600E-08 | 1.732E-09 | 1.400E-08 | 1.700E-08 |
| 17984 | 900 | F900D | C | 9e-09 | 7.5e-09 | 1.3e-08 | 9.833E-09 | 2.843E-09 | 7.500E-09 | 1.300E-08 |
| 17985 | 900 | F900C | C | 9.9e-09 | 1.1e-08 | 1.8e-09 | 7.567E-09 | 5.024E-09 | 1.800E-09 | 1.100E-08 |
| 17986 | 900 | F900E | C | 1.1e-08 | 1.3e-08 | 4e-09 | 9.333E-09 | 4.726E-09 | 4.000E-09 | 1.300E-08 |
| 17987 | 900 | F900Q | C | 2.1e-08 | 5.3e-09 | 7.1e-09 | 1.113E-08 | 8.592E-09 | 5.300E-09 | 2.100E-08 |
| 17988 | 900 | F900G | C | 2.4e-09 | 1.5e-09 | 4.9e-09 | 2.933E-09 | 1.762E-09 | 1.500E-09 | 4.900E-09 |
| 17989 | 900 | F900H | C | 7.1e-09 | 1.8e-08 | 2e-08 | 1.503E-08 | 6.943E-09 | 7.100E-09 | 2.000E-08 |
| 17990 | 900 | F900I | C | 1e-08 | 8.3e-09 | 3.7e-09 | 7.333E-09 | 3.259E-09 | 3.700E-09 | 1.000E-08 |
| 17991 | 900 | F900L | C | 1.1e-08 | 4e-09 | 4.6e-09 | 6.533E-09 | 3.880E-09 | 4.000E-09 | 1.100E-08 |
| 17992 | 900 | F900K | C | 7.3e-09 | 4.7e-09 | 1e-08 | 7.333E-09 | 2.650E-09 | 4.700E-09 | 1.000E-08 |
| 17993 | 900 | F900M | C | 2.3e-09 | 1.6e-08 | 8e-09 | 8.767E-09 | 6.882E-09 | 2.300E-09 | 1.600E-08 |
| 17994 | 900 | F900F | C | 6.7e-09 | 8.6e-09 | 1.3e-08 | 9.433E-09 | 3.232E-09 | 6.700E-09 | 1.300E-08 |
| 17995 | 900 | F900P | C | 7.8e-09 | 1.2e-08 | 4.5e-09 | 8.100E-09 | 3.759E-09 | 4.500E-09 | 1.200E-08 |
| 17996 | 900 | F900S | C | 5.3e-09 | 7.2e-09 | 6.5e-09 | 6.333E-09 | 9.609E-10 | 5.300E-09 | 7.200E-09 |
| 17997 | 900 | F900T | C | 3.3e-09 | 9.1e-09 | 3.2e-08 | 1.480E-08 | 1.518E-08 | 3.300E-09 | 3.200E-08 |
| 17998 | 900 | F900W | C | 1.7e-08 | 6e-09 | 3.4e-09 | 8.800E-09 | 7.219E-09 | 3.400E-09 | 1.700E-08 |
| 17999 | 900 | F900Y | C | 8.9e-09 | 2.5e-08 | 1.2e-08 | 1.530E-08 | 8.542E-09 | 8.900E-09 | 2.500E-08 |
| 18000 | 900 | F900V | C | 3.6e-09 | 1.3e-08 | 7e-09 | 7.867E-09 | 4.760E-09 | 3.600E-09 | 1.300E-08 |
| 18001 | 901 | K901A | C | 5.8e-09 | 8.3e-09 | 6.1e-09 | 6.733E-09 | 1.365E-09 | 5.800E-09 | 8.300E-09 |
| 18002 | 901 | K901R | C | 7e-09 | 1.3e-08 | 3.7e-09 | 7.900E-09 | 4.715E-09 | 3.700E-09 | 1.300E-08 |
| 18003 | 901 | K901N | C | 5.5e-09 | 5.5e-09 | 5.8e-09 | 5.600E-09 | 1.732E-10 | 5.500E-09 | 5.800E-09 |
| 18004 | 901 | K901D | C | 2.4e-08 | 1.1e-08 | 9.5e-09 | 1.483E-08 | 7.974E-09 | 9.500E-09 | 2.400E-08 |
| 18005 | 901 | K901C | C | 1.1e-08 | 6e-09 | 2.7e-09 | 6.567E-09 | 4.179E-09 | 2.700E-09 | 1.100E-08 |
| 18006 | 901 | K901E | C | 7.6e-09 | 9e-09 | 1.3e-08 | 9.867E-09 | 2.802E-09 | 7.600E-09 | 1.300E-08 |
| 18007 | 901 | K901Q | C | 8.9e-09 | 9.8e-09 | 7.9e-09 | 8.867E-09 | 9.504E-10 | 7.900E-09 | 9.800E-09 |
| 18008 | 901 | K901G | C | 8.7e-09 | 5e-09 | 3.5e-09 | 5.733E-09 | 2.676E-09 | 3.500E-09 | 8.700E-09 |
| 18009 | 901 | K901H | C | 9.7e-09 | 3.5e-09 | 1.1e-08 | 8.067E-09 | 4.008E-09 | 3.500E-09 | 1.100E-08 |
| 18010 | 901 | K901I | C | 4.4e-09 | 1.2e-08 | 2.5e-09 | 6.300E-09 | 5.027E-09 | 2.500E-09 | 1.200E-08 |
| 18011 | 901 | K901L | C | 7.4e-09 | 4.2e-09 | 6.6e-09 | 6.067E-09 | 1.665E-09 | 4.200E-09 | 7.400E-09 |
| 18012 | 901 | K901K | C | 8.4e-09 | 5.7e-09 | 2.6e-08 | 1.337E-08 | 1.102E-08 | 5.700E-09 | 2.600E-08 |
| 18013 | 901 | K901M | C | 3e-09 | 4.9e-09 | 1.1e-08 | 6.300E-09 | 4.180E-09 | 3.000E-09 | 1.100E-08 |
| 18014 | 901 | K901F | C | 5.3e-09 | 6e-09 | 1.9e-09 | 4.400E-09 | 2.193E-09 | 1.900E-09 | 6.000E-09 |
| 18015 | 901 | K901P | C | 9.1e-09 | 1.4e-08 | 8e-09 | 1.037E-08 | 3.194E-09 | 8.000E-09 | 1.400E-08 |
| 18016 | 901 | K901S | C | 8.3e-09 | 1.7e-08 | 2e-08 | 1.510E-08 | 6.077E-09 | 8.300E-09 | 2.000E-08 |
| 18017 | 901 | K901T | C | 8.7e-09 | 5e-09 | 1.1e-08 | 8.233E-09 | 3.027E-09 | 5.000E-09 | 1.100E-08 |
| 18018 | 901 | K901W | C | 6.3e-09 | 5.2e-09 | 6e-09 | 5.833E-09 | 5.686E-10 | 5.200E-09 | 6.300E-09 |
| 18019 | 901 | K901Y | C | 8.8e-09 | 5.7e-09 | 2.2e-09 | 5.567E-09 | 3.302E-09 | 2.200E-09 | 8.800E-09 |

| | | | | | | | | | | |
|---|---|---|---|---|---|---|---|---|---|---|
| 18020 | 901 | K901V | C | 1.1e-08 | 3.9e-09 | 4e-09 | 6.300E-09 | 4.071E-09 | 3.900E-09 | 1.100E-08 |
| 18021 | 902 | G902A | C | 4.6e-09 | 5.5e-09 | 5.4e-09 | 5.167E-09 | 4.933E-10 | 4.600E-09 | 5.500E-09 |
| 18022 | 902 | G902R | C | 2.2e-08 | 7.2e-09 | 6.5e-09 | 1.190E-08 | 8.754E-09 | 6.500E-09 | 2.200E-08 |
| 18023 | 902 | G902N | C | 4.9e-09 | 4.7e-09 | 9.5e-09 | 6.367E-09 | 2.715E-09 | 4.700E-09 | 9.500E-09 |
| 18024 | 902 | G902D | C | 7.3e-09 | 9.7e-09 | 5.4e-09 | 7.467E-09 | 2.155E-09 | 5.400E-09 | 9.700E-09 |
| 18025 | 902 | G902C | C | 1.9e-08 | 9.3e-09 | 6.5e-09 | 1.160E-08 | 6.560E-09 | 6.500E-09 | 1.900E-08 |
| 18026 | 902 | G902E | C | 1.1e-08 | 4.9e-09 | 9.2e-09 | 8.367E-09 | 3.134E-09 | 4.900E-09 | 1.100E-08 |
| 18027 | 902 | G902Q | C | 4.8e-09 | 7.6e-09 | 7.1e-09 | 6.500E-09 | 1.493E-09 | 4.800E-09 | 7.600E-09 |
| 18028 | 902 | G902G | C | 6.2e-09 | 1.4e-08 | 9.9e-09 | 1.003E-08 | 3.902E-09 | 6.200E-09 | 1.400E-08 |
| 18029 | 902 | G902H | C | 2.6e-08 | 9.5e-09 | 2.6e-08 | 2.050E-08 | 9.526E-09 | 9.500E-09 | 2.600E-08 |
| 18030 | 902 | G902I | C | 3.2e-09 | 1.2e-08 | 1e-08 | 8.400E-09 | 4.613E-09 | 3.200E-09 | 1.200E-08 |
| 18031 | 902 | G902L | C | 5.5e-09 | 7.4e-09 | 1.8e-08 | 1.030E-08 | 6.736E-09 | 5.500E-09 | 1.800E-08 |
| 18032 | 902 | G902K | C | 6.9e-09 | 4.2e-09 | 9.9e-09 | 7.000E-09 | 2.851E-09 | 4.200E-09 | 9.900E-09 |
| 18033 | 902 | G902M | C | 1.9e-08 | 2.1e-09 | 7.2e-09 | 9.433E-09 | 8.669E-09 | 2.100E-09 | 1.900E-08 |
| 18034 | 902 | G902F | C | 8.2e-09 | 1.1e-08 | 3.1e-09 | 7.433E-09 | 4.005E-09 | 3.100E-09 | 1.100E-08 |
| 18035 | 902 | G902P | C | 7.1e-09 | 2.2e-08 | 9.8e-09 | 1.297E-08 | 7.939E-09 | 7.100E-09 | 2.200E-08 |
| 18036 | 902 | G902S | C | 1.5e-08 | 4.6e-09 | 1.6e-08 | 1.187E-08 | 6.313E-09 | 4.600E-09 | 1.600E-08 |
| 18037 | 902 | G902T | C | 1.5e-08 | 1.1e-08 | 8.1e-09 | 1.137E-08 | 3.465E-09 | 8.100E-09 | 1.500E-08 |
| 18038 | 902 | G902W | C | 1.2e-08 | 6.5e-09 | 4.3e-09 | 7.600E-09 | 3.966E-09 | 4.300E-09 | 1.200E-08 |
| 18039 | 902 | G902Y | C | 1.8e-08 | 7e-09 | 3.2e-09 | 9.400E-09 | 7.686E-09 | 3.200E-09 | 1.800E-08 |
| 18040 | 902 | G902V | C | 6.3e-09 | 7.2e-09 | 6.4e-09 | 6.633E-09 | 4.933E-10 | 6.300E-09 | 7.200E-09 |
| 18041 | 903 | R903A | C | 1.2e-08 | 8.8e-09 | 3.5e-09 | 8.100E-09 | 4.293E-09 | 3.500E-09 | 1.200E-08 |
| 18042 | 903 | R903R | C | 1.5e-08 | 1.4e-08 | 8.9e-09 | 1.263E-08 | 3.272E-09 | 8.900E-09 | 1.500E-08 |
| 18043 | 903 | R903N | C | 7.4e-09 | 2.6e-09 | 6e-09 | 5.333E-09 | 2.468E-09 | 2.600E-09 | 7.400E-09 |
| 18044 | 903 | R903D | C | 9.7e-09 | 3.7e-09 | 9.2e-09 | 7.533E-09 | 3.329E-09 | 3.700E-09 | 9.700E-09 |
| 18045 | 903 | R903C | C | 4.1e-09 | 7e-09 | 2.4e-09 | 4.500E-09 | 2.326E-09 | 2.400E-09 | 7.000E-09 |
| 18046 | 903 | R903E | C | 3.1e-08 | 3.2e-09 | 5.5e-08 | 2.973E-08 | 2.592E-08 | 3.200E-09 | 5.500E-08 |
| 18047 | 903 | R903Q | C | 3.4e-09 | 4e-09 | 1e-08 | 5.800E-09 | 3.650E-09 | 3.400E-09 | 1.000E-08 |
| 18048 | 903 | R903G | C | 1.1e-08 | 7.7e-09 | 2.9e-09 | 7.200E-09 | 4.073E-09 | 2.900E-09 | 1.100E-08 |
| 18049 | 903 | R903H | C | 8.2e-09 | 1.6e-08 | 3.9e-09 | 9.367E-09 | 6.134E-09 | 3.900E-09 | 1.600E-08 |
| 18050 | 903 | R903I | C | 1e-08 | 3.4e-09 | 1.1e-08 | 8.133E-09 | 4.130E-09 | 3.400E-09 | 1.100E-08 |
| 18051 | 903 | R903L | C | 2.2e-08 | 7.9e-09 | 5.4e-09 | 1.177E-08 | 8.950E-09 | 5.400E-09 | 2.200E-08 |
| 18052 | 903 | R903K | C | 1.9e-09 | 6.7e-09 | 3.1e-08 | 1.320E-08 | 1.560E-08 | 1.900E-09 | 3.100E-08 |
| 18053 | 903 | R903M | C | 2.7e-09 | 5.4e-09 | 3e-09 | 3.700E-09 | 1.480E-09 | 2.700E-09 | 5.400E-09 |
| 18054 | 903 | R903F | C | 4.6e-09 | 7.3e-09 | 4.9e-09 | 5.600E-09 | 1.480E-09 | 4.600E-09 | 7.300E-09 |
| 18055 | 903 | R903P | C | 4.4e-09 | 3.7e-09 | 8.1e-09 | 5.400E-09 | 2.364E-09 | 3.700E-09 | 8.100E-09 |
| 18056 | 903 | R903S | C | 1.6e-08 | 3.9e-09 | 3.1e-09 | 7.667E-09 | 7.228E-09 | 3.100E-09 | 1.600E-08 |
| 18057 | 903 | R903T | C | 8e-09 | 4.2e-09 | 1.2e-08 | 8.067E-09 | 3.900E-09 | 4.200E-09 | 1.200E-08 |
| 18058 | 903 | R903W | C | 9.8e-09 | 1.8e-08 | 1.2e-08 | 1.327E-08 | 4.244E-09 | 9.800E-09 | 1.800E-08 |
| 18059 | 903 | R903Y | C | 1e-08 | 3e-09 | 5.9e-09 | 6.300E-09 | 3.517E-09 | 3.000E-09 | 1.000E-08 |
| 18060 | 903 | R903V | C | 9.4e-09 | 5.6e-09 | 7.4e-09 | 7.467E-09 | 1.901E-09 | 5.600E-09 | 9.400E-09 |
| 18061 | 904 | F904A | C | 7.6e-09 | 4.3e-09 | 3e-09 | 4.967E-09 | 2.371E-09 | 3.000E-09 | 7.600E-09 |
| 18062 | 904 | F904R | C | 5.1e-09 | 9e-09 | 6.1e-09 | 6.733E-09 | 2.026E-09 | 5.100E-09 | 9.000E-09 |
| 18063 | 904 | F904N | C | 6.8e-09 | 4.5e-09 | 3.5e-09 | 4.933E-09 | 1.692E-09 | 3.500E-09 | 6.800E-09 |
| 18064 | 904 | F904D | C | 1.3e-08 | 5.4e-09 | 2.9e-09 | 7.100E-09 | 5.260E-09 | 2.900E-09 | 1.300E-08 |
| 18065 | 904 | F904C | C | 1.9e-08 | 4.6e-09 | 6.4e-09 | 1.000E-08 | 7.846E-09 | 4.600E-09 | 1.900E-08 |
| 18066 | 904 | F904E | C | 4.7e-09 | 6.5e-09 | 6.2e-09 | 5.800E-09 | 9.644E-10 | 4.700E-09 | 6.500E-09 |
| 18067 | 904 | F904Q | C | 1.1e-08 | 3.3e-09 | 9e-09 | 7.767E-09 | 3.995E-09 | 3.300E-09 | 1.100E-08 |
| 18068 | 904 | F904G | C | 7.1e-09 | 3e-09 | 1e-08 | 6.700E-09 | 3.517E-09 | 3.000E-09 | 1.000E-08 |
| 18069 | 904 | F904H | C | 1.3e-08 | 3.3e-09 | 8.1e-09 | 8.133E-09 | 4.850E-09 | 3.300E-09 | 1.300E-08 |
| 18070 | 904 | F904I | C | 4.4e-09 | 2.9e-09 | 3.3e-09 | 3.533E-09 | 7.767E-10 | 2.900E-09 | 4.400E-09 |
| 18071 | 904 | F904L | C | 9.7e-09 | 6.6e-09 | 7e-09 | 7.767E-09 | 1.686E-09 | 6.600E-09 | 9.700E-09 |
| 18072 | 904 | F904K | C | 2.8e-09 | 2.9e-09 | 6.3e-09 | 4.000E-09 | 1.992E-09 | 2.800E-09 | 6.300E-09 |
| 18073 | 904 | F904M | C | 4.9e-09 | 4.5e-09 | 6.2e-09 | 5.200E-09 | 8.888E-10 | 4.500E-09 | 6.200E-09 |
| 18074 | 904 | F904F | C | 6.1e-09 | 4.8e-09 | 8.9e-09 | 6.600E-09 | 2.095E-09 | 4.800E-09 | 8.900E-09 |
| 18075 | 904 | F904P | C | 2.9e-09 | 4.2e-09 | 9.7e-09 | 5.600E-09 | 3.610E-09 | 2.900E-09 | 9.700E-09 |
| 18076 | 904 | F904S | C | 4.3e-09 | 3.4e-09 | 9.3e-09 | 5.667E-09 | 3.179E-09 | 3.400E-09 | 9.300E-09 |
| 18077 | 904 | F904T | C | 9.5e-09 | 1e-08 | 7.5e-09 | 9.000E-09 | 1.323E-09 | 7.500E-09 | 1.000E-08 |
| 18078 | 904 | F904W | C | 4.9e-09 | 9.3e-09 | 8.4e-09 | 7.533E-09 | 2.325E-09 | 4.900E-09 | 9.300E-09 |
| 18079 | 904 | F904Y | C | 7.5e-09 | 6.4e-09 | 1.6e-08 | 9.967E-09 | 5.254E-09 | 6.400E-09 | 1.600E-08 |
| 18080 | 904 | F904V | C | 6.8e-09 | 6.3e-09 | 1.1e-08 | 8.033E-09 | 2.581E-09 | 6.300E-09 | 1.100E-08 |
| 18081 | 905 | T905A | C | 1.4e-08 | 4.3e-09 | 6.4e-09 | 8.233E-09 | 5.103E-09 | 4.300E-09 | 1.400E-08 |
| 18082 | 905 | T905R | C | 3.5e-09 | 1.5e-08 | 5.5e-09 | 8.000E-09 | 6.144E-09 | 3.500E-09 | 1.500E-08 |
| 18083 | 905 | T905N | C | 1.8e-09 | 3.7e-09 | 3e-09 | 2.833E-09 | 9.609E-10 | 1.800E-09 | 3.700E-09 |
| 18084 | 905 | T905D | C | 7.8e-09 | 8.9e-09 | 1.9e-08 | 1.190E-08 | 6.173E-09 | 7.800E-09 | 1.900E-08 |
| 18085 | 905 | T905C | C | 4.9e-09 | 5.5e-09 | 1.6e-08 | 8.800E-09 | 6.243E-09 | 4.900E-09 | 1.600E-08 |
| 18086 | 905 | T905E | C | 5e-09 | 7e-09 | 4.4e-09 | 5.467E-09 | 1.361E-09 | 4.400E-09 | 7.000E-09 |
| 18087 | 905 | T905Q | C | 1.1e-08 | 8.7e-09 | 1.3e-08 | 1.090E-08 | 2.152E-09 | 8.700E-09 | 1.300E-08 |
| 18088 | 905 | T905G | C | 7.9e-09 | 7.5e-09 | 3.5e-09 | 6.300E-09 | 2.433E-09 | 3.500E-09 | 7.900E-09 |
| 18089 | 905 | T905H | C | 6.3e-09 | 5.9e-09 | 5.8e-09 | 6.000E-09 | 2.646E-10 | 5.800E-09 | 6.300E-09 |
| 18090 | 905 | T905I | C | 6.6e-09 | 5.1e-09 | 3.4e-09 | 5.033E-09 | 1.601E-09 | 3.400E-09 | 6.600E-09 |
| 18091 | 905 | T905L | C | 8.1e-09 | 1.1e-08 | 2e-09 | 7.033E-09 | 4.594E-09 | 2.000E-09 | 1.100E-08 |
| 18092 | 905 | T905K | C | 3.3e-09 | 3.1e-09 | 4.7e-09 | 3.700E-09 | 8.718E-10 | 3.100E-09 | 4.700E-09 |
| 18093 | 905 | T905M | C | 2e-09 | 2.5e-09 | 4.8e-09 | 3.100E-09 | 1.493E-09 | 2.000E-09 | 4.800E-09 |
| 18094 | 905 | T905F | C | 2.2e-08 | 1.5e-08 | 6.5e-09 | 1.450E-08 | 7.762E-09 | 6.500E-09 | 2.200E-08 |
| 18095 | 905 | T905P | C | 1.5e-08 | 7.2e-09 | 3.6e-09 | 8.600E-09 | 5.828E-09 | 3.600E-09 | 1.500E-08 |
| 18096 | 905 | T905S | C | 5.3e-09 | 2.6e-09 | 9.8e-09 | 5.900E-09 | 3.637E-09 | 2.600E-09 | 9.800E-09 |
| 18097 | 905 | T905T | C | 7.2e-09 | 4.2e-09 | 7.7e-09 | 6.367E-09 | 1.893E-09 | 4.200E-09 | 7.700E-09 |
| 18098 | 905 | T905W | C | 3.9e-09 | 6.1e-09 | 5e-09 | 5.000E-09 | 1.100E-09 | 3.900E-09 | 6.100E-09 |
| 18099 | 905 | T905Y | C | 6.8e-09 | 3.1e-09 | 7.7e-09 | 5.867E-09 | 2.438E-09 | 3.100E-09 | 7.700E-09 |
| 18100 | 905 | T905V | C | 1.4e-08 | 1.1e-08 | 6.5e-09 | 1.050E-08 | 3.775E-09 | 6.500E-09 | 1.400E-08 |
| 18101 | 906 | L906A | C | 5.9e-09 | 3e-09 | 4.1e-09 | 4.333E-09 | 1.464E-09 | 3.000E-09 | 5.900E-09 |
| 18102 | 906 | L906R | C | 2.3e-09 | 5.7e-09 | 2.2e-08 | 1.000E-08 | 1.053E-08 | 2.300E-09 | 2.200E-08 |
| 18103 | 906 | L906N | C | 1.3e-08 | 8.4e-09 | 2.7e-09 | 8.033E-09 | 5.160E-09 | 2.700E-09 | 1.300E-08 |
| 18104 | 906 | L906D | C | 2.4e-09 | 1.2e-08 | 2.8e-09 | 5.733E-09 | 5.431E-09 | 2.400E-09 | 1.200E-08 |

| | | | | | | | | | | |
|---|---|---|---|---|---|---|---|---|---|---|
| 18105 | 906 | L906C | C | 3.5e-09 | 1e-08 | 5.3e-09 | 6.267E-09 | 3.356E-09 | 3.500E-09 | 1.000E-08 |
| 18106 | 906 | L906E | C | 7.6e-09 | 3.6e-09 | 2.4e-09 | 4.533E-09 | 2.723E-09 | 2.400E-09 | 7.600E-09 |
| 18107 | 906 | L906Q | C | 6.5e-09 | 1e-08 | 1.4e-09 | 5.967E-09 | 4.325E-09 | 1.400E-09 | 1.000E-08 |
| 18108 | 906 | L906G | C | 2.3e-09 | 4.1e-09 | 5.5e-09 | 3.967E-09 | 1.604E-09 | 2.300E-09 | 5.500E-09 |
| 18109 | 906 | L906H | C | 3.1e-08 | 8.7e-09 | 6e-09 | 1.523E-08 | 1.372E-08 | 6.000E-09 | 3.100E-08 |
| 18110 | 906 | L906I | C | 9.5e-09 | 1.3e-08 | 6.1e-09 | 9.533E-09 | 3.450E-09 | 6.100E-09 | 1.300E-08 |
| 18111 | 906 | L906L | C | 2.6e-09 | 3.4e-09 | 7.1e-09 | 4.367E-09 | 2.401E-09 | 2.600E-09 | 7.100E-09 |
| 18112 | 906 | L906K | C | 8e-09 | 9.2e-09 | 4e-09 | 7.067E-09 | 2.723E-09 | 4.000E-09 | 9.200E-09 |
| 18113 | 906 | L906M | C | 6.5e-09 | 7.3e-09 | 8e-09 | 7.267E-09 | 7.506E-10 | 6.500E-09 | 8.000E-09 |
| 18114 | 906 | L906F | C | 6.2e-09 | 7.7e-09 | 1.1e-08 | 8.300E-09 | 2.456E-09 | 6.200E-09 | 1.100E-08 |
| 18115 | 906 | L906P | C | 1.5e-08 | 3.2e-09 | 6.3e-09 | 8.167E-09 | 6.117E-09 | 3.200E-09 | 1.500E-08 |
| 18116 | 906 | L906S | C | 2e-08 | 9.2e-09 | 6e-09 | 1.173E-08 | 7.336E-09 | 6.000E-09 | 2.000E-08 |
| 18117 | 906 | L906T | C | 6.8e-09 | 2.4e-09 | 1.3e-08 | 7.400E-09 | 5.325E-09 | 2.400E-09 | 1.300E-08 |
| 18118 | 906 | L906W | C | 7.4e-09 | 4.5e-09 | 1.2e-08 | 7.967E-09 | 3.782E-09 | 4.500E-09 | 1.200E-08 |
| 18119 | 906 | L906Y | C | 1.5e-08 | 6.3e-09 | 1.7e-08 | 1.277E-08 | 5.689E-09 | 6.300E-09 | 1.700E-08 |
| 18120 | 906 | L906V | C | 1.8e-08 | 1.1e-08 | 1.5e-08 | 1.467E-08 | 3.512E-09 | 1.100E-08 | 1.800E-08 |
| 18121 | 907 | S907A | C | 1.2e-08 | 6.4e-09 | 6.9e-09 | 8.433E-09 | 3.099E-09 | 6.400E-09 | 1.200E-08 |
| 18122 | 907 | S907R | C | 5e-09 | 1.7e-09 | 2.7e-09 | 3.133E-09 | 1.692E-09 | 1.700E-09 | 5.000E-09 |
| 18123 | 907 | S907N | C | 3e-09 | 4.3e-09 | 3.6e-09 | 3.633E-09 | 6.506E-10 | 3.000E-09 | 4.300E-09 |
| 18124 | 907 | S907D | C | 8.7e-09 | 3.2e-09 | 5.8e-09 | 5.900E-09 | 2.751E-09 | 3.200E-09 | 8.700E-09 |
| 18125 | 907 | S907C | C | 8.5e-09 | 7.8e-09 | 1.2e-09 | 5.833E-09 | 4.028E-09 | 1.200E-09 | 8.500E-09 |
| 18126 | 907 | S907E | C | 3.7e-09 | 4.5e-09 | 1.7e-08 | 8.400E-09 | 7.459E-09 | 3.700E-09 | 1.700E-08 |
| 18127 | 907 | S907Q | C | 8e-09 | 3.7e-09 | 1.4e-08 | 8.567E-09 | 5.173E-09 | 3.700E-09 | 1.400E-08 |
| 18128 | 907 | S907G | C | 4.7e-09 | 3.5e-09 | 1.1e-08 | 6.400E-09 | 4.029E-09 | 3.500E-09 | 1.100E-08 |
| 18129 | 907 | S907H | C | 6.1e-09 | 5.6e-09 | 7.2e-09 | 6.300E-09 | 8.185E-10 | 5.600E-09 | 7.200E-09 |
| 18130 | 907 | S907I | C | 8.5e-09 | 2.1e-09 | 1.3e-08 | 7.867E-09 | 5.478E-09 | 2.100E-09 | 1.300E-08 |
| 18131 | 907 | S907L | C | 1.2e-08 | 4.5e-09 | 6.2e-09 | 7.567E-09 | 3.932E-09 | 4.500E-09 | 1.200E-08 |
| 18132 | 907 | S907K | C | 2.8e-09 | 1.2e-09 | 3e-09 | 2.333E-09 | 9.866E-10 | 1.200E-09 | 3.000E-09 |
| 18133 | 907 | S907M | C | 9.1e-09 | 1.7e-08 | 5.2e-09 | 1.043E-08 | 6.012E-09 | 5.200E-09 | 1.700E-08 |
| 18134 | 907 | S907F | C | 5.9e-09 | 8.9e-09 | 3.7e-09 | 6.167E-09 | 2.610E-09 | 3.700E-09 | 8.900E-09 |
| 18135 | 907 | S907P | C | 1.2e-08 | 3.6e-09 | 8.4e-09 | 8.000E-09 | 4.214E-09 | 3.600E-09 | 1.200E-08 |
| 18136 | 907 | S907S | C | 6.7e-09 | 1.2e-08 | 5.9e-09 | 8.200E-09 | 3.315E-09 | 5.900E-09 | 1.200E-08 |
| 18137 | 907 | S907T | C | 1e-08 | 5.1e-09 | 4.2e-09 | 6.433E-09 | 3.121E-09 | 4.200E-09 | 1.000E-08 |
| 18138 | 907 | S907W | C | 8.2e-09 | 1e-08 | 1.5e-08 | 1.107E-08 | 3.523E-09 | 8.200E-09 | 1.500E-08 |
| 18139 | 907 | S907Y | C | 1.2e-08 | 4.1e-09 | 3.3e-09 | 6.467E-09 | 4.809E-09 | 3.300E-09 | 1.200E-08 |
| 18140 | 907 | S907V | C | 3.5e-09 | 5.1e-09 | 1.2e-08 | 6.867E-09 | 4.517E-09 | 3.500E-09 | 1.200E-08 |
| 18141 | 908 | V908A | C | 8.1e-09 | 8.8e-09 | 7.7e-09 | 8.200E-09 | 5.568E-10 | 7.700E-09 | 8.800E-09 |
| 18142 | 908 | V908R | C | 1.2e-08 | 7.1e-09 | 3.5e-09 | 7.533E-09 | 4.267E-09 | 3.500E-09 | 1.200E-08 |
| 18143 | 908 | V908N | C | 6.7e-09 | 1.4e-08 | 1.2e-08 | 1.090E-08 | 3.772E-09 | 6.700E-09 | 1.400E-08 |
| 18144 | 908 | V908D | C | 6.5e-09 | 1.2e-08 | 5.2e-09 | 7.900E-09 | 3.610E-09 | 5.200E-09 | 1.200E-08 |
| 18145 | 908 | V908C | C | 8.3e-09 | 3.6e-09 | 4.7e-09 | 5.533E-09 | 2.458E-09 | 3.600E-09 | 8.300E-09 |
| 18146 | 908 | V908E | C | 5.3e-09 | 9e-09 | 4.3e-09 | 6.200E-09 | 2.476E-09 | 4.300E-09 | 9.000E-09 |
| 18147 | 908 | V908Q | C | 3e-09 | 2.2e-09 | 7.4e-09 | 4.200E-09 | 2.800E-09 | 2.200E-09 | 7.400E-09 |
| 18148 | 908 | V908G | C | 6.4e-09 | 4e-09 | 4.7e-09 | 5.033E-09 | 1.234E-09 | 4.000E-09 | 6.400E-09 |
| 18149 | 908 | V908H | C | 1.2e-08 | 3e-09 | 2.9e-09 | 5.967E-09 | 5.225E-09 | 2.900E-09 | 1.200E-08 |
| 18150 | 908 | V908I | C | 1.8e-08 | 3e-08 | 5.2e-09 | 1.773E-08 | 1.240E-08 | 5.200E-09 | 3.000E-08 |
| 18151 | 908 | V908L | C | 5.8e-09 | 4.5e-09 | 4.8e-09 | 5.033E-09 | 6.807E-10 | 4.500E-09 | 5.800E-09 |
| 18152 | 908 | V908K | C | 8.4e-09 | 5e-09 | 9.2e-09 | 7.533E-09 | 2.230E-09 | 5.000E-09 | 9.200E-09 |
| 18153 | 908 | V908M | C | 1.7e-08 | 6.4e-09 | 9.6e-09 | 1.100E-08 | 5.437E-09 | 6.400E-09 | 1.700E-08 |
| 18154 | 908 | V908F | C | 9.8e-09 | 8.7e-09 | 4.3e-08 | 2.050E-08 | 1.949E-08 | 8.700E-09 | 4.300E-08 |
| 18155 | 908 | V908P | C | 9.8e-09 | 5.8e-09 | 6e-09 | 7.200E-09 | 2.254E-09 | 5.800E-09 | 9.800E-09 |
| 18156 | 908 | V908S | C | 1.2e-08 | 3.6e-09 | 1.5e-08 | 1.020E-08 | 5.909E-09 | 3.600E-09 | 1.500E-08 |
| 18157 | 908 | V908T | C | 1.6e-08 | 1.3e-08 | 4.3e-09 | 1.110E-08 | 6.077E-09 | 4.300E-09 | 1.600E-08 |
| 18158 | 908 | V908W | C | 6.9e-09 | 1.1e-08 | 4.4e-09 | 7.433E-09 | 3.332E-09 | 4.400E-09 | 1.100E-08 |
| 18159 | 908 | V908Y | C | 5.6e-09 | 3.6e-09 | 5.3e-09 | 4.833E-09 | 1.079E-09 | 3.600E-09 | 5.600E-09 |
| 18160 | 908 | V908V | C | 7.1e-09 | 4.9e-09 | 3.7e-09 | 5.233E-09 | 1.724E-09 | 3.700E-09 | 7.100E-09 |
| 18161 | 909 | D909A | C | 5.6e-09 | 4.1e-09 | 4.1e-09 | 4.600E-09 | 8.660E-10 | 4.100E-09 | 5.600E-09 |
| 18162 | 909 | D909R | C | 6.4e-09 | 2.3e-09 | 1e-08 | 6.233E-09 | 3.853E-09 | 2.300E-09 | 1.000E-08 |
| 18163 | 909 | D909N | C | 8.3e-09 | 1.2e-08 | 2.1e-08 | 1.377E-08 | 6.532E-09 | 8.300E-09 | 2.100E-08 |
| 18164 | 909 | D909D | C | 5.4e-09 | 3.6e-09 | 5.5e-09 | 4.833E-09 | 1.069E-09 | 3.600E-09 | 5.500E-09 |
| 18165 | 909 | D909C | C | 9.1e-09 | 2.3e-08 | 5e-09 | 1.237E-08 | 9.434E-09 | 5.000E-09 | 2.300E-08 |
| 18166 | 909 | D909E | C | 2.2e-09 | 8.5e-09 | 4.3e-09 | 5.000E-09 | 3.208E-09 | 2.200E-09 | 8.500E-09 |
| 18167 | 909 | D909Q | C | 3.3e-09 | 1.1e-08 | 6.6e-09 | 6.967E-09 | 3.863E-09 | 3.300E-09 | 1.100E-08 |
| 18168 | 909 | D909G | C | 9.4e-09 | 1.4e-08 | 3.1e-09 | 8.833E-09 | 5.472E-09 | 3.100E-09 | 1.400E-08 |
| 18169 | 909 | D909H | C | 6.1e-09 | 2e-08 | 1e-08 | 1.203E-08 | 7.170E-09 | 6.100E-09 | 2.000E-08 |
| 18170 | 909 | D909I | C | 1.1e-08 | 1.3e-08 | 2.1e-09 | 8.700E-09 | 5.803E-09 | 2.100E-09 | 1.300E-08 |
| 18171 | 909 | D909L | C | 2.1e-09 | 6.2e-09 | 1.8e-08 | 8.767E-09 | 8.255E-09 | 2.100E-09 | 1.800E-08 |
| 18172 | 909 | D909K | C | 2.3e-09 | 6.2e-09 | 4e-09 | 4.167E-09 | 1.955E-09 | 2.300E-09 | 6.200E-09 |
| 18173 | 909 | D909M | C | 2.3e-08 | 1.5e-08 | 6.9e-09 | 1.497E-08 | 8.050E-09 | 6.900E-09 | 2.300E-08 |
| 18174 | 909 | D909F | C | 6.2e-09 | 2.5e-08 | 1.9e-09 | 1.103E-08 | 1.229E-08 | 1.900E-09 | 2.500E-08 |
| 18175 | 909 | D909P | C | 1.1e-08 | 2.2e-09 | 3.1e-09 | 5.433E-09 | 4.842E-09 | 2.200E-09 | 1.100E-08 |
| 18176 | 909 | D909S | C | 1.2e-08 | 5.7e-09 | 5.6e-09 | 7.767E-09 | 3.667E-09 | 5.600E-09 | 1.200E-08 |
| 18177 | 909 | D909T | C | 6.4e-09 | 9.9e-09 | 8.9e-09 | 8.400E-09 | 1.803E-09 | 6.400E-09 | 9.900E-09 |
| 18178 | 909 | D909W | C | 1.3e-08 | 8.6e-09 | 5.8e-09 | 9.133E-09 | 3.630E-09 | 5.800E-09 | 1.300E-08 |
| 18179 | 909 | D909Y | C | 2.7e-09 | 1.3e-08 | 8.6e-09 | 8.100E-09 | 5.168E-09 | 2.700E-09 | 1.300E-08 |
| 18180 | 909 | D909V | C | 1.1e-08 | 8.6e-09 | 3.5e-09 | 7.700E-09 | 3.830E-09 | 3.500E-09 | 1.100E-08 |
| 18181 | 910 | R910A | C | 1.2e-08 | 7.3e-09 | 6.2e-09 | 8.500E-09 | 3.081E-09 | 6.200E-09 | 1.200E-08 |
| 18182 | 910 | R910R | C | 1.3e-09 | 7.6e-09 | 1.1e-08 | 6.633E-09 | 4.922E-09 | 1.300E-09 | 1.100E-08 |
| 18183 | 910 | R910N | C | 8.1e-09 | 6e-09 | 5.1e-09 | 6.400E-09 | 1.539E-09 | 5.100E-09 | 8.100E-09 |
| 18184 | 910 | R910D | C | 1.4e-08 | 5.6e-09 | 6.9e-09 | 8.833E-09 | 4.521E-09 | 5.600E-09 | 1.400E-08 |
| 18185 | 910 | R910C | C | 2.1e-08 | 5.6e-09 | 1.2e-08 | 1.287E-08 | 7.736E-09 | 5.600E-09 | 2.100E-08 |
| 18186 | 910 | R910E | C | 4.1e-09 | 6.7e-09 | 1.1e-08 | 7.267E-09 | 3.485E-09 | 4.100E-09 | 1.100E-08 |
| 18187 | 910 | R910Q | C | 9.3e-09 | 3.9e-09 | 4.8e-09 | 6.000E-09 | 2.893E-09 | 3.900E-09 | 9.300E-09 |
| 18188 | 910 | R910G | C | 8.4e-09 | 4.4e-09 | 1.4e-08 | 8.933E-09 | 4.822E-09 | 4.400E-09 | 1.400E-08 |
| 18189 | 910 | R910H | C | 1.1e-08 | 6.2e-09 | 5.9e-09 | 7.700E-09 | 2.862E-09 | 5.900E-09 | 1.100E-08 |

| | | | | | | | | | | |
|---|---|---|---|---|---|---|---|---|---|---|
| 18190 | 910 | R910I | C | 1.1e-08 | 4.8e-09 | 2.2e-08 | 1.260E-08 | 8.711E-09 | 4.800E-09 | 2.200E-08 |
| 18191 | 910 | R910L | C | 5.7e-09 | 7.6e-09 | 3.6e-09 | 5.633E-09 | 2.001E-09 | 3.600E-09 | 7.600E-09 |
| 18192 | 910 | R910K | C | 7.6e-09 | 9.3e-09 | 1.2e-08 | 9.633E-09 | 2.219E-09 | 7.600E-09 | 1.200E-08 |
| 18193 | 910 | R910M | C | 3.2e-08 | 1.7e-08 | 9.6e-09 | 1.953E-08 | 1.141E-08 | 9.600E-09 | 3.200E-08 |
| 18194 | 910 | R910F | C | 4e-09 | 9.1e-09 | 7.7e-09 | 6.933E-09 | 2.635E-09 | 4.000E-09 | 9.100E-09 |
| 18195 | 910 | R910P | C | 2e-08 | 2e-08 | 9.9e-09 | 1.663E-08 | 5.831E-09 | 9.900E-09 | 2.000E-08 |
| 18196 | 910 | R910S | C | 4.8e-09 | 4.7e-09 | 1.5e-08 | 8.167E-09 | 5.918E-09 | 4.700E-09 | 1.500E-08 |
| 18197 | 910 | R910T | C | 5.5e-09 | 6.8e-09 | 2e-08 | 1.077E-08 | 8.023E-09 | 5.500E-09 | 2.000E-08 |
| 18198 | 910 | R910W | C | 1e-08 | 1e-08 | 3.6e-09 | 7.867E-09 | 3.695E-09 | 3.600E-09 | 1.000E-08 |
| 18199 | 910 | R910Y | C | 1.6e-08 | 1.4e-08 | 6.9e-09 | 1.230E-08 | 4.782E-09 | 6.900E-09 | 1.600E-08 |
| 18200 | 910 | R910V | C | 7.9e-09 | 8.6e-09 | 1.8e-08 | 1.150E-08 | 5.640E-09 | 7.900E-09 | 1.800E-08 |
| 18201 | 911 | S911A | C | 1.8e-09 | 9.8e-09 | 1.7e-08 | 9.533E-09 | 7.604E-09 | 1.800E-09 | 1.700E-08 |
| 18202 | 911 | S911R | C | 2.9e-09 | 3e-09 | 9.9e-10 | 2.297E-09 | 1.133E-09 | 9.900E-10 | 3.000E-09 |
| 18203 | 911 | S911N | C | 6.7e-09 | 3.8e-09 | 2.9e-09 | 4.467E-09 | 1.986E-09 | 2.900E-09 | 6.700E-09 |
| 18204 | 911 | S911D | C | 2e-09 | 7.2e-09 | 1.4e-08 | 7.733E-09 | 6.018E-09 | 2.000E-09 | 1.400E-08 |
| 18205 | 911 | S911C | C | 2.5e-09 | 1.3e-08 | 3.8e-09 | 6.433E-09 | 5.724E-09 | 2.500E-09 | 1.300E-08 |
| 18206 | 911 | S911E | C | 4.7e-09 | 9.3e-09 | 4.6e-09 | 6.200E-09 | 2.685E-09 | 4.600E-09 | 9.300E-09 |
| 18207 | 911 | S911Q | C | 7.4e-09 | 1.5e-09 | 6e-09 | 4.967E-09 | 3.083E-09 | 1.500E-09 | 7.400E-09 |
| 18208 | 911 | S911G | C | 2.1e-09 | 3.2e-09 | 5.8e-09 | 3.700E-09 | 1.900E-09 | 2.100E-09 | 5.800E-09 |
| 18209 | 911 | S911H | C | 5e-09 | 9.6e-09 | 3.6e-09 | 6.067E-09 | 3.139E-09 | 3.600E-09 | 9.600E-09 |
| 18210 | 911 | S911I | C | 1.8e-08 | 1.2e-09 | 1.1e-08 | 1.007E-08 | 8.439E-09 | 1.200E-09 | 1.800E-08 |
| 18211 | 911 | S911L | C | 2.8e-09 | 2.7e-09 | 3.1e-09 | 2.867E-09 | 2.082E-10 | 2.700E-09 | 3.100E-09 |
| 18212 | 911 | S911K | C | 3.1e-09 | 2.4e-09 | 3.6e-09 | 3.033E-09 | 6.028E-10 | 2.400E-09 | 3.600E-09 |
| 18213 | 911 | S911M | C | 3e-09 | 2.1e-09 | 3.8e-09 | 2.967E-09 | 8.505E-10 | 2.100E-09 | 3.800E-09 |
| 18214 | 911 | S911F | C | 3.5e-09 | 2.9e-10 | 2.7e-09 | 2.163E-09 | 1.671E-09 | 2.900E-10 | 3.500E-09 |
| 18215 | 911 | S911P | C | 1e-08 | 5.4e-09 | 5.5e-09 | 6.967E-09 | 2.627E-09 | 5.400E-09 | 1.000E-08 |
| 18216 | 911 | S911S | C | 5.3e-09 | 3e-09 | 4.6e-09 | 4.300E-09 | 1.179E-09 | 3.000E-09 | 5.300E-09 |
| 18217 | 911 | S911T | C | 7.4e-09 | 3.6e-09 | 8.6e-09 | 6.533E-09 | 2.610E-09 | 3.600E-09 | 8.600E-09 |
| 18218 | 911 | S911W | C | 1.7e-09 | 1.4e-09 | 2.4e-09 | 1.833E-09 | 5.132E-10 | 1.400E-09 | 2.400E-09 |
| 18219 | 911 | S911Y | C | 6.9e-09 | 1.2e-09 | 1.5e-09 | 3.200E-09 | 3.208E-09 | 1.200E-09 | 6.900E-09 |
| 18220 | 911 | S911V | C | 4e-09 | 9e-10 | 4.6e-09 | 3.167E-09 | 1.986E-09 | 9.000E-10 | 4.600E-09 |
| 18221 | 912 | K912A | C | 3.3e-08 | 4.4e-09 | 7.3e-09 | 1.490E-08 | 1.574E-08 | 4.400E-09 | 3.300E-08 |
| 18222 | 912 | K912R | C | 1.3e-09 | 9e-09 | 7.1e-09 | 5.800E-09 | 4.011E-09 | 1.300E-09 | 9.000E-09 |
| 18223 | 912 | K912N | C | 6e-09 | 8.3e-09 | 1e-08 | 8.100E-09 | 2.007E-09 | 6.000E-09 | 1.000E-08 |
| 18224 | 912 | K912D | C | 8.2e-09 | 1.9e-09 | 7.9e-09 | 6.000E-09 | 3.554E-09 | 1.900E-09 | 8.200E-09 |
| 18225 | 912 | K912C | C | 9.6e-09 | 1.3e-08 | 3.5e-09 | 8.700E-09 | 4.814E-09 | 3.500E-09 | 1.300E-08 |
| 18226 | 912 | K912E | C | 2.5e-08 | 1.1e-08 | 1e-08 | 1.533E-08 | 8.386E-09 | 1.000E-08 | 2.500E-08 |
| 18227 | 912 | K912Q | C | 6.3e-09 | 1e-08 | 1.4e-08 | 1.010E-08 | 3.851E-09 | 6.300E-09 | 1.400E-08 |
| 18228 | 912 | K912G | C | 2e-09 | 6.9e-09 | 3.9e-09 | 4.267E-09 | 2.470E-09 | 2.000E-09 | 6.900E-09 |
| 18229 | 912 | K912H | C | 2.5e-08 | 7.5e-09 | 6.4e-09 | 1.297E-08 | 1.044E-08 | 6.400E-09 | 2.500E-08 |
| 18230 | 912 | K912I | C | 4.8e-09 | 2.6e-08 | 1.1e-08 | 1.393E-08 | 1.090E-08 | 4.800E-09 | 2.600E-08 |
| 18231 | 912 | K912L | C | 7.6e-09 | 7.2e-09 | 1.6e-09 | 5.467E-09 | 3.355E-09 | 1.600E-09 | 7.600E-09 |
| 18232 | 912 | K912K | C | 4.1e-09 | 2.8e-09 | 1.1e-09 | 2.667E-09 | 1.504E-09 | 1.100E-09 | 4.100E-09 |
| 18233 | 912 | K912M | C | 1.4e-08 | 8.2e-09 | 2.4e-09 | 8.200E-09 | 5.800E-09 | 2.400E-09 | 1.400E-08 |
| 18234 | 912 | K912F | C | 8.3e-09 | 8.9e-09 | 8.6e-09 | 8.600E-09 | 3.000E-10 | 8.300E-09 | 8.900E-09 |
| 18235 | 912 | K912P | C | 5.6e-09 | 1.2e-08 | 9.9e-09 | 9.167E-09 | 3.262E-09 | 5.600E-09 | 1.200E-08 |
| 18236 | 912 | K912S | C | 1.8e-08 | 5.5e-09 | 1.1e-08 | 1.150E-08 | 6.265E-09 | 5.500E-09 | 1.800E-08 |
| 18237 | 912 | K912T | C | 2.3e-08 | 1.4e-08 | 1.3e-08 | 1.667E-08 | 5.508E-09 | 1.300E-08 | 2.300E-08 |
| 18238 | 912 | K912W | C | 1.1e-08 | 1.7e-09 | 1.2e-08 | 8.233E-09 | 5.680E-09 | 1.700E-09 | 1.200E-08 |
| 18239 | 912 | K912Y | C | 3.8e-09 | 2.8e-09 | 8.8e-10 | 2.493E-09 | 1.484E-09 | 8.800E-10 | 3.800E-09 |
| 18240 | 912 | K912V | C | 2.1e-09 | 8.8e-09 | 5.9e-09 | 5.600E-09 | 3.360E-09 | 2.100E-09 | 8.800E-09 |
| 18241 | 913 | N913A | C | 2.2e-09 | 5.5e-09 | 6.8e-09 | 4.833E-09 | 2.371E-09 | 2.200E-09 | 6.800E-09 |
| 18242 | 913 | N913R | C | 4.8e-09 | 2.9e-09 | 1.8e-08 | 8.567E-09 | 8.225E-09 | 2.900E-09 | 1.800E-08 |
| 18243 | 913 | N913N | C | 5.7e-09 | 6.3e-09 | 1e-08 | 7.333E-09 | 2.329E-09 | 5.700E-09 | 1.000E-08 |
| 18244 | 913 | N913D | C | 6.9e-09 | 9.5e-09 | 2.3e-09 | 6.233E-09 | 3.646E-09 | 2.300E-09 | 9.500E-09 |
| 18245 | 913 | N913C | C | 7.2e-09 | 1.2e-08 | 7.4e-09 | 8.867E-09 | 2.715E-09 | 7.200E-09 | 1.200E-08 |
| 18246 | 913 | N913E | C | 4.8e-09 | 7.3e-09 | 5.6e-09 | 5.900E-09 | 1.277E-09 | 4.800E-09 | 7.300E-09 |
| 18247 | 913 | N913Q | C | 9e-09 | 2.1e-09 | 1.6e-08 | 9.033E-09 | 6.950E-09 | 2.100E-09 | 1.600E-08 |
| 18248 | 913 | N913G | C | 5.7e-09 | 8.5e-09 | 7.5e-09 | 7.233E-09 | 1.419E-09 | 5.700E-09 | 8.500E-09 |
| 18249 | 913 | N913H | C | 4.9e-09 | 8.3e-09 | 1.9e-09 | 5.033E-09 | 3.202E-09 | 1.900E-09 | 8.300E-09 |
| 18250 | 913 | N913I | C | 3.8e-09 | 4e-09 | 1.6e-08 | 7.933E-09 | 6.987E-09 | 3.800E-09 | 1.600E-08 |
| 18251 | 913 | N913L | C | 5e-09 | 1.2e-08 | 6.8e-09 | 7.933E-09 | 3.635E-09 | 5.000E-09 | 1.200E-08 |
| 18252 | 913 | N913K | C | 5.9e-09 | 1e-08 | 8.8e-09 | 8.233E-09 | 2.108E-09 | 5.900E-09 | 1.000E-08 |
| 18253 | 913 | N913M | C | 4.4e-09 | 2e-09 | 1.7e-08 | 7.800E-09 | 8.057E-09 | 2.000E-09 | 1.700E-08 |
| 18254 | 913 | N913F | C | 2.7e-09 | 3.8e-09 | 3.8e-09 | 3.433E-09 | 6.351E-10 | 2.700E-09 | 3.800E-09 |
| 18255 | 913 | N913P | C | 1.2e-08 | 4e-09 | 6.4e-09 | 7.467E-09 | 4.105E-09 | 4.000E-09 | 1.200E-08 |
| 18256 | 913 | N913S | C | 7.6e-09 | 8.1e-09 | 7.9e-09 | 7.867E-09 | 2.517E-10 | 7.600E-09 | 8.100E-09 |
| 18257 | 913 | N913T | C | 5.8e-09 | 9.3e-09 | 1.3e-08 | 9.367E-09 | 3.600E-09 | 5.800E-09 | 1.300E-08 |
| 18258 | 913 | N913W | C | 5.4e-09 | 9.8e-10 | 4.4e-09 | 3.593E-09 | 2.318E-09 | 9.800E-10 | 5.400E-09 |
| 18259 | 913 | N913Y | C | 2.7e-09 | 6.5e-09 | 1.2e-08 | 7.067E-09 | 4.676E-09 | 2.700E-09 | 1.200E-08 |
| 18260 | 913 | N913V | C | 1.3e-08 | 1.1e-08 | 3.8e-09 | 9.267E-09 | 4.839E-09 | 3.800E-09 | 1.300E-08 |
| 18261 | 914 | T914A | C | 6.6e-09 | 7.8e-09 | 2.2e-08 | 1.213E-08 | 8.566E-09 | 6.600E-09 | 2.200E-08 |
| 18262 | 914 | T914R | C | 7.5e-09 | 1e-08 | 5.4e-09 | 7.633E-09 | 2.303E-09 | 5.400E-09 | 1.000E-08 |
| 18263 | 914 | T914N | C | 2.4e-09 | 3.4e-09 | 3e-09 | 2.933E-09 | 5.033E-10 | 2.400E-09 | 3.400E-09 |
| 18264 | 914 | T914D | C | 5.6e-09 | 7.9e-09 | 7e-09 | 6.833E-09 | 1.159E-09 | 5.600E-09 | 7.900E-09 |
| 18265 | 914 | T914C | C | 6e-09 | 4.1e-09 | 2e-08 | 1.003E-08 | 8.684E-09 | 4.100E-09 | 2.000E-08 |
| 18266 | 914 | T914E | C | 1.3e-08 | 8.4e-09 | 1.4e-08 | 1.180E-08 | 2.987E-09 | 8.400E-09 | 1.400E-08 |
| 18267 | 914 | T914Q | C | 3.8e-09 | 5.4e-09 | 4.9e-09 | 4.700E-09 | 8.185E-10 | 3.800E-09 | 5.400E-09 |
| 18268 | 914 | T914G | C | 1.4e-08 | 1e-08 | 2.5e-09 | 8.833E-09 | 5.838E-09 | 2.500E-09 | 1.400E-08 |
| 18269 | 914 | T914H | C | 6.5e-09 | 3.6e-09 | 9.5e-09 | 6.533E-09 | 2.950E-09 | 3.600E-09 | 9.500E-09 |
| 18270 | 914 | T914I | C | 6.6e-09 | 5.3e-09 | 3.2e-09 | 5.033E-09 | 1.716E-09 | 3.200E-09 | 6.600E-09 |
| 18271 | 914 | T914L | C | 6.7e-09 | 1e-08 | 3.8e-09 | 6.833E-09 | 3.102E-09 | 3.800E-09 | 1.000E-08 |
| 18272 | 914 | T914K | C | 1e-08 | 7e-09 | 5.1e-09 | 7.367E-09 | 2.470E-09 | 5.100E-09 | 1.000E-08 |
| 18273 | 914 | T914M | C | 1.6e-08 | 2.5e-09 | 1e-08 | 9.500E-09 | 6.764E-09 | 2.500E-09 | 1.600E-08 |
| 18274 | 914 | T914F | C | 1e-08 | 6.2e-09 | 7.3e-09 | 7.833E-09 | 1.955E-09 | 6.200E-09 | 1.000E-08 |

| | | | | | | | | | | |
|---|---|---|---|---|---|---|---|---|---|---|
| 18275 | 914 | T914P | C | 9.1e-09 | 8.5e-09 | 8.4e-09 | 8.667E-09 | 3.786E-10 | 8.400E-09 | 9.100E-09 |
| 18276 | 914 | T914S | C | 9.4e-09 | 3.5e-09 | 1.9e-08 | 1.063E-08 | 7.823E-09 | 3.500E-09 | 1.900E-08 |
| 18277 | 914 | T914T | C | 2.5e-08 | 9.2e-09 | 1.7e-08 | 1.707E-08 | 7.900E-09 | 9.200E-09 | 2.500E-08 |
| 18278 | 914 | T914W | C | 2.4e-09 | 2e-09 | 6.8e-09 | 3.733E-09 | 2.663E-09 | 2.000E-09 | 6.800E-09 |
| 18279 | 914 | T914Y | C | 1.8e-08 | 5.2e-09 | 7.8e-09 | 1.033E-08 | 6.766E-09 | 5.200E-09 | 1.800E-08 |
| 18280 | 914 | T914V | C | 9.9e-09 | 6.2e-09 | 1e-08 | 8.700E-09 | 2.166E-09 | 6.200E-09 | 1.000E-08 |
| 18281 | 915 | L915A | C | 3.2e-09 | 2e-08 | 9.3e-09 | 1.083E-08 | 8.504E-09 | 3.200E-09 | 2.000E-08 |
| 18282 | 915 | L915R | C | 8.5e-09 | 1.2e-08 | 5.7e-09 | 8.733E-09 | 3.156E-09 | 5.700E-09 | 1.200E-08 |
| 18283 | 915 | L915N | C | 8.6e-09 | 1.1e-08 | 6.5e-09 | 8.700E-09 | 2.252E-09 | 6.500E-09 | 1.100E-08 |
| 18284 | 915 | L915D | C | 1.3e-08 | 1.4e-08 | 4.5e-09 | 1.050E-08 | 5.220E-09 | 4.500E-09 | 1.400E-08 |
| 18285 | 915 | L915C | C | 6.8e-09 | 1.1e-08 | 7.1e-09 | 8.300E-09 | 2.343E-09 | 6.800E-09 | 1.100E-08 |
| 18286 | 915 | L915E | C | 8.8e-09 | 1.7e-09 | 9.3e-09 | 6.600E-09 | 4.251E-09 | 1.700E-09 | 9.300E-09 |
| 18287 | 915 | L915Q | C | 1.5e-08 | 3.6e-09 | 9.1e-09 | 9.233E-09 | 5.701E-09 | 3.600E-09 | 1.500E-08 |
| 18288 | 915 | L915G | C | 1.9e-09 | 4.4e-09 | 4.6e-09 | 3.633E-09 | 1.504E-09 | 1.900E-09 | 4.600E-09 |
| 18289 | 915 | L915H | C | 5.3e-09 | 5.4e-09 | 8.9e-09 | 6.533E-09 | 2.050E-09 | 5.300E-09 | 8.900E-09 |
| 18290 | 915 | L915I | C | 8e-09 | 3.4e-09 | 5.5e-09 | 5.633E-09 | 2.303E-09 | 3.400E-09 | 8.000E-09 |
| 18291 | 915 | L915L | C | 2e-08 | 4.7e-09 | 8.4e-09 | 1.103E-08 | 7.983E-09 | 4.700E-09 | 2.000E-08 |
| 18292 | 915 | L915K | C | 8.9e-09 | 5.4e-09 | 7.7e-09 | 7.333E-09 | 1.779E-09 | 5.400E-09 | 8.900E-09 |
| 18293 | 915 | L915M | C | 2.8e-08 | 2.3e-09 | 8.5e-09 | 1.293E-08 | 1.341E-08 | 2.300E-09 | 2.800E-08 |
| 18294 | 915 | L915F | C | 8e-09 | 1.9e-09 | 1.3e-08 | 7.633E-09 | 5.559E-09 | 1.900E-09 | 1.300E-08 |
| 18295 | 915 | L915P | C | 1e-08 | 2.1e-09 | 7.4e-09 | 6.500E-09 | 4.026E-09 | 2.100E-09 | 1.000E-08 |
| 18296 | 915 | L915S | C | 1e-08 | 5.6e-09 | 4.1e-09 | 6.567E-09 | 3.066E-09 | 4.100E-09 | 1.000E-08 |
| 18297 | 915 | L915T | C | 1.6e-08 | 4.2e-09 | 5.5e-09 | 8.567E-09 | 6.470E-09 | 4.200E-09 | 1.600E-08 |
| 18298 | 915 | L915W | C | 7.1e-09 | 9.4e-09 | 8e-09 | 8.167E-09 | 1.159E-09 | 7.100E-09 | 9.400E-09 |
| 18299 | 915 | L915Y | C | 1.7e-08 | 2.1e-08 | 6.6e-09 | 1.487E-08 | 7.433E-09 | 6.600E-09 | 2.100E-08 |
| 18300 | 915 | L915V | C | 4.9e-09 | 5.7e-09 | 1.4e-09 | 4.000E-09 | 2.287E-09 | 1.400E-09 | 5.700E-09 |
| 18301 | 916 | Y916A | C | 4.7e-09 | 2.3e-08 | 1.6e-08 | 1.457E-08 | 9.234E-09 | 4.700E-09 | 2.300E-08 |
| 18302 | 916 | Y916R | C | 5.8e-09 | 1.9e-08 | 9.2e-09 | 1.133E-08 | 6.854E-09 | 5.800E-09 | 1.900E-08 |
| 18303 | 916 | Y916N | C | 6.5e-09 | 7e-09 | 5.9e-09 | 6.467E-09 | 5.508E-10 | 5.900E-09 | 7.000E-09 |
| 18304 | 916 | Y916D | C | 3.8e-09 | 1.7e-08 | 2.3e-09 | 7.700E-09 | 8.089E-09 | 2.300E-09 | 1.700E-08 |
| 18305 | 916 | Y916C | C | 5.7e-09 | 2.2e-08 | 1.1e-08 | 1.290E-08 | 8.314E-09 | 5.700E-09 | 2.200E-08 |
| 18306 | 916 | Y916E | C | 5.1e-09 | 8e-09 | 1e-08 | 7.700E-09 | 2.464E-09 | 5.100E-09 | 1.000E-08 |
| 18307 | 916 | Y916Q | C | 1e-08 | 8.9e-09 | 8.2e-09 | 9.033E-09 | 9.074E-10 | 8.200E-09 | 1.000E-08 |
| 18308 | 916 | Y916G | C | 2.7e-08 | 1.1e-08 | 9.1e-09 | 1.570E-08 | 9.832E-09 | 9.100E-09 | 2.700E-08 |
| 18309 | 916 | Y916H | C | 1.5e-09 | 9.5e-09 | 8.7e-09 | 6.567E-09 | 4.406E-09 | 1.500E-09 | 9.500E-09 |
| 18310 | 916 | Y916I | C | 1.8e-08 | 1.6e-08 | 5.8e-09 | 1.327E-08 | 6.543E-09 | 5.800E-09 | 1.800E-08 |
| 18311 | 916 | Y916L | C | 4.5e-09 | 8.2e-09 | 9.5e-09 | 7.400E-09 | 2.594E-09 | 4.500E-09 | 9.500E-09 |
| 18312 | 916 | Y916K | C | 6.8e-09 | 2.6e-08 | 9.1e-09 | 1.397E-08 | 1.048E-08 | 6.800E-09 | 2.600E-08 |
| 18313 | 916 | Y916M | C | 4e-09 | 4e-09 | 1.3e-08 | 7.000E-09 | 5.196E-09 | 4.000E-09 | 1.300E-08 |
| 18314 | 916 | Y916F | C | 1.8e-08 | 2.1e-08 | 9.3e-09 | 1.610E-08 | 6.077E-09 | 9.300E-09 | 2.100E-08 |
| 18315 | 916 | Y916P | C | 1.2e-08 | 2.8e-09 | 1.6e-08 | 1.027E-08 | 6.769E-09 | 2.800E-09 | 1.600E-08 |
| 18316 | 916 | Y916S | C | 6.7e-09 | 4.6e-09 | 4.3e-09 | 5.200E-09 | 1.308E-09 | 4.300E-09 | 6.700E-09 |
| 18317 | 916 | Y916T | C | 5e-09 | 1.5e-08 | 8.3e-09 | 9.433E-09 | 5.095E-09 | 5.000E-09 | 1.500E-08 |
| 18318 | 916 | Y916W | C | 1.6e-08 | 1e-08 | 7.6e-09 | 1.120E-08 | 4.327E-09 | 7.600E-09 | 1.600E-08 |
| 18319 | 916 | Y916Y | C | 3.5e-09 | 8.3e-09 | 7.7e-09 | 6.500E-09 | 2.615E-09 | 3.500E-09 | 8.300E-09 |
| 18320 | 916 | Y916V | C | 1.1e-08 | 3.6e-09 | 8.6e-09 | 7.733E-09 | 3.775E-09 | 3.600E-09 | 1.100E-08 |
| 18321 | 917 | L917A | C | 9.3e-09 | 1.1e-08 | 1.3e-08 | 1.110E-08 | 1.852E-09 | 9.300E-09 | 1.300E-08 |
| 18322 | 917 | L917R | C | 8.5e-09 | 6.8e-09 | 3.2e-09 | 6.167E-09 | 2.706E-09 | 3.200E-09 | 8.500E-09 |
| 18323 | 917 | L917N | C | 7.1e-09 | 1.5e-08 | 6e-09 | 9.367E-09 | 4.910E-09 | 6.000E-09 | 1.500E-08 |
| 18324 | 917 | L917D | C | 4.6e-09 | 3.7e-09 | 8.9e-09 | 5.733E-09 | 2.779E-09 | 3.700E-09 | 8.900E-09 |
| 18325 | 917 | L917C | C | 1.5e-08 | 4.9e-09 | 4.2e-09 | 8.033E-09 | 6.043E-09 | 4.200E-09 | 1.500E-08 |
| 18326 | 917 | L917E | C | 1.2e-08 | 2.5e-09 | 5e-09 | 6.500E-09 | 4.924E-09 | 2.500E-09 | 1.200E-08 |
| 18327 | 917 | L917Q | C | 6.8e-09 | 3.3e-09 | 8.7e-09 | 6.267E-09 | 2.739E-09 | 3.300E-09 | 8.700E-09 |
| 18328 | 917 | L917G | C | 5.8e-09 | 1.4e-08 | 3.4e-08 | 1.793E-08 | 1.451E-08 | 5.800E-09 | 3.400E-08 |
| 18329 | 917 | L917H | C | 1.4e-08 | 1.6e-08 | 1.1e-08 | 1.367E-08 | 2.517E-09 | 1.100E-08 | 1.600E-08 |
| 18330 | 917 | L917I | C | 1e-08 | 1.9e-08 | 4.1e-09 | 1.103E-08 | 7.504E-09 | 4.100E-09 | 1.900E-08 |
| 18331 | 917 | L917L | C | 6e-09 | 5.1e-09 | 6.7e-09 | 5.933E-09 | 8.021E-10 | 5.100E-09 | 6.700E-09 |
| 18332 | 917 | L917K | C | 1.1e-08 | 6.1e-09 | 6.7e-09 | 7.933E-09 | 2.673E-09 | 6.100E-09 | 1.100E-08 |
| 18333 | 917 | L917M | C | 3.6e-09 | 4.9e-09 | 2.9e-09 | 3.800E-09 | 1.015E-09 | 2.900E-09 | 4.900E-09 |
| 18334 | 917 | L917F | C | 2.5e-08 | 3.8e-09 | 1e-08 | 1.293E-08 | 1.090E-08 | 3.800E-09 | 2.500E-08 |
| 18335 | 917 | L917P | C | 1.6e-08 | 4.9e-09 | 6.6e-09 | 9.167E-09 | 5.979E-09 | 4.900E-09 | 1.600E-08 |
| 18336 | 917 | L917S | C | 2.3e-08 | 1.1e-08 | 8.3e-09 | 1.410E-08 | 7.825E-09 | 8.300E-09 | 2.300E-08 |
| 18337 | 917 | L917T | C | 6e-09 | 3.3e-09 | 8.8e-09 | 6.033E-09 | 2.750E-09 | 3.300E-09 | 8.800E-09 |
| 18338 | 917 | L917W | C | 4.4e-09 | 2.2e-08 | 4.6e-09 | 1.033E-08 | 1.010E-08 | 4.400E-09 | 2.200E-08 |
| 18339 | 917 | L917Y | C | 5.1e-09 | 2.7e-09 | 2.7e-09 | 3.500E-09 | 1.386E-09 | 2.700E-09 | 5.100E-09 |
| 18340 | 917 | L917V | C | 5.3e-09 | 3.8e-09 | 8.7e-09 | 5.933E-09 | 2.511E-09 | 3.800E-09 | 8.700E-09 |
| 18341 | 918 | Q918A | C | 6.7e-09 | 2.7e-09 | 6.1e-09 | 5.167E-09 | 2.157E-09 | 2.700E-09 | 6.700E-09 |
| 18342 | 918 | Q918R | C | 1e-08 | 2.4e-09 | 6.7e-09 | 6.367E-09 | 3.811E-09 | 2.400E-09 | 1.000E-08 |
| 18343 | 918 | Q918N | C | 4.3e-09 | 2.7e-08 | 8.7e-09 | 1.333E-08 | 1.204E-08 | 4.300E-09 | 2.700E-08 |
| 18344 | 918 | Q918D | C | 5.4e-09 | 7.1e-09 | 1.9e-08 | 1.050E-08 | 7.410E-09 | 5.400E-09 | 1.900E-08 |
| 18345 | 918 | Q918C | C | 7.3e-09 | 9.7e-09 | 4.9e-09 | 7.300E-09 | 2.400E-09 | 4.900E-09 | 9.700E-09 |
| 18346 | 918 | Q918E | C | 1.4e-09 | 1.1e-08 | 3.5e-09 | 5.300E-09 | 5.047E-09 | 1.400E-09 | 1.100E-08 |
| 18347 | 918 | Q918Q | C | 1.2e-08 | 3.5e-09 | 7.2e-09 | 7.567E-09 | 4.262E-09 | 3.500E-09 | 1.200E-08 |
| 18348 | 918 | Q918G | C | 6.6e-09 | 1.7e-08 | 9e-09 | 1.087E-08 | 5.445E-09 | 6.600E-09 | 1.700E-08 |
| 18349 | 918 | Q918H | C | 7e-09 | 1.1e-08 | 4.7e-09 | 7.567E-09 | 3.188E-09 | 4.700E-09 | 1.100E-08 |
| 18350 | 918 | Q918I | C | 6.6e-09 | 1e-08 | 8.3e-09 | 8.300E-09 | 1.700E-09 | 6.600E-09 | 1.000E-08 |
| 18351 | 918 | Q918L | C | 1.2e-08 | 4.7e-09 | 5.9e-09 | 7.533E-09 | 3.915E-09 | 4.700E-09 | 1.200E-08 |
| 18352 | 918 | Q918K | C | 2.4e-09 | 4.7e-09 | 5.4e-09 | 4.167E-09 | 1.570E-09 | 2.400E-09 | 5.400E-09 |
| 18353 | 918 | Q918M | C | 3.3e-09 | 4.1e-09 | 1.3e-08 | 6.800E-09 | 5.384E-09 | 3.300E-09 | 1.300E-08 |
| 18354 | 918 | Q918F | C | 9.9e-09 | 3.1e-08 | 8.3e-09 | 1.640E-08 | 1.267E-08 | 8.300E-09 | 3.100E-08 |
| 18355 | 918 | Q918P | C | 6.6e-09 | 1.3e-08 | 1.8e-08 | 1.253E-08 | 5.714E-09 | 6.600E-09 | 1.800E-08 |
| 18356 | 918 | Q918S | C | 5.5e-09 | 8.2e-09 | 9.7e-09 | 7.800E-09 | 2.128E-09 | 5.500E-09 | 9.700E-09 |
| 18357 | 918 | Q918T | C | 3.9e-09 | 1.3e-09 | 1.2e-08 | 5.733E-09 | 5.581E-09 | 1.300E-09 | 1.200E-08 |
| 18358 | 918 | Q918W | C | 2.1e-09 | 1.1e-08 | 1.1e-08 | 8.033E-09 | 5.138E-09 | 2.100E-09 | 1.100E-08 |
| 18359 | 918 | Q918Y | C | 7.3e-09 | 2.5e-09 | 1e-08 | 6.600E-09 | 3.799E-09 | 2.500E-09 | 1.000E-08 |

| | | | | | | | | | | |
|---|---|---|---|---|---|---|---|---|---|---|
| 18360 | 918 | Q918V | C | 8.7e-09 | 1.1e-08 | 1.7e-08 | 1.223E-08 | 4.285E-09 | 8.700E-09 | 1.700E-08 |
| 18361 | 919 | M919A | C | 1.8e-08 | 3.1e-09 | 7.5e-09 | 9.533E-09 | 7.655E-09 | 3.100E-09 | 1.800E-08 |
| 18362 | 919 | M919R | C | 8.9e-09 | 4.9e-09 | 2.7e-09 | 5.500E-09 | 3.143E-09 | 2.700E-09 | 8.900E-09 |
| 18363 | 919 | M919N | C | 1.8e-08 | 2.8e-09 | 3.5e-09 | 8.100E-09 | 8.581E-09 | 2.800E-09 | 1.800E-08 |
| 18364 | 919 | M919D | C | 7e-09 | 8.7e-09 | 1.2e-08 | 9.233E-09 | 2.542E-09 | 7.000E-09 | 1.200E-08 |
| 18365 | 919 | M919C | C | 1.1e-08 | 6.9e-09 | 1.4e-08 | 1.063E-08 | 3.564E-09 | 6.900E-09 | 1.400E-08 |
| 18366 | 919 | M919E | C | 5.8e-09 | 9.6e-09 | 2.2e-09 | 5.867E-09 | 3.700E-09 | 2.200E-09 | 9.600E-09 |
| 18367 | 919 | M919Q | C | 9e-09 | 4.4e-09 | 7.6e-09 | 7.000E-09 | 2.358E-09 | 4.400E-09 | 9.000E-09 |
| 18368 | 919 | M919G | C | 3.2e-09 | 4.5e-09 | 4.1e-09 | 3.933E-09 | 6.658E-10 | 3.200E-09 | 4.500E-09 |
| 18369 | 919 | M919H | C | 1.9e-08 | 3.6e-09 | 4.5e-09 | 9.033E-09 | 8.643E-09 | 3.600E-09 | 1.900E-08 |
| 18370 | 919 | M919I | C | 1.1e-08 | 9.6e-09 | 2.3e-08 | 1.453E-08 | 7.366E-09 | 9.600E-09 | 2.300E-08 |
| 18371 | 919 | M919L | C | 4.5e-09 | 2.3e-09 | 9e-09 | 5.267E-09 | 3.415E-09 | 2.300E-09 | 9.000E-09 |
| 18372 | 919 | M919K | C | 6e-09 | 4.6e-09 | 1.4e-08 | 8.200E-09 | 5.071E-09 | 4.600E-09 | 1.400E-08 |
| 18373 | 919 | M919M | C | 3.3e-09 | 7.7e-09 | 2e-09 | 4.333E-09 | 2.987E-09 | 2.000E-09 | 7.700E-09 |
| 18374 | 919 | M919F | C | 7.7e-09 | 3.1e-09 | 1.1e-08 | 7.267E-09 | 3.968E-09 | 3.100E-09 | 1.100E-08 |
| 18375 | 919 | M919P | C | 8.2e-09 | 7.1e-09 | 1.1e-08 | 8.767E-09 | 2.011E-09 | 7.100E-09 | 1.100E-08 |
| 18376 | 919 | M919S | C | 1.3e-08 | 1.2e-08 | 9.9e-09 | 1.163E-08 | 1.582E-09 | 9.900E-09 | 1.300E-08 |
| 18377 | 919 | M919T | C | 7.5e-09 | 1.2e-08 | 5.4e-09 | 8.300E-09 | 3.372E-09 | 5.400E-09 | 1.200E-08 |
| 18378 | 919 | M919W | C | 5.5e-09 | 5.1e-09 | 3.6e-09 | 4.733E-09 | 1.002E-09 | 3.600E-09 | 5.500E-09 |
| 18379 | 919 | M919Y | C | 1.1e-08 | 1.2e-08 | 7.7e-09 | 1.023E-08 | 2.250E-09 | 7.700E-09 | 1.200E-08 |
| 18380 | 919 | M919V | C | 1.4e-08 | 9.8e-09 | 5.4e-09 | 9.733E-09 | 4.300E-09 | 5.400E-09 | 1.400E-08 |
| 18381 | 920 | N920A | C | 5e-09 | 2.5e-09 | 7.3e-09 | 4.933E-09 | 2.401E-09 | 2.500E-09 | 7.300E-09 |
| 18382 | 920 | N920R | C | 1.3e-08 | 1.7e-08 | 1.5e-08 | 1.500E-08 | 2.000E-09 | 1.300E-08 | 1.700E-08 |
| 18383 | 920 | N920N | C | 2.6e-09 | 1.7e-08 | 5.5e-09 | 8.367E-09 | 7.616E-09 | 2.600E-09 | 1.700E-08 |
| 18384 | 920 | N920D | C | 1.1e-08 | 3.7e-09 | 1.2e-08 | 8.900E-09 | 4.531E-09 | 3.700E-09 | 1.200E-08 |
| 18385 | 920 | N920C | C | 8e-09 | 6.3e-09 | 6e-09 | 6.767E-09 | 1.079E-09 | 6.000E-09 | 8.000E-09 |
| 18386 | 920 | N920E | C | 1.3e-08 | 9.4e-09 | 6.9e-09 | 9.767E-09 | 3.066E-09 | 6.900E-09 | 1.300E-08 |
| 18387 | 920 | N920Q | C | 4e-09 | 5.2e-09 | 5.2e-09 | 4.800E-09 | 6.928E-10 | 4.000E-09 | 5.200E-09 |
| 18388 | 920 | N920G | C | 5e-09 | 2.3e-08 | 1.2e-08 | 1.333E-08 | 9.074E-09 | 5.000E-09 | 2.300E-08 |
| 18389 | 920 | N920H | C | 1.2e-08 | 6.1e-09 | 5.3e-09 | 7.800E-09 | 3.659E-09 | 5.300E-09 | 1.200E-08 |
| 18390 | 920 | N920I | C | 2.7e-09 | 3e-09 | 3.5e-09 | 3.067E-09 | 4.041E-10 | 2.700E-09 | 3.500E-09 |
| 18391 | 920 | N920L | C | 8e-09 | 5.8e-09 | 9.1e-09 | 7.633E-09 | 1.680E-09 | 5.800E-09 | 9.100E-09 |
| 18392 | 920 | N920K | C | 5.9e-09 | 1e-08 | 5.3e-09 | 7.067E-09 | 2.558E-09 | 5.300E-09 | 1.000E-08 |
| 18393 | 920 | N920M | C | 2.8e-09 | 9.5e-09 | 5.7e-09 | 6.000E-09 | 3.360E-09 | 2.800E-09 | 9.500E-09 |
| 18394 | 920 | N920F | C | 1.1e-08 | 3.4e-09 | 3.6e-09 | 6.000E-09 | 4.331E-09 | 3.400E-09 | 1.100E-08 |
| 18395 | 920 | N920P | C | 4e-09 | 3e-08 | 5.6e-09 | 1.320E-08 | 1.457E-08 | 4.000E-09 | 3.000E-08 |
| 18396 | 920 | N920S | C | 1.9e-08 | 1.4e-08 | 9.1e-09 | 1.403E-08 | 4.950E-09 | 9.100E-09 | 1.900E-08 |
| 18397 | 920 | N920T | C | 6e-09 | 4e-09 | 6.3e-09 | 5.433E-09 | 1.250E-09 | 4.000E-09 | 6.300E-09 |
| 18398 | 920 | N920W | C | 7.2e-09 | 6.8e-09 | 7.9e-09 | 7.300E-09 | 5.568E-10 | 6.800E-09 | 7.900E-09 |
| 18399 | 920 | N920Y | C | 4e-09 | 2.8e-09 | 5.4e-09 | 4.067E-09 | 1.301E-09 | 2.800E-09 | 5.400E-09 |
| 18400 | 920 | N920V | C | 4.5e-09 | 3.8e-08 | 8.7e-09 | 1.707E-08 | 1.825E-08 | 4.500E-09 | 3.800E-08 |
| 18401 | 921 | S921A | C | 4.7e-09 | 1.1e-08 | 9.6e-09 | 8.433E-09 | 3.308E-09 | 4.700E-09 | 1.100E-08 |
| 18402 | 921 | S921R | C | 6.2e-09 | 8.4e-09 | 8e-09 | 7.533E-09 | 1.172E-09 | 6.200E-09 | 8.400E-09 |
| 18403 | 921 | S921N | C | 5.3e-09 | 2.5e-09 | 3.6e-09 | 3.800E-09 | 1.411E-09 | 2.500E-09 | 5.300E-09 |
| 18404 | 921 | S921D | C | 8.4e-09 | 1.5e-08 | 1.4e-08 | 1.247E-08 | 3.557E-09 | 8.400E-09 | 1.500E-08 |
| 18405 | 921 | S921C | C | 1.2e-08 | 1.2e-08 | 2e-08 | 1.467E-08 | 4.619E-09 | 1.200E-08 | 2.000E-08 |
| 18406 | 921 | S921E | C | 2e-09 | 1.6e-08 | 1.6e-08 | 1.133E-08 | 8.083E-09 | 2.000E-09 | 1.600E-08 |
| 18407 | 921 | S921Q | C | 1.1e-08 | 1e-08 | 1.7e-08 | 1.267E-08 | 3.786E-09 | 1.000E-08 | 1.700E-08 |
| 18408 | 921 | S921G | C | 1e-08 | 1.5e-08 | 1.5e-08 | 1.333E-08 | 2.887E-09 | 1.000E-08 | 1.500E-08 |
| 18409 | 921 | S921H | C | 4.4e-09 | 7.5e-09 | 1.6e-08 | 9.300E-09 | 6.006E-09 | 4.400E-09 | 1.600E-08 |
| 18410 | 921 | S921I | C | 4e-08 | 5e-09 | 6.7e-09 | 1.723E-08 | 1.973E-08 | 5.000E-09 | 4.000E-08 |
| 18411 | 921 | S921L | C | 5.6e-09 | 1.6e-08 | 2e-08 | 1.387E-08 | 7.433E-09 | 5.600E-09 | 2.000E-08 |
| 18412 | 921 | S921K | C | 4.8e-09 | 3.1e-09 | 9.7e-09 | 5.867E-09 | 3.427E-09 | 3.100E-09 | 9.700E-09 |
| 18413 | 921 | S921M | C | 8.2e-09 | 8e-09 | 2.2e-09 | 6.133E-09 | 3.408E-09 | 2.200E-09 | 8.200E-09 |
| 18414 | 921 | S921F | C | 4.2e-09 | 8.8e-09 | 3.2e-09 | 5.400E-09 | 2.987E-09 | 3.200E-09 | 8.800E-09 |
| 18415 | 921 | S921P | C | 6.7e-09 | 1.3e-08 | 2.7e-09 | 7.467E-09 | 5.193E-09 | 2.700E-09 | 1.300E-08 |
| 18416 | 921 | S921S | C | 3.6e-09 | 8.7e-09 | 2.7e-08 | 1.310E-08 | 1.230E-08 | 3.600E-09 | 2.700E-08 |
| 18417 | 921 | S921T | C | 2.5e-09 | 1.6e-09 | 1e-08 | 4.700E-09 | 4.612E-09 | 1.600E-09 | 1.000E-08 |
| 18418 | 921 | S921W | C | 7e-09 | 1.1e-08 | 4.4e-09 | 7.467E-09 | 3.325E-09 | 4.400E-09 | 1.100E-08 |
| 18419 | 921 | S921Y | C | 1.7e-08 | 3.9e-09 | 1.1e-08 | 1.063E-08 | 6.558E-09 | 3.900E-09 | 1.700E-08 |
| 18420 | 921 | S921V | C | 2.7e-09 | 1.2e-08 | 5.6e-09 | 6.767E-09 | 4.759E-09 | 2.700E-09 | 1.200E-08 |
| 18421 | 922 | L922A | C | 7.8e-09 | 7.1e-09 | 4.5e-09 | 6.467E-09 | 1.739E-09 | 4.500E-09 | 7.800E-09 |
| 18422 | 922 | L922R | C | 9.1e-09 | 2.4e-08 | 1.1e-08 | 1.470E-08 | 8.110E-09 | 9.100E-09 | 2.400E-08 |
| 18423 | 922 | L922N | C | 6e-09 | 6.8e-09 | 6.3e-09 | 6.367E-09 | 4.041E-10 | 6.000E-09 | 6.800E-09 |
| 18424 | 922 | L922D | C | 3.5e-09 | 1.8e-08 | 1.2e-08 | 1.117E-08 | 7.286E-09 | 3.500E-09 | 1.800E-08 |
| 18425 | 922 | L922C | C | 4.3e-09 | 1.3e-08 | 1.5e-08 | 1.077E-08 | 5.689E-09 | 4.300E-09 | 1.500E-08 |
| 18426 | 922 | L922E | C | 7.7e-09 | 8.9e-09 | 2e-08 | 1.220E-08 | 6.782E-09 | 7.700E-09 | 2.000E-08 |
| 18427 | 922 | L922Q | C | 1.9e-08 | 7.5e-09 | 5.4e-09 | 1.063E-08 | 7.321E-09 | 5.400E-09 | 1.900E-08 |
| 18428 | 922 | L922G | C | 1.8e-08 | 4.7e-09 | 1.1e-08 | 1.123E-08 | 6.653E-09 | 4.700E-09 | 1.800E-08 |
| 18429 | 922 | L922H | C | 4.7e-09 | 4.7e-09 | 1.3e-08 | 7.467E-09 | 4.792E-09 | 4.700E-09 | 1.300E-08 |
| 18430 | 922 | L922I | C | 6.2e-09 | 6.5e-09 | 1.1e-08 | 7.900E-09 | 2.689E-09 | 6.200E-09 | 1.100E-08 |
| 18431 | 922 | L922L | C | 2.3e-09 | 1.3e-08 | 1e-08 | 8.433E-09 | 5.519E-09 | 2.300E-09 | 1.300E-08 |
| 18432 | 922 | L922K | C | 1.8e-08 | 4.2e-09 | 9e-09 | 1.040E-08 | 7.006E-09 | 4.200E-09 | 1.800E-08 |
| 18433 | 922 | L922M | C | 4e-09 | 4.9e-09 | 7.6e-09 | 5.500E-09 | 1.873E-09 | 4.000E-09 | 7.600E-09 |
| 18434 | 922 | L922F | C | 1.9e-08 | 2.8e-09 | 1.2e-08 | 1.127E-08 | 8.125E-09 | 2.800E-09 | 1.900E-08 |
| 18435 | 922 | L922P | C | 3.7e-09 | 1.4e-08 | 5.4e-09 | 7.700E-09 | 5.522E-09 | 3.700E-09 | 1.400E-08 |
| 18436 | 922 | L922S | C | 1.1e-08 | 5.3e-09 | 4.3e-09 | 6.867E-09 | 3.614E-09 | 4.300E-09 | 1.100E-08 |
| 18437 | 922 | L922T | C | 7.9e-09 | 2.7e-09 | 1.2e-08 | 7.533E-09 | 4.661E-09 | 2.700E-09 | 1.200E-08 |
| 18438 | 922 | L922W | C | 6.8e-09 | 2.3e-09 | 9.2e-09 | 6.100E-09 | 3.503E-09 | 2.300E-09 | 9.200E-09 |
| 18439 | 922 | L922Y | C | 1.7e-08 | 5.6e-09 | 4.7e-09 | 9.100E-09 | 6.856E-09 | 4.700E-09 | 1.700E-08 |
| 18440 | 922 | L922V | C | 1.6e-08 | 6.1e-09 | 7.9e-09 | 1.000E-08 | 5.274E-09 | 6.100E-09 | 1.600E-08 |
| 18441 | 923 | R923A | C | 6.9e-09 | 2.4e-09 | 8.6e-09 | 5.967E-09 | 3.204E-09 | 2.400E-09 | 8.600E-09 |
| 18442 | 923 | R923R | C | 9.9e-09 | 4.7e-09 | 7.1e-09 | 7.233E-09 | 2.603E-09 | 4.700E-09 | 9.900E-09 |
| 18443 | 923 | R923N | C | 4.2e-09 | 4.1e-09 | 2.3e-09 | 3.533E-09 | 1.069E-09 | 2.300E-09 | 4.200E-09 |
| 18444 | 923 | R923D | C | 5e-09 | 2e-09 | 9.8e-09 | 5.600E-09 | 3.934E-09 | 2.000E-09 | 9.800E-09 |

| | | | | | | | | | | |
|---|---|---|---|---|---|---|---|---|---|---|
| 18445 | 923 | R923C | C | 9.7e-09 | 3.1e-09 | 9.1e-09 | 7.300E-09 | 3.650E-09 | 3.100E-09 | 9.700E-09 |
| 18446 | 923 | R923E | C | 1.3e-08 | 6.5e-09 | 2.6e-09 | 7.367E-09 | 5.254E-09 | 2.600E-09 | 1.300E-08 |
| 18447 | 923 | R923Q | C | 1.2e-08 | 1.2e-08 | 1.1e-08 | 1.167E-08 | 5.774E-10 | 1.100E-08 | 1.200E-08 |
| 18448 | 923 | R923G | C | 6.4e-09 | 4.8e-09 | 1.3e-08 | 8.067E-09 | 4.347E-09 | 4.800E-09 | 1.300E-08 |
| 18449 | 923 | R923H | C | 8.3e-09 | 6.9e-09 | 8.6e-09 | 7.933E-09 | 9.074E-10 | 6.900E-09 | 8.600E-09 |
| 18450 | 923 | R923I | C | 5.9e-09 | 6.7e-09 | 1.2e-08 | 8.200E-09 | 3.315E-09 | 5.900E-09 | 1.200E-08 |
| 18451 | 923 | R923L | C | 8.2e-09 | 1e-08 | 7.2e-09 | 8.467E-09 | 1.419E-09 | 7.200E-09 | 1.000E-08 |
| 18452 | 923 | R923K | C | 5.7e-09 | 4.2e-09 | 8.9e-09 | 6.267E-09 | 2.401E-09 | 4.200E-09 | 8.900E-09 |
| 18453 | 923 | R923M | C | 6.9e-09 | 1.3e-08 | 1.7e-08 | 1.230E-08 | 5.086E-09 | 6.900E-09 | 1.700E-08 |
| 18454 | 923 | R923F | C | 1e-08 | 5.5e-09 | 5.8e-09 | 7.100E-09 | 2.516E-09 | 5.500E-09 | 1.000E-08 |
| 18455 | 923 | R923P | C | 6.3e-09 | 6.8e-09 | 3.9e-09 | 5.667E-09 | 1.550E-09 | 3.900E-09 | 6.800E-09 |
| 18456 | 923 | R923S | C | 1e-09 | 1e-08 | 1e-08 | 7.000E-09 | 5.196E-09 | 1.000E-09 | 1.000E-08 |
| 18457 | 923 | R923T | C | 1.3e-08 | 8.2e-09 | 4.6e-09 | 8.600E-09 | 4.214E-09 | 4.600E-09 | 1.300E-08 |
| 18458 | 923 | R923W | C | 4e-09 | 4e-09 | 5.2e-09 | 4.400E-09 | 6.928E-10 | 4.000E-09 | 5.200E-09 |
| 18459 | 923 | R923Y | C | 3.6e-09 | 2e-08 | 2e-08 | 1.453E-08 | 9.469E-09 | 3.600E-09 | 2.000E-08 |
| 18460 | 923 | R923V | C | 3.1e-09 | 5.6e-09 | 2.2e-08 | 1.023E-08 | 1.027E-08 | 3.100E-09 | 2.200E-08 |
| 18461 | 924 | A924A | C | 6.2e-09 | 1.8e-08 | 6.5e-09 | 1.023E-08 | 6.728E-09 | 6.200E-09 | 1.800E-08 |
| 18462 | 924 | A924R | C | 1.6e-08 | 4.8e-09 | 1.8e-08 | 1.293E-08 | 7.114E-09 | 4.800E-09 | 1.800E-08 |
| 18463 | 924 | A924N | C | 2.4e-09 | 1e-08 | 6.7e-09 | 6.367E-09 | 3.811E-09 | 2.400E-09 | 1.000E-08 |
| 18464 | 924 | A924D | C | 8.6e-09 | 2.8e-09 | 8.4e-09 | 6.600E-09 | 3.292E-09 | 2.800E-09 | 8.600E-09 |
| 18465 | 924 | A924C | C | 2.6e-09 | 5.4e-09 | 1.8e-08 | 8.667E-09 | 8.203E-09 | 2.600E-09 | 1.800E-08 |
| 18466 | 924 | A924E | C | 6.8e-09 | 1.1e-08 | 9.5e-09 | 9.100E-09 | 2.128E-09 | 6.800E-09 | 1.100E-08 |
| 18467 | 924 | A924Q | C | 1e-08 | 9.4e-09 | 9.9e-09 | 9.767E-09 | 3.215E-10 | 9.400E-09 | 1.000E-08 |
| 18468 | 924 | A924G | C | 5e-09 | 3.3e-09 | 1.2e-08 | 6.767E-09 | 4.611E-09 | 3.300E-09 | 1.200E-08 |
| 18469 | 924 | A924H | C | 1.1e-08 | 2.7e-09 | 2.8e-09 | 5.500E-09 | 4.763E-09 | 2.700E-09 | 1.100E-08 |
| 18470 | 924 | A924I | C | 3.6e-09 | 7e-09 | 9.6e-09 | 6.733E-09 | 3.009E-09 | 3.600E-09 | 9.600E-09 |
| 18471 | 924 | A924L | C | 2.5e-08 | 1.6e-08 | 4.1e-09 | 1.503E-08 | 1.048E-08 | 4.100E-09 | 2.500E-08 |
| 18472 | 924 | A924K | C | 4.8e-09 | 3.6e-09 | 5.9e-09 | 4.767E-09 | 1.150E-09 | 3.600E-09 | 5.900E-09 |
| 18473 | 924 | A924M | C | 6.4e-09 | 4.1e-09 | 3e-09 | 4.500E-09 | 1.735E-09 | 3.000E-09 | 6.400E-09 |
| 18474 | 924 | A924F | C | 3.2e-09 | 6.3e-09 | 1.1e-08 | 6.833E-09 | 3.927E-09 | 3.200E-09 | 1.100E-08 |
| 18475 | 924 | A924P | C | 1.4e-08 | 6.1e-09 | 3e-09 | 7.700E-09 | 5.672E-09 | 3.000E-09 | 1.400E-08 |
| 18476 | 924 | A924S | C | 7.9e-09 | 5.2e-09 | 7.7e-09 | 6.933E-09 | 1.504E-09 | 5.200E-09 | 7.900E-09 |
| 18477 | 924 | A924T | C | 5.3e-09 | 2e-09 | 2.4e-09 | 3.233E-09 | 1.801E-09 | 2.000E-09 | 5.300E-09 |
| 18478 | 924 | A924W | C | 8.4e-09 | 4.4e-09 | 8.3e-09 | 7.033E-09 | 2.281E-09 | 4.400E-09 | 8.400E-09 |
| 18479 | 924 | A924Y | C | 3.8e-09 | 4.9e-09 | 1e-08 | 6.233E-09 | 3.308E-09 | 3.800E-09 | 1.000E-08 |
| 18480 | 924 | A924V | C | 1.4e-08 | 5.7e-09 | 3.5e-09 | 7.733E-09 | 5.537E-09 | 3.500E-09 | 1.400E-08 |
| 18481 | 925 | E925A | C | 7.4e-09 | 1e-08 | 9.5e-09 | 8.967E-09 | 1.380E-09 | 7.400E-09 | 1.000E-08 |
| 18482 | 925 | E925R | C | 3.3e-09 | 3.2e-09 | 5.3e-09 | 3.933E-09 | 1.185E-09 | 3.200E-09 | 5.300E-09 |
| 18483 | 925 | E925N | C | 6.5e-09 | 3.9e-09 | 3.9e-09 | 4.767E-09 | 1.501E-09 | 3.900E-09 | 6.500E-09 |
| 18484 | 925 | E925D | C | 7.6e-09 | 6.6e-09 | 5e-09 | 6.400E-09 | 1.311E-09 | 5.000E-09 | 7.600E-09 |
| 18485 | 925 | E925C | C | 1.6e-08 | 1.7e-08 | 7.3e-09 | 1.343E-08 | 5.335E-09 | 7.300E-09 | 1.700E-08 |
| 18486 | 925 | E925E | C | 2.9e-09 | 5.2e-09 | 1.1e-08 | 6.367E-09 | 4.174E-09 | 2.900E-09 | 1.100E-08 |
| 18487 | 925 | E925Q | C | 5.3e-09 | 2.8e-08 | 4.2e-09 | 1.250E-08 | 1.343E-08 | 4.200E-09 | 2.800E-08 |
| 18488 | 925 | E925G | C | 8.7e-09 | 9.8e-09 | 5.5e-09 | 8.000E-09 | 2.234E-09 | 5.500E-09 | 9.800E-09 |
| 18489 | 925 | E925H | C | 9.8e-09 | 1e-08 | 5.3e-09 | 8.367E-09 | 2.658E-09 | 5.300E-09 | 1.000E-08 |
| 18490 | 925 | E925I | C | 2.4e-08 | 9.9e-09 | 5.1e-09 | 1.300E-08 | 9.824E-09 | 5.100E-09 | 2.400E-08 |
| 18491 | 925 | E925L | C | 5.2e-09 | 2.6e-09 | 2e-08 | 9.267E-09 | 9.386E-09 | 2.600E-09 | 2.000E-08 |
| 18492 | 925 | E925K | C | 9.2e-09 | 1.4e-08 | 2.9e-09 | 8.700E-09 | 5.567E-09 | 2.900E-09 | 1.400E-08 |
| 18493 | 925 | E925M | C | 8.1e-09 | 5.3e-09 | 7.7e-09 | 7.033E-09 | 1.514E-09 | 5.300E-09 | 8.100E-09 |
| 18494 | 925 | E925F | C | 3.7e-09 | 1.2e-08 | 5.2e-09 | 6.967E-09 | 4.423E-09 | 3.700E-09 | 1.200E-08 |
| 18495 | 925 | E925P | C | 2.1e-08 | 1.4e-09 | 9e-09 | 1.047E-08 | 9.882E-09 | 1.400E-09 | 2.100E-08 |
| 18496 | 925 | E925S | C | 7.3e-09 | 5.6e-09 | 7.9e-09 | 6.933E-09 | 1.193E-09 | 5.600E-09 | 7.900E-09 |
| 18497 | 925 | E925T | C | 2.5e-09 | 1e-08 | 1.1e-08 | 7.833E-09 | 4.646E-09 | 2.500E-09 | 1.100E-08 |
| 18498 | 925 | E925W | C | 4.3e-09 | 1.2e-08 | 5.2e-09 | 7.167E-09 | 4.210E-09 | 4.300E-09 | 1.200E-08 |
| 18499 | 925 | E925Y | C | 7.3e-09 | 7.2e-09 | 8.6e-09 | 7.700E-09 | 7.810E-10 | 7.200E-09 | 8.600E-09 |
| 18500 | 925 | E925V | C | 6.1e-09 | 2.2e-09 | 4.8e-09 | 4.367E-09 | 1.986E-09 | 2.200E-09 | 6.100E-09 |
| 18501 | 926 | D926A | C | 3.1e-09 | 3.3e-09 | 5.7e-09 | 4.033E-09 | 1.447E-09 | 3.100E-09 | 5.700E-09 |
| 18502 | 926 | D926R | C | 6e-09 | 5.2e-09 | 8.3e-09 | 6.500E-09 | 1.609E-09 | 5.200E-09 | 8.300E-09 |
| 18503 | 926 | D926N | C | 2.6e-09 | 1.2e-09 | 9.3e-09 | 4.367E-09 | 4.329E-09 | 1.200E-09 | 9.300E-09 |
| 18504 | 926 | D926D | C | 4.6e-09 | 4.3e-09 | 4.7e-09 | 4.533E-09 | 2.082E-10 | 4.300E-09 | 4.700E-09 |
| 18505 | 926 | D926C | C | 1.1e-08 | 2.4e-09 | 5.7e-09 | 6.367E-09 | 4.339E-09 | 2.400E-09 | 1.100E-08 |
| 18506 | 926 | D926E | C | 5.8e-09 | 1.2e-08 | 4.7e-09 | 7.500E-09 | 3.936E-09 | 4.700E-09 | 1.200E-08 |
| 18507 | 926 | D926Q | C | 8.8e-09 | 4.1e-09 | 6.4e-09 | 6.433E-09 | 2.350E-09 | 4.100E-09 | 8.800E-09 |
| 18508 | 926 | D926G | C | 6.1e-09 | 1.8e-09 | 8.2e-09 | 5.367E-09 | 3.262E-09 | 1.800E-09 | 8.200E-09 |
| 18509 | 926 | D926H | C | 8.7e-09 | 4.6e-09 | 4.4e-09 | 5.900E-09 | 2.427E-09 | 4.400E-09 | 8.700E-09 |
| 18510 | 926 | D926I | C | 5.8e-09 | 1.1e-08 | 4.8e-09 | 7.200E-09 | 3.329E-09 | 4.800E-09 | 1.100E-08 |
| 18511 | 926 | D926L | C | 1.9e-08 | 1e-08 | 1e-08 | 1.300E-08 | 5.196E-09 | 1.000E-08 | 1.900E-08 |
| 18512 | 926 | D926K | C | 1.4e-08 | 6.2e-09 | 1e-08 | 1.007E-08 | 3.900E-09 | 6.200E-09 | 1.400E-08 |
| 18513 | 926 | D926M | C | 3.9e-09 | 2e-09 | 4.7e-09 | 3.533E-09 | 1.387E-09 | 2.000E-09 | 4.700E-09 |
| 18514 | 926 | D926F | C | 5.4e-09 | 3.4e-09 | 4e-09 | 4.267E-09 | 1.026E-09 | 3.400E-09 | 5.400E-09 |
| 18515 | 926 | D926P | C | 5.3e-09 | 6.2e-09 | 4.8e-09 | 5.433E-09 | 7.095E-10 | 4.800E-09 | 6.200E-09 |
| 18516 | 926 | D926S | C | 1.1e-08 | 6.2e-09 | 3.3e-09 | 6.833E-09 | 3.889E-09 | 3.300E-09 | 1.100E-08 |
| 18517 | 926 | D926T | C | 1.8e-09 | 5e-09 | 6.5e-09 | 4.433E-09 | 2.401E-09 | 1.800E-09 | 6.500E-09 |
| 18518 | 926 | D926W | C | 3.6e-09 | 2.9e-09 | 2.9e-09 | 3.133E-09 | 4.041E-10 | 2.900E-09 | 3.600E-09 |
| 18519 | 926 | D926Y | C | 1.7e-08 | 6.9e-09 | 2.5e-09 | 8.800E-09 | 7.434E-09 | 2.500E-09 | 1.700E-08 |
| 18520 | 926 | D926V | C | 4.5e-09 | 1.4e-08 | 1.6e-08 | 1.150E-08 | 6.144E-09 | 4.500E-09 | 1.600E-08 |
| 18521 | 927 | T927A | C | 1.3e-08 | 1e-08 | 1.8e-08 | 1.367E-08 | 4.041E-09 | 1.000E-08 | 1.800E-08 |
| 18522 | 927 | T927R | C | 8.5e-09 | 5.6e-09 | 9.8e-09 | 7.967E-09 | 2.150E-09 | 5.600E-09 | 9.800E-09 |
| 18523 | 927 | T927N | C | 2.1e-09 | 7.5e-09 | 4.1e-09 | 4.567E-09 | 2.730E-09 | 2.100E-09 | 7.500E-09 |
| 18524 | 927 | T927D | C | 5.2e-09 | 8.4e-09 | 5e-09 | 6.200E-09 | 1.908E-09 | 5.000E-09 | 8.400E-09 |
| 18525 | 927 | T927C | C | 8.4e-09 | 1.6e-08 | 2e-09 | 8.800E-09 | 7.009E-09 | 2.000E-09 | 1.600E-08 |
| 18526 | 927 | T927E | C | 6.9e-09 | 5.7e-09 | 6.5e-09 | 6.367E-09 | 6.110E-10 | 5.700E-09 | 6.900E-09 |
| 18527 | 927 | T927Q | C | 2.8e-09 | 5.7e-09 | 2e-08 | 9.500E-09 | 9.208E-09 | 2.800E-09 | 2.000E-08 |
| 18528 | 927 | T927G | C | 5.2e-09 | 3e-09 | 8e-09 | 5.400E-09 | 2.506E-09 | 3.000E-09 | 8.000E-09 |
| 18529 | 927 | T927H | C | 3.5e-09 | 5.4e-09 | 9.7e-09 | 6.200E-09 | 3.176E-09 | 3.500E-09 | 9.700E-09 |

| | | | | | | | | | | |
|---|---|---|---|---|---|---|---|---|---|---|
| 18530 | 927 | T927I | C | 1.4e-08 | 4.4e-09 | 6.5e-09 | 8.300E-09 | 5.047E-09 | 4.400E-09 | 1.400E-08 |
| 18531 | 927 | T927L | C | 1.8e-09 | 5.7e-09 | 2.6e-09 | 3.367E-09 | 2.060E-09 | 1.800E-09 | 5.700E-09 |
| 18532 | 927 | T927K | C | 1.4e-08 | 4.7e-09 | 8.5e-09 | 9.067E-09 | 4.676E-09 | 4.700E-09 | 1.400E-08 |
| 18533 | 927 | T927M | C | 3.9e-09 | 9.1e-09 | 1.8e-08 | 1.033E-08 | 7.130E-09 | 3.900E-09 | 1.800E-08 |
| 18534 | 927 | T927F | C | 1e-08 | 6.8e-09 | 1.7e-08 | 1.127E-08 | 5.217E-09 | 6.800E-09 | 1.700E-08 |
| 18535 | 927 | T927P | C | 4.9e-09 | 1.7e-08 | 2.9e-09 | 8.267E-09 | 7.629E-09 | 2.900E-09 | 1.700E-08 |
| 18536 | 927 | T927S | C | 7.3e-09 | 4.2e-09 | 1.4e-08 | 8.500E-09 | 5.009E-09 | 4.200E-09 | 1.400E-08 |
| 18537 | 927 | T927T | C | 9.5e-09 | 8.6e-09 | 2.4e-09 | 6.833E-09 | 3.866E-09 | 2.400E-09 | 9.500E-09 |
| 18538 | 927 | T927W | C | 6.5e-09 | 1.4e-08 | 4e-09 | 8.167E-09 | 5.204E-09 | 4.000E-09 | 1.400E-08 |
| 18539 | 927 | T927Y | C | 1.3e-08 | 6.1e-09 | 2.3e-09 | 7.133E-09 | 5.424E-09 | 2.300E-09 | 1.300E-08 |
| 18540 | 927 | T927V | C | 4e-09 | 8.8e-09 | 6.1e-09 | 6.300E-09 | 2.406E-09 | 4.000E-09 | 8.800E-09 |
| 18541 | 928 | A928A | C | 6.6e-09 | 1.2e-08 | 9e-09 | 9.200E-09 | 2.706E-09 | 6.600E-09 | 1.200E-08 |
| 18542 | 928 | A928R | C | 1e-08 | 5.2e-09 | 4.2e-09 | 6.467E-09 | 3.101E-09 | 4.200E-09 | 1.000E-08 |
| 18543 | 928 | A928N | C | 2.2e-09 | 4.3e-09 | 6.3e-09 | 4.267E-09 | 2.050E-09 | 2.200E-09 | 6.300E-09 |
| 18544 | 928 | A928D | C | 4.6e-09 | 3.6e-09 | 1.6e-09 | 3.267E-09 | 1.528E-09 | 1.600E-09 | 4.600E-09 |
| 18545 | 928 | A928C | C | 4.3e-09 | 1e-08 | 1.2e-08 | 8.767E-09 | 3.995E-09 | 4.300E-09 | 1.200E-08 |
| 18546 | 928 | A928E | C | 8.9e-09 | 3.4e-09 | 6.9e-09 | 6.400E-09 | 2.784E-09 | 3.400E-09 | 8.900E-09 |
| 18547 | 928 | A928Q | C | 3.3e-09 | 1e-08 | 4e-09 | 5.767E-09 | 3.683E-09 | 3.300E-09 | 1.000E-08 |
| 18548 | 928 | A928G | C | 9.1e-09 | 6.7e-09 | 5.5e-09 | 7.100E-09 | 1.833E-09 | 5.500E-09 | 9.100E-09 |
| 18549 | 928 | A928H | C | 1.7e-09 | 2.6e-09 | 2.4e-09 | 2.233E-09 | 4.726E-10 | 1.700E-09 | 2.600E-09 |
| 18550 | 928 | A928I | C | 1.4e-08 | 3.2e-09 | 1.3e-08 | 1.007E-08 | 5.968E-09 | 3.200E-09 | 1.400E-08 |
| 18551 | 928 | A928L | C | 7.9e-09 | 4.4e-09 | 9.5e-09 | 7.267E-09 | 2.608E-09 | 4.400E-09 | 9.500E-09 |
| 18552 | 928 | A928K | C | 4.4e-09 | 2.7e-09 | 5.8e-09 | 4.300E-09 | 1.552E-09 | 2.700E-09 | 5.800E-09 |
| 18553 | 928 | A928M | C | 4.3e-09 | 1e-08 | 3.6e-09 | 5.967E-09 | 3.510E-09 | 3.600E-09 | 1.000E-08 |
| 18554 | 928 | A928F | C | 1.4e-08 | 2.5e-09 | 5.4e-09 | 7.300E-09 | 5.981E-09 | 2.500E-09 | 1.400E-08 |
| 18555 | 928 | A928P | C | 1.4e-08 | 5.4e-09 | 1.5e-08 | 1.147E-08 | 5.278E-09 | 5.400E-09 | 1.500E-08 |
| 18556 | 928 | A928S | C | 2.7e-08 | 4.8e-09 | 3.8e-09 | 1.187E-08 | 1.312E-08 | 3.800E-09 | 2.700E-08 |
| 18557 | 928 | A928T | C | 7.4e-09 | 4e-09 | 4.1e-09 | 5.167E-09 | 1.935E-09 | 4.000E-09 | 7.400E-09 |
| 18558 | 928 | A928W | C | 3.7e-09 | 1e-08 | 2.3e-09 | 5.333E-09 | 4.102E-09 | 2.300E-09 | 1.000E-08 |
| 18559 | 928 | A928Y | C | 1.7e-08 | 3.3e-09 | 7.8e-09 | 9.367E-09 | 6.983E-09 | 3.300E-09 | 1.700E-08 |
| 18560 | 928 | A928V | C | 1.4e-08 | 1.5e-08 | 7e-09 | 1.200E-08 | 4.359E-09 | 7.000E-09 | 1.500E-08 |
| 18561 | 929 | V929A | C | 4.1e-09 | 7.7e-09 | 5.5e-09 | 5.767E-09 | 1.815E-09 | 4.100E-09 | 7.700E-09 |
| 18562 | 929 | V929R | C | 5.6e-09 | 9.3e-09 | 3.3e-09 | 6.067E-09 | 3.027E-09 | 3.300E-09 | 9.300E-09 |
| 18563 | 929 | V929N | C | 4.2e-09 | 7.4e-09 | 2.9e-09 | 4.833E-09 | 2.316E-09 | 2.900E-09 | 7.400E-09 |
| 18564 | 929 | V929D | C | 4.1e-09 | 1.4e-08 | 6.4e-09 | 8.167E-09 | 5.181E-09 | 4.100E-09 | 1.400E-08 |
| 18565 | 929 | V929C | C | 6.4e-09 | 1.5e-08 | 4.5e-09 | 8.633E-09 | 5.595E-09 | 4.500E-09 | 1.500E-08 |
| 18566 | 929 | V929E | C | 2.2e-08 | 1e-08 | 1.1e-08 | 1.433E-08 | 6.658E-09 | 1.000E-08 | 2.200E-08 |
| 18567 | 929 | V929Q | C | 6.6e-09 | 6.4e-09 | 1.5e-08 | 9.333E-09 | 4.908E-09 | 6.400E-09 | 1.500E-08 |
| 18568 | 929 | V929G | C | 1.2e-08 | 4.3e-09 | 3.3e-09 | 6.533E-09 | 4.761E-09 | 3.300E-09 | 1.200E-08 |
| 18569 | 929 | V929H | C | 4e-09 | 3.6e-09 | 6.6e-09 | 4.733E-09 | 1.629E-09 | 3.600E-09 | 6.600E-09 |
| 18570 | 929 | V929I | C | 4.7e-09 | 1.6e-08 | 4.8e-09 | 8.500E-09 | 6.495E-09 | 4.700E-09 | 1.600E-08 |
| 18571 | 929 | V929L | C | 7.4e-09 | 6.1e-09 | 9e-09 | 7.500E-09 | 1.453E-09 | 6.100E-09 | 9.000E-09 |
| 18572 | 929 | V929K | C | 4.1e-09 | 2.1e-08 | 4.4e-09 | 9.833E-09 | 9.672E-09 | 4.100E-09 | 2.100E-08 |
| 18573 | 929 | V929M | C | 6.4e-09 | 9.5e-09 | 9.6e-09 | 8.500E-09 | 1.819E-09 | 6.400E-09 | 9.600E-09 |
| 18574 | 929 | V929F | C | 7.5e-09 | 1.8e-08 | 4.3e-09 | 9.933E-09 | 7.167E-09 | 4.300E-09 | 1.800E-08 |
| 18575 | 929 | V929P | C | 3.1e-09 | 7.3e-09 | 3.4e-09 | 4.600E-09 | 2.343E-09 | 3.100E-09 | 7.300E-09 |
| 18576 | 929 | V929S | C | 7.6e-09 | 8.5e-09 | 1.4e-08 | 1.003E-08 | 3.465E-09 | 7.600E-09 | 1.400E-08 |
| 18577 | 929 | V929T | C | 1.8e-09 | 2.2e-08 | 1.1e-08 | 1.160E-08 | 1.011E-08 | 1.800E-09 | 2.200E-08 |
| 18578 | 929 | V929W | C | 1.1e-08 | 3.1e-09 | 1.7e-09 | 5.267E-09 | 5.014E-09 | 1.700E-09 | 1.100E-08 |
| 18579 | 929 | V929Y | C | 5.6e-09 | 1.9e-09 | 6.4e-09 | 4.633E-09 | 2.401E-09 | 1.900E-09 | 6.400E-09 |
| 18580 | 929 | V929V | C | 9.8e-09 | 8.4e-09 | 3.2e-09 | 7.133E-09 | 3.478E-09 | 3.200E-09 | 9.800E-09 |
| 18581 | 930 | Y930A | C | 2.6e-09 | 2.3e-08 | 1.1e-08 | 1.220E-08 | 1.025E-08 | 2.600E-09 | 2.300E-08 |
| 18582 | 930 | Y930R | C | 1.2e-08 | 1.2e-08 | 5.5e-09 | 9.833E-09 | 3.753E-09 | 5.500E-09 | 1.200E-08 |
| 18583 | 930 | Y930N | C | 6.7e-09 | 9e-09 | 4.2e-09 | 6.633E-09 | 2.401E-09 | 4.200E-09 | 9.000E-09 |
| 18584 | 930 | Y930D | C | 6.3e-09 | 8.4e-09 | 2.5e-09 | 5.733E-09 | 2.991E-09 | 2.500E-09 | 8.400E-09 |
| 18585 | 930 | Y930C | C | 1.4e-08 | 5.1e-09 | 6.5e-09 | 8.533E-09 | 4.786E-09 | 5.100E-09 | 1.400E-08 |
| 18586 | 930 | Y930E | C | 6.2e-09 | 6e-09 | 1.9e-08 | 1.040E-08 | 7.448E-09 | 6.000E-09 | 1.900E-08 |
| 18587 | 930 | Y930Q | C | 1.4e-08 | 5.7e-09 | 3.8e-09 | 7.833E-09 | 5.424E-09 | 3.800E-09 | 1.400E-08 |
| 18588 | 930 | Y930G | C | 5.7e-09 | 2.9e-09 | 3.8e-09 | 4.133E-09 | 1.429E-09 | 2.900E-09 | 5.700E-09 |
| 18589 | 930 | Y930H | C | 3.6e-09 | 4e-09 | 1e-08 | 5.867E-09 | 3.585E-09 | 3.600E-09 | 1.000E-08 |
| 18590 | 930 | Y930I | C | 3.9e-09 | 6.1e-09 | 7.2e-09 | 5.733E-09 | 1.680E-09 | 3.900E-09 | 7.200E-09 |
| 18591 | 930 | Y930L | C | 4.3e-09 | 1.8e-08 | 1.5e-08 | 1.243E-08 | 7.202E-09 | 4.300E-09 | 1.800E-08 |
| 18592 | 930 | Y930K | C | 7.9e-09 | 9.3e-09 | 8.4e-09 | 8.533E-09 | 7.095E-10 | 7.900E-09 | 9.300E-09 |
| 18593 | 930 | Y930M | C | 1.3e-08 | 4.7e-09 | 5.7e-09 | 7.800E-09 | 4.531E-09 | 4.700E-09 | 1.300E-08 |
| 18594 | 930 | Y930F | C | 6.3e-09 | 8.5e-09 | 2e-09 | 5.600E-09 | 3.306E-09 | 2.000E-09 | 8.500E-09 |
| 18595 | 930 | Y930P | C | 4.7e-09 | 8.1e-09 | 2e-08 | 1.093E-08 | 8.034E-09 | 4.700E-09 | 2.000E-08 |
| 18596 | 930 | Y930S | C | 1.2e-08 | 6.5e-09 | 5.5e-09 | 8.000E-09 | 3.500E-09 | 5.500E-09 | 1.200E-08 |
| 18597 | 930 | Y930T | C | 1.5e-08 | 5e-09 | 2e-08 | 1.333E-08 | 7.638E-09 | 5.000E-09 | 2.000E-08 |
| 18598 | 930 | Y930W | C | 1.1e-08 | 6.8e-09 | 1.1e-08 | 9.600E-09 | 2.425E-09 | 6.800E-09 | 1.100E-08 |
| 18599 | 930 | Y930Y | C | 1.1e-08 | 3.1e-09 | 2.8e-09 | 5.633E-09 | 4.650E-09 | 2.800E-09 | 1.100E-08 |
| 18600 | 930 | Y930V | C | 1.5e-08 | 5.4e-09 | 2.2e-09 | 7.533E-09 | 6.661E-09 | 2.200E-09 | 1.500E-08 |
| 18601 | 931 | Y931A | C | 1.8e-08 | 1.2e-08 | 2.8e-09 | 1.093E-08 | 7.656E-09 | 2.800E-09 | 1.800E-08 |
| 18602 | 931 | Y931R | C | 1.4e-08 | 1.5e-08 | 1.9e-09 | 1.030E-08 | 7.292E-09 | 1.900E-09 | 1.500E-08 |
| 18603 | 931 | Y931N | C | 4.5e-09 | 5.2e-09 | 1.8e-08 | 9.233E-09 | 7.600E-09 | 4.500E-09 | 1.800E-08 |
| 18604 | 931 | Y931D | C | 6.8e-10 | 4.3e-09 | 3.1e-09 | 2.693E-09 | 1.844E-09 | 6.800E-10 | 4.300E-09 |
| 18605 | 931 | Y931C | C | 1.2e-08 | 1.3e-08 | 9.8e-09 | 1.160E-08 | 1.637E-09 | 9.800E-09 | 1.300E-08 |
| 18606 | 931 | Y931E | C | 9.1e-09 | 1.1e-08 | 1.4e-08 | 1.137E-08 | 2.470E-09 | 9.100E-09 | 1.400E-08 |
| 18607 | 931 | Y931Q | C | 9.2e-09 | 4.3e-09 | 7.6e-09 | 7.033E-09 | 2.499E-09 | 4.300E-09 | 9.200E-09 |
| 18608 | 931 | Y931G | C | 5.7e-09 | 1.2e-08 | 6.2e-09 | 7.967E-09 | 3.502E-09 | 5.700E-09 | 1.200E-08 |
| 18609 | 931 | Y931H | C | 2.7e-09 | 1.6e-08 | 3e-08 | 1.623E-08 | 1.365E-08 | 2.700E-09 | 3.000E-08 |
| 18610 | 931 | Y931I | C | 9.1e-09 | 8e-09 | 1e-08 | 9.033E-09 | 1.002E-09 | 8.000E-09 | 1.000E-08 |
| 18611 | 931 | Y931L | C | 3.9e-09 | 1.1e-08 | 2.4e-09 | 5.767E-09 | 4.594E-09 | 2.400E-09 | 1.100E-08 |
| 18612 | 931 | Y931K | C | 4.1e-09 | 3.2e-09 | 3.6e-09 | 3.633E-09 | 4.509E-10 | 3.200E-09 | 4.100E-09 |
| 18613 | 931 | Y931M | C | 7e-09 | 9.5e-09 | 1.7e-08 | 1.117E-08 | 5.204E-09 | 7.000E-09 | 1.700E-08 |
| 18614 | 931 | Y931F | C | 3.7e-09 | 1.7e-08 | 3.2e-09 | 7.967E-09 | 7.827E-09 | 3.200E-09 | 1.700E-08 |

| | | | | | | | | | | |
|---|---|---|---|---|---|---|---|---|---|---|
| 18615 | 931 | Y931P | C | 2.9e-09 | 1.3e-08 | 7.1e-09 | 7.667E-09 | 5.074E-09 | 2.900E-09 | 1.300E-08 |
| 18616 | 931 | Y931S | C | 7.7e-09 | 1e-08 | 1.4e-08 | 1.057E-08 | 3.188E-09 | 7.700E-09 | 1.400E-08 |
| 18617 | 931 | Y931T | C | 1.9e-08 | 7.2e-09 | 6.2e-09 | 1.080E-08 | 7.119E-09 | 6.200E-09 | 1.900E-08 |
| 18618 | 931 | Y931W | C | 6e-09 | 1.1e-08 | 7.9e-09 | 8.300E-09 | 2.524E-09 | 6.000E-09 | 1.100E-08 |
| 18619 | 931 | Y931Y | C | 1.4e-08 | 1.2e-08 | 1.6e-08 | 1.400E-08 | 2.000E-09 | 1.200E-08 | 1.600E-08 |
| 18620 | 931 | Y931V | C | 1e-08 | 7.4e-09 | 7.4e-09 | 8.267E-09 | 1.501E-09 | 7.400E-09 | 1.000E-08 |
| 18621 | 932 | C932A | C | 6.2e-09 | 9.6e-09 | 4e-09 | 6.600E-09 | 2.821E-09 | 4.000E-09 | 9.600E-09 |
| 18622 | 932 | C932R | C | 1.4e-08 | 1.3e-08 | 5.4e-09 | 1.080E-08 | 4.703E-09 | 5.400E-09 | 1.400E-08 |
| 18623 | 932 | C932N | C | 4.1e-09 | 4.6e-09 | 4e-09 | 4.233E-09 | 3.215E-10 | 4.000E-09 | 4.600E-09 |
| 18624 | 932 | C932D | C | 4.6e-09 | 1.6e-09 | 1.5e-08 | 7.067E-09 | 7.032E-09 | 1.600E-09 | 1.500E-08 |
| 18625 | 932 | C932C | C | 5.8e-09 | 2.6e-09 | 3.6e-09 | 4.000E-09 | 1.637E-09 | 2.600E-09 | 5.800E-09 |
| 18626 | 932 | C932E | C | 9.3e-09 | 6.9e-09 | 1.2e-09 | 5.800E-09 | 4.161E-09 | 1.200E-09 | 9.300E-09 |
| 18627 | 932 | C932Q | C | 4e-09 | 1.9e-08 | 1.1e-08 | 1.133E-08 | 7.506E-09 | 4.000E-09 | 1.900E-08 |
| 18628 | 932 | C932G | C | 2.2e-09 | 1.3e-08 | 2.1e-09 | 5.767E-09 | 6.264E-09 | 2.100E-09 | 1.300E-08 |
| 18629 | 932 | C932H | C | 1.3e-08 | 8.4e-09 | 5.2e-09 | 8.867E-09 | 3.921E-09 | 5.200E-09 | 1.300E-08 |
| 18630 | 932 | C932I | C | 5.6e-09 | 1.7e-08 | 5.3e-09 | 9.300E-09 | 6.670E-09 | 5.300E-09 | 1.700E-08 |
| 18631 | 932 | C932L | C | 1.9e-08 | 7.1e-09 | 1.3e-08 | 1.303E-08 | 5.950E-09 | 7.100E-09 | 1.900E-08 |
| 18632 | 932 | C932K | C | 7.5e-09 | 8.2e-09 | 2.2e-09 | 5.967E-09 | 3.281E-09 | 2.200E-09 | 8.200E-09 |
| 18633 | 932 | C932M | C | 1.5e-08 | 6.9e-09 | 2.2e-08 | 1.463E-08 | 7.557E-09 | 6.900E-09 | 2.200E-08 |
| 18634 | 932 | C932F | C | 2.2e-09 | 6.2e-09 | 8.2e-09 | 5.533E-09 | 3.055E-09 | 2.200E-09 | 8.200E-09 |
| 18635 | 932 | C932P | C | 8.5e-09 | 1.2e-08 | 1.3e-09 | 7.267E-09 | 5.456E-09 | 1.300E-09 | 1.200E-08 |
| 18636 | 932 | C932S | C | 8.3e-09 | 6.3e-09 | 1.2e-08 | 8.867E-09 | 2.892E-09 | 6.300E-09 | 1.200E-08 |
| 18637 | 932 | C932T | C | 6.6e-09 | 9.9e-09 | 6.8e-09 | 7.767E-09 | 1.850E-09 | 6.600E-09 | 9.900E-09 |
| 18638 | 932 | C932W | C | 3.8e-09 | 2.7e-09 | 2.3e-09 | 2.933E-09 | 7.767E-10 | 2.300E-09 | 3.800E-09 |
| 18639 | 932 | C932Y | C | 1e-08 | 5.9e-09 | 2e-09 | 5.967E-09 | 4.000E-09 | 2.000E-09 | 1.000E-08 |
| 18640 | 932 | C932V | C | 4.9e-09 | 7e-09 | 8.6e-09 | 6.833E-09 | 1.856E-09 | 4.900E-09 | 8.600E-09 |
| 18641 | 933 | A933A | C | 4.1e-09 | 1.7e-08 | 2.8e-09 | 7.967E-09 | 7.850E-09 | 2.800E-09 | 1.700E-08 |
| 18642 | 933 | A933R | C | 6.5e-09 | 1.5e-08 | 8e-09 | 9.833E-09 | 4.537E-09 | 6.500E-09 | 1.500E-08 |
| 18643 | 933 | A933N | C | 1.1e-08 | 7.8e-09 | 7.3e-09 | 8.700E-09 | 2.007E-09 | 7.300E-09 | 1.100E-08 |
| 18644 | 933 | A933D | C | 7.5e-09 | 7.4e-09 | 1e-08 | 8.300E-09 | 1.473E-09 | 7.400E-09 | 1.000E-08 |
| 18645 | 933 | A933C | C | 1.1e-08 | 6e-09 | 8e-09 | 8.333E-09 | 2.517E-09 | 6.000E-09 | 1.100E-08 |
| 18646 | 933 | A933E | C | 1.8e-08 | 1e-08 | 9.1e-09 | 1.237E-08 | 4.899E-09 | 9.100E-09 | 1.800E-08 |
| 18647 | 933 | A933Q | C | 3.3e-09 | 7.2e-09 | 5.2e-09 | 5.233E-09 | 1.950E-09 | 3.300E-09 | 7.200E-09 |
| 18648 | 933 | A933G | C | 7.9e-09 | 8.4e-09 | 3.9e-09 | 6.733E-09 | 2.466E-09 | 3.900E-09 | 8.400E-09 |
| 18649 | 933 | A933H | C | 2e-08 | 6.8e-09 | 9e-09 | 1.193E-08 | 7.072E-09 | 6.800E-09 | 2.000E-08 |
| 18650 | 933 | A933I | C | 7.4e-09 | 7.1e-09 | 1.8e-08 | 1.083E-08 | 6.208E-09 | 7.100E-09 | 1.800E-08 |
| 18651 | 933 | A933L | C | 1e-08 | 3.6e-09 | 6e-09 | 6.533E-09 | 3.233E-09 | 3.600E-09 | 1.000E-08 |
| 18652 | 933 | A933K | C | 4.1e-09 | 3.6e-09 | 4.1e-09 | 3.933E-09 | 2.887E-10 | 3.600E-09 | 4.100E-09 |
| 18653 | 933 | A933M | C | 9.4e-09 | 5.9e-09 | 8.3e-09 | 7.867E-09 | 1.790E-09 | 5.900E-09 | 9.400E-09 |
| 18654 | 933 | A933F | C | 7.7e-09 | 1.2e-08 | 6.1e-09 | 8.600E-09 | 3.051E-09 | 6.100E-09 | 1.200E-08 |
| 18655 | 933 | A933P | C | 1.1e-08 | 4.8e-09 | 5.8e-09 | 7.200E-09 | 3.329E-09 | 4.800E-09 | 1.100E-08 |
| 18656 | 933 | A933S | C | 3.4e-09 | 4.8e-09 | 7.8e-09 | 5.333E-09 | 2.248E-09 | 3.400E-09 | 7.800E-09 |
| 18657 | 933 | A933T | C | 1.6e-08 | 1.2e-08 | 9e-09 | 1.233E-08 | 3.512E-09 | 9.000E-09 | 1.600E-08 |
| 18658 | 933 | A933W | C | 3.8e-09 | 5.2e-09 | 4.6e-09 | 4.533E-09 | 7.024E-10 | 3.800E-09 | 5.200E-09 |
| 18659 | 933 | A933Y | C | 2.2e-08 | 5.7e-09 | 9.2e-09 | 1.230E-08 | 8.581E-09 | 5.700E-09 | 2.200E-08 |
| 18660 | 933 | A933V | C | 8.2e-09 | 6.1e-09 | 5.5e-09 | 6.600E-09 | 1.418E-09 | 5.500E-09 | 8.200E-09 |
| 18661 | 934 | R934A | C | 2e-09 | 1.2e-09 | 1.2e-08 | 5.067E-09 | 6.018E-09 | 1.200E-09 | 1.200E-08 |
| 18662 | 934 | R934R | C | 1.2e-08 | 8.9e-09 | 9.8e-09 | 1.023E-08 | 1.595E-09 | 8.900E-09 | 1.200E-08 |
| 18663 | 934 | R934N | C | 1e-08 | 7.8e-09 | 9.9e-09 | 9.233E-09 | 1.242E-09 | 7.800E-09 | 1.000E-08 |
| 18664 | 934 | R934D | C | 5.3e-09 | 4e-09 | 4.6e-09 | 4.633E-09 | 6.506E-10 | 4.000E-09 | 5.300E-09 |
| 18665 | 934 | R934C | C | 7.6e-09 | 5.9e-09 | 5e-09 | 6.167E-09 | 1.320E-09 | 5.000E-09 | 7.600E-09 |
| 18666 | 934 | R934E | C | 9.3e-09 | 8.5e-09 | 5.8e-09 | 7.867E-09 | 1.834E-09 | 5.800E-09 | 9.300E-09 |
| 18667 | 934 | R934Q | C | 2e-08 | 1.4e-08 | 5.6e-09 | 1.320E-08 | 7.233E-09 | 5.600E-09 | 2.000E-08 |
| 18668 | 934 | R934G | C | 5.7e-09 | 1.3e-08 | 6.7e-09 | 8.467E-09 | 3.958E-09 | 5.700E-09 | 1.300E-08 |
| 18669 | 934 | R934H | C | 7e-09 | 5.2e-09 | 1.5e-08 | 9.067E-09 | 5.217E-09 | 5.200E-09 | 1.500E-08 |
| 18670 | 934 | R934I | C | 5.7e-09 | 1.4e-08 | 3.3e-08 | 1.757E-08 | 1.400E-08 | 5.700E-09 | 3.300E-08 |
| 18671 | 934 | R934L | C | 5.3e-09 | 1e-08 | 6e-09 | 7.100E-09 | 2.536E-09 | 5.300E-09 | 1.000E-08 |
| 18672 | 934 | R934K | C | 1.4e-08 | 4.3e-09 | 3.2e-09 | 7.167E-09 | 5.943E-09 | 3.200E-09 | 1.400E-08 |
| 18673 | 934 | R934M | C | 4.8e-09 | 1.1e-08 | 8.9e-09 | 8.233E-09 | 3.153E-09 | 4.800E-09 | 1.100E-08 |
| 18674 | 934 | R934F | C | 4e-09 | 8.3e-09 | 7.3e-09 | 6.533E-09 | 2.250E-09 | 4.000E-09 | 8.300E-09 |
| 18675 | 934 | R934P | C | 1.5e-08 | 7.6e-09 | 1.5e-08 | 1.253E-08 | 4.272E-09 | 7.600E-09 | 1.500E-08 |
| 18676 | 934 | R934S | C | 5.7e-09 | 6.4e-09 | 1.5e-08 | 9.033E-09 | 5.179E-09 | 5.700E-09 | 1.500E-08 |
| 18677 | 934 | R934T | C | 9.5e-09 | 2.3e-08 | 4.7e-09 | 1.240E-08 | 9.488E-09 | 4.700E-09 | 2.300E-08 |
| 18678 | 934 | R934W | C | 6.6e-09 | 3.3e-09 | 8.2e-09 | 6.033E-09 | 2.499E-09 | 3.300E-09 | 8.200E-09 |
| 18679 | 934 | R934Y | C | 1.7e-08 | 7.4e-09 | 1.1e-08 | 1.180E-08 | 4.850E-09 | 7.400E-09 | 1.700E-08 |
| 18680 | 934 | R934V | C | 6.9e-09 | 1.4e-08 | 4.9e-09 | 8.600E-09 | 4.782E-09 | 4.900E-09 | 1.400E-08 |
| 18681 | 935 | N935A | C | 6e-09 | 4.5e-09 | 7.4e-09 | 5.967E-09 | 1.450E-09 | 4.500E-09 | 7.400E-09 |
| 18682 | 935 | N935R | C | 1.2e-08 | 1.7e-09 | 6.6e-09 | 6.767E-09 | 5.152E-09 | 1.700E-09 | 1.200E-08 |
| 18683 | 935 | N935N | C | 4.9e-09 | 6.8e-09 | 7.1e-09 | 6.267E-09 | 1.193E-09 | 4.900E-09 | 7.100E-09 |
| 18684 | 935 | N935D | C | 2.4e-09 | 7.8e-09 | 8.4e-09 | 6.200E-09 | 3.305E-09 | 2.400E-09 | 8.400E-09 |
| 18685 | 935 | N935C | C | 3.8e-09 | 5.5e-09 | 3.4e-09 | 4.233E-09 | 1.115E-09 | 3.400E-09 | 5.500E-09 |
| 18686 | 935 | N935E | C | 2e-09 | 2.3e-09 | 7.6e-09 | 3.967E-09 | 3.150E-09 | 2.000E-09 | 7.600E-09 |
| 18687 | 935 | N935Q | C | 9.7e-09 | 7.3e-09 | 6.5e-09 | 7.833E-09 | 1.665E-09 | 6.500E-09 | 9.700E-09 |
| 18688 | 935 | N935G | C | 5.6e-09 | 1.8e-08 | 1.1e-08 | 1.153E-08 | 6.217E-09 | 5.600E-09 | 1.800E-08 |
| 18689 | 935 | N935H | C | 5.8e-09 | 3.6e-09 | 2.8e-09 | 4.067E-09 | 1.553E-09 | 2.800E-09 | 5.800E-09 |
| 18690 | 935 | N935I | C | 8.4e-09 | 4.7e-09 | 5.6e-09 | 6.233E-09 | 1.930E-09 | 4.700E-09 | 8.400E-09 |
| 18691 | 935 | N935L | C | 5.1e-09 | 1e-08 | 2.2e-09 | 5.767E-09 | 3.943E-09 | 2.200E-09 | 1.000E-08 |
| 18692 | 935 | N935K | C | 1.5e-08 | 5.4e-09 | 4.4e-09 | 8.267E-09 | 5.853E-09 | 4.400E-09 | 1.500E-08 |
| 18693 | 935 | N935M | C | 1.6e-08 | 1.4e-08 | 6.7e-09 | 1.223E-08 | 4.895E-09 | 6.700E-09 | 1.600E-08 |
| 18694 | 935 | N935F | C | 5.7e-09 | 3.2e-09 | 6.9e-09 | 5.267E-09 | 1.888E-09 | 3.200E-09 | 6.900E-09 |
| 18695 | 935 | N935P | C | 4.2e-09 | 2.2e-08 | 8.7e-09 | 1.163E-08 | 9.255E-09 | 4.200E-09 | 2.200E-08 |
| 18696 | 935 | N935S | C | 3.5e-09 | 7e-09 | 2.8e-08 | 1.283E-08 | 1.325E-08 | 3.500E-09 | 2.800E-08 |
| 18697 | 935 | N935T | C | 1.9e-08 | 2.6e-09 | 8.6e-09 | 1.007E-08 | 8.298E-09 | 2.600E-09 | 1.900E-08 |
| 18698 | 935 | N935W | C | 5.1e-09 | 2.9e-09 | 5.2e-09 | 4.400E-09 | 1.300E-09 | 2.900E-09 | 5.200E-09 |
| 18699 | 935 | N935Y | C | 3.5e-09 | 5.2e-09 | 6.5e-09 | 5.067E-09 | 1.504E-09 | 3.500E-09 | 6.500E-09 |

| | | | | | | | | | | |
|---|---|---|---|---|---|---|---|---|---|---|
| 18700 | 935 | N935V | C | 4.5e-09 | 7.2e-09 | 7.7e-09 | 6.467E-09 | 1.721E-09 | 4.500E-09 | 7.700E-09 |
| 18701 | 936 | L936A | C | 7.1e-09 | 2.6e-09 | 4.9e-09 | 4.867E-09 | 2.250E-09 | 2.600E-09 | 7.100E-09 |
| 18702 | 936 | L936R | C | 1.2e-08 | 7.1e-09 | 5.8e-09 | 8.300E-09 | 3.270E-09 | 5.800E-09 | 1.200E-08 |
| 18703 | 936 | L936N | C | 3.3e-08 | 1.1e-08 | 1.9e-08 | 2.100E-08 | 1.114E-08 | 1.100E-08 | 3.300E-08 |
| 18704 | 936 | L936D | C | 8.4e-09 | 6.8e-09 | 1.4e-08 | 9.733E-09 | 3.781E-09 | 6.800E-09 | 1.400E-08 |
| 18705 | 936 | L936C | C | 7.1e-09 | 7.7e-09 | 1.1e-08 | 8.600E-09 | 2.100E-09 | 7.100E-09 | 1.100E-08 |
| 18706 | 936 | L936E | C | 8.3e-09 | 1.1e-08 | 1.9e-08 | 1.277E-08 | 5.564E-09 | 8.300E-09 | 1.900E-08 |
| 18707 | 936 | L936Q | C | 1e-08 | 1.9e-08 | 1.3e-08 | 1.400E-08 | 4.583E-09 | 1.000E-08 | 1.900E-08 |
| 18708 | 936 | L936G | C | 7.4e-09 | 1.5e-08 | 8.2e-09 | 1.020E-08 | 4.176E-09 | 7.400E-09 | 1.500E-08 |
| 18709 | 936 | L936H | C | 2.8e-09 | 7.3e-09 | 2.3e-08 | 1.103E-08 | 1.060E-08 | 2.800E-09 | 2.300E-08 |
| 18710 | 936 | L936I | C | 5.7e-09 | 5.9e-09 | 1.2e-08 | 7.867E-09 | 3.581E-09 | 5.700E-09 | 1.200E-08 |
| 18711 | 936 | L936L | C | 1.1e-08 | 1.6e-08 | 5.4e-09 | 1.080E-08 | 5.303E-09 | 5.400E-09 | 1.600E-08 |
| 18712 | 936 | L936K | C | 3.7e-09 | 5.8e-09 | 1.1e-08 | 6.833E-09 | 3.758E-09 | 3.700E-09 | 1.100E-08 |
| 18713 | 936 | L936M | C | 1.5e-08 | 5.9e-09 | 9.3e-09 | 1.007E-08 | 4.598E-09 | 5.900E-09 | 1.500E-08 |
| 18714 | 936 | L936F | C | 4.1e-09 | 3.9e-09 | 2e-08 | 9.333E-09 | 9.238E-09 | 3.900E-09 | 2.000E-08 |
| 18715 | 936 | L936P | C | 1.5e-08 | 3.7e-09 | 4.5e-09 | 7.733E-09 | 6.306E-09 | 3.700E-09 | 1.500E-08 |
| 18716 | 936 | L936S | C | 6.1e-09 | 5e-09 | 1e-08 | 7.033E-09 | 2.627E-09 | 5.000E-09 | 1.000E-08 |
| 18717 | 936 | L936T | C | 1.8e-08 | 9.2e-09 | 1.8e-08 | 1.507E-08 | 5.081E-09 | 9.200E-09 | 1.800E-08 |
| 18718 | 936 | L936W | C | 4.2e-08 | 1.7e-08 | 1.3e-08 | 2.400E-08 | 1.572E-08 | 1.300E-08 | 4.200E-08 |
| 18719 | 936 | L936Y | C | 5.2e-09 | 3.7e-09 | 1.8e-08 | 8.967E-09 | 7.859E-09 | 3.700E-09 | 1.800E-08 |
| 18720 | 936 | L936V | C | 2.8e-09 | 2e-08 | 8e-09 | 1.027E-08 | 8.821E-09 | 2.800E-09 | 2.000E-08 |
| 18721 | 937 | G937A | C | 6.8e-09 | 3e-09 | 6.9e-09 | 5.567E-09 | 2.223E-09 | 3.000E-09 | 6.900E-09 |
| 18722 | 937 | G937R | C | 5.5e-09 | 4.7e-09 | 6.5e-09 | 5.567E-09 | 9.018E-10 | 4.700E-09 | 6.500E-09 |
| 18723 | 937 | G937N | C | 4.6e-09 | 5.5e-09 | 5.6e-09 | 5.233E-09 | 5.508E-10 | 4.600E-09 | 5.600E-09 |
| 18724 | 937 | G937D | C | 7.3e-09 | 5.5e-09 | 1.7e-09 | 4.833E-09 | 2.859E-09 | 1.700E-09 | 7.300E-09 |
| 18725 | 937 | G937C | C | 9.1e-09 | 1.7e-08 | 6e-09 | 1.070E-08 | 5.672E-09 | 6.000E-09 | 1.700E-08 |
| 18726 | 937 | G937E | C | 5.7e-09 | 1.4e-08 | 2.7e-09 | 7.467E-09 | 5.853E-09 | 2.700E-09 | 1.400E-08 |
| 18727 | 937 | G937Q | C | 1.7e-08 | 1.4e-08 | 6.1e-09 | 1.237E-08 | 5.631E-09 | 6.100E-09 | 1.700E-08 |
| 18728 | 937 | G937G | C | 2.5e-09 | 5.7e-09 | 1.1e-08 | 6.400E-09 | 4.293E-09 | 2.500E-09 | 1.100E-08 |
| 18729 | 937 | G937H | C | 2.9e-09 | 4.5e-09 | 3.2e-09 | 3.533E-09 | 8.505E-10 | 2.900E-09 | 4.500E-09 |
| 18730 | 937 | G937I | C | 5.1e-09 | 4.9e-09 | 4e-09 | 4.667E-09 | 5.859E-10 | 4.000E-09 | 5.100E-09 |
| 18731 | 937 | G937L | C | 7.6e-09 | 5.4e-09 | 4.2e-09 | 5.733E-09 | 1.724E-09 | 4.200E-09 | 7.600E-09 |
| 18732 | 937 | G937K | C | 2.8e-09 | 1.1e-08 | 3.1e-09 | 5.633E-09 | 4.650E-09 | 2.800E-09 | 1.100E-08 |
| 18733 | 937 | G937M | C | 9.1e-09 | 1.1e-08 | 1.4e-09 | 7.167E-09 | 5.084E-09 | 1.400E-09 | 1.100E-08 |
| 18734 | 937 | G937F | C | 4.3e-09 | 9.2e-09 | 6e-09 | 6.500E-09 | 2.488E-09 | 4.300E-09 | 9.200E-09 |
| 18735 | 937 | G937P | C | 5.2e-09 | 1.3e-08 | 1.3e-08 | 1.040E-08 | 4.503E-09 | 5.200E-09 | 1.300E-08 |
| 18736 | 937 | G937S | C | 2.9e-09 | 6.6e-09 | 4.1e-09 | 4.533E-09 | 1.888E-09 | 2.900E-09 | 6.600E-09 |
| 18737 | 937 | G937T | C | 2.6e-09 | 6.8e-09 | 8.2e-09 | 5.867E-09 | 2.914E-09 | 2.600E-09 | 8.200E-09 |
| 18738 | 937 | G937W | C | 2.1e-09 | 3e-09 | 3.3e-09 | 2.800E-09 | 6.245E-10 | 2.100E-09 | 3.300E-09 |
| 18739 | 937 | G937Y | C | 7.4e-09 | 1.9e-08 | 2.1e-09 | 9.500E-09 | 8.643E-09 | 2.100E-09 | 1.900E-08 |
| 18740 | 937 | G937V | C | 1.3e-08 | 4.2e-09 | 6.6e-09 | 7.933E-09 | 4.549E-09 | 4.200E-09 | 1.300E-08 |
| 18741 | 938 | P938A | C | 2e-08 | 8.5e-09 | 1.9e-08 | 1.583E-08 | 6.371E-09 | 8.500E-09 | 2.000E-08 |
| 18742 | 938 | P938R | C | 3.7e-09 | 9.3e-09 | 1.3e-08 | 8.667E-09 | 4.682E-09 | 3.700E-09 | 1.300E-08 |
| 18743 | 938 | P938N | C | 1.7e-08 | 7.3e-09 | 2.3e-08 | 1.577E-08 | 7.922E-09 | 7.300E-09 | 2.300E-08 |
| 18744 | 938 | P938D | C | 5.3e-09 | 1.3e-08 | 1.2e-08 | 1.010E-08 | 4.187E-09 | 5.300E-09 | 1.300E-08 |
| 18745 | 938 | P938C | C | 4.9e-09 | 9.3e-09 | 6.9e-09 | 7.033E-09 | 2.203E-09 | 4.900E-09 | 9.300E-09 |
| 18746 | 938 | P938E | C | 1.8e-08 | 4.4e-09 | 8.6e-09 | 1.033E-08 | 6.964E-09 | 4.400E-09 | 1.800E-08 |
| 18747 | 938 | P938Q | C | 1.2e-08 | 6.1e-09 | 3.4e-08 | 1.737E-08 | 1.470E-08 | 6.100E-09 | 3.400E-08 |
| 18748 | 938 | P938G | C | 1.3e-08 | 9.8e-09 | 1.2e-08 | 1.160E-08 | 1.637E-09 | 9.800E-09 | 1.300E-08 |
| 18749 | 938 | P938H | C | 5.3e-09 | 7.5e-09 | 4.4e-09 | 5.733E-09 | 1.595E-09 | 4.400E-09 | 7.500E-09 |
| 18750 | 938 | P938I | C | 9e-09 | 1.5e-08 | 4.6e-09 | 9.533E-09 | 5.220E-09 | 4.600E-09 | 1.500E-08 |
| 18751 | 938 | P938L | C | 5.4e-09 | 8.8e-09 | 6.7e-09 | 6.967E-09 | 1.716E-09 | 5.400E-09 | 8.800E-09 |
| 18752 | 938 | P938K | C | 1.3e-08 | 7.7e-09 | 2.3e-08 | 1.457E-08 | 7.769E-09 | 7.700E-09 | 2.300E-08 |
| 18753 | 938 | P938M | C | 2.3e-09 | 4.6e-09 | 7e-09 | 4.633E-09 | 2.350E-09 | 2.300E-09 | 7.000E-09 |
| 18754 | 938 | P938F | C | 1.9e-09 | 8.3e-09 | 1.5e-08 | 8.400E-09 | 6.551E-09 | 1.900E-09 | 1.500E-08 |
| 18755 | 938 | P938P | C | 1.3e-08 | 4.3e-09 | 7.3e-09 | 8.200E-09 | 4.419E-09 | 4.300E-09 | 1.300E-08 |
| 18756 | 938 | P938S | C | 2.2e-08 | 1.9e-08 | 1.7e-08 | 1.933E-08 | 2.517E-09 | 1.700E-08 | 2.200E-08 |
| 18757 | 938 | P938T | C | 7.3e-09 | 2.1e-08 | 2.2e-08 | 1.677E-08 | 8.214E-09 | 7.300E-09 | 2.200E-08 |
| 18758 | 938 | P938W | C | 1.2e-08 | 1.2e-08 | 6e-09 | 1.000E-08 | 3.464E-09 | 6.000E-09 | 1.200E-08 |
| 18759 | 938 | P938Y | C | 4.1e-09 | 1.4e-08 | 3.1e-09 | 7.067E-09 | 6.025E-09 | 3.100E-09 | 1.400E-08 |
| 18760 | 938 | P938V | C | 1.2e-08 | 1.8e-09 | 4.3e-09 | 6.033E-09 | 5.316E-09 | 1.800E-09 | 1.200E-08 |
| 18761 | 939 | S939A | C | 4.3e-09 | 1.1e-08 | 9.1e-09 | 8.133E-09 | 3.453E-09 | 4.300E-09 | 1.100E-08 |
| 18762 | 939 | S939R | C | 2e-08 | 1e-08 | 6.6e-09 | 1.220E-08 | 6.966E-09 | 6.600E-09 | 2.000E-08 |
| 18763 | 939 | S939N | C | 3.6e-09 | 3.6e-09 | 6.3e-09 | 4.500E-09 | 1.559E-09 | 3.600E-09 | 6.300E-09 |
| 18764 | 939 | S939D | C | 1.1e-08 | 1.1e-08 | 7.6e-09 | 9.867E-09 | 1.963E-09 | 7.600E-09 | 1.100E-08 |
| 18765 | 939 | S939C | C | 2.3e-09 | 8.1e-09 | 6.7e-09 | 5.700E-09 | 3.027E-09 | 2.300E-09 | 8.100E-09 |
| 18766 | 939 | S939E | C | 8.8e-09 | 3.4e-09 | 2.5e-09 | 4.900E-09 | 3.407E-09 | 2.500E-09 | 8.800E-09 |
| 18767 | 939 | S939Q | C | 7.9e-09 | 8.8e-09 | 6e-09 | 7.567E-09 | 1.429E-09 | 6.000E-09 | 8.800E-09 |
| 18768 | 939 | S939G | C | 2.5e-09 | 8.8e-09 | 7.5e-09 | 6.267E-09 | 3.326E-09 | 2.500E-09 | 8.800E-09 |
| 18769 | 939 | S939H | C | 1.1e-08 | 8e-09 | 3.7e-09 | 7.567E-09 | 3.669E-09 | 3.700E-09 | 1.100E-08 |
| 18770 | 939 | S939I | C | 4e-09 | 5.5e-09 | 3.1e-09 | 4.200E-09 | 1.212E-09 | 3.100E-09 | 5.500E-09 |
| 18771 | 939 | S939L | C | 7.9e-09 | 2.8e-09 | 1e-08 | 6.900E-09 | 3.703E-09 | 2.800E-09 | 1.000E-08 |
| 18772 | 939 | S939K | C | 8.1e-09 | 8.4e-09 | 7.1e-09 | 7.867E-09 | 6.807E-10 | 7.100E-09 | 8.400E-09 |
| 18773 | 939 | S939M | C | 1.8e-08 | 3.5e-09 | 7.1e-09 | 9.533E-09 | 7.550E-09 | 3.500E-09 | 1.800E-08 |
| 18774 | 939 | S939F | C | 3.8e-09 | 5.2e-09 | 4.9e-09 | 4.633E-09 | 7.371E-10 | 3.800E-09 | 5.200E-09 |
| 18775 | 939 | S939P | C | 3.3e-09 | 2.5e-08 | 2.1e-09 | 1.013E-08 | 1.289E-08 | 2.100E-09 | 2.500E-08 |
| 18776 | 939 | S939S | C | 1.1e-08 | 2.3e-09 | 4.3e-09 | 5.867E-09 | 4.557E-09 | 2.300E-09 | 1.100E-08 |
| 18777 | 939 | S939T | C | 6.9e-09 | 2.2e-08 | 7.2e-09 | 1.203E-08 | 8.633E-09 | 6.900E-09 | 2.200E-08 |
| 18778 | 939 | S939W | C | 8.6e-09 | 7.3e-09 | 2.9e-09 | 6.267E-09 | 2.987E-09 | 2.900E-09 | 8.600E-09 |
| 18779 | 939 | S939Y | C | 9.5e-09 | 1.8e-09 | 7e-09 | 6.100E-09 | 3.928E-09 | 1.800E-09 | 9.500E-09 |
| 18780 | 939 | S939V | C | 1.8e-08 | 1.2e-08 | 3.5e-09 | 1.117E-08 | 7.286E-09 | 3.500E-09 | 1.800E-08 |
| 18781 | 940 | F940A | C | 5.7e-09 | 5.8e-09 | 4.2e-09 | 5.233E-09 | 8.963E-10 | 4.200E-09 | 5.800E-09 |
| 18782 | 940 | F940R | C | 1e-08 | 1.6e-08 | 5.8e-09 | 1.060E-08 | 5.126E-09 | 5.800E-09 | 1.600E-08 |
| 18783 | 940 | F940N | C | 6.6e-09 | 3e-09 | 6.9e-09 | 5.500E-09 | 2.170E-09 | 3.000E-09 | 6.900E-09 |
| 18784 | 940 | F940D | C | 1.1e-08 | 5.9e-09 | 4.5e-09 | 7.133E-09 | 3.421E-09 | 4.500E-09 | 1.100E-08 |

| | | | | | | | | | | |
|---|---|---|---|---|---|---|---|---|---|---|
| 18785 | 940 | F940C | C | 1.4e-08 | 6.2e-09 | 7.3e-09 | 9.167E-09 | 4.222E-09 | 6.200E-09 | 1.400E-08 |
| 18786 | 940 | F940E | C | 4.1e-09 | 1.5e-09 | 7.6e-09 | 4.400E-09 | 3.061E-09 | 1.500E-09 | 7.600E-09 |
| 18787 | 940 | F940Q | C | 7.9e-09 | 2.1e-09 | 3.9e-09 | 4.633E-09 | 2.969E-09 | 2.100E-09 | 7.900E-09 |
| 18788 | 940 | F940G | C | 4.9e-09 | 8.5e-09 | 8.7e-09 | 7.367E-09 | 2.139E-09 | 4.900E-09 | 8.700E-09 |
| 18789 | 940 | F940H | C | 6.6e-09 | 1.3e-08 | 7.5e-09 | 9.033E-09 | 3.465E-09 | 6.600E-09 | 1.300E-08 |
| 18790 | 940 | F940I | C | 1.2e-08 | 4.8e-09 | 9.8e-09 | 8.867E-09 | 3.690E-09 | 4.800E-09 | 1.200E-08 |
| 18791 | 940 | F940L | C | 6e-09 | 3.1e-09 | 7.5e-09 | 5.533E-09 | 2.237E-09 | 3.100E-09 | 7.500E-09 |
| 18792 | 940 | F940K | C | 5.3e-09 | 1.8e-08 | 7.2e-09 | 1.017E-08 | 6.850E-09 | 5.300E-09 | 1.800E-08 |
| 18793 | 940 | F940M | C | 1.8e-08 | 6.1e-09 | 3.3e-09 | 9.133E-09 | 7.805E-09 | 3.300E-09 | 1.800E-08 |
| 18794 | 940 | F940F | C | 6.5e-09 | 3.2e-09 | 3.1e-09 | 4.267E-09 | 1.935E-09 | 3.100E-09 | 6.500E-09 |
| 18795 | 940 | F940P | C | 6.8e-09 | 4.3e-09 | 3.8e-09 | 4.967E-09 | 1.607E-09 | 3.800E-09 | 6.800E-09 |
| 18796 | 940 | F940S | C | 6.3e-09 | 4.9e-09 | 6.4e-09 | 5.867E-09 | 8.386E-10 | 4.900E-09 | 6.400E-09 |
| 18797 | 940 | F940T | C | 3e-09 | 6.5e-09 | 1.8e-08 | 9.167E-09 | 7.848E-09 | 3.000E-09 | 1.800E-08 |
| 18798 | 940 | F940W | C | 4.1e-09 | 9.4e-09 | 1e-08 | 7.833E-09 | 3.247E-09 | 4.100E-09 | 1.000E-08 |
| 18799 | 940 | F940Y | C | 1.6e-08 | 4.5e-09 | 2.7e-09 | 7.733E-09 | 7.215E-09 | 2.700E-09 | 1.600E-08 |
| 18800 | 940 | F940V | C | 1e-08 | 9e-09 | 1.6e-08 | 1.167E-08 | 3.786E-09 | 9.000E-09 | 1.600E-08 |
| 18801 | 941 | Y941A | C | 2.8e-08 | 3.1e-09 | 7.8e-09 | 1.297E-08 | 1.323E-08 | 3.100E-09 | 2.800E-08 |
| 18802 | 941 | Y941R | C | 2.2e-08 | 1e-08 | 6.6e-09 | 1.287E-08 | 8.090E-09 | 6.600E-09 | 2.200E-08 |
| 18803 | 941 | Y941N | C | 7.1e-09 | 1.8e-08 | 7.3e-09 | 1.080E-08 | 6.236E-09 | 7.100E-09 | 1.800E-08 |
| 18804 | 941 | Y941D | C | 1.4e-08 | 5.1e-09 | 5.6e-09 | 8.233E-09 | 5.000E-09 | 5.100E-09 | 1.400E-08 |
| 18805 | 941 | Y941C | C | 7.3e-09 | 3.3e-09 | 1.2e-08 | 7.533E-09 | 4.355E-09 | 3.300E-09 | 1.200E-08 |
| 18806 | 941 | Y941E | C | 3.8e-09 | 7.6e-09 | 1.1e-08 | 7.467E-09 | 3.602E-09 | 3.800E-09 | 1.100E-08 |
| 18807 | 941 | Y941Q | C | 1.1e-08 | 8.4e-09 | 3.7e-09 | 7.700E-09 | 3.700E-09 | 3.700E-09 | 1.100E-08 |
| 18808 | 941 | Y941G | C | 3.6e-09 | 7.7e-09 | 8.5e-09 | 6.600E-09 | 2.629E-09 | 3.600E-09 | 8.500E-09 |
| 18809 | 941 | Y941H | C | 7.9e-09 | 5.2e-09 | 1.3e-08 | 8.700E-09 | 3.961E-09 | 5.200E-09 | 1.300E-08 |
| 18810 | 941 | Y941I | C | 1.3e-08 | 1e-08 | 1.8e-09 | 8.267E-09 | 5.798E-09 | 1.800E-09 | 1.300E-08 |
| 18811 | 941 | Y941L | C | 7.4e-09 | 1.5e-08 | 9.8e-09 | 1.073E-08 | 3.885E-09 | 7.400E-09 | 1.500E-08 |
| 18812 | 941 | Y941K | C | 5.4e-09 | 6.1e-09 | 3.3e-09 | 4.933E-09 | 1.457E-09 | 3.300E-09 | 6.100E-09 |
| 18813 | 941 | Y941M | C | 8.7e-09 | 8.7e-09 | 3.8e-09 | 7.067E-09 | 2.829E-09 | 3.800E-09 | 8.700E-09 |
| 18814 | 941 | Y941F | C | 2.1e-08 | 7.1e-09 | 1.1e-08 | 1.303E-08 | 7.170E-09 | 7.100E-09 | 2.100E-08 |
| 18815 | 941 | Y941P | C | 1.3e-08 | 5.8e-09 | 3.6e-09 | 7.467E-09 | 4.917E-09 | 3.600E-09 | 1.300E-08 |
| 18816 | 941 | Y941S | C | 1.2e-08 | 1.3e-08 | 4.5e-09 | 9.833E-09 | 4.646E-09 | 4.500E-09 | 1.300E-08 |
| 18817 | 941 | Y941T | C | 2.2e-08 | 1.8e-08 | 8.2e-09 | 1.607E-08 | 7.100E-09 | 8.200E-09 | 2.200E-08 |
| 18818 | 941 | Y941W | C | 4.4e-09 | 1.6e-08 | 7.7e-09 | 9.367E-09 | 5.977E-09 | 4.400E-09 | 1.600E-08 |
| 18819 | 941 | Y941Y | C | 4.3e-09 | 1.1e-08 | 5.6e-09 | 6.967E-09 | 3.553E-09 | 4.300E-09 | 1.100E-08 |
| 18820 | 941 | Y941V | C | 7.8e-09 | 1.2e-08 | 2.9e-09 | 7.567E-09 | 4.554E-09 | 2.900E-09 | 1.200E-08 |
| 18821 | 942 | F942A | C | 7.1e-09 | 5e-09 | 3.8e-09 | 5.300E-09 | 1.670E-09 | 3.800E-09 | 7.100E-09 |
| 18822 | 942 | F942R | C | 3.4e-09 | 3e-09 | 1.9e-08 | 8.467E-09 | 9.124E-09 | 3.000E-09 | 1.900E-08 |
| 18823 | 942 | F942N | C | 6.1e-09 | 9.3e-09 | 4.9e-09 | 6.767E-09 | 2.274E-09 | 4.900E-09 | 9.300E-09 |
| 18824 | 942 | F942D | C | 1.2e-08 | 7.7e-09 | 3.9e-09 | 7.867E-09 | 4.053E-09 | 3.900E-09 | 1.200E-08 |
| 18825 | 942 | F942C | C | 1e-08 | 3.3e-09 | 7.8e-09 | 7.033E-09 | 3.415E-09 | 3.300E-09 | 1.000E-08 |
| 18826 | 942 | F942E | C | 6.6e-09 | 1.2e-08 | 4.8e-09 | 7.800E-09 | 3.747E-09 | 4.800E-09 | 1.200E-08 |
| 18827 | 942 | F942Q | C | 3.6e-09 | 7.7e-09 | 2e-08 | 1.043E-08 | 8.535E-09 | 3.600E-09 | 2.000E-08 |
| 18828 | 942 | F942G | C | 6.4e-09 | 1.2e-08 | 7.3e-09 | 8.567E-09 | 3.007E-09 | 6.400E-09 | 1.200E-08 |
| 18829 | 942 | F942H | C | 4e-09 | 1.4e-08 | 4.5e-09 | 7.500E-09 | 5.635E-09 | 4.000E-09 | 1.400E-08 |
| 18830 | 942 | F942I | C | 5.2e-09 | 4.2e-09 | 9.5e-09 | 6.300E-09 | 2.816E-09 | 4.200E-09 | 9.500E-09 |
| 18831 | 942 | F942L | C | 6.2e-09 | 5e-09 | 6.2e-09 | 5.800E-09 | 6.928E-10 | 5.000E-09 | 6.200E-09 |
| 18832 | 942 | F942K | C | 8.1e-09 | 7.4e-09 | 7.3e-09 | 7.600E-09 | 4.359E-10 | 7.300E-09 | 8.100E-09 |
| 18833 | 942 | F942M | C | 6.5e-09 | 4.9e-09 | 1e-08 | 7.133E-09 | 2.608E-09 | 4.900E-09 | 1.000E-08 |
| 18834 | 942 | F942F | C | 1.2e-08 | 4.2e-09 | 5e-09 | 7.067E-09 | 4.291E-09 | 4.200E-09 | 1.200E-08 |
| 18835 | 942 | F942P | C | 1.3e-08 | 8.4e-09 | 4.2e-09 | 8.533E-09 | 4.402E-09 | 4.200E-09 | 1.300E-08 |
| 18836 | 942 | F942S | C | 9.6e-09 | 1.7e-08 | 6.1e-09 | 1.090E-08 | 5.565E-09 | 6.100E-09 | 1.700E-08 |
| 18837 | 942 | F942T | C | 7.5e-09 | 4.2e-09 | 5.6e-09 | 5.767E-09 | 1.656E-09 | 4.200E-09 | 7.500E-09 |
| 18838 | 942 | F942W | C | 2.9e-09 | 4.3e-09 | 6.5e-09 | 4.567E-09 | 1.815E-09 | 2.900E-09 | 6.500E-09 |
| 18839 | 942 | F942Y | C | 5.4e-09 | 3.4e-09 | 2.9e-09 | 3.900E-09 | 1.323E-09 | 2.900E-09 | 5.400E-09 |
| 18840 | 942 | F942V | C | 9.5e-09 | 3.1e-09 | 9.9e-09 | 7.500E-09 | 3.816E-09 | 3.100E-09 | 9.900E-09 |
| 18841 | 943 | D943A | C | 2.9e-08 | 5.8e-09 | 1.2e-08 | 1.560E-08 | 1.201E-08 | 5.800E-09 | 2.900E-08 |
| 18842 | 943 | D943R | C | 8.6e-09 | 8.9e-09 | 4.2e-09 | 7.233E-09 | 2.631E-09 | 4.200E-09 | 8.900E-09 |
| 18843 | 943 | D943N | C | 1.4e-08 | 9.7e-09 | 5.1e-09 | 9.600E-09 | 4.451E-09 | 5.100E-09 | 1.400E-08 |
| 18844 | 943 | D943D | C | 1.1e-08 | 1.4e-08 | 7.4e-10 | 8.580E-09 | 6.953E-09 | 7.400E-10 | 1.400E-08 |
| 18845 | 943 | D943C | C | 8.6e-09 | 3.4e-09 | 4.5e-09 | 5.500E-09 | 2.740E-09 | 3.400E-09 | 8.600E-09 |
| 18846 | 943 | D943E | C | 1.2e-08 | 1.4e-08 | 2.2e-09 | 9.400E-09 | 6.315E-09 | 2.200E-09 | 1.400E-08 |
| 18847 | 943 | D943Q | C | 3.6e-09 | 1.3e-08 | 6.8e-09 | 7.800E-09 | 4.779E-09 | 3.600E-09 | 1.300E-08 |
| 18848 | 943 | D943G | C | 7.1e-09 | 4.9e-09 | 3.2e-08 | 1.467E-08 | 1.505E-08 | 4.900E-09 | 3.200E-08 |
| 18849 | 943 | D943H | C | 6.7e-09 | 1.6e-08 | 8e-09 | 1.023E-08 | 5.036E-09 | 6.700E-09 | 1.600E-08 |
| 18850 | 943 | D943I | C | 1.4e-08 | 2.1e-08 | 1.8e-08 | 1.767E-08 | 3.512E-09 | 1.400E-08 | 2.100E-08 |
| 18851 | 943 | D943L | C | 8.7e-09 | 3.8e-09 | 4.4e-09 | 5.633E-09 | 2.673E-09 | 3.800E-09 | 8.700E-09 |
| 18852 | 943 | D943K | C | 1.6e-08 | 1e-08 | 1e-08 | 1.200E-08 | 3.464E-09 | 1.000E-08 | 1.600E-08 |
| 18853 | 943 | D943M | C | 1.5e-08 | 9.8e-09 | 5.4e-09 | 1.007E-08 | 4.806E-09 | 5.400E-09 | 1.500E-08 |
| 18854 | 943 | D943F | C | 1.4e-08 | 4.8e-09 | 8.7e-09 | 9.167E-09 | 4.618E-09 | 4.800E-09 | 1.400E-08 |
| 18855 | 943 | D943P | C | 1e-08 | 9e-09 | 1.1e-08 | 1.000E-08 | 1.000E-09 | 9.000E-09 | 1.100E-08 |
| 18856 | 943 | D943S | C | 8.4e-09 | 9.5e-09 | 1.1e-08 | 9.633E-09 | 1.305E-09 | 8.400E-09 | 1.100E-08 |
| 18857 | 943 | D943T | C | 4.6e-09 | 7.6e-09 | 5.5e-09 | 5.900E-09 | 1.539E-09 | 4.600E-09 | 7.600E-09 |
| 18858 | 943 | D943W | C | 5.1e-09 | 6.5e-09 | 5.3e-09 | 5.633E-09 | 7.572E-10 | 5.100E-09 | 6.500E-09 |
| 18859 | 943 | D943Y | C | 1.1e-08 | 1.6e-08 | 1.1e-08 | 1.267E-08 | 2.887E-09 | 1.100E-08 | 1.600E-08 |
| 18860 | 943 | D943V | C | 2.7e-09 | 5.7e-09 | 9.2e-09 | 5.867E-09 | 3.253E-09 | 2.700E-09 | 9.200E-09 |
| 18861 | 944 | Y944A | C | 1.8e-08 | 3.4e-09 | 3e-09 | 8.133E-09 | 8.547E-09 | 3.000E-09 | 1.800E-08 |
| 18862 | 944 | Y944R | C | 8.2e-09 | 1.7e-08 | 1.4e-08 | 1.307E-08 | 4.474E-09 | 8.200E-09 | 1.700E-08 |
| 18863 | 944 | Y944N | C | 9.6e-09 | 1.1e-08 | 5e-09 | 8.533E-09 | 3.139E-09 | 5.000E-09 | 1.100E-08 |
| 18864 | 944 | Y944D | C | 5.4e-09 | 6.8e-09 | 2.4e-08 | 1.207E-08 | 1.036E-08 | 5.400E-09 | 2.400E-08 |
| 18865 | 944 | Y944C | C | 7.6e-09 | 1.8e-09 | 1.9e-08 | 9.467E-09 | 8.751E-09 | 1.800E-09 | 1.900E-08 |
| 18866 | 944 | Y944E | C | 2.4e-09 | 1.3e-09 | 7.2e-09 | 3.633E-09 | 3.137E-09 | 1.300E-09 | 7.200E-09 |
| 18867 | 944 | Y944Q | C | 5.4e-09 | 9.8e-09 | 1.9e-09 | 5.700E-09 | 3.959E-09 | 1.900E-09 | 9.800E-09 |
| 18868 | 944 | Y944G | C | 2.5e-08 | 7.7e-09 | 1.4e-08 | 1.557E-08 | 8.756E-09 | 7.700E-09 | 2.500E-08 |
| 18869 | 944 | Y944H | C | 5.9e-09 | 1.1e-08 | 5.7e-09 | 7.533E-09 | 3.004E-09 | 5.700E-09 | 1.100E-08 |

| | | | | | | | | | | |
|---|---|---|---|---|---|---|---|---|---|---|
| 18870 | 944 | Y944I | C | 2.1e-09 | 1.3e-08 | 6.9e-09 | 7.333E-09 | 5.463E-09 | 2.100E-09 | 1.300E-08 |
| 18871 | 944 | Y944L | C | 1.3e-08 | 3.4e-09 | 1.4e-08 | 1.013E-08 | 5.853E-09 | 3.400E-09 | 1.400E-08 |
| 18872 | 944 | Y944K | C | 1.8e-09 | 6.8e-09 | 6.5e-09 | 5.033E-09 | 2.804E-09 | 1.800E-09 | 6.800E-09 |
| 18873 | 944 | Y944M | C | 2.1e-09 | 6.6e-09 | 3.2e-08 | 1.357E-08 | 1.612E-08 | 2.100E-09 | 3.200E-08 |
| 18874 | 944 | Y944F | C | 3.1e-09 | 7.5e-09 | 7.8e-10 | 3.793E-09 | 3.413E-09 | 7.800E-10 | 7.500E-09 |
| 18875 | 944 | Y944P | C | 4.6e-09 | 3.9e-09 | 1.5e-08 | 7.833E-09 | 6.216E-09 | 3.900E-09 | 1.500E-08 |
| 18876 | 944 | Y944S | C | 1.1e-08 | 1.2e-08 | 6.3e-09 | 9.767E-09 | 3.044E-09 | 6.300E-09 | 1.200E-08 |
| 18877 | 944 | Y944T | C | 6.8e-09 | 2.6e-08 | 1.2e-08 | 1.493E-08 | 9.930E-09 | 6.800E-09 | 2.600E-08 |
| 18878 | 944 | Y944W | C | 6.2e-09 | 4.3e-09 | 7.6e-09 | 6.033E-09 | 1.656E-09 | 4.300E-09 | 7.600E-09 |
| 18879 | 944 | Y944Y | C | 6.4e-09 | 9.6e-09 | 7.7e-09 | 7.900E-09 | 1.609E-09 | 6.400E-09 | 9.600E-09 |
| 18880 | 944 | Y944V | C | 2e-08 | 3.5e-09 | 8.8e-09 | 1.077E-08 | 8.424E-09 | 3.500E-09 | 2.000E-08 |
| 18881 | 945 | W945A | C | 7.1e-09 | 3.2e-09 | 5.9e-09 | 5.400E-09 | 1.997E-09 | 3.200E-09 | 7.100E-09 |
| 18882 | 945 | W945R | C | 8.8e-09 | 9.6e-09 | 3.9e-09 | 7.433E-09 | 3.086E-09 | 3.900E-09 | 9.600E-09 |
| 18883 | 945 | W945N | C | 1.4e-08 | 2.2e-09 | 2.2e-08 | 1.273E-08 | 9.961E-09 | 2.200E-09 | 2.200E-08 |
| 18884 | 945 | W945D | C | 6e-09 | 4.3e-09 | 4.1e-09 | 4.800E-09 | 1.044E-09 | 4.100E-09 | 6.000E-09 |
| 18885 | 945 | W945C | C | 3.4e-09 | 4.6e-09 | 7e-09 | 5.000E-09 | 1.833E-09 | 3.400E-09 | 7.000E-09 |
| 18886 | 945 | W945E | C | 1.2e-08 | 4.2e-09 | 1.1e-08 | 9.067E-09 | 4.244E-09 | 4.200E-09 | 1.200E-08 |
| 18887 | 945 | W945Q | C | 4e-09 | 5.7e-09 | 6.3e-09 | 5.333E-09 | 1.193E-09 | 4.000E-09 | 6.300E-09 |
| 18888 | 945 | W945G | C | 5e-09 | 5.7e-09 | 1.4e-08 | 8.233E-09 | 5.006E-09 | 5.000E-09 | 1.400E-08 |
| 18889 | 945 | W945H | C | 9e-09 | 2.5e-09 | 2.1e-08 | 1.083E-08 | 9.385E-09 | 2.500E-09 | 2.100E-08 |
| 18890 | 945 | W945I | C | 9.3e-09 | 7.5e-09 | 7.3e-09 | 8.033E-09 | 1.102E-09 | 7.300E-09 | 9.300E-09 |
| 18891 | 945 | W945L | C | 4.2e-09 | 1.1e-08 | 6.2e-09 | 7.133E-09 | 3.495E-09 | 4.200E-09 | 1.100E-08 |
| 18892 | 945 | W945K | C | 7.5e-09 | 1e-08 | 8.5e-09 | 8.667E-09 | 1.258E-09 | 7.500E-09 | 1.000E-08 |
| 18893 | 945 | W945M | C | 9.4e-09 | 2.8e-09 | 2.1e-08 | 1.107E-08 | 9.214E-09 | 2.800E-09 | 2.100E-08 |
| 18894 | 945 | W945F | C | 1.5e-08 | 3e-09 | 7e-09 | 8.333E-09 | 6.110E-09 | 3.000E-09 | 1.500E-08 |
| 18895 | 945 | W945P | C | 2.6e-09 | 5.7e-09 | 1.1e-08 | 6.433E-09 | 4.248E-09 | 2.600E-09 | 1.100E-08 |
| 18896 | 945 | W945S | C | 3.6e-09 | 4.4e-09 | 1.3e-09 | 3.100E-09 | 1.609E-09 | 1.300E-09 | 4.400E-09 |
| 18897 | 945 | W945T | C | 2e-09 | 5.6e-09 | 8.1e-09 | 5.233E-09 | 3.066E-09 | 2.000E-09 | 8.100E-09 |
| 18898 | 945 | W945W | C | 3.7e-09 | 5.4e-09 | 6.8e-09 | 5.300E-09 | 1.552E-09 | 3.700E-09 | 6.800E-09 |
| 18899 | 945 | W945Y | C | 4.8e-09 | 1e-08 | 5.5e-09 | 6.767E-09 | 2.822E-09 | 4.800E-09 | 1.000E-08 |
| 18900 | 945 | W945V | C | 8.7e-09 | 5.4e-09 | 2.1e-08 | 1.170E-08 | 8.221E-09 | 5.400E-09 | 2.100E-08 |
| 18901 | 946 | G946A | C | 8.8e-09 | 1.4e-08 | 4.4e-09 | 9.067E-09 | 4.806E-09 | 4.400E-09 | 1.400E-08 |
| 18902 | 946 | G946R | C | 2.4e-08 | 2.5e-08 | 6.5e-09 | 1.850E-08 | 1.040E-08 | 6.500E-09 | 2.500E-08 |
| 18903 | 946 | G946N | C | 1e-08 | 2.9e-09 | 3.4e-08 | 1.563E-08 | 1.630E-08 | 2.900E-09 | 3.400E-08 |
| 18904 | 946 | G946D | C | 7.2e-09 | 5.3e-09 | 1.1e-08 | 7.833E-09 | 2.902E-09 | 5.300E-09 | 1.100E-08 |
| 18905 | 946 | G946C | C | 8.1e-09 | 2.1e-08 | 4e-09 | 1.103E-08 | 8.871E-09 | 4.000E-09 | 2.100E-08 |
| 18906 | 946 | G946E | C | 8.2e-09 | 1.2e-08 | 1.4e-08 | 1.140E-08 | 2.946E-09 | 8.200E-09 | 1.400E-08 |
| 18907 | 946 | G946Q | C | 8.4e-09 | 8.1e-09 | 9e-09 | 8.500E-09 | 4.583E-10 | 8.100E-09 | 9.000E-09 |
| 18908 | 946 | G946G | C | 6.2e-09 | 1.6e-08 | 4.4e-09 | 8.867E-09 | 6.243E-09 | 4.400E-09 | 1.600E-08 |
| 18909 | 946 | G946H | C | 6.4e-09 | 1.9e-08 | 8.2e-09 | 1.120E-08 | 6.815E-09 | 6.400E-09 | 1.900E-08 |
| 18910 | 946 | G946I | C | 7.5e-09 | 2.6e-09 | 9.2e-09 | 6.433E-09 | 3.427E-09 | 2.600E-09 | 9.200E-09 |
| 18911 | 946 | G946L | C | 5.5e-09 | 7.7e-09 | 1.3e-08 | 8.733E-09 | 3.855E-09 | 5.500E-09 | 1.300E-08 |
| 18912 | 946 | G946K | C | 8.5e-09 | 1.7e-09 | 9.6e-09 | 6.600E-09 | 4.279E-09 | 1.700E-09 | 9.600E-09 |
| 18913 | 946 | G946M | C | 1.9e-09 | 1.3e-08 | 8.3e-09 | 7.733E-09 | 5.572E-09 | 1.900E-09 | 1.300E-08 |
| 18914 | 946 | G946F | C | 8.7e-09 | 8.6e-09 | 2.3e-09 | 6.533E-09 | 3.667E-09 | 2.300E-09 | 8.700E-09 |
| 18915 | 946 | G946P | C | 8.7e-09 | 6.4e-09 | 1.3e-08 | 9.367E-09 | 3.350E-09 | 6.400E-09 | 1.300E-08 |
| 18916 | 946 | G946S | C | 3.5e-09 | 7.6e-09 | 3.9e-09 | 5.000E-09 | 2.261E-09 | 3.500E-09 | 7.600E-09 |
| 18917 | 946 | G946T | C | 7.2e-09 | 5.5e-09 | 5.4e-09 | 6.033E-09 | 1.012E-09 | 5.400E-09 | 7.200E-09 |
| 18918 | 946 | G946W | C | 1.8e-08 | 3e-09 | 4.8e-09 | 8.600E-09 | 8.190E-09 | 3.000E-09 | 1.800E-08 |
| 18919 | 946 | G946Y | C | 5.9e-09 | 1.3e-08 | 5.3e-09 | 8.067E-09 | 4.283E-09 | 5.300E-09 | 1.300E-08 |
| 18920 | 946 | G946V | C | 1.6e-08 | 9e-09 | 7e-09 | 1.067E-08 | 4.726E-09 | 7.000E-09 | 1.600E-08 |
| 18921 | 947 | Q947A | C | 1.1e-08 | 5.7e-09 | 8.5e-09 | 8.400E-09 | 2.651E-09 | 5.700E-09 | 1.100E-08 |
| 18922 | 947 | Q947R | C | 5.5e-09 | 9.5e-09 | 5.9e-09 | 6.967E-09 | 2.203E-09 | 5.500E-09 | 9.500E-09 |
| 18923 | 947 | Q947N | C | 7e-09 | 2.5e-09 | 6.5e-09 | 5.333E-09 | 2.466E-09 | 2.500E-09 | 7.000E-09 |
| 18924 | 947 | Q947D | C | 4.1e-09 | 5.4e-09 | 5.3e-09 | 4.933E-09 | 7.234E-10 | 4.100E-09 | 5.400E-09 |
| 18925 | 947 | Q947C | C | 1e-08 | 4.5e-09 | 3.9e-09 | 6.133E-09 | 3.362E-09 | 3.900E-09 | 1.000E-08 |
| 18926 | 947 | Q947E | C | 6.3e-09 | 6.4e-09 | 7.4e-09 | 6.700E-09 | 6.083E-10 | 6.300E-09 | 7.400E-09 |
| 18927 | 947 | Q947Q | C | 5.4e-09 | 3.6e-09 | 3e-09 | 4.000E-09 | 1.249E-09 | 3.000E-09 | 5.400E-09 |
| 18928 | 947 | Q947G | C | 1.5e-08 | 3.8e-09 | 4.7e-09 | 7.833E-09 | 6.223E-09 | 3.800E-09 | 1.500E-08 |
| 18929 | 947 | Q947H | C | 8.2e-09 | 3.4e-08 | 1.1e-08 | 1.773E-08 | 1.416E-08 | 8.200E-09 | 3.400E-08 |
| 18930 | 947 | Q947I | C | 1.1e-08 | 1.1e-08 | 3.6e-09 | 8.533E-09 | 4.272E-09 | 3.600E-09 | 1.100E-08 |
| 18931 | 947 | Q947L | C | 5.4e-09 | 9.2e-09 | 3.9e-09 | 6.167E-09 | 2.732E-09 | 3.900E-09 | 9.200E-09 |
| 18932 | 947 | Q947K | C | 4.3e-08 | 6.2e-09 | 2.5e-09 | 1.723E-08 | 2.239E-08 | 2.500E-09 | 4.300E-08 |
| 18933 | 947 | Q947M | C | 2.9e-09 | 6.1e-09 | 1.2e-08 | 7.000E-09 | 4.616E-09 | 2.900E-09 | 1.200E-08 |
| 18934 | 947 | Q947F | C | 9.6e-09 | 9.3e-09 | 1.1e-08 | 9.967E-09 | 9.074E-10 | 9.300E-09 | 1.100E-08 |
| 18935 | 947 | Q947P | C | 5.4e-09 | 1.1e-08 | 2.9e-09 | 6.433E-09 | 4.148E-09 | 2.900E-09 | 1.100E-08 |
| 18936 | 947 | Q947S | C | 4.5e-09 | 2.7e-09 | 1.5e-08 | 7.400E-09 | 6.643E-09 | 2.700E-09 | 1.500E-08 |
| 18937 | 947 | Q947T | C | 5.1e-09 | 7.8e-09 | 1.1e-08 | 7.967E-09 | 2.954E-09 | 5.100E-09 | 1.100E-08 |
| 18938 | 947 | Q947W | C | 8.7e-09 | 8.8e-09 | 5.3e-09 | 7.600E-09 | 1.992E-09 | 5.300E-09 | 8.800E-09 |
| 18939 | 947 | Q947Y | C | 4.9e-09 | 1.5e-08 | 6e-09 | 8.633E-09 | 5.541E-09 | 4.900E-09 | 1.500E-08 |
| 18940 | 947 | Q947V | C | 4.9e-09 | 1.6e-09 | 7.1e-09 | 4.533E-09 | 2.768E-09 | 1.600E-09 | 7.100E-09 |
| 18941 | 948 | G948A | C | 2e-09 | 1.5e-08 | 1.7e-08 | 1.133E-08 | 8.145E-09 | 2.000E-09 | 1.700E-08 |
| 18942 | 948 | G948R | C | 2.6e-09 | 5.6e-09 | 3.7e-09 | 3.967E-09 | 1.518E-09 | 2.600E-09 | 5.600E-09 |
| 18943 | 948 | G948N | C | 1e-08 | 5.2e-09 | 6.7e-09 | 7.300E-09 | 2.456E-09 | 5.200E-09 | 1.000E-08 |
| 18944 | 948 | G948D | C | 3e-09 | 1e-08 | 1.2e-08 | 8.333E-09 | 4.726E-09 | 3.000E-09 | 1.200E-08 |
| 18945 | 948 | G948C | C | 1.2e-08 | 5.5e-09 | 8e-09 | 8.500E-09 | 3.279E-09 | 5.500E-09 | 1.200E-08 |
| 18946 | 948 | G948E | C | 8.3e-09 | 6.9e-09 | 1.1e-08 | 8.733E-09 | 2.084E-09 | 6.900E-09 | 1.100E-08 |
| 18947 | 948 | G948Q | C | 1.7e-08 | 9.6e-09 | 1.2e-08 | 1.287E-08 | 3.775E-09 | 9.600E-09 | 1.700E-08 |
| 18948 | 948 | G948G | C | 1.4e-08 | 7.9e-09 | 7.4e-09 | 9.767E-09 | 3.675E-09 | 7.400E-09 | 1.400E-08 |
| 18949 | 948 | G948H | C | 1.5e-08 | 2.3e-09 | 8.2e-09 | 8.500E-09 | 6.355E-09 | 2.300E-09 | 1.500E-08 |
| 18950 | 948 | G948I | C | 7.6e-09 | 2e-09 | 8e-09 | 5.867E-09 | 3.355E-09 | 2.000E-09 | 8.000E-09 |
| 18951 | 948 | G948L | C | 1.7e-09 | 3.5e-09 | 2.1e-09 | 2.433E-09 | 9.452E-10 | 1.700E-09 | 3.500E-09 |
| 18952 | 948 | G948K | C | 1e-08 | 1.1e-08 | 5.6e-09 | 8.867E-09 | 2.873E-09 | 5.600E-09 | 1.100E-08 |
| 18953 | 948 | G948M | C | 2.2e-09 | 2.4e-09 | 8.1e-09 | 4.233E-09 | 3.350E-09 | 2.200E-09 | 8.100E-09 |
| 18954 | 948 | G948F | C | 8e-09 | 1.3e-08 | 1.3e-08 | 1.133E-08 | 2.887E-09 | 8.000E-09 | 1.300E-08 |

| | | | | | | | | | | |
|---|---|---|---|---|---|---|---|---|---|---|
| 18955 | 948 | G948P | C | 1.3e-08 | 1.5e-08 | 3.5e-09 | 1.050E-08 | 6.144E-09 | 3.500E-09 | 1.500E-08 |
| 18956 | 948 | G948S | C | 1.2e-08 | 1.5e-08 | 1.2e-08 | 1.300E-08 | 1.732E-09 | 1.200E-08 | 1.500E-08 |
| 18957 | 948 | G948T | C | 1.1e-08 | 3.5e-09 | 5.1e-09 | 6.533E-09 | 3.950E-09 | 3.500E-09 | 1.100E-08 |
| 18958 | 948 | G948W | C | 1.8e-09 | 5.7e-09 | 8.9e-09 | 5.467E-09 | 3.556E-09 | 1.800E-09 | 8.900E-09 |
| 18959 | 948 | G948Y | C | 4.9e-09 | 2.4e-08 | 1.3e-08 | 1.397E-08 | 9.587E-09 | 4.900E-09 | 2.400E-08 |
| 18960 | 948 | G948V | C | 1.2e-08 | 3e-09 | 4.6e-09 | 6.533E-09 | 4.801E-09 | 3.000E-09 | 1.200E-08 |
| 18961 | 949 | T949A | C | 6.2e-09 | 3.7e-09 | 4.2e-09 | 4.700E-09 | 1.323E-09 | 3.700E-09 | 6.200E-09 |
| 18962 | 949 | T949R | C | 1.6e-08 | 4.3e-09 | 4.5e-09 | 8.267E-09 | 6.698E-09 | 4.300E-09 | 1.600E-08 |
| 18963 | 949 | T949N | C | 5.6e-09 | 7.4e-09 | 1.2e-08 | 8.333E-09 | 3.301E-09 | 5.600E-09 | 1.200E-08 |
| 18964 | 949 | T949D | C | 6.2e-09 | 1.3e-08 | 9.2e-09 | 9.467E-09 | 3.408E-09 | 6.200E-09 | 1.300E-08 |
| 18965 | 949 | T949C | C | 6e-09 | 6.8e-09 | 4.5e-09 | 5.767E-09 | 1.168E-09 | 4.500E-09 | 6.800E-09 |
| 18966 | 949 | T949E | C | 1.2e-08 | 8.3e-09 | 9.1e-09 | 9.800E-09 | 1.947E-09 | 8.300E-09 | 1.200E-08 |
| 18967 | 949 | T949Q | C | 6.1e-09 | 1.5e-08 | 9.4e-09 | 1.017E-08 | 4.499E-09 | 6.100E-09 | 1.500E-08 |
| 18968 | 949 | T949G | C | 1.2e-08 | 6.8e-09 | 7.4e-09 | 8.733E-09 | 2.845E-09 | 6.800E-09 | 1.200E-08 |
| 18969 | 949 | T949H | C | 1.1e-08 | 1.4e-08 | 7.4e-09 | 1.080E-08 | 3.305E-09 | 7.400E-09 | 1.400E-08 |
| 18970 | 949 | T949I | C | 1.1e-08 | 1.5e-08 | 1.1e-08 | 1.233E-08 | 2.309E-09 | 1.100E-08 | 1.500E-08 |
| 18971 | 949 | T949L | C | 5.6e-09 | 9.3e-09 | 6.6e-09 | 7.167E-09 | 1.914E-09 | 5.600E-09 | 9.300E-09 |
| 18972 | 949 | T949K | C | 9.4e-09 | 1.7e-08 | 2.6e-09 | 9.667E-09 | 7.204E-09 | 2.600E-09 | 1.700E-08 |
| 18973 | 949 | T949M | C | 7.9e-09 | 1.3e-08 | 8.9e-09 | 9.933E-09 | 2.702E-09 | 7.900E-09 | 1.300E-08 |
| 18974 | 949 | T949F | C | 6.1e-09 | 7.2e-09 | 8.4e-09 | 7.233E-09 | 1.150E-09 | 6.100E-09 | 8.400E-09 |
| 18975 | 949 | T949P | C | 5.4e-09 | 1.6e-08 | 6.6e-09 | 9.333E-09 | 5.805E-09 | 5.400E-09 | 1.600E-08 |
| 18976 | 949 | T949S | C | 2.3e-09 | 3.4e-09 | 1.6e-08 | 7.233E-09 | 7.612E-09 | 2.300E-09 | 1.600E-08 |
| 18977 | 949 | T949T | C | 7.7e-09 | 5.7e-09 | 2.7e-08 | 1.347E-08 | 1.176E-08 | 5.700E-09 | 2.700E-08 |
| 18978 | 949 | T949W | C | 5.5e-09 | 7e-09 | 3.5e-09 | 5.333E-09 | 1.756E-09 | 3.500E-09 | 7.000E-09 |
| 18979 | 949 | T949Y | C | 6.2e-09 | 7.4e-09 | 3e-09 | 5.533E-09 | 2.274E-09 | 3.000E-09 | 7.400E-09 |
| 18980 | 949 | T949V | C | 9.6e-09 | 4.8e-09 | 6.4e-09 | 6.933E-09 | 2.444E-09 | 4.800E-09 | 9.600E-09 |
| 18981 | 950 | L950A | C | 1e-08 | 1.7e-08 | 4.4e-09 | 1.047E-08 | 6.313E-09 | 4.400E-09 | 1.700E-08 |
| 18982 | 950 | L950R | C | 1e-08 | 8.2e-09 | 3.7e-09 | 7.300E-09 | 3.245E-09 | 3.700E-09 | 1.000E-08 |
| 18983 | 950 | L950N | C | 5.7e-09 | 2.4e-09 | 2.8e-09 | 3.633E-09 | 1.801E-09 | 2.400E-09 | 5.700E-09 |
| 18984 | 950 | L950D | C | 2.2e-09 | 1.8e-09 | 8e-09 | 4.000E-09 | 3.470E-09 | 1.800E-09 | 8.000E-09 |
| 18985 | 950 | L950C | C | 2.1e-09 | 6e-09 | 4.2e-09 | 4.100E-09 | 1.952E-09 | 2.100E-09 | 6.000E-09 |
| 18986 | 950 | L950E | C | 4e-09 | 7.7e-09 | 1.1e-08 | 7.567E-09 | 3.502E-09 | 4.000E-09 | 1.100E-08 |
| 18987 | 950 | L950Q | C | 9.4e-09 | 6.2e-09 | 1.5e-09 | 5.700E-09 | 3.974E-09 | 1.500E-09 | 9.400E-09 |
| 18988 | 950 | L950G | C | 1.2e-08 | 3.4e-09 | 7.3e-09 | 7.567E-09 | 4.306E-09 | 3.400E-09 | 1.200E-08 |
| 18989 | 950 | L950H | C | 3.6e-09 | 7.8e-09 | 7.6e-09 | 6.333E-09 | 2.369E-09 | 3.600E-09 | 7.800E-09 |
| 18990 | 950 | L950I | C | 1.5e-08 | 9e-09 | 1.7e-08 | 1.367E-08 | 4.163E-09 | 9.000E-09 | 1.700E-08 |
| 18991 | 950 | L950L | C | 6.2e-09 | 8e-09 | 7.7e-09 | 7.300E-09 | 9.644E-10 | 6.200E-09 | 8.000E-09 |
| 18992 | 950 | L950K | C | 6.7e-09 | 1.4e-08 | 7.7e-09 | 9.467E-09 | 3.958E-09 | 6.700E-09 | 1.400E-08 |
| 18993 | 950 | L950M | C | 2.9e-08 | 8.1e-09 | 5.9e-09 | 1.433E-08 | 1.275E-08 | 5.900E-09 | 2.900E-08 |
| 18994 | 950 | L950F | C | 1.9e-09 | 1.6e-08 | 8.9e-09 | 8.933E-09 | 7.050E-09 | 1.900E-09 | 1.600E-08 |
| 18995 | 950 | L950P | C | 8.1e-09 | 1.7e-08 | 1.2e-08 | 1.237E-08 | 4.461E-09 | 8.100E-09 | 1.700E-08 |
| 18996 | 950 | L950S | C | 1.7e-08 | 1.2e-08 | 8.2e-09 | 1.240E-08 | 4.414E-09 | 8.200E-09 | 1.700E-08 |
| 18997 | 950 | L950T | C | 4.1e-09 | 3.4e-09 | 7.5e-09 | 5.000E-09 | 2.193E-09 | 3.400E-09 | 7.500E-09 |
| 18998 | 950 | L950W | C | 8e-09 | 6.2e-09 | 1.2e-08 | 8.733E-09 | 2.969E-09 | 6.200E-09 | 1.200E-08 |
| 18999 | 950 | L950Y | C | 4e-09 | 3.6e-09 | 5.1e-09 | 4.233E-09 | 7.767E-10 | 3.600E-09 | 5.100E-09 |
| 19000 | 950 | L950V | C | 8.8e-09 | 3e-09 | 2.4e-08 | 1.193E-08 | 1.084E-08 | 3.000E-09 | 2.400E-08 |
| 19001 | 951 | V951A | C | 7.5e-09 | 5.6e-09 | 3.5e-09 | 5.533E-09 | 2.001E-09 | 3.500E-09 | 7.500E-09 |
| 19002 | 951 | V951R | C | 4.3e-09 | 1.1e-08 | 1.1e-08 | 8.767E-09 | 3.868E-09 | 4.300E-09 | 1.100E-08 |
| 19003 | 951 | V951N | C | 3.5e-09 | 7.5e-09 | 7e-09 | 6.000E-09 | 2.179E-09 | 3.500E-09 | 7.500E-09 |
| 19004 | 951 | V951D | C | 6.4e-09 | 5e-09 | 2.5e-09 | 4.633E-09 | 1.976E-09 | 2.500E-09 | 6.400E-09 |
| 19005 | 951 | V951C | C | 6e-09 | 3e-09 | 8.9e-09 | 5.967E-09 | 2.950E-09 | 3.000E-09 | 8.900E-09 |
| 19006 | 951 | V951E | C | 1.6e-09 | 6.8e-09 | 6.8e-09 | 5.067E-09 | 3.002E-09 | 1.600E-09 | 6.800E-09 |
| 19007 | 951 | V951Q | C | 1.1e-08 | 1e-08 | 2.5e-09 | 7.833E-09 | 4.646E-09 | 2.500E-09 | 1.100E-08 |
| 19008 | 951 | V951G | C | 3.2e-09 | 3.9e-09 | 1.3e-08 | 6.700E-09 | 5.467E-09 | 3.200E-09 | 1.300E-08 |
| 19009 | 951 | V951H | C | 7.5e-09 | 1.3e-08 | 1.4e-08 | 1.150E-08 | 3.500E-09 | 7.500E-09 | 1.400E-08 |
| 19010 | 951 | V951I | C | 2.2e-09 | 8.8e-09 | 3e-09 | 4.667E-09 | 3.602E-09 | 2.200E-09 | 8.800E-09 |
| 19011 | 951 | V951L | C | 4.6e-09 | 3.2e-09 | 8.7e-09 | 5.500E-09 | 2.858E-09 | 3.200E-09 | 8.700E-09 |
| 19012 | 951 | V951K | C | 6.7e-09 | 1.4e-08 | 1e-08 | 1.023E-08 | 3.656E-09 | 6.700E-09 | 1.400E-08 |
| 19013 | 951 | V951M | C | 4e-09 | 6.6e-09 | 9.1e-09 | 6.567E-09 | 2.550E-09 | 4.000E-09 | 9.100E-09 |
| 19014 | 951 | V951F | C | 7.6e-09 | 1.5e-08 | 8.8e-09 | 1.047E-08 | 3.972E-09 | 7.600E-09 | 1.500E-08 |
| 19015 | 951 | V951P | C | 2.1e-09 | 1.8e-09 | 1.5e-08 | 6.300E-09 | 7.536E-09 | 1.800E-09 | 1.500E-08 |
| 19016 | 951 | V951S | C | 1.3e-08 | 8.5e-09 | 1.2e-08 | 1.117E-08 | 2.363E-09 | 8.500E-09 | 1.300E-08 |
| 19017 | 951 | V951T | C | 5.6e-09 | 5.5e-09 | 6.3e-09 | 5.800E-09 | 4.359E-10 | 5.500E-09 | 6.300E-09 |
| 19018 | 951 | V951W | C | 1.4e-08 | 6.7e-09 | 6.6e-09 | 9.100E-09 | 4.244E-09 | 6.600E-09 | 1.400E-08 |
| 19019 | 951 | V951Y | C | 9.4e-09 | 6.5e-09 | 9.1e-09 | 8.333E-09 | 1.595E-09 | 6.500E-09 | 9.400E-09 |
| 19020 | 951 | V951V | C | 8.7e-09 | 7.5e-09 | 2.8e-09 | 6.333E-09 | 3.118E-09 | 2.800E-09 | 8.700E-09 |
| 19021 | 952 | T952A | C | 5.8e-09 | 3.6e-09 | 1.2e-08 | 7.133E-09 | 4.356E-09 | 3.600E-09 | 1.200E-08 |
| 19022 | 952 | T952R | C | 2.2e-09 | 5e-09 | 2.4e-08 | 1.040E-08 | 1.186E-08 | 2.200E-09 | 2.400E-08 |
| 19023 | 952 | T952N | C | 4.3e-09 | 1.2e-08 | 1.8e-08 | 1.143E-08 | 6.868E-09 | 4.300E-09 | 1.800E-08 |
| 19024 | 952 | T952D | C | 7.9e-09 | 5.7e-09 | 1.5e-09 | 5.033E-09 | 3.252E-09 | 1.500E-09 | 7.900E-09 |
| 19025 | 952 | T952C | C | 1.1e-08 | 1e-08 | 2.5e-08 | 1.533E-08 | 8.386E-09 | 1.000E-08 | 2.500E-08 |
| 19026 | 952 | T952E | C | 4.9e-09 | 7.5e-09 | 2.3e-09 | 4.900E-09 | 2.600E-09 | 2.300E-09 | 7.500E-09 |
| 19027 | 952 | T952Q | C | 4.5e-09 | 3.7e-09 | 6.4e-09 | 4.867E-09 | 1.387E-09 | 3.700E-09 | 6.400E-09 |
| 19028 | 952 | T952G | C | 1e-08 | 9.7e-09 | 8.7e-09 | 9.467E-09 | 6.807E-10 | 8.700E-09 | 1.000E-08 |
| 19029 | 952 | T952H | C | 7.7e-09 | 1.7e-08 | 1.1e-08 | 1.190E-08 | 4.715E-09 | 7.700E-09 | 1.700E-08 |
| 19030 | 952 | T952I | C | 1.1e-08 | 7.8e-09 | 2.4e-09 | 7.067E-09 | 4.347E-09 | 2.400E-09 | 1.100E-08 |
| 19031 | 952 | T952L | C | 1.3e-08 | 1e-08 | 1.3e-08 | 1.200E-08 | 1.732E-09 | 1.000E-08 | 1.300E-08 |
| 19032 | 952 | T952K | C | 1.2e-08 | 2.3e-09 | 1.5e-08 | 9.767E-09 | 6.638E-09 | 2.300E-09 | 1.500E-08 |
| 19033 | 952 | T952M | C | 1.1e-08 | 1.2e-08 | 1e-08 | 1.100E-08 | 1.000E-09 | 1.000E-08 | 1.200E-08 |
| 19034 | 952 | T952F | C | 7.8e-09 | 6.7e-09 | 7.3e-09 | 7.267E-09 | 5.508E-10 | 6.700E-09 | 7.800E-09 |
| 19035 | 952 | T952P | C | 4.2e-09 | 1.7e-08 | 2.3e-08 | 1.473E-08 | 9.603E-09 | 4.200E-09 | 2.300E-08 |
| 19036 | 952 | T952S | C | 9.3e-09 | 4.9e-09 | 9.9e-09 | 8.033E-09 | 2.730E-09 | 4.900E-09 | 9.900E-09 |
| 19037 | 952 | T952T | C | 6e-09 | 3.7e-09 | 1.5e-08 | 8.233E-09 | 5.972E-09 | 3.700E-09 | 1.500E-08 |
| 19038 | 952 | T952W | C | 6.4e-09 | 5.2e-09 | 3.1e-09 | 4.900E-09 | 1.670E-09 | 3.100E-09 | 6.400E-09 |
| 19039 | 952 | T952Y | C | 3.2e-09 | 7.6e-09 | 3e-09 | 4.600E-09 | 2.600E-09 | 3.000E-09 | 7.600E-09 |

| | | | | | | | | | | |
|---|---|---|---|---|---|---|---|---|---|---|
| 19040 | 952 | T952V | C | 6.8e-09 | 2.2e-09 | 4e-09 | 4.333E-09 | 2.318E-09 | 2.200E-09 | 6.800E-09 |
| 19041 | 953 | V953A | C | 5.1e-09 | 5.8e-09 | 4.3e-09 | 5.067E-09 | 7.506E-10 | 4.300E-09 | 5.800E-09 |
| 19042 | 953 | V953R | C | 4.2e-09 | 4.5e-09 | 1.7e-08 | 8.567E-09 | 7.305E-09 | 4.200E-09 | 1.700E-08 |
| 19043 | 953 | V953N | C | 8.5e-09 | 1.2e-08 | 2.4e-09 | 7.633E-09 | 4.858E-09 | 2.400E-09 | 1.200E-08 |
| 19044 | 953 | V953D | C | 5.7e-09 | 2.7e-09 | 4.3e-09 | 4.233E-09 | 1.501E-09 | 2.700E-09 | 5.700E-09 |
| 19045 | 953 | V953C | C | 3.7e-09 | 1.9e-09 | 4.3e-09 | 3.300E-09 | 1.249E-09 | 1.900E-09 | 4.300E-09 |
| 19046 | 953 | V953E | C | 8.9e-09 | 7.6e-09 | 6.5e-09 | 7.667E-09 | 1.201E-09 | 6.500E-09 | 8.900E-09 |
| 19047 | 953 | V953Q | C | 8e-09 | 2.6e-09 | 6.5e-09 | 5.700E-09 | 2.787E-09 | 2.600E-09 | 8.000E-09 |
| 19048 | 953 | V953G | C | 3.7e-09 | 1.3e-08 | 2.2e-09 | 6.300E-09 | 5.851E-09 | 2.200E-09 | 1.300E-08 |
| 19049 | 953 | V953H | C | 6.7e-09 | 5e-09 | 4.1e-09 | 5.267E-09 | 1.320E-09 | 4.100E-09 | 6.700E-09 |
| 19050 | 953 | V953I | C | 7.9e-09 | 1.1e-08 | 3.6e-09 | 7.500E-09 | 3.716E-09 | 3.600E-09 | 1.100E-08 |
| 19051 | 953 | V953L | C | 4e-09 | 2.9e-09 | 9.1e-09 | 5.333E-09 | 3.308E-09 | 2.900E-09 | 9.100E-09 |
| 19052 | 953 | V953K | C | 2.3e-09 | 2.1e-09 | 4e-09 | 2.800E-09 | 1.044E-09 | 2.100E-09 | 4.000E-09 |
| 19053 | 953 | V953M | C | 8.6e-09 | 1.7e-08 | 1.4e-08 | 1.320E-08 | 4.257E-09 | 8.600E-09 | 1.700E-08 |
| 19054 | 953 | V953F | C | 6.4e-09 | 2e-08 | 6.5e-09 | 1.097E-08 | 7.823E-09 | 6.400E-09 | 2.000E-08 |
| 19055 | 953 | V953P | C | 9e-09 | 6.3e-09 | 7.2e-09 | 7.500E-09 | 1.375E-09 | 6.300E-09 | 9.000E-09 |
| 19056 | 953 | V953S | C | 4.3e-09 | 1.4e-08 | 9.9e-09 | 9.400E-09 | 4.869E-09 | 4.300E-09 | 1.400E-08 |
| 19057 | 953 | V953T | C | 5.3e-09 | 3.2e-09 | 1.1e-08 | 6.500E-09 | 4.036E-09 | 3.200E-09 | 1.100E-08 |
| 19058 | 953 | V953W | C | 5.8e-09 | 1.7e-08 | 6.8e-09 | 9.867E-09 | 6.198E-09 | 5.800E-09 | 1.700E-08 |
| 19059 | 953 | V953Y | C | 5.3e-09 | 3.4e-09 | 5.6e-09 | 4.767E-09 | 1.193E-09 | 3.400E-09 | 5.600E-09 |
| 19060 | 953 | V953V | C | 2.9e-09 | 3.7e-09 | 9.1e-09 | 5.233E-09 | 3.372E-09 | 2.900E-09 | 9.100E-09 |
| 19061 | 954 | S954A | C | 3.2e-09 | 4.3e-09 | 1.1e-08 | 6.167E-09 | 4.222E-09 | 3.200E-09 | 1.100E-08 |
| 19062 | 954 | S954R | C | 3.7e-09 | 1.2e-08 | 3.7e-09 | 6.467E-09 | 4.792E-09 | 3.700E-09 | 1.200E-08 |
| 19063 | 954 | S954N | C | 7.8e-09 | 7.3e-09 | 6.2e-09 | 7.100E-09 | 8.185E-10 | 6.200E-09 | 7.800E-09 |
| 19064 | 954 | S954D | C | 3.8e-09 | 6.2e-09 | 4.9e-09 | 4.967E-09 | 1.201E-09 | 3.800E-09 | 6.200E-09 |
| 19065 | 954 | S954C | C | 3.9e-09 | 5.7e-09 | 4.7e-09 | 4.767E-09 | 9.018E-10 | 3.900E-09 | 5.700E-09 |
| 19066 | 954 | S954E | C | 3.4e-08 | 1.1e-08 | 3.5e-09 | 1.617E-08 | 1.589E-08 | 3.500E-09 | 3.400E-08 |
| 19067 | 954 | S954Q | C | 5.5e-09 | 8.4e-09 | 1.2e-08 | 8.633E-09 | 3.256E-09 | 5.500E-09 | 1.200E-08 |
| 19068 | 954 | S954G | C | 8.1e-09 | 1.7e-08 | 9.2e-09 | 1.143E-08 | 4.852E-09 | 8.100E-09 | 1.700E-08 |
| 19069 | 954 | S954H | C | 6.7e-09 | 8.2e-09 | 1.4e-08 | 9.633E-09 | 3.855E-09 | 6.700E-09 | 1.400E-08 |
| 19070 | 954 | S954I | C | 3.2e-09 | 2.5e-09 | 6.6e-09 | 4.100E-09 | 2.193E-09 | 2.500E-09 | 6.600E-09 |
| 19071 | 954 | S954L | C | 5.5e-09 | 6.8e-09 | 3e-09 | 5.100E-09 | 1.931E-09 | 3.000E-09 | 6.800E-09 |
| 19072 | 954 | S954K | C | 1.4e-08 | 6.9e-09 | 2.4e-09 | 7.767E-09 | 5.848E-09 | 2.400E-09 | 1.400E-08 |
| 19073 | 954 | S954M | C | 2.8e-08 | 1.8e-08 | 4e-09 | 1.667E-08 | 1.206E-08 | 4.000E-09 | 2.800E-08 |
| 19074 | 954 | S954F | C | 8.4e-09 | 8.5e-09 | 5.5e-09 | 7.467E-09 | 1.704E-09 | 5.500E-09 | 8.500E-09 |
| 19075 | 954 | S954P | C | 9e-09 | 8.2e-09 | 9.9e-09 | 9.033E-09 | 8.505E-10 | 8.200E-09 | 9.900E-09 |
| 19076 | 954 | S954S | C | 7.6e-09 | 3.6e-09 | 7.8e-09 | 6.333E-09 | 2.369E-09 | 3.600E-09 | 7.800E-09 |
| 19077 | 954 | S954T | C | 9.9e-09 | 6.7e-09 | 1e-08 | 8.867E-09 | 1.877E-09 | 6.700E-09 | 1.000E-08 |
| 19078 | 954 | S954W | C | 9.4e-09 | 7.3e-09 | 1.1e-08 | 9.233E-09 | 1.856E-09 | 7.300E-09 | 1.100E-08 |
| 19079 | 954 | S954Y | C | 4e-09 | 9e-09 | 1.1e-08 | 8.000E-09 | 3.606E-09 | 4.000E-09 | 1.100E-08 |
| 19080 | 954 | S954V | C | 1.9e-08 | 6.4e-09 | 9.5e-09 | 1.163E-08 | 6.565E-09 | 6.400E-09 | 1.900E-08 |
| 19081 | 955 | S955A | C | 4.3e-09 | 2.2e-08 | 7.7e-09 | 1.133E-08 | 9.393E-09 | 4.300E-09 | 2.200E-08 |
| 19082 | 955 | S955R | C | 5.3e-09 | 1.4e-08 | 1.5e-08 | 1.143E-08 | 5.335E-09 | 5.300E-09 | 1.500E-08 |
| 19083 | 955 | S955N | C | 4.3e-09 | 1.2e-08 | 8.6e-09 | 8.300E-09 | 3.859E-09 | 4.300E-09 | 1.200E-08 |
| 19084 | 955 | S955D | C | 8.4e-09 | 1.3e-08 | 7.9e-10 | 7.397E-09 | 6.167E-09 | 7.900E-10 | 1.300E-08 |
| 19085 | 955 | S955C | C | 2.4e-09 | 2.5e-09 | 9.3e-09 | 4.733E-09 | 3.955E-09 | 2.400E-09 | 9.300E-09 |
| 19086 | 955 | S955E | C | 7.7e-09 | 6.2e-09 | 1.2e-08 | 8.633E-09 | 3.011E-09 | 6.200E-09 | 1.200E-08 |
| 19087 | 955 | S955Q | C | 1e-08 | 1.3e-09 | 1.2e-09 | 4.167E-09 | 5.052E-09 | 1.200E-09 | 1.000E-08 |
| 19088 | 955 | S955G | C | 6.5e-09 | 6.3e-09 | 5.2e-09 | 6.000E-09 | 7.000E-10 | 5.200E-09 | 6.500E-09 |
| 19089 | 955 | S955H | C | 2.7e-09 | 1.2e-08 | 1.1e-08 | 8.567E-09 | 5.105E-09 | 2.700E-09 | 1.200E-08 |
| 19090 | 955 | S955I | C | 5.2e-09 | 9.4e-09 | 1.4e-08 | 9.533E-09 | 4.402E-09 | 5.200E-09 | 1.400E-08 |
| 19091 | 955 | S955L | C | 4.5e-09 | 2.7e-09 | 2.9e-08 | 1.207E-08 | 1.469E-08 | 2.700E-09 | 2.900E-08 |
| 19092 | 955 | S955K | C | 6.4e-09 | 1.1e-08 | 7.1e-09 | 8.167E-09 | 2.479E-09 | 6.400E-09 | 1.100E-08 |
| 19093 | 955 | S955M | C | 3e-09 | 1.2e-08 | 5.2e-09 | 6.733E-09 | 4.692E-09 | 3.000E-09 | 1.200E-08 |
| 19094 | 955 | S955F | C | 7.1e-09 | 4.1e-09 | 1.3e-08 | 8.067E-09 | 4.528E-09 | 4.100E-09 | 1.300E-08 |
| 19095 | 955 | S955P | C | 5.9e-09 | 5.2e-09 | 1.6e-08 | 9.033E-09 | 6.043E-09 | 5.200E-09 | 1.600E-08 |
| 19096 | 955 | S955S | C | 8.9e-09 | 9.1e-09 | 4.8e-09 | 7.600E-09 | 2.427E-09 | 4.800E-09 | 9.100E-09 |
| 19097 | 955 | S955T | C | 5.2e-09 | 5.1e-09 | 5.7e-09 | 5.333E-09 | 3.215E-10 | 5.100E-09 | 5.700E-09 |
| 19098 | 955 | S955W | C | 1.1e-08 | 1.5e-08 | 3.2e-09 | 9.733E-09 | 6.001E-09 | 3.200E-09 | 1.500E-08 |
| 19099 | 955 | S955Y | C | 8.3e-09 | 6.2e-09 | 6.7e-09 | 7.067E-09 | 1.097E-09 | 6.200E-09 | 8.300E-09 |
| 19100 | 955 | S955V | C | 1.9e-09 | 1e-08 | 1e-08 | 7.300E-09 | 4.677E-09 | 1.900E-09 | 1.000E-08 |
| 19101 | 956 | A956A | C | 1.4e-08 | 4.1e-09 | 1.2e-08 | 1.003E-08 | 5.235E-09 | 4.100E-09 | 1.400E-08 |
| 19102 | 956 | A956R | C | 7.9e-09 | 3.2e-09 | 1.2e-08 | 7.700E-09 | 4.403E-09 | 3.200E-09 | 1.200E-08 |
| 19103 | 956 | A956N | C | 4e-09 | 5.3e-09 | 2.3e-09 | 3.867E-09 | 1.504E-09 | 2.300E-09 | 5.300E-09 |
| 19104 | 956 | A956D | C | 2.5e-09 | 6.6e-09 | 5e-09 | 4.700E-09 | 2.066E-09 | 2.500E-09 | 6.600E-09 |
| 19105 | 956 | A956C | C | 3.2e-09 | 6.3e-09 | 9e-09 | 6.167E-09 | 2.902E-09 | 3.200E-09 | 9.000E-09 |
| 19106 | 956 | A956E | C | 2.1e-09 | 4e-09 | 7.2e-09 | 4.433E-09 | 2.577E-09 | 2.100E-09 | 7.200E-09 |
| 19107 | 956 | A956Q | C | 1.2e-08 | 5.3e-09 | 5.4e-09 | 7.567E-09 | 3.840E-09 | 5.300E-09 | 1.200E-08 |
| 19108 | 956 | A956G | C | 4.7e-09 | 2.5e-08 | 5.4e-09 | 1.170E-08 | 1.152E-08 | 4.700E-09 | 2.500E-08 |
| 19109 | 956 | A956H | C | 7.9e-09 | 8.6e-09 | 9.8e-09 | 8.767E-09 | 9.609E-10 | 7.900E-09 | 9.800E-09 |
| 19110 | 956 | A956I | C | 4.3e-09 | 5.9e-09 | 7.6e-09 | 5.933E-09 | 1.650E-09 | 4.300E-09 | 7.600E-09 |
| 19111 | 956 | A956L | C | 3.5e-09 | 2.6e-09 | 7.9e-09 | 4.667E-09 | 2.836E-09 | 2.600E-09 | 7.900E-09 |
| 19112 | 956 | A956K | C | 1.3e-08 | 2.6e-09 | 2.6e-08 | 1.387E-08 | 1.172E-08 | 2.600E-09 | 2.600E-08 |
| 19113 | 956 | A956M | C | 2e-08 | 3.5e-08 | 5.7e-09 | 2.023E-08 | 1.465E-08 | 5.700E-09 | 3.500E-08 |
| 19114 | 956 | A956F | C | 1.2e-08 | 1.2e-08 | 3.2e-09 | 9.067E-09 | 5.081E-09 | 3.200E-09 | 1.200E-08 |
| 19115 | 956 | A956P | C | 1.1e-08 | 2e-09 | 7.9e-09 | 6.967E-09 | 4.572E-09 | 2.000E-09 | 1.100E-08 |
| 19116 | 956 | A956S | C | 8.2e-09 | 5.7e-09 | 3e-09 | 5.633E-09 | 2.601E-09 | 3.000E-09 | 8.200E-09 |
| 19117 | 956 | A956T | C | 5.2e-09 | 1.2e-08 | 8.1e-09 | 8.433E-09 | 3.412E-09 | 5.200E-09 | 1.200E-08 |
| 19118 | 956 | A956W | C | 1.4e-09 | 1.9e-09 | 5.7e-09 | 3.000E-09 | 2.352E-09 | 1.400E-09 | 5.700E-09 |
| 19119 | 956 | A956Y | C | 4.9e-09 | 7.9e-09 | 7.6e-09 | 6.800E-09 | 1.652E-09 | 4.900E-09 | 7.900E-09 |
| 19120 | 956 | A956V | C | 5.7e-09 | 5.4e-09 | 2.8e-09 | 4.633E-09 | 1.595E-09 | 2.800E-09 | 5.700E-09 |
| 19121 | 957 | S957A | C | 8e-09 | 6e-09 | 9.2e-09 | 7.733E-09 | 1.617E-09 | 6.000E-09 | 9.200E-09 |
| 19122 | 957 | S957R | C | 5e-09 | 2.4e-08 | 7.6e-09 | 1.220E-08 | 1.030E-08 | 5.000E-09 | 2.400E-08 |
| 19123 | 957 | S957N | C | 3.5e-09 | 1.1e-08 | 1.8e-09 | 5.433E-09 | 4.895E-09 | 1.800E-09 | 1.100E-08 |
| 19124 | 957 | S957D | C | 7.1e-09 | 4e-09 | 2.2e-09 | 4.433E-09 | 2.479E-09 | 2.200E-09 | 7.100E-09 |

| | | | | | | | | | | |
|---|---|---|---|---|---|---|---|---|---|---|
| 19125 | 957 | S957C | C | 1.2e-08 | 1.2e-08 | 4.6e-09 | 9.533E-09 | 4.272E-09 | 4.600E-09 | 1.200E-08 |
| 19126 | 957 | S957E | C | 8.7e-09 | 8.4e-09 | 8.5e-09 | 8.533E-09 | 1.528E-10 | 8.400E-09 | 8.700E-09 |
| 19127 | 957 | S957Q | C | 9.6e-09 | 7.7e-09 | 6.6e-09 | 7.967E-09 | 1.518E-09 | 6.600E-09 | 9.600E-09 |
| 19128 | 957 | S957G | C | 2.3e-09 | 2.5e-09 | 7.7e-09 | 4.167E-09 | 3.062E-09 | 2.300E-09 | 7.700E-09 |
| 19129 | 957 | S957H | C | 6.3e-09 | 2e-09 | 6.1e-09 | 4.800E-09 | 2.427E-09 | 2.000E-09 | 6.300E-09 |
| 19130 | 957 | S957I | C | 7.1e-09 | 1.6e-08 | 7.4e-09 | 1.017E-08 | 5.054E-09 | 7.100E-09 | 1.600E-08 |
| 19131 | 957 | S957L | C | 9.8e-09 | 1.7e-08 | 7.3e-09 | 1.137E-08 | 5.036E-09 | 7.300E-09 | 1.700E-08 |
| 19132 | 957 | S957K | C | 5.1e-09 | 1.1e-08 | 1.4e-09 | 5.833E-09 | 4.842E-09 | 1.400E-09 | 1.100E-08 |
| 19133 | 957 | S957M | C | 4.8e-09 | 2.9e-09 | 5.3e-09 | 4.333E-09 | 1.266E-09 | 2.900E-09 | 5.300E-09 |
| 19134 | 957 | S957F | C | 3.5e-08 | 8.2e-09 | 2.1e-09 | 1.510E-08 | 1.750E-08 | 2.100E-09 | 3.500E-08 |
| 19135 | 957 | S957P | C | 5.9e-09 | 9.3e-09 | 6.2e-09 | 7.133E-09 | 1.882E-09 | 5.900E-09 | 9.300E-09 |
| 19136 | 957 | S957S | C | 9.7e-09 | 6.3e-09 | 9.6e-09 | 8.533E-09 | 1.935E-09 | 6.300E-09 | 9.700E-09 |
| 19137 | 957 | S957T | C | 6.7e-09 | 6.4e-09 | 1.4e-09 | 4.833E-09 | 2.977E-09 | 1.400E-09 | 6.700E-09 |
| 19138 | 957 | S957W | C | 1.3e-08 | 1.3e-08 | 5.2e-09 | 1.040E-08 | 4.503E-09 | 5.200E-09 | 1.300E-08 |
| 19139 | 957 | S957Y | C | 4.8e-09 | 5e-09 | 7.1e-09 | 5.633E-09 | 1.274E-09 | 4.800E-09 | 7.100E-09 |
| 19140 | 957 | S957V | C | 5.2e-09 | 1.1e-08 | 1.5e-08 | 1.040E-08 | 4.927E-09 | 5.200E-09 | 1.500E-08 |
| 19141 | 958 | T958A | C | 3.6e-09 | 1.5e-08 | 2e-09 | 6.867E-09 | 7.089E-09 | 2.000E-09 | 1.500E-08 |
| 19142 | 958 | T958R | C | 2.9e-08 | 1.1e-08 | 2.2e-08 | 2.067E-08 | 9.074E-09 | 1.100E-08 | 2.900E-08 |
| 19143 | 958 | T958N | C | 2.3e-08 | 9.1e-09 | 1.1e-08 | 1.437E-08 | 7.537E-09 | 9.100E-09 | 2.300E-08 |
| 19144 | 958 | T958D | C | 9.9e-09 | 1.7e-08 | 8.7e-09 | 1.187E-08 | 4.486E-09 | 8.700E-09 | 1.700E-08 |
| 19145 | 958 | T958C | C | 1.4e-08 | 3.6e-09 | 4.9e-09 | 7.500E-09 | 5.667E-09 | 3.600E-09 | 1.400E-08 |
| 19146 | 958 | T958E | C | 1.1e-08 | 4.3e-09 | 2.9e-09 | 6.067E-09 | 4.329E-09 | 2.900E-09 | 1.100E-08 |
| 19147 | 958 | T958Q | C | 5.7e-09 | 1.6e-08 | 5.4e-09 | 9.033E-09 | 6.035E-09 | 5.400E-09 | 1.600E-08 |
| 19148 | 958 | T958G | C | 9.7e-09 | 2e-08 | 2.8e-09 | 1.083E-08 | 8.656E-09 | 2.800E-09 | 2.000E-08 |
| 19149 | 958 | T958H | C | 6.3e-09 | 9.6e-09 | 4.8e-09 | 6.900E-09 | 2.456E-09 | 4.800E-09 | 9.600E-09 |
| 19150 | 958 | T958I | C | 1.3e-08 | 7.8e-09 | 9e-09 | 9.933E-09 | 2.723E-09 | 7.800E-09 | 1.300E-08 |
| 19151 | 958 | T958L | C | 7.2e-09 | 3.9e-09 | 8.8e-09 | 6.633E-09 | 2.499E-09 | 3.900E-09 | 8.800E-09 |
| 19152 | 958 | T958K | C | 1.3e-08 | 7.6e-09 | 1.6e-08 | 1.220E-08 | 4.257E-09 | 7.600E-09 | 1.600E-08 |
| 19153 | 958 | T958M | C | 1.2e-08 | 3e-09 | 3.8e-09 | 6.267E-09 | 4.981E-09 | 3.000E-09 | 1.200E-08 |
| 19154 | 958 | T958F | C | 6.2e-09 | 1.2e-08 | 2.1e-08 | 1.307E-08 | 7.457E-09 | 6.200E-09 | 2.100E-08 |
| 19155 | 958 | T958P | C | 1.1e-08 | 4.9e-09 | 1.2e-08 | 9.300E-09 | 3.843E-09 | 4.900E-09 | 1.200E-08 |
| 19156 | 958 | T958S | C | 2.9e-09 | 5.4e-09 | 3.5e-09 | 3.933E-09 | 1.305E-09 | 2.900E-09 | 5.400E-09 |
| 19157 | 958 | T958T | C | 1.7e-09 | 1.1e-08 | 2e-09 | 4.900E-09 | 5.285E-09 | 1.700E-09 | 1.100E-08 |
| 19158 | 958 | T958W | C | 5.7e-09 | 2.1e-08 | 4.9e-09 | 1.053E-08 | 9.073E-09 | 4.900E-09 | 2.100E-08 |
| 19159 | 958 | T958Y | C | 4.4e-09 | 9.8e-09 | 6.9e-09 | 7.033E-09 | 2.702E-09 | 4.400E-09 | 9.800E-09 |
| 19160 | 958 | T958V | C | 5e-09 | 7.5e-09 | 5.1e-09 | 5.867E-09 | 1.415E-09 | 5.000E-09 | 7.500E-09 |
| 19161 | 959 | K959A | C | 4e-09 | 1.5e-08 | 7.7e-09 | 8.900E-09 | 5.597E-09 | 4.000E-09 | 1.500E-08 |
| 19162 | 959 | K959R | C | 8.3e-09 | 3.7e-09 | 2.8e-09 | 4.933E-09 | 2.950E-09 | 2.800E-09 | 8.300E-09 |
| 19163 | 959 | K959N | C | 3.6e-09 | 6.8e-09 | 8.2e-09 | 6.200E-09 | 2.358E-09 | 3.600E-09 | 8.200E-09 |
| 19164 | 959 | K959D | C | 1.7e-08 | 6.4e-09 | 1.9e-08 | 1.413E-08 | 6.772E-09 | 6.400E-09 | 1.900E-08 |
| 19165 | 959 | K959C | C | 9.3e-09 | 8.7e-09 | 8.7e-09 | 8.900E-09 | 3.464E-10 | 8.700E-09 | 9.300E-09 |
| 19166 | 959 | K959E | C | 4.2e-09 | 9.2e-09 | 2.5e-09 | 5.300E-09 | 3.483E-09 | 2.500E-09 | 9.200E-09 |
| 19167 | 959 | K959Q | C | 1e-09 | 4.5e-09 | 4e-09 | 3.167E-09 | 1.893E-09 | 1.000E-09 | 4.500E-09 |
| 19168 | 959 | K959G | C | 8.1e-09 | 5.9e-09 | 3.7e-09 | 5.900E-09 | 2.200E-09 | 3.700E-09 | 8.100E-09 |
| 19169 | 959 | K959H | C | 3.8e-09 | 1e-08 | 3.9e-09 | 5.900E-09 | 3.551E-09 | 3.800E-09 | 1.000E-08 |
| 19170 | 959 | K959I | C | 8.4e-09 | 3.7e-09 | 5.1e-09 | 5.733E-09 | 2.413E-09 | 3.700E-09 | 8.400E-09 |
| 19171 | 959 | K959L | C | 4.2e-09 | 5.7e-09 | 1.1e-08 | 6.967E-09 | 3.573E-09 | 4.200E-09 | 1.100E-08 |
| 19172 | 959 | K959K | C | 2.6e-09 | 1.1e-08 | 7.9e-09 | 7.167E-09 | 4.248E-09 | 2.600E-09 | 1.100E-08 |
| 19173 | 959 | K959M | C | 3.4e-09 | 4.3e-09 | 1e-08 | 5.900E-09 | 3.579E-09 | 3.400E-09 | 1.000E-08 |
| 19174 | 959 | K959F | C | 5.1e-08 | 3.9e-09 | 1.7e-08 | 2.397E-08 | 2.431E-08 | 3.900E-09 | 5.100E-08 |
| 19175 | 959 | K959P | C | 2.6e-09 | 4.4e-09 | 4.2e-09 | 3.733E-09 | 9.866E-10 | 2.600E-09 | 4.400E-09 |
| 19176 | 959 | K959S | C | 9.1e-09 | 9.7e-09 | 6.1e-09 | 8.300E-09 | 1.929E-09 | 6.100E-09 | 9.700E-09 |
| 19177 | 959 | K959T | C | 1e-08 | 6.2e-09 | 4.6e-09 | 6.933E-09 | 2.774E-09 | 4.600E-09 | 1.000E-08 |
| 19178 | 959 | K959W | C | 1.1e-08 | 5.7e-09 | 7.2e-09 | 7.967E-09 | 2.732E-09 | 5.700E-09 | 1.100E-08 |
| 19179 | 959 | K959Y | C | 1.2e-08 | 5e-09 | 4.6e-09 | 7.200E-09 | 4.162E-09 | 4.600E-09 | 1.200E-08 |
| 19180 | 959 | K959V | C | 4.1e-09 | 1.7e-09 | 1.4e-08 | 6.600E-09 | 6.520E-09 | 1.700E-09 | 1.400E-08 |
| 19181 | 960 | G960A | C | 4.9e-09 | 5.3e-09 | 1.3e-08 | 7.733E-09 | 4.565E-09 | 4.900E-09 | 1.300E-08 |
| 19182 | 960 | G960R | C | 6.1e-09 | 3.4e-09 | 7e-09 | 5.500E-09 | 1.873E-09 | 3.400E-09 | 7.000E-09 |
| 19183 | 960 | G960N | C | 3.3e-09 | 6.8e-09 | 4.6e-09 | 4.900E-09 | 1.769E-09 | 3.300E-09 | 6.800E-09 |
| 19184 | 960 | G960D | C | 1.2e-08 | 9.6e-09 | 7.8e-09 | 9.800E-09 | 2.107E-09 | 7.800E-09 | 1.200E-08 |
| 19185 | 960 | G960C | C | 9.5e-09 | 3.1e-09 | 5.3e-09 | 5.967E-09 | 3.252E-09 | 3.100E-09 | 9.500E-09 |
| 19186 | 960 | G960E | C | 2.1e-09 | 3.3e-09 | 1.6e-08 | 7.133E-09 | 7.702E-09 | 2.100E-09 | 1.600E-08 |
| 19187 | 960 | G960Q | C | 8e-09 | 6.7e-09 | 7.1e-09 | 7.267E-09 | 6.658E-10 | 6.700E-09 | 8.000E-09 |
| 19188 | 960 | G960G | C | 1.1e-08 | 6.2e-09 | 6.9e-09 | 8.033E-09 | 2.593E-09 | 6.200E-09 | 1.100E-08 |
| 19189 | 960 | G960H | C | 1.8e-09 | 3.8e-09 | 1.7e-08 | 7.533E-09 | 8.259E-09 | 1.800E-09 | 1.700E-08 |
| 19190 | 960 | G960I | C | 5.5e-09 | 3.4e-09 | 1e-08 | 6.300E-09 | 3.372E-09 | 3.400E-09 | 1.000E-08 |
| 19191 | 960 | G960L | C | 2.6e-09 | 5.1e-09 | 5e-09 | 4.233E-09 | 1.415E-09 | 2.600E-09 | 5.100E-09 |
| 19192 | 960 | G960K | C | 7.2e-09 | 4.4e-09 | 3.6e-09 | 5.067E-09 | 1.890E-09 | 3.600E-09 | 7.200E-09 |
| 19193 | 960 | G960M | C | 9.7e-09 | 7.8e-09 | 3.2e-09 | 6.900E-09 | 3.342E-09 | 3.200E-09 | 9.700E-09 |
| 19194 | 960 | G960F | C | 3e-09 | 2.7e-09 | 6.7e-09 | 4.133E-09 | 2.228E-09 | 2.700E-09 | 6.700E-09 |
| 19195 | 960 | G960P | C | 2.4e-09 | 4.8e-09 | 6.5e-09 | 4.567E-09 | 2.060E-09 | 2.400E-09 | 6.500E-09 |
| 19196 | 960 | G960S | C | 6.1e-09 | 1.4e-08 | 3.3e-09 | 7.800E-09 | 5.549E-09 | 3.300E-09 | 1.400E-08 |
| 19197 | 960 | G960T | C | 8.2e-09 | 5.6e-09 | 7e-09 | 6.933E-09 | 1.301E-09 | 5.600E-09 | 8.200E-09 |
| 19198 | 960 | G960W | C | 1.1e-08 | 7.9e-09 | 2.8e-09 | 7.233E-09 | 4.140E-09 | 2.800E-09 | 1.100E-08 |
| 19199 | 960 | G960Y | C | 1.2e-08 | 7.7e-09 | 4.6e-09 | 8.100E-09 | 3.716E-09 | 4.600E-09 | 1.200E-08 |
| 19200 | 960 | G960V | C | 4.1e-09 | 2.4e-09 | 1.4e-08 | 6.833E-09 | 6.264E-09 | 2.400E-09 | 1.400E-08 |
| 19201 | 961 | P961A | C | 8e-09 | 1.5e-08 | 9.6e-09 | 1.087E-08 | 3.668E-09 | 8.000E-09 | 1.500E-08 |
| 19202 | 961 | P961R | C | 8.9e-09 | 1.5e-08 | 6.9e-09 | 1.027E-08 | 4.219E-09 | 6.900E-09 | 1.500E-08 |
| 19203 | 961 | P961N | C | 8.8e-09 | 8e-09 | 6.3e-09 | 7.700E-09 | 1.277E-09 | 6.300E-09 | 8.800E-09 |
| 19204 | 961 | P961D | C | 1.5e-08 | 1.5e-08 | 2.6e-09 | 1.087E-08 | 7.159E-09 | 2.600E-09 | 1.500E-08 |
| 19205 | 961 | P961C | C | 2e-09 | 4.8e-09 | 1.3e-08 | 6.600E-09 | 5.717E-09 | 2.000E-09 | 1.300E-08 |
| 19206 | 961 | P961E | C | 8.5e-09 | 6.9e-09 | 6.1e-09 | 7.167E-09 | 1.222E-09 | 6.100E-09 | 8.500E-09 |
| 19207 | 961 | P961Q | C | 6.7e-09 | 1.3e-08 | 7.2e-09 | 8.967E-09 | 3.502E-09 | 6.700E-09 | 1.300E-08 |
| 19208 | 961 | P961G | C | 2e-09 | 5.5e-09 | 6.7e-09 | 4.733E-09 | 2.442E-09 | 2.000E-09 | 6.700E-09 |
| 19209 | 961 | P961H | C | 1.6e-08 | 1.1e-08 | 9.2e-09 | 1.207E-08 | 3.523E-09 | 9.200E-09 | 1.600E-08 |

| | | | | | | | | | | |
|---|---|---|---|---|---|---|---|---|---|---|
| 19210 | 961 | P961I | C | 7.7e-09 | 4.5e-09 | 4.7e-09 | 5.633E-09 | 1.793E-09 | 4.500E-09 | 7.700E-09 |
| 19211 | 961 | P961L | C | 7.3e-09 | 2e-09 | 9.5e-09 | 6.267E-09 | 3.855E-09 | 2.000E-09 | 9.500E-09 |
| 19212 | 961 | P961K | C | 1.4e-08 | 3.8e-09 | 9.5e-09 | 9.100E-09 | 5.112E-09 | 3.800E-09 | 1.400E-08 |
| 19213 | 961 | P961M | C | 6.9e-09 | 1.3e-08 | 1.2e-08 | 1.063E-08 | 3.272E-09 | 6.900E-09 | 1.300E-08 |
| 19214 | 961 | P961F | C | 6.8e-09 | 1.4e-08 | 3.7e-09 | 8.167E-09 | 5.284E-09 | 3.700E-09 | 1.400E-08 |
| 19215 | 961 | P961P | C | 9.5e-09 | 2.2e-09 | 5.4e-09 | 5.700E-09 | 3.659E-09 | 2.200E-09 | 9.500E-09 |
| 19216 | 961 | P961S | C | 7e-09 | 8.5e-09 | 6.1e-09 | 7.200E-09 | 1.212E-09 | 6.100E-09 | 8.500E-09 |
| 19217 | 961 | P961T | C | 6e-09 | 7.4e-09 | 4.2e-09 | 5.867E-09 | 1.604E-09 | 4.200E-09 | 7.400E-09 |
| 19218 | 961 | P961W | C | 1.4e-08 | 7e-09 | 1.2e-08 | 1.100E-08 | 3.606E-09 | 7.000E-09 | 1.400E-08 |
| 19219 | 961 | P961Y | C | 9.5e-09 | 6.4e-09 | 1.1e-08 | 8.967E-09 | 2.346E-09 | 6.400E-09 | 1.100E-08 |
| 19220 | 961 | P961V | C | 3.2e-09 | 2.2e-08 | 2.4e-09 | 9.200E-09 | 1.109E-08 | 2.400E-09 | 2.200E-08 |
| 19221 | 962 | S962A | C | 6.3e-10 | 9.6e-09 | 2.3e-08 | 1.108E-08 | 1.126E-08 | 6.300E-10 | 2.300E-08 |
| 19222 | 962 | S962R | C | 4.4e-09 | 4.4e-09 | 6.5e-09 | 5.100E-09 | 1.212E-09 | 4.400E-09 | 6.500E-09 |
| 19223 | 962 | S962N | C | 9.5e-09 | 5e-09 | 7.5e-09 | 7.333E-09 | 2.255E-09 | 5.000E-09 | 9.500E-09 |
| 19224 | 962 | S962D | C | 8.9e-09 | 4.4e-09 | 2.5e-09 | 5.267E-09 | 3.287E-09 | 2.500E-09 | 8.900E-09 |
| 19225 | 962 | S962C | C | 6.6e-09 | 2.1e-08 | 3.7e-09 | 1.043E-08 | 9.265E-09 | 3.700E-09 | 2.100E-08 |
| 19226 | 962 | S962E | C | 2.3e-09 | 8.9e-09 | 1.5e-08 | 8.733E-09 | 6.352E-09 | 2.300E-09 | 1.500E-08 |
| 19227 | 962 | S962Q | C | 8e-09 | 5.2e-09 | 1.9e-08 | 1.073E-08 | 7.295E-09 | 5.200E-09 | 1.900E-08 |
| 19228 | 962 | S962G | C | 8.8e-09 | 2.3e-09 | 8e-09 | 6.367E-09 | 3.544E-09 | 2.300E-09 | 8.800E-09 |
| 19229 | 962 | S962H | C | 1.6e-08 | 3e-08 | 8.6e-09 | 1.820E-08 | 1.087E-08 | 8.600E-09 | 3.000E-08 |
| 19230 | 962 | S962I | C | 1.2e-08 | 4.8e-09 | 6.5e-09 | 7.767E-09 | 3.763E-09 | 4.800E-09 | 1.200E-08 |
| 19231 | 962 | S962L | C | 1.1e-08 | 3.5e-09 | 5.4e-09 | 6.633E-09 | 3.899E-09 | 3.500E-09 | 1.100E-08 |
| 19232 | 962 | S962K | C | 2.5e-09 | 8.3e-09 | 9.9e-09 | 6.900E-09 | 3.894E-09 | 2.500E-09 | 9.900E-09 |
| 19233 | 962 | S962M | C | 1.2e-08 | 3.5e-09 | 8.7e-09 | 8.067E-09 | 4.285E-09 | 3.500E-09 | 1.200E-08 |
| 19234 | 962 | S962F | C | 1.4e-08 | 5.3e-09 | 3.9e-09 | 7.733E-09 | 5.472E-09 | 3.900E-09 | 1.400E-08 |
| 19235 | 962 | S962P | C | 7.6e-09 | 4.1e-09 | 5.9e-09 | 5.867E-09 | 1.750E-09 | 4.100E-09 | 7.600E-09 |
| 19236 | 962 | S962S | C | 1.8e-08 | 5.8e-09 | 1.6e-08 | 1.327E-08 | 6.543E-09 | 5.800E-09 | 1.800E-08 |
| 19237 | 962 | S962T | C | 4.7e-09 | 1.1e-08 | 7.7e-09 | 7.800E-09 | 3.151E-09 | 4.700E-09 | 1.100E-08 |
| 19238 | 962 | S962W | C | 2.6e-08 | 6.5e-09 | 1.8e-08 | 1.683E-08 | 9.802E-09 | 6.500E-09 | 2.600E-08 |
| 19239 | 962 | S962Y | C | 7.2e-09 | 4.9e-09 | 5e-09 | 5.700E-09 | 1.300E-09 | 4.900E-09 | 7.200E-09 |
| 19240 | 962 | S962V | C | 1e-08 | 3.4e-09 | 5.9e-09 | 6.433E-09 | 3.332E-09 | 3.400E-09 | 1.000E-08 |
| 19241 | 963 | V963A | C | 1.9e-08 | 9.5e-09 | 6e-09 | 1.150E-08 | 6.727E-09 | 6.000E-09 | 1.900E-08 |
| 19242 | 963 | V963R | C | 4.5e-09 | 2.2e-08 | 1.5e-08 | 1.383E-08 | 8.808E-09 | 4.500E-09 | 2.200E-08 |
| 19243 | 963 | V963N | C | 8.1e-09 | 9.2e-09 | 1.3e-08 | 1.010E-08 | 2.571E-09 | 8.100E-09 | 1.300E-08 |
| 19244 | 963 | V963D | C | 7.5e-09 | 7.9e-09 | 1.3e-08 | 9.467E-09 | 3.066E-09 | 7.500E-09 | 1.300E-08 |
| 19245 | 963 | V963C | C | 1.2e-08 | 2.1e-08 | 6.1e-09 | 1.303E-08 | 7.504E-09 | 6.100E-09 | 2.100E-08 |
| 19246 | 963 | V963E | C | 3.7e-09 | 1.8e-08 | 1.8e-09 | 7.833E-09 | 8.856E-09 | 1.800E-09 | 1.800E-08 |
| 19247 | 963 | V963Q | C | 1.4e-08 | 7.1e-09 | 4.9e-09 | 8.667E-09 | 4.748E-09 | 4.900E-09 | 1.400E-08 |
| 19248 | 963 | V963G | C | 1.1e-08 | 8.7e-09 | 4.3e-09 | 8.000E-09 | 3.404E-09 | 4.300E-09 | 1.100E-08 |
| 19249 | 963 | V963H | C | 1.1e-08 | 1e-08 | 1e-08 | 1.033E-08 | 5.774E-10 | 1.000E-08 | 1.100E-08 |
| 19250 | 963 | V963I | C | 1.2e-09 | 9.4e-09 | 2.4e-09 | 4.333E-09 | 4.429E-09 | 1.200E-09 | 9.400E-09 |
| 19251 | 963 | V963L | C | 1.3e-08 | 1.8e-09 | 1.5e-08 | 9.933E-09 | 7.114E-09 | 1.800E-09 | 1.500E-08 |
| 19252 | 963 | V963K | C | 5.2e-09 | 7.3e-09 | 3.7e-09 | 5.400E-09 | 1.808E-09 | 3.700E-09 | 7.300E-09 |
| 19253 | 963 | V963M | C | 1.4e-08 | 4.3e-09 | 3.4e-09 | 7.233E-09 | 5.877E-09 | 3.400E-09 | 1.400E-08 |
| 19254 | 963 | V963F | C | 5.4e-09 | 3.7e-09 | 1.5e-08 | 8.033E-09 | 6.093E-09 | 3.700E-09 | 1.500E-08 |
| 19255 | 963 | V963P | C | 1.2e-08 | 3.4e-09 | 1.1e-08 | 8.800E-09 | 4.703E-09 | 3.400E-09 | 1.200E-08 |
| 19256 | 963 | V963S | C | 1.4e-08 | 1.9e-08 | 3.6e-09 | 1.220E-08 | 7.856E-09 | 3.600E-09 | 1.900E-08 |
| 19257 | 963 | V963T | C | 1.2e-08 | 1e-08 | 1e-08 | 1.067E-08 | 1.155E-09 | 1.000E-08 | 1.200E-08 |
| 19258 | 963 | V963W | C | 2.1e-08 | 5.4e-09 | 3.4e-09 | 9.933E-09 | 9.636E-09 | 3.400E-09 | 2.100E-08 |
| 19259 | 963 | V963Y | C | 3.9e-09 | 7.5e-09 | 6.4e-09 | 5.933E-09 | 1.845E-09 | 3.900E-09 | 7.500E-09 |
| 19260 | 963 | V963V | C | 3.5e-09 | 5.4e-09 | 6.2e-09 | 5.033E-09 | 1.387E-09 | 3.500E-09 | 6.200E-09 |
| 19261 | 964 | F964A | C | 4.4e-09 | 4.9e-09 | 7.2e-09 | 5.500E-09 | 1.493E-09 | 4.400E-09 | 7.200E-09 |
| 19262 | 964 | F964R | C | 3e-09 | 6.3e-09 | 1.4e-08 | 7.767E-09 | 5.645E-09 | 3.000E-09 | 1.400E-08 |
| 19263 | 964 | F964N | C | 6.2e-09 | 8.4e-09 | 1.9e-09 | 5.500E-09 | 3.306E-09 | 1.900E-09 | 8.400E-09 |
| 19264 | 964 | F964D | C | 8e-09 | 1.4e-08 | 6.9e-09 | 9.633E-09 | 3.821E-09 | 6.900E-09 | 1.400E-08 |
| 19265 | 964 | F964C | C | 1.2e-08 | 5.8e-09 | 5.8e-09 | 7.867E-09 | 3.580E-09 | 5.800E-09 | 1.200E-08 |
| 19266 | 964 | F964E | C | 1.5e-08 | 4.4e-09 | 1.4e-08 | 1.113E-08 | 5.853E-09 | 4.400E-09 | 1.500E-08 |
| 19267 | 964 | F964Q | C | 9.7e-09 | 1.3e-08 | 1.5e-08 | 1.257E-08 | 2.676E-09 | 9.700E-09 | 1.500E-08 |
| 19268 | 964 | F964G | C | 6.9e-09 | 4.3e-09 | 3.2e-09 | 4.800E-09 | 1.900E-09 | 3.200E-09 | 6.900E-09 |
| 19269 | 964 | F964H | C | 5.1e-09 | 2.1e-09 | 4.6e-09 | 3.933E-09 | 1.607E-09 | 2.100E-09 | 5.100E-09 |
| 19270 | 964 | F964I | C | 6.6e-09 | 4.8e-09 | 4.6e-09 | 5.333E-09 | 1.102E-09 | 4.600E-09 | 6.600E-09 |
| 19271 | 964 | F964L | C | 1.4e-08 | 7.6e-09 | 3.4e-09 | 8.333E-09 | 5.338E-09 | 3.400E-09 | 1.400E-08 |
| 19272 | 964 | F964K | C | 4.7e-09 | 2.1e-08 | 1.2e-08 | 1.257E-08 | 8.165E-09 | 4.700E-09 | 2.100E-08 |
| 19273 | 964 | F964M | C | 2.5e-09 | 1.2e-08 | 1.1e-08 | 8.500E-09 | 5.220E-09 | 2.500E-09 | 1.200E-08 |
| 19274 | 964 | F964F | C | 1.2e-08 | 3.1e-08 | 3.1e-09 | 1.537E-08 | 1.425E-08 | 3.100E-09 | 3.100E-08 |
| 19275 | 964 | F964P | C | 2.9e-09 | 8.3e-09 | 6.4e-09 | 5.867E-09 | 2.739E-09 | 2.900E-09 | 8.300E-09 |
| 19276 | 964 | F964S | C | 5.8e-09 | 4.9e-09 | 1.2e-08 | 7.567E-09 | 3.866E-09 | 4.900E-09 | 1.200E-08 |
| 19277 | 964 | F964T | C | 8.2e-09 | 1.3e-08 | 3.2e-09 | 8.133E-09 | 4.900E-09 | 3.200E-09 | 1.300E-08 |
| 19278 | 964 | F964W | C | 7.1e-09 | 1.4e-08 | 6.9e-09 | 9.333E-09 | 4.043E-09 | 6.900E-09 | 1.400E-08 |
| 19279 | 964 | F964Y | C | 1.4e-08 | 9.3e-09 | 9.6e-09 | 1.097E-08 | 2.631E-09 | 9.300E-09 | 1.400E-08 |
| 19280 | 964 | F964V | C | 3.6e-09 | 1.7e-08 | 4.8e-09 | 8.467E-09 | 7.414E-09 | 3.600E-09 | 1.700E-08 |
| 19281 | 965 | P965A | C | 1.7e-08 | 8e-09 | 6.7e-09 | 1.057E-08 | 5.609E-09 | 6.700E-09 | 1.700E-08 |
| 19282 | 965 | P965R | C | 8.3e-09 | 4.9e-09 | 1.2e-08 | 8.400E-09 | 3.551E-09 | 4.900E-09 | 1.200E-08 |
| 19283 | 965 | P965N | C | 7.3e-09 | 9.7e-09 | 7.3e-09 | 8.100E-09 | 1.386E-09 | 7.300E-09 | 9.700E-09 |
| 19284 | 965 | P965D | C | 2.9e-09 | 7.4e-09 | 5.2e-09 | 5.167E-09 | 2.250E-09 | 2.900E-09 | 7.400E-09 |
| 19285 | 965 | P965C | C | 8.9e-09 | 4e-09 | 2.5e-09 | 5.133E-09 | 3.347E-09 | 2.500E-09 | 8.900E-09 |
| 19286 | 965 | P965E | C | 5.7e-09 | 3.6e-09 | 9e-09 | 6.100E-09 | 2.722E-09 | 3.600E-09 | 9.000E-09 |
| 19287 | 965 | P965Q | C | 1.3e-08 | 4.5e-09 | 3.6e-08 | 1.783E-08 | 1.630E-08 | 4.500E-09 | 3.600E-08 |
| 19288 | 965 | P965G | C | 5.4e-09 | 6.6e-09 | 9.7e-09 | 7.233E-09 | 2.219E-09 | 5.400E-09 | 9.700E-09 |
| 19289 | 965 | P965H | C | 1.1e-08 | 1.7e-09 | 9.9e-09 | 7.533E-09 | 5.082E-09 | 1.700E-09 | 1.100E-08 |
| 19290 | 965 | P965I | C | 6.1e-09 | 1.5e-08 | 1.1e-08 | 1.070E-08 | 4.458E-09 | 6.100E-09 | 1.500E-08 |
| 19291 | 965 | P965L | C | 4.6e-09 | 1.1e-08 | 9.2e-09 | 8.267E-09 | 3.301E-09 | 4.600E-09 | 1.100E-08 |
| 19292 | 965 | P965K | C | 4.6e-09 | 2.8e-09 | 6.9e-09 | 4.767E-09 | 2.055E-09 | 2.800E-09 | 6.900E-09 |
| 19293 | 965 | P965M | C | 8.4e-09 | 9.6e-09 | 7.9e-09 | 8.633E-09 | 8.737E-10 | 7.900E-09 | 9.600E-09 |
| 19294 | 965 | P965F | C | 1.1e-08 | 4.8e-09 | 2.5e-09 | 6.100E-09 | 4.397E-09 | 2.500E-09 | 1.100E-08 |

| | | | | | | | | | | |
|---|---|---|---|---|---|---|---|---|---|---|
| 19295 | 965 | P965P | C | 2.9e-09 | 9.6e-09 | 8.2e-09 | 6.900E-09 | 3.534E-09 | 2.900E-09 | 9.600E-09 |
| 19296 | 965 | P965S | C | 2.2e-08 | 1.2e-08 | 6.2e-09 | 1.340E-08 | 7.992E-09 | 6.200E-09 | 2.200E-08 |
| 19297 | 965 | P965T | C | 1.3e-08 | 2.5e-09 | 1e-08 | 8.500E-09 | 5.408E-09 | 2.500E-09 | 1.300E-08 |
| 19298 | 965 | P965W | C | 6.8e-09 | 1.7e-08 | 1.6e-08 | 1.327E-08 | 5.623E-09 | 6.800E-09 | 1.700E-08 |
| 19299 | 965 | P965Y | C | 1e-08 | 6.9e-09 | 3.5e-09 | 6.800E-09 | 3.251E-09 | 3.500E-09 | 1.000E-08 |
| 19300 | 965 | P965V | C | 7.4e-09 | 1.2e-08 | 9e-09 | 9.467E-09 | 2.335E-09 | 7.400E-09 | 1.200E-08 |
| 19301 | 966 | L966A | C | 5.9e-09 | 3.5e-09 | 1.5e-08 | 8.133E-09 | 6.067E-09 | 3.500E-09 | 1.500E-08 |
| 19302 | 966 | L966R | C | 3.2e-09 | 3.9e-09 | 2.7e-09 | 3.267E-09 | 6.028E-10 | 2.700E-09 | 3.900E-09 |
| 19303 | 966 | L966N | C | 3.1e-09 | 5.4e-09 | 5.6e-09 | 4.700E-09 | 1.389E-09 | 3.100E-09 | 5.600E-09 |
| 19304 | 966 | L966D | C | 4.2e-08 | 1.9e-09 | 9.2e-09 | 1.770E-08 | 2.136E-08 | 1.900E-09 | 4.200E-08 |
| 19305 | 966 | L966C | C | 9e-09 | 7.7e-09 | 5.9e-09 | 7.533E-09 | 1.557E-09 | 5.900E-09 | 9.000E-09 |
| 19306 | 966 | L966E | C | 7.7e-09 | 9e-09 | 7.7e-09 | 8.133E-09 | 7.506E-10 | 7.700E-09 | 9.000E-09 |
| 19307 | 966 | L966Q | C | 6.1e-09 | 3.6e-09 | 1.3e-08 | 7.567E-09 | 4.869E-09 | 3.600E-09 | 1.300E-08 |
| 19308 | 966 | L966G | C | 8.6e-09 | 1.8e-08 | 3e-09 | 9.867E-09 | 7.580E-09 | 3.000E-09 | 1.800E-08 |
| 19309 | 966 | L966H | C | 6.9e-09 | 8e-09 | 6.3e-09 | 7.067E-09 | 8.622E-10 | 6.300E-09 | 8.000E-09 |
| 19310 | 966 | L966I | C | 1.1e-08 | 5.4e-09 | 6.8e-09 | 7.733E-09 | 2.914E-09 | 5.400E-09 | 1.100E-08 |
| 19311 | 966 | L966L | C | 4e-09 | 1.1e-08 | 1.9e-08 | 1.133E-08 | 7.506E-09 | 4.000E-09 | 1.900E-08 |
| 19312 | 966 | L966K | C | 5.1e-09 | 3e-09 | 1.2e-08 | 6.700E-09 | 4.709E-09 | 3.000E-09 | 1.200E-08 |
| 19313 | 966 | L966M | C | 1e-08 | 6.6e-09 | 9.9e-09 | 8.833E-09 | 1.935E-09 | 6.600E-09 | 1.000E-08 |
| 19314 | 966 | L966F | C | 7.1e-09 | 1.6e-08 | 9.6e-09 | 1.090E-08 | 4.590E-09 | 7.100E-09 | 1.600E-08 |
| 19315 | 966 | L966P | C | 2.7e-09 | 6.7e-09 | 5.6e-09 | 5.000E-09 | 2.066E-09 | 2.700E-09 | 6.700E-09 |
| 19316 | 966 | L966S | C | 6.8e-09 | 7.6e-09 | 5.4e-09 | 6.600E-09 | 1.114E-09 | 5.400E-09 | 7.600E-09 |
| 19317 | 966 | L966T | C | 1e-08 | 1.3e-08 | 1.1e-08 | 1.133E-08 | 1.528E-09 | 1.000E-08 | 1.300E-08 |
| 19318 | 966 | L966W | C | 7.5e-09 | 1.9e-08 | 3.6e-09 | 1.003E-08 | 8.006E-09 | 3.600E-09 | 1.900E-08 |
| 19319 | 966 | L966Y | C | 5.1e-09 | 2.7e-09 | 4.6e-09 | 4.133E-09 | 1.266E-09 | 2.700E-09 | 5.100E-09 |
| 19320 | 966 | L966V | C | 7.6e-09 | 8.3e-09 | 3.9e-09 | 6.600E-09 | 2.364E-09 | 3.900E-09 | 8.300E-09 |
| 19321 | 967 | A967A | C | 8.2e-09 | 8.5e-09 | 8.1e-09 | 8.267E-09 | 2.082E-10 | 8.100E-09 | 8.500E-09 |
| 19322 | 967 | A967R | C | 7e-09 | 7.3e-09 | 5.7e-09 | 6.667E-09 | 8.505E-10 | 5.700E-09 | 7.300E-09 |
| 19323 | 967 | A967N | C | 7.3e-09 | 2.2e-09 | 2.4e-08 | 1.117E-08 | 1.140E-08 | 2.200E-09 | 2.400E-08 |
| 19324 | 967 | A967D | C | 1.2e-08 | 7e-09 | 1e-08 | 9.667E-09 | 2.517E-09 | 7.000E-09 | 1.200E-08 |
| 19325 | 967 | A967C | C | 2.4e-09 | 4.4e-09 | 7.7e-09 | 4.833E-09 | 2.676E-09 | 2.400E-09 | 7.700E-09 |
| 19326 | 967 | A967E | C | 9.4e-09 | 1e-08 | 1.2e-08 | 1.047E-08 | 1.361E-09 | 9.400E-09 | 1.200E-08 |
| 19327 | 967 | A967Q | C | 2.2e-09 | 5.4e-09 | 5.1e-09 | 4.233E-09 | 1.767E-09 | 2.200E-09 | 5.400E-09 |
| 19328 | 967 | A967G | C | 1e-08 | 2.3e-09 | 1.3e-08 | 8.433E-09 | 5.519E-09 | 2.300E-09 | 1.300E-08 |
| 19329 | 967 | A967H | C | 1e-08 | 1e-08 | 4.8e-09 | 8.267E-09 | 3.002E-09 | 4.800E-09 | 1.000E-08 |
| 19330 | 967 | A967I | C | 1.3e-08 | 8.3e-09 | 5.9e-09 | 9.067E-09 | 3.612E-09 | 5.900E-09 | 1.300E-08 |
| 19331 | 967 | A967L | C | 9e-09 | 1.2e-08 | 5.9e-09 | 8.967E-09 | 3.050E-09 | 5.900E-09 | 1.200E-08 |
| 19332 | 967 | A967K | C | 9.6e-09 | 2.8e-09 | 2.7e-09 | 5.033E-09 | 3.955E-09 | 2.700E-09 | 9.600E-09 |
| 19333 | 967 | A967M | C | 7e-09 | 7e-09 | 4.2e-09 | 6.067E-09 | 1.617E-09 | 4.200E-09 | 7.000E-09 |
| 19334 | 967 | A967F | C | 2e-09 | 9.9e-09 | 5.6e-09 | 5.833E-09 | 3.955E-09 | 2.000E-09 | 9.900E-09 |
| 19335 | 967 | A967P | C | 3.2e-09 | 6.1e-09 | 6.9e-09 | 5.400E-09 | 1.947E-09 | 3.200E-09 | 6.900E-09 |
| 19336 | 967 | A967S | C | 9.8e-09 | 2.9e-09 | 7e-09 | 6.567E-09 | 3.470E-09 | 2.900E-09 | 9.800E-09 |
| 19337 | 967 | A967T | C | 8.6e-09 | 7.2e-09 | 9.9e-09 | 8.567E-09 | 1.350E-09 | 7.200E-09 | 9.900E-09 |
| 19338 | 967 | A967W | C | 6.5e-09 | 1.3e-09 | 7.5e-09 | 5.100E-09 | 3.329E-09 | 1.300E-09 | 7.500E-09 |
| 19339 | 967 | A967Y | C | 9.3e-09 | 4.8e-09 | 2e-08 | 1.137E-08 | 7.808E-09 | 4.800E-09 | 2.000E-08 |
| 19340 | 967 | A967V | C | 2.3e-09 | 8.1e-09 | 2.6e-09 | 4.333E-09 | 3.265E-09 | 2.300E-09 | 8.100E-09 |
| 19341 | 968 | P968A | C | 6.1e-09 | 4.7e-09 | 5.7e-09 | 5.500E-09 | 7.211E-10 | 4.700E-09 | 6.100E-09 |
| 19342 | 968 | P968R | C | 8e-09 | 9.9e-09 | 6.5e-09 | 8.133E-09 | 1.704E-09 | 6.500E-09 | 9.900E-09 |
| 19343 | 968 | P968N | C | 3e-09 | 1.4e-08 | 2.9e-09 | 6.633E-09 | 6.380E-09 | 2.900E-09 | 1.400E-08 |
| 19344 | 968 | P968D | C | 6.9e-09 | 6e-09 | 2.6e-08 | 1.297E-08 | 1.130E-08 | 6.000E-09 | 2.600E-08 |
| 19345 | 968 | P968C | C | 2e-08 | 7.5e-09 | 9.6e-09 | 1.237E-08 | 6.694E-09 | 7.500E-09 | 2.000E-08 |
| 19346 | 968 | P968E | C | 7.7e-09 | 6e-09 | 6.6e-09 | 6.767E-09 | 8.622E-10 | 6.000E-09 | 7.700E-09 |
| 19347 | 968 | P968Q | C | 8.2e-09 | 8.5e-09 | 7.1e-09 | 7.933E-09 | 7.371E-10 | 7.100E-09 | 8.500E-09 |
| 19348 | 968 | P968G | C | 7.3e-09 | 1.4e-08 | 1.7e-08 | 1.277E-08 | 4.966E-09 | 7.300E-09 | 1.700E-08 |
| 19349 | 968 | P968H | C | 3.6e-09 | 4.4e-09 | 6e-09 | 4.667E-09 | 1.222E-09 | 3.600E-09 | 6.000E-09 |
| 19350 | 968 | P968I | C | 7.8e-09 | 4.2e-09 | 2.3e-09 | 4.767E-09 | 2.793E-09 | 2.300E-09 | 7.800E-09 |
| 19351 | 968 | P968L | C | 4e-09 | 8.5e-09 | 9.9e-09 | 7.467E-09 | 3.083E-09 | 4.000E-09 | 9.900E-09 |
| 19352 | 968 | P968K | C | 8.6e-09 | 7e-09 | 1.2e-08 | 9.200E-09 | 2.553E-09 | 7.000E-09 | 1.200E-08 |
| 19353 | 968 | P968M | C | 5.7e-09 | 1.6e-09 | 5.9e-09 | 4.400E-09 | 2.427E-09 | 1.600E-09 | 5.900E-09 |
| 19354 | 968 | P968F | C | 7.2e-09 | 6.5e-09 | 1.6e-08 | 9.900E-09 | 5.294E-09 | 6.500E-09 | 1.600E-08 |
| 19355 | 968 | P968P | C | 9.7e-09 | 7.9e-09 | 8.2e-09 | 8.600E-09 | 9.644E-10 | 7.900E-09 | 9.700E-09 |
| 19356 | 968 | P968S | C | 8.6e-09 | 4.7e-09 | 3.4e-09 | 5.567E-09 | 2.706E-09 | 3.400E-09 | 8.600E-09 |
| 19357 | 968 | P968T | C | 2.2e-09 | 2.3e-09 | 5.7e-09 | 3.400E-09 | 1.992E-09 | 2.200E-09 | 5.700E-09 |
| 19358 | 968 | P968W | C | 5.4e-09 | 5.6e-09 | 8.7e-09 | 6.567E-09 | 1.850E-09 | 5.400E-09 | 8.700E-09 |
| 19359 | 968 | P968Y | C | 1.2e-08 | 4.6e-09 | 2.7e-09 | 6.433E-09 | 4.914E-09 | 2.700E-09 | 1.200E-08 |
| 19360 | 968 | P968V | C | 4.7e-09 | 9.2e-09 | 3.8e-09 | 5.900E-09 | 2.893E-09 | 3.800E-09 | 9.200E-09 |
| 19361 | 969 | S969A | C | 7.7e-09 | 9.2e-09 | 6.7e-09 | 7.867E-09 | 1.258E-09 | 6.700E-09 | 9.200E-09 |
| 19362 | 969 | S969R | C | 5.7e-09 | 9e-09 | 6.9e-09 | 7.200E-09 | 1.670E-09 | 5.700E-09 | 9.000E-09 |
| 19363 | 969 | S969N | C | 7.4e-09 | 4.4e-09 | 7.5e-09 | 6.433E-09 | 1.762E-09 | 4.400E-09 | 7.500E-09 |
| 19364 | 969 | S969D | C | 7.3e-09 | 1e-08 | 2.2e-08 | 1.310E-08 | 7.825E-09 | 7.300E-09 | 2.200E-08 |
| 19365 | 969 | S969C | C | 7.6e-09 | 3.5e-09 | 4.8e-09 | 5.300E-09 | 2.095E-09 | 3.500E-09 | 7.600E-09 |
| 19366 | 969 | S969E | C | 6.2e-09 | 9.5e-09 | 4.5e-09 | 6.733E-09 | 2.542E-09 | 4.500E-09 | 9.500E-09 |
| 19367 | 969 | S969Q | C | 9.8e-09 | 8.2e-09 | 1.7e-08 | 1.167E-08 | 4.688E-09 | 8.200E-09 | 1.700E-08 |
| 19368 | 969 | S969G | C | 1.5e-08 | 1.3e-08 | 7.5e-09 | 1.183E-08 | 3.884E-09 | 7.500E-09 | 1.500E-08 |
| 19369 | 969 | S969H | C | 1.5e-08 | 2.5e-08 | 9.3e-09 | 1.643E-08 | 7.948E-09 | 9.300E-09 | 2.500E-08 |
| 19370 | 969 | S969I | C | 1.4e-08 | 8e-09 | 9.2e-09 | 1.040E-08 | 3.175E-09 | 8.000E-09 | 1.400E-08 |
| 19371 | 969 | S969L | C | 6.7e-09 | 7.3e-09 | 4e-09 | 6.000E-09 | 1.758E-09 | 4.000E-09 | 7.300E-09 |
| 19372 | 969 | S969K | C | 1.7e-08 | 1.6e-08 | 4.2e-09 | 1.240E-08 | 7.119E-09 | 4.200E-09 | 1.700E-08 |
| 19373 | 969 | S969M | C | 1.7e-08 | 1e-08 | 9.3e-09 | 1.210E-08 | 4.258E-09 | 9.300E-09 | 1.700E-08 |
| 19374 | 969 | S969F | C | 3.2e-09 | 1.9e-08 | 5.9e-09 | 9.367E-09 | 8.451E-09 | 3.200E-09 | 1.900E-08 |
| 19375 | 969 | S969P | C | 1.1e-08 | 4.7e-09 | 1.2e-08 | 9.233E-09 | 3.958E-09 | 4.700E-09 | 1.200E-08 |
| 19376 | 969 | S969S | C | 3.4e-09 | 6.1e-09 | 1.3e-08 | 7.500E-09 | 4.951E-09 | 3.400E-09 | 1.300E-08 |
| 19377 | 969 | S969T | C | 1.1e-08 | 3e-09 | 1.1e-08 | 8.333E-09 | 4.619E-09 | 3.000E-09 | 1.100E-08 |
| 19378 | 969 | S969W | C | 1.5e-08 | 8.2e-09 | 6.1e-09 | 9.767E-09 | 4.652E-09 | 6.100E-09 | 1.500E-08 |
| 19379 | 969 | S969Y | C | 6.5e-09 | 1.2e-08 | 5.2e-09 | 7.900E-09 | 3.610E-09 | 5.200E-09 | 1.200E-08 |

| | | | | | | | | | | |
|---|---|---|---|---|---|---|---|---|---|---|
| 19380 | 969 | S969V | C | 5.3e-09 | 9.3e-09 | 6.8e-09 | 7.133E-09 | 2.021E-09 | 5.300E-09 | 9.300E-09 |
| 19381 | 970 | S970A | C | 1e-08 | 1.7e-09 | 2.6e-08 | 1.257E-08 | 1.235E-08 | 1.700E-09 | 2.600E-08 |
| 19382 | 970 | S970R | C | 4.4e-09 | 1.8e-08 | 1.5e-08 | 1.247E-08 | 7.145E-09 | 4.400E-09 | 1.800E-08 |
| 19383 | 970 | S970N | C | 6.3e-09 | 6e-09 | 1.3e-08 | 8.433E-09 | 3.958E-09 | 6.000E-09 | 1.300E-08 |
| 19384 | 970 | S970D | C | 5.4e-09 | 5.9e-09 | 6.9e-09 | 6.067E-09 | 7.638E-10 | 5.400E-09 | 6.900E-09 |
| 19385 | 970 | S970C | C | 1.3e-08 | 4.7e-09 | 3.4e-09 | 7.033E-09 | 5.208E-09 | 3.400E-09 | 1.300E-08 |
| 19386 | 970 | S970E | C | 2.6e-08 | 1.2e-08 | 2e-09 | 1.333E-08 | 1.206E-08 | 2.000E-09 | 2.600E-08 |
| 19387 | 970 | S970Q | C | 8e-09 | 4.6e-09 | 2.3e-08 | 1.187E-08 | 9.790E-09 | 4.600E-09 | 2.300E-08 |
| 19388 | 970 | S970G | C | 9.9e-09 | 1.4e-08 | 5.3e-09 | 9.733E-09 | 4.352E-09 | 5.300E-09 | 1.400E-08 |
| 19389 | 970 | S970H | C | 5.2e-09 | 4.9e-09 | 1.2e-08 | 7.367E-09 | 4.015E-09 | 4.900E-09 | 1.200E-08 |
| 19390 | 970 | S970I | C | 2.8e-09 | 3.8e-09 | 8.1e-09 | 4.900E-09 | 2.816E-09 | 2.800E-09 | 8.100E-09 |
| 19391 | 970 | S970L | C | 5.3e-09 | 8e-09 | 9e-09 | 7.433E-09 | 1.914E-09 | 5.300E-09 | 9.000E-09 |
| 19392 | 970 | S970K | C | 6.2e-09 | 4.9e-09 | 5e-09 | 5.367E-09 | 7.234E-10 | 4.900E-09 | 6.200E-09 |
| 19393 | 970 | S970M | C | 9.6e-09 | 4.4e-09 | 5.7e-09 | 6.567E-09 | 2.706E-09 | 4.400E-09 | 9.600E-09 |
| 19394 | 970 | S970F | C | 1.2e-08 | 9.6e-09 | 4.2e-09 | 8.600E-09 | 3.995E-09 | 4.200E-09 | 1.200E-08 |
| 19395 | 970 | S970P | C | 1.4e-08 | 8e-09 | 9.5e-09 | 1.050E-08 | 3.122E-09 | 8.000E-09 | 1.400E-08 |
| 19396 | 970 | S970S | C | 5.1e-09 | 9.1e-09 | 2.3e-08 | 1.240E-08 | 9.395E-09 | 5.100E-09 | 2.300E-08 |
| 19397 | 970 | S970T | C | 1.8e-08 | 6.7e-09 | 1.2e-08 | 1.223E-08 | 5.654E-09 | 6.700E-09 | 1.800E-08 |
| 19398 | 970 | S970W | C | 3.7e-09 | 7.9e-09 | 8e-09 | 6.533E-09 | 2.454E-09 | 3.700E-09 | 8.000E-09 |
| 19399 | 970 | S970Y | C | 5.6e-09 | 6.5e-09 | 4.7e-09 | 5.600E-09 | 9.000E-10 | 4.700E-09 | 6.500E-09 |
| 19400 | 970 | S970V | C | 7.3e-09 | 2.3e-09 | 1.6e-08 | 8.533E-09 | 6.933E-09 | 2.300E-09 | 1.600E-08 |
| 19401 | 971 | K971A | C | 1.2e-08 | 1.1e-08 | 4.1e-09 | 9.033E-09 | 4.302E-09 | 4.100E-09 | 1.200E-08 |
| 19402 | 971 | K971R | C | 6.2e-09 | 1.1e-08 | 3.2e-09 | 6.800E-09 | 3.934E-09 | 3.200E-09 | 1.100E-08 |
| 19403 | 971 | K971N | C | 5.6e-09 | 3.8e-09 | 4.7e-09 | 4.700E-09 | 9.000E-10 | 3.800E-09 | 5.600E-09 |
| 19404 | 971 | K971D | C | 3.2e-09 | 4.2e-09 | 7.2e-09 | 4.867E-09 | 2.082E-09 | 3.200E-09 | 7.200E-09 |
| 19405 | 971 | K971C | C | 5.6e-09 | 8.3e-09 | 4.4e-09 | 6.100E-09 | 1.997E-09 | 4.400E-09 | 8.300E-09 |
| 19406 | 971 | K971E | C | 5.7e-09 | 6.6e-09 | 1.3e-08 | 8.433E-09 | 3.980E-09 | 5.700E-09 | 1.300E-08 |
| 19407 | 971 | K971Q | C | 5.3e-09 | 3.2e-09 | 4.3e-09 | 4.267E-09 | 1.050E-09 | 3.200E-09 | 5.300E-09 |
| 19408 | 971 | K971G | C | 9.5e-09 | 6.3e-09 | 8e-09 | 7.933E-09 | 1.601E-09 | 6.300E-09 | 9.500E-09 |
| 19409 | 971 | K971H | C | 2.8e-09 | 5.1e-09 | 1.1e-08 | 6.300E-09 | 4.230E-09 | 2.800E-09 | 1.100E-08 |
| 19410 | 971 | K971I | C | 1.3e-08 | 8.1e-09 | 2.5e-09 | 7.867E-09 | 5.254E-09 | 2.500E-09 | 1.300E-08 |
| 19411 | 971 | K971L | C | 1.2e-08 | 1.9e-08 | 1e-08 | 1.367E-08 | 4.726E-09 | 1.000E-08 | 1.900E-08 |
| 19412 | 971 | K971K | C | 2.6e-09 | 1.6e-08 | 8.5e-09 | 9.033E-09 | 6.716E-09 | 2.600E-09 | 1.600E-08 |
| 19413 | 971 | K971M | C | 3e-08 | 1e-08 | 1.5e-08 | 1.833E-08 | 1.041E-08 | 1.000E-08 | 3.000E-08 |
| 19414 | 971 | K971F | C | 9.3e-09 | 6.6e-09 | 5.7e-09 | 7.200E-09 | 1.873E-09 | 5.700E-09 | 9.300E-09 |
| 19415 | 971 | K971P | C | 4.1e-09 | 4.7e-09 | 5.8e-09 | 4.867E-09 | 8.622E-10 | 4.100E-09 | 5.800E-09 |
| 19416 | 971 | K971S | C | 1.4e-08 | 3.7e-09 | 1.3e-08 | 1.023E-08 | 5.680E-09 | 3.700E-09 | 1.400E-08 |
| 19417 | 971 | K971T | C | 9.5e-09 | 7.8e-09 | 1.1e-08 | 9.433E-09 | 1.601E-09 | 7.800E-09 | 1.100E-08 |
| 19418 | 971 | K971W | C | 9.7e-09 | 1.5e-08 | 1e-08 | 1.157E-08 | 2.977E-09 | 9.700E-09 | 1.500E-08 |
| 19419 | 971 | K971Y | C | 1.4e-08 | 1.7e-08 | 6.3e-09 | 1.243E-08 | 5.519E-09 | 6.300E-09 | 1.700E-08 |
| 19420 | 971 | K971V | C | 1.5e-08 | 1e-08 | 6.9e-09 | 1.063E-08 | 4.087E-09 | 6.900E-09 | 1.500E-08 |
| 19421 | 972 | S972A | C | 3.1e-09 | 8.7e-09 | 2.3e-08 | 1.160E-08 | 1.026E-08 | 3.100E-09 | 2.300E-08 |
| 19422 | 972 | S972R | C | 3.8e-09 | 5.6e-09 | 1.2e-08 | 7.133E-09 | 4.310E-09 | 3.800E-09 | 1.200E-08 |
| 19423 | 972 | S972N | C | 6.8e-09 | 4.7e-09 | 7e-09 | 6.167E-09 | 1.274E-09 | 4.700E-09 | 7.000E-09 |
| 19424 | 972 | S972D | C | 8.3e-09 | 9.6e-09 | 1.5e-08 | 1.097E-08 | 3.553E-09 | 8.300E-09 | 1.500E-08 |
| 19425 | 972 | S972C | C | 1.8e-08 | 6e-09 | 3.4e-08 | 1.933E-08 | 1.405E-08 | 6.000E-09 | 3.400E-08 |
| 19426 | 972 | S972E | C | 2.2e-08 | 1.4e-08 | 4.4e-09 | 1.347E-08 | 8.812E-09 | 4.400E-09 | 2.200E-08 |
| 19427 | 972 | S972Q | C | 6.4e-09 | 2.3e-08 | 4.1e-09 | 1.117E-08 | 1.031E-08 | 4.100E-09 | 2.300E-08 |
| 19428 | 972 | S972G | C | 4.4e-09 | 3.7e-09 | 6.7e-09 | 4.933E-09 | 1.570E-09 | 3.700E-09 | 6.700E-09 |
| 19429 | 972 | S972H | C | 7.7e-09 | 1.7e-09 | 8.1e-09 | 5.833E-09 | 3.585E-09 | 1.700E-09 | 8.100E-09 |
| 19430 | 972 | S972I | C | 6.5e-09 | 9.2e-09 | 7.5e-09 | 7.733E-09 | 1.365E-09 | 6.500E-09 | 9.200E-09 |
| 19431 | 972 | S972L | C | 5e-09 | 1e-08 | 1.5e-08 | 1.000E-08 | 5.000E-09 | 5.000E-09 | 1.500E-08 |
| 19432 | 972 | S972K | C | 5.7e-09 | 7.9e-09 | 6.7e-09 | 6.767E-09 | 1.102E-09 | 5.700E-09 | 7.900E-09 |
| 19433 | 972 | S972M | C | 1.4e-09 | 7.5e-09 | 1.9e-08 | 9.300E-09 | 8.937E-09 | 1.400E-09 | 1.900E-08 |
| 19434 | 972 | S972F | C | 2.1e-09 | 5.4e-09 | 4.8e-09 | 4.100E-09 | 1.758E-09 | 2.100E-09 | 5.400E-09 |
| 19435 | 972 | S972P | C | 7.3e-09 | 8.1e-09 | 4.8e-09 | 6.733E-09 | 1.721E-09 | 4.800E-09 | 8.100E-09 |
| 19436 | 972 | S972S | C | 3.1e-09 | 4.1e-09 | 1.5e-08 | 7.400E-09 | 6.601E-09 | 3.100E-09 | 1.500E-08 |
| 19437 | 972 | S972T | C | 3.3e-09 | 9.5e-09 | 6.9e-09 | 6.567E-09 | 3.113E-09 | 3.300E-09 | 9.500E-09 |
| 19438 | 972 | S972W | C | 2e-08 | 6.4e-09 | 1e-08 | 1.213E-08 | 7.047E-09 | 6.400E-09 | 2.000E-08 |
| 19439 | 972 | S972Y | C | 4.2e-09 | 2.8e-09 | 5.1e-09 | 4.033E-09 | 1.159E-09 | 2.800E-09 | 5.100E-09 |
| 19440 | 972 | S972V | C | 8.8e-09 | 5.4e-09 | 1.2e-08 | 8.733E-09 | 3.301E-09 | 5.400E-09 | 1.200E-08 |
| 19441 | 973 | T973A | C | 8.9e-09 | 6.1e-09 | 7.8e-09 | 7.600E-09 | 1.411E-09 | 6.100E-09 | 8.900E-09 |
| 19442 | 973 | T973R | C | 5.3e-09 | 5.3e-09 | 1.3e-08 | 7.867E-09 | 4.446E-09 | 5.300E-09 | 1.300E-08 |
| 19443 | 973 | T973N | C | 3.5e-08 | 1.8e-08 | 1.4e-08 | 2.233E-08 | 1.115E-08 | 1.400E-08 | 3.500E-08 |
| 19444 | 973 | T973D | C | 8.1e-09 | 2.3e-08 | 6.5e-09 | 1.253E-08 | 9.100E-09 | 6.500E-09 | 2.300E-08 |
| 19445 | 973 | T973C | C | 2.6e-09 | 6.7e-09 | 1.1e-08 | 6.767E-09 | 4.200E-09 | 2.600E-09 | 1.100E-08 |
| 19446 | 973 | T973E | C | 1.2e-08 | 7.4e-09 | 9.7e-09 | 9.700E-09 | 2.300E-09 | 7.400E-09 | 1.200E-08 |
| 19447 | 973 | T973Q | C | 5.1e-09 | 1e-08 | 2.7e-09 | 5.933E-09 | 3.721E-09 | 2.700E-09 | 1.000E-08 |
| 19448 | 973 | T973G | C | 6e-09 | 1.9e-08 | 8.6e-09 | 1.120E-08 | 6.879E-09 | 6.000E-09 | 1.900E-08 |
| 19449 | 973 | T973H | C | 8.2e-09 | 7.6e-09 | 3.2e-09 | 6.333E-09 | 2.730E-09 | 3.200E-09 | 8.200E-09 |
| 19450 | 973 | T973I | C | 2.1e-08 | 4.7e-09 | 1e-08 | 1.190E-08 | 8.314E-09 | 4.700E-09 | 2.100E-08 |
| 19451 | 973 | T973L | C | 3e-09 | 2.5e-09 | 1.5e-08 | 6.833E-09 | 7.077E-09 | 2.500E-09 | 1.500E-08 |
| 19452 | 973 | T973K | C | 3.1e-08 | 9.8e-09 | 8e-09 | 1.627E-08 | 1.279E-08 | 8.000E-09 | 3.100E-08 |
| 19453 | 973 | T973M | C | 3.7e-09 | 7e-09 | 2.7e-09 | 4.467E-09 | 2.250E-09 | 2.700E-09 | 7.000E-09 |
| 19454 | 973 | T973F | C | 2.9e-09 | 6.1e-09 | 2e-09 | 3.667E-09 | 2.155E-09 | 2.000E-09 | 6.100E-09 |
| 19455 | 973 | T973P | C | 1.7e-09 | 6.3e-09 | 1.1e-08 | 6.333E-09 | 4.650E-09 | 1.700E-09 | 1.100E-08 |
| 19456 | 973 | T973S | C | 3.7e-09 | 4.6e-09 | 1.1e-08 | 6.433E-09 | 3.980E-09 | 3.700E-09 | 1.100E-08 |
| 19457 | 973 | T973T | C | 3.1e-09 | 7.3e-09 | 7.9e-09 | 6.100E-09 | 2.615E-09 | 3.100E-09 | 7.900E-09 |
| 19458 | 973 | T973W | C | 1.2e-08 | 7.3e-09 | 1e-08 | 9.767E-09 | 2.359E-09 | 7.300E-09 | 1.200E-08 |
| 19459 | 973 | T973Y | C | 1.4e-08 | 7.2e-09 | 5.5e-09 | 8.900E-09 | 4.498E-09 | 5.500E-09 | 1.400E-08 |
| 19460 | 973 | T973V | C | 7.2e-09 | 7.4e-09 | 1e-08 | 8.200E-09 | 1.562E-09 | 7.200E-09 | 1.000E-08 |
| 19461 | 974 | S974A | C | 3.7e-09 | 1.7e-08 | 1.3e-08 | 1.123E-08 | 6.824E-09 | 3.700E-09 | 1.700E-08 |
| 19462 | 974 | S974R | C | 7e-09 | 7.5e-09 | 1.8e-08 | 1.083E-08 | 6.212E-09 | 7.000E-09 | 1.800E-08 |
| 19463 | 974 | S974N | C | 9.1e-09 | 5.6e-09 | 4e-09 | 6.233E-09 | 2.608E-09 | 4.000E-09 | 9.100E-09 |
| 19464 | 974 | S974D | C | 1.6e-08 | 9.2e-09 | 7.4e-09 | 1.087E-08 | 4.536E-09 | 7.400E-09 | 1.600E-08 |

| | | | | | | | | | | |
|---|---|---|---|---|---|---|---|---|---|---|
| 19465 | 974 | S974C | C | 1.2e-08 | 1.2e-08 | 5.4e-09 | 9.800E-09 | 3.811E-09 | 5.400E-09 | 1.200E-08 |
| 19466 | 974 | S974E | C | 7e-09 | 1.7e-08 | 1.7e-08 | 1.367E-08 | 5.774E-09 | 7.000E-09 | 1.700E-08 |
| 19467 | 974 | S974Q | C | 4.4e-09 | 9.9e-09 | 5.7e-09 | 6.667E-09 | 2.875E-09 | 4.400E-09 | 9.900E-09 |
| 19468 | 974 | S974G | C | 6.1e-09 | 5e-09 | 2e-08 | 1.037E-08 | 8.361E-09 | 5.000E-09 | 2.000E-08 |
| 19469 | 974 | S974H | C | 2.4e-09 | 1.5e-08 | 8.7e-09 | 8.700E-09 | 6.300E-09 | 2.400E-09 | 1.500E-08 |
| 19470 | 974 | S974I | C | 4.5e-09 | 2e-08 | 3e-09 | 9.167E-09 | 9.412E-09 | 3.000E-09 | 2.000E-08 |
| 19471 | 974 | S974L | C | 1.1e-08 | 8.6e-09 | 1.5e-08 | 1.153E-08 | 3.233E-09 | 8.600E-09 | 1.500E-08 |
| 19472 | 974 | S974K | C | 6.2e-09 | 5.2e-09 | 7.9e-09 | 6.433E-09 | 1.365E-09 | 5.200E-09 | 7.900E-09 |
| 19473 | 974 | S974M | C | 1.6e-08 | 3.8e-09 | 1.4e-08 | 1.127E-08 | 6.543E-09 | 3.800E-09 | 1.600E-08 |
| 19474 | 974 | S974F | C | 2.6e-09 | 1.6e-08 | 8e-09 | 8.867E-09 | 6.742E-09 | 2.600E-09 | 1.600E-08 |
| 19475 | 974 | S974P | C | 1.3e-08 | 1.5e-09 | 8e-09 | 7.500E-09 | 5.766E-09 | 1.500E-09 | 1.300E-08 |
| 19476 | 974 | S974S | C | 5.2e-09 | 1.1e-08 | 1.1e-08 | 9.067E-09 | 3.349E-09 | 5.200E-09 | 1.100E-08 |
| 19477 | 974 | S974T | C | 4.9e-09 | 6.4e-09 | 2.4e-09 | 4.567E-09 | 2.021E-09 | 2.400E-09 | 6.400E-09 |
| 19478 | 974 | S974W | C | 1e-08 | 3.3e-09 | 1.2e-08 | 8.433E-09 | 4.557E-09 | 3.300E-09 | 1.200E-08 |
| 19479 | 974 | S974Y | C | 8.7e-09 | 2.9e-08 | 3.7e-09 | 1.380E-08 | 1.340E-08 | 3.700E-09 | 2.900E-08 |
| 19480 | 974 | S974V | C | 2.5e-09 | 3.5e-09 | 4.8e-09 | 3.600E-09 | 1.153E-09 | 2.500E-09 | 4.800E-09 |
| 19481 | 975 | G975A | C | 2.1e-09 | 8.7e-09 | 2.3e-09 | 4.367E-09 | 3.754E-09 | 2.100E-09 | 8.700E-09 |
| 19482 | 975 | G975R | C | 1.3e-08 | 9.1e-09 | 5.5e-09 | 9.200E-09 | 3.751E-09 | 5.500E-09 | 1.300E-08 |
| 19483 | 975 | G975N | C | 8.5e-09 | 6.8e-09 | 7.3e-09 | 7.533E-09 | 8.737E-10 | 6.800E-09 | 8.500E-09 |
| 19484 | 975 | G975D | C | 5.5e-09 | 1.9e-08 | 1.2e-08 | 1.217E-08 | 6.752E-09 | 5.500E-09 | 1.900E-08 |
| 19485 | 975 | G975C | C | 3.4e-09 | 3.1e-09 | 1.3e-08 | 6.500E-09 | 5.631E-09 | 3.100E-09 | 1.300E-08 |
| 19486 | 975 | G975E | C | 1.4e-08 | 6e-09 | 4.8e-09 | 8.267E-09 | 5.001E-09 | 4.800E-09 | 1.400E-08 |
| 19487 | 975 | G975Q | C | 4.5e-09 | 3.8e-09 | 8.7e-09 | 5.667E-09 | 2.650E-09 | 3.800E-09 | 8.700E-09 |
| 19488 | 975 | G975G | C | 2e-08 | 6.2e-09 | 1.5e-08 | 1.373E-08 | 6.987E-09 | 6.200E-09 | 2.000E-08 |
| 19489 | 975 | G975H | C | 8.3e-09 | 2.4e-09 | 6.2e-09 | 5.633E-09 | 2.991E-09 | 2.400E-09 | 8.300E-09 |
| 19490 | 975 | G975I | C | 4.9e-09 | 1.5e-08 | 4.4e-09 | 8.100E-09 | 5.981E-09 | 4.400E-09 | 1.500E-08 |
| 19491 | 975 | G975L | C | 3e-08 | 6.6e-09 | 3.6e-09 | 1.340E-08 | 1.445E-08 | 3.600E-09 | 3.000E-08 |
| 19492 | 975 | G975K | C | 1.4e-08 | 2.4e-09 | 8.6e-09 | 8.333E-09 | 5.805E-09 | 2.400E-09 | 1.400E-08 |
| 19493 | 975 | G975M | C | 5.9e-09 | 8.3e-09 | 5.1e-09 | 6.433E-09 | 1.665E-09 | 5.100E-09 | 8.300E-09 |
| 19494 | 975 | G975F | C | 7.1e-09 | 7.2e-09 | 3.2e-09 | 5.833E-09 | 2.281E-09 | 3.200E-09 | 7.200E-09 |
| 19495 | 975 | G975P | C | 6.2e-09 | 4.3e-09 | 1.7e-09 | 4.067E-09 | 2.259E-09 | 1.700E-09 | 6.200E-09 |
| 19496 | 975 | G975S | C | 1.9e-08 | 1.2e-08 | 1e-08 | 1.367E-08 | 4.726E-09 | 1.000E-08 | 1.900E-08 |
| 19497 | 975 | G975T | C | 1.3e-08 | 5.8e-09 | 5.9e-09 | 8.233E-09 | 4.128E-09 | 5.800E-09 | 1.300E-08 |
| 19498 | 975 | G975W | C | 6.5e-09 | 7e-09 | 7.4e-09 | 6.967E-09 | 4.509E-10 | 6.500E-09 | 7.400E-09 |
| 19499 | 975 | G975Y | C | 7.2e-09 | 8.4e-09 | 6.3e-09 | 7.300E-09 | 1.054E-09 | 6.300E-09 | 8.400E-09 |
| 19500 | 975 | G975V | C | 7e-09 | 4.4e-09 | 4.2e-09 | 5.200E-09 | 1.562E-09 | 4.200E-09 | 7.000E-09 |
| 19501 | 976 | G976A | C | 5.3e-09 | 4.2e-09 | 4.4e-09 | 4.633E-09 | 5.859E-10 | 4.200E-09 | 5.300E-09 |
| 19502 | 976 | G976R | C | 6.5e-09 | 4.1e-09 | 5.3e-09 | 5.300E-09 | 1.200E-09 | 4.100E-09 | 6.500E-09 |
| 19503 | 976 | G976N | C | 3.2e-09 | 8e-09 | 7e-09 | 6.067E-09 | 2.532E-09 | 3.200E-09 | 8.000E-09 |
| 19504 | 976 | G976D | C | 8.8e-09 | 1.5e-08 | 1e-08 | 1.127E-08 | 3.288E-09 | 8.800E-09 | 1.500E-08 |
| 19505 | 976 | G976C | C | 3e-09 | 1.6e-08 | 1.6e-08 | 1.167E-08 | 7.506E-09 | 3.000E-09 | 1.600E-08 |
| 19506 | 976 | G976E | C | 5.3e-09 | 6e-09 | 2.8e-09 | 4.700E-09 | 1.682E-09 | 2.800E-09 | 6.000E-09 |
| 19507 | 976 | G976Q | C | 2.1e-09 | 5e-09 | 9.1e-09 | 5.400E-09 | 3.517E-09 | 2.100E-09 | 9.100E-09 |
| 19508 | 976 | G976G | C | 5.8e-09 | 5.5e-09 | 3e-08 | 1.377E-08 | 1.406E-08 | 5.500E-09 | 3.000E-08 |
| 19509 | 976 | G976H | C | 5.3e-09 | 1.5e-08 | 6.6e-09 | 8.967E-09 | 5.265E-09 | 5.300E-09 | 1.500E-08 |
| 19510 | 976 | G976I | C | 4.4e-09 | 2.8e-09 | 5.9e-09 | 4.367E-09 | 1.550E-09 | 2.800E-09 | 5.900E-09 |
| 19511 | 976 | G976L | C | 7e-09 | 6.7e-09 | 6.8e-09 | 6.833E-09 | 1.528E-10 | 6.700E-09 | 7.000E-09 |
| 19512 | 976 | G976K | C | 5.1e-09 | 1.5e-08 | 1.2e-08 | 1.070E-08 | 5.076E-09 | 5.100E-09 | 1.500E-08 |
| 19513 | 976 | G976M | C | 8.2e-09 | 6e-09 | 1.1e-08 | 8.400E-09 | 2.506E-09 | 6.000E-09 | 1.100E-08 |
| 19514 | 976 | G976F | C | 4.6e-09 | 2.7e-08 | 4e-09 | 1.187E-08 | 1.311E-08 | 4.000E-09 | 2.700E-08 |
| 19515 | 976 | G976P | C | 4.8e-09 | 1.2e-08 | 4.1e-09 | 6.967E-09 | 4.373E-09 | 4.100E-09 | 1.200E-08 |
| 19516 | 976 | G976S | C | 6.4e-09 | 7e-09 | 2e-08 | 1.113E-08 | 7.685E-09 | 6.400E-09 | 2.000E-08 |
| 19517 | 976 | G976T | C | 5.4e-09 | 6.5e-09 | 6.4e-09 | 6.100E-09 | 6.083E-10 | 5.400E-09 | 6.500E-09 |
| 19518 | 976 | G976W | C | 1.6e-08 | 6.7e-09 | 8.9e-09 | 1.053E-08 | 4.860E-09 | 6.700E-09 | 1.600E-08 |
| 19519 | 976 | G976Y | C | 3.4e-09 | 4.4e-09 | 6.7e-09 | 4.833E-09 | 1.692E-09 | 3.400E-09 | 6.700E-09 |
| 19520 | 976 | G976V | C | 4.1e-09 | 3.2e-09 | 4.3e-09 | 3.867E-09 | 5.859E-10 | 3.200E-09 | 4.300E-09 |
| 19521 | 977 | T977A | C | 5.1e-09 | 3.4e-09 | 1.4e-08 | 7.500E-09 | 5.693E-09 | 3.400E-09 | 1.400E-08 |
| 19522 | 977 | T977R | C | 7.2e-09 | 5.2e-09 | 8.7e-09 | 7.033E-09 | 1.756E-09 | 5.200E-09 | 8.700E-09 |
| 19523 | 977 | T977N | C | 4.9e-09 | 6.4e-09 | 8.7e-09 | 6.667E-09 | 1.914E-09 | 4.900E-09 | 8.700E-09 |
| 19524 | 977 | T977D | C | 6.2e-09 | 6.5e-09 | 8.5e-09 | 7.067E-09 | 1.250E-09 | 6.200E-09 | 8.500E-09 |
| 19525 | 977 | T977C | C | 8.5e-09 | 1.2e-08 | 4.8e-09 | 8.433E-09 | 3.600E-09 | 4.800E-09 | 1.200E-08 |
| 19526 | 977 | T977E | C | 2.2e-09 | 5.4e-09 | 7e-09 | 4.867E-09 | 2.444E-09 | 2.200E-09 | 7.000E-09 |
| 19527 | 977 | T977Q | C | 9.3e-09 | 4e-09 | 9.8e-09 | 7.700E-09 | 3.214E-09 | 4.000E-09 | 9.800E-09 |
| 19528 | 977 | T977G | C | 7.4e-09 | 2e-08 | 8.6e-09 | 1.200E-08 | 6.954E-09 | 7.400E-09 | 2.000E-08 |
| 19529 | 977 | T977H | C | 9.6e-09 | 6.8e-09 | 7.7e-09 | 8.033E-09 | 1.429E-09 | 6.800E-09 | 9.600E-09 |
| 19530 | 977 | T977I | C | 2.6e-08 | 5e-09 | 9e-09 | 1.333E-08 | 1.115E-08 | 5.000E-09 | 2.600E-08 |
| 19531 | 977 | T977L | C | 6.4e-09 | 7.8e-09 | 7.2e-09 | 7.133E-09 | 7.024E-10 | 6.400E-09 | 7.800E-09 |
| 19532 | 977 | T977K | C | 9.9e-09 | 4.7e-09 | 2.8e-09 | 5.800E-09 | 3.676E-09 | 2.800E-09 | 9.900E-09 |
| 19533 | 977 | T977M | C | 1.1e-08 | 5.3e-09 | 8.5e-09 | 8.267E-09 | 2.857E-09 | 5.300E-09 | 1.100E-08 |
| 19534 | 977 | T977F | C | 1.1e-08 | 6.7e-09 | 1.8e-08 | 1.190E-08 | 5.704E-09 | 6.700E-09 | 1.800E-08 |
| 19535 | 977 | T977P | C | 9.4e-09 | 1.1e-08 | 6.6e-09 | 9.000E-09 | 2.227E-09 | 6.600E-09 | 1.100E-08 |
| 19536 | 977 | T977S | C | 2.7e-09 | 5.6e-09 | 1.6e-08 | 8.100E-09 | 6.994E-09 | 2.700E-09 | 1.600E-08 |
| 19537 | 977 | T977T | C | 1.2e-08 | 4.6e-09 | 5.3e-09 | 7.300E-09 | 4.085E-09 | 4.600E-09 | 1.200E-08 |
| 19538 | 977 | T977W | C | 8e-09 | 4e-09 | 8.6e-09 | 6.867E-09 | 2.501E-09 | 4.000E-09 | 8.600E-09 |
| 19539 | 977 | T977Y | C | 3.3e-09 | 1.3e-08 | 7e-09 | 7.767E-09 | 4.895E-09 | 3.300E-09 | 1.300E-08 |
| 19540 | 977 | T977V | C | 5.1e-09 | 6.6e-09 | 6.5e-09 | 6.067E-09 | 8.386E-10 | 5.100E-09 | 6.600E-09 |
| 19541 | 978 | A978A | C | 5e-09 | 4.4e-09 | 1.4e-08 | 7.800E-09 | 5.378E-09 | 4.400E-09 | 1.400E-08 |
| 19542 | 978 | A978R | C | 6.4e-09 | 9.5e-09 | 3.4e-09 | 6.433E-09 | 3.050E-09 | 3.400E-09 | 9.500E-09 |
| 19543 | 978 | A978N | C | 8.5e-09 | 2.8e-09 | 7.4e-09 | 6.233E-09 | 3.024E-09 | 2.800E-09 | 8.500E-09 |
| 19544 | 978 | A978D | C | 1.1e-08 | 5.3e-09 | 9.3e-09 | 8.533E-09 | 2.926E-09 | 5.300E-09 | 1.100E-08 |
| 19545 | 978 | A978C | C | 2.7e-09 | 3e-09 | 1.2e-08 | 5.900E-09 | 5.285E-09 | 2.700E-09 | 1.200E-08 |
| 19546 | 978 | A978E | C | 1.5e-08 | 8e-09 | 8.1e-09 | 1.037E-08 | 4.013E-09 | 8.000E-09 | 1.500E-08 |
| 19547 | 978 | A978Q | C | 1.9e-08 | 4.3e-09 | 1.5e-08 | 1.277E-08 | 7.600E-09 | 4.300E-09 | 1.900E-08 |
| 19548 | 978 | A978G | C | 8.3e-09 | 3.9e-09 | 5.4e-09 | 5.867E-09 | 2.237E-09 | 3.900E-09 | 8.300E-09 |
| 19549 | 978 | A978H | C | 9.6e-09 | 2e-08 | 4e-09 | 1.120E-08 | 8.119E-09 | 4.000E-09 | 2.000E-08 |

| | | | | | | | | | | |
|---|---|---|---|---|---|---|---|---|---|---|
| 19550 | 978 | A978I | C | 1.1e-08 | 4.5e-09 | 8.3e-09 | 7.933E-09 | 3.265E-09 | 4.500E-09 | 1.100E-08 |
| 19551 | 978 | A978L | C | 3.5e-09 | 9.2e-09 | 2e-09 | 4.900E-09 | 3.799E-09 | 2.000E-09 | 9.200E-09 |
| 19552 | 978 | A978K | C | 1.2e-08 | 9.3e-09 | 2.1e-08 | 1.410E-08 | 6.126E-09 | 9.300E-09 | 2.100E-08 |
| 19553 | 978 | A978M | C | 3.9e-09 | 8.8e-09 | 4.3e-09 | 5.667E-09 | 2.721E-09 | 3.900E-09 | 8.800E-09 |
| 19554 | 978 | A978F | C | 1.4e-08 | 1.8e-09 | 1.3e-08 | 9.600E-09 | 6.773E-09 | 1.800E-09 | 1.400E-08 |
| 19555 | 978 | A978P | C | 3.7e-09 | 5e-09 | 2.2e-09 | 3.633E-09 | 1.401E-09 | 2.200E-09 | 5.000E-09 |
| 19556 | 978 | A978S | C | 7e-09 | 3.3e-09 | 1.3e-09 | 3.867E-09 | 2.892E-09 | 1.300E-09 | 7.000E-09 |
| 19557 | 978 | A978T | C | 7.9e-09 | 5.5e-09 | 1e-08 | 7.800E-09 | 2.252E-09 | 5.500E-09 | 1.000E-08 |
| 19558 | 978 | A978W | C | 6.2e-09 | 8.6e-09 | 5.9e-09 | 6.900E-09 | 1.480E-09 | 5.900E-09 | 8.600E-09 |
| 19559 | 978 | A978Y | C | 4e-09 | 7.8e-09 | 9.4e-09 | 7.067E-09 | 2.774E-09 | 4.000E-09 | 9.400E-09 |
| 19560 | 978 | A978V | C | 3.6e-09 | 9.2e-09 | 7e-09 | 6.600E-09 | 2.821E-09 | 3.600E-09 | 9.200E-09 |
| 19561 | 979 | A979A | C | 7.8e-09 | 1.8e-08 | 8.6e-09 | 1.147E-08 | 5.672E-09 | 7.800E-09 | 1.800E-08 |
| 19562 | 979 | A979R | C | 1e-08 | 9.3e-09 | 3.2e-09 | 7.500E-09 | 3.740E-09 | 3.200E-09 | 1.000E-08 |
| 19563 | 979 | A979N | C | 7.3e-09 | 2.4e-09 | 4.7e-09 | 4.800E-09 | 2.452E-09 | 2.400E-09 | 7.300E-09 |
| 19564 | 979 | A979D | C | 8.4e-09 | 5.6e-09 | 4.3e-09 | 6.100E-09 | 2.095E-09 | 4.300E-09 | 8.400E-09 |
| 19565 | 979 | A979C | C | 1.7e-08 | 7e-09 | 1e-08 | 1.133E-08 | 5.132E-09 | 7.000E-09 | 1.700E-08 |
| 19566 | 979 | A979E | C | 7.3e-09 | 4.8e-09 | 1.6e-08 | 9.367E-09 | 5.879E-09 | 4.800E-09 | 1.600E-08 |
| 19567 | 979 | A979Q | C | 1.7e-08 | 2e-09 | 1.1e-08 | 1.000E-08 | 7.550E-09 | 2.000E-09 | 1.700E-08 |
| 19568 | 979 | A979G | C | 6.8e-09 | 3.5e-09 | 5.6e-09 | 5.300E-09 | 1.670E-09 | 3.500E-09 | 6.800E-09 |
| 19569 | 979 | A979H | C | 2.6e-09 | 1.2e-08 | 9.8e-09 | 8.133E-09 | 4.917E-09 | 2.600E-09 | 1.200E-08 |
| 19570 | 979 | A979I | C | 7.6e-09 | 7.5e-09 | 4.6e-09 | 6.567E-09 | 1.704E-09 | 4.600E-09 | 7.600E-09 |
| 19571 | 979 | A979L | C | 4e-09 | 7.2e-09 | 1.7e-08 | 9.400E-09 | 6.773E-09 | 4.000E-09 | 1.700E-08 |
| 19572 | 979 | A979K | C | 9.7e-09 | 6.6e-09 | 3.9e-09 | 6.733E-09 | 2.902E-09 | 3.900E-09 | 9.700E-09 |
| 19573 | 979 | A979M | C | 1.4e-08 | 6.6e-09 | 5.9e-09 | 8.833E-09 | 4.488E-09 | 5.900E-09 | 1.400E-08 |
| 19574 | 979 | A979F | C | 6.8e-09 | 2.5e-09 | 1.1e-08 | 6.767E-09 | 4.250E-09 | 2.500E-09 | 1.100E-08 |
| 19575 | 979 | A979P | C | 8.8e-09 | 8e-09 | 7.5e-09 | 8.100E-09 | 6.557E-10 | 7.500E-09 | 8.800E-09 |
| 19576 | 979 | A979S | C | 3.2e-09 | 1.1e-08 | 3e-09 | 5.733E-09 | 4.562E-09 | 3.000E-09 | 1.100E-08 |
| 19577 | 979 | A979T | C | 3.9e-09 | 4.3e-09 | 8.1e-09 | 5.433E-09 | 2.318E-09 | 3.900E-09 | 8.100E-09 |
| 19578 | 979 | A979W | C | 4.7e-09 | 3.6e-09 | 5.8e-09 | 4.700E-09 | 1.100E-09 | 3.600E-09 | 5.800E-09 |
| 19579 | 979 | A979Y | C | 7.6e-09 | 7.8e-09 | 2.7e-09 | 6.033E-09 | 2.888E-09 | 2.700E-09 | 7.800E-09 |
| 19580 | 979 | A979V | C | 4.5e-09 | 1.9e-09 | 2.2e-09 | 2.867E-09 | 1.422E-09 | 1.900E-09 | 4.500E-09 |
| 19581 | 980 | L980A | C | 2.2e-09 | 8.9e-09 | 1.8e-08 | 9.700E-09 | 7.930E-09 | 2.200E-09 | 1.800E-08 |
| 19582 | 980 | L980R | C | 3.3e-09 | 9.6e-09 | 6.8e-09 | 6.567E-09 | 3.156E-09 | 3.300E-09 | 9.600E-09 |
| 19583 | 980 | L980N | C | 5.3e-09 | 9.5e-09 | 9.7e-09 | 8.167E-09 | 2.485E-09 | 5.300E-09 | 9.700E-09 |
| 19584 | 980 | L980D | C | 8.7e-09 | 1.2e-08 | 1.7e-08 | 1.257E-08 | 4.179E-09 | 8.700E-09 | 1.700E-08 |
| 19585 | 980 | L980C | C | 5.5e-09 | 7.8e-09 | 4.6e-09 | 5.967E-09 | 1.650E-09 | 4.600E-09 | 7.800E-09 |
| 19586 | 980 | L980E | C | 1.1e-08 | 4.2e-09 | 7.9e-09 | 7.700E-09 | 3.404E-09 | 4.200E-09 | 1.100E-08 |
| 19587 | 980 | L980Q | C | 6.4e-09 | 8.7e-09 | 5.2e-09 | 6.767E-09 | 1.779E-09 | 5.200E-09 | 8.700E-09 |
| 19588 | 980 | L980G | C | 1.2e-08 | 1.4e-08 | 8.6e-09 | 1.153E-08 | 2.730E-09 | 8.600E-09 | 1.400E-08 |
| 19589 | 980 | L980H | C | 1e-08 | 6.1e-09 | 5.9e-09 | 7.333E-09 | 2.312E-09 | 5.900E-09 | 1.000E-08 |
| 19590 | 980 | L980I | C | 7.7e-09 | 1.7e-08 | 1.3e-08 | 1.257E-08 | 4.665E-09 | 7.700E-09 | 1.700E-08 |
| 19591 | 980 | L980L | C | 8.2e-09 | 8.5e-09 | 1.2e-08 | 9.567E-09 | 2.113E-09 | 8.200E-09 | 1.200E-08 |
| 19592 | 980 | L980K | C | 4.1e-09 | 8e-09 | 6.3e-09 | 6.133E-09 | 1.955E-09 | 4.100E-09 | 8.000E-09 |
| 19593 | 980 | L980M | C | 4.9e-09 | 7e-09 | 1.1e-08 | 7.633E-09 | 3.099E-09 | 4.900E-09 | 1.100E-08 |
| 19594 | 980 | L980F | C | 1.7e-08 | 1.7e-08 | 6.8e-09 | 1.360E-08 | 5.889E-09 | 6.800E-09 | 1.700E-08 |
| 19595 | 980 | L980P | C | 7.6e-09 | 7.5e-09 | 6.4e-09 | 7.167E-09 | 6.658E-10 | 6.400E-09 | 7.600E-09 |
| 19596 | 980 | L980S | C | 2e-09 | 2.8e-09 | 1.5e-08 | 6.600E-09 | 7.286E-09 | 2.000E-09 | 1.500E-08 |
| 19597 | 980 | L980T | C | 7.6e-09 | 3.3e-09 | 5.2e-09 | 5.367E-09 | 2.155E-09 | 3.300E-09 | 7.600E-09 |
| 19598 | 980 | L980W | C | 6.3e-09 | 7.2e-09 | 9.3e-09 | 7.600E-09 | 1.539E-09 | 6.300E-09 | 9.300E-09 |
| 19599 | 980 | L980Y | C | 7.7e-09 | 3.8e-09 | 4.9e-09 | 5.467E-09 | 2.011E-09 | 3.800E-09 | 7.700E-09 |
| 19600 | 980 | L980V | C | 9.3e-09 | 5.7e-09 | 4.1e-09 | 6.367E-09 | 2.663E-09 | 4.100E-09 | 9.300E-09 |
| 19601 | 981 | G981A | C | 1e-08 | 2.9e-08 | 1e-08 | 1.633E-08 | 1.097E-08 | 1.000E-08 | 2.900E-08 |
| 19602 | 981 | G981R | C | 4.2e-09 | 2.9e-09 | 1.1e-09 | 2.733E-09 | 1.557E-09 | 1.100E-09 | 4.200E-09 |
| 19603 | 981 | G981N | C | 5.9e-09 | 8.4e-09 | 1.6e-09 | 5.300E-09 | 3.439E-09 | 1.600E-09 | 8.400E-09 |
| 19604 | 981 | G981D | C | 1e-08 | 1.6e-08 | 4.5e-09 | 1.017E-08 | 5.752E-09 | 4.500E-09 | 1.600E-08 |
| 19605 | 981 | G981C | C | 2e-08 | 2.1e-08 | 3.4e-09 | 1.480E-08 | 9.885E-09 | 3.400E-09 | 2.100E-08 |
| 19606 | 981 | G981E | C | 4.9e-09 | 1e-08 | 7.4e-09 | 7.433E-09 | 2.550E-09 | 4.900E-09 | 1.000E-08 |
| 19607 | 981 | G981Q | C | 7.3e-09 | 5.1e-09 | 4.7e-09 | 5.700E-09 | 1.400E-09 | 4.700E-09 | 7.300E-09 |
| 19608 | 981 | G981G | C | 5.3e-09 | 1.4e-08 | 4.4e-09 | 7.900E-09 | 5.302E-09 | 4.400E-09 | 1.400E-08 |
| 19609 | 981 | G981H | C | 6e-09 | 4.8e-09 | 1.1e-08 | 7.267E-09 | 3.288E-09 | 4.800E-09 | 1.100E-08 |
| 19610 | 981 | G981I | C | 8.4e-09 | 6.5e-09 | 6.9e-09 | 7.267E-09 | 1.002E-09 | 6.500E-09 | 8.400E-09 |
| 19611 | 981 | G981L | C | 5.5e-09 | 5.7e-09 | 4.1e-09 | 5.100E-09 | 8.718E-10 | 4.100E-09 | 5.700E-09 |
| 19612 | 981 | G981K | C | 8.1e-09 | 6.9e-09 | 7.7e-09 | 7.567E-09 | 6.110E-10 | 6.900E-09 | 8.100E-09 |
| 19613 | 981 | G981M | C | 4.6e-09 | 5.6e-09 | 2.1e-08 | 1.040E-08 | 9.193E-09 | 4.600E-09 | 2.100E-08 |
| 19614 | 981 | G981F | C | 2.6e-08 | 9.1e-09 | 1.6e-08 | 1.703E-08 | 8.497E-09 | 9.100E-09 | 2.600E-08 |
| 19615 | 981 | G981P | C | 6.2e-09 | 1.3e-08 | 1.9e-09 | 7.033E-09 | 5.597E-09 | 1.900E-09 | 1.300E-08 |
| 19616 | 981 | G981S | C | 6.8e-09 | 4.9e-09 | 1.9e-08 | 1.023E-08 | 7.651E-09 | 4.900E-09 | 1.900E-08 |
| 19617 | 981 | G981T | C | 5.1e-09 | 1.4e-08 | 4.8e-09 | 7.967E-09 | 5.227E-09 | 4.800E-09 | 1.400E-08 |
| 19618 | 981 | G981W | C | 1.2e-08 | 3.6e-09 | 4.3e-09 | 6.633E-09 | 4.661E-09 | 3.600E-09 | 1.200E-08 |
| 19619 | 981 | G981Y | C | 5.5e-09 | 1.1e-08 | 6.1e-09 | 7.533E-09 | 3.017E-09 | 5.500E-09 | 1.100E-08 |
| 19620 | 981 | G981V | C | 5e-09 | 9.2e-09 | 1e-08 | 8.067E-09 | 2.686E-09 | 5.000E-09 | 1.000E-08 |
| 19621 | 982 | C982A | C | 2.8e-09 | 5.5e-09 | 1.9e-08 | 9.100E-09 | 8.679E-09 | 2.800E-09 | 1.900E-08 |
| 19622 | 982 | C982R | C | 1e-08 | 4.6e-09 | 5.4e-09 | 6.667E-09 | 2.914E-09 | 4.600E-09 | 1.000E-08 |
| 19623 | 982 | C982N | C | 5.2e-09 | 6.8e-09 | 4.6e-09 | 5.533E-09 | 1.137E-09 | 4.600E-09 | 6.800E-09 |
| 19624 | 982 | C982D | C | 9.4e-09 | 3.8e-09 | 3.1e-08 | 1.473E-08 | 1.436E-08 | 3.800E-09 | 3.100E-08 |
| 19625 | 982 | C982C | C | 8.2e-09 | 9.1e-09 | 1.3e-08 | 1.010E-08 | 2.551E-09 | 8.200E-09 | 1.300E-08 |
| 19626 | 982 | C982E | C | 1.8e-08 | 6.2e-09 | 8.5e-09 | 1.090E-08 | 6.255E-09 | 6.200E-09 | 1.800E-08 |
| 19627 | 982 | C982Q | C | 2.7e-09 | 2.5e-09 | 1.5e-08 | 6.733E-09 | 7.160E-09 | 2.500E-09 | 1.500E-08 |
| 19628 | 982 | C982G | C | 5.1e-09 | 4.4e-09 | 5.9e-09 | 5.133E-09 | 7.506E-10 | 4.400E-09 | 5.900E-09 |
| 19629 | 982 | C982H | C | 2.2e-09 | 8.4e-09 | 6.7e-09 | 5.767E-09 | 3.204E-09 | 2.200E-09 | 8.400E-09 |
| 19630 | 982 | C982I | C | 2e-08 | 6.4e-09 | 7.6e-09 | 1.133E-08 | 7.529E-09 | 6.400E-09 | 2.000E-08 |
| 19631 | 982 | C982L | C | 1.3e-08 | 1.2e-08 | 1.2e-08 | 1.233E-08 | 5.774E-10 | 1.200E-08 | 1.300E-08 |
| 19632 | 982 | C982K | C | 7.6e-09 | 4.3e-09 | 9.2e-09 | 7.033E-09 | 2.499E-09 | 4.300E-09 | 9.200E-09 |
| 19633 | 982 | C982M | C | 1e-08 | 7.4e-09 | 5.3e-09 | 7.567E-09 | 2.354E-09 | 5.300E-09 | 1.000E-08 |
| 19634 | 982 | C982F | C | 1.9e-08 | 8.4e-09 | 2.5e-09 | 9.967E-09 | 8.361E-09 | 2.500E-09 | 1.900E-08 |

| | | | | | | | | | | |
|---|---|---|---|---|---|---|---|---|---|---|
| 19635 | 982 | C982P | C | 4.1e-09 | 6e-09 | 9.8e-09 | 6.633E-09 | 2.902E-09 | 4.100E-09 | 9.800E-09 |
| 19636 | 982 | C982S | C | 7.5e-09 | 4.3e-09 | 7e-09 | 6.267E-09 | 1.721E-09 | 4.300E-09 | 7.500E-09 |
| 19637 | 982 | C982T | C | 3.4e-08 | 2.7e-09 | 1.2e-08 | 1.623E-08 | 1.607E-08 | 2.700E-09 | 3.400E-08 |
| 19638 | 982 | C982W | C | 8.5e-09 | 1.1e-08 | 9.4e-09 | 9.633E-09 | 1.266E-09 | 8.500E-09 | 1.100E-08 |
| 19639 | 982 | C982Y | C | 4.6e-09 | 4.2e-09 | 8.5e-09 | 5.767E-09 | 2.376E-09 | 4.200E-09 | 8.500E-09 |
| 19640 | 982 | C982V | C | 2.9e-08 | 9.4e-09 | 2.4e-08 | 2.080E-08 | 1.018E-08 | 9.400E-09 | 2.900E-08 |
| 19641 | 983 | L983A | C | 4.9e-09 | 5.2e-09 | 1e-08 | 6.700E-09 | 2.862E-09 | 4.900E-09 | 1.000E-08 |
| 19642 | 983 | L983R | C | 6.5e-09 | 1.1e-08 | 8e-09 | 8.500E-09 | 2.291E-09 | 6.500E-09 | 1.100E-08 |
| 19643 | 983 | L983N | C | 1.2e-08 | 1e-08 | 1.4e-08 | 1.200E-08 | 2.000E-09 | 1.000E-08 | 1.400E-08 |
| 19644 | 983 | L983D | C | 4e-09 | 9.3e-09 | 2.3e-09 | 5.200E-09 | 3.651E-09 | 2.300E-09 | 9.300E-09 |
| 19645 | 983 | L983C | C | 5.4e-09 | 1.3e-08 | 3.5e-09 | 7.300E-09 | 5.027E-09 | 3.500E-09 | 1.300E-08 |
| 19646 | 983 | L983E | C | 3.2e-09 | 5e-09 | 1.3e-08 | 7.067E-09 | 5.217E-09 | 3.200E-09 | 1.300E-08 |
| 19647 | 983 | L983Q | C | 1.7e-08 | 5.8e-09 | 4.7e-09 | 9.167E-09 | 6.806E-09 | 4.700E-09 | 1.700E-08 |
| 19648 | 983 | L983G | C | 4.3e-09 | 1.5e-08 | 4.7e-09 | 8.000E-09 | 6.065E-09 | 4.300E-09 | 1.500E-08 |
| 19649 | 983 | L983H | C | 1.1e-08 | 5.2e-09 | 2.9e-09 | 6.367E-09 | 4.174E-09 | 2.900E-09 | 1.100E-08 |
| 19650 | 983 | L983I | C | 3.7e-09 | 4.7e-09 | 7.3e-09 | 5.233E-09 | 1.858E-09 | 3.700E-09 | 7.300E-09 |
| 19651 | 983 | L983L | C | 8.5e-09 | 1.5e-08 | 2.4e-09 | 8.633E-09 | 6.301E-09 | 2.400E-09 | 1.500E-08 |
| 19652 | 983 | L983K | C | 4.1e-09 | 6.8e-09 | 7.1e-09 | 6.000E-09 | 1.652E-09 | 4.100E-09 | 7.100E-09 |
| 19653 | 983 | L983M | C | 3.4e-09 | 9.6e-09 | 7e-09 | 6.667E-09 | 3.113E-09 | 3.400E-09 | 9.600E-09 |
| 19654 | 983 | L983F | C | 1.4e-08 | 3e-09 | 2.4e-09 | 6.467E-09 | 6.531E-09 | 2.400E-09 | 1.400E-08 |
| 19655 | 983 | L983P | C | 3e-09 | 6.6e-09 | 9e-09 | 6.200E-09 | 3.020E-09 | 3.000E-09 | 9.000E-09 |
| 19656 | 983 | L983S | C | 9.4e-09 | 1.6e-08 | 2e-08 | 1.513E-08 | 5.353E-09 | 9.400E-09 | 2.000E-08 |
| 19657 | 983 | L983T | C | 1.5e-08 | 2.1e-08 | 1.2e-08 | 1.600E-08 | 4.583E-09 | 1.200E-08 | 2.100E-08 |
| 19658 | 983 | L983W | C | 2.7e-09 | 1.9e-08 | 2.6e-09 | 8.100E-09 | 9.440E-09 | 2.600E-09 | 1.900E-08 |
| 19659 | 983 | L983Y | C | 8.8e-09 | 2.4e-09 | 6.5e-09 | 5.900E-09 | 3.242E-09 | 2.400E-09 | 8.800E-09 |
| 19660 | 983 | L983V | C | 6.3e-09 | 9e-09 | 5.8e-09 | 7.033E-09 | 1.721E-09 | 5.800E-09 | 9.000E-09 |
| 19661 | 984 | V984A | C | 1.6e-08 | 5.5e-09 | 1.7e-08 | 1.283E-08 | 6.371E-09 | 5.500E-09 | 1.700E-08 |
| 19662 | 984 | V984R | C | 4.3e-09 | 1.1e-08 | 3.5e-09 | 6.267E-09 | 4.119E-09 | 3.500E-09 | 1.100E-08 |
| 19663 | 984 | V984N | C | 6.8e-09 | 5.9e-09 | 1.1e-08 | 7.900E-09 | 2.722E-09 | 5.900E-09 | 1.100E-08 |
| 19664 | 984 | V984D | C | 1e-08 | 5.4e-09 | 1e-08 | 8.467E-09 | 2.656E-09 | 5.400E-09 | 1.000E-08 |
| 19665 | 984 | V984C | C | 1.2e-08 | 3.3e-09 | 5.2e-09 | 6.833E-09 | 4.574E-09 | 3.300E-09 | 1.200E-08 |
| 19666 | 984 | V984E | C | 6.4e-09 | 6.1e-09 | 7.6e-09 | 6.700E-09 | 7.937E-10 | 6.100E-09 | 7.600E-09 |
| 19667 | 984 | V984Q | C | 1e-08 | 8.7e-09 | 3.7e-09 | 7.467E-09 | 3.326E-09 | 3.700E-09 | 1.000E-08 |
| 19668 | 984 | V984G | C | 6.9e-09 | 7.8e-09 | 9.1e-09 | 7.933E-09 | 1.106E-09 | 6.900E-09 | 9.100E-09 |
| 19669 | 984 | V984H | C | 1e-08 | 5.8e-09 | 2.6e-09 | 6.133E-09 | 3.711E-09 | 2.600E-09 | 1.000E-08 |
| 19670 | 984 | V984I | C | 4.7e-09 | 7.9e-09 | 6.8e-09 | 6.467E-09 | 1.626E-09 | 4.700E-09 | 7.900E-09 |
| 19671 | 984 | V984L | C | 1.1e-08 | 6.8e-09 | 1.7e-08 | 1.160E-08 | 5.126E-09 | 6.800E-09 | 1.700E-08 |
| 19672 | 984 | V984K | C | 1.2e-08 | 6.6e-09 | 4.3e-09 | 7.633E-09 | 3.953E-09 | 4.300E-09 | 1.200E-08 |
| 19673 | 984 | V984M | C | 1.2e-08 | 3.4e-09 | 2.1e-08 | 1.213E-08 | 8.801E-09 | 3.400E-09 | 2.100E-08 |
| 19674 | 984 | V984F | C | 1.4e-08 | 4.4e-09 | 8.6e-09 | 9.000E-09 | 4.812E-09 | 4.400E-09 | 1.400E-08 |
| 19675 | 984 | V984P | C | 4.9e-09 | 4.6e-09 | 6.9e-09 | 5.467E-09 | 1.250E-09 | 4.600E-09 | 6.900E-09 |
| 19676 | 984 | V984S | C | 4.3e-09 | 6.5e-09 | 6.2e-09 | 5.667E-09 | 1.193E-09 | 4.300E-09 | 6.500E-09 |
| 19677 | 984 | V984T | C | 4.7e-09 | 5.8e-09 | 5.4e-09 | 5.300E-09 | 5.568E-10 | 4.700E-09 | 5.800E-09 |
| 19678 | 984 | V984W | C | 3e-08 | 2.1e-08 | 9.9e-09 | 2.030E-08 | 1.007E-08 | 9.900E-09 | 3.000E-08 |
| 19679 | 984 | V984Y | C | 2.1e-08 | 4.1e-09 | 6e-09 | 1.037E-08 | 9.258E-09 | 4.100E-09 | 2.100E-08 |
| 19680 | 984 | V984V | C | 6.7e-09 | 6.2e-09 | 5.2e-09 | 6.033E-09 | 7.638E-10 | 5.200E-09 | 6.700E-09 |
| 19681 | 985 | K985A | C | 6e-09 | 1e-08 | 6.1e-09 | 7.367E-09 | 2.281E-09 | 6.000E-09 | 1.000E-08 |
| 19682 | 985 | K985R | C | 6.6e-09 | 7.8e-09 | 8.4e-09 | 7.600E-09 | 9.165E-10 | 6.600E-09 | 8.400E-09 |
| 19683 | 985 | K985N | C | 1.4e-08 | 6.8e-09 | 5e-09 | 8.600E-09 | 4.762E-09 | 5.000E-09 | 1.400E-08 |
| 19684 | 985 | K985D | C | 6.3e-09 | 7.7e-09 | 5.1e-09 | 6.367E-09 | 1.301E-09 | 5.100E-09 | 7.700E-09 |
| 19685 | 985 | K985C | C | 3.3e-09 | 4.7e-09 | 3.3e-09 | 3.767E-09 | 8.083E-10 | 3.300E-09 | 4.700E-09 |
| 19686 | 985 | K985E | C | 7.9e-09 | 8.1e-09 | 6.5e-09 | 7.500E-09 | 8.718E-10 | 6.500E-09 | 8.100E-09 |
| 19687 | 985 | K985Q | C | 5e-09 | 5.4e-09 | 4.3e-09 | 4.900E-09 | 5.568E-10 | 4.300E-09 | 5.400E-09 |
| 19688 | 985 | K985G | C | 2.2e-08 | 1.6e-08 | 3.2e-09 | 1.373E-08 | 9.603E-09 | 3.200E-09 | 2.200E-08 |
| 19689 | 985 | K985H | C | 1.1e-08 | 1.3e-08 | 3.1e-09 | 9.033E-09 | 5.235E-09 | 3.100E-09 | 1.300E-08 |
| 19690 | 985 | K985I | C | 6.3e-09 | 4.5e-09 | 3.8e-09 | 4.867E-09 | 1.290E-09 | 3.800E-09 | 6.300E-09 |
| 19691 | 985 | K985L | C | 2.2e-08 | 2.6e-09 | 6.4e-09 | 1.033E-08 | 1.028E-08 | 2.600E-09 | 2.200E-08 |
| 19692 | 985 | K985K | C | 1.1e-08 | 7.9e-09 | 2.2e-08 | 1.363E-08 | 7.410E-09 | 7.900E-09 | 2.200E-08 |
| 19693 | 985 | K985M | C | 4.8e-09 | 5.3e-09 | 1.1e-08 | 7.033E-09 | 3.444E-09 | 4.800E-09 | 1.100E-08 |
| 19694 | 985 | K985F | C | 1.1e-08 | 1.3e-08 | 9.7e-09 | 1.123E-08 | 1.662E-09 | 9.700E-09 | 1.300E-08 |
| 19695 | 985 | K985P | C | 2.8e-09 | 4.2e-09 | 4.6e-09 | 3.867E-09 | 9.452E-10 | 2.800E-09 | 4.600E-09 |
| 19696 | 985 | K985S | C | 4.3e-09 | 1.1e-08 | 4.1e-09 | 6.467E-09 | 3.927E-09 | 4.100E-09 | 1.100E-08 |
| 19697 | 985 | K985T | C | 8.4e-09 | 5.2e-09 | 2.5e-09 | 5.367E-09 | 2.954E-09 | 2.500E-09 | 8.400E-09 |
| 19698 | 985 | K985W | C | 2e-09 | 4.2e-09 | 1.9e-09 | 2.700E-09 | 1.300E-09 | 1.900E-09 | 4.200E-09 |
| 19699 | 985 | K985Y | C | 1.5e-08 | 2.8e-09 | 1e-08 | 9.267E-09 | 6.133E-09 | 2.800E-09 | 1.500E-08 |
| 19700 | 985 | K985V | C | 4.7e-09 | 4.9e-09 | 1.7e-08 | 8.867E-09 | 7.044E-09 | 4.700E-09 | 1.700E-08 |
| 19701 | 986 | D986A | C | 6e-09 | 1.1e-08 | 7.8e-09 | 8.267E-09 | 2.532E-09 | 6.000E-09 | 1.100E-08 |
| 19702 | 986 | D986R | C | 6e-09 | 4.4e-09 | 1e-08 | 6.800E-09 | 2.884E-09 | 4.400E-09 | 1.000E-08 |
| 19703 | 986 | D986N | C | 1.6e-08 | 2.5e-09 | 1.3e-08 | 1.050E-08 | 7.089E-09 | 2.500E-09 | 1.600E-08 |
| 19704 | 986 | D986D | C | 4.4e-09 | 1.9e-08 | 1.8e-08 | 1.380E-08 | 8.156E-09 | 4.400E-09 | 1.900E-08 |
| 19705 | 986 | D986C | C | 2.4e-09 | 4e-09 | 7.5e-09 | 4.633E-09 | 2.608E-09 | 2.400E-09 | 7.500E-09 |
| 19706 | 986 | D986E | C | 1.2e-08 | 1.3e-08 | 1.4e-08 | 1.300E-08 | 1.000E-09 | 1.200E-08 | 1.400E-08 |
| 19707 | 986 | D986Q | C | 9.1e-09 | 6.3e-09 | 9.3e-09 | 8.233E-09 | 1.677E-09 | 6.300E-09 | 9.300E-09 |
| 19708 | 986 | D986G | C | 1.1e-08 | 2e-09 | 4.7e-09 | 5.900E-09 | 4.618E-09 | 2.000E-09 | 1.100E-08 |
| 19709 | 986 | D986H | C | 2.4e-08 | 1.1e-08 | 5.7e-09 | 1.357E-08 | 9.416E-09 | 5.700E-09 | 2.400E-08 |
| 19710 | 986 | D986I | C | 9.7e-09 | 5.3e-09 | 2.5e-09 | 5.833E-09 | 3.630E-09 | 2.500E-09 | 9.700E-09 |
| 19711 | 986 | D986L | C | 5.8e-09 | 2.2e-08 | 8.6e-09 | 1.213E-08 | 8.659E-09 | 5.800E-09 | 2.200E-08 |
| 19712 | 986 | D986K | C | 5e-09 | 7.4e-09 | 9.9e-09 | 7.433E-09 | 2.450E-09 | 5.000E-09 | 9.900E-09 |
| 19713 | 986 | D986M | C | 2.9e-09 | 4.4e-09 | 5.2e-09 | 4.167E-09 | 1.168E-09 | 2.900E-09 | 5.200E-09 |
| 19714 | 986 | D986F | C | 5.2e-09 | 3.9e-09 | 7.3e-09 | 5.467E-09 | 1.716E-09 | 3.900E-09 | 7.300E-09 |
| 19715 | 986 | D986P | C | 1.1e-08 | 1.7e-08 | 5.9e-09 | 1.130E-08 | 5.556E-09 | 5.900E-09 | 1.700E-08 |
| 19716 | 986 | D986S | C | 9.8e-09 | 3.7e-09 | 6.5e-09 | 6.667E-09 | 3.053E-09 | 3.700E-09 | 9.800E-09 |
| 19717 | 986 | D986T | C | 6.5e-09 | 1.3e-08 | 4.3e-09 | 7.933E-09 | 4.524E-09 | 4.300E-09 | 1.300E-08 |
| 19718 | 986 | D986W | C | 9.1e-09 | 7e-09 | 6.7e-09 | 7.600E-09 | 1.308E-09 | 6.700E-09 | 9.100E-09 |
| 19719 | 986 | D986Y | C | 1.5e-08 | 2.4e-08 | 1.1e-08 | 1.667E-08 | 6.658E-09 | 1.100E-08 | 2.400E-08 |

| | | | | | | | | | | |
|---|---|---|---|---|---|---|---|---|---|---|
| 19720 | 986 | D986V | C | 7.2e-09 | 3.8e-09 | 6.6e-09 | 5.867E-09 | 1.815E-09 | 3.800E-09 | 7.200E-09 |
| 19721 | 987 | Y987A | C | 2.3e-09 | 8.9e-09 | 5.4e-09 | 5.533E-09 | 3.302E-09 | 2.300E-09 | 8.900E-09 |
| 19722 | 987 | Y987R | C | 9.5e-09 | 2.2e-08 | 7.2e-09 | 1.290E-08 | 7.964E-09 | 7.200E-09 | 2.200E-08 |
| 19723 | 987 | Y987N | C | 5.1e-09 | 6e-09 | 8.4e-09 | 6.500E-09 | 1.706E-09 | 5.100E-09 | 8.400E-09 |
| 19724 | 987 | Y987D | C | 4.4e-09 | 4.2e-09 | 1.5e-09 | 3.367E-09 | 1.620E-09 | 1.500E-09 | 4.400E-09 |
| 19725 | 987 | Y987C | C | 4.9e-09 | 2.7e-09 | 1.2e-08 | 6.533E-09 | 4.860E-09 | 2.700E-09 | 1.200E-08 |
| 19726 | 987 | Y987E | C | 1.5e-09 | 5.1e-09 | 4.8e-09 | 3.800E-09 | 1.997E-09 | 1.500E-09 | 5.100E-09 |
| 19727 | 987 | Y987Q | C | 2.5e-08 | 1.1e-08 | 4.5e-09 | 1.350E-08 | 1.048E-08 | 4.500E-09 | 2.500E-08 |
| 19728 | 987 | Y987G | C | 1.5e-08 | 1e-08 | 1e-08 | 1.167E-08 | 2.887E-09 | 1.000E-08 | 1.500E-08 |
| 19729 | 987 | Y987H | C | 7.1e-09 | 2.1e-08 | 1.5e-08 | 1.437E-08 | 6.972E-09 | 7.100E-09 | 2.100E-08 |
| 19730 | 987 | Y987I | C | 1.8e-08 | 2.1e-09 | 9.7e-09 | 9.933E-09 | 7.953E-09 | 2.100E-09 | 1.800E-08 |
| 19731 | 987 | Y987L | C | 3.5e-09 | 1.7e-08 | 7.1e-09 | 9.200E-09 | 6.991E-09 | 3.500E-09 | 1.700E-08 |
| 19732 | 987 | Y987K | C | 3.1e-09 | 6.1e-09 | 1.1e-08 | 6.733E-09 | 3.988E-09 | 3.100E-09 | 1.100E-08 |
| 19733 | 987 | Y987M | C | 8.4e-09 | 1.5e-09 | 2.5e-08 | 1.163E-08 | 1.208E-08 | 1.500E-09 | 2.500E-08 |
| 19734 | 987 | Y987F | C | 7.9e-09 | 7.9e-09 | 6.6e-09 | 7.467E-09 | 7.506E-10 | 6.600E-09 | 7.900E-09 |
| 19735 | 987 | Y987P | C | 1.1e-08 | 1.4e-08 | 4.5e-09 | 9.833E-09 | 4.856E-09 | 4.500E-09 | 1.400E-08 |
| 19736 | 987 | Y987S | C | 1.2e-08 | 2.5e-09 | 1.7e-09 | 5.400E-09 | 5.730E-09 | 1.700E-09 | 1.200E-08 |
| 19737 | 987 | Y987T | C | 9.6e-09 | 5e-09 | 1.5e-08 | 9.867E-09 | 5.005E-09 | 5.000E-09 | 1.500E-08 |
| 19738 | 987 | Y987W | C | 1.4e-08 | 3.7e-09 | 5.8e-09 | 7.833E-09 | 5.443E-09 | 3.700E-09 | 1.400E-08 |
| 19739 | 987 | Y987Y | C | 2.6e-09 | 2.1e-08 | 1e-08 | 1.120E-08 | 9.259E-09 | 2.600E-09 | 2.100E-08 |
| 19740 | 987 | Y987V | C | 2.8e-09 | 4e-09 | 1e-08 | 5.600E-09 | 3.857E-09 | 2.800E-09 | 1.000E-08 |
| 19741 | 988 | F988A | C | 3.7e-09 | 1.3e-08 | 3.3e-09 | 6.667E-09 | 5.488E-09 | 3.300E-09 | 1.300E-08 |
| 19742 | 988 | F988R | C | 8.2e-09 | 5.3e-09 | 9.3e-09 | 7.600E-09 | 2.066E-09 | 5.300E-09 | 9.300E-09 |
| 19743 | 988 | F988N | C | 7.1e-09 | 2.8e-09 | 3.4e-09 | 4.433E-09 | 2.329E-09 | 2.800E-09 | 7.100E-09 |
| 19744 | 988 | F988D | C | 4.8e-09 | 5.7e-09 | 6.4e-09 | 5.633E-09 | 8.021E-10 | 4.800E-09 | 6.400E-09 |
| 19745 | 988 | F988C | C | 3.6e-09 | 9.5e-09 | 3e-09 | 5.367E-09 | 3.592E-09 | 3.000E-09 | 9.500E-09 |
| 19746 | 988 | F988E | C | 3.7e-09 | 6.7e-09 | 3.4e-09 | 4.600E-09 | 1.825E-09 | 3.400E-09 | 6.700E-09 |
| 19747 | 988 | F988Q | C | 1.1e-08 | 3.4e-09 | 1.7e-08 | 1.047E-08 | 6.816E-09 | 3.400E-09 | 1.700E-08 |
| 19748 | 988 | F988G | C | 4.4e-09 | 4.1e-09 | 1.2e-08 | 6.833E-09 | 4.477E-09 | 4.100E-09 | 1.200E-08 |
| 19749 | 988 | F988H | C | 2.9e-09 | 1.7e-08 | 4.8e-09 | 8.233E-09 | 7.651E-09 | 2.900E-09 | 1.700E-08 |
| 19750 | 988 | F988I | C | 2e-08 | 9.6e-09 | 8e-09 | 1.253E-08 | 6.516E-09 | 8.000E-09 | 2.000E-08 |
| 19751 | 988 | F988L | C | 8e-09 | 6.4e-09 | 1e-08 | 8.133E-09 | 1.804E-09 | 6.400E-09 | 1.000E-08 |
| 19752 | 988 | F988K | C | 1.6e-08 | 2.1e-09 | 1.4e-08 | 1.070E-08 | 7.515E-09 | 2.100E-09 | 1.600E-08 |
| 19753 | 988 | F988M | C | 1.1e-08 | 9.8e-09 | 5.9e-09 | 8.900E-09 | 2.666E-09 | 5.900E-09 | 1.100E-08 |
| 19754 | 988 | F988F | C | 1e-08 | 3.9e-09 | 1.2e-08 | 8.633E-09 | 4.219E-09 | 3.900E-09 | 1.200E-08 |
| 19755 | 988 | F988P | C | 5.5e-09 | 7.1e-09 | 7.3e-09 | 6.633E-09 | 9.866E-10 | 5.500E-09 | 7.300E-09 |
| 19756 | 988 | F988S | C | 1.3e-08 | 8.5e-09 | 3.5e-09 | 8.333E-09 | 4.752E-09 | 3.500E-09 | 1.300E-08 |
| 19757 | 988 | F988T | C | 3.3e-09 | 1.7e-08 | 4.7e-09 | 8.333E-09 | 7.538E-09 | 3.300E-09 | 1.700E-08 |
| 19758 | 988 | F988W | C | 5e-09 | 2.3e-08 | 5.5e-09 | 1.117E-08 | 1.025E-08 | 5.000E-09 | 2.300E-08 |
| 19759 | 988 | F988Y | C | 6.2e-09 | 8.4e-09 | 2.5e-08 | 1.320E-08 | 1.028E-08 | 6.200E-09 | 2.500E-08 |
| 19760 | 988 | F988V | C | 3.9e-09 | 5.8e-09 | 7.8e-09 | 5.833E-09 | 1.950E-09 | 3.900E-09 | 7.800E-09 |
| 19761 | 989 | P989A | C | 1.2e-08 | 7.1e-09 | 2.6e-09 | 7.233E-09 | 4.701E-09 | 2.600E-09 | 1.200E-08 |
| 19762 | 989 | P989R | C | 6.5e-09 | 5.9e-09 | 9e-09 | 7.133E-09 | 1.644E-09 | 5.900E-09 | 9.000E-09 |
| 19763 | 989 | P989N | C | 8.4e-10 | 2.6e-09 | 4.7e-09 | 2.713E-09 | 1.932E-09 | 8.400E-10 | 4.700E-09 |
| 19764 | 989 | P989D | C | 5.1e-09 | 4.1e-09 | 1.6e-08 | 8.400E-09 | 6.601E-09 | 4.100E-09 | 1.600E-08 |
| 19765 | 989 | P989C | C | 5.9e-09 | 1.9e-08 | 1e-08 | 1.163E-08 | 6.701E-09 | 5.900E-09 | 1.900E-08 |
| 19766 | 989 | P989E | C | 1.7e-08 | 1.4e-08 | 1e-08 | 1.367E-08 | 3.512E-09 | 1.000E-08 | 1.700E-08 |
| 19767 | 989 | P989Q | C | 2.5e-09 | 1.5e-08 | 8.5e-09 | 8.667E-09 | 6.252E-09 | 2.500E-09 | 1.500E-08 |
| 19768 | 989 | P989G | C | 9.2e-09 | 7.4e-09 | 1.5e-09 | 6.033E-09 | 4.028E-09 | 1.500E-09 | 9.200E-09 |
| 19769 | 989 | P989H | C | 1.1e-08 | 1.5e-09 | 1e-08 | 7.500E-09 | 5.220E-09 | 1.500E-09 | 1.100E-08 |
| 19770 | 989 | P989I | C | 4.6e-09 | 5e-09 | 7.9e-09 | 5.833E-09 | 1.801E-09 | 4.600E-09 | 7.900E-09 |
| 19771 | 989 | P989L | C | 6.2e-09 | 6.7e-09 | 3.4e-09 | 5.433E-09 | 1.779E-09 | 3.400E-09 | 6.700E-09 |
| 19772 | 989 | P989K | C | 2e-08 | 3.5e-09 | 1.2e-08 | 1.183E-08 | 8.251E-09 | 3.500E-09 | 2.000E-08 |
| 19773 | 989 | P989M | C | 5.2e-09 | 2.9e-08 | 4.5e-09 | 1.290E-08 | 1.395E-08 | 4.500E-09 | 2.900E-08 |
| 19774 | 989 | P989F | C | 8e-09 | 4.8e-09 | 9.9e-09 | 7.567E-09 | 2.577E-09 | 4.800E-09 | 9.900E-09 |
| 19775 | 989 | P989P | C | 5.4e-09 | 2.9e-09 | 8.9e-09 | 5.733E-09 | 3.014E-09 | 2.900E-09 | 8.900E-09 |
| 19776 | 989 | P989S | C | 1.2e-08 | 3.5e-09 | 8e-09 | 7.833E-09 | 4.252E-09 | 3.500E-09 | 1.200E-08 |
| 19777 | 989 | P989T | C | 3.9e-09 | 7e-09 | 1.7e-08 | 9.300E-09 | 6.846E-09 | 3.900E-09 | 1.700E-08 |
| 19778 | 989 | P989W | C | 3.6e-09 | 2.3e-08 | 2.4e-08 | 1.687E-08 | 1.150E-08 | 3.600E-09 | 2.400E-08 |
| 19779 | 989 | P989Y | C | 8.7e-09 | 3.8e-09 | 2.7e-09 | 5.067E-09 | 3.194E-09 | 2.700E-09 | 8.700E-09 |
| 19780 | 989 | P989V | C | 2.3e-08 | 8.4e-09 | 8.3e-09 | 1.323E-08 | 8.458E-09 | 8.300E-09 | 2.300E-08 |
| 19781 | 990 | E990A | C | 5.5e-09 | 1.9e-09 | 3.3e-09 | 3.567E-09 | 1.815E-09 | 1.900E-09 | 5.500E-09 |
| 19782 | 990 | E990R | C | 7.9e-09 | 6.5e-09 | 3.7e-09 | 6.033E-09 | 2.139E-09 | 3.700E-09 | 7.900E-09 |
| 19783 | 990 | E990N | C | 8.4e-09 | 9.2e-09 | 4.6e-09 | 7.400E-09 | 2.458E-09 | 4.600E-09 | 9.200E-09 |
| 19784 | 990 | E990D | C | 4.5e-09 | 1e-08 | 6.8e-09 | 7.100E-09 | 2.762E-09 | 4.500E-09 | 1.000E-08 |
| 19785 | 990 | E990C | C | 3.1e-09 | 6e-09 | 5.7e-09 | 4.933E-09 | 1.595E-09 | 3.100E-09 | 6.000E-09 |
| 19786 | 990 | E990E | C | 8.1e-09 | 1.4e-08 | 1.8e-09 | 7.967E-09 | 6.101E-09 | 1.800E-09 | 1.400E-08 |
| 19787 | 990 | E990Q | C | 1e-08 | 6.4e-09 | 1.4e-08 | 1.013E-08 | 3.802E-09 | 6.400E-09 | 1.400E-08 |
| 19788 | 990 | E990G | C | 4.7e-09 | 6.1e-09 | 5e-09 | 5.267E-09 | 7.371E-10 | 4.700E-09 | 6.100E-09 |
| 19789 | 990 | E990H | C | 9.2e-09 | 3.4e-09 | 6e-09 | 6.200E-09 | 2.905E-09 | 3.400E-09 | 9.200E-09 |
| 19790 | 990 | E990I | C | 7.9e-09 | 2.7e-09 | 1e-08 | 6.867E-09 | 3.758E-09 | 2.700E-09 | 1.000E-08 |
| 19791 | 990 | E990L | C | 9.3e-09 | 8.1e-09 | 1.2e-08 | 9.800E-09 | 1.997E-09 | 8.100E-09 | 1.200E-08 |
| 19792 | 990 | E990K | C | 7.1e-09 | 6.1e-09 | 2.2e-08 | 1.173E-08 | 8.905E-09 | 6.100E-09 | 2.200E-08 |
| 19793 | 990 | E990M | C | 7.5e-09 | 2.4e-09 | 9.1e-09 | 6.333E-09 | 3.499E-09 | 2.400E-09 | 9.100E-09 |
| 19794 | 990 | E990F | C | 3.4e-09 | 6.8e-09 | 3.6e-09 | 4.600E-09 | 1.908E-09 | 3.400E-09 | 6.800E-09 |
| 19795 | 990 | E990P | C | 2.3e-09 | 3.9e-09 | 2.9e-09 | 3.033E-09 | 8.083E-10 | 2.300E-09 | 3.900E-09 |
| 19796 | 990 | E990S | C | 3.4e-09 | 5.3e-09 | 2.7e-09 | 3.800E-09 | 1.345E-09 | 2.700E-09 | 5.300E-09 |
| 19797 | 990 | E990T | C | 2.6e-09 | 8.7e-09 | 9.2e-09 | 6.833E-09 | 3.675E-09 | 2.600E-09 | 9.200E-09 |
| 19798 | 990 | E990W | C | 2.9e-09 | 7.8e-09 | 1.2e-08 | 7.567E-09 | 4.554E-09 | 2.900E-09 | 1.200E-08 |
| 19799 | 990 | E990Y | C | 9.3e-09 | 9e-09 | 3.6e-09 | 7.300E-09 | 3.208E-09 | 3.600E-09 | 9.300E-09 |
| 19800 | 990 | E990V | C | 2.8e-09 | 1.9e-09 | 1.1e-08 | 5.233E-09 | 5.014E-09 | 1.900E-09 | 1.100E-08 |
| 19801 | 991 | P991A | C | 4.6e-09 | 3.8e-09 | 4.3e-09 | 4.233E-09 | 4.041E-10 | 3.800E-09 | 4.600E-09 |
| 19802 | 991 | P991R | C | 1.8e-08 | 9.4e-09 | 7.7e-09 | 1.170E-08 | 5.522E-09 | 7.700E-09 | 1.800E-08 |
| 19803 | 991 | P991N | C | 5.4e-09 | 1.6e-08 | 3.8e-09 | 8.400E-09 | 6.630E-09 | 3.800E-09 | 1.600E-08 |
| 19804 | 991 | P991D | C | 6.1e-09 | 2.9e-09 | 6.2e-09 | 5.067E-09 | 1.877E-09 | 2.900E-09 | 6.200E-09 |

| | | | | | | | | | | |
|---|---|---|---|---|---|---|---|---|---|---|
| 19805 | 991 | P991C | C | 9.7e-09 | 4.9e-09 | 4.4e-09 | 6.333E-09 | 2.926E-09 | 4.400E-09 | 9.700E-09 |
| 19806 | 991 | P991E | C | 5.4e-09 | 3.3e-09 | 4.2e-09 | 4.300E-09 | 1.054E-09 | 3.300E-09 | 5.400E-09 |
| 19807 | 991 | P991Q | C | 6.6e-09 | 1.3e-08 | 7.7e-09 | 9.100E-09 | 3.422E-09 | 6.600E-09 | 1.300E-08 |
| 19808 | 991 | P991G | C | 1.4e-08 | 3.8e-09 | 1.7e-08 | 1.160E-08 | 6.920E-09 | 3.800E-09 | 1.700E-08 |
| 19809 | 991 | P991H | C | 1.3e-08 | 4.7e-09 | 3.8e-09 | 7.167E-09 | 5.072E-09 | 3.800E-09 | 1.300E-08 |
| 19810 | 991 | P991I | C | 6.8e-09 | 6.4e-09 | 1e-08 | 7.733E-09 | 1.973E-09 | 6.400E-09 | 1.000E-08 |
| 19811 | 991 | P991L | C | 4.9e-09 | 1.3e-08 | 6.8e-09 | 8.233E-09 | 4.236E-09 | 4.900E-09 | 1.300E-08 |
| 19812 | 991 | P991K | C | 5.4e-09 | 5e-09 | 1.4e-08 | 8.133E-09 | 5.085E-09 | 5.000E-09 | 1.400E-08 |
| 19813 | 991 | P991M | C | 2.8e-09 | 2e-08 | 2.2e-08 | 1.493E-08 | 1.056E-08 | 2.800E-09 | 2.200E-08 |
| 19814 | 991 | P991F | C | 1.1e-08 | 5.1e-09 | 8.4e-09 | 8.167E-09 | 2.957E-09 | 5.100E-09 | 1.100E-08 |
| 19815 | 991 | P991P | C | 7.9e-09 | 7.9e-09 | 5.5e-09 | 7.100E-09 | 1.386E-09 | 5.500E-09 | 7.900E-09 |
| 19816 | 991 | P991S | C | 1.3e-08 | 5.2e-09 | 3.3e-09 | 7.167E-09 | 5.140E-09 | 3.300E-09 | 1.300E-08 |
| 19817 | 991 | P991T | C | 5.7e-09 | 1.4e-08 | 1.3e-08 | 1.090E-08 | 4.531E-09 | 5.700E-09 | 1.400E-08 |
| 19818 | 991 | P991W | C | 8.4e-09 | 9.9e-09 | 9.8e-09 | 9.367E-09 | 8.386E-10 | 8.400E-09 | 9.900E-09 |
| 19819 | 991 | P991Y | C | 3.5e-09 | 3.1e-09 | 7.9e-09 | 4.833E-09 | 2.663E-09 | 3.100E-09 | 7.900E-09 |
| 19820 | 991 | P991V | C | 8.2e-09 | 1.1e-08 | 1.2e-08 | 1.040E-08 | 1.970E-09 | 8.200E-09 | 1.200E-08 |
| 19821 | 992 | V992A | C | 2.8e-09 | 3.5e-09 | 4.1e-09 | 3.467E-09 | 6.506E-10 | 2.800E-09 | 4.100E-09 |
| 19822 | 992 | V992R | C | 2.4e-09 | 4e-09 | 2.6e-09 | 3.000E-09 | 8.718E-10 | 2.400E-09 | 4.000E-09 |
| 19823 | 992 | V992N | C | 4e-09 | 3.3e-09 | 1.4e-08 | 7.100E-09 | 5.986E-09 | 3.300E-09 | 1.400E-08 |
| 19824 | 992 | V992D | C | 5e-09 | 4.9e-09 | 6.8e-09 | 5.567E-09 | 1.069E-09 | 4.900E-09 | 6.800E-09 |
| 19825 | 992 | V992C | C | 9.7e-09 | 1.2e-08 | 9e-09 | 1.023E-08 | 1.570E-09 | 9.000E-09 | 1.200E-08 |
| 19826 | 992 | V992E | C | 5.3e-09 | 4.2e-09 | 6.7e-09 | 5.400E-09 | 1.253E-09 | 4.200E-09 | 6.700E-09 |
| 19827 | 992 | V992Q | C | 3.3e-09 | 2.1e-09 | 9.7e-09 | 5.033E-09 | 4.086E-09 | 2.100E-09 | 9.700E-09 |
| 19828 | 992 | V992G | C | 3e-09 | 4.3e-09 | 8.1e-09 | 5.133E-09 | 2.650E-09 | 3.000E-09 | 8.100E-09 |
| 19829 | 992 | V992H | C | 5.7e-09 | 4.1e-09 | 1.1e-08 | 6.933E-09 | 3.612E-09 | 4.100E-09 | 1.100E-08 |
| 19830 | 992 | V992I | C | 3.8e-09 | 6.1e-09 | 4.3e-09 | 4.733E-09 | 1.210E-09 | 3.800E-09 | 6.100E-09 |
| 19831 | 992 | V992L | C | 1.3e-08 | 5.4e-09 | 6.4e-09 | 8.267E-09 | 4.130E-09 | 5.400E-09 | 1.300E-08 |
| 19832 | 992 | V992K | C | 4.3e-09 | 5.3e-09 | 5.1e-09 | 4.900E-09 | 5.292E-10 | 4.300E-09 | 5.300E-09 |
| 19833 | 992 | V992M | C | 7.4e-09 | 1.7e-08 | 6.9e-09 | 1.043E-08 | 5.692E-09 | 6.900E-09 | 1.700E-08 |
| 19834 | 992 | V992F | C | 4e-09 | 3.7e-09 | 2.3e-08 | 1.023E-08 | 1.106E-08 | 3.700E-09 | 2.300E-08 |
| 19835 | 992 | V992P | C | 9.7e-09 | 5.1e-09 | 6.9e-09 | 7.233E-09 | 2.318E-09 | 5.100E-09 | 9.700E-09 |
| 19836 | 992 | V992S | C | 4.6e-09 | 3.8e-09 | 1.2e-08 | 6.800E-09 | 4.521E-09 | 3.800E-09 | 1.200E-08 |
| 19837 | 992 | V992T | C | 6.4e-09 | 1.3e-08 | 3.1e-09 | 7.500E-09 | 5.041E-09 | 3.100E-09 | 1.300E-08 |
| 19838 | 992 | V992W | C | 6.6e-09 | 5.5e-09 | 3.7e-09 | 5.267E-09 | 1.464E-09 | 3.700E-09 | 6.600E-09 |
| 19839 | 992 | V992Y | C | 6.3e-09 | 4e-09 | 3.4e-09 | 4.567E-09 | 1.531E-09 | 3.400E-09 | 6.300E-09 |
| 19840 | 992 | V992V | C | 7e-09 | 4.1e-09 | 2.5e-08 | 1.203E-08 | 1.132E-08 | 4.100E-09 | 2.500E-08 |
| 19841 | 993 | T993A | C | 3.3e-09 | 1.5e-08 | 7.4e-09 | 8.567E-09 | 5.937E-09 | 3.300E-09 | 1.500E-08 |
| 19842 | 993 | T993R | C | 5.7e-09 | 8.8e-09 | 1.3e-08 | 9.167E-09 | 3.664E-09 | 5.700E-09 | 1.300E-08 |
| 19843 | 993 | T993N | C | 1.5e-08 | 1e-08 | 9.4e-09 | 1.147E-08 | 3.075E-09 | 9.400E-09 | 1.500E-08 |
| 19844 | 993 | T993D | C | 6.5e-09 | 9.1e-09 | 9.6e-09 | 8.400E-09 | 1.664E-09 | 6.500E-09 | 9.600E-09 |
| 19845 | 993 | T993C | C | 2.8e-09 | 7.7e-09 | 7.5e-09 | 6.000E-09 | 2.773E-09 | 2.800E-09 | 7.700E-09 |
| 19846 | 993 | T993E | C | 7.6e-09 | 2.2e-08 | 5.5e-09 | 1.170E-08 | 8.982E-09 | 5.500E-09 | 2.200E-08 |
| 19847 | 993 | T993Q | C | 8.4e-09 | 9.7e-09 | 1.2e-08 | 1.003E-08 | 1.823E-09 | 8.400E-09 | 1.200E-08 |
| 19848 | 993 | T993G | C | 2.7e-09 | 1.1e-08 | 1.7e-08 | 1.023E-08 | 7.181E-09 | 2.700E-09 | 1.700E-08 |
| 19849 | 993 | T993H | C | 4.1e-09 | 3.8e-09 | 9.3e-09 | 5.733E-09 | 3.092E-09 | 3.800E-09 | 9.300E-09 |
| 19850 | 993 | T993I | C | 1e-08 | 2.4e-09 | 1.9e-08 | 1.047E-08 | 8.310E-09 | 2.400E-09 | 1.900E-08 |
| 19851 | 993 | T993L | C | 1.3e-08 | 1.2e-08 | 7.3e-09 | 1.077E-08 | 3.044E-09 | 7.300E-09 | 1.300E-08 |
| 19852 | 993 | T993K | C | 3.9e-09 | 1e-08 | 3.2e-09 | 5.700E-09 | 3.740E-09 | 3.200E-09 | 1.000E-08 |
| 19853 | 993 | T993M | C | 5.9e-09 | 9.2e-09 | 6.9e-09 | 7.333E-09 | 1.692E-09 | 5.900E-09 | 9.200E-09 |
| 19854 | 993 | T993F | C | 1.6e-09 | 5.4e-09 | 5.5e-09 | 4.167E-09 | 2.223E-09 | 1.600E-09 | 5.500E-09 |
| 19855 | 993 | T993P | C | 8.2e-09 | 6.4e-09 | 1.4e-08 | 9.533E-09 | 3.972E-09 | 6.400E-09 | 1.400E-08 |
| 19856 | 993 | T993S | C | 4.7e-09 | 3.9e-09 | 7.4e-09 | 5.333E-09 | 1.834E-09 | 3.900E-09 | 7.400E-09 |
| 19857 | 993 | T993T | C | 8e-09 | 4.3e-09 | 1.4e-08 | 8.767E-09 | 4.895E-09 | 4.300E-09 | 1.400E-08 |
| 19858 | 993 | T993W | C | 6.8e-09 | 9.9e-09 | 5.8e-09 | 7.500E-09 | 2.138E-09 | 5.800E-09 | 9.900E-09 |
| 19859 | 993 | T993Y | C | 9e-09 | 2e-08 | 1.4e-08 | 1.433E-08 | 5.508E-09 | 9.000E-09 | 2.000E-08 |
| 19860 | 993 | T993V | C | 4.4e-09 | 1.6e-08 | 5.5e-09 | 8.633E-09 | 6.403E-09 | 4.400E-09 | 1.600E-08 |
| 19861 | 994 | V994A | C | 5.9e-09 | 1.1e-08 | 5.9e-09 | 7.600E-09 | 2.944E-09 | 5.900E-09 | 1.100E-08 |
| 19862 | 994 | V994R | C | 1.3e-08 | 6e-09 | 3.3e-09 | 7.433E-09 | 5.006E-09 | 3.300E-09 | 1.300E-08 |
| 19863 | 994 | V994N | C | 3.4e-09 | 1.4e-08 | 1.1e-08 | 9.467E-09 | 5.464E-09 | 3.400E-09 | 1.400E-08 |
| 19864 | 994 | V994D | C | 6.4e-09 | 8e-09 | 6.4e-09 | 6.933E-09 | 9.238E-10 | 6.400E-09 | 8.000E-09 |
| 19865 | 994 | V994C | C | 3.9e-09 | 8.5e-09 | 9.8e-09 | 7.400E-09 | 3.100E-09 | 3.900E-09 | 9.800E-09 |
| 19866 | 994 | V994E | C | 3.8e-09 | 5.5e-09 | 7.4e-09 | 5.567E-09 | 1.801E-09 | 3.800E-09 | 7.400E-09 |
| 19867 | 994 | V994Q | C | 6.9e-09 | 6.9e-09 | 3.9e-09 | 5.900E-09 | 1.732E-09 | 3.900E-09 | 6.900E-09 |
| 19868 | 994 | V994G | C | 6.4e-09 | 7.7e-09 | 5.5e-09 | 6.533E-09 | 1.106E-09 | 5.500E-09 | 7.700E-09 |
| 19869 | 994 | V994H | C | 6e-09 | 1.3e-08 | 6.1e-09 | 8.367E-09 | 4.013E-09 | 6.000E-09 | 1.300E-08 |
| 19870 | 994 | V994I | C | 4.7e-09 | 1.8e-09 | 3.7e-09 | 3.400E-09 | 1.473E-09 | 1.800E-09 | 4.700E-09 |
| 19871 | 994 | V994L | C | 9.3e-09 | 2.4e-09 | 2.8e-09 | 4.833E-09 | 3.873E-09 | 2.400E-09 | 9.300E-09 |
| 19872 | 994 | V994K | C | 5.2e-09 | 4.4e-09 | 4e-09 | 4.533E-09 | 6.110E-10 | 4.000E-09 | 5.200E-09 |
| 19873 | 994 | V994M | C | 7.9e-09 | 2.2e-08 | 7.5e-09 | 1.247E-08 | 8.259E-09 | 7.500E-09 | 2.200E-08 |
| 19874 | 994 | V994F | C | 8.8e-09 | 7.3e-09 | 9.9e-10 | 5.697E-09 | 4.145E-09 | 9.900E-10 | 8.800E-09 |
| 19875 | 994 | V994P | C | 4e-09 | 6.9e-09 | 3.7e-09 | 4.867E-09 | 1.767E-09 | 3.700E-09 | 6.900E-09 |
| 19876 | 994 | V994S | C | 2.3e-09 | 2.1e-08 | 6.2e-09 | 9.833E-09 | 9.865E-09 | 2.300E-09 | 2.100E-08 |
| 19877 | 994 | V994T | C | 4e-09 | 6.1e-09 | 6.2e-09 | 5.433E-09 | 1.242E-09 | 4.000E-09 | 6.200E-09 |
| 19878 | 994 | V994W | C | 5.7e-09 | 1.1e-08 | 3e-08 | 1.557E-08 | 1.278E-08 | 5.700E-09 | 3.000E-08 |
| 19879 | 994 | V994Y | C | 3.6e-09 | 1.7e-08 | 2.7e-09 | 7.767E-09 | 8.009E-09 | 2.700E-09 | 1.700E-08 |
| 19880 | 994 | V994V | C | 2.8e-09 | 8.3e-09 | 9.1e-09 | 6.733E-09 | 3.430E-09 | 2.800E-09 | 9.100E-09 |
| 19881 | 995 | S995A | C | 6.5e-09 | 1e-08 | 6e-09 | 7.500E-09 | 2.179E-09 | 6.000E-09 | 1.000E-08 |
| 19882 | 995 | S995R | C | 9.8e-09 | 8.7e-09 | 3.9e-09 | 7.467E-09 | 3.137E-09 | 3.900E-09 | 9.800E-09 |
| 19883 | 995 | S995N | C | 2.7e-09 | 1.2e-08 | 1.1e-08 | 8.567E-09 | 5.105E-09 | 2.700E-09 | 1.200E-08 |
| 19884 | 995 | S995D | C | 4.1e-09 | 1e-08 | 4.9e-09 | 6.333E-09 | 3.201E-09 | 4.100E-09 | 1.000E-08 |
| 19885 | 995 | S995C | C | 8.9e-09 | 5.2e-09 | 5.2e-09 | 6.433E-09 | 2.136E-09 | 5.200E-09 | 8.900E-09 |
| 19886 | 995 | S995E | C | 2.9e-09 | 9.7e-09 | 3.4e-09 | 5.333E-09 | 3.790E-09 | 2.900E-09 | 9.700E-09 |
| 19887 | 995 | S995Q | C | 5.4e-09 | 9.3e-09 | 4.1e-09 | 6.267E-09 | 2.706E-09 | 4.100E-09 | 9.300E-09 |
| 19888 | 995 | S995G | C | 1.7e-09 | 5.6e-09 | 3.8e-09 | 3.700E-09 | 1.952E-09 | 1.700E-09 | 5.600E-09 |
| 19889 | 995 | S995H | C | 4.8e-09 | 1.1e-08 | 7e-09 | 7.600E-09 | 3.143E-09 | 4.800E-09 | 1.100E-08 |

| | | | | | | | | | | |
|---|---|---|---|---|---|---|---|---|---|---|
| 19890 | 995 | S995I | C | 6.5e-09 | 5.8e-09 | 1.7e-09 | 4.667E-09 | 2.593E-09 | 1.700E-09 | 6.500E-09 |
| 19891 | 995 | S995L | C | 4.3e-09 | 6.2e-09 | 5.9e-09 | 5.467E-09 | 1.021E-09 | 4.300E-09 | 6.200E-09 |
| 19892 | 995 | S995K | C | 8.4e-09 | 3.2e-09 | 1.9e-09 | 4.500E-09 | 3.439E-09 | 1.900E-09 | 8.400E-09 |
| 19893 | 995 | S995M | C | 1.5e-08 | 1.8e-08 | 4.3e-09 | 1.243E-08 | 7.202E-09 | 4.300E-09 | 1.800E-08 |
| 19894 | 995 | S995F | C | 4.4e-09 | 3.2e-09 | 8.7e-09 | 5.433E-09 | 2.892E-09 | 3.200E-09 | 8.700E-09 |
| 19895 | 995 | S995P | C | 1.9e-08 | 9.9e-09 | 3.4e-09 | 1.077E-08 | 7.836E-09 | 3.400E-09 | 1.900E-08 |
| 19896 | 995 | S995S | C | 4.1e-09 | 7.6e-09 | 1.3e-08 | 8.233E-09 | 4.484E-09 | 4.100E-09 | 1.300E-08 |
| 19897 | 995 | S995T | C | 4.5e-09 | 1.8e-08 | 1.1e-08 | 1.117E-08 | 6.752E-09 | 4.500E-09 | 1.800E-08 |
| 19898 | 995 | S995W | C | 6.4e-09 | 9e-09 | 4.7e-09 | 6.700E-09 | 2.166E-09 | 4.700E-09 | 9.000E-09 |
| 19899 | 995 | S995Y | C | 5e-09 | 4.8e-09 | 9e-09 | 6.267E-09 | 2.369E-09 | 4.800E-09 | 9.000E-09 |
| 19900 | 995 | S995V | C | 3.4e-09 | 5.5e-09 | 7.6e-09 | 5.500E-09 | 2.100E-09 | 3.400E-09 | 7.600E-09 |
| 19901 | 996 | W996A | C | 5.1e-09 | 1.5e-08 | 5.8e-09 | 8.633E-09 | 5.525E-09 | 5.100E-09 | 1.500E-08 |
| 19902 | 996 | W996R | C | 5.6e-09 | 8.6e-09 | 6.7e-09 | 6.967E-09 | 1.518E-09 | 5.600E-09 | 8.600E-09 |
| 19903 | 996 | W996N | C | 1.1e-08 | 1.4e-08 | 6.5e-09 | 1.050E-08 | 3.775E-09 | 6.500E-09 | 1.400E-08 |
| 19904 | 996 | W996D | C | 3e-09 | 1e-08 | 5.1e-09 | 6.033E-09 | 3.592E-09 | 3.000E-09 | 1.000E-08 |
| 19905 | 996 | W996C | C | 1.2e-08 | 7.3e-09 | 8.7e-09 | 9.333E-09 | 2.413E-09 | 7.300E-09 | 1.200E-08 |
| 19906 | 996 | W996E | C | 1.2e-08 | 6e-09 | 9.6e-09 | 9.200E-09 | 3.020E-09 | 6.000E-09 | 1.200E-08 |
| 19907 | 996 | W996Q | C | 1.2e-08 | 2e-08 | 9.7e-09 | 1.390E-08 | 5.406E-09 | 9.700E-09 | 2.000E-08 |
| 19908 | 996 | W996G | C | 1.4e-08 | 3.2e-09 | 1.2e-08 | 9.733E-09 | 5.746E-09 | 3.200E-09 | 1.400E-08 |
| 19909 | 996 | W996H | C | 2.9e-09 | 6.1e-09 | 1.3e-08 | 7.333E-09 | 5.162E-09 | 2.900E-09 | 1.300E-08 |
| 19910 | 996 | W996I | C | 1.1e-08 | 3.4e-09 | 3.7e-09 | 6.033E-09 | 4.304E-09 | 3.400E-09 | 1.100E-08 |
| 19911 | 996 | W996L | C | 4.3e-09 | 7.8e-09 | 1.2e-08 | 8.033E-09 | 3.855E-09 | 4.300E-09 | 1.200E-08 |
| 19912 | 996 | W996K | C | 2.5e-09 | 1.5e-09 | 5.9e-09 | 3.300E-09 | 2.307E-09 | 1.500E-09 | 5.900E-09 |
| 19913 | 996 | W996M | C | 3.3e-08 | 1.1e-08 | 3.5e-09 | 1.583E-08 | 1.533E-08 | 3.500E-09 | 3.300E-08 |
| 19914 | 996 | W996F | C | 2.9e-09 | 8e-09 | 9.3e-09 | 6.733E-09 | 3.383E-09 | 2.900E-09 | 9.300E-09 |
| 19915 | 996 | W996P | C | 5.8e-09 | 8.8e-09 | 7.3e-09 | 7.300E-09 | 1.500E-09 | 5.800E-09 | 8.800E-09 |
| 19916 | 996 | W996S | C | 7.1e-09 | 5.4e-09 | 5.4e-09 | 5.967E-09 | 9.815E-10 | 5.400E-09 | 7.100E-09 |
| 19917 | 996 | W996T | C | 1.2e-08 | 3.3e-09 | 2.1e-08 | 1.210E-08 | 8.850E-09 | 3.300E-09 | 2.100E-08 |
| 19918 | 996 | W996W | C | 1.3e-08 | 3.5e-09 | 1.8e-08 | 1.150E-08 | 7.365E-09 | 3.500E-09 | 1.800E-08 |
| 19919 | 996 | W996Y | C | 1.4e-08 | 1.3e-08 | 8.1e-09 | 1.170E-08 | 3.158E-09 | 8.100E-09 | 1.400E-08 |
| 19920 | 996 | W996V | C | 1.9e-09 | 3.5e-09 | 4.4e-09 | 3.267E-09 | 1.266E-09 | 1.900E-09 | 4.400E-09 |
| 19921 | 997 | N997A | C | 1.1e-08 | 5.4e-09 | 1.4e-08 | 1.013E-08 | 4.365E-09 | 5.400E-09 | 1.400E-08 |
| 19922 | 997 | N997R | C | 1.3e-08 | 6.5e-09 | 1.8e-08 | 1.250E-08 | 5.766E-09 | 6.500E-09 | 1.800E-08 |
| 19923 | 997 | N997N | C | 4e-09 | 4.9e-09 | 7.4e-09 | 5.433E-09 | 1.762E-09 | 4.000E-09 | 7.400E-09 |
| 19924 | 997 | N997D | C | 3.8e-09 | 1.1e-08 | 7.4e-09 | 7.400E-09 | 3.600E-09 | 3.800E-09 | 1.100E-08 |
| 19925 | 997 | N997C | C | 1.8e-08 | 1.5e-08 | 7.7e-09 | 1.357E-08 | 5.297E-09 | 7.700E-09 | 1.800E-08 |
| 19926 | 997 | N997E | C | 6.1e-09 | 8e-09 | 1.2e-08 | 8.700E-09 | 3.012E-09 | 6.100E-09 | 1.200E-08 |
| 19927 | 997 | N997Q | C | 3.9e-09 | 4e-09 | 2.1e-08 | 9.633E-09 | 9.844E-09 | 3.900E-09 | 2.100E-08 |
| 19928 | 997 | N997G | C | 3.4e-09 | 7.1e-09 | 1.7e-08 | 9.167E-09 | 7.032E-09 | 3.400E-09 | 1.700E-08 |
| 19929 | 997 | N997H | C | 3.3e-09 | 4.9e-09 | 4.5e-09 | 4.233E-09 | 8.327E-10 | 3.300E-09 | 4.900E-09 |
| 19930 | 997 | N997I | C | 1.7e-08 | 5.9e-09 | 5.3e-09 | 9.400E-09 | 6.589E-09 | 5.300E-09 | 1.700E-08 |
| 19931 | 997 | N997L | C | 1e-08 | 1.9e-08 | 2.1e-08 | 1.667E-08 | 5.859E-09 | 1.000E-08 | 2.100E-08 |
| 19932 | 997 | N997K | C | 9.1e-09 | 9.5e-09 | 1.2e-08 | 1.020E-08 | 1.572E-09 | 9.100E-09 | 1.200E-08 |
| 19933 | 997 | N997M | C | 6.6e-09 | 7.6e-09 | 8.7e-09 | 7.633E-09 | 1.050E-09 | 6.600E-09 | 8.700E-09 |
| 19934 | 997 | N997F | C | 6.5e-09 | 2.1e-08 | 1.8e-09 | 9.767E-09 | 1.001E-08 | 1.800E-09 | 2.100E-08 |
| 19935 | 997 | N997P | C | 4.4e-09 | 4.5e-09 | 7.7e-09 | 5.533E-09 | 1.877E-09 | 4.400E-09 | 7.700E-09 |
| 19936 | 997 | N997S | C | 7.1e-09 | 6.6e-09 | 2.1e-09 | 5.267E-09 | 2.754E-09 | 2.100E-09 | 7.100E-09 |
| 19937 | 997 | N997T | C | 1.4e-08 | 4.8e-09 | 1e-08 | 9.600E-09 | 4.613E-09 | 4.800E-09 | 1.400E-08 |
| 19938 | 997 | N997W | C | 6e-09 | 4.9e-09 | 3.3e-09 | 4.733E-09 | 1.358E-09 | 3.300E-09 | 6.000E-09 |
| 19939 | 997 | N997Y | C | 3.2e-08 | 3.8e-09 | 6.5e-09 | 1.410E-08 | 1.556E-08 | 3.800E-09 | 3.200E-08 |
| 19940 | 997 | N997V | C | 9.1e-09 | 5.2e-09 | 4.3e-09 | 6.200E-09 | 2.551E-09 | 4.300E-09 | 9.100E-09 |
| 19941 | 998 | S998A | C | 1.5e-08 | 8.3e-09 | 1.9e-09 | 8.400E-09 | 6.551E-09 | 1.900E-09 | 1.500E-08 |
| 19942 | 998 | S998R | C | 6.9e-09 | 6.8e-09 | 8e-09 | 7.233E-09 | 6.658E-10 | 6.800E-09 | 8.000E-09 |
| 19943 | 998 | S998N | C | 4.1e-09 | 5.7e-09 | 1.3e-08 | 7.600E-09 | 4.744E-09 | 4.100E-09 | 1.300E-08 |
| 19944 | 998 | S998D | C | 1.3e-08 | 5.3e-09 | 1.3e-09 | 6.533E-09 | 5.947E-09 | 1.300E-09 | 1.300E-08 |
| 19945 | 998 | S998C | C | 2.5e-08 | 1.1e-08 | 8.6e-09 | 1.487E-08 | 8.857E-09 | 8.600E-09 | 2.500E-08 |
| 19946 | 998 | S998E | C | 6.7e-09 | 4.2e-09 | 9.6e-09 | 6.833E-09 | 2.702E-09 | 4.200E-09 | 9.600E-09 |
| 19947 | 998 | S998Q | C | 3.5e-09 | 5.9e-09 | 3e-09 | 4.133E-09 | 1.550E-09 | 3.000E-09 | 5.900E-09 |
| 19948 | 998 | S998G | C | 1.3e-08 | 6.8e-09 | 2e-08 | 1.327E-08 | 6.604E-09 | 6.800E-09 | 2.000E-08 |
| 19949 | 998 | S998H | C | 6.7e-09 | 4.8e-09 | 2.5e-09 | 4.667E-09 | 2.103E-09 | 2.500E-09 | 6.700E-09 |
| 19950 | 998 | S998I | C | 6.2e-09 | 3.3e-08 | 7.4e-09 | 1.553E-08 | 1.514E-08 | 6.200E-09 | 3.300E-08 |
| 19951 | 998 | S998L | C | 1.4e-08 | 8e-09 | 2.5e-08 | 1.567E-08 | 8.622E-09 | 8.000E-09 | 2.500E-08 |
| 19952 | 998 | S998K | C | 1e-08 | 3.4e-09 | 4e-09 | 5.800E-09 | 3.650E-09 | 3.400E-09 | 1.000E-08 |
| 19953 | 998 | S998M | C | 3.8e-09 | 7.7e-09 | 6.6e-09 | 6.033E-09 | 2.011E-09 | 3.800E-09 | 7.700E-09 |
| 19954 | 998 | S998F | C | 3.1e-09 | 1.7e-08 | 7.3e-09 | 9.133E-09 | 7.129E-09 | 3.100E-09 | 1.700E-08 |
| 19955 | 998 | S998P | C | 1.1e-08 | 3.2e-09 | 7.8e-09 | 7.333E-09 | 3.921E-09 | 3.200E-09 | 1.100E-08 |
| 19956 | 998 | S998S | C | 3.2e-09 | 2.6e-08 | 1.3e-08 | 1.407E-08 | 1.144E-08 | 3.200E-09 | 2.600E-08 |
| 19957 | 998 | S998T | C | 6.8e-09 | 8.2e-09 | 4.2e-09 | 6.400E-09 | 2.030E-09 | 4.200E-09 | 8.200E-09 |
| 19958 | 998 | S998W | C | 2.8e-09 | 4.7e-09 | 1.7e-08 | 8.167E-09 | 7.709E-09 | 2.800E-09 | 1.700E-08 |
| 19959 | 998 | S998Y | C | 1e-08 | 1e-08 | 9.4e-09 | 9.800E-09 | 3.464E-10 | 9.400E-09 | 1.000E-08 |
| 19960 | 998 | S998V | C | 2.5e-08 | 1.2e-08 | 7.1e-09 | 1.470E-08 | 9.250E-09 | 7.100E-09 | 2.500E-08 |
| 19961 | 999 | G999A | C | 9e-09 | 8.9e-09 | 5.4e-09 | 7.767E-09 | 2.050E-09 | 5.400E-09 | 9.000E-09 |
| 19962 | 999 | G999R | C | 6.5e-09 | 2.7e-09 | 3.1e-09 | 4.100E-09 | 2.088E-09 | 2.700E-09 | 6.500E-09 |
| 19963 | 999 | G999N | C | 6.5e-09 | 1.1e-08 | 5.6e-09 | 7.700E-09 | 2.893E-09 | 5.600E-09 | 1.100E-08 |
| 19964 | 999 | G999D | C | 6.6e-09 | 8.8e-09 | 2.6e-09 | 6.000E-09 | 3.143E-09 | 2.600E-09 | 8.800E-09 |
| 19965 | 999 | G999C | C | 3.8e-09 | 5.5e-09 | 1.1e-08 | 6.767E-09 | 3.763E-09 | 3.800E-09 | 1.100E-08 |
| 19966 | 999 | G999E | C | 9.2e-09 | 1.2e-08 | 6.9e-09 | 9.367E-09 | 2.554E-09 | 6.900E-09 | 1.200E-08 |
| 19967 | 999 | G999Q | C | 3.5e-09 | 9.2e-09 | 1.3e-08 | 8.567E-09 | 4.782E-09 | 3.500E-09 | 1.300E-08 |
| 19968 | 999 | G999G | C | 1.2e-08 | 8.7e-09 | 9.1e-09 | 9.933E-09 | 1.801E-09 | 8.700E-09 | 1.200E-08 |
| 19969 | 999 | G999H | C | 1.3e-08 | 1.1e-08 | 7.7e-09 | 1.057E-08 | 2.676E-09 | 7.700E-09 | 1.300E-08 |
| 19970 | 999 | G999I | C | 5.1e-09 | 7e-09 | 6.5e-09 | 6.200E-09 | 9.849E-10 | 5.100E-09 | 7.000E-09 |
| 19971 | 999 | G999L | C | 6.2e-09 | 9.6e-09 | 9.2e-09 | 8.333E-09 | 1.858E-09 | 6.200E-09 | 9.600E-09 |
| 19972 | 999 | G999K | C | 6e-09 | 1.2e-08 | 5.2e-09 | 7.733E-09 | 3.717E-09 | 5.200E-09 | 1.200E-08 |
| 19973 | 999 | G999M | C | 2.8e-08 | 5.2e-09 | 8.6e-09 | 1.393E-08 | 1.230E-08 | 5.200E-09 | 2.800E-08 |
| 19974 | 999 | G999F | C | 1.9e-08 | 1e-08 | 8e-09 | 1.233E-08 | 5.859E-09 | 8.000E-09 | 1.900E-08 |

| | | | | | | | | | | |
|---|---|---|---|---|---|---|---|---|---|---|
| 19975 | 999 | G999P | C | 4.4e-09 | 6.1e-09 | 4.3e-09 | 4.933E-09 | 1.012E-09 | 4.300E-09 | 6.100E-09 |
| 19976 | 999 | G999S | C | 7e-09 | 4.2e-09 | 1.6e-08 | 9.067E-09 | 6.165E-09 | 4.200E-09 | 1.600E-08 |
| 19977 | 999 | G999T | C | 3.4e-09 | 3.2e-09 | 9.9e-09 | 5.500E-09 | 3.812E-09 | 3.200E-09 | 9.900E-09 |
| 19978 | 999 | G999W | C | 1.1e-08 | 5.3e-09 | 8.7e-09 | 8.333E-09 | 2.868E-09 | 5.300E-09 | 1.100E-08 |
| 19979 | 999 | G999Y | C | 5e-09 | 3.7e-09 | 8.6e-09 | 5.767E-09 | 2.538E-09 | 3.700E-09 | 8.600E-09 |
| 19980 | 999 | G999V | C | 7.4e-09 | 8.2e-09 | 3.2e-09 | 6.267E-09 | 2.686E-09 | 3.200E-09 | 8.200E-09 |
| 19981 | 1000 | A1000A | C | 1.2e-08 | 2.3e-09 | 5.7e-09 | 6.667E-09 | 4.922E-09 | 2.300E-09 | 1.200E-08 |
| 19982 | 1000 | A1000R | C | 9.2e-09 | 1.5e-08 | 5.3e-09 | 9.833E-09 | 4.881E-09 | 5.300E-09 | 1.500E-08 |
| 19983 | 1000 | A1000N | C | 8.9e-09 | 5.4e-09 | 7.4e-09 | 7.233E-09 | 1.756E-09 | 5.400E-09 | 8.900E-09 |
| 19984 | 1000 | A1000D | C | 3e-09 | 5.9e-09 | 4.5e-09 | 4.467E-09 | 1.450E-09 | 3.000E-09 | 5.900E-09 |
| 19985 | 1000 | A1000C | C | 6.5e-09 | 1.4e-08 | 4.9e-09 | 8.467E-09 | 4.858E-09 | 4.900E-09 | 1.400E-08 |
| 19986 | 1000 | A1000E | C | 7.6e-09 | 2.4e-09 | 6.4e-09 | 5.467E-09 | 2.723E-09 | 2.400E-09 | 7.600E-09 |
| 19987 | 1000 | A1000Q | C | 6.6e-09 | 1.4e-08 | 3e-09 | 7.867E-09 | 5.608E-09 | 3.000E-09 | 1.400E-08 |
| 19988 | 1000 | A1000G | C | 5.3e-09 | 1.8e-08 | 6.1e-09 | 9.800E-09 | 7.113E-09 | 5.300E-09 | 1.800E-08 |
| 19989 | 1000 | A1000H | C | 3.7e-09 | 5.7e-09 | 1.9e-08 | 9.467E-09 | 8.316E-09 | 3.700E-09 | 1.900E-08 |
| 19990 | 1000 | A1000I | C | 7.7e-09 | 1.5e-08 | 3e-09 | 8.567E-09 | 6.047E-09 | 3.000E-09 | 1.500E-08 |
| 19991 | 1000 | A1000L | C | 1e-08 | 2.3e-09 | 3e-09 | 5.100E-09 | 4.258E-09 | 2.300E-09 | 1.000E-08 |
| 19992 | 1000 | A1000K | C | 5.7e-09 | 4.5e-09 | 1.2e-08 | 7.400E-09 | 4.029E-09 | 4.500E-09 | 1.200E-08 |
| 19993 | 1000 | A1000M | C | 6e-09 | 5.9e-09 | 1.9e-08 | 1.030E-08 | 7.535E-09 | 5.900E-09 | 1.900E-08 |
| 19994 | 1000 | A1000F | C | 4.8e-09 | 1.3e-08 | 7e-09 | 8.267E-09 | 4.244E-09 | 4.800E-09 | 1.300E-08 |
| 19995 | 1000 | A1000P | C | 7.8e-09 | 4.5e-09 | 1.8e-09 | 4.700E-09 | 3.005E-09 | 1.800E-09 | 7.800E-09 |
| 19996 | 1000 | A1000S | C | 3.9e-09 | 1.5e-09 | 1.3e-08 | 6.133E-09 | 6.067E-09 | 1.500E-09 | 1.300E-08 |
| 19997 | 1000 | A1000T | C | 1.2e-08 | 3.5e-09 | 3.7e-09 | 6.400E-09 | 4.851E-09 | 3.500E-09 | 1.200E-08 |
| 19998 | 1000 | A1000W | C | 4.7e-09 | 6.7e-09 | 4.1e-09 | 5.167E-09 | 1.361E-09 | 4.100E-09 | 6.700E-09 |
| 19999 | 1000 | A1000Y | C | 3.3e-09 | 1.8e-09 | 4.1e-09 | 3.067E-09 | 1.168E-09 | 1.800E-09 | 4.100E-09 |
| 20000 | 1000 | A1000V | C | 1e-08 | 6.6e-09 | 4.6e-09 | 7.067E-09 | 2.730E-09 | 4.600E-09 | 1.000E-08 |
| 20001 | 1001 | L1001A | C | 1.5e-08 | 7.9e-09 | 5.6e-09 | 9.500E-09 | 4.900E-09 | 5.600E-09 | 1.500E-08 |
| 20002 | 1001 | L1001R | C | 4.6e-09 | 1.2e-08 | 2.3e-09 | 6.300E-09 | 5.069E-09 | 2.300E-09 | 1.200E-08 |
| 20003 | 1001 | L1001N | C | 2.4e-09 | 4.7e-09 | 4.5e-09 | 3.867E-09 | 1.274E-09 | 2.400E-09 | 4.700E-09 |
| 20004 | 1001 | L1001D | C | 8.9e-09 | 4.5e-09 | 7.7e-09 | 7.033E-09 | 2.274E-09 | 4.500E-09 | 8.900E-09 |
| 20005 | 1001 | L1001C | C | 2.9e-09 | 2.7e-09 | 1e-08 | 5.200E-09 | 4.158E-09 | 2.700E-09 | 1.000E-08 |
| 20006 | 1001 | L1001E | C | 4.2e-09 | 3e-09 | 7.5e-09 | 4.900E-09 | 2.330E-09 | 3.000E-09 | 7.500E-09 |
| 20007 | 1001 | L1001Q | C | 5.2e-09 | 1.1e-08 | 1.4e-08 | 1.007E-08 | 4.474E-09 | 5.200E-09 | 1.400E-08 |
| 20008 | 1001 | L1001G | C | 5.6e-09 | 1.8e-08 | 2e-08 | 1.453E-08 | 7.801E-09 | 5.600E-09 | 2.000E-08 |
| 20009 | 1001 | L1001H | C | 3.2e-09 | 9.3e-09 | 4.8e-09 | 5.767E-09 | 3.163E-09 | 3.200E-09 | 9.300E-09 |
| 20010 | 1001 | L1001I | C | 6.8e-09 | 8.2e-09 | 3.2e-09 | 6.067E-09 | 2.579E-09 | 3.200E-09 | 8.200E-09 |
| 20011 | 1001 | L1001L | C | 9e-09 | 1.1e-08 | 1.3e-08 | 1.100E-08 | 2.000E-09 | 9.000E-09 | 1.300E-08 |
| 20012 | 1001 | L1001K | C | 1.1e-08 | 4.2e-09 | 5e-09 | 6.733E-09 | 3.717E-09 | 4.200E-09 | 1.100E-08 |
| 20013 | 1001 | L1001M | C | 1.9e-08 | 7e-09 | 3.7e-09 | 9.900E-09 | 8.052E-09 | 3.700E-09 | 1.900E-08 |
| 20014 | 1001 | L1001F | C | 1.2e-08 | 6.2e-09 | 5.4e-09 | 7.867E-09 | 3.602E-09 | 5.400E-09 | 1.200E-08 |
| 20015 | 1001 | L1001P | C | 9e-09 | 4.4e-09 | 3.8e-09 | 5.733E-09 | 2.845E-09 | 3.800E-09 | 9.000E-09 |
| 20016 | 1001 | L1001S | C | 4.9e-09 | 3.8e-09 | 1.1e-08 | 6.567E-09 | 3.879E-09 | 3.800E-09 | 1.100E-08 |
| 20017 | 1001 | L1001T | C | 3.4e-09 | 1.1e-08 | 1.2e-08 | 8.800E-09 | 4.703E-09 | 3.400E-09 | 1.200E-08 |
| 20018 | 1001 | L1001W | C | 6.8e-09 | 7.5e-09 | 1.8e-08 | 1.077E-08 | 6.274E-09 | 6.800E-09 | 1.800E-08 |
| 20019 | 1001 | L1001Y | C | 4.9e-09 | 1.1e-08 | 1.7e-08 | 1.097E-08 | 6.050E-09 | 4.900E-09 | 1.700E-08 |
| 20020 | 1001 | L1001V | C | 1.8e-08 | 2.4e-09 | 8.4e-09 | 9.600E-09 | 7.869E-09 | 2.400E-09 | 1.800E-08 |
| 20021 | 1002 | T1002A | C | 1e-08 | 9.4e-09 | 5e-09 | 8.133E-09 | 2.730E-09 | 5.000E-09 | 1.000E-08 |
| 20022 | 1002 | T1002R | C | 6.9e-09 | 5.4e-09 | 6.2e-09 | 6.167E-09 | 7.506E-10 | 5.400E-09 | 6.900E-09 |
| 20023 | 1002 | T1002N | C | 6.5e-09 | 7.2e-09 | 9.4e-09 | 7.700E-09 | 1.513E-09 | 6.500E-09 | 9.400E-09 |
| 20024 | 1002 | T1002D | C | 1e-08 | 4.6e-09 | 2.3e-08 | 1.253E-08 | 9.458E-09 | 4.600E-09 | 2.300E-08 |
| 20025 | 1002 | T1002C | C | 6e-09 | 2.5e-08 | 7.6e-09 | 1.287E-08 | 1.054E-08 | 6.000E-09 | 2.500E-08 |
| 20026 | 1002 | T1002E | C | 7.7e-09 | 3.4e-09 | 2.4e-09 | 4.500E-09 | 2.816E-09 | 2.400E-09 | 7.700E-09 |
| 20027 | 1002 | T1002Q | C | 7.2e-09 | 8.2e-09 | 8.6e-09 | 8.000E-09 | 7.211E-10 | 7.200E-09 | 8.600E-09 |
| 20028 | 1002 | T1002G | C | 2.7e-09 | 4.2e-09 | 6.4e-09 | 4.433E-09 | 1.861E-09 | 2.700E-09 | 6.400E-09 |
| 20029 | 1002 | T1002H | C | 3e-09 | 4.5e-09 | 1.2e-08 | 6.500E-09 | 4.822E-09 | 3.000E-09 | 1.200E-08 |
| 20030 | 1002 | T1002I | C | 2.1e-09 | 4.1e-09 | 7.5e-09 | 4.567E-09 | 2.730E-09 | 2.100E-09 | 7.500E-09 |
| 20031 | 1002 | T1002L | C | 8.7e-09 | 6.9e-09 | 3.3e-09 | 6.300E-09 | 2.750E-09 | 3.300E-09 | 8.700E-09 |
| 20032 | 1002 | T1002K | C | 6.5e-09 | 6.2e-09 | 8.4e-09 | 7.033E-09 | 1.193E-09 | 6.200E-09 | 8.400E-09 |
| 20033 | 1002 | T1002M | C | 8.6e-09 | 5.8e-09 | 9.2e-09 | 7.867E-09 | 1.815E-09 | 5.800E-09 | 9.200E-09 |
| 20034 | 1002 | T1002F | C | 2.8e-09 | 5.2e-09 | 4.1e-09 | 4.033E-09 | 1.201E-09 | 2.800E-09 | 5.200E-09 |
| 20035 | 1002 | T1002P | C | 6e-09 | 7.9e-09 | 7.5e-09 | 7.133E-09 | 1.002E-09 | 6.000E-09 | 7.900E-09 |
| 20036 | 1002 | T1002S | C | 3e-09 | 1.4e-08 | 1.1e-08 | 9.333E-09 | 5.686E-09 | 3.000E-09 | 1.400E-08 |
| 20037 | 1002 | T1002T | C | 9.5e-09 | 1.2e-08 | 5e-09 | 8.833E-09 | 3.547E-09 | 5.000E-09 | 1.200E-08 |
| 20038 | 1002 | T1002W | C | 8.8e-09 | 7.6e-09 | 4.3e-09 | 6.900E-09 | 2.330E-09 | 4.300E-09 | 8.800E-09 |
| 20039 | 1002 | T1002Y | C | 9.2e-09 | 6.6e-09 | 6.2e-09 | 7.333E-09 | 1.629E-09 | 6.200E-09 | 9.200E-09 |
| 20040 | 1002 | T1002V | C | 3.6e-09 | 6.9e-09 | 3.8e-09 | 4.767E-09 | 1.850E-09 | 3.600E-09 | 6.900E-09 |
| 20041 | 1003 | S1003A | C | 3.3e-09 | 4.1e-09 | 8.3e-09 | 5.233E-09 | 2.686E-09 | 3.300E-09 | 8.300E-09 |
| 20042 | 1003 | S1003R | C | 2.9e-09 | 2.2e-08 | 4.6e-09 | 9.833E-09 | 1.057E-08 | 2.900E-09 | 2.200E-08 |
| 20043 | 1003 | S1003N | C | 1.5e-08 | 6.4e-09 | 4.4e-09 | 8.600E-09 | 5.632E-09 | 4.400E-09 | 1.500E-08 |
| 20044 | 1003 | S1003D | C | 1.4e-08 | 5e-09 | 5.8e-09 | 8.267E-09 | 4.981E-09 | 5.000E-09 | 1.400E-08 |
| 20045 | 1003 | S1003C | C | 1.9e-08 | 1.5e-08 | 7.3e-09 | 1.377E-08 | 5.947E-09 | 7.300E-09 | 1.900E-08 |
| 20046 | 1003 | S1003E | C | 2.2e-09 | 6.9e-09 | 2e-08 | 9.700E-09 | 9.224E-09 | 2.200E-09 | 2.000E-08 |
| 20047 | 1003 | S1003Q | C | 7.5e-09 | 8.3e-09 | 1.5e-08 | 1.027E-08 | 4.119E-09 | 7.500E-09 | 1.500E-08 |
| 20048 | 1003 | S1003G | C | 7.5e-09 | 3.3e-09 | 3.8e-09 | 4.867E-09 | 2.294E-09 | 3.300E-09 | 7.500E-09 |
| 20049 | 1003 | S1003H | C | 3e-09 | 3.7e-09 | 9.4e-09 | 5.367E-09 | 3.510E-09 | 3.000E-09 | 9.400E-09 |
| 20050 | 1003 | S1003I | C | 5.5e-09 | 5.3e-09 | 1.6e-08 | 8.933E-09 | 6.121E-09 | 5.300E-09 | 1.600E-08 |
| 20051 | 1003 | S1003L | C | 1.5e-08 | 1.1e-08 | 5e-09 | 1.033E-08 | 5.033E-09 | 5.000E-09 | 1.500E-08 |
| 20052 | 1003 | S1003K | C | 1.6e-08 | 7.7e-09 | 1.1e-08 | 1.157E-08 | 4.179E-09 | 7.700E-09 | 1.600E-08 |
| 20053 | 1003 | S1003M | C | 5.1e-09 | 1.1e-08 | 1e-08 | 8.700E-09 | 3.158E-09 | 5.100E-09 | 1.100E-08 |
| 20054 | 1003 | S1003F | C | 1.4e-08 | 2.3e-08 | 5.1e-09 | 1.403E-08 | 8.950E-09 | 5.100E-09 | 2.300E-08 |
| 20055 | 1003 | S1003P | C | 7.7e-09 | 6.6e-09 | 2.4e-09 | 5.567E-09 | 2.797E-09 | 2.400E-09 | 7.700E-09 |
| 20056 | 1003 | S1003S | C | 9.6e-09 | 2.6e-09 | 1.6e-08 | 9.400E-09 | 6.702E-09 | 2.600E-09 | 1.600E-08 |
| 20057 | 1003 | S1003T | C | 4.8e-09 | 1.3e-08 | 2.8e-08 | 1.527E-08 | 1.176E-08 | 4.800E-09 | 2.800E-08 |
| 20058 | 1003 | S1003W | C | 1.1e-08 | 5.3e-09 | 1e-09 | 5.767E-09 | 5.016E-09 | 1.000E-09 | 1.100E-08 |
| 20059 | 1003 | S1003Y | C | 7.7e-09 | 9.5e-09 | 1e-08 | 9.067E-09 | 1.210E-09 | 7.700E-09 | 1.000E-08 |

| | | | | | | | | | | |
|---|---|---|---|---|---|---|---|---|---|---|
| 20060 | 1003 | S1003V | C | 2.8e-09 | 9.2e-09 | 6.2e-09 | 6.067E-09 | 3.202E-09 | 2.800E-09 | 9.200E-09 |
| 20061 | 1004 | G1004A | C | 8.5e-09 | 1.6e-08 | 2.9e-08 | 1.783E-08 | 1.037E-08 | 8.500E-09 | 2.900E-08 |
| 20062 | 1004 | G1004R | C | 2.1e-09 | 7.4e-09 | 1.3e-08 | 7.500E-09 | 5.451E-09 | 2.100E-09 | 1.300E-08 |
| 20063 | 1004 | G1004N | C | 5.4e-09 | 2.8e-09 | 1.2e-08 | 6.733E-09 | 4.743E-09 | 2.800E-09 | 1.200E-08 |
| 20064 | 1004 | G1004D | C | 1.7e-08 | 4.1e-09 | 9.3e-09 | 1.013E-08 | 6.490E-09 | 4.100E-09 | 1.700E-08 |
| 20065 | 1004 | G1004C | C | 1e-09 | 1.3e-08 | 6e-09 | 6.667E-09 | 6.028E-09 | 1.000E-09 | 1.300E-08 |
| 20066 | 1004 | G1004E | C | 2e-08 | 3.2e-09 | 1.7e-08 | 1.340E-08 | 8.960E-09 | 3.200E-09 | 2.000E-08 |
| 20067 | 1004 | G1004Q | C | 2.9e-09 | 1.2e-08 | 5.9e-09 | 6.933E-09 | 4.637E-09 | 2.900E-09 | 1.200E-08 |
| 20068 | 1004 | G1004G | C | 3.3e-09 | 1.3e-08 | 6.9e-09 | 7.733E-09 | 4.903E-09 | 3.300E-09 | 1.300E-08 |
| 20069 | 1004 | G1004H | C | 3.4e-09 | 8.3e-09 | 6e-09 | 5.900E-09 | 2.452E-09 | 3.400E-09 | 8.300E-09 |
| 20070 | 1004 | G1004I | C | 4.8e-09 | 5.3e-09 | 1.9e-09 | 4.000E-09 | 1.836E-09 | 1.900E-09 | 5.300E-09 |
| 20071 | 1004 | G1004L | C | 1e-08 | 1.2e-08 | 3.8e-09 | 8.600E-09 | 4.276E-09 | 3.800E-09 | 1.200E-08 |
| 20072 | 1004 | G1004K | C | 6.7e-09 | 6e-09 | 5.8e-09 | 6.167E-09 | 4.726E-10 | 5.800E-09 | 6.700E-09 |
| 20073 | 1004 | G1004M | C | 1.4e-08 | 9.9e-09 | 5.2e-09 | 9.700E-09 | 4.403E-09 | 5.200E-09 | 1.400E-08 |
| 20074 | 1004 | G1004F | C | 6.1e-09 | 1.8e-09 | 2.3e-08 | 1.030E-08 | 1.121E-08 | 1.800E-09 | 2.300E-08 |
| 20075 | 1004 | G1004P | C | 9.8e-09 | 5.3e-09 | 3.9e-09 | 6.333E-09 | 3.083E-09 | 3.900E-09 | 9.800E-09 |
| 20076 | 1004 | G1004S | C | 6e-09 | 6.3e-09 | 1.7e-09 | 4.667E-09 | 2.574E-09 | 1.700E-09 | 6.300E-09 |
| 20077 | 1004 | G1004T | C | 6.2e-09 | 2e-08 | 1.2e-08 | 1.273E-08 | 6.929E-09 | 6.200E-09 | 2.000E-08 |
| 20078 | 1004 | G1004W | C | 8.9e-09 | 4.9e-09 | 8.1e-09 | 7.300E-09 | 2.117E-09 | 4.900E-09 | 8.900E-09 |
| 20079 | 1004 | G1004Y | C | 7.4e-09 | 1e-08 | 5.2e-09 | 7.533E-09 | 2.403E-09 | 5.200E-09 | 1.000E-08 |
| 20080 | 1004 | G1004V | C | 9.3e-09 | 9.2e-09 | 1.7e-09 | 6.733E-09 | 4.359E-09 | 1.700E-09 | 9.300E-09 |
| 20081 | 1005 | V1005A | C | 1.1e-08 | 1.3e-08 | 1.1e-08 | 1.167E-08 | 1.155E-09 | 1.100E-08 | 1.300E-08 |
| 20082 | 1005 | V1005R | C | 8.5e-09 | 2.7e-08 | 1.4e-08 | 1.650E-08 | 9.500E-09 | 8.500E-09 | 2.700E-08 |
| 20083 | 1005 | V1005N | C | 3.2e-09 | 2.2e-08 | 4.9e-09 | 1.003E-08 | 1.040E-08 | 3.200E-09 | 2.200E-08 |
| 20084 | 1005 | V1005D | C | 6.3e-09 | 1e-08 | 1.1e-08 | 9.100E-09 | 2.476E-09 | 6.300E-09 | 1.100E-08 |
| 20085 | 1005 | V1005C | C | 3.8e-09 | 9.7e-09 | 7.6e-09 | 7.033E-09 | 2.991E-09 | 3.800E-09 | 9.700E-09 |
| 20086 | 1005 | V1005E | C | 9.8e-09 | 7.8e-09 | 6.9e-09 | 8.167E-09 | 1.484E-09 | 6.900E-09 | 9.800E-09 |
| 20087 | 1005 | V1005Q | C | 8.5e-09 | 6.4e-09 | 2.4e-09 | 5.767E-09 | 3.099E-09 | 2.400E-09 | 8.500E-09 |
| 20088 | 1005 | V1005G | C | 2.1e-08 | 1.1e-08 | 1e-08 | 1.400E-08 | 6.083E-09 | 1.000E-08 | 2.100E-08 |
| 20089 | 1005 | V1005H | C | 4.5e-09 | 3.7e-09 | 2.6e-09 | 3.600E-09 | 9.539E-10 | 2.600E-09 | 4.500E-09 |
| 20090 | 1005 | V1005I | C | 3.8e-09 | 4e-09 | 1.6e-08 | 7.933E-09 | 6.987E-09 | 3.800E-09 | 1.600E-08 |
| 20091 | 1005 | V1005L | C | 6.1e-09 | 1.1e-08 | 1e-08 | 9.033E-09 | 2.589E-09 | 6.100E-09 | 1.100E-08 |
| 20092 | 1005 | V1005K | C | 4.4e-09 | 7.4e-09 | 1e-08 | 7.267E-09 | 2.802E-09 | 4.400E-09 | 1.000E-08 |
| 20093 | 1005 | V1005M | C | 5.2e-09 | 2.8e-09 | 5e-09 | 4.333E-09 | 1.332E-09 | 2.800E-09 | 5.200E-09 |
| 20094 | 1005 | V1005F | C | 5e-09 | 3.7e-09 | 1.2e-08 | 6.900E-09 | 4.464E-09 | 3.700E-09 | 1.200E-08 |
| 20095 | 1005 | V1005P | C | 1.2e-08 | 1.6e-08 | 1.5e-08 | 1.433E-08 | 2.082E-09 | 1.200E-08 | 1.600E-08 |
| 20096 | 1005 | V1005S | C | 1.5e-08 | 7.4e-09 | 6.4e-09 | 9.600E-09 | 4.703E-09 | 6.400E-09 | 1.500E-08 |
| 20097 | 1005 | V1005T | C | 1.2e-08 | 1.2e-08 | 2.3e-08 | 1.567E-08 | 6.351E-09 | 1.200E-08 | 2.300E-08 |
| 20098 | 1005 | V1005W | C | 4.5e-09 | 8.1e-09 | 6e-09 | 6.200E-09 | 1.808E-09 | 4.500E-09 | 8.100E-09 |
| 20099 | 1005 | V1005Y | C | 1.1e-08 | 2e-09 | 6.7e-09 | 6.567E-09 | 4.501E-09 | 2.000E-09 | 1.100E-08 |
| 20100 | 1005 | V1005V | C | 8e-09 | 4.9e-09 | 2.2e-08 | 1.163E-08 | 9.111E-09 | 4.900E-09 | 2.200E-08 |
| 20101 | 1006 | H1006A | C | 5.5e-09 | 6.3e-09 | 1.3e-08 | 8.267E-09 | 4.119E-09 | 5.500E-09 | 1.300E-08 |
| 20102 | 1006 | H1006R | C | 8.1e-09 | 8.5e-09 | 1.3e-08 | 9.867E-09 | 2.721E-09 | 8.100E-09 | 1.300E-08 |
| 20103 | 1006 | H1006N | C | 4.6e-09 | 4.6e-09 | 5.1e-09 | 4.767E-09 | 2.887E-10 | 4.600E-09 | 5.100E-09 |
| 20104 | 1006 | H1006D | C | 1.9e-08 | 1.2e-08 | 1.5e-08 | 1.533E-08 | 3.512E-09 | 1.200E-08 | 1.900E-08 |
| 20105 | 1006 | H1006C | C | 4.3e-09 | 1.4e-08 | 7.6e-09 | 8.633E-09 | 4.932E-09 | 4.300E-09 | 1.400E-08 |
| 20106 | 1006 | H1006E | C | 6e-09 | 1.1e-08 | 5.6e-09 | 7.533E-09 | 3.009E-09 | 5.600E-09 | 1.100E-08 |
| 20107 | 1006 | H1006Q | C | 2.9e-08 | 1.2e-08 | 8.8e-09 | 1.660E-08 | 1.086E-08 | 8.800E-09 | 2.900E-08 |
| 20108 | 1006 | H1006G | C | 1.4e-08 | 2.1e-08 | 1e-08 | 1.500E-08 | 5.568E-09 | 1.000E-08 | 2.100E-08 |
| 20109 | 1006 | H1006H | C | 1.4e-08 | 5.5e-09 | 4.4e-09 | 7.967E-09 | 5.254E-09 | 4.400E-09 | 1.400E-08 |
| 20110 | 1006 | H1006I | C | 6.4e-09 | 2.3e-09 | 5.3e-09 | 4.667E-09 | 2.122E-09 | 2.300E-09 | 6.400E-09 |
| 20111 | 1006 | H1006L | C | 1.1e-08 | 9.2e-09 | 7.3e-10 | 6.977E-09 | 5.484E-09 | 7.300E-10 | 1.100E-08 |
| 20112 | 1006 | H1006K | C | 6.1e-09 | 6.1e-09 | 8.1e-09 | 6.767E-09 | 1.155E-09 | 6.100E-09 | 8.100E-09 |
| 20113 | 1006 | H1006M | C | 4.2e-09 | 2.8e-09 | 1.3e-08 | 6.667E-09 | 5.529E-09 | 2.800E-09 | 1.300E-08 |
| 20114 | 1006 | H1006F | C | 7.3e-09 | 4.9e-09 | 2e-08 | 1.073E-08 | 8.114E-09 | 4.900E-09 | 2.000E-08 |
| 20115 | 1006 | H1006P | C | 6.5e-09 | 1e-08 | 6.4e-09 | 7.633E-09 | 2.050E-09 | 6.400E-09 | 1.000E-08 |
| 20116 | 1006 | H1006S | C | 4.3e-09 | 1.9e-09 | 6.4e-09 | 4.200E-09 | 2.252E-09 | 1.900E-09 | 6.400E-09 |
| 20117 | 1006 | H1006T | C | 5.6e-09 | 6.6e-09 | 6e-09 | 6.067E-09 | 5.033E-10 | 5.600E-09 | 6.600E-09 |
| 20118 | 1006 | H1006W | C | 7.6e-09 | 6e-09 | 3.8e-09 | 5.800E-09 | 1.908E-09 | 3.800E-09 | 7.600E-09 |
| 20119 | 1006 | H1006Y | C | 5e-09 | 7.5e-09 | 7.1e-09 | 6.533E-09 | 1.343E-09 | 5.000E-09 | 7.500E-09 |
| 20120 | 1006 | H1006V | C | 4.9e-09 | 2.1e-08 | 7.1e-09 | 1.100E-08 | 8.730E-09 | 4.900E-09 | 2.100E-08 |
| 20121 | 1007 | T1007A | C | 6e-09 | 2.4e-09 | 3.3e-09 | 3.900E-09 | 1.873E-09 | 2.400E-09 | 6.000E-09 |
| 20122 | 1007 | T1007R | C | 9.9e-09 | 4.6e-09 | 1.2e-08 | 8.833E-09 | 3.814E-09 | 4.600E-09 | 1.200E-08 |
| 20123 | 1007 | T1007N | C | 1.3e-08 | 5.3e-09 | 1.4e-08 | 1.077E-08 | 4.761E-09 | 5.300E-09 | 1.400E-08 |
| 20124 | 1007 | T1007D | C | 1.1e-08 | 4.7e-09 | 1.1e-08 | 8.900E-09 | 3.637E-09 | 4.700E-09 | 1.100E-08 |
| 20125 | 1007 | T1007C | C | 2.5e-09 | 1.2e-08 | 1.2e-08 | 8.833E-09 | 5.485E-09 | 2.500E-09 | 1.200E-08 |
| 20126 | 1007 | T1007E | C | 6.4e-09 | 7.1e-09 | 4.6e-09 | 6.033E-09 | 1.290E-09 | 4.600E-09 | 7.100E-09 |
| 20127 | 1007 | T1007Q | C | 4.7e-09 | 3.7e-09 | 3.8e-09 | 4.067E-09 | 5.508E-10 | 3.700E-09 | 4.700E-09 |
| 20128 | 1007 | T1007G | C | 4e-09 | 2.1e-08 | 2.9e-09 | 9.300E-09 | 1.015E-08 | 2.900E-09 | 2.100E-08 |
| 20129 | 1007 | T1007H | C | 6.8e-09 | 7.8e-09 | 6.4e-09 | 7.000E-09 | 7.211E-10 | 6.400E-09 | 7.800E-09 |
| 20130 | 1007 | T1007I | C | 1.2e-08 | 7.8e-09 | 1.5e-08 | 1.160E-08 | 3.617E-09 | 7.800E-09 | 1.500E-08 |
| 20131 | 1007 | T1007L | C | 3.7e-09 | 4.8e-09 | 3.4e-09 | 3.967E-09 | 7.371E-10 | 3.400E-09 | 4.800E-09 |
| 20132 | 1007 | T1007K | C | 1.1e-08 | 4.2e-09 | 2.4e-08 | 1.307E-08 | 1.006E-08 | 4.200E-09 | 2.400E-08 |
| 20133 | 1007 | T1007M | C | 7.2e-09 | 2.7e-09 | 6.6e-09 | 5.500E-09 | 2.443E-09 | 2.700E-09 | 7.200E-09 |
| 20134 | 1007 | T1007F | C | 9.8e-09 | 1.1e-08 | 3.8e-09 | 8.200E-09 | 3.857E-09 | 3.800E-09 | 1.100E-08 |
| 20135 | 1007 | T1007P | C | 5.1e-09 | 8.2e-09 | 3e-09 | 5.433E-09 | 2.616E-09 | 3.000E-09 | 8.200E-09 |
| 20136 | 1007 | T1007S | C | 4.4e-09 | 4.9e-09 | 7.1e-09 | 5.467E-09 | 1.436E-09 | 4.400E-09 | 7.100E-09 |
| 20137 | 1007 | T1007T | C | 7.5e-09 | 1.1e-08 | 1.6e-08 | 1.150E-08 | 4.272E-09 | 7.500E-09 | 1.600E-08 |
| 20138 | 1007 | T1007W | C | 5.5e-09 | 1.1e-08 | 7e-09 | 7.833E-09 | 2.843E-09 | 5.500E-09 | 1.100E-08 |
| 20139 | 1007 | T1007Y | C | 6.8e-09 | 1.4e-08 | 2.5e-09 | 7.767E-09 | 5.811E-09 | 2.500E-09 | 1.400E-08 |
| 20140 | 1007 | T1007V | C | 3.9e-09 | 1e-08 | 1.3e-08 | 8.967E-09 | 4.637E-09 | 3.900E-09 | 1.300E-08 |
| 20141 | 1008 | F1008A | C | 6.9e-09 | 1.1e-08 | 6.6e-09 | 8.167E-09 | 2.458E-09 | 6.600E-09 | 1.100E-08 |
| 20142 | 1008 | F1008R | C | 4.5e-09 | 4.5e-09 | 4.4e-09 | 4.467E-09 | 5.774E-11 | 4.400E-09 | 4.500E-09 |
| 20143 | 1008 | F1008N | C | 5.2e-09 | 3.6e-09 | 6.4e-09 | 5.067E-09 | 1.405E-09 | 3.600E-09 | 6.400E-09 |
| 20144 | 1008 | F1008D | C | 2.4e-08 | 6.4e-09 | 8.7e-09 | 1.303E-08 | 9.567E-09 | 6.400E-09 | 2.400E-08 |

| | | | | | | | | | | |
|---|---|---|---|---|---|---|---|---|---|---|
| 20145 | 1008 | F1008C | C | 1e-08 | 5.3e-09 | 5.6e-09 | 6.967E-09 | 2.631E-09 | 5.300E-09 | 1.000E-08 |
| 20146 | 1008 | F1008E | C | 4.5e-09 | 7.5e-09 | 2.5e-09 | 4.833E-09 | 2.517E-09 | 2.500E-09 | 7.500E-09 |
| 20147 | 1008 | F1008Q | C | 7.9e-09 | 2.2e-09 | 4.4e-09 | 4.833E-09 | 2.875E-09 | 2.200E-09 | 7.900E-09 |
| 20148 | 1008 | F1008G | C | 5.9e-09 | 5.3e-09 | 3.5e-09 | 4.900E-09 | 1.249E-09 | 3.500E-09 | 5.900E-09 |
| 20149 | 1008 | F1008H | C | 2.6e-09 | 2.1e-09 | 1.2e-08 | 5.567E-09 | 5.577E-09 | 2.100E-09 | 1.200E-08 |
| 20150 | 1008 | F1008I | C | 6.1e-09 | 5.1e-09 | 1e-08 | 7.067E-09 | 2.589E-09 | 5.100E-09 | 1.000E-08 |
| 20151 | 1008 | F1008L | C | 5.7e-09 | 1.4e-08 | 1.2e-08 | 1.057E-08 | 4.332E-09 | 5.700E-09 | 1.400E-08 |
| 20152 | 1008 | F1008K | C | 1.2e-08 | 2.1e-09 | 7.3e-09 | 7.133E-09 | 4.952E-09 | 2.100E-09 | 1.200E-08 |
| 20153 | 1008 | F1008M | C | 4.4e-09 | 4.6e-09 | 4.2e-09 | 4.400E-09 | 2.000E-10 | 4.200E-09 | 4.600E-09 |
| 20154 | 1008 | F1008F | C | 6.8e-09 | 4.2e-09 | 8.2e-09 | 6.400E-09 | 2.030E-09 | 4.200E-09 | 8.200E-09 |
| 20155 | 1008 | F1008P | C | 1.4e-08 | 2.8e-09 | 1.6e-08 | 1.093E-08 | 7.114E-09 | 2.800E-09 | 1.600E-08 |
| 20156 | 1008 | F1008S | C | 1.4e-08 | 9.5e-09 | 3.8e-09 | 9.100E-09 | 5.112E-09 | 3.800E-09 | 1.400E-08 |
| 20157 | 1008 | F1008T | C | 1e-08 | 2.6e-08 | 1.5e-08 | 1.700E-08 | 8.185E-09 | 1.000E-08 | 2.600E-08 |
| 20158 | 1008 | F1008W | C | 2.2e-08 | 7.2e-09 | 5.8e-09 | 1.167E-08 | 8.976E-09 | 5.800E-09 | 2.200E-08 |
| 20159 | 1008 | F1008Y | C | 2.2e-09 | 6e-09 | 4.5e-09 | 4.233E-09 | 1.914E-09 | 2.200E-09 | 6.000E-09 |
| 20160 | 1008 | F1008V | C | 2.4e-08 | 5.7e-09 | 4e-09 | 1.123E-08 | 1.109E-08 | 4.000E-09 | 2.400E-08 |
| 20161 | 1009 | P1009A | C | 7.8e-09 | 7.8e-09 | 5e-09 | 6.867E-09 | 1.617E-09 | 5.000E-09 | 7.800E-09 |
| 20162 | 1009 | P1009R | C | 2.1e-08 | 4.7e-09 | 9.7e-09 | 1.180E-08 | 8.350E-09 | 4.700E-09 | 2.100E-08 |
| 20163 | 1009 | P1009N | C | 3.1e-09 | 3.9e-09 | 7.4e-09 | 4.800E-09 | 2.287E-09 | 3.100E-09 | 7.400E-09 |
| 20164 | 1009 | P1009D | C | 2.7e-09 | 3.7e-09 | 6.2e-09 | 4.200E-09 | 1.803E-09 | 2.700E-09 | 6.200E-09 |
| 20165 | 1009 | P1009C | C | 1.6e-08 | 3.5e-09 | 6.2e-09 | 8.567E-09 | 6.577E-09 | 3.500E-09 | 1.600E-08 |
| 20166 | 1009 | P1009E | C | 3.9e-09 | 8.8e-09 | 5.7e-09 | 6.133E-09 | 2.479E-09 | 3.900E-09 | 8.800E-09 |
| 20167 | 1009 | P1009Q | C | 3e-09 | 1.1e-08 | 4.8e-09 | 6.267E-09 | 4.197E-09 | 3.000E-09 | 1.100E-08 |
| 20168 | 1009 | P1009G | C | 1.1e-08 | 5.3e-09 | 6.9e-09 | 7.733E-09 | 2.940E-09 | 5.300E-09 | 1.100E-08 |
| 20169 | 1009 | P1009H | C | 2.5e-09 | 1.4e-08 | 6.2e-09 | 7.567E-09 | 5.871E-09 | 2.500E-09 | 1.400E-08 |
| 20170 | 1009 | P1009I | C | 6e-09 | 1.6e-08 | 4.5e-09 | 8.833E-09 | 6.252E-09 | 4.500E-09 | 1.600E-08 |
| 20171 | 1009 | P1009L | C | 6.3e-09 | 7.4e-09 | 6e-09 | 6.567E-09 | 7.371E-10 | 6.000E-09 | 7.400E-09 |
| 20172 | 1009 | P1009K | C | 1e-08 | 3.7e-09 | 6.3e-09 | 6.667E-09 | 3.166E-09 | 3.700E-09 | 1.000E-08 |
| 20173 | 1009 | P1009M | C | 4.5e-09 | 8.2e-09 | 1.2e-08 | 8.233E-09 | 3.750E-09 | 4.500E-09 | 1.200E-08 |
| 20174 | 1009 | P1009F | C | 3.7e-09 | 7.8e-09 | 1.5e-08 | 8.833E-09 | 5.720E-09 | 3.700E-09 | 1.500E-08 |
| 20175 | 1009 | P1009P | C | 2.9e-08 | 4.6e-09 | 9e-09 | 1.420E-08 | 1.300E-08 | 4.600E-09 | 2.900E-08 |
| 20176 | 1009 | P1009S | C | 1e-08 | 3.8e-09 | 2.7e-09 | 5.500E-09 | 3.936E-09 | 2.700E-09 | 1.000E-08 |
| 20177 | 1009 | P1009T | C | 8.2e-09 | 8.8e-09 | 1e-08 | 9.000E-09 | 9.165E-10 | 8.200E-09 | 1.000E-08 |
| 20178 | 1009 | P1009W | C | 6.4e-09 | 1.3e-08 | 1.4e-08 | 1.113E-08 | 4.130E-09 | 6.400E-09 | 1.400E-08 |
| 20179 | 1009 | P1009Y | C | 3.5e-09 | 1.8e-08 | 3.4e-09 | 8.300E-09 | 8.401E-09 | 3.400E-09 | 1.800E-08 |
| 20180 | 1009 | P1009V | C | 2.5e-09 | 8.2e-09 | 9.1e-09 | 6.600E-09 | 3.579E-09 | 2.500E-09 | 9.100E-09 |
| 20181 | 1010 | A1010A | C | 8.7e-10 | 7.7e-09 | 3.9e-09 | 4.157E-09 | 3.422E-09 | 8.700E-10 | 7.700E-09 |
| 20182 | 1010 | A1010R | C | 8.9e-09 | 3.6e-09 | 5.5e-09 | 6.000E-09 | 2.685E-09 | 3.600E-09 | 8.900E-09 |
| 20183 | 1010 | A1010N | C | 1.1e-08 | 2.6e-09 | 6.4e-09 | 6.667E-09 | 4.206E-09 | 2.600E-09 | 1.100E-08 |
| 20184 | 1010 | A1010D | C | 7.9e-09 | 8.2e-09 | 9.5e-09 | 8.533E-09 | 8.505E-10 | 7.900E-09 | 9.500E-09 |
| 20185 | 1010 | A1010C | C | 1.3e-08 | 3.7e-09 | 1.1e-08 | 9.233E-09 | 4.895E-09 | 3.700E-09 | 1.300E-08 |
| 20186 | 1010 | A1010E | C | 9.1e-09 | 9.9e-09 | 7.1e-09 | 8.700E-09 | 1.442E-09 | 7.100E-09 | 9.900E-09 |
| 20187 | 1010 | A1010Q | C | 6.8e-09 | 1.7e-08 | 5.6e-09 | 9.800E-09 | 6.264E-09 | 5.600E-09 | 1.700E-08 |
| 20188 | 1010 | A1010G | C | 3.5e-09 | 8.6e-09 | 2e-08 | 1.070E-08 | 8.448E-09 | 3.500E-09 | 2.000E-08 |
| 20189 | 1010 | A1010H | C | 3.4e-09 | 8.8e-09 | 1.5e-08 | 9.067E-09 | 5.805E-09 | 3.400E-09 | 1.500E-08 |
| 20190 | 1010 | A1010I | C | 4e-09 | 4.3e-09 | 1.1e-08 | 6.433E-09 | 3.958E-09 | 4.000E-09 | 1.100E-08 |
| 20191 | 1010 | A1010L | C | 2.6e-09 | 5e-09 | 1.2e-08 | 6.533E-09 | 4.884E-09 | 2.600E-09 | 1.200E-08 |
| 20192 | 1010 | A1010K | C | 6e-09 | 1.1e-08 | 2e-08 | 1.233E-08 | 7.095E-09 | 6.000E-09 | 2.000E-08 |
| 20193 | 1010 | A1010M | C | 7.6e-09 | 2.1e-08 | 3.3e-09 | 1.063E-08 | 9.232E-09 | 3.300E-09 | 2.100E-08 |
| 20194 | 1010 | A1010F | C | 9.7e-09 | 7.6e-09 | 4.5e-09 | 7.267E-09 | 2.616E-09 | 4.500E-09 | 9.700E-09 |
| 20195 | 1010 | A1010P | C | 1.4e-08 | 9.3e-09 | 1.3e-08 | 1.210E-08 | 2.476E-09 | 9.300E-09 | 1.400E-08 |
| 20196 | 1010 | A1010S | C | 1.1e-08 | 1.8e-08 | 4.9e-09 | 1.130E-08 | 6.555E-09 | 4.900E-09 | 1.800E-08 |
| 20197 | 1010 | A1010T | C | 5.3e-09 | 8.1e-09 | 2.7e-09 | 5.367E-09 | 2.701E-09 | 2.700E-09 | 8.100E-09 |
| 20198 | 1010 | A1010W | C | 7.4e-09 | 7.7e-09 | 5e-09 | 6.700E-09 | 1.480E-09 | 5.000E-09 | 7.700E-09 |
| 20199 | 1010 | A1010Y | C | 6.8e-09 | 4.7e-09 | 1.9e-08 | 1.017E-08 | 7.722E-09 | 4.700E-09 | 1.900E-08 |
| 20200 | 1010 | A1010V | C | 6.5e-09 | 1.3e-08 | 4.9e-09 | 8.133E-09 | 4.290E-09 | 4.900E-09 | 1.300E-08 |
| 20201 | 1011 | V1011A | C | 4.1e-09 | 4.1e-09 | 4.1e-08 | 1.640E-08 | 2.130E-08 | 4.100E-09 | 4.100E-08 |
| 20202 | 1011 | V1011R | C | 5.9e-09 | 7.6e-09 | 1.6e-08 | 9.833E-09 | 5.408E-09 | 5.900E-09 | 1.600E-08 |
| 20203 | 1011 | V1011N | C | 1.2e-08 | 6e-09 | 1.8e-08 | 1.200E-08 | 6.000E-09 | 6.000E-09 | 1.800E-08 |
| 20204 | 1011 | V1011D | C | 8.9e-09 | 1.1e-08 | 3.6e-09 | 7.833E-09 | 3.814E-09 | 3.600E-09 | 1.100E-08 |
| 20205 | 1011 | V1011C | C | 6.6e-09 | 2.4e-09 | 5.5e-09 | 4.833E-09 | 2.178E-09 | 2.400E-09 | 6.600E-09 |
| 20206 | 1011 | V1011E | C | 5.9e-09 | 5.3e-09 | 1.1e-08 | 7.400E-09 | 3.132E-09 | 5.300E-09 | 1.100E-08 |
| 20207 | 1011 | V1011Q | C | 1.4e-08 | 4.8e-09 | 1.7e-08 | 1.193E-08 | 6.357E-09 | 4.800E-09 | 1.700E-08 |
| 20208 | 1011 | V1011G | C | 6.1e-09 | 9.4e-09 | 6.6e-09 | 7.367E-09 | 1.779E-09 | 6.100E-09 | 9.400E-09 |
| 20209 | 1011 | V1011H | C | 6.1e-09 | 6.9e-09 | 1.1e-08 | 8.000E-09 | 2.629E-09 | 6.100E-09 | 1.100E-08 |
| 20210 | 1011 | V1011I | C | 2.6e-09 | 1.2e-08 | 2.5e-09 | 5.700E-09 | 5.456E-09 | 2.500E-09 | 1.200E-08 |
| 20211 | 1011 | V1011L | C | 1.8e-09 | 1.8e-08 | 1.2e-08 | 1.060E-08 | 8.190E-09 | 1.800E-09 | 1.800E-08 |
| 20212 | 1011 | V1011K | C | 4.3e-09 | 1.9e-09 | 1.4e-08 | 6.733E-09 | 6.407E-09 | 1.900E-09 | 1.400E-08 |
| 20213 | 1011 | V1011M | C | 8.5e-09 | 7.3e-09 | 5.9e-09 | 7.233E-09 | 1.301E-09 | 5.900E-09 | 8.500E-09 |
| 20214 | 1011 | V1011F | C | 9.1e-09 | 1.9e-09 | 2.2e-08 | 1.100E-08 | 1.018E-08 | 1.900E-09 | 2.200E-08 |
| 20215 | 1011 | V1011P | C | 3.1e-09 | 4.1e-09 | 4.1e-09 | 3.767E-09 | 5.774E-10 | 3.100E-09 | 4.100E-09 |
| 20216 | 1011 | V1011S | C | 2.7e-09 | 1.5e-08 | 1.4e-08 | 1.057E-08 | 6.831E-09 | 2.700E-09 | 1.500E-08 |
| 20217 | 1011 | V1011T | C | 6.7e-09 | 6.6e-09 | 4.8e-09 | 6.033E-09 | 1.069E-09 | 4.800E-09 | 6.700E-09 |
| 20218 | 1011 | V1011W | C | 3.7e-09 | 1.6e-08 | 1.9e-08 | 1.290E-08 | 8.107E-09 | 3.700E-09 | 1.900E-08 |
| 20219 | 1011 | V1011Y | C | 1e-08 | 3.6e-09 | 3e-09 | 5.533E-09 | 3.880E-09 | 3.000E-09 | 1.000E-08 |
| 20220 | 1011 | V1011V | C | 7.2e-09 | 8.8e-09 | 4.1e-09 | 6.700E-09 | 2.390E-09 | 4.100E-09 | 8.800E-09 |
| 20221 | 1012 | L1012A | C | 4.8e-09 | 9.4e-09 | 4.5e-09 | 6.233E-09 | 2.747E-09 | 4.500E-09 | 9.400E-09 |
| 20222 | 1012 | L1012R | C | 7.4e-09 | 1.8e-08 | 3e-09 | 9.467E-09 | 7.711E-09 | 3.000E-09 | 1.800E-08 |
| 20223 | 1012 | L1012N | C | 3.7e-09 | 3.1e-09 | 1.1e-08 | 5.933E-09 | 4.398E-09 | 3.100E-09 | 1.100E-08 |
| 20224 | 1012 | L1012D | C | 3.5e-09 | 9.7e-09 | 5.7e-09 | 6.300E-09 | 3.143E-09 | 3.500E-09 | 9.700E-09 |
| 20225 | 1012 | L1012C | C | 2.2e-08 | 4.1e-09 | 6.6e-09 | 1.090E-08 | 9.694E-09 | 4.100E-09 | 2.200E-08 |
| 20226 | 1012 | L1012E | C | 1.9e-08 | 1.3e-08 | 1.1e-08 | 1.433E-08 | 4.163E-09 | 1.100E-08 | 1.900E-08 |
| 20227 | 1012 | L1012Q | C | 2.1e-08 | 1.2e-08 | 4.3e-09 | 1.243E-08 | 8.358E-09 | 4.300E-09 | 2.100E-08 |
| 20228 | 1012 | L1012G | C | 3.3e-09 | 4.9e-09 | 1.1e-08 | 6.400E-09 | 4.063E-09 | 3.300E-09 | 1.100E-08 |
| 20229 | 1012 | L1012H | C | 9.1e-09 | 1e-08 | 1.4e-08 | 1.103E-08 | 2.608E-09 | 9.100E-09 | 1.400E-08 |

| | | | | | | | | | | |
|---|---|---|---|---|---|---|---|---|---|---|
| 20230 | 1012 | L1012I | C | 1.8e-08 | 2.7e-09 | 5.1e-09 | 8.600E-09 | 8.229E-09 | 2.700E-09 | 1.800E-08 |
| 20231 | 1012 | L1012L | C | 4.2e-09 | 6.9e-09 | 1e-08 | 7.033E-09 | 2.902E-09 | 4.200E-09 | 1.000E-08 |
| 20232 | 1012 | L1012K | C | 1.9e-08 | 1.3e-08 | 9.4e-09 | 1.380E-08 | 4.850E-09 | 9.400E-09 | 1.900E-08 |
| 20233 | 1012 | L1012M | C | 1e-08 | 1.1e-08 | 2.5e-09 | 7.833E-09 | 4.646E-09 | 2.500E-09 | 1.100E-08 |
| 20234 | 1012 | L1012F | C | 4e-09 | 4.3e-09 | 1.5e-09 | 3.267E-09 | 1.537E-09 | 1.500E-09 | 4.300E-09 |
| 20235 | 1012 | L1012P | C | 1.7e-09 | 1.4e-08 | 9.1e-09 | 8.267E-09 | 6.192E-09 | 1.700E-09 | 1.400E-08 |
| 20236 | 1012 | L1012S | C | 6.3e-09 | 1.5e-08 | 3.8e-09 | 8.367E-09 | 5.879E-09 | 3.800E-09 | 1.500E-08 |
| 20237 | 1012 | L1012T | C | 2e-08 | 6.8e-09 | 1e-08 | 1.227E-08 | 6.886E-09 | 6.800E-09 | 2.000E-08 |
| 20238 | 1012 | L1012W | C | 7.8e-09 | 5.3e-09 | 1.8e-09 | 4.967E-09 | 3.014E-09 | 1.800E-09 | 7.800E-09 |
| 20239 | 1012 | L1012Y | C | 7e-09 | 1.8e-08 | 2.5e-09 | 9.167E-09 | 7.974E-09 | 2.500E-09 | 1.800E-08 |
| 20240 | 1012 | L1012V | C | 3.6e-09 | 9.7e-09 | 4.8e-09 | 6.033E-09 | 3.232E-09 | 3.600E-09 | 9.700E-09 |
| 20241 | 1013 | Q1013A | C | 4e-09 | 3.5e-09 | 5.3e-09 | 4.267E-09 | 9.292E-10 | 3.500E-09 | 5.300E-09 |
| 20242 | 1013 | Q1013R | C | 3.4e-09 | 4.4e-09 | 7.2e-09 | 5.000E-09 | 1.970E-09 | 3.400E-09 | 7.200E-09 |
| 20243 | 1013 | Q1013N | C | 3e-09 | 4.4e-09 | 4.5e-09 | 3.967E-09 | 8.386E-10 | 3.000E-09 | 4.500E-09 |
| 20244 | 1013 | Q1013D | C | 3.4e-09 | 5.1e-09 | 7.9e-09 | 5.467E-09 | 2.272E-09 | 3.400E-09 | 7.900E-09 |
| 20245 | 1013 | Q1013C | C | 9.1e-09 | 7.9e-09 | 6.3e-09 | 7.767E-09 | 1.405E-09 | 6.300E-09 | 9.100E-09 |
| 20246 | 1013 | Q1013E | C | 8.6e-09 | 7.5e-09 | 1.2e-08 | 9.367E-09 | 2.346E-09 | 7.500E-09 | 1.200E-08 |
| 20247 | 1013 | Q1013Q | C | 3.1e-08 | 1.1e-08 | 4.1e-09 | 1.537E-08 | 1.397E-08 | 4.100E-09 | 3.100E-08 |
| 20248 | 1013 | Q1013G | C | 1.2e-08 | 8.1e-09 | 1.2e-08 | 1.070E-08 | 2.252E-09 | 8.100E-09 | 1.200E-08 |
| 20249 | 1013 | Q1013H | C | 5.5e-09 | 4.6e-09 | 3.5e-09 | 4.533E-09 | 1.002E-09 | 3.500E-09 | 5.500E-09 |
| 20250 | 1013 | Q1013I | C | 3.3e-09 | 5.2e-09 | 5.9e-09 | 4.800E-09 | 1.345E-09 | 3.300E-09 | 5.900E-09 |
| 20251 | 1013 | Q1013L | C | 1.2e-08 | 1e-08 | 8.2e-09 | 1.007E-08 | 1.901E-09 | 8.200E-09 | 1.200E-08 |
| 20252 | 1013 | Q1013K | C | 1.2e-08 | 4.8e-09 | 3.5e-09 | 6.767E-09 | 4.579E-09 | 3.500E-09 | 1.200E-08 |
| 20253 | 1013 | Q1013M | C | 1.6e-09 | 7e-09 | 5e-09 | 4.533E-09 | 2.730E-09 | 1.600E-09 | 7.000E-09 |
| 20254 | 1013 | Q1013F | C | 6.7e-09 | 4.2e-09 | 1.3e-08 | 7.967E-09 | 4.535E-09 | 4.200E-09 | 1.300E-08 |
| 20255 | 1013 | Q1013P | C | 4.4e-09 | 3.2e-09 | 9.1e-09 | 5.567E-09 | 3.118E-09 | 3.200E-09 | 9.100E-09 |
| 20256 | 1013 | Q1013S | C | 8.1e-09 | 5.1e-09 | 1.8e-08 | 1.040E-08 | 6.751E-09 | 5.100E-09 | 1.800E-08 |
| 20257 | 1013 | Q1013T | C | 4.5e-09 | 5.5e-09 | 8.2e-09 | 6.067E-09 | 1.914E-09 | 4.500E-09 | 8.200E-09 |
| 20258 | 1013 | Q1013W | C | 1.6e-08 | 3.9e-09 | 5.7e-09 | 8.533E-09 | 6.529E-09 | 3.900E-09 | 1.600E-08 |
| 20259 | 1013 | Q1013Y | C | 3.6e-09 | 1.1e-08 | 1.1e-08 | 8.533E-09 | 4.272E-09 | 3.600E-09 | 1.100E-08 |
| 20260 | 1013 | Q1013V | C | 7.9e-09 | 4e-09 | 3.8e-09 | 5.233E-09 | 2.312E-09 | 3.800E-09 | 7.900E-09 |
| 20261 | 1014 | S1014A | C | 5.7e-09 | 2.9e-09 | 8.2e-09 | 5.600E-09 | 2.651E-09 | 2.900E-09 | 8.200E-09 |
| 20262 | 1014 | S1014R | C | 6.4e-09 | 6.1e-09 | 4.3e-09 | 5.600E-09 | 1.136E-09 | 4.300E-09 | 6.400E-09 |
| 20263 | 1014 | S1014N | C | 5.4e-09 | 1.1e-08 | 7.8e-09 | 8.067E-09 | 2.810E-09 | 5.400E-09 | 1.100E-08 |
| 20264 | 1014 | S1014D | C | 5.4e-09 | 1.5e-08 | 8e-09 | 9.467E-09 | 4.965E-09 | 5.400E-09 | 1.500E-08 |
| 20265 | 1014 | S1014C | C | 1.6e-08 | 1.5e-08 | 5.3e-09 | 1.210E-08 | 5.910E-09 | 5.300E-09 | 1.600E-08 |
| 20266 | 1014 | S1014E | C | 6.2e-09 | 1.3e-08 | 3.5e-09 | 7.567E-09 | 4.895E-09 | 3.500E-09 | 1.300E-08 |
| 20267 | 1014 | S1014Q | C | 4.7e-09 | 7.6e-09 | 5.5e-09 | 5.933E-09 | 1.498E-09 | 4.700E-09 | 7.600E-09 |
| 20268 | 1014 | S1014G | C | 5.8e-09 | 8.9e-09 | 9e-09 | 7.900E-09 | 1.819E-09 | 5.800E-09 | 9.000E-09 |
| 20269 | 1014 | S1014H | C | 1e-08 | 5.8e-09 | 1.5e-08 | 1.027E-08 | 4.606E-09 | 5.800E-09 | 1.500E-08 |
| 20270 | 1014 | S1014I | C | 8.9e-09 | 7.4e-09 | 1.4e-08 | 1.010E-08 | 3.460E-09 | 7.400E-09 | 1.400E-08 |
| 20271 | 1014 | S1014L | C | 1.1e-08 | 2.7e-09 | 7.7e-09 | 7.133E-09 | 4.179E-09 | 2.700E-09 | 1.100E-08 |
| 20272 | 1014 | S1014K | C | 2.5e-08 | 1.4e-08 | 8e-09 | 1.567E-08 | 8.622E-09 | 8.000E-09 | 2.500E-08 |
| 20273 | 1014 | S1014M | C | 4.1e-09 | 2.5e-09 | 6.8e-09 | 4.467E-09 | 2.173E-09 | 2.500E-09 | 6.800E-09 |
| 20274 | 1014 | S1014F | C | 1.6e-08 | 9.1e-09 | 4.8e-09 | 9.967E-09 | 5.650E-09 | 4.800E-09 | 1.600E-08 |
| 20275 | 1014 | S1014P | C | 9.2e-09 | 7.9e-09 | 5.3e-09 | 7.467E-09 | 1.986E-09 | 5.300E-09 | 9.200E-09 |
| 20276 | 1014 | S1014S | C | 5.5e-09 | 1.6e-08 | 4.5e-09 | 8.667E-09 | 6.371E-09 | 4.500E-09 | 1.600E-08 |
| 20277 | 1014 | S1014T | C | 5.3e-09 | 8.8e-09 | 4.6e-09 | 6.233E-09 | 2.250E-09 | 4.600E-09 | 8.800E-09 |
| 20278 | 1014 | S1014W | C | 1.2e-08 | 4.4e-09 | 8.9e-09 | 8.433E-09 | 3.821E-09 | 4.400E-09 | 1.200E-08 |
| 20279 | 1014 | S1014Y | C | 7.3e-09 | 6.7e-09 | 4.1e-09 | 6.033E-09 | 1.701E-09 | 4.100E-09 | 7.300E-09 |
| 20280 | 1014 | S1014V | C | 2.5e-09 | 4.2e-09 | 1.1e-09 | 2.600E-09 | 1.552E-09 | 1.100E-09 | 4.200E-09 |
| 20281 | 1015 | S1015A | C | 9.5e-09 | 6.6e-09 | 5.1e-09 | 7.067E-09 | 2.237E-09 | 5.100E-09 | 9.500E-09 |
| 20282 | 1015 | S1015R | C | 1.5e-08 | 6.7e-09 | 1.8e-08 | 1.323E-08 | 5.853E-09 | 6.700E-09 | 1.800E-08 |
| 20283 | 1015 | S1015N | C | 1.3e-08 | 1e-08 | 1.5e-08 | 1.267E-08 | 2.517E-09 | 1.000E-08 | 1.500E-08 |
| 20284 | 1015 | S1015D | C | 4.7e-09 | 4.3e-09 | 1.1e-08 | 6.667E-09 | 3.758E-09 | 4.300E-09 | 1.100E-08 |
| 20285 | 1015 | S1015C | C | 1.8e-08 | 1.7e-08 | 1.3e-08 | 1.600E-08 | 2.646E-09 | 1.300E-08 | 1.800E-08 |
| 20286 | 1015 | S1015E | C | 6.9e-09 | 3.7e-09 | 9.3e-09 | 6.633E-09 | 2.810E-09 | 3.700E-09 | 9.300E-09 |
| 20287 | 1015 | S1015Q | C | 4.8e-09 | 1.8e-08 | 1.4e-08 | 1.227E-08 | 6.769E-09 | 4.800E-09 | 1.800E-08 |
| 20288 | 1015 | S1015G | C | 1.2e-08 | 8.9e-09 | 1.7e-08 | 1.263E-08 | 4.087E-09 | 8.900E-09 | 1.700E-08 |
| 20289 | 1015 | S1015H | C | 3.9e-09 | 2.9e-09 | 2.2e-08 | 9.600E-09 | 1.075E-08 | 2.900E-09 | 2.200E-08 |
| 20290 | 1015 | S1015I | C | 7.7e-09 | 8.6e-09 | 1.2e-08 | 9.433E-09 | 2.268E-09 | 7.700E-09 | 1.200E-08 |
| 20291 | 1015 | S1015L | C | 6e-09 | 3.4e-09 | 5.7e-09 | 5.033E-09 | 1.422E-09 | 3.400E-09 | 6.000E-09 |
| 20292 | 1015 | S1015K | C | 1.7e-08 | 9.2e-09 | 1.5e-08 | 1.373E-08 | 4.051E-09 | 9.200E-09 | 1.700E-08 |
| 20293 | 1015 | S1015M | C | 5e-09 | 6.4e-09 | 6e-09 | 5.800E-09 | 7.211E-10 | 5.000E-09 | 6.400E-09 |
| 20294 | 1015 | S1015F | C | 2e-08 | 8.9e-09 | 4.3e-09 | 1.107E-08 | 8.071E-09 | 4.300E-09 | 2.000E-08 |
| 20295 | 1015 | S1015P | C | 8.3e-09 | 2.4e-09 | 1.2e-08 | 7.567E-09 | 4.842E-09 | 2.400E-09 | 1.200E-08 |
| 20296 | 1015 | S1015S | C | 4.4e-09 | 1e-08 | 2.5e-09 | 5.633E-09 | 3.899E-09 | 2.500E-09 | 1.000E-08 |
| 20297 | 1015 | S1015T | C | 3.5e-09 | 6.9e-09 | 7.9e-09 | 6.100E-09 | 2.307E-09 | 3.500E-09 | 7.900E-09 |
| 20298 | 1015 | S1015W | C | 7.3e-09 | 1.7e-08 | 2.1e-08 | 1.510E-08 | 7.045E-09 | 7.300E-09 | 2.100E-08 |
| 20299 | 1015 | S1015Y | C | 4.1e-09 | 7.2e-09 | 7.5e-09 | 6.267E-09 | 1.882E-09 | 4.100E-09 | 7.500E-09 |
| 20300 | 1015 | S1015V | C | 7.8e-09 | 7.7e-09 | 2.2e-08 | 1.250E-08 | 8.227E-09 | 7.700E-09 | 2.200E-08 |
| 20301 | 1016 | G1016A | C | 2.1e-08 | 7.6e-09 | 1.8e-08 | 1.553E-08 | 7.032E-09 | 7.600E-09 | 2.100E-08 |
| 20302 | 1016 | G1016R | C | 8.4e-09 | 1.1e-08 | 1.1e-08 | 1.013E-08 | 1.501E-09 | 8.400E-09 | 1.100E-08 |
| 20303 | 1016 | G1016N | C | 8e-09 | 4.3e-09 | 6e-09 | 6.100E-09 | 1.852E-09 | 4.300E-09 | 8.000E-09 |
| 20304 | 1016 | G1016D | C | 7.7e-09 | 1.1e-08 | 9.8e-09 | 9.500E-09 | 1.670E-09 | 7.700E-09 | 1.100E-08 |
| 20305 | 1016 | G1016C | C | 3e-09 | 1e-08 | 4.7e-09 | 5.900E-09 | 3.651E-09 | 3.000E-09 | 1.000E-08 |
| 20306 | 1016 | G1016E | C | 6.6e-09 | 3e-09 | 9.4e-09 | 6.333E-09 | 3.208E-09 | 3.000E-09 | 9.400E-09 |
| 20307 | 1016 | G1016Q | C | 4.4e-09 | 6.6e-09 | 5.8e-09 | 5.600E-09 | 1.114E-09 | 4.400E-09 | 6.600E-09 |
| 20308 | 1016 | G1016G | C | 7.1e-09 | 6.9e-09 | 2.7e-09 | 5.567E-09 | 2.485E-09 | 2.700E-09 | 7.100E-09 |
| 20309 | 1016 | G1016H | C | 7.8e-09 | 4.3e-09 | 6.3e-09 | 6.133E-09 | 1.756E-09 | 4.300E-09 | 7.800E-09 |
| 20310 | 1016 | G1016I | C | 2.8e-09 | 4.5e-09 | 6.5e-09 | 4.600E-09 | 1.852E-09 | 2.800E-09 | 6.500E-09 |
| 20311 | 1016 | G1016L | C | 3.6e-09 | 5.6e-09 | 3.8e-09 | 4.333E-09 | 1.102E-09 | 3.600E-09 | 5.600E-09 |
| 20312 | 1016 | G1016K | C | 5.1e-09 | 8e-09 | 3e-09 | 5.367E-09 | 2.511E-09 | 3.000E-09 | 8.000E-09 |
| 20313 | 1016 | G1016M | C | 5.5e-09 | 2.4e-08 | 6.6e-09 | 1.203E-08 | 1.038E-08 | 5.500E-09 | 2.400E-08 |
| 20314 | 1016 | G1016F | C | 2.4e-08 | 7.8e-09 | 1.5e-08 | 1.560E-08 | 8.117E-09 | 7.800E-09 | 2.400E-08 |

| | | | | | | | | | | |
|---|---|---|---|---|---|---|---|---|---|---|
| 20315 | 1016 | G1016P | C | 5e-09 | 5.9e-09 | 1.2e-08 | 7.633E-09 | 3.808E-09 | 5.000E-09 | 1.200E-08 |
| 20316 | 1016 | G1016S | C | 9.1e-10 | 5.5e-09 | 2.4e-09 | 2.937E-09 | 2.342E-09 | 9.100E-10 | 5.500E-09 |
| 20317 | 1016 | G1016T | C | 2.4e-09 | 6.5e-09 | 7.5e-09 | 5.467E-09 | 2.702E-09 | 2.400E-09 | 7.500E-09 |
| 20318 | 1016 | G1016W | C | 8.7e-09 | 7.1e-09 | 3.2e-09 | 6.333E-09 | 2.829E-09 | 3.200E-09 | 8.700E-09 |
| 20319 | 1016 | G1016Y | C | 8e-09 | 1.8e-08 | 6.4e-09 | 1.080E-08 | 6.286E-09 | 6.400E-09 | 1.800E-08 |
| 20320 | 1016 | G1016V | C | 5e-09 | 9.3e-09 | 7.6e-09 | 7.300E-09 | 2.166E-09 | 5.000E-09 | 9.300E-09 |
| 20321 | 1017 | L1017A | C | 2.8e-09 | 4.5e-08 | 1.5e-08 | 2.093E-08 | 2.172E-08 | 2.800E-09 | 4.500E-08 |
| 20322 | 1017 | L1017R | C | 3.7e-09 | 2.5e-08 | 5.4e-09 | 1.137E-08 | 1.184E-08 | 3.700E-09 | 2.500E-08 |
| 20323 | 1017 | L1017N | C | 2e-09 | 5.7e-09 | 8.1e-09 | 5.267E-09 | 3.073E-09 | 2.000E-09 | 8.100E-09 |
| 20324 | 1017 | L1017D | C | 1.5e-08 | 1.8e-08 | 2.4e-09 | 1.180E-08 | 8.278E-09 | 2.400E-09 | 1.800E-08 |
| 20325 | 1017 | L1017C | C | 1.2e-08 | 3e-09 | 4.9e-09 | 6.633E-09 | 4.744E-09 | 3.000E-09 | 1.200E-08 |
| 20326 | 1017 | L1017E | C | 6.6e-09 | 2.2e-09 | 7.5e-09 | 5.433E-09 | 2.836E-09 | 2.200E-09 | 7.500E-09 |
| 20327 | 1017 | L1017Q | C | 6.4e-09 | 9.3e-09 | 2.1e-09 | 5.933E-09 | 3.623E-09 | 2.100E-09 | 9.300E-09 |
| 20328 | 1017 | L1017G | C | 5.3e-09 | 6.8e-09 | 6.9e-09 | 6.333E-09 | 8.963E-10 | 5.300E-09 | 6.900E-09 |
| 20329 | 1017 | L1017H | C | 5.7e-09 | 5.7e-09 | 7.2e-09 | 6.200E-09 | 8.660E-10 | 5.700E-09 | 7.200E-09 |
| 20330 | 1017 | L1017I | C | 7.1e-09 | 6.8e-09 | 4.4e-09 | 6.100E-09 | 1.480E-09 | 4.400E-09 | 7.100E-09 |
| 20331 | 1017 | L1017L | C | 3e-09 | 1.3e-08 | 8e-09 | 8.000E-09 | 5.000E-09 | 3.000E-09 | 1.300E-08 |
| 20332 | 1017 | L1017K | C | 3.5e-09 | 1.1e-08 | 5.5e-09 | 6.667E-09 | 3.884E-09 | 3.500E-09 | 1.100E-08 |
| 20333 | 1017 | L1017M | C | 1.2e-08 | 8.3e-09 | 7.1e-09 | 9.133E-09 | 2.554E-09 | 7.100E-09 | 1.200E-08 |
| 20334 | 1017 | L1017F | C | 9.9e-09 | 1.2e-08 | 9.2e-09 | 1.037E-08 | 1.457E-09 | 9.200E-09 | 1.200E-08 |
| 20335 | 1017 | L1017P | C | 3.8e-09 | 1.6e-08 | 1.1e-08 | 1.027E-08 | 6.133E-09 | 3.800E-09 | 1.600E-08 |
| 20336 | 1017 | L1017S | C | 8.4e-09 | 4.4e-09 | 8.2e-09 | 7.000E-09 | 2.254E-09 | 4.400E-09 | 8.400E-09 |
| 20337 | 1017 | L1017T | C | 8.8e-09 | 1.4e-08 | 4.2e-09 | 9.000E-09 | 4.903E-09 | 4.200E-09 | 1.400E-08 |
| 20338 | 1017 | L1017W | C | 1.2e-08 | 2.7e-08 | 5.8e-09 | 1.493E-08 | 1.090E-08 | 5.800E-09 | 2.700E-08 |
| 20339 | 1017 | L1017Y | C | 1.8e-09 | 1.8e-08 | 1.2e-08 | 1.060E-08 | 8.190E-09 | 1.800E-09 | 1.800E-08 |
| 20340 | 1017 | L1017V | C | 2.4e-09 | 1.2e-08 | 8.4e-09 | 7.600E-09 | 4.850E-09 | 2.400E-09 | 1.200E-08 |
| 20341 | 1018 | Y1018A | C | 3.4e-09 | 5.6e-09 | 4.3e-09 | 4.433E-09 | 1.106E-09 | 3.400E-09 | 5.600E-09 |
| 20342 | 1018 | Y1018R | C | 4.7e-09 | 6.8e-09 | 4.3e-09 | 5.267E-09 | 1.343E-09 | 4.300E-09 | 6.800E-09 |
| 20343 | 1018 | Y1018N | C | 6.2e-09 | 3.5e-09 | 2.9e-09 | 4.200E-09 | 1.758E-09 | 2.900E-09 | 6.200E-09 |
| 20344 | 1018 | Y1018D | C | 3.2e-09 | 1.7e-08 | 2.3e-09 | 7.500E-09 | 8.240E-09 | 2.300E-09 | 1.700E-08 |
| 20345 | 1018 | Y1018C | C | 4.9e-09 | 7.7e-09 | 3e-09 | 5.200E-09 | 2.364E-09 | 3.000E-09 | 7.700E-09 |
| 20346 | 1018 | Y1018E | C | 4.3e-09 | 1.3e-08 | 4.2e-09 | 7.167E-09 | 5.052E-09 | 4.200E-09 | 1.300E-08 |
| 20347 | 1018 | Y1018Q | C | 3.8e-09 | 6.9e-09 | 1.1e-08 | 7.233E-09 | 3.612E-09 | 3.800E-09 | 1.100E-08 |
| 20348 | 1018 | Y1018G | C | 9.7e-09 | 1e-08 | 9e-09 | 9.567E-09 | 5.132E-10 | 9.000E-09 | 1.000E-08 |
| 20349 | 1018 | Y1018H | C | 6.6e-09 | 5.4e-09 | 1.5e-09 | 4.500E-09 | 2.666E-09 | 1.500E-09 | 6.600E-09 |
| 20350 | 1018 | Y1018I | C | 6.7e-09 | 5e-09 | 5.1e-09 | 5.600E-09 | 9.539E-10 | 5.000E-09 | 6.700E-09 |
| 20351 | 1018 | Y1018L | C | 3.5e-09 | 3.5e-08 | 6e-09 | 1.483E-08 | 1.751E-08 | 3.500E-09 | 3.500E-08 |
| 20352 | 1018 | Y1018K | C | 5.5e-09 | 2.2e-08 | 9.8e-09 | 1.243E-08 | 8.559E-09 | 5.500E-09 | 2.200E-08 |
| 20353 | 1018 | Y1018M | C | 9.9e-09 | 6.3e-09 | 6.5e-09 | 7.567E-09 | 2.023E-09 | 6.300E-09 | 9.900E-09 |
| 20354 | 1018 | Y1018F | C | 3.5e-09 | 7.9e-09 | 1.1e-09 | 4.167E-09 | 3.449E-09 | 1.100E-09 | 7.900E-09 |
| 20355 | 1018 | Y1018P | C | 7.1e-09 | 8.1e-09 | 1.3e-08 | 9.400E-09 | 3.158E-09 | 7.100E-09 | 1.300E-08 |
| 20356 | 1018 | Y1018S | C | 5e-09 | 6.7e-09 | 9.9e-09 | 7.200E-09 | 2.488E-09 | 5.000E-09 | 9.900E-09 |
| 20357 | 1018 | Y1018T | C | 8.6e-09 | 5.3e-09 | 1.4e-08 | 9.300E-09 | 4.392E-09 | 5.300E-09 | 1.400E-08 |
| 20358 | 1018 | Y1018W | C | 5.5e-09 | 1.9e-09 | 3e-09 | 3.467E-09 | 1.845E-09 | 1.900E-09 | 5.500E-09 |
| 20359 | 1018 | Y1018Y | C | 8.6e-09 | 2.9e-09 | 7e-09 | 6.167E-09 | 2.940E-09 | 2.900E-09 | 8.600E-09 |
| 20360 | 1018 | Y1018V | C | 5.9e-09 | 5.1e-09 | 9e-09 | 6.667E-09 | 2.060E-09 | 5.100E-09 | 9.000E-09 |
| 20361 | 1019 | S1019A | C | 4.9e-09 | 1.2e-08 | 1e-08 | 8.967E-09 | 3.661E-09 | 4.900E-09 | 1.200E-08 |
| 20362 | 1019 | S1019R | C | 5.4e-09 | 7.3e-09 | 2.5e-09 | 5.067E-09 | 2.417E-09 | 2.500E-09 | 7.300E-09 |
| 20363 | 1019 | S1019N | C | 3.4e-09 | 3.6e-09 | 6.3e-09 | 4.433E-09 | 1.620E-09 | 3.400E-09 | 6.300E-09 |
| 20364 | 1019 | S1019D | C | 5.8e-09 | 6.9e-09 | 5.5e-09 | 6.067E-09 | 7.371E-10 | 5.500E-09 | 6.900E-09 |
| 20365 | 1019 | S1019C | C | 6.4e-09 | 2.1e-08 | 1.1e-08 | 1.280E-08 | 7.465E-09 | 6.400E-09 | 2.100E-08 |
| 20366 | 1019 | S1019E | C | 2.1e-08 | 5.2e-09 | 1.4e-08 | 1.340E-08 | 7.917E-09 | 5.200E-09 | 2.100E-08 |
| 20367 | 1019 | S1019Q | C | 8.6e-09 | 8.8e-09 | 2.7e-09 | 6.700E-09 | 3.466E-09 | 2.700E-09 | 8.800E-09 |
| 20368 | 1019 | S1019G | C | 4.3e-09 | 1.7e-08 | 4.3e-09 | 8.533E-09 | 7.332E-09 | 4.300E-09 | 1.700E-08 |
| 20369 | 1019 | S1019H | C | 1.2e-08 | 1.7e-08 | 1.4e-08 | 1.433E-08 | 2.517E-09 | 1.200E-08 | 1.700E-08 |
| 20370 | 1019 | S1019I | C | 7.1e-09 | 1.3e-08 | 4.6e-09 | 8.233E-09 | 4.313E-09 | 4.600E-09 | 1.300E-08 |
| 20371 | 1019 | S1019L | C | 4.5e-09 | 2.6e-09 | 2.4e-09 | 3.167E-09 | 1.159E-09 | 2.400E-09 | 4.500E-09 |
| 20372 | 1019 | S1019K | C | 1.9e-08 | 1.2e-08 | 1.6e-09 | 1.087E-08 | 8.755E-09 | 1.600E-09 | 1.900E-08 |
| 20373 | 1019 | S1019M | C | 9.7e-09 | 4.4e-09 | 5.4e-09 | 6.500E-09 | 2.816E-09 | 4.400E-09 | 9.700E-09 |
| 20374 | 1019 | S1019F | C | 1.4e-08 | 7.6e-09 | 9.4e-09 | 1.033E-08 | 3.301E-09 | 7.600E-09 | 1.400E-08 |
| 20375 | 1019 | S1019P | C | 9.9e-09 | 3.3e-09 | 1e-08 | 7.733E-09 | 3.840E-09 | 3.300E-09 | 1.000E-08 |
| 20376 | 1019 | S1019S | C | 1.1e-08 | 1.3e-08 | 1.7e-08 | 1.367E-08 | 3.055E-09 | 1.100E-08 | 1.700E-08 |
| 20377 | 1019 | S1019T | C | 3.7e-09 | 1.1e-08 | 1.2e-08 | 8.900E-09 | 4.531E-09 | 3.700E-09 | 1.200E-08 |
| 20378 | 1019 | S1019W | C | 4.6e-09 | 6.3e-09 | 6e-09 | 5.633E-09 | 9.074E-10 | 4.600E-09 | 6.300E-09 |
| 20379 | 1019 | S1019Y | C | 1.2e-08 | 4.8e-09 | 7.8e-09 | 8.200E-09 | 3.617E-09 | 4.800E-09 | 1.200E-08 |
| 20380 | 1019 | S1019V | C | 5.8e-09 | 8.3e-09 | 4.5e-09 | 6.200E-09 | 1.931E-09 | 4.500E-09 | 8.300E-09 |
| 20381 | 1020 | L1020A | C | 5.7e-09 | 4.3e-09 | 2.6e-09 | 4.200E-09 | 1.552E-09 | 2.600E-09 | 5.700E-09 |
| 20382 | 1020 | L1020R | C | 2.1e-09 | 7.8e-09 | 3.1e-09 | 4.333E-09 | 3.044E-09 | 2.100E-09 | 7.800E-09 |
| 20383 | 1020 | L1020N | C | 7e-09 | 9.1e-09 | 3.2e-09 | 6.433E-09 | 2.991E-09 | 3.200E-09 | 9.100E-09 |
| 20384 | 1020 | L1020D | C | 2.1e-09 | 4.9e-09 | 5.5e-09 | 4.167E-09 | 1.815E-09 | 2.100E-09 | 5.500E-09 |
| 20385 | 1020 | L1020C | C | 3.9e-09 | 7.1e-09 | 1.4e-08 | 8.333E-09 | 5.162E-09 | 3.900E-09 | 1.400E-08 |
| 20386 | 1020 | L1020E | C | 6.3e-09 | 6.3e-09 | 3.5e-09 | 5.367E-09 | 1.617E-09 | 3.500E-09 | 6.300E-09 |
| 20387 | 1020 | L1020Q | C | 7.4e-09 | 9.4e-09 | 1.8e-08 | 1.160E-08 | 5.632E-09 | 7.400E-09 | 1.800E-08 |
| 20388 | 1020 | L1020G | C | 7e-09 | 6.3e-09 | 6.3e-09 | 6.533E-09 | 4.041E-10 | 6.300E-09 | 7.000E-09 |
| 20389 | 1020 | L1020H | C | 1.6e-08 | 5.4e-09 | 1e-08 | 1.047E-08 | 5.315E-09 | 5.400E-09 | 1.600E-08 |
| 20390 | 1020 | L1020I | C | 7.7e-09 | 6.5e-09 | 8e-09 | 7.400E-09 | 7.937E-10 | 6.500E-09 | 8.000E-09 |
| 20391 | 1020 | L1020L | C | 5.8e-09 | 1.7e-08 | 6.5e-09 | 9.767E-09 | 6.274E-09 | 5.800E-09 | 1.700E-08 |
| 20392 | 1020 | L1020K | C | 1.7e-08 | 3.9e-09 | 2.2e-08 | 1.430E-08 | 9.347E-09 | 3.900E-09 | 2.200E-08 |
| 20393 | 1020 | L1020M | C | 4.8e-09 | 1e-08 | 1.3e-08 | 9.267E-09 | 4.149E-09 | 4.800E-09 | 1.300E-08 |
| 20394 | 1020 | L1020F | C | 1.1e-08 | 1e-08 | 8.8e-09 | 9.933E-09 | 1.102E-09 | 8.800E-09 | 1.100E-08 |
| 20395 | 1020 | L1020P | C | 9.1e-09 | 1.2e-09 | 6.5e-09 | 5.600E-09 | 4.026E-09 | 1.200E-09 | 9.100E-09 |
| 20396 | 1020 | L1020S | C | 1.8e-09 | 3.5e-09 | 1.2e-08 | 5.767E-09 | 5.465E-09 | 1.800E-09 | 1.200E-08 |
| 20397 | 1020 | L1020T | C | 2.5e-09 | 9.6e-09 | 5.4e-09 | 5.833E-09 | 3.570E-09 | 2.500E-09 | 9.600E-09 |
| 20398 | 1020 | L1020W | C | 5.7e-09 | 8.4e-09 | 1.1e-08 | 8.367E-09 | 2.650E-09 | 5.700E-09 | 1.100E-08 |
| 20399 | 1020 | L1020Y | C | 8.7e-09 | 6.2e-09 | 9.9e-09 | 8.267E-09 | 1.888E-09 | 6.200E-09 | 9.900E-09 |

| | | | | | | | | | | |
|---|---|---|---|---|---|---|---|---|---|---|
| 20400 | 1020 | L1020V | C | 3.8e-09 | 3.9e-09 | 3.9e-09 | 3.867E-09 | 5.774E-11 | 3.800E-09 | 3.900E-09 |
| 20401 | 1021 | S1021A | C | 7.3e-09 | 7.1e-09 | 1.6e-08 | 1.013E-08 | 5.082E-09 | 7.100E-09 | 1.600E-08 |
| 20402 | 1021 | S1021R | C | 8.9e-09 | 5.3e-09 | 4.9e-09 | 6.367E-09 | 2.203E-09 | 4.900E-09 | 8.900E-09 |
| 20403 | 1021 | S1021N | C | 5e-09 | 1.2e-08 | 2.1e-08 | 1.267E-08 | 8.021E-09 | 5.000E-09 | 2.100E-08 |
| 20404 | 1021 | S1021D | C | 5.8e-09 | 8.3e-09 | 3.7e-09 | 5.933E-09 | 2.303E-09 | 3.700E-09 | 8.300E-09 |
| 20405 | 1021 | S1021C | C | 2.5e-09 | 7e-09 | 8e-09 | 5.833E-09 | 2.930E-09 | 2.500E-09 | 8.000E-09 |
| 20406 | 1021 | S1021E | C | 8e-09 | 5.8e-09 | 7e-09 | 6.933E-09 | 1.102E-09 | 5.800E-09 | 8.000E-09 |
| 20407 | 1021 | S1021Q | C | 2.4e-09 | 8.1e-09 | 1.1e-08 | 7.167E-09 | 4.375E-09 | 2.400E-09 | 1.100E-08 |
| 20408 | 1021 | S1021G | C | 3.6e-09 | 8e-09 | 4.4e-09 | 5.333E-09 | 2.344E-09 | 3.600E-09 | 8.000E-09 |
| 20409 | 1021 | S1021H | C | 8e-09 | 8e-09 | 2.2e-08 | 1.267E-08 | 8.083E-09 | 8.000E-09 | 2.200E-08 |
| 20410 | 1021 | S1021I | C | 7.8e-09 | 2.7e-09 | 3.8e-09 | 4.767E-09 | 2.684E-09 | 2.700E-09 | 7.800E-09 |
| 20411 | 1021 | S1021L | C | 6.1e-09 | 1.1e-08 | 1.4e-08 | 1.037E-08 | 3.988E-09 | 6.100E-09 | 1.400E-08 |
| 20412 | 1021 | S1021K | C | 9.2e-09 | 5.6e-09 | 1.5e-08 | 9.933E-09 | 4.743E-09 | 5.600E-09 | 1.500E-08 |
| 20413 | 1021 | S1021M | C | 6.7e-09 | 4.2e-09 | 5.1e-09 | 5.333E-09 | 1.266E-09 | 4.200E-09 | 6.700E-09 |
| 20414 | 1021 | S1021F | C | 5e-09 | 8.5e-09 | 3.1e-09 | 5.533E-09 | 2.739E-09 | 3.100E-09 | 8.500E-09 |
| 20415 | 1021 | S1021P | C | 3.1e-09 | 3.5e-09 | 4.8e-09 | 3.800E-09 | 8.888E-10 | 3.100E-09 | 4.800E-09 |
| 20416 | 1021 | S1021S | C | 1.2e-08 | 1.2e-08 | 3.4e-09 | 9.133E-09 | 4.965E-09 | 3.400E-09 | 1.200E-08 |
| 20417 | 1021 | S1021T | C | 9.6e-09 | 3.2e-09 | 4.3e-09 | 5.700E-09 | 3.422E-09 | 3.200E-09 | 9.600E-09 |
| 20418 | 1021 | S1021W | C | 6.5e-09 | 1.6e-08 | 5.4e-09 | 9.300E-09 | 5.828E-09 | 5.400E-09 | 1.600E-08 |
| 20419 | 1021 | S1021Y | C | 1.3e-08 | 9.8e-09 | 3.4e-09 | 8.733E-09 | 4.888E-09 | 3.400E-09 | 1.300E-08 |
| 20420 | 1021 | S1021V | C | 3.3e-09 | 6.3e-09 | 1.8e-08 | 9.200E-09 | 7.767E-09 | 3.300E-09 | 1.800E-08 |
| 20421 | 1022 | S1022A | C | 4.2e-09 | 7.4e-09 | 2.7e-09 | 4.767E-09 | 2.401E-09 | 2.700E-09 | 7.400E-09 |
| 20422 | 1022 | S1022R | C | 1.2e-08 | 1.6e-08 | 5.6e-09 | 1.120E-08 | 5.246E-09 | 5.600E-09 | 1.600E-08 |
| 20423 | 1022 | S1022N | C | 3.9e-09 | 1.1e-08 | 1.3e-08 | 9.300E-09 | 4.782E-09 | 3.900E-09 | 1.300E-08 |
| 20424 | 1022 | S1022D | C | 5.6e-09 | 5.9e-09 | 3e-09 | 4.833E-09 | 1.595E-09 | 3.000E-09 | 5.900E-09 |
| 20425 | 1022 | S1022C | C | 6.6e-09 | 1.1e-08 | 6.7e-09 | 8.100E-09 | 2.512E-09 | 6.600E-09 | 1.100E-08 |
| 20426 | 1022 | S1022E | C | 4e-09 | 1.1e-08 | 9.5e-09 | 8.167E-09 | 3.686E-09 | 4.000E-09 | 1.100E-08 |
| 20427 | 1022 | S1022Q | C | 8.4e-09 | 3.6e-09 | 3e-09 | 5.000E-09 | 2.960E-09 | 3.000E-09 | 8.400E-09 |
| 20428 | 1022 | S1022G | C | 1.1e-08 | 1.1e-08 | 3.1e-09 | 8.367E-09 | 4.561E-09 | 3.100E-09 | 1.100E-08 |
| 20429 | 1022 | S1022H | C | 4.4e-09 | 7.1e-09 | 2.5e-09 | 4.667E-09 | 2.312E-09 | 2.500E-09 | 7.100E-09 |
| 20430 | 1022 | S1022I | C | 6.1e-09 | 5e-09 | 3.7e-09 | 4.933E-09 | 1.201E-09 | 3.700E-09 | 6.100E-09 |
| 20431 | 1022 | S1022L | C | 1e-08 | 1.4e-08 | 4.5e-09 | 9.500E-09 | 4.770E-09 | 4.500E-09 | 1.400E-08 |
| 20432 | 1022 | S1022K | C | 2.4e-09 | 6.1e-09 | 5.2e-09 | 4.567E-09 | 1.930E-09 | 2.400E-09 | 6.100E-09 |
| 20433 | 1022 | S1022M | C | 1e-08 | 7.7e-09 | 3.5e-08 | 1.757E-08 | 1.514E-08 | 7.700E-09 | 3.500E-08 |
| 20434 | 1022 | S1022F | C | 1.3e-08 | 9e-09 | 7.3e-09 | 9.767E-09 | 2.926E-09 | 7.300E-09 | 1.300E-08 |
| 20435 | 1022 | S1022P | C | 6.8e-09 | 1e-08 | 1.9e-09 | 6.233E-09 | 4.080E-09 | 1.900E-09 | 1.000E-08 |
| 20436 | 1022 | S1022S | C | 1.1e-08 | 5.1e-09 | 1.3e-08 | 9.700E-09 | 4.107E-09 | 5.100E-09 | 1.300E-08 |
| 20437 | 1022 | S1022T | C | 9.3e-09 | 6.5e-09 | 6.1e-09 | 7.300E-09 | 1.744E-09 | 6.100E-09 | 9.300E-09 |
| 20438 | 1022 | S1022W | C | 1.4e-08 | 8.8e-09 | 4.9e-09 | 9.233E-09 | 4.565E-09 | 4.900E-09 | 1.400E-08 |
| 20439 | 1022 | S1022Y | C | 1.9e-09 | 7.6e-09 | 2.1e-09 | 3.867E-09 | 3.235E-09 | 1.900E-09 | 7.600E-09 |
| 20440 | 1022 | S1022V | C | 1.2e-08 | 6.2e-09 | 1.3e-08 | 1.040E-08 | 3.672E-09 | 6.200E-09 | 1.300E-08 |
| 20441 | 1023 | V1023A | C | 7.4e-09 | 2.3e-08 | 4.5e-09 | 1.163E-08 | 9.950E-09 | 4.500E-09 | 2.300E-08 |
| 20442 | 1023 | V1023R | C | 9.3e-09 | 8.9e-09 | 7.5e-09 | 8.567E-09 | 9.452E-10 | 7.500E-09 | 9.300E-09 |
| 20443 | 1023 | V1023N | C | 5.2e-09 | 5.6e-09 | 5.3e-09 | 5.367E-09 | 2.082E-10 | 5.200E-09 | 5.600E-09 |
| 20444 | 1023 | V1023D | C | 1.2e-08 | 8e-09 | 8.4e-09 | 9.467E-09 | 2.203E-09 | 8.000E-09 | 1.200E-08 |
| 20445 | 1023 | V1023C | C | 5.8e-09 | 1.4e-08 | 6.4e-09 | 8.733E-09 | 4.571E-09 | 5.800E-09 | 1.400E-08 |
| 20446 | 1023 | V1023E | C | 1.6e-09 | 1.6e-09 | 8.6e-09 | 3.933E-09 | 4.041E-09 | 1.600E-09 | 8.600E-09 |
| 20447 | 1023 | V1023Q | C | 1.2e-08 | 8.3e-09 | 8.9e-09 | 9.733E-09 | 1.986E-09 | 8.300E-09 | 1.200E-08 |
| 20448 | 1023 | V1023G | C | 7.6e-09 | 4.7e-09 | 7.7e-09 | 6.667E-09 | 1.704E-09 | 4.700E-09 | 7.700E-09 |
| 20449 | 1023 | V1023H | C | 2e-09 | 5.1e-09 | 5.2e-09 | 4.100E-09 | 1.819E-09 | 2.000E-09 | 5.200E-09 |
| 20450 | 1023 | V1023I | C | 1.4e-08 | 6.6e-09 | 1.4e-08 | 1.153E-08 | 4.272E-09 | 6.600E-09 | 1.400E-08 |
| 20451 | 1023 | V1023L | C | 1.6e-09 | 3e-09 | 1.1e-08 | 5.200E-09 | 5.071E-09 | 1.600E-09 | 1.100E-08 |
| 20452 | 1023 | V1023K | C | 5.5e-09 | 2.4e-09 | 7.6e-10 | 2.887E-09 | 2.407E-09 | 7.600E-10 | 5.500E-09 |
| 20453 | 1023 | V1023M | C | 5.7e-09 | 8.1e-09 | 1.3e-08 | 8.933E-09 | 3.721E-09 | 5.700E-09 | 1.300E-08 |
| 20454 | 1023 | V1023F | C | 8.5e-09 | 3.8e-09 | 5.9e-09 | 6.067E-09 | 2.354E-09 | 3.800E-09 | 8.500E-09 |
| 20455 | 1023 | V1023P | C | 1.1e-08 | 4.6e-09 | 7.5e-09 | 7.700E-09 | 3.205E-09 | 4.600E-09 | 1.100E-08 |
| 20456 | 1023 | V1023S | C | 7.8e-09 | 7.3e-09 | 7.8e-09 | 7.633E-09 | 2.887E-10 | 7.300E-09 | 7.800E-09 |
| 20457 | 1023 | V1023T | C | 5.9e-09 | 1.1e-08 | 7e-09 | 7.967E-09 | 2.684E-09 | 5.900E-09 | 1.100E-08 |
| 20458 | 1023 | V1023W | C | 1.5e-08 | 6.2e-09 | 7.1e-09 | 9.433E-09 | 4.842E-09 | 6.200E-09 | 1.500E-08 |
| 20459 | 1023 | V1023Y | C | 6.4e-09 | 1.6e-08 | 8.1e-09 | 1.017E-08 | 5.123E-09 | 6.400E-09 | 1.600E-08 |
| 20460 | 1023 | V1023V | C | 9.3e-09 | 1.8e-08 | 7.9e-09 | 1.173E-08 | 5.472E-09 | 7.900E-09 | 1.800E-08 |
| 20461 | 1024 | V1024A | C | 4e-09 | 1e-08 | 1e-08 | 8.000E-09 | 3.464E-09 | 4.000E-09 | 1.000E-08 |
| 20462 | 1024 | V1024R | C | 5.1e-09 | 3e-09 | 1.3e-08 | 7.033E-09 | 5.273E-09 | 3.000E-09 | 1.300E-08 |
| 20463 | 1024 | V1024N | C | 1.5e-08 | 4.7e-09 | 3.4e-09 | 7.700E-09 | 6.355E-09 | 3.400E-09 | 1.500E-08 |
| 20464 | 1024 | V1024D | C | 1e-08 | 8.6e-09 | 5.5e-09 | 8.033E-09 | 2.303E-09 | 5.500E-09 | 1.000E-08 |
| 20465 | 1024 | V1024C | C | 4.9e-09 | 2.5e-09 | 6.2e-09 | 4.533E-09 | 1.877E-09 | 2.500E-09 | 6.200E-09 |
| 20466 | 1024 | V1024E | C | 6.9e-09 | 1.3e-08 | 1.3e-08 | 1.097E-08 | 3.522E-09 | 6.900E-09 | 1.300E-08 |
| 20467 | 1024 | V1024Q | C | 1.3e-08 | 1.4e-08 | 6.4e-09 | 1.113E-08 | 4.130E-09 | 6.400E-09 | 1.400E-08 |
| 20468 | 1024 | V1024G | C | 9.3e-09 | 1.5e-08 | 1.5e-08 | 1.310E-08 | 3.291E-09 | 9.300E-09 | 1.500E-08 |
| 20469 | 1024 | V1024H | C | 1e-08 | 4.4e-09 | 9.6e-09 | 8.000E-09 | 3.124E-09 | 4.400E-09 | 1.000E-08 |
| 20470 | 1024 | V1024I | C | 1.3e-08 | 5.7e-09 | 7.4e-09 | 8.700E-09 | 3.820E-09 | 5.700E-09 | 1.300E-08 |
| 20471 | 1024 | V1024L | C | 4.6e-09 | 4.1e-08 | 1e-08 | 1.853E-08 | 1.964E-08 | 4.600E-09 | 4.100E-08 |
| 20472 | 1024 | V1024K | C | 1e-09 | 9.5e-09 | 2.8e-09 | 4.433E-09 | 4.479E-09 | 1.000E-09 | 9.500E-09 |
| 20473 | 1024 | V1024M | C | 8.5e-09 | 2.2e-09 | 1.6e-08 | 8.900E-09 | 6.909E-09 | 2.200E-09 | 1.600E-08 |
| 20474 | 1024 | V1024F | C | 1.7e-08 | 8e-09 | 3.6e-09 | 9.533E-09 | 6.830E-09 | 3.600E-09 | 1.700E-08 |
| 20475 | 1024 | V1024P | C | 3.3e-09 | 1.4e-08 | 3.3e-09 | 6.867E-09 | 6.178E-09 | 3.300E-09 | 1.400E-08 |
| 20476 | 1024 | V1024S | C | 5.3e-09 | 3.5e-09 | 9.6e-09 | 6.133E-09 | 3.134E-09 | 3.500E-09 | 9.600E-09 |
| 20477 | 1024 | V1024T | C | 8.9e-09 | 1.2e-08 | 3.2e-08 | 1.763E-08 | 1.254E-08 | 8.900E-09 | 3.200E-08 |
| 20478 | 1024 | V1024W | C | 8.4e-09 | 3.4e-09 | 1e-08 | 7.267E-09 | 3.443E-09 | 3.400E-09 | 1.000E-08 |
| 20479 | 1024 | V1024Y | C | 1.5e-08 | 4.2e-09 | 5.4e-09 | 8.200E-09 | 5.919E-09 | 4.200E-09 | 1.500E-08 |
| 20480 | 1024 | V1024V | C | 1.7e-08 | 4.1e-09 | 7.3e-09 | 9.467E-09 | 6.717E-09 | 4.100E-09 | 1.700E-08 |
| 20481 | 1025 | T1025A | C | 2.1e-08 | 7.6e-09 | 8.8e-09 | 1.247E-08 | 7.414E-09 | 7.600E-09 | 2.100E-08 |
| 20482 | 1025 | T1025R | C | 8.6e-09 | 4.5e-09 | 3.9e-09 | 5.667E-09 | 2.558E-09 | 3.900E-09 | 8.600E-09 |
| 20483 | 1025 | T1025N | C | 9.4e-09 | 1.3e-08 | 5.3e-09 | 9.233E-09 | 3.853E-09 | 5.300E-09 | 1.300E-08 |
| 20484 | 1025 | T1025D | C | 3.2e-08 | 2.6e-08 | 9.3e-09 | 2.243E-08 | 1.176E-08 | 9.300E-09 | 3.200E-08 |

| | | | | | | | | | | |
|---|---|---|---|---|---|---|---|---|---|---|
| 20485 | 1025 | T1025C | C | 6e-09 | 1.2e-08 | 3.2e-09 | 7.067E-09 | 4.496E-09 | 3.200E-09 | 1.200E-08 |
| 20486 | 1025 | T1025E | C | 6.8e-09 | 1.4e-08 | 2.6e-08 | 1.560E-08 | 9.699E-09 | 6.800E-09 | 2.600E-08 |
| 20487 | 1025 | T1025Q | C | 1.1e-08 | 8.9e-09 | 4.2e-09 | 8.033E-09 | 3.482E-09 | 4.200E-09 | 1.100E-08 |
| 20488 | 1025 | T1025G | C | 2.9e-09 | 2e-08 | 9.5e-09 | 1.080E-08 | 8.624E-09 | 2.900E-09 | 2.000E-08 |
| 20489 | 1025 | T1025H | C | 2.1e-09 | 7.5e-09 | 9.6e-09 | 6.400E-09 | 3.869E-09 | 2.100E-09 | 9.600E-09 |
| 20490 | 1025 | T1025I | C | 7.3e-09 | 6.3e-09 | 2.2e-08 | 1.187E-08 | 8.790E-09 | 6.300E-09 | 2.200E-08 |
| 20491 | 1025 | T1025L | C | 1.6e-08 | 6.1e-09 | 1.2e-08 | 1.137E-08 | 4.980E-09 | 6.100E-09 | 1.600E-08 |
| 20492 | 1025 | T1025K | C | 2e-08 | 5e-09 | 1.5e-08 | 1.333E-08 | 7.638E-09 | 5.000E-09 | 2.000E-08 |
| 20493 | 1025 | T1025M | C | 1.1e-08 | 4.8e-09 | 6.1e-09 | 7.300E-09 | 3.270E-09 | 4.800E-09 | 1.100E-08 |
| 20494 | 1025 | T1025F | C | 4.6e-09 | 5.2e-09 | 1.1e-08 | 6.933E-09 | 3.535E-09 | 4.600E-09 | 1.100E-08 |
| 20495 | 1025 | T1025P | C | 7.9e-09 | 5.3e-09 | 6.3e-09 | 6.500E-09 | 1.311E-09 | 5.300E-09 | 7.900E-09 |
| 20496 | 1025 | T1025S | C | 1.1e-08 | 2.5e-09 | 4.2e-09 | 5.900E-09 | 4.498E-09 | 2.500E-09 | 1.100E-08 |
| 20497 | 1025 | T1025T | C | 1.5e-08 | 2.4e-08 | 3.3e-09 | 1.410E-08 | 1.038E-08 | 3.300E-09 | 2.400E-08 |
| 20498 | 1025 | T1025W | C | 2.8e-09 | 1.3e-08 | 1e-08 | 8.600E-09 | 5.242E-09 | 2.800E-09 | 1.300E-08 |
| 20499 | 1025 | T1025Y | C | 1.4e-08 | 4.3e-09 | 7.1e-09 | 8.467E-09 | 4.992E-09 | 4.300E-09 | 1.400E-08 |
| 20500 | 1025 | T1025V | C | 6.7e-09 | 3.9e-09 | 1.7e-08 | 9.200E-09 | 6.899E-09 | 3.900E-09 | 1.700E-08 |
| 20501 | 1026 | V1026A | C | 2.6e-08 | 1.2e-08 | 7.7e-09 | 1.523E-08 | 9.569E-09 | 7.700E-09 | 2.600E-08 |
| 20502 | 1026 | V1026R | C | 1.1e-08 | 2.6e-09 | 1e-08 | 7.867E-09 | 4.588E-09 | 2.600E-09 | 1.100E-08 |
| 20503 | 1026 | V1026N | C | 7.5e-10 | 1e-08 | 1.7e-09 | 4.150E-09 | 5.088E-09 | 7.500E-10 | 1.000E-08 |
| 20504 | 1026 | V1026D | C | 4.2e-09 | 5.8e-09 | 8.1e-09 | 6.033E-09 | 1.960E-09 | 4.200E-09 | 8.100E-09 |
| 20505 | 1026 | V1026C | C | 1.8e-09 | 1.8e-09 | 1.1e-08 | 4.867E-09 | 5.312E-09 | 1.800E-09 | 1.100E-08 |
| 20506 | 1026 | V1026E | C | 5.7e-09 | 2e-08 | 9.5e-09 | 1.173E-08 | 7.407E-09 | 5.700E-09 | 2.000E-08 |
| 20507 | 1026 | V1026Q | C | 3.5e-09 | 4.2e-09 | 4.7e-09 | 4.133E-09 | 6.028E-10 | 3.500E-09 | 4.700E-09 |
| 20508 | 1026 | V1026G | C | 5e-09 | 1.1e-08 | 1e-08 | 8.667E-09 | 3.215E-09 | 5.000E-09 | 1.100E-08 |
| 20509 | 1026 | V1026H | C | 8.2e-09 | 9e-09 | 5.5e-09 | 7.567E-09 | 1.834E-09 | 5.500E-09 | 9.000E-09 |
| 20510 | 1026 | V1026I | C | 4.4e-09 | 2.8e-09 | 8e-09 | 5.067E-09 | 2.663E-09 | 2.800E-09 | 8.000E-09 |
| 20511 | 1026 | V1026L | C | 4.8e-09 | 6.1e-09 | 2.2e-09 | 4.367E-09 | 1.986E-09 | 2.200E-09 | 6.100E-09 |
| 20512 | 1026 | V1026K | C | 4.1e-09 | 7.1e-09 | 6.3e-09 | 5.833E-09 | 1.553E-09 | 4.100E-09 | 7.100E-09 |
| 20513 | 1026 | V1026M | C | 1.9e-08 | 1.2e-08 | 2.8e-09 | 1.127E-08 | 8.125E-09 | 2.800E-09 | 1.900E-08 |
| 20514 | 1026 | V1026F | C | 6.2e-09 | 2e-08 | 1.4e-08 | 1.340E-08 | 6.920E-09 | 6.200E-09 | 2.000E-08 |
| 20515 | 1026 | V1026P | C | 2.3e-08 | 9.2e-09 | 2.6e-09 | 1.160E-08 | 1.041E-08 | 2.600E-09 | 2.300E-08 |
| 20516 | 1026 | V1026S | C | 1.1e-08 | 6e-09 | 1.9e-09 | 6.300E-09 | 4.557E-09 | 1.900E-09 | 1.100E-08 |
| 20517 | 1026 | V1026T | C | 7e-09 | 6e-09 | 4e-09 | 5.667E-09 | 1.528E-09 | 4.000E-09 | 7.000E-09 |
| 20518 | 1026 | V1026W | C | 1.3e-08 | 2.5e-09 | 6.2e-09 | 7.233E-09 | 5.326E-09 | 2.500E-09 | 1.300E-08 |
| 20519 | 1026 | V1026Y | C | 4.8e-09 | 5.9e-09 | 2.2e-09 | 4.300E-09 | 1.900E-09 | 2.200E-09 | 5.900E-09 |
| 20520 | 1026 | V1026V | C | 1.1e-08 | 1e-08 | 4.6e-09 | 8.533E-09 | 3.443E-09 | 4.600E-09 | 1.100E-08 |
| 20521 | 1027 | P1027A | C | 9.9e-09 | 1.5e-08 | 9.3e-09 | 1.140E-08 | 3.132E-09 | 9.300E-09 | 1.500E-08 |
| 20522 | 1027 | P1027R | C | 5.9e-09 | 2.3e-09 | 1.3e-08 | 7.067E-09 | 5.445E-09 | 2.300E-09 | 1.300E-08 |
| 20523 | 1027 | P1027N | C | 1.6e-08 | 4.4e-09 | 6.3e-09 | 8.900E-09 | 6.222E-09 | 4.400E-09 | 1.600E-08 |
| 20524 | 1027 | P1027D | C | 5.7e-09 | 7.4e-09 | 8.6e-09 | 7.233E-09 | 1.457E-09 | 5.700E-09 | 8.600E-09 |
| 20525 | 1027 | P1027C | C | 8.5e-09 | 5.3e-09 | 4.1e-09 | 5.967E-09 | 2.274E-09 | 4.100E-09 | 8.500E-09 |
| 20526 | 1027 | P1027E | C | 1.1e-08 | 4.8e-09 | 5.2e-09 | 7.000E-09 | 3.470E-09 | 4.800E-09 | 1.100E-08 |
| 20527 | 1027 | P1027Q | C | 7.9e-09 | 6.1e-09 | 7.6e-09 | 7.200E-09 | 9.644E-10 | 6.100E-09 | 7.900E-09 |
| 20528 | 1027 | P1027G | C | 8e-09 | 1.7e-09 | 4.1e-09 | 4.600E-09 | 3.180E-09 | 1.700E-09 | 8.000E-09 |
| 20529 | 1027 | P1027H | C | 2.4e-09 | 4.3e-09 | 1.5e-08 | 7.233E-09 | 6.793E-09 | 2.400E-09 | 1.500E-08 |
| 20530 | 1027 | P1027I | C | 7.5e-09 | 4.1e-09 | 4.8e-09 | 5.467E-09 | 1.795E-09 | 4.100E-09 | 7.500E-09 |
| 20531 | 1027 | P1027L | C | 8.2e-09 | 1.1e-08 | 6.4e-09 | 8.533E-09 | 2.318E-09 | 6.400E-09 | 1.100E-08 |
| 20532 | 1027 | P1027K | C | 2.1e-08 | 5.4e-09 | 2.9e-09 | 9.767E-09 | 9.808E-09 | 2.900E-09 | 2.100E-08 |
| 20533 | 1027 | P1027M | C | 6.8e-09 | 7.1e-09 | 6.5e-09 | 6.800E-09 | 3.000E-10 | 6.500E-09 | 7.100E-09 |
| 20534 | 1027 | P1027F | C | 5.8e-09 | 5.5e-09 | 5.9e-09 | 5.733E-09 | 2.082E-10 | 5.500E-09 | 5.900E-09 |
| 20535 | 1027 | P1027P | C | 9.2e-09 | 1e-09 | 7.7e-09 | 5.967E-09 | 4.366E-09 | 1.000E-09 | 9.200E-09 |
| 20536 | 1027 | P1027S | C | 3.8e-09 | 6.3e-09 | 4.8e-09 | 4.967E-09 | 1.258E-09 | 3.800E-09 | 6.300E-09 |
| 20537 | 1027 | P1027T | C | 5.4e-09 | 1.2e-08 | 2.5e-09 | 6.633E-09 | 4.869E-09 | 2.500E-09 | 1.200E-08 |
| 20538 | 1027 | P1027W | C | 6.3e-09 | 1.6e-08 | 5.4e-09 | 9.233E-09 | 5.877E-09 | 5.400E-09 | 1.600E-08 |
| 20539 | 1027 | P1027Y | C | 3.6e-09 | 3.9e-09 | 1.4e-08 | 7.167E-09 | 5.920E-09 | 3.600E-09 | 1.400E-08 |
| 20540 | 1027 | P1027V | C | 7.4e-09 | 1.3e-08 | 5.8e-09 | 8.733E-09 | 3.781E-09 | 5.800E-09 | 1.300E-08 |
| 20541 | 1028 | S1028A | C | 2.2e-09 | 8.6e-09 | 1.4e-08 | 8.267E-09 | 5.907E-09 | 2.200E-09 | 1.400E-08 |
| 20542 | 1028 | S1028R | C | 6.5e-09 | 5e-09 | 6.4e-09 | 5.967E-09 | 8.386E-10 | 5.000E-09 | 6.500E-09 |
| 20543 | 1028 | S1028N | C | 1.3e-08 | 5.6e-09 | 1.7e-08 | 1.187E-08 | 5.784E-09 | 5.600E-09 | 1.700E-08 |
| 20544 | 1028 | S1028D | C | 3.9e-09 | 9e-09 | 1.3e-08 | 8.633E-09 | 4.561E-09 | 3.900E-09 | 1.300E-08 |
| 20545 | 1028 | S1028C | C | 1.1e-08 | 1e-08 | 2.8e-09 | 7.933E-09 | 4.474E-09 | 2.800E-09 | 1.100E-08 |
| 20546 | 1028 | S1028E | C | 7.3e-09 | 1e-08 | 9.6e-09 | 8.967E-09 | 1.457E-09 | 7.300E-09 | 1.000E-08 |
| 20547 | 1028 | S1028Q | C | 9.7e-09 | 9.1e-09 | 2.6e-08 | 1.493E-08 | 9.589E-09 | 9.100E-09 | 2.600E-08 |
| 20548 | 1028 | S1028G | C | 9.8e-09 | 3.9e-09 | 1.3e-08 | 8.900E-09 | 4.616E-09 | 3.900E-09 | 1.300E-08 |
| 20549 | 1028 | S1028H | C | 2e-08 | 6.4e-09 | 7.3e-09 | 1.123E-08 | 7.605E-09 | 6.400E-09 | 2.000E-08 |
| 20550 | 1028 | S1028I | C | 8.8e-09 | 5.1e-09 | 6e-09 | 6.633E-09 | 1.930E-09 | 5.100E-09 | 8.800E-09 |
| 20551 | 1028 | S1028L | C | 6.5e-09 | 2.5e-08 | 4.4e-09 | 1.197E-08 | 1.134E-08 | 4.400E-09 | 2.500E-08 |
| 20552 | 1028 | S1028K | C | 7.4e-09 | 6.7e-09 | 4.4e-09 | 6.167E-09 | 1.570E-09 | 4.400E-09 | 7.400E-09 |
| 20553 | 1028 | S1028M | C | 3.6e-09 | 2.8e-09 | 3.6e-09 | 3.333E-09 | 4.619E-10 | 2.800E-09 | 3.600E-09 |
| 20554 | 1028 | S1028F | C | 8.4e-09 | 3.5e-09 | 1.4e-08 | 8.633E-09 | 5.254E-09 | 3.500E-09 | 1.400E-08 |
| 20555 | 1028 | S1028P | C | 5.4e-08 | 2.1e-09 | 1.5e-08 | 2.370E-08 | 2.702E-08 | 2.100E-09 | 5.400E-08 |
| 20556 | 1028 | S1028S | C | 3.9e-09 | 2.7e-09 | 1.1e-08 | 5.867E-09 | 4.486E-09 | 2.700E-09 | 1.100E-08 |
| 20557 | 1028 | S1028T | C | 2.4e-09 | 9.4e-09 | 1.3e-08 | 8.267E-09 | 5.390E-09 | 2.400E-09 | 1.300E-08 |
| 20558 | 1028 | S1028W | C | 9e-09 | 1.4e-08 | 8.7e-09 | 1.057E-08 | 2.977E-09 | 8.700E-09 | 1.400E-08 |
| 20559 | 1028 | S1028Y | C | 1.2e-08 | 6.7e-09 | 6.6e-09 | 8.433E-09 | 3.089E-09 | 6.600E-09 | 1.200E-08 |
| 20560 | 1028 | S1028V | C | 1e-08 | 5.1e-09 | 4.6e-09 | 6.567E-09 | 2.984E-09 | 4.600E-09 | 1.000E-08 |
| 20561 | 1029 | S1029A | C | 7.4e-09 | 8.7e-09 | 6.5e-09 | 7.533E-09 | 1.106E-09 | 6.500E-09 | 8.700E-09 |
| 20562 | 1029 | S1029R | C | 1.5e-08 | 4.6e-09 | 3.4e-09 | 7.667E-09 | 6.379E-09 | 3.400E-09 | 1.500E-08 |
| 20563 | 1029 | S1029N | C | 2.3e-09 | 2.3e-09 | 4.3e-09 | 2.967E-09 | 1.155E-09 | 2.300E-09 | 4.300E-09 |
| 20564 | 1029 | S1029D | C | 1.3e-08 | 1.6e-08 | 9e-09 | 1.267E-08 | 3.512E-09 | 9.000E-09 | 1.600E-08 |
| 20565 | 1029 | S1029C | C | 3e-09 | 3.8e-09 | 6.4e-09 | 4.400E-09 | 1.778E-09 | 3.000E-09 | 6.400E-09 |
| 20566 | 1029 | S1029E | C | 3.2e-09 | 4.9e-09 | 5.6e-09 | 4.567E-09 | 1.234E-09 | 3.200E-09 | 5.600E-09 |
| 20567 | 1029 | S1029Q | C | 2e-08 | 7.9e-09 | 1.3e-08 | 1.363E-08 | 6.075E-09 | 7.900E-09 | 2.000E-08 |
| 20568 | 1029 | S1029G | C | 6.5e-09 | 1.7e-08 | 5.6e-09 | 9.700E-09 | 6.338E-09 | 5.600E-09 | 1.700E-08 |
| 20569 | 1029 | S1029H | C | 1e-08 | 4.3e-09 | 7.8e-09 | 7.367E-09 | 2.875E-09 | 4.300E-09 | 1.000E-08 |

| | | | | | | | | | | |
|---|---|---|---|---|---|---|---|---|---|---|
| 20570 | 1029 | S1029I | C | 9.4e-09 | 7.2e-09 | 8e-09 | 8.200E-09 | 1.114E-09 | 7.200E-09 | 9.400E-09 |
| 20571 | 1029 | S1029L | C | 7.5e-09 | 4.2e-09 | 1.3e-08 | 8.233E-09 | 4.446E-09 | 4.200E-09 | 1.300E-08 |
| 20572 | 1029 | S1029K | C | 6.9e-09 | 7e-09 | 2.3e-09 | 5.400E-09 | 2.685E-09 | 2.300E-09 | 7.000E-09 |
| 20573 | 1029 | S1029M | C | 4.2e-09 | 1.1e-08 | 1.1e-08 | 8.733E-09 | 3.926E-09 | 4.200E-09 | 1.100E-08 |
| 20574 | 1029 | S1029F | C | 5.5e-09 | 1e-09 | 6.6e-09 | 4.367E-09 | 2.967E-09 | 1.000E-09 | 6.600E-09 |
| 20575 | 1029 | S1029P | C | 3.3e-09 | 5.4e-09 | 5.6e-09 | 4.767E-09 | 1.274E-09 | 3.300E-09 | 5.600E-09 |
| 20576 | 1029 | S1029S | C | 2.1e-09 | 5.9e-09 | 1.3e-08 | 7.000E-09 | 5.533E-09 | 2.100E-09 | 1.300E-08 |
| 20577 | 1029 | S1029T | C | 5.2e-09 | 7.7e-09 | 4.4e-09 | 5.767E-09 | 1.721E-09 | 4.400E-09 | 7.700E-09 |
| 20578 | 1029 | S1029W | C | 2.6e-09 | 1.8e-08 | 9.9e-09 | 1.017E-08 | 7.703E-09 | 2.600E-09 | 1.800E-08 |
| 20579 | 1029 | S1029Y | C | 3e-09 | 5e-09 | 5.3e-09 | 4.433E-09 | 1.250E-09 | 3.000E-09 | 5.300E-09 |
| 20580 | 1029 | S1029V | C | 1.1e-08 | 1.4e-08 | 1.9e-08 | 1.467E-08 | 4.041E-09 | 1.100E-08 | 1.900E-08 |
| 20581 | 1030 | S1030A | C | 9.7e-09 | 7e-09 | 5.3e-09 | 7.333E-09 | 2.219E-09 | 5.300E-09 | 9.700E-09 |
| 20582 | 1030 | S1030R | C | 8.9e-09 | 7.4e-09 | 1e-08 | 8.767E-09 | 1.305E-09 | 7.400E-09 | 1.000E-08 |
| 20583 | 1030 | S1030N | C | 6.2e-09 | 4.5e-09 | 7.3e-09 | 6.000E-09 | 1.411E-09 | 4.500E-09 | 7.300E-09 |
| 20584 | 1030 | S1030D | C | 8.5e-09 | 6.8e-09 | 2e-08 | 1.177E-08 | 7.181E-09 | 6.800E-09 | 2.000E-08 |
| 20585 | 1030 | S1030C | C | 5.7e-09 | 5.5e-09 | 1.2e-08 | 7.733E-09 | 3.696E-09 | 5.500E-09 | 1.200E-08 |
| 20586 | 1030 | S1030E | C | 2.2e-08 | 9e-09 | 7.5e-09 | 1.283E-08 | 7.974E-09 | 7.500E-09 | 2.200E-08 |
| 20587 | 1030 | S1030Q | C | 4.4e-09 | 4.5e-09 | 1.3e-08 | 7.300E-09 | 4.937E-09 | 4.400E-09 | 1.300E-08 |
| 20588 | 1030 | S1030G | C | 5.6e-09 | 1.1e-08 | 1.1e-08 | 9.200E-09 | 3.118E-09 | 5.600E-09 | 1.100E-08 |
| 20589 | 1030 | S1030H | C | 1.2e-08 | 9.4e-09 | 5e-09 | 8.800E-09 | 3.538E-09 | 5.000E-09 | 1.200E-08 |
| 20590 | 1030 | S1030I | C | 4.5e-09 | 7.4e-09 | 1.2e-08 | 7.967E-09 | 3.782E-09 | 4.500E-09 | 1.200E-08 |
| 20591 | 1030 | S1030L | C | 9.7e-09 | 4.5e-09 | 5e-09 | 6.400E-09 | 2.869E-09 | 4.500E-09 | 9.700E-09 |
| 20592 | 1030 | S1030K | C | 9.2e-09 | 1.4e-09 | 6.8e-09 | 5.800E-09 | 3.995E-09 | 1.400E-09 | 9.200E-09 |
| 20593 | 1030 | S1030M | C | 6.7e-09 | 4.2e-09 | 1.5e-09 | 4.133E-09 | 2.601E-09 | 1.500E-09 | 6.700E-09 |
| 20594 | 1030 | S1030F | C | 1e-08 | 8.8e-09 | 7.4e-09 | 8.733E-09 | 1.301E-09 | 7.400E-09 | 1.000E-08 |
| 20595 | 1030 | S1030P | C | 6.2e-09 | 1.1e-08 | 5.4e-09 | 7.533E-09 | 3.029E-09 | 5.400E-09 | 1.100E-08 |
| 20596 | 1030 | S1030S | C | 4.3e-09 | 1.2e-08 | 4e-09 | 6.767E-09 | 4.535E-09 | 4.000E-09 | 1.200E-08 |
| 20597 | 1030 | S1030T | C | 6e-09 | 1.6e-08 | 5.8e-09 | 9.267E-09 | 5.832E-09 | 5.800E-09 | 1.600E-08 |
| 20598 | 1030 | S1030W | C | 4.4e-09 | 3e-09 | 5.4e-09 | 4.267E-09 | 1.206E-09 | 3.000E-09 | 5.400E-09 |
| 20599 | 1030 | S1030Y | C | 8.8e-09 | 1.7e-08 | 2.1e-08 | 1.560E-08 | 6.219E-09 | 8.800E-09 | 2.100E-08 |
| 20600 | 1030 | S1030V | C | 7.4e-09 | 2.1e-08 | 1.7e-08 | 1.513E-08 | 6.990E-09 | 7.400E-09 | 2.100E-08 |
| 20601 | 1031 | L1031A | C | 6.9e-09 | 6.1e-09 | 8.4e-09 | 7.133E-09 | 1.168E-09 | 6.100E-09 | 8.400E-09 |
| 20602 | 1031 | L1031R | C | 2.6e-09 | 1.3e-08 | 3.7e-09 | 6.433E-09 | 5.713E-09 | 2.600E-09 | 1.300E-08 |
| 20603 | 1031 | L1031N | C | 7.2e-09 | 5.8e-09 | 3.7e-09 | 5.567E-09 | 1.762E-09 | 3.700E-09 | 7.200E-09 |
| 20604 | 1031 | L1031D | C | 4.8e-09 | 2.6e-08 | 3.6e-09 | 1.147E-08 | 1.260E-08 | 3.600E-09 | 2.600E-08 |
| 20605 | 1031 | L1031C | C | 7.1e-09 | 2.3e-08 | 5.6e-09 | 1.190E-08 | 9.642E-09 | 5.600E-09 | 2.300E-08 |
| 20606 | 1031 | L1031E | C | 5.1e-09 | 2e-09 | 6.9e-09 | 4.667E-09 | 2.479E-09 | 2.000E-09 | 6.900E-09 |
| 20607 | 1031 | L1031Q | C | 6.8e-09 | 1e-08 | 4.1e-09 | 6.967E-09 | 2.954E-09 | 4.100E-09 | 1.000E-08 |
| 20608 | 1031 | L1031G | C | 5e-09 | 3.1e-09 | 3.1e-09 | 3.733E-09 | 1.097E-09 | 3.100E-09 | 5.000E-09 |
| 20609 | 1031 | L1031H | C | 2.1e-08 | 1e-08 | 9.9e-09 | 1.363E-08 | 6.380E-09 | 9.900E-09 | 2.100E-08 |
| 20610 | 1031 | L1031I | C | 4.9e-09 | 6.2e-09 | 3.8e-09 | 4.967E-09 | 1.201E-09 | 3.800E-09 | 6.200E-09 |
| 20611 | 1031 | L1031L | C | 8.3e-09 | 8.9e-09 | 5.7e-09 | 7.633E-09 | 1.701E-09 | 5.700E-09 | 8.900E-09 |
| 20612 | 1031 | L1031K | C | 5.1e-09 | 1.8e-08 | 8.1e-09 | 1.040E-08 | 6.751E-09 | 5.100E-09 | 1.800E-08 |
| 20613 | 1031 | L1031M | C | 1.1e-08 | 6.2e-09 | 1.2e-08 | 9.733E-09 | 3.101E-09 | 6.200E-09 | 1.200E-08 |
| 20614 | 1031 | L1031F | C | 3.9e-09 | 7.1e-09 | 5.7e-09 | 5.567E-09 | 1.604E-09 | 3.900E-09 | 7.100E-09 |
| 20615 | 1031 | L1031P | C | 5.1e-09 | 1.9e-09 | 8.1e-09 | 5.033E-09 | 3.101E-09 | 1.900E-09 | 8.100E-09 |
| 20616 | 1031 | L1031S | C | 1.3e-08 | 1.2e-08 | 3.5e-09 | 9.500E-09 | 5.220E-09 | 3.500E-09 | 1.300E-08 |
| 20617 | 1031 | L1031T | C | 1.1e-08 | 1.4e-09 | 1.5e-08 | 9.133E-09 | 6.990E-09 | 1.400E-09 | 1.500E-08 |
| 20618 | 1031 | L1031W | C | 1.6e-08 | 7.3e-09 | 3.7e-09 | 9.000E-09 | 6.324E-09 | 3.700E-09 | 1.600E-08 |
| 20619 | 1031 | L1031Y | C | 4.8e-09 | 4.2e-09 | 1.1e-08 | 6.667E-09 | 3.765E-09 | 4.200E-09 | 1.100E-08 |
| 20620 | 1031 | L1031V | C | 1.5e-08 | 3.7e-09 | 8.1e-10 | 6.503E-09 | 7.499E-09 | 8.100E-10 | 1.500E-08 |
| 20621 | 1032 | G1032A | C | 2.6e-09 | 5.7e-09 | 5.1e-09 | 4.467E-09 | 1.644E-09 | 2.600E-09 | 5.700E-09 |
| 20622 | 1032 | G1032R | C | 1.8e-08 | 8.8e-09 | 4.1e-09 | 1.030E-08 | 7.070E-09 | 4.100E-09 | 1.800E-08 |
| 20623 | 1032 | G1032N | C | 6.3e-09 | 1.1e-08 | 1e-08 | 9.100E-09 | 2.476E-09 | 6.300E-09 | 1.100E-08 |
| 20624 | 1032 | G1032D | C | 2.1e-08 | 8.2e-09 | 8.7e-09 | 1.263E-08 | 7.250E-09 | 8.200E-09 | 2.100E-08 |
| 20625 | 1032 | G1032C | C | 1.9e-09 | 4e-09 | 9.8e-09 | 5.233E-09 | 4.092E-09 | 1.900E-09 | 9.800E-09 |
| 20626 | 1032 | G1032E | C | 1.2e-08 | 4.4e-09 | 5e-09 | 7.133E-09 | 4.225E-09 | 4.400E-09 | 1.200E-08 |
| 20627 | 1032 | G1032Q | C | 1.2e-08 | 7.2e-09 | 6.8e-09 | 8.667E-09 | 2.894E-09 | 6.800E-09 | 1.200E-08 |
| 20628 | 1032 | G1032G | C | 6.5e-09 | 3e-09 | 6.9e-09 | 5.467E-09 | 2.146E-09 | 3.000E-09 | 6.900E-09 |
| 20629 | 1032 | G1032H | C | 1.2e-08 | 8.4e-09 | 2.3e-09 | 7.567E-09 | 4.903E-09 | 2.300E-09 | 1.200E-08 |
| 20630 | 1032 | G1032I | C | 5.3e-09 | 5.3e-09 | 1e-08 | 6.867E-09 | 2.714E-09 | 5.300E-09 | 1.000E-08 |
| 20631 | 1032 | G1032L | C | 6.1e-09 | 1.7e-08 | 5e-09 | 9.367E-09 | 6.634E-09 | 5.000E-09 | 1.700E-08 |
| 20632 | 1032 | G1032K | C | 1e-08 | 4.5e-09 | 8.3e-09 | 7.600E-09 | 2.816E-09 | 4.500E-09 | 1.000E-08 |
| 20633 | 1032 | G1032M | C | 7.3e-09 | 5.2e-09 | 8.8e-09 | 7.100E-09 | 1.808E-09 | 5.200E-09 | 8.800E-09 |
| 20634 | 1032 | G1032F | C | 2e-08 | 1.5e-08 | 4.2e-09 | 1.307E-08 | 8.075E-09 | 4.200E-09 | 2.000E-08 |
| 20635 | 1032 | G1032P | C | 5.2e-09 | 1e-08 | 8.7e-09 | 7.967E-09 | 2.483E-09 | 5.200E-09 | 1.000E-08 |
| 20636 | 1032 | G1032S | C | 1.1e-08 | 6.2e-09 | 1.4e-08 | 1.040E-08 | 3.934E-09 | 6.200E-09 | 1.400E-08 |
| 20637 | 1032 | G1032T | C | 1.1e-08 | 6.3e-09 | 7.1e-09 | 8.133E-09 | 2.515E-09 | 6.300E-09 | 1.100E-08 |
| 20638 | 1032 | G1032W | C | 3e-09 | 4.5e-09 | 1.2e-08 | 6.500E-09 | 4.822E-09 | 3.000E-09 | 1.200E-08 |
| 20639 | 1032 | G1032Y | C | 3.9e-09 | 2.1e-08 | 1.9e-09 | 8.933E-09 | 1.050E-08 | 1.900E-09 | 2.100E-08 |
| 20640 | 1032 | G1032V | C | 1.3e-08 | 1.1e-08 | 7.4e-09 | 1.047E-08 | 2.838E-09 | 7.400E-09 | 1.300E-08 |
| 20641 | 1033 | T1033A | C | 2.8e-09 | 3e-09 | 1.4e-08 | 6.600E-09 | 6.409E-09 | 2.800E-09 | 1.400E-08 |
| 20642 | 1033 | T1033R | C | 3.1e-09 | 1.2e-08 | 8.9e-09 | 8.000E-09 | 4.518E-09 | 3.100E-09 | 1.200E-08 |
| 20643 | 1033 | T1033N | C | 4.2e-09 | 7.4e-09 | 1.3e-08 | 8.200E-09 | 4.454E-09 | 4.200E-09 | 1.300E-08 |
| 20644 | 1033 | T1033D | C | 3.9e-09 | 7.1e-09 | 1.1e-08 | 7.333E-09 | 3.556E-09 | 3.900E-09 | 1.100E-08 |
| 20645 | 1033 | T1033C | C | 3.7e-09 | 1e-08 | 9.9e-09 | 7.867E-09 | 3.609E-09 | 3.700E-09 | 1.000E-08 |
| 20646 | 1033 | T1033E | C | 6.1e-09 | 1.1e-08 | 1.7e-08 | 1.137E-08 | 5.459E-09 | 6.100E-09 | 1.700E-08 |
| 20647 | 1033 | T1033Q | C | 7.5e-09 | 2.3e-08 | 7.2e-09 | 1.257E-08 | 9.037E-09 | 7.200E-09 | 2.300E-08 |
| 20648 | 1033 | T1033G | C | 1.6e-08 | 8.2e-09 | 3.3e-09 | 9.167E-09 | 6.405E-09 | 3.300E-09 | 1.600E-08 |
| 20649 | 1033 | T1033H | C | 6.1e-09 | 2.7e-09 | 9.1e-09 | 5.967E-09 | 3.202E-09 | 2.700E-09 | 9.100E-09 |
| 20650 | 1033 | T1033I | C | 5.6e-09 | 8e-09 | 8.9e-09 | 7.500E-09 | 1.706E-09 | 5.600E-09 | 8.900E-09 |
| 20651 | 1033 | T1033L | C | 6.4e-09 | 2.5e-09 | 9.5e-09 | 6.133E-09 | 3.508E-09 | 2.500E-09 | 9.500E-09 |
| 20652 | 1033 | T1033K | C | 3.5e-09 | 6e-09 | 6.6e-09 | 5.367E-09 | 1.644E-09 | 3.500E-09 | 6.600E-09 |
| 20653 | 1033 | T1033M | C | 4e-09 | 5.7e-09 | 7.6e-09 | 5.767E-09 | 1.801E-09 | 4.000E-09 | 7.600E-09 |
| 20654 | 1033 | T1033F | C | 8.4e-09 | 1.3e-08 | 3e-09 | 8.133E-09 | 5.005E-09 | 3.000E-09 | 1.300E-08 |

| | | | | | | | | | | |
|---|---|---|---|---|---|---|---|---|---|---|
| 20655 | 1033 | T1033P | C | 1e-08 | 8.3e-09 | 9.2e-09 | 9.167E-09 | 8.505E-10 | 8.300E-09 | 1.000E-08 |
| 20656 | 1033 | T1033S | C | 5.7e-09 | 7.7e-09 | 6e-09 | 6.467E-09 | 1.079E-09 | 5.700E-09 | 7.700E-09 |
| 20657 | 1033 | T1033T | C | 3.1e-09 | 5.8e-09 | 1e-08 | 6.300E-09 | 3.477E-09 | 3.100E-09 | 1.000E-08 |
| 20658 | 1033 | T1033W | C | 1e-08 | 1e-08 | 9.5e-09 | 9.833E-09 | 2.887E-10 | 9.500E-09 | 1.000E-08 |
| 20659 | 1033 | T1033Y | C | 1.5e-08 | 2e-08 | 7.2e-09 | 1.407E-08 | 6.451E-09 | 7.200E-09 | 2.000E-08 |
| 20660 | 1033 | T1033V | C | 4.4e-09 | 2.7e-09 | 5.2e-09 | 4.100E-09 | 1.277E-09 | 2.700E-09 | 5.200E-09 |
| 20661 | 1034 | Q1034A | C | 7.9e-09 | 1.2e-08 | 1.3e-08 | 1.097E-08 | 2.702E-09 | 7.900E-09 | 1.300E-08 |
| 20662 | 1034 | Q1034R | C | 3.9e-09 | 2.6e-09 | 1.2e-08 | 6.167E-09 | 5.093E-09 | 2.600E-09 | 1.200E-08 |
| 20663 | 1034 | Q1034N | C | 2.4e-09 | 6.1e-09 | 6.3e-09 | 4.933E-09 | 2.196E-09 | 2.400E-09 | 6.300E-09 |
| 20664 | 1034 | Q1034D | C | 8.5e-09 | 1e-08 | 3.4e-08 | 1.750E-08 | 1.431E-08 | 8.500E-09 | 3.400E-08 |
| 20665 | 1034 | Q1034C | C | 6.9e-09 | 3.8e-09 | 6e-09 | 5.567E-09 | 1.595E-09 | 3.800E-09 | 6.900E-09 |
| 20666 | 1034 | Q1034E | C | 6.3e-09 | 9.2e-09 | 4.4e-09 | 6.633E-09 | 2.417E-09 | 4.400E-09 | 9.200E-09 |
| 20667 | 1034 | Q1034Q | C | 1.3e-08 | 4.8e-09 | 1e-08 | 9.267E-09 | 4.149E-09 | 4.800E-09 | 1.300E-08 |
| 20668 | 1034 | Q1034G | C | 5.5e-09 | 1e-08 | 3.8e-09 | 6.433E-09 | 3.204E-09 | 3.800E-09 | 1.000E-08 |
| 20669 | 1034 | Q1034H | C | 5.8e-09 | 5.5e-09 | 4e-09 | 5.100E-09 | 9.644E-10 | 4.000E-09 | 5.800E-09 |
| 20670 | 1034 | Q1034I | C | 3.1e-09 | 1.7e-08 | 3.3e-09 | 7.800E-09 | 7.968E-09 | 3.100E-09 | 1.700E-08 |
| 20671 | 1034 | Q1034L | C | 1.4e-08 | 1.1e-08 | 8.8e-09 | 1.127E-08 | 2.610E-09 | 8.800E-09 | 1.400E-08 |
| 20672 | 1034 | Q1034K | C | 6.1e-09 | 3.7e-09 | 7e-09 | 5.600E-09 | 1.706E-09 | 3.700E-09 | 7.000E-09 |
| 20673 | 1034 | Q1034M | C | 7.9e-09 | 1.4e-08 | 8.3e-09 | 1.007E-08 | 3.412E-09 | 7.900E-09 | 1.400E-08 |
| 20674 | 1034 | Q1034F | C | 9.2e-09 | 3.3e-09 | 3.9e-09 | 5.467E-09 | 3.247E-09 | 3.300E-09 | 9.200E-09 |
| 20675 | 1034 | Q1034P | C | 1e-08 | 1.4e-08 | 1.2e-08 | 1.200E-08 | 2.000E-09 | 1.000E-08 | 1.400E-08 |
| 20676 | 1034 | Q1034S | C | 1.1e-08 | 4e-09 | 1.6e-08 | 1.033E-08 | 6.028E-09 | 4.000E-09 | 1.600E-08 |
| 20677 | 1034 | Q1034T | C | 1.2e-08 | 1.6e-08 | 5.4e-09 | 1.113E-08 | 5.353E-09 | 5.400E-09 | 1.600E-08 |
| 20678 | 1034 | Q1034W | C | 4.7e-09 | 7.6e-09 | 2.2e-09 | 4.833E-09 | 2.702E-09 | 2.200E-09 | 7.600E-09 |
| 20679 | 1034 | Q1034Y | C | 3.6e-08 | 4.5e-09 | 2.7e-09 | 1.440E-08 | 1.873E-08 | 2.700E-09 | 3.600E-08 |
| 20680 | 1034 | Q1034V | C | 8.4e-09 | 1.1e-08 | 2.8e-09 | 7.400E-09 | 4.190E-09 | 2.800E-09 | 1.100E-08 |
| 20681 | 1035 | T1035A | C | 1.8e-08 | 7.8e-09 | 1.1e-08 | 1.227E-08 | 5.217E-09 | 7.800E-09 | 1.800E-08 |
| 20682 | 1035 | T1035R | C | 3.5e-09 | 6.3e-09 | 5.2e-09 | 5.000E-09 | 1.411E-09 | 3.500E-09 | 6.300E-09 |
| 20683 | 1035 | T1035N | C | 1.3e-08 | 5.6e-09 | 6.1e-09 | 8.233E-09 | 4.136E-09 | 5.600E-09 | 1.300E-08 |
| 20684 | 1035 | T1035D | C | 4.9e-09 | 8.9e-09 | 1.3e-08 | 8.933E-09 | 4.050E-09 | 4.900E-09 | 1.300E-08 |
| 20685 | 1035 | T1035C | C | 3.9e-09 | 1.3e-08 | 9.1e-09 | 8.667E-09 | 4.565E-09 | 3.900E-09 | 1.300E-08 |
| 20686 | 1035 | T1035E | C | 4e-09 | 1.1e-08 | 4.7e-09 | 6.567E-09 | 3.855E-09 | 4.000E-09 | 1.100E-08 |
| 20687 | 1035 | T1035Q | C | 6.4e-09 | 1.7e-08 | 1.6e-08 | 1.313E-08 | 5.853E-09 | 6.400E-09 | 1.700E-08 |
| 20688 | 1035 | T1035G | C | 1.3e-08 | 7.9e-09 | 1.8e-08 | 1.297E-08 | 5.050E-09 | 7.900E-09 | 1.800E-08 |
| 20689 | 1035 | T1035H | C | 3e-09 | 7.2e-09 | 8.3e-09 | 6.167E-09 | 2.797E-09 | 3.000E-09 | 8.300E-09 |
| 20690 | 1035 | T1035I | C | 7.9e-09 | 8.8e-09 | 6.6e-09 | 7.767E-09 | 1.106E-09 | 6.600E-09 | 8.800E-09 |
| 20691 | 1035 | T1035L | C | 6.2e-09 | 7.9e-09 | 3.9e-09 | 6.000E-09 | 2.007E-09 | 3.900E-09 | 7.900E-09 |
| 20692 | 1035 | T1035K | C | 1e-08 | 1.4e-08 | 1.7e-08 | 1.367E-08 | 3.512E-09 | 1.000E-08 | 1.700E-08 |
| 20693 | 1035 | T1035M | C | 1.2e-08 | 7e-09 | 1e-08 | 9.667E-09 | 2.517E-09 | 7.000E-09 | 1.200E-08 |
| 20694 | 1035 | T1035F | C | 5.9e-09 | 1.3e-08 | 9.9e-09 | 9.600E-09 | 3.559E-09 | 5.900E-09 | 1.300E-08 |
| 20695 | 1035 | T1035P | C | 2.2e-08 | 1.2e-08 | 6.4e-09 | 1.347E-08 | 7.903E-09 | 6.400E-09 | 2.200E-08 |
| 20696 | 1035 | T1035S | C | 8.5e-09 | 1e-08 | 8.3e-09 | 8.933E-09 | 9.292E-10 | 8.300E-09 | 1.000E-08 |
| 20697 | 1035 | T1035T | C | 6.1e-08 | 6.4e-09 | 4.1e-09 | 2.383E-08 | 3.221E-08 | 4.100E-09 | 6.100E-08 |
| 20698 | 1035 | T1035W | C | 3.3e-09 | 5.1e-09 | 5.6e-09 | 4.667E-09 | 1.210E-09 | 3.300E-09 | 5.600E-09 |
| 20699 | 1035 | T1035Y | C | 3.9e-09 | 1.7e-08 | 2.8e-08 | 1.630E-08 | 1.207E-08 | 3.900E-09 | 2.800E-08 |
| 20700 | 1035 | T1035V | C | 7.8e-09 | 1.3e-08 | 9.4e-09 | 1.007E-08 | 2.663E-09 | 7.800E-09 | 1.300E-08 |
| 20701 | 1036 | Y1036A | C | 5e-09 | 8.9e-09 | 1.5e-08 | 9.633E-09 | 5.040E-09 | 5.000E-09 | 1.500E-08 |
| 20702 | 1036 | Y1036R | C | 1.3e-08 | 9.2e-10 | 9.8e-09 | 7.907E-09 | 6.259E-09 | 9.200E-10 | 1.300E-08 |
| 20703 | 1036 | Y1036N | C | 6.5e-09 | 8.6e-09 | 7.8e-09 | 7.633E-09 | 1.060E-09 | 6.500E-09 | 8.600E-09 |
| 20704 | 1036 | Y1036D | C | 1.1e-08 | 3.5e-09 | 9e-09 | 7.833E-09 | 3.884E-09 | 3.500E-09 | 1.100E-08 |
| 20705 | 1036 | Y1036C | C | 2.4e-09 | 9.5e-09 | 8e-09 | 6.633E-09 | 3.742E-09 | 2.400E-09 | 9.500E-09 |
| 20706 | 1036 | Y1036E | C | 5.9e-09 | 7.3e-09 | 7.9e-09 | 7.033E-09 | 1.026E-09 | 5.900E-09 | 7.900E-09 |
| 20707 | 1036 | Y1036Q | C | 9.6e-09 | 3.4e-09 | 4.6e-09 | 5.867E-09 | 3.288E-09 | 3.400E-09 | 9.600E-09 |
| 20708 | 1036 | Y1036G | C | 7.1e-09 | 4.8e-09 | 2.9e-08 | 1.363E-08 | 1.336E-08 | 4.800E-09 | 2.900E-08 |
| 20709 | 1036 | Y1036H | C | 2e-08 | 4.6e-09 | 7.3e-09 | 1.063E-08 | 8.223E-09 | 4.600E-09 | 2.000E-08 |
| 20710 | 1036 | Y1036I | C | 3.4e-09 | 5.2e-09 | 5.9e-09 | 4.833E-09 | 1.290E-09 | 3.400E-09 | 5.900E-09 |
| 20711 | 1036 | Y1036L | C | 6.5e-09 | 6.8e-09 | 6.4e-09 | 6.567E-09 | 2.082E-10 | 6.400E-09 | 6.800E-09 |
| 20712 | 1036 | Y1036K | C | 6.6e-09 | 8.4e-09 | 4.5e-09 | 6.500E-09 | 1.952E-09 | 4.500E-09 | 8.400E-09 |
| 20713 | 1036 | Y1036M | C | 1.3e-08 | 5e-09 | 4.7e-09 | 7.567E-09 | 4.708E-09 | 4.700E-09 | 1.300E-08 |
| 20714 | 1036 | Y1036F | C | 1.2e-08 | 7.4e-09 | 1.1e-08 | 1.013E-08 | 2.419E-09 | 7.400E-09 | 1.200E-08 |
| 20715 | 1036 | Y1036P | C | 1.5e-08 | 1.3e-08 | 1.2e-08 | 1.333E-08 | 1.528E-09 | 1.200E-08 | 1.500E-08 |
| 20716 | 1036 | Y1036S | C | 6.3e-09 | 5.3e-09 | 4.7e-09 | 5.433E-09 | 8.083E-10 | 4.700E-09 | 6.300E-09 |
| 20717 | 1036 | Y1036T | C | 3.1e-09 | 7.3e-09 | 3.7e-08 | 1.580E-08 | 1.848E-08 | 3.100E-09 | 3.700E-08 |
| 20718 | 1036 | Y1036W | C | 3.1e-08 | 1.2e-08 | 1.6e-08 | 1.967E-08 | 1.002E-08 | 1.200E-08 | 3.100E-08 |
| 20719 | 1036 | Y1036Y | C | 8.4e-09 | 5.5e-09 | 8.6e-09 | 7.500E-09 | 1.735E-09 | 5.500E-09 | 8.600E-09 |
| 20720 | 1036 | Y1036V | C | 8.3e-09 | 4.9e-09 | 1.2e-08 | 8.400E-09 | 3.551E-09 | 4.900E-09 | 1.200E-08 |
| 20721 | 1037 | I1037A | C | 1.2e-08 | 6e-09 | 6e-09 | 8.000E-09 | 3.464E-09 | 6.000E-09 | 1.200E-08 |
| 20722 | 1037 | I1037R | C | 7.8e-09 | 1.2e-08 | 7.4e-09 | 9.067E-09 | 2.548E-09 | 7.400E-09 | 1.200E-08 |
| 20723 | 1037 | I1037N | C | 5e-09 | 1.4e-08 | 4.4e-09 | 7.800E-09 | 5.378E-09 | 4.400E-09 | 1.400E-08 |
| 20724 | 1037 | I1037D | C | 9e-09 | 1.2e-08 | 3.9e-09 | 8.300E-09 | 4.095E-09 | 3.900E-09 | 1.200E-08 |
| 20725 | 1037 | I1037C | C | 5.4e-09 | 9.7e-09 | 6e-09 | 7.033E-09 | 2.329E-09 | 5.400E-09 | 9.700E-09 |
| 20726 | 1037 | I1037E | C | 1.6e-08 | 6.1e-09 | 4.6e-09 | 8.900E-09 | 6.194E-09 | 4.600E-09 | 1.600E-08 |
| 20727 | 1037 | I1037Q | C | 1.5e-08 | 6.8e-09 | 7.9e-09 | 9.900E-09 | 4.451E-09 | 6.800E-09 | 1.500E-08 |
| 20728 | 1037 | I1037G | C | 5.7e-09 | 6.5e-09 | 5.2e-09 | 5.800E-09 | 6.557E-10 | 5.200E-09 | 6.500E-09 |
| 20729 | 1037 | I1037H | C | 3.1e-09 | 1.3e-08 | 7.8e-09 | 7.967E-09 | 4.952E-09 | 3.100E-09 | 1.300E-08 |
| 20730 | 1037 | I1037I | C | 8.9e-09 | 1.1e-08 | 1.9e-08 | 1.297E-08 | 5.329E-09 | 8.900E-09 | 1.900E-08 |
| 20731 | 1037 | I1037L | C | 7.6e-09 | 1.9e-09 | 4.1e-09 | 4.533E-09 | 2.875E-09 | 1.900E-09 | 7.600E-09 |
| 20732 | 1037 | I1037K | C | 6.6e-09 | 3e-09 | 5e-09 | 4.867E-09 | 1.804E-09 | 3.000E-09 | 6.600E-09 |
| 20733 | 1037 | I1037M | C | 3.4e-09 | 2.1e-08 | 4.4e-09 | 9.600E-09 | 9.885E-09 | 3.400E-09 | 2.100E-08 |
| 20734 | 1037 | I1037F | C | 4.5e-09 | 5.5e-09 | 5.1e-09 | 5.033E-09 | 5.033E-10 | 4.500E-09 | 5.500E-09 |
| 20735 | 1037 | I1037P | C | 3.3e-09 | 9.4e-09 | 4.1e-09 | 5.600E-09 | 3.315E-09 | 3.300E-09 | 9.400E-09 |
| 20736 | 1037 | I1037S | C | 6.5e-09 | 4.8e-09 | 6.4e-09 | 5.900E-09 | 9.539E-10 | 4.800E-09 | 6.500E-09 |
| 20737 | 1037 | I1037T | C | 5.6e-09 | 9.3e-09 | 4.2e-09 | 6.367E-09 | 2.635E-09 | 4.200E-09 | 9.300E-09 |
| 20738 | 1037 | I1037W | C | 3.9e-09 | 2.1e-08 | 1.5e-08 | 1.330E-08 | 8.676E-09 | 3.900E-09 | 2.100E-08 |
| 20739 | 1037 | I1037Y | C | 7.7e-09 | 5.3e-09 | 1.1e-08 | 8.000E-09 | 2.862E-09 | 5.300E-09 | 1.100E-08 |

| | | | | | | | | | | |
|---|---|---|---|---|---|---|---|---|---|---|
| 20740 | 1037 | I1037V | C | 6.6e-09 | 7.7e-09 | 1.4e-08 | 9.433E-09 | 3.993E-09 | 6.600E-09 | 1.400E-08 |
| 20741 | 1038 | C1038A | C | 5.2e-09 | 4.6e-09 | 5.9e-09 | 5.233E-09 | 6.506E-10 | 4.600E-09 | 5.900E-09 |
| 20742 | 1038 | C1038R | C | 1.5e-08 | 9.3e-09 | 9.7e-09 | 1.133E-08 | 3.182E-09 | 9.300E-09 | 1.500E-08 |
| 20743 | 1038 | C1038N | C | 1.8e-08 | 3.8e-09 | 7.2e-09 | 9.667E-09 | 7.414E-09 | 3.800E-09 | 1.800E-08 |
| 20744 | 1038 | C1038D | C | 3.4e-09 | 1.2e-08 | 2.5e-09 | 5.967E-09 | 5.244E-09 | 2.500E-09 | 1.200E-08 |
| 20745 | 1038 | C1038C | C | 1.2e-08 | 1.2e-08 | 6.8e-09 | 1.027E-08 | 3.002E-09 | 6.800E-09 | 1.200E-08 |
| 20746 | 1038 | C1038E | C | 1.8e-09 | 8.6e-09 | 3e-09 | 4.467E-09 | 3.630E-09 | 1.800E-09 | 8.600E-09 |
| 20747 | 1038 | C1038Q | C | 1.1e-08 | 7.3e-09 | 6.5e-09 | 8.267E-09 | 2.401E-09 | 6.500E-09 | 1.100E-08 |
| 20748 | 1038 | C1038G | C | 6.1e-09 | 5.3e-09 | 1.4e-08 | 8.467E-09 | 4.809E-09 | 5.300E-09 | 1.400E-08 |
| 20749 | 1038 | C1038H | C | 1.4e-08 | 3e-09 | 3.8e-09 | 6.933E-09 | 6.133E-09 | 3.000E-09 | 1.400E-08 |
| 20750 | 1038 | C1038I | C | 8.8e-09 | 6.3e-09 | 2e-09 | 5.700E-09 | 3.439E-09 | 2.000E-09 | 8.800E-09 |
| 20751 | 1038 | C1038L | C | 1.6e-08 | 6.6e-09 | 1.1e-08 | 1.120E-08 | 4.703E-09 | 6.600E-09 | 1.600E-08 |
| 20752 | 1038 | C1038K | C | 9.6e-09 | 1.6e-08 | 8.9e-09 | 1.150E-08 | 3.913E-09 | 8.900E-09 | 1.600E-08 |
| 20753 | 1038 | C1038M | C | 1.3e-08 | 1.8e-09 | 6.9e-09 | 7.233E-09 | 5.607E-09 | 1.800E-09 | 1.300E-08 |
| 20754 | 1038 | C1038F | C | 8.8e-09 | 1.5e-08 | 7.2e-09 | 1.033E-08 | 4.120E-09 | 7.200E-09 | 1.500E-08 |
| 20755 | 1038 | C1038P | C | 1.1e-08 | 5.8e-09 | 9.2e-09 | 8.667E-09 | 2.641E-09 | 5.800E-09 | 1.100E-08 |
| 20756 | 1038 | C1038S | C | 1.2e-08 | 1.4e-08 | 7.4e-09 | 1.113E-08 | 3.384E-09 | 7.400E-09 | 1.400E-08 |
| 20757 | 1038 | C1038T | C | 1.5e-08 | 1.7e-08 | 6.5e-09 | 1.283E-08 | 5.575E-09 | 6.500E-09 | 1.700E-08 |
| 20758 | 1038 | C1038W | C | 8.1e-09 | 3.1e-09 | 3.3e-09 | 4.833E-09 | 2.831E-09 | 3.100E-09 | 8.100E-09 |
| 20759 | 1038 | C1038Y | C | 6.4e-09 | 4.8e-09 | 5.1e-09 | 5.433E-09 | 8.505E-10 | 4.800E-09 | 6.400E-09 |
| 20760 | 1038 | C1038V | C | 1e-08 | 9.7e-09 | 7.2e-09 | 8.967E-09 | 1.537E-09 | 7.200E-09 | 1.000E-08 |
| 20761 | 1039 | N1039A | C | 2.5e-08 | 7.9e-09 | 8.8e-09 | 1.390E-08 | 9.623E-09 | 7.900E-09 | 2.500E-08 |
| 20762 | 1039 | N1039R | C | 3.6e-09 | 1.1e-08 | 6.4e-09 | 7.000E-09 | 3.736E-09 | 3.600E-09 | 1.100E-08 |
| 20763 | 1039 | N1039N | C | 1.6e-08 | 8.2e-09 | 1.6e-09 | 8.600E-09 | 7.208E-09 | 1.600E-09 | 1.600E-08 |
| 20764 | 1039 | N1039D | C | 5.2e-09 | 1e-08 | 4.8e-09 | 6.667E-09 | 2.894E-09 | 4.800E-09 | 1.000E-08 |
| 20765 | 1039 | N1039C | C | 3.7e-09 | 8.1e-09 | 4.2e-09 | 5.333E-09 | 2.409E-09 | 3.700E-09 | 8.100E-09 |
| 20766 | 1039 | N1039E | C | 8.3e-09 | 8.6e-09 | 3.7e-09 | 6.867E-09 | 2.747E-09 | 3.700E-09 | 8.600E-09 |
| 20767 | 1039 | N1039Q | C | 7.6e-09 | 1.7e-08 | 1.1e-08 | 1.187E-08 | 4.760E-09 | 7.600E-09 | 1.700E-08 |
| 20768 | 1039 | N1039G | C | 3.2e-09 | 2.1e-08 | 1.1e-08 | 1.173E-08 | 8.923E-09 | 3.200E-09 | 2.100E-08 |
| 20769 | 1039 | N1039H | C | 3e-09 | 1.1e-08 | 4.3e-09 | 6.100E-09 | 4.293E-09 | 3.000E-09 | 1.100E-08 |
| 20770 | 1039 | N1039I | C | 4.8e-09 | 9.6e-09 | 6.3e-09 | 6.900E-09 | 2.456E-09 | 4.800E-09 | 9.600E-09 |
| 20771 | 1039 | N1039L | C | 2.4e-08 | 6.8e-09 | 6.7e-09 | 1.250E-08 | 9.959E-09 | 6.700E-09 | 2.400E-08 |
| 20772 | 1039 | N1039K | C | 2.2e-08 | 3e-09 | 1.3e-08 | 1.267E-08 | 9.504E-09 | 3.000E-09 | 2.200E-08 |
| 20773 | 1039 | N1039M | C | 5.8e-09 | 4.3e-09 | 9.4e-09 | 6.500E-09 | 2.621E-09 | 4.300E-09 | 9.400E-09 |
| 20774 | 1039 | N1039F | C | 2.6e-09 | 3.3e-09 | 1.3e-08 | 6.300E-09 | 5.813E-09 | 2.600E-09 | 1.300E-08 |
| 20775 | 1039 | N1039P | C | 1.5e-08 | 3.6e-09 | 3.8e-09 | 7.467E-09 | 6.525E-09 | 3.600E-09 | 1.500E-08 |
| 20776 | 1039 | N1039S | C | 3.3e-09 | 7e-09 | 6.8e-09 | 5.700E-09 | 2.081E-09 | 3.300E-09 | 7.000E-09 |
| 20777 | 1039 | N1039T | C | 2e-08 | 5e-09 | 7.1e-09 | 1.070E-08 | 8.122E-09 | 5.000E-09 | 2.000E-08 |
| 20778 | 1039 | N1039W | C | 3.1e-09 | 6.5e-09 | 9.8e-09 | 6.467E-09 | 3.350E-09 | 3.100E-09 | 9.800E-09 |
| 20779 | 1039 | N1039Y | C | 8e-09 | 7.4e-09 | 9.3e-09 | 8.233E-09 | 9.713E-10 | 7.400E-09 | 9.300E-09 |
| 20780 | 1039 | N1039V | C | 1e-08 | 1.6e-08 | 4.1e-09 | 1.003E-08 | 5.950E-09 | 4.100E-09 | 1.600E-08 |
| 20781 | 1040 | V1040A | C | 1.7e-08 | 4e-09 | 4.8e-09 | 8.600E-09 | 7.286E-09 | 4.000E-09 | 1.700E-08 |
| 20782 | 1040 | V1040R | C | 7.1e-09 | 6.2e-09 | 9.6e-09 | 7.633E-09 | 1.762E-09 | 6.200E-09 | 9.600E-09 |
| 20783 | 1040 | V1040N | C | 4.5e-09 | 5e-09 | 1.3e-08 | 7.500E-09 | 4.770E-09 | 4.500E-09 | 1.300E-08 |
| 20784 | 1040 | V1040D | C | 5.3e-09 | 9e-09 | 4.9e-09 | 6.400E-09 | 2.261E-09 | 4.900E-09 | 9.000E-09 |
| 20785 | 1040 | V1040C | C | 1.7e-08 | 1e-08 | 4e-09 | 1.033E-08 | 6.506E-09 | 4.000E-09 | 1.700E-08 |
| 20786 | 1040 | V1040E | C | 9e-09 | 2e-08 | 6.9e-09 | 1.197E-08 | 7.036E-09 | 6.900E-09 | 2.000E-08 |
| 20787 | 1040 | V1040Q | C | 6.7e-09 | 6.2e-09 | 1.1e-08 | 7.967E-09 | 2.639E-09 | 6.200E-09 | 1.100E-08 |
| 20788 | 1040 | V1040G | C | 5.5e-09 | 1.5e-08 | 7e-09 | 9.167E-09 | 5.107E-09 | 5.500E-09 | 1.500E-08 |
| 20789 | 1040 | V1040H | C | 8.2e-09 | 1.4e-08 | 2.8e-09 | 8.333E-09 | 5.601E-09 | 2.800E-09 | 1.400E-08 |
| 20790 | 1040 | V1040I | C | 1.7e-08 | 6.9e-09 | 1.9e-08 | 1.430E-08 | 6.486E-09 | 6.900E-09 | 1.900E-08 |
| 20791 | 1040 | V1040L | C | 1.8e-08 | 1.4e-08 | 3.7e-09 | 1.190E-08 | 7.378E-09 | 3.700E-09 | 1.800E-08 |
| 20792 | 1040 | V1040K | C | 1.4e-08 | 6.6e-09 | 1e-08 | 1.020E-08 | 3.704E-09 | 6.600E-09 | 1.400E-08 |
| 20793 | 1040 | V1040M | C | 2e-08 | 6.9e-09 | 5.4e-09 | 1.077E-08 | 8.031E-09 | 5.400E-09 | 2.000E-08 |
| 20794 | 1040 | V1040F | C | 4.4e-09 | 2.2e-09 | 2.8e-09 | 3.133E-09 | 1.137E-09 | 2.200E-09 | 4.400E-09 |
| 20795 | 1040 | V1040P | C | 7.2e-09 | 1.1e-08 | 3.7e-09 | 7.300E-09 | 3.651E-09 | 3.700E-09 | 1.100E-08 |
| 20796 | 1040 | V1040S | C | 4.6e-09 | 1.3e-08 | 1.8e-08 | 1.187E-08 | 6.772E-09 | 4.600E-09 | 1.800E-08 |
| 20797 | 1040 | V1040T | C | 1.3e-08 | 6e-09 | 5.2e-09 | 8.067E-09 | 4.291E-09 | 5.200E-09 | 1.300E-08 |
| 20798 | 1040 | V1040W | C | 7.1e-09 | 4.3e-09 | 6.3e-09 | 5.900E-09 | 1.442E-09 | 4.300E-09 | 7.100E-09 |
| 20799 | 1040 | V1040Y | C | 2.9e-09 | 1.1e-08 | 5.7e-09 | 6.533E-09 | 4.114E-09 | 2.900E-09 | 1.100E-08 |
| 20800 | 1040 | V1040V | C | 1.8e-09 | 6.5e-09 | 1.8e-08 | 8.767E-09 | 8.334E-09 | 1.800E-09 | 1.800E-08 |
| 20801 | 1041 | N1041A | C | 3.4e-09 | 5.3e-09 | 1.1e-08 | 6.567E-09 | 3.955E-09 | 3.400E-09 | 1.100E-08 |
| 20802 | 1041 | N1041R | C | 4.9e-09 | 1.3e-08 | 4.6e-09 | 7.500E-09 | 4.766E-09 | 4.600E-09 | 1.300E-08 |
| 20803 | 1041 | N1041N | C | 1.4e-08 | 3e-09 | 7.6e-09 | 8.200E-09 | 5.524E-09 | 3.000E-09 | 1.400E-08 |
| 20804 | 1041 | N1041D | C | 8.8e-09 | 2.3e-09 | 3.7e-09 | 4.933E-09 | 3.421E-09 | 2.300E-09 | 8.800E-09 |
| 20805 | 1041 | N1041C | C | 1.3e-08 | 1.2e-08 | 9.8e-09 | 1.160E-08 | 1.637E-09 | 9.800E-09 | 1.300E-08 |
| 20806 | 1041 | N1041E | C | 4.5e-09 | 1.9e-09 | 8e-09 | 4.800E-09 | 3.061E-09 | 1.900E-09 | 8.000E-09 |
| 20807 | 1041 | N1041Q | C | 7.8e-09 | 5.3e-09 | 3.3e-09 | 5.467E-09 | 2.255E-09 | 3.300E-09 | 7.800E-09 |
| 20808 | 1041 | N1041G | C | 6.5e-09 | 3.6e-09 | 6e-09 | 5.367E-09 | 1.550E-09 | 3.600E-09 | 6.500E-09 |
| 20809 | 1041 | N1041H | C | 6.5e-09 | 1.3e-08 | 1.1e-08 | 1.017E-08 | 3.329E-09 | 6.500E-09 | 1.300E-08 |
| 20810 | 1041 | N1041I | C | 5.6e-09 | 1.4e-08 | 4.5e-09 | 8.033E-09 | 5.196E-09 | 4.500E-09 | 1.400E-08 |
| 20811 | 1041 | N1041L | C | 8e-09 | 4.5e-09 | 6.5e-09 | 6.333E-09 | 1.756E-09 | 4.500E-09 | 8.000E-09 |
| 20812 | 1041 | N1041K | C | 1.4e-08 | 9.4e-09 | 1.1e-08 | 1.147E-08 | 2.335E-09 | 9.400E-09 | 1.400E-08 |
| 20813 | 1041 | N1041M | C | 1.9e-08 | 1e-08 | 1.1e-08 | 1.333E-08 | 4.933E-09 | 1.000E-08 | 1.900E-08 |
| 20814 | 1041 | N1041F | C | 7.3e-09 | 1.2e-08 | 1e-08 | 9.767E-09 | 2.359E-09 | 7.300E-09 | 1.200E-08 |
| 20815 | 1041 | N1041P | C | 3.8e-09 | 6.1e-09 | 1.3e-08 | 7.633E-09 | 4.788E-09 | 3.800E-09 | 1.300E-08 |
| 20816 | 1041 | N1041S | C | 8.4e-09 | 6.6e-09 | 8.5e-09 | 7.833E-09 | 1.069E-09 | 6.600E-09 | 8.500E-09 |
| 20817 | 1041 | N1041T | C | 5.7e-09 | 2e-09 | 1.8e-08 | 8.567E-09 | 8.376E-09 | 2.000E-09 | 1.800E-08 |
| 20818 | 1041 | N1041W | C | 1e-08 | 4.4e-09 | 1.2e-08 | 8.800E-09 | 3.940E-09 | 4.400E-09 | 1.200E-08 |
| 20819 | 1041 | N1041Y | C | 6.4e-09 | 4.3e-09 | 2.8e-09 | 4.500E-09 | 1.808E-09 | 2.800E-09 | 6.400E-09 |
| 20820 | 1041 | N1041V | C | 3e-09 | 5.3e-09 | 1.8e-08 | 8.767E-09 | 8.079E-09 | 3.000E-09 | 1.800E-08 |
| 20821 | 1042 | H1042A | C | 1.1e-08 | 1.3e-08 | 4.3e-09 | 9.433E-09 | 4.557E-09 | 4.300E-09 | 1.300E-08 |
| 20822 | 1042 | H1042R | C | 5e-09 | 6.5e-09 | 9e-09 | 6.833E-09 | 2.021E-09 | 5.000E-09 | 9.000E-09 |
| 20823 | 1042 | H1042N | C | 4.2e-09 | 2.5e-09 | 8e-09 | 4.900E-09 | 2.816E-09 | 2.500E-09 | 8.000E-09 |
| 20824 | 1042 | H1042D | C | 9.2e-09 | 4.9e-09 | 6.8e-09 | 6.967E-09 | 2.155E-09 | 4.900E-09 | 9.200E-09 |

| | | | | | | | | | | |
|---|---|---|---|---|---|---|---|---|---|---|
| 20825 | 1042 | H1042C | C | 6.3e-09 | 7.9e-09 | 6.4e-09 | 6.867E-09 | 8.963E-10 | 6.300E-09 | 7.900E-09 |
| 20826 | 1042 | H1042E | C | 1.1e-08 | 7e-09 | 7.7e-09 | 8.567E-09 | 2.136E-09 | 7.000E-09 | 1.100E-08 |
| 20827 | 1042 | H1042Q | C | 1.2e-08 | 1e-08 | 6.8e-09 | 9.600E-09 | 2.623E-09 | 6.800E-09 | 1.200E-08 |
| 20828 | 1042 | H1042G | C | 5.1e-09 | 7.7e-09 | 1.3e-08 | 8.600E-09 | 4.026E-09 | 5.100E-09 | 1.300E-08 |
| 20829 | 1042 | H1042H | C | 1.1e-08 | 1.9e-08 | 3.7e-09 | 1.123E-08 | 7.653E-09 | 3.700E-09 | 1.900E-08 |
| 20830 | 1042 | H1042I | C | 4.6e-09 | 6e-09 | 2.2e-08 | 1.087E-08 | 9.667E-09 | 4.600E-09 | 2.200E-08 |
| 20831 | 1042 | H1042L | C | 8.1e-09 | 9.8e-09 | 1.6e-08 | 1.130E-08 | 4.158E-09 | 8.100E-09 | 1.600E-08 |
| 20832 | 1042 | H1042K | C | 2.1e-09 | 4.8e-09 | 1.3e-08 | 6.633E-09 | 5.677E-09 | 2.100E-09 | 1.300E-08 |
| 20833 | 1042 | H1042M | C | 7.1e-09 | 6.7e-09 | 7.4e-09 | 7.067E-09 | 3.512E-10 | 6.700E-09 | 7.400E-09 |
| 20834 | 1042 | H1042F | C | 5.9e-09 | 7.1e-09 | 5e-09 | 6.000E-09 | 1.054E-09 | 5.000E-09 | 7.100E-09 |
| 20835 | 1042 | H1042P | C | 4.1e-09 | 1e-08 | 4.8e-09 | 6.300E-09 | 3.223E-09 | 4.100E-09 | 1.000E-08 |
| 20836 | 1042 | H1042S | C | 2e-09 | 1.5e-08 | 2.8e-09 | 6.600E-09 | 7.286E-09 | 2.000E-09 | 1.500E-08 |
| 20837 | 1042 | H1042T | C | 8.6e-09 | 1.8e-08 | 6e-09 | 1.087E-08 | 6.313E-09 | 6.000E-09 | 1.800E-08 |
| 20838 | 1042 | H1042W | C | 1.6e-09 | 2.3e-09 | 4.8e-09 | 2.900E-09 | 1.682E-09 | 1.600E-09 | 4.800E-09 |
| 20839 | 1042 | H1042Y | C | 1e-08 | 3.2e-09 | 1.1e-08 | 8.067E-09 | 4.244E-09 | 3.200E-09 | 1.100E-08 |
| 20840 | 1042 | H1042V | C | 9.7e-09 | 4.2e-09 | 7.5e-09 | 7.133E-09 | 2.768E-09 | 4.200E-09 | 9.700E-09 |
| 20841 | 1043 | K1043A | C | 4.2e-09 | 8.5e-09 | 4e-09 | 5.567E-09 | 2.542E-09 | 4.000E-09 | 8.500E-09 |
| 20842 | 1043 | K1043R | C | 4.5e-09 | 1.1e-08 | 1.3e-09 | 5.600E-09 | 4.943E-09 | 1.300E-09 | 1.100E-08 |
| 20843 | 1043 | K1043N | C | 3.8e-09 | 2.7e-09 | 4.3e-08 | 1.650E-08 | 2.296E-08 | 2.700E-09 | 4.300E-08 |
| 20844 | 1043 | K1043D | C | 1.1e-08 | 8.1e-09 | 1.6e-08 | 1.170E-08 | 3.996E-09 | 8.100E-09 | 1.600E-08 |
| 20845 | 1043 | K1043C | C | 7.7e-09 | 7.2e-09 | 1.4e-08 | 9.633E-09 | 3.790E-09 | 7.200E-09 | 1.400E-08 |
| 20846 | 1043 | K1043E | C | 2.3e-09 | 5.1e-09 | 1.3e-08 | 6.800E-09 | 5.549E-09 | 2.300E-09 | 1.300E-08 |
| 20847 | 1043 | K1043Q | C | 8.8e-09 | 6.3e-09 | 1.1e-08 | 8.700E-09 | 2.352E-09 | 6.300E-09 | 1.100E-08 |
| 20848 | 1043 | K1043G | C | 3.1e-08 | 5.4e-09 | 8.4e-09 | 1.493E-08 | 1.399E-08 | 5.400E-09 | 3.100E-08 |
| 20849 | 1043 | K1043H | C | 1e-08 | 1.7e-08 | 3.8e-09 | 1.027E-08 | 6.604E-09 | 3.800E-09 | 1.700E-08 |
| 20850 | 1043 | K1043I | C | 1.4e-08 | 6e-09 | 1.1e-08 | 1.033E-08 | 4.041E-09 | 6.000E-09 | 1.400E-08 |
| 20851 | 1043 | K1043L | C | 4.4e-09 | 7.2e-09 | 4.6e-09 | 5.400E-09 | 1.562E-09 | 4.400E-09 | 7.200E-09 |
| 20852 | 1043 | K1043K | C | 1.4e-08 | 6e-09 | 4e-09 | 8.000E-09 | 5.292E-09 | 4.000E-09 | 1.400E-08 |
| 20853 | 1043 | K1043M | C | 1.1e-08 | 4.5e-09 | 2.3e-09 | 5.933E-09 | 4.524E-09 | 2.300E-09 | 1.100E-08 |
| 20854 | 1043 | K1043F | C | 5.6e-09 | 1.6e-08 | 6.9e-09 | 9.500E-09 | 5.667E-09 | 5.600E-09 | 1.600E-08 |
| 20855 | 1043 | K1043P | C | 4.4e-09 | 1.2e-08 | 2.7e-09 | 6.367E-09 | 4.952E-09 | 2.700E-09 | 1.200E-08 |
| 20856 | 1043 | K1043S | C | 2.6e-09 | 4.8e-09 | 8.1e-09 | 5.167E-09 | 2.768E-09 | 2.600E-09 | 8.100E-09 |
| 20857 | 1043 | K1043T | C | 7.4e-09 | 7.7e-09 | 2.2e-08 | 1.237E-08 | 8.344E-09 | 7.400E-09 | 2.200E-08 |
| 20858 | 1043 | K1043W | C | 7.7e-09 | 3.4e-09 | 1.8e-08 | 9.700E-09 | 7.503E-09 | 3.400E-09 | 1.800E-08 |
| 20859 | 1043 | K1043Y | C | 5.5e-09 | 9e-09 | 5.9e-09 | 6.800E-09 | 1.916E-09 | 5.500E-09 | 9.000E-09 |
| 20860 | 1043 | K1043V | C | 4.3e-09 | 1.4e-08 | 1.3e-08 | 1.043E-08 | 5.335E-09 | 4.300E-09 | 1.400E-08 |
| 20861 | 1044 | P1044A | C | 2.6e-08 | 1.1e-08 | 1.2e-08 | 1.633E-08 | 8.386E-09 | 1.100E-08 | 2.600E-08 |
| 20862 | 1044 | P1044R | C | 3.2e-09 | 3.7e-09 | 3.1e-09 | 3.333E-09 | 3.215E-10 | 3.100E-09 | 3.700E-09 |
| 20863 | 1044 | P1044N | C | 1.3e-08 | 1.1e-08 | 1.1e-08 | 1.167E-08 | 1.155E-09 | 1.100E-08 | 1.300E-08 |
| 20864 | 1044 | P1044D | C | 5.6e-09 | 8.8e-09 | 5.3e-09 | 6.567E-09 | 1.940E-09 | 5.300E-09 | 8.800E-09 |
| 20865 | 1044 | P1044C | C | 1e-08 | 4e-09 | 4.8e-09 | 6.267E-09 | 3.258E-09 | 4.000E-09 | 1.000E-08 |
| 20866 | 1044 | P1044E | C | 1.5e-09 | 1.1e-08 | 5.3e-09 | 5.933E-09 | 4.782E-09 | 1.500E-09 | 1.100E-08 |
| 20867 | 1044 | P1044Q | C | 5.9e-09 | 9.6e-09 | 4.4e-09 | 6.633E-09 | 2.676E-09 | 4.400E-09 | 9.600E-09 |
| 20868 | 1044 | P1044G | C | 2.4e-09 | 8.8e-09 | 3.7e-09 | 4.967E-09 | 3.383E-09 | 2.400E-09 | 8.800E-09 |
| 20869 | 1044 | P1044H | C | 3.2e-09 | 3.4e-09 | 5.9e-09 | 4.167E-09 | 1.504E-09 | 3.200E-09 | 5.900E-09 |
| 20870 | 1044 | P1044I | C | 6e-09 | 1e-08 | 4.3e-09 | 6.767E-09 | 2.926E-09 | 4.300E-09 | 1.000E-08 |
| 20871 | 1044 | P1044L | C | 4.8e-09 | 5.6e-09 | 7.4e-09 | 5.933E-09 | 1.332E-09 | 4.800E-09 | 7.400E-09 |
| 20872 | 1044 | P1044K | C | 6.7e-09 | 6.8e-09 | 2.9e-09 | 5.467E-09 | 2.223E-09 | 2.900E-09 | 6.800E-09 |
| 20873 | 1044 | P1044M | C | 3.6e-09 | 1e-08 | 8.8e-09 | 7.467E-09 | 3.402E-09 | 3.600E-09 | 1.000E-08 |
| 20874 | 1044 | P1044F | C | 6.3e-09 | 4.7e-09 | 1.1e-08 | 7.333E-09 | 3.275E-09 | 4.700E-09 | 1.100E-08 |
| 20875 | 1044 | P1044P | C | 3e-09 | 7.8e-09 | 1.5e-08 | 8.600E-09 | 6.040E-09 | 3.000E-09 | 1.500E-08 |
| 20876 | 1044 | P1044S | C | 6.3e-09 | 1e-08 | 1.7e-08 | 1.110E-08 | 5.434E-09 | 6.300E-09 | 1.700E-08 |
| 20877 | 1044 | P1044T | C | 4.7e-09 | 1e-08 | 6.1e-09 | 6.933E-09 | 2.747E-09 | 4.700E-09 | 1.000E-08 |
| 20878 | 1044 | P1044W | C | 8.9e-09 | 2e-09 | 1e-08 | 6.967E-09 | 4.336E-09 | 2.000E-09 | 1.000E-08 |
| 20879 | 1044 | P1044Y | C | 2e-09 | 1.3e-08 | 3.7e-09 | 6.233E-09 | 5.921E-09 | 2.000E-09 | 1.300E-08 |
| 20880 | 1044 | P1044V | C | 4.1e-09 | 4.1e-09 | 7.5e-09 | 5.233E-09 | 1.963E-09 | 4.100E-09 | 7.500E-09 |
| 20881 | 1045 | S1045A | C | 1.1e-08 | 5.2e-09 | 1.1e-08 | 9.067E-09 | 3.349E-09 | 5.200E-09 | 1.100E-08 |
| 20882 | 1045 | S1045R | C | 7.2e-09 | 3.5e-09 | 7e-09 | 5.900E-09 | 2.081E-09 | 3.500E-09 | 7.200E-09 |
| 20883 | 1045 | S1045N | C | 8.4e-09 | 9.9e-09 | 3.4e-09 | 7.233E-09 | 3.403E-09 | 3.400E-09 | 9.900E-09 |
| 20884 | 1045 | S1045D | C | 7e-09 | 3.1e-09 | 5.7e-09 | 5.267E-09 | 1.986E-09 | 3.100E-09 | 7.000E-09 |
| 20885 | 1045 | S1045C | C | 8.4e-09 | 7.8e-09 | 4.8e-09 | 7.000E-09 | 1.929E-09 | 4.800E-09 | 8.400E-09 |
| 20886 | 1045 | S1045E | C | 1e-08 | 1.1e-08 | 8.7e-09 | 9.900E-09 | 1.153E-09 | 8.700E-09 | 1.100E-08 |
| 20887 | 1045 | S1045Q | C | 8.7e-09 | 7.5e-09 | 6.2e-09 | 7.467E-09 | 1.250E-09 | 6.200E-09 | 8.700E-09 |
| 20888 | 1045 | S1045G | C | 6.9e-09 | 5.3e-09 | 8.8e-09 | 7.000E-09 | 1.752E-09 | 5.300E-09 | 8.800E-09 |
| 20889 | 1045 | S1045H | C | 8.8e-09 | 7.6e-09 | 9.2e-09 | 8.533E-09 | 8.327E-10 | 7.600E-09 | 9.200E-09 |
| 20890 | 1045 | S1045I | C | 1.3e-08 | 5.2e-09 | 6.4e-09 | 8.200E-09 | 4.200E-09 | 5.200E-09 | 1.300E-08 |
| 20891 | 1045 | S1045L | C | 8.6e-09 | 9.7e-09 | 3.4e-09 | 7.233E-09 | 3.365E-09 | 3.400E-09 | 9.700E-09 |
| 20892 | 1045 | S1045K | C | 9.4e-09 | 5.6e-09 | 8.3e-09 | 7.767E-09 | 1.955E-09 | 5.600E-09 | 9.400E-09 |
| 20893 | 1045 | S1045M | C | 5.8e-09 | 1.2e-08 | 9.6e-09 | 9.133E-09 | 3.126E-09 | 5.800E-09 | 1.200E-08 |
| 20894 | 1045 | S1045F | C | 7.2e-09 | 7.3e-09 | 3.7e-09 | 6.067E-09 | 2.050E-09 | 3.700E-09 | 7.300E-09 |
| 20895 | 1045 | S1045P | C | 9.1e-09 | 7.3e-09 | 4.3e-09 | 6.900E-09 | 2.425E-09 | 4.300E-09 | 9.100E-09 |
| 20896 | 1045 | S1045S | C | 5.5e-09 | 6.1e-09 | 6.4e-09 | 6.000E-09 | 4.583E-10 | 5.500E-09 | 6.400E-09 |
| 20897 | 1045 | S1045T | C | 6.4e-09 | 6.7e-09 | 2.3e-08 | 1.203E-08 | 9.499E-09 | 6.400E-09 | 2.300E-08 |
| 20898 | 1045 | S1045W | C | 9.4e-09 | 3.1e-09 | 4e-09 | 5.500E-09 | 3.407E-09 | 3.100E-09 | 9.400E-09 |
| 20899 | 1045 | S1045Y | C | 8.8e-09 | 2.8e-09 | 4.4e-09 | 5.333E-09 | 3.107E-09 | 2.800E-09 | 8.800E-09 |
| 20900 | 1045 | S1045V | C | 8.2e-09 | 7.7e-09 | 1.1e-08 | 8.967E-09 | 1.779E-09 | 7.700E-09 | 1.100E-08 |
| 20901 | 1046 | N1046A | C | 5.7e-09 | 5e-09 | 1.1e-08 | 7.233E-09 | 3.281E-09 | 5.000E-09 | 1.100E-08 |
| 20902 | 1046 | N1046R | C | 5.7e-09 | 1.1e-08 | 7.4e-09 | 8.033E-09 | 2.706E-09 | 5.700E-09 | 1.100E-08 |
| 20903 | 1046 | N1046N | C | 4.1e-09 | 6.7e-09 | 2.5e-09 | 4.433E-09 | 2.120E-09 | 2.500E-09 | 6.700E-09 |
| 20904 | 1046 | N1046D | C | 6.1e-09 | 1.4e-08 | 5.4e-09 | 8.500E-09 | 4.776E-09 | 5.400E-09 | 1.400E-08 |
| 20905 | 1046 | N1046C | C | 4.9e-09 | 1.5e-08 | 1.3e-08 | 1.097E-08 | 5.348E-09 | 4.900E-09 | 1.500E-08 |
| 20906 | 1046 | N1046E | C | 1.3e-08 | 3.5e-09 | 6.6e-09 | 7.700E-09 | 4.845E-09 | 3.500E-09 | 1.300E-08 |
| 20907 | 1046 | N1046Q | C | 2.5e-08 | 2.9e-09 | 6.3e-09 | 1.140E-08 | 1.190E-08 | 2.900E-09 | 2.500E-08 |
| 20908 | 1046 | N1046G | C | 1.8e-08 | 9.4e-09 | 2.2e-09 | 9.867E-09 | 7.910E-09 | 2.200E-09 | 1.800E-08 |
| 20909 | 1046 | N1046H | C | 5.8e-09 | 5.2e-09 | 1e-08 | 7.000E-09 | 2.615E-09 | 5.200E-09 | 1.000E-08 |

| | | | | | | | | | | |
|---|---|---|---|---|---|---|---|---|---|---|
| 20910 | 1046 | N1046I | C | 2e-08 | 5.9e-09 | 6.2e-09 | 1.070E-08 | 8.055E-09 | 5.900E-09 | 2.000E-08 |
| 20911 | 1046 | N1046L | C | 5.1e-09 | 7.9e-09 | 9.2e-09 | 7.400E-09 | 2.095E-09 | 5.100E-09 | 9.200E-09 |
| 20912 | 1046 | N1046K | C | 8.6e-09 | 3.6e-08 | 6.6e-09 | 1.707E-08 | 1.643E-08 | 6.600E-09 | 3.600E-08 |
| 20913 | 1046 | N1046M | C | 1.6e-08 | 2.8e-09 | 1e-08 | 9.600E-09 | 6.609E-09 | 2.800E-09 | 1.600E-08 |
| 20914 | 1046 | N1046F | C | 6.9e-09 | 7e-09 | 5.4e-09 | 6.433E-09 | 8.963E-10 | 5.400E-09 | 7.000E-09 |
| 20915 | 1046 | N1046P | C | 2.9e-09 | 2.8e-09 | 9.6e-09 | 5.100E-09 | 3.897E-09 | 2.800E-09 | 9.600E-09 |
| 20916 | 1046 | N1046S | C | 8.1e-10 | 5.4e-09 | 5.4e-09 | 3.870E-09 | 2.650E-09 | 8.100E-10 | 5.400E-09 |
| 20917 | 1046 | N1046T | C | 6e-09 | 7.9e-09 | 9.5e-09 | 7.800E-09 | 1.752E-09 | 6.000E-09 | 9.500E-09 |
| 20918 | 1046 | N1046W | C | 1.1e-08 | 6.9e-09 | 9.4e-09 | 9.100E-09 | 2.066E-09 | 6.900E-09 | 1.100E-08 |
| 20919 | 1046 | N1046Y | C | 5.8e-09 | 1.3e-08 | 2e-08 | 1.293E-08 | 7.100E-09 | 5.800E-09 | 2.000E-08 |
| 20920 | 1046 | N1046V | C | 8.7e-09 | 3.3e-09 | 9.2e-09 | 7.067E-09 | 3.272E-09 | 3.300E-09 | 9.200E-09 |
| 20921 | 1047 | T1047A | C | 5.7e-09 | 7.9e-09 | 3.6e-09 | 5.733E-09 | 2.150E-09 | 3.600E-09 | 7.900E-09 |
| 20922 | 1047 | T1047R | C | 4.8e-09 | 1.4e-08 | 8.7e-09 | 9.167E-09 | 4.618E-09 | 4.800E-09 | 1.400E-08 |
| 20923 | 1047 | T1047N | C | 7.1e-09 | 8.8e-09 | 7.2e-09 | 7.700E-09 | 9.539E-10 | 7.100E-09 | 8.800E-09 |
| 20924 | 1047 | T1047D | C | 3.6e-09 | 5.6e-09 | 4.6e-09 | 4.600E-09 | 1.000E-09 | 3.600E-09 | 5.600E-09 |
| 20925 | 1047 | T1047C | C | 8.3e-09 | 1.4e-09 | 1.8e-08 | 9.233E-09 | 8.339E-09 | 1.400E-09 | 1.800E-08 |
| 20926 | 1047 | T1047E | C | 6.9e-09 | 7.6e-09 | 4.7e-09 | 6.400E-09 | 1.513E-09 | 4.700E-09 | 7.600E-09 |
| 20927 | 1047 | T1047Q | C | 1.1e-08 | 2.7e-09 | 4.8e-09 | 6.167E-09 | 4.315E-09 | 2.700E-09 | 1.100E-08 |
| 20928 | 1047 | T1047G | C | 5.5e-09 | 1.6e-08 | 8.1e-09 | 9.867E-09 | 5.468E-09 | 5.500E-09 | 1.600E-08 |
| 20929 | 1047 | T1047H | C | 5.6e-09 | 6.2e-09 | 4.3e-09 | 5.367E-09 | 9.713E-10 | 4.300E-09 | 6.200E-09 |
| 20930 | 1047 | T1047I | C | 4.4e-09 | 2e-08 | 3.5e-09 | 9.300E-09 | 9.277E-09 | 3.500E-09 | 2.000E-08 |
| 20931 | 1047 | T1047L | C | 7.1e-09 | 8e-09 | 1.8e-08 | 1.103E-08 | 6.050E-09 | 7.100E-09 | 1.800E-08 |
| 20932 | 1047 | T1047K | C | 3.8e-09 | 5.7e-09 | 6.2e-09 | 5.233E-09 | 1.266E-09 | 3.800E-09 | 6.200E-09 |
| 20933 | 1047 | T1047M | C | 1.3e-08 | 1.1e-08 | 1.6e-08 | 1.333E-08 | 2.517E-09 | 1.100E-08 | 1.600E-08 |
| 20934 | 1047 | T1047F | C | 5e-09 | 4.4e-09 | 2.7e-09 | 4.033E-09 | 1.193E-09 | 2.700E-09 | 5.000E-09 |
| 20935 | 1047 | T1047P | C | 6.3e-09 | 2.8e-09 | 6.5e-09 | 5.200E-09 | 2.081E-09 | 2.800E-09 | 6.500E-09 |
| 20936 | 1047 | T1047S | C | 6.1e-09 | 1e-08 | 5.1e-09 | 7.067E-09 | 2.589E-09 | 5.100E-09 | 1.000E-08 |
| 20937 | 1047 | T1047T | C | 4.4e-09 | 1e-08 | 2e-08 | 1.147E-08 | 7.903E-09 | 4.400E-09 | 2.000E-08 |
| 20938 | 1047 | T1047W | C | 7.4e-09 | 5.8e-09 | 4.4e-09 | 5.867E-09 | 1.501E-09 | 4.400E-09 | 7.400E-09 |
| 20939 | 1047 | T1047Y | C | 2.1e-08 | 4e-09 | 1.1e-08 | 1.200E-08 | 8.544E-09 | 4.000E-09 | 2.100E-08 |
| 20940 | 1047 | T1047V | C | 3e-09 | 1.2e-08 | 3.2e-09 | 6.067E-09 | 5.139E-09 | 3.000E-09 | 1.200E-08 |
| 20941 | 1048 | K1048A | C | 6.1e-09 | 5.4e-09 | 1.8e-08 | 9.833E-09 | 7.081E-09 | 5.400E-09 | 1.800E-08 |
| 20942 | 1048 | K1048R | C | 2e-08 | 1.5e-08 | 9.8e-09 | 1.493E-08 | 5.100E-09 | 9.800E-09 | 2.000E-08 |
| 20943 | 1048 | K1048N | C | 5.3e-09 | 2.1e-09 | 4.7e-09 | 4.033E-09 | 1.701E-09 | 2.100E-09 | 5.300E-09 |
| 20944 | 1048 | K1048D | C | 8e-09 | 3.1e-09 | 1.2e-08 | 7.700E-09 | 4.458E-09 | 3.100E-09 | 1.200E-08 |
| 20945 | 1048 | K1048C | C | 2.3e-08 | 4.5e-09 | 9.1e-09 | 1.220E-08 | 9.632E-09 | 4.500E-09 | 2.300E-08 |
| 20946 | 1048 | K1048E | C | 5.2e-09 | 5.5e-09 | 1.2e-08 | 7.567E-09 | 3.842E-09 | 5.200E-09 | 1.200E-08 |
| 20947 | 1048 | K1048Q | C | 1.4e-08 | 7e-09 | 8.2e-09 | 9.733E-09 | 3.743E-09 | 7.000E-09 | 1.400E-08 |
| 20948 | 1048 | K1048G | C | 6.6e-09 | 1.8e-08 | 7.3e-09 | 1.063E-08 | 6.389E-09 | 6.600E-09 | 1.800E-08 |
| 20949 | 1048 | K1048H | C | 9.4e-09 | 7.4e-09 | 4.8e-09 | 7.200E-09 | 2.307E-09 | 4.800E-09 | 9.400E-09 |
| 20950 | 1048 | K1048I | C | 1.1e-08 | 3.5e-09 | 1.1e-08 | 8.500E-09 | 4.330E-09 | 3.500E-09 | 1.100E-08 |
| 20951 | 1048 | K1048L | C | 1.1e-09 | 7e-09 | 1.3e-08 | 7.033E-09 | 5.950E-09 | 1.100E-09 | 1.300E-08 |
| 20952 | 1048 | K1048K | C | 4.8e-09 | 4.9e-09 | 4.6e-09 | 4.767E-09 | 1.528E-10 | 4.600E-09 | 4.900E-09 |
| 20953 | 1048 | K1048M | C | 9.5e-09 | 4.5e-09 | 3.8e-09 | 5.933E-09 | 3.109E-09 | 3.800E-09 | 9.500E-09 |
| 20954 | 1048 | K1048F | C | 3.8e-09 | 4.4e-09 | 7.7e-09 | 5.300E-09 | 2.100E-09 | 3.800E-09 | 7.700E-09 |
| 20955 | 1048 | K1048P | C | 7.3e-09 | 1.1e-09 | 5.8e-09 | 4.733E-09 | 3.235E-09 | 1.100E-09 | 7.300E-09 |
| 20956 | 1048 | K1048S | C | 1.6e-08 | 1.1e-08 | 8e-09 | 1.167E-08 | 4.041E-09 | 8.000E-09 | 1.600E-08 |
| 20957 | 1048 | K1048T | C | 8.5e-09 | 1.1e-08 | 7.4e-09 | 8.967E-09 | 1.845E-09 | 7.400E-09 | 1.100E-08 |
| 20958 | 1048 | K1048W | C | 3.1e-09 | 3e-09 | 6.9e-09 | 4.333E-09 | 2.223E-09 | 3.000E-09 | 6.900E-09 |
| 20959 | 1048 | K1048Y | C | 1e-08 | 4.1e-09 | 1.9e-09 | 5.333E-09 | 4.188E-09 | 1.900E-09 | 1.000E-08 |
| 20960 | 1048 | K1048V | C | 4.1e-09 | 5.2e-09 | 7.2e-09 | 5.500E-09 | 1.572E-09 | 4.100E-09 | 7.200E-09 |
| 20961 | 1049 | V1049A | C | 1.2e-08 | 1.4e-08 | 1.6e-08 | 1.400E-08 | 2.000E-09 | 1.200E-08 | 1.600E-08 |
| 20962 | 1049 | V1049R | C | 1.3e-08 | 8e-09 | 6.7e-09 | 9.233E-09 | 3.326E-09 | 6.700E-09 | 1.300E-08 |
| 20963 | 1049 | V1049N | C | 8.5e-09 | 1.3e-08 | 2.7e-09 | 8.067E-09 | 5.164E-09 | 2.700E-09 | 1.300E-08 |
| 20964 | 1049 | V1049D | C | 5.1e-09 | 7.5e-09 | 6.2e-09 | 6.267E-09 | 1.201E-09 | 5.100E-09 | 7.500E-09 |
| 20965 | 1049 | V1049C | C | 1.5e-08 | 2.6e-09 | 5.7e-09 | 7.767E-09 | 6.453E-09 | 2.600E-09 | 1.500E-08 |
| 20966 | 1049 | V1049E | C | 4.2e-09 | 1.4e-08 | 7.8e-09 | 8.667E-09 | 4.957E-09 | 4.200E-09 | 1.400E-08 |
| 20967 | 1049 | V1049Q | C | 1e-08 | 3.6e-09 | 7.8e-09 | 7.133E-09 | 3.252E-09 | 3.600E-09 | 1.000E-08 |
| 20968 | 1049 | V1049G | C | 8.2e-09 | 4e-09 | 3.3e-09 | 5.167E-09 | 2.650E-09 | 3.300E-09 | 8.200E-09 |
| 20969 | 1049 | V1049H | C | 5e-09 | 8.5e-09 | 5e-09 | 6.167E-09 | 2.021E-09 | 5.000E-09 | 8.500E-09 |
| 20970 | 1049 | V1049I | C | 4.4e-09 | 9.1e-09 | 4.6e-09 | 6.033E-09 | 2.658E-09 | 4.400E-09 | 9.100E-09 |
| 20971 | 1049 | V1049L | C | 5e-09 | 1.3e-08 | 3.6e-09 | 7.200E-09 | 5.071E-09 | 3.600E-09 | 1.300E-08 |
| 20972 | 1049 | V1049K | C | 8.8e-09 | 1e-08 | 2.5e-09 | 7.100E-09 | 4.029E-09 | 2.500E-09 | 1.000E-08 |
| 20973 | 1049 | V1049M | C | 8.1e-09 | 9.8e-09 | 1.5e-08 | 1.097E-08 | 3.595E-09 | 8.100E-09 | 1.500E-08 |
| 20974 | 1049 | V1049F | C | 9.8e-09 | 1.9e-08 | 2.7e-08 | 1.860E-08 | 8.607E-09 | 9.800E-09 | 2.700E-08 |
| 20975 | 1049 | V1049P | C | 1.7e-08 | 4.8e-09 | 3.4e-09 | 8.400E-09 | 7.481E-09 | 3.400E-09 | 1.700E-08 |
| 20976 | 1049 | V1049S | C | 9.5e-09 | 2.7e-08 | 4.1e-09 | 1.353E-08 | 1.197E-08 | 4.100E-09 | 2.700E-08 |
| 20977 | 1049 | V1049T | C | 7e-09 | 8.1e-09 | 7.1e-09 | 7.400E-09 | 6.083E-10 | 7.000E-09 | 8.100E-09 |
| 20978 | 1049 | V1049W | C | 6.1e-09 | 5.2e-09 | 8.9e-09 | 6.733E-09 | 1.930E-09 | 5.200E-09 | 8.900E-09 |
| 20979 | 1049 | V1049Y | C | 4.6e-09 | 2.8e-09 | 1e-08 | 5.800E-09 | 3.747E-09 | 2.800E-09 | 1.000E-08 |
| 20980 | 1049 | V1049V | C | 2.3e-09 | 3.4e-09 | 7.4e-09 | 4.367E-09 | 2.684E-09 | 2.300E-09 | 7.400E-09 |
| 20981 | 1050 | D1050A | C | 6.4e-09 | 8.3e-09 | 5.7e-09 | 6.800E-09 | 1.345E-09 | 5.700E-09 | 8.300E-09 |
| 20982 | 1050 | D1050R | C | 1.3e-08 | 1.3e-08 | 4.3e-09 | 1.010E-08 | 5.023E-09 | 4.300E-09 | 1.300E-08 |
| 20983 | 1050 | D1050N | C | 1.7e-08 | 5.7e-09 | 8.5e-09 | 1.040E-08 | 5.885E-09 | 5.700E-09 | 1.700E-08 |
| 20984 | 1050 | D1050D | C | 6.1e-09 | 6.4e-09 | 7.4e-09 | 6.633E-09 | 6.807E-10 | 6.100E-09 | 7.400E-09 |
| 20985 | 1050 | D1050C | C | 9e-09 | 2.8e-08 | 7e-09 | 1.467E-08 | 1.159E-08 | 7.000E-09 | 2.800E-08 |
| 20986 | 1050 | D1050E | C | 1.2e-08 | 1.1e-08 | 3.8e-09 | 8.933E-09 | 4.474E-09 | 3.800E-09 | 1.200E-08 |
| 20987 | 1050 | D1050Q | C | 1.2e-08 | 2.3e-09 | 8.2e-09 | 7.500E-09 | 4.888E-09 | 2.300E-09 | 1.200E-08 |
| 20988 | 1050 | D1050G | C | 1.6e-08 | 2.5e-08 | 2.5e-08 | 2.200E-08 | 5.196E-09 | 1.600E-08 | 2.500E-08 |
| 20989 | 1050 | D1050H | C | 5.5e-09 | 3e-09 | 9.5e-09 | 6.000E-09 | 3.279E-09 | 3.000E-09 | 9.500E-09 |
| 20990 | 1050 | D1050I | C | 8.2e-09 | 3.6e-09 | 7.8e-09 | 6.533E-09 | 2.548E-09 | 3.600E-09 | 8.200E-09 |
| 20991 | 1050 | D1050L | C | 2.8e-09 | 3.1e-09 | 5.7e-09 | 3.867E-09 | 1.595E-09 | 2.800E-09 | 5.700E-09 |
| 20992 | 1050 | D1050K | C | 2.3e-09 | 7.2e-09 | 1.2e-08 | 7.167E-09 | 4.850E-09 | 2.300E-09 | 1.200E-08 |
| 20993 | 1050 | D1050M | C | 7.1e-09 | 1.4e-08 | 1.3e-08 | 1.137E-08 | 3.729E-09 | 7.100E-09 | 1.400E-08 |
| 20994 | 1050 | D1050F | C | 6e-09 | 4e-09 | 2.8e-09 | 4.267E-09 | 1.617E-09 | 2.800E-09 | 6.000E-09 |

| | | | | | | | | | | |
|---|---|---|---|---|---|---|---|---|---|---|
| 20995 | 1050 | D1050P | C | 1.1e-08 | 8.1e-09 | 9e-09 | 9.367E-09 | 1.484E-09 | 8.100E-09 | 1.100E-08 |
| 20996 | 1050 | D1050S | C | 5.6e-09 | 1.1e-08 | 1.1e-08 | 9.200E-09 | 3.118E-09 | 5.600E-09 | 1.100E-08 |
| 20997 | 1050 | D1050T | C | 9.4e-09 | 1.4e-08 | 7.3e-09 | 1.023E-08 | 3.427E-09 | 7.300E-09 | 1.400E-08 |
| 20998 | 1050 | D1050W | C | 5.6e-09 | 6.9e-09 | 3.3e-09 | 5.267E-09 | 1.823E-09 | 3.300E-09 | 6.900E-09 |
| 20999 | 1050 | D1050Y | C | 9e-09 | 1e-08 | 3.7e-09 | 7.567E-09 | 3.386E-09 | 3.700E-09 | 1.000E-08 |
| 21000 | 1050 | D1050V | C | 2.7e-08 | 5.4e-09 | 4e-09 | 1.213E-08 | 1.289E-08 | 4.000E-09 | 2.700E-08 |
| 21001 | 1051 | K1051A | C | 1.9e-08 | 1.5e-08 | 1.5e-08 | 1.633E-08 | 2.309E-09 | 1.500E-08 | 1.900E-08 |
| 21002 | 1051 | K1051R | C | 1.1e-08 | 2.9e-09 | 5.3e-09 | 6.400E-09 | 4.161E-09 | 2.900E-09 | 1.100E-08 |
| 21003 | 1051 | K1051N | C | 3.2e-09 | 4.5e-09 | 7.1e-09 | 4.933E-09 | 1.986E-09 | 3.200E-09 | 7.100E-09 |
| 21004 | 1051 | K1051D | C | 3.5e-09 | 9.6e-09 | 2.1e-09 | 5.067E-09 | 3.988E-09 | 2.100E-09 | 9.600E-09 |
| 21005 | 1051 | K1051C | C | 1.9e-08 | 5e-09 | 1.5e-08 | 1.300E-08 | 7.211E-09 | 5.000E-09 | 1.900E-08 |
| 21006 | 1051 | K1051E | C | 5.8e-09 | 2e-09 | 1.4e-08 | 7.267E-09 | 6.133E-09 | 2.000E-09 | 1.400E-08 |
| 21007 | 1051 | K1051Q | C | 4.8e-09 | 1.1e-08 | 9.8e-09 | 8.533E-09 | 3.288E-09 | 4.800E-09 | 1.100E-08 |
| 21008 | 1051 | K1051G | C | 1.7e-09 | 6.4e-09 | 3.3e-09 | 3.800E-09 | 2.390E-09 | 1.700E-09 | 6.400E-09 |
| 21009 | 1051 | K1051H | C | 1.3e-08 | 1.2e-08 | 3e-09 | 9.333E-09 | 5.508E-09 | 3.000E-09 | 1.300E-08 |
| 21010 | 1051 | K1051I | C | 5.2e-09 | 1.8e-08 | 1.9e-08 | 1.407E-08 | 7.695E-09 | 5.200E-09 | 1.900E-08 |
| 21011 | 1051 | K1051L | C | 2.5e-09 | 6e-09 | 1.1e-08 | 6.500E-09 | 4.272E-09 | 2.500E-09 | 1.100E-08 |
| 21012 | 1051 | K1051K | C | 3.2e-09 | 5.3e-09 | 2.8e-09 | 3.767E-09 | 1.343E-09 | 2.800E-09 | 5.300E-09 |
| 21013 | 1051 | K1051M | C | 1.3e-08 | 9.7e-09 | 2e-09 | 8.233E-09 | 5.645E-09 | 2.000E-09 | 1.300E-08 |
| 21014 | 1051 | K1051F | C | 6.9e-09 | 1.4e-08 | 8.1e-09 | 9.667E-09 | 3.800E-09 | 6.900E-09 | 1.400E-08 |
| 21015 | 1051 | K1051P | C | 7.1e-09 | 4.3e-09 | 1.4e-08 | 8.467E-09 | 4.992E-09 | 4.300E-09 | 1.400E-08 |
| 21016 | 1051 | K1051S | C | 1.6e-08 | 4.3e-09 | 5.2e-09 | 8.500E-09 | 6.511E-09 | 4.300E-09 | 1.600E-08 |
| 21017 | 1051 | K1051T | C | 9.3e-09 | 5e-09 | 1.3e-08 | 9.100E-09 | 4.004E-09 | 5.000E-09 | 1.300E-08 |
| 21018 | 1051 | K1051W | C | 1.1e-08 | 8.2e-09 | 5.1e-09 | 8.100E-09 | 2.951E-09 | 5.100E-09 | 1.100E-08 |
| 21019 | 1051 | K1051Y | C | 6.9e-09 | 3.2e-09 | 9.1e-09 | 6.400E-09 | 2.982E-09 | 3.200E-09 | 9.100E-09 |
| 21020 | 1051 | K1051V | C | 5.5e-09 | 1.4e-08 | 4.1e-09 | 7.867E-09 | 5.358E-09 | 4.100E-09 | 1.400E-08 |
| 21021 | 1052 | K1052A | C | 8.5e-09 | 2.6e-09 | 9.3e-09 | 6.800E-09 | 3.659E-09 | 2.600E-09 | 9.300E-09 |
| 21022 | 1052 | K1052R | C | 9.7e-09 | 6.9e-09 | 1.2e-08 | 9.533E-09 | 2.554E-09 | 6.900E-09 | 1.200E-08 |
| 21023 | 1052 | K1052N | C | 5.3e-09 | 8.1e-09 | 1.2e-08 | 8.467E-09 | 3.365E-09 | 5.300E-09 | 1.200E-08 |
| 21024 | 1052 | K1052D | C | 1.6e-09 | 4e-09 | 9.9e-09 | 5.167E-09 | 4.271E-09 | 1.600E-09 | 9.900E-09 |
| 21025 | 1052 | K1052C | C | 4.6e-09 | 1.3e-08 | 4.2e-09 | 7.267E-09 | 4.969E-09 | 4.200E-09 | 1.300E-08 |
| 21026 | 1052 | K1052E | C | 2.9e-09 | 1.1e-08 | 5.8e-09 | 6.567E-09 | 4.104E-09 | 2.900E-09 | 1.100E-08 |
| 21027 | 1052 | K1052Q | C | 1.2e-08 | 1.5e-09 | 9.8e-09 | 7.767E-09 | 5.537E-09 | 1.500E-09 | 1.200E-08 |
| 21028 | 1052 | K1052G | C | 1.7e-09 | 3.5e-09 | 3e-08 | 1.173E-08 | 1.584E-08 | 1.700E-09 | 3.000E-08 |
| 21029 | 1052 | K1052H | C | 6.2e-09 | 1.1e-08 | 5.8e-09 | 7.667E-09 | 2.894E-09 | 5.800E-09 | 1.100E-08 |
| 21030 | 1052 | K1052I | C | 1.1e-08 | 4.8e-09 | 7.3e-09 | 7.700E-09 | 3.119E-09 | 4.800E-09 | 1.100E-08 |
| 21031 | 1052 | K1052L | C | 1.8e-08 | 5.4e-09 | 9.1e-09 | 1.083E-08 | 6.476E-09 | 5.400E-09 | 1.800E-08 |
| 21032 | 1052 | K1052K | C | 4.5e-09 | 3.3e-09 | 1.8e-08 | 8.600E-09 | 8.163E-09 | 3.300E-09 | 1.800E-08 |
| 21033 | 1052 | K1052M | C | 4.1e-09 | 3.2e-09 | 5.4e-09 | 4.233E-09 | 1.106E-09 | 3.200E-09 | 5.400E-09 |
| 21034 | 1052 | K1052F | C | 8.4e-09 | 9.2e-09 | 1.3e-08 | 1.020E-08 | 2.458E-09 | 8.400E-09 | 1.300E-08 |
| 21035 | 1052 | K1052P | C | 1.1e-08 | 3.3e-09 | 8.4e-09 | 7.567E-09 | 3.917E-09 | 3.300E-09 | 1.100E-08 |
| 21036 | 1052 | K1052S | C | 8.1e-09 | 1e-08 | 5.2e-09 | 7.767E-09 | 2.417E-09 | 5.200E-09 | 1.000E-08 |
| 21037 | 1052 | K1052T | C | 9e-09 | 1.2e-08 | 5.2e-09 | 8.733E-09 | 3.408E-09 | 5.200E-09 | 1.200E-08 |
| 21038 | 1052 | K1052W | C | 4e-09 | 9.3e-09 | 1.2e-08 | 8.433E-09 | 4.070E-09 | 4.000E-09 | 1.200E-08 |
| 21039 | 1052 | K1052Y | C | 6.2e-09 | 5.6e-09 | 9.3e-09 | 7.033E-09 | 1.986E-09 | 5.600E-09 | 9.300E-09 |
| 21040 | 1052 | K1052V | C | 3.5e-09 | 3.9e-09 | 4.3e-09 | 3.900E-09 | 4.000E-10 | 3.500E-09 | 4.300E-09 |
| 21041 | 1053 | V1053A | C | 1.1e-08 | 7.7e-09 | 5.5e-09 | 8.067E-09 | 2.768E-09 | 5.500E-09 | 1.100E-08 |
| 21042 | 1053 | V1053R | C | 7e-09 | 1.4e-08 | 5.6e-09 | 8.867E-09 | 4.500E-09 | 5.600E-09 | 1.400E-08 |
| 21043 | 1053 | V1053N | C | 6.5e-09 | 1.3e-08 | 7.4e-09 | 8.967E-09 | 3.522E-09 | 6.500E-09 | 1.300E-08 |
| 21044 | 1053 | V1053D | C | 7.1e-09 | 3.4e-09 | 4.6e-09 | 5.033E-09 | 1.888E-09 | 3.400E-09 | 7.100E-09 |
| 21045 | 1053 | V1053C | C | 4.7e-09 | 4.4e-09 | 1e-08 | 6.367E-09 | 3.150E-09 | 4.400E-09 | 1.000E-08 |
| 21046 | 1053 | V1053E | C | 6.3e-09 | 6.4e-09 | 5.7e-09 | 6.133E-09 | 3.786E-10 | 5.700E-09 | 6.400E-09 |
| 21047 | 1053 | V1053Q | C | 5.6e-09 | 6.3e-09 | 1.1e-08 | 7.633E-09 | 2.937E-09 | 5.600E-09 | 1.100E-08 |
| 21048 | 1053 | V1053G | C | 1.9e-09 | 1.1e-08 | 7.1e-09 | 6.667E-09 | 4.565E-09 | 1.900E-09 | 1.100E-08 |
| 21049 | 1053 | V1053H | C | 5e-09 | 6.3e-09 | 1.5e-08 | 8.767E-09 | 5.437E-09 | 5.000E-09 | 1.500E-08 |
| 21050 | 1053 | V1053I | C | 9.5e-09 | 7e-09 | 1e-08 | 8.833E-09 | 1.607E-09 | 7.000E-09 | 1.000E-08 |
| 21051 | 1053 | V1053L | C | 2.3e-09 | 3.2e-09 | 1.8e-09 | 2.433E-09 | 7.095E-10 | 1.800E-09 | 3.200E-09 |
| 21052 | 1053 | V1053K | C | 7.8e-09 | 1.6e-09 | 1.3e-08 | 7.467E-09 | 5.707E-09 | 1.600E-09 | 1.300E-08 |
| 21053 | 1053 | V1053M | C | 3.6e-09 | 8.3e-09 | 9.1e-09 | 7.000E-09 | 2.972E-09 | 3.600E-09 | 9.100E-09 |
| 21054 | 1053 | V1053F | C | 6.4e-09 | 2.5e-08 | 4.5e-09 | 1.197E-08 | 1.133E-08 | 4.500E-09 | 2.500E-08 |
| 21055 | 1053 | V1053P | C | 8.1e-09 | 5.2e-09 | 6.6e-09 | 6.633E-09 | 1.450E-09 | 5.200E-09 | 8.100E-09 |
| 21056 | 1053 | V1053S | C | 6.1e-09 | 1.6e-09 | 2.3e-09 | 3.333E-09 | 2.421E-09 | 1.600E-09 | 6.100E-09 |
| 21057 | 1053 | V1053T | C | 7e-09 | 1.2e-08 | 4.9e-09 | 7.967E-09 | 3.647E-09 | 4.900E-09 | 1.200E-08 |
| 21058 | 1053 | V1053W | C | 1.1e-08 | 5.9e-09 | 8.7e-09 | 8.533E-09 | 2.554E-09 | 5.900E-09 | 1.100E-08 |
| 21059 | 1053 | V1053Y | C | 3.7e-09 | 9.6e-09 | 2.7e-09 | 5.333E-09 | 3.729E-09 | 2.700E-09 | 9.600E-09 |
| 21060 | 1053 | V1053V | C | 8.3e-09 | 6.4e-09 | 1.3e-08 | 9.233E-09 | 3.398E-09 | 6.400E-09 | 1.300E-08 |
| 21061 | 1054 | E1054A | C | 1.5e-08 | 1.7e-08 | 1.3e-08 | 1.500E-08 | 2.000E-09 | 1.300E-08 | 1.700E-08 |
| 21062 | 1054 | E1054R | C | 4.4e-09 | 7.3e-09 | 6.5e-09 | 6.067E-09 | 1.498E-09 | 4.400E-09 | 7.300E-09 |
| 21063 | 1054 | E1054N | C | 6.7e-09 | 7.7e-09 | 1.8e-08 | 1.080E-08 | 6.255E-09 | 6.700E-09 | 1.800E-08 |
| 21064 | 1054 | E1054D | C | 1.4e-08 | 4.5e-09 | 2.3e-09 | 6.933E-09 | 6.218E-09 | 2.300E-09 | 1.400E-08 |
| 21065 | 1054 | E1054C | C | 2e-08 | 1e-08 | 6.7e-09 | 1.223E-08 | 6.926E-09 | 6.700E-09 | 2.000E-08 |
| 21066 | 1054 | E1054E | C | 3.2e-09 | 4.7e-09 | 8.1e-09 | 5.333E-09 | 2.511E-09 | 3.200E-09 | 8.100E-09 |
| 21067 | 1054 | E1054Q | C | 5.6e-09 | 4.1e-09 | 8.6e-09 | 6.100E-09 | 2.291E-09 | 4.100E-09 | 8.600E-09 |
| 21068 | 1054 | E1054G | C | 6.1e-09 | 7.1e-09 | 1.7e-08 | 1.007E-08 | 6.025E-09 | 6.100E-09 | 1.700E-08 |
| 21069 | 1054 | E1054H | C | 3.7e-09 | 9.9e-09 | 8.6e-09 | 7.400E-09 | 3.270E-09 | 3.700E-09 | 9.900E-09 |
| 21070 | 1054 | E1054I | C | 7.5e-09 | 7e-09 | 8.3e-09 | 7.600E-09 | 6.557E-10 | 7.000E-09 | 8.300E-09 |
| 21071 | 1054 | E1054L | C | 8.5e-09 | 4.4e-09 | 5.9e-09 | 6.267E-09 | 2.074E-09 | 4.400E-09 | 8.500E-09 |
| 21072 | 1054 | E1054K | C | 1.1e-08 | 1.1e-08 | 1.2e-08 | 1.133E-08 | 5.774E-10 | 1.100E-08 | 1.200E-08 |
| 21073 | 1054 | E1054M | C | 1.7e-09 | 1.1e-08 | 3e-09 | 5.233E-09 | 5.036E-09 | 1.700E-09 | 1.100E-08 |
| 21074 | 1054 | E1054F | C | 2.6e-09 | 7.6e-09 | 5.9e-09 | 5.367E-09 | 2.542E-09 | 2.600E-09 | 7.600E-09 |
| 21075 | 1054 | E1054P | C | 2e-09 | 6e-09 | 4.2e-09 | 4.067E-09 | 2.003E-09 | 2.000E-09 | 6.000E-09 |
| 21076 | 1054 | E1054S | C | 2.2e-09 | 3.8e-09 | 2e-08 | 8.667E-09 | 9.848E-09 | 2.200E-09 | 2.000E-08 |
| 21077 | 1054 | E1054T | C | 1.6e-08 | 7.5e-09 | 6.6e-09 | 1.003E-08 | 5.187E-09 | 6.600E-09 | 1.600E-08 |
| 21078 | 1054 | E1054W | C | 9.2e-09 | 8.2e-09 | 8e-09 | 8.467E-09 | 6.429E-10 | 8.000E-09 | 9.200E-09 |
| 21079 | 1054 | E1054Y | C | 1.8e-08 | 1.2e-09 | 6.3e-09 | 8.500E-09 | 8.613E-09 | 1.200E-09 | 1.800E-08 |

| | | | | | | | | | | |
|---|---|---|---|---|---|---|---|---|---|---|
| 21080 | 1054 | E1054V | C | 1.8e-08 | 1.5e-08 | 1.2e-08 | 1.500E-08 | 3.000E-09 | 1.200E-08 | 1.800E-08 |
| 21081 | 1055 | P1055A | C | 6.9e-09 | 9.4e-09 | 7.6e-09 | 7.967E-09 | 1.290E-09 | 6.900E-09 | 9.400E-09 |
| 21082 | 1055 | P1055R | C | 3.4e-09 | 6.7e-09 | 8.8e-09 | 6.300E-09 | 2.722E-09 | 3.400E-09 | 8.800E-09 |
| 21083 | 1055 | P1055N | C | 1.2e-08 | 3.6e-09 | 3.5e-09 | 6.367E-09 | 4.879E-09 | 3.500E-09 | 1.200E-08 |
| 21084 | 1055 | P1055D | C | 4.4e-09 | 7.3e-09 | 6.8e-09 | 6.167E-09 | 1.550E-09 | 4.400E-09 | 7.300E-09 |
| 21085 | 1055 | P1055C | C | 1.4e-08 | 8.9e-09 | 1.1e-08 | 1.130E-08 | 2.563E-09 | 8.900E-09 | 1.400E-08 |
| 21086 | 1055 | P1055E | C | 5.9e-09 | 1.8e-09 | 1e-08 | 5.900E-09 | 4.100E-09 | 1.800E-09 | 1.000E-08 |
| 21087 | 1055 | P1055Q | C | 4.3e-09 | 1.1e-08 | 1.7e-09 | 5.667E-09 | 4.798E-09 | 1.700E-09 | 1.100E-08 |
| 21088 | 1055 | P1055G | C | 8.3e-09 | 3.8e-09 | 6.2e-09 | 6.100E-09 | 2.252E-09 | 3.800E-09 | 8.300E-09 |
| 21089 | 1055 | P1055H | C | 1.1e-08 | 8e-09 | 5.7e-09 | 8.233E-09 | 2.658E-09 | 5.700E-09 | 1.100E-08 |
| 21090 | 1055 | P1055I | C | 7e-09 | 2.1e-08 | 6.2e-09 | 1.140E-08 | 8.323E-09 | 6.200E-09 | 2.100E-08 |
| 21091 | 1055 | P1055L | C | 3.7e-09 | 5.9e-09 | 2.9e-09 | 4.167E-09 | 1.553E-09 | 2.900E-09 | 5.900E-09 |
| 21092 | 1055 | P1055K | C | 6.7e-09 | 2.2e-09 | 1.8e-08 | 8.967E-09 | 8.140E-09 | 2.200E-09 | 1.800E-08 |
| 21093 | 1055 | P1055M | C | 4e-09 | 8.4e-09 | 8.6e-09 | 7.000E-09 | 2.600E-09 | 4.000E-09 | 8.600E-09 |
| 21094 | 1055 | P1055F | C | 7.7e-09 | 2.2e-08 | 1.6e-08 | 1.523E-08 | 7.181E-09 | 7.700E-09 | 2.200E-08 |
| 21095 | 1055 | P1055P | C | 7.8e-09 | 5.7e-09 | 1.3e-08 | 8.833E-09 | 3.758E-09 | 5.700E-09 | 1.300E-08 |
| 21096 | 1055 | P1055S | C | 1.3e-08 | 8.4e-09 | 4.1e-09 | 8.500E-09 | 4.451E-09 | 4.100E-09 | 1.300E-08 |
| 21097 | 1055 | P1055T | C | 2e-08 | 7.3e-09 | 8.2e-09 | 1.183E-08 | 7.087E-09 | 7.300E-09 | 2.000E-08 |
| 21098 | 1055 | P1055W | C | 4.5e-09 | 4.2e-09 | 4.8e-09 | 4.500E-09 | 3.000E-10 | 4.200E-09 | 4.800E-09 |
| 21099 | 1055 | P1055Y | C | 3.7e-09 | 6.8e-09 | 8.3e-09 | 6.267E-09 | 2.346E-09 | 3.700E-09 | 8.300E-09 |
| 21100 | 1055 | P1055V | C | 2.3e-09 | 9e-09 | 9.8e-09 | 7.033E-09 | 4.119E-09 | 2.300E-09 | 9.800E-09 |
| 21101 | 1056 | K1056A | C | 1.1e-08 | 3.6e-09 | 1.5e-08 | 9.867E-09 | 5.784E-09 | 3.600E-09 | 1.500E-08 |
| 21102 | 1056 | K1056R | C | 7.6e-09 | 5.5e-09 | 6.1e-09 | 6.400E-09 | 1.082E-09 | 5.500E-09 | 7.600E-09 |
| 21103 | 1056 | K1056N | C | 7.7e-09 | 6.1e-09 | 4.5e-09 | 6.100E-09 | 1.600E-09 | 4.500E-09 | 7.700E-09 |
| 21104 | 1056 | K1056D | C | 6.5e-09 | 1.1e-08 | 6.4e-09 | 7.967E-09 | 2.627E-09 | 6.400E-09 | 1.100E-08 |
| 21105 | 1056 | K1056C | C | 6e-09 | 1e-08 | 5.3e-09 | 7.100E-09 | 2.536E-09 | 5.300E-09 | 1.000E-08 |
| 21106 | 1056 | K1056E | C | 5.9e-09 | 2.6e-08 | 8.1e-09 | 1.333E-08 | 1.102E-08 | 5.900E-09 | 2.600E-08 |
| 21107 | 1056 | K1056Q | C | 8.9e-09 | 5.3e-09 | 3.8e-09 | 6.000E-09 | 2.621E-09 | 3.800E-09 | 8.900E-09 |
| 21108 | 1056 | K1056G | C | 6e-09 | 6.6e-09 | 1.4e-08 | 8.867E-09 | 4.456E-09 | 6.000E-09 | 1.400E-08 |
| 21109 | 1056 | K1056H | C | 5e-09 | 1.6e-08 | 2.1e-08 | 1.400E-08 | 8.185E-09 | 5.000E-09 | 2.100E-08 |
| 21110 | 1056 | K1056I | C | 4.3e-09 | 1.2e-08 | 6.8e-09 | 7.700E-09 | 3.928E-09 | 4.300E-09 | 1.200E-08 |
| 21111 | 1056 | K1056L | C | 3.6e-09 | 9.8e-09 | 2e-09 | 5.133E-09 | 4.120E-09 | 2.000E-09 | 9.800E-09 |
| 21112 | 1056 | K1056K | C | 5.3e-09 | 9e-09 | 7e-09 | 7.100E-09 | 1.852E-09 | 5.300E-09 | 9.000E-09 |
| 21113 | 1056 | K1056M | C | 6.6e-09 | 5.5e-09 | 8.9e-09 | 7.000E-09 | 1.735E-09 | 5.500E-09 | 8.900E-09 |
| 21114 | 1056 | K1056F | C | 1.6e-08 | 1.2e-08 | 4.8e-09 | 1.093E-08 | 5.676E-09 | 4.800E-09 | 1.600E-08 |
| 21115 | 1056 | K1056P | C | 7.8e-09 | 3.5e-09 | 3.6e-09 | 4.967E-09 | 2.454E-09 | 3.500E-09 | 7.800E-09 |
| 21116 | 1056 | K1056S | C | 9.5e-09 | 1.1e-08 | 1.6e-08 | 1.217E-08 | 3.403E-09 | 9.500E-09 | 1.600E-08 |
| 21117 | 1056 | K1056T | C | 6.5e-09 | 9.3e-09 | 5.3e-09 | 7.033E-09 | 2.053E-09 | 5.300E-09 | 9.300E-09 |
| 21118 | 1056 | K1056W | C | 6.2e-09 | 9.4e-09 | 8.6e-09 | 8.067E-09 | 1.665E-09 | 6.200E-09 | 9.400E-09 |
| 21119 | 1056 | K1056Y | C | 5.7e-09 | 6.2e-09 | 1e-08 | 7.300E-09 | 2.352E-09 | 5.700E-09 | 1.000E-08 |
| 21120 | 1056 | K1056V | C | 1e-08 | 1.5e-09 | 5.1e-09 | 5.533E-09 | 4.267E-09 | 1.500E-09 | 1.000E-08 |
| 21121 | 1057 | S1057A | C | 1.1e-09 | 1.2e-08 | 1.3e-08 | 8.700E-09 | 6.601E-09 | 1.100E-09 | 1.300E-08 |
| 21122 | 1057 | S1057R | C | 2.8e-08 | 4.8e-09 | 5.7e-09 | 1.283E-08 | 1.314E-08 | 4.800E-09 | 2.800E-08 |
| 21123 | 1057 | S1057N | C | 5.2e-09 | 3.2e-08 | 7.2e-09 | 1.480E-08 | 1.493E-08 | 5.200E-09 | 3.200E-08 |
| 21124 | 1057 | S1057D | C | 6.9e-09 | 1.6e-08 | 2.6e-09 | 8.500E-09 | 6.842E-09 | 2.600E-09 | 1.600E-08 |
| 21125 | 1057 | S1057C | C | 2.1e-08 | 3.5e-09 | 6.9e-09 | 1.047E-08 | 9.279E-09 | 3.500E-09 | 2.100E-08 |
| 21126 | 1057 | S1057E | C | 3.2e-09 | 3.9e-09 | 5.3e-09 | 4.133E-09 | 1.069E-09 | 3.200E-09 | 5.300E-09 |
| 21127 | 1057 | S1057Q | C | 6.6e-09 | 1.4e-09 | 5e-09 | 4.333E-09 | 2.663E-09 | 1.400E-09 | 6.600E-09 |
| 21128 | 1057 | S1057G | C | 5.7e-09 | 1.4e-08 | 7.1e-09 | 8.933E-09 | 4.443E-09 | 5.700E-09 | 1.400E-08 |
| 21129 | 1057 | S1057H | C | 7.6e-09 | 8.1e-09 | 7.2e-09 | 7.633E-09 | 4.509E-10 | 7.200E-09 | 8.100E-09 |
| 21130 | 1057 | S1057I | C | 2.8e-09 | 2.1e-08 | 3.3e-09 | 9.033E-09 | 1.037E-08 | 2.800E-09 | 2.100E-08 |
| 21131 | 1057 | S1057L | C | 9.6e-09 | 7e-09 | 4.3e-09 | 6.967E-09 | 2.650E-09 | 4.300E-09 | 9.600E-09 |
| 21132 | 1057 | S1057K | C | 6.6e-09 | 1.9e-08 | 3.4e-09 | 9.667E-09 | 8.240E-09 | 3.400E-09 | 1.900E-08 |
| 21133 | 1057 | S1057M | C | 2.9e-08 | 1.7e-08 | 5e-09 | 1.700E-08 | 1.200E-08 | 5.000E-09 | 2.900E-08 |
| 21134 | 1057 | S1057F | C | 6.1e-09 | 1.3e-08 | 4.4e-09 | 7.833E-09 | 4.554E-09 | 4.400E-09 | 1.300E-08 |
| 21135 | 1057 | S1057P | C | 3.4e-09 | 4.7e-09 | 3.1e-09 | 3.733E-09 | 8.505E-10 | 3.100E-09 | 4.700E-09 |
| 21136 | 1057 | S1057S | C | 8.6e-09 | 6.4e-09 | 9.5e-09 | 8.167E-09 | 1.595E-09 | 6.400E-09 | 9.500E-09 |
| 21137 | 1057 | S1057T | C | 1.1e-08 | 8.2e-09 | 9.5e-09 | 9.567E-09 | 1.401E-09 | 8.200E-09 | 1.100E-08 |
| 21138 | 1057 | S1057W | C | 6.1e-09 | 1.5e-08 | 1.4e-08 | 1.170E-08 | 4.875E-09 | 6.100E-09 | 1.500E-08 |
| 21139 | 1057 | S1057Y | C | 1e-08 | 3.5e-08 | 1.7e-08 | 2.067E-08 | 1.290E-08 | 1.000E-08 | 3.500E-08 |
| 21140 | 1057 | S1057V | C | 1.1e-08 | 5.8e-09 | 3.5e-09 | 6.767E-09 | 3.842E-09 | 3.500E-09 | 1.100E-08 |
| 21141 | 1058 | C1058A | C | 3.7e-09 | 4.4e-09 | 1.1e-08 | 6.367E-09 | 4.028E-09 | 3.700E-09 | 1.100E-08 |
| 21142 | 1058 | C1058R | C | 1.1e-08 | 1.3e-09 | 1.9e-08 | 1.043E-08 | 8.864E-09 | 1.300E-09 | 1.900E-08 |
| 21143 | 1058 | C1058N | C | 4.2e-09 | 2.9e-09 | 9.2e-09 | 5.433E-09 | 3.326E-09 | 2.900E-09 | 9.200E-09 |
| 21144 | 1058 | C1058D | C | 1.3e-08 | 1.3e-08 | 1.4e-09 | 9.133E-09 | 6.697E-09 | 1.400E-09 | 1.300E-08 |
| 21145 | 1058 | C1058C | C | 7.5e-09 | 4.7e-09 | 5.6e-09 | 5.933E-09 | 1.429E-09 | 4.700E-09 | 7.500E-09 |
| 21146 | 1058 | C1058E | C | 1.2e-08 | 4.3e-09 | 1.2e-08 | 9.433E-09 | 4.446E-09 | 4.300E-09 | 1.200E-08 |
| 21147 | 1058 | C1058Q | C | 3.2e-09 | 7e-09 | 1.1e-08 | 7.067E-09 | 3.900E-09 | 3.200E-09 | 1.100E-08 |
| 21148 | 1058 | C1058G | C | 5.2e-09 | 4.3e-09 | 7.3e-09 | 5.600E-09 | 1.539E-09 | 4.300E-09 | 7.300E-09 |
| 21149 | 1058 | C1058H | C | 1.5e-08 | 6.3e-09 | 2.5e-09 | 7.933E-09 | 6.408E-09 | 2.500E-09 | 1.500E-08 |
| 21150 | 1058 | C1058I | C | 1.4e-08 | 8.3e-09 | 1e-08 | 1.077E-08 | 2.926E-09 | 8.300E-09 | 1.400E-08 |
| 21151 | 1058 | C1058L | C | 3.6e-09 | 6.9e-09 | 3.5e-09 | 4.667E-09 | 1.935E-09 | 3.500E-09 | 6.900E-09 |
| 21152 | 1058 | C1058K | C | 6.6e-09 | 3.2e-09 | 6.1e-09 | 5.300E-09 | 1.836E-09 | 3.200E-09 | 6.600E-09 |
| 21153 | 1058 | C1058M | C | 2.1e-08 | 1.2e-08 | 1.4e-08 | 1.567E-08 | 4.726E-09 | 1.200E-08 | 2.100E-08 |
| 21154 | 1058 | C1058F | C | 3.9e-09 | 6e-09 | 6.7e-09 | 5.533E-09 | 1.457E-09 | 3.900E-09 | 6.700E-09 |
| 21155 | 1058 | C1058P | C | 1.2e-08 | 7.6e-09 | 6e-09 | 8.533E-09 | 3.107E-09 | 6.000E-09 | 1.200E-08 |
| 21156 | 1058 | C1058S | C | 1.7e-08 | 5.2e-09 | 3.8e-09 | 8.667E-09 | 7.251E-09 | 3.800E-09 | 1.700E-08 |
| 21157 | 1058 | C1058T | C | 1.3e-09 | 6.9e-09 | 5.2e-09 | 4.467E-09 | 2.871E-09 | 1.300E-09 | 6.900E-09 |
| 21158 | 1058 | C1058W | C | 6.5e-09 | 4.6e-09 | 8.5e-09 | 6.533E-09 | 1.950E-09 | 4.600E-09 | 8.500E-09 |
| 21159 | 1058 | C1058Y | C | 5.2e-09 | 6.8e-09 | 1.7e-08 | 9.667E-09 | 6.401E-09 | 5.200E-09 | 1.700E-08 |
| 21160 | 1058 | C1058V | C | 7.3e-09 | 5e-09 | 6.7e-09 | 6.333E-09 | 1.193E-09 | 5.000E-09 | 7.300E-09 |
| 21161 | 1059 | D1059A | D | 1.4e-08 | 6.9e-09 | 4.9e-09 | 8.600E-09 | 4.782E-09 | 4.900E-09 | 1.400E-08 |
| 21162 | 1059 | D1059R | D | 8.9e-09 | 3e-09 | 7e-09 | 6.300E-09 | 3.012E-09 | 3.000E-09 | 8.900E-09 |
| 21163 | 1059 | D1059N | D | 1.1e-08 | 1.3e-08 | 7.3e-09 | 1.043E-08 | 2.892E-09 | 7.300E-09 | 1.300E-08 |
| 21164 | 1059 | D1059D | D | 7.8e-09 | 5.2e-09 | 6.8e-09 | 6.600E-09 | 1.311E-09 | 5.200E-09 | 7.800E-09 |

| | | | | | | | | | | |
|---|---|---|---|---|---|---|---|---|---|---|
| 21165 | 1059 | D1059C | D | 7.7e-09 | 2.3e-09 | 4.1e-09 | 4.700E-09 | 2.750E-09 | 2.300E-09 | 7.700E-09 |
| 21166 | 1059 | D1059E | D | 4.7e-09 | 2.2e-09 | 4.1e-09 | 3.667E-09 | 1.305E-09 | 2.200E-09 | 4.700E-09 |
| 21167 | 1059 | D1059Q | D | 1.8e-08 | 1.6e-08 | 5.6e-09 | 1.320E-08 | 6.657E-09 | 5.600E-09 | 1.800E-08 |
| 21168 | 1059 | D1059G | D | 2.2e-09 | 1e-08 | 5.8e-09 | 6.000E-09 | 3.904E-09 | 2.200E-09 | 1.000E-08 |
| 21169 | 1059 | D1059H | D | 1.4e-08 | 5.2e-09 | 1.1e-08 | 1.007E-08 | 4.474E-09 | 5.200E-09 | 1.400E-08 |
| 21170 | 1059 | D1059I | D | 1.4e-08 | 8.6e-09 | 1.6e-08 | 1.287E-08 | 3.828E-09 | 8.600E-09 | 1.600E-08 |
| 21171 | 1059 | D1059L | D | 5.2e-09 | 4.4e-09 | 6.8e-09 | 5.467E-09 | 1.222E-09 | 4.400E-09 | 6.800E-09 |
| 21172 | 1059 | D1059K | D | 2.3e-09 | 9.3e-09 | 1e-08 | 7.200E-09 | 4.258E-09 | 2.300E-09 | 1.000E-08 |
| 21173 | 1059 | D1059M | D | 1.5e-09 | 2.4e-08 | 1.6e-08 | 1.383E-08 | 1.141E-08 | 1.500E-09 | 2.400E-08 |
| 21174 | 1059 | D1059F | D | 8.8e-09 | 1.4e-08 | 2.1e-08 | 1.460E-08 | 6.122E-09 | 8.800E-09 | 2.100E-08 |
| 21175 | 1059 | D1059P | D | 5.4e-09 | 8.4e-09 | 4.8e-09 | 6.200E-09 | 1.929E-09 | 4.800E-09 | 8.400E-09 |
| 21176 | 1059 | D1059S | D | 9.6e-09 | 4.7e-09 | 1e-08 | 8.100E-09 | 2.951E-09 | 4.700E-09 | 1.000E-08 |
| 21177 | 1059 | D1059T | D | 7.7e-09 | 5.6e-09 | 2.1e-08 | 1.143E-08 | 8.351E-09 | 5.600E-09 | 2.100E-08 |
| 21178 | 1059 | D1059W | D | 2.1e-08 | 6.7e-09 | 1.1e-08 | 1.290E-08 | 7.337E-09 | 6.700E-09 | 2.100E-08 |
| 21179 | 1059 | D1059Y | D | 1e-08 | 1.7e-08 | 1e-08 | 1.233E-08 | 4.041E-09 | 1.000E-08 | 1.700E-08 |
| 21180 | 1059 | D1059V | D | 2.3e-09 | 4e-09 | 8.9e-09 | 5.067E-09 | 3.427E-09 | 2.300E-09 | 8.900E-09 |
| 21181 | 1060 | I1060A | D | 8.9e-09 | 1.6e-08 | 4.2e-09 | 9.700E-09 | 5.941E-09 | 4.200E-09 | 1.600E-08 |
| 21182 | 1060 | I1060R | D | 1.2e-08 | 3.2e-09 | 6.7e-09 | 7.300E-09 | 4.431E-09 | 3.200E-09 | 1.200E-08 |
| 21183 | 1060 | I1060N | D | 2.3e-08 | 3.7e-09 | 6.3e-09 | 1.100E-08 | 1.047E-08 | 3.700E-09 | 2.300E-08 |
| 21184 | 1060 | I1060D | D | 7e-09 | 1.1e-08 | 7.3e-09 | 8.433E-09 | 2.228E-09 | 7.000E-09 | 1.100E-08 |
| 21185 | 1060 | I1060C | D | 7.6e-09 | 1.7e-09 | 8.6e-09 | 5.967E-09 | 3.729E-09 | 1.700E-09 | 8.600E-09 |
| 21186 | 1060 | I1060E | D | 1.2e-08 | 8.9e-09 | 8.5e-09 | 9.800E-09 | 1.916E-09 | 8.500E-09 | 1.200E-08 |
| 21187 | 1060 | I1060Q | D | 1.8e-09 | 9.9e-09 | 7e-09 | 6.233E-09 | 4.104E-09 | 1.800E-09 | 9.900E-09 |
| 21188 | 1060 | I1060G | D | 4.5e-09 | 2e-08 | 9.8e-09 | 1.143E-08 | 7.878E-09 | 4.500E-09 | 2.000E-08 |
| 21189 | 1060 | I1060H | D | 5.1e-09 | 7.6e-09 | 4.9e-09 | 5.867E-09 | 1.504E-09 | 4.900E-09 | 7.600E-09 |
| 21190 | 1060 | I1060I | D | 4.3e-09 | 2.5e-09 | 7.7e-09 | 4.833E-09 | 2.641E-09 | 2.500E-09 | 7.700E-09 |
| 21191 | 1060 | I1060L | D | 8.2e-09 | 8.8e-09 | 1.2e-08 | 9.667E-09 | 2.043E-09 | 8.200E-09 | 1.200E-08 |
| 21192 | 1060 | I1060K | D | 4.8e-09 | 5.2e-09 | 1.1e-08 | 7.000E-09 | 3.470E-09 | 4.800E-09 | 1.100E-08 |
| 21193 | 1060 | I1060M | D | 2.8e-09 | 4.8e-09 | 7.5e-09 | 5.033E-09 | 2.359E-09 | 2.800E-09 | 7.500E-09 |
| 21194 | 1060 | I1060F | D | 5.8e-09 | 4.7e-09 | 3.9e-09 | 4.800E-09 | 9.539E-10 | 3.900E-09 | 5.800E-09 |
| 21195 | 1060 | I1060P | D | 3.1e-08 | 3.2e-08 | 3.9e-09 | 2.230E-08 | 1.594E-08 | 3.900E-09 | 3.200E-08 |
| 21196 | 1060 | I1060S | D | 1.8e-08 | 2.3e-08 | 4.6e-09 | 1.520E-08 | 9.514E-09 | 4.600E-09 | 2.300E-08 |
| 21197 | 1060 | I1060T | D | 1.5e-08 | 9.8e-09 | 6.7e-09 | 1.050E-08 | 4.194E-09 | 6.700E-09 | 1.500E-08 |
| 21198 | 1060 | I1060W | D | 2.7e-09 | 5.8e-09 | 7.9e-09 | 5.467E-09 | 2.616E-09 | 2.700E-09 | 7.900E-09 |
| 21199 | 1060 | I1060Y | D | 7.4e-09 | 1.6e-08 | 1.2e-08 | 1.180E-08 | 4.303E-09 | 7.400E-09 | 1.600E-08 |
| 21200 | 1060 | I1060V | D | 2.1e-09 | 5.6e-09 | 4e-09 | 3.900E-09 | 1.752E-09 | 2.100E-09 | 5.600E-09 |
| 21201 | 1061 | Q1061A | D | 1.2e-08 | 4.8e-09 | 5e-09 | 7.267E-09 | 4.100E-09 | 4.800E-09 | 1.200E-08 |
| 21202 | 1061 | Q1061R | D | 1.3e-08 | 9.7e-09 | 1.3e-08 | 1.190E-08 | 1.905E-09 | 9.700E-09 | 1.300E-08 |
| 21203 | 1061 | Q1061N | D | 2e-09 | 7.4e-09 | 1.5e-08 | 8.133E-09 | 6.531E-09 | 2.000E-09 | 1.500E-08 |
| 21204 | 1061 | Q1061D | D | 1.4e-09 | 4e-09 | 4.1e-09 | 3.167E-09 | 1.531E-09 | 1.400E-09 | 4.100E-09 |
| 21205 | 1061 | Q1061C | D | 1e-08 | 2.3e-09 | 5.2e-09 | 5.833E-09 | 3.889E-09 | 2.300E-09 | 1.000E-08 |
| 21206 | 1061 | Q1061E | D | 6.4e-09 | 5.9e-09 | 8.8e-09 | 7.033E-09 | 1.550E-09 | 5.900E-09 | 8.800E-09 |
| 21207 | 1061 | Q1061Q | D | 9.5e-09 | 4.9e-09 | 4.2e-09 | 6.200E-09 | 2.879E-09 | 4.200E-09 | 9.500E-09 |
| 21208 | 1061 | Q1061G | D | 1.3e-08 | 6.9e-09 | 2e-08 | 1.330E-08 | 6.555E-09 | 6.900E-09 | 2.000E-08 |
| 21209 | 1061 | Q1061H | D | 8.1e-09 | 5.8e-09 | 1.1e-08 | 8.300E-09 | 2.606E-09 | 5.800E-09 | 1.100E-08 |
| 21210 | 1061 | Q1061I | D | 7.8e-09 | 5.7e-09 | 1.1e-08 | 8.167E-09 | 2.669E-09 | 5.700E-09 | 1.100E-08 |
| 21211 | 1061 | Q1061L | D | 3.5e-09 | 3.3e-09 | 2.9e-09 | 3.233E-09 | 3.055E-10 | 2.900E-09 | 3.500E-09 |
| 21212 | 1061 | Q1061K | D | 6.3e-09 | 7.2e-09 | 5.7e-09 | 6.400E-09 | 7.550E-10 | 5.700E-09 | 7.200E-09 |
| 21213 | 1061 | Q1061M | D | 4.7e-09 | 2.1e-09 | 3.4e-09 | 3.400E-09 | 1.300E-09 | 2.100E-09 | 4.700E-09 |
| 21214 | 1061 | Q1061F | D | 7.6e-09 | 1.4e-08 | 1.5e-08 | 1.220E-08 | 4.015E-09 | 7.600E-09 | 1.500E-08 |
| 21215 | 1061 | Q1061P | D | 4.1e-09 | 3.5e-09 | 9.8e-09 | 5.800E-09 | 3.477E-09 | 3.500E-09 | 9.800E-09 |
| 21216 | 1061 | Q1061S | D | 5.6e-09 | 1.9e-08 | 1.3e-08 | 1.253E-08 | 6.712E-09 | 5.600E-09 | 1.900E-08 |
| 21217 | 1061 | Q1061T | D | 1.6e-08 | 3.1e-09 | 8e-09 | 9.033E-09 | 6.512E-09 | 3.100E-09 | 1.600E-08 |
| 21218 | 1061 | Q1061W | D | 8.4e-09 | 4.5e-09 | 4.4e-09 | 5.767E-09 | 2.281E-09 | 4.400E-09 | 8.400E-09 |
| 21219 | 1061 | Q1061Y | D | 9.8e-09 | 3.5e-09 | 5.5e-09 | 6.267E-09 | 3.219E-09 | 3.500E-09 | 9.800E-09 |
| 21220 | 1061 | Q1061V | D | 2.3e-08 | 7.6e-09 | 1.1e-08 | 1.387E-08 | 8.090E-09 | 7.600E-09 | 2.300E-08 |
| 21221 | 1062 | M1062A | D | 6.3e-09 | 2.7e-09 | 1.1e-08 | 6.667E-09 | 4.162E-09 | 2.700E-09 | 1.100E-08 |
| 21222 | 1062 | M1062R | D | 7.2e-09 | 6.6e-09 | 2.2e-09 | 5.333E-09 | 2.730E-09 | 2.200E-09 | 7.200E-09 |
| 21223 | 1062 | M1062N | D | 1.9e-08 | 1.2e-08 | 5e-09 | 1.200E-08 | 7.000E-09 | 5.000E-09 | 1.900E-08 |
| 21224 | 1062 | M1062D | D | 7.4e-09 | 5.2e-09 | 9e-09 | 7.200E-09 | 1.908E-09 | 5.200E-09 | 9.000E-09 |
| 21225 | 1062 | M1062C | D | 3.5e-09 | 6.1e-09 | 2.1e-09 | 3.900E-09 | 2.030E-09 | 2.100E-09 | 6.100E-09 |
| 21226 | 1062 | M1062E | D | 4.9e-09 | 2.9e-09 | 6.5e-09 | 4.767E-09 | 1.804E-09 | 2.900E-09 | 6.500E-09 |
| 21227 | 1062 | M1062Q | D | 3.3e-09 | 8.9e-09 | 1.6e-08 | 9.400E-09 | 6.365E-09 | 3.300E-09 | 1.600E-08 |
| 21228 | 1062 | M1062G | D | 4e-09 | 7.3e-09 | 6e-09 | 5.767E-09 | 1.662E-09 | 4.000E-09 | 7.300E-09 |
| 21229 | 1062 | M1062H | D | 1e-08 | 5.8e-09 | 3.5e-09 | 6.433E-09 | 3.296E-09 | 3.500E-09 | 1.000E-08 |
| 21230 | 1062 | M1062I | D | 4.8e-09 | 8.6e-09 | 6.5e-09 | 6.633E-09 | 1.904E-09 | 4.800E-09 | 8.600E-09 |
| 21231 | 1062 | M1062L | D | 5.3e-09 | 3.8e-09 | 9.4e-09 | 6.167E-09 | 2.899E-09 | 3.800E-09 | 9.400E-09 |
| 21232 | 1062 | M1062K | D | 7.4e-09 | 9.4e-09 | 3e-08 | 1.560E-08 | 1.251E-08 | 7.400E-09 | 3.000E-08 |
| 21233 | 1062 | M1062M | D | 5.9e-09 | 1.4e-08 | 1.8e-08 | 1.263E-08 | 6.165E-09 | 5.900E-09 | 1.800E-08 |
| 21234 | 1062 | M1062F | D | 3.4e-09 | 2.2e-08 | 3e-09 | 9.467E-09 | 1.086E-08 | 3.000E-09 | 2.200E-08 |
| 21235 | 1062 | M1062P | D | 3.7e-09 | 7.9e-09 | 7.4e-09 | 6.333E-09 | 2.294E-09 | 3.700E-09 | 7.900E-09 |
| 21236 | 1062 | M1062S | D | 6.2e-09 | 2.6e-09 | 4.1e-09 | 4.300E-09 | 1.808E-09 | 2.600E-09 | 6.200E-09 |
| 21237 | 1062 | M1062T | D | 8.7e-09 | 4.8e-09 | 6.5e-09 | 6.667E-09 | 1.955E-09 | 4.800E-09 | 8.700E-09 |
| 21238 | 1062 | M1062W | D | 3e-09 | 5.4e-09 | 5.6e-09 | 4.667E-09 | 1.447E-09 | 3.000E-09 | 5.600E-09 |
| 21239 | 1062 | M1062Y | D | 1.3e-08 | 1.4e-08 | 5.5e-09 | 1.083E-08 | 4.646E-09 | 5.500E-09 | 1.400E-08 |
| 21240 | 1062 | M1062V | D | 4.5e-09 | 1.1e-08 | 1.9e-08 | 1.150E-08 | 7.263E-09 | 4.500E-09 | 1.900E-08 |
| 21241 | 1063 | T1063A | D | 1.2e-08 | 2.3e-09 | 8.3e-09 | 7.533E-09 | 4.895E-09 | 2.300E-09 | 1.200E-08 |
| 21242 | 1063 | T1063R | D | 4.3e-09 | 1.6e-08 | 3.2e-09 | 7.833E-09 | 7.094E-09 | 3.200E-09 | 1.600E-08 |
| 21243 | 1063 | T1063N | D | 1.3e-08 | 3.6e-09 | 1.2e-08 | 9.533E-09 | 5.163E-09 | 3.600E-09 | 1.300E-08 |
| 21244 | 1063 | T1063D | D | 9.2e-09 | 8.4e-09 | 1e-08 | 9.200E-09 | 8.000E-10 | 8.400E-09 | 1.000E-08 |
| 21245 | 1063 | T1063C | D | 6.4e-09 | 2.2e-09 | 7.6e-09 | 5.400E-09 | 2.835E-09 | 2.200E-09 | 7.600E-09 |
| 21246 | 1063 | T1063E | D | 4.8e-09 | 9.5e-09 | 4.6e-09 | 6.300E-09 | 2.773E-09 | 4.600E-09 | 9.500E-09 |
| 21247 | 1063 | T1063Q | D | 4.4e-09 | 9.8e-09 | 8.9e-09 | 7.700E-09 | 2.893E-09 | 4.400E-09 | 9.800E-09 |
| 21248 | 1063 | T1063G | D | 2.1e-09 | 6.9e-09 | 6.6e-09 | 5.200E-09 | 2.689E-09 | 2.100E-09 | 6.900E-09 |
| 21249 | 1063 | T1063H | D | 1.9e-08 | 1.4e-08 | 7.3e-09 | 1.343E-08 | 5.871E-09 | 7.300E-09 | 1.900E-08 |

| | | | | | | | | | | |
|---|---|---|---|---|---|---|---|---|---|---|
| 21250 | 1063 | T1063I | D | 2.6e-08 | 5.4e-09 | 2.1e-08 | 1.747E-08 | 1.074E-08 | 5.400E-09 | 2.600E-08 |
| 21251 | 1063 | T1063L | D | 2.7e-09 | 1.4e-08 | 1.8e-08 | 1.157E-08 | 7.935E-09 | 2.700E-09 | 1.800E-08 |
| 21252 | 1063 | T1063K | D | 3.6e-09 | 3.3e-09 | 8.4e-09 | 5.100E-09 | 2.862E-09 | 3.300E-09 | 8.400E-09 |
| 21253 | 1063 | T1063M | D | 7.1e-09 | 6.2e-09 | 1.4e-08 | 9.100E-09 | 4.267E-09 | 6.200E-09 | 1.400E-08 |
| 21254 | 1063 | T1063F | D | 3.2e-09 | 1.2e-08 | 9.3e-09 | 8.167E-09 | 4.508E-09 | 3.200E-09 | 1.200E-08 |
| 21255 | 1063 | T1063P | D | 2.5e-09 | 2e-08 | 3.9e-09 | 8.800E-09 | 9.725E-09 | 2.500E-09 | 2.000E-08 |
| 21256 | 1063 | T1063S | D | 6.6e-09 | 1.8e-09 | 1.5e-08 | 7.800E-09 | 6.681E-09 | 1.800E-09 | 1.500E-08 |
| 21257 | 1063 | T1063T | D | 6.1e-09 | 6e-09 | 1.5e-08 | 9.033E-09 | 5.168E-09 | 6.000E-09 | 1.500E-08 |
| 21258 | 1063 | T1063W | D | 3.2e-09 | 6.1e-09 | 6.9e-09 | 5.400E-09 | 1.947E-09 | 3.200E-09 | 6.900E-09 |
| 21259 | 1063 | T1063Y | D | 8.1e-09 | 1.2e-08 | 3.8e-09 | 7.967E-09 | 4.102E-09 | 3.800E-09 | 1.200E-08 |
| 21260 | 1063 | T1063V | D | 3.9e-09 | 2.2e-09 | 8.3e-09 | 4.800E-09 | 3.148E-09 | 2.200E-09 | 8.300E-09 |
| 21261 | 1064 | Q1064A | D | 2e-08 | 1.2e-09 | 5.3e-09 | 8.833E-09 | 9.886E-09 | 1.200E-09 | 2.000E-08 |
| 21262 | 1064 | Q1064R | D | 6.9e-09 | 1.8e-08 | 3e-09 | 9.300E-09 | 7.783E-09 | 3.000E-09 | 1.800E-08 |
| 21263 | 1064 | Q1064N | D | 1e-08 | 2.7e-09 | 5.6e-09 | 6.100E-09 | 3.676E-09 | 2.700E-09 | 1.000E-08 |
| 21264 | 1064 | Q1064D | D | 7.2e-09 | 1.6e-08 | 9.1e-09 | 1.077E-08 | 4.631E-09 | 7.200E-09 | 1.600E-08 |
| 21265 | 1064 | Q1064C | D | 3.5e-09 | 8.4e-09 | 1.3e-08 | 8.300E-09 | 4.751E-09 | 3.500E-09 | 1.300E-08 |
| 21266 | 1064 | Q1064E | D | 2.7e-09 | 1.2e-08 | 8.2e-09 | 7.633E-09 | 4.676E-09 | 2.700E-09 | 1.200E-08 |
| 21267 | 1064 | Q1064Q | D | 3.7e-09 | 7.1e-09 | 4.3e-09 | 5.033E-09 | 1.815E-09 | 3.700E-09 | 7.100E-09 |
| 21268 | 1064 | Q1064G | D | 3.2e-09 | 7.8e-09 | 5.4e-09 | 5.467E-09 | 2.301E-09 | 3.200E-09 | 7.800E-09 |
| 21269 | 1064 | Q1064H | D | 5.5e-09 | 2e-09 | 2e-08 | 9.167E-09 | 9.544E-09 | 2.000E-09 | 2.000E-08 |
| 21270 | 1064 | Q1064I | D | 4.8e-09 | 7.1e-09 | 1.3e-08 | 8.300E-09 | 4.230E-09 | 4.800E-09 | 1.300E-08 |
| 21271 | 1064 | Q1064L | D | 2.9e-09 | 1.3e-08 | 4.3e-09 | 6.733E-09 | 5.472E-09 | 2.900E-09 | 1.300E-08 |
| 21272 | 1064 | Q1064K | D | 8.5e-09 | 4.1e-09 | 6.6e-09 | 6.400E-09 | 2.207E-09 | 4.100E-09 | 8.500E-09 |
| 21273 | 1064 | Q1064M | D | 3.2e-09 | 8.7e-09 | 5.3e-09 | 5.733E-09 | 2.775E-09 | 3.200E-09 | 8.700E-09 |
| 21274 | 1064 | Q1064F | D | 5.9e-09 | 1.4e-08 | 4.2e-09 | 8.033E-09 | 5.237E-09 | 4.200E-09 | 1.400E-08 |
| 21275 | 1064 | Q1064P | D | 1.2e-08 | 3.3e-09 | 5.1e-09 | 6.800E-09 | 4.592E-09 | 3.300E-09 | 1.200E-08 |
| 21276 | 1064 | Q1064S | D | 6e-09 | 9.2e-09 | 2.5e-09 | 5.900E-09 | 3.351E-09 | 2.500E-09 | 9.200E-09 |
| 21277 | 1064 | Q1064T | D | 1e-08 | 5.1e-09 | 6.2e-09 | 7.100E-09 | 2.571E-09 | 5.100E-09 | 1.000E-08 |
| 21278 | 1064 | Q1064W | D | 1.9e-08 | 5.8e-08 | 1.1e-08 | 2.933E-08 | 2.515E-08 | 1.100E-08 | 5.800E-08 |
| 21279 | 1064 | Q1064Y | D | 1.2e-08 | 5.2e-09 | 1.5e-08 | 1.073E-08 | 5.021E-09 | 5.200E-09 | 1.500E-08 |
| 21280 | 1064 | Q1064V | D | 4.2e-09 | 7.3e-09 | 1.4e-08 | 8.500E-09 | 5.009E-09 | 4.200E-09 | 1.400E-08 |
| 21281 | 1065 | S1065A | D | 3.7e-09 | 1.2e-08 | 9.3e-09 | 8.333E-09 | 4.234E-09 | 3.700E-09 | 1.200E-08 |
| 21282 | 1065 | S1065R | D | 5.6e-09 | 2.9e-09 | 1.6e-09 | 3.367E-09 | 2.040E-09 | 1.600E-09 | 5.600E-09 |
| 21283 | 1065 | S1065N | D | 3.1e-09 | 9.8e-09 | 1.6e-08 | 9.633E-09 | 6.452E-09 | 3.100E-09 | 1.600E-08 |
| 21284 | 1065 | S1065D | D | 6e-09 | 8.7e-09 | 4.6e-09 | 6.433E-09 | 2.084E-09 | 4.600E-09 | 8.700E-09 |
| 21285 | 1065 | S1065C | D | 6.8e-09 | 1.9e-08 | 1.3e-08 | 1.293E-08 | 6.100E-09 | 6.800E-09 | 1.900E-08 |
| 21286 | 1065 | S1065E | D | 4.8e-09 | 2.1e-09 | 1.1e-08 | 5.967E-09 | 4.563E-09 | 2.100E-09 | 1.100E-08 |
| 21287 | 1065 | S1065Q | D | 1.1e-08 | 2.9e-09 | 6.3e-09 | 6.733E-09 | 4.067E-09 | 2.900E-09 | 1.100E-08 |
| 21288 | 1065 | S1065G | D | 4.4e-09 | 2e-09 | 1.1e-08 | 5.800E-09 | 4.660E-09 | 2.000E-09 | 1.100E-08 |
| 21289 | 1065 | S1065H | D | 2.3e-09 | 4.3e-09 | 2.7e-08 | 1.120E-08 | 1.372E-08 | 2.300E-09 | 2.700E-08 |
| 21290 | 1065 | S1065I | D | 5.6e-09 | 1.1e-08 | 2.9e-08 | 1.520E-08 | 1.225E-08 | 5.600E-09 | 2.900E-08 |
| 21291 | 1065 | S1065L | D | 1.8e-08 | 8.9e-09 | 4.8e-09 | 1.057E-08 | 6.756E-09 | 4.800E-09 | 1.800E-08 |
| 21292 | 1065 | S1065K | D | 5.1e-09 | 1.2e-08 | 6.7e-09 | 7.933E-09 | 3.612E-09 | 5.100E-09 | 1.200E-08 |
| 21293 | 1065 | S1065M | D | 6.8e-09 | 4.4e-09 | 5.6e-09 | 5.600E-09 | 1.200E-09 | 4.400E-09 | 6.800E-09 |
| 21294 | 1065 | S1065F | D | 7e-09 | 7.5e-09 | 1.8e-08 | 1.083E-08 | 6.212E-09 | 7.000E-09 | 1.800E-08 |
| 21295 | 1065 | S1065P | D | 2.2e-09 | 4.4e-09 | 3.1e-09 | 3.233E-09 | 1.106E-09 | 2.200E-09 | 4.400E-09 |
| 21296 | 1065 | S1065S | D | 6.1e-09 | 2.8e-09 | 8.1e-09 | 5.667E-09 | 2.676E-09 | 2.800E-09 | 8.100E-09 |
| 21297 | 1065 | S1065T | D | 4.1e-09 | 1.1e-08 | 3.5e-09 | 6.200E-09 | 4.168E-09 | 3.500E-09 | 1.100E-08 |
| 21298 | 1065 | S1065W | D | 7.3e-09 | 6.3e-09 | 1e-08 | 7.867E-09 | 1.914E-09 | 6.300E-09 | 1.000E-08 |
| 21299 | 1065 | S1065Y | D | 6.3e-09 | 1e-08 | 5.1e-09 | 7.133E-09 | 2.554E-09 | 5.100E-09 | 1.000E-08 |
| 21300 | 1065 | S1065V | D | 1.3e-08 | 2.3e-08 | 1.2e-08 | 1.600E-08 | 6.083E-09 | 1.200E-08 | 2.300E-08 |
| 21301 | 1066 | P1066A | D | 3.7e-09 | 6.6e-09 | 7.3e-09 | 5.867E-09 | 1.909E-09 | 3.700E-09 | 7.300E-09 |
| 21302 | 1066 | P1066R | D | 1.4e-08 | 8.9e-09 | 7.5e-09 | 1.013E-08 | 3.421E-09 | 7.500E-09 | 1.400E-08 |
| 21303 | 1066 | P1066N | D | 1.4e-08 | 1.1e-08 | 2.8e-08 | 1.767E-08 | 9.074E-09 | 1.100E-08 | 2.800E-08 |
| 21304 | 1066 | P1066D | D | 6.6e-09 | 3e-09 | 2e-09 | 3.867E-09 | 2.419E-09 | 2.000E-09 | 6.600E-09 |
| 21305 | 1066 | P1066C | D | 2e-09 | 1.5e-08 | 5.9e-09 | 7.633E-09 | 6.671E-09 | 2.000E-09 | 1.500E-08 |
| 21306 | 1066 | P1066E | D | 8.3e-09 | 5.2e-09 | 8.2e-09 | 7.233E-09 | 1.762E-09 | 5.200E-09 | 8.300E-09 |
| 21307 | 1066 | P1066Q | D | 2.7e-09 | 3.3e-08 | 1.1e-08 | 1.557E-08 | 1.566E-08 | 2.700E-09 | 3.300E-08 |
| 21308 | 1066 | P1066G | D | 9.6e-09 | 6.3e-09 | 6.8e-09 | 7.567E-09 | 1.779E-09 | 6.300E-09 | 9.600E-09 |
| 21309 | 1066 | P1066H | D | 3.9e-09 | 1.2e-08 | 1.7e-08 | 1.097E-08 | 6.611E-09 | 3.900E-09 | 1.700E-08 |
| 21310 | 1066 | P1066I | D | 9.7e-09 | 8.7e-09 | 8.4e-09 | 8.933E-09 | 6.807E-10 | 8.400E-09 | 9.700E-09 |
| 21311 | 1066 | P1066L | D | 2.8e-09 | 1.5e-08 | 1.6e-08 | 1.127E-08 | 7.349E-09 | 2.800E-09 | 1.600E-08 |
| 21312 | 1066 | P1066K | D | 3.5e-09 | 4.5e-09 | 3.5e-09 | 3.833E-09 | 5.774E-10 | 3.500E-09 | 4.500E-09 |
| 21313 | 1066 | P1066M | D | 2.8e-09 | 1.8e-08 | 4.4e-09 | 8.400E-09 | 8.352E-09 | 2.800E-09 | 1.800E-08 |
| 21314 | 1066 | P1066F | D | 2.2e-08 | 9.3e-09 | 2.8e-09 | 1.137E-08 | 9.765E-09 | 2.800E-09 | 2.200E-08 |
| 21315 | 1066 | P1066P | D | 5e-09 | 1.7e-08 | 4.7e-09 | 8.900E-09 | 7.016E-09 | 4.700E-09 | 1.700E-08 |
| 21316 | 1066 | P1066S | D | 2.1e-09 | 1e-08 | 1.2e-08 | 8.033E-09 | 5.235E-09 | 2.100E-09 | 1.200E-08 |
| 21317 | 1066 | P1066T | D | 8.4e-09 | 1.2e-08 | 5.8e-09 | 8.733E-09 | 3.113E-09 | 5.800E-09 | 1.200E-08 |
| 21318 | 1066 | P1066W | D | 7.3e-09 | 2.2e-09 | 2e-09 | 3.833E-09 | 3.004E-09 | 2.000E-09 | 7.300E-09 |
| 21319 | 1066 | P1066Y | D | 1.5e-08 | 6.1e-09 | 4.2e-09 | 8.433E-09 | 5.766E-09 | 4.200E-09 | 1.500E-08 |
| 21320 | 1066 | P1066V | D | 9.6e-09 | 1.6e-08 | 7.2e-09 | 1.093E-08 | 4.549E-09 | 7.200E-09 | 1.600E-08 |
| 21321 | 1067 | S1067A | D | 4.3e-09 | 1.3e-08 | 3.4e-09 | 6.900E-09 | 5.302E-09 | 3.400E-09 | 1.300E-08 |
| 21322 | 1067 | S1067R | D | 9.8e-09 | 1.2e-08 | 3.1e-09 | 8.300E-09 | 4.636E-09 | 3.100E-09 | 1.200E-08 |
| 21323 | 1067 | S1067N | D | 8.3e-09 | 5.6e-09 | 8e-09 | 7.300E-09 | 1.480E-09 | 5.600E-09 | 8.300E-09 |
| 21324 | 1067 | S1067D | D | 4.9e-09 | 6.8e-09 | 3.7e-08 | 1.623E-08 | 1.801E-08 | 4.900E-09 | 3.700E-08 |
| 21325 | 1067 | S1067C | D | 6.2e-09 | 9.8e-09 | 1.2e-08 | 9.333E-09 | 2.928E-09 | 6.200E-09 | 1.200E-08 |
| 21326 | 1067 | S1067E | D | 1.2e-08 | 4.9e-09 | 5.3e-09 | 7.400E-09 | 3.989E-09 | 4.900E-09 | 1.200E-08 |
| 21327 | 1067 | S1067Q | D | 4.7e-09 | 4.7e-09 | 9.5e-09 | 6.300E-09 | 2.771E-09 | 4.700E-09 | 9.500E-09 |
| 21328 | 1067 | S1067G | D | 8.9e-09 | 1.1e-08 | 1.1e-08 | 1.030E-08 | 1.212E-09 | 8.900E-09 | 1.100E-08 |
| 21329 | 1067 | S1067H | D | 9.3e-09 | 3.3e-09 | 2.2e-09 | 4.933E-09 | 3.821E-09 | 2.200E-09 | 9.300E-09 |
| 21330 | 1067 | S1067I | D | 2.1e-08 | 2.2e-09 | 6.1e-09 | 9.767E-09 | 9.922E-09 | 2.200E-09 | 2.100E-08 |
| 21331 | 1067 | S1067L | D | 4.6e-09 | 8.3e-09 | 8.6e-09 | 7.167E-09 | 2.228E-09 | 4.600E-09 | 8.600E-09 |
| 21332 | 1067 | S1067K | D | 1.3e-08 | 1.1e-08 | 4.2e-09 | 9.400E-09 | 4.613E-09 | 4.200E-09 | 1.300E-08 |
| 21333 | 1067 | S1067M | D | 7e-09 | 1e-08 | 1e-08 | 9.000E-09 | 1.732E-09 | 7.000E-09 | 1.000E-08 |
| 21334 | 1067 | S1067F | D | 6.3e-09 | 1e-08 | 1.8e-08 | 1.143E-08 | 5.980E-09 | 6.300E-09 | 1.800E-08 |

| | | | | | | | | | | |
|---|---|---|---|---|---|---|---|---|---|---|
| 21335 | 1067 | S1067P | D | 1.1e-08 | 1.1e-08 | 1.6e-08 | 1.267E-08 | 2.887E-09 | 1.100E-08 | 1.600E-08 |
| 21336 | 1067 | S1067S | D | 9.1e-09 | 4e-09 | 7.8e-09 | 6.967E-09 | 2.650E-09 | 4.000E-09 | 9.100E-09 |
| 21337 | 1067 | S1067T | D | 3.6e-09 | 1.5e-08 | 1.8e-08 | 1.220E-08 | 7.597E-09 | 3.600E-09 | 1.800E-08 |
| 21338 | 1067 | S1067W | D | 2.5e-09 | 9e-09 | 1.5e-08 | 8.833E-09 | 6.252E-09 | 2.500E-09 | 1.500E-08 |
| 21339 | 1067 | S1067Y | D | 5.6e-09 | 1.4e-08 | 2.3e-09 | 7.300E-09 | 6.032E-09 | 2.300E-09 | 1.400E-08 |
| 21340 | 1067 | S1067V | D | 8.3e-09 | 8.5e-09 | 1.2e-09 | 6.000E-09 | 4.158E-09 | 1.200E-09 | 8.500E-09 |
| 21341 | 1068 | S1068A | D | 1.4e-08 | 9.5e-09 | 6.7e-09 | 1.007E-08 | 3.683E-09 | 6.700E-09 | 1.400E-08 |
| 21342 | 1068 | S1068R | D | 1.1e-08 | 8.9e-09 | 3.8e-09 | 7.900E-09 | 3.703E-09 | 3.800E-09 | 1.100E-08 |
| 21343 | 1068 | S1068N | D | 3e-08 | 6.6e-09 | 1.7e-08 | 1.787E-08 | 1.172E-08 | 6.600E-09 | 3.000E-08 |
| 21344 | 1068 | S1068D | D | 5.7e-09 | 7.3e-09 | 2.7e-09 | 5.233E-09 | 2.335E-09 | 2.700E-09 | 7.300E-09 |
| 21345 | 1068 | S1068C | D | 4.2e-09 | 1.1e-08 | 2e-08 | 1.173E-08 | 7.925E-09 | 4.200E-09 | 2.000E-08 |
| 21346 | 1068 | S1068E | D | 8.6e-09 | 3.7e-09 | 6.7e-09 | 6.333E-09 | 2.470E-09 | 3.700E-09 | 8.600E-09 |
| 21347 | 1068 | S1068Q | D | 5.5e-09 | 6e-09 | 1e-08 | 7.167E-09 | 2.466E-09 | 5.500E-09 | 1.000E-08 |
| 21348 | 1068 | S1068G | D | 1.4e-08 | 1.3e-08 | 1.1e-08 | 1.267E-08 | 1.528E-09 | 1.100E-08 | 1.400E-08 |
| 21349 | 1068 | S1068H | D | 2.4e-08 | 5.3e-09 | 1.3e-08 | 1.410E-08 | 9.398E-09 | 5.300E-09 | 2.400E-08 |
| 21350 | 1068 | S1068I | D | 1.1e-08 | 1.1e-08 | 3.3e-08 | 1.833E-08 | 1.270E-08 | 1.100E-08 | 3.300E-08 |
| 21351 | 1068 | S1068L | D | 6.5e-09 | 7.4e-09 | 1.3e-08 | 8.967E-09 | 3.522E-09 | 6.500E-09 | 1.300E-08 |
| 21352 | 1068 | S1068K | D | 1e-08 | 5.5e-09 | 8.2e-09 | 7.900E-09 | 2.265E-09 | 5.500E-09 | 1.000E-08 |
| 21353 | 1068 | S1068M | D | 5.3e-09 | 3.8e-09 | 9.9e-09 | 6.333E-09 | 3.179E-09 | 3.800E-09 | 9.900E-09 |
| 21354 | 1068 | S1068F | D | 2.1e-08 | 7.2e-09 | 1.8e-09 | 1.000E-08 | 9.902E-09 | 1.800E-09 | 2.100E-08 |
| 21355 | 1068 | S1068P | D | 1.1e-08 | 3.7e-09 | 7.4e-09 | 7.367E-09 | 3.650E-09 | 3.700E-09 | 1.100E-08 |
| 21356 | 1068 | S1068S | D | 1e-08 | 1.3e-08 | 4.6e-09 | 9.200E-09 | 4.257E-09 | 4.600E-09 | 1.300E-08 |
| 21357 | 1068 | S1068T | D | 1.2e-08 | 1.3e-08 | 1.3e-08 | 1.267E-08 | 5.774E-10 | 1.200E-08 | 1.300E-08 |
| 21358 | 1068 | S1068W | D | 2.6e-08 | 8.8e-09 | 2.6e-08 | 2.027E-08 | 9.930E-09 | 8.800E-09 | 2.600E-08 |
| 21359 | 1068 | S1068Y | D | 2.7e-09 | 4.8e-09 | 6.3e-09 | 4.600E-09 | 1.808E-09 | 2.700E-09 | 6.300E-09 |
| 21360 | 1068 | S1068V | D | 1.3e-08 | 1.2e-08 | 4.7e-09 | 9.900E-09 | 4.531E-09 | 4.700E-09 | 1.300E-08 |
| 21361 | 1069 | L1069A | D | 7.3e-09 | 1.3e-08 | 1.1e-08 | 1.043E-08 | 2.892E-09 | 7.300E-09 | 1.300E-08 |
| 21362 | 1069 | L1069R | D | 1.6e-09 | 7.5e-09 | 1.7e-08 | 8.700E-09 | 7.770E-09 | 1.600E-09 | 1.700E-08 |
| 21363 | 1069 | L1069N | D | 8.6e-09 | 1.2e-08 | 3.1e-09 | 7.900E-09 | 4.491E-09 | 3.100E-09 | 1.200E-08 |
| 21364 | 1069 | L1069D | D | 7.6e-09 | 7.4e-09 | 5.5e-09 | 6.833E-09 | 1.159E-09 | 5.500E-09 | 7.600E-09 |
| 21365 | 1069 | L1069C | D | 4.3e-09 | 5.7e-09 | 3.4e-09 | 4.467E-09 | 1.159E-09 | 3.400E-09 | 5.700E-09 |
| 21366 | 1069 | L1069E | D | 6.4e-09 | 1.6e-09 | 3.1e-09 | 3.700E-09 | 2.456E-09 | 1.600E-09 | 6.400E-09 |
| 21367 | 1069 | L1069Q | D | 8.5e-09 | 1.6e-09 | 4.2e-09 | 4.767E-09 | 3.485E-09 | 1.600E-09 | 8.500E-09 |
| 21368 | 1069 | L1069G | D | 1.2e-08 | 1.5e-08 | 3.7e-09 | 1.023E-08 | 5.853E-09 | 3.700E-09 | 1.500E-08 |
| 21369 | 1069 | L1069H | D | 1e-08 | 5.9e-09 | 3.5e-09 | 6.467E-09 | 3.287E-09 | 3.500E-09 | 1.000E-08 |
| 21370 | 1069 | L1069I | D | 5.6e-09 | 1.7e-08 | 6.3e-09 | 9.633E-09 | 6.389E-09 | 5.600E-09 | 1.700E-08 |
| 21371 | 1069 | L1069L | D | 4.3e-09 | 5.1e-09 | 6.7e-09 | 5.367E-09 | 1.222E-09 | 4.300E-09 | 6.700E-09 |
| 21372 | 1069 | L1069K | D | 1.1e-08 | 7.3e-09 | 4.8e-09 | 7.700E-09 | 3.119E-09 | 4.800E-09 | 1.100E-08 |
| 21373 | 1069 | L1069M | D | 6.8e-09 | 5.7e-09 | 6.9e-09 | 6.467E-09 | 6.658E-10 | 5.700E-09 | 6.900E-09 |
| 21374 | 1069 | L1069F | D | 1.3e-08 | 1.1e-08 | 2.5e-09 | 8.833E-09 | 5.575E-09 | 2.500E-09 | 1.300E-08 |
| 21375 | 1069 | L1069P | D | 1.3e-08 | 3.3e-09 | 5.8e-09 | 7.367E-09 | 5.036E-09 | 3.300E-09 | 1.300E-08 |
| 21376 | 1069 | L1069S | D | 1.2e-08 | 5.2e-09 | 3.8e-09 | 7.000E-09 | 4.386E-09 | 3.800E-09 | 1.200E-08 |
| 21377 | 1069 | L1069T | D | 1.1e-08 | 9.1e-09 | 1.8e-09 | 7.300E-09 | 4.857E-09 | 1.800E-09 | 1.100E-08 |
| 21378 | 1069 | L1069W | D | 8.4e-09 | 3.3e-09 | 3e-09 | 4.900E-09 | 3.035E-09 | 3.000E-09 | 8.400E-09 |
| 21379 | 1069 | L1069Y | D | 8.5e-09 | 4.6e-09 | 1.5e-08 | 9.367E-09 | 5.254E-09 | 4.600E-09 | 1.500E-08 |
| 21380 | 1069 | L1069V | D | 9.9e-09 | 3.4e-09 | 1.5e-08 | 9.433E-09 | 5.814E-09 | 3.400E-09 | 1.500E-08 |
| 21381 | 1070 | S1070A | D | 3e-09 | 8.7e-09 | 1.7e-08 | 9.567E-09 | 7.040E-09 | 3.000E-09 | 1.700E-08 |
| 21382 | 1070 | S1070R | D | 8.4e-09 | 1.1e-08 | 9.5e-09 | 9.633E-09 | 1.305E-09 | 8.400E-09 | 1.100E-08 |
| 21383 | 1070 | S1070N | D | 1.5e-08 | 1.6e-08 | 1.6e-08 | 1.567E-08 | 5.774E-10 | 1.500E-08 | 1.600E-08 |
| 21384 | 1070 | S1070D | D | 4.9e-09 | 4.7e-09 | 8.9e-09 | 6.167E-09 | 2.369E-09 | 4.700E-09 | 8.900E-09 |
| 21385 | 1070 | S1070C | D | 4.6e-09 | 8e-09 | 3.3e-09 | 5.300E-09 | 2.427E-09 | 3.300E-09 | 8.000E-09 |
| 21386 | 1070 | S1070E | D | 5.9e-09 | 1.1e-08 | 8e-09 | 8.300E-09 | 2.563E-09 | 5.900E-09 | 1.100E-08 |
| 21387 | 1070 | S1070Q | D | 1.9e-08 | 1.1e-08 | 9.9e-09 | 1.330E-08 | 4.967E-09 | 9.900E-09 | 1.900E-08 |
| 21388 | 1070 | S1070G | D | 5.9e-09 | 1.8e-08 | 5.5e-09 | 9.800E-09 | 7.104E-09 | 5.500E-09 | 1.800E-08 |
| 21389 | 1070 | S1070H | D | 5.8e-09 | 5e-09 | 4.8e-09 | 5.200E-09 | 5.292E-10 | 4.800E-09 | 5.800E-09 |
| 21390 | 1070 | S1070I | D | 4.4e-09 | 8.6e-09 | 2.7e-09 | 5.233E-09 | 3.037E-09 | 2.700E-09 | 8.600E-09 |
| 21391 | 1070 | S1070L | D | 6.9e-09 | 3.4e-08 | 1.5e-08 | 1.863E-08 | 1.391E-08 | 6.900E-09 | 3.400E-08 |
| 21392 | 1070 | S1070K | D | 1.1e-08 | 7.1e-09 | 8.1e-09 | 8.733E-09 | 2.026E-09 | 7.100E-09 | 1.100E-08 |
| 21393 | 1070 | S1070M | D | 1.1e-08 | 6.8e-09 | 6.5e-09 | 8.100E-09 | 2.516E-09 | 6.500E-09 | 1.100E-08 |
| 21394 | 1070 | S1070F | D | 5.5e-09 | 5.7e-09 | 5.5e-09 | 5.567E-09 | 1.155E-10 | 5.500E-09 | 5.700E-09 |
| 21395 | 1070 | S1070P | D | 1.7e-08 | 5.9e-09 | 6.7e-09 | 9.867E-09 | 6.191E-09 | 5.900E-09 | 1.700E-08 |
| 21396 | 1070 | S1070S | D | 3.2e-09 | 2.5e-09 | 1.7e-09 | 2.467E-09 | 7.506E-10 | 1.700E-09 | 3.200E-09 |
| 21397 | 1070 | S1070T | D | 4.5e-09 | 8.3e-09 | 6.2e-09 | 6.333E-09 | 1.904E-09 | 4.500E-09 | 8.300E-09 |
| 21398 | 1070 | S1070W | D | 3e-09 | 1.1e-08 | 1.1e-08 | 8.333E-09 | 4.619E-09 | 3.000E-09 | 1.100E-08 |
| 21399 | 1070 | S1070Y | D | 5.1e-09 | 7.1e-09 | 5e-09 | 5.733E-09 | 1.185E-09 | 5.000E-09 | 7.100E-09 |
| 21400 | 1070 | S1070V | D | 3.2e-09 | 8e-09 | 1.4e-08 | 8.400E-09 | 5.411E-09 | 3.200E-09 | 1.400E-08 |
| 21401 | 1071 | A1071A | D | 2.5e-09 | 5.5e-09 | 6.6e-09 | 4.867E-09 | 2.122E-09 | 2.500E-09 | 6.600E-09 |
| 21402 | 1071 | A1071R | D | 1.8e-08 | 1.2e-09 | 7.9e-09 | 9.033E-09 | 8.457E-09 | 1.200E-09 | 1.800E-08 |
| 21403 | 1071 | A1071N | D | 5.9e-09 | 4.4e-09 | 4.7e-09 | 5.000E-09 | 7.937E-10 | 4.400E-09 | 5.900E-09 |
| 21404 | 1071 | A1071D | D | 1.4e-08 | 1.8e-08 | 8.5e-09 | 1.350E-08 | 4.770E-09 | 8.500E-09 | 1.800E-08 |
| 21405 | 1071 | A1071C | D | 9.6e-09 | 9.9e-09 | 3.6e-09 | 7.700E-09 | 3.554E-09 | 3.600E-09 | 9.900E-09 |
| 21406 | 1071 | A1071E | D | 1.1e-08 | 2.9e-09 | 1.3e-08 | 8.967E-09 | 5.348E-09 | 2.900E-09 | 1.300E-08 |
| 21407 | 1071 | A1071Q | D | 2.2e-09 | 4e-09 | 4.7e-09 | 3.633E-09 | 1.290E-09 | 2.200E-09 | 4.700E-09 |
| 21408 | 1071 | A1071G | D | 2e-08 | 8.8e-09 | 8.9e-09 | 1.257E-08 | 6.438E-09 | 8.800E-09 | 2.000E-08 |
| 21409 | 1071 | A1071H | D | 3.1e-09 | 5.8e-09 | 4.4e-09 | 4.433E-09 | 1.350E-09 | 3.100E-09 | 5.800E-09 |
| 21410 | 1071 | A1071I | D | 4.9e-09 | 6.5e-09 | 1.4e-08 | 8.467E-09 | 4.858E-09 | 4.900E-09 | 1.400E-08 |
| 21411 | 1071 | A1071L | D | 6.5e-09 | 1.1e-08 | 7.8e-09 | 8.433E-09 | 2.316E-09 | 6.500E-09 | 1.100E-08 |
| 21412 | 1071 | A1071K | D | 2.1e-08 | 1.4e-08 | 6.5e-09 | 1.383E-08 | 7.251E-09 | 6.500E-09 | 2.100E-08 |
| 21413 | 1071 | A1071M | D | 8.9e-09 | 6e-09 | 5.3e-09 | 6.733E-09 | 1.909E-09 | 5.300E-09 | 8.900E-09 |
| 21414 | 1071 | A1071F | D | 2.9e-09 | 5.6e-09 | 3.5e-09 | 4.000E-09 | 1.418E-09 | 2.900E-09 | 5.600E-09 |
| 21415 | 1071 | A1071P | D | 1.1e-08 | 6.7e-09 | 6.3e-09 | 8.000E-09 | 2.606E-09 | 6.300E-09 | 1.100E-08 |
| 21416 | 1071 | A1071S | D | 1.1e-08 | 1e-08 | 1.5e-08 | 1.200E-08 | 2.646E-09 | 1.000E-08 | 1.500E-08 |
| 21417 | 1071 | A1071T | D | 6.1e-09 | 4.1e-09 | 1.3e-08 | 7.733E-09 | 4.669E-09 | 4.100E-09 | 1.300E-08 |
| 21418 | 1071 | A1071W | D | 1.2e-08 | 3.4e-09 | 9.2e-09 | 8.200E-09 | 4.386E-09 | 3.400E-09 | 1.200E-08 |
| 21419 | 1071 | A1071Y | D | 4.8e-09 | 3.4e-09 | 2.6e-09 | 3.600E-09 | 1.114E-09 | 2.600E-09 | 4.800E-09 |

| | | | | | | | | | | |
|---|---|---|---|---|---|---|---|---|---|---|
| 21420 | 1071 | A1071V | D | 5.1e-09 | 4.7e-09 | 1.9e-08 | 9.600E-09 | 8.143E-09 | 4.700E-09 | 1.900E-08 |
| 21421 | 1072 | S1072A | D | 6.5e-09 | 1.4e-08 | 7.1e-09 | 9.200E-09 | 4.168E-09 | 6.500E-09 | 1.400E-08 |
| 21422 | 1072 | S1072R | D | 9.8e-09 | 5.2e-09 | 4.7e-09 | 6.567E-09 | 2.811E-09 | 4.700E-09 | 9.800E-09 |
| 21423 | 1072 | S1072N | D | 2.3e-08 | 1.1e-08 | 3.8e-09 | 1.260E-08 | 9.699E-09 | 3.800E-09 | 2.300E-08 |
| 21424 | 1072 | S1072D | D | 8.8e-09 | 4.6e-09 | 6.3e-09 | 6.567E-09 | 2.113E-09 | 4.600E-09 | 8.800E-09 |
| 21425 | 1072 | S1072C | D | 7.1e-09 | 1.5e-08 | 3.6e-09 | 8.567E-09 | 5.840E-09 | 3.600E-09 | 1.500E-08 |
| 21426 | 1072 | S1072E | D | 1.1e-08 | 6.5e-09 | 1.2e-08 | 9.833E-09 | 2.930E-09 | 6.500E-09 | 1.200E-08 |
| 21427 | 1072 | S1072Q | D | 5.6e-09 | 1.1e-08 | 1.2e-08 | 9.533E-09 | 3.443E-09 | 5.600E-09 | 1.200E-08 |
| 21428 | 1072 | S1072G | D | 1e-08 | 1.9e-08 | 4.7e-09 | 1.123E-08 | 7.229E-09 | 4.700E-09 | 1.900E-08 |
| 21429 | 1072 | S1072H | D | 1e-08 | 6.1e-09 | 1.6e-08 | 1.070E-08 | 4.987E-09 | 6.100E-09 | 1.600E-08 |
| 21430 | 1072 | S1072I | D | 9.6e-09 | 5.1e-09 | 1.3e-08 | 9.233E-09 | 3.963E-09 | 5.100E-09 | 1.300E-08 |
| 21431 | 1072 | S1072L | D | 1.1e-08 | 1.3e-08 | 3.2e-09 | 9.067E-09 | 5.178E-09 | 3.200E-09 | 1.300E-08 |
| 21432 | 1072 | S1072K | D | 5.2e-09 | 6.5e-09 | 6e-09 | 5.900E-09 | 6.557E-10 | 5.200E-09 | 6.500E-09 |
| 21433 | 1072 | S1072M | D | 5.4e-09 | 2.2e-08 | 7e-09 | 1.147E-08 | 9.157E-09 | 5.400E-09 | 2.200E-08 |
| 21434 | 1072 | S1072F | D | 9.7e-09 | 1.5e-08 | 1.2e-08 | 1.223E-08 | 2.658E-09 | 9.700E-09 | 1.500E-08 |
| 21435 | 1072 | S1072P | D | 1.2e-08 | 1.3e-08 | 4.9e-09 | 9.967E-09 | 4.416E-09 | 4.900E-09 | 1.300E-08 |
| 21436 | 1072 | S1072S | D | 1.6e-08 | 7.3e-09 | 1.3e-08 | 1.210E-08 | 4.419E-09 | 7.300E-09 | 1.600E-08 |
| 21437 | 1072 | S1072T | D | 2.4e-09 | 4.8e-09 | 5.6e-09 | 4.267E-09 | 1.665E-09 | 2.400E-09 | 5.600E-09 |
| 21438 | 1072 | S1072W | D | 1.3e-08 | 1e-08 | 6.9e-09 | 9.967E-09 | 3.050E-09 | 6.900E-09 | 1.300E-08 |
| 21439 | 1072 | S1072Y | D | 2.2e-09 | 1.2e-08 | 3.1e-09 | 5.767E-09 | 5.417E-09 | 2.200E-09 | 1.200E-08 |
| 21440 | 1072 | S1072V | D | 1.7e-08 | 5.2e-09 | 5e-09 | 9.067E-09 | 6.871E-09 | 5.000E-09 | 1.700E-08 |
| 21441 | 1073 | V1073A | D | 3.1e-09 | 4.3e-08 | 1.2e-09 | 1.577E-08 | 2.360E-08 | 1.200E-09 | 4.300E-08 |
| 21442 | 1073 | V1073R | D | 1.3e-08 | 1.1e-08 | 3e-09 | 9.000E-09 | 5.292E-09 | 3.000E-09 | 1.300E-08 |
| 21443 | 1073 | V1073N | D | 2.5e-09 | 4.8e-09 | 6e-09 | 4.433E-09 | 1.779E-09 | 2.500E-09 | 6.000E-09 |
| 21444 | 1073 | V1073D | D | 4.9e-09 | 1.2e-08 | 2.5e-08 | 1.397E-08 | 1.019E-08 | 4.900E-09 | 2.500E-08 |
| 21445 | 1073 | V1073C | D | 1.8e-08 | 4.1e-09 | 9.1e-09 | 1.040E-08 | 7.041E-09 | 4.100E-09 | 1.800E-08 |
| 21446 | 1073 | V1073E | D | 1.5e-08 | 8.5e-09 | 1.3e-08 | 1.217E-08 | 3.329E-09 | 8.500E-09 | 1.500E-08 |
| 21447 | 1073 | V1073Q | D | 3.8e-09 | 8.9e-09 | 2.2e-09 | 4.967E-09 | 3.499E-09 | 2.200E-09 | 8.900E-09 |
| 21448 | 1073 | V1073G | D | 1.2e-08 | 2.7e-09 | 2.8e-09 | 5.833E-09 | 5.341E-09 | 2.700E-09 | 1.200E-08 |
| 21449 | 1073 | V1073H | D | 1.1e-08 | 5.3e-09 | 9.7e-09 | 8.667E-09 | 2.987E-09 | 5.300E-09 | 1.100E-08 |
| 21450 | 1073 | V1073I | D | 6.8e-09 | 7.1e-09 | 3.8e-09 | 5.900E-09 | 1.825E-09 | 3.800E-09 | 7.100E-09 |
| 21451 | 1073 | V1073L | D | 8.2e-09 | 1e-08 | 6.2e-09 | 8.133E-09 | 1.901E-09 | 6.200E-09 | 1.000E-08 |
| 21452 | 1073 | V1073K | D | 5.4e-09 | 3.9e-09 | 8.7e-09 | 6.000E-09 | 2.456E-09 | 3.900E-09 | 8.700E-09 |
| 21453 | 1073 | V1073M | D | 7.9e-09 | 1.2e-08 | 8.3e-09 | 9.400E-09 | 2.261E-09 | 7.900E-09 | 1.200E-08 |
| 21454 | 1073 | V1073F | D | 5.6e-09 | 8.6e-09 | 3.6e-09 | 5.933E-09 | 2.517E-09 | 3.600E-09 | 8.600E-09 |
| 21455 | 1073 | V1073P | D | 1e-08 | 1.4e-08 | 1.1e-08 | 1.167E-08 | 2.082E-09 | 1.000E-08 | 1.400E-08 |
| 21456 | 1073 | V1073S | D | 4.3e-09 | 4.9e-09 | 3.9e-09 | 4.367E-09 | 5.033E-10 | 3.900E-09 | 4.900E-09 |
| 21457 | 1073 | V1073T | D | 6.2e-09 | 2.6e-09 | 9.2e-09 | 6.000E-09 | 3.305E-09 | 2.600E-09 | 9.200E-09 |
| 21458 | 1073 | V1073W | D | 4.6e-09 | 7.7e-09 | 3.5e-09 | 5.267E-09 | 2.178E-09 | 3.500E-09 | 7.700E-09 |
| 21459 | 1073 | V1073Y | D | 6.7e-09 | 7e-09 | 6.3e-09 | 6.667E-09 | 3.512E-10 | 6.300E-09 | 7.000E-09 |
| 21460 | 1073 | V1073V | D | 3.5e-09 | 7.1e-09 | 1.7e-08 | 9.200E-09 | 6.991E-09 | 3.500E-09 | 1.700E-08 |
| 21461 | 1074 | G1074A | D | 1e-08 | 4.7e-09 | 8.6e-09 | 7.767E-09 | 2.747E-09 | 4.700E-09 | 1.000E-08 |
| 21462 | 1074 | G1074R | D | 6.6e-09 | 2.9e-09 | 4.3e-09 | 4.600E-09 | 1.868E-09 | 2.900E-09 | 6.600E-09 |
| 21463 | 1074 | G1074N | D | 6.6e-09 | 5.7e-09 | 3.7e-09 | 5.333E-09 | 1.484E-09 | 3.700E-09 | 6.600E-09 |
| 21464 | 1074 | G1074D | D | 8.3e-09 | 5.3e-09 | 9.8e-09 | 7.800E-09 | 2.291E-09 | 5.300E-09 | 9.800E-09 |
| 21465 | 1074 | G1074C | D | 2.3e-09 | 8e-09 | 5.7e-09 | 5.333E-09 | 2.868E-09 | 2.300E-09 | 8.000E-09 |
| 21466 | 1074 | G1074E | D | 1.2e-08 | 1.8e-09 | 4.7e-09 | 6.167E-09 | 5.256E-09 | 1.800E-09 | 1.200E-08 |
| 21467 | 1074 | G1074Q | D | 6.6e-09 | 2.1e-08 | 1.1e-08 | 1.287E-08 | 7.379E-09 | 6.600E-09 | 2.100E-08 |
| 21468 | 1074 | G1074G | D | 2.4e-09 | 4.1e-09 | 3.1e-09 | 3.200E-09 | 8.544E-10 | 2.400E-09 | 4.100E-09 |
| 21469 | 1074 | G1074H | D | 7.6e-09 | 6.5e-09 | 2.7e-09 | 5.600E-09 | 2.571E-09 | 2.700E-09 | 7.600E-09 |
| 21470 | 1074 | G1074I | D | 1.5e-08 | 7.8e-09 | 1.3e-08 | 1.193E-08 | 3.717E-09 | 7.800E-09 | 1.500E-08 |
| 21471 | 1074 | G1074L | D | 3.6e-09 | 5.9e-09 | 7.1e-09 | 5.533E-09 | 1.779E-09 | 3.600E-09 | 7.100E-09 |
| 21472 | 1074 | G1074K | D | 4.4e-09 | 4.7e-09 | 1e-08 | 6.367E-09 | 3.150E-09 | 4.400E-09 | 1.000E-08 |
| 21473 | 1074 | G1074M | D | 5.6e-09 | 5.3e-09 | 6.9e-09 | 5.933E-09 | 8.505E-10 | 5.300E-09 | 6.900E-09 |
| 21474 | 1074 | G1074F | D | 1.8e-09 | 4.7e-09 | 4.8e-09 | 3.767E-09 | 1.704E-09 | 1.800E-09 | 4.800E-09 |
| 21475 | 1074 | G1074P | D | 3.5e-09 | 1.7e-08 | 1.2e-08 | 1.083E-08 | 6.825E-09 | 3.500E-09 | 1.700E-08 |
| 21476 | 1074 | G1074S | D | 6.1e-09 | 9.4e-09 | 4.9e-09 | 6.800E-09 | 2.330E-09 | 4.900E-09 | 9.400E-09 |
| 21477 | 1074 | G1074T | D | 9.8e-09 | 2.3e-09 | 1.5e-08 | 9.033E-09 | 6.385E-09 | 2.300E-09 | 1.500E-08 |
| 21478 | 1074 | G1074W | D | 1.3e-09 | 4.6e-09 | 1.1e-08 | 5.633E-09 | 4.932E-09 | 1.300E-09 | 1.100E-08 |
| 21479 | 1074 | G1074Y | D | 6.9e-09 | 7.2e-09 | 7.6e-09 | 7.233E-09 | 3.512E-10 | 6.900E-09 | 7.600E-09 |
| 21480 | 1074 | G1074V | D | 5e-09 | 4.1e-09 | 5.5e-09 | 4.867E-09 | 7.095E-10 | 4.100E-09 | 5.500E-09 |
| 21481 | 1075 | D1075A | D | 1.3e-08 | 5.4e-09 | 2.8e-09 | 7.067E-09 | 5.300E-09 | 2.800E-09 | 1.300E-08 |
| 21482 | 1075 | D1075R | D | 1.7e-08 | 7.1e-09 | 1.1e-08 | 1.170E-08 | 4.987E-09 | 7.100E-09 | 1.700E-08 |
| 21483 | 1075 | D1075N | D | 5e-09 | 2.4e-08 | 8.3e-09 | 1.243E-08 | 1.015E-08 | 5.000E-09 | 2.400E-08 |
| 21484 | 1075 | D1075D | D | 5.7e-09 | 3.8e-08 | 7.7e-09 | 1.713E-08 | 1.810E-08 | 5.700E-09 | 3.800E-08 |
| 21485 | 1075 | D1075C | D | 1.2e-08 | 3e-09 | 5.1e-09 | 6.700E-09 | 4.709E-09 | 3.000E-09 | 1.200E-08 |
| 21486 | 1075 | D1075E | D | 5.8e-09 | 5.5e-09 | 1.8e-08 | 9.767E-09 | 7.132E-09 | 5.500E-09 | 1.800E-08 |
| 21487 | 1075 | D1075Q | D | 7.8e-09 | 1e-08 | 4.8e-09 | 7.533E-09 | 2.610E-09 | 4.800E-09 | 1.000E-08 |
| 21488 | 1075 | D1075G | D | 4.9e-09 | 1.6e-08 | 3.8e-09 | 8.233E-09 | 6.749E-09 | 3.800E-09 | 1.600E-08 |
| 21489 | 1075 | D1075H | D | 1.8e-08 | 9.6e-09 | 9.5e-09 | 1.237E-08 | 4.879E-09 | 9.500E-09 | 1.800E-08 |
| 21490 | 1075 | D1075I | D | 1.2e-08 | 5.4e-09 | 6.4e-09 | 7.933E-09 | 3.557E-09 | 5.400E-09 | 1.200E-08 |
| 21491 | 1075 | D1075L | D | 6.7e-09 | 8.8e-09 | 1.1e-08 | 8.833E-09 | 2.150E-09 | 6.700E-09 | 1.100E-08 |
| 21492 | 1075 | D1075K | D | 1.5e-08 | 1.5e-08 | 3.8e-09 | 1.127E-08 | 6.466E-09 | 3.800E-09 | 1.500E-08 |
| 21493 | 1075 | D1075M | D | 7.2e-09 | 8.2e-09 | 1.1e-08 | 8.800E-09 | 1.970E-09 | 7.200E-09 | 1.100E-08 |
| 21494 | 1075 | D1075F | D | 1.3e-08 | 5.8e-09 | 4.6e-09 | 7.800E-09 | 4.543E-09 | 4.600E-09 | 1.300E-08 |
| 21495 | 1075 | D1075P | D | 4.1e-09 | 1e-08 | 1.2e-08 | 8.700E-09 | 4.107E-09 | 4.100E-09 | 1.200E-08 |
| 21496 | 1075 | D1075S | D | 1.5e-09 | 1.1e-08 | 6.7e-09 | 6.400E-09 | 4.757E-09 | 1.500E-09 | 1.100E-08 |
| 21497 | 1075 | D1075T | D | 2.9e-08 | 4.3e-09 | 5.9e-09 | 1.307E-08 | 1.382E-08 | 4.300E-09 | 2.900E-08 |
| 21498 | 1075 | D1075W | D | 6e-09 | 3.3e-09 | 2.5e-09 | 3.933E-09 | 1.834E-09 | 2.500E-09 | 6.000E-09 |
| 21499 | 1075 | D1075Y | D | 4e-09 | 4.9e-09 | 5.3e-09 | 4.733E-09 | 6.658E-10 | 4.000E-09 | 5.300E-09 |
| 21500 | 1075 | D1075V | D | 8.1e-09 | 5.3e-09 | 2.1e-09 | 5.167E-09 | 3.002E-09 | 2.100E-09 | 8.100E-09 |
| 21501 | 1076 | R1076A | D | 2.5e-09 | 7.8e-09 | 4.5e-09 | 4.933E-09 | 2.676E-09 | 2.500E-09 | 7.800E-09 |
| 21502 | 1076 | R1076R | D | 9.5e-09 | 4.2e-09 | 1e-09 | 4.900E-09 | 4.293E-09 | 1.000E-09 | 9.500E-09 |
| 21503 | 1076 | R1076N | D | 6.4e-09 | 9.4e-09 | 4.6e-08 | 2.060E-08 | 2.205E-08 | 6.400E-09 | 4.600E-08 |
| 21504 | 1076 | R1076D | D | 7.8e-09 | 7.5e-09 | 3.6e-09 | 6.300E-09 | 2.343E-09 | 3.600E-09 | 7.800E-09 |

| | | | | | | | | | | |
|---|---|---|---|---|---|---|---|---|---|---|
| 21505 | 1076 | R1076C | D | 6.9e-09 | 4.8e-09 | 3e-09 | 4.900E-09 | 1.952E-09 | 3.000E-09 | 6.900E-09 |
| 21506 | 1076 | R1076E | D | 3.7e-09 | 2.7e-09 | 1.8e-09 | 2.733E-09 | 9.504E-10 | 1.800E-09 | 3.700E-09 |
| 21507 | 1076 | R1076Q | D | 4.6e-09 | 1.2e-08 | 1.9e-09 | 6.167E-09 | 5.229E-09 | 1.900E-09 | 1.200E-08 |
| 21508 | 1076 | R1076G | D | 7e-09 | 1e-08 | 1.7e-09 | 6.233E-09 | 4.203E-09 | 1.700E-09 | 1.000E-08 |
| 21509 | 1076 | R1076H | D | 4.9e-09 | 9.3e-09 | 3.8e-09 | 6.000E-09 | 2.910E-09 | 3.800E-09 | 9.300E-09 |
| 21510 | 1076 | R1076I | D | 3.6e-09 | 3.8e-09 | 4.5e-09 | 3.967E-09 | 4.726E-10 | 3.600E-09 | 4.500E-09 |
| 21511 | 1076 | R1076L | D | 1.5e-09 | 1.8e-09 | 8.4e-09 | 3.900E-09 | 3.900E-09 | 1.500E-09 | 8.400E-09 |
| 21512 | 1076 | R1076K | D | 2e-09 | 2.1e-08 | 1.6e-08 | 1.300E-08 | 9.849E-09 | 2.000E-09 | 2.100E-08 |
| 21513 | 1076 | R1076M | D | 5.8e-09 | 9.4e-09 | 2.1e-09 | 5.767E-09 | 3.650E-09 | 2.100E-09 | 9.400E-09 |
| 21514 | 1076 | R1076F | D | 3.1e-09 | 2.3e-08 | 4.2e-09 | 1.010E-08 | 1.119E-08 | 3.100E-09 | 2.300E-08 |
| 21515 | 1076 | R1076P | D | 4.5e-09 | 6.5e-09 | 2e-08 | 1.033E-08 | 8.431E-09 | 4.500E-09 | 2.000E-08 |
| 21516 | 1076 | R1076S | D | 1e-08 | 1.6e-08 | 1.2e-08 | 1.267E-08 | 3.055E-09 | 1.000E-08 | 1.600E-08 |
| 21517 | 1076 | R1076T | D | 3.8e-09 | 8.1e-09 | 1.8e-08 | 9.967E-09 | 7.282E-09 | 3.800E-09 | 1.800E-08 |
| 21518 | 1076 | R1076W | D | 2.6e-09 | 1e-08 | 3.5e-09 | 5.367E-09 | 4.038E-09 | 2.600E-09 | 1.000E-08 |
| 21519 | 1076 | R1076Y | D | 1.6e-08 | 4.5e-09 | 1.1e-08 | 1.050E-08 | 5.766E-09 | 4.500E-09 | 1.600E-08 |
| 21520 | 1076 | R1076V | D | 6.3e-09 | 9.6e-09 | 5.4e-09 | 7.100E-09 | 2.211E-09 | 5.400E-09 | 9.600E-09 |
| 21521 | 1077 | V1077A | D | 5e-09 | 1.7e-08 | 7.5e-09 | 9.833E-09 | 6.331E-09 | 5.000E-09 | 1.700E-08 |
| 21522 | 1077 | V1077R | D | 6e-09 | 3.4e-09 | 1.9e-08 | 9.467E-09 | 8.358E-09 | 3.400E-09 | 1.900E-08 |
| 21523 | 1077 | V1077N | D | 6.2e-09 | 6.2e-09 | 2.5e-08 | 1.247E-08 | 1.085E-08 | 6.200E-09 | 2.500E-08 |
| 21524 | 1077 | V1077D | D | 3.2e-09 | 2.6e-08 | 2.9e-09 | 1.070E-08 | 1.325E-08 | 2.900E-09 | 2.600E-08 |
| 21525 | 1077 | V1077C | D | 6.6e-09 | 5.9e-09 | 6.5e-09 | 6.333E-09 | 3.786E-10 | 5.900E-09 | 6.600E-09 |
| 21526 | 1077 | V1077E | D | 3.3e-09 | 7.8e-09 | 8.5e-09 | 6.533E-09 | 2.822E-09 | 3.300E-09 | 8.500E-09 |
| 21527 | 1077 | V1077Q | D | 5.6e-09 | 5.9e-09 | 3.6e-09 | 5.033E-09 | 1.250E-09 | 3.600E-09 | 5.900E-09 |
| 21528 | 1077 | V1077G | D | 6.8e-09 | 6.7e-09 | 9.9e-09 | 7.800E-09 | 1.819E-09 | 6.700E-09 | 9.900E-09 |
| 21529 | 1077 | V1077H | D | 5.2e-08 | 6.1e-09 | 5.8e-09 | 2.130E-08 | 2.659E-08 | 5.800E-09 | 5.200E-08 |
| 21530 | 1077 | V1077I | D | 7.8e-09 | 3.5e-09 | 7.6e-09 | 6.300E-09 | 2.427E-09 | 3.500E-09 | 7.800E-09 |
| 21531 | 1077 | V1077L | D | 4.6e-09 | 4.3e-09 | 7e-09 | 5.300E-09 | 1.480E-09 | 4.300E-09 | 7.000E-09 |
| 21532 | 1077 | V1077K | D | 3.9e-09 | 6.7e-09 | 5.1e-09 | 5.233E-09 | 1.405E-09 | 3.900E-09 | 6.700E-09 |
| 21533 | 1077 | V1077M | D | 2.9e-09 | 9.5e-09 | 6.4e-09 | 6.267E-09 | 3.302E-09 | 2.900E-09 | 9.500E-09 |
| 21534 | 1077 | V1077F | D | 4.6e-09 | 5.9e-09 | 4.4e-09 | 4.967E-09 | 8.145E-10 | 4.400E-09 | 5.900E-09 |
| 21535 | 1077 | V1077P | D | 8.8e-09 | 4.3e-09 | 6e-09 | 6.367E-09 | 2.272E-09 | 4.300E-09 | 8.800E-09 |
| 21536 | 1077 | V1077S | D | 1.3e-08 | 1.1e-08 | 7.1e-09 | 1.037E-08 | 3.001E-09 | 7.100E-09 | 1.300E-08 |
| 21537 | 1077 | V1077T | D | 1.3e-08 | 8.7e-09 | 6.4e-09 | 9.367E-09 | 3.350E-09 | 6.400E-09 | 1.300E-08 |
| 21538 | 1077 | V1077W | D | 7.6e-09 | 7.9e-09 | 3.2e-09 | 6.233E-09 | 2.631E-09 | 3.200E-09 | 7.900E-09 |
| 21539 | 1077 | V1077Y | D | 5.7e-09 | 1.6e-09 | 6e-09 | 4.433E-09 | 2.458E-09 | 1.600E-09 | 6.000E-09 |
| 21540 | 1077 | V1077V | D | 3.5e-09 | 2e-09 | 4.3e-09 | 3.267E-09 | 1.168E-09 | 2.000E-09 | 4.300E-09 |
| 21541 | 1078 | T1078A | D | 9.2e-09 | 8.8e-09 | 4.2e-09 | 7.400E-09 | 2.778E-09 | 4.200E-09 | 9.200E-09 |
| 21542 | 1078 | T1078R | D | 2e-08 | 3.6e-09 | 4.4e-09 | 9.333E-09 | 9.246E-09 | 3.600E-09 | 2.000E-08 |
| 21543 | 1078 | T1078N | D | 2.6e-09 | 7.1e-09 | 7.7e-09 | 5.800E-09 | 2.787E-09 | 2.600E-09 | 7.700E-09 |
| 21544 | 1078 | T1078D | D | 5.6e-09 | 8.1e-09 | 8.3e-09 | 7.333E-09 | 1.504E-09 | 5.600E-09 | 8.300E-09 |
| 21545 | 1078 | T1078C | D | 8.9e-09 | 6.5e-09 | 5.6e-09 | 7.000E-09 | 1.706E-09 | 5.600E-09 | 8.900E-09 |
| 21546 | 1078 | T1078E | D | 1.3e-08 | 1.3e-08 | 3.4e-09 | 9.800E-09 | 5.543E-09 | 3.400E-09 | 1.300E-08 |
| 21547 | 1078 | T1078Q | D | 8.1e-09 | 3.8e-09 | 5.2e-09 | 5.700E-09 | 2.193E-09 | 3.800E-09 | 8.100E-09 |
| 21548 | 1078 | T1078G | D | 5e-09 | 2.4e-09 | 5.2e-09 | 4.200E-09 | 1.562E-09 | 2.400E-09 | 5.200E-09 |
| 21549 | 1078 | T1078H | D | 3e-09 | 6e-09 | 1.1e-08 | 6.667E-09 | 4.041E-09 | 3.000E-09 | 1.100E-08 |
| 21550 | 1078 | T1078I | D | 4e-09 | 3e-09 | 1.1e-08 | 6.000E-09 | 4.359E-09 | 3.000E-09 | 1.100E-08 |
| 21551 | 1078 | T1078L | D | 7.7e-09 | 1.3e-08 | 1.2e-08 | 1.090E-08 | 2.816E-09 | 7.700E-09 | 1.300E-08 |
| 21552 | 1078 | T1078K | D | 7.8e-09 | 1.5e-08 | 7.3e-09 | 1.003E-08 | 4.309E-09 | 7.300E-09 | 1.500E-08 |
| 21553 | 1078 | T1078M | D | 6.1e-09 | 1.6e-08 | 9.3e-09 | 1.047E-08 | 5.052E-09 | 6.100E-09 | 1.600E-08 |
| 21554 | 1078 | T1078F | D | 6.9e-09 | 6e-09 | 7.4e-09 | 6.767E-09 | 7.095E-10 | 6.000E-09 | 7.400E-09 |
| 21555 | 1078 | T1078P | D | 8.4e-09 | 4.4e-09 | 8.4e-09 | 7.067E-09 | 2.309E-09 | 4.400E-09 | 8.400E-09 |
| 21556 | 1078 | T1078S | D | 8.9e-09 | 1.3e-08 | 5.7e-09 | 9.200E-09 | 3.659E-09 | 5.700E-09 | 1.300E-08 |
| 21557 | 1078 | T1078T | D | 5.3e-09 | 5.8e-09 | 8.6e-09 | 6.567E-09 | 1.779E-09 | 5.300E-09 | 8.600E-09 |
| 21558 | 1078 | T1078W | D | 1e-08 | 1e-08 | 8.6e-09 | 9.533E-09 | 8.083E-10 | 8.600E-09 | 1.000E-08 |
| 21559 | 1078 | T1078Y | D | 9e-09 | 7.2e-09 | 4.9e-09 | 7.033E-09 | 2.055E-09 | 4.900E-09 | 9.000E-09 |
| 21560 | 1078 | T1078V | D | 9.1e-09 | 4.8e-09 | 7.6e-09 | 7.167E-09 | 2.183E-09 | 4.800E-09 | 9.100E-09 |
| 21561 | 1079 | I1079A | D | 6.1e-09 | 1.4e-08 | 3.8e-09 | 7.967E-09 | 5.350E-09 | 3.800E-09 | 1.400E-08 |
| 21562 | 1079 | I1079R | D | 9.5e-09 | 2.8e-09 | 1.3e-08 | 8.433E-09 | 5.183E-09 | 2.800E-09 | 1.300E-08 |
| 21563 | 1079 | I1079N | D | 4.8e-09 | 8.9e-09 | 6.2e-09 | 6.633E-09 | 2.084E-09 | 4.800E-09 | 8.900E-09 |
| 21564 | 1079 | I1079D | D | 5.7e-09 | 3.4e-09 | 4e-09 | 4.367E-09 | 1.193E-09 | 3.400E-09 | 5.700E-09 |
| 21565 | 1079 | I1079C | D | 1.2e-08 | 4.2e-09 | 4e-09 | 6.733E-09 | 4.562E-09 | 4.000E-09 | 1.200E-08 |
| 21566 | 1079 | I1079E | D | 1.6e-09 | 4.3e-09 | 6.3e-09 | 4.067E-09 | 2.359E-09 | 1.600E-09 | 6.300E-09 |
| 21567 | 1079 | I1079Q | D | 2.7e-09 | 5.1e-09 | 3.1e-09 | 3.633E-09 | 1.286E-09 | 2.700E-09 | 5.100E-09 |
| 21568 | 1079 | I1079G | D | 8e-09 | 9e-09 | 7.5e-09 | 8.167E-09 | 7.638E-10 | 7.500E-09 | 9.000E-09 |
| 21569 | 1079 | I1079H | D | 4.5e-09 | 8.2e-09 | 5.2e-09 | 5.967E-09 | 1.966E-09 | 4.500E-09 | 8.200E-09 |
| 21570 | 1079 | I1079I | D | 1.1e-08 | 5.3e-09 | 1.4e-08 | 1.010E-08 | 4.419E-09 | 5.300E-09 | 1.400E-08 |
| 21571 | 1079 | I1079L | D | 1.4e-08 | 1.1e-08 | 1.1e-08 | 1.200E-08 | 1.732E-09 | 1.100E-08 | 1.400E-08 |
| 21572 | 1079 | I1079K | D | 9.1e-09 | 2e-08 | 1.6e-08 | 1.503E-08 | 5.514E-09 | 9.100E-09 | 2.000E-08 |
| 21573 | 1079 | I1079M | D | 5e-09 | 4e-09 | 5.4e-09 | 4.800E-09 | 7.211E-10 | 4.000E-09 | 5.400E-09 |
| 21574 | 1079 | I1079F | D | 7.6e-09 | 5.8e-09 | 4e-09 | 5.800E-09 | 1.800E-09 | 4.000E-09 | 7.600E-09 |
| 21575 | 1079 | I1079P | D | 2e-08 | 7.9e-09 | 5.5e-09 | 1.113E-08 | 7.772E-09 | 5.500E-09 | 2.000E-08 |
| 21576 | 1079 | I1079S | D | 1.8e-08 | 9.5e-09 | 3.2e-09 | 1.023E-08 | 7.427E-09 | 3.200E-09 | 1.800E-08 |
| 21577 | 1079 | I1079T | D | 7.1e-09 | 2.8e-09 | 5.6e-09 | 5.167E-09 | 2.183E-09 | 2.800E-09 | 7.100E-09 |
| 21578 | 1079 | I1079W | D | 8.7e-09 | 6.1e-09 | 1.2e-08 | 8.933E-09 | 2.957E-09 | 6.100E-09 | 1.200E-08 |
| 21579 | 1079 | I1079Y | D | 4.1e-09 | 1.3e-08 | 4.4e-09 | 7.167E-09 | 5.054E-09 | 4.100E-09 | 1.300E-08 |
| 21580 | 1079 | I1079V | D | 7.2e-09 | 1.2e-08 | 7.4e-09 | 8.867E-09 | 2.715E-09 | 7.200E-09 | 1.200E-08 |
| 21581 | 1080 | T1080A | D | 7e-09 | 2e-08 | 6.1e-09 | 1.103E-08 | 7.778E-09 | 6.100E-09 | 2.000E-08 |
| 21582 | 1080 | T1080R | D | 7.1e-09 | 3.5e-09 | 5.1e-09 | 5.233E-09 | 1.804E-09 | 3.500E-09 | 7.100E-09 |
| 21583 | 1080 | T1080N | D | 2.1e-08 | 3.7e-09 | 5.4e-08 | 2.623E-08 | 2.556E-08 | 3.700E-09 | 5.400E-08 |
| 21584 | 1080 | T1080D | D | 6.6e-09 | 2.8e-09 | 1.8e-08 | 9.133E-09 | 7.910E-09 | 2.800E-09 | 1.800E-08 |
| 21585 | 1080 | T1080C | D | 3.2e-09 | 3.9e-09 | 5e-09 | 4.033E-09 | 9.074E-10 | 3.200E-09 | 5.000E-09 |
| 21586 | 1080 | T1080E | D | 8.2e-09 | 8.9e-09 | 8.5e-09 | 8.533E-09 | 3.512E-10 | 8.200E-09 | 8.900E-09 |
| 21587 | 1080 | T1080Q | D | 1.1e-08 | 5.9e-09 | 1.3e-08 | 9.967E-09 | 3.661E-09 | 5.900E-09 | 1.300E-08 |
| 21588 | 1080 | T1080G | D | 4e-09 | 1.7e-08 | 2.8e-09 | 7.933E-09 | 7.875E-09 | 2.800E-09 | 1.700E-08 |
| 21589 | 1080 | T1080H | D | 1.8e-08 | 9.9e-09 | 8.2e-09 | 1.203E-08 | 5.237E-09 | 8.200E-09 | 1.800E-08 |

| | | | | | | | | | | |
|---|---|---|---|---|---|---|---|---|---|---|
| 21590 | 1080 | T1080I | D | 6e-09 | 1.1e-08 | 1.1e-08 | 9.333E-09 | 2.887E-09 | 6.000E-09 | 1.100E-08 |
| 21591 | 1080 | T1080L | D | 1.1e-08 | 8.8e-09 | 1.5e-08 | 1.160E-08 | 3.143E-09 | 8.800E-09 | 1.500E-08 |
| 21592 | 1080 | T1080K | D | 6.6e-09 | 1.5e-08 | 9e-09 | 1.020E-08 | 4.327E-09 | 6.600E-09 | 1.500E-08 |
| 21593 | 1080 | T1080M | D | 3.9e-09 | 5.3e-09 | 4.7e-09 | 4.633E-09 | 7.024E-10 | 3.900E-09 | 5.300E-09 |
| 21594 | 1080 | T1080F | D | 1.1e-08 | 6.4e-09 | 2.7e-08 | 1.480E-08 | 1.081E-08 | 6.400E-09 | 2.700E-08 |
| 21595 | 1080 | T1080P | D | 4.1e-09 | 1.9e-09 | 8.6e-09 | 4.867E-09 | 3.415E-09 | 1.900E-09 | 8.600E-09 |
| 21596 | 1080 | T1080S | D | 4.7e-09 | 5.3e-09 | 3.7e-09 | 4.567E-09 | 8.083E-10 | 3.700E-09 | 5.300E-09 |
| 21597 | 1080 | T1080T | D | 2.2e-09 | 4e-09 | 1.3e-08 | 6.400E-09 | 5.786E-09 | 2.200E-09 | 1.300E-08 |
| 21598 | 1080 | T1080W | D | 7.1e-09 | 8.8e-09 | 1.2e-08 | 9.300E-09 | 2.488E-09 | 7.100E-09 | 1.200E-08 |
| 21599 | 1080 | T1080Y | D | 5.6e-09 | 9.8e-09 | 3.4e-09 | 6.267E-09 | 3.252E-09 | 3.400E-09 | 9.800E-09 |
| 21600 | 1080 | T1080V | D | 5.8e-09 | 9e-09 | 1.7e-08 | 1.060E-08 | 5.769E-09 | 5.800E-09 | 1.700E-08 |
| 21601 | 1081 | C1081A | D | 1.3e-08 | 1e-08 | 7.3e-09 | 1.010E-08 | 2.851E-09 | 7.300E-09 | 1.300E-08 |
| 21602 | 1081 | C1081R | D | 5.9e-09 | 8.8e-09 | 4.2e-09 | 6.300E-09 | 2.326E-09 | 4.200E-09 | 8.800E-09 |
| 21603 | 1081 | C1081N | D | 6.5e-09 | 5.9e-09 | 4.7e-09 | 5.700E-09 | 9.165E-10 | 4.700E-09 | 6.500E-09 |
| 21604 | 1081 | C1081D | D | 2.1e-08 | 1.3e-08 | 3e-09 | 1.233E-08 | 9.018E-09 | 3.000E-09 | 2.100E-08 |
| 21605 | 1081 | C1081C | D | 4e-09 | 7e-09 | 6.3e-09 | 5.767E-09 | 1.570E-09 | 4.000E-09 | 7.000E-09 |
| 21606 | 1081 | C1081E | D | 8.7e-09 | 5.9e-09 | 3.5e-08 | 1.653E-08 | 1.605E-08 | 5.900E-09 | 3.500E-08 |
| 21607 | 1081 | C1081Q | D | 3.3e-09 | 1.1e-08 | 2.6e-08 | 1.343E-08 | 1.154E-08 | 3.300E-09 | 2.600E-08 |
| 21608 | 1081 | C1081G | D | 3.7e-09 | 6.4e-09 | 3.6e-09 | 4.567E-09 | 1.589E-09 | 3.600E-09 | 6.400E-09 |
| 21609 | 1081 | C1081H | D | 1.4e-08 | 4.5e-09 | 6.2e-09 | 8.233E-09 | 5.066E-09 | 4.500E-09 | 1.400E-08 |
| 21610 | 1081 | C1081I | D | 6.9e-09 | 1.3e-08 | 9.3e-09 | 9.733E-09 | 3.073E-09 | 6.900E-09 | 1.300E-08 |
| 21611 | 1081 | C1081L | D | 7.3e-09 | 1.1e-08 | 9.2e-09 | 9.167E-09 | 1.850E-09 | 7.300E-09 | 1.100E-08 |
| 21612 | 1081 | C1081K | D | 1e-08 | 1.3e-08 | 1e-08 | 1.100E-08 | 1.732E-09 | 1.000E-08 | 1.300E-08 |
| 21613 | 1081 | C1081M | D | 1.5e-08 | 6.2e-09 | 6.1e-09 | 9.100E-09 | 5.110E-09 | 6.100E-09 | 1.500E-08 |
| 21614 | 1081 | C1081F | D | 1.9e-08 | 5.5e-09 | 4.5e-09 | 9.667E-09 | 8.098E-09 | 4.500E-09 | 1.900E-08 |
| 21615 | 1081 | C1081P | D | 5.3e-09 | 2e-08 | 3.4e-09 | 9.567E-09 | 9.085E-09 | 3.400E-09 | 2.000E-08 |
| 21616 | 1081 | C1081S | D | 2.4e-08 | 1e-08 | 7.1e-09 | 1.370E-08 | 9.037E-09 | 7.100E-09 | 2.400E-08 |
| 21617 | 1081 | C1081T | D | 1.6e-08 | 8.1e-09 | 8.7e-09 | 1.093E-08 | 4.398E-09 | 8.100E-09 | 1.600E-08 |
| 21618 | 1081 | C1081W | D | 4.9e-09 | 1.1e-08 | 6.7e-09 | 7.533E-09 | 3.134E-09 | 4.900E-09 | 1.100E-08 |
| 21619 | 1081 | C1081Y | D | 6.5e-09 | 1.9e-09 | 7.9e-09 | 5.433E-09 | 3.139E-09 | 1.900E-09 | 7.900E-09 |
| 21620 | 1081 | C1081V | D | 2.1e-08 | 6.1e-09 | 2e-09 | 9.700E-09 | 9.998E-09 | 2.000E-09 | 2.100E-08 |
| 21621 | 1082 | R1082A | D | 3e-09 | 6.7e-09 | 4.8e-09 | 4.833E-09 | 1.850E-09 | 3.000E-09 | 6.700E-09 |
| 21622 | 1082 | R1082R | D | 2e-08 | 2.4e-09 | 1.5e-08 | 1.247E-08 | 9.069E-09 | 2.400E-09 | 2.000E-08 |
| 21623 | 1082 | R1082N | D | 1.4e-09 | 7.5e-09 | 9.9e-09 | 6.267E-09 | 4.382E-09 | 1.400E-09 | 9.900E-09 |
| 21624 | 1082 | R1082D | D | 8.7e-09 | 5.3e-09 | 8.4e-09 | 7.467E-09 | 1.882E-09 | 5.300E-09 | 8.700E-09 |
| 21625 | 1082 | R1082C | D | 8.6e-09 | 9.8e-09 | 7.8e-09 | 8.733E-09 | 1.007E-09 | 7.800E-09 | 9.800E-09 |
| 21626 | 1082 | R1082E | D | 4.5e-09 | 6.6e-09 | 9.9e-09 | 7.000E-09 | 2.722E-09 | 4.500E-09 | 9.900E-09 |
| 21627 | 1082 | R1082Q | D | 3.5e-09 | 1.6e-08 | 6.9e-09 | 8.800E-09 | 6.463E-09 | 3.500E-09 | 1.600E-08 |
| 21628 | 1082 | R1082G | D | 2e-09 | 4.9e-09 | 9.1e-09 | 5.333E-09 | 3.570E-09 | 2.000E-09 | 9.100E-09 |
| 21629 | 1082 | R1082H | D | 4.9e-09 | 1.3e-08 | 1.6e-08 | 1.130E-08 | 5.742E-09 | 4.900E-09 | 1.600E-08 |
| 21630 | 1082 | R1082I | D | 2e-08 | 5.9e-09 | 3.6e-09 | 9.833E-09 | 8.879E-09 | 3.600E-09 | 2.000E-08 |
| 21631 | 1082 | R1082L | D | 1.3e-08 | 3.3e-09 | 1.3e-08 | 9.767E-09 | 5.600E-09 | 3.300E-09 | 1.300E-08 |
| 21632 | 1082 | R1082K | D | 7e-09 | 9.4e-09 | 5.7e-09 | 7.367E-09 | 1.877E-09 | 5.700E-09 | 9.400E-09 |
| 21633 | 1082 | R1082M | D | 2e-08 | 1.1e-08 | 4.4e-09 | 1.180E-08 | 7.831E-09 | 4.400E-09 | 2.000E-08 |
| 21634 | 1082 | R1082F | D | 7.6e-09 | 3.5e-09 | 5.9e-09 | 5.667E-09 | 2.060E-09 | 3.500E-09 | 7.600E-09 |
| 21635 | 1082 | R1082P | D | 6.9e-09 | 2.2e-09 | 5e-09 | 4.700E-09 | 2.364E-09 | 2.200E-09 | 6.900E-09 |
| 21636 | 1082 | R1082S | D | 1.3e-08 | 8e-09 | 7.5e-09 | 9.500E-09 | 3.041E-09 | 7.500E-09 | 1.300E-08 |
| 21637 | 1082 | R1082T | D | 5.8e-09 | 3.1e-09 | 1.1e-08 | 6.633E-09 | 4.015E-09 | 3.100E-09 | 1.100E-08 |
| 21638 | 1082 | R1082W | D | 4.2e-09 | 1.6e-08 | 8.5e-09 | 9.567E-09 | 5.972E-09 | 4.200E-09 | 1.600E-08 |
| 21639 | 1082 | R1082Y | D | 7.8e-09 | 1.4e-08 | 3.1e-09 | 8.300E-09 | 5.467E-09 | 3.100E-09 | 1.400E-08 |
| 21640 | 1082 | R1082V | D | 1.3e-08 | 1.3e-09 | 4.8e-09 | 6.367E-09 | 6.005E-09 | 1.300E-09 | 1.300E-08 |
| 21641 | 1083 | A1083A | D | 3.8e-09 | 3.9e-09 | 3.7e-09 | 3.800E-09 | 1.000E-10 | 3.700E-09 | 3.900E-09 |
| 21642 | 1083 | A1083R | D | 9.7e-09 | 6.3e-09 | 1.3e-08 | 9.667E-09 | 3.350E-09 | 6.300E-09 | 1.300E-08 |
| 21643 | 1083 | A1083N | D | 1.3e-08 | 7e-09 | 5.4e-09 | 8.467E-09 | 4.007E-09 | 5.400E-09 | 1.300E-08 |
| 21644 | 1083 | A1083D | D | 5.3e-09 | 6.4e-09 | 2.1e-08 | 1.090E-08 | 8.764E-09 | 5.300E-09 | 2.100E-08 |
| 21645 | 1083 | A1083C | D | 5.3e-09 | 1.1e-08 | 1.5e-08 | 1.043E-08 | 4.875E-09 | 5.300E-09 | 1.500E-08 |
| 21646 | 1083 | A1083E | D | 1.2e-08 | 3.5e-09 | 9.3e-09 | 8.267E-09 | 4.343E-09 | 3.500E-09 | 1.200E-08 |
| 21647 | 1083 | A1083Q | D | 4.1e-09 | 1.9e-08 | 1.1e-08 | 1.137E-08 | 7.457E-09 | 4.100E-09 | 1.900E-08 |
| 21648 | 1083 | A1083G | D | 9.6e-09 | 6.7e-09 | 9.3e-09 | 8.533E-09 | 1.595E-09 | 6.700E-09 | 9.600E-09 |
| 21649 | 1083 | A1083H | D | 7e-09 | 1.8e-09 | 5.4e-09 | 4.733E-09 | 2.663E-09 | 1.800E-09 | 7.000E-09 |
| 21650 | 1083 | A1083I | D | 3.3e-09 | 1e-08 | 1.1e-08 | 8.100E-09 | 4.187E-09 | 3.300E-09 | 1.100E-08 |
| 21651 | 1083 | A1083L | D | 1.2e-08 | 6.1e-09 | 1.5e-08 | 1.103E-08 | 4.528E-09 | 6.100E-09 | 1.500E-08 |
| 21652 | 1083 | A1083K | D | 1.6e-08 | 1.5e-08 | 7.6e-09 | 1.287E-08 | 4.588E-09 | 7.600E-09 | 1.600E-08 |
| 21653 | 1083 | A1083M | D | 6.7e-09 | 3.4e-09 | 1.1e-08 | 7.033E-09 | 3.811E-09 | 3.400E-09 | 1.100E-08 |
| 21654 | 1083 | A1083F | D | 8.6e-09 | 6.8e-09 | 1.8e-08 | 1.113E-08 | 6.014E-09 | 6.800E-09 | 1.800E-08 |
| 21655 | 1083 | A1083P | D | 6.9e-09 | 4.6e-09 | 9e-09 | 6.833E-09 | 2.201E-09 | 4.600E-09 | 9.000E-09 |
| 21656 | 1083 | A1083S | D | 1e-08 | 1.3e-08 | 4.9e-09 | 9.300E-09 | 4.095E-09 | 4.900E-09 | 1.300E-08 |
| 21657 | 1083 | A1083T | D | 4.7e-09 | 2.1e-08 | 1.3e-08 | 1.290E-08 | 8.150E-09 | 4.700E-09 | 2.100E-08 |
| 21658 | 1083 | A1083W | D | 1.6e-08 | 1.3e-08 | 7e-09 | 1.200E-08 | 4.583E-09 | 7.000E-09 | 1.600E-08 |
| 21659 | 1083 | A1083Y | D | 7.6e-09 | 5.3e-09 | 3.5e-09 | 5.467E-09 | 2.055E-09 | 3.500E-09 | 7.600E-09 |
| 21660 | 1083 | A1083V | D | 1.9e-08 | 4.1e-09 | 1.1e-08 | 1.137E-08 | 7.457E-09 | 4.100E-09 | 1.900E-08 |
| 21661 | 1084 | S1084A | D | 1.3e-08 | 2.4e-08 | 1.1e-08 | 1.600E-08 | 7.000E-09 | 1.100E-08 | 2.400E-08 |
| 21662 | 1084 | S1084R | D | 6.8e-09 | 4.1e-09 | 1.1e-08 | 7.300E-09 | 3.477E-09 | 4.100E-09 | 1.100E-08 |
| 21663 | 1084 | S1084N | D | 7.1e-09 | 1.1e-08 | 9.8e-09 | 9.300E-09 | 1.997E-09 | 7.100E-09 | 1.100E-08 |
| 21664 | 1084 | S1084D | D | 4.1e-09 | 4.8e-09 | 7.5e-09 | 5.467E-09 | 1.795E-09 | 4.100E-09 | 7.500E-09 |
| 21665 | 1084 | S1084C | D | 1.1e-08 | 2e-08 | 3.6e-09 | 1.153E-08 | 8.213E-09 | 3.600E-09 | 2.000E-08 |
| 21666 | 1084 | S1084E | D | 7.3e-09 | 8.1e-09 | 5.3e-09 | 6.900E-09 | 1.442E-09 | 5.300E-09 | 8.100E-09 |
| 21667 | 1084 | S1084Q | D | 6.2e-09 | 3.2e-09 | 1.1e-08 | 6.800E-09 | 3.934E-09 | 3.200E-09 | 1.100E-08 |
| 21668 | 1084 | S1084G | D | 8.2e-09 | 2.7e-09 | 5.2e-09 | 5.367E-09 | 2.754E-09 | 2.700E-09 | 8.200E-09 |
| 21669 | 1084 | S1084H | D | 1.2e-08 | 8.9e-09 | 6.7e-09 | 9.200E-09 | 2.663E-09 | 6.700E-09 | 1.200E-08 |
| 21670 | 1084 | S1084I | D | 6.8e-09 | 2.6e-08 | 1.2e-08 | 1.493E-08 | 9.930E-09 | 6.800E-09 | 2.600E-08 |
| 21671 | 1084 | S1084L | D | 1e-08 | 2.1e-09 | 9.4e-09 | 7.167E-09 | 4.398E-09 | 2.100E-09 | 1.000E-08 |
| 21672 | 1084 | S1084K | D | 1.5e-08 | 4.1e-09 | 7.1e-09 | 8.733E-09 | 5.631E-09 | 4.100E-09 | 1.500E-08 |
| 21673 | 1084 | S1084M | D | 1.7e-08 | 4.4e-09 | 1.5e-08 | 1.213E-08 | 6.772E-09 | 4.400E-09 | 1.700E-08 |
| 21674 | 1084 | S1084F | D | 3.6e-09 | 1.4e-08 | 2e-09 | 6.533E-09 | 6.516E-09 | 2.000E-09 | 1.400E-08 |

| | | | | | | | | | | |
|---|---|---|---|---|---|---|---|---|---|---|
| 21675 | 1084 | S1084P | D | 2.6e-09 | 1.6e-08 | 9e-09 | 9.200E-09 | 6.702E-09 | 2.600E-09 | 1.600E-08 |
| 21676 | 1084 | S1084S | D | 1.6e-08 | 1.5e-08 | 2.1e-08 | 1.733E-08 | 3.215E-09 | 1.500E-08 | 2.100E-08 |
| 21677 | 1084 | S1084T | D | 3.4e-09 | 2.4e-09 | 1.2e-08 | 5.933E-09 | 5.278E-09 | 2.400E-09 | 1.200E-08 |
| 21678 | 1084 | S1084W | D | 5.3e-09 | 1.9e-08 | 5.6e-09 | 9.967E-09 | 7.825E-09 | 5.300E-09 | 1.900E-08 |
| 21679 | 1084 | S1084Y | D | 1.5e-08 | 1.5e-08 | 1.5e-08 | 1.500E-08 | 0.000E+00 | 1.500E-08 | 1.500E-08 |
| 21680 | 1084 | S1084V | D | 2.2e-08 | 1.1e-08 | 3.8e-08 | 2.367E-08 | 1.358E-08 | 1.100E-08 | 3.800E-08 |
| 21681 | 1085 | Q1085A | D | 1.1e-08 | 5e-09 | 4.3e-09 | 6.767E-09 | 3.683E-09 | 4.300E-09 | 1.100E-08 |
| 21682 | 1085 | Q1085R | D | 1.4e-08 | 8.9e-09 | 6.5e-09 | 9.800E-09 | 3.830E-09 | 6.500E-09 | 1.400E-08 |
| 21683 | 1085 | Q1085N | D | 4.5e-09 | 6.4e-09 | 1.3e-08 | 7.967E-09 | 4.461E-09 | 4.500E-09 | 1.300E-08 |
| 21684 | 1085 | Q1085D | D | 8.1e-09 | 9e-09 | 4.8e-09 | 7.300E-09 | 2.211E-09 | 4.800E-09 | 9.000E-09 |
| 21685 | 1085 | Q1085C | D | 4.6e-09 | 7.7e-09 | 8.1e-09 | 6.800E-09 | 1.916E-09 | 4.600E-09 | 8.100E-09 |
| 21686 | 1085 | Q1085E | D | 6.2e-09 | 7e-09 | 6.1e-09 | 6.433E-09 | 4.933E-10 | 6.100E-09 | 7.000E-09 |
| 21687 | 1085 | Q1085Q | D | 1.2e-08 | 5.8e-09 | 2.1e-08 | 1.293E-08 | 7.643E-09 | 5.800E-09 | 2.100E-08 |
| 21688 | 1085 | Q1085G | D | 1.8e-08 | 6.6e-09 | 5.4e-09 | 1.000E-08 | 6.954E-09 | 5.400E-09 | 1.800E-08 |
| 21689 | 1085 | Q1085H | D | 1.1e-08 | 3.1e-09 | 2.6e-09 | 5.567E-09 | 4.712E-09 | 2.600E-09 | 1.100E-08 |
| 21690 | 1085 | Q1085I | D | 4.5e-09 | 7.7e-09 | 1e-08 | 7.400E-09 | 2.762E-09 | 4.500E-09 | 1.000E-08 |
| 21691 | 1085 | Q1085L | D | 3.4e-09 | 1e-08 | 5.7e-09 | 6.367E-09 | 3.350E-09 | 3.400E-09 | 1.000E-08 |
| 21692 | 1085 | Q1085K | D | 5.3e-09 | 7.7e-09 | 2.3e-09 | 5.100E-09 | 2.706E-09 | 2.300E-09 | 7.700E-09 |
| 21693 | 1085 | Q1085M | D | 1.9e-08 | 1.6e-09 | 8.3e-09 | 9.633E-09 | 8.776E-09 | 1.600E-09 | 1.900E-08 |
| 21694 | 1085 | Q1085F | D | 4.3e-09 | 1.1e-08 | 1.3e-09 | 5.533E-09 | 4.966E-09 | 1.300E-09 | 1.100E-08 |
| 21695 | 1085 | Q1085P | D | 2e-08 | 3.1e-08 | 8e-09 | 1.967E-08 | 1.150E-08 | 8.000E-09 | 3.100E-08 |
| 21696 | 1085 | Q1085S | D | 3e-09 | 1.4e-08 | 2.8e-09 | 6.600E-09 | 6.409E-09 | 2.800E-09 | 1.400E-08 |
| 21697 | 1085 | Q1085T | D | 1e-08 | 6.7e-09 | 7.5e-09 | 8.067E-09 | 1.721E-09 | 6.700E-09 | 1.000E-08 |
| 21698 | 1085 | Q1085W | D | 9.7e-09 | 1.9e-08 | 5.8e-09 | 1.150E-08 | 6.782E-09 | 5.800E-09 | 1.900E-08 |
| 21699 | 1085 | Q1085Y | D | 1.2e-08 | 5.5e-09 | 3.6e-09 | 7.033E-09 | 4.405E-09 | 3.600E-09 | 1.200E-08 |
| 21700 | 1085 | Q1085V | D | 1.9e-08 | 4.7e-09 | 5.2e-09 | 9.633E-09 | 8.116E-09 | 4.700E-09 | 1.900E-08 |
| 21701 | 1086 | D1086A | D | 1.3e-08 | 1.6e-08 | 8.6e-09 | 1.253E-08 | 3.722E-09 | 8.600E-09 | 1.600E-08 |
| 21702 | 1086 | D1086R | D | 1.3e-08 | 3.3e-09 | 4.6e-09 | 6.967E-09 | 5.265E-09 | 3.300E-09 | 1.300E-08 |
| 21703 | 1086 | D1086N | D | 1.2e-08 | 1.4e-08 | 1.3e-08 | 1.300E-08 | 1.000E-09 | 1.200E-08 | 1.400E-08 |
| 21704 | 1086 | D1086D | D | 5e-09 | 1.4e-08 | 6.3e-09 | 8.433E-09 | 4.864E-09 | 5.000E-09 | 1.400E-08 |
| 21705 | 1086 | D1086C | D | 2.5e-08 | 6.4e-09 | 7.6e-09 | 1.300E-08 | 1.041E-08 | 6.400E-09 | 2.500E-08 |
| 21706 | 1086 | D1086E | D | 7.7e-09 | 1.1e-08 | 9.8e-09 | 9.500E-09 | 1.670E-09 | 7.700E-09 | 1.100E-08 |
| 21707 | 1086 | D1086Q | D | 9e-09 | 3.5e-09 | 4.3e-09 | 5.600E-09 | 2.972E-09 | 3.500E-09 | 9.000E-09 |
| 21708 | 1086 | D1086G | D | 6e-09 | 8.9e-09 | 2.1e-09 | 5.667E-09 | 3.412E-09 | 2.100E-09 | 8.900E-09 |
| 21709 | 1086 | D1086H | D | 1.1e-08 | 7.8e-09 | 4.8e-09 | 7.867E-09 | 3.101E-09 | 4.800E-09 | 1.100E-08 |
| 21710 | 1086 | D1086I | D | 6.2e-09 | 9.8e-09 | 4.3e-09 | 6.767E-09 | 2.793E-09 | 4.300E-09 | 9.800E-09 |
| 21711 | 1086 | D1086L | D | 5.8e-09 | 6.6e-09 | 8.4e-09 | 6.933E-09 | 1.332E-09 | 5.800E-09 | 8.400E-09 |
| 21712 | 1086 | D1086K | D | 3.6e-09 | 8.8e-09 | 5.1e-09 | 5.833E-09 | 2.676E-09 | 3.600E-09 | 8.800E-09 |
| 21713 | 1086 | D1086M | D | 1.2e-08 | 1.2e-08 | 1.4e-08 | 1.267E-08 | 1.155E-09 | 1.200E-08 | 1.400E-08 |
| 21714 | 1086 | D1086F | D | 6.5e-09 | 1e-08 | 7.7e-09 | 8.067E-09 | 1.779E-09 | 6.500E-09 | 1.000E-08 |
| 21715 | 1086 | D1086P | D | 1.1e-08 | 8.4e-09 | 1.3e-08 | 1.080E-08 | 2.307E-09 | 8.400E-09 | 1.300E-08 |
| 21716 | 1086 | D1086S | D | 5.2e-09 | 7.4e-09 | 4.4e-09 | 5.667E-09 | 1.553E-09 | 4.400E-09 | 7.400E-09 |
| 21717 | 1086 | D1086T | D | 3e-09 | 1.5e-08 | 2.1e-08 | 1.300E-08 | 9.165E-09 | 3.000E-09 | 2.100E-08 |
| 21718 | 1086 | D1086W | D | 7.2e-09 | 6.1e-09 | 1e-08 | 7.767E-09 | 2.011E-09 | 6.100E-09 | 1.000E-08 |
| 21719 | 1086 | D1086Y | D | 2e-08 | 5.4e-09 | 6.7e-09 | 1.070E-08 | 8.080E-09 | 5.400E-09 | 2.000E-08 |
| 21720 | 1086 | D1086V | D | 1.5e-08 | 2.2e-08 | 8.3e-09 | 1.510E-08 | 6.851E-09 | 8.300E-09 | 2.200E-08 |
| 21721 | 1087 | V1087A | D | 1.1e-08 | 1.1e-08 | 5.7e-09 | 9.233E-09 | 3.060E-09 | 5.700E-09 | 1.100E-08 |
| 21722 | 1087 | V1087R | D | 9.7e-09 | 4e-09 | 7.3e-09 | 7.000E-09 | 2.862E-09 | 4.000E-09 | 9.700E-09 |
| 21723 | 1087 | V1087N | D | 1e-08 | 5e-09 | 1.4e-08 | 9.667E-09 | 4.509E-09 | 5.000E-09 | 1.400E-08 |
| 21724 | 1087 | V1087D | D | 5.9e-09 | 5.8e-09 | 5.2e-09 | 5.633E-09 | 3.786E-10 | 5.200E-09 | 5.900E-09 |
| 21725 | 1087 | V1087C | D | 1.2e-08 | 5.2e-09 | 3.5e-09 | 6.900E-09 | 4.498E-09 | 3.500E-09 | 1.200E-08 |
| 21726 | 1087 | V1087E | D | 4.1e-09 | 1e-08 | 3.9e-09 | 6.000E-09 | 3.466E-09 | 3.900E-09 | 1.000E-08 |
| 21727 | 1087 | V1087Q | D | 2e-08 | 2.7e-09 | 7e-10 | 7.800E-09 | 1.061E-08 | 7.000E-10 | 2.000E-08 |
| 21728 | 1087 | V1087G | D | 6.9e-09 | 7.7e-09 | 1.7e-08 | 1.053E-08 | 5.615E-09 | 6.900E-09 | 1.700E-08 |
| 21729 | 1087 | V1087H | D | 1.2e-08 | 3.9e-09 | 5.7e-09 | 7.200E-09 | 4.253E-09 | 3.900E-09 | 1.200E-08 |
| 21730 | 1087 | V1087I | D | 5.7e-09 | 1.2e-08 | 3.7e-09 | 7.133E-09 | 4.332E-09 | 3.700E-09 | 1.200E-08 |
| 21731 | 1087 | V1087L | D | 1.2e-08 | 2.1e-09 | 5.9e-09 | 6.667E-09 | 4.994E-09 | 2.100E-09 | 1.200E-08 |
| 21732 | 1087 | V1087K | D | 6.8e-09 | 6.6e-09 | 4.8e-09 | 6.067E-09 | 1.102E-09 | 4.800E-09 | 6.800E-09 |
| 21733 | 1087 | V1087M | D | 4.1e-09 | 1.2e-08 | 5.7e-09 | 7.267E-09 | 4.177E-09 | 4.100E-09 | 1.200E-08 |
| 21734 | 1087 | V1087F | D | 4e-09 | 8.3e-09 | 2.7e-09 | 5.000E-09 | 2.931E-09 | 2.700E-09 | 8.300E-09 |
| 21735 | 1087 | V1087P | D | 4.8e-09 | 6e-09 | 7.1e-09 | 5.967E-09 | 1.150E-09 | 4.800E-09 | 7.100E-09 |
| 21736 | 1087 | V1087S | D | 2.7e-09 | 7.3e-09 | 8.3e-09 | 6.100E-09 | 2.987E-09 | 2.700E-09 | 8.300E-09 |
| 21737 | 1087 | V1087T | D | 2e-09 | 1.1e-08 | 3.3e-09 | 5.433E-09 | 4.864E-09 | 2.000E-09 | 1.100E-08 |
| 21738 | 1087 | V1087W | D | 9.8e-09 | 4.2e-09 | 1.2e-08 | 8.667E-09 | 4.022E-09 | 4.200E-09 | 1.200E-08 |
| 21739 | 1087 | V1087Y | D | 6.3e-09 | 1.5e-08 | 1.2e-08 | 1.110E-08 | 4.419E-09 | 6.300E-09 | 1.500E-08 |
| 21740 | 1087 | V1087V | D | 7e-09 | 3.7e-08 | 2.3e-09 | 1.543E-08 | 1.882E-08 | 2.300E-09 | 3.700E-08 |
| 21741 | 1088 | N1088A | D | 1.5e-08 | 9.1e-09 | 4.5e-09 | 9.533E-09 | 5.263E-09 | 4.500E-09 | 1.500E-08 |
| 21742 | 1088 | N1088R | D | 1e-08 | 9.7e-09 | 8.5e-09 | 9.400E-09 | 7.937E-10 | 8.500E-09 | 1.000E-08 |
| 21743 | 1088 | N1088N | D | 6.3e-09 | 9.1e-09 | 1e-08 | 8.467E-09 | 1.930E-09 | 6.300E-09 | 1.000E-08 |
| 21744 | 1088 | N1088D | D | 1.4e-08 | 1.5e-08 | 6.3e-09 | 1.177E-08 | 4.761E-09 | 6.300E-09 | 1.500E-08 |
| 21745 | 1088 | N1088C | D | 4.1e-09 | 1.3e-08 | 1.3e-08 | 1.003E-08 | 5.138E-09 | 4.100E-09 | 1.300E-08 |
| 21746 | 1088 | N1088E | D | 4.4e-09 | 1.8e-08 | 9.9e-09 | 1.077E-08 | 6.841E-09 | 4.400E-09 | 1.800E-08 |
| 21747 | 1088 | N1088Q | D | 7.7e-09 | 1.4e-08 | 2.6e-08 | 1.590E-08 | 9.297E-09 | 7.700E-09 | 2.600E-08 |
| 21748 | 1088 | N1088G | D | 6.1e-09 | 5.9e-09 | 1.6e-08 | 9.333E-09 | 5.774E-09 | 5.900E-09 | 1.600E-08 |
| 21749 | 1088 | N1088H | D | 3.4e-09 | 4.6e-09 | 1.1e-08 | 6.333E-09 | 4.086E-09 | 3.400E-09 | 1.100E-08 |
| 21750 | 1088 | N1088I | D | 7.2e-09 | 1e-08 | 4.6e-09 | 7.267E-09 | 2.701E-09 | 4.600E-09 | 1.000E-08 |
| 21751 | 1088 | N1088L | D | 3.6e-09 | 4.7e-09 | 1.3e-08 | 7.100E-09 | 5.139E-09 | 3.600E-09 | 1.300E-08 |
| 21752 | 1088 | N1088K | D | 1.5e-09 | 1e-08 | 7.7e-09 | 6.400E-09 | 4.397E-09 | 1.500E-09 | 1.000E-08 |
| 21753 | 1088 | N1088M | D | 7.1e-09 | 6.8e-09 | 1.3e-08 | 8.967E-09 | 3.496E-09 | 6.800E-09 | 1.300E-08 |
| 21754 | 1088 | N1088F | D | 7.1e-09 | 3.6e-09 | 8.5e-09 | 6.400E-09 | 2.524E-09 | 3.600E-09 | 8.500E-09 |
| 21755 | 1088 | N1088P | D | 2.6e-09 | 9.4e-09 | 1e-08 | 7.333E-09 | 4.110E-09 | 2.600E-09 | 1.000E-08 |
| 21756 | 1088 | N1088S | D | 5.1e-09 | 1.1e-08 | 6.8e-09 | 7.633E-09 | 3.037E-09 | 5.100E-09 | 1.100E-08 |
| 21757 | 1088 | N1088T | D | 6e-09 | 1.1e-08 | 1.5e-08 | 1.067E-08 | 4.509E-09 | 6.000E-09 | 1.500E-08 |
| 21758 | 1088 | N1088W | D | 4.8e-09 | 1.4e-08 | 8.7e-09 | 9.167E-09 | 4.618E-09 | 4.800E-09 | 1.400E-08 |
| 21759 | 1088 | N1088Y | D | 9.6e-09 | 2.1e-08 | 1.1e-08 | 1.387E-08 | 6.217E-09 | 9.600E-09 | 2.100E-08 |

| | | | | | | | | | | |
|---|---|---|---|---|---|---|---|---|---|---|
| 21760 | 1088 | N1088V | D | 1.6e-08 | 1.3e-08 | 8.1e-09 | 1.237E-08 | 3.988E-09 | 8.100E-09 | 1.600E-08 |
| 21761 | 1089 | T1089A | D | 7.9e-09 | 1.3e-08 | 8.5e-09 | 9.800E-09 | 2.787E-09 | 7.900E-09 | 1.300E-08 |
| 21762 | 1089 | T1089R | D | 7.3e-09 | 2.6e-09 | 3.1e-09 | 4.333E-09 | 2.581E-09 | 2.600E-09 | 7.300E-09 |
| 21763 | 1089 | T1089N | D | 8.1e-09 | 5.1e-09 | 9.9e-09 | 7.700E-09 | 2.425E-09 | 5.100E-09 | 9.900E-09 |
| 21764 | 1089 | T1089D | D | 1.3e-08 | 1.1e-08 | 3.7e-09 | 9.233E-09 | 4.895E-09 | 3.700E-09 | 1.300E-08 |
| 21765 | 1089 | T1089C | D | 3.3e-09 | 9.8e-09 | 2.9e-09 | 5.333E-09 | 3.873E-09 | 2.900E-09 | 9.800E-09 |
| 21766 | 1089 | T1089E | D | 7.8e-09 | 1e-08 | 3.6e-09 | 7.133E-09 | 3.252E-09 | 3.600E-09 | 1.000E-08 |
| 21767 | 1089 | T1089Q | D | 1.1e-08 | 3.6e-09 | 5.6e-09 | 6.733E-09 | 3.828E-09 | 3.600E-09 | 1.100E-08 |
| 21768 | 1089 | T1089G | D | 7.6e-09 | 2.2e-09 | 8.4e-09 | 6.067E-09 | 3.372E-09 | 2.200E-09 | 8.400E-09 |
| 21769 | 1089 | T1089H | D | 1.1e-08 | 2.2e-09 | 2.6e-09 | 5.267E-09 | 4.969E-09 | 2.200E-09 | 1.100E-08 |
| 21770 | 1089 | T1089I | D | 3.4e-09 | 4e-09 | 1.4e-08 | 7.133E-09 | 5.954E-09 | 3.400E-09 | 1.400E-08 |
| 21771 | 1089 | T1089L | D | 3.3e-09 | 6.7e-09 | 8.9e-09 | 6.300E-09 | 2.821E-09 | 3.300E-09 | 8.900E-09 |
| 21772 | 1089 | T1089K | D | 4.9e-09 | 2e-08 | 4.9e-09 | 9.933E-09 | 8.718E-09 | 4.900E-09 | 2.000E-08 |
| 21773 | 1089 | T1089M | D | 4.4e-09 | 2.3e-08 | 9e-09 | 1.213E-08 | 9.688E-09 | 4.400E-09 | 2.300E-08 |
| 21774 | 1089 | T1089F | D | 6.5e-09 | 7e-09 | 3.3e-09 | 5.600E-09 | 2.007E-09 | 3.300E-09 | 7.000E-09 |
| 21775 | 1089 | T1089P | D | 5.7e-09 | 5e-09 | 2.5e-08 | 1.190E-08 | 1.135E-08 | 5.000E-09 | 2.500E-08 |
| 21776 | 1089 | T1089S | D | 2.4e-09 | 4.7e-09 | 1.3e-08 | 6.700E-09 | 5.576E-09 | 2.400E-09 | 1.300E-08 |
| 21777 | 1089 | T1089T | D | 2.4e-08 | 4.6e-09 | 1.9e-08 | 1.587E-08 | 1.007E-08 | 4.600E-09 | 2.400E-08 |
| 21778 | 1089 | T1089W | D | 1.1e-08 | 1e-08 | 5.2e-09 | 8.733E-09 | 3.101E-09 | 5.200E-09 | 1.100E-08 |
| 21779 | 1089 | T1089Y | D | 7.1e-09 | 1.5e-08 | 4.5e-09 | 8.867E-09 | 5.468E-09 | 4.500E-09 | 1.500E-08 |
| 21780 | 1089 | T1089V | D | 7.8e-09 | 7.4e-09 | 4.7e-09 | 6.633E-09 | 1.686E-09 | 4.700E-09 | 7.800E-09 |
| 21781 | 1090 | A1090A | D | 4.5e-09 | 4.9e-09 | 9.2e-09 | 6.200E-09 | 2.606E-09 | 4.500E-09 | 9.200E-09 |
| 21782 | 1090 | A1090R | D | 1.4e-08 | 1.9e-09 | 1.4e-08 | 9.967E-09 | 6.986E-09 | 1.900E-09 | 1.400E-08 |
| 21783 | 1090 | A1090N | D | 3.4e-09 | 1.8e-08 | 6.2e-09 | 9.200E-09 | 7.749E-09 | 3.400E-09 | 1.800E-08 |
| 21784 | 1090 | A1090D | D | 5.6e-09 | 3.8e-09 | 4.6e-09 | 4.667E-09 | 9.018E-10 | 3.800E-09 | 5.600E-09 |
| 21785 | 1090 | A1090C | D | 6.4e-09 | 1.5e-08 | 5.6e-09 | 9.000E-09 | 5.212E-09 | 5.600E-09 | 1.500E-08 |
| 21786 | 1090 | A1090E | D | 1.1e-08 | 1.6e-08 | 4.4e-09 | 1.047E-08 | 5.818E-09 | 4.400E-09 | 1.600E-08 |
| 21787 | 1090 | A1090Q | D | 1.5e-08 | 6.6e-09 | 1.7e-08 | 1.287E-08 | 5.518E-09 | 6.600E-09 | 1.700E-08 |
| 21788 | 1090 | A1090G | D | 1.1e-08 | 5.2e-09 | 2.2e-08 | 1.273E-08 | 8.533E-09 | 5.200E-09 | 2.200E-08 |
| 21789 | 1090 | A1090H | D | 7.9e-09 | 3.4e-09 | 2.7e-09 | 4.667E-09 | 2.822E-09 | 2.700E-09 | 7.900E-09 |
| 21790 | 1090 | A1090I | D | 3.5e-09 | 5.3e-09 | 1.4e-08 | 7.600E-09 | 5.615E-09 | 3.500E-09 | 1.400E-08 |
| 21791 | 1090 | A1090L | D | 1.4e-08 | 1.2e-08 | 6.5e-09 | 1.083E-08 | 3.884E-09 | 6.500E-09 | 1.400E-08 |
| 21792 | 1090 | A1090K | D | 1e-08 | 1.1e-08 | 3.5e-09 | 8.167E-09 | 4.072E-09 | 3.500E-09 | 1.100E-08 |
| 21793 | 1090 | A1090M | D | 3e-09 | 7.6e-09 | 1.6e-08 | 8.867E-09 | 6.592E-09 | 3.000E-09 | 1.600E-08 |
| 21794 | 1090 | A1090F | D | 1.3e-08 | 5.2e-09 | 1.7e-09 | 6.633E-09 | 5.785E-09 | 1.700E-09 | 1.300E-08 |
| 21795 | 1090 | A1090P | D | 3.9e-09 | 8.7e-09 | 1.3e-08 | 8.533E-09 | 4.552E-09 | 3.900E-09 | 1.300E-08 |
| 21796 | 1090 | A1090S | D | 3.1e-09 | 5.1e-09 | 1.7e-08 | 8.400E-09 | 7.515E-09 | 3.100E-09 | 1.700E-08 |
| 21797 | 1090 | A1090T | D | 7.5e-09 | 1.9e-08 | 9.5e-09 | 1.200E-08 | 6.144E-09 | 7.500E-09 | 1.900E-08 |
| 21798 | 1090 | A1090W | D | 7.8e-09 | 1.3e-08 | 4.9e-09 | 8.567E-09 | 4.104E-09 | 4.900E-09 | 1.300E-08 |
| 21799 | 1090 | A1090Y | D | 1e-08 | 1.3e-08 | 1.1e-08 | 1.133E-08 | 1.528E-09 | 1.000E-08 | 1.300E-08 |
| 21800 | 1090 | A1090V | D | 9e-09 | 1.5e-08 | 3.9e-09 | 9.300E-09 | 5.556E-09 | 3.900E-09 | 1.500E-08 |
| 21801 | 1091 | V1091A | D | 6.1e-09 | 1.1e-08 | 6e-09 | 7.700E-09 | 2.858E-09 | 6.000E-09 | 1.100E-08 |
| 21802 | 1091 | V1091R | D | 2.4e-09 | 2.3e-08 | 2.2e-08 | 1.580E-08 | 1.162E-08 | 2.400E-09 | 2.300E-08 |
| 21803 | 1091 | V1091N | D | 5e-09 | 6.8e-09 | 8.6e-09 | 6.800E-09 | 1.800E-09 | 5.000E-09 | 8.600E-09 |
| 21804 | 1091 | V1091D | D | 5e-09 | 2.5e-09 | 4.5e-09 | 4.000E-09 | 1.323E-09 | 2.500E-09 | 5.000E-09 |
| 21805 | 1091 | V1091C | D | 5.1e-09 | 7e-09 | 9.8e-09 | 7.300E-09 | 2.364E-09 | 5.100E-09 | 9.800E-09 |
| 21806 | 1091 | V1091E | D | 5.5e-09 | 4.7e-09 | 1.1e-08 | 7.067E-09 | 3.430E-09 | 4.700E-09 | 1.100E-08 |
| 21807 | 1091 | V1091Q | D | 8.6e-09 | 1.3e-08 | 1.2e-08 | 1.120E-08 | 2.307E-09 | 8.600E-09 | 1.300E-08 |
| 21808 | 1091 | V1091G | D | 4.6e-09 | 2.2e-09 | 3.6e-09 | 3.467E-09 | 1.206E-09 | 2.200E-09 | 4.600E-09 |
| 21809 | 1091 | V1091H | D | 1.4e-08 | 7.9e-09 | 6.2e-09 | 9.367E-09 | 4.102E-09 | 6.200E-09 | 1.400E-08 |
| 21810 | 1091 | V1091I | D | 2e-08 | 5.4e-09 | 1.4e-08 | 1.313E-08 | 7.338E-09 | 5.400E-09 | 2.000E-08 |
| 21811 | 1091 | V1091L | D | 5.1e-09 | 2.3e-08 | 1.4e-08 | 1.403E-08 | 8.950E-09 | 5.100E-09 | 2.300E-08 |
| 21812 | 1091 | V1091K | D | 1.4e-08 | 1.5e-08 | 6.2e-09 | 1.173E-08 | 4.818E-09 | 6.200E-09 | 1.500E-08 |
| 21813 | 1091 | V1091M | D | 4.3e-09 | 8.9e-09 | 4.2e-09 | 5.800E-09 | 2.685E-09 | 4.200E-09 | 8.900E-09 |
| 21814 | 1091 | V1091F | D | 1.6e-08 | 2.2e-08 | 6.1e-09 | 1.470E-08 | 8.029E-09 | 6.100E-09 | 2.200E-08 |
| 21815 | 1091 | V1091P | D | 1.6e-08 | 1.6e-08 | 5.7e-09 | 1.257E-08 | 5.947E-09 | 5.700E-09 | 1.600E-08 |
| 21816 | 1091 | V1091S | D | 1.3e-08 | 1.1e-08 | 4.8e-09 | 9.600E-09 | 4.276E-09 | 4.800E-09 | 1.300E-08 |
| 21817 | 1091 | V1091T | D | 6.3e-09 | 2.7e-09 | 5.4e-09 | 4.800E-09 | 1.873E-09 | 2.700E-09 | 6.300E-09 |
| 21818 | 1091 | V1091W | D | 7.7e-09 | 3.5e-09 | 6.7e-09 | 5.967E-09 | 2.194E-09 | 3.500E-09 | 7.700E-09 |
| 21819 | 1091 | V1091Y | D | 1.1e-08 | 8.2e-09 | 5.5e-09 | 8.233E-09 | 2.750E-09 | 5.500E-09 | 1.100E-08 |
| 21820 | 1091 | V1091V | D | 7e-09 | 4e-09 | 3.1e-09 | 4.700E-09 | 2.042E-09 | 3.100E-09 | 7.000E-09 |
| 21821 | 1092 | A1092A | D | 8.2e-09 | 5.6e-09 | 6.7e-09 | 6.833E-09 | 1.305E-09 | 5.600E-09 | 8.200E-09 |
| 21822 | 1092 | A1092R | D | 4.4e-09 | 7.3e-09 | 1.5e-08 | 8.900E-09 | 5.478E-09 | 4.400E-09 | 1.500E-08 |
| 21823 | 1092 | A1092N | D | 2.3e-09 | 3.4e-09 | 1.1e-08 | 5.567E-09 | 4.737E-09 | 2.300E-09 | 1.100E-08 |
| 21824 | 1092 | A1092D | D | 5.1e-09 | 6.5e-09 | 3.5e-09 | 5.033E-09 | 1.501E-09 | 3.500E-09 | 6.500E-09 |
| 21825 | 1092 | A1092C | D | 1.4e-08 | 8.5e-09 | 4.2e-09 | 8.900E-09 | 4.912E-09 | 4.200E-09 | 1.400E-08 |
| 21826 | 1092 | A1092E | D | 1e-08 | 1.3e-08 | 4e-08 | 2.100E-08 | 1.652E-08 | 1.000E-08 | 4.000E-08 |
| 21827 | 1092 | A1092Q | D | 4.2e-09 | 2.1e-09 | 3.9e-09 | 3.400E-09 | 1.136E-09 | 2.100E-09 | 4.200E-09 |
| 21828 | 1092 | A1092G | D | 4.6e-09 | 8.3e-09 | 9.4e-09 | 7.433E-09 | 2.515E-09 | 4.600E-09 | 9.400E-09 |
| 21829 | 1092 | A1092H | D | 2.7e-09 | 5.3e-09 | 1.8e-08 | 8.667E-09 | 8.187E-09 | 2.700E-09 | 1.800E-08 |
| 21830 | 1092 | A1092I | D | 5.5e-09 | 3.9e-09 | 4.7e-09 | 4.700E-09 | 8.000E-10 | 3.900E-09 | 5.500E-09 |
| 21831 | 1092 | A1092L | D | 1.1e-08 | 6.6e-09 | 3.9e-09 | 7.167E-09 | 3.584E-09 | 3.900E-09 | 1.100E-08 |
| 21832 | 1092 | A1092K | D | 1.1e-08 | 2.4e-08 | 1.3e-08 | 1.600E-08 | 7.000E-09 | 1.100E-08 | 2.400E-08 |
| 21833 | 1092 | A1092M | D | 1.4e-08 | 9.6e-09 | 5.4e-09 | 9.667E-09 | 4.300E-09 | 5.400E-09 | 1.400E-08 |
| 21834 | 1092 | A1092F | D | 3.8e-09 | 9.1e-09 | 2.7e-08 | 1.330E-08 | 1.216E-08 | 3.800E-09 | 2.700E-08 |
| 21835 | 1092 | A1092P | D | 5.8e-09 | 6.5e-09 | 5.9e-09 | 6.067E-09 | 3.786E-10 | 5.800E-09 | 6.500E-09 |
| 21836 | 1092 | A1092S | D | 8e-09 | 2.7e-09 | 1e-08 | 6.900E-09 | 3.772E-09 | 2.700E-09 | 1.000E-08 |
| 21837 | 1092 | A1092T | D | 1e-08 | 7.2e-09 | 7e-09 | 8.067E-09 | 1.677E-09 | 7.000E-09 | 1.000E-08 |
| 21838 | 1092 | A1092W | D | 1.1e-08 | 6.1e-09 | 4.7e-09 | 7.267E-09 | 3.308E-09 | 4.700E-09 | 1.100E-08 |
| 21839 | 1092 | A1092Y | D | 2.9e-08 | 3.3e-09 | 3.2e-09 | 1.183E-08 | 1.487E-08 | 3.200E-09 | 2.900E-08 |
| 21840 | 1092 | A1092V | D | 6.1e-09 | 1.7e-08 | 1e-08 | 1.103E-08 | 5.523E-09 | 6.100E-09 | 1.700E-08 |
| 21841 | 1093 | W1093A | D | 2.2e-08 | 2e-08 | 9.2e-09 | 1.707E-08 | 6.886E-09 | 9.200E-09 | 2.200E-08 |
| 21842 | 1093 | W1093R | D | 6.4e-09 | 5.7e-09 | 1.6e-08 | 9.367E-09 | 5.755E-09 | 5.700E-09 | 1.600E-08 |
| 21843 | 1093 | W1093N | D | 1.9e-09 | 3.9e-09 | 6.1e-09 | 3.967E-09 | 2.101E-09 | 1.900E-09 | 6.100E-09 |
| 21844 | 1093 | W1093D | D | 3.4e-09 | 5e-09 | 4.9e-09 | 4.433E-09 | 8.963E-10 | 3.400E-09 | 5.000E-09 |

| | | | | | | | | | | |
|---|---|---|---|---|---|---|---|---|---|---|
| 21845 | 1093 | W1093C | D | 8.1e-09 | 3.7e-09 | 5.8e-09 | 5.867E-09 | 2.201E-09 | 3.700E-09 | 8.100E-09 |
| 21846 | 1093 | W1093E | D | 2.3e-09 | 1.5e-08 | 1.4e-08 | 1.043E-08 | 7.061E-09 | 2.300E-09 | 1.500E-08 |
| 21847 | 1093 | W1093Q | D | 8.6e-09 | 7.3e-09 | 9.8e-09 | 8.567E-09 | 1.250E-09 | 7.300E-09 | 9.800E-09 |
| 21848 | 1093 | W1093G | D | 8.9e-09 | 5.2e-09 | 1.8e-08 | 1.070E-08 | 6.587E-09 | 5.200E-09 | 1.800E-08 |
| 21849 | 1093 | W1093H | D | 9.6e-09 | 6.4e-09 | 9.9e-09 | 8.633E-09 | 1.940E-09 | 6.400E-09 | 9.900E-09 |
| 21850 | 1093 | W1093I | D | 5e-09 | 5.1e-09 | 3.2e-09 | 4.433E-09 | 1.069E-09 | 3.200E-09 | 5.100E-09 |
| 21851 | 1093 | W1093L | D | 6.8e-09 | 7.7e-09 | 1.1e-08 | 8.500E-09 | 2.211E-09 | 6.800E-09 | 1.100E-08 |
| 21852 | 1093 | W1093K | D | 5.3e-09 | 6.2e-09 | 5.5e-09 | 5.667E-09 | 4.726E-10 | 5.300E-09 | 6.200E-09 |
| 21853 | 1093 | W1093M | D | 6.3e-09 | 4.1e-09 | 9.4e-09 | 6.600E-09 | 2.663E-09 | 4.100E-09 | 9.400E-09 |
| 21854 | 1093 | W1093F | D | 6.6e-09 | 7.1e-09 | 9.7e-09 | 7.800E-09 | 1.664E-09 | 6.600E-09 | 9.700E-09 |
| 21855 | 1093 | W1093P | D | 7.4e-09 | 1e-08 | 4e-09 | 7.133E-09 | 3.009E-09 | 4.000E-09 | 1.000E-08 |
| 21856 | 1093 | W1093S | D | 3e-09 | 9.6e-09 | 1e-08 | 7.533E-09 | 3.931E-09 | 3.000E-09 | 1.000E-08 |
| 21857 | 1093 | W1093T | D | 1.2e-08 | 7.8e-09 | 5.1e-09 | 8.300E-09 | 3.477E-09 | 5.100E-09 | 1.200E-08 |
| 21858 | 1093 | W1093W | D | 1.1e-08 | 1.8e-08 | 7.3e-09 | 1.210E-08 | 5.434E-09 | 7.300E-09 | 1.800E-08 |
| 21859 | 1093 | W1093Y | D | 7.4e-09 | 7.1e-09 | 7.2e-09 | 7.233E-09 | 1.528E-10 | 7.100E-09 | 7.400E-09 |
| 21860 | 1093 | W1093V | D | 1.5e-08 | 1.5e-08 | 8.4e-09 | 1.280E-08 | 3.811E-09 | 8.400E-09 | 1.500E-08 |
| 21861 | 1094 | Y1094A | D | 8.8e-09 | 7.6e-09 | 2.2e-09 | 6.200E-09 | 3.516E-09 | 2.200E-09 | 8.800E-09 |
| 21862 | 1094 | Y1094R | D | 8.2e-09 | 1e-08 | 3.3e-09 | 7.167E-09 | 3.467E-09 | 3.300E-09 | 1.000E-08 |
| 21863 | 1094 | Y1094N | D | 8.6e-09 | 3e-08 | 3.8e-09 | 1.413E-08 | 1.395E-08 | 3.800E-09 | 3.000E-08 |
| 21864 | 1094 | Y1094D | D | 1.2e-08 | 5.7e-09 | 4.1e-09 | 7.267E-09 | 4.177E-09 | 4.100E-09 | 1.200E-08 |
| 21865 | 1094 | Y1094C | D | 7e-09 | 6.7e-09 | 4.4e-09 | 6.033E-09 | 1.422E-09 | 4.400E-09 | 7.000E-09 |
| 21866 | 1094 | Y1094E | D | 8.3e-09 | 5.2e-09 | 1.4e-08 | 9.167E-09 | 4.464E-09 | 5.200E-09 | 1.400E-08 |
| 21867 | 1094 | Y1094Q | D | 1.3e-08 | 1.1e-08 | 3.1e-09 | 9.033E-09 | 5.235E-09 | 3.100E-09 | 1.300E-08 |
| 21868 | 1094 | Y1094G | D | 5.4e-09 | 1.1e-08 | 8.6e-09 | 8.333E-09 | 2.810E-09 | 5.400E-09 | 1.100E-08 |
| 21869 | 1094 | Y1094H | D | 9.7e-09 | 5.4e-09 | 8.3e-09 | 7.800E-09 | 2.193E-09 | 5.400E-09 | 9.700E-09 |
| 21870 | 1094 | Y1094I | D | 4.2e-09 | 2.8e-08 | 1e-08 | 1.407E-08 | 1.241E-08 | 4.200E-09 | 2.800E-08 |
| 21871 | 1094 | Y1094L | D | 9.3e-09 | 7.7e-09 | 1.8e-09 | 6.267E-09 | 3.950E-09 | 1.800E-09 | 9.300E-09 |
| 21872 | 1094 | Y1094K | D | 1.3e-08 | 5.6e-09 | 6.7e-09 | 8.433E-09 | 3.993E-09 | 5.600E-09 | 1.300E-08 |
| 21873 | 1094 | Y1094M | D | 2.4e-09 | 9e-09 | 6.4e-09 | 5.933E-09 | 3.325E-09 | 2.400E-09 | 9.000E-09 |
| 21874 | 1094 | Y1094F | D | 4e-09 | 1.5e-09 | 4.9e-09 | 3.467E-09 | 1.762E-09 | 1.500E-09 | 4.900E-09 |
| 21875 | 1094 | Y1094P | D | 8.2e-09 | 9.1e-09 | 1.1e-08 | 9.433E-09 | 1.429E-09 | 8.200E-09 | 1.100E-08 |
| 21876 | 1094 | Y1094S | D | 2e-09 | 8.9e-09 | 1.2e-08 | 7.633E-09 | 5.119E-09 | 2.000E-09 | 1.200E-08 |
| 21877 | 1094 | Y1094T | D | 3.1e-09 | 4.2e-09 | 2e-09 | 3.100E-09 | 1.100E-09 | 2.000E-09 | 4.200E-09 |
| 21878 | 1094 | Y1094W | D | 1.4e-08 | 6.1e-09 | 1.2e-08 | 1.070E-08 | 4.107E-09 | 6.100E-09 | 1.400E-08 |
| 21879 | 1094 | Y1094Y | D | 1.5e-09 | 5.5e-09 | 1.2e-08 | 6.333E-09 | 5.299E-09 | 1.500E-09 | 1.200E-08 |
| 21880 | 1094 | Y1094V | D | 5.6e-09 | 1.5e-08 | 6.6e-09 | 9.067E-09 | 5.163E-09 | 5.600E-09 | 1.500E-08 |
| 21881 | 1095 | Q1095A | D | 1.3e-08 | 7.1e-10 | 6.1e-09 | 6.603E-09 | 6.160E-09 | 7.100E-10 | 1.300E-08 |
| 21882 | 1095 | Q1095R | D | 6.4e-09 | 1.1e-08 | 6.1e-09 | 7.833E-09 | 2.747E-09 | 6.100E-09 | 1.100E-08 |
| 21883 | 1095 | Q1095N | D | 1.4e-08 | 8.7e-09 | 6.3e-09 | 9.667E-09 | 3.940E-09 | 6.300E-09 | 1.400E-08 |
| 21884 | 1095 | Q1095D | D | 4.3e-09 | 2.7e-09 | 4.7e-09 | 3.900E-09 | 1.058E-09 | 2.700E-09 | 4.700E-09 |
| 21885 | 1095 | Q1095C | D | 8.5e-09 | 9e-09 | 1.2e-08 | 9.833E-09 | 1.893E-09 | 8.500E-09 | 1.200E-08 |
| 21886 | 1095 | Q1095E | D | 6.9e-09 | 6.8e-09 | 1.2e-08 | 8.567E-09 | 2.974E-09 | 6.800E-09 | 1.200E-08 |
| 21887 | 1095 | Q1095Q | D | 3.3e-09 | 6.3e-09 | 7.3e-09 | 5.633E-09 | 2.082E-09 | 3.300E-09 | 7.300E-09 |
| 21888 | 1095 | Q1095G | D | 6e-09 | 8.2e-09 | 1.1e-08 | 8.400E-09 | 2.506E-09 | 6.000E-09 | 1.100E-08 |
| 21889 | 1095 | Q1095H | D | 8.1e-09 | 7.1e-09 | 4.6e-09 | 6.600E-09 | 1.803E-09 | 4.600E-09 | 8.100E-09 |
| 21890 | 1095 | Q1095I | D | 6.4e-09 | 1.2e-08 | 1.1e-08 | 9.800E-09 | 2.987E-09 | 6.400E-09 | 1.200E-08 |
| 21891 | 1095 | Q1095L | D | 6.3e-09 | 7e-09 | 2.6e-09 | 5.300E-09 | 2.364E-09 | 2.600E-09 | 7.000E-09 |
| 21892 | 1095 | Q1095K | D | 5.6e-09 | 4.7e-09 | 2.1e-08 | 1.043E-08 | 9.162E-09 | 4.700E-09 | 2.100E-08 |
| 21893 | 1095 | Q1095M | D | 8.1e-09 | 6.3e-09 | 3.5e-09 | 5.967E-09 | 2.318E-09 | 3.500E-09 | 8.100E-09 |
| 21894 | 1095 | Q1095F | D | 4.6e-09 | 9.2e-09 | 5.7e-09 | 6.500E-09 | 2.402E-09 | 4.600E-09 | 9.200E-09 |
| 21895 | 1095 | Q1095P | D | 1.5e-08 | 1.4e-08 | 3.2e-09 | 1.073E-08 | 6.543E-09 | 3.200E-09 | 1.500E-08 |
| 21896 | 1095 | Q1095S | D | 1.1e-08 | 3.1e-09 | 1.2e-08 | 8.700E-09 | 4.875E-09 | 3.100E-09 | 1.200E-08 |
| 21897 | 1095 | Q1095T | D | 6.2e-09 | 4.4e-09 | 7.4e-09 | 6.000E-09 | 1.510E-09 | 4.400E-09 | 7.400E-09 |
| 21898 | 1095 | Q1095W | D | 1.2e-08 | 1.2e-08 | 6.9e-09 | 1.030E-08 | 2.944E-09 | 6.900E-09 | 1.200E-08 |
| 21899 | 1095 | Q1095Y | D | 1.3e-08 | 8.2e-09 | 9.8e-09 | 1.033E-08 | 2.444E-09 | 8.200E-09 | 1.300E-08 |
| 21900 | 1095 | Q1095V | D | 5.6e-09 | 1.6e-08 | 2.9e-08 | 1.687E-08 | 1.172E-08 | 5.600E-09 | 2.900E-08 |
| 21901 | 1096 | Q1096A | D | 1.1e-08 | 7.4e-09 | 5.9e-09 | 8.100E-09 | 2.621E-09 | 5.900E-09 | 1.100E-08 |
| 21902 | 1096 | Q1096R | D | 1.1e-08 | 1e-08 | 1.3e-08 | 1.133E-08 | 1.528E-09 | 1.000E-08 | 1.300E-08 |
| 21903 | 1096 | Q1096N | D | 3.7e-09 | 5.9e-09 | 1.6e-08 | 8.533E-09 | 6.559E-09 | 3.700E-09 | 1.600E-08 |
| 21904 | 1096 | Q1096D | D | 1e-08 | 7e-09 | 8.7e-09 | 8.567E-09 | 1.504E-09 | 7.000E-09 | 1.000E-08 |
| 21905 | 1096 | Q1096C | D | 9.2e-09 | 1.2e-08 | 7.6e-09 | 9.600E-09 | 2.227E-09 | 7.600E-09 | 1.200E-08 |
| 21906 | 1096 | Q1096E | D | 6.3e-09 | 1.1e-08 | 1.8e-08 | 1.177E-08 | 5.888E-09 | 6.300E-09 | 1.800E-08 |
| 21907 | 1096 | Q1096Q | D | 5.6e-09 | 8e-09 | 1.1e-08 | 8.200E-09 | 2.706E-09 | 5.600E-09 | 1.100E-08 |
| 21908 | 1096 | Q1096G | D | 1.3e-08 | 5.5e-09 | 3e-09 | 7.167E-09 | 5.204E-09 | 3.000E-09 | 1.300E-08 |
| 21909 | 1096 | Q1096H | D | 6.5e-09 | 1.4e-09 | 2.7e-08 | 1.163E-08 | 1.355E-08 | 1.400E-09 | 2.700E-08 |
| 21910 | 1096 | Q1096I | D | 1.8e-08 | 7.4e-09 | 1.1e-08 | 1.213E-08 | 5.390E-09 | 7.400E-09 | 1.800E-08 |
| 21911 | 1096 | Q1096L | D | 5.7e-09 | 2.4e-08 | 1.6e-08 | 1.523E-08 | 9.174E-09 | 5.700E-09 | 2.400E-08 |
| 21912 | 1096 | Q1096K | D | 2.3e-08 | 3.3e-09 | 1.7e-09 | 9.333E-09 | 1.186E-08 | 1.700E-09 | 2.300E-08 |
| 21913 | 1096 | Q1096M | D | 7.4e-09 | 1.4e-08 | 1.9e-08 | 1.347E-08 | 5.818E-09 | 7.400E-09 | 1.900E-08 |
| 21914 | 1096 | Q1096F | D | 7.5e-09 | 6.3e-09 | 6.6e-09 | 6.800E-09 | 6.245E-10 | 6.300E-09 | 7.500E-09 |
| 21915 | 1096 | Q1096P | D | 7.4e-09 | 1.3e-08 | 8e-09 | 9.467E-09 | 3.075E-09 | 7.400E-09 | 1.300E-08 |
| 21916 | 1096 | Q1096S | D | 4.5e-09 | 9.7e-09 | 1.3e-08 | 9.067E-09 | 4.285E-09 | 4.500E-09 | 1.300E-08 |
| 21917 | 1096 | Q1096T | D | 1.1e-08 | 1.2e-08 | 4.6e-09 | 9.200E-09 | 4.015E-09 | 4.600E-09 | 1.200E-08 |
| 21918 | 1096 | Q1096W | D | 4e-09 | 1.2e-08 | 6.4e-09 | 7.467E-09 | 4.105E-09 | 4.000E-09 | 1.200E-08 |
| 21919 | 1096 | Q1096Y | D | 1.2e-08 | 7.8e-09 | 1.1e-08 | 1.027E-08 | 2.194E-09 | 7.800E-09 | 1.200E-08 |
| 21920 | 1096 | Q1096V | D | 1.7e-08 | 5.9e-09 | 1.8e-09 | 8.233E-09 | 7.864E-09 | 1.800E-09 | 1.700E-08 |
| 21921 | 1097 | K1097A | D | 4.2e-09 | 8.4e-09 | 3e-09 | 5.200E-09 | 2.835E-09 | 3.000E-09 | 8.400E-09 |
| 21922 | 1097 | K1097R | D | 1.2e-08 | 5.4e-09 | 6.3e-09 | 7.900E-09 | 3.579E-09 | 5.400E-09 | 1.200E-08 |
| 21923 | 1097 | K1097N | D | 4.8e-09 | 6.6e-09 | 6.8e-09 | 6.067E-09 | 1.102E-09 | 4.800E-09 | 6.800E-09 |
| 21924 | 1097 | K1097D | D | 2e-09 | 7.1e-09 | 4e-09 | 4.367E-09 | 2.570E-09 | 2.000E-09 | 7.100E-09 |
| 21925 | 1097 | K1097C | D | 2.9e-09 | 1.5e-08 | 2.6e-09 | 6.833E-09 | 7.074E-09 | 2.600E-09 | 1.500E-08 |
| 21926 | 1097 | K1097E | D | 2.4e-09 | 6.2e-09 | 5.5e-09 | 4.700E-09 | 2.022E-09 | 2.400E-09 | 6.200E-09 |
| 21927 | 1097 | K1097Q | D | 3.4e-09 | 1.1e-08 | 4.5e-09 | 6.300E-09 | 4.107E-09 | 3.400E-09 | 1.100E-08 |
| 21928 | 1097 | K1097G | D | 2.6e-08 | 9.1e-09 | 5.5e-09 | 1.353E-08 | 1.095E-08 | 5.500E-09 | 2.600E-08 |
| 21929 | 1097 | K1097H | D | 3.5e-09 | 1.3e-08 | 2e-09 | 6.167E-09 | 5.965E-09 | 2.000E-09 | 1.300E-08 |

| | | | | | | | | | | |
|---|---|---|---|---|---|---|---|---|---|---|
| 21930 | 1097 | K1097I | D | 6.3e-09 | 1.1e-08 | 6.9e-09 | 8.067E-09 | 2.558E-09 | 6.300E-09 | 1.100E-08 |
| 21931 | 1097 | K1097L | D | 1.1e-08 | 1.7e-08 | 9.2e-09 | 1.240E-08 | 4.084E-09 | 9.200E-09 | 1.700E-08 |
| 21932 | 1097 | K1097K | D | 9.7e-09 | 5.7e-09 | 6.9e-09 | 7.433E-09 | 2.053E-09 | 5.700E-09 | 9.700E-09 |
| 21933 | 1097 | K1097M | D | 1e-08 | 6.8e-09 | 6.2e-09 | 7.667E-09 | 2.043E-09 | 6.200E-09 | 1.000E-08 |
| 21934 | 1097 | K1097F | D | 7e-09 | 2.9e-09 | 7.3e-09 | 5.733E-09 | 2.458E-09 | 2.900E-09 | 7.300E-09 |
| 21935 | 1097 | K1097P | D | 4.2e-09 | 3.8e-09 | 5.2e-09 | 4.400E-09 | 7.211E-10 | 3.800E-09 | 5.200E-09 |
| 21936 | 1097 | K1097S | D | 1.1e-08 | 1.1e-08 | 1e-08 | 1.067E-08 | 5.774E-10 | 1.000E-08 | 1.100E-08 |
| 21937 | 1097 | K1097T | D | 8.3e-09 | 5.3e-09 | 7.6e-09 | 7.067E-09 | 1.570E-09 | 5.300E-09 | 8.300E-09 |
| 21938 | 1097 | K1097W | D | 1.6e-08 | 2.3e-08 | 4.9e-09 | 1.463E-08 | 9.127E-09 | 4.900E-09 | 2.300E-08 |
| 21939 | 1097 | K1097Y | D | 7.8e-09 | 1.3e-08 | 6e-09 | 8.933E-09 | 3.635E-09 | 6.000E-09 | 1.300E-08 |
| 21940 | 1097 | K1097V | D | 4.8e-09 | 9e-09 | 1.7e-08 | 1.027E-08 | 6.198E-09 | 4.800E-09 | 1.700E-08 |
| 21941 | 1098 | P1098A | D | 5.5e-09 | 1.3e-08 | 8e-09 | 8.833E-09 | 3.819E-09 | 5.500E-09 | 1.300E-08 |
| 21942 | 1098 | P1098R | D | 1.1e-08 | 1e-08 | 3.2e-09 | 8.067E-09 | 4.244E-09 | 3.200E-09 | 1.100E-08 |
| 21943 | 1098 | P1098N | D | 5.9e-09 | 4.8e-09 | 3.2e-09 | 4.633E-09 | 1.358E-09 | 3.200E-09 | 5.900E-09 |
| 21944 | 1098 | P1098D | D | 7.8e-09 | 4.5e-09 | 8.8e-09 | 7.033E-09 | 2.250E-09 | 4.500E-09 | 8.800E-09 |
| 21945 | 1098 | P1098C | D | 9.1e-09 | 7.5e-09 | 3.5e-09 | 6.700E-09 | 2.884E-09 | 3.500E-09 | 9.100E-09 |
| 21946 | 1098 | P1098E | D | 2.9e-09 | 6.2e-09 | 9.5e-09 | 6.200E-09 | 3.300E-09 | 2.900E-09 | 9.500E-09 |
| 21947 | 1098 | P1098Q | D | 6e-09 | 5.6e-09 | 2.2e-09 | 4.600E-09 | 2.088E-09 | 2.200E-09 | 6.000E-09 |
| 21948 | 1098 | P1098G | D | 9.9e-09 | 2.1e-08 | 6.2e-09 | 1.237E-08 | 7.702E-09 | 6.200E-09 | 2.100E-08 |
| 21949 | 1098 | P1098H | D | 4.6e-09 | 7.1e-09 | 4.3e-09 | 5.333E-09 | 1.537E-09 | 4.300E-09 | 7.100E-09 |
| 21950 | 1098 | P1098I | D | 2.4e-08 | 1.6e-08 | 4.4e-09 | 1.480E-08 | 9.855E-09 | 4.400E-09 | 2.400E-08 |
| 21951 | 1098 | P1098L | D | 1.6e-09 | 8.9e-09 | 8.3e-09 | 6.267E-09 | 4.053E-09 | 1.600E-09 | 8.900E-09 |
| 21952 | 1098 | P1098K | D | 5.3e-09 | 5.3e-09 | 2.1e-08 | 1.053E-08 | 9.064E-09 | 5.300E-09 | 2.100E-08 |
| 21953 | 1098 | P1098M | D | 5.9e-09 | 1e-08 | 1.6e-08 | 1.063E-08 | 5.080E-09 | 5.900E-09 | 1.600E-08 |
| 21954 | 1098 | P1098F | D | 2.3e-09 | 5e-09 | 1.3e-08 | 6.767E-09 | 5.564E-09 | 2.300E-09 | 1.300E-08 |
| 21955 | 1098 | P1098P | D | 1.5e-08 | 1.1e-08 | 4.2e-09 | 1.007E-08 | 5.460E-09 | 4.200E-09 | 1.500E-08 |
| 21956 | 1098 | P1098S | D | 1.2e-08 | 1.5e-08 | 1e-08 | 1.233E-08 | 2.517E-09 | 1.000E-08 | 1.500E-08 |
| 21957 | 1098 | P1098T | D | 3.2e-09 | 4.8e-09 | 4.7e-09 | 4.233E-09 | 8.963E-10 | 3.200E-09 | 4.800E-09 |
| 21958 | 1098 | P1098W | D | 6.3e-09 | 1.6e-08 | 6.5e-09 | 9.600E-09 | 5.543E-09 | 6.300E-09 | 1.600E-08 |
| 21959 | 1098 | P1098Y | D | 7.1e-09 | 1e-08 | 7.9e-09 | 8.333E-09 | 1.498E-09 | 7.100E-09 | 1.000E-08 |
| 21960 | 1098 | P1098V | D | 1.5e-09 | 1.3e-09 | 7.4e-09 | 3.400E-09 | 3.466E-09 | 1.300E-09 | 7.400E-09 |
| 21961 | 1099 | G1099A | D | 6.2e-09 | 6.3e-09 | 6.9e-09 | 6.467E-09 | 3.786E-10 | 6.200E-09 | 6.900E-09 |
| 21962 | 1099 | G1099R | D | 3.3e-09 | 4e-09 | 8.8e-09 | 5.367E-09 | 2.994E-09 | 3.300E-09 | 8.800E-09 |
| 21963 | 1099 | G1099N | D | 4e-09 | 5e-09 | 1.3e-08 | 7.333E-09 | 4.933E-09 | 4.000E-09 | 1.300E-08 |
| 21964 | 1099 | G1099D | D | 4.2e-09 | 7.8e-09 | 3.6e-09 | 5.200E-09 | 2.272E-09 | 3.600E-09 | 7.800E-09 |
| 21965 | 1099 | G1099C | D | 2.7e-09 | 1.1e-08 | 2.6e-08 | 1.323E-08 | 1.181E-08 | 2.700E-09 | 2.600E-08 |
| 21966 | 1099 | G1099E | D | 6.3e-09 | 6.9e-09 | 5.9e-09 | 6.367E-09 | 5.033E-10 | 5.900E-09 | 6.900E-09 |
| 21967 | 1099 | G1099Q | D | 1.2e-08 | 3.6e-09 | 1e-08 | 8.533E-09 | 4.388E-09 | 3.600E-09 | 1.200E-08 |
| 21968 | 1099 | G1099G | D | 1.5e-08 | 9.4e-09 | 7e-09 | 1.047E-08 | 4.105E-09 | 7.000E-09 | 1.500E-08 |
| 21969 | 1099 | G1099H | D | 5.6e-09 | 9.1e-09 | 3.9e-09 | 6.200E-09 | 2.651E-09 | 3.900E-09 | 9.100E-09 |
| 21970 | 1099 | G1099I | D | 7.3e-09 | 5.7e-09 | 5.8e-09 | 6.267E-09 | 8.963E-10 | 5.700E-09 | 7.300E-09 |
| 21971 | 1099 | G1099L | D | 1.2e-08 | 5.7e-09 | 3.6e-09 | 7.100E-09 | 4.371E-09 | 3.600E-09 | 1.200E-08 |
| 21972 | 1099 | G1099K | D | 9.8e-09 | 1.9e-08 | 3.7e-09 | 1.083E-08 | 7.702E-09 | 3.700E-09 | 1.900E-08 |
| 21973 | 1099 | G1099M | D | 4.9e-09 | 1.3e-08 | 6.8e-09 | 8.233E-09 | 4.236E-09 | 4.900E-09 | 1.300E-08 |
| 21974 | 1099 | G1099F | D | 4.8e-09 | 4.9e-09 | 3.6e-09 | 4.433E-09 | 7.234E-10 | 3.600E-09 | 4.900E-09 |
| 21975 | 1099 | G1099P | D | 1.2e-08 | 2.8e-09 | 9.1e-09 | 7.967E-09 | 4.704E-09 | 2.800E-09 | 1.200E-08 |
| 21976 | 1099 | G1099S | D | 4.1e-09 | 3.9e-09 | 8.8e-09 | 5.600E-09 | 2.773E-09 | 3.900E-09 | 8.800E-09 |
| 21977 | 1099 | G1099T | D | 1.6e-08 | 2.2e-09 | 3.8e-09 | 7.333E-09 | 7.548E-09 | 2.200E-09 | 1.600E-08 |
| 21978 | 1099 | G1099W | D | 6.2e-09 | 5.8e-09 | 7.8e-09 | 6.600E-09 | 1.058E-09 | 5.800E-09 | 7.800E-09 |
| 21979 | 1099 | G1099Y | D | 3.8e-09 | 2.9e-09 | 2.2e-09 | 2.967E-09 | 8.021E-10 | 2.200E-09 | 3.800E-09 |
| 21980 | 1099 | G1099V | D | 1.8e-08 | 5.5e-09 | 4.7e-09 | 9.400E-09 | 7.459E-09 | 4.700E-09 | 1.800E-08 |
| 21981 | 1100 | K1100A | D | 7.9e-09 | 4.7e-09 | 6.9e-09 | 6.500E-09 | 1.637E-09 | 4.700E-09 | 7.900E-09 |
| 21982 | 1100 | K1100R | D | 1.4e-08 | 4.5e-09 | 4.2e-09 | 7.567E-09 | 5.573E-09 | 4.200E-09 | 1.400E-08 |
| 21983 | 1100 | K1100N | D | 1.8e-08 | 1.7e-08 | 1.5e-08 | 1.667E-08 | 1.528E-09 | 1.500E-08 | 1.800E-08 |
| 21984 | 1100 | K1100D | D | 3.2e-09 | 8.1e-09 | 4.7e-09 | 5.333E-09 | 2.511E-09 | 3.200E-09 | 8.100E-09 |
| 21985 | 1100 | K1100C | D | 7.6e-09 | 1.9e-08 | 7.4e-09 | 1.133E-08 | 6.640E-09 | 7.400E-09 | 1.900E-08 |
| 21986 | 1100 | K1100E | D | 4.8e-09 | 8.3e-09 | 1.6e-08 | 9.700E-09 | 5.730E-09 | 4.800E-09 | 1.600E-08 |
| 21987 | 1100 | K1100Q | D | 7.7e-09 | 4.6e-09 | 3.9e-09 | 5.400E-09 | 2.022E-09 | 3.900E-09 | 7.700E-09 |
| 21988 | 1100 | K1100G | D | 2.4e-09 | 7.2e-09 | 1.9e-08 | 9.533E-09 | 8.542E-09 | 2.400E-09 | 1.900E-08 |
| 21989 | 1100 | K1100H | D | 8.2e-09 | 1.3e-09 | 2e-09 | 3.833E-09 | 3.798E-09 | 1.300E-09 | 8.200E-09 |
| 21990 | 1100 | K1100I | D | 4.2e-09 | 4.2e-09 | 3.9e-09 | 4.100E-09 | 1.732E-10 | 3.900E-09 | 4.200E-09 |
| 21991 | 1100 | K1100L | D | 2.4e-09 | 3.1e-09 | 1.8e-08 | 7.833E-09 | 8.812E-09 | 2.400E-09 | 1.800E-08 |
| 21992 | 1100 | K1100K | D | 5.5e-09 | 2.9e-09 | 9.1e-09 | 5.833E-09 | 3.113E-09 | 2.900E-09 | 9.100E-09 |
| 21993 | 1100 | K1100M | D | 7.5e-09 | 1.2e-08 | 1.4e-08 | 1.117E-08 | 3.329E-09 | 7.500E-09 | 1.400E-08 |
| 21994 | 1100 | K1100F | D | 1.4e-08 | 7.5e-09 | 1.3e-09 | 7.600E-09 | 6.351E-09 | 1.300E-09 | 1.400E-08 |
| 21995 | 1100 | K1100P | D | 1.5e-09 | 9.5e-09 | 7.3e-09 | 6.100E-09 | 4.133E-09 | 1.500E-09 | 9.500E-09 |
| 21996 | 1100 | K1100S | D | 1.1e-08 | 5.9e-09 | 6.2e-09 | 7.700E-09 | 2.862E-09 | 5.900E-09 | 1.100E-08 |
| 21997 | 1100 | K1100T | D | 8.2e-09 | 6e-09 | 3.8e-09 | 6.000E-09 | 2.200E-09 | 3.800E-09 | 8.200E-09 |
| 21998 | 1100 | K1100W | D | 6e-09 | 1e-08 | 6.7e-09 | 7.567E-09 | 2.136E-09 | 6.000E-09 | 1.000E-08 |
| 21999 | 1100 | K1100Y | D | 3.5e-08 | 7.3e-09 | 3.1e-09 | 1.513E-08 | 1.733E-08 | 3.100E-09 | 3.500E-08 |
| 22000 | 1100 | K1100V | D | 5.2e-09 | 6.3e-09 | 9.3e-09 | 6.933E-09 | 2.122E-09 | 5.200E-09 | 9.300E-09 |
| 22001 | 1101 | A1101A | D | 8.1e-09 | 2.1e-09 | 1.1e-08 | 7.067E-09 | 4.539E-09 | 2.100E-09 | 1.100E-08 |
| 22002 | 1101 | A1101R | D | 5.8e-09 | 8.4e-09 | 8.8e-09 | 7.667E-09 | 1.629E-09 | 5.800E-09 | 8.800E-09 |
| 22003 | 1101 | A1101N | D | 1.6e-08 | 1.1e-08 | 1.1e-08 | 1.267E-08 | 2.887E-09 | 1.100E-08 | 1.600E-08 |
| 22004 | 1101 | A1101D | D | 1.5e-08 | 1.4e-08 | 6e-09 | 1.167E-08 | 4.933E-09 | 6.000E-09 | 1.500E-08 |
| 22005 | 1101 | A1101C | D | 4.8e-09 | 3.9e-09 | 2.4e-09 | 3.700E-09 | 1.212E-09 | 2.400E-09 | 4.800E-09 |
| 22006 | 1101 | A1101E | D | 5.6e-09 | 1e-08 | 9.6e-09 | 8.400E-09 | 2.433E-09 | 5.600E-09 | 1.000E-08 |
| 22007 | 1101 | A1101Q | D | 8.9e-09 | 1.2e-08 | 2.7e-09 | 7.867E-09 | 4.735E-09 | 2.700E-09 | 1.200E-08 |
| 22008 | 1101 | A1101G | D | 1.5e-08 | 8.4e-09 | 1.2e-08 | 1.180E-08 | 3.305E-09 | 8.400E-09 | 1.500E-08 |
| 22009 | 1101 | A1101H | D | 8.4e-09 | 4.3e-09 | 1e-08 | 7.567E-09 | 2.940E-09 | 4.300E-09 | 1.000E-08 |
| 22010 | 1101 | A1101I | D | 1.8e-08 | 5.8e-09 | 5.7e-09 | 9.833E-09 | 7.073E-09 | 5.700E-09 | 1.800E-08 |
| 22011 | 1101 | A1101L | D | 4.6e-09 | 2.3e-08 | 1.1e-08 | 1.287E-08 | 9.341E-09 | 4.600E-09 | 2.300E-08 |
| 22012 | 1101 | A1101K | D | 1.1e-08 | 3.4e-09 | 8.5e-09 | 7.633E-09 | 3.873E-09 | 3.400E-09 | 1.100E-08 |
| 22013 | 1101 | A1101M | D | 6.2e-09 | 9.9e-09 | 5.5e-09 | 7.200E-09 | 2.364E-09 | 5.500E-09 | 9.900E-09 |
| 22014 | 1101 | A1101F | D | 8.4e-09 | 7.9e-09 | 4.4e-09 | 6.900E-09 | 2.179E-09 | 4.400E-09 | 8.400E-09 |

| | | | | | | | | | | |
|---|---|---|---|---|---|---|---|---|---|---|
| 22015 | 1101 | A1101P | D | 1.5e-08 | 6.8e-09 | 7.3e-09 | 9.700E-09 | 4.597E-09 | 6.800E-09 | 1.500E-08 |
| 22016 | 1101 | A1101S | D | 2.1e-09 | 1.4e-08 | 1.1e-08 | 9.033E-09 | 6.189E-09 | 2.100E-09 | 1.400E-08 |
| 22017 | 1101 | A1101T | D | 1.2e-08 | 4.6e-09 | 1.2e-08 | 9.533E-09 | 4.272E-09 | 4.600E-09 | 1.200E-08 |
| 22018 | 1101 | A1101W | D | 6.2e-09 | 8.5e-09 | 9.5e-09 | 8.067E-09 | 1.692E-09 | 6.200E-09 | 9.500E-09 |
| 22019 | 1101 | A1101Y | D | 1.2e-08 | 4.9e-09 | 3.9e-09 | 6.933E-09 | 4.416E-09 | 3.900E-09 | 1.200E-08 |
| 22020 | 1101 | A1101V | D | 4.4e-08 | 9.3e-09 | 6.8e-09 | 2.003E-08 | 2.079E-08 | 6.800E-09 | 4.400E-08 |
| 22021 | 1102 | P1102A | D | 1.2e-08 | 4.3e-08 | 9.8e-09 | 2.160E-08 | 1.857E-08 | 9.800E-09 | 4.300E-08 |
| 22022 | 1102 | P1102R | D | 4.1e-09 | 3.5e-09 | 5.5e-09 | 4.367E-09 | 1.026E-09 | 3.500E-09 | 5.500E-09 |
| 22023 | 1102 | P1102N | D | 2.9e-09 | 1.3e-08 | 5.2e-09 | 7.033E-09 | 5.294E-09 | 2.900E-09 | 1.300E-08 |
| 22024 | 1102 | P1102D | D | 8e-09 | 5.1e-09 | 8.7e-09 | 7.267E-09 | 1.909E-09 | 5.100E-09 | 8.700E-09 |
| 22025 | 1102 | P1102C | D | 3.6e-09 | 1.3e-08 | 9e-09 | 8.533E-09 | 4.717E-09 | 3.600E-09 | 1.300E-08 |
| 22026 | 1102 | P1102E | D | 7.7e-09 | 7.8e-09 | 1e-08 | 8.500E-09 | 1.300E-09 | 7.700E-09 | 1.000E-08 |
| 22027 | 1102 | P1102Q | D | 5.4e-09 | 7.9e-09 | 1.1e-09 | 4.800E-09 | 3.439E-09 | 1.100E-09 | 7.900E-09 |
| 22028 | 1102 | P1102G | D | 2.5e-09 | 3e-09 | 6.1e-09 | 3.867E-09 | 1.950E-09 | 2.500E-09 | 6.100E-09 |
| 22029 | 1102 | P1102H | D | 2e-08 | 7.7e-09 | 2.2e-08 | 1.657E-08 | 7.744E-09 | 7.700E-09 | 2.200E-08 |
| 22030 | 1102 | P1102I | D | 6.8e-09 | 1.1e-08 | 4.4e-09 | 7.400E-09 | 3.341E-09 | 4.400E-09 | 1.100E-08 |
| 22031 | 1102 | P1102L | D | 1.8e-08 | 4.1e-09 | 9.5e-09 | 1.053E-08 | 7.007E-09 | 4.100E-09 | 1.800E-08 |
| 22032 | 1102 | P1102K | D | 2.4e-09 | 1.7e-09 | 9.3e-09 | 4.467E-09 | 4.200E-09 | 1.700E-09 | 9.300E-09 |
| 22033 | 1102 | P1102M | D | 6.9e-09 | 7e-09 | 2.6e-09 | 5.500E-09 | 2.512E-09 | 2.600E-09 | 7.000E-09 |
| 22034 | 1102 | P1102F | D | 9.8e-09 | 2.3e-08 | 7.8e-09 | 1.353E-08 | 8.259E-09 | 7.800E-09 | 2.300E-08 |
| 22035 | 1102 | P1102P | D | 5.8e-09 | 9e-09 | 5.3e-09 | 6.700E-09 | 2.007E-09 | 5.300E-09 | 9.000E-09 |
| 22036 | 1102 | P1102S | D | 6.3e-09 | 1.2e-08 | 5e-09 | 7.767E-09 | 3.723E-09 | 5.000E-09 | 1.200E-08 |
| 22037 | 1102 | P1102T | D | 1.2e-08 | 8.6e-09 | 3.3e-09 | 7.967E-09 | 4.384E-09 | 3.300E-09 | 1.200E-08 |
| 22038 | 1102 | P1102W | D | 5.5e-09 | 7.8e-09 | 3e-09 | 5.433E-09 | 2.401E-09 | 3.000E-09 | 7.800E-09 |
| 22039 | 1102 | P1102Y | D | 4.4e-09 | 1.1e-08 | 4.7e-09 | 6.700E-09 | 3.727E-09 | 4.400E-09 | 1.100E-08 |
| 22040 | 1102 | P1102V | D | 1e-08 | 1.5e-08 | 1.8e-09 | 8.933E-09 | 6.664E-09 | 1.800E-09 | 1.500E-08 |
| 22041 | 1103 | K1103A | D | 1.7e-08 | 6.1e-09 | 6.6e-09 | 9.900E-09 | 6.154E-09 | 6.100E-09 | 1.700E-08 |
| 22042 | 1103 | K1103R | D | 5.9e-09 | 7.3e-09 | 3.5e-08 | 1.607E-08 | 1.641E-08 | 5.900E-09 | 3.500E-08 |
| 22043 | 1103 | K1103N | D | 8.1e-09 | 4.9e-09 | 4.5e-09 | 5.833E-09 | 1.973E-09 | 4.500E-09 | 8.100E-09 |
| 22044 | 1103 | K1103D | D | 3.1e-09 | 1.2e-08 | 6e-09 | 7.033E-09 | 4.539E-09 | 3.100E-09 | 1.200E-08 |
| 22045 | 1103 | K1103C | D | 3.5e-09 | 6e-09 | 2.8e-09 | 4.100E-09 | 1.682E-09 | 2.800E-09 | 6.000E-09 |
| 22046 | 1103 | K1103E | D | 3.9e-09 | 4.4e-09 | 1.4e-08 | 7.433E-09 | 5.692E-09 | 3.900E-09 | 1.400E-08 |
| 22047 | 1103 | K1103Q | D | 5.2e-09 | 2.6e-09 | 4.9e-09 | 4.233E-09 | 1.422E-09 | 2.600E-09 | 5.200E-09 |
| 22048 | 1103 | K1103G | D | 1.3e-08 | 1.1e-08 | 1.7e-08 | 1.367E-08 | 3.055E-09 | 1.100E-08 | 1.700E-08 |
| 22049 | 1103 | K1103H | D | 1.5e-08 | 7.2e-09 | 1.3e-08 | 1.173E-08 | 4.051E-09 | 7.200E-09 | 1.500E-08 |
| 22050 | 1103 | K1103I | D | 2.7e-09 | 1.8e-08 | 1.9e-09 | 7.533E-09 | 9.073E-09 | 1.900E-09 | 1.800E-08 |
| 22051 | 1103 | K1103L | D | 1.3e-08 | 1.5e-08 | 2.4e-08 | 1.733E-08 | 5.859E-09 | 1.300E-08 | 2.400E-08 |
| 22052 | 1103 | K1103K | D | 1.5e-08 | 3.7e-09 | 5.7e-09 | 8.133E-09 | 6.030E-09 | 3.700E-09 | 1.500E-08 |
| 22053 | 1103 | K1103M | D | 1.8e-08 | 6.1e-09 | 2.6e-09 | 8.900E-09 | 8.073E-09 | 2.600E-09 | 1.800E-08 |
| 22054 | 1103 | K1103F | D | 2.3e-08 | 9.2e-09 | 5.6e-09 | 1.260E-08 | 9.185E-09 | 5.600E-09 | 2.300E-08 |
| 22055 | 1103 | K1103P | D | 1.1e-08 | 1.1e-08 | 5e-09 | 9.000E-09 | 3.464E-09 | 5.000E-09 | 1.100E-08 |
| 22056 | 1103 | K1103S | D | 1.5e-08 | 5.4e-09 | 8.6e-09 | 9.667E-09 | 4.888E-09 | 5.400E-09 | 1.500E-08 |
| 22057 | 1103 | K1103T | D | 7.9e-09 | 5.9e-09 | 1.4e-08 | 9.267E-09 | 4.219E-09 | 5.900E-09 | 1.400E-08 |
| 22058 | 1103 | K1103W | D | 2.3e-09 | 1.9e-08 | 4.6e-09 | 8.633E-09 | 9.051E-09 | 2.300E-09 | 1.900E-08 |
| 22059 | 1103 | K1103Y | D | 1.2e-08 | 3.1e-09 | 3e-08 | 1.503E-08 | 1.370E-08 | 3.100E-09 | 3.000E-08 |
| 22060 | 1103 | K1103V | D | 1.1e-08 | 3.6e-09 | 2.8e-08 | 1.420E-08 | 1.251E-08 | 3.600E-09 | 2.800E-08 |
| 22061 | 1104 | L1104A | D | 4.4e-09 | 1.6e-09 | 9.4e-09 | 5.133E-09 | 3.951E-09 | 1.600E-09 | 9.400E-09 |
| 22062 | 1104 | L1104R | D | 1.3e-08 | 9.8e-09 | 1.2e-08 | 1.160E-08 | 1.637E-09 | 9.800E-09 | 1.300E-08 |
| 22063 | 1104 | L1104N | D | 1.6e-08 | 9.6e-09 | 1.6e-08 | 1.387E-08 | 3.695E-09 | 9.600E-09 | 1.600E-08 |
| 22064 | 1104 | L1104D | D | 1.4e-08 | 1.3e-08 | 7e-09 | 1.133E-08 | 3.786E-09 | 7.000E-09 | 1.400E-08 |
| 22065 | 1104 | L1104C | D | 3.6e-09 | 4.5e-09 | 1.1e-08 | 6.367E-09 | 4.038E-09 | 3.600E-09 | 1.100E-08 |
| 22066 | 1104 | L1104E | D | 5.8e-09 | 8.6e-09 | 7.5e-09 | 7.300E-09 | 1.411E-09 | 5.800E-09 | 8.600E-09 |
| 22067 | 1104 | L1104Q | D | 2e-09 | 6.4e-09 | 9.7e-09 | 6.033E-09 | 3.863E-09 | 2.000E-09 | 9.700E-09 |
| 22068 | 1104 | L1104G | D | 3.4e-09 | 6.5e-09 | 6.5e-09 | 5.467E-09 | 1.790E-09 | 3.400E-09 | 6.500E-09 |
| 22069 | 1104 | L1104H | D | 4e-09 | 1.4e-08 | 9.5e-09 | 9.167E-09 | 5.008E-09 | 4.000E-09 | 1.400E-08 |
| 22070 | 1104 | L1104I | D | 5.5e-09 | 1.7e-08 | 2.4e-09 | 8.300E-09 | 7.692E-09 | 2.400E-09 | 1.700E-08 |
| 22071 | 1104 | L1104L | D | 4.4e-09 | 8.9e-09 | 2.9e-09 | 5.400E-09 | 3.122E-09 | 2.900E-09 | 8.900E-09 |
| 22072 | 1104 | L1104K | D | 4.4e-09 | 5.4e-09 | 9e-09 | 6.267E-09 | 2.419E-09 | 4.400E-09 | 9.000E-09 |
| 22073 | 1104 | L1104M | D | 1.3e-08 | 3.6e-08 | 4.3e-09 | 1.777E-08 | 1.638E-08 | 4.300E-09 | 3.600E-08 |
| 22074 | 1104 | L1104F | D | 1.3e-08 | 8.9e-09 | 1e-08 | 1.063E-08 | 2.122E-09 | 8.900E-09 | 1.300E-08 |
| 22075 | 1104 | L1104P | D | 4.8e-09 | 6e-09 | 2.4e-09 | 4.400E-09 | 1.833E-09 | 2.400E-09 | 6.000E-09 |
| 22076 | 1104 | L1104S | D | 7.7e-09 | 4.9e-09 | 1.5e-08 | 9.200E-09 | 5.214E-09 | 4.900E-09 | 1.500E-08 |
| 22077 | 1104 | L1104T | D | 5.9e-09 | 1.1e-08 | 5.8e-09 | 7.567E-09 | 2.974E-09 | 5.800E-09 | 1.100E-08 |
| 22078 | 1104 | L1104W | D | 2.8e-09 | 8.3e-09 | 7.1e-09 | 6.067E-09 | 2.892E-09 | 2.800E-09 | 8.300E-09 |
| 22079 | 1104 | L1104Y | D | 4.9e-09 | 3.3e-09 | 6.3e-09 | 4.833E-09 | 1.501E-09 | 3.300E-09 | 6.300E-09 |
| 22080 | 1104 | L1104V | D | 3.9e-09 | 4.8e-09 | 9.8e-09 | 6.167E-09 | 3.179E-09 | 3.900E-09 | 9.800E-09 |
| 22081 | 1105 | L1105A | D | 3.5e-09 | 8.1e-09 | 8.3e-09 | 6.633E-09 | 2.715E-09 | 3.500E-09 | 8.300E-09 |
| 22082 | 1105 | L1105R | D | 1.5e-08 | 1.1e-08 | 7.4e-09 | 1.113E-08 | 3.802E-09 | 7.400E-09 | 1.500E-08 |
| 22083 | 1105 | L1105N | D | 3.9e-09 | 4.2e-09 | 1.7e-08 | 8.367E-09 | 7.478E-09 | 3.900E-09 | 1.700E-08 |
| 22084 | 1105 | L1105D | D | 4e-09 | 7.6e-09 | 3e-09 | 4.867E-09 | 2.419E-09 | 3.000E-09 | 7.600E-09 |
| 22085 | 1105 | L1105C | D | 1.4e-08 | 3.4e-09 | 4.6e-09 | 7.333E-09 | 5.805E-09 | 3.400E-09 | 1.400E-08 |
| 22086 | 1105 | L1105E | D | 1e-08 | 6.7e-09 | 2.6e-09 | 6.433E-09 | 3.707E-09 | 2.600E-09 | 1.000E-08 |
| 22087 | 1105 | L1105Q | D | 6.9e-09 | 2e-08 | 7e-09 | 1.130E-08 | 7.535E-09 | 6.900E-09 | 2.000E-08 |
| 22088 | 1105 | L1105G | D | 9.3e-09 | 1.3e-08 | 7.2e-09 | 9.833E-09 | 2.937E-09 | 7.200E-09 | 1.300E-08 |
| 22089 | 1105 | L1105H | D | 6.5e-09 | 1.2e-08 | 6.2e-09 | 8.233E-09 | 3.265E-09 | 6.200E-09 | 1.200E-08 |
| 22090 | 1105 | L1105I | D | 1.2e-08 | 1.7e-09 | 2.5e-09 | 5.400E-09 | 5.730E-09 | 1.700E-09 | 1.200E-08 |
| 22091 | 1105 | L1105L | D | 4e-09 | 3.9e-09 | 6.4e-09 | 4.767E-09 | 1.415E-09 | 3.900E-09 | 6.400E-09 |
| 22092 | 1105 | L1105K | D | 6.2e-09 | 7.3e-09 | 1.9e-09 | 5.133E-09 | 2.854E-09 | 1.900E-09 | 7.300E-09 |
| 22093 | 1105 | L1105M | D | 5.4e-09 | 1.2e-08 | 3.2e-09 | 6.867E-09 | 4.580E-09 | 3.200E-09 | 1.200E-08 |
| 22094 | 1105 | L1105F | D | 4.7e-09 | 5.7e-09 | 9.3e-09 | 6.567E-09 | 2.419E-09 | 4.700E-09 | 9.300E-09 |
| 22095 | 1105 | L1105P | D | 5.1e-09 | 1.1e-08 | 6.6e-09 | 7.567E-09 | 3.066E-09 | 5.100E-09 | 1.100E-08 |
| 22096 | 1105 | L1105S | D | 1.8e-08 | 5.6e-09 | 2e-09 | 8.533E-09 | 8.394E-09 | 2.000E-09 | 1.800E-08 |
| 22097 | 1105 | L1105T | D | 3.9e-09 | 8.8e-09 | 8.5e-09 | 7.067E-09 | 2.747E-09 | 3.900E-09 | 8.800E-09 |
| 22098 | 1105 | L1105W | D | 9.6e-09 | 2.8e-09 | 2.3e-08 | 1.180E-08 | 1.028E-08 | 2.800E-09 | 2.300E-08 |
| 22099 | 1105 | L1105Y | D | 2.9e-09 | 3.5e-09 | 2.1e-08 | 9.133E-09 | 1.028E-08 | 2.900E-09 | 2.100E-08 |

| | | | | | | | | | | |
|---|---|---|---|---|---|---|---|---|---|---|
| 22100 | 1105 | L1105V | D | 1.1e-08 | 1.6e-09 | 5.7e-09 | 6.100E-09 | 4.713E-09 | 1.600E-09 | 1.100E-08 |
| 22101 | 1106 | I1106A | D | 6.8e-09 | 5e-09 | 8.7e-09 | 6.833E-09 | 1.850E-09 | 5.000E-09 | 8.700E-09 |
| 22102 | 1106 | I1106R | D | 7.4e-09 | 1.5e-08 | 4.6e-09 | 9.000E-09 | 5.381E-09 | 4.600E-09 | 1.500E-08 |
| 22103 | 1106 | I1106N | D | 6.5e-09 | 1.1e-08 | 5.1e-09 | 7.533E-09 | 3.083E-09 | 5.100E-09 | 1.100E-08 |
| 22104 | 1106 | I1106D | D | 1.4e-08 | 4.7e-09 | 2.3e-09 | 7.000E-09 | 6.180E-09 | 2.300E-09 | 1.400E-08 |
| 22105 | 1106 | I1106C | D | 3.1e-09 | 1.7e-09 | 8.4e-09 | 4.400E-09 | 3.534E-09 | 1.700E-09 | 8.400E-09 |
| 22106 | 1106 | I1106E | D | 9.6e-09 | 5e-09 | 8.7e-09 | 7.767E-09 | 2.438E-09 | 5.000E-09 | 9.600E-09 |
| 22107 | 1106 | I1106Q | D | 4.9e-09 | 1.1e-08 | 6.5e-09 | 7.467E-09 | 3.163E-09 | 4.900E-09 | 1.100E-08 |
| 22108 | 1106 | I1106G | D | 1.5e-08 | 4.1e-09 | 8e-09 | 9.033E-09 | 5.523E-09 | 4.100E-09 | 1.500E-08 |
| 22109 | 1106 | I1106H | D | 2.4e-09 | 1.1e-08 | 5.1e-09 | 6.167E-09 | 4.398E-09 | 2.400E-09 | 1.100E-08 |
| 22110 | 1106 | I1106I | D | 1e-08 | 5.6e-09 | 8.1e-09 | 7.900E-09 | 2.207E-09 | 5.600E-09 | 1.000E-08 |
| 22111 | 1106 | I1106L | D | 1.8e-08 | 1.3e-08 | 5.1e-09 | 1.203E-08 | 6.504E-09 | 5.100E-09 | 1.800E-08 |
| 22112 | 1106 | I1106K | D | 6.8e-09 | 3.1e-08 | 6.7e-09 | 1.483E-08 | 1.400E-08 | 6.700E-09 | 3.100E-08 |
| 22113 | 1106 | I1106M | D | 5.3e-09 | 1.5e-08 | 4.9e-09 | 8.400E-09 | 5.719E-09 | 4.900E-09 | 1.500E-08 |
| 22114 | 1106 | I1106F | D | 1e-08 | 8.6e-09 | 3.6e-08 | 1.820E-08 | 1.543E-08 | 8.600E-09 | 3.600E-08 |
| 22115 | 1106 | I1106P | D | 6e-09 | 9.5e-09 | 4.6e-09 | 6.700E-09 | 2.524E-09 | 4.600E-09 | 9.500E-09 |
| 22116 | 1106 | I1106S | D | 7.1e-09 | 1.2e-08 | 5.5e-09 | 8.200E-09 | 3.387E-09 | 5.500E-09 | 1.200E-08 |
| 22117 | 1106 | I1106T | D | 4.8e-09 | 1.3e-08 | 2.8e-09 | 6.867E-09 | 5.405E-09 | 2.800E-09 | 1.300E-08 |
| 22118 | 1106 | I1106W | D | 8.2e-09 | 2.1e-08 | 4.3e-09 | 1.117E-08 | 8.736E-09 | 4.300E-09 | 2.100E-08 |
| 22119 | 1106 | I1106Y | D | 5e-09 | 5.5e-09 | 3e-08 | 1.350E-08 | 1.429E-08 | 5.000E-09 | 3.000E-08 |
| 22120 | 1106 | I1106V | D | 1.1e-08 | 3.9e-09 | 2.1e-08 | 1.197E-08 | 8.591E-09 | 3.900E-09 | 2.100E-08 |
| 22121 | 1107 | Y1107A | D | 1.5e-08 | 3.3e-09 | 2.4e-09 | 6.900E-09 | 7.029E-09 | 2.400E-09 | 1.500E-08 |
| 22122 | 1107 | Y1107R | D | 5.8e-09 | 3e-09 | 9.6e-09 | 6.133E-09 | 3.313E-09 | 3.000E-09 | 9.600E-09 |
| 22123 | 1107 | Y1107N | D | 6.1e-09 | 1e-08 | 1.2e-08 | 9.367E-09 | 3.001E-09 | 6.100E-09 | 1.200E-08 |
| 22124 | 1107 | Y1107D | D | 2.7e-09 | 1.3e-08 | 9.1e-09 | 8.267E-09 | 5.200E-09 | 2.700E-09 | 1.300E-08 |
| 22125 | 1107 | Y1107C | D | 3e-09 | 1.1e-08 | 9e-09 | 7.667E-09 | 4.163E-09 | 3.000E-09 | 1.100E-08 |
| 22126 | 1107 | Y1107E | D | 3.7e-09 | 5.1e-09 | 5.2e-09 | 4.667E-09 | 8.386E-10 | 3.700E-09 | 5.200E-09 |
| 22127 | 1107 | Y1107Q | D | 9.5e-09 | 5.1e-09 | 4.9e-09 | 6.500E-09 | 2.600E-09 | 4.900E-09 | 9.500E-09 |
| 22128 | 1107 | Y1107G | D | 3.2e-09 | 5.7e-09 | 2.7e-08 | 1.197E-08 | 1.308E-08 | 3.200E-09 | 2.700E-08 |
| 22129 | 1107 | Y1107H | D | 4e-09 | 4.8e-09 | 2.7e-09 | 3.833E-09 | 1.060E-09 | 2.700E-09 | 4.800E-09 |
| 22130 | 1107 | Y1107I | D | 1.1e-08 | 3.5e-08 | 7.2e-09 | 1.773E-08 | 1.507E-08 | 7.200E-09 | 3.500E-08 |
| 22131 | 1107 | Y1107L | D | 6.5e-09 | 2.5e-08 | 1.9e-08 | 1.683E-08 | 9.438E-09 | 6.500E-09 | 2.500E-08 |
| 22132 | 1107 | Y1107K | D | 1.3e-08 | 6.1e-09 | 1e-08 | 9.700E-09 | 3.460E-09 | 6.100E-09 | 1.300E-08 |
| 22133 | 1107 | Y1107M | D | 1.3e-08 | 4.7e-09 | 4.5e-09 | 7.400E-09 | 4.851E-09 | 4.500E-09 | 1.300E-08 |
| 22134 | 1107 | Y1107F | D | 4.3e-09 | 2.5e-09 | 1e-08 | 5.600E-09 | 3.915E-09 | 2.500E-09 | 1.000E-08 |
| 22135 | 1107 | Y1107P | D | 1e-08 | 1.3e-08 | 1.3e-08 | 1.200E-08 | 1.732E-09 | 1.000E-08 | 1.300E-08 |
| 22136 | 1107 | Y1107S | D | 1.2e-09 | 2.8e-09 | 5.7e-09 | 3.233E-09 | 2.281E-09 | 1.200E-09 | 5.700E-09 |
| 22137 | 1107 | Y1107T | D | 6.6e-09 | 1.1e-08 | 1e-08 | 9.200E-09 | 2.307E-09 | 6.600E-09 | 1.100E-08 |
| 22138 | 1107 | Y1107W | D | 6.5e-09 | 1.1e-08 | 6.3e-09 | 7.933E-09 | 2.658E-09 | 6.300E-09 | 1.100E-08 |
| 22139 | 1107 | Y1107Y | D | 6.5e-09 | 1e-09 | 1e-08 | 5.833E-09 | 4.537E-09 | 1.000E-09 | 1.000E-08 |
| 22140 | 1107 | Y1107V | D | 6.2e-09 | 1.3e-09 | 4.3e-09 | 3.933E-09 | 2.470E-09 | 1.300E-09 | 6.200E-09 |
| 22141 | 1108 | S1108A | D | 1.4e-08 | 4.5e-09 | 5.9e-09 | 8.133E-09 | 5.129E-09 | 4.500E-09 | 1.400E-08 |
| 22142 | 1108 | S1108R | D | 2e-09 | 2.8e-09 | 7.2e-09 | 4.000E-09 | 2.800E-09 | 2.000E-09 | 7.200E-09 |
| 22143 | 1108 | S1108N | D | 7.3e-09 | 3.5e-09 | 4.9e-09 | 5.233E-09 | 1.922E-09 | 3.500E-09 | 7.300E-09 |
| 22144 | 1108 | S1108D | D | 5.4e-09 | 7.5e-09 | 5.3e-09 | 6.067E-09 | 1.242E-09 | 5.300E-09 | 7.500E-09 |
| 22145 | 1108 | S1108C | D | 2.8e-09 | 2.4e-09 | 1.5e-08 | 6.733E-09 | 7.162E-09 | 2.400E-09 | 1.500E-08 |
| 22146 | 1108 | S1108E | D | 9.6e-09 | 1.3e-08 | 3.9e-09 | 8.833E-09 | 4.598E-09 | 3.900E-09 | 1.300E-08 |
| 22147 | 1108 | S1108Q | D | 2e-08 | 1.2e-08 | 7.8e-09 | 1.327E-08 | 6.198E-09 | 7.800E-09 | 2.000E-08 |
| 22148 | 1108 | S1108G | D | 6.9e-09 | 9.3e-09 | 4.8e-09 | 7.000E-09 | 2.252E-09 | 4.800E-09 | 9.300E-09 |
| 22149 | 1108 | S1108H | D | 1.5e-08 | 2.8e-09 | 5.9e-09 | 7.900E-09 | 6.341E-09 | 2.800E-09 | 1.500E-08 |
| 22150 | 1108 | S1108I | D | 8.2e-09 | 5e-09 | 6.1e-09 | 6.433E-09 | 1.626E-09 | 5.000E-09 | 8.200E-09 |
| 22151 | 1108 | S1108L | D | 4.9e-09 | 2.8e-08 | 8.2e-09 | 1.370E-08 | 1.249E-08 | 4.900E-09 | 2.800E-08 |
| 22152 | 1108 | S1108K | D | 9.6e-09 | 1.1e-08 | 4.2e-09 | 8.267E-09 | 3.591E-09 | 4.200E-09 | 1.100E-08 |
| 22153 | 1108 | S1108M | D | 3.3e-09 | 1.4e-08 | 6.3e-09 | 7.867E-09 | 5.519E-09 | 3.300E-09 | 1.400E-08 |
| 22154 | 1108 | S1108F | D | 1.5e-08 | 4.5e-09 | 4.1e-09 | 7.867E-09 | 6.181E-09 | 4.100E-09 | 1.500E-08 |
| 22155 | 1108 | S1108P | D | 7.2e-09 | 8.8e-09 | 6.1e-09 | 7.367E-09 | 1.358E-09 | 6.100E-09 | 8.800E-09 |
| 22156 | 1108 | S1108S | D | 6.8e-09 | 8.9e-09 | 5.1e-09 | 6.933E-09 | 1.904E-09 | 5.100E-09 | 8.900E-09 |
| 22157 | 1108 | S1108T | D | 2.7e-08 | 7.6e-09 | 7.9e-09 | 1.417E-08 | 1.112E-08 | 7.600E-09 | 2.700E-08 |
| 22158 | 1108 | S1108W | D | 4.9e-09 | 2.7e-09 | 8.9e-09 | 5.500E-09 | 3.143E-09 | 2.700E-09 | 8.900E-09 |
| 22159 | 1108 | S1108Y | D | 3.1e-09 | 5.1e-09 | 2.1e-08 | 9.733E-09 | 9.808E-09 | 3.100E-09 | 2.100E-08 |
| 22160 | 1108 | S1108V | D | 7.7e-09 | 3.1e-09 | 1.2e-08 | 7.600E-09 | 4.451E-09 | 3.100E-09 | 1.200E-08 |
| 22161 | 1109 | A1109A | D | 1.1e-08 | 8.4e-09 | 3.7e-09 | 7.700E-09 | 3.700E-09 | 3.700E-09 | 1.100E-08 |
| 22162 | 1109 | A1109R | D | 2.5e-09 | 5e-09 | 2.8e-09 | 3.433E-09 | 1.365E-09 | 2.500E-09 | 5.000E-09 |
| 22163 | 1109 | A1109N | D | 8.5e-09 | 8.7e-09 | 3.4e-09 | 6.867E-09 | 3.004E-09 | 3.400E-09 | 8.700E-09 |
| 22164 | 1109 | A1109D | D | 1.4e-08 | 3.5e-09 | 5.9e-09 | 7.800E-09 | 5.502E-09 | 3.500E-09 | 1.400E-08 |
| 22165 | 1109 | A1109C | D | 6.2e-09 | 4.9e-09 | 4.5e-09 | 5.200E-09 | 8.888E-10 | 4.500E-09 | 6.200E-09 |
| 22166 | 1109 | A1109E | D | 4.4e-09 | 2e-09 | 7.9e-09 | 4.767E-09 | 2.967E-09 | 2.000E-09 | 7.900E-09 |
| 22167 | 1109 | A1109Q | D | 5e-09 | 1e-08 | 8.3e-09 | 7.767E-09 | 2.542E-09 | 5.000E-09 | 1.000E-08 |
| 22168 | 1109 | A1109G | D | 1e-08 | 1.1e-08 | 7.3e-09 | 9.433E-09 | 1.914E-09 | 7.300E-09 | 1.100E-08 |
| 22169 | 1109 | A1109H | D | 9.7e-09 | 2.3e-09 | 1.1e-08 | 7.667E-09 | 4.693E-09 | 2.300E-09 | 1.100E-08 |
| 22170 | 1109 | A1109I | D | 7.1e-09 | 5.2e-09 | 8.6e-09 | 6.967E-09 | 1.704E-09 | 5.200E-09 | 8.600E-09 |
| 22171 | 1109 | A1109L | D | 3.9e-09 | 1.2e-08 | 8.6e-09 | 8.167E-09 | 4.067E-09 | 3.900E-09 | 1.200E-08 |
| 22172 | 1109 | A1109K | D | 2.5e-08 | 4.3e-09 | 4.2e-09 | 1.117E-08 | 1.198E-08 | 4.200E-09 | 2.500E-08 |
| 22173 | 1109 | A1109M | D | 8.5e-09 | 1.3e-08 | 9.6e-09 | 1.037E-08 | 2.346E-09 | 8.500E-09 | 1.300E-08 |
| 22174 | 1109 | A1109F | D | 5.9e-09 | 7.2e-09 | 3.4e-09 | 5.500E-09 | 1.931E-09 | 3.400E-09 | 7.200E-09 |
| 22175 | 1109 | A1109P | D | 1.3e-08 | 1.9e-08 | 9.3e-09 | 1.377E-08 | 4.895E-09 | 9.300E-09 | 1.900E-08 |
| 22176 | 1109 | A1109S | D | 3e-09 | 8e-09 | 2.2e-09 | 4.400E-09 | 3.143E-09 | 2.200E-09 | 8.000E-09 |
| 22177 | 1109 | A1109T | D | 1.1e-08 | 1.2e-08 | 1.2e-08 | 1.167E-08 | 5.774E-10 | 1.100E-08 | 1.200E-08 |
| 22178 | 1109 | A1109W | D | 5.6e-09 | 9.9e-09 | 2.1e-09 | 5.867E-09 | 3.907E-09 | 2.100E-09 | 9.900E-09 |
| 22179 | 1109 | A1109Y | D | 2e-09 | 2e-08 | 5.8e-09 | 9.267E-09 | 9.488E-09 | 2.000E-09 | 2.000E-08 |
| 22180 | 1109 | A1109V | D | 3e-09 | 1.4e-08 | 3.1e-09 | 6.700E-09 | 6.322E-09 | 3.000E-09 | 1.400E-08 |
| 22181 | 1110 | S1110A | D | 6.9e-09 | 8.1e-09 | 4.6e-09 | 6.533E-09 | 1.779E-09 | 4.600E-09 | 8.100E-09 |
| 22182 | 1110 | S1110R | D | 3.7e-09 | 1.3e-09 | 1.6e-08 | 7.000E-09 | 7.886E-09 | 1.300E-09 | 1.600E-08 |
| 22183 | 1110 | S1110N | D | 6.7e-09 | 1.2e-08 | 4.1e-09 | 7.600E-09 | 4.026E-09 | 4.100E-09 | 1.200E-08 |
| 22184 | 1110 | S1110D | D | 2.5e-08 | 7.6e-09 | 3.4e-08 | 2.220E-08 | 1.342E-08 | 7.600E-09 | 3.400E-08 |

| | | | | | | | | | | |
|---|---|---|---|---|---|---|---|---|---|---|
| 22185 | 1110 | S1110C | D | 6.3e-09 | 4.7e-09 | 5.7e-09 | 5.567E-09 | 8.083E-10 | 4.700E-09 | 6.300E-09 |
| 22186 | 1110 | S1110E | D | 5.9e-09 | 2.5e-09 | 4.4e-09 | 4.267E-09 | 1.704E-09 | 2.500E-09 | 5.900E-09 |
| 22187 | 1110 | S1110Q | D | 1e-08 | 1.2e-08 | 4.4e-09 | 8.800E-09 | 3.940E-09 | 4.400E-09 | 1.200E-08 |
| 22188 | 1110 | S1110G | D | 9.2e-09 | 8.1e-09 | 3.3e-09 | 6.867E-09 | 3.137E-09 | 3.300E-09 | 9.200E-09 |
| 22189 | 1110 | S1110H | D | 5.6e-09 | 1.3e-08 | 8.7e-09 | 9.100E-09 | 3.716E-09 | 5.600E-09 | 1.300E-08 |
| 22190 | 1110 | S1110I | D | 5.1e-09 | 1.7e-08 | 4.6e-09 | 8.900E-09 | 7.019E-09 | 4.600E-09 | 1.700E-08 |
| 22191 | 1110 | S1110L | D | 1.9e-08 | 5.2e-09 | 1.2e-08 | 1.207E-08 | 6.900E-09 | 5.200E-09 | 1.900E-08 |
| 22192 | 1110 | S1110K | D | 7.4e-09 | 1.3e-08 | 4.6e-09 | 8.333E-09 | 4.277E-09 | 4.600E-09 | 1.300E-08 |
| 22193 | 1110 | S1110M | D | 1.2e-08 | 9.8e-09 | 6.9e-09 | 9.567E-09 | 2.558E-09 | 6.900E-09 | 1.200E-08 |
| 22194 | 1110 | S1110F | D | 5.4e-09 | 9e-09 | 9e-09 | 7.800E-09 | 2.078E-09 | 5.400E-09 | 9.000E-09 |
| 22195 | 1110 | S1110P | D | 7.5e-09 | 1.2e-08 | 3.7e-09 | 7.733E-09 | 4.155E-09 | 3.700E-09 | 1.200E-08 |
| 22196 | 1110 | S1110S | D | 3.2e-09 | 7.9e-09 | 7.2e-09 | 6.100E-09 | 2.536E-09 | 3.200E-09 | 7.900E-09 |
| 22197 | 1110 | S1110T | D | 7.6e-09 | 1e-08 | 3.3e-09 | 6.967E-09 | 3.395E-09 | 3.300E-09 | 1.000E-08 |
| 22198 | 1110 | S1110W | D | 1.3e-08 | 1.7e-08 | 8.3e-09 | 1.277E-08 | 4.355E-09 | 8.300E-09 | 1.700E-08 |
| 22199 | 1110 | S1110Y | D | 5e-09 | 4.9e-09 | 4.6e-09 | 4.833E-09 | 2.082E-10 | 4.600E-09 | 5.000E-09 |
| 22200 | 1110 | S1110V | D | 1.7e-09 | 4.8e-09 | 6.8e-09 | 4.433E-09 | 2.570E-09 | 1.700E-09 | 6.800E-09 |
| 22201 | 1111 | F1111A | D | 4e-09 | 1.3e-08 | 4.4e-09 | 7.133E-09 | 5.085E-09 | 4.000E-09 | 1.300E-08 |
| 22202 | 1111 | F1111R | D | 7e-09 | 1.4e-08 | 1.9e-09 | 7.633E-09 | 6.075E-09 | 1.900E-09 | 1.400E-08 |
| 22203 | 1111 | F1111N | D | 2.6e-08 | 5.6e-09 | 3.3e-09 | 1.163E-08 | 1.249E-08 | 3.300E-09 | 2.600E-08 |
| 22204 | 1111 | F1111D | D | 2.7e-09 | 2.3e-09 | 1.2e-08 | 5.667E-09 | 5.488E-09 | 2.300E-09 | 1.200E-08 |
| 22205 | 1111 | F1111C | D | 6.2e-09 | 3.9e-09 | 1.7e-08 | 9.033E-09 | 6.995E-09 | 3.900E-09 | 1.700E-08 |
| 22206 | 1111 | F1111E | D | 3.9e-09 | 4.4e-09 | 3.9e-08 | 1.577E-08 | 2.012E-08 | 3.900E-09 | 3.900E-08 |
| 22207 | 1111 | F1111Q | D | 7.2e-09 | 3.8e-09 | 1.3e-08 | 8.000E-09 | 4.652E-09 | 3.800E-09 | 1.300E-08 |
| 22208 | 1111 | F1111G | D | 6.1e-09 | 7.6e-09 | 7.7e-09 | 7.133E-09 | 8.963E-10 | 6.100E-09 | 7.700E-09 |
| 22209 | 1111 | F1111H | D | 9.5e-09 | 6.2e-09 | 2.3e-08 | 1.290E-08 | 8.901E-09 | 6.200E-09 | 2.300E-08 |
| 22210 | 1111 | F1111I | D | 1.7e-08 | 6e-09 | 3.9e-09 | 8.967E-09 | 7.036E-09 | 3.900E-09 | 1.700E-08 |
| 22211 | 1111 | F1111L | D | 1.1e-08 | 7.8e-09 | 8.1e-09 | 8.967E-09 | 1.767E-09 | 7.800E-09 | 1.100E-08 |
| 22212 | 1111 | F1111K | D | 1.3e-08 | 4.7e-09 | 5.4e-09 | 7.700E-09 | 4.603E-09 | 4.700E-09 | 1.300E-08 |
| 22213 | 1111 | F1111M | D | 9.5e-09 | 1.2e-08 | 1.2e-08 | 1.117E-08 | 1.443E-09 | 9.500E-09 | 1.200E-08 |
| 22214 | 1111 | F1111F | D | 6.5e-09 | 1.7e-08 | 6e-09 | 9.833E-09 | 6.212E-09 | 6.000E-09 | 1.700E-08 |
| 22215 | 1111 | F1111P | D | 8.9e-09 | 6.4e-09 | 1.1e-08 | 8.767E-09 | 2.303E-09 | 6.400E-09 | 1.100E-08 |
| 22216 | 1111 | F1111S | D | 1e-08 | 7.8e-09 | 4.2e-09 | 7.333E-09 | 2.928E-09 | 4.200E-09 | 1.000E-08 |
| 22217 | 1111 | F1111T | D | 4.8e-09 | 6.7e-09 | 6.9e-09 | 6.133E-09 | 1.159E-09 | 4.800E-09 | 6.900E-09 |
| 22218 | 1111 | F1111W | D | 8e-09 | 4.3e-09 | 5.1e-09 | 5.800E-09 | 1.947E-09 | 4.300E-09 | 8.000E-09 |
| 22219 | 1111 | F1111Y | D | 4.5e-09 | 7.3e-09 | 4.1e-09 | 5.300E-09 | 1.744E-09 | 4.100E-09 | 7.300E-09 |
| 22220 | 1111 | F1111V | D | 9.7e-09 | 5.5e-09 | 3e-09 | 6.067E-09 | 3.386E-09 | 3.000E-09 | 9.700E-09 |
| 22221 | 1112 | L1112A | D | 4.8e-09 | 1.6e-08 | 1.7e-08 | 1.260E-08 | 6.773E-09 | 4.800E-09 | 1.700E-08 |
| 22222 | 1112 | L1112R | D | 5.3e-09 | 7.6e-09 | 6.6e-09 | 6.500E-09 | 1.153E-09 | 5.300E-09 | 7.600E-09 |
| 22223 | 1112 | L1112N | D | 7.1e-09 | 4.2e-09 | 9.5e-09 | 6.933E-09 | 2.654E-09 | 4.200E-09 | 9.500E-09 |
| 22224 | 1112 | L1112D | D | 6.2e-09 | 2.8e-09 | 4.2e-09 | 4.400E-09 | 1.709E-09 | 2.800E-09 | 6.200E-09 |
| 22225 | 1112 | L1112C | D | 3.4e-09 | 6.4e-09 | 1.2e-08 | 7.267E-09 | 4.365E-09 | 3.400E-09 | 1.200E-08 |
| 22226 | 1112 | L1112E | D | 3.6e-09 | 7.9e-09 | 7.9e-09 | 6.467E-09 | 2.483E-09 | 3.600E-09 | 7.900E-09 |
| 22227 | 1112 | L1112Q | D | 8.8e-09 | 1.2e-08 | 3.3e-09 | 8.033E-09 | 4.400E-09 | 3.300E-09 | 1.200E-08 |
| 22228 | 1112 | L1112G | D | 7.5e-09 | 3.2e-08 | 1.9e-08 | 1.950E-08 | 1.226E-08 | 7.500E-09 | 3.200E-08 |
| 22229 | 1112 | L1112H | D | 9.4e-09 | 2.8e-09 | 2.2e-09 | 4.800E-09 | 3.995E-09 | 2.200E-09 | 9.400E-09 |
| 22230 | 1112 | L1112I | D | 9.6e-09 | 1.5e-08 | 1.4e-08 | 1.287E-08 | 2.873E-09 | 9.600E-09 | 1.500E-08 |
| 22231 | 1112 | L1112L | D | 7.1e-09 | 4.8e-09 | 5.5e-09 | 5.800E-09 | 1.179E-09 | 4.800E-09 | 7.100E-09 |
| 22232 | 1112 | L1112K | D | 7.5e-09 | 3.2e-09 | 2.9e-09 | 4.533E-09 | 2.574E-09 | 2.900E-09 | 7.500E-09 |
| 22233 | 1112 | L1112M | D | 2.5e-09 | 1.7e-08 | 5.9e-09 | 8.467E-09 | 7.583E-09 | 2.500E-09 | 1.700E-08 |
| 22234 | 1112 | L1112F | D | 2.1e-08 | 6.5e-09 | 6.8e-09 | 1.143E-08 | 8.286E-09 | 6.500E-09 | 2.100E-08 |
| 22235 | 1112 | L1112P | D | 4.5e-09 | 9.3e-09 | 1e-08 | 7.933E-09 | 2.994E-09 | 4.500E-09 | 1.000E-08 |
| 22236 | 1112 | L1112S | D | 3.8e-09 | 8.6e-09 | 6.2e-09 | 6.200E-09 | 2.400E-09 | 3.800E-09 | 8.600E-09 |
| 22237 | 1112 | L1112T | D | 1.8e-08 | 1.7e-08 | 1.3e-08 | 1.600E-08 | 2.646E-09 | 1.300E-08 | 1.800E-08 |
| 22238 | 1112 | L1112W | D | 6.4e-09 | 8.5e-09 | 9.1e-09 | 8.000E-09 | 1.418E-09 | 6.400E-09 | 9.100E-09 |
| 22239 | 1112 | L1112Y | D | 4.1e-09 | 3.7e-09 | 2.7e-09 | 3.500E-09 | 7.211E-10 | 2.700E-09 | 4.100E-09 |
| 22240 | 1112 | L1112V | D | 1.6e-08 | 2.7e-09 | 4.2e-09 | 7.633E-09 | 7.284E-09 | 2.700E-09 | 1.600E-08 |
| 22241 | 1113 | Y1113A | D | 9.2e-09 | 5e-09 | 4.5e-09 | 6.233E-09 | 2.581E-09 | 4.500E-09 | 9.200E-09 |
| 22242 | 1113 | Y1113R | D | 1.1e-08 | 1.3e-08 | 4.3e-09 | 9.433E-09 | 4.557E-09 | 4.300E-09 | 1.300E-08 |
| 22243 | 1113 | Y1113N | D | 3.7e-09 | 4.6e-09 | 4.3e-09 | 4.200E-09 | 4.583E-10 | 3.700E-09 | 4.600E-09 |
| 22244 | 1113 | Y1113D | D | 9.4e-09 | 2.6e-09 | 3.9e-09 | 5.300E-09 | 3.610E-09 | 2.600E-09 | 9.400E-09 |
| 22245 | 1113 | Y1113C | D | 1.3e-08 | 9.5e-09 | 2.5e-09 | 8.333E-09 | 5.346E-09 | 2.500E-09 | 1.300E-08 |
| 22246 | 1113 | Y1113E | D | 2.1e-08 | 8.2e-09 | 1.3e-08 | 1.407E-08 | 6.466E-09 | 8.200E-09 | 2.100E-08 |
| 22247 | 1113 | Y1113Q | D | 3.2e-09 | 7e-09 | 5.3e-09 | 5.167E-09 | 1.904E-09 | 3.200E-09 | 7.000E-09 |
| 22248 | 1113 | Y1113G | D | 6.2e-09 | 8e-09 | 4.5e-09 | 6.233E-09 | 1.750E-09 | 4.500E-09 | 8.000E-09 |
| 22249 | 1113 | Y1113H | D | 7e-09 | 6.1e-09 | 5.8e-09 | 6.300E-09 | 6.245E-10 | 5.800E-09 | 7.000E-09 |
| 22250 | 1113 | Y1113I | D | 1e-08 | 9.7e-09 | 4.5e-09 | 8.067E-09 | 3.092E-09 | 4.500E-09 | 1.000E-08 |
| 22251 | 1113 | Y1113L | D | 1.5e-08 | 1.4e-08 | 8.5e-09 | 1.250E-08 | 3.500E-09 | 8.500E-09 | 1.500E-08 |
| 22252 | 1113 | Y1113K | D | 2.1e-08 | 1.2e-09 | 2.5e-09 | 8.233E-09 | 1.108E-08 | 1.200E-09 | 2.100E-08 |
| 22253 | 1113 | Y1113M | D | 9.7e-09 | 5e-09 | 3.8e-09 | 6.167E-09 | 3.118E-09 | 3.800E-09 | 9.700E-09 |
| 22254 | 1113 | Y1113F | D | 7.7e-09 | 7.1e-09 | 6.2e-09 | 7.000E-09 | 7.550E-10 | 6.200E-09 | 7.700E-09 |
| 22255 | 1113 | Y1113P | D | 1.3e-08 | 6.8e-09 | 6.8e-09 | 8.867E-09 | 3.580E-09 | 6.800E-09 | 1.300E-08 |
| 22256 | 1113 | Y1113S | D | 4.6e-09 | 3.2e-09 | 2.2e-08 | 9.933E-09 | 1.047E-08 | 3.200E-09 | 2.200E-08 |
| 22257 | 1113 | Y1113T | D | 4.8e-09 | 8.5e-09 | 7.3e-09 | 6.867E-09 | 1.888E-09 | 4.800E-09 | 8.500E-09 |
| 22258 | 1113 | Y1113W | D | 8.7e-09 | 4.1e-09 | 5.7e-10 | 4.457E-09 | 4.077E-09 | 5.700E-10 | 8.700E-09 |
| 22259 | 1113 | Y1113Y | D | 1.1e-08 | 6.7e-09 | 1.1e-08 | 9.567E-09 | 2.483E-09 | 6.700E-09 | 1.100E-08 |
| 22260 | 1113 | Y1113V | D | 6.3e-09 | 4.8e-09 | 6.3e-09 | 5.800E-09 | 8.660E-10 | 4.800E-09 | 6.300E-09 |
| 22261 | 1114 | S1114A | D | 1.6e-08 | 7.5e-09 | 1.5e-08 | 1.283E-08 | 4.646E-09 | 7.500E-09 | 1.600E-08 |
| 22262 | 1114 | S1114R | D | 4.8e-09 | 6.7e-09 | 1.1e-08 | 7.500E-09 | 3.176E-09 | 4.800E-09 | 1.100E-08 |
| 22263 | 1114 | S1114N | D | 5.8e-09 | 2.5e-08 | 7.7e-09 | 1.283E-08 | 1.058E-08 | 5.800E-09 | 2.500E-08 |
| 22264 | 1114 | S1114D | D | 7.7e-09 | 1.4e-09 | 9.5e-09 | 6.200E-09 | 4.253E-09 | 1.400E-09 | 9.500E-09 |
| 22265 | 1114 | S1114C | D | 9.5e-09 | 1.1e-08 | 7.7e-09 | 9.400E-09 | 1.652E-09 | 7.700E-09 | 1.100E-08 |
| 22266 | 1114 | S1114E | D | 1.4e-08 | 3.6e-09 | 7.3e-09 | 8.300E-09 | 5.272E-09 | 3.600E-09 | 1.400E-08 |
| 22267 | 1114 | S1114Q | D | 5.9e-09 | 5.8e-09 | 2e-08 | 1.057E-08 | 8.170E-09 | 5.800E-09 | 2.000E-08 |
| 22268 | 1114 | S1114G | D | 3.9e-09 | 4.7e-09 | 5.9e-09 | 4.833E-09 | 1.007E-09 | 3.900E-09 | 5.900E-09 |
| 22269 | 1114 | S1114H | D | 1.1e-08 | 9e-09 | 4.9e-09 | 8.300E-09 | 3.110E-09 | 4.900E-09 | 1.100E-08 |

| | | | | | | | | | | |
|---|---|---|---|---|---|---|---|---|---|---|
| 22270 | 1114 | S1114I | D | 6e-09 | 9.5e-09 | 4.1e-09 | 6.533E-09 | 2.739E-09 | 4.100E-09 | 9.500E-09 |
| 22271 | 1114 | S1114L | D | 1.8e-08 | 2.9e-09 | 3.9e-09 | 8.267E-09 | 8.444E-09 | 2.900E-09 | 1.800E-08 |
| 22272 | 1114 | S1114K | D | 1.6e-08 | 1e-08 | 1e-08 | 1.200E-08 | 3.464E-09 | 1.000E-08 | 1.600E-08 |
| 22273 | 1114 | S1114M | D | 5.6e-09 | 1.2e-08 | 1.5e-08 | 1.087E-08 | 4.801E-09 | 5.600E-09 | 1.500E-08 |
| 22274 | 1114 | S1114F | D | 6.6e-09 | 6.2e-09 | 5.3e-09 | 6.033E-09 | 6.658E-10 | 5.300E-09 | 6.600E-09 |
| 22275 | 1114 | S1114P | D | 1.3e-08 | 1.7e-09 | 8.5e-09 | 7.733E-09 | 5.689E-09 | 1.700E-09 | 1.300E-08 |
| 22276 | 1114 | S1114S | D | 5e-09 | 4.9e-09 | 2e-08 | 9.967E-09 | 8.689E-09 | 4.900E-09 | 2.000E-08 |
| 22277 | 1114 | S1114T | D | 1.3e-08 | 5.9e-09 | 7.3e-09 | 8.733E-09 | 3.761E-09 | 5.900E-09 | 1.300E-08 |
| 22278 | 1114 | S1114W | D | 3e-09 | 5.7e-09 | 1.7e-08 | 8.567E-09 | 7.427E-09 | 3.000E-09 | 1.700E-08 |
| 22279 | 1114 | S1114Y | D | 6.4e-09 | 1.4e-08 | 5.8e-09 | 8.733E-09 | 4.571E-09 | 5.800E-09 | 1.400E-08 |
| 22280 | 1114 | S1114V | D | 1e-08 | 4.6e-09 | 7.1e-09 | 7.233E-09 | 2.702E-09 | 4.600E-09 | 1.000E-08 |
| 22281 | 1115 | G1115A | D | 9.7e-09 | 1e-08 | 4.6e-09 | 8.100E-09 | 3.035E-09 | 4.600E-09 | 1.000E-08 |
| 22282 | 1115 | G1115R | D | 3.7e-09 | 8.8e-09 | 4e-09 | 5.500E-09 | 2.862E-09 | 3.700E-09 | 8.800E-09 |
| 22283 | 1115 | G1115N | D | 4.2e-09 | 3.9e-09 | 3.3e-09 | 3.800E-09 | 4.583E-10 | 3.300E-09 | 4.200E-09 |
| 22284 | 1115 | G1115D | D | 1.1e-08 | 8.8e-09 | 1.8e-08 | 1.260E-08 | 4.804E-09 | 8.800E-09 | 1.800E-08 |
| 22285 | 1115 | G1115C | D | 1.3e-08 | 6.9e-09 | 1.1e-08 | 1.030E-08 | 3.110E-09 | 6.900E-09 | 1.300E-08 |
| 22286 | 1115 | G1115E | D | 1.4e-08 | 7.2e-09 | 7.9e-09 | 9.700E-09 | 3.740E-09 | 7.200E-09 | 1.400E-08 |
| 22287 | 1115 | G1115Q | D | 9.4e-09 | 7.1e-09 | 5.9e-09 | 7.467E-09 | 1.779E-09 | 5.900E-09 | 9.400E-09 |
| 22288 | 1115 | G1115G | D | 6e-09 | 2.3e-09 | 9.8e-09 | 6.033E-09 | 3.750E-09 | 2.300E-09 | 9.800E-09 |
| 22289 | 1115 | G1115H | D | 1e-08 | 3.6e-09 | 7.4e-09 | 7.000E-09 | 3.219E-09 | 3.600E-09 | 1.000E-08 |
| 22290 | 1115 | G1115I | D | 4.8e-09 | 7.7e-09 | 1.1e-08 | 7.833E-09 | 3.102E-09 | 4.800E-09 | 1.100E-08 |
| 22291 | 1115 | G1115L | D | 3.3e-09 | 6.8e-09 | 7.2e-09 | 5.767E-09 | 2.146E-09 | 3.300E-09 | 7.200E-09 |
| 22292 | 1115 | G1115K | D | 1.5e-08 | 1.2e-08 | 7.2e-09 | 1.140E-08 | 3.934E-09 | 7.200E-09 | 1.500E-08 |
| 22293 | 1115 | G1115M | D | 1.5e-08 | 6.7e-09 | 8.9e-09 | 1.020E-08 | 4.300E-09 | 6.700E-09 | 1.500E-08 |
| 22294 | 1115 | G1115F | D | 3e-09 | 8.2e-09 | 8.8e-09 | 6.667E-09 | 3.190E-09 | 3.000E-09 | 8.800E-09 |
| 22295 | 1115 | G1115P | D | 1e-08 | 4.2e-09 | 9.5e-09 | 7.900E-09 | 3.214E-09 | 4.200E-09 | 1.000E-08 |
| 22296 | 1115 | G1115S | D | 1.3e-08 | 1e-08 | 5.2e-09 | 9.400E-09 | 3.934E-09 | 5.200E-09 | 1.300E-08 |
| 22297 | 1115 | G1115T | D | 7.8e-09 | 9e-09 | 6.3e-09 | 7.700E-09 | 1.353E-09 | 6.300E-09 | 9.000E-09 |
| 22298 | 1115 | G1115W | D | 1.1e-08 | 5.7e-09 | 3.9e-09 | 6.867E-09 | 3.691E-09 | 3.900E-09 | 1.100E-08 |
| 22299 | 1115 | G1115Y | D | 6.9e-09 | 1.1e-08 | 4.1e-09 | 7.333E-09 | 3.470E-09 | 4.100E-09 | 1.100E-08 |
| 22300 | 1115 | G1115V | D | 2.2e-08 | 1.3e-08 | 8.9e-09 | 1.463E-08 | 6.701E-09 | 8.900E-09 | 2.200E-08 |
| 22301 | 1116 | V1116A | D | 1.1e-08 | 6.9e-09 | 1.1e-08 | 9.633E-09 | 2.367E-09 | 6.900E-09 | 1.100E-08 |
| 22302 | 1116 | V1116R | D | 3.7e-09 | 2.4e-09 | 4e-09 | 3.367E-09 | 8.505E-10 | 2.400E-09 | 4.000E-09 |
| 22303 | 1116 | V1116N | D | 8e-09 | 6e-09 | 4.1e-09 | 6.033E-09 | 1.950E-09 | 4.100E-09 | 8.000E-09 |
| 22304 | 1116 | V1116D | D | 4.4e-09 | 5.2e-09 | 1.3e-08 | 7.533E-09 | 4.751E-09 | 4.400E-09 | 1.300E-08 |
| 22305 | 1116 | V1116C | D | 1.1e-08 | 6.3e-09 | 1e-08 | 9.100E-09 | 2.476E-09 | 6.300E-09 | 1.100E-08 |
| 22306 | 1116 | V1116E | D | 2.6e-09 | 9.1e-09 | 1.4e-08 | 8.567E-09 | 5.719E-09 | 2.600E-09 | 1.400E-08 |
| 22307 | 1116 | V1116Q | D | 6.3e-09 | 9e-09 | 7.8e-09 | 7.700E-09 | 1.353E-09 | 6.300E-09 | 9.000E-09 |
| 22308 | 1116 | V1116G | D | 9.1e-09 | 1.8e-09 | 2.8e-08 | 1.297E-08 | 1.352E-08 | 1.800E-09 | 2.800E-08 |
| 22309 | 1116 | V1116H | D | 6.6e-09 | 5.9e-09 | 1.9e-08 | 1.050E-08 | 7.370E-09 | 5.900E-09 | 1.900E-08 |
| 22310 | 1116 | V1116I | D | 6.8e-09 | 4.6e-09 | 4.6e-09 | 5.333E-09 | 1.270E-09 | 4.600E-09 | 6.800E-09 |
| 22311 | 1116 | V1116L | D | 9.6e-09 | 2.7e-09 | 9e-09 | 7.100E-09 | 3.822E-09 | 2.700E-09 | 9.600E-09 |
| 22312 | 1116 | V1116K | D | 5e-09 | 3.2e-09 | 1.8e-08 | 8.733E-09 | 8.075E-09 | 3.200E-09 | 1.800E-08 |
| 22313 | 1116 | V1116M | D | 2.4e-09 | 2.4e-09 | 2.1e-08 | 8.600E-09 | 1.074E-08 | 2.400E-09 | 2.100E-08 |
| 22314 | 1116 | V1116F | D | 4.9e-09 | 1e-08 | 6.1e-09 | 7.000E-09 | 2.666E-09 | 4.900E-09 | 1.000E-08 |
| 22315 | 1116 | V1116P | D | 5.5e-09 | 1.2e-08 | 3.9e-09 | 7.133E-09 | 4.290E-09 | 3.900E-09 | 1.200E-08 |
| 22316 | 1116 | V1116S | D | 4.2e-09 | 5.1e-09 | 7.2e-09 | 5.500E-09 | 1.539E-09 | 4.200E-09 | 7.200E-09 |
| 22317 | 1116 | V1116T | D | 1.4e-08 | 5.7e-09 | 2.3e-09 | 7.333E-09 | 6.019E-09 | 2.300E-09 | 1.400E-08 |
| 22318 | 1116 | V1116W | D | 6e-09 | 2.4e-09 | 1.3e-08 | 7.133E-09 | 5.390E-09 | 2.400E-09 | 1.300E-08 |
| 22319 | 1116 | V1116Y | D | 3.9e-09 | 3.2e-09 | 1e-08 | 5.700E-09 | 3.740E-09 | 3.200E-09 | 1.000E-08 |
| 22320 | 1116 | V1116V | D | 9e-09 | 6.2e-09 | 8.5e-09 | 7.900E-09 | 1.493E-09 | 6.200E-09 | 9.000E-09 |
| 22321 | 1117 | P1117A | D | 9.3e-09 | 2.6e-09 | 6.4e-09 | 6.100E-09 | 3.360E-09 | 2.600E-09 | 9.300E-09 |
| 22322 | 1117 | P1117R | D | 7.1e-09 | 3.7e-09 | 7.7e-09 | 6.167E-09 | 2.157E-09 | 3.700E-09 | 7.700E-09 |
| 22323 | 1117 | P1117N | D | 5.7e-09 | 8.6e-09 | 5.3e-09 | 6.533E-09 | 1.801E-09 | 5.300E-09 | 8.600E-09 |
| 22324 | 1117 | P1117D | D | 9.5e-09 | 5.5e-09 | 8e-09 | 7.667E-09 | 2.021E-09 | 5.500E-09 | 9.500E-09 |
| 22325 | 1117 | P1117C | D | 3.5e-09 | 4.2e-09 | 7.2e-09 | 4.967E-09 | 1.966E-09 | 3.500E-09 | 7.200E-09 |
| 22326 | 1117 | P1117E | D | 1e-08 | 1.4e-08 | 1.9e-08 | 1.433E-08 | 4.509E-09 | 1.000E-08 | 1.900E-08 |
| 22327 | 1117 | P1117Q | D | 8.4e-09 | 6.3e-09 | 1.4e-08 | 9.567E-09 | 3.980E-09 | 6.300E-09 | 1.400E-08 |
| 22328 | 1117 | P1117G | D | 3.1e-08 | 2.2e-09 | 5.5e-09 | 1.290E-08 | 1.576E-08 | 2.200E-09 | 3.100E-08 |
| 22329 | 1117 | P1117H | D | 1e-08 | 3.2e-09 | 1.4e-08 | 9.067E-09 | 5.460E-09 | 3.200E-09 | 1.400E-08 |
| 22330 | 1117 | P1117I | D | 5.8e-09 | 8.5e-09 | 5.3e-09 | 6.533E-09 | 1.721E-09 | 5.300E-09 | 8.500E-09 |
| 22331 | 1117 | P1117L | D | 9.6e-09 | 7.5e-09 | 5.1e-09 | 7.400E-09 | 2.252E-09 | 5.100E-09 | 9.600E-09 |
| 22332 | 1117 | P1117K | D | 6.2e-09 | 1.7e-08 | 2.1e-08 | 1.473E-08 | 7.656E-09 | 6.200E-09 | 2.100E-08 |
| 22333 | 1117 | P1117M | D | 4.6e-09 | 1.2e-08 | 1.8e-08 | 1.153E-08 | 6.712E-09 | 4.600E-09 | 1.800E-08 |
| 22334 | 1117 | P1117F | D | 2.1e-09 | 4.5e-09 | 5.5e-09 | 4.033E-09 | 1.747E-09 | 2.100E-09 | 5.500E-09 |
| 22335 | 1117 | P1117P | D | 5.1e-09 | 9.6e-09 | 4.7e-09 | 6.467E-09 | 2.721E-09 | 4.700E-09 | 9.600E-09 |
| 22336 | 1117 | P1117S | D | 3.3e-09 | 7.3e-09 | 4.8e-09 | 5.133E-09 | 2.021E-09 | 3.300E-09 | 7.300E-09 |
| 22337 | 1117 | P1117T | D | 5.5e-09 | 5.3e-09 | 6.2e-09 | 5.667E-09 | 4.726E-10 | 5.300E-09 | 6.200E-09 |
| 22338 | 1117 | P1117W | D | 1.2e-08 | 4.2e-09 | 2.1e-09 | 6.100E-09 | 5.216E-09 | 2.100E-09 | 1.200E-08 |
| 22339 | 1117 | P1117Y | D | 8.5e-09 | 5.5e-09 | 6.5e-09 | 6.833E-09 | 1.528E-09 | 5.500E-09 | 8.500E-09 |
| 22340 | 1117 | P1117V | D | 1.4e-08 | 9.4e-09 | 5.5e-09 | 9.633E-09 | 4.255E-09 | 5.500E-09 | 1.400E-08 |
| 22341 | 1118 | S1118A | D | 3.7e-09 | 6.6e-09 | 2.4e-09 | 4.233E-09 | 2.150E-09 | 2.400E-09 | 6.600E-09 |
| 22342 | 1118 | S1118R | D | 1.3e-08 | 2.6e-09 | 3.4e-09 | 6.333E-09 | 5.787E-09 | 2.600E-09 | 1.300E-08 |
| 22343 | 1118 | S1118N | D | 7.6e-09 | 4.2e-09 | 1.4e-09 | 4.400E-09 | 3.105E-09 | 1.400E-09 | 7.600E-09 |
| 22344 | 1118 | S1118D | D | 3.8e-09 | 1.6e-08 | 9.2e-09 | 9.667E-09 | 6.113E-09 | 3.800E-09 | 1.600E-08 |
| 22345 | 1118 | S1118C | D | 3.3e-09 | 5.6e-09 | 2.9e-09 | 3.933E-09 | 1.457E-09 | 2.900E-09 | 5.600E-09 |
| 22346 | 1118 | S1118E | D | 6.1e-09 | 3.2e-09 | 2.2e-08 | 1.043E-08 | 1.012E-08 | 3.200E-09 | 2.200E-08 |
| 22347 | 1118 | S1118Q | D | 2e-08 | 4.7e-09 | 9.6e-09 | 1.143E-08 | 7.813E-09 | 4.700E-09 | 2.000E-08 |
| 22348 | 1118 | S1118G | D | 1.3e-08 | 1.1e-08 | 7.7e-09 | 1.057E-08 | 2.676E-09 | 7.700E-09 | 1.300E-08 |
| 22349 | 1118 | S1118H | D | 2.7e-09 | 1.6e-08 | 9.7e-09 | 9.467E-09 | 6.653E-09 | 2.700E-09 | 1.600E-08 |
| 22350 | 1118 | S1118I | D | 3.9e-09 | 2.8e-09 | 1e-08 | 5.567E-09 | 3.879E-09 | 2.800E-09 | 1.000E-08 |
| 22351 | 1118 | S1118L | D | 5.4e-09 | 3.6e-09 | 1e-08 | 6.333E-09 | 3.301E-09 | 3.600E-09 | 1.000E-08 |
| 22352 | 1118 | S1118K | D | 5e-09 | 8.7e-09 | 6.8e-09 | 6.833E-09 | 1.850E-09 | 5.000E-09 | 8.700E-09 |
| 22353 | 1118 | S1118M | D | 1.3e-08 | 3.1e-09 | 8.3e-09 | 8.133E-09 | 4.952E-09 | 3.100E-09 | 1.300E-08 |
| 22354 | 1118 | S1118F | D | 2.5e-09 | 4.5e-09 | 8.9e-09 | 5.300E-09 | 3.274E-09 | 2.500E-09 | 8.900E-09 |

| | | | | | | | | | | |
|---|---|---|---|---|---|---|---|---|---|---|
| 22355 | 1118 | S1118P | D | 1.9e-08 | 1.3e-08 | 6e-09 | 1.267E-08 | 6.506E-09 | 6.000E-09 | 1.900E-08 |
| 22356 | 1118 | S1118S | D | 4.4e-09 | 5.5e-09 | 4e-09 | 4.633E-09 | 7.767E-10 | 4.000E-09 | 5.500E-09 |
| 22357 | 1118 | S1118T | D | 1.4e-08 | 3.4e-09 | 8.7e-09 | 8.700E-09 | 5.300E-09 | 3.400E-09 | 1.400E-08 |
| 22358 | 1118 | S1118W | D | 8.8e-09 | 4.7e-09 | 1e-08 | 7.833E-09 | 2.779E-09 | 4.700E-09 | 1.000E-08 |
| 22359 | 1118 | S1118Y | D | 4.8e-09 | 4e-09 | 6.1e-09 | 4.967E-09 | 1.060E-09 | 4.000E-09 | 6.100E-09 |
| 22360 | 1118 | S1118V | D | 7.1e-09 | 6.7e-09 | 6.1e-09 | 6.633E-09 | 5.033E-10 | 6.100E-09 | 7.100E-09 |
| 22361 | 1119 | R1119A | D | 6.4e-09 | 7.9e-09 | 6.2e-09 | 6.833E-09 | 9.292E-10 | 6.200E-09 | 7.900E-09 |
| 22362 | 1119 | R1119R | D | 6.4e-09 | 1.8e-08 | 1.2e-08 | 1.213E-08 | 5.801E-09 | 6.400E-09 | 1.800E-08 |
| 22363 | 1119 | R1119N | D | 8.5e-09 | 1.3e-08 | 7.3e-09 | 9.600E-09 | 3.005E-09 | 7.300E-09 | 1.300E-08 |
| 22364 | 1119 | R1119D | D | 9.8e-09 | 5.3e-09 | 4.5e-09 | 6.533E-09 | 2.857E-09 | 4.500E-09 | 9.800E-09 |
| 22365 | 1119 | R1119C | D | 3.5e-09 | 7.2e-09 | 6.5e-09 | 5.733E-09 | 1.966E-09 | 3.500E-09 | 7.200E-09 |
| 22366 | 1119 | R1119E | D | 1.5e-09 | 8.4e-09 | 7.1e-09 | 5.667E-09 | 3.667E-09 | 1.500E-09 | 8.400E-09 |
| 22367 | 1119 | R1119Q | D | 1.4e-08 | 1e-08 | 7.2e-09 | 1.040E-08 | 3.418E-09 | 7.200E-09 | 1.400E-08 |
| 22368 | 1119 | R1119G | D | 5e-09 | 7.3e-09 | 2.2e-09 | 4.833E-09 | 2.554E-09 | 2.200E-09 | 7.300E-09 |
| 22369 | 1119 | R1119H | D | 3.4e-09 | 5.9e-09 | 7e-09 | 5.433E-09 | 1.845E-09 | 3.400E-09 | 7.000E-09 |
| 22370 | 1119 | R1119I | D | 6.4e-09 | 5.2e-09 | 1.6e-08 | 9.200E-09 | 5.919E-09 | 5.200E-09 | 1.600E-08 |
| 22371 | 1119 | R1119L | D | 3e-08 | 1.4e-08 | 6.7e-09 | 1.690E-08 | 1.192E-08 | 6.700E-09 | 3.000E-08 |
| 22372 | 1119 | R1119K | D | 6.9e-09 | 7.5e-09 | 6.1e-09 | 6.833E-09 | 7.024E-10 | 6.100E-09 | 7.500E-09 |
| 22373 | 1119 | R1119M | D | 8.1e-09 | 5.1e-09 | 1e-08 | 7.733E-09 | 2.470E-09 | 5.100E-09 | 1.000E-08 |
| 22374 | 1119 | R1119F | D | 9.8e-09 | 1e-08 | 7e-09 | 8.933E-09 | 1.677E-09 | 7.000E-09 | 1.000E-08 |
| 22375 | 1119 | R1119P | D | 4.7e-09 | 1.2e-08 | 1.3e-08 | 9.900E-09 | 4.531E-09 | 4.700E-09 | 1.300E-08 |
| 22376 | 1119 | R1119S | D | 7.5e-09 | 8.4e-09 | 2.9e-09 | 6.267E-09 | 2.950E-09 | 2.900E-09 | 8.400E-09 |
| 22377 | 1119 | R1119T | D | 6.3e-09 | 5.3e-09 | 2.5e-08 | 1.220E-08 | 1.110E-08 | 5.300E-09 | 2.500E-08 |
| 22378 | 1119 | R1119W | D | 8.4e-09 | 8e-09 | 1.2e-08 | 9.467E-09 | 2.203E-09 | 8.000E-09 | 1.200E-08 |
| 22379 | 1119 | R1119Y | D | 5.4e-09 | 4.1e-09 | 8e-09 | 5.833E-09 | 1.986E-09 | 4.100E-09 | 8.000E-09 |
| 22380 | 1119 | R1119V | D | 6e-09 | 5.6e-09 | 5.1e-09 | 5.567E-09 | 4.509E-10 | 5.100E-09 | 6.000E-09 |
| 22381 | 1120 | F1120A | D | 8.8e-09 | 1.2e-08 | 1.2e-08 | 1.093E-08 | 1.848E-09 | 8.800E-09 | 1.200E-08 |
| 22382 | 1120 | F1120R | D | 6.6e-09 | 1.5e-08 | 4.5e-09 | 8.700E-09 | 5.556E-09 | 4.500E-09 | 1.500E-08 |
| 22383 | 1120 | F1120N | D | 6e-09 | 3.7e-09 | 1.5e-08 | 8.233E-09 | 5.972E-09 | 3.700E-09 | 1.500E-08 |
| 22384 | 1120 | F1120D | D | 5.9e-09 | 5e-09 | 7.5e-09 | 6.133E-09 | 1.266E-09 | 5.000E-09 | 7.500E-09 |
| 22385 | 1120 | F1120C | D | 5.5e-09 | 7.4e-09 | 9.7e-09 | 7.533E-09 | 2.103E-09 | 5.500E-09 | 9.700E-09 |
| 22386 | 1120 | F1120E | D | 1.1e-08 | 9.3e-09 | 6.1e-09 | 8.800E-09 | 2.488E-09 | 6.100E-09 | 1.100E-08 |
| 22387 | 1120 | F1120Q | D | 3.5e-09 | 1.3e-08 | 3.3e-09 | 6.600E-09 | 5.543E-09 | 3.300E-09 | 1.300E-08 |
| 22388 | 1120 | F1120G | D | 1.8e-08 | 4.7e-08 | 3.6e-09 | 2.287E-08 | 2.211E-08 | 3.600E-09 | 4.700E-08 |
| 22389 | 1120 | F1120H | D | 2.1e-08 | 4.4e-09 | 2e-09 | 9.133E-09 | 1.035E-08 | 2.000E-09 | 2.100E-08 |
| 22390 | 1120 | F1120I | D | 1.7e-08 | 8e-09 | 1.1e-08 | 1.200E-08 | 4.583E-09 | 8.000E-09 | 1.700E-08 |
| 22391 | 1120 | F1120L | D | 4.4e-09 | 2.7e-09 | 6.2e-09 | 4.433E-09 | 1.750E-09 | 2.700E-09 | 6.200E-09 |
| 22392 | 1120 | F1120K | D | 8.2e-09 | 5.2e-09 | 1e-08 | 7.800E-09 | 2.425E-09 | 5.200E-09 | 1.000E-08 |
| 22393 | 1120 | F1120M | D | 1.2e-08 | 6.6e-09 | 1.8e-08 | 1.220E-08 | 5.703E-09 | 6.600E-09 | 1.800E-08 |
| 22394 | 1120 | F1120F | D | 4.6e-09 | 3.4e-09 | 5.5e-09 | 4.500E-09 | 1.054E-09 | 3.400E-09 | 5.500E-09 |
| 22395 | 1120 | F1120P | D | 7.6e-09 | 5.1e-09 | 1.9e-08 | 1.057E-08 | 7.410E-09 | 5.100E-09 | 1.900E-08 |
| 22396 | 1120 | F1120S | D | 1.5e-09 | 7e-09 | 2.9e-09 | 3.800E-09 | 2.858E-09 | 1.500E-09 | 7.000E-09 |
| 22397 | 1120 | F1120T | D | 8.3e-09 | 9.8e-09 | 5.3e-09 | 7.800E-09 | 2.291E-09 | 5.300E-09 | 9.800E-09 |
| 22398 | 1120 | F1120W | D | 1.5e-08 | 9.8e-09 | 5.8e-09 | 1.020E-08 | 4.613E-09 | 5.800E-09 | 1.500E-08 |
| 22399 | 1120 | F1120Y | D | 1.1e-08 | 8e-09 | 5.2e-09 | 8.067E-09 | 2.901E-09 | 5.200E-09 | 1.100E-08 |
| 22400 | 1120 | F1120V | D | 1.2e-08 | 6.8e-09 | 1.1e-08 | 9.933E-09 | 2.759E-09 | 6.800E-09 | 1.200E-08 |
| 22401 | 1121 | S1121A | D | 4.7e-09 | 6.4e-09 | 7e-09 | 6.033E-09 | 1.193E-09 | 4.700E-09 | 7.000E-09 |
| 22402 | 1121 | S1121R | D | 8.3e-09 | 8.3e-09 | 1.6e-08 | 1.087E-08 | 4.446E-09 | 8.300E-09 | 1.600E-08 |
| 22403 | 1121 | S1121N | D | 5.1e-09 | 3.3e-09 | 3.3e-09 | 3.900E-09 | 1.039E-09 | 3.300E-09 | 5.100E-09 |
| 22404 | 1121 | S1121D | D | 6.6e-09 | 1.9e-08 | 1.7e-09 | 9.100E-09 | 8.917E-09 | 1.700E-09 | 1.900E-08 |
| 22405 | 1121 | S1121C | D | 1.9e-08 | 1.1e-08 | 4.1e-09 | 1.137E-08 | 7.457E-09 | 4.100E-09 | 1.900E-08 |
| 22406 | 1121 | S1121E | D | 3.4e-09 | 7.4e-09 | 5.2e-09 | 5.333E-09 | 2.003E-09 | 3.400E-09 | 7.400E-09 |
| 22407 | 1121 | S1121Q | D | 3.4e-09 | 8.5e-09 | 2.9e-08 | 1.363E-08 | 1.355E-08 | 3.400E-09 | 2.900E-08 |
| 22408 | 1121 | S1121G | D | 1.9e-08 | 4.3e-09 | 3.1e-09 | 8.800E-09 | 8.854E-09 | 3.100E-09 | 1.900E-08 |
| 22409 | 1121 | S1121H | D | 9.2e-09 | 6.4e-09 | 3.8e-09 | 6.467E-09 | 2.701E-09 | 3.800E-09 | 9.200E-09 |
| 22410 | 1121 | S1121I | D | 2.1e-09 | 3e-09 | 1.7e-08 | 7.367E-09 | 8.355E-09 | 2.100E-09 | 1.700E-08 |
| 22411 | 1121 | S1121L | D | 5.7e-09 | 2.1e-08 | 6.5e-09 | 1.107E-08 | 8.612E-09 | 5.700E-09 | 2.100E-08 |
| 22412 | 1121 | S1121K | D | 3.8e-09 | 1.1e-08 | 7.8e-09 | 7.533E-09 | 3.607E-09 | 3.800E-09 | 1.100E-08 |
| 22413 | 1121 | S1121M | D | 2e-08 | 1.2e-08 | 1.4e-08 | 1.533E-08 | 4.163E-09 | 1.200E-08 | 2.000E-08 |
| 22414 | 1121 | S1121F | D | 7.3e-09 | 1.2e-08 | 2e-08 | 1.310E-08 | 6.421E-09 | 7.300E-09 | 2.000E-08 |
| 22415 | 1121 | S1121P | D | 1.4e-08 | 3.6e-09 | 5.6e-09 | 7.733E-09 | 5.518E-09 | 3.600E-09 | 1.400E-08 |
| 22416 | 1121 | S1121S | D | 7.5e-09 | 9.1e-09 | 2e-08 | 1.220E-08 | 6.802E-09 | 7.500E-09 | 2.000E-08 |
| 22417 | 1121 | S1121T | D | 6.5e-09 | 4.1e-09 | 4.7e-09 | 5.100E-09 | 1.249E-09 | 4.100E-09 | 6.500E-09 |
| 22418 | 1121 | S1121W | D | 9.4e-09 | 5.2e-09 | 6.1e-09 | 6.900E-09 | 2.211E-09 | 5.200E-09 | 9.400E-09 |
| 22419 | 1121 | S1121Y | D | 1.8e-08 | 1.2e-08 | 5.7e-09 | 1.190E-08 | 6.151E-09 | 5.700E-09 | 1.800E-08 |
| 22420 | 1121 | S1121V | D | 6e-09 | 3.9e-09 | 6.4e-09 | 5.433E-09 | 1.343E-09 | 3.900E-09 | 6.400E-09 |
| 22421 | 1122 | G1122A | D | 7.7e-09 | 2.2e-09 | 3.4e-09 | 4.433E-09 | 2.892E-09 | 2.200E-09 | 7.700E-09 |
| 22422 | 1122 | G1122R | D | 7.3e-09 | 9.2e-09 | 7e-09 | 7.833E-09 | 1.193E-09 | 7.000E-09 | 9.200E-09 |
| 22423 | 1122 | G1122N | D | 4.3e-09 | 1.8e-09 | 6.7e-09 | 4.267E-09 | 2.450E-09 | 1.800E-09 | 6.700E-09 |
| 22424 | 1122 | G1122D | D | 7.5e-09 | 8.8e-09 | 1e-08 | 8.767E-09 | 1.250E-09 | 7.500E-09 | 1.000E-08 |
| 22425 | 1122 | G1122C | D | 7.8e-09 | 3.3e-09 | 5.4e-09 | 5.500E-09 | 2.252E-09 | 3.300E-09 | 7.800E-09 |
| 22426 | 1122 | G1122E | D | 7.3e-09 | 1.4e-08 | 6.9e-09 | 9.400E-09 | 3.989E-09 | 6.900E-09 | 1.400E-08 |
| 22427 | 1122 | G1122Q | D | 3.2e-09 | 5.8e-09 | 3.7e-09 | 4.233E-09 | 1.380E-09 | 3.200E-09 | 5.800E-09 |
| 22428 | 1122 | G1122G | D | 5.1e-09 | 1.4e-08 | 1.4e-08 | 1.103E-08 | 5.138E-09 | 5.100E-09 | 1.400E-08 |
| 22429 | 1122 | G1122H | D | 5.3e-09 | 9.3e-09 | 6.3e-09 | 6.967E-09 | 2.082E-09 | 5.300E-09 | 9.300E-09 |
| 22430 | 1122 | G1122I | D | 4.7e-09 | 4.9e-09 | 1.1e-08 | 6.867E-09 | 3.581E-09 | 4.700E-09 | 1.100E-08 |
| 22431 | 1122 | G1122L | D | 5e-09 | 2.5e-08 | 1.2e-08 | 1.400E-08 | 1.015E-08 | 5.000E-09 | 2.500E-08 |
| 22432 | 1122 | G1122K | D | 4.9e-09 | 9.3e-09 | 9.4e-09 | 7.867E-09 | 2.570E-09 | 4.900E-09 | 9.400E-09 |
| 22433 | 1122 | G1122M | D | 1.5e-08 | 4.9e-09 | 8.8e-09 | 9.567E-09 | 5.093E-09 | 4.900E-09 | 1.500E-08 |
| 22434 | 1122 | G1122F | D | 8.7e-09 | 8.7e-09 | 2.3e-09 | 6.567E-09 | 3.695E-09 | 2.300E-09 | 8.700E-09 |
| 22435 | 1122 | G1122P | D | 6.7e-09 | 4.3e-09 | 1.6e-08 | 9.000E-09 | 6.180E-09 | 4.300E-09 | 1.600E-08 |
| 22436 | 1122 | G1122S | D | 2e-09 | 4.1e-09 | 9.1e-09 | 5.067E-09 | 3.647E-09 | 2.000E-09 | 9.100E-09 |
| 22437 | 1122 | G1122T | D | 4e-09 | 4.9e-09 | 3.1e-08 | 1.330E-08 | 1.534E-08 | 4.000E-09 | 3.100E-08 |
| 22438 | 1122 | G1122W | D | 8.3e-09 | 6.5e-09 | 6.6e-09 | 7.133E-09 | 1.012E-09 | 6.500E-09 | 8.300E-09 |
| 22439 | 1122 | G1122Y | D | 8.8e-09 | 6e-09 | 4.3e-09 | 6.367E-09 | 2.272E-09 | 4.300E-09 | 8.800E-09 |

| | | | | | | | | | | |
|---|---|---|---|---|---|---|---|---|---|---|
| 22440 | 1122 | G1122V | D | 6.2e-09 | 4.9e-09 | 2.3e-08 | 1.137E-08 | 1.010E-08 | 4.900E-09 | 2.300E-08 |
| 22441 | 1123 | S1123A | D | 3.1e-09 | 3.3e-09 | 6e-09 | 4.133E-09 | 1.620E-09 | 3.100E-09 | 6.000E-09 |
| 22442 | 1123 | S1123R | D | 3.3e-09 | 6.7e-09 | 2.5e-09 | 4.167E-09 | 2.230E-09 | 2.500E-09 | 6.700E-09 |
| 22443 | 1123 | S1123N | D | 5.1e-09 | 7.9e-09 | 7.8e-09 | 6.933E-09 | 1.589E-09 | 5.100E-09 | 7.900E-09 |
| 22444 | 1123 | S1123D | D | 9.2e-09 | 1.2e-09 | 3.2e-09 | 4.533E-09 | 4.163E-09 | 1.200E-09 | 9.200E-09 |
| 22445 | 1123 | S1123C | D | 9.7e-09 | 8.4e-09 | 1.9e-09 | 6.667E-09 | 4.179E-09 | 1.900E-09 | 9.700E-09 |
| 22446 | 1123 | S1123E | D | 9.5e-09 | 1.3e-08 | 9e-09 | 1.050E-08 | 2.179E-09 | 9.000E-09 | 1.300E-08 |
| 22447 | 1123 | S1123Q | D | 1.1e-08 | 5.8e-09 | 4.7e-09 | 7.167E-09 | 3.365E-09 | 4.700E-09 | 1.100E-08 |
| 22448 | 1123 | S1123G | D | 1.4e-08 | 8.5e-09 | 5.7e-09 | 9.400E-09 | 4.223E-09 | 5.700E-09 | 1.400E-08 |
| 22449 | 1123 | S1123H | D | 9.8e-09 | 5.7e-09 | 5.1e-09 | 6.867E-09 | 2.558E-09 | 5.100E-09 | 9.800E-09 |
| 22450 | 1123 | S1123I | D | 2.2e-09 | 6.5e-09 | 4.7e-09 | 4.467E-09 | 2.159E-09 | 2.200E-09 | 6.500E-09 |
| 22451 | 1123 | S1123L | D | 3.8e-09 | 4.4e-09 | 5.4e-09 | 4.533E-09 | 8.083E-10 | 3.800E-09 | 5.400E-09 |
| 22452 | 1123 | S1123K | D | 9e-09 | 5.2e-09 | 6.5e-09 | 6.900E-09 | 1.931E-09 | 5.200E-09 | 9.000E-09 |
| 22453 | 1123 | S1123M | D | 7.7e-09 | 1.3e-08 | 1.8e-08 | 1.290E-08 | 5.151E-09 | 7.700E-09 | 1.800E-08 |
| 22454 | 1123 | S1123F | D | 8.1e-09 | 9.5e-09 | 4.1e-09 | 7.233E-09 | 2.802E-09 | 4.100E-09 | 9.500E-09 |
| 22455 | 1123 | S1123P | D | 9.6e-09 | 7.6e-09 | 8.3e-09 | 8.500E-09 | 1.015E-09 | 7.600E-09 | 9.600E-09 |
| 22456 | 1123 | S1123S | D | 5.8e-09 | 2.6e-09 | 9.1e-09 | 5.833E-09 | 3.250E-09 | 2.600E-09 | 9.100E-09 |
| 22457 | 1123 | S1123T | D | 7.6e-09 | 3.5e-09 | 3.8e-09 | 4.967E-09 | 2.285E-09 | 3.500E-09 | 7.600E-09 |
| 22458 | 1123 | S1123W | D | 2.1e-09 | 4.5e-10 | 7.8e-09 | 3.450E-09 | 3.856E-09 | 4.500E-10 | 7.800E-09 |
| 22459 | 1123 | S1123Y | D | 3e-09 | 2.7e-09 | 4.5e-09 | 3.400E-09 | 9.644E-10 | 2.700E-09 | 4.500E-09 |
| 22460 | 1123 | S1123V | D | 4e-09 | 1.8e-08 | 6.2e-09 | 9.400E-09 | 7.529E-09 | 4.000E-09 | 1.800E-08 |
| 22461 | 1124 | R1124A | D | 1.7e-08 | 1.9e-08 | 4.1e-09 | 1.337E-08 | 8.087E-09 | 4.100E-09 | 1.900E-08 |
| 22462 | 1124 | R1124R | D | 9.2e-09 | 5.3e-09 | 3.6e-09 | 6.033E-09 | 2.871E-09 | 3.600E-09 | 9.200E-09 |
| 22463 | 1124 | R1124N | D | 5.3e-09 | 1.1e-08 | 5e-09 | 7.100E-09 | 3.381E-09 | 5.000E-09 | 1.100E-08 |
| 22464 | 1124 | R1124D | D | 8.1e-09 | 1.6e-08 | 6.5e-09 | 1.020E-08 | 5.086E-09 | 6.500E-09 | 1.600E-08 |
| 22465 | 1124 | R1124C | D | 1e-08 | 2.8e-09 | 8.3e-09 | 7.033E-09 | 3.763E-09 | 2.800E-09 | 1.000E-08 |
| 22466 | 1124 | R1124E | D | 2.1e-08 | 6.2e-09 | 6.6e-09 | 1.127E-08 | 8.432E-09 | 6.200E-09 | 2.100E-08 |
| 22467 | 1124 | R1124Q | D | 1.9e-08 | 1.8e-08 | 8.9e-09 | 1.530E-08 | 5.565E-09 | 8.900E-09 | 1.900E-08 |
| 22468 | 1124 | R1124G | D | 3.4e-09 | 1e-08 | 1e-08 | 7.800E-09 | 3.811E-09 | 3.400E-09 | 1.000E-08 |
| 22469 | 1124 | R1124H | D | 4.6e-09 | 1.7e-08 | 2.8e-09 | 8.133E-09 | 7.731E-09 | 2.800E-09 | 1.700E-08 |
| 22470 | 1124 | R1124I | D | 9.4e-09 | 8.1e-09 | 1.6e-08 | 1.117E-08 | 4.236E-09 | 8.100E-09 | 1.600E-08 |
| 22471 | 1124 | R1124L | D | 8.6e-09 | 1.7e-08 | 1.1e-08 | 1.220E-08 | 4.327E-09 | 8.600E-09 | 1.700E-08 |
| 22472 | 1124 | R1124K | D | 1.1e-08 | 6.5e-09 | 5.4e-09 | 7.633E-09 | 2.967E-09 | 5.400E-09 | 1.100E-08 |
| 22473 | 1124 | R1124M | D | 3.9e-09 | 1.4e-08 | 4.7e-09 | 7.533E-09 | 5.615E-09 | 3.900E-09 | 1.400E-08 |
| 22474 | 1124 | R1124F | D | 4e-09 | 1.7e-08 | 4.4e-09 | 8.467E-09 | 7.393E-09 | 4.000E-09 | 1.700E-08 |
| 22475 | 1124 | R1124P | D | 1.1e-08 | 4.7e-09 | 1.1e-08 | 8.900E-09 | 3.637E-09 | 4.700E-09 | 1.100E-08 |
| 22476 | 1124 | R1124S | D | 5.7e-09 | 1.2e-08 | 4.7e-09 | 7.467E-09 | 3.958E-09 | 4.700E-09 | 1.200E-08 |
| 22477 | 1124 | R1124T | D | 6.5e-09 | 5.8e-09 | 6.5e-09 | 6.267E-09 | 4.041E-10 | 5.800E-09 | 6.500E-09 |
| 22478 | 1124 | R1124W | D | 9.2e-09 | 4e-09 | 1e-08 | 7.733E-09 | 3.258E-09 | 4.000E-09 | 1.000E-08 |
| 22479 | 1124 | R1124Y | D | 3e-09 | 2.2e-08 | 5.4e-09 | 1.013E-08 | 1.035E-08 | 3.000E-09 | 2.200E-08 |
| 22480 | 1124 | R1124V | D | 1e-08 | 4.6e-09 | 8.6e-09 | 7.733E-09 | 2.802E-09 | 4.600E-09 | 1.000E-08 |
| 22481 | 1125 | S1125A | D | 8.2e-09 | 7e-09 | 3e-09 | 6.067E-09 | 2.723E-09 | 3.000E-09 | 8.200E-09 |
| 22482 | 1125 | S1125R | D | 2.2e-08 | 2.7e-08 | 4.2e-09 | 1.773E-08 | 1.198E-08 | 4.200E-09 | 2.700E-08 |
| 22483 | 1125 | S1125N | D | 1.8e-08 | 1.5e-08 | 1.2e-08 | 1.500E-08 | 3.000E-09 | 1.200E-08 | 1.800E-08 |
| 22484 | 1125 | S1125D | D | 1.7e-09 | 2.8e-08 | 3.6e-09 | 1.110E-08 | 1.467E-08 | 1.700E-09 | 2.800E-08 |
| 22485 | 1125 | S1125C | D | 3.3e-08 | 4.7e-09 | 1.2e-08 | 1.657E-08 | 1.469E-08 | 4.700E-09 | 3.300E-08 |
| 22486 | 1125 | S1125E | D | 1.2e-08 | 7.1e-09 | 2.9e-09 | 7.333E-09 | 4.554E-09 | 2.900E-09 | 1.200E-08 |
| 22487 | 1125 | S1125Q | D | 8.4e-09 | 6.3e-09 | 1.1e-08 | 8.567E-09 | 2.354E-09 | 6.300E-09 | 1.100E-08 |
| 22488 | 1125 | S1125G | D | 1.5e-08 | 4.6e-09 | 1.6e-08 | 1.187E-08 | 6.313E-09 | 4.600E-09 | 1.600E-08 |
| 22489 | 1125 | S1125H | D | 4.1e-09 | 6.5e-09 | 1.7e-09 | 4.100E-09 | 2.400E-09 | 1.700E-09 | 6.500E-09 |
| 22490 | 1125 | S1125I | D | 9.4e-09 | 4e-09 | 7.8e-09 | 7.067E-09 | 2.774E-09 | 4.000E-09 | 9.400E-09 |
| 22491 | 1125 | S1125L | D | 1.4e-08 | 9.4e-09 | 7.6e-09 | 1.033E-08 | 3.301E-09 | 7.600E-09 | 1.400E-08 |
| 22492 | 1125 | S1125K | D | 8.2e-09 | 5.5e-09 | 6.9e-09 | 6.867E-09 | 1.350E-09 | 5.500E-09 | 8.200E-09 |
| 22493 | 1125 | S1125M | D | 8.3e-09 | 1.8e-09 | 3.1e-09 | 4.400E-09 | 3.439E-09 | 1.800E-09 | 8.300E-09 |
| 22494 | 1125 | S1125F | D | 5e-09 | 1.3e-08 | 5.7e-09 | 7.900E-09 | 4.431E-09 | 5.000E-09 | 1.300E-08 |
| 22495 | 1125 | S1125P | D | 3.1e-09 | 6.4e-09 | 6.8e-09 | 5.433E-09 | 2.031E-09 | 3.100E-09 | 6.800E-09 |
| 22496 | 1125 | S1125S | D | 5e-09 | 2.7e-09 | 8.9e-09 | 5.533E-09 | 3.134E-09 | 2.700E-09 | 8.900E-09 |
| 22497 | 1125 | S1125T | D | 8.6e-09 | 9.8e-09 | 1.7e-08 | 1.180E-08 | 4.543E-09 | 8.600E-09 | 1.700E-08 |
| 22498 | 1125 | S1125W | D | 2e-08 | 5.1e-09 | 5e-09 | 1.003E-08 | 8.632E-09 | 5.000E-09 | 2.000E-08 |
| 22499 | 1125 | S1125Y | D | 2.6e-09 | 4.9e-09 | 1.3e-08 | 6.833E-09 | 5.463E-09 | 2.600E-09 | 1.300E-08 |
| 22500 | 1125 | S1125V | D | 7.4e-09 | 5.6e-09 | 6.2e-09 | 6.400E-09 | 9.165E-10 | 5.600E-09 | 7.400E-09 |
| 22501 | 1126 | G1126A | D | 1.2e-08 | 1.1e-08 | 5.1e-09 | 9.367E-09 | 3.729E-09 | 5.100E-09 | 1.200E-08 |
| 22502 | 1126 | G1126R | D | 1e-08 | 1.8e-08 | 3.3e-09 | 1.043E-08 | 7.360E-09 | 3.300E-09 | 1.800E-08 |
| 22503 | 1126 | G1126N | D | 1.3e-08 | 7e-09 | 1.3e-08 | 1.100E-08 | 3.464E-09 | 7.000E-09 | 1.300E-08 |
| 22504 | 1126 | G1126D | D | 7.7e-09 | 5.9e-09 | 9.3e-09 | 7.633E-09 | 1.701E-09 | 5.900E-09 | 9.300E-09 |
| 22505 | 1126 | G1126C | D | 9.8e-09 | 1e-08 | 1.2e-08 | 1.060E-08 | 1.217E-09 | 9.800E-09 | 1.200E-08 |
| 22506 | 1126 | G1126E | D | 1.2e-08 | 1e-08 | 5.1e-09 | 9.033E-09 | 3.550E-09 | 5.100E-09 | 1.200E-08 |
| 22507 | 1126 | G1126Q | D | 3.2e-09 | 2.3e-09 | 1.6e-08 | 7.167E-09 | 7.663E-09 | 2.300E-09 | 1.600E-08 |
| 22508 | 1126 | G1126G | D | 1.8e-08 | 5.7e-09 | 3.6e-09 | 9.100E-09 | 7.779E-09 | 3.600E-09 | 1.800E-08 |
| 22509 | 1126 | G1126H | D | 2.5e-09 | 6.3e-09 | 6.9e-09 | 5.233E-09 | 2.386E-09 | 2.500E-09 | 6.900E-09 |
| 22510 | 1126 | G1126I | D | 1e-08 | 1e-08 | 4.9e-09 | 8.300E-09 | 2.944E-09 | 4.900E-09 | 1.000E-08 |
| 22511 | 1126 | G1126L | D | 1.4e-08 | 1.1e-08 | 4.5e-09 | 9.833E-09 | 4.856E-09 | 4.500E-09 | 1.400E-08 |
| 22512 | 1126 | G1126K | D | 8.2e-09 | 1.3e-08 | 6.3e-09 | 9.167E-09 | 3.453E-09 | 6.300E-09 | 1.300E-08 |
| 22513 | 1126 | G1126M | D | 6.8e-09 | 5.5e-09 | 5.3e-09 | 5.867E-09 | 8.145E-10 | 5.300E-09 | 6.800E-09 |
| 22514 | 1126 | G1126F | D | 2.1e-09 | 6.6e-09 | 9.4e-09 | 6.033E-09 | 3.683E-09 | 2.100E-09 | 9.400E-09 |
| 22515 | 1126 | G1126P | D | 4.8e-09 | 8.4e-09 | 3.8e-09 | 5.667E-09 | 2.419E-09 | 3.800E-09 | 8.400E-09 |
| 22516 | 1126 | G1126S | D | 4.8e-09 | 5.9e-09 | 9.3e-09 | 6.667E-09 | 2.346E-09 | 4.800E-09 | 9.300E-09 |
| 22517 | 1126 | G1126T | D | 7.6e-09 | 3e-09 | 5.1e-09 | 5.233E-09 | 2.303E-09 | 3.000E-09 | 7.600E-09 |
| 22518 | 1126 | G1126W | D | 2.3e-09 | 4.4e-09 | 4.8e-09 | 3.833E-09 | 1.343E-09 | 2.300E-09 | 4.800E-09 |
| 22519 | 1126 | G1126Y | D | 1.3e-08 | 2.8e-08 | 2.7e-09 | 1.457E-08 | 1.272E-08 | 2.700E-09 | 2.800E-08 |
| 22520 | 1126 | G1126V | D | 1.1e-08 | 6.7e-09 | 2.6e-09 | 6.767E-09 | 4.200E-09 | 2.600E-09 | 1.100E-08 |
| 22521 | 1127 | T1127A | D | 7.2e-09 | 2.7e-09 | 1.1e-08 | 6.967E-09 | 4.155E-09 | 2.700E-09 | 1.100E-08 |
| 22522 | 1127 | T1127R | D | 1.4e-08 | 3.8e-09 | 2.8e-09 | 6.867E-09 | 6.198E-09 | 2.800E-09 | 1.400E-08 |
| 22523 | 1127 | T1127N | D | 2.5e-09 | 6.3e-09 | 4.3e-09 | 4.367E-09 | 1.901E-09 | 2.500E-09 | 6.300E-09 |
| 22524 | 1127 | T1127D | D | 1.5e-08 | 6.8e-09 | 5.4e-09 | 9.067E-09 | 5.186E-09 | 5.400E-09 | 1.500E-08 |

| | | | | | | | | | | |
|---|---|---|---|---|---|---|---|---|---|---|
| 22525 | 1127 | T1127C | D | 7.5e-09 | 5.2e-09 | 2.1e-09 | 4.933E-09 | 2.710E-09 | 2.100E-09 | 7.500E-09 |
| 22526 | 1127 | T1127E | D | 5.6e-09 | 1.4e-08 | 3.2e-09 | 7.600E-09 | 5.671E-09 | 3.200E-09 | 1.400E-08 |
| 22527 | 1127 | T1127Q | D | 3.9e-09 | 5.6e-09 | 4.8e-09 | 4.767E-09 | 8.505E-10 | 3.900E-09 | 5.600E-09 |
| 22528 | 1127 | T1127G | D | 8.3e-09 | 1.4e-08 | 9.9e-09 | 1.073E-08 | 2.940E-09 | 8.300E-09 | 1.400E-08 |
| 22529 | 1127 | T1127H | D | 3.2e-09 | 3.4e-09 | 6e-09 | 4.200E-09 | 1.562E-09 | 3.200E-09 | 6.000E-09 |
| 22530 | 1127 | T1127I | D | 2e-09 | 8.7e-09 | 4.3e-09 | 5.000E-09 | 3.404E-09 | 2.000E-09 | 8.700E-09 |
| 22531 | 1127 | T1127L | D | 2.7e-09 | 3e-09 | 7e-09 | 4.233E-09 | 2.401E-09 | 2.700E-09 | 7.000E-09 |
| 22532 | 1127 | T1127K | D | 1.4e-08 | 1.5e-08 | 6.4e-09 | 1.180E-08 | 4.703E-09 | 6.400E-09 | 1.500E-08 |
| 22533 | 1127 | T1127M | D | 8.7e-09 | 1.6e-08 | 3.6e-09 | 9.433E-09 | 6.232E-09 | 3.600E-09 | 1.600E-08 |
| 22534 | 1127 | T1127F | D | 7.4e-09 | 2.5e-09 | 1.2e-08 | 7.300E-09 | 4.751E-09 | 2.500E-09 | 1.200E-08 |
| 22535 | 1127 | T1127P | D | 3.4e-09 | 7e-09 | 1.7e-08 | 9.133E-09 | 7.047E-09 | 3.400E-09 | 1.700E-08 |
| 22536 | 1127 | T1127S | D | 5.5e-09 | 8.5e-09 | 1.3e-08 | 9.000E-09 | 3.775E-09 | 5.500E-09 | 1.300E-08 |
| 22537 | 1127 | T1127T | D | 1.4e-08 | 5.5e-09 | 1.1e-08 | 1.017E-08 | 4.311E-09 | 5.500E-09 | 1.400E-08 |
| 22538 | 1127 | T1127W | D | 7e-09 | 1e-08 | 2.5e-09 | 6.500E-09 | 3.775E-09 | 2.500E-09 | 1.000E-08 |
| 22539 | 1127 | T1127Y | D | 4.5e-09 | 6.1e-09 | 1.8e-08 | 9.533E-09 | 7.376E-09 | 4.500E-09 | 1.800E-08 |
| 22540 | 1127 | T1127V | D | 4.1e-09 | 1.8e-08 | 5.3e-09 | 9.133E-09 | 7.702E-09 | 4.100E-09 | 1.800E-08 |
| 22541 | 1128 | D1128A | D | 2.1e-08 | 3.5e-09 | 5.9e-09 | 1.013E-08 | 9.487E-09 | 3.500E-09 | 2.100E-08 |
| 22542 | 1128 | D1128R | D | 1.3e-08 | 6.3e-09 | 1.4e-08 | 1.110E-08 | 4.187E-09 | 6.300E-09 | 1.400E-08 |
| 22543 | 1128 | D1128N | D | 1.5e-08 | 1.5e-08 | 1.2e-08 | 1.400E-08 | 1.732E-09 | 1.200E-08 | 1.500E-08 |
| 22544 | 1128 | D1128D | D | 3.4e-09 | 1.7e-09 | 3.8e-09 | 2.967E-09 | 1.115E-09 | 1.700E-09 | 3.800E-09 |
| 22545 | 1128 | D1128C | D | 1.7e-08 | 2.5e-08 | 7.5e-09 | 1.650E-08 | 8.761E-09 | 7.500E-09 | 2.500E-08 |
| 22546 | 1128 | D1128E | D | 8.1e-09 | 4.2e-09 | 1e-08 | 7.433E-09 | 2.957E-09 | 4.200E-09 | 1.000E-08 |
| 22547 | 1128 | D1128Q | D | 4.3e-09 | 1.5e-08 | 1.4e-08 | 1.110E-08 | 5.910E-09 | 4.300E-09 | 1.500E-08 |
| 22548 | 1128 | D1128G | D | 1.8e-09 | 1e-08 | 1.8e-08 | 9.933E-09 | 8.100E-09 | 1.800E-09 | 1.800E-08 |
| 22549 | 1128 | D1128H | D | 1.9e-09 | 3.4e-09 | 4.7e-09 | 3.333E-09 | 1.401E-09 | 1.900E-09 | 4.700E-09 |
| 22550 | 1128 | D1128I | D | 1.1e-08 | 7.7e-09 | 4.7e-09 | 7.800E-09 | 3.151E-09 | 4.700E-09 | 1.100E-08 |
| 22551 | 1128 | D1128L | D | 9.1e-09 | 3.8e-08 | 5.6e-09 | 1.757E-08 | 1.778E-08 | 5.600E-09 | 3.800E-08 |
| 22552 | 1128 | D1128K | D | 1.8e-08 | 1.1e-08 | 4.3e-09 | 1.110E-08 | 6.851E-09 | 4.300E-09 | 1.800E-08 |
| 22553 | 1128 | D1128M | D | 8.7e-09 | 4.3e-09 | 8.3e-09 | 7.100E-09 | 2.433E-09 | 4.300E-09 | 8.700E-09 |
| 22554 | 1128 | D1128F | D | 7e-09 | 5.4e-09 | 5.6e-09 | 6.000E-09 | 8.718E-10 | 5.400E-09 | 7.000E-09 |
| 22555 | 1128 | D1128P | D | 3.2e-09 | 4e-09 | 4e-09 | 3.733E-09 | 4.619E-10 | 3.200E-09 | 4.000E-09 |
| 22556 | 1128 | D1128S | D | 9.7e-09 | 1.3e-08 | 1.1e-08 | 1.123E-08 | 1.662E-09 | 9.700E-09 | 1.300E-08 |
| 22557 | 1128 | D1128T | D | 1.5e-08 | 7.1e-09 | 6.3e-09 | 9.467E-09 | 4.809E-09 | 6.300E-09 | 1.500E-08 |
| 22558 | 1128 | D1128W | D | 9.1e-09 | 2.2e-08 | 1.4e-08 | 1.503E-08 | 6.512E-09 | 9.100E-09 | 2.200E-08 |
| 22559 | 1128 | D1128Y | D | 9.2e-09 | 4.1e-09 | 9.1e-09 | 7.467E-09 | 2.916E-09 | 4.100E-09 | 9.200E-09 |
| 22560 | 1128 | D1128V | D | 5.7e-09 | 1e-08 | 1.4e-08 | 9.900E-09 | 4.151E-09 | 5.700E-09 | 1.400E-08 |
| 22561 | 1129 | F1129A | D | 6e-09 | 4.7e-09 | 1.1e-08 | 7.233E-09 | 3.326E-09 | 4.700E-09 | 1.100E-08 |
| 22562 | 1129 | F1129R | D | 7.8e-09 | 1.4e-08 | 6.5e-09 | 9.433E-09 | 4.008E-09 | 6.500E-09 | 1.400E-08 |
| 22563 | 1129 | F1129N | D | 1.2e-08 | 6.1e-09 | 6.7e-09 | 8.267E-09 | 3.247E-09 | 6.100E-09 | 1.200E-08 |
| 22564 | 1129 | F1129D | D | 1e-08 | 1.5e-08 | 5.7e-09 | 1.023E-08 | 4.654E-09 | 5.700E-09 | 1.500E-08 |
| 22565 | 1129 | F1129C | D | 1.4e-08 | 1.3e-08 | 2.1e-08 | 1.600E-08 | 4.359E-09 | 1.300E-08 | 2.100E-08 |
| 22566 | 1129 | F1129E | D | 3.5e-08 | 4e-09 | 5.4e-09 | 1.480E-08 | 1.751E-08 | 4.000E-09 | 3.500E-08 |
| 22567 | 1129 | F1129Q | D | 9.8e-09 | 1.1e-09 | 4.3e-09 | 5.067E-09 | 4.400E-09 | 1.100E-09 | 9.800E-09 |
| 22568 | 1129 | F1129G | D | 3.2e-09 | 2.7e-09 | 4.5e-08 | 1.697E-08 | 2.428E-08 | 2.700E-09 | 4.500E-08 |
| 22569 | 1129 | F1129H | D | 3.2e-09 | 1e-08 | 9e-09 | 7.400E-09 | 3.672E-09 | 3.200E-09 | 1.000E-08 |
| 22570 | 1129 | F1129I | D | 8.8e-09 | 4.7e-09 | 1.3e-08 | 8.833E-09 | 4.150E-09 | 4.700E-09 | 1.300E-08 |
| 22571 | 1129 | F1129L | D | 2.2e-09 | 5.9e-09 | 8.1e-09 | 5.400E-09 | 2.982E-09 | 2.200E-09 | 8.100E-09 |
| 22572 | 1129 | F1129K | D | 6.7e-09 | 2.9e-09 | 9.5e-09 | 6.367E-09 | 3.313E-09 | 2.900E-09 | 9.500E-09 |
| 22573 | 1129 | F1129M | D | 1.4e-08 | 7.2e-09 | 9.7e-09 | 1.030E-08 | 3.439E-09 | 7.200E-09 | 1.400E-08 |
| 22574 | 1129 | F1129F | D | 1.1e-08 | 1.7e-09 | 6.2e-09 | 6.300E-09 | 4.651E-09 | 1.700E-09 | 1.100E-08 |
| 22575 | 1129 | F1129P | D | 1.3e-08 | 7.7e-09 | 1.6e-09 | 7.433E-09 | 5.705E-09 | 1.600E-09 | 1.300E-08 |
| 22576 | 1129 | F1129S | D | 4.5e-09 | 3.4e-09 | 7.4e-09 | 5.100E-09 | 2.066E-09 | 3.400E-09 | 7.400E-09 |
| 22577 | 1129 | F1129T | D | 9e-09 | 4.1e-09 | 5.8e-09 | 6.300E-09 | 2.488E-09 | 4.100E-09 | 9.000E-09 |
| 22578 | 1129 | F1129W | D | 4.1e-08 | 2.7e-09 | 2.1e-08 | 2.157E-08 | 1.916E-08 | 2.700E-09 | 4.100E-08 |
| 22579 | 1129 | F1129Y | D | 1.1e-08 | 7.5e-09 | 1e-08 | 9.500E-09 | 1.803E-09 | 7.500E-09 | 1.100E-08 |
| 22580 | 1129 | F1129V | D | 1.2e-08 | 4.2e-09 | 7.8e-09 | 8.000E-09 | 3.904E-09 | 4.200E-09 | 1.200E-08 |
| 22581 | 1130 | T1130A | D | 3.9e-09 | 5.9e-09 | 9.3e-09 | 6.367E-09 | 2.730E-09 | 3.900E-09 | 9.300E-09 |
| 22582 | 1130 | T1130R | D | 1.2e-08 | 1.6e-08 | 6e-09 | 1.133E-08 | 5.033E-09 | 6.000E-09 | 1.600E-08 |
| 22583 | 1130 | T1130N | D | 7.8e-09 | 4.3e-09 | 3.8e-09 | 5.300E-09 | 2.179E-09 | 3.800E-09 | 7.800E-09 |
| 22584 | 1130 | T1130D | D | 1.3e-08 | 5.2e-09 | 4e-09 | 7.400E-09 | 4.887E-09 | 4.000E-09 | 1.300E-08 |
| 22585 | 1130 | T1130C | D | 4.2e-09 | 1.2e-08 | 9.1e-09 | 8.433E-09 | 3.943E-09 | 4.200E-09 | 1.200E-08 |
| 22586 | 1130 | T1130E | D | 2.2e-09 | 9e-09 | 1.2e-08 | 7.733E-09 | 5.021E-09 | 2.200E-09 | 1.200E-08 |
| 22587 | 1130 | T1130Q | D | 8.8e-09 | 5.4e-09 | 7.7e-09 | 7.300E-09 | 1.735E-09 | 5.400E-09 | 8.800E-09 |
| 22588 | 1130 | T1130G | D | 2e-08 | 7.8e-09 | 5.7e-09 | 1.117E-08 | 7.722E-09 | 5.700E-09 | 2.000E-08 |
| 22589 | 1130 | T1130H | D | 1e-08 | 1.1e-08 | 1.6e-08 | 1.233E-08 | 3.215E-09 | 1.000E-08 | 1.600E-08 |
| 22590 | 1130 | T1130I | D | 8e-09 | 1.5e-08 | 9e-09 | 1.067E-08 | 3.786E-09 | 8.000E-09 | 1.500E-08 |
| 22591 | 1130 | T1130L | D | 1.5e-08 | 7.1e-09 | 7.2e-09 | 9.767E-09 | 4.532E-09 | 7.100E-09 | 1.500E-08 |
| 22592 | 1130 | T1130K | D | 4.5e-08 | 5.4e-09 | 1.5e-08 | 2.180E-08 | 2.066E-08 | 5.400E-09 | 4.500E-08 |
| 22593 | 1130 | T1130M | D | 4.1e-09 | 6.5e-09 | 1.8e-08 | 9.533E-09 | 7.430E-09 | 4.100E-09 | 1.800E-08 |
| 22594 | 1130 | T1130F | D | 5.4e-09 | 8.5e-10 | 6.4e-09 | 4.217E-09 | 2.958E-09 | 8.500E-10 | 6.400E-09 |
| 22595 | 1130 | T1130P | D | 3.4e-09 | 5.4e-09 | 6.6e-09 | 5.133E-09 | 1.617E-09 | 3.400E-09 | 6.600E-09 |
| 22596 | 1130 | T1130S | D | 1.2e-08 | 2.2e-09 | 4.9e-09 | 6.367E-09 | 5.062E-09 | 2.200E-09 | 1.200E-08 |
| 22597 | 1130 | T1130T | D | 1.2e-08 | 1e-08 | 8.9e-09 | 1.030E-08 | 1.572E-09 | 8.900E-09 | 1.200E-08 |
| 22598 | 1130 | T1130W | D | 4.5e-09 | 6.4e-09 | 1e-08 | 6.967E-09 | 2.793E-09 | 4.500E-09 | 1.000E-08 |
| 22599 | 1130 | T1130Y | D | 1.2e-08 | 2.7e-09 | 3.9e-09 | 6.200E-09 | 5.059E-09 | 2.700E-09 | 1.200E-08 |
| 22600 | 1130 | T1130V | D | 1.5e-08 | 1.8e-08 | 8.6e-09 | 1.387E-08 | 4.801E-09 | 8.600E-09 | 1.800E-08 |
| 22601 | 1131 | L1131A | D | 9.2e-09 | 9.9e-09 | 5e-09 | 8.033E-09 | 2.650E-09 | 5.000E-09 | 9.900E-09 |
| 22602 | 1131 | L1131R | D | 8e-09 | 1.6e-08 | 1.1e-08 | 1.167E-08 | 4.041E-09 | 8.000E-09 | 1.600E-08 |
| 22603 | 1131 | L1131N | D | 1.2e-08 | 5.8e-09 | 7.4e-09 | 8.400E-09 | 3.219E-09 | 5.800E-09 | 1.200E-08 |
| 22604 | 1131 | L1131D | D | 5.4e-09 | 1.3e-08 | 5e-09 | 7.800E-09 | 4.508E-09 | 5.000E-09 | 1.300E-08 |
| 22605 | 1131 | L1131C | D | 5e-09 | 8.8e-09 | 2.9e-09 | 5.567E-09 | 2.991E-09 | 2.900E-09 | 8.800E-09 |
| 22606 | 1131 | L1131E | D | 3.4e-09 | 9.1e-09 | 6.3e-09 | 6.267E-09 | 2.850E-09 | 3.400E-09 | 9.100E-09 |
| 22607 | 1131 | L1131Q | D | 6.9e-09 | 9.7e-09 | 8.4e-09 | 8.333E-09 | 1.401E-09 | 6.900E-09 | 9.700E-09 |
| 22608 | 1131 | L1131G | D | 8.6e-09 | 2.1e-08 | 4.5e-09 | 1.137E-08 | 8.591E-09 | 4.500E-09 | 2.100E-08 |
| 22609 | 1131 | L1131H | D | 1.6e-08 | 3.8e-09 | 4.9e-09 | 8.233E-09 | 6.749E-09 | 3.800E-09 | 1.600E-08 |

| | | | | | | | | | | |
|---|---|---|---|---|---|---|---|---|---|---|
| 22610 | 1131 | L1131I | D | 4.7e-09 | 1.2e-08 | 3.1e-09 | 6.600E-09 | 4.744E-09 | 3.100E-09 | 1.200E-08 |
| 22611 | 1131 | L1131L | D | 2e-09 | 8.9e-09 | 4.9e-09 | 5.267E-09 | 3.465E-09 | 2.000E-09 | 8.900E-09 |
| 22612 | 1131 | L1131K | D | 4.6e-09 | 7.5e-09 | 9.7e-09 | 7.267E-09 | 2.558E-09 | 4.600E-09 | 9.700E-09 |
| 22613 | 1131 | L1131M | D | 6.6e-09 | 7.7e-09 | 1.2e-08 | 8.767E-09 | 2.854E-09 | 6.600E-09 | 1.200E-08 |
| 22614 | 1131 | L1131F | D | 1.3e-08 | 7.8e-09 | 3.9e-09 | 8.233E-09 | 4.565E-09 | 3.900E-09 | 1.300E-08 |
| 22615 | 1131 | L1131P | D | 7.5e-09 | 2.5e-09 | 6e-09 | 5.333E-09 | 2.566E-09 | 2.500E-09 | 7.500E-09 |
| 22616 | 1131 | L1131S | D | 5.2e-09 | 3.9e-09 | 8.8e-09 | 5.967E-09 | 2.538E-09 | 3.900E-09 | 8.800E-09 |
| 22617 | 1131 | L1131T | D | 5e-09 | 6.6e-09 | 2.1e-08 | 1.087E-08 | 8.812E-09 | 5.000E-09 | 2.100E-08 |
| 22618 | 1131 | L1131W | D | 1.2e-08 | 9.5e-09 | 1.1e-08 | 1.083E-08 | 1.258E-09 | 9.500E-09 | 1.200E-08 |
| 22619 | 1131 | L1131Y | D | 4.8e-09 | 9.1e-09 | 9.2e-09 | 7.700E-09 | 2.512E-09 | 4.800E-09 | 9.200E-09 |
| 22620 | 1131 | L1131V | D | 1.3e-08 | 5.7e-09 | 6.1e-09 | 8.267E-09 | 4.104E-09 | 5.700E-09 | 1.300E-08 |
| 22621 | 1132 | T1132A | D | 9.8e-09 | 1.1e-08 | 2.3e-09 | 7.700E-09 | 4.715E-09 | 2.300E-09 | 1.100E-08 |
| 22622 | 1132 | T1132R | D | 3e-09 | 5.8e-09 | 1e-08 | 6.267E-09 | 3.523E-09 | 3.000E-09 | 1.000E-08 |
| 22623 | 1132 | T1132N | D | 1.8e-08 | 8e-09 | 2.5e-08 | 1.700E-08 | 8.544E-09 | 8.000E-09 | 2.500E-08 |
| 22624 | 1132 | T1132D | D | 7.6e-09 | 1.2e-08 | 7e-09 | 8.867E-09 | 2.730E-09 | 7.000E-09 | 1.200E-08 |
| 22625 | 1132 | T1132C | D | 1.3e-08 | 7.3e-09 | 2.5e-09 | 7.600E-09 | 5.256E-09 | 2.500E-09 | 1.300E-08 |
| 22626 | 1132 | T1132E | D | 1.3e-08 | 1.1e-08 | 6.4e-09 | 1.013E-08 | 3.384E-09 | 6.400E-09 | 1.300E-08 |
| 22627 | 1132 | T1132Q | D | 1.5e-08 | 1.5e-08 | 9e-09 | 1.300E-08 | 3.464E-09 | 9.000E-09 | 1.500E-08 |
| 22628 | 1132 | T1132G | D | 8.7e-09 | 1.9e-08 | 1.1e-08 | 1.290E-08 | 5.406E-09 | 8.700E-09 | 1.900E-08 |
| 22629 | 1132 | T1132H | D | 1.1e-08 | 6.2e-09 | 3.8e-09 | 7.000E-09 | 3.666E-09 | 3.800E-09 | 1.100E-08 |
| 22630 | 1132 | T1132I | D | 3.9e-09 | 9.2e-09 | 5.6e-09 | 6.233E-09 | 2.706E-09 | 3.900E-09 | 9.200E-09 |
| 22631 | 1132 | T1132L | D | 8.7e-09 | 7.6e-09 | 8.7e-09 | 8.333E-09 | 6.351E-10 | 7.600E-09 | 8.700E-09 |
| 22632 | 1132 | T1132K | D | 4.1e-09 | 6.5e-09 | 7.9e-09 | 6.167E-09 | 1.922E-09 | 4.100E-09 | 7.900E-09 |
| 22633 | 1132 | T1132M | D | 9.2e-09 | 7.7e-09 | 1.8e-08 | 1.163E-08 | 5.564E-09 | 7.700E-09 | 1.800E-08 |
| 22634 | 1132 | T1132F | D | 4.8e-09 | 4.4e-09 | 5.8e-09 | 5.000E-09 | 7.211E-10 | 4.400E-09 | 5.800E-09 |
| 22635 | 1132 | T1132P | D | 3.8e-09 | 6.9e-09 | 2.1e-08 | 1.057E-08 | 9.168E-09 | 3.800E-09 | 2.100E-08 |
| 22636 | 1132 | T1132S | D | 1.7e-09 | 1.2e-09 | 2e-08 | 7.633E-09 | 1.071E-08 | 1.200E-09 | 2.000E-08 |
| 22637 | 1132 | T1132T | D | 8.8e-09 | 1.4e-08 | 6.7e-09 | 9.833E-09 | 3.758E-09 | 6.700E-09 | 1.400E-08 |
| 22638 | 1132 | T1132W | D | 1e-08 | 7.4e-09 | 8.9e-09 | 8.767E-09 | 1.305E-09 | 7.400E-09 | 1.000E-08 |
| 22639 | 1132 | T1132Y | D | 9.3e-09 | 4.8e-09 | 7e-09 | 7.033E-09 | 2.250E-09 | 4.800E-09 | 9.300E-09 |
| 22640 | 1132 | T1132V | D | 3.6e-09 | 1.5e-08 | 6.7e-09 | 8.433E-09 | 5.894E-09 | 3.600E-09 | 1.500E-08 |
| 22641 | 1133 | I1133A | D | 5.9e-09 | 1.9e-08 | 9.4e-09 | 1.143E-08 | 6.783E-09 | 5.900E-09 | 1.900E-08 |
| 22642 | 1133 | I1133R | D | 5.7e-09 | 3.9e-09 | 7.6e-09 | 5.733E-09 | 1.850E-09 | 3.900E-09 | 7.600E-09 |
| 22643 | 1133 | I1133N | D | 4.5e-09 | 4e-09 | 5.5e-09 | 4.667E-09 | 7.638E-10 | 4.000E-09 | 5.500E-09 |
| 22644 | 1133 | I1133D | D | 4.3e-09 | 5.4e-09 | 1.3e-08 | 7.567E-09 | 4.737E-09 | 4.300E-09 | 1.300E-08 |
| 22645 | 1133 | I1133C | D | 8.1e-09 | 7.9e-09 | 1.2e-08 | 9.333E-09 | 2.312E-09 | 7.900E-09 | 1.200E-08 |
| 22646 | 1133 | I1133E | D | 7.7e-09 | 1e-08 | 7.9e-09 | 8.533E-09 | 1.274E-09 | 7.700E-09 | 1.000E-08 |
| 22647 | 1133 | I1133Q | D | 2.6e-08 | 4.6e-09 | 9.7e-09 | 1.343E-08 | 1.118E-08 | 4.600E-09 | 2.600E-08 |
| 22648 | 1133 | I1133G | D | 5.9e-09 | 3.3e-09 | 7.4e-09 | 5.533E-09 | 2.074E-09 | 3.300E-09 | 7.400E-09 |
| 22649 | 1133 | I1133H | D | 8.5e-09 | 6.2e-09 | 1.1e-08 | 8.567E-09 | 2.401E-09 | 6.200E-09 | 1.100E-08 |
| 22650 | 1133 | I1133I | D | 6.8e-09 | 5.4e-09 | 1.1e-08 | 7.733E-09 | 2.914E-09 | 5.400E-09 | 1.100E-08 |
| 22651 | 1133 | I1133L | D | 1.1e-08 | 9.2e-09 | 3.3e-09 | 7.833E-09 | 4.028E-09 | 3.300E-09 | 1.100E-08 |
| 22652 | 1133 | I1133K | D | 6.8e-09 | 1.5e-08 | 8.8e-09 | 1.020E-08 | 4.276E-09 | 6.800E-09 | 1.500E-08 |
| 22653 | 1133 | I1133M | D | 1.4e-08 | 1.1e-08 | 7.4e-09 | 1.080E-08 | 3.305E-09 | 7.400E-09 | 1.400E-08 |
| 22654 | 1133 | I1133F | D | 1.5e-08 | 9.9e-09 | 9e-09 | 1.130E-08 | 3.236E-09 | 9.000E-09 | 1.500E-08 |
| 22655 | 1133 | I1133P | D | 9e-09 | 5.6e-09 | 8.6e-09 | 7.733E-09 | 1.858E-09 | 5.600E-09 | 9.000E-09 |
| 22656 | 1133 | I1133S | D | 7e-09 | 2.9e-09 | 2.9e-09 | 4.267E-09 | 2.367E-09 | 2.900E-09 | 7.000E-09 |
| 22657 | 1133 | I1133T | D | 2.5e-09 | 3.9e-09 | 7e-09 | 4.467E-09 | 2.303E-09 | 2.500E-09 | 7.000E-09 |
| 22658 | 1133 | I1133W | D | 5.3e-09 | 3.5e-09 | 3.4e-09 | 4.067E-09 | 1.069E-09 | 3.400E-09 | 5.300E-09 |
| 22659 | 1133 | I1133Y | D | 2.4e-09 | 2.6e-09 | 9e-09 | 4.667E-09 | 3.754E-09 | 2.400E-09 | 9.000E-09 |
| 22660 | 1133 | I1133V | D | 7.8e-09 | 6.8e-09 | 6.1e-09 | 6.900E-09 | 8.544E-10 | 6.100E-09 | 7.800E-09 |
| 22661 | 1134 | S1134A | D | 9e-09 | 2.1e-09 | 7.3e-09 | 6.133E-09 | 3.595E-09 | 2.100E-09 | 9.000E-09 |
| 22662 | 1134 | S1134R | D | 5.2e-09 | 3.7e-09 | 4.7e-09 | 4.533E-09 | 7.638E-10 | 3.700E-09 | 5.200E-09 |
| 22663 | 1134 | S1134N | D | 4.7e-09 | 7.9e-09 | 9e-09 | 7.200E-09 | 2.234E-09 | 4.700E-09 | 9.000E-09 |
| 22664 | 1134 | S1134D | D | 4.7e-09 | 1.3e-08 | 4.1e-09 | 7.267E-09 | 4.974E-09 | 4.100E-09 | 1.300E-08 |
| 22665 | 1134 | S1134C | D | 1.4e-09 | 2.1e-08 | 5.9e-09 | 9.433E-09 | 1.027E-08 | 1.400E-09 | 2.100E-08 |
| 22666 | 1134 | S1134E | D | 4.1e-09 | 1.6e-08 | 1.1e-08 | 1.037E-08 | 5.975E-09 | 4.100E-09 | 1.600E-08 |
| 22667 | 1134 | S1134Q | D | 1.2e-08 | 1.3e-08 | 8.9e-09 | 1.130E-08 | 2.138E-09 | 8.900E-09 | 1.300E-08 |
| 22668 | 1134 | S1134G | D | 7.9e-09 | 7e-09 | 6.4e-09 | 7.100E-09 | 7.550E-10 | 6.400E-09 | 7.900E-09 |
| 22669 | 1134 | S1134H | D | 3.9e-09 | 1.3e-08 | 4e-09 | 6.967E-09 | 5.225E-09 | 3.900E-09 | 1.300E-08 |
| 22670 | 1134 | S1134I | D | 3e-09 | 5.6e-09 | 2.2e-08 | 1.020E-08 | 1.030E-08 | 3.000E-09 | 2.200E-08 |
| 22671 | 1134 | S1134L | D | 2.9e-09 | 5.9e-09 | 1.3e-08 | 7.267E-09 | 5.187E-09 | 2.900E-09 | 1.300E-08 |
| 22672 | 1134 | S1134K | D | 2.6e-09 | 8e-09 | 2.9e-09 | 4.500E-09 | 3.035E-09 | 2.600E-09 | 8.000E-09 |
| 22673 | 1134 | S1134M | D | 2.7e-09 | 6.1e-09 | 5.4e-09 | 4.733E-09 | 1.795E-09 | 2.700E-09 | 6.100E-09 |
| 22674 | 1134 | S1134F | D | 5.2e-09 | 9.5e-09 | 7.1e-09 | 7.267E-09 | 2.155E-09 | 5.200E-09 | 9.500E-09 |
| 22675 | 1134 | S1134P | D | 6.2e-09 | 2e-09 | 4.8e-09 | 4.333E-09 | 2.139E-09 | 2.000E-09 | 6.200E-09 |
| 22676 | 1134 | S1134S | D | 8.5e-09 | 3.8e-09 | 2.8e-09 | 5.033E-09 | 3.044E-09 | 2.800E-09 | 8.500E-09 |
| 22677 | 1134 | S1134T | D | 4.2e-09 | 4.2e-09 | 2.8e-09 | 3.733E-09 | 8.083E-10 | 2.800E-09 | 4.200E-09 |
| 22678 | 1134 | S1134W | D | 2.4e-08 | 4.8e-09 | 1.2e-08 | 1.360E-08 | 9.699E-09 | 4.800E-09 | 2.400E-08 |
| 22679 | 1134 | S1134Y | D | 7e-09 | 7.8e-09 | 6.3e-09 | 7.033E-09 | 7.506E-10 | 6.300E-09 | 7.800E-09 |
| 22680 | 1134 | S1134V | D | 5.3e-09 | 1.1e-08 | 1.4e-08 | 1.010E-08 | 4.419E-09 | 5.300E-09 | 1.400E-08 |
| 22681 | 1135 | S1135A | D | 1e-08 | 6.9e-09 | 1.1e-08 | 9.300E-09 | 2.138E-09 | 6.900E-09 | 1.100E-08 |
| 22682 | 1135 | S1135R | D | 6.8e-09 | 8.2e-09 | 1.1e-08 | 8.667E-09 | 2.139E-09 | 6.800E-09 | 1.100E-08 |
| 22683 | 1135 | S1135N | D | 1.2e-08 | 8.9e-09 | 2.8e-08 | 1.630E-08 | 1.025E-08 | 8.900E-09 | 2.800E-08 |
| 22684 | 1135 | S1135D | D | 1e-08 | 1.1e-08 | 6.3e-09 | 9.100E-09 | 2.476E-09 | 6.300E-09 | 1.100E-08 |
| 22685 | 1135 | S1135C | D | 7.2e-09 | 7.6e-09 | 7.6e-09 | 7.467E-09 | 2.309E-10 | 7.200E-09 | 7.600E-09 |
| 22686 | 1135 | S1135E | D | 2.5e-09 | 6.9e-09 | 9.4e-09 | 6.267E-09 | 3.493E-09 | 2.500E-09 | 9.400E-09 |
| 22687 | 1135 | S1135Q | D | 4.9e-09 | 7.8e-09 | 5.5e-09 | 6.067E-09 | 1.531E-09 | 4.900E-09 | 7.800E-09 |
| 22688 | 1135 | S1135G | D | 5.7e-09 | 7.4e-09 | 7e-09 | 6.700E-09 | 8.888E-10 | 5.700E-09 | 7.400E-09 |
| 22689 | 1135 | S1135H | D | 2.5e-09 | 3.6e-09 | 8.4e-09 | 4.833E-09 | 3.137E-09 | 2.500E-09 | 8.400E-09 |
| 22690 | 1135 | S1135I | D | 7.3e-09 | 7e-09 | 7.7e-09 | 7.333E-09 | 3.512E-10 | 7.000E-09 | 7.700E-09 |
| 22691 | 1135 | S1135L | D | 8.2e-09 | 1.5e-08 | 8e-09 | 1.040E-08 | 3.985E-09 | 8.000E-09 | 1.500E-08 |
| 22692 | 1135 | S1135K | D | 8.3e-09 | 1.4e-08 | 3.5e-09 | 8.600E-09 | 5.256E-09 | 3.500E-09 | 1.400E-08 |
| 22693 | 1135 | S1135M | D | 6.7e-09 | 8.2e-09 | 8e-09 | 7.633E-09 | 8.145E-10 | 6.700E-09 | 8.200E-09 |
| 22694 | 1135 | S1135F | D | 8.4e-09 | 8.5e-09 | 9e-09 | 8.633E-09 | 3.215E-10 | 8.400E-09 | 9.000E-09 |

| | | | | | | | | | | |
|---|---|---|---|---|---|---|---|---|---|---|
| 22695 | 1135 | S1135P | D | 6.7e-09 | 8.7e-09 | 4.2e-09 | 6.533E-09 | 2.255E-09 | 4.200E-09 | 8.700E-09 |
| 22696 | 1135 | S1135S | D | 5.9e-09 | 7.4e-09 | 3.2e-09 | 5.500E-09 | 2.128E-09 | 3.200E-09 | 7.400E-09 |
| 22697 | 1135 | S1135T | D | 4.6e-09 | 6.2e-09 | 1.1e-08 | 7.267E-09 | 3.331E-09 | 4.600E-09 | 1.100E-08 |
| 22698 | 1135 | S1135W | D | 1.7e-08 | 4.6e-09 | 9.8e-09 | 1.047E-08 | 6.227E-09 | 4.600E-09 | 1.700E-08 |
| 22699 | 1135 | S1135Y | D | 7.5e-09 | 1.8e-09 | 4e-09 | 4.433E-09 | 2.875E-09 | 1.800E-09 | 7.500E-09 |
| 22700 | 1135 | S1135V | D | 2.4e-09 | 1.8e-08 | 1.4e-08 | 1.147E-08 | 8.103E-09 | 2.400E-09 | 1.800E-08 |
| 22701 | 1136 | L1136A | D | 1.3e-08 | 3.3e-08 | 3.7e-09 | 1.657E-08 | 1.497E-08 | 3.700E-09 | 3.300E-08 |
| 22702 | 1136 | L1136R | D | 2e-09 | 3.8e-09 | 1.9e-08 | 8.267E-09 | 9.339E-09 | 2.000E-09 | 1.900E-08 |
| 22703 | 1136 | L1136N | D | 1.1e-08 | 1.2e-08 | 6.6e-09 | 9.867E-09 | 2.873E-09 | 6.600E-09 | 1.200E-08 |
| 22704 | 1136 | L1136D | D | 7e-09 | 2.3e-09 | 1.1e-08 | 6.767E-09 | 4.355E-09 | 2.300E-09 | 1.100E-08 |
| 22705 | 1136 | L1136C | D | 4.5e-09 | 6.3e-09 | 2.7e-09 | 4.500E-09 | 1.800E-09 | 2.700E-09 | 6.300E-09 |
| 22706 | 1136 | L1136E | D | 5.5e-09 | 5.5e-09 | 1.1e-08 | 7.333E-09 | 3.175E-09 | 5.500E-09 | 1.100E-08 |
| 22707 | 1136 | L1136Q | D | 8.6e-09 | 1.3e-08 | 4.1e-09 | 8.567E-09 | 4.450E-09 | 4.100E-09 | 1.300E-08 |
| 22708 | 1136 | L1136G | D | 5.9e-09 | 5.9e-09 | 1.1e-08 | 7.600E-09 | 2.944E-09 | 5.900E-09 | 1.100E-08 |
| 22709 | 1136 | L1136H | D | 4e-09 | 5.2e-09 | 5e-09 | 4.733E-09 | 6.429E-10 | 4.000E-09 | 5.200E-09 |
| 22710 | 1136 | L1136I | D | 3e-09 | 1.9e-09 | 1.7e-08 | 7.300E-09 | 8.418E-09 | 1.900E-09 | 1.700E-08 |
| 22711 | 1136 | L1136L | D | 1.2e-08 | 1e-08 | 5.4e-09 | 9.133E-09 | 3.384E-09 | 5.400E-09 | 1.200E-08 |
| 22712 | 1136 | L1136K | D | 1e-08 | 5.3e-09 | 4.8e-09 | 6.700E-09 | 2.869E-09 | 4.800E-09 | 1.000E-08 |
| 22713 | 1136 | L1136M | D | 1.6e-08 | 5.6e-09 | 6e-09 | 9.200E-09 | 5.892E-09 | 5.600E-09 | 1.600E-08 |
| 22714 | 1136 | L1136F | D | 9.5e-09 | 1.3e-08 | 5.1e-09 | 9.200E-09 | 3.959E-09 | 5.100E-09 | 1.300E-08 |
| 22715 | 1136 | L1136P | D | 5.6e-09 | 5.5e-09 | 6.4e-09 | 5.833E-09 | 4.933E-10 | 5.500E-09 | 6.400E-09 |
| 22716 | 1136 | L1136S | D | 8.2e-09 | 1.5e-08 | 2.3e-09 | 8.500E-09 | 6.355E-09 | 2.300E-09 | 1.500E-08 |
| 22717 | 1136 | L1136T | D | 7.8e-09 | 6.5e-09 | 2.1e-08 | 1.177E-08 | 8.023E-09 | 6.500E-09 | 2.100E-08 |
| 22718 | 1136 | L1136W | D | 7e-09 | 3.8e-09 | 4.8e-09 | 5.200E-09 | 1.637E-09 | 3.800E-09 | 7.000E-09 |
| 22719 | 1136 | L1136Y | D | 2.1e-09 | 2.6e-09 | 7.8e-09 | 4.167E-09 | 3.156E-09 | 2.100E-09 | 7.800E-09 |
| 22720 | 1136 | L1136V | D | 7.3e-09 | 9.6e-09 | 5.6e-09 | 7.500E-09 | 2.007E-09 | 5.600E-09 | 9.600E-09 |
| 22721 | 1137 | Q1137A | D | 1.1e-08 | 4.8e-09 | 7.2e-09 | 7.667E-09 | 3.126E-09 | 4.800E-09 | 1.100E-08 |
| 22722 | 1137 | Q1137R | D | 6.4e-09 | 9.9e-09 | 9.2e-09 | 8.500E-09 | 1.852E-09 | 6.400E-09 | 9.900E-09 |
| 22723 | 1137 | Q1137N | D | 1.7e-08 | 4.2e-09 | 6.4e-09 | 9.200E-09 | 6.844E-09 | 4.200E-09 | 1.700E-08 |
| 22724 | 1137 | Q1137D | D | 1.1e-08 | 1e-08 | 7.2e-09 | 9.400E-09 | 1.970E-09 | 7.200E-09 | 1.100E-08 |
| 22725 | 1137 | Q1137C | D | 3.6e-09 | 5.9e-09 | 5.8e-09 | 5.100E-09 | 1.300E-09 | 3.600E-09 | 5.900E-09 |
| 22726 | 1137 | Q1137E | D | 7.1e-09 | 6.7e-09 | 8e-09 | 7.267E-09 | 6.658E-10 | 6.700E-09 | 8.000E-09 |
| 22727 | 1137 | Q1137Q | D | 8.9e-09 | 4.4e-09 | 3.3e-09 | 5.533E-09 | 2.967E-09 | 3.300E-09 | 8.900E-09 |
| 22728 | 1137 | Q1137G | D | 1.4e-08 | 5.8e-09 | 1.2e-08 | 1.060E-08 | 4.276E-09 | 5.800E-09 | 1.400E-08 |
| 22729 | 1137 | Q1137H | D | 7.1e-09 | 5.8e-09 | 7.1e-09 | 6.667E-09 | 7.506E-10 | 5.800E-09 | 7.100E-09 |
| 22730 | 1137 | Q1137I | D | 9.2e-09 | 3.9e-09 | 1.1e-08 | 8.033E-09 | 3.691E-09 | 3.900E-09 | 1.100E-08 |
| 22731 | 1137 | Q1137L | D | 3.8e-09 | 1.1e-08 | 1.1e-08 | 8.600E-09 | 4.157E-09 | 3.800E-09 | 1.100E-08 |
| 22732 | 1137 | Q1137K | D | 4.3e-09 | 9.7e-09 | 2.3e-09 | 5.433E-09 | 3.828E-09 | 2.300E-09 | 9.700E-09 |
| 22733 | 1137 | Q1137M | D | 5.8e-09 | 2.8e-09 | 4.3e-09 | 4.300E-09 | 1.500E-09 | 2.800E-09 | 5.800E-09 |
| 22734 | 1137 | Q1137F | D | 6.1e-09 | 1.1e-08 | 4.2e-09 | 7.100E-09 | 3.509E-09 | 4.200E-09 | 1.100E-08 |
| 22735 | 1137 | Q1137P | D | 6.6e-09 | 1.1e-08 | 8.5e-09 | 8.700E-09 | 2.207E-09 | 6.600E-09 | 1.100E-08 |
| 22736 | 1137 | Q1137S | D | 8.1e-09 | 5.8e-09 | 3.7e-09 | 5.867E-09 | 2.201E-09 | 3.700E-09 | 8.100E-09 |
| 22737 | 1137 | Q1137T | D | 1.1e-08 | 8.5e-09 | 7.3e-09 | 8.933E-09 | 1.888E-09 | 7.300E-09 | 1.100E-08 |
| 22738 | 1137 | Q1137W | D | 4.1e-09 | 8.1e-09 | 9.1e-09 | 7.100E-09 | 2.646E-09 | 4.100E-09 | 9.100E-09 |
| 22739 | 1137 | Q1137Y | D | 1.7e-08 | 2.9e-09 | 7.5e-09 | 9.133E-09 | 7.191E-09 | 2.900E-09 | 1.700E-08 |
| 22740 | 1137 | Q1137V | D | 9.8e-09 | 6.1e-09 | 5.5e-09 | 7.133E-09 | 2.329E-09 | 5.500E-09 | 9.800E-09 |
| 22741 | 1138 | P1138A | D | 2.7e-09 | 1.1e-08 | 1.5e-09 | 5.067E-09 | 5.173E-09 | 1.500E-09 | 1.100E-08 |
| 22742 | 1138 | P1138R | D | 4.4e-08 | 6.5e-09 | 3.7e-09 | 1.807E-08 | 2.250E-08 | 3.700E-09 | 4.400E-08 |
| 22743 | 1138 | P1138N | D | 6.5e-09 | 4e-09 | 6.1e-09 | 5.533E-09 | 1.343E-09 | 4.000E-09 | 6.500E-09 |
| 22744 | 1138 | P1138D | D | 6.4e-09 | 1.4e-08 | 1.5e-08 | 1.180E-08 | 4.703E-09 | 6.400E-09 | 1.500E-08 |
| 22745 | 1138 | P1138C | D | 6.5e-09 | 5.2e-09 | 7.3e-09 | 6.333E-09 | 1.060E-09 | 5.200E-09 | 7.300E-09 |
| 22746 | 1138 | P1138E | D | 5.4e-09 | 1.7e-09 | 2.8e-08 | 1.170E-08 | 1.424E-08 | 1.700E-09 | 2.800E-08 |
| 22747 | 1138 | P1138Q | D | 7.2e-09 | 9.2e-09 | 1.8e-09 | 6.067E-09 | 3.828E-09 | 1.800E-09 | 9.200E-09 |
| 22748 | 1138 | P1138G | D | 4.7e-09 | 7.2e-09 | 3.5e-09 | 5.133E-09 | 1.888E-09 | 3.500E-09 | 7.200E-09 |
| 22749 | 1138 | P1138H | D | 6.2e-09 | 6.3e-09 | 3.3e-09 | 5.267E-09 | 1.704E-09 | 3.300E-09 | 6.300E-09 |
| 22750 | 1138 | P1138I | D | 5.3e-09 | 5.5e-09 | 6.3e-09 | 5.700E-09 | 5.292E-10 | 5.300E-09 | 6.300E-09 |
| 22751 | 1138 | P1138L | D | 2.3e-09 | 5.4e-09 | 1.1e-08 | 6.233E-09 | 4.409E-09 | 2.300E-09 | 1.100E-08 |
| 22752 | 1138 | P1138K | D | 1.1e-08 | 1.1e-08 | 6.5e-09 | 9.500E-09 | 2.598E-09 | 6.500E-09 | 1.100E-08 |
| 22753 | 1138 | P1138M | D | 6.1e-09 | 5.2e-09 | 1.8e-09 | 4.367E-09 | 2.268E-09 | 1.800E-09 | 6.100E-09 |
| 22754 | 1138 | P1138F | D | 1.1e-08 | 2.9e-09 | 5.8e-09 | 6.567E-09 | 4.104E-09 | 2.900E-09 | 1.100E-08 |
| 22755 | 1138 | P1138P | D | 4.3e-09 | 5.5e-09 | 6.5e-09 | 5.433E-09 | 1.102E-09 | 4.300E-09 | 6.500E-09 |
| 22756 | 1138 | P1138S | D | 7.8e-09 | 5.5e-09 | 6.2e-09 | 6.500E-09 | 1.179E-09 | 5.500E-09 | 7.800E-09 |
| 22757 | 1138 | P1138T | D | 5.5e-09 | 2.1e-09 | 8e-09 | 5.200E-09 | 2.961E-09 | 2.100E-09 | 8.000E-09 |
| 22758 | 1138 | P1138W | D | 1.3e-08 | 5.4e-09 | 1.3e-08 | 1.047E-08 | 4.388E-09 | 5.400E-09 | 1.300E-08 |
| 22759 | 1138 | P1138Y | D | 5.6e-09 | 4.9e-09 | 7e-09 | 5.833E-09 | 1.069E-09 | 4.900E-09 | 7.000E-09 |
| 22760 | 1138 | P1138V | D | 1.2e-08 | 2.8e-09 | 2.3e-09 | 5.700E-09 | 5.462E-09 | 2.300E-09 | 1.200E-08 |
| 22761 | 1139 | E1139A | D | 1.5e-08 | 1.1e-08 | 1.4e-08 | 1.333E-08 | 2.082E-09 | 1.100E-08 | 1.500E-08 |
| 22762 | 1139 | E1139R | D | 2.6e-09 | 1.9e-08 | 9.2e-09 | 1.027E-08 | 8.252E-09 | 2.600E-09 | 1.900E-08 |
| 22763 | 1139 | E1139N | D | 1.2e-08 | 3.3e-09 | 1.6e-08 | 1.043E-08 | 6.493E-09 | 3.300E-09 | 1.600E-08 |
| 22764 | 1139 | E1139D | D | 7.4e-09 | 8.2e-09 | 7.1e-09 | 7.567E-09 | 5.686E-10 | 7.100E-09 | 8.200E-09 |
| 22765 | 1139 | E1139C | D | 7.3e-09 | 3.4e-09 | 3.6e-09 | 4.767E-09 | 2.196E-09 | 3.400E-09 | 7.300E-09 |
| 22766 | 1139 | E1139E | D | 8.5e-09 | 1.8e-08 | 8.9e-09 | 1.180E-08 | 5.373E-09 | 8.500E-09 | 1.800E-08 |
| 22767 | 1139 | E1139Q | D | 6.7e-09 | 1.2e-08 | 5.4e-09 | 8.033E-09 | 3.496E-09 | 5.400E-09 | 1.200E-08 |
| 22768 | 1139 | E1139G | D | 1.2e-08 | 1.7e-08 | 5.1e-09 | 1.137E-08 | 5.975E-09 | 5.100E-09 | 1.700E-08 |
| 22769 | 1139 | E1139H | D | 7.2e-09 | 3.2e-09 | 1.4e-08 | 8.133E-09 | 5.460E-09 | 3.200E-09 | 1.400E-08 |
| 22770 | 1139 | E1139I | D | 5.3e-09 | 4.7e-09 | 7.5e-09 | 5.833E-09 | 1.474E-09 | 4.700E-09 | 7.500E-09 |
| 22771 | 1139 | E1139L | D | 2.6e-09 | 2.8e-09 | 9.1e-09 | 4.833E-09 | 3.696E-09 | 2.600E-09 | 9.100E-09 |
| 22772 | 1139 | E1139K | D | 1.5e-08 | 3.8e-09 | 2.5e-08 | 1.460E-08 | 1.061E-08 | 3.800E-09 | 2.500E-08 |
| 22773 | 1139 | E1139M | D | 2.3e-09 | 1.1e-08 | 4e-09 | 5.767E-09 | 4.611E-09 | 2.300E-09 | 1.100E-08 |
| 22774 | 1139 | E1139F | D | 8e-09 | 1.2e-08 | 7.4e-09 | 9.133E-09 | 2.501E-09 | 7.400E-09 | 1.200E-08 |
| 22775 | 1139 | E1139P | D | 3.7e-09 | 1.2e-08 | 4.8e-09 | 6.833E-09 | 4.508E-09 | 3.700E-09 | 1.200E-08 |
| 22776 | 1139 | E1139S | D | 5.3e-09 | 1.4e-08 | 5e-09 | 8.100E-09 | 5.112E-09 | 5.000E-09 | 1.400E-08 |
| 22777 | 1139 | E1139T | D | 9.9e-09 | 5e-09 | 2.7e-09 | 5.867E-09 | 3.677E-09 | 2.700E-09 | 9.900E-09 |
| 22778 | 1139 | E1139W | D | 3.2e-09 | 2.3e-08 | 2.6e-09 | 9.600E-09 | 1.161E-08 | 2.600E-09 | 2.300E-08 |
| 22779 | 1139 | E1139Y | D | 4.4e-09 | 7.5e-09 | 6.7e-09 | 6.200E-09 | 1.609E-09 | 4.400E-09 | 7.500E-09 |

| | | | | | | | | | | |
|---|---|---|---|---|---|---|---|---|---|---|
| 22780 | 1139 | E1139V | D | 9.6e-09 | 5.1e-09 | 8.2e-10 | 5.173E-09 | 4.390E-09 | 8.200E-10 | 9.600E-09 |
| 22781 | 1140 | D1140A | D | 1.2e-08 | 9.1e-09 | 2.2e-08 | 1.437E-08 | 6.768E-09 | 9.100E-09 | 2.200E-08 |
| 22782 | 1140 | D1140R | D | 5.2e-09 | 5.2e-09 | 1.2e-08 | 7.467E-09 | 3.926E-09 | 5.200E-09 | 1.200E-08 |
| 22783 | 1140 | D1140N | D | 4.2e-09 | 1.1e-09 | 4.2e-09 | 3.167E-09 | 1.790E-09 | 1.100E-09 | 4.200E-09 |
| 22784 | 1140 | D1140D | D | 5.7e-09 | 8e-09 | 9.3e-09 | 7.667E-09 | 1.823E-09 | 5.700E-09 | 9.300E-09 |
| 22785 | 1140 | D1140C | D | 2.7e-09 | 9.3e-09 | 9.6e-09 | 7.200E-09 | 3.900E-09 | 2.700E-09 | 9.600E-09 |
| 22786 | 1140 | D1140E | D | 3.3e-09 | 1.5e-08 | 1.1e-08 | 9.767E-09 | 5.947E-09 | 3.300E-09 | 1.500E-08 |
| 22787 | 1140 | D1140Q | D | 3.8e-09 | 7.1e-09 | 8.6e-09 | 6.500E-09 | 2.456E-09 | 3.800E-09 | 8.600E-09 |
| 22788 | 1140 | D1140G | D | 6.2e-09 | 3.9e-09 | 7.6e-09 | 5.900E-09 | 1.868E-09 | 3.900E-09 | 7.600E-09 |
| 22789 | 1140 | D1140H | D | 3.9e-09 | 3.5e-09 | 4.5e-09 | 3.967E-09 | 5.033E-10 | 3.500E-09 | 4.500E-09 |
| 22790 | 1140 | D1140I | D | 5.7e-09 | 2.5e-09 | 1.1e-08 | 6.400E-09 | 4.293E-09 | 2.500E-09 | 1.100E-08 |
| 22791 | 1140 | D1140L | D | 8e-09 | 9.3e-09 | 8.2e-09 | 8.500E-09 | 7.000E-10 | 8.000E-09 | 9.300E-09 |
| 22792 | 1140 | D1140K | D | 9e-09 | 8.8e-09 | 1.2e-08 | 9.933E-09 | 1.793E-09 | 8.800E-09 | 1.200E-08 |
| 22793 | 1140 | D1140M | D | 4.6e-09 | 9.5e-09 | 3.3e-09 | 5.800E-09 | 3.270E-09 | 3.300E-09 | 9.500E-09 |
| 22794 | 1140 | D1140F | D | 4.8e-09 | 1.5e-08 | 7e-09 | 8.933E-09 | 5.368E-09 | 4.800E-09 | 1.500E-08 |
| 22795 | 1140 | D1140P | D | 4.2e-09 | 1.4e-08 | 6.2e-09 | 8.133E-09 | 5.178E-09 | 4.200E-09 | 1.400E-08 |
| 22796 | 1140 | D1140S | D | 7.4e-09 | 3.7e-09 | 7.8e-09 | 6.300E-09 | 2.261E-09 | 3.700E-09 | 7.800E-09 |
| 22797 | 1140 | D1140T | D | 1.4e-08 | 9.3e-09 | 9.2e-09 | 1.083E-08 | 2.743E-09 | 9.200E-09 | 1.400E-08 |
| 22798 | 1140 | D1140W | D | 1e-08 | 4.8e-09 | 1.6e-08 | 1.027E-08 | 5.605E-09 | 4.800E-09 | 1.600E-08 |
| 22799 | 1140 | D1140Y | D | 9.1e-09 | 2.3e-09 | 8.1e-09 | 6.500E-09 | 3.672E-09 | 2.300E-09 | 9.100E-09 |
| 22800 | 1140 | D1140V | D | 4.3e-09 | 5.4e-09 | 1.1e-08 | 6.900E-09 | 3.593E-09 | 4.300E-09 | 1.100E-08 |
| 22801 | 1141 | F1141A | D | 8.6e-09 | 1.3e-08 | 3.2e-09 | 8.267E-09 | 4.908E-09 | 3.200E-09 | 1.300E-08 |
| 22802 | 1141 | F1141R | D | 7.3e-09 | 2e-09 | 3.1e-09 | 4.133E-09 | 2.797E-09 | 2.000E-09 | 7.300E-09 |
| 22803 | 1141 | F1141N | D | 6.4e-09 | 3.1e-09 | 2.1e-09 | 3.867E-09 | 2.250E-09 | 2.100E-09 | 6.400E-09 |
| 22804 | 1141 | F1141D | D | 2.9e-08 | 8e-09 | 1.6e-08 | 1.767E-08 | 1.060E-08 | 8.000E-09 | 2.900E-08 |
| 22805 | 1141 | F1141C | D | 9.5e-09 | 2e-08 | 1.4e-08 | 1.450E-08 | 5.268E-09 | 9.500E-09 | 2.000E-08 |
| 22806 | 1141 | F1141E | D | 3.1e-09 | 3.4e-09 | 5.3e-09 | 3.933E-09 | 1.193E-09 | 3.100E-09 | 5.300E-09 |
| 22807 | 1141 | F1141Q | D | 5.7e-09 | 9.9e-09 | 5.3e-09 | 6.967E-09 | 2.548E-09 | 5.300E-09 | 9.900E-09 |
| 22808 | 1141 | F1141G | D | 1.4e-08 | 3.9e-09 | 6.6e-09 | 8.167E-09 | 5.229E-09 | 3.900E-09 | 1.400E-08 |
| 22809 | 1141 | F1141H | D | 7.1e-09 | 8.2e-09 | 1.1e-08 | 8.767E-09 | 2.011E-09 | 7.100E-09 | 1.100E-08 |
| 22810 | 1141 | F1141I | D | 6e-09 | 1.3e-08 | 1.2e-08 | 1.033E-08 | 3.786E-09 | 6.000E-09 | 1.300E-08 |
| 22811 | 1141 | F1141L | D | 7e-09 | 7.3e-09 | 4.6e-09 | 6.300E-09 | 1.480E-09 | 4.600E-09 | 7.300E-09 |
| 22812 | 1141 | F1141K | D | 5.5e-09 | 1.4e-08 | 3.9e-09 | 7.800E-09 | 5.429E-09 | 3.900E-09 | 1.400E-08 |
| 22813 | 1141 | F1141M | D | 5.6e-09 | 6.7e-09 | 4e-09 | 5.433E-09 | 1.358E-09 | 4.000E-09 | 6.700E-09 |
| 22814 | 1141 | F1141F | D | 6.7e-09 | 4e-09 | 4.7e-09 | 5.133E-09 | 1.401E-09 | 4.000E-09 | 6.700E-09 |
| 22815 | 1141 | F1141P | D | 2.4e-09 | 1.1e-08 | 1.5e-08 | 9.467E-09 | 6.438E-09 | 2.400E-09 | 1.500E-08 |
| 22816 | 1141 | F1141S | D | 1.4e-08 | 4.8e-09 | 4.9e-09 | 7.900E-09 | 5.283E-09 | 4.800E-09 | 1.400E-08 |
| 22817 | 1141 | F1141T | D | 7.5e-09 | 7e-09 | 3.8e-08 | 1.750E-08 | 1.776E-08 | 7.000E-09 | 3.800E-08 |
| 22818 | 1141 | F1141W | D | 1.3e-08 | 7.1e-09 | 9.5e-09 | 9.867E-09 | 2.967E-09 | 7.100E-09 | 1.300E-08 |
| 22819 | 1141 | F1141Y | D | 1.2e-08 | 3.9e-09 | 7.2e-09 | 7.700E-09 | 4.073E-09 | 3.900E-09 | 1.200E-08 |
| 22820 | 1141 | F1141V | D | 2.8e-08 | 1e-08 | 6.1e-09 | 1.470E-08 | 1.168E-08 | 6.100E-09 | 2.800E-08 |
| 22821 | 1142 | A1142A | D | 9.5e-09 | 1.4e-08 | 9.1e-09 | 1.087E-08 | 2.721E-09 | 9.100E-09 | 1.400E-08 |
| 22822 | 1142 | A1142R | D | 1.5e-08 | 4.1e-09 | 1e-08 | 9.700E-09 | 5.456E-09 | 4.100E-09 | 1.500E-08 |
| 22823 | 1142 | A1142N | D | 8.6e-09 | 6.1e-09 | 3e-09 | 5.900E-09 | 2.805E-09 | 3.000E-09 | 8.600E-09 |
| 22824 | 1142 | A1142D | D | 1.2e-08 | 1.1e-08 | 4.3e-09 | 9.100E-09 | 4.187E-09 | 4.300E-09 | 1.200E-08 |
| 22825 | 1142 | A1142C | D | 4.1e-09 | 1.8e-08 | 9.9e-09 | 1.067E-08 | 6.982E-09 | 4.100E-09 | 1.800E-08 |
| 22826 | 1142 | A1142E | D | 2.6e-09 | 6e-09 | 9.6e-09 | 6.067E-09 | 3.500E-09 | 2.600E-09 | 9.600E-09 |
| 22827 | 1142 | A1142Q | D | 4.2e-09 | 8.4e-09 | 8.7e-09 | 7.100E-09 | 2.516E-09 | 4.200E-09 | 8.700E-09 |
| 22828 | 1142 | A1142G | D | 8.4e-09 | 2.5e-09 | 1.5e-08 | 8.633E-09 | 6.253E-09 | 2.500E-09 | 1.500E-08 |
| 22829 | 1142 | A1142H | D | 6.6e-09 | 1.4e-09 | 9.5e-09 | 5.833E-09 | 4.104E-09 | 1.400E-09 | 9.500E-09 |
| 22830 | 1142 | A1142I | D | 2.2e-08 | 1e-08 | 7e-09 | 1.300E-08 | 7.937E-09 | 7.000E-09 | 2.200E-08 |
| 22831 | 1142 | A1142L | D | 5.2e-09 | 5.3e-09 | 6e-09 | 5.500E-09 | 4.359E-10 | 5.200E-09 | 6.000E-09 |
| 22832 | 1142 | A1142K | D | 1e-08 | 5.1e-09 | 5.3e-09 | 6.800E-09 | 2.773E-09 | 5.100E-09 | 1.000E-08 |
| 22833 | 1142 | A1142M | D | 1.6e-08 | 1e-08 | 6.6e-09 | 1.087E-08 | 4.760E-09 | 6.600E-09 | 1.600E-08 |
| 22834 | 1142 | A1142F | D | 1.6e-08 | 1.3e-08 | 2.7e-08 | 1.867E-08 | 7.371E-09 | 1.300E-08 | 2.700E-08 |
| 22835 | 1142 | A1142P | D | 3.4e-09 | 2.3e-09 | 3.6e-09 | 3.100E-09 | 7.000E-10 | 2.300E-09 | 3.600E-09 |
| 22836 | 1142 | A1142S | D | 4.6e-09 | 6.3e-09 | 5.9e-09 | 5.600E-09 | 8.888E-10 | 4.600E-09 | 6.300E-09 |
| 22837 | 1142 | A1142T | D | 2.9e-09 | 4.6e-09 | 1.1e-08 | 6.167E-09 | 4.271E-09 | 2.900E-09 | 1.100E-08 |
| 22838 | 1142 | A1142W | D | 5.6e-09 | 2.1e-09 | 3.3e-08 | 1.357E-08 | 1.692E-08 | 2.100E-09 | 3.300E-08 |
| 22839 | 1142 | A1142Y | D | 9.4e-09 | 3.9e-09 | 1.5e-08 | 9.433E-09 | 5.550E-09 | 3.900E-09 | 1.500E-08 |
| 22840 | 1142 | A1142V | D | 4.6e-09 | 4.8e-09 | 2e-08 | 9.800E-09 | 8.834E-09 | 4.600E-09 | 2.000E-08 |
| 22841 | 1143 | T1143A | D | 2.6e-08 | 2.4e-09 | 8.2e-09 | 1.220E-08 | 1.230E-08 | 2.400E-09 | 2.600E-08 |
| 22842 | 1143 | T1143R | D | 6.4e-09 | 5.9e-09 | 3.6e-09 | 5.300E-09 | 1.493E-09 | 3.600E-09 | 6.400E-09 |
| 22843 | 1143 | T1143N | D | 6.8e-09 | 3.4e-09 | 8.4e-09 | 6.200E-09 | 2.553E-09 | 3.400E-09 | 8.400E-09 |
| 22844 | 1143 | T1143D | D | 1.3e-08 | 8.9e-09 | 1.4e-08 | 1.197E-08 | 2.702E-09 | 8.900E-09 | 1.400E-08 |
| 22845 | 1143 | T1143C | D | 2.9e-09 | 7.3e-09 | 1.2e-08 | 7.400E-09 | 4.551E-09 | 2.900E-09 | 1.200E-08 |
| 22846 | 1143 | T1143E | D | 8.7e-09 | 1.2e-08 | 2e-09 | 7.567E-09 | 5.095E-09 | 2.000E-09 | 1.200E-08 |
| 22847 | 1143 | T1143Q | D | 4.4e-09 | 1.9e-08 | 8.4e-09 | 1.060E-08 | 7.545E-09 | 4.400E-09 | 1.900E-08 |
| 22848 | 1143 | T1143G | D | 1.5e-08 | 1.4e-08 | 1.2e-08 | 1.367E-08 | 1.528E-09 | 1.200E-08 | 1.500E-08 |
| 22849 | 1143 | T1143H | D | 3.6e-09 | 1.3e-08 | 4.8e-09 | 7.133E-09 | 5.116E-09 | 3.600E-09 | 1.300E-08 |
| 22850 | 1143 | T1143I | D | 1.8e-08 | 1.5e-08 | 2.9e-09 | 1.197E-08 | 7.994E-09 | 2.900E-09 | 1.800E-08 |
| 22851 | 1143 | T1143L | D | 1.1e-08 | 8.8e-10 | 1.1e-08 | 7.627E-09 | 5.843E-09 | 8.800E-10 | 1.100E-08 |
| 22852 | 1143 | T1143K | D | 2.2e-09 | 2.3e-09 | 4.9e-09 | 3.133E-09 | 1.531E-09 | 2.200E-09 | 4.900E-09 |
| 22853 | 1143 | T1143M | D | 2.6e-08 | 5.3e-09 | 2.3e-09 | 1.120E-08 | 1.290E-08 | 2.300E-09 | 2.600E-08 |
| 22854 | 1143 | T1143F | D | 4.2e-09 | 1.2e-08 | 4.7e-09 | 6.967E-09 | 4.366E-09 | 4.200E-09 | 1.200E-08 |
| 22855 | 1143 | T1143P | D | 1.1e-08 | 2.5e-09 | 3.9e-09 | 5.800E-09 | 4.557E-09 | 2.500E-09 | 1.100E-08 |
| 22856 | 1143 | T1143S | D | 2.9e-09 | 4.5e-09 | 2.6e-09 | 3.333E-09 | 1.021E-09 | 2.600E-09 | 4.500E-09 |
| 22857 | 1143 | T1143T | D | 9.8e-09 | 1.3e-08 | 1.2e-08 | 1.160E-08 | 1.637E-09 | 9.800E-09 | 1.300E-08 |
| 22858 | 1143 | T1143W | D | 2e-08 | 2.5e-09 | 1e-08 | 1.083E-08 | 8.780E-09 | 2.500E-09 | 2.000E-08 |
| 22859 | 1143 | T1143Y | D | 8.2e-09 | 1.3e-08 | 7.4e-09 | 9.533E-09 | 3.029E-09 | 7.400E-09 | 1.300E-08 |
| 22860 | 1143 | T1143V | D | 4.9e-09 | 2.1e-08 | 9.5e-09 | 1.180E-08 | 8.293E-09 | 4.900E-09 | 2.100E-08 |
| 22861 | 1144 | Y1144A | D | 5.5e-09 | 1.9e-08 | 6e-09 | 1.017E-08 | 7.654E-09 | 5.500E-09 | 1.900E-08 |
| 22862 | 1144 | Y1144R | D | 4.6e-09 | 3.1e-08 | 1.3e-08 | 1.620E-08 | 1.349E-08 | 4.600E-09 | 3.100E-08 |
| 22863 | 1144 | Y1144N | D | 8.7e-09 | 1.3e-08 | 4.3e-09 | 8.667E-09 | 4.350E-09 | 4.300E-09 | 1.300E-08 |
| 22864 | 1144 | Y1144D | D | 1.9e-09 | 4.7e-09 | 9.1e-09 | 5.233E-09 | 3.630E-09 | 1.900E-09 | 9.100E-09 |

| | | | | | | | | | | |
|---|---|---|---|---|---|---|---|---|---|---|
| 22865 | 1144 | Y1144C | D | 8.6e-09 | 7.1e-09 | 1.1e-08 | 8.900E-09 | 1.967E-09 | 7.100E-09 | 1.100E-08 |
| 22866 | 1144 | Y1144E | D | 1.5e-09 | 7e-09 | 2.2e-08 | 1.017E-08 | 1.061E-08 | 1.500E-09 | 2.200E-08 |
| 22867 | 1144 | Y1144Q | D | 1.1e-08 | 1e-08 | 1.2e-08 | 1.100E-08 | 1.000E-09 | 1.000E-08 | 1.200E-08 |
| 22868 | 1144 | Y1144G | D | 1.8e-08 | 1.4e-08 | 5.2e-09 | 1.240E-08 | 6.548E-09 | 5.200E-09 | 1.800E-08 |
| 22869 | 1144 | Y1144H | D | 7.1e-09 | 1.8e-08 | 6.1e-09 | 1.040E-08 | 6.601E-09 | 6.100E-09 | 1.800E-08 |
| 22870 | 1144 | Y1144I | D | 2.6e-09 | 5.9e-09 | 5.9e-09 | 4.800E-09 | 1.905E-09 | 2.600E-09 | 5.900E-09 |
| 22871 | 1144 | Y1144L | D | 5.9e-09 | 5.6e-09 | 2.8e-09 | 4.767E-09 | 1.710E-09 | 2.800E-09 | 5.900E-09 |
| 22872 | 1144 | Y1144K | D | 1.2e-09 | 3.2e-09 | 1.2e-08 | 5.467E-09 | 5.746E-09 | 1.200E-09 | 1.200E-08 |
| 22873 | 1144 | Y1144M | D | 2.2e-08 | 1.2e-08 | 1.8e-08 | 1.733E-08 | 5.033E-09 | 1.200E-08 | 2.200E-08 |
| 22874 | 1144 | Y1144F | D | 2.4e-08 | 2.8e-09 | 1.1e-08 | 1.260E-08 | 1.069E-08 | 2.800E-09 | 2.400E-08 |
| 22875 | 1144 | Y1144P | D | 1e-08 | 4.5e-09 | 8e-09 | 7.500E-09 | 2.784E-09 | 4.500E-09 | 1.000E-08 |
| 22876 | 1144 | Y1144S | D | 1.6e-08 | 1.9e-08 | 3.6e-09 | 1.287E-08 | 8.164E-09 | 3.600E-09 | 1.900E-08 |
| 22877 | 1144 | Y1144T | D | 1.6e-08 | 2.2e-09 | 2.2e-08 | 1.340E-08 | 1.015E-08 | 2.200E-09 | 2.200E-08 |
| 22878 | 1144 | Y1144W | D | 8.3e-09 | 1.4e-08 | 1.5e-08 | 1.243E-08 | 3.614E-09 | 8.300E-09 | 1.500E-08 |
| 22879 | 1144 | Y1144Y | D | 1.6e-09 | 4e-09 | 8.8e-09 | 4.800E-09 | 3.666E-09 | 1.600E-09 | 8.800E-09 |
| 22880 | 1144 | Y1144V | D | 7e-09 | 3.8e-09 | 5e-09 | 5.267E-09 | 1.617E-09 | 3.800E-09 | 7.000E-09 |
| 22881 | 1145 | Y1145A | D | 7e-09 | 4.8e-09 | 5.9e-09 | 5.900E-09 | 1.100E-09 | 4.800E-09 | 7.000E-09 |
| 22882 | 1145 | Y1145R | D | 7.7e-09 | 3e-09 | 1.1e-08 | 7.233E-09 | 4.020E-09 | 3.000E-09 | 1.100E-08 |
| 22883 | 1145 | Y1145N | D | 9.3e-09 | 2.9e-09 | 6e-09 | 6.067E-09 | 3.201E-09 | 2.900E-09 | 9.300E-09 |
| 22884 | 1145 | Y1145D | D | 1.7e-08 | 6.2e-09 | 1.7e-08 | 1.340E-08 | 6.235E-09 | 6.200E-09 | 1.700E-08 |
| 22885 | 1145 | Y1145C | D | 5.1e-09 | 6.7e-09 | 2.8e-09 | 4.867E-09 | 1.960E-09 | 2.800E-09 | 6.700E-09 |
| 22886 | 1145 | Y1145E | D | 8.1e-09 | 1.2e-08 | 5.5e-09 | 8.533E-09 | 3.272E-09 | 5.500E-09 | 1.200E-08 |
| 22887 | 1145 | Y1145Q | D | 5.3e-09 | 1.1e-08 | 1.3e-08 | 9.767E-09 | 3.995E-09 | 5.300E-09 | 1.300E-08 |
| 22888 | 1145 | Y1145G | D | 3e-09 | 6.8e-09 | 1.5e-08 | 8.267E-09 | 6.133E-09 | 3.000E-09 | 1.500E-08 |
| 22889 | 1145 | Y1145H | D | 6.5e-09 | 5.3e-09 | 1e-08 | 7.267E-09 | 2.442E-09 | 5.300E-09 | 1.000E-08 |
| 22890 | 1145 | Y1145I | D | 1.4e-08 | 7.1e-09 | 8.7e-09 | 9.933E-09 | 3.612E-09 | 7.100E-09 | 1.400E-08 |
| 22891 | 1145 | Y1145L | D | 3.6e-09 | 2.1e-09 | 1.3e-08 | 6.233E-09 | 5.908E-09 | 2.100E-09 | 1.300E-08 |
| 22892 | 1145 | Y1145K | D | 3e-09 | 7.3e-09 | 9.8e-09 | 6.700E-09 | 3.439E-09 | 3.000E-09 | 9.800E-09 |
| 22893 | 1145 | Y1145M | D | 5.5e-09 | 6.5e-09 | 1.4e-08 | 8.667E-09 | 4.646E-09 | 5.500E-09 | 1.400E-08 |
| 22894 | 1145 | Y1145F | D | 1.3e-08 | 5.6e-09 | 3.7e-09 | 7.433E-09 | 4.914E-09 | 3.700E-09 | 1.300E-08 |
| 22895 | 1145 | Y1145P | D | 3e-09 | 1e-08 | 1.3e-08 | 8.667E-09 | 5.132E-09 | 3.000E-09 | 1.300E-08 |
| 22896 | 1145 | Y1145S | D | 7.2e-09 | 3.9e-09 | 5.4e-09 | 5.500E-09 | 1.652E-09 | 3.900E-09 | 7.200E-09 |
| 22897 | 1145 | Y1145T | D | 3.2e-09 | 5.1e-09 | 9.3e-09 | 5.867E-09 | 3.121E-09 | 3.200E-09 | 9.300E-09 |
| 22898 | 1145 | Y1145W | D | 9.5e-09 | 4.8e-09 | 1e-08 | 8.100E-09 | 2.869E-09 | 4.800E-09 | 1.000E-08 |
| 22899 | 1145 | Y1145Y | D | 1.6e-09 | 3.3e-09 | 2.7e-09 | 2.533E-09 | 8.622E-10 | 1.600E-09 | 3.300E-09 |
| 22900 | 1145 | Y1145V | D | 4.5e-09 | 5.5e-09 | 2e-08 | 1.000E-08 | 8.675E-09 | 4.500E-09 | 2.000E-08 |
| 22901 | 1146 | C1146A | D | 4.5e-09 | 5.9e-09 | 3.7e-09 | 4.700E-09 | 1.114E-09 | 3.700E-09 | 5.900E-09 |
| 22902 | 1146 | C1146R | D | 5.6e-09 | 8.5e-09 | 1e-08 | 8.033E-09 | 2.237E-09 | 5.600E-09 | 1.000E-08 |
| 22903 | 1146 | C1146N | D | 6.5e-09 | 1.5e-08 | 2e-08 | 1.383E-08 | 6.825E-09 | 6.500E-09 | 2.000E-08 |
| 22904 | 1146 | C1146D | D | 4.8e-09 | 3.7e-09 | 8.3e-09 | 5.600E-09 | 2.402E-09 | 3.700E-09 | 8.300E-09 |
| 22905 | 1146 | C1146C | D | 1.3e-08 | 1.2e-08 | 5.1e-09 | 1.003E-08 | 4.302E-09 | 5.100E-09 | 1.300E-08 |
| 22906 | 1146 | C1146E | D | 5.4e-09 | 7.4e-09 | 5.7e-09 | 6.167E-09 | 1.079E-09 | 5.400E-09 | 7.400E-09 |
| 22907 | 1146 | C1146Q | D | 6.2e-09 | 9.3e-09 | 8.8e-09 | 8.100E-09 | 1.664E-09 | 6.200E-09 | 9.300E-09 |
| 22908 | 1146 | C1146G | D | 4.2e-09 | 6.1e-09 | 2e-08 | 1.010E-08 | 8.626E-09 | 4.200E-09 | 2.000E-08 |
| 22909 | 1146 | C1146H | D | 5e-09 | 3.6e-09 | 6.6e-09 | 5.067E-09 | 1.501E-09 | 3.600E-09 | 6.600E-09 |
| 22910 | 1146 | C1146I | D | 8e-09 | 9.6e-09 | 4.5e-09 | 7.367E-09 | 2.608E-09 | 4.500E-09 | 9.600E-09 |
| 22911 | 1146 | C1146L | D | 4.4e-09 | 5.1e-09 | 3.4e-09 | 4.300E-09 | 8.544E-10 | 3.400E-09 | 5.100E-09 |
| 22912 | 1146 | C1146K | D | 5.2e-09 | 7.2e-09 | 8.9e-09 | 7.100E-09 | 1.852E-09 | 5.200E-09 | 8.900E-09 |
| 22913 | 1146 | C1146M | D | 4.5e-09 | 7.2e-09 | 3.9e-09 | 5.200E-09 | 1.758E-09 | 3.900E-09 | 7.200E-09 |
| 22914 | 1146 | C1146F | D | 1.4e-08 | 2.1e-08 | 7.7e-09 | 1.423E-08 | 6.653E-09 | 7.700E-09 | 2.100E-08 |
| 22915 | 1146 | C1146P | D | 6.3e-09 | 4.7e-09 | 8.1e-09 | 6.367E-09 | 1.701E-09 | 4.700E-09 | 8.100E-09 |
| 22916 | 1146 | C1146S | D | 1.2e-08 | 8.5e-09 | 4.6e-09 | 8.367E-09 | 3.702E-09 | 4.600E-09 | 1.200E-08 |
| 22917 | 1146 | C1146T | D | 3.4e-09 | 1.2e-08 | 7e-09 | 7.467E-09 | 4.319E-09 | 3.400E-09 | 1.200E-08 |
| 22918 | 1146 | C1146W | D | 1e-08 | 1.2e-08 | 8.2e-09 | 1.007E-08 | 1.901E-09 | 8.200E-09 | 1.200E-08 |
| 22919 | 1146 | C1146Y | D | 9.2e-09 | 1.1e-08 | 4.1e-09 | 8.100E-09 | 3.579E-09 | 4.100E-09 | 1.100E-08 |
| 22920 | 1146 | C1146V | D | 6.9e-09 | 2.8e-08 | 3.6e-09 | 1.283E-08 | 1.324E-08 | 3.600E-09 | 2.800E-08 |
| 22921 | 1147 | Q1147A | D | 5.9e-09 | 5.4e-09 | 1.1e-08 | 7.433E-09 | 3.099E-09 | 5.400E-09 | 1.100E-08 |
| 22922 | 1147 | Q1147R | D | 8.6e-09 | 8.3e-09 | 1.2e-08 | 9.633E-09 | 2.055E-09 | 8.300E-09 | 1.200E-08 |
| 22923 | 1147 | Q1147N | D | 9e-09 | 2.6e-09 | 4.6e-09 | 5.400E-09 | 3.274E-09 | 2.600E-09 | 9.000E-09 |
| 22924 | 1147 | Q1147D | D | 3.1e-09 | 6.7e-09 | 5.1e-09 | 4.967E-09 | 1.804E-09 | 3.100E-09 | 6.700E-09 |
| 22925 | 1147 | Q1147C | D | 5.5e-09 | 1.1e-08 | 8.1e-09 | 8.200E-09 | 2.751E-09 | 5.500E-09 | 1.100E-08 |
| 22926 | 1147 | Q1147E | D | 9.1e-09 | 2e-09 | 7.2e-09 | 6.100E-09 | 3.676E-09 | 2.000E-09 | 9.100E-09 |
| 22927 | 1147 | Q1147Q | D | 9.3e-09 | 6.9e-09 | 8.9e-09 | 8.367E-09 | 1.286E-09 | 6.900E-09 | 9.300E-09 |
| 22928 | 1147 | Q1147G | D | 1.7e-08 | 1.6e-08 | 1.4e-08 | 1.567E-08 | 1.528E-09 | 1.400E-08 | 1.700E-08 |
| 22929 | 1147 | Q1147H | D | 5.9e-09 | 7.1e-09 | 3e-09 | 5.333E-09 | 2.108E-09 | 3.000E-09 | 7.100E-09 |
| 22930 | 1147 | Q1147I | D | 8.3e-09 | 5.9e-09 | 4e-09 | 6.067E-09 | 2.155E-09 | 4.000E-09 | 8.300E-09 |
| 22931 | 1147 | Q1147L | D | 3.1e-09 | 2e-08 | 7.5e-09 | 1.020E-08 | 8.768E-09 | 3.100E-09 | 2.000E-08 |
| 22932 | 1147 | Q1147K | D | 1.2e-08 | 1.7e-08 | 1e-08 | 1.300E-08 | 3.606E-09 | 1.000E-08 | 1.700E-08 |
| 22933 | 1147 | Q1147M | D | 1.3e-08 | 1.7e-08 | 1.3e-08 | 1.433E-08 | 2.309E-09 | 1.300E-08 | 1.700E-08 |
| 22934 | 1147 | Q1147F | D | 8.3e-09 | 7.3e-09 | 1.6e-08 | 1.053E-08 | 4.761E-09 | 7.300E-09 | 1.600E-08 |
| 22935 | 1147 | Q1147P | D | 1e-08 | 4.6e-09 | 4.2e-09 | 6.267E-09 | 3.239E-09 | 4.200E-09 | 1.000E-08 |
| 22936 | 1147 | Q1147S | D | 1.9e-09 | 8.2e-09 | 1e-08 | 6.700E-09 | 4.253E-09 | 1.900E-09 | 1.000E-08 |
| 22937 | 1147 | Q1147T | D | 1.5e-08 | 1.7e-08 | 6.7e-09 | 1.290E-08 | 5.462E-09 | 6.700E-09 | 1.700E-08 |
| 22938 | 1147 | Q1147W | D | 4.4e-09 | 6e-09 | 7.7e-09 | 6.033E-09 | 1.650E-09 | 4.400E-09 | 7.700E-09 |
| 22939 | 1147 | Q1147Y | D | 6.9e-09 | 7.2e-09 | 1.1e-08 | 8.367E-09 | 2.285E-09 | 6.900E-09 | 1.100E-08 |
| 22940 | 1147 | Q1147V | D | 1.4e-08 | 3.1e-09 | 6e-09 | 7.700E-09 | 5.645E-09 | 3.100E-09 | 1.400E-08 |
| 22941 | 1148 | Q1148A | D | 1.5e-08 | 5.4e-09 | 2e-08 | 1.347E-08 | 7.420E-09 | 5.400E-09 | 2.000E-08 |
| 22942 | 1148 | Q1148R | D | 1.1e-08 | 7e-09 | 4.2e-09 | 7.400E-09 | 3.418E-09 | 4.200E-09 | 1.100E-08 |
| 22943 | 1148 | Q1148N | D | 6.3e-09 | 8.1e-09 | 1.3e-08 | 9.133E-09 | 3.467E-09 | 6.300E-09 | 1.300E-08 |
| 22944 | 1148 | Q1148D | D | 6.2e-09 | 2.5e-09 | 4.8e-09 | 4.500E-09 | 1.868E-09 | 2.500E-09 | 6.200E-09 |
| 22945 | 1148 | Q1148C | D | 1e-08 | 9.8e-09 | 4.5e-09 | 8.100E-09 | 3.119E-09 | 4.500E-09 | 1.000E-08 |
| 22946 | 1148 | Q1148E | D | 6.4e-09 | 7.7e-09 | 1.8e-09 | 5.300E-09 | 3.100E-09 | 1.800E-09 | 7.700E-09 |
| 22947 | 1148 | Q1148Q | D | 8.5e-09 | 5.2e-09 | 1.8e-09 | 5.167E-09 | 3.350E-09 | 1.800E-09 | 8.500E-09 |
| 22948 | 1148 | Q1148G | D | 6e-09 | 1.3e-08 | 1.1e-08 | 1.000E-08 | 3.606E-09 | 6.000E-09 | 1.300E-08 |
| 22949 | 1148 | Q1148H | D | 8e-09 | 2.1e-09 | 9e-09 | 6.367E-09 | 3.729E-09 | 2.100E-09 | 9.000E-09 |

| | | | | | | | | | | |
|---|---|---|---|---|---|---|---|---|---|---|
| 22950 | 1148 | Q1148I | D | 1.5e-08 | 1.2e-08 | 1.4e-08 | 1.367E-08 | 1.528E-09 | 1.200E-08 | 1.500E-08 |
| 22951 | 1148 | Q1148L | D | 9.3e-09 | 8.3e-09 | 7.5e-09 | 8.367E-09 | 9.018E-10 | 7.500E-09 | 9.300E-09 |
| 22952 | 1148 | Q1148K | D | 4.5e-09 | 5e-09 | 3.7e-09 | 4.400E-09 | 6.557E-10 | 3.700E-09 | 5.000E-09 |
| 22953 | 1148 | Q1148M | D | 7e-09 | 9e-09 | 2.6e-09 | 6.200E-09 | 3.274E-09 | 2.600E-09 | 9.000E-09 |
| 22954 | 1148 | Q1148F | D | 8.6e-09 | 9.1e-09 | 5.8e-09 | 7.833E-09 | 1.779E-09 | 5.800E-09 | 9.100E-09 |
| 22955 | 1148 | Q1148P | D | 1.6e-09 | 6.9e-09 | 1.3e-08 | 7.167E-09 | 5.705E-09 | 1.600E-09 | 1.300E-08 |
| 22956 | 1148 | Q1148S | D | 1e-08 | 5.8e-09 | 5.7e-09 | 7.167E-09 | 2.454E-09 | 5.700E-09 | 1.000E-08 |
| 22957 | 1148 | Q1148T | D | 6.5e-09 | 1.1e-08 | 8e-09 | 8.500E-09 | 2.291E-09 | 6.500E-09 | 1.100E-08 |
| 22958 | 1148 | Q1148W | D | 1.4e-08 | 8.7e-09 | 1.4e-08 | 1.223E-08 | 3.060E-09 | 8.700E-09 | 1.400E-08 |
| 22959 | 1148 | Q1148Y | D | 4.2e-09 | 1.2e-08 | 8.3e-09 | 8.167E-09 | 3.902E-09 | 4.200E-09 | 1.200E-08 |
| 22960 | 1148 | Q1148V | D | 1.2e-08 | 3.9e-09 | 2.1e-08 | 1.230E-08 | 8.554E-09 | 3.900E-09 | 2.100E-08 |
| 22961 | 1149 | H1149A | D | 1.1e-08 | 1.9e-08 | 3.2e-09 | 1.107E-08 | 7.900E-09 | 3.200E-09 | 1.900E-08 |
| 22962 | 1149 | H1149R | D | 7.2e-09 | 8e-09 | 6.2e-09 | 7.133E-09 | 9.018E-10 | 6.200E-09 | 8.000E-09 |
| 22963 | 1149 | H1149N | D | 1.3e-08 | 2e-08 | 8.9e-09 | 1.397E-08 | 5.613E-09 | 8.900E-09 | 2.000E-08 |
| 22964 | 1149 | H1149D | D | 1.2e-08 | 4.8e-09 | 7.9e-09 | 8.233E-09 | 3.612E-09 | 4.800E-09 | 1.200E-08 |
| 22965 | 1149 | H1149C | D | 5.9e-09 | 1.3e-08 | 5.3e-09 | 8.067E-09 | 4.283E-09 | 5.300E-09 | 1.300E-08 |
| 22966 | 1149 | H1149E | D | 3.3e-09 | 8.3e-09 | 7.3e-09 | 6.300E-09 | 2.646E-09 | 3.300E-09 | 8.300E-09 |
| 22967 | 1149 | H1149Q | D | 4e-09 | 5.8e-09 | 7.5e-09 | 5.767E-09 | 1.750E-09 | 4.000E-09 | 7.500E-09 |
| 22968 | 1149 | H1149G | D | 3.3e-09 | 1.6e-08 | 2.7e-09 | 7.333E-09 | 7.512E-09 | 2.700E-09 | 1.600E-08 |
| 22969 | 1149 | H1149H | D | 6.7e-09 | 9.7e-09 | 1.3e-08 | 9.800E-09 | 3.151E-09 | 6.700E-09 | 1.300E-08 |
| 22970 | 1149 | H1149I | D | 1.3e-08 | 3.7e-09 | 2.3e-09 | 6.333E-09 | 5.816E-09 | 2.300E-09 | 1.300E-08 |
| 22971 | 1149 | H1149L | D | 3.4e-09 | 8.3e-09 | 5.4e-09 | 5.700E-09 | 2.464E-09 | 3.400E-09 | 8.300E-09 |
| 22972 | 1149 | H1149K | D | 2.8e-09 | 3.6e-09 | 6.9e-09 | 4.433E-09 | 2.173E-09 | 2.800E-09 | 6.900E-09 |
| 22973 | 1149 | H1149M | D | 9e-09 | 9.3e-09 | 9.2e-09 | 9.167E-09 | 1.528E-10 | 9.000E-09 | 9.300E-09 |
| 22974 | 1149 | H1149F | D | 7.6e-09 | 8.7e-09 | 3e-09 | 6.433E-09 | 3.024E-09 | 3.000E-09 | 8.700E-09 |
| 22975 | 1149 | H1149P | D | 8.3e-09 | 4.2e-09 | 1.1e-08 | 7.833E-09 | 3.424E-09 | 4.200E-09 | 1.100E-08 |
| 22976 | 1149 | H1149S | D | 7.6e-09 | 8.3e-09 | 4.2e-09 | 6.700E-09 | 2.193E-09 | 4.200E-09 | 8.300E-09 |
| 22977 | 1149 | H1149T | D | 6e-09 | 1.1e-08 | 4.6e-09 | 7.200E-09 | 3.365E-09 | 4.600E-09 | 1.100E-08 |
| 22978 | 1149 | H1149W | D | 1e-08 | 6.8e-09 | 2e-09 | 6.267E-09 | 4.027E-09 | 2.000E-09 | 1.000E-08 |
| 22979 | 1149 | H1149Y | D | 4.4e-09 | 2.6e-09 | 6.8e-09 | 4.600E-09 | 2.107E-09 | 2.600E-09 | 6.800E-09 |
| 22980 | 1149 | H1149V | D | 2.8e-09 | 9.3e-09 | 1.4e-08 | 8.700E-09 | 5.624E-09 | 2.800E-09 | 1.400E-08 |
| 22981 | 1150 | Y1150A | D | 1.3e-08 | 5.4e-09 | 1.1e-08 | 9.800E-09 | 3.940E-09 | 5.400E-09 | 1.300E-08 |
| 22982 | 1150 | Y1150R | D | 7.2e-09 | 7.7e-09 | 5.3e-09 | 6.733E-09 | 1.266E-09 | 5.300E-09 | 7.700E-09 |
| 22983 | 1150 | Y1150N | D | 3.5e-09 | 6.7e-09 | 1.1e-08 | 7.067E-09 | 3.763E-09 | 3.500E-09 | 1.100E-08 |
| 22984 | 1150 | Y1150D | D | 7.1e-09 | 6.8e-09 | 2e-08 | 1.130E-08 | 7.536E-09 | 6.800E-09 | 2.000E-08 |
| 22985 | 1150 | Y1150C | D | 5.7e-09 | 8.7e-09 | 2e-09 | 5.467E-09 | 3.356E-09 | 2.000E-09 | 8.700E-09 |
| 22986 | 1150 | Y1150E | D | 1.1e-08 | 7.8e-09 | 3.6e-09 | 7.467E-09 | 3.711E-09 | 3.600E-09 | 1.100E-08 |
| 22987 | 1150 | Y1150Q | D | 6.4e-09 | 2e-08 | 7.3e-09 | 1.123E-08 | 7.605E-09 | 6.400E-09 | 2.000E-08 |
| 22988 | 1150 | Y1150G | D | 1.7e-08 | 7.4e-09 | 1.4e-08 | 1.280E-08 | 4.911E-09 | 7.400E-09 | 1.700E-08 |
| 22989 | 1150 | Y1150H | D | 3.2e-09 | 9.5e-09 | 5.8e-09 | 6.167E-09 | 3.166E-09 | 3.200E-09 | 9.500E-09 |
| 22990 | 1150 | Y1150I | D | 1.1e-08 | 7.2e-09 | 4.1e-09 | 7.433E-09 | 3.456E-09 | 4.100E-09 | 1.100E-08 |
| 22991 | 1150 | Y1150L | D | 5.3e-09 | 2.2e-09 | 6.5e-09 | 4.667E-09 | 2.219E-09 | 2.200E-09 | 6.500E-09 |
| 22992 | 1150 | Y1150K | D | 1.6e-08 | 4.7e-09 | 4.4e-09 | 8.367E-09 | 6.612E-09 | 4.400E-09 | 1.600E-08 |
| 22993 | 1150 | Y1150M | D | 6e-09 | 4e-09 | 6.6e-09 | 5.533E-09 | 1.361E-09 | 4.000E-09 | 6.600E-09 |
| 22994 | 1150 | Y1150F | D | 1.1e-08 | 1.2e-08 | 1.4e-08 | 1.233E-08 | 1.528E-09 | 1.100E-08 | 1.400E-08 |
| 22995 | 1150 | Y1150P | D | 5.5e-09 | 5.7e-09 | 5.6e-09 | 5.600E-09 | 1.000E-10 | 5.500E-09 | 5.700E-09 |
| 22996 | 1150 | Y1150S | D | 1e-08 | 5.6e-09 | 8.4e-09 | 8.000E-09 | 2.227E-09 | 5.600E-09 | 1.000E-08 |
| 22997 | 1150 | Y1150T | D | 1.2e-08 | 1.1e-08 | 3.9e-09 | 8.967E-09 | 4.416E-09 | 3.900E-09 | 1.200E-08 |
| 22998 | 1150 | Y1150W | D | 6.2e-09 | 7.6e-09 | 7.3e-09 | 7.033E-09 | 7.371E-10 | 6.200E-09 | 7.600E-09 |
| 22999 | 1150 | Y1150Y | D | 8.7e-09 | 9e-09 | 5.5e-09 | 7.733E-09 | 1.940E-09 | 5.500E-09 | 9.000E-09 |
| 23000 | 1150 | Y1150V | D | 1.7e-08 | 6.5e-09 | 3.6e-09 | 9.033E-09 | 7.050E-09 | 3.600E-09 | 1.700E-08 |
| 23001 | 1151 | T1151A | D | 1.4e-08 | 7.9e-09 | 1.5e-08 | 1.230E-08 | 3.843E-09 | 7.900E-09 | 1.500E-08 |
| 23002 | 1151 | T1151R | D | 4e-09 | 3.2e-09 | 1.7e-08 | 8.067E-09 | 7.747E-09 | 3.200E-09 | 1.700E-08 |
| 23003 | 1151 | T1151N | D | 1.1e-08 | 8.5e-09 | 9.8e-09 | 9.767E-09 | 1.250E-09 | 8.500E-09 | 1.100E-08 |
| 23004 | 1151 | T1151D | D | 4.1e-09 | 4.9e-09 | 1.1e-08 | 6.667E-09 | 3.774E-09 | 4.100E-09 | 1.100E-08 |
| 23005 | 1151 | T1151C | D | 1.9e-08 | 6.4e-09 | 1.2e-09 | 8.867E-09 | 9.153E-09 | 1.200E-09 | 1.900E-08 |
| 23006 | 1151 | T1151E | D | 1.4e-08 | 7e-09 | 1.4e-08 | 1.167E-08 | 4.041E-09 | 7.000E-09 | 1.400E-08 |
| 23007 | 1151 | T1151Q | D | 1.2e-08 | 7.5e-09 | 6.2e-09 | 8.567E-09 | 3.044E-09 | 6.200E-09 | 1.200E-08 |
| 23008 | 1151 | T1151G | D | 3e-09 | 2.9e-09 | 1.4e-08 | 6.633E-09 | 6.380E-09 | 2.900E-09 | 1.400E-08 |
| 23009 | 1151 | T1151H | D | 9.7e-09 | 1.3e-08 | 1.2e-08 | 1.157E-08 | 1.692E-09 | 9.700E-09 | 1.300E-08 |
| 23010 | 1151 | T1151I | D | 5.2e-09 | 1.3e-08 | 5.6e-09 | 7.933E-09 | 4.392E-09 | 5.200E-09 | 1.300E-08 |
| 23011 | 1151 | T1151L | D | 8.9e-09 | 8.8e-09 | 9e-09 | 8.900E-09 | 1.000E-10 | 8.800E-09 | 9.000E-09 |
| 23012 | 1151 | T1151K | D | 1.1e-08 | 7.9e-09 | 1e-08 | 9.633E-09 | 1.582E-09 | 7.900E-09 | 1.100E-08 |
| 23013 | 1151 | T1151M | D | 4e-09 | 8e-09 | 4.5e-09 | 5.500E-09 | 2.179E-09 | 4.000E-09 | 8.000E-09 |
| 23014 | 1151 | T1151F | D | 8.3e-09 | 1.3e-08 | 6.6e-09 | 9.300E-09 | 3.315E-09 | 6.600E-09 | 1.300E-08 |
| 23015 | 1151 | T1151P | D | 5.3e-09 | 5.6e-09 | 3.7e-09 | 4.867E-09 | 1.021E-09 | 3.700E-09 | 5.600E-09 |
| 23016 | 1151 | T1151S | D | 1.5e-08 | 8.3e-09 | 9.1e-09 | 1.080E-08 | 3.659E-09 | 8.300E-09 | 1.500E-08 |
| 23017 | 1151 | T1151T | D | 1.3e-08 | 9.2e-09 | 1.4e-08 | 1.207E-08 | 2.532E-09 | 9.200E-09 | 1.400E-08 |
| 23018 | 1151 | T1151W | D | 7.3e-09 | 9.4e-09 | 7.2e-09 | 7.967E-09 | 1.242E-09 | 7.200E-09 | 9.400E-09 |
| 23019 | 1151 | T1151Y | D | 3.2e-09 | 1.4e-08 | 4.1e-09 | 7.100E-09 | 5.992E-09 | 3.200E-09 | 1.400E-08 |
| 23020 | 1151 | T1151V | D | 7.8e-09 | 4e-09 | 6.7e-09 | 6.167E-09 | 1.955E-09 | 4.000E-09 | 7.800E-09 |
| 23021 | 1152 | T1152A | D | 4.8e-09 | 7e-09 | 8.7e-09 | 6.833E-09 | 1.955E-09 | 4.800E-09 | 8.700E-09 |
| 23022 | 1152 | T1152R | D | 1.1e-08 | 5.4e-09 | 1.2e-08 | 9.467E-09 | 3.557E-09 | 5.400E-09 | 1.200E-08 |
| 23023 | 1152 | T1152N | D | 1e-08 | 1.1e-08 | 2.7e-09 | 7.900E-09 | 4.531E-09 | 2.700E-09 | 1.100E-08 |
| 23024 | 1152 | T1152D | D | 1.5e-08 | 1.2e-08 | 4.3e-09 | 1.043E-08 | 5.519E-09 | 4.300E-09 | 1.500E-08 |
| 23025 | 1152 | T1152C | D | 7.1e-09 | 1.2e-09 | 3.8e-09 | 4.033E-09 | 2.957E-09 | 1.200E-09 | 7.100E-09 |
| 23026 | 1152 | T1152E | D | 7.2e-09 | 8.3e-09 | 1.9e-08 | 1.150E-08 | 6.518E-09 | 7.200E-09 | 1.900E-08 |
| 23027 | 1152 | T1152Q | D | 6.2e-09 | 3.4e-09 | 4.2e-09 | 4.600E-09 | 1.442E-09 | 3.400E-09 | 6.200E-09 |
| 23028 | 1152 | T1152G | D | 1.2e-08 | 7.3e-09 | 3.1e-08 | 1.677E-08 | 1.255E-08 | 7.300E-09 | 3.100E-08 |
| 23029 | 1152 | T1152H | D | 7.2e-09 | 4.7e-09 | 4.8e-09 | 5.567E-09 | 1.415E-09 | 4.700E-09 | 7.200E-09 |
| 23030 | 1152 | T1152I | D | 6.3e-09 | 2.5e-09 | 3.4e-09 | 4.067E-09 | 1.986E-09 | 2.500E-09 | 6.300E-09 |
| 23031 | 1152 | T1152L | D | 1.1e-08 | 5.9e-09 | 9.7e-09 | 8.867E-09 | 2.650E-09 | 5.900E-09 | 1.100E-08 |
| 23032 | 1152 | T1152K | D | 9.2e-09 | 6.5e-09 | 5.9e-09 | 7.200E-09 | 1.758E-09 | 5.900E-09 | 9.200E-09 |
| 23033 | 1152 | T1152M | D | 5.3e-09 | 9e-09 | 3.7e-09 | 6.000E-09 | 2.718E-09 | 3.700E-09 | 9.000E-09 |
| 23034 | 1152 | T1152F | D | 7.1e-09 | 6.2e-09 | 7e-09 | 6.767E-09 | 4.933E-10 | 6.200E-09 | 7.100E-09 |

| | | | | | | | | | | |
|---|---|---|---|---|---|---|---|---|---|---|
| 23035 | 1152 | T1152P | D | 1.7e-09 | 7.6e-09 | 6.2e-09 | 5.167E-09 | 3.083E-09 | 1.700E-09 | 7.600E-09 |
| 23036 | 1152 | T1152S | D | 2.4e-09 | 5.6e-09 | 6.1e-09 | 4.700E-09 | 2.007E-09 | 2.400E-09 | 6.100E-09 |
| 23037 | 1152 | T1152T | D | 7.2e-09 | 3.9e-09 | 9.9e-09 | 7.000E-09 | 3.005E-09 | 3.900E-09 | 9.900E-09 |
| 23038 | 1152 | T1152W | D | 5.7e-09 | 1.1e-08 | 5.7e-09 | 7.467E-09 | 3.060E-09 | 5.700E-09 | 1.100E-08 |
| 23039 | 1152 | T1152Y | D | 6.4e-09 | 7.1e-09 | 7.8e-09 | 7.100E-09 | 7.000E-10 | 6.400E-09 | 7.800E-09 |
| 23040 | 1152 | T1152V | D | 4.4e-09 | 6.2e-09 | 4.1e-09 | 4.900E-09 | 1.136E-09 | 4.100E-09 | 6.200E-09 |
| 23041 | 1153 | P1153A | D | 1.3e-08 | 6e-09 | 1.2e-08 | 1.033E-08 | 3.786E-09 | 6.000E-09 | 1.300E-08 |
| 23042 | 1153 | P1153R | D | 1e-08 | 4.1e-09 | 5.5e-09 | 6.533E-09 | 3.083E-09 | 4.100E-09 | 1.000E-08 |
| 23043 | 1153 | P1153N | D | 5.8e-09 | 3.5e-09 | 1e-08 | 6.433E-09 | 3.296E-09 | 3.500E-09 | 1.000E-08 |
| 23044 | 1153 | P1153D | D | 1e-08 | 1e-08 | 3.9e-09 | 7.967E-09 | 3.522E-09 | 3.900E-09 | 1.000E-08 |
| 23045 | 1153 | P1153C | D | 2.9e-09 | 6.3e-09 | 7.5e-09 | 5.567E-09 | 2.386E-09 | 2.900E-09 | 7.500E-09 |
| 23046 | 1153 | P1153E | D | 3e-09 | 2.2e-08 | 9.9e-09 | 1.163E-08 | 9.618E-09 | 3.000E-09 | 2.200E-08 |
| 23047 | 1153 | P1153Q | D | 2.4e-09 | 3.7e-09 | 8.7e-09 | 4.933E-09 | 3.326E-09 | 2.400E-09 | 8.700E-09 |
| 23048 | 1153 | P1153G | D | 1.8e-08 | 2.8e-09 | 1.4e-08 | 1.160E-08 | 7.879E-09 | 2.800E-09 | 1.800E-08 |
| 23049 | 1153 | P1153H | D | 1.9e-08 | 7.4e-09 | 3.9e-09 | 1.010E-08 | 7.904E-09 | 3.900E-09 | 1.900E-08 |
| 23050 | 1153 | P1153I | D | 5.5e-09 | 1e-08 | 1.2e-08 | 9.167E-09 | 3.329E-09 | 5.500E-09 | 1.200E-08 |
| 23051 | 1153 | P1153L | D | 5.2e-09 | 1.3e-08 | 7.4e-09 | 8.533E-09 | 4.022E-09 | 5.200E-09 | 1.300E-08 |
| 23052 | 1153 | P1153K | D | 8e-09 | 3.6e-09 | 1.2e-08 | 7.867E-09 | 4.202E-09 | 3.600E-09 | 1.200E-08 |
| 23053 | 1153 | P1153M | D | 8.1e-09 | 1.6e-08 | 4.7e-09 | 9.600E-09 | 5.797E-09 | 4.700E-09 | 1.600E-08 |
| 23054 | 1153 | P1153F | D | 1.4e-08 | 4e-09 | 3.1e-08 | 1.633E-08 | 1.365E-08 | 4.000E-09 | 3.100E-08 |
| 23055 | 1153 | P1153P | D | 3.5e-09 | 6e-09 | 3.7e-08 | 1.550E-08 | 1.866E-08 | 3.500E-09 | 3.700E-08 |
| 23056 | 1153 | P1153S | D | 3.3e-09 | 9.4e-09 | 1.5e-08 | 9.233E-09 | 5.852E-09 | 3.300E-09 | 1.500E-08 |
| 23057 | 1153 | P1153T | D | 4.4e-09 | 5e-09 | 6.2e-09 | 5.200E-09 | 9.165E-10 | 4.400E-09 | 6.200E-09 |
| 23058 | 1153 | P1153W | D | 1.2e-08 | 1.7e-08 | 7.7e-09 | 1.223E-08 | 4.654E-09 | 7.700E-09 | 1.700E-08 |
| 23059 | 1153 | P1153Y | D | 7.2e-09 | 2.7e-09 | 6.3e-09 | 5.400E-09 | 2.381E-09 | 2.700E-09 | 7.200E-09 |
| 23060 | 1153 | P1153V | D | 7.3e-09 | 8.6e-09 | 1.4e-08 | 9.967E-09 | 3.553E-09 | 7.300E-09 | 1.400E-08 |
| 23061 | 1154 | P1154A | D | 1.8e-09 | 7.4e-09 | 4.1e-09 | 4.433E-09 | 2.815E-09 | 1.800E-09 | 7.400E-09 |
| 23062 | 1154 | P1154R | D | 4.7e-09 | 1.4e-08 | 7.3e-09 | 8.667E-09 | 4.798E-09 | 4.700E-09 | 1.400E-08 |
| 23063 | 1154 | P1154N | D | 6.1e-09 | 8.9e-09 | 5e-09 | 6.667E-09 | 2.011E-09 | 5.000E-09 | 8.900E-09 |
| 23064 | 1154 | P1154D | D | 2.3e-09 | 1.1e-08 | 5.1e-09 | 6.133E-09 | 4.441E-09 | 2.300E-09 | 1.100E-08 |
| 23065 | 1154 | P1154C | D | 5.7e-09 | 5.6e-09 | 1.5e-08 | 8.767E-09 | 5.398E-09 | 5.600E-09 | 1.500E-08 |
| 23066 | 1154 | P1154E | D | 3.1e-09 | 3.7e-09 | 1.3e-08 | 6.600E-09 | 5.551E-09 | 3.100E-09 | 1.300E-08 |
| 23067 | 1154 | P1154Q | D | 9.6e-09 | 1.2e-08 | 5.3e-09 | 8.967E-09 | 3.395E-09 | 5.300E-09 | 1.200E-08 |
| 23068 | 1154 | P1154G | D | 1.6e-08 | 1.1e-08 | 2.7e-09 | 9.900E-09 | 6.718E-09 | 2.700E-09 | 1.600E-08 |
| 23069 | 1154 | P1154H | D | 5.8e-09 | 2.1e-08 | 7.7e-09 | 1.150E-08 | 8.282E-09 | 5.800E-09 | 2.100E-08 |
| 23070 | 1154 | P1154I | D | 8.1e-09 | 2.7e-08 | 3.1e-09 | 1.273E-08 | 1.261E-08 | 3.100E-09 | 2.700E-08 |
| 23071 | 1154 | P1154L | D | 4.2e-09 | 7.6e-09 | 1.2e-08 | 7.933E-09 | 3.911E-09 | 4.200E-09 | 1.200E-08 |
| 23072 | 1154 | P1154K | D | 3.6e-09 | 5.5e-09 | 3.5e-09 | 4.200E-09 | 1.127E-09 | 3.500E-09 | 5.500E-09 |
| 23073 | 1154 | P1154M | D | 1.9e-08 | 1.3e-08 | 8.3e-09 | 1.343E-08 | 5.363E-09 | 8.300E-09 | 1.900E-08 |
| 23074 | 1154 | P1154F | D | 1e-08 | 1.2e-08 | 4.9e-09 | 8.967E-09 | 3.661E-09 | 4.900E-09 | 1.200E-08 |
| 23075 | 1154 | P1154P | D | 1.1e-08 | 2.9e-09 | 1.9e-08 | 1.097E-08 | 8.050E-09 | 2.900E-09 | 1.900E-08 |
| 23076 | 1154 | P1154S | D | 4.7e-09 | 1.7e-08 | 5.1e-09 | 8.933E-09 | 6.989E-09 | 4.700E-09 | 1.700E-08 |
| 23077 | 1154 | P1154T | D | 5.4e-09 | 2e-09 | 6.5e-09 | 4.633E-09 | 2.346E-09 | 2.000E-09 | 6.500E-09 |
| 23078 | 1154 | P1154W | D | 1.8e-08 | 1.1e-08 | 4.1e-09 | 1.103E-08 | 6.950E-09 | 4.100E-09 | 1.800E-08 |
| 23079 | 1154 | P1154Y | D | 8.3e-09 | 1.8e-08 | 4.8e-09 | 1.037E-08 | 6.838E-09 | 4.800E-09 | 1.800E-08 |
| 23080 | 1154 | P1154V | D | 6.7e-09 | 6.7e-09 | 7e-09 | 6.800E-09 | 1.732E-10 | 6.700E-09 | 7.000E-09 |
| 23081 | 1155 | T1155A | D | 2e-09 | 1.6e-08 | 1.2e-08 | 1.000E-08 | 7.211E-09 | 2.000E-09 | 1.600E-08 |
| 23082 | 1155 | T1155R | D | 1e-08 | 1.3e-08 | 1.5e-08 | 1.267E-08 | 2.517E-09 | 1.000E-08 | 1.500E-08 |
| 23083 | 1155 | T1155N | D | 2e-09 | 1.7e-08 | 1.4e-08 | 1.100E-08 | 7.937E-09 | 2.000E-09 | 1.700E-08 |
| 23084 | 1155 | T1155D | D | 5.2e-09 | 2.6e-08 | 7.6e-09 | 1.293E-08 | 1.138E-08 | 5.200E-09 | 2.600E-08 |
| 23085 | 1155 | T1155C | D | 9.2e-09 | 2.3e-08 | 2e-09 | 1.140E-08 | 1.067E-08 | 2.000E-09 | 2.300E-08 |
| 23086 | 1155 | T1155E | D | 7.8e-09 | 1.5e-09 | 4e-09 | 4.433E-09 | 3.172E-09 | 1.500E-09 | 7.800E-09 |
| 23087 | 1155 | T1155Q | D | 1.1e-08 | 4.3e-09 | 1.2e-08 | 9.100E-09 | 4.187E-09 | 4.300E-09 | 1.200E-08 |
| 23088 | 1155 | T1155G | D | 1.3e-08 | 1.7e-08 | 7e-09 | 1.233E-08 | 5.033E-09 | 7.000E-09 | 1.700E-08 |
| 23089 | 1155 | T1155H | D | 2.9e-08 | 8.4e-09 | 9.4e-09 | 1.560E-08 | 1.162E-08 | 8.400E-09 | 2.900E-08 |
| 23090 | 1155 | T1155I | D | 7.9e-09 | 1.5e-08 | 3.7e-09 | 8.867E-09 | 5.712E-09 | 3.700E-09 | 1.500E-08 |
| 23091 | 1155 | T1155L | D | 1.1e-08 | 5.3e-09 | 6.5e-09 | 7.600E-09 | 3.005E-09 | 5.300E-09 | 1.100E-08 |
| 23092 | 1155 | T1155K | D | 1.2e-08 | 4.9e-09 | 2.6e-09 | 6.500E-09 | 4.900E-09 | 2.600E-09 | 1.200E-08 |
| 23093 | 1155 | T1155M | D | 1.2e-08 | 5.4e-09 | 6.2e-09 | 7.867E-09 | 3.602E-09 | 5.400E-09 | 1.200E-08 |
| 23094 | 1155 | T1155F | D | 7.6e-09 | 1e-08 | 1.1e-09 | 6.233E-09 | 4.605E-09 | 1.100E-09 | 1.000E-08 |
| 23095 | 1155 | T1155P | D | 4e-09 | 4e-09 | 1.2e-08 | 6.667E-09 | 4.619E-09 | 4.000E-09 | 1.200E-08 |
| 23096 | 1155 | T1155S | D | 8.2e-09 | 5.3e-09 | 4.9e-09 | 6.133E-09 | 1.801E-09 | 4.900E-09 | 8.200E-09 |
| 23097 | 1155 | T1155T | D | 7.3e-09 | 4.7e-09 | 5.2e-09 | 5.733E-09 | 1.380E-09 | 4.700E-09 | 7.300E-09 |
| 23098 | 1155 | T1155W | D | 4.8e-09 | 6.1e-09 | 2e-08 | 1.030E-08 | 8.426E-09 | 4.800E-09 | 2.000E-08 |
| 23099 | 1155 | T1155Y | D | 2.3e-09 | 4e-09 | 9.8e-09 | 5.367E-09 | 3.932E-09 | 2.300E-09 | 9.800E-09 |
| 23100 | 1155 | T1155V | D | 3.7e-09 | 3.5e-09 | 1.6e-08 | 7.733E-09 | 7.160E-09 | 3.500E-09 | 1.600E-08 |
| 23101 | 1156 | F1156A | D | 5.7e-09 | 3.3e-09 | 2e-09 | 3.667E-09 | 1.877E-09 | 2.000E-09 | 5.700E-09 |
| 23102 | 1156 | F1156R | D | 4.4e-09 | 6.4e-09 | 9.9e-09 | 6.900E-09 | 2.784E-09 | 4.400E-09 | 9.900E-09 |
| 23103 | 1156 | F1156N | D | 7.6e-09 | 2.3e-09 | 3.6e-09 | 4.500E-09 | 2.762E-09 | 2.300E-09 | 7.600E-09 |
| 23104 | 1156 | F1156D | D | 3.3e-09 | 1.4e-08 | 6.7e-09 | 8.000E-09 | 5.467E-09 | 3.300E-09 | 1.400E-08 |
| 23105 | 1156 | F1156C | D | 5.4e-09 | 1.4e-08 | 1.2e-08 | 1.047E-08 | 4.500E-09 | 5.400E-09 | 1.400E-08 |
| 23106 | 1156 | F1156E | D | 1.6e-08 | 6.4e-09 | 2.8e-09 | 8.400E-09 | 6.823E-09 | 2.800E-09 | 1.600E-08 |
| 23107 | 1156 | F1156Q | D | 1.4e-08 | 6.3e-09 | 1.1e-08 | 1.043E-08 | 3.881E-09 | 6.300E-09 | 1.400E-08 |
| 23108 | 1156 | F1156G | D | 7e-09 | 8.5e-09 | 2e-09 | 5.833E-09 | 3.403E-09 | 2.000E-09 | 8.500E-09 |
| 23109 | 1156 | F1156H | D | 9.5e-09 | 4.6e-09 | 8.2e-09 | 7.433E-09 | 2.538E-09 | 4.600E-09 | 9.500E-09 |
| 23110 | 1156 | F1156I | D | 1.1e-08 | 1.1e-08 | 4.2e-09 | 8.733E-09 | 3.926E-09 | 4.200E-09 | 1.100E-08 |
| 23111 | 1156 | F1156L | D | 1e-08 | 5.1e-09 | 4.4e-09 | 6.500E-09 | 3.051E-09 | 4.400E-09 | 1.000E-08 |
| 23112 | 1156 | F1156K | D | 6.4e-09 | 5.4e-09 | 3.3e-09 | 5.033E-09 | 1.582E-09 | 3.300E-09 | 6.400E-09 |
| 23113 | 1156 | F1156M | D | 1.7e-08 | 2.9e-09 | 2.7e-09 | 7.533E-09 | 8.199E-09 | 2.700E-09 | 1.700E-08 |
| 23114 | 1156 | F1156F | D | 8.5e-09 | 1.1e-08 | 3.9e-09 | 7.800E-09 | 3.601E-09 | 3.900E-09 | 1.100E-08 |
| 23115 | 1156 | F1156P | D | 5.6e-09 | 1.2e-08 | 9e-09 | 8.867E-09 | 3.202E-09 | 5.600E-09 | 1.200E-08 |
| 23116 | 1156 | F1156S | D | 7.3e-09 | 2.4e-09 | 6.7e-09 | 5.467E-09 | 2.673E-09 | 2.400E-09 | 7.300E-09 |
| 23117 | 1156 | F1156T | D | 1.1e-08 | 4.3e-09 | 8.6e-10 | 5.387E-09 | 5.157E-09 | 8.600E-10 | 1.100E-08 |
| 23118 | 1156 | F1156W | D | 3.1e-09 | 1.4e-08 | 9.3e-09 | 8.800E-09 | 5.467E-09 | 3.100E-09 | 1.400E-08 |
| 23119 | 1156 | F1156Y | D | 4.2e-09 | 1.5e-08 | 6e-09 | 8.400E-09 | 5.786E-09 | 4.200E-09 | 1.500E-08 |

| | | | | | | | | | | |
|---|---|---|---|---|---|---|---|---|---|---|
| 23120 | 1156 | F1156V | D | 1e-08 | 8.5e-09 | 4.4e-09 | 7.633E-09 | 2.899E-09 | 4.400E-09 | 1.000E-08 |
| 23121 | 1157 | G1157A | D | 5e-09 | 1.8e-08 | 3.4e-09 | 8.800E-09 | 8.007E-09 | 3.400E-09 | 1.800E-08 |
| 23122 | 1157 | G1157R | D | 1e-08 | 9.7e-09 | 1.3e-08 | 1.090E-08 | 1.825E-09 | 9.700E-09 | 1.300E-08 |
| 23123 | 1157 | G1157N | D | 9.4e-09 | 9.2e-09 | 6.7e-09 | 8.433E-09 | 1.504E-09 | 6.700E-09 | 9.400E-09 |
| 23124 | 1157 | G1157D | D | 1.3e-08 | 8.9e-10 | 8.4e-09 | 7.430E-09 | 6.113E-09 | 8.900E-10 | 1.300E-08 |
| 23125 | 1157 | G1157C | D | 2.4e-08 | 3.5e-09 | 9.4e-09 | 1.230E-08 | 1.055E-08 | 3.500E-09 | 2.400E-08 |
| 23126 | 1157 | G1157E | D | 2.4e-08 | 5.4e-09 | 1.8e-08 | 1.580E-08 | 9.493E-09 | 5.400E-09 | 2.400E-08 |
| 23127 | 1157 | G1157Q | D | 5.6e-09 | 2.7e-09 | 5.9e-09 | 4.733E-09 | 1.767E-09 | 2.700E-09 | 5.900E-09 |
| 23128 | 1157 | G1157G | D | 5.3e-09 | 5.4e-09 | 5.3e-09 | 5.333E-09 | 5.774E-11 | 5.300E-09 | 5.400E-09 |
| 23129 | 1157 | G1157H | D | 6.1e-09 | 2.4e-09 | 6.9e-09 | 5.133E-09 | 2.401E-09 | 2.400E-09 | 6.900E-09 |
| 23130 | 1157 | G1157I | D | 6.9e-09 | 4.3e-09 | 1e-08 | 7.067E-09 | 2.854E-09 | 4.300E-09 | 1.000E-08 |
| 23131 | 1157 | G1157L | D | 7.6e-09 | 1.1e-08 | 7.1e-09 | 8.567E-09 | 2.122E-09 | 7.100E-09 | 1.100E-08 |
| 23132 | 1157 | G1157K | D | 5.1e-09 | 5.6e-09 | 3.8e-09 | 4.833E-09 | 9.292E-10 | 3.800E-09 | 5.600E-09 |
| 23133 | 1157 | G1157M | D | 1.1e-08 | 8e-09 | 1.1e-08 | 1.000E-08 | 1.732E-09 | 8.000E-09 | 1.100E-08 |
| 23134 | 1157 | G1157F | D | 1.4e-08 | 7.4e-09 | 1.1e-08 | 1.080E-08 | 3.305E-09 | 7.400E-09 | 1.400E-08 |
| 23135 | 1157 | G1157P | D | 1.7e-08 | 7.4e-09 | 6.3e-09 | 1.023E-08 | 5.886E-09 | 6.300E-09 | 1.700E-08 |
| 23136 | 1157 | G1157S | D | 9.7e-09 | 2.1e-09 | 1.3e-08 | 8.267E-09 | 5.590E-09 | 2.100E-09 | 1.300E-08 |
| 23137 | 1157 | G1157T | D | 6.8e-09 | 4.1e-09 | 6.7e-09 | 5.867E-09 | 1.531E-09 | 4.100E-09 | 6.800E-09 |
| 23138 | 1157 | G1157W | D | 7.4e-09 | 1.2e-08 | 1.6e-08 | 1.180E-08 | 4.303E-09 | 7.400E-09 | 1.600E-08 |
| 23139 | 1157 | G1157Y | D | 6.5e-09 | 7.3e-09 | 4.7e-09 | 6.167E-09 | 1.332E-09 | 4.700E-09 | 7.300E-09 |
| 23140 | 1157 | G1157V | D | 8.9e-09 | 3.2e-08 | 8.9e-09 | 1.660E-08 | 1.334E-08 | 8.900E-09 | 3.200E-08 |
| 23141 | 1158 | Q1158A | D | 1.7e-09 | 6.4e-09 | 6.3e-09 | 4.800E-09 | 2.685E-09 | 1.700E-09 | 6.400E-09 |
| 23142 | 1158 | Q1158R | D | 8.8e-09 | 1.8e-08 | 1.3e-08 | 1.327E-08 | 4.606E-09 | 8.800E-09 | 1.800E-08 |
| 23143 | 1158 | Q1158N | D | 9.5e-09 | 6.7e-09 | 9.4e-09 | 8.533E-09 | 1.589E-09 | 6.700E-09 | 9.500E-09 |
| 23144 | 1158 | Q1158D | D | 4e-09 | 6.8e-09 | 7.7e-09 | 6.167E-09 | 1.930E-09 | 4.000E-09 | 7.700E-09 |
| 23145 | 1158 | Q1158C | D | 6e-09 | 5.8e-09 | 1.8e-08 | 9.933E-09 | 6.987E-09 | 5.800E-09 | 1.800E-08 |
| 23146 | 1158 | Q1158E | D | 1.3e-08 | 8.2e-09 | 1.3e-08 | 1.140E-08 | 2.771E-09 | 8.200E-09 | 1.300E-08 |
| 23147 | 1158 | Q1158Q | D | 8.4e-09 | 1.5e-08 | 9.9e-09 | 1.110E-08 | 3.460E-09 | 8.400E-09 | 1.500E-08 |
| 23148 | 1158 | Q1158G | D | 5e-09 | 2.6e-09 | 1.5e-08 | 7.533E-09 | 6.577E-09 | 2.600E-09 | 1.500E-08 |
| 23149 | 1158 | Q1158H | D | 2.7e-09 | 6.8e-09 | 1.2e-08 | 7.167E-09 | 4.661E-09 | 2.700E-09 | 1.200E-08 |
| 23150 | 1158 | Q1158I | D | 1e-08 | 9.1e-09 | 6.2e-09 | 8.433E-09 | 1.986E-09 | 6.200E-09 | 1.000E-08 |
| 23151 | 1158 | Q1158L | D | 9.8e-09 | 4.6e-09 | 3.5e-09 | 5.967E-09 | 3.365E-09 | 3.500E-09 | 9.800E-09 |
| 23152 | 1158 | Q1158K | D | 3.7e-09 | 6.4e-09 | 1.1e-08 | 7.033E-09 | 3.691E-09 | 3.700E-09 | 1.100E-08 |
| 23153 | 1158 | Q1158M | D | 2.7e-09 | 1.3e-08 | 2.3e-09 | 6.000E-09 | 6.065E-09 | 2.300E-09 | 1.300E-08 |
| 23154 | 1158 | Q1158F | D | 2e-08 | 8e-09 | 3.1e-09 | 1.037E-08 | 8.695E-09 | 3.100E-09 | 2.000E-08 |
| 23155 | 1158 | Q1158P | D | 1.4e-08 | 2.3e-09 | 4.8e-09 | 7.033E-09 | 6.161E-09 | 2.300E-09 | 1.400E-08 |
| 23156 | 1158 | Q1158S | D | 8.3e-09 | 9.3e-09 | 2.6e-08 | 1.453E-08 | 9.943E-09 | 8.300E-09 | 2.600E-08 |
| 23157 | 1158 | Q1158T | D | 1.3e-08 | 2.5e-09 | 9.3e-09 | 8.267E-09 | 5.326E-09 | 2.500E-09 | 1.300E-08 |
| 23158 | 1158 | Q1158W | D | 6.1e-09 | 7e-09 | 1.5e-08 | 9.367E-09 | 4.899E-09 | 6.100E-09 | 1.500E-08 |
| 23159 | 1158 | Q1158Y | D | 5.4e-09 | 4e-09 | 3.9e-09 | 4.433E-09 | 8.386E-10 | 3.900E-09 | 5.400E-09 |
| 23160 | 1158 | Q1158V | D | 2.3e-08 | 2e-09 | 3.2e-09 | 9.400E-09 | 1.179E-08 | 2.000E-09 | 2.300E-08 |
| 23161 | 1159 | G1159A | D | 3.8e-09 | 1.6e-08 | 1.9e-08 | 1.293E-08 | 8.051E-09 | 3.800E-09 | 1.900E-08 |
| 23162 | 1159 | G1159R | D | 7.4e-09 | 1.3e-08 | 3.6e-09 | 8.000E-09 | 4.729E-09 | 3.600E-09 | 1.300E-08 |
| 23163 | 1159 | G1159N | D | 7.1e-09 | 7.6e-09 | 1.7e-08 | 1.057E-08 | 5.577E-09 | 7.100E-09 | 1.700E-08 |
| 23164 | 1159 | G1159D | D | 4.8e-09 | 5.6e-09 | 6.8e-09 | 5.733E-09 | 1.007E-09 | 4.800E-09 | 6.800E-09 |
| 23165 | 1159 | G1159C | D | 1.4e-08 | 1.3e-08 | 6.3e-09 | 1.110E-08 | 4.187E-09 | 6.300E-09 | 1.400E-08 |
| 23166 | 1159 | G1159E | D | 1.3e-08 | 2e-09 | 7.6e-09 | 7.533E-09 | 5.500E-09 | 2.000E-09 | 1.300E-08 |
| 23167 | 1159 | G1159Q | D | 1.9e-09 | 9e-09 | 1.1e-08 | 7.300E-09 | 4.782E-09 | 1.900E-09 | 1.100E-08 |
| 23168 | 1159 | G1159G | D | 8.8e-09 | 5e-09 | 8.9e-09 | 7.567E-09 | 2.223E-09 | 5.000E-09 | 8.900E-09 |
| 23169 | 1159 | G1159H | D | 8.6e-09 | 6.7e-09 | 2.6e-09 | 5.967E-09 | 3.066E-09 | 2.600E-09 | 8.600E-09 |
| 23170 | 1159 | G1159I | D | 8e-09 | 7e-09 | 3.7e-09 | 6.233E-09 | 2.250E-09 | 3.700E-09 | 8.000E-09 |
| 23171 | 1159 | G1159L | D | 1.1e-08 | 1.7e-09 | 1e-08 | 7.567E-09 | 5.105E-09 | 1.700E-09 | 1.100E-08 |
| 23172 | 1159 | G1159K | D | 9.2e-09 | 1e-08 | 5.2e-09 | 8.133E-09 | 2.572E-09 | 5.200E-09 | 1.000E-08 |
| 23173 | 1159 | G1159M | D | 1.5e-09 | 1.4e-08 | 3e-09 | 6.167E-09 | 6.825E-09 | 1.500E-09 | 1.400E-08 |
| 23174 | 1159 | G1159F | D | 2.4e-08 | 5.3e-09 | 2.1e-08 | 1.677E-08 | 1.004E-08 | 5.300E-09 | 2.400E-08 |
| 23175 | 1159 | G1159P | D | 1.4e-08 | 1.1e-08 | 1.9e-09 | 8.967E-09 | 6.301E-09 | 1.900E-09 | 1.400E-08 |
| 23176 | 1159 | G1159S | D | 6.4e-09 | 1.9e-08 | 5.7e-09 | 1.037E-08 | 7.485E-09 | 5.700E-09 | 1.900E-08 |
| 23177 | 1159 | G1159T | D | 9.1e-09 | 3.1e-09 | 1e-08 | 7.400E-09 | 3.751E-09 | 3.100E-09 | 1.000E-08 |
| 23178 | 1159 | G1159W | D | 4.9e-09 | 4.5e-09 | 9.6e-09 | 6.333E-09 | 2.836E-09 | 4.500E-09 | 9.600E-09 |
| 23179 | 1159 | G1159Y | D | 3.5e-09 | 8.9e-09 | 7e-09 | 6.467E-09 | 2.739E-09 | 3.500E-09 | 8.900E-09 |
| 23180 | 1159 | G1159V | D | 3.7e-09 | 3.5e-09 | 5.8e-09 | 4.333E-09 | 1.274E-09 | 3.500E-09 | 5.800E-09 |
| 23181 | 1160 | T1160A | D | 3.8e-09 | 1.2e-08 | 1.5e-08 | 1.027E-08 | 5.798E-09 | 3.800E-09 | 1.500E-08 |
| 23182 | 1160 | T1160R | D | 7e-09 | 1.9e-08 | 2.4e-09 | 9.467E-09 | 8.570E-09 | 2.400E-09 | 1.900E-08 |
| 23183 | 1160 | T1160N | D | 1.9e-08 | 5.6e-09 | 1.1e-08 | 1.187E-08 | 6.742E-09 | 5.600E-09 | 1.900E-08 |
| 23184 | 1160 | T1160D | D | 4.9e-09 | 4.5e-09 | 7.2e-09 | 5.533E-09 | 1.457E-09 | 4.500E-09 | 7.200E-09 |
| 23185 | 1160 | T1160C | D | 1.4e-08 | 5e-09 | 2.8e-09 | 7.267E-09 | 5.934E-09 | 2.800E-09 | 1.400E-08 |
| 23186 | 1160 | T1160E | D | 1.7e-08 | 2.4e-08 | 2.7e-08 | 2.267E-08 | 5.132E-09 | 1.700E-08 | 2.700E-08 |
| 23187 | 1160 | T1160Q | D | 3.5e-09 | 6e-09 | 2.6e-09 | 4.033E-09 | 1.762E-09 | 2.600E-09 | 6.000E-09 |
| 23188 | 1160 | T1160G | D | 5.7e-09 | 9.6e-09 | 1.1e-08 | 8.767E-09 | 2.747E-09 | 5.700E-09 | 1.100E-08 |
| 23189 | 1160 | T1160H | D | 9.1e-09 | 9.4e-09 | 2.8e-09 | 7.100E-09 | 3.727E-09 | 2.800E-09 | 9.400E-09 |
| 23190 | 1160 | T1160I | D | 2.1e-08 | 7.5e-09 | 2e-08 | 1.617E-08 | 7.522E-09 | 7.500E-09 | 2.100E-08 |
| 23191 | 1160 | T1160L | D | 2e-09 | 6.7e-09 | 6.3e-09 | 5.000E-09 | 2.606E-09 | 2.000E-09 | 6.700E-09 |
| 23192 | 1160 | T1160K | D | 1.1e-08 | 8.1e-09 | 5.8e-09 | 8.300E-09 | 2.606E-09 | 5.800E-09 | 1.100E-08 |
| 23193 | 1160 | T1160M | D | 7.6e-09 | 1.3e-08 | 3.7e-09 | 8.100E-09 | 4.670E-09 | 3.700E-09 | 1.300E-08 |
| 23194 | 1160 | T1160F | D | 5.1e-09 | 5.7e-09 | 4.7e-09 | 5.167E-09 | 5.033E-10 | 4.700E-09 | 5.700E-09 |
| 23195 | 1160 | T1160P | D | 6.9e-09 | 1.5e-08 | 1.5e-08 | 1.230E-08 | 4.677E-09 | 6.900E-09 | 1.500E-08 |
| 23196 | 1160 | T1160S | D | 7.2e-09 | 2.5e-09 | 1.3e-08 | 7.567E-09 | 5.260E-09 | 2.500E-09 | 1.300E-08 |
| 23197 | 1160 | T1160T | D | 1.3e-08 | 9.4e-09 | 1.4e-08 | 1.213E-08 | 2.419E-09 | 9.400E-09 | 1.400E-08 |
| 23198 | 1160 | T1160W | D | 6.6e-09 | 4.3e-09 | 1.8e-09 | 4.233E-09 | 2.401E-09 | 1.800E-09 | 6.600E-09 |
| 23199 | 1160 | T1160Y | D | 1.2e-08 | 1.7e-09 | 2.7e-09 | 5.467E-09 | 5.680E-09 | 1.700E-09 | 1.200E-08 |
| 23200 | 1160 | T1160V | D | 6.6e-09 | 7.9e-09 | 4.2e-09 | 6.233E-09 | 1.877E-09 | 4.200E-09 | 7.900E-09 |
| 23201 | 1161 | K1161A | D | 1.1e-08 | 5.8e-09 | 4.1e-09 | 6.967E-09 | 3.595E-09 | 4.100E-09 | 1.100E-08 |
| 23202 | 1161 | K1161R | D | 9.7e-09 | 9.2e-09 | 9.3e-09 | 9.400E-09 | 2.646E-10 | 9.200E-09 | 9.700E-09 |
| 23203 | 1161 | K1161N | D | 6e-09 | 1.2e-08 | 7.6e-09 | 8.533E-09 | 3.107E-09 | 6.000E-09 | 1.200E-08 |
| 23204 | 1161 | K1161D | D | 1.1e-08 | 7.4e-09 | 9.2e-09 | 9.200E-09 | 1.800E-09 | 7.400E-09 | 1.100E-08 |

| | | | | | | | | | | |
|---|---|---|---|---|---|---|---|---|---|---|
| 23205 | 1161 | K1161C | D | 6.2e-09 | 1.1e-08 | 3.5e-09 | 6.900E-09 | 3.799E-09 | 3.500E-09 | 1.100E-08 |
| 23206 | 1161 | K1161E | D | 5.2e-09 | 4.3e-09 | 3.2e-09 | 4.233E-09 | 1.002E-09 | 3.200E-09 | 5.200E-09 |
| 23207 | 1161 | K1161Q | D | 2.2e-08 | 1.4e-08 | 1.2e-08 | 1.600E-08 | 5.292E-09 | 1.200E-08 | 2.200E-08 |
| 23208 | 1161 | K1161G | D | 1.3e-08 | 6.7e-09 | 7.8e-09 | 9.167E-09 | 3.365E-09 | 6.700E-09 | 1.300E-08 |
| 23209 | 1161 | K1161H | D | 1e-08 | 1.5e-08 | 6.6e-09 | 1.053E-08 | 4.225E-09 | 6.600E-09 | 1.500E-08 |
| 23210 | 1161 | K1161I | D | 2.2e-09 | 6.1e-08 | 1.8e-08 | 2.707E-08 | 3.043E-08 | 2.200E-09 | 6.100E-08 |
| 23211 | 1161 | K1161L | D | 1.7e-08 | 8.4e-09 | 3e-09 | 9.467E-09 | 7.061E-09 | 3.000E-09 | 1.700E-08 |
| 23212 | 1161 | K1161K | D | 7.8e-09 | 1.6e-08 | 7.6e-09 | 1.047E-08 | 4.793E-09 | 7.600E-09 | 1.600E-08 |
| 23213 | 1161 | K1161M | D | 7.6e-09 | 1.2e-08 | 4.2e-09 | 7.933E-09 | 3.911E-09 | 4.200E-09 | 1.200E-08 |
| 23214 | 1161 | K1161F | D | 3.8e-09 | 2.7e-09 | 1.9e-08 | 8.500E-09 | 9.110E-09 | 2.700E-09 | 1.900E-08 |
| 23215 | 1161 | K1161P | D | 4.6e-09 | 1.5e-08 | 9.1e-09 | 9.567E-09 | 5.216E-09 | 4.600E-09 | 1.500E-08 |
| 23216 | 1161 | K1161S | D | 8.2e-09 | 4.8e-09 | 1.3e-08 | 8.667E-09 | 4.120E-09 | 4.800E-09 | 1.300E-08 |
| 23217 | 1161 | K1161T | D | 9.5e-09 | 5.6e-09 | 1.6e-08 | 1.037E-08 | 5.254E-09 | 5.600E-09 | 1.600E-08 |
| 23218 | 1161 | K1161W | D | 1.8e-08 | 1.3e-08 | 1.1e-08 | 1.400E-08 | 3.606E-09 | 1.100E-08 | 1.800E-08 |
| 23219 | 1161 | K1161Y | D | 2.3e-09 | 2.3e-09 | 5.2e-09 | 3.267E-09 | 1.674E-09 | 2.300E-09 | 5.200E-09 |
| 23220 | 1161 | K1161V | D | 3.6e-09 | 7.9e-09 | 1.8e-09 | 4.433E-09 | 3.134E-09 | 1.800E-09 | 7.900E-09 |
| 23221 | 1162 | V1162A | D | 1.6e-08 | 1.5e-08 | 3.4e-09 | 1.147E-08 | 7.004E-09 | 3.400E-09 | 1.600E-08 |
| 23222 | 1162 | V1162R | D | 9e-09 | 1.4e-08 | 4.4e-09 | 9.133E-09 | 4.801E-09 | 4.400E-09 | 1.400E-08 |
| 23223 | 1162 | V1162N | D | 1.4e-08 | 6e-09 | 3.5e-09 | 7.833E-09 | 5.485E-09 | 3.500E-09 | 1.400E-08 |
| 23224 | 1162 | V1162D | D | 1.2e-08 | 7.4e-09 | 7e-09 | 8.800E-09 | 2.778E-09 | 7.000E-09 | 1.200E-08 |
| 23225 | 1162 | V1162C | D | 5.4e-09 | 1.4e-08 | 4.4e-09 | 7.933E-09 | 5.278E-09 | 4.400E-09 | 1.400E-08 |
| 23226 | 1162 | V1162E | D | 1.4e-08 | 2.9e-08 | 4.8e-09 | 1.593E-08 | 1.222E-08 | 4.800E-09 | 2.900E-08 |
| 23227 | 1162 | V1162Q | D | 6.1e-09 | 6.2e-09 | 1.3e-08 | 8.433E-09 | 3.955E-09 | 6.100E-09 | 1.300E-08 |
| 23228 | 1162 | V1162G | D | 1.3e-08 | 1.1e-08 | 5.9e-09 | 9.967E-09 | 3.661E-09 | 5.900E-09 | 1.300E-08 |
| 23229 | 1162 | V1162H | D | 4.4e-09 | 2.1e-08 | 6.6e-09 | 1.067E-08 | 9.016E-09 | 4.400E-09 | 2.100E-08 |
| 23230 | 1162 | V1162I | D | 1e-08 | 8.9e-09 | 4.1e-09 | 7.667E-09 | 3.137E-09 | 4.100E-09 | 1.000E-08 |
| 23231 | 1162 | V1162L | D | 1.4e-08 | 3.4e-09 | 6.6e-09 | 8.000E-09 | 5.437E-09 | 3.400E-09 | 1.400E-08 |
| 23232 | 1162 | V1162K | D | 9.2e-09 | 4.1e-09 | 1.8e-08 | 1.043E-08 | 7.032E-09 | 4.100E-09 | 1.800E-08 |
| 23233 | 1162 | V1162M | D | 1e-08 | 2.1e-09 | 5.2e-09 | 5.767E-09 | 3.980E-09 | 2.100E-09 | 1.000E-08 |
| 23234 | 1162 | V1162F | D | 4.7e-09 | 2.1e-08 | 6.8e-09 | 1.083E-08 | 8.867E-09 | 4.700E-09 | 2.100E-08 |
| 23235 | 1162 | V1162P | D | 2.4e-09 | 8e-09 | 6.7e-09 | 5.700E-09 | 2.931E-09 | 2.400E-09 | 8.000E-09 |
| 23236 | 1162 | V1162S | D | 5.6e-09 | 1.1e-08 | 5.7e-09 | 7.433E-09 | 3.089E-09 | 5.600E-09 | 1.100E-08 |
| 23237 | 1162 | V1162T | D | 9e-09 | 9.7e-09 | 5e-09 | 7.900E-09 | 2.536E-09 | 5.000E-09 | 9.700E-09 |
| 23238 | 1162 | V1162W | D | 9e-09 | 3.9e-09 | 5e-09 | 5.967E-09 | 2.684E-09 | 3.900E-09 | 9.000E-09 |
| 23239 | 1162 | V1162Y | D | 6e-09 | 2.6e-09 | 4.3e-09 | 4.300E-09 | 1.700E-09 | 2.600E-09 | 6.000E-09 |
| 23240 | 1162 | V1162V | D | 1e-08 | 5.4e-09 | 8.8e-09 | 8.067E-09 | 2.386E-09 | 5.400E-09 | 1.000E-08 |
| 23241 | 1163 | E1163A | D | 3.1e-09 | 1e-08 | 7.2e-09 | 6.767E-09 | 3.470E-09 | 3.100E-09 | 1.000E-08 |
| 23242 | 1163 | E1163R | D | 1.2e-08 | 6.8e-09 | 8.4e-09 | 9.067E-09 | 2.663E-09 | 6.800E-09 | 1.200E-08 |
| 23243 | 1163 | E1163N | D | 5e-09 | 1.9e-08 | 2.7e-08 | 1.700E-08 | 1.114E-08 | 5.000E-09 | 2.700E-08 |
| 23244 | 1163 | E1163D | D | 4.7e-09 | 5.9e-09 | 1e-08 | 6.867E-09 | 2.779E-09 | 4.700E-09 | 1.000E-08 |
| 23245 | 1163 | E1163C | D | 3.3e-09 | 1.3e-08 | 3.6e-09 | 6.633E-09 | 5.516E-09 | 3.300E-09 | 1.300E-08 |
| 23246 | 1163 | E1163E | D | 4.9e-09 | 5.2e-09 | 1.2e-08 | 7.367E-09 | 4.015E-09 | 4.900E-09 | 1.200E-08 |
| 23247 | 1163 | E1163Q | D | 9.8e-09 | 8.8e-09 | 7.9e-09 | 8.833E-09 | 9.504E-10 | 7.900E-09 | 9.800E-09 |
| 23248 | 1163 | E1163G | D | 2.5e-08 | 1.2e-08 | 3e-09 | 1.333E-08 | 1.106E-08 | 3.000E-09 | 2.500E-08 |
| 23249 | 1163 | E1163H | D | 1.2e-08 | 6.1e-09 | 4.7e-09 | 7.600E-09 | 3.874E-09 | 4.700E-09 | 1.200E-08 |
| 23250 | 1163 | E1163I | D | 5.3e-09 | 2.4e-09 | 6.4e-09 | 4.700E-09 | 2.066E-09 | 2.400E-09 | 6.400E-09 |
| 23251 | 1163 | E1163L | D | 8.8e-09 | 3.9e-09 | 7.8e-09 | 6.833E-09 | 2.589E-09 | 3.900E-09 | 8.800E-09 |
| 23252 | 1163 | E1163K | D | 3.1e-08 | 1.3e-08 | 1.9e-08 | 2.100E-08 | 9.165E-09 | 1.300E-08 | 3.100E-08 |
| 23253 | 1163 | E1163M | D | 8.8e-09 | 7.9e-09 | 1.1e-08 | 9.233E-09 | 1.595E-09 | 7.900E-09 | 1.100E-08 |
| 23254 | 1163 | E1163F | D | 3.1e-09 | 5.1e-09 | 8.8e-09 | 5.667E-09 | 2.892E-09 | 3.100E-09 | 8.800E-09 |
| 23255 | 1163 | E1163P | D | 5e-09 | 8.1e-09 | 8.7e-09 | 7.267E-09 | 1.986E-09 | 5.000E-09 | 8.700E-09 |
| 23256 | 1163 | E1163S | D | 5.2e-09 | 4.2e-09 | 3.5e-09 | 4.300E-09 | 8.544E-10 | 3.500E-09 | 5.200E-09 |
| 23257 | 1163 | E1163T | D | 4.5e-09 | 5.6e-09 | 5.5e-09 | 5.200E-09 | 6.083E-10 | 4.500E-09 | 5.600E-09 |
| 23258 | 1163 | E1163W | D | 1.4e-08 | 4.6e-09 | 4.1e-09 | 7.567E-09 | 5.577E-09 | 4.100E-09 | 1.400E-08 |
| 23259 | 1163 | E1163Y | D | 4.5e-09 | 6e-09 | 4.6e-09 | 5.033E-09 | 8.386E-10 | 4.500E-09 | 6.000E-09 |
| 23260 | 1163 | E1163V | D | 1.8e-09 | 8.9e-09 | 8.3e-09 | 6.333E-09 | 3.937E-09 | 1.800E-09 | 8.900E-09 |
| 23261 | 1164 | I1164A | D | 7.5e-09 | 9.1e-09 | 8.4e-09 | 8.333E-09 | 8.021E-10 | 7.500E-09 | 9.100E-09 |
| 23262 | 1164 | I1164R | D | 5.2e-09 | 6.1e-09 | 2.8e-09 | 4.700E-09 | 1.706E-09 | 2.800E-09 | 6.100E-09 |
| 23263 | 1164 | I1164N | D | 1.3e-08 | 3.3e-08 | 9.2e-09 | 1.840E-08 | 1.279E-08 | 9.200E-09 | 3.300E-08 |
| 23264 | 1164 | I1164D | D | 4.8e-09 | 6.5e-09 | 1e-08 | 7.100E-09 | 2.651E-09 | 4.800E-09 | 1.000E-08 |
| 23265 | 1164 | I1164C | D | 5.1e-09 | 6.7e-09 | 6e-09 | 5.933E-09 | 8.021E-10 | 5.100E-09 | 6.700E-09 |
| 23266 | 1164 | I1164E | D | 3.8e-09 | 7.8e-09 | 1.1e-08 | 7.533E-09 | 3.607E-09 | 3.800E-09 | 1.100E-08 |
| 23267 | 1164 | I1164Q | D | 2.8e-09 | 3.7e-09 | 3.3e-09 | 3.267E-09 | 4.509E-10 | 2.800E-09 | 3.700E-09 |
| 23268 | 1164 | I1164G | D | 2.7e-09 | 1.9e-08 | 6.4e-09 | 9.367E-09 | 8.545E-09 | 2.700E-09 | 1.900E-08 |
| 23269 | 1164 | I1164H | D | 5.9e-09 | 1.8e-09 | 1.1e-08 | 6.233E-09 | 4.609E-09 | 1.800E-09 | 1.100E-08 |
| 23270 | 1164 | I1164I | D | 7e-09 | 7.3e-09 | 5.2e-09 | 6.500E-09 | 1.136E-09 | 5.200E-09 | 7.300E-09 |
| 23271 | 1164 | I1164L | D | 9.4e-09 | 3.7e-09 | 1.7e-08 | 1.003E-08 | 6.673E-09 | 3.700E-09 | 1.700E-08 |
| 23272 | 1164 | I1164K | D | 9.9e-09 | 8.5e-09 | 5.9e-09 | 8.100E-09 | 2.030E-09 | 5.900E-09 | 9.900E-09 |
| 23273 | 1164 | I1164M | D | 5e-09 | 9.5e-09 | 5.1e-09 | 6.533E-09 | 2.570E-09 | 5.000E-09 | 9.500E-09 |
| 23274 | 1164 | I1164F | D | 2.8e-09 | 6.6e-09 | 3.7e-09 | 4.367E-09 | 1.986E-09 | 2.800E-09 | 6.600E-09 |
| 23275 | 1164 | I1164P | D | 1.1e-08 | 2.1e-09 | 5.2e-09 | 6.100E-09 | 4.518E-09 | 2.100E-09 | 1.100E-08 |
| 23276 | 1164 | I1164S | D | 2.7e-08 | 2.3e-09 | 9.4e-09 | 1.290E-08 | 1.272E-08 | 2.300E-09 | 2.700E-08 |
| 23277 | 1164 | I1164T | D | 5.5e-09 | 1.5e-08 | 6.8e-09 | 9.100E-09 | 5.151E-09 | 5.500E-09 | 1.500E-08 |
| 23278 | 1164 | I1164W | D | 5.3e-09 | 5.6e-09 | 1.7e-09 | 4.200E-09 | 2.170E-09 | 1.700E-09 | 5.600E-09 |
| 23279 | 1164 | I1164Y | D | 6.3e-09 | 5.4e-09 | 5.1e-09 | 5.600E-09 | 6.245E-10 | 5.100E-09 | 6.300E-09 |
| 23280 | 1164 | I1164V | D | 2.1e-08 | 7.4e-09 | 2.2e-09 | 1.020E-08 | 9.708E-09 | 2.200E-09 | 2.100E-08 |
| 23281 | 1165 | K1165A | D | 3.5e-09 | 8.4e-09 | 1.2e-08 | 7.967E-09 | 4.267E-09 | 3.500E-09 | 1.200E-08 |
| 23282 | 1165 | K1165R | D | 1.9e-08 | 7.5e-09 | 1.8e-08 | 1.483E-08 | 6.371E-09 | 7.500E-09 | 1.900E-08 |
| 23283 | 1165 | K1165N | D | 4.8e-09 | 7.7e-09 | 1.1e-08 | 7.833E-09 | 3.102E-09 | 4.800E-09 | 1.100E-08 |
| 23284 | 1165 | K1165D | D | 2.8e-09 | 2.4e-08 | 6e-09 | 1.093E-08 | 1.143E-08 | 2.800E-09 | 2.400E-08 |
| 23285 | 1165 | K1165C | D | 6.8e-09 | 7.8e-09 | 4.8e-09 | 6.467E-09 | 1.528E-09 | 4.800E-09 | 7.800E-09 |
| 23286 | 1165 | K1165E | D | 3.7e-08 | 9e-09 | 2.9e-09 | 1.630E-08 | 1.818E-08 | 2.900E-09 | 3.700E-08 |
| 23287 | 1165 | K1165Q | D | 1.1e-08 | 1e-08 | 7.4e-09 | 9.467E-09 | 1.858E-09 | 7.400E-09 | 1.100E-08 |
| 23288 | 1165 | K1165G | D | 1.5e-08 | 1.8e-08 | 5.4e-09 | 1.280E-08 | 6.582E-09 | 5.400E-09 | 1.800E-08 |
| 23289 | 1165 | K1165H | D | 9.7e-09 | 1.1e-08 | 1.7e-08 | 1.257E-08 | 3.894E-09 | 9.700E-09 | 1.700E-08 |

| | | | | | | | | | | |
|---|---|---|---|---|---|---|---|---|---|---|
| 23290 | 1165 | K1165I | D | 3e-09 | 2.2e-09 | 8.9e-09 | 4.700E-09 | 3.659E-09 | 2.200E-09 | 8.900E-09 |
| 23291 | 1165 | K1165L | D | 7.1e-09 | 6.8e-09 | 1.8e-08 | 1.063E-08 | 6.381E-09 | 6.800E-09 | 1.800E-08 |
| 23292 | 1165 | K1165K | D | 4.3e-09 | 6.5e-09 | 4.9e-09 | 5.233E-09 | 1.137E-09 | 4.300E-09 | 6.500E-09 |
| 23293 | 1165 | K1165M | D | 7.9e-09 | 2.2e-08 | 4.6e-09 | 1.150E-08 | 9.242E-09 | 4.600E-09 | 2.200E-08 |
| 23294 | 1165 | K1165F | D | 1.2e-08 | 8e-09 | 1.7e-08 | 1.233E-08 | 4.509E-09 | 8.000E-09 | 1.700E-08 |
| 23295 | 1165 | K1165P | D | 2.4e-09 | 6.9e-09 | 1.3e-08 | 7.433E-09 | 5.320E-09 | 2.400E-09 | 1.300E-08 |
| 23296 | 1165 | K1165S | D | 6e-09 | 5.9e-09 | 1.4e-08 | 8.633E-09 | 4.648E-09 | 5.900E-09 | 1.400E-08 |
| 23297 | 1165 | K1165T | D | 2.9e-09 | 7.5e-09 | 2.3e-08 | 1.113E-08 | 1.053E-08 | 2.900E-09 | 2.300E-08 |
| 23298 | 1165 | K1165W | D | 3.7e-09 | 1.3e-08 | 4.5e-09 | 7.067E-09 | 5.154E-09 | 3.700E-09 | 1.300E-08 |
| 23299 | 1165 | K1165Y | D | 9.6e-09 | 9.1e-09 | 8.6e-09 | 9.100E-09 | 5.000E-10 | 8.600E-09 | 9.600E-09 |
| 23300 | 1165 | K1165V | D | 3.8e-09 | 6e-09 | 8.8e-09 | 6.200E-09 | 2.506E-09 | 3.800E-09 | 8.800E-09 |
| 23301 | 1166 | R1166A | D | 1.3e-08 | 6.6e-09 | 2.1e-09 | 7.233E-09 | 5.478E-09 | 2.100E-09 | 1.300E-08 |
| 23302 | 1166 | R1166R | D | 1.3e-08 | 2.8e-08 | 5.9e-09 | 1.563E-08 | 1.128E-08 | 5.900E-09 | 2.800E-08 |
| 23303 | 1166 | R1166N | D | 5.8e-09 | 8.6e-09 | 1.5e-08 | 9.800E-09 | 4.716E-09 | 5.800E-09 | 1.500E-08 |
| 23304 | 1166 | R1166D | D | 4.8e-09 | 8.5e-09 | 3e-09 | 5.433E-09 | 2.804E-09 | 3.000E-09 | 8.500E-09 |
| 23305 | 1166 | R1166C | D | 1.4e-08 | 4.7e-09 | 1.2e-08 | 1.023E-08 | 4.895E-09 | 4.700E-09 | 1.400E-08 |
| 23306 | 1166 | R1166E | D | 8.3e-09 | 6.2e-09 | 5.2e-09 | 6.567E-09 | 1.582E-09 | 5.200E-09 | 8.300E-09 |
| 23307 | 1166 | R1166Q | D | 7.8e-09 | 7.2e-09 | 5.2e-09 | 6.733E-09 | 1.361E-09 | 5.200E-09 | 7.800E-09 |
| 23308 | 1166 | R1166G | D | 1.1e-08 | 4.5e-09 | 5.6e-09 | 7.033E-09 | 3.479E-09 | 4.500E-09 | 1.100E-08 |
| 23309 | 1166 | R1166H | D | 9.6e-09 | 8.7e-09 | 8.4e-09 | 8.900E-09 | 6.245E-10 | 8.400E-09 | 9.600E-09 |
| 23310 | 1166 | R1166I | D | 5.5e-09 | 3.8e-09 | 9.7e-09 | 6.333E-09 | 3.037E-09 | 3.800E-09 | 9.700E-09 |
| 23311 | 1166 | R1166L | D | 1.2e-08 | 2.4e-09 | 2.2e-09 | 5.533E-09 | 5.601E-09 | 2.200E-09 | 1.200E-08 |
| 23312 | 1166 | R1166K | D | 4.2e-09 | 1.5e-08 | 1.1e-08 | 1.007E-08 | 5.460E-09 | 4.200E-09 | 1.500E-08 |
| 23313 | 1166 | R1166M | D | 4.6e-09 | 5.2e-09 | 9.5e-09 | 6.433E-09 | 2.673E-09 | 4.600E-09 | 9.500E-09 |
| 23314 | 1166 | R1166F | D | 3.1e-09 | 5.8e-09 | 1e-08 | 6.300E-09 | 3.477E-09 | 3.100E-09 | 1.000E-08 |
| 23315 | 1166 | R1166P | D | 9.5e-09 | 8.5e-09 | 1.5e-08 | 1.100E-08 | 3.500E-09 | 8.500E-09 | 1.500E-08 |
| 23316 | 1166 | R1166S | D | 7.5e-09 | 6.7e-09 | 7e-09 | 7.067E-09 | 4.041E-10 | 6.700E-09 | 7.500E-09 |
| 23317 | 1166 | R1166T | D | 7.1e-09 | 1.5e-08 | 5.1e-09 | 9.067E-09 | 5.235E-09 | 5.100E-09 | 1.500E-08 |
| 23318 | 1166 | R1166W | D | 1.2e-08 | 8.6e-09 | 5.1e-09 | 8.567E-09 | 3.450E-09 | 5.100E-09 | 1.200E-08 |
| 23319 | 1166 | R1166Y | D | 7.5e-09 | 2.9e-09 | 1.2e-08 | 7.467E-09 | 4.550E-09 | 2.900E-09 | 1.200E-08 |
| 23320 | 1166 | R1166V | D | 1.6e-08 | 1.9e-09 | 9.1e-09 | 9.000E-09 | 7.051E-09 | 1.900E-09 | 1.600E-08 |
| 23321 | 1167 | T1167A | D | 1.2e-08 | 3.9e-09 | 5.2e-09 | 7.033E-09 | 4.350E-09 | 3.900E-09 | 1.200E-08 |
| 23322 | 1167 | T1167R | D | 6.1e-09 | 4.6e-09 | 1.2e-08 | 7.567E-09 | 3.912E-09 | 4.600E-09 | 1.200E-08 |
| 23323 | 1167 | T1167N | D | 8.6e-09 | 7.4e-09 | 7.7e-09 | 7.900E-09 | 6.245E-10 | 7.400E-09 | 8.600E-09 |
| 23324 | 1167 | T1167D | D | 9.6e-09 | 9.1e-09 | 8.8e-09 | 9.167E-09 | 4.041E-10 | 8.800E-09 | 9.600E-09 |
| 23325 | 1167 | T1167C | D | 1.4e-08 | 1e-08 | 5.9e-09 | 9.967E-09 | 4.050E-09 | 5.900E-09 | 1.400E-08 |
| 23326 | 1167 | T1167E | D | 7.7e-09 | 7.5e-09 | 8.8e-10 | 5.360E-09 | 3.881E-09 | 8.800E-10 | 7.700E-09 |
| 23327 | 1167 | T1167Q | D | 1.8e-08 | 8.4e-09 | 3.8e-09 | 1.007E-08 | 7.245E-09 | 3.800E-09 | 1.800E-08 |
| 23328 | 1167 | T1167G | D | 9.4e-09 | 1.2e-08 | 5.6e-09 | 9.000E-09 | 3.219E-09 | 5.600E-09 | 1.200E-08 |
| 23329 | 1167 | T1167H | D | 6.5e-09 | 4.7e-09 | 6e-09 | 5.733E-09 | 9.292E-10 | 4.700E-09 | 6.500E-09 |
| 23330 | 1167 | T1167I | D | 9.2e-09 | 6.7e-09 | 2.5e-08 | 1.363E-08 | 9.923E-09 | 6.700E-09 | 2.500E-08 |
| 23331 | 1167 | T1167L | D | 2.6e-09 | 7.8e-09 | 5.8e-09 | 5.400E-09 | 2.623E-09 | 2.600E-09 | 7.800E-09 |
| 23332 | 1167 | T1167K | D | 2e-08 | 8.7e-09 | 1.2e-08 | 1.357E-08 | 5.811E-09 | 8.700E-09 | 2.000E-08 |
| 23333 | 1167 | T1167M | D | 6.9e-09 | 5.4e-09 | 2.4e-08 | 1.210E-08 | 1.033E-08 | 5.400E-09 | 2.400E-08 |
| 23334 | 1167 | T1167F | D | 1e-08 | 7.5e-09 | 8.6e-09 | 8.700E-09 | 1.253E-09 | 7.500E-09 | 1.000E-08 |
| 23335 | 1167 | T1167P | D | 5.1e-09 | 7e-09 | 1.2e-08 | 8.033E-09 | 3.564E-09 | 5.100E-09 | 1.200E-08 |
| 23336 | 1167 | T1167S | D | 1.5e-08 | 7.9e-09 | 1.4e-08 | 1.230E-08 | 3.843E-09 | 7.900E-09 | 1.500E-08 |
| 23337 | 1167 | T1167T | D | 1.2e-08 | 6.7e-09 | 6.3e-09 | 8.333E-09 | 3.182E-09 | 6.300E-09 | 1.200E-08 |
| 23338 | 1167 | T1167W | D | 3e-09 | 2.5e-09 | 9.6e-09 | 5.033E-09 | 3.963E-09 | 2.500E-09 | 9.600E-09 |
| 23339 | 1167 | T1167Y | D | 4.4e-09 | 1.4e-08 | 6.2e-09 | 8.200E-09 | 5.103E-09 | 4.400E-09 | 1.400E-08 |
| 23340 | 1167 | T1167V | D | 1.6e-08 | 4.9e-09 | 1.5e-08 | 1.197E-08 | 6.140E-09 | 4.900E-09 | 1.600E-08 |
| 23341 | 1168 | V1168A | D | 2.6e-09 | 7.9e-09 | 7.8e-09 | 6.100E-09 | 3.032E-09 | 2.600E-09 | 7.900E-09 |
| 23342 | 1168 | V1168R | D | 1.3e-08 | 1.1e-08 | 1.1e-08 | 1.167E-08 | 1.155E-09 | 1.100E-08 | 1.300E-08 |
| 23343 | 1168 | V1168N | D | 4.9e-09 | 1e-08 | 2.4e-08 | 1.297E-08 | 9.890E-09 | 4.900E-09 | 2.400E-08 |
| 23344 | 1168 | V1168D | D | 7.9e-09 | 6.7e-09 | 9.7e-09 | 8.100E-09 | 1.510E-09 | 6.700E-09 | 9.700E-09 |
| 23345 | 1168 | V1168C | D | 9.5e-09 | 5e-09 | 1.6e-08 | 1.017E-08 | 5.530E-09 | 5.000E-09 | 1.600E-08 |
| 23346 | 1168 | V1168E | D | 6.4e-09 | 7.4e-09 | 3.5e-09 | 5.767E-09 | 2.026E-09 | 3.500E-09 | 7.400E-09 |
| 23347 | 1168 | V1168Q | D | 1.7e-08 | 3.3e-09 | 6.6e-09 | 8.967E-09 | 7.150E-09 | 3.300E-09 | 1.700E-08 |
| 23348 | 1168 | V1168G | D | 4.4e-09 | 1.5e-08 | 2.6e-09 | 7.333E-09 | 6.700E-09 | 2.600E-09 | 1.500E-08 |
| 23349 | 1168 | V1168H | D | 8e-09 | 3.2e-09 | 2.2e-08 | 1.107E-08 | 9.768E-09 | 3.200E-09 | 2.200E-08 |
| 23350 | 1168 | V1168I | D | 1.2e-08 | 2.6e-09 | 1e-08 | 8.200E-09 | 4.952E-09 | 2.600E-09 | 1.200E-08 |
| 23351 | 1168 | V1168L | D | 6.7e-09 | 8e-09 | 4e-09 | 6.233E-09 | 2.040E-09 | 4.000E-09 | 8.000E-09 |
| 23352 | 1168 | V1168K | D | 3.5e-09 | 1.6e-08 | 1.3e-08 | 1.083E-08 | 6.526E-09 | 3.500E-09 | 1.600E-08 |
| 23353 | 1168 | V1168M | D | 5.7e-09 | 1e-08 | 1.5e-08 | 1.023E-08 | 4.654E-09 | 5.700E-09 | 1.500E-08 |
| 23354 | 1168 | V1168F | D | 1.3e-08 | 1.2e-08 | 4.2e-09 | 9.733E-09 | 4.818E-09 | 4.200E-09 | 1.300E-08 |
| 23355 | 1168 | V1168P | D | 1.1e-08 | 5.4e-09 | 7.7e-09 | 8.033E-09 | 2.815E-09 | 5.400E-09 | 1.100E-08 |
| 23356 | 1168 | V1168S | D | 3.4e-09 | 1.7e-08 | 6.6e-09 | 9.000E-09 | 7.111E-09 | 3.400E-09 | 1.700E-08 |
| 23357 | 1168 | V1168T | D | 1.5e-09 | 7.8e-09 | 8.3e-09 | 5.867E-09 | 3.790E-09 | 1.500E-09 | 8.300E-09 |
| 23358 | 1168 | V1168W | D | 1.1e-08 | 7.1e-09 | 5.2e-09 | 7.767E-09 | 2.957E-09 | 5.200E-09 | 1.100E-08 |
| 23359 | 1168 | V1168Y | D | 3e-08 | 1.2e-08 | 6e-09 | 1.600E-08 | 1.249E-08 | 6.000E-09 | 3.000E-08 |
| 23360 | 1168 | V1168V | D | 4.5e-09 | 7e-09 | 1.1e-08 | 7.500E-09 | 3.279E-09 | 4.500E-09 | 1.100E-08 |
| 23361 | 1169 | A1169A | D | 1e-08 | 7e-09 | 5.2e-09 | 7.400E-09 | 2.425E-09 | 5.200E-09 | 1.000E-08 |
| 23362 | 1169 | A1169R | D | 9.7e-09 | 7.6e-09 | 2.3e-08 | 1.343E-08 | 8.351E-09 | 7.600E-09 | 2.300E-08 |
| 23363 | 1169 | A1169N | D | 1.6e-09 | 1.1e-08 | 9.6e-09 | 7.400E-09 | 5.071E-09 | 1.600E-09 | 1.100E-08 |
| 23364 | 1169 | A1169D | D | 4.7e-09 | 9.5e-09 | 3.1e-09 | 5.767E-09 | 3.331E-09 | 3.100E-09 | 9.500E-09 |
| 23365 | 1169 | A1169C | D | 9.9e-09 | 5.9e-09 | 2.3e-08 | 1.293E-08 | 8.944E-09 | 5.900E-09 | 2.300E-08 |
| 23366 | 1169 | A1169E | D | 1.1e-08 | 2.2e-09 | 4.5e-09 | 5.900E-09 | 4.564E-09 | 2.200E-09 | 1.100E-08 |
| 23367 | 1169 | A1169Q | D | 1.2e-08 | 9.9e-09 | 1.2e-08 | 1.130E-08 | 1.212E-09 | 9.900E-09 | 1.200E-08 |
| 23368 | 1169 | A1169G | D | 5.1e-09 | 3.3e-09 | 1.1e-08 | 6.467E-09 | 4.028E-09 | 3.300E-09 | 1.100E-08 |
| 23369 | 1169 | A1169H | D | 9.2e-09 | 8.9e-09 | 9.9e-09 | 9.333E-09 | 5.132E-10 | 8.900E-09 | 9.900E-09 |
| 23370 | 1169 | A1169I | D | 6.6e-09 | 2.3e-09 | 3.8e-09 | 4.233E-09 | 2.183E-09 | 2.300E-09 | 6.600E-09 |
| 23371 | 1169 | A1169L | D | 2.1e-08 | 4.1e-09 | 4.3e-09 | 9.800E-09 | 9.700E-09 | 4.100E-09 | 2.100E-08 |
| 23372 | 1169 | A1169K | D | 4.3e-09 | 5.2e-09 | 6.6e-09 | 5.367E-09 | 1.159E-09 | 4.300E-09 | 6.600E-09 |
| 23373 | 1169 | A1169M | D | 3.7e-09 | 2.8e-09 | 7e-09 | 4.500E-09 | 2.211E-09 | 2.800E-09 | 7.000E-09 |
| 23374 | 1169 | A1169F | D | 4.6e-09 | 2.2e-09 | 2.7e-09 | 3.167E-09 | 1.266E-09 | 2.200E-09 | 4.600E-09 |

| | | | | | | | | | | |
|---|---|---|---|---|---|---|---|---|---|---|
| 23375 | 1169 | A1169P | D | 1.6e-08 | 9.7e-09 | 4.9e-09 | 1.020E-08 | 5.567E-09 | 4.900E-09 | 1.600E-08 |
| 23376 | 1169 | A1169S | D | 5.8e-09 | 1.2e-08 | 6.2e-09 | 8.000E-09 | 3.470E-09 | 5.800E-09 | 1.200E-08 |
| 23377 | 1169 | A1169T | D | 1.4e-08 | 2.8e-09 | 6.2e-09 | 7.667E-09 | 5.742E-09 | 2.800E-09 | 1.400E-08 |
| 23378 | 1169 | A1169W | D | 3.7e-09 | 1.3e-08 | 8.9e-09 | 8.533E-09 | 4.661E-09 | 3.700E-09 | 1.300E-08 |
| 23379 | 1169 | A1169Y | D | 4.3e-09 | 5.2e-09 | 6.8e-09 | 5.433E-09 | 1.266E-09 | 4.300E-09 | 6.800E-09 |
| 23380 | 1169 | A1169V | D | 4.5e-09 | 1.8e-08 | 9.5e-09 | 1.067E-08 | 6.825E-09 | 4.500E-09 | 1.800E-08 |
| 23381 | 1170 | A1170A | D | 6.7e-09 | 9.5e-09 | 9.9e-09 | 8.700E-09 | 1.744E-09 | 6.700E-09 | 9.900E-09 |
| 23382 | 1170 | A1170R | D | 1.4e-08 | 4.7e-09 | 1.2e-08 | 1.023E-08 | 4.895E-09 | 4.700E-09 | 1.400E-08 |
| 23383 | 1170 | A1170N | D | 1.8e-08 | 8.6e-09 | 1.3e-08 | 1.320E-08 | 4.703E-09 | 8.600E-09 | 1.800E-08 |
| 23384 | 1170 | A1170D | D | 5.1e-09 | 2.4e-08 | 1.1e-08 | 1.337E-08 | 9.670E-09 | 5.100E-09 | 2.400E-08 |
| 23385 | 1170 | A1170C | D | 7.2e-09 | 1.5e-08 | 6.2e-09 | 9.467E-09 | 4.818E-09 | 6.200E-09 | 1.500E-08 |
| 23386 | 1170 | A1170E | D | 1.6e-09 | 1.3e-08 | 2.5e-09 | 5.700E-09 | 6.338E-09 | 1.600E-09 | 1.300E-08 |
| 23387 | 1170 | A1170Q | D | 2e-08 | 6.7e-09 | 8.5e-09 | 1.173E-08 | 7.215E-09 | 6.700E-09 | 2.000E-08 |
| 23388 | 1170 | A1170G | D | 4.3e-09 | 9.2e-09 | 2e-08 | 1.117E-08 | 8.033E-09 | 4.300E-09 | 2.000E-08 |
| 23389 | 1170 | A1170H | D | 3.3e-09 | 8.7e-09 | 2.5e-09 | 4.833E-09 | 3.372E-09 | 2.500E-09 | 8.700E-09 |
| 23390 | 1170 | A1170I | D | 5.8e-09 | 9.9e-09 | 4e-09 | 6.567E-09 | 3.024E-09 | 4.000E-09 | 9.900E-09 |
| 23391 | 1170 | A1170L | D | 2.5e-09 | 5.4e-09 | 1.1e-08 | 6.300E-09 | 4.321E-09 | 2.500E-09 | 1.100E-08 |
| 23392 | 1170 | A1170K | D | 1.7e-08 | 1.6e-09 | 8.9e-09 | 9.167E-09 | 7.703E-09 | 1.600E-09 | 1.700E-08 |
| 23393 | 1170 | A1170M | D | 3.9e-09 | 1.8e-08 | 3.3e-09 | 8.400E-09 | 8.319E-09 | 3.300E-09 | 1.800E-08 |
| 23394 | 1170 | A1170F | D | 1.3e-08 | 5e-09 | 7.5e-09 | 8.500E-09 | 4.093E-09 | 5.000E-09 | 1.300E-08 |
| 23395 | 1170 | A1170P | D | 7.9e-09 | 5.2e-09 | 1e-08 | 7.700E-09 | 2.406E-09 | 5.200E-09 | 1.000E-08 |
| 23396 | 1170 | A1170S | D | 6.4e-09 | 2e-08 | 6e-09 | 1.080E-08 | 7.970E-09 | 6.000E-09 | 2.000E-08 |
| 23397 | 1170 | A1170T | D | 2.1e-09 | 9.1e-09 | 2.5e-08 | 1.207E-08 | 1.173E-08 | 2.100E-09 | 2.500E-08 |
| 23398 | 1170 | A1170W | D | 2e-08 | 1.2e-08 | 2e-08 | 1.733E-08 | 4.619E-09 | 1.200E-08 | 2.000E-08 |
| 23399 | 1170 | A1170Y | D | 3.8e-09 | 9.7e-09 | 5.4e-09 | 6.300E-09 | 3.051E-09 | 3.800E-09 | 9.700E-09 |
| 23400 | 1170 | A1170V | D | 8.9e-09 | 6.2e-09 | 6.6e-09 | 7.233E-09 | 1.457E-09 | 6.200E-09 | 8.900E-09 |
| 23401 | 1171 | P1171A | D | 4.8e-09 | 8.6e-09 | 2.1e-09 | 5.167E-09 | 3.265E-09 | 2.100E-09 | 8.600E-09 |
| 23402 | 1171 | P1171R | D | 1.2e-09 | 9.4e-09 | 6.8e-09 | 5.800E-09 | 4.190E-09 | 1.200E-09 | 9.400E-09 |
| 23403 | 1171 | P1171N | D | 5.6e-09 | 4.5e-09 | 1.1e-08 | 7.033E-09 | 3.479E-09 | 4.500E-09 | 1.100E-08 |
| 23404 | 1171 | P1171D | D | 1.1e-08 | 2.5e-09 | 6.5e-09 | 6.667E-09 | 4.252E-09 | 2.500E-09 | 1.100E-08 |
| 23405 | 1171 | P1171C | D | 5.5e-09 | 2.1e-08 | 3.4e-09 | 9.967E-09 | 9.613E-09 | 3.400E-09 | 2.100E-08 |
| 23406 | 1171 | P1171E | D | 3e-09 | 1.2e-08 | 4.7e-09 | 6.567E-09 | 4.782E-09 | 3.000E-09 | 1.200E-08 |
| 23407 | 1171 | P1171Q | D | 1.3e-08 | 6.3e-09 | 4.3e-09 | 7.867E-09 | 4.557E-09 | 4.300E-09 | 1.300E-08 |
| 23408 | 1171 | P1171G | D | 6.9e-09 | 8e-09 | 7.8e-09 | 7.567E-09 | 5.859E-10 | 6.900E-09 | 8.000E-09 |
| 23409 | 1171 | P1171H | D | 3.8e-09 | 3.8e-09 | 6.9e-09 | 4.833E-09 | 1.790E-09 | 3.800E-09 | 6.900E-09 |
| 23410 | 1171 | P1171I | D | 8.9e-09 | 6.2e-09 | 7.8e-09 | 7.633E-09 | 1.358E-09 | 6.200E-09 | 8.900E-09 |
| 23411 | 1171 | P1171L | D | 1e-08 | 2.8e-09 | 3.5e-09 | 5.433E-09 | 3.970E-09 | 2.800E-09 | 1.000E-08 |
| 23412 | 1171 | P1171K | D | 6.8e-09 | 1.4e-08 | 9.6e-09 | 1.013E-08 | 3.630E-09 | 6.800E-09 | 1.400E-08 |
| 23413 | 1171 | P1171M | D | 8.8e-09 | 2.8e-09 | 5.2e-09 | 5.600E-09 | 3.020E-09 | 2.800E-09 | 8.800E-09 |
| 23414 | 1171 | P1171F | D | 5.1e-09 | 3e-09 | 1.8e-08 | 8.700E-09 | 8.122E-09 | 3.000E-09 | 1.800E-08 |
| 23415 | 1171 | P1171P | D | 1.4e-08 | 1.2e-08 | 1.1e-08 | 1.233E-08 | 1.528E-09 | 1.100E-08 | 1.400E-08 |
| 23416 | 1171 | P1171S | D | 1.1e-09 | 4.4e-09 | 7.6e-09 | 4.367E-09 | 3.250E-09 | 1.100E-09 | 7.600E-09 |
| 23417 | 1171 | P1171T | D | 1e-08 | 4e-09 | 1.1e-08 | 8.333E-09 | 3.786E-09 | 4.000E-09 | 1.100E-08 |
| 23418 | 1171 | P1171W | D | 1e-08 | 9.2e-09 | 7.8e-09 | 9.000E-09 | 1.114E-09 | 7.800E-09 | 1.000E-08 |
| 23419 | 1171 | P1171Y | D | 6.3e-09 | 1.8e-08 | 4.4e-09 | 9.567E-09 | 7.365E-09 | 4.400E-09 | 1.800E-08 |
| 23420 | 1171 | P1171V | D | 8.5e-09 | 2.6e-09 | 3.3e-09 | 4.800E-09 | 3.223E-09 | 2.600E-09 | 8.500E-09 |
| 23421 | 1172 | S1172A | D | 2.3e-09 | 6.5e-09 | 1.1e-08 | 6.600E-09 | 4.351E-09 | 2.300E-09 | 1.100E-08 |
| 23422 | 1172 | S1172R | D | 2.6e-09 | 3.4e-09 | 6.3e-09 | 4.100E-09 | 1.947E-09 | 2.600E-09 | 6.300E-09 |
| 23423 | 1172 | S1172N | D | 1.4e-08 | 8.3e-09 | 7.3e-09 | 9.867E-09 | 3.614E-09 | 7.300E-09 | 1.400E-08 |
| 23424 | 1172 | S1172D | D | 1.1e-08 | 5e-09 | 4.4e-09 | 6.800E-09 | 3.650E-09 | 4.400E-09 | 1.100E-08 |
| 23425 | 1172 | S1172C | D | 5.7e-09 | 6e-09 | 2.3e-09 | 4.667E-09 | 2.055E-09 | 2.300E-09 | 6.000E-09 |
| 23426 | 1172 | S1172E | D | 1e-08 | 1.9e-08 | 1.6e-08 | 1.500E-08 | 4.583E-09 | 1.000E-08 | 1.900E-08 |
| 23427 | 1172 | S1172Q | D | 4.3e-09 | 1.3e-08 | 5.7e-09 | 7.667E-09 | 4.672E-09 | 4.300E-09 | 1.300E-08 |
| 23428 | 1172 | S1172G | D | 5.2e-09 | 6.4e-09 | 1.4e-08 | 8.533E-09 | 4.772E-09 | 5.200E-09 | 1.400E-08 |
| 23429 | 1172 | S1172H | D | 3.1e-09 | 5.4e-09 | 2.7e-09 | 3.733E-09 | 1.457E-09 | 2.700E-09 | 5.400E-09 |
| 23430 | 1172 | S1172I | D | 4.6e-09 | 5.5e-09 | 1.7e-08 | 9.033E-09 | 6.914E-09 | 4.600E-09 | 1.700E-08 |
| 23431 | 1172 | S1172L | D | 4.2e-09 | 6e-09 | 8.5e-09 | 6.233E-09 | 2.159E-09 | 4.200E-09 | 8.500E-09 |
| 23432 | 1172 | S1172K | D | 7e-09 | 1.2e-08 | 3.4e-09 | 7.467E-09 | 4.319E-09 | 3.400E-09 | 1.200E-08 |
| 23433 | 1172 | S1172M | D | 6e-09 | 1.9e-08 | 7.5e-09 | 1.083E-08 | 7.112E-09 | 6.000E-09 | 1.900E-08 |
| 23434 | 1172 | S1172F | D | 8.2e-09 | 1.5e-08 | 1.2e-08 | 1.173E-08 | 3.408E-09 | 8.200E-09 | 1.500E-08 |
| 23435 | 1172 | S1172P | D | 2e-09 | 6.9e-09 | 1e-08 | 6.300E-09 | 4.034E-09 | 2.000E-09 | 1.000E-08 |
| 23436 | 1172 | S1172S | D | 1.9e-08 | 3.9e-09 | 4.7e-09 | 9.200E-09 | 8.496E-09 | 3.900E-09 | 1.900E-08 |
| 23437 | 1172 | S1172T | D | 6.9e-09 | 4.5e-09 | 3.7e-09 | 5.033E-09 | 1.665E-09 | 3.700E-09 | 6.900E-09 |
| 23438 | 1172 | S1172W | D | 1.1e-09 | 5.8e-09 | 1.2e-08 | 6.300E-09 | 5.467E-09 | 1.100E-09 | 1.200E-08 |
| 23439 | 1172 | S1172Y | D | 4.9e-09 | 1.1e-08 | 6.5e-09 | 7.467E-09 | 3.163E-09 | 4.900E-09 | 1.100E-08 |
| 23440 | 1172 | S1172V | D | 7e-09 | 4.8e-09 | 5e-09 | 5.600E-09 | 1.217E-09 | 4.800E-09 | 7.000E-09 |
| 23441 | 1173 | V1173A | D | 6.4e-09 | 1.4e-09 | 4.8e-09 | 4.200E-09 | 2.553E-09 | 1.400E-09 | 6.400E-09 |
| 23442 | 1173 | V1173R | D | 9.2e-09 | 3.8e-09 | 4.2e-09 | 5.733E-09 | 3.009E-09 | 3.800E-09 | 9.200E-09 |
| 23443 | 1173 | V1173N | D | 1.1e-08 | 1.1e-08 | 7.4e-09 | 9.800E-09 | 2.078E-09 | 7.400E-09 | 1.100E-08 |
| 23444 | 1173 | V1173D | D | 1e-08 | 5.4e-09 | 3.9e-09 | 6.433E-09 | 3.179E-09 | 3.900E-09 | 1.000E-08 |
| 23445 | 1173 | V1173C | D | 6e-09 | 1.3e-08 | 1.5e-08 | 1.133E-08 | 4.726E-09 | 6.000E-09 | 1.500E-08 |
| 23446 | 1173 | V1173E | D | 5e-09 | 7.1e-09 | 6.2e-09 | 6.100E-09 | 1.054E-09 | 5.000E-09 | 7.100E-09 |
| 23447 | 1173 | V1173Q | D | 4.1e-09 | 5.6e-09 | 5e-09 | 4.900E-09 | 7.550E-10 | 4.100E-09 | 5.600E-09 |
| 23448 | 1173 | V1173G | D | 3.7e-09 | 6.1e-09 | 3.3e-09 | 4.367E-09 | 1.514E-09 | 3.300E-09 | 6.100E-09 |
| 23449 | 1173 | V1173H | D | 1.3e-08 | 6.8e-09 | 2.1e-08 | 1.360E-08 | 7.119E-09 | 6.800E-09 | 2.100E-08 |
| 23450 | 1173 | V1173I | D | 5.6e-09 | 1.6e-08 | 1.6e-08 | 1.253E-08 | 6.004E-09 | 5.600E-09 | 1.600E-08 |
| 23451 | 1173 | V1173L | D | 6.8e-09 | 7.8e-09 | 7.4e-09 | 7.333E-09 | 5.033E-10 | 6.800E-09 | 7.800E-09 |
| 23452 | 1173 | V1173K | D | 1.5e-08 | 8.3e-09 | 5.3e-09 | 9.533E-09 | 4.966E-09 | 5.300E-09 | 1.500E-08 |
| 23453 | 1173 | V1173M | D | 1.1e-08 | 2.8e-09 | 5.7e-09 | 6.500E-09 | 4.158E-09 | 2.800E-09 | 1.100E-08 |
| 23454 | 1173 | V1173F | D | 2.6e-09 | 4.6e-09 | 1.7e-08 | 8.067E-09 | 7.801E-09 | 2.600E-09 | 1.700E-08 |
| 23455 | 1173 | V1173P | D | 1.4e-08 | 8.7e-09 | 4.5e-09 | 9.067E-09 | 4.761E-09 | 4.500E-09 | 1.400E-08 |
| 23456 | 1173 | V1173S | D | 2.6e-08 | 2.3e-08 | 1.4e-08 | 2.100E-08 | 6.245E-09 | 1.400E-08 | 2.600E-08 |
| 23457 | 1173 | V1173T | D | 5.9e-09 | 3.8e-09 | 1.1e-08 | 6.900E-09 | 3.703E-09 | 3.800E-09 | 1.100E-08 |
| 23458 | 1173 | V1173W | D | 3.8e-09 | 3.4e-09 | 5.6e-09 | 4.267E-09 | 1.172E-09 | 3.400E-09 | 5.600E-09 |
| 23459 | 1173 | V1173Y | D | 1.6e-08 | 7.1e-09 | 9.7e-09 | 1.093E-08 | 4.576E-09 | 7.100E-09 | 1.600E-08 |

| | | | | | | | | | | |
|---|---|---|---|---|---|---|---|---|---|---|
| 23460 | 1173 | V1173V | D | 6.3e-09 | 1.4e-08 | 8.1e-09 | 9.467E-09 | 4.028E-09 | 6.300E-09 | 1.400E-08 |
| 23461 | 1174 | F1174A | D | 2e-08 | 1.5e-08 | 1e-08 | 1.500E-08 | 5.000E-09 | 1.000E-08 | 2.000E-08 |
| 23462 | 1174 | F1174R | D | 8.7e-09 | 2.2e-08 | 1e-08 | 1.357E-08 | 7.332E-09 | 8.700E-09 | 2.200E-08 |
| 23463 | 1174 | F1174N | D | 9.3e-09 | 4.3e-09 | 1.1e-08 | 8.200E-09 | 3.483E-09 | 4.300E-09 | 1.100E-08 |
| 23464 | 1174 | F1174D | D | 6.3e-09 | 9.3e-09 | 2e-08 | 1.187E-08 | 7.202E-09 | 6.300E-09 | 2.000E-08 |
| 23465 | 1174 | F1174C | D | 1.8e-08 | 4.1e-09 | 3.4e-09 | 8.500E-09 | 8.235E-09 | 3.400E-09 | 1.800E-08 |
| 23466 | 1174 | F1174E | D | 5.9e-09 | 5e-09 | 1e-08 | 6.967E-09 | 2.665E-09 | 5.000E-09 | 1.000E-08 |
| 23467 | 1174 | F1174Q | D | 3.9e-09 | 1.9e-09 | 6.1e-09 | 3.967E-09 | 2.101E-09 | 1.900E-09 | 6.100E-09 |
| 23468 | 1174 | F1174G | D | 7e-09 | 1.2e-08 | 5.4e-09 | 8.133E-09 | 3.443E-09 | 5.400E-09 | 1.200E-08 |
| 23469 | 1174 | F1174H | D | 5.2e-09 | 3.8e-09 | 4.8e-09 | 4.600E-09 | 7.211E-10 | 3.800E-09 | 5.200E-09 |
| 23470 | 1174 | F1174I | D | 4.7e-09 | 3.4e-09 | 1.1e-08 | 6.367E-09 | 4.065E-09 | 3.400E-09 | 1.100E-08 |
| 23471 | 1174 | F1174L | D | 1.3e-08 | 5.7e-09 | 4.9e-09 | 7.867E-09 | 4.464E-09 | 4.900E-09 | 1.300E-08 |
| 23472 | 1174 | F1174K | D | 3.4e-09 | 1.3e-08 | 4.1e-09 | 6.833E-09 | 5.352E-09 | 3.400E-09 | 1.300E-08 |
| 23473 | 1174 | F1174M | D | 6.6e-09 | 9.8e-09 | 6.6e-09 | 7.667E-09 | 1.848E-09 | 6.600E-09 | 9.800E-09 |
| 23474 | 1174 | F1174F | D | 1.4e-08 | 5.5e-09 | 5.2e-09 | 8.233E-09 | 4.996E-09 | 5.200E-09 | 1.400E-08 |
| 23475 | 1174 | F1174P | D | 1.6e-08 | 1.5e-09 | 5.2e-09 | 7.567E-09 | 7.534E-09 | 1.500E-09 | 1.600E-08 |
| 23476 | 1174 | F1174S | D | 1.9e-09 | 3.6e-08 | 1.3e-08 | 1.697E-08 | 1.739E-08 | 1.900E-09 | 3.600E-08 |
| 23477 | 1174 | F1174T | D | 8.5e-09 | 8.1e-09 | 4.7e-09 | 7.100E-09 | 2.088E-09 | 4.700E-09 | 8.500E-09 |
| 23478 | 1174 | F1174W | D | 6.1e-09 | 5.3e-09 | 8.8e-09 | 6.733E-09 | 1.834E-09 | 5.300E-09 | 8.800E-09 |
| 23479 | 1174 | F1174Y | D | 1e-08 | 3.4e-09 | 2.6e-08 | 1.313E-08 | 1.162E-08 | 3.400E-09 | 2.600E-08 |
| 23480 | 1174 | F1174V | D | 4.3e-09 | 8.1e-09 | 8.5e-09 | 6.967E-09 | 2.318E-09 | 4.300E-09 | 8.500E-09 |
| 23481 | 1175 | I1175A | D | 8.6e-09 | 6.9e-09 | 8.2e-09 | 7.900E-09 | 8.888E-10 | 6.900E-09 | 8.600E-09 |
| 23482 | 1175 | I1175R | D | 7e-09 | 3e-09 | 9.8e-09 | 6.600E-09 | 3.418E-09 | 3.000E-09 | 9.800E-09 |
| 23483 | 1175 | I1175N | D | 8.6e-09 | 4.1e-09 | 6.2e-09 | 6.300E-09 | 2.252E-09 | 4.100E-09 | 8.600E-09 |
| 23484 | 1175 | I1175D | D | 1.9e-09 | 1.8e-09 | 3.7e-09 | 2.467E-09 | 1.069E-09 | 1.800E-09 | 3.700E-09 |
| 23485 | 1175 | I1175C | D | 5.2e-09 | 5.5e-09 | 7e-09 | 5.900E-09 | 9.644E-10 | 5.200E-09 | 7.000E-09 |
| 23486 | 1175 | I1175E | D | 4.2e-09 | 1.2e-08 | 9.9e-09 | 8.700E-09 | 4.036E-09 | 4.200E-09 | 1.200E-08 |
| 23487 | 1175 | I1175Q | D | 1.2e-09 | 1.6e-08 | 5.7e-09 | 7.633E-09 | 7.587E-09 | 1.200E-09 | 1.600E-08 |
| 23488 | 1175 | I1175G | D | 4.5e-09 | 4.6e-09 | 7.4e-09 | 5.500E-09 | 1.646E-09 | 4.500E-09 | 7.400E-09 |
| 23489 | 1175 | I1175H | D | 5.1e-09 | 4.5e-09 | 8.9e-09 | 6.167E-09 | 2.386E-09 | 4.500E-09 | 8.900E-09 |
| 23490 | 1175 | I1175I | D | 1.4e-09 | 1.4e-08 | 1e-08 | 8.467E-09 | 6.438E-09 | 1.400E-09 | 1.400E-08 |
| 23491 | 1175 | I1175L | D | 2e-09 | 1.4e-08 | 1.2e-08 | 9.333E-09 | 6.429E-09 | 2.000E-09 | 1.400E-08 |
| 23492 | 1175 | I1175K | D | 7.2e-09 | 4.1e-09 | 3.1e-09 | 4.800E-09 | 2.138E-09 | 3.100E-09 | 7.200E-09 |
| 23493 | 1175 | I1175M | D | 1.4e-08 | 9.5e-09 | 6.7e-09 | 1.007E-08 | 3.683E-09 | 6.700E-09 | 1.400E-08 |
| 23494 | 1175 | I1175F | D | 6.8e-09 | 3.9e-09 | 4.8e-09 | 5.167E-09 | 1.484E-09 | 3.900E-09 | 6.800E-09 |
| 23495 | 1175 | I1175P | D | 1e-08 | 1.1e-08 | 4e-09 | 8.333E-09 | 3.786E-09 | 4.000E-09 | 1.100E-08 |
| 23496 | 1175 | I1175S | D | 1.7e-09 | 2.3e-08 | 1e-08 | 1.157E-08 | 1.074E-08 | 1.700E-09 | 2.300E-08 |
| 23497 | 1175 | I1175T | D | 1.6e-08 | 1.9e-08 | 9.6e-09 | 1.487E-08 | 4.801E-09 | 9.600E-09 | 1.900E-08 |
| 23498 | 1175 | I1175W | D | 4.6e-09 | 4.5e-09 | 1.4e-08 | 7.700E-09 | 5.456E-09 | 4.500E-09 | 1.400E-08 |
| 23499 | 1175 | I1175Y | D | 8.9e-09 | 1e-08 | 3.5e-09 | 7.467E-09 | 3.479E-09 | 3.500E-09 | 1.000E-08 |
| 23500 | 1175 | I1175V | D | 3.8e-09 | 1.7e-09 | 7.4e-09 | 4.300E-09 | 2.883E-09 | 1.700E-09 | 7.400E-09 |
| 23501 | 1176 | F1176A | D | 6.8e-09 | 1.5e-08 | 1.5e-08 | 1.227E-08 | 4.734E-09 | 6.800E-09 | 1.500E-08 |
| 23502 | 1176 | F1176R | D | 8.6e-09 | 4.7e-09 | 1.6e-08 | 9.767E-09 | 5.740E-09 | 4.700E-09 | 1.600E-08 |
| 23503 | 1176 | F1176N | D | 3e-09 | 7.1e-09 | 6.2e-09 | 5.433E-09 | 2.155E-09 | 3.000E-09 | 7.100E-09 |
| 23504 | 1176 | F1176D | D | 1.4e-08 | 1.2e-08 | 1.3e-08 | 1.300E-08 | 1.000E-09 | 1.200E-08 | 1.400E-08 |
| 23505 | 1176 | F1176C | D | 6.8e-09 | 4.1e-09 | 1.5e-08 | 8.633E-09 | 5.677E-09 | 4.100E-09 | 1.500E-08 |
| 23506 | 1176 | F1176E | D | 7e-09 | 7.7e-09 | 6.2e-09 | 6.967E-09 | 7.506E-10 | 6.200E-09 | 7.700E-09 |
| 23507 | 1176 | F1176Q | D | 5.3e-09 | 9.2e-09 | 1.3e-08 | 9.167E-09 | 3.850E-09 | 5.300E-09 | 1.300E-08 |
| 23508 | 1176 | F1176G | D | 1.4e-08 | 1.9e-08 | 1e-08 | 1.433E-08 | 4.509E-09 | 1.000E-08 | 1.900E-08 |
| 23509 | 1176 | F1176H | D | 4.6e-09 | 5.5e-09 | 7.3e-09 | 5.800E-09 | 1.375E-09 | 4.600E-09 | 7.300E-09 |
| 23510 | 1176 | F1176I | D | 6.9e-09 | 3.9e-09 | 6.4e-09 | 5.733E-09 | 1.607E-09 | 3.900E-09 | 6.900E-09 |
| 23511 | 1176 | F1176L | D | 5e-09 | 4e-09 | 2.7e-09 | 3.900E-09 | 1.153E-09 | 2.700E-09 | 5.000E-09 |
| 23512 | 1176 | F1176K | D | 4.5e-09 | 6e-09 | 5.9e-09 | 5.467E-09 | 8.386E-10 | 4.500E-09 | 6.000E-09 |
| 23513 | 1176 | F1176M | D | 1.6e-08 | 3.3e-09 | 8.1e-09 | 9.133E-09 | 6.413E-09 | 3.300E-09 | 1.600E-08 |
| 23514 | 1176 | F1176F | D | 4.2e-09 | 7.5e-09 | 1.6e-09 | 4.433E-09 | 2.957E-09 | 1.600E-09 | 7.500E-09 |
| 23515 | 1176 | F1176P | D | 3.1e-09 | 1.7e-08 | 1e-08 | 1.003E-08 | 6.950E-09 | 3.100E-09 | 1.700E-08 |
| 23516 | 1176 | F1176S | D | 1.1e-08 | 5.8e-09 | 1.1e-08 | 9.267E-09 | 3.002E-09 | 5.800E-09 | 1.100E-08 |
| 23517 | 1176 | F1176T | D | 3.4e-09 | 1.1e-08 | 7.5e-09 | 7.300E-09 | 3.804E-09 | 3.400E-09 | 1.100E-08 |
| 23518 | 1176 | F1176W | D | 6.8e-09 | 6.4e-09 | 6e-09 | 6.400E-09 | 4.000E-10 | 6.000E-09 | 6.800E-09 |
| 23519 | 1176 | F1176Y | D | 5.5e-09 | 4.9e-09 | 1e-08 | 6.800E-09 | 2.787E-09 | 4.900E-09 | 1.000E-08 |
| 23520 | 1176 | F1176V | D | 3.4e-09 | 9e-09 | 2.4e-08 | 1.213E-08 | 1.065E-08 | 3.400E-09 | 2.400E-08 |
| 23521 | 1177 | P1177A | D | 7.8e-09 | 2.4e-08 | 1.2e-08 | 1.460E-08 | 8.407E-09 | 7.800E-09 | 2.400E-08 |
| 23522 | 1177 | P1177R | D | 7.2e-09 | 3.9e-09 | 1.5e-09 | 4.200E-09 | 2.862E-09 | 1.500E-09 | 7.200E-09 |
| 23523 | 1177 | P1177N | D | 1.3e-08 | 7.3e-09 | 2.9e-09 | 7.733E-09 | 5.064E-09 | 2.900E-09 | 1.300E-08 |
| 23524 | 1177 | P1177D | D | 8.7e-09 | 9.7e-09 | 7.5e-09 | 8.633E-09 | 1.102E-09 | 7.500E-09 | 9.700E-09 |
| 23525 | 1177 | P1177C | D | 1.1e-08 | 4.6e-09 | 7.2e-09 | 7.600E-09 | 3.219E-09 | 4.600E-09 | 1.100E-08 |
| 23526 | 1177 | P1177E | D | 8.7e-09 | 3e-09 | 8.2e-09 | 6.633E-09 | 3.156E-09 | 3.000E-09 | 8.700E-09 |
| 23527 | 1177 | P1177Q | D | 2.4e-09 | 2.7e-09 | 7.7e-09 | 4.267E-09 | 2.977E-09 | 2.400E-09 | 7.700E-09 |
| 23528 | 1177 | P1177G | D | 4.6e-09 | 1.2e-08 | 2.2e-09 | 6.267E-09 | 5.108E-09 | 2.200E-09 | 1.200E-08 |
| 23529 | 1177 | P1177H | D | 5.2e-09 | 4.8e-09 | 3.4e-09 | 4.467E-09 | 9.452E-10 | 3.400E-09 | 5.200E-09 |
| 23530 | 1177 | P1177I | D | 1e-08 | 3.9e-09 | 5.6e-09 | 6.500E-09 | 3.148E-09 | 3.900E-09 | 1.000E-08 |
| 23531 | 1177 | P1177L | D | 8.6e-09 | 1.9e-08 | 9.3e-09 | 1.230E-08 | 5.813E-09 | 8.600E-09 | 1.900E-08 |
| 23532 | 1177 | P1177K | D | 4.2e-09 | 1.2e-08 | 6.6e-09 | 7.600E-09 | 3.995E-09 | 4.200E-09 | 1.200E-08 |
| 23533 | 1177 | P1177M | D | 1.5e-08 | 1.1e-08 | 9.2e-09 | 1.173E-08 | 2.969E-09 | 9.200E-09 | 1.500E-08 |
| 23534 | 1177 | P1177F | D | 5.7e-09 | 7.4e-09 | 5.5e-09 | 6.200E-09 | 1.044E-09 | 5.500E-09 | 7.400E-09 |
| 23535 | 1177 | P1177P | D | 2.5e-09 | 7.2e-09 | 6.5e-09 | 5.400E-09 | 2.536E-09 | 2.500E-09 | 7.200E-09 |
| 23536 | 1177 | P1177S | D | 1.3e-08 | 1.1e-08 | 2.6e-09 | 8.867E-09 | 5.518E-09 | 2.600E-09 | 1.300E-08 |
| 23537 | 1177 | P1177T | D | 3.4e-09 | 5.5e-09 | 9.9e-09 | 6.267E-09 | 3.317E-09 | 3.400E-09 | 9.900E-09 |
| 23538 | 1177 | P1177W | D | 9.7e-09 | 3.4e-09 | 7.5e-09 | 6.867E-09 | 3.197E-09 | 3.400E-09 | 9.700E-09 |
| 23539 | 1177 | P1177Y | D | 8.1e-09 | 4.8e-09 | 1.4e-08 | 8.967E-09 | 4.661E-09 | 4.800E-09 | 1.400E-08 |
| 23540 | 1177 | P1177V | D | 5.1e-09 | 2.1e-08 | 1.2e-08 | 1.270E-08 | 7.973E-09 | 5.100E-09 | 2.100E-08 |
| 23541 | 1178 | P1178A | D | 9e-09 | 8.7e-09 | 9.8e-09 | 9.167E-09 | 5.686E-10 | 8.700E-09 | 9.800E-09 |
| 23542 | 1178 | P1178R | D | 2.4e-08 | 5.6e-09 | 3.1e-09 | 1.090E-08 | 1.141E-08 | 3.100E-09 | 2.400E-08 |
| 23543 | 1178 | P1178N | D | 6.4e-10 | 6.1e-09 | 9.4e-09 | 5.380E-09 | 4.424E-09 | 6.400E-10 | 9.400E-09 |
| 23544 | 1178 | P1178D | D | 1e-08 | 1.6e-08 | 3.4e-09 | 9.800E-09 | 6.302E-09 | 3.400E-09 | 1.600E-08 |

| | | | | | | | | | | |
|---|---|---|---|---|---|---|---|---|---|---|
| 23545 | 1178 | P1178C | D | 5.4e-09 | 7e-09 | 3.6e-08 | 1.613E-08 | 1.722E-08 | 5.400E-09 | 3.600E-08 |
| 23546 | 1178 | P1178E | D | 4e-09 | 9.8e-09 | 3.1e-09 | 5.633E-09 | 3.636E-09 | 3.100E-09 | 9.800E-09 |
| 23547 | 1178 | P1178Q | D | 2.3e-08 | 5e-09 | 9.7e-09 | 1.257E-08 | 9.336E-09 | 5.000E-09 | 2.300E-08 |
| 23548 | 1178 | P1178G | D | 7.6e-09 | 1.6e-08 | 3.1e-08 | 1.820E-08 | 1.185E-08 | 7.600E-09 | 3.100E-08 |
| 23549 | 1178 | P1178H | D | 5.3e-09 | 2.3e-08 | 1.4e-08 | 1.410E-08 | 8.850E-09 | 5.300E-09 | 2.300E-08 |
| 23550 | 1178 | P1178I | D | 1.4e-08 | 4.8e-09 | 1.7e-08 | 1.193E-08 | 6.357E-09 | 4.800E-09 | 1.700E-08 |
| 23551 | 1178 | P1178L | D | 9.7e-09 | 4.7e-09 | 8.3e-09 | 7.567E-09 | 2.579E-09 | 4.700E-09 | 9.700E-09 |
| 23552 | 1178 | P1178K | D | 6.1e-09 | 1.3e-08 | 1.3e-08 | 1.070E-08 | 3.984E-09 | 6.100E-09 | 1.300E-08 |
| 23553 | 1178 | P1178M | D | 3.6e-08 | 9.4e-09 | 1.2e-08 | 1.913E-08 | 1.466E-08 | 9.400E-09 | 3.600E-08 |
| 23554 | 1178 | P1178F | D | 1.5e-08 | 5e-09 | 1.7e-08 | 1.233E-08 | 6.429E-09 | 5.000E-09 | 1.700E-08 |
| 23555 | 1178 | P1178P | D | 3.5e-09 | 1e-08 | 7.9e-09 | 7.133E-09 | 3.317E-09 | 3.500E-09 | 1.000E-08 |
| 23556 | 1178 | P1178S | D | 7.9e-09 | 7.8e-10 | 2.4e-08 | 1.089E-08 | 1.190E-08 | 7.800E-10 | 2.400E-08 |
| 23557 | 1178 | P1178T | D | 5.6e-09 | 7e-09 | 1e-08 | 7.533E-09 | 2.248E-09 | 5.600E-09 | 1.000E-08 |
| 23558 | 1178 | P1178W | D | 1.1e-08 | 1.7e-08 | 1.9e-09 | 9.967E-09 | 7.603E-09 | 1.900E-09 | 1.700E-08 |
| 23559 | 1178 | P1178Y | D | 4.9e-09 | 2.4e-09 | 4.3e-09 | 3.867E-09 | 1.305E-09 | 2.400E-09 | 4.900E-09 |
| 23560 | 1178 | P1178V | D | 9.4e-09 | 9.3e-09 | 2.3e-09 | 7.000E-09 | 4.071E-09 | 2.300E-09 | 9.400E-09 |
| 23561 | 1179 | S1179A | D | 9.6e-09 | 6.9e-09 | 1.8e-08 | 1.150E-08 | 5.789E-09 | 6.900E-09 | 1.800E-08 |
| 23562 | 1179 | S1179R | D | 1e-08 | 4.2e-09 | 3.6e-09 | 5.933E-09 | 3.535E-09 | 3.600E-09 | 1.000E-08 |
| 23563 | 1179 | S1179N | D | 1.9e-08 | 8.8e-09 | 6.8e-09 | 1.153E-08 | 6.543E-09 | 6.800E-09 | 1.900E-08 |
| 23564 | 1179 | S1179D | D | 8.8e-09 | 4.1e-09 | 2e-09 | 4.967E-09 | 3.482E-09 | 2.000E-09 | 8.800E-09 |
| 23565 | 1179 | S1179C | D | 9.7e-09 | 6.1e-09 | 2.3e-09 | 6.033E-09 | 3.700E-09 | 2.300E-09 | 9.700E-09 |
| 23566 | 1179 | S1179E | D | 1.9e-08 | 3.5e-09 | 9.1e-09 | 1.053E-08 | 7.849E-09 | 3.500E-09 | 1.900E-08 |
| 23567 | 1179 | S1179Q | D | 1.4e-08 | 1e-08 | 6.4e-09 | 1.013E-08 | 3.802E-09 | 6.400E-09 | 1.400E-08 |
| 23568 | 1179 | S1179G | D | 5.5e-09 | 5.2e-09 | 5.4e-09 | 5.367E-09 | 1.528E-10 | 5.200E-09 | 5.500E-09 |
| 23569 | 1179 | S1179H | D | 5.1e-09 | 1e-08 | 1.9e-08 | 1.137E-08 | 7.050E-09 | 5.100E-09 | 1.900E-08 |
| 23570 | 1179 | S1179I | D | 2.1e-08 | 4.1e-09 | 3.6e-09 | 9.567E-09 | 9.905E-09 | 3.600E-09 | 2.100E-08 |
| 23571 | 1179 | S1179L | D | 8.5e-09 | 6.3e-09 | 2.1e-08 | 1.193E-08 | 7.929E-09 | 6.300E-09 | 2.100E-08 |
| 23572 | 1179 | S1179K | D | 1.1e-08 | 7.1e-09 | 1.1e-08 | 9.700E-09 | 2.252E-09 | 7.100E-09 | 1.100E-08 |
| 23573 | 1179 | S1179M | D | 4.1e-09 | 1e-08 | 4.2e-09 | 6.100E-09 | 3.378E-09 | 4.100E-09 | 1.000E-08 |
| 23574 | 1179 | S1179F | D | 1.7e-08 | 1.1e-08 | 6.2e-09 | 1.140E-08 | 5.411E-09 | 6.200E-09 | 1.700E-08 |
| 23575 | 1179 | S1179P | D | 6.9e-09 | 2.6e-08 | 3.5e-09 | 1.213E-08 | 1.213E-08 | 3.500E-09 | 2.600E-08 |
| 23576 | 1179 | S1179S | D | 3.5e-09 | 6e-09 | 9.4e-09 | 6.300E-09 | 2.961E-09 | 3.500E-09 | 9.400E-09 |
| 23577 | 1179 | S1179T | D | 2.1e-08 | 1.3e-08 | 6.3e-09 | 1.343E-08 | 7.360E-09 | 6.300E-09 | 2.100E-08 |
| 23578 | 1179 | S1179W | D | 1.5e-09 | 2.2e-08 | 4.4e-09 | 9.300E-09 | 1.109E-08 | 1.500E-09 | 2.200E-08 |
| 23579 | 1179 | S1179Y | D | 2.6e-09 | 1.2e-08 | 1.2e-08 | 8.867E-09 | 5.427E-09 | 2.600E-09 | 1.200E-08 |
| 23580 | 1179 | S1179V | D | 5.2e-09 | 6.7e-09 | 5e-09 | 5.633E-09 | 9.292E-10 | 5.000E-09 | 6.700E-09 |
| 23581 | 1180 | D1180A | D | 1.9e-09 | 9.3e-09 | 3.6e-09 | 4.933E-09 | 3.876E-09 | 1.900E-09 | 9.300E-09 |
| 23582 | 1180 | D1180R | D | 4.3e-08 | 5.1e-09 | 1.1e-08 | 1.970E-08 | 2.039E-08 | 5.100E-09 | 4.300E-08 |
| 23583 | 1180 | D1180N | D | 1e-08 | 8.1e-09 | 6.9e-09 | 8.333E-09 | 1.563E-09 | 6.900E-09 | 1.000E-08 |
| 23584 | 1180 | D1180D | D | 7.2e-09 | 4.9e-09 | 3.4e-08 | 1.537E-08 | 1.618E-08 | 4.900E-09 | 3.400E-08 |
| 23585 | 1180 | D1180C | D | 3.2e-09 | 2.5e-08 | 2.4e-08 | 1.740E-08 | 1.231E-08 | 3.200E-09 | 2.500E-08 |
| 23586 | 1180 | D1180E | D | 6.3e-09 | 9.9e-09 | 4.8e-09 | 7.000E-09 | 2.621E-09 | 4.800E-09 | 9.900E-09 |
| 23587 | 1180 | D1180Q | D | 6.3e-09 | 1.8e-08 | 9.6e-09 | 1.130E-08 | 6.032E-09 | 6.300E-09 | 1.800E-08 |
| 23588 | 1180 | D1180G | D | 6.7e-09 | 6.4e-09 | 1.2e-08 | 8.367E-09 | 3.150E-09 | 6.400E-09 | 1.200E-08 |
| 23589 | 1180 | D1180H | D | 8.2e-09 | 9.9e-09 | 5.9e-09 | 8.000E-09 | 2.007E-09 | 5.900E-09 | 9.900E-09 |
| 23590 | 1180 | D1180I | D | 8.2e-09 | 6.1e-09 | 1e-08 | 8.100E-09 | 1.952E-09 | 6.100E-09 | 1.000E-08 |
| 23591 | 1180 | D1180L | D | 3.6e-09 | 1.8e-08 | 1.1e-08 | 1.087E-08 | 7.201E-09 | 3.600E-09 | 1.800E-08 |
| 23592 | 1180 | D1180K | D | 2.6e-08 | 3.6e-09 | 5.8e-09 | 1.180E-08 | 1.235E-08 | 3.600E-09 | 2.600E-08 |
| 23593 | 1180 | D1180M | D | 1.7e-08 | 5.9e-09 | 5.5e-09 | 9.467E-09 | 6.527E-09 | 5.500E-09 | 1.700E-08 |
| 23594 | 1180 | D1180F | D | 3.5e-09 | 1.2e-08 | 4.7e-09 | 6.733E-09 | 4.600E-09 | 3.500E-09 | 1.200E-08 |
| 23595 | 1180 | D1180P | D | 4e-09 | 3.8e-09 | 8.8e-09 | 5.533E-09 | 2.831E-09 | 3.800E-09 | 8.800E-09 |
| 23596 | 1180 | D1180S | D | 7.3e-09 | 1.5e-08 | 1.2e-08 | 1.143E-08 | 3.881E-09 | 7.300E-09 | 1.500E-08 |
| 23597 | 1180 | D1180T | D | 8.2e-09 | 6.9e-09 | 1e-08 | 8.367E-09 | 1.557E-09 | 6.900E-09 | 1.000E-08 |
| 23598 | 1180 | D1180W | D | 4.2e-09 | 2.4e-08 | 1.1e-08 | 1.307E-08 | 1.006E-08 | 4.200E-09 | 2.400E-08 |
| 23599 | 1180 | D1180Y | D | 2.3e-09 | 3.2e-09 | 1.5e-08 | 6.833E-09 | 7.087E-09 | 2.300E-09 | 1.500E-08 |
| 23600 | 1180 | D1180V | D | 8.1e-10 | 1e-08 | 4.7e-09 | 5.170E-09 | 4.613E-09 | 8.100E-10 | 1.000E-08 |
| 23601 | 1181 | E1181A | D | 6.5e-09 | 2.6e-09 | 5.7e-09 | 4.933E-09 | 2.060E-09 | 2.600E-09 | 6.500E-09 |
| 23602 | 1181 | E1181R | D | 2.7e-09 | 3.9e-09 | 4.4e-09 | 3.667E-09 | 8.737E-10 | 2.700E-09 | 4.400E-09 |
| 23603 | 1181 | E1181N | D | 2.3e-09 | 4.5e-09 | 1e-08 | 5.600E-09 | 3.966E-09 | 2.300E-09 | 1.000E-08 |
| 23604 | 1181 | E1181D | D | 5.3e-09 | 1e-08 | 4.3e-09 | 6.533E-09 | 3.044E-09 | 4.300E-09 | 1.000E-08 |
| 23605 | 1181 | E1181C | D | 9.5e-09 | 9.9e-09 | 8.5e-10 | 6.750E-09 | 5.113E-09 | 8.500E-10 | 9.900E-09 |
| 23606 | 1181 | E1181E | D | 5.7e-09 | 1.3e-08 | 1e-08 | 9.567E-09 | 3.669E-09 | 5.700E-09 | 1.300E-08 |
| 23607 | 1181 | E1181Q | D | 6.4e-09 | 6.1e-09 | 4.8e-09 | 5.767E-09 | 8.505E-10 | 4.800E-09 | 6.400E-09 |
| 23608 | 1181 | E1181G | D | 4.7e-09 | 1e-08 | 2.6e-08 | 1.357E-08 | 1.109E-08 | 4.700E-09 | 2.600E-08 |
| 23609 | 1181 | E1181H | D | 1.5e-08 | 7.2e-09 | 3.5e-09 | 8.567E-09 | 5.871E-09 | 3.500E-09 | 1.500E-08 |
| 23610 | 1181 | E1181I | D | 5.6e-09 | 1.1e-08 | 5.3e-09 | 7.300E-09 | 3.208E-09 | 5.300E-09 | 1.100E-08 |
| 23611 | 1181 | E1181L | D | 6.8e-09 | 7.9e-09 | 6.2e-09 | 6.967E-09 | 8.622E-10 | 6.200E-09 | 7.900E-09 |
| 23612 | 1181 | E1181K | D | 1.9e-08 | 5.8e-09 | 7.6e-09 | 1.080E-08 | 7.158E-09 | 5.800E-09 | 1.900E-08 |
| 23613 | 1181 | E1181M | D | 1.2e-08 | 5.2e-09 | 3.4e-09 | 6.867E-09 | 4.536E-09 | 3.400E-09 | 1.200E-08 |
| 23614 | 1181 | E1181F | D | 2.6e-08 | 9.5e-09 | 1.1e-08 | 1.550E-08 | 9.124E-09 | 9.500E-09 | 2.600E-08 |
| 23615 | 1181 | E1181P | D | 1.2e-08 | 4.1e-09 | 4.3e-09 | 6.800E-09 | 4.504E-09 | 4.100E-09 | 1.200E-08 |
| 23616 | 1181 | E1181S | D | 4.9e-09 | 7.7e-09 | 6.8e-09 | 6.467E-09 | 1.429E-09 | 4.900E-09 | 7.700E-09 |
| 23617 | 1181 | E1181T | D | 3.1e-08 | 5.6e-09 | 8e-09 | 1.487E-08 | 1.402E-08 | 5.600E-09 | 3.100E-08 |
| 23618 | 1181 | E1181W | D | 1.7e-09 | 1.3e-08 | 1.3e-08 | 9.233E-09 | 6.524E-09 | 1.700E-09 | 1.300E-08 |
| 23619 | 1181 | E1181Y | D | 1.2e-08 | 1.7e-08 | 2.1e-09 | 1.037E-08 | 7.583E-09 | 2.100E-09 | 1.700E-08 |
| 23620 | 1181 | E1181V | D | 7.8e-09 | 1.2e-08 | 6.4e-09 | 8.733E-09 | 2.914E-09 | 6.400E-09 | 1.200E-08 |
| 23621 | 1182 | Q1182A | D | 3.6e-08 | 4e-09 | 1.1e-08 | 1.700E-08 | 1.682E-08 | 4.000E-09 | 3.600E-08 |
| 23622 | 1182 | Q1182R | D | 2.8e-08 | 8.9e-09 | 5.4e-09 | 1.410E-08 | 1.216E-08 | 5.400E-09 | 2.800E-08 |
| 23623 | 1182 | Q1182N | D | 5.7e-09 | 2.4e-09 | 5.4e-09 | 4.500E-09 | 1.825E-09 | 2.400E-09 | 5.700E-09 |
| 23624 | 1182 | Q1182D | D | 7.3e-09 | 7.2e-09 | 2.7e-09 | 5.733E-09 | 2.627E-09 | 2.700E-09 | 7.300E-09 |
| 23625 | 1182 | Q1182C | D | 6.3e-09 | 1e-08 | 1.3e-08 | 9.767E-09 | 3.356E-09 | 6.300E-09 | 1.300E-08 |
| 23626 | 1182 | Q1182E | D | 2e-08 | 5.2e-09 | 2.3e-08 | 1.607E-08 | 9.530E-09 | 5.200E-09 | 2.300E-08 |
| 23627 | 1182 | Q1182Q | D | 5.1e-09 | 6.9e-09 | 2e-09 | 4.667E-09 | 2.479E-09 | 2.000E-09 | 6.900E-09 |
| 23628 | 1182 | Q1182G | D | 7.6e-09 | 4.6e-09 | 5.5e-09 | 5.900E-09 | 1.539E-09 | 4.600E-09 | 7.600E-09 |
| 23629 | 1182 | Q1182H | D | 1.5e-08 | 9.1e-09 | 3.1e-09 | 9.067E-09 | 5.950E-09 | 3.100E-09 | 1.500E-08 |

| | | | | | | | | | | |
|---|---|---|---|---|---|---|---|---|---|---|
| 23630 | 1182 | Q1182I | D | 1.1e-08 | 5e-09 | 1.3e-08 | 9.667E-09 | 4.163E-09 | 5.000E-09 | 1.300E-08 |
| 23631 | 1182 | Q1182L | D | 1e-08 | 6.2e-09 | 2.1e-08 | 1.240E-08 | 7.686E-09 | 6.200E-09 | 2.100E-08 |
| 23632 | 1182 | Q1182K | D | 2.9e-09 | 4.9e-09 | 8.9e-09 | 5.567E-09 | 3.055E-09 | 2.900E-09 | 8.900E-09 |
| 23633 | 1182 | Q1182M | D | 3.4e-09 | 1e-08 | 9.5e-09 | 7.633E-09 | 3.675E-09 | 3.400E-09 | 1.000E-08 |
| 23634 | 1182 | Q1182F | D | 2.4e-08 | 6.6e-09 | 8.3e-09 | 1.297E-08 | 9.593E-09 | 6.600E-09 | 2.400E-08 |
| 23635 | 1182 | Q1182P | D | 1.1e-08 | 6.6e-09 | 6.2e-09 | 7.933E-09 | 2.663E-09 | 6.200E-09 | 1.100E-08 |
| 23636 | 1182 | Q1182S | D | 6.5e-09 | 8.5e-09 | 7.3e-09 | 7.433E-09 | 1.007E-09 | 6.500E-09 | 8.500E-09 |
| 23637 | 1182 | Q1182T | D | 1.2e-08 | 5.9e-09 | 8.7e-09 | 8.867E-09 | 3.053E-09 | 5.900E-09 | 1.200E-08 |
| 23638 | 1182 | Q1182W | D | 6.3e-09 | 9.8e-09 | 7e-09 | 7.700E-09 | 1.852E-09 | 6.300E-09 | 9.800E-09 |
| 23639 | 1182 | Q1182Y | D | 7.7e-09 | 2.4e-09 | 5e-09 | 5.033E-09 | 2.650E-09 | 2.400E-09 | 7.700E-09 |
| 23640 | 1182 | Q1182V | D | 6.6e-09 | 4.5e-09 | 7.6e-09 | 6.233E-09 | 1.582E-09 | 4.500E-09 | 7.600E-09 |
| 23641 | 1183 | L1183A | D | 5.7e-09 | 1e-08 | 8e-09 | 7.900E-09 | 2.152E-09 | 5.700E-09 | 1.000E-08 |
| 23642 | 1183 | L1183R | D | 8.5e-09 | 8.7e-09 | 3.5e-09 | 6.900E-09 | 2.946E-09 | 3.500E-09 | 8.700E-09 |
| 23643 | 1183 | L1183N | D | 9.4e-09 | 4.2e-09 | 1.7e-08 | 1.020E-08 | 6.437E-09 | 4.200E-09 | 1.700E-08 |
| 23644 | 1183 | L1183D | D | 1.3e-08 | 9.4e-09 | 1.5e-08 | 1.247E-08 | 2.838E-09 | 9.400E-09 | 1.500E-08 |
| 23645 | 1183 | L1183C | D | 4.6e-09 | 1.4e-08 | 2e-08 | 1.287E-08 | 7.762E-09 | 4.600E-09 | 2.000E-08 |
| 23646 | 1183 | L1183E | D | 1.2e-08 | 3.3e-08 | 5.2e-09 | 1.673E-08 | 1.449E-08 | 5.200E-09 | 3.300E-08 |
| 23647 | 1183 | L1183Q | D | 1.6e-08 | 1.1e-08 | 9.1e-09 | 1.203E-08 | 3.564E-09 | 9.100E-09 | 1.600E-08 |
| 23648 | 1183 | L1183G | D | 7.6e-09 | 2.1e-09 | 7.2e-09 | 5.633E-09 | 3.066E-09 | 2.100E-09 | 7.600E-09 |
| 23649 | 1183 | L1183H | D | 3.6e-09 | 7.1e-09 | 2.7e-08 | 1.257E-08 | 1.262E-08 | 3.600E-09 | 2.700E-08 |
| 23650 | 1183 | L1183I | D | 3.9e-09 | 5e-09 | 1.2e-08 | 6.967E-09 | 4.394E-09 | 3.900E-09 | 1.200E-08 |
| 23651 | 1183 | L1183L | D | 1.7e-08 | 2.3e-09 | 2.7e-09 | 7.333E-09 | 8.374E-09 | 2.300E-09 | 1.700E-08 |
| 23652 | 1183 | L1183K | D | 5.7e-09 | 5.4e-09 | 1.3e-08 | 8.033E-09 | 4.304E-09 | 5.400E-09 | 1.300E-08 |
| 23653 | 1183 | L1183M | D | 4.1e-09 | 8.2e-09 | 4.5e-09 | 5.600E-09 | 2.261E-09 | 4.100E-09 | 8.200E-09 |
| 23654 | 1183 | L1183F | D | 1.1e-08 | 8.8e-09 | 5.7e-09 | 8.500E-09 | 2.663E-09 | 5.700E-09 | 1.100E-08 |
| 23655 | 1183 | L1183P | D | 7e-09 | 8e-09 | 2.9e-09 | 5.967E-09 | 2.702E-09 | 2.900E-09 | 8.000E-09 |
| 23656 | 1183 | L1183S | D | 2.5e-09 | 5.4e-09 | 1e-08 | 5.967E-09 | 3.782E-09 | 2.500E-09 | 1.000E-08 |
| 23657 | 1183 | L1183T | D | 4.6e-09 | 9e-09 | 8.5e-09 | 7.367E-09 | 2.409E-09 | 4.600E-09 | 9.000E-09 |
| 23658 | 1183 | L1183W | D | 6.7e-09 | 4.3e-09 | 9.6e-09 | 6.867E-09 | 2.654E-09 | 4.300E-09 | 9.600E-09 |
| 23659 | 1183 | L1183Y | D | 1.3e-08 | 4.5e-09 | 2.8e-09 | 6.767E-09 | 5.465E-09 | 2.800E-09 | 1.300E-08 |
| 23660 | 1183 | L1183V | D | 1.8e-08 | 7.8e-09 | 1.6e-08 | 1.393E-08 | 5.405E-09 | 7.800E-09 | 1.800E-08 |
| 23661 | 1184 | K1184A | D | 6.2e-09 | 1.6e-08 | 1.1e-08 | 1.107E-08 | 4.900E-09 | 6.200E-09 | 1.600E-08 |
| 23662 | 1184 | K1184R | D | 4.1e-09 | 1.5e-08 | 1.7e-08 | 1.203E-08 | 6.943E-09 | 4.100E-09 | 1.700E-08 |
| 23663 | 1184 | K1184N | D | 3.7e-09 | 4.6e-09 | 1.9e-08 | 9.100E-09 | 8.585E-09 | 3.700E-09 | 1.900E-08 |
| 23664 | 1184 | K1184D | D | 2.2e-09 | 7.1e-09 | 6.2e-09 | 5.167E-09 | 2.608E-09 | 2.200E-09 | 7.100E-09 |
| 23665 | 1184 | K1184C | D | 1.2e-08 | 6.8e-09 | 2e-08 | 1.293E-08 | 6.649E-09 | 6.800E-09 | 2.000E-08 |
| 23666 | 1184 | K1184E | D | 2e-09 | 7.9e-09 | 6.6e-09 | 5.500E-09 | 3.100E-09 | 2.000E-09 | 7.900E-09 |
| 23667 | 1184 | K1184Q | D | 3.2e-08 | 8.4e-09 | 5.1e-09 | 1.517E-08 | 1.467E-08 | 5.100E-09 | 3.200E-08 |
| 23668 | 1184 | K1184G | D | 9.9e-09 | 5.6e-09 | 8e-09 | 7.833E-09 | 2.155E-09 | 5.600E-09 | 9.900E-09 |
| 23669 | 1184 | K1184H | D | 3.9e-09 | 3.9e-09 | 4.4e-09 | 4.067E-09 | 2.887E-10 | 3.900E-09 | 4.400E-09 |
| 23670 | 1184 | K1184I | D | 4.3e-09 | 8.9e-09 | 1.4e-08 | 9.067E-09 | 4.852E-09 | 4.300E-09 | 1.400E-08 |
| 23671 | 1184 | K1184L | D | 1.5e-08 | 6.1e-09 | 7.3e-09 | 9.467E-09 | 4.829E-09 | 6.100E-09 | 1.500E-08 |
| 23672 | 1184 | K1184K | D | 5.1e-09 | 2.1e-09 | 7.7e-09 | 4.967E-09 | 2.802E-09 | 2.100E-09 | 7.700E-09 |
| 23673 | 1184 | K1184M | D | 6.2e-09 | 6.7e-09 | 1.3e-08 | 8.633E-09 | 3.790E-09 | 6.200E-09 | 1.300E-08 |
| 23674 | 1184 | K1184F | D | 1.3e-08 | 5.8e-09 | 8.5e-09 | 9.100E-09 | 3.637E-09 | 5.800E-09 | 1.300E-08 |
| 23675 | 1184 | K1184P | D | 1.6e-08 | 7.7e-09 | 6e-09 | 9.900E-09 | 5.351E-09 | 6.000E-09 | 1.600E-08 |
| 23676 | 1184 | K1184S | D | 2.1e-08 | 1e-08 | 9.3e-09 | 1.343E-08 | 6.562E-09 | 9.300E-09 | 2.100E-08 |
| 23677 | 1184 | K1184T | D | 2.1e-08 | 3.8e-09 | 9.4e-09 | 1.140E-08 | 8.773E-09 | 3.800E-09 | 2.100E-08 |
| 23678 | 1184 | K1184W | D | 7.4e-09 | 1e-08 | 1.2e-08 | 9.800E-09 | 2.307E-09 | 7.400E-09 | 1.200E-08 |
| 23679 | 1184 | K1184Y | D | 6.8e-09 | 4.2e-09 | 4.2e-09 | 5.067E-09 | 1.501E-09 | 4.200E-09 | 6.800E-09 |
| 23680 | 1184 | K1184V | D | 2.5e-08 | 2e-09 | 2.6e-09 | 9.867E-09 | 1.311E-08 | 2.000E-09 | 2.500E-08 |
| 23681 | 1185 | S1185A | D | 9.4e-09 | 1e-08 | 6.6e-09 | 8.667E-09 | 1.815E-09 | 6.600E-09 | 1.000E-08 |
| 23682 | 1185 | S1185R | D | 6.4e-09 | 9e-09 | 3.7e-09 | 6.367E-09 | 2.650E-09 | 3.700E-09 | 9.000E-09 |
| 23683 | 1185 | S1185N | D | 8.3e-09 | 1.3e-08 | 9.5e-09 | 1.027E-08 | 2.442E-09 | 8.300E-09 | 1.300E-08 |
| 23684 | 1185 | S1185D | D | 3.2e-09 | 1.2e-08 | 4.8e-09 | 6.667E-09 | 4.688E-09 | 3.200E-09 | 1.200E-08 |
| 23685 | 1185 | S1185C | D | 8.8e-09 | 8.7e-09 | 2.3e-09 | 6.600E-09 | 3.724E-09 | 2.300E-09 | 8.800E-09 |
| 23686 | 1185 | S1185E | D | 1.8e-08 | 1.5e-08 | 5e-09 | 1.267E-08 | 6.807E-09 | 5.000E-09 | 1.800E-08 |
| 23687 | 1185 | S1185Q | D | 5e-09 | 7.6e-09 | 1e-08 | 7.533E-09 | 2.501E-09 | 5.000E-09 | 1.000E-08 |
| 23688 | 1185 | S1185G | D | 1.8e-08 | 5.9e-09 | 3.2e-08 | 1.863E-08 | 1.306E-08 | 5.900E-09 | 3.200E-08 |
| 23689 | 1185 | S1185H | D | 8e-09 | 8.2e-09 | 2.9e-09 | 6.367E-09 | 3.004E-09 | 2.900E-09 | 8.200E-09 |
| 23690 | 1185 | S1185I | D | 1.8e-08 | 6.8e-09 | 1.2e-08 | 1.227E-08 | 5.605E-09 | 6.800E-09 | 1.800E-08 |
| 23691 | 1185 | S1185L | D | 7.7e-09 | 1.4e-08 | 8.1e-09 | 9.933E-09 | 3.528E-09 | 7.700E-09 | 1.400E-08 |
| 23692 | 1185 | S1185K | D | 6e-09 | 4.7e-09 | 1.1e-08 | 7.233E-09 | 3.326E-09 | 4.700E-09 | 1.100E-08 |
| 23693 | 1185 | S1185M | D | 8e-09 | 2e-09 | 4.3e-09 | 4.767E-09 | 3.027E-09 | 2.000E-09 | 8.000E-09 |
| 23694 | 1185 | S1185F | D | 5.1e-09 | 4.9e-09 | 3.5e-09 | 4.500E-09 | 8.718E-10 | 3.500E-09 | 5.100E-09 |
| 23695 | 1185 | S1185P | D | 6.4e-09 | 1.2e-08 | 1.9e-09 | 6.767E-09 | 5.060E-09 | 1.900E-09 | 1.200E-08 |
| 23696 | 1185 | S1185S | D | 1e-08 | 6.7e-09 | 8.1e-09 | 8.267E-09 | 1.656E-09 | 6.700E-09 | 1.000E-08 |
| 23697 | 1185 | S1185T | D | 4.2e-09 | 1.6e-08 | 9.3e-09 | 9.833E-09 | 5.918E-09 | 4.200E-09 | 1.600E-08 |
| 23698 | 1185 | S1185W | D | 1.9e-08 | 9e-09 | 5.7e-09 | 1.123E-08 | 6.926E-09 | 5.700E-09 | 1.900E-08 |
| 23699 | 1185 | S1185Y | D | 3.6e-09 | 3.3e-09 | 1.1e-08 | 5.967E-09 | 4.362E-09 | 3.300E-09 | 1.100E-08 |
| 23700 | 1185 | S1185V | D | 1.3e-08 | 2.6e-09 | 1.1e-08 | 8.867E-09 | 5.518E-09 | 2.600E-09 | 1.300E-08 |
| 23701 | 1186 | G1186A | D | 1.1e-08 | 7.2e-09 | 3.5e-09 | 7.233E-09 | 3.750E-09 | 3.500E-09 | 1.100E-08 |
| 23702 | 1186 | G1186R | D | 1.1e-08 | 8e-09 | 7.8e-09 | 8.933E-09 | 1.793E-09 | 7.800E-09 | 1.100E-08 |
| 23703 | 1186 | G1186N | D | 7.5e-09 | 8.8e-09 | 1.2e-08 | 9.433E-09 | 2.316E-09 | 7.500E-09 | 1.200E-08 |
| 23704 | 1186 | G1186D | D | 9.4e-09 | 1.7e-09 | 8.8e-09 | 6.633E-09 | 4.283E-09 | 1.700E-09 | 9.400E-09 |
| 23705 | 1186 | G1186C | D | 4.4e-09 | 5.8e-09 | 8.9e-09 | 6.367E-09 | 2.303E-09 | 4.400E-09 | 8.900E-09 |
| 23706 | 1186 | G1186E | D | 7.6e-09 | 5.2e-09 | 3.1e-08 | 1.460E-08 | 1.425E-08 | 5.200E-09 | 3.100E-08 |
| 23707 | 1186 | G1186Q | D | 3.5e-09 | 2.7e-09 | 1e-08 | 5.400E-09 | 4.004E-09 | 2.700E-09 | 1.000E-08 |
| 23708 | 1186 | G1186G | D | 7.4e-09 | 1.2e-08 | 9.7e-09 | 9.700E-09 | 2.300E-09 | 7.400E-09 | 1.200E-08 |
| 23709 | 1186 | G1186H | D | 1.6e-08 | 9e-09 | 1.5e-08 | 1.333E-08 | 3.786E-09 | 9.000E-09 | 1.600E-08 |
| 23710 | 1186 | G1186I | D | 7.1e-09 | 1.7e-08 | 5.9e-09 | 1.000E-08 | 6.092E-09 | 5.900E-09 | 1.700E-08 |
| 23711 | 1186 | G1186L | D | 4.1e-09 | 8.7e-09 | 6.9e-09 | 6.567E-09 | 2.318E-09 | 4.100E-09 | 8.700E-09 |
| 23712 | 1186 | G1186K | D | 9.9e-09 | 1.1e-08 | 6.4e-09 | 9.100E-09 | 2.402E-09 | 6.400E-09 | 1.100E-08 |
| 23713 | 1186 | G1186M | D | 4.5e-09 | 4.7e-09 | 4.9e-09 | 4.700E-09 | 2.000E-10 | 4.500E-09 | 4.900E-09 |
| 23714 | 1186 | G1186F | D | 5.8e-09 | 1.1e-08 | 8.9e-09 | 8.567E-09 | 2.616E-09 | 5.800E-09 | 1.100E-08 |

| | | | | | | | | | | |
|---|---|---|---|---|---|---|---|---|---|---|
| 23715 | 1186 | G1186P | D | 1.1e-08 | 7.6e-09 | 8.9e-09 | 9.167E-09 | 1.716E-09 | 7.600E-09 | 1.100E-08 |
| 23716 | 1186 | G1186S | D | 4.3e-09 | 1.5e-08 | 6e-09 | 8.433E-09 | 5.750E-09 | 4.300E-09 | 1.500E-08 |
| 23717 | 1186 | G1186T | D | 1.2e-08 | 2.4e-09 | 1.2e-08 | 8.800E-09 | 5.543E-09 | 2.400E-09 | 1.200E-08 |
| 23718 | 1186 | G1186W | D | 1.6e-09 | 3.3e-09 | 9.3e-09 | 4.733E-09 | 4.045E-09 | 1.600E-09 | 9.300E-09 |
| 23719 | 1186 | G1186Y | D | 1e-08 | 9.1e-09 | 1.8e-08 | 1.237E-08 | 4.899E-09 | 9.100E-09 | 1.800E-08 |
| 23720 | 1186 | G1186V | D | 1e-08 | 1.7e-08 | 6.1e-09 | 1.103E-08 | 5.523E-09 | 6.100E-09 | 1.700E-08 |
| 23721 | 1187 | T1187A | D | 1.1e-08 | 1.4e-09 | 2.9e-09 | 5.100E-09 | 5.164E-09 | 1.400E-09 | 1.100E-08 |
| 23722 | 1187 | T1187R | D | 6.8e-09 | 2.9e-09 | 7e-09 | 5.567E-09 | 2.312E-09 | 2.900E-09 | 7.000E-09 |
| 23723 | 1187 | T1187N | D | 7.6e-09 | 3.6e-09 | 5.1e-09 | 5.433E-09 | 2.021E-09 | 3.600E-09 | 7.600E-09 |
| 23724 | 1187 | T1187D | D | 4.2e-09 | 5.1e-09 | 1.2e-08 | 7.100E-09 | 4.267E-09 | 4.200E-09 | 1.200E-08 |
| 23725 | 1187 | T1187C | D | 4e-09 | 1.5e-08 | 7.3e-09 | 8.767E-09 | 5.645E-09 | 4.000E-09 | 1.500E-08 |
| 23726 | 1187 | T1187E | D | 1.4e-08 | 7.4e-09 | 1.3e-08 | 1.147E-08 | 3.557E-09 | 7.400E-09 | 1.400E-08 |
| 23727 | 1187 | T1187Q | D | 4.6e-09 | 2.5e-09 | 1.1e-08 | 6.033E-09 | 4.428E-09 | 2.500E-09 | 1.100E-08 |
| 23728 | 1187 | T1187G | D | 1.5e-09 | 2.3e-09 | 6.6e-09 | 3.467E-09 | 2.743E-09 | 1.500E-09 | 6.600E-09 |
| 23729 | 1187 | T1187H | D | 5.5e-09 | 1.2e-08 | 3.6e-09 | 7.033E-09 | 4.405E-09 | 3.600E-09 | 1.200E-08 |
| 23730 | 1187 | T1187I | D | 1.6e-08 | 4.3e-09 | 1.6e-08 | 1.210E-08 | 6.755E-09 | 4.300E-09 | 1.600E-08 |
| 23731 | 1187 | T1187L | D | 9.4e-09 | 1.1e-08 | 7.2e-09 | 9.200E-09 | 1.908E-09 | 7.200E-09 | 1.100E-08 |
| 23732 | 1187 | T1187K | D | 2.1e-08 | 6.4e-09 | 1.9e-08 | 1.547E-08 | 7.915E-09 | 6.400E-09 | 2.100E-08 |
| 23733 | 1187 | T1187M | D | 1.9e-09 | 4.7e-09 | 9.4e-09 | 5.333E-09 | 3.790E-09 | 1.900E-09 | 9.400E-09 |
| 23734 | 1187 | T1187F | D | 3.7e-09 | 2e-09 | 8.2e-09 | 4.633E-09 | 3.204E-09 | 2.000E-09 | 8.200E-09 |
| 23735 | 1187 | T1187P | D | 8.5e-09 | 4e-09 | 7.4e-09 | 6.633E-09 | 2.346E-09 | 4.000E-09 | 8.500E-09 |
| 23736 | 1187 | T1187S | D | 1.1e-08 | 1.2e-08 | 1.8e-08 | 1.367E-08 | 3.786E-09 | 1.100E-08 | 1.800E-08 |
| 23737 | 1187 | T1187T | D | 6.5e-09 | 3.2e-09 | 1.3e-08 | 7.567E-09 | 4.986E-09 | 3.200E-09 | 1.300E-08 |
| 23738 | 1187 | T1187W | D | 1.4e-08 | 6.3e-09 | 6.9e-09 | 9.067E-09 | 4.283E-09 | 6.300E-09 | 1.400E-08 |
| 23739 | 1187 | T1187Y | D | 9.1e-09 | 3.2e-09 | 1.2e-08 | 8.100E-09 | 4.484E-09 | 3.200E-09 | 1.200E-08 |
| 23740 | 1187 | T1187V | D | 2.8e-09 | 5.9e-09 | 2.5e-09 | 3.733E-09 | 1.882E-09 | 2.500E-09 | 5.900E-09 |
| 23741 | 1188 | A1188A | D | 4.6e-09 | 1e-08 | 6.4e-09 | 7.000E-09 | 2.750E-09 | 4.600E-09 | 1.000E-08 |
| 23742 | 1188 | A1188R | D | 2.9e-09 | 6.9e-09 | 1.6e-08 | 8.600E-09 | 6.713E-09 | 2.900E-09 | 1.600E-08 |
| 23743 | 1188 | A1188N | D | 6e-09 | 1.4e-08 | 9.4e-09 | 9.800E-09 | 4.015E-09 | 6.000E-09 | 1.400E-08 |
| 23744 | 1188 | A1188D | D | 4.6e-09 | 1.7e-08 | 3.5e-09 | 8.367E-09 | 7.497E-09 | 3.500E-09 | 1.700E-08 |
| 23745 | 1188 | A1188C | D | 4.5e-09 | 1.2e-08 | 1.2e-08 | 9.500E-09 | 4.330E-09 | 4.500E-09 | 1.200E-08 |
| 23746 | 1188 | A1188E | D | 5.3e-09 | 3.9e-09 | 2.9e-09 | 4.033E-09 | 1.206E-09 | 2.900E-09 | 5.300E-09 |
| 23747 | 1188 | A1188Q | D | 1.3e-08 | 3.1e-09 | 5e-09 | 7.033E-09 | 5.254E-09 | 3.100E-09 | 1.300E-08 |
| 23748 | 1188 | A1188G | D | 9.1e-09 | 4.6e-09 | 9e-09 | 7.567E-09 | 2.570E-09 | 4.600E-09 | 9.100E-09 |
| 23749 | 1188 | A1188H | D | 1.1e-08 | 3.2e-08 | 5.7e-09 | 1.623E-08 | 1.391E-08 | 5.700E-09 | 3.200E-08 |
| 23750 | 1188 | A1188I | D | 6.3e-09 | 3.5e-09 | 8.5e-09 | 6.100E-09 | 2.506E-09 | 3.500E-09 | 8.500E-09 |
| 23751 | 1188 | A1188L | D | 7.2e-09 | 3.7e-09 | 6.6e-09 | 5.833E-09 | 1.872E-09 | 3.700E-09 | 7.200E-09 |
| 23752 | 1188 | A1188K | D | 7.9e-09 | 9e-09 | 5.3e-09 | 7.400E-09 | 1.900E-09 | 5.300E-09 | 9.000E-09 |
| 23753 | 1188 | A1188M | D | 5.8e-09 | 7.7e-09 | 1.7e-08 | 1.017E-08 | 5.994E-09 | 5.800E-09 | 1.700E-08 |
| 23754 | 1188 | A1188F | D | 1.4e-08 | 1.1e-08 | 1.4e-08 | 1.300E-08 | 1.732E-09 | 1.100E-08 | 1.400E-08 |
| 23755 | 1188 | A1188P | D | 5.3e-09 | 3.5e-09 | 7.4e-09 | 5.400E-09 | 1.952E-09 | 3.500E-09 | 7.400E-09 |
| 23756 | 1188 | A1188S | D | 3.9e-09 | 9.1e-09 | 6.6e-09 | 6.533E-09 | 2.601E-09 | 3.900E-09 | 9.100E-09 |
| 23757 | 1188 | A1188T | D | 7.7e-09 | 8.7e-09 | 3.4e-09 | 6.600E-09 | 2.816E-09 | 3.400E-09 | 8.700E-09 |
| 23758 | 1188 | A1188W | D | 9.5e-09 | 8.3e-09 | 6.9e-09 | 8.233E-09 | 1.301E-09 | 6.900E-09 | 9.500E-09 |
| 23759 | 1188 | A1188Y | D | 8.4e-09 | 4e-09 | 4.1e-09 | 5.500E-09 | 2.512E-09 | 4.000E-09 | 8.400E-09 |
| 23760 | 1188 | A1188V | D | 3.7e-09 | 2.6e-08 | 5.9e-09 | 1.187E-08 | 1.229E-08 | 3.700E-09 | 2.600E-08 |
| 23761 | 1189 | S1189A | D | 7.3e-09 | 2.3e-09 | 1.1e-08 | 6.867E-09 | 4.366E-09 | 2.300E-09 | 1.100E-08 |
| 23762 | 1189 | S1189R | D | 7.1e-09 | 1.5e-08 | 1.1e-08 | 1.103E-08 | 3.950E-09 | 7.100E-09 | 1.500E-08 |
| 23763 | 1189 | S1189N | D | 6.9e-09 | 3.1e-09 | 1.7e-08 | 9.000E-09 | 7.184E-09 | 3.100E-09 | 1.700E-08 |
| 23764 | 1189 | S1189D | D | 4.6e-09 | 3.1e-09 | 1.1e-08 | 6.233E-09 | 4.196E-09 | 3.100E-09 | 1.100E-08 |
| 23765 | 1189 | S1189C | D | 2.2e-08 | 1.4e-08 | 9e-09 | 1.500E-08 | 6.557E-09 | 9.000E-09 | 2.200E-08 |
| 23766 | 1189 | S1189E | D | 1.3e-08 | 2.5e-08 | 9.4e-09 | 1.580E-08 | 8.168E-09 | 9.400E-09 | 2.500E-08 |
| 23767 | 1189 | S1189Q | D | 7.8e-09 | 9.7e-09 | 2.7e-09 | 6.733E-09 | 3.620E-09 | 2.700E-09 | 9.700E-09 |
| 23768 | 1189 | S1189G | D | 1.1e-08 | 6.4e-09 | 2.5e-08 | 1.413E-08 | 9.688E-09 | 6.400E-09 | 2.500E-08 |
| 23769 | 1189 | S1189H | D | 1.6e-08 | 7.3e-09 | 4e-09 | 9.100E-09 | 6.199E-09 | 4.000E-09 | 1.600E-08 |
| 23770 | 1189 | S1189I | D | 5.8e-09 | 4.8e-09 | 6.9e-09 | 5.833E-09 | 1.050E-09 | 4.800E-09 | 6.900E-09 |
| 23771 | 1189 | S1189L | D | 5.8e-09 | 2.6e-08 | 6.4e-09 | 1.273E-08 | 1.149E-08 | 5.800E-09 | 2.600E-08 |
| 23772 | 1189 | S1189K | D | 4.6e-09 | 2e-08 | 1.7e-08 | 1.387E-08 | 8.164E-09 | 4.600E-09 | 2.000E-08 |
| 23773 | 1189 | S1189M | D | 1.5e-09 | 7.1e-09 | 2.1e-08 | 9.867E-09 | 1.004E-08 | 1.500E-09 | 2.100E-08 |
| 23774 | 1189 | S1189F | D | 6.2e-09 | 2e-09 | 2.5e-08 | 1.107E-08 | 1.225E-08 | 2.000E-09 | 2.500E-08 |
| 23775 | 1189 | S1189P | D | 1.2e-08 | 1.4e-08 | 2e-08 | 1.533E-08 | 4.163E-09 | 1.200E-08 | 2.000E-08 |
| 23776 | 1189 | S1189S | D | 4e-09 | 9.8e-09 | 8.4e-09 | 7.400E-09 | 3.027E-09 | 4.000E-09 | 9.800E-09 |
| 23777 | 1189 | S1189T | D | 3e-08 | 2.4e-09 | 8.2e-09 | 1.353E-08 | 1.455E-08 | 2.400E-09 | 3.000E-08 |
| 23778 | 1189 | S1189W | D | 1.1e-08 | 4.1e-09 | 9e-09 | 8.033E-09 | 3.550E-09 | 4.100E-09 | 1.100E-08 |
| 23779 | 1189 | S1189Y | D | 4.2e-09 | 8.9e-09 | 3.2e-09 | 5.433E-09 | 3.044E-09 | 3.200E-09 | 8.900E-09 |
| 23780 | 1189 | S1189V | D | 3.3e-09 | 7.2e-09 | 6.9e-09 | 5.800E-09 | 2.170E-09 | 3.300E-09 | 7.200E-09 |
| 23781 | 1190 | V1190A | D | 7.6e-09 | 6.4e-09 | 1.1e-08 | 8.333E-09 | 2.386E-09 | 6.400E-09 | 1.100E-08 |
| 23782 | 1190 | V1190R | D | 1.3e-08 | 1.1e-08 | 5.1e-09 | 9.700E-09 | 4.107E-09 | 5.100E-09 | 1.300E-08 |
| 23783 | 1190 | V1190N | D | 4.8e-09 | 1.1e-08 | 7.5e-09 | 7.767E-09 | 3.109E-09 | 4.800E-09 | 1.100E-08 |
| 23784 | 1190 | V1190D | D | 1.8e-08 | 6e-09 | 9.4e-09 | 1.113E-08 | 6.185E-09 | 6.000E-09 | 1.800E-08 |
| 23785 | 1190 | V1190C | D | 6.6e-09 | 1e-08 | 3.9e-09 | 6.833E-09 | 3.057E-09 | 3.900E-09 | 1.000E-08 |
| 23786 | 1190 | V1190E | D | 5.6e-09 | 6.3e-09 | 6.8e-09 | 6.233E-09 | 6.028E-10 | 5.600E-09 | 6.800E-09 |
| 23787 | 1190 | V1190Q | D | 2.6e-09 | 8.8e-09 | 8.3e-09 | 6.567E-09 | 3.444E-09 | 2.600E-09 | 8.800E-09 |
| 23788 | 1190 | V1190G | D | 8.5e-09 | 2.2e-08 | 4.8e-09 | 1.177E-08 | 9.053E-09 | 4.800E-09 | 2.200E-08 |
| 23789 | 1190 | V1190H | D | 1.1e-08 | 7e-09 | 1.1e-08 | 9.667E-09 | 2.309E-09 | 7.000E-09 | 1.100E-08 |
| 23790 | 1190 | V1190I | D | 5.9e-09 | 1.2e-08 | 4.9e-09 | 7.600E-09 | 3.843E-09 | 4.900E-09 | 1.200E-08 |
| 23791 | 1190 | V1190L | D | 1.6e-08 | 1.4e-08 | 9.8e-09 | 1.327E-08 | 3.164E-09 | 9.800E-09 | 1.600E-08 |
| 23792 | 1190 | V1190K | D | 6.2e-09 | 6.5e-09 | 7.4e-09 | 6.700E-09 | 6.245E-10 | 6.200E-09 | 7.400E-09 |
| 23793 | 1190 | V1190M | D | 5.6e-09 | 5.9e-09 | 1.7e-08 | 9.500E-09 | 6.497E-09 | 5.600E-09 | 1.700E-08 |
| 23794 | 1190 | V1190F | D | 7.5e-09 | 6.3e-09 | 3.1e-09 | 5.633E-09 | 2.274E-09 | 3.100E-09 | 7.500E-09 |
| 23795 | 1190 | V1190P | D | 7.6e-09 | 5.4e-09 | 1.2e-08 | 8.333E-09 | 3.361E-09 | 5.400E-09 | 1.200E-08 |
| 23796 | 1190 | V1190S | D | 1.9e-09 | 1.3e-08 | 3.1e-09 | 6.000E-09 | 6.092E-09 | 1.900E-09 | 1.300E-08 |
| 23797 | 1190 | V1190T | D | 9.2e-09 | 7.7e-09 | 6e-09 | 7.633E-09 | 1.601E-09 | 6.000E-09 | 9.200E-09 |
| 23798 | 1190 | V1190W | D | 4.4e-09 | 1.1e-08 | 9.3e-09 | 8.233E-09 | 3.427E-09 | 4.400E-09 | 1.100E-08 |
| 23799 | 1190 | V1190Y | D | 6.5e-09 | 1.7e-08 | 7.6e-09 | 1.037E-08 | 5.771E-09 | 6.500E-09 | 1.700E-08 |

| | | | | | | | | | | |
|---|---|---|---|---|---|---|---|---|---|---|
| 23800 | 1190 | V1190V | D | 5.4e-09 | 2.5e-09 | 1.4e-08 | 7.300E-09 | 5.981E-09 | 2.500E-09 | 1.400E-08 |
| 23801 | 1191 | V1191A | D | 8.9e-09 | 2.7e-09 | 8.4e-09 | 6.667E-09 | 3.444E-09 | 2.700E-09 | 8.900E-09 |
| 23802 | 1191 | V1191R | D | 9.3e-09 | 2.9e-08 | 9.9e-09 | 1.607E-08 | 1.120E-08 | 9.300E-09 | 2.900E-08 |
| 23803 | 1191 | V1191N | D | 7.2e-09 | 8.5e-09 | 7.2e-09 | 7.633E-09 | 7.506E-10 | 7.200E-09 | 8.500E-09 |
| 23804 | 1191 | V1191D | D | 5.1e-09 | 1e-08 | 9.4e-09 | 8.167E-09 | 2.673E-09 | 5.100E-09 | 1.000E-08 |
| 23805 | 1191 | V1191C | D | 6.8e-09 | 6.8e-09 | 1.7e-08 | 1.020E-08 | 5.889E-09 | 6.800E-09 | 1.700E-08 |
| 23806 | 1191 | V1191E | D | 4e-09 | 8e-09 | 5.2e-09 | 5.733E-09 | 2.053E-09 | 4.000E-09 | 8.000E-09 |
| 23807 | 1191 | V1191Q | D | 1.8e-08 | 1e-08 | 9.3e-09 | 1.243E-08 | 4.834E-09 | 9.300E-09 | 1.800E-08 |
| 23808 | 1191 | V1191G | D | 2.2e-08 | 9.4e-09 | 7.1e-09 | 1.283E-08 | 8.021E-09 | 7.100E-09 | 2.200E-08 |
| 23809 | 1191 | V1191H | D | 6e-09 | 3.6e-09 | 7.3e-09 | 5.633E-09 | 1.877E-09 | 3.600E-09 | 7.300E-09 |
| 23810 | 1191 | V1191I | D | 1.5e-08 | 5e-09 | 3.2e-09 | 7.733E-09 | 6.357E-09 | 3.200E-09 | 1.500E-08 |
| 23811 | 1191 | V1191L | D | 4e-09 | 7.6e-09 | 3.9e-09 | 5.167E-09 | 2.108E-09 | 3.900E-09 | 7.600E-09 |
| 23812 | 1191 | V1191K | D | 2.4e-08 | 7.1e-09 | 3.3e-09 | 1.147E-08 | 1.102E-08 | 3.300E-09 | 2.400E-08 |
| 23813 | 1191 | V1191M | D | 8.4e-09 | 1.4e-09 | 4.3e-09 | 4.700E-09 | 3.517E-09 | 1.400E-09 | 8.400E-09 |
| 23814 | 1191 | V1191F | D | 7.8e-09 | 4.1e-09 | 1.2e-08 | 7.967E-09 | 3.953E-09 | 4.100E-09 | 1.200E-08 |
| 23815 | 1191 | V1191P | D | 1e-08 | 7.5e-09 | 2.6e-09 | 6.700E-09 | 3.764E-09 | 2.600E-09 | 1.000E-08 |
| 23816 | 1191 | V1191S | D | 2.7e-09 | 8.7e-09 | 7.7e-09 | 6.367E-09 | 3.215E-09 | 2.700E-09 | 8.700E-09 |
| 23817 | 1191 | V1191T | D | 3.8e-08 | 5e-09 | 3.8e-09 | 1.560E-08 | 1.941E-08 | 3.800E-09 | 3.800E-08 |
| 23818 | 1191 | V1191W | D | 5.1e-09 | 1e-08 | 4.5e-09 | 6.533E-09 | 3.017E-09 | 4.500E-09 | 1.000E-08 |
| 23819 | 1191 | V1191Y | D | 3.8e-09 | 1e-08 | 6.2e-09 | 6.667E-09 | 3.126E-09 | 3.800E-09 | 1.000E-08 |
| 23820 | 1191 | V1191V | D | 3.6e-09 | 2.9e-09 | 1.1e-08 | 5.833E-09 | 4.488E-09 | 2.900E-09 | 1.100E-08 |
| 23821 | 1192 | C1192A | D | 5.7e-09 | 3.2e-09 | 1.2e-08 | 6.967E-09 | 4.535E-09 | 3.200E-09 | 1.200E-08 |
| 23822 | 1192 | C1192R | D | 1.5e-08 | 5.4e-09 | 7.5e-09 | 9.300E-09 | 5.047E-09 | 5.400E-09 | 1.500E-08 |
| 23823 | 1192 | C1192N | D | 4.2e-09 | 3.1e-09 | 1.4e-08 | 7.100E-09 | 6.001E-09 | 3.100E-09 | 1.400E-08 |
| 23824 | 1192 | C1192D | D | 2.1e-09 | 5.5e-09 | 1.2e-08 | 6.533E-09 | 5.030E-09 | 2.100E-09 | 1.200E-08 |
| 23825 | 1192 | C1192C | D | 4.8e-09 | 7.7e-09 | 1.4e-08 | 8.833E-09 | 4.704E-09 | 4.800E-09 | 1.400E-08 |
| 23826 | 1192 | C1192E | D | 5.2e-09 | 1.8e-09 | 4.4e-09 | 3.800E-09 | 1.778E-09 | 1.800E-09 | 5.200E-09 |
| 23827 | 1192 | C1192Q | D | 6e-09 | 6.1e-09 | 4.3e-09 | 5.467E-09 | 1.012E-09 | 4.300E-09 | 6.100E-09 |
| 23828 | 1192 | C1192G | D | 8.2e-09 | 6.8e-09 | 9.2e-09 | 8.067E-09 | 1.206E-09 | 6.800E-09 | 9.200E-09 |
| 23829 | 1192 | C1192H | D | 2.2e-08 | 9.1e-09 | 3.8e-09 | 1.163E-08 | 9.361E-09 | 3.800E-09 | 2.200E-08 |
| 23830 | 1192 | C1192I | D | 7.6e-09 | 9.2e-09 | 1.5e-08 | 1.060E-08 | 3.894E-09 | 7.600E-09 | 1.500E-08 |
| 23831 | 1192 | C1192L | D | 6.3e-09 | 6.7e-09 | 2.5e-09 | 5.167E-09 | 2.318E-09 | 2.500E-09 | 6.700E-09 |
| 23832 | 1192 | C1192K | D | 3.3e-09 | 1.2e-08 | 3.2e-09 | 6.167E-09 | 5.052E-09 | 3.200E-09 | 1.200E-08 |
| 23833 | 1192 | C1192M | D | 3.8e-09 | 1.2e-08 | 2.3e-08 | 1.293E-08 | 9.634E-09 | 3.800E-09 | 2.300E-08 |
| 23834 | 1192 | C1192F | D | 1.1e-08 | 5.3e-09 | 5e-09 | 7.100E-09 | 3.381E-09 | 5.000E-09 | 1.100E-08 |
| 23835 | 1192 | C1192P | D | 8.1e-09 | 1.2e-08 | 6.3e-09 | 8.800E-09 | 2.914E-09 | 6.300E-09 | 1.200E-08 |
| 23836 | 1192 | C1192S | D | 1e-08 | 2.5e-08 | 2.7e-08 | 2.067E-08 | 9.292E-09 | 1.000E-08 | 2.700E-08 |
| 23837 | 1192 | C1192T | D | 8.4e-09 | 1.2e-08 | 5.5e-09 | 8.633E-09 | 3.256E-09 | 5.500E-09 | 1.200E-08 |
| 23838 | 1192 | C1192W | D | 1.2e-08 | 7.8e-09 | 3.8e-09 | 7.867E-09 | 4.100E-09 | 3.800E-09 | 1.200E-08 |
| 23839 | 1192 | C1192Y | D | 4.9e-09 | 1.3e-08 | 1.4e-08 | 1.063E-08 | 4.990E-09 | 4.900E-09 | 1.400E-08 |
| 23840 | 1192 | C1192V | D | 6.7e-09 | 5.1e-09 | 5.5e-09 | 5.767E-09 | 8.327E-10 | 5.100E-09 | 6.700E-09 |
| 23841 | 1193 | L1193A | D | 1.4e-08 | 7.2e-09 | 1.2e-08 | 1.107E-08 | 3.495E-09 | 7.200E-09 | 1.400E-08 |
| 23842 | 1193 | L1193R | D | 6.6e-09 | 4.4e-09 | 1.4e-08 | 8.333E-09 | 5.029E-09 | 4.400E-09 | 1.400E-08 |
| 23843 | 1193 | L1193N | D | 1e-08 | 1.6e-08 | 4.6e-09 | 1.020E-08 | 5.703E-09 | 4.600E-09 | 1.600E-08 |
| 23844 | 1193 | L1193D | D | 4.1e-09 | 1e-08 | 1.2e-08 | 8.700E-09 | 4.107E-09 | 4.100E-09 | 1.200E-08 |
| 23845 | 1193 | L1193C | D | 3.5e-09 | 3.2e-09 | 5.1e-09 | 3.933E-09 | 1.021E-09 | 3.200E-09 | 5.100E-09 |
| 23846 | 1193 | L1193E | D | 4.7e-09 | 2e-08 | 1.9e-09 | 8.867E-09 | 9.743E-09 | 1.900E-09 | 2.000E-08 |
| 23847 | 1193 | L1193Q | D | 2.4e-09 | 1.4e-08 | 9.5e-09 | 8.633E-09 | 5.848E-09 | 2.400E-09 | 1.400E-08 |
| 23848 | 1193 | L1193G | D | 1.4e-08 | 1.9e-08 | 9.7e-09 | 1.423E-08 | 4.654E-09 | 9.700E-09 | 1.900E-08 |
| 23849 | 1193 | L1193H | D | 4.5e-09 | 1.3e-08 | 3.7e-09 | 7.067E-09 | 5.154E-09 | 3.700E-09 | 1.300E-08 |
| 23850 | 1193 | L1193I | D | 9.8e-09 | 6.3e-09 | 6.7e-09 | 7.600E-09 | 1.916E-09 | 6.300E-09 | 9.800E-09 |
| 23851 | 1193 | L1193L | D | 6.8e-09 | 4.8e-09 | 4.5e-09 | 5.367E-09 | 1.250E-09 | 4.500E-09 | 6.800E-09 |
| 23852 | 1193 | L1193K | D | 4.1e-09 | 3.3e-09 | 1.1e-08 | 6.133E-09 | 4.234E-09 | 3.300E-09 | 1.100E-08 |
| 23853 | 1193 | L1193M | D | 1.5e-08 | 1.1e-08 | 4e-09 | 1.000E-08 | 5.568E-09 | 4.000E-09 | 1.500E-08 |
| 23854 | 1193 | L1193F | D | 7.9e-09 | 7.6e-09 | 1e-08 | 8.500E-09 | 1.308E-09 | 7.600E-09 | 1.000E-08 |
| 23855 | 1193 | L1193P | D | 1.3e-08 | 6.3e-09 | 3.1e-08 | 1.677E-08 | 1.277E-08 | 6.300E-09 | 3.100E-08 |
| 23856 | 1193 | L1193S | D | 5.7e-09 | 7.4e-09 | 7.4e-09 | 6.833E-09 | 9.815E-10 | 5.700E-09 | 7.400E-09 |
| 23857 | 1193 | L1193T | D | 5.7e-09 | 6.2e-09 | 6.9e-09 | 6.267E-09 | 6.028E-10 | 5.700E-09 | 6.900E-09 |
| 23858 | 1193 | L1193W | D | 6.1e-09 | 4.5e-09 | 1.2e-08 | 7.533E-09 | 3.950E-09 | 4.500E-09 | 1.200E-08 |
| 23859 | 1193 | L1193Y | D | 1e-08 | 5.6e-09 | 1e-08 | 8.533E-09 | 2.540E-09 | 5.600E-09 | 1.000E-08 |
| 23860 | 1193 | L1193V | D | 6.2e-09 | 5.6e-09 | 5.3e-09 | 5.700E-09 | 4.583E-10 | 5.300E-09 | 6.200E-09 |
| 23861 | 1194 | L1194A | D | 1.1e-08 | 5.3e-09 | 2.9e-09 | 6.400E-09 | 4.161E-09 | 2.900E-09 | 1.100E-08 |
| 23862 | 1194 | L1194R | D | 2.3e-08 | 1.4e-08 | 1.8e-08 | 1.833E-08 | 4.509E-09 | 1.400E-08 | 2.300E-08 |
| 23863 | 1194 | L1194N | D | 6.3e-09 | 3e-09 | 2.8e-09 | 4.033E-09 | 1.966E-09 | 2.800E-09 | 6.300E-09 |
| 23864 | 1194 | L1194D | D | 8e-09 | 7.7e-09 | 1.2e-08 | 9.233E-09 | 2.401E-09 | 7.700E-09 | 1.200E-08 |
| 23865 | 1194 | L1194C | D | 4e-09 | 5e-09 | 3.3e-09 | 4.100E-09 | 8.544E-10 | 3.300E-09 | 5.000E-09 |
| 23866 | 1194 | L1194E | D | 7e-09 | 5.1e-09 | 8e-09 | 6.700E-09 | 1.473E-09 | 5.100E-09 | 8.000E-09 |
| 23867 | 1194 | L1194Q | D | 5.7e-09 | 6.8e-09 | 7.1e-09 | 6.533E-09 | 7.371E-10 | 5.700E-09 | 7.100E-09 |
| 23868 | 1194 | L1194G | D | 5.5e-09 | 2e-08 | 1.7e-08 | 1.417E-08 | 7.654E-09 | 5.500E-09 | 2.000E-08 |
| 23869 | 1194 | L1194H | D | 1.8e-08 | 1.6e-08 | 1.3e-08 | 1.567E-08 | 2.517E-09 | 1.300E-08 | 1.800E-08 |
| 23870 | 1194 | L1194I | D | 3.8e-09 | 1.4e-08 | 1e-08 | 9.267E-09 | 5.139E-09 | 3.800E-09 | 1.400E-08 |
| 23871 | 1194 | L1194L | D | 8.1e-09 | 1e-08 | 4.3e-09 | 7.467E-09 | 2.902E-09 | 4.300E-09 | 1.000E-08 |
| 23872 | 1194 | L1194K | D | 9.2e-09 | 7.2e-09 | 5.2e-09 | 7.200E-09 | 2.000E-09 | 5.200E-09 | 9.200E-09 |
| 23873 | 1194 | L1194M | D | 1.2e-08 | 9.8e-09 | 2.4e-09 | 8.067E-09 | 5.029E-09 | 2.400E-09 | 1.200E-08 |
| 23874 | 1194 | L1194F | D | 1.5e-08 | 4.4e-09 | 1e-08 | 9.800E-09 | 5.303E-09 | 4.400E-09 | 1.500E-08 |
| 23875 | 1194 | L1194P | D | 9e-09 | 8.5e-09 | 8.2e-09 | 8.567E-09 | 4.041E-10 | 8.200E-09 | 9.000E-09 |
| 23876 | 1194 | L1194S | D | 5e-09 | 3.6e-09 | 3.1e-08 | 1.320E-08 | 1.543E-08 | 3.600E-09 | 3.100E-08 |
| 23877 | 1194 | L1194T | D | 4.1e-09 | 4.7e-09 | 2e-08 | 9.600E-09 | 9.012E-09 | 4.100E-09 | 2.000E-08 |
| 23878 | 1194 | L1194W | D | 1.2e-08 | 8.7e-09 | 6.4e-09 | 9.033E-09 | 2.815E-09 | 6.400E-09 | 1.200E-08 |
| 23879 | 1194 | L1194Y | D | 1.1e-08 | 1.2e-08 | 6.9e-09 | 9.967E-09 | 2.702E-09 | 6.900E-09 | 1.200E-08 |
| 23880 | 1194 | L1194V | D | 9.5e-09 | 1.6e-08 | 1.1e-08 | 1.217E-08 | 3.403E-09 | 9.500E-09 | 1.600E-08 |
| 23881 | 1195 | N1195A | D | 4.2e-09 | 7.2e-09 | 1.2e-08 | 7.800E-09 | 3.934E-09 | 4.200E-09 | 1.200E-08 |
| 23882 | 1195 | N1195R | D | 6.2e-09 | 4e-09 | 9.6e-09 | 6.600E-09 | 2.821E-09 | 4.000E-09 | 9.600E-09 |
| 23883 | 1195 | N1195N | D | 1.2e-08 | 1.7e-08 | 2.9e-09 | 1.063E-08 | 7.149E-09 | 2.900E-09 | 1.700E-08 |
| 23884 | 1195 | N1195D | D | 5.1e-09 | 1.3e-08 | 6.3e-09 | 8.133E-09 | 4.257E-09 | 5.100E-09 | 1.300E-08 |

| | | | | | | | | | | |
|---|---|---|---|---|---|---|---|---|---|---|
| 23885 | 1195 | N1195C | D | 1.4e-08 | 7.3e-09 | 6.4e-09 | 9.233E-09 | 4.153E-09 | 6.400E-09 | 1.400E-08 |
| 23886 | 1195 | N1195E | D | 6.7e-09 | 5.1e-09 | 1e-08 | 7.267E-09 | 2.499E-09 | 5.100E-09 | 1.000E-08 |
| 23887 | 1195 | N1195Q | D | 7.8e-09 | 8.2e-09 | 5.4e-09 | 7.133E-09 | 1.514E-09 | 5.400E-09 | 8.200E-09 |
| 23888 | 1195 | N1195G | D | 2.7e-09 | 5.5e-09 | 7e-09 | 5.067E-09 | 2.183E-09 | 2.700E-09 | 7.000E-09 |
| 23889 | 1195 | N1195H | D | 4.5e-09 | 1.6e-09 | 9.4e-09 | 5.167E-09 | 3.943E-09 | 1.600E-09 | 9.400E-09 |
| 23890 | 1195 | N1195I | D | 3.8e-09 | 8.7e-09 | 6.7e-09 | 6.400E-09 | 2.464E-09 | 3.800E-09 | 8.700E-09 |
| 23891 | 1195 | N1195L | D | 1.2e-08 | 1.5e-08 | 1.6e-08 | 1.433E-08 | 2.082E-09 | 1.200E-08 | 1.600E-08 |
| 23892 | 1195 | N1195K | D | 6e-09 | 1.9e-09 | 1.9e-08 | 8.967E-09 | 8.928E-09 | 1.900E-09 | 1.900E-08 |
| 23893 | 1195 | N1195M | D | 5.1e-09 | 4e-09 | 1.3e-08 | 7.367E-09 | 4.910E-09 | 4.000E-09 | 1.300E-08 |
| 23894 | 1195 | N1195F | D | 8.8e-09 | 2.2e-08 | 1.6e-08 | 1.560E-08 | 6.609E-09 | 8.800E-09 | 2.200E-08 |
| 23895 | 1195 | N1195P | D | 2.1e-09 | 3.1e-09 | 6e-09 | 3.733E-09 | 2.026E-09 | 2.100E-09 | 6.000E-09 |
| 23896 | 1195 | N1195S | D | 5.7e-09 | 1.1e-08 | 8.6e-09 | 8.433E-09 | 2.654E-09 | 5.700E-09 | 1.100E-08 |
| 23897 | 1195 | N1195T | D | 5.7e-09 | 6.6e-09 | 1.3e-08 | 8.433E-09 | 3.980E-09 | 5.700E-09 | 1.300E-08 |
| 23898 | 1195 | N1195W | D | 5.4e-09 | 6.7e-09 | 1.5e-08 | 9.033E-09 | 5.208E-09 | 5.400E-09 | 1.500E-08 |
| 23899 | 1195 | N1195Y | D | 4.1e-09 | 1.1e-08 | 1.5e-08 | 1.003E-08 | 5.514E-09 | 4.100E-09 | 1.500E-08 |
| 23900 | 1195 | N1195V | D | 4.3e-09 | 3e-09 | 6e-09 | 4.433E-09 | 1.504E-09 | 3.000E-09 | 6.000E-09 |
| 23901 | 1196 | N1196A | D | 1.2e-08 | 7.5e-09 | 2.9e-09 | 7.467E-09 | 4.550E-09 | 2.900E-09 | 1.200E-08 |
| 23902 | 1196 | N1196R | D | 1.6e-08 | 3.4e-09 | 5.9e-09 | 8.433E-09 | 6.671E-09 | 3.400E-09 | 1.600E-08 |
| 23903 | 1196 | N1196N | D | 1.1e-08 | 9.7e-09 | 6.4e-09 | 9.033E-09 | 2.371E-09 | 6.400E-09 | 1.100E-08 |
| 23904 | 1196 | N1196D | D | 5.7e-09 | 1.8e-09 | 1.1e-08 | 6.167E-09 | 4.618E-09 | 1.800E-09 | 1.100E-08 |
| 23905 | 1196 | N1196C | D | 7e-09 | 6.3e-09 | 4.4e-09 | 5.900E-09 | 1.345E-09 | 4.400E-09 | 7.000E-09 |
| 23906 | 1196 | N1196E | D | 4.8e-09 | 3.2e-09 | 8e-09 | 5.333E-09 | 2.444E-09 | 3.200E-09 | 8.000E-09 |
| 23907 | 1196 | N1196Q | D | 1.4e-08 | 4.1e-09 | 6.1e-09 | 8.067E-09 | 5.235E-09 | 4.100E-09 | 1.400E-08 |
| 23908 | 1196 | N1196G | D | 5.8e-09 | 1e-08 | 4.9e-09 | 6.900E-09 | 2.722E-09 | 4.900E-09 | 1.000E-08 |
| 23909 | 1196 | N1196H | D | 1.5e-08 | 6.4e-09 | 2.7e-09 | 8.033E-09 | 6.311E-09 | 2.700E-09 | 1.500E-08 |
| 23910 | 1196 | N1196I | D | 7.7e-09 | 6.4e-09 | 3.7e-09 | 5.933E-09 | 2.040E-09 | 3.700E-09 | 7.700E-09 |
| 23911 | 1196 | N1196L | D | 6.9e-09 | 1.1e-08 | 8.2e-09 | 8.700E-09 | 2.095E-09 | 6.900E-09 | 1.100E-08 |
| 23912 | 1196 | N1196K | D | 4.8e-09 | 1.1e-08 | 9e-09 | 8.267E-09 | 3.164E-09 | 4.800E-09 | 1.100E-08 |
| 23913 | 1196 | N1196M | D | 6.9e-09 | 4.1e-09 | 7.3e-09 | 6.100E-09 | 1.744E-09 | 4.100E-09 | 7.300E-09 |
| 23914 | 1196 | N1196F | D | 4.3e-09 | 1.3e-08 | 1.4e-08 | 1.043E-08 | 5.335E-09 | 4.300E-09 | 1.400E-08 |
| 23915 | 1196 | N1196P | D | 8.8e-09 | 5.2e-09 | 7.2e-09 | 7.067E-09 | 1.804E-09 | 5.200E-09 | 8.800E-09 |
| 23916 | 1196 | N1196S | D | 9.5e-09 | 5e-09 | 8.6e-09 | 7.700E-09 | 2.381E-09 | 5.000E-09 | 9.500E-09 |
| 23917 | 1196 | N1196T | D | 3.7e-09 | 3.9e-09 | 1.2e-08 | 6.533E-09 | 4.735E-09 | 3.700E-09 | 1.200E-08 |
| 23918 | 1196 | N1196W | D | 2.1e-08 | 1.2e-08 | 4.2e-09 | 1.240E-08 | 8.407E-09 | 4.200E-09 | 2.100E-08 |
| 23919 | 1196 | N1196Y | D | 3.2e-09 | 3.7e-09 | 4.4e-09 | 3.767E-09 | 6.028E-10 | 3.200E-09 | 4.400E-09 |
| 23920 | 1196 | N1196V | D | 7e-09 | 7.3e-09 | 3.2e-09 | 5.833E-09 | 2.285E-09 | 3.200E-09 | 7.300E-09 |
| 23921 | 1197 | F1197A | D | 7.9e-09 | 6.4e-09 | 3.2e-09 | 5.833E-09 | 2.401E-09 | 3.200E-09 | 7.900E-09 |
| 23922 | 1197 | F1197R | D | 1.3e-08 | 2e-08 | 1.3e-08 | 1.533E-08 | 4.041E-09 | 1.300E-08 | 2.000E-08 |
| 23923 | 1197 | F1197N | D | 4.7e-09 | 2.4e-08 | 9.2e-09 | 1.263E-08 | 1.010E-08 | 4.700E-09 | 2.400E-08 |
| 23924 | 1197 | F1197D | D | 6.6e-09 | 6e-09 | 5e-09 | 5.867E-09 | 8.083E-10 | 5.000E-09 | 6.600E-09 |
| 23925 | 1197 | F1197C | D | 1.6e-09 | 4.6e-09 | 7.4e-09 | 4.533E-09 | 2.901E-09 | 1.600E-09 | 7.400E-09 |
| 23926 | 1197 | F1197E | D | 6.6e-09 | 7.1e-09 | 4.7e-09 | 6.133E-09 | 1.266E-09 | 4.700E-09 | 7.100E-09 |
| 23927 | 1197 | F1197Q | D | 4.6e-09 | 1e-08 | 2.4e-09 | 5.667E-09 | 3.911E-09 | 2.400E-09 | 1.000E-08 |
| 23928 | 1197 | F1197G | D | 7.9e-09 | 2e-09 | 8.4e-09 | 6.100E-09 | 3.559E-09 | 2.000E-09 | 8.400E-09 |
| 23929 | 1197 | F1197H | D | 5.4e-09 | 6.4e-09 | 6.5e-09 | 6.100E-09 | 6.083E-10 | 5.400E-09 | 6.500E-09 |
| 23930 | 1197 | F1197I | D | 8e-09 | 2.5e-08 | 5.7e-09 | 1.290E-08 | 1.054E-08 | 5.700E-09 | 2.500E-08 |
| 23931 | 1197 | F1197L | D | 8.4e-09 | 3.5e-09 | 8e-09 | 6.633E-09 | 2.721E-09 | 3.500E-09 | 8.400E-09 |
| 23932 | 1197 | F1197K | D | 1.4e-08 | 4.1e-09 | 1.6e-08 | 1.137E-08 | 6.372E-09 | 4.100E-09 | 1.600E-08 |
| 23933 | 1197 | F1197M | D | 7.3e-09 | 8.1e-09 | 3.8e-09 | 6.400E-09 | 2.287E-09 | 3.800E-09 | 8.100E-09 |
| 23934 | 1197 | F1197F | D | 4.7e-09 | 4.3e-09 | 7.6e-09 | 5.533E-09 | 1.801E-09 | 4.300E-09 | 7.600E-09 |
| 23935 | 1197 | F1197P | D | 8.4e-09 | 6.4e-09 | 4.6e-09 | 6.467E-09 | 1.901E-09 | 4.600E-09 | 8.400E-09 |
| 23936 | 1197 | F1197S | D | 9.9e-09 | 1.4e-08 | 2.9e-09 | 8.933E-09 | 5.613E-09 | 2.900E-09 | 1.400E-08 |
| 23937 | 1197 | F1197T | D | 6e-09 | 4.2e-09 | 1.5e-08 | 8.400E-09 | 5.786E-09 | 4.200E-09 | 1.500E-08 |
| 23938 | 1197 | F1197W | D | 8.4e-09 | 8.5e-09 | 7.2e-09 | 8.033E-09 | 7.234E-10 | 7.200E-09 | 8.500E-09 |
| 23939 | 1197 | F1197Y | D | 9.7e-09 | 1.6e-08 | 4.7e-09 | 1.013E-08 | 5.662E-09 | 4.700E-09 | 1.600E-08 |
| 23940 | 1197 | F1197V | D | 4.5e-09 | 1.6e-08 | 1.3e-08 | 1.117E-08 | 5.965E-09 | 4.500E-09 | 1.600E-08 |
| 23941 | 1198 | Y1198A | D | 6e-09 | 6.6e-09 | 1.2e-08 | 8.200E-09 | 3.305E-09 | 6.000E-09 | 1.200E-08 |
| 23942 | 1198 | Y1198R | D | 8e-09 | 9.1e-09 | 3.9e-09 | 7.000E-09 | 2.740E-09 | 3.900E-09 | 9.100E-09 |
| 23943 | 1198 | Y1198N | D | 1.7e-08 | 3.3e-09 | 8.5e-09 | 9.600E-09 | 6.916E-09 | 3.300E-09 | 1.700E-08 |
| 23944 | 1198 | Y1198D | D | 5.1e-09 | 7e-09 | 2.4e-09 | 4.833E-09 | 2.312E-09 | 2.400E-09 | 7.000E-09 |
| 23945 | 1198 | Y1198C | D | 4.5e-09 | 1.8e-08 | 5.6e-09 | 9.367E-09 | 7.497E-09 | 4.500E-09 | 1.800E-08 |
| 23946 | 1198 | Y1198E | D | 5e-09 | 2.7e-09 | 4.1e-09 | 3.933E-09 | 1.159E-09 | 2.700E-09 | 5.000E-09 |
| 23947 | 1198 | Y1198Q | D | 1.9e-08 | 5.4e-09 | 1.1e-08 | 1.180E-08 | 6.835E-09 | 5.400E-09 | 1.900E-08 |
| 23948 | 1198 | Y1198G | D | 2.8e-09 | 1.2e-08 | 8.8e-09 | 7.867E-09 | 4.670E-09 | 2.800E-09 | 1.200E-08 |
| 23949 | 1198 | Y1198H | D | 7.1e-09 | 1.5e-08 | 8.7e-09 | 1.027E-08 | 4.177E-09 | 7.100E-09 | 1.500E-08 |
| 23950 | 1198 | Y1198I | D | 2.6e-09 | 1.2e-08 | 6.8e-09 | 7.133E-09 | 4.709E-09 | 2.600E-09 | 1.200E-08 |
| 23951 | 1198 | Y1198L | D | 8.5e-09 | 4.8e-09 | 1.7e-08 | 1.010E-08 | 6.255E-09 | 4.800E-09 | 1.700E-08 |
| 23952 | 1198 | Y1198K | D | 1.4e-08 | 3.4e-09 | 7.9e-09 | 8.433E-09 | 5.320E-09 | 3.400E-09 | 1.400E-08 |
| 23953 | 1198 | Y1198M | D | 6.4e-09 | 5.3e-09 | 9.1e-09 | 6.933E-09 | 1.955E-09 | 5.300E-09 | 9.100E-09 |
| 23954 | 1198 | Y1198F | D | 1.2e-08 | 2.1e-08 | 7.5e-09 | 1.350E-08 | 6.874E-09 | 7.500E-09 | 2.100E-08 |
| 23955 | 1198 | Y1198P | D | 5e-09 | 1.1e-08 | 3.7e-09 | 6.567E-09 | 3.894E-09 | 3.700E-09 | 1.100E-08 |
| 23956 | 1198 | Y1198S | D | 7.2e-09 | 4.2e-09 | 4.8e-09 | 5.400E-09 | 1.587E-09 | 4.200E-09 | 7.200E-09 |
| 23957 | 1198 | Y1198T | D | 2.1e-08 | 8.5e-09 | 5.8e-09 | 1.177E-08 | 8.109E-09 | 5.800E-09 | 2.100E-08 |
| 23958 | 1198 | Y1198W | D | 6.1e-09 | 4.7e-09 | 1.2e-08 | 7.600E-09 | 3.874E-09 | 4.700E-09 | 1.200E-08 |
| 23959 | 1198 | Y1198Y | D | 1.2e-08 | 1.2e-08 | 4.5e-09 | 9.500E-09 | 4.330E-09 | 4.500E-09 | 1.200E-08 |
| 23960 | 1198 | Y1198V | D | 1.4e-08 | 3.8e-09 | 1e-08 | 9.267E-09 | 5.139E-09 | 3.800E-09 | 1.400E-08 |
| 23961 | 1199 | P1199A | D | 7.9e-09 | 8.3e-09 | 1.5e-08 | 1.040E-08 | 3.989E-09 | 7.900E-09 | 1.500E-08 |
| 23962 | 1199 | P1199R | D | 6.2e-09 | 1.9e-08 | 4.3e-09 | 9.833E-09 | 7.995E-09 | 4.300E-09 | 1.900E-08 |
| 23963 | 1199 | P1199N | D | 2.6e-09 | 1.1e-08 | 1.5e-09 | 5.033E-09 | 5.196E-09 | 1.500E-09 | 1.100E-08 |
| 23964 | 1199 | P1199D | D | 1.1e-08 | 1.1e-08 | 3.6e-09 | 8.533E-09 | 4.272E-09 | 3.600E-09 | 1.100E-08 |
| 23965 | 1199 | P1199C | D | 1.1e-08 | 1.3e-08 | 6.8e-09 | 1.027E-08 | 3.164E-09 | 6.800E-09 | 1.300E-08 |
| 23966 | 1199 | P1199E | D | 1.7e-08 | 1.2e-08 | 4e-09 | 1.100E-08 | 6.557E-09 | 4.000E-09 | 1.700E-08 |
| 23967 | 1199 | P1199Q | D | 1.2e-08 | 1.8e-08 | 1.1e-08 | 1.367E-08 | 3.786E-09 | 1.100E-08 | 1.800E-08 |
| 23968 | 1199 | P1199G | D | 2e-08 | 9.3e-09 | 9.7e-09 | 1.300E-08 | 6.065E-09 | 9.300E-09 | 2.000E-08 |
| 23969 | 1199 | P1199H | D | 3.5e-09 | 8.4e-09 | 8.6e-09 | 6.833E-09 | 2.888E-09 | 3.500E-09 | 8.600E-09 |

| | | | | | | | | | | |
|---|---|---|---|---|---|---|---|---|---|---|
| 23970 | 1199 | P1199I | D | 3.1e-09 | 1.2e-09 | 4.4e-08 | 1.610E-08 | 2.418E-08 | 1.200E-09 | 4.400E-08 |
| 23971 | 1199 | P1199L | D | 3.7e-09 | 2.7e-08 | 1.7e-08 | 1.590E-08 | 1.169E-08 | 3.700E-09 | 2.700E-08 |
| 23972 | 1199 | P1199K | D | 6e-09 | 1.1e-08 | 6e-09 | 7.667E-09 | 2.887E-09 | 6.000E-09 | 1.100E-08 |
| 23973 | 1199 | P1199M | D | 3.3e-09 | 2.6e-08 | 9e-09 | 1.277E-08 | 1.181E-08 | 3.300E-09 | 2.600E-08 |
| 23974 | 1199 | P1199F | D | 7.7e-09 | 8.5e-09 | 1.3e-08 | 9.733E-09 | 2.857E-09 | 7.700E-09 | 1.300E-08 |
| 23975 | 1199 | P1199P | D | 1.2e-08 | 2.8e-09 | 5.9e-09 | 6.900E-09 | 4.681E-09 | 2.800E-09 | 1.200E-08 |
| 23976 | 1199 | P1199S | D | 6.9e-09 | 1.5e-08 | 3e-09 | 8.300E-09 | 6.121E-09 | 3.000E-09 | 1.500E-08 |
| 23977 | 1199 | P1199T | D | 8.3e-09 | 1e-08 | 1.3e-08 | 1.043E-08 | 2.380E-09 | 8.300E-09 | 1.300E-08 |
| 23978 | 1199 | P1199W | D | 7.9e-09 | 6e-09 | 9.5e-09 | 7.800E-09 | 1.752E-09 | 6.000E-09 | 9.500E-09 |
| 23979 | 1199 | P1199Y | D | 7.5e-09 | 8.9e-09 | 1.3e-08 | 9.800E-09 | 2.858E-09 | 7.500E-09 | 1.300E-08 |
| 23980 | 1199 | P1199V | D | 1.1e-08 | 7.1e-09 | 1.2e-08 | 1.003E-08 | 2.589E-09 | 7.100E-09 | 1.200E-08 |
| 23981 | 1200 | R1200A | D | 4.7e-09 | 4.7e-09 | 8.3e-09 | 5.900E-09 | 2.078E-09 | 4.700E-09 | 8.300E-09 |
| 23982 | 1200 | R1200R | D | 6.4e-09 | 3.2e-09 | 5.1e-09 | 4.900E-09 | 1.609E-09 | 3.200E-09 | 6.400E-09 |
| 23983 | 1200 | R1200N | D | 5.3e-09 | 8.4e-09 | 1.1e-08 | 8.233E-09 | 2.854E-09 | 5.300E-09 | 1.100E-08 |
| 23984 | 1200 | R1200D | D | 7.1e-09 | 4.4e-09 | 1.1e-08 | 7.500E-09 | 3.318E-09 | 4.400E-09 | 1.100E-08 |
| 23985 | 1200 | R1200C | D | 5.8e-09 | 6.9e-09 | 1.6e-08 | 9.567E-09 | 5.599E-09 | 5.800E-09 | 1.600E-08 |
| 23986 | 1200 | R1200E | D | 1.2e-08 | 2.2e-09 | 3.3e-09 | 5.833E-09 | 5.369E-09 | 2.200E-09 | 1.200E-08 |
| 23987 | 1200 | R1200Q | D | 3e-09 | 4.7e-09 | 4.5e-09 | 4.067E-09 | 9.292E-10 | 3.000E-09 | 4.700E-09 |
| 23988 | 1200 | R1200G | D | 1.8e-08 | 1.2e-08 | 1.9e-08 | 1.633E-08 | 3.786E-09 | 1.200E-08 | 1.900E-08 |
| 23989 | 1200 | R1200H | D | 3.9e-09 | 5.3e-09 | 3.6e-09 | 4.267E-09 | 9.074E-10 | 3.600E-09 | 5.300E-09 |
| 23990 | 1200 | R1200I | D | 6.4e-09 | 1.1e-08 | 4.4e-09 | 7.267E-09 | 3.384E-09 | 4.400E-09 | 1.100E-08 |
| 23991 | 1200 | R1200L | D | 3.1e-09 | 4.5e-09 | 2e-08 | 9.200E-09 | 9.379E-09 | 3.100E-09 | 2.000E-08 |
| 23992 | 1200 | R1200K | D | 3.1e-09 | 4.1e-09 | 1.1e-08 | 6.067E-09 | 4.302E-09 | 3.100E-09 | 1.100E-08 |
| 23993 | 1200 | R1200M | D | 3.9e-09 | 4e-09 | 4.4e-09 | 4.100E-09 | 2.646E-10 | 3.900E-09 | 4.400E-09 |
| 23994 | 1200 | R1200F | D | 4.9e-09 | 1.2e-08 | 1e-08 | 8.967E-09 | 3.661E-09 | 4.900E-09 | 1.200E-08 |
| 23995 | 1200 | R1200P | D | 5.4e-09 | 9.8e-09 | 5.9e-09 | 7.033E-09 | 2.409E-09 | 5.400E-09 | 9.800E-09 |
| 23996 | 1200 | R1200S | D | 5.8e-09 | 5.6e-09 | 8.1e-09 | 6.500E-09 | 1.389E-09 | 5.600E-09 | 8.100E-09 |
| 23997 | 1200 | R1200T | D | 5.2e-09 | 4.1e-09 | 5.6e-09 | 4.967E-09 | 7.767E-10 | 4.100E-09 | 5.600E-09 |
| 23998 | 1200 | R1200W | D | 1.3e-08 | 4.6e-09 | 4e-09 | 7.200E-09 | 5.032E-09 | 4.000E-09 | 1.300E-08 |
| 23999 | 1200 | R1200Y | D | 1.7e-08 | 4.1e-09 | 5.4e-09 | 8.833E-09 | 7.102E-09 | 4.100E-09 | 1.700E-08 |
| 24000 | 1200 | R1200V | D | 1.5e-08 | 4.7e-09 | 8.9e-09 | 9.533E-09 | 5.179E-09 | 4.700E-09 | 1.500E-08 |
| 24001 | 1201 | E1201A | D | 4.3e-09 | 4e-09 | 2.6e-08 | 1.143E-08 | 1.262E-08 | 4.000E-09 | 2.600E-08 |
| 24002 | 1201 | E1201R | D | 1.4e-08 | 5.2e-09 | 6.8e-09 | 8.667E-09 | 4.688E-09 | 5.200E-09 | 1.400E-08 |
| 24003 | 1201 | E1201N | D | 7e-09 | 1.5e-08 | 4.9e-09 | 8.967E-09 | 5.329E-09 | 4.900E-09 | 1.500E-08 |
| 24004 | 1201 | E1201D | D | 3.1e-09 | 7.8e-09 | 7.6e-09 | 6.167E-09 | 2.658E-09 | 3.100E-09 | 7.800E-09 |
| 24005 | 1201 | E1201C | D | 5.6e-09 | 2.8e-09 | 1.9e-08 | 9.133E-09 | 8.659E-09 | 2.800E-09 | 1.900E-08 |
| 24006 | 1201 | E1201E | D | 3.8e-09 | 1.4e-08 | 4.7e-09 | 7.500E-09 | 5.647E-09 | 3.800E-09 | 1.400E-08 |
| 24007 | 1201 | E1201Q | D | 6.9e-09 | 6.2e-09 | 7.8e-09 | 6.967E-09 | 8.021E-10 | 6.200E-09 | 7.800E-09 |
| 24008 | 1201 | E1201G | D | 8.5e-09 | 8.4e-09 | 7.9e-09 | 8.267E-09 | 3.215E-10 | 7.900E-09 | 8.500E-09 |
| 24009 | 1201 | E1201H | D | 9.3e-09 | 1.5e-08 | 7.9e-09 | 1.073E-08 | 3.761E-09 | 7.900E-09 | 1.500E-08 |
| 24010 | 1201 | E1201I | D | 8.1e-09 | 3.7e-09 | 8.7e-09 | 6.833E-09 | 2.730E-09 | 3.700E-09 | 8.700E-09 |
| 24011 | 1201 | E1201L | D | 2.2e-08 | 6.8e-09 | 8.4e-09 | 1.240E-08 | 8.352E-09 | 6.800E-09 | 2.200E-08 |
| 24012 | 1201 | E1201K | D | 9.1e-09 | 3.3e-09 | 1.2e-09 | 4.533E-09 | 4.092E-09 | 1.200E-09 | 9.100E-09 |
| 24013 | 1201 | E1201M | D | 4.6e-09 | 5.8e-09 | 6.9e-09 | 5.767E-09 | 1.150E-09 | 4.600E-09 | 6.900E-09 |
| 24014 | 1201 | E1201F | D | 3.5e-09 | 1.3e-08 | 1.1e-08 | 9.167E-09 | 5.008E-09 | 3.500E-09 | 1.300E-08 |
| 24015 | 1201 | E1201P | D | 1.7e-08 | 7.6e-09 | 6.9e-09 | 1.050E-08 | 5.640E-09 | 6.900E-09 | 1.700E-08 |
| 24016 | 1201 | E1201S | D | 3.1e-09 | 4.8e-09 | 8.5e-09 | 5.467E-09 | 2.761E-09 | 3.100E-09 | 8.500E-09 |
| 24017 | 1201 | E1201T | D | 8.5e-09 | 1.1e-08 | 2.8e-08 | 1.583E-08 | 1.061E-08 | 8.500E-09 | 2.800E-08 |
| 24018 | 1201 | E1201W | D | 2.8e-09 | 6.8e-09 | 2.5e-08 | 1.153E-08 | 1.183E-08 | 2.800E-09 | 2.500E-08 |
| 24019 | 1201 | E1201Y | D | 2.8e-08 | 4.7e-09 | 9e-09 | 1.390E-08 | 1.240E-08 | 4.700E-09 | 2.800E-08 |
| 24020 | 1201 | E1201V | D | 1.9e-08 | 5.9e-09 | 1.7e-08 | 1.397E-08 | 7.057E-09 | 5.900E-09 | 1.900E-08 |
| 24021 | 1202 | A1202A | D | 1e-08 | 1.1e-08 | 8.7e-09 | 9.900E-09 | 1.153E-09 | 8.700E-09 | 1.100E-08 |
| 24022 | 1202 | A1202R | D | 5.9e-09 | 2.1e-09 | 5.6e-09 | 4.533E-09 | 2.113E-09 | 2.100E-09 | 5.900E-09 |
| 24023 | 1202 | A1202N | D | 2.5e-09 | 3.1e-09 | 3e-09 | 2.867E-09 | 3.215E-10 | 2.500E-09 | 3.100E-09 |
| 24024 | 1202 | A1202D | D | 7e-09 | 2.6e-09 | 1e-08 | 6.533E-09 | 3.722E-09 | 2.600E-09 | 1.000E-08 |
| 24025 | 1202 | A1202C | D | 4.6e-09 | 6.3e-09 | 2.8e-09 | 4.567E-09 | 1.750E-09 | 2.800E-09 | 6.300E-09 |
| 24026 | 1202 | A1202E | D | 3.1e-09 | 6.4e-09 | 1.3e-08 | 7.500E-09 | 5.041E-09 | 3.100E-09 | 1.300E-08 |
| 24027 | 1202 | A1202Q | D | 1.1e-08 | 1.4e-08 | 1.2e-08 | 1.233E-08 | 1.528E-09 | 1.100E-08 | 1.400E-08 |
| 24028 | 1202 | A1202G | D | 8.6e-09 | 6.6e-09 | 3.4e-09 | 6.200E-09 | 2.623E-09 | 3.400E-09 | 8.600E-09 |
| 24029 | 1202 | A1202H | D | 2.7e-09 | 2.7e-09 | 7.8e-09 | 4.400E-09 | 2.944E-09 | 2.700E-09 | 7.800E-09 |
| 24030 | 1202 | A1202I | D | 3.7e-09 | 2.5e-09 | 4.5e-09 | 3.567E-09 | 1.007E-09 | 2.500E-09 | 4.500E-09 |
| 24031 | 1202 | A1202L | D | 9.9e-09 | 8.1e-09 | 4.9e-09 | 7.633E-09 | 2.532E-09 | 4.900E-09 | 9.900E-09 |
| 24032 | 1202 | A1202K | D | 3.4e-09 | 5.9e-09 | 8.2e-09 | 5.833E-09 | 2.401E-09 | 3.400E-09 | 8.200E-09 |
| 24033 | 1202 | A1202M | D | 5.8e-09 | 7.1e-09 | 2.7e-09 | 5.200E-09 | 2.261E-09 | 2.700E-09 | 7.100E-09 |
| 24034 | 1202 | A1202F | D | 5.2e-09 | 4.1e-09 | 9.5e-09 | 6.267E-09 | 2.854E-09 | 4.100E-09 | 9.500E-09 |
| 24035 | 1202 | A1202P | D | 4.5e-09 | 7.7e-09 | 4.3e-09 | 5.500E-09 | 1.908E-09 | 4.300E-09 | 7.700E-09 |
| 24036 | 1202 | A1202S | D | 5.5e-09 | 4.3e-09 | 1.9e-09 | 3.900E-09 | 1.833E-09 | 1.900E-09 | 5.500E-09 |
| 24037 | 1202 | A1202T | D | 8.4e-09 | 6.5e-09 | 4.8e-09 | 6.567E-09 | 1.801E-09 | 4.800E-09 | 8.400E-09 |
| 24038 | 1202 | A1202W | D | 3.2e-09 | 5.1e-09 | 2.4e-09 | 3.567E-09 | 1.387E-09 | 2.400E-09 | 5.100E-09 |
| 24039 | 1202 | A1202Y | D | 1.2e-08 | 6.2e-09 | 5.2e-09 | 7.800E-09 | 3.672E-09 | 5.200E-09 | 1.200E-08 |
| 24040 | 1202 | A1202V | D | 1.2e-08 | 8.5e-09 | 5.8e-09 | 8.767E-09 | 3.109E-09 | 5.800E-09 | 1.200E-08 |
| 24041 | 1203 | K1203A | D | 4.5e-08 | 4.7e-09 | 6.4e-09 | 1.870E-08 | 2.279E-08 | 4.700E-09 | 4.500E-08 |
| 24042 | 1203 | K1203R | D | 2.9e-09 | 6.4e-09 | 8e-09 | 5.767E-09 | 2.608E-09 | 2.900E-09 | 8.000E-09 |
| 24043 | 1203 | K1203N | D | 3.1e-09 | 6.4e-09 | 2.2e-09 | 3.900E-09 | 2.211E-09 | 2.200E-09 | 6.400E-09 |
| 24044 | 1203 | K1203D | D | 6.9e-09 | 2.8e-09 | 5.2e-09 | 4.967E-09 | 2.060E-09 | 2.800E-09 | 6.900E-09 |
| 24045 | 1203 | K1203C | D | 7e-09 | 5.1e-09 | 2.7e-09 | 4.933E-09 | 2.155E-09 | 2.700E-09 | 7.000E-09 |
| 24046 | 1203 | K1203E | D | 1.7e-08 | 2.6e-09 | 5.8e-09 | 8.467E-09 | 7.561E-09 | 2.600E-09 | 1.700E-08 |
| 24047 | 1203 | K1203Q | D | 5.2e-09 | 8.8e-09 | 9.5e-09 | 7.833E-09 | 2.307E-09 | 5.200E-09 | 9.500E-09 |
| 24048 | 1203 | K1203G | D | 1.2e-08 | 5.9e-09 | 3.3e-09 | 7.067E-09 | 4.466E-09 | 3.300E-09 | 1.200E-08 |
| 24049 | 1203 | K1203H | D | 4.3e-09 | 8.6e-09 | 3.5e-09 | 5.467E-09 | 2.743E-09 | 3.500E-09 | 8.600E-09 |
| 24050 | 1203 | K1203I | D | 6.9e-09 | 3.2e-09 | 2.8e-08 | 1.270E-08 | 1.338E-08 | 3.200E-09 | 2.800E-08 |
| 24051 | 1203 | K1203L | D | 1.4e-08 | 1.1e-08 | 1e-08 | 1.167E-08 | 2.082E-09 | 1.000E-08 | 1.400E-08 |
| 24052 | 1203 | K1203K | D | 1.4e-08 | 1.3e-08 | 5e-09 | 1.067E-08 | 4.933E-09 | 5.000E-09 | 1.400E-08 |
| 24053 | 1203 | K1203M | D | 2.3e-09 | 4.1e-09 | 4.3e-09 | 3.567E-09 | 1.102E-09 | 2.300E-09 | 4.300E-09 |
| 24054 | 1203 | K1203F | D | 1.1e-08 | 9.9e-09 | 1.2e-08 | 1.097E-08 | 1.050E-09 | 9.900E-09 | 1.200E-08 |

| | | | | | | | | | | |
|---|---|---|---|---|---|---|---|---|---|---|
| 24055 | 1203 | K1203P | D | 2.7e-09 | 6.7e-09 | 4.3e-09 | 4.567E-09 | 2.013E-09 | 2.700E-09 | 6.700E-09 |
| 24056 | 1203 | K1203S | D | 1.7e-08 | 5.3e-09 | 3.1e-09 | 8.467E-09 | 7.472E-09 | 3.100E-09 | 1.700E-08 |
| 24057 | 1203 | K1203T | D | 5.2e-09 | 8.9e-09 | 6.3e-09 | 6.800E-09 | 1.900E-09 | 5.200E-09 | 8.900E-09 |
| 24058 | 1203 | K1203W | D | 9.4e-09 | 7e-09 | 1.8e-08 | 1.147E-08 | 5.784E-09 | 7.000E-09 | 1.800E-08 |
| 24059 | 1203 | K1203Y | D | 8e-09 | 1.3e-08 | 6.8e-09 | 9.267E-09 | 3.288E-09 | 6.800E-09 | 1.300E-08 |
| 24060 | 1203 | K1203V | D | 7.4e-09 | 1.3e-08 | 8e-09 | 9.467E-09 | 3.075E-09 | 7.400E-09 | 1.300E-08 |
| 24061 | 1204 | V1204A | D | 3.8e-09 | 1.1e-08 | 2.2e-09 | 5.667E-09 | 4.688E-09 | 2.200E-09 | 1.100E-08 |
| 24062 | 1204 | V1204R | D | 5.9e-09 | 1.2e-08 | 4.1e-09 | 7.333E-09 | 4.140E-09 | 4.100E-09 | 1.200E-08 |
| 24063 | 1204 | V1204N | D | 1.4e-08 | 3.4e-09 | 6e-09 | 7.800E-09 | 5.524E-09 | 3.400E-09 | 1.400E-08 |
| 24064 | 1204 | V1204D | D | 1.3e-08 | 5.3e-09 | 1.6e-08 | 1.143E-08 | 5.519E-09 | 5.300E-09 | 1.600E-08 |
| 24065 | 1204 | V1204C | D | 7.9e-09 | 7.6e-09 | 6e-09 | 7.167E-09 | 1.021E-09 | 6.000E-09 | 7.900E-09 |
| 24066 | 1204 | V1204E | D | 6.1e-09 | 4.7e-09 | 8.4e-09 | 6.400E-09 | 1.868E-09 | 4.700E-09 | 8.400E-09 |
| 24067 | 1204 | V1204Q | D | 1.7e-08 | 8.7e-09 | 9.1e-09 | 1.160E-08 | 4.681E-09 | 8.700E-09 | 1.700E-08 |
| 24068 | 1204 | V1204G | D | 1.6e-08 | 7.5e-09 | 1.8e-08 | 1.383E-08 | 5.575E-09 | 7.500E-09 | 1.800E-08 |
| 24069 | 1204 | V1204H | D | 3.6e-09 | 6e-09 | 4.4e-09 | 4.667E-09 | 1.222E-09 | 3.600E-09 | 6.000E-09 |
| 24070 | 1204 | V1204I | D | 3.3e-09 | 4.5e-09 | 4.2e-09 | 4.000E-09 | 6.245E-10 | 3.300E-09 | 4.500E-09 |
| 24071 | 1204 | V1204L | D | 5.6e-09 | 8.1e-09 | 2.4e-09 | 5.367E-09 | 2.857E-09 | 2.400E-09 | 8.100E-09 |
| 24072 | 1204 | V1204K | D | 5.5e-09 | 1.8e-08 | 8.2e-09 | 1.057E-08 | 6.577E-09 | 5.500E-09 | 1.800E-08 |
| 24073 | 1204 | V1204M | D | 9.4e-09 | 1.1e-08 | 3.9e-09 | 8.100E-09 | 3.724E-09 | 3.900E-09 | 1.100E-08 |
| 24074 | 1204 | V1204F | D | 3.5e-09 | 3.4e-08 | 4.7e-09 | 1.407E-08 | 1.727E-08 | 3.500E-09 | 3.400E-08 |
| 24075 | 1204 | V1204P | D | 4.5e-09 | 7.5e-09 | 4.5e-09 | 5.500E-09 | 1.732E-09 | 4.500E-09 | 7.500E-09 |
| 24076 | 1204 | V1204S | D | 6.4e-09 | 8.3e-09 | 8.3e-09 | 7.667E-09 | 1.097E-09 | 6.400E-09 | 8.300E-09 |
| 24077 | 1204 | V1204T | D | 1.2e-08 | 1.1e-08 | 5.1e-09 | 9.367E-09 | 3.729E-09 | 5.100E-09 | 1.200E-08 |
| 24078 | 1204 | V1204W | D | 7.7e-09 | 5.9e-09 | 1.3e-08 | 8.867E-09 | 3.691E-09 | 5.900E-09 | 1.300E-08 |
| 24079 | 1204 | V1204Y | D | 4.1e-09 | 8.2e-09 | 6.5e-09 | 6.267E-09 | 2.060E-09 | 4.100E-09 | 8.200E-09 |
| 24080 | 1204 | V1204V | D | 3.1e-09 | 1.4e-08 | 4.4e-09 | 7.167E-09 | 5.953E-09 | 3.100E-09 | 1.400E-08 |
| 24081 | 1205 | Q1205A | D | 3.7e-09 | 2.5e-08 | 1.2e-08 | 1.357E-08 | 1.074E-08 | 3.700E-09 | 2.500E-08 |
| 24082 | 1205 | Q1205R | D | 2.2e-08 | 5.5e-09 | 8.8e-09 | 1.210E-08 | 8.731E-09 | 5.500E-09 | 2.200E-08 |
| 24083 | 1205 | Q1205N | D | 1.2e-08 | 6.9e-09 | 1.6e-08 | 1.163E-08 | 4.561E-09 | 6.900E-09 | 1.600E-08 |
| 24084 | 1205 | Q1205D | D | 5e-09 | 8e-09 | 6.4e-09 | 6.467E-09 | 1.501E-09 | 5.000E-09 | 8.000E-09 |
| 24085 | 1205 | Q1205C | D | 3.4e-09 | 1.2e-08 | 1.3e-08 | 9.467E-09 | 5.278E-09 | 3.400E-09 | 1.300E-08 |
| 24086 | 1205 | Q1205E | D | 3.5e-09 | 5.1e-09 | 1.6e-09 | 3.400E-09 | 1.752E-09 | 1.600E-09 | 5.100E-09 |
| 24087 | 1205 | Q1205Q | D | 9.8e-09 | 6.4e-09 | 1.7e-08 | 1.107E-08 | 5.412E-09 | 6.400E-09 | 1.700E-08 |
| 24088 | 1205 | Q1205G | D | 4.3e-09 | 7.5e-09 | 7.7e-09 | 6.500E-09 | 1.908E-09 | 4.300E-09 | 7.700E-09 |
| 24089 | 1205 | Q1205H | D | 6.1e-09 | 1.7e-08 | 6.8e-09 | 9.967E-09 | 6.101E-09 | 6.100E-09 | 1.700E-08 |
| 24090 | 1205 | Q1205I | D | 3e-08 | 9.5e-09 | 6e-09 | 1.517E-08 | 1.296E-08 | 6.000E-09 | 3.000E-08 |
| 24091 | 1205 | Q1205L | D | 8.6e-09 | 1.3e-08 | 7.8e-09 | 9.800E-09 | 2.800E-09 | 7.800E-09 | 1.300E-08 |
| 24092 | 1205 | Q1205K | D | 1.3e-08 | 7.1e-09 | 2.9e-09 | 7.667E-09 | 5.074E-09 | 2.900E-09 | 1.300E-08 |
| 24093 | 1205 | Q1205M | D | 1.1e-08 | 1e-08 | 1.3e-08 | 1.133E-08 | 1.528E-09 | 1.000E-08 | 1.300E-08 |
| 24094 | 1205 | Q1205F | D | 3e-09 | 1.6e-09 | 3.3e-09 | 2.633E-09 | 9.074E-10 | 1.600E-09 | 3.300E-09 |
| 24095 | 1205 | Q1205P | D | 5.9e-09 | 7.1e-09 | 1.5e-08 | 9.333E-09 | 4.944E-09 | 5.900E-09 | 1.500E-08 |
| 24096 | 1205 | Q1205S | D | 6.4e-09 | 1.5e-08 | 1.7e-09 | 7.700E-09 | 6.745E-09 | 1.700E-09 | 1.500E-08 |
| 24097 | 1205 | Q1205T | D | 5.6e-09 | 7.7e-09 | 1.4e-08 | 9.100E-09 | 4.371E-09 | 5.600E-09 | 1.400E-08 |
| 24098 | 1205 | Q1205W | D | 6.1e-09 | 6.8e-09 | 4.1e-09 | 5.667E-09 | 1.401E-09 | 4.100E-09 | 6.800E-09 |
| 24099 | 1205 | Q1205Y | D | 2e-08 | 1.2e-08 | 3.2e-09 | 1.173E-08 | 8.403E-09 | 3.200E-09 | 2.000E-08 |
| 24100 | 1205 | Q1205V | D | 8.9e-09 | 1.4e-08 | 1.8e-08 | 1.363E-08 | 4.561E-09 | 8.900E-09 | 1.800E-08 |
| 24101 | 1206 | W1206A | D | 5e-09 | 2e-09 | 9.3e-09 | 5.433E-09 | 3.669E-09 | 2.000E-09 | 9.300E-09 |
| 24102 | 1206 | W1206R | D | 1.5e-08 | 4.8e-09 | 1.5e-08 | 1.160E-08 | 5.889E-09 | 4.800E-09 | 1.500E-08 |
| 24103 | 1206 | W1206N | D | 7.7e-09 | 7.9e-09 | 1.8e-08 | 1.120E-08 | 5.890E-09 | 7.700E-09 | 1.800E-08 |
| 24104 | 1206 | W1206D | D | 1.2e-08 | 3.5e-09 | 8.4e-09 | 7.967E-09 | 4.267E-09 | 3.500E-09 | 1.200E-08 |
| 24105 | 1206 | W1206C | D | 1e-08 | 6.5e-09 | 9.7e-09 | 8.733E-09 | 1.940E-09 | 6.500E-09 | 1.000E-08 |
| 24106 | 1206 | W1206E | D | 1.5e-08 | 5.2e-09 | 4e-09 | 8.067E-09 | 6.034E-09 | 4.000E-09 | 1.500E-08 |
| 24107 | 1206 | W1206Q | D | 3.2e-09 | 2.6e-09 | 5e-09 | 3.600E-09 | 1.249E-09 | 2.600E-09 | 5.000E-09 |
| 24108 | 1206 | W1206G | D | 1e-08 | 1.2e-08 | 2.8e-09 | 8.267E-09 | 4.839E-09 | 2.800E-09 | 1.200E-08 |
| 24109 | 1206 | W1206H | D | 8.5e-09 | 7.1e-09 | 1.2e-08 | 9.200E-09 | 2.524E-09 | 7.100E-09 | 1.200E-08 |
| 24110 | 1206 | W1206I | D | 5.3e-09 | 9.3e-09 | 8.7e-09 | 7.767E-09 | 2.157E-09 | 5.300E-09 | 9.300E-09 |
| 24111 | 1206 | W1206L | D | 1.5e-08 | 3.9e-09 | 8.1e-09 | 9.000E-09 | 5.604E-09 | 3.900E-09 | 1.500E-08 |
| 24112 | 1206 | W1206K | D | 7.6e-09 | 1e-08 | 1.9e-09 | 6.500E-09 | 4.161E-09 | 1.900E-09 | 1.000E-08 |
| 24113 | 1206 | W1206M | D | 7.4e-09 | 1.2e-08 | 5e-09 | 8.133E-09 | 3.557E-09 | 5.000E-09 | 1.200E-08 |
| 24114 | 1206 | W1206F | D | 4.9e-09 | 2e-09 | 6.4e-09 | 4.433E-09 | 2.237E-09 | 2.000E-09 | 6.400E-09 |
| 24115 | 1206 | W1206P | D | 2.2e-09 | 2.8e-09 | 4.6e-09 | 3.200E-09 | 1.249E-09 | 2.200E-09 | 4.600E-09 |
| 24116 | 1206 | W1206S | D | 3.8e-09 | 4.5e-09 | 8.4e-09 | 5.567E-09 | 2.479E-09 | 3.800E-09 | 8.400E-09 |
| 24117 | 1206 | W1206T | D | 1.5e-08 | 1.3e-08 | 3.5e-09 | 1.050E-08 | 6.144E-09 | 3.500E-09 | 1.500E-08 |
| 24118 | 1206 | W1206W | D | 4.3e-09 | 3.5e-09 | 1.8e-08 | 8.600E-09 | 8.150E-09 | 3.500E-09 | 1.800E-08 |
| 24119 | 1206 | W1206Y | D | 1.4e-08 | 9.4e-09 | 2.7e-09 | 8.700E-09 | 5.682E-09 | 2.700E-09 | 1.400E-08 |
| 24120 | 1206 | W1206V | D | 6.7e-09 | 7.8e-09 | 1.7e-08 | 1.050E-08 | 5.656E-09 | 6.700E-09 | 1.700E-08 |
| 24121 | 1207 | K1207A | D | 8.3e-09 | 3.2e-09 | 1.2e-08 | 7.833E-09 | 4.419E-09 | 3.200E-09 | 1.200E-08 |
| 24122 | 1207 | K1207R | D | 1.7e-08 | 4.2e-09 | 2.9e-09 | 8.033E-09 | 7.793E-09 | 2.900E-09 | 1.700E-08 |
| 24123 | 1207 | K1207N | D | 3.9e-09 | 4.4e-09 | 8.7e-09 | 5.667E-09 | 2.639E-09 | 3.900E-09 | 8.700E-09 |
| 24124 | 1207 | K1207D | D | 1.3e-08 | 3.7e-09 | 3.9e-09 | 6.867E-09 | 5.313E-09 | 3.700E-09 | 1.300E-08 |
| 24125 | 1207 | K1207C | D | 5e-09 | 1.2e-08 | 8.4e-09 | 8.467E-09 | 3.500E-09 | 5.000E-09 | 1.200E-08 |
| 24126 | 1207 | K1207E | D | 8.6e-09 | 4.7e-09 | 7.3e-09 | 6.867E-09 | 1.986E-09 | 4.700E-09 | 8.600E-09 |
| 24127 | 1207 | K1207Q | D | 4.4e-09 | 6.4e-09 | 7.3e-09 | 6.033E-09 | 1.484E-09 | 4.400E-09 | 7.300E-09 |
| 24128 | 1207 | K1207G | D | 1.1e-08 | 5.9e-09 | 6.5e-09 | 7.800E-09 | 2.787E-09 | 5.900E-09 | 1.100E-08 |
| 24129 | 1207 | K1207H | D | 1.9e-08 | 6.4e-09 | 7.5e-09 | 1.097E-08 | 6.979E-09 | 6.400E-09 | 1.900E-08 |
| 24130 | 1207 | K1207I | D | 1.4e-08 | 1.1e-08 | 5.6e-09 | 1.020E-08 | 4.257E-09 | 5.600E-09 | 1.400E-08 |
| 24131 | 1207 | K1207L | D | 1.2e-08 | 4.5e-09 | 7.6e-09 | 8.033E-09 | 3.769E-09 | 4.500E-09 | 1.200E-08 |
| 24132 | 1207 | K1207K | D | 1.4e-08 | 7.8e-09 | 5.1e-09 | 8.967E-09 | 4.563E-09 | 5.100E-09 | 1.400E-08 |
| 24133 | 1207 | K1207M | D | 7.9e-09 | 4.7e-09 | 4.8e-09 | 5.800E-09 | 1.819E-09 | 4.700E-09 | 7.900E-09 |
| 24134 | 1207 | K1207F | D | 4.2e-09 | 4.1e-09 | 2.1e-09 | 3.467E-09 | 1.185E-09 | 2.100E-09 | 4.200E-09 |
| 24135 | 1207 | K1207P | D | 8.6e-09 | 3.7e-09 | 1.1e-08 | 7.767E-09 | 3.721E-09 | 3.700E-09 | 1.100E-08 |
| 24136 | 1207 | K1207S | D | 6.2e-09 | 7.2e-09 | 5.4e-09 | 6.267E-09 | 9.018E-10 | 5.400E-09 | 7.200E-09 |
| 24137 | 1207 | K1207T | D | 2e-09 | 5.1e-09 | 6.2e-09 | 4.433E-09 | 2.178E-09 | 2.000E-09 | 6.200E-09 |
| 24138 | 1207 | K1207W | D | 1.5e-08 | 6.5e-09 | 4.5e-08 | 2.217E-08 | 2.023E-08 | 6.500E-09 | 4.500E-08 |
| 24139 | 1207 | K1207Y | D | 7.9e-09 | 7e-09 | 9.6e-09 | 8.167E-09 | 1.320E-09 | 7.000E-09 | 9.600E-09 |

| | | | | | | | | | | |
|---|---|---|---|---|---|---|---|---|---|---|
| 24140 | 1207 | K1207V | D | 1.1e-08 | 1.6e-08 | 8.3e-09 | 1.177E-08 | 3.907E-09 | 8.300E-09 | 1.600E-08 |
| 24141 | 1208 | V1208A | D | 1.1e-08 | 3e-08 | 3.1e-09 | 1.470E-08 | 1.383E-08 | 3.100E-09 | 3.000E-08 |
| 24142 | 1208 | V1208R | D | 1.3e-08 | 2.9e-09 | 2.5e-09 | 6.133E-09 | 5.950E-09 | 2.500E-09 | 1.300E-08 |
| 24143 | 1208 | V1208N | D | 3.4e-09 | 5.8e-09 | 5.8e-09 | 5.000E-09 | 1.386E-09 | 3.400E-09 | 5.800E-09 |
| 24144 | 1208 | V1208D | D | 1e-08 | 3.5e-09 | 1.4e-08 | 9.167E-09 | 5.299E-09 | 3.500E-09 | 1.400E-08 |
| 24145 | 1208 | V1208C | D | 2e-08 | 9.2e-09 | 2.4e-09 | 1.053E-08 | 8.875E-09 | 2.400E-09 | 2.000E-08 |
| 24146 | 1208 | V1208E | D | 5.9e-09 | 9e-09 | 5.5e-09 | 6.800E-09 | 1.916E-09 | 5.500E-09 | 9.000E-09 |
| 24147 | 1208 | V1208Q | D | 6.9e-09 | 8.8e-09 | 2.3e-09 | 6.000E-09 | 3.342E-09 | 2.300E-09 | 8.800E-09 |
| 24148 | 1208 | V1208G | D | 1.1e-08 | 1.9e-09 | 1.2e-08 | 8.300E-09 | 5.565E-09 | 1.900E-09 | 1.200E-08 |
| 24149 | 1208 | V1208H | D | 3.3e-09 | 4.2e-09 | 8.1e-09 | 5.200E-09 | 2.551E-09 | 3.300E-09 | 8.100E-09 |
| 24150 | 1208 | V1208I | D | 1.3e-08 | 1e-08 | 5.7e-09 | 9.567E-09 | 3.669E-09 | 5.700E-09 | 1.300E-08 |
| 24151 | 1208 | V1208L | D | 5.5e-09 | 1.5e-08 | 1.3e-09 | 7.267E-09 | 7.019E-09 | 1.300E-09 | 1.500E-08 |
| 24152 | 1208 | V1208K | D | 4.5e-09 | 2.4e-08 | 5.4e-09 | 1.130E-08 | 1.101E-08 | 4.500E-09 | 2.400E-08 |
| 24153 | 1208 | V1208M | D | 8e-09 | 3.1e-09 | 4.2e-09 | 5.100E-09 | 2.571E-09 | 3.100E-09 | 8.000E-09 |
| 24154 | 1208 | V1208F | D | 3.8e-09 | 3e-09 | 3.4e-09 | 3.400E-09 | 4.000E-10 | 3.000E-09 | 3.800E-09 |
| 24155 | 1208 | V1208P | D | 5.3e-09 | 5.8e-09 | 7.3e-09 | 6.133E-09 | 1.041E-09 | 5.300E-09 | 7.300E-09 |
| 24156 | 1208 | V1208S | D | 8.4e-09 | 8.3e-09 | 4.2e-09 | 6.967E-09 | 2.397E-09 | 4.200E-09 | 8.400E-09 |
| 24157 | 1208 | V1208T | D | 5.6e-09 | 7.8e-09 | 9.1e-09 | 7.500E-09 | 1.769E-09 | 5.600E-09 | 9.100E-09 |
| 24158 | 1208 | V1208W | D | 2e-08 | 1.1e-08 | 7e-09 | 1.267E-08 | 6.658E-09 | 7.000E-09 | 2.000E-08 |
| 24159 | 1208 | V1208Y | D | 1.3e-08 | 7.6e-09 | 2e-08 | 1.353E-08 | 6.217E-09 | 7.600E-09 | 2.000E-08 |
| 24160 | 1208 | V1208V | D | 1.2e-08 | 2.8e-09 | 1.3e-08 | 9.267E-09 | 5.623E-09 | 2.800E-09 | 1.300E-08 |
| 24161 | 1209 | D1209A | D | 8e-09 | 1.6e-08 | 1.3e-08 | 1.233E-08 | 4.041E-09 | 8.000E-09 | 1.600E-08 |
| 24162 | 1209 | D1209R | D | 6.3e-09 | 6e-09 | 2.1e-08 | 1.110E-08 | 8.575E-09 | 6.000E-09 | 2.100E-08 |
| 24163 | 1209 | D1209N | D | 1.2e-08 | 3.8e-09 | 3.9e-08 | 1.827E-08 | 1.842E-08 | 3.800E-09 | 3.900E-08 |
| 24164 | 1209 | D1209D | D | 1.1e-08 | 8.6e-09 | 9.1e-09 | 9.567E-09 | 1.266E-09 | 8.600E-09 | 1.100E-08 |
| 24165 | 1209 | D1209C | D | 7.7e-09 | 2.3e-09 | 5.6e-09 | 5.200E-09 | 2.722E-09 | 2.300E-09 | 7.700E-09 |
| 24166 | 1209 | D1209E | D | 1e-08 | 8.6e-09 | 7.3e-09 | 8.633E-09 | 1.350E-09 | 7.300E-09 | 1.000E-08 |
| 24167 | 1209 | D1209Q | D | 7.6e-09 | 3.3e-09 | 1.4e-08 | 8.300E-09 | 5.384E-09 | 3.300E-09 | 1.400E-08 |
| 24168 | 1209 | D1209G | D | 3.9e-09 | 4.4e-09 | 3.7e-09 | 4.000E-09 | 3.606E-10 | 3.700E-09 | 4.400E-09 |
| 24169 | 1209 | D1209H | D | 5.4e-09 | 4.5e-09 | 8.3e-09 | 6.067E-09 | 1.986E-09 | 4.500E-09 | 8.300E-09 |
| 24170 | 1209 | D1209I | D | 8.3e-09 | 1e-08 | 4.6e-09 | 7.633E-09 | 2.761E-09 | 4.600E-09 | 1.000E-08 |
| 24171 | 1209 | D1209L | D | 3e-09 | 9.6e-09 | 1.6e-08 | 9.533E-09 | 6.500E-09 | 3.000E-09 | 1.600E-08 |
| 24172 | 1209 | D1209K | D | 4.2e-09 | 8.2e-09 | 2.7e-08 | 1.313E-08 | 1.217E-08 | 4.200E-09 | 2.700E-08 |
| 24173 | 1209 | D1209M | D | 4.4e-09 | 4.6e-09 | 6.3e-09 | 5.100E-09 | 1.044E-09 | 4.400E-09 | 6.300E-09 |
| 24174 | 1209 | D1209F | D | 1.2e-08 | 4.5e-09 | 8.5e-09 | 8.333E-09 | 3.753E-09 | 4.500E-09 | 1.200E-08 |
| 24175 | 1209 | D1209P | D | 8.2e-09 | 9.3e-09 | 1.4e-08 | 1.050E-08 | 3.081E-09 | 8.200E-09 | 1.400E-08 |
| 24176 | 1209 | D1209S | D | 4.8e-09 | 2.2e-09 | 4.8e-09 | 3.933E-09 | 1.501E-09 | 2.200E-09 | 4.800E-09 |
| 24177 | 1209 | D1209T | D | 1.8e-09 | 8e-09 | 5.1e-09 | 4.967E-09 | 3.102E-09 | 1.800E-09 | 8.000E-09 |
| 24178 | 1209 | D1209W | D | 9.2e-09 | 4.8e-09 | 1.8e-08 | 1.067E-08 | 6.721E-09 | 4.800E-09 | 1.800E-08 |
| 24179 | 1209 | D1209Y | D | 6.6e-09 | 9.8e-09 | 3.5e-09 | 6.633E-09 | 3.150E-09 | 3.500E-09 | 9.800E-09 |
| 24180 | 1209 | D1209V | D | 1e-08 | 1.5e-08 | 5.9e-09 | 1.030E-08 | 4.557E-09 | 5.900E-09 | 1.500E-08 |
| 24181 | 1210 | N1210A | D | 2.7e-09 | 9.3e-09 | 9e-09 | 7.000E-09 | 3.727E-09 | 2.700E-09 | 9.300E-09 |
| 24182 | 1210 | N1210R | D | 1.8e-09 | 4.3e-09 | 1.3e-08 | 6.367E-09 | 5.879E-09 | 1.800E-09 | 1.300E-08 |
| 24183 | 1210 | N1210N | D | 7.2e-09 | 5.5e-09 | 3.4e-09 | 5.367E-09 | 1.904E-09 | 3.400E-09 | 7.200E-09 |
| 24184 | 1210 | N1210D | D | 4.6e-09 | 5.4e-09 | 1.8e-08 | 9.333E-09 | 7.516E-09 | 4.600E-09 | 1.800E-08 |
| 24185 | 1210 | N1210C | D | 1.8e-08 | 7.3e-09 | 5.9e-09 | 1.040E-08 | 6.619E-09 | 5.900E-09 | 1.800E-08 |
| 24186 | 1210 | N1210E | D | 6.3e-09 | 6.9e-09 | 5.3e-09 | 6.167E-09 | 8.083E-10 | 5.300E-09 | 6.900E-09 |
| 24187 | 1210 | N1210Q | D | 1.3e-08 | 1.6e-08 | 3.8e-09 | 1.093E-08 | 6.357E-09 | 3.800E-09 | 1.600E-08 |
| 24188 | 1210 | N1210G | D | 1.1e-08 | 1.1e-09 | 2.6e-08 | 1.270E-08 | 1.254E-08 | 1.100E-09 | 2.600E-08 |
| 24189 | 1210 | N1210H | D | 2.5e-08 | 3.2e-09 | 1.3e-08 | 1.373E-08 | 1.092E-08 | 3.200E-09 | 2.500E-08 |
| 24190 | 1210 | N1210I | D | 3.7e-09 | 6e-09 | 1.7e-08 | 8.900E-09 | 7.108E-09 | 3.700E-09 | 1.700E-08 |
| 24191 | 1210 | N1210L | D | 7.5e-09 | 2.9e-09 | 1e-08 | 6.800E-09 | 3.601E-09 | 2.900E-09 | 1.000E-08 |
| 24192 | 1210 | N1210K | D | 1.4e-08 | 8e-09 | 6.7e-09 | 9.567E-09 | 3.894E-09 | 6.700E-09 | 1.400E-08 |
| 24193 | 1210 | N1210M | D | 1.4e-08 | 3.9e-09 | 1.5e-08 | 1.097E-08 | 6.140E-09 | 3.900E-09 | 1.500E-08 |
| 24194 | 1210 | N1210F | D | 2e-09 | 8.1e-09 | 5.9e-09 | 5.333E-09 | 3.089E-09 | 2.000E-09 | 8.100E-09 |
| 24195 | 1210 | N1210P | D | 5.4e-09 | 2.7e-09 | 7.6e-09 | 5.233E-09 | 2.454E-09 | 2.700E-09 | 7.600E-09 |
| 24196 | 1210 | N1210S | D | 1.3e-08 | 8.7e-09 | 3.9e-09 | 8.533E-09 | 4.552E-09 | 3.900E-09 | 1.300E-08 |
| 24197 | 1210 | N1210T | D | 8.7e-09 | 1.9e-08 | 6.8e-09 | 1.150E-08 | 6.564E-09 | 6.800E-09 | 1.900E-08 |
| 24198 | 1210 | N1210W | D | 4.3e-09 | 7.5e-09 | 6.2e-09 | 6.000E-09 | 1.609E-09 | 4.300E-09 | 7.500E-09 |
| 24199 | 1210 | N1210Y | D | 1.3e-08 | 1e-08 | 5.6e-09 | 9.533E-09 | 3.722E-09 | 5.600E-09 | 1.300E-08 |
| 24200 | 1210 | N1210V | D | 8.8e-09 | 5e-09 | 3.5e-09 | 5.767E-09 | 2.732E-09 | 3.500E-09 | 8.800E-09 |
| 24201 | 1211 | A1211A | D | 1.9e-08 | 1.2e-08 | 7.9e-09 | 1.297E-08 | 5.613E-09 | 7.900E-09 | 1.900E-08 |
| 24202 | 1211 | A1211R | D | 9e-09 | 1.5e-08 | 9.4e-09 | 1.113E-08 | 3.355E-09 | 9.000E-09 | 1.500E-08 |
| 24203 | 1211 | A1211N | D | 4.8e-09 | 6.1e-09 | 8e-09 | 6.300E-09 | 1.609E-09 | 4.800E-09 | 8.000E-09 |
| 24204 | 1211 | A1211D | D | 3.6e-09 | 8e-09 | 3.2e-09 | 4.933E-09 | 2.663E-09 | 3.200E-09 | 8.000E-09 |
| 24205 | 1211 | A1211C | D | 9.2e-09 | 1.5e-08 | 1.4e-08 | 1.273E-08 | 3.101E-09 | 9.200E-09 | 1.500E-08 |
| 24206 | 1211 | A1211E | D | 4.9e-09 | 1.1e-08 | 3e-09 | 6.300E-09 | 4.180E-09 | 3.000E-09 | 1.100E-08 |
| 24207 | 1211 | A1211Q | D | 1.4e-08 | 7.8e-09 | 6.1e-09 | 9.300E-09 | 4.158E-09 | 6.100E-09 | 1.400E-08 |
| 24208 | 1211 | A1211G | D | 5.8e-09 | 2.8e-09 | 5.1e-09 | 4.567E-09 | 1.570E-09 | 2.800E-09 | 5.800E-09 |
| 24209 | 1211 | A1211H | D | 3.2e-09 | 1.4e-08 | 7.9e-09 | 8.367E-09 | 5.415E-09 | 3.200E-09 | 1.400E-08 |
| 24210 | 1211 | A1211I | D | 2.3e-09 | 6e-09 | 7.7e-09 | 5.333E-09 | 2.761E-09 | 2.300E-09 | 7.700E-09 |
| 24211 | 1211 | A1211L | D | 2.6e-08 | 7.1e-09 | 8.4e-09 | 1.383E-08 | 1.056E-08 | 7.100E-09 | 2.600E-08 |
| 24212 | 1211 | A1211K | D | 3.2e-09 | 6.4e-09 | 4.7e-09 | 4.767E-09 | 1.601E-09 | 3.200E-09 | 6.400E-09 |
| 24213 | 1211 | A1211M | D | 5.1e-09 | 7.7e-09 | 7.6e-09 | 6.800E-09 | 1.473E-09 | 5.100E-09 | 7.700E-09 |
| 24214 | 1211 | A1211F | D | 8.7e-09 | 5.1e-09 | 1.4e-09 | 5.067E-09 | 3.650E-09 | 1.400E-09 | 8.700E-09 |
| 24215 | 1211 | A1211P | D | 5.3e-09 | 5e-09 | 3.7e-08 | 1.577E-08 | 1.839E-08 | 5.000E-09 | 3.700E-08 |
| 24216 | 1211 | A1211S | D | 6.1e-09 | 5.4e-09 | 5e-09 | 5.500E-09 | 5.568E-10 | 5.000E-09 | 6.100E-09 |
| 24217 | 1211 | A1211T | D | 1.1e-08 | 1.2e-08 | 1e-08 | 1.100E-08 | 1.000E-09 | 1.000E-08 | 1.200E-08 |
| 24218 | 1211 | A1211W | D | 1.2e-08 | 1.5e-08 | 3.8e-09 | 1.027E-08 | 5.798E-09 | 3.800E-09 | 1.500E-08 |
| 24219 | 1211 | A1211Y | D | 4.1e-09 | 5.8e-09 | 5.2e-09 | 5.033E-09 | 8.622E-10 | 4.100E-09 | 5.800E-09 |
| 24220 | 1211 | A1211V | D | 1.7e-08 | 4.3e-09 | 2.7e-09 | 8.000E-09 | 7.835E-09 | 2.700E-09 | 1.700E-08 |
| 24221 | 1212 | L1212A | D | 2e-09 | 1.4e-08 | 3.4e-09 | 6.467E-09 | 6.562E-09 | 2.000E-09 | 1.400E-08 |
| 24222 | 1212 | L1212R | D | 6.2e-09 | 8e-09 | 3.4e-09 | 5.867E-09 | 2.318E-09 | 3.400E-09 | 8.000E-09 |
| 24223 | 1212 | L1212N | D | 1.8e-08 | 7.5e-09 | 1e-08 | 1.183E-08 | 5.485E-09 | 7.500E-09 | 1.800E-08 |
| 24224 | 1212 | L1212D | D | 4.4e-09 | 7e-09 | 3.2e-09 | 4.867E-09 | 1.943E-09 | 3.200E-09 | 7.000E-09 |

| | | | | | | | | | | |
|---|---|---|---|---|---|---|---|---|---|---|
| 24225 | 1212 | L1212C | D | 3.5e-09 | 7.8e-09 | 1.5e-09 | 4.267E-09 | 3.219E-09 | 1.500E-09 | 7.800E-09 |
| 24226 | 1212 | L1212E | D | 6.1e-09 | 1.2e-08 | 1e-08 | 9.367E-09 | 3.001E-09 | 6.100E-09 | 1.200E-08 |
| 24227 | 1212 | L1212Q | D | 1.1e-08 | 1.2e-08 | 2.9e-08 | 1.733E-08 | 1.012E-08 | 1.100E-08 | 2.900E-08 |
| 24228 | 1212 | L1212G | D | 1e-08 | 4.6e-09 | 3.3e-09 | 5.967E-09 | 3.553E-09 | 3.300E-09 | 1.000E-08 |
| 24229 | 1212 | L1212H | D | 4.9e-09 | 1.3e-08 | 4.1e-09 | 7.333E-09 | 4.924E-09 | 4.100E-09 | 1.300E-08 |
| 24230 | 1212 | L1212I | D | 4.8e-09 | 3.3e-09 | 2.5e-09 | 3.533E-09 | 1.168E-09 | 2.500E-09 | 4.800E-09 |
| 24231 | 1212 | L1212L | D | 7.2e-09 | 1.5e-08 | 1.2e-08 | 1.140E-08 | 3.934E-09 | 7.200E-09 | 1.500E-08 |
| 24232 | 1212 | L1212K | D | 1.5e-08 | 1.8e-08 | 1.3e-08 | 1.533E-08 | 2.517E-09 | 1.300E-08 | 1.800E-08 |
| 24233 | 1212 | L1212M | D | 5e-09 | 7.7e-09 | 4.6e-09 | 5.767E-09 | 1.686E-09 | 4.600E-09 | 7.700E-09 |
| 24234 | 1212 | L1212F | D | 3e-09 | 1.8e-09 | 4.2e-09 | 3.000E-09 | 1.200E-09 | 1.800E-09 | 4.200E-09 |
| 24235 | 1212 | L1212P | D | 1.3e-08 | 1.7e-08 | 5.7e-09 | 1.190E-08 | 5.730E-09 | 5.700E-09 | 1.700E-08 |
| 24236 | 1212 | L1212S | D | 2.1e-09 | 2.5e-09 | 3.1e-09 | 2.567E-09 | 5.033E-10 | 2.100E-09 | 3.100E-09 |
| 24237 | 1212 | L1212T | D | 5.8e-09 | 2.4e-08 | 9.4e-09 | 1.307E-08 | 9.638E-09 | 5.800E-09 | 2.400E-08 |
| 24238 | 1212 | L1212W | D | 1e-08 | 2.7e-09 | 6.1e-09 | 6.267E-09 | 3.653E-09 | 2.700E-09 | 1.000E-08 |
| 24239 | 1212 | L1212Y | D | 5.8e-09 | 6.5e-09 | 1.9e-09 | 4.733E-09 | 2.479E-09 | 1.900E-09 | 6.500E-09 |
| 24240 | 1212 | L1212V | D | 3.2e-08 | 1.1e-08 | 3e-09 | 1.533E-08 | 1.498E-08 | 3.000E-09 | 3.200E-08 |
| 24241 | 1213 | Q1213A | D | 9.1e-09 | 3.3e-09 | 7.8e-09 | 6.733E-09 | 3.044E-09 | 3.300E-09 | 9.100E-09 |
| 24242 | 1213 | Q1213R | D | 1.5e-09 | 1.2e-08 | 2e-08 | 1.117E-08 | 9.278E-09 | 1.500E-09 | 2.000E-08 |
| 24243 | 1213 | Q1213N | D | 8e-09 | 2e-08 | 2.7e-09 | 1.023E-08 | 8.864E-09 | 2.700E-09 | 2.000E-08 |
| 24244 | 1213 | Q1213D | D | 5.3e-09 | 5.9e-09 | 5e-09 | 5.400E-09 | 4.583E-10 | 5.000E-09 | 5.900E-09 |
| 24245 | 1213 | Q1213C | D | 7.3e-09 | 9.4e-09 | 2e-08 | 1.223E-08 | 6.808E-09 | 7.300E-09 | 2.000E-08 |
| 24246 | 1213 | Q1213E | D | 1e-08 | 4.2e-09 | 9.4e-09 | 7.867E-09 | 3.190E-09 | 4.200E-09 | 1.000E-08 |
| 24247 | 1213 | Q1213Q | D | 1.1e-08 | 3.7e-09 | 3.5e-09 | 6.067E-09 | 4.274E-09 | 3.500E-09 | 1.100E-08 |
| 24248 | 1213 | Q1213G | D | 1.2e-08 | 1.2e-08 | 7.4e-09 | 1.047E-08 | 2.656E-09 | 7.400E-09 | 1.200E-08 |
| 24249 | 1213 | Q1213H | D | 9.4e-09 | 6.7e-09 | 3.8e-09 | 6.633E-09 | 2.801E-09 | 3.800E-09 | 9.400E-09 |
| 24250 | 1213 | Q1213I | D | 1.4e-08 | 4.5e-09 | 1e-08 | 9.500E-09 | 4.770E-09 | 4.500E-09 | 1.400E-08 |
| 24251 | 1213 | Q1213L | D | 1.4e-08 | 3.2e-09 | 1.6e-08 | 1.107E-08 | 6.886E-09 | 3.200E-09 | 1.600E-08 |
| 24252 | 1213 | Q1213K | D | 5.6e-09 | 2.8e-09 | 1.1e-08 | 6.467E-09 | 4.168E-09 | 2.800E-09 | 1.100E-08 |
| 24253 | 1213 | Q1213M | D | 1.2e-08 | 1.1e-08 | 6.1e-09 | 9.700E-09 | 3.158E-09 | 6.100E-09 | 1.200E-08 |
| 24254 | 1213 | Q1213F | D | 4.7e-09 | 8.5e-09 | 6.6e-09 | 6.600E-09 | 1.900E-09 | 4.700E-09 | 8.500E-09 |
| 24255 | 1213 | Q1213P | D | 4.3e-09 | 4.3e-09 | 4.8e-09 | 4.467E-09 | 2.887E-10 | 4.300E-09 | 4.800E-09 |
| 24256 | 1213 | Q1213S | D | 1e-08 | 8.8e-09 | 7.6e-09 | 8.800E-09 | 1.200E-09 | 7.600E-09 | 1.000E-08 |
| 24257 | 1213 | Q1213T | D | 1.6e-08 | 8e-09 | 1.2e-08 | 1.200E-08 | 4.000E-09 | 8.000E-09 | 1.600E-08 |
| 24258 | 1213 | Q1213W | D | 1.1e-08 | 1.2e-09 | 4.4e-09 | 5.533E-09 | 4.997E-09 | 1.200E-09 | 1.100E-08 |
| 24259 | 1213 | Q1213Y | D | 2e-09 | 1.1e-08 | 1.9e-08 | 1.067E-08 | 8.505E-09 | 2.000E-09 | 1.900E-08 |
| 24260 | 1213 | Q1213V | D | 1.1e-08 | 6.5e-09 | 9.6e-09 | 9.033E-09 | 2.303E-09 | 6.500E-09 | 1.100E-08 |
| 24261 | 1214 | S1214A | D | 1e-08 | 7e-09 | 6.3e-09 | 7.767E-09 | 1.966E-09 | 6.300E-09 | 1.000E-08 |
| 24262 | 1214 | S1214R | D | 1e-08 | 2e-08 | 5.2e-09 | 1.173E-08 | 7.551E-09 | 5.200E-09 | 2.000E-08 |
| 24263 | 1214 | S1214N | D | 3.8e-09 | 1.1e-08 | 1.5e-08 | 9.933E-09 | 5.676E-09 | 3.800E-09 | 1.500E-08 |
| 24264 | 1214 | S1214D | D | 1.4e-08 | 3.6e-09 | 4.3e-09 | 7.300E-09 | 5.813E-09 | 3.600E-09 | 1.400E-08 |
| 24265 | 1214 | S1214C | D | 6.8e-09 | 1e-08 | 8.1e-09 | 8.300E-09 | 1.609E-09 | 6.800E-09 | 1.000E-08 |
| 24266 | 1214 | S1214E | D | 7e-09 | 4.5e-09 | 5.3e-09 | 5.600E-09 | 1.277E-09 | 4.500E-09 | 7.000E-09 |
| 24267 | 1214 | S1214Q | D | 7.2e-09 | 6.1e-09 | 9.7e-09 | 7.667E-09 | 1.845E-09 | 6.100E-09 | 9.700E-09 |
| 24268 | 1214 | S1214G | D | 9.9e-09 | 1.1e-08 | 2.2e-09 | 7.700E-09 | 4.795E-09 | 2.200E-09 | 1.100E-08 |
| 24269 | 1214 | S1214H | D | 6.7e-09 | 3.9e-09 | 6.3e-09 | 5.633E-09 | 1.514E-09 | 3.900E-09 | 6.700E-09 |
| 24270 | 1214 | S1214I | D | 8.2e-09 | 1.2e-08 | 5.4e-09 | 8.533E-09 | 3.313E-09 | 5.400E-09 | 1.200E-08 |
| 24271 | 1214 | S1214L | D | 1.3e-08 | 1.6e-08 | 6.5e-09 | 1.183E-08 | 4.856E-09 | 6.500E-09 | 1.600E-08 |
| 24272 | 1214 | S1214K | D | 4.5e-09 | 2.7e-08 | 1.3e-08 | 1.483E-08 | 1.136E-08 | 4.500E-09 | 2.700E-08 |
| 24273 | 1214 | S1214M | D | 1.4e-08 | 6.4e-09 | 1.4e-08 | 1.147E-08 | 4.388E-09 | 6.400E-09 | 1.400E-08 |
| 24274 | 1214 | S1214F | D | 2.9e-09 | 7.4e-09 | 6.9e-09 | 5.733E-09 | 2.466E-09 | 2.900E-09 | 7.400E-09 |
| 24275 | 1214 | S1214P | D | 1.5e-08 | 8.3e-09 | 3.5e-09 | 8.933E-09 | 5.776E-09 | 3.500E-09 | 1.500E-08 |
| 24276 | 1214 | S1214S | D | 1.7e-09 | 3.9e-09 | 3.4e-09 | 3.000E-09 | 1.153E-09 | 1.700E-09 | 3.900E-09 |
| 24277 | 1214 | S1214T | D | 5.4e-09 | 7.2e-09 | 5.1e-09 | 5.900E-09 | 1.136E-09 | 5.100E-09 | 7.200E-09 |
| 24278 | 1214 | S1214W | D | 6.5e-09 | 5e-09 | 1.7e-09 | 4.400E-09 | 2.456E-09 | 1.700E-09 | 6.500E-09 |
| 24279 | 1214 | S1214Y | D | 5e-09 | 5.3e-09 | 6.5e-09 | 5.600E-09 | 7.937E-10 | 5.000E-09 | 6.500E-09 |
| 24280 | 1214 | S1214V | D | 7.1e-09 | 4e-09 | 9e-09 | 6.700E-09 | 2.524E-09 | 4.000E-09 | 9.000E-09 |
| 24281 | 1215 | G1215A | D | 1.7e-08 | 8.7e-09 | 1.2e-08 | 1.257E-08 | 4.179E-09 | 8.700E-09 | 1.700E-08 |
| 24282 | 1215 | G1215R | D | 4.8e-09 | 1.2e-08 | 3.4e-09 | 6.733E-09 | 4.614E-09 | 3.400E-09 | 1.200E-08 |
| 24283 | 1215 | G1215N | D | 7.9e-09 | 3.9e-09 | 1.9e-09 | 4.567E-09 | 3.055E-09 | 1.900E-09 | 7.900E-09 |
| 24284 | 1215 | G1215D | D | 5.6e-09 | 1e-08 | 1.1e-08 | 8.867E-09 | 2.873E-09 | 5.600E-09 | 1.100E-08 |
| 24285 | 1215 | G1215C | D | 8.9e-09 | 4.5e-09 | 1e-08 | 7.800E-09 | 2.910E-09 | 4.500E-09 | 1.000E-08 |
| 24286 | 1215 | G1215E | D | 5.8e-09 | 1e-08 | 8.1e-09 | 7.967E-09 | 2.103E-09 | 5.800E-09 | 1.000E-08 |
| 24287 | 1215 | G1215Q | D | 1.4e-09 | 1.2e-08 | 1.6e-08 | 9.800E-09 | 7.545E-09 | 1.400E-09 | 1.600E-08 |
| 24288 | 1215 | G1215G | D | 7.5e-09 | 9.3e-09 | 1.1e-08 | 9.267E-09 | 1.750E-09 | 7.500E-09 | 1.100E-08 |
| 24289 | 1215 | G1215H | D | 1.6e-09 | 5e-09 | 8.7e-09 | 5.100E-09 | 3.551E-09 | 1.600E-09 | 8.700E-09 |
| 24290 | 1215 | G1215I | D | 5.5e-09 | 1.3e-08 | 2.1e-09 | 6.867E-09 | 5.577E-09 | 2.100E-09 | 1.300E-08 |
| 24291 | 1215 | G1215L | D | 4.1e-09 | 1.3e-08 | 3.6e-09 | 6.900E-09 | 5.289E-09 | 3.600E-09 | 1.300E-08 |
| 24292 | 1215 | G1215K | D | 4e-09 | 1.2e-08 | 1e-08 | 8.667E-09 | 4.163E-09 | 4.000E-09 | 1.200E-08 |
| 24293 | 1215 | G1215M | D | 2.9e-09 | 6.7e-09 | 5.6e-09 | 5.067E-09 | 1.955E-09 | 2.900E-09 | 6.700E-09 |
| 24294 | 1215 | G1215F | D | 5.8e-09 | 6.2e-09 | 4.4e-09 | 5.467E-09 | 9.452E-10 | 4.400E-09 | 6.200E-09 |
| 24295 | 1215 | G1215P | D | 4.2e-09 | 1e-08 | 7.8e-09 | 7.333E-09 | 2.928E-09 | 4.200E-09 | 1.000E-08 |
| 24296 | 1215 | G1215S | D | 2.2e-08 | 1.6e-08 | 4.8e-09 | 1.427E-08 | 8.730E-09 | 4.800E-09 | 2.200E-08 |
| 24297 | 1215 | G1215T | D | 1.1e-08 | 9.2e-09 | 1.6e-08 | 1.207E-08 | 3.523E-09 | 9.200E-09 | 1.600E-08 |
| 24298 | 1215 | G1215W | D | 1e-08 | 5.3e-09 | 6.3e-09 | 7.200E-09 | 2.476E-09 | 5.300E-09 | 1.000E-08 |
| 24299 | 1215 | G1215Y | D | 9.7e-09 | 2e-08 | 2.1e-09 | 1.060E-08 | 8.984E-09 | 2.100E-09 | 2.000E-08 |
| 24300 | 1215 | G1215V | D | 5.6e-09 | 2.7e-09 | 3e-09 | 3.767E-09 | 1.595E-09 | 2.700E-09 | 5.600E-09 |
| 24301 | 1216 | N1216A | D | 3.9e-09 | 5.8e-09 | 4.3e-09 | 4.667E-09 | 1.002E-09 | 3.900E-09 | 5.800E-09 |
| 24302 | 1216 | N1216R | D | 1.2e-08 | 2.7e-08 | 9.8e-09 | 1.627E-08 | 9.360E-09 | 9.800E-09 | 2.700E-08 |
| 24303 | 1216 | N1216N | D | 6.6e-09 | 6.3e-09 | 6.5e-09 | 6.467E-09 | 1.528E-10 | 6.300E-09 | 6.600E-09 |
| 24304 | 1216 | N1216D | D | 7.1e-09 | 1.1e-08 | 7.5e-09 | 8.533E-09 | 2.146E-09 | 7.100E-09 | 1.100E-08 |
| 24305 | 1216 | N1216C | D | 9e-09 | 5e-09 | 1.8e-08 | 1.067E-08 | 6.658E-09 | 5.000E-09 | 1.800E-08 |
| 24306 | 1216 | N1216E | D | 7.5e-09 | 9.7e-09 | 3.8e-09 | 7.000E-09 | 2.982E-09 | 3.800E-09 | 9.700E-09 |
| 24307 | 1216 | N1216Q | D | 3.5e-09 | 4.1e-09 | 1.2e-08 | 6.533E-09 | 4.744E-09 | 3.500E-09 | 1.200E-08 |
| 24308 | 1216 | N1216G | D | 3.1e-09 | 1.9e-08 | 1.2e-08 | 1.137E-08 | 7.969E-09 | 3.100E-09 | 1.900E-08 |
| 24309 | 1216 | N1216H | D | 6.2e-09 | 6.5e-09 | 9e-09 | 7.233E-09 | 1.537E-09 | 6.200E-09 | 9.000E-09 |

| | | | | | | | | | | |
|---|---|---|---|---|---|---|---|---|---|---|
| 24310 | 1216 | N1216I | D | 5e-09 | 5.6e-09 | 1.8e-09 | 4.133E-09 | 2.043E-09 | 1.800E-09 | 5.600E-09 |
| 24311 | 1216 | N1216L | D | 2.3e-09 | 3.5e-09 | 3.2e-09 | 3.000E-09 | 6.245E-10 | 2.300E-09 | 3.500E-09 |
| 24312 | 1216 | N1216K | D | 1.1e-08 | 1e-08 | 6.8e-09 | 9.267E-09 | 2.194E-09 | 6.800E-09 | 1.100E-08 |
| 24313 | 1216 | N1216M | D | 1.1e-08 | 1.5e-08 | 4.6e-09 | 1.020E-08 | 5.246E-09 | 4.600E-09 | 1.500E-08 |
| 24314 | 1216 | N1216F | D | 4.3e-09 | 4.9e-09 | 7.5e-09 | 5.567E-09 | 1.701E-09 | 4.300E-09 | 7.500E-09 |
| 24315 | 1216 | N1216P | D | 3.5e-09 | 1.5e-08 | 8.6e-09 | 9.033E-09 | 5.762E-09 | 3.500E-09 | 1.500E-08 |
| 24316 | 1216 | N1216S | D | 6.8e-09 | 1.9e-08 | 8.4e-09 | 1.140E-08 | 6.630E-09 | 6.800E-09 | 1.900E-08 |
| 24317 | 1216 | N1216T | D | 3.2e-09 | 1.2e-08 | 3.7e-09 | 6.300E-09 | 4.943E-09 | 3.200E-09 | 1.200E-08 |
| 24318 | 1216 | N1216W | D | 7e-09 | 1.2e-08 | 1e-08 | 9.667E-09 | 2.517E-09 | 7.000E-09 | 1.200E-08 |
| 24319 | 1216 | N1216Y | D | 9.7e-09 | 3.3e-09 | 1.2e-09 | 4.733E-09 | 4.428E-09 | 1.200E-09 | 9.700E-09 |
| 24320 | 1216 | N1216V | D | 8.3e-09 | 3.1e-08 | 1.1e-08 | 1.677E-08 | 1.240E-08 | 8.300E-09 | 3.100E-08 |
| 24321 | 1217 | S1217A | D | 9e-09 | 6.8e-09 | 3.9e-09 | 6.567E-09 | 2.558E-09 | 3.900E-09 | 9.000E-09 |
| 24322 | 1217 | S1217R | D | 8.9e-09 | 1.5e-08 | 1.5e-08 | 1.297E-08 | 3.522E-09 | 8.900E-09 | 1.500E-08 |
| 24323 | 1217 | S1217N | D | 6.3e-09 | 7.7e-09 | 1.2e-08 | 8.667E-09 | 2.970E-09 | 6.300E-09 | 1.200E-08 |
| 24324 | 1217 | S1217D | D | 6.5e-09 | 2e-08 | 7.7e-09 | 1.140E-08 | 7.472E-09 | 6.500E-09 | 2.000E-08 |
| 24325 | 1217 | S1217C | D | 1e-08 | 7.1e-09 | 3.9e-09 | 7.000E-09 | 3.051E-09 | 3.900E-09 | 1.000E-08 |
| 24326 | 1217 | S1217E | D | 7.5e-09 | 4.2e-09 | 1.1e-08 | 7.567E-09 | 3.400E-09 | 4.200E-09 | 1.100E-08 |
| 24327 | 1217 | S1217Q | D | 2.9e-09 | 9.2e-09 | 9.3e-09 | 7.133E-09 | 3.667E-09 | 2.900E-09 | 9.300E-09 |
| 24328 | 1217 | S1217G | D | 6e-09 | 3e-09 | 2.3e-09 | 3.767E-09 | 1.966E-09 | 2.300E-09 | 6.000E-09 |
| 24329 | 1217 | S1217H | D | 4.4e-09 | 1e-08 | 4.8e-09 | 6.400E-09 | 3.124E-09 | 4.400E-09 | 1.000E-08 |
| 24330 | 1217 | S1217I | D | 1.1e-08 | 6.7e-09 | 1.5e-08 | 1.090E-08 | 4.151E-09 | 6.700E-09 | 1.500E-08 |
| 24331 | 1217 | S1217L | D | 1.1e-08 | 8.5e-09 | 4.2e-09 | 7.900E-09 | 3.439E-09 | 4.200E-09 | 1.100E-08 |
| 24332 | 1217 | S1217K | D | 2.7e-08 | 3e-09 | 1.8e-08 | 1.600E-08 | 1.212E-08 | 3.000E-09 | 2.700E-08 |
| 24333 | 1217 | S1217M | D | 2e-09 | 1.9e-08 | 6.6e-09 | 9.200E-09 | 8.793E-09 | 2.000E-09 | 1.900E-08 |
| 24334 | 1217 | S1217F | D | 9.2e-09 | 3.2e-09 | 2.6e-09 | 5.000E-09 | 3.650E-09 | 2.600E-09 | 9.200E-09 |
| 24335 | 1217 | S1217P | D | 6.9e-09 | 5.4e-09 | 3.3e-09 | 5.200E-09 | 1.808E-09 | 3.300E-09 | 6.900E-09 |
| 24336 | 1217 | S1217S | D | 5.9e-09 | 1.1e-08 | 7.6e-09 | 8.167E-09 | 2.597E-09 | 5.900E-09 | 1.100E-08 |
| 24337 | 1217 | S1217T | D | 7.5e-09 | 2.6e-09 | 8.1e-09 | 6.067E-09 | 3.017E-09 | 2.600E-09 | 8.100E-09 |
| 24338 | 1217 | S1217W | D | 1.4e-08 | 3.6e-09 | 2.9e-09 | 6.833E-09 | 6.216E-09 | 2.900E-09 | 1.400E-08 |
| 24339 | 1217 | S1217Y | D | 4.4e-09 | 5.4e-09 | 3.8e-09 | 4.533E-09 | 8.083E-10 | 3.800E-09 | 5.400E-09 |
| 24340 | 1217 | S1217V | D | 1.1e-08 | 8e-09 | 8.2e-09 | 9.067E-09 | 1.677E-09 | 8.000E-09 | 1.100E-08 |
| 24341 | 1218 | Q1218A | D | 2.4e-08 | 8.4e-09 | 7.8e-09 | 1.340E-08 | 9.185E-09 | 7.800E-09 | 2.400E-08 |
| 24342 | 1218 | Q1218R | D | 1e-08 | 7.2e-09 | 4e-09 | 7.067E-09 | 3.002E-09 | 4.000E-09 | 1.000E-08 |
| 24343 | 1218 | Q1218N | D | 9.2e-09 | 8.2e-09 | 4.8e-09 | 7.400E-09 | 2.307E-09 | 4.800E-09 | 9.200E-09 |
| 24344 | 1218 | Q1218D | D | 4.5e-09 | 7.6e-09 | 1.3e-08 | 8.367E-09 | 4.302E-09 | 4.500E-09 | 1.300E-08 |
| 24345 | 1218 | Q1218C | D | 6.1e-09 | 8e-09 | 1e-08 | 8.033E-09 | 1.950E-09 | 6.100E-09 | 1.000E-08 |
| 24346 | 1218 | Q1218E | D | 1.3e-08 | 3.4e-09 | 1.3e-08 | 9.800E-09 | 5.543E-09 | 3.400E-09 | 1.300E-08 |
| 24347 | 1218 | Q1218Q | D | 1.3e-08 | 6e-09 | 1.2e-08 | 1.033E-08 | 3.786E-09 | 6.000E-09 | 1.300E-08 |
| 24348 | 1218 | Q1218G | D | 1.2e-08 | 1e-08 | 6.7e-09 | 9.567E-09 | 2.676E-09 | 6.700E-09 | 1.200E-08 |
| 24349 | 1218 | Q1218H | D | 9.4e-09 | 3.8e-09 | 6.7e-09 | 6.633E-09 | 2.801E-09 | 3.800E-09 | 9.400E-09 |
| 24350 | 1218 | Q1218I | D | 4e-09 | 1.1e-08 | 1.3e-08 | 9.333E-09 | 4.726E-09 | 4.000E-09 | 1.300E-08 |
| 24351 | 1218 | Q1218L | D | 2.6e-09 | 8.7e-09 | 3.8e-09 | 5.033E-09 | 3.232E-09 | 2.600E-09 | 8.700E-09 |
| 24352 | 1218 | Q1218K | D | 5.9e-09 | 8.6e-09 | 4.5e-09 | 6.333E-09 | 2.084E-09 | 4.500E-09 | 8.600E-09 |
| 24353 | 1218 | Q1218M | D | 2.5e-09 | 7e-09 | 1.1e-08 | 6.833E-09 | 4.252E-09 | 2.500E-09 | 1.100E-08 |
| 24354 | 1218 | Q1218F | D | 7.3e-09 | 6.5e-09 | 8.5e-09 | 7.433E-09 | 1.007E-09 | 6.500E-09 | 8.500E-09 |
| 24355 | 1218 | Q1218P | D | 9.4e-09 | 7.2e-09 | 1.1e-08 | 9.200E-09 | 1.908E-09 | 7.200E-09 | 1.100E-08 |
| 24356 | 1218 | Q1218S | D | 1e-08 | 1.2e-08 | 1e-08 | 1.067E-08 | 1.155E-09 | 1.000E-08 | 1.200E-08 |
| 24357 | 1218 | Q1218T | D | 2.3e-09 | 9.8e-09 | 4.8e-09 | 5.633E-09 | 3.819E-09 | 2.300E-09 | 9.800E-09 |
| 24358 | 1218 | Q1218W | D | 7.9e-09 | 2.1e-08 | 5.7e-09 | 1.153E-08 | 8.272E-09 | 5.700E-09 | 2.100E-08 |
| 24359 | 1218 | Q1218Y | D | 3.7e-09 | 3.2e-09 | 7e-09 | 4.633E-09 | 2.065E-09 | 3.200E-09 | 7.000E-09 |
| 24360 | 1218 | Q1218V | D | 1.2e-08 | 1.1e-08 | 4.5e-09 | 9.167E-09 | 4.072E-09 | 4.500E-09 | 1.200E-08 |
| 24361 | 1219 | E1219A | D | 1.2e-08 | 3.6e-09 | 1.7e-08 | 1.087E-08 | 6.772E-09 | 3.600E-09 | 1.700E-08 |
| 24362 | 1219 | E1219R | D | 4.9e-09 | 5.3e-09 | 2.1e-08 | 1.040E-08 | 9.182E-09 | 4.900E-09 | 2.100E-08 |
| 24363 | 1219 | E1219N | D | 4.9e-09 | 8.1e-09 | 1.6e-08 | 9.667E-09 | 5.713E-09 | 4.900E-09 | 1.600E-08 |
| 24364 | 1219 | E1219D | D | 1.5e-08 | 7.1e-09 | 6.1e-09 | 9.400E-09 | 4.875E-09 | 6.100E-09 | 1.500E-08 |
| 24365 | 1219 | E1219C | D | 5.5e-09 | 1.5e-08 | 8.5e-09 | 9.667E-09 | 4.856E-09 | 5.500E-09 | 1.500E-08 |
| 24366 | 1219 | E1219E | D | 2.1e-09 | 9.9e-09 | 7.7e-09 | 6.567E-09 | 4.022E-09 | 2.100E-09 | 9.900E-09 |
| 24367 | 1219 | E1219Q | D | 4e-09 | 1.9e-08 | 1.6e-08 | 1.300E-08 | 7.937E-09 | 4.000E-09 | 1.900E-08 |
| 24368 | 1219 | E1219G | D | 1e-08 | 3.3e-09 | 2.6e-09 | 5.300E-09 | 4.085E-09 | 2.600E-09 | 1.000E-08 |
| 24369 | 1219 | E1219H | D | 5.3e-09 | 9.3e-09 | 1.5e-08 | 9.867E-09 | 4.875E-09 | 5.300E-09 | 1.500E-08 |
| 24370 | 1219 | E1219I | D | 7.3e-09 | 5.1e-09 | 9.5e-09 | 7.300E-09 | 2.200E-09 | 5.100E-09 | 9.500E-09 |
| 24371 | 1219 | E1219L | D | 8.2e-09 | 1.7e-08 | 6e-09 | 1.040E-08 | 5.821E-09 | 6.000E-09 | 1.700E-08 |
| 24372 | 1219 | E1219K | D | 9.4e-09 | 5.3e-09 | 7.2e-09 | 7.300E-09 | 2.052E-09 | 5.300E-09 | 9.400E-09 |
| 24373 | 1219 | E1219M | D | 4.8e-09 | 7.5e-09 | 7.7e-09 | 6.667E-09 | 1.620E-09 | 4.800E-09 | 7.700E-09 |
| 24374 | 1219 | E1219F | D | 7.4e-09 | 2.1e-08 | 1.9e-08 | 1.580E-08 | 7.343E-09 | 7.400E-09 | 2.100E-08 |
| 24375 | 1219 | E1219P | D | 1.2e-08 | 3.5e-09 | 6e-09 | 7.167E-09 | 4.368E-09 | 3.500E-09 | 1.200E-08 |
| 24376 | 1219 | E1219S | D | 5e-09 | 7.1e-09 | 5.9e-09 | 6.000E-09 | 1.054E-09 | 5.000E-09 | 7.100E-09 |
| 24377 | 1219 | E1219T | D | 1.8e-09 | 2.7e-08 | 8.7e-09 | 1.250E-08 | 1.302E-08 | 1.800E-09 | 2.700E-08 |
| 24378 | 1219 | E1219W | D | 8.6e-09 | 1.5e-08 | 2.6e-09 | 8.733E-09 | 6.201E-09 | 2.600E-09 | 1.500E-08 |
| 24379 | 1219 | E1219Y | D | 7.5e-09 | 1e-08 | 2e-09 | 6.500E-09 | 4.093E-09 | 2.000E-09 | 1.000E-08 |
| 24380 | 1219 | E1219V | D | 1.2e-08 | 1.5e-09 | 1.1e-08 | 8.167E-09 | 5.795E-09 | 1.500E-09 | 1.200E-08 |
| 24381 | 1220 | S1220A | D | 1e-08 | 4.2e-09 | 3.5e-09 | 5.900E-09 | 3.568E-09 | 3.500E-09 | 1.000E-08 |
| 24382 | 1220 | S1220R | D | 5.7e-09 | 7.9e-09 | 1.3e-08 | 8.867E-09 | 3.745E-09 | 5.700E-09 | 1.300E-08 |
| 24383 | 1220 | S1220N | D | 2.6e-09 | 8e-09 | 1e-08 | 6.867E-09 | 3.828E-09 | 2.600E-09 | 1.000E-08 |
| 24384 | 1220 | S1220D | D | 2.6e-09 | 2.3e-09 | 3.9e-09 | 2.933E-09 | 8.505E-10 | 2.300E-09 | 3.900E-09 |
| 24385 | 1220 | S1220C | D | 1.1e-08 | 9.5e-09 | 1.5e-09 | 7.333E-09 | 5.107E-09 | 1.500E-09 | 1.100E-08 |
| 24386 | 1220 | S1220E | D | 1.6e-08 | 4.7e-09 | 3e-09 | 7.900E-09 | 7.066E-09 | 3.000E-09 | 1.600E-08 |
| 24387 | 1220 | S1220Q | D | 1e-08 | 9.2e-09 | 3.3e-08 | 1.740E-08 | 1.352E-08 | 9.200E-09 | 3.300E-08 |
| 24388 | 1220 | S1220G | D | 4.6e-09 | 5.8e-09 | 4.2e-09 | 4.867E-09 | 8.327E-10 | 4.200E-09 | 5.800E-09 |
| 24389 | 1220 | S1220H | D | 2.6e-09 | 8.3e-09 | 1.1e-08 | 7.300E-09 | 4.288E-09 | 2.600E-09 | 1.100E-08 |
| 24390 | 1220 | S1220I | D | 7.4e-09 | 4e-09 | 9.6e-09 | 7.000E-09 | 2.821E-09 | 4.000E-09 | 9.600E-09 |
| 24391 | 1220 | S1220L | D | 1.7e-08 | 2.9e-09 | 6.9e-09 | 8.933E-09 | 7.267E-09 | 2.900E-09 | 1.700E-08 |
| 24392 | 1220 | S1220K | D | 4.2e-09 | 4.9e-09 | 1.1e-08 | 6.700E-09 | 3.740E-09 | 4.200E-09 | 1.100E-08 |
| 24393 | 1220 | S1220M | D | 1.3e-08 | 3.8e-09 | 7.2e-09 | 8.000E-09 | 4.652E-09 | 3.800E-09 | 1.300E-08 |
| 24394 | 1220 | S1220F | D | 6.1e-09 | 6e-09 | 4.6e-09 | 5.567E-09 | 8.386E-10 | 4.600E-09 | 6.100E-09 |

| | | | | | | | | | | |
|---|---|---|---|---|---|---|---|---|---|---|
| 24395 | 1220 | S1220P | D | 1.2e-08 | 7.8e-09 | 6.5e-09 | 8.767E-09 | 2.875E-09 | 6.500E-09 | 1.200E-08 |
| 24396 | 1220 | S1220S | D | 6.6e-09 | 7.9e-09 | 7e-09 | 7.167E-09 | 6.658E-10 | 6.600E-09 | 7.900E-09 |
| 24397 | 1220 | S1220T | D | 2.7e-09 | 9.4e-09 | 9.5e-09 | 7.200E-09 | 3.897E-09 | 2.700E-09 | 9.500E-09 |
| 24398 | 1220 | S1220W | D | 1.1e-08 | 8.6e-09 | 2.5e-09 | 7.367E-09 | 4.382E-09 | 2.500E-09 | 1.100E-08 |
| 24399 | 1220 | S1220Y | D | 6.6e-09 | 1.8e-08 | 1.1e-08 | 1.187E-08 | 5.749E-09 | 6.600E-09 | 1.800E-08 |
| 24400 | 1220 | S1220V | D | 9e-09 | 1.3e-08 | 1.2e-08 | 1.133E-08 | 2.082E-09 | 9.000E-09 | 1.300E-08 |
| 24401 | 1221 | V1221A | D | 3.8e-09 | 1.1e-08 | 1e-08 | 8.267E-09 | 3.900E-09 | 3.800E-09 | 1.100E-08 |
| 24402 | 1221 | V1221R | D | 8.9e-09 | 1e-08 | 9.4e-09 | 9.433E-09 | 5.508E-10 | 8.900E-09 | 1.000E-08 |
| 24403 | 1221 | V1221N | D | 1.3e-08 | 4.6e-09 | 1.1e-08 | 9.533E-09 | 4.388E-09 | 4.600E-09 | 1.300E-08 |
| 24404 | 1221 | V1221D | D | 1e-08 | 7.4e-09 | 4.8e-09 | 7.400E-09 | 2.600E-09 | 4.800E-09 | 1.000E-08 |
| 24405 | 1221 | V1221C | D | 3.1e-09 | 7.7e-09 | 3.5e-09 | 4.767E-09 | 2.548E-09 | 3.100E-09 | 7.700E-09 |
| 24406 | 1221 | V1221E | D | 7.6e-09 | 6.7e-09 | 1.2e-08 | 8.767E-09 | 2.836E-09 | 6.700E-09 | 1.200E-08 |
| 24407 | 1221 | V1221Q | D | 5.1e-09 | 6.4e-09 | 6.7e-09 | 6.067E-09 | 8.505E-10 | 5.100E-09 | 6.700E-09 |
| 24408 | 1221 | V1221G | D | 5.5e-09 | 6.7e-09 | 4.3e-09 | 5.500E-09 | 1.200E-09 | 4.300E-09 | 6.700E-09 |
| 24409 | 1221 | V1221H | D | 4.6e-09 | 1.7e-08 | 5.7e-09 | 9.100E-09 | 6.864E-09 | 4.600E-09 | 1.700E-08 |
| 24410 | 1221 | V1221I | D | 7.1e-09 | 8.3e-09 | 8.6e-09 | 8.000E-09 | 7.937E-10 | 7.100E-09 | 8.600E-09 |
| 24411 | 1221 | V1221L | D | 6.7e-09 | 1.2e-08 | 5e-09 | 7.900E-09 | 3.651E-09 | 5.000E-09 | 1.200E-08 |
| 24412 | 1221 | V1221K | D | 5.2e-09 | 2.8e-09 | 7e-09 | 5.000E-09 | 2.107E-09 | 2.800E-09 | 7.000E-09 |
| 24413 | 1221 | V1221M | D | 1.8e-08 | 1.7e-08 | 1.1e-08 | 1.533E-08 | 3.786E-09 | 1.100E-08 | 1.800E-08 |
| 24414 | 1221 | V1221F | D | 6.4e-09 | 2.8e-09 | 8.2e-09 | 5.800E-09 | 2.750E-09 | 2.800E-09 | 8.200E-09 |
| 24415 | 1221 | V1221P | D | 2.8e-08 | 3.1e-09 | 1.7e-08 | 1.603E-08 | 1.248E-08 | 3.100E-09 | 2.800E-08 |
| 24416 | 1221 | V1221S | D | 1.1e-08 | 1.2e-08 | 7.9e-09 | 1.030E-08 | 2.138E-09 | 7.900E-09 | 1.200E-08 |
| 24417 | 1221 | V1221T | D | 5.3e-09 | 2.6e-09 | 3.3e-09 | 3.733E-09 | 1.401E-09 | 2.600E-09 | 5.300E-09 |
| 24418 | 1221 | V1221W | D | 1.1e-08 | 9e-09 | 2.2e-09 | 7.400E-09 | 4.613E-09 | 2.200E-09 | 1.100E-08 |
| 24419 | 1221 | V1221Y | D | 5.3e-09 | 6.1e-09 | 1e-08 | 7.133E-09 | 2.515E-09 | 5.300E-09 | 1.000E-08 |
| 24420 | 1221 | V1221V | D | 4e-09 | 8.2e-09 | 2.4e-09 | 4.867E-09 | 2.996E-09 | 2.400E-09 | 8.200E-09 |
| 24421 | 1222 | T1222A | D | 4e-09 | 1.1e-08 | 1.2e-08 | 9.000E-09 | 4.359E-09 | 4.000E-09 | 1.200E-08 |
| 24422 | 1222 | T1222R | D | 1e-08 | 1.4e-08 | 9.2e-09 | 1.107E-08 | 2.572E-09 | 9.200E-09 | 1.400E-08 |
| 24423 | 1222 | T1222N | D | 8e-09 | 2e-08 | 3.6e-09 | 1.053E-08 | 8.488E-09 | 3.600E-09 | 2.000E-08 |
| 24424 | 1222 | T1222D | D | 5.9e-09 | 3.4e-09 | 5.8e-09 | 5.033E-09 | 1.415E-09 | 3.400E-09 | 5.900E-09 |
| 24425 | 1222 | T1222C | D | 3.4e-08 | 1.6e-08 | 4.1e-09 | 1.803E-08 | 1.505E-08 | 4.100E-09 | 3.400E-08 |
| 24426 | 1222 | T1222E | D | 4.3e-09 | 1.1e-08 | 3.7e-09 | 6.333E-09 | 4.053E-09 | 3.700E-09 | 1.100E-08 |
| 24427 | 1222 | T1222Q | D | 9.7e-09 | 5.3e-09 | 8.6e-09 | 7.867E-09 | 2.290E-09 | 5.300E-09 | 9.700E-09 |
| 24428 | 1222 | T1222G | D | 1.3e-08 | 6.3e-09 | 1e-08 | 9.767E-09 | 3.356E-09 | 6.300E-09 | 1.300E-08 |
| 24429 | 1222 | T1222H | D | 1e-08 | 4e-09 | 6e-09 | 6.667E-09 | 3.055E-09 | 4.000E-09 | 1.000E-08 |
| 24430 | 1222 | T1222I | D | 6.1e-09 | 1.3e-08 | 1e-08 | 9.700E-09 | 3.460E-09 | 6.100E-09 | 1.300E-08 |
| 24431 | 1222 | T1222L | D | 1.2e-08 | 1e-08 | 5.1e-09 | 9.033E-09 | 3.550E-09 | 5.100E-09 | 1.200E-08 |
| 24432 | 1222 | T1222K | D | 2.6e-09 | 4.9e-09 | 6.6e-09 | 4.700E-09 | 2.007E-09 | 2.600E-09 | 6.600E-09 |
| 24433 | 1222 | T1222M | D | 5.8e-09 | 2.8e-09 | 8e-09 | 5.533E-09 | 2.610E-09 | 2.800E-09 | 8.000E-09 |
| 24434 | 1222 | T1222F | D | 1.3e-08 | 2.6e-09 | 1.1e-08 | 8.867E-09 | 5.518E-09 | 2.600E-09 | 1.300E-08 |
| 24435 | 1222 | T1222P | D | 9.7e-09 | 7.9e-09 | 4.9e-09 | 7.500E-09 | 2.425E-09 | 4.900E-09 | 9.700E-09 |
| 24436 | 1222 | T1222S | D | 8.2e-09 | 1e-08 | 5.5e-09 | 7.900E-09 | 2.265E-09 | 5.500E-09 | 1.000E-08 |
| 24437 | 1222 | T1222T | D | 1.2e-08 | 1.2e-09 | 7.6e-09 | 6.933E-09 | 5.431E-09 | 1.200E-09 | 1.200E-08 |
| 24438 | 1222 | T1222W | D | 2.2e-09 | 5.4e-09 | 6.3e-09 | 4.633E-09 | 2.155E-09 | 2.200E-09 | 6.300E-09 |
| 24439 | 1222 | T1222Y | D | 5.8e-09 | 1.8e-08 | 3.4e-09 | 9.067E-09 | 7.829E-09 | 3.400E-09 | 1.800E-08 |
| 24440 | 1222 | T1222V | D | 3.8e-09 | 3.4e-09 | 7e-09 | 4.733E-09 | 1.973E-09 | 3.400E-09 | 7.000E-09 |
| 24441 | 1223 | E1223A | D | 1.4e-08 | 3.6e-09 | 9.3e-09 | 8.967E-09 | 5.208E-09 | 3.600E-09 | 1.400E-08 |
| 24442 | 1223 | E1223R | D | 4.3e-09 | 6.3e-09 | 6.1e-09 | 5.567E-09 | 1.102E-09 | 4.300E-09 | 6.300E-09 |
| 24443 | 1223 | E1223N | D | 6.6e-09 | 2.2e-08 | 1.9e-09 | 1.017E-08 | 1.051E-08 | 1.900E-09 | 2.200E-08 |
| 24444 | 1223 | E1223D | D | 4.2e-09 | 7.2e-09 | 4.6e-09 | 5.333E-09 | 1.629E-09 | 4.200E-09 | 7.200E-09 |
| 24445 | 1223 | E1223C | D | 8.6e-09 | 3.9e-09 | 1.6e-08 | 9.500E-09 | 6.100E-09 | 3.900E-09 | 1.600E-08 |
| 24446 | 1223 | E1223E | D | 4.5e-09 | 9.8e-09 | 1e-08 | 8.100E-09 | 3.119E-09 | 4.500E-09 | 1.000E-08 |
| 24447 | 1223 | E1223Q | D | 1.8e-08 | 8.9e-09 | 8.9e-09 | 1.193E-08 | 5.254E-09 | 8.900E-09 | 1.800E-08 |
| 24448 | 1223 | E1223G | D | 3.6e-09 | 1.4e-08 | 3.6e-09 | 7.067E-09 | 6.004E-09 | 3.600E-09 | 1.400E-08 |
| 24449 | 1223 | E1223H | D | 7.8e-09 | 2.4e-08 | 4.5e-09 | 1.210E-08 | 1.044E-08 | 4.500E-09 | 2.400E-08 |
| 24450 | 1223 | E1223I | D | 9.3e-09 | 6e-09 | 7.2e-09 | 7.500E-09 | 1.670E-09 | 6.000E-09 | 9.300E-09 |
| 24451 | 1223 | E1223L | D | 6.3e-09 | 1.6e-08 | 5.4e-10 | 7.613E-09 | 7.813E-09 | 5.400E-10 | 1.600E-08 |
| 24452 | 1223 | E1223K | D | 5.7e-09 | 1.4e-08 | 1.3e-08 | 1.090E-08 | 4.531E-09 | 5.700E-09 | 1.400E-08 |
| 24453 | 1223 | E1223M | D | 5.5e-09 | 3.6e-09 | 8e-09 | 5.700E-09 | 2.207E-09 | 3.600E-09 | 8.000E-09 |
| 24454 | 1223 | E1223F | D | 1.1e-08 | 7.6e-09 | 6e-09 | 8.200E-09 | 2.553E-09 | 6.000E-09 | 1.100E-08 |
| 24455 | 1223 | E1223P | D | 2.9e-09 | 2.2e-09 | 1.1e-08 | 5.367E-09 | 4.891E-09 | 2.200E-09 | 1.100E-08 |
| 24456 | 1223 | E1223S | D | 5.6e-09 | 4.7e-09 | 8.4e-09 | 6.233E-09 | 1.930E-09 | 4.700E-09 | 8.400E-09 |
| 24457 | 1223 | E1223T | D | 5.3e-09 | 1.7e-08 | 1.3e-08 | 1.177E-08 | 5.947E-09 | 5.300E-09 | 1.700E-08 |
| 24458 | 1223 | E1223W | D | 8.6e-09 | 6.5e-09 | 1.3e-08 | 9.367E-09 | 3.317E-09 | 6.500E-09 | 1.300E-08 |
| 24459 | 1223 | E1223Y | D | 8.5e-09 | 1.4e-08 | 1.2e-08 | 1.150E-08 | 2.784E-09 | 8.500E-09 | 1.400E-08 |
| 24460 | 1223 | E1223V | D | 7.6e-09 | 1.1e-08 | 6.9e-09 | 8.500E-09 | 2.193E-09 | 6.900E-09 | 1.100E-08 |
| 24461 | 1224 | Q1224A | D | 7e-09 | 8.2e-09 | 1.3e-08 | 9.400E-09 | 3.175E-09 | 7.000E-09 | 1.300E-08 |
| 24462 | 1224 | Q1224R | D | 1.8e-08 | 8.6e-09 | 1.1e-08 | 1.253E-08 | 4.884E-09 | 8.600E-09 | 1.800E-08 |
| 24463 | 1224 | Q1224N | D | 2.3e-09 | 9e-09 | 1.2e-08 | 7.767E-09 | 4.966E-09 | 2.300E-09 | 1.200E-08 |
| 24464 | 1224 | Q1224D | D | 6.7e-09 | 4e-09 | 6.4e-09 | 5.700E-09 | 1.480E-09 | 4.000E-09 | 6.700E-09 |
| 24465 | 1224 | Q1224C | D | 6.8e-09 | 1.2e-08 | 2.4e-09 | 7.067E-09 | 4.806E-09 | 2.400E-09 | 1.200E-08 |
| 24466 | 1224 | Q1224E | D | 1e-08 | 7.8e-09 | 6.9e-09 | 8.233E-09 | 1.595E-09 | 6.900E-09 | 1.000E-08 |
| 24467 | 1224 | Q1224Q | D | 3.5e-09 | 4.1e-09 | 6.5e-09 | 4.700E-09 | 1.587E-09 | 3.500E-09 | 6.500E-09 |
| 24468 | 1224 | Q1224G | D | 8.3e-09 | 6.6e-09 | 6.2e-09 | 7.033E-09 | 1.115E-09 | 6.200E-09 | 8.300E-09 |
| 24469 | 1224 | Q1224H | D | 6.1e-09 | 9.7e-09 | 8.4e-09 | 8.067E-09 | 1.823E-09 | 6.100E-09 | 9.700E-09 |
| 24470 | 1224 | Q1224I | D | 7.2e-09 | 4.4e-09 | 1.7e-08 | 9.533E-09 | 6.616E-09 | 4.400E-09 | 1.700E-08 |
| 24471 | 1224 | Q1224L | D | 8.8e-09 | 5.1e-09 | 7.7e-09 | 7.200E-09 | 1.900E-09 | 5.100E-09 | 8.800E-09 |
| 24472 | 1224 | Q1224K | D | 1.4e-08 | 6.9e-09 | 4.8e-09 | 8.567E-09 | 4.821E-09 | 4.800E-09 | 1.400E-08 |
| 24473 | 1224 | Q1224M | D | 3.4e-09 | 6.2e-09 | 5.2e-09 | 4.933E-09 | 1.419E-09 | 3.400E-09 | 6.200E-09 |
| 24474 | 1224 | Q1224F | D | 2e-08 | 1.7e-09 | 2.6e-08 | 1.590E-08 | 1.266E-08 | 1.700E-09 | 2.600E-08 |
| 24475 | 1224 | Q1224P | D | 3.6e-09 | 4.4e-09 | 3e-09 | 3.667E-09 | 7.024E-10 | 3.000E-09 | 4.400E-09 |
| 24476 | 1224 | Q1224S | D | 2e-09 | 3.4e-09 | 3.5e-09 | 2.967E-09 | 8.386E-10 | 2.000E-09 | 3.500E-09 |
| 24477 | 1224 | Q1224T | D | 9.8e-09 | 5.1e-09 | 4.5e-09 | 6.467E-09 | 2.902E-09 | 4.500E-09 | 9.800E-09 |
| 24478 | 1224 | Q1224W | D | 1.2e-08 | 4.5e-09 | 4.8e-09 | 7.100E-09 | 4.246E-09 | 4.500E-09 | 1.200E-08 |
| 24479 | 1224 | Q1224Y | D | 9.9e-09 | 4.4e-09 | 5.7e-09 | 6.667E-09 | 2.875E-09 | 4.400E-09 | 9.900E-09 |

| | | | | | | | | | | |
|---|---|---|---|---|---|---|---|---|---|---|
| 24480 | 1224 | Q1224V | D | 1.5e-08 | 3.8e-09 | 4.6e-09 | 7.800E-09 | 6.248E-09 | 3.800E-09 | 1.500E-08 |
| 24481 | 1225 | D1225A | D | 1.1e-08 | 1.3e-08 | 9.2e-09 | 1.107E-08 | 1.901E-09 | 9.200E-09 | 1.300E-08 |
| 24482 | 1225 | D1225R | D | 1.4e-08 | 5.5e-09 | 3.2e-09 | 7.567E-09 | 5.689E-09 | 3.200E-09 | 1.400E-08 |
| 24483 | 1225 | D1225N | D | 4.4e-08 | 3e-09 | 1.7e-08 | 2.133E-08 | 2.084E-08 | 3.000E-09 | 4.400E-08 |
| 24484 | 1225 | D1225D | D | 5.5e-09 | 3.8e-09 | 5.6e-09 | 4.967E-09 | 1.012E-09 | 3.800E-09 | 5.600E-09 |
| 24485 | 1225 | D1225C | D | 1e-08 | 3.9e-09 | 1.4e-08 | 9.300E-09 | 5.086E-09 | 3.900E-09 | 1.400E-08 |
| 24486 | 1225 | D1225E | D | 1.5e-08 | 8.1e-09 | 1e-08 | 1.103E-08 | 3.564E-09 | 8.100E-09 | 1.500E-08 |
| 24487 | 1225 | D1225Q | D | 1.4e-08 | 4.5e-09 | 5.6e-09 | 8.033E-09 | 5.196E-09 | 4.500E-09 | 1.400E-08 |
| 24488 | 1225 | D1225G | D | 6.6e-09 | 1.6e-08 | 5.7e-09 | 9.433E-09 | 5.705E-09 | 5.700E-09 | 1.600E-08 |
| 24489 | 1225 | D1225H | D | 4.6e-09 | 1.7e-08 | 1.2e-09 | 7.600E-09 | 8.316E-09 | 1.200E-09 | 1.700E-08 |
| 24490 | 1225 | D1225I | D | 9.7e-09 | 1.7e-08 | 1.4e-08 | 1.357E-08 | 3.669E-09 | 9.700E-09 | 1.700E-08 |
| 24491 | 1225 | D1225L | D | 2.3e-09 | 2.4e-08 | 4.3e-09 | 1.020E-08 | 1.199E-08 | 2.300E-09 | 2.400E-08 |
| 24492 | 1225 | D1225K | D | 6.7e-09 | 5.1e-09 | 2.2e-08 | 1.127E-08 | 9.330E-09 | 5.100E-09 | 2.200E-08 |
| 24493 | 1225 | D1225M | D | 6.9e-09 | 1.8e-08 | 3.8e-09 | 9.567E-09 | 7.466E-09 | 3.800E-09 | 1.800E-08 |
| 24494 | 1225 | D1225F | D | 5.2e-09 | 4.8e-09 | 4.3e-09 | 4.767E-09 | 4.509E-10 | 4.300E-09 | 5.200E-09 |
| 24495 | 1225 | D1225P | D | 1.1e-08 | 8.2e-09 | 1e-08 | 9.733E-09 | 1.419E-09 | 8.200E-09 | 1.100E-08 |
| 24496 | 1225 | D1225S | D | 1.7e-08 | 6.5e-09 | 7.8e-09 | 1.043E-08 | 5.724E-09 | 6.500E-09 | 1.700E-08 |
| 24497 | 1225 | D1225T | D | 6.2e-09 | 5.1e-09 | 3e-09 | 4.767E-09 | 1.626E-09 | 3.000E-09 | 6.200E-09 |
| 24498 | 1225 | D1225W | D | 7.6e-09 | 7e-09 | 1.6e-08 | 1.020E-08 | 5.032E-09 | 7.000E-09 | 1.600E-08 |
| 24499 | 1225 | D1225Y | D | 2e-09 | 4.3e-09 | 6.3e-09 | 4.200E-09 | 2.152E-09 | 2.000E-09 | 6.300E-09 |
| 24500 | 1225 | D1225V | D | 1.4e-08 | 5.1e-09 | 1.8e-08 | 1.237E-08 | 6.603E-09 | 5.100E-09 | 1.800E-08 |
| 24501 | 1226 | S1226A | D | 8.7e-09 | 5.5e-09 | 1.5e-08 | 9.733E-09 | 4.834E-09 | 5.500E-09 | 1.500E-08 |
| 24502 | 1226 | S1226R | D | 7.1e-09 | 7.1e-09 | 6.3e-09 | 6.833E-09 | 4.619E-10 | 6.300E-09 | 7.100E-09 |
| 24503 | 1226 | S1226N | D | 7.5e-09 | 8.9e-09 | 7.9e-09 | 8.100E-09 | 7.211E-10 | 7.500E-09 | 8.900E-09 |
| 24504 | 1226 | S1226D | D | 1.1e-08 | 1.8e-08 | 4e-08 | 2.300E-08 | 1.513E-08 | 1.100E-08 | 4.000E-08 |
| 24505 | 1226 | S1226C | D | 5.3e-09 | 4.4e-09 | 4e-09 | 4.567E-09 | 6.658E-10 | 4.000E-09 | 5.300E-09 |
| 24506 | 1226 | S1226E | D | 6e-09 | 2.4e-08 | 3.3e-09 | 1.110E-08 | 1.125E-08 | 3.300E-09 | 2.400E-08 |
| 24507 | 1226 | S1226Q | D | 3.9e-09 | 1.6e-08 | 1.2e-08 | 1.063E-08 | 6.165E-09 | 3.900E-09 | 1.600E-08 |
| 24508 | 1226 | S1226G | D | 3.8e-09 | 4.8e-09 | 4e-09 | 4.200E-09 | 5.292E-10 | 3.800E-09 | 4.800E-09 |
| 24509 | 1226 | S1226H | D | 1.9e-08 | 6.8e-09 | 1.2e-08 | 1.260E-08 | 6.122E-09 | 6.800E-09 | 1.900E-08 |
| 24510 | 1226 | S1226I | D | 1.1e-08 | 7e-09 | 4.8e-09 | 7.600E-09 | 3.143E-09 | 4.800E-09 | 1.100E-08 |
| 24511 | 1226 | S1226L | D | 1e-08 | 6.8e-09 | 4e-09 | 6.933E-09 | 3.002E-09 | 4.000E-09 | 1.000E-08 |
| 24512 | 1226 | S1226K | D | 1.3e-09 | 7.5e-09 | 2.6e-09 | 3.800E-09 | 3.270E-09 | 1.300E-09 | 7.500E-09 |
| 24513 | 1226 | S1226M | D | 6.8e-09 | 1.7e-09 | 3.3e-09 | 3.933E-09 | 2.608E-09 | 1.700E-09 | 6.800E-09 |
| 24514 | 1226 | S1226F | D | 1e-08 | 4.4e-09 | 7e-09 | 7.133E-09 | 2.802E-09 | 4.400E-09 | 1.000E-08 |
| 24515 | 1226 | S1226P | D | 4.6e-09 | 2.2e-08 | 6.5e-09 | 1.103E-08 | 9.545E-09 | 4.600E-09 | 2.200E-08 |
| 24516 | 1226 | S1226S | D | 4.2e-09 | 7.5e-09 | 6.6e-09 | 6.100E-09 | 1.706E-09 | 4.200E-09 | 7.500E-09 |
| 24517 | 1226 | S1226T | D | 5e-09 | 6.9e-09 | 1.8e-08 | 9.967E-09 | 7.022E-09 | 5.000E-09 | 1.800E-08 |
| 24518 | 1226 | S1226W | D | 7.5e-09 | 1.1e-08 | 8.6e-09 | 9.033E-09 | 1.790E-09 | 7.500E-09 | 1.100E-08 |
| 24519 | 1226 | S1226Y | D | 1.1e-08 | 6.5e-09 | 2.5e-08 | 1.417E-08 | 9.648E-09 | 6.500E-09 | 2.500E-08 |
| 24520 | 1226 | S1226V | D | 9.8e-09 | 1.9e-08 | 7.2e-09 | 1.200E-08 | 6.200E-09 | 7.200E-09 | 1.900E-08 |
| 24521 | 1227 | K1227A | D | 3.2e-09 | 3.9e-09 | 3.8e-09 | 3.633E-09 | 3.786E-10 | 3.200E-09 | 3.900E-09 |
| 24522 | 1227 | K1227R | D | 8.4e-09 | 7.5e-09 | 9.6e-09 | 8.500E-09 | 1.054E-09 | 7.500E-09 | 9.600E-09 |
| 24523 | 1227 | K1227N | D | 6.6e-09 | 1e-09 | 5e-09 | 4.200E-09 | 2.884E-09 | 1.000E-09 | 6.600E-09 |
| 24524 | 1227 | K1227D | D | 6.1e-09 | 8.4e-09 | 7.7e-09 | 7.400E-09 | 1.179E-09 | 6.100E-09 | 8.400E-09 |
| 24525 | 1227 | K1227C | D | 9.9e-09 | 9.7e-09 | 3.9e-09 | 7.833E-09 | 3.408E-09 | 3.900E-09 | 9.900E-09 |
| 24526 | 1227 | K1227E | D | 1.7e-09 | 7.2e-09 | 8.8e-09 | 5.900E-09 | 3.724E-09 | 1.700E-09 | 8.800E-09 |
| 24527 | 1227 | K1227Q | D | 1.6e-08 | 3.2e-09 | 5.5e-09 | 8.233E-09 | 6.824E-09 | 3.200E-09 | 1.600E-08 |
| 24528 | 1227 | K1227G | D | 1e-08 | 1.1e-08 | 6e-09 | 9.000E-09 | 2.646E-09 | 6.000E-09 | 1.100E-08 |
| 24529 | 1227 | K1227H | D | 2.1e-09 | 1.3e-08 | 1.1e-08 | 8.700E-09 | 5.803E-09 | 2.100E-09 | 1.300E-08 |
| 24530 | 1227 | K1227I | D | 1.4e-08 | 5.7e-09 | 8.5e-09 | 9.400E-09 | 4.223E-09 | 5.700E-09 | 1.400E-08 |
| 24531 | 1227 | K1227L | D | 7.4e-09 | 1.1e-08 | 2.2e-08 | 1.347E-08 | 7.606E-09 | 7.400E-09 | 2.200E-08 |
| 24532 | 1227 | K1227K | D | 5.8e-09 | 9e-09 | 3.6e-09 | 6.133E-09 | 2.715E-09 | 3.600E-09 | 9.000E-09 |
| 24533 | 1227 | K1227M | D | 1.5e-08 | 4.8e-09 | 5.8e-09 | 8.533E-09 | 5.623E-09 | 4.800E-09 | 1.500E-08 |
| 24534 | 1227 | K1227F | D | 7.7e-09 | 9.5e-09 | 4.8e-09 | 7.333E-09 | 2.371E-09 | 4.800E-09 | 9.500E-09 |
| 24535 | 1227 | K1227P | D | 6.7e-09 | 6.8e-09 | 8.6e-09 | 7.367E-09 | 1.069E-09 | 6.700E-09 | 8.600E-09 |
| 24536 | 1227 | K1227S | D | 3.3e-09 | 7.3e-09 | 1e-08 | 6.867E-09 | 3.371E-09 | 3.300E-09 | 1.000E-08 |
| 24537 | 1227 | K1227T | D | 4.3e-09 | 4.1e-09 | 2.7e-09 | 3.700E-09 | 8.718E-10 | 2.700E-09 | 4.300E-09 |
| 24538 | 1227 | K1227W | D | 3.7e-09 | 7.1e-09 | 1.1e-08 | 7.267E-09 | 3.653E-09 | 3.700E-09 | 1.100E-08 |
| 24539 | 1227 | K1227Y | D | 1.3e-08 | 4.3e-09 | 5.6e-09 | 7.633E-09 | 4.693E-09 | 4.300E-09 | 1.300E-08 |
| 24540 | 1227 | K1227V | D | 2.4e-08 | 4.2e-09 | 2.6e-09 | 1.027E-08 | 1.192E-08 | 2.600E-09 | 2.400E-08 |
| 24541 | 1228 | D1228A | D | 5.2e-09 | 8.4e-09 | 9e-09 | 7.533E-09 | 2.043E-09 | 5.200E-09 | 9.000E-09 |
| 24542 | 1228 | D1228R | D | 6e-09 | 3.8e-09 | 1.9e-08 | 9.600E-09 | 8.215E-09 | 3.800E-09 | 1.900E-08 |
| 24543 | 1228 | D1228N | D | 6e-09 | 6.8e-09 | 4.9e-09 | 5.900E-09 | 9.539E-10 | 4.900E-09 | 6.800E-09 |
| 24544 | 1228 | D1228D | D | 1.3e-08 | 5.1e-09 | 1.5e-08 | 1.103E-08 | 5.235E-09 | 5.100E-09 | 1.500E-08 |
| 24545 | 1228 | D1228C | D | 8.7e-09 | 1.2e-08 | 7.9e-09 | 9.533E-09 | 2.173E-09 | 7.900E-09 | 1.200E-08 |
| 24546 | 1228 | D1228E | D | 9.4e-09 | 7.4e-09 | 7e-09 | 7.933E-09 | 1.286E-09 | 7.000E-09 | 9.400E-09 |
| 24547 | 1228 | D1228Q | D | 1.3e-09 | 9.1e-09 | 1.2e-08 | 7.467E-09 | 5.534E-09 | 1.300E-09 | 1.200E-08 |
| 24548 | 1228 | D1228G | D | 1.3e-08 | 5.2e-09 | 1.7e-08 | 1.173E-08 | 6.001E-09 | 5.200E-09 | 1.700E-08 |
| 24549 | 1228 | D1228H | D | 4.4e-09 | 4.7e-09 | 1.3e-08 | 7.367E-09 | 4.881E-09 | 4.400E-09 | 1.300E-08 |
| 24550 | 1228 | D1228I | D | 6.5e-09 | 1e-08 | 1.3e-08 | 9.833E-09 | 3.253E-09 | 6.500E-09 | 1.300E-08 |
| 24551 | 1228 | D1228L | D | 6.4e-09 | 1.1e-08 | 3.8e-09 | 7.067E-09 | 3.646E-09 | 3.800E-09 | 1.100E-08 |
| 24552 | 1228 | D1228K | D | 4e-09 | 9.3e-09 | 1.2e-08 | 8.433E-09 | 4.070E-09 | 4.000E-09 | 1.200E-08 |
| 24553 | 1228 | D1228M | D | 1.3e-08 | 1.6e-08 | 8.9e-09 | 1.263E-08 | 3.564E-09 | 8.900E-09 | 1.600E-08 |
| 24554 | 1228 | D1228F | D | 6e-09 | 7e-09 | 4.4e-09 | 5.800E-09 | 1.311E-09 | 4.400E-09 | 7.000E-09 |
| 24555 | 1228 | D1228P | D | 8.5e-09 | 7e-09 | 9.2e-09 | 8.233E-09 | 1.124E-09 | 7.000E-09 | 9.200E-09 |
| 24556 | 1228 | D1228S | D | 5.7e-09 | 6.1e-09 | 3.1e-09 | 4.967E-09 | 1.629E-09 | 3.100E-09 | 6.100E-09 |
| 24557 | 1228 | D1228T | D | 7.8e-09 | 7.2e-09 | 7.2e-09 | 7.400E-09 | 3.464E-10 | 7.200E-09 | 7.800E-09 |
| 24558 | 1228 | D1228W | D | 1e-08 | 4e-09 | 4.8e-09 | 6.267E-09 | 3.258E-09 | 4.000E-09 | 1.000E-08 |
| 24559 | 1228 | D1228Y | D | 1.8e-08 | 1.2e-08 | 6.1e-09 | 1.203E-08 | 5.950E-09 | 6.100E-09 | 1.800E-08 |
| 24560 | 1228 | D1228V | D | 1.7e-08 | 1.2e-08 | 3.7e-09 | 1.090E-08 | 6.718E-09 | 3.700E-09 | 1.700E-08 |
| 24561 | 1229 | S1229A | D | 1.2e-08 | 7.9e-09 | 5.2e-08 | 2.397E-08 | 2.436E-08 | 7.900E-09 | 5.200E-08 |
| 24562 | 1229 | S1229R | D | 4.5e-09 | 6.1e-09 | 7.9e-09 | 6.167E-09 | 1.701E-09 | 4.500E-09 | 7.900E-09 |
| 24563 | 1229 | S1229N | D | 1.1e-08 | 3.9e-09 | 4.5e-09 | 6.467E-09 | 3.937E-09 | 3.900E-09 | 1.100E-08 |
| 24564 | 1229 | S1229D | D | 1.2e-08 | 1.1e-08 | 4.8e-09 | 9.267E-09 | 3.900E-09 | 4.800E-09 | 1.200E-08 |

| | | | | | | | | | | |
|---|---|---|---|---|---|---|---|---|---|---|
| 24565 | 1229 | S1229C | D | 3.2e-09 | 2.1e-08 | 1.2e-08 | 1.207E-08 | 8.900E-09 | 3.200E-09 | 2.100E-08 |
| 24566 | 1229 | S1229E | D | 5.4e-09 | 8.6e-09 | 5.3e-09 | 6.433E-09 | 1.877E-09 | 5.300E-09 | 8.600E-09 |
| 24567 | 1229 | S1229Q | D | 3.4e-09 | 1.4e-08 | 7.1e-09 | 8.167E-09 | 5.380E-09 | 3.400E-09 | 1.400E-08 |
| 24568 | 1229 | S1229G | D | 1.2e-08 | 8.4e-09 | 6.9e-09 | 9.100E-09 | 2.621E-09 | 6.900E-09 | 1.200E-08 |
| 24569 | 1229 | S1229H | D | 9.2e-09 | 9.9e-09 | 1.2e-08 | 1.037E-08 | 1.457E-09 | 9.200E-09 | 1.200E-08 |
| 24570 | 1229 | S1229I | D | 9.1e-09 | 5.1e-09 | 8.4e-09 | 7.533E-09 | 2.136E-09 | 5.100E-09 | 9.100E-09 |
| 24571 | 1229 | S1229L | D | 5.7e-09 | 9.6e-09 | 3.4e-08 | 1.643E-08 | 1.534E-08 | 5.700E-09 | 3.400E-08 |
| 24572 | 1229 | S1229K | D | 2.3e-08 | 1.5e-08 | 5.4e-09 | 1.447E-08 | 8.812E-09 | 5.400E-09 | 2.300E-08 |
| 24573 | 1229 | S1229M | D | 4.3e-09 | 1.2e-08 | 2e-09 | 6.100E-09 | 5.237E-09 | 2.000E-09 | 1.200E-08 |
| 24574 | 1229 | S1229F | D | 1.2e-08 | 3.2e-09 | 1.3e-08 | 9.400E-09 | 5.393E-09 | 3.200E-09 | 1.300E-08 |
| 24575 | 1229 | S1229P | D | 3.7e-09 | 7.7e-09 | 5e-09 | 5.467E-09 | 2.040E-09 | 3.700E-09 | 7.700E-09 |
| 24576 | 1229 | S1229S | D | 5.6e-09 | 8.5e-09 | 7.8e-09 | 7.300E-09 | 1.513E-09 | 5.600E-09 | 8.500E-09 |
| 24577 | 1229 | S1229T | D | 8.4e-09 | 7.2e-09 | 7.4e-09 | 7.667E-09 | 6.429E-10 | 7.200E-09 | 8.400E-09 |
| 24578 | 1229 | S1229W | D | 2.5e-09 | 4.6e-09 | 8.3e-09 | 5.133E-09 | 2.937E-09 | 2.500E-09 | 8.300E-09 |
| 24579 | 1229 | S1229Y | D | 1.6e-08 | 1.2e-08 | 1.1e-08 | 1.300E-08 | 2.646E-09 | 1.100E-08 | 1.600E-08 |
| 24580 | 1229 | S1229V | D | 3.4e-09 | 5.9e-09 | 7.1e-09 | 5.467E-09 | 1.888E-09 | 3.400E-09 | 7.100E-09 |
| 24581 | 1230 | T1230A | D | 1.5e-08 | 3.4e-09 | 1.7e-08 | 1.180E-08 | 7.343E-09 | 3.400E-09 | 1.700E-08 |
| 24582 | 1230 | T1230R | D | 5.5e-09 | 7.4e-09 | 1.2e-08 | 8.300E-09 | 3.342E-09 | 5.500E-09 | 1.200E-08 |
| 24583 | 1230 | T1230N | D | 3.5e-09 | 1.5e-08 | 1.1e-08 | 9.833E-09 | 5.838E-09 | 3.500E-09 | 1.500E-08 |
| 24584 | 1230 | T1230D | D | 4.7e-09 | 1.2e-08 | 7.4e-09 | 8.033E-09 | 3.691E-09 | 4.700E-09 | 1.200E-08 |
| 24585 | 1230 | T1230C | D | 7.6e-09 | 1.6e-08 | 8.1e-09 | 1.057E-08 | 4.712E-09 | 7.600E-09 | 1.600E-08 |
| 24586 | 1230 | T1230E | D | 7.8e-09 | 1.6e-08 | 4.7e-09 | 9.500E-09 | 5.839E-09 | 4.700E-09 | 1.600E-08 |
| 24587 | 1230 | T1230Q | D | 1.1e-08 | 5.4e-09 | 4.6e-09 | 7.000E-09 | 3.487E-09 | 4.600E-09 | 1.100E-08 |
| 24588 | 1230 | T1230G | D | 6.2e-09 | 1.1e-08 | 1.1e-08 | 9.400E-09 | 2.771E-09 | 6.200E-09 | 1.100E-08 |
| 24589 | 1230 | T1230H | D | 6.6e-09 | 1.2e-08 | 6.1e-09 | 8.233E-09 | 3.272E-09 | 6.100E-09 | 1.200E-08 |
| 24590 | 1230 | T1230I | D | 1.3e-08 | 7e-09 | 7.8e-09 | 9.267E-09 | 3.258E-09 | 7.000E-09 | 1.300E-08 |
| 24591 | 1230 | T1230L | D | 1.4e-08 | 7.7e-09 | 5e-09 | 8.900E-09 | 4.618E-09 | 5.000E-09 | 1.400E-08 |
| 24592 | 1230 | T1230K | D | 3.8e-09 | 8.4e-09 | 4.3e-09 | 5.500E-09 | 2.524E-09 | 3.800E-09 | 8.400E-09 |
| 24593 | 1230 | T1230M | D | 7.1e-09 | 8.5e-09 | 4.8e-09 | 6.800E-09 | 1.868E-09 | 4.800E-09 | 8.500E-09 |
| 24594 | 1230 | T1230F | D | 2e-08 | 7.8e-09 | 1e-08 | 1.260E-08 | 6.502E-09 | 7.800E-09 | 2.000E-08 |
| 24595 | 1230 | T1230P | D | 3.9e-09 | 1e-08 | 1.1e-08 | 8.300E-09 | 3.843E-09 | 3.900E-09 | 1.100E-08 |
| 24596 | 1230 | T1230S | D | 9.7e-09 | 3.2e-09 | 6.5e-09 | 6.467E-09 | 3.250E-09 | 3.200E-09 | 9.700E-09 |
| 24597 | 1230 | T1230T | D | 6.4e-09 | 8.9e-09 | 6.3e-09 | 7.200E-09 | 1.473E-09 | 6.300E-09 | 8.900E-09 |
| 24598 | 1230 | T1230W | D | 8.5e-09 | 5e-09 | 3.2e-09 | 5.567E-09 | 2.695E-09 | 3.200E-09 | 8.500E-09 |
| 24599 | 1230 | T1230Y | D | 6.6e-09 | 5.6e-09 | 1.9e-08 | 1.040E-08 | 7.465E-09 | 5.600E-09 | 1.900E-08 |
| 24600 | 1230 | T1230V | D | 5.7e-09 | 4e-09 | 1.7e-09 | 3.800E-09 | 2.007E-09 | 1.700E-09 | 5.700E-09 |
| 24601 | 1231 | Y1231A | D | 3.2e-09 | 6.3e-09 | 3.7e-09 | 4.400E-09 | 1.664E-09 | 3.200E-09 | 6.300E-09 |
| 24602 | 1231 | Y1231R | D | 1.8e-08 | 1.5e-08 | 6.1e-09 | 1.303E-08 | 6.189E-09 | 6.100E-09 | 1.800E-08 |
| 24603 | 1231 | Y1231N | D | 6e-09 | 2.6e-09 | 5e-09 | 4.533E-09 | 1.747E-09 | 2.600E-09 | 6.000E-09 |
| 24604 | 1231 | Y1231D | D | 6.1e-09 | 6.3e-09 | 9.3e-09 | 7.233E-09 | 1.793E-09 | 6.100E-09 | 9.300E-09 |
| 24605 | 1231 | Y1231C | D | 5.1e-09 | 1.5e-08 | 3.7e-09 | 7.933E-09 | 6.160E-09 | 3.700E-09 | 1.500E-08 |
| 24606 | 1231 | Y1231E | D | 1.3e-08 | 9.4e-09 | 7.1e-09 | 9.833E-09 | 2.974E-09 | 7.100E-09 | 1.300E-08 |
| 24607 | 1231 | Y1231Q | D | 6.2e-09 | 4.3e-09 | 1.5e-08 | 8.500E-09 | 5.709E-09 | 4.300E-09 | 1.500E-08 |
| 24608 | 1231 | Y1231G | D | 3.8e-09 | 4.4e-09 | 4.3e-09 | 4.167E-09 | 3.215E-10 | 3.800E-09 | 4.400E-09 |
| 24609 | 1231 | Y1231H | D | 3e-09 | 1.3e-09 | 7.6e-09 | 3.967E-09 | 3.259E-09 | 1.300E-09 | 7.600E-09 |
| 24610 | 1231 | Y1231I | D | 2.9e-08 | 3.2e-09 | 3.3e-09 | 1.183E-08 | 1.487E-08 | 3.200E-09 | 2.900E-08 |
| 24611 | 1231 | Y1231L | D | 9.2e-09 | 1.9e-09 | 2.3e-09 | 4.467E-09 | 4.104E-09 | 1.900E-09 | 9.200E-09 |
| 24612 | 1231 | Y1231K | D | 6e-09 | 2.5e-08 | 1e-08 | 1.367E-08 | 1.002E-08 | 6.000E-09 | 2.500E-08 |
| 24613 | 1231 | Y1231M | D | 9.5e-09 | 4.8e-09 | 5.3e-09 | 6.533E-09 | 2.581E-09 | 4.800E-09 | 9.500E-09 |
| 24614 | 1231 | Y1231F | D | 1.3e-08 | 2.9e-09 | 6.2e-09 | 7.367E-09 | 5.150E-09 | 2.900E-09 | 1.300E-08 |
| 24615 | 1231 | Y1231P | D | 5.9e-09 | 8.9e-09 | 1.1e-08 | 8.600E-09 | 2.563E-09 | 5.900E-09 | 1.100E-08 |
| 24616 | 1231 | Y1231S | D | 2.4e-08 | 5.4e-09 | 4e-09 | 1.113E-08 | 1.116E-08 | 4.000E-09 | 2.400E-08 |
| 24617 | 1231 | Y1231T | D | 4e-09 | 8.6e-09 | 4.9e-09 | 5.833E-09 | 2.438E-09 | 4.000E-09 | 8.600E-09 |
| 24618 | 1231 | Y1231W | D | 1.1e-08 | 5.6e-09 | 1.8e-08 | 1.153E-08 | 6.217E-09 | 5.600E-09 | 1.800E-08 |
| 24619 | 1231 | Y1231Y | D | 1.4e-08 | 7.4e-09 | 9.6e-09 | 1.033E-08 | 3.361E-09 | 7.400E-09 | 1.400E-08 |
| 24620 | 1231 | Y1231V | D | 9.3e-09 | 1.4e-08 | 3.3e-09 | 8.867E-09 | 5.363E-09 | 3.300E-09 | 1.400E-08 |
| 24621 | 1232 | S1232A | D | 3.5e-09 | 7.3e-09 | 6.3e-09 | 5.700E-09 | 1.970E-09 | 3.500E-09 | 7.300E-09 |
| 24622 | 1232 | S1232R | D | 1e-08 | 1.5e-08 | 4.3e-09 | 9.767E-09 | 5.354E-09 | 4.300E-09 | 1.500E-08 |
| 24623 | 1232 | S1232N | D | 4.6e-09 | 1.1e-08 | 9.9e-09 | 8.500E-09 | 3.422E-09 | 4.600E-09 | 1.100E-08 |
| 24624 | 1232 | S1232D | D | 9.9e-09 | 6.1e-09 | 1.3e-08 | 9.667E-09 | 3.456E-09 | 6.100E-09 | 1.300E-08 |
| 24625 | 1232 | S1232C | D | 2.5e-08 | 1.6e-08 | 6.2e-09 | 1.573E-08 | 9.403E-09 | 6.200E-09 | 2.500E-08 |
| 24626 | 1232 | S1232E | D | 2e-09 | 6.8e-09 | 5.4e-09 | 4.733E-09 | 2.468E-09 | 2.000E-09 | 6.800E-09 |
| 24627 | 1232 | S1232Q | D | 6e-09 | 2.6e-09 | 1.2e-08 | 6.867E-09 | 4.760E-09 | 2.600E-09 | 1.200E-08 |
| 24628 | 1232 | S1232G | D | 8.4e-09 | 5.1e-09 | 3.6e-09 | 5.700E-09 | 2.456E-09 | 3.600E-09 | 8.400E-09 |
| 24629 | 1232 | S1232H | D | 8.8e-09 | 1.1e-08 | 8.6e-09 | 9.467E-09 | 1.332E-09 | 8.600E-09 | 1.100E-08 |
| 24630 | 1232 | S1232I | D | 3.5e-09 | 7.3e-09 | 6.4e-09 | 5.733E-09 | 1.986E-09 | 3.500E-09 | 7.300E-09 |
| 24631 | 1232 | S1232L | D | 6.7e-09 | 1.1e-08 | 1.3e-08 | 1.023E-08 | 3.219E-09 | 6.700E-09 | 1.300E-08 |
| 24632 | 1232 | S1232K | D | 7.6e-09 | 1.5e-08 | 2.6e-09 | 8.400E-09 | 6.239E-09 | 2.600E-09 | 1.500E-08 |
| 24633 | 1232 | S1232M | D | 1.4e-08 | 1.3e-08 | 8e-09 | 1.167E-08 | 3.215E-09 | 8.000E-09 | 1.400E-08 |
| 24634 | 1232 | S1232F | D | 1.2e-08 | 7.2e-09 | 1e-08 | 9.733E-09 | 2.411E-09 | 7.200E-09 | 1.200E-08 |
| 24635 | 1232 | S1232P | D | 1.8e-08 | 1.9e-08 | 7e-09 | 1.467E-08 | 6.658E-09 | 7.000E-09 | 1.900E-08 |
| 24636 | 1232 | S1232S | D | 5.3e-09 | 7.4e-09 | 7e-09 | 6.567E-09 | 1.115E-09 | 5.300E-09 | 7.400E-09 |
| 24637 | 1232 | S1232T | D | 4.2e-09 | 9e-09 | 3.5e-09 | 5.567E-09 | 2.994E-09 | 3.500E-09 | 9.000E-09 |
| 24638 | 1232 | S1232W | D | 1.1e-08 | 6.9e-09 | 6.6e-09 | 8.167E-09 | 2.458E-09 | 6.600E-09 | 1.100E-08 |
| 24639 | 1232 | S1232Y | D | 8.1e-09 | 1.7e-08 | 7.8e-09 | 1.097E-08 | 5.227E-09 | 7.800E-09 | 1.700E-08 |
| 24640 | 1232 | S1232V | D | 1.1e-08 | 1.4e-09 | 1.3e-08 | 8.467E-09 | 6.201E-09 | 1.400E-09 | 1.300E-08 |
| 24641 | 1233 | L1233A | D | 1.1e-08 | 4e-09 | 3.4e-09 | 6.133E-09 | 4.225E-09 | 3.400E-09 | 1.100E-08 |
| 24642 | 1233 | L1233R | D | 2.2e-08 | 7.1e-09 | 9.2e-09 | 1.277E-08 | 8.065E-09 | 7.100E-09 | 2.200E-08 |
| 24643 | 1233 | L1233N | D | 2.6e-09 | 1.9e-09 | 2.8e-09 | 2.433E-09 | 4.726E-10 | 1.900E-09 | 2.800E-09 |
| 24644 | 1233 | L1233D | D | 1e-08 | 3.9e-09 | 9.9e-09 | 7.933E-09 | 3.493E-09 | 3.900E-09 | 1.000E-08 |
| 24645 | 1233 | L1233C | D | 5.2e-09 | 7.1e-09 | 4.4e-09 | 5.567E-09 | 1.387E-09 | 4.400E-09 | 7.100E-09 |
| 24646 | 1233 | L1233E | D | 9.2e-09 | 9.9e-09 | 7.9e-09 | 9.000E-09 | 1.015E-09 | 7.900E-09 | 9.900E-09 |
| 24647 | 1233 | L1233Q | D | 5.6e-09 | 1.6e-08 | 1.4e-08 | 1.187E-08 | 5.518E-09 | 5.600E-09 | 1.600E-08 |
| 24648 | 1233 | L1233G | D | 6.1e-09 | 1.2e-08 | 7e-09 | 8.367E-09 | 3.179E-09 | 6.100E-09 | 1.200E-08 |
| 24649 | 1233 | L1233H | D | 6.7e-09 | 1e-08 | 1.7e-08 | 1.123E-08 | 5.260E-09 | 6.700E-09 | 1.700E-08 |

| | | | | | | | | | | |
|---|---|---|---|---|---|---|---|---|---|---|
| 24650 | 1233 | L1233I | D | 4.4e-09 | 1.1e-08 | 6.4e-09 | 7.267E-09 | 3.384E-09 | 4.400E-09 | 1.100E-08 |
| 24651 | 1233 | L1233L | D | 7.8e-09 | 5.8e-09 | 7.6e-09 | 7.067E-09 | 1.102E-09 | 5.800E-09 | 7.800E-09 |
| 24652 | 1233 | L1233K | D | 1.1e-08 | 9.4e-09 | 7.9e-09 | 9.433E-09 | 1.550E-09 | 7.900E-09 | 1.100E-08 |
| 24653 | 1233 | L1233M | D | 8e-09 | 1.1e-08 | 6.5e-09 | 8.500E-09 | 2.291E-09 | 6.500E-09 | 1.100E-08 |
| 24654 | 1233 | L1233F | D | 1.6e-08 | 2.3e-09 | 1.5e-08 | 1.110E-08 | 7.637E-09 | 2.300E-09 | 1.600E-08 |
| 24655 | 1233 | L1233P | D | 8.4e-09 | 3.9e-09 | 1.3e-08 | 8.433E-09 | 4.550E-09 | 3.900E-09 | 1.300E-08 |
| 24656 | 1233 | L1233S | D | 1.9e-08 | 1.3e-08 | 6.2e-09 | 1.273E-08 | 6.404E-09 | 6.200E-09 | 1.900E-08 |
| 24657 | 1233 | L1233T | D | 2.8e-09 | 8.8e-09 | 9.8e-09 | 7.133E-09 | 3.786E-09 | 2.800E-09 | 9.800E-09 |
| 24658 | 1233 | L1233W | D | 1.9e-09 | 1.2e-08 | 1.2e-08 | 8.633E-09 | 5.831E-09 | 1.900E-09 | 1.200E-08 |
| 24659 | 1233 | L1233Y | D | 9.2e-09 | 7.4e-09 | 9.4e-09 | 8.667E-09 | 1.102E-09 | 7.400E-09 | 9.400E-09 |
| 24660 | 1233 | L1233V | D | 5.6e-09 | 3e-09 | 3.7e-09 | 4.100E-09 | 1.345E-09 | 3.000E-09 | 5.600E-09 |
| 24661 | 1234 | S1234A | D | 1.1e-08 | 6.6e-09 | 1.6e-08 | 1.120E-08 | 4.703E-09 | 6.600E-09 | 1.600E-08 |
| 24662 | 1234 | S1234R | D | 1.2e-08 | 5.5e-09 | 5e-09 | 7.500E-09 | 3.905E-09 | 5.000E-09 | 1.200E-08 |
| 24663 | 1234 | S1234N | D | 6.4e-09 | 7e-09 | 8.3e-09 | 7.233E-09 | 9.713E-10 | 6.400E-09 | 8.300E-09 |
| 24664 | 1234 | S1234D | D | 1e-08 | 1.3e-08 | 7.6e-09 | 1.020E-08 | 2.706E-09 | 7.600E-09 | 1.300E-08 |
| 24665 | 1234 | S1234C | D | 1.1e-08 | 8.6e-09 | 1.5e-08 | 1.153E-08 | 3.233E-09 | 8.600E-09 | 1.500E-08 |
| 24666 | 1234 | S1234E | D | 1.6e-09 | 6.3e-09 | 4.7e-09 | 4.200E-09 | 2.390E-09 | 1.600E-09 | 6.300E-09 |
| 24667 | 1234 | S1234Q | D | 1.1e-08 | 1.5e-08 | 4.5e-09 | 1.017E-08 | 5.299E-09 | 4.500E-09 | 1.500E-08 |
| 24668 | 1234 | S1234G | D | 4.3e-09 | 1.1e-08 | 5.2e-09 | 6.833E-09 | 3.636E-09 | 4.300E-09 | 1.100E-08 |
| 24669 | 1234 | S1234H | D | 1.1e-08 | 3.8e-09 | 6.7e-09 | 7.167E-09 | 3.623E-09 | 3.800E-09 | 1.100E-08 |
| 24670 | 1234 | S1234I | D | 3.8e-09 | 2.9e-09 | 2.8e-09 | 3.167E-09 | 5.508E-10 | 2.800E-09 | 3.800E-09 |
| 24671 | 1234 | S1234L | D | 9.2e-09 | 3.6e-09 | 1.2e-08 | 8.267E-09 | 4.277E-09 | 3.600E-09 | 1.200E-08 |
| 24672 | 1234 | S1234K | D | 4.6e-09 | 5.9e-09 | 4.2e-09 | 4.900E-09 | 8.888E-10 | 4.200E-09 | 5.900E-09 |
| 24673 | 1234 | S1234M | D | 1e-08 | 3.5e-09 | 4e-08 | 1.783E-08 | 1.947E-08 | 3.500E-09 | 4.000E-08 |
| 24674 | 1234 | S1234F | D | 3.4e-09 | 1e-08 | 6e-09 | 6.467E-09 | 3.325E-09 | 3.400E-09 | 1.000E-08 |
| 24675 | 1234 | S1234P | D | 1.7e-08 | 1e-08 | 7.1e-09 | 1.137E-08 | 5.090E-09 | 7.100E-09 | 1.700E-08 |
| 24676 | 1234 | S1234S | D | 7.5e-09 | 6.6e-09 | 5.5e-09 | 6.533E-09 | 1.002E-09 | 5.500E-09 | 7.500E-09 |
| 24677 | 1234 | S1234T | D | 8.8e-09 | 2.2e-08 | 9.6e-09 | 1.347E-08 | 7.401E-09 | 8.800E-09 | 2.200E-08 |
| 24678 | 1234 | S1234W | D | 6.5e-09 | 1e-08 | 4.1e-09 | 6.867E-09 | 2.967E-09 | 4.100E-09 | 1.000E-08 |
| 24679 | 1234 | S1234Y | D | 7.4e-09 | 1.6e-08 | 1.5e-08 | 1.280E-08 | 4.703E-09 | 7.400E-09 | 1.600E-08 |
| 24680 | 1234 | S1234V | D | 1.1e-08 | 1e-08 | 1.2e-08 | 1.100E-08 | 1.000E-09 | 1.000E-08 | 1.200E-08 |
| 24681 | 1235 | S1235A | D | 9.5e-09 | 1e-08 | 5.8e-09 | 8.433E-09 | 2.294E-09 | 5.800E-09 | 1.000E-08 |
| 24682 | 1235 | S1235R | D | 9.4e-09 | 8.9e-09 | 5.9e-09 | 8.067E-09 | 1.893E-09 | 5.900E-09 | 9.400E-09 |
| 24683 | 1235 | S1235N | D | 4.7e-09 | 3.1e-09 | 4e-09 | 3.933E-09 | 8.021E-10 | 3.100E-09 | 4.700E-09 |
| 24684 | 1235 | S1235D | D | 3.7e-09 | 4.6e-09 | 3.2e-09 | 3.833E-09 | 7.095E-10 | 3.200E-09 | 4.600E-09 |
| 24685 | 1235 | S1235C | D | 6.1e-09 | 4.4e-09 | 6.3e-09 | 5.600E-09 | 1.044E-09 | 4.400E-09 | 6.300E-09 |
| 24686 | 1235 | S1235E | D | 1e-08 | 6.4e-09 | 1.7e-09 | 6.033E-09 | 4.162E-09 | 1.700E-09 | 1.000E-08 |
| 24687 | 1235 | S1235Q | D | 5.8e-09 | 5.3e-09 | 3.8e-09 | 4.967E-09 | 1.041E-09 | 3.800E-09 | 5.800E-09 |
| 24688 | 1235 | S1235G | D | 1.2e-08 | 4.2e-09 | 1.1e-08 | 9.067E-09 | 4.244E-09 | 4.200E-09 | 1.200E-08 |
| 24689 | 1235 | S1235H | D | 4.9e-09 | 1.1e-08 | 3.2e-09 | 6.367E-09 | 4.102E-09 | 3.200E-09 | 1.100E-08 |
| 24690 | 1235 | S1235I | D | 1.1e-08 | 3e-09 | 1.4e-08 | 9.333E-09 | 5.686E-09 | 3.000E-09 | 1.400E-08 |
| 24691 | 1235 | S1235L | D | 4.2e-09 | 1.2e-08 | 2.6e-09 | 6.267E-09 | 5.029E-09 | 2.600E-09 | 1.200E-08 |
| 24692 | 1235 | S1235K | D | 2.4e-09 | 2.4e-08 | 6.7e-09 | 1.103E-08 | 1.143E-08 | 2.400E-09 | 2.400E-08 |
| 24693 | 1235 | S1235M | D | 6.1e-09 | 1.2e-08 | 3.1e-09 | 7.067E-09 | 4.528E-09 | 3.100E-09 | 1.200E-08 |
| 24694 | 1235 | S1235F | D | 2.3e-08 | 4.8e-09 | 2.1e-09 | 9.967E-09 | 1.137E-08 | 2.100E-09 | 2.300E-08 |
| 24695 | 1235 | S1235P | D | 1.2e-08 | 7.2e-09 | 4.7e-09 | 7.967E-09 | 3.710E-09 | 4.700E-09 | 1.200E-08 |
| 24696 | 1235 | S1235S | D | 1.9e-08 | 9.9e-09 | 2.3e-09 | 1.040E-08 | 8.361E-09 | 2.300E-09 | 1.900E-08 |
| 24697 | 1235 | S1235T | D | 5.7e-09 | 1.2e-08 | 4.5e-09 | 7.400E-09 | 4.029E-09 | 4.500E-09 | 1.200E-08 |
| 24698 | 1235 | S1235W | D | 1.5e-08 | 3.7e-09 | 7.8e-09 | 8.833E-09 | 5.720E-09 | 3.700E-09 | 1.500E-08 |
| 24699 | 1235 | S1235Y | D | 1.6e-08 | 9.9e-09 | 6.2e-09 | 1.070E-08 | 4.949E-09 | 6.200E-09 | 1.600E-08 |
| 24700 | 1235 | S1235V | D | 3.9e-09 | 5.8e-09 | 4e-09 | 4.567E-09 | 1.069E-09 | 3.900E-09 | 5.800E-09 |
| 24701 | 1236 | T1236A | D | 8.7e-09 | 4.2e-09 | 9.8e-09 | 7.567E-09 | 2.967E-09 | 4.200E-09 | 9.800E-09 |
| 24702 | 1236 | T1236R | D | 6.5e-09 | 5.5e-09 | 6.7e-09 | 6.233E-09 | 6.429E-10 | 5.500E-09 | 6.700E-09 |
| 24703 | 1236 | T1236N | D | 6.5e-09 | 4.5e-09 | 8.3e-09 | 6.433E-09 | 1.901E-09 | 4.500E-09 | 8.300E-09 |
| 24704 | 1236 | T1236D | D | 4.5e-09 | 1.8e-08 | 1.6e-08 | 1.283E-08 | 7.286E-09 | 4.500E-09 | 1.800E-08 |
| 24705 | 1236 | T1236C | D | 1.7e-09 | 4.7e-09 | 5.2e-09 | 3.867E-09 | 1.893E-09 | 1.700E-09 | 5.200E-09 |
| 24706 | 1236 | T1236E | D | 6.9e-09 | 4e-09 | 1.2e-08 | 7.633E-09 | 4.050E-09 | 4.000E-09 | 1.200E-08 |
| 24707 | 1236 | T1236Q | D | 6.1e-09 | 4.6e-09 | 7.6e-09 | 6.100E-09 | 1.500E-09 | 4.600E-09 | 7.600E-09 |
| 24708 | 1236 | T1236G | D | 3.1e-09 | 2.6e-09 | 1.5e-08 | 6.900E-09 | 7.019E-09 | 2.600E-09 | 1.500E-08 |
| 24709 | 1236 | T1236H | D | 7.3e-09 | 4.9e-09 | 5.4e-09 | 5.867E-09 | 1.266E-09 | 4.900E-09 | 7.300E-09 |
| 24710 | 1236 | T1236I | D | 1.1e-08 | 1.5e-09 | 8e-09 | 6.833E-09 | 4.856E-09 | 1.500E-09 | 1.100E-08 |
| 24711 | 1236 | T1236L | D | 5.5e-09 | 9.5e-09 | 2.3e-09 | 5.767E-09 | 3.607E-09 | 2.300E-09 | 9.500E-09 |
| 24712 | 1236 | T1236K | D | 5.9e-09 | 7.7e-09 | 2.9e-09 | 5.500E-09 | 2.425E-09 | 2.900E-09 | 7.700E-09 |
| 24713 | 1236 | T1236M | D | 3.5e-09 | 1.5e-08 | 7.1e-09 | 8.533E-09 | 5.882E-09 | 3.500E-09 | 1.500E-08 |
| 24714 | 1236 | T1236F | D | 4.7e-09 | 1.9e-09 | 9.5e-09 | 5.367E-09 | 3.844E-09 | 1.900E-09 | 9.500E-09 |
| 24715 | 1236 | T1236P | D | 3.4e-09 | 8.6e-09 | 8.8e-09 | 6.933E-09 | 3.062E-09 | 3.400E-09 | 8.800E-09 |
| 24716 | 1236 | T1236S | D | 8.4e-09 | 2.3e-08 | 1.1e-08 | 1.413E-08 | 7.788E-09 | 8.400E-09 | 2.300E-08 |
| 24717 | 1236 | T1236T | D | 2.4e-09 | 1.2e-08 | 1.6e-08 | 1.013E-08 | 6.990E-09 | 2.400E-09 | 1.600E-08 |
| 24718 | 1236 | T1236W | D | 8.7e-09 | 8.2e-09 | 1.5e-08 | 1.063E-08 | 3.790E-09 | 8.200E-09 | 1.500E-08 |
| 24719 | 1236 | T1236Y | D | 2.9e-08 | 3.6e-09 | 7e-09 | 1.320E-08 | 1.379E-08 | 3.600E-09 | 2.900E-08 |
| 24720 | 1236 | T1236V | D | 6.6e-09 | 3.8e-09 | 8.5e-09 | 6.300E-09 | 2.364E-09 | 3.800E-09 | 8.500E-09 |
| 24721 | 1237 | L1237A | D | 6.2e-09 | 1.3e-08 | 7.2e-09 | 8.800E-09 | 3.672E-09 | 6.200E-09 | 1.300E-08 |
| 24722 | 1237 | L1237R | D | 1.6e-08 | 7.7e-09 | 4.4e-09 | 9.367E-09 | 5.977E-09 | 4.400E-09 | 1.600E-08 |
| 24723 | 1237 | L1237N | D | 6.8e-09 | 1.8e-08 | 7e-09 | 1.060E-08 | 6.409E-09 | 6.800E-09 | 1.800E-08 |
| 24724 | 1237 | L1237D | D | 4.7e-09 | 7.8e-09 | 5.7e-09 | 6.067E-09 | 1.582E-09 | 4.700E-09 | 7.800E-09 |
| 24725 | 1237 | L1237C | D | 3.1e-09 | 1e-09 | 7.1e-09 | 3.733E-09 | 3.099E-09 | 1.000E-09 | 7.100E-09 |
| 24726 | 1237 | L1237E | D | 3.1e-09 | 5.6e-09 | 7.2e-09 | 5.300E-09 | 2.066E-09 | 3.100E-09 | 7.200E-09 |
| 24727 | 1237 | L1237Q | D | 3.4e-09 | 4.3e-09 | 8.1e-09 | 5.267E-09 | 2.495E-09 | 3.400E-09 | 8.100E-09 |
| 24728 | 1237 | L1237G | D | 5.8e-09 | 2.4e-08 | 8e-09 | 1.260E-08 | 9.934E-09 | 5.800E-09 | 2.400E-08 |
| 24729 | 1237 | L1237H | D | 1.2e-08 | 4.8e-09 | 1.9e-08 | 1.193E-08 | 7.100E-09 | 4.800E-09 | 1.900E-08 |
| 24730 | 1237 | L1237I | D | 6.1e-09 | 8.6e-09 | 3.4e-09 | 6.033E-09 | 2.601E-09 | 3.400E-09 | 8.600E-09 |
| 24731 | 1237 | L1237L | D | 9.7e-09 | 8.9e-09 | 5.3e-09 | 7.967E-09 | 2.344E-09 | 5.300E-09 | 9.700E-09 |
| 24732 | 1237 | L1237K | D | 3.8e-09 | 5.9e-09 | 9.8e-09 | 6.500E-09 | 3.045E-09 | 3.800E-09 | 9.800E-09 |
| 24733 | 1237 | L1237M | D | 3.9e-09 | 8e-09 | 3e-09 | 4.967E-09 | 2.665E-09 | 3.000E-09 | 8.000E-09 |
| 24734 | 1237 | L1237F | D | 1.7e-08 | 1.9e-08 | 1.5e-08 | 1.700E-08 | 2.000E-09 | 1.500E-08 | 1.900E-08 |

| | | | | | | | | | | |
|---|---|---|---|---|---|---|---|---|---|---|
| 24735 | 1237 | L1237P | D | 1.7e-08 | 6.5e-09 | 1.4e-08 | 1.250E-08 | 5.408E-09 | 6.500E-09 | 1.700E-08 |
| 24736 | 1237 | L1237S | D | 5.1e-09 | 5.3e-09 | 7.7e-09 | 6.033E-09 | 1.447E-09 | 5.100E-09 | 7.700E-09 |
| 24737 | 1237 | L1237T | D | 1.1e-08 | 4.5e-09 | 6.3e-09 | 7.267E-09 | 3.356E-09 | 4.500E-09 | 1.100E-08 |
| 24738 | 1237 | L1237W | D | 9.7e-09 | 1.1e-08 | 6.2e-09 | 8.967E-09 | 2.483E-09 | 6.200E-09 | 1.100E-08 |
| 24739 | 1237 | L1237Y | D | 2.4e-09 | 3.9e-09 | 3.9e-09 | 3.400E-09 | 8.660E-10 | 2.400E-09 | 3.900E-09 |
| 24740 | 1237 | L1237V | D | 4.7e-09 | 5.5e-09 | 6.5e-09 | 5.567E-09 | 9.018E-10 | 4.700E-09 | 6.500E-09 |
| 24741 | 1238 | T1238A | D | 3.4e-09 | 2.1e-08 | 7.2e-09 | 1.053E-08 | 9.261E-09 | 3.400E-09 | 2.100E-08 |
| 24742 | 1238 | T1238R | D | 1.8e-09 | 1.1e-08 | 9.1e-09 | 7.300E-09 | 4.857E-09 | 1.800E-09 | 1.100E-08 |
| 24743 | 1238 | T1238N | D | 9.9e-09 | 1.4e-08 | 8.3e-09 | 1.073E-08 | 2.940E-09 | 8.300E-09 | 1.400E-08 |
| 24744 | 1238 | T1238D | D | 7.6e-09 | 1.4e-08 | 3.7e-09 | 8.433E-09 | 5.200E-09 | 3.700E-09 | 1.400E-08 |
| 24745 | 1238 | T1238C | D | 9.8e-09 | 6.2e-09 | 6e-09 | 7.333E-09 | 2.139E-09 | 6.000E-09 | 9.800E-09 |
| 24746 | 1238 | T1238E | D | 8.6e-09 | 5.4e-09 | 1e-08 | 8.000E-09 | 2.358E-09 | 5.400E-09 | 1.000E-08 |
| 24747 | 1238 | T1238Q | D | 8.7e-09 | 1.1e-08 | 2e-08 | 1.323E-08 | 5.972E-09 | 8.700E-09 | 2.000E-08 |
| 24748 | 1238 | T1238G | D | 9e-09 | 4.4e-09 | 2.4e-09 | 5.267E-09 | 3.384E-09 | 2.400E-09 | 9.000E-09 |
| 24749 | 1238 | T1238H | D | 6.3e-09 | 5.5e-09 | 3.1e-09 | 4.967E-09 | 1.665E-09 | 3.100E-09 | 6.300E-09 |
| 24750 | 1238 | T1238I | D | 7.9e-09 | 7.4e-09 | 6.1e-09 | 7.133E-09 | 9.292E-10 | 6.100E-09 | 7.900E-09 |
| 24751 | 1238 | T1238L | D | 1.4e-09 | 8.2e-09 | 6.2e-09 | 5.267E-09 | 3.495E-09 | 1.400E-09 | 8.200E-09 |
| 24752 | 1238 | T1238K | D | 1.6e-09 | 6.2e-09 | 5.2e-09 | 4.333E-09 | 2.419E-09 | 1.600E-09 | 6.200E-09 |
| 24753 | 1238 | T1238M | D | 9e-09 | 6.7e-09 | 1.2e-08 | 9.233E-09 | 2.658E-09 | 6.700E-09 | 1.200E-08 |
| 24754 | 1238 | T1238F | D | 6.5e-09 | 3.6e-09 | 6.5e-09 | 5.533E-09 | 1.674E-09 | 3.600E-09 | 6.500E-09 |
| 24755 | 1238 | T1238P | D | 5.7e-09 | 6.7e-09 | 1.2e-08 | 8.133E-09 | 3.386E-09 | 5.700E-09 | 1.200E-08 |
| 24756 | 1238 | T1238S | D | 2.8e-09 | 4.8e-09 | 1.3e-08 | 6.867E-09 | 5.405E-09 | 2.800E-09 | 1.300E-08 |
| 24757 | 1238 | T1238T | D | 8.3e-09 | 3.9e-09 | 6.1e-09 | 6.100E-09 | 2.200E-09 | 3.900E-09 | 8.300E-09 |
| 24758 | 1238 | T1238W | D | 7.9e-09 | 1.9e-09 | 9.9e-09 | 6.567E-09 | 4.163E-09 | 1.900E-09 | 9.900E-09 |
| 24759 | 1238 | T1238Y | D | 1.6e-08 | 1e-08 | 4.4e-09 | 1.013E-08 | 5.801E-09 | 4.400E-09 | 1.600E-08 |
| 24760 | 1238 | T1238V | D | 4.8e-09 | 5.6e-09 | 1.9e-09 | 4.100E-09 | 1.947E-09 | 1.900E-09 | 5.600E-09 |
| 24761 | 1239 | L1239A | D | 8.7e-09 | 7.1e-09 | 1.1e-08 | 8.933E-09 | 1.960E-09 | 7.100E-09 | 1.100E-08 |
| 24762 | 1239 | L1239R | D | 4.3e-09 | 3.9e-09 | 1.2e-08 | 6.733E-09 | 4.565E-09 | 3.900E-09 | 1.200E-08 |
| 24763 | 1239 | L1239N | D | 1.2e-08 | 6.9e-09 | 7.3e-09 | 8.733E-09 | 2.836E-09 | 6.900E-09 | 1.200E-08 |
| 24764 | 1239 | L1239D | D | 8.8e-09 | 6.9e-09 | 9e-09 | 8.233E-09 | 1.159E-09 | 6.900E-09 | 9.000E-09 |
| 24765 | 1239 | L1239C | D | 9.7e-09 | 6.2e-09 | 2.9e-09 | 6.267E-09 | 3.400E-09 | 2.900E-09 | 9.700E-09 |
| 24766 | 1239 | L1239E | D | 8.7e-09 | 2.7e-09 | 2.7e-09 | 4.700E-09 | 3.464E-09 | 2.700E-09 | 8.700E-09 |
| 24767 | 1239 | L1239Q | D | 3.9e-09 | 6e-09 | 8.2e-09 | 6.033E-09 | 2.150E-09 | 3.900E-09 | 8.200E-09 |
| 24768 | 1239 | L1239G | D | 1.6e-08 | 6.4e-09 | 7.7e-09 | 1.003E-08 | 5.208E-09 | 6.400E-09 | 1.600E-08 |
| 24769 | 1239 | L1239H | D | 2.5e-08 | 6.2e-09 | 1.2e-08 | 1.440E-08 | 9.627E-09 | 6.200E-09 | 2.500E-08 |
| 24770 | 1239 | L1239I | D | 4.7e-09 | 3.5e-09 | 3e-09 | 3.733E-09 | 8.737E-10 | 3.000E-09 | 4.700E-09 |
| 24771 | 1239 | L1239L | D | 3.3e-09 | 4e-09 | 3.3e-09 | 3.533E-09 | 4.041E-10 | 3.300E-09 | 4.000E-09 |
| 24772 | 1239 | L1239K | D | 1e-08 | 6.4e-09 | 4.8e-09 | 7.067E-09 | 2.663E-09 | 4.800E-09 | 1.000E-08 |
| 24773 | 1239 | L1239M | D | 2.4e-09 | 2e-08 | 2.8e-09 | 8.400E-09 | 1.005E-08 | 2.400E-09 | 2.000E-08 |
| 24774 | 1239 | L1239F | D | 8.2e-09 | 1.1e-08 | 1.2e-08 | 1.040E-08 | 1.970E-09 | 8.200E-09 | 1.200E-08 |
| 24775 | 1239 | L1239P | D | 5e-09 | 8e-09 | 4.3e-09 | 5.767E-09 | 1.966E-09 | 4.300E-09 | 8.000E-09 |
| 24776 | 1239 | L1239S | D | 1.3e-08 | 5.4e-09 | 4.2e-09 | 7.533E-09 | 4.772E-09 | 4.200E-09 | 1.300E-08 |
| 24777 | 1239 | L1239T | D | 5.1e-09 | 8.8e-09 | 3.2e-09 | 5.700E-09 | 2.848E-09 | 3.200E-09 | 8.800E-09 |
| 24778 | 1239 | L1239W | D | 8.5e-09 | 1.9e-08 | 1.1e-08 | 1.283E-08 | 5.485E-09 | 8.500E-09 | 1.900E-08 |
| 24779 | 1239 | L1239Y | D | 4.4e-09 | 4.3e-09 | 9.2e-09 | 5.967E-09 | 2.801E-09 | 4.300E-09 | 9.200E-09 |
| 24780 | 1239 | L1239V | D | 2.2e-09 | 1.4e-08 | 5.5e-09 | 7.233E-09 | 6.088E-09 | 2.200E-09 | 1.400E-08 |
| 24781 | 1240 | S1240A | D | 7.2e-09 | 5e-09 | 9.5e-09 | 7.233E-09 | 2.250E-09 | 5.000E-09 | 9.500E-09 |
| 24782 | 1240 | S1240R | D | 6.5e-09 | 3.1e-09 | 8.5e-09 | 6.033E-09 | 2.730E-09 | 3.100E-09 | 8.500E-09 |
| 24783 | 1240 | S1240N | D | 4.7e-09 | 6.1e-09 | 3.2e-09 | 4.667E-09 | 1.450E-09 | 3.200E-09 | 6.100E-09 |
| 24784 | 1240 | S1240D | D | 5.8e-09 | 8.2e-09 | 2.3e-09 | 5.433E-09 | 2.967E-09 | 2.300E-09 | 8.200E-09 |
| 24785 | 1240 | S1240C | D | 5.6e-09 | 5.8e-09 | 1.2e-09 | 4.200E-09 | 2.600E-09 | 1.200E-09 | 5.800E-09 |
| 24786 | 1240 | S1240E | D | 3.6e-09 | 8.9e-09 | 1.5e-08 | 9.167E-09 | 5.705E-09 | 3.600E-09 | 1.500E-08 |
| 24787 | 1240 | S1240Q | D | 3.5e-09 | 6.3e-09 | 6.4e-09 | 5.400E-09 | 1.646E-09 | 3.500E-09 | 6.400E-09 |
| 24788 | 1240 | S1240G | D | 3.2e-09 | 9.9e-09 | 8e-09 | 7.033E-09 | 3.453E-09 | 3.200E-09 | 9.900E-09 |
| 24789 | 1240 | S1240H | D | 1e-08 | 9.2e-09 | 1.3e-08 | 1.073E-08 | 2.003E-09 | 9.200E-09 | 1.300E-08 |
| 24790 | 1240 | S1240I | D | 1.1e-08 | 3e-09 | 9.8e-09 | 7.933E-09 | 4.314E-09 | 3.000E-09 | 1.100E-08 |
| 24791 | 1240 | S1240L | D | 1.1e-08 | 8.4e-09 | 7.6e-09 | 9.000E-09 | 1.778E-09 | 7.600E-09 | 1.100E-08 |
| 24792 | 1240 | S1240K | D | 2.1e-08 | 3.7e-09 | 8e-09 | 1.090E-08 | 9.007E-09 | 3.700E-09 | 2.100E-08 |
| 24793 | 1240 | S1240M | D | 8.6e-09 | 7.6e-09 | 5.1e-09 | 7.100E-09 | 1.803E-09 | 5.100E-09 | 8.600E-09 |
| 24794 | 1240 | S1240F | D | 1.1e-08 | 6.7e-09 | 1.4e-09 | 6.367E-09 | 4.809E-09 | 1.400E-09 | 1.100E-08 |
| 24795 | 1240 | S1240P | D | 2.3e-09 | 4.2e-09 | 1.2e-08 | 6.167E-09 | 5.140E-09 | 2.300E-09 | 1.200E-08 |
| 24796 | 1240 | S1240S | D | 9.3e-09 | 4.1e-09 | 4.1e-09 | 5.833E-09 | 3.002E-09 | 4.100E-09 | 9.300E-09 |
| 24797 | 1240 | S1240T | D | 5e-09 | 9.2e-09 | 8.8e-09 | 7.667E-09 | 2.318E-09 | 5.000E-09 | 9.200E-09 |
| 24798 | 1240 | S1240W | D | 3.5e-09 | 5e-09 | 4.6e-09 | 4.367E-09 | 7.767E-10 | 3.500E-09 | 5.000E-09 |
| 24799 | 1240 | S1240Y | D | 4.4e-09 | 6.5e-09 | 3.4e-09 | 4.767E-09 | 1.582E-09 | 3.400E-09 | 6.500E-09 |
| 24800 | 1240 | S1240V | D | 2.5e-09 | 1.4e-08 | 5.4e-09 | 7.300E-09 | 5.981E-09 | 2.500E-09 | 1.400E-08 |
| 24801 | 1241 | K1241A | D | 2.3e-09 | 6.9e-09 | 1.1e-08 | 6.733E-09 | 4.352E-09 | 2.300E-09 | 1.100E-08 |
| 24802 | 1241 | K1241R | D | 2.6e-09 | 7.6e-09 | 6.6e-09 | 5.600E-09 | 2.646E-09 | 2.600E-09 | 7.600E-09 |
| 24803 | 1241 | K1241N | D | 7.1e-09 | 2.2e-09 | 5.9e-09 | 5.067E-09 | 2.554E-09 | 2.200E-09 | 7.100E-09 |
| 24804 | 1241 | K1241D | D | 3e-09 | 2.6e-09 | 1.5e-08 | 6.867E-09 | 7.047E-09 | 2.600E-09 | 1.500E-08 |
| 24805 | 1241 | K1241C | D | 4.4e-09 | 4.4e-09 | 1.7e-08 | 8.600E-09 | 7.275E-09 | 4.400E-09 | 1.700E-08 |
| 24806 | 1241 | K1241E | D | 3.4e-09 | 8e-09 | 5.9e-09 | 5.767E-09 | 2.303E-09 | 3.400E-09 | 8.000E-09 |
| 24807 | 1241 | K1241Q | D | 6.9e-09 | 6.6e-09 | 7.2e-09 | 6.900E-09 | 3.000E-10 | 6.600E-09 | 7.200E-09 |
| 24808 | 1241 | K1241G | D | 3e-09 | 1.4e-08 | 8.2e-09 | 8.400E-09 | 5.503E-09 | 3.000E-09 | 1.400E-08 |
| 24809 | 1241 | K1241H | D | 5.8e-09 | 1.2e-08 | 1.2e-08 | 9.933E-09 | 3.580E-09 | 5.800E-09 | 1.200E-08 |
| 24810 | 1241 | K1241I | D | 1.4e-08 | 1.7e-08 | 1.5e-09 | 1.083E-08 | 8.221E-09 | 1.500E-09 | 1.700E-08 |
| 24811 | 1241 | K1241L | D | 3.9e-09 | 2.2e-08 | 2.4e-08 | 1.663E-08 | 1.107E-08 | 3.900E-09 | 2.400E-08 |
| 24812 | 1241 | K1241K | D | 4.3e-09 | 4.8e-09 | 4.2e-09 | 4.433E-09 | 3.215E-10 | 4.200E-09 | 4.800E-09 |
| 24813 | 1241 | K1241M | D | 1.2e-08 | 6.6e-09 | 1.8e-08 | 1.220E-08 | 5.703E-09 | 6.600E-09 | 1.800E-08 |
| 24814 | 1241 | K1241F | D | 1.4e-08 | 4.5e-09 | 8.1e-09 | 8.867E-09 | 4.796E-09 | 4.500E-09 | 1.400E-08 |
| 24815 | 1241 | K1241P | D | 7.8e-09 | 1.6e-08 | 2.1e-09 | 8.633E-09 | 6.987E-09 | 2.100E-09 | 1.600E-08 |
| 24816 | 1241 | K1241S | D | 2e-08 | 1.4e-08 | 3.3e-09 | 1.243E-08 | 8.460E-09 | 3.300E-09 | 2.000E-08 |
| 24817 | 1241 | K1241T | D | 3.2e-09 | 2e-08 | 1.4e-08 | 1.240E-08 | 8.514E-09 | 3.200E-09 | 2.000E-08 |
| 24818 | 1241 | K1241W | D | 3.2e-09 | 9.2e-09 | 8.5e-09 | 6.967E-09 | 3.281E-09 | 3.200E-09 | 9.200E-09 |
| 24819 | 1241 | K1241Y | D | 1.1e-08 | 5.4e-09 | 7.3e-09 | 7.900E-09 | 2.848E-09 | 5.400E-09 | 1.100E-08 |

| | | | | | | | | | | |
|---|---|---|---|---|---|---|---|---|---|---|
| 24820 | 1241 | K1241V | D | 7e-09 | 4.4e-09 | 9.2e-09 | 6.867E-09 | 2.403E-09 | 4.400E-09 | 9.200E-09 |
| 24821 | 1242 | A1242A | D | 1.4e-08 | 1.9e-08 | 1e-08 | 1.433E-08 | 4.509E-09 | 1.000E-08 | 1.900E-08 |
| 24822 | 1242 | A1242R | D | 3.5e-09 | 9e-09 | 6.3e-09 | 6.267E-09 | 2.750E-09 | 3.500E-09 | 9.000E-09 |
| 24823 | 1242 | A1242N | D | 2.4e-09 | 1.9e-09 | 1.3e-09 | 1.867E-09 | 5.508E-10 | 1.300E-09 | 2.400E-09 |
| 24824 | 1242 | A1242D | D | 1.3e-08 | 1.1e-08 | 7.3e-09 | 1.043E-08 | 2.892E-09 | 7.300E-09 | 1.300E-08 |
| 24825 | 1242 | A1242C | D | 2.5e-08 | 1.2e-09 | 2.1e-09 | 9.433E-09 | 1.349E-08 | 1.200E-09 | 2.500E-08 |
| 24826 | 1242 | A1242E | D | 3.9e-09 | 9.3e-09 | 1.4e-08 | 9.067E-09 | 5.054E-09 | 3.900E-09 | 1.400E-08 |
| 24827 | 1242 | A1242Q | D | 1e-08 | 9.5e-09 | 1.7e-08 | 1.217E-08 | 4.193E-09 | 9.500E-09 | 1.700E-08 |
| 24828 | 1242 | A1242G | D | 1.4e-08 | 8.8e-09 | 3.4e-09 | 8.733E-09 | 5.300E-09 | 3.400E-09 | 1.400E-08 |
| 24829 | 1242 | A1242H | D | 6.3e-09 | 1.1e-08 | 1.4e-08 | 1.043E-08 | 3.881E-09 | 6.300E-09 | 1.400E-08 |
| 24830 | 1242 | A1242I | D | 4e-09 | 9.5e-09 | 5.3e-08 | 2.217E-08 | 2.684E-08 | 4.000E-09 | 5.300E-08 |
| 24831 | 1242 | A1242L | D | 1.1e-08 | 6.8e-09 | 7e-09 | 8.267E-09 | 2.369E-09 | 6.800E-09 | 1.100E-08 |
| 24832 | 1242 | A1242K | D | 7e-09 | 9e-09 | 1.4e-08 | 1.000E-08 | 3.606E-09 | 7.000E-09 | 1.400E-08 |
| 24833 | 1242 | A1242M | D | 7e-09 | 1.7e-08 | 4.1e-09 | 9.367E-09 | 6.768E-09 | 4.100E-09 | 1.700E-08 |
| 24834 | 1242 | A1242F | D | 9.3e-09 | 8.3e-09 | 1.8e-08 | 1.187E-08 | 5.335E-09 | 8.300E-09 | 1.800E-08 |
| 24835 | 1242 | A1242P | D | 3.8e-09 | 1.9e-08 | 5.2e-09 | 9.333E-09 | 8.401E-09 | 3.800E-09 | 1.900E-08 |
| 24836 | 1242 | A1242S | D | 1.3e-08 | 2.1e-08 | 2.9e-09 | 1.230E-08 | 9.070E-09 | 2.900E-09 | 2.100E-08 |
| 24837 | 1242 | A1242T | D | 2.1e-09 | 2e-09 | 3.9e-09 | 2.667E-09 | 1.069E-09 | 2.000E-09 | 3.900E-09 |
| 24838 | 1242 | A1242W | D | 2.2e-09 | 7e-09 | 5.8e-09 | 5.000E-09 | 2.498E-09 | 2.200E-09 | 7.000E-09 |
| 24839 | 1242 | A1242Y | D | 6.8e-09 | 4.5e-09 | 4.1e-09 | 5.133E-09 | 1.457E-09 | 4.100E-09 | 6.800E-09 |
| 24840 | 1242 | A1242V | D | 2.2e-08 | 1.6e-08 | 2.3e-09 | 1.343E-08 | 1.010E-08 | 2.300E-09 | 2.200E-08 |
| 24841 | 1243 | D1243A | D | 4.4e-09 | 1.4e-08 | 6e-09 | 8.133E-09 | 5.143E-09 | 4.400E-09 | 1.400E-08 |
| 24842 | 1243 | D1243R | D | 7.2e-09 | 9.2e-09 | 1.3e-08 | 9.800E-09 | 2.946E-09 | 7.200E-09 | 1.300E-08 |
| 24843 | 1243 | D1243N | D | 2e-09 | 1.2e-08 | 1.5e-08 | 9.667E-09 | 6.807E-09 | 2.000E-09 | 1.500E-08 |
| 24844 | 1243 | D1243D | D | 5.6e-09 | 1.8e-08 | 7.8e-09 | 1.047E-08 | 6.616E-09 | 5.600E-09 | 1.800E-08 |
| 24845 | 1243 | D1243C | D | 1.5e-08 | 6e-09 | 1.4e-08 | 1.167E-08 | 4.933E-09 | 6.000E-09 | 1.500E-08 |
| 24846 | 1243 | D1243E | D | 1.2e-08 | 4.1e-09 | 4e-09 | 6.700E-09 | 4.590E-09 | 4.000E-09 | 1.200E-08 |
| 24847 | 1243 | D1243Q | D | 6e-09 | 7.2e-09 | 1.6e-08 | 9.733E-09 | 5.460E-09 | 6.000E-09 | 1.600E-08 |
| 24848 | 1243 | D1243G | D | 3e-09 | 7e-09 | 1.4e-08 | 8.000E-09 | 5.568E-09 | 3.000E-09 | 1.400E-08 |
| 24849 | 1243 | D1243H | D | 1.1e-08 | 1.4e-08 | 1.2e-08 | 1.233E-08 | 1.528E-09 | 1.100E-08 | 1.400E-08 |
| 24850 | 1243 | D1243I | D | 9.5e-09 | 1.7e-09 | 1e-08 | 7.067E-09 | 4.654E-09 | 1.700E-09 | 1.000E-08 |
| 24851 | 1243 | D1243L | D | 5.3e-09 | 7.5e-09 | 6.2e-09 | 6.333E-09 | 1.106E-09 | 5.300E-09 | 7.500E-09 |
| 24852 | 1243 | D1243K | D | 2.9e-08 | 1e-08 | 2e-09 | 1.367E-08 | 1.387E-08 | 2.000E-09 | 2.900E-08 |
| 24853 | 1243 | D1243M | D | 2.4e-08 | 1.6e-08 | 7.9e-09 | 1.597E-08 | 8.050E-09 | 7.900E-09 | 2.400E-08 |
| 24854 | 1243 | D1243F | D | 1.3e-08 | 7.1e-09 | 1.1e-08 | 1.037E-08 | 3.001E-09 | 7.100E-09 | 1.300E-08 |
| 24855 | 1243 | D1243P | D | 1e-08 | 7.2e-09 | 1.7e-08 | 1.140E-08 | 5.048E-09 | 7.200E-09 | 1.700E-08 |
| 24856 | 1243 | D1243S | D | 1.2e-08 | 2.8e-09 | 6.8e-09 | 7.200E-09 | 4.613E-09 | 2.800E-09 | 1.200E-08 |
| 24857 | 1243 | D1243T | D | 4.5e-09 | 2.7e-09 | 3.6e-09 | 3.600E-09 | 9.000E-10 | 2.700E-09 | 4.500E-09 |
| 24858 | 1243 | D1243W | D | 6.9e-09 | 5.9e-09 | 4.1e-09 | 5.633E-09 | 1.419E-09 | 4.100E-09 | 6.900E-09 |
| 24859 | 1243 | D1243Y | D | 2e-08 | 1.6e-08 | 1.5e-08 | 1.700E-08 | 2.646E-09 | 1.500E-08 | 2.000E-08 |
| 24860 | 1243 | D1243V | D | 3.9e-09 | 1.1e-08 | 5e-09 | 6.633E-09 | 3.821E-09 | 3.900E-09 | 1.100E-08 |
| 24861 | 1244 | Y1244A | D | 8.5e-09 | 7.9e-09 | 1.5e-08 | 1.047E-08 | 3.937E-09 | 7.900E-09 | 1.500E-08 |
| 24862 | 1244 | Y1244R | D | 5e-09 | 4.2e-09 | 1e-08 | 6.400E-09 | 3.143E-09 | 4.200E-09 | 1.000E-08 |
| 24863 | 1244 | Y1244N | D | 5.4e-09 | 3.7e-09 | 5.8e-09 | 4.967E-09 | 1.115E-09 | 3.700E-09 | 5.800E-09 |
| 24864 | 1244 | Y1244D | D | 1.4e-08 | 1.2e-08 | 1.4e-08 | 1.333E-08 | 1.155E-09 | 1.200E-08 | 1.400E-08 |
| 24865 | 1244 | Y1244C | D | 7.5e-09 | 1e-08 | 9.4e-09 | 8.967E-09 | 1.305E-09 | 7.500E-09 | 1.000E-08 |
| 24866 | 1244 | Y1244E | D | 6.3e-09 | 2.1e-09 | 5.4e-09 | 4.600E-09 | 2.211E-09 | 2.100E-09 | 6.300E-09 |
| 24867 | 1244 | Y1244Q | D | 1.7e-09 | 4.5e-09 | 3.4e-09 | 3.200E-09 | 1.411E-09 | 1.700E-09 | 4.500E-09 |
| 24868 | 1244 | Y1244G | D | 3.7e-09 | 6.7e-09 | 1e-08 | 6.800E-09 | 3.151E-09 | 3.700E-09 | 1.000E-08 |
| 24869 | 1244 | Y1244H | D | 1.1e-08 | 2.1e-08 | 4.6e-09 | 1.220E-08 | 8.266E-09 | 4.600E-09 | 2.100E-08 |
| 24870 | 1244 | Y1244I | D | 4.8e-09 | 1.2e-08 | 7.6e-09 | 8.133E-09 | 3.630E-09 | 4.800E-09 | 1.200E-08 |
| 24871 | 1244 | Y1244L | D | 6.2e-09 | 1.4e-09 | 1.9e-08 | 8.867E-09 | 9.098E-09 | 1.400E-09 | 1.900E-08 |
| 24872 | 1244 | Y1244K | D | 1.9e-08 | 6.7e-09 | 4.9e-09 | 1.020E-08 | 7.674E-09 | 4.900E-09 | 1.900E-08 |
| 24873 | 1244 | Y1244M | D | 4.5e-08 | 4.5e-09 | 6e-09 | 1.850E-08 | 2.296E-08 | 4.500E-09 | 4.500E-08 |
| 24874 | 1244 | Y1244F | D | 3.7e-09 | 3.7e-09 | 5.3e-09 | 4.233E-09 | 9.238E-10 | 3.700E-09 | 5.300E-09 |
| 24875 | 1244 | Y1244P | D | 4.1e-09 | 6.6e-09 | 2.8e-09 | 4.500E-09 | 1.931E-09 | 2.800E-09 | 6.600E-09 |
| 24876 | 1244 | Y1244S | D | 6.9e-09 | 1.2e-08 | 2.3e-08 | 1.397E-08 | 8.228E-09 | 6.900E-09 | 2.300E-08 |
| 24877 | 1244 | Y1244T | D | 5.6e-09 | 4.3e-09 | 5.1e-09 | 5.000E-09 | 6.557E-10 | 4.300E-09 | 5.600E-09 |
| 24878 | 1244 | Y1244W | D | 8.9e-09 | 7.3e-09 | 7.5e-09 | 7.900E-09 | 8.718E-10 | 7.300E-09 | 8.900E-09 |
| 24879 | 1244 | Y1244Y | D | 9.1e-09 | 4.9e-09 | 1.3e-08 | 9.000E-09 | 4.051E-09 | 4.900E-09 | 1.300E-08 |
| 24880 | 1244 | Y1244V | D | 8.9e-09 | 1.1e-08 | 3e-09 | 7.633E-09 | 4.148E-09 | 3.000E-09 | 1.100E-08 |
| 24881 | 1245 | E1245A | D | 7.6e-09 | 8.9e-09 | 1e-08 | 8.833E-09 | 1.201E-09 | 7.600E-09 | 1.000E-08 |
| 24882 | 1245 | E1245R | D | 6.1e-09 | 7.3e-09 | 7e-09 | 6.800E-09 | 6.245E-10 | 6.100E-09 | 7.300E-09 |
| 24883 | 1245 | E1245N | D | 9.3e-09 | 2.3e-09 | 4.4e-09 | 5.333E-09 | 3.592E-09 | 2.300E-09 | 9.300E-09 |
| 24884 | 1245 | E1245D | D | 6.6e-09 | 3.8e-09 | 3.9e-09 | 4.767E-09 | 1.589E-09 | 3.800E-09 | 6.600E-09 |
| 24885 | 1245 | E1245C | D | 4.6e-09 | 5.9e-09 | 3.6e-09 | 4.700E-09 | 1.153E-09 | 3.600E-09 | 5.900E-09 |
| 24886 | 1245 | E1245E | D | 3.7e-09 | 2.3e-08 | 6.2e-09 | 1.097E-08 | 1.050E-08 | 3.700E-09 | 2.300E-08 |
| 24887 | 1245 | E1245Q | D | 4.5e-09 | 9.4e-09 | 7.9e-10 | 4.897E-09 | 4.319E-09 | 7.900E-10 | 9.400E-09 |
| 24888 | 1245 | E1245G | D | 1.6e-08 | 2.9e-09 | 6.8e-09 | 8.567E-09 | 6.726E-09 | 2.900E-09 | 1.600E-08 |
| 24889 | 1245 | E1245H | D | 1.6e-08 | 2.8e-08 | 4.5e-09 | 1.617E-08 | 1.175E-08 | 4.500E-09 | 2.800E-08 |
| 24890 | 1245 | E1245I | D | 3.9e-09 | 1.4e-09 | 6.9e-09 | 4.067E-09 | 2.754E-09 | 1.400E-09 | 6.900E-09 |
| 24891 | 1245 | E1245L | D | 6.5e-09 | 4.8e-09 | 3.6e-09 | 4.967E-09 | 1.457E-09 | 3.600E-09 | 6.500E-09 |
| 24892 | 1245 | E1245K | D | 1.1e-08 | 5.6e-09 | 1e-08 | 8.867E-09 | 2.873E-09 | 5.600E-09 | 1.100E-08 |
| 24893 | 1245 | E1245M | D | 9.1e-09 | 2.2e-08 | 3.6e-09 | 1.157E-08 | 9.445E-09 | 3.600E-09 | 2.200E-08 |
| 24894 | 1245 | E1245F | D | 5.2e-09 | 1.5e-08 | 7.3e-09 | 9.167E-09 | 5.160E-09 | 5.200E-09 | 1.500E-08 |
| 24895 | 1245 | E1245P | D | 5.4e-09 | 6.5e-09 | 5e-09 | 5.633E-09 | 7.767E-10 | 5.000E-09 | 6.500E-09 |
| 24896 | 1245 | E1245S | D | 8.9e-09 | 6.9e-09 | 6.9e-09 | 7.567E-09 | 1.155E-09 | 6.900E-09 | 8.900E-09 |
| 24897 | 1245 | E1245T | D | 4.4e-09 | 9e-09 | 2e-08 | 1.113E-08 | 8.016E-09 | 4.400E-09 | 2.000E-08 |
| 24898 | 1245 | E1245W | D | 3.5e-09 | 1.1e-08 | 1.5e-08 | 9.833E-09 | 5.838E-09 | 3.500E-09 | 1.500E-08 |
| 24899 | 1245 | E1245Y | D | 1.8e-08 | 4.2e-09 | 5.7e-09 | 9.300E-09 | 7.572E-09 | 4.200E-09 | 1.800E-08 |
| 24900 | 1245 | E1245V | D | 1.1e-08 | 7.9e-09 | 7.2e-09 | 8.700E-09 | 2.022E-09 | 7.200E-09 | 1.100E-08 |
| 24901 | 1246 | K1246A | D | 6.7e-09 | 5.4e-09 | 8.7e-09 | 6.933E-09 | 1.662E-09 | 5.400E-09 | 8.700E-09 |
| 24902 | 1246 | K1246R | D | 9.2e-09 | 5.2e-09 | 6.3e-09 | 6.900E-09 | 2.066E-09 | 5.200E-09 | 9.200E-09 |
| 24903 | 1246 | K1246N | D | 9.2e-09 | 1.1e-08 | 5.8e-09 | 8.667E-09 | 2.641E-09 | 5.800E-09 | 1.100E-08 |
| 24904 | 1246 | K1246D | D | 2.9e-08 | 6.8e-09 | 2e-08 | 1.860E-08 | 1.117E-08 | 6.800E-09 | 2.900E-08 |

| | | | | | | | | | | |
|---|---|---|---|---|---|---|---|---|---|---|
| 24905 | 1246 | K1246C | D | 5.7e-09 | 6.4e-09 | 1.3e-08 | 8.367E-09 | 4.028E-09 | 5.700E-09 | 1.300E-08 |
| 24906 | 1246 | K1246E | D | 9.5e-09 | 1.2e-08 | 7.8e-09 | 9.767E-09 | 2.113E-09 | 7.800E-09 | 1.200E-08 |
| 24907 | 1246 | K1246Q | D | 7.1e-09 | 1.8e-08 | 5.1e-09 | 1.007E-08 | 6.943E-09 | 5.100E-09 | 1.800E-08 |
| 24908 | 1246 | K1246G | D | 7.8e-09 | 6.1e-09 | 1.5e-08 | 9.633E-09 | 4.725E-09 | 6.100E-09 | 1.500E-08 |
| 24909 | 1246 | K1246H | D | 1e-08 | 1.2e-08 | 3.1e-09 | 8.367E-09 | 4.669E-09 | 3.100E-09 | 1.200E-08 |
| 24910 | 1246 | K1246I | D | 9.6e-09 | 7.5e-09 | 5.9e-09 | 7.667E-09 | 1.856E-09 | 5.900E-09 | 9.600E-09 |
| 24911 | 1246 | K1246L | D | 6.2e-09 | 5.9e-09 | 1e-08 | 7.367E-09 | 2.285E-09 | 5.900E-09 | 1.000E-08 |
| 24912 | 1246 | K1246K | D | 8.2e-09 | 1.1e-08 | 5e-09 | 8.067E-09 | 3.002E-09 | 5.000E-09 | 1.100E-08 |
| 24913 | 1246 | K1246M | D | 3.5e-09 | 6.8e-09 | 4.5e-09 | 4.933E-09 | 1.692E-09 | 3.500E-09 | 6.800E-09 |
| 24914 | 1246 | K1246F | D | 4.4e-09 | 1.2e-08 | 5.9e-09 | 7.433E-09 | 4.025E-09 | 4.400E-09 | 1.200E-08 |
| 24915 | 1246 | K1246P | D | 7.1e-09 | 2.9e-09 | 6.1e-09 | 5.367E-09 | 2.194E-09 | 2.900E-09 | 7.100E-09 |
| 24916 | 1246 | K1246S | D | 2.4e-09 | 5.9e-09 | 9.1e-09 | 5.800E-09 | 3.351E-09 | 2.400E-09 | 9.100E-09 |
| 24917 | 1246 | K1246T | D | 5.9e-09 | 5.7e-09 | 1.1e-08 | 7.533E-09 | 3.004E-09 | 5.700E-09 | 1.100E-08 |
| 24918 | 1246 | K1246W | D | 5.9e-09 | 1.5e-08 | 1e-08 | 1.030E-08 | 4.557E-09 | 5.900E-09 | 1.500E-08 |
| 24919 | 1246 | K1246Y | D | 5.9e-09 | 2e-09 | 2.8e-08 | 1.197E-08 | 1.402E-08 | 2.000E-09 | 2.800E-08 |
| 24920 | 1246 | K1246V | D | 3e-09 | 1.1e-08 | 4e-09 | 6.000E-09 | 4.359E-09 | 3.000E-09 | 1.100E-08 |
| 24921 | 1247 | H1247A | D | 2.8e-09 | 3e-09 | 4.9e-09 | 3.567E-09 | 1.159E-09 | 2.800E-09 | 4.900E-09 |
| 24922 | 1247 | H1247R | D | 5.4e-09 | 2.7e-08 | 1.2e-08 | 1.480E-08 | 1.107E-08 | 5.400E-09 | 2.700E-08 |
| 24923 | 1247 | H1247N | D | 4.7e-09 | 1.2e-08 | 4.5e-09 | 7.067E-09 | 4.274E-09 | 4.500E-09 | 1.200E-08 |
| 24924 | 1247 | H1247D | D | 8.7e-09 | 1.8e-08 | 2.4e-09 | 9.700E-09 | 7.848E-09 | 2.400E-09 | 1.800E-08 |
| 24925 | 1247 | H1247C | D | 1.1e-09 | 6.2e-09 | 6.9e-09 | 4.733E-09 | 3.166E-09 | 1.100E-09 | 6.900E-09 |
| 24926 | 1247 | H1247E | D | 1e-08 | 9.2e-09 | 4.5e-09 | 7.900E-09 | 2.972E-09 | 4.500E-09 | 1.000E-08 |
| 24927 | 1247 | H1247Q | D | 9.4e-09 | 9.5e-09 | 2.4e-08 | 1.430E-08 | 8.401E-09 | 9.400E-09 | 2.400E-08 |
| 24928 | 1247 | H1247G | D | 1.3e-08 | 3.3e-09 | 7.5e-09 | 7.933E-09 | 4.864E-09 | 3.300E-09 | 1.300E-08 |
| 24929 | 1247 | H1247H | D | 1.9e-08 | 7.3e-09 | 4.9e-09 | 1.040E-08 | 7.544E-09 | 4.900E-09 | 1.900E-08 |
| 24930 | 1247 | H1247I | D | 8.9e-09 | 6.3e-09 | 8.4e-09 | 7.867E-09 | 1.380E-09 | 6.300E-09 | 8.900E-09 |
| 24931 | 1247 | H1247L | D | 2.2e-08 | 3.6e-08 | 6.1e-09 | 2.137E-08 | 1.496E-08 | 6.100E-09 | 3.600E-08 |
| 24932 | 1247 | H1247K | D | 5.5e-09 | 2.2e-09 | 5.4e-09 | 4.367E-09 | 1.877E-09 | 2.200E-09 | 5.500E-09 |
| 24933 | 1247 | H1247M | D | 6e-09 | 6.7e-09 | 2.5e-09 | 5.067E-09 | 2.250E-09 | 2.500E-09 | 6.700E-09 |
| 24934 | 1247 | H1247F | D | 4.8e-09 | 1.8e-09 | 1.4e-08 | 6.867E-09 | 6.357E-09 | 1.800E-09 | 1.400E-08 |
| 24935 | 1247 | H1247P | D | 1e-08 | 1.1e-08 | 4.5e-09 | 8.500E-09 | 3.500E-09 | 4.500E-09 | 1.100E-08 |
| 24936 | 1247 | H1247S | D | 4.5e-09 | 1.6e-08 | 8.6e-09 | 9.700E-09 | 5.828E-09 | 4.500E-09 | 1.600E-08 |
| 24937 | 1247 | H1247T | D | 9.8e-09 | 1.1e-08 | 1.7e-08 | 1.260E-08 | 3.857E-09 | 9.800E-09 | 1.700E-08 |
| 24938 | 1247 | H1247W | D | 1.4e-08 | 3.5e-09 | 2.5e-09 | 6.667E-09 | 6.371E-09 | 2.500E-09 | 1.400E-08 |
| 24939 | 1247 | H1247Y | D | 9.2e-09 | 9.5e-09 | 9.7e-09 | 9.467E-09 | 2.517E-10 | 9.200E-09 | 9.700E-09 |
| 24940 | 1247 | H1247V | D | 2.3e-09 | 2.1e-09 | 1.4e-08 | 6.133E-09 | 6.813E-09 | 2.100E-09 | 1.400E-08 |
| 24941 | 1248 | K1248A | D | 1.8e-08 | 1e-08 | 4.7e-09 | 1.090E-08 | 6.696E-09 | 4.700E-09 | 1.800E-08 |
| 24942 | 1248 | K1248R | D | 6e-09 | 8.5e-09 | 4.8e-09 | 6.433E-09 | 1.888E-09 | 4.800E-09 | 8.500E-09 |
| 24943 | 1248 | K1248N | D | 6e-09 | 1.5e-08 | 4.7e-09 | 8.567E-09 | 5.609E-09 | 4.700E-09 | 1.500E-08 |
| 24944 | 1248 | K1248D | D | 1.1e-08 | 1e-08 | 8.5e-09 | 9.833E-09 | 1.258E-09 | 8.500E-09 | 1.100E-08 |
| 24945 | 1248 | K1248C | D | 1.5e-08 | 6.7e-09 | 6.8e-09 | 9.500E-09 | 4.763E-09 | 6.700E-09 | 1.500E-08 |
| 24946 | 1248 | K1248E | D | 1.1e-08 | 3.7e-09 | 4.2e-09 | 6.300E-09 | 4.078E-09 | 3.700E-09 | 1.100E-08 |
| 24947 | 1248 | K1248Q | D | 1.1e-08 | 2.4e-08 | 9e-09 | 1.467E-08 | 8.145E-09 | 9.000E-09 | 2.400E-08 |
| 24948 | 1248 | K1248G | D | 7.8e-09 | 8.8e-09 | 2.1e-09 | 6.233E-09 | 3.614E-09 | 2.100E-09 | 8.800E-09 |
| 24949 | 1248 | K1248H | D | 7e-09 | 5.3e-09 | 8.5e-09 | 6.933E-09 | 1.601E-09 | 5.300E-09 | 8.500E-09 |
| 24950 | 1248 | K1248I | D | 4.5e-09 | 8.6e-10 | 2.7e-09 | 2.687E-09 | 1.820E-09 | 8.600E-10 | 4.500E-09 |
| 24951 | 1248 | K1248L | D | 1.9e-08 | 3.9e-09 | 6.8e-09 | 9.900E-09 | 8.013E-09 | 3.900E-09 | 1.900E-08 |
| 24952 | 1248 | K1248K | D | 1.2e-08 | 1.2e-08 | 9.4e-09 | 1.113E-08 | 1.501E-09 | 9.400E-09 | 1.200E-08 |
| 24953 | 1248 | K1248M | D | 7.1e-09 | 1e-08 | 5.8e-09 | 7.633E-09 | 2.150E-09 | 5.800E-09 | 1.000E-08 |
| 24954 | 1248 | K1248F | D | 8.1e-09 | 6.1e-09 | 6.5e-09 | 6.900E-09 | 1.058E-09 | 6.100E-09 | 8.100E-09 |
| 24955 | 1248 | K1248P | D | 1.3e-08 | 5.2e-09 | 1.3e-08 | 1.040E-08 | 4.503E-09 | 5.200E-09 | 1.300E-08 |
| 24956 | 1248 | K1248S | D | 1.6e-08 | 4e-09 | 1.1e-08 | 1.033E-08 | 6.028E-09 | 4.000E-09 | 1.600E-08 |
| 24957 | 1248 | K1248T | D | 5.4e-09 | 6.4e-09 | 6.7e-09 | 6.167E-09 | 6.807E-10 | 5.400E-09 | 6.700E-09 |
| 24958 | 1248 | K1248W | D | 8.2e-09 | 1.7e-08 | 1.4e-08 | 1.307E-08 | 4.474E-09 | 8.200E-09 | 1.700E-08 |
| 24959 | 1248 | K1248Y | D | 4.3e-09 | 7.7e-09 | 2.4e-09 | 4.800E-09 | 2.685E-09 | 2.400E-09 | 7.700E-09 |
| 24960 | 1248 | K1248V | D | 1.7e-08 | 6.4e-09 | 2.2e-08 | 1.513E-08 | 7.966E-09 | 6.400E-09 | 2.200E-08 |
| 24961 | 1249 | V1249A | D | 3.5e-09 | 6.7e-09 | 3.3e-09 | 4.500E-09 | 1.908E-09 | 3.300E-09 | 6.700E-09 |
| 24962 | 1249 | V1249R | D | 5.5e-09 | 8.1e-09 | 8.7e-09 | 7.433E-09 | 1.701E-09 | 5.500E-09 | 8.700E-09 |
| 24963 | 1249 | V1249N | D | 2.1e-08 | 3.4e-09 | 7.9e-09 | 1.077E-08 | 9.143E-09 | 3.400E-09 | 2.100E-08 |
| 24964 | 1249 | V1249D | D | 5.2e-09 | 8.7e-09 | 5.7e-09 | 6.533E-09 | 1.893E-09 | 5.200E-09 | 8.700E-09 |
| 24965 | 1249 | V1249C | D | 9.6e-09 | 3.9e-09 | 2.9e-09 | 5.467E-09 | 3.614E-09 | 2.900E-09 | 9.600E-09 |
| 24966 | 1249 | V1249E | D | 5.5e-09 | 9.3e-09 | 9.6e-09 | 8.133E-09 | 2.285E-09 | 5.500E-09 | 9.600E-09 |
| 24967 | 1249 | V1249Q | D | 1.3e-08 | 5.5e-09 | 1.3e-08 | 1.050E-08 | 4.330E-09 | 5.500E-09 | 1.300E-08 |
| 24968 | 1249 | V1249G | D | 1.1e-08 | 8.7e-09 | 6.1e-09 | 8.600E-09 | 2.452E-09 | 6.100E-09 | 1.100E-08 |
| 24969 | 1249 | V1249H | D | 1.2e-08 | 8.2e-09 | 5.7e-09 | 8.633E-09 | 3.172E-09 | 5.700E-09 | 1.200E-08 |
| 24970 | 1249 | V1249I | D | 9.9e-09 | 6.4e-09 | 2.4e-09 | 6.233E-09 | 3.753E-09 | 2.400E-09 | 9.900E-09 |
| 24971 | 1249 | V1249L | D | 1.5e-08 | 2.8e-09 | 7.9e-09 | 8.567E-09 | 6.127E-09 | 2.800E-09 | 1.500E-08 |
| 24972 | 1249 | V1249K | D | 9.6e-09 | 1.8e-09 | 1.2e-08 | 7.800E-09 | 5.333E-09 | 1.800E-09 | 1.200E-08 |
| 24973 | 1249 | V1249M | D | 3.1e-08 | 3.9e-09 | 3.1e-08 | 2.197E-08 | 1.565E-08 | 3.900E-09 | 3.100E-08 |
| 24974 | 1249 | V1249F | D | 2.8e-09 | 4.1e-09 | 1.1e-08 | 5.967E-09 | 4.407E-09 | 2.800E-09 | 1.100E-08 |
| 24975 | 1249 | V1249P | D | 6.3e-09 | 3.8e-09 | 2.1e-09 | 4.067E-09 | 2.113E-09 | 2.100E-09 | 6.300E-09 |
| 24976 | 1249 | V1249S | D | 2e-08 | 1.3e-08 | 7.3e-09 | 1.343E-08 | 6.361E-09 | 7.300E-09 | 2.000E-08 |
| 24977 | 1249 | V1249T | D | 7.1e-09 | 8.4e-09 | 8.4e-09 | 7.967E-09 | 7.506E-10 | 7.100E-09 | 8.400E-09 |
| 24978 | 1249 | V1249W | D | 6.8e-09 | 7.9e-09 | 4.6e-09 | 6.433E-09 | 1.680E-09 | 4.600E-09 | 7.900E-09 |
| 24979 | 1249 | V1249Y | D | 1.6e-08 | 4.5e-09 | 6.3e-09 | 8.933E-09 | 6.186E-09 | 4.500E-09 | 1.600E-08 |
| 24980 | 1249 | V1249V | D | 6.8e-09 | 2.3e-08 | 7.7e-09 | 1.250E-08 | 9.104E-09 | 6.800E-09 | 2.300E-08 |
| 24981 | 1250 | Y1250A | D | 3.8e-09 | 8.4e-09 | 1.5e-08 | 9.067E-09 | 5.630E-09 | 3.800E-09 | 1.500E-08 |
| 24982 | 1250 | Y1250R | D | 1.1e-08 | 8.2e-09 | 1e-08 | 9.733E-09 | 1.419E-09 | 8.200E-09 | 1.100E-08 |
| 24983 | 1250 | Y1250N | D | 1e-08 | 4.3e-09 | 4.4e-09 | 6.233E-09 | 3.262E-09 | 4.300E-09 | 1.000E-08 |
| 24984 | 1250 | Y1250D | D | 4.5e-09 | 1.2e-08 | 1.5e-08 | 1.050E-08 | 5.408E-09 | 4.500E-09 | 1.500E-08 |
| 24985 | 1250 | Y1250C | D | 4.7e-09 | 2.4e-08 | 5e-09 | 1.123E-08 | 1.106E-08 | 4.700E-09 | 2.400E-08 |
| 24986 | 1250 | Y1250E | D | 1.1e-08 | 2.8e-09 | 2.2e-09 | 5.333E-09 | 4.917E-09 | 2.200E-09 | 1.100E-08 |
| 24987 | 1250 | Y1250Q | D | 4.7e-09 | 5.1e-09 | 6.3e-09 | 5.367E-09 | 8.327E-10 | 4.700E-09 | 6.300E-09 |
| 24988 | 1250 | Y1250G | D | 4.7e-09 | 7.4e-09 | 1.6e-09 | 4.567E-09 | 2.902E-09 | 1.600E-09 | 7.400E-09 |
| 24989 | 1250 | Y1250H | D | 9.4e-09 | 1.6e-08 | 3.1e-09 | 9.500E-09 | 6.451E-09 | 3.100E-09 | 1.600E-08 |

| | | | | | | | | | | |
|---|---|---|---|---|---|---|---|---|---|---|
| 24990 | 1250 | Y1250I | D | 6e-09 | 5.6e-09 | 5.3e-09 | 5.633E-09 | 3.512E-10 | 5.300E-09 | 6.000E-09 |
| 24991 | 1250 | Y1250L | D | 3.1e-09 | 9.2e-09 | 5.9e-09 | 6.067E-09 | 3.053E-09 | 3.100E-09 | 9.200E-09 |
| 24992 | 1250 | Y1250K | D | 9.8e-09 | 5.8e-09 | 5.9e-09 | 7.167E-09 | 2.281E-09 | 5.800E-09 | 9.800E-09 |
| 24993 | 1250 | Y1250M | D | 8.5e-09 | 5.7e-09 | 1.5e-08 | 9.733E-09 | 4.771E-09 | 5.700E-09 | 1.500E-08 |
| 24994 | 1250 | Y1250F | D | 3.8e-09 | 1.1e-08 | 1.6e-08 | 1.027E-08 | 6.133E-09 | 3.800E-09 | 1.600E-08 |
| 24995 | 1250 | Y1250P | D | 2.7e-09 | 1.3e-08 | 4e-09 | 6.567E-09 | 5.609E-09 | 2.700E-09 | 1.300E-08 |
| 24996 | 1250 | Y1250S | D | 4.7e-09 | 4.2e-09 | 1.5e-08 | 7.967E-09 | 6.096E-09 | 4.200E-09 | 1.500E-08 |
| 24997 | 1250 | Y1250T | D | 8.6e-09 | 9.6e-09 | 2.9e-09 | 7.033E-09 | 3.614E-09 | 2.900E-09 | 9.600E-09 |
| 24998 | 1250 | Y1250W | D | 8.3e-09 | 3.7e-09 | 7.4e-09 | 6.467E-09 | 2.438E-09 | 3.700E-09 | 8.300E-09 |
| 24999 | 1250 | Y1250Y | D | 1e-08 | 7.4e-09 | 3.4e-09 | 6.933E-09 | 3.325E-09 | 3.400E-09 | 1.000E-08 |
| 25000 | 1250 | Y1250V | D | 4.7e-09 | 1e-08 | 1.2e-08 | 8.900E-09 | 3.772E-09 | 4.700E-09 | 1.200E-08 |
| 25001 | 1251 | A1251A | D | 5.2e-09 | 8.1e-09 | 8e-09 | 7.100E-09 | 1.646E-09 | 5.200E-09 | 8.100E-09 |
| 25002 | 1251 | A1251R | D | 1e-08 | 1.4e-08 | 1.6e-08 | 1.333E-08 | 3.055E-09 | 1.000E-08 | 1.600E-08 |
| 25003 | 1251 | A1251N | D | 1e-08 | 7.9e-09 | 4.7e-09 | 7.533E-09 | 2.669E-09 | 4.700E-09 | 1.000E-08 |
| 25004 | 1251 | A1251D | D | 2.1e-09 | 7.7e-09 | 8.2e-09 | 6.000E-09 | 3.387E-09 | 2.100E-09 | 8.200E-09 |
| 25005 | 1251 | A1251C | D | 4.2e-09 | 2.3e-09 | 4.9e-09 | 3.800E-09 | 1.345E-09 | 2.300E-09 | 4.900E-09 |
| 25006 | 1251 | A1251E | D | 9.4e-09 | 3.2e-09 | 6e-09 | 6.200E-09 | 3.105E-09 | 3.200E-09 | 9.400E-09 |
| 25007 | 1251 | A1251Q | D | 1.3e-08 | 1.1e-08 | 1.5e-08 | 1.300E-08 | 2.000E-09 | 1.100E-08 | 1.500E-08 |
| 25008 | 1251 | A1251G | D | 3.2e-09 | 7.6e-09 | 3.2e-09 | 4.667E-09 | 2.540E-09 | 3.200E-09 | 7.600E-09 |
| 25009 | 1251 | A1251H | D | 1.3e-08 | 1.2e-08 | 6.4e-09 | 1.047E-08 | 3.557E-09 | 6.400E-09 | 1.300E-08 |
| 25010 | 1251 | A1251I | D | 1.7e-09 | 2.9e-09 | 1.6e-08 | 6.867E-09 | 7.932E-09 | 1.700E-09 | 1.600E-08 |
| 25011 | 1251 | A1251L | D | 5e-09 | 4.9e-09 | 1.5e-08 | 8.300E-09 | 5.803E-09 | 4.900E-09 | 1.500E-08 |
| 25012 | 1251 | A1251K | D | 6.6e-09 | 4.5e-09 | 5.9e-09 | 5.667E-09 | 1.069E-09 | 4.500E-09 | 6.600E-09 |
| 25013 | 1251 | A1251M | D | 8.7e-09 | 7.9e-09 | 4.7e-09 | 7.100E-09 | 2.117E-09 | 4.700E-09 | 8.700E-09 |
| 25014 | 1251 | A1251F | D | 7.7e-09 | 4e-09 | 2.7e-09 | 4.800E-09 | 2.594E-09 | 2.700E-09 | 7.700E-09 |
| 25015 | 1251 | A1251P | D | 3.4e-09 | 3.6e-09 | 1.4e-08 | 7.000E-09 | 6.063E-09 | 3.400E-09 | 1.400E-08 |
| 25016 | 1251 | A1251S | D | 7.2e-09 | 6.1e-09 | 8.3e-09 | 7.200E-09 | 1.100E-09 | 6.100E-09 | 8.300E-09 |
| 25017 | 1251 | A1251T | D | 2.7e-09 | 1.5e-08 | 3.1e-09 | 6.933E-09 | 6.989E-09 | 2.700E-09 | 1.500E-08 |
| 25018 | 1251 | A1251W | D | 8.1e-09 | 3.8e-09 | 9.8e-09 | 7.233E-09 | 3.092E-09 | 3.800E-09 | 9.800E-09 |
| 25019 | 1251 | A1251Y | D | 5.9e-09 | 3.4e-09 | 1.1e-08 | 6.767E-09 | 3.873E-09 | 3.400E-09 | 1.100E-08 |
| 25020 | 1251 | A1251V | D | 9e-09 | 4e-09 | 1.3e-08 | 8.667E-09 | 4.509E-09 | 4.000E-09 | 1.300E-08 |
| 25021 | 1252 | C1252A | D | 1e-08 | 9.4e-09 | 4.7e-09 | 8.033E-09 | 2.902E-09 | 4.700E-09 | 1.000E-08 |
| 25022 | 1252 | C1252R | D | 5.2e-09 | 1.2e-08 | 3.3e-09 | 6.833E-09 | 4.574E-09 | 3.300E-09 | 1.200E-08 |
| 25023 | 1252 | C1252N | D | 1.1e-08 | 7.7e-09 | 5.6e-09 | 8.100E-09 | 2.722E-09 | 5.600E-09 | 1.100E-08 |
| 25024 | 1252 | C1252D | D | 4.9e-09 | 3e-09 | 1.9e-08 | 8.967E-09 | 8.741E-09 | 3.000E-09 | 1.900E-08 |
| 25025 | 1252 | C1252C | D | 7.5e-09 | 1.3e-08 | 8.6e-09 | 9.700E-09 | 2.910E-09 | 7.500E-09 | 1.300E-08 |
| 25026 | 1252 | C1252E | D | 3.5e-09 | 1.5e-08 | 5.5e-09 | 8.000E-09 | 6.144E-09 | 3.500E-09 | 1.500E-08 |
| 25027 | 1252 | C1252Q | D | 1.2e-08 | 1.7e-08 | 4.6e-09 | 1.120E-08 | 6.239E-09 | 4.600E-09 | 1.700E-08 |
| 25028 | 1252 | C1252G | D | 3e-09 | 8.4e-09 | 6.5e-09 | 5.967E-09 | 2.739E-09 | 3.000E-09 | 8.400E-09 |
| 25029 | 1252 | C1252H | D | 2.2e-09 | 3.5e-09 | 3.8e-09 | 3.167E-09 | 8.505E-10 | 2.200E-09 | 3.800E-09 |
| 25030 | 1252 | C1252I | D | 6.6e-09 | 8.9e-09 | 1.4e-08 | 9.833E-09 | 3.787E-09 | 6.600E-09 | 1.400E-08 |
| 25031 | 1252 | C1252L | D | 4.4e-09 | 1.9e-08 | 5.3e-09 | 9.567E-09 | 8.182E-09 | 4.400E-09 | 1.900E-08 |
| 25032 | 1252 | C1252K | D | 5.3e-09 | 1.1e-08 | 6.8e-09 | 7.700E-09 | 2.955E-09 | 5.300E-09 | 1.100E-08 |
| 25033 | 1252 | C1252M | D | 7.5e-09 | 3.8e-09 | 5e-09 | 5.433E-09 | 1.888E-09 | 3.800E-09 | 7.500E-09 |
| 25034 | 1252 | C1252F | D | 2.3e-09 | 1.5e-08 | 9.7e-09 | 9.000E-09 | 6.379E-09 | 2.300E-09 | 1.500E-08 |
| 25035 | 1252 | C1252P | D | 4e-09 | 2.1e-09 | 5.8e-09 | 3.967E-09 | 1.850E-09 | 2.100E-09 | 5.800E-09 |
| 25036 | 1252 | C1252S | D | 6.1e-09 | 4.4e-09 | 7.6e-09 | 6.033E-09 | 1.601E-09 | 4.400E-09 | 7.600E-09 |
| 25037 | 1252 | C1252T | D | 4.8e-09 | 9.4e-09 | 1.8e-08 | 1.073E-08 | 6.700E-09 | 4.800E-09 | 1.800E-08 |
| 25038 | 1252 | C1252W | D | 2.5e-08 | 7e-09 | 2.5e-09 | 1.150E-08 | 1.191E-08 | 2.500E-09 | 2.500E-08 |
| 25039 | 1252 | C1252Y | D | 7.5e-09 | 1.7e-08 | 6.6e-09 | 1.037E-08 | 5.762E-09 | 6.600E-09 | 1.700E-08 |
| 25040 | 1252 | C1252V | D | 2.5e-08 | 8.6e-09 | 4.6e-09 | 1.273E-08 | 1.081E-08 | 4.600E-09 | 2.500E-08 |
| 25041 | 1253 | E1253A | D | 8.5e-09 | 1.2e-08 | 6.5e-09 | 9.000E-09 | 2.784E-09 | 6.500E-09 | 1.200E-08 |
| 25042 | 1253 | E1253R | D | 1.1e-08 | 7.7e-09 | 3.6e-09 | 7.433E-09 | 3.707E-09 | 3.600E-09 | 1.100E-08 |
| 25043 | 1253 | E1253N | D | 4.9e-09 | 8.9e-09 | 7.6e-09 | 7.133E-09 | 2.040E-09 | 4.900E-09 | 8.900E-09 |
| 25044 | 1253 | E1253D | D | 6.7e-09 | 6.9e-09 | 1.5e-08 | 9.533E-09 | 4.735E-09 | 6.700E-09 | 1.500E-08 |
| 25045 | 1253 | E1253C | D | 6e-09 | 1.2e-08 | 1.5e-09 | 6.500E-09 | 5.268E-09 | 1.500E-09 | 1.200E-08 |
| 25046 | 1253 | E1253E | D | 1.3e-08 | 8.4e-09 | 6.7e-09 | 9.367E-09 | 3.259E-09 | 6.700E-09 | 1.300E-08 |
| 25047 | 1253 | E1253Q | D | 1.4e-08 | 5.2e-09 | 1.4e-08 | 1.107E-08 | 5.081E-09 | 5.200E-09 | 1.400E-08 |
| 25048 | 1253 | E1253G | D | 6.3e-09 | 3.9e-09 | 5.6e-09 | 5.267E-09 | 1.234E-09 | 3.900E-09 | 6.300E-09 |
| 25049 | 1253 | E1253H | D | 7.6e-09 | 3.9e-09 | 4.7e-09 | 5.400E-09 | 1.947E-09 | 3.900E-09 | 7.600E-09 |
| 25050 | 1253 | E1253I | D | 7.7e-09 | 9.4e-09 | 4.3e-09 | 7.133E-09 | 2.597E-09 | 4.300E-09 | 9.400E-09 |
| 25051 | 1253 | E1253L | D | 9.7e-09 | 2.9e-09 | 1.2e-08 | 8.200E-09 | 4.732E-09 | 2.900E-09 | 1.200E-08 |
| 25052 | 1253 | E1253K | D | 1.6e-08 | 3.6e-09 | 8e-09 | 9.200E-09 | 6.286E-09 | 3.600E-09 | 1.600E-08 |
| 25053 | 1253 | E1253M | D | 7.2e-09 | 4.3e-09 | 3e-09 | 4.833E-09 | 2.150E-09 | 3.000E-09 | 7.200E-09 |
| 25054 | 1253 | E1253F | D | 1e-08 | 8.9e-09 | 7.4e-09 | 8.767E-09 | 1.305E-09 | 7.400E-09 | 1.000E-08 |
| 25055 | 1253 | E1253P | D | 3e-09 | 6.1e-09 | 6.2e-09 | 5.100E-09 | 1.819E-09 | 3.000E-09 | 6.200E-09 |
| 25056 | 1253 | E1253S | D | 8.6e-09 | 1.4e-08 | 1.5e-08 | 1.253E-08 | 3.443E-09 | 8.600E-09 | 1.500E-08 |
| 25057 | 1253 | E1253T | D | 7.7e-09 | 7.1e-09 | 1.5e-08 | 9.933E-09 | 4.398E-09 | 7.100E-09 | 1.500E-08 |
| 25058 | 1253 | E1253W | D | 2.4e-09 | 1.3e-08 | 8.9e-09 | 8.100E-09 | 5.345E-09 | 2.400E-09 | 1.300E-08 |
| 25059 | 1253 | E1253Y | D | 7.6e-09 | 3.2e-09 | 7.1e-09 | 5.967E-09 | 2.409E-09 | 3.200E-09 | 7.600E-09 |
| 25060 | 1253 | E1253V | D | 1.8e-08 | 7.1e-09 | 7.4e-09 | 1.083E-08 | 6.208E-09 | 7.100E-09 | 1.800E-08 |
| 25061 | 1254 | V1254A | D | 6.1e-09 | 2.5e-09 | 2.2e-08 | 1.020E-08 | 1.038E-08 | 2.500E-09 | 2.200E-08 |
| 25062 | 1254 | V1254R | D | 3.8e-09 | 8.7e-09 | 6.3e-09 | 6.267E-09 | 2.450E-09 | 3.800E-09 | 8.700E-09 |
| 25063 | 1254 | V1254N | D | 9.1e-09 | 6.5e-09 | 2.2e-08 | 1.253E-08 | 8.301E-09 | 6.500E-09 | 2.200E-08 |
| 25064 | 1254 | V1254D | D | 5.5e-09 | 3.7e-09 | 7.8e-09 | 5.667E-09 | 2.055E-09 | 3.700E-09 | 7.800E-09 |
| 25065 | 1254 | V1254C | D | 5.8e-09 | 9e-09 | 3.1e-09 | 5.967E-09 | 2.954E-09 | 3.100E-09 | 9.000E-09 |
| 25066 | 1254 | V1254E | D | 6e-09 | 3.4e-09 | 4.9e-09 | 4.767E-09 | 1.305E-09 | 3.400E-09 | 6.000E-09 |
| 25067 | 1254 | V1254Q | D | 5.2e-09 | 4.6e-09 | 4.3e-09 | 4.700E-09 | 4.583E-10 | 4.300E-09 | 5.200E-09 |
| 25068 | 1254 | V1254G | D | 1.5e-08 | 2.5e-08 | 4.6e-09 | 1.487E-08 | 1.020E-08 | 4.600E-09 | 2.500E-08 |
| 25069 | 1254 | V1254H | D | 3.7e-09 | 4.7e-09 | 3.8e-09 | 4.067E-09 | 5.508E-10 | 3.700E-09 | 4.700E-09 |
| 25070 | 1254 | V1254I | D | 3.8e-09 | 1.3e-08 | 1.5e-08 | 1.060E-08 | 5.973E-09 | 3.800E-09 | 1.500E-08 |
| 25071 | 1254 | V1254L | D | 2e-08 | 1.1e-08 | 6.5e-09 | 1.250E-08 | 6.874E-09 | 6.500E-09 | 2.000E-08 |
| 25072 | 1254 | V1254K | D | 1.3e-08 | 1.1e-08 | 9.3e-09 | 1.110E-08 | 1.852E-09 | 9.300E-09 | 1.300E-08 |
| 25073 | 1254 | V1254M | D | 9e-09 | 6.4e-09 | 3.2e-09 | 6.200E-09 | 2.905E-09 | 3.200E-09 | 9.000E-09 |
| 25074 | 1254 | V1254F | D | 5.8e-09 | 7.8e-09 | 1e-08 | 7.867E-09 | 2.101E-09 | 5.800E-09 | 1.000E-08 |

| | | | | | | | | | | |
|---|---|---|---|---|---|---|---|---|---|---|
| 25075 | 1254 | V1254P | D | 7.4e-09 | 4.2e-09 | 9.6e-09 | 7.067E-09 | 2.715E-09 | 4.200E-09 | 9.600E-09 |
| 25076 | 1254 | V1254S | D | 6.8e-09 | 6.1e-09 | 7.6e-09 | 6.833E-09 | 7.506E-10 | 6.100E-09 | 7.600E-09 |
| 25077 | 1254 | V1254T | D | 7.8e-09 | 1e-08 | 9.2e-09 | 9.000E-09 | 1.114E-09 | 7.800E-09 | 1.000E-08 |
| 25078 | 1254 | V1254W | D | 1.1e-08 | 1.4e-09 | 1.3e-08 | 8.467E-09 | 6.201E-09 | 1.400E-09 | 1.300E-08 |
| 25079 | 1254 | V1254Y | D | 8.5e-09 | 4.3e-09 | 1.9e-08 | 1.060E-08 | 7.572E-09 | 4.300E-09 | 1.900E-08 |
| 25080 | 1254 | V1254V | D | 5.5e-09 | 1e-08 | 8.2e-09 | 7.900E-09 | 2.265E-09 | 5.500E-09 | 1.000E-08 |
| 25081 | 1255 | T1255A | D | 9.4e-09 | 1.3e-08 | 4e-09 | 8.800E-09 | 4.530E-09 | 4.000E-09 | 1.300E-08 |
| 25082 | 1255 | T1255R | D | 2.5e-09 | 8.1e-09 | 7.7e-09 | 6.100E-09 | 3.124E-09 | 2.500E-09 | 8.100E-09 |
| 25083 | 1255 | T1255N | D | 7.7e-09 | 1.1e-08 | 3.1e-08 | 1.657E-08 | 1.261E-08 | 7.700E-09 | 3.100E-08 |
| 25084 | 1255 | T1255D | D | 8.9e-09 | 1.2e-08 | 5e-09 | 8.633E-09 | 3.508E-09 | 5.000E-09 | 1.200E-08 |
| 25085 | 1255 | T1255C | D | 9e-09 | 4.1e-09 | 1.5e-08 | 9.367E-09 | 5.459E-09 | 4.100E-09 | 1.500E-08 |
| 25086 | 1255 | T1255E | D | 1.6e-08 | 9.2e-09 | 1.4e-08 | 1.307E-08 | 3.495E-09 | 9.200E-09 | 1.600E-08 |
| 25087 | 1255 | T1255Q | D | 5e-09 | 2.9e-09 | 9.3e-09 | 5.733E-09 | 3.262E-09 | 2.900E-09 | 9.300E-09 |
| 25088 | 1255 | T1255G | D | 1.1e-09 | 6.7e-09 | 3.6e-08 | 1.460E-08 | 1.874E-08 | 1.100E-09 | 3.600E-08 |
| 25089 | 1255 | T1255H | D | 4.2e-09 | 1.6e-09 | 6.7e-09 | 4.167E-09 | 2.550E-09 | 1.600E-09 | 6.700E-09 |
| 25090 | 1255 | T1255I | D | 9.3e-09 | 1.1e-08 | 1.1e-08 | 1.043E-08 | 9.815E-10 | 9.300E-09 | 1.100E-08 |
| 25091 | 1255 | T1255L | D | 1.6e-08 | 4.9e-09 | 8.5e-09 | 9.800E-09 | 5.663E-09 | 4.900E-09 | 1.600E-08 |
| 25092 | 1255 | T1255K | D | 5.8e-09 | 7.1e-09 | 9.6e-09 | 7.500E-09 | 1.931E-09 | 5.800E-09 | 9.600E-09 |
| 25093 | 1255 | T1255M | D | 1.3e-09 | 2.3e-08 | 1.6e-08 | 1.343E-08 | 1.108E-08 | 1.300E-09 | 2.300E-08 |
| 25094 | 1255 | T1255F | D | 5.4e-09 | 1.1e-08 | 2.3e-09 | 6.233E-09 | 4.409E-09 | 2.300E-09 | 1.100E-08 |
| 25095 | 1255 | T1255P | D | 7.8e-09 | 1.7e-09 | 8e-09 | 5.833E-09 | 3.581E-09 | 1.700E-09 | 8.000E-09 |
| 25096 | 1255 | T1255S | D | 3.9e-09 | 4.8e-09 | 8.1e-09 | 5.600E-09 | 2.211E-09 | 3.900E-09 | 8.100E-09 |
| 25097 | 1255 | T1255T | D | 1e-08 | 5.5e-09 | 1.1e-08 | 8.833E-09 | 2.930E-09 | 5.500E-09 | 1.100E-08 |
| 25098 | 1255 | T1255W | D | 7.4e-09 | 1.2e-08 | 5.8e-09 | 8.400E-09 | 3.219E-09 | 5.800E-09 | 1.200E-08 |
| 25099 | 1255 | T1255Y | D | 3.7e-09 | 5.8e-09 | 1.6e-08 | 8.500E-09 | 6.580E-09 | 3.700E-09 | 1.600E-08 |
| 25100 | 1255 | T1255V | D | 9.7e-09 | 7.7e-09 | 1.4e-08 | 1.047E-08 | 3.219E-09 | 7.700E-09 | 1.400E-08 |
| 25101 | 1256 | H1256A | D | 3.5e-09 | 1.3e-08 | 8.8e-09 | 8.433E-09 | 4.761E-09 | 3.500E-09 | 1.300E-08 |
| 25102 | 1256 | H1256R | D | 3.6e-09 | 6.7e-09 | 8.2e-09 | 6.167E-09 | 2.346E-09 | 3.600E-09 | 8.200E-09 |
| 25103 | 1256 | H1256N | D | 1.1e-08 | 1e-08 | 1.8e-08 | 1.300E-08 | 4.359E-09 | 1.000E-08 | 1.800E-08 |
| 25104 | 1256 | H1256D | D | 9.3e-09 | 4.2e-09 | 6.7e-09 | 6.733E-09 | 2.550E-09 | 4.200E-09 | 9.300E-09 |
| 25105 | 1256 | H1256C | D | 6.6e-09 | 2.9e-09 | 1.6e-08 | 8.500E-09 | 6.754E-09 | 2.900E-09 | 1.600E-08 |
| 25106 | 1256 | H1256E | D | 6.2e-09 | 4.7e-09 | 3.5e-09 | 4.800E-09 | 1.353E-09 | 3.500E-09 | 6.200E-09 |
| 25107 | 1256 | H1256Q | D | 9.2e-09 | 8.2e-09 | 9.5e-09 | 8.967E-09 | 6.807E-10 | 8.200E-09 | 9.500E-09 |
| 25108 | 1256 | H1256G | D | 2.2e-08 | 1.2e-08 | 1.1e-08 | 1.500E-08 | 6.083E-09 | 1.100E-08 | 2.200E-08 |
| 25109 | 1256 | H1256H | D | 1e-08 | 4.1e-09 | 2.7e-09 | 5.600E-09 | 3.874E-09 | 2.700E-09 | 1.000E-08 |
| 25110 | 1256 | H1256I | D | 1.2e-08 | 5.1e-09 | 5.7e-09 | 7.600E-09 | 3.822E-09 | 5.100E-09 | 1.200E-08 |
| 25111 | 1256 | H1256L | D | 9.8e-09 | 6.8e-09 | 5.2e-09 | 7.267E-09 | 2.335E-09 | 5.200E-09 | 9.800E-09 |
| 25112 | 1256 | H1256K | D | 6.9e-09 | 1.1e-08 | 1.1e-08 | 9.633E-09 | 2.367E-09 | 6.900E-09 | 1.100E-08 |
| 25113 | 1256 | H1256M | D | 9.6e-09 | 1e-08 | 1.2e-08 | 1.053E-08 | 1.286E-09 | 9.600E-09 | 1.200E-08 |
| 25114 | 1256 | H1256F | D | 6.2e-09 | 1.8e-08 | 1.1e-08 | 1.173E-08 | 5.934E-09 | 6.200E-09 | 1.800E-08 |
| 25115 | 1256 | H1256P | D | 1.1e-08 | 4.7e-09 | 1.2e-08 | 9.233E-09 | 3.958E-09 | 4.700E-09 | 1.200E-08 |
| 25116 | 1256 | H1256S | D | 1e-08 | 7.7e-09 | 9.7e-09 | 9.133E-09 | 1.250E-09 | 7.700E-09 | 1.000E-08 |
| 25117 | 1256 | H1256T | D | 1.7e-08 | 7.5e-09 | 6.9e-09 | 1.047E-08 | 5.666E-09 | 6.900E-09 | 1.700E-08 |
| 25118 | 1256 | H1256W | D | 1.7e-08 | 8.5e-09 | 3.7e-09 | 9.733E-09 | 6.735E-09 | 3.700E-09 | 1.700E-08 |
| 25119 | 1256 | H1256Y | D | 3.1e-09 | 1.5e-08 | 9.8e-09 | 9.300E-09 | 5.966E-09 | 3.100E-09 | 1.500E-08 |
| 25120 | 1256 | H1256V | D | 8.4e-09 | 7.1e-09 | 8.6e-09 | 8.033E-09 | 8.145E-10 | 7.100E-09 | 8.600E-09 |
| 25121 | 1257 | Q1257A | D | 2.5e-08 | 8.3e-09 | 8.1e-09 | 1.380E-08 | 9.700E-09 | 8.100E-09 | 2.500E-08 |
| 25122 | 1257 | Q1257R | D | 1.6e-08 | 5.2e-09 | 2.9e-09 | 8.033E-09 | 6.995E-09 | 2.900E-09 | 1.600E-08 |
| 25123 | 1257 | Q1257N | D | 5.3e-09 | 2.2e-08 | 6.3e-09 | 1.120E-08 | 9.366E-09 | 5.300E-09 | 2.200E-08 |
| 25124 | 1257 | Q1257D | D | 2.2e-09 | 7.1e-09 | 5.9e-09 | 5.067E-09 | 2.554E-09 | 2.200E-09 | 7.100E-09 |
| 25125 | 1257 | Q1257C | D | 3e-09 | 1.5e-08 | 1.1e-08 | 9.667E-09 | 6.110E-09 | 3.000E-09 | 1.500E-08 |
| 25126 | 1257 | Q1257E | D | 9.6e-09 | 2.9e-09 | 4.1e-09 | 5.533E-09 | 3.573E-09 | 2.900E-09 | 9.600E-09 |
| 25127 | 1257 | Q1257Q | D | 1.3e-08 | 6.3e-09 | 1.9e-09 | 7.067E-09 | 5.590E-09 | 1.900E-09 | 1.300E-08 |
| 25128 | 1257 | Q1257G | D | 9.8e-09 | 2.2e-08 | 2.5e-08 | 1.893E-08 | 8.051E-09 | 9.800E-09 | 2.500E-08 |
| 25129 | 1257 | Q1257H | D | 2.5e-08 | 9.1e-09 | 6.4e-09 | 1.350E-08 | 1.005E-08 | 6.400E-09 | 2.500E-08 |
| 25130 | 1257 | Q1257I | D | 1.6e-08 | 1.4e-09 | 3.5e-09 | 6.967E-09 | 7.893E-09 | 1.400E-09 | 1.600E-08 |
| 25131 | 1257 | Q1257L | D | 5.1e-09 | 8.9e-09 | 1.8e-08 | 1.067E-08 | 6.629E-09 | 5.100E-09 | 1.800E-08 |
| 25132 | 1257 | Q1257K | D | 7.1e-09 | 5.9e-09 | 6.6e-09 | 6.533E-09 | 6.028E-10 | 5.900E-09 | 7.100E-09 |
| 25133 | 1257 | Q1257M | D | 5.2e-09 | 2.8e-09 | 3.1e-09 | 3.700E-09 | 1.308E-09 | 2.800E-09 | 5.200E-09 |
| 25134 | 1257 | Q1257F | D | 7.7e-09 | 7.1e-09 | 5.4e-09 | 6.733E-09 | 1.193E-09 | 5.400E-09 | 7.700E-09 |
| 25135 | 1257 | Q1257P | D | 8.4e-09 | 1.7e-08 | 1.7e-08 | 1.413E-08 | 4.965E-09 | 8.400E-09 | 1.700E-08 |
| 25136 | 1257 | Q1257S | D | 8e-09 | 4.1e-09 | 7.3e-09 | 6.467E-09 | 2.079E-09 | 4.100E-09 | 8.000E-09 |
| 25137 | 1257 | Q1257T | D | 6.3e-09 | 4.9e-09 | 4.2e-09 | 5.133E-09 | 1.069E-09 | 4.200E-09 | 6.300E-09 |
| 25138 | 1257 | Q1257W | D | 8.6e-09 | 5.9e-09 | 6.3e-09 | 6.933E-09 | 1.457E-09 | 5.900E-09 | 8.600E-09 |
| 25139 | 1257 | Q1257Y | D | 1.7e-08 | 6.3e-09 | 3.4e-09 | 8.900E-09 | 7.163E-09 | 3.400E-09 | 1.700E-08 |
| 25140 | 1257 | Q1257V | D | 1.3e-08 | 1.7e-08 | 1.5e-09 | 1.050E-08 | 8.047E-09 | 1.500E-09 | 1.700E-08 |
| 25141 | 1258 | G1258A | D | 3.6e-09 | 5.9e-09 | 8.3e-09 | 5.933E-09 | 2.350E-09 | 3.600E-09 | 8.300E-09 |
| 25142 | 1258 | G1258R | D | 8.2e-09 | 1.3e-08 | 3.4e-09 | 8.200E-09 | 4.800E-09 | 3.400E-09 | 1.300E-08 |
| 25143 | 1258 | G1258N | D | 5.2e-09 | 8.1e-09 | 6.1e-09 | 6.467E-09 | 1.484E-09 | 5.200E-09 | 8.100E-09 |
| 25144 | 1258 | G1258D | D | 3.5e-09 | 3.6e-09 | 7.9e-09 | 5.000E-09 | 2.512E-09 | 3.500E-09 | 7.900E-09 |
| 25145 | 1258 | G1258C | D | 3.8e-09 | 2.6e-09 | 5.3e-09 | 3.900E-09 | 1.353E-09 | 2.600E-09 | 5.300E-09 |
| 25146 | 1258 | G1258E | D | 6.7e-09 | 1.3e-08 | 7.1e-09 | 8.933E-09 | 3.528E-09 | 6.700E-09 | 1.300E-08 |
| 25147 | 1258 | G1258Q | D | 3.5e-09 | 3.9e-09 | 7.3e-09 | 4.900E-09 | 2.088E-09 | 3.500E-09 | 7.300E-09 |
| 25148 | 1258 | G1258G | D | 5.2e-09 | 6.8e-09 | 8.8e-09 | 6.933E-09 | 1.804E-09 | 5.200E-09 | 8.800E-09 |
| 25149 | 1258 | G1258H | D | 6.5e-09 | 4.2e-09 | 5.4e-09 | 5.367E-09 | 1.150E-09 | 4.200E-09 | 6.500E-09 |
| 25150 | 1258 | G1258I | D | 6.7e-09 | 1.2e-08 | 9.5e-09 | 9.400E-09 | 2.651E-09 | 6.700E-09 | 1.200E-08 |
| 25151 | 1258 | G1258L | D | 2.5e-08 | 2.6e-09 | 5.5e-09 | 1.103E-08 | 1.218E-08 | 2.600E-09 | 2.500E-08 |
| 25152 | 1258 | G1258K | D | 1.3e-08 | 2.4e-09 | 1.8e-08 | 1.113E-08 | 7.966E-09 | 2.400E-09 | 1.800E-08 |
| 25153 | 1258 | G1258M | D | 4.9e-09 | 1.3e-08 | 5.5e-09 | 7.800E-09 | 4.513E-09 | 4.900E-09 | 1.300E-08 |
| 25154 | 1258 | G1258F | D | 4.6e-09 | 9.7e-09 | 1.1e-08 | 8.433E-09 | 3.383E-09 | 4.600E-09 | 1.100E-08 |
| 25155 | 1258 | G1258P | D | 7.2e-09 | 1.2e-08 | 1.3e-08 | 1.073E-08 | 3.101E-09 | 7.200E-09 | 1.300E-08 |
| 25156 | 1258 | G1258S | D | 5.9e-09 | 1.1e-08 | 7.3e-09 | 8.067E-09 | 2.635E-09 | 5.900E-09 | 1.100E-08 |
| 25157 | 1258 | G1258T | D | 2.7e-09 | 4.3e-09 | 7.1e-09 | 4.700E-09 | 2.227E-09 | 2.700E-09 | 7.100E-09 |
| 25158 | 1258 | G1258W | D | 1.3e-08 | 4.8e-09 | 5.6e-09 | 7.800E-09 | 4.521E-09 | 4.800E-09 | 1.300E-08 |
| 25159 | 1258 | G1258Y | D | 6.8e-09 | 4.9e-09 | 5.9e-09 | 5.867E-09 | 9.504E-10 | 4.900E-09 | 6.800E-09 |

| | | | | | | | | | | |
|---|---|---|---|---|---|---|---|---|---|---|
| 25160 | 1258 | G1258V | D | 1.2e-08 | 1.6e-08 | 1.2e-08 | 1.333E-08 | 2.309E-09 | 1.200E-08 | 1.600E-08 |
| 25161 | 1259 | L1259A | D | 1.5e-08 | 3.2e-09 | 2.6e-08 | 1.473E-08 | 1.140E-08 | 3.200E-09 | 2.600E-08 |
| 25162 | 1259 | L1259R | D | 6.3e-09 | 8.6e-09 | 1.3e-08 | 9.300E-09 | 3.404E-09 | 6.300E-09 | 1.300E-08 |
| 25163 | 1259 | L1259N | D | 2.4e-09 | 2.2e-09 | 4.8e-09 | 3.133E-09 | 1.447E-09 | 2.200E-09 | 4.800E-09 |
| 25164 | 1259 | L1259D | D | 6.7e-09 | 1.2e-09 | 3.7e-09 | 3.867E-09 | 2.754E-09 | 1.200E-09 | 6.700E-09 |
| 25165 | 1259 | L1259C | D | 5.4e-09 | 1.3e-08 | 7.5e-09 | 8.633E-09 | 3.925E-09 | 5.400E-09 | 1.300E-08 |
| 25166 | 1259 | L1259E | D | 2.6e-09 | 6e-09 | 5.4e-09 | 4.667E-09 | 1.815E-09 | 2.600E-09 | 6.000E-09 |
| 25167 | 1259 | L1259Q | D | 5.4e-09 | 6.7e-09 | 7.6e-09 | 6.567E-09 | 1.106E-09 | 5.400E-09 | 7.600E-09 |
| 25168 | 1259 | L1259G | D | 6.8e-09 | 4.3e-09 | 5.7e-09 | 5.600E-09 | 1.253E-09 | 4.300E-09 | 6.800E-09 |
| 25169 | 1259 | L1259H | D | 4.8e-09 | 8.9e-09 | 6.8e-09 | 6.833E-09 | 2.050E-09 | 4.800E-09 | 8.900E-09 |
| 25170 | 1259 | L1259I | D | 3.7e-09 | 1.1e-08 | 1.8e-08 | 1.090E-08 | 7.151E-09 | 3.700E-09 | 1.800E-08 |
| 25171 | 1259 | L1259L | D | 8e-09 | 1.9e-09 | 8.8e-09 | 6.233E-09 | 3.774E-09 | 1.900E-09 | 8.800E-09 |
| 25172 | 1259 | L1259K | D | 8.6e-09 | 7.8e-09 | 2.4e-09 | 6.267E-09 | 3.372E-09 | 2.400E-09 | 8.600E-09 |
| 25173 | 1259 | L1259M | D | 1e-08 | 4.2e-09 | 8.7e-09 | 7.633E-09 | 3.044E-09 | 4.200E-09 | 1.000E-08 |
| 25174 | 1259 | L1259F | D | 4.4e-09 | 3.8e-09 | 7.6e-09 | 5.267E-09 | 2.043E-09 | 3.800E-09 | 7.600E-09 |
| 25175 | 1259 | L1259P | D | 5.8e-09 | 3e-09 | 1.4e-08 | 7.600E-09 | 5.717E-09 | 3.000E-09 | 1.400E-08 |
| 25176 | 1259 | L1259S | D | 7.1e-09 | 4.4e-09 | 3e-09 | 4.833E-09 | 2.084E-09 | 3.000E-09 | 7.100E-09 |
| 25177 | 1259 | L1259T | D | 5e-09 | 3.1e-09 | 5.4e-09 | 4.500E-09 | 1.229E-09 | 3.100E-09 | 5.400E-09 |
| 25178 | 1259 | L1259W | D | 5.3e-09 | 2.1e-08 | 7.3e-09 | 1.120E-08 | 8.546E-09 | 5.300E-09 | 2.100E-08 |
| 25179 | 1259 | L1259Y | D | 4.1e-09 | 5.4e-09 | 4.9e-09 | 4.800E-09 | 6.557E-10 | 4.100E-09 | 5.400E-09 |
| 25180 | 1259 | L1259V | D | 1.1e-08 | 5.7e-09 | 4.3e-09 | 7.000E-09 | 3.534E-09 | 4.300E-09 | 1.100E-08 |
| 25181 | 1260 | S1260A | D | 6.9e-09 | 7.4e-09 | 6.8e-09 | 7.033E-09 | 3.215E-10 | 6.800E-09 | 7.400E-09 |
| 25182 | 1260 | S1260R | D | 1.1e-08 | 1.6e-08 | 4.7e-09 | 1.057E-08 | 5.662E-09 | 4.700E-09 | 1.600E-08 |
| 25183 | 1260 | S1260N | D | 3.2e-09 | 9.4e-09 | 5.7e-09 | 6.100E-09 | 3.119E-09 | 3.200E-09 | 9.400E-09 |
| 25184 | 1260 | S1260D | D | 5.2e-09 | 2.4e-09 | 1.1e-08 | 6.200E-09 | 4.386E-09 | 2.400E-09 | 1.100E-08 |
| 25185 | 1260 | S1260C | D | 6.9e-09 | 1.8e-09 | 3.2e-09 | 3.967E-09 | 2.635E-09 | 1.800E-09 | 6.900E-09 |
| 25186 | 1260 | S1260E | D | 1.2e-08 | 9.3e-09 | 6.5e-09 | 9.267E-09 | 2.750E-09 | 6.500E-09 | 1.200E-08 |
| 25187 | 1260 | S1260Q | D | 1e-08 | 3.3e-09 | 4.7e-09 | 6.000E-09 | 3.534E-09 | 3.300E-09 | 1.000E-08 |
| 25188 | 1260 | S1260G | D | 1.3e-08 | 3.7e-09 | 3.1e-09 | 6.600E-09 | 5.551E-09 | 3.100E-09 | 1.300E-08 |
| 25189 | 1260 | S1260H | D | 4.1e-09 | 5.9e-09 | 1.2e-08 | 7.333E-09 | 4.140E-09 | 4.100E-09 | 1.200E-08 |
| 25190 | 1260 | S1260I | D | 7.5e-09 | 1.4e-08 | 5.1e-09 | 8.867E-09 | 4.605E-09 | 5.100E-09 | 1.400E-08 |
| 25191 | 1260 | S1260L | D | 1e-08 | 6.1e-09 | 3.8e-09 | 6.633E-09 | 3.134E-09 | 3.800E-09 | 1.000E-08 |
| 25192 | 1260 | S1260K | D | 7.6e-09 | 7.6e-09 | 1.3e-08 | 9.400E-09 | 3.118E-09 | 7.600E-09 | 1.300E-08 |
| 25193 | 1260 | S1260M | D | 1.1e-08 | 4.3e-09 | 3.4e-09 | 6.233E-09 | 4.153E-09 | 3.400E-09 | 1.100E-08 |
| 25194 | 1260 | S1260F | D | 1.1e-08 | 7.5e-09 | 2.8e-09 | 7.100E-09 | 4.115E-09 | 2.800E-09 | 1.100E-08 |
| 25195 | 1260 | S1260P | D | 1.2e-08 | 4.4e-09 | 8.3e-09 | 8.233E-09 | 3.800E-09 | 4.400E-09 | 1.200E-08 |
| 25196 | 1260 | S1260S | D | 2.4e-09 | 1.3e-08 | 1.4e-08 | 9.800E-09 | 6.428E-09 | 2.400E-09 | 1.400E-08 |
| 25197 | 1260 | S1260T | D | 9.2e-09 | 4.8e-09 | 2.7e-08 | 1.367E-08 | 1.175E-08 | 4.800E-09 | 2.700E-08 |
| 25198 | 1260 | S1260W | D | 3.5e-09 | 1.1e-08 | 4.5e-09 | 6.333E-09 | 4.072E-09 | 3.500E-09 | 1.100E-08 |
| 25199 | 1260 | S1260Y | D | 7.5e-09 | 6.3e-09 | 5.8e-09 | 6.533E-09 | 8.737E-10 | 5.800E-09 | 7.500E-09 |
| 25200 | 1260 | S1260V | D | 8e-09 | 1.3e-08 | 3.5e-09 | 8.167E-09 | 4.752E-09 | 3.500E-09 | 1.300E-08 |
| 25201 | 1261 | S1261A | D | 8.4e-09 | 1e-08 | 1.5e-08 | 1.113E-08 | 3.443E-09 | 8.400E-09 | 1.500E-08 |
| 25202 | 1261 | S1261R | D | 9.1e-09 | 6.3e-09 | 1.1e-08 | 8.800E-09 | 2.364E-09 | 6.300E-09 | 1.100E-08 |
| 25203 | 1261 | S1261N | D | 6.6e-09 | 5.4e-09 | 8.3e-09 | 6.767E-09 | 1.457E-09 | 5.400E-09 | 8.300E-09 |
| 25204 | 1261 | S1261D | D | 3.1e-09 | 1.1e-08 | 5.3e-09 | 6.467E-09 | 4.077E-09 | 3.100E-09 | 1.100E-08 |
| 25205 | 1261 | S1261C | D | 7.1e-09 | 5.7e-09 | 1.7e-08 | 9.933E-09 | 6.160E-09 | 5.700E-09 | 1.700E-08 |
| 25206 | 1261 | S1261E | D | 6.3e-09 | 1.1e-08 | 1.1e-08 | 9.433E-09 | 2.714E-09 | 6.300E-09 | 1.100E-08 |
| 25207 | 1261 | S1261Q | D | 8.6e-09 | 2.1e-09 | 6e-09 | 5.567E-09 | 3.272E-09 | 2.100E-09 | 8.600E-09 |
| 25208 | 1261 | S1261G | D | 1e-08 | 4.3e-09 | 3.4e-09 | 5.900E-09 | 3.579E-09 | 3.400E-09 | 1.000E-08 |
| 25209 | 1261 | S1261H | D | 1.8e-08 | 3.3e-08 | 6.7e-09 | 1.923E-08 | 1.319E-08 | 6.700E-09 | 3.300E-08 |
| 25210 | 1261 | S1261I | D | 5.1e-09 | 5.1e-09 | 1.1e-08 | 7.067E-09 | 3.406E-09 | 5.100E-09 | 1.100E-08 |
| 25211 | 1261 | S1261L | D | 8.5e-09 | 1.9e-08 | 3.1e-08 | 1.950E-08 | 1.126E-08 | 8.500E-09 | 3.100E-08 |
| 25212 | 1261 | S1261K | D | 1.3e-08 | 9.4e-09 | 1.5e-08 | 1.247E-08 | 2.838E-09 | 9.400E-09 | 1.500E-08 |
| 25213 | 1261 | S1261M | D | 3.4e-09 | 1.7e-08 | 8.8e-09 | 9.733E-09 | 6.848E-09 | 3.400E-09 | 1.700E-08 |
| 25214 | 1261 | S1261F | D | 4.3e-09 | 2e-08 | 6.1e-09 | 1.013E-08 | 8.592E-09 | 4.300E-09 | 2.000E-08 |
| 25215 | 1261 | S1261P | D | 5.1e-09 | 6.9e-09 | 4.7e-09 | 5.567E-09 | 1.172E-09 | 4.700E-09 | 6.900E-09 |
| 25216 | 1261 | S1261S | D | 5.1e-09 | 7.4e-09 | 4.5e-09 | 5.667E-09 | 1.531E-09 | 4.500E-09 | 7.400E-09 |
| 25217 | 1261 | S1261T | D | 5.7e-09 | 9.5e-09 | 6.1e-09 | 7.100E-09 | 2.088E-09 | 5.700E-09 | 9.500E-09 |
| 25218 | 1261 | S1261W | D | 8.4e-09 | 9.8e-09 | 4.9e-09 | 7.700E-09 | 2.524E-09 | 4.900E-09 | 9.800E-09 |
| 25219 | 1261 | S1261Y | D | 4e-09 | 8.1e-09 | 3.9e-09 | 5.333E-09 | 2.397E-09 | 3.900E-09 | 8.100E-09 |
| 25220 | 1261 | S1261V | D | 6.8e-09 | 3.4e-09 | 4.5e-09 | 4.900E-09 | 1.735E-09 | 3.400E-09 | 6.800E-09 |
| 25221 | 1262 | P1262A | D | 1e-08 | 4.2e-09 | 3.8e-09 | 6.000E-09 | 3.470E-09 | 3.800E-09 | 1.000E-08 |
| 25222 | 1262 | P1262R | D | 3.2e-09 | 4.9e-09 | 2.5e-09 | 3.533E-09 | 1.234E-09 | 2.500E-09 | 4.900E-09 |
| 25223 | 1262 | P1262N | D | 1.3e-08 | 5.8e-09 | 3.5e-08 | 1.793E-08 | 1.521E-08 | 5.800E-09 | 3.500E-08 |
| 25224 | 1262 | P1262D | D | 1.2e-08 | 1.8e-08 | 6.6e-09 | 1.220E-08 | 5.703E-09 | 6.600E-09 | 1.800E-08 |
| 25225 | 1262 | P1262C | D | 9e-09 | 6e-09 | 5.8e-09 | 6.933E-09 | 1.793E-09 | 5.800E-09 | 9.000E-09 |
| 25226 | 1262 | P1262E | D | 8.5e-09 | 1.3e-08 | 7e-09 | 9.500E-09 | 3.122E-09 | 7.000E-09 | 1.300E-08 |
| 25227 | 1262 | P1262Q | D | 5.2e-09 | 2.6e-09 | 1.1e-08 | 6.267E-09 | 4.300E-09 | 2.600E-09 | 1.100E-08 |
| 25228 | 1262 | P1262G | D | 5.5e-09 | 2.5e-09 | 2e-09 | 3.333E-09 | 1.893E-09 | 2.000E-09 | 5.500E-09 |
| 25229 | 1262 | P1262H | D | 5.3e-09 | 6.4e-09 | 4.8e-09 | 5.500E-09 | 8.185E-10 | 4.800E-09 | 6.400E-09 |
| 25230 | 1262 | P1262I | D | 1e-08 | 7e-09 | 8.9e-09 | 8.633E-09 | 1.518E-09 | 7.000E-09 | 1.000E-08 |
| 25231 | 1262 | P1262L | D | 5.9e-09 | 1.3e-08 | 2.3e-08 | 1.397E-08 | 8.591E-09 | 5.900E-09 | 2.300E-08 |
| 25232 | 1262 | P1262K | D | 4.6e-09 | 7.1e-09 | 2.9e-09 | 4.867E-09 | 2.113E-09 | 2.900E-09 | 7.100E-09 |
| 25233 | 1262 | P1262M | D | 2.6e-09 | 1.6e-08 | 1.3e-08 | 1.053E-08 | 7.032E-09 | 2.600E-09 | 1.600E-08 |
| 25234 | 1262 | P1262F | D | 8e-09 | 1.1e-08 | 4.8e-09 | 7.933E-09 | 3.101E-09 | 4.800E-09 | 1.100E-08 |
| 25235 | 1262 | P1262P | D | 6e-09 | 3.6e-09 | 3.6e-09 | 4.400E-09 | 1.386E-09 | 3.600E-09 | 6.000E-09 |
| 25236 | 1262 | P1262S | D | 5e-09 | 6.7e-09 | 8.2e-09 | 6.633E-09 | 1.601E-09 | 5.000E-09 | 8.200E-09 |
| 25237 | 1262 | P1262T | D | 6.3e-09 | 6.4e-09 | 9.8e-09 | 7.500E-09 | 1.992E-09 | 6.300E-09 | 9.800E-09 |
| 25238 | 1262 | P1262W | D | 6e-09 | 4.8e-09 | 6.8e-09 | 5.867E-09 | 1.007E-09 | 4.800E-09 | 6.800E-09 |
| 25239 | 1262 | P1262Y | D | 1.9e-08 | 1.1e-08 | 4.8e-09 | 1.160E-08 | 7.119E-09 | 4.800E-09 | 1.900E-08 |
| 25240 | 1262 | P1262V | D | 8.2e-09 | 1.1e-08 | 7.1e-09 | 8.767E-09 | 2.011E-09 | 7.100E-09 | 1.100E-08 |
| 25241 | 1263 | V1263A | D | 4.1e-09 | 7.1e-09 | 1.8e-08 | 9.733E-09 | 7.315E-09 | 4.100E-09 | 1.800E-08 |
| 25242 | 1263 | V1263R | D | 6.5e-09 | 5.7e-09 | 5.4e-09 | 5.867E-09 | 5.686E-10 | 5.400E-09 | 6.500E-09 |
| 25243 | 1263 | V1263N | D | 9.6e-09 | 7.9e-09 | 4e-09 | 7.167E-09 | 2.871E-09 | 4.000E-09 | 9.600E-09 |
| 25244 | 1263 | V1263D | D | 1e-08 | 4.3e-09 | 4.1e-09 | 6.133E-09 | 3.350E-09 | 4.100E-09 | 1.000E-08 |

| | | | | | | | | | | |
|---|---|---|---|---|---|---|---|---|---|---|
| 25245 | 1263 | V1263C | D | 1.9e-09 | 5.8e-09 | 7.5e-09 | 5.067E-09 | 2.871E-09 | 1.900E-09 | 7.500E-09 |
| 25246 | 1263 | V1263E | D | 1.7e-08 | 7.1e-09 | 7e-09 | 1.037E-08 | 5.745E-09 | 7.000E-09 | 1.700E-08 |
| 25247 | 1263 | V1263Q | D | 6.7e-09 | 1.9e-08 | 9.2e-09 | 1.163E-08 | 6.501E-09 | 6.700E-09 | 1.900E-08 |
| 25248 | 1263 | V1263G | D | 1.3e-08 | 3e-09 | 9.1e-09 | 8.367E-09 | 5.040E-09 | 3.000E-09 | 1.300E-08 |
| 25249 | 1263 | V1263H | D | 3.7e-09 | 1.7e-08 | 2.1e-08 | 1.390E-08 | 9.057E-09 | 3.700E-09 | 2.100E-08 |
| 25250 | 1263 | V1263I | D | 4.8e-09 | 9.9e-09 | 7.2e-09 | 7.300E-09 | 2.551E-09 | 4.800E-09 | 9.900E-09 |
| 25251 | 1263 | V1263L | D | 4.1e-09 | 5.8e-09 | 8.6e-09 | 6.167E-09 | 2.272E-09 | 4.100E-09 | 8.600E-09 |
| 25252 | 1263 | V1263K | D | 5.8e-09 | 5.8e-09 | 1.4e-08 | 8.533E-09 | 4.734E-09 | 5.800E-09 | 1.400E-08 |
| 25253 | 1263 | V1263M | D | 4.5e-09 | 8.1e-09 | 1.2e-08 | 8.200E-09 | 3.751E-09 | 4.500E-09 | 1.200E-08 |
| 25254 | 1263 | V1263F | D | 7.3e-09 | 2e-08 | 6.9e-09 | 1.140E-08 | 7.451E-09 | 6.900E-09 | 2.000E-08 |
| 25255 | 1263 | V1263P | D | 1.1e-08 | 5.8e-09 | 2.7e-09 | 6.500E-09 | 4.194E-09 | 2.700E-09 | 1.100E-08 |
| 25256 | 1263 | V1263S | D | 7e-09 | 6.6e-09 | 4.9e-09 | 6.167E-09 | 1.115E-09 | 4.900E-09 | 7.000E-09 |
| 25257 | 1263 | V1263T | D | 6.4e-09 | 8.5e-09 | 4.8e-09 | 6.567E-09 | 1.856E-09 | 4.800E-09 | 8.500E-09 |
| 25258 | 1263 | V1263W | D | 5.5e-09 | 1e-08 | 7.1e-09 | 7.533E-09 | 2.281E-09 | 5.500E-09 | 1.000E-08 |
| 25259 | 1263 | V1263Y | D | 1.2e-08 | 4e-09 | 1.1e-08 | 9.000E-09 | 4.359E-09 | 4.000E-09 | 1.200E-08 |
| 25260 | 1263 | V1263V | D | 6.5e-09 | 5.4e-09 | 3.1e-09 | 5.000E-09 | 1.735E-09 | 3.100E-09 | 6.500E-09 |
| 25261 | 1264 | T1264A | D | 1e-08 | 6.6e-09 | 5.3e-09 | 7.300E-09 | 2.427E-09 | 5.300E-09 | 1.000E-08 |
| 25262 | 1264 | T1264R | D | 2e-09 | 3.7e-09 | 1.9e-09 | 2.533E-09 | 1.012E-09 | 1.900E-09 | 3.700E-09 |
| 25263 | 1264 | T1264N | D | 2.2e-09 | 2.1e-09 | 6.6e-09 | 3.633E-09 | 2.570E-09 | 2.100E-09 | 6.600E-09 |
| 25264 | 1264 | T1264D | D | 3.8e-09 | 2.6e-08 | 1.2e-08 | 1.393E-08 | 1.123E-08 | 3.800E-09 | 2.600E-08 |
| 25265 | 1264 | T1264C | D | 1.9e-09 | 1e-08 | 1e-08 | 7.300E-09 | 4.677E-09 | 1.900E-09 | 1.000E-08 |
| 25266 | 1264 | T1264E | D | 1.2e-08 | 1.1e-08 | 5.5e-09 | 9.500E-09 | 3.500E-09 | 5.500E-09 | 1.200E-08 |
| 25267 | 1264 | T1264Q | D | 4.9e-09 | 1e-08 | 2.1e-09 | 5.667E-09 | 4.005E-09 | 2.100E-09 | 1.000E-08 |
| 25268 | 1264 | T1264G | D | 1.6e-08 | 4e-09 | 3.4e-08 | 1.800E-08 | 1.510E-08 | 4.000E-09 | 3.400E-08 |
| 25269 | 1264 | T1264H | D | 1.3e-08 | 8.3e-09 | 4.2e-09 | 8.500E-09 | 4.403E-09 | 4.200E-09 | 1.300E-08 |
| 25270 | 1264 | T1264I | D | 8.6e-09 | 2.3e-08 | 9e-09 | 1.353E-08 | 8.201E-09 | 8.600E-09 | 2.300E-08 |
| 25271 | 1264 | T1264L | D | 1.7e-08 | 4.8e-09 | 1e-08 | 1.060E-08 | 6.122E-09 | 4.800E-09 | 1.700E-08 |
| 25272 | 1264 | T1264K | D | 6.4e-09 | 6.8e-09 | 8e-09 | 7.067E-09 | 8.327E-10 | 6.400E-09 | 8.000E-09 |
| 25273 | 1264 | T1264M | D | 7.6e-09 | 2.2e-08 | 6.2e-09 | 1.193E-08 | 8.746E-09 | 6.200E-09 | 2.200E-08 |
| 25274 | 1264 | T1264F | D | 1.2e-08 | 6.2e-09 | 9.5e-09 | 9.233E-09 | 2.909E-09 | 6.200E-09 | 1.200E-08 |
| 25275 | 1264 | T1264P | D | 1.7e-08 | 8.3e-09 | 1.1e-08 | 1.210E-08 | 4.453E-09 | 8.300E-09 | 1.700E-08 |
| 25276 | 1264 | T1264S | D | 6.3e-09 | 3.3e-09 | 7.6e-09 | 5.733E-09 | 2.205E-09 | 3.300E-09 | 7.600E-09 |
| 25277 | 1264 | T1264T | D | 1.1e-08 | 8.8e-09 | 8.1e-09 | 9.300E-09 | 1.513E-09 | 8.100E-09 | 1.100E-08 |
| 25278 | 1264 | T1264W | D | 7.4e-09 | 5.8e-09 | 1.3e-08 | 8.733E-09 | 3.781E-09 | 5.800E-09 | 1.300E-08 |
| 25279 | 1264 | T1264Y | D | 1.4e-09 | 7.8e-09 | 2.6e-08 | 1.173E-08 | 1.276E-08 | 1.400E-09 | 2.600E-08 |
| 25280 | 1264 | T1264V | D | 6.8e-09 | 6.6e-09 | 6.8e-09 | 6.733E-09 | 1.155E-10 | 6.600E-09 | 6.800E-09 |
| 25281 | 1265 | K1265A | D | 7.9e-09 | 3.3e-09 | 1.2e-08 | 7.733E-09 | 4.352E-09 | 3.300E-09 | 1.200E-08 |
| 25282 | 1265 | K1265R | D | 3.7e-09 | 4.4e-09 | 2.2e-08 | 1.003E-08 | 1.037E-08 | 3.700E-09 | 2.200E-08 |
| 25283 | 1265 | K1265N | D | 8.2e-09 | 5e-09 | 1.1e-08 | 8.067E-09 | 3.002E-09 | 5.000E-09 | 1.100E-08 |
| 25284 | 1265 | K1265D | D | 6.4e-09 | 3e-09 | 6.1e-09 | 5.167E-09 | 1.882E-09 | 3.000E-09 | 6.400E-09 |
| 25285 | 1265 | K1265C | D | 7.8e-09 | 3.3e-09 | 4e-09 | 5.033E-09 | 2.421E-09 | 3.300E-09 | 7.800E-09 |
| 25286 | 1265 | K1265E | D | 5.4e-09 | 7.7e-09 | 6.8e-09 | 6.633E-09 | 1.159E-09 | 5.400E-09 | 7.700E-09 |
| 25287 | 1265 | K1265Q | D | 4.2e-09 | 3.1e-09 | 4.8e-09 | 4.033E-09 | 8.622E-10 | 3.100E-09 | 4.800E-09 |
| 25288 | 1265 | K1265G | D | 1.1e-08 | 2e-09 | 7.9e-09 | 6.967E-09 | 4.572E-09 | 2.000E-09 | 1.100E-08 |
| 25289 | 1265 | K1265H | D | 1.1e-08 | 5.8e-09 | 3e-08 | 1.560E-08 | 1.274E-08 | 5.800E-09 | 3.000E-08 |
| 25290 | 1265 | K1265I | D | 6.3e-09 | 1.3e-08 | 1e-08 | 9.767E-09 | 3.356E-09 | 6.300E-09 | 1.300E-08 |
| 25291 | 1265 | K1265L | D | 6e-09 | 7.1e-09 | 1.5e-08 | 9.367E-09 | 4.910E-09 | 6.000E-09 | 1.500E-08 |
| 25292 | 1265 | K1265K | D | 1.1e-08 | 5.6e-09 | 4.9e-09 | 7.167E-09 | 3.338E-09 | 4.900E-09 | 1.100E-08 |
| 25293 | 1265 | K1265M | D | 4.5e-09 | 1.4e-08 | 4.5e-09 | 7.667E-09 | 5.485E-09 | 4.500E-09 | 1.400E-08 |
| 25294 | 1265 | K1265F | D | 4.8e-09 | 8.8e-09 | 7.6e-09 | 7.067E-09 | 2.053E-09 | 4.800E-09 | 8.800E-09 |
| 25295 | 1265 | K1265P | D | 1.2e-08 | 1.6e-08 | 5.6e-09 | 1.120E-08 | 5.246E-09 | 5.600E-09 | 1.600E-08 |
| 25296 | 1265 | K1265S | D | 1.5e-08 | 5.8e-09 | 3.6e-09 | 8.133E-09 | 6.048E-09 | 3.600E-09 | 1.500E-08 |
| 25297 | 1265 | K1265T | D | 1.2e-08 | 2.9e-09 | 5e-09 | 6.633E-09 | 4.765E-09 | 2.900E-09 | 1.200E-08 |
| 25298 | 1265 | K1265W | D | 2.4e-09 | 1.2e-08 | 9.1e-09 | 7.833E-09 | 4.924E-09 | 2.400E-09 | 1.200E-08 |
| 25299 | 1265 | K1265Y | D | 7.8e-09 | 1e-08 | 1.2e-08 | 9.933E-09 | 2.101E-09 | 7.800E-09 | 1.200E-08 |
| 25300 | 1265 | K1265V | D | 2.3e-08 | 4.7e-09 | 8e-09 | 1.190E-08 | 9.753E-09 | 4.700E-09 | 2.300E-08 |
| 25301 | 1266 | S1266A | D | 1.2e-08 | 7.9e-09 | 8.1e-09 | 9.333E-09 | 2.312E-09 | 7.900E-09 | 1.200E-08 |
| 25302 | 1266 | S1266R | D | 1.1e-08 | 4.3e-09 | 3.5e-09 | 6.267E-09 | 4.119E-09 | 3.500E-09 | 1.100E-08 |
| 25303 | 1266 | S1266N | D | 3.5e-09 | 1.8e-08 | 1.2e-08 | 1.117E-08 | 7.286E-09 | 3.500E-09 | 1.800E-08 |
| 25304 | 1266 | S1266D | D | 1.1e-08 | 6.7e-09 | 4.4e-09 | 7.367E-09 | 3.350E-09 | 4.400E-09 | 1.100E-08 |
| 25305 | 1266 | S1266C | D | 6.1e-09 | 5e-09 | 5.1e-09 | 5.400E-09 | 6.083E-10 | 5.000E-09 | 6.100E-09 |
| 25306 | 1266 | S1266E | D | 5.9e-09 | 3.9e-09 | 7.1e-09 | 5.633E-09 | 1.617E-09 | 3.900E-09 | 7.100E-09 |
| 25307 | 1266 | S1266Q | D | 3.1e-09 | 7.3e-09 | 1.5e-08 | 8.467E-09 | 6.035E-09 | 3.100E-09 | 1.500E-08 |
| 25308 | 1266 | S1266G | D | 1.3e-08 | 3.5e-09 | 2.5e-09 | 6.333E-09 | 5.795E-09 | 2.500E-09 | 1.300E-08 |
| 25309 | 1266 | S1266H | D | 3.8e-09 | 4.1e-08 | 3.5e-09 | 1.610E-08 | 2.156E-08 | 3.500E-09 | 4.100E-08 |
| 25310 | 1266 | S1266I | D | 4.8e-09 | 1.7e-08 | 4.8e-09 | 8.867E-09 | 7.044E-09 | 4.800E-09 | 1.700E-08 |
| 25311 | 1266 | S1266L | D | 8.3e-09 | 3.5e-09 | 2.3e-09 | 4.700E-09 | 3.175E-09 | 2.300E-09 | 8.300E-09 |
| 25312 | 1266 | S1266K | D | 1.2e-08 | 2.8e-09 | 7e-09 | 7.267E-09 | 4.606E-09 | 2.800E-09 | 1.200E-08 |
| 25313 | 1266 | S1266M | D | 1.2e-08 | 3e-09 | 1.4e-08 | 9.667E-09 | 5.859E-09 | 3.000E-09 | 1.400E-08 |
| 25314 | 1266 | S1266F | D | 3.1e-09 | 8.9e-09 | 1.4e-08 | 8.667E-09 | 5.454E-09 | 3.100E-09 | 1.400E-08 |
| 25315 | 1266 | S1266P | D | 3.6e-09 | 8.9e-09 | 3.9e-09 | 5.467E-09 | 2.977E-09 | 3.600E-09 | 8.900E-09 |
| 25316 | 1266 | S1266S | D | 7.8e-09 | 5.6e-09 | 6.3e-09 | 6.567E-09 | 1.124E-09 | 5.600E-09 | 7.800E-09 |
| 25317 | 1266 | S1266T | D | 6e-09 | 1e-08 | 1.3e-09 | 5.767E-09 | 4.355E-09 | 1.300E-09 | 1.000E-08 |
| 25318 | 1266 | S1266W | D | 6.5e-09 | 3.7e-09 | 7.5e-09 | 5.900E-09 | 1.970E-09 | 3.700E-09 | 7.500E-09 |
| 25319 | 1266 | S1266Y | D | 4.8e-09 | 3.5e-09 | 3.4e-09 | 3.900E-09 | 7.810E-10 | 3.400E-09 | 4.800E-09 |
| 25320 | 1266 | S1266V | D | 8.4e-09 | 1.6e-08 | 3.6e-09 | 9.333E-09 | 6.252E-09 | 3.600E-09 | 1.600E-08 |
| 25321 | 1267 | F1267A | D | 1.9e-08 | 3.1e-09 | 6.5e-09 | 9.533E-09 | 8.373E-09 | 3.100E-09 | 1.900E-08 |
| 25322 | 1267 | F1267R | D | 6.2e-09 | 1.3e-08 | 3.5e-09 | 7.567E-09 | 4.895E-09 | 3.500E-09 | 1.300E-08 |
| 25323 | 1267 | F1267N | D | 5.2e-09 | 6.4e-09 | 6.1e-09 | 5.900E-09 | 6.245E-10 | 5.200E-09 | 6.400E-09 |
| 25324 | 1267 | F1267D | D | 7.7e-09 | 1.1e-08 | 8.3e-09 | 9.000E-09 | 1.758E-09 | 7.700E-09 | 1.100E-08 |
| 25325 | 1267 | F1267C | D | 2.8e-08 | 1.7e-08 | 6.2e-09 | 1.707E-08 | 1.090E-08 | 6.200E-09 | 2.800E-08 |
| 25326 | 1267 | F1267E | D | 5.9e-09 | 1.5e-08 | 1.2e-08 | 1.097E-08 | 4.637E-09 | 5.900E-09 | 1.500E-08 |
| 25327 | 1267 | F1267Q | D | 9.6e-09 | 6.9e-09 | 1.3e-08 | 9.833E-09 | 3.057E-09 | 6.900E-09 | 1.300E-08 |
| 25328 | 1267 | F1267G | D | 1.2e-08 | 2.8e-09 | 3.4e-09 | 6.067E-09 | 5.147E-09 | 2.800E-09 | 1.200E-08 |
| 25329 | 1267 | F1267H | D | 5.9e-09 | 2.9e-09 | 1.7e-08 | 8.600E-09 | 7.428E-09 | 2.900E-09 | 1.700E-08 |

| | | | | | | | | | | |
|---|---|---|---|---|---|---|---|---|---|---|
| 25330 | 1267 | F1267I | D | 1.5e-08 | 7.1e-09 | 5.5e-09 | 9.200E-09 | 5.086E-09 | 5.500E-09 | 1.500E-08 |
| 25331 | 1267 | F1267L | D | 1.2e-08 | 1.1e-08 | 8.5e-09 | 1.050E-08 | 1.803E-09 | 8.500E-09 | 1.200E-08 |
| 25332 | 1267 | F1267K | D | 7.8e-09 | 1.3e-08 | 5.1e-09 | 8.633E-09 | 4.015E-09 | 5.100E-09 | 1.300E-08 |
| 25333 | 1267 | F1267M | D | 8.3e-09 | 3.6e-09 | 3.8e-09 | 5.233E-09 | 2.658E-09 | 3.600E-09 | 8.300E-09 |
| 25334 | 1267 | F1267F | D | 1.1e-08 | 8e-09 | 1.1e-08 | 1.000E-08 | 1.732E-09 | 8.000E-09 | 1.100E-08 |
| 25335 | 1267 | F1267P | D | 4.2e-09 | 4.7e-09 | 5.2e-09 | 4.700E-09 | 5.000E-10 | 4.200E-09 | 5.200E-09 |
| 25336 | 1267 | F1267S | D | 5.5e-09 | 1.4e-08 | 4.2e-09 | 7.900E-09 | 5.323E-09 | 4.200E-09 | 1.400E-08 |
| 25337 | 1267 | F1267T | D | 9e-09 | 1.1e-08 | 3.1e-09 | 7.700E-09 | 4.107E-09 | 3.100E-09 | 1.100E-08 |
| 25338 | 1267 | F1267W | D | 1.9e-09 | 1e-08 | 1.3e-08 | 8.300E-09 | 5.742E-09 | 1.900E-09 | 1.300E-08 |
| 25339 | 1267 | F1267Y | D | 3.5e-09 | 5e-09 | 6.2e-09 | 4.900E-09 | 1.353E-09 | 3.500E-09 | 6.200E-09 |
| 25340 | 1267 | F1267V | D | 3.2e-09 | 5.6e-09 | 3.5e-09 | 4.100E-09 | 1.308E-09 | 3.200E-09 | 5.600E-09 |
| 25341 | 1268 | N1268A | D | 6.9e-09 | 1.1e-08 | 2.4e-09 | 6.767E-09 | 4.302E-09 | 2.400E-09 | 1.100E-08 |
| 25342 | 1268 | N1268R | D | 1.1e-08 | 9.9e-09 | 1.4e-08 | 1.163E-08 | 2.122E-09 | 9.900E-09 | 1.400E-08 |
| 25343 | 1268 | N1268N | D | 6.2e-09 | 3.8e-09 | 1.2e-08 | 7.333E-09 | 4.216E-09 | 3.800E-09 | 1.200E-08 |
| 25344 | 1268 | N1268D | D | 1.1e-09 | 7.1e-09 | 5.3e-09 | 4.500E-09 | 3.079E-09 | 1.100E-09 | 7.100E-09 |
| 25345 | 1268 | N1268C | D | 1e-08 | 3e-08 | 4.5e-09 | 1.483E-08 | 1.342E-08 | 4.500E-09 | 3.000E-08 |
| 25346 | 1268 | N1268E | D | 2.1e-08 | 2e-09 | 1.2e-08 | 1.167E-08 | 9.504E-09 | 2.000E-09 | 2.100E-08 |
| 25347 | 1268 | N1268Q | D | 1.2e-08 | 9.9e-09 | 3.1e-09 | 8.333E-09 | 4.652E-09 | 3.100E-09 | 1.200E-08 |
| 25348 | 1268 | N1268G | D | 3.1e-09 | 8.3e-09 | 1.2e-08 | 7.800E-09 | 4.471E-09 | 3.100E-09 | 1.200E-08 |
| 25349 | 1268 | N1268H | D | 5.5e-09 | 1.1e-08 | 1.4e-08 | 1.017E-08 | 4.311E-09 | 5.500E-09 | 1.400E-08 |
| 25350 | 1268 | N1268I | D | 5.6e-09 | 1.1e-08 | 5e-09 | 7.200E-09 | 3.305E-09 | 5.000E-09 | 1.100E-08 |
| 25351 | 1268 | N1268L | D | 3.7e-09 | 1.6e-08 | 7.6e-09 | 9.100E-09 | 6.286E-09 | 3.700E-09 | 1.600E-08 |
| 25352 | 1268 | N1268K | D | 3.5e-09 | 7.3e-09 | 2e-09 | 4.267E-09 | 2.732E-09 | 2.000E-09 | 7.300E-09 |
| 25353 | 1268 | N1268M | D | 1.1e-08 | 9.6e-09 | 5.8e-09 | 8.800E-09 | 2.691E-09 | 5.800E-09 | 1.100E-08 |
| 25354 | 1268 | N1268F | D | 2.9e-09 | 5.7e-09 | 2.1e-09 | 3.567E-09 | 1.890E-09 | 2.100E-09 | 5.700E-09 |
| 25355 | 1268 | N1268P | D | 1.4e-08 | 8.8e-09 | 1.9e-08 | 1.393E-08 | 5.100E-09 | 8.800E-09 | 1.900E-08 |
| 25356 | 1268 | N1268S | D | 3.9e-09 | 8.6e-09 | 5.8e-09 | 6.100E-09 | 2.364E-09 | 3.900E-09 | 8.600E-09 |
| 25357 | 1268 | N1268T | D | 1.8e-08 | 1e-08 | 1.6e-08 | 1.467E-08 | 4.163E-09 | 1.000E-08 | 1.800E-08 |
| 25358 | 1268 | N1268W | D | 5.8e-09 | 5.3e-09 | 7.9e-09 | 6.333E-09 | 1.380E-09 | 5.300E-09 | 7.900E-09 |
| 25359 | 1268 | N1268Y | D | 1.2e-08 | 3.5e-09 | 3.3e-08 | 1.617E-08 | 1.518E-08 | 3.500E-09 | 3.300E-08 |
| 25360 | 1268 | N1268V | D | 8.7e-09 | 8.5e-09 | 1.9e-08 | 1.207E-08 | 6.005E-09 | 8.500E-09 | 1.900E-08 |
| 25361 | 1269 | R1269A | D | 1e-08 | 8.2e-09 | 9.2e-09 | 9.133E-09 | 9.018E-10 | 8.200E-09 | 1.000E-08 |
| 25362 | 1269 | R1269R | D | 9.9e-09 | 5.5e-09 | 5.9e-09 | 7.100E-09 | 2.433E-09 | 5.500E-09 | 9.900E-09 |
| 25363 | 1269 | R1269N | D | 1.7e-08 | 7e-09 | 6.9e-09 | 1.030E-08 | 5.803E-09 | 6.900E-09 | 1.700E-08 |
| 25364 | 1269 | R1269D | D | 3.4e-09 | 5.3e-09 | 1.2e-08 | 6.900E-09 | 4.518E-09 | 3.400E-09 | 1.200E-08 |
| 25365 | 1269 | R1269C | D | 2.9e-09 | 1.4e-08 | 6.5e-09 | 7.800E-09 | 5.663E-09 | 2.900E-09 | 1.400E-08 |
| 25366 | 1269 | R1269E | D | 9.4e-09 | 3.3e-09 | 1.4e-08 | 8.900E-09 | 5.367E-09 | 3.300E-09 | 1.400E-08 |
| 25367 | 1269 | R1269Q | D | 3.9e-09 | 2.5e-09 | 1.9e-09 | 2.767E-09 | 1.026E-09 | 1.900E-09 | 3.900E-09 |
| 25368 | 1269 | R1269G | D | 3.2e-09 | 2.7e-09 | 1e-08 | 5.300E-09 | 4.078E-09 | 2.700E-09 | 1.000E-08 |
| 25369 | 1269 | R1269H | D | 6.6e-09 | 7.8e-09 | 5.7e-09 | 6.700E-09 | 1.054E-09 | 5.700E-09 | 7.800E-09 |
| 25370 | 1269 | R1269I | D | 8.3e-09 | 2.9e-09 | 8.9e-09 | 6.700E-09 | 3.305E-09 | 2.900E-09 | 8.900E-09 |
| 25371 | 1269 | R1269L | D | 1.2e-08 | 6.2e-09 | 8.5e-09 | 8.900E-09 | 2.921E-09 | 6.200E-09 | 1.200E-08 |
| 25372 | 1269 | R1269K | D | 7.6e-09 | 5e-09 | 6.5e-09 | 6.367E-09 | 1.305E-09 | 5.000E-09 | 7.600E-09 |
| 25373 | 1269 | R1269M | D | 6.7e-09 | 7.3e-09 | 8.8e-09 | 7.600E-09 | 1.082E-09 | 6.700E-09 | 8.800E-09 |
| 25374 | 1269 | R1269F | D | 5.7e-09 | 7.8e-09 | 8e-09 | 7.167E-09 | 1.274E-09 | 5.700E-09 | 8.000E-09 |
| 25375 | 1269 | R1269P | D | 5.5e-09 | 6.2e-09 | 1.1e-08 | 7.567E-09 | 2.994E-09 | 5.500E-09 | 1.100E-08 |
| 25376 | 1269 | R1269S | D | 1.8e-08 | 1.9e-09 | 7.1e-09 | 9.000E-09 | 8.216E-09 | 1.900E-09 | 1.800E-08 |
| 25377 | 1269 | R1269T | D | 1.8e-08 | 1.7e-08 | 6.3e-09 | 1.377E-08 | 6.486E-09 | 6.300E-09 | 1.800E-08 |
| 25378 | 1269 | R1269W | D | 2.6e-09 | 1.2e-08 | 1e-08 | 8.200E-09 | 4.952E-09 | 2.600E-09 | 1.200E-08 |
| 25379 | 1269 | R1269Y | D | 1.3e-08 | 7.5e-09 | 5.3e-09 | 8.600E-09 | 3.966E-09 | 5.300E-09 | 1.300E-08 |
| 25380 | 1269 | R1269V | D | 4.6e-09 | 4.4e-09 | 5.8e-09 | 4.933E-09 | 7.572E-10 | 4.400E-09 | 5.800E-09 |
| 25381 | 1270 | G1270A | D | 6.2e-09 | 7e-09 | 8.9e-09 | 7.367E-09 | 1.387E-09 | 6.200E-09 | 8.900E-09 |
| 25382 | 1270 | G1270R | D | 5.5e-09 | 8e-09 | 7.1e-09 | 6.867E-09 | 1.266E-09 | 5.500E-09 | 8.000E-09 |
| 25383 | 1270 | G1270N | D | 3.5e-08 | 7.2e-09 | 1.2e-08 | 1.807E-08 | 1.486E-08 | 7.200E-09 | 3.500E-08 |
| 25384 | 1270 | G1270D | D | 1.2e-08 | 9.3e-09 | 1.2e-08 | 1.110E-08 | 1.559E-09 | 9.300E-09 | 1.200E-08 |
| 25385 | 1270 | G1270C | D | 2.8e-09 | 5e-09 | 6.3e-09 | 4.700E-09 | 1.769E-09 | 2.800E-09 | 6.300E-09 |
| 25386 | 1270 | G1270E | D | 3.4e-09 | 2.3e-09 | 3.5e-09 | 3.067E-09 | 6.658E-10 | 2.300E-09 | 3.500E-09 |
| 25387 | 1270 | G1270Q | D | 5.3e-09 | 2.4e-09 | 2.4e-08 | 1.057E-08 | 1.172E-08 | 2.400E-09 | 2.400E-08 |
| 25388 | 1270 | G1270G | D | 9.3e-09 | 7.1e-09 | 4.9e-09 | 7.100E-09 | 2.200E-09 | 4.900E-09 | 9.300E-09 |
| 25389 | 1270 | G1270H | D | 4.5e-09 | 4.9e-09 | 4e-09 | 4.467E-09 | 4.509E-10 | 4.000E-09 | 4.900E-09 |
| 25390 | 1270 | G1270I | D | 3.2e-09 | 8.5e-09 | 3.6e-09 | 5.100E-09 | 2.951E-09 | 3.200E-09 | 8.500E-09 |
| 25391 | 1270 | G1270L | D | 3.5e-08 | 1.1e-08 | 4e-09 | 1.667E-08 | 1.626E-08 | 4.000E-09 | 3.500E-08 |
| 25392 | 1270 | G1270K | D | 5.7e-09 | 1.5e-08 | 6.4e-09 | 9.033E-09 | 5.179E-09 | 5.700E-09 | 1.500E-08 |
| 25393 | 1270 | G1270M | D | 4.4e-09 | 9.5e-09 | 7.1e-09 | 7.000E-09 | 2.551E-09 | 4.400E-09 | 9.500E-09 |
| 25394 | 1270 | G1270F | D | 7.9e-09 | 2.9e-09 | 2.2e-08 | 1.093E-08 | 9.905E-09 | 2.900E-09 | 2.200E-08 |
| 25395 | 1270 | G1270P | D | 4.2e-09 | 6.9e-09 | 6e-09 | 5.700E-09 | 1.375E-09 | 4.200E-09 | 6.900E-09 |
| 25396 | 1270 | G1270S | D | 2.2e-08 | 6.3e-09 | 8e-09 | 1.210E-08 | 8.616E-09 | 6.300E-09 | 2.200E-08 |
| 25397 | 1270 | G1270T | D | 1.8e-08 | 1.2e-08 | 1.7e-08 | 1.567E-08 | 3.215E-09 | 1.200E-08 | 1.800E-08 |
| 25398 | 1270 | G1270W | D | 3.9e-09 | 4.8e-09 | 4.3e-09 | 4.333E-09 | 4.509E-10 | 3.900E-09 | 4.800E-09 |
| 25399 | 1270 | G1270Y | D | 1e-08 | 1.5e-08 | 2.5e-08 | 1.667E-08 | 7.638E-09 | 1.000E-08 | 2.500E-08 |
| 25400 | 1270 | G1270V | D | 9.7e-09 | 4.6e-09 | 1.2e-08 | 8.767E-09 | 3.787E-09 | 4.600E-09 | 1.200E-08 |
| 25401 | 1271 | E1271A | D | 1.1e-08 | 7.9e-09 | 4.3e-09 | 7.733E-09 | 3.353E-09 | 4.300E-09 | 1.100E-08 |
| 25402 | 1271 | E1271R | D | 1.4e-08 | 7.1e-09 | 3.1e-09 | 8.067E-09 | 5.514E-09 | 3.100E-09 | 1.400E-08 |
| 25403 | 1271 | E1271N | D | 3.1e-09 | 5e-09 | 1.8e-09 | 3.300E-09 | 1.609E-09 | 1.800E-09 | 5.000E-09 |
| 25404 | 1271 | E1271D | D | 6.9e-09 | 8.3e-09 | 7.6e-09 | 7.600E-09 | 7.000E-10 | 6.900E-09 | 8.300E-09 |
| 25405 | 1271 | E1271C | D | 5.3e-09 | 6.3e-09 | 6.9e-09 | 6.167E-09 | 8.083E-10 | 5.300E-09 | 6.900E-09 |
| 25406 | 1271 | E1271E | D | 7.3e-09 | 5.4e-09 | 2.1e-08 | 1.123E-08 | 8.511E-09 | 5.400E-09 | 2.100E-08 |
| 25407 | 1271 | E1271Q | D | 1.9e-08 | 1.3e-08 | 1.1e-08 | 1.433E-08 | 4.163E-09 | 1.100E-08 | 1.900E-08 |
| 25408 | 1271 | E1271G | D | 2e-09 | 5.3e-09 | 4e-09 | 3.767E-09 | 1.662E-09 | 2.000E-09 | 5.300E-09 |
| 25409 | 1271 | E1271H | D | 1.7e-08 | 1e-08 | 1.9e-08 | 1.533E-08 | 4.726E-09 | 1.000E-08 | 1.900E-08 |
| 25410 | 1271 | E1271I | D | 1.3e-09 | 1.1e-08 | 2.5e-08 | 1.243E-08 | 1.191E-08 | 1.300E-09 | 2.500E-08 |
| 25411 | 1271 | E1271L | D | 5e-09 | 7.2e-09 | 1.1e-08 | 7.733E-09 | 3.035E-09 | 5.000E-09 | 1.100E-08 |
| 25412 | 1271 | E1271K | D | 2.9e-09 | 1.3e-08 | 1.5e-08 | 1.030E-08 | 6.486E-09 | 2.900E-09 | 1.500E-08 |
| 25413 | 1271 | E1271M | D | 1e-08 | 6.3e-09 | 8.3e-09 | 8.200E-09 | 1.852E-09 | 6.300E-09 | 1.000E-08 |
| 25414 | 1271 | E1271F | D | 5e-09 | 1.1e-08 | 5.9e-09 | 7.300E-09 | 3.236E-09 | 5.000E-09 | 1.100E-08 |

| | | | | | | | | | | |
|---|---|---|---|---|---|---|---|---|---|---|
| 25415 | 1271 | E1271P | D | 1.5e-08 | 1.8e-08 | 6.3e-09 | 1.310E-08 | 6.077E-09 | 6.300E-09 | 1.800E-08 |
| 25416 | 1271 | E1271S | D | 9.6e-09 | 6.9e-09 | 6.1e-09 | 7.533E-09 | 1.834E-09 | 6.100E-09 | 9.600E-09 |
| 25417 | 1271 | E1271T | D | 5.5e-09 | 6.9e-09 | 4.5e-09 | 5.633E-09 | 1.206E-09 | 4.500E-09 | 6.900E-09 |
| 25418 | 1271 | E1271W | D | 1e-08 | 1.1e-08 | 8.2e-09 | 9.733E-09 | 1.419E-09 | 8.200E-09 | 1.100E-08 |
| 25419 | 1271 | E1271Y | D | 4.4e-09 | 7e-09 | 7.9e-09 | 6.433E-09 | 1.818E-09 | 4.400E-09 | 7.900E-09 |
| 25420 | 1271 | E1271V | D | 2.1e-08 | 4.5e-09 | 6.1e-09 | 1.053E-08 | 9.100E-09 | 4.500E-09 | 2.100E-08 |
| 25421 | 1272 | C1272A | D | 3.3e-09 | 1.5e-08 | 5.8e-09 | 8.033E-09 | 6.161E-09 | 3.300E-09 | 1.500E-08 |
| 25422 | 1272 | C1272R | D | 6.1e-09 | 2e-09 | 9.5e-09 | 5.867E-09 | 3.755E-09 | 2.000E-09 | 9.500E-09 |
| 25423 | 1272 | C1272N | D | 1.3e-08 | 9.2e-09 | 1.2e-08 | 1.140E-08 | 1.970E-09 | 9.200E-09 | 1.300E-08 |
| 25424 | 1272 | C1272D | D | 4.5e-09 | 6.9e-09 | 5.5e-09 | 5.633E-09 | 1.206E-09 | 4.500E-09 | 6.900E-09 |
| 25425 | 1272 | C1272C | D | 6.7e-09 | 7.9e-09 | 1.2e-08 | 8.867E-09 | 2.779E-09 | 6.700E-09 | 1.200E-08 |
| 25426 | 1272 | C1272E | D | 5.2e-09 | 3.7e-09 | 2.9e-09 | 3.933E-09 | 1.168E-09 | 2.900E-09 | 5.200E-09 |
| 25427 | 1272 | C1272Q | D | 7.2e-09 | 1e-08 | 6.5e-09 | 7.900E-09 | 1.852E-09 | 6.500E-09 | 1.000E-08 |
| 25428 | 1272 | C1272G | D | 6.9e-09 | 7e-09 | 8.9e-09 | 7.600E-09 | 1.127E-09 | 6.900E-09 | 8.900E-09 |
| 25429 | 1272 | C1272H | D | 3.6e-09 | 2.7e-09 | 1.3e-08 | 6.433E-09 | 5.705E-09 | 2.700E-09 | 1.300E-08 |
| 25430 | 1272 | C1272I | D | 3.8e-09 | 6.8e-09 | 2.6e-09 | 4.400E-09 | 2.163E-09 | 2.600E-09 | 6.800E-09 |
| 25431 | 1272 | C1272L | D | 4.7e-09 | 7e-09 | 5.5e-09 | 5.733E-09 | 1.168E-09 | 4.700E-09 | 7.000E-09 |
| 25432 | 1272 | C1272K | D | 4.8e-09 | 7.3e-09 | 2e-08 | 1.070E-08 | 8.150E-09 | 4.800E-09 | 2.000E-08 |
| 25433 | 1272 | C1272M | D | 1.1e-08 | 6.1e-09 | 4.1e-09 | 7.067E-09 | 3.550E-09 | 4.100E-09 | 1.100E-08 |
| 25434 | 1272 | C1272F | D | 2.7e-09 | 1.1e-08 | 7.9e-09 | 7.200E-09 | 4.194E-09 | 2.700E-09 | 1.100E-08 |
| 25435 | 1272 | C1272P | D | 5.3e-09 | 4.8e-09 | 3.1e-09 | 4.400E-09 | 1.153E-09 | 3.100E-09 | 5.300E-09 |
| 25436 | 1272 | C1272S | D | 2.5e-08 | 7.3e-09 | 2.8e-09 | 1.170E-08 | 1.174E-08 | 2.800E-09 | 2.500E-08 |
| 25437 | 1272 | C1272T | D | 3.6e-09 | 3.7e-09 | 4.2e-09 | 3.833E-09 | 3.215E-10 | 3.600E-09 | 4.200E-09 |
| 25438 | 1272 | C1272W | D | 7.3e-09 | 6.6e-09 | 3.4e-09 | 5.767E-09 | 2.079E-09 | 3.400E-09 | 7.300E-09 |
| 25439 | 1272 | C1272Y | D | 8.9e-09 | 2.2e-08 | 4e-09 | 1.163E-08 | 9.306E-09 | 4.000E-09 | 2.200E-08 |
| 25440 | 1272 | C1272V | D | 8.3e-09 | 3.5e-09 | 1.3e-08 | 8.267E-09 | 4.750E-09 | 3.500E-09 | 1.300E-08 |
| 25441 | 1273 | E1273A | E | 5.9e-09 | 5.7e-09 | 3.4e-09 | 5.000E-09 | 1.389E-09 | 3.400E-09 | 5.900E-09 |
| 25442 | 1273 | E1273R | E | 5.3e-09 | 1.1e-08 | 4.8e-09 | 7.033E-09 | 3.444E-09 | 4.800E-09 | 1.100E-08 |
| 25443 | 1273 | E1273N | E | 1.9e-08 | 1.7e-08 | 9.7e-09 | 1.523E-08 | 4.895E-09 | 9.700E-09 | 1.900E-08 |
| 25444 | 1273 | E1273D | E | 3.6e-09 | 8.2e-09 | 5.1e-09 | 5.633E-09 | 2.346E-09 | 3.600E-09 | 8.200E-09 |
| 25445 | 1273 | E1273C | E | 9.1e-09 | 1.7e-08 | 9.3e-09 | 1.180E-08 | 4.504E-09 | 9.100E-09 | 1.700E-08 |
| 25446 | 1273 | E1273E | E | 7.1e-09 | 5.2e-09 | 1.1e-08 | 7.767E-09 | 2.957E-09 | 5.200E-09 | 1.100E-08 |
| 25447 | 1273 | E1273Q | E | 3.9e-09 | 9.8e-09 | 1.3e-08 | 8.900E-09 | 4.616E-09 | 3.900E-09 | 1.300E-08 |
| 25448 | 1273 | E1273G | E | 1.1e-08 | 4.6e-09 | 2.1e-09 | 5.900E-09 | 4.590E-09 | 2.100E-09 | 1.100E-08 |
| 25449 | 1273 | E1273H | E | 6.7e-09 | 7.1e-09 | 8e-09 | 7.267E-09 | 6.658E-10 | 6.700E-09 | 8.000E-09 |
| 25450 | 1273 | E1273I | E | 5.7e-09 | 3.7e-09 | 2.3e-09 | 3.900E-09 | 1.709E-09 | 2.300E-09 | 5.700E-09 |
| 25451 | 1273 | E1273L | E | 9.5e-09 | 7.7e-09 | 8.8e-09 | 8.667E-09 | 9.074E-10 | 7.700E-09 | 9.500E-09 |
| 25452 | 1273 | E1273K | E | 3.7e-09 | 5.8e-09 | 1.6e-08 | 8.500E-09 | 6.580E-09 | 3.700E-09 | 1.600E-08 |
| 25453 | 1273 | E1273M | E | 9.1e-09 | 6.6e-09 | 7.7e-09 | 7.800E-09 | 1.253E-09 | 6.600E-09 | 9.100E-09 |
| 25454 | 1273 | E1273F | E | 9.3e-09 | 1e-08 | 2.5e-08 | 1.477E-08 | 8.869E-09 | 9.300E-09 | 2.500E-08 |
| 25455 | 1273 | E1273P | E | 1.2e-08 | 3.8e-09 | 5.5e-09 | 7.100E-09 | 4.328E-09 | 3.800E-09 | 1.200E-08 |
| 25456 | 1273 | E1273S | E | 1.4e-08 | 5.6e-09 | 5.6e-09 | 8.400E-09 | 4.850E-09 | 5.600E-09 | 1.400E-08 |
| 25457 | 1273 | E1273T | E | 1.2e-08 | 5.3e-09 | 6.3e-09 | 7.867E-09 | 3.614E-09 | 5.300E-09 | 1.200E-08 |
| 25458 | 1273 | E1273W | E | 8.3e-09 | 1.9e-08 | 1.3e-08 | 1.343E-08 | 5.363E-09 | 8.300E-09 | 1.900E-08 |
| 25459 | 1273 | E1273Y | E | 2.4e-09 | 6.2e-09 | 1.4e-08 | 7.533E-09 | 5.914E-09 | 2.400E-09 | 1.400E-08 |
| 25460 | 1273 | E1273V | E | 1.3e-08 | 4.6e-09 | 1.7e-08 | 1.153E-08 | 6.329E-09 | 4.600E-09 | 1.700E-08 |
| 25461 | 1274 | V1274A | E | 2.6e-09 | 1.7e-08 | 7.3e-09 | 8.967E-09 | 7.343E-09 | 2.600E-09 | 1.700E-08 |
| 25462 | 1274 | V1274R | E | 1.3e-08 | 1.6e-09 | 9e-09 | 7.867E-09 | 5.784E-09 | 1.600E-09 | 1.300E-08 |
| 25463 | 1274 | V1274N | E | 3.3e-09 | 8.6e-09 | 4.1e-09 | 5.333E-09 | 2.857E-09 | 3.300E-09 | 8.600E-09 |
| 25464 | 1274 | V1274D | E | 9.9e-09 | 9.6e-09 | 9.6e-09 | 9.700E-09 | 1.732E-10 | 9.600E-09 | 9.900E-09 |
| 25465 | 1274 | V1274C | E | 1.1e-08 | 8.6e-09 | 7.8e-09 | 9.133E-09 | 1.665E-09 | 7.800E-09 | 1.100E-08 |
| 25466 | 1274 | V1274E | E | 8.5e-09 | 1e-09 | 4.5e-09 | 4.667E-09 | 3.753E-09 | 1.000E-09 | 8.500E-09 |
| 25467 | 1274 | V1274Q | E | 1e-08 | 5.9e-09 | 6.2e-09 | 7.367E-09 | 2.285E-09 | 5.900E-09 | 1.000E-08 |
| 25468 | 1274 | V1274G | E | 5.6e-09 | 3.1e-09 | 1.1e-08 | 6.567E-09 | 4.038E-09 | 3.100E-09 | 1.100E-08 |
| 25469 | 1274 | V1274H | E | 5.9e-09 | 5.5e-09 | 6.7e-09 | 6.033E-09 | 6.110E-10 | 5.500E-09 | 6.700E-09 |
| 25470 | 1274 | V1274I | E | 4.8e-09 | 4.1e-09 | 1.5e-08 | 7.967E-09 | 6.101E-09 | 4.100E-09 | 1.500E-08 |
| 25471 | 1274 | V1274L | E | 4.5e-09 | 6.5e-09 | 5e-09 | 5.333E-09 | 1.041E-09 | 4.500E-09 | 6.500E-09 |
| 25472 | 1274 | V1274K | E | 6.1e-09 | 1.2e-09 | 1.3e-08 | 6.767E-09 | 5.928E-09 | 1.200E-09 | 1.300E-08 |
| 25473 | 1274 | V1274M | E | 9.1e-09 | 1.1e-08 | 1.3e-08 | 1.103E-08 | 1.950E-09 | 9.100E-09 | 1.300E-08 |
| 25474 | 1274 | V1274F | E | 4.7e-09 | 7.3e-09 | 8.6e-09 | 6.867E-09 | 1.986E-09 | 4.700E-09 | 8.600E-09 |
| 25475 | 1274 | V1274P | E | 4.3e-09 | 7.1e-09 | 1.2e-09 | 4.200E-09 | 2.951E-09 | 1.200E-09 | 7.100E-09 |
| 25476 | 1274 | V1274S | E | 1e-08 | 1.6e-08 | 5.6e-09 | 1.053E-08 | 5.220E-09 | 5.600E-09 | 1.600E-08 |
| 25477 | 1274 | V1274T | E | 6.4e-09 | 1.7e-08 | 6.1e-09 | 9.833E-09 | 6.208E-09 | 6.100E-09 | 1.700E-08 |
| 25478 | 1274 | V1274W | E | 9.4e-09 | 3.3e-09 | 1.3e-08 | 8.567E-09 | 4.903E-09 | 3.300E-09 | 1.300E-08 |
| 25479 | 1274 | V1274Y | E | 1.1e-08 | 2e-09 | 3.3e-09 | 5.433E-09 | 4.864E-09 | 2.000E-09 | 1.100E-08 |
| 25480 | 1274 | V1274V | E | 5.5e-09 | 8.7e-09 | 7.8e-09 | 7.333E-09 | 1.650E-09 | 5.500E-09 | 8.700E-09 |
| 25481 | 1275 | Q1275A | E | 3.9e-09 | 8.5e-09 | 9.1e-09 | 7.167E-09 | 2.845E-09 | 3.900E-09 | 9.100E-09 |
| 25482 | 1275 | Q1275R | E | 6.5e-09 | 1.1e-08 | 1.4e-08 | 1.050E-08 | 3.775E-09 | 6.500E-09 | 1.400E-08 |
| 25483 | 1275 | Q1275N | E | 1.6e-08 | 8.9e-09 | 6.6e-09 | 1.050E-08 | 4.900E-09 | 6.600E-09 | 1.600E-08 |
| 25484 | 1275 | Q1275D | E | 7e-09 | 1.2e-08 | 1.8e-08 | 1.233E-08 | 5.508E-09 | 7.000E-09 | 1.800E-08 |
| 25485 | 1275 | Q1275C | E | 1.7e-08 | 2e-08 | 5e-09 | 1.400E-08 | 7.937E-09 | 5.000E-09 | 2.000E-08 |
| 25486 | 1275 | Q1275E | E | 6.6e-09 | 9e-09 | 1.2e-08 | 9.200E-09 | 2.706E-09 | 6.600E-09 | 1.200E-08 |
| 25487 | 1275 | Q1275Q | E | 8.1e-10 | 9.6e-09 | 7.2e-09 | 5.870E-09 | 4.543E-09 | 8.100E-10 | 9.600E-09 |
| 25488 | 1275 | Q1275G | E | 2.5e-08 | 1.3e-08 | 2.7e-09 | 1.357E-08 | 1.116E-08 | 2.700E-09 | 2.500E-08 |
| 25489 | 1275 | Q1275H | E | 3e-09 | 6.1e-09 | 5.2e-09 | 4.767E-09 | 1.595E-09 | 3.000E-09 | 6.100E-09 |
| 25490 | 1275 | Q1275I | E | 9.7e-09 | 7.3e-09 | 6e-09 | 7.667E-09 | 1.877E-09 | 6.000E-09 | 9.700E-09 |
| 25491 | 1275 | Q1275L | E | 4.6e-09 | 6.4e-09 | 2.3e-09 | 4.433E-09 | 2.055E-09 | 2.300E-09 | 6.400E-09 |
| 25492 | 1275 | Q1275K | E | 5e-09 | 4.8e-09 | 1e-08 | 6.600E-09 | 2.946E-09 | 4.800E-09 | 1.000E-08 |
| 25493 | 1275 | Q1275M | E | 1.3e-08 | 4.8e-09 | 1.9e-08 | 1.227E-08 | 7.128E-09 | 4.800E-09 | 1.900E-08 |
| 25494 | 1275 | Q1275F | E | 4.2e-09 | 1.6e-08 | 3.8e-09 | 8.000E-09 | 6.931E-09 | 3.800E-09 | 1.600E-08 |
| 25495 | 1275 | Q1275P | E | 4.2e-09 | 7.3e-09 | 1.2e-08 | 7.833E-09 | 3.927E-09 | 4.200E-09 | 1.200E-08 |
| 25496 | 1275 | Q1275S | E | 1.3e-08 | 5.9e-09 | 1.4e-08 | 1.097E-08 | 4.416E-09 | 5.900E-09 | 1.400E-08 |
| 25497 | 1275 | Q1275T | E | 8.2e-09 | 4.3e-09 | 1.1e-09 | 4.533E-09 | 3.556E-09 | 1.100E-09 | 8.200E-09 |
| 25498 | 1275 | Q1275W | E | 3e-08 | 1.8e-08 | 9.6e-09 | 1.920E-08 | 1.025E-08 | 9.600E-09 | 3.000E-08 |
| 25499 | 1275 | Q1275Y | E | 5.7e-09 | 7.8e-09 | 6.1e-09 | 6.533E-09 | 1.115E-09 | 5.700E-09 | 7.800E-09 |

| | | | | | | | | | | |
|---|---|---|---|---|---|---|---|---|---|---|
| 25500 | 1275 | Q1275V | E | 1.7e-08 | 9.9e-09 | 5.5e-08 | 2.730E-08 | 2.425E-08 | 9.900E-09 | 5.500E-08 |
| 25501 | 1276 | L1276A | E | 5.6e-09 | 5.7e-09 | 3e-09 | 4.767E-09 | 1.531E-09 | 3.000E-09 | 5.700E-09 |
| 25502 | 1276 | L1276R | E | 5e-09 | 2.5e-08 | 1.1e-08 | 1.367E-08 | 1.026E-08 | 5.000E-09 | 2.500E-08 |
| 25503 | 1276 | L1276N | E | 5.7e-09 | 3.7e-09 | 8.1e-09 | 5.833E-09 | 2.203E-09 | 3.700E-09 | 8.100E-09 |
| 25504 | 1276 | L1276D | E | 7.6e-09 | 2.2e-08 | 1.7e-08 | 1.553E-08 | 7.311E-09 | 7.600E-09 | 2.200E-08 |
| 25505 | 1276 | L1276C | E | 4.1e-09 | 1.3e-08 | 2.4e-09 | 6.500E-09 | 5.693E-09 | 2.400E-09 | 1.300E-08 |
| 25506 | 1276 | L1276E | E | 2.3e-09 | 2.3e-08 | 5.7e-09 | 1.033E-08 | 1.110E-08 | 2.300E-09 | 2.300E-08 |
| 25507 | 1276 | L1276Q | E | 1.7e-09 | 1.4e-08 | 1.6e-08 | 1.057E-08 | 7.744E-09 | 1.700E-09 | 1.600E-08 |
| 25508 | 1276 | L1276G | E | 2.5e-09 | 1.3e-08 | 1.7e-09 | 5.733E-09 | 6.306E-09 | 1.700E-09 | 1.300E-08 |
| 25509 | 1276 | L1276H | E | 6.5e-09 | 6.3e-09 | 6.9e-09 | 6.567E-09 | 3.055E-10 | 6.300E-09 | 6.900E-09 |
| 25510 | 1276 | L1276I | E | 8.5e-09 | 1.3e-08 | 9.2e-09 | 1.023E-08 | 2.421E-09 | 8.500E-09 | 1.300E-08 |
| 25511 | 1276 | L1276L | E | 9.4e-09 | 6.3e-09 | 4.8e-09 | 6.833E-09 | 2.346E-09 | 4.800E-09 | 9.400E-09 |
| 25512 | 1276 | L1276K | E | 5e-09 | 8.9e-09 | 5.6e-09 | 6.500E-09 | 2.100E-09 | 5.000E-09 | 8.900E-09 |
| 25513 | 1276 | L1276M | E | 1e-08 | 4.1e-09 | 7.6e-09 | 7.233E-09 | 2.967E-09 | 4.100E-09 | 1.000E-08 |
| 25514 | 1276 | L1276F | E | 2.6e-09 | 6.8e-09 | 8.3e-09 | 5.900E-09 | 2.955E-09 | 2.600E-09 | 8.300E-09 |
| 25515 | 1276 | L1276P | E | 1.8e-08 | 3.2e-08 | 1.1e-08 | 2.033E-08 | 1.069E-08 | 1.100E-08 | 3.200E-08 |
| 25516 | 1276 | L1276S | E | 5.4e-09 | 6.9e-09 | 4.5e-09 | 5.600E-09 | 1.212E-09 | 4.500E-09 | 6.900E-09 |
| 25517 | 1276 | L1276T | E | 3.9e-09 | 3.4e-09 | 1.9e-08 | 8.767E-09 | 8.866E-09 | 3.400E-09 | 1.900E-08 |
| 25518 | 1276 | L1276W | E | 1.1e-08 | 5.9e-09 | 6.3e-09 | 7.733E-09 | 2.836E-09 | 5.900E-09 | 1.100E-08 |
| 25519 | 1276 | L1276Y | E | 1.3e-08 | 8.8e-09 | 2.8e-09 | 8.200E-09 | 5.126E-09 | 2.800E-09 | 1.300E-08 |
| 25520 | 1276 | L1276V | E | 6.2e-09 | 4e-09 | 7.1e-09 | 5.767E-09 | 1.595E-09 | 4.000E-09 | 7.100E-09 |
| 25521 | 1277 | V1277A | E | 8e-09 | 5.4e-09 | 2.1e-09 | 5.167E-09 | 2.957E-09 | 2.100E-09 | 8.000E-09 |
| 25522 | 1277 | V1277R | E | 7.7e-09 | 1.4e-08 | 4e-09 | 8.567E-09 | 5.056E-09 | 4.000E-09 | 1.400E-08 |
| 25523 | 1277 | V1277N | E | 5.7e-09 | 1.4e-08 | 3.7e-09 | 7.800E-09 | 5.462E-09 | 3.700E-09 | 1.400E-08 |
| 25524 | 1277 | V1277D | E | 1.7e-08 | 7.1e-09 | 1.1e-08 | 1.170E-08 | 4.987E-09 | 7.100E-09 | 1.700E-08 |
| 25525 | 1277 | V1277C | E | 7.4e-09 | 5.8e-09 | 8.1e-09 | 7.100E-09 | 1.179E-09 | 5.800E-09 | 8.100E-09 |
| 25526 | 1277 | V1277E | E | 4e-09 | 2.6e-09 | 1.8e-08 | 8.200E-09 | 8.516E-09 | 2.600E-09 | 1.800E-08 |
| 25527 | 1277 | V1277Q | E | 9e-09 | 1.7e-09 | 3.2e-09 | 4.633E-09 | 3.855E-09 | 1.700E-09 | 9.000E-09 |
| 25528 | 1277 | V1277G | E | 6e-09 | 2.8e-09 | 6.1e-09 | 4.967E-09 | 1.877E-09 | 2.800E-09 | 6.100E-09 |
| 25529 | 1277 | V1277H | E | 1e-08 | 3.8e-09 | 9.4e-09 | 7.733E-09 | 3.420E-09 | 3.800E-09 | 1.000E-08 |
| 25530 | 1277 | V1277I | E | 7e-09 | 4.6e-09 | 1e-08 | 7.200E-09 | 2.706E-09 | 4.600E-09 | 1.000E-08 |
| 25531 | 1277 | V1277L | E | 6.5e-09 | 1.3e-08 | 3e-09 | 7.500E-09 | 5.074E-09 | 3.000E-09 | 1.300E-08 |
| 25532 | 1277 | V1277K | E | 1.3e-08 | 3.7e-09 | 5.1e-09 | 7.267E-09 | 5.014E-09 | 3.700E-09 | 1.300E-08 |
| 25533 | 1277 | V1277M | E | 1.9e-08 | 1.2e-08 | 6.9e-09 | 1.263E-08 | 6.075E-09 | 6.900E-09 | 1.900E-08 |
| 25534 | 1277 | V1277F | E | 1.5e-08 | 8.6e-09 | 2.9e-09 | 8.833E-09 | 6.053E-09 | 2.900E-09 | 1.500E-08 |
| 25535 | 1277 | V1277P | E | 4.5e-09 | 3.2e-09 | 8e-09 | 5.233E-09 | 2.483E-09 | 3.200E-09 | 8.000E-09 |
| 25536 | 1277 | V1277S | E | 7.7e-09 | 5.8e-09 | 3.2e-09 | 5.567E-09 | 2.259E-09 | 3.200E-09 | 7.700E-09 |
| 25537 | 1277 | V1277T | E | 1.2e-08 | 9.2e-09 | 3.7e-09 | 8.300E-09 | 4.223E-09 | 3.700E-09 | 1.200E-08 |
| 25538 | 1277 | V1277W | E | 1.5e-08 | 4.2e-09 | 5.4e-09 | 8.200E-09 | 5.919E-09 | 4.200E-09 | 1.500E-08 |
| 25539 | 1277 | V1277Y | E | 9.2e-09 | 8.4e-09 | 8.3e-09 | 8.633E-09 | 4.933E-10 | 8.300E-09 | 9.200E-09 |
| 25540 | 1277 | V1277V | E | 1.6e-08 | 5e-09 | 5.4e-09 | 8.800E-09 | 6.239E-09 | 5.000E-09 | 1.600E-08 |
| 25541 | 1278 | E1278A | E | 1.2e-08 | 1.2e-08 | 8.1e-09 | 1.070E-08 | 2.252E-09 | 8.100E-09 | 1.200E-08 |
| 25542 | 1278 | E1278R | E | 8.9e-09 | 5e-09 | 1.1e-08 | 8.300E-09 | 3.045E-09 | 5.000E-09 | 1.100E-08 |
| 25543 | 1278 | E1278N | E | 5.4e-09 | 1.2e-08 | 2e-08 | 1.247E-08 | 7.311E-09 | 5.400E-09 | 2.000E-08 |
| 25544 | 1278 | E1278D | E | 1.4e-08 | 3.9e-09 | 8.1e-09 | 8.667E-09 | 5.074E-09 | 3.900E-09 | 1.400E-08 |
| 25545 | 1278 | E1278C | E | 2e-08 | 1e-08 | 1.1e-08 | 1.367E-08 | 5.508E-09 | 1.000E-08 | 2.000E-08 |
| 25546 | 1278 | E1278E | E | 5.7e-09 | 9.4e-09 | 2.1e-08 | 1.203E-08 | 7.983E-09 | 5.700E-09 | 2.100E-08 |
| 25547 | 1278 | E1278Q | E | 4.4e-09 | 4.8e-09 | 9.5e-09 | 6.233E-09 | 2.836E-09 | 4.400E-09 | 9.500E-09 |
| 25548 | 1278 | E1278G | E | 1.9e-08 | 5.6e-09 | 3.1e-09 | 9.233E-09 | 8.550E-09 | 3.100E-09 | 1.900E-08 |
| 25549 | 1278 | E1278H | E | 1.4e-08 | 8.3e-09 | 1.7e-08 | 1.310E-08 | 4.419E-09 | 8.300E-09 | 1.700E-08 |
| 25550 | 1278 | E1278I | E | 5e-09 | 1.6e-08 | 4.8e-09 | 8.600E-09 | 6.409E-09 | 4.800E-09 | 1.600E-08 |
| 25551 | 1278 | E1278L | E | 1.5e-08 | 1e-08 | 1.2e-08 | 1.233E-08 | 2.517E-09 | 1.000E-08 | 1.500E-08 |
| 25552 | 1278 | E1278K | E | 9.5e-09 | 6.9e-09 | 3.6e-09 | 6.667E-09 | 2.957E-09 | 3.600E-09 | 9.500E-09 |
| 25553 | 1278 | E1278M | E | 1.9e-09 | 1.1e-08 | 1.5e-08 | 9.300E-09 | 6.713E-09 | 1.900E-09 | 1.500E-08 |
| 25554 | 1278 | E1278F | E | 2.6e-09 | 1.2e-09 | 2.4e-08 | 9.267E-09 | 1.278E-08 | 1.200E-09 | 2.400E-08 |
| 25555 | 1278 | E1278P | E | 4e-09 | 1.1e-08 | 1.1e-08 | 8.667E-09 | 4.041E-09 | 4.000E-09 | 1.100E-08 |
| 25556 | 1278 | E1278S | E | 3.5e-09 | 1.9e-08 | 3.6e-09 | 8.700E-09 | 8.920E-09 | 3.500E-09 | 1.900E-08 |
| 25557 | 1278 | E1278T | E | 5e-09 | 2e-08 | 9.6e-09 | 1.153E-08 | 7.685E-09 | 5.000E-09 | 2.000E-08 |
| 25558 | 1278 | E1278W | E | 6.4e-09 | 7.3e-09 | 2.2e-08 | 1.190E-08 | 8.758E-09 | 6.400E-09 | 2.200E-08 |
| 25559 | 1278 | E1278Y | E | 1.3e-08 | 1.4e-08 | 1.1e-08 | 1.267E-08 | 1.528E-09 | 1.100E-08 | 1.400E-08 |
| 25560 | 1278 | E1278V | E | 1.2e-08 | 1.3e-08 | 5.9e-09 | 1.030E-08 | 3.843E-09 | 5.900E-09 | 1.300E-08 |
| 25561 | 1279 | S1279A | E | 1.1e-08 | 1.1e-08 | 4.7e-09 | 8.900E-09 | 3.637E-09 | 4.700E-09 | 1.100E-08 |
| 25562 | 1279 | S1279R | E | 4.6e-09 | 1.4e-08 | 4.6e-09 | 7.733E-09 | 5.427E-09 | 4.600E-09 | 1.400E-08 |
| 25563 | 1279 | S1279N | E | 9.3e-09 | 1.2e-08 | 1.1e-08 | 1.077E-08 | 1.365E-09 | 9.300E-09 | 1.200E-08 |
| 25564 | 1279 | S1279D | E | 2.1e-09 | 8.9e-09 | 2e-09 | 4.333E-09 | 3.955E-09 | 2.000E-09 | 8.900E-09 |
| 25565 | 1279 | S1279C | E | 6.9e-09 | 1.7e-08 | 5e-09 | 9.633E-09 | 6.450E-09 | 5.000E-09 | 1.700E-08 |
| 25566 | 1279 | S1279E | E | 7e-09 | 1.3e-08 | 6.1e-09 | 8.700E-09 | 3.751E-09 | 6.100E-09 | 1.300E-08 |
| 25567 | 1279 | S1279Q | E | 7.7e-09 | 6.3e-09 | 1.1e-08 | 8.333E-09 | 2.413E-09 | 6.300E-09 | 1.100E-08 |
| 25568 | 1279 | S1279G | E | 5.4e-09 | 5.7e-09 | 5.7e-09 | 5.600E-09 | 1.732E-10 | 5.400E-09 | 5.700E-09 |
| 25569 | 1279 | S1279H | E | 8.6e-09 | 9.3e-09 | 2.4e-08 | 1.397E-08 | 8.696E-09 | 8.600E-09 | 2.400E-08 |
| 25570 | 1279 | S1279I | E | 3.3e-09 | 1e-09 | 1.1e-08 | 5.100E-09 | 5.237E-09 | 1.000E-09 | 1.100E-08 |
| 25571 | 1279 | S1279L | E | 2.8e-09 | 4.2e-09 | 3.4e-09 | 3.467E-09 | 7.024E-10 | 2.800E-09 | 4.200E-09 |
| 25572 | 1279 | S1279K | E | 7.2e-09 | 4e-09 | 3.3e-09 | 4.833E-09 | 2.079E-09 | 3.300E-09 | 7.200E-09 |
| 25573 | 1279 | S1279M | E | 3.7e-09 | 1.7e-09 | 6.9e-09 | 4.100E-09 | 2.623E-09 | 1.700E-09 | 6.900E-09 |
| 25574 | 1279 | S1279F | E | 1.1e-08 | 1.1e-08 | 8.3e-09 | 1.010E-08 | 1.559E-09 | 8.300E-09 | 1.100E-08 |
| 25575 | 1279 | S1279P | E | 7e-09 | 1.2e-08 | 8.4e-09 | 9.133E-09 | 2.579E-09 | 7.000E-09 | 1.200E-08 |
| 25576 | 1279 | S1279S | E | 8.6e-09 | 6.8e-09 | 7.7e-09 | 7.700E-09 | 9.000E-10 | 6.800E-09 | 8.600E-09 |
| 25577 | 1279 | S1279T | E | 3.5e-09 | 5.3e-09 | 1.7e-08 | 8.600E-09 | 7.330E-09 | 3.500E-09 | 1.700E-08 |
| 25578 | 1279 | S1279W | E | 6.4e-09 | 5.5e-09 | 1.4e-08 | 8.633E-09 | 4.669E-09 | 5.500E-09 | 1.400E-08 |
| 25579 | 1279 | S1279Y | E | 5.6e-09 | 1.1e-08 | 7.9e-09 | 8.167E-09 | 2.710E-09 | 5.600E-09 | 1.100E-08 |
| 25580 | 1279 | S1279V | E | 1.2e-08 | 1.1e-08 | 1.5e-08 | 1.267E-08 | 2.082E-09 | 1.100E-08 | 1.500E-08 |
| 25581 | 1280 | G1280A | E | 4.3e-09 | 1.1e-08 | 5e-09 | 6.767E-09 | 3.683E-09 | 4.300E-09 | 1.100E-08 |
| 25582 | 1280 | G1280R | E | 7e-09 | 9.2e-09 | 9.4e-09 | 8.533E-09 | 1.332E-09 | 7.000E-09 | 9.400E-09 |
| 25583 | 1280 | G1280N | E | 9.7e-09 | 6.4e-09 | 1.5e-08 | 1.037E-08 | 4.339E-09 | 6.400E-09 | 1.500E-08 |
| 25584 | 1280 | G1280D | E | 5e-09 | 2.8e-08 | 8.9e-09 | 1.397E-08 | 1.231E-08 | 5.000E-09 | 2.800E-08 |

| | | | | | | | | | | |
|---|---|---|---|---|---|---|---|---|---|---|
| 25585 | 1280 | G1280C | E | 6.1e-09 | 1.5e-08 | 3.2e-09 | 8.100E-09 | 6.149E-09 | 3.200E-09 | 1.500E-08 |
| 25586 | 1280 | G1280E | E | 4.8e-09 | 6.5e-09 | 1.3e-09 | 4.200E-09 | 2.651E-09 | 1.300E-09 | 6.500E-09 |
| 25587 | 1280 | G1280Q | E | 5e-09 | 7.1e-09 | 1e-08 | 7.367E-09 | 2.511E-09 | 5.000E-09 | 1.000E-08 |
| 25588 | 1280 | G1280G | E | 4.3e-09 | 6.1e-09 | 6.1e-09 | 5.500E-09 | 1.039E-09 | 4.300E-09 | 6.100E-09 |
| 25589 | 1280 | G1280H | E | 7.8e-09 | 5.9e-09 | 1.2e-08 | 8.567E-09 | 3.121E-09 | 5.900E-09 | 1.200E-08 |
| 25590 | 1280 | G1280I | E | 2.1e-08 | 9.5e-09 | 4e-09 | 1.150E-08 | 8.675E-09 | 4.000E-09 | 2.100E-08 |
| 25591 | 1280 | G1280L | E | 1.2e-08 | 3.3e-09 | 1.2e-08 | 9.100E-09 | 5.023E-09 | 3.300E-09 | 1.200E-08 |
| 25592 | 1280 | G1280K | E | 4.6e-09 | 4.6e-09 | 7.6e-09 | 5.600E-09 | 1.732E-09 | 4.600E-09 | 7.600E-09 |
| 25593 | 1280 | G1280M | E | 5.5e-09 | 7.7e-09 | 5e-09 | 6.067E-09 | 1.436E-09 | 5.000E-09 | 7.700E-09 |
| 25594 | 1280 | G1280F | E | 4.2e-09 | 3.4e-09 | 1.8e-08 | 8.533E-09 | 8.208E-09 | 3.400E-09 | 1.800E-08 |
| 25595 | 1280 | G1280P | E | 4.5e-09 | 3.7e-09 | 4.7e-09 | 4.300E-09 | 5.292E-10 | 3.700E-09 | 4.700E-09 |
| 25596 | 1280 | G1280S | E | 7.8e-09 | 5.4e-09 | 6.1e-09 | 6.433E-09 | 1.234E-09 | 5.400E-09 | 7.800E-09 |
| 25597 | 1280 | G1280T | E | 1.5e-08 | 1.3e-08 | 1.3e-08 | 1.367E-08 | 1.155E-09 | 1.300E-08 | 1.500E-08 |
| 25598 | 1280 | G1280W | E | 1.8e-08 | 1.1e-08 | 2.9e-09 | 1.063E-08 | 7.557E-09 | 2.900E-09 | 1.800E-08 |
| 25599 | 1280 | G1280Y | E | 4.2e-09 | 3.7e-09 | 8.8e-09 | 5.567E-09 | 2.811E-09 | 3.700E-09 | 8.800E-09 |
| 25600 | 1280 | G1280V | E | 5.3e-09 | 1.3e-08 | 6.3e-09 | 8.200E-09 | 4.187E-09 | 5.300E-09 | 1.300E-08 |
| 25601 | 1281 | G1281A | E | 4.8e-09 | 2.9e-09 | 5.4e-09 | 4.367E-09 | 1.305E-09 | 2.900E-09 | 5.400E-09 |
| 25602 | 1281 | G1281R | E | 1.8e-08 | 9.1e-09 | 5.6e-09 | 1.090E-08 | 6.393E-09 | 5.600E-09 | 1.800E-08 |
| 25603 | 1281 | G1281N | E | 9.2e-09 | 5.7e-09 | 5.9e-09 | 6.933E-09 | 1.966E-09 | 5.700E-09 | 9.200E-09 |
| 25604 | 1281 | G1281D | E | 7.8e-09 | 6.1e-09 | 5.9e-09 | 6.600E-09 | 1.044E-09 | 5.900E-09 | 7.800E-09 |
| 25605 | 1281 | G1281C | E | 5.1e-09 | 7.9e-09 | 5.9e-09 | 6.300E-09 | 1.442E-09 | 5.100E-09 | 7.900E-09 |
| 25606 | 1281 | G1281E | E | 7e-09 | 5.6e-09 | 7e-09 | 6.533E-09 | 8.083E-10 | 5.600E-09 | 7.000E-09 |
| 25607 | 1281 | G1281Q | E | 2.5e-09 | 1.1e-08 | 7.7e-09 | 7.067E-09 | 4.285E-09 | 2.500E-09 | 1.100E-08 |
| 25608 | 1281 | G1281G | E | 2.7e-09 | 8.7e-09 | 8.2e-09 | 6.533E-09 | 3.329E-09 | 2.700E-09 | 8.700E-09 |
| 25609 | 1281 | G1281H | E | 9.1e-09 | 5.8e-09 | 4e-09 | 6.300E-09 | 2.587E-09 | 4.000E-09 | 9.100E-09 |
| 25610 | 1281 | G1281I | E | 2.1e-09 | 1e-08 | 7.1e-09 | 6.400E-09 | 3.996E-09 | 2.100E-09 | 1.000E-08 |
| 25611 | 1281 | G1281L | E | 5.8e-09 | 1.5e-08 | 1.1e-08 | 1.060E-08 | 4.613E-09 | 5.800E-09 | 1.500E-08 |
| 25612 | 1281 | G1281K | E | 5.8e-09 | 1.3e-08 | 5.4e-09 | 8.067E-09 | 4.277E-09 | 5.400E-09 | 1.300E-08 |
| 25613 | 1281 | G1281M | E | 1.9e-08 | 8.2e-09 | 1.4e-08 | 1.373E-08 | 5.405E-09 | 8.200E-09 | 1.900E-08 |
| 25614 | 1281 | G1281F | E | 5.4e-09 | 1e-08 | 7.8e-09 | 7.733E-09 | 2.301E-09 | 5.400E-09 | 1.000E-08 |
| 25615 | 1281 | G1281P | E | 3.8e-09 | 3.9e-09 | 1e-08 | 5.900E-09 | 3.551E-09 | 3.800E-09 | 1.000E-08 |
| 25616 | 1281 | G1281S | E | 8e-09 | 4.7e-09 | 6.8e-09 | 6.500E-09 | 1.670E-09 | 4.700E-09 | 8.000E-09 |
| 25617 | 1281 | G1281T | E | 1.1e-08 | 7.3e-09 | 3.6e-09 | 7.300E-09 | 3.700E-09 | 3.600E-09 | 1.100E-08 |
| 25618 | 1281 | G1281W | E | 5.6e-09 | 1.6e-08 | 3.9e-09 | 8.500E-09 | 6.551E-09 | 3.900E-09 | 1.600E-08 |
| 25619 | 1281 | G1281Y | E | 4.3e-09 | 8.9e-09 | 8.7e-09 | 7.300E-09 | 2.600E-09 | 4.300E-09 | 8.900E-09 |
| 25620 | 1281 | G1281V | E | 5e-09 | 2.2e-09 | 6.1e-09 | 4.433E-09 | 2.011E-09 | 2.200E-09 | 6.100E-09 |
| 25621 | 1282 | G1282A | E | 1.4e-08 | 1.1e-08 | 9e-09 | 1.133E-08 | 2.517E-09 | 9.000E-09 | 1.400E-08 |
| 25622 | 1282 | G1282R | E | 8.3e-09 | 2.5e-09 | 4.7e-09 | 5.167E-09 | 2.928E-09 | 2.500E-09 | 8.300E-09 |
| 25623 | 1282 | G1282N | E | 1.3e-08 | 4.5e-09 | 2.7e-09 | 6.733E-09 | 5.501E-09 | 2.700E-09 | 1.300E-08 |
| 25624 | 1282 | G1282D | E | 7.5e-09 | 4.7e-09 | 9.8e-09 | 7.333E-09 | 2.554E-09 | 4.700E-09 | 9.800E-09 |
| 25625 | 1282 | G1282C | E | 1.6e-08 | 1e-08 | 1.9e-08 | 1.500E-08 | 4.583E-09 | 1.000E-08 | 1.900E-08 |
| 25626 | 1282 | G1282E | E | 3e-09 | 4.2e-09 | 3.5e-09 | 3.567E-09 | 6.028E-10 | 3.000E-09 | 4.200E-09 |
| 25627 | 1282 | G1282Q | E | 4.9e-09 | 6.5e-09 | 6e-09 | 5.800E-09 | 8.185E-10 | 4.900E-09 | 6.500E-09 |
| 25628 | 1282 | G1282G | E | 6.6e-09 | 5.1e-09 | 5.7e-09 | 5.800E-09 | 7.550E-10 | 5.100E-09 | 6.600E-09 |
| 25629 | 1282 | G1282H | E | 5.5e-09 | 7e-09 | 4.1e-09 | 5.533E-09 | 1.450E-09 | 4.100E-09 | 7.000E-09 |
| 25630 | 1282 | G1282I | E | 4.6e-09 | 5.6e-09 | 3.2e-09 | 4.467E-09 | 1.206E-09 | 3.200E-09 | 5.600E-09 |
| 25631 | 1282 | G1282L | E | 1e-08 | 2e-08 | 5.8e-09 | 1.193E-08 | 7.295E-09 | 5.800E-09 | 2.000E-08 |
| 25632 | 1282 | G1282K | E | 1.4e-08 | 2.5e-09 | 1.1e-08 | 9.167E-09 | 5.965E-09 | 2.500E-09 | 1.400E-08 |
| 25633 | 1282 | G1282M | E | 8.6e-09 | 1.2e-08 | 4.6e-09 | 8.400E-09 | 3.704E-09 | 4.600E-09 | 1.200E-08 |
| 25634 | 1282 | G1282F | E | 4.3e-09 | 6.1e-09 | 9.4e-09 | 6.600E-09 | 2.587E-09 | 4.300E-09 | 9.400E-09 |
| 25635 | 1282 | G1282P | E | 5.2e-09 | 2.2e-09 | 1.5e-08 | 7.467E-09 | 6.694E-09 | 2.200E-09 | 1.500E-08 |
| 25636 | 1282 | G1282S | E | 4.2e-09 | 5.9e-09 | 7.9e-09 | 6.000E-09 | 1.852E-09 | 4.200E-09 | 7.900E-09 |
| 25637 | 1282 | G1282T | E | 8.4e-09 | 4.6e-09 | 1.2e-08 | 8.333E-09 | 3.700E-09 | 4.600E-09 | 1.200E-08 |
| 25638 | 1282 | G1282W | E | 2.7e-09 | 4.9e-09 | 8.4e-09 | 5.333E-09 | 2.875E-09 | 2.700E-09 | 8.400E-09 |
| 25639 | 1282 | G1282Y | E | 1.1e-08 | 1e-08 | 5.8e-09 | 8.933E-09 | 2.759E-09 | 5.800E-09 | 1.100E-08 |
| 25640 | 1282 | G1282V | E | 1.8e-09 | 1e-08 | 1.2e-08 | 7.933E-09 | 5.405E-09 | 1.800E-09 | 1.200E-08 |
| 25641 | 1283 | L1283A | E | 6e-09 | 5e-09 | 4.8e-09 | 5.267E-09 | 6.429E-10 | 4.800E-09 | 6.000E-09 |
| 25642 | 1283 | L1283R | E | 1.3e-08 | 6.1e-09 | 3.4e-09 | 7.500E-09 | 4.951E-09 | 3.400E-09 | 1.300E-08 |
| 25643 | 1283 | L1283N | E | 3.3e-09 | 9.9e-09 | 8.6e-09 | 7.267E-09 | 3.496E-09 | 3.300E-09 | 9.900E-09 |
| 25644 | 1283 | L1283D | E | 5.9e-09 | 1.3e-08 | 1.3e-08 | 1.063E-08 | 4.099E-09 | 5.900E-09 | 1.300E-08 |
| 25645 | 1283 | L1283C | E | 4.5e-09 | 6.1e-09 | 6.7e-09 | 5.767E-09 | 1.137E-09 | 4.500E-09 | 6.700E-09 |
| 25646 | 1283 | L1283E | E | 4.6e-09 | 1e-08 | 5.5e-09 | 6.700E-09 | 2.893E-09 | 4.600E-09 | 1.000E-08 |
| 25647 | 1283 | L1283Q | E | 9.9e-09 | 9.9e-09 | 3e-09 | 7.600E-09 | 3.984E-09 | 3.000E-09 | 9.900E-09 |
| 25648 | 1283 | L1283G | E | 1.3e-08 | 8.6e-09 | 1.1e-08 | 1.087E-08 | 2.203E-09 | 8.600E-09 | 1.300E-08 |
| 25649 | 1283 | L1283H | E | 1e-08 | 1.4e-08 | 9.8e-09 | 1.127E-08 | 2.369E-09 | 9.800E-09 | 1.400E-08 |
| 25650 | 1283 | L1283I | E | 5.4e-09 | 1.8e-08 | 2.2e-08 | 1.513E-08 | 8.663E-09 | 5.400E-09 | 2.200E-08 |
| 25651 | 1283 | L1283L | E | 1.2e-08 | 8.3e-09 | 6.6e-09 | 8.967E-09 | 2.761E-09 | 6.600E-09 | 1.200E-08 |
| 25652 | 1283 | L1283K | E | 1e-08 | 7e-09 | 9.6e-09 | 8.867E-09 | 1.629E-09 | 7.000E-09 | 1.000E-08 |
| 25653 | 1283 | L1283M | E | 5.3e-09 | 3.3e-09 | 5.8e-09 | 4.800E-09 | 1.323E-09 | 3.300E-09 | 5.800E-09 |
| 25654 | 1283 | L1283F | E | 9.5e-09 | 5e-09 | 1.4e-08 | 9.500E-09 | 4.500E-09 | 5.000E-09 | 1.400E-08 |
| 25655 | 1283 | L1283P | E | 1.6e-09 | 6.2e-09 | 7e-09 | 4.933E-09 | 2.914E-09 | 1.600E-09 | 7.000E-09 |
| 25656 | 1283 | L1283S | E | 4.3e-09 | 7.5e-09 | 6e-09 | 5.933E-09 | 1.601E-09 | 4.300E-09 | 7.500E-09 |
| 25657 | 1283 | L1283T | E | 7.2e-09 | 4.8e-09 | 7.6e-09 | 6.533E-09 | 1.514E-09 | 4.800E-09 | 7.600E-09 |
| 25658 | 1283 | L1283W | E | 1.1e-08 | 9.3e-09 | 1.5e-08 | 1.177E-08 | 2.926E-09 | 9.300E-09 | 1.500E-08 |
| 25659 | 1283 | L1283Y | E | 7.4e-09 | 8.2e-09 | 9.2e-09 | 8.267E-09 | 9.018E-10 | 7.400E-09 | 9.200E-09 |
| 25660 | 1283 | L1283V | E | 7.8e-09 | 3.4e-09 | 4.3e-09 | 5.167E-09 | 2.325E-09 | 3.400E-09 | 7.800E-09 |
| 25661 | 1284 | V1284A | E | 5.8e-09 | 5.7e-09 | 8.4e-09 | 6.633E-09 | 1.531E-09 | 5.700E-09 | 8.400E-09 |
| 25662 | 1284 | V1284R | E | 9.7e-09 | 4.9e-09 | 1.2e-08 | 8.867E-09 | 3.623E-09 | 4.900E-09 | 1.200E-08 |
| 25663 | 1284 | V1284N | E | 1.1e-08 | 7.6e-09 | 1.1e-08 | 9.867E-09 | 1.963E-09 | 7.600E-09 | 1.100E-08 |
| 25664 | 1284 | V1284D | E | 1.4e-08 | 1.3e-08 | 3.9e-09 | 1.030E-08 | 5.565E-09 | 3.900E-09 | 1.400E-08 |
| 25665 | 1284 | V1284C | E | 6.4e-09 | 9.5e-09 | 5.7e-09 | 7.200E-09 | 2.022E-09 | 5.700E-09 | 9.500E-09 |
| 25666 | 1284 | V1284E | E | 1.3e-08 | 4.9e-09 | 4.2e-09 | 7.367E-09 | 4.891E-09 | 4.200E-09 | 1.300E-08 |
| 25667 | 1284 | V1284Q | E | 4.2e-09 | 1.2e-08 | 8.4e-09 | 8.200E-09 | 3.904E-09 | 4.200E-09 | 1.200E-08 |
| 25668 | 1284 | V1284G | E | 1.3e-08 | 1e-08 | 3.7e-09 | 8.900E-09 | 4.747E-09 | 3.700E-09 | 1.300E-08 |
| 25669 | 1284 | V1284H | E | 1.4e-08 | 3.3e-09 | 1.2e-08 | 9.767E-09 | 5.689E-09 | 3.300E-09 | 1.400E-08 |

| | | | | | | | | | | |
|---|---|---|---|---|---|---|---|---|---|---|
| 25670 | 1284 | V1284I | E | 5.1e-09 | 1e-08 | 1.2e-08 | 9.033E-09 | 3.550E-09 | 5.100E-09 | 1.200E-08 |
| 25671 | 1284 | V1284L | E | 3.9e-09 | 4.6e-09 | 7e-09 | 5.167E-09 | 1.626E-09 | 3.900E-09 | 7.000E-09 |
| 25672 | 1284 | V1284K | E | 3.3e-09 | 9e-09 | 3.7e-09 | 5.333E-09 | 3.182E-09 | 3.300E-09 | 9.000E-09 |
| 25673 | 1284 | V1284M | E | 5e-09 | 9.2e-09 | 1.1e-08 | 8.400E-09 | 3.079E-09 | 5.000E-09 | 1.100E-08 |
| 25674 | 1284 | V1284F | E | 1.5e-08 | 4e-09 | 2e-08 | 1.300E-08 | 8.185E-09 | 4.000E-09 | 2.000E-08 |
| 25675 | 1284 | V1284P | E | 1.2e-08 | 4.1e-09 | 1.2e-08 | 9.367E-09 | 4.561E-09 | 4.100E-09 | 1.200E-08 |
| 25676 | 1284 | V1284S | E | 4.9e-09 | 8.5e-09 | 9.7e-09 | 7.700E-09 | 2.498E-09 | 4.900E-09 | 9.700E-09 |
| 25677 | 1284 | V1284T | E | 5.4e-09 | 1.6e-08 | 3.6e-09 | 8.333E-09 | 6.700E-09 | 3.600E-09 | 1.600E-08 |
| 25678 | 1284 | V1284W | E | 1.8e-08 | 9.3e-09 | 6e-09 | 1.110E-08 | 6.199E-09 | 6.000E-09 | 1.800E-08 |
| 25679 | 1284 | V1284Y | E | 2.2e-08 | 1.8e-08 | 8.9e-09 | 1.630E-08 | 6.713E-09 | 8.900E-09 | 2.200E-08 |
| 25680 | 1284 | V1284V | E | 6.4e-09 | 9.9e-09 | 9.3e-09 | 8.533E-09 | 1.872E-09 | 6.400E-09 | 9.900E-09 |
| 25681 | 1285 | Q1285A | E | 1.4e-09 | 1.1e-08 | 1e-08 | 7.467E-09 | 5.278E-09 | 1.400E-09 | 1.100E-08 |
| 25682 | 1285 | Q1285R | E | 1.1e-08 | 5.9e-09 | 1.4e-09 | 6.100E-09 | 4.803E-09 | 1.400E-09 | 1.100E-08 |
| 25683 | 1285 | Q1285N | E | 4.9e-09 | 2.6e-09 | 5.6e-09 | 4.367E-09 | 1.570E-09 | 2.600E-09 | 5.600E-09 |
| 25684 | 1285 | Q1285D | E | 1.1e-08 | 5.3e-09 | 4.6e-09 | 6.967E-09 | 3.510E-09 | 4.600E-09 | 1.100E-08 |
| 25685 | 1285 | Q1285C | E | 8.2e-09 | 9.4e-09 | 4.7e-09 | 7.433E-09 | 2.442E-09 | 4.700E-09 | 9.400E-09 |
| 25686 | 1285 | Q1285E | E | 1.3e-08 | 5.2e-09 | 5.2e-09 | 7.800E-09 | 4.503E-09 | 5.200E-09 | 1.300E-08 |
| 25687 | 1285 | Q1285Q | E | 7.6e-09 | 2.1e-09 | 1e-08 | 6.567E-09 | 4.050E-09 | 2.100E-09 | 1.000E-08 |
| 25688 | 1285 | Q1285G | E | 1.7e-08 | 4.9e-09 | 1.3e-08 | 1.163E-08 | 6.165E-09 | 4.900E-09 | 1.700E-08 |
| 25689 | 1285 | Q1285H | E | 1.7e-08 | 9.3e-09 | 1.7e-09 | 9.333E-09 | 7.650E-09 | 1.700E-09 | 1.700E-08 |
| 25690 | 1285 | Q1285I | E | 6.1e-09 | 9.8e-09 | 1.4e-08 | 9.967E-09 | 3.953E-09 | 6.100E-09 | 1.400E-08 |
| 25691 | 1285 | Q1285L | E | 2e-08 | 2.5e-09 | 3.1e-09 | 8.533E-09 | 9.935E-09 | 2.500E-09 | 2.000E-08 |
| 25692 | 1285 | Q1285K | E | 5e-09 | 1.4e-09 | 4.9e-09 | 3.767E-09 | 2.050E-09 | 1.400E-09 | 5.000E-09 |
| 25693 | 1285 | Q1285M | E | 4.8e-09 | 1.9e-08 | 4.1e-09 | 9.300E-09 | 8.408E-09 | 4.100E-09 | 1.900E-08 |
| 25694 | 1285 | Q1285F | E | 6.1e-09 | 5.6e-09 | 1.2e-08 | 7.900E-09 | 3.559E-09 | 5.600E-09 | 1.200E-08 |
| 25695 | 1285 | Q1285P | E | 6.6e-09 | 2.8e-09 | 1.4e-08 | 7.800E-09 | 5.696E-09 | 2.800E-09 | 1.400E-08 |
| 25696 | 1285 | Q1285S | E | 1.1e-08 | 1.2e-08 | 4.8e-09 | 9.267E-09 | 3.900E-09 | 4.800E-09 | 1.200E-08 |
| 25697 | 1285 | Q1285T | E | 8.1e-09 | 3.4e-09 | 7.4e-09 | 6.300E-09 | 2.536E-09 | 3.400E-09 | 8.100E-09 |
| 25698 | 1285 | Q1285W | E | 7.5e-09 | 5.5e-09 | 4.2e-09 | 5.733E-09 | 1.662E-09 | 4.200E-09 | 7.500E-09 |
| 25699 | 1285 | Q1285Y | E | 2.9e-09 | 8.9e-09 | 6.9e-09 | 6.233E-09 | 3.055E-09 | 2.900E-09 | 8.900E-09 |
| 25700 | 1285 | Q1285V | E | 1.7e-08 | 7.2e-09 | 5.4e-09 | 9.867E-09 | 6.243E-09 | 5.400E-09 | 1.700E-08 |
| 25701 | 1286 | P1286A | E | 5.1e-09 | 5.9e-09 | 2.9e-09 | 4.633E-09 | 1.553E-09 | 2.900E-09 | 5.900E-09 |
| 25702 | 1286 | P1286R | E | 2.4e-09 | 5e-09 | 4.4e-09 | 3.933E-09 | 1.361E-09 | 2.400E-09 | 5.000E-09 |
| 25703 | 1286 | P1286N | E | 2.3e-09 | 7.8e-09 | 2.1e-09 | 4.067E-09 | 3.235E-09 | 2.100E-09 | 7.800E-09 |
| 25704 | 1286 | P1286D | E | 1.1e-08 | 3.7e-09 | 8.4e-09 | 7.700E-09 | 3.700E-09 | 3.700E-09 | 1.100E-08 |
| 25705 | 1286 | P1286C | E | 6.3e-09 | 1.2e-08 | 6.5e-09 | 8.267E-09 | 3.235E-09 | 6.300E-09 | 1.200E-08 |
| 25706 | 1286 | P1286E | E | 1.5e-08 | 1.1e-08 | 1.2e-08 | 1.267E-08 | 2.082E-09 | 1.100E-08 | 1.500E-08 |
| 25707 | 1286 | P1286Q | E | 8.2e-09 | 7.4e-09 | 1.3e-08 | 9.533E-09 | 3.029E-09 | 7.400E-09 | 1.300E-08 |
| 25708 | 1286 | P1286G | E | 4.1e-09 | 1.5e-08 | 2.9e-09 | 7.333E-09 | 6.667E-09 | 2.900E-09 | 1.500E-08 |
| 25709 | 1286 | P1286H | E | 1.6e-08 | 5.2e-09 | 5e-09 | 8.733E-09 | 6.294E-09 | 5.000E-09 | 1.600E-08 |
| 25710 | 1286 | P1286I | E | 8e-09 | 1.1e-08 | 5.9e-09 | 8.300E-09 | 2.563E-09 | 5.900E-09 | 1.100E-08 |
| 25711 | 1286 | P1286L | E | 1.3e-08 | 8.7e-09 | 2e-08 | 1.390E-08 | 5.704E-09 | 8.700E-09 | 2.000E-08 |
| 25712 | 1286 | P1286K | E | 3.8e-09 | 4.9e-09 | 4.2e-09 | 4.300E-09 | 5.568E-10 | 3.800E-09 | 4.900E-09 |
| 25713 | 1286 | P1286M | E | 3.6e-09 | 2.2e-09 | 2.4e-09 | 2.733E-09 | 7.572E-10 | 2.200E-09 | 3.600E-09 |
| 25714 | 1286 | P1286F | E | 3.2e-08 | 6.8e-09 | 7.3e-09 | 1.537E-08 | 1.441E-08 | 6.800E-09 | 3.200E-08 |
| 25715 | 1286 | P1286P | E | 1.7e-09 | 4.6e-09 | 5.5e-09 | 3.933E-09 | 1.986E-09 | 1.700E-09 | 5.500E-09 |
| 25716 | 1286 | P1286S | E | 5e-09 | 7.5e-09 | 7e-09 | 6.500E-09 | 1.323E-09 | 5.000E-09 | 7.500E-09 |
| 25717 | 1286 | P1286T | E | 7.4e-09 | 4e-09 | 3.6e-09 | 5.000E-09 | 2.088E-09 | 3.600E-09 | 7.400E-09 |
| 25718 | 1286 | P1286W | E | 1.1e-08 | 2.3e-09 | 4.9e-09 | 6.067E-09 | 4.466E-09 | 2.300E-09 | 1.100E-08 |
| 25719 | 1286 | P1286Y | E | 6e-09 | 3.8e-09 | 1e-08 | 6.600E-09 | 3.143E-09 | 3.800E-09 | 1.000E-08 |
| 25720 | 1286 | P1286V | E | 6.1e-09 | 8.4e-09 | 4.6e-09 | 6.367E-09 | 1.914E-09 | 4.600E-09 | 8.400E-09 |
| 25721 | 1287 | G1287A | E | 4.8e-09 | 1.5e-08 | 6e-09 | 8.600E-09 | 5.575E-09 | 4.800E-09 | 1.500E-08 |
| 25722 | 1287 | G1287R | E | 4.5e-09 | 9.6e-09 | 8.3e-09 | 7.467E-09 | 2.650E-09 | 4.500E-09 | 9.600E-09 |
| 25723 | 1287 | G1287N | E | 7.6e-09 | 9.6e-09 | 4.1e-09 | 7.100E-09 | 2.784E-09 | 4.100E-09 | 9.600E-09 |
| 25724 | 1287 | G1287D | E | 3.9e-09 | 9.2e-09 | 1e-08 | 7.700E-09 | 3.315E-09 | 3.900E-09 | 1.000E-08 |
| 25725 | 1287 | G1287C | E | 7.2e-09 | 8.9e-09 | 2.5e-09 | 6.200E-09 | 3.315E-09 | 2.500E-09 | 8.900E-09 |
| 25726 | 1287 | G1287E | E | 8.4e-09 | 6.2e-09 | 3.5e-09 | 6.033E-09 | 2.454E-09 | 3.500E-09 | 8.400E-09 |
| 25727 | 1287 | G1287Q | E | 1.6e-09 | 6.6e-10 | 9.2e-09 | 3.820E-09 | 4.683E-09 | 6.600E-10 | 9.200E-09 |
| 25728 | 1287 | G1287G | E | 1.2e-08 | 2.5e-08 | 3.6e-09 | 1.353E-08 | 1.078E-08 | 3.600E-09 | 2.500E-08 |
| 25729 | 1287 | G1287H | E | 4.8e-09 | 6.2e-09 | 1.8e-09 | 4.267E-09 | 2.248E-09 | 1.800E-09 | 6.200E-09 |
| 25730 | 1287 | G1287I | E | 1.3e-08 | 1.4e-08 | 9.2e-09 | 1.207E-08 | 2.532E-09 | 9.200E-09 | 1.400E-08 |
| 25731 | 1287 | G1287L | E | 4.4e-09 | 1.1e-08 | 7.2e-09 | 7.533E-09 | 3.313E-09 | 4.400E-09 | 1.100E-08 |
| 25732 | 1287 | G1287K | E | 1.7e-08 | 1.8e-08 | 1.7e-08 | 1.733E-08 | 5.774E-10 | 1.700E-08 | 1.800E-08 |
| 25733 | 1287 | G1287M | E | 2.9e-09 | 2.8e-09 | 1e-08 | 5.233E-09 | 4.128E-09 | 2.800E-09 | 1.000E-08 |
| 25734 | 1287 | G1287F | E | 5.1e-09 | 8.4e-09 | 5e-09 | 6.167E-09 | 1.935E-09 | 5.000E-09 | 8.400E-09 |
| 25735 | 1287 | G1287P | E | 8.8e-09 | 2.4e-09 | 2.2e-09 | 4.467E-09 | 3.754E-09 | 2.200E-09 | 8.800E-09 |
| 25736 | 1287 | G1287S | E | 7.6e-09 | 1.8e-09 | 9.7e-09 | 6.367E-09 | 4.092E-09 | 1.800E-09 | 9.700E-09 |
| 25737 | 1287 | G1287T | E | 3.1e-08 | 5.9e-09 | 5.8e-09 | 1.423E-08 | 1.452E-08 | 5.800E-09 | 3.100E-08 |
| 25738 | 1287 | G1287W | E | 1.4e-08 | 9.8e-09 | 2.5e-09 | 8.767E-09 | 5.819E-09 | 2.500E-09 | 1.400E-08 |
| 25739 | 1287 | G1287Y | E | 3.2e-09 | 8.9e-09 | 3e-09 | 5.033E-09 | 3.350E-09 | 3.000E-09 | 8.900E-09 |
| 25740 | 1287 | G1287V | E | 1.2e-08 | 6.2e-09 | 9.5e-09 | 9.233E-09 | 2.909E-09 | 6.200E-09 | 1.200E-08 |
| 25741 | 1288 | G1288A | E | 1.2e-08 | 1.4e-08 | 1.9e-09 | 9.300E-09 | 6.486E-09 | 1.900E-09 | 1.400E-08 |
| 25742 | 1288 | G1288R | E | 6.2e-09 | 4.2e-09 | 1.2e-08 | 7.467E-09 | 4.051E-09 | 4.200E-09 | 1.200E-08 |
| 25743 | 1288 | G1288N | E | 1.2e-09 | 4.4e-09 | 1.6e-08 | 7.200E-09 | 7.787E-09 | 1.200E-09 | 1.600E-08 |
| 25744 | 1288 | G1288D | E | 7.3e-09 | 1.2e-08 | 1.5e-08 | 1.143E-08 | 3.881E-09 | 7.300E-09 | 1.500E-08 |
| 25745 | 1288 | G1288C | E | 8.4e-09 | 4.9e-09 | 1.4e-08 | 9.100E-09 | 4.590E-09 | 4.900E-09 | 1.400E-08 |
| 25746 | 1288 | G1288E | E | 4.7e-09 | 6.8e-09 | 5.6e-09 | 5.700E-09 | 1.054E-09 | 4.700E-09 | 6.800E-09 |
| 25747 | 1288 | G1288Q | E | 1.2e-08 | 7.1e-09 | 1.9e-08 | 1.270E-08 | 5.981E-09 | 7.100E-09 | 1.900E-08 |
| 25748 | 1288 | G1288G | E | 3.2e-09 | 7.9e-09 | 8.2e-09 | 6.433E-09 | 2.804E-09 | 3.200E-09 | 8.200E-09 |
| 25749 | 1288 | G1288H | E | 2.2e-09 | 8.5e-09 | 7.4e-09 | 6.033E-09 | 3.365E-09 | 2.200E-09 | 8.500E-09 |
| 25750 | 1288 | G1288I | E | 7.6e-09 | 2.7e-09 | 1e-08 | 6.767E-09 | 3.721E-09 | 2.700E-09 | 1.000E-08 |
| 25751 | 1288 | G1288L | E | 8.5e-09 | 2.7e-09 | 1.1e-08 | 7.400E-09 | 4.258E-09 | 2.700E-09 | 1.100E-08 |
| 25752 | 1288 | G1288K | E | 1.5e-08 | 7.6e-09 | 1.5e-08 | 1.253E-08 | 4.272E-09 | 7.600E-09 | 1.500E-08 |
| 25753 | 1288 | G1288M | E | 6.4e-09 | 1.4e-08 | 3e-08 | 1.680E-08 | 1.205E-08 | 6.400E-09 | 3.000E-08 |
| 25754 | 1288 | G1288F | E | 3.3e-09 | 9e-09 | 7.5e-09 | 6.600E-09 | 2.955E-09 | 3.300E-09 | 9.000E-09 |

| | | | | | | | | | | |
|---|---|---|---|---|---|---|---|---|---|---|
| 25755 | 1288 | G1288P | E | 7.3e-09 | 5.5e-09 | 1e-08 | 7.600E-09 | 2.265E-09 | 5.500E-09 | 1.000E-08 |
| 25756 | 1288 | G1288S | E | 7.5e-09 | 9.1e-09 | 8.8e-09 | 8.467E-09 | 8.505E-10 | 7.500E-09 | 9.100E-09 |
| 25757 | 1288 | G1288T | E | 8.8e-09 | 5.6e-09 | 2.1e-08 | 1.180E-08 | 8.126E-09 | 5.600E-09 | 2.100E-08 |
| 25758 | 1288 | G1288W | E | 2.9e-08 | 1.6e-08 | 5.3e-09 | 1.677E-08 | 1.187E-08 | 5.300E-09 | 2.900E-08 |
| 25759 | 1288 | G1288Y | E | 1.2e-08 | 8.6e-09 | 1.1e-08 | 1.053E-08 | 1.747E-09 | 8.600E-09 | 1.200E-08 |
| 25760 | 1288 | G1288V | E | 4.1e-09 | 1e-08 | 9.2e-09 | 7.767E-09 | 3.201E-09 | 4.100E-09 | 1.000E-08 |
| 25761 | 1289 | S1289A | E | 5.1e-09 | 8.5e-09 | 1.6e-08 | 9.867E-09 | 5.577E-09 | 5.100E-09 | 1.600E-08 |
| 25762 | 1289 | S1289R | E | 2.3e-08 | 1.2e-08 | 3e-09 | 1.267E-08 | 1.002E-08 | 3.000E-09 | 2.300E-08 |
| 25763 | 1289 | S1289N | E | 5.9e-09 | 4e-09 | 5.6e-09 | 5.167E-09 | 1.021E-09 | 4.000E-09 | 5.900E-09 |
| 25764 | 1289 | S1289D | E | 1.1e-08 | 2.7e-09 | 9.8e-09 | 7.833E-09 | 4.486E-09 | 2.700E-09 | 1.100E-08 |
| 25765 | 1289 | S1289C | E | 3.9e-09 | 1.6e-08 | 5.9e-09 | 8.600E-09 | 6.486E-09 | 3.900E-09 | 1.600E-08 |
| 25766 | 1289 | S1289E | E | 2.7e-09 | 6.3e-09 | 4e-09 | 4.333E-09 | 1.823E-09 | 2.700E-09 | 6.300E-09 |
| 25767 | 1289 | S1289Q | E | 1.2e-08 | 1.8e-08 | 9.9e-09 | 1.330E-08 | 4.204E-09 | 9.900E-09 | 1.800E-08 |
| 25768 | 1289 | S1289G | E | 5.5e-09 | 7.1e-09 | 1.1e-08 | 7.867E-09 | 2.829E-09 | 5.500E-09 | 1.100E-08 |
| 25769 | 1289 | S1289H | E | 6.1e-09 | 3.3e-09 | 5.4e-09 | 4.933E-09 | 1.457E-09 | 3.300E-09 | 6.100E-09 |
| 25770 | 1289 | S1289I | E | 8.8e-09 | 6.1e-09 | 8.2e-09 | 7.700E-09 | 1.418E-09 | 6.100E-09 | 8.800E-09 |
| 25771 | 1289 | S1289L | E | 6e-09 | 8.4e-09 | 4.6e-09 | 6.333E-09 | 1.922E-09 | 4.600E-09 | 8.400E-09 |
| 25772 | 1289 | S1289K | E | 7.8e-09 | 4.6e-09 | 1.1e-08 | 7.800E-09 | 3.200E-09 | 4.600E-09 | 1.100E-08 |
| 25773 | 1289 | S1289M | E | 1.6e-08 | 5.1e-09 | 1.5e-08 | 1.203E-08 | 6.025E-09 | 5.100E-09 | 1.600E-08 |
| 25774 | 1289 | S1289F | E | 1.5e-08 | 4.5e-09 | 6.3e-09 | 8.600E-09 | 5.615E-09 | 4.500E-09 | 1.500E-08 |
| 25775 | 1289 | S1289P | E | 4.9e-09 | 9.2e-09 | 1.4e-08 | 9.367E-09 | 4.552E-09 | 4.900E-09 | 1.400E-08 |
| 25776 | 1289 | S1289S | E | 2e-09 | 1.7e-09 | 1.4e-08 | 5.900E-09 | 7.016E-09 | 1.700E-09 | 1.400E-08 |
| 25777 | 1289 | S1289T | E | 1.3e-08 | 1.5e-08 | 5.6e-09 | 1.120E-08 | 4.952E-09 | 5.600E-09 | 1.500E-08 |
| 25778 | 1289 | S1289W | E | 3.7e-09 | 1.1e-08 | 2.6e-09 | 5.767E-09 | 4.565E-09 | 2.600E-09 | 1.100E-08 |
| 25779 | 1289 | S1289Y | E | 1.1e-08 | 6e-09 | 1.5e-08 | 1.067E-08 | 4.509E-09 | 6.000E-09 | 1.500E-08 |
| 25780 | 1289 | S1289V | E | 5e-09 | 1.3e-08 | 3.1e-09 | 7.033E-09 | 5.254E-09 | 3.100E-09 | 1.300E-08 |
| 25781 | 1290 | L1290A | E | 4.7e-09 | 3.8e-09 | 3.5e-09 | 4.000E-09 | 6.245E-10 | 3.500E-09 | 4.700E-09 |
| 25782 | 1290 | L1290R | E | 7.8e-09 | 5.8e-09 | 5.7e-09 | 6.433E-09 | 1.185E-09 | 5.700E-09 | 7.800E-09 |
| 25783 | 1290 | L1290N | E | 8.7e-09 | 1.4e-08 | 1.6e-08 | 1.290E-08 | 3.772E-09 | 8.700E-09 | 1.600E-08 |
| 25784 | 1290 | L1290D | E | 4.7e-09 | 8.8e-09 | 5.4e-09 | 6.300E-09 | 2.193E-09 | 4.700E-09 | 8.800E-09 |
| 25785 | 1290 | L1290C | E | 1.4e-08 | 1.4e-08 | 7.6e-09 | 1.187E-08 | 3.695E-09 | 7.600E-09 | 1.400E-08 |
| 25786 | 1290 | L1290E | E | 1.9e-08 | 3.5e-09 | 5.7e-09 | 9.400E-09 | 8.386E-09 | 3.500E-09 | 1.900E-08 |
| 25787 | 1290 | L1290Q | E | 1.1e-08 | 7.9e-09 | 9.9e-09 | 9.600E-09 | 1.572E-09 | 7.900E-09 | 1.100E-08 |
| 25788 | 1290 | L1290G | E | 5.3e-09 | 1.3e-08 | 8.7e-09 | 9.000E-09 | 3.859E-09 | 5.300E-09 | 1.300E-08 |
| 25789 | 1290 | L1290H | E | 2e-08 | 6e-09 | 4.4e-09 | 1.013E-08 | 8.582E-09 | 4.400E-09 | 2.000E-08 |
| 25790 | 1290 | L1290I | E | 3.5e-09 | 5.2e-09 | 4.6e-09 | 4.433E-09 | 8.622E-10 | 3.500E-09 | 5.200E-09 |
| 25791 | 1290 | L1290L | E | 8.3e-09 | 6.9e-09 | 9.9e-09 | 8.367E-09 | 1.501E-09 | 6.900E-09 | 9.900E-09 |
| 25792 | 1290 | L1290K | E | 1.6e-08 | 5.8e-09 | 9.2e-09 | 1.033E-08 | 5.194E-09 | 5.800E-09 | 1.600E-08 |
| 25793 | 1290 | L1290M | E | 1.6e-08 | 4.2e-09 | 6e-09 | 8.733E-09 | 6.357E-09 | 4.200E-09 | 1.600E-08 |
| 25794 | 1290 | L1290F | E | 8.6e-09 | 1.4e-08 | 1.2e-08 | 1.153E-08 | 2.730E-09 | 8.600E-09 | 1.400E-08 |
| 25795 | 1290 | L1290P | E | 9.7e-09 | 3e-08 | 4.7e-09 | 1.480E-08 | 1.340E-08 | 4.700E-09 | 3.000E-08 |
| 25796 | 1290 | L1290S | E | 1.2e-08 | 6.3e-09 | 3.5e-09 | 7.267E-09 | 4.332E-09 | 3.500E-09 | 1.200E-08 |
| 25797 | 1290 | L1290T | E | 1e-08 | 2.2e-08 | 6.4e-09 | 1.280E-08 | 8.168E-09 | 6.400E-09 | 2.200E-08 |
| 25798 | 1290 | L1290W | E | 9.6e-09 | 6.6e-09 | 8.5e-09 | 8.233E-09 | 1.518E-09 | 6.600E-09 | 9.600E-09 |
| 25799 | 1290 | L1290Y | E | 4.9e-09 | 3.9e-09 | 7.7e-09 | 5.500E-09 | 1.970E-09 | 3.900E-09 | 7.700E-09 |
| 25800 | 1290 | L1290V | E | 7.5e-09 | 1.1e-08 | 2.8e-09 | 7.100E-09 | 4.115E-09 | 2.800E-09 | 1.100E-08 |
| 25801 | 1291 | R1291A | E | 1.1e-08 | 9.4e-09 | 1.4e-08 | 1.147E-08 | 2.335E-09 | 9.400E-09 | 1.400E-08 |
| 25802 | 1291 | R1291R | E | 5.3e-09 | 1.3e-08 | 1.8e-08 | 1.210E-08 | 6.398E-09 | 5.300E-09 | 1.800E-08 |
| 25803 | 1291 | R1291N | E | 5.1e-09 | 5.1e-09 | 9e-09 | 6.400E-09 | 2.252E-09 | 5.100E-09 | 9.000E-09 |
| 25804 | 1291 | R1291D | E | 5.8e-09 | 9.7e-09 | 5.9e-09 | 7.133E-09 | 2.223E-09 | 5.800E-09 | 9.700E-09 |
| 25805 | 1291 | R1291C | E | 6.4e-09 | 1.5e-08 | 8.9e-09 | 1.010E-08 | 4.424E-09 | 6.400E-09 | 1.500E-08 |
| 25806 | 1291 | R1291E | E | 1.1e-09 | 1.3e-09 | 5.9e-09 | 2.767E-09 | 2.715E-09 | 1.100E-09 | 5.900E-09 |
| 25807 | 1291 | R1291Q | E | 5.9e-09 | 7.7e-09 | 5.9e-09 | 6.500E-09 | 1.039E-09 | 5.900E-09 | 7.700E-09 |
| 25808 | 1291 | R1291G | E | 1.4e-08 | 4.1e-09 | 2.7e-09 | 6.933E-09 | 6.160E-09 | 2.700E-09 | 1.400E-08 |
| 25809 | 1291 | R1291H | E | 4.1e-09 | 5.6e-09 | 3.3e-09 | 4.333E-09 | 1.168E-09 | 3.300E-09 | 5.600E-09 |
| 25810 | 1291 | R1291I | E | 9.6e-09 | 1.1e-08 | 4.1e-09 | 8.233E-09 | 3.647E-09 | 4.100E-09 | 1.100E-08 |
| 25811 | 1291 | R1291L | E | 1.3e-08 | 1e-08 | 1.5e-08 | 1.267E-08 | 2.517E-09 | 1.000E-08 | 1.500E-08 |
| 25812 | 1291 | R1291K | E | 3.9e-09 | 7.1e-09 | 7.9e-09 | 6.300E-09 | 2.117E-09 | 3.900E-09 | 7.900E-09 |
| 25813 | 1291 | R1291M | E | 2.9e-08 | 4.8e-09 | 3.4e-09 | 1.240E-08 | 1.439E-08 | 3.400E-09 | 2.900E-08 |
| 25814 | 1291 | R1291F | E | 1.1e-08 | 4.3e-09 | 7.7e-09 | 7.667E-09 | 3.350E-09 | 4.300E-09 | 1.100E-08 |
| 25815 | 1291 | R1291P | E | 8.6e-09 | 1.7e-09 | 2e-09 | 4.100E-09 | 3.900E-09 | 1.700E-09 | 8.600E-09 |
| 25816 | 1291 | R1291S | E | 1.2e-08 | 1.5e-08 | 1.5e-08 | 1.400E-08 | 1.732E-09 | 1.200E-08 | 1.500E-08 |
| 25817 | 1291 | R1291T | E | 4.1e-09 | 4.6e-09 | 6.4e-09 | 5.033E-09 | 1.210E-09 | 4.100E-09 | 6.400E-09 |
| 25818 | 1291 | R1291W | E | 7.1e-09 | 6.6e-09 | 2.6e-09 | 5.433E-09 | 2.466E-09 | 2.600E-09 | 7.100E-09 |
| 25819 | 1291 | R1291Y | E | 1.2e-08 | 1.6e-09 | 1.9e-08 | 1.087E-08 | 8.755E-09 | 1.600E-09 | 1.900E-08 |
| 25820 | 1291 | R1291V | E | 1.2e-08 | 4.8e-09 | 1.1e-08 | 9.267E-09 | 3.900E-09 | 4.800E-09 | 1.200E-08 |
| 25821 | 1292 | L1292A | E | 7.4e-09 | 7.2e-09 | 3.9e-09 | 6.167E-09 | 1.966E-09 | 3.900E-09 | 7.400E-09 |
| 25822 | 1292 | L1292R | E | 5.3e-09 | 1e-08 | 4.5e-09 | 6.600E-09 | 2.972E-09 | 4.500E-09 | 1.000E-08 |
| 25823 | 1292 | L1292N | E | 2.8e-09 | 5.9e-09 | 2.3e-09 | 3.667E-09 | 1.950E-09 | 2.300E-09 | 5.900E-09 |
| 25824 | 1292 | L1292D | E | 7.3e-09 | 1.3e-08 | 1.5e-08 | 1.177E-08 | 3.995E-09 | 7.300E-09 | 1.500E-08 |
| 25825 | 1292 | L1292C | E | 6.3e-09 | 5.6e-09 | 5.6e-09 | 5.833E-09 | 4.041E-10 | 5.600E-09 | 6.300E-09 |
| 25826 | 1292 | L1292E | E | 2.4e-08 | 4.8e-09 | 3.1e-09 | 1.063E-08 | 1.161E-08 | 3.100E-09 | 2.400E-08 |
| 25827 | 1292 | L1292Q | E | 1.4e-08 | 3.8e-09 | 1.2e-08 | 9.933E-09 | 5.405E-09 | 3.800E-09 | 1.400E-08 |
| 25828 | 1292 | L1292G | E | 4.4e-09 | 8.9e-09 | 6.5e-09 | 6.600E-09 | 2.252E-09 | 4.400E-09 | 8.900E-09 |
| 25829 | 1292 | L1292H | E | 8.8e-09 | 8.8e-09 | 2.4e-08 | 1.387E-08 | 8.776E-09 | 8.800E-09 | 2.400E-08 |
| 25830 | 1292 | L1292I | E | 6.6e-09 | 2.1e-09 | 7e-09 | 5.233E-09 | 2.721E-09 | 2.100E-09 | 7.000E-09 |
| 25831 | 1292 | L1292L | E | 4.6e-09 | 1.4e-08 | 3.4e-09 | 7.333E-09 | 5.805E-09 | 3.400E-09 | 1.400E-08 |
| 25832 | 1292 | L1292K | E | 8.5e-09 | 4.2e-09 | 6.3e-09 | 6.333E-09 | 2.150E-09 | 4.200E-09 | 8.500E-09 |
| 25833 | 1292 | L1292M | E | 1.2e-08 | 1.7e-09 | 4.7e-09 | 6.133E-09 | 5.297E-09 | 1.700E-09 | 1.200E-08 |
| 25834 | 1292 | L1292F | E | 8.4e-09 | 2.9e-09 | 1.2e-08 | 7.767E-09 | 4.583E-09 | 2.900E-09 | 1.200E-08 |
| 25835 | 1292 | L1292P | E | 1.6e-08 | 8.7e-09 | 8.7e-09 | 1.113E-08 | 4.215E-09 | 8.700E-09 | 1.600E-08 |
| 25836 | 1292 | L1292S | E | 9.1e-09 | 1e-08 | 6.6e-09 | 8.567E-09 | 1.762E-09 | 6.600E-09 | 1.000E-08 |
| 25837 | 1292 | L1292T | E | 8.5e-09 | 1e-08 | 3.9e-09 | 7.467E-09 | 3.179E-09 | 3.900E-09 | 1.000E-08 |
| 25838 | 1292 | L1292W | E | 1.1e-08 | 2.3e-09 | 9.5e-09 | 7.600E-09 | 4.651E-09 | 2.300E-09 | 1.100E-08 |
| 25839 | 1292 | L1292Y | E | 8.7e-09 | 4.8e-09 | 1.4e-08 | 9.167E-09 | 4.618E-09 | 4.800E-09 | 1.400E-08 |

| | | | | | | | | | | |
|---|---|---|---|---|---|---|---|---|---|---|
| 25840 | 1292 | L1292V | E | 6.1e-09 | 1.3e-08 | 6.5e-09 | 8.533E-09 | 3.873E-09 | 6.100E-09 | 1.300E-08 |
| 25841 | 1293 | S1293A | E | 1.3e-08 | 4.8e-09 | 1.5e-08 | 1.093E-08 | 5.405E-09 | 4.800E-09 | 1.500E-08 |
| 25842 | 1293 | S1293R | E | 1.4e-09 | 1.1e-08 | 1.3e-08 | 8.467E-09 | 6.201E-09 | 1.400E-09 | 1.300E-08 |
| 25843 | 1293 | S1293N | E | 4.4e-09 | 8e-09 | 1.3e-09 | 4.567E-09 | 3.353E-09 | 1.300E-09 | 8.000E-09 |
| 25844 | 1293 | S1293D | E | 6.9e-09 | 6.7e-09 | 2.3e-09 | 5.300E-09 | 2.600E-09 | 2.300E-09 | 6.900E-09 |
| 25845 | 1293 | S1293C | E | 4.2e-09 | 5e-09 | 7.5e-09 | 5.567E-09 | 1.721E-09 | 4.200E-09 | 7.500E-09 |
| 25846 | 1293 | S1293E | E | 3.7e-09 | 9.6e-09 | 1e-08 | 7.767E-09 | 3.528E-09 | 3.700E-09 | 1.000E-08 |
| 25847 | 1293 | S1293Q | E | 1.2e-08 | 6e-09 | 4.6e-09 | 7.533E-09 | 3.931E-09 | 4.600E-09 | 1.200E-08 |
| 25848 | 1293 | S1293G | E | 9.7e-09 | 9.8e-09 | 6.3e-09 | 8.600E-09 | 1.992E-09 | 6.300E-09 | 9.800E-09 |
| 25849 | 1293 | S1293H | E | 6.5e-09 | 9.3e-09 | 1.1e-08 | 8.933E-09 | 2.272E-09 | 6.500E-09 | 1.100E-08 |
| 25850 | 1293 | S1293I | E | 4.4e-09 | 3.6e-09 | 2.7e-09 | 3.567E-09 | 8.505E-10 | 2.700E-09 | 4.400E-09 |
| 25851 | 1293 | S1293L | E | 2.1e-09 | 6.7e-09 | 7.3e-09 | 5.367E-09 | 2.845E-09 | 2.100E-09 | 7.300E-09 |
| 25852 | 1293 | S1293K | E | 7.6e-09 | 6.3e-09 | 1.1e-08 | 8.300E-09 | 2.427E-09 | 6.300E-09 | 1.100E-08 |
| 25853 | 1293 | S1293M | E | 8e-09 | 6e-09 | 8.1e-09 | 7.367E-09 | 1.185E-09 | 6.000E-09 | 8.100E-09 |
| 25854 | 1293 | S1293F | E | 8.9e-09 | 1e-08 | 9.1e-09 | 9.333E-09 | 5.859E-10 | 8.900E-09 | 1.000E-08 |
| 25855 | 1293 | S1293P | E | 8.2e-09 | 6.4e-09 | 4.2e-09 | 6.267E-09 | 2.003E-09 | 4.200E-09 | 8.200E-09 |
| 25856 | 1293 | S1293S | E | 6.7e-09 | 6.1e-09 | 1.1e-08 | 7.933E-09 | 2.673E-09 | 6.100E-09 | 1.100E-08 |
| 25857 | 1293 | S1293T | E | 4.9e-09 | 8e-09 | 1.5e-08 | 9.300E-09 | 5.174E-09 | 4.900E-09 | 1.500E-08 |
| 25858 | 1293 | S1293W | E | 3.3e-09 | 4.8e-09 | 4.2e-09 | 4.100E-09 | 7.550E-10 | 3.300E-09 | 4.800E-09 |
| 25859 | 1293 | S1293Y | E | 2.6e-09 | 9.8e-09 | 4.4e-09 | 5.600E-09 | 3.747E-09 | 2.600E-09 | 9.800E-09 |
| 25860 | 1293 | S1293V | E | 1.1e-08 | 5.9e-09 | 4.5e-09 | 7.133E-09 | 3.421E-09 | 4.500E-09 | 1.100E-08 |
| 25861 | 1294 | C1294A | E | 6.6e-09 | 7.7e-09 | 6.2e-09 | 6.833E-09 | 7.767E-10 | 6.200E-09 | 7.700E-09 |
| 25862 | 1294 | C1294R | E | 1.1e-08 | 5.4e-09 | 1.5e-08 | 1.047E-08 | 4.822E-09 | 5.400E-09 | 1.500E-08 |
| 25863 | 1294 | C1294N | E | 1.2e-09 | 1.1e-08 | 4.4e-09 | 5.533E-09 | 4.997E-09 | 1.200E-09 | 1.100E-08 |
| 25864 | 1294 | C1294D | E | 1.5e-08 | 7.4e-09 | 7.6e-09 | 1.000E-08 | 4.331E-09 | 7.400E-09 | 1.500E-08 |
| 25865 | 1294 | C1294C | E | 2.2e-09 | 4.8e-09 | 7e-09 | 4.667E-09 | 2.403E-09 | 2.200E-09 | 7.000E-09 |
| 25866 | 1294 | C1294E | E | 1.6e-08 | 1.1e-08 | 6.8e-09 | 1.127E-08 | 4.606E-09 | 6.800E-09 | 1.600E-08 |
| 25867 | 1294 | C1294Q | E | 3.8e-09 | 7e-09 | 8.9e-09 | 6.567E-09 | 2.577E-09 | 3.800E-09 | 8.900E-09 |
| 25868 | 1294 | C1294G | E | 1.6e-08 | 6e-09 | 1.1e-08 | 1.100E-08 | 5.000E-09 | 6.000E-09 | 1.600E-08 |
| 25869 | 1294 | C1294H | E | 1.1e-08 | 4.4e-09 | 8.5e-09 | 7.967E-09 | 3.332E-09 | 4.400E-09 | 1.100E-08 |
| 25870 | 1294 | C1294I | E | 1.2e-08 | 2.2e-08 | 5.4e-09 | 1.313E-08 | 8.358E-09 | 5.400E-09 | 2.200E-08 |
| 25871 | 1294 | C1294L | E | 1.5e-08 | 4.7e-09 | 1.2e-08 | 1.057E-08 | 5.297E-09 | 4.700E-09 | 1.500E-08 |
| 25872 | 1294 | C1294K | E | 8.1e-09 | 6.6e-09 | 6.7e-09 | 7.133E-09 | 8.386E-10 | 6.600E-09 | 8.100E-09 |
| 25873 | 1294 | C1294M | E | 5.9e-09 | 3.3e-09 | 8.3e-09 | 5.833E-09 | 2.501E-09 | 3.300E-09 | 8.300E-09 |
| 25874 | 1294 | C1294F | E | 7.5e-09 | 1.1e-08 | 4e-09 | 7.500E-09 | 3.500E-09 | 4.000E-09 | 1.100E-08 |
| 25875 | 1294 | C1294P | E | 1.8e-08 | 3.5e-09 | 1.8e-09 | 7.767E-09 | 8.903E-09 | 1.800E-09 | 1.800E-08 |
| 25876 | 1294 | C1294S | E | 4.3e-09 | 3.3e-09 | 3.8e-09 | 3.800E-09 | 5.000E-10 | 3.300E-09 | 4.300E-09 |
| 25877 | 1294 | C1294T | E | 1e-08 | 5.2e-09 | 8.1e-09 | 7.767E-09 | 2.417E-09 | 5.200E-09 | 1.000E-08 |
| 25878 | 1294 | C1294W | E | 1e-08 | 1.4e-08 | 8.7e-09 | 1.090E-08 | 2.762E-09 | 8.700E-09 | 1.400E-08 |
| 25879 | 1294 | C1294Y | E | 2.3e-09 | 6.4e-09 | 1.1e-08 | 6.567E-09 | 4.352E-09 | 2.300E-09 | 1.100E-08 |
| 25880 | 1294 | C1294V | E | 8.4e-09 | 1.2e-08 | 8.5e-09 | 9.633E-09 | 2.050E-09 | 8.400E-09 | 1.200E-08 |
| 25881 | 1295 | A1295A | E | 8.8e-09 | 1e-08 | 7.1e-09 | 8.633E-09 | 1.457E-09 | 7.100E-09 | 1.000E-08 |
| 25882 | 1295 | A1295R | E | 3.2e-09 | 1.4e-08 | 1.1e-08 | 9.400E-09 | 5.575E-09 | 3.200E-09 | 1.400E-08 |
| 25883 | 1295 | A1295N | E | 5.9e-09 | 3.5e-08 | 3.7e-09 | 1.487E-08 | 1.747E-08 | 3.700E-09 | 3.500E-08 |
| 25884 | 1295 | A1295D | E | 8.9e-09 | 1.1e-08 | 7.6e-09 | 9.167E-09 | 1.716E-09 | 7.600E-09 | 1.100E-08 |
| 25885 | 1295 | A1295C | E | 4.1e-08 | 1.1e-08 | 9.5e-10 | 1.765E-08 | 2.084E-08 | 9.500E-10 | 4.100E-08 |
| 25886 | 1295 | A1295E | E | 4.1e-09 | 2.9e-09 | 2.1e-08 | 9.333E-09 | 1.012E-08 | 2.900E-09 | 2.100E-08 |
| 25887 | 1295 | A1295Q | E | 8.6e-09 | 7.5e-09 | 5e-09 | 7.033E-09 | 1.845E-09 | 5.000E-09 | 8.600E-09 |
| 25888 | 1295 | A1295G | E | 1e-08 | 4e-09 | 2.7e-09 | 5.567E-09 | 3.894E-09 | 2.700E-09 | 1.000E-08 |
| 25889 | 1295 | A1295H | E | 6.1e-09 | 1.5e-09 | 1.3e-08 | 6.867E-09 | 5.788E-09 | 1.500E-09 | 1.300E-08 |
| 25890 | 1295 | A1295I | E | 6.2e-09 | 3.9e-09 | 3.5e-09 | 4.533E-09 | 1.457E-09 | 3.500E-09 | 6.200E-09 |
| 25891 | 1295 | A1295L | E | 8.6e-09 | 6.5e-09 | 9.7e-09 | 8.267E-09 | 1.626E-09 | 6.500E-09 | 9.700E-09 |
| 25892 | 1295 | A1295K | E | 4.2e-09 | 1.7e-08 | 3.3e-09 | 8.167E-09 | 7.663E-09 | 3.300E-09 | 1.700E-08 |
| 25893 | 1295 | A1295M | E | 5.6e-08 | 2.4e-08 | 8.9e-09 | 2.963E-08 | 2.405E-08 | 8.900E-09 | 5.600E-08 |
| 25894 | 1295 | A1295F | E | 7.9e-09 | 5.3e-09 | 6.7e-09 | 6.633E-09 | 1.301E-09 | 5.300E-09 | 7.900E-09 |
| 25895 | 1295 | A1295P | E | 1.4e-08 | 7.3e-09 | 7.6e-09 | 9.633E-09 | 3.785E-09 | 7.300E-09 | 1.400E-08 |
| 25896 | 1295 | A1295S | E | 6.5e-09 | 1.5e-08 | 8.9e-09 | 1.013E-08 | 4.382E-09 | 6.500E-09 | 1.500E-08 |
| 25897 | 1295 | A1295T | E | 2.6e-09 | 1.4e-08 | 5.8e-09 | 7.467E-09 | 5.880E-09 | 2.600E-09 | 1.400E-08 |
| 25898 | 1295 | A1295W | E | 6.8e-09 | 6.1e-09 | 5.7e-09 | 6.200E-09 | 5.568E-10 | 5.700E-09 | 6.800E-09 |
| 25899 | 1295 | A1295Y | E | 5.6e-09 | 5.8e-09 | 2.3e-09 | 4.567E-09 | 1.966E-09 | 2.300E-09 | 5.800E-09 |
| 25900 | 1295 | A1295V | E | 3.5e-09 | 6.7e-09 | 7.2e-09 | 5.800E-09 | 2.007E-09 | 3.500E-09 | 7.200E-09 |
| 25901 | 1296 | A1296A | E | 2e-09 | 1.1e-08 | 9.3e-09 | 7.433E-09 | 4.782E-09 | 2.000E-09 | 1.100E-08 |
| 25902 | 1296 | A1296R | E | 3.1e-09 | 7.9e-09 | 2.9e-09 | 4.633E-09 | 2.831E-09 | 2.900E-09 | 7.900E-09 |
| 25903 | 1296 | A1296N | E | 2.6e-08 | 8.5e-09 | 1.5e-08 | 1.650E-08 | 8.846E-09 | 8.500E-09 | 2.600E-08 |
| 25904 | 1296 | A1296D | E | 2.8e-09 | 6.9e-09 | 1.9e-09 | 3.867E-09 | 2.665E-09 | 1.900E-09 | 6.900E-09 |
| 25905 | 1296 | A1296C | E | 1.4e-08 | 5.6e-09 | 4e-09 | 7.867E-09 | 5.372E-09 | 4.000E-09 | 1.400E-08 |
| 25906 | 1296 | A1296E | E | 2.7e-09 | 1.6e-09 | 1.6e-08 | 6.767E-09 | 8.015E-09 | 1.600E-09 | 1.600E-08 |
| 25907 | 1296 | A1296Q | E | 7.8e-09 | 5e-09 | 1.3e-08 | 8.600E-09 | 4.060E-09 | 5.000E-09 | 1.300E-08 |
| 25908 | 1296 | A1296G | E | 9.4e-09 | 5.9e-09 | 9.6e-09 | 8.300E-09 | 2.081E-09 | 5.900E-09 | 9.600E-09 |
| 25909 | 1296 | A1296H | E | 1e-08 | 6e-09 | 8.3e-09 | 8.100E-09 | 2.007E-09 | 6.000E-09 | 1.000E-08 |
| 25910 | 1296 | A1296I | E | 7.4e-09 | 3.3e-08 | 6.5e-09 | 1.563E-08 | 1.505E-08 | 6.500E-09 | 3.300E-08 |
| 25911 | 1296 | A1296L | E | 8.9e-09 | 5.8e-09 | 2.2e-09 | 5.633E-09 | 3.353E-09 | 2.200E-09 | 8.900E-09 |
| 25912 | 1296 | A1296K | E | 5.8e-09 | 1.2e-08 | 8.2e-09 | 8.667E-09 | 3.126E-09 | 5.800E-09 | 1.200E-08 |
| 25913 | 1296 | A1296M | E | 4.1e-09 | 7.2e-09 | 7.7e-09 | 6.333E-09 | 1.950E-09 | 4.100E-09 | 7.700E-09 |
| 25914 | 1296 | A1296F | E | 9e-09 | 8.7e-09 | 6.9e-09 | 8.200E-09 | 1.136E-09 | 6.900E-09 | 9.000E-09 |
| 25915 | 1296 | A1296P | E | 5.4e-09 | 4e-09 | 2.6e-08 | 1.180E-08 | 1.232E-08 | 4.000E-09 | 2.600E-08 |
| 25916 | 1296 | A1296S | E | 7.1e-09 | 7e-09 | 7.9e-09 | 7.333E-09 | 4.933E-10 | 7.000E-09 | 7.900E-09 |
| 25917 | 1296 | A1296T | E | 3.6e-09 | 4.8e-09 | 9.8e-09 | 6.067E-09 | 3.288E-09 | 3.600E-09 | 9.800E-09 |
| 25918 | 1296 | A1296W | E | 5.2e-09 | 3.4e-09 | 9.1e-09 | 5.900E-09 | 2.914E-09 | 3.400E-09 | 9.100E-09 |
| 25919 | 1296 | A1296Y | E | 3.2e-09 | 9e-09 | 5.2e-09 | 5.800E-09 | 2.946E-09 | 3.200E-09 | 9.000E-09 |
| 25920 | 1296 | A1296V | E | 3.5e-09 | 7.1e-09 | 1.3e-08 | 7.867E-09 | 4.796E-09 | 3.500E-09 | 1.300E-08 |
| 25921 | 1297 | S1297A | E | 1.7e-08 | 6.9e-09 | 4.7e-09 | 9.533E-09 | 6.559E-09 | 4.700E-09 | 1.700E-08 |
| 25922 | 1297 | S1297R | E | 4.8e-09 | 2.7e-09 | 1.1e-08 | 6.167E-09 | 4.315E-09 | 2.700E-09 | 1.100E-08 |
| 25923 | 1297 | S1297N | E | 4.9e-09 | 6.1e-09 | 5.1e-09 | 5.367E-09 | 6.429E-10 | 4.900E-09 | 6.100E-09 |
| 25924 | 1297 | S1297D | E | 1e-08 | 1.5e-08 | 3.2e-09 | 9.400E-09 | 5.923E-09 | 3.200E-09 | 1.500E-08 |

| | | | | | | | | | | |
|---|---|---|---|---|---|---|---|---|---|---|
| 25925 | 1297 | S1297C | E | 2.3e-08 | 1.6e-08 | 8.8e-09 | 1.593E-08 | 7.100E-09 | 8.800E-09 | 2.300E-08 |
| 25926 | 1297 | S1297E | E | 2e-08 | 6.2e-09 | 9.3e-09 | 1.183E-08 | 7.240E-09 | 6.200E-09 | 2.000E-08 |
| 25927 | 1297 | S1297Q | E | 3.6e-09 | 7.9e-09 | 7.9e-09 | 6.467E-09 | 2.483E-09 | 3.600E-09 | 7.900E-09 |
| 25928 | 1297 | S1297G | E | 4.3e-09 | 4.6e-09 | 7.7e-09 | 5.533E-09 | 1.882E-09 | 4.300E-09 | 7.700E-09 |
| 25929 | 1297 | S1297H | E | 9e-09 | 3.9e-09 | 6.7e-09 | 6.533E-09 | 2.554E-09 | 3.900E-09 | 9.000E-09 |
| 25930 | 1297 | S1297I | E | 1.1e-08 | 4.5e-09 | 1.1e-08 | 8.833E-09 | 3.753E-09 | 4.500E-09 | 1.100E-08 |
| 25931 | 1297 | S1297L | E | 2.4e-09 | 3.3e-09 | 9.9e-09 | 5.200E-09 | 4.095E-09 | 2.400E-09 | 9.900E-09 |
| 25932 | 1297 | S1297K | E | 6.8e-09 | 2.5e-08 | 6.6e-09 | 1.280E-08 | 1.057E-08 | 6.600E-09 | 2.500E-08 |
| 25933 | 1297 | S1297M | E | 9.3e-09 | 2e-09 | 5.9e-09 | 5.733E-09 | 3.653E-09 | 2.000E-09 | 9.300E-09 |
| 25934 | 1297 | S1297F | E | 4.6e-09 | 6.4e-09 | 4.5e-09 | 5.167E-09 | 1.069E-09 | 4.500E-09 | 6.400E-09 |
| 25935 | 1297 | S1297P | E | 4.9e-09 | 4.8e-09 | 2.5e-09 | 4.067E-09 | 1.358E-09 | 2.500E-09 | 4.900E-09 |
| 25936 | 1297 | S1297S | E | 1.6e-08 | 1.7e-08 | 1.1e-08 | 1.467E-08 | 3.215E-09 | 1.100E-08 | 1.700E-08 |
| 25937 | 1297 | S1297T | E | 1.8e-09 | 5.5e-09 | 1.2e-08 | 6.433E-09 | 5.164E-09 | 1.800E-09 | 1.200E-08 |
| 25938 | 1297 | S1297W | E | 5.5e-09 | 7.1e-09 | 8e-09 | 6.867E-09 | 1.266E-09 | 5.500E-09 | 8.000E-09 |
| 25939 | 1297 | S1297Y | E | 5.3e-09 | 8.7e-09 | 1.1e-08 | 8.333E-09 | 2.868E-09 | 5.300E-09 | 1.100E-08 |
| 25940 | 1297 | S1297V | E | 2.4e-09 | 3.4e-09 | 1.1e-08 | 5.600E-09 | 4.703E-09 | 2.400E-09 | 1.100E-08 |
| 25941 | 1298 | G1298A | E | 7.2e-09 | 2.1e-08 | 7.3e-09 | 1.183E-08 | 7.939E-09 | 7.200E-09 | 2.100E-08 |
| 25942 | 1298 | G1298R | E | 1.2e-08 | 3.2e-09 | 4.6e-09 | 6.600E-09 | 4.729E-09 | 3.200E-09 | 1.200E-08 |
| 25943 | 1298 | G1298N | E | 9.1e-09 | 5.8e-09 | 2e-09 | 5.633E-09 | 3.553E-09 | 2.000E-09 | 9.100E-09 |
| 25944 | 1298 | G1298D | E | 1.4e-08 | 4.3e-09 | 2.4e-09 | 6.900E-09 | 6.222E-09 | 2.400E-09 | 1.400E-08 |
| 25945 | 1298 | G1298C | E | 3.4e-09 | 7e-09 | 9.4e-09 | 6.600E-09 | 3.020E-09 | 3.400E-09 | 9.400E-09 |
| 25946 | 1298 | G1298E | E | 1.6e-08 | 8.6e-09 | 1.8e-09 | 8.800E-09 | 7.102E-09 | 1.800E-09 | 1.600E-08 |
| 25947 | 1298 | G1298Q | E | 8.4e-09 | 2.2e-08 | 3.3e-09 | 1.123E-08 | 9.667E-09 | 3.300E-09 | 2.200E-08 |
| 25948 | 1298 | G1298G | E | 6.3e-09 | 1.1e-08 | 4e-09 | 7.100E-09 | 3.568E-09 | 4.000E-09 | 1.100E-08 |
| 25949 | 1298 | G1298H | E | 1.9e-08 | 8.5e-09 | 2.6e-09 | 1.003E-08 | 8.307E-09 | 2.600E-09 | 1.900E-08 |
| 25950 | 1298 | G1298I | E | 1.7e-08 | 6e-09 | 3.7e-09 | 8.900E-09 | 7.108E-09 | 3.700E-09 | 1.700E-08 |
| 25951 | 1298 | G1298L | E | 9e-09 | 1e-08 | 9.7e-09 | 9.567E-09 | 5.132E-10 | 9.000E-09 | 1.000E-08 |
| 25952 | 1298 | G1298K | E | 7.9e-09 | 5e-09 | 5.7e-09 | 6.200E-09 | 1.513E-09 | 5.000E-09 | 7.900E-09 |
| 25953 | 1298 | G1298M | E | 1.4e-08 | 5.7e-09 | 5.7e-09 | 8.467E-09 | 4.792E-09 | 5.700E-09 | 1.400E-08 |
| 25954 | 1298 | G1298F | E | 7.8e-09 | 2.1e-08 | 1.6e-08 | 1.493E-08 | 6.664E-09 | 7.800E-09 | 2.100E-08 |
| 25955 | 1298 | G1298P | E | 9.2e-09 | 8.7e-09 | 8.8e-09 | 8.900E-09 | 2.646E-10 | 8.700E-09 | 9.200E-09 |
| 25956 | 1298 | G1298S | E | 3.6e-09 | 6.9e-09 | 6e-09 | 5.500E-09 | 1.706E-09 | 3.600E-09 | 6.900E-09 |
| 25957 | 1298 | G1298T | E | 3.9e-09 | 7.4e-09 | 4.9e-09 | 5.400E-09 | 1.803E-09 | 3.900E-09 | 7.400E-09 |
| 25958 | 1298 | G1298W | E | 1.4e-08 | 5.9e-09 | 5.7e-09 | 8.533E-09 | 4.735E-09 | 5.700E-09 | 1.400E-08 |
| 25959 | 1298 | G1298Y | E | 2.8e-09 | 2.4e-09 | 2.3e-09 | 2.500E-09 | 2.646E-10 | 2.300E-09 | 2.800E-09 |
| 25960 | 1298 | G1298V | E | 8e-09 | 9.9e-09 | 2e-08 | 1.263E-08 | 6.450E-09 | 8.000E-09 | 2.000E-08 |
| 25961 | 1299 | F1299A | E | 6.4e-09 | 1.1e-08 | 6.6e-09 | 8.000E-09 | 2.600E-09 | 6.400E-09 | 1.100E-08 |
| 25962 | 1299 | F1299R | E | 9e-09 | 1e-08 | 1.2e-08 | 1.033E-08 | 1.528E-09 | 9.000E-09 | 1.200E-08 |
| 25963 | 1299 | F1299N | E | 6.2e-09 | 4.8e-09 | 1.4e-09 | 4.133E-09 | 2.468E-09 | 1.400E-09 | 6.200E-09 |
| 25964 | 1299 | F1299D | E | 4.5e-09 | 1e-08 | 1.4e-08 | 9.500E-09 | 4.770E-09 | 4.500E-09 | 1.400E-08 |
| 25965 | 1299 | F1299C | E | 7.2e-09 | 6.1e-09 | 6.8e-09 | 6.700E-09 | 5.568E-10 | 6.100E-09 | 7.200E-09 |
| 25966 | 1299 | F1299E | E | 1.7e-08 | 4.1e-09 | 1.7e-08 | 1.270E-08 | 7.448E-09 | 4.100E-09 | 1.700E-08 |
| 25967 | 1299 | F1299Q | E | 9.3e-09 | 5e-09 | 6.2e-09 | 6.833E-09 | 2.219E-09 | 5.000E-09 | 9.300E-09 |
| 25968 | 1299 | F1299G | E | 3.6e-09 | 6.7e-09 | 2.8e-08 | 1.277E-08 | 1.328E-08 | 3.600E-09 | 2.800E-08 |
| 25969 | 1299 | F1299H | E | 2.7e-09 | 4e-09 | 4.3e-09 | 3.667E-09 | 8.505E-10 | 2.700E-09 | 4.300E-09 |
| 25970 | 1299 | F1299I | E | 4.9e-09 | 1.8e-08 | 3.5e-09 | 8.800E-09 | 7.998E-09 | 3.500E-09 | 1.800E-08 |
| 25971 | 1299 | F1299L | E | 2.7e-09 | 5e-09 | 1.5e-08 | 7.567E-09 | 6.539E-09 | 2.700E-09 | 1.500E-08 |
| 25972 | 1299 | F1299K | E | 1.2e-08 | 8.9e-09 | 6.3e-09 | 9.067E-09 | 2.854E-09 | 6.300E-09 | 1.200E-08 |
| 25973 | 1299 | F1299M | E | 2.4e-08 | 1.3e-08 | 3.6e-09 | 1.353E-08 | 1.021E-08 | 3.600E-09 | 2.400E-08 |
| 25974 | 1299 | F1299F | E | 1.4e-08 | 6.1e-09 | 5.3e-09 | 8.467E-09 | 4.809E-09 | 5.300E-09 | 1.400E-08 |
| 25975 | 1299 | F1299P | E | 4.2e-09 | 1e-08 | 1.1e-08 | 8.400E-09 | 3.672E-09 | 4.200E-09 | 1.100E-08 |
| 25976 | 1299 | F1299S | E | 1.7e-08 | 2.9e-09 | 1e-08 | 9.967E-09 | 7.050E-09 | 2.900E-09 | 1.700E-08 |
| 25977 | 1299 | F1299T | E | 1.4e-08 | 4.2e-09 | 6.3e-09 | 8.167E-09 | 5.160E-09 | 4.200E-09 | 1.400E-08 |
| 25978 | 1299 | F1299W | E | 8.9e-09 | 2.1e-09 | 1.4e-08 | 8.333E-09 | 5.970E-09 | 2.100E-09 | 1.400E-08 |
| 25979 | 1299 | F1299Y | E | 3.2e-09 | 5.6e-09 | 9e-09 | 5.933E-09 | 2.914E-09 | 3.200E-09 | 9.000E-09 |
| 25980 | 1299 | F1299V | E | 8.6e-09 | 6.7e-09 | 6.9e-09 | 7.400E-09 | 1.044E-09 | 6.700E-09 | 8.600E-09 |
| 25981 | 1300 | N1300A | E | 7.2e-09 | 6.2e-09 | 9.2e-09 | 7.533E-09 | 1.528E-09 | 6.200E-09 | 9.200E-09 |
| 25982 | 1300 | N1300R | E | 6.2e-09 | 3.8e-09 | 5.1e-09 | 5.033E-09 | 1.201E-09 | 3.800E-09 | 6.200E-09 |
| 25983 | 1300 | N1300N | E | 2.9e-09 | 7.6e-09 | 4e-09 | 4.833E-09 | 2.458E-09 | 2.900E-09 | 7.600E-09 |
| 25984 | 1300 | N1300D | E | 2.5e-09 | 3.4e-09 | 3.5e-09 | 3.133E-09 | 5.508E-10 | 2.500E-09 | 3.500E-09 |
| 25985 | 1300 | N1300C | E | 1.1e-08 | 2.8e-09 | 1e-08 | 7.933E-09 | 4.474E-09 | 2.800E-09 | 1.100E-08 |
| 25986 | 1300 | N1300E | E | 1.1e-08 | 2.7e-09 | 3.9e-09 | 5.867E-09 | 4.486E-09 | 2.700E-09 | 1.100E-08 |
| 25987 | 1300 | N1300Q | E | 6.4e-09 | 4.2e-09 | 5.5e-09 | 5.367E-09 | 1.106E-09 | 4.200E-09 | 6.400E-09 |
| 25988 | 1300 | N1300G | E | 7.5e-09 | 8.7e-09 | 7.4e-09 | 7.867E-09 | 7.234E-10 | 7.400E-09 | 8.700E-09 |
| 25989 | 1300 | N1300H | E | 8.8e-09 | 6.7e-09 | 9e-09 | 8.167E-09 | 1.274E-09 | 6.700E-09 | 9.000E-09 |
| 25990 | 1300 | N1300I | E | 1.7e-08 | 3.9e-09 | 6.1e-09 | 9.000E-09 | 7.015E-09 | 3.900E-09 | 1.700E-08 |
| 25991 | 1300 | N1300L | E | 7.4e-09 | 9.7e-09 | 7.6e-09 | 8.233E-09 | 1.274E-09 | 7.400E-09 | 9.700E-09 |
| 25992 | 1300 | N1300K | E | 9.1e-09 | 5.6e-09 | 1e-08 | 8.233E-09 | 2.325E-09 | 5.600E-09 | 1.000E-08 |
| 25993 | 1300 | N1300M | E | 6.2e-09 | 2e-09 | 1.6e-08 | 8.067E-09 | 7.184E-09 | 2.000E-09 | 1.600E-08 |
| 25994 | 1300 | N1300F | E | 1e-08 | 5.3e-09 | 2.5e-09 | 5.933E-09 | 3.790E-09 | 2.500E-09 | 1.000E-08 |
| 25995 | 1300 | N1300P | E | 3.1e-08 | 3.6e-09 | 7.4e-09 | 1.400E-08 | 1.484E-08 | 3.600E-09 | 3.100E-08 |
| 25996 | 1300 | N1300S | E | 7.5e-09 | 7.4e-09 | 3.6e-09 | 6.167E-09 | 2.223E-09 | 3.600E-09 | 7.500E-09 |
| 25997 | 1300 | N1300T | E | 5.3e-09 | 6.9e-09 | 4.3e-09 | 5.500E-09 | 1.311E-09 | 4.300E-09 | 6.900E-09 |
| 25998 | 1300 | N1300W | E | 5e-09 | 5.9e-09 | 3.9e-09 | 4.933E-09 | 1.002E-09 | 3.900E-09 | 5.900E-09 |
| 25999 | 1300 | N1300Y | E | 4.2e-09 | 3.2e-09 | 9.9e-10 | 2.797E-09 | 1.643E-09 | 9.900E-10 | 4.200E-09 |
| 26000 | 1300 | N1300V | E | 2.9e-09 | 7.9e-09 | 3.8e-09 | 4.867E-09 | 2.665E-09 | 2.900E-09 | 7.900E-09 |
| 26001 | 1301 | I1301A | E | 3.6e-09 | 7.1e-09 | 4.6e-09 | 5.100E-09 | 1.803E-09 | 3.600E-09 | 7.100E-09 |
| 26002 | 1301 | I1301R | E | 4.5e-09 | 1e-08 | 4.6e-08 | 2.017E-08 | 2.254E-08 | 4.500E-09 | 4.600E-08 |
| 26003 | 1301 | I1301N | E | 1.4e-08 | 9.5e-09 | 2.9e-09 | 8.800E-09 | 5.583E-09 | 2.900E-09 | 1.400E-08 |
| 26004 | 1301 | I1301D | E | 4.6e-09 | 3.6e-09 | 7e-09 | 5.067E-09 | 1.747E-09 | 3.600E-09 | 7.000E-09 |
| 26005 | 1301 | I1301C | E | 2.6e-09 | 2.9e-09 | 2.4e-08 | 9.833E-09 | 1.227E-08 | 2.600E-09 | 2.400E-08 |
| 26006 | 1301 | I1301E | E | 1.1e-08 | 6.9e-09 | 1.1e-08 | 9.633E-09 | 2.367E-09 | 6.900E-09 | 1.100E-08 |
| 26007 | 1301 | I1301Q | E | 2.6e-09 | 4.8e-09 | 9.5e-09 | 5.633E-09 | 3.525E-09 | 2.600E-09 | 9.500E-09 |
| 26008 | 1301 | I1301G | E | 4.4e-09 | 1.4e-08 | 7.6e-09 | 8.667E-09 | 4.888E-09 | 4.400E-09 | 1.400E-08 |
| 26009 | 1301 | I1301H | E | 6.4e-09 | 8.3e-09 | 1e-08 | 8.233E-09 | 1.801E-09 | 6.400E-09 | 1.000E-08 |

| | | | | | | | | | | |
|---|---|---|---|---|---|---|---|---|---|---|
| 26010 | 1301 | I1301I | E | 5.6e-09 | 1.6e-09 | 3e-09 | 3.400E-09 | 2.030E-09 | 1.600E-09 | 5.600E-09 |
| 26011 | 1301 | I1301L | E | 2e-09 | 4.7e-09 | 2.3e-09 | 3.000E-09 | 1.480E-09 | 2.000E-09 | 4.700E-09 |
| 26012 | 1301 | I1301K | E | 7.8e-09 | 1.6e-08 | 4.9e-09 | 9.567E-09 | 5.757E-09 | 4.900E-09 | 1.600E-08 |
| 26013 | 1301 | I1301M | E | 4.2e-09 | 8.2e-09 | 7e-09 | 6.467E-09 | 2.053E-09 | 4.200E-09 | 8.200E-09 |
| 26014 | 1301 | I1301F | E | 6.3e-09 | 5.2e-09 | 6.3e-09 | 5.933E-09 | 6.351E-10 | 5.200E-09 | 6.300E-09 |
| 26015 | 1301 | I1301P | E | 1.9e-08 | 1.8e-08 | 4.4e-09 | 1.380E-08 | 8.156E-09 | 4.400E-09 | 1.900E-08 |
| 26016 | 1301 | I1301S | E | 3.8e-09 | 1.4e-08 | 5.2e-09 | 7.667E-09 | 5.529E-09 | 3.800E-09 | 1.400E-08 |
| 26017 | 1301 | I1301T | E | 4.6e-09 | 1.1e-08 | 6.4e-09 | 7.333E-09 | 3.301E-09 | 4.600E-09 | 1.100E-08 |
| 26018 | 1301 | I1301W | E | 7.4e-09 | 1.8e-08 | 2.2e-09 | 9.200E-09 | 8.052E-09 | 2.200E-09 | 1.800E-08 |
| 26019 | 1301 | I1301Y | E | 1.3e-08 | 4.1e-09 | 2.1e-08 | 1.270E-08 | 8.454E-09 | 4.100E-09 | 2.100E-08 |
| 26020 | 1301 | I1301V | E | 9.3e-10 | 1.1e-08 | 1.3e-08 | 8.310E-09 | 6.469E-09 | 9.300E-10 | 1.300E-08 |
| 26021 | 1302 | K1302A | E | 6.5e-09 | 4.8e-09 | 7.3e-09 | 6.200E-09 | 1.277E-09 | 4.800E-09 | 7.300E-09 |
| 26022 | 1302 | K1302R | E | 2.8e-09 | 7.3e-09 | 9.1e-09 | 6.400E-09 | 3.245E-09 | 2.800E-09 | 9.100E-09 |
| 26023 | 1302 | K1302N | E | 3.8e-09 | 4.4e-09 | 4.7e-09 | 4.300E-09 | 4.583E-10 | 3.800E-09 | 4.700E-09 |
| 26024 | 1302 | K1302D | E | 7.4e-09 | 3.5e-09 | 8.8e-09 | 6.567E-09 | 2.747E-09 | 3.500E-09 | 8.800E-09 |
| 26025 | 1302 | K1302C | E | 1.5e-08 | 1.7e-09 | 9.9e-09 | 8.867E-09 | 6.710E-09 | 1.700E-09 | 1.500E-08 |
| 26026 | 1302 | K1302E | E | 4.4e-09 | 1.7e-08 | 2.8e-09 | 8.067E-09 | 7.778E-09 | 2.800E-09 | 1.700E-08 |
| 26027 | 1302 | K1302Q | E | 1.1e-08 | 1.7e-09 | 6.8e-09 | 6.500E-09 | 4.657E-09 | 1.700E-09 | 1.100E-08 |
| 26028 | 1302 | K1302G | E | 9.1e-09 | 2e-08 | 5.5e-09 | 1.153E-08 | 7.550E-09 | 5.500E-09 | 2.000E-08 |
| 26029 | 1302 | K1302H | E | 1.3e-08 | 1.4e-08 | 5.2e-09 | 1.073E-08 | 4.818E-09 | 5.200E-09 | 1.400E-08 |
| 26030 | 1302 | K1302I | E | 7.8e-09 | 3.5e-09 | 4.4e-09 | 5.233E-09 | 2.268E-09 | 3.500E-09 | 7.800E-09 |
| 26031 | 1302 | K1302L | E | 5.6e-09 | 1.3e-08 | 7.4e-09 | 8.667E-09 | 3.859E-09 | 5.600E-09 | 1.300E-08 |
| 26032 | 1302 | K1302K | E | 4.9e-09 | 1.6e-08 | 9.9e-09 | 1.027E-08 | 5.559E-09 | 4.900E-09 | 1.600E-08 |
| 26033 | 1302 | K1302M | E | 5.1e-09 | 1.2e-08 | 1.1e-08 | 9.367E-09 | 3.729E-09 | 5.100E-09 | 1.200E-08 |
| 26034 | 1302 | K1302F | E | 8.1e-09 | 5.1e-09 | 1.1e-08 | 8.067E-09 | 2.950E-09 | 5.100E-09 | 1.100E-08 |
| 26035 | 1302 | K1302P | E | 4.6e-09 | 1.8e-08 | 5.5e-09 | 9.367E-09 | 7.490E-09 | 4.600E-09 | 1.800E-08 |
| 26036 | 1302 | K1302S | E | 7.4e-09 | 8.6e-09 | 5.7e-09 | 7.233E-09 | 1.457E-09 | 5.700E-09 | 8.600E-09 |
| 26037 | 1302 | K1302T | E | 1.9e-09 | 1.1e-08 | 5.4e-09 | 6.100E-09 | 4.590E-09 | 1.900E-09 | 1.100E-08 |
| 26038 | 1302 | K1302W | E | 6.5e-09 | 8.3e-09 | 2e-08 | 1.160E-08 | 7.330E-09 | 6.500E-09 | 2.000E-08 |
| 26039 | 1302 | K1302Y | E | 4.5e-09 | 9.2e-09 | 1.2e-08 | 8.567E-09 | 3.790E-09 | 4.500E-09 | 1.200E-08 |
| 26040 | 1302 | K1302V | E | 1.6e-08 | 2.8e-09 | 7.7e-09 | 8.833E-09 | 6.673E-09 | 2.800E-09 | 1.600E-08 |
| 26041 | 1303 | D1303A | E | 9e-09 | 3.4e-09 | 3.5e-09 | 5.300E-09 | 3.205E-09 | 3.400E-09 | 9.000E-09 |
| 26042 | 1303 | D1303R | E | 1.1e-08 | 6e-09 | 1.1e-08 | 9.333E-09 | 2.887E-09 | 6.000E-09 | 1.100E-08 |
| 26043 | 1303 | D1303N | E | 9.8e-09 | 5.1e-09 | 8.9e-09 | 7.933E-09 | 2.495E-09 | 5.100E-09 | 9.800E-09 |
| 26044 | 1303 | D1303D | E | 5.8e-09 | 6.5e-09 | 1e-08 | 7.433E-09 | 2.250E-09 | 5.800E-09 | 1.000E-08 |
| 26045 | 1303 | D1303C | E | 6.3e-09 | 5.5e-09 | 9.7e-09 | 7.167E-09 | 2.230E-09 | 5.500E-09 | 9.700E-09 |
| 26046 | 1303 | D1303E | E | 6.9e-09 | 7.9e-09 | 6.9e-09 | 7.233E-09 | 5.774E-10 | 6.900E-09 | 7.900E-09 |
| 26047 | 1303 | D1303Q | E | 9.7e-09 | 1.7e-08 | 1.2e-08 | 1.290E-08 | 3.732E-09 | 9.700E-09 | 1.700E-08 |
| 26048 | 1303 | D1303G | E | 7.3e-09 | 1e-08 | 1.1e-08 | 9.433E-09 | 1.914E-09 | 7.300E-09 | 1.100E-08 |
| 26049 | 1303 | D1303H | E | 6e-09 | 5.5e-09 | 4.7e-09 | 5.400E-09 | 6.557E-10 | 4.700E-09 | 6.000E-09 |
| 26050 | 1303 | D1303I | E | 4.2e-09 | 1.3e-08 | 9.3e-09 | 8.833E-09 | 4.419E-09 | 4.200E-09 | 1.300E-08 |
| 26051 | 1303 | D1303L | E | 3e-09 | 1.8e-08 | 7.9e-09 | 9.633E-09 | 7.649E-09 | 3.000E-09 | 1.800E-08 |
| 26052 | 1303 | D1303K | E | 2.2e-09 | 3.1e-09 | 3.9e-09 | 3.067E-09 | 8.505E-10 | 2.200E-09 | 3.900E-09 |
| 26053 | 1303 | D1303M | E | 1.1e-08 | 1.2e-08 | 3.7e-09 | 8.900E-09 | 4.531E-09 | 3.700E-09 | 1.200E-08 |
| 26054 | 1303 | D1303F | E | 2.4e-09 | 3.6e-09 | 5.8e-09 | 3.933E-09 | 1.724E-09 | 2.400E-09 | 5.800E-09 |
| 26055 | 1303 | D1303P | E | 7.2e-10 | 1.9e-08 | 1.9e-09 | 7.207E-09 | 1.023E-08 | 7.200E-10 | 1.900E-08 |
| 26056 | 1303 | D1303S | E | 3.8e-09 | 5.5e-09 | 4.4e-09 | 4.567E-09 | 8.622E-10 | 3.800E-09 | 5.500E-09 |
| 26057 | 1303 | D1303T | E | 7.9e-09 | 1.1e-08 | 6.4e-09 | 8.433E-09 | 2.346E-09 | 6.400E-09 | 1.100E-08 |
| 26058 | 1303 | D1303W | E | 1e-09 | 6.9e-09 | 6.3e-09 | 4.733E-09 | 3.247E-09 | 1.000E-09 | 6.900E-09 |
| 26059 | 1303 | D1303Y | E | 9.6e-09 | 7.7e-09 | 8.7e-09 | 8.667E-09 | 9.504E-10 | 7.700E-09 | 9.600E-09 |
| 26060 | 1303 | D1303V | E | 9.6e-09 | 4.8e-09 | 8.1e-09 | 7.500E-09 | 2.456E-09 | 4.800E-09 | 9.600E-09 |
| 26061 | 1304 | T1304A | E | 5.5e-09 | 5.9e-09 | 6.2e-09 | 5.867E-09 | 3.512E-10 | 5.500E-09 | 6.200E-09 |
| 26062 | 1304 | T1304R | E | 4.5e-09 | 6.1e-09 | 4.6e-09 | 5.067E-09 | 8.963E-10 | 4.500E-09 | 6.100E-09 |
| 26063 | 1304 | T1304N | E | 7.1e-09 | 1.2e-08 | 6.3e-09 | 8.467E-09 | 3.086E-09 | 6.300E-09 | 1.200E-08 |
| 26064 | 1304 | T1304D | E | 3.2e-09 | 6.4e-09 | 7.9e-09 | 5.833E-09 | 2.401E-09 | 3.200E-09 | 7.900E-09 |
| 26065 | 1304 | T1304C | E | 1.2e-08 | 1.1e-08 | 1.9e-08 | 1.400E-08 | 4.359E-09 | 1.100E-08 | 1.900E-08 |
| 26066 | 1304 | T1304E | E | 1.9e-09 | 6.6e-09 | 8.5e-09 | 5.667E-09 | 3.398E-09 | 1.900E-09 | 8.500E-09 |
| 26067 | 1304 | T1304Q | E | 7.5e-09 | 1.4e-08 | 6.6e-09 | 9.367E-09 | 4.038E-09 | 6.600E-09 | 1.400E-08 |
| 26068 | 1304 | T1304G | E | 2.5e-09 | 4.2e-08 | 6.2e-09 | 1.690E-08 | 2.182E-08 | 2.500E-09 | 4.200E-08 |
| 26069 | 1304 | T1304H | E | 9.1e-09 | 5.3e-09 | 1e-08 | 8.133E-09 | 2.495E-09 | 5.300E-09 | 1.000E-08 |
| 26070 | 1304 | T1304I | E | 3.4e-09 | 3.3e-09 | 1.1e-08 | 5.900E-09 | 4.417E-09 | 3.300E-09 | 1.100E-08 |
| 26071 | 1304 | T1304L | E | 8.2e-09 | 6.5e-09 | 2.2e-08 | 1.223E-08 | 8.501E-09 | 6.500E-09 | 2.200E-08 |
| 26072 | 1304 | T1304K | E | 8.1e-09 | 1.2e-08 | 1e-08 | 1.003E-08 | 1.950E-09 | 8.100E-09 | 1.200E-08 |
| 26073 | 1304 | T1304M | E | 1.1e-08 | 8.7e-09 | 4.6e-09 | 8.100E-09 | 3.242E-09 | 4.600E-09 | 1.100E-08 |
| 26074 | 1304 | T1304F | E | 7.3e-09 | 6.7e-09 | 1.8e-08 | 1.067E-08 | 6.358E-09 | 6.700E-09 | 1.800E-08 |
| 26075 | 1304 | T1304P | E | 7e-09 | 8.9e-09 | 6.4e-09 | 7.433E-09 | 1.305E-09 | 6.400E-09 | 8.900E-09 |
| 26076 | 1304 | T1304S | E | 1.2e-08 | 7e-09 | 5.9e-09 | 8.300E-09 | 3.251E-09 | 5.900E-09 | 1.200E-08 |
| 26077 | 1304 | T1304T | E | 8.7e-09 | 1.6e-08 | 4.3e-09 | 9.667E-09 | 5.910E-09 | 4.300E-09 | 1.600E-08 |
| 26078 | 1304 | T1304W | E | 8.7e-09 | 5e-09 | 8.7e-09 | 7.467E-09 | 2.136E-09 | 5.000E-09 | 8.700E-09 |
| 26079 | 1304 | T1304Y | E | 3e-09 | 2.4e-09 | 6.3e-09 | 3.900E-09 | 2.100E-09 | 2.400E-09 | 6.300E-09 |
| 26080 | 1304 | T1304V | E | 1.5e-08 | 6.7e-09 | 1.3e-08 | 1.157E-08 | 4.332E-09 | 6.700E-09 | 1.500E-08 |
| 26081 | 1305 | Y1305A | E | 1.7e-08 | 1.4e-08 | 8.5e-09 | 1.317E-08 | 4.311E-09 | 8.500E-09 | 1.700E-08 |
| 26082 | 1305 | Y1305R | E | 1.8e-08 | 2.6e-09 | 6.9e-09 | 9.167E-09 | 7.946E-09 | 2.600E-09 | 1.800E-08 |
| 26083 | 1305 | Y1305N | E | 5e-09 | 8.8e-09 | 2e-08 | 1.127E-08 | 7.798E-09 | 5.000E-09 | 2.000E-08 |
| 26084 | 1305 | Y1305D | E | 8.6e-09 | 6.1e-09 | 1.2e-08 | 8.900E-09 | 2.961E-09 | 6.100E-09 | 1.200E-08 |
| 26085 | 1305 | Y1305C | E | 1.5e-08 | 2.1e-09 | 1.1e-08 | 9.367E-09 | 6.603E-09 | 2.100E-09 | 1.500E-08 |
| 26086 | 1305 | Y1305E | E | 1.8e-08 | 4.5e-09 | 4.6e-09 | 9.033E-09 | 7.766E-09 | 4.500E-09 | 1.800E-08 |
| 26087 | 1305 | Y1305Q | E | 7.5e-09 | 9.7e-09 | 6e-09 | 7.733E-09 | 1.861E-09 | 6.000E-09 | 9.700E-09 |
| 26088 | 1305 | Y1305G | E | 7e-09 | 9.2e-09 | 1.2e-08 | 9.400E-09 | 2.506E-09 | 7.000E-09 | 1.200E-08 |
| 26089 | 1305 | Y1305H | E | 4.1e-09 | 1.7e-08 | 3.4e-09 | 8.167E-09 | 7.658E-09 | 3.400E-09 | 1.700E-08 |
| 26090 | 1305 | Y1305I | E | 3.6e-09 | 3.6e-09 | 4.3e-09 | 3.833E-09 | 4.041E-10 | 3.600E-09 | 4.300E-09 |
| 26091 | 1305 | Y1305L | E | 3.6e-09 | 9.2e-09 | 1.3e-08 | 8.600E-09 | 4.729E-09 | 3.600E-09 | 1.300E-08 |
| 26092 | 1305 | Y1305K | E | 8.3e-09 | 2.9e-09 | 8.3e-09 | 6.500E-09 | 3.118E-09 | 2.900E-09 | 8.300E-09 |
| 26093 | 1305 | Y1305M | E | 1.8e-09 | 1.2e-08 | 6.4e-09 | 6.733E-09 | 5.108E-09 | 1.800E-09 | 1.200E-08 |
| 26094 | 1305 | Y1305F | E | 1.2e-08 | 6.5e-09 | 2.1e-09 | 6.867E-09 | 4.960E-09 | 2.100E-09 | 1.200E-08 |

| | | | | | | | | | | |
|---|---|---|---|---|---|---|---|---|---|---|
| 26095 | 1305 | Y1305P | E | 1e-08 | 4.3e-09 | 7.1e-09 | 7.133E-09 | 2.850E-09 | 4.300E-09 | 1.000E-08 |
| 26096 | 1305 | Y1305S | E | 3.1e-09 | 7.9e-09 | 1.4e-08 | 8.333E-09 | 5.463E-09 | 3.100E-09 | 1.400E-08 |
| 26097 | 1305 | Y1305T | E | 7.5e-09 | 4.7e-09 | 1.6e-08 | 9.400E-09 | 5.885E-09 | 4.700E-09 | 1.600E-08 |
| 26098 | 1305 | Y1305W | E | 6.4e-09 | 2.2e-09 | 1.2e-08 | 6.867E-09 | 4.917E-09 | 2.200E-09 | 1.200E-08 |
| 26099 | 1305 | Y1305Y | E | 4.5e-09 | 6.5e-09 | 4e-09 | 5.000E-09 | 1.323E-09 | 4.000E-09 | 6.500E-09 |
| 26100 | 1305 | Y1305V | E | 7e-09 | 3.8e-08 | 1e-08 | 1.833E-08 | 1.710E-08 | 7.000E-09 | 3.800E-08 |
| 26101 | 1306 | I1306A | E | 6.2e-09 | 6e-09 | 4.9e-09 | 5.700E-09 | 7.000E-10 | 4.900E-09 | 6.200E-09 |
| 26102 | 1306 | I1306R | E | 5.6e-09 | 1.7e-08 | 1.6e-08 | 1.287E-08 | 6.313E-09 | 5.600E-09 | 1.700E-08 |
| 26103 | 1306 | I1306N | E | 2.5e-09 | 2e-08 | 7.9e-09 | 1.013E-08 | 8.961E-09 | 2.500E-09 | 2.000E-08 |
| 26104 | 1306 | I1306D | E | 5e-09 | 2.6e-09 | 2.2e-09 | 3.267E-09 | 1.514E-09 | 2.200E-09 | 5.000E-09 |
| 26105 | 1306 | I1306C | E | 2.9e-09 | 8.4e-09 | 1.2e-08 | 7.767E-09 | 4.583E-09 | 2.900E-09 | 1.200E-08 |
| 26106 | 1306 | I1306E | E | 1.2e-08 | 1.8e-08 | 5.7e-09 | 1.190E-08 | 6.151E-09 | 5.700E-09 | 1.800E-08 |
| 26107 | 1306 | I1306Q | E | 4.6e-09 | 1.7e-09 | 4.4e-09 | 3.567E-09 | 1.620E-09 | 1.700E-09 | 4.600E-09 |
| 26108 | 1306 | I1306G | E | 2.5e-08 | 2.9e-09 | 5.6e-09 | 1.117E-08 | 1.206E-08 | 2.900E-09 | 2.500E-08 |
| 26109 | 1306 | I1306H | E | 1.5e-08 | 5.8e-09 | 7.6e-09 | 9.467E-09 | 4.876E-09 | 5.800E-09 | 1.500E-08 |
| 26110 | 1306 | I1306I | E | 6.9e-09 | 2.1e-08 | 9.5e-09 | 1.247E-08 | 7.504E-09 | 6.900E-09 | 2.100E-08 |
| 26111 | 1306 | I1306L | E | 4.2e-09 | 1.9e-09 | 1.1e-08 | 5.700E-09 | 4.732E-09 | 1.900E-09 | 1.100E-08 |
| 26112 | 1306 | I1306K | E | 5.5e-09 | 8.4e-09 | 6e-09 | 6.633E-09 | 1.550E-09 | 5.500E-09 | 8.400E-09 |
| 26113 | 1306 | I1306M | E | 1.2e-08 | 6.3e-09 | 1e-09 | 6.433E-09 | 5.501E-09 | 1.000E-09 | 1.200E-08 |
| 26114 | 1306 | I1306F | E | 3.1e-08 | 2.7e-09 | 7.1e-09 | 1.360E-08 | 1.523E-08 | 2.700E-09 | 3.100E-08 |
| 26115 | 1306 | I1306P | E | 4.9e-09 | 6.6e-09 | 6.4e-09 | 5.967E-09 | 9.292E-10 | 4.900E-09 | 6.600E-09 |
| 26116 | 1306 | I1306S | E | 6.5e-09 | 7.6e-09 | 2.2e-09 | 5.433E-09 | 2.854E-09 | 2.200E-09 | 7.600E-09 |
| 26117 | 1306 | I1306T | E | 5.9e-09 | 2.1e-09 | 4e-09 | 4.000E-09 | 1.900E-09 | 2.100E-09 | 5.900E-09 |
| 26118 | 1306 | I1306W | E | 1.4e-08 | 7.3e-09 | 4.2e-09 | 8.500E-09 | 5.009E-09 | 4.200E-09 | 1.400E-08 |
| 26119 | 1306 | I1306Y | E | 2.5e-09 | 6.3e-09 | 3.4e-09 | 4.067E-09 | 1.986E-09 | 2.500E-09 | 6.300E-09 |
| 26120 | 1306 | I1306V | E | 8.9e-09 | 1.5e-09 | 7.8e-09 | 6.067E-09 | 3.993E-09 | 1.500E-09 | 8.900E-09 |
| 26121 | 1307 | H1307A | E | 5.3e-09 | 7.5e-09 | 8.5e-09 | 7.100E-09 | 1.637E-09 | 5.300E-09 | 8.500E-09 |
| 26122 | 1307 | H1307R | E | 1.1e-08 | 8.4e-09 | 1.9e-09 | 7.100E-09 | 4.687E-09 | 1.900E-09 | 1.100E-08 |
| 26123 | 1307 | H1307N | E | 1.2e-08 | 9.2e-09 | 7.8e-09 | 9.667E-09 | 2.139E-09 | 7.800E-09 | 1.200E-08 |
| 26124 | 1307 | H1307D | E | 5.2e-09 | 6.9e-09 | 1.3e-08 | 8.367E-09 | 4.102E-09 | 5.200E-09 | 1.300E-08 |
| 26125 | 1307 | H1307C | E | 2.8e-09 | 1e-08 | 8.8e-09 | 7.200E-09 | 3.857E-09 | 2.800E-09 | 1.000E-08 |
| 26126 | 1307 | H1307E | E | 4.7e-09 | 5.7e-09 | 3.6e-09 | 4.667E-09 | 1.050E-09 | 3.600E-09 | 5.700E-09 |
| 26127 | 1307 | H1307Q | E | 3.3e-09 | 8.5e-09 | 3e-09 | 4.933E-09 | 3.092E-09 | 3.000E-09 | 8.500E-09 |
| 26128 | 1307 | H1307G | E | 5.6e-09 | 1.5e-08 | 8.1e-09 | 9.567E-09 | 4.869E-09 | 5.600E-09 | 1.500E-08 |
| 26129 | 1307 | H1307H | E | 8.1e-09 | 5.1e-09 | 7e-09 | 6.733E-09 | 1.518E-09 | 5.100E-09 | 8.100E-09 |
| 26130 | 1307 | H1307I | E | 9.7e-09 | 8.9e-09 | 2e-08 | 1.287E-08 | 6.191E-09 | 8.900E-09 | 2.000E-08 |
| 26131 | 1307 | H1307L | E | 4.9e-09 | 1.1e-08 | 4.9e-09 | 6.933E-09 | 3.522E-09 | 4.900E-09 | 1.100E-08 |
| 26132 | 1307 | H1307K | E | 1.8e-08 | 9.6e-09 | 6.3e-09 | 1.130E-08 | 6.032E-09 | 6.300E-09 | 1.800E-08 |
| 26133 | 1307 | H1307M | E | 4.6e-09 | 7.8e-09 | 9.3e-09 | 7.233E-09 | 2.401E-09 | 4.600E-09 | 9.300E-09 |
| 26134 | 1307 | H1307F | E | 1.7e-08 | 5.1e-09 | 2.1e-08 | 1.437E-08 | 8.271E-09 | 5.100E-09 | 2.100E-08 |
| 26135 | 1307 | H1307P | E | 2.2e-09 | 1.1e-08 | 7.3e-09 | 6.833E-09 | 4.419E-09 | 2.200E-09 | 1.100E-08 |
| 26136 | 1307 | H1307S | E | 1.2e-08 | 6.1e-09 | 2.2e-08 | 1.337E-08 | 8.038E-09 | 6.100E-09 | 2.200E-08 |
| 26137 | 1307 | H1307T | E | 4.2e-09 | 1.9e-08 | 2.6e-08 | 1.640E-08 | 1.113E-08 | 4.200E-09 | 2.600E-08 |
| 26138 | 1307 | H1307W | E | 7.9e-09 | 4.7e-09 | 3.6e-09 | 5.400E-09 | 2.234E-09 | 3.600E-09 | 7.900E-09 |
| 26139 | 1307 | H1307Y | E | 2.1e-08 | 9e-09 | 8.6e-09 | 1.287E-08 | 7.047E-09 | 8.600E-09 | 2.100E-08 |
| 26140 | 1307 | H1307V | E | 4.7e-09 | 3.6e-09 | 2.7e-09 | 3.667E-09 | 1.002E-09 | 2.700E-09 | 4.700E-09 |
| 26141 | 1308 | W1308A | E | 2e-09 | 8.8e-09 | 9.8e-09 | 6.867E-09 | 4.244E-09 | 2.000E-09 | 9.800E-09 |
| 26142 | 1308 | W1308R | E | 5.1e-09 | 3.1e-09 | 1.3e-08 | 7.067E-09 | 5.235E-09 | 3.100E-09 | 1.300E-08 |
| 26143 | 1308 | W1308N | E | 7.5e-09 | 3.4e-09 | 4.1e-09 | 5.000E-09 | 2.193E-09 | 3.400E-09 | 7.500E-09 |
| 26144 | 1308 | W1308D | E | 2.9e-09 | 1.3e-08 | 3.7e-09 | 6.533E-09 | 5.615E-09 | 2.900E-09 | 1.300E-08 |
| 26145 | 1308 | W1308C | E | 9e-09 | 8.6e-09 | 1e-08 | 9.200E-09 | 7.211E-10 | 8.600E-09 | 1.000E-08 |
| 26146 | 1308 | W1308E | E | 3.4e-09 | 6.4e-09 | 3.1e-09 | 4.300E-09 | 1.825E-09 | 3.100E-09 | 6.400E-09 |
| 26147 | 1308 | W1308Q | E | 6.7e-09 | 1.9e-09 | 1.3e-08 | 7.200E-09 | 5.567E-09 | 1.900E-09 | 1.300E-08 |
| 26148 | 1308 | W1308G | E | 2.6e-09 | 5.1e-09 | 8.1e-09 | 5.267E-09 | 2.754E-09 | 2.600E-09 | 8.100E-09 |
| 26149 | 1308 | W1308H | E | 4.4e-09 | 4e-09 | 2.2e-09 | 3.533E-09 | 1.172E-09 | 2.200E-09 | 4.400E-09 |
| 26150 | 1308 | W1308I | E | 1.7e-08 | 1.1e-08 | 3.9e-09 | 1.063E-08 | 6.558E-09 | 3.900E-09 | 1.700E-08 |
| 26151 | 1308 | W1308L | E | 1.2e-08 | 1.8e-09 | 1.5e-08 | 9.600E-09 | 6.920E-09 | 1.800E-09 | 1.500E-08 |
| 26152 | 1308 | W1308K | E | 1.1e-08 | 6.8e-09 | 7.1e-09 | 8.300E-09 | 2.343E-09 | 6.800E-09 | 1.100E-08 |
| 26153 | 1308 | W1308M | E | 3.2e-09 | 4.9e-09 | 2.1e-09 | 3.400E-09 | 1.411E-09 | 2.100E-09 | 4.900E-09 |
| 26154 | 1308 | W1308F | E | 5.5e-09 | 1.4e-08 | 2.1e-09 | 7.200E-09 | 6.129E-09 | 2.100E-09 | 1.400E-08 |
| 26155 | 1308 | W1308P | E | 7.7e-09 | 2.5e-09 | 1.1e-08 | 7.067E-09 | 4.285E-09 | 2.500E-09 | 1.100E-08 |
| 26156 | 1308 | W1308S | E | 9.9e-09 | 1.7e-09 | 4.6e-09 | 5.400E-09 | 4.158E-09 | 1.700E-09 | 9.900E-09 |
| 26157 | 1308 | W1308T | E | 1.7e-09 | 1.2e-08 | 3.6e-09 | 5.767E-09 | 5.481E-09 | 1.700E-09 | 1.200E-08 |
| 26158 | 1308 | W1308W | E | 5.2e-09 | 1.1e-08 | 1.4e-08 | 1.007E-08 | 4.474E-09 | 5.200E-09 | 1.400E-08 |
| 26159 | 1308 | W1308Y | E | 4.7e-09 | 1.7e-08 | 2.6e-09 | 8.100E-09 | 7.779E-09 | 2.600E-09 | 1.700E-08 |
| 26160 | 1308 | W1308V | E | 4.8e-09 | 1.2e-08 | 8.8e-09 | 8.533E-09 | 3.607E-09 | 4.800E-09 | 1.200E-08 |
| 26161 | 1309 | V1309A | E | 1.3e-08 | 1.3e-08 | 5.9e-09 | 1.063E-08 | 4.099E-09 | 5.900E-09 | 1.300E-08 |
| 26162 | 1309 | V1309R | E | 1.1e-08 | 2.5e-09 | 1.3e-08 | 8.833E-09 | 5.575E-09 | 2.500E-09 | 1.300E-08 |
| 26163 | 1309 | V1309N | E | 8.4e-09 | 1e-08 | 1.6e-08 | 1.147E-08 | 4.007E-09 | 8.400E-09 | 1.600E-08 |
| 26164 | 1309 | V1309D | E | 5.5e-09 | 9.5e-09 | 1.9e-08 | 1.133E-08 | 6.934E-09 | 5.500E-09 | 1.900E-08 |
| 26165 | 1309 | V1309C | E | 7.1e-09 | 8.2e-09 | 7e-09 | 7.433E-09 | 6.658E-10 | 7.000E-09 | 8.200E-09 |
| 26166 | 1309 | V1309E | E | 4.8e-09 | 5.5e-09 | 3.8e-09 | 4.700E-09 | 8.544E-10 | 3.800E-09 | 5.500E-09 |
| 26167 | 1309 | V1309Q | E | 3.3e-09 | 7e-09 | 9e-09 | 6.433E-09 | 2.892E-09 | 3.300E-09 | 9.000E-09 |
| 26168 | 1309 | V1309G | E | 6.4e-09 | 8e-09 | 5.5e-09 | 6.633E-09 | 1.266E-09 | 5.500E-09 | 8.000E-09 |
| 26169 | 1309 | V1309H | E | 6.9e-09 | 6e-09 | 4.4e-09 | 5.767E-09 | 1.266E-09 | 4.400E-09 | 6.900E-09 |
| 26170 | 1309 | V1309I | E | 4.6e-09 | 2.3e-09 | 1e-08 | 5.633E-09 | 3.953E-09 | 2.300E-09 | 1.000E-08 |
| 26171 | 1309 | V1309L | E | 1.6e-08 | 3.8e-09 | 1e-08 | 9.933E-09 | 6.100E-09 | 3.800E-09 | 1.600E-08 |
| 26172 | 1309 | V1309K | E | 7.1e-09 | 8.8e-09 | 2.6e-09 | 6.167E-09 | 3.204E-09 | 2.600E-09 | 8.800E-09 |
| 26173 | 1309 | V1309M | E | 4.1e-09 | 4e-09 | 2.8e-09 | 3.633E-09 | 7.234E-10 | 2.800E-09 | 4.100E-09 |
| 26174 | 1309 | V1309F | E | 1.8e-08 | 5.9e-09 | 7.4e-09 | 1.043E-08 | 6.596E-09 | 5.900E-09 | 1.800E-08 |
| 26175 | 1309 | V1309P | E | 2.5e-09 | 5.8e-09 | 5.8e-09 | 4.700E-09 | 1.905E-09 | 2.500E-09 | 5.800E-09 |
| 26176 | 1309 | V1309S | E | 9.3e-09 | 3.7e-09 | 8.7e-09 | 7.233E-09 | 3.075E-09 | 3.700E-09 | 9.300E-09 |
| 26177 | 1309 | V1309T | E | 2.8e-09 | 9.2e-09 | 5.5e-09 | 5.833E-09 | 3.213E-09 | 2.800E-09 | 9.200E-09 |
| 26178 | 1309 | V1309W | E | 3.5e-09 | 6.1e-09 | 1e-08 | 6.533E-09 | 3.272E-09 | 3.500E-09 | 1.000E-08 |
| 26179 | 1309 | V1309Y | E | 7e-09 | 6e-09 | 7.4e-09 | 6.800E-09 | 7.211E-10 | 6.000E-09 | 7.400E-09 |

| | | | | | | | | | | |
|---|---|---|---|---|---|---|---|---|---|---|
| 26180 | 1309 | V1309V | E | 2.6e-09 | 4.9e-09 | 8e-09 | 5.167E-09 | 2.710E-09 | 2.600E-09 | 8.000E-09 |
| 26181 | 1310 | R1310A | E | 7.1e-09 | 1e-08 | 6.5e-09 | 7.867E-09 | 1.872E-09 | 6.500E-09 | 1.000E-08 |
| 26182 | 1310 | R1310R | E | 1.3e-08 | 4.6e-09 | 1.6e-08 | 1.120E-08 | 5.909E-09 | 4.600E-09 | 1.600E-08 |
| 26183 | 1310 | R1310N | E | 2.8e-09 | 8.2e-09 | 4.3e-09 | 5.100E-09 | 2.787E-09 | 2.800E-09 | 8.200E-09 |
| 26184 | 1310 | R1310D | E | 4e-09 | 1.3e-08 | 7.4e-09 | 8.133E-09 | 4.545E-09 | 4.000E-09 | 1.300E-08 |
| 26185 | 1310 | R1310C | E | 5.2e-09 | 1.3e-08 | 3e-09 | 7.067E-09 | 5.255E-09 | 3.000E-09 | 1.300E-08 |
| 26186 | 1310 | R1310E | E | 1.2e-08 | 9.5e-09 | 2.8e-09 | 8.100E-09 | 4.757E-09 | 2.800E-09 | 1.200E-08 |
| 26187 | 1310 | R1310Q | E | 7.6e-09 | 2.4e-08 | 7.5e-09 | 1.303E-08 | 9.498E-09 | 7.500E-09 | 2.400E-08 |
| 26188 | 1310 | R1310G | E | 7e-09 | 8e-09 | 1.8e-08 | 1.100E-08 | 6.083E-09 | 7.000E-09 | 1.800E-08 |
| 26189 | 1310 | R1310H | E | 6.3e-09 | 6.4e-09 | 7.1e-09 | 6.600E-09 | 4.359E-10 | 6.300E-09 | 7.100E-09 |
| 26190 | 1310 | R1310I | E | 5.8e-09 | 5.7e-09 | 2.6e-08 | 1.250E-08 | 1.169E-08 | 5.700E-09 | 2.600E-08 |
| 26191 | 1310 | R1310L | E | 7.3e-09 | 8.3e-09 | 7.1e-09 | 7.567E-09 | 6.429E-10 | 7.100E-09 | 8.300E-09 |
| 26192 | 1310 | R1310K | E | 3.3e-09 | 1e-08 | 2.9e-09 | 5.400E-09 | 3.989E-09 | 2.900E-09 | 1.000E-08 |
| 26193 | 1310 | R1310M | E | 1.6e-08 | 2.1e-09 | 5.2e-09 | 7.767E-09 | 7.297E-09 | 2.100E-09 | 1.600E-08 |
| 26194 | 1310 | R1310F | E | 6.4e-09 | 5.9e-09 | 1.3e-08 | 8.433E-09 | 3.963E-09 | 5.900E-09 | 1.300E-08 |
| 26195 | 1310 | R1310P | E | 7.6e-09 | 5.5e-09 | 7e-09 | 6.700E-09 | 1.082E-09 | 5.500E-09 | 7.600E-09 |
| 26196 | 1310 | R1310S | E | 6e-09 | 7.1e-09 | 6.7e-09 | 6.600E-09 | 5.568E-10 | 6.000E-09 | 7.100E-09 |
| 26197 | 1310 | R1310T | E | 6.4e-09 | 1.1e-08 | 1e-08 | 9.133E-09 | 2.419E-09 | 6.400E-09 | 1.100E-08 |
| 26198 | 1310 | R1310W | E | 1.3e-08 | 4.4e-09 | 6.9e-09 | 8.100E-09 | 4.424E-09 | 4.400E-09 | 1.300E-08 |
| 26199 | 1310 | R1310Y | E | 4.5e-09 | 1.9e-09 | 6.2e-09 | 4.200E-09 | 2.166E-09 | 1.900E-09 | 6.200E-09 |
| 26200 | 1310 | R1310V | E | 1.5e-08 | 1.4e-08 | 6e-09 | 1.167E-08 | 4.933E-09 | 6.000E-09 | 1.500E-08 |
| 26201 | 1311 | Q1311A | E | 3.7e-09 | 5.6e-09 | 6.1e-09 | 5.133E-09 | 1.266E-09 | 3.700E-09 | 6.100E-09 |
| 26202 | 1311 | Q1311R | E | 5.6e-09 | 6.5e-09 | 4.1e-09 | 5.400E-09 | 1.212E-09 | 4.100E-09 | 6.500E-09 |
| 26203 | 1311 | Q1311N | E | 1.4e-08 | 1.7e-08 | 6.2e-09 | 1.240E-08 | 5.575E-09 | 6.200E-09 | 1.700E-08 |
| 26204 | 1311 | Q1311D | E | 4.7e-09 | 4.7e-09 | 1.9e-08 | 9.467E-09 | 8.256E-09 | 4.700E-09 | 1.900E-08 |
| 26205 | 1311 | Q1311C | E | 1.3e-08 | 1.7e-09 | 1.3e-08 | 9.233E-09 | 6.524E-09 | 1.700E-09 | 1.300E-08 |
| 26206 | 1311 | Q1311E | E | 2.6e-08 | 5.5e-09 | 9.4e-09 | 1.363E-08 | 1.089E-08 | 5.500E-09 | 2.600E-08 |
| 26207 | 1311 | Q1311Q | E | 2.8e-08 | 8.8e-09 | 5.5e-09 | 1.410E-08 | 1.215E-08 | 5.500E-09 | 2.800E-08 |
| 26208 | 1311 | Q1311G | E | 3e-09 | 7.8e-09 | 6.7e-09 | 5.833E-09 | 2.515E-09 | 3.000E-09 | 7.800E-09 |
| 26209 | 1311 | Q1311H | E | 3.9e-09 | 9.2e-09 | 6.9e-09 | 6.667E-09 | 2.658E-09 | 3.900E-09 | 9.200E-09 |
| 26210 | 1311 | Q1311I | E | 4.5e-09 | 6.4e-09 | 4.8e-09 | 5.233E-09 | 1.021E-09 | 4.500E-09 | 6.400E-09 |
| 26211 | 1311 | Q1311L | E | 1.6e-08 | 1.2e-08 | 5.6e-09 | 1.120E-08 | 5.246E-09 | 5.600E-09 | 1.600E-08 |
| 26212 | 1311 | Q1311K | E | 6.3e-09 | 3.8e-09 | 4.4e-09 | 4.833E-09 | 1.305E-09 | 3.800E-09 | 6.300E-09 |
| 26213 | 1311 | Q1311M | E | 5.3e-09 | 7.7e-09 | 2e-09 | 5.000E-09 | 2.862E-09 | 2.000E-09 | 7.700E-09 |
| 26214 | 1311 | Q1311F | E | 1.5e-08 | 1.7e-08 | 3.4e-09 | 1.180E-08 | 7.343E-09 | 3.400E-09 | 1.700E-08 |
| 26215 | 1311 | Q1311P | E | 3.3e-09 | 7.4e-09 | 5.9e-09 | 5.533E-09 | 2.074E-09 | 3.300E-09 | 7.400E-09 |
| 26216 | 1311 | Q1311S | E | 2e-08 | 8.1e-09 | 1e-08 | 1.270E-08 | 6.393E-09 | 8.100E-09 | 2.000E-08 |
| 26217 | 1311 | Q1311T | E | 4.9e-09 | 7.5e-09 | 1.2e-08 | 8.133E-09 | 3.592E-09 | 4.900E-09 | 1.200E-08 |
| 26218 | 1311 | Q1311W | E | 1.5e-08 | 9.8e-09 | 3.7e-09 | 9.500E-09 | 5.656E-09 | 3.700E-09 | 1.500E-08 |
| 26219 | 1311 | Q1311Y | E | 3e-08 | 1.3e-08 | 7e-09 | 1.667E-08 | 1.193E-08 | 7.000E-09 | 3.000E-08 |
| 26220 | 1311 | Q1311V | E | 7.2e-09 | 2.8e-09 | 4.7e-09 | 4.900E-09 | 2.207E-09 | 2.800E-09 | 7.200E-09 |
| 26221 | 1312 | A1312A | E | 6.5e-09 | 6e-09 | 3e-09 | 5.167E-09 | 1.893E-09 | 3.000E-09 | 6.500E-09 |
| 26222 | 1312 | A1312R | E | 7.3e-09 | 6e-09 | 1.8e-08 | 1.043E-08 | 6.585E-09 | 6.000E-09 | 1.800E-08 |
| 26223 | 1312 | A1312N | E | 1.1e-08 | 3.4e-09 | 6.6e-09 | 7.000E-09 | 3.816E-09 | 3.400E-09 | 1.100E-08 |
| 26224 | 1312 | A1312D | E | 1.8e-08 | 6.1e-09 | 1.8e-08 | 1.403E-08 | 6.870E-09 | 6.100E-09 | 1.800E-08 |
| 26225 | 1312 | A1312C | E | 1e-08 | 2.6e-09 | 7e-09 | 6.533E-09 | 3.722E-09 | 2.600E-09 | 1.000E-08 |
| 26226 | 1312 | A1312E | E | 1.6e-08 | 8.5e-09 | 1.8e-08 | 1.417E-08 | 5.008E-09 | 8.500E-09 | 1.800E-08 |
| 26227 | 1312 | A1312Q | E | 1.1e-08 | 4.7e-09 | 2.6e-09 | 6.100E-09 | 4.371E-09 | 2.600E-09 | 1.100E-08 |
| 26228 | 1312 | A1312G | E | 5.4e-09 | 1.5e-08 | 1.3e-09 | 7.233E-09 | 7.032E-09 | 1.300E-09 | 1.500E-08 |
| 26229 | 1312 | A1312H | E | 7.8e-09 | 1.1e-08 | 5e-09 | 7.933E-09 | 3.002E-09 | 5.000E-09 | 1.100E-08 |
| 26230 | 1312 | A1312I | E | 7.2e-09 | 6.2e-09 | 8.1e-09 | 7.167E-09 | 9.504E-10 | 6.200E-09 | 8.100E-09 |
| 26231 | 1312 | A1312L | E | 7.5e-09 | 1.4e-08 | 9.8e-09 | 1.043E-08 | 3.296E-09 | 7.500E-09 | 1.400E-08 |
| 26232 | 1312 | A1312K | E | 8.9e-09 | 2.6e-09 | 1e-08 | 7.167E-09 | 3.993E-09 | 2.600E-09 | 1.000E-08 |
| 26233 | 1312 | A1312M | E | 4.9e-09 | 8.3e-09 | 2.1e-08 | 1.140E-08 | 8.486E-09 | 4.900E-09 | 2.100E-08 |
| 26234 | 1312 | A1312F | E | 1e-08 | 2e-09 | 7.3e-09 | 6.433E-09 | 4.070E-09 | 2.000E-09 | 1.000E-08 |
| 26235 | 1312 | A1312P | E | 7.8e-09 | 7.7e-09 | 1.5e-08 | 1.017E-08 | 4.186E-09 | 7.700E-09 | 1.500E-08 |
| 26236 | 1312 | A1312S | E | 1.6e-08 | 1.7e-08 | 9.3e-09 | 1.410E-08 | 4.187E-09 | 9.300E-09 | 1.700E-08 |
| 26237 | 1312 | A1312T | E | 8e-09 | 1.6e-08 | 1.3e-08 | 1.233E-08 | 4.041E-09 | 8.000E-09 | 1.600E-08 |
| 26238 | 1312 | A1312W | E | 5.2e-09 | 4.7e-09 | 1e-08 | 6.633E-09 | 2.926E-09 | 4.700E-09 | 1.000E-08 |
| 26239 | 1312 | A1312Y | E | 7e-09 | 1.6e-09 | 1.3e-08 | 7.200E-09 | 5.703E-09 | 1.600E-09 | 1.300E-08 |
| 26240 | 1312 | A1312V | E | 7.2e-09 | 3.6e-09 | 9.5e-09 | 6.767E-09 | 2.974E-09 | 3.600E-09 | 9.500E-09 |
| 26241 | 1313 | P1313A | E | 2.3e-09 | 5.4e-09 | 6.1e-09 | 4.600E-09 | 2.022E-09 | 2.300E-09 | 6.100E-09 |
| 26242 | 1313 | P1313R | E | 1.2e-08 | 1.1e-08 | 7.1e-09 | 1.003E-08 | 2.589E-09 | 7.100E-09 | 1.200E-08 |
| 26243 | 1313 | P1313N | E | 2.9e-09 | 2.7e-09 | 1.3e-08 | 6.200E-09 | 5.890E-09 | 2.700E-09 | 1.300E-08 |
| 26244 | 1313 | P1313D | E | 4.4e-09 | 9.2e-09 | 2.1e-08 | 1.153E-08 | 8.542E-09 | 4.400E-09 | 2.100E-08 |
| 26245 | 1313 | P1313C | E | 6.9e-09 | 1e-08 | 6.9e-09 | 7.933E-09 | 1.790E-09 | 6.900E-09 | 1.000E-08 |
| 26246 | 1313 | P1313E | E | 8e-09 | 6.8e-09 | 7.7e-09 | 7.500E-09 | 6.245E-10 | 6.800E-09 | 8.000E-09 |
| 26247 | 1313 | P1313Q | E | 8.7e-09 | 1.2e-09 | 4.3e-09 | 4.733E-09 | 3.769E-09 | 1.200E-09 | 8.700E-09 |
| 26248 | 1313 | P1313G | E | 6.3e-09 | 3.8e-09 | 9.2e-09 | 6.433E-09 | 2.702E-09 | 3.800E-09 | 9.200E-09 |
| 26249 | 1313 | P1313H | E | 9.1e-09 | 1.7e-08 | 9.3e-09 | 1.180E-08 | 4.504E-09 | 9.100E-09 | 1.700E-08 |
| 26250 | 1313 | P1313I | E | 5.8e-09 | 3.9e-09 | 3.7e-09 | 4.467E-09 | 1.159E-09 | 3.700E-09 | 5.800E-09 |
| 26251 | 1313 | P1313L | E | 6.2e-09 | 1.3e-08 | 7.6e-09 | 8.933E-09 | 3.591E-09 | 6.200E-09 | 1.300E-08 |
| 26252 | 1313 | P1313K | E | 8.4e-09 | 1.6e-08 | 1.2e-08 | 1.213E-08 | 3.802E-09 | 8.400E-09 | 1.600E-08 |
| 26253 | 1313 | P1313M | E | 1e-08 | 5.9e-09 | 3.4e-09 | 6.433E-09 | 3.332E-09 | 3.400E-09 | 1.000E-08 |
| 26254 | 1313 | P1313F | E | 2.3e-09 | 2.4e-08 | 5.2e-09 | 1.050E-08 | 1.178E-08 | 2.300E-09 | 2.400E-08 |
| 26255 | 1313 | P1313P | E | 7.1e-09 | 5.8e-09 | 8.6e-09 | 7.167E-09 | 1.401E-09 | 5.800E-09 | 8.600E-09 |
| 26256 | 1313 | P1313S | E | 1e-08 | 1.5e-08 | 7.1e-09 | 1.070E-08 | 3.996E-09 | 7.100E-09 | 1.500E-08 |
| 26257 | 1313 | P1313T | E | 1.9e-09 | 1.2e-08 | 1.2e-08 | 8.633E-09 | 5.831E-09 | 1.900E-09 | 1.200E-08 |
| 26258 | 1313 | P1313W | E | 6.4e-09 | 1.1e-08 | 1.4e-08 | 1.047E-08 | 3.828E-09 | 6.400E-09 | 1.400E-08 |
| 26259 | 1313 | P1313Y | E | 7.3e-09 | 3.2e-09 | 1.2e-08 | 7.500E-09 | 4.403E-09 | 3.200E-09 | 1.200E-08 |
| 26260 | 1313 | P1313V | E | 1.1e-08 | 1.6e-08 | 1.3e-08 | 1.333E-08 | 2.517E-09 | 1.100E-08 | 1.600E-08 |
| 26261 | 1314 | G1314A | E | 1.7e-08 | 2.4e-09 | 6.8e-09 | 8.733E-09 | 7.490E-09 | 2.400E-09 | 1.700E-08 |
| 26262 | 1314 | G1314R | E | 3.6e-09 | 6.4e-09 | 7.7e-09 | 5.900E-09 | 2.095E-09 | 3.600E-09 | 7.700E-09 |
| 26263 | 1314 | G1314N | E | 4.5e-09 | 2.1e-08 | 5.4e-09 | 1.030E-08 | 9.277E-09 | 4.500E-09 | 2.100E-08 |
| 26264 | 1314 | G1314D | E | 1.7e-08 | 1e-08 | 4e-09 | 1.033E-08 | 6.506E-09 | 4.000E-09 | 1.700E-08 |

| | | | | | | | | | | |
|---|---|---|---|---|---|---|---|---|---|---|
| 26265 | 1314 | G1314C | E | 1e-08 | 1.5e-08 | 3.5e-09 | 9.500E-09 | 5.766E-09 | 3.500E-09 | 1.500E-08 |
| 26266 | 1314 | G1314E | E | 3.4e-09 | 3.3e-09 | 7.2e-09 | 4.633E-09 | 2.223E-09 | 3.300E-09 | 7.200E-09 |
| 26267 | 1314 | G1314Q | E | 2.6e-09 | 4.8e-09 | 8.2e-09 | 5.200E-09 | 2.821E-09 | 2.600E-09 | 8.200E-09 |
| 26268 | 1314 | G1314G | E | 4.7e-09 | 8.4e-09 | 5.7e-09 | 6.267E-09 | 1.914E-09 | 4.700E-09 | 8.400E-09 |
| 26269 | 1314 | G1314H | E | 9.9e-09 | 9.9e-09 | 6.6e-09 | 8.800E-09 | 1.905E-09 | 6.600E-09 | 9.900E-09 |
| 26270 | 1314 | G1314I | E | 1e-08 | 2.8e-09 | 1.8e-09 | 4.867E-09 | 4.474E-09 | 1.800E-09 | 1.000E-08 |
| 26271 | 1314 | G1314L | E | 5.3e-09 | 8e-09 | 5e-09 | 6.100E-09 | 1.652E-09 | 5.000E-09 | 8.000E-09 |
| 26272 | 1314 | G1314K | E | 1.3e-08 | 2.8e-09 | 1.2e-08 | 9.267E-09 | 5.623E-09 | 2.800E-09 | 1.300E-08 |
| 26273 | 1314 | G1314M | E | 1.2e-08 | 9.3e-09 | 7.5e-09 | 9.600E-09 | 2.265E-09 | 7.500E-09 | 1.200E-08 |
| 26274 | 1314 | G1314F | E | 7.5e-09 | 4.7e-09 | 4e-09 | 5.400E-09 | 1.852E-09 | 4.000E-09 | 7.500E-09 |
| 26275 | 1314 | G1314P | E | 5.1e-09 | 7e-09 | 1e-08 | 7.367E-09 | 2.470E-09 | 5.100E-09 | 1.000E-08 |
| 26276 | 1314 | G1314S | E | 1.4e-08 | 2.4e-09 | 8.1e-09 | 8.167E-09 | 5.800E-09 | 2.400E-09 | 1.400E-08 |
| 26277 | 1314 | G1314T | E | 4.9e-09 | 3.8e-09 | 6.2e-09 | 4.967E-09 | 1.201E-09 | 3.800E-09 | 6.200E-09 |
| 26278 | 1314 | G1314W | E | 2.5e-09 | 5.9e-09 | 5.5e-09 | 4.633E-09 | 1.858E-09 | 2.500E-09 | 5.900E-09 |
| 26279 | 1314 | G1314Y | E | 6.2e-09 | 6.9e-09 | 7.9e-09 | 7.000E-09 | 8.544E-10 | 6.200E-09 | 7.900E-09 |
| 26280 | 1314 | G1314V | E | 4.3e-09 | 4.8e-09 | 3.3e-09 | 4.133E-09 | 7.638E-10 | 3.300E-09 | 4.800E-09 |
| 26281 | 1315 | K1315A | E | 9.1e-09 | 8.1e-09 | 9.1e-09 | 8.767E-09 | 5.774E-10 | 8.100E-09 | 9.100E-09 |
| 26282 | 1315 | K1315R | E | 3e-09 | 2.8e-09 | 9.6e-09 | 5.133E-09 | 3.870E-09 | 2.800E-09 | 9.600E-09 |
| 26283 | 1315 | K1315N | E | 6e-09 | 2.8e-08 | 9.3e-09 | 1.443E-08 | 1.186E-08 | 6.000E-09 | 2.800E-08 |
| 26284 | 1315 | K1315D | E | 1.6e-08 | 6.2e-09 | 1.8e-08 | 1.340E-08 | 6.315E-09 | 6.200E-09 | 1.800E-08 |
| 26285 | 1315 | K1315C | E | 1.1e-08 | 5.7e-09 | 4.6e-09 | 7.100E-09 | 3.422E-09 | 4.600E-09 | 1.100E-08 |
| 26286 | 1315 | K1315E | E | 1.2e-08 | 8.7e-09 | 3.6e-09 | 8.100E-09 | 4.232E-09 | 3.600E-09 | 1.200E-08 |
| 26287 | 1315 | K1315Q | E | 4.8e-09 | 9.9e-09 | 4.2e-09 | 6.300E-09 | 3.132E-09 | 4.200E-09 | 9.900E-09 |
| 26288 | 1315 | K1315G | E | 3.7e-09 | 4.7e-09 | 2.2e-09 | 3.533E-09 | 1.258E-09 | 2.200E-09 | 4.700E-09 |
| 26289 | 1315 | K1315H | E | 1.3e-08 | 8.5e-09 | 3.6e-09 | 8.367E-09 | 4.701E-09 | 3.600E-09 | 1.300E-08 |
| 26290 | 1315 | K1315I | E | 1.7e-08 | 4.9e-09 | 2.3e-09 | 8.067E-09 | 7.845E-09 | 2.300E-09 | 1.700E-08 |
| 26291 | 1315 | K1315L | E | 9e-09 | 6e-09 | 1.7e-08 | 1.067E-08 | 5.686E-09 | 6.000E-09 | 1.700E-08 |
| 26292 | 1315 | K1315K | E | 7.6e-09 | 1.2e-08 | 2.5e-09 | 7.367E-09 | 4.754E-09 | 2.500E-09 | 1.200E-08 |
| 26293 | 1315 | K1315M | E | 4.3e-09 | 4.5e-09 | 3.4e-09 | 4.067E-09 | 5.859E-10 | 3.400E-09 | 4.500E-09 |
| 26294 | 1315 | K1315F | E | 5.6e-09 | 9e-09 | 4.3e-09 | 6.300E-09 | 2.427E-09 | 4.300E-09 | 9.000E-09 |
| 26295 | 1315 | K1315P | E | 7.8e-09 | 1.4e-08 | 3.7e-09 | 8.500E-09 | 5.186E-09 | 3.700E-09 | 1.400E-08 |
| 26296 | 1315 | K1315S | E | 1.7e-08 | 1.8e-08 | 1.2e-08 | 1.567E-08 | 3.215E-09 | 1.200E-08 | 1.800E-08 |
| 26297 | 1315 | K1315T | E | 3.7e-09 | 2.1e-08 | 5.7e-09 | 1.013E-08 | 9.464E-09 | 3.700E-09 | 2.100E-08 |
| 26298 | 1315 | K1315W | E | 1.2e-09 | 6.2e-09 | 8.8e-09 | 5.400E-09 | 3.863E-09 | 1.200E-09 | 8.800E-09 |
| 26299 | 1315 | K1315Y | E | 6.1e-09 | 6.9e-09 | 1.8e-08 | 1.033E-08 | 6.652E-09 | 6.100E-09 | 1.800E-08 |
| 26300 | 1315 | K1315V | E | 6.6e-09 | 6.4e-09 | 4.6e-09 | 5.867E-09 | 1.102E-09 | 4.600E-09 | 6.600E-09 |
| 26301 | 1316 | G1316A | E | 1.5e-08 | 5.3e-09 | 8.1e-09 | 9.467E-09 | 4.992E-09 | 5.300E-09 | 1.500E-08 |
| 26302 | 1316 | G1316R | E | 7.5e-09 | 4.2e-09 | 5.1e-09 | 5.600E-09 | 1.706E-09 | 4.200E-09 | 7.500E-09 |
| 26303 | 1316 | G1316N | E | 1e-08 | 4.6e-09 | 2.4e-09 | 5.667E-09 | 3.911E-09 | 2.400E-09 | 1.000E-08 |
| 26304 | 1316 | G1316D | E | 3.6e-09 | 1.5e-08 | 7.1e-09 | 8.567E-09 | 5.840E-09 | 3.600E-09 | 1.500E-08 |
| 26305 | 1316 | G1316C | E | 3.2e-09 | 5.2e-09 | 2.7e-08 | 1.180E-08 | 1.320E-08 | 3.200E-09 | 2.700E-08 |
| 26306 | 1316 | G1316E | E | 2.7e-08 | 3.7e-09 | 3.3e-09 | 1.133E-08 | 1.357E-08 | 3.300E-09 | 2.700E-08 |
| 26307 | 1316 | G1316Q | E | 1.4e-08 | 9e-09 | 6.5e-09 | 9.833E-09 | 3.819E-09 | 6.500E-09 | 1.400E-08 |
| 26308 | 1316 | G1316G | E | 7.2e-09 | 4.9e-09 | 2.1e-09 | 4.733E-09 | 2.554E-09 | 2.100E-09 | 7.200E-09 |
| 26309 | 1316 | G1316H | E | 2.6e-09 | 1.1e-08 | 6.5e-09 | 6.700E-09 | 4.204E-09 | 2.600E-09 | 1.100E-08 |
| 26310 | 1316 | G1316I | E | 1.1e-08 | 1.4e-08 | 8.1e-09 | 1.103E-08 | 2.950E-09 | 8.100E-09 | 1.400E-08 |
| 26311 | 1316 | G1316L | E | 9.9e-09 | 6e-09 | 6.5e-09 | 7.467E-09 | 2.122E-09 | 6.000E-09 | 9.900E-09 |
| 26312 | 1316 | G1316K | E | 1.4e-08 | 9.5e-09 | 1.7e-09 | 8.400E-09 | 6.223E-09 | 1.700E-09 | 1.400E-08 |
| 26313 | 1316 | G1316M | E | 1.3e-08 | 1.6e-08 | 5.3e-09 | 1.143E-08 | 5.519E-09 | 5.300E-09 | 1.600E-08 |
| 26314 | 1316 | G1316F | E | 5.2e-09 | 4.1e-09 | 8.4e-10 | 3.380E-09 | 2.267E-09 | 8.400E-10 | 5.200E-09 |
| 26315 | 1316 | G1316P | E | 5.6e-09 | 1.1e-07 | 4.8e-09 | 4.013E-08 | 6.051E-08 | 4.800E-09 | 1.100E-07 |
| 26316 | 1316 | G1316S | E | 1.1e-08 | 7.1e-09 | 1.2e-08 | 1.003E-08 | 2.589E-09 | 7.100E-09 | 1.200E-08 |
| 26317 | 1316 | G1316T | E | 6.5e-09 | 9.8e-09 | 8.8e-09 | 8.367E-09 | 1.692E-09 | 6.500E-09 | 9.800E-09 |
| 26318 | 1316 | G1316W | E | 1e-08 | 4e-09 | 4.6e-09 | 6.200E-09 | 3.305E-09 | 4.000E-09 | 1.000E-08 |
| 26319 | 1316 | G1316Y | E | 1.4e-08 | 3.3e-08 | 4e-08 | 2.900E-08 | 1.345E-08 | 1.400E-08 | 4.000E-08 |
| 26320 | 1316 | G1316V | E | 9.7e-09 | 1.7e-08 | 7e-09 | 1.123E-08 | 5.173E-09 | 7.000E-09 | 1.700E-08 |
| 26321 | 1317 | L1317A | E | 8.9e-09 | 9.3e-09 | 2.7e-09 | 6.967E-09 | 3.700E-09 | 2.700E-09 | 9.300E-09 |
| 26322 | 1317 | L1317R | E | 1.5e-08 | 1.6e-08 | 8.2e-09 | 1.307E-08 | 4.244E-09 | 8.200E-09 | 1.600E-08 |
| 26323 | 1317 | L1317N | E | 7.2e-09 | 1.4e-08 | 7.5e-09 | 9.567E-09 | 3.842E-09 | 7.200E-09 | 1.400E-08 |
| 26324 | 1317 | L1317D | E | 7.6e-09 | 2.9e-09 | 6.3e-09 | 5.600E-09 | 2.427E-09 | 2.900E-09 | 7.600E-09 |
| 26325 | 1317 | L1317C | E | 6.8e-09 | 1.6e-08 | 1.2e-08 | 1.160E-08 | 4.613E-09 | 6.800E-09 | 1.600E-08 |
| 26326 | 1317 | L1317E | E | 1.4e-08 | 5.5e-09 | 7.3e-09 | 8.933E-09 | 4.479E-09 | 5.500E-09 | 1.400E-08 |
| 26327 | 1317 | L1317Q | E | 1.3e-08 | 8e-09 | 1.9e-08 | 1.333E-08 | 5.508E-09 | 8.000E-09 | 1.900E-08 |
| 26328 | 1317 | L1317G | E | 3.1e-09 | 5.6e-09 | 9.2e-09 | 5.967E-09 | 3.066E-09 | 3.100E-09 | 9.200E-09 |
| 26329 | 1317 | L1317H | E | 3.8e-09 | 8.4e-09 | 8.9e-09 | 7.033E-09 | 2.811E-09 | 3.800E-09 | 8.900E-09 |
| 26330 | 1317 | L1317I | E | 8.2e-09 | 1.2e-08 | 7.4e-09 | 9.200E-09 | 2.458E-09 | 7.400E-09 | 1.200E-08 |
| 26331 | 1317 | L1317L | E | 6.5e-09 | 1.4e-08 | 6.1e-09 | 8.867E-09 | 4.450E-09 | 6.100E-09 | 1.400E-08 |
| 26332 | 1317 | L1317K | E | 5.5e-09 | 5.3e-09 | 1.2e-08 | 7.600E-09 | 3.812E-09 | 5.300E-09 | 1.200E-08 |
| 26333 | 1317 | L1317M | E | 1.4e-08 | 1.7e-08 | 5.6e-09 | 1.220E-08 | 5.909E-09 | 5.600E-09 | 1.700E-08 |
| 26334 | 1317 | L1317F | E | 1e-08 | 3e-08 | 1.3e-08 | 1.767E-08 | 1.079E-08 | 1.000E-08 | 3.000E-08 |
| 26335 | 1317 | L1317P | E | 1.8e-08 | 1.6e-08 | 1.7e-08 | 1.700E-08 | 1.000E-09 | 1.600E-08 | 1.800E-08 |
| 26336 | 1317 | L1317S | E | 8.9e-09 | 1.1e-08 | 5.6e-09 | 8.500E-09 | 2.722E-09 | 5.600E-09 | 1.100E-08 |
| 26337 | 1317 | L1317T | E | 5.7e-09 | 2.9e-09 | 1e-08 | 6.200E-09 | 3.576E-09 | 2.900E-09 | 1.000E-08 |
| 26338 | 1317 | L1317W | E | 1.6e-08 | 8.2e-09 | 9.7e-09 | 1.130E-08 | 4.139E-09 | 8.200E-09 | 1.600E-08 |
| 26339 | 1317 | L1317Y | E | 8.3e-09 | 7e-09 | 1.8e-08 | 1.110E-08 | 6.011E-09 | 7.000E-09 | 1.800E-08 |
| 26340 | 1317 | L1317V | E | 5.6e-09 | 5.1e-09 | 1.2e-08 | 7.567E-09 | 3.848E-09 | 5.100E-09 | 1.200E-08 |
| 26341 | 1318 | E1318A | E | 6.3e-09 | 3.9e-09 | 9e-09 | 6.400E-09 | 2.551E-09 | 3.900E-09 | 9.000E-09 |
| 26342 | 1318 | E1318R | E | 3.7e-09 | 6.9e-09 | 3.3e-09 | 4.633E-09 | 1.973E-09 | 3.300E-09 | 6.900E-09 |
| 26343 | 1318 | E1318N | E | 5.4e-09 | 3.8e-09 | 6.8e-09 | 5.333E-09 | 1.501E-09 | 3.800E-09 | 6.800E-09 |
| 26344 | 1318 | E1318D | E | 4.8e-09 | 1.2e-08 | 4.3e-09 | 7.033E-09 | 4.309E-09 | 4.300E-09 | 1.200E-08 |
| 26345 | 1318 | E1318C | E | 4.8e-09 | 6e-09 | 3.4e-09 | 4.733E-09 | 1.301E-09 | 3.400E-09 | 6.000E-09 |
| 26346 | 1318 | E1318E | E | 1.2e-08 | 1.1e-08 | 1.3e-08 | 1.200E-08 | 1.000E-09 | 1.100E-08 | 1.300E-08 |
| 26347 | 1318 | E1318Q | E | 5.1e-09 | 4.7e-09 | 5e-09 | 4.933E-09 | 2.082E-10 | 4.700E-09 | 5.100E-09 |
| 26348 | 1318 | E1318G | E | 7.8e-09 | 4.8e-09 | 1.1e-08 | 7.867E-09 | 3.101E-09 | 4.800E-09 | 1.100E-08 |
| 26349 | 1318 | E1318H | E | 9.9e-09 | 3.9e-09 | 6.1e-09 | 6.633E-09 | 3.035E-09 | 3.900E-09 | 9.900E-09 |

| | | | | | | | | | | |
|---|---|---|---|---|---|---|---|---|---|---|
| 26350 | 1318 | E1318I | E | 4e-09 | 5.4e-09 | 6.6e-09 | 5.333E-09 | 1.301E-09 | 4.000E-09 | 6.600E-09 |
| 26351 | 1318 | E1318L | E | 4.9e-09 | 1.1e-08 | 2.4e-08 | 1.330E-08 | 9.756E-09 | 4.900E-09 | 2.400E-08 |
| 26352 | 1318 | E1318K | E | 1.4e-08 | 5.7e-09 | 7.3e-09 | 9.000E-09 | 4.403E-09 | 5.700E-09 | 1.400E-08 |
| 26353 | 1318 | E1318M | E | 1e-08 | 5.7e-09 | 1.4e-08 | 9.900E-09 | 4.151E-09 | 5.700E-09 | 1.400E-08 |
| 26354 | 1318 | E1318F | E | 1.4e-08 | 1.5e-08 | 1.4e-08 | 1.433E-08 | 5.774E-10 | 1.400E-08 | 1.500E-08 |
| 26355 | 1318 | E1318P | E | 4e-09 | 4.6e-09 | 6.9e-09 | 5.167E-09 | 1.531E-09 | 4.000E-09 | 6.900E-09 |
| 26356 | 1318 | E1318S | E | 7e-09 | 2.1e-08 | 1e-09 | 9.667E-09 | 1.026E-08 | 1.000E-09 | 2.100E-08 |
| 26357 | 1318 | E1318T | E | 1.1e-08 | 5.4e-09 | 5.1e-09 | 7.167E-09 | 3.323E-09 | 5.100E-09 | 1.100E-08 |
| 26358 | 1318 | E1318W | E | 8.9e-09 | 6.3e-09 | 5e-09 | 6.733E-09 | 1.986E-09 | 5.000E-09 | 8.900E-09 |
| 26359 | 1318 | E1318Y | E | 4.4e-09 | 4.4e-09 | 6.2e-09 | 5.000E-09 | 1.039E-09 | 4.400E-09 | 6.200E-09 |
| 26360 | 1318 | E1318V | E | 2.5e-09 | 2.1e-09 | 1.2e-08 | 5.533E-09 | 5.604E-09 | 2.100E-09 | 1.200E-08 |
| 26361 | 1319 | W1319A | E | 6.4e-09 | 5.4e-09 | 6.4e-09 | 6.067E-09 | 5.774E-10 | 5.400E-09 | 6.400E-09 |
| 26362 | 1319 | W1319R | E | 1.7e-08 | 1.1e-08 | 3.7e-09 | 1.057E-08 | 6.661E-09 | 3.700E-09 | 1.700E-08 |
| 26363 | 1319 | W1319N | E | 9.8e-09 | 8.1e-09 | 9.5e-09 | 9.133E-09 | 9.074E-10 | 8.100E-09 | 9.800E-09 |
| 26364 | 1319 | W1319D | E | 4.8e-09 | 2.9e-09 | 5.3e-08 | 2.023E-08 | 2.839E-08 | 2.900E-09 | 5.300E-08 |
| 26365 | 1319 | W1319C | E | 1.9e-08 | 7.6e-09 | 6.9e-09 | 1.117E-08 | 6.793E-09 | 6.900E-09 | 1.900E-08 |
| 26366 | 1319 | W1319E | E | 1.3e-08 | 7.7e-09 | 1e-08 | 1.023E-08 | 2.658E-09 | 7.700E-09 | 1.300E-08 |
| 26367 | 1319 | W1319Q | E | 7.9e-09 | 1e-08 | 3.3e-09 | 7.067E-09 | 3.427E-09 | 3.300E-09 | 1.000E-08 |
| 26368 | 1319 | W1319G | E | 6.3e-09 | 7.4e-09 | 4e-09 | 5.900E-09 | 1.735E-09 | 4.000E-09 | 7.400E-09 |
| 26369 | 1319 | W1319H | E | 2.3e-09 | 7.1e-09 | 9.7e-09 | 6.367E-09 | 3.754E-09 | 2.300E-09 | 9.700E-09 |
| 26370 | 1319 | W1319I | E | 9.4e-09 | 3.8e-09 | 8e-09 | 7.067E-09 | 2.914E-09 | 3.800E-09 | 9.400E-09 |
| 26371 | 1319 | W1319L | E | 5.8e-09 | 5.7e-09 | 1.8e-08 | 9.833E-09 | 7.073E-09 | 5.700E-09 | 1.800E-08 |
| 26372 | 1319 | W1319K | E | 6.5e-09 | 3.1e-09 | 2.5e-09 | 4.033E-09 | 2.157E-09 | 2.500E-09 | 6.500E-09 |
| 26373 | 1319 | W1319M | E | 6.8e-09 | 2.5e-09 | 5.5e-09 | 4.933E-09 | 2.205E-09 | 2.500E-09 | 6.800E-09 |
| 26374 | 1319 | W1319F | E | 2.8e-09 | 9.8e-09 | 4.4e-09 | 5.667E-09 | 3.668E-09 | 2.800E-09 | 9.800E-09 |
| 26375 | 1319 | W1319P | E | 9.1e-09 | 7.6e-09 | 5.9e-09 | 7.533E-09 | 1.601E-09 | 5.900E-09 | 9.100E-09 |
| 26376 | 1319 | W1319S | E | 5.8e-09 | 1.5e-08 | 8.9e-09 | 9.900E-09 | 4.681E-09 | 5.800E-09 | 1.500E-08 |
| 26377 | 1319 | W1319T | E | 7.9e-09 | 2.6e-09 | 6.6e-09 | 5.700E-09 | 2.762E-09 | 2.600E-09 | 7.900E-09 |
| 26378 | 1319 | W1319W | E | 1.1e-08 | 6.8e-09 | 1.3e-09 | 6.367E-09 | 4.864E-09 | 1.300E-09 | 1.100E-08 |
| 26379 | 1319 | W1319Y | E | 9.1e-09 | 1.2e-08 | 1.4e-08 | 1.170E-08 | 2.464E-09 | 9.100E-09 | 1.400E-08 |
| 26380 | 1319 | W1319V | E | 7.1e-09 | 7.7e-09 | 3.2e-09 | 6.000E-09 | 2.443E-09 | 3.200E-09 | 7.700E-09 |
| 26381 | 1320 | V1320A | E | 5.2e-09 | 3.6e-09 | 6.3e-09 | 5.033E-09 | 1.358E-09 | 3.600E-09 | 6.300E-09 |
| 26382 | 1320 | V1320R | E | 1.1e-08 | 9.3e-09 | 4e-09 | 8.100E-09 | 3.651E-09 | 4.000E-09 | 1.100E-08 |
| 26383 | 1320 | V1320N | E | 3.2e-09 | 2.6e-08 | 9.5e-09 | 1.290E-08 | 1.177E-08 | 3.200E-09 | 2.600E-08 |
| 26384 | 1320 | V1320D | E | 6.1e-09 | 6.5e-09 | 3e-09 | 5.200E-09 | 1.916E-09 | 3.000E-09 | 6.500E-09 |
| 26385 | 1320 | V1320C | E | 8.9e-09 | 6.9e-09 | 2.8e-09 | 6.200E-09 | 3.110E-09 | 2.800E-09 | 8.900E-09 |
| 26386 | 1320 | V1320E | E | 6e-09 | 5.7e-09 | 1.3e-08 | 8.233E-09 | 4.131E-09 | 5.700E-09 | 1.300E-08 |
| 26387 | 1320 | V1320Q | E | 7.4e-09 | 8.9e-09 | 5.2e-09 | 7.167E-09 | 1.861E-09 | 5.200E-09 | 8.900E-09 |
| 26388 | 1320 | V1320G | E | 4.7e-09 | 4.3e-09 | 5.1e-09 | 4.700E-09 | 4.000E-10 | 4.300E-09 | 5.100E-09 |
| 26389 | 1320 | V1320H | E | 4.5e-09 | 3.4e-09 | 9.7e-09 | 5.867E-09 | 3.365E-09 | 3.400E-09 | 9.700E-09 |
| 26390 | 1320 | V1320I | E | 4.1e-09 | 8.4e-09 | 1.4e-08 | 8.833E-09 | 4.964E-09 | 4.100E-09 | 1.400E-08 |
| 26391 | 1320 | V1320L | E | 7.6e-09 | 3.9e-09 | 6.8e-09 | 6.100E-09 | 1.947E-09 | 3.900E-09 | 7.600E-09 |
| 26392 | 1320 | V1320K | E | 4.3e-09 | 2.9e-09 | 4.6e-09 | 3.933E-09 | 9.074E-10 | 2.900E-09 | 4.600E-09 |
| 26393 | 1320 | V1320M | E | 6.3e-09 | 1.9e-09 | 2.8e-09 | 3.667E-09 | 2.325E-09 | 1.900E-09 | 6.300E-09 |
| 26394 | 1320 | V1320F | E | 1.3e-08 | 1.3e-08 | 5.1e-09 | 1.037E-08 | 4.561E-09 | 5.100E-09 | 1.300E-08 |
| 26395 | 1320 | V1320P | E | 7.2e-09 | 4.5e-09 | 7.2e-09 | 6.300E-09 | 1.559E-09 | 4.500E-09 | 7.200E-09 |
| 26396 | 1320 | V1320S | E | 2.4e-09 | 4.1e-09 | 1.5e-08 | 7.167E-09 | 6.837E-09 | 2.400E-09 | 1.500E-08 |
| 26397 | 1320 | V1320T | E | 1.1e-08 | 3.1e-09 | 3.7e-09 | 5.933E-09 | 4.398E-09 | 3.100E-09 | 1.100E-08 |
| 26398 | 1320 | V1320W | E | 1e-08 | 3.2e-09 | 3.7e-09 | 5.633E-09 | 3.790E-09 | 3.200E-09 | 1.000E-08 |
| 26399 | 1320 | V1320Y | E | 4.4e-09 | 8.2e-09 | 7.5e-09 | 6.700E-09 | 2.022E-09 | 4.400E-09 | 8.200E-09 |
| 26400 | 1320 | V1320V | E | 4.6e-09 | 9.4e-09 | 5.7e-09 | 6.567E-09 | 2.515E-09 | 4.600E-09 | 9.400E-09 |
| 26401 | 1321 | A1321A | E | 1e-08 | 6.6e-09 | 2.6e-09 | 6.400E-09 | 3.704E-09 | 2.600E-09 | 1.000E-08 |
| 26402 | 1321 | A1321R | E | 7.6e-09 | 4.9e-09 | 2.7e-09 | 5.067E-09 | 2.454E-09 | 2.700E-09 | 7.600E-09 |
| 26403 | 1321 | A1321N | E | 9.8e-09 | 1.1e-08 | 1.1e-08 | 1.060E-08 | 6.928E-10 | 9.800E-09 | 1.100E-08 |
| 26404 | 1321 | A1321D | E | 9.1e-09 | 9.2e-09 | 3e-09 | 7.100E-09 | 3.551E-09 | 3.000E-09 | 9.200E-09 |
| 26405 | 1321 | A1321C | E | 8.8e-09 | 2.8e-09 | 3.2e-09 | 4.933E-09 | 3.355E-09 | 2.800E-09 | 8.800E-09 |
| 26406 | 1321 | A1321E | E | 5.3e-09 | 2.9e-09 | 7.7e-09 | 5.300E-09 | 2.400E-09 | 2.900E-09 | 7.700E-09 |
| 26407 | 1321 | A1321Q | E | 6.9e-09 | 5.2e-09 | 4.2e-09 | 5.433E-09 | 1.365E-09 | 4.200E-09 | 6.900E-09 |
| 26408 | 1321 | A1321G | E | 2.7e-09 | 2e-09 | 4.9e-09 | 3.200E-09 | 1.513E-09 | 2.000E-09 | 4.900E-09 |
| 26409 | 1321 | A1321H | E | 1.8e-08 | 8.5e-09 | 3.7e-09 | 1.007E-08 | 7.278E-09 | 3.700E-09 | 1.800E-08 |
| 26410 | 1321 | A1321I | E | 9.2e-09 | 5.9e-09 | 5.5e-09 | 6.867E-09 | 2.031E-09 | 5.500E-09 | 9.200E-09 |
| 26411 | 1321 | A1321L | E | 3.2e-09 | 8.1e-09 | 7.1e-09 | 6.133E-09 | 2.589E-09 | 3.200E-09 | 8.100E-09 |
| 26412 | 1321 | A1321K | E | 5.6e-09 | 2.3e-09 | 6.7e-09 | 4.867E-09 | 2.290E-09 | 2.300E-09 | 6.700E-09 |
| 26413 | 1321 | A1321M | E | 1.1e-08 | 8.8e-09 | 9.8e-09 | 9.867E-09 | 1.102E-09 | 8.800E-09 | 1.100E-08 |
| 26414 | 1321 | A1321F | E | 1.5e-08 | 6.6e-09 | 4.2e-09 | 8.600E-09 | 5.671E-09 | 4.200E-09 | 1.500E-08 |
| 26415 | 1321 | A1321P | E | 4.9e-09 | 9.9e-09 | 1.7e-09 | 5.500E-09 | 4.133E-09 | 1.700E-09 | 9.900E-09 |
| 26416 | 1321 | A1321S | E | 1.4e-08 | 8.1e-09 | 8.7e-09 | 1.027E-08 | 3.247E-09 | 8.100E-09 | 1.400E-08 |
| 26417 | 1321 | A1321T | E | 2.3e-08 | 2.9e-09 | 1.1e-08 | 1.230E-08 | 1.011E-08 | 2.900E-09 | 2.300E-08 |
| 26418 | 1321 | A1321W | E | 1.1e-08 | 3.8e-09 | 1.4e-08 | 9.600E-09 | 5.242E-09 | 3.800E-09 | 1.400E-08 |
| 26419 | 1321 | A1321Y | E | 3.2e-09 | 8.1e-09 | 5.5e-09 | 5.600E-09 | 2.452E-09 | 3.200E-09 | 8.100E-09 |
| 26420 | 1321 | A1321V | E | 5e-09 | 1.5e-08 | 6.4e-09 | 8.800E-09 | 5.415E-09 | 5.000E-09 | 1.500E-08 |
| 26421 | 1322 | R1322A | E | 5.6e-09 | 6.9e-09 | 1.6e-08 | 9.500E-09 | 5.667E-09 | 5.600E-09 | 1.600E-08 |
| 26422 | 1322 | R1322R | E | 1.1e-08 | 2.1e-08 | 9.7e-09 | 1.390E-08 | 6.183E-09 | 9.700E-09 | 2.100E-08 |
| 26423 | 1322 | R1322N | E | 6.1e-09 | 6.4e-09 | 6.7e-09 | 6.400E-09 | 3.000E-10 | 6.100E-09 | 6.700E-09 |
| 26424 | 1322 | R1322D | E | 1.9e-09 | 1.1e-08 | 6.8e-09 | 6.567E-09 | 4.554E-09 | 1.900E-09 | 1.100E-08 |
| 26425 | 1322 | R1322C | E | 6e-09 | 2.1e-08 | 1.2e-08 | 1.300E-08 | 7.550E-09 | 6.000E-09 | 2.100E-08 |
| 26426 | 1322 | R1322E | E | 4.6e-09 | 8e-09 | 6.4e-09 | 6.333E-09 | 1.701E-09 | 4.600E-09 | 8.000E-09 |
| 26427 | 1322 | R1322Q | E | 5e-09 | 9.6e-09 | 8.1e-09 | 7.567E-09 | 2.346E-09 | 5.000E-09 | 9.600E-09 |
| 26428 | 1322 | R1322G | E | 1.7e-08 | 3.9e-09 | 7e-09 | 9.300E-09 | 6.846E-09 | 3.900E-09 | 1.700E-08 |
| 26429 | 1322 | R1322H | E | 5.8e-09 | 3.2e-09 | 7.4e-09 | 5.467E-09 | 2.120E-09 | 3.200E-09 | 7.400E-09 |
| 26430 | 1322 | R1322I | E | 7.3e-09 | 1.5e-08 | 2.4e-09 | 8.233E-09 | 6.352E-09 | 2.400E-09 | 1.500E-08 |
| 26431 | 1322 | R1322L | E | 4.8e-09 | 6.9e-09 | 4.8e-09 | 5.500E-09 | 1.212E-09 | 4.800E-09 | 6.900E-09 |
| 26432 | 1322 | R1322K | E | 8.2e-09 | 6.5e-09 | 3e-09 | 5.900E-09 | 2.651E-09 | 3.000E-09 | 8.200E-09 |
| 26433 | 1322 | R1322M | E | 5e-09 | 1.4e-08 | 5.3e-09 | 8.100E-09 | 5.112E-09 | 5.000E-09 | 1.400E-08 |
| 26434 | 1322 | R1322F | E | 5.1e-09 | 3.5e-09 | 1e-08 | 6.200E-09 | 3.387E-09 | 3.500E-09 | 1.000E-08 |

| | | | | | | | | | | |
|---|---|---|---|---|---|---|---|---|---|---|
| 26435 | 1322 | R1322P | E | 2.5e-09 | 4.1e-09 | 1.4e-08 | 6.867E-09 | 6.229E-09 | 2.500E-09 | 1.400E-08 |
| 26436 | 1322 | R1322S | E | 4.2e-09 | 8.9e-09 | 1.1e-08 | 8.033E-09 | 3.482E-09 | 4.200E-09 | 1.100E-08 |
| 26437 | 1322 | R1322T | E | 1.4e-08 | 2.4e-09 | 4.3e-09 | 6.900E-09 | 6.222E-09 | 2.400E-09 | 1.400E-08 |
| 26438 | 1322 | R1322W | E | 4.8e-09 | 1e-08 | 6.4e-09 | 7.067E-09 | 2.663E-09 | 4.800E-09 | 1.000E-08 |
| 26439 | 1322 | R1322Y | E | 1.2e-08 | 8.7e-09 | 1.6e-08 | 1.223E-08 | 3.656E-09 | 8.700E-09 | 1.600E-08 |
| 26440 | 1322 | R1322V | E | 4.7e-09 | 1.3e-08 | 2.9e-09 | 6.867E-09 | 5.387E-09 | 2.900E-09 | 1.300E-08 |
| 26441 | 1323 | I1323A | E | 7.1e-09 | 2.9e-08 | 1.7e-08 | 1.770E-08 | 1.097E-08 | 7.100E-09 | 2.900E-08 |
| 26442 | 1323 | I1323R | E | 1.3e-08 | 3.7e-09 | 1.2e-08 | 9.567E-09 | 5.105E-09 | 3.700E-09 | 1.300E-08 |
| 26443 | 1323 | I1323N | E | 1.1e-08 | 2.1e-09 | 3.3e-09 | 5.467E-09 | 4.829E-09 | 2.100E-09 | 1.100E-08 |
| 26444 | 1323 | I1323D | E | 3.1e-08 | 9.4e-09 | 1.4e-08 | 1.813E-08 | 1.138E-08 | 9.400E-09 | 3.100E-08 |
| 26445 | 1323 | I1323C | E | 5.7e-09 | 8.5e-09 | 2e-08 | 1.140E-08 | 7.578E-09 | 5.700E-09 | 2.000E-08 |
| 26446 | 1323 | I1323E | E | 1.4e-08 | 7.4e-09 | 2.1e-09 | 7.833E-09 | 5.962E-09 | 2.100E-09 | 1.400E-08 |
| 26447 | 1323 | I1323Q | E | 3.2e-09 | 6.3e-09 | 1.4e-08 | 7.833E-09 | 5.561E-09 | 3.200E-09 | 1.400E-08 |
| 26448 | 1323 | I1323G | E | 6.3e-09 | 3.8e-09 | 9.8e-09 | 6.633E-09 | 3.014E-09 | 3.800E-09 | 9.800E-09 |
| 26449 | 1323 | I1323H | E | 1.3e-08 | 7e-09 | 6.6e-09 | 8.867E-09 | 3.585E-09 | 6.600E-09 | 1.300E-08 |
| 26450 | 1323 | I1323I | E | 5.3e-09 | 8.3e-09 | 5e-09 | 6.200E-09 | 1.825E-09 | 5.000E-09 | 8.300E-09 |
| 26451 | 1323 | I1323L | E | 1.4e-09 | 1.6e-08 | 6.5e-09 | 7.967E-09 | 7.410E-09 | 1.400E-09 | 1.600E-08 |
| 26452 | 1323 | I1323K | E | 5.3e-09 | 1.7e-08 | 1.1e-08 | 1.110E-08 | 5.851E-09 | 5.300E-09 | 1.700E-08 |
| 26453 | 1323 | I1323M | E | 7.6e-09 | 5.1e-09 | 1.6e-08 | 9.567E-09 | 5.710E-09 | 5.100E-09 | 1.600E-08 |
| 26454 | 1323 | I1323F | E | 1.5e-08 | 8.6e-09 | 5.5e-09 | 9.700E-09 | 4.845E-09 | 5.500E-09 | 1.500E-08 |
| 26455 | 1323 | I1323P | E | 1.5e-08 | 7.8e-09 | 9.6e-09 | 1.080E-08 | 3.747E-09 | 7.800E-09 | 1.500E-08 |
| 26456 | 1323 | I1323S | E | 9.1e-09 | 4.3e-09 | 5.1e-09 | 6.167E-09 | 2.572E-09 | 4.300E-09 | 9.100E-09 |
| 26457 | 1323 | I1323T | E | 9.2e-09 | 8.1e-09 | 6.8e-09 | 8.033E-09 | 1.201E-09 | 6.800E-09 | 9.200E-09 |
| 26458 | 1323 | I1323W | E | 3.2e-09 | 6.9e-09 | 7.4e-09 | 5.833E-09 | 2.294E-09 | 3.200E-09 | 7.400E-09 |
| 26459 | 1323 | I1323Y | E | 3.8e-09 | 7.9e-09 | 3.5e-09 | 5.067E-09 | 2.458E-09 | 3.500E-09 | 7.900E-09 |
| 26460 | 1323 | I1323V | E | 1.1e-08 | 1e-08 | 1e-08 | 1.033E-08 | 5.774E-10 | 1.000E-08 | 1.100E-08 |
| 26461 | 1324 | Y1324A | E | 1.3e-08 | 1.3e-08 | 6.3e-09 | 1.077E-08 | 3.868E-09 | 6.300E-09 | 1.300E-08 |
| 26462 | 1324 | Y1324R | E | 8.4e-09 | 1.3e-08 | 1.2e-08 | 1.113E-08 | 2.419E-09 | 8.400E-09 | 1.300E-08 |
| 26463 | 1324 | Y1324N | E | 3.5e-09 | 6.1e-09 | 8.1e-09 | 5.900E-09 | 2.307E-09 | 3.500E-09 | 8.100E-09 |
| 26464 | 1324 | Y1324D | E | 4.9e-09 | 8.8e-09 | 8e-09 | 7.233E-09 | 2.060E-09 | 4.900E-09 | 8.800E-09 |
| 26465 | 1324 | Y1324C | E | 2.5e-08 | 1.3e-08 | 7.3e-09 | 1.510E-08 | 9.035E-09 | 7.300E-09 | 2.500E-08 |
| 26466 | 1324 | Y1324E | E | 5.7e-09 | 1e-08 | 5.2e-09 | 6.967E-09 | 2.639E-09 | 5.200E-09 | 1.000E-08 |
| 26467 | 1324 | Y1324Q | E | 7.9e-09 | 4.6e-09 | 8.2e-09 | 6.900E-09 | 1.997E-09 | 4.600E-09 | 8.200E-09 |
| 26468 | 1324 | Y1324G | E | 8.2e-09 | 2e-08 | 7.3e-09 | 1.183E-08 | 7.087E-09 | 7.300E-09 | 2.000E-08 |
| 26469 | 1324 | Y1324H | E | 1.8e-08 | 8.5e-09 | 3.7e-09 | 1.007E-08 | 7.278E-09 | 3.700E-09 | 1.800E-08 |
| 26470 | 1324 | Y1324I | E | 5.6e-09 | 9.6e-09 | 7e-09 | 7.400E-09 | 2.030E-09 | 5.600E-09 | 9.600E-09 |
| 26471 | 1324 | Y1324L | E | 1.3e-08 | 9.9e-09 | 6.7e-09 | 9.867E-09 | 3.150E-09 | 6.700E-09 | 1.300E-08 |
| 26472 | 1324 | Y1324K | E | 5.4e-09 | 1.2e-08 | 4.2e-09 | 7.200E-09 | 4.200E-09 | 4.200E-09 | 1.200E-08 |
| 26473 | 1324 | Y1324M | E | 2.7e-08 | 7.7e-09 | 5.6e-09 | 1.343E-08 | 1.180E-08 | 5.600E-09 | 2.700E-08 |
| 26474 | 1324 | Y1324F | E | 6.2e-09 | 3.6e-09 | 3.4e-09 | 4.400E-09 | 1.562E-09 | 3.400E-09 | 6.200E-09 |
| 26475 | 1324 | Y1324P | E | 9.6e-09 | 5.2e-08 | 1.1e-08 | 2.420E-08 | 2.409E-08 | 9.600E-09 | 5.200E-08 |
| 26476 | 1324 | Y1324S | E | 1.9e-08 | 3.6e-09 | 3.5e-09 | 8.700E-09 | 8.920E-09 | 3.500E-09 | 1.900E-08 |
| 26477 | 1324 | Y1324T | E | 3.3e-08 | 5.3e-09 | 6e-09 | 1.477E-08 | 1.579E-08 | 5.300E-09 | 3.300E-08 |
| 26478 | 1324 | Y1324W | E | 1.4e-08 | 8.2e-09 | 1.5e-08 | 1.240E-08 | 3.672E-09 | 8.200E-09 | 1.500E-08 |
| 26479 | 1324 | Y1324Y | E | 1.4e-08 | 8.3e-09 | 8.7e-09 | 1.033E-08 | 3.182E-09 | 8.300E-09 | 1.400E-08 |
| 26480 | 1324 | Y1324V | E | 5.4e-09 | 4.4e-09 | 4.9e-09 | 4.900E-09 | 5.000E-10 | 4.400E-09 | 5.400E-09 |
| 26481 | 1325 | P1325A | E | 4.3e-09 | 1.4e-08 | 1.8e-08 | 1.210E-08 | 7.045E-09 | 4.300E-09 | 1.800E-08 |
| 26482 | 1325 | P1325R | E | 5.4e-09 | 1.6e-08 | 5.3e-09 | 8.900E-09 | 6.149E-09 | 5.300E-09 | 1.600E-08 |
| 26483 | 1325 | P1325N | E | 2.5e-09 | 1.4e-09 | 1.1e-08 | 4.967E-09 | 5.254E-09 | 1.400E-09 | 1.100E-08 |
| 26484 | 1325 | P1325D | E | 1.2e-08 | 1.1e-08 | 8.4e-09 | 1.047E-08 | 1.858E-09 | 8.400E-09 | 1.200E-08 |
| 26485 | 1325 | P1325C | E | 4.1e-09 | 2.1e-09 | 5.9e-09 | 4.033E-09 | 1.901E-09 | 2.100E-09 | 5.900E-09 |
| 26486 | 1325 | P1325E | E | 6.9e-09 | 7.2e-09 | 3e-09 | 5.700E-09 | 2.343E-09 | 3.000E-09 | 7.200E-09 |
| 26487 | 1325 | P1325Q | E | 1.3e-08 | 6.1e-09 | 7.2e-09 | 8.767E-09 | 3.707E-09 | 6.100E-09 | 1.300E-08 |
| 26488 | 1325 | P1325G | E | 1.1e-08 | 4.7e-09 | 6.6e-09 | 7.433E-09 | 3.232E-09 | 4.700E-09 | 1.100E-08 |
| 26489 | 1325 | P1325H | E | 6.3e-09 | 2.3e-09 | 5.4e-09 | 4.667E-09 | 2.098E-09 | 2.300E-09 | 6.300E-09 |
| 26490 | 1325 | P1325I | E | 1.3e-08 | 8.5e-09 | 6.2e-09 | 9.233E-09 | 3.459E-09 | 6.200E-09 | 1.300E-08 |
| 26491 | 1325 | P1325L | E | 2.3e-09 | 3.2e-09 | 7.3e-09 | 4.267E-09 | 2.665E-09 | 2.300E-09 | 7.300E-09 |
| 26492 | 1325 | P1325K | E | 3.4e-09 | 7.3e-09 | 6e-09 | 5.567E-09 | 1.986E-09 | 3.400E-09 | 7.300E-09 |
| 26493 | 1325 | P1325M | E | 1.6e-08 | 6.7e-09 | 1.2e-08 | 1.157E-08 | 4.665E-09 | 6.700E-09 | 1.600E-08 |
| 26494 | 1325 | P1325F | E | 3.1e-09 | 6.3e-09 | 8.5e-09 | 5.967E-09 | 2.715E-09 | 3.100E-09 | 8.500E-09 |
| 26495 | 1325 | P1325P | E | 8.7e-09 | 2.2e-09 | 4.1e-09 | 5.000E-09 | 3.342E-09 | 2.200E-09 | 8.700E-09 |
| 26496 | 1325 | P1325S | E | 3e-09 | 5.1e-09 | 1.3e-08 | 7.033E-09 | 5.273E-09 | 3.000E-09 | 1.300E-08 |
| 26497 | 1325 | P1325T | E | 3.1e-09 | 6.5e-09 | 4e-09 | 4.533E-09 | 1.762E-09 | 3.100E-09 | 6.500E-09 |
| 26498 | 1325 | P1325W | E | 9.2e-09 | 2.8e-09 | 4.6e-09 | 5.533E-09 | 3.301E-09 | 2.800E-09 | 9.200E-09 |
| 26499 | 1325 | P1325Y | E | 1.4e-08 | 2.2e-08 | 4.4e-09 | 1.347E-08 | 8.812E-09 | 4.400E-09 | 2.200E-08 |
| 26500 | 1325 | P1325V | E | 8.7e-09 | 1.2e-08 | 6.6e-09 | 9.100E-09 | 2.722E-09 | 6.600E-09 | 1.200E-08 |
| 26501 | 1326 | T1326A | E | 5.4e-09 | 1.4e-08 | 4.8e-09 | 8.067E-09 | 5.147E-09 | 4.800E-09 | 1.400E-08 |
| 26502 | 1326 | T1326R | E | 4.2e-09 | 1.6e-08 | 1.4e-09 | 7.200E-09 | 7.749E-09 | 1.400E-09 | 1.600E-08 |
| 26503 | 1326 | T1326N | E | 4.9e-09 | 1.2e-08 | 5.3e-09 | 7.400E-09 | 3.989E-09 | 4.900E-09 | 1.200E-08 |
| 26504 | 1326 | T1326D | E | 1.1e-08 | 6.6e-09 | 1.1e-08 | 9.533E-09 | 2.540E-09 | 6.600E-09 | 1.100E-08 |
| 26505 | 1326 | T1326C | E | 3.3e-09 | 4.4e-08 | 2.6e-09 | 1.663E-08 | 2.370E-08 | 2.600E-09 | 4.400E-08 |
| 26506 | 1326 | T1326E | E | 9.5e-09 | 5.9e-09 | 3.2e-08 | 1.580E-08 | 1.414E-08 | 5.900E-09 | 3.200E-08 |
| 26507 | 1326 | T1326Q | E | 3.5e-09 | 5.1e-09 | 1.3e-08 | 7.200E-09 | 5.086E-09 | 3.500E-09 | 1.300E-08 |
| 26508 | 1326 | T1326G | E | 1e-08 | 5.4e-09 | 7.4e-09 | 7.600E-09 | 2.307E-09 | 5.400E-09 | 1.000E-08 |
| 26509 | 1326 | T1326H | E | 4.6e-09 | 7.6e-09 | 1e-08 | 7.400E-09 | 2.706E-09 | 4.600E-09 | 1.000E-08 |
| 26510 | 1326 | T1326I | E | 9.2e-09 | 7.1e-09 | 1.9e-08 | 1.177E-08 | 6.352E-09 | 7.100E-09 | 1.900E-08 |
| 26511 | 1326 | T1326L | E | 5.9e-09 | 6.1e-09 | 2.1e-09 | 4.700E-09 | 2.254E-09 | 2.100E-09 | 6.100E-09 |
| 26512 | 1326 | T1326K | E | 3.8e-09 | 5.2e-09 | 3.2e-09 | 4.067E-09 | 1.026E-09 | 3.200E-09 | 5.200E-09 |
| 26513 | 1326 | T1326M | E | 4.9e-09 | 7.4e-09 | 7.7e-09 | 6.667E-09 | 1.537E-09 | 4.900E-09 | 7.700E-09 |
| 26514 | 1326 | T1326F | E | 3.7e-09 | 7.3e-09 | 1.5e-08 | 8.667E-09 | 5.773E-09 | 3.700E-09 | 1.500E-08 |
| 26515 | 1326 | T1326P | E | 1.5e-08 | 4.2e-09 | 9.2e-09 | 9.467E-09 | 5.405E-09 | 4.200E-09 | 1.500E-08 |
| 26516 | 1326 | T1326S | E | 5.7e-09 | 3.7e-09 | 2.4e-08 | 1.113E-08 | 1.119E-08 | 3.700E-09 | 2.400E-08 |
| 26517 | 1326 | T1326T | E | 8.6e-09 | 2.7e-08 | 3.9e-09 | 1.317E-08 | 1.221E-08 | 3.900E-09 | 2.700E-08 |
| 26518 | 1326 | T1326W | E | 1.3e-08 | 1e-08 | 4.1e-09 | 9.033E-09 | 4.528E-09 | 4.100E-09 | 1.300E-08 |
| 26519 | 1326 | T1326Y | E | 6.6e-09 | 2.7e-09 | 1.7e-08 | 8.767E-09 | 7.392E-09 | 2.700E-09 | 1.700E-08 |

| | | | | | | | | | | |
|---|---|---|---|---|---|---|---|---|---|---|
| 26520 | 1326 | T1326V | E | 4.8e-09 | 3.1e-09 | 3.3e-09 | 3.733E-09 | 9.292E-10 | 3.100E-09 | 4.800E-09 |
| 26521 | 1327 | N1327A | E | 4.6e-09 | 1.1e-08 | 4.4e-09 | 6.667E-09 | 3.754E-09 | 4.400E-09 | 1.100E-08 |
| 26522 | 1327 | N1327R | E | 1.9e-08 | 2.6e-08 | 8.9e-09 | 1.797E-08 | 8.597E-09 | 8.900E-09 | 2.600E-08 |
| 26523 | 1327 | N1327N | E | 4e-09 | 6.6e-09 | 1.5e-08 | 8.533E-09 | 5.749E-09 | 4.000E-09 | 1.500E-08 |
| 26524 | 1327 | N1327D | E | 5.5e-09 | 5.9e-09 | 5.5e-09 | 5.633E-09 | 2.309E-10 | 5.500E-09 | 5.900E-09 |
| 26525 | 1327 | N1327C | E | 1.2e-08 | 1.5e-08 | 9.8e-09 | 1.227E-08 | 2.610E-09 | 9.800E-09 | 1.500E-08 |
| 26526 | 1327 | N1327E | E | 3.6e-09 | 9e-09 | 6.3e-09 | 6.300E-09 | 2.700E-09 | 3.600E-09 | 9.000E-09 |
| 26527 | 1327 | N1327Q | E | 3.1e-09 | 2.1e-09 | 1.1e-08 | 5.400E-09 | 4.875E-09 | 2.100E-09 | 1.100E-08 |
| 26528 | 1327 | N1327G | E | 1.4e-08 | 7.3e-09 | 7.9e-09 | 9.733E-09 | 3.707E-09 | 7.300E-09 | 1.400E-08 |
| 26529 | 1327 | N1327H | E | 6.8e-09 | 3.4e-09 | 6.9e-09 | 5.700E-09 | 1.992E-09 | 3.400E-09 | 6.900E-09 |
| 26530 | 1327 | N1327I | E | 5e-09 | 5.2e-09 | 6.8e-09 | 5.667E-09 | 9.866E-10 | 5.000E-09 | 6.800E-09 |
| 26531 | 1327 | N1327L | E | 8.1e-09 | 3.4e-09 | 5.1e-09 | 5.533E-09 | 2.380E-09 | 3.400E-09 | 8.100E-09 |
| 26532 | 1327 | N1327K | E | 6e-09 | 7.6e-09 | 5.4e-09 | 6.333E-09 | 1.137E-09 | 5.400E-09 | 7.600E-09 |
| 26533 | 1327 | N1327M | E | 3.1e-09 | 1e-08 | 1e-08 | 7.700E-09 | 3.984E-09 | 3.100E-09 | 1.000E-08 |
| 26534 | 1327 | N1327F | E | 1.1e-08 | 7.9e-09 | 1.9e-09 | 6.933E-09 | 4.626E-09 | 1.900E-09 | 1.100E-08 |
| 26535 | 1327 | N1327P | E | 1.8e-08 | 4.4e-09 | 8.3e-09 | 1.023E-08 | 7.003E-09 | 4.400E-09 | 1.800E-08 |
| 26536 | 1327 | N1327S | E | 8.6e-09 | 7.5e-09 | 9.3e-09 | 8.467E-09 | 9.074E-10 | 7.500E-09 | 9.300E-09 |
| 26537 | 1327 | N1327T | E | 2.5e-09 | 4e-09 | 6.4e-09 | 4.300E-09 | 1.967E-09 | 2.500E-09 | 6.400E-09 |
| 26538 | 1327 | N1327W | E | 6.5e-09 | 7.1e-09 | 3.9e-09 | 5.833E-09 | 1.701E-09 | 3.900E-09 | 7.100E-09 |
| 26539 | 1327 | N1327Y | E | 2.4e-09 | 4.4e-09 | 1.4e-08 | 6.933E-09 | 6.201E-09 | 2.400E-09 | 1.400E-08 |
| 26540 | 1327 | N1327V | E | 5.6e-09 | 8.9e-09 | 3e-09 | 5.833E-09 | 2.957E-09 | 3.000E-09 | 8.900E-09 |
| 26541 | 1328 | G1328A | E | 1.5e-08 | 2.6e-09 | 2.1e-08 | 1.287E-08 | 9.384E-09 | 2.600E-09 | 2.100E-08 |
| 26542 | 1328 | G1328R | E | 1.8e-08 | 6.9e-09 | 1.8e-08 | 1.430E-08 | 6.409E-09 | 6.900E-09 | 1.800E-08 |
| 26543 | 1328 | G1328N | E | 8.5e-09 | 6.1e-09 | 7.3e-09 | 7.300E-09 | 1.200E-09 | 6.100E-09 | 8.500E-09 |
| 26544 | 1328 | G1328D | E | 5.8e-09 | 8.4e-09 | 1.8e-08 | 1.073E-08 | 6.426E-09 | 5.800E-09 | 1.800E-08 |
| 26545 | 1328 | G1328C | E | 4.2e-09 | 4.4e-09 | 7.3e-09 | 5.300E-09 | 1.735E-09 | 4.200E-09 | 7.300E-09 |
| 26546 | 1328 | G1328E | E | 5.6e-09 | 6.6e-09 | 6e-09 | 6.067E-09 | 5.033E-10 | 5.600E-09 | 6.600E-09 |
| 26547 | 1328 | G1328Q | E | 9.5e-09 | 1.3e-09 | 8.1e-09 | 6.300E-09 | 4.386E-09 | 1.300E-09 | 9.500E-09 |
| 26548 | 1328 | G1328G | E | 4.8e-09 | 1.2e-08 | 6.6e-09 | 7.800E-09 | 3.747E-09 | 4.800E-09 | 1.200E-08 |
| 26549 | 1328 | G1328H | E | 5.3e-09 | 9.2e-09 | 8.6e-09 | 7.700E-09 | 2.100E-09 | 5.300E-09 | 9.200E-09 |
| 26550 | 1328 | G1328I | E | 8.1e-09 | 1.3e-08 | 2e-08 | 1.370E-08 | 5.981E-09 | 8.100E-09 | 2.000E-08 |
| 26551 | 1328 | G1328L | E | 8.6e-09 | 2.4e-09 | 1.5e-08 | 8.667E-09 | 6.300E-09 | 2.400E-09 | 1.500E-08 |
| 26552 | 1328 | G1328K | E | 2.9e-08 | 6.5e-09 | 4e-09 | 1.317E-08 | 1.377E-08 | 4.000E-09 | 2.900E-08 |
| 26553 | 1328 | G1328M | E | 9.7e-09 | 5.3e-09 | 1.2e-08 | 9.000E-09 | 3.404E-09 | 5.300E-09 | 1.200E-08 |
| 26554 | 1328 | G1328F | E | 9.6e-09 | 1.5e-08 | 2.7e-09 | 9.100E-09 | 6.165E-09 | 2.700E-09 | 1.500E-08 |
| 26555 | 1328 | G1328P | E | 5e-09 | 9.9e-09 | 6.2e-09 | 7.033E-09 | 2.554E-09 | 5.000E-09 | 9.900E-09 |
| 26556 | 1328 | G1328S | E | 1.5e-08 | 4.7e-09 | 5.9e-09 | 8.533E-09 | 5.632E-09 | 4.700E-09 | 1.500E-08 |
| 26557 | 1328 | G1328T | E | 6.2e-09 | 9.2e-09 | 3e-09 | 6.133E-09 | 3.101E-09 | 3.000E-09 | 9.200E-09 |
| 26558 | 1328 | G1328W | E | 8.5e-09 | 1.3e-09 | 1.3e-08 | 7.600E-09 | 5.902E-09 | 1.300E-09 | 1.300E-08 |
| 26559 | 1328 | G1328Y | E | 1.2e-09 | 5.1e-09 | 1.1e-08 | 5.767E-09 | 4.934E-09 | 1.200E-09 | 1.100E-08 |
| 26560 | 1328 | G1328V | E | 5.9e-09 | 6.6e-09 | 2.6e-08 | 1.283E-08 | 1.141E-08 | 5.900E-09 | 2.600E-08 |
| 26561 | 1329 | Y1329A | E | 6.5e-09 | 5.8e-09 | 4e-09 | 5.433E-09 | 1.290E-09 | 4.000E-09 | 6.500E-09 |
| 26562 | 1329 | Y1329R | E | 4.2e-09 | 4.8e-09 | 1.3e-09 | 3.433E-09 | 1.872E-09 | 1.300E-09 | 4.800E-09 |
| 26563 | 1329 | Y1329N | E | 4.5e-09 | 1.4e-08 | 6.7e-09 | 8.400E-09 | 4.973E-09 | 4.500E-09 | 1.400E-08 |
| 26564 | 1329 | Y1329D | E | 2.7e-09 | 7.9e-09 | 1.4e-08 | 8.200E-09 | 5.656E-09 | 2.700E-09 | 1.400E-08 |
| 26565 | 1329 | Y1329C | E | 4.6e-09 | 5.6e-09 | 6.1e-09 | 5.433E-09 | 7.638E-10 | 4.600E-09 | 6.100E-09 |
| 26566 | 1329 | Y1329E | E | 2e-08 | 2e-08 | 1.3e-08 | 1.767E-08 | 4.041E-09 | 1.300E-08 | 2.000E-08 |
| 26567 | 1329 | Y1329Q | E | 8.3e-09 | 1.5e-08 | 2.3e-09 | 8.533E-09 | 6.353E-09 | 2.300E-09 | 1.500E-08 |
| 26568 | 1329 | Y1329G | E | 7.3e-09 | 7.8e-09 | 1.5e-08 | 1.003E-08 | 4.309E-09 | 7.300E-09 | 1.500E-08 |
| 26569 | 1329 | Y1329H | E | 9.2e-09 | 7.1e-09 | 2.7e-09 | 6.333E-09 | 3.317E-09 | 2.700E-09 | 9.200E-09 |
| 26570 | 1329 | Y1329I | E | 1.8e-08 | 3.9e-09 | 7.3e-09 | 9.733E-09 | 7.358E-09 | 3.900E-09 | 1.800E-08 |
| 26571 | 1329 | Y1329L | E | 4.9e-09 | 4.6e-09 | 1.3e-08 | 7.500E-09 | 4.766E-09 | 4.600E-09 | 1.300E-08 |
| 26572 | 1329 | Y1329K | E | 1.1e-08 | 3.6e-09 | 2.9e-08 | 1.453E-08 | 1.306E-08 | 3.600E-09 | 2.900E-08 |
| 26573 | 1329 | Y1329M | E | 6.6e-09 | 8.1e-09 | 1.6e-08 | 1.023E-08 | 5.050E-09 | 6.600E-09 | 1.600E-08 |
| 26574 | 1329 | Y1329F | E | 3.2e-09 | 2.6e-08 | 7.3e-09 | 1.217E-08 | 1.215E-08 | 3.200E-09 | 2.600E-08 |
| 26575 | 1329 | Y1329P | E | 5.9e-09 | 7.1e-09 | 4.9e-09 | 5.967E-09 | 1.102E-09 | 4.900E-09 | 7.100E-09 |
| 26576 | 1329 | Y1329S | E | 2.2e-09 | 4.2e-09 | 7.9e-09 | 4.767E-09 | 2.892E-09 | 2.200E-09 | 7.900E-09 |
| 26577 | 1329 | Y1329T | E | 3.3e-08 | 7.9e-09 | 3.1e-09 | 1.467E-08 | 1.606E-08 | 3.100E-09 | 3.300E-08 |
| 26578 | 1329 | Y1329W | E | 3e-09 | 6.9e-09 | 3e-09 | 4.300E-09 | 2.252E-09 | 3.000E-09 | 6.900E-09 |
| 26579 | 1329 | Y1329Y | E | 1e-08 | 4.3e-09 | 8.2e-09 | 7.500E-09 | 2.914E-09 | 4.300E-09 | 1.000E-08 |
| 26580 | 1329 | Y1329V | E | 5.8e-09 | 6.9e-09 | 7.8e-09 | 6.833E-09 | 1.002E-09 | 5.800E-09 | 7.800E-09 |
| 26581 | 1330 | T1330A | E | 4.5e-09 | 7.4e-09 | 1.2e-08 | 7.967E-09 | 3.782E-09 | 4.500E-09 | 1.200E-08 |
| 26582 | 1330 | T1330R | E | 8.1e-09 | 1.1e-08 | 1.1e-08 | 1.003E-08 | 1.674E-09 | 8.100E-09 | 1.100E-08 |
| 26583 | 1330 | T1330N | E | 1.3e-08 | 9.3e-09 | 6.9e-09 | 9.733E-09 | 3.073E-09 | 6.900E-09 | 1.300E-08 |
| 26584 | 1330 | T1330D | E | 8.2e-09 | 8e-09 | 3.8e-09 | 6.667E-09 | 2.485E-09 | 3.800E-09 | 8.200E-09 |
| 26585 | 1330 | T1330C | E | 6.7e-09 | 2.2e-09 | 8.6e-09 | 5.833E-09 | 3.287E-09 | 2.200E-09 | 8.600E-09 |
| 26586 | 1330 | T1330E | E | 1.4e-08 | 5.7e-09 | 7.1e-09 | 8.933E-09 | 4.443E-09 | 5.700E-09 | 1.400E-08 |
| 26587 | 1330 | T1330Q | E | 7.3e-09 | 1.4e-08 | 1.1e-08 | 1.077E-08 | 3.356E-09 | 7.300E-09 | 1.400E-08 |
| 26588 | 1330 | T1330G | E | 6.3e-09 | 9.1e-09 | 7.3e-09 | 7.567E-09 | 1.419E-09 | 6.300E-09 | 9.100E-09 |
| 26589 | 1330 | T1330H | E | 1.3e-08 | 3.1e-09 | 8.2e-09 | 8.100E-09 | 4.951E-09 | 3.100E-09 | 1.300E-08 |
| 26590 | 1330 | T1330I | E | 1.4e-08 | 4.4e-09 | 1.7e-09 | 6.700E-09 | 6.465E-09 | 1.700E-09 | 1.400E-08 |
| 26591 | 1330 | T1330L | E | 1.1e-08 | 8.8e-09 | 1.1e-08 | 1.027E-08 | 1.270E-09 | 8.800E-09 | 1.100E-08 |
| 26592 | 1330 | T1330K | E | 1.5e-08 | 3.2e-09 | 4.4e-09 | 7.533E-09 | 6.494E-09 | 3.200E-09 | 1.500E-08 |
| 26593 | 1330 | T1330M | E | 8.9e-09 | 1.5e-08 | 6.3e-09 | 1.007E-08 | 4.466E-09 | 6.300E-09 | 1.500E-08 |
| 26594 | 1330 | T1330F | E | 4.5e-09 | 5e-09 | 4.3e-09 | 4.600E-09 | 3.606E-10 | 4.300E-09 | 5.000E-09 |
| 26595 | 1330 | T1330P | E | 1.2e-08 | 1.3e-08 | 3.6e-09 | 9.533E-09 | 5.163E-09 | 3.600E-09 | 1.300E-08 |
| 26596 | 1330 | T1330S | E | 1.1e-08 | 7.9e-09 | 1.4e-08 | 1.097E-08 | 3.050E-09 | 7.900E-09 | 1.400E-08 |
| 26597 | 1330 | T1330T | E | 4.2e-09 | 1.2e-08 | 9.8e-09 | 8.667E-09 | 4.022E-09 | 4.200E-09 | 1.200E-08 |
| 26598 | 1330 | T1330W | E | 2.9e-09 | 4.6e-09 | 4e-09 | 3.833E-09 | 8.622E-10 | 2.900E-09 | 4.600E-09 |
| 26599 | 1330 | T1330Y | E | 9.4e-09 | 8.5e-09 | 2.5e-08 | 1.430E-08 | 9.277E-09 | 8.500E-09 | 2.500E-08 |
| 26600 | 1330 | T1330V | E | 1.4e-08 | 4e-09 | 5.1e-09 | 7.700E-09 | 5.484E-09 | 4.000E-09 | 1.400E-08 |
| 26601 | 1331 | R1331A | E | 1e-08 | 7.2e-09 | 2.1e-08 | 1.273E-08 | 7.295E-09 | 7.200E-09 | 2.100E-08 |
| 26602 | 1331 | R1331R | E | 5.5e-09 | 4.8e-09 | 4e-09 | 4.767E-09 | 7.506E-10 | 4.000E-09 | 5.500E-09 |
| 26603 | 1331 | R1331N | E | 1.1e-08 | 9.4e-09 | 3.6e-09 | 8.000E-09 | 3.894E-09 | 3.600E-09 | 1.100E-08 |
| 26604 | 1331 | R1331D | E | 3.7e-09 | 1.6e-08 | 9.1e-09 | 9.600E-09 | 6.165E-09 | 3.700E-09 | 1.600E-08 |

| | | | | | | | | | | |
|---|---|---|---|---|---|---|---|---|---|---|
| 26605 | 1331 | R1331C | E | 7.4e-09 | 1e-08 | 9.9e-09 | 9.100E-09 | 1.473E-09 | 7.400E-09 | 1.000E-08 |
| 26606 | 1331 | R1331E | E | 8.1e-09 | 3.2e-09 | 1e-08 | 7.100E-09 | 3.509E-09 | 3.200E-09 | 1.000E-08 |
| 26607 | 1331 | R1331Q | E | 2.1e-08 | 6.5e-09 | 8.1e-09 | 1.187E-08 | 7.950E-09 | 6.500E-09 | 2.100E-08 |
| 26608 | 1331 | R1331G | E | 8.5e-09 | 6.4e-09 | 2.9e-09 | 5.933E-09 | 2.829E-09 | 2.900E-09 | 8.500E-09 |
| 26609 | 1331 | R1331H | E | 6e-09 | 8.6e-09 | 2.7e-09 | 5.767E-09 | 2.957E-09 | 2.700E-09 | 8.600E-09 |
| 26610 | 1331 | R1331I | E | 7.2e-09 | 1.7e-08 | 1.6e-08 | 1.340E-08 | 5.393E-09 | 7.200E-09 | 1.700E-08 |
| 26611 | 1331 | R1331L | E | 1.4e-09 | 6.2e-09 | 7e-09 | 4.867E-09 | 3.029E-09 | 1.400E-09 | 7.000E-09 |
| 26612 | 1331 | R1331K | E | 7e-09 | 6.2e-09 | 1.1e-08 | 8.067E-09 | 2.572E-09 | 6.200E-09 | 1.100E-08 |
| 26613 | 1331 | R1331M | E | 4.1e-08 | 1.3e-08 | 1.1e-08 | 2.167E-08 | 1.677E-08 | 1.100E-08 | 4.100E-08 |
| 26614 | 1331 | R1331F | E | 7.1e-09 | 5.5e-09 | 4e-09 | 5.533E-09 | 1.550E-09 | 4.000E-09 | 7.100E-09 |
| 26615 | 1331 | R1331P | E | 1.9e-08 | 3.2e-08 | 1.5e-08 | 2.200E-08 | 8.888E-09 | 1.500E-08 | 3.200E-08 |
| 26616 | 1331 | R1331S | E | 4.3e-09 | 1.9e-08 | 1.2e-08 | 1.177E-08 | 7.353E-09 | 4.300E-09 | 1.900E-08 |
| 26617 | 1331 | R1331T | E | 5.6e-09 | 8.4e-09 | 1.8e-08 | 1.067E-08 | 6.503E-09 | 5.600E-09 | 1.800E-08 |
| 26618 | 1331 | R1331W | E | 8.8e-09 | 2.7e-09 | 1.3e-08 | 8.167E-09 | 5.179E-09 | 2.700E-09 | 1.300E-08 |
| 26619 | 1331 | R1331Y | E | 7.1e-09 | 5.8e-09 | 8.5e-09 | 7.133E-09 | 1.350E-09 | 5.800E-09 | 8.500E-09 |
| 26620 | 1331 | R1331V | E | 9.1e-09 | 1.3e-08 | 1.4e-08 | 1.203E-08 | 2.589E-09 | 9.100E-09 | 1.400E-08 |
| 26621 | 1332 | Y1332A | E | 6e-09 | 9.5e-09 | 3.8e-09 | 6.433E-09 | 2.875E-09 | 3.800E-09 | 9.500E-09 |
| 26622 | 1332 | Y1332R | E | 9.3e-09 | 6.9e-09 | 8e-09 | 8.067E-09 | 1.201E-09 | 6.900E-09 | 9.300E-09 |
| 26623 | 1332 | Y1332N | E | 8.3e-09 | 1.2e-08 | 5.3e-09 | 8.533E-09 | 3.356E-09 | 5.300E-09 | 1.200E-08 |
| 26624 | 1332 | Y1332D | E | 4.5e-09 | 6.8e-09 | 1.1e-08 | 7.433E-09 | 3.296E-09 | 4.500E-09 | 1.100E-08 |
| 26625 | 1332 | Y1332C | E | 8.4e-09 | 9.6e-09 | 1.2e-09 | 6.400E-09 | 4.543E-09 | 1.200E-09 | 9.600E-09 |
| 26626 | 1332 | Y1332E | E | 4.2e-09 | 1.2e-08 | 5.5e-09 | 7.233E-09 | 4.179E-09 | 4.200E-09 | 1.200E-08 |
| 26627 | 1332 | Y1332Q | E | 2.7e-09 | 4.1e-09 | 1.1e-08 | 5.933E-09 | 4.443E-09 | 2.700E-09 | 1.100E-08 |
| 26628 | 1332 | Y1332G | E | 6.6e-09 | 1.6e-09 | 1.4e-08 | 7.400E-09 | 6.239E-09 | 1.600E-09 | 1.400E-08 |
| 26629 | 1332 | Y1332H | E | 1.2e-08 | 5.1e-09 | 8.5e-09 | 8.533E-09 | 3.450E-09 | 5.100E-09 | 1.200E-08 |
| 26630 | 1332 | Y1332I | E | 6e-09 | 1.5e-08 | 3.5e-09 | 8.167E-09 | 6.048E-09 | 3.500E-09 | 1.500E-08 |
| 26631 | 1332 | Y1332L | E | 1.2e-08 | 1e-08 | 4.3e-09 | 8.767E-09 | 3.995E-09 | 4.300E-09 | 1.200E-08 |
| 26632 | 1332 | Y1332K | E | 5.7e-09 | 6.7e-09 | 6.1e-09 | 6.167E-09 | 5.033E-10 | 5.700E-09 | 6.700E-09 |
| 26633 | 1332 | Y1332M | E | 1.9e-08 | 1.6e-08 | 4.3e-09 | 1.310E-08 | 7.767E-09 | 4.300E-09 | 1.900E-08 |
| 26634 | 1332 | Y1332F | E | 5.8e-09 | 4.5e-09 | 3.8e-09 | 4.700E-09 | 1.015E-09 | 3.800E-09 | 5.800E-09 |
| 26635 | 1332 | Y1332P | E | 6.3e-09 | 3.9e-09 | 5.4e-09 | 5.200E-09 | 1.212E-09 | 3.900E-09 | 6.300E-09 |
| 26636 | 1332 | Y1332S | E | 7.3e-09 | 8.8e-09 | 1.2e-08 | 9.367E-09 | 2.401E-09 | 7.300E-09 | 1.200E-08 |
| 26637 | 1332 | Y1332T | E | 1.4e-08 | 1e-08 | 1.1e-09 | 8.367E-09 | 6.603E-09 | 1.100E-09 | 1.400E-08 |
| 26638 | 1332 | Y1332W | E | 7.2e-09 | 7.2e-09 | 1.8e-08 | 1.080E-08 | 6.235E-09 | 7.200E-09 | 1.800E-08 |
| 26639 | 1332 | Y1332Y | E | 1.1e-08 | 4.8e-09 | 3.4e-09 | 6.400E-09 | 4.045E-09 | 3.400E-09 | 1.100E-08 |
| 26640 | 1332 | Y1332V | E | 6.7e-09 | 1.4e-08 | 4.8e-09 | 8.500E-09 | 4.857E-09 | 4.800E-09 | 1.400E-08 |
| 26641 | 1333 | A1333A | E | 1.1e-08 | 4.9e-09 | 2.5e-09 | 6.133E-09 | 4.382E-09 | 2.500E-09 | 1.100E-08 |
| 26642 | 1333 | A1333R | E | 4.3e-09 | 4.2e-09 | 8.1e-09 | 5.533E-09 | 2.223E-09 | 4.200E-09 | 8.100E-09 |
| 26643 | 1333 | A1333N | E | 2e-09 | 2.2e-08 | 5.5e-09 | 9.833E-09 | 1.068E-08 | 2.000E-09 | 2.200E-08 |
| 26644 | 1333 | A1333D | E | 8.5e-09 | 1.9e-08 | 4.7e-09 | 1.073E-08 | 7.407E-09 | 4.700E-09 | 1.900E-08 |
| 26645 | 1333 | A1333C | E | 3.1e-08 | 1.5e-08 | 2.9e-09 | 1.630E-08 | 1.410E-08 | 2.900E-09 | 3.100E-08 |
| 26646 | 1333 | A1333E | E | 4.8e-09 | 1.3e-08 | 4.2e-09 | 7.333E-09 | 4.917E-09 | 4.200E-09 | 1.300E-08 |
| 26647 | 1333 | A1333Q | E | 6.6e-09 | 3.9e-09 | 1.6e-08 | 8.833E-09 | 6.352E-09 | 3.900E-09 | 1.600E-08 |
| 26648 | 1333 | A1333G | E | 6.6e-09 | 7.7e-09 | 1.4e-08 | 9.433E-09 | 3.993E-09 | 6.600E-09 | 1.400E-08 |
| 26649 | 1333 | A1333H | E | 9.8e-09 | 9.8e-09 | 9.3e-09 | 9.633E-09 | 2.887E-10 | 9.300E-09 | 9.800E-09 |
| 26650 | 1333 | A1333I | E | 2.5e-09 | 5.3e-09 | 7.2e-09 | 5.000E-09 | 2.364E-09 | 2.500E-09 | 7.200E-09 |
| 26651 | 1333 | A1333L | E | 7.4e-09 | 9.9e-09 | 1.1e-08 | 9.433E-09 | 1.845E-09 | 7.400E-09 | 1.100E-08 |
| 26652 | 1333 | A1333K | E | 5e-09 | 4.6e-09 | 7.2e-09 | 5.600E-09 | 1.400E-09 | 4.600E-09 | 7.200E-09 |
| 26653 | 1333 | A1333M | E | 1.4e-08 | 1.6e-09 | 5.3e-09 | 6.967E-09 | 6.366E-09 | 1.600E-09 | 1.400E-08 |
| 26654 | 1333 | A1333F | E | 5.4e-09 | 1.2e-08 | 5.6e-09 | 7.667E-09 | 3.754E-09 | 5.400E-09 | 1.200E-08 |
| 26655 | 1333 | A1333P | E | 4.1e-09 | 6e-09 | 6.1e-09 | 5.400E-09 | 1.127E-09 | 4.100E-09 | 6.100E-09 |
| 26656 | 1333 | A1333S | E | 4.7e-09 | 4e-09 | 1.6e-08 | 8.233E-09 | 6.735E-09 | 4.000E-09 | 1.600E-08 |
| 26657 | 1333 | A1333T | E | 8.1e-09 | 8e-09 | 5.2e-09 | 7.100E-09 | 1.646E-09 | 5.200E-09 | 8.100E-09 |
| 26658 | 1333 | A1333W | E | 1.7e-08 | 8.6e-09 | 6.9e-09 | 1.083E-08 | 5.408E-09 | 6.900E-09 | 1.700E-08 |
| 26659 | 1333 | A1333Y | E | 1.1e-08 | 8.8e-09 | 7.1e-09 | 8.967E-09 | 1.955E-09 | 7.100E-09 | 1.100E-08 |
| 26660 | 1333 | A1333V | E | 3.7e-09 | 7.5e-09 | 9.6e-09 | 6.933E-09 | 2.991E-09 | 3.700E-09 | 9.600E-09 |
| 26661 | 1334 | D1334A | E | 1.2e-08 | 6.8e-09 | 1.1e-08 | 9.933E-09 | 2.759E-09 | 6.800E-09 | 1.200E-08 |
| 26662 | 1334 | D1334R | E | 1.2e-08 | 4.3e-09 | 8.8e-09 | 8.367E-09 | 3.868E-09 | 4.300E-09 | 1.200E-08 |
| 26663 | 1334 | D1334N | E | 2.5e-08 | 3.9e-08 | 6.2e-09 | 2.340E-08 | 1.646E-08 | 6.200E-09 | 3.900E-08 |
| 26664 | 1334 | D1334D | E | 3.9e-09 | 4e-09 | 6.7e-09 | 4.867E-09 | 1.589E-09 | 3.900E-09 | 6.700E-09 |
| 26665 | 1334 | D1334C | E | 4.6e-09 | 2.4e-09 | 1.5e-08 | 7.333E-09 | 6.730E-09 | 2.400E-09 | 1.500E-08 |
| 26666 | 1334 | D1334E | E | 5.9e-09 | 1.2e-08 | 1.8e-08 | 1.197E-08 | 6.050E-09 | 5.900E-09 | 1.800E-08 |
| 26667 | 1334 | D1334Q | E | 9.1e-09 | 1.2e-08 | 5.3e-09 | 8.800E-09 | 3.360E-09 | 5.300E-09 | 1.200E-08 |
| 26668 | 1334 | D1334G | E | 5.1e-09 | 8.3e-09 | 7.6e-09 | 7.000E-09 | 1.682E-09 | 5.100E-09 | 8.300E-09 |
| 26669 | 1334 | D1334H | E | 8.9e-09 | 6.2e-09 | 5.9e-09 | 7.000E-09 | 1.652E-09 | 5.900E-09 | 8.900E-09 |
| 26670 | 1334 | D1334I | E | 2.3e-09 | 1.3e-08 | 6.7e-09 | 7.333E-09 | 5.378E-09 | 2.300E-09 | 1.300E-08 |
| 26671 | 1334 | D1334L | E | 1e-08 | 1.4e-08 | 4.1e-09 | 9.367E-09 | 4.980E-09 | 4.100E-09 | 1.400E-08 |
| 26672 | 1334 | D1334K | E | 1.6e-08 | 4.4e-09 | 1.6e-08 | 1.213E-08 | 6.697E-09 | 4.400E-09 | 1.600E-08 |
| 26673 | 1334 | D1334M | E | 4.8e-09 | 1.6e-08 | 5.2e-09 | 8.667E-09 | 6.354E-09 | 4.800E-09 | 1.600E-08 |
| 26674 | 1334 | D1334F | E | 1.7e-08 | 1.6e-08 | 8.5e-09 | 1.383E-08 | 4.646E-09 | 8.500E-09 | 1.700E-08 |
| 26675 | 1334 | D1334P | E | 3.8e-09 | 5.9e-09 | 1.1e-08 | 6.900E-09 | 3.703E-09 | 3.800E-09 | 1.100E-08 |
| 26676 | 1334 | D1334S | E | 1.1e-08 | 5.1e-09 | 2e-09 | 6.033E-09 | 4.572E-09 | 2.000E-09 | 1.100E-08 |
| 26677 | 1334 | D1334T | E | 4.9e-09 | 8.1e-09 | 1.4e-08 | 9.000E-09 | 4.616E-09 | 4.900E-09 | 1.400E-08 |
| 26678 | 1334 | D1334W | E | 3.3e-09 | 7.6e-09 | 7.8e-09 | 6.233E-09 | 2.542E-09 | 3.300E-09 | 7.800E-09 |
| 26679 | 1334 | D1334Y | E | 1.2e-08 | 1.4e-08 | 4.9e-09 | 1.030E-08 | 4.782E-09 | 4.900E-09 | 1.400E-08 |
| 26680 | 1334 | D1334V | E | 1.3e-08 | 5.1e-09 | 2.9e-08 | 1.570E-08 | 1.218E-08 | 5.100E-09 | 2.900E-08 |
| 26681 | 1335 | S1335A | E | 5.8e-09 | 9e-09 | 2.9e-09 | 5.900E-09 | 3.051E-09 | 2.900E-09 | 9.000E-09 |
| 26682 | 1335 | S1335R | E | 6.2e-09 | 1.1e-08 | 5e-09 | 7.400E-09 | 3.175E-09 | 5.000E-09 | 1.100E-08 |
| 26683 | 1335 | S1335N | E | 7.6e-09 | 4e-09 | 9.7e-09 | 7.100E-09 | 2.883E-09 | 4.000E-09 | 9.700E-09 |
| 26684 | 1335 | S1335D | E | 7.5e-09 | 7.9e-09 | 3.7e-09 | 6.367E-09 | 2.318E-09 | 3.700E-09 | 7.900E-09 |
| 26685 | 1335 | S1335C | E | 2.8e-08 | 4.3e-09 | 3.7e-09 | 1.200E-08 | 1.386E-08 | 3.700E-09 | 2.800E-08 |
| 26686 | 1335 | S1335E | E | 3.7e-09 | 6.8e-09 | 1.8e-08 | 9.500E-09 | 7.523E-09 | 3.700E-09 | 1.800E-08 |
| 26687 | 1335 | S1335Q | E | 8.5e-09 | 1.1e-08 | 8.3e-09 | 9.267E-09 | 1.504E-09 | 8.300E-09 | 1.100E-08 |
| 26688 | 1335 | S1335G | E | 1.4e-08 | 4.8e-09 | 6.7e-09 | 8.500E-09 | 4.857E-09 | 4.800E-09 | 1.400E-08 |
| 26689 | 1335 | S1335H | E | 1.6e-08 | 9e-09 | 5.2e-09 | 1.007E-08 | 5.478E-09 | 5.200E-09 | 1.600E-08 |

| | | | | | | | | | | |
|---|---|---|---|---|---|---|---|---|---|---|
| 26690 | 1335 | S1335I | E | 3.6e-09 | 7e-09 | 4.9e-09 | 5.167E-09 | 1.716E-09 | 3.600E-09 | 7.000E-09 |
| 26691 | 1335 | S1335L | E | 5.7e-09 | 1.1e-08 | 4.6e-09 | 7.100E-09 | 3.422E-09 | 4.600E-09 | 1.100E-08 |
| 26692 | 1335 | S1335K | E | 1.2e-08 | 1.2e-08 | 2.9e-09 | 8.967E-09 | 5.254E-09 | 2.900E-09 | 1.200E-08 |
| 26693 | 1335 | S1335M | E | 1.2e-08 | 5.1e-09 | 9.7e-09 | 8.933E-09 | 3.513E-09 | 5.100E-09 | 1.200E-08 |
| 26694 | 1335 | S1335F | E | 1.8e-08 | 3.8e-09 | 5.8e-09 | 9.200E-09 | 7.686E-09 | 3.800E-09 | 1.800E-08 |
| 26695 | 1335 | S1335P | E | 1.8e-08 | 4.9e-08 | 7.8e-09 | 2.493E-08 | 2.146E-08 | 7.800E-09 | 4.900E-08 |
| 26696 | 1335 | S1335S | E | 1.5e-08 | 7.1e-09 | 1.6e-08 | 1.270E-08 | 4.875E-09 | 7.100E-09 | 1.600E-08 |
| 26697 | 1335 | S1335T | E | 3.4e-09 | 9e-09 | 1.9e-09 | 4.767E-09 | 3.742E-09 | 1.900E-09 | 9.000E-09 |
| 26698 | 1335 | S1335W | E | 4e-09 | 5.1e-09 | 6.7e-09 | 5.267E-09 | 1.358E-09 | 4.000E-09 | 6.700E-09 |
| 26699 | 1335 | S1335Y | E | 5e-09 | 1.6e-08 | 4.3e-09 | 8.433E-09 | 6.562E-09 | 4.300E-09 | 1.600E-08 |
| 26700 | 1335 | S1335V | E | 9.6e-09 | 6.2e-09 | 3.5e-09 | 6.433E-09 | 3.057E-09 | 3.500E-09 | 9.600E-09 |
| 26701 | 1336 | V1336A | E | 5.8e-09 | 5.5e-09 | 8.8e-09 | 6.700E-09 | 1.825E-09 | 5.500E-09 | 8.800E-09 |
| 26702 | 1336 | V1336R | E | 6e-09 | 3.7e-09 | 3.4e-09 | 4.367E-09 | 1.422E-09 | 3.400E-09 | 6.000E-09 |
| 26703 | 1336 | V1336N | E | 1e-08 | 1.4e-08 | 8.2e-09 | 1.073E-08 | 2.969E-09 | 8.200E-09 | 1.400E-08 |
| 26704 | 1336 | V1336D | E | 9.8e-09 | 4.2e-09 | 4.9e-09 | 6.300E-09 | 3.051E-09 | 4.200E-09 | 9.800E-09 |
| 26705 | 1336 | V1336C | E | 2.3e-09 | 2.9e-09 | 3.5e-09 | 2.900E-09 | 6.000E-10 | 2.300E-09 | 3.500E-09 |
| 26706 | 1336 | V1336E | E | 5.5e-09 | 6.1e-09 | 1e-08 | 7.200E-09 | 2.443E-09 | 5.500E-09 | 1.000E-08 |
| 26707 | 1336 | V1336Q | E | 1.4e-08 | 5.5e-09 | 7.6e-09 | 9.033E-09 | 4.428E-09 | 5.500E-09 | 1.400E-08 |
| 26708 | 1336 | V1336G | E | 9.9e-09 | 1.2e-08 | 4.1e-08 | 2.097E-08 | 1.738E-08 | 9.900E-09 | 4.100E-08 |
| 26709 | 1336 | V1336H | E | 8.6e-09 | 5.9e-09 | 1.6e-09 | 5.367E-09 | 3.530E-09 | 1.600E-09 | 8.600E-09 |
| 26710 | 1336 | V1336I | E | 9.9e-09 | 1.9e-08 | 3.7e-09 | 1.087E-08 | 7.696E-09 | 3.700E-09 | 1.900E-08 |
| 26711 | 1336 | V1336L | E | 5.7e-09 | 7.6e-09 | 8.5e-09 | 7.267E-09 | 1.429E-09 | 5.700E-09 | 8.500E-09 |
| 26712 | 1336 | V1336K | E | 7.8e-09 | 1.8e-08 | 6.8e-09 | 1.087E-08 | 6.198E-09 | 6.800E-09 | 1.800E-08 |
| 26713 | 1336 | V1336M | E | 4e-09 | 5.5e-09 | 4.5e-09 | 4.667E-09 | 7.638E-10 | 4.000E-09 | 5.500E-09 |
| 26714 | 1336 | V1336F | E | 8.9e-09 | 9.8e-09 | 7.3e-09 | 8.667E-09 | 1.266E-09 | 7.300E-09 | 9.800E-09 |
| 26715 | 1336 | V1336P | E | 1.4e-08 | 2.2e-08 | 6.2e-09 | 1.407E-08 | 7.900E-09 | 6.200E-09 | 2.200E-08 |
| 26716 | 1336 | V1336S | E | 3.8e-09 | 6.5e-09 | 4.9e-09 | 5.067E-09 | 1.358E-09 | 3.800E-09 | 6.500E-09 |
| 26717 | 1336 | V1336T | E | 4.7e-09 | 1.3e-08 | 2.1e-09 | 6.600E-09 | 5.693E-09 | 2.100E-09 | 1.300E-08 |
| 26718 | 1336 | V1336W | E | 1.2e-08 | 1.3e-09 | 1.1e-08 | 8.100E-09 | 5.910E-09 | 1.300E-09 | 1.200E-08 |
| 26719 | 1336 | V1336Y | E | 1.2e-08 | 4.1e-09 | 6.4e-09 | 7.500E-09 | 4.063E-09 | 4.100E-09 | 1.200E-08 |
| 26720 | 1336 | V1336V | E | 2.1e-08 | 2.2e-08 | 9.6e-09 | 1.753E-08 | 6.889E-09 | 9.600E-09 | 2.200E-08 |
| 26721 | 1337 | K1337A | E | 4e-09 | 1.2e-08 | 1.6e-08 | 1.067E-08 | 6.110E-09 | 4.000E-09 | 1.600E-08 |
| 26722 | 1337 | K1337R | E | 4.9e-09 | 5e-09 | 1.3e-08 | 7.633E-09 | 4.648E-09 | 4.900E-09 | 1.300E-08 |
| 26723 | 1337 | K1337N | E | 1.3e-08 | 7e-09 | 4.8e-09 | 8.267E-09 | 4.244E-09 | 4.800E-09 | 1.300E-08 |
| 26724 | 1337 | K1337D | E | 8e-09 | 5.9e-09 | 3.9e-09 | 5.933E-09 | 2.050E-09 | 3.900E-09 | 8.000E-09 |
| 26725 | 1337 | K1337C | E | 1.8e-08 | 5.6e-09 | 5.7e-09 | 9.767E-09 | 7.130E-09 | 5.600E-09 | 1.800E-08 |
| 26726 | 1337 | K1337E | E | 8.2e-09 | 1.3e-08 | 4e-09 | 8.400E-09 | 4.503E-09 | 4.000E-09 | 1.300E-08 |
| 26727 | 1337 | K1337Q | E | 7.2e-09 | 4.6e-09 | 3e-08 | 1.393E-08 | 1.397E-08 | 4.600E-09 | 3.000E-08 |
| 26728 | 1337 | K1337G | E | 7.7e-09 | 7.1e-09 | 2.4e-08 | 1.293E-08 | 9.589E-09 | 7.100E-09 | 2.400E-08 |
| 26729 | 1337 | K1337H | E | 4.9e-09 | 1.1e-08 | 5.8e-09 | 7.233E-09 | 3.293E-09 | 4.900E-09 | 1.100E-08 |
| 26730 | 1337 | K1337I | E | 5.8e-09 | 3.7e-09 | 1.1e-08 | 6.833E-09 | 3.758E-09 | 3.700E-09 | 1.100E-08 |
| 26731 | 1337 | K1337L | E | 4.5e-09 | 1.8e-08 | 5.6e-09 | 9.367E-09 | 7.497E-09 | 4.500E-09 | 1.800E-08 |
| 26732 | 1337 | K1337K | E | 6.4e-09 | 6.2e-09 | 9.9e-09 | 7.500E-09 | 2.081E-09 | 6.200E-09 | 9.900E-09 |
| 26733 | 1337 | K1337M | E | 5e-09 | 6.1e-09 | 3.2e-09 | 4.767E-09 | 1.464E-09 | 3.200E-09 | 6.100E-09 |
| 26734 | 1337 | K1337F | E | 9.9e-09 | 1e-08 | 2.2e-08 | 1.397E-08 | 6.957E-09 | 9.900E-09 | 2.200E-08 |
| 26735 | 1337 | K1337P | E | 9.7e-09 | 4.7e-09 | 1e-08 | 8.133E-09 | 2.977E-09 | 4.700E-09 | 1.000E-08 |
| 26736 | 1337 | K1337S | E | 1.7e-09 | 2.7e-09 | 1.2e-08 | 5.467E-09 | 5.680E-09 | 1.700E-09 | 1.200E-08 |
| 26737 | 1337 | K1337T | E | 3.2e-09 | 1.2e-08 | 7.2e-09 | 7.467E-09 | 4.406E-09 | 3.200E-09 | 1.200E-08 |
| 26738 | 1337 | K1337W | E | 1.3e-08 | 6.8e-09 | 4.7e-09 | 8.167E-09 | 4.315E-09 | 4.700E-09 | 1.300E-08 |
| 26739 | 1337 | K1337Y | E | 2.4e-09 | 1.9e-09 | 6.3e-09 | 3.533E-09 | 2.409E-09 | 1.900E-09 | 6.300E-09 |
| 26740 | 1337 | K1337V | E | 7e-09 | 3.1e-09 | 4.1e-09 | 4.733E-09 | 2.026E-09 | 3.100E-09 | 7.000E-09 |
| 26741 | 1338 | G1338A | E | 4.8e-09 | 4.7e-09 | 5.1e-09 | 4.867E-09 | 2.082E-10 | 4.700E-09 | 5.100E-09 |
| 26742 | 1338 | G1338R | E | 5.9e-09 | 2.5e-08 | 8e-09 | 1.297E-08 | 1.047E-08 | 5.900E-09 | 2.500E-08 |
| 26743 | 1338 | G1338N | E | 1.1e-08 | 4.4e-09 | 8e-09 | 7.800E-09 | 3.305E-09 | 4.400E-09 | 1.100E-08 |
| 26744 | 1338 | G1338D | E | 7.5e-09 | 6.3e-09 | 1.3e-08 | 8.933E-09 | 3.573E-09 | 6.300E-09 | 1.300E-08 |
| 26745 | 1338 | G1338C | E | 1.9e-09 | 8.7e-09 | 1e-08 | 6.867E-09 | 4.350E-09 | 1.900E-09 | 1.000E-08 |
| 26746 | 1338 | G1338E | E | 4.4e-09 | 6.7e-09 | 1.5e-08 | 8.700E-09 | 5.576E-09 | 4.400E-09 | 1.500E-08 |
| 26747 | 1338 | G1338Q | E | 7.4e-09 | 6.1e-09 | 8.7e-09 | 7.400E-09 | 1.300E-09 | 6.100E-09 | 8.700E-09 |
| 26748 | 1338 | G1338G | E | 9.5e-09 | 6.6e-09 | 3.2e-09 | 6.433E-09 | 3.153E-09 | 3.200E-09 | 9.500E-09 |
| 26749 | 1338 | G1338H | E | 2.3e-09 | 2.6e-09 | 2.8e-09 | 2.567E-09 | 2.517E-10 | 2.300E-09 | 2.800E-09 |
| 26750 | 1338 | G1338I | E | 5.8e-09 | 7.1e-09 | 1.8e-08 | 1.030E-08 | 6.700E-09 | 5.800E-09 | 1.800E-08 |
| 26751 | 1338 | G1338L | E | 6.7e-09 | 9.7e-09 | 1.1e-08 | 9.133E-09 | 2.205E-09 | 6.700E-09 | 1.100E-08 |
| 26752 | 1338 | G1338K | E | 7.6e-09 | 2.3e-09 | 1.4e-08 | 7.967E-09 | 5.859E-09 | 2.300E-09 | 1.400E-08 |
| 26753 | 1338 | G1338M | E | 2.6e-09 | 2.8e-09 | 2e-08 | 8.467E-09 | 9.989E-09 | 2.600E-09 | 2.000E-08 |
| 26754 | 1338 | G1338F | E | 2.9e-09 | 6.1e-09 | 2.3e-09 | 3.767E-09 | 2.043E-09 | 2.300E-09 | 6.100E-09 |
| 26755 | 1338 | G1338P | E | 2.1e-08 | 2e-08 | 8.7e-09 | 1.657E-08 | 6.831E-09 | 8.700E-09 | 2.100E-08 |
| 26756 | 1338 | G1338S | E | 7.2e-09 | 4.1e-09 | 4e-09 | 5.100E-09 | 1.819E-09 | 4.000E-09 | 7.200E-09 |
| 26757 | 1338 | G1338T | E | 5.5e-09 | 8.5e-09 | 8.7e-09 | 7.567E-09 | 1.793E-09 | 5.500E-09 | 8.700E-09 |
| 26758 | 1338 | G1338W | E | 8.1e-09 | 3.6e-09 | 7e-09 | 6.233E-09 | 2.346E-09 | 3.600E-09 | 8.100E-09 |
| 26759 | 1338 | G1338Y | E | 1.2e-08 | 8.1e-09 | 1.5e-08 | 1.170E-08 | 3.460E-09 | 8.100E-09 | 1.500E-08 |
| 26760 | 1338 | G1338V | E | 8.9e-09 | 2.1e-08 | 4.1e-09 | 1.133E-08 | 8.709E-09 | 4.100E-09 | 2.100E-08 |
| 26761 | 1339 | R1339A | E | 4.7e-09 | 5.1e-09 | 4.8e-09 | 4.867E-09 | 2.082E-10 | 4.700E-09 | 5.100E-09 |
| 26762 | 1339 | R1339R | E | 3.2e-09 | 1.4e-08 | 1.1e-08 | 9.400E-09 | 5.575E-09 | 3.200E-09 | 1.400E-08 |
| 26763 | 1339 | R1339N | E | 9.4e-09 | 4.7e-09 | 8.1e-09 | 7.400E-09 | 2.427E-09 | 4.700E-09 | 9.400E-09 |
| 26764 | 1339 | R1339D | E | 9.5e-09 | 3.4e-08 | 3.2e-08 | 2.517E-08 | 1.360E-08 | 9.500E-09 | 3.400E-08 |
| 26765 | 1339 | R1339C | E | 8e-09 | 9.1e-09 | 4.2e-09 | 7.100E-09 | 2.571E-09 | 4.200E-09 | 9.100E-09 |
| 26766 | 1339 | R1339E | E | 1.5e-08 | 9.1e-09 | 1e-08 | 1.137E-08 | 3.179E-09 | 9.100E-09 | 1.500E-08 |
| 26767 | 1339 | R1339Q | E | 4.3e-09 | 7.6e-09 | 1.8e-08 | 9.967E-09 | 7.150E-09 | 4.300E-09 | 1.800E-08 |
| 26768 | 1339 | R1339G | E | 1.4e-08 | 3.3e-09 | 7.4e-09 | 8.233E-09 | 5.398E-09 | 3.300E-09 | 1.400E-08 |
| 26769 | 1339 | R1339H | E | 1e-08 | 7.6e-09 | 8.6e-09 | 8.733E-09 | 1.206E-09 | 7.600E-09 | 1.000E-08 |
| 26770 | 1339 | R1339I | E | 2.1e-08 | 9e-09 | 6.2e-09 | 1.207E-08 | 7.862E-09 | 6.200E-09 | 2.100E-08 |
| 26771 | 1339 | R1339L | E | 5.5e-09 | 1.1e-08 | 3.8e-09 | 6.767E-09 | 3.763E-09 | 3.800E-09 | 1.100E-08 |
| 26772 | 1339 | R1339K | E | 5.1e-09 | 4.8e-09 | 2.2e-08 | 1.063E-08 | 9.845E-09 | 4.800E-09 | 2.200E-08 |
| 26773 | 1339 | R1339M | E | 3.4e-09 | 1.1e-08 | 7.7e-09 | 7.367E-09 | 3.811E-09 | 3.400E-09 | 1.100E-08 |
| 26774 | 1339 | R1339F | E | 7.2e-09 | 5.4e-09 | 7.3e-09 | 6.633E-09 | 1.069E-09 | 5.400E-09 | 7.300E-09 |

| | | | | | | | | | | |
|---|---|---|---|---|---|---|---|---|---|---|
| 26775 | 1339 | R1339P | E | 5.5e-09 | 3.9e-09 | 1.2e-08 | 7.133E-09 | 4.290E-09 | 3.900E-09 | 1.200E-08 |
| 26776 | 1339 | R1339S | E | 8.6e-09 | 5.9e-09 | 5.1e-09 | 6.533E-09 | 1.834E-09 | 5.100E-09 | 8.600E-09 |
| 26777 | 1339 | R1339T | E | 3.3e-09 | 6.7e-09 | 4.9e-09 | 4.967E-09 | 1.701E-09 | 3.300E-09 | 6.700E-09 |
| 26778 | 1339 | R1339W | E | 3.7e-09 | 1.1e-08 | 1.6e-08 | 1.023E-08 | 6.186E-09 | 3.700E-09 | 1.600E-08 |
| 26779 | 1339 | R1339Y | E | 1.1e-08 | 6.9e-09 | 5.6e-09 | 7.833E-09 | 2.818E-09 | 5.600E-09 | 1.100E-08 |
| 26780 | 1339 | R1339V | E | 6.7e-09 | 6.1e-09 | 5.8e-09 | 6.200E-09 | 4.583E-10 | 5.800E-09 | 6.700E-09 |
| 26781 | 1340 | F1340A | E | 8.6e-09 | 5.8e-09 | 1.1e-08 | 8.467E-09 | 2.603E-09 | 5.800E-09 | 1.100E-08 |
| 26782 | 1340 | F1340R | E | 7.9e-09 | 7.3e-09 | 7.9e-09 | 7.700E-09 | 3.464E-10 | 7.300E-09 | 7.900E-09 |
| 26783 | 1340 | F1340N | E | 3.9e-09 | 1.2e-08 | 5e-09 | 6.967E-09 | 4.394E-09 | 3.900E-09 | 1.200E-08 |
| 26784 | 1340 | F1340D | E | 5.8e-09 | 7.8e-09 | 1e-08 | 7.867E-09 | 2.101E-09 | 5.800E-09 | 1.000E-08 |
| 26785 | 1340 | F1340C | E | 6.7e-09 | 7.5e-09 | 1.1e-08 | 8.400E-09 | 2.287E-09 | 6.700E-09 | 1.100E-08 |
| 26786 | 1340 | F1340E | E | 9.9e-09 | 9.4e-09 | 4.1e-09 | 7.800E-09 | 3.214E-09 | 4.100E-09 | 9.900E-09 |
| 26787 | 1340 | F1340Q | E | 7.7e-09 | 8.8e-09 | 7.4e-09 | 7.967E-09 | 7.371E-10 | 7.400E-09 | 8.800E-09 |
| 26788 | 1340 | F1340G | E | 7.8e-09 | 1.8e-08 | 4.8e-09 | 1.020E-08 | 6.920E-09 | 4.800E-09 | 1.800E-08 |
| 26789 | 1340 | F1340H | E | 1.5e-08 | 5.7e-09 | 1.7e-08 | 1.257E-08 | 6.030E-09 | 5.700E-09 | 1.700E-08 |
| 26790 | 1340 | F1340I | E | 5.2e-09 | 3.7e-09 | 7.4e-09 | 5.433E-09 | 1.861E-09 | 3.700E-09 | 7.400E-09 |
| 26791 | 1340 | F1340L | E | 9.1e-09 | 8.4e-09 | 4.3e-09 | 7.267E-09 | 2.593E-09 | 4.300E-09 | 9.100E-09 |
| 26792 | 1340 | F1340K | E | 9.3e-09 | 2.5e-08 | 1.1e-08 | 1.510E-08 | 8.616E-09 | 9.300E-09 | 2.500E-08 |
| 26793 | 1340 | F1340M | E | 5.6e-09 | 1.8e-08 | 1.3e-08 | 1.220E-08 | 6.239E-09 | 5.600E-09 | 1.800E-08 |
| 26794 | 1340 | F1340F | E | 7.3e-09 | 1.1e-08 | 7.1e-09 | 8.467E-09 | 2.196E-09 | 7.100E-09 | 1.100E-08 |
| 26795 | 1340 | F1340P | E | 6.6e-09 | 5.7e-09 | 5.1e-09 | 5.800E-09 | 7.550E-10 | 5.100E-09 | 6.600E-09 |
| 26796 | 1340 | F1340S | E | 1.5e-08 | 3.3e-08 | 2e-08 | 2.267E-08 | 9.292E-09 | 1.500E-08 | 3.300E-08 |
| 26797 | 1340 | F1340T | E | 1.9e-08 | 9.2e-09 | 5.1e-09 | 1.110E-08 | 7.142E-09 | 5.100E-09 | 1.900E-08 |
| 26798 | 1340 | F1340W | E | 3.1e-09 | 2.5e-08 | 8.6e-09 | 1.223E-08 | 1.139E-08 | 3.100E-09 | 2.500E-08 |
| 26799 | 1340 | F1340Y | E | 4.5e-09 | 1.1e-08 | 3.8e-09 | 6.433E-09 | 3.970E-09 | 3.800E-09 | 1.100E-08 |
| 26800 | 1340 | F1340V | E | 1.3e-08 | 3.8e-09 | 5.5e-09 | 7.433E-09 | 4.895E-09 | 3.800E-09 | 1.300E-08 |
| 26801 | 1341 | T1341A | E | 7.5e-09 | 5.9e-09 | 5.2e-09 | 6.200E-09 | 1.179E-09 | 5.200E-09 | 7.500E-09 |
| 26802 | 1341 | T1341R | E | 4.1e-09 | 5.9e-09 | 2.4e-09 | 4.133E-09 | 1.750E-09 | 2.400E-09 | 5.900E-09 |
| 26803 | 1341 | T1341N | E | 1.3e-08 | 5.6e-09 | 5.3e-09 | 7.967E-09 | 4.362E-09 | 5.300E-09 | 1.300E-08 |
| 26804 | 1341 | T1341D | E | 9.7e-09 | 2.6e-08 | 2.3e-09 | 1.267E-08 | 1.213E-08 | 2.300E-09 | 2.600E-08 |
| 26805 | 1341 | T1341C | E | 6.6e-09 | 1.2e-08 | 6.8e-09 | 8.467E-09 | 3.062E-09 | 6.600E-09 | 1.200E-08 |
| 26806 | 1341 | T1341E | E | 9.2e-09 | 3.2e-09 | 7.5e-09 | 6.633E-09 | 3.092E-09 | 3.200E-09 | 9.200E-09 |
| 26807 | 1341 | T1341Q | E | 8.5e-09 | 1.1e-08 | 9.9e-09 | 9.800E-09 | 1.253E-09 | 8.500E-09 | 1.100E-08 |
| 26808 | 1341 | T1341G | E | 8.4e-09 | 9.5e-09 | 2.2e-09 | 6.700E-09 | 3.936E-09 | 2.200E-09 | 9.500E-09 |
| 26809 | 1341 | T1341H | E | 6.6e-09 | 2.7e-09 | 1.5e-09 | 3.600E-09 | 2.666E-09 | 1.500E-09 | 6.600E-09 |
| 26810 | 1341 | T1341I | E | 1.1e-08 | 1.9e-08 | 2.8e-09 | 1.093E-08 | 8.100E-09 | 2.800E-09 | 1.900E-08 |
| 26811 | 1341 | T1341L | E | 7.2e-09 | 4.9e-09 | 2.6e-08 | 1.270E-08 | 1.158E-08 | 4.900E-09 | 2.600E-08 |
| 26812 | 1341 | T1341K | E | 1.3e-08 | 3e-09 | 3.7e-09 | 6.567E-09 | 5.582E-09 | 3.000E-09 | 1.300E-08 |
| 26813 | 1341 | T1341M | E | 3.2e-09 | 9e-09 | 5e-09 | 5.733E-09 | 2.969E-09 | 3.200E-09 | 9.000E-09 |
| 26814 | 1341 | T1341F | E | 2e-08 | 3.2e-09 | 1.2e-08 | 1.173E-08 | 8.403E-09 | 3.200E-09 | 2.000E-08 |
| 26815 | 1341 | T1341P | E | 8.3e-09 | 9.8e-09 | 2.2e-09 | 6.767E-09 | 4.025E-09 | 2.200E-09 | 9.800E-09 |
| 26816 | 1341 | T1341S | E | 1.2e-08 | 1.5e-08 | 4.7e-09 | 1.057E-08 | 5.297E-09 | 4.700E-09 | 1.500E-08 |
| 26817 | 1341 | T1341T | E | 9.6e-09 | 3e-09 | 9.1e-09 | 7.233E-09 | 3.675E-09 | 3.000E-09 | 9.600E-09 |
| 26818 | 1341 | T1341W | E | 1e-08 | 3.4e-09 | 6.7e-09 | 6.700E-09 | 3.300E-09 | 3.400E-09 | 1.000E-08 |
| 26819 | 1341 | T1341Y | E | 1.1e-08 | 5.7e-09 | 2.7e-08 | 1.457E-08 | 1.109E-08 | 5.700E-09 | 2.700E-08 |
| 26820 | 1341 | T1341V | E | 2.1e-08 | 1.9e-09 | 7.2e-09 | 1.003E-08 | 9.860E-09 | 1.900E-09 | 2.100E-08 |
| 26821 | 1342 | I1342A | E | 1.4e-08 | 3.6e-09 | 1.2e-08 | 9.867E-09 | 5.518E-09 | 3.600E-09 | 1.400E-08 |
| 26822 | 1342 | I1342R | E | 8.7e-09 | 1.2e-08 | 2.6e-09 | 7.767E-09 | 4.769E-09 | 2.600E-09 | 1.200E-08 |
| 26823 | 1342 | I1342N | E | 2.8e-09 | 1.3e-08 | 1.5e-08 | 1.027E-08 | 6.543E-09 | 2.800E-09 | 1.500E-08 |
| 26824 | 1342 | I1342D | E | 1.2e-08 | 7e-09 | 1.6e-08 | 1.167E-08 | 4.509E-09 | 7.000E-09 | 1.600E-08 |
| 26825 | 1342 | I1342C | E | 7.8e-09 | 4.7e-09 | 6e-09 | 6.167E-09 | 1.557E-09 | 4.700E-09 | 7.800E-09 |
| 26826 | 1342 | I1342E | E | 5e-09 | 7.3e-09 | 2.1e-09 | 4.800E-09 | 2.606E-09 | 2.100E-09 | 7.300E-09 |
| 26827 | 1342 | I1342Q | E | 3.6e-09 | 1.2e-08 | 3.1e-09 | 6.233E-09 | 5.000E-09 | 3.100E-09 | 1.200E-08 |
| 26828 | 1342 | I1342G | E | 8e-09 | 4.3e-09 | 2.2e-08 | 1.143E-08 | 9.336E-09 | 4.300E-09 | 2.200E-08 |
| 26829 | 1342 | I1342H | E | 2.5e-09 | 2.8e-08 | 9.3e-09 | 1.327E-08 | 1.320E-08 | 2.500E-09 | 2.800E-08 |
| 26830 | 1342 | I1342I | E | 2e-08 | 1.4e-08 | 9.2e-09 | 1.440E-08 | 5.411E-09 | 9.200E-09 | 2.000E-08 |
| 26831 | 1342 | I1342L | E | 2e-09 | 4.1e-09 | 2.1e-09 | 2.733E-09 | 1.185E-09 | 2.000E-09 | 4.100E-09 |
| 26832 | 1342 | I1342K | E | 1.2e-08 | 2.8e-08 | 3e-09 | 1.433E-08 | 1.266E-08 | 3.000E-09 | 2.800E-08 |
| 26833 | 1342 | I1342M | E | 8.2e-09 | 5.8e-09 | 1.8e-09 | 5.267E-09 | 3.233E-09 | 1.800E-09 | 8.200E-09 |
| 26834 | 1342 | I1342F | E | 6.3e-09 | 1.8e-08 | 1.3e-08 | 1.243E-08 | 5.871E-09 | 6.300E-09 | 1.800E-08 |
| 26835 | 1342 | I1342P | E | 9e-09 | 8.7e-09 | 2e-08 | 1.257E-08 | 6.439E-09 | 8.700E-09 | 2.000E-08 |
| 26836 | 1342 | I1342S | E | 5.3e-09 | 1.2e-08 | 5.7e-09 | 7.667E-09 | 3.758E-09 | 5.300E-09 | 1.200E-08 |
| 26837 | 1342 | I1342T | E | 5.2e-09 | 5.2e-09 | 1.1e-08 | 7.133E-09 | 3.349E-09 | 5.200E-09 | 1.100E-08 |
| 26838 | 1342 | I1342W | E | 3.6e-09 | 1e-08 | 1.3e-08 | 8.867E-09 | 4.801E-09 | 3.600E-09 | 1.300E-08 |
| 26839 | 1342 | I1342Y | E | 5.3e-09 | 4.5e-09 | 1.6e-08 | 8.600E-09 | 6.421E-09 | 4.500E-09 | 1.600E-08 |
| 26840 | 1342 | I1342V | E | 6.1e-09 | 4.8e-09 | 6.1e-09 | 5.667E-09 | 7.506E-10 | 4.800E-09 | 6.100E-09 |
| 26841 | 1343 | S1343A | E | 3.7e-09 | 1.8e-08 | 1e-08 | 1.057E-08 | 7.167E-09 | 3.700E-09 | 1.800E-08 |
| 26842 | 1343 | S1343R | E | 3.2e-09 | 3.8e-09 | 8.6e-09 | 5.200E-09 | 2.960E-09 | 3.200E-09 | 8.600E-09 |
| 26843 | 1343 | S1343N | E | 9.2e-09 | 1.1e-08 | 3.9e-09 | 8.033E-09 | 3.691E-09 | 3.900E-09 | 1.100E-08 |
| 26844 | 1343 | S1343D | E | 1.4e-08 | 5.2e-09 | 3.5e-09 | 7.567E-09 | 5.636E-09 | 3.500E-09 | 1.400E-08 |
| 26845 | 1343 | S1343C | E | 4.1e-09 | 1.1e-08 | 1.4e-08 | 9.700E-09 | 5.076E-09 | 4.100E-09 | 1.400E-08 |
| 26846 | 1343 | S1343E | E | 6.2e-09 | 2.8e-08 | 3.7e-09 | 1.263E-08 | 1.337E-08 | 3.700E-09 | 2.800E-08 |
| 26847 | 1343 | S1343Q | E | 4.7e-09 | 1.1e-08 | 9.9e-09 | 8.533E-09 | 3.365E-09 | 4.700E-09 | 1.100E-08 |
| 26848 | 1343 | S1343G | E | 1.2e-08 | 6.5e-09 | 3.6e-09 | 7.367E-09 | 4.267E-09 | 3.600E-09 | 1.200E-08 |
| 26849 | 1343 | S1343H | E | 3.8e-09 | 5.5e-09 | 5.3e-09 | 4.867E-09 | 9.292E-10 | 3.800E-09 | 5.500E-09 |
| 26850 | 1343 | S1343I | E | 9.9e-09 | 9.3e-09 | 1.4e-08 | 1.107E-08 | 2.558E-09 | 9.300E-09 | 1.400E-08 |
| 26851 | 1343 | S1343L | E | 6.9e-09 | 3e-09 | 5.7e-09 | 5.200E-09 | 1.997E-09 | 3.000E-09 | 6.900E-09 |
| 26852 | 1343 | S1343K | E | 7.9e-09 | 1.9e-08 | 4.8e-09 | 1.057E-08 | 7.466E-09 | 4.800E-09 | 1.900E-08 |
| 26853 | 1343 | S1343M | E | 4.8e-09 | 4.4e-09 | 7.9e-09 | 5.700E-09 | 1.916E-09 | 4.400E-09 | 7.900E-09 |
| 26854 | 1343 | S1343F | E | 4.3e-09 | 6.3e-09 | 3e-08 | 1.353E-08 | 1.430E-08 | 4.300E-09 | 3.000E-08 |
| 26855 | 1343 | S1343P | E | 2.3e-08 | 6.2e-09 | 1.5e-08 | 1.473E-08 | 8.403E-09 | 6.200E-09 | 2.300E-08 |
| 26856 | 1343 | S1343S | E | 9.3e-09 | 1e-08 | 9.5e-10 | 6.750E-09 | 5.035E-09 | 9.500E-10 | 1.000E-08 |
| 26857 | 1343 | S1343T | E | 1.6e-08 | 9.2e-09 | 5.2e-09 | 1.013E-08 | 5.460E-09 | 5.200E-09 | 1.600E-08 |
| 26858 | 1343 | S1343W | E | 1.2e-09 | 2.2e-09 | 4.5e-09 | 2.633E-09 | 1.692E-09 | 1.200E-09 | 4.500E-09 |
| 26859 | 1343 | S1343Y | E | 1.1e-08 | 8.7e-09 | 1e-08 | 9.900E-09 | 1.153E-09 | 8.700E-09 | 1.100E-08 |

| | | | | | | | | | | |
|---|---|---|---|---|---|---|---|---|---|---|
| 26860 | 1343 | S1343V | E | 6.7e-09 | 3.1e-09 | 3.5e-08 | 1.493E-08 | 1.747E-08 | 3.100E-09 | 3.500E-08 |
| 26861 | 1344 | A1344A | E | 5.8e-09 | 1.4e-08 | 6.6e-09 | 8.800E-09 | 4.521E-09 | 5.800E-09 | 1.400E-08 |
| 26862 | 1344 | A1344R | E | 8.5e-09 | 1.7e-09 | 4.8e-09 | 5.000E-09 | 3.404E-09 | 1.700E-09 | 8.500E-09 |
| 26863 | 1344 | A1344N | E | 1.1e-08 | 9.2e-09 | 2.8e-09 | 7.667E-09 | 4.310E-09 | 2.800E-09 | 1.100E-08 |
| 26864 | 1344 | A1344D | E | 1.2e-08 | 4.3e-09 | 1.3e-09 | 5.867E-09 | 5.519E-09 | 1.300E-09 | 1.200E-08 |
| 26865 | 1344 | A1344C | E | 4.9e-09 | 3.6e-08 | 3.8e-09 | 1.490E-08 | 1.828E-08 | 3.800E-09 | 3.600E-08 |
| 26866 | 1344 | A1344E | E | 9.9e-09 | 1.8e-08 | 1.1e-08 | 1.297E-08 | 4.394E-09 | 9.900E-09 | 1.800E-08 |
| 26867 | 1344 | A1344Q | E | 9.2e-09 | 4.8e-09 | 7.8e-09 | 7.267E-09 | 2.248E-09 | 4.800E-09 | 9.200E-09 |
| 26868 | 1344 | A1344G | E | 1e-08 | 2.5e-08 | 6.1e-09 | 1.370E-08 | 9.978E-09 | 6.100E-09 | 2.500E-08 |
| 26869 | 1344 | A1344H | E | 7.3e-09 | 1.2e-08 | 5.7e-09 | 8.333E-09 | 3.275E-09 | 5.700E-09 | 1.200E-08 |
| 26870 | 1344 | A1344I | E | 9e-09 | 3e-09 | 6.9e-09 | 6.300E-09 | 3.045E-09 | 3.000E-09 | 9.000E-09 |
| 26871 | 1344 | A1344L | E | 9.1e-09 | 1.5e-08 | 1e-08 | 1.137E-08 | 3.179E-09 | 9.100E-09 | 1.500E-08 |
| 26872 | 1344 | A1344K | E | 5.8e-09 | 5.7e-09 | 1.5e-08 | 8.833E-09 | 5.341E-09 | 5.700E-09 | 1.500E-08 |
| 26873 | 1344 | A1344M | E | 9.3e-09 | 5.1e-09 | 9.3e-09 | 7.900E-09 | 2.425E-09 | 5.100E-09 | 9.300E-09 |
| 26874 | 1344 | A1344F | E | 4.3e-09 | 6.1e-09 | 1.5e-08 | 8.467E-09 | 5.729E-09 | 4.300E-09 | 1.500E-08 |
| 26875 | 1344 | A1344P | E | 4.2e-09 | 8.6e-09 | 2.1e-08 | 1.127E-08 | 8.712E-09 | 4.200E-09 | 2.100E-08 |
| 26876 | 1344 | A1344S | E | 5.8e-09 | 7.7e-09 | 9.8e-09 | 7.767E-09 | 2.001E-09 | 5.800E-09 | 9.800E-09 |
| 26877 | 1344 | A1344T | E | 1.3e-09 | 6.4e-09 | 4.7e-09 | 4.133E-09 | 2.597E-09 | 1.300E-09 | 6.400E-09 |
| 26878 | 1344 | A1344W | E | 2.7e-08 | 7e-09 | 1.8e-08 | 1.733E-08 | 1.002E-08 | 7.000E-09 | 2.700E-08 |
| 26879 | 1344 | A1344Y | E | 5.9e-09 | 1.4e-08 | 6.1e-09 | 8.667E-09 | 4.620E-09 | 5.900E-09 | 1.400E-08 |
| 26880 | 1344 | A1344V | E | 1.2e-08 | 3.2e-09 | 1.4e-08 | 9.733E-09 | 5.746E-09 | 3.200E-09 | 1.400E-08 |
| 26881 | 1345 | D1345A | E | 8.9e-09 | 7.2e-09 | 5.5e-09 | 7.200E-09 | 1.700E-09 | 5.500E-09 | 8.900E-09 |
| 26882 | 1345 | D1345R | E | 3.9e-09 | 1.7e-08 | 7.1e-09 | 9.333E-09 | 6.830E-09 | 3.900E-09 | 1.700E-08 |
| 26883 | 1345 | D1345N | E | 1.6e-08 | 2e-08 | 8.4e-09 | 1.480E-08 | 5.892E-09 | 8.400E-09 | 2.000E-08 |
| 26884 | 1345 | D1345D | E | 2e-09 | 1.3e-08 | 3.3e-08 | 1.600E-08 | 1.572E-08 | 2.000E-09 | 3.300E-08 |
| 26885 | 1345 | D1345C | E | 4.8e-09 | 8.2e-09 | 2.4e-09 | 5.133E-09 | 2.914E-09 | 2.400E-09 | 8.200E-09 |
| 26886 | 1345 | D1345E | E | 1.2e-08 | 1.5e-09 | 6.3e-09 | 6.600E-09 | 5.256E-09 | 1.500E-09 | 1.200E-08 |
| 26887 | 1345 | D1345Q | E | 8e-09 | 6e-09 | 5.9e-09 | 6.633E-09 | 1.185E-09 | 5.900E-09 | 8.000E-09 |
| 26888 | 1345 | D1345G | E | 1.9e-08 | 7.7e-09 | 4.3e-09 | 1.033E-08 | 7.696E-09 | 4.300E-09 | 1.900E-08 |
| 26889 | 1345 | D1345H | E | 1.7e-09 | 3e-09 | 5.1e-09 | 3.267E-09 | 1.716E-09 | 1.700E-09 | 5.100E-09 |
| 26890 | 1345 | D1345I | E | 5.6e-09 | 9.6e-09 | 4.4e-09 | 6.533E-09 | 2.723E-09 | 4.400E-09 | 9.600E-09 |
| 26891 | 1345 | D1345L | E | 2.1e-09 | 3.4e-09 | 1e-08 | 5.167E-09 | 4.236E-09 | 2.100E-09 | 1.000E-08 |
| 26892 | 1345 | D1345K | E | 1.5e-08 | 4.2e-09 | 1.4e-08 | 1.107E-08 | 5.968E-09 | 4.200E-09 | 1.500E-08 |
| 26893 | 1345 | D1345M | E | 8.8e-09 | 3.5e-09 | 3.3e-09 | 5.200E-09 | 3.119E-09 | 3.300E-09 | 8.800E-09 |
| 26894 | 1345 | D1345F | E | 8.5e-09 | 4.4e-09 | 2.8e-09 | 5.233E-09 | 2.940E-09 | 2.800E-09 | 8.500E-09 |
| 26895 | 1345 | D1345P | E | 3.5e-09 | 6.1e-09 | 1.9e-08 | 9.533E-09 | 8.301E-09 | 3.500E-09 | 1.900E-08 |
| 26896 | 1345 | D1345S | E | 3.9e-09 | 5.6e-09 | 4.4e-09 | 4.633E-09 | 8.737E-10 | 3.900E-09 | 5.600E-09 |
| 26897 | 1345 | D1345T | E | 1.9e-08 | 6.8e-10 | 1.2e-08 | 1.056E-08 | 9.245E-09 | 6.800E-10 | 1.900E-08 |
| 26898 | 1345 | D1345W | E | 6.2e-09 | 5.6e-09 | 2.9e-09 | 4.900E-09 | 1.758E-09 | 2.900E-09 | 6.200E-09 |
| 26899 | 1345 | D1345Y | E | 1.3e-08 | 7.9e-09 | 9.6e-09 | 1.017E-08 | 2.597E-09 | 7.900E-09 | 1.300E-08 |
| 26900 | 1345 | D1345V | E | 6.2e-09 | 2e-08 | 4e-09 | 1.007E-08 | 8.673E-09 | 4.000E-09 | 2.000E-08 |
| 26901 | 1346 | T1346A | E | 1.2e-08 | 1.6e-08 | 1.5e-09 | 9.833E-09 | 7.489E-09 | 1.500E-09 | 1.600E-08 |
| 26902 | 1346 | T1346R | E | 7.9e-09 | 3.6e-09 | 1.1e-08 | 7.500E-09 | 3.716E-09 | 3.600E-09 | 1.100E-08 |
| 26903 | 1346 | T1346N | E | 1.1e-08 | 1e-08 | 6e-09 | 9.000E-09 | 2.646E-09 | 6.000E-09 | 1.100E-08 |
| 26904 | 1346 | T1346D | E | 5.1e-09 | 6.6e-09 | 1.6e-08 | 9.233E-09 | 5.908E-09 | 5.100E-09 | 1.600E-08 |
| 26905 | 1346 | T1346C | E | 6.1e-09 | 6.1e-09 | 4.9e-09 | 5.700E-09 | 6.928E-10 | 4.900E-09 | 6.100E-09 |
| 26906 | 1346 | T1346E | E | 7.6e-09 | 3.4e-09 | 1.3e-08 | 8.000E-09 | 4.812E-09 | 3.400E-09 | 1.300E-08 |
| 26907 | 1346 | T1346Q | E | 5e-09 | 6.8e-09 | 6.5e-09 | 6.100E-09 | 9.644E-10 | 5.000E-09 | 6.800E-09 |
| 26908 | 1346 | T1346G | E | 6.6e-09 | 1e-08 | 6.2e-09 | 7.600E-09 | 2.088E-09 | 6.200E-09 | 1.000E-08 |
| 26909 | 1346 | T1346H | E | 7.8e-09 | 8e-09 | 1.5e-08 | 1.027E-08 | 4.100E-09 | 7.800E-09 | 1.500E-08 |
| 26910 | 1346 | T1346I | E | 6e-09 | 8.8e-09 | 1.4e-08 | 9.600E-09 | 4.060E-09 | 6.000E-09 | 1.400E-08 |
| 26911 | 1346 | T1346L | E | 8.5e-09 | 2.8e-09 | 1.3e-08 | 8.100E-09 | 5.112E-09 | 2.800E-09 | 1.300E-08 |
| 26912 | 1346 | T1346K | E | 7.9e-09 | 3.4e-09 | 1.5e-08 | 8.767E-09 | 5.848E-09 | 3.400E-09 | 1.500E-08 |
| 26913 | 1346 | T1346M | E | 1.2e-08 | 8.4e-09 | 8.8e-09 | 9.733E-09 | 1.973E-09 | 8.400E-09 | 1.200E-08 |
| 26914 | 1346 | T1346F | E | 4.1e-09 | 5.8e-09 | 2.1e-08 | 1.030E-08 | 9.305E-09 | 4.100E-09 | 2.100E-08 |
| 26915 | 1346 | T1346P | E | 8.6e-09 | 7.5e-09 | 3.2e-09 | 6.433E-09 | 2.854E-09 | 3.200E-09 | 8.600E-09 |
| 26916 | 1346 | T1346S | E | 3.5e-09 | 6.3e-09 | 2.7e-09 | 4.167E-09 | 1.890E-09 | 2.700E-09 | 6.300E-09 |
| 26917 | 1346 | T1346T | E | 3.6e-09 | 1.5e-08 | 1e-08 | 9.533E-09 | 5.714E-09 | 3.600E-09 | 1.500E-08 |
| 26918 | 1346 | T1346W | E | 1.5e-08 | 2.4e-08 | 1.3e-08 | 1.733E-08 | 5.859E-09 | 1.300E-08 | 2.400E-08 |
| 26919 | 1346 | T1346Y | E | 9.8e-09 | 1.1e-08 | 7.5e-09 | 9.433E-09 | 1.779E-09 | 7.500E-09 | 1.100E-08 |
| 26920 | 1346 | T1346V | E | 1.9e-08 | 9e-09 | 9.1e-09 | 1.237E-08 | 5.745E-09 | 9.000E-09 | 1.900E-08 |
| 26921 | 1347 | S1347A | E | 6.8e-09 | 1e-08 | 3.6e-08 | 1.760E-08 | 1.601E-08 | 6.800E-09 | 3.600E-08 |
| 26922 | 1347 | S1347R | E | 4.9e-09 | 6.4e-09 | 3.3e-09 | 4.867E-09 | 1.550E-09 | 3.300E-09 | 6.400E-09 |
| 26923 | 1347 | S1347N | E | 1.3e-08 | 3.4e-09 | 2.4e-08 | 1.347E-08 | 1.031E-08 | 3.400E-09 | 2.400E-08 |
| 26924 | 1347 | S1347D | E | 6.1e-09 | 7.2e-09 | 5.2e-09 | 6.167E-09 | 1.002E-09 | 5.200E-09 | 7.200E-09 |
| 26925 | 1347 | S1347C | E | 6.3e-09 | 2.2e-08 | 4.3e-09 | 1.087E-08 | 9.693E-09 | 4.300E-09 | 2.200E-08 |
| 26926 | 1347 | S1347E | E | 3.9e-09 | 5.8e-09 | 1e-08 | 6.567E-09 | 3.121E-09 | 3.900E-09 | 1.000E-08 |
| 26927 | 1347 | S1347Q | E | 4.1e-09 | 2.8e-09 | 1.4e-08 | 6.967E-09 | 6.126E-09 | 2.800E-09 | 1.400E-08 |
| 26928 | 1347 | S1347G | E | 9.3e-09 | 4.3e-09 | 7.3e-09 | 6.967E-09 | 2.517E-09 | 4.300E-09 | 9.300E-09 |
| 26929 | 1347 | S1347H | E | 5.2e-09 | 5.1e-09 | 1.2e-08 | 7.433E-09 | 3.955E-09 | 5.100E-09 | 1.200E-08 |
| 26930 | 1347 | S1347I | E | 1.1e-08 | 1.8e-09 | 9e-09 | 7.267E-09 | 4.839E-09 | 1.800E-09 | 1.100E-08 |
| 26931 | 1347 | S1347L | E | 5.7e-09 | 5.1e-09 | 5.8e-09 | 5.533E-09 | 3.786E-10 | 5.100E-09 | 5.800E-09 |
| 26932 | 1347 | S1347K | E | 1e-08 | 3.6e-09 | 5.4e-09 | 6.333E-09 | 3.301E-09 | 3.600E-09 | 1.000E-08 |
| 26933 | 1347 | S1347M | E | 1.1e-08 | 9.1e-09 | 4.5e-09 | 8.200E-09 | 3.342E-09 | 4.500E-09 | 1.100E-08 |
| 26934 | 1347 | S1347F | E | 1.8e-08 | 1.2e-08 | 3.8e-09 | 1.127E-08 | 7.128E-09 | 3.800E-09 | 1.800E-08 |
| 26935 | 1347 | S1347P | E | 2.2e-09 | 7e-09 | 1.1e-08 | 6.733E-09 | 4.406E-09 | 2.200E-09 | 1.100E-08 |
| 26936 | 1347 | S1347S | E | 1.1e-08 | 2.5e-08 | 3.9e-09 | 1.330E-08 | 1.074E-08 | 3.900E-09 | 2.500E-08 |
| 26937 | 1347 | S1347T | E | 3.3e-09 | 4.4e-09 | 1.3e-08 | 6.900E-09 | 5.311E-09 | 3.300E-09 | 1.300E-08 |
| 26938 | 1347 | S1347W | E | 8.8e-09 | 1.2e-08 | 4.3e-09 | 8.367E-09 | 3.868E-09 | 4.300E-09 | 1.200E-08 |
| 26939 | 1347 | S1347Y | E | 1e-08 | 6.7e-09 | 9.3e-09 | 8.667E-09 | 1.739E-09 | 6.700E-09 | 1.000E-08 |
| 26940 | 1347 | S1347V | E | 8.4e-09 | 8e-09 | 3.5e-09 | 6.633E-09 | 2.721E-09 | 3.500E-09 | 8.400E-09 |
| 26941 | 1348 | K1348A | E | 1.3e-09 | 9.8e-09 | 4.2e-09 | 5.100E-09 | 4.321E-09 | 1.300E-09 | 9.800E-09 |
| 26942 | 1348 | K1348R | E | 1.9e-09 | 1.3e-08 | 6.3e-09 | 7.067E-09 | 5.590E-09 | 1.900E-09 | 1.300E-08 |
| 26943 | 1348 | K1348N | E | 5e-09 | 6.6e-09 | 2.8e-09 | 4.800E-09 | 1.908E-09 | 2.800E-09 | 6.600E-09 |
| 26944 | 1348 | K1348D | E | 2e-09 | 4.2e-09 | 1.1e-08 | 5.733E-09 | 4.692E-09 | 2.000E-09 | 1.100E-08 |

| | | | | | | | | | | |
|---|---|---|---|---|---|---|---|---|---|---|
| 26945 | 1348 | K1348C | E | 3.7e-08 | 1.4e-08 | 2.2e-08 | 2.433E-08 | 1.168E-08 | 1.400E-08 | 3.700E-08 |
| 26946 | 1348 | K1348E | E | 3.7e-09 | 1.1e-08 | 9e-09 | 7.900E-09 | 3.772E-09 | 3.700E-09 | 1.100E-08 |
| 26947 | 1348 | K1348Q | E | 5.8e-09 | 5.4e-09 | 3.5e-09 | 4.900E-09 | 1.229E-09 | 3.500E-09 | 5.800E-09 |
| 26948 | 1348 | K1348G | E | 5.2e-09 | 6.2e-09 | 5.1e-09 | 5.500E-09 | 6.083E-10 | 5.100E-09 | 6.200E-09 |
| 26949 | 1348 | K1348H | E | 4.9e-09 | 4e-09 | 8.4e-09 | 5.767E-09 | 2.325E-09 | 4.000E-09 | 8.400E-09 |
| 26950 | 1348 | K1348I | E | 1.3e-08 | 7.4e-09 | 6.1e-09 | 8.833E-09 | 3.667E-09 | 6.100E-09 | 1.300E-08 |
| 26951 | 1348 | K1348L | E | 4.9e-09 | 5.8e-09 | 5.9e-09 | 5.533E-09 | 5.508E-10 | 4.900E-09 | 5.900E-09 |
| 26952 | 1348 | K1348K | E | 8.2e-09 | 1.3e-08 | 8.4e-09 | 9.867E-09 | 2.715E-09 | 8.200E-09 | 1.300E-08 |
| 26953 | 1348 | K1348M | E | 1.7e-08 | 5.4e-09 | 9.6e-09 | 1.067E-08 | 5.873E-09 | 5.400E-09 | 1.700E-08 |
| 26954 | 1348 | K1348F | E | 3.3e-09 | 7.9e-09 | 8.6e-09 | 6.600E-09 | 2.879E-09 | 3.300E-09 | 8.600E-09 |
| 26955 | 1348 | K1348P | E | 1.2e-08 | 5.6e-09 | 4.3e-09 | 7.300E-09 | 4.122E-09 | 4.300E-09 | 1.200E-08 |
| 26956 | 1348 | K1348S | E | 5.9e-09 | 4.9e-09 | 5.1e-09 | 5.300E-09 | 5.292E-10 | 4.900E-09 | 5.900E-09 |
| 26957 | 1348 | K1348T | E | 3.1e-09 | 6.2e-09 | 2.2e-08 | 1.043E-08 | 1.014E-08 | 3.100E-09 | 2.200E-08 |
| 26958 | 1348 | K1348W | E | 4.4e-09 | 5.9e-09 | 8e-09 | 6.100E-09 | 1.808E-09 | 4.400E-09 | 8.000E-09 |
| 26959 | 1348 | K1348Y | E | 2.9e-09 | 5.2e-09 | 1.3e-08 | 7.033E-09 | 5.294E-09 | 2.900E-09 | 1.300E-08 |
| 26960 | 1348 | K1348V | E | 7.3e-09 | 6.5e-09 | 7.6e-09 | 7.133E-09 | 5.686E-10 | 6.500E-09 | 7.600E-09 |
| 26961 | 1349 | N1349A | E | 9.8e-09 | 4.5e-09 | 8.7e-09 | 7.667E-09 | 2.797E-09 | 4.500E-09 | 9.800E-09 |
| 26962 | 1349 | N1349R | E | 1.9e-08 | 9.6e-09 | 8.1e-09 | 1.223E-08 | 5.908E-09 | 8.100E-09 | 1.900E-08 |
| 26963 | 1349 | N1349N | E | 1e-08 | 8e-09 | 8.1e-09 | 8.700E-09 | 1.127E-09 | 8.000E-09 | 1.000E-08 |
| 26964 | 1349 | N1349D | E | 2.7e-09 | 5.1e-09 | 1e-08 | 5.933E-09 | 3.721E-09 | 2.700E-09 | 1.000E-08 |
| 26965 | 1349 | N1349C | E | 1.4e-08 | 1.3e-08 | 6.4e-09 | 1.113E-08 | 4.130E-09 | 6.400E-09 | 1.400E-08 |
| 26966 | 1349 | N1349E | E | 1.4e-08 | 3.5e-09 | 2.7e-08 | 1.483E-08 | 1.177E-08 | 3.500E-09 | 2.700E-08 |
| 26967 | 1349 | N1349Q | E | 8.3e-09 | 8.9e-09 | 1.5e-08 | 1.073E-08 | 3.707E-09 | 8.300E-09 | 1.500E-08 |
| 26968 | 1349 | N1349G | E | 1.2e-08 | 5.9e-09 | 1e-08 | 9.300E-09 | 3.110E-09 | 5.900E-09 | 1.200E-08 |
| 26969 | 1349 | N1349H | E | 1.2e-08 | 4.6e-09 | 4.8e-09 | 7.133E-09 | 4.216E-09 | 4.600E-09 | 1.200E-08 |
| 26970 | 1349 | N1349I | E | 1.5e-09 | 3.4e-09 | 6.1e-09 | 3.667E-09 | 2.312E-09 | 1.500E-09 | 6.100E-09 |
| 26971 | 1349 | N1349L | E | 1.7e-08 | 7.9e-09 | 1.3e-08 | 1.263E-08 | 4.561E-09 | 7.900E-09 | 1.700E-08 |
| 26972 | 1349 | N1349K | E | 7.6e-09 | 8.4e-09 | 6.1e-09 | 7.367E-09 | 1.168E-09 | 6.100E-09 | 8.400E-09 |
| 26973 | 1349 | N1349M | E | 5.3e-09 | 1.2e-08 | 2.7e-08 | 1.477E-08 | 1.111E-08 | 5.300E-09 | 2.700E-08 |
| 26974 | 1349 | N1349F | E | 1.1e-08 | 4e-09 | 3.1e-09 | 6.033E-09 | 4.325E-09 | 3.100E-09 | 1.100E-08 |
| 26975 | 1349 | N1349P | E | 9e-09 | 4.6e-09 | 3e-08 | 1.453E-08 | 1.357E-08 | 4.600E-09 | 3.000E-08 |
| 26976 | 1349 | N1349S | E | 4.6e-09 | 7.9e-09 | 1.1e-08 | 7.833E-09 | 3.201E-09 | 4.600E-09 | 1.100E-08 |
| 26977 | 1349 | N1349T | E | 1.3e-08 | 8.1e-09 | 1.3e-08 | 1.137E-08 | 2.829E-09 | 8.100E-09 | 1.300E-08 |
| 26978 | 1349 | N1349W | E | 6.7e-09 | 5.3e-09 | 1.6e-08 | 9.333E-09 | 5.816E-09 | 5.300E-09 | 1.600E-08 |
| 26979 | 1349 | N1349Y | E | 4.3e-09 | 1.9e-08 | 8.2e-09 | 1.050E-08 | 7.615E-09 | 4.300E-09 | 1.900E-08 |
| 26980 | 1349 | N1349V | E | 3.3e-08 | 7.6e-09 | 6.7e-09 | 1.577E-08 | 1.493E-08 | 6.700E-09 | 3.300E-08 |
| 26981 | 1350 | T1350A | E | 6.8e-09 | 4.8e-09 | 8.8e-09 | 6.800E-09 | 2.000E-09 | 4.800E-09 | 8.800E-09 |
| 26982 | 1350 | T1350R | E | 8.5e-09 | 6.2e-09 | 4.4e-09 | 6.367E-09 | 2.055E-09 | 4.400E-09 | 8.500E-09 |
| 26983 | 1350 | T1350N | E | 1.4e-08 | 9.1e-09 | 2.6e-09 | 8.567E-09 | 5.719E-09 | 2.600E-09 | 1.400E-08 |
| 26984 | 1350 | T1350D | E | 4.7e-09 | 6.4e-09 | 3.2e-09 | 4.767E-09 | 1.601E-09 | 3.200E-09 | 6.400E-09 |
| 26985 | 1350 | T1350C | E | 6.7e-09 | 9.2e-09 | 8.8e-09 | 8.233E-09 | 1.343E-09 | 6.700E-09 | 9.200E-09 |
| 26986 | 1350 | T1350E | E | 1.6e-08 | 5.7e-09 | 8.3e-09 | 1.000E-08 | 5.356E-09 | 5.700E-09 | 1.600E-08 |
| 26987 | 1350 | T1350Q | E | 5.4e-09 | 2.1e-08 | 6.4e-09 | 1.093E-08 | 8.732E-09 | 5.400E-09 | 2.100E-08 |
| 26988 | 1350 | T1350G | E | 7.3e-09 | 1.2e-08 | 6.5e-09 | 8.600E-09 | 2.972E-09 | 6.500E-09 | 1.200E-08 |
| 26989 | 1350 | T1350H | E | 1e-08 | 1.1e-08 | 9e-09 | 1.000E-08 | 1.000E-09 | 9.000E-09 | 1.100E-08 |
| 26990 | 1350 | T1350I | E | 3.4e-09 | 6.7e-09 | 2e-08 | 1.003E-08 | 8.788E-09 | 3.400E-09 | 2.000E-08 |
| 26991 | 1350 | T1350L | E | 1.3e-08 | 1e-08 | 8.9e-09 | 1.063E-08 | 2.122E-09 | 8.900E-09 | 1.300E-08 |
| 26992 | 1350 | T1350K | E | 1.3e-08 | 1.2e-08 | 2.7e-09 | 9.233E-09 | 5.680E-09 | 2.700E-09 | 1.300E-08 |
| 26993 | 1350 | T1350M | E | 1.7e-08 | 7.3e-09 | 8.5e-09 | 1.093E-08 | 5.288E-09 | 7.300E-09 | 1.700E-08 |
| 26994 | 1350 | T1350F | E | 6.2e-09 | 5.5e-09 | 1.1e-08 | 7.567E-09 | 2.994E-09 | 5.500E-09 | 1.100E-08 |
| 26995 | 1350 | T1350P | E | 1.7e-08 | 1e-08 | 7.5e-09 | 1.150E-08 | 4.924E-09 | 7.500E-09 | 1.700E-08 |
| 26996 | 1350 | T1350S | E | 9.8e-09 | 5.4e-09 | 6.7e-09 | 7.300E-09 | 2.261E-09 | 5.400E-09 | 9.800E-09 |
| 26997 | 1350 | T1350T | E | 4.5e-09 | 4e-09 | 2.2e-08 | 1.017E-08 | 1.025E-08 | 4.000E-09 | 2.200E-08 |
| 26998 | 1350 | T1350W | E | 9.7e-09 | 2.3e-08 | 6.6e-09 | 1.310E-08 | 8.713E-09 | 6.600E-09 | 2.300E-08 |
| 26999 | 1350 | T1350Y | E | 1.2e-08 | 1.7e-08 | 2.9e-09 | 1.063E-08 | 7.149E-09 | 2.900E-09 | 1.700E-08 |
| 27000 | 1350 | T1350V | E | 3.3e-08 | 1.2e-08 | 1.7e-08 | 2.067E-08 | 1.097E-08 | 1.200E-08 | 3.300E-08 |
| 27001 | 1351 | A1351A | E | 3.4e-09 | 3.2e-09 | 5.2e-09 | 3.933E-09 | 1.102E-09 | 3.200E-09 | 5.200E-09 |
| 27002 | 1351 | A1351R | E | 7.8e-09 | 4.6e-09 | 8.6e-09 | 7.000E-09 | 2.117E-09 | 4.600E-09 | 8.600E-09 |
| 27003 | 1351 | A1351N | E | 4.4e-09 | 6.2e-09 | 6.1e-09 | 5.567E-09 | 1.012E-09 | 4.400E-09 | 6.200E-09 |
| 27004 | 1351 | A1351D | E | 8e-09 | 7.6e-09 | 3.4e-09 | 6.333E-09 | 2.548E-09 | 3.400E-09 | 8.000E-09 |
| 27005 | 1351 | A1351C | E | 7.7e-09 | 7.4e-09 | 9.9e-09 | 8.333E-09 | 1.365E-09 | 7.400E-09 | 9.900E-09 |
| 27006 | 1351 | A1351E | E | 1.3e-08 | 4.7e-09 | 1.1e-08 | 9.567E-09 | 4.332E-09 | 4.700E-09 | 1.300E-08 |
| 27007 | 1351 | A1351Q | E | 1.1e-08 | 7.1e-09 | 1.4e-08 | 1.070E-08 | 3.460E-09 | 7.100E-09 | 1.400E-08 |
| 27008 | 1351 | A1351G | E | 9.4e-09 | 5.7e-09 | 9.8e-09 | 8.300E-09 | 2.261E-09 | 5.700E-09 | 9.800E-09 |
| 27009 | 1351 | A1351H | E | 1.2e-08 | 6.6e-09 | 1.1e-08 | 9.867E-09 | 2.873E-09 | 6.600E-09 | 1.200E-08 |
| 27010 | 1351 | A1351I | E | 1.6e-09 | 2.9e-08 | 1.2e-08 | 1.420E-08 | 1.383E-08 | 1.600E-09 | 2.900E-08 |
| 27011 | 1351 | A1351L | E | 6.5e-09 | 1.1e-08 | 7.7e-09 | 8.400E-09 | 2.330E-09 | 6.500E-09 | 1.100E-08 |
| 27012 | 1351 | A1351K | E | 8e-09 | 6.7e-09 | 1.5e-08 | 9.900E-09 | 4.464E-09 | 6.700E-09 | 1.500E-08 |
| 27013 | 1351 | A1351M | E | 2.3e-08 | 7.6e-09 | 5.5e-09 | 1.203E-08 | 9.555E-09 | 5.500E-09 | 2.300E-08 |
| 27014 | 1351 | A1351F | E | 1.4e-08 | 9.1e-09 | 5.6e-09 | 9.567E-09 | 4.219E-09 | 5.600E-09 | 1.400E-08 |
| 27015 | 1351 | A1351P | E | 1.1e-08 | 5.6e-09 | 6.3e-09 | 7.633E-09 | 2.937E-09 | 5.600E-09 | 1.100E-08 |
| 27016 | 1351 | A1351S | E | 4.1e-09 | 5.7e-09 | 6.5e-09 | 5.433E-09 | 1.222E-09 | 4.100E-09 | 6.500E-09 |
| 27017 | 1351 | A1351T | E | 1.5e-08 | 1.3e-08 | 7.5e-09 | 1.183E-08 | 3.884E-09 | 7.500E-09 | 1.500E-08 |
| 27018 | 1351 | A1351W | E | 6.4e-09 | 9.7e-09 | 6e-09 | 7.367E-09 | 2.031E-09 | 6.000E-09 | 9.700E-09 |
| 27019 | 1351 | A1351Y | E | 9.7e-09 | 4.3e-09 | 3e-09 | 5.667E-09 | 3.553E-09 | 3.000E-09 | 9.700E-09 |
| 27020 | 1351 | A1351V | E | 4.6e-09 | 4.4e-09 | 1.2e-08 | 7.000E-09 | 4.331E-09 | 4.400E-09 | 1.200E-08 |
| 27021 | 1352 | Y1352A | E | 6.3e-09 | 1.4e-08 | 5.2e-09 | 8.500E-09 | 4.795E-09 | 5.200E-09 | 1.400E-08 |
| 27022 | 1352 | Y1352R | E | 4.9e-09 | 9e-09 | 1.8e-08 | 1.063E-08 | 6.701E-09 | 4.900E-09 | 1.800E-08 |
| 27023 | 1352 | Y1352N | E | 7.3e-09 | 1.3e-08 | 1.9e-08 | 1.310E-08 | 5.851E-09 | 7.300E-09 | 1.900E-08 |
| 27024 | 1352 | Y1352D | E | 1.9e-08 | 3e-09 | 6.4e-09 | 9.467E-09 | 8.429E-09 | 3.000E-09 | 1.900E-08 |
| 27025 | 1352 | Y1352C | E | 1.9e-08 | 1.1e-08 | 6.2e-09 | 1.207E-08 | 6.466E-09 | 6.200E-09 | 1.900E-08 |
| 27026 | 1352 | Y1352E | E | 3e-09 | 1.1e-08 | 2.7e-09 | 5.567E-09 | 4.708E-09 | 2.700E-09 | 1.100E-08 |
| 27027 | 1352 | Y1352Q | E | 6e-09 | 9.2e-09 | 8.8e-09 | 8.000E-09 | 1.744E-09 | 6.000E-09 | 9.200E-09 |
| 27028 | 1352 | Y1352G | E | 1e-08 | 2.2e-08 | 9.3e-09 | 1.377E-08 | 7.139E-09 | 9.300E-09 | 2.200E-08 |
| 27029 | 1352 | Y1352H | E | 7.1e-09 | 2.8e-09 | 6.1e-09 | 5.333E-09 | 2.250E-09 | 2.800E-09 | 7.100E-09 |

| | | | | | | | | | | |
|---|---|---|---|---|---|---|---|---|---|---|
| 27030 | 1352 | Y1352I | E | 7.8e-09 | 3e-08 | 9.3e-09 | 1.570E-08 | 1.241E-08 | 7.800E-09 | 3.000E-08 |
| 27031 | 1352 | Y1352L | E | 6.3e-09 | 1.1e-08 | 9.4e-09 | 8.900E-09 | 2.390E-09 | 6.300E-09 | 1.100E-08 |
| 27032 | 1352 | Y1352K | E | 1.9e-08 | 6.6e-09 | 1.7e-08 | 1.420E-08 | 6.657E-09 | 6.600E-09 | 1.900E-08 |
| 27033 | 1352 | Y1352M | E | 1.5e-08 | 8.2e-09 | 1.1e-08 | 1.140E-08 | 3.418E-09 | 8.200E-09 | 1.500E-08 |
| 27034 | 1352 | Y1352F | E | 1.3e-08 | 7.7e-09 | 5.4e-09 | 8.700E-09 | 3.897E-09 | 5.400E-09 | 1.300E-08 |
| 27035 | 1352 | Y1352P | E | 3.5e-09 | 4.6e-09 | 9.1e-09 | 5.733E-09 | 2.967E-09 | 3.500E-09 | 9.100E-09 |
| 27036 | 1352 | Y1352S | E | 4.5e-09 | 1.7e-08 | 3.2e-08 | 1.783E-08 | 1.377E-08 | 4.500E-09 | 3.200E-08 |
| 27037 | 1352 | Y1352T | E | 1e-08 | 3.9e-09 | 2.9e-09 | 5.600E-09 | 3.843E-09 | 2.900E-09 | 1.000E-08 |
| 27038 | 1352 | Y1352W | E | 9e-09 | 5.5e-09 | 8e-09 | 7.500E-09 | 1.803E-09 | 5.500E-09 | 9.000E-09 |
| 27039 | 1352 | Y1352Y | E | 1.4e-08 | 1.2e-08 | 7.1e-09 | 1.103E-08 | 3.550E-09 | 7.100E-09 | 1.400E-08 |
| 27040 | 1352 | Y1352V | E | 5.8e-09 | 7.3e-09 | 4.2e-09 | 5.767E-09 | 1.550E-09 | 4.200E-09 | 7.300E-09 |
| 27041 | 1353 | L1353A | E | 5.2e-09 | 5e-09 | 8.6e-09 | 6.267E-09 | 2.023E-09 | 5.000E-09 | 8.600E-09 |
| 27042 | 1353 | L1353R | E | 7.9e-09 | 4.4e-09 | 6.7e-09 | 6.333E-09 | 1.779E-09 | 4.400E-09 | 7.900E-09 |
| 27043 | 1353 | L1353N | E | 1.8e-08 | 1.2e-08 | 5.7e-09 | 1.190E-08 | 6.151E-09 | 5.700E-09 | 1.800E-08 |
| 27044 | 1353 | L1353D | E | 4.5e-09 | 4.3e-09 | 8.8e-09 | 5.867E-09 | 2.542E-09 | 4.300E-09 | 8.800E-09 |
| 27045 | 1353 | L1353C | E | 2.1e-09 | 1.2e-08 | 8e-09 | 7.367E-09 | 4.980E-09 | 2.100E-09 | 1.200E-08 |
| 27046 | 1353 | L1353E | E | 8.9e-09 | 1e-08 | 1.1e-08 | 9.967E-09 | 1.050E-09 | 8.900E-09 | 1.100E-08 |
| 27047 | 1353 | L1353Q | E | 5.5e-09 | 3.9e-09 | 6e-09 | 5.133E-09 | 1.097E-09 | 3.900E-09 | 6.000E-09 |
| 27048 | 1353 | L1353G | E | 8.3e-09 | 1.2e-08 | 7.2e-09 | 9.167E-09 | 2.515E-09 | 7.200E-09 | 1.200E-08 |
| 27049 | 1353 | L1353H | E | 6.7e-09 | 2.7e-09 | 2.8e-09 | 4.067E-09 | 2.281E-09 | 2.700E-09 | 6.700E-09 |
| 27050 | 1353 | L1353I | E | 1.9e-09 | 7.4e-09 | 3e-09 | 4.100E-09 | 2.910E-09 | 1.900E-09 | 7.400E-09 |
| 27051 | 1353 | L1353L | E | 6.7e-09 | 1.2e-08 | 3.3e-09 | 7.333E-09 | 4.384E-09 | 3.300E-09 | 1.200E-08 |
| 27052 | 1353 | L1353K | E | 6.3e-09 | 1.1e-08 | 3.8e-09 | 7.033E-09 | 3.656E-09 | 3.800E-09 | 1.100E-08 |
| 27053 | 1353 | L1353M | E | 5.2e-09 | 4.5e-09 | 5.3e-09 | 5.000E-09 | 4.359E-10 | 4.500E-09 | 5.300E-09 |
| 27054 | 1353 | L1353F | E | 9.2e-09 | 1.2e-08 | 3.4e-09 | 8.200E-09 | 4.386E-09 | 3.400E-09 | 1.200E-08 |
| 27055 | 1353 | L1353P | E | 5.4e-09 | 8.2e-09 | 6.6e-09 | 6.733E-09 | 1.405E-09 | 5.400E-09 | 8.200E-09 |
| 27056 | 1353 | L1353S | E | 8.6e-09 | 1.1e-09 | 6.2e-09 | 5.300E-09 | 3.830E-09 | 1.100E-09 | 8.600E-09 |
| 27057 | 1353 | L1353T | E | 2.2e-08 | 2.1e-09 | 3.7e-09 | 9.267E-09 | 1.106E-08 | 2.100E-09 | 2.200E-08 |
| 27058 | 1353 | L1353W | E | 5.3e-09 | 1.3e-08 | 7.4e-09 | 8.567E-09 | 3.980E-09 | 5.300E-09 | 1.300E-08 |
| 27059 | 1353 | L1353Y | E | 2.3e-09 | 9.6e-09 | 7.1e-09 | 6.333E-09 | 3.710E-09 | 2.300E-09 | 9.600E-09 |
| 27060 | 1353 | L1353V | E | 1.6e-08 | 1.3e-08 | 2.6e-08 | 1.833E-08 | 6.807E-09 | 1.300E-08 | 2.600E-08 |
| 27061 | 1354 | Q1354A | E | 1.6e-08 | 8.7e-09 | 9.9e-09 | 1.153E-08 | 3.915E-09 | 8.700E-09 | 1.600E-08 |
| 27062 | 1354 | Q1354R | E | 3.3e-09 | 3.3e-09 | 5.5e-09 | 4.033E-09 | 1.270E-09 | 3.300E-09 | 5.500E-09 |
| 27063 | 1354 | Q1354N | E | 8.6e-09 | 3.9e-09 | 4.2e-09 | 5.567E-09 | 2.631E-09 | 3.900E-09 | 8.600E-09 |
| 27064 | 1354 | Q1354D | E | 7.5e-09 | 4.2e-09 | 6.3e-09 | 6.000E-09 | 1.670E-09 | 4.200E-09 | 7.500E-09 |
| 27065 | 1354 | Q1354C | E | 1.6e-08 | 1.3e-08 | 1.3e-08 | 1.400E-08 | 1.732E-09 | 1.300E-08 | 1.600E-08 |
| 27066 | 1354 | Q1354E | E | 9.4e-09 | 6.9e-09 | 1.4e-08 | 1.010E-08 | 3.601E-09 | 6.900E-09 | 1.400E-08 |
| 27067 | 1354 | Q1354Q | E | 4.3e-09 | 1.3e-08 | 1.2e-08 | 9.767E-09 | 4.761E-09 | 4.300E-09 | 1.300E-08 |
| 27068 | 1354 | Q1354G | E | 2e-08 | 5.9e-09 | 1.1e-08 | 1.230E-08 | 7.139E-09 | 5.900E-09 | 2.000E-08 |
| 27069 | 1354 | Q1354H | E | 6.1e-09 | 1e-08 | 4.3e-09 | 6.800E-09 | 2.914E-09 | 4.300E-09 | 1.000E-08 |
| 27070 | 1354 | Q1354I | E | 1.7e-08 | 4.7e-09 | 7.1e-09 | 9.600E-09 | 6.520E-09 | 4.700E-09 | 1.700E-08 |
| 27071 | 1354 | Q1354L | E | 8.3e-09 | 6.3e-09 | 2.1e-08 | 1.187E-08 | 7.973E-09 | 6.300E-09 | 2.100E-08 |
| 27072 | 1354 | Q1354K | E | 7.8e-09 | 5.9e-09 | 5.9e-09 | 6.533E-09 | 1.097E-09 | 5.900E-09 | 7.800E-09 |
| 27073 | 1354 | Q1354M | E | 5.6e-09 | 1.8e-08 | 1.1e-08 | 1.153E-08 | 6.217E-09 | 5.600E-09 | 1.800E-08 |
| 27074 | 1354 | Q1354F | E | 4e-09 | 8.2e-09 | 5.3e-09 | 5.833E-09 | 2.150E-09 | 4.000E-09 | 8.200E-09 |
| 27075 | 1354 | Q1354P | E | 2.6e-09 | 8.2e-09 | 9.3e-09 | 6.700E-09 | 3.593E-09 | 2.600E-09 | 9.300E-09 |
| 27076 | 1354 | Q1354S | E | 6.6e-09 | 8e-09 | 3.5e-09 | 6.033E-09 | 2.303E-09 | 3.500E-09 | 8.000E-09 |
| 27077 | 1354 | Q1354T | E | 7.3e-09 | 1.4e-08 | 3.5e-09 | 8.267E-09 | 5.316E-09 | 3.500E-09 | 1.400E-08 |
| 27078 | 1354 | Q1354W | E | 7.1e-09 | 1.3e-08 | 4e-09 | 8.033E-09 | 4.572E-09 | 4.000E-09 | 1.300E-08 |
| 27079 | 1354 | Q1354Y | E | 1.8e-09 | 3.6e-09 | 3.3e-09 | 2.900E-09 | 9.644E-10 | 1.800E-09 | 3.600E-09 |
| 27080 | 1354 | Q1354V | E | 2.2e-09 | 1.2e-08 | 7.8e-09 | 7.333E-09 | 4.917E-09 | 2.200E-09 | 1.200E-08 |
| 27081 | 1355 | M1355A | E | 8.9e-09 | 6.2e-09 | 7.7e-09 | 7.600E-09 | 1.353E-09 | 6.200E-09 | 8.900E-09 |
| 27082 | 1355 | M1355R | E | 7.8e-09 | 3.5e-09 | 8e-09 | 6.433E-09 | 2.542E-09 | 3.500E-09 | 8.000E-09 |
| 27083 | 1355 | M1355N | E | 6.3e-09 | 5.9e-09 | 1.6e-08 | 9.400E-09 | 5.719E-09 | 5.900E-09 | 1.600E-08 |
| 27084 | 1355 | M1355D | E | 5.9e-09 | 3.4e-09 | 8.8e-09 | 6.033E-09 | 2.702E-09 | 3.400E-09 | 8.800E-09 |
| 27085 | 1355 | M1355C | E | 3.9e-09 | 4.3e-09 | 8.4e-09 | 5.533E-09 | 2.491E-09 | 3.900E-09 | 8.400E-09 |
| 27086 | 1355 | M1355E | E | 1.1e-08 | 8.2e-09 | 9.9e-09 | 9.700E-09 | 1.411E-09 | 8.200E-09 | 1.100E-08 |
| 27087 | 1355 | M1355Q | E | 6.3e-09 | 1.4e-08 | 1.3e-08 | 1.110E-08 | 4.187E-09 | 6.300E-09 | 1.400E-08 |
| 27088 | 1355 | M1355G | E | 1.2e-08 | 4.3e-09 | 1.6e-08 | 1.077E-08 | 5.947E-09 | 4.300E-09 | 1.600E-08 |
| 27089 | 1355 | M1355H | E | 4.6e-09 | 9.1e-09 | 1.1e-08 | 8.233E-09 | 3.287E-09 | 4.600E-09 | 1.100E-08 |
| 27090 | 1355 | M1355I | E | 1.8e-08 | 9.5e-09 | 8.7e-09 | 1.207E-08 | 5.154E-09 | 8.700E-09 | 1.800E-08 |
| 27091 | 1355 | M1355L | E | 4.6e-09 | 2.7e-09 | 3.6e-09 | 3.633E-09 | 9.504E-10 | 2.700E-09 | 4.600E-09 |
| 27092 | 1355 | M1355K | E | 1.1e-08 | 6.5e-09 | 2.3e-08 | 1.350E-08 | 8.529E-09 | 6.500E-09 | 2.300E-08 |
| 27093 | 1355 | M1355M | E | 7.6e-09 | 9.9e-09 | 5.6e-09 | 7.700E-09 | 2.152E-09 | 5.600E-09 | 9.900E-09 |
| 27094 | 1355 | M1355F | E | 5.6e-09 | 9.8e-09 | 1.5e-08 | 1.013E-08 | 4.709E-09 | 5.600E-09 | 1.500E-08 |
| 27095 | 1355 | M1355P | E | 1.3e-08 | 9.6e-09 | 3.9e-09 | 8.833E-09 | 4.598E-09 | 3.900E-09 | 1.300E-08 |
| 27096 | 1355 | M1355S | E | 3.1e-09 | 1e-08 | 7.2e-09 | 6.767E-09 | 3.470E-09 | 3.100E-09 | 1.000E-08 |
| 27097 | 1355 | M1355T | E | 7.3e-09 | 8.9e-09 | 7.8e-09 | 8.000E-09 | 8.185E-10 | 7.300E-09 | 8.900E-09 |
| 27098 | 1355 | M1355W | E | 8.6e-09 | 1e-08 | 2.5e-09 | 7.033E-09 | 3.988E-09 | 2.500E-09 | 1.000E-08 |
| 27099 | 1355 | M1355Y | E | 5.7e-09 | 3e-09 | 5.6e-09 | 4.767E-09 | 1.531E-09 | 3.000E-09 | 5.700E-09 |
| 27100 | 1355 | M1355V | E | 8.3e-09 | 3.7e-09 | 6.8e-09 | 6.267E-09 | 2.346E-09 | 3.700E-09 | 8.300E-09 |
| 27101 | 1356 | N1356A | E | 6.1e-09 | 5.9e-09 | 9.5e-09 | 7.167E-09 | 2.023E-09 | 5.900E-09 | 9.500E-09 |
| 27102 | 1356 | N1356R | E | 4.3e-09 | 4.3e-09 | 2.3e-08 | 1.053E-08 | 1.080E-08 | 4.300E-09 | 2.300E-08 |
| 27103 | 1356 | N1356N | E | 9.5e-09 | 2.1e-09 | 4e-09 | 5.200E-09 | 3.843E-09 | 2.100E-09 | 9.500E-09 |
| 27104 | 1356 | N1356D | E | 6.6e-09 | 1.1e-08 | 1.4e-08 | 1.053E-08 | 3.722E-09 | 6.600E-09 | 1.400E-08 |
| 27105 | 1356 | N1356C | E | 1.1e-08 | 5.5e-09 | 9e-09 | 8.500E-09 | 2.784E-09 | 5.500E-09 | 1.100E-08 |
| 27106 | 1356 | N1356E | E | 2e-08 | 1.4e-08 | 4.1e-09 | 1.270E-08 | 8.029E-09 | 4.100E-09 | 2.000E-08 |
| 27107 | 1356 | N1356Q | E | 1.4e-08 | 4.7e-09 | 8e-09 | 8.900E-09 | 4.715E-09 | 4.700E-09 | 1.400E-08 |
| 27108 | 1356 | N1356G | E | 7.8e-09 | 5.1e-09 | 6e-09 | 6.300E-09 | 1.375E-09 | 5.100E-09 | 7.800E-09 |
| 27109 | 1356 | N1356H | E | 6e-09 | 5.6e-09 | 8e-09 | 6.533E-09 | 1.286E-09 | 5.600E-09 | 8.000E-09 |
| 27110 | 1356 | N1356I | E | 7.4e-09 | 8.8e-09 | 5.6e-09 | 7.267E-09 | 1.604E-09 | 5.600E-09 | 8.800E-09 |
| 27111 | 1356 | N1356L | E | 4.8e-09 | 6.2e-09 | 9e-09 | 6.667E-09 | 2.139E-09 | 4.800E-09 | 9.000E-09 |
| 27112 | 1356 | N1356K | E | 1.3e-08 | 9e-09 | 2.7e-08 | 1.633E-08 | 9.452E-09 | 9.000E-09 | 2.700E-08 |
| 27113 | 1356 | N1356M | E | 7e-09 | 6.5e-09 | 6.5e-09 | 6.667E-09 | 2.887E-10 | 6.500E-09 | 7.000E-09 |
| 27114 | 1356 | N1356F | E | 4.2e-09 | 2.8e-09 | 1e-08 | 5.667E-09 | 3.818E-09 | 2.800E-09 | 1.000E-08 |

| | | | | | | | | | | |
|---|---|---|---|---|---|---|---|---|---|---|
| 27115 | 1356 | N1356P | E | 5.2e-09 | 4e-09 | 1.1e-08 | 6.733E-09 | 3.743E-09 | 4.000E-09 | 1.100E-08 |
| 27116 | 1356 | N1356S | E | 6.5e-09 | 5.4e-09 | 5.1e-09 | 5.667E-09 | 7.371E-10 | 5.100E-09 | 6.500E-09 |
| 27117 | 1356 | N1356T | E | 1.9e-09 | 6.8e-09 | 1.6e-08 | 8.233E-09 | 7.158E-09 | 1.900E-09 | 1.600E-08 |
| 27118 | 1356 | N1356W | E | 2.8e-09 | 1.2e-08 | 3e-09 | 5.933E-09 | 5.255E-09 | 2.800E-09 | 1.200E-08 |
| 27119 | 1356 | N1356Y | E | 2.6e-09 | 2.8e-09 | 1.1e-08 | 5.467E-09 | 4.793E-09 | 2.600E-09 | 1.100E-08 |
| 27120 | 1356 | N1356V | E | 5.5e-09 | 1e-08 | 3.3e-09 | 6.267E-09 | 3.415E-09 | 3.300E-09 | 1.000E-08 |
| 27121 | 1357 | S1357A | E | 6.2e-09 | 2.4e-09 | 4.7e-09 | 4.433E-09 | 1.914E-09 | 2.400E-09 | 6.200E-09 |
| 27122 | 1357 | S1357R | E | 5.7e-09 | 1.8e-08 | 1.1e-08 | 1.157E-08 | 6.170E-09 | 5.700E-09 | 1.800E-08 |
| 27123 | 1357 | S1357N | E | 3.8e-09 | 1.2e-08 | 5.3e-09 | 7.033E-09 | 4.366E-09 | 3.800E-09 | 1.200E-08 |
| 27124 | 1357 | S1357D | E | 1.5e-08 | 5e-08 | 6.9e-09 | 2.397E-08 | 2.291E-08 | 6.900E-09 | 5.000E-08 |
| 27125 | 1357 | S1357C | E | 6.7e-09 | 6.7e-09 | 9.7e-09 | 7.700E-09 | 1.732E-09 | 6.700E-09 | 9.700E-09 |
| 27126 | 1357 | S1357E | E | 5.8e-09 | 1.6e-08 | 3.7e-09 | 8.500E-09 | 6.580E-09 | 3.700E-09 | 1.600E-08 |
| 27127 | 1357 | S1357Q | E | 5e-09 | 5e-09 | 1.4e-08 | 8.000E-09 | 5.196E-09 | 5.000E-09 | 1.400E-08 |
| 27128 | 1357 | S1357G | E | 1.3e-08 | 1.6e-08 | 1.2e-08 | 1.367E-08 | 2.082E-09 | 1.200E-08 | 1.600E-08 |
| 27129 | 1357 | S1357H | E | 1.1e-08 | 4.5e-09 | 8.3e-09 | 7.933E-09 | 3.265E-09 | 4.500E-09 | 1.100E-08 |
| 27130 | 1357 | S1357I | E | 2.8e-09 | 3.1e-09 | 5.7e-09 | 3.867E-09 | 1.595E-09 | 2.800E-09 | 5.700E-09 |
| 27131 | 1357 | S1357L | E | 1.3e-08 | 1.5e-08 | 8.1e-09 | 1.203E-08 | 3.550E-09 | 8.100E-09 | 1.500E-08 |
| 27132 | 1357 | S1357K | E | 4.4e-09 | 1.4e-08 | 8.8e-09 | 9.067E-09 | 4.806E-09 | 4.400E-09 | 1.400E-08 |
| 27133 | 1357 | S1357M | E | 3.9e-09 | 1.4e-08 | 5.1e-09 | 7.667E-09 | 5.518E-09 | 3.900E-09 | 1.400E-08 |
| 27134 | 1357 | S1357F | E | 1.5e-08 | 1.2e-08 | 6.3e-09 | 1.110E-08 | 4.419E-09 | 6.300E-09 | 1.500E-08 |
| 27135 | 1357 | S1357P | E | 2.1e-09 | 1.2e-08 | 2.4e-08 | 1.270E-08 | 1.097E-08 | 2.100E-09 | 2.400E-08 |
| 27136 | 1357 | S1357S | E | 6.4e-09 | 1.7e-08 | 8.7e-09 | 1.070E-08 | 5.576E-09 | 6.400E-09 | 1.700E-08 |
| 27137 | 1357 | S1357T | E | 1e-08 | 2.5e-09 | 3.6e-09 | 5.367E-09 | 4.050E-09 | 2.500E-09 | 1.000E-08 |
| 27138 | 1357 | S1357W | E | 2.6e-09 | 8.5e-09 | 8.2e-09 | 6.433E-09 | 3.323E-09 | 2.600E-09 | 8.500E-09 |
| 27139 | 1357 | S1357Y | E | 6.9e-09 | 6.1e-09 | 7.9e-09 | 6.967E-09 | 9.018E-10 | 6.100E-09 | 7.900E-09 |
| 27140 | 1357 | S1357V | E | 9e-09 | 1e-08 | 9e-09 | 9.333E-09 | 5.774E-10 | 9.000E-09 | 1.000E-08 |
| 27141 | 1358 | L1358A | E | 7.5e-09 | 7.4e-09 | 5.4e-09 | 6.767E-09 | 1.185E-09 | 5.400E-09 | 7.500E-09 |
| 27142 | 1358 | L1358R | E | 1.5e-08 | 2.2e-08 | 9.5e-09 | 1.550E-08 | 6.265E-09 | 9.500E-09 | 2.200E-08 |
| 27143 | 1358 | L1358N | E | 1.2e-08 | 6.9e-09 | 6.6e-09 | 8.500E-09 | 3.035E-09 | 6.600E-09 | 1.200E-08 |
| 27144 | 1358 | L1358D | E | 9e-09 | 8e-09 | 7.4e-09 | 8.133E-09 | 8.083E-10 | 7.400E-09 | 9.000E-09 |
| 27145 | 1358 | L1358C | E | 3.3e-09 | 1.3e-08 | 6.3e-09 | 7.533E-09 | 4.966E-09 | 3.300E-09 | 1.300E-08 |
| 27146 | 1358 | L1358E | E | 5.2e-09 | 1.3e-08 | 8.8e-09 | 9.000E-09 | 3.904E-09 | 5.200E-09 | 1.300E-08 |
| 27147 | 1358 | L1358Q | E | 2e-09 | 1.6e-08 | 3.4e-09 | 7.133E-09 | 7.711E-09 | 2.000E-09 | 1.600E-08 |
| 27148 | 1358 | L1358G | E | 6.1e-09 | 6.8e-09 | 7.3e-09 | 6.733E-09 | 6.028E-10 | 6.100E-09 | 7.300E-09 |
| 27149 | 1358 | L1358H | E | 4.4e-09 | 4.8e-09 | 1.4e-08 | 7.733E-09 | 5.431E-09 | 4.400E-09 | 1.400E-08 |
| 27150 | 1358 | L1358I | E | 1.5e-08 | 6.6e-09 | 1.8e-08 | 1.320E-08 | 5.909E-09 | 6.600E-09 | 1.800E-08 |
| 27151 | 1358 | L1358L | E | 6.1e-09 | 1.6e-08 | 1.5e-08 | 1.237E-08 | 5.450E-09 | 6.100E-09 | 1.600E-08 |
| 27152 | 1358 | L1358K | E | 8.6e-09 | 4.2e-09 | 1.1e-08 | 7.933E-09 | 3.449E-09 | 4.200E-09 | 1.100E-08 |
| 27153 | 1358 | L1358M | E | 4.5e-09 | 3.3e-08 | 8.9e-09 | 1.547E-08 | 1.534E-08 | 4.500E-09 | 3.300E-08 |
| 27154 | 1358 | L1358F | E | 6.5e-09 | 6.3e-09 | 1.2e-08 | 8.267E-09 | 3.235E-09 | 6.300E-09 | 1.200E-08 |
| 27155 | 1358 | L1358P | E | 2.4e-09 | 3.6e-09 | 8.7e-09 | 4.900E-09 | 3.345E-09 | 2.400E-09 | 8.700E-09 |
| 27156 | 1358 | L1358S | E | 6.1e-09 | 2.3e-08 | 3e-09 | 1.070E-08 | 1.076E-08 | 3.000E-09 | 2.300E-08 |
| 27157 | 1358 | L1358T | E | 1.2e-08 | 9.2e-09 | 2e-08 | 1.373E-08 | 5.605E-09 | 9.200E-09 | 2.000E-08 |
| 27158 | 1358 | L1358W | E | 1.4e-08 | 3.8e-09 | 6e-09 | 7.933E-09 | 5.368E-09 | 3.800E-09 | 1.400E-08 |
| 27159 | 1358 | L1358Y | E | 5.9e-09 | 1e-08 | 5.4e-09 | 7.100E-09 | 2.524E-09 | 5.400E-09 | 1.000E-08 |
| 27160 | 1358 | L1358V | E | 5.9e-09 | 5.6e-09 | 5.1e-09 | 5.533E-09 | 4.041E-10 | 5.100E-09 | 5.900E-09 |
| 27161 | 1359 | R1359A | E | 9.6e-09 | 5.9e-09 | 1.2e-08 | 9.167E-09 | 3.073E-09 | 5.900E-09 | 1.200E-08 |
| 27162 | 1359 | R1359R | E | 7e-09 | 4e-09 | 1.2e-08 | 7.667E-09 | 4.041E-09 | 4.000E-09 | 1.200E-08 |
| 27163 | 1359 | R1359N | E | 1.6e-08 | 7.6e-09 | 7.1e-09 | 1.023E-08 | 5.000E-09 | 7.100E-09 | 1.600E-08 |
| 27164 | 1359 | R1359D | E | 8e-09 | 1.1e-08 | 5.1e-09 | 8.033E-09 | 2.950E-09 | 5.100E-09 | 1.100E-08 |
| 27165 | 1359 | R1359C | E | 9.2e-09 | 3.2e-09 | 3.5e-09 | 5.300E-09 | 3.381E-09 | 3.200E-09 | 9.200E-09 |
| 27166 | 1359 | R1359E | E | 4.6e-09 | 3.1e-09 | 2.3e-09 | 3.333E-09 | 1.168E-09 | 2.300E-09 | 4.600E-09 |
| 27167 | 1359 | R1359Q | E | 3.1e-09 | 1.6e-08 | 5.2e-09 | 8.100E-09 | 6.922E-09 | 3.100E-09 | 1.600E-08 |
| 27168 | 1359 | R1359G | E | 4.6e-09 | 3.3e-09 | 4.1e-09 | 4.000E-09 | 6.557E-10 | 3.300E-09 | 4.600E-09 |
| 27169 | 1359 | R1359H | E | 6.2e-09 | 8.7e-09 | 3.5e-09 | 6.133E-09 | 2.601E-09 | 3.500E-09 | 8.700E-09 |
| 27170 | 1359 | R1359I | E | 4.5e-09 | 1.8e-08 | 9.8e-09 | 1.077E-08 | 6.802E-09 | 4.500E-09 | 1.800E-08 |
| 27171 | 1359 | R1359L | E | 1.9e-08 | 8.4e-09 | 6.5e-09 | 1.130E-08 | 6.736E-09 | 6.500E-09 | 1.900E-08 |
| 27172 | 1359 | R1359K | E | 1.1e-08 | 4.9e-09 | 8.4e-09 | 8.100E-09 | 3.061E-09 | 4.900E-09 | 1.100E-08 |
| 27173 | 1359 | R1359M | E | 3.7e-09 | 4.2e-09 | 1.2e-08 | 6.633E-09 | 4.654E-09 | 3.700E-09 | 1.200E-08 |
| 27174 | 1359 | R1359F | E | 5.3e-09 | 5.3e-09 | 1.9e-09 | 4.167E-09 | 1.963E-09 | 1.900E-09 | 5.300E-09 |
| 27175 | 1359 | R1359P | E | 9.9e-09 | 5.1e-09 | 1.4e-08 | 9.667E-09 | 4.455E-09 | 5.100E-09 | 1.400E-08 |
| 27176 | 1359 | R1359S | E | 1.7e-08 | 4.8e-09 | 1.3e-08 | 1.160E-08 | 6.219E-09 | 4.800E-09 | 1.700E-08 |
| 27177 | 1359 | R1359T | E | 4.3e-09 | 1.1e-08 | 9.7e-09 | 8.333E-09 | 3.553E-09 | 4.300E-09 | 1.100E-08 |
| 27178 | 1359 | R1359W | E | 6.2e-09 | 9.2e-09 | 7.2e-09 | 7.533E-09 | 1.528E-09 | 6.200E-09 | 9.200E-09 |
| 27179 | 1359 | R1359Y | E | 4.2e-08 | 5.6e-09 | 1.7e-08 | 2.153E-08 | 1.862E-08 | 5.600E-09 | 4.200E-08 |
| 27180 | 1359 | R1359V | E | 4.4e-09 | 1.1e-08 | 2.2e-08 | 1.247E-08 | 8.891E-09 | 4.400E-09 | 2.200E-08 |
| 27181 | 1360 | A1360A | E | 1.1e-08 | 1.1e-08 | 3.9e-09 | 8.633E-09 | 4.099E-09 | 3.900E-09 | 1.100E-08 |
| 27182 | 1360 | A1360R | E | 2.4e-09 | 7.4e-09 | 1.3e-08 | 7.600E-09 | 5.303E-09 | 2.400E-09 | 1.300E-08 |
| 27183 | 1360 | A1360N | E | 3.5e-09 | 7.4e-09 | 9.1e-09 | 6.667E-09 | 2.871E-09 | 3.500E-09 | 9.100E-09 |
| 27184 | 1360 | A1360D | E | 1.2e-08 | 4.9e-09 | 1.1e-08 | 9.300E-09 | 3.843E-09 | 4.900E-09 | 1.200E-08 |
| 27185 | 1360 | A1360C | E | 3.8e-09 | 1.3e-08 | 8.5e-09 | 8.433E-09 | 4.600E-09 | 3.800E-09 | 1.300E-08 |
| 27186 | 1360 | A1360E | E | 1.1e-08 | 3.4e-09 | 6.8e-09 | 7.067E-09 | 3.807E-09 | 3.400E-09 | 1.100E-08 |
| 27187 | 1360 | A1360Q | E | 5.9e-09 | 1.4e-08 | 3.8e-09 | 7.900E-09 | 5.386E-09 | 3.800E-09 | 1.400E-08 |
| 27188 | 1360 | A1360G | E | 9.4e-09 | 5.3e-09 | 1.5e-08 | 9.900E-09 | 4.869E-09 | 5.300E-09 | 1.500E-08 |
| 27189 | 1360 | A1360H | E | 5.7e-09 | 3.4e-09 | 1.6e-08 | 8.367E-09 | 6.710E-09 | 3.400E-09 | 1.600E-08 |
| 27190 | 1360 | A1360I | E | 5.9e-09 | 2.2e-08 | 6.8e-09 | 1.157E-08 | 9.047E-09 | 5.900E-09 | 2.200E-08 |
| 27191 | 1360 | A1360L | E | 6.8e-09 | 1.2e-08 | 5.8e-09 | 8.200E-09 | 3.329E-09 | 5.800E-09 | 1.200E-08 |
| 27192 | 1360 | A1360K | E | 1.2e-08 | 7.5e-09 | 4.2e-09 | 7.900E-09 | 3.915E-09 | 4.200E-09 | 1.200E-08 |
| 27193 | 1360 | A1360M | E | 6.5e-09 | 1e-08 | 2e-08 | 1.217E-08 | 7.006E-09 | 6.500E-09 | 2.000E-08 |
| 27194 | 1360 | A1360F | E | 8.7e-09 | 1.1e-08 | 5.1e-09 | 8.267E-09 | 2.974E-09 | 5.100E-09 | 1.100E-08 |
| 27195 | 1360 | A1360P | E | 7.8e-09 | 1.3e-08 | 6.8e-09 | 9.200E-09 | 3.329E-09 | 6.800E-09 | 1.300E-08 |
| 27196 | 1360 | A1360S | E | 1.1e-08 | 7.8e-09 | 6.9e-09 | 8.567E-09 | 2.155E-09 | 6.900E-09 | 1.100E-08 |
| 27197 | 1360 | A1360T | E | 4.8e-09 | 1.9e-08 | 2e-08 | 1.460E-08 | 8.502E-09 | 4.800E-09 | 2.000E-08 |
| 27198 | 1360 | A1360W | E | 1.4e-08 | 7.4e-09 | 5.6e-09 | 9.000E-09 | 4.423E-09 | 5.600E-09 | 1.400E-08 |
| 27199 | 1360 | A1360Y | E | 3.1e-09 | 1.1e-08 | 4e-09 | 6.033E-09 | 4.325E-09 | 3.100E-09 | 1.100E-08 |

| | | | | | | | | | | |
|---|---|---|---|---|---|---|---|---|---|---|
| 27200 | 1360 | A1360V | E | 3.9e-09 | 1.5e-08 | 2.4e-08 | 1.430E-08 | 1.007E-08 | 3.900E-09 | 2.400E-08 |
| 27201 | 1361 | E1361A | E | 2.2e-09 | 6.8e-09 | 7.1e-09 | 5.367E-09 | 2.747E-09 | 2.200E-09 | 7.100E-09 |
| 27202 | 1361 | E1361R | E | 6.6e-09 | 2.6e-09 | 1.5e-08 | 8.067E-09 | 6.329E-09 | 2.600E-09 | 1.500E-08 |
| 27203 | 1361 | E1361N | E | 6.6e-09 | 7.8e-09 | 9.2e-09 | 7.867E-09 | 1.301E-09 | 6.600E-09 | 9.200E-09 |
| 27204 | 1361 | E1361D | E | 1.5e-08 | 7e-09 | 5.9e-09 | 9.300E-09 | 4.967E-09 | 5.900E-09 | 1.500E-08 |
| 27205 | 1361 | E1361C | E | 7.6e-09 | 2.6e-09 | 9.9e-10 | 3.730E-09 | 3.447E-09 | 9.900E-10 | 7.600E-09 |
| 27206 | 1361 | E1361E | E | 1.5e-08 | 5.8e-09 | 8.2e-09 | 9.667E-09 | 4.772E-09 | 5.800E-09 | 1.500E-08 |
| 27207 | 1361 | E1361Q | E | 2.2e-08 | 6.8e-09 | 1e-08 | 1.293E-08 | 8.013E-09 | 6.800E-09 | 2.200E-08 |
| 27208 | 1361 | E1361G | E | 1.6e-09 | 6.7e-09 | 5.2e-09 | 4.500E-09 | 2.621E-09 | 1.600E-09 | 6.700E-09 |
| 27209 | 1361 | E1361H | E | 7.3e-09 | 6e-09 | 7.3e-09 | 6.867E-09 | 7.506E-10 | 6.000E-09 | 7.300E-09 |
| 27210 | 1361 | E1361I | E | 2.1e-08 | 2.4e-08 | 8.1e-09 | 1.770E-08 | 8.448E-09 | 8.100E-09 | 2.400E-08 |
| 27211 | 1361 | E1361L | E | 9.6e-09 | 8.7e-09 | 9.4e-09 | 9.233E-09 | 4.726E-10 | 8.700E-09 | 9.600E-09 |
| 27212 | 1361 | E1361K | E | 5.4e-09 | 1.4e-08 | 3e-08 | 1.647E-08 | 1.248E-08 | 5.400E-09 | 3.000E-08 |
| 27213 | 1361 | E1361M | E | 8.3e-09 | 6.6e-09 | 1.1e-08 | 8.633E-09 | 2.219E-09 | 6.600E-09 | 1.100E-08 |
| 27214 | 1361 | E1361F | E | 5.9e-09 | 8e-09 | 7.9e-09 | 7.267E-09 | 1.185E-09 | 5.900E-09 | 8.000E-09 |
| 27215 | 1361 | E1361P | E | 4.3e-09 | 2e-09 | 2.4e-09 | 2.900E-09 | 1.229E-09 | 2.000E-09 | 4.300E-09 |
| 27216 | 1361 | E1361S | E | 5.9e-09 | 2.5e-09 | 6.7e-09 | 5.033E-09 | 2.230E-09 | 2.500E-09 | 6.700E-09 |
| 27217 | 1361 | E1361T | E | 7.1e-09 | 1.2e-08 | 3.8e-09 | 7.633E-09 | 4.126E-09 | 3.800E-09 | 1.200E-08 |
| 27218 | 1361 | E1361W | E | 6.8e-09 | 4.5e-09 | 1.5e-08 | 8.767E-09 | 5.519E-09 | 4.500E-09 | 1.500E-08 |
| 27219 | 1361 | E1361Y | E | 2.3e-08 | 1e-08 | 4.4e-09 | 1.247E-08 | 9.542E-09 | 4.400E-09 | 2.300E-08 |
| 27220 | 1361 | E1361V | E | 6.8e-09 | 5.9e-09 | 6.2e-09 | 6.300E-09 | 4.583E-10 | 5.900E-09 | 6.800E-09 |
| 27221 | 1362 | D1362A | E | 6e-09 | 1.2e-08 | 8.2e-09 | 8.733E-09 | 3.035E-09 | 6.000E-09 | 1.200E-08 |
| 27222 | 1362 | D1362R | E | 8.4e-09 | 1.7e-08 | 5.7e-09 | 1.037E-08 | 5.901E-09 | 5.700E-09 | 1.700E-08 |
| 27223 | 1362 | D1362N | E | 9.8e-09 | 2.9e-09 | 1.6e-08 | 9.567E-09 | 6.553E-09 | 2.900E-09 | 1.600E-08 |
| 27224 | 1362 | D1362D | E | 1.7e-08 | 5e-09 | 1.6e-08 | 1.267E-08 | 6.658E-09 | 5.000E-09 | 1.700E-08 |
| 27225 | 1362 | D1362C | E | 4.2e-09 | 1.1e-08 | 1.2e-08 | 9.067E-09 | 4.244E-09 | 4.200E-09 | 1.200E-08 |
| 27226 | 1362 | D1362E | E | 7.4e-09 | 1.1e-08 | 7.3e-09 | 8.567E-09 | 2.108E-09 | 7.300E-09 | 1.100E-08 |
| 27227 | 1362 | D1362Q | E | 9e-09 | 9.2e-09 | 5.4e-09 | 7.867E-09 | 2.139E-09 | 5.400E-09 | 9.200E-09 |
| 27228 | 1362 | D1362G | E | 9.8e-09 | 1.5e-08 | 1.2e-08 | 1.227E-08 | 2.610E-09 | 9.800E-09 | 1.500E-08 |
| 27229 | 1362 | D1362H | E | 6.3e-09 | 5.7e-09 | 4.7e-09 | 5.567E-09 | 8.083E-10 | 4.700E-09 | 6.300E-09 |
| 27230 | 1362 | D1362I | E | 1e-08 | 5.6e-09 | 1.1e-08 | 8.867E-09 | 2.873E-09 | 5.600E-09 | 1.100E-08 |
| 27231 | 1362 | D1362L | E | 3.1e-08 | 1.4e-08 | 2.9e-08 | 2.467E-08 | 9.292E-09 | 1.400E-08 | 3.100E-08 |
| 27232 | 1362 | D1362K | E | 7.4e-09 | 1.8e-09 | 7.9e-09 | 5.700E-09 | 3.387E-09 | 1.800E-09 | 7.900E-09 |
| 27233 | 1362 | D1362M | E | 7.5e-09 | 3.8e-09 | 3.1e-09 | 4.800E-09 | 2.364E-09 | 3.100E-09 | 7.500E-09 |
| 27234 | 1362 | D1362F | E | 4.8e-09 | 1.1e-08 | 9.3e-09 | 8.367E-09 | 3.204E-09 | 4.800E-09 | 1.100E-08 |
| 27235 | 1362 | D1362P | E | 7.1e-09 | 1.4e-08 | 4.6e-09 | 8.567E-09 | 4.869E-09 | 4.600E-09 | 1.400E-08 |
| 27236 | 1362 | D1362S | E | 2.9e-09 | 8.9e-09 | 1.5e-08 | 8.933E-09 | 6.050E-09 | 2.900E-09 | 1.500E-08 |
| 27237 | 1362 | D1362T | E | 2.4e-08 | 4.5e-09 | 4.2e-09 | 1.090E-08 | 1.135E-08 | 4.200E-09 | 2.400E-08 |
| 27238 | 1362 | D1362W | E | 1.2e-08 | 9.3e-09 | 7.8e-09 | 9.700E-09 | 2.128E-09 | 7.800E-09 | 1.200E-08 |
| 27239 | 1362 | D1362Y | E | 1.1e-08 | 4.5e-09 | 2.6e-08 | 1.383E-08 | 1.103E-08 | 4.500E-09 | 2.600E-08 |
| 27240 | 1362 | D1362V | E | 2.7e-09 | 5.8e-09 | 8.2e-09 | 5.567E-09 | 2.757E-09 | 2.700E-09 | 8.200E-09 |
| 27241 | 1363 | T1363A | E | 2.4e-08 | 6.9e-09 | 1.2e-08 | 1.430E-08 | 8.779E-09 | 6.900E-09 | 2.400E-08 |
| 27242 | 1363 | T1363R | E | 4.3e-09 | 3.8e-09 | 1.3e-08 | 7.033E-09 | 5.173E-09 | 3.800E-09 | 1.300E-08 |
| 27243 | 1363 | T1363N | E | 1.2e-08 | 1.5e-09 | 6.7e-09 | 6.733E-09 | 5.250E-09 | 1.500E-09 | 1.200E-08 |
| 27244 | 1363 | T1363D | E | 5.2e-09 | 7.3e-09 | 8.3e-09 | 6.933E-09 | 1.582E-09 | 5.200E-09 | 8.300E-09 |
| 27245 | 1363 | T1363C | E | 1.1e-08 | 4e-09 | 3.6e-09 | 6.200E-09 | 4.162E-09 | 3.600E-09 | 1.100E-08 |
| 27246 | 1363 | T1363E | E | 1.5e-08 | 5e-09 | 2e-08 | 1.333E-08 | 7.638E-09 | 5.000E-09 | 2.000E-08 |
| 27247 | 1363 | T1363Q | E | 5.2e-09 | 2.3e-08 | 4.9e-09 | 1.103E-08 | 1.036E-08 | 4.900E-09 | 2.300E-08 |
| 27248 | 1363 | T1363G | E | 1.2e-08 | 8.3e-09 | 9.6e-09 | 9.967E-09 | 1.877E-09 | 8.300E-09 | 1.200E-08 |
| 27249 | 1363 | T1363H | E | 1.5e-08 | 8.4e-09 | 6.8e-09 | 1.007E-08 | 4.347E-09 | 6.800E-09 | 1.500E-08 |
| 27250 | 1363 | T1363I | E | 7.2e-09 | 6.9e-09 | 7.7e-09 | 7.267E-09 | 4.041E-10 | 6.900E-09 | 7.700E-09 |
| 27251 | 1363 | T1363L | E | 1.3e-08 | 1.5e-09 | 4.1e-09 | 6.200E-09 | 6.031E-09 | 1.500E-09 | 1.300E-08 |
| 27252 | 1363 | T1363K | E | 3.5e-09 | 4.6e-09 | 1.3e-08 | 7.033E-09 | 5.196E-09 | 3.500E-09 | 1.300E-08 |
| 27253 | 1363 | T1363M | E | 6.1e-09 | 4.6e-09 | 5.3e-09 | 5.333E-09 | 7.506E-10 | 4.600E-09 | 6.100E-09 |
| 27254 | 1363 | T1363F | E | 2.8e-08 | 2e-08 | 7.1e-09 | 1.837E-08 | 1.055E-08 | 7.100E-09 | 2.800E-08 |
| 27255 | 1363 | T1363P | E | 7.4e-09 | 6.6e-09 | 1.2e-08 | 8.667E-09 | 2.914E-09 | 6.600E-09 | 1.200E-08 |
| 27256 | 1363 | T1363S | E | 8.4e-09 | 4.5e-09 | 7.4e-09 | 6.767E-09 | 2.026E-09 | 4.500E-09 | 8.400E-09 |
| 27257 | 1363 | T1363T | E | 5.6e-09 | 8.4e-09 | 1.4e-08 | 9.333E-09 | 4.277E-09 | 5.600E-09 | 1.400E-08 |
| 27258 | 1363 | T1363W | E | 1.1e-08 | 1.5e-08 | 8.7e-09 | 1.157E-08 | 3.188E-09 | 8.700E-09 | 1.500E-08 |
| 27259 | 1363 | T1363Y | E | 1.1e-08 | 5.1e-09 | 5.6e-09 | 7.233E-09 | 3.272E-09 | 5.100E-09 | 1.100E-08 |
| 27260 | 1363 | T1363V | E | 7.6e-09 | 8.2e-09 | 5.8e-09 | 7.200E-09 | 1.249E-09 | 5.800E-09 | 8.200E-09 |
| 27261 | 1364 | A1364A | E | 1.1e-08 | 8.9e-09 | 6.4e-09 | 8.767E-09 | 2.303E-09 | 6.400E-09 | 1.100E-08 |
| 27262 | 1364 | A1364R | E | 1e-09 | 7.7e-09 | 3.2e-08 | 1.357E-08 | 1.631E-08 | 1.000E-09 | 3.200E-08 |
| 27263 | 1364 | A1364N | E | 1.6e-09 | 8e-09 | 1e-08 | 6.533E-09 | 4.388E-09 | 1.600E-09 | 1.000E-08 |
| 27264 | 1364 | A1364D | E | 7.7e-09 | 3e-09 | 6.4e-09 | 5.700E-09 | 2.427E-09 | 3.000E-09 | 7.700E-09 |
| 27265 | 1364 | A1364C | E | 6.7e-09 | 7.4e-09 | 6e-09 | 6.700E-09 | 7.000E-10 | 6.000E-09 | 7.400E-09 |
| 27266 | 1364 | A1364E | E | 6e-09 | 3e-09 | 1.2e-08 | 7.000E-09 | 4.583E-09 | 3.000E-09 | 1.200E-08 |
| 27267 | 1364 | A1364Q | E | 4.1e-09 | 1.7e-09 | 5.8e-09 | 3.867E-09 | 2.060E-09 | 1.700E-09 | 5.800E-09 |
| 27268 | 1364 | A1364G | E | 3.8e-09 | 1.6e-09 | 1.3e-08 | 6.133E-09 | 6.048E-09 | 1.600E-09 | 1.300E-08 |
| 27269 | 1364 | A1364H | E | 8.1e-09 | 4.2e-09 | 1.1e-08 | 7.767E-09 | 3.412E-09 | 4.200E-09 | 1.100E-08 |
| 27270 | 1364 | A1364I | E | 6.4e-09 | 8e-09 | 5.9e-09 | 6.767E-09 | 1.097E-09 | 5.900E-09 | 8.000E-09 |
| 27271 | 1364 | A1364L | E | 1.2e-08 | 1.2e-08 | 1.1e-08 | 1.167E-08 | 5.774E-10 | 1.100E-08 | 1.200E-08 |
| 27272 | 1364 | A1364K | E | 7e-09 | 3.2e-08 | 6.8e-09 | 1.527E-08 | 1.449E-08 | 6.800E-09 | 3.200E-08 |
| 27273 | 1364 | A1364M | E | 3.3e-09 | 6.2e-09 | 1e-08 | 6.500E-09 | 3.360E-09 | 3.300E-09 | 1.000E-08 |
| 27274 | 1364 | A1364F | E | 6e-09 | 1.6e-09 | 6.1e-09 | 4.567E-09 | 2.570E-09 | 1.600E-09 | 6.100E-09 |
| 27275 | 1364 | A1364P | E | 9e-09 | 7.8e-09 | 3.7e-09 | 6.833E-09 | 2.779E-09 | 3.700E-09 | 9.000E-09 |
| 27276 | 1364 | A1364S | E | 2.5e-08 | 1.5e-08 | 2.1e-09 | 1.403E-08 | 1.148E-08 | 2.100E-09 | 2.500E-08 |
| 27277 | 1364 | A1364T | E | 6.6e-09 | 1.1e-08 | 5.8e-09 | 7.800E-09 | 2.800E-09 | 5.800E-09 | 1.100E-08 |
| 27278 | 1364 | A1364W | E | 6e-09 | 2e-08 | 1.8e-08 | 1.467E-08 | 7.572E-09 | 6.000E-09 | 2.000E-08 |
| 27279 | 1364 | A1364Y | E | 6.5e-09 | 4.7e-09 | 3.7e-09 | 4.967E-09 | 1.419E-09 | 3.700E-09 | 6.500E-09 |
| 27280 | 1364 | A1364V | E | 4.8e-09 | 2.6e-09 | 6.5e-09 | 4.633E-09 | 1.955E-09 | 2.600E-09 | 6.500E-09 |
| 27281 | 1365 | V1365A | E | 1.1e-08 | 4.4e-09 | 1.6e-08 | 1.047E-08 | 5.818E-09 | 4.400E-09 | 1.600E-08 |
| 27282 | 1365 | V1365R | E | 4.6e-09 | 1.7e-08 | 6.2e-09 | 9.267E-09 | 6.745E-09 | 4.600E-09 | 1.700E-08 |
| 27283 | 1365 | V1365N | E | 4.5e-09 | 6.9e-09 | 3.6e-09 | 5.000E-09 | 1.706E-09 | 3.600E-09 | 6.900E-09 |
| 27284 | 1365 | V1365D | E | 2e-09 | 2.5e-09 | 8.2e-09 | 4.233E-09 | 3.444E-09 | 2.000E-09 | 8.200E-09 |

| | | | | | | | | | | |
|---|---|---|---|---|---|---|---|---|---|---|
| 27285 | 1365 | V1365C | E | 8.8e-09 | 1.3e-08 | 6e-09 | 9.267E-09 | 3.523E-09 | 6.000E-09 | 1.300E-08 |
| 27286 | 1365 | V1365E | E | 5.7e-09 | 1.1e-09 | 2e-09 | 2.933E-09 | 2.438E-09 | 1.100E-09 | 5.700E-09 |
| 27287 | 1365 | V1365Q | E | 8.2e-09 | 1.9e-09 | 9.5e-09 | 6.533E-09 | 4.065E-09 | 1.900E-09 | 9.500E-09 |
| 27288 | 1365 | V1365G | E | 2.1e-09 | 7.9e-09 | 4.7e-09 | 4.900E-09 | 2.905E-09 | 2.100E-09 | 7.900E-09 |
| 27289 | 1365 | V1365H | E | 8.4e-09 | 1e-08 | 8.5e-09 | 8.967E-09 | 8.963E-10 | 8.400E-09 | 1.000E-08 |
| 27290 | 1365 | V1365I | E | 4.8e-09 | 7.1e-09 | 1.5e-08 | 8.967E-09 | 5.350E-09 | 4.800E-09 | 1.500E-08 |
| 27291 | 1365 | V1365L | E | 1.1e-08 | 5.9e-09 | 2.5e-09 | 6.467E-09 | 4.278E-09 | 2.500E-09 | 1.100E-08 |
| 27292 | 1365 | V1365K | E | 4.8e-09 | 4.5e-09 | 9.4e-09 | 6.233E-09 | 2.747E-09 | 4.500E-09 | 9.400E-09 |
| 27293 | 1365 | V1365M | E | 1.5e-09 | 5.9e-09 | 5.6e-09 | 4.333E-09 | 2.458E-09 | 1.500E-09 | 5.900E-09 |
| 27294 | 1365 | V1365F | E | 3.5e-09 | 5.8e-09 | 6.6e-09 | 5.300E-09 | 1.609E-09 | 3.500E-09 | 6.600E-09 |
| 27295 | 1365 | V1365P | E | 1e-08 | 7.1e-09 | 1.3e-09 | 6.133E-09 | 4.430E-09 | 1.300E-09 | 1.000E-08 |
| 27296 | 1365 | V1365S | E | 3e-09 | 1.1e-08 | 2.8e-09 | 5.600E-09 | 4.678E-09 | 2.800E-09 | 1.100E-08 |
| 27297 | 1365 | V1365T | E | 2e-08 | 5.5e-09 | 5.1e-09 | 1.020E-08 | 8.489E-09 | 5.100E-09 | 2.000E-08 |
| 27298 | 1365 | V1365W | E | 3.6e-09 | 3.5e-09 | 3.3e-09 | 3.467E-09 | 1.528E-10 | 3.300E-09 | 3.600E-09 |
| 27299 | 1365 | V1365Y | E | 3.9e-09 | 1.3e-08 | 5.2e-09 | 7.367E-09 | 4.922E-09 | 3.900E-09 | 1.300E-08 |
| 27300 | 1365 | V1365V | E | 1.3e-08 | 1.2e-08 | 7.1e-09 | 1.070E-08 | 3.158E-09 | 7.100E-09 | 1.300E-08 |
| 27301 | 1366 | Y1366A | E | 8.5e-09 | 1.2e-08 | 1.7e-08 | 1.250E-08 | 4.272E-09 | 8.500E-09 | 1.700E-08 |
| 27302 | 1366 | Y1366R | E | 9.8e-09 | 9.2e-09 | 1.8e-08 | 1.233E-08 | 4.917E-09 | 9.200E-09 | 1.800E-08 |
| 27303 | 1366 | Y1366N | E | 2e-09 | 6.4e-09 | 1.4e-08 | 7.467E-09 | 6.071E-09 | 2.000E-09 | 1.400E-08 |
| 27304 | 1366 | Y1366D | E | 3.9e-09 | 6.6e-09 | 7e-09 | 5.833E-09 | 1.686E-09 | 3.900E-09 | 7.000E-09 |
| 27305 | 1366 | Y1366C | E | 1.4e-08 | 1.2e-08 | 5.8e-09 | 1.060E-08 | 4.276E-09 | 5.800E-09 | 1.400E-08 |
| 27306 | 1366 | Y1366E | E | 1.3e-08 | 1.8e-09 | 5.2e-09 | 6.667E-09 | 5.742E-09 | 1.800E-09 | 1.300E-08 |
| 27307 | 1366 | Y1366Q | E | 1.3e-08 | 1.1e-08 | 3.2e-09 | 9.067E-09 | 5.178E-09 | 3.200E-09 | 1.300E-08 |
| 27308 | 1366 | Y1366G | E | 3.1e-09 | 1.2e-08 | 1.3e-08 | 9.367E-09 | 5.450E-09 | 3.100E-09 | 1.300E-08 |
| 27309 | 1366 | Y1366H | E | 5.4e-09 | 1.3e-08 | 2.4e-09 | 6.933E-09 | 5.464E-09 | 2.400E-09 | 1.300E-08 |
| 27310 | 1366 | Y1366I | E | 4.3e-09 | 1.1e-08 | 2.9e-09 | 6.067E-09 | 4.329E-09 | 2.900E-09 | 1.100E-08 |
| 27311 | 1366 | Y1366L | E | 7.4e-09 | 6e-08 | 6.6e-09 | 2.467E-08 | 3.060E-08 | 6.600E-09 | 6.000E-08 |
| 27312 | 1366 | Y1366K | E | 4.6e-09 | 6.5e-09 | 1.3e-08 | 8.033E-09 | 4.405E-09 | 4.600E-09 | 1.300E-08 |
| 27313 | 1366 | Y1366M | E | 3.7e-09 | 7.2e-09 | 1.2e-08 | 7.633E-09 | 4.167E-09 | 3.700E-09 | 1.200E-08 |
| 27314 | 1366 | Y1366F | E | 7.7e-09 | 3.1e-09 | 5.7e-09 | 5.500E-09 | 2.307E-09 | 3.100E-09 | 7.700E-09 |
| 27315 | 1366 | Y1366P | E | 6.2e-09 | 7.5e-09 | 1.2e-08 | 8.567E-09 | 3.044E-09 | 6.200E-09 | 1.200E-08 |
| 27316 | 1366 | Y1366S | E | 1.6e-08 | 9.4e-09 | 1.2e-08 | 1.247E-08 | 3.325E-09 | 9.400E-09 | 1.600E-08 |
| 27317 | 1366 | Y1366T | E | 4.3e-09 | 1e-08 | 3.8e-09 | 6.033E-09 | 3.444E-09 | 3.800E-09 | 1.000E-08 |
| 27318 | 1366 | Y1366W | E | 6.6e-09 | 1.1e-08 | 4.5e-08 | 2.087E-08 | 2.102E-08 | 6.600E-09 | 4.500E-08 |
| 27319 | 1366 | Y1366Y | E | 7.1e-09 | 9e-09 | 6.5e-09 | 7.533E-09 | 1.305E-09 | 6.500E-09 | 9.000E-09 |
| 27320 | 1366 | Y1366V | E | 3e-09 | 3.2e-08 | 1.6e-08 | 1.700E-08 | 1.453E-08 | 3.000E-09 | 3.200E-08 |
| 27321 | 1367 | Y1367A | E | 5.1e-09 | 8.3e-09 | 5.9e-09 | 6.433E-09 | 1.665E-09 | 5.100E-09 | 8.300E-09 |
| 27322 | 1367 | Y1367R | E | 3.4e-09 | 9.8e-09 | 6.4e-09 | 6.533E-09 | 3.202E-09 | 3.400E-09 | 9.800E-09 |
| 27323 | 1367 | Y1367N | E | 1.1e-08 | 2.1e-09 | 6.1e-09 | 6.400E-09 | 4.458E-09 | 2.100E-09 | 1.100E-08 |
| 27324 | 1367 | Y1367D | E | 5.5e-09 | 9.3e-09 | 8.1e-09 | 7.633E-09 | 1.943E-09 | 5.500E-09 | 9.300E-09 |
| 27325 | 1367 | Y1367C | E | 4.1e-09 | 3.9e-09 | 6.6e-09 | 4.867E-09 | 1.504E-09 | 3.900E-09 | 6.600E-09 |
| 27326 | 1367 | Y1367E | E | 1.3e-08 | 1e-08 | 5.9e-09 | 9.633E-09 | 3.564E-09 | 5.900E-09 | 1.300E-08 |
| 27327 | 1367 | Y1367Q | E | 7.5e-09 | 1.3e-08 | 2.5e-09 | 7.667E-09 | 5.252E-09 | 2.500E-09 | 1.300E-08 |
| 27328 | 1367 | Y1367G | E | 1.7e-08 | 6.4e-09 | 8.3e-09 | 1.057E-08 | 5.652E-09 | 6.400E-09 | 1.700E-08 |
| 27329 | 1367 | Y1367H | E | 1.6e-08 | 1.1e-09 | 1.7e-08 | 1.137E-08 | 8.905E-09 | 1.100E-09 | 1.700E-08 |
| 27330 | 1367 | Y1367I | E | 8.6e-09 | 1.1e-08 | 1.2e-08 | 1.053E-08 | 1.747E-09 | 8.600E-09 | 1.200E-08 |
| 27331 | 1367 | Y1367L | E | 8e-09 | 5.1e-09 | 9e-09 | 7.367E-09 | 2.026E-09 | 5.100E-09 | 9.000E-09 |
| 27332 | 1367 | Y1367K | E | 1.8e-09 | 5.4e-09 | 7.1e-09 | 4.767E-09 | 2.706E-09 | 1.800E-09 | 7.100E-09 |
| 27333 | 1367 | Y1367M | E | 4.2e-09 | 3.4e-09 | 5.4e-09 | 4.333E-09 | 1.007E-09 | 3.400E-09 | 5.400E-09 |
| 27334 | 1367 | Y1367F | E | 3.6e-09 | 1.3e-08 | 1.4e-08 | 1.020E-08 | 5.738E-09 | 3.600E-09 | 1.400E-08 |
| 27335 | 1367 | Y1367P | E | 5.3e-09 | 1.4e-08 | 4.3e-09 | 7.867E-09 | 5.335E-09 | 4.300E-09 | 1.400E-08 |
| 27336 | 1367 | Y1367S | E | 2.8e-08 | 4.9e-09 | 1.1e-08 | 1.463E-08 | 1.197E-08 | 4.900E-09 | 2.800E-08 |
| 27337 | 1367 | Y1367T | E | 2.7e-09 | 2.3e-08 | 1.9e-09 | 9.200E-09 | 1.196E-08 | 1.900E-09 | 2.300E-08 |
| 27338 | 1367 | Y1367W | E | 8e-09 | 5.7e-09 | 1e-08 | 7.900E-09 | 2.152E-09 | 5.700E-09 | 1.000E-08 |
| 27339 | 1367 | Y1367Y | E | 1.5e-08 | 5.6e-09 | 4e-09 | 8.200E-09 | 5.943E-09 | 4.000E-09 | 1.500E-08 |
| 27340 | 1367 | Y1367V | E | 1.4e-08 | 9.6e-09 | 9.6e-09 | 1.107E-08 | 2.540E-09 | 9.600E-09 | 1.400E-08 |
| 27341 | 1368 | C1368A | E | 1.4e-08 | 4.9e-09 | 6e-09 | 8.300E-09 | 4.967E-09 | 4.900E-09 | 1.400E-08 |
| 27342 | 1368 | C1368R | E | 7.2e-09 | 3.8e-09 | 1.4e-08 | 8.333E-09 | 5.194E-09 | 3.800E-09 | 1.400E-08 |
| 27343 | 1368 | C1368N | E | 1.5e-08 | 4.9e-09 | 9.1e-09 | 9.667E-09 | 5.074E-09 | 4.900E-09 | 1.500E-08 |
| 27344 | 1368 | C1368D | E | 7.5e-09 | 2.9e-08 | 6.6e-09 | 1.437E-08 | 1.268E-08 | 6.600E-09 | 2.900E-08 |
| 27345 | 1368 | C1368C | E | 1.3e-09 | 4e-08 | 5.2e-09 | 1.550E-08 | 2.131E-08 | 1.300E-09 | 4.000E-08 |
| 27346 | 1368 | C1368E | E | 2.5e-09 | 2.2e-09 | 7.6e-09 | 4.100E-09 | 3.035E-09 | 2.200E-09 | 7.600E-09 |
| 27347 | 1368 | C1368Q | E | 8e-09 | 8e-09 | 6.8e-09 | 7.600E-09 | 6.928E-10 | 6.800E-09 | 8.000E-09 |
| 27348 | 1368 | C1368G | E | 8.4e-09 | 1.4e-08 | 1.4e-08 | 1.213E-08 | 3.233E-09 | 8.400E-09 | 1.400E-08 |
| 27349 | 1368 | C1368H | E | 9.6e-09 | 3.9e-09 | 5.9e-09 | 6.467E-09 | 2.892E-09 | 3.900E-09 | 9.600E-09 |
| 27350 | 1368 | C1368I | E | 6.2e-09 | 4.4e-09 | 1.5e-08 | 8.533E-09 | 5.672E-09 | 4.400E-09 | 1.500E-08 |
| 27351 | 1368 | C1368L | E | 8.8e-09 | 7.5e-09 | 4.4e-09 | 6.900E-09 | 2.261E-09 | 4.400E-09 | 8.800E-09 |
| 27352 | 1368 | C1368K | E | 9.9e-09 | 8.5e-09 | 9.3e-09 | 9.233E-09 | 7.024E-10 | 8.500E-09 | 9.900E-09 |
| 27353 | 1368 | C1368M | E | 2.2e-09 | 1.8e-08 | 7.3e-09 | 9.167E-09 | 8.064E-09 | 2.200E-09 | 1.800E-08 |
| 27354 | 1368 | C1368F | E | 4.5e-09 | 3.8e-09 | 1.1e-08 | 6.433E-09 | 3.970E-09 | 3.800E-09 | 1.100E-08 |
| 27355 | 1368 | C1368P | E | 5.1e-09 | 5.2e-09 | 2.1e-09 | 4.133E-09 | 1.762E-09 | 2.100E-09 | 5.200E-09 |
| 27356 | 1368 | C1368S | E | 2.1e-08 | 1.1e-08 | 7.6e-09 | 1.320E-08 | 6.966E-09 | 7.600E-09 | 2.100E-08 |
| 27357 | 1368 | C1368T | E | 2.2e-09 | 1.6e-08 | 2.6e-08 | 1.473E-08 | 1.195E-08 | 2.200E-09 | 2.600E-08 |
| 27358 | 1368 | C1368W | E | 7.6e-09 | 6.7e-09 | 1.6e-08 | 1.010E-08 | 5.129E-09 | 6.700E-09 | 1.600E-08 |
| 27359 | 1368 | C1368Y | E | 7.1e-09 | 8.5e-09 | 8.3e-09 | 7.967E-09 | 7.572E-10 | 7.100E-09 | 8.500E-09 |
| 27360 | 1368 | C1368V | E | 2.5e-09 | 4.2e-09 | 3.4e-09 | 3.367E-09 | 8.505E-10 | 2.500E-09 | 4.200E-09 |
| 27361 | 1369 | S1369A | E | 5.6e-09 | 3.3e-09 | 6.1e-09 | 5.000E-09 | 1.493E-09 | 3.300E-09 | 6.100E-09 |
| 27362 | 1369 | S1369R | E | 6.6e-09 | 7.9e-09 | 4.9e-09 | 6.467E-09 | 1.504E-09 | 4.900E-09 | 7.900E-09 |
| 27363 | 1369 | S1369N | E | 3.9e-09 | 1.7e-08 | 6.1e-09 | 9.000E-09 | 7.015E-09 | 3.900E-09 | 1.700E-08 |
| 27364 | 1369 | S1369D | E | 9.7e-09 | 7.5e-09 | 9.4e-09 | 8.867E-09 | 1.193E-09 | 7.500E-09 | 9.700E-09 |
| 27365 | 1369 | S1369C | E | 4.8e-09 | 4e-09 | 8.9e-09 | 5.900E-09 | 2.629E-09 | 4.000E-09 | 8.900E-09 |
| 27366 | 1369 | S1369E | E | 4.7e-09 | 2e-09 | 8.2e-09 | 4.967E-09 | 3.109E-09 | 2.000E-09 | 8.200E-09 |
| 27367 | 1369 | S1369Q | E | 7.8e-09 | 9.6e-09 | 9.2e-09 | 8.867E-09 | 9.452E-10 | 7.800E-09 | 9.600E-09 |
| 27368 | 1369 | S1369G | E | 6.5e-09 | 9.7e-09 | 6.8e-09 | 7.667E-09 | 1.767E-09 | 6.500E-09 | 9.700E-09 |
| 27369 | 1369 | S1369H | E | 4.7e-09 | 1.2e-08 | 1.4e-08 | 1.023E-08 | 4.895E-09 | 4.700E-09 | 1.400E-08 |

| | | | | | | | | | | |
|---|---|---|---|---|---|---|---|---|---|---|
| 27370 | 1369 | S1369I | E | 4.2e-09 | 4.2e-09 | 6.1e-09 | 4.833E-09 | 1.097E-09 | 4.200E-09 | 6.100E-09 |
| 27371 | 1369 | S1369L | E | 8.6e-09 | 9.2e-09 | 7e-09 | 8.267E-09 | 1.137E-09 | 7.000E-09 | 9.200E-09 |
| 27372 | 1369 | S1369K | E | 2.3e-09 | 7.5e-09 | 9e-09 | 6.267E-09 | 3.516E-09 | 2.300E-09 | 9.000E-09 |
| 27373 | 1369 | S1369M | E | 2.2e-08 | 6.1e-09 | 7.9e-09 | 1.200E-08 | 8.707E-09 | 6.100E-09 | 2.200E-08 |
| 27374 | 1369 | S1369F | E | 1.2e-08 | 1.9e-09 | 5.5e-09 | 6.467E-09 | 5.119E-09 | 1.900E-09 | 1.200E-08 |
| 27375 | 1369 | S1369P | E | 1.6e-08 | 1.6e-08 | 2e-08 | 1.733E-08 | 2.309E-09 | 1.600E-08 | 2.000E-08 |
| 27376 | 1369 | S1369S | E | 6.7e-09 | 2e-08 | 7.7e-09 | 1.147E-08 | 7.407E-09 | 6.700E-09 | 2.000E-08 |
| 27377 | 1369 | S1369T | E | 7.7e-09 | 8.6e-09 | 1e-08 | 8.767E-09 | 1.159E-09 | 7.700E-09 | 1.000E-08 |
| 27378 | 1369 | S1369W | E | 1.1e-08 | 5.2e-09 | 1.3e-08 | 9.733E-09 | 4.051E-09 | 5.200E-09 | 1.300E-08 |
| 27379 | 1369 | S1369Y | E | 9.7e-09 | 1e-08 | 1.2e-08 | 1.057E-08 | 1.250E-09 | 9.700E-09 | 1.200E-08 |
| 27380 | 1369 | S1369V | E | 3.3e-09 | 5.7e-09 | 1.1e-08 | 6.667E-09 | 3.940E-09 | 3.300E-09 | 1.100E-08 |
| 27381 | 1370 | R1370A | E | 4.5e-09 | 5.6e-09 | 2.9e-09 | 4.333E-09 | 1.358E-09 | 2.900E-09 | 5.600E-09 |
| 27382 | 1370 | R1370R | E | 6.2e-09 | 9.5e-09 | 6.8e-09 | 7.500E-09 | 1.758E-09 | 6.200E-09 | 9.500E-09 |
| 27383 | 1370 | R1370N | E | 5.2e-09 | 1e-08 | 5.4e-09 | 6.867E-09 | 2.715E-09 | 5.200E-09 | 1.000E-08 |
| 27384 | 1370 | R1370D | E | 1e-08 | 4.8e-09 | 3.5e-09 | 6.100E-09 | 3.439E-09 | 3.500E-09 | 1.000E-08 |
| 27385 | 1370 | R1370C | E | 4e-09 | 6.6e-09 | 1e-08 | 6.867E-09 | 3.009E-09 | 4.000E-09 | 1.000E-08 |
| 27386 | 1370 | R1370E | E | 1.1e-08 | 1.5e-08 | 5.1e-09 | 1.037E-08 | 4.980E-09 | 5.100E-09 | 1.500E-08 |
| 27387 | 1370 | R1370Q | E | 9.9e-09 | 1.2e-08 | 8.6e-09 | 1.017E-08 | 1.716E-09 | 8.600E-09 | 1.200E-08 |
| 27388 | 1370 | R1370G | E | 3e-09 | 3.6e-09 | 5.7e-09 | 4.100E-09 | 1.418E-09 | 3.000E-09 | 5.700E-09 |
| 27389 | 1370 | R1370H | E | 4.9e-09 | 4.1e-09 | 5.6e-09 | 4.867E-09 | 7.506E-10 | 4.100E-09 | 5.600E-09 |
| 27390 | 1370 | R1370I | E | 9.2e-09 | 9e-09 | 4.8e-09 | 7.667E-09 | 2.485E-09 | 4.800E-09 | 9.200E-09 |
| 27391 | 1370 | R1370L | E | 1e-08 | 8.6e-09 | 5.5e-09 | 8.033E-09 | 2.303E-09 | 5.500E-09 | 1.000E-08 |
| 27392 | 1370 | R1370K | E | 1e-08 | 6.2e-09 | 1.4e-08 | 1.007E-08 | 3.900E-09 | 6.200E-09 | 1.400E-08 |
| 27393 | 1370 | R1370M | E | 1.1e-08 | 1e-08 | 1.3e-08 | 1.133E-08 | 1.528E-09 | 1.000E-08 | 1.300E-08 |
| 27394 | 1370 | R1370F | E | 4e-09 | 2.6e-09 | 5.6e-09 | 4.067E-09 | 1.501E-09 | 2.600E-09 | 5.600E-09 |
| 27395 | 1370 | R1370P | E | 1.2e-08 | 6.8e-09 | 4.4e-09 | 7.733E-09 | 3.885E-09 | 4.400E-09 | 1.200E-08 |
| 27396 | 1370 | R1370S | E | 1e-08 | 1.5e-08 | 1.5e-09 | 8.833E-09 | 6.825E-09 | 1.500E-09 | 1.500E-08 |
| 27397 | 1370 | R1370T | E | 6.4e-09 | 7.1e-09 | 8.3e-09 | 7.267E-09 | 9.609E-10 | 6.400E-09 | 8.300E-09 |
| 27398 | 1370 | R1370W | E | 6e-09 | 5.6e-09 | 3.9e-09 | 5.167E-09 | 1.115E-09 | 3.900E-09 | 6.000E-09 |
| 27399 | 1370 | R1370Y | E | 2.1e-08 | 1.1e-08 | 6.6e-09 | 1.287E-08 | 7.379E-09 | 6.600E-09 | 2.100E-08 |
| 27400 | 1370 | R1370V | E | 4e-09 | 5.8e-09 | 7.2e-09 | 5.667E-09 | 1.604E-09 | 4.000E-09 | 7.200E-09 |
| 27401 | 1371 | W1371A | E | 4.8e-09 | 7.8e-09 | 1.2e-09 | 4.600E-09 | 3.305E-09 | 1.200E-09 | 7.800E-09 |
| 27402 | 1371 | W1371R | E | 1.3e-08 | 9.1e-09 | 1.8e-08 | 1.337E-08 | 4.461E-09 | 9.100E-09 | 1.800E-08 |
| 27403 | 1371 | W1371N | E | 2.1e-09 | 5.6e-09 | 3.7e-09 | 3.800E-09 | 1.752E-09 | 2.100E-09 | 5.600E-09 |
| 27404 | 1371 | W1371D | E | 1.7e-08 | 5.9e-09 | 4.3e-09 | 9.067E-09 | 6.917E-09 | 4.300E-09 | 1.700E-08 |
| 27405 | 1371 | W1371C | E | 9.9e-09 | 9.8e-09 | 1e-08 | 9.900E-09 | 1.000E-10 | 9.800E-09 | 1.000E-08 |
| 27406 | 1371 | W1371E | E | 1.1e-08 | 4.5e-09 | 2.8e-08 | 1.450E-08 | 1.213E-08 | 4.500E-09 | 2.800E-08 |
| 27407 | 1371 | W1371Q | E | 1e-08 | 1e-08 | 6.2e-09 | 8.733E-09 | 2.194E-09 | 6.200E-09 | 1.000E-08 |
| 27408 | 1371 | W1371G | E | 1.4e-08 | 9.1e-09 | 3.8e-09 | 8.967E-09 | 5.101E-09 | 3.800E-09 | 1.400E-08 |
| 27409 | 1371 | W1371H | E | 4.8e-09 | 5.4e-09 | 5.4e-09 | 5.200E-09 | 3.464E-10 | 4.800E-09 | 5.400E-09 |
| 27410 | 1371 | W1371I | E | 8.1e-09 | 6.4e-09 | 7.9e-09 | 7.467E-09 | 9.292E-10 | 6.400E-09 | 8.100E-09 |
| 27411 | 1371 | W1371L | E | 2.9e-09 | 6.7e-09 | 6e-09 | 5.200E-09 | 2.022E-09 | 2.900E-09 | 6.700E-09 |
| 27412 | 1371 | W1371K | E | 1.3e-08 | 5.7e-09 | 2.9e-09 | 7.200E-09 | 5.214E-09 | 2.900E-09 | 1.300E-08 |
| 27413 | 1371 | W1371M | E | 1.2e-08 | 6.4e-09 | 4.4e-09 | 7.600E-09 | 3.940E-09 | 4.400E-09 | 1.200E-08 |
| 27414 | 1371 | W1371F | E | 4e-09 | 1.5e-08 | 7.3e-09 | 8.767E-09 | 5.645E-09 | 4.000E-09 | 1.500E-08 |
| 27415 | 1371 | W1371P | E | 6.1e-09 | 4.6e-09 | 1.7e-09 | 4.133E-09 | 2.237E-09 | 1.700E-09 | 6.100E-09 |
| 27416 | 1371 | W1371S | E | 5.4e-09 | 9.1e-09 | 6.4e-09 | 6.967E-09 | 1.914E-09 | 5.400E-09 | 9.100E-09 |
| 27417 | 1371 | W1371T | E | 8.6e-09 | 8e-09 | 6.1e-09 | 7.567E-09 | 1.305E-09 | 6.100E-09 | 8.600E-09 |
| 27418 | 1371 | W1371W | E | 1.1e-08 | 5.3e-09 | 1.4e-08 | 1.010E-08 | 4.419E-09 | 5.300E-09 | 1.400E-08 |
| 27419 | 1371 | W1371Y | E | 3.3e-09 | 9.3e-09 | 8.5e-09 | 7.033E-09 | 3.258E-09 | 3.300E-09 | 9.300E-09 |
| 27420 | 1371 | W1371V | E | 2.6e-09 | 1e-08 | 6.4e-09 | 6.333E-09 | 3.700E-09 | 2.600E-09 | 1.000E-08 |
| 27421 | 1372 | G1372A | E | 5e-09 | 1.3e-08 | 6.4e-09 | 8.133E-09 | 4.272E-09 | 5.000E-09 | 1.300E-08 |
| 27422 | 1372 | G1372R | E | 6.3e-09 | 7.6e-09 | 7.4e-09 | 7.100E-09 | 7.000E-10 | 6.300E-09 | 7.600E-09 |
| 27423 | 1372 | G1372N | E | 1.9e-08 | 7.9e-09 | 6.5e-09 | 1.113E-08 | 6.849E-09 | 6.500E-09 | 1.900E-08 |
| 27424 | 1372 | G1372D | E | 8.1e-09 | 1.9e-09 | 3.6e-09 | 4.533E-09 | 3.204E-09 | 1.900E-09 | 8.100E-09 |
| 27425 | 1372 | G1372C | E | 4e-08 | 8.2e-09 | 6e-09 | 1.807E-08 | 1.903E-08 | 6.000E-09 | 4.000E-08 |
| 27426 | 1372 | G1372E | E | 4.4e-09 | 9.8e-09 | 5.5e-09 | 6.567E-09 | 2.854E-09 | 4.400E-09 | 9.800E-09 |
| 27427 | 1372 | G1372Q | E | 5.6e-09 | 2.8e-09 | 7e-09 | 5.133E-09 | 2.139E-09 | 2.800E-09 | 7.000E-09 |
| 27428 | 1372 | G1372G | E | 1.1e-08 | 5.8e-09 | 1.7e-08 | 1.127E-08 | 5.605E-09 | 5.800E-09 | 1.700E-08 |
| 27429 | 1372 | G1372H | E | 2e-08 | 4e-09 | 6.5e-09 | 1.017E-08 | 8.607E-09 | 4.000E-09 | 2.000E-08 |
| 27430 | 1372 | G1372I | E | 4.2e-09 | 2.9e-09 | 9.5e-09 | 5.533E-09 | 3.496E-09 | 2.900E-09 | 9.500E-09 |
| 27431 | 1372 | G1372L | E | 7.1e-09 | 2.8e-09 | 1e-08 | 6.633E-09 | 3.623E-09 | 2.800E-09 | 1.000E-08 |
| 27432 | 1372 | G1372K | E | 5e-09 | 7.2e-09 | 9.3e-09 | 7.167E-09 | 2.150E-09 | 5.000E-09 | 9.300E-09 |
| 27433 | 1372 | G1372M | E | 4.7e-09 | 3.9e-09 | 8.7e-09 | 5.767E-09 | 2.572E-09 | 3.900E-09 | 8.700E-09 |
| 27434 | 1372 | G1372F | E | 6.6e-09 | 9.1e-09 | 3.8e-09 | 6.500E-09 | 2.651E-09 | 3.800E-09 | 9.100E-09 |
| 27435 | 1372 | G1372P | E | 1.3e-09 | 6.6e-09 | 8.3e-09 | 5.400E-09 | 3.651E-09 | 1.300E-09 | 8.300E-09 |
| 27436 | 1372 | G1372S | E | 7.1e-09 | 3.7e-09 | 7e-09 | 5.933E-09 | 1.935E-09 | 3.700E-09 | 7.100E-09 |
| 27437 | 1372 | G1372T | E | 2.7e-08 | 1.2e-08 | 1.5e-08 | 1.800E-08 | 7.937E-09 | 1.200E-08 | 2.700E-08 |
| 27438 | 1372 | G1372W | E | 8.2e-09 | 1.1e-08 | 6.7e-09 | 8.633E-09 | 2.183E-09 | 6.700E-09 | 1.100E-08 |
| 27439 | 1372 | G1372Y | E | 3.9e-09 | 5.8e-09 | 2.7e-08 | 1.223E-08 | 1.282E-08 | 3.900E-09 | 2.700E-08 |
| 27440 | 1372 | G1372V | E | 7.8e-09 | 8.1e-09 | 3e-09 | 6.300E-09 | 2.862E-09 | 3.000E-09 | 8.100E-09 |
| 27441 | 1373 | G1373A | E | 4.8e-09 | 8.8e-09 | 2.9e-09 | 5.500E-09 | 3.012E-09 | 2.900E-09 | 8.800E-09 |
| 27442 | 1373 | G1373R | E | 6.4e-09 | 1.9e-08 | 5.4e-09 | 1.027E-08 | 7.580E-09 | 5.400E-09 | 1.900E-08 |
| 27443 | 1373 | G1373N | E | 7.1e-09 | 4.9e-09 | 3.6e-09 | 5.200E-09 | 1.769E-09 | 3.600E-09 | 7.100E-09 |
| 27444 | 1373 | G1373D | E | 9.5e-09 | 1.1e-08 | 5.6e-09 | 8.700E-09 | 2.787E-09 | 5.600E-09 | 1.100E-08 |
| 27445 | 1373 | G1373C | E | 5.5e-09 | 8e-09 | 3.7e-09 | 5.733E-09 | 2.159E-09 | 3.700E-09 | 8.000E-09 |
| 27446 | 1373 | G1373E | E | 7.5e-09 | 9e-09 | 6.8e-09 | 7.767E-09 | 1.124E-09 | 6.800E-09 | 9.000E-09 |
| 27447 | 1373 | G1373Q | E | 6.1e-09 | 7.9e-09 | 4.5e-09 | 6.167E-09 | 1.701E-09 | 4.500E-09 | 7.900E-09 |
| 27448 | 1373 | G1373G | E | 3.6e-09 | 3.7e-09 | 1.4e-09 | 2.900E-09 | 1.300E-09 | 1.400E-09 | 3.700E-09 |
| 27449 | 1373 | G1373H | E | 9.7e-09 | 8.6e-09 | 1.2e-08 | 1.010E-08 | 1.735E-09 | 8.600E-09 | 1.200E-08 |
| 27450 | 1373 | G1373I | E | 6.9e-09 | 7.6e-09 | 4.5e-09 | 6.333E-09 | 1.626E-09 | 4.500E-09 | 7.600E-09 |
| 27451 | 1373 | G1373L | E | 1.2e-08 | 8e-09 | 1.1e-08 | 1.033E-08 | 2.082E-09 | 8.000E-09 | 1.200E-08 |
| 27452 | 1373 | G1373K | E | 4.8e-09 | 6.9e-09 | 1.1e-08 | 7.567E-09 | 3.153E-09 | 4.800E-09 | 1.100E-08 |
| 27453 | 1373 | G1373M | E | 6.4e-09 | 2.5e-08 | 4.8e-09 | 1.207E-08 | 1.123E-08 | 4.800E-09 | 2.500E-08 |
| 27454 | 1373 | G1373F | E | 2.5e-09 | 1.7e-08 | 5e-09 | 8.167E-09 | 7.751E-09 | 2.500E-09 | 1.700E-08 |

| | | | | | | | | | | |
|---|---|---|---|---|---|---|---|---|---|---|
| 27455 | 1373 | G1373P | E | 9.5e-09 | 4e-09 | 8.4e-09 | 7.300E-09 | 2.910E-09 | 4.000E-09 | 9.500E-09 |
| 27456 | 1373 | G1373S | E | 3.1e-09 | 7.2e-09 | 9.6e-09 | 6.633E-09 | 3.287E-09 | 3.100E-09 | 9.600E-09 |
| 27457 | 1373 | G1373T | E | 1.4e-08 | 6.5e-09 | 5e-09 | 8.500E-09 | 4.822E-09 | 5.000E-09 | 1.400E-08 |
| 27458 | 1373 | G1373W | E | 1.1e-08 | 7.7e-09 | 1.1e-08 | 9.900E-09 | 1.905E-09 | 7.700E-09 | 1.100E-08 |
| 27459 | 1373 | G1373Y | E | 1e-08 | 3.9e-09 | 6.2e-09 | 6.700E-09 | 3.081E-09 | 3.900E-09 | 1.000E-08 |
| 27460 | 1373 | G1373V | E | 1.9e-08 | 4.8e-09 | 6.6e-09 | 1.013E-08 | 7.731E-09 | 4.800E-09 | 1.900E-08 |
| 27461 | 1374 | D1374A | E | 6.5e-09 | 5.7e-09 | 5.1e-09 | 5.767E-09 | 7.024E-10 | 5.100E-09 | 6.500E-09 |
| 27462 | 1374 | D1374R | E | 2.1e-08 | 1.5e-08 | 5e-09 | 1.367E-08 | 8.083E-09 | 5.000E-09 | 2.100E-08 |
| 27463 | 1374 | D1374N | E | 3.1e-09 | 6.6e-09 | 4.8e-09 | 4.833E-09 | 1.750E-09 | 3.100E-09 | 6.600E-09 |
| 27464 | 1374 | D1374D | E | 1e-08 | 7.9e-09 | 8.6e-09 | 8.833E-09 | 1.069E-09 | 7.900E-09 | 1.000E-08 |
| 27465 | 1374 | D1374C | E | 1.7e-08 | 2.4e-09 | 4.8e-09 | 8.067E-09 | 7.829E-09 | 2.400E-09 | 1.700E-08 |
| 27466 | 1374 | D1374E | E | 5.9e-09 | 1.5e-08 | 4.7e-09 | 8.533E-09 | 5.632E-09 | 4.700E-09 | 1.500E-08 |
| 27467 | 1374 | D1374Q | E | 1.8e-09 | 1.2e-08 | 7.5e-09 | 7.100E-09 | 5.112E-09 | 1.800E-09 | 1.200E-08 |
| 27468 | 1374 | D1374G | E | 6.1e-09 | 1e-08 | 3.8e-09 | 6.633E-09 | 3.134E-09 | 3.800E-09 | 1.000E-08 |
| 27469 | 1374 | D1374H | E | 4.2e-09 | 7e-09 | 1.2e-08 | 7.733E-09 | 3.951E-09 | 4.200E-09 | 1.200E-08 |
| 27470 | 1374 | D1374I | E | 5.3e-09 | 1.6e-08 | 1.2e-09 | 7.500E-09 | 7.641E-09 | 1.200E-09 | 1.600E-08 |
| 27471 | 1374 | D1374L | E | 1.8e-08 | 1.8e-08 | 3.8e-09 | 1.327E-08 | 8.198E-09 | 3.800E-09 | 1.800E-08 |
| 27472 | 1374 | D1374K | E | 8.2e-09 | 8.5e-09 | 7.3e-09 | 8.000E-09 | 6.245E-10 | 7.300E-09 | 8.500E-09 |
| 27473 | 1374 | D1374M | E | 1.1e-08 | 7.2e-09 | 5.6e-09 | 7.933E-09 | 2.774E-09 | 5.600E-09 | 1.100E-08 |
| 27474 | 1374 | D1374F | E | 9.9e-09 | 1.6e-08 | 4.2e-09 | 1.003E-08 | 5.901E-09 | 4.200E-09 | 1.600E-08 |
| 27475 | 1374 | D1374P | E | 6.2e-09 | 4e-09 | 8.5e-09 | 6.233E-09 | 2.250E-09 | 4.000E-09 | 8.500E-09 |
| 27476 | 1374 | D1374S | E | 5e-09 | 1.8e-08 | 6.2e-09 | 9.733E-09 | 7.184E-09 | 5.000E-09 | 1.800E-08 |
| 27477 | 1374 | D1374T | E | 1.4e-08 | 1.9e-08 | 1.2e-08 | 1.500E-08 | 3.606E-09 | 1.200E-08 | 1.900E-08 |
| 27478 | 1374 | D1374W | E | 6.7e-09 | 6.3e-09 | 7.3e-09 | 6.767E-09 | 5.033E-10 | 6.300E-09 | 7.300E-09 |
| 27479 | 1374 | D1374Y | E | 1.4e-08 | 8.9e-09 | 9.9e-09 | 1.093E-08 | 2.702E-09 | 8.900E-09 | 1.400E-08 |
| 27480 | 1374 | D1374V | E | 7.3e-09 | 1.3e-08 | 5.5e-09 | 8.600E-09 | 3.915E-09 | 5.500E-09 | 1.300E-08 |
| 27481 | 1375 | G1375A | E | 4e-09 | 6.4e-09 | 9.4e-09 | 6.600E-09 | 2.706E-09 | 4.000E-09 | 9.400E-09 |
| 27482 | 1375 | G1375R | E | 7.6e-09 | 1.4e-08 | 8.3e-09 | 9.967E-09 | 3.510E-09 | 7.600E-09 | 1.400E-08 |
| 27483 | 1375 | G1375N | E | 2e-09 | 2e-08 | 3.2e-09 | 8.400E-09 | 1.006E-08 | 2.000E-09 | 2.000E-08 |
| 27484 | 1375 | G1375D | E | 8.7e-09 | 1.8e-09 | 4e-09 | 4.833E-09 | 3.525E-09 | 1.800E-09 | 8.700E-09 |
| 27485 | 1375 | G1375C | E | 4.2e-09 | 4.8e-09 | 4.7e-09 | 4.567E-09 | 3.215E-10 | 4.200E-09 | 4.800E-09 |
| 27486 | 1375 | G1375E | E | 1.3e-08 | 2.8e-09 | 2.9e-09 | 6.233E-09 | 5.860E-09 | 2.800E-09 | 1.300E-08 |
| 27487 | 1375 | G1375Q | E | 4e-09 | 4.5e-09 | 1.2e-08 | 6.833E-09 | 4.481E-09 | 4.000E-09 | 1.200E-08 |
| 27488 | 1375 | G1375G | E | 9.6e-09 | 1e-08 | 1.3e-08 | 1.087E-08 | 1.858E-09 | 9.600E-09 | 1.300E-08 |
| 27489 | 1375 | G1375H | E | 6.9e-09 | 1.2e-08 | 7.6e-09 | 8.833E-09 | 2.765E-09 | 6.900E-09 | 1.200E-08 |
| 27490 | 1375 | G1375I | E | 1.4e-08 | 1e-08 | 2.2e-08 | 1.533E-08 | 6.110E-09 | 1.000E-08 | 2.200E-08 |
| 27491 | 1375 | G1375L | E | 6.6e-09 | 5.3e-09 | 2.1e-09 | 4.667E-09 | 2.316E-09 | 2.100E-09 | 6.600E-09 |
| 27492 | 1375 | G1375K | E | 1.3e-08 | 8.7e-09 | 1.4e-08 | 1.190E-08 | 2.816E-09 | 8.700E-09 | 1.400E-08 |
| 27493 | 1375 | G1375M | E | 5.4e-09 | 9.8e-09 | 8.8e-09 | 8.000E-09 | 2.307E-09 | 5.400E-09 | 9.800E-09 |
| 27494 | 1375 | G1375F | E | 4.7e-09 | 8.5e-09 | 7.4e-09 | 6.867E-09 | 1.955E-09 | 4.700E-09 | 8.500E-09 |
| 27495 | 1375 | G1375P | E | 6.1e-09 | 1.4e-08 | 1e-08 | 1.003E-08 | 3.950E-09 | 6.100E-09 | 1.400E-08 |
| 27496 | 1375 | G1375S | E | 3.5e-09 | 4.7e-09 | 8.2e-09 | 5.467E-09 | 2.442E-09 | 3.500E-09 | 8.200E-09 |
| 27497 | 1375 | G1375T | E | 3.2e-09 | 2.6e-09 | 3.8e-09 | 3.200E-09 | 6.000E-10 | 2.600E-09 | 3.800E-09 |
| 27498 | 1375 | G1375W | E | 2.7e-09 | 5.7e-09 | 6.9e-09 | 5.100E-09 | 2.163E-09 | 2.700E-09 | 6.900E-09 |
| 27499 | 1375 | G1375Y | E | 7e-09 | 1.2e-08 | 3.2e-09 | 7.400E-09 | 4.414E-09 | 3.200E-09 | 1.200E-08 |
| 27500 | 1375 | G1375V | E | 4.6e-09 | 8.2e-09 | 1.7e-08 | 9.933E-09 | 6.379E-09 | 4.600E-09 | 1.700E-08 |
| 27501 | 1376 | F1376A | E | 3.5e-09 | 1.3e-09 | 9.3e-09 | 4.700E-09 | 4.133E-09 | 1.300E-09 | 9.300E-09 |
| 27502 | 1376 | F1376R | E | 7.3e-09 | 1.2e-08 | 1e-08 | 9.767E-09 | 2.359E-09 | 7.300E-09 | 1.200E-08 |
| 27503 | 1376 | F1376N | E | 3.9e-09 | 6.6e-09 | 1.3e-08 | 7.833E-09 | 4.674E-09 | 3.900E-09 | 1.300E-08 |
| 27504 | 1376 | F1376D | E | 1.6e-08 | 1.6e-08 | 5.2e-09 | 1.240E-08 | 6.235E-09 | 5.200E-09 | 1.600E-08 |
| 27505 | 1376 | F1376C | E | 4.3e-09 | 4.3e-09 | 8.8e-09 | 5.800E-09 | 2.598E-09 | 4.300E-09 | 8.800E-09 |
| 27506 | 1376 | F1376E | E | 6.3e-09 | 1.6e-08 | 3.5e-09 | 8.600E-09 | 6.560E-09 | 3.500E-09 | 1.600E-08 |
| 27507 | 1376 | F1376Q | E | 3.3e-09 | 1.3e-08 | 6.8e-09 | 7.700E-09 | 4.912E-09 | 3.300E-09 | 1.300E-08 |
| 27508 | 1376 | F1376G | E | 1e-08 | 9.7e-09 | 9.5e-09 | 9.733E-09 | 2.517E-10 | 9.500E-09 | 1.000E-08 |
| 27509 | 1376 | F1376H | E | 9.1e-09 | 8e-09 | 1.1e-08 | 9.367E-09 | 1.518E-09 | 8.000E-09 | 1.100E-08 |
| 27510 | 1376 | F1376I | E | 1.1e-08 | 1e-08 | 1.7e-08 | 1.267E-08 | 3.786E-09 | 1.000E-08 | 1.700E-08 |
| 27511 | 1376 | F1376L | E | 4.3e-09 | 6.9e-09 | 3.3e-08 | 1.473E-08 | 1.587E-08 | 4.300E-09 | 3.300E-08 |
| 27512 | 1376 | F1376K | E | 7.3e-09 | 1.3e-08 | 7.6e-09 | 9.300E-09 | 3.208E-09 | 7.300E-09 | 1.300E-08 |
| 27513 | 1376 | F1376M | E | 1.8e-08 | 5.7e-09 | 3.6e-09 | 9.100E-09 | 7.779E-09 | 3.600E-09 | 1.800E-08 |
| 27514 | 1376 | F1376F | E | 4.6e-09 | 1.1e-08 | 4e-09 | 6.533E-09 | 3.880E-09 | 4.000E-09 | 1.100E-08 |
| 27515 | 1376 | F1376P | E | 1.2e-08 | 1e-08 | 4.1e-09 | 8.700E-09 | 4.107E-09 | 4.100E-09 | 1.200E-08 |
| 27516 | 1376 | F1376S | E | 3.2e-08 | 5.7e-08 | 8.7e-09 | 3.257E-08 | 2.415E-08 | 8.700E-09 | 5.700E-08 |
| 27517 | 1376 | F1376T | E | 6.9e-09 | 9.7e-09 | 9.5e-09 | 8.700E-09 | 1.562E-09 | 6.900E-09 | 9.700E-09 |
| 27518 | 1376 | F1376W | E | 2.9e-09 | 1.2e-08 | 6.6e-09 | 7.167E-09 | 4.576E-09 | 2.900E-09 | 1.200E-08 |
| 27519 | 1376 | F1376Y | E | 2e-09 | 6.8e-09 | 1.3e-08 | 7.267E-09 | 5.515E-09 | 2.000E-09 | 1.300E-08 |
| 27520 | 1376 | F1376V | E | 1.5e-08 | 8.4e-09 | 8.2e-09 | 1.053E-08 | 3.870E-09 | 8.200E-09 | 1.500E-08 |
| 27521 | 1377 | Y1377A | E | 4e-09 | 6.7e-09 | 2.6e-09 | 4.433E-09 | 2.084E-09 | 2.600E-09 | 6.700E-09 |
| 27522 | 1377 | Y1377R | E | 5.3e-09 | 4.8e-09 | 3.6e-08 | 1.537E-08 | 1.787E-08 | 4.800E-09 | 3.600E-08 |
| 27523 | 1377 | Y1377N | E | 6.7e-09 | 4.2e-09 | 4.5e-09 | 5.133E-09 | 1.365E-09 | 4.200E-09 | 6.700E-09 |
| 27524 | 1377 | Y1377D | E | 3.2e-09 | 2e-09 | 7.7e-09 | 4.300E-09 | 3.005E-09 | 2.000E-09 | 7.700E-09 |
| 27525 | 1377 | Y1377C | E | 3.6e-09 | 5.4e-09 | 6.7e-09 | 5.233E-09 | 1.557E-09 | 3.600E-09 | 6.700E-09 |
| 27526 | 1377 | Y1377E | E | 1.8e-08 | 6.8e-09 | 1.1e-08 | 1.193E-08 | 5.658E-09 | 6.800E-09 | 1.800E-08 |
| 27527 | 1377 | Y1377Q | E | 3e-08 | 5.5e-09 | 6.6e-09 | 1.403E-08 | 1.384E-08 | 5.500E-09 | 3.000E-08 |
| 27528 | 1377 | Y1377G | E | 8.3e-09 | 3.9e-09 | 5.8e-09 | 6.000E-09 | 2.207E-09 | 3.900E-09 | 8.300E-09 |
| 27529 | 1377 | Y1377H | E | 1.8e-08 | 1.1e-08 | 1.2e-08 | 1.367E-08 | 3.786E-09 | 1.100E-08 | 1.800E-08 |
| 27530 | 1377 | Y1377I | E | 5.1e-09 | 2.9e-09 | 5.8e-09 | 4.600E-09 | 1.513E-09 | 2.900E-09 | 5.800E-09 |
| 27531 | 1377 | Y1377L | E | 5.2e-09 | 4.3e-09 | 4.6e-09 | 4.700E-09 | 4.583E-10 | 4.300E-09 | 5.200E-09 |
| 27532 | 1377 | Y1377K | E | 6.7e-09 | 3.2e-09 | 1.2e-08 | 7.300E-09 | 4.431E-09 | 3.200E-09 | 1.200E-08 |
| 27533 | 1377 | Y1377M | E | 6.2e-09 | 1.2e-08 | 8.6e-09 | 8.933E-09 | 2.914E-09 | 6.200E-09 | 1.200E-08 |
| 27534 | 1377 | Y1377F | E | 8.7e-09 | 5.1e-09 | 4.9e-09 | 6.233E-09 | 2.139E-09 | 4.900E-09 | 8.700E-09 |
| 27535 | 1377 | Y1377P | E | 8.8e-09 | 2.9e-09 | 1.2e-08 | 7.900E-09 | 4.616E-09 | 2.900E-09 | 1.200E-08 |
| 27536 | 1377 | Y1377S | E | 8.3e-09 | 9e-09 | 2.4e-09 | 6.567E-09 | 3.625E-09 | 2.400E-09 | 9.000E-09 |
| 27537 | 1377 | Y1377T | E | 9.3e-09 | 1.6e-08 | 1e-08 | 1.177E-08 | 3.683E-09 | 9.300E-09 | 1.600E-08 |
| 27538 | 1377 | Y1377W | E | 9.1e-09 | 7.4e-09 | 7.7e-09 | 8.067E-09 | 9.074E-10 | 7.400E-09 | 9.100E-09 |
| 27539 | 1377 | Y1377Y | E | 6.1e-09 | 1.1e-08 | 2.8e-09 | 6.633E-09 | 4.126E-09 | 2.800E-09 | 1.100E-08 |

| | | | | | | | | | | |
|---|---|---|---|---|---|---|---|---|---|---|
| 27540 | 1377 | Y1377V | E | 6.8e-09 | 6.4e-09 | 5.2e-09 | 6.133E-09 | 8.327E-10 | 5.200E-09 | 6.800E-09 |
| 27541 | 1378 | A1378A | E | 4.8e-09 | 6.9e-09 | 3.4e-09 | 5.033E-09 | 1.762E-09 | 3.400E-09 | 6.900E-09 |
| 27542 | 1378 | A1378R | E | 3.7e-09 | 4.5e-09 | 3.1e-09 | 3.767E-09 | 7.024E-10 | 3.100E-09 | 4.500E-09 |
| 27543 | 1378 | A1378N | E | 7.4e-09 | 4.7e-09 | 1.3e-08 | 8.367E-09 | 4.234E-09 | 4.700E-09 | 1.300E-08 |
| 27544 | 1378 | A1378D | E | 2.1e-08 | 1e-08 | 3.2e-09 | 1.140E-08 | 8.982E-09 | 3.200E-09 | 2.100E-08 |
| 27545 | 1378 | A1378C | E | 5.2e-09 | 1.7e-09 | 2.7e-09 | 3.200E-09 | 1.803E-09 | 1.700E-09 | 5.200E-09 |
| 27546 | 1378 | A1378E | E | 6e-09 | 4.7e-09 | 9.8e-09 | 6.833E-09 | 2.650E-09 | 4.700E-09 | 9.800E-09 |
| 27547 | 1378 | A1378Q | E | 2.5e-08 | 3.9e-09 | 9.7e-09 | 1.287E-08 | 1.090E-08 | 3.900E-09 | 2.500E-08 |
| 27548 | 1378 | A1378G | E | 5e-09 | 1.4e-08 | 6.6e-09 | 8.533E-09 | 4.801E-09 | 5.000E-09 | 1.400E-08 |
| 27549 | 1378 | A1378H | E | 1.8e-08 | 6e-09 | 1e-08 | 1.133E-08 | 6.110E-09 | 6.000E-09 | 1.800E-08 |
| 27550 | 1378 | A1378I | E | 9.8e-09 | 1.1e-08 | 2e-09 | 7.600E-09 | 4.887E-09 | 2.000E-09 | 1.100E-08 |
| 27551 | 1378 | A1378L | E | 5.7e-09 | 1.8e-08 | 9.1e-10 | 8.203E-09 | 8.816E-09 | 9.100E-10 | 1.800E-08 |
| 27552 | 1378 | A1378K | E | 1.2e-08 | 8.6e-09 | 8e-09 | 9.533E-09 | 2.157E-09 | 8.000E-09 | 1.200E-08 |
| 27553 | 1378 | A1378M | E | 1.7e-08 | 3.6e-09 | 3.6e-09 | 8.067E-09 | 7.736E-09 | 3.600E-09 | 1.700E-08 |
| 27554 | 1378 | A1378F | E | 6.2e-09 | 6.6e-09 | 1.3e-08 | 8.600E-09 | 3.816E-09 | 6.200E-09 | 1.300E-08 |
| 27555 | 1378 | A1378P | E | 3.6e-09 | 5e-09 | 4.8e-09 | 4.467E-09 | 7.572E-10 | 3.600E-09 | 5.000E-09 |
| 27556 | 1378 | A1378S | E | 8.5e-09 | 1.4e-08 | 5.1e-09 | 9.200E-09 | 4.491E-09 | 5.100E-09 | 1.400E-08 |
| 27557 | 1378 | A1378T | E | 1e-08 | 1e-08 | 6.3e-09 | 8.767E-09 | 2.136E-09 | 6.300E-09 | 1.000E-08 |
| 27558 | 1378 | A1378W | E | 7.7e-09 | 5.9e-09 | 1e-08 | 7.867E-09 | 2.055E-09 | 5.900E-09 | 1.000E-08 |
| 27559 | 1378 | A1378Y | E | 8.8e-09 | 1.9e-08 | 1.3e-08 | 1.360E-08 | 5.126E-09 | 8.800E-09 | 1.900E-08 |
| 27560 | 1378 | A1378V | E | 3.3e-09 | 5.9e-09 | 7.3e-09 | 5.500E-09 | 2.030E-09 | 3.300E-09 | 7.300E-09 |
| 27561 | 1379 | M1379A | E | 9e-09 | 5.8e-09 | 4.2e-09 | 6.333E-09 | 2.444E-09 | 4.200E-09 | 9.000E-09 |
| 27562 | 1379 | M1379R | E | 8.8e-09 | 8.8e-09 | 9.5e-09 | 9.033E-09 | 4.041E-10 | 8.800E-09 | 9.500E-09 |
| 27563 | 1379 | M1379N | E | 1.1e-08 | 7.9e-09 | 1.3e-08 | 1.063E-08 | 2.570E-09 | 7.900E-09 | 1.300E-08 |
| 27564 | 1379 | M1379D | E | 5.6e-09 | 2e-09 | 6.8e-09 | 4.800E-09 | 2.498E-09 | 2.000E-09 | 6.800E-09 |
| 27565 | 1379 | M1379C | E | 1e-08 | 1.6e-08 | 6.6e-09 | 1.087E-08 | 4.760E-09 | 6.600E-09 | 1.600E-08 |
| 27566 | 1379 | M1379E | E | 5.6e-09 | 1.3e-08 | 4.3e-09 | 7.633E-09 | 4.693E-09 | 4.300E-09 | 1.300E-08 |
| 27567 | 1379 | M1379Q | E | 4.9e-09 | 1e-08 | 4.6e-09 | 6.500E-09 | 3.035E-09 | 4.600E-09 | 1.000E-08 |
| 27568 | 1379 | M1379G | E | 4.8e-09 | 6.4e-09 | 6.1e-09 | 5.767E-09 | 8.505E-10 | 4.800E-09 | 6.400E-09 |
| 27569 | 1379 | M1379H | E | 6.3e-09 | 1.7e-09 | 4.8e-09 | 4.267E-09 | 2.346E-09 | 1.700E-09 | 6.300E-09 |
| 27570 | 1379 | M1379I | E | 5.1e-09 | 1.1e-08 | 4.1e-09 | 6.733E-09 | 3.729E-09 | 4.100E-09 | 1.100E-08 |
| 27571 | 1379 | M1379L | E | 7.3e-09 | 1e-08 | 5.1e-09 | 7.467E-09 | 2.454E-09 | 5.100E-09 | 1.000E-08 |
| 27572 | 1379 | M1379K | E | 8.3e-09 | 3.9e-09 | 6.7e-09 | 6.300E-09 | 2.227E-09 | 3.900E-09 | 8.300E-09 |
| 27573 | 1379 | M1379M | E | 1.7e-08 | 6e-09 | 5.3e-09 | 9.433E-09 | 6.562E-09 | 5.300E-09 | 1.700E-08 |
| 27574 | 1379 | M1379F | E | 8.7e-09 | 3.4e-09 | 4.1e-09 | 5.400E-09 | 2.879E-09 | 3.400E-09 | 8.700E-09 |
| 27575 | 1379 | M1379P | E | 7.4e-09 | 8.9e-09 | 3.2e-09 | 6.500E-09 | 2.955E-09 | 3.200E-09 | 8.900E-09 |
| 27576 | 1379 | M1379S | E | 1e-08 | 1.2e-08 | 1.1e-08 | 1.100E-08 | 1.000E-09 | 1.000E-08 | 1.200E-08 |
| 27577 | 1379 | M1379T | E | 3.4e-09 | 1.3e-08 | 3.8e-09 | 6.733E-09 | 5.431E-09 | 3.400E-09 | 1.300E-08 |
| 27578 | 1379 | M1379W | E | 9.1e-09 | 1.1e-08 | 1.8e-08 | 1.270E-08 | 4.687E-09 | 9.100E-09 | 1.800E-08 |
| 27579 | 1379 | M1379Y | E | 3.4e-09 | 9.4e-09 | 1.3e-08 | 8.600E-09 | 4.850E-09 | 3.400E-09 | 1.300E-08 |
| 27580 | 1379 | M1379V | E | 5.3e-09 | 8.5e-09 | 4.7e-09 | 6.167E-09 | 2.043E-09 | 4.700E-09 | 8.500E-09 |
| 27581 | 1380 | D1380A | E | 1.2e-08 | 8.6e-09 | 5e-09 | 8.533E-09 | 3.500E-09 | 5.000E-09 | 1.200E-08 |
| 27582 | 1380 | D1380R | E | 1.5e-08 | 7e-09 | 2.6e-09 | 8.200E-09 | 6.286E-09 | 2.600E-09 | 1.500E-08 |
| 27583 | 1380 | D1380N | E | 2e-08 | 1.6e-08 | 1.1e-08 | 1.567E-08 | 4.509E-09 | 1.100E-08 | 2.000E-08 |
| 27584 | 1380 | D1380D | E | 8.8e-09 | 8.7e-09 | 5.5e-09 | 7.667E-09 | 1.877E-09 | 5.500E-09 | 8.800E-09 |
| 27585 | 1380 | D1380C | E | 1.5e-08 | 1.5e-09 | 2.2e-09 | 6.233E-09 | 7.600E-09 | 1.500E-09 | 1.500E-08 |
| 27586 | 1380 | D1380E | E | 1.3e-08 | 2.5e-08 | 4.5e-09 | 1.417E-08 | 1.030E-08 | 4.500E-09 | 2.500E-08 |
| 27587 | 1380 | D1380Q | E | 5.1e-09 | 2.8e-09 | 1.3e-08 | 6.967E-09 | 5.350E-09 | 2.800E-09 | 1.300E-08 |
| 27588 | 1380 | D1380G | E | 1.5e-08 | 9.6e-09 | 7.4e-09 | 1.067E-08 | 3.911E-09 | 7.400E-09 | 1.500E-08 |
| 27589 | 1380 | D1380H | E | 4e-09 | 5.3e-09 | 4.2e-09 | 4.500E-09 | 7.000E-10 | 4.000E-09 | 5.300E-09 |
| 27590 | 1380 | D1380I | E | 1.5e-08 | 8.5e-09 | 3.2e-08 | 1.850E-08 | 1.213E-08 | 8.500E-09 | 3.200E-08 |
| 27591 | 1380 | D1380L | E | 1.9e-08 | 7.3e-09 | 7.2e-09 | 1.117E-08 | 6.784E-09 | 7.200E-09 | 1.900E-08 |
| 27592 | 1380 | D1380K | E | 4.4e-09 | 6.2e-09 | 3e-09 | 4.533E-09 | 1.604E-09 | 3.000E-09 | 6.200E-09 |
| 27593 | 1380 | D1380M | E | 8.6e-09 | 1.5e-08 | 8e-09 | 1.053E-08 | 3.880E-09 | 8.000E-09 | 1.500E-08 |
| 27594 | 1380 | D1380F | E | 5.5e-09 | 1.1e-08 | 3.3e-09 | 6.600E-09 | 3.966E-09 | 3.300E-09 | 1.100E-08 |
| 27595 | 1380 | D1380P | E | 7e-09 | 1.2e-08 | 1e-08 | 9.667E-09 | 2.517E-09 | 7.000E-09 | 1.200E-08 |
| 27596 | 1380 | D1380S | E | 5.8e-09 | 1.2e-08 | 5.9e-09 | 7.900E-09 | 3.551E-09 | 5.800E-09 | 1.200E-08 |
| 27597 | 1380 | D1380T | E | 6.2e-09 | 3.9e-09 | 3.5e-09 | 4.533E-09 | 1.457E-09 | 3.500E-09 | 6.200E-09 |
| 27598 | 1380 | D1380W | E | 1.5e-08 | 8.6e-09 | 2.4e-09 | 8.667E-09 | 6.300E-09 | 2.400E-09 | 1.500E-08 |
| 27599 | 1380 | D1380Y | E | 8e-10 | 8.2e-09 | 6.3e-09 | 5.100E-09 | 3.843E-09 | 8.000E-10 | 8.200E-09 |
| 27600 | 1380 | D1380V | E | 4.8e-09 | 1.4e-09 | 4.8e-09 | 3.667E-09 | 1.963E-09 | 1.400E-09 | 4.800E-09 |
| 27601 | 1381 | Y1381A | E | 4.9e-09 | 1e-08 | 6.3e-09 | 7.067E-09 | 2.635E-09 | 4.900E-09 | 1.000E-08 |
| 27602 | 1381 | Y1381R | E | 1e-08 | 9.1e-09 | 6.1e-09 | 8.400E-09 | 2.042E-09 | 6.100E-09 | 1.000E-08 |
| 27603 | 1381 | Y1381N | E | 2.3e-09 | 3.7e-09 | 4e-09 | 3.333E-09 | 9.074E-10 | 2.300E-09 | 4.000E-09 |
| 27604 | 1381 | Y1381D | E | 6.4e-09 | 1.2e-08 | 6.3e-09 | 8.233E-09 | 3.262E-09 | 6.300E-09 | 1.200E-08 |
| 27605 | 1381 | Y1381C | E | 6.4e-09 | 1.2e-08 | 4e-09 | 7.467E-09 | 4.105E-09 | 4.000E-09 | 1.200E-08 |
| 27606 | 1381 | Y1381E | E | 6.7e-09 | 6.8e-09 | 8.8e-09 | 7.433E-09 | 1.185E-09 | 6.700E-09 | 8.800E-09 |
| 27607 | 1381 | Y1381Q | E | 8.3e-09 | 1.2e-08 | 1e-08 | 1.010E-08 | 1.852E-09 | 8.300E-09 | 1.200E-08 |
| 27608 | 1381 | Y1381G | E | 1.3e-08 | 6.1e-09 | 3.2e-09 | 7.433E-09 | 5.034E-09 | 3.200E-09 | 1.300E-08 |
| 27609 | 1381 | Y1381H | E | 9.3e-09 | 1.1e-08 | 5.2e-09 | 8.500E-09 | 2.982E-09 | 5.200E-09 | 1.100E-08 |
| 27610 | 1381 | Y1381I | E | 8.7e-09 | 8.3e-09 | 6.4e-09 | 7.800E-09 | 1.229E-09 | 6.400E-09 | 8.700E-09 |
| 27611 | 1381 | Y1381L | E | 5.6e-09 | 3e-09 | 9e-09 | 5.867E-09 | 3.009E-09 | 3.000E-09 | 9.000E-09 |
| 27612 | 1381 | Y1381K | E | 2.2e-08 | 7.8e-09 | 5.3e-09 | 1.170E-08 | 9.007E-09 | 5.300E-09 | 2.200E-08 |
| 27613 | 1381 | Y1381M | E | 2.6e-09 | 6.8e-09 | 2e-09 | 3.800E-09 | 2.615E-09 | 2.000E-09 | 6.800E-09 |
| 27614 | 1381 | Y1381F | E | 1.6e-09 | 6.6e-09 | 8.2e-09 | 5.467E-09 | 3.443E-09 | 1.600E-09 | 8.200E-09 |
| 27615 | 1381 | Y1381P | E | 4.2e-09 | 4e-09 | 8.2e-09 | 5.467E-09 | 2.369E-09 | 4.000E-09 | 8.200E-09 |
| 27616 | 1381 | Y1381S | E | 2.6e-09 | 6.2e-09 | 5.5e-09 | 4.767E-09 | 1.909E-09 | 2.600E-09 | 6.200E-09 |
| 27617 | 1381 | Y1381T | E | 2.3e-08 | 5e-09 | 1.1e-08 | 1.300E-08 | 9.165E-09 | 5.000E-09 | 2.300E-08 |
| 27618 | 1381 | Y1381W | E | 2.3e-08 | 4.8e-09 | 8.1e-09 | 1.197E-08 | 9.697E-09 | 4.800E-09 | 2.300E-08 |
| 27619 | 1381 | Y1381Y | E | 3.3e-09 | 1.4e-08 | 1.6e-08 | 1.110E-08 | 6.829E-09 | 3.300E-09 | 1.600E-08 |
| 27620 | 1381 | Y1381V | E | 3.4e-09 | 3.3e-09 | 7.5e-09 | 4.733E-09 | 2.397E-09 | 3.300E-09 | 7.500E-09 |
| 27621 | 1382 | W1382A | E | 6.7e-09 | 2.9e-09 | 3.3e-09 | 4.300E-09 | 2.088E-09 | 2.900E-09 | 6.700E-09 |
| 27622 | 1382 | W1382R | E | 2.6e-09 | 1.4e-08 | 1.1e-08 | 9.200E-09 | 5.909E-09 | 2.600E-09 | 1.400E-08 |
| 27623 | 1382 | W1382N | E | 8.3e-09 | 8.8e-09 | 2.8e-08 | 1.503E-08 | 1.123E-08 | 8.300E-09 | 2.800E-08 |
| 27624 | 1382 | W1382D | E | 9.1e-09 | 1.9e-08 | 8.6e-09 | 1.223E-08 | 5.865E-09 | 8.600E-09 | 1.900E-08 |

| | | | | | | | | | | |
|---|---|---|---|---|---|---|---|---|---|---|
| 27625 | 1382 | W1382C | E | 9.4e-09 | 1.1e-08 | 2.2e-08 | 1.413E-08 | 6.860E-09 | 9.400E-09 | 2.200E-08 |
| 27626 | 1382 | W1382E | E | 1.3e-08 | 1.7e-08 | 6.7e-09 | 1.223E-08 | 5.193E-09 | 6.700E-09 | 1.700E-08 |
| 27627 | 1382 | W1382Q | E | 1.3e-08 | 5.4e-09 | 3e-09 | 7.133E-09 | 5.220E-09 | 3.000E-09 | 1.300E-08 |
| 27628 | 1382 | W1382G | E | 1.3e-09 | 1.8e-08 | 1.2e-08 | 1.043E-08 | 8.460E-09 | 1.300E-09 | 1.800E-08 |
| 27629 | 1382 | W1382H | E | 3.4e-09 | 7.8e-09 | 4.9e-09 | 5.367E-09 | 2.237E-09 | 3.400E-09 | 7.800E-09 |
| 27630 | 1382 | W1382I | E | 6.3e-09 | 1.1e-08 | 6.3e-09 | 7.867E-09 | 2.714E-09 | 6.300E-09 | 1.100E-08 |
| 27631 | 1382 | W1382L | E | 6e-09 | 4.2e-09 | 6.3e-09 | 5.500E-09 | 1.136E-09 | 4.200E-09 | 6.300E-09 |
| 27632 | 1382 | W1382K | E | 6.3e-09 | 1.8e-08 | 7.8e-09 | 1.070E-08 | 6.366E-09 | 6.300E-09 | 1.800E-08 |
| 27633 | 1382 | W1382M | E | 5.6e-09 | 4.7e-09 | 2.5e-09 | 4.267E-09 | 1.595E-09 | 2.500E-09 | 5.600E-09 |
| 27634 | 1382 | W1382F | E | 1.2e-08 | 1.4e-08 | 5.3e-09 | 1.043E-08 | 4.557E-09 | 5.300E-09 | 1.400E-08 |
| 27635 | 1382 | W1382P | E | 4.9e-09 | 9.3e-09 | 5.5e-09 | 6.567E-09 | 2.386E-09 | 4.900E-09 | 9.300E-09 |
| 27636 | 1382 | W1382S | E | 4.2e-09 | 6.4e-09 | 7e-09 | 5.867E-09 | 1.474E-09 | 4.200E-09 | 7.000E-09 |
| 27637 | 1382 | W1382T | E | 5.1e-09 | 1.4e-08 | 9.5e-09 | 9.533E-09 | 4.450E-09 | 5.100E-09 | 1.400E-08 |
| 27638 | 1382 | W1382W | E | 5e-09 | 3e-09 | 3.9e-09 | 3.967E-09 | 1.002E-09 | 3.000E-09 | 5.000E-09 |
| 27639 | 1382 | W1382Y | E | 2.3e-09 | 7.5e-09 | 8.6e-09 | 6.133E-09 | 3.365E-09 | 2.300E-09 | 8.600E-09 |
| 27640 | 1382 | W1382V | E | 5.9e-09 | 5.1e-09 | 1.3e-08 | 8.000E-09 | 4.349E-09 | 5.100E-09 | 1.300E-08 |
| 27641 | 1383 | G1383A | E | 3.7e-09 | 7.5e-09 | 3.6e-09 | 4.933E-09 | 2.223E-09 | 3.600E-09 | 7.500E-09 |
| 27642 | 1383 | G1383R | E | 4.5e-09 | 1.2e-08 | 9.5e-09 | 8.667E-09 | 3.819E-09 | 4.500E-09 | 1.200E-08 |
| 27643 | 1383 | G1383N | E | 4.6e-09 | 8.6e-09 | 1.5e-08 | 9.400E-09 | 5.246E-09 | 4.600E-09 | 1.500E-08 |
| 27644 | 1383 | G1383D | E | 4.3e-09 | 7.5e-09 | 4.2e-09 | 5.333E-09 | 1.877E-09 | 4.200E-09 | 7.500E-09 |
| 27645 | 1383 | G1383C | E | 6.9e-09 | 1.8e-08 | 3.2e-09 | 9.367E-09 | 7.702E-09 | 3.200E-09 | 1.800E-08 |
| 27646 | 1383 | G1383E | E | 5.4e-09 | 6.6e-09 | 9.5e-09 | 7.167E-09 | 2.108E-09 | 5.400E-09 | 9.500E-09 |
| 27647 | 1383 | G1383Q | E | 6.3e-09 | 4.8e-09 | 1.4e-08 | 8.367E-09 | 4.936E-09 | 4.800E-09 | 1.400E-08 |
| 27648 | 1383 | G1383G | E | 8.7e-09 | 1.3e-08 | 5.4e-09 | 9.033E-09 | 3.811E-09 | 5.400E-09 | 1.300E-08 |
| 27649 | 1383 | G1383H | E | 9.1e-09 | 4e-09 | 2.4e-09 | 5.167E-09 | 3.499E-09 | 2.400E-09 | 9.100E-09 |
| 27650 | 1383 | G1383I | E | 4.6e-09 | 9.9e-09 | 2.9e-09 | 5.800E-09 | 3.651E-09 | 2.900E-09 | 9.900E-09 |
| 27651 | 1383 | G1383L | E | 1.9e-08 | 7.5e-09 | 1.4e-08 | 1.350E-08 | 5.766E-09 | 7.500E-09 | 1.900E-08 |
| 27652 | 1383 | G1383K | E | 1.7e-08 | 3.2e-09 | 6e-09 | 8.733E-09 | 7.295E-09 | 3.200E-09 | 1.700E-08 |
| 27653 | 1383 | G1383M | E | 8.1e-09 | 2.2e-09 | 1e-08 | 6.767E-09 | 4.067E-09 | 2.200E-09 | 1.000E-08 |
| 27654 | 1383 | G1383F | E | 1.2e-08 | 6e-09 | 5.9e-09 | 7.967E-09 | 3.493E-09 | 5.900E-09 | 1.200E-08 |
| 27655 | 1383 | G1383P | E | 2.1e-09 | 1e-08 | 1.4e-08 | 8.700E-09 | 6.056E-09 | 2.100E-09 | 1.400E-08 |
| 27656 | 1383 | G1383S | E | 1.1e-08 | 5.6e-09 | 9.4e-09 | 8.667E-09 | 2.774E-09 | 5.600E-09 | 1.100E-08 |
| 27657 | 1383 | G1383T | E | 1.5e-08 | 1.1e-08 | 5e-09 | 1.033E-08 | 5.033E-09 | 5.000E-09 | 1.500E-08 |
| 27658 | 1383 | G1383W | E | 1.5e-08 | 5.4e-09 | 3e-09 | 7.800E-09 | 6.350E-09 | 3.000E-09 | 1.500E-08 |
| 27659 | 1383 | G1383Y | E | 8.1e-09 | 5.7e-09 | 7.7e-09 | 7.167E-09 | 1.286E-09 | 5.700E-09 | 8.100E-09 |
| 27660 | 1383 | G1383V | E | 1.3e-08 | 1.2e-08 | 3.8e-08 | 2.100E-08 | 1.473E-08 | 1.200E-08 | 3.800E-08 |
| 27661 | 1384 | Q1384A | E | 4.9e-09 | 5.7e-09 | 4e-09 | 4.867E-09 | 8.505E-10 | 4.000E-09 | 5.700E-09 |
| 27662 | 1384 | Q1384R | E | 7.1e-09 | 1.4e-08 | 7.3e-09 | 9.467E-09 | 3.927E-09 | 7.100E-09 | 1.400E-08 |
| 27663 | 1384 | Q1384N | E | 1.2e-08 | 1.1e-08 | 1.6e-08 | 1.300E-08 | 2.646E-09 | 1.100E-08 | 1.600E-08 |
| 27664 | 1384 | Q1384D | E | 2.9e-09 | 7.2e-09 | 7.3e-09 | 5.800E-09 | 2.512E-09 | 2.900E-09 | 7.300E-09 |
| 27665 | 1384 | Q1384C | E | 8.9e-09 | 4e-09 | 1e-08 | 7.633E-09 | 3.194E-09 | 4.000E-09 | 1.000E-08 |
| 27666 | 1384 | Q1384E | E | 3.2e-09 | 2.9e-09 | 1.3e-08 | 6.367E-09 | 5.747E-09 | 2.900E-09 | 1.300E-08 |
| 27667 | 1384 | Q1384Q | E | 3.1e-09 | 4.2e-09 | 1.2e-08 | 6.433E-09 | 4.852E-09 | 3.100E-09 | 1.200E-08 |
| 27668 | 1384 | Q1384G | E | 2e-09 | 7.1e-09 | 7.7e-09 | 5.600E-09 | 3.132E-09 | 2.000E-09 | 7.700E-09 |
| 27669 | 1384 | Q1384H | E | 5.9e-09 | 1.1e-08 | 3.5e-09 | 6.800E-09 | 3.830E-09 | 3.500E-09 | 1.100E-08 |
| 27670 | 1384 | Q1384I | E | 5.7e-09 | 1.4e-08 | 5.7e-09 | 8.467E-09 | 4.792E-09 | 5.700E-09 | 1.400E-08 |
| 27671 | 1384 | Q1384L | E | 5.8e-09 | 8e-09 | 1.1e-08 | 8.267E-09 | 2.610E-09 | 5.800E-09 | 1.100E-08 |
| 27672 | 1384 | Q1384K | E | 6e-09 | 7.1e-09 | 1.1e-08 | 8.033E-09 | 2.627E-09 | 6.000E-09 | 1.100E-08 |
| 27673 | 1384 | Q1384M | E | 8.9e-09 | 9.9e-09 | 8.3e-09 | 9.033E-09 | 8.083E-10 | 8.300E-09 | 9.900E-09 |
| 27674 | 1384 | Q1384F | E | 6.9e-09 | 1.1e-08 | 5.4e-09 | 7.767E-09 | 2.899E-09 | 5.400E-09 | 1.100E-08 |
| 27675 | 1384 | Q1384P | E | 8.2e-09 | 2e-09 | 9.2e-09 | 6.467E-09 | 3.900E-09 | 2.000E-09 | 9.200E-09 |
| 27676 | 1384 | Q1384S | E | 1.1e-08 | 5.8e-09 | 3.8e-09 | 6.867E-09 | 3.717E-09 | 3.800E-09 | 1.100E-08 |
| 27677 | 1384 | Q1384T | E | 8.9e-09 | 4e-09 | 9.2e-09 | 7.367E-09 | 2.919E-09 | 4.000E-09 | 9.200E-09 |
| 27678 | 1384 | Q1384W | E | 7.7e-09 | 5.9e-09 | 3.7e-09 | 5.767E-09 | 2.003E-09 | 3.700E-09 | 7.700E-09 |
| 27679 | 1384 | Q1384Y | E | 1.4e-08 | 1.2e-08 | 1.2e-08 | 1.267E-08 | 1.155E-09 | 1.200E-08 | 1.400E-08 |
| 27680 | 1384 | Q1384V | E | 1.2e-08 | 5.5e-09 | 6.1e-09 | 7.867E-09 | 3.592E-09 | 5.500E-09 | 1.200E-08 |
| 27681 | 1385 | G1385A | E | 1.2e-08 | 9.7e-09 | 1.3e-08 | 1.157E-08 | 1.692E-09 | 9.700E-09 | 1.300E-08 |
| 27682 | 1385 | G1385R | E | 6.3e-09 | 3.6e-09 | 1.5e-08 | 8.300E-09 | 5.957E-09 | 3.600E-09 | 1.500E-08 |
| 27683 | 1385 | G1385N | E | 8.8e-09 | 9.8e-09 | 1.1e-08 | 9.867E-09 | 1.102E-09 | 8.800E-09 | 1.100E-08 |
| 27684 | 1385 | G1385D | E | 5.9e-09 | 9.4e-09 | 4.5e-09 | 6.600E-09 | 2.524E-09 | 4.500E-09 | 9.400E-09 |
| 27685 | 1385 | G1385C | E | 5.9e-09 | 6e-09 | 4.4e-09 | 5.433E-09 | 8.963E-10 | 4.400E-09 | 6.000E-09 |
| 27686 | 1385 | G1385E | E | 7.4e-09 | 9.2e-09 | 6e-09 | 7.533E-09 | 1.604E-09 | 6.000E-09 | 9.200E-09 |
| 27687 | 1385 | G1385Q | E | 6e-09 | 9.4e-09 | 7.5e-09 | 7.633E-09 | 1.704E-09 | 6.000E-09 | 9.400E-09 |
| 27688 | 1385 | G1385G | E | 5.9e-09 | 4.5e-09 | 1.2e-08 | 7.467E-09 | 3.988E-09 | 4.500E-09 | 1.200E-08 |
| 27689 | 1385 | G1385H | E | 8.1e-09 | 1.4e-08 | 1.1e-08 | 1.103E-08 | 2.950E-09 | 8.100E-09 | 1.400E-08 |
| 27690 | 1385 | G1385I | E | 1.4e-08 | 3.5e-09 | 1.2e-08 | 9.833E-09 | 5.575E-09 | 3.500E-09 | 1.400E-08 |
| 27691 | 1385 | G1385L | E | 7e-09 | 9e-09 | 1e-08 | 8.667E-09 | 1.528E-09 | 7.000E-09 | 1.000E-08 |
| 27692 | 1385 | G1385K | E | 1.5e-08 | 9e-09 | 5.1e-09 | 9.700E-09 | 4.987E-09 | 5.100E-09 | 1.500E-08 |
| 27693 | 1385 | G1385M | E | 5e-09 | 6.7e-09 | 1.2e-08 | 7.900E-09 | 3.651E-09 | 5.000E-09 | 1.200E-08 |
| 27694 | 1385 | G1385F | E | 8.9e-09 | 5.7e-09 | 1.3e-08 | 9.200E-09 | 3.659E-09 | 5.700E-09 | 1.300E-08 |
| 27695 | 1385 | G1385P | E | 5.6e-09 | 8.6e-09 | 1.5e-08 | 9.733E-09 | 4.801E-09 | 5.600E-09 | 1.500E-08 |
| 27696 | 1385 | G1385S | E | 9.4e-09 | 1.9e-08 | 1.8e-08 | 1.547E-08 | 5.278E-09 | 9.400E-09 | 1.900E-08 |
| 27697 | 1385 | G1385T | E | 4.4e-09 | 2.8e-09 | 8.4e-09 | 5.200E-09 | 2.884E-09 | 2.800E-09 | 8.400E-09 |
| 27698 | 1385 | G1385W | E | 8.8e-09 | 2.5e-08 | 2.6e-09 | 1.213E-08 | 1.157E-08 | 2.600E-09 | 2.500E-08 |
| 27699 | 1385 | G1385Y | E | 1e-08 | 2.3e-09 | 5.3e-09 | 5.867E-09 | 3.881E-09 | 2.300E-09 | 1.000E-08 |
| 27700 | 1385 | G1385V | E | 1e-08 | 2.1e-09 | 3.6e-09 | 5.233E-09 | 4.196E-09 | 2.100E-09 | 1.000E-08 |
| 27701 | 1386 | T1386A | E | 2.1e-08 | 1.1e-08 | 1e-08 | 1.400E-08 | 6.083E-09 | 1.000E-08 | 2.100E-08 |
| 27702 | 1386 | T1386R | E | 9.1e-09 | 6.9e-09 | 1.2e-08 | 9.333E-09 | 2.558E-09 | 6.900E-09 | 1.200E-08 |
| 27703 | 1386 | T1386N | E | 4.9e-09 | 3.7e-08 | 7.6e-09 | 1.650E-08 | 1.780E-08 | 4.900E-09 | 3.700E-08 |
| 27704 | 1386 | T1386D | E | 7.6e-09 | 3.1e-09 | 1.7e-08 | 9.233E-09 | 7.092E-09 | 3.100E-09 | 1.700E-08 |
| 27705 | 1386 | T1386C | E | 9.1e-09 | 9.5e-09 | 5.7e-09 | 8.100E-09 | 2.088E-09 | 5.700E-09 | 9.500E-09 |
| 27706 | 1386 | T1386E | E | 7.7e-09 | 6.5e-09 | 1.5e-08 | 9.733E-09 | 4.600E-09 | 6.500E-09 | 1.500E-08 |
| 27707 | 1386 | T1386Q | E | 6.8e-09 | 3.1e-08 | 6.9e-09 | 1.490E-08 | 1.394E-08 | 6.800E-09 | 3.100E-08 |
| 27708 | 1386 | T1386G | E | 7.6e-09 | 3e-09 | 1.4e-08 | 8.200E-09 | 5.524E-09 | 3.000E-09 | 1.400E-08 |
| 27709 | 1386 | T1386H | E | 1.5e-08 | 1.5e-09 | 4.7e-09 | 7.067E-09 | 7.054E-09 | 1.500E-09 | 1.500E-08 |

| | | | | | | | | | | |
|---|---|---|---|---|---|---|---|---|---|---|
| 27710 | 1386 | T1386I | E | 1.4e-08 | 1.1e-08 | 6.6e-09 | 1.053E-08 | 3.722E-09 | 6.600E-09 | 1.400E-08 |
| 27711 | 1386 | T1386L | E | 5.9e-09 | 2e-09 | 4.4e-09 | 4.100E-09 | 1.967E-09 | 2.000E-09 | 5.900E-09 |
| 27712 | 1386 | T1386K | E | 5.3e-09 | 5.7e-09 | 3.8e-09 | 4.933E-09 | 1.002E-09 | 3.800E-09 | 5.700E-09 |
| 27713 | 1386 | T1386M | E | 6.1e-09 | 3.4e-09 | 1.1e-08 | 6.833E-09 | 3.853E-09 | 3.400E-09 | 1.100E-08 |
| 27714 | 1386 | T1386F | E | 7.9e-09 | 7.5e-09 | 5.3e-09 | 6.900E-09 | 1.400E-09 | 5.300E-09 | 7.900E-09 |
| 27715 | 1386 | T1386P | E | 1.5e-08 | 7.3e-09 | 1.5e-08 | 1.243E-08 | 4.446E-09 | 7.300E-09 | 1.500E-08 |
| 27716 | 1386 | T1386S | E | 9.6e-09 | 1.1e-08 | 9.9e-09 | 1.017E-08 | 7.371E-10 | 9.600E-09 | 1.100E-08 |
| 27717 | 1386 | T1386T | E | 4.3e-09 | 8.8e-09 | 7.9e-09 | 7.000E-09 | 2.381E-09 | 4.300E-09 | 8.800E-09 |
| 27718 | 1386 | T1386W | E | 2.7e-09 | 2.2e-09 | 4.5e-09 | 3.133E-09 | 1.210E-09 | 2.200E-09 | 4.500E-09 |
| 27719 | 1386 | T1386Y | E | 1.5e-08 | 6.2e-09 | 5.3e-09 | 8.833E-09 | 5.359E-09 | 5.300E-09 | 1.500E-08 |
| 27720 | 1386 | T1386V | E | 9.1e-09 | 7.8e-09 | 4.5e-09 | 7.133E-09 | 2.371E-09 | 4.500E-09 | 9.100E-09 |
| 27721 | 1387 | L1387A | E | 2.8e-09 | 5.9e-09 | 4e-09 | 4.233E-09 | 1.563E-09 | 2.800E-09 | 5.900E-09 |
| 27722 | 1387 | L1387R | E | 5.5e-09 | 1e-08 | 4.6e-09 | 6.700E-09 | 2.893E-09 | 4.600E-09 | 1.000E-08 |
| 27723 | 1387 | L1387N | E | 7.7e-09 | 3.5e-09 | 3.5e-08 | 1.540E-08 | 1.710E-08 | 3.500E-09 | 3.500E-08 |
| 27724 | 1387 | L1387D | E | 4.8e-09 | 9.4e-09 | 3.5e-09 | 5.900E-09 | 3.100E-09 | 3.500E-09 | 9.400E-09 |
| 27725 | 1387 | L1387C | E | 2.6e-09 | 9.4e-09 | 5.8e-09 | 5.933E-09 | 3.402E-09 | 2.600E-09 | 9.400E-09 |
| 27726 | 1387 | L1387E | E | 5.1e-09 | 9.8e-09 | 1.2e-08 | 8.967E-09 | 3.525E-09 | 5.100E-09 | 1.200E-08 |
| 27727 | 1387 | L1387Q | E | 3.7e-09 | 4.1e-09 | 4.9e-09 | 4.233E-09 | 6.110E-10 | 3.700E-09 | 4.900E-09 |
| 27728 | 1387 | L1387G | E | 1.6e-08 | 7e-09 | 5.8e-09 | 9.600E-09 | 5.575E-09 | 5.800E-09 | 1.600E-08 |
| 27729 | 1387 | L1387H | E | 2.3e-09 | 5.4e-09 | 4.3e-09 | 4.000E-09 | 1.572E-09 | 2.300E-09 | 5.400E-09 |
| 27730 | 1387 | L1387I | E | 2.3e-08 | 8.9e-09 | 2.4e-09 | 1.143E-08 | 1.053E-08 | 2.400E-09 | 2.300E-08 |
| 27731 | 1387 | L1387L | E | 1.2e-08 | 4.5e-09 | 4.5e-09 | 7.000E-09 | 4.330E-09 | 4.500E-09 | 1.200E-08 |
| 27732 | 1387 | L1387K | E | 7.1e-09 | 6e-09 | 6.1e-09 | 6.400E-09 | 6.083E-10 | 6.000E-09 | 7.100E-09 |
| 27733 | 1387 | L1387M | E | 6.3e-09 | 1e-08 | 4.4e-09 | 6.900E-09 | 2.848E-09 | 4.400E-09 | 1.000E-08 |
| 27734 | 1387 | L1387F | E | 3.4e-09 | 1.4e-08 | 9.4e-09 | 8.933E-09 | 5.315E-09 | 3.400E-09 | 1.400E-08 |
| 27735 | 1387 | L1387P | E | 8.3e-09 | 1.9e-09 | 1.4e-08 | 8.067E-09 | 6.053E-09 | 1.900E-09 | 1.400E-08 |
| 27736 | 1387 | L1387S | E | 5.2e-09 | 1.3e-08 | 1.9e-08 | 1.240E-08 | 6.920E-09 | 5.200E-09 | 1.900E-08 |
| 27737 | 1387 | L1387T | E | 1.7e-08 | 1.6e-08 | 1.1e-08 | 1.467E-08 | 3.215E-09 | 1.100E-08 | 1.700E-08 |
| 27738 | 1387 | L1387W | E | 7.6e-09 | 5.9e-09 | 2e-09 | 5.167E-09 | 2.871E-09 | 2.000E-09 | 7.600E-09 |
| 27739 | 1387 | L1387Y | E | 8.4e-09 | 6.3e-09 | 1.4e-09 | 5.367E-09 | 3.592E-09 | 1.400E-09 | 8.400E-09 |
| 27740 | 1387 | L1387V | E | 1.9e-09 | 5.5e-09 | 6.5e-09 | 4.633E-09 | 2.419E-09 | 1.900E-09 | 6.500E-09 |
| 27741 | 1388 | V1388A | E | 3e-08 | 5e-09 | 5e-09 | 1.333E-08 | 1.443E-08 | 5.000E-09 | 3.000E-08 |
| 27742 | 1388 | V1388R | E | 1.1e-08 | 1.5e-09 | 6.4e-09 | 6.300E-09 | 4.751E-09 | 1.500E-09 | 1.100E-08 |
| 27743 | 1388 | V1388N | E | 1.2e-08 | 4.9e-09 | 5.7e-09 | 7.533E-09 | 3.889E-09 | 4.900E-09 | 1.200E-08 |
| 27744 | 1388 | V1388D | E | 1.7e-08 | 1.1e-08 | 9.1e-09 | 1.237E-08 | 4.124E-09 | 9.100E-09 | 1.700E-08 |
| 27745 | 1388 | V1388C | E | 9.5e-09 | 1e-08 | 1e-08 | 9.833E-09 | 2.887E-10 | 9.500E-09 | 1.000E-08 |
| 27746 | 1388 | V1388E | E | 1e-08 | 2.1e-08 | 4.4e-09 | 1.180E-08 | 8.445E-09 | 4.400E-09 | 2.100E-08 |
| 27747 | 1388 | V1388Q | E | 9e-09 | 1.9e-09 | 1.7e-08 | 9.300E-09 | 7.554E-09 | 1.900E-09 | 1.700E-08 |
| 27748 | 1388 | V1388G | E | 1.8e-08 | 8.4e-09 | 3.1e-09 | 9.833E-09 | 7.553E-09 | 3.100E-09 | 1.800E-08 |
| 27749 | 1388 | V1388H | E | 1.2e-08 | 7.5e-09 | 4.5e-09 | 8.000E-09 | 3.775E-09 | 4.500E-09 | 1.200E-08 |
| 27750 | 1388 | V1388I | E | 7.7e-09 | 9.4e-09 | 5.2e-09 | 7.433E-09 | 2.113E-09 | 5.200E-09 | 9.400E-09 |
| 27751 | 1388 | V1388L | E | 2.9e-09 | 6.5e-09 | 1.2e-08 | 7.133E-09 | 4.583E-09 | 2.900E-09 | 1.200E-08 |
| 27752 | 1388 | V1388K | E | 1.4e-08 | 1.1e-08 | 8.3e-09 | 1.110E-08 | 2.851E-09 | 8.300E-09 | 1.400E-08 |
| 27753 | 1388 | V1388M | E | 7.5e-09 | 2.6e-09 | 5.4e-09 | 5.167E-09 | 2.458E-09 | 2.600E-09 | 7.500E-09 |
| 27754 | 1388 | V1388F | E | 7.8e-09 | 7.3e-09 | 9.3e-09 | 8.133E-09 | 1.041E-09 | 7.300E-09 | 9.300E-09 |
| 27755 | 1388 | V1388P | E | 8.8e-09 | 9.4e-09 | 6.3e-09 | 8.167E-09 | 1.644E-09 | 6.300E-09 | 9.400E-09 |
| 27756 | 1388 | V1388S | E | 9.2e-09 | 8.8e-09 | 1.7e-08 | 1.167E-08 | 4.623E-09 | 8.800E-09 | 1.700E-08 |
| 27757 | 1388 | V1388T | E | 3e-09 | 2.2e-08 | 6.6e-09 | 1.053E-08 | 1.009E-08 | 3.000E-09 | 2.200E-08 |
| 27758 | 1388 | V1388W | E | 5.7e-09 | 3.5e-09 | 2.7e-09 | 3.967E-09 | 1.553E-09 | 2.700E-09 | 5.700E-09 |
| 27759 | 1388 | V1388Y | E | 3.3e-09 | 1.3e-08 | 7.9e-09 | 8.067E-09 | 4.852E-09 | 3.300E-09 | 1.300E-08 |
| 27760 | 1388 | V1388V | E | 5.1e-09 | 5.7e-09 | 1.6e-09 | 4.133E-09 | 2.214E-09 | 1.600E-09 | 5.700E-09 |
| 27761 | 1389 | T1389A | E | 1.3e-08 | 1.4e-08 | 9.9e-09 | 1.230E-08 | 2.138E-09 | 9.900E-09 | 1.400E-08 |
| 27762 | 1389 | T1389R | E | 3.1e-09 | 5.4e-09 | 7.9e-09 | 5.467E-09 | 2.401E-09 | 3.100E-09 | 7.900E-09 |
| 27763 | 1389 | T1389N | E | 2.9e-09 | 3.1e-09 | 5e-09 | 3.667E-09 | 1.159E-09 | 2.900E-09 | 5.000E-09 |
| 27764 | 1389 | T1389D | E | 9.3e-09 | 8.7e-09 | 7.2e-09 | 8.400E-09 | 1.082E-09 | 7.200E-09 | 9.300E-09 |
| 27765 | 1389 | T1389C | E | 9e-09 | 3.3e-09 | 3.9e-09 | 5.400E-09 | 3.132E-09 | 3.300E-09 | 9.000E-09 |
| 27766 | 1389 | T1389E | E | 3.9e-09 | 7e-09 | 6.3e-09 | 5.733E-09 | 1.626E-09 | 3.900E-09 | 7.000E-09 |
| 27767 | 1389 | T1389Q | E | 1.8e-08 | 1.2e-08 | 4.7e-09 | 1.157E-08 | 6.661E-09 | 4.700E-09 | 1.800E-08 |
| 27768 | 1389 | T1389G | E | 6.6e-09 | 8.1e-09 | 6.4e-09 | 7.033E-09 | 9.292E-10 | 6.400E-09 | 8.100E-09 |
| 27769 | 1389 | T1389H | E | 5.6e-09 | 7e-09 | 5.7e-09 | 6.100E-09 | 7.810E-10 | 5.600E-09 | 7.000E-09 |
| 27770 | 1389 | T1389I | E | 3.4e-08 | 6.8e-09 | 5.3e-09 | 1.537E-08 | 1.615E-08 | 5.300E-09 | 3.400E-08 |
| 27771 | 1389 | T1389L | E | 3.9e-09 | 3.3e-09 | 1.2e-08 | 6.400E-09 | 4.859E-09 | 3.300E-09 | 1.200E-08 |
| 27772 | 1389 | T1389K | E | 8.5e-09 | 3.2e-09 | 8.7e-09 | 6.800E-09 | 3.119E-09 | 3.200E-09 | 8.700E-09 |
| 27773 | 1389 | T1389M | E | 1.5e-08 | 1.4e-08 | 9.7e-09 | 1.290E-08 | 2.816E-09 | 9.700E-09 | 1.500E-08 |
| 27774 | 1389 | T1389F | E | 1.3e-08 | 2.2e-09 | 4.3e-09 | 6.500E-09 | 5.726E-09 | 2.200E-09 | 1.300E-08 |
| 27775 | 1389 | T1389P | E | 7.5e-09 | 2.7e-09 | 1.1e-08 | 7.067E-09 | 4.167E-09 | 2.700E-09 | 1.100E-08 |
| 27776 | 1389 | T1389S | E | 2.9e-09 | 8.9e-09 | 2.6e-09 | 4.800E-09 | 3.554E-09 | 2.600E-09 | 8.900E-09 |
| 27777 | 1389 | T1389T | E | 6.2e-09 | 1.1e-08 | 7.6e-09 | 8.267E-09 | 2.468E-09 | 6.200E-09 | 1.100E-08 |
| 27778 | 1389 | T1389W | E | 5.6e-09 | 1.3e-08 | 1.2e-08 | 1.020E-08 | 4.015E-09 | 5.600E-09 | 1.300E-08 |
| 27779 | 1389 | T1389Y | E | 6.5e-09 | 8.2e-09 | 5.8e-09 | 6.833E-09 | 1.234E-09 | 5.800E-09 | 8.200E-09 |
| 27780 | 1389 | T1389V | E | 1.3e-08 | 1e-08 | 9.2e-09 | 1.073E-08 | 2.003E-09 | 9.200E-09 | 1.300E-08 |
| 27781 | 1390 | V1390A | E | 5.4e-09 | 4.1e-09 | 1.4e-08 | 7.833E-09 | 5.380E-09 | 4.100E-09 | 1.400E-08 |
| 27782 | 1390 | V1390R | E | 1.3e-08 | 1.3e-08 | 7.1e-09 | 1.103E-08 | 3.406E-09 | 7.100E-09 | 1.300E-08 |
| 27783 | 1390 | V1390N | E | 1.3e-08 | 5.8e-09 | 4.6e-09 | 7.800E-09 | 4.543E-09 | 4.600E-09 | 1.300E-08 |
| 27784 | 1390 | V1390D | E | 9.6e-09 | 2.9e-09 | 5e-09 | 5.833E-09 | 3.427E-09 | 2.900E-09 | 9.600E-09 |
| 27785 | 1390 | V1390C | E | 4.4e-09 | 5e-09 | 3.6e-09 | 4.333E-09 | 7.024E-10 | 3.600E-09 | 5.000E-09 |
| 27786 | 1390 | V1390E | E | 9.6e-09 | 1.2e-08 | 3e-09 | 8.200E-09 | 4.660E-09 | 3.000E-09 | 1.200E-08 |
| 27787 | 1390 | V1390Q | E | 3.9e-09 | 7.5e-09 | 8.9e-09 | 6.767E-09 | 2.579E-09 | 3.900E-09 | 8.900E-09 |
| 27788 | 1390 | V1390G | E | 1.2e-09 | 6.6e-09 | 8.1e-09 | 5.300E-09 | 3.629E-09 | 1.200E-09 | 8.100E-09 |
| 27789 | 1390 | V1390H | E | 9.8e-09 | 4.5e-09 | 5.4e-09 | 6.567E-09 | 2.836E-09 | 4.500E-09 | 9.800E-09 |
| 27790 | 1390 | V1390I | E | 1.3e-08 | 1.6e-09 | 7.7e-09 | 7.433E-09 | 5.705E-09 | 1.600E-09 | 1.300E-08 |
| 27791 | 1390 | V1390L | E | 2.3e-09 | 8.1e-09 | 3.3e-08 | 1.447E-08 | 1.631E-08 | 2.300E-09 | 3.300E-08 |
| 27792 | 1390 | V1390K | E | 1.4e-08 | 3.9e-09 | 6.5e-09 | 8.133E-09 | 5.244E-09 | 3.900E-09 | 1.400E-08 |
| 27793 | 1390 | V1390M | E | 4.9e-09 | 1.1e-08 | 8.1e-09 | 8.000E-09 | 3.051E-09 | 4.900E-09 | 1.100E-08 |
| 27794 | 1390 | V1390F | E | 4.8e-09 | 1.5e-08 | 4e-09 | 7.933E-09 | 6.133E-09 | 4.000E-09 | 1.500E-08 |

| | | | | | | | | | | |
|---|---|---|---|---|---|---|---|---|---|---|
| 27795 | 1390 | V1390P | E | 3.9e-09 | 8.1e-09 | 4.7e-09 | 5.567E-09 | 2.230E-09 | 3.900E-09 | 8.100E-09 |
| 27796 | 1390 | V1390S | E | 8.1e-09 | 1.3e-08 | 8.4e-09 | 9.833E-09 | 2.747E-09 | 8.100E-09 | 1.300E-08 |
| 27797 | 1390 | V1390T | E | 2.6e-09 | 1.3e-08 | 1.6e-08 | 1.053E-08 | 7.032E-09 | 2.600E-09 | 1.600E-08 |
| 27798 | 1390 | V1390W | E | 2.9e-09 | 1.1e-08 | 7.3e-09 | 7.067E-09 | 4.055E-09 | 2.900E-09 | 1.100E-08 |
| 27799 | 1390 | V1390Y | E | 1.8e-08 | 1.5e-08 | 5.3e-09 | 1.277E-08 | 6.638E-09 | 5.300E-09 | 1.800E-08 |
| 27800 | 1390 | V1390V | E | 1.8e-08 | 2.2e-08 | 7.4e-09 | 1.580E-08 | 7.545E-09 | 7.400E-09 | 2.200E-08 |
| 27801 | 1391 | S1391A | E | 6.9e-09 | 3e-09 | 1.7e-09 | 3.867E-09 | 2.706E-09 | 1.700E-09 | 6.900E-09 |
| 27802 | 1391 | S1391R | E | 4.2e-09 | 3.9e-09 | 2.2e-08 | 1.003E-08 | 1.036E-08 | 3.900E-09 | 2.200E-08 |
| 27803 | 1391 | S1391N | E | 8.8e-09 | 1.7e-08 | 9.2e-09 | 1.167E-08 | 4.623E-09 | 8.800E-09 | 1.700E-08 |
| 27804 | 1391 | S1391D | E | 8.8e-09 | 8.4e-09 | 8.8e-09 | 8.667E-09 | 2.309E-10 | 8.400E-09 | 8.800E-09 |
| 27805 | 1391 | S1391C | E | 1.2e-08 | 6.4e-09 | 8.2e-09 | 8.867E-09 | 2.859E-09 | 6.400E-09 | 1.200E-08 |
| 27806 | 1391 | S1391E | E | 3e-09 | 9.3e-09 | 1.4e-08 | 8.767E-09 | 5.519E-09 | 3.000E-09 | 1.400E-08 |
| 27807 | 1391 | S1391Q | E | 4.1e-09 | 7.1e-09 | 8.1e-09 | 6.433E-09 | 2.082E-09 | 4.100E-09 | 8.100E-09 |
| 27808 | 1391 | S1391G | E | 1.2e-08 | 1.8e-08 | 2.5e-08 | 1.833E-08 | 6.506E-09 | 1.200E-08 | 2.500E-08 |
| 27809 | 1391 | S1391H | E | 1e-08 | 7.8e-09 | 3.4e-09 | 7.067E-09 | 3.361E-09 | 3.400E-09 | 1.000E-08 |
| 27810 | 1391 | S1391I | E | 5.7e-09 | 7.4e-09 | 2e-09 | 5.033E-09 | 2.761E-09 | 2.000E-09 | 7.400E-09 |
| 27811 | 1391 | S1391L | E | 4.6e-09 | 1e-08 | 3.9e-09 | 6.167E-09 | 3.338E-09 | 3.900E-09 | 1.000E-08 |
| 27812 | 1391 | S1391K | E | 2e-08 | 1.3e-08 | 1.7e-08 | 1.667E-08 | 3.512E-09 | 1.300E-08 | 2.000E-08 |
| 27813 | 1391 | S1391M | E | 7e-09 | 5.9e-09 | 8.9e-09 | 7.267E-09 | 1.518E-09 | 5.900E-09 | 8.900E-09 |
| 27814 | 1391 | S1391F | E | 3.8e-08 | 1.1e-08 | 7.8e-09 | 1.893E-08 | 1.659E-08 | 7.800E-09 | 3.800E-08 |
| 27815 | 1391 | S1391P | E | 2.4e-09 | 3.8e-09 | 3e-09 | 3.067E-09 | 7.024E-10 | 2.400E-09 | 3.800E-09 |
| 27816 | 1391 | S1391S | E | 7.2e-09 | 1.8e-09 | 1.1e-08 | 6.667E-09 | 4.623E-09 | 1.800E-09 | 1.100E-08 |
| 27817 | 1391 | S1391T | E | 3.8e-09 | 9.1e-09 | 8.4e-09 | 7.100E-09 | 2.879E-09 | 3.800E-09 | 9.100E-09 |
| 27818 | 1391 | S1391W | E | 2.8e-08 | 4.2e-09 | 2.1e-09 | 1.143E-08 | 1.439E-08 | 2.100E-09 | 2.800E-08 |
| 27819 | 1391 | S1391Y | E | 1e-08 | 1.4e-08 | 1.9e-08 | 1.433E-08 | 4.509E-09 | 1.000E-08 | 1.900E-08 |
| 27820 | 1391 | S1391V | E | 3.5e-09 | 2.6e-09 | 1.6e-08 | 7.367E-09 | 7.490E-09 | 2.600E-09 | 1.600E-08 |
| 27821 | 1392 | S1392A | E | 7.6e-09 | 8.9e-09 | 1.3e-08 | 9.833E-09 | 2.818E-09 | 7.600E-09 | 1.300E-08 |
| 27822 | 1392 | S1392R | E | 1e-08 | 4.8e-09 | 4.5e-09 | 6.433E-09 | 3.092E-09 | 4.500E-09 | 1.000E-08 |
| 27823 | 1392 | S1392N | E | 1.1e-08 | 7.3e-09 | 9.3e-09 | 9.200E-09 | 1.852E-09 | 7.300E-09 | 1.100E-08 |
| 27824 | 1392 | S1392D | E | 1.6e-08 | 3.3e-09 | 3.4e-09 | 7.567E-09 | 7.304E-09 | 3.300E-09 | 1.600E-08 |
| 27825 | 1392 | S1392C | E | 4.9e-09 | 9.7e-09 | 6.6e-09 | 7.067E-09 | 2.434E-09 | 4.900E-09 | 9.700E-09 |
| 27826 | 1392 | S1392E | E | 8.7e-09 | 2.7e-09 | 1.5e-08 | 8.800E-09 | 6.151E-09 | 2.700E-09 | 1.500E-08 |
| 27827 | 1392 | S1392Q | E | 3.8e-09 | 8e-09 | 5e-09 | 5.600E-09 | 2.163E-09 | 3.800E-09 | 8.000E-09 |
| 27828 | 1392 | S1392G | E | 2.3e-08 | 2.2e-08 | 8e-09 | 1.767E-08 | 8.386E-09 | 8.000E-09 | 2.300E-08 |
| 27829 | 1392 | S1392H | E | 9.5e-09 | 8.9e-09 | 4.8e-09 | 7.733E-09 | 2.558E-09 | 4.800E-09 | 9.500E-09 |
| 27830 | 1392 | S1392I | E | 1.2e-08 | 5.8e-09 | 7.9e-09 | 8.567E-09 | 3.153E-09 | 5.800E-09 | 1.200E-08 |
| 27831 | 1392 | S1392L | E | 8.8e-09 | 2.1e-09 | 1.7e-08 | 9.300E-09 | 7.463E-09 | 2.100E-09 | 1.700E-08 |
| 27832 | 1392 | S1392K | E | 3.1e-09 | 1.6e-08 | 1.8e-09 | 6.967E-09 | 7.850E-09 | 1.800E-09 | 1.600E-08 |
| 27833 | 1392 | S1392M | E | 1.2e-08 | 7.4e-09 | 2.8e-08 | 1.580E-08 | 1.081E-08 | 7.400E-09 | 2.800E-08 |
| 27834 | 1392 | S1392F | E | 2.6e-09 | 1.8e-09 | 1.2e-08 | 5.467E-09 | 5.672E-09 | 1.800E-09 | 1.200E-08 |
| 27835 | 1392 | S1392P | E | 6.2e-09 | 7.4e-09 | 5.4e-09 | 6.333E-09 | 1.007E-09 | 5.400E-09 | 7.400E-09 |
| 27836 | 1392 | S1392S | E | 7.3e-09 | 4.4e-09 | 9e-09 | 6.900E-09 | 2.326E-09 | 4.400E-09 | 9.000E-09 |
| 27837 | 1392 | S1392T | E | 1.1e-08 | 2.1e-09 | 1.4e-08 | 9.033E-09 | 6.189E-09 | 2.100E-09 | 1.400E-08 |
| 27838 | 1392 | S1392W | E | 1.4e-08 | 1.5e-08 | 1.1e-08 | 1.333E-08 | 2.082E-09 | 1.100E-08 | 1.500E-08 |
| 27839 | 1392 | S1392Y | E | 2.8e-09 | 6.4e-09 | 1.1e-08 | 6.733E-09 | 4.110E-09 | 2.800E-09 | 1.100E-08 |
| 27840 | 1392 | S1392V | E | 3e-09 | 5.1e-09 | 1e-08 | 6.033E-09 | 3.592E-09 | 3.000E-09 | 1.000E-08 |
| 27841 | 1393 | A1393A | E | 7.9e-09 | 2.9e-09 | 8.2e-09 | 6.333E-09 | 2.977E-09 | 2.900E-09 | 8.200E-09 |
| 27842 | 1393 | A1393R | E | 3e-09 | 1e-09 | 4.8e-09 | 2.933E-09 | 1.901E-09 | 1.000E-09 | 4.800E-09 |
| 27843 | 1393 | A1393N | E | 3.4e-09 | 4.1e-09 | 8.7e-09 | 5.400E-09 | 2.879E-09 | 3.400E-09 | 8.700E-09 |
| 27844 | 1393 | A1393D | E | 4.7e-09 | 4.2e-09 | 2.2e-08 | 1.030E-08 | 1.014E-08 | 4.200E-09 | 2.200E-08 |
| 27845 | 1393 | A1393C | E | 6.7e-09 | 4.3e-09 | 8.6e-09 | 6.533E-09 | 2.155E-09 | 4.300E-09 | 8.600E-09 |
| 27846 | 1393 | A1393E | E | 1.3e-08 | 4.7e-09 | 1.1e-08 | 9.567E-09 | 4.332E-09 | 4.700E-09 | 1.300E-08 |
| 27847 | 1393 | A1393Q | E | 6e-09 | 4.5e-09 | 9.1e-09 | 6.533E-09 | 2.346E-09 | 4.500E-09 | 9.100E-09 |
| 27848 | 1393 | A1393G | E | 8.7e-09 | 4.3e-09 | 8.9e-09 | 7.300E-09 | 2.600E-09 | 4.300E-09 | 8.900E-09 |
| 27849 | 1393 | A1393H | E | 4.2e-09 | 6.3e-09 | 3.2e-09 | 4.567E-09 | 1.582E-09 | 3.200E-09 | 6.300E-09 |
| 27850 | 1393 | A1393I | E | 1.5e-08 | 1.3e-08 | 6.8e-09 | 1.160E-08 | 4.276E-09 | 6.800E-09 | 1.500E-08 |
| 27851 | 1393 | A1393L | E | 1e-08 | 1.6e-08 | 6.3e-09 | 1.077E-08 | 4.895E-09 | 6.300E-09 | 1.600E-08 |
| 27852 | 1393 | A1393K | E | 5.6e-09 | 6.4e-09 | 7.9e-09 | 6.633E-09 | 1.168E-09 | 5.600E-09 | 7.900E-09 |
| 27853 | 1393 | A1393M | E | 7.2e-09 | 2.8e-09 | 6.3e-09 | 5.433E-09 | 2.325E-09 | 2.800E-09 | 7.200E-09 |
| 27854 | 1393 | A1393F | E | 1.8e-09 | 9.7e-09 | 1.9e-08 | 1.017E-08 | 8.609E-09 | 1.800E-09 | 1.900E-08 |
| 27855 | 1393 | A1393P | E | 3.8e-09 | 3.6e-09 | 6e-09 | 4.467E-09 | 1.332E-09 | 3.600E-09 | 6.000E-09 |
| 27856 | 1393 | A1393S | E | 8.9e-09 | 6.2e-09 | 1.3e-08 | 9.367E-09 | 3.424E-09 | 6.200E-09 | 1.300E-08 |
| 27857 | 1393 | A1393T | E | 1.1e-08 | 7.7e-09 | 1.6e-08 | 1.157E-08 | 4.179E-09 | 7.700E-09 | 1.600E-08 |
| 27858 | 1393 | A1393W | E | 5.3e-09 | 3.6e-09 | 5.8e-09 | 4.900E-09 | 1.153E-09 | 3.600E-09 | 5.800E-09 |
| 27859 | 1393 | A1393Y | E | 1.2e-08 | 2.8e-09 | 1.2e-08 | 8.933E-09 | 5.312E-09 | 2.800E-09 | 1.200E-08 |
| 27860 | 1393 | A1393V | E | 2.5e-09 | 1.6e-08 | 2.3e-09 | 6.933E-09 | 7.853E-09 | 2.300E-09 | 1.600E-08 |
| 27861 | 1394 | S1394A | E | 3e-09 | 4.7e-09 | 1.1e-08 | 6.233E-09 | 4.215E-09 | 3.000E-09 | 1.100E-08 |
| 27862 | 1394 | S1394R | E | 7.4e-09 | 6.8e-09 | 8.8e-09 | 7.667E-09 | 1.026E-09 | 6.800E-09 | 8.800E-09 |
| 27863 | 1394 | S1394N | E | 1.3e-08 | 3e-09 | 5e-09 | 7.000E-09 | 5.292E-09 | 3.000E-09 | 1.300E-08 |
| 27864 | 1394 | S1394D | E | 1.7e-08 | 1.4e-09 | 6.5e-09 | 8.300E-09 | 7.954E-09 | 1.400E-09 | 1.700E-08 |
| 27865 | 1394 | S1394C | E | 6.4e-09 | 3.5e-09 | 1.4e-08 | 7.967E-09 | 5.422E-09 | 3.500E-09 | 1.400E-08 |
| 27866 | 1394 | S1394E | E | 1.5e-08 | 4.8e-09 | 1.1e-08 | 1.027E-08 | 5.139E-09 | 4.800E-09 | 1.500E-08 |
| 27867 | 1394 | S1394Q | E | 8.3e-09 | 4.9e-09 | 7.1e-09 | 6.767E-09 | 1.724E-09 | 4.900E-09 | 8.300E-09 |
| 27868 | 1394 | S1394G | E | 1.3e-08 | 2.7e-09 | 6.8e-09 | 7.500E-09 | 5.186E-09 | 2.700E-09 | 1.300E-08 |
| 27869 | 1394 | S1394H | E | 2.5e-09 | 3.3e-09 | 4.3e-09 | 3.367E-09 | 9.018E-10 | 2.500E-09 | 4.300E-09 |
| 27870 | 1394 | S1394I | E | 1e-08 | 9e-09 | 5.8e-09 | 8.267E-09 | 2.194E-09 | 5.800E-09 | 1.000E-08 |
| 27871 | 1394 | S1394L | E | 4.4e-09 | 6e-09 | 8.7e-09 | 6.367E-09 | 2.173E-09 | 4.400E-09 | 8.700E-09 |
| 27872 | 1394 | S1394K | E | 1.7e-08 | 9.8e-09 | 7.5e-09 | 1.143E-08 | 4.956E-09 | 7.500E-09 | 1.700E-08 |
| 27873 | 1394 | S1394M | E | 4.2e-09 | 8.7e-09 | 7.8e-09 | 6.900E-09 | 2.381E-09 | 4.200E-09 | 8.700E-09 |
| 27874 | 1394 | S1394F | E | 1e-08 | 5.2e-09 | 1.4e-08 | 9.733E-09 | 4.406E-09 | 5.200E-09 | 1.400E-08 |
| 27875 | 1394 | S1394P | E | 9.5e-09 | 3.1e-09 | 2.3e-09 | 4.967E-09 | 3.946E-09 | 2.300E-09 | 9.500E-09 |
| 27876 | 1394 | S1394S | E | 6.1e-09 | 3.1e-09 | 1.1e-08 | 6.733E-09 | 3.988E-09 | 3.100E-09 | 1.100E-08 |
| 27877 | 1394 | S1394T | E | 6.2e-09 | 4.4e-09 | 3e-09 | 4.533E-09 | 1.604E-09 | 3.000E-09 | 6.200E-09 |
| 27878 | 1394 | S1394W | E | 5.2e-09 | 7.6e-09 | 1.1e-08 | 7.933E-09 | 2.914E-09 | 5.200E-09 | 1.100E-08 |
| 27879 | 1394 | S1394Y | E | 7.2e-09 | 1.6e-08 | 2.2e-08 | 1.507E-08 | 7.444E-09 | 7.200E-09 | 2.200E-08 |

| | | | | | | | | | | |
|---|---|---|---|---|---|---|---|---|---|---|
| 27880 | 1394 | S1394V | E | 1.4e-08 | 5.4e-09 | 5.3e-09 | 8.233E-09 | 4.994E-09 | 5.300E-09 | 1.400E-08 |
| 27881 | 1395 | T1395A | E | 6.6e-09 | 6.5e-09 | 5.1e-09 | 6.067E-09 | 8.386E-10 | 5.100E-09 | 6.600E-09 |
| 27882 | 1395 | T1395R | E | 7.9e-09 | 1.5e-08 | 3.8e-09 | 8.900E-09 | 5.667E-09 | 3.800E-09 | 1.500E-08 |
| 27883 | 1395 | T1395N | E | 7.8e-09 | 2.1e-08 | 1.3e-08 | 1.393E-08 | 6.649E-09 | 7.800E-09 | 2.100E-08 |
| 27884 | 1395 | T1395D | E | 6.1e-09 | 6.9e-09 | 1.2e-08 | 8.333E-09 | 3.201E-09 | 6.100E-09 | 1.200E-08 |
| 27885 | 1395 | T1395C | E | 4.4e-09 | 4.1e-09 | 9.9e-09 | 6.133E-09 | 3.265E-09 | 4.100E-09 | 9.900E-09 |
| 27886 | 1395 | T1395E | E | 8.3e-09 | 9.1e-09 | 1.2e-08 | 9.800E-09 | 1.947E-09 | 8.300E-09 | 1.200E-08 |
| 27887 | 1395 | T1395Q | E | 1.5e-08 | 1.2e-08 | 1.2e-08 | 1.300E-08 | 1.732E-09 | 1.200E-08 | 1.500E-08 |
| 27888 | 1395 | T1395G | E | 7.3e-09 | 1.1e-08 | 7.2e-09 | 8.500E-09 | 2.166E-09 | 7.200E-09 | 1.100E-08 |
| 27889 | 1395 | T1395H | E | 9.3e-09 | 2.9e-08 | 5.5e-09 | 1.460E-08 | 1.261E-08 | 5.500E-09 | 2.900E-08 |
| 27890 | 1395 | T1395I | E | 5.7e-09 | 9.6e-09 | 5.1e-09 | 6.800E-09 | 2.443E-09 | 5.100E-09 | 9.600E-09 |
| 27891 | 1395 | T1395L | E | 3.6e-09 | 1.4e-08 | 1.3e-08 | 1.020E-08 | 5.738E-09 | 3.600E-09 | 1.400E-08 |
| 27892 | 1395 | T1395K | E | 4.1e-09 | 1.7e-08 | 7.1e-09 | 9.400E-09 | 6.751E-09 | 4.100E-09 | 1.700E-08 |
| 27893 | 1395 | T1395M | E | 1.2e-08 | 3.4e-09 | 8.9e-09 | 8.100E-09 | 4.355E-09 | 3.400E-09 | 1.200E-08 |
| 27894 | 1395 | T1395F | E | 1.1e-08 | 3.7e-09 | 8e-09 | 7.567E-09 | 3.669E-09 | 3.700E-09 | 1.100E-08 |
| 27895 | 1395 | T1395P | E | 1.6e-08 | 8.9e-09 | 4.8e-09 | 9.900E-09 | 5.667E-09 | 4.800E-09 | 1.600E-08 |
| 27896 | 1395 | T1395S | E | 2.1e-09 | 4.8e-09 | 7e-09 | 4.633E-09 | 2.454E-09 | 2.100E-09 | 7.000E-09 |
| 27897 | 1395 | T1395T | E | 5.2e-09 | 9.1e-09 | 1.5e-08 | 9.767E-09 | 4.934E-09 | 5.200E-09 | 1.500E-08 |
| 27898 | 1395 | T1395W | E | 4.7e-09 | 9.6e-09 | 7e-09 | 7.100E-09 | 2.452E-09 | 4.700E-09 | 9.600E-09 |
| 27899 | 1395 | T1395Y | E | 5.1e-09 | 4.7e-09 | 1.7e-08 | 8.933E-09 | 6.989E-09 | 4.700E-09 | 1.700E-08 |
| 27900 | 1395 | T1395V | E | 6.1e-09 | 1.7e-08 | 9.8e-09 | 1.097E-08 | 5.543E-09 | 6.100E-09 | 1.700E-08 |
| 27901 | 1396 | K1396A | E | 6e-09 | 6.8e-09 | 9.8e-09 | 7.533E-09 | 2.003E-09 | 6.000E-09 | 9.800E-09 |
| 27902 | 1396 | K1396R | E | 1.1e-08 | 1.4e-08 | 1.4e-08 | 1.300E-08 | 1.732E-09 | 1.100E-08 | 1.400E-08 |
| 27903 | 1396 | K1396N | E | 4.5e-09 | 1.2e-08 | 5.4e-09 | 7.300E-09 | 4.095E-09 | 4.500E-09 | 1.200E-08 |
| 27904 | 1396 | K1396D | E | 3.9e-09 | 1.4e-08 | 2.4e-08 | 1.397E-08 | 1.005E-08 | 3.900E-09 | 2.400E-08 |
| 27905 | 1396 | K1396C | E | 1.5e-08 | 1.1e-08 | 4.4e-09 | 1.013E-08 | 5.353E-09 | 4.400E-09 | 1.500E-08 |
| 27906 | 1396 | K1396E | E | 5e-09 | 5.3e-09 | 1.2e-08 | 7.433E-09 | 3.958E-09 | 5.000E-09 | 1.200E-08 |
| 27907 | 1396 | K1396Q | E | 1.3e-08 | 2.5e-09 | 1.6e-08 | 1.050E-08 | 7.089E-09 | 2.500E-09 | 1.600E-08 |
| 27908 | 1396 | K1396G | E | 3.9e-09 | 3.5e-09 | 5.9e-09 | 4.433E-09 | 1.286E-09 | 3.500E-09 | 5.900E-09 |
| 27909 | 1396 | K1396H | E | 1.1e-08 | 3.8e-09 | 4.4e-09 | 6.400E-09 | 3.995E-09 | 3.800E-09 | 1.100E-08 |
| 27910 | 1396 | K1396I | E | 8e-09 | 1.1e-08 | 1.8e-08 | 1.233E-08 | 5.132E-09 | 8.000E-09 | 1.800E-08 |
| 27911 | 1396 | K1396L | E | 9.6e-09 | 7.8e-09 | 2.5e-08 | 1.413E-08 | 9.454E-09 | 7.800E-09 | 2.500E-08 |
| 27912 | 1396 | K1396K | E | 5.3e-09 | 5.8e-09 | 6.3e-09 | 5.800E-09 | 5.000E-10 | 5.300E-09 | 6.300E-09 |
| 27913 | 1396 | K1396M | E | 7.5e-09 | 2.6e-09 | 5.6e-09 | 5.233E-09 | 2.470E-09 | 2.600E-09 | 7.500E-09 |
| 27914 | 1396 | K1396F | E | 9.5e-09 | 6.1e-09 | 5.7e-09 | 7.100E-09 | 2.088E-09 | 5.700E-09 | 9.500E-09 |
| 27915 | 1396 | K1396P | E | 9.6e-09 | 7.1e-09 | 1.3e-08 | 9.900E-09 | 2.961E-09 | 7.100E-09 | 1.300E-08 |
| 27916 | 1396 | K1396S | E | 6.7e-09 | 9.1e-09 | 2e-08 | 1.193E-08 | 7.088E-09 | 6.700E-09 | 2.000E-08 |
| 27917 | 1396 | K1396T | E | 5.6e-09 | 1.5e-08 | 2.5e-09 | 7.700E-09 | 6.509E-09 | 2.500E-09 | 1.500E-08 |
| 27918 | 1396 | K1396W | E | 6.4e-09 | 2.9e-09 | 7.6e-09 | 5.633E-09 | 2.442E-09 | 2.900E-09 | 7.600E-09 |
| 27919 | 1396 | K1396Y | E | 2.1e-09 | 3.7e-09 | 4.5e-09 | 3.433E-09 | 1.222E-09 | 2.100E-09 | 4.500E-09 |
| 27920 | 1396 | K1396V | E | 3.1e-09 | 3.3e-09 | 9.8e-09 | 5.400E-09 | 3.812E-09 | 3.100E-09 | 9.800E-09 |
| 27921 | 1397 | G1397A | E | 4.2e-09 | 4.3e-09 | 2.9e-09 | 3.800E-09 | 7.810E-10 | 2.900E-09 | 4.300E-09 |
| 27922 | 1397 | G1397R | E | 4.4e-09 | 8.1e-09 | 5e-09 | 5.833E-09 | 1.986E-09 | 4.400E-09 | 8.100E-09 |
| 27923 | 1397 | G1397N | E | 6.5e-09 | 4.3e-09 | 5.9e-09 | 5.567E-09 | 1.137E-09 | 4.300E-09 | 6.500E-09 |
| 27924 | 1397 | G1397D | E | 8.4e-09 | 1.8e-09 | 1.3e-08 | 7.733E-09 | 5.630E-09 | 1.800E-09 | 1.300E-08 |
| 27925 | 1397 | G1397C | E | 1.4e-08 | 1.5e-08 | 9.6e-09 | 1.287E-08 | 2.873E-09 | 9.600E-09 | 1.500E-08 |
| 27926 | 1397 | G1397E | E | 6e-09 | 6.1e-09 | 3.9e-09 | 5.333E-09 | 1.242E-09 | 3.900E-09 | 6.100E-09 |
| 27927 | 1397 | G1397Q | E | 2.6e-09 | 9.1e-09 | 4.8e-09 | 5.500E-09 | 3.306E-09 | 2.600E-09 | 9.100E-09 |
| 27928 | 1397 | G1397G | E | 4.7e-09 | 2.4e-09 | 1.3e-08 | 6.700E-09 | 5.576E-09 | 2.400E-09 | 1.300E-08 |
| 27929 | 1397 | G1397H | E | 1.5e-08 | 5.6e-09 | 1.6e-08 | 1.220E-08 | 5.738E-09 | 5.600E-09 | 1.600E-08 |
| 27930 | 1397 | G1397I | E | 1.1e-08 | 7.2e-09 | 6.7e-09 | 8.300E-09 | 2.352E-09 | 6.700E-09 | 1.100E-08 |
| 27931 | 1397 | G1397L | E | 6.3e-09 | 4.9e-09 | 2.6e-08 | 1.240E-08 | 1.180E-08 | 4.900E-09 | 2.600E-08 |
| 27932 | 1397 | G1397K | E | 1.3e-08 | 5.1e-09 | 7.2e-09 | 8.433E-09 | 4.092E-09 | 5.100E-09 | 1.300E-08 |
| 27933 | 1397 | G1397M | E | 1.3e-08 | 6.2e-09 | 2e-09 | 7.067E-09 | 5.551E-09 | 2.000E-09 | 1.300E-08 |
| 27934 | 1397 | G1397F | E | 4.2e-09 | 6.4e-09 | 1e-08 | 6.867E-09 | 2.928E-09 | 4.200E-09 | 1.000E-08 |
| 27935 | 1397 | G1397P | E | 6.7e-09 | 1.2e-08 | 3.7e-09 | 7.467E-09 | 4.203E-09 | 3.700E-09 | 1.200E-08 |
| 27936 | 1397 | G1397S | E | 7.5e-09 | 1.2e-08 | 3.9e-09 | 7.800E-09 | 4.058E-09 | 3.900E-09 | 1.200E-08 |
| 27937 | 1397 | G1397T | E | 3.7e-09 | 6e-09 | 1.2e-08 | 7.233E-09 | 4.285E-09 | 3.700E-09 | 1.200E-08 |
| 27938 | 1397 | G1397W | E | 1.4e-08 | 7.7e-09 | 7.6e-09 | 9.767E-09 | 3.667E-09 | 7.600E-09 | 1.400E-08 |
| 27939 | 1397 | G1397Y | E | 8.2e-09 | 5.3e-09 | 7.9e-09 | 7.133E-09 | 1.595E-09 | 5.300E-09 | 8.200E-09 |
| 27940 | 1397 | G1397V | E | 9e-09 | 2.3e-09 | 1.3e-08 | 8.100E-09 | 5.406E-09 | 2.300E-09 | 1.300E-08 |
| 27941 | 1398 | P1398A | E | 8.2e-09 | 4.2e-09 | 4.3e-09 | 5.567E-09 | 2.281E-09 | 4.200E-09 | 8.200E-09 |
| 27942 | 1398 | P1398R | E | 1.6e-08 | 7.9e-09 | 9.7e-09 | 1.120E-08 | 4.253E-09 | 7.900E-09 | 1.600E-08 |
| 27943 | 1398 | P1398N | E | 5e-09 | 4.7e-09 | 6.1e-09 | 5.267E-09 | 7.371E-10 | 4.700E-09 | 6.100E-09 |
| 27944 | 1398 | P1398D | E | 2.4e-08 | 3.3e-09 | 4.3e-09 | 1.053E-08 | 1.167E-08 | 3.300E-09 | 2.400E-08 |
| 27945 | 1398 | P1398C | E | 5.2e-09 | 2.7e-09 | 1.4e-08 | 7.300E-09 | 5.935E-09 | 2.700E-09 | 1.400E-08 |
| 27946 | 1398 | P1398E | E | 7.7e-09 | 5.9e-09 | 1.6e-08 | 9.867E-09 | 5.387E-09 | 5.900E-09 | 1.600E-08 |
| 27947 | 1398 | P1398Q | E | 2.6e-09 | 2.3e-08 | 8.1e-09 | 1.123E-08 | 1.055E-08 | 2.600E-09 | 2.300E-08 |
| 27948 | 1398 | P1398G | E | 2.7e-09 | 1.7e-08 | 2.5e-09 | 7.400E-09 | 8.314E-09 | 2.500E-09 | 1.700E-08 |
| 27949 | 1398 | P1398H | E | 8.2e-09 | 2.5e-09 | 6.4e-09 | 5.700E-09 | 2.914E-09 | 2.500E-09 | 8.200E-09 |
| 27950 | 1398 | P1398I | E | 3.8e-09 | 8.6e-09 | 3.7e-09 | 5.367E-09 | 2.801E-09 | 3.700E-09 | 8.600E-09 |
| 27951 | 1398 | P1398L | E | 2.3e-08 | 4.3e-09 | 7.9e-09 | 1.173E-08 | 9.922E-09 | 4.300E-09 | 2.300E-08 |
| 27952 | 1398 | P1398K | E | 6.4e-09 | 1.2e-08 | 8.7e-09 | 9.033E-09 | 2.815E-09 | 6.400E-09 | 1.200E-08 |
| 27953 | 1398 | P1398M | E | 2.5e-09 | 3.8e-09 | 4.9e-09 | 3.733E-09 | 1.201E-09 | 2.500E-09 | 4.900E-09 |
| 27954 | 1398 | P1398F | E | 2e-08 | 3e-09 | 4.3e-09 | 9.100E-09 | 9.462E-09 | 3.000E-09 | 2.000E-08 |
| 27955 | 1398 | P1398P | E | 1e-08 | 1e-08 | 8.9e-09 | 9.633E-09 | 6.351E-10 | 8.900E-09 | 1.000E-08 |
| 27956 | 1398 | P1398S | E | 1.9e-08 | 2.9e-09 | 8.8e-09 | 1.023E-08 | 8.145E-09 | 2.900E-09 | 1.900E-08 |
| 27957 | 1398 | P1398T | E | 1.3e-08 | 8.3e-09 | 6.3e-09 | 9.200E-09 | 3.439E-09 | 6.300E-09 | 1.300E-08 |
| 27958 | 1398 | P1398W | E | 5.8e-09 | 9.1e-09 | 6.7e-09 | 7.200E-09 | 1.706E-09 | 5.800E-09 | 9.100E-09 |
| 27959 | 1398 | P1398Y | E | 3.1e-09 | 1.3e-08 | 7.1e-09 | 7.733E-09 | 4.980E-09 | 3.100E-09 | 1.300E-08 |
| 27960 | 1398 | P1398V | E | 5.8e-09 | 1.1e-08 | 1.1e-08 | 9.267E-09 | 3.002E-09 | 5.800E-09 | 1.100E-08 |
| 27961 | 1399 | S1399A | E | 1.4e-08 | 2.8e-09 | 4.2e-08 | 1.960E-08 | 2.019E-08 | 2.800E-09 | 4.200E-08 |
| 27962 | 1399 | S1399R | E | 5.2e-09 | 2.1e-08 | 3e-09 | 9.733E-09 | 9.819E-09 | 3.000E-09 | 2.100E-08 |
| 27963 | 1399 | S1399N | E | 2.4e-09 | 7.2e-09 | 8.5e-09 | 6.033E-09 | 3.213E-09 | 2.400E-09 | 8.500E-09 |
| 27964 | 1399 | S1399D | E | 1.6e-08 | 4.2e-09 | 3.7e-09 | 7.967E-09 | 6.962E-09 | 3.700E-09 | 1.600E-08 |

| | | | | | | | | | | |
|---|---|---|---|---|---|---|---|---|---|---|
| 27965 | 1399 | S1399C | E | 3.7e-09 | 5.4e-09 | 9.8e-09 | 6.300E-09 | 3.148E-09 | 3.700E-09 | 9.800E-09 |
| 27966 | 1399 | S1399E | E | 7.7e-09 | 3.6e-09 | 2.5e-09 | 4.600E-09 | 2.740E-09 | 2.500E-09 | 7.700E-09 |
| 27967 | 1399 | S1399Q | E | 6e-09 | 6.8e-09 | 1.4e-08 | 8.933E-09 | 4.406E-09 | 6.000E-09 | 1.400E-08 |
| 27968 | 1399 | S1399G | E | 1.7e-08 | 2e-08 | 4.8e-09 | 1.393E-08 | 8.051E-09 | 4.800E-09 | 2.000E-08 |
| 27969 | 1399 | S1399H | E | 5e-09 | 1.8e-08 | 9.9e-09 | 1.097E-08 | 6.565E-09 | 5.000E-09 | 1.800E-08 |
| 27970 | 1399 | S1399I | E | 1.1e-08 | 2.8e-09 | 4.2e-09 | 6.000E-09 | 4.386E-09 | 2.800E-09 | 1.100E-08 |
| 27971 | 1399 | S1399L | E | 3.5e-09 | 1.8e-08 | 1.1e-08 | 1.083E-08 | 7.251E-09 | 3.500E-09 | 1.800E-08 |
| 27972 | 1399 | S1399K | E | 8.6e-09 | 9.6e-09 | 1e-08 | 9.400E-09 | 7.211E-10 | 8.600E-09 | 1.000E-08 |
| 27973 | 1399 | S1399M | E | 6.1e-09 | 3.5e-09 | 5.3e-09 | 4.967E-09 | 1.332E-09 | 3.500E-09 | 6.100E-09 |
| 27974 | 1399 | S1399F | E | 7.4e-09 | 3e-09 | 1.1e-08 | 7.133E-09 | 4.007E-09 | 3.000E-09 | 1.100E-08 |
| 27975 | 1399 | S1399P | E | 6e-09 | 2.7e-09 | 4.6e-09 | 4.433E-09 | 1.656E-09 | 2.700E-09 | 6.000E-09 |
| 27976 | 1399 | S1399S | E | 1.1e-08 | 3.8e-09 | 3.5e-09 | 6.100E-09 | 4.246E-09 | 3.500E-09 | 1.100E-08 |
| 27977 | 1399 | S1399T | E | 3e-09 | 3.6e-09 | 7.6e-09 | 4.733E-09 | 2.501E-09 | 3.000E-09 | 7.600E-09 |
| 27978 | 1399 | S1399W | E | 5.3e-09 | 4.2e-09 | 5.3e-09 | 4.933E-09 | 6.351E-10 | 4.200E-09 | 5.300E-09 |
| 27979 | 1399 | S1399Y | E | 5.4e-09 | 8.6e-09 | 5.8e-09 | 6.600E-09 | 1.744E-09 | 5.400E-09 | 8.600E-09 |
| 27980 | 1399 | S1399V | E | 9.1e-09 | 1.1e-08 | 1.4e-08 | 1.137E-08 | 2.470E-09 | 9.100E-09 | 1.400E-08 |
| 27981 | 1400 | V1400A | E | 1.1e-08 | 7.1e-09 | 5.1e-09 | 7.733E-09 | 3.001E-09 | 5.100E-09 | 1.100E-08 |
| 27982 | 1400 | V1400R | E | 8.3e-09 | 1.3e-08 | 1.8e-08 | 1.310E-08 | 4.851E-09 | 8.300E-09 | 1.800E-08 |
| 27983 | 1400 | V1400N | E | 1.3e-08 | 7.6e-09 | 1.2e-08 | 1.087E-08 | 2.873E-09 | 7.600E-09 | 1.300E-08 |
| 27984 | 1400 | V1400D | E | 2.9e-09 | 5.4e-09 | 2.5e-08 | 1.110E-08 | 1.210E-08 | 2.900E-09 | 2.500E-08 |
| 27985 | 1400 | V1400C | E | 7.6e-09 | 1.2e-08 | 8.4e-09 | 9.333E-09 | 2.344E-09 | 7.600E-09 | 1.200E-08 |
| 27986 | 1400 | V1400E | E | 4.7e-09 | 2.8e-09 | 3.7e-09 | 3.733E-09 | 9.504E-10 | 2.800E-09 | 4.700E-09 |
| 27987 | 1400 | V1400Q | E | 4.1e-09 | 7.5e-09 | 2.4e-08 | 1.187E-08 | 1.064E-08 | 4.100E-09 | 2.400E-08 |
| 27988 | 1400 | V1400G | E | 9.2e-09 | 6.4e-09 | 6.7e-09 | 7.433E-09 | 1.537E-09 | 6.400E-09 | 9.200E-09 |
| 27989 | 1400 | V1400H | E | 8.6e-09 | 5.7e-09 | 8.6e-09 | 7.633E-09 | 1.674E-09 | 5.700E-09 | 8.600E-09 |
| 27990 | 1400 | V1400I | E | 9.4e-09 | 5e-09 | 6.8e-09 | 7.067E-09 | 2.212E-09 | 5.000E-09 | 9.400E-09 |
| 27991 | 1400 | V1400L | E | 5.5e-09 | 6.3e-09 | 3.1e-09 | 4.967E-09 | 1.665E-09 | 3.100E-09 | 6.300E-09 |
| 27992 | 1400 | V1400K | E | 4.1e-09 | 5.8e-09 | 1.8e-08 | 9.300E-09 | 7.582E-09 | 4.100E-09 | 1.800E-08 |
| 27993 | 1400 | V1400M | E | 8.2e-09 | 5.8e-09 | 9.6e-09 | 7.867E-09 | 1.922E-09 | 5.800E-09 | 9.600E-09 |
| 27994 | 1400 | V1400F | E | 9.5e-09 | 1.1e-08 | 8e-09 | 9.500E-09 | 1.500E-09 | 8.000E-09 | 1.100E-08 |
| 27995 | 1400 | V1400P | E | 4.3e-09 | 1.2e-08 | 1.1e-08 | 9.100E-09 | 4.187E-09 | 4.300E-09 | 1.200E-08 |
| 27996 | 1400 | V1400S | E | 2.4e-09 | 9.7e-09 | 1e-08 | 7.367E-09 | 4.304E-09 | 2.400E-09 | 1.000E-08 |
| 27997 | 1400 | V1400T | E | 4e-09 | 5.8e-09 | 1.6e-08 | 8.600E-09 | 6.471E-09 | 4.000E-09 | 1.600E-08 |
| 27998 | 1400 | V1400W | E | 2.5e-08 | 1.6e-08 | 1.4e-08 | 1.833E-08 | 5.859E-09 | 1.400E-08 | 2.500E-08 |
| 27999 | 1400 | V1400Y | E | 6.9e-09 | 5.8e-09 | 3.3e-09 | 5.333E-09 | 1.845E-09 | 3.300E-09 | 6.900E-09 |
| 28000 | 1400 | V1400V | E | 1.1e-08 | 1.5e-08 | 9.4e-09 | 1.180E-08 | 2.884E-09 | 9.400E-09 | 1.500E-08 |
| 28001 | 1401 | F1401A | E | 7.9e-09 | 7.4e-09 | 1.2e-08 | 9.100E-09 | 2.524E-09 | 7.400E-09 | 1.200E-08 |
| 28002 | 1401 | F1401R | E | 1.1e-08 | 6.3e-09 | 7e-09 | 8.100E-09 | 2.536E-09 | 6.300E-09 | 1.100E-08 |
| 28003 | 1401 | F1401N | E | 4.6e-09 | 6.5e-09 | 9.3e-09 | 6.800E-09 | 2.364E-09 | 4.600E-09 | 9.300E-09 |
| 28004 | 1401 | F1401D | E | 1e-08 | 4.9e-09 | 3.9e-09 | 6.267E-09 | 3.272E-09 | 3.900E-09 | 1.000E-08 |
| 28005 | 1401 | F1401C | E | 7e-09 | 3.6e-09 | 1.1e-08 | 7.200E-09 | 3.704E-09 | 3.600E-09 | 1.100E-08 |
| 28006 | 1401 | F1401E | E | 3.1e-09 | 6.2e-09 | 1.9e-09 | 3.733E-09 | 2.219E-09 | 1.900E-09 | 6.200E-09 |
| 28007 | 1401 | F1401Q | E | 3.3e-09 | 1.5e-08 | 3.3e-09 | 7.200E-09 | 6.755E-09 | 3.300E-09 | 1.500E-08 |
| 28008 | 1401 | F1401G | E | 8.2e-09 | 1.3e-08 | 9.1e-09 | 1.010E-08 | 2.551E-09 | 8.200E-09 | 1.300E-08 |
| 28009 | 1401 | F1401H | E | 1.1e-08 | 2.9e-09 | 1.4e-08 | 9.300E-09 | 5.742E-09 | 2.900E-09 | 1.400E-08 |
| 28010 | 1401 | F1401I | E | 9.1e-10 | 8.4e-09 | 5.5e-09 | 4.937E-09 | 3.777E-09 | 9.100E-10 | 8.400E-09 |
| 28011 | 1401 | F1401L | E | 1.4e-09 | 3.8e-09 | 3.5e-09 | 2.900E-09 | 1.308E-09 | 1.400E-09 | 3.800E-09 |
| 28012 | 1401 | F1401K | E | 1.4e-08 | 1.9e-09 | 1.1e-08 | 8.967E-09 | 6.301E-09 | 1.900E-09 | 1.400E-08 |
| 28013 | 1401 | F1401M | E | 5.3e-09 | 9.7e-09 | 4.6e-09 | 6.533E-09 | 2.765E-09 | 4.600E-09 | 9.700E-09 |
| 28014 | 1401 | F1401F | E | 1.3e-08 | 3.1e-09 | 7.3e-09 | 7.800E-09 | 4.969E-09 | 3.100E-09 | 1.300E-08 |
| 28015 | 1401 | F1401P | E | 5e-09 | 6e-09 | 8.8e-09 | 6.600E-09 | 1.970E-09 | 5.000E-09 | 8.800E-09 |
| 28016 | 1401 | F1401S | E | 7.1e-09 | 5.4e-09 | 1.4e-08 | 8.833E-09 | 4.554E-09 | 5.400E-09 | 1.400E-08 |
| 28017 | 1401 | F1401T | E | 7.8e-09 | 4.9e-09 | 4e-09 | 5.567E-09 | 1.986E-09 | 4.000E-09 | 7.800E-09 |
| 28018 | 1401 | F1401W | E | 9.7e-09 | 5.4e-09 | 2.5e-09 | 5.867E-09 | 3.623E-09 | 2.500E-09 | 9.700E-09 |
| 28019 | 1401 | F1401Y | E | 6.2e-09 | 1.8e-08 | 1.4e-08 | 1.273E-08 | 6.001E-09 | 6.200E-09 | 1.800E-08 |
| 28020 | 1401 | F1401V | E | 5.4e-09 | 6.1e-09 | 2.6e-09 | 4.700E-09 | 1.852E-09 | 2.600E-09 | 6.100E-09 |
| 28021 | 1402 | P1402A | E | 6.1e-09 | 1.1e-08 | 4.7e-09 | 7.267E-09 | 3.308E-09 | 4.700E-09 | 1.100E-08 |
| 28022 | 1402 | P1402R | E | 1.4e-08 | 2.1e-08 | 1e-08 | 1.500E-08 | 5.568E-09 | 1.000E-08 | 2.100E-08 |
| 28023 | 1402 | P1402N | E | 4.5e-09 | 5.3e-09 | 1.2e-08 | 7.267E-09 | 4.119E-09 | 4.500E-09 | 1.200E-08 |
| 28024 | 1402 | P1402D | E | 1.2e-08 | 5.2e-09 | 4.8e-09 | 7.333E-09 | 4.046E-09 | 4.800E-09 | 1.200E-08 |
| 28025 | 1402 | P1402C | E | 7.6e-09 | 6.8e-09 | 7.1e-09 | 7.167E-09 | 4.041E-10 | 6.800E-09 | 7.600E-09 |
| 28026 | 1402 | P1402E | E | 5.7e-09 | 6.1e-09 | 4e-09 | 5.267E-09 | 1.115E-09 | 4.000E-09 | 6.100E-09 |
| 28027 | 1402 | P1402Q | E | 2.2e-09 | 3.3e-09 | 5.2e-09 | 3.567E-09 | 1.518E-09 | 2.200E-09 | 5.200E-09 |
| 28028 | 1402 | P1402G | E | 9.9e-09 | 4.9e-09 | 8.7e-09 | 7.833E-09 | 2.610E-09 | 4.900E-09 | 9.900E-09 |
| 28029 | 1402 | P1402H | E | 6.6e-09 | 8.4e-09 | 1.8e-08 | 1.100E-08 | 6.129E-09 | 6.600E-09 | 1.800E-08 |
| 28030 | 1402 | P1402I | E | 3.5e-09 | 5.8e-09 | 3.3e-09 | 4.200E-09 | 1.389E-09 | 3.300E-09 | 5.800E-09 |
| 28031 | 1402 | P1402L | E | 1.2e-08 | 7e-09 | 8.2e-09 | 9.067E-09 | 2.610E-09 | 7.000E-09 | 1.200E-08 |
| 28032 | 1402 | P1402K | E | 2.8e-09 | 7.3e-09 | 9.3e-09 | 6.467E-09 | 3.329E-09 | 2.800E-09 | 9.300E-09 |
| 28033 | 1402 | P1402M | E | 9.9e-09 | 4.5e-09 | 8.6e-09 | 7.667E-09 | 2.818E-09 | 4.500E-09 | 9.900E-09 |
| 28034 | 1402 | P1402F | E | 1.3e-08 | 1.6e-08 | 6.3e-09 | 1.177E-08 | 4.966E-09 | 6.300E-09 | 1.600E-08 |
| 28035 | 1402 | P1402P | E | 5.7e-09 | 6.5e-09 | 2.5e-09 | 4.900E-09 | 2.117E-09 | 2.500E-09 | 6.500E-09 |
| 28036 | 1402 | P1402S | E | 2.9e-09 | 1e-08 | 3.8e-09 | 5.567E-09 | 3.866E-09 | 2.900E-09 | 1.000E-08 |
| 28037 | 1402 | P1402T | E | 8.7e-09 | 4.4e-09 | 2.8e-09 | 5.300E-09 | 3.051E-09 | 2.800E-09 | 8.700E-09 |
| 28038 | 1402 | P1402W | E | 5.6e-09 | 6.4e-09 | 4e-09 | 5.333E-09 | 1.222E-09 | 4.000E-09 | 6.400E-09 |
| 28039 | 1402 | P1402Y | E | 4.6e-09 | 6.3e-09 | 7e-09 | 5.967E-09 | 1.234E-09 | 4.600E-09 | 7.000E-09 |
| 28040 | 1402 | P1402V | E | 1.1e-08 | 1.6e-08 | 8.5e-09 | 1.183E-08 | 3.819E-09 | 8.500E-09 | 1.600E-08 |
| 28041 | 1403 | L1403A | E | 5.8e-09 | 6.2e-09 | 1e-08 | 7.333E-09 | 2.318E-09 | 5.800E-09 | 1.000E-08 |
| 28042 | 1403 | L1403R | E | 5.1e-09 | 1.3e-08 | 3.4e-09 | 7.167E-09 | 5.123E-09 | 3.400E-09 | 1.300E-08 |
| 28043 | 1403 | L1403N | E | 1.9e-08 | 1.6e-08 | 2.5e-09 | 1.250E-08 | 8.789E-09 | 2.500E-09 | 1.900E-08 |
| 28044 | 1403 | L1403D | E | 7.7e-09 | 1.4e-08 | 1.6e-08 | 1.257E-08 | 4.332E-09 | 7.700E-09 | 1.600E-08 |
| 28045 | 1403 | L1403C | E | 9.6e-09 | 1.5e-08 | 6.2e-09 | 1.027E-08 | 4.438E-09 | 6.200E-09 | 1.500E-08 |
| 28046 | 1403 | L1403E | E | 1.3e-08 | 2.2e-08 | 5.2e-09 | 1.340E-08 | 8.407E-09 | 5.200E-09 | 2.200E-08 |
| 28047 | 1403 | L1403Q | E | 5.8e-09 | 1.1e-08 | 2.9e-09 | 6.567E-09 | 4.104E-09 | 2.900E-09 | 1.100E-08 |
| 28048 | 1403 | L1403G | E | 6.5e-09 | 8.1e-09 | 4.4e-09 | 6.333E-09 | 1.856E-09 | 4.400E-09 | 8.100E-09 |
| 28049 | 1403 | L1403H | E | 6.6e-09 | 1.4e-08 | 4e-09 | 8.200E-09 | 5.188E-09 | 4.000E-09 | 1.400E-08 |

| | | | | | | | | | | |
|---|---|---|---|---|---|---|---|---|---|---|
| 28050 | 1403 | L1403I | E | 5.9e-09 | 1.5e-08 | 1.1e-08 | 1.063E-08 | 4.561E-09 | 5.900E-09 | 1.500E-08 |
| 28051 | 1403 | L1403L | E | 1.1e-08 | 9.7e-09 | 5.8e-09 | 8.833E-09 | 2.706E-09 | 5.800E-09 | 1.100E-08 |
| 28052 | 1403 | L1403K | E | 3.8e-09 | 7.9e-09 | 4.6e-09 | 5.433E-09 | 2.173E-09 | 3.800E-09 | 7.900E-09 |
| 28053 | 1403 | L1403M | E | 1.4e-08 | 1.1e-08 | 6.8e-09 | 1.060E-08 | 3.617E-09 | 6.800E-09 | 1.400E-08 |
| 28054 | 1403 | L1403F | E | 3.1e-09 | 7.9e-09 | 8.1e-09 | 6.367E-09 | 2.831E-09 | 3.100E-09 | 8.100E-09 |
| 28055 | 1403 | L1403P | E | 2.8e-08 | 1.3e-08 | 9.4e-09 | 1.680E-08 | 9.865E-09 | 9.400E-09 | 2.800E-08 |
| 28056 | 1403 | L1403S | E | 1.8e-08 | 7.1e-09 | 1.2e-08 | 1.237E-08 | 5.459E-09 | 7.100E-09 | 1.800E-08 |
| 28057 | 1403 | L1403T | E | 3.1e-09 | 8.5e-09 | 4.3e-09 | 5.300E-09 | 2.835E-09 | 3.100E-09 | 8.500E-09 |
| 28058 | 1403 | L1403W | E | 6e-09 | 1.1e-08 | 1.2e-08 | 9.667E-09 | 3.215E-09 | 6.000E-09 | 1.200E-08 |
| 28059 | 1403 | L1403Y | E | 1.1e-08 | 1.5e-08 | 9.8e-09 | 1.193E-08 | 2.723E-09 | 9.800E-09 | 1.500E-08 |
| 28060 | 1403 | L1403V | E | 1.3e-08 | 2e-08 | 1.7e-08 | 1.667E-08 | 3.512E-09 | 1.300E-08 | 2.000E-08 |
| 28061 | 1404 | A1404A | E | 6.3e-09 | 2e-09 | 6.2e-09 | 4.833E-09 | 2.454E-09 | 2.000E-09 | 6.300E-09 |
| 28062 | 1404 | A1404R | E | 1.4e-08 | 4.3e-09 | 6.2e-09 | 8.167E-09 | 5.140E-09 | 4.300E-09 | 1.400E-08 |
| 28063 | 1404 | A1404N | E | 6.6e-09 | 2e-08 | 8.3e-09 | 1.163E-08 | 7.295E-09 | 6.600E-09 | 2.000E-08 |
| 28064 | 1404 | A1404D | E | 1.4e-08 | 7.1e-09 | 8.1e-09 | 9.733E-09 | 3.729E-09 | 7.100E-09 | 1.400E-08 |
| 28065 | 1404 | A1404C | E | 1.2e-08 | 9.9e-09 | 1e-08 | 1.063E-08 | 1.185E-09 | 9.900E-09 | 1.200E-08 |
| 28066 | 1404 | A1404E | E | 1.2e-09 | 6.7e-09 | 5.2e-09 | 4.367E-09 | 2.843E-09 | 1.200E-09 | 6.700E-09 |
| 28067 | 1404 | A1404Q | E | 1.1e-08 | 1.2e-08 | 4.7e-09 | 9.233E-09 | 3.958E-09 | 4.700E-09 | 1.200E-08 |
| 28068 | 1404 | A1404G | E | 5e-09 | 1.1e-08 | 1.4e-08 | 1.000E-08 | 4.583E-09 | 5.000E-09 | 1.400E-08 |
| 28069 | 1404 | A1404H | E | 4.2e-09 | 2.4e-08 | 1.3e-08 | 1.373E-08 | 9.920E-09 | 4.200E-09 | 2.400E-08 |
| 28070 | 1404 | A1404I | E | 6.5e-09 | 3.5e-09 | 3.4e-09 | 4.467E-09 | 1.762E-09 | 3.400E-09 | 6.500E-09 |
| 28071 | 1404 | A1404L | E | 5.2e-09 | 1.7e-08 | 5.4e-09 | 9.200E-09 | 6.756E-09 | 5.200E-09 | 1.700E-08 |
| 28072 | 1404 | A1404K | E | 9.3e-09 | 6.3e-09 | 1.8e-08 | 1.120E-08 | 6.077E-09 | 6.300E-09 | 1.800E-08 |
| 28073 | 1404 | A1404M | E | 2.3e-08 | 1.7e-08 | 5.1e-09 | 1.503E-08 | 9.111E-09 | 5.100E-09 | 2.300E-08 |
| 28074 | 1404 | A1404F | E | 8.7e-09 | 4.9e-09 | 1.9e-08 | 1.087E-08 | 7.295E-09 | 4.900E-09 | 1.900E-08 |
| 28075 | 1404 | A1404P | E | 8.7e-09 | 6.5e-09 | 5.7e-09 | 6.967E-09 | 1.553E-09 | 5.700E-09 | 8.700E-09 |
| 28076 | 1404 | A1404S | E | 1.7e-09 | 9.1e-09 | 1.5e-08 | 8.600E-09 | 6.664E-09 | 1.700E-09 | 1.500E-08 |
| 28077 | 1404 | A1404T | E | 1.4e-09 | 6.8e-09 | 1.8e-08 | 8.733E-09 | 8.467E-09 | 1.400E-09 | 1.800E-08 |
| 28078 | 1404 | A1404W | E | 4.7e-09 | 2.8e-09 | 6.8e-09 | 4.767E-09 | 2.001E-09 | 2.800E-09 | 6.800E-09 |
| 28079 | 1404 | A1404Y | E | 8e-09 | 5.3e-09 | 1e-08 | 7.767E-09 | 2.359E-09 | 5.300E-09 | 1.000E-08 |
| 28080 | 1404 | A1404V | E | 3.6e-09 | 4.5e-09 | 1.6e-08 | 8.033E-09 | 6.914E-09 | 3.600E-09 | 1.600E-08 |
| 28081 | 1405 | P1405A | E | 1.5e-08 | 4.5e-09 | 2.7e-09 | 7.400E-09 | 6.643E-09 | 2.700E-09 | 1.500E-08 |
| 28082 | 1405 | P1405R | E | 7.3e-09 | 7.7e-09 | 3.7e-09 | 6.233E-09 | 2.203E-09 | 3.700E-09 | 7.700E-09 |
| 28083 | 1405 | P1405N | E | 7.1e-09 | 1.5e-08 | 8.9e-09 | 1.033E-08 | 4.140E-09 | 7.100E-09 | 1.500E-08 |
| 28084 | 1405 | P1405D | E | 1.9e-08 | 3.7e-09 | 1.7e-08 | 1.323E-08 | 8.316E-09 | 3.700E-09 | 1.900E-08 |
| 28085 | 1405 | P1405C | E | 1.2e-08 | 8.8e-09 | 7.5e-09 | 9.433E-09 | 2.316E-09 | 7.500E-09 | 1.200E-08 |
| 28086 | 1405 | P1405E | E | 3.7e-09 | 4.7e-09 | 1.2e-08 | 6.800E-09 | 4.531E-09 | 3.700E-09 | 1.200E-08 |
| 28087 | 1405 | P1405Q | E | 7e-09 | 1.9e-08 | 3.5e-09 | 9.833E-09 | 8.129E-09 | 3.500E-09 | 1.900E-08 |
| 28088 | 1405 | P1405G | E | 4.6e-09 | 2.3e-08 | 6.5e-09 | 1.137E-08 | 1.012E-08 | 4.600E-09 | 2.300E-08 |
| 28089 | 1405 | P1405H | E | 2.3e-09 | 4.3e-09 | 9e-09 | 5.200E-09 | 3.439E-09 | 2.300E-09 | 9.000E-09 |
| 28090 | 1405 | P1405I | E | 5.9e-09 | 8.8e-09 | 1e-08 | 8.233E-09 | 2.108E-09 | 5.900E-09 | 1.000E-08 |
| 28091 | 1405 | P1405L | E | 4.5e-09 | 6e-09 | 6.8e-09 | 5.767E-09 | 1.168E-09 | 4.500E-09 | 6.800E-09 |
| 28092 | 1405 | P1405K | E | 1e-08 | 1.9e-08 | 7.9e-09 | 1.230E-08 | 5.897E-09 | 7.900E-09 | 1.900E-08 |
| 28093 | 1405 | P1405M | E | 6.4e-09 | 4.7e-09 | 4.7e-09 | 5.267E-09 | 9.815E-10 | 4.700E-09 | 6.400E-09 |
| 28094 | 1405 | P1405F | E | 9.4e-09 | 1e-08 | 5.2e-09 | 8.200E-09 | 2.615E-09 | 5.200E-09 | 1.000E-08 |
| 28095 | 1405 | P1405P | E | 7.5e-09 | 8.1e-09 | 1.7e-08 | 1.087E-08 | 5.320E-09 | 7.500E-09 | 1.700E-08 |
| 28096 | 1405 | P1405S | E | 5.1e-09 | 4.7e-09 | 8.7e-09 | 6.167E-09 | 2.203E-09 | 4.700E-09 | 8.700E-09 |
| 28097 | 1405 | P1405T | E | 5.1e-09 | 5.9e-09 | 8.2e-09 | 6.400E-09 | 1.609E-09 | 5.100E-09 | 8.200E-09 |
| 28098 | 1405 | P1405W | E | 1.5e-08 | 8.7e-09 | 9.1e-09 | 1.093E-08 | 3.528E-09 | 8.700E-09 | 1.500E-08 |
| 28099 | 1405 | P1405Y | E | 1.2e-08 | 1e-08 | 9.2e-09 | 1.040E-08 | 1.442E-09 | 9.200E-09 | 1.200E-08 |
| 28100 | 1405 | P1405V | E | 3.9e-09 | 2.2e-08 | 1.1e-08 | 1.230E-08 | 9.120E-09 | 3.900E-09 | 2.200E-08 |
| 28101 | 1406 | S1406A | E | 9.4e-09 | 1.1e-08 | 1.7e-08 | 1.247E-08 | 4.007E-09 | 9.400E-09 | 1.700E-08 |
| 28102 | 1406 | S1406R | E | 1.1e-08 | 3e-08 | 6.4e-09 | 1.580E-08 | 1.251E-08 | 6.400E-09 | 3.000E-08 |
| 28103 | 1406 | S1406N | E | 1.1e-08 | 3e-09 | 1.2e-08 | 8.667E-09 | 4.933E-09 | 3.000E-09 | 1.200E-08 |
| 28104 | 1406 | S1406D | E | 8.7e-09 | 9.2e-09 | 5.5e-09 | 7.800E-09 | 2.007E-09 | 5.500E-09 | 9.200E-09 |
| 28105 | 1406 | S1406C | E | 6.3e-09 | 1.3e-08 | 1.6e-08 | 1.177E-08 | 4.966E-09 | 6.300E-09 | 1.600E-08 |
| 28106 | 1406 | S1406E | E | 8.1e-09 | 7.9e-09 | 1.5e-09 | 5.833E-09 | 3.754E-09 | 1.500E-09 | 8.100E-09 |
| 28107 | 1406 | S1406Q | E | 1.7e-08 | 8.7e-09 | 9.7e-09 | 1.180E-08 | 4.531E-09 | 8.700E-09 | 1.700E-08 |
| 28108 | 1406 | S1406G | E | 1.4e-08 | 2.6e-09 | 4.2e-09 | 6.933E-09 | 6.172E-09 | 2.600E-09 | 1.400E-08 |
| 28109 | 1406 | S1406H | E | 6.3e-09 | 1e-08 | 1.1e-08 | 9.100E-09 | 2.476E-09 | 6.300E-09 | 1.100E-08 |
| 28110 | 1406 | S1406I | E | 1.4e-08 | 2e-08 | 5.8e-09 | 1.327E-08 | 7.128E-09 | 5.800E-09 | 2.000E-08 |
| 28111 | 1406 | S1406L | E | 7.8e-09 | 9.5e-09 | 2.4e-08 | 1.377E-08 | 8.903E-09 | 7.800E-09 | 2.400E-08 |
| 28112 | 1406 | S1406K | E | 1.1e-08 | 1.1e-08 | 1.2e-08 | 1.133E-08 | 5.774E-10 | 1.100E-08 | 1.200E-08 |
| 28113 | 1406 | S1406M | E | 5.3e-09 | 8.5e-09 | 7.9e-09 | 7.233E-09 | 1.701E-09 | 5.300E-09 | 8.500E-09 |
| 28114 | 1406 | S1406F | E | 3.7e-09 | 1.1e-08 | 8.6e-09 | 7.767E-09 | 3.721E-09 | 3.700E-09 | 1.100E-08 |
| 28115 | 1406 | S1406P | E | 1.4e-08 | 2.5e-09 | 1.2e-08 | 9.500E-09 | 6.144E-09 | 2.500E-09 | 1.400E-08 |
| 28116 | 1406 | S1406S | E | 3e-09 | 4.3e-09 | 2.1e-09 | 3.133E-09 | 1.106E-09 | 2.100E-09 | 4.300E-09 |
| 28117 | 1406 | S1406T | E | 6.3e-09 | 1.7e-08 | 5.8e-09 | 9.700E-09 | 6.327E-09 | 5.800E-09 | 1.700E-08 |
| 28118 | 1406 | S1406W | E | 1.5e-08 | 5.5e-09 | 2.4e-08 | 1.483E-08 | 9.251E-09 | 5.500E-09 | 2.400E-08 |
| 28119 | 1406 | S1406Y | E | 2.3e-08 | 8.3e-09 | 3.9e-09 | 1.173E-08 | 1.000E-08 | 3.900E-09 | 2.300E-08 |
| 28120 | 1406 | S1406V | E | 7.7e-09 | 1.7e-08 | 2.9e-09 | 9.200E-09 | 7.169E-09 | 2.900E-09 | 1.700E-08 |
| 28121 | 1407 | S1407A | E | 8.9e-09 | 2.2e-09 | 3.8e-09 | 4.967E-09 | 3.499E-09 | 2.200E-09 | 8.900E-09 |
| 28122 | 1407 | S1407R | E | 7.9e-09 | 7.2e-09 | 1.3e-08 | 9.367E-09 | 3.166E-09 | 7.200E-09 | 1.300E-08 |
| 28123 | 1407 | S1407N | E | 3.6e-09 | 3.4e-09 | 1.3e-08 | 6.667E-09 | 5.486E-09 | 3.400E-09 | 1.300E-08 |
| 28124 | 1407 | S1407D | E | 6.2e-09 | 5.7e-09 | 1.4e-08 | 8.633E-09 | 4.654E-09 | 5.700E-09 | 1.400E-08 |
| 28125 | 1407 | S1407C | E | 1.5e-08 | 2.8e-09 | 6.2e-09 | 8.000E-09 | 6.296E-09 | 2.800E-09 | 1.500E-08 |
| 28126 | 1407 | S1407E | E | 6e-09 | 9.1e-09 | 4.7e-09 | 6.600E-09 | 2.261E-09 | 4.700E-09 | 9.100E-09 |
| 28127 | 1407 | S1407Q | E | 1.2e-08 | 3.2e-09 | 1e-08 | 8.400E-09 | 4.613E-09 | 3.200E-09 | 1.200E-08 |
| 28128 | 1407 | S1407G | E | 8.3e-09 | 2.2e-08 | 4.2e-09 | 1.150E-08 | 9.321E-09 | 4.200E-09 | 2.200E-08 |
| 28129 | 1407 | S1407H | E | 8.2e-09 | 1.7e-08 | 1.2e-08 | 1.240E-08 | 4.414E-09 | 8.200E-09 | 1.700E-08 |
| 28130 | 1407 | S1407I | E | 1.2e-08 | 1.1e-08 | 9.7e-09 | 1.090E-08 | 1.153E-09 | 9.700E-09 | 1.200E-08 |
| 28131 | 1407 | S1407L | E | 1.6e-08 | 8.3e-09 | 5.4e-09 | 9.900E-09 | 5.478E-09 | 5.400E-09 | 1.600E-08 |
| 28132 | 1407 | S1407K | E | 1.2e-08 | 1.1e-08 | 1.5e-08 | 1.267E-08 | 2.082E-09 | 1.100E-08 | 1.500E-08 |
| 28133 | 1407 | S1407M | E | 1.2e-08 | 3.5e-09 | 7.7e-09 | 7.733E-09 | 4.250E-09 | 3.500E-09 | 1.200E-08 |
| 28134 | 1407 | S1407F | E | 1e-08 | 4.7e-09 | 7.4e-09 | 7.367E-09 | 2.650E-09 | 4.700E-09 | 1.000E-08 |

| | | | | | | | | | | |
|---|---|---|---|---|---|---|---|---|---|---|
| 28135 | 1407 | S1407P | E | 2.6e-09 | 3e-09 | 4.2e-09 | 3.267E-09 | 8.327E-10 | 2.600E-09 | 4.200E-09 |
| 28136 | 1407 | S1407S | E | 6.5e-09 | 3.1e-08 | 3.1e-09 | 1.353E-08 | 1.522E-08 | 3.100E-09 | 3.100E-08 |
| 28137 | 1407 | S1407T | E | 3.8e-09 | 6.9e-09 | 5.3e-09 | 5.333E-09 | 1.550E-09 | 3.800E-09 | 6.900E-09 |
| 28138 | 1407 | S1407W | E | 2.1e-09 | 6.3e-09 | 1.6e-08 | 8.133E-09 | 7.129E-09 | 2.100E-09 | 1.600E-08 |
| 28139 | 1407 | S1407Y | E | 5.7e-09 | 1.6e-08 | 2.7e-09 | 8.133E-09 | 6.976E-09 | 2.700E-09 | 1.600E-08 |
| 28140 | 1407 | S1407V | E | 3.7e-09 | 1.2e-08 | 1.2e-08 | 9.233E-09 | 4.792E-09 | 3.700E-09 | 1.200E-08 |
| 28141 | 1408 | K1408A | E | 4.9e-09 | 1.1e-08 | 5.1e-09 | 7.000E-09 | 3.466E-09 | 4.900E-09 | 1.100E-08 |
| 28142 | 1408 | K1408R | E | 9.7e-09 | 9.3e-09 | 7.2e-09 | 8.733E-09 | 1.343E-09 | 7.200E-09 | 9.700E-09 |
| 28143 | 1408 | K1408N | E | 7.4e-09 | 9.2e-09 | 2.7e-09 | 6.433E-09 | 3.356E-09 | 2.700E-09 | 9.200E-09 |
| 28144 | 1408 | K1408D | E | 4.5e-09 | 1.1e-08 | 7.4e-09 | 7.633E-09 | 3.256E-09 | 4.500E-09 | 1.100E-08 |
| 28145 | 1408 | K1408C | E | 6e-09 | 1.3e-08 | 5.1e-09 | 8.033E-09 | 4.325E-09 | 5.100E-09 | 1.300E-08 |
| 28146 | 1408 | K1408E | E | 6.4e-09 | 5.5e-09 | 6.2e-09 | 6.033E-09 | 4.726E-10 | 5.500E-09 | 6.400E-09 |
| 28147 | 1408 | K1408Q | E | 3.2e-09 | 5.4e-09 | 1e-08 | 6.200E-09 | 3.470E-09 | 3.200E-09 | 1.000E-08 |
| 28148 | 1408 | K1408G | E | 7.7e-09 | 8.7e-09 | 6.4e-09 | 7.600E-09 | 1.153E-09 | 6.400E-09 | 8.700E-09 |
| 28149 | 1408 | K1408H | E | 5.8e-09 | 1.4e-08 | 4.7e-09 | 8.167E-09 | 5.082E-09 | 4.700E-09 | 1.400E-08 |
| 28150 | 1408 | K1408I | E | 8.2e-09 | 1.3e-08 | 4.4e-09 | 8.533E-09 | 4.310E-09 | 4.400E-09 | 1.300E-08 |
| 28151 | 1408 | K1408L | E | 3.8e-08 | 8e-09 | 4.1e-09 | 1.670E-08 | 1.855E-08 | 4.100E-09 | 3.800E-08 |
| 28152 | 1408 | K1408K | E | 6.9e-09 | 3e-09 | 8.8e-09 | 6.233E-09 | 2.957E-09 | 3.000E-09 | 8.800E-09 |
| 28153 | 1408 | K1408M | E | 8e-09 | 4.5e-09 | 7.2e-09 | 6.567E-09 | 1.834E-09 | 4.500E-09 | 8.000E-09 |
| 28154 | 1408 | K1408F | E | 6.9e-09 | 5.3e-09 | 4.8e-09 | 5.667E-09 | 1.097E-09 | 4.800E-09 | 6.900E-09 |
| 28155 | 1408 | K1408P | E | 5.8e-09 | 4.2e-09 | 8.9e-09 | 6.300E-09 | 2.390E-09 | 4.200E-09 | 8.900E-09 |
| 28156 | 1408 | K1408S | E | 1.5e-08 | 4.8e-09 | 1.2e-08 | 1.060E-08 | 5.242E-09 | 4.800E-09 | 1.500E-08 |
| 28157 | 1408 | K1408T | E | 2.3e-09 | 5.1e-09 | 4.8e-09 | 4.067E-09 | 1.537E-09 | 2.300E-09 | 5.100E-09 |
| 28158 | 1408 | K1408W | E | 5.8e-09 | 4.7e-09 | 2.7e-09 | 4.400E-09 | 1.572E-09 | 2.700E-09 | 5.800E-09 |
| 28159 | 1408 | K1408Y | E | 8e-09 | 1.6e-08 | 7.3e-09 | 1.043E-08 | 4.834E-09 | 7.300E-09 | 1.600E-08 |
| 28160 | 1408 | K1408V | E | 1.4e-08 | 9.9e-09 | 5.2e-09 | 9.700E-09 | 4.403E-09 | 5.200E-09 | 1.400E-08 |
| 28161 | 1409 | S1409A | E | 6e-09 | 1.2e-08 | 8.5e-09 | 8.833E-09 | 3.014E-09 | 6.000E-09 | 1.200E-08 |
| 28162 | 1409 | S1409R | E | 1.4e-08 | 6.2e-09 | 1.2e-08 | 1.073E-08 | 4.051E-09 | 6.200E-09 | 1.400E-08 |
| 28163 | 1409 | S1409N | E | 3.8e-09 | 3.4e-09 | 1.6e-08 | 7.733E-09 | 7.162E-09 | 3.400E-09 | 1.600E-08 |
| 28164 | 1409 | S1409D | E | 6.7e-09 | 4.6e-09 | 2.1e-09 | 4.467E-09 | 2.303E-09 | 2.100E-09 | 6.700E-09 |
| 28165 | 1409 | S1409C | E | 4.5e-09 | 4.2e-09 | 9.1e-09 | 5.933E-09 | 2.747E-09 | 4.200E-09 | 9.100E-09 |
| 28166 | 1409 | S1409E | E | 3.3e-09 | 2.4e-08 | 5.3e-09 | 1.087E-08 | 1.142E-08 | 3.300E-09 | 2.400E-08 |
| 28167 | 1409 | S1409Q | E | 3.5e-09 | 9.8e-09 | 1.2e-09 | 4.833E-09 | 4.452E-09 | 1.200E-09 | 9.800E-09 |
| 28168 | 1409 | S1409G | E | 1.3e-08 | 1.7e-08 | 4.6e-09 | 1.153E-08 | 6.329E-09 | 4.600E-09 | 1.700E-08 |
| 28169 | 1409 | S1409H | E | 8.1e-09 | 8.5e-09 | 5.7e-09 | 7.433E-09 | 1.514E-09 | 5.700E-09 | 8.500E-09 |
| 28170 | 1409 | S1409I | E | 4.2e-09 | 7.3e-09 | 8.5e-09 | 6.667E-09 | 2.219E-09 | 4.200E-09 | 8.500E-09 |
| 28171 | 1409 | S1409L | E | 3.1e-08 | 6.2e-09 | 5.5e-09 | 1.423E-08 | 1.452E-08 | 5.500E-09 | 3.100E-08 |
| 28172 | 1409 | S1409K | E | 6.3e-09 | 8.9e-09 | 1.3e-08 | 9.400E-09 | 3.378E-09 | 6.300E-09 | 1.300E-08 |
| 28173 | 1409 | S1409M | E | 5.8e-09 | 8.8e-09 | 1.3e-08 | 9.200E-09 | 3.617E-09 | 5.800E-09 | 1.300E-08 |
| 28174 | 1409 | S1409F | E | 2.9e-09 | 1.9e-09 | 5.6e-09 | 3.467E-09 | 1.914E-09 | 1.900E-09 | 5.600E-09 |
| 28175 | 1409 | S1409P | E | 2.3e-09 | 1.1e-08 | 1e-08 | 7.767E-09 | 4.761E-09 | 2.300E-09 | 1.100E-08 |
| 28176 | 1409 | S1409S | E | 7.4e-09 | 3.5e-09 | 1.3e-08 | 7.967E-09 | 4.775E-09 | 3.500E-09 | 1.300E-08 |
| 28177 | 1409 | S1409T | E | 2.6e-09 | 1.1e-08 | 2.8e-09 | 5.467E-09 | 4.793E-09 | 2.600E-09 | 1.100E-08 |
| 28178 | 1409 | S1409W | E | 1.1e-08 | 6.6e-09 | 1e-08 | 9.200E-09 | 2.307E-09 | 6.600E-09 | 1.100E-08 |
| 28179 | 1409 | S1409Y | E | 1.2e-08 | 2.1e-09 | 1.3e-08 | 9.033E-09 | 6.025E-09 | 2.100E-09 | 1.300E-08 |
| 28180 | 1409 | S1409V | E | 3.7e-09 | 9.6e-09 | 1.5e-08 | 9.433E-09 | 5.652E-09 | 3.700E-09 | 1.500E-08 |
| 28181 | 1410 | T1410A | E | 5e-09 | 4.9e-09 | 8.9e-09 | 6.267E-09 | 2.281E-09 | 4.900E-09 | 8.900E-09 |
| 28182 | 1410 | T1410R | E | 8.8e-09 | 4.1e-09 | 2.5e-09 | 5.133E-09 | 3.275E-09 | 2.500E-09 | 8.800E-09 |
| 28183 | 1410 | T1410N | E | 1.8e-08 | 1.5e-08 | 2.4e-09 | 1.180E-08 | 8.278E-09 | 2.400E-09 | 1.800E-08 |
| 28184 | 1410 | T1410D | E | 9.5e-09 | 3.6e-09 | 5.8e-09 | 6.300E-09 | 2.982E-09 | 3.600E-09 | 9.500E-09 |
| 28185 | 1410 | T1410C | E | 2.3e-09 | 8.7e-09 | 9.1e-09 | 6.700E-09 | 3.816E-09 | 2.300E-09 | 9.100E-09 |
| 28186 | 1410 | T1410E | E | 1.2e-08 | 2.4e-09 | 6.1e-09 | 6.833E-09 | 4.842E-09 | 2.400E-09 | 1.200E-08 |
| 28187 | 1410 | T1410Q | E | 7.6e-09 | 3.3e-09 | 8e-09 | 6.300E-09 | 2.606E-09 | 3.300E-09 | 8.000E-09 |
| 28188 | 1410 | T1410G | E | 1.4e-08 | 4.1e-09 | 4e-09 | 7.367E-09 | 5.745E-09 | 4.000E-09 | 1.400E-08 |
| 28189 | 1410 | T1410H | E | 3.3e-09 | 5.9e-09 | 5.2e-09 | 4.800E-09 | 1.345E-09 | 3.300E-09 | 5.900E-09 |
| 28190 | 1410 | T1410I | E | 2.5e-08 | 8.7e-09 | 4.3e-09 | 1.267E-08 | 1.091E-08 | 4.300E-09 | 2.500E-08 |
| 28191 | 1410 | T1410L | E | 1.8e-08 | 4.2e-09 | 8.8e-09 | 1.033E-08 | 7.027E-09 | 4.200E-09 | 1.800E-08 |
| 28192 | 1410 | T1410K | E | 2.6e-09 | 5.5e-09 | 1.1e-08 | 6.367E-09 | 4.267E-09 | 2.600E-09 | 1.100E-08 |
| 28193 | 1410 | T1410M | E | 1.3e-08 | 5.2e-09 | 1.2e-08 | 1.007E-08 | 4.244E-09 | 5.200E-09 | 1.300E-08 |
| 28194 | 1410 | T1410F | E | 4e-09 | 3.7e-09 | 1.1e-08 | 6.233E-09 | 4.131E-09 | 3.700E-09 | 1.100E-08 |
| 28195 | 1410 | T1410P | E | 3.5e-09 | 9.7e-09 | 4.7e-09 | 5.967E-09 | 3.288E-09 | 3.500E-09 | 9.700E-09 |
| 28196 | 1410 | T1410S | E | 4.3e-09 | 6.7e-09 | 3.2e-09 | 4.733E-09 | 1.790E-09 | 3.200E-09 | 6.700E-09 |
| 28197 | 1410 | T1410T | E | 7.3e-09 | 8.3e-09 | 5.7e-09 | 7.100E-09 | 1.311E-09 | 5.700E-09 | 8.300E-09 |
| 28198 | 1410 | T1410W | E | 2e-09 | 1e-08 | 1e-08 | 7.333E-09 | 4.619E-09 | 2.000E-09 | 1.000E-08 |
| 28199 | 1410 | T1410Y | E | 7.5e-09 | 4.5e-09 | 1.2e-08 | 8.000E-09 | 3.775E-09 | 4.500E-09 | 1.200E-08 |
| 28200 | 1410 | T1410V | E | 7.6e-09 | 1.2e-08 | 8.8e-09 | 9.467E-09 | 2.274E-09 | 7.600E-09 | 1.200E-08 |
| 28201 | 1411 | S1411A | E | 1.6e-09 | 4.8e-09 | 5.2e-09 | 3.867E-09 | 1.973E-09 | 1.600E-09 | 5.200E-09 |
| 28202 | 1411 | S1411R | E | 1.9e-08 | 9.2e-09 | 1.1e-08 | 1.307E-08 | 5.217E-09 | 9.200E-09 | 1.900E-08 |
| 28203 | 1411 | S1411N | E | 9.4e-09 | 5.9e-09 | 6.6e-09 | 7.300E-09 | 1.852E-09 | 5.900E-09 | 9.400E-09 |
| 28204 | 1411 | S1411D | E | 2.1e-09 | 6.4e-09 | 6.1e-09 | 4.867E-09 | 2.401E-09 | 2.100E-09 | 6.400E-09 |
| 28205 | 1411 | S1411C | E | 8.1e-09 | 8.1e-09 | 5.6e-09 | 7.267E-09 | 1.443E-09 | 5.600E-09 | 8.100E-09 |
| 28206 | 1411 | S1411E | E | 4.3e-09 | 1.2e-08 | 2.8e-08 | 1.477E-08 | 1.209E-08 | 4.300E-09 | 2.800E-08 |
| 28207 | 1411 | S1411Q | E | 3e-09 | 9.8e-09 | 4.3e-09 | 5.700E-09 | 3.610E-09 | 3.000E-09 | 9.800E-09 |
| 28208 | 1411 | S1411G | E | 4.1e-09 | 1.4e-08 | 4.8e-09 | 7.633E-09 | 5.525E-09 | 4.100E-09 | 1.400E-08 |
| 28209 | 1411 | S1411H | E | 4.9e-09 | 2e-09 | 7.4e-09 | 4.767E-09 | 2.702E-09 | 2.000E-09 | 7.400E-09 |
| 28210 | 1411 | S1411I | E | 1.1e-08 | 1.3e-08 | 5.4e-09 | 9.800E-09 | 3.940E-09 | 5.400E-09 | 1.300E-08 |
| 28211 | 1411 | S1411L | E | 3.5e-09 | 8e-09 | 1.2e-08 | 7.833E-09 | 4.252E-09 | 3.500E-09 | 1.200E-08 |
| 28212 | 1411 | S1411K | E | 2.5e-09 | 7.2e-09 | 6.8e-09 | 5.500E-09 | 2.606E-09 | 2.500E-09 | 7.200E-09 |
| 28213 | 1411 | S1411M | E | 2.6e-09 | 1.2e-08 | 8.7e-09 | 7.767E-09 | 4.769E-09 | 2.600E-09 | 1.200E-08 |
| 28214 | 1411 | S1411F | E | 2.6e-08 | 9.7e-09 | 5.6e-09 | 1.377E-08 | 1.079E-08 | 5.600E-09 | 2.600E-08 |
| 28215 | 1411 | S1411P | E | 6.6e-09 | 1.2e-08 | 5.8e-09 | 8.133E-09 | 3.372E-09 | 5.800E-09 | 1.200E-08 |
| 28216 | 1411 | S1411S | E | 1.3e-08 | 8.8e-09 | 1.6e-08 | 1.260E-08 | 3.617E-09 | 8.800E-09 | 1.600E-08 |
| 28217 | 1411 | S1411T | E | 1.1e-08 | 3.5e-09 | 1.1e-08 | 8.500E-09 | 4.330E-09 | 3.500E-09 | 1.100E-08 |
| 28218 | 1411 | S1411W | E | 1.5e-08 | 4.5e-09 | 8.1e-09 | 9.200E-09 | 5.336E-09 | 4.500E-09 | 1.500E-08 |
| 28219 | 1411 | S1411Y | E | 7.1e-09 | 1.3e-08 | 8e-09 | 9.367E-09 | 3.179E-09 | 7.100E-09 | 1.300E-08 |

| | | | | | | | | | | |
|---|---|---|---|---|---|---|---|---|---|---|
| 28220 | 1411 | S1411V | E | 1.9e-08 | 5.7e-09 | 8.1e-09 | 1.093E-08 | 7.088E-09 | 5.700E-09 | 1.900E-08 |
| 28221 | 1412 | G1412A | E | 4.2e-09 | 7.3e-09 | 3.7e-09 | 5.067E-09 | 1.950E-09 | 3.700E-09 | 7.300E-09 |
| 28222 | 1412 | G1412R | E | 4.2e-09 | 7e-09 | 9.8e-09 | 7.000E-09 | 2.800E-09 | 4.200E-09 | 9.800E-09 |
| 28223 | 1412 | G1412N | E | 4.8e-09 | 5e-09 | 6.3e-09 | 5.367E-09 | 8.145E-10 | 4.800E-09 | 6.300E-09 |
| 28224 | 1412 | G1412D | E | 6.9e-09 | 6.2e-09 | 4.1e-09 | 5.733E-09 | 1.457E-09 | 4.100E-09 | 6.900E-09 |
| 28225 | 1412 | G1412C | E | 4.9e-09 | 4.3e-09 | 5.2e-09 | 4.800E-09 | 4.583E-10 | 4.300E-09 | 5.200E-09 |
| 28226 | 1412 | G1412E | E | 1.2e-08 | 1.3e-08 | 6.2e-09 | 1.040E-08 | 3.672E-09 | 6.200E-09 | 1.300E-08 |
| 28227 | 1412 | G1412Q | E | 1e-08 | 7.3e-09 | 7.4e-09 | 8.233E-09 | 1.531E-09 | 7.300E-09 | 1.000E-08 |
| 28228 | 1412 | G1412G | E | 1.1e-08 | 9.5e-09 | 3e-09 | 7.833E-09 | 4.252E-09 | 3.000E-09 | 1.100E-08 |
| 28229 | 1412 | G1412H | E | 9.9e-09 | 1.4e-08 | 2.4e-09 | 8.767E-09 | 5.882E-09 | 2.400E-09 | 1.400E-08 |
| 28230 | 1412 | G1412I | E | 4.9e-09 | 8e-09 | 6.7e-09 | 6.533E-09 | 1.557E-09 | 4.900E-09 | 8.000E-09 |
| 28231 | 1412 | G1412L | E | 1.1e-08 | 1.3e-08 | 3.3e-09 | 9.100E-09 | 5.122E-09 | 3.300E-09 | 1.300E-08 |
| 28232 | 1412 | G1412K | E | 9.6e-09 | 7e-09 | 1.1e-08 | 9.200E-09 | 2.030E-09 | 7.000E-09 | 1.100E-08 |
| 28233 | 1412 | G1412M | E | 5e-09 | 3e-09 | 9.5e-09 | 5.833E-09 | 3.329E-09 | 3.000E-09 | 9.500E-09 |
| 28234 | 1412 | G1412F | E | 1.5e-08 | 5.7e-09 | 3.7e-09 | 8.133E-09 | 6.030E-09 | 3.700E-09 | 1.500E-08 |
| 28235 | 1412 | G1412P | E | 1.2e-08 | 6.9e-09 | 8.5e-09 | 9.133E-09 | 2.608E-09 | 6.900E-09 | 1.200E-08 |
| 28236 | 1412 | G1412S | E | 7.7e-09 | 1.8e-09 | 6.6e-09 | 5.367E-09 | 3.137E-09 | 1.800E-09 | 7.700E-09 |
| 28237 | 1412 | G1412T | E | 3.6e-09 | 1e-08 | 2e-08 | 1.120E-08 | 8.266E-09 | 3.600E-09 | 2.000E-08 |
| 28238 | 1412 | G1412W | E | 3.4e-09 | 5.2e-09 | 1.2e-08 | 6.867E-09 | 4.536E-09 | 3.400E-09 | 1.200E-08 |
| 28239 | 1412 | G1412Y | E | 8.9e-09 | 5.7e-09 | 1.2e-08 | 8.867E-09 | 3.150E-09 | 5.700E-09 | 1.200E-08 |
| 28240 | 1412 | G1412V | E | 4.1e-09 | 2e-08 | 4.2e-08 | 2.203E-08 | 1.903E-08 | 4.100E-09 | 4.200E-08 |
| 28241 | 1413 | G1413A | E | 2.4e-09 | 1.4e-08 | 3.9e-09 | 6.767E-09 | 6.309E-09 | 2.400E-09 | 1.400E-08 |
| 28242 | 1413 | G1413R | E | 7e-09 | 9.2e-09 | 8.9e-09 | 8.367E-09 | 1.193E-09 | 7.000E-09 | 9.200E-09 |
| 28243 | 1413 | G1413N | E | 1.7e-08 | 7e-09 | 9.9e-09 | 1.130E-08 | 5.145E-09 | 7.000E-09 | 1.700E-08 |
| 28244 | 1413 | G1413D | E | 1.5e-08 | 1.6e-08 | 1.3e-08 | 1.467E-08 | 1.528E-09 | 1.300E-08 | 1.600E-08 |
| 28245 | 1413 | G1413C | E | 4e-09 | 5.5e-09 | 8.1e-09 | 5.867E-09 | 2.074E-09 | 4.000E-09 | 8.100E-09 |
| 28246 | 1413 | G1413E | E | 8.5e-09 | 5.1e-09 | 1.6e-08 | 9.867E-09 | 5.577E-09 | 5.100E-09 | 1.600E-08 |
| 28247 | 1413 | G1413Q | E | 1.4e-08 | 4.6e-09 | 6.9e-09 | 8.500E-09 | 4.900E-09 | 4.600E-09 | 1.400E-08 |
| 28248 | 1413 | G1413G | E | 6.2e-09 | 2e-08 | 1.3e-08 | 1.307E-08 | 6.900E-09 | 6.200E-09 | 2.000E-08 |
| 28249 | 1413 | G1413H | E | 4.1e-09 | 9.6e-09 | 1.5e-08 | 9.567E-09 | 5.450E-09 | 4.100E-09 | 1.500E-08 |
| 28250 | 1413 | G1413I | E | 6.6e-09 | 1.1e-08 | 6.6e-09 | 8.067E-09 | 2.540E-09 | 6.600E-09 | 1.100E-08 |
| 28251 | 1413 | G1413L | E | 4.8e-09 | 1.3e-08 | 1.2e-08 | 9.933E-09 | 4.474E-09 | 4.800E-09 | 1.300E-08 |
| 28252 | 1413 | G1413K | E | 2.5e-09 | 7.2e-09 | 2.9e-09 | 4.200E-09 | 2.606E-09 | 2.500E-09 | 7.200E-09 |
| 28253 | 1413 | G1413M | E | 3.1e-09 | 6.6e-09 | 1.3e-08 | 7.567E-09 | 5.020E-09 | 3.100E-09 | 1.300E-08 |
| 28254 | 1413 | G1413F | E | 6.3e-09 | 3.5e-09 | 3.4e-09 | 4.400E-09 | 1.646E-09 | 3.400E-09 | 6.300E-09 |
| 28255 | 1413 | G1413P | E | 3.8e-09 | 8.5e-09 | 4.9e-09 | 5.733E-09 | 2.458E-09 | 3.800E-09 | 8.500E-09 |
| 28256 | 1413 | G1413S | E | 8.6e-09 | 5e-09 | 9.6e-09 | 7.733E-09 | 2.419E-09 | 5.000E-09 | 9.600E-09 |
| 28257 | 1413 | G1413T | E | 5.1e-09 | 5.3e-09 | 8.2e-09 | 6.200E-09 | 1.735E-09 | 5.100E-09 | 8.200E-09 |
| 28258 | 1413 | G1413W | E | 8e-09 | 1.3e-08 | 6.5e-09 | 9.167E-09 | 3.403E-09 | 6.500E-09 | 1.300E-08 |
| 28259 | 1413 | G1413Y | E | 2.1e-09 | 9.6e-09 | 1.1e-08 | 7.567E-09 | 4.786E-09 | 2.100E-09 | 1.100E-08 |
| 28260 | 1413 | G1413V | E | 7e-09 | 6.4e-09 | 6.3e-09 | 6.567E-09 | 3.786E-10 | 6.300E-09 | 7.000E-09 |
| 28261 | 1414 | T1414A | E | 1.4e-08 | 3.2e-09 | 2.4e-09 | 6.533E-09 | 6.479E-09 | 2.400E-09 | 1.400E-08 |
| 28262 | 1414 | T1414R | E | 7.7e-09 | 5.1e-09 | 3.4e-09 | 5.400E-09 | 2.166E-09 | 3.400E-09 | 7.700E-09 |
| 28263 | 1414 | T1414N | E | 6.8e-09 | 2e-08 | 2.7e-09 | 9.833E-09 | 9.040E-09 | 2.700E-09 | 2.000E-08 |
| 28264 | 1414 | T1414D | E | 4.1e-09 | 2.8e-08 | 3.9e-09 | 1.200E-08 | 1.386E-08 | 3.900E-09 | 2.800E-08 |
| 28265 | 1414 | T1414C | E | 6.2e-09 | 1.5e-09 | 2.2e-09 | 3.300E-09 | 2.536E-09 | 1.500E-09 | 6.200E-09 |
| 28266 | 1414 | T1414E | E | 3.5e-09 | 4e-09 | 3.5e-09 | 3.667E-09 | 2.887E-10 | 3.500E-09 | 4.000E-09 |
| 28267 | 1414 | T1414Q | E | 2.7e-09 | 2.7e-09 | 1.3e-08 | 6.133E-09 | 5.947E-09 | 2.700E-09 | 1.300E-08 |
| 28268 | 1414 | T1414G | E | 3.4e-09 | 1.4e-08 | 6.4e-09 | 7.933E-09 | 5.464E-09 | 3.400E-09 | 1.400E-08 |
| 28269 | 1414 | T1414H | E | 7e-09 | 3.4e-09 | 1.3e-08 | 7.800E-09 | 4.850E-09 | 3.400E-09 | 1.300E-08 |
| 28270 | 1414 | T1414I | E | 7.3e-09 | 3.6e-09 | 6.2e-09 | 5.700E-09 | 1.900E-09 | 3.600E-09 | 7.300E-09 |
| 28271 | 1414 | T1414L | E | 5.9e-09 | 3.3e-09 | 7.4e-09 | 5.533E-09 | 2.074E-09 | 3.300E-09 | 7.400E-09 |
| 28272 | 1414 | T1414K | E | 1.5e-09 | 4.7e-09 | 4.7e-09 | 3.633E-09 | 1.848E-09 | 1.500E-09 | 4.700E-09 |
| 28273 | 1414 | T1414M | E | 2.5e-09 | 3.5e-09 | 4.6e-09 | 3.533E-09 | 1.050E-09 | 2.500E-09 | 4.600E-09 |
| 28274 | 1414 | T1414F | E | 4.6e-09 | 5.8e-09 | 2.6e-09 | 4.333E-09 | 1.617E-09 | 2.600E-09 | 5.800E-09 |
| 28275 | 1414 | T1414P | E | 1.1e-08 | 3.7e-09 | 6.4e-09 | 7.033E-09 | 3.691E-09 | 3.700E-09 | 1.100E-08 |
| 28276 | 1414 | T1414S | E | 3.5e-09 | 9.2e-09 | 2.7e-09 | 5.133E-09 | 3.544E-09 | 2.700E-09 | 9.200E-09 |
| 28277 | 1414 | T1414T | E | 4e-09 | 8.1e-09 | 1e-08 | 7.367E-09 | 3.066E-09 | 4.000E-09 | 1.000E-08 |
| 28278 | 1414 | T1414W | E | 1.7e-08 | 8.4e-09 | 1e-08 | 1.180E-08 | 4.574E-09 | 8.400E-09 | 1.700E-08 |
| 28279 | 1414 | T1414Y | E | 3.8e-09 | 4.6e-09 | 7.3e-09 | 5.233E-09 | 1.834E-09 | 3.800E-09 | 7.300E-09 |
| 28280 | 1414 | T1414V | E | 4.8e-09 | 7.7e-09 | 1.2e-08 | 8.167E-09 | 3.623E-09 | 4.800E-09 | 1.200E-08 |
| 28281 | 1415 | A1415A | E | 5.7e-09 | 8.6e-09 | 7.7e-09 | 7.333E-09 | 1.484E-09 | 5.700E-09 | 8.600E-09 |
| 28282 | 1415 | A1415R | E | 5.9e-09 | 1.1e-08 | 4.6e-09 | 7.167E-09 | 3.383E-09 | 4.600E-09 | 1.100E-08 |
| 28283 | 1415 | A1415N | E | 1.2e-08 | 6.8e-09 | 1.8e-08 | 1.227E-08 | 5.605E-09 | 6.800E-09 | 1.800E-08 |
| 28284 | 1415 | A1415D | E | 6e-10 | 2e-08 | 7.4e-09 | 9.333E-09 | 9.843E-09 | 6.000E-10 | 2.000E-08 |
| 28285 | 1415 | A1415C | E | 2e-09 | 1.1e-08 | 5.8e-09 | 6.267E-09 | 4.518E-09 | 2.000E-09 | 1.100E-08 |
| 28286 | 1415 | A1415E | E | 8.3e-09 | 4.8e-09 | 8e-09 | 7.033E-09 | 1.940E-09 | 4.800E-09 | 8.300E-09 |
| 28287 | 1415 | A1415Q | E | 4.5e-09 | 6.8e-09 | 7.8e-09 | 6.367E-09 | 1.692E-09 | 4.500E-09 | 7.800E-09 |
| 28288 | 1415 | A1415G | E | 1.5e-08 | 3.3e-08 | 5.6e-09 | 1.787E-08 | 1.392E-08 | 5.600E-09 | 3.300E-08 |
| 28289 | 1415 | A1415H | E | 1.9e-08 | 4.7e-09 | 9e-09 | 1.090E-08 | 7.337E-09 | 4.700E-09 | 1.900E-08 |
| 28290 | 1415 | A1415I | E | 4.5e-09 | 3e-09 | 3e-09 | 3.500E-09 | 8.660E-10 | 3.000E-09 | 4.500E-09 |
| 28291 | 1415 | A1415L | E | 9e-09 | 6.9e-09 | 7.5e-09 | 7.800E-09 | 1.082E-09 | 6.900E-09 | 9.000E-09 |
| 28292 | 1415 | A1415K | E | 6.3e-09 | 9.5e-09 | 2.8e-09 | 6.200E-09 | 3.351E-09 | 2.800E-09 | 9.500E-09 |
| 28293 | 1415 | A1415M | E | 7.3e-09 | 3.7e-09 | 6.1e-09 | 5.700E-09 | 1.833E-09 | 3.700E-09 | 7.300E-09 |
| 28294 | 1415 | A1415F | E | 3.8e-09 | 4.3e-09 | 1e-08 | 6.033E-09 | 3.444E-09 | 3.800E-09 | 1.000E-08 |
| 28295 | 1415 | A1415P | E | 1.2e-09 | 7.4e-09 | 4.3e-09 | 4.300E-09 | 3.100E-09 | 1.200E-09 | 7.400E-09 |
| 28296 | 1415 | A1415S | E | 3.9e-09 | 8.8e-09 | 4.2e-09 | 5.633E-09 | 2.747E-09 | 3.900E-09 | 8.800E-09 |
| 28297 | 1415 | A1415T | E | 7.1e-09 | 1.2e-08 | 5.2e-09 | 8.100E-09 | 3.509E-09 | 5.200E-09 | 1.200E-08 |
| 28298 | 1415 | A1415W | E | 6.8e-09 | 6.3e-09 | 1.2e-08 | 8.367E-09 | 3.156E-09 | 6.300E-09 | 1.200E-08 |
| 28299 | 1415 | A1415Y | E | 7.5e-09 | 1.8e-08 | 4.7e-09 | 1.007E-08 | 7.012E-09 | 4.700E-09 | 1.800E-08 |
| 28300 | 1415 | A1415V | E | 1.2e-08 | 1.5e-08 | 1.1e-08 | 1.267E-08 | 2.082E-09 | 1.100E-08 | 1.500E-08 |
| 28301 | 1416 | A1416A | E | 7.2e-09 | 2.2e-08 | 2.4e-09 | 1.053E-08 | 1.022E-08 | 2.400E-09 | 2.200E-08 |
| 28302 | 1416 | A1416R | E | 8.8e-09 | 1.2e-08 | 7.3e-09 | 9.367E-09 | 2.401E-09 | 7.300E-09 | 1.200E-08 |
| 28303 | 1416 | A1416N | E | 1.1e-08 | 2.3e-09 | 9.2e-09 | 7.500E-09 | 4.592E-09 | 2.300E-09 | 1.100E-08 |
| 28304 | 1416 | A1416D | E | 7.1e-09 | 7.5e-09 | 7e-09 | 7.200E-09 | 2.646E-10 | 7.000E-09 | 7.500E-09 |

| | | | | | | | | | | |
|---|---|---|---|---|---|---|---|---|---|---|
| 28305 | 1416 | A1416C | E | 8.1e-09 | 1e-08 | 1.1e-08 | 9.700E-09 | 1.473E-09 | 8.100E-09 | 1.100E-08 |
| 28306 | 1416 | A1416E | E | 1e-08 | 7.5e-09 | 1.3e-08 | 1.017E-08 | 2.754E-09 | 7.500E-09 | 1.300E-08 |
| 28307 | 1416 | A1416Q | E | 7.5e-09 | 9e-09 | 7.8e-09 | 8.100E-09 | 7.937E-10 | 7.500E-09 | 9.000E-09 |
| 28308 | 1416 | A1416G | E | 8.2e-09 | 6.9e-09 | 2.9e-09 | 6.000E-09 | 2.762E-09 | 2.900E-09 | 8.200E-09 |
| 28309 | 1416 | A1416H | E | 6.9e-09 | 1.1e-08 | 8.9e-09 | 8.933E-09 | 2.050E-09 | 6.900E-09 | 1.100E-08 |
| 28310 | 1416 | A1416I | E | 8.5e-09 | 3.6e-09 | 6.2e-09 | 6.100E-09 | 2.452E-09 | 3.600E-09 | 8.500E-09 |
| 28311 | 1416 | A1416L | E | 5.2e-09 | 5e-09 | 1.3e-08 | 7.733E-09 | 4.562E-09 | 5.000E-09 | 1.300E-08 |
| 28312 | 1416 | A1416K | E | 4.1e-09 | 8.7e-09 | 9.9e-09 | 7.567E-09 | 3.062E-09 | 4.100E-09 | 9.900E-09 |
| 28313 | 1416 | A1416M | E | 1.1e-08 | 8.7e-09 | 3.3e-09 | 7.667E-09 | 3.953E-09 | 3.300E-09 | 1.100E-08 |
| 28314 | 1416 | A1416F | E | 1.1e-08 | 4.9e-09 | 7.5e-09 | 7.800E-09 | 3.061E-09 | 4.900E-09 | 1.100E-08 |
| 28315 | 1416 | A1416P | E | 1e-08 | 3.3e-09 | 1.6e-08 | 9.767E-09 | 6.353E-09 | 3.300E-09 | 1.600E-08 |
| 28316 | 1416 | A1416S | E | 5e-09 | 5.5e-09 | 5.4e-09 | 5.300E-09 | 2.646E-10 | 5.000E-09 | 5.500E-09 |
| 28317 | 1416 | A1416T | E | 2e-08 | 4e-09 | 3.8e-09 | 9.267E-09 | 9.296E-09 | 3.800E-09 | 2.000E-08 |
| 28318 | 1416 | A1416W | E | 3.7e-09 | 2.4e-09 | 4e-09 | 3.367E-09 | 8.505E-10 | 2.400E-09 | 4.000E-09 |
| 28319 | 1416 | A1416Y | E | 5.4e-09 | 9.8e-09 | 6.2e-09 | 7.133E-09 | 2.344E-09 | 5.400E-09 | 9.800E-09 |
| 28320 | 1416 | A1416V | E | 1.2e-08 | 3.1e-08 | 7.7e-09 | 1.690E-08 | 1.240E-08 | 7.700E-09 | 3.100E-08 |
| 28321 | 1417 | L1417A | E | 1.8e-09 | 1.4e-08 | 1.8e-08 | 1.127E-08 | 8.439E-09 | 1.800E-09 | 1.800E-08 |
| 28322 | 1417 | L1417R | E | 5.6e-09 | 6e-09 | 1.2e-08 | 7.867E-09 | 3.585E-09 | 5.600E-09 | 1.200E-08 |
| 28323 | 1417 | L1417N | E | 3.1e-08 | 1e-08 | 7.8e-09 | 1.627E-08 | 1.281E-08 | 7.800E-09 | 3.100E-08 |
| 28324 | 1417 | L1417D | E | 2.7e-08 | 3.6e-09 | 5.5e-09 | 1.203E-08 | 1.300E-08 | 3.600E-09 | 2.700E-08 |
| 28325 | 1417 | L1417C | E | 9.5e-09 | 6e-09 | 9.9e-09 | 8.467E-09 | 2.146E-09 | 6.000E-09 | 9.900E-09 |
| 28326 | 1417 | L1417E | E | 8.8e-09 | 3.6e-09 | 4.9e-09 | 5.767E-09 | 2.706E-09 | 3.600E-09 | 8.800E-09 |
| 28327 | 1417 | L1417Q | E | 1.2e-08 | 2.6e-09 | 7.6e-09 | 7.400E-09 | 4.703E-09 | 2.600E-09 | 1.200E-08 |
| 28328 | 1417 | L1417G | E | 1.3e-08 | 7.5e-09 | 6.5e-09 | 9.000E-09 | 3.500E-09 | 6.500E-09 | 1.300E-08 |
| 28329 | 1417 | L1417H | E | 5.1e-09 | 6.7e-09 | 8.3e-09 | 6.700E-09 | 1.600E-09 | 5.100E-09 | 8.300E-09 |
| 28330 | 1417 | L1417I | E | 8.8e-09 | 1.8e-09 | 8.6e-09 | 6.400E-09 | 3.985E-09 | 1.800E-09 | 8.800E-09 |
| 28331 | 1417 | L1417L | E | 3.3e-09 | 1.7e-09 | 9.5e-09 | 4.833E-09 | 4.120E-09 | 1.700E-09 | 9.500E-09 |
| 28332 | 1417 | L1417K | E | 1.1e-08 | 1.6e-08 | 1e-08 | 1.233E-08 | 3.215E-09 | 1.000E-08 | 1.600E-08 |
| 28333 | 1417 | L1417M | E | 5.5e-09 | 6.2e-09 | 5.5e-09 | 5.733E-09 | 4.041E-10 | 5.500E-09 | 6.200E-09 |
| 28334 | 1417 | L1417F | E | 4.5e-09 | 4.3e-09 | 5.1e-09 | 4.633E-09 | 4.163E-10 | 4.300E-09 | 5.100E-09 |
| 28335 | 1417 | L1417P | E | 3.3e-09 | 1.5e-08 | 9.6e-09 | 9.300E-09 | 5.856E-09 | 3.300E-09 | 1.500E-08 |
| 28336 | 1417 | L1417S | E | 3.7e-08 | 4.8e-09 | 1.4e-08 | 1.860E-08 | 1.659E-08 | 4.800E-09 | 3.700E-08 |
| 28337 | 1417 | L1417T | E | 1e-08 | 8.6e-09 | 1.2e-09 | 6.600E-09 | 4.729E-09 | 1.200E-09 | 1.000E-08 |
| 28338 | 1417 | L1417W | E | 6.9e-09 | 1.9e-08 | 5.1e-09 | 1.033E-08 | 7.559E-09 | 5.100E-09 | 1.900E-08 |
| 28339 | 1417 | L1417Y | E | 3.7e-09 | 6.7e-09 | 6.4e-09 | 5.600E-09 | 1.652E-09 | 3.700E-09 | 6.700E-09 |
| 28340 | 1417 | L1417V | E | 6.9e-09 | 2.2e-08 | 1.5e-08 | 1.463E-08 | 7.557E-09 | 6.900E-09 | 2.200E-08 |
| 28341 | 1418 | G1418A | E | 7e-09 | 6.9e-09 | 7.4e-09 | 7.100E-09 | 2.646E-10 | 6.900E-09 | 7.400E-09 |
| 28342 | 1418 | G1418R | E | 1.6e-08 | 8e-09 | 7.2e-09 | 1.040E-08 | 4.866E-09 | 7.200E-09 | 1.600E-08 |
| 28343 | 1418 | G1418N | E | 4.2e-09 | 4.7e-09 | 1.2e-08 | 6.967E-09 | 4.366E-09 | 4.200E-09 | 1.200E-08 |
| 28344 | 1418 | G1418D | E | 2.8e-09 | 6.3e-09 | 1.2e-08 | 7.033E-09 | 4.644E-09 | 2.800E-09 | 1.200E-08 |
| 28345 | 1418 | G1418C | E | 4.2e-09 | 3.5e-09 | 1.1e-08 | 6.233E-09 | 4.143E-09 | 3.500E-09 | 1.100E-08 |
| 28346 | 1418 | G1418E | E | 5.9e-09 | 2.3e-09 | 5.7e-09 | 4.633E-09 | 2.023E-09 | 2.300E-09 | 5.900E-09 |
| 28347 | 1418 | G1418Q | E | 1.3e-08 | 7.2e-09 | 5.3e-09 | 8.500E-09 | 4.011E-09 | 5.300E-09 | 1.300E-08 |
| 28348 | 1418 | G1418G | E | 6.5e-09 | 1.7e-09 | 4.6e-09 | 4.267E-09 | 2.417E-09 | 1.700E-09 | 6.500E-09 |
| 28349 | 1418 | G1418H | E | 3e-09 | 5.5e-09 | 7.1e-09 | 5.200E-09 | 2.066E-09 | 3.000E-09 | 7.100E-09 |
| 28350 | 1418 | G1418I | E | 1.1e-08 | 4.5e-09 | 1.6e-08 | 1.050E-08 | 5.766E-09 | 4.500E-09 | 1.600E-08 |
| 28351 | 1418 | G1418L | E | 5.1e-09 | 1.4e-08 | 4.7e-09 | 7.933E-09 | 5.258E-09 | 4.700E-09 | 1.400E-08 |
| 28352 | 1418 | G1418K | E | 1.6e-08 | 7.5e-09 | 5.4e-09 | 9.633E-09 | 5.613E-09 | 5.400E-09 | 1.600E-08 |
| 28353 | 1418 | G1418M | E | 8.4e-09 | 3.4e-09 | 3.7e-09 | 5.167E-09 | 2.804E-09 | 3.400E-09 | 8.400E-09 |
| 28354 | 1418 | G1418F | E | 1e-08 | 1.1e-08 | 4e-09 | 8.333E-09 | 3.786E-09 | 4.000E-09 | 1.100E-08 |
| 28355 | 1418 | G1418P | E | 1.2e-08 | 2.8e-09 | 7.4e-09 | 7.400E-09 | 4.600E-09 | 2.800E-09 | 1.200E-08 |
| 28356 | 1418 | G1418S | E | 6.3e-09 | 1.7e-09 | 1.2e-08 | 6.667E-09 | 5.160E-09 | 1.700E-09 | 1.200E-08 |
| 28357 | 1418 | G1418T | E | 4.5e-09 | 6.2e-09 | 1.2e-08 | 7.567E-09 | 3.932E-09 | 4.500E-09 | 1.200E-08 |
| 28358 | 1418 | G1418W | E | 5.3e-09 | 1e-08 | 4.7e-09 | 6.667E-09 | 2.902E-09 | 4.700E-09 | 1.000E-08 |
| 28359 | 1418 | G1418Y | E | 4.7e-09 | 1.1e-08 | 5.3e-09 | 7.000E-09 | 3.477E-09 | 4.700E-09 | 1.100E-08 |
| 28360 | 1418 | G1418V | E | 9.4e-09 | 9.3e-09 | 1.2e-08 | 1.023E-08 | 1.531E-09 | 9.300E-09 | 1.200E-08 |
| 28361 | 1419 | C1419A | E | 9.7e-09 | 1.1e-08 | 8.7e-09 | 9.800E-09 | 1.153E-09 | 8.700E-09 | 1.100E-08 |
| 28362 | 1419 | C1419R | E | 3.4e-09 | 1.1e-08 | 1e-08 | 8.133E-09 | 4.130E-09 | 3.400E-09 | 1.100E-08 |
| 28363 | 1419 | C1419N | E | 6.1e-09 | 2.8e-09 | 6.6e-09 | 5.167E-09 | 2.065E-09 | 2.800E-09 | 6.600E-09 |
| 28364 | 1419 | C1419D | E | 4.7e-09 | 2.8e-09 | 4.3e-09 | 3.933E-09 | 1.002E-09 | 2.800E-09 | 4.700E-09 |
| 28365 | 1419 | C1419C | E | 4.3e-09 | 1.6e-08 | 4.2e-09 | 8.167E-09 | 6.784E-09 | 4.200E-09 | 1.600E-08 |
| 28366 | 1419 | C1419E | E | 2.2e-08 | 4.6e-09 | 1.8e-09 | 9.467E-09 | 1.094E-08 | 1.800E-09 | 2.200E-08 |
| 28367 | 1419 | C1419Q | E | 4e-09 | 1.1e-08 | 8.5e-09 | 7.833E-09 | 3.547E-09 | 4.000E-09 | 1.100E-08 |
| 28368 | 1419 | C1419G | E | 9.7e-09 | 1.3e-08 | 4.1e-09 | 8.933E-09 | 4.499E-09 | 4.100E-09 | 1.300E-08 |
| 28369 | 1419 | C1419H | E | 6.7e-09 | 1.2e-08 | 7e-09 | 8.567E-09 | 2.977E-09 | 6.700E-09 | 1.200E-08 |
| 28370 | 1419 | C1419I | E | 1.9e-09 | 1.2e-08 | 3.3e-09 | 5.733E-09 | 5.472E-09 | 1.900E-09 | 1.200E-08 |
| 28371 | 1419 | C1419L | E | 1.9e-08 | 3.8e-09 | 2.7e-09 | 8.500E-09 | 9.110E-09 | 2.700E-09 | 1.900E-08 |
| 28372 | 1419 | C1419K | E | 2.5e-08 | 9.7e-09 | 4.6e-09 | 1.310E-08 | 1.062E-08 | 4.600E-09 | 2.500E-08 |
| 28373 | 1419 | C1419M | E | 3.2e-09 | 5.9e-09 | 3.6e-09 | 4.233E-09 | 1.457E-09 | 3.200E-09 | 5.900E-09 |
| 28374 | 1419 | C1419F | E | 2.7e-08 | 7.6e-09 | 8.9e-09 | 1.450E-08 | 1.084E-08 | 7.600E-09 | 2.700E-08 |
| 28375 | 1419 | C1419P | E | 5.4e-09 | 4.5e-08 | 1.3e-08 | 2.113E-08 | 2.102E-08 | 5.400E-09 | 4.500E-08 |
| 28376 | 1419 | C1419S | E | 9.2e-09 | 4.8e-09 | 1.3e-08 | 9.000E-09 | 4.104E-09 | 4.800E-09 | 1.300E-08 |
| 28377 | 1419 | C1419T | E | 1.9e-08 | 4.2e-09 | 6.9e-09 | 1.003E-08 | 7.882E-09 | 4.200E-09 | 1.900E-08 |
| 28378 | 1419 | C1419W | E | 8e-09 | 6.2e-09 | 7.7e-09 | 7.300E-09 | 9.644E-10 | 6.200E-09 | 8.000E-09 |
| 28379 | 1419 | C1419Y | E | 4.8e-09 | 9.1e-09 | 3.5e-09 | 5.800E-09 | 2.931E-09 | 3.500E-09 | 9.100E-09 |
| 28380 | 1419 | C1419V | E | 1.2e-08 | 3.4e-09 | 7.2e-09 | 7.533E-09 | 4.310E-09 | 3.400E-09 | 1.200E-08 |
| 28381 | 1420 | L1420A | E | 8.3e-09 | 1e-08 | 9.1e-09 | 9.133E-09 | 8.505E-10 | 8.300E-09 | 1.000E-08 |
| 28382 | 1420 | L1420R | E | 1.3e-08 | 5.7e-09 | 1.2e-08 | 1.023E-08 | 3.958E-09 | 5.700E-09 | 1.300E-08 |
| 28383 | 1420 | L1420N | E | 7.7e-09 | 7.2e-09 | 7.6e-09 | 7.500E-09 | 2.646E-10 | 7.200E-09 | 7.700E-09 |
| 28384 | 1420 | L1420D | E | 5.7e-09 | 1.3e-08 | 1.4e-08 | 1.090E-08 | 4.531E-09 | 5.700E-09 | 1.400E-08 |
| 28385 | 1420 | L1420C | E | 1.5e-08 | 9.4e-09 | 1e-08 | 1.147E-08 | 3.075E-09 | 9.400E-09 | 1.500E-08 |
| 28386 | 1420 | L1420E | E | 8.6e-09 | 1.7e-08 | 3.9e-09 | 9.833E-09 | 6.637E-09 | 3.900E-09 | 1.700E-08 |
| 28387 | 1420 | L1420Q | E | 6.7e-09 | 4.4e-09 | 1.1e-08 | 7.367E-09 | 3.350E-09 | 4.400E-09 | 1.100E-08 |
| 28388 | 1420 | L1420G | E | 8.9e-09 | 9.6e-09 | 1.2e-08 | 1.017E-08 | 1.626E-09 | 8.900E-09 | 1.200E-08 |
| 28389 | 1420 | L1420H | E | 5.7e-09 | 8.9e-09 | 8.9e-09 | 7.833E-09 | 1.848E-09 | 5.700E-09 | 8.900E-09 |

| | | | | | | | | | | |
|---|---|---|---|---|---|---|---|---|---|---|
| 28390 | 1420 | L1420I | E | 1.1e-08 | 5.5e-09 | 2.3e-08 | 1.317E-08 | 8.949E-09 | 5.500E-09 | 2.300E-08 |
| 28391 | 1420 | L1420L | E | 1.2e-08 | 4.3e-09 | 5.4e-09 | 7.233E-09 | 4.165E-09 | 4.300E-09 | 1.200E-08 |
| 28392 | 1420 | L1420K | E | 2.9e-09 | 1.6e-08 | 1.1e-08 | 9.967E-09 | 6.611E-09 | 2.900E-09 | 1.600E-08 |
| 28393 | 1420 | L1420M | E | 1.3e-08 | 4.2e-09 | 1.5e-08 | 1.073E-08 | 5.746E-09 | 4.200E-09 | 1.500E-08 |
| 28394 | 1420 | L1420F | E | 4.2e-09 | 5e-09 | 5.9e-09 | 5.033E-09 | 8.505E-10 | 4.200E-09 | 5.900E-09 |
| 28395 | 1420 | L1420P | E | 3.8e-09 | 1.2e-08 | 7.7e-09 | 7.833E-09 | 4.102E-09 | 3.800E-09 | 1.200E-08 |
| 28396 | 1420 | L1420S | E | 5.2e-09 | 2.1e-08 | 2.5e-09 | 9.567E-09 | 9.993E-09 | 2.500E-09 | 2.100E-08 |
| 28397 | 1420 | L1420T | E | 9e-09 | 1.2e-08 | 8.3e-09 | 9.767E-09 | 1.966E-09 | 8.300E-09 | 1.200E-08 |
| 28398 | 1420 | L1420W | E | 4.1e-09 | 3.6e-09 | 4.1e-09 | 3.933E-09 | 2.887E-10 | 3.600E-09 | 4.100E-09 |
| 28399 | 1420 | L1420Y | E | 1.3e-08 | 1.6e-08 | 6.3e-09 | 1.177E-08 | 4.966E-09 | 6.300E-09 | 1.600E-08 |
| 28400 | 1420 | L1420V | E | 2.5e-09 | 9.1e-09 | 5.9e-09 | 5.833E-09 | 3.301E-09 | 2.500E-09 | 9.100E-09 |
| 28401 | 1421 | V1421A | E | 1.6e-08 | 1.2e-08 | 9.6e-09 | 1.253E-08 | 3.233E-09 | 9.600E-09 | 1.600E-08 |
| 28402 | 1421 | V1421R | E | 1.2e-08 | 6.2e-09 | 8.2e-09 | 8.800E-09 | 2.946E-09 | 6.200E-09 | 1.200E-08 |
| 28403 | 1421 | V1421N | E | 5.3e-09 | 5.1e-09 | 1.1e-08 | 7.133E-09 | 3.350E-09 | 5.100E-09 | 1.100E-08 |
| 28404 | 1421 | V1421D | E | 1.1e-08 | 5.7e-09 | 3.7e-09 | 6.800E-09 | 3.772E-09 | 3.700E-09 | 1.100E-08 |
| 28405 | 1421 | V1421C | E | 7.9e-09 | 5.6e-09 | 2.1e-08 | 1.150E-08 | 8.307E-09 | 5.600E-09 | 2.100E-08 |
| 28406 | 1421 | V1421E | E | 4.1e-09 | 8.9e-09 | 7.9e-09 | 6.967E-09 | 2.532E-09 | 4.100E-09 | 8.900E-09 |
| 28407 | 1421 | V1421Q | E | 1e-08 | 7e-09 | 9.5e-09 | 8.833E-09 | 1.607E-09 | 7.000E-09 | 1.000E-08 |
| 28408 | 1421 | V1421G | E | 8.7e-09 | 8.2e-09 | 8.4e-09 | 8.433E-09 | 2.517E-10 | 8.200E-09 | 8.700E-09 |
| 28409 | 1421 | V1421H | E | 7.2e-09 | 9.3e-09 | 1.4e-08 | 1.017E-08 | 3.482E-09 | 7.200E-09 | 1.400E-08 |
| 28410 | 1421 | V1421I | E | 1.2e-08 | 1.1e-08 | 1e-08 | 1.100E-08 | 1.000E-09 | 1.000E-08 | 1.200E-08 |
| 28411 | 1421 | V1421L | E | 6.8e-09 | 1e-08 | 3.7e-09 | 6.833E-09 | 3.150E-09 | 3.700E-09 | 1.000E-08 |
| 28412 | 1421 | V1421K | E | 3.9e-09 | 9e-09 | 4e-09 | 5.633E-09 | 2.916E-09 | 3.900E-09 | 9.000E-09 |
| 28413 | 1421 | V1421M | E | 2.1e-09 | 5.5e-09 | 4e-09 | 3.867E-09 | 1.704E-09 | 2.100E-09 | 5.500E-09 |
| 28414 | 1421 | V1421F | E | 5e-09 | 1.2e-08 | 6.8e-09 | 7.933E-09 | 3.635E-09 | 5.000E-09 | 1.200E-08 |
| 28415 | 1421 | V1421P | E | 7.5e-09 | 1e-08 | 7.8e-09 | 8.433E-09 | 1.365E-09 | 7.500E-09 | 1.000E-08 |
| 28416 | 1421 | V1421S | E | 2.8e-08 | 1.2e-08 | 1.1e-08 | 1.700E-08 | 9.539E-09 | 1.100E-08 | 2.800E-08 |
| 28417 | 1421 | V1421T | E | 1.4e-08 | 8.2e-09 | 9.3e-09 | 1.050E-08 | 3.081E-09 | 8.200E-09 | 1.400E-08 |
| 28418 | 1421 | V1421W | E | 1.2e-09 | 3.5e-09 | 2e-09 | 2.233E-09 | 1.168E-09 | 1.200E-09 | 3.500E-09 |
| 28419 | 1421 | V1421Y | E | 1.9e-08 | 6.3e-09 | 1.2e-08 | 1.243E-08 | 6.361E-09 | 6.300E-09 | 1.900E-08 |
| 28420 | 1421 | V1421V | E | 1.9e-09 | 1.4e-08 | 1.3e-08 | 9.633E-09 | 6.716E-09 | 1.900E-09 | 1.400E-08 |
| 28421 | 1422 | K1422A | E | 1.7e-08 | 4.6e-09 | 5.2e-09 | 8.933E-09 | 6.992E-09 | 4.600E-09 | 1.700E-08 |
| 28422 | 1422 | K1422R | E | 4.4e-09 | 2.1e-08 | 7.2e-09 | 1.087E-08 | 8.887E-09 | 4.400E-09 | 2.100E-08 |
| 28423 | 1422 | K1422N | E | 5.4e-09 | 1.6e-09 | 1.2e-08 | 6.333E-09 | 5.262E-09 | 1.600E-09 | 1.200E-08 |
| 28424 | 1422 | K1422D | E | 1.2e-08 | 1e-08 | 1.8e-08 | 1.333E-08 | 4.163E-09 | 1.000E-08 | 1.800E-08 |
| 28425 | 1422 | K1422C | E | 6.1e-09 | 6.3e-09 | 2.7e-09 | 5.033E-09 | 2.023E-09 | 2.700E-09 | 6.300E-09 |
| 28426 | 1422 | K1422E | E | 1.3e-08 | 5.3e-09 | 4.2e-09 | 7.500E-09 | 4.795E-09 | 4.200E-09 | 1.300E-08 |
| 28427 | 1422 | K1422Q | E | 9.2e-09 | 7.7e-09 | 5.3e-09 | 7.400E-09 | 1.967E-09 | 5.300E-09 | 9.200E-09 |
| 28428 | 1422 | K1422G | E | 4.2e-09 | 4.1e-09 | 1.2e-08 | 6.767E-09 | 4.532E-09 | 4.100E-09 | 1.200E-08 |
| 28429 | 1422 | K1422H | E | 3.6e-08 | 1.8e-08 | 9.7e-09 | 2.123E-08 | 1.344E-08 | 9.700E-09 | 3.600E-08 |
| 28430 | 1422 | K1422I | E | 4.5e-09 | 7.3e-09 | 9e-09 | 6.933E-09 | 2.272E-09 | 4.500E-09 | 9.000E-09 |
| 28431 | 1422 | K1422L | E | 7.7e-09 | 1.1e-08 | 8.3e-09 | 9.000E-09 | 1.758E-09 | 7.700E-09 | 1.100E-08 |
| 28432 | 1422 | K1422K | E | 9.5e-09 | 7.7e-09 | 7.2e-09 | 8.133E-09 | 1.210E-09 | 7.200E-09 | 9.500E-09 |
| 28433 | 1422 | K1422M | E | 1.3e-08 | 3.2e-09 | 6.4e-09 | 7.533E-09 | 4.997E-09 | 3.200E-09 | 1.300E-08 |
| 28434 | 1422 | K1422F | E | 7.8e-09 | 2.7e-09 | 1.2e-08 | 7.500E-09 | 4.657E-09 | 2.700E-09 | 1.200E-08 |
| 28435 | 1422 | K1422P | E | 6.9e-09 | 3.5e-09 | 7.4e-09 | 5.933E-09 | 2.122E-09 | 3.500E-09 | 7.400E-09 |
| 28436 | 1422 | K1422S | E | 9.7e-09 | 3e-09 | 6.7e-09 | 6.467E-09 | 3.356E-09 | 3.000E-09 | 9.700E-09 |
| 28437 | 1422 | K1422T | E | 1.8e-09 | 4.7e-09 | 5.9e-09 | 4.133E-09 | 2.108E-09 | 1.800E-09 | 5.900E-09 |
| 28438 | 1422 | K1422W | E | 5.3e-09 | 9.2e-09 | 4e-09 | 6.167E-09 | 2.706E-09 | 4.000E-09 | 9.200E-09 |
| 28439 | 1422 | K1422Y | E | 9.2e-09 | 1.5e-08 | 8.2e-09 | 1.080E-08 | 3.672E-09 | 8.200E-09 | 1.500E-08 |
| 28440 | 1422 | K1422V | E | 1.2e-08 | 2e-09 | 6.5e-09 | 6.833E-09 | 5.008E-09 | 2.000E-09 | 1.200E-08 |
| 28441 | 1423 | D1423A | E | 3.9e-09 | 8.5e-09 | 2.9e-09 | 5.100E-09 | 2.987E-09 | 2.900E-09 | 8.500E-09 |
| 28442 | 1423 | D1423R | E | 1.3e-08 | 4.4e-09 | 8.4e-09 | 8.600E-09 | 4.303E-09 | 4.400E-09 | 1.300E-08 |
| 28443 | 1423 | D1423N | E | 4.5e-09 | 4.4e-09 | 1.6e-08 | 8.300E-09 | 6.669E-09 | 4.400E-09 | 1.600E-08 |
| 28444 | 1423 | D1423D | E | 6.2e-09 | 7.5e-09 | 9.2e-09 | 7.633E-09 | 1.504E-09 | 6.200E-09 | 9.200E-09 |
| 28445 | 1423 | D1423C | E | 6.4e-09 | 8.4e-09 | 7.5e-09 | 7.433E-09 | 1.002E-09 | 6.400E-09 | 8.400E-09 |
| 28446 | 1423 | D1423E | E | 3e-09 | 6e-09 | 5.6e-09 | 4.867E-09 | 1.629E-09 | 3.000E-09 | 6.000E-09 |
| 28447 | 1423 | D1423Q | E | 8.1e-09 | 1.1e-08 | 4.8e-09 | 7.967E-09 | 3.102E-09 | 4.800E-09 | 1.100E-08 |
| 28448 | 1423 | D1423G | E | 4.4e-09 | 5.9e-09 | 3.7e-09 | 4.667E-09 | 1.124E-09 | 3.700E-09 | 5.900E-09 |
| 28449 | 1423 | D1423H | E | 9.4e-09 | 1e-08 | 4.8e-09 | 8.067E-09 | 2.845E-09 | 4.800E-09 | 1.000E-08 |
| 28450 | 1423 | D1423I | E | 6.3e-09 | 6.7e-09 | 8.7e-09 | 7.233E-09 | 1.286E-09 | 6.300E-09 | 8.700E-09 |
| 28451 | 1423 | D1423L | E | 4.6e-09 | 2.6e-08 | 7.3e-09 | 1.263E-08 | 1.165E-08 | 4.600E-09 | 2.600E-08 |
| 28452 | 1423 | D1423K | E | 4.7e-09 | 1.6e-08 | 7.8e-09 | 9.500E-09 | 5.839E-09 | 4.700E-09 | 1.600E-08 |
| 28453 | 1423 | D1423M | E | 2.4e-08 | 1e-08 | 9.3e-09 | 1.443E-08 | 8.292E-09 | 9.300E-09 | 2.400E-08 |
| 28454 | 1423 | D1423F | E | 5.7e-09 | 1.5e-08 | 1.4e-08 | 1.157E-08 | 5.105E-09 | 5.700E-09 | 1.500E-08 |
| 28455 | 1423 | D1423P | E | 1.4e-08 | 2.6e-08 | 3.2e-09 | 1.440E-08 | 1.141E-08 | 3.200E-09 | 2.600E-08 |
| 28456 | 1423 | D1423S | E | 5.8e-09 | 7.7e-09 | 5.3e-09 | 6.267E-09 | 1.266E-09 | 5.300E-09 | 7.700E-09 |
| 28457 | 1423 | D1423T | E | 2.4e-09 | 5.3e-09 | 5.1e-09 | 4.267E-09 | 1.620E-09 | 2.400E-09 | 5.300E-09 |
| 28458 | 1423 | D1423W | E | 2.1e-08 | 4.2e-09 | 4.6e-09 | 9.933E-09 | 9.586E-09 | 4.200E-09 | 2.100E-08 |
| 28459 | 1423 | D1423Y | E | 6.5e-09 | 2.6e-09 | 1e-08 | 6.367E-09 | 3.702E-09 | 2.600E-09 | 1.000E-08 |
| 28460 | 1423 | D1423V | E | 2.2e-09 | 1.4e-08 | 1e-08 | 8.733E-09 | 6.001E-09 | 2.200E-09 | 1.400E-08 |
| 28461 | 1424 | Y1424A | E | 1.7e-08 | 5.4e-09 | 1e-08 | 1.080E-08 | 5.841E-09 | 5.400E-09 | 1.700E-08 |
| 28462 | 1424 | Y1424R | E | 7.1e-09 | 1.1e-08 | 1.3e-08 | 1.037E-08 | 3.001E-09 | 7.100E-09 | 1.300E-08 |
| 28463 | 1424 | Y1424N | E | 2.5e-09 | 6e-09 | 9.6e-09 | 6.033E-09 | 3.550E-09 | 2.500E-09 | 9.600E-09 |
| 28464 | 1424 | Y1424D | E | 1.2e-08 | 3.3e-09 | 6.2e-09 | 7.167E-09 | 4.430E-09 | 3.300E-09 | 1.200E-08 |
| 28465 | 1424 | Y1424C | E | 5.6e-09 | 1.4e-08 | 3.7e-09 | 7.767E-09 | 5.481E-09 | 3.700E-09 | 1.400E-08 |
| 28466 | 1424 | Y1424E | E | 4.3e-09 | 4.5e-09 | 1.3e-08 | 7.267E-09 | 4.966E-09 | 4.300E-09 | 1.300E-08 |
| 28467 | 1424 | Y1424Q | E | 8.1e-09 | 3.9e-09 | 1.2e-08 | 8.000E-09 | 4.051E-09 | 3.900E-09 | 1.200E-08 |
| 28468 | 1424 | Y1424G | E | 1.2e-08 | 4.9e-09 | 9.9e-09 | 8.933E-09 | 3.647E-09 | 4.900E-09 | 1.200E-08 |
| 28469 | 1424 | Y1424H | E | 1e-08 | 4.3e-09 | 7.4e-09 | 7.233E-09 | 2.854E-09 | 4.300E-09 | 1.000E-08 |
| 28470 | 1424 | Y1424I | E | 2.5e-09 | 1.5e-08 | 6.8e-09 | 8.100E-09 | 6.351E-09 | 2.500E-09 | 1.500E-08 |
| 28471 | 1424 | Y1424L | E | 5.7e-09 | 4.4e-09 | 1.4e-08 | 8.033E-09 | 5.208E-09 | 4.400E-09 | 1.400E-08 |
| 28472 | 1424 | Y1424K | E | 6e-09 | 4.9e-09 | 5.9e-09 | 5.600E-09 | 6.083E-10 | 4.900E-09 | 6.000E-09 |
| 28473 | 1424 | Y1424M | E | 8e-09 | 1.3e-08 | 1.5e-08 | 1.200E-08 | 3.606E-09 | 8.000E-09 | 1.500E-08 |
| 28474 | 1424 | Y1424F | E | 4.9e-09 | 8.8e-09 | 1e-08 | 7.900E-09 | 2.666E-09 | 4.900E-09 | 1.000E-08 |

| | | | | | | | | | | |
|---|---|---|---|---|---|---|---|---|---|---|
| 28475 | 1424 | Y1424P | E | 3.4e-09 | 1.6e-09 | 1.4e-08 | 6.333E-09 | 6.700E-09 | 1.600E-09 | 1.400E-08 |
| 28476 | 1424 | Y1424S | E | 2e-09 | 8.1e-09 | 3.1e-09 | 4.400E-09 | 3.251E-09 | 2.000E-09 | 8.100E-09 |
| 28477 | 1424 | Y1424T | E | 1.2e-08 | 1.3e-08 | 4.1e-09 | 9.700E-09 | 4.875E-09 | 4.100E-09 | 1.300E-08 |
| 28478 | 1424 | Y1424W | E | 4.3e-09 | 6.1e-09 | 1.7e-08 | 9.133E-09 | 6.872E-09 | 4.300E-09 | 1.700E-08 |
| 28479 | 1424 | Y1424Y | E | 5e-09 | 1e-08 | 3.6e-09 | 6.200E-09 | 3.365E-09 | 3.600E-09 | 1.000E-08 |
| 28480 | 1424 | Y1424V | E | 7.7e-09 | 8.4e-09 | 7.5e-09 | 7.867E-09 | 4.726E-10 | 7.500E-09 | 8.400E-09 |
| 28481 | 1425 | F1425A | E | 1.6e-08 | 3.1e-09 | 5.6e-09 | 8.233E-09 | 6.841E-09 | 3.100E-09 | 1.600E-08 |
| 28482 | 1425 | F1425R | E | 3e-09 | 8.5e-09 | 7.7e-09 | 6.400E-09 | 2.972E-09 | 3.000E-09 | 8.500E-09 |
| 28483 | 1425 | F1425N | E | 6e-09 | 3.5e-09 | 5.9e-09 | 5.133E-09 | 1.415E-09 | 3.500E-09 | 6.000E-09 |
| 28484 | 1425 | F1425D | E | 2.4e-09 | 1.1e-08 | 5.6e-09 | 6.333E-09 | 4.347E-09 | 2.400E-09 | 1.100E-08 |
| 28485 | 1425 | F1425C | E | 8e-09 | 1e-08 | 1.3e-08 | 1.033E-08 | 2.517E-09 | 8.000E-09 | 1.300E-08 |
| 28486 | 1425 | F1425E | E | 6.9e-09 | 1e-08 | 7.7e-09 | 8.200E-09 | 1.609E-09 | 6.900E-09 | 1.000E-08 |
| 28487 | 1425 | F1425Q | E | 4.8e-09 | 8.4e-09 | 2.8e-09 | 5.333E-09 | 2.838E-09 | 2.800E-09 | 8.400E-09 |
| 28488 | 1425 | F1425G | E | 1.1e-08 | 2.1e-08 | 4.7e-09 | 1.223E-08 | 8.220E-09 | 4.700E-09 | 2.100E-08 |
| 28489 | 1425 | F1425H | E | 6.7e-09 | 1e-08 | 1.5e-09 | 6.067E-09 | 4.285E-09 | 1.500E-09 | 1.000E-08 |
| 28490 | 1425 | F1425I | E | 8.1e-09 | 6.1e-09 | 1.1e-08 | 8.400E-09 | 2.464E-09 | 6.100E-09 | 1.100E-08 |
| 28491 | 1425 | F1425L | E | 1.5e-08 | 1.6e-08 | 4.4e-09 | 1.180E-08 | 6.428E-09 | 4.400E-09 | 1.600E-08 |
| 28492 | 1425 | F1425K | E | 5.7e-09 | 7.1e-09 | 6.7e-09 | 6.500E-09 | 7.211E-10 | 5.700E-09 | 7.100E-09 |
| 28493 | 1425 | F1425M | E | 4.8e-09 | 7.7e-09 | 1.9e-08 | 1.050E-08 | 7.503E-09 | 4.800E-09 | 1.900E-08 |
| 28494 | 1425 | F1425F | E | 2.1e-08 | 1.3e-08 | 1.2e-08 | 1.533E-08 | 4.933E-09 | 1.200E-08 | 2.100E-08 |
| 28495 | 1425 | F1425P | E | 1.8e-08 | 7.2e-09 | 1.8e-09 | 9.000E-09 | 8.249E-09 | 1.800E-09 | 1.800E-08 |
| 28496 | 1425 | F1425S | E | 2.8e-09 | 3.9e-09 | 4.8e-09 | 3.833E-09 | 1.002E-09 | 2.800E-09 | 4.800E-09 |
| 28497 | 1425 | F1425T | E | 2.4e-09 | 1.4e-08 | 3.3e-09 | 6.567E-09 | 6.453E-09 | 2.400E-09 | 1.400E-08 |
| 28498 | 1425 | F1425W | E | 7.1e-09 | 3.8e-09 | 1.2e-08 | 7.633E-09 | 4.126E-09 | 3.800E-09 | 1.200E-08 |
| 28499 | 1425 | F1425Y | E | 3.2e-09 | 4.4e-09 | 6.9e-09 | 4.833E-09 | 1.888E-09 | 3.200E-09 | 6.900E-09 |
| 28500 | 1425 | F1425V | E | 1.1e-08 | 3.9e-09 | 9.2e-09 | 8.033E-09 | 3.691E-09 | 3.900E-09 | 1.100E-08 |
| 28501 | 1426 | P1426A | E | 4.3e-09 | 3.9e-09 | 2.7e-09 | 3.633E-09 | 8.327E-10 | 2.700E-09 | 4.300E-09 |
| 28502 | 1426 | P1426R | E | 1.3e-08 | 2.6e-09 | 9.4e-09 | 8.333E-09 | 5.281E-09 | 2.600E-09 | 1.300E-08 |
| 28503 | 1426 | P1426N | E | 3.8e-09 | 1e-08 | 7.6e-09 | 7.133E-09 | 3.126E-09 | 3.800E-09 | 1.000E-08 |
| 28504 | 1426 | P1426D | E | 6.6e-09 | 1e-08 | 5.9e-09 | 7.500E-09 | 2.193E-09 | 5.900E-09 | 1.000E-08 |
| 28505 | 1426 | P1426C | E | 3.6e-09 | 7.6e-09 | 6.9e-09 | 6.033E-09 | 2.136E-09 | 3.600E-09 | 7.600E-09 |
| 28506 | 1426 | P1426E | E | 1.1e-08 | 2.6e-09 | 1e-08 | 7.867E-09 | 4.588E-09 | 2.600E-09 | 1.100E-08 |
| 28507 | 1426 | P1426Q | E | 6.7e-09 | 1.3e-08 | 1e-08 | 9.900E-09 | 3.151E-09 | 6.700E-09 | 1.300E-08 |
| 28508 | 1426 | P1426G | E | 8.8e-09 | 4.1e-09 | 3.1e-09 | 5.333E-09 | 3.044E-09 | 3.100E-09 | 8.800E-09 |
| 28509 | 1426 | P1426H | E | 9.7e-09 | 3.5e-09 | 7.8e-09 | 7.000E-09 | 3.176E-09 | 3.500E-09 | 9.700E-09 |
| 28510 | 1426 | P1426I | E | 7.2e-09 | 3.6e-09 | 1.7e-09 | 4.167E-09 | 2.793E-09 | 1.700E-09 | 7.200E-09 |
| 28511 | 1426 | P1426L | E | 5.9e-09 | 1.1e-08 | 4.3e-09 | 7.067E-09 | 3.499E-09 | 4.300E-09 | 1.100E-08 |
| 28512 | 1426 | P1426K | E | 4.2e-09 | 6.9e-09 | 7.9e-09 | 6.333E-09 | 1.914E-09 | 4.200E-09 | 7.900E-09 |
| 28513 | 1426 | P1426M | E | 4.1e-09 | 4.4e-09 | 1.3e-08 | 7.167E-09 | 5.054E-09 | 4.100E-09 | 1.300E-08 |
| 28514 | 1426 | P1426F | E | 1.2e-08 | 9.3e-09 | 3.7e-09 | 8.333E-09 | 4.234E-09 | 3.700E-09 | 1.200E-08 |
| 28515 | 1426 | P1426P | E | 3.2e-09 | 9.8e-09 | 7.1e-09 | 6.700E-09 | 3.318E-09 | 3.200E-09 | 9.800E-09 |
| 28516 | 1426 | P1426S | E | 1e-08 | 1.2e-08 | 1e-08 | 1.067E-08 | 1.155E-09 | 1.000E-08 | 1.200E-08 |
| 28517 | 1426 | P1426T | E | 1.4e-08 | 4.8e-09 | 8.8e-09 | 9.200E-09 | 4.613E-09 | 4.800E-09 | 1.400E-08 |
| 28518 | 1426 | P1426W | E | 2e-09 | 4.8e-09 | 8.9e-09 | 5.233E-09 | 3.470E-09 | 2.000E-09 | 8.900E-09 |
| 28519 | 1426 | P1426Y | E | 1.2e-08 | 8.3e-09 | 1.5e-08 | 1.177E-08 | 3.356E-09 | 8.300E-09 | 1.500E-08 |
| 28520 | 1426 | P1426V | E | 4.7e-09 | 6.2e-09 | 8.4e-09 | 6.433E-09 | 1.861E-09 | 4.700E-09 | 8.400E-09 |
| 28521 | 1427 | E1427A | E | 6.1e-09 | 7e-09 | 1.3e-08 | 8.700E-09 | 3.751E-09 | 6.100E-09 | 1.300E-08 |
| 28522 | 1427 | E1427R | E | 3.5e-09 | 6.5e-09 | 4.9e-09 | 4.967E-09 | 1.501E-09 | 3.500E-09 | 6.500E-09 |
| 28523 | 1427 | E1427N | E | 5.1e-09 | 4.2e-09 | 2e-09 | 3.767E-09 | 1.595E-09 | 2.000E-09 | 5.100E-09 |
| 28524 | 1427 | E1427D | E | 6.9e-09 | 8.6e-09 | 1.2e-08 | 9.167E-09 | 2.597E-09 | 6.900E-09 | 1.200E-08 |
| 28525 | 1427 | E1427C | E | 8.7e-09 | 3.1e-09 | 5.9e-09 | 5.900E-09 | 2.800E-09 | 3.100E-09 | 8.700E-09 |
| 28526 | 1427 | E1427E | E | 3.1e-09 | 1.1e-08 | 1.7e-08 | 1.037E-08 | 6.972E-09 | 3.100E-09 | 1.700E-08 |
| 28527 | 1427 | E1427Q | E | 3.1e-09 | 7.7e-09 | 8.8e-09 | 6.533E-09 | 3.024E-09 | 3.100E-09 | 8.800E-09 |
| 28528 | 1427 | E1427G | E | 8.5e-09 | 6.7e-09 | 1.5e-08 | 1.007E-08 | 4.366E-09 | 6.700E-09 | 1.500E-08 |
| 28529 | 1427 | E1427H | E | 7.1e-09 | 4.6e-09 | 3.3e-09 | 5.000E-09 | 1.931E-09 | 3.300E-09 | 7.100E-09 |
| 28530 | 1427 | E1427I | E | 9.1e-09 | 2.3e-09 | 4.6e-09 | 5.333E-09 | 3.459E-09 | 2.300E-09 | 9.100E-09 |
| 28531 | 1427 | E1427L | E | 7.8e-09 | 1.2e-08 | 6.8e-09 | 8.867E-09 | 2.759E-09 | 6.800E-09 | 1.200E-08 |
| 28532 | 1427 | E1427K | E | 9.4e-09 | 3e-09 | 2.6e-09 | 5.000E-09 | 3.816E-09 | 2.600E-09 | 9.400E-09 |
| 28533 | 1427 | E1427M | E | 9.6e-09 | 1.3e-08 | 6.3e-09 | 9.633E-09 | 3.350E-09 | 6.300E-09 | 1.300E-08 |
| 28534 | 1427 | E1427F | E | 1.2e-08 | 1.6e-08 | 5e-09 | 1.100E-08 | 5.568E-09 | 5.000E-09 | 1.600E-08 |
| 28535 | 1427 | E1427P | E | 8.1e-09 | 8.4e-09 | 5.7e-09 | 7.400E-09 | 1.480E-09 | 5.700E-09 | 8.400E-09 |
| 28536 | 1427 | E1427S | E | 3.8e-09 | 5.4e-09 | 8.3e-09 | 5.833E-09 | 2.281E-09 | 3.800E-09 | 8.300E-09 |
| 28537 | 1427 | E1427T | E | 5.2e-09 | 1.1e-08 | 5.8e-09 | 7.333E-09 | 3.190E-09 | 5.200E-09 | 1.100E-08 |
| 28538 | 1427 | E1427W | E | 4.6e-09 | 1.9e-08 | 8.5e-09 | 1.070E-08 | 7.448E-09 | 4.600E-09 | 1.900E-08 |
| 28539 | 1427 | E1427Y | E | 4.3e-09 | 4.6e-09 | 6.9e-09 | 5.267E-09 | 1.422E-09 | 4.300E-09 | 6.900E-09 |
| 28540 | 1427 | E1427V | E | 1.1e-08 | 1.5e-08 | 1.4e-08 | 1.333E-08 | 2.082E-09 | 1.100E-08 | 1.500E-08 |
| 28541 | 1428 | P1428A | E | 9.7e-09 | 5.4e-09 | 2.7e-08 | 1.403E-08 | 1.143E-08 | 5.400E-09 | 2.700E-08 |
| 28542 | 1428 | P1428R | E | 8e-09 | 4.7e-09 | 8.3e-09 | 7.000E-09 | 1.997E-09 | 4.700E-09 | 8.300E-09 |
| 28543 | 1428 | P1428N | E | 1.1e-09 | 1.5e-08 | 5.2e-09 | 7.100E-09 | 7.142E-09 | 1.100E-09 | 1.500E-08 |
| 28544 | 1428 | P1428D | E | 5.1e-09 | 3e-09 | 4.1e-08 | 1.637E-08 | 2.136E-08 | 3.000E-09 | 4.100E-08 |
| 28545 | 1428 | P1428C | E | 1.3e-08 | 1.8e-08 | 6.6e-09 | 1.253E-08 | 5.714E-09 | 6.600E-09 | 1.800E-08 |
| 28546 | 1428 | P1428E | E | 4.8e-09 | 9e-09 | 1.1e-08 | 8.267E-09 | 3.164E-09 | 4.800E-09 | 1.100E-08 |
| 28547 | 1428 | P1428Q | E | 5.1e-09 | 3.7e-09 | 1.2e-08 | 6.933E-09 | 4.443E-09 | 3.700E-09 | 1.200E-08 |
| 28548 | 1428 | P1428G | E | 4e-09 | 7.4e-09 | 4.4e-09 | 5.267E-09 | 1.858E-09 | 4.000E-09 | 7.400E-09 |
| 28549 | 1428 | P1428H | E | 7.6e-09 | 6.8e-09 | 5.6e-09 | 6.667E-09 | 1.007E-09 | 5.600E-09 | 7.600E-09 |
| 28550 | 1428 | P1428I | E | 5e-09 | 1.3e-08 | 8.2e-09 | 8.733E-09 | 4.027E-09 | 5.000E-09 | 1.300E-08 |
| 28551 | 1428 | P1428L | E | 7.5e-09 | 4.8e-09 | 8.6e-09 | 6.967E-09 | 1.955E-09 | 4.800E-09 | 8.600E-09 |
| 28552 | 1428 | P1428K | E | 7e-09 | 7.4e-09 | 4.5e-09 | 6.300E-09 | 1.572E-09 | 4.500E-09 | 7.400E-09 |
| 28553 | 1428 | P1428M | E | 3.7e-09 | 9.3e-09 | 4.7e-09 | 5.900E-09 | 2.987E-09 | 3.700E-09 | 9.300E-09 |
| 28554 | 1428 | P1428F | E | 1e-08 | 7e-09 | 8.7e-09 | 8.567E-09 | 1.504E-09 | 7.000E-09 | 1.000E-08 |
| 28555 | 1428 | P1428P | E | 3.2e-09 | 9.5e-09 | 7.6e-09 | 6.767E-09 | 3.232E-09 | 3.200E-09 | 9.500E-09 |
| 28556 | 1428 | P1428S | E | 4.9e-09 | 6.6e-09 | 5e-09 | 5.500E-09 | 9.539E-10 | 4.900E-09 | 6.600E-09 |
| 28557 | 1428 | P1428T | E | 2.3e-09 | 1.4e-08 | 3.2e-09 | 6.500E-09 | 6.511E-09 | 2.300E-09 | 1.400E-08 |
| 28558 | 1428 | P1428W | E | 2.8e-08 | 6.8e-09 | 5.5e-09 | 1.343E-08 | 1.263E-08 | 5.500E-09 | 2.800E-08 |
| 28559 | 1428 | P1428Y | E | 3.9e-09 | 6.9e-09 | 9e-09 | 6.600E-09 | 2.563E-09 | 3.900E-09 | 9.000E-09 |

| | | | | | | | | | | |
|---|---|---|---|---|---|---|---|---|---|---|
| 28560 | 1428 | P1428V | E | 5.8e-09 | 2.2e-09 | 4.2e-09 | 4.067E-09 | 1.804E-09 | 2.200E-09 | 5.800E-09 |
| 28561 | 1429 | V1429A | E | 4.4e-09 | 7.7e-09 | 8.4e-09 | 6.833E-09 | 2.136E-09 | 4.400E-09 | 8.400E-09 |
| 28562 | 1429 | V1429R | E | 1.5e-08 | 4.2e-09 | 2.9e-09 | 7.367E-09 | 6.643E-09 | 2.900E-09 | 1.500E-08 |
| 28563 | 1429 | V1429N | E | 3.4e-09 | 1.3e-08 | 8.7e-09 | 8.367E-09 | 4.809E-09 | 3.400E-09 | 1.300E-08 |
| 28564 | 1429 | V1429D | E | 9.8e-09 | 1.5e-08 | 1e-08 | 1.160E-08 | 2.946E-09 | 9.800E-09 | 1.500E-08 |
| 28565 | 1429 | V1429C | E | 2.6e-08 | 9.3e-09 | 1.2e-08 | 1.577E-08 | 8.965E-09 | 9.300E-09 | 2.600E-08 |
| 28566 | 1429 | V1429E | E | 1.1e-08 | 1e-08 | 5.1e-09 | 8.700E-09 | 3.158E-09 | 5.100E-09 | 1.100E-08 |
| 28567 | 1429 | V1429Q | E | 2.3e-09 | 7.8e-09 | 3.2e-09 | 4.433E-09 | 2.950E-09 | 2.300E-09 | 7.800E-09 |
| 28568 | 1429 | V1429G | E | 3.3e-09 | 5.1e-09 | 6.4e-09 | 4.933E-09 | 1.557E-09 | 3.300E-09 | 6.400E-09 |
| 28569 | 1429 | V1429H | E | 4.7e-09 | 3.7e-09 | 7.9e-09 | 5.433E-09 | 2.194E-09 | 3.700E-09 | 7.900E-09 |
| 28570 | 1429 | V1429I | E | 4.4e-08 | 8.2e-09 | 1.5e-08 | 2.240E-08 | 1.901E-08 | 8.200E-09 | 4.400E-08 |
| 28571 | 1429 | V1429L | E | 6.1e-09 | 1.1e-08 | 8.8e-09 | 8.633E-09 | 2.454E-09 | 6.100E-09 | 1.100E-08 |
| 28572 | 1429 | V1429K | E | 1.8e-08 | 1.2e-08 | 3.9e-09 | 1.130E-08 | 7.076E-09 | 3.900E-09 | 1.800E-08 |
| 28573 | 1429 | V1429M | E | 6.2e-09 | 9.3e-09 | 6.7e-09 | 7.400E-09 | 1.664E-09 | 6.200E-09 | 9.300E-09 |
| 28574 | 1429 | V1429F | E | 7.6e-09 | 3.6e-09 | 9.1e-09 | 6.767E-09 | 2.843E-09 | 3.600E-09 | 9.100E-09 |
| 28575 | 1429 | V1429P | E | 3.2e-09 | 7.6e-09 | 2.3e-09 | 4.367E-09 | 2.836E-09 | 2.300E-09 | 7.600E-09 |
| 28576 | 1429 | V1429S | E | 1.3e-08 | 2.5e-09 | 8.7e-09 | 8.067E-09 | 5.279E-09 | 2.500E-09 | 1.300E-08 |
| 28577 | 1429 | V1429T | E | 3.6e-09 | 9.1e-09 | 2.8e-09 | 5.167E-09 | 3.430E-09 | 2.800E-09 | 9.100E-09 |
| 28578 | 1429 | V1429W | E | 1e-08 | 1.6e-08 | 4.7e-09 | 1.023E-08 | 5.654E-09 | 4.700E-09 | 1.600E-08 |
| 28579 | 1429 | V1429Y | E | 6e-09 | 3.7e-09 | 1.1e-08 | 6.900E-09 | 3.732E-09 | 3.700E-09 | 1.100E-08 |
| 28580 | 1429 | V1429V | E | 1.1e-08 | 1.6e-08 | 5e-09 | 1.067E-08 | 5.508E-09 | 5.000E-09 | 1.600E-08 |
| 28581 | 1430 | T1430A | E | 2.7e-09 | 5.1e-08 | 1.5e-08 | 2.290E-08 | 2.510E-08 | 2.700E-09 | 5.100E-08 |
| 28582 | 1430 | T1430R | E | 3.2e-09 | 7.7e-09 | 3.2e-09 | 4.700E-09 | 2.598E-09 | 3.200E-09 | 7.700E-09 |
| 28583 | 1430 | T1430N | E | 1e-08 | 2.6e-09 | 4.5e-09 | 5.700E-09 | 3.843E-09 | 2.600E-09 | 1.000E-08 |
| 28584 | 1430 | T1430D | E | 4.3e-09 | 3.5e-09 | 2.6e-08 | 1.127E-08 | 1.277E-08 | 3.500E-09 | 2.600E-08 |
| 28585 | 1430 | T1430C | E | 9.1e-09 | 5.4e-09 | 4.5e-09 | 6.333E-09 | 2.438E-09 | 4.500E-09 | 9.100E-09 |
| 28586 | 1430 | T1430E | E | 4.5e-09 | 1.7e-08 | 3.8e-09 | 8.433E-09 | 7.427E-09 | 3.800E-09 | 1.700E-08 |
| 28587 | 1430 | T1430Q | E | 6.9e-09 | 3.1e-09 | 3.5e-09 | 4.500E-09 | 2.088E-09 | 3.100E-09 | 6.900E-09 |
| 28588 | 1430 | T1430G | E | 6.4e-09 | 2.6e-09 | 6.3e-09 | 5.100E-09 | 2.166E-09 | 2.600E-09 | 6.400E-09 |
| 28589 | 1430 | T1430H | E | 3.8e-09 | 8.8e-09 | 6.8e-09 | 6.467E-09 | 2.517E-09 | 3.800E-09 | 8.800E-09 |
| 28590 | 1430 | T1430I | E | 8.9e-09 | 3.8e-09 | 1e-08 | 7.567E-09 | 3.308E-09 | 3.800E-09 | 1.000E-08 |
| 28591 | 1430 | T1430L | E | 1.3e-08 | 1.4e-08 | 7.8e-09 | 1.160E-08 | 3.329E-09 | 7.800E-09 | 1.400E-08 |
| 28592 | 1430 | T1430K | E | 1.6e-08 | 8.2e-09 | 3.5e-09 | 9.233E-09 | 6.314E-09 | 3.500E-09 | 1.600E-08 |
| 28593 | 1430 | T1430M | E | 3.7e-09 | 4.5e-09 | 1.8e-08 | 8.733E-09 | 8.035E-09 | 3.700E-09 | 1.800E-08 |
| 28594 | 1430 | T1430F | E | 1.7e-08 | 1.1e-08 | 6.2e-09 | 1.140E-08 | 5.411E-09 | 6.200E-09 | 1.700E-08 |
| 28595 | 1430 | T1430P | E | 5.3e-09 | 3e-09 | 1.5e-08 | 7.767E-09 | 6.369E-09 | 3.000E-09 | 1.500E-08 |
| 28596 | 1430 | T1430S | E | 7.1e-09 | 1.4e-08 | 7.5e-09 | 9.533E-09 | 3.873E-09 | 7.100E-09 | 1.400E-08 |
| 28597 | 1430 | T1430T | E | 1e-08 | 6.3e-09 | 3e-09 | 6.433E-09 | 3.502E-09 | 3.000E-09 | 1.000E-08 |
| 28598 | 1430 | T1430W | E | 4.2e-09 | 9.5e-09 | 4.2e-09 | 5.967E-09 | 3.060E-09 | 4.200E-09 | 9.500E-09 |
| 28599 | 1430 | T1430Y | E | 7.7e-09 | 6.7e-09 | 6.6e-09 | 7.000E-09 | 6.083E-10 | 6.600E-09 | 7.700E-09 |
| 28600 | 1430 | T1430V | E | 1.5e-08 | 1.8e-08 | 1.3e-08 | 1.533E-08 | 2.517E-09 | 1.300E-08 | 1.800E-08 |
| 28601 | 1431 | V1431A | E | 8.7e-09 | 3.7e-09 | 2.4e-09 | 4.933E-09 | 3.326E-09 | 2.400E-09 | 8.700E-09 |
| 28602 | 1431 | V1431R | E | 4.5e-09 | 5.4e-09 | 8.1e-09 | 6.000E-09 | 1.873E-09 | 4.500E-09 | 8.100E-09 |
| 28603 | 1431 | V1431N | E | 1.9e-08 | 9.4e-09 | 5.8e-09 | 1.140E-08 | 6.823E-09 | 5.800E-09 | 1.900E-08 |
| 28604 | 1431 | V1431D | E | 5.1e-09 | 8.2e-09 | 5.2e-09 | 6.167E-09 | 1.762E-09 | 5.100E-09 | 8.200E-09 |
| 28605 | 1431 | V1431C | E | 9.4e-09 | 1.7e-08 | 1.7e-09 | 9.367E-09 | 7.650E-09 | 1.700E-09 | 1.700E-08 |
| 28606 | 1431 | V1431E | E | 6.8e-09 | 2.5e-09 | 9.8e-09 | 6.367E-09 | 3.669E-09 | 2.500E-09 | 9.800E-09 |
| 28607 | 1431 | V1431Q | E | 1.3e-08 | 1.1e-08 | 6e-09 | 1.000E-08 | 3.606E-09 | 6.000E-09 | 1.300E-08 |
| 28608 | 1431 | V1431G | E | 4.1e-09 | 4.2e-09 | 1.6e-08 | 8.100E-09 | 6.842E-09 | 4.100E-09 | 1.600E-08 |
| 28609 | 1431 | V1431H | E | 9.9e-09 | 4.5e-09 | 1.5e-08 | 9.800E-09 | 5.251E-09 | 4.500E-09 | 1.500E-08 |
| 28610 | 1431 | V1431I | E | 1.2e-09 | 2.3e-08 | 1.2e-08 | 1.207E-08 | 1.090E-08 | 1.200E-09 | 2.300E-08 |
| 28611 | 1431 | V1431L | E | 9.1e-09 | 8.7e-09 | 6.1e-09 | 7.967E-09 | 1.629E-09 | 6.100E-09 | 9.100E-09 |
| 28612 | 1431 | V1431K | E | 3.5e-09 | 2.1e-08 | 5.8e-09 | 1.010E-08 | 9.509E-09 | 3.500E-09 | 2.100E-08 |
| 28613 | 1431 | V1431M | E | 7.1e-09 | 1.1e-08 | 5.9e-09 | 8.000E-09 | 2.666E-09 | 5.900E-09 | 1.100E-08 |
| 28614 | 1431 | V1431F | E | 1.4e-08 | 1.2e-08 | 3.2e-09 | 9.733E-09 | 5.746E-09 | 3.200E-09 | 1.400E-08 |
| 28615 | 1431 | V1431P | E | 7e-09 | 1.2e-08 | 1e-08 | 9.667E-09 | 2.517E-09 | 7.000E-09 | 1.200E-08 |
| 28616 | 1431 | V1431S | E | 4.2e-09 | 5.7e-09 | 7.9e-09 | 5.933E-09 | 1.861E-09 | 4.200E-09 | 7.900E-09 |
| 28617 | 1431 | V1431T | E | 8.9e-09 | 8.5e-09 | 5.6e-09 | 7.667E-09 | 1.801E-09 | 5.600E-09 | 8.900E-09 |
| 28618 | 1431 | V1431W | E | 3.7e-09 | 5e-09 | 8.1e-09 | 5.600E-09 | 2.261E-09 | 3.700E-09 | 8.100E-09 |
| 28619 | 1431 | V1431Y | E | 1.4e-08 | 2.2e-09 | 3.7e-09 | 6.633E-09 | 6.424E-09 | 2.200E-09 | 1.400E-08 |
| 28620 | 1431 | V1431V | E | 7.8e-09 | 1e-08 | 3.4e-09 | 7.067E-09 | 3.361E-09 | 3.400E-09 | 1.000E-08 |
| 28621 | 1432 | S1432A | E | 8e-09 | 2.4e-08 | 1.2e-08 | 1.467E-08 | 8.327E-09 | 8.000E-09 | 2.400E-08 |
| 28622 | 1432 | S1432R | E | 4.6e-09 | 8.4e-09 | 9.8e-09 | 7.600E-09 | 2.691E-09 | 4.600E-09 | 9.800E-09 |
| 28623 | 1432 | S1432N | E | 3.4e-09 | 6e-09 | 1.1e-08 | 6.800E-09 | 3.863E-09 | 3.400E-09 | 1.100E-08 |
| 28624 | 1432 | S1432D | E | 5.3e-09 | 1e-08 | 8.4e-09 | 7.900E-09 | 2.390E-09 | 5.300E-09 | 1.000E-08 |
| 28625 | 1432 | S1432C | E | 5.6e-09 | 4.5e-09 | 8.2e-09 | 6.100E-09 | 1.900E-09 | 4.500E-09 | 8.200E-09 |
| 28626 | 1432 | S1432E | E | 1.4e-08 | 4.2e-09 | 1.3e-08 | 1.040E-08 | 5.393E-09 | 4.200E-09 | 1.400E-08 |
| 28627 | 1432 | S1432Q | E | 1.4e-08 | 6.5e-09 | 8.4e-09 | 9.633E-09 | 3.899E-09 | 6.500E-09 | 1.400E-08 |
| 28628 | 1432 | S1432G | E | 2.7e-09 | 5.9e-09 | 7.6e-09 | 5.400E-09 | 2.488E-09 | 2.700E-09 | 7.600E-09 |
| 28629 | 1432 | S1432H | E | 1.1e-08 | 4e-09 | 5.2e-09 | 6.733E-09 | 3.743E-09 | 4.000E-09 | 1.100E-08 |
| 28630 | 1432 | S1432I | E | 1e-08 | 6.7e-09 | 1.5e-08 | 1.057E-08 | 4.179E-09 | 6.700E-09 | 1.500E-08 |
| 28631 | 1432 | S1432L | E | 2.9e-09 | 6.7e-09 | 8.2e-09 | 5.933E-09 | 2.732E-09 | 2.900E-09 | 8.200E-09 |
| 28632 | 1432 | S1432K | E | 8.4e-09 | 6.5e-09 | 1.5e-08 | 9.967E-09 | 4.461E-09 | 6.500E-09 | 1.500E-08 |
| 28633 | 1432 | S1432M | E | 4.9e-09 | 8.4e-09 | 5.3e-09 | 6.200E-09 | 1.916E-09 | 4.900E-09 | 8.400E-09 |
| 28634 | 1432 | S1432F | E | 1.7e-08 | 1.2e-08 | 1.9e-08 | 1.600E-08 | 3.606E-09 | 1.200E-08 | 1.900E-08 |
| 28635 | 1432 | S1432P | E | 3.6e-09 | 1.4e-08 | 5.1e-09 | 7.567E-09 | 5.622E-09 | 3.600E-09 | 1.400E-08 |
| 28636 | 1432 | S1432S | E | 1e-08 | 2.2e-09 | 6.2e-09 | 6.133E-09 | 3.900E-09 | 2.200E-09 | 1.000E-08 |
| 28637 | 1432 | S1432T | E | 2.1e-08 | 1.1e-08 | 4.2e-09 | 1.207E-08 | 8.451E-09 | 4.200E-09 | 2.100E-08 |
| 28638 | 1432 | S1432W | E | 7.3e-09 | 5.8e-09 | 6.3e-09 | 6.467E-09 | 7.638E-10 | 5.800E-09 | 7.300E-09 |
| 28639 | 1432 | S1432Y | E | 8e-09 | 2.1e-09 | 6.2e-09 | 5.433E-09 | 3.024E-09 | 2.100E-09 | 8.000E-09 |
| 28640 | 1432 | S1432V | E | 5.5e-09 | 9.1e-09 | 8.6e-09 | 7.733E-09 | 1.950E-09 | 5.500E-09 | 9.100E-09 |
| 28641 | 1433 | W1433A | E | 2.7e-09 | 2.1e-08 | 3.5e-09 | 9.067E-09 | 1.034E-08 | 2.700E-09 | 2.100E-08 |
| 28642 | 1433 | W1433R | E | 6.5e-09 | 8.8e-09 | 1.4e-08 | 9.767E-09 | 3.842E-09 | 6.500E-09 | 1.400E-08 |
| 28643 | 1433 | W1433N | E | 4.4e-09 | 7.7e-09 | 8.7e-09 | 6.933E-09 | 2.250E-09 | 4.400E-09 | 8.700E-09 |
| 28644 | 1433 | W1433D | E | 2.3e-08 | 1.3e-08 | 1.6e-08 | 1.733E-08 | 5.132E-09 | 1.300E-08 | 2.300E-08 |

| | | | | | | | | | | |
|---|---|---|---|---|---|---|---|---|---|---|
| 28645 | 1433 | W1433C | E | 9e-09 | 5.8e-09 | 5.6e-09 | 6.800E-09 | 1.908E-09 | 5.600E-09 | 9.000E-09 |
| 28646 | 1433 | W1433E | E | 7.5e-09 | 1.1e-08 | 1.3e-08 | 1.050E-08 | 2.784E-09 | 7.500E-09 | 1.300E-08 |
| 28647 | 1433 | W1433Q | E | 4.9e-09 | 1.2e-08 | 3.5e-09 | 6.800E-09 | 4.557E-09 | 3.500E-09 | 1.200E-08 |
| 28648 | 1433 | W1433G | E | 2.1e-09 | 8.9e-09 | 7.4e-09 | 6.133E-09 | 3.573E-09 | 2.100E-09 | 8.900E-09 |
| 28649 | 1433 | W1433H | E | 4.5e-09 | 6.1e-09 | 1.1e-09 | 3.900E-09 | 2.553E-09 | 1.100E-09 | 6.100E-09 |
| 28650 | 1433 | W1433I | E | 4.4e-09 | 7.2e-09 | 5.5e-09 | 5.700E-09 | 1.411E-09 | 4.400E-09 | 7.200E-09 |
| 28651 | 1433 | W1433L | E | 1.7e-08 | 2.9e-08 | 1.8e-08 | 2.133E-08 | 6.658E-09 | 1.700E-08 | 2.900E-08 |
| 28652 | 1433 | W1433K | E | 7.1e-09 | 1e-08 | 2.7e-09 | 6.600E-09 | 3.676E-09 | 2.700E-09 | 1.000E-08 |
| 28653 | 1433 | W1433M | E | 7.4e-09 | 4.2e-09 | 9e-09 | 6.867E-09 | 2.444E-09 | 4.200E-09 | 9.000E-09 |
| 28654 | 1433 | W1433F | E | 2.7e-09 | 1.3e-08 | 1.3e-08 | 9.567E-09 | 5.947E-09 | 2.700E-09 | 1.300E-08 |
| 28655 | 1433 | W1433P | E | 1.1e-08 | 5.4e-09 | 2.5e-09 | 6.300E-09 | 4.321E-09 | 2.500E-09 | 1.100E-08 |
| 28656 | 1433 | W1433S | E | 4.1e-09 | 3e-09 | 9.5e-09 | 5.533E-09 | 3.479E-09 | 3.000E-09 | 9.500E-09 |
| 28657 | 1433 | W1433T | E | 6e-09 | 1.2e-08 | 1e-08 | 9.333E-09 | 3.055E-09 | 6.000E-09 | 1.200E-08 |
| 28658 | 1433 | W1433W | E | 7e-09 | 4.7e-09 | 1.3e-08 | 8.233E-09 | 4.285E-09 | 4.700E-09 | 1.300E-08 |
| 28659 | 1433 | W1433Y | E | 1.3e-08 | 8.9e-09 | 1.1e-08 | 1.097E-08 | 2.050E-09 | 8.900E-09 | 1.300E-08 |
| 28660 | 1433 | W1433V | E | 6.5e-09 | 1.1e-08 | 1.4e-08 | 1.050E-08 | 3.775E-09 | 6.500E-09 | 1.400E-08 |
| 28661 | 1434 | N1434A | E | 8.1e-09 | 2.3e-09 | 1.3e-08 | 7.800E-09 | 5.356E-09 | 2.300E-09 | 1.300E-08 |
| 28662 | 1434 | N1434R | E | 2.7e-09 | 7.9e-09 | 1.1e-08 | 7.200E-09 | 4.194E-09 | 2.700E-09 | 1.100E-08 |
| 28663 | 1434 | N1434N | E | 1e-08 | 4.8e-09 | 3.8e-09 | 6.200E-09 | 3.329E-09 | 3.800E-09 | 1.000E-08 |
| 28664 | 1434 | N1434D | E | 5.7e-09 | 2.7e-09 | 2.6e-09 | 3.667E-09 | 1.762E-09 | 2.600E-09 | 5.700E-09 |
| 28665 | 1434 | N1434C | E | 2.5e-09 | 3.2e-09 | 4.4e-09 | 3.367E-09 | 9.609E-10 | 2.500E-09 | 4.400E-09 |
| 28666 | 1434 | N1434E | E | 7.2e-09 | 1.2e-08 | 2.4e-09 | 7.200E-09 | 4.800E-09 | 2.400E-09 | 1.200E-08 |
| 28667 | 1434 | N1434Q | E | 1e-08 | 5.9e-09 | 1.2e-08 | 9.300E-09 | 3.110E-09 | 5.900E-09 | 1.200E-08 |
| 28668 | 1434 | N1434G | E | 2.3e-08 | 6e-09 | 1.4e-08 | 1.433E-08 | 8.505E-09 | 6.000E-09 | 2.300E-08 |
| 28669 | 1434 | N1434H | E | 6.7e-09 | 4.2e-09 | 4.3e-09 | 5.067E-09 | 1.415E-09 | 4.200E-09 | 6.700E-09 |
| 28670 | 1434 | N1434I | E | 1.1e-08 | 5.6e-09 | 1.3e-08 | 9.867E-09 | 3.828E-09 | 5.600E-09 | 1.300E-08 |
| 28671 | 1434 | N1434L | E | 8.5e-09 | 2.7e-08 | 8.4e-09 | 1.463E-08 | 1.071E-08 | 8.400E-09 | 2.700E-08 |
| 28672 | 1434 | N1434K | E | 1.1e-08 | 1.3e-08 | 8.2e-09 | 1.073E-08 | 2.411E-09 | 8.200E-09 | 1.300E-08 |
| 28673 | 1434 | N1434M | E | 7.1e-09 | 1.2e-08 | 6.1e-09 | 8.400E-09 | 3.158E-09 | 6.100E-09 | 1.200E-08 |
| 28674 | 1434 | N1434F | E | 6.1e-09 | 4.6e-09 | 5.7e-09 | 5.467E-09 | 7.767E-10 | 4.600E-09 | 6.100E-09 |
| 28675 | 1434 | N1434P | E | 7.2e-09 | 2e-09 | 8.3e-09 | 5.833E-09 | 3.365E-09 | 2.000E-09 | 8.300E-09 |
| 28676 | 1434 | N1434S | E | 1.4e-08 | 8.2e-09 | 1.1e-08 | 1.107E-08 | 2.901E-09 | 8.200E-09 | 1.400E-08 |
| 28677 | 1434 | N1434T | E | 8e-09 | 8.1e-09 | 4.7e-09 | 6.933E-09 | 1.935E-09 | 4.700E-09 | 8.100E-09 |
| 28678 | 1434 | N1434W | E | 9.7e-09 | 1.2e-08 | 3.8e-09 | 8.500E-09 | 4.230E-09 | 3.800E-09 | 1.200E-08 |
| 28679 | 1434 | N1434Y | E | 7.1e-09 | 1.1e-08 | 6.2e-09 | 8.100E-09 | 2.551E-09 | 6.200E-09 | 1.100E-08 |
| 28680 | 1434 | N1434V | E | 5e-09 | 5.3e-09 | 7.4e-09 | 5.900E-09 | 1.308E-09 | 5.000E-09 | 7.400E-09 |
| 28681 | 1435 | S1435A | E | 5.3e-09 | 1e-08 | 1.8e-08 | 1.110E-08 | 6.421E-09 | 5.300E-09 | 1.800E-08 |
| 28682 | 1435 | S1435R | E | 9.5e-09 | 1.1e-08 | 4.4e-09 | 8.300E-09 | 3.460E-09 | 4.400E-09 | 1.100E-08 |
| 28683 | 1435 | S1435N | E | 6.4e-09 | 3.3e-09 | 6.8e-09 | 5.500E-09 | 1.916E-09 | 3.300E-09 | 6.800E-09 |
| 28684 | 1435 | S1435D | E | 8e-09 | 1.3e-08 | 5e-09 | 8.667E-09 | 4.041E-09 | 5.000E-09 | 1.300E-08 |
| 28685 | 1435 | S1435C | E | 5.6e-09 | 2.3e-09 | 6.6e-09 | 4.833E-09 | 2.250E-09 | 2.300E-09 | 6.600E-09 |
| 28686 | 1435 | S1435E | E | 2.8e-09 | 8.7e-10 | 1.4e-08 | 5.890E-09 | 7.089E-09 | 8.700E-10 | 1.400E-08 |
| 28687 | 1435 | S1435Q | E | 3.9e-09 | 9e-09 | 8e-09 | 6.967E-09 | 2.702E-09 | 3.900E-09 | 9.000E-09 |
| 28688 | 1435 | S1435G | E | 8.1e-09 | 9.9e-09 | 1e-08 | 9.333E-09 | 1.069E-09 | 8.100E-09 | 1.000E-08 |
| 28689 | 1435 | S1435H | E | 1.3e-08 | 8.8e-09 | 4.9e-09 | 8.900E-09 | 4.051E-09 | 4.900E-09 | 1.300E-08 |
| 28690 | 1435 | S1435I | E | 5.6e-09 | 1.5e-08 | 3.2e-09 | 7.933E-09 | 6.236E-09 | 3.200E-09 | 1.500E-08 |
| 28691 | 1435 | S1435L | E | 2.8e-09 | 7.5e-09 | 7.7e-09 | 6.000E-09 | 2.773E-09 | 2.800E-09 | 7.700E-09 |
| 28692 | 1435 | S1435K | E | 5.3e-09 | 8.3e-09 | 6.8e-09 | 6.800E-09 | 1.500E-09 | 5.300E-09 | 8.300E-09 |
| 28693 | 1435 | S1435M | E | 9.2e-09 | 5e-09 | 1.6e-08 | 1.007E-08 | 5.551E-09 | 5.000E-09 | 1.600E-08 |
| 28694 | 1435 | S1435F | E | 6.2e-09 | 3.4e-09 | 3.9e-09 | 4.500E-09 | 1.493E-09 | 3.400E-09 | 6.200E-09 |
| 28695 | 1435 | S1435P | E | 1.5e-08 | 9.9e-09 | 8.3e-09 | 1.107E-08 | 3.499E-09 | 8.300E-09 | 1.500E-08 |
| 28696 | 1435 | S1435S | E | 2.9e-09 | 6.5e-09 | 7e-09 | 5.467E-09 | 2.237E-09 | 2.900E-09 | 7.000E-09 |
| 28697 | 1435 | S1435T | E | 1.6e-08 | 1.2e-08 | 4.2e-09 | 1.073E-08 | 6.001E-09 | 4.200E-09 | 1.600E-08 |
| 28698 | 1435 | S1435W | E | 1e-08 | 1.4e-08 | 3.3e-09 | 9.100E-09 | 5.406E-09 | 3.300E-09 | 1.400E-08 |
| 28699 | 1435 | S1435Y | E | 5.5e-09 | 2.9e-09 | 2.5e-09 | 3.633E-09 | 1.629E-09 | 2.500E-09 | 5.500E-09 |
| 28700 | 1435 | S1435V | E | 9.9e-09 | 7.8e-09 | 6.3e-09 | 8.000E-09 | 1.808E-09 | 6.300E-09 | 9.900E-09 |
| 28701 | 1436 | G1436A | E | 4.7e-09 | 1.3e-08 | 9.7e-09 | 9.133E-09 | 4.179E-09 | 4.700E-09 | 1.300E-08 |
| 28702 | 1436 | G1436R | E | 7.3e-09 | 5.4e-09 | 2.1e-09 | 4.933E-09 | 2.631E-09 | 2.100E-09 | 7.300E-09 |
| 28703 | 1436 | G1436N | E | 6.2e-09 | 5e-09 | 8.9e-09 | 6.700E-09 | 1.997E-09 | 5.000E-09 | 8.900E-09 |
| 28704 | 1436 | G1436D | E | 1.3e-08 | 4.1e-09 | 4e-09 | 7.033E-09 | 5.168E-09 | 4.000E-09 | 1.300E-08 |
| 28705 | 1436 | G1436C | E | 1.4e-08 | 1.1e-08 | 1.3e-08 | 1.267E-08 | 1.528E-09 | 1.100E-08 | 1.400E-08 |
| 28706 | 1436 | G1436E | E | 1.1e-08 | 5.9e-09 | 1.1e-08 | 9.300E-09 | 2.944E-09 | 5.900E-09 | 1.100E-08 |
| 28707 | 1436 | G1436Q | E | 8.6e-09 | 1.6e-09 | 1e-08 | 6.733E-09 | 4.500E-09 | 1.600E-09 | 1.000E-08 |
| 28708 | 1436 | G1436G | E | 4.1e-09 | 1e-08 | 5.8e-09 | 6.633E-09 | 3.037E-09 | 4.100E-09 | 1.000E-08 |
| 28709 | 1436 | G1436H | E | 6.6e-09 | 1e-08 | 9.7e-09 | 8.767E-09 | 1.882E-09 | 6.600E-09 | 1.000E-08 |
| 28710 | 1436 | G1436I | E | 3.7e-09 | 5.1e-09 | 3.6e-09 | 4.133E-09 | 8.386E-10 | 3.600E-09 | 5.100E-09 |
| 28711 | 1436 | G1436L | E | 2.3e-09 | 2.9e-09 | 5.6e-09 | 3.600E-09 | 1.758E-09 | 2.300E-09 | 5.600E-09 |
| 28712 | 1436 | G1436K | E | 1.5e-08 | 6.7e-09 | 3.4e-09 | 8.367E-09 | 5.977E-09 | 3.400E-09 | 1.500E-08 |
| 28713 | 1436 | G1436M | E | 1.4e-08 | 8.6e-09 | 5.5e-09 | 9.367E-09 | 4.302E-09 | 5.500E-09 | 1.400E-08 |
| 28714 | 1436 | G1436F | E | 6.8e-09 | 8.6e-09 | 7.9e-09 | 7.767E-09 | 9.074E-10 | 6.800E-09 | 8.600E-09 |
| 28715 | 1436 | G1436P | E | 9.4e-09 | 1.3e-08 | 6.5e-09 | 9.633E-09 | 3.256E-09 | 6.500E-09 | 1.300E-08 |
| 28716 | 1436 | G1436S | E | 2.3e-09 | 1.5e-08 | 1.2e-08 | 9.767E-09 | 6.638E-09 | 2.300E-09 | 1.500E-08 |
| 28717 | 1436 | G1436T | E | 8.1e-09 | 1.7e-09 | 9.2e-09 | 6.333E-09 | 4.050E-09 | 1.700E-09 | 9.200E-09 |
| 28718 | 1436 | G1436W | E | 1.1e-08 | 5.1e-09 | 1.3e-08 | 9.700E-09 | 4.107E-09 | 5.100E-09 | 1.300E-08 |
| 28719 | 1436 | G1436Y | E | 7.2e-09 | 3.8e-09 | 3.9e-09 | 4.967E-09 | 1.935E-09 | 3.800E-09 | 7.200E-09 |
| 28720 | 1436 | G1436V | E | 1.1e-08 | 5.2e-09 | 9.5e-09 | 8.567E-09 | 3.011E-09 | 5.200E-09 | 1.100E-08 |
| 28721 | 1437 | A1437A | E | 1.2e-08 | 4.4e-09 | 6.7e-09 | 7.700E-09 | 3.897E-09 | 4.400E-09 | 1.200E-08 |
| 28722 | 1437 | A1437R | E | 4.8e-09 | 7.1e-09 | 4.3e-09 | 5.400E-09 | 1.493E-09 | 4.300E-09 | 7.100E-09 |
| 28723 | 1437 | A1437N | E | 2.1e-08 | 9.9e-09 | 1e-08 | 1.363E-08 | 6.380E-09 | 9.900E-09 | 2.100E-08 |
| 28724 | 1437 | A1437D | E | 1.4e-08 | 1.4e-08 | 2.6e-09 | 1.020E-08 | 6.582E-09 | 2.600E-09 | 1.400E-08 |
| 28725 | 1437 | A1437C | E | 3e-09 | 7e-09 | 4.7e-09 | 4.900E-09 | 2.007E-09 | 3.000E-09 | 7.000E-09 |
| 28726 | 1437 | A1437E | E | 3e-09 | 1.2e-08 | 4.9e-09 | 6.633E-09 | 4.744E-09 | 3.000E-09 | 1.200E-08 |
| 28727 | 1437 | A1437Q | E | 4.7e-09 | 8.3e-09 | 1.6e-08 | 9.667E-09 | 5.773E-09 | 4.700E-09 | 1.600E-08 |
| 28728 | 1437 | A1437G | E | 6e-09 | 5.5e-09 | 3.8e-09 | 5.100E-09 | 1.153E-09 | 3.800E-09 | 6.000E-09 |
| 28729 | 1437 | A1437H | E | 1e-08 | 7.9e-09 | 2e-08 | 1.263E-08 | 6.466E-09 | 7.900E-09 | 2.000E-08 |

| | | | | | | | | | | |
|---|---|---|---|---|---|---|---|---|---|---|
| 28730 | 1437 | A1437I | E | 2.1e-09 | 5.2e-09 | 1.6e-08 | 7.767E-09 | 7.297E-09 | 2.100E-09 | 1.600E-08 |
| 28731 | 1437 | A1437L | E | 1.6e-08 | 3.1e-09 | 1.4e-08 | 1.103E-08 | 6.943E-09 | 3.100E-09 | 1.600E-08 |
| 28732 | 1437 | A1437K | E | 5.8e-09 | 8e-09 | 9.2e-09 | 7.667E-09 | 1.724E-09 | 5.800E-09 | 9.200E-09 |
| 28733 | 1437 | A1437M | E | 2.9e-09 | 5.2e-09 | 7.6e-09 | 5.233E-09 | 2.350E-09 | 2.900E-09 | 7.600E-09 |
| 28734 | 1437 | A1437F | E | 1.5e-08 | 1e-08 | 1.1e-08 | 1.200E-08 | 2.646E-09 | 1.000E-08 | 1.500E-08 |
| 28735 | 1437 | A1437P | E | 2.3e-09 | 9.5e-09 | 1.2e-08 | 7.933E-09 | 5.036E-09 | 2.300E-09 | 1.200E-08 |
| 28736 | 1437 | A1437S | E | 7.7e-09 | 1.6e-08 | 9e-09 | 1.090E-08 | 4.464E-09 | 7.700E-09 | 1.600E-08 |
| 28737 | 1437 | A1437T | E | 1.1e-08 | 7.4e-09 | 7.3e-09 | 8.567E-09 | 2.108E-09 | 7.300E-09 | 1.100E-08 |
| 28738 | 1437 | A1437W | E | 6.4e-09 | 3.9e-09 | 1.3e-08 | 7.767E-09 | 4.701E-09 | 3.900E-09 | 1.300E-08 |
| 28739 | 1437 | A1437Y | E | 1.5e-08 | 1.4e-08 | 5.3e-09 | 1.143E-08 | 5.335E-09 | 5.300E-09 | 1.500E-08 |
| 28740 | 1437 | A1437V | E | 9e-09 | 1.8e-09 | 1.1e-08 | 7.267E-09 | 4.839E-09 | 1.800E-09 | 1.100E-08 |
| 28741 | 1438 | L1438A | E | 7.5e-09 | 2.3e-09 | 1e-08 | 6.600E-09 | 3.928E-09 | 2.300E-09 | 1.000E-08 |
| 28742 | 1438 | L1438R | E | 5.1e-09 | 1.8e-09 | 5.8e-09 | 4.233E-09 | 2.136E-09 | 1.800E-09 | 5.800E-09 |
| 28743 | 1438 | L1438N | E | 7.3e-09 | 6.7e-09 | 7.7e-09 | 7.233E-09 | 5.033E-10 | 6.700E-09 | 7.700E-09 |
| 28744 | 1438 | L1438D | E | 3.2e-09 | 2.2e-08 | 5.6e-09 | 1.027E-08 | 1.023E-08 | 3.200E-09 | 2.200E-08 |
| 28745 | 1438 | L1438C | E | 9.5e-09 | 1.3e-08 | 6.7e-09 | 9.733E-09 | 3.156E-09 | 6.700E-09 | 1.300E-08 |
| 28746 | 1438 | L1438E | E | 2.9e-09 | 1.5e-08 | 8.9e-09 | 8.933E-09 | 6.050E-09 | 2.900E-09 | 1.500E-08 |
| 28747 | 1438 | L1438Q | E | 1.1e-08 | 5.7e-09 | 5.5e-09 | 7.400E-09 | 3.119E-09 | 5.500E-09 | 1.100E-08 |
| 28748 | 1438 | L1438G | E | 1.4e-08 | 9.9e-09 | 1e-08 | 1.130E-08 | 2.339E-09 | 9.900E-09 | 1.400E-08 |
| 28749 | 1438 | L1438H | E | 3.1e-08 | 1.7e-08 | 5.5e-09 | 1.783E-08 | 1.277E-08 | 5.500E-09 | 3.100E-08 |
| 28750 | 1438 | L1438I | E | 1.1e-08 | 3.7e-09 | 8.4e-09 | 7.700E-09 | 3.700E-09 | 3.700E-09 | 1.100E-08 |
| 28751 | 1438 | L1438L | E | 1.3e-08 | 1e-08 | 6e-09 | 9.667E-09 | 3.512E-09 | 6.000E-09 | 1.300E-08 |
| 28752 | 1438 | L1438K | E | 2.1e-08 | 1.7e-08 | 1.4e-08 | 1.733E-08 | 3.512E-09 | 1.400E-08 | 2.100E-08 |
| 28753 | 1438 | L1438M | E | 3.7e-08 | 1.3e-08 | 3.8e-09 | 1.793E-08 | 1.714E-08 | 3.800E-09 | 3.700E-08 |
| 28754 | 1438 | L1438F | E | 3.7e-09 | 1.7e-08 | 1.8e-08 | 1.290E-08 | 7.983E-09 | 3.700E-09 | 1.800E-08 |
| 28755 | 1438 | L1438P | E | 1.3e-08 | 7e-09 | 4.6e-09 | 8.200E-09 | 4.327E-09 | 4.600E-09 | 1.300E-08 |
| 28756 | 1438 | L1438S | E | 9.9e-09 | 7.7e-09 | 4.9e-09 | 7.500E-09 | 2.506E-09 | 4.900E-09 | 9.900E-09 |
| 28757 | 1438 | L1438T | E | 5.3e-09 | 1e-08 | 7.7e-09 | 7.667E-09 | 2.350E-09 | 5.300E-09 | 1.000E-08 |
| 28758 | 1438 | L1438W | E | 7.4e-09 | 6.9e-09 | 7.9e-09 | 7.400E-09 | 5.000E-10 | 6.900E-09 | 7.900E-09 |
| 28759 | 1438 | L1438Y | E | 1.4e-08 | 1e-08 | 6e-09 | 1.000E-08 | 4.000E-09 | 6.000E-09 | 1.400E-08 |
| 28760 | 1438 | L1438V | E | 1.4e-08 | 8.8e-09 | 2.5e-09 | 8.433E-09 | 5.759E-09 | 2.500E-09 | 1.400E-08 |
| 28761 | 1439 | T1439A | E | 3.5e-09 | 6.6e-09 | 8.2e-09 | 6.100E-09 | 2.390E-09 | 3.500E-09 | 8.200E-09 |
| 28762 | 1439 | T1439R | E | 3.8e-09 | 4.3e-09 | 1.1e-08 | 6.367E-09 | 4.020E-09 | 3.800E-09 | 1.100E-08 |
| 28763 | 1439 | T1439N | E | 4.4e-09 | 3.6e-08 | 1.1e-08 | 1.713E-08 | 1.667E-08 | 4.400E-09 | 3.600E-08 |
| 28764 | 1439 | T1439D | E | 2.3e-09 | 1.2e-08 | 6.5e-09 | 6.933E-09 | 4.864E-09 | 2.300E-09 | 1.200E-08 |
| 28765 | 1439 | T1439C | E | 9.9e-09 | 3.7e-09 | 3.9e-09 | 5.833E-09 | 3.523E-09 | 3.700E-09 | 9.900E-09 |
| 28766 | 1439 | T1439E | E | 9.9e-09 | 2.2e-08 | 7.9e-09 | 1.327E-08 | 7.629E-09 | 7.900E-09 | 2.200E-08 |
| 28767 | 1439 | T1439Q | E | 6.9e-09 | 1.8e-08 | 6.4e-09 | 1.043E-08 | 6.558E-09 | 6.400E-09 | 1.800E-08 |
| 28768 | 1439 | T1439G | E | 1.1e-08 | 1.4e-08 | 4.3e-09 | 9.767E-09 | 4.966E-09 | 4.300E-09 | 1.400E-08 |
| 28769 | 1439 | T1439H | E | 6.1e-09 | 2.9e-09 | 9.1e-09 | 6.033E-09 | 3.101E-09 | 2.900E-09 | 9.100E-09 |
| 28770 | 1439 | T1439I | E | 1.4e-08 | 3.7e-09 | 5.2e-09 | 7.633E-09 | 5.564E-09 | 3.700E-09 | 1.400E-08 |
| 28771 | 1439 | T1439L | E | 2.3e-08 | 4.3e-09 | 4.5e-08 | 2.410E-08 | 2.037E-08 | 4.300E-09 | 4.500E-08 |
| 28772 | 1439 | T1439K | E | 4.2e-09 | 3.2e-09 | 2.1e-08 | 9.467E-09 | 1.000E-08 | 3.200E-09 | 2.100E-08 |
| 28773 | 1439 | T1439M | E | 4.4e-09 | 1.2e-08 | 4.8e-09 | 7.067E-09 | 4.277E-09 | 4.400E-09 | 1.200E-08 |
| 28774 | 1439 | T1439F | E | 5.7e-09 | 8.4e-09 | 6.4e-09 | 6.833E-09 | 1.401E-09 | 5.700E-09 | 8.400E-09 |
| 28775 | 1439 | T1439P | E | 6.4e-09 | 3.5e-09 | 8.1e-09 | 6.000E-09 | 2.326E-09 | 3.500E-09 | 8.100E-09 |
| 28776 | 1439 | T1439S | E | 8.8e-09 | 6.5e-09 | 1.1e-08 | 8.767E-09 | 2.250E-09 | 6.500E-09 | 1.100E-08 |
| 28777 | 1439 | T1439T | E | 7.1e-09 | 1.9e-09 | 8.2e-09 | 5.733E-09 | 3.365E-09 | 1.900E-09 | 8.200E-09 |
| 28778 | 1439 | T1439W | E | 3e-09 | 8.7e-09 | 1.3e-08 | 8.233E-09 | 5.016E-09 | 3.000E-09 | 1.300E-08 |
| 28779 | 1439 | T1439Y | E | 5.8e-09 | 1.3e-08 | 7.7e-09 | 8.833E-09 | 3.731E-09 | 5.800E-09 | 1.300E-08 |
| 28780 | 1439 | T1439V | E | 4.7e-09 | 1.5e-09 | 2e-08 | 8.733E-09 | 9.888E-09 | 1.500E-09 | 2.000E-08 |
| 28781 | 1440 | S1440A | E | 8.7e-09 | 1.6e-08 | 4.7e-09 | 9.800E-09 | 5.730E-09 | 4.700E-09 | 1.600E-08 |
| 28782 | 1440 | S1440R | E | 1e-08 | 2.5e-09 | 6.6e-09 | 6.367E-09 | 3.755E-09 | 2.500E-09 | 1.000E-08 |
| 28783 | 1440 | S1440N | E | 1.4e-08 | 5.8e-09 | 7.3e-09 | 9.033E-09 | 4.366E-09 | 5.800E-09 | 1.400E-08 |
| 28784 | 1440 | S1440D | E | 2.8e-09 | 5.7e-09 | 5.4e-09 | 4.633E-09 | 1.595E-09 | 2.800E-09 | 5.700E-09 |
| 28785 | 1440 | S1440C | E | 7.4e-09 | 3.7e-09 | 1.2e-08 | 7.700E-09 | 4.158E-09 | 3.700E-09 | 1.200E-08 |
| 28786 | 1440 | S1440E | E | 7.7e-09 | 3.4e-09 | 7.2e-09 | 6.100E-09 | 2.352E-09 | 3.400E-09 | 7.700E-09 |
| 28787 | 1440 | S1440Q | E | 8.2e-09 | 3.6e-09 | 4.8e-09 | 5.533E-09 | 2.386E-09 | 3.600E-09 | 8.200E-09 |
| 28788 | 1440 | S1440G | E | 6.2e-09 | 7.8e-09 | 8.2e-09 | 7.400E-09 | 1.058E-09 | 6.200E-09 | 8.200E-09 |
| 28789 | 1440 | S1440H | E | 3.1e-09 | 1.3e-08 | 1.3e-08 | 9.700E-09 | 5.716E-09 | 3.100E-09 | 1.300E-08 |
| 28790 | 1440 | S1440I | E | 3.1e-09 | 4.7e-09 | 9e-09 | 5.600E-09 | 3.051E-09 | 3.100E-09 | 9.000E-09 |
| 28791 | 1440 | S1440L | E | 7.7e-09 | 5.1e-09 | 1.6e-08 | 9.600E-09 | 5.693E-09 | 5.100E-09 | 1.600E-08 |
| 28792 | 1440 | S1440K | E | 1.2e-08 | 9.6e-09 | 1.3e-08 | 1.153E-08 | 1.747E-09 | 9.600E-09 | 1.300E-08 |
| 28793 | 1440 | S1440M | E | 3.3e-09 | 1.6e-08 | 5.8e-09 | 8.367E-09 | 6.728E-09 | 3.300E-09 | 1.600E-08 |
| 28794 | 1440 | S1440F | E | 7.5e-09 | 3.7e-09 | 4.3e-09 | 5.167E-09 | 2.043E-09 | 3.700E-09 | 7.500E-09 |
| 28795 | 1440 | S1440P | E | 8.5e-09 | 3.6e-09 | 6e-09 | 6.033E-09 | 2.450E-09 | 3.600E-09 | 8.500E-09 |
| 28796 | 1440 | S1440S | E | 2.9e-09 | 2.1e-08 | 1.3e-08 | 1.230E-08 | 9.070E-09 | 2.900E-09 | 2.100E-08 |
| 28797 | 1440 | S1440T | E | 1.6e-08 | 2.3e-09 | 4.9e-09 | 7.733E-09 | 7.276E-09 | 2.300E-09 | 1.600E-08 |
| 28798 | 1440 | S1440W | E | 1.5e-08 | 7.9e-09 | 2.4e-09 | 8.433E-09 | 6.317E-09 | 2.400E-09 | 1.500E-08 |
| 28799 | 1440 | S1440Y | E | 7.4e-09 | 1.7e-08 | 1.5e-08 | 1.313E-08 | 5.065E-09 | 7.400E-09 | 1.700E-08 |
| 28800 | 1440 | S1440V | E | 2.6e-09 | 1.6e-08 | 1.6e-08 | 1.153E-08 | 7.736E-09 | 2.600E-09 | 1.600E-08 |
| 28801 | 1441 | G1441A | E | 4.5e-09 | 1.2e-08 | 1.4e-08 | 1.017E-08 | 5.008E-09 | 4.500E-09 | 1.400E-08 |
| 28802 | 1441 | G1441R | E | 2.4e-08 | 2.7e-08 | 1.8e-08 | 2.300E-08 | 4.583E-09 | 1.800E-08 | 2.700E-08 |
| 28803 | 1441 | G1441N | E | 5.8e-09 | 4.8e-09 | 6.9e-09 | 5.833E-09 | 1.050E-09 | 4.800E-09 | 6.900E-09 |
| 28804 | 1441 | G1441D | E | 6.2e-09 | 5e-09 | 1.7e-08 | 9.400E-09 | 6.609E-09 | 5.000E-09 | 1.700E-08 |
| 28805 | 1441 | G1441C | E | 6.7e-09 | 1.2e-08 | 2.6e-09 | 7.100E-09 | 4.713E-09 | 2.600E-09 | 1.200E-08 |
| 28806 | 1441 | G1441E | E | 6.8e-09 | 6.5e-09 | 1.2e-08 | 8.433E-09 | 3.092E-09 | 6.500E-09 | 1.200E-08 |
| 28807 | 1441 | G1441Q | E | 6.6e-09 | 3.2e-08 | 3e-09 | 1.387E-08 | 1.581E-08 | 3.000E-09 | 3.200E-08 |
| 28808 | 1441 | G1441G | E | 4.3e-09 | 4.9e-09 | 1.1e-08 | 6.733E-09 | 3.707E-09 | 4.300E-09 | 1.100E-08 |
| 28809 | 1441 | G1441H | E | 8e-09 | 8.7e-09 | 1.1e-08 | 9.233E-09 | 1.570E-09 | 8.000E-09 | 1.100E-08 |
| 28810 | 1441 | G1441I | E | 2.6e-08 | 4.1e-09 | 6.6e-09 | 1.223E-08 | 1.199E-08 | 4.100E-09 | 2.600E-08 |
| 28811 | 1441 | G1441L | E | 1.7e-08 | 1.4e-08 | 1.2e-08 | 1.433E-08 | 2.517E-09 | 1.200E-08 | 1.700E-08 |
| 28812 | 1441 | G1441K | E | 6.4e-09 | 1.2e-08 | 4.9e-09 | 7.767E-09 | 3.742E-09 | 4.900E-09 | 1.200E-08 |
| 28813 | 1441 | G1441M | E | 6.2e-09 | 1.1e-08 | 9.7e-09 | 8.967E-09 | 2.483E-09 | 6.200E-09 | 1.100E-08 |
| 28814 | 1441 | G1441F | E | 1.2e-08 | 6.1e-09 | 3.8e-09 | 7.300E-09 | 4.230E-09 | 3.800E-09 | 1.200E-08 |

| | | | | | | | | | | |
|---|---|---|---|---|---|---|---|---|---|---|
| 28815 | 1441 | G1441P | E | 9.4e-09 | 5.8e-09 | 1e-08 | 8.400E-09 | 2.272E-09 | 5.800E-09 | 1.000E-08 |
| 28816 | 1441 | G1441S | E | 7.2e-09 | 1.1e-08 | 4.9e-09 | 7.700E-09 | 3.081E-09 | 4.900E-09 | 1.100E-08 |
| 28817 | 1441 | G1441T | E | 2.2e-09 | 8.9e-09 | 7.7e-09 | 6.267E-09 | 3.573E-09 | 2.200E-09 | 8.900E-09 |
| 28818 | 1441 | G1441W | E | 8.6e-09 | 6.3e-09 | 3.8e-09 | 6.233E-09 | 2.401E-09 | 3.800E-09 | 8.600E-09 |
| 28819 | 1441 | G1441Y | E | 7.7e-09 | 6.2e-09 | 1.7e-08 | 1.030E-08 | 5.851E-09 | 6.200E-09 | 1.700E-08 |
| 28820 | 1441 | G1441V | E | 1.9e-09 | 3.8e-09 | 1.6e-08 | 7.233E-09 | 7.651E-09 | 1.900E-09 | 1.600E-08 |
| 28821 | 1442 | V1442A | E | 3.1e-09 | 1.5e-09 | 7.5e-09 | 4.033E-09 | 3.107E-09 | 1.500E-09 | 7.500E-09 |
| 28822 | 1442 | V1442R | E | 4.6e-09 | 3.2e-09 | 1.2e-08 | 6.600E-09 | 4.729E-09 | 3.200E-09 | 1.200E-08 |
| 28823 | 1442 | V1442N | E | 8.3e-09 | 2.1e-08 | 1.1e-08 | 1.343E-08 | 6.691E-09 | 8.300E-09 | 2.100E-08 |
| 28824 | 1442 | V1442D | E | 1.2e-08 | 6.8e-09 | 3.5e-09 | 7.433E-09 | 4.285E-09 | 3.500E-09 | 1.200E-08 |
| 28825 | 1442 | V1442C | E | 3.5e-09 | 7.3e-09 | 1.1e-08 | 7.267E-09 | 3.750E-09 | 3.500E-09 | 1.100E-08 |
| 28826 | 1442 | V1442E | E | 3.7e-09 | 2.1e-09 | 4.6e-09 | 3.467E-09 | 1.266E-09 | 2.100E-09 | 4.600E-09 |
| 28827 | 1442 | V1442Q | E | 6.1e-09 | 8.7e-09 | 4.5e-09 | 6.433E-09 | 2.120E-09 | 4.500E-09 | 8.700E-09 |
| 28828 | 1442 | V1442G | E | 2.1e-09 | 3.1e-09 | 1e-08 | 5.067E-09 | 4.302E-09 | 2.100E-09 | 1.000E-08 |
| 28829 | 1442 | V1442H | E | 5.4e-09 | 4.6e-09 | 1.2e-08 | 7.333E-09 | 4.061E-09 | 4.600E-09 | 1.200E-08 |
| 28830 | 1442 | V1442I | E | 6.6e-09 | 9e-09 | 1.3e-08 | 9.533E-09 | 3.233E-09 | 6.600E-09 | 1.300E-08 |
| 28831 | 1442 | V1442L | E | 9.6e-09 | 3.9e-09 | 8.5e-09 | 7.333E-09 | 3.024E-09 | 3.900E-09 | 9.600E-09 |
| 28832 | 1442 | V1442K | E | 5.1e-09 | 2.2e-08 | 1.1e-09 | 9.400E-09 | 1.109E-08 | 1.100E-09 | 2.200E-08 |
| 28833 | 1442 | V1442M | E | 8.8e-09 | 3e-09 | 8.4e-09 | 6.733E-09 | 3.239E-09 | 3.000E-09 | 8.800E-09 |
| 28834 | 1442 | V1442F | E | 8.1e-09 | 3.2e-09 | 5.6e-09 | 5.633E-09 | 2.450E-09 | 3.200E-09 | 8.100E-09 |
| 28835 | 1442 | V1442P | E | 9.3e-09 | 8e-09 | 2.3e-09 | 6.533E-09 | 3.723E-09 | 2.300E-09 | 9.300E-09 |
| 28836 | 1442 | V1442S | E | 4.3e-09 | 5.8e-09 | 5.7e-09 | 5.267E-09 | 8.386E-10 | 4.300E-09 | 5.800E-09 |
| 28837 | 1442 | V1442T | E | 6.6e-09 | 9e-09 | 6.6e-09 | 7.400E-09 | 1.386E-09 | 6.600E-09 | 9.000E-09 |
| 28838 | 1442 | V1442W | E | 1.5e-09 | 5.2e-09 | 8.6e-09 | 5.100E-09 | 3.551E-09 | 1.500E-09 | 8.600E-09 |
| 28839 | 1442 | V1442Y | E | 1.3e-08 | 1.1e-08 | 9.7e-09 | 1.123E-08 | 1.662E-09 | 9.700E-09 | 1.300E-08 |
| 28840 | 1442 | V1442V | E | 6.5e-09 | 6e-09 | 3.2e-09 | 5.233E-09 | 1.779E-09 | 3.200E-09 | 6.500E-09 |
| 28841 | 1443 | H1443A | E | 4.8e-09 | 1.8e-08 | 5.9e-09 | 9.567E-09 | 7.324E-09 | 4.800E-09 | 1.800E-08 |
| 28842 | 1443 | H1443R | E | 6.2e-09 | 1.2e-08 | 1.4e-09 | 6.533E-09 | 5.308E-09 | 1.400E-09 | 1.200E-08 |
| 28843 | 1443 | H1443N | E | 1.3e-08 | 1.5e-08 | 6.2e-09 | 1.140E-08 | 4.613E-09 | 6.200E-09 | 1.500E-08 |
| 28844 | 1443 | H1443D | E | 5e-09 | 1.9e-09 | 1.1e-08 | 5.967E-09 | 4.626E-09 | 1.900E-09 | 1.100E-08 |
| 28845 | 1443 | H1443C | E | 1.2e-08 | 4e-09 | 7.4e-09 | 7.800E-09 | 4.015E-09 | 4.000E-09 | 1.200E-08 |
| 28846 | 1443 | H1443E | E | 1.3e-08 | 1.2e-08 | 1.2e-08 | 1.233E-08 | 5.774E-10 | 1.200E-08 | 1.300E-08 |
| 28847 | 1443 | H1443Q | E | 1.3e-08 | 8.5e-09 | 8.2e-09 | 9.900E-09 | 2.689E-09 | 8.200E-09 | 1.300E-08 |
| 28848 | 1443 | H1443G | E | 6.4e-09 | 7.7e-09 | 2.6e-08 | 1.337E-08 | 1.096E-08 | 6.400E-09 | 2.600E-08 |
| 28849 | 1443 | H1443H | E | 7.6e-09 | 6.6e-09 | 4.2e-09 | 6.133E-09 | 1.747E-09 | 4.200E-09 | 7.600E-09 |
| 28850 | 1443 | H1443I | E | 8.5e-09 | 6.4e-09 | 3.3e-09 | 6.067E-09 | 2.616E-09 | 3.300E-09 | 8.500E-09 |
| 28851 | 1443 | H1443L | E | 2e-08 | 1e-08 | 4.9e-09 | 1.163E-08 | 7.681E-09 | 4.900E-09 | 2.000E-08 |
| 28852 | 1443 | H1443K | E | 9.7e-09 | 5.3e-09 | 4e-09 | 6.333E-09 | 2.987E-09 | 4.000E-09 | 9.700E-09 |
| 28853 | 1443 | H1443M | E | 9.6e-09 | 4.9e-09 | 4e-09 | 6.167E-09 | 3.007E-09 | 4.000E-09 | 9.600E-09 |
| 28854 | 1443 | H1443F | E | 3.6e-09 | 1.2e-08 | 2.8e-08 | 1.453E-08 | 1.240E-08 | 3.600E-09 | 2.800E-08 |
| 28855 | 1443 | H1443P | E | 5.7e-09 | 7.9e-09 | 1.7e-08 | 1.020E-08 | 5.991E-09 | 5.700E-09 | 1.700E-08 |
| 28856 | 1443 | H1443S | E | 1.1e-08 | 5.4e-09 | 1.6e-08 | 1.080E-08 | 5.303E-09 | 5.400E-09 | 1.600E-08 |
| 28857 | 1443 | H1443T | E | 1.4e-08 | 4e-09 | 1e-08 | 9.333E-09 | 5.033E-09 | 4.000E-09 | 1.400E-08 |
| 28858 | 1443 | H1443W | E | 5.3e-09 | 7e-09 | 3.3e-09 | 5.200E-09 | 1.852E-09 | 3.300E-09 | 7.000E-09 |
| 28859 | 1443 | H1443Y | E | 2.8e-09 | 1.1e-08 | 1.3e-08 | 8.933E-09 | 5.405E-09 | 2.800E-09 | 1.300E-08 |
| 28860 | 1443 | H1443V | E | 5.9e-09 | 7.4e-09 | 5.3e-09 | 6.200E-09 | 1.082E-09 | 5.300E-09 | 7.400E-09 |
| 28861 | 1444 | T1444A | E | 3.9e-09 | 5.4e-09 | 1.2e-08 | 7.100E-09 | 4.309E-09 | 3.900E-09 | 1.200E-08 |
| 28862 | 1444 | T1444R | E | 8.1e-09 | 9e-09 | 1.6e-08 | 1.103E-08 | 4.325E-09 | 8.100E-09 | 1.600E-08 |
| 28863 | 1444 | T1444N | E | 3.5e-09 | 7.5e-09 | 2.7e-09 | 4.567E-09 | 2.572E-09 | 2.700E-09 | 7.500E-09 |
| 28864 | 1444 | T1444D | E | 2.2e-09 | 1.3e-08 | 5e-09 | 6.733E-09 | 5.605E-09 | 2.200E-09 | 1.300E-08 |
| 28865 | 1444 | T1444C | E | 4e-09 | 4.1e-09 | 5.6e-09 | 4.567E-09 | 8.963E-10 | 4.000E-09 | 5.600E-09 |
| 28866 | 1444 | T1444E | E | 8.2e-09 | 1.6e-08 | 4.8e-09 | 9.667E-09 | 5.742E-09 | 4.800E-09 | 1.600E-08 |
| 28867 | 1444 | T1444Q | E | 2.1e-09 | 6e-09 | 6e-09 | 4.700E-09 | 2.252E-09 | 2.100E-09 | 6.000E-09 |
| 28868 | 1444 | T1444G | E | 1e-08 | 5.5e-09 | 1e-08 | 8.500E-09 | 2.598E-09 | 5.500E-09 | 1.000E-08 |
| 28869 | 1444 | T1444H | E | 1.2e-08 | 6.9e-09 | 4.3e-09 | 7.733E-09 | 3.917E-09 | 4.300E-09 | 1.200E-08 |
| 28870 | 1444 | T1444I | E | 6.1e-09 | 3.1e-08 | 1.1e-08 | 1.603E-08 | 1.319E-08 | 6.100E-09 | 3.100E-08 |
| 28871 | 1444 | T1444L | E | 1.1e-08 | 3.5e-08 | 1.1e-08 | 1.900E-08 | 1.386E-08 | 1.100E-08 | 3.500E-08 |
| 28872 | 1444 | T1444K | E | 7.4e-09 | 1.2e-08 | 5.4e-09 | 8.267E-09 | 3.384E-09 | 5.400E-09 | 1.200E-08 |
| 28873 | 1444 | T1444M | E | 7.3e-09 | 3.8e-09 | 8.4e-09 | 6.500E-09 | 2.402E-09 | 3.800E-09 | 8.400E-09 |
| 28874 | 1444 | T1444F | E | 1.1e-08 | 5.2e-09 | 8.8e-09 | 8.333E-09 | 2.928E-09 | 5.200E-09 | 1.100E-08 |
| 28875 | 1444 | T1444P | E | 9.9e-09 | 6.7e-09 | 5.1e-09 | 7.233E-09 | 2.444E-09 | 5.100E-09 | 9.900E-09 |
| 28876 | 1444 | T1444S | E | 9.4e-09 | 1.7e-08 | 9e-09 | 1.180E-08 | 4.508E-09 | 9.000E-09 | 1.700E-08 |
| 28877 | 1444 | T1444T | E | 1.2e-08 | 5.2e-09 | 4.2e-09 | 7.133E-09 | 4.244E-09 | 4.200E-09 | 1.200E-08 |
| 28878 | 1444 | T1444W | E | 1.4e-08 | 5.7e-09 | 4.6e-09 | 8.100E-09 | 5.139E-09 | 4.600E-09 | 1.400E-08 |
| 28879 | 1444 | T1444Y | E | 6.5e-09 | 8.2e-09 | 3.3e-08 | 1.590E-08 | 1.483E-08 | 6.500E-09 | 3.300E-08 |
| 28880 | 1444 | T1444V | E | 1.5e-09 | 4.2e-09 | 9.5e-09 | 5.067E-09 | 4.070E-09 | 1.500E-09 | 9.500E-09 |
| 28881 | 1445 | F1445A | E | 7.2e-09 | 9.2e-09 | 1e-08 | 8.800E-09 | 1.442E-09 | 7.200E-09 | 1.000E-08 |
| 28882 | 1445 | F1445R | E | 1.8e-09 | 4.7e-09 | 5.3e-09 | 3.933E-09 | 1.872E-09 | 1.800E-09 | 5.300E-09 |
| 28883 | 1445 | F1445N | E | 4.7e-09 | 5e-09 | 4.9e-09 | 4.867E-09 | 1.528E-10 | 4.700E-09 | 5.000E-09 |
| 28884 | 1445 | F1445D | E | 2.8e-08 | 4.9e-09 | 7.7e-09 | 1.353E-08 | 1.261E-08 | 4.900E-09 | 2.800E-08 |
| 28885 | 1445 | F1445C | E | 2.8e-09 | 4.6e-09 | 1.2e-08 | 6.467E-09 | 4.876E-09 | 2.800E-09 | 1.200E-08 |
| 28886 | 1445 | F1445E | E | 4.9e-09 | 1.5e-08 | 1.1e-08 | 1.030E-08 | 5.086E-09 | 4.900E-09 | 1.500E-08 |
| 28887 | 1445 | F1445Q | E | 2e-08 | 4e-09 | 3.5e-09 | 9.167E-09 | 9.385E-09 | 3.500E-09 | 2.000E-08 |
| 28888 | 1445 | F1445G | E | 2.7e-08 | 6.5e-09 | 5.2e-09 | 1.290E-08 | 1.223E-08 | 5.200E-09 | 2.700E-08 |
| 28889 | 1445 | F1445H | E | 6e-09 | 2.9e-09 | 5.5e-09 | 4.800E-09 | 1.664E-09 | 2.900E-09 | 6.000E-09 |
| 28890 | 1445 | F1445I | E | 5.3e-09 | 2.3e-09 | 1.2e-08 | 6.533E-09 | 4.966E-09 | 2.300E-09 | 1.200E-08 |
| 28891 | 1445 | F1445L | E | 6.8e-09 | 3.2e-08 | 7e-09 | 1.527E-08 | 1.449E-08 | 6.800E-09 | 3.200E-08 |
| 28892 | 1445 | F1445K | E | 2.4e-09 | 5.5e-09 | 8.1e-09 | 5.333E-09 | 2.854E-09 | 2.400E-09 | 8.100E-09 |
| 28893 | 1445 | F1445M | E | 4.2e-09 | 1.2e-08 | 6e-09 | 7.400E-09 | 4.084E-09 | 4.200E-09 | 1.200E-08 |
| 28894 | 1445 | F1445F | E | 4.3e-09 | 2.4e-09 | 3.1e-09 | 3.267E-09 | 9.609E-10 | 2.400E-09 | 4.300E-09 |
| 28895 | 1445 | F1445P | E | 6e-09 | 9.8e-09 | 1.8e-09 | 5.867E-09 | 4.002E-09 | 1.800E-09 | 9.800E-09 |
| 28896 | 1445 | F1445S | E | 8.2e-09 | 4.5e-09 | 7.1e-09 | 6.600E-09 | 1.900E-09 | 4.500E-09 | 8.200E-09 |
| 28897 | 1445 | F1445T | E | 6.2e-09 | 1.6e-08 | 5.3e-09 | 9.167E-09 | 5.935E-09 | 5.300E-09 | 1.600E-08 |
| 28898 | 1445 | F1445W | E | 8.4e-09 | 4.9e-09 | 5.1e-09 | 6.133E-09 | 1.966E-09 | 4.900E-09 | 8.400E-09 |
| 28899 | 1445 | F1445Y | E | 4.3e-09 | 1.2e-08 | 6.1e-09 | 7.467E-09 | 4.028E-09 | 4.300E-09 | 1.200E-08 |

| | | | | | | | | | | |
|---|---|---|---|---|---|---|---|---|---|---|
| 28900 | 1445 | F1445V | E | 7.3e-09 | 5.6e-09 | 5.7e-09 | 6.200E-09 | 9.539E-10 | 5.600E-09 | 7.300E-09 |
| 28901 | 1446 | P1446A | E | 1.8e-08 | 6.2e-09 | 2.3e-09 | 8.833E-09 | 8.175E-09 | 2.300E-09 | 1.800E-08 |
| 28902 | 1446 | P1446R | E | 1.1e-08 | 2.9e-09 | 1.7e-08 | 1.030E-08 | 7.076E-09 | 2.900E-09 | 1.700E-08 |
| 28903 | 1446 | P1446N | E | 9.6e-09 | 6.2e-09 | 6.9e-09 | 7.567E-09 | 1.795E-09 | 6.200E-09 | 9.600E-09 |
| 28904 | 1446 | P1446D | E | 4.3e-09 | 5.1e-09 | 4.9e-08 | 1.947E-08 | 2.558E-08 | 4.300E-09 | 4.900E-08 |
| 28905 | 1446 | P1446C | E | 3e-09 | 8.2e-09 | 5.7e-09 | 5.633E-09 | 2.601E-09 | 3.000E-09 | 8.200E-09 |
| 28906 | 1446 | P1446E | E | 6.6e-09 | 1.1e-08 | 5.8e-09 | 7.800E-09 | 2.800E-09 | 5.800E-09 | 1.100E-08 |
| 28907 | 1446 | P1446Q | E | 1.4e-08 | 7.6e-09 | 4.1e-09 | 8.567E-09 | 5.020E-09 | 4.100E-09 | 1.400E-08 |
| 28908 | 1446 | P1446G | E | 6.6e-09 | 9.7e-09 | 2.7e-09 | 6.333E-09 | 3.508E-09 | 2.700E-09 | 9.700E-09 |
| 28909 | 1446 | P1446H | E | 8e-09 | 4e-09 | 2.2e-09 | 4.733E-09 | 2.969E-09 | 2.200E-09 | 8.000E-09 |
| 28910 | 1446 | P1446I | E | 2.8e-09 | 2.5e-08 | 1e-08 | 1.260E-08 | 1.133E-08 | 2.800E-09 | 2.500E-08 |
| 28911 | 1446 | P1446L | E | 8e-09 | 8.2e-09 | 1.3e-08 | 9.733E-09 | 2.831E-09 | 8.000E-09 | 1.300E-08 |
| 28912 | 1446 | P1446K | E | 1.3e-08 | 5.6e-09 | 4.6e-09 | 7.733E-09 | 4.588E-09 | 4.600E-09 | 1.300E-08 |
| 28913 | 1446 | P1446M | E | 1e-08 | 2.4e-09 | 1.1e-09 | 4.500E-09 | 4.807E-09 | 1.100E-09 | 1.000E-08 |
| 28914 | 1446 | P1446F | E | 1.6e-08 | 8e-09 | 4.9e-09 | 9.633E-09 | 5.727E-09 | 4.900E-09 | 1.600E-08 |
| 28915 | 1446 | P1446P | E | 8.3e-09 | 2.5e-09 | 1.2e-08 | 7.600E-09 | 4.789E-09 | 2.500E-09 | 1.200E-08 |
| 28916 | 1446 | P1446S | E | 1e-08 | 5.5e-09 | 1e-08 | 8.500E-09 | 2.598E-09 | 5.500E-09 | 1.000E-08 |
| 28917 | 1446 | P1446T | E | 9.1e-09 | 4.1e-09 | 3e-09 | 5.400E-09 | 3.251E-09 | 3.000E-09 | 9.100E-09 |
| 28918 | 1446 | P1446W | E | 8e-09 | 1.3e-08 | 1.2e-08 | 1.100E-08 | 2.646E-09 | 8.000E-09 | 1.300E-08 |
| 28919 | 1446 | P1446Y | E | 4.5e-09 | 2e-08 | 5e-09 | 9.833E-09 | 8.808E-09 | 4.500E-09 | 2.000E-08 |
| 28920 | 1446 | P1446V | E | 1.5e-08 | 3.8e-09 | 3e-09 | 7.267E-09 | 6.709E-09 | 3.000E-09 | 1.500E-08 |
| 28921 | 1447 | A1447A | E | 1.2e-08 | 8e-09 | 5.7e-09 | 8.567E-09 | 3.188E-09 | 5.700E-09 | 1.200E-08 |
| 28922 | 1447 | A1447R | E | 2.9e-09 | 6.1e-09 | 4.2e-09 | 4.400E-09 | 1.609E-09 | 2.900E-09 | 6.100E-09 |
| 28923 | 1447 | A1447N | E | 5.3e-09 | 1.4e-08 | 9.6e-09 | 9.633E-09 | 4.350E-09 | 5.300E-09 | 1.400E-08 |
| 28924 | 1447 | A1447D | E | 5.5e-09 | 1.5e-08 | 7.7e-09 | 9.400E-09 | 4.973E-09 | 5.500E-09 | 1.500E-08 |
| 28925 | 1447 | A1447C | E | 1.1e-08 | 1.4e-09 | 1.5e-08 | 9.133E-09 | 6.990E-09 | 1.400E-09 | 1.500E-08 |
| 28926 | 1447 | A1447E | E | 3.5e-09 | 1.5e-08 | 1.1e-08 | 9.833E-09 | 5.838E-09 | 3.500E-09 | 1.500E-08 |
| 28927 | 1447 | A1447Q | E | 3.5e-09 | 1.9e-09 | 1.1e-08 | 5.467E-09 | 4.858E-09 | 1.900E-09 | 1.100E-08 |
| 28928 | 1447 | A1447G | E | 3.2e-09 | 4.6e-09 | 9.5e-09 | 5.767E-09 | 3.308E-09 | 3.200E-09 | 9.500E-09 |
| 28929 | 1447 | A1447H | E | 7.2e-09 | 5.9e-09 | 6.2e-09 | 6.433E-09 | 6.807E-10 | 5.900E-09 | 7.200E-09 |
| 28930 | 1447 | A1447I | E | 1e-08 | 4.5e-09 | 5.1e-09 | 6.533E-09 | 3.017E-09 | 4.500E-09 | 1.000E-08 |
| 28931 | 1447 | A1447L | E | 9.4e-09 | 6.8e-09 | 3.3e-09 | 6.500E-09 | 3.061E-09 | 3.300E-09 | 9.400E-09 |
| 28932 | 1447 | A1447K | E | 1.5e-08 | 1.8e-08 | 7.6e-09 | 1.353E-08 | 5.353E-09 | 7.600E-09 | 1.800E-08 |
| 28933 | 1447 | A1447M | E | 7.4e-09 | 6.8e-09 | 7e-09 | 7.067E-09 | 3.055E-10 | 6.800E-09 | 7.400E-09 |
| 28934 | 1447 | A1447F | E | 5.4e-09 | 1.3e-08 | 5.2e-09 | 7.867E-09 | 4.447E-09 | 5.200E-09 | 1.300E-08 |
| 28935 | 1447 | A1447P | E | 1.5e-08 | 5e-09 | 3.3e-09 | 7.767E-09 | 6.322E-09 | 3.300E-09 | 1.500E-08 |
| 28936 | 1447 | A1447S | E | 7.8e-09 | 1.2e-08 | 1.5e-08 | 1.160E-08 | 3.617E-09 | 7.800E-09 | 1.500E-08 |
| 28937 | 1447 | A1447T | E | 1.9e-09 | 4.4e-09 | 1.4e-08 | 6.767E-09 | 6.388E-09 | 1.900E-09 | 1.400E-08 |
| 28938 | 1447 | A1447W | E | 7e-09 | 1.1e-08 | 3.2e-09 | 7.067E-09 | 3.900E-09 | 3.200E-09 | 1.100E-08 |
| 28939 | 1447 | A1447Y | E | 9.4e-09 | 7.1e-09 | 9.4e-09 | 8.633E-09 | 1.328E-09 | 7.100E-09 | 9.400E-09 |
| 28940 | 1447 | A1447V | E | 3e-09 | 1.3e-08 | 7.9e-09 | 7.967E-09 | 5.000E-09 | 3.000E-09 | 1.300E-08 |
| 28941 | 1448 | V1448A | E | 1.6e-08 | 5e-09 | 5.6e-09 | 8.867E-09 | 6.185E-09 | 5.000E-09 | 1.600E-08 |
| 28942 | 1448 | V1448R | E | 2.8e-09 | 9.8e-09 | 5.3e-09 | 5.967E-09 | 3.547E-09 | 2.800E-09 | 9.800E-09 |
| 28943 | 1448 | V1448N | E | 4.7e-09 | 4.7e-09 | 2.1e-08 | 1.013E-08 | 9.411E-09 | 4.700E-09 | 2.100E-08 |
| 28944 | 1448 | V1448D | E | 6.2e-09 | 9.2e-09 | 1.5e-08 | 1.013E-08 | 4.474E-09 | 6.200E-09 | 1.500E-08 |
| 28945 | 1448 | V1448C | E | 8.4e-09 | 3.9e-09 | 6.6e-09 | 6.300E-09 | 2.265E-09 | 3.900E-09 | 8.400E-09 |
| 28946 | 1448 | V1448E | E | 2.1e-09 | 9.2e-09 | 8.9e-09 | 6.733E-09 | 4.015E-09 | 2.100E-09 | 9.200E-09 |
| 28947 | 1448 | V1448Q | E | 5.3e-09 | 1.9e-09 | 1.5e-08 | 7.400E-09 | 6.798E-09 | 1.900E-09 | 1.500E-08 |
| 28948 | 1448 | V1448G | E | 1.1e-08 | 6.5e-09 | 4.4e-09 | 7.300E-09 | 3.372E-09 | 4.400E-09 | 1.100E-08 |
| 28949 | 1448 | V1448H | E | 5.9e-09 | 2.6e-08 | 6.2e-09 | 1.270E-08 | 1.152E-08 | 5.900E-09 | 2.600E-08 |
| 28950 | 1448 | V1448I | E | 7.2e-09 | 7.5e-09 | 4.8e-09 | 6.500E-09 | 1.480E-09 | 4.800E-09 | 7.500E-09 |
| 28951 | 1448 | V1448L | E | 4.4e-09 | 1.3e-08 | 2.6e-08 | 1.447E-08 | 1.087E-08 | 4.400E-09 | 2.600E-08 |
| 28952 | 1448 | V1448K | E | 4.1e-09 | 2.2e-09 | 1.1e-08 | 5.767E-09 | 4.631E-09 | 2.200E-09 | 1.100E-08 |
| 28953 | 1448 | V1448M | E | 2.3e-08 | 6.6e-09 | 1.1e-08 | 1.353E-08 | 8.488E-09 | 6.600E-09 | 2.300E-08 |
| 28954 | 1448 | V1448F | E | 1e-08 | 7.1e-09 | 1e-08 | 9.033E-09 | 1.674E-09 | 7.100E-09 | 1.000E-08 |
| 28955 | 1448 | V1448P | E | 1.4e-09 | 5.6e-09 | 7.7e-09 | 4.900E-09 | 3.208E-09 | 1.400E-09 | 7.700E-09 |
| 28956 | 1448 | V1448S | E | 2.2e-09 | 4.8e-09 | 3.4e-09 | 3.467E-09 | 1.301E-09 | 2.200E-09 | 4.800E-09 |
| 28957 | 1448 | V1448T | E | 4.8e-09 | 6.5e-09 | 2.3e-09 | 4.533E-09 | 2.113E-09 | 2.300E-09 | 6.500E-09 |
| 28958 | 1448 | V1448W | E | 3.4e-09 | 5.3e-09 | 3.4e-09 | 4.033E-09 | 1.097E-09 | 3.400E-09 | 5.300E-09 |
| 28959 | 1448 | V1448Y | E | 4.8e-09 | 3.3e-09 | 6.7e-09 | 4.933E-09 | 1.704E-09 | 3.300E-09 | 6.700E-09 |
| 28960 | 1448 | V1448V | E | 1.2e-08 | 9.3e-09 | 1.3e-08 | 1.143E-08 | 1.914E-09 | 9.300E-09 | 1.300E-08 |
| 28961 | 1449 | L1449A | E | 8.1e-09 | 5.7e-09 | 1.2e-08 | 8.600E-09 | 3.180E-09 | 5.700E-09 | 1.200E-08 |
| 28962 | 1449 | L1449R | E | 6.6e-09 | 3.6e-09 | 5.1e-09 | 5.100E-09 | 1.500E-09 | 3.600E-09 | 6.600E-09 |
| 28963 | 1449 | L1449N | E | 1.6e-08 | 3.1e-08 | 1.6e-08 | 2.100E-08 | 8.660E-09 | 1.600E-08 | 3.100E-08 |
| 28964 | 1449 | L1449D | E | 1.2e-08 | 3.5e-09 | 4.5e-09 | 6.667E-09 | 4.646E-09 | 3.500E-09 | 1.200E-08 |
| 28965 | 1449 | L1449C | E | 6e-09 | 6.4e-09 | 5.1e-09 | 5.833E-09 | 6.658E-10 | 5.100E-09 | 6.400E-09 |
| 28966 | 1449 | L1449E | E | 3.7e-09 | 5.8e-09 | 5.6e-09 | 5.033E-09 | 1.159E-09 | 3.700E-09 | 5.800E-09 |
| 28967 | 1449 | L1449Q | E | 5.8e-09 | 9.4e-09 | 1.1e-08 | 8.733E-09 | 2.663E-09 | 5.800E-09 | 1.100E-08 |
| 28968 | 1449 | L1449G | E | 9.2e-09 | 5.8e-09 | 3.5e-09 | 6.167E-09 | 2.868E-09 | 3.500E-09 | 9.200E-09 |
| 28969 | 1449 | L1449H | E | 2.5e-08 | 1.9e-09 | 1.7e-08 | 1.463E-08 | 1.173E-08 | 1.900E-09 | 2.500E-08 |
| 28970 | 1449 | L1449I | E | 4.9e-09 | 1.1e-08 | 7e-09 | 7.633E-09 | 3.099E-09 | 4.900E-09 | 1.100E-08 |
| 28971 | 1449 | L1449L | E | 5.4e-09 | 2.5e-08 | 7.3e-09 | 1.257E-08 | 1.081E-08 | 5.400E-09 | 2.500E-08 |
| 28972 | 1449 | L1449K | E | 1e-08 | 6.9e-09 | 1.1e-08 | 9.300E-09 | 2.138E-09 | 6.900E-09 | 1.100E-08 |
| 28973 | 1449 | L1449M | E | 3.1e-09 | 1e-08 | 1.3e-08 | 8.700E-09 | 5.076E-09 | 3.100E-09 | 1.300E-08 |
| 28974 | 1449 | L1449F | E | 3.3e-09 | 8.1e-09 | 1e-08 | 7.133E-09 | 3.453E-09 | 3.300E-09 | 1.000E-08 |
| 28975 | 1449 | L1449P | E | 4.1e-09 | 7.1e-09 | 1e-08 | 7.067E-09 | 2.950E-09 | 4.100E-09 | 1.000E-08 |
| 28976 | 1449 | L1449S | E | 1.7e-08 | 8.7e-09 | 5.8e-09 | 1.050E-08 | 5.813E-09 | 5.800E-09 | 1.700E-08 |
| 28977 | 1449 | L1449T | E | 9.2e-09 | 9.5e-09 | 1.5e-08 | 1.123E-08 | 3.265E-09 | 9.200E-09 | 1.500E-08 |
| 28978 | 1449 | L1449W | E | 4.2e-09 | 5.3e-09 | 8.7e-09 | 6.067E-09 | 2.346E-09 | 4.200E-09 | 8.700E-09 |
| 28979 | 1449 | L1449Y | E | 1.5e-09 | 1.1e-08 | 5.8e-09 | 6.100E-09 | 4.757E-09 | 1.500E-09 | 1.100E-08 |
| 28980 | 1449 | L1449V | E | 7.2e-09 | 2.5e-08 | 1.6e-08 | 1.607E-08 | 8.900E-09 | 7.200E-09 | 2.500E-08 |
| 28981 | 1450 | Q1450A | E | 9.7e-09 | 1e-08 | 3.9e-09 | 7.867E-09 | 3.439E-09 | 3.900E-09 | 1.000E-08 |
| 28982 | 1450 | Q1450R | E | 3.8e-09 | 1.4e-08 | 8.1e-09 | 8.633E-09 | 5.121E-09 | 3.800E-09 | 1.400E-08 |
| 28983 | 1450 | Q1450N | E | 7.1e-09 | 6.7e-09 | 7.4e-09 | 7.067E-09 | 3.512E-10 | 6.700E-09 | 7.400E-09 |
| 28984 | 1450 | Q1450D | E | 7.9e-09 | 2.2e-08 | 4.1e-09 | 1.133E-08 | 9.431E-09 | 4.100E-09 | 2.200E-08 |

| | | | | | | | | | | |
|---|---|---|---|---|---|---|---|---|---|---|
| 28985 | 1450 | Q1450C | E | 1.2e-08 | 2.9e-09 | 5.1e-09 | 6.667E-09 | 4.748E-09 | 2.900E-09 | 1.200E-08 |
| 28986 | 1450 | Q1450E | E | 1.3e-09 | 1e-08 | 1.1e-08 | 7.433E-09 | 5.335E-09 | 1.300E-09 | 1.100E-08 |
| 28987 | 1450 | Q1450Q | E | 3.4e-09 | 7.6e-09 | 1.9e-09 | 4.300E-09 | 2.955E-09 | 1.900E-09 | 7.600E-09 |
| 28988 | 1450 | Q1450G | E | 1.1e-08 | 7.2e-09 | 6.2e-09 | 8.133E-09 | 2.532E-09 | 6.200E-09 | 1.100E-08 |
| 28989 | 1450 | Q1450H | E | 4.1e-09 | 5.8e-09 | 4.3e-09 | 4.733E-09 | 9.292E-10 | 4.100E-09 | 5.800E-09 |
| 28990 | 1450 | Q1450I | E | 1.1e-08 | 1.4e-08 | 1.2e-08 | 1.233E-08 | 1.528E-09 | 1.100E-08 | 1.400E-08 |
| 28991 | 1450 | Q1450L | E | 4.3e-09 | 1.5e-08 | 5.1e-09 | 8.133E-09 | 5.960E-09 | 4.300E-09 | 1.500E-08 |
| 28992 | 1450 | Q1450K | E | 9.9e-09 | 3.6e-09 | 6.1e-09 | 6.533E-09 | 3.172E-09 | 3.600E-09 | 9.900E-09 |
| 28993 | 1450 | Q1450M | E | 6.3e-09 | 9.5e-09 | 7.1e-09 | 7.633E-09 | 1.665E-09 | 6.300E-09 | 9.500E-09 |
| 28994 | 1450 | Q1450F | E | 6.7e-09 | 1.1e-08 | 8.4e-09 | 8.700E-09 | 2.166E-09 | 6.700E-09 | 1.100E-08 |
| 28995 | 1450 | Q1450P | E | 1e-08 | 5.4e-09 | 1.8e-09 | 5.733E-09 | 4.110E-09 | 1.800E-09 | 1.000E-08 |
| 28996 | 1450 | Q1450S | E | 9.9e-09 | 4e-09 | 5.5e-09 | 6.467E-09 | 3.066E-09 | 4.000E-09 | 9.900E-09 |
| 28997 | 1450 | Q1450T | E | 2.3e-08 | 6.3e-09 | 9.1e-09 | 1.280E-08 | 8.944E-09 | 6.300E-09 | 2.300E-08 |
| 28998 | 1450 | Q1450W | E | 2.7e-09 | 7.3e-09 | 5.4e-09 | 5.133E-09 | 2.312E-09 | 2.700E-09 | 7.300E-09 |
| 28999 | 1450 | Q1450Y | E | 5.9e-09 | 8.8e-09 | 1.1e-08 | 8.567E-09 | 2.558E-09 | 5.900E-09 | 1.100E-08 |
| 29000 | 1450 | Q1450V | E | 1.5e-08 | 5.1e-09 | 3e-09 | 7.700E-09 | 6.409E-09 | 3.000E-09 | 1.500E-08 |
| 29001 | 1451 | S1451A | E | 3.5e-09 | 4.4e-09 | 9.5e-09 | 5.800E-09 | 3.236E-09 | 3.500E-09 | 9.500E-09 |
| 29002 | 1451 | S1451R | E | 4.3e-09 | 2.2e-09 | 6.2e-09 | 4.233E-09 | 2.001E-09 | 2.200E-09 | 6.200E-09 |
| 29003 | 1451 | S1451N | E | 9.5e-09 | 2.8e-09 | 4.2e-09 | 5.500E-09 | 3.534E-09 | 2.800E-09 | 9.500E-09 |
| 29004 | 1451 | S1451D | E | 2.6e-08 | 6.1e-09 | 8.2e-09 | 1.343E-08 | 1.093E-08 | 6.100E-09 | 2.600E-08 |
| 29005 | 1451 | S1451C | E | 1.4e-08 | 4.8e-09 | 1.5e-08 | 1.127E-08 | 5.623E-09 | 4.800E-09 | 1.500E-08 |
| 29006 | 1451 | S1451E | E | 9.3e-09 | 8.3e-09 | 1e-08 | 9.200E-09 | 8.544E-10 | 8.300E-09 | 1.000E-08 |
| 29007 | 1451 | S1451Q | E | 3.6e-09 | 9.4e-09 | 9.1e-09 | 7.367E-09 | 3.265E-09 | 3.600E-09 | 9.400E-09 |
| 29008 | 1451 | S1451G | E | 4.3e-09 | 1e-08 | 1.4e-08 | 9.433E-09 | 4.875E-09 | 4.300E-09 | 1.400E-08 |
| 29009 | 1451 | S1451H | E | 7.8e-09 | 3.5e-09 | 5e-09 | 5.433E-09 | 2.183E-09 | 3.500E-09 | 7.800E-09 |
| 29010 | 1451 | S1451I | E | 1.5e-08 | 1.1e-08 | 5.9e-09 | 1.063E-08 | 4.561E-09 | 5.900E-09 | 1.500E-08 |
| 29011 | 1451 | S1451L | E | 7.7e-09 | 5.7e-09 | 1.9e-08 | 1.080E-08 | 7.171E-09 | 5.700E-09 | 1.900E-08 |
| 29012 | 1451 | S1451K | E | 5.1e-09 | 5.9e-09 | 8.1e-09 | 6.367E-09 | 1.553E-09 | 5.100E-09 | 8.100E-09 |
| 29013 | 1451 | S1451M | E | 5.8e-09 | 1.2e-08 | 7.2e-09 | 8.333E-09 | 3.252E-09 | 5.800E-09 | 1.200E-08 |
| 29014 | 1451 | S1451F | E | 6e-08 | 3.8e-09 | 6.5e-09 | 2.343E-08 | 3.170E-08 | 3.800E-09 | 6.000E-08 |
| 29015 | 1451 | S1451P | E | 4.8e-09 | 4.7e-09 | 8.6e-09 | 6.033E-09 | 2.223E-09 | 4.700E-09 | 8.600E-09 |
| 29016 | 1451 | S1451S | E | 1.1e-08 | 2.6e-09 | 6e-09 | 6.533E-09 | 4.225E-09 | 2.600E-09 | 1.100E-08 |
| 29017 | 1451 | S1451T | E | 8.5e-09 | 4.8e-09 | 5.5e-09 | 6.267E-09 | 1.966E-09 | 4.800E-09 | 8.500E-09 |
| 29018 | 1451 | S1451W | E | 1.1e-08 | 9.2e-09 | 4.5e-09 | 8.233E-09 | 3.356E-09 | 4.500E-09 | 1.100E-08 |
| 29019 | 1451 | S1451Y | E | 6.8e-09 | 9.9e-09 | 7.6e-09 | 8.100E-09 | 1.609E-09 | 6.800E-09 | 9.900E-09 |
| 29020 | 1451 | S1451V | E | 8.3e-09 | 1.4e-08 | 3.3e-09 | 8.533E-09 | 5.354E-09 | 3.300E-09 | 1.400E-08 |
| 29021 | 1452 | S1452A | E | 6e-09 | 4.1e-09 | 6e-09 | 5.367E-09 | 1.097E-09 | 4.100E-09 | 6.000E-09 |
| 29022 | 1452 | S1452R | E | 8.9e-09 | 9.6e-09 | 9.3e-09 | 9.267E-09 | 3.512E-10 | 8.900E-09 | 9.600E-09 |
| 29023 | 1452 | S1452N | E | 2e-09 | 5e-09 | 6.5e-09 | 4.500E-09 | 2.291E-09 | 2.000E-09 | 6.500E-09 |
| 29024 | 1452 | S1452D | E | 6.5e-09 | 9.5e-09 | 2.9e-09 | 6.300E-09 | 3.305E-09 | 2.900E-09 | 9.500E-09 |
| 29025 | 1452 | S1452C | E | 9.2e-09 | 8.8e-09 | 5.2e-09 | 7.733E-09 | 2.203E-09 | 5.200E-09 | 9.200E-09 |
| 29026 | 1452 | S1452E | E | 2.6e-09 | 4.7e-09 | 5.4e-09 | 4.233E-09 | 1.457E-09 | 2.600E-09 | 5.400E-09 |
| 29027 | 1452 | S1452Q | E | 4.3e-09 | 4.5e-09 | 4.1e-09 | 4.300E-09 | 2.000E-10 | 4.100E-09 | 4.500E-09 |
| 29028 | 1452 | S1452G | E | 8.3e-09 | 5.9e-09 | 5.1e-09 | 6.433E-09 | 1.665E-09 | 5.100E-09 | 8.300E-09 |
| 29029 | 1452 | S1452H | E | 5.4e-09 | 1.1e-08 | 8.8e-09 | 8.400E-09 | 2.821E-09 | 5.400E-09 | 1.100E-08 |
| 29030 | 1452 | S1452I | E | 4.2e-08 | 8.6e-09 | 1.3e-08 | 2.120E-08 | 1.815E-08 | 8.600E-09 | 4.200E-08 |
| 29031 | 1452 | S1452L | E | 2.2e-09 | 2.8e-09 | 8.9e-09 | 4.633E-09 | 3.707E-09 | 2.200E-09 | 8.900E-09 |
| 29032 | 1452 | S1452K | E | 7.8e-09 | 7.6e-09 | 1.1e-08 | 8.800E-09 | 1.908E-09 | 7.600E-09 | 1.100E-08 |
| 29033 | 1452 | S1452M | E | 1.7e-08 | 2.1e-09 | 8.5e-09 | 9.200E-09 | 7.475E-09 | 2.100E-09 | 1.700E-08 |
| 29034 | 1452 | S1452F | E | 3.2e-09 | 2.8e-09 | 4.9e-09 | 3.633E-09 | 1.115E-09 | 2.800E-09 | 4.900E-09 |
| 29035 | 1452 | S1452P | E | 5.4e-09 | 1.6e-08 | 9e-09 | 1.013E-08 | 5.390E-09 | 5.400E-09 | 1.600E-08 |
| 29036 | 1452 | S1452S | E | 3.5e-09 | 2.4e-09 | 2.1e-08 | 8.967E-09 | 1.044E-08 | 2.400E-09 | 2.100E-08 |
| 29037 | 1452 | S1452T | E | 3.8e-09 | 8.2e-09 | 1.1e-08 | 7.667E-09 | 3.630E-09 | 3.800E-09 | 1.100E-08 |
| 29038 | 1452 | S1452W | E | 3.1e-09 | 7.5e-09 | 1e-08 | 6.867E-09 | 3.493E-09 | 3.100E-09 | 1.000E-08 |
| 29039 | 1452 | S1452Y | E | 5.9e-09 | 1.5e-08 | 1.2e-08 | 1.097E-08 | 4.637E-09 | 5.900E-09 | 1.500E-08 |
| 29040 | 1452 | S1452V | E | 1.2e-08 | 3.7e-09 | 3e-09 | 6.233E-09 | 5.006E-09 | 3.000E-09 | 1.200E-08 |
| 29041 | 1453 | G1453A | E | 1.5e-08 | 1e-08 | 6e-09 | 1.033E-08 | 4.509E-09 | 6.000E-09 | 1.500E-08 |
| 29042 | 1453 | G1453R | E | 1.1e-08 | 7.1e-09 | 7.3e-09 | 8.467E-09 | 2.196E-09 | 7.100E-09 | 1.100E-08 |
| 29043 | 1453 | G1453N | E | 9.4e-09 | 5.7e-09 | 9e-09 | 8.033E-09 | 2.031E-09 | 5.700E-09 | 9.400E-09 |
| 29044 | 1453 | G1453D | E | 6.6e-09 | 1.5e-08 | 6.9e-09 | 9.500E-09 | 4.766E-09 | 6.600E-09 | 1.500E-08 |
| 29045 | 1453 | G1453C | E | 9.1e-09 | 7.1e-09 | 8.4e-09 | 8.200E-09 | 1.015E-09 | 7.100E-09 | 9.100E-09 |
| 29046 | 1453 | G1453E | E | 4.8e-09 | 8.2e-09 | 5.8e-09 | 6.267E-09 | 1.747E-09 | 4.800E-09 | 8.200E-09 |
| 29047 | 1453 | G1453Q | E | 6e-09 | 9.5e-09 | 1.1e-08 | 8.833E-09 | 2.566E-09 | 6.000E-09 | 1.100E-08 |
| 29048 | 1453 | G1453G | E | 6.7e-09 | 7e-09 | 1.7e-08 | 1.023E-08 | 5.862E-09 | 6.700E-09 | 1.700E-08 |
| 29049 | 1453 | G1453H | E | 1.2e-08 | 1.8e-09 | 4.6e-09 | 6.133E-09 | 5.270E-09 | 1.800E-09 | 1.200E-08 |
| 29050 | 1453 | G1453I | E | 6.2e-09 | 3e-09 | 6.8e-09 | 5.333E-09 | 2.043E-09 | 3.000E-09 | 6.800E-09 |
| 29051 | 1453 | G1453L | E | 1.5e-08 | 8.1e-09 | 3.9e-09 | 9.000E-09 | 5.604E-09 | 3.900E-09 | 1.500E-08 |
| 29052 | 1453 | G1453K | E | 5.8e-09 | 1e-08 | 1.5e-08 | 1.027E-08 | 4.606E-09 | 5.800E-09 | 1.500E-08 |
| 29053 | 1453 | G1453M | E | 1.8e-09 | 2.2e-08 | 4e-09 | 9.267E-09 | 1.108E-08 | 1.800E-09 | 2.200E-08 |
| 29054 | 1453 | G1453F | E | 8.4e-09 | 6.9e-09 | 6.8e-09 | 7.367E-09 | 8.963E-10 | 6.800E-09 | 8.400E-09 |
| 29055 | 1453 | G1453P | E | 5.6e-09 | 1.1e-08 | 2.2e-08 | 1.287E-08 | 8.358E-09 | 5.600E-09 | 2.200E-08 |
| 29056 | 1453 | G1453S | E | 1.8e-09 | 5e-09 | 9.6e-09 | 5.467E-09 | 3.921E-09 | 1.800E-09 | 9.600E-09 |
| 29057 | 1453 | G1453T | E | 4.3e-09 | 9.7e-10 | 1.5e-08 | 6.757E-09 | 7.331E-09 | 9.700E-10 | 1.500E-08 |
| 29058 | 1453 | G1453W | E | 5.1e-09 | 9.4e-09 | 1.4e-08 | 9.500E-09 | 4.451E-09 | 5.100E-09 | 1.400E-08 |
| 29059 | 1453 | G1453Y | E | 1.9e-08 | 5.1e-09 | 4.3e-09 | 9.467E-09 | 8.266E-09 | 4.300E-09 | 1.900E-08 |
| 29060 | 1453 | G1453V | E | 5.6e-09 | 1.2e-08 | 1.4e-08 | 1.053E-08 | 4.388E-09 | 5.600E-09 | 1.400E-08 |
| 29061 | 1454 | L1454A | E | 4e-09 | 1.4e-08 | 4.7e-09 | 7.567E-09 | 5.582E-09 | 4.000E-09 | 1.400E-08 |
| 29062 | 1454 | L1454R | E | 6.8e-09 | 1.2e-08 | 3.8e-09 | 7.533E-09 | 4.149E-09 | 3.800E-09 | 1.200E-08 |
| 29063 | 1454 | L1454N | E | 1e-08 | 6.6e-09 | 4.3e-09 | 6.967E-09 | 2.868E-09 | 4.300E-09 | 1.000E-08 |
| 29064 | 1454 | L1454D | E | 7.1e-09 | 4.3e-09 | 7e-09 | 6.133E-09 | 1.589E-09 | 4.300E-09 | 7.100E-09 |
| 29065 | 1454 | L1454C | E | 7.2e-09 | 1.7e-08 | 5.1e-09 | 9.767E-09 | 6.352E-09 | 5.100E-09 | 1.700E-08 |
| 29066 | 1454 | L1454E | E | 4.9e-09 | 2e-09 | 1.8e-08 | 8.300E-09 | 8.525E-09 | 2.000E-09 | 1.800E-08 |
| 29067 | 1454 | L1454Q | E | 4.2e-09 | 7.1e-09 | 1.1e-08 | 7.433E-09 | 3.412E-09 | 4.200E-09 | 1.100E-08 |
| 29068 | 1454 | L1454G | E | 3.7e-08 | 1.2e-08 | 9.5e-09 | 1.950E-08 | 1.521E-08 | 9.500E-09 | 3.700E-08 |
| 29069 | 1454 | L1454H | E | 7.5e-09 | 1.4e-08 | 8.2e-09 | 9.900E-09 | 3.568E-09 | 7.500E-09 | 1.400E-08 |

| | | | | | | | | | | |
|---|---|---|---|---|---|---|---|---|---|---|
| 29070 | 1454 | L1454I | E | 6.5e-09 | 1.9e-08 | 6.1e-09 | 1.053E-08 | 7.335E-09 | 6.100E-09 | 1.900E-08 |
| 29071 | 1454 | L1454L | E | 2.2e-09 | 2.9e-09 | 1.9e-08 | 8.033E-09 | 9.504E-09 | 2.200E-09 | 1.900E-08 |
| 29072 | 1454 | L1454K | E | 1.2e-08 | 6.6e-09 | 8.9e-09 | 9.167E-09 | 2.710E-09 | 6.600E-09 | 1.200E-08 |
| 29073 | 1454 | L1454M | E | 8.8e-09 | 4.6e-09 | 5e-09 | 6.133E-09 | 2.318E-09 | 4.600E-09 | 8.800E-09 |
| 29074 | 1454 | L1454F | E | 6.5e-09 | 1.6e-08 | 9.5e-09 | 1.067E-08 | 4.856E-09 | 6.500E-09 | 1.600E-08 |
| 29075 | 1454 | L1454P | E | 4.4e-09 | 5.4e-09 | 1.9e-08 | 9.600E-09 | 8.156E-09 | 4.400E-09 | 1.900E-08 |
| 29076 | 1454 | L1454S | E | 1.3e-08 | 6.8e-09 | 3e-09 | 7.600E-09 | 5.048E-09 | 3.000E-09 | 1.300E-08 |
| 29077 | 1454 | L1454T | E | 7e-09 | 1.6e-08 | 1.2e-08 | 1.167E-08 | 4.509E-09 | 7.000E-09 | 1.600E-08 |
| 29078 | 1454 | L1454W | E | 1.3e-08 | 1.7e-09 | 5e-09 | 6.567E-09 | 5.811E-09 | 1.700E-09 | 1.300E-08 |
| 29079 | 1454 | L1454Y | E | 3.9e-09 | 5.9e-09 | 4e-09 | 4.600E-09 | 1.127E-09 | 3.900E-09 | 5.900E-09 |
| 29080 | 1454 | L1454V | E | 5.5e-09 | 2.8e-09 | 1.3e-08 | 7.100E-09 | 5.285E-09 | 2.800E-09 | 1.300E-08 |
| 29081 | 1455 | Y1455A | E | 5.6e-09 | 8.3e-09 | 1.3e-08 | 8.967E-09 | 3.745E-09 | 5.600E-09 | 1.300E-08 |
| 29082 | 1455 | Y1455R | E | 5.5e-09 | 3e-09 | 3.5e-08 | 1.450E-08 | 1.780E-08 | 3.000E-09 | 3.500E-08 |
| 29083 | 1455 | Y1455N | E | 6.2e-09 | 8.3e-09 | 1.1e-08 | 8.500E-09 | 2.406E-09 | 6.200E-09 | 1.100E-08 |
| 29084 | 1455 | Y1455D | E | 3.8e-09 | 5.7e-09 | 4.2e-09 | 4.567E-09 | 1.002E-09 | 3.800E-09 | 5.700E-09 |
| 29085 | 1455 | Y1455C | E | 3.4e-09 | 2e-08 | 2.7e-09 | 8.700E-09 | 9.792E-09 | 2.700E-09 | 2.000E-08 |
| 29086 | 1455 | Y1455E | E | 1.2e-08 | 5.4e-09 | 2.9e-09 | 6.767E-09 | 4.701E-09 | 2.900E-09 | 1.200E-08 |
| 29087 | 1455 | Y1455Q | E | 5.8e-09 | 7.6e-09 | 1.2e-09 | 4.867E-09 | 3.301E-09 | 1.200E-09 | 7.600E-09 |
| 29088 | 1455 | Y1455G | E | 1.3e-08 | 5.8e-09 | 5.2e-09 | 8.000E-09 | 4.341E-09 | 5.200E-09 | 1.300E-08 |
| 29089 | 1455 | Y1455H | E | 8e-09 | 5.6e-09 | 8e-09 | 7.200E-09 | 1.386E-09 | 5.600E-09 | 8.000E-09 |
| 29090 | 1455 | Y1455I | E | 4.9e-09 | 9.9e-09 | 2.6e-08 | 1.360E-08 | 1.103E-08 | 4.900E-09 | 2.600E-08 |
| 29091 | 1455 | Y1455L | E | 2.7e-09 | 8.8e-09 | 4.9e-09 | 5.467E-09 | 3.089E-09 | 2.700E-09 | 8.800E-09 |
| 29092 | 1455 | Y1455K | E | 1.4e-08 | 5.2e-09 | 1.3e-08 | 1.073E-08 | 4.818E-09 | 5.200E-09 | 1.400E-08 |
| 29093 | 1455 | Y1455M | E | 1.5e-08 | 3e-09 | 2.5e-08 | 1.433E-08 | 1.102E-08 | 3.000E-09 | 2.500E-08 |
| 29094 | 1455 | Y1455F | E | 6.4e-09 | 6.1e-09 | 8.6e-09 | 7.033E-09 | 1.365E-09 | 6.100E-09 | 8.600E-09 |
| 29095 | 1455 | Y1455P | E | 1.4e-08 | 6.7e-09 | 8.1e-09 | 9.600E-09 | 3.874E-09 | 6.700E-09 | 1.400E-08 |
| 29096 | 1455 | Y1455S | E | 7.6e-09 | 7.2e-09 | 1e-08 | 8.267E-09 | 1.514E-09 | 7.200E-09 | 1.000E-08 |
| 29097 | 1455 | Y1455T | E | 1.1e-08 | 8.4e-09 | 5.5e-09 | 8.300E-09 | 2.751E-09 | 5.500E-09 | 1.100E-08 |
| 29098 | 1455 | Y1455W | E | 4.2e-09 | 6.3e-09 | 5.7e-09 | 5.400E-09 | 1.082E-09 | 4.200E-09 | 6.300E-09 |
| 29099 | 1455 | Y1455Y | E | 7.9e-09 | 2.5e-09 | 4.2e-09 | 4.867E-09 | 2.761E-09 | 2.500E-09 | 7.900E-09 |
| 29100 | 1455 | Y1455V | E | 1.2e-08 | 6.4e-09 | 2.7e-09 | 7.033E-09 | 4.682E-09 | 2.700E-09 | 1.200E-08 |
| 29101 | 1456 | S1456A | E | 5.7e-09 | 7.3e-09 | 5.8e-09 | 6.267E-09 | 8.963E-10 | 5.700E-09 | 7.300E-09 |
| 29102 | 1456 | S1456R | E | 1e-08 | 5.1e-09 | 1.1e-08 | 8.700E-09 | 3.158E-09 | 5.100E-09 | 1.100E-08 |
| 29103 | 1456 | S1456N | E | 2.2e-08 | 1.7e-08 | 3.1e-09 | 1.403E-08 | 9.793E-09 | 3.100E-09 | 2.200E-08 |
| 29104 | 1456 | S1456D | E | 1.1e-08 | 5.8e-09 | 4.9e-09 | 7.233E-09 | 3.293E-09 | 4.900E-09 | 1.100E-08 |
| 29105 | 1456 | S1456C | E | 1e-08 | 3.2e-09 | 2.7e-09 | 5.300E-09 | 4.078E-09 | 2.700E-09 | 1.000E-08 |
| 29106 | 1456 | S1456E | E | 6.2e-09 | 1.4e-09 | 8.1e-09 | 5.233E-09 | 3.453E-09 | 1.400E-09 | 8.100E-09 |
| 29107 | 1456 | S1456Q | E | 7.8e-09 | 4.4e-09 | 1.2e-08 | 8.067E-09 | 3.807E-09 | 4.400E-09 | 1.200E-08 |
| 29108 | 1456 | S1456G | E | 1.2e-08 | 9e-09 | 1.1e-08 | 1.067E-08 | 1.528E-09 | 9.000E-09 | 1.200E-08 |
| 29109 | 1456 | S1456H | E | 2.6e-09 | 4e-09 | 7.7e-09 | 4.767E-09 | 2.635E-09 | 2.600E-09 | 7.700E-09 |
| 29110 | 1456 | S1456I | E | 5.6e-09 | 2e-08 | 6.9e-09 | 1.083E-08 | 7.965E-09 | 5.600E-09 | 2.000E-08 |
| 29111 | 1456 | S1456L | E | 5.9e-09 | 6.4e-09 | 7.8e-09 | 6.700E-09 | 9.849E-10 | 5.900E-09 | 7.800E-09 |
| 29112 | 1456 | S1456K | E | 1.3e-08 | 1.6e-08 | 4.3e-09 | 1.110E-08 | 6.077E-09 | 4.300E-09 | 1.600E-08 |
| 29113 | 1456 | S1456M | E | 1.6e-08 | 1.6e-08 | 1.2e-08 | 1.467E-08 | 2.309E-09 | 1.200E-08 | 1.600E-08 |
| 29114 | 1456 | S1456F | E | 9.4e-09 | 1.1e-08 | 1.4e-08 | 1.147E-08 | 2.335E-09 | 9.400E-09 | 1.400E-08 |
| 29115 | 1456 | S1456P | E | 2.9e-09 | 5.7e-09 | 4.5e-09 | 4.367E-09 | 1.405E-09 | 2.900E-09 | 5.700E-09 |
| 29116 | 1456 | S1456S | E | 6e-09 | 7.8e-09 | 5.7e-09 | 6.500E-09 | 1.136E-09 | 5.700E-09 | 7.800E-09 |
| 29117 | 1456 | S1456T | E | 4e-09 | 8.6e-09 | 4.4e-09 | 5.667E-09 | 2.548E-09 | 4.000E-09 | 8.600E-09 |
| 29118 | 1456 | S1456W | E | 7.6e-09 | 7.8e-09 | 1.1e-08 | 8.800E-09 | 1.908E-09 | 7.600E-09 | 1.100E-08 |
| 29119 | 1456 | S1456Y | E | 1.2e-08 | 6.2e-09 | 2.2e-09 | 6.800E-09 | 4.927E-09 | 2.200E-09 | 1.200E-08 |
| 29120 | 1456 | S1456V | E | 5.1e-09 | 1.4e-08 | 4.7e-09 | 7.933E-09 | 5.258E-09 | 4.700E-09 | 1.400E-08 |
| 29121 | 1457 | L1457A | E | 3.6e-09 | 6e-09 | 1.6e-08 | 8.533E-09 | 6.577E-09 | 3.600E-09 | 1.600E-08 |
| 29122 | 1457 | L1457R | E | 5.2e-09 | 5.7e-09 | 1.4e-08 | 8.300E-09 | 4.943E-09 | 5.200E-09 | 1.400E-08 |
| 29123 | 1457 | L1457N | E | 2.5e-09 | 1.5e-08 | 3.5e-09 | 7.000E-09 | 6.946E-09 | 2.500E-09 | 1.500E-08 |
| 29124 | 1457 | L1457D | E | 4e-09 | 8e-09 | 6.3e-09 | 6.100E-09 | 2.007E-09 | 4.000E-09 | 8.000E-09 |
| 29125 | 1457 | L1457C | E | 8.5e-09 | 8.5e-10 | 5.8e-09 | 5.050E-09 | 3.880E-09 | 8.500E-10 | 8.500E-09 |
| 29126 | 1457 | L1457E | E | 1.2e-08 | 3.4e-09 | 5.4e-09 | 6.933E-09 | 4.500E-09 | 3.400E-09 | 1.200E-08 |
| 29127 | 1457 | L1457Q | E | 9.5e-09 | 3.4e-09 | 7e-09 | 6.633E-09 | 3.066E-09 | 3.400E-09 | 9.500E-09 |
| 29128 | 1457 | L1457G | E | 2.5e-08 | 3.5e-09 | 6.3e-09 | 1.160E-08 | 1.169E-08 | 3.500E-09 | 2.500E-08 |
| 29129 | 1457 | L1457H | E | 3e-09 | 3.9e-09 | 1.4e-08 | 6.967E-09 | 6.108E-09 | 3.000E-09 | 1.400E-08 |
| 29130 | 1457 | L1457I | E | 1.3e-08 | 6.5e-09 | 1e-08 | 9.833E-09 | 3.253E-09 | 6.500E-09 | 1.300E-08 |
| 29131 | 1457 | L1457L | E | 3.1e-09 | 3.1e-09 | 8.8e-10 | 2.360E-09 | 1.282E-09 | 8.800E-10 | 3.100E-09 |
| 29132 | 1457 | L1457K | E | 1.5e-08 | 5.2e-09 | 1e-08 | 1.007E-08 | 4.900E-09 | 5.200E-09 | 1.500E-08 |
| 29133 | 1457 | L1457M | E | 4.3e-09 | 1.2e-08 | 5.7e-09 | 7.333E-09 | 4.102E-09 | 4.300E-09 | 1.200E-08 |
| 29134 | 1457 | L1457F | E | 1.1e-08 | 4.9e-09 | 8.6e-09 | 8.167E-09 | 3.073E-09 | 4.900E-09 | 1.100E-08 |
| 29135 | 1457 | L1457P | E | 7.1e-09 | 5e-09 | 6e-09 | 6.033E-09 | 1.050E-09 | 5.000E-09 | 7.100E-09 |
| 29136 | 1457 | L1457S | E | 7.1e-09 | 1.9e-08 | 3.6e-09 | 9.900E-09 | 8.073E-09 | 3.600E-09 | 1.900E-08 |
| 29137 | 1457 | L1457T | E | 1.7e-08 | 1e-08 | 4.8e-09 | 1.060E-08 | 6.122E-09 | 4.800E-09 | 1.700E-08 |
| 29138 | 1457 | L1457W | E | 4.5e-09 | 6.6e-09 | 8e-09 | 6.367E-09 | 1.762E-09 | 4.500E-09 | 8.000E-09 |
| 29139 | 1457 | L1457Y | E | 6.2e-09 | 1.8e-08 | 1.8e-08 | 1.407E-08 | 6.813E-09 | 6.200E-09 | 1.800E-08 |
| 29140 | 1457 | L1457V | E | 8.7e-09 | 2.4e-08 | 1e-08 | 1.423E-08 | 8.483E-09 | 8.700E-09 | 2.400E-08 |
| 29141 | 1458 | S1458A | E | 1.9e-08 | 3.2e-09 | 1.5e-08 | 1.240E-08 | 8.215E-09 | 3.200E-09 | 1.900E-08 |
| 29142 | 1458 | S1458R | E | 6.7e-09 | 1.9e-08 | 1e-08 | 1.190E-08 | 6.366E-09 | 6.700E-09 | 1.900E-08 |
| 29143 | 1458 | S1458N | E | 5.2e-09 | 1.3e-08 | 5.3e-09 | 7.833E-09 | 4.475E-09 | 5.200E-09 | 1.300E-08 |
| 29144 | 1458 | S1458D | E | 9.1e-09 | 7.3e-09 | 1.4e-08 | 1.013E-08 | 3.467E-09 | 7.300E-09 | 1.400E-08 |
| 29145 | 1458 | S1458C | E | 2.2e-09 | 4.5e-09 | 7.5e-09 | 4.733E-09 | 2.658E-09 | 2.200E-09 | 7.500E-09 |
| 29146 | 1458 | S1458E | E | 3.5e-09 | 5.7e-09 | 3.3e-09 | 4.167E-09 | 1.332E-09 | 3.300E-09 | 5.700E-09 |
| 29147 | 1458 | S1458Q | E | 6.9e-09 | 8.6e-09 | 4.3e-09 | 6.600E-09 | 2.166E-09 | 4.300E-09 | 8.600E-09 |
| 29148 | 1458 | S1458G | E | 1.7e-08 | 8.7e-09 | 4.4e-09 | 1.003E-08 | 6.405E-09 | 4.400E-09 | 1.700E-08 |
| 29149 | 1458 | S1458H | E | 3.5e-09 | 1.7e-08 | 3.4e-09 | 7.967E-09 | 7.823E-09 | 3.400E-09 | 1.700E-08 |
| 29150 | 1458 | S1458I | E | 7.1e-09 | 5.1e-09 | 6e-09 | 6.067E-09 | 1.002E-09 | 5.100E-09 | 7.100E-09 |
| 29151 | 1458 | S1458L | E | 1.4e-08 | 3.5e-09 | 1.8e-08 | 1.183E-08 | 7.489E-09 | 3.500E-09 | 1.800E-08 |
| 29152 | 1458 | S1458K | E | 2.3e-08 | 5.6e-09 | 4e-09 | 1.087E-08 | 1.054E-08 | 4.000E-09 | 2.300E-08 |
| 29153 | 1458 | S1458M | E | 6.3e-09 | 7.7e-09 | 5.1e-09 | 6.367E-09 | 1.301E-09 | 5.100E-09 | 7.700E-09 |
| 29154 | 1458 | S1458F | E | 1.2e-08 | 1.7e-08 | 8.9e-09 | 1.263E-08 | 4.087E-09 | 8.900E-09 | 1.700E-08 |

| | | | | | | | | | | |
|---|---|---|---|---|---|---|---|---|---|---|
| 29155 | 1458 | S1458P | E | 6.3e-09 | 1.3e-08 | 1.5e-08 | 1.143E-08 | 4.557E-09 | 6.300E-09 | 1.500E-08 |
| 29156 | 1458 | S1458S | E | 1.3e-08 | 3.1e-09 | 4.9e-09 | 7.000E-09 | 5.274E-09 | 3.100E-09 | 1.300E-08 |
| 29157 | 1458 | S1458T | E | 3e-09 | 2e-08 | 4.1e-09 | 9.033E-09 | 9.513E-09 | 3.000E-09 | 2.000E-08 |
| 29158 | 1458 | S1458W | E | 4.2e-09 | 1.3e-08 | 3.9e-09 | 7.033E-09 | 5.169E-09 | 3.900E-09 | 1.300E-08 |
| 29159 | 1458 | S1458Y | E | 6.2e-09 | 8.8e-09 | 2.1e-09 | 5.700E-09 | 3.378E-09 | 2.100E-09 | 8.800E-09 |
| 29160 | 1458 | S1458V | E | 1.7e-08 | 7e-09 | 1.1e-08 | 1.167E-08 | 5.033E-09 | 7.000E-09 | 1.700E-08 |
| 29161 | 1459 | S1459A | E | 2.7e-09 | 4.3e-09 | 2.2e-09 | 3.067E-09 | 1.097E-09 | 2.200E-09 | 4.300E-09 |
| 29162 | 1459 | S1459R | E | 1.2e-08 | 7.3e-09 | 2.6e-08 | 1.510E-08 | 9.728E-09 | 7.300E-09 | 2.600E-08 |
| 29163 | 1459 | S1459N | E | 3.6e-09 | 1.2e-08 | 3.1e-09 | 6.233E-09 | 5.000E-09 | 3.100E-09 | 1.200E-08 |
| 29164 | 1459 | S1459D | E | 1.9e-08 | 4.6e-09 | 2.9e-09 | 8.833E-09 | 8.846E-09 | 2.900E-09 | 1.900E-08 |
| 29165 | 1459 | S1459C | E | 6.6e-09 | 1.9e-09 | 1.3e-08 | 7.167E-09 | 5.572E-09 | 1.900E-09 | 1.300E-08 |
| 29166 | 1459 | S1459E | E | 1.2e-08 | 5.1e-09 | 7.6e-09 | 8.233E-09 | 3.493E-09 | 5.100E-09 | 1.200E-08 |
| 29167 | 1459 | S1459Q | E | 5.5e-09 | 5.7e-09 | 4.4e-09 | 5.200E-09 | 7.000E-10 | 4.400E-09 | 5.700E-09 |
| 29168 | 1459 | S1459G | E | 4.1e-09 | 1.1e-08 | 3.5e-09 | 6.200E-09 | 4.168E-09 | 3.500E-09 | 1.100E-08 |
| 29169 | 1459 | S1459H | E | 4.4e-09 | 5.1e-09 | 7.5e-09 | 5.667E-09 | 1.626E-09 | 4.400E-09 | 7.500E-09 |
| 29170 | 1459 | S1459I | E | 5.8e-09 | 7.1e-09 | 7.7e-09 | 6.867E-09 | 9.713E-10 | 5.800E-09 | 7.700E-09 |
| 29171 | 1459 | S1459L | E | 3.4e-09 | 2.9e-09 | 9.5e-09 | 5.267E-09 | 3.675E-09 | 2.900E-09 | 9.500E-09 |
| 29172 | 1459 | S1459K | E | 4.5e-09 | 1.3e-08 | 1.5e-08 | 1.083E-08 | 5.575E-09 | 4.500E-09 | 1.500E-08 |
| 29173 | 1459 | S1459M | E | 1.8e-09 | 8e-09 | 6.6e-09 | 5.467E-09 | 3.252E-09 | 1.800E-09 | 8.000E-09 |
| 29174 | 1459 | S1459F | E | 4.8e-09 | 2.3e-09 | 7.5e-09 | 4.867E-09 | 2.601E-09 | 2.300E-09 | 7.500E-09 |
| 29175 | 1459 | S1459P | E | 6.1e-09 | 7.5e-09 | 1.2e-08 | 8.533E-09 | 3.083E-09 | 6.100E-09 | 1.200E-08 |
| 29176 | 1459 | S1459S | E | 8.6e-09 | 6.1e-09 | 7.6e-09 | 7.433E-09 | 1.258E-09 | 6.100E-09 | 8.600E-09 |
| 29177 | 1459 | S1459T | E | 1.6e-08 | 8.8e-09 | 3.4e-09 | 9.400E-09 | 6.321E-09 | 3.400E-09 | 1.600E-08 |
| 29178 | 1459 | S1459W | E | 1.2e-08 | 6.8e-09 | 5.4e-09 | 8.067E-09 | 3.478E-09 | 5.400E-09 | 1.200E-08 |
| 29179 | 1459 | S1459Y | E | 1.1e-08 | 8.2e-09 | 2.2e-09 | 7.133E-09 | 4.496E-09 | 2.200E-09 | 1.100E-08 |
| 29180 | 1459 | S1459V | E | 7.7e-09 | 8.5e-09 | 4.5e-09 | 6.900E-09 | 2.117E-09 | 4.500E-09 | 8.500E-09 |
| 29181 | 1460 | V1460A | E | 5.2e-09 | 3.1e-09 | 1.2e-08 | 6.767E-09 | 4.652E-09 | 3.100E-09 | 1.200E-08 |
| 29182 | 1460 | V1460R | E | 1.7e-08 | 2e-08 | 1.3e-08 | 1.667E-08 | 3.512E-09 | 1.300E-08 | 2.000E-08 |
| 29183 | 1460 | V1460N | E | 6e-09 | 8.4e-09 | 6.3e-09 | 6.900E-09 | 1.308E-09 | 6.000E-09 | 8.400E-09 |
| 29184 | 1460 | V1460D | E | 4.4e-09 | 2.6e-08 | 3.6e-09 | 1.133E-08 | 1.271E-08 | 3.600E-09 | 2.600E-08 |
| 29185 | 1460 | V1460C | E | 1.8e-09 | 1.3e-08 | 7.4e-09 | 7.400E-09 | 5.600E-09 | 1.800E-09 | 1.300E-08 |
| 29186 | 1460 | V1460E | E | 9.8e-09 | 2.1e-08 | 2e-08 | 1.693E-08 | 6.198E-09 | 9.800E-09 | 2.100E-08 |
| 29187 | 1460 | V1460Q | E | 1e-09 | 6.8e-09 | 6.7e-09 | 4.833E-09 | 3.320E-09 | 1.000E-09 | 6.800E-09 |
| 29188 | 1460 | V1460G | E | 9.5e-09 | 4.7e-09 | 3e-09 | 5.733E-09 | 3.371E-09 | 3.000E-09 | 9.500E-09 |
| 29189 | 1460 | V1460H | E | 4.2e-09 | 1.4e-08 | 4.7e-09 | 7.633E-09 | 5.519E-09 | 4.200E-09 | 1.400E-08 |
| 29190 | 1460 | V1460I | E | 1.1e-08 | 4.8e-09 | 3.6e-09 | 6.467E-09 | 3.972E-09 | 3.600E-09 | 1.100E-08 |
| 29191 | 1460 | V1460L | E | 1.7e-08 | 1.2e-08 | 5.8e-09 | 1.160E-08 | 5.611E-09 | 5.800E-09 | 1.700E-08 |
| 29192 | 1460 | V1460K | E | 4.7e-09 | 4.6e-09 | 8.2e-09 | 5.833E-09 | 2.050E-09 | 4.600E-09 | 8.200E-09 |
| 29193 | 1460 | V1460M | E | 5.9e-09 | 1.2e-08 | 6.4e-09 | 8.100E-09 | 3.387E-09 | 5.900E-09 | 1.200E-08 |
| 29194 | 1460 | V1460F | E | 4.4e-09 | 2.4e-09 | 1.5e-08 | 7.267E-09 | 6.772E-09 | 2.400E-09 | 1.500E-08 |
| 29195 | 1460 | V1460P | E | 8.8e-09 | 6.7e-09 | 1.1e-08 | 8.833E-09 | 2.150E-09 | 6.700E-09 | 1.100E-08 |
| 29196 | 1460 | V1460S | E | 6.8e-09 | 2.8e-09 | 3e-09 | 4.200E-09 | 2.254E-09 | 2.800E-09 | 6.800E-09 |
| 29197 | 1460 | V1460T | E | 3.3e-09 | 1e-08 | 2.4e-09 | 5.233E-09 | 4.153E-09 | 2.400E-09 | 1.000E-08 |
| 29198 | 1460 | V1460W | E | 1.3e-08 | 1.1e-08 | 5e-09 | 9.667E-09 | 4.163E-09 | 5.000E-09 | 1.300E-08 |
| 29199 | 1460 | V1460Y | E | 3.4e-09 | 1.1e-08 | 1.4e-08 | 9.467E-09 | 5.464E-09 | 3.400E-09 | 1.400E-08 |
| 29200 | 1460 | V1460V | E | 5.6e-09 | 3.2e-09 | 5.8e-09 | 4.867E-09 | 1.447E-09 | 3.200E-09 | 5.800E-09 |
| 29201 | 1461 | V1461A | E | 5.2e-09 | 2.9e-09 | 5.5e-09 | 4.533E-09 | 1.422E-09 | 2.900E-09 | 5.500E-09 |
| 29202 | 1461 | V1461R | E | 6.1e-09 | 4.1e-09 | 2.8e-09 | 4.333E-09 | 1.662E-09 | 2.800E-09 | 6.100E-09 |
| 29203 | 1461 | V1461N | E | 1.3e-08 | 1.1e-08 | 7.8e-09 | 1.060E-08 | 2.623E-09 | 7.800E-09 | 1.300E-08 |
| 29204 | 1461 | V1461D | E | 2.8e-09 | 1.1e-08 | 8.7e-09 | 7.500E-09 | 4.230E-09 | 2.800E-09 | 1.100E-08 |
| 29205 | 1461 | V1461C | E | 7.7e-09 | 4.2e-09 | 1.4e-08 | 8.633E-09 | 4.966E-09 | 4.200E-09 | 1.400E-08 |
| 29206 | 1461 | V1461E | E | 6.6e-09 | 1e-08 | 1.2e-08 | 9.533E-09 | 2.730E-09 | 6.600E-09 | 1.200E-08 |
| 29207 | 1461 | V1461Q | E | 1.9e-08 | 7.2e-09 | 7e-09 | 1.107E-08 | 6.871E-09 | 7.000E-09 | 1.900E-08 |
| 29208 | 1461 | V1461G | E | 4.9e-09 | 5.6e-09 | 5.4e-09 | 5.300E-09 | 3.606E-10 | 4.900E-09 | 5.600E-09 |
| 29209 | 1461 | V1461H | E | 9.8e-09 | 1.9e-08 | 3.5e-09 | 1.077E-08 | 7.795E-09 | 3.500E-09 | 1.900E-08 |
| 29210 | 1461 | V1461I | E | 7.3e-09 | 1.9e-08 | 1e-08 | 1.210E-08 | 6.126E-09 | 7.300E-09 | 1.900E-08 |
| 29211 | 1461 | V1461L | E | 5.3e-09 | 7.1e-09 | 1.2e-08 | 8.133E-09 | 3.467E-09 | 5.300E-09 | 1.200E-08 |
| 29212 | 1461 | V1461K | E | 4.2e-09 | 4.9e-09 | 8.6e-09 | 5.900E-09 | 2.364E-09 | 4.200E-09 | 8.600E-09 |
| 29213 | 1461 | V1461M | E | 1.8e-08 | 7.8e-09 | 3.7e-09 | 9.833E-09 | 7.364E-09 | 3.700E-09 | 1.800E-08 |
| 29214 | 1461 | V1461F | E | 5.8e-09 | 6.3e-09 | 5.5e-09 | 5.867E-09 | 4.041E-10 | 5.500E-09 | 6.300E-09 |
| 29215 | 1461 | V1461P | E | 9.3e-09 | 3e-09 | 1.2e-09 | 4.500E-09 | 4.253E-09 | 1.200E-09 | 9.300E-09 |
| 29216 | 1461 | V1461S | E | 3.6e-09 | 1.1e-08 | 8.1e-09 | 7.567E-09 | 3.729E-09 | 3.600E-09 | 1.100E-08 |
| 29217 | 1461 | V1461T | E | 2.3e-09 | 2.8e-09 | 9.4e-09 | 4.833E-09 | 3.963E-09 | 2.300E-09 | 9.400E-09 |
| 29218 | 1461 | V1461W | E | 8.1e-09 | 5e-09 | 2.6e-09 | 5.233E-09 | 2.757E-09 | 2.600E-09 | 8.100E-09 |
| 29219 | 1461 | V1461Y | E | 1.5e-08 | 4.9e-09 | 6.9e-09 | 8.933E-09 | 5.348E-09 | 4.900E-09 | 1.500E-08 |
| 29220 | 1461 | V1461V | E | 1.2e-08 | 1.3e-08 | 3.6e-09 | 9.533E-09 | 5.163E-09 | 3.600E-09 | 1.300E-08 |
| 29221 | 1462 | T1462A | E | 3.9e-09 | 8.2e-09 | 2.4e-09 | 4.833E-09 | 3.011E-09 | 2.400E-09 | 8.200E-09 |
| 29222 | 1462 | T1462R | E | 1e-08 | 2.7e-09 | 5.5e-09 | 6.067E-09 | 3.683E-09 | 2.700E-09 | 1.000E-08 |
| 29223 | 1462 | T1462N | E | 2e-08 | 3.5e-09 | 3.1e-09 | 8.867E-09 | 9.644E-09 | 3.100E-09 | 2.000E-08 |
| 29224 | 1462 | T1462D | E | 1.5e-08 | 7.2e-09 | 9.5e-09 | 1.057E-08 | 4.008E-09 | 7.200E-09 | 1.500E-08 |
| 29225 | 1462 | T1462C | E | 4e-09 | 5e-09 | 9.9e-09 | 6.300E-09 | 3.158E-09 | 4.000E-09 | 9.900E-09 |
| 29226 | 1462 | T1462E | E | 2.2e-08 | 1.1e-08 | 5.7e-09 | 1.290E-08 | 8.314E-09 | 5.700E-09 | 2.200E-08 |
| 29227 | 1462 | T1462Q | E | 7.1e-09 | 1.1e-08 | 2.1e-09 | 6.733E-09 | 4.461E-09 | 2.100E-09 | 1.100E-08 |
| 29228 | 1462 | T1462G | E | 1e-08 | 6.8e-09 | 1.7e-08 | 1.127E-08 | 5.217E-09 | 6.800E-09 | 1.700E-08 |
| 29229 | 1462 | T1462H | E | 5.3e-09 | 9.6e-09 | 8.4e-09 | 7.767E-09 | 2.219E-09 | 5.300E-09 | 9.600E-09 |
| 29230 | 1462 | T1462I | E | 7e-09 | 5.6e-09 | 8.7e-09 | 7.100E-09 | 1.552E-09 | 5.600E-09 | 8.700E-09 |
| 29231 | 1462 | T1462L | E | 8e-09 | 5.2e-09 | 3.8e-09 | 5.667E-09 | 2.139E-09 | 3.800E-09 | 8.000E-09 |
| 29232 | 1462 | T1462K | E | 6.9e-09 | 9.5e-09 | 8e-09 | 8.133E-09 | 1.305E-09 | 6.900E-09 | 9.500E-09 |
| 29233 | 1462 | T1462M | E | 1.7e-08 | 5.8e-09 | 5.9e-09 | 9.567E-09 | 6.438E-09 | 5.800E-09 | 1.700E-08 |
| 29234 | 1462 | T1462F | E | 4.9e-09 | 9.2e-09 | 2e-08 | 1.137E-08 | 7.780E-09 | 4.900E-09 | 2.000E-08 |
| 29235 | 1462 | T1462P | E | 2.2e-09 | 7.8e-09 | 7.1e-09 | 5.700E-09 | 3.051E-09 | 2.200E-09 | 7.800E-09 |
| 29236 | 1462 | T1462S | E | 5.7e-09 | 3.5e-09 | 8.6e-09 | 5.933E-09 | 2.558E-09 | 3.500E-09 | 8.600E-09 |
| 29237 | 1462 | T1462T | E | 3.9e-09 | 1.2e-08 | 2e-08 | 1.197E-08 | 8.050E-09 | 3.900E-09 | 2.000E-08 |
| 29238 | 1462 | T1462W | E | 3.2e-09 | 4e-09 | 1.2e-08 | 6.400E-09 | 4.866E-09 | 3.200E-09 | 1.200E-08 |
| 29239 | 1462 | T1462Y | E | 7.1e-09 | 1e-08 | 5.5e-09 | 7.533E-09 | 2.281E-09 | 5.500E-09 | 1.000E-08 |

| | | | | | | | | | | |
|---|---|---|---|---|---|---|---|---|---|---|
| 29240 | 1462 | T1462V | E | 6.9e-09 | 1.4e-09 | 1.4e-08 | 7.433E-09 | 6.317E-09 | 1.400E-09 | 1.400E-08 |
| 29241 | 1463 | V1463A | E | 1.4e-08 | 1.3e-08 | 9.5e-09 | 1.217E-08 | 2.363E-09 | 9.500E-09 | 1.400E-08 |
| 29242 | 1463 | V1463R | E | 2e-09 | 3.6e-09 | 4.7e-09 | 3.433E-09 | 1.358E-09 | 2.000E-09 | 4.700E-09 |
| 29243 | 1463 | V1463N | E | 2.8e-09 | 6.7e-09 | 1.8e-08 | 9.167E-09 | 7.895E-09 | 2.800E-09 | 1.800E-08 |
| 29244 | 1463 | V1463D | E | 7.7e-09 | 4.5e-09 | 8.7e-09 | 6.967E-09 | 2.194E-09 | 4.500E-09 | 8.700E-09 |
| 29245 | 1463 | V1463C | E | 5e-09 | 6.5e-09 | 2e-09 | 4.500E-09 | 2.291E-09 | 2.000E-09 | 6.500E-09 |
| 29246 | 1463 | V1463E | E | 6.5e-09 | 8.7e-09 | 6e-09 | 7.067E-09 | 1.436E-09 | 6.000E-09 | 8.700E-09 |
| 29247 | 1463 | V1463Q | E | 6e-09 | 5.6e-09 | 1.1e-08 | 7.533E-09 | 3.009E-09 | 5.600E-09 | 1.100E-08 |
| 29248 | 1463 | V1463G | E | 4.6e-09 | 3.7e-09 | 2.9e-09 | 3.733E-09 | 8.505E-10 | 2.900E-09 | 4.600E-09 |
| 29249 | 1463 | V1463H | E | 1.3e-08 | 1e-08 | 2.9e-09 | 8.633E-09 | 5.187E-09 | 2.900E-09 | 1.300E-08 |
| 29250 | 1463 | V1463I | E | 7.8e-09 | 1e-08 | 1.1e-08 | 9.600E-09 | 1.637E-09 | 7.800E-09 | 1.100E-08 |
| 29251 | 1463 | V1463L | E | 4.2e-09 | 5.1e-09 | 7.5e-09 | 5.600E-09 | 1.706E-09 | 4.200E-09 | 7.500E-09 |
| 29252 | 1463 | V1463K | E | 9.1e-09 | 6.5e-09 | 1.2e-08 | 9.200E-09 | 2.751E-09 | 6.500E-09 | 1.200E-08 |
| 29253 | 1463 | V1463M | E | 7e-09 | 4.5e-09 | 3.1e-08 | 1.417E-08 | 1.463E-08 | 4.500E-09 | 3.100E-08 |
| 29254 | 1463 | V1463F | E | 3.4e-09 | 1.4e-08 | 1.7e-08 | 1.147E-08 | 7.145E-09 | 3.400E-09 | 1.700E-08 |
| 29255 | 1463 | V1463P | E | 7e-09 | 2.3e-08 | 8.1e-09 | 1.270E-08 | 8.937E-09 | 7.000E-09 | 2.300E-08 |
| 29256 | 1463 | V1463S | E | 8.8e-09 | 7.1e-09 | 7.7e-09 | 7.867E-09 | 8.622E-10 | 7.100E-09 | 8.800E-09 |
| 29257 | 1463 | V1463T | E | 1e-08 | 5.4e-09 | 8.5e-09 | 7.967E-09 | 2.346E-09 | 5.400E-09 | 1.000E-08 |
| 29258 | 1463 | V1463W | E | 6.9e-09 | 9.7e-09 | 9.2e-09 | 8.600E-09 | 1.493E-09 | 6.900E-09 | 9.700E-09 |
| 29259 | 1463 | V1463Y | E | 9.1e-09 | 6.7e-09 | 1.8e-08 | 1.127E-08 | 5.953E-09 | 6.700E-09 | 1.800E-08 |
| 29260 | 1463 | V1463V | E | 4.1e-08 | 3.8e-09 | 1.8e-09 | 1.553E-08 | 2.208E-08 | 1.800E-09 | 4.100E-08 |
| 29261 | 1464 | P1464A | E | 5.4e-09 | 1.4e-08 | 9.4e-09 | 9.600E-09 | 4.303E-09 | 5.400E-09 | 1.400E-08 |
| 29262 | 1464 | P1464R | E | 2e-08 | 6.4e-09 | 7.9e-09 | 1.143E-08 | 7.457E-09 | 6.400E-09 | 2.000E-08 |
| 29263 | 1464 | P1464N | E | 7e-09 | 7.3e-09 | 1.3e-08 | 9.100E-09 | 3.381E-09 | 7.000E-09 | 1.300E-08 |
| 29264 | 1464 | P1464D | E | 2.2e-09 | 7.2e-09 | 7.4e-09 | 5.600E-09 | 2.946E-09 | 2.200E-09 | 7.400E-09 |
| 29265 | 1464 | P1464C | E | 2.9e-09 | 7.6e-09 | 5.4e-09 | 5.300E-09 | 2.352E-09 | 2.900E-09 | 7.600E-09 |
| 29266 | 1464 | P1464E | E | 1.2e-08 | 7e-09 | 2.7e-09 | 7.233E-09 | 4.654E-09 | 2.700E-09 | 1.200E-08 |
| 29267 | 1464 | P1464Q | E | 3.5e-09 | 1e-08 | 1.1e-08 | 8.167E-09 | 4.072E-09 | 3.500E-09 | 1.100E-08 |
| 29268 | 1464 | P1464G | E | 2.6e-09 | 1e-08 | 1.3e-08 | 8.533E-09 | 5.353E-09 | 2.600E-09 | 1.300E-08 |
| 29269 | 1464 | P1464H | E | 5.4e-09 | 5.9e-09 | 1.1e-08 | 7.433E-09 | 3.099E-09 | 5.400E-09 | 1.100E-08 |
| 29270 | 1464 | P1464I | E | 1.4e-08 | 1.9e-08 | 8.5e-09 | 1.383E-08 | 5.252E-09 | 8.500E-09 | 1.900E-08 |
| 29271 | 1464 | P1464L | E | 2.4e-09 | 6.3e-09 | 3.2e-09 | 3.967E-09 | 2.060E-09 | 2.400E-09 | 6.300E-09 |
| 29272 | 1464 | P1464K | E | 2.8e-08 | 5.9e-09 | 8.4e-09 | 1.410E-08 | 1.210E-08 | 5.900E-09 | 2.800E-08 |
| 29273 | 1464 | P1464M | E | 3.7e-09 | 4.8e-09 | 6.4e-09 | 4.967E-09 | 1.358E-09 | 3.700E-09 | 6.400E-09 |
| 29274 | 1464 | P1464F | E | 9.7e-09 | 1.1e-08 | 9.5e-09 | 1.007E-08 | 8.145E-10 | 9.500E-09 | 1.100E-08 |
| 29275 | 1464 | P1464P | E | 1.6e-08 | 2e-08 | 1.4e-08 | 1.667E-08 | 3.055E-09 | 1.400E-08 | 2.000E-08 |
| 29276 | 1464 | P1464S | E | 2.4e-09 | 1.7e-08 | 1.9e-08 | 1.280E-08 | 9.062E-09 | 2.400E-09 | 1.900E-08 |
| 29277 | 1464 | P1464T | E | 6.1e-09 | 4.7e-09 | 8.8e-09 | 6.533E-09 | 2.084E-09 | 4.700E-09 | 8.800E-09 |
| 29278 | 1464 | P1464W | E | 6.2e-09 | 2.4e-09 | 7.6e-09 | 5.400E-09 | 2.691E-09 | 2.400E-09 | 7.600E-09 |
| 29279 | 1464 | P1464Y | E | 3.5e-09 | 6.6e-09 | 6.8e-09 | 5.633E-09 | 1.850E-09 | 3.500E-09 | 6.800E-09 |
| 29280 | 1464 | P1464V | E | 8.2e-09 | 6.5e-09 | 6.6e-09 | 7.100E-09 | 9.539E-10 | 6.500E-09 | 8.200E-09 |
| 29281 | 1465 | S1465A | E | 1.3e-08 | 8.5e-09 | 7.5e-09 | 9.667E-09 | 2.930E-09 | 7.500E-09 | 1.300E-08 |
| 29282 | 1465 | S1465R | E | 5.2e-09 | 1.1e-08 | 2.2e-08 | 1.273E-08 | 8.533E-09 | 5.200E-09 | 2.200E-08 |
| 29283 | 1465 | S1465N | E | 3.8e-09 | 4.6e-09 | 1.5e-08 | 7.800E-09 | 6.248E-09 | 3.800E-09 | 1.500E-08 |
| 29284 | 1465 | S1465D | E | 8.9e-09 | 1.5e-08 | 5.9e-09 | 9.933E-09 | 4.637E-09 | 5.900E-09 | 1.500E-08 |
| 29285 | 1465 | S1465C | E | 1.2e-08 | 1.3e-08 | 4.1e-09 | 9.700E-09 | 4.875E-09 | 4.100E-09 | 1.300E-08 |
| 29286 | 1465 | S1465E | E | 6.9e-09 | 1.1e-08 | 7.7e-09 | 8.533E-09 | 2.173E-09 | 6.900E-09 | 1.100E-08 |
| 29287 | 1465 | S1465Q | E | 1.2e-08 | 5.9e-09 | 8.7e-09 | 8.867E-09 | 3.053E-09 | 5.900E-09 | 1.200E-08 |
| 29288 | 1465 | S1465G | E | 1.3e-08 | 9.4e-09 | 8.2e-09 | 1.020E-08 | 2.498E-09 | 8.200E-09 | 1.300E-08 |
| 29289 | 1465 | S1465H | E | 9.8e-09 | 3.4e-09 | 8.8e-09 | 7.333E-09 | 3.443E-09 | 3.400E-09 | 9.800E-09 |
| 29290 | 1465 | S1465I | E | 5.7e-09 | 2.5e-09 | 1.8e-08 | 8.733E-09 | 8.183E-09 | 2.500E-09 | 1.800E-08 |
| 29291 | 1465 | S1465L | E | 8.7e-09 | 7.5e-09 | 1.2e-08 | 9.400E-09 | 2.330E-09 | 7.500E-09 | 1.200E-08 |
| 29292 | 1465 | S1465K | E | 3.7e-09 | 6.7e-09 | 6.8e-09 | 5.733E-09 | 1.762E-09 | 3.700E-09 | 6.800E-09 |
| 29293 | 1465 | S1465M | E | 6.3e-09 | 1e-08 | 6.4e-09 | 7.567E-09 | 2.108E-09 | 6.300E-09 | 1.000E-08 |
| 29294 | 1465 | S1465F | E | 4.2e-09 | 4.3e-09 | 6.6e-09 | 5.033E-09 | 1.358E-09 | 4.200E-09 | 6.600E-09 |
| 29295 | 1465 | S1465P | E | 2.8e-09 | 1e-08 | 3e-09 | 5.267E-09 | 4.100E-09 | 2.800E-09 | 1.000E-08 |
| 29296 | 1465 | S1465S | E | 8.4e-09 | 1.5e-08 | 5.3e-09 | 9.567E-09 | 4.954E-09 | 5.300E-09 | 1.500E-08 |
| 29297 | 1465 | S1465T | E | 7.4e-09 | 6.8e-09 | 5e-09 | 6.400E-09 | 1.249E-09 | 5.000E-09 | 7.400E-09 |
| 29298 | 1465 | S1465W | E | 1.1e-08 | 2e-08 | 9.2e-09 | 1.340E-08 | 5.786E-09 | 9.200E-09 | 2.000E-08 |
| 29299 | 1465 | S1465Y | E | 4.6e-09 | 4.3e-09 | 8.6e-09 | 5.833E-09 | 2.401E-09 | 4.300E-09 | 8.600E-09 |
| 29300 | 1465 | S1465V | E | 5e-09 | 6.5e-09 | 2.2e-08 | 1.117E-08 | 9.412E-09 | 5.000E-09 | 2.200E-08 |
| 29301 | 1466 | S1466A | E | 1e-08 | 8.1e-09 | 6.2e-09 | 8.100E-09 | 1.900E-09 | 6.200E-09 | 1.000E-08 |
| 29302 | 1466 | S1466R | E | 8.6e-09 | 1.4e-08 | 9.6e-09 | 1.073E-08 | 2.873E-09 | 8.600E-09 | 1.400E-08 |
| 29303 | 1466 | S1466N | E | 6.3e-09 | 1.6e-08 | 6.7e-09 | 9.667E-09 | 5.488E-09 | 6.300E-09 | 1.600E-08 |
| 29304 | 1466 | S1466D | E | 3.5e-09 | 7.8e-09 | 9.2e-09 | 6.833E-09 | 2.970E-09 | 3.500E-09 | 9.200E-09 |
| 29305 | 1466 | S1466C | E | 2.7e-09 | 1.1e-08 | 8.6e-09 | 7.433E-09 | 4.271E-09 | 2.700E-09 | 1.100E-08 |
| 29306 | 1466 | S1466E | E | 6.8e-09 | 6.4e-09 | 1.2e-08 | 8.400E-09 | 3.124E-09 | 6.400E-09 | 1.200E-08 |
| 29307 | 1466 | S1466Q | E | 4.6e-09 | 5.3e-09 | 9e-09 | 6.300E-09 | 2.364E-09 | 4.600E-09 | 9.000E-09 |
| 29308 | 1466 | S1466G | E | 6.7e-09 | 1.1e-08 | 3.3e-08 | 1.690E-08 | 1.411E-08 | 6.700E-09 | 3.300E-08 |
| 29309 | 1466 | S1466H | E | 2.5e-08 | 3.7e-09 | 1.2e-08 | 1.357E-08 | 1.074E-08 | 3.700E-09 | 2.500E-08 |
| 29310 | 1466 | S1466I | E | 1.6e-08 | 1e-08 | 4.6e-09 | 1.020E-08 | 5.703E-09 | 4.600E-09 | 1.600E-08 |
| 29311 | 1466 | S1466L | E | 5.4e-09 | 3.2e-09 | 9.2e-09 | 5.933E-09 | 3.035E-09 | 3.200E-09 | 9.200E-09 |
| 29312 | 1466 | S1466K | E | 1.8e-09 | 5.6e-09 | 8.9e-09 | 5.433E-09 | 3.553E-09 | 1.800E-09 | 8.900E-09 |
| 29313 | 1466 | S1466M | E | 4.5e-09 | 4.9e-09 | 7.2e-09 | 5.533E-09 | 1.457E-09 | 4.500E-09 | 7.200E-09 |
| 29314 | 1466 | S1466F | E | 8.5e-09 | 2.6e-09 | 2.3e-09 | 4.467E-09 | 3.496E-09 | 2.300E-09 | 8.500E-09 |
| 29315 | 1466 | S1466P | E | 1.2e-08 | 4.4e-09 | 8.6e-09 | 8.333E-09 | 3.807E-09 | 4.400E-09 | 1.200E-08 |
| 29316 | 1466 | S1466S | E | 6.1e-09 | 7.4e-09 | 7.1e-09 | 6.867E-09 | 6.807E-10 | 6.100E-09 | 7.400E-09 |
| 29317 | 1466 | S1466T | E | 1.5e-08 | 9e-09 | 6e-09 | 1.000E-08 | 4.583E-09 | 6.000E-09 | 1.500E-08 |
| 29318 | 1466 | S1466W | E | 3.4e-09 | 1.2e-08 | 1.3e-08 | 9.467E-09 | 5.278E-09 | 3.400E-09 | 1.300E-08 |
| 29319 | 1466 | S1466Y | E | 1e-08 | 8.7e-09 | 9.7e-09 | 9.467E-09 | 6.807E-10 | 8.700E-09 | 1.000E-08 |
| 29320 | 1466 | S1466V | E | 1.6e-08 | 4.3e-09 | 5.6e-09 | 8.633E-09 | 6.413E-09 | 4.300E-09 | 1.600E-08 |
| 29321 | 1467 | S1467A | E | 1e-08 | 6e-09 | 2.5e-09 | 6.167E-09 | 3.753E-09 | 2.500E-09 | 1.000E-08 |
| 29322 | 1467 | S1467R | E | 4.8e-09 | 4.3e-09 | 1e-08 | 6.367E-09 | 3.156E-09 | 4.300E-09 | 1.000E-08 |
| 29323 | 1467 | S1467N | E | 1.1e-08 | 1.8e-08 | 4.3e-08 | 2.400E-08 | 1.682E-08 | 1.100E-08 | 4.300E-08 |
| 29324 | 1467 | S1467D | E | 6.6e-09 | 6.1e-09 | 1.3e-08 | 8.567E-09 | 3.848E-09 | 6.100E-09 | 1.300E-08 |

| | | | | | | | | | | |
|---|---|---|---|---|---|---|---|---|---|---|
| 29325 | 1467 | S1467C | E | 6.1e-09 | 2e-08 | 1e-08 | 1.203E-08 | 7.170E-09 | 6.100E-09 | 2.000E-08 |
| 29326 | 1467 | S1467E | E | 1e-08 | 3.3e-09 | 4.4e-09 | 5.900E-09 | 3.593E-09 | 3.300E-09 | 1.000E-08 |
| 29327 | 1467 | S1467Q | E | 1.2e-08 | 3.6e-09 | 7.1e-09 | 7.567E-09 | 4.219E-09 | 3.600E-09 | 1.200E-08 |
| 29328 | 1467 | S1467G | E | 3.5e-09 | 9.2e-09 | 3.8e-09 | 5.500E-09 | 3.208E-09 | 3.500E-09 | 9.200E-09 |
| 29329 | 1467 | S1467H | E | 1.5e-08 | 5.4e-09 | 1.6e-09 | 7.333E-09 | 6.906E-09 | 1.600E-09 | 1.500E-08 |
| 29330 | 1467 | S1467I | E | 6e-09 | 1.2e-08 | 1.3e-08 | 1.033E-08 | 3.786E-09 | 6.000E-09 | 1.300E-08 |
| 29331 | 1467 | S1467L | E | 7.2e-09 | 2.7e-09 | 5.1e-09 | 5.000E-09 | 2.252E-09 | 2.700E-09 | 7.200E-09 |
| 29332 | 1467 | S1467K | E | 5.3e-09 | 1.1e-08 | 8.4e-09 | 8.233E-09 | 2.854E-09 | 5.300E-09 | 1.100E-08 |
| 29333 | 1467 | S1467M | E | 2.8e-09 | 1.4e-09 | 1.4e-08 | 6.067E-09 | 6.906E-09 | 1.400E-09 | 1.400E-08 |
| 29334 | 1467 | S1467F | E | 2.2e-08 | 3.4e-09 | 6.2e-09 | 1.053E-08 | 1.003E-08 | 3.400E-09 | 2.200E-08 |
| 29335 | 1467 | S1467P | E | 5.1e-09 | 7.6e-09 | 7.7e-09 | 6.800E-09 | 1.473E-09 | 5.100E-09 | 7.700E-09 |
| 29336 | 1467 | S1467S | E | 1.5e-08 | 4.7e-09 | 2.4e-09 | 7.367E-09 | 6.710E-09 | 2.400E-09 | 1.500E-08 |
| 29337 | 1467 | S1467T | E | 3.4e-09 | 1e-08 | 9.1e-09 | 7.500E-09 | 3.579E-09 | 3.400E-09 | 1.000E-08 |
| 29338 | 1467 | S1467W | E | 6.1e-09 | 9.5e-09 | 9.4e-09 | 8.333E-09 | 1.935E-09 | 6.100E-09 | 9.500E-09 |
| 29339 | 1467 | S1467Y | E | 1.1e-08 | 1.9e-08 | 1.5e-08 | 1.500E-08 | 4.000E-09 | 1.100E-08 | 1.900E-08 |
| 29340 | 1467 | S1467V | E | 9.6e-09 | 5.1e-09 | 1.3e-08 | 9.233E-09 | 3.963E-09 | 5.100E-09 | 1.300E-08 |
| 29341 | 1468 | L1468A | E | 2e-09 | 6.1e-09 | 1.7e-09 | 3.267E-09 | 2.458E-09 | 1.700E-09 | 6.100E-09 |
| 29342 | 1468 | L1468R | E | 4.4e-09 | 7.1e-09 | 1.7e-08 | 9.500E-09 | 6.634E-09 | 4.400E-09 | 1.700E-08 |
| 29343 | 1468 | L1468N | E | 8.5e-09 | 1.1e-08 | 1.1e-08 | 1.017E-08 | 1.443E-09 | 8.500E-09 | 1.100E-08 |
| 29344 | 1468 | L1468D | E | 7.5e-09 | 8.8e-09 | 5.6e-09 | 7.300E-09 | 1.609E-09 | 5.600E-09 | 8.800E-09 |
| 29345 | 1468 | L1468C | E | 4.6e-09 | 1.8e-08 | 5.2e-09 | 9.267E-09 | 7.569E-09 | 4.600E-09 | 1.800E-08 |
| 29346 | 1468 | L1468E | E | 5.5e-09 | 9.3e-09 | 1.1e-08 | 8.600E-09 | 2.816E-09 | 5.500E-09 | 1.100E-08 |
| 29347 | 1468 | L1468Q | E | 7.5e-09 | 2.8e-09 | 2.1e-08 | 1.043E-08 | 9.448E-09 | 2.800E-09 | 2.100E-08 |
| 29348 | 1468 | L1468G | E | 7.4e-09 | 2.6e-09 | 8.7e-09 | 6.233E-09 | 3.213E-09 | 2.600E-09 | 8.700E-09 |
| 29349 | 1468 | L1468H | E | 2.5e-09 | 2.9e-09 | 4.8e-09 | 3.400E-09 | 1.229E-09 | 2.500E-09 | 4.800E-09 |
| 29350 | 1468 | L1468I | E | 5.4e-09 | 2.3e-09 | 7.7e-09 | 5.133E-09 | 2.710E-09 | 2.300E-09 | 7.700E-09 |
| 29351 | 1468 | L1468L | E | 6.3e-09 | 1.2e-09 | 2.2e-08 | 9.833E-09 | 1.084E-08 | 1.200E-09 | 2.200E-08 |
| 29352 | 1468 | L1468K | E | 1.4e-09 | 2e-09 | 1.4e-08 | 5.800E-09 | 7.108E-09 | 1.400E-09 | 1.400E-08 |
| 29353 | 1468 | L1468M | E | 5.9e-09 | 8.8e-09 | 1.1e-08 | 8.567E-09 | 2.558E-09 | 5.900E-09 | 1.100E-08 |
| 29354 | 1468 | L1468F | E | 5.5e-09 | 4.6e-09 | 7.8e-09 | 5.967E-09 | 1.650E-09 | 4.600E-09 | 7.800E-09 |
| 29355 | 1468 | L1468P | E | 7.8e-09 | 1e-08 | 4.5e-09 | 7.433E-09 | 2.768E-09 | 4.500E-09 | 1.000E-08 |
| 29356 | 1468 | L1468S | E | 6.1e-09 | 8.6e-09 | 5.2e-09 | 6.633E-09 | 1.762E-09 | 5.200E-09 | 8.600E-09 |
| 29357 | 1468 | L1468T | E | 5.2e-09 | 9.6e-09 | 6.6e-09 | 7.133E-09 | 2.248E-09 | 5.200E-09 | 9.600E-09 |
| 29358 | 1468 | L1468W | E | 7.9e-09 | 1.3e-08 | 8.1e-09 | 9.667E-09 | 2.888E-09 | 7.900E-09 | 1.300E-08 |
| 29359 | 1468 | L1468Y | E | 3.1e-09 | 4.3e-09 | 2.1e-09 | 3.167E-09 | 1.102E-09 | 2.100E-09 | 4.300E-09 |
| 29360 | 1468 | L1468V | E | 1.1e-08 | 6.4e-09 | 1.5e-08 | 1.080E-08 | 4.303E-09 | 6.400E-09 | 1.500E-08 |
| 29361 | 1469 | G1469A | E | 5.2e-09 | 1.1e-08 | 1.6e-09 | 5.933E-09 | 4.743E-09 | 1.600E-09 | 1.100E-08 |
| 29362 | 1469 | G1469R | E | 6.6e-09 | 8.7e-09 | 5.8e-09 | 7.033E-09 | 1.498E-09 | 5.800E-09 | 8.700E-09 |
| 29363 | 1469 | G1469N | E | 1.5e-08 | 1.5e-08 | 6.2e-09 | 1.207E-08 | 5.081E-09 | 6.200E-09 | 1.500E-08 |
| 29364 | 1469 | G1469D | E | 8.7e-09 | 4.8e-09 | 2.1e-08 | 1.150E-08 | 8.455E-09 | 4.800E-09 | 2.100E-08 |
| 29365 | 1469 | G1469C | E | 1.4e-08 | 5.1e-09 | 1.7e-08 | 1.203E-08 | 6.189E-09 | 5.100E-09 | 1.700E-08 |
| 29366 | 1469 | G1469E | E | 1.4e-08 | 7e-09 | 7.9e-09 | 9.633E-09 | 3.808E-09 | 7.000E-09 | 1.400E-08 |
| 29367 | 1469 | G1469Q | E | 1.2e-08 | 1.2e-08 | 1.4e-09 | 8.467E-09 | 6.120E-09 | 1.400E-09 | 1.200E-08 |
| 29368 | 1469 | G1469G | E | 9.3e-09 | 3.3e-09 | 4.4e-09 | 5.667E-09 | 3.194E-09 | 3.300E-09 | 9.300E-09 |
| 29369 | 1469 | G1469H | E | 4.2e-09 | 9.3e-09 | 7.7e-09 | 7.067E-09 | 2.608E-09 | 4.200E-09 | 9.300E-09 |
| 29370 | 1469 | G1469I | E | 1.2e-08 | 4e-09 | 1.4e-08 | 1.000E-08 | 5.292E-09 | 4.000E-09 | 1.400E-08 |
| 29371 | 1469 | G1469L | E | 1.2e-08 | 4.6e-09 | 4.4e-09 | 7.000E-09 | 4.331E-09 | 4.400E-09 | 1.200E-08 |
| 29372 | 1469 | G1469K | E | 3.5e-09 | 1e-08 | 3e-09 | 5.500E-09 | 3.905E-09 | 3.000E-09 | 1.000E-08 |
| 29373 | 1469 | G1469M | E | 8.6e-09 | 3.4e-09 | 1.6e-08 | 9.333E-09 | 6.332E-09 | 3.400E-09 | 1.600E-08 |
| 29374 | 1469 | G1469F | E | 3.6e-09 | 3.4e-09 | 1.6e-08 | 7.667E-09 | 7.218E-09 | 3.400E-09 | 1.600E-08 |
| 29375 | 1469 | G1469P | E | 1.2e-08 | 7.7e-09 | 4.3e-09 | 8.000E-09 | 3.859E-09 | 4.300E-09 | 1.200E-08 |
| 29376 | 1469 | G1469S | E | 2.6e-09 | 9.5e-09 | 6e-09 | 6.033E-09 | 3.450E-09 | 2.600E-09 | 9.500E-09 |
| 29377 | 1469 | G1469T | E | 4.2e-09 | 9.1e-09 | 4.6e-09 | 5.967E-09 | 2.721E-09 | 4.200E-09 | 9.100E-09 |
| 29378 | 1469 | G1469W | E | 1.3e-09 | 3e-09 | 5.5e-09 | 3.267E-09 | 2.113E-09 | 1.300E-09 | 5.500E-09 |
| 29379 | 1469 | G1469Y | E | 2.4e-09 | 1e-08 | 6.8e-09 | 6.400E-09 | 3.816E-09 | 2.400E-09 | 1.000E-08 |
| 29380 | 1469 | G1469V | E | 8.4e-09 | 3.5e-09 | 6e-09 | 5.967E-09 | 2.450E-09 | 3.500E-09 | 8.400E-09 |
| 29381 | 1470 | T1470A | E | 7.6e-09 | 5.4e-09 | 7.5e-09 | 6.833E-09 | 1.242E-09 | 5.400E-09 | 7.600E-09 |
| 29382 | 1470 | T1470R | E | 5.2e-09 | 1.6e-08 | 2.8e-09 | 8.000E-09 | 7.031E-09 | 2.800E-09 | 1.600E-08 |
| 29383 | 1470 | T1470N | E | 3.4e-09 | 1.4e-08 | 4.1e-09 | 7.167E-09 | 5.928E-09 | 3.400E-09 | 1.400E-08 |
| 29384 | 1470 | T1470D | E | 8.7e-09 | 1.4e-08 | 5.7e-09 | 9.467E-09 | 4.203E-09 | 5.700E-09 | 1.400E-08 |
| 29385 | 1470 | T1470C | E | 7e-09 | 5e-09 | 1.2e-08 | 8.000E-09 | 3.606E-09 | 5.000E-09 | 1.200E-08 |
| 29386 | 1470 | T1470E | E | 8.6e-09 | 5.9e-09 | 1.1e-08 | 8.500E-09 | 2.551E-09 | 5.900E-09 | 1.100E-08 |
| 29387 | 1470 | T1470Q | E | 1.4e-08 | 8.7e-09 | 7.8e-09 | 1.017E-08 | 3.350E-09 | 7.800E-09 | 1.400E-08 |
| 29388 | 1470 | T1470G | E | 1.2e-08 | 1.5e-08 | 5.5e-09 | 1.083E-08 | 4.856E-09 | 5.500E-09 | 1.500E-08 |
| 29389 | 1470 | T1470H | E | 8.9e-09 | 6e-09 | 3.4e-09 | 6.100E-09 | 2.751E-09 | 3.400E-09 | 8.900E-09 |
| 29390 | 1470 | T1470I | E | 3.8e-09 | 7.9e-09 | 4.8e-09 | 5.500E-09 | 2.138E-09 | 3.800E-09 | 7.900E-09 |
| 29391 | 1470 | T1470L | E | 9.5e-09 | 4.5e-09 | 6.7e-09 | 6.900E-09 | 2.506E-09 | 4.500E-09 | 9.500E-09 |
| 29392 | 1470 | T1470K | E | 1e-08 | 7.3e-09 | 4.9e-09 | 7.400E-09 | 2.551E-09 | 4.900E-09 | 1.000E-08 |
| 29393 | 1470 | T1470M | E | 6.5e-09 | 8.7e-09 | 1.8e-08 | 1.107E-08 | 6.104E-09 | 6.500E-09 | 1.800E-08 |
| 29394 | 1470 | T1470F | E | 8.2e-09 | 7.9e-09 | 9.2e-09 | 8.433E-09 | 6.807E-10 | 7.900E-09 | 9.200E-09 |
| 29395 | 1470 | T1470P | E | 6.1e-09 | 4.4e-09 | 6.8e-09 | 5.767E-09 | 1.234E-09 | 4.400E-09 | 6.800E-09 |
| 29396 | 1470 | T1470S | E | 1.3e-08 | 6.9e-09 | 1e-08 | 9.967E-09 | 3.050E-09 | 6.900E-09 | 1.300E-08 |
| 29397 | 1470 | T1470T | E | 2.9e-09 | 8.6e-09 | 8.3e-09 | 6.600E-09 | 3.208E-09 | 2.900E-09 | 8.600E-09 |
| 29398 | 1470 | T1470W | E | 5.3e-09 | 3.3e-09 | 4.3e-09 | 4.300E-09 | 1.000E-09 | 3.300E-09 | 5.300E-09 |
| 29399 | 1470 | T1470Y | E | 3.2e-09 | 6.4e-09 | 1.4e-08 | 7.867E-09 | 5.547E-09 | 3.200E-09 | 1.400E-08 |
| 29400 | 1470 | T1470V | E | 1.2e-08 | 1e-08 | 4.5e-09 | 8.833E-09 | 3.884E-09 | 4.500E-09 | 1.200E-08 |
| 29401 | 1471 | Q1471A | E | 6e-09 | 4.7e-09 | 1.6e-08 | 8.900E-09 | 6.183E-09 | 4.700E-09 | 1.600E-08 |
| 29402 | 1471 | Q1471R | E | 1.1e-08 | 5.2e-09 | 2.5e-08 | 1.373E-08 | 1.018E-08 | 5.200E-09 | 2.500E-08 |
| 29403 | 1471 | Q1471N | E | 1.2e-08 | 4.6e-09 | 6.1e-09 | 7.567E-09 | 3.912E-09 | 4.600E-09 | 1.200E-08 |
| 29404 | 1471 | Q1471D | E | 5.9e-09 | 1.1e-08 | 2.2e-08 | 1.297E-08 | 8.228E-09 | 5.900E-09 | 2.200E-08 |
| 29405 | 1471 | Q1471C | E | 1e-08 | 6e-09 | 7.6e-09 | 7.867E-09 | 2.013E-09 | 6.000E-09 | 1.000E-08 |
| 29406 | 1471 | Q1471E | E | 5.7e-09 | 1.1e-08 | 8e-09 | 8.233E-09 | 2.658E-09 | 5.700E-09 | 1.100E-08 |
| 29407 | 1471 | Q1471Q | E | 1.1e-08 | 4.5e-09 | 5.8e-09 | 7.100E-09 | 3.439E-09 | 4.500E-09 | 1.100E-08 |
| 29408 | 1471 | Q1471G | E | 5.1e-09 | 1.8e-08 | 8.1e-09 | 1.040E-08 | 6.751E-09 | 5.100E-09 | 1.800E-08 |
| 29409 | 1471 | Q1471H | E | 1.3e-08 | 1.4e-08 | 1.1e-08 | 1.267E-08 | 1.528E-09 | 1.100E-08 | 1.400E-08 |

| | | | | | | | | | | |
|---|---|---|---|---|---|---|---|---|---|---|
| 29410 | 1471 | Q1471I | E | 4.6e-09 | 4.4e-09 | 1.2e-08 | 7.000E-09 | 4.331E-09 | 4.400E-09 | 1.200E-08 |
| 29411 | 1471 | Q1471L | E | 2.5e-09 | 8.3e-09 | 1.1e-08 | 7.267E-09 | 4.343E-09 | 2.500E-09 | 1.100E-08 |
| 29412 | 1471 | Q1471K | E | 4.4e-09 | 4.1e-09 | 2.8e-09 | 3.767E-09 | 8.505E-10 | 2.800E-09 | 4.400E-09 |
| 29413 | 1471 | Q1471M | E | 1.4e-08 | 3.7e-09 | 1.8e-09 | 6.500E-09 | 6.564E-09 | 1.800E-09 | 1.400E-08 |
| 29414 | 1471 | Q1471F | E | 8.1e-09 | 2.1e-08 | 3.6e-09 | 1.090E-08 | 9.032E-09 | 3.600E-09 | 2.100E-08 |
| 29415 | 1471 | Q1471P | E | 2e-08 | 8e-09 | 6.1e-09 | 1.137E-08 | 7.537E-09 | 6.100E-09 | 2.000E-08 |
| 29416 | 1471 | Q1471S | E | 3.1e-09 | 1.6e-08 | 1.5e-08 | 1.137E-08 | 7.177E-09 | 3.100E-09 | 1.600E-08 |
| 29417 | 1471 | Q1471T | E | 5e-09 | 1.6e-08 | 7e-09 | 9.333E-09 | 5.859E-09 | 5.000E-09 | 1.600E-08 |
| 29418 | 1471 | Q1471W | E | 2e-08 | 1.3e-08 | 1.2e-08 | 1.500E-08 | 4.359E-09 | 1.200E-08 | 2.000E-08 |
| 29419 | 1471 | Q1471Y | E | 8.8e-09 | 1.9e-09 | 1.4e-08 | 8.233E-09 | 6.070E-09 | 1.900E-09 | 1.400E-08 |
| 29420 | 1471 | Q1471V | E | 5.1e-09 | 8.5e-09 | 3e-08 | 1.453E-08 | 1.350E-08 | 5.100E-09 | 3.000E-08 |
| 29421 | 1472 | T1472A | E | 2.7e-08 | 8.6e-09 | 2.2e-08 | 1.920E-08 | 9.514E-09 | 8.600E-09 | 2.700E-08 |
| 29422 | 1472 | T1472R | E | 2.7e-09 | 8.4e-09 | 5e-09 | 5.367E-09 | 2.868E-09 | 2.700E-09 | 8.400E-09 |
| 29423 | 1472 | T1472N | E | 2.5e-09 | 5.6e-09 | 6.5e-09 | 4.867E-09 | 2.098E-09 | 2.500E-09 | 6.500E-09 |
| 29424 | 1472 | T1472D | E | 9.7e-09 | 9.8e-09 | 3.1e-09 | 7.533E-09 | 3.840E-09 | 3.100E-09 | 9.800E-09 |
| 29425 | 1472 | T1472C | E | 4.6e-09 | 2.5e-08 | 4.8e-09 | 1.147E-08 | 1.172E-08 | 4.600E-09 | 2.500E-08 |
| 29426 | 1472 | T1472E | E | 5.3e-09 | 8e-09 | 8e-09 | 7.100E-09 | 1.559E-09 | 5.300E-09 | 8.000E-09 |
| 29427 | 1472 | T1472Q | E | 2e-09 | 2.3e-08 | 7.3e-09 | 1.077E-08 | 1.092E-08 | 2.000E-09 | 2.300E-08 |
| 29428 | 1472 | T1472G | E | 7.8e-09 | 1e-08 | 5.4e-09 | 7.733E-09 | 2.301E-09 | 5.400E-09 | 1.000E-08 |
| 29429 | 1472 | T1472H | E | 3.2e-09 | 4.8e-09 | 6e-09 | 4.667E-09 | 1.405E-09 | 3.200E-09 | 6.000E-09 |
| 29430 | 1472 | T1472I | E | 1.3e-09 | 2.2e-09 | 1.2e-08 | 5.167E-09 | 5.935E-09 | 1.300E-09 | 1.200E-08 |
| 29431 | 1472 | T1472L | E | 9.5e-09 | 4.3e-09 | 4.3e-09 | 6.033E-09 | 3.002E-09 | 4.300E-09 | 9.500E-09 |
| 29432 | 1472 | T1472K | E | 7.3e-09 | 1.9e-08 | 1.5e-08 | 1.377E-08 | 5.947E-09 | 7.300E-09 | 1.900E-08 |
| 29433 | 1472 | T1472M | E | 1.2e-08 | 6.7e-09 | 5.9e-09 | 8.200E-09 | 3.315E-09 | 5.900E-09 | 1.200E-08 |
| 29434 | 1472 | T1472F | E | 2.5e-09 | 3.2e-09 | 4.3e-09 | 3.333E-09 | 9.074E-10 | 2.500E-09 | 4.300E-09 |
| 29435 | 1472 | T1472P | E | 2.2e-08 | 1.4e-08 | 1e-08 | 1.533E-08 | 6.110E-09 | 1.000E-08 | 2.200E-08 |
| 29436 | 1472 | T1472S | E | 2.2e-09 | 8e-09 | 8.4e-09 | 6.200E-09 | 3.470E-09 | 2.200E-09 | 8.400E-09 |
| 29437 | 1472 | T1472T | E | 1.3e-08 | 6.5e-09 | 1.2e-08 | 1.050E-08 | 3.500E-09 | 6.500E-09 | 1.300E-08 |
| 29438 | 1472 | T1472W | E | 4e-08 | 2.1e-09 | 9.4e-09 | 1.717E-08 | 2.011E-08 | 2.100E-09 | 4.000E-08 |
| 29439 | 1472 | T1472Y | E | 1.1e-08 | 7.3e-09 | 1.9e-08 | 1.243E-08 | 5.980E-09 | 7.300E-09 | 1.900E-08 |
| 29440 | 1472 | T1472V | E | 3e-08 | 5.9e-09 | 2.5e-09 | 1.280E-08 | 1.499E-08 | 2.500E-09 | 3.000E-08 |
| 29441 | 1473 | Y1473A | E | 7.9e-09 | 1.4e-08 | 5.7e-09 | 9.200E-09 | 4.300E-09 | 5.700E-09 | 1.400E-08 |
| 29442 | 1473 | Y1473R | E | 9e-09 | 6.9e-09 | 4.6e-09 | 6.833E-09 | 2.201E-09 | 4.600E-09 | 9.000E-09 |
| 29443 | 1473 | Y1473N | E | 6.4e-09 | 7.7e-09 | 1.5e-08 | 9.700E-09 | 4.636E-09 | 6.400E-09 | 1.500E-08 |
| 29444 | 1473 | Y1473D | E | 8.1e-09 | 6.3e-09 | 9.9e-09 | 8.100E-09 | 1.800E-09 | 6.300E-09 | 9.900E-09 |
| 29445 | 1473 | Y1473C | E | 6.8e-09 | 5.1e-09 | 1.6e-08 | 9.300E-09 | 5.864E-09 | 5.100E-09 | 1.600E-08 |
| 29446 | 1473 | Y1473E | E | 1.4e-08 | 9.5e-09 | 1.9e-08 | 1.417E-08 | 4.752E-09 | 9.500E-09 | 1.900E-08 |
| 29447 | 1473 | Y1473Q | E | 2e-08 | 1.6e-09 | 1.1e-08 | 1.087E-08 | 9.201E-09 | 1.600E-09 | 2.000E-08 |
| 29448 | 1473 | Y1473G | E | 8.1e-09 | 6.5e-09 | 6.5e-09 | 7.033E-09 | 9.238E-10 | 6.500E-09 | 8.100E-09 |
| 29449 | 1473 | Y1473H | E | 8e-09 | 2e-08 | 4.7e-09 | 1.090E-08 | 8.052E-09 | 4.700E-09 | 2.000E-08 |
| 29450 | 1473 | Y1473I | E | 1.4e-08 | 4e-09 | 1.1e-08 | 9.667E-09 | 5.132E-09 | 4.000E-09 | 1.400E-08 |
| 29451 | 1473 | Y1473L | E | 6.5e-09 | 3.2e-09 | 1.5e-08 | 8.233E-09 | 6.088E-09 | 3.200E-09 | 1.500E-08 |
| 29452 | 1473 | Y1473K | E | 1.1e-08 | 1.9e-08 | 1.1e-08 | 1.367E-08 | 4.619E-09 | 1.100E-08 | 1.900E-08 |
| 29453 | 1473 | Y1473M | E | 5.2e-09 | 1.8e-08 | 1.6e-08 | 1.307E-08 | 6.886E-09 | 5.200E-09 | 1.800E-08 |
| 29454 | 1473 | Y1473F | E | 1.4e-08 | 5.9e-09 | 2.7e-09 | 7.533E-09 | 5.824E-09 | 2.700E-09 | 1.400E-08 |
| 29455 | 1473 | Y1473P | E | 2.8e-09 | 3.6e-09 | 2.5e-08 | 1.047E-08 | 1.259E-08 | 2.800E-09 | 2.500E-08 |
| 29456 | 1473 | Y1473S | E | 7.1e-09 | 1.5e-08 | 5.3e-09 | 9.133E-09 | 5.160E-09 | 5.300E-09 | 1.500E-08 |
| 29457 | 1473 | Y1473T | E | 3.6e-09 | 5.9e-09 | 8e-09 | 5.833E-09 | 2.201E-09 | 3.600E-09 | 8.000E-09 |
| 29458 | 1473 | Y1473W | E | 7.3e-09 | 4.2e-09 | 9.8e-09 | 7.100E-09 | 2.805E-09 | 4.200E-09 | 9.800E-09 |
| 29459 | 1473 | Y1473Y | E | 7.4e-09 | 3.6e-09 | 1.3e-08 | 8.000E-09 | 4.729E-09 | 3.600E-09 | 1.300E-08 |
| 29460 | 1473 | Y1473V | E | 3.7e-09 | 9.6e-09 | 8.1e-09 | 7.133E-09 | 3.066E-09 | 3.700E-09 | 9.600E-09 |
| 29461 | 1474 | I1474A | E | 6.3e-09 | 2.7e-09 | 5.7e-09 | 4.900E-09 | 1.929E-09 | 2.700E-09 | 6.300E-09 |
| 29462 | 1474 | I1474R | E | 5.6e-09 | 5e-09 | 1.6e-08 | 8.867E-09 | 6.185E-09 | 5.000E-09 | 1.600E-08 |
| 29463 | 1474 | I1474N | E | 4.2e-09 | 1.6e-08 | 3.6e-09 | 7.933E-09 | 6.992E-09 | 3.600E-09 | 1.600E-08 |
| 29464 | 1474 | I1474D | E | 5.7e-09 | 7.3e-09 | 6.2e-09 | 6.400E-09 | 8.185E-10 | 5.700E-09 | 7.300E-09 |
| 29465 | 1474 | I1474C | E | 1.1e-08 | 1.2e-08 | 8.1e-09 | 1.037E-08 | 2.026E-09 | 8.100E-09 | 1.200E-08 |
| 29466 | 1474 | I1474E | E | 3.8e-09 | 3e-09 | 3.2e-09 | 3.333E-09 | 4.163E-10 | 3.000E-09 | 3.800E-09 |
| 29467 | 1474 | I1474Q | E | 2.7e-09 | 4.5e-08 | 5.1e-09 | 1.760E-08 | 2.376E-08 | 2.700E-09 | 4.500E-08 |
| 29468 | 1474 | I1474G | E | 1.4e-08 | 9.5e-09 | 3e-09 | 8.833E-09 | 5.530E-09 | 3.000E-09 | 1.400E-08 |
| 29469 | 1474 | I1474H | E | 2.7e-09 | 2e-08 | 7.4e-09 | 1.003E-08 | 8.946E-09 | 2.700E-09 | 2.000E-08 |
| 29470 | 1474 | I1474I | E | 3.3e-09 | 5.5e-09 | 1.7e-08 | 8.600E-09 | 7.357E-09 | 3.300E-09 | 1.700E-08 |
| 29471 | 1474 | I1474L | E | 1.8e-08 | 6.7e-09 | 1.3e-08 | 1.257E-08 | 5.662E-09 | 6.700E-09 | 1.800E-08 |
| 29472 | 1474 | I1474K | E | 6.4e-09 | 1.9e-08 | 6.3e-09 | 1.057E-08 | 7.304E-09 | 6.300E-09 | 1.900E-08 |
| 29473 | 1474 | I1474M | E | 4.9e-09 | 4.2e-09 | 3e-09 | 4.033E-09 | 9.609E-10 | 3.000E-09 | 4.900E-09 |
| 29474 | 1474 | I1474F | E | 8.3e-09 | 7.2e-09 | 4.5e-09 | 6.667E-09 | 1.955E-09 | 4.500E-09 | 8.300E-09 |
| 29475 | 1474 | I1474P | E | 2.4e-08 | 4.4e-09 | 3.3e-09 | 1.057E-08 | 1.165E-08 | 3.300E-09 | 2.400E-08 |
| 29476 | 1474 | I1474S | E | 2.6e-09 | 1.6e-08 | 5.6e-09 | 8.067E-09 | 7.032E-09 | 2.600E-09 | 1.600E-08 |
| 29477 | 1474 | I1474T | E | 7.9e-09 | 3.7e-09 | 4.1e-09 | 5.233E-09 | 2.318E-09 | 3.700E-09 | 7.900E-09 |
| 29478 | 1474 | I1474W | E | 1.3e-08 | 7.1e-09 | 9.4e-09 | 9.833E-09 | 2.974E-09 | 7.100E-09 | 1.300E-08 |
| 29479 | 1474 | I1474Y | E | 5.8e-09 | 7.1e-09 | 6.7e-09 | 6.533E-09 | 6.658E-10 | 5.800E-09 | 7.100E-09 |
| 29480 | 1474 | I1474V | E | 7.8e-09 | 3.1e-09 | 7.9e-09 | 6.267E-09 | 2.743E-09 | 3.100E-09 | 7.900E-09 |
| 29481 | 1475 | C1475A | E | 8e-09 | 1e-08 | 4.8e-09 | 7.600E-09 | 2.623E-09 | 4.800E-09 | 1.000E-08 |
| 29482 | 1475 | C1475R | E | 2.7e-09 | 6.5e-09 | 1.8e-08 | 9.067E-09 | 7.966E-09 | 2.700E-09 | 1.800E-08 |
| 29483 | 1475 | C1475N | E | 7.1e-09 | 9.4e-09 | 6.8e-09 | 7.767E-09 | 1.422E-09 | 6.800E-09 | 9.400E-09 |
| 29484 | 1475 | C1475D | E | 5.6e-09 | 1.6e-08 | 2.7e-09 | 8.100E-09 | 6.994E-09 | 2.700E-09 | 1.600E-08 |
| 29485 | 1475 | C1475C | E | 1.8e-08 | 5.1e-09 | 8.9e-09 | 1.067E-08 | 6.629E-09 | 5.100E-09 | 1.800E-08 |
| 29486 | 1475 | C1475E | E | 1.6e-08 | 3.5e-09 | 6.6e-09 | 8.700E-09 | 6.509E-09 | 3.500E-09 | 1.600E-08 |
| 29487 | 1475 | C1475Q | E | 9.1e-09 | 8.4e-09 | 3.6e-09 | 7.033E-09 | 2.994E-09 | 3.600E-09 | 9.100E-09 |
| 29488 | 1475 | C1475G | E | 8e-09 | 9.9e-09 | 4.5e-09 | 7.467E-09 | 2.739E-09 | 4.500E-09 | 9.900E-09 |
| 29489 | 1475 | C1475H | E | 1.1e-09 | 7.2e-09 | 1.5e-08 | 7.767E-09 | 6.967E-09 | 1.100E-09 | 1.500E-08 |
| 29490 | 1475 | C1475I | E | 5e-09 | 3.4e-09 | 4.4e-09 | 4.267E-09 | 8.083E-10 | 3.400E-09 | 5.000E-09 |
| 29491 | 1475 | C1475L | E | 2.7e-09 | 8.6e-09 | 6.2e-09 | 5.833E-09 | 2.967E-09 | 2.700E-09 | 8.600E-09 |
| 29492 | 1475 | C1475K | E | 5.5e-09 | 6.3e-09 | 3e-09 | 4.933E-09 | 1.721E-09 | 3.000E-09 | 6.300E-09 |
| 29493 | 1475 | C1475M | E | 9.3e-09 | 6.1e-09 | 3.6e-09 | 6.333E-09 | 2.857E-09 | 3.600E-09 | 9.300E-09 |
| 29494 | 1475 | C1475F | E | 7.1e-09 | 2.7e-09 | 7.5e-09 | 5.767E-09 | 2.663E-09 | 2.700E-09 | 7.500E-09 |

| | | | | | | | | | | |
|---|---|---|---|---|---|---|---|---|---|---|
| 29495 | 1475 | C1475P | E | 9.8e-09 | 2.9e-09 | 5.9e-09 | 6.200E-09 | 3.460E-09 | 2.900E-09 | 9.800E-09 |
| 29496 | 1475 | C1475S | E | 7.3e-09 | 6.1e-09 | 7.5e-09 | 6.967E-09 | 7.572E-10 | 6.100E-09 | 7.500E-09 |
| 29497 | 1475 | C1475T | E | 1.7e-08 | 1e-08 | 9.1e-09 | 1.203E-08 | 4.325E-09 | 9.100E-09 | 1.700E-08 |
| 29498 | 1475 | C1475W | E | 3e-09 | 5.2e-09 | 2.5e-08 | 1.107E-08 | 1.212E-08 | 3.000E-09 | 2.500E-08 |
| 29499 | 1475 | C1475Y | E | 7.9e-09 | 1.8e-08 | 1.5e-08 | 1.363E-08 | 5.187E-09 | 7.900E-09 | 1.800E-08 |
| 29500 | 1475 | C1475V | E | 1.3e-09 | 1.3e-08 | 9.1e-09 | 7.800E-09 | 5.957E-09 | 1.300E-09 | 1.300E-08 |
| 29501 | 1476 | N1476A | E | 8.1e-09 | 6.8e-09 | 9.6e-09 | 8.167E-09 | 1.401E-09 | 6.800E-09 | 9.600E-09 |
| 29502 | 1476 | N1476R | E | 8.2e-09 | 1.5e-08 | 2.8e-09 | 8.667E-09 | 6.113E-09 | 2.800E-09 | 1.500E-08 |
| 29503 | 1476 | N1476N | E | 4.8e-09 | 5.4e-09 | 4.5e-09 | 4.900E-09 | 4.583E-10 | 4.500E-09 | 5.400E-09 |
| 29504 | 1476 | N1476D | E | 6.9e-09 | 9.3e-09 | 4.6e-09 | 6.933E-09 | 2.350E-09 | 4.600E-09 | 9.300E-09 |
| 29505 | 1476 | N1476C | E | 6.3e-09 | 3.8e-09 | 9.5e-09 | 6.533E-09 | 2.857E-09 | 3.800E-09 | 9.500E-09 |
| 29506 | 1476 | N1476E | E | 1.3e-08 | 1.9e-08 | 1.4e-08 | 1.533E-08 | 3.215E-09 | 1.300E-08 | 1.900E-08 |
| 29507 | 1476 | N1476Q | E | 1e-08 | 1.8e-09 | 7.9e-09 | 6.567E-09 | 4.259E-09 | 1.800E-09 | 1.000E-08 |
| 29508 | 1476 | N1476G | E | 8.4e-09 | 8.7e-09 | 6.1e-09 | 7.733E-09 | 1.422E-09 | 6.100E-09 | 8.700E-09 |
| 29509 | 1476 | N1476H | E | 6.1e-09 | 5.2e-09 | 6.5e-09 | 5.933E-09 | 6.658E-10 | 5.200E-09 | 6.500E-09 |
| 29510 | 1476 | N1476I | E | 1.2e-08 | 7.5e-09 | 2.8e-09 | 7.433E-09 | 4.600E-09 | 2.800E-09 | 1.200E-08 |
| 29511 | 1476 | N1476L | E | 1e-08 | 3e-09 | 7.4e-09 | 6.800E-09 | 3.538E-09 | 3.000E-09 | 1.000E-08 |
| 29512 | 1476 | N1476K | E | 1.3e-08 | 4.3e-09 | 1.4e-08 | 1.043E-08 | 5.335E-09 | 4.300E-09 | 1.400E-08 |
| 29513 | 1476 | N1476M | E | 2.9e-09 | 1.6e-08 | 9.5e-09 | 9.467E-09 | 6.550E-09 | 2.900E-09 | 1.600E-08 |
| 29514 | 1476 | N1476F | E | 9e-09 | 9.1e-09 | 1.1e-08 | 9.700E-09 | 1.127E-09 | 9.000E-09 | 1.100E-08 |
| 29515 | 1476 | N1476P | E | 6.5e-09 | 4.7e-09 | 6.7e-09 | 5.967E-09 | 1.102E-09 | 4.700E-09 | 6.700E-09 |
| 29516 | 1476 | N1476S | E | 2.6e-09 | 1.8e-08 | 6.5e-09 | 9.033E-09 | 8.006E-09 | 2.600E-09 | 1.800E-08 |
| 29517 | 1476 | N1476T | E | 5.9e-09 | 3.6e-09 | 1e-08 | 6.500E-09 | 3.242E-09 | 3.600E-09 | 1.000E-08 |
| 29518 | 1476 | N1476W | E | 7.2e-09 | 1.4e-08 | 5.8e-09 | 9.000E-09 | 4.386E-09 | 5.800E-09 | 1.400E-08 |
| 29519 | 1476 | N1476Y | E | 1e-08 | 5e-09 | 5.1e-09 | 6.700E-09 | 2.858E-09 | 5.000E-09 | 1.000E-08 |
| 29520 | 1476 | N1476V | E | 9.7e-09 | 4.3e-09 | 2.5e-09 | 5.500E-09 | 3.747E-09 | 2.500E-09 | 9.700E-09 |
| 29521 | 1477 | V1477A | E | 1.1e-08 | 2.6e-09 | 3.2e-09 | 5.600E-09 | 4.686E-09 | 2.600E-09 | 1.100E-08 |
| 29522 | 1477 | V1477R | E | 1.7e-08 | 9.5e-09 | 8.9e-09 | 1.180E-08 | 4.513E-09 | 8.900E-09 | 1.700E-08 |
| 29523 | 1477 | V1477N | E | 1.5e-08 | 1e-08 | 6.7e-09 | 1.057E-08 | 4.179E-09 | 6.700E-09 | 1.500E-08 |
| 29524 | 1477 | V1477D | E | 9.1e-09 | 3.5e-09 | 7.7e-09 | 6.767E-09 | 2.914E-09 | 3.500E-09 | 9.100E-09 |
| 29525 | 1477 | V1477C | E | 2.8e-09 | 4.4e-09 | 2.9e-09 | 3.367E-09 | 8.963E-10 | 2.800E-09 | 4.400E-09 |
| 29526 | 1477 | V1477E | E | 3.2e-09 | 4.6e-09 | 5.2e-09 | 4.333E-09 | 1.026E-09 | 3.200E-09 | 5.200E-09 |
| 29527 | 1477 | V1477Q | E | 6e-09 | 2.4e-09 | 4.1e-09 | 4.167E-09 | 1.801E-09 | 2.400E-09 | 6.000E-09 |
| 29528 | 1477 | V1477G | E | 7e-09 | 9.7e-09 | 4e-09 | 6.900E-09 | 2.851E-09 | 4.000E-09 | 9.700E-09 |
| 29529 | 1477 | V1477H | E | 3.8e-09 | 1.1e-08 | 5.6e-09 | 6.800E-09 | 3.747E-09 | 3.800E-09 | 1.100E-08 |
| 29530 | 1477 | V1477I | E | 7.4e-09 | 4.7e-09 | 1.7e-08 | 9.700E-09 | 6.465E-09 | 4.700E-09 | 1.700E-08 |
| 29531 | 1477 | V1477L | E | 8.5e-09 | 6.2e-09 | 6.1e-09 | 6.933E-09 | 1.358E-09 | 6.100E-09 | 8.500E-09 |
| 29532 | 1477 | V1477K | E | 5.9e-09 | 7.3e-09 | 1.6e-08 | 9.733E-09 | 5.472E-09 | 5.900E-09 | 1.600E-08 |
| 29533 | 1477 | V1477M | E | 7.8e-09 | 4.3e-09 | 2.1e-08 | 1.103E-08 | 8.807E-09 | 4.300E-09 | 2.100E-08 |
| 29534 | 1477 | V1477F | E | 1.6e-09 | 2.8e-08 | 9.2e-09 | 1.293E-08 | 1.359E-08 | 1.600E-09 | 2.800E-08 |
| 29535 | 1477 | V1477P | E | 3.4e-09 | 1.2e-08 | 6.6e-09 | 7.333E-09 | 4.347E-09 | 3.400E-09 | 1.200E-08 |
| 29536 | 1477 | V1477S | E | 2e-08 | 1.1e-08 | 2.3e-09 | 1.110E-08 | 8.850E-09 | 2.300E-09 | 2.000E-08 |
| 29537 | 1477 | V1477T | E | 6.2e-09 | 9.8e-09 | 1.1e-08 | 9.000E-09 | 2.498E-09 | 6.200E-09 | 1.100E-08 |
| 29538 | 1477 | V1477W | E | 5.3e-09 | 2.4e-09 | 1.1e-08 | 6.233E-09 | 4.375E-09 | 2.400E-09 | 1.100E-08 |
| 29539 | 1477 | V1477Y | E | 5.7e-09 | 8.9e-09 | 5.1e-09 | 6.567E-09 | 2.043E-09 | 5.100E-09 | 8.900E-09 |
| 29540 | 1477 | V1477V | E | 7.8e-09 | 1e-08 | 8e-09 | 8.600E-09 | 1.217E-09 | 7.800E-09 | 1.000E-08 |
| 29541 | 1478 | N1478A | E | 1.9e-08 | 9.1e-09 | 2.7e-09 | 1.027E-08 | 8.212E-09 | 2.700E-09 | 1.900E-08 |
| 29542 | 1478 | N1478R | E | 5.7e-09 | 3.6e-09 | 3.9e-09 | 4.400E-09 | 1.136E-09 | 3.600E-09 | 5.700E-09 |
| 29543 | 1478 | N1478N | E | 2.7e-09 | 6.3e-09 | 8.4e-09 | 5.800E-09 | 2.883E-09 | 2.700E-09 | 8.400E-09 |
| 29544 | 1478 | N1478D | E | 6.3e-09 | 2.1e-09 | 4.9e-09 | 4.433E-09 | 2.139E-09 | 2.100E-09 | 6.300E-09 |
| 29545 | 1478 | N1478C | E | 9.7e-09 | 8.1e-09 | 6.3e-09 | 8.033E-09 | 1.701E-09 | 6.300E-09 | 9.700E-09 |
| 29546 | 1478 | N1478E | E | 2.6e-09 | 5.1e-09 | 3e-09 | 3.567E-09 | 1.343E-09 | 2.600E-09 | 5.100E-09 |
| 29547 | 1478 | N1478Q | E | 7.8e-09 | 5.2e-09 | 9.2e-09 | 7.400E-09 | 2.030E-09 | 5.200E-09 | 9.200E-09 |
| 29548 | 1478 | N1478G | E | 9.7e-09 | 5e-09 | 8.1e-09 | 7.600E-09 | 2.390E-09 | 5.000E-09 | 9.700E-09 |
| 29549 | 1478 | N1478H | E | 7e-09 | 3.7e-09 | 3e-09 | 4.567E-09 | 2.136E-09 | 3.000E-09 | 7.000E-09 |
| 29550 | 1478 | N1478I | E | 4e-09 | 1.3e-08 | 1.1e-08 | 9.333E-09 | 4.726E-09 | 4.000E-09 | 1.300E-08 |
| 29551 | 1478 | N1478L | E | 7.5e-09 | 7.8e-09 | 2.1e-08 | 1.210E-08 | 7.709E-09 | 7.500E-09 | 2.100E-08 |
| 29552 | 1478 | N1478K | E | 1.9e-09 | 4.3e-09 | 1.7e-08 | 7.733E-09 | 8.114E-09 | 1.900E-09 | 1.700E-08 |
| 29553 | 1478 | N1478M | E | 8.5e-09 | 1e-08 | 3.9e-09 | 7.467E-09 | 3.179E-09 | 3.900E-09 | 1.000E-08 |
| 29554 | 1478 | N1478F | E | 1.2e-08 | 4.7e-09 | 7e-09 | 7.900E-09 | 3.732E-09 | 4.700E-09 | 1.200E-08 |
| 29555 | 1478 | N1478P | E | 4.1e-09 | 8.4e-09 | 2.2e-09 | 4.900E-09 | 3.176E-09 | 2.200E-09 | 8.400E-09 |
| 29556 | 1478 | N1478S | E | 5.6e-09 | 1.4e-08 | 8.7e-09 | 9.433E-09 | 4.248E-09 | 5.600E-09 | 1.400E-08 |
| 29557 | 1478 | N1478T | E | 3.9e-09 | 2.5e-08 | 4e-09 | 1.097E-08 | 1.215E-08 | 3.900E-09 | 2.500E-08 |
| 29558 | 1478 | N1478W | E | 8.1e-09 | 1.3e-08 | 6.2e-09 | 9.100E-09 | 3.509E-09 | 6.200E-09 | 1.300E-08 |
| 29559 | 1478 | N1478Y | E | 2.9e-09 | 1.6e-08 | 8.5e-09 | 9.133E-09 | 6.573E-09 | 2.900E-09 | 1.600E-08 |
| 29560 | 1478 | N1478V | E | 1.5e-08 | 2.9e-09 | 1.1e-08 | 9.633E-09 | 6.165E-09 | 2.900E-09 | 1.500E-08 |
| 29561 | 1479 | H1479A | E | 5.1e-09 | 1.5e-08 | 3.3e-09 | 7.800E-09 | 6.300E-09 | 3.300E-09 | 1.500E-08 |
| 29562 | 1479 | H1479R | E | 7.2e-09 | 5.9e-09 | 4.1e-09 | 5.733E-09 | 1.557E-09 | 4.100E-09 | 7.200E-09 |
| 29563 | 1479 | H1479N | E | 7.3e-09 | 2.5e-09 | 8.1e-09 | 5.967E-09 | 3.029E-09 | 2.500E-09 | 8.100E-09 |
| 29564 | 1479 | H1479D | E | 3.6e-09 | 5.1e-09 | 1.4e-08 | 7.567E-09 | 5.622E-09 | 3.600E-09 | 1.400E-08 |
| 29565 | 1479 | H1479C | E | 6.1e-09 | 9.8e-09 | 1.8e-08 | 1.130E-08 | 6.090E-09 | 6.100E-09 | 1.800E-08 |
| 29566 | 1479 | H1479E | E | 1e-08 | 7.4e-09 | 3.8e-09 | 7.067E-09 | 3.113E-09 | 3.800E-09 | 1.000E-08 |
| 29567 | 1479 | H1479Q | E | 1.2e-08 | 6.6e-09 | 9.9e-09 | 9.500E-09 | 2.722E-09 | 6.600E-09 | 1.200E-08 |
| 29568 | 1479 | H1479G | E | 4.9e-09 | 2e-08 | 3.9e-09 | 9.600E-09 | 9.021E-09 | 3.900E-09 | 2.000E-08 |
| 29569 | 1479 | H1479H | E | 5.1e-09 | 3.8e-09 | 5.2e-09 | 4.700E-09 | 7.810E-10 | 3.800E-09 | 5.200E-09 |
| 29570 | 1479 | H1479I | E | 6e-09 | 8.9e-09 | 9.5e-09 | 8.133E-09 | 1.872E-09 | 6.000E-09 | 9.500E-09 |
| 29571 | 1479 | H1479L | E | 4.1e-09 | 5.8e-09 | 8.4e-09 | 6.100E-09 | 2.166E-09 | 4.100E-09 | 8.400E-09 |
| 29572 | 1479 | H1479K | E | 7.3e-09 | 6.7e-09 | 7.7e-09 | 7.233E-09 | 5.033E-10 | 6.700E-09 | 7.700E-09 |
| 29573 | 1479 | H1479M | E | 9e-09 | 5.1e-09 | 5.5e-09 | 6.533E-09 | 2.146E-09 | 5.100E-09 | 9.000E-09 |
| 29574 | 1479 | H1479F | E | 1.5e-08 | 4.3e-09 | 1e-08 | 9.767E-09 | 5.354E-09 | 4.300E-09 | 1.500E-08 |
| 29575 | 1479 | H1479P | E | 5.7e-09 | 6.9e-09 | 1.3e-08 | 8.533E-09 | 3.915E-09 | 5.700E-09 | 1.300E-08 |
| 29576 | 1479 | H1479S | E | 5.7e-09 | 6.4e-09 | 6.4e-09 | 6.167E-09 | 4.041E-10 | 5.700E-09 | 6.400E-09 |
| 29577 | 1479 | H1479T | E | 1.2e-08 | 7.3e-09 | 4.6e-09 | 7.967E-09 | 3.745E-09 | 4.600E-09 | 1.200E-08 |
| 29578 | 1479 | H1479W | E | 1.1e-08 | 2.8e-08 | 1.1e-08 | 1.667E-08 | 9.815E-09 | 1.100E-08 | 2.800E-08 |
| 29579 | 1479 | H1479Y | E | 1.7e-09 | 1.3e-08 | 1.7e-08 | 1.057E-08 | 7.935E-09 | 1.700E-09 | 1.700E-08 |

| | | | | | | | | | | |
|---|---|---|---|---|---|---|---|---|---|---|
| 29580 | 1479 | H1479V | E | 1.3e-08 | 2.5e-09 | 6.4e-09 | 7.300E-09 | 5.308E-09 | 2.500E-09 | 1.300E-08 |
| 29581 | 1480 | K1480A | E | 1e-08 | 1.3e-08 | 1.8e-08 | 1.367E-08 | 4.041E-09 | 1.000E-08 | 1.800E-08 |
| 29582 | 1480 | K1480R | E | 6.9e-09 | 1.4e-08 | 1.5e-08 | 1.197E-08 | 4.416E-09 | 6.900E-09 | 1.500E-08 |
| 29583 | 1480 | K1480N | E | 7.4e-09 | 4.7e-09 | 1.5e-08 | 9.033E-09 | 5.341E-09 | 4.700E-09 | 1.500E-08 |
| 29584 | 1480 | K1480D | E | 4.7e-09 | 3.4e-09 | 1.2e-08 | 6.700E-09 | 4.636E-09 | 3.400E-09 | 1.200E-08 |
| 29585 | 1480 | K1480C | E | 7.7e-09 | 4e-09 | 1.6e-08 | 9.233E-09 | 6.145E-09 | 4.000E-09 | 1.600E-08 |
| 29586 | 1480 | K1480E | E | 9e-09 | 4.1e-09 | 6.7e-09 | 6.600E-09 | 2.452E-09 | 4.100E-09 | 9.000E-09 |
| 29587 | 1480 | K1480Q | E | 5.8e-09 | 3.7e-09 | 3.9e-09 | 4.467E-09 | 1.159E-09 | 3.700E-09 | 5.800E-09 |
| 29588 | 1480 | K1480G | E | 1.2e-08 | 2.2e-08 | 1e-08 | 1.467E-08 | 6.429E-09 | 1.000E-08 | 2.200E-08 |
| 29589 | 1480 | K1480H | E | 2.5e-08 | 9.7e-09 | 1.1e-08 | 1.523E-08 | 8.483E-09 | 9.700E-09 | 2.500E-08 |
| 29590 | 1480 | K1480I | E | 4e-09 | 5.5e-09 | 6.1e-09 | 5.200E-09 | 1.082E-09 | 4.000E-09 | 6.100E-09 |
| 29591 | 1480 | K1480L | E | 9.9e-09 | 7.1e-09 | 3.1e-09 | 6.700E-09 | 3.418E-09 | 3.100E-09 | 9.900E-09 |
| 29592 | 1480 | K1480K | E | 3.8e-09 | 6.5e-09 | 1.4e-08 | 8.100E-09 | 5.285E-09 | 3.800E-09 | 1.400E-08 |
| 29593 | 1480 | K1480M | E | 7.3e-09 | 1.2e-08 | 9.3e-09 | 9.533E-09 | 2.359E-09 | 7.300E-09 | 1.200E-08 |
| 29594 | 1480 | K1480F | E | 4.5e-09 | 8.7e-09 | 1.2e-08 | 8.400E-09 | 3.759E-09 | 4.500E-09 | 1.200E-08 |
| 29595 | 1480 | K1480P | E | 2e-09 | 5.4e-09 | 4.3e-09 | 3.900E-09 | 1.735E-09 | 2.000E-09 | 5.400E-09 |
| 29596 | 1480 | K1480S | E | 2.6e-09 | 4e-09 | 4.8e-09 | 3.800E-09 | 1.114E-09 | 2.600E-09 | 4.800E-09 |
| 29597 | 1480 | K1480T | E | 1.4e-08 | 2.5e-09 | 4.6e-09 | 7.033E-09 | 6.124E-09 | 2.500E-09 | 1.400E-08 |
| 29598 | 1480 | K1480W | E | 6.5e-09 | 1.5e-08 | 6.9e-09 | 9.467E-09 | 4.796E-09 | 6.500E-09 | 1.500E-08 |
| 29599 | 1480 | K1480Y | E | 1.1e-08 | 3e-09 | 2.3e-08 | 1.233E-08 | 1.007E-08 | 3.000E-09 | 2.300E-08 |
| 29600 | 1480 | K1480V | E | 9.1e-09 | 5.9e-09 | 9e-09 | 8.000E-09 | 1.819E-09 | 5.900E-09 | 9.100E-09 |
| 29601 | 1481 | P1481A | E | 5.6e-09 | 2.5e-09 | 4.1e-09 | 4.067E-09 | 1.550E-09 | 2.500E-09 | 5.600E-09 |
| 29602 | 1481 | P1481R | E | 2e-09 | 7.3e-09 | 9.4e-09 | 6.233E-09 | 3.814E-09 | 2.000E-09 | 9.400E-09 |
| 29603 | 1481 | P1481N | E | 1.5e-08 | 3.1e-09 | 1.1e-08 | 9.700E-09 | 6.056E-09 | 3.100E-09 | 1.500E-08 |
| 29604 | 1481 | P1481D | E | 4.1e-09 | 1.1e-08 | 6.4e-09 | 7.167E-09 | 3.513E-09 | 4.100E-09 | 1.100E-08 |
| 29605 | 1481 | P1481C | E | 7e-09 | 1.1e-08 | 3.2e-09 | 7.067E-09 | 3.900E-09 | 3.200E-09 | 1.100E-08 |
| 29606 | 1481 | P1481E | E | 1.6e-09 | 8.8e-09 | 6.8e-09 | 5.733E-09 | 3.717E-09 | 1.600E-09 | 8.800E-09 |
| 29607 | 1481 | P1481Q | E | 8.7e-09 | 4.1e-09 | 2.7e-09 | 5.167E-09 | 3.139E-09 | 2.700E-09 | 8.700E-09 |
| 29608 | 1481 | P1481G | E | 5.6e-09 | 1.2e-08 | 7.3e-09 | 8.300E-09 | 3.315E-09 | 5.600E-09 | 1.200E-08 |
| 29609 | 1481 | P1481H | E | 1.3e-08 | 2.6e-09 | 7.7e-09 | 7.767E-09 | 5.200E-09 | 2.600E-09 | 1.300E-08 |
| 29610 | 1481 | P1481I | E | 6.7e-09 | 1.5e-08 | 2e-08 | 1.390E-08 | 6.718E-09 | 6.700E-09 | 2.000E-08 |
| 29611 | 1481 | P1481L | E | 4.1e-09 | 5.5e-09 | 7.6e-09 | 5.733E-09 | 1.762E-09 | 4.100E-09 | 7.600E-09 |
| 29612 | 1481 | P1481K | E | 7.4e-09 | 7.2e-09 | 1.2e-08 | 8.867E-09 | 2.715E-09 | 7.200E-09 | 1.200E-08 |
| 29613 | 1481 | P1481M | E | 5.9e-09 | 2e-08 | 1.3e-09 | 9.067E-09 | 9.744E-09 | 1.300E-09 | 2.000E-08 |
| 29614 | 1481 | P1481F | E | 6e-09 | 2.8e-09 | 6.4e-09 | 5.067E-09 | 1.973E-09 | 2.800E-09 | 6.400E-09 |
| 29615 | 1481 | P1481P | E | 1.7e-08 | 2.4e-08 | 2.9e-09 | 1.463E-08 | 1.075E-08 | 2.900E-09 | 2.400E-08 |
| 29616 | 1481 | P1481S | E | 3.7e-09 | 4.2e-09 | 6.1e-09 | 4.667E-09 | 1.266E-09 | 3.700E-09 | 6.100E-09 |
| 29617 | 1481 | P1481T | E | 8.7e-09 | 1.3e-08 | 4.3e-09 | 8.667E-09 | 4.350E-09 | 4.300E-09 | 1.300E-08 |
| 29618 | 1481 | P1481W | E | 5.6e-09 | 4.7e-09 | 1.8e-08 | 9.433E-09 | 7.433E-09 | 4.700E-09 | 1.800E-08 |
| 29619 | 1481 | P1481Y | E | 2.4e-09 | 1.2e-08 | 1.8e-08 | 1.080E-08 | 7.869E-09 | 2.400E-09 | 1.800E-08 |
| 29620 | 1481 | P1481V | E | 5.1e-09 | 6.9e-09 | 6.3e-09 | 6.100E-09 | 9.165E-10 | 5.100E-09 | 6.900E-09 |
| 29621 | 1482 | S1482A | E | 6.7e-09 | 1e-08 | 4.3e-09 | 7.000E-09 | 2.862E-09 | 4.300E-09 | 1.000E-08 |
| 29622 | 1482 | S1482R | E | 5.8e-09 | 1.3e-08 | 4.3e-09 | 7.700E-09 | 4.651E-09 | 4.300E-09 | 1.300E-08 |
| 29623 | 1482 | S1482N | E | 1.9e-09 | 4.1e-09 | 6e-09 | 4.000E-09 | 2.052E-09 | 1.900E-09 | 6.000E-09 |
| 29624 | 1482 | S1482D | E | 2.5e-09 | 8.3e-09 | 6.4e-09 | 5.733E-09 | 2.957E-09 | 2.500E-09 | 8.300E-09 |
| 29625 | 1482 | S1482C | E | 5e-09 | 8.3e-09 | 4.7e-09 | 6.000E-09 | 1.997E-09 | 4.700E-09 | 8.300E-09 |
| 29626 | 1482 | S1482E | E | 5.4e-09 | 2.1e-09 | 1.1e-08 | 6.167E-09 | 4.499E-09 | 2.100E-09 | 1.100E-08 |
| 29627 | 1482 | S1482Q | E | 4.1e-09 | 3e-09 | 2.1e-08 | 9.367E-09 | 1.009E-08 | 3.000E-09 | 2.100E-08 |
| 29628 | 1482 | S1482G | E | 5.6e-09 | 9.9e-09 | 2.2e-09 | 5.900E-09 | 3.859E-09 | 2.200E-09 | 9.900E-09 |
| 29629 | 1482 | S1482H | E | 8.8e-09 | 7.8e-09 | 3.5e-09 | 6.700E-09 | 2.816E-09 | 3.500E-09 | 8.800E-09 |
| 29630 | 1482 | S1482I | E | 5.8e-09 | 4.7e-09 | 3.3e-09 | 4.600E-09 | 1.253E-09 | 3.300E-09 | 5.800E-09 |
| 29631 | 1482 | S1482L | E | 9.8e-09 | 5.5e-09 | 4.8e-09 | 6.700E-09 | 2.707E-09 | 4.800E-09 | 9.800E-09 |
| 29632 | 1482 | S1482K | E | 1.3e-08 | 4.5e-09 | 3.5e-09 | 7.000E-09 | 5.220E-09 | 3.500E-09 | 1.300E-08 |
| 29633 | 1482 | S1482M | E | 2.6e-09 | 1e-08 | 3.1e-09 | 5.233E-09 | 4.136E-09 | 2.600E-09 | 1.000E-08 |
| 29634 | 1482 | S1482F | E | 9.5e-09 | 2.1e-08 | 4.7e-09 | 1.173E-08 | 8.376E-09 | 4.700E-09 | 2.100E-08 |
| 29635 | 1482 | S1482P | E | 4.3e-09 | 1.3e-08 | 2.5e-08 | 1.410E-08 | 1.039E-08 | 4.300E-09 | 2.500E-08 |
| 29636 | 1482 | S1482S | E | 3.1e-09 | 1e-08 | 1.4e-08 | 9.033E-09 | 5.514E-09 | 3.100E-09 | 1.400E-08 |
| 29637 | 1482 | S1482T | E | 3.3e-09 | 1.2e-08 | 5.3e-09 | 6.867E-09 | 4.557E-09 | 3.300E-09 | 1.200E-08 |
| 29638 | 1482 | S1482W | E | 6.3e-09 | 5.2e-09 | 3.8e-09 | 5.100E-09 | 1.253E-09 | 3.800E-09 | 6.300E-09 |
| 29639 | 1482 | S1482Y | E | 8e-09 | 2.6e-09 | 3.1e-09 | 4.567E-09 | 2.984E-09 | 2.600E-09 | 8.000E-09 |
| 29640 | 1482 | S1482V | E | 2.9e-09 | 8.2e-09 | 7.4e-09 | 6.167E-09 | 2.857E-09 | 2.900E-09 | 8.200E-09 |
| 29641 | 1483 | N1483A | E | 1.1e-08 | 9.2e-09 | 5.3e-09 | 8.500E-09 | 2.914E-09 | 5.300E-09 | 1.100E-08 |
| 29642 | 1483 | N1483R | E | 1.1e-08 | 4.8e-09 | 1.1e-08 | 8.933E-09 | 3.580E-09 | 4.800E-09 | 1.100E-08 |
| 29643 | 1483 | N1483N | E | 7.6e-09 | 3.6e-09 | 1.8e-08 | 9.733E-09 | 7.433E-09 | 3.600E-09 | 1.800E-08 |
| 29644 | 1483 | N1483D | E | 9.6e-09 | 1.3e-08 | 9.8e-09 | 1.080E-08 | 1.908E-09 | 9.600E-09 | 1.300E-08 |
| 29645 | 1483 | N1483C | E | 2.9e-09 | 8.3e-09 | 9.2e-09 | 6.800E-09 | 3.407E-09 | 2.900E-09 | 9.200E-09 |
| 29646 | 1483 | N1483E | E | 2.7e-09 | 2.8e-09 | 7.7e-09 | 4.400E-09 | 2.858E-09 | 2.700E-09 | 7.700E-09 |
| 29647 | 1483 | N1483Q | E | 1.2e-08 | 7.8e-09 | 1e-08 | 9.933E-09 | 2.101E-09 | 7.800E-09 | 1.200E-08 |
| 29648 | 1483 | N1483G | E | 5.3e-09 | 4.3e-09 | 6.7e-09 | 5.433E-09 | 1.206E-09 | 4.300E-09 | 6.700E-09 |
| 29649 | 1483 | N1483H | E | 8.5e-09 | 4.8e-09 | 1.4e-08 | 9.100E-09 | 4.629E-09 | 4.800E-09 | 1.400E-08 |
| 29650 | 1483 | N1483I | E | 4e-09 | 4.8e-09 | 1.1e-08 | 6.600E-09 | 3.831E-09 | 4.000E-09 | 1.100E-08 |
| 29651 | 1483 | N1483L | E | 6.1e-09 | 5.2e-09 | 7.9e-09 | 6.400E-09 | 1.375E-09 | 5.200E-09 | 7.900E-09 |
| 29652 | 1483 | N1483K | E | 1.2e-08 | 1.2e-08 | 3.4e-09 | 9.133E-09 | 4.965E-09 | 3.400E-09 | 1.200E-08 |
| 29653 | 1483 | N1483M | E | 7.4e-09 | 3.7e-09 | 1.3e-08 | 8.033E-09 | 4.682E-09 | 3.700E-09 | 1.300E-08 |
| 29654 | 1483 | N1483F | E | 7.7e-09 | 2.7e-08 | 5.3e-09 | 1.333E-08 | 1.190E-08 | 5.300E-09 | 2.700E-08 |
| 29655 | 1483 | N1483P | E | 8.3e-09 | 6e-09 | 1.4e-08 | 9.433E-09 | 4.119E-09 | 6.000E-09 | 1.400E-08 |
| 29656 | 1483 | N1483S | E | 1.6e-08 | 2.7e-09 | 1.9e-09 | 6.867E-09 | 7.920E-09 | 1.900E-09 | 1.600E-08 |
| 29657 | 1483 | N1483T | E | 1.1e-08 | 6.6e-09 | 3.8e-09 | 7.133E-09 | 3.630E-09 | 3.800E-09 | 1.100E-08 |
| 29658 | 1483 | N1483W | E | 5.9e-09 | 8.9e-09 | 8.2e-09 | 7.667E-09 | 1.570E-09 | 5.900E-09 | 8.900E-09 |
| 29659 | 1483 | N1483Y | E | 1.8e-08 | 8.2e-09 | 4.6e-09 | 1.027E-08 | 6.935E-09 | 4.600E-09 | 1.800E-08 |
| 29660 | 1483 | N1483V | E | 6.8e-09 | 2.7e-08 | 1.1e-08 | 1.493E-08 | 1.066E-08 | 6.800E-09 | 2.700E-08 |
| 29661 | 1484 | T1484A | E | 9.5e-09 | 8.8e-09 | 6.1e-09 | 8.133E-09 | 1.795E-09 | 6.100E-09 | 9.500E-09 |
| 29662 | 1484 | T1484R | E | 8.2e-09 | 6.8e-09 | 1.1e-08 | 8.667E-09 | 2.139E-09 | 6.800E-09 | 1.100E-08 |
| 29663 | 1484 | T1484N | E | 1.2e-08 | 8.2e-09 | 6e-09 | 8.733E-09 | 3.035E-09 | 6.000E-09 | 1.200E-08 |
| 29664 | 1484 | T1484D | E | 3.8e-09 | 3.1e-09 | 8.1e-09 | 5.000E-09 | 2.707E-09 | 3.100E-09 | 8.100E-09 |

| | | | | | | | | | | |
|---|---|---|---|---|---|---|---|---|---|---|
| 29665 | 1484 | T1484C | E | 9.5e-09 | 6.2e-09 | 1.2e-08 | 9.233E-09 | 2.909E-09 | 6.200E-09 | 1.200E-08 |
| 29666 | 1484 | T1484E | E | 9.2e-09 | 5.7e-09 | 3.4e-09 | 6.100E-09 | 2.921E-09 | 3.400E-09 | 9.200E-09 |
| 29667 | 1484 | T1484Q | E | 6.4e-09 | 1.2e-08 | 1.3e-08 | 1.047E-08 | 3.557E-09 | 6.400E-09 | 1.300E-08 |
| 29668 | 1484 | T1484G | E | 7.7e-09 | 1.3e-08 | 7.8e-09 | 9.500E-09 | 3.032E-09 | 7.700E-09 | 1.300E-08 |
| 29669 | 1484 | T1484H | E | 1.1e-08 | 3.2e-09 | 7e-09 | 7.067E-09 | 3.900E-09 | 3.200E-09 | 1.100E-08 |
| 29670 | 1484 | T1484I | E | 5.1e-09 | 9.2e-09 | 2.4e-09 | 5.567E-09 | 3.424E-09 | 2.400E-09 | 9.200E-09 |
| 29671 | 1484 | T1484L | E | 3.5e-09 | 1e-08 | 1.5e-08 | 9.500E-09 | 5.766E-09 | 3.500E-09 | 1.500E-08 |
| 29672 | 1484 | T1484K | E | 7e-09 | 1.4e-08 | 8e-09 | 9.667E-09 | 3.786E-09 | 7.000E-09 | 1.400E-08 |
| 29673 | 1484 | T1484M | E | 1.2e-08 | 9.4e-09 | 9.9e-09 | 1.043E-08 | 1.380E-09 | 9.400E-09 | 1.200E-08 |
| 29674 | 1484 | T1484F | E | 5.4e-09 | 8.8e-09 | 1.5e-09 | 5.233E-09 | 3.653E-09 | 1.500E-09 | 8.800E-09 |
| 29675 | 1484 | T1484P | E | 8.6e-09 | 6.3e-09 | 1.7e-08 | 1.063E-08 | 5.632E-09 | 6.300E-09 | 1.700E-08 |
| 29676 | 1484 | T1484S | E | 8.2e-09 | 2.3e-09 | 9.3e-09 | 6.600E-09 | 3.764E-09 | 2.300E-09 | 9.300E-09 |
| 29677 | 1484 | T1484T | E | 8.6e-10 | 2.5e-08 | 6.9e-09 | 1.092E-08 | 1.256E-08 | 8.600E-10 | 2.500E-08 |
| 29678 | 1484 | T1484W | E | 1e-08 | 7.9e-09 | 6.5e-09 | 8.133E-09 | 1.762E-09 | 6.500E-09 | 1.000E-08 |
| 29679 | 1484 | T1484Y | E | 4.5e-09 | 1e-08 | 5.5e-09 | 6.667E-09 | 2.930E-09 | 4.500E-09 | 1.000E-08 |
| 29680 | 1484 | T1484V | E | 2.9e-09 | 6.9e-09 | 4e-09 | 4.600E-09 | 2.066E-09 | 2.900E-09 | 6.900E-09 |
| 29681 | 1485 | K1485A | E | 1.5e-08 | 2.2e-09 | 7.2e-09 | 8.133E-09 | 6.451E-09 | 2.200E-09 | 1.500E-08 |
| 29682 | 1485 | K1485R | E | 4.6e-09 | 1.2e-08 | 5.7e-09 | 7.433E-09 | 3.993E-09 | 4.600E-09 | 1.200E-08 |
| 29683 | 1485 | K1485N | E | 8.2e-09 | 6.8e-09 | 5e-09 | 6.667E-09 | 1.604E-09 | 5.000E-09 | 8.200E-09 |
| 29684 | 1485 | K1485D | E | 1.3e-08 | 3.6e-09 | 1.6e-08 | 1.087E-08 | 6.469E-09 | 3.600E-09 | 1.600E-08 |
| 29685 | 1485 | K1485C | E | 1.3e-08 | 1.7e-08 | 9.8e-09 | 1.327E-08 | 3.607E-09 | 9.800E-09 | 1.700E-08 |
| 29686 | 1485 | K1485E | E | 5.9e-09 | 8.7e-09 | 2.5e-09 | 5.700E-09 | 3.105E-09 | 2.500E-09 | 8.700E-09 |
| 29687 | 1485 | K1485Q | E | 7.1e-09 | 3.3e-09 | 2.4e-09 | 4.267E-09 | 2.495E-09 | 2.400E-09 | 7.100E-09 |
| 29688 | 1485 | K1485G | E | 3.4e-09 | 3.7e-09 | 2.3e-09 | 3.133E-09 | 7.371E-10 | 2.300E-09 | 3.700E-09 |
| 29689 | 1485 | K1485H | E | 3.4e-09 | 3.6e-08 | 7e-09 | 1.547E-08 | 1.787E-08 | 3.400E-09 | 3.600E-08 |
| 29690 | 1485 | K1485I | E | 6.7e-09 | 7.6e-09 | 5.2e-09 | 6.500E-09 | 1.212E-09 | 5.200E-09 | 7.600E-09 |
| 29691 | 1485 | K1485L | E | 4.3e-09 | 5.2e-09 | 1.8e-08 | 9.167E-09 | 7.663E-09 | 4.300E-09 | 1.800E-08 |
| 29692 | 1485 | K1485K | E | 7.5e-09 | 1e-08 | 8.9e-09 | 8.800E-09 | 1.253E-09 | 7.500E-09 | 1.000E-08 |
| 29693 | 1485 | K1485M | E | 1.6e-08 | 4.8e-09 | 6.9e-09 | 9.233E-09 | 5.953E-09 | 4.800E-09 | 1.600E-08 |
| 29694 | 1485 | K1485F | E | 8.9e-09 | 1.8e-08 | 6.5e-09 | 1.113E-08 | 6.067E-09 | 6.500E-09 | 1.800E-08 |
| 29695 | 1485 | K1485P | E | 9.2e-09 | 3e-09 | 6.4e-09 | 6.200E-09 | 3.105E-09 | 3.000E-09 | 9.200E-09 |
| 29696 | 1485 | K1485S | E | 6.4e-09 | 1.2e-08 | 4.4e-09 | 7.600E-09 | 3.940E-09 | 4.400E-09 | 1.200E-08 |
| 29697 | 1485 | K1485T | E | 1.5e-08 | 4.2e-09 | 2.4e-08 | 1.440E-08 | 9.914E-09 | 4.200E-09 | 2.400E-08 |
| 29698 | 1485 | K1485W | E | 6.2e-09 | 3.4e-09 | 5.9e-09 | 5.167E-09 | 1.537E-09 | 3.400E-09 | 6.200E-09 |
| 29699 | 1485 | K1485Y | E | 4.7e-09 | 1.1e-08 | 1.8e-08 | 1.123E-08 | 6.653E-09 | 4.700E-09 | 1.800E-08 |
| 29700 | 1485 | K1485V | E | 5.6e-09 | 1.1e-08 | 4.5e-09 | 7.033E-09 | 3.479E-09 | 4.500E-09 | 1.100E-08 |
| 29701 | 1486 | V1486A | E | 5.6e-09 | 1.1e-08 | 9.4e-09 | 8.667E-09 | 2.774E-09 | 5.600E-09 | 1.100E-08 |
| 29702 | 1486 | V1486R | E | 5.7e-09 | 5.3e-09 | 2.1e-08 | 1.067E-08 | 8.951E-09 | 5.300E-09 | 2.100E-08 |
| 29703 | 1486 | V1486N | E | 2.1e-09 | 1.2e-08 | 5.9e-09 | 6.667E-09 | 4.994E-09 | 2.100E-09 | 1.200E-08 |
| 29704 | 1486 | V1486D | E | 8.1e-09 | 1.3e-08 | 4e-09 | 8.367E-09 | 4.506E-09 | 4.000E-09 | 1.300E-08 |
| 29705 | 1486 | V1486C | E | 1.1e-08 | 6.2e-09 | 2.9e-09 | 6.700E-09 | 4.073E-09 | 2.900E-09 | 1.100E-08 |
| 29706 | 1486 | V1486E | E | 1.5e-08 | 1.3e-08 | 7.2e-09 | 1.173E-08 | 4.051E-09 | 7.200E-09 | 1.500E-08 |
| 29707 | 1486 | V1486Q | E | 3.9e-09 | 8.9e-09 | 3.4e-09 | 5.400E-09 | 3.041E-09 | 3.400E-09 | 8.900E-09 |
| 29708 | 1486 | V1486G | E | 9.7e-09 | 5.8e-09 | 1e-08 | 8.500E-09 | 2.343E-09 | 5.800E-09 | 1.000E-08 |
| 29709 | 1486 | V1486H | E | 4.1e-09 | 4e-09 | 5e-09 | 4.367E-09 | 5.508E-10 | 4.000E-09 | 5.000E-09 |
| 29710 | 1486 | V1486I | E | 6.7e-09 | 1.1e-09 | 4.9e-09 | 4.233E-09 | 2.859E-09 | 1.100E-09 | 6.700E-09 |
| 29711 | 1486 | V1486L | E | 9.9e-09 | 3.8e-09 | 1.2e-08 | 8.567E-09 | 4.259E-09 | 3.800E-09 | 1.200E-08 |
| 29712 | 1486 | V1486K | E | 7.3e-09 | 8.2e-09 | 4.2e-09 | 6.567E-09 | 2.098E-09 | 4.200E-09 | 8.200E-09 |
| 29713 | 1486 | V1486M | E | 2.8e-09 | 7.1e-09 | 1.9e-08 | 9.633E-09 | 8.392E-09 | 2.800E-09 | 1.900E-08 |
| 29714 | 1486 | V1486F | E | 6.6e-09 | 3.3e-09 | 8.7e-09 | 6.200E-09 | 2.722E-09 | 3.300E-09 | 8.700E-09 |
| 29715 | 1486 | V1486P | E | 7.4e-09 | 4.5e-09 | 9.7e-09 | 7.200E-09 | 2.606E-09 | 4.500E-09 | 9.700E-09 |
| 29716 | 1486 | V1486S | E | 3.5e-09 | 1.9e-09 | 4.6e-09 | 3.333E-09 | 1.358E-09 | 1.900E-09 | 4.600E-09 |
| 29717 | 1486 | V1486T | E | 3.5e-08 | 1.2e-08 | 1.4e-08 | 2.033E-08 | 1.274E-08 | 1.200E-08 | 3.500E-08 |
| 29718 | 1486 | V1486W | E | 5.9e-09 | 7.9e-09 | 3.7e-08 | 1.693E-08 | 1.741E-08 | 5.900E-09 | 3.700E-08 |
| 29719 | 1486 | V1486Y | E | 1.5e-08 | 6.3e-09 | 5.9e-09 | 9.067E-09 | 5.142E-09 | 5.900E-09 | 1.500E-08 |
| 29720 | 1486 | V1486V | E | 5.7e-09 | 9.4e-09 | 5.5e-09 | 6.867E-09 | 2.196E-09 | 5.500E-09 | 9.400E-09 |
| 29721 | 1487 | D1487A | E | 1.1e-08 | 3.8e-09 | 8.5e-09 | 7.767E-09 | 3.656E-09 | 3.800E-09 | 1.100E-08 |
| 29722 | 1487 | D1487R | E | 1.1e-08 | 9.9e-09 | 6.6e-09 | 9.167E-09 | 2.290E-09 | 6.600E-09 | 1.100E-08 |
| 29723 | 1487 | D1487N | E | 7.4e-09 | 6.1e-09 | 2.1e-08 | 1.150E-08 | 8.253E-09 | 6.100E-09 | 2.100E-08 |
| 29724 | 1487 | D1487D | E | 7.4e-09 | 8.2e-09 | 6.2e-09 | 7.267E-09 | 1.007E-09 | 6.200E-09 | 8.200E-09 |
| 29725 | 1487 | D1487C | E | 6.1e-09 | 1.3e-08 | 8.3e-09 | 9.133E-09 | 3.525E-09 | 6.100E-09 | 1.300E-08 |
| 29726 | 1487 | D1487E | E | 1.1e-08 | 1.6e-08 | 6.4e-09 | 1.113E-08 | 4.801E-09 | 6.400E-09 | 1.600E-08 |
| 29727 | 1487 | D1487Q | E | 8.7e-09 | 3.2e-09 | 5.9e-09 | 5.933E-09 | 2.750E-09 | 3.200E-09 | 8.700E-09 |
| 29728 | 1487 | D1487G | E | 1.9e-08 | 8.6e-09 | 4.5e-09 | 1.070E-08 | 7.475E-09 | 4.500E-09 | 1.900E-08 |
| 29729 | 1487 | D1487H | E | 4.8e-09 | 4e-09 | 9.7e-09 | 6.167E-09 | 3.086E-09 | 4.000E-09 | 9.700E-09 |
| 29730 | 1487 | D1487I | E | 2.6e-09 | 8e-09 | 1.5e-08 | 8.533E-09 | 6.217E-09 | 2.600E-09 | 1.500E-08 |
| 29731 | 1487 | D1487L | E | 2.3e-08 | 5.4e-09 | 7.5e-09 | 1.197E-08 | 9.613E-09 | 5.400E-09 | 2.300E-08 |
| 29732 | 1487 | D1487K | E | 7.2e-09 | 3.7e-09 | 3.7e-09 | 4.867E-09 | 2.021E-09 | 3.700E-09 | 7.200E-09 |
| 29733 | 1487 | D1487M | E | 6e-09 | 2.1e-09 | 5.8e-09 | 4.633E-09 | 2.196E-09 | 2.100E-09 | 6.000E-09 |
| 29734 | 1487 | D1487F | E | 1.1e-08 | 6.7e-09 | 8.1e-09 | 8.600E-09 | 2.193E-09 | 6.700E-09 | 1.100E-08 |
| 29735 | 1487 | D1487P | E | 4.9e-09 | 7.4e-09 | 3.4e-09 | 5.233E-09 | 2.021E-09 | 3.400E-09 | 7.400E-09 |
| 29736 | 1487 | D1487S | E | 3.3e-09 | 9.8e-09 | 8.8e-09 | 7.300E-09 | 3.500E-09 | 3.300E-09 | 9.800E-09 |
| 29737 | 1487 | D1487T | E | 3.9e-09 | 5.2e-09 | 6.2e-09 | 5.100E-09 | 1.153E-09 | 3.900E-09 | 6.200E-09 |
| 29738 | 1487 | D1487W | E | 5.4e-09 | 1.9e-08 | 8.6e-09 | 1.100E-08 | 7.111E-09 | 5.400E-09 | 1.900E-08 |
| 29739 | 1487 | D1487Y | E | 7.2e-09 | 4.1e-08 | 9.1e-09 | 1.910E-08 | 1.899E-08 | 7.200E-09 | 4.100E-08 |
| 29740 | 1487 | D1487V | E | 1.4e-08 | 1.2e-08 | 3.5e-09 | 9.833E-09 | 5.575E-09 | 3.500E-09 | 1.400E-08 |
| 29741 | 1488 | K1488A | E | 3.4e-09 | 2.7e-09 | 5.7e-09 | 3.933E-09 | 1.570E-09 | 2.700E-09 | 5.700E-09 |
| 29742 | 1488 | K1488R | E | 5.9e-09 | 1.2e-08 | 1e-08 | 9.300E-09 | 3.110E-09 | 5.900E-09 | 1.200E-08 |
| 29743 | 1488 | K1488N | E | 6.9e-09 | 5.4e-09 | 1.8e-09 | 4.700E-09 | 2.621E-09 | 1.800E-09 | 6.900E-09 |
| 29744 | 1488 | K1488D | E | 4e-09 | 2.8e-09 | 7.6e-09 | 4.800E-09 | 2.498E-09 | 2.800E-09 | 7.600E-09 |
| 29745 | 1488 | K1488C | E | 6.6e-09 | 5.2e-09 | 5.6e-09 | 5.800E-09 | 7.211E-10 | 5.200E-09 | 6.600E-09 |
| 29746 | 1488 | K1488E | E | 7.6e-09 | 6.7e-09 | 1e-08 | 8.100E-09 | 1.706E-09 | 6.700E-09 | 1.000E-08 |
| 29747 | 1488 | K1488Q | E | 2e-08 | 5.1e-09 | 5.1e-09 | 1.007E-08 | 8.603E-09 | 5.100E-09 | 2.000E-08 |
| 29748 | 1488 | K1488G | E | 3.9e-09 | 6.4e-09 | 1.4e-08 | 8.100E-09 | 5.260E-09 | 3.900E-09 | 1.400E-08 |
| 29749 | 1488 | K1488H | E | 3.6e-09 | 4.2e-09 | 3.8e-09 | 3.867E-09 | 3.055E-10 | 3.600E-09 | 4.200E-09 |

| | | | | | | | | | | |
|---|---|---|---|---|---|---|---|---|---|---|
| 29750 | 1488 | K1488I | E | 2.2e-09 | 5.6e-09 | 6.3e-09 | 4.700E-09 | 2.193E-09 | 2.200E-09 | 6.300E-09 |
| 29751 | 1488 | K1488L | E | 1e-08 | 5.6e-09 | 9.1e-09 | 8.233E-09 | 2.325E-09 | 5.600E-09 | 1.000E-08 |
| 29752 | 1488 | K1488K | E | 2.8e-09 | 4.1e-09 | 5.7e-09 | 4.200E-09 | 1.453E-09 | 2.800E-09 | 5.700E-09 |
| 29753 | 1488 | K1488M | E | 1.2e-08 | 9.1e-09 | 9.3e-09 | 1.013E-08 | 1.620E-09 | 9.100E-09 | 1.200E-08 |
| 29754 | 1488 | K1488F | E | 1.6e-08 | 1.1e-08 | 1.1e-08 | 1.267E-08 | 2.887E-09 | 1.100E-08 | 1.600E-08 |
| 29755 | 1488 | K1488P | E | 7e-09 | 2e-08 | 8.9e-09 | 1.197E-08 | 7.022E-09 | 7.000E-09 | 2.000E-08 |
| 29756 | 1488 | K1488S | E | 5.9e-09 | 3e-09 | 1.1e-08 | 6.633E-09 | 4.050E-09 | 3.000E-09 | 1.100E-08 |
| 29757 | 1488 | K1488T | E | 7.7e-09 | 1.7e-09 | 1.2e-08 | 7.133E-09 | 5.173E-09 | 1.700E-09 | 1.200E-08 |
| 29758 | 1488 | K1488W | E | 5.6e-09 | 8.6e-09 | 4.4e-09 | 6.200E-09 | 2.163E-09 | 4.400E-09 | 8.600E-09 |
| 29759 | 1488 | K1488Y | E | 5.4e-09 | 3e-09 | 3.6e-09 | 4.000E-09 | 1.249E-09 | 3.000E-09 | 5.400E-09 |
| 29760 | 1488 | K1488V | E | 3.5e-09 | 2.6e-08 | 5.9e-09 | 1.180E-08 | 1.236E-08 | 3.500E-09 | 2.600E-08 |
| 29761 | 1489 | K1489A | E | 5.7e-09 | 1.1e-08 | 8.9e-09 | 8.533E-09 | 2.669E-09 | 5.700E-09 | 1.100E-08 |
| 29762 | 1489 | K1489R | E | 4.6e-09 | 1.1e-08 | 2e-09 | 5.867E-09 | 4.632E-09 | 2.000E-09 | 1.100E-08 |
| 29763 | 1489 | K1489N | E | 7.2e-09 | 9.3e-09 | 1.6e-09 | 6.033E-09 | 3.980E-09 | 1.600E-09 | 9.300E-09 |
| 29764 | 1489 | K1489D | E | 8.3e-09 | 3.8e-08 | 1.8e-09 | 1.603E-08 | 1.930E-08 | 1.800E-09 | 3.800E-08 |
| 29765 | 1489 | K1489C | E | 3.8e-09 | 1.3e-08 | 1e-08 | 8.933E-09 | 4.692E-09 | 3.800E-09 | 1.300E-08 |
| 29766 | 1489 | K1489E | E | 6.3e-09 | 3.1e-09 | 2.3e-09 | 3.900E-09 | 2.117E-09 | 2.300E-09 | 6.300E-09 |
| 29767 | 1489 | K1489Q | E | 5.3e-09 | 3.2e-09 | 4.8e-09 | 4.433E-09 | 1.097E-09 | 3.200E-09 | 5.300E-09 |
| 29768 | 1489 | K1489G | E | 6.8e-09 | 1.1e-08 | 6.4e-09 | 8.067E-09 | 2.548E-09 | 6.400E-09 | 1.100E-08 |
| 29769 | 1489 | K1489H | E | 3.7e-09 | 3.2e-09 | 8.1e-09 | 5.000E-09 | 2.696E-09 | 3.200E-09 | 8.100E-09 |
| 29770 | 1489 | K1489I | E | 1.1e-08 | 3.1e-08 | 2.1e-09 | 1.470E-08 | 1.480E-08 | 2.100E-09 | 3.100E-08 |
| 29771 | 1489 | K1489L | E | 3.3e-09 | 4.3e-09 | 1.1e-08 | 6.200E-09 | 4.187E-09 | 3.300E-09 | 1.100E-08 |
| 29772 | 1489 | K1489K | E | 6.6e-09 | 3.4e-09 | 8.2e-09 | 6.067E-09 | 2.444E-09 | 3.400E-09 | 8.200E-09 |
| 29773 | 1489 | K1489M | E | 1.7e-08 | 6.4e-09 | 1.1e-08 | 1.147E-08 | 5.315E-09 | 6.400E-09 | 1.700E-08 |
| 29774 | 1489 | K1489F | E | 6.2e-09 | 6.2e-09 | 1.6e-08 | 9.467E-09 | 5.658E-09 | 6.200E-09 | 1.600E-08 |
| 29775 | 1489 | K1489P | E | 2.7e-08 | 9.3e-09 | 6.4e-09 | 1.423E-08 | 1.115E-08 | 6.400E-09 | 2.700E-08 |
| 29776 | 1489 | K1489S | E | 8.4e-09 | 8e-09 | 3.5e-09 | 6.633E-09 | 2.721E-09 | 3.500E-09 | 8.400E-09 |
| 29777 | 1489 | K1489T | E | 9.1e-09 | 1.9e-08 | 4.5e-09 | 1.087E-08 | 7.410E-09 | 4.500E-09 | 1.900E-08 |
| 29778 | 1489 | K1489W | E | 3.4e-09 | 8.7e-09 | 2.2e-09 | 4.767E-09 | 3.459E-09 | 2.200E-09 | 8.700E-09 |
| 29779 | 1489 | K1489Y | E | 6.5e-09 | 3.3e-09 | 1.3e-08 | 7.600E-09 | 4.943E-09 | 3.300E-09 | 1.300E-08 |
| 29780 | 1489 | K1489V | E | 6.6e-09 | 6.9e-09 | 4.6e-09 | 6.033E-09 | 1.250E-09 | 4.600E-09 | 6.900E-09 |
| 29781 | 1490 | V1490A | E | 1.6e-09 | 4.9e-09 | 8.2e-09 | 4.900E-09 | 3.300E-09 | 1.600E-09 | 8.200E-09 |
| 29782 | 1490 | V1490R | E | 6.1e-09 | 1.4e-09 | 4.1e-09 | 3.867E-09 | 2.359E-09 | 1.400E-09 | 6.100E-09 |
| 29783 | 1490 | V1490N | E | 2.3e-08 | 7.4e-09 | 1.9e-08 | 1.647E-08 | 8.103E-09 | 7.400E-09 | 2.300E-08 |
| 29784 | 1490 | V1490D | E | 5e-09 | 1.2e-08 | 1e-08 | 9.000E-09 | 3.606E-09 | 5.000E-09 | 1.200E-08 |
| 29785 | 1490 | V1490C | E | 2.3e-09 | 6e-09 | 6.6e-09 | 4.967E-09 | 2.329E-09 | 2.300E-09 | 6.600E-09 |
| 29786 | 1490 | V1490E | E | 4.3e-09 | 2.2e-08 | 5.2e-09 | 1.050E-08 | 9.969E-09 | 4.300E-09 | 2.200E-08 |
| 29787 | 1490 | V1490Q | E | 9.7e-09 | 4.9e-09 | 4.1e-09 | 6.233E-09 | 3.029E-09 | 4.100E-09 | 9.700E-09 |
| 29788 | 1490 | V1490G | E | 9.9e-09 | 1.9e-08 | 2.6e-09 | 1.050E-08 | 8.216E-09 | 2.600E-09 | 1.900E-08 |
| 29789 | 1490 | V1490H | E | 1.7e-09 | 1.5e-08 | 8.4e-09 | 8.367E-09 | 6.650E-09 | 1.700E-09 | 1.500E-08 |
| 29790 | 1490 | V1490I | E | 1e-08 | 5.3e-09 | 1e-08 | 8.433E-09 | 2.714E-09 | 5.300E-09 | 1.000E-08 |
| 29791 | 1490 | V1490L | E | 6.7e-09 | 5.6e-09 | 1e-08 | 7.433E-09 | 2.290E-09 | 5.600E-09 | 1.000E-08 |
| 29792 | 1490 | V1490K | E | 1.7e-08 | 6.7e-09 | 7.1e-09 | 1.027E-08 | 5.835E-09 | 6.700E-09 | 1.700E-08 |
| 29793 | 1490 | V1490M | E | 1.2e-08 | 1.7e-09 | 3.6e-09 | 5.767E-09 | 5.481E-09 | 1.700E-09 | 1.200E-08 |
| 29794 | 1490 | V1490F | E | 2.7e-09 | 7.3e-09 | 1.4e-08 | 8.000E-09 | 5.682E-09 | 2.700E-09 | 1.400E-08 |
| 29795 | 1490 | V1490P | E | 2.7e-09 | 5.7e-09 | 5.2e-09 | 4.533E-09 | 1.607E-09 | 2.700E-09 | 5.700E-09 |
| 29796 | 1490 | V1490S | E | 1.2e-08 | 6.6e-09 | 1.1e-08 | 9.867E-09 | 2.873E-09 | 6.600E-09 | 1.200E-08 |
| 29797 | 1490 | V1490T | E | 3.3e-09 | 3.7e-09 | 1e-08 | 5.667E-09 | 3.758E-09 | 3.300E-09 | 1.000E-08 |
| 29798 | 1490 | V1490W | E | 3.5e-09 | 1.1e-08 | 4.2e-08 | 1.883E-08 | 2.041E-08 | 3.500E-09 | 4.200E-08 |
| 29799 | 1490 | V1490Y | E | 8.2e-09 | 6.1e-09 | 8.2e-09 | 7.500E-09 | 1.212E-09 | 6.100E-09 | 8.200E-09 |
| 29800 | 1490 | V1490V | E | 1.5e-08 | 2.5e-09 | 6.6e-09 | 8.033E-09 | 6.372E-09 | 2.500E-09 | 1.500E-08 |
| 29801 | 1491 | E1491A | E | 7.7e-09 | 1.1e-08 | 9e-09 | 9.233E-09 | 1.662E-09 | 7.700E-09 | 1.100E-08 |
| 29802 | 1491 | E1491R | E | 8.7e-09 | 6e-09 | 2.4e-09 | 5.700E-09 | 3.161E-09 | 2.400E-09 | 8.700E-09 |
| 29803 | 1491 | E1491N | E | 1.1e-08 | 6.5e-09 | 8.5e-09 | 8.667E-09 | 2.255E-09 | 6.500E-09 | 1.100E-08 |
| 29804 | 1491 | E1491D | E | 2.6e-09 | 4.2e-09 | 1.9e-09 | 2.900E-09 | 1.179E-09 | 1.900E-09 | 4.200E-09 |
| 29805 | 1491 | E1491C | E | 6.8e-09 | 1.5e-08 | 9.9e-09 | 1.057E-08 | 4.140E-09 | 6.800E-09 | 1.500E-08 |
| 29806 | 1491 | E1491E | E | 1.1e-08 | 6.9e-09 | 3.5e-09 | 7.133E-09 | 3.755E-09 | 3.500E-09 | 1.100E-08 |
| 29807 | 1491 | E1491Q | E | 7.3e-09 | 1.4e-08 | 4e-09 | 8.433E-09 | 5.095E-09 | 4.000E-09 | 1.400E-08 |
| 29808 | 1491 | E1491G | E | 1.1e-08 | 7.9e-09 | 4.4e-09 | 7.767E-09 | 3.302E-09 | 4.400E-09 | 1.100E-08 |
| 29809 | 1491 | E1491H | E | 8.6e-09 | 8.2e-09 | 1.7e-09 | 6.167E-09 | 3.873E-09 | 1.700E-09 | 8.600E-09 |
| 29810 | 1491 | E1491I | E | 6.6e-09 | 3.4e-09 | 1.4e-08 | 8.000E-09 | 5.437E-09 | 3.400E-09 | 1.400E-08 |
| 29811 | 1491 | E1491L | E | 6.3e-09 | 4.8e-09 | 9.1e-09 | 6.733E-09 | 2.183E-09 | 4.800E-09 | 9.100E-09 |
| 29812 | 1491 | E1491K | E | 5.4e-09 | 1.2e-08 | 2.2e-09 | 6.533E-09 | 4.997E-09 | 2.200E-09 | 1.200E-08 |
| 29813 | 1491 | E1491M | E | 3.7e-08 | 8.4e-09 | 9e-09 | 1.813E-08 | 1.634E-08 | 8.400E-09 | 3.700E-08 |
| 29814 | 1491 | E1491F | E | 6.9e-09 | 3e-09 | 1.2e-08 | 7.300E-09 | 4.513E-09 | 3.000E-09 | 1.200E-08 |
| 29815 | 1491 | E1491P | E | 5.3e-09 | 6.6e-09 | 7.4e-09 | 6.433E-09 | 1.060E-09 | 5.300E-09 | 7.400E-09 |
| 29816 | 1491 | E1491S | E | 1.1e-08 | 4.8e-09 | 2.2e-08 | 1.260E-08 | 8.711E-09 | 4.800E-09 | 2.200E-08 |
| 29817 | 1491 | E1491T | E | 1.8e-08 | 7.4e-09 | 1.7e-08 | 1.413E-08 | 5.853E-09 | 7.400E-09 | 1.800E-08 |
| 29818 | 1491 | E1491W | E | 6.4e-09 | 8.3e-09 | 6.4e-09 | 7.033E-09 | 1.097E-09 | 6.400E-09 | 8.300E-09 |
| 29819 | 1491 | E1491Y | E | 2.9e-09 | 4.6e-09 | 1.4e-08 | 7.167E-09 | 5.979E-09 | 2.900E-09 | 1.400E-08 |
| 29820 | 1491 | E1491V | E | 1.3e-08 | 7.3e-09 | 7.5e-09 | 9.267E-09 | 3.235E-09 | 7.300E-09 | 1.300E-08 |
| 29821 | 1492 | P1492A | E | 4.2e-09 | 1.2e-08 | 5.6e-09 | 7.267E-09 | 4.159E-09 | 4.200E-09 | 1.200E-08 |
| 29822 | 1492 | P1492R | E | 3.2e-09 | 4.7e-09 | 4.4e-09 | 4.100E-09 | 7.937E-10 | 3.200E-09 | 4.700E-09 |
| 29823 | 1492 | P1492N | E | 1.2e-08 | 6e-09 | 8.5e-09 | 8.833E-09 | 3.014E-09 | 6.000E-09 | 1.200E-08 |
| 29824 | 1492 | P1492D | E | 1.5e-08 | 1.4e-09 | 4e-09 | 6.800E-09 | 7.219E-09 | 1.400E-09 | 1.500E-08 |
| 29825 | 1492 | P1492C | E | 5.7e-09 | 3.5e-09 | 1.4e-08 | 7.733E-09 | 5.537E-09 | 3.500E-09 | 1.400E-08 |
| 29826 | 1492 | P1492E | E | 9e-09 | 4.3e-09 | 6.6e-09 | 6.633E-09 | 2.350E-09 | 4.300E-09 | 9.000E-09 |
| 29827 | 1492 | P1492Q | E | 6e-09 | 7.9e-09 | 1.2e-08 | 8.633E-09 | 3.066E-09 | 6.000E-09 | 1.200E-08 |
| 29828 | 1492 | P1492G | E | 1.5e-08 | 1e-08 | 8.7e-09 | 1.123E-08 | 3.326E-09 | 8.700E-09 | 1.500E-08 |
| 29829 | 1492 | P1492H | E | 5.4e-09 | 7.5e-09 | 7.9e-09 | 6.933E-09 | 1.343E-09 | 5.400E-09 | 7.900E-09 |
| 29830 | 1492 | P1492I | E | 1.9e-08 | 2.4e-08 | 5.4e-09 | 1.613E-08 | 9.626E-09 | 5.400E-09 | 2.400E-08 |
| 29831 | 1492 | P1492L | E | 2.5e-09 | 1.1e-08 | 7.3e-09 | 6.933E-09 | 4.262E-09 | 2.500E-09 | 1.100E-08 |
| 29832 | 1492 | P1492K | E | 1.2e-08 | 7.2e-09 | 8.8e-09 | 9.333E-09 | 2.444E-09 | 7.200E-09 | 1.200E-08 |
| 29833 | 1492 | P1492M | E | 1.3e-08 | 2.2e-09 | 6.6e-09 | 7.267E-09 | 5.431E-09 | 2.200E-09 | 1.300E-08 |
| 29834 | 1492 | P1492F | E | 6.5e-09 | 2.7e-08 | 5.1e-09 | 1.287E-08 | 1.226E-08 | 5.100E-09 | 2.700E-08 |

| | | | | | | | | | | |
|---|---|---|---|---|---|---|---|---|---|---|
| 29835 | 1492 | P1492P | E | 4.3e-09 | 1.3e-08 | 1e-08 | 9.100E-09 | 4.419E-09 | 4.300E-09 | 1.300E-08 |
| 29836 | 1492 | P1492S | E | 1.1e-08 | 1.4e-08 | 1.6e-09 | 8.867E-09 | 6.469E-09 | 1.600E-09 | 1.400E-08 |
| 29837 | 1492 | P1492T | E | 2.9e-09 | 5.7e-09 | 8e-09 | 5.533E-09 | 2.554E-09 | 2.900E-09 | 8.000E-09 |
| 29838 | 1492 | P1492W | E | 8.7e-09 | 9.4e-09 | 9.5e-09 | 9.200E-09 | 4.359E-10 | 8.700E-09 | 9.500E-09 |
| 29839 | 1492 | P1492Y | E | 8e-09 | 8.7e-09 | 1.8e-09 | 6.167E-09 | 3.798E-09 | 1.800E-09 | 8.700E-09 |
| 29840 | 1492 | P1492V | E | 2.1e-08 | 3.7e-09 | 6.6e-09 | 1.043E-08 | 9.265E-09 | 3.700E-09 | 2.100E-08 |

Table 7: A scalable Her2-Trastuzumab-Pertuzumab binding affinity landscape. In this table, **ResID** represents the amino acid residue number of PDB entry 6OGE, while **Muta** represents the site-specific missense mutation introduced into the 1492 amino acid residues in PDB entry 6OGE, **Min**, **Max**, **Mean** and **Std** represent the minimum, the maximum, the average and the standard deviation of the $K_d$ values calculated using Prodigy, respectively.

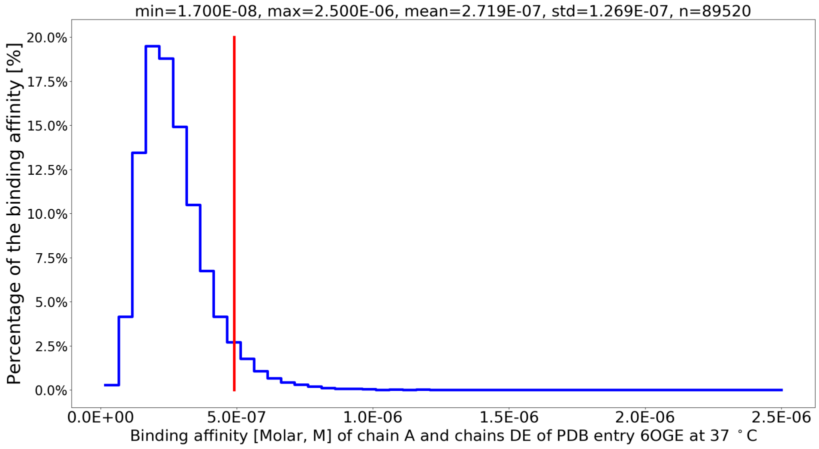


Figure 6: A histogram of the distribution pattern of the Her2-Trastuzumab-Pertuzumab binding affinities between chain A and chain DE (listed in Table 1) of PDB entry 6OGE (with one site-specific missense mutation). In this figure, the vertical red line represents the $K_d$ between chain A (Her2) and chain DE (Trastuzumab FAB, Table 1) of PDB entry 6OGE, i.e., native complex structure of Her2-Trastuzumab-Pertuzumab.

| No. | ResID | Muta | ChainID | $K_{d1}$ (M) | $K_{d2}$ (M) | $K_{d3}$ (M) | Mean (M) | Std (M) | Min (M) | Max (M) |
|---|---|---|---|---|---|---|---|---|---|---|
| 1 | 1 | T1A | A | 3.2e-07 | 3.2e-07 | 3.2e-07 | 3.200E-07 | 6.484E-23 | 3.200E-07 | 3.200E-07 |
| 2 | 1 | T1R | A | 2.7e-07 | 2.7e-07 | 2.7e-07 | 2.700E-07 | 0.000E+00 | 2.700E-07 | 2.700E-07 |
| 3 | 1 | T1N | A | 1.7e-07 | 1.7e-07 | 1.7e-07 | 1.700E-07 | 0.000E+00 | 1.700E-07 | 1.700E-07 |
| 4 | 1 | T1D | A | 4.4e-07 | 4.4e-07 | 4.4e-07 | 4.400E-07 | 0.000E+00 | 4.400E-07 | 4.400E-07 |
| 5 | 1 | T1C | A | 1.4e-07 | 1.4e-07 | 1.4e-07 | 1.400E-07 | 0.000E+00 | 1.400E-07 | 1.400E-07 |
| 6 | 1 | T1E | A | 1.7e-07 | 1.7e-07 | 1.7e-07 | 1.700E-07 | 0.000E+00 | 1.700E-07 | 1.700E-07 |
| 7 | 1 | T1Q | A | 2.5e-07 | 2.5e-07 | 2.5e-07 | 2.500E-07 | 0.000E+00 | 2.500E-07 | 2.500E-07 |
| 8 | 1 | T1G | A | 2.7e-07 | 2.7e-07 | 2.7e-07 | 2.700E-07 | 0.000E+00 | 2.700E-07 | 2.700E-07 |
| 9 | 1 | T1H | A | 2.2e-07 | 2.2e-07 | 2.2e-07 | 2.200E-07 | 0.000E+00 | 2.200E-07 | 2.200E-07 |
| 10 | 1 | T1I | A | 4.9e-07 | 4.9e-07 | 4.9e-07 | 4.900E-07 | 0.000E+00 | 4.900E-07 | 4.900E-07 |
| 11 | 1 | T1L | A | 3.9e-07 | 3.9e-07 | 3.9e-07 | 3.900E-07 | 0.000E+00 | 3.900E-07 | 3.900E-07 |
| 12 | 1 | T1K | A | 1.8e-07 | 1.8e-07 | 1.8e-07 | 1.800E-07 | 0.000E+00 | 1.800E-07 | 1.800E-07 |
| 13 | 1 | T1M | A | 2.5e-07 | 2.5e-07 | 2.5e-07 | 2.500E-07 | 0.000E+00 | 2.500E-07 | 2.500E-07 |
| 14 | 1 | T1F | A | 8.2e-07 | 8.2e-07 | 8.2e-07 | 8.200E-07 | 1.297E-22 | 8.200E-07 | 8.200E-07 |
| 15 | 1 | T1P | A | 2.2e-07 | 2.2e-07 | 2.2e-07 | 2.200E-07 | 0.000E+00 | 2.200E-07 | 2.200E-07 |
| 16 | 1 | T1S | A | 2e-07 | 2e-07 | 2e-07 | 2.000E-07 | 0.000E+00 | 2.000E-07 | 2.000E-07 |
| 17 | 1 | T1T | A | 2e-07 | 2e-07 | 2e-07 | 2.000E-07 | 0.000E+00 | 2.000E-07 | 2.000E-07 |
| 18 | 1 | T1W | A | 2.1e-07 | 2.1e-07 | 2.1e-07 | 2.100E-07 | 0.000E+00 | 2.100E-07 | 2.100E-07 |
| 19 | 1 | T1Y | A | 4.8e-07 | 4.8e-07 | 4.8e-07 | 4.800E-07 | 0.000E+00 | 4.800E-07 | 4.800E-07 |
| 20 | 1 | T1V | A | 2e-07 | 2e-07 | 2e-07 | 2.000E-07 | 0.000E+00 | 2.000E-07 | 2.000E-07 |
| 21 | 2 | Q2A | A | 3.8e-07 | 3.8e-07 | 3.8e-07 | 3.800E-07 | 0.000E+00 | 3.800E-07 | 3.800E-07 |
| 22 | 2 | Q2R | A | 4.6e-07 | 4.6e-07 | 4.6e-07 | 4.600E-07 | 0.000E+00 | 4.600E-07 | 4.600E-07 |
| 23 | 2 | Q2N | A | 2.7e-07 | 2.7e-07 | 2.7e-07 | 2.700E-07 | 0.000E+00 | 2.700E-07 | 2.700E-07 |
| 24 | 2 | Q2D | A | 5.2e-07 | 5.2e-07 | 5.2e-07 | 5.200E-07 | 0.000E+00 | 5.200E-07 | 5.200E-07 |
| 25 | 2 | Q2C | A | 1.9e-07 | 5.6e-07 | 5.6e-07 | 4.367E-07 | 2.136E-07 | 1.900E-07 | 5.600E-07 |
| 26 | 2 | Q2E | A | 1.4e-07 | 4.7e-07 | 4e-07 | 3.367E-07 | 1.739E-07 | 1.400E-07 | 4.700E-07 |
| 27 | 2 | Q2Q | A | 5.3e-07 | 8.4e-08 | 3.3e-07 | 3.147E-07 | 2.234E-07 | 8.400E-08 | 5.300E-07 |
| 28 | 2 | Q2G | A | 3e-07 | 4.1e-07 | 2e-07 | 3.033E-07 | 1.050E-07 | 2.000E-07 | 4.100E-07 |
| 29 | 2 | Q2H | A | 1.8e-07 | 5.7e-07 | 5.7e-07 | 4.400E-07 | 2.252E-07 | 1.800E-07 | 5.700E-07 |
| 30 | 2 | Q2I | A | 9.3e-08 | 1.8e-07 | 9.3e-08 | 1.220E-07 | 5.023E-08 | 9.300E-08 | 1.800E-07 |
| 31 | 2 | Q2L | A | 3.8e-07 | 2.7e-07 | 3.8e-07 | 3.433E-07 | 6.351E-08 | 2.700E-07 | 3.800E-07 |
| 32 | 2 | Q2K | A | 2.2e-07 | 2.2e-07 | 3.7e-07 | 2.700E-07 | 8.660E-08 | 2.200E-07 | 3.700E-07 |
| 33 | 2 | Q2M | A | 4.7e-07 | 4.7e-07 | 2.4e-07 | 3.933E-07 | 1.328E-07 | 2.400E-07 | 4.700E-07 |
| 34 | 2 | Q2F | A | 1.6e-07 | 3.7e-07 | 1.6e-07 | 2.300E-07 | 1.212E-07 | 1.600E-07 | 3.700E-07 |
| 35 | 2 | Q2P | A | 4.2e-07 | 2.6e-07 | 4.2e-07 | 3.667E-07 | 9.238E-08 | 2.600E-07 | 4.200E-07 |
| 36 | 2 | Q2S | A | 4.6e-07 | 4.6e-07 | 4.6e-07 | 4.600E-07 | 0.000E+00 | 4.600E-07 | 4.600E-07 |
| 37 | 2 | Q2T | A | 3.5e-07 | 1.8e-07 | 1.3e-07 | 2.200E-07 | 1.153E-07 | 1.300E-07 | 3.500E-07 |
| 38 | 2 | Q2W | A | 3.7e-07 | 2.2e-07 | 2.3e-07 | 2.733E-07 | 8.386E-08 | 2.200E-07 | 3.700E-07 |
| 39 | 2 | Q2Y | A | 1.6e-07 | 1.8e-07 | 5.8e-07 | 3.067E-07 | 2.369E-07 | 1.600E-07 | 5.800E-07 |
| 40 | 2 | Q2V | A | 2.4e-07 | 1.3e-07 | 1.3e-07 | 1.667E-07 | 6.351E-08 | 1.300E-07 | 2.400E-07 |
| 41 | 3 | V3A | A | 2.6e-07 | 2.6e-07 | 3.3e-07 | 2.833E-07 | 4.041E-08 | 2.600E-07 | 3.300E-07 |
| 42 | 3 | V3R | A | 2.9e-07 | 4.7e-07 | 4.7e-07 | 4.100E-07 | 1.039E-07 | 2.900E-07 | 4.700E-07 |
| 43 | 3 | V3N | A | 2.9e-07 | 2.9e-07 | 2.9e-07 | 2.900E-07 | 0.000E+00 | 2.900E-07 | 2.900E-07 |
| 44 | 3 | V3D | A | 2e-07 | 2e-07 | 2e-07 | 2.000E-07 | 0.000E+00 | 2.000E-07 | 2.000E-07 |
| 45 | 3 | V3C | A | 4.4e-07 | 4.4e-07 | 2e-07 | 3.600E-07 | 1.386E-07 | 2.000E-07 | 4.400E-07 |
| 46 | 3 | V3E | A | 1.5e-07 | 1.5e-07 | 2.2e-07 | 1.733E-07 | 4.041E-08 | 1.500E-07 | 2.200E-07 |
| 47 | 3 | V3Q | A | 3.7e-07 | 3.7e-07 | 4.1e-07 | 3.833E-07 | 2.309E-08 | 3.700E-07 | 4.100E-07 |
| 48 | 3 | V3G | A | 1.5e-07 | 1.5e-07 | 1.5e-07 | 1.500E-07 | 0.000E+00 | 1.500E-07 | 1.500E-07 |
| 49 | 3 | V3H | A | 2.3e-07 | 2.5e-07 | 2.1e-07 | 2.300E-07 | 2.000E-08 | 2.100E-07 | 2.500E-07 |
| 50 | 3 | V3I | A | 1.4e-07 | 3.2e-07 | 1.4e-07 | 2.000E-07 | 1.039E-07 | 1.400E-07 | 3.200E-07 |
| 51 | 3 | V3L | A | 3.3e-07 | 1.9e-07 | 1.8e-07 | 2.333E-07 | 8.386E-08 | 1.800E-07 | 3.300E-07 |
| 52 | 3 | V3K | A | 4.8e-07 | 2.3e-07 | 2.4e-07 | 3.167E-07 | 1.415E-07 | 2.300E-07 | 4.800E-07 |
| 53 | 3 | V3M | A | 2.7e-07 | 9.8e-08 | 1.8e-07 | 1.827E-07 | 8.603E-08 | 9.800E-08 | 2.700E-07 |
| 54 | 3 | V3F | A | 3.3e-07 | 1.1e-07 | 3.6e-07 | 2.667E-07 | 1.365E-07 | 1.100E-07 | 3.600E-07 |
| 55 | 3 | V3P | A | 2.6e-07 | 1.5e-07 | 1.5e-07 | 1.867E-07 | 6.351E-08 | 1.500E-07 | 2.600E-07 |
| 56 | 3 | V3S | A | 2.5e-07 | 2.5e-07 | 1.4e-07 | 2.133E-07 | 6.351E-08 | 1.400E-07 | 2.500E-07 |
| 57 | 3 | V3T | A | 1.1e-07 | 2.1e-07 | 1.8e-07 | 1.667E-07 | 5.132E-08 | 1.100E-07 | 2.100E-07 |
| 58 | 3 | V3W | A | 3.1e-07 | 3.1e-07 | 1.7e-07 | 2.633E-07 | 8.083E-08 | 1.700E-07 | 3.100E-07 |
| 59 | 3 | V3Y | A | 2.1e-07 | 2.6e-07 | 2.8e-07 | 2.500E-07 | 3.606E-08 | 2.100E-07 | 2.800E-07 |
| 60 | 3 | V3V | A | 2.7e-07 | 2.4e-07 | 2.8e-07 | 2.633E-07 | 2.082E-08 | 2.400E-07 | 2.800E-07 |
| 61 | 4 | C4A | A | 2.9e-07 | 6.1e-07 | 2.2e-07 | 3.733E-07 | 2.079E-07 | 2.200E-07 | 6.100E-07 |
| 62 | 4 | C4R | A | 1.9e-07 | 1.5e-07 | 1.6e-07 | 1.667E-07 | 2.082E-08 | 1.500E-07 | 1.900E-07 |
| 63 | 4 | C4N | A | 2.9e-07 | 3.1e-07 | 3.1e-07 | 3.033E-07 | 1.155E-08 | 2.900E-07 | 3.100E-07 |
| 64 | 4 | C4D | A | 2.1e-07 | 2.9e-07 | 1.6e-07 | 2.200E-07 | 6.557E-08 | 1.600E-07 | 2.900E-07 |
| 65 | 4 | C4C | A | 3.2e-07 | 3.2e-07 | 3.2e-07 | 3.200E-07 | 6.484E-23 | 3.200E-07 | 3.200E-07 |
| 66 | 4 | C4E | A | 2.6e-07 | 2.1e-07 | 1.6e-07 | 2.100E-07 | 5.000E-08 | 1.600E-07 | 2.600E-07 |
| 67 | 4 | C4Q | A | 1.2e-07 | 2.7e-07 | 1.3e-07 | 1.733E-07 | 8.386E-08 | 1.200E-07 | 2.700E-07 |
| 68 | 4 | C4G | A | 4.5e-07 | 3.2e-07 | 2.5e-07 | 3.400E-07 | 1.015E-07 | 2.500E-07 | 4.500E-07 |
| 69 | 4 | C4H | A | 4.2e-07 | 2.5e-07 | 1.6e-07 | 2.767E-07 | 1.320E-07 | 1.600E-07 | 4.200E-07 |
| 70 | 4 | C4I | A | 1.7e-07 | 1.7e-07 | 3.8e-07 | 2.400E-07 | 1.212E-07 | 1.700E-07 | 3.800E-07 |
| 71 | 4 | C4L | A | 1.3e-07 | 2e-07 | 2.4e-07 | 1.900E-07 | 5.568E-08 | 1.300E-07 | 2.400E-07 |
| 72 | 4 | C4K | A | 2.6e-07 | 2.6e-07 | 3.9e-07 | 3.033E-07 | 7.506E-08 | 2.600E-07 | 3.900E-07 |
| 73 | 4 | C4M | A | 3.9e-07 | 1.8e-07 | 1.7e-07 | 2.467E-07 | 1.242E-07 | 1.700E-07 | 3.900E-07 |
| 74 | 4 | C4F | A | 2.6e-07 | 2.2e-07 | 3.1e-07 | 2.633E-07 | 4.509E-08 | 2.200E-07 | 3.100E-07 |
| 75 | 4 | C4P | A | 2e-07 | 1.8e-07 | 3.2e-07 | 2.333E-07 | 7.572E-08 | 1.800E-07 | 3.200E-07 |
| 76 | 4 | C4S | A | 3.9e-07 | 8.7e-08 | 2.4e-07 | 2.390E-07 | 1.515E-07 | 8.700E-08 | 3.900E-07 |
| 77 | 4 | C4T | A | 1.9e-07 | 2.5e-07 | 4e-07 | 2.800E-07 | 1.082E-07 | 1.900E-07 | 4.000E-07 |
| 78 | 4 | C4W | A | 2.8e-07 | 1.8e-07 | 2e-07 | 2.200E-07 | 5.292E-08 | 1.800E-07 | 2.800E-07 |
| 79 | 4 | C4Y | A | 1.9e-07 | 4.1e-07 | 1.4e-07 | 2.467E-07 | 1.436E-07 | 1.400E-07 | 4.100E-07 |
| 80 | 4 | C4V | A | 4.5e-07 | 2.3e-07 | 2.3e-07 | 3.033E-07 | 1.270E-07 | 2.300E-07 | 4.500E-07 |
| 81 | 5 | T5A | A | 2.3e-07 | 3.1e-07 | 3.6e-07 | 3.000E-07 | 6.557E-08 | 2.300E-07 | 3.600E-07 |
| 82 | 5 | T5R | A | 1.6e-07 | 3.2e-07 | 6.6e-07 | 3.800E-07 | 2.553E-07 | 1.600E-07 | 6.600E-07 |
| 83 | 5 | T5N | A | 1.4e-07 | 4.3e-07 | 1.6e-07 | 2.433E-07 | 1.620E-07 | 1.400E-07 | 4.300E-07 |
| 84 | 5 | T5D | A | 4.1e-07 | 1.3e-07 | 1.3e-07 | 2.233E-07 | 1.617E-07 | 1.300E-07 | 4.100E-07 |

| | | | | | | | | | | |
|---|---|---|---|---|---|---|---|---|---|---|
| 85 | 5 | T5C | A | 1.9e-07 | 3.4e-07 | 1.3e-07 | 2.200E-07 | 1.082E-07 | 1.300E-07 | 3.400E-07 |
| 86 | 5 | T5E | A | 5.3e-07 | 3.1e-07 | 4.6e-07 | 4.333E-07 | 1.124E-07 | 3.100E-07 | 5.300E-07 |
| 87 | 5 | T5Q | A | 2e-07 | 1.2e-07 | 1.9e-07 | 1.700E-07 | 4.359E-08 | 1.200E-07 | 2.000E-07 |
| 88 | 5 | T5G | A | 2.6e-07 | 2e-07 | 2.5e-07 | 2.367E-07 | 3.215E-08 | 2.000E-07 | 2.600E-07 |
| 89 | 5 | T5H | A | 2.2e-07 | 2.3e-07 | 7.3e-07 | 3.933E-07 | 2.916E-07 | 2.200E-07 | 7.300E-07 |
| 90 | 5 | T5I | A | 3.6e-07 | 2.4e-07 | 2e-07 | 2.667E-07 | 8.327E-08 | 2.000E-07 | 3.600E-07 |
| 91 | 5 | T5L | A | 3.2e-07 | 3.2e-07 | 1.9e-07 | 2.767E-07 | 7.506E-08 | 1.900E-07 | 3.200E-07 |
| 92 | 5 | T5K | A | 1.4e-07 | 4e-07 | 3.1e-07 | 2.833E-07 | 1.320E-07 | 1.400E-07 | 4.000E-07 |
| 93 | 5 | T5M | A | 2.6e-07 | 1.7e-07 | 3e-07 | 2.433E-07 | 6.658E-08 | 1.700E-07 | 3.000E-07 |
| 94 | 5 | T5F | A | 2.2e-07 | 2.9e-07 | 2.9e-07 | 2.667E-07 | 4.041E-08 | 2.200E-07 | 2.900E-07 |
| 95 | 5 | T5P | A | 3e-07 | 2.5e-07 | 5.5e-07 | 3.667E-07 | 1.607E-07 | 2.500E-07 | 5.500E-07 |
| 96 | 5 | T5S | A | 2.7e-07 | 1.2e-07 | 3.5e-07 | 2.467E-07 | 1.168E-07 | 1.200E-07 | 3.500E-07 |
| 97 | 5 | T5T | A | 1.5e-07 | 3.7e-07 | 3.6e-07 | 2.933E-07 | 1.242E-07 | 1.500E-07 | 3.700E-07 |
| 98 | 5 | T5W | A | 2.3e-07 | 3e-07 | 2.8e-07 | 2.700E-07 | 3.606E-08 | 2.300E-07 | 3.000E-07 |
| 99 | 5 | T5Y | A | 1.6e-07 | 2.2e-07 | 3.3e-07 | 2.367E-07 | 8.622E-08 | 1.600E-07 | 3.300E-07 |
| 100 | 5 | T5V | A | 1.9e-07 | 2.9e-07 | 4.2e-07 | 3.000E-07 | 1.153E-07 | 1.900E-07 | 4.200E-07 |
| 101 | 6 | G6A | A | 2.2e-07 | 2.5e-07 | 1.3e-07 | 2.000E-07 | 6.245E-08 | 1.300E-07 | 2.500E-07 |
| 102 | 6 | G6R | A | 4.2e-07 | 1.6e-07 | 3.2e-07 | 3.000E-07 | 1.311E-07 | 1.600E-07 | 4.200E-07 |
| 103 | 6 | G6N | A | 3.3e-07 | 1.9e-07 | 4.4e-07 | 3.200E-07 | 1.253E-07 | 1.900E-07 | 4.400E-07 |
| 104 | 6 | G6D | A | 3e-07 | 2e-07 | 3.4e-07 | 2.800E-07 | 7.211E-08 | 2.000E-07 | 3.400E-07 |
| 105 | 6 | G6C | A | 2e-07 | 7.9e-08 | 2.9e-07 | 1.897E-07 | 1.059E-07 | 7.900E-08 | 2.900E-07 |
| 106 | 6 | G6E | A | 1.1e-07 | 4.1e-07 | 3.3e-07 | 2.833E-07 | 1.553E-07 | 1.100E-07 | 4.100E-07 |
| 107 | 6 | G6Q | A | 2.1e-07 | 4.6e-07 | 2.1e-07 | 2.933E-07 | 1.443E-07 | 2.100E-07 | 4.600E-07 |
| 108 | 6 | G6G | A | 3.5e-07 | 8.7e-08 | 5.6e-08 | 1.643E-07 | 1.615E-07 | 5.600E-08 | 3.500E-07 |
| 109 | 6 | G6H | A | 3.6e-07 | 3.2e-07 | 1.5e-07 | 2.767E-07 | 1.115E-07 | 1.500E-07 | 3.600E-07 |
| 110 | 6 | G6I | A | 3e-07 | 1.3e-07 | 3.2e-07 | 2.500E-07 | 1.044E-07 | 1.300E-07 | 3.200E-07 |
| 111 | 6 | G6L | A | 4.1e-07 | 3.5e-07 | 3.8e-07 | 3.800E-07 | 3.000E-08 | 3.500E-07 | 4.100E-07 |
| 112 | 6 | G6K | A | 1.1e-07 | 4e-07 | 1.3e-07 | 2.133E-07 | 1.620E-07 | 1.100E-07 | 4.000E-07 |
| 113 | 6 | G6M | A | 3.2e-07 | 1.5e-07 | 4.1e-07 | 2.933E-07 | 1.320E-07 | 1.500E-07 | 4.100E-07 |
| 114 | 6 | G6F | A | 3.9e-07 | 3.3e-07 | 2.7e-07 | 3.300E-07 | 6.000E-08 | 2.700E-07 | 3.900E-07 |
| 115 | 6 | G6P | A | 2.5e-07 | 3.6e-07 | 1.3e-07 | 2.467E-07 | 1.150E-07 | 1.300E-07 | 3.600E-07 |
| 116 | 6 | G6S | A | 4.2e-07 | 3e-07 | 1.9e-07 | 3.033E-07 | 1.150E-07 | 1.900E-07 | 4.200E-07 |
| 117 | 6 | G6T | A | 4.5e-07 | 3.7e-07 | 5e-07 | 4.400E-07 | 6.557E-08 | 3.700E-07 | 5.000E-07 |
| 118 | 6 | G6W | A | 2.6e-07 | 4.3e-07 | 2.7e-07 | 3.200E-07 | 9.539E-08 | 2.600E-07 | 4.300E-07 |
| 119 | 6 | G6Y | A | 3.6e-07 | 3.3e-07 | 2.7e-07 | 3.200E-07 | 4.583E-08 | 2.700E-07 | 3.600E-07 |
| 120 | 6 | G6V | A | 1.5e-07 | 1.9e-07 | 2.2e-07 | 1.867E-07 | 3.512E-08 | 1.500E-07 | 2.200E-07 |
| 121 | 7 | T7A | A | 2.8e-07 | 2.2e-07 | 3.9e-07 | 2.967E-07 | 8.622E-08 | 2.200E-07 | 3.900E-07 |
| 122 | 7 | T7R | A | 1.6e-07 | 4e-07 | 2.3e-07 | 2.633E-07 | 1.234E-07 | 1.600E-07 | 4.000E-07 |
| 123 | 7 | T7N | A | 1.5e-07 | 1.3e-07 | 1.5e-07 | 1.433E-07 | 1.155E-08 | 1.300E-07 | 1.500E-07 |
| 124 | 7 | T7D | A | 3e-07 | 2.7e-07 | 1.4e-07 | 2.367E-07 | 8.505E-08 | 1.400E-07 | 3.000E-07 |
| 125 | 7 | T7C | A | 3e-07 | 4.6e-07 | 2.2e-07 | 3.267E-07 | 1.222E-07 | 2.200E-07 | 4.600E-07 |
| 126 | 7 | T7E | A | 1.7e-07 | 2.9e-07 | 1.9e-07 | 2.167E-07 | 6.429E-08 | 1.700E-07 | 2.900E-07 |
| 127 | 7 | T7Q | A | 2.9e-07 | 1.5e-07 | 5.6e-07 | 3.333E-07 | 2.084E-07 | 1.500E-07 | 5.600E-07 |
| 128 | 7 | T7G | A | 1.6e-07 | 1.9e-07 | 1.9e-07 | 1.800E-07 | 1.732E-08 | 1.600E-07 | 1.900E-07 |
| 129 | 7 | T7H | A | 6.5e-07 | 2.5e-07 | 2.5e-07 | 3.833E-07 | 2.309E-07 | 2.500E-07 | 6.500E-07 |
| 130 | 7 | T7I | A | 4.1e-07 | 2.6e-07 | 3.2e-07 | 3.300E-07 | 7.550E-08 | 2.600E-07 | 4.100E-07 |
| 131 | 7 | T7L | A | 3.6e-07 | 1.6e-07 | 2.1e-07 | 2.433E-07 | 1.041E-07 | 1.600E-07 | 3.600E-07 |
| 132 | 7 | T7K | A | 1.6e-07 | 1.2e-07 | 1.8e-07 | 1.533E-07 | 3.055E-08 | 1.200E-07 | 1.800E-07 |
| 133 | 7 | T7M | A | 2.6e-07 | 2.3e-07 | 1.7e-07 | 2.200E-07 | 4.583E-08 | 1.700E-07 | 2.600E-07 |
| 134 | 7 | T7F | A | 1.1e-07 | 1.5e-07 | 2.5e-07 | 1.700E-07 | 7.211E-08 | 1.100E-07 | 2.500E-07 |
| 135 | 7 | T7P | A | 3.2e-07 | 2e-07 | 1.2e-07 | 2.133E-07 | 1.007E-07 | 1.200E-07 | 3.200E-07 |
| 136 | 7 | T7S | A | 1.4e-07 | 3.3e-07 | 2.3e-07 | 2.333E-07 | 9.504E-08 | 1.400E-07 | 3.300E-07 |
| 137 | 7 | T7T | A | 2.4e-07 | 4.1e-07 | 2.9e-07 | 3.133E-07 | 8.737E-08 | 2.400E-07 | 4.100E-07 |
| 138 | 7 | T7W | A | 2.3e-07 | 3e-07 | 2.5e-07 | 2.600E-07 | 3.606E-08 | 2.300E-07 | 3.000E-07 |
| 139 | 7 | T7Y | A | 2.2e-07 | 1.5e-07 | 2.7e-07 | 2.133E-07 | 6.028E-08 | 1.500E-07 | 2.700E-07 |
| 140 | 7 | T7V | A | 2e-07 | 2.9e-07 | 1.1e-07 | 2.000E-07 | 9.000E-08 | 1.100E-07 | 2.900E-07 |
| 141 | 8 | D8A | A | 9.3e-08 | 3.2e-07 | 1.6e-07 | 1.910E-07 | 1.166E-07 | 9.300E-08 | 3.200E-07 |
| 142 | 8 | D8R | A | 2.7e-07 | 4.3e-07 | 4.3e-07 | 3.767E-07 | 9.238E-08 | 2.700E-07 | 4.300E-07 |
| 143 | 8 | D8N | A | 4.3e-07 | 1.3e-07 | 2.3e-07 | 2.633E-07 | 1.528E-07 | 1.300E-07 | 4.300E-07 |
| 144 | 8 | D8D | A | 2.3e-07 | 1.5e-07 | 3.3e-07 | 2.367E-07 | 9.018E-08 | 1.500E-07 | 3.300E-07 |
| 145 | 8 | D8C | A | 3.6e-07 | 4.2e-07 | 2.3e-07 | 3.367E-07 | 9.713E-08 | 2.300E-07 | 4.200E-07 |
| 146 | 8 | D8E | A | 2e-07 | 1.9e-07 | 3.2e-07 | 2.367E-07 | 7.234E-08 | 1.900E-07 | 3.200E-07 |
| 147 | 8 | D8Q | A | 2.2e-07 | 2.6e-07 | 1.7e-07 | 2.167E-07 | 4.509E-08 | 1.700E-07 | 2.600E-07 |
| 148 | 8 | D8G | A | 5.4e-07 | 3.4e-07 | 4.7e-07 | 4.500E-07 | 1.015E-07 | 3.400E-07 | 5.400E-07 |
| 149 | 8 | D8H | A | 6.7e-08 | 3.6e-07 | 2.7e-07 | 2.323E-07 | 1.501E-07 | 6.700E-08 | 3.600E-07 |
| 150 | 8 | D8I | A | 2.1e-07 | 4.9e-07 | 1.8e-07 | 2.933E-07 | 1.710E-07 | 1.800E-07 | 4.900E-07 |
| 151 | 8 | D8L | A | 2.2e-07 | 3.4e-07 | 2.8e-07 | 2.800E-07 | 6.000E-08 | 2.200E-07 | 3.400E-07 |
| 152 | 8 | D8K | A | 1.3e-07 | 1.9e-07 | 3.4e-07 | 2.200E-07 | 1.082E-07 | 1.300E-07 | 3.400E-07 |
| 153 | 8 | D8M | A | 1.7e-07 | 2.3e-07 | 1.4e-07 | 1.800E-07 | 4.583E-08 | 1.400E-07 | 2.300E-07 |
| 154 | 8 | D8F | A | 2.3e-07 | 9.8e-08 | 2.3e-07 | 1.860E-07 | 7.621E-08 | 9.800E-08 | 2.300E-07 |
| 155 | 8 | D8P | A | 3e-07 | 2e-07 | 3.4e-07 | 2.800E-07 | 7.211E-08 | 2.000E-07 | 3.400E-07 |
| 156 | 8 | D8S | A | 2.6e-07 | 3.5e-07 | 3.1e-07 | 3.067E-07 | 4.509E-08 | 2.600E-07 | 3.500E-07 |
| 157 | 8 | D8T | A | 3.4e-07 | 1.9e-07 | 2.4e-07 | 2.567E-07 | 7.638E-08 | 1.900E-07 | 3.400E-07 |
| 158 | 8 | D8W | A | 1.7e-07 | 2.9e-07 | 3.9e-07 | 2.833E-07 | 1.102E-07 | 1.700E-07 | 3.900E-07 |
| 159 | 8 | D8Y | A | 2.4e-07 | 3e-07 | 3.2e-07 | 2.867E-07 | 4.163E-08 | 2.400E-07 | 3.200E-07 |
| 160 | 8 | D8V | A | 2.6e-07 | 1.9e-07 | 3.7e-07 | 2.733E-07 | 9.074E-08 | 1.900E-07 | 3.700E-07 |
| 161 | 9 | M9A | A | 3.4e-07 | 8.9e-08 | 2e-07 | 2.097E-07 | 1.258E-07 | 8.900E-08 | 3.400E-07 |
| 162 | 9 | M9R | A | 3.9e-07 | 3.7e-07 | 2.4e-07 | 3.333E-07 | 8.145E-08 | 2.400E-07 | 3.900E-07 |
| 163 | 9 | M9N | A | 3e-07 | 2.3e-07 | 1.6e-07 | 2.300E-07 | 7.000E-08 | 1.600E-07 | 3.000E-07 |
| 164 | 9 | M9D | A | 1.8e-07 | 2.1e-07 | 3.4e-07 | 2.433E-07 | 8.505E-08 | 1.800E-07 | 3.400E-07 |
| 165 | 9 | M9C | A | 3.8e-07 | 2.9e-07 | 3.1e-07 | 3.267E-07 | 4.726E-08 | 2.900E-07 | 3.800E-07 |
| 166 | 9 | M9E | A | 5e-07 | 1.4e-07 | 2.4e-07 | 2.933E-07 | 1.858E-07 | 1.400E-07 | 5.000E-07 |
| 167 | 9 | M9Q | A | 1.3e-07 | 2.6e-07 | 1.4e-07 | 1.767E-07 | 7.234E-08 | 1.300E-07 | 2.600E-07 |
| 168 | 9 | M9G | A | 3.3e-07 | 8.1e-08 | 2.4e-07 | 2.170E-07 | 1.261E-07 | 8.100E-08 | 3.300E-07 |
| 169 | 9 | M9H | A | 1.5e-07 | 1.5e-07 | 1.4e-07 | 1.467E-07 | 5.774E-09 | 1.400E-07 | 1.500E-07 |

| | | | | | | | | | | |
|---|---|---|---|---|---|---|---|---|---|---|
| 170 | 9 | M9I | A | 2.1e-07 | 1.9e-07 | 1.9e-07 | 1.967E-07 | 1.155E-08 | 1.900E-07 | 2.100E-07 |
| 171 | 9 | M9L | A | 2.4e-07 | 5.9e-07 | 2.2e-07 | 3.500E-07 | 2.081E-07 | 2.200E-07 | 5.900E-07 |
| 172 | 9 | M9K | A | 1.1e-07 | 1.7e-07 | 2.8e-07 | 1.867E-07 | 8.622E-08 | 1.100E-07 | 2.800E-07 |
| 173 | 9 | M9M | A | 2.9e-07 | 2.5e-07 | 2e-07 | 2.467E-07 | 4.509E-08 | 2.000E-07 | 2.900E-07 |
| 174 | 9 | M9F | A | 1.5e-07 | 4e-07 | 2.1e-07 | 2.533E-07 | 1.305E-07 | 1.500E-07 | 4.000E-07 |
| 175 | 9 | M9P | A | 1.4e-07 | 2.5e-07 | 2.7e-07 | 2.200E-07 | 7.000E-08 | 1.400E-07 | 2.700E-07 |
| 176 | 9 | M9S | A | 2.4e-07 | 1.7e-07 | 6.8e-08 | 1.593E-07 | 8.649E-08 | 6.800E-08 | 2.400E-07 |
| 177 | 9 | M9T | A | 4.9e-07 | 2.4e-07 | 2.2e-07 | 3.167E-07 | 1.504E-07 | 2.200E-07 | 4.900E-07 |
| 178 | 9 | M9W | A | 1.4e-07 | 1.7e-07 | 1.9e-07 | 1.667E-07 | 2.517E-08 | 1.400E-07 | 1.900E-07 |
| 179 | 9 | M9Y | A | 3.6e-07 | 2e-07 | 2.2e-07 | 2.600E-07 | 8.718E-08 | 2.000E-07 | 3.600E-07 |
| 180 | 9 | M9V | A | 2.9e-07 | 2e-07 | 3.4e-07 | 2.767E-07 | 7.095E-08 | 2.000E-07 | 3.400E-07 |
| 181 | 10 | K10A | A | 3e-07 | 1.1e-07 | 1.5e-07 | 1.867E-07 | 1.002E-07 | 1.100E-07 | 3.000E-07 |
| 182 | 10 | K10R | A | 4.5e-07 | 2.3e-07 | 3.8e-07 | 3.533E-07 | 1.124E-07 | 2.300E-07 | 4.500E-07 |
| 183 | 10 | K10N | A | 1.7e-07 | 2.1e-07 | 2.4e-07 | 2.067E-07 | 3.512E-08 | 1.700E-07 | 2.400E-07 |
| 184 | 10 | K10D | A | 1e-07 | 3.5e-07 | 1.6e-07 | 2.033E-07 | 1.305E-07 | 1.000E-07 | 3.500E-07 |
| 185 | 10 | K10C | A | 1.3e-07 | 8.9e-08 | 1.1e-07 | 1.097E-07 | 2.050E-08 | 8.900E-08 | 1.300E-07 |
| 186 | 10 | K10E | A | 4e-07 | 1.9e-07 | 1.8e-07 | 2.567E-07 | 1.242E-07 | 1.800E-07 | 4.000E-07 |
| 187 | 10 | K10Q | A | 2.2e-07 | 2.7e-07 | 4.4e-07 | 3.100E-07 | 1.153E-07 | 2.200E-07 | 4.400E-07 |
| 188 | 10 | K10G | A | 3e-07 | 2e-07 | 3.1e-07 | 2.700E-07 | 6.083E-08 | 2.000E-07 | 3.100E-07 |
| 189 | 10 | K10H | A | 1.8e-07 | 2.4e-07 | 3.8e-07 | 2.667E-07 | 1.026E-07 | 1.800E-07 | 3.800E-07 |
| 190 | 10 | K10I | A | 4.1e-07 | 2.9e-07 | 9e-08 | 2.633E-07 | 1.617E-07 | 9.000E-08 | 4.100E-07 |
| 191 | 10 | K10L | A | 1.9e-07 | 3.3e-07 | 2.5e-07 | 2.567E-07 | 7.024E-08 | 1.900E-07 | 3.300E-07 |
| 192 | 10 | K10K | A | 2.2e-07 | 5e-07 | 1.5e-07 | 2.900E-07 | 1.852E-07 | 1.500E-07 | 5.000E-07 |
| 193 | 10 | K10M | A | 3.1e-07 | 1.1e-07 | 1.9e-07 | 2.033E-07 | 1.007E-07 | 1.100E-07 | 3.100E-07 |
| 194 | 10 | K10F | A | 3e-07 | 3e-07 | 2.7e-07 | 2.900E-07 | 1.732E-08 | 2.700E-07 | 3.000E-07 |
| 195 | 10 | K10P | A | 3.2e-07 | 3.3e-07 | 1.5e-07 | 2.667E-07 | 1.012E-07 | 1.500E-07 | 3.300E-07 |
| 196 | 10 | K10S | A | 1.1e-07 | 3e-07 | 1.9e-07 | 2.000E-07 | 9.539E-08 | 1.100E-07 | 3.000E-07 |
| 197 | 10 | K10T | A | 2.8e-07 | 3.6e-07 | 4.7e-07 | 3.700E-07 | 9.539E-08 | 2.800E-07 | 4.700E-07 |
| 198 | 10 | K10W | A | 2.4e-07 | 1.5e-07 | 3.2e-07 | 2.367E-07 | 8.505E-08 | 1.500E-07 | 3.200E-07 |
| 199 | 10 | K10Y | A | 3e-07 | 2.1e-07 | 1.3e-07 | 2.133E-07 | 8.505E-08 | 1.300E-07 | 3.000E-07 |
| 200 | 10 | K10V | A | 1.4e-07 | 2.4e-07 | 2.7e-07 | 2.167E-07 | 6.807E-08 | 1.400E-07 | 2.700E-07 |
| 201 | 11 | L11A | A | 4e-07 | 4e-07 | 3.5e-07 | 3.833E-07 | 2.887E-08 | 3.500E-07 | 4.000E-07 |
| 202 | 11 | L11R | A | 1.1e-07 | 1.7e-07 | 3.3e-07 | 2.033E-07 | 1.137E-07 | 1.100E-07 | 3.300E-07 |
| 203 | 11 | L11N | A | 2.2e-07 | 5e-07 | 2e-07 | 3.067E-07 | 1.677E-07 | 2.000E-07 | 5.000E-07 |
| 204 | 11 | L11D | A | 2.3e-07 | 2.9e-07 | 3.5e-07 | 2.900E-07 | 6.000E-08 | 2.300E-07 | 3.500E-07 |
| 205 | 11 | L11C | A | 3.9e-07 | 2.6e-07 | 1.3e-07 | 2.600E-07 | 1.300E-07 | 1.300E-07 | 3.900E-07 |
| 206 | 11 | L11E | A | 1.9e-07 | 1.9e-07 | 2.2e-07 | 2.000E-07 | 1.732E-08 | 1.900E-07 | 2.200E-07 |
| 207 | 11 | L11Q | A | 2.7e-07 | 1.6e-07 | 3.9e-07 | 2.733E-07 | 1.150E-07 | 1.600E-07 | 3.900E-07 |
| 208 | 11 | L11G | A | 3.1e-07 | 2.3e-07 | 2.4e-07 | 2.600E-07 | 4.359E-08 | 2.300E-07 | 3.100E-07 |
| 209 | 11 | L11H | A | 1.9e-07 | 2.6e-07 | 3.5e-07 | 2.667E-07 | 8.021E-08 | 1.900E-07 | 3.500E-07 |
| 210 | 11 | L11I | A | 1.9e-07 | 1.8e-07 | 1.8e-07 | 1.833E-07 | 5.774E-09 | 1.800E-07 | 1.900E-07 |
| 211 | 11 | L11L | A | 2.8e-07 | 3.3e-07 | 4.1e-07 | 3.400E-07 | 6.557E-08 | 2.800E-07 | 4.100E-07 |
| 212 | 11 | L11K | A | 5.1e-07 | 3e-07 | 1.5e-07 | 3.200E-07 | 1.808E-07 | 1.500E-07 | 5.100E-07 |
| 213 | 11 | L11M | A | 2.9e-07 | 4.2e-07 | 3.3e-07 | 3.467E-07 | 6.658E-08 | 2.900E-07 | 4.200E-07 |
| 214 | 11 | L11F | A | 2.9e-07 | 1.8e-07 | 3.1e-07 | 2.600E-07 | 7.000E-08 | 1.800E-07 | 3.100E-07 |
| 215 | 11 | L11P | A | 3.4e-07 | 2.3e-07 | 2.4e-07 | 2.700E-07 | 6.083E-08 | 2.300E-07 | 3.400E-07 |
| 216 | 11 | L11S | A | 3.5e-07 | 2.5e-07 | 2.8e-07 | 2.933E-07 | 5.132E-08 | 2.500E-07 | 3.500E-07 |
| 217 | 11 | L11T | A | 1.7e-07 | 5.6e-07 | 1.5e-07 | 2.933E-07 | 2.312E-07 | 1.500E-07 | 5.600E-07 |
| 218 | 11 | L11W | A | 2.3e-07 | 1.1e-07 | 5.2e-07 | 2.867E-07 | 2.108E-07 | 1.100E-07 | 5.200E-07 |
| 219 | 11 | L11Y | A | 2.1e-07 | 2e-07 | 4.7e-07 | 2.933E-07 | 1.531E-07 | 2.000E-07 | 4.700E-07 |
| 220 | 11 | L11V | A | 1.3e-07 | 2.4e-07 | 3.6e-07 | 2.433E-07 | 1.150E-07 | 1.300E-07 | 3.600E-07 |
| 221 | 12 | R12A | A | 9.9e-08 | 2.1e-07 | 1e-07 | 1.363E-07 | 6.380E-08 | 9.900E-08 | 2.100E-07 |
| 222 | 12 | R12R | A | 1.7e-07 | 2.4e-07 | 2.7e-07 | 2.267E-07 | 5.132E-08 | 1.700E-07 | 2.700E-07 |
| 223 | 12 | R12N | A | 2.7e-07 | 1.3e-07 | 2.6e-07 | 2.200E-07 | 7.810E-08 | 1.300E-07 | 2.700E-07 |
| 224 | 12 | R12D | A | 2.4e-07 | 3.7e-07 | 1.7e-07 | 2.600E-07 | 1.015E-07 | 1.700E-07 | 3.700E-07 |
| 225 | 12 | R12C | A | 1.5e-07 | 3e-07 | 3.7e-07 | 2.733E-07 | 1.124E-07 | 1.500E-07 | 3.700E-07 |
| 226 | 12 | R12E | A | 1.4e-07 | 2e-07 | 3e-07 | 2.133E-07 | 8.083E-08 | 1.400E-07 | 3.000E-07 |
| 227 | 12 | R12Q | A | 1.9e-07 | 3.2e-07 | 2.4e-07 | 2.500E-07 | 6.557E-08 | 1.900E-07 | 3.200E-07 |
| 228 | 12 | R12G | A | 1.9e-07 | 2.6e-07 | 4.2e-07 | 2.900E-07 | 1.179E-07 | 1.900E-07 | 4.200E-07 |
| 229 | 12 | R12H | A | 3.5e-07 | 2.9e-07 | 3e-07 | 3.133E-07 | 3.215E-08 | 2.900E-07 | 3.500E-07 |
| 230 | 12 | R12I | A | 2.6e-07 | 2.9e-07 | 2.2e-07 | 2.567E-07 | 3.512E-08 | 2.200E-07 | 2.900E-07 |
| 231 | 12 | R12L | A | 3.4e-07 | 1.8e-07 | 2.3e-07 | 2.500E-07 | 8.185E-08 | 1.800E-07 | 3.400E-07 |
| 232 | 12 | R12K | A | 4.2e-07 | 1.6e-07 | 2.1e-07 | 2.633E-07 | 1.380E-07 | 1.600E-07 | 4.200E-07 |
| 233 | 12 | R12M | A | 1.7e-07 | 4.5e-07 | 2.4e-07 | 2.867E-07 | 1.457E-07 | 1.700E-07 | 4.500E-07 |
| 234 | 12 | R12F | A | 2.4e-07 | 2.9e-07 | 3.5e-07 | 2.933E-07 | 5.508E-08 | 2.400E-07 | 3.500E-07 |
| 235 | 12 | R12P | A | 4.2e-07 | 1.7e-07 | 2.8e-07 | 2.900E-07 | 1.253E-07 | 1.700E-07 | 4.200E-07 |
| 236 | 12 | R12S | A | 2.6e-07 | 1.9e-07 | 1.6e-07 | 2.033E-07 | 5.132E-08 | 1.600E-07 | 2.600E-07 |
| 237 | 12 | R12T | A | 1.6e-07 | 1.4e-07 | 4e-07 | 2.333E-07 | 1.447E-07 | 1.400E-07 | 4.000E-07 |
| 238 | 12 | R12W | A | 1.9e-07 | 2.4e-07 | 2.7e-07 | 2.333E-07 | 4.041E-08 | 1.900E-07 | 2.700E-07 |
| 239 | 12 | R12Y | A | 4.1e-07 | 3.3e-07 | 3.1e-07 | 3.500E-07 | 5.292E-08 | 3.100E-07 | 4.100E-07 |
| 240 | 12 | R12V | A | 1.7e-07 | 3.1e-07 | 2.8e-07 | 2.533E-07 | 7.371E-08 | 1.700E-07 | 3.100E-07 |
| 241 | 13 | L13A | A | 3.6e-07 | 1.5e-07 | 2.8e-07 | 2.633E-07 | 1.060E-07 | 1.500E-07 | 3.600E-07 |
| 242 | 13 | L13R | A | 2.2e-07 | 3.1e-07 | 1.5e-07 | 2.267E-07 | 8.021E-08 | 1.500E-07 | 3.100E-07 |
| 243 | 13 | L13N | A | 2.5e-07 | 1.5e-07 | 2.7e-07 | 2.233E-07 | 6.429E-08 | 1.500E-07 | 2.700E-07 |
| 244 | 13 | L13D | A | 4.4e-07 | 2.1e-07 | 2.4e-07 | 2.967E-07 | 1.250E-07 | 2.100E-07 | 4.400E-07 |
| 245 | 13 | L13C | A | 2e-07 | 1.5e-07 | 2.8e-07 | 2.100E-07 | 6.557E-08 | 1.500E-07 | 2.800E-07 |
| 246 | 13 | L13E | A | 2.5e-07 | 3.3e-07 | 3.6e-07 | 3.133E-07 | 5.686E-08 | 2.500E-07 | 3.600E-07 |
| 247 | 13 | L13Q | A | 2.3e-07 | 3.1e-07 | 1.8e-07 | 2.400E-07 | 6.557E-08 | 1.800E-07 | 3.100E-07 |
| 248 | 13 | L13G | A | 2.1e-07 | 1.2e-07 | 2.1e-07 | 1.800E-07 | 5.196E-08 | 1.200E-07 | 2.100E-07 |
| 249 | 13 | L13H | A | 2.6e-07 | 1.5e-07 | 2.1e-07 | 2.067E-07 | 5.508E-08 | 1.500E-07 | 2.600E-07 |
| 250 | 13 | L13I | A | 2.9e-07 | 4.2e-07 | 3e-07 | 3.367E-07 | 7.234E-08 | 2.900E-07 | 4.200E-07 |
| 251 | 13 | L13L | A | 5.9e-07 | 3e-07 | 4e-07 | 4.300E-07 | 1.473E-07 | 3.000E-07 | 5.900E-07 |
| 252 | 13 | L13K | A | 7.2e-07 | 3.4e-07 | 1.9e-07 | 4.167E-07 | 2.732E-07 | 1.900E-07 | 7.200E-07 |
| 253 | 13 | L13M | A | 1.4e-07 | 6.1e-07 | 3.6e-07 | 3.700E-07 | 2.352E-07 | 1.400E-07 | 6.100E-07 |
| 254 | 13 | L13F | A | 4.5e-07 | 2.5e-07 | 3.4e-07 | 3.467E-07 | 1.002E-07 | 2.500E-07 | 4.500E-07 |

| | | | | | | | | | | |
|---|---|---|---|---|---|---|---|---|---|---|
| 255 | 13 | L13P | A | 1.2e-07 | 4.9e-07 | 4.1e-07 | 3.400E-07 | 1.947E-07 | 1.200E-07 | 4.900E-07 |
| 256 | 13 | L13S | A | 1.5e-07 | 3.4e-07 | 2.6e-07 | 2.500E-07 | 9.539E-08 | 1.500E-07 | 3.400E-07 |
| 257 | 13 | L13T | A | 2.1e-07 | 9.7e-08 | 1.1e-07 | 1.390E-07 | 6.183E-08 | 9.700E-08 | 2.100E-07 |
| 258 | 13 | L13W | A | 3.4e-07 | 3.7e-07 | 2.2e-07 | 3.100E-07 | 7.937E-08 | 2.200E-07 | 3.700E-07 |
| 259 | 13 | L13Y | A | 3.3e-07 | 2.8e-07 | 2.4e-07 | 2.833E-07 | 4.509E-08 | 2.400E-07 | 3.300E-07 |
| 260 | 13 | L13V | A | 1.1e-07 | 4.2e-07 | 3.4e-07 | 2.900E-07 | 1.609E-07 | 1.100E-07 | 4.200E-07 |
| 261 | 14 | P14A | A | 3e-07 | 1.7e-07 | 2.3e-07 | 2.333E-07 | 6.506E-08 | 1.700E-07 | 3.000E-07 |
| 262 | 14 | P14R | A | 1.5e-07 | 2.3e-07 | 2.3e-07 | 2.033E-07 | 4.619E-08 | 1.500E-07 | 2.300E-07 |
| 263 | 14 | P14N | A | 3.9e-07 | 3.2e-07 | 2.7e-07 | 3.267E-07 | 6.028E-08 | 2.700E-07 | 3.900E-07 |
| 264 | 14 | P14D | A | 2.6e-07 | 2.6e-07 | 3.4e-07 | 2.867E-07 | 4.619E-08 | 2.600E-07 | 3.400E-07 |
| 265 | 14 | P14C | A | 5.4e-08 | 2.2e-07 | 3e-07 | 1.913E-07 | 1.255E-07 | 5.400E-08 | 3.000E-07 |
| 266 | 14 | P14E | A | 2e-07 | 3e-07 | 2.8e-07 | 2.600E-07 | 5.292E-08 | 2.000E-07 | 3.000E-07 |
| 267 | 14 | P14Q | A | 1.5e-07 | 2.6e-07 | 2.1e-07 | 2.067E-07 | 5.508E-08 | 1.500E-07 | 2.600E-07 |
| 268 | 14 | P14G | A | 1.9e-07 | 3.5e-07 | 1.6e-07 | 2.333E-07 | 1.021E-07 | 1.600E-07 | 3.500E-07 |
| 269 | 14 | P14H | A | 3.3e-07 | 1.2e-07 | 2.4e-07 | 2.300E-07 | 1.054E-07 | 1.200E-07 | 3.300E-07 |
| 270 | 14 | P14I | A | 2.3e-07 | 2.9e-07 | 6.4e-07 | 3.867E-07 | 2.214E-07 | 2.300E-07 | 6.400E-07 |
| 271 | 14 | P14L | A | 2.5e-07 | 2.3e-07 | 1.6e-07 | 2.133E-07 | 4.726E-08 | 1.600E-07 | 2.500E-07 |
| 272 | 14 | P14K | A | 2.3e-07 | 2.9e-07 | 3.5e-07 | 2.900E-07 | 6.000E-08 | 2.300E-07 | 3.500E-07 |
| 273 | 14 | P14M | A | 3.2e-07 | 2.6e-07 | 2.5e-07 | 2.767E-07 | 3.786E-08 | 2.500E-07 | 3.200E-07 |
| 274 | 14 | P14F | A | 1.8e-07 | 3.3e-07 | 1.6e-07 | 2.233E-07 | 9.292E-08 | 1.600E-07 | 3.300E-07 |
| 275 | 14 | P14P | A | 3.9e-07 | 1.6e-07 | 9.5e-08 | 2.150E-07 | 1.550E-07 | 9.500E-08 | 3.900E-07 |
| 276 | 14 | P14S | A | 3.6e-07 | 3.2e-07 | 1.4e-07 | 2.733E-07 | 1.172E-07 | 1.400E-07 | 3.600E-07 |
| 277 | 14 | P14T | A | 1.7e-07 | 3e-07 | 3.6e-07 | 2.767E-07 | 9.713E-08 | 1.700E-07 | 3.600E-07 |
| 278 | 14 | P14W | A | 2.8e-07 | 2.3e-07 | 8.5e-08 | 1.983E-07 | 1.013E-07 | 8.500E-08 | 2.800E-07 |
| 279 | 14 | P14Y | A | 2e-07 | 1.2e-07 | 2e-07 | 1.733E-07 | 4.619E-08 | 1.200E-07 | 2.000E-07 |
| 280 | 14 | P14V | A | 2.5e-07 | 2.3e-07 | 3.7e-07 | 2.833E-07 | 7.572E-08 | 2.300E-07 | 3.700E-07 |
| 281 | 15 | A15A | A | 3.8e-07 | 4.7e-07 | 1.8e-07 | 3.433E-07 | 1.484E-07 | 1.800E-07 | 4.700E-07 |
| 282 | 15 | A15R | A | 1.7e-07 | 3.2e-07 | 2.3e-07 | 2.400E-07 | 7.550E-08 | 1.700E-07 | 3.200E-07 |
| 283 | 15 | A15N | A | 2.9e-07 | 3.3e-07 | 2.2e-07 | 2.800E-07 | 5.568E-08 | 2.200E-07 | 3.300E-07 |
| 284 | 15 | A15D | A | 1.9e-07 | 1.8e-07 | 1.8e-07 | 1.833E-07 | 5.774E-09 | 1.800E-07 | 1.900E-07 |
| 285 | 15 | A15C | A | 1.9e-07 | 2.2e-07 | 4.1e-07 | 2.733E-07 | 1.193E-07 | 1.900E-07 | 4.100E-07 |
| 286 | 15 | A15E | A | 2.2e-07 | 1.5e-07 | 3.1e-07 | 2.267E-07 | 8.021E-08 | 1.500E-07 | 3.100E-07 |
| 287 | 15 | A15Q | A | 2e-07 | 2.9e-07 | 1.9e-07 | 2.267E-07 | 5.508E-08 | 1.900E-07 | 2.900E-07 |
| 288 | 15 | A15G | A | 3.4e-07 | 2.8e-07 | 5.2e-07 | 3.800E-07 | 1.249E-07 | 2.800E-07 | 5.200E-07 |
| 289 | 15 | A15H | A | 2.9e-07 | 2.2e-07 | 3e-07 | 2.700E-07 | 4.359E-08 | 2.200E-07 | 3.000E-07 |
| 290 | 15 | A15I | A | 1.1e-07 | 3.1e-07 | 4.8e-07 | 3.000E-07 | 1.852E-07 | 1.100E-07 | 4.800E-07 |
| 291 | 15 | A15L | A | 2e-07 | 3.2e-07 | 2.8e-07 | 2.667E-07 | 6.110E-08 | 2.000E-07 | 3.200E-07 |
| 292 | 15 | A15K | A | 2.2e-07 | 3.3e-07 | 3.1e-07 | 2.867E-07 | 5.859E-08 | 2.200E-07 | 3.300E-07 |
| 293 | 15 | A15M | A | 3.1e-07 | 3.1e-07 | 2.9e-07 | 3.033E-07 | 1.155E-08 | 2.900E-07 | 3.100E-07 |
| 294 | 15 | A15F | A | 5.6e-07 | 3.8e-07 | 1.1e-07 | 3.500E-07 | 2.265E-07 | 1.100E-07 | 5.600E-07 |
| 295 | 15 | A15P | A | 5.4e-07 | 3.4e-07 | 1.7e-07 | 3.500E-07 | 1.852E-07 | 1.700E-07 | 5.400E-07 |
| 296 | 15 | A15S | A | 2.4e-07 | 3.2e-07 | 2.7e-07 | 2.767E-07 | 4.041E-08 | 2.400E-07 | 3.200E-07 |
| 297 | 15 | A15T | A | 3.9e-07 | 2.4e-07 | 1.5e-07 | 2.600E-07 | 1.212E-07 | 1.500E-07 | 3.900E-07 |
| 298 | 15 | A15W | A | 1.9e-07 | 2.2e-07 | 4.1e-07 | 2.733E-07 | 1.193E-07 | 1.900E-07 | 4.100E-07 |
| 299 | 15 | A15Y | A | 2.4e-07 | 1.1e-07 | 2.8e-07 | 2.100E-07 | 8.888E-08 | 1.100E-07 | 2.800E-07 |
| 300 | 15 | A15V | A | 2.1e-07 | 4.4e-07 | 1.9e-07 | 2.800E-07 | 1.389E-07 | 1.900E-07 | 4.400E-07 |
| 301 | 16 | S16A | A | 1.6e-07 | 1.9e-07 | 3.5e-07 | 2.333E-07 | 1.021E-07 | 1.600E-07 | 3.500E-07 |
| 302 | 16 | S16R | A | 2.7e-07 | 1.2e-07 | 4.9e-07 | 2.933E-07 | 1.861E-07 | 1.200E-07 | 4.900E-07 |
| 303 | 16 | S16N | A | 1.9e-07 | 2.1e-07 | 1.7e-07 | 1.900E-07 | 2.000E-08 | 1.700E-07 | 2.100E-07 |
| 304 | 16 | S16D | A | 1.8e-07 | 1.9e-07 | 2.2e-07 | 1.967E-07 | 2.082E-08 | 1.800E-07 | 2.200E-07 |
| 305 | 16 | S16C | A | 1.5e-07 | 1.5e-07 | 4.7e-07 | 2.567E-07 | 1.848E-07 | 1.500E-07 | 4.700E-07 |
| 306 | 16 | S16E | A | 1.7e-07 | 6.8e-07 | 1.9e-07 | 3.467E-07 | 2.888E-07 | 1.700E-07 | 6.800E-07 |
| 307 | 16 | S16Q | A | 1.4e-07 | 1.9e-07 | 3.6e-07 | 2.300E-07 | 1.153E-07 | 1.400E-07 | 3.600E-07 |
| 308 | 16 | S16G | A | 3.8e-07 | 1.9e-07 | 2.1e-07 | 2.600E-07 | 1.044E-07 | 1.900E-07 | 3.800E-07 |
| 309 | 16 | S16H | A | 6.9e-07 | 4.2e-07 | 2.5e-07 | 4.533E-07 | 2.219E-07 | 2.500E-07 | 6.900E-07 |
| 310 | 16 | S16I | A | 5.5e-07 | 3.7e-07 | 1.6e-07 | 3.600E-07 | 1.952E-07 | 1.600E-07 | 5.500E-07 |
| 311 | 16 | S16L | A | 2.9e-07 | 2.5e-07 | 2.9e-07 | 2.767E-07 | 2.309E-08 | 2.500E-07 | 2.900E-07 |
| 312 | 16 | S16K | A | 2e-07 | 2.4e-07 | 2.6e-07 | 2.333E-07 | 3.055E-08 | 2.000E-07 | 2.600E-07 |
| 313 | 16 | S16M | A | 3.6e-07 | 2.8e-07 | 3e-07 | 3.133E-07 | 4.163E-08 | 2.800E-07 | 3.600E-07 |
| 314 | 16 | S16F | A | 4.4e-07 | 2.5e-07 | 1.2e-07 | 2.700E-07 | 1.609E-07 | 1.200E-07 | 4.400E-07 |
| 315 | 16 | S16P | A | 1.2e-07 | 2.3e-07 | 1.6e-07 | 1.700E-07 | 5.568E-08 | 1.200E-07 | 2.300E-07 |
| 316 | 16 | S16S | A | 3.4e-07 | 1.1e-07 | 2.5e-07 | 2.333E-07 | 1.159E-07 | 1.100E-07 | 3.400E-07 |
| 317 | 16 | S16T | A | 1.7e-07 | 2.9e-07 | 3.2e-07 | 2.600E-07 | 7.937E-08 | 1.700E-07 | 3.200E-07 |
| 318 | 16 | S16W | A | 1.4e-07 | 3.4e-07 | 2e-07 | 2.267E-07 | 1.026E-07 | 1.400E-07 | 3.400E-07 |
| 319 | 16 | S16Y | A | 1.7e-07 | 1.1e-07 | 1.7e-07 | 1.500E-07 | 3.464E-08 | 1.100E-07 | 1.700E-07 |
| 320 | 16 | S16V | A | 2.6e-07 | 3.1e-07 | 1.4e-07 | 2.367E-07 | 8.737E-08 | 1.400E-07 | 3.100E-07 |
| 321 | 17 | P17A | A | 2.4e-07 | 4.1e-07 | 3.7e-07 | 3.400E-07 | 8.888E-08 | 2.400E-07 | 4.100E-07 |
| 322 | 17 | P17R | A | 2.4e-07 | 2.7e-07 | 3.3e-07 | 2.800E-07 | 4.583E-08 | 2.400E-07 | 3.300E-07 |
| 323 | 17 | P17N | A | 5.5e-07 | 2.3e-07 | 1.6e-07 | 3.133E-07 | 2.079E-07 | 1.600E-07 | 5.500E-07 |
| 324 | 17 | P17D | A | 2.5e-07 | 3.3e-07 | 3.3e-07 | 3.033E-07 | 4.619E-08 | 2.500E-07 | 3.300E-07 |
| 325 | 17 | P17C | A | 3.2e-07 | 3.2e-07 | 2.2e-07 | 2.867E-07 | 5.774E-08 | 2.200E-07 | 3.200E-07 |
| 326 | 17 | P17E | A | 2.6e-07 | 2.4e-07 | 2.5e-07 | 2.500E-07 | 1.000E-08 | 2.400E-07 | 2.600E-07 |
| 327 | 17 | P17Q | A | 4.3e-07 | 2e-07 | 2.5e-07 | 2.933E-07 | 1.210E-07 | 2.000E-07 | 4.300E-07 |
| 328 | 17 | P17G | A | 5.1e-07 | 2e-07 | 2.8e-07 | 3.300E-07 | 1.609E-07 | 2.000E-07 | 5.100E-07 |
| 329 | 17 | P17H | A | 2.3e-07 | 2.9e-07 | 2.5e-07 | 2.567E-07 | 3.055E-08 | 2.300E-07 | 2.900E-07 |
| 330 | 17 | P17I | A | 1.7e-07 | 2e-07 | 2.2e-07 | 1.967E-07 | 2.517E-08 | 1.700E-07 | 2.200E-07 |
| 331 | 17 | P17L | A | 2.2e-07 | 2.9e-07 | 2.5e-07 | 2.533E-07 | 3.512E-08 | 2.200E-07 | 2.900E-07 |
| 332 | 17 | P17K | A | 2.9e-07 | 2.6e-07 | 2.2e-07 | 2.567E-07 | 3.512E-08 | 2.200E-07 | 2.900E-07 |
| 333 | 17 | P17M | A | 1.9e-07 | 3.6e-07 | 2.9e-07 | 2.800E-07 | 8.544E-08 | 1.900E-07 | 3.600E-07 |
| 334 | 17 | P17F | A | 5e-07 | 3.3e-07 | 2.5e-07 | 3.600E-07 | 1.277E-07 | 2.500E-07 | 5.000E-07 |
| 335 | 17 | P17P | A | 6.3e-07 | 1.6e-07 | 2.7e-07 | 3.533E-07 | 2.458E-07 | 1.600E-07 | 6.300E-07 |
| 336 | 17 | P17S | A | 4e-07 | 2e-07 | 2.9e-07 | 2.967E-07 | 1.002E-07 | 2.000E-07 | 4.000E-07 |
| 337 | 17 | P17T | A | 2.5e-07 | 2.8e-07 | 1.6e-07 | 2.300E-07 | 6.245E-08 | 1.600E-07 | 2.800E-07 |
| 338 | 17 | P17W | A | 3.5e-07 | 1.6e-07 | 1.1e-07 | 2.067E-07 | 1.266E-07 | 1.100E-07 | 3.500E-07 |
| 339 | 17 | P17Y | A | 2.2e-07 | 1.7e-07 | 2.2e-07 | 2.033E-07 | 2.887E-08 | 1.700E-07 | 2.200E-07 |

| | | | | | | | | | | |
|---|---|---|---|---|---|---|---|---|---|---|
| 340 | 17 | P17V | A | 4.6e-07 | 3.2e-07 | 2.9e-07 | 3.567E-07 | 9.074E-08 | 2.900E-07 | 4.600E-07 |
| 341 | 18 | E18A | A | 1.8e-07 | 3.4e-07 | 2.3e-07 | 2.500E-07 | 8.185E-08 | 1.800E-07 | 3.400E-07 |
| 342 | 18 | E18R | A | 2.1e-07 | 1.3e-07 | 3.7e-07 | 2.367E-07 | 1.222E-07 | 1.300E-07 | 3.700E-07 |
| 343 | 18 | E18N | A | 2.4e-07 | 3.5e-07 | 1.5e-07 | 2.467E-07 | 1.002E-07 | 1.500E-07 | 3.500E-07 |
| 344 | 18 | E18D | A | 3.8e-07 | 1.6e-07 | 2.9e-07 | 2.767E-07 | 1.106E-07 | 1.600E-07 | 3.800E-07 |
| 345 | 18 | E18C | A | 4.3e-07 | 3.6e-07 | 1.7e-07 | 3.200E-07 | 1.345E-07 | 1.700E-07 | 4.300E-07 |
| 346 | 18 | E18E | A | 3.8e-07 | 1.7e-07 | 2.2e-07 | 2.567E-07 | 1.097E-07 | 1.700E-07 | 3.800E-07 |
| 347 | 18 | E18Q | A | 2.2e-07 | 9.7e-08 | 2.5e-07 | 1.890E-07 | 8.107E-08 | 9.700E-08 | 2.500E-07 |
| 348 | 18 | E18G | A | 2.3e-07 | 2.6e-07 | 1e-07 | 1.967E-07 | 8.505E-08 | 1.000E-07 | 2.600E-07 |
| 349 | 18 | E18H | A | 2.2e-07 | 2e-07 | 1.5e-07 | 1.900E-07 | 3.606E-08 | 1.500E-07 | 2.200E-07 |
| 350 | 18 | E18I | A | 5.8e-07 | 1e-07 | 2.4e-07 | 3.067E-07 | 2.468E-07 | 1.000E-07 | 5.800E-07 |
| 351 | 18 | E18L | A | 2.8e-07 | 2.3e-07 | 2.3e-07 | 2.467E-07 | 2.887E-08 | 2.300E-07 | 2.800E-07 |
| 352 | 18 | E18K | A | 1e-07 | 3.1e-07 | 1.9e-07 | 2.000E-07 | 1.054E-07 | 1.000E-07 | 3.100E-07 |
| 353 | 18 | E18M | A | 2e-07 | 1.8e-07 | 2.1e-07 | 1.967E-07 | 1.528E-08 | 1.800E-07 | 2.100E-07 |
| 354 | 18 | E18F | A | 3.7e-07 | 2.5e-07 | 3.3e-07 | 3.167E-07 | 6.110E-08 | 2.500E-07 | 3.700E-07 |
| 355 | 18 | E18P | A | 1.4e-07 | 3.2e-07 | 4.2e-07 | 2.933E-07 | 1.419E-07 | 1.400E-07 | 4.200E-07 |
| 356 | 18 | E18S | A | 5.6e-07 | 1.2e-07 | 3.6e-07 | 3.467E-07 | 2.203E-07 | 1.200E-07 | 5.600E-07 |
| 357 | 18 | E18T | A | 2.6e-07 | 1.9e-07 | 4.4e-07 | 2.967E-07 | 1.290E-07 | 1.900E-07 | 4.400E-07 |
| 358 | 18 | E18W | A | 1.7e-07 | 3.1e-07 | 1.5e-07 | 2.100E-07 | 8.718E-08 | 1.500E-07 | 3.100E-07 |
| 359 | 18 | E18Y | A | 1.1e-07 | 4.6e-07 | 3.1e-07 | 2.933E-07 | 1.756E-07 | 1.100E-07 | 4.600E-07 |
| 360 | 18 | E18V | A | 2.6e-07 | 7.6e-07 | 2.3e-07 | 4.167E-07 | 2.977E-07 | 2.300E-07 | 7.600E-07 |
| 361 | 19 | T19A | A | 1.7e-07 | 3.7e-07 | 6.1e-07 | 3.833E-07 | 2.203E-07 | 1.700E-07 | 6.100E-07 |
| 362 | 19 | T19R | A | 2.2e-07 | 2e-07 | 3.7e-07 | 2.633E-07 | 9.292E-08 | 2.000E-07 | 3.700E-07 |
| 363 | 19 | T19N | A | 2.2e-07 | 2.2e-07 | 2.3e-07 | 2.233E-07 | 5.774E-09 | 2.200E-07 | 2.300E-07 |
| 364 | 19 | T19D | A | 2.8e-07 | 2.2e-07 | 2.7e-07 | 2.567E-07 | 3.215E-08 | 2.200E-07 | 2.800E-07 |
| 365 | 19 | T19C | A | 3.1e-07 | 2.6e-07 | 1.9e-07 | 2.533E-07 | 6.028E-08 | 1.900E-07 | 3.100E-07 |
| 366 | 19 | T19E | A | 2.2e-07 | 4.5e-07 | 2.2e-07 | 2.967E-07 | 1.328E-07 | 2.200E-07 | 4.500E-07 |
| 367 | 19 | T19Q | A | 3.8e-07 | 1.8e-07 | 4.7e-07 | 3.433E-07 | 1.484E-07 | 1.800E-07 | 4.700E-07 |
| 368 | 19 | T19G | A | 1.7e-07 | 2e-07 | 2.9e-07 | 2.200E-07 | 6.245E-08 | 1.700E-07 | 2.900E-07 |
| 369 | 19 | T19H | A | 2.3e-07 | 1.5e-07 | 2.8e-07 | 2.200E-07 | 6.557E-08 | 1.500E-07 | 2.800E-07 |
| 370 | 19 | T19I | A | 3e-07 | 2.9e-07 | 2.1e-07 | 2.667E-07 | 4.933E-08 | 2.100E-07 | 3.000E-07 |
| 371 | 19 | T19L | A | 3.1e-07 | 5.2e-07 | 2.1e-07 | 3.467E-07 | 1.582E-07 | 2.100E-07 | 5.200E-07 |
| 372 | 19 | T19K | A | 2.5e-07 | 1.2e-07 | 2.7e-07 | 2.133E-07 | 8.145E-08 | 1.200E-07 | 2.700E-07 |
| 373 | 19 | T19M | A | 1.5e-07 | 2.3e-07 | 1.9e-07 | 1.900E-07 | 4.000E-08 | 1.500E-07 | 2.300E-07 |
| 374 | 19 | T19F | A | 3.6e-07 | 3.9e-07 | 2.1e-07 | 3.200E-07 | 9.644E-08 | 2.100E-07 | 3.900E-07 |
| 375 | 19 | T19P | A | 2e-07 | 3.1e-07 | 3.7e-07 | 2.933E-07 | 8.622E-08 | 2.000E-07 | 3.700E-07 |
| 376 | 19 | T19S | A | 2.4e-07 | 1.7e-07 | 3e-07 | 2.367E-07 | 6.506E-08 | 1.700E-07 | 3.000E-07 |
| 377 | 19 | T19T | A | 2.8e-07 | 9.9e-08 | 6.2e-07 | 3.330E-07 | 2.645E-07 | 9.900E-08 | 6.200E-07 |
| 378 | 19 | T19W | A | 7e-08 | 4.6e-07 | 1.6e-07 | 2.300E-07 | 2.042E-07 | 7.000E-08 | 4.600E-07 |
| 379 | 19 | T19Y | A | 4e-07 | 3.7e-07 | 2.6e-07 | 3.433E-07 | 7.371E-08 | 2.600E-07 | 4.000E-07 |
| 380 | 19 | T19V | A | 2.8e-07 | 2.9e-07 | 3.3e-07 | 3.000E-07 | 2.646E-08 | 2.800E-07 | 3.300E-07 |
| 381 | 20 | H20A | A | 2.3e-07 | 1.9e-07 | 3.6e-07 | 2.600E-07 | 8.888E-08 | 1.900E-07 | 3.600E-07 |
| 382 | 20 | H20R | A | 2.9e-07 | 1.9e-07 | 3.6e-07 | 2.800E-07 | 8.544E-08 | 1.900E-07 | 3.600E-07 |
| 383 | 20 | H20N | A | 1.9e-07 | 3.3e-07 | 2.8e-07 | 2.667E-07 | 7.095E-08 | 1.900E-07 | 3.300E-07 |
| 384 | 20 | H20D | A | 3.6e-07 | 1.9e-07 | 2.9e-07 | 2.800E-07 | 8.544E-08 | 1.900E-07 | 3.600E-07 |
| 385 | 20 | H20C | A | 3.2e-07 | 3.6e-07 | 3.5e-07 | 3.433E-07 | 2.082E-08 | 3.200E-07 | 3.600E-07 |
| 386 | 20 | H20E | A | 2.6e-07 | 3.8e-07 | 9.2e-07 | 5.200E-07 | 3.516E-07 | 2.600E-07 | 9.200E-07 |
| 387 | 20 | H20Q | A | 1.2e-07 | 5.8e-07 | 3.9e-07 | 3.633E-07 | 2.312E-07 | 1.200E-07 | 5.800E-07 |
| 388 | 20 | H20G | A | 3.2e-07 | 4.9e-07 | 4.2e-07 | 4.100E-07 | 8.544E-08 | 3.200E-07 | 4.900E-07 |
| 389 | 20 | H20H | A | 5.2e-07 | 2.4e-07 | 5e-07 | 4.200E-07 | 1.562E-07 | 2.400E-07 | 5.200E-07 |
| 390 | 20 | H20I | A | 1.9e-07 | 1.6e-07 | 1.9e-07 | 1.800E-07 | 1.732E-08 | 1.600E-07 | 1.900E-07 |
| 391 | 20 | H20L | A | 1.9e-07 | 4.3e-07 | 3.1e-07 | 3.100E-07 | 1.200E-07 | 1.900E-07 | 4.300E-07 |
| 392 | 20 | H20K | A | 5.1e-07 | 2.3e-07 | 2.1e-07 | 3.167E-07 | 1.677E-07 | 2.100E-07 | 5.100E-07 |
| 393 | 20 | H20M | A | 1.9e-07 | 3.7e-07 | 1.9e-07 | 2.500E-07 | 1.039E-07 | 1.900E-07 | 3.700E-07 |
| 394 | 20 | H20F | A | 3.5e-07 | 1.3e-07 | 5.1e-07 | 3.300E-07 | 1.908E-07 | 1.300E-07 | 5.100E-07 |
| 395 | 20 | H20P | A | 4.2e-07 | 2.2e-07 | 2.3e-07 | 2.900E-07 | 1.127E-07 | 2.200E-07 | 4.200E-07 |
| 396 | 20 | H20S | A | 3.4e-07 | 2.6e-07 | 4.5e-07 | 3.500E-07 | 9.539E-08 | 2.600E-07 | 4.500E-07 |
| 397 | 20 | H20T | A | 2e-07 | 1.3e-07 | 2.1e-07 | 1.800E-07 | 4.359E-08 | 1.300E-07 | 2.100E-07 |
| 398 | 20 | H20W | A | 2.1e-07 | 3e-07 | 2.5e-07 | 2.533E-07 | 4.509E-08 | 2.100E-07 | 3.000E-07 |
| 399 | 20 | H20Y | A | 2.7e-07 | 4e-07 | 1.5e-07 | 2.733E-07 | 1.250E-07 | 1.500E-07 | 4.000E-07 |
| 400 | 20 | H20V | A | 2.2e-07 | 5.5e-07 | 4e-07 | 3.900E-07 | 1.652E-07 | 2.200E-07 | 5.500E-07 |
| 401 | 21 | L21A | A | 2.6e-07 | 5e-07 | 1.9e-07 | 3.167E-07 | 1.626E-07 | 1.900E-07 | 5.000E-07 |
| 402 | 21 | L21R | A | 3.2e-07 | 2.3e-07 | 2.9e-07 | 2.800E-07 | 4.583E-08 | 2.300E-07 | 3.200E-07 |
| 403 | 21 | L21N | A | 1.1e-07 | 2.1e-07 | 1.4e-07 | 1.533E-07 | 5.132E-08 | 1.100E-07 | 2.100E-07 |
| 404 | 21 | L21D | A | 3.1e-07 | 3.2e-07 | 3.7e-07 | 3.333E-07 | 3.215E-08 | 3.100E-07 | 3.700E-07 |
| 405 | 21 | L21C | A | 3.3e-07 | 3.4e-07 | 2.9e-07 | 3.200E-07 | 2.646E-08 | 2.900E-07 | 3.400E-07 |
| 406 | 21 | L21E | A | 2.8e-07 | 1.3e-07 | 2.6e-07 | 2.233E-07 | 8.145E-08 | 1.300E-07 | 2.800E-07 |
| 407 | 21 | L21Q | A | 2.5e-07 | 2.2e-07 | 1.4e-07 | 2.033E-07 | 5.686E-08 | 1.400E-07 | 2.500E-07 |
| 408 | 21 | L21G | A | 1.6e-07 | 2.7e-07 | 1.2e-07 | 1.833E-07 | 7.767E-08 | 1.200E-07 | 2.700E-07 |
| 409 | 21 | L21H | A | 2.6e-07 | 2.8e-07 | 8.7e-08 | 2.090E-07 | 1.061E-07 | 8.700E-08 | 2.800E-07 |
| 410 | 21 | L21I | A | 2.1e-07 | 2.3e-07 | 2.8e-07 | 2.400E-07 | 3.606E-08 | 2.100E-07 | 2.800E-07 |
| 411 | 21 | L21L | A | 2.5e-07 | 3e-07 | 2.8e-07 | 2.767E-07 | 2.517E-08 | 2.500E-07 | 3.000E-07 |
| 412 | 21 | L21K | A | 5.2e-07 | 2.9e-07 | 4.6e-07 | 4.233E-07 | 1.193E-07 | 2.900E-07 | 5.200E-07 |
| 413 | 21 | L21M | A | 1.8e-07 | 4.5e-07 | 2.6e-07 | 2.967E-07 | 1.387E-07 | 1.800E-07 | 4.500E-07 |
| 414 | 21 | L21F | A | 2.2e-07 | 3.2e-07 | 3.4e-07 | 2.933E-07 | 6.429E-08 | 2.200E-07 | 3.400E-07 |
| 415 | 21 | L21P | A | 2.8e-07 | 2.9e-07 | 3.1e-07 | 2.933E-07 | 1.528E-08 | 2.800E-07 | 3.100E-07 |
| 416 | 21 | L21S | A | 1.1e-07 | 6.4e-07 | 8.6e-08 | 2.787E-07 | 3.132E-07 | 8.600E-08 | 6.400E-07 |
| 417 | 21 | L21T | A | 2.7e-07 | 9.6e-08 | 3.7e-07 | 2.453E-07 | 1.387E-07 | 9.600E-08 | 3.700E-07 |
| 418 | 21 | L21W | A | 1.5e-07 | 2.7e-07 | 1.1e-07 | 1.767E-07 | 8.327E-08 | 1.100E-07 | 2.700E-07 |
| 419 | 21 | L21Y | A | 3.1e-07 | 2.3e-07 | 1.4e-07 | 2.267E-07 | 8.505E-08 | 1.400E-07 | 3.100E-07 |
| 420 | 21 | L21V | A | 3.8e-07 | 1.4e-07 | 4.1e-07 | 3.100E-07 | 1.480E-07 | 1.400E-07 | 4.100E-07 |
| 421 | 22 | D22A | A | 2.4e-07 | 3.5e-07 | 3.8e-07 | 3.233E-07 | 7.371E-08 | 2.400E-07 | 3.800E-07 |
| 422 | 22 | D22R | A | 1.1e-07 | 4.2e-07 | 2.7e-07 | 2.667E-07 | 1.550E-07 | 1.100E-07 | 4.200E-07 |
| 423 | 22 | D22N | A | 1.4e-07 | 2.6e-07 | 4e-07 | 2.667E-07 | 1.301E-07 | 1.400E-07 | 4.000E-07 |
| 424 | 22 | D22D | A | 2.2e-07 | 1.7e-07 | 1.7e-07 | 1.867E-07 | 2.887E-08 | 1.700E-07 | 2.200E-07 |

| | | | | | | | | | | |
|---|---|---|---|---|---|---|---|---|---|---|
| 425 | 22 | D22C | A | 2.4e-07 | 3e-07 | 3.1e-07 | 2.833E-07 | 3.786E-08 | 2.400E-07 | 3.100E-07 |
| 426 | 22 | D22E | A | 1.6e-07 | 2e-07 | 2.3e-07 | 1.967E-07 | 3.512E-08 | 1.600E-07 | 2.300E-07 |
| 427 | 22 | D22Q | A | 2.6e-07 | 1.8e-07 | 2.8e-07 | 2.400E-07 | 5.292E-08 | 1.800E-07 | 2.800E-07 |
| 428 | 22 | D22G | A | 3.3e-07 | 2.6e-07 | 4.2e-07 | 3.367E-07 | 8.021E-08 | 2.600E-07 | 4.200E-07 |
| 429 | 22 | D22H | A | 2.6e-07 | 1e-06 | 2.7e-07 | 5.100E-07 | 4.244E-07 | 2.600E-07 | 1.000E-06 |
| 430 | 22 | D22I | A | 1.6e-07 | 3.2e-07 | 3.6e-07 | 2.800E-07 | 1.058E-07 | 1.600E-07 | 3.600E-07 |
| 431 | 22 | D22L | A | 1.2e-07 | 9.1e-08 | 1.4e-07 | 1.170E-07 | 2.464E-08 | 9.100E-08 | 1.400E-07 |
| 432 | 22 | D22K | A | 2.6e-07 | 3.2e-07 | 1.9e-07 | 2.567E-07 | 6.506E-08 | 1.900E-07 | 3.200E-07 |
| 433 | 22 | D22M | A | 3e-07 | 2.2e-07 | 4.1e-07 | 3.100E-07 | 9.539E-08 | 2.200E-07 | 4.100E-07 |
| 434 | 22 | D22F | A | 5.1e-07 | 1.7e-07 | 2.2e-07 | 3.000E-07 | 1.836E-07 | 1.700E-07 | 5.100E-07 |
| 435 | 22 | D22P | A | 2.6e-07 | 2.4e-07 | 2.9e-07 | 2.633E-07 | 2.517E-08 | 2.400E-07 | 2.900E-07 |
| 436 | 22 | D22S | A | 1.1e-07 | 4.5e-07 | 3.3e-07 | 2.967E-07 | 1.724E-07 | 1.100E-07 | 4.500E-07 |
| 437 | 22 | D22T | A | 3.9e-07 | 2.6e-07 | 2.5e-07 | 3.000E-07 | 7.810E-08 | 2.500E-07 | 3.900E-07 |
| 438 | 22 | D22W | A | 1.1e-07 | 1.8e-07 | 2.1e-07 | 1.667E-07 | 5.132E-08 | 1.100E-07 | 2.100E-07 |
| 439 | 22 | D22Y | A | 3e-07 | 2.3e-07 | 5.4e-07 | 3.567E-07 | 1.626E-07 | 2.300E-07 | 5.400E-07 |
| 440 | 22 | D22V | A | 1.5e-07 | 3.4e-07 | 3.6e-07 | 2.833E-07 | 1.159E-07 | 1.500E-07 | 3.600E-07 |
| 441 | 23 | M23A | A | 2.8e-07 | 2.1e-07 | 3.1e-07 | 2.667E-07 | 5.132E-08 | 2.100E-07 | 3.100E-07 |
| 442 | 23 | M23R | A | 2.9e-07 | 2.1e-07 | 2.1e-07 | 2.367E-07 | 4.619E-08 | 2.100E-07 | 2.900E-07 |
| 443 | 23 | M23N | A | 1.4e-07 | 1.8e-07 | 1.4e-07 | 1.533E-07 | 2.309E-08 | 1.400E-07 | 1.800E-07 |
| 444 | 23 | M23D | A | 1.8e-07 | 2.1e-07 | 2.8e-07 | 2.233E-07 | 5.132E-08 | 1.800E-07 | 2.800E-07 |
| 445 | 23 | M23C | A | 3.3e-07 | 7e-07 | 1.3e-07 | 3.867E-07 | 2.892E-07 | 1.300E-07 | 7.000E-07 |
| 446 | 23 | M23E | A | 3e-07 | 2e-07 | 2.6e-07 | 2.533E-07 | 5.033E-08 | 2.000E-07 | 3.000E-07 |
| 447 | 23 | M23Q | A | 3.7e-07 | 3.7e-07 | 3.7e-07 | 3.700E-07 | 0.000E+00 | 3.700E-07 | 3.700E-07 |
| 448 | 23 | M23G | A | 5.3e-07 | 4.2e-07 | 2.5e-07 | 4.000E-07 | 1.411E-07 | 2.500E-07 | 5.300E-07 |
| 449 | 23 | M23H | A | 6e-07 | 3.8e-07 | 1.8e-07 | 3.867E-07 | 2.101E-07 | 1.800E-07 | 6.000E-07 |
| 450 | 23 | M23I | A | 3.7e-07 | 3.6e-07 | 4e-07 | 3.767E-07 | 2.082E-08 | 3.600E-07 | 4.000E-07 |
| 451 | 23 | M23L | A | 2.6e-07 | 2.7e-07 | 1.8e-07 | 2.367E-07 | 4.933E-08 | 1.800E-07 | 2.700E-07 |
| 452 | 23 | M23K | A | 3.6e-07 | 2.4e-07 | 2.3e-07 | 2.767E-07 | 7.234E-08 | 2.300E-07 | 3.600E-07 |
| 453 | 23 | M23M | A | 4.9e-07 | 1.2e-07 | 2.1e-07 | 2.733E-07 | 1.930E-07 | 1.200E-07 | 4.900E-07 |
| 454 | 23 | M23F | A | 4.9e-07 | 1.5e-07 | 2.8e-07 | 3.067E-07 | 1.716E-07 | 1.500E-07 | 4.900E-07 |
| 455 | 23 | M23P | A | 2.1e-07 | 2.1e-07 | 2.3e-07 | 2.167E-07 | 1.155E-08 | 2.100E-07 | 2.300E-07 |
| 456 | 23 | M23S | A | 5.9e-07 | 3.1e-07 | 3.5e-07 | 4.167E-07 | 1.514E-07 | 3.100E-07 | 5.900E-07 |
| 457 | 23 | M23T | A | 1.4e-07 | 2.1e-07 | 3.2e-07 | 2.233E-07 | 9.074E-08 | 1.400E-07 | 3.200E-07 |
| 458 | 23 | M23W | A | 4.3e-07 | 3.3e-07 | 2.3e-07 | 3.300E-07 | 1.000E-07 | 2.300E-07 | 4.300E-07 |
| 459 | 23 | M23Y | A | 6.4e-08 | 1.7e-07 | 3.5e-07 | 1.947E-07 | 1.446E-07 | 6.400E-08 | 3.500E-07 |
| 460 | 23 | M23V | A | 4e-07 | 2.1e-07 | 2.1e-07 | 2.733E-07 | 1.097E-07 | 2.100E-07 | 4.000E-07 |
| 461 | 24 | L24A | A | 3.4e-07 | 1.4e-07 | 2.5e-07 | 2.433E-07 | 1.002E-07 | 1.400E-07 | 3.400E-07 |
| 462 | 24 | L24R | A | 2.4e-07 | 1.9e-07 | 1.3e-07 | 1.867E-07 | 5.508E-08 | 1.300E-07 | 2.400E-07 |
| 463 | 24 | L24N | A | 2.5e-07 | 2.6e-07 | 5.5e-07 | 3.533E-07 | 1.704E-07 | 2.500E-07 | 5.500E-07 |
| 464 | 24 | L24D | A | 9.5e-08 | 2.1e-07 | 4.2e-07 | 2.417E-07 | 1.648E-07 | 9.500E-08 | 4.200E-07 |
| 465 | 24 | L24C | A | 1.6e-07 | 8.8e-07 | 3.5e-07 | 4.633E-07 | 3.731E-07 | 1.600E-07 | 8.800E-07 |
| 466 | 24 | L24E | A | 4.6e-07 | 2.7e-07 | 2.4e-07 | 3.233E-07 | 1.193E-07 | 2.400E-07 | 4.600E-07 |
| 467 | 24 | L24Q | A | 3.3e-07 | 2.1e-07 | 2.5e-07 | 2.633E-07 | 6.110E-08 | 2.100E-07 | 3.300E-07 |
| 468 | 24 | L24G | A | 1.6e-07 | 2.7e-07 | 2.6e-07 | 2.300E-07 | 6.083E-08 | 1.600E-07 | 2.700E-07 |
| 469 | 24 | L24H | A | 7e-07 | 3.5e-07 | 3.5e-07 | 4.667E-07 | 2.021E-07 | 3.500E-07 | 7.000E-07 |
| 470 | 24 | L24I | A | 2.3e-07 | 1.4e-07 | 5.9e-07 | 3.200E-07 | 2.381E-07 | 1.400E-07 | 5.900E-07 |
| 471 | 24 | L24L | A | 8.4e-08 | 1.3e-07 | 2e-07 | 1.380E-07 | 5.841E-08 | 8.400E-08 | 2.000E-07 |
| 472 | 24 | L24K | A | 1.7e-07 | 1.6e-07 | 1.7e-07 | 1.667E-07 | 5.774E-09 | 1.600E-07 | 1.700E-07 |
| 473 | 24 | L24M | A | 1.2e-07 | 3.4e-07 | 3.7e-07 | 2.767E-07 | 1.365E-07 | 1.200E-07 | 3.700E-07 |
| 474 | 24 | L24F | A | 1.8e-07 | 3e-07 | 1.6e-07 | 2.133E-07 | 7.572E-08 | 1.600E-07 | 3.000E-07 |
| 475 | 24 | L24P | A | 2.2e-07 | 3.4e-07 | 2.6e-07 | 2.733E-07 | 6.110E-08 | 2.200E-07 | 3.400E-07 |
| 476 | 24 | L24S | A | 2.1e-07 | 3.3e-07 | 2.8e-07 | 2.733E-07 | 6.028E-08 | 2.100E-07 | 3.300E-07 |
| 477 | 24 | L24T | A | 4.5e-07 | 4.2e-07 | 3.2e-07 | 3.967E-07 | 6.807E-08 | 3.200E-07 | 4.500E-07 |
| 478 | 24 | L24W | A | 2.8e-07 | 2.7e-07 | 3.6e-07 | 3.033E-07 | 4.933E-08 | 2.700E-07 | 3.600E-07 |
| 479 | 24 | L24Y | A | 2.3e-07 | 4e-07 | 4.1e-07 | 3.467E-07 | 1.012E-07 | 2.300E-07 | 4.100E-07 |
| 480 | 24 | L24V | A | 4e-07 | 2e-07 | 1.1e-06 | 5.667E-07 | 4.726E-07 | 2.000E-07 | 1.100E-06 |
| 481 | 25 | R25A | A | 1.7e-07 | 1.4e-07 | 3.3e-07 | 2.133E-07 | 1.021E-07 | 1.400E-07 | 3.300E-07 |
| 482 | 25 | R25R | A | 3.2e-07 | 1.8e-07 | 1.4e-07 | 2.133E-07 | 9.452E-08 | 1.400E-07 | 3.200E-07 |
| 483 | 25 | R25N | A | 3.4e-07 | 2.7e-07 | 1.6e-07 | 2.567E-07 | 9.074E-08 | 1.600E-07 | 3.400E-07 |
| 484 | 25 | R25D | A | 1.5e-07 | 2.8e-07 | 1.9e-07 | 2.067E-07 | 6.658E-08 | 1.500E-07 | 2.800E-07 |
| 485 | 25 | R25C | A | 1.8e-07 | 2.5e-07 | 1.2e-07 | 1.833E-07 | 6.506E-08 | 1.200E-07 | 2.500E-07 |
| 486 | 25 | R25E | A | 4.3e-07 | 1.2e-07 | 2.2e-07 | 2.567E-07 | 1.582E-07 | 1.200E-07 | 4.300E-07 |
| 487 | 25 | R25Q | A | 1.4e-07 | 1.2e-07 | 4.9e-07 | 2.500E-07 | 2.081E-07 | 1.200E-07 | 4.900E-07 |
| 488 | 25 | R25G | A | 1.7e-07 | 1.8e-07 | 1.6e-07 | 1.700E-07 | 1.000E-08 | 1.600E-07 | 1.800E-07 |
| 489 | 25 | R25H | A | 1.6e-07 | 1.8e-07 | 2.8e-07 | 2.067E-07 | 6.429E-08 | 1.600E-07 | 2.800E-07 |
| 490 | 25 | R25I | A | 2e-07 | 3.5e-07 | 1.7e-07 | 2.400E-07 | 9.644E-08 | 1.700E-07 | 3.500E-07 |
| 491 | 25 | R25L | A | 1.5e-07 | 1.9e-07 | 3.8e-07 | 2.400E-07 | 1.229E-07 | 1.500E-07 | 3.800E-07 |
| 492 | 25 | R25K | A | 1.2e-07 | 3.1e-07 | 3e-07 | 2.433E-07 | 1.069E-07 | 1.200E-07 | 3.100E-07 |
| 493 | 25 | R25M | A | 1.4e-07 | 3.1e-07 | 2.1e-07 | 2.200E-07 | 8.544E-08 | 1.400E-07 | 3.100E-07 |
| 494 | 25 | R25F | A | 2.3e-07 | 3.5e-07 | 3.3e-07 | 3.033E-07 | 6.429E-08 | 2.300E-07 | 3.500E-07 |
| 495 | 25 | R25P | A | 2e-07 | 3.5e-07 | 4.6e-07 | 3.367E-07 | 1.305E-07 | 2.000E-07 | 4.600E-07 |
| 496 | 25 | R25S | A | 2e-07 | 6e-08 | 3.4e-07 | 2.000E-07 | 1.400E-07 | 6.000E-08 | 3.400E-07 |
| 497 | 25 | R25T | A | 2.4e-07 | 5.4e-07 | 2e-07 | 3.267E-07 | 1.858E-07 | 2.000E-07 | 5.400E-07 |
| 498 | 25 | R25W | A | 3.7e-07 | 2.9e-07 | 2.6e-07 | 3.067E-07 | 5.686E-08 | 2.600E-07 | 3.700E-07 |
| 499 | 25 | R25Y | A | 1.5e-07 | 2.4e-07 | 2.3e-07 | 2.067E-07 | 4.933E-08 | 1.500E-07 | 2.400E-07 |
| 500 | 25 | R25V | A | 2.5e-07 | 2.5e-07 | 2.6e-07 | 2.533E-07 | 5.774E-09 | 2.500E-07 | 2.600E-07 |
| 501 | 26 | H26A | A | 5.6e-07 | 1.4e-07 | 2.6e-07 | 3.200E-07 | 2.163E-07 | 1.400E-07 | 5.600E-07 |
| 502 | 26 | H26R | A | 1.9e-07 | 3e-07 | 2.2e-07 | 2.367E-07 | 5.686E-08 | 1.900E-07 | 3.000E-07 |
| 503 | 26 | H26N | A | 2.2e-07 | 4.6e-07 | 1.1e-07 | 2.633E-07 | 1.790E-07 | 1.100E-07 | 4.600E-07 |
| 504 | 26 | H26D | A | 3.1e-07 | 2.1e-07 | 5.1e-07 | 3.433E-07 | 1.528E-07 | 2.100E-07 | 5.100E-07 |
| 505 | 26 | H26C | A | 5.8e-07 | 3.5e-07 | 3.5e-07 | 4.267E-07 | 1.328E-07 | 3.500E-07 | 5.800E-07 |
| 506 | 26 | H26E | A | 2.6e-07 | 3.5e-07 | 2.2e-07 | 2.767E-07 | 6.658E-08 | 2.200E-07 | 3.500E-07 |
| 507 | 26 | H26Q | A | 2e-07 | 1.3e-07 | 1.4e-07 | 1.567E-07 | 3.786E-08 | 1.300E-07 | 2.000E-07 |
| 508 | 26 | H26G | A | 3.4e-07 | 2.8e-07 | 4e-07 | 3.400E-07 | 6.000E-08 | 2.800E-07 | 4.000E-07 |
| 509 | 26 | H26H | A | 2.2e-07 | 6.4e-07 | 2.3e-07 | 3.633E-07 | 2.397E-07 | 2.200E-07 | 6.400E-07 |

| | | | | | | | | | | |
|---|---|---|---|---|---|---|---|---|---|---|
| 510 | 26 | H26I | A | 3e-07 | 1.6e-07 | 3.4e-07 | 2.667E-07 | 9.452E-08 | 1.600E-07 | 3.400E-07 |
| 511 | 26 | H26L | A | 2.7e-07 | 3.4e-07 | 1.5e-07 | 2.533E-07 | 9.609E-08 | 1.500E-07 | 3.400E-07 |
| 512 | 26 | H26K | A | 2.4e-07 | 3e-07 | 3.4e-07 | 2.933E-07 | 5.033E-08 | 2.400E-07 | 3.400E-07 |
| 513 | 26 | H26M | A | 1.4e-07 | 3.2e-07 | 9.5e-08 | 1.850E-07 | 1.191E-07 | 9.500E-08 | 3.200E-07 |
| 514 | 26 | H26F | A | 2.5e-07 | 3.7e-07 | 1.4e-07 | 2.533E-07 | 1.150E-07 | 1.400E-07 | 3.700E-07 |
| 515 | 26 | H26P | A | 4.7e-07 | 4e-07 | 1.9e-07 | 3.533E-07 | 1.457E-07 | 1.900E-07 | 4.700E-07 |
| 516 | 26 | H26S | A | 2.2e-07 | 7.8e-07 | 3.1e-07 | 4.367E-07 | 3.007E-07 | 2.200E-07 | 7.800E-07 |
| 517 | 26 | H26T | A | 1.6e-07 | 1.3e-07 | 1.5e-07 | 1.467E-07 | 1.528E-08 | 1.300E-07 | 1.600E-07 |
| 518 | 26 | H26W | A | 3.9e-07 | 3.1e-07 | 5.6e-07 | 4.200E-07 | 1.277E-07 | 3.100E-07 | 5.600E-07 |
| 519 | 26 | H26Y | A | 1.8e-07 | 3e-07 | 2.5e-07 | 2.433E-07 | 6.028E-08 | 1.800E-07 | 3.000E-07 |
| 520 | 26 | H26V | A | 3.4e-07 | 3.9e-07 | 2.6e-07 | 3.300E-07 | 6.557E-08 | 2.600E-07 | 3.900E-07 |
| 521 | 27 | L27A | A | 2.4e-07 | 1.2e-07 | 3.6e-07 | 2.400E-07 | 1.200E-07 | 1.200E-07 | 3.600E-07 |
| 522 | 27 | L27R | A | 1.7e-07 | 6.7e-07 | 2.8e-07 | 3.733E-07 | 2.627E-07 | 1.700E-07 | 6.700E-07 |
| 523 | 27 | L27N | A | 2e-07 | 1.7e-07 | 5.4e-07 | 3.033E-07 | 2.055E-07 | 1.700E-07 | 5.400E-07 |
| 524 | 27 | L27D | A | 2.5e-07 | 4.8e-07 | 2.9e-07 | 3.400E-07 | 1.229E-07 | 2.500E-07 | 4.800E-07 |
| 525 | 27 | L27C | A | 3.5e-07 | 5e-07 | 2.3e-07 | 3.600E-07 | 1.353E-07 | 2.300E-07 | 5.000E-07 |
| 526 | 27 | L27E | A | 3.6e-07 | 2.5e-07 | 5e-07 | 3.700E-07 | 1.253E-07 | 2.500E-07 | 5.000E-07 |
| 527 | 27 | L27Q | A | 3.5e-07 | 2.5e-07 | 1.8e-07 | 2.600E-07 | 8.544E-08 | 1.800E-07 | 3.500E-07 |
| 528 | 27 | L27G | A | 2.5e-07 | 2.3e-07 | 3.3e-07 | 2.700E-07 | 5.292E-08 | 2.300E-07 | 3.300E-07 |
| 529 | 27 | L27H | A | 1.6e-07 | 1.9e-07 | 1.8e-07 | 1.767E-07 | 1.528E-08 | 1.600E-07 | 1.900E-07 |
| 530 | 27 | L27I | A | 3.2e-07 | 2.1e-07 | 1.8e-07 | 2.367E-07 | 7.371E-08 | 1.800E-07 | 3.200E-07 |
| 531 | 27 | L27L | A | 3.1e-07 | 3.4e-07 | 2.2e-07 | 2.900E-07 | 6.245E-08 | 2.200E-07 | 3.400E-07 |
| 532 | 27 | L27K | A | 1.2e-07 | 1.8e-07 | 1.8e-07 | 1.600E-07 | 3.464E-08 | 1.200E-07 | 1.800E-07 |
| 533 | 27 | L27M | A | 2.4e-07 | 2.1e-07 | 3.8e-07 | 2.767E-07 | 9.074E-08 | 2.100E-07 | 3.800E-07 |
| 534 | 27 | L27F | A | 3.3e-07 | 2.4e-07 | 3.9e-07 | 3.200E-07 | 7.550E-08 | 2.400E-07 | 3.900E-07 |
| 535 | 27 | L27P | A | 6.2e-07 | 1.9e-07 | 1.3e-07 | 3.133E-07 | 2.673E-07 | 1.300E-07 | 6.200E-07 |
| 536 | 27 | L27S | A | 3.9e-07 | 2.3e-07 | 2e-07 | 2.733E-07 | 1.021E-07 | 2.000E-07 | 3.900E-07 |
| 537 | 27 | L27T | A | 3.2e-07 | 3.6e-07 | 3.1e-07 | 3.300E-07 | 2.646E-08 | 3.100E-07 | 3.600E-07 |
| 538 | 27 | L27W | A | 2.4e-07 | 3.2e-07 | 2.2e-07 | 2.600E-07 | 5.292E-08 | 2.200E-07 | 3.200E-07 |
| 539 | 27 | L27Y | A | 1.2e-07 | 1.8e-07 | 4.3e-07 | 2.433E-07 | 1.644E-07 | 1.200E-07 | 4.300E-07 |
| 540 | 27 | L27V | A | 2.5e-07 | 3.1e-07 | 2.3e-07 | 2.633E-07 | 4.163E-08 | 2.300E-07 | 3.100E-07 |
| 541 | 28 | Y28A | A | 5.6e-07 | 4.6e-07 | 4.5e-07 | 4.900E-07 | 6.083E-08 | 4.500E-07 | 5.600E-07 |
| 542 | 28 | Y28R | A | 3.3e-07 | 2.8e-07 | 4.3e-07 | 3.467E-07 | 7.638E-08 | 2.800E-07 | 4.300E-07 |
| 543 | 28 | Y28N | A | 2e-07 | 1.8e-07 | 4.2e-07 | 2.667E-07 | 1.332E-07 | 1.800E-07 | 4.200E-07 |
| 544 | 28 | Y28D | A | 2.5e-07 | 4.3e-07 | 4.7e-07 | 3.833E-07 | 1.172E-07 | 2.500E-07 | 4.700E-07 |
| 545 | 28 | Y28C | A | 4.2e-07 | 2.5e-07 | 1.6e-07 | 2.767E-07 | 1.320E-07 | 1.600E-07 | 4.200E-07 |
| 546 | 28 | Y28E | A | 4.1e-07 | 3.8e-07 | 2.1e-07 | 3.333E-07 | 1.079E-07 | 2.100E-07 | 4.100E-07 |
| 547 | 28 | Y28Q | A | 2.5e-07 | 2.5e-07 | 2.6e-07 | 2.533E-07 | 5.774E-09 | 2.500E-07 | 2.600E-07 |
| 548 | 28 | Y28G | A | 2.2e-07 | 1.3e-07 | 2e-07 | 1.833E-07 | 4.726E-08 | 1.300E-07 | 2.200E-07 |
| 549 | 28 | Y28H | A | 2.1e-07 | 2.1e-07 | 2.1e-07 | 2.100E-07 | 0.000E+00 | 2.100E-07 | 2.100E-07 |
| 550 | 28 | Y28I | A | 3.7e-07 | 4.5e-07 | 4.5e-07 | 4.233E-07 | 4.619E-08 | 3.700E-07 | 4.500E-07 |
| 551 | 28 | Y28L | A | 3.1e-07 | 3.2e-07 | 2.9e-07 | 3.067E-07 | 1.528E-08 | 2.900E-07 | 3.200E-07 |
| 552 | 28 | Y28K | A | 4e-07 | 1.3e-07 | 2.2e-07 | 2.500E-07 | 1.375E-07 | 1.300E-07 | 4.000E-07 |
| 553 | 28 | Y28M | A | 2.2e-07 | 2.7e-07 | 4e-07 | 2.967E-07 | 9.292E-08 | 2.200E-07 | 4.000E-07 |
| 554 | 28 | Y28F | A | 1.8e-07 | 3.6e-07 | 2.7e-07 | 2.700E-07 | 9.000E-08 | 1.800E-07 | 3.600E-07 |
| 555 | 28 | Y28P | A | 6e-07 | 1.1e-07 | 3.4e-07 | 3.500E-07 | 2.452E-07 | 1.100E-07 | 6.000E-07 |
| 556 | 28 | Y28S | A | 1.1e-07 | 1.7e-07 | 8.7e-08 | 1.223E-07 | 4.285E-08 | 8.700E-08 | 1.700E-07 |
| 557 | 28 | Y28T | A | 2.9e-07 | 3.6e-07 | 3.7e-07 | 3.400E-07 | 4.359E-08 | 2.900E-07 | 3.700E-07 |
| 558 | 28 | Y28W | A | 1.4e-07 | 2.7e-07 | 3.2e-07 | 2.433E-07 | 9.292E-08 | 1.400E-07 | 3.200E-07 |
| 559 | 28 | Y28Y | A | 2.8e-07 | 2.7e-07 | 1e-07 | 2.167E-07 | 1.012E-07 | 1.000E-07 | 2.800E-07 |
| 560 | 28 | Y28V | A | 2.9e-07 | 3.2e-07 | 3.4e-07 | 3.167E-07 | 2.517E-08 | 2.900E-07 | 3.400E-07 |
| 561 | 29 | Q29A | A | 2.7e-07 | 1.5e-07 | 4.9e-07 | 3.033E-07 | 1.724E-07 | 1.500E-07 | 4.900E-07 |
| 562 | 29 | Q29R | A | 4.5e-07 | 2.4e-07 | 1.4e-07 | 2.767E-07 | 1.582E-07 | 1.400E-07 | 4.500E-07 |
| 563 | 29 | Q29N | A | 4.8e-07 | 2e-07 | 2.5e-07 | 3.100E-07 | 1.493E-07 | 2.000E-07 | 4.800E-07 |
| 564 | 29 | Q29D | A | 2.1e-07 | 3.7e-07 | 1.7e-07 | 2.500E-07 | 1.058E-07 | 1.700E-07 | 3.700E-07 |
| 565 | 29 | Q29C | A | 3.3e-07 | 2.3e-07 | 1.4e-07 | 2.333E-07 | 9.504E-08 | 1.400E-07 | 3.300E-07 |
| 566 | 29 | Q29E | A | 3.4e-07 | 2.9e-07 | 2.2e-07 | 2.833E-07 | 6.028E-08 | 2.200E-07 | 3.400E-07 |
| 567 | 29 | Q29Q | A | 1.4e-07 | 2.4e-07 | 2.2e-07 | 2.000E-07 | 5.292E-08 | 1.400E-07 | 2.400E-07 |
| 568 | 29 | Q29G | A | 1.4e-07 | 4.2e-07 | 2.8e-07 | 2.800E-07 | 1.400E-07 | 1.400E-07 | 4.200E-07 |
| 569 | 29 | Q29H | A | 3.3e-07 | 2.9e-07 | 3.7e-07 | 3.300E-07 | 4.000E-08 | 2.900E-07 | 3.700E-07 |
| 570 | 29 | Q29I | A | 2.8e-07 | 2e-07 | 2.7e-07 | 2.500E-07 | 4.359E-08 | 2.000E-07 | 2.800E-07 |
| 571 | 29 | Q29L | A | 5.1e-07 | 2.9e-07 | 6.4e-07 | 4.800E-07 | 1.769E-07 | 2.900E-07 | 6.400E-07 |
| 572 | 29 | Q29K | A | 1.3e-07 | 9.6e-08 | 2.8e-07 | 1.687E-07 | 9.790E-08 | 9.600E-08 | 2.800E-07 |
| 573 | 29 | Q29M | A | 1.9e-06 | 9.4e-08 | 3.5e-07 | 7.813E-07 | 9.772E-07 | 9.400E-08 | 1.900E-06 |
| 574 | 29 | Q29F | A | 3e-07 | 1.7e-07 | 2.1e-07 | 2.267E-07 | 6.658E-08 | 1.700E-07 | 3.000E-07 |
| 575 | 29 | Q29P | A | 3e-07 | 4.2e-07 | 2.7e-07 | 3.300E-07 | 7.937E-08 | 2.700E-07 | 4.200E-07 |
| 576 | 29 | Q29S | A | 1.6e-07 | 4.9e-07 | 1.9e-07 | 2.800E-07 | 1.825E-07 | 1.600E-07 | 4.900E-07 |
| 577 | 29 | Q29T | A | 2.3e-07 | 2.6e-07 | 2e-07 | 2.300E-07 | 3.000E-08 | 2.000E-07 | 2.600E-07 |
| 578 | 29 | Q29W | A | 3.6e-07 | 4.5e-07 | 1.8e-07 | 3.300E-07 | 1.375E-07 | 1.800E-07 | 4.500E-07 |
| 579 | 29 | Q29Y | A | 2.1e-07 | 2.4e-07 | 2.6e-07 | 2.367E-07 | 2.517E-08 | 2.100E-07 | 2.600E-07 |
| 580 | 29 | Q29V | A | 3.1e-07 | 1.2e-07 | 4.3e-07 | 2.867E-07 | 1.563E-07 | 1.200E-07 | 4.300E-07 |
| 581 | 30 | G30A | A | 4.6e-07 | 9.9e-08 | 2.6e-07 | 2.730E-07 | 1.809E-07 | 9.900E-08 | 4.600E-07 |
| 582 | 30 | G30R | A | 2.2e-07 | 3.2e-07 | 1.8e-07 | 2.400E-07 | 7.211E-08 | 1.800E-07 | 3.200E-07 |
| 583 | 30 | G30N | A | 3.7e-07 | 2.1e-07 | 2.9e-07 | 2.900E-07 | 8.000E-08 | 2.100E-07 | 3.700E-07 |
| 584 | 30 | G30D | A | 2.3e-07 | 2e-07 | 4.1e-07 | 2.800E-07 | 1.136E-07 | 2.000E-07 | 4.100E-07 |
| 585 | 30 | G30C | A | 1.7e-07 | 2.6e-07 | 1.2e-07 | 1.833E-07 | 7.095E-08 | 1.200E-07 | 2.600E-07 |
| 586 | 30 | G30E | A | 1.3e-07 | 4.1e-07 | 2.3e-07 | 2.567E-07 | 1.419E-07 | 1.300E-07 | 4.100E-07 |
| 587 | 30 | G30Q | A | 1.5e-07 | 1.9e-07 | 2e-07 | 1.800E-07 | 2.646E-08 | 1.500E-07 | 2.000E-07 |
| 588 | 30 | G30G | A | 3.2e-07 | 2.8e-07 | 4e-07 | 3.333E-07 | 6.110E-08 | 2.800E-07 | 4.000E-07 |
| 589 | 30 | G30H | A | 2.5e-07 | 2.7e-07 | 3.2e-07 | 2.800E-07 | 3.606E-08 | 2.500E-07 | 3.200E-07 |
| 590 | 30 | G30I | A | 2.7e-07 | 2e-07 | 3.3e-07 | 2.667E-07 | 6.506E-08 | 2.000E-07 | 3.300E-07 |
| 591 | 30 | G30L | A | 2.5e-07 | 5.2e-07 | 2e-07 | 3.233E-07 | 1.721E-07 | 2.000E-07 | 5.200E-07 |
| 592 | 30 | G30K | A | 3.3e-07 | 2.7e-07 | 3.5e-07 | 3.167E-07 | 4.163E-08 | 2.700E-07 | 3.500E-07 |
| 593 | 30 | G30M | A | 2.2e-07 | 2.6e-07 | 3.3e-07 | 2.700E-07 | 5.568E-08 | 2.200E-07 | 3.300E-07 |
| 594 | 30 | G30F | A | 4.1e-07 | 1.3e-07 | 4.1e-07 | 3.167E-07 | 1.617E-07 | 1.300E-07 | 4.100E-07 |

| | | | | | | | | | | |
|---|---|---|---|---|---|---|---|---|---|---|
| 595 | 30 | G30P | A | 2.5e-07 | 2e-07 | 1.7e-07 | 2.067E-07 | 4.041E-08 | 1.700E-07 | 2.500E-07 |
| 596 | 30 | G30S | A | 1.6e-07 | 3.2e-07 | 3.5e-07 | 2.767E-07 | 1.021E-07 | 1.600E-07 | 3.500E-07 |
| 597 | 30 | G30T | A | 3.9e-07 | 2.6e-07 | 3.6e-07 | 3.367E-07 | 6.807E-08 | 2.600E-07 | 3.900E-07 |
| 598 | 30 | G30W | A | 3.3e-07 | 2.1e-07 | 2.4e-07 | 2.600E-07 | 6.245E-08 | 2.100E-07 | 3.300E-07 |
| 599 | 30 | G30Y | A | 3.2e-07 | 3e-07 | 9e-08 | 2.367E-07 | 1.274E-07 | 9.000E-08 | 3.200E-07 |
| 600 | 30 | G30V | A | 4.2e-07 | 2.3e-07 | 2.3e-07 | 2.933E-07 | 1.097E-07 | 2.300E-07 | 4.200E-07 |
| 601 | 31 | C31A | A | 2.1e-07 | 4.4e-07 | 2.8e-07 | 3.100E-07 | 1.179E-07 | 2.100E-07 | 4.400E-07 |
| 602 | 31 | C31R | A | 4.6e-07 | 1.7e-07 | 3.9e-07 | 3.400E-07 | 1.513E-07 | 1.700E-07 | 4.600E-07 |
| 603 | 31 | C31N | A | 3.2e-07 | 2.8e-07 | 2.8e-07 | 2.933E-07 | 2.309E-08 | 2.800E-07 | 3.200E-07 |
| 604 | 31 | C31D | A | 1.6e-07 | 1.4e-07 | 3.1e-07 | 2.033E-07 | 9.292E-08 | 1.400E-07 | 3.100E-07 |
| 605 | 31 | C31C | A | 2.6e-07 | 1.7e-07 | 3.6e-07 | 2.633E-07 | 9.504E-08 | 1.700E-07 | 3.600E-07 |
| 606 | 31 | C31E | A | 2.4e-07 | 3.3e-07 | 3.9e-07 | 3.200E-07 | 7.550E-08 | 2.400E-07 | 3.900E-07 |
| 607 | 31 | C31Q | A | 1.6e-07 | 2.8e-07 | 4e-07 | 2.800E-07 | 1.200E-07 | 1.600E-07 | 4.000E-07 |
| 608 | 31 | C31G | A | 1.8e-07 | 3.1e-07 | 2.2e-07 | 2.367E-07 | 6.658E-08 | 1.800E-07 | 3.100E-07 |
| 609 | 31 | C31H | A | 1.9e-07 | 2.6e-07 | 4e-07 | 2.833E-07 | 1.069E-07 | 1.900E-07 | 4.000E-07 |
| 610 | 31 | C31I | A | 6.6e-07 | 2.3e-07 | 1.7e-07 | 3.533E-07 | 2.673E-07 | 1.700E-07 | 6.600E-07 |
| 611 | 31 | C31L | A | 3.6e-07 | 4.7e-07 | 1.6e-07 | 3.300E-07 | 1.572E-07 | 1.600E-07 | 4.700E-07 |
| 612 | 31 | C31K | A | 3.2e-07 | 2e-07 | 1.2e-07 | 2.133E-07 | 1.007E-07 | 1.200E-07 | 3.200E-07 |
| 613 | 31 | C31M | A | 2.2e-07 | 4e-07 | 2.2e-07 | 2.800E-07 | 1.039E-07 | 2.200E-07 | 4.000E-07 |
| 614 | 31 | C31F | A | 3.7e-07 | 9e-08 | 3.4e-07 | 2.667E-07 | 1.537E-07 | 9.000E-08 | 3.700E-07 |
| 615 | 31 | C31P | A | 2.7e-07 | 2e-07 | 2.7e-07 | 2.467E-07 | 4.041E-08 | 2.000E-07 | 2.700E-07 |
| 616 | 31 | C31S | A | 1.2e-07 | 2.6e-07 | 1.7e-07 | 1.833E-07 | 7.095E-08 | 1.200E-07 | 2.600E-07 |
| 617 | 31 | C31T | A | 3.5e-07 | 1.5e-07 | 1.6e-07 | 2.200E-07 | 1.127E-07 | 1.500E-07 | 3.500E-07 |
| 618 | 31 | C31W | A | 3.6e-07 | 2.6e-07 | 3.3e-07 | 3.167E-07 | 5.132E-08 | 2.600E-07 | 3.600E-07 |
| 619 | 31 | C31Y | A | 2e-07 | 1.4e-07 | 1.5e-07 | 1.633E-07 | 3.215E-08 | 1.400E-07 | 2.000E-07 |
| 620 | 31 | C31V | A | 2.1e-07 | 2.5e-07 | 3.2e-07 | 2.600E-07 | 5.568E-08 | 2.100E-07 | 3.200E-07 |
| 621 | 32 | Q32A | A | 1.8e-07 | 4.9e-07 | 2.2e-07 | 2.967E-07 | 1.686E-07 | 1.800E-07 | 4.900E-07 |
| 622 | 32 | Q32R | A | 1.8e-07 | 1.3e-07 | 1.6e-07 | 1.567E-07 | 2.517E-08 | 1.300E-07 | 1.800E-07 |
| 623 | 32 | Q32N | A | 1.4e-07 | 1.6e-07 | 3.4e-07 | 2.133E-07 | 1.102E-07 | 1.400E-07 | 3.400E-07 |
| 624 | 32 | Q32D | A | 6.8e-07 | 2.9e-07 | 5.2e-07 | 4.967E-07 | 1.960E-07 | 2.900E-07 | 6.800E-07 |
| 625 | 32 | Q32C | A | 1.1e-07 | 1.8e-07 | 1.4e-07 | 1.433E-07 | 3.512E-08 | 1.100E-07 | 1.800E-07 |
| 626 | 32 | Q32E | A | 2.3e-07 | 4.7e-07 | 2.3e-07 | 3.100E-07 | 1.386E-07 | 2.300E-07 | 4.700E-07 |
| 627 | 32 | Q32Q | A | 1.7e-07 | 2.8e-07 | 3.8e-07 | 2.767E-07 | 1.050E-07 | 1.700E-07 | 3.800E-07 |
| 628 | 32 | Q32G | A | 3.6e-07 | 1.3e-07 | 5.7e-07 | 3.533E-07 | 2.201E-07 | 1.300E-07 | 5.700E-07 |
| 629 | 32 | Q32H | A | 2.4e-07 | 2.2e-07 | 1.9e-07 | 2.167E-07 | 2.517E-08 | 1.900E-07 | 2.400E-07 |
| 630 | 32 | Q32I | A | 1.8e-07 | 2.2e-07 | 3e-07 | 2.333E-07 | 6.110E-08 | 1.800E-07 | 3.000E-07 |
| 631 | 32 | Q32L | A | 2.7e-07 | 2.1e-07 | 2e-07 | 2.267E-07 | 3.786E-08 | 2.000E-07 | 2.700E-07 |
| 632 | 32 | Q32K | A | 1.4e-07 | 2.6e-07 | 3.1e-07 | 2.367E-07 | 8.737E-08 | 1.400E-07 | 3.100E-07 |
| 633 | 32 | Q32M | A | 2.2e-07 | 1.6e-07 | 1.6e-07 | 1.800E-07 | 3.464E-08 | 1.600E-07 | 2.200E-07 |
| 634 | 32 | Q32F | A | 2.6e-07 | 2.4e-07 | 3.7e-07 | 2.900E-07 | 7.000E-08 | 2.400E-07 | 3.700E-07 |
| 635 | 32 | Q32P | A | 1.5e-07 | 4.1e-07 | 3.2e-07 | 2.933E-07 | 1.320E-07 | 1.500E-07 | 4.100E-07 |
| 636 | 32 | Q32S | A | 1.8e-07 | 1.9e-07 | 1.1e-07 | 1.600E-07 | 4.359E-08 | 1.100E-07 | 1.900E-07 |
| 637 | 32 | Q32T | A | 1.7e-07 | 2.8e-07 | 2.8e-07 | 2.433E-07 | 6.351E-08 | 1.700E-07 | 2.800E-07 |
| 638 | 32 | Q32W | A | 3.1e-07 | 4.3e-07 | 1.1e-07 | 2.833E-07 | 1.617E-07 | 1.100E-07 | 4.300E-07 |
| 639 | 32 | Q32Y | A | 1.9e-07 | 1.5e-07 | 1.9e-07 | 1.767E-07 | 2.309E-08 | 1.500E-07 | 1.900E-07 |
| 640 | 32 | Q32V | A | 2.9e-07 | 1.3e-07 | 2.4e-07 | 2.200E-07 | 8.185E-08 | 1.300E-07 | 2.900E-07 |
| 641 | 33 | V33A | A | 1.3e-07 | 5.4e-07 | 3.3e-07 | 3.333E-07 | 2.050E-07 | 1.300E-07 | 5.400E-07 |
| 642 | 33 | V33R | A | 1.8e-07 | 1.7e-07 | 3.3e-07 | 2.267E-07 | 8.963E-08 | 1.700E-07 | 3.300E-07 |
| 643 | 33 | V33N | A | 1.4e-07 | 3.8e-07 | 4.1e-07 | 3.100E-07 | 1.480E-07 | 1.400E-07 | 4.100E-07 |
| 644 | 33 | V33D | A | 3.2e-07 | 2.7e-07 | 1e-07 | 2.300E-07 | 1.153E-07 | 1.000E-07 | 3.200E-07 |
| 645 | 33 | V33C | A | 4.4e-07 | 1.8e-07 | 9e-08 | 2.367E-07 | 1.818E-07 | 9.000E-08 | 4.400E-07 |
| 646 | 33 | V33E | A | 1.9e-07 | 3.9e-07 | 1e-07 | 2.267E-07 | 1.484E-07 | 1.000E-07 | 3.900E-07 |
| 647 | 33 | V33Q | A | 3.9e-07 | 3.1e-07 | 3.1e-07 | 3.367E-07 | 4.619E-08 | 3.100E-07 | 3.900E-07 |
| 648 | 33 | V33G | A | 1.4e-07 | 1.3e-07 | 4.4e-07 | 2.367E-07 | 1.762E-07 | 1.300E-07 | 4.400E-07 |
| 649 | 33 | V33H | A | 4.4e-07 | 1.6e-07 | 2.2e-07 | 2.733E-07 | 1.474E-07 | 1.600E-07 | 4.400E-07 |
| 650 | 33 | V33I | A | 2.1e-07 | 2.1e-07 | 2e-07 | 2.067E-07 | 5.774E-09 | 2.000E-07 | 2.100E-07 |
| 651 | 33 | V33L | A | 1.4e-07 | 2.7e-07 | 4.6e-07 | 2.900E-07 | 1.609E-07 | 1.400E-07 | 4.600E-07 |
| 652 | 33 | V33K | A | 1.6e-07 | 4.6e-07 | 1.5e-07 | 2.567E-07 | 1.762E-07 | 1.500E-07 | 4.600E-07 |
| 653 | 33 | V33M | A | 2.1e-07 | 2.4e-07 | 1.3e-07 | 1.933E-07 | 5.686E-08 | 1.300E-07 | 2.400E-07 |
| 654 | 33 | V33F | A | 2.5e-07 | 4.6e-07 | 1.7e-07 | 2.933E-07 | 1.498E-07 | 1.700E-07 | 4.600E-07 |
| 655 | 33 | V33P | A | 1.7e-07 | 1.8e-07 | 5e-07 | 2.833E-07 | 1.877E-07 | 1.700E-07 | 5.000E-07 |
| 656 | 33 | V33S | A | 1.6e-07 | 2.1e-07 | 2.3e-07 | 2.000E-07 | 3.606E-08 | 1.600E-07 | 2.300E-07 |
| 657 | 33 | V33T | A | 2.2e-07 | 3e-07 | 1.6e-07 | 2.267E-07 | 7.024E-08 | 1.600E-07 | 3.000E-07 |
| 658 | 33 | V33W | A | 1.9e-07 | 1.9e-07 | 4.1e-07 | 2.633E-07 | 1.270E-07 | 1.900E-07 | 4.100E-07 |
| 659 | 33 | V33Y | A | 2.3e-07 | 1.6e-07 | 1.4e-07 | 1.767E-07 | 4.726E-08 | 1.400E-07 | 2.300E-07 |
| 660 | 33 | V33V | A | 2.4e-07 | 2.7e-07 | 2.4e-07 | 2.500E-07 | 1.732E-08 | 2.400E-07 | 2.700E-07 |
| 661 | 34 | V34A | A | 3.8e-07 | 3.4e-07 | 3.7e-07 | 3.633E-07 | 2.082E-08 | 3.400E-07 | 3.800E-07 |
| 662 | 34 | V34R | A | 7.3e-07 | 2.2e-07 | 4.6e-07 | 4.700E-07 | 2.551E-07 | 2.200E-07 | 7.300E-07 |
| 663 | 34 | V34N | A | 6.1e-07 | 2.5e-07 | 1.1e-07 | 3.233E-07 | 2.579E-07 | 1.100E-07 | 6.100E-07 |
| 664 | 34 | V34D | A | 2.8e-07 | 1.4e-07 | 2.2e-07 | 2.133E-07 | 7.024E-08 | 1.400E-07 | 2.800E-07 |
| 665 | 34 | V34C | A | 3.4e-07 | 2.9e-07 | 1.4e-07 | 2.567E-07 | 1.041E-07 | 1.400E-07 | 3.400E-07 |
| 666 | 34 | V34E | A | 1.3e-07 | 3.3e-07 | 4.5e-07 | 3.033E-07 | 1.617E-07 | 1.300E-07 | 4.500E-07 |
| 667 | 34 | V34Q | A | 2.5e-07 | 2.4e-07 | 1.9e-07 | 2.267E-07 | 3.215E-08 | 1.900E-07 | 2.500E-07 |
| 668 | 34 | V34G | A | 2.5e-07 | 1e-07 | 3.8e-07 | 2.433E-07 | 1.401E-07 | 1.000E-07 | 3.800E-07 |
| 669 | 34 | V34H | A | 1.7e-07 | 3.5e-07 | 3.4e-07 | 2.867E-07 | 1.012E-07 | 1.700E-07 | 3.500E-07 |
| 670 | 34 | V34I | A | 3e-07 | 2.2e-07 | 1.8e-07 | 2.333E-07 | 6.110E-08 | 1.800E-07 | 3.000E-07 |
| 671 | 34 | V34L | A | 1.5e-07 | 1.9e-07 | 4.8e-07 | 2.733E-07 | 1.801E-07 | 1.500E-07 | 4.800E-07 |
| 672 | 34 | V34K | A | 2.3e-07 | 2.6e-07 | 4.9e-07 | 3.267E-07 | 1.422E-07 | 2.300E-07 | 4.900E-07 |
| 673 | 34 | V34M | A | 2.8e-07 | 2.2e-07 | 2.8e-07 | 2.600E-07 | 3.464E-08 | 2.200E-07 | 2.800E-07 |
| 674 | 34 | V34F | A | 5.8e-07 | 2.2e-07 | 3.3e-07 | 3.767E-07 | 1.845E-07 | 2.200E-07 | 5.800E-07 |
| 675 | 34 | V34P | A | 9.5e-08 | 2.4e-07 | 2.5e-07 | 1.950E-07 | 8.675E-08 | 9.500E-08 | 2.500E-07 |
| 676 | 34 | V34S | A | 2.6e-07 | 2.5e-07 | 3.6e-07 | 2.900E-07 | 6.083E-08 | 2.500E-07 | 3.600E-07 |
| 677 | 34 | V34T | A | 2.9e-07 | 5.1e-07 | 2.1e-07 | 3.367E-07 | 1.553E-07 | 2.100E-07 | 5.100E-07 |
| 678 | 34 | V34W | A | 1.9e-07 | 2.1e-07 | 1.5e-07 | 1.833E-07 | 3.055E-08 | 1.500E-07 | 2.100E-07 |
| 679 | 34 | V34Y | A | 7.9e-08 | 4.5e-07 | 2.7e-07 | 2.663E-07 | 1.855E-07 | 7.900E-08 | 4.500E-07 |

| | | | | | | | | | | |
|---|---|---|---|---|---|---|---|---|---|---|
| 680 | 34 | V34V | A | 2.2e-07 | 1.5e-07 | 5.3e-07 | 3.000E-07 | 2.022E-07 | 1.500E-07 | 5.300E-07 |
| 681 | 35 | Q35A | A | 5.1e-07 | 3.5e-07 | 4e-07 | 4.200E-07 | 8.185E-08 | 3.500E-07 | 5.100E-07 |
| 682 | 35 | Q35R | A | 2.3e-07 | 3.7e-07 | 1.6e-07 | 2.533E-07 | 1.069E-07 | 1.600E-07 | 3.700E-07 |
| 683 | 35 | Q35N | A | 2.5e-07 | 3.4e-07 | 2.9e-07 | 2.933E-07 | 4.509E-08 | 2.500E-07 | 3.400E-07 |
| 684 | 35 | Q35D | A | 2.4e-07 | 3.3e-07 | 6.4e-07 | 4.033E-07 | 2.098E-07 | 2.400E-07 | 6.400E-07 |
| 685 | 35 | Q35C | A | 4.4e-07 | 4.7e-07 | 6.8e-07 | 5.300E-07 | 1.308E-07 | 4.400E-07 | 6.800E-07 |
| 686 | 35 | Q35E | A | 1.1e-07 | 2.6e-07 | 2.3e-07 | 2.000E-07 | 7.937E-08 | 1.100E-07 | 2.600E-07 |
| 687 | 35 | Q35Q | A | 1.9e-07 | 3.2e-07 | 1.7e-07 | 2.267E-07 | 8.145E-08 | 1.700E-07 | 3.200E-07 |
| 688 | 35 | Q35G | A | 3.9e-07 | 2.4e-07 | 2.8e-07 | 3.033E-07 | 7.767E-08 | 2.400E-07 | 3.900E-07 |
| 689 | 35 | Q35H | A | 2.4e-07 | 5e-07 | 9.1e-08 | 2.770E-07 | 2.070E-07 | 9.100E-08 | 5.000E-07 |
| 690 | 35 | Q35I | A | 4.2e-07 | 2.5e-07 | 1.4e-07 | 2.700E-07 | 1.411E-07 | 1.400E-07 | 4.200E-07 |
| 691 | 35 | Q35L | A | 3.4e-07 | 1.4e-07 | 1.3e-07 | 2.033E-07 | 1.185E-07 | 1.300E-07 | 3.400E-07 |
| 692 | 35 | Q35K | A | 1.3e-07 | 1.5e-07 | 2.9e-07 | 1.900E-07 | 8.718E-08 | 1.300E-07 | 2.900E-07 |
| 693 | 35 | Q35M | A | 3.2e-07 | 1.2e-07 | 1.4e-07 | 1.933E-07 | 1.102E-07 | 1.200E-07 | 3.200E-07 |
| 694 | 35 | Q35F | A | 2.5e-07 | 4.4e-07 | 3.1e-07 | 3.333E-07 | 9.713E-08 | 2.500E-07 | 4.400E-07 |
| 695 | 35 | Q35P | A | 5.5e-07 | 1.7e-07 | 2.1e-07 | 3.100E-07 | 2.088E-07 | 1.700E-07 | 5.500E-07 |
| 696 | 35 | Q35S | A | 2.1e-07 | 2.8e-07 | 2.1e-07 | 2.333E-07 | 4.041E-08 | 2.100E-07 | 2.800E-07 |
| 697 | 35 | Q35T | A | 3.4e-07 | 2.8e-07 | 1.9e-07 | 2.700E-07 | 7.550E-08 | 1.900E-07 | 3.400E-07 |
| 698 | 35 | Q35W | A | 2.5e-07 | 2.1e-07 | 4.6e-07 | 3.067E-07 | 1.343E-07 | 2.100E-07 | 4.600E-07 |
| 699 | 35 | Q35Y | A | 3.8e-07 | 2.5e-07 | 2.5e-07 | 2.933E-07 | 7.506E-08 | 2.500E-07 | 3.800E-07 |
| 700 | 35 | Q35V | A | 4.3e-07 | 2.8e-07 | 1.4e-07 | 2.833E-07 | 1.450E-07 | 1.400E-07 | 4.300E-07 |
| 701 | 36 | G36A | A | 2.1e-07 | 3.7e-07 | 7.2e-07 | 4.333E-07 | 2.608E-07 | 2.100E-07 | 7.200E-07 |
| 702 | 36 | G36R | A | 8.5e-08 | 1.8e-07 | 5.3e-07 | 2.650E-07 | 2.344E-07 | 8.500E-08 | 5.300E-07 |
| 703 | 36 | G36N | A | 2.5e-07 | 2.9e-07 | 2e-07 | 2.467E-07 | 4.509E-08 | 2.000E-07 | 2.900E-07 |
| 704 | 36 | G36D | A | 1.5e-07 | 1.9e-07 | 1.9e-07 | 1.767E-07 | 2.309E-08 | 1.500E-07 | 1.900E-07 |
| 705 | 36 | G36C | A | 3.7e-07 | 1.8e-07 | 2.9e-07 | 2.800E-07 | 9.539E-08 | 1.800E-07 | 3.700E-07 |
| 706 | 36 | G36E | A | 4.5e-07 | 3.2e-07 | 2.6e-07 | 3.433E-07 | 9.713E-08 | 2.600E-07 | 4.500E-07 |
| 707 | 36 | G36Q | A | 1.8e-07 | 5.3e-07 | 2.4e-07 | 3.167E-07 | 1.872E-07 | 1.800E-07 | 5.300E-07 |
| 708 | 36 | G36G | A | 3e-07 | 1.7e-07 | 2.1e-07 | 2.267E-07 | 6.658E-08 | 1.700E-07 | 3.000E-07 |
| 709 | 36 | G36H | A | 2.5e-07 | 2.2e-07 | 1.4e-07 | 2.033E-07 | 5.686E-08 | 1.400E-07 | 2.500E-07 |
| 710 | 36 | G36I | A | 3.9e-07 | 3.1e-07 | 2.5e-07 | 3.167E-07 | 7.024E-08 | 2.500E-07 | 3.900E-07 |
| 711 | 36 | G36L | A | 3.9e-07 | 1.8e-07 | 2.1e-07 | 2.600E-07 | 1.136E-07 | 1.800E-07 | 3.900E-07 |
| 712 | 36 | G36K | A | 4.7e-07 | 4.1e-07 | 2.8e-07 | 3.867E-07 | 9.713E-08 | 2.800E-07 | 4.700E-07 |
| 713 | 36 | G36M | A | 4.7e-07 | 1.4e-07 | 1.8e-07 | 2.633E-07 | 1.801E-07 | 1.400E-07 | 4.700E-07 |
| 714 | 36 | G36F | A | 5.4e-07 | 2e-07 | 2.3e-07 | 3.233E-07 | 1.882E-07 | 2.000E-07 | 5.400E-07 |
| 715 | 36 | G36P | A | 4.1e-07 | 2.5e-07 | 2.1e-07 | 2.900E-07 | 1.058E-07 | 2.100E-07 | 4.100E-07 |
| 716 | 36 | G36S | A | 1.3e-07 | 3.6e-07 | 1.6e-07 | 2.167E-07 | 1.250E-07 | 1.300E-07 | 3.600E-07 |
| 717 | 36 | G36T | A | 1.6e-07 | 1.2e-07 | 2.4e-07 | 1.733E-07 | 6.110E-08 | 1.200E-07 | 2.400E-07 |
| 718 | 36 | G36W | A | 2.1e-07 | 1.6e-07 | 1.7e-07 | 1.800E-07 | 2.646E-08 | 1.600E-07 | 2.100E-07 |
| 719 | 36 | G36Y | A | 5.6e-07 | 2.8e-07 | 1.6e-07 | 3.333E-07 | 2.053E-07 | 1.600E-07 | 5.600E-07 |
| 720 | 36 | G36V | A | 4e-07 | 2.2e-07 | 2.9e-07 | 3.033E-07 | 9.074E-08 | 2.200E-07 | 4.000E-07 |
| 721 | 37 | N37A | A | 2.8e-07 | 6.1e-07 | 2.4e-07 | 3.767E-07 | 2.031E-07 | 2.400E-07 | 6.100E-07 |
| 722 | 37 | N37R | A | 2.8e-07 | 9.4e-08 | 2.9e-07 | 2.213E-07 | 1.104E-07 | 9.400E-08 | 2.900E-07 |
| 723 | 37 | N37N | A | 1.6e-07 | 2.3e-07 | 2.5e-07 | 2.133E-07 | 4.726E-08 | 1.600E-07 | 2.500E-07 |
| 724 | 37 | N37D | A | 1.9e-07 | 3.2e-07 | 1.3e-07 | 2.133E-07 | 9.713E-08 | 1.300E-07 | 3.200E-07 |
| 725 | 37 | N37C | A | 2e-07 | 3e-07 | 2.4e-07 | 2.467E-07 | 5.033E-08 | 2.000E-07 | 3.000E-07 |
| 726 | 37 | N37E | A | 3.8e-07 | 1.9e-07 | 2e-07 | 2.567E-07 | 1.069E-07 | 1.900E-07 | 3.800E-07 |
| 727 | 37 | N37Q | A | 1.7e-07 | 4e-07 | 7.1e-08 | 2.137E-07 | 1.688E-07 | 7.100E-08 | 4.000E-07 |
| 728 | 37 | N37G | A | 1.6e-07 | 1.6e-07 | 2.4e-07 | 1.867E-07 | 4.619E-08 | 1.600E-07 | 2.400E-07 |
| 729 | 37 | N37H | A | 3.1e-07 | 2e-07 | 3.1e-07 | 2.733E-07 | 6.351E-08 | 2.000E-07 | 3.100E-07 |
| 730 | 37 | N37I | A | 1.6e-07 | 2e-07 | 2e-07 | 1.867E-07 | 2.309E-08 | 1.600E-07 | 2.000E-07 |
| 731 | 37 | N37L | A | 1.5e-07 | 3.2e-07 | 2e-07 | 2.233E-07 | 8.737E-08 | 1.500E-07 | 3.200E-07 |
| 732 | 37 | N37K | A | 4.4e-07 | 1.6e-07 | 2.8e-07 | 2.933E-07 | 1.405E-07 | 1.600E-07 | 4.400E-07 |
| 733 | 37 | N37M | A | 4.1e-07 | 2.7e-07 | 3.3e-07 | 3.367E-07 | 7.024E-08 | 2.700E-07 | 4.100E-07 |
| 734 | 37 | N37F | A | 3.8e-07 | 2e-07 | 2.8e-07 | 2.867E-07 | 9.018E-08 | 2.000E-07 | 3.800E-07 |
| 735 | 37 | N37P | A | 1.9e-07 | 8.6e-07 | 1.4e-07 | 3.967E-07 | 4.020E-07 | 1.400E-07 | 8.600E-07 |
| 736 | 37 | N37S | A | 5.1e-07 | 1.9e-07 | 3.6e-07 | 3.533E-07 | 1.601E-07 | 1.900E-07 | 5.100E-07 |
| 737 | 37 | N37T | A | 3.7e-07 | 1.8e-07 | 3e-07 | 2.833E-07 | 9.609E-08 | 1.800E-07 | 3.700E-07 |
| 738 | 37 | N37W | A | 2.3e-07 | 1.9e-07 | 2.6e-07 | 2.267E-07 | 3.512E-08 | 1.900E-07 | 2.600E-07 |
| 739 | 37 | N37Y | A | 1.5e-07 | 1.5e-07 | 3.3e-07 | 2.100E-07 | 1.039E-07 | 1.500E-07 | 3.300E-07 |
| 740 | 37 | N37V | A | 2.4e-07 | 8.9e-08 | 4.6e-07 | 2.630E-07 | 1.866E-07 | 8.900E-08 | 4.600E-07 |
| 741 | 38 | L38A | A | 1.3e-07 | 1.2e-07 | 2.9e-07 | 1.800E-07 | 9.539E-08 | 1.200E-07 | 2.900E-07 |
| 742 | 38 | L38R | A | 2.3e-07 | 4.3e-07 | 2.3e-07 | 2.967E-07 | 1.155E-07 | 2.300E-07 | 4.300E-07 |
| 743 | 38 | L38N | A | 2.8e-07 | 3.4e-07 | 2.6e-07 | 2.933E-07 | 4.163E-08 | 2.600E-07 | 3.400E-07 |
| 744 | 38 | L38D | A | 1.3e-07 | 2e-07 | 3.2e-07 | 2.167E-07 | 9.609E-08 | 1.300E-07 | 3.200E-07 |
| 745 | 38 | L38C | A | 3.8e-07 | 6.1e-07 | 2.8e-07 | 4.233E-07 | 1.692E-07 | 2.800E-07 | 6.100E-07 |
| 746 | 38 | L38E | A | 1e-07 | 1.8e-07 | 4.8e-07 | 2.533E-07 | 2.003E-07 | 1.000E-07 | 4.800E-07 |
| 747 | 38 | L38Q | A | 4.5e-07 | 2.4e-07 | 2.2e-07 | 3.033E-07 | 1.274E-07 | 2.200E-07 | 4.500E-07 |
| 748 | 38 | L38G | A | 3.4e-07 | 2.5e-07 | 2.7e-07 | 2.867E-07 | 4.726E-08 | 2.500E-07 | 3.400E-07 |
| 749 | 38 | L38H | A | 1.1e-07 | 2.2e-07 | 1.8e-07 | 1.700E-07 | 5.568E-08 | 1.100E-07 | 2.200E-07 |
| 750 | 38 | L38I | A | 2.5e-07 | 1.3e-07 | 3e-07 | 2.267E-07 | 8.737E-08 | 1.300E-07 | 3.000E-07 |
| 751 | 38 | L38L | A | 4.6e-07 | 3.5e-07 | 1.2e-07 | 3.100E-07 | 1.735E-07 | 1.200E-07 | 4.600E-07 |
| 752 | 38 | L38K | A | 2.8e-07 | 2e-07 | 9e-08 | 1.900E-07 | 9.539E-08 | 9.000E-08 | 2.800E-07 |
| 753 | 38 | L38M | A | 1.9e-07 | 7.7e-08 | 5.2e-07 | 2.623E-07 | 2.302E-07 | 7.700E-08 | 5.200E-07 |
| 754 | 38 | L38F | A | 4.1e-07 | 1.9e-07 | 1.6e-07 | 2.533E-07 | 1.365E-07 | 1.600E-07 | 4.100E-07 |
| 755 | 38 | L38P | A | 1.9e-07 | 1.4e-07 | 2.3e-07 | 1.867E-07 | 4.509E-08 | 1.400E-07 | 2.300E-07 |
| 756 | 38 | L38S | A | 2.7e-07 | 7.9e-07 | 2.2e-07 | 4.267E-07 | 3.156E-07 | 2.200E-07 | 7.900E-07 |
| 757 | 38 | L38T | A | 2.8e-07 | 1.9e-07 | 2e-07 | 2.233E-07 | 4.933E-08 | 1.900E-07 | 2.800E-07 |
| 758 | 38 | L38W | A | 4e-07 | 2.1e-07 | 3.8e-07 | 3.300E-07 | 1.044E-07 | 2.100E-07 | 4.000E-07 |
| 759 | 38 | L38Y | A | 2.4e-07 | 2.2e-07 | 2.9e-07 | 2.500E-07 | 3.606E-08 | 2.200E-07 | 2.900E-07 |
| 760 | 38 | L38V | A | 1.8e-07 | 2.7e-07 | 3.3e-07 | 2.600E-07 | 7.550E-08 | 1.800E-07 | 3.300E-07 |
| 761 | 39 | E39A | A | 2.6e-07 | 1.9e-07 | 1.9e-07 | 2.133E-07 | 4.041E-08 | 1.900E-07 | 2.600E-07 |
| 762 | 39 | E39R | A | 3.8e-07 | 2.2e-07 | 2.1e-07 | 2.700E-07 | 9.539E-08 | 2.100E-07 | 3.800E-07 |
| 763 | 39 | E39N | A | 3.1e-07 | 3.3e-07 | 3.5e-07 | 3.300E-07 | 2.000E-08 | 3.100E-07 | 3.500E-07 |
| 764 | 39 | E39D | A | 7.2e-07 | 5.8e-07 | 2.6e-07 | 5.200E-07 | 2.358E-07 | 2.600E-07 | 7.200E-07 |

| | | | | | | | | | | |
|---|---|---|---|---|---|---|---|---|---|---|
| 765 | 39 | E39C | A | 2.9e-07 | 2.7e-07 | 1.1e-07 | 2.233E-07 | 9.866E-08 | 1.100E-07 | 2.900E-07 |
| 766 | 39 | E39E | A | 1.6e-07 | 2.5e-07 | 1.6e-07 | 1.900E-07 | 5.196E-08 | 1.600E-07 | 2.500E-07 |
| 767 | 39 | E39Q | A | 3.2e-07 | 3.2e-07 | 2.3e-07 | 2.900E-07 | 5.196E-08 | 2.300E-07 | 3.200E-07 |
| 768 | 39 | E39G | A | 3e-07 | 1.6e-07 | 5.2e-07 | 3.267E-07 | 1.815E-07 | 1.600E-07 | 5.200E-07 |
| 769 | 39 | E39H | A | 3.1e-07 | 2.7e-07 | 3.2e-07 | 3.000E-07 | 2.646E-08 | 2.700E-07 | 3.200E-07 |
| 770 | 39 | E39I | A | 5.6e-07 | 2.3e-07 | 2.6e-07 | 3.500E-07 | 1.825E-07 | 2.300E-07 | 5.600E-07 |
| 771 | 39 | E39L | A | 1.9e-07 | 2e-07 | 2.1e-07 | 2.000E-07 | 1.000E-08 | 1.900E-07 | 2.100E-07 |
| 772 | 39 | E39K | A | 4.7e-07 | 1.5e-07 | 2.9e-07 | 3.033E-07 | 1.604E-07 | 1.500E-07 | 4.700E-07 |
| 773 | 39 | E39M | A | 1.7e-07 | 2.5e-07 | 2.5e-07 | 2.233E-07 | 4.619E-08 | 1.700E-07 | 2.500E-07 |
| 774 | 39 | E39F | A | 1.9e-07 | 2.6e-07 | 5.9e-07 | 3.467E-07 | 2.136E-07 | 1.900E-07 | 5.900E-07 |
| 775 | 39 | E39P | A | 4.8e-07 | 2.2e-07 | 1.8e-07 | 2.933E-07 | 1.629E-07 | 1.800E-07 | 4.800E-07 |
| 776 | 39 | E39S | A | 2.4e-07 | 1.6e-07 | 1.6e-07 | 1.867E-07 | 4.619E-08 | 1.600E-07 | 2.400E-07 |
| 777 | 39 | E39T | A | 2.8e-07 | 1.1e-07 | 1.6e-07 | 1.833E-07 | 8.737E-08 | 1.100E-07 | 2.800E-07 |
| 778 | 39 | E39W | A | 1.5e-07 | 3e-07 | 2.8e-07 | 2.433E-07 | 8.145E-08 | 1.500E-07 | 3.000E-07 |
| 779 | 39 | E39Y | A | 1.8e-07 | 1.5e-07 | 9.3e-08 | 1.410E-07 | 4.419E-08 | 9.300E-08 | 1.800E-07 |
| 780 | 39 | E39V | A | 2.6e-07 | 8.9e-08 | 2e-07 | 1.830E-07 | 8.676E-08 | 8.900E-08 | 2.600E-07 |
| 781 | 40 | L40A | A | 2.6e-07 | 3.1e-07 | 1.7e-07 | 2.467E-07 | 7.095E-08 | 1.700E-07 | 3.100E-07 |
| 782 | 40 | L40R | A | 1.9e-07 | 4.2e-07 | 2.3e-07 | 2.800E-07 | 1.229E-07 | 1.900E-07 | 4.200E-07 |
| 783 | 40 | L40N | A | 1.2e-07 | 1.7e-07 | 3.1e-07 | 2.000E-07 | 9.849E-08 | 1.200E-07 | 3.100E-07 |
| 784 | 40 | L40D | A | 3e-07 | 2e-07 | 2e-07 | 2.333E-07 | 5.774E-08 | 2.000E-07 | 3.000E-07 |
| 785 | 40 | L40C | A | 2.8e-07 | 9.7e-08 | 2.7e-07 | 2.157E-07 | 1.029E-07 | 9.700E-08 | 2.800E-07 |
| 786 | 40 | L40E | A | 1.2e-07 | 1.7e-07 | 2.7e-07 | 1.867E-07 | 7.638E-08 | 1.200E-07 | 2.700E-07 |
| 787 | 40 | L40Q | A | 4.7e-07 | 6.4e-07 | 3.8e-07 | 4.967E-07 | 1.320E-07 | 3.800E-07 | 6.400E-07 |
| 788 | 40 | L40G | A | 2.9e-07 | 2.2e-07 | 1.8e-07 | 2.300E-07 | 5.568E-08 | 1.800E-07 | 2.900E-07 |
| 789 | 40 | L40H | A | 4.9e-07 | 1.6e-07 | 3.7e-07 | 3.400E-07 | 1.670E-07 | 1.600E-07 | 4.900E-07 |
| 790 | 40 | L40I | A | 3.7e-07 | 1.8e-07 | 3.6e-07 | 3.033E-07 | 1.069E-07 | 1.800E-07 | 3.700E-07 |
| 791 | 40 | L40L | A | 1.9e-07 | 3.3e-07 | 1.6e-07 | 2.267E-07 | 9.074E-08 | 1.600E-07 | 3.300E-07 |
| 792 | 40 | L40K | A | 2.2e-07 | 2.8e-07 | 2e-07 | 2.333E-07 | 4.163E-08 | 2.000E-07 | 2.800E-07 |
| 793 | 40 | L40M | A | 1.5e-07 | 3e-07 | 2.4e-07 | 2.300E-07 | 7.550E-08 | 1.500E-07 | 3.000E-07 |
| 794 | 40 | L40F | A | 2.5e-07 | 2e-07 | 2.2e-07 | 2.233E-07 | 2.517E-08 | 2.000E-07 | 2.500E-07 |
| 795 | 40 | L40P | A | 2.1e-07 | 3.8e-07 | 3.1e-07 | 3.000E-07 | 8.544E-08 | 2.100E-07 | 3.800E-07 |
| 796 | 40 | L40S | A | 1.2e-07 | 1.5e-07 | 1.7e-07 | 1.467E-07 | 2.517E-08 | 1.200E-07 | 1.700E-07 |
| 797 | 40 | L40T | A | 2.1e-07 | 2.9e-07 | 4.2e-07 | 3.067E-07 | 1.060E-07 | 2.100E-07 | 4.200E-07 |
| 798 | 40 | L40W | A | 2.3e-07 | 3.6e-07 | 3.4e-07 | 3.100E-07 | 7.000E-08 | 2.300E-07 | 3.600E-07 |
| 799 | 40 | L40Y | A | 1.6e-07 | 2e-07 | 2.4e-07 | 2.000E-07 | 4.000E-08 | 1.600E-07 | 2.400E-07 |
| 800 | 40 | L40V | A | 4.5e-07 | 1.9e-07 | 2e-07 | 2.800E-07 | 1.473E-07 | 1.900E-07 | 4.500E-07 |
| 801 | 41 | T41A | A | 3e-07 | 4.8e-07 | 2.2e-07 | 3.333E-07 | 1.332E-07 | 2.200E-07 | 4.800E-07 |
| 802 | 41 | T41R | A | 2.6e-07 | 2e-07 | 2.1e-07 | 2.233E-07 | 3.215E-08 | 2.000E-07 | 2.600E-07 |
| 803 | 41 | T41N | A | 3.4e-07 | 1.6e-07 | 2.4e-07 | 2.467E-07 | 9.018E-08 | 1.600E-07 | 3.400E-07 |
| 804 | 41 | T41D | A | 1.5e-07 | 1.2e-07 | 1.3e-07 | 1.333E-07 | 1.528E-08 | 1.200E-07 | 1.500E-07 |
| 805 | 41 | T41C | A | 2.2e-07 | 3.2e-07 | 3.1e-07 | 2.833E-07 | 5.508E-08 | 2.200E-07 | 3.200E-07 |
| 806 | 41 | T41E | A | 4.3e-07 | 1.1e-07 | 3.1e-07 | 2.833E-07 | 1.617E-07 | 1.100E-07 | 4.300E-07 |
| 807 | 41 | T41Q | A | 2.9e-07 | 5e-07 | 1.6e-07 | 3.167E-07 | 1.716E-07 | 1.600E-07 | 5.000E-07 |
| 808 | 41 | T41G | A | 2.6e-07 | 2.6e-07 | 3.2e-07 | 2.800E-07 | 3.464E-08 | 2.600E-07 | 3.200E-07 |
| 809 | 41 | T41H | A | 2.5e-07 | 2.8e-07 | 4.2e-07 | 3.167E-07 | 9.074E-08 | 2.500E-07 | 4.200E-07 |
| 810 | 41 | T41I | A | 1.9e-07 | 3.3e-07 | 4e-07 | 3.067E-07 | 1.069E-07 | 1.900E-07 | 4.000E-07 |
| 811 | 41 | T41L | A | 1.4e-07 | 4.2e-07 | 2.2e-07 | 2.600E-07 | 1.442E-07 | 1.400E-07 | 4.200E-07 |
| 812 | 41 | T41K | A | 2.9e-07 | 1.7e-07 | 1.2e-06 | 5.533E-07 | 5.632E-07 | 1.700E-07 | 1.200E-06 |
| 813 | 41 | T41M | A | 2.7e-07 | 1.3e-07 | 2.8e-07 | 2.267E-07 | 8.386E-08 | 1.300E-07 | 2.800E-07 |
| 814 | 41 | T41F | A | 1.9e-07 | 2e-07 | 2.4e-07 | 2.100E-07 | 2.646E-08 | 1.900E-07 | 2.400E-07 |
| 815 | 41 | T41P | A | 2.5e-07 | 2e-07 | 2.5e-07 | 2.333E-07 | 2.887E-08 | 2.000E-07 | 2.500E-07 |
| 816 | 41 | T41S | A | 1.3e-07 | 8.2e-07 | 1.4e-07 | 3.633E-07 | 3.955E-07 | 1.300E-07 | 8.200E-07 |
| 817 | 41 | T41T | A | 1.2e-07 | 8.4e-08 | 1.9e-07 | 1.313E-07 | 5.390E-08 | 8.400E-08 | 1.900E-07 |
| 818 | 41 | T41W | A | 3.1e-07 | 3e-07 | 1.9e-07 | 2.667E-07 | 6.658E-08 | 1.900E-07 | 3.100E-07 |
| 819 | 41 | T41Y | A | 4.3e-07 | 1.3e-07 | 2.2e-07 | 2.600E-07 | 1.539E-07 | 1.300E-07 | 4.300E-07 |
| 820 | 41 | T41V | A | 2.6e-07 | 3.7e-07 | 4.6e-07 | 3.633E-07 | 1.002E-07 | 2.600E-07 | 4.600E-07 |
| 821 | 42 | Y42A | A | 2.7e-07 | 5.9e-07 | 4.4e-07 | 4.333E-07 | 1.601E-07 | 2.700E-07 | 5.900E-07 |
| 822 | 42 | Y42R | A | 3.3e-07 | 1.6e-07 | 3.2e-07 | 2.700E-07 | 9.539E-08 | 1.600E-07 | 3.300E-07 |
| 823 | 42 | Y42N | A | 2.2e-07 | 2.6e-07 | 1.9e-07 | 2.233E-07 | 3.512E-08 | 1.900E-07 | 2.600E-07 |
| 824 | 42 | Y42D | A | 1.7e-07 | 2.6e-07 | 1.5e-07 | 1.933E-07 | 5.859E-08 | 1.500E-07 | 2.600E-07 |
| 825 | 42 | Y42C | A | 4.7e-07 | 1.6e-07 | 3.6e-07 | 3.300E-07 | 1.572E-07 | 1.600E-07 | 4.700E-07 |
| 826 | 42 | Y42E | A | 1.7e-07 | 2.5e-07 | 2.8e-07 | 2.333E-07 | 5.686E-08 | 1.700E-07 | 2.800E-07 |
| 827 | 42 | Y42Q | A | 3.8e-07 | 3.6e-07 | 1.7e-07 | 3.033E-07 | 1.159E-07 | 1.700E-07 | 3.800E-07 |
| 828 | 42 | Y42G | A | 3.1e-07 | 2.3e-07 | 5.6e-07 | 3.667E-07 | 1.721E-07 | 2.300E-07 | 5.600E-07 |
| 829 | 42 | Y42H | A | 5e-07 | 2.6e-07 | 1.8e-07 | 3.133E-07 | 1.665E-07 | 1.800E-07 | 5.000E-07 |
| 830 | 42 | Y42I | A | 3.3e-07 | 2.5e-07 | 1.9e-07 | 2.567E-07 | 7.024E-08 | 1.900E-07 | 3.300E-07 |
| 831 | 42 | Y42L | A | 1.9e-07 | 1.9e-07 | 2.7e-07 | 2.167E-07 | 4.619E-08 | 1.900E-07 | 2.700E-07 |
| 832 | 42 | Y42K | A | 1.5e-07 | 1.7e-07 | 2.4e-07 | 1.867E-07 | 4.726E-08 | 1.500E-07 | 2.400E-07 |
| 833 | 42 | Y42M | A | 2e-07 | 1.7e-07 | 2.1e-07 | 1.933E-07 | 2.082E-08 | 1.700E-07 | 2.100E-07 |
| 834 | 42 | Y42F | A | 4.4e-07 | 1.8e-07 | 8.7e-08 | 2.357E-07 | 1.830E-07 | 8.700E-08 | 4.400E-07 |
| 835 | 42 | Y42P | A | 1.4e-07 | 7.7e-08 | 2.2e-07 | 1.457E-07 | 7.167E-08 | 7.700E-08 | 2.200E-07 |
| 836 | 42 | Y42S | A | 2.2e-07 | 2.8e-07 | 1.2e-07 | 2.067E-07 | 8.083E-08 | 1.200E-07 | 2.800E-07 |
| 837 | 42 | Y42T | A | 2.5e-07 | 1e-07 | 1.6e-07 | 1.700E-07 | 7.550E-08 | 1.000E-07 | 2.500E-07 |
| 838 | 42 | Y42W | A | 3.5e-07 | 2.5e-07 | 2.7e-07 | 2.900E-07 | 5.292E-08 | 2.500E-07 | 3.500E-07 |
| 839 | 42 | Y42Y | A | 2.6e-07 | 1.7e-07 | 4.3e-07 | 2.867E-07 | 1.320E-07 | 1.700E-07 | 4.300E-07 |
| 840 | 42 | Y42V | A | 2.6e-07 | 1.9e-07 | 3.5e-07 | 2.667E-07 | 8.021E-08 | 1.900E-07 | 3.500E-07 |
| 841 | 43 | L43A | A | 1.9e-07 | 2.9e-07 | 3.5e-07 | 2.767E-07 | 8.083E-08 | 1.900E-07 | 3.500E-07 |
| 842 | 43 | L43R | A | 2.4e-07 | 1.6e-07 | 2.4e-07 | 2.133E-07 | 4.619E-08 | 1.600E-07 | 2.400E-07 |
| 843 | 43 | L43N | A | 5.8e-07 | 5.3e-07 | 2.2e-07 | 4.433E-07 | 1.950E-07 | 2.200E-07 | 5.800E-07 |
| 844 | 43 | L43D | A | 2.4e-07 | 3.7e-07 | 2e-07 | 2.700E-07 | 8.888E-08 | 2.000E-07 | 3.700E-07 |
| 845 | 43 | L43C | A | 2.5e-07 | 3e-07 | 8.9e-07 | 4.800E-07 | 3.559E-07 | 2.500E-07 | 8.900E-07 |
| 846 | 43 | L43E | A | 2.9e-07 | 5.6e-07 | 5.3e-07 | 4.600E-07 | 1.480E-07 | 2.900E-07 | 5.600E-07 |
| 847 | 43 | L43Q | A | 2.1e-07 | 1.9e-07 | 1.7e-07 | 1.900E-07 | 2.000E-08 | 1.700E-07 | 2.100E-07 |
| 848 | 43 | L43G | A | 6.9e-07 | 1.2e-07 | 5e-07 | 4.367E-07 | 2.902E-07 | 1.200E-07 | 6.900E-07 |
| 849 | 43 | L43H | A | 2e-07 | 2.5e-07 | 2.4e-07 | 2.300E-07 | 2.646E-08 | 2.000E-07 | 2.500E-07 |

| | | | | | | | | | | |
|---|---|---|---|---|---|---|---|---|---|---|
| 850 | 43 | L43I | A | 4.1e-07 | 3.5e-07 | 2e-07 | 3.200E-07 | 1.082E-07 | 2.000E-07 | 4.100E-07 |
| 851 | 43 | L43L | A | 3.4e-07 | 3.5e-07 | 2.9e-07 | 3.267E-07 | 3.215E-08 | 2.900E-07 | 3.500E-07 |
| 852 | 43 | L43K | A | 1.6e-07 | 1.7e-07 | 2.9e-07 | 2.067E-07 | 7.234E-08 | 1.600E-07 | 2.900E-07 |
| 853 | 43 | L43M | A | 2.9e-07 | 4e-07 | 3.6e-07 | 3.500E-07 | 5.568E-08 | 2.900E-07 | 4.000E-07 |
| 854 | 43 | L43F | A | 1.7e-07 | 3e-07 | 6.4e-07 | 3.700E-07 | 2.427E-07 | 1.700E-07 | 6.400E-07 |
| 855 | 43 | L43P | A | 2.3e-07 | 2.3e-07 | 1.4e-07 | 2.000E-07 | 5.196E-08 | 1.400E-07 | 2.300E-07 |
| 856 | 43 | L43S | A | 3e-07 | 2.3e-07 | 1.9e-07 | 2.400E-07 | 5.568E-08 | 1.900E-07 | 3.000E-07 |
| 857 | 43 | L43T | A | 2.4e-07 | 2.2e-07 | 3.1e-07 | 2.567E-07 | 4.726E-08 | 2.200E-07 | 3.100E-07 |
| 858 | 43 | L43W | A | 1.6e-07 | 2e-07 | 3.1e-07 | 2.233E-07 | 7.767E-08 | 1.600E-07 | 3.100E-07 |
| 859 | 43 | L43Y | A | 4.2e-07 | 2.8e-07 | 7.7e-07 | 4.900E-07 | 2.524E-07 | 2.800E-07 | 7.700E-07 |
| 860 | 43 | L43V | A | 1.7e-07 | 2.5e-07 | 7.9e-07 | 4.033E-07 | 3.372E-07 | 1.700E-07 | 7.900E-07 |
| 861 | 44 | P44A | A | 6e-07 | 1.1e-07 | 1.5e-07 | 2.867E-07 | 2.721E-07 | 1.100E-07 | 6.000E-07 |
| 862 | 44 | P44R | A | 1.6e-07 | 2.2e-07 | 1.9e-07 | 1.900E-07 | 3.000E-08 | 1.600E-07 | 2.200E-07 |
| 863 | 44 | P44N | A | 2.1e-07 | 1.8e-07 | 1.6e-07 | 1.833E-07 | 2.517E-08 | 1.600E-07 | 2.100E-07 |
| 864 | 44 | P44D | A | 2e-07 | 2.1e-07 | 2.8e-07 | 2.300E-07 | 4.359E-08 | 2.000E-07 | 2.800E-07 |
| 865 | 44 | P44C | A | 3.3e-07 | 4.6e-07 | 1.7e-07 | 3.200E-07 | 1.453E-07 | 1.700E-07 | 4.600E-07 |
| 866 | 44 | P44E | A | 2.2e-07 | 1.5e-07 | 1.1e-07 | 1.600E-07 | 5.568E-08 | 1.100E-07 | 2.200E-07 |
| 867 | 44 | P44Q | A | 1.8e-07 | 3.7e-07 | 2.5e-07 | 2.667E-07 | 9.609E-08 | 1.800E-07 | 3.700E-07 |
| 868 | 44 | P44G | A | 1.7e-07 | 2.3e-07 | 3.4e-07 | 2.467E-07 | 8.622E-08 | 1.700E-07 | 3.400E-07 |
| 869 | 44 | P44H | A | 1.9e-07 | 3.1e-07 | 2.4e-07 | 2.467E-07 | 6.028E-08 | 1.900E-07 | 3.100E-07 |
| 870 | 44 | P44I | A | 2.1e-07 | 2.2e-07 | 2.3e-07 | 2.200E-07 | 1.000E-08 | 2.100E-07 | 2.300E-07 |
| 871 | 44 | P44L | A | 5.1e-07 | 2.5e-07 | 1.8e-07 | 3.133E-07 | 1.739E-07 | 1.800E-07 | 5.100E-07 |
| 872 | 44 | P44K | A | 2.9e-07 | 3.7e-07 | 1.5e-07 | 2.700E-07 | 1.114E-07 | 1.500E-07 | 3.700E-07 |
| 873 | 44 | P44M | A | 3.2e-07 | 3e-07 | 4.2e-07 | 3.467E-07 | 6.429E-08 | 3.000E-07 | 4.200E-07 |
| 874 | 44 | P44F | A | 1.4e-07 | 2.3e-07 | 6.8e-07 | 3.500E-07 | 2.893E-07 | 1.400E-07 | 6.800E-07 |
| 875 | 44 | P44P | A | 7.2e-08 | 1.9e-07 | 1.9e-07 | 1.507E-07 | 6.813E-08 | 7.200E-08 | 1.900E-07 |
| 876 | 44 | P44S | A | 2.2e-07 | 1.8e-07 | 1.9e-07 | 1.967E-07 | 2.082E-08 | 1.800E-07 | 2.200E-07 |
| 877 | 44 | P44T | A | 1.5e-07 | 2.1e-07 | 2.5e-07 | 2.033E-07 | 5.033E-08 | 1.500E-07 | 2.500E-07 |
| 878 | 44 | P44W | A | 1.3e-07 | 1.2e-07 | 8.3e-08 | 1.110E-07 | 2.476E-08 | 8.300E-08 | 1.300E-07 |
| 879 | 44 | P44Y | A | 7.8e-07 | 4e-07 | 2.9e-07 | 4.900E-07 | 2.571E-07 | 2.900E-07 | 7.800E-07 |
| 880 | 44 | P44V | A | 2.4e-07 | 1.5e-07 | 1.6e-07 | 1.833E-07 | 4.933E-08 | 1.500E-07 | 2.400E-07 |
| 881 | 45 | T45A | A | 1.8e-07 | 2.6e-07 | 3e-07 | 2.467E-07 | 6.110E-08 | 1.800E-07 | 3.000E-07 |
| 882 | 45 | T45R | A | 3.4e-07 | 3e-07 | 3e-07 | 3.133E-07 | 2.309E-08 | 3.000E-07 | 3.400E-07 |
| 883 | 45 | T45N | A | 1.8e-07 | 2.7e-07 | 3e-07 | 2.500E-07 | 6.245E-08 | 1.800E-07 | 3.000E-07 |
| 884 | 45 | T45D | A | 2.8e-07 | 2.7e-07 | 3.4e-07 | 2.967E-07 | 3.786E-08 | 2.700E-07 | 3.400E-07 |
| 885 | 45 | T45C | A | 1.4e-07 | 2.4e-07 | 1.4e-07 | 1.733E-07 | 5.774E-08 | 1.400E-07 | 2.400E-07 |
| 886 | 45 | T45E | A | 2.9e-07 | 1.4e-07 | 2.7e-07 | 2.333E-07 | 8.145E-08 | 1.400E-07 | 2.900E-07 |
| 887 | 45 | T45Q | A | 2.7e-07 | 1.4e-07 | 2.5e-07 | 2.200E-07 | 7.000E-08 | 1.400E-07 | 2.700E-07 |
| 888 | 45 | T45G | A | 3.1e-07 | 5.8e-07 | 3.6e-07 | 4.167E-07 | 1.436E-07 | 3.100E-07 | 5.800E-07 |
| 889 | 45 | T45H | A | 1.4e-07 | 3.2e-07 | 2.3e-07 | 2.300E-07 | 9.000E-08 | 1.400E-07 | 3.200E-07 |
| 890 | 45 | T45I | A | 3.7e-07 | 3.2e-07 | 3.7e-07 | 3.533E-07 | 2.887E-08 | 3.200E-07 | 3.700E-07 |
| 891 | 45 | T45L | A | 1.8e-07 | 2e-07 | 2.4e-07 | 2.067E-07 | 3.055E-08 | 1.800E-07 | 2.400E-07 |
| 892 | 45 | T45K | A | 2.9e-07 | 2.3e-07 | 4.1e-07 | 3.100E-07 | 9.165E-08 | 2.300E-07 | 4.100E-07 |
| 893 | 45 | T45M | A | 6.5e-07 | 4.4e-07 | 2.3e-07 | 4.400E-07 | 2.100E-07 | 2.300E-07 | 6.500E-07 |
| 894 | 45 | T45F | A | 2e-07 | 3.7e-07 | 1.6e-07 | 2.433E-07 | 1.115E-07 | 1.600E-07 | 3.700E-07 |
| 895 | 45 | T45P | A | 2.9e-07 | 2e-07 | 3e-07 | 2.633E-07 | 5.508E-08 | 2.000E-07 | 3.000E-07 |
| 896 | 45 | T45S | A | 1.7e-07 | 1.5e-07 | 3.1e-07 | 2.100E-07 | 8.718E-08 | 1.500E-07 | 3.100E-07 |
| 897 | 45 | T45T | A | 3.2e-07 | 2e-07 | 1.4e-07 | 2.200E-07 | 9.165E-08 | 1.400E-07 | 3.200E-07 |
| 898 | 45 | T45W | A | 9.3e-08 | 1.9e-07 | 2.3e-07 | 1.710E-07 | 7.045E-08 | 9.300E-08 | 2.300E-07 |
| 899 | 45 | T45Y | A | 2.3e-07 | 1.3e-07 | 1.8e-07 | 1.800E-07 | 5.000E-08 | 1.300E-07 | 2.300E-07 |
| 900 | 45 | T45V | A | 5.6e-08 | 4.5e-07 | 3.6e-07 | 2.887E-07 | 2.065E-07 | 5.600E-08 | 4.500E-07 |
| 901 | 46 | N46A | A | 2e-07 | 7.2e-07 | 3.4e-07 | 4.200E-07 | 2.691E-07 | 2.000E-07 | 7.200E-07 |
| 902 | 46 | N46R | A | 4.9e-07 | 3.3e-07 | 2.4e-07 | 3.533E-07 | 1.266E-07 | 2.400E-07 | 4.900E-07 |
| 903 | 46 | N46N | A | 3.8e-07 | 3.6e-07 | 3.7e-07 | 3.700E-07 | 1.000E-08 | 3.600E-07 | 3.800E-07 |
| 904 | 46 | N46D | A | 2.3e-07 | 2.4e-07 | 9.2e-08 | 1.873E-07 | 8.271E-08 | 9.200E-08 | 2.400E-07 |
| 905 | 46 | N46C | A | 4.7e-07 | 4.1e-07 | 1.7e-07 | 3.500E-07 | 1.587E-07 | 1.700E-07 | 4.700E-07 |
| 906 | 46 | N46E | A | 1.4e-07 | 2.8e-07 | 3.1e-07 | 2.433E-07 | 9.074E-08 | 1.400E-07 | 3.100E-07 |
| 907 | 46 | N46Q | A | 3.1e-07 | 4.6e-07 | 2.1e-07 | 3.267E-07 | 1.258E-07 | 2.100E-07 | 4.600E-07 |
| 908 | 46 | N46G | A | 4.1e-07 | 3.9e-07 | 1.8e-07 | 3.267E-07 | 1.274E-07 | 1.800E-07 | 4.100E-07 |
| 909 | 46 | N46H | A | 1.2e-07 | 2.2e-07 | 2.5e-07 | 1.967E-07 | 6.807E-08 | 1.200E-07 | 2.500E-07 |
| 910 | 46 | N46I | A | 3.1e-07 | 3e-07 | 2e-07 | 2.700E-07 | 6.083E-08 | 2.000E-07 | 3.100E-07 |
| 911 | 46 | N46L | A | 3.2e-07 | 3.7e-07 | 8.1e-08 | 2.570E-07 | 1.545E-07 | 8.100E-08 | 3.700E-07 |
| 912 | 46 | N46K | A | 3.9e-07 | 4.2e-07 | 2.5e-07 | 3.533E-07 | 9.074E-08 | 2.500E-07 | 4.200E-07 |
| 913 | 46 | N46M | A | 6.1e-07 | 1.4e-07 | 2.4e-07 | 3.300E-07 | 2.476E-07 | 1.400E-07 | 6.100E-07 |
| 914 | 46 | N46F | A | 2.2e-07 | 3.9e-07 | 6.3e-07 | 4.133E-07 | 2.060E-07 | 2.200E-07 | 6.300E-07 |
| 915 | 46 | N46P | A | 2.2e-07 | 3e-07 | 1.7e-07 | 2.300E-07 | 6.557E-08 | 1.700E-07 | 3.000E-07 |
| 916 | 46 | N46S | A | 1.5e-07 | 1.9e-07 | 1.8e-07 | 1.733E-07 | 2.082E-08 | 1.500E-07 | 1.900E-07 |
| 917 | 46 | N46T | A | 3.6e-07 | 1.7e-07 | 3.6e-07 | 2.967E-07 | 1.097E-07 | 1.700E-07 | 3.600E-07 |
| 918 | 46 | N46W | A | 2.9e-07 | 3.3e-07 | 6.5e-08 | 2.283E-07 | 1.429E-07 | 6.500E-08 | 3.300E-07 |
| 919 | 46 | N46Y | A | 3.3e-07 | 1.4e-07 | 1e-07 | 1.900E-07 | 1.229E-07 | 1.000E-07 | 3.300E-07 |
| 920 | 46 | N46V | A | 1.7e-07 | 3.2e-07 | 3.2e-07 | 2.700E-07 | 8.660E-08 | 1.700E-07 | 3.200E-07 |
| 921 | 47 | A47A | A | 2.8e-07 | 3.8e-07 | 2e-07 | 2.867E-07 | 9.018E-08 | 2.000E-07 | 3.800E-07 |
| 922 | 47 | A47R | A | 3.9e-07 | 1.4e-07 | 1.7e-07 | 2.333E-07 | 1.365E-07 | 1.400E-07 | 3.900E-07 |
| 923 | 47 | A47N | A | 2.9e-07 | 2.2e-07 | 2.2e-07 | 2.433E-07 | 4.041E-08 | 2.200E-07 | 2.900E-07 |
| 924 | 47 | A47D | A | 2.7e-07 | 4.7e-07 | 3.1e-07 | 3.500E-07 | 1.058E-07 | 2.700E-07 | 4.700E-07 |
| 925 | 47 | A47C | A | 2.3e-07 | 9e-08 | 2.3e-07 | 1.833E-07 | 8.083E-08 | 9.000E-08 | 2.300E-07 |
| 926 | 47 | A47E | A | 1.3e-07 | 1.4e-07 | 1.6e-07 | 1.433E-07 | 1.528E-08 | 1.300E-07 | 1.600E-07 |
| 927 | 47 | A47Q | A | 2.1e-07 | 4.8e-07 | 2.1e-07 | 3.000E-07 | 1.559E-07 | 2.100E-07 | 4.800E-07 |
| 928 | 47 | A47G | A | 2e-07 | 4.2e-07 | 2.7e-07 | 2.967E-07 | 1.124E-07 | 2.000E-07 | 4.200E-07 |
| 929 | 47 | A47H | A | 4e-07 | 4.6e-07 | 1.7e-07 | 3.433E-07 | 1.531E-07 | 1.700E-07 | 4.600E-07 |
| 930 | 47 | A47I | A | 2.4e-07 | 4e-07 | 1.8e-07 | 2.733E-07 | 1.137E-07 | 1.800E-07 | 4.000E-07 |
| 931 | 47 | A47L | A | 7e-07 | 3.3e-07 | 2.5e-07 | 4.267E-07 | 2.401E-07 | 2.500E-07 | 7.000E-07 |
| 932 | 47 | A47K | A | 1.5e-07 | 2.2e-07 | 2e-07 | 1.900E-07 | 3.606E-08 | 1.500E-07 | 2.200E-07 |
| 933 | 47 | A47M | A | 5.6e-07 | 2.3e-07 | 3.8e-07 | 3.900E-07 | 1.652E-07 | 2.300E-07 | 5.600E-07 |
| 934 | 47 | A47F | A | 1.2e-07 | 3.9e-07 | 7.1e-08 | 1.937E-07 | 1.718E-07 | 7.100E-08 | 3.900E-07 |

| | | | | | | | | | | |
|---|---|---|---|---|---|---|---|---|---|---|
| 935 | 47 | A47P | A | 2.8e-07 | 1.2e-07 | 4.4e-07 | 2.800E-07 | 1.600E-07 | 1.200E-07 | 4.400E-07 |
| 936 | 47 | A47S | A | 1.9e-07 | 3.2e-07 | 2.4e-07 | 2.500E-07 | 6.557E-08 | 1.900E-07 | 3.200E-07 |
| 937 | 47 | A47T | A | 3.2e-07 | 2.3e-07 | 4.1e-07 | 3.200E-07 | 9.000E-08 | 2.300E-07 | 4.100E-07 |
| 938 | 47 | A47W | A | 5.7e-07 | 2e-07 | 1.4e-07 | 3.033E-07 | 2.329E-07 | 1.400E-07 | 5.700E-07 |
| 939 | 47 | A47Y | A | 2.3e-07 | 9.8e-08 | 2.4e-07 | 1.893E-07 | 7.925E-08 | 9.800E-08 | 2.400E-07 |
| 940 | 47 | A47V | A | 2.5e-07 | 3.7e-07 | 3.2e-07 | 3.133E-07 | 6.028E-08 | 2.500E-07 | 3.700E-07 |
| 941 | 48 | S48A | A | 1.6e-07 | 2.4e-07 | 1.9e-07 | 1.967E-07 | 4.041E-08 | 1.600E-07 | 2.400E-07 |
| 942 | 48 | S48R | A | 1.6e-07 | 3.1e-07 | 6.5e-07 | 3.733E-07 | 2.511E-07 | 1.600E-07 | 6.500E-07 |
| 943 | 48 | S48N | A | 1.9e-07 | 1.6e-07 | 2.4e-07 | 1.967E-07 | 4.041E-08 | 1.600E-07 | 2.400E-07 |
| 944 | 48 | S48D | A | 9e-07 | 2.8e-07 | 1.7e-07 | 4.500E-07 | 3.936E-07 | 1.700E-07 | 9.000E-07 |
| 945 | 48 | S48C | A | 2e-07 | 2.4e-07 | 1.6e-07 | 2.000E-07 | 4.000E-08 | 1.600E-07 | 2.400E-07 |
| 946 | 48 | S48E | A | 2.4e-07 | 8.2e-07 | 2.8e-07 | 4.467E-07 | 3.239E-07 | 2.400E-07 | 8.200E-07 |
| 947 | 48 | S48Q | A | 4.3e-07 | 4.5e-07 | 3.3e-07 | 4.033E-07 | 6.429E-08 | 3.300E-07 | 4.500E-07 |
| 948 | 48 | S48G | A | 1.8e-07 | 1.7e-07 | 2.8e-07 | 2.100E-07 | 6.083E-08 | 1.700E-07 | 2.800E-07 |
| 949 | 48 | S48H | A | 3.9e-07 | 2.1e-07 | 1.4e-07 | 2.467E-07 | 1.290E-07 | 1.400E-07 | 3.900E-07 |
| 950 | 48 | S48I | A | 1.5e-07 | 2.1e-07 | 1.8e-07 | 1.800E-07 | 3.000E-08 | 1.500E-07 | 2.100E-07 |
| 951 | 48 | S48L | A | 5e-07 | 3.9e-07 | 6.5e-07 | 5.133E-07 | 1.305E-07 | 3.900E-07 | 6.500E-07 |
| 952 | 48 | S48K | A | 1.4e-07 | 2.3e-07 | 2.7e-07 | 2.133E-07 | 6.658E-08 | 1.400E-07 | 2.700E-07 |
| 953 | 48 | S48M | A | 2.4e-07 | 3.9e-07 | 3.4e-07 | 3.233E-07 | 7.638E-08 | 2.400E-07 | 3.900E-07 |
| 954 | 48 | S48F | A | 1.7e-07 | 2.5e-07 | 1.6e-07 | 1.933E-07 | 4.933E-08 | 1.600E-07 | 2.500E-07 |
| 955 | 48 | S48P | A | 3.7e-07 | 2.7e-07 | 3.4e-07 | 3.267E-07 | 5.132E-08 | 2.700E-07 | 3.700E-07 |
| 956 | 48 | S48S | A | 1.7e-07 | 2.6e-07 | 1.7e-07 | 2.000E-07 | 5.196E-08 | 1.700E-07 | 2.600E-07 |
| 957 | 48 | S48T | A | 2.4e-07 | 1.9e-07 | 2.5e-07 | 2.267E-07 | 3.215E-08 | 1.900E-07 | 2.500E-07 |
| 958 | 48 | S48W | A | 1.9e-07 | 2.7e-07 | 3.1e-07 | 2.567E-07 | 6.110E-08 | 1.900E-07 | 3.100E-07 |
| 959 | 48 | S48Y | A | 3.3e-07 | 3.8e-07 | 2.6e-07 | 3.233E-07 | 6.028E-08 | 2.600E-07 | 3.800E-07 |
| 960 | 48 | S48V | A | 1.4e-07 | 2.4e-07 | 1e-07 | 1.600E-07 | 7.211E-08 | 1.000E-07 | 2.400E-07 |
| 961 | 49 | L49A | A | 2.3e-07 | 1.4e-07 | 2.9e-07 | 2.200E-07 | 7.550E-08 | 1.400E-07 | 2.900E-07 |
| 962 | 49 | L49R | A | 1.1e-07 | 1.2e-07 | 2.2e-07 | 1.500E-07 | 6.083E-08 | 1.100E-07 | 2.200E-07 |
| 963 | 49 | L49N | A | 1.5e-07 | 4.4e-07 | 4.8e-07 | 3.567E-07 | 1.801E-07 | 1.500E-07 | 4.800E-07 |
| 964 | 49 | L49D | A | 1.6e-07 | 1.9e-07 | 2.3e-07 | 1.933E-07 | 3.512E-08 | 1.600E-07 | 2.300E-07 |
| 965 | 49 | L49C | A | 2.8e-07 | 3e-07 | 3e-07 | 2.933E-07 | 1.155E-08 | 2.800E-07 | 3.000E-07 |
| 966 | 49 | L49E | A | 1.3e-07 | 1.7e-07 | 3.3e-07 | 2.100E-07 | 1.058E-07 | 1.300E-07 | 3.300E-07 |
| 967 | 49 | L49Q | A | 1.8e-07 | 3.9e-07 | 1.5e-07 | 2.400E-07 | 1.308E-07 | 1.500E-07 | 3.900E-07 |
| 968 | 49 | L49G | A | 1.9e-07 | 8.7e-08 | 2.1e-07 | 1.623E-07 | 6.600E-08 | 8.700E-08 | 2.100E-07 |
| 969 | 49 | L49H | A | 1.4e-07 | 2.4e-07 | 1.8e-07 | 1.867E-07 | 5.033E-08 | 1.400E-07 | 2.400E-07 |
| 970 | 49 | L49I | A | 1.8e-07 | 1.8e-07 | 1.6e-07 | 1.733E-07 | 1.155E-08 | 1.600E-07 | 1.800E-07 |
| 971 | 49 | L49L | A | 3.8e-07 | 2.6e-07 | 5.3e-07 | 3.900E-07 | 1.353E-07 | 2.600E-07 | 5.300E-07 |
| 972 | 49 | L49K | A | 2.4e-07 | 6.4e-07 | 3.1e-07 | 3.967E-07 | 2.136E-07 | 2.400E-07 | 6.400E-07 |
| 973 | 49 | L49M | A | 5.4e-07 | 1.3e-07 | 1.7e-07 | 2.800E-07 | 2.261E-07 | 1.300E-07 | 5.400E-07 |
| 974 | 49 | L49F | A | 3.3e-07 | 2e-07 | 4.2e-07 | 3.167E-07 | 1.106E-07 | 2.000E-07 | 4.200E-07 |
| 975 | 49 | L49P | A | 1.6e-07 | 4e-07 | 3.7e-07 | 3.100E-07 | 1.308E-07 | 1.600E-07 | 4.000E-07 |
| 976 | 49 | L49S | A | 2.1e-07 | 3.2e-07 | 2.4e-07 | 2.567E-07 | 5.686E-08 | 2.100E-07 | 3.200E-07 |
| 977 | 49 | L49T | A | 2.7e-07 | 2.3e-07 | 1.3e-07 | 2.100E-07 | 7.211E-08 | 1.300E-07 | 2.700E-07 |
| 978 | 49 | L49W | A | 1.8e-07 | 1.5e-07 | 5.7e-07 | 3.000E-07 | 2.343E-07 | 1.500E-07 | 5.700E-07 |
| 979 | 49 | L49Y | A | 1.8e-07 | 2e-07 | 3.1e-07 | 2.300E-07 | 7.000E-08 | 1.800E-07 | 3.100E-07 |
| 980 | 49 | L49V | A | 6.5e-07 | 5.4e-07 | 1.4e-07 | 4.433E-07 | 2.684E-07 | 1.400E-07 | 6.500E-07 |
| 981 | 50 | S50A | A | 2.8e-07 | 3.8e-07 | 3.3e-07 | 3.300E-07 | 5.000E-08 | 2.800E-07 | 3.800E-07 |
| 982 | 50 | S50R | A | 8.7e-08 | 1.8e-07 | 4.6e-07 | 2.423E-07 | 1.942E-07 | 8.700E-08 | 4.600E-07 |
| 983 | 50 | S50N | A | 3.4e-07 | 1.3e-07 | 4.8e-07 | 3.167E-07 | 1.762E-07 | 1.300E-07 | 4.800E-07 |
| 984 | 50 | S50D | A | 2e-07 | 3.7e-07 | 1.9e-07 | 2.533E-07 | 1.012E-07 | 1.900E-07 | 3.700E-07 |
| 985 | 50 | S50C | A | 2.9e-07 | 4.2e-07 | 2.9e-07 | 3.333E-07 | 7.506E-08 | 2.900E-07 | 4.200E-07 |
| 986 | 50 | S50E | A | 2.6e-07 | 1.6e-07 | 3.7e-07 | 2.633E-07 | 1.050E-07 | 1.600E-07 | 3.700E-07 |
| 987 | 50 | S50Q | A | 2.6e-07 | 7.1e-07 | 2.4e-07 | 4.033E-07 | 2.658E-07 | 2.400E-07 | 7.100E-07 |
| 988 | 50 | S50G | A | 2.8e-07 | 2.4e-07 | 3e-07 | 2.733E-07 | 3.055E-08 | 2.400E-07 | 3.000E-07 |
| 989 | 50 | S50H | A | 3.4e-07 | 1.2e-07 | 5.7e-07 | 3.433E-07 | 2.250E-07 | 1.200E-07 | 5.700E-07 |
| 990 | 50 | S50I | A | 3.9e-07 | 2e-07 | 1.8e-07 | 2.567E-07 | 1.159E-07 | 1.800E-07 | 3.900E-07 |
| 991 | 50 | S50L | A | 2.4e-07 | 2.4e-07 | 2.2e-07 | 2.333E-07 | 1.155E-08 | 2.200E-07 | 2.400E-07 |
| 992 | 50 | S50K | A | 2.3e-07 | 3.8e-07 | 3.7e-07 | 3.267E-07 | 8.386E-08 | 2.300E-07 | 3.800E-07 |
| 993 | 50 | S50M | A | 3.3e-07 | 2.1e-07 | 3.8e-07 | 3.067E-07 | 8.737E-08 | 2.100E-07 | 3.800E-07 |
| 994 | 50 | S50F | A | 1.8e-07 | 3.9e-07 | 3.5e-07 | 3.067E-07 | 1.115E-07 | 1.800E-07 | 3.900E-07 |
| 995 | 50 | S50P | A | 1.4e-07 | 4.9e-07 | 4.9e-07 | 3.733E-07 | 2.021E-07 | 1.400E-07 | 4.900E-07 |
| 996 | 50 | S50S | A | 2.1e-07 | 1.6e-07 | 2.3e-07 | 2.000E-07 | 3.606E-08 | 1.600E-07 | 2.300E-07 |
| 997 | 50 | S50T | A | 4.2e-07 | 2.8e-07 | 1.4e-07 | 2.800E-07 | 1.400E-07 | 1.400E-07 | 4.200E-07 |
| 998 | 50 | S50W | A | 2.3e-07 | 1.7e-07 | 2.6e-07 | 2.200E-07 | 4.583E-08 | 1.700E-07 | 2.600E-07 |
| 999 | 50 | S50Y | A | 2.5e-07 | 3.8e-07 | 3e-07 | 3.100E-07 | 6.557E-08 | 2.500E-07 | 3.800E-07 |
| 1000 | 50 | S50V | A | 2.4e-07 | 2.4e-07 | 2e-07 | 2.267E-07 | 2.309E-08 | 2.000E-07 | 2.400E-07 |
| 1001 | 51 | F51A | A | 1.7e-07 | 2.7e-07 | 1.5e-07 | 1.967E-07 | 6.429E-08 | 1.500E-07 | 2.700E-07 |
| 1002 | 51 | F51R | A | 3.4e-07 | 1.1e-07 | 2.3e-07 | 2.267E-07 | 1.150E-07 | 1.100E-07 | 3.400E-07 |
| 1003 | 51 | F51N | A | 5.8e-07 | 1.4e-07 | 4.1e-07 | 3.767E-07 | 2.219E-07 | 1.400E-07 | 5.800E-07 |
| 1004 | 51 | F51D | A | 4e-07 | 2.8e-07 | 5.7e-07 | 4.167E-07 | 1.457E-07 | 2.800E-07 | 5.700E-07 |
| 1005 | 51 | F51C | A | 4.2e-07 | 4e-07 | 3e-07 | 3.733E-07 | 6.429E-08 | 3.000E-07 | 4.200E-07 |
| 1006 | 51 | F51E | A | 2.7e-07 | 8.4e-08 | 1.6e-07 | 1.713E-07 | 9.352E-08 | 8.400E-08 | 2.700E-07 |
| 1007 | 51 | F51Q | A | 1.8e-07 | 1.2e-07 | 3e-07 | 2.000E-07 | 9.165E-08 | 1.200E-07 | 3.000E-07 |
| 1008 | 51 | F51G | A | 3.3e-07 | 3.3e-07 | 1.6e-07 | 2.733E-07 | 9.815E-08 | 1.600E-07 | 3.300E-07 |
| 1009 | 51 | F51H | A | 2.8e-07 | 3.1e-07 | 1.6e-07 | 2.500E-07 | 7.937E-08 | 1.600E-07 | 3.100E-07 |
| 1010 | 51 | F51I | A | 2e-07 | 4.1e-07 | 4.1e-07 | 3.400E-07 | 1.212E-07 | 2.000E-07 | 4.100E-07 |
| 1011 | 51 | F51L | A | 4.2e-07 | 3.1e-07 | 3.9e-07 | 3.733E-07 | 5.686E-08 | 3.100E-07 | 4.200E-07 |
| 1012 | 51 | F51K | A | 5.3e-07 | 5.1e-07 | 5.9e-07 | 5.433E-07 | 4.163E-08 | 5.100E-07 | 5.900E-07 |
| 1013 | 51 | F51M | A | 2e-07 | 8.9e-08 | 2.1e-07 | 1.663E-07 | 6.716E-08 | 8.900E-08 | 2.100E-07 |
| 1014 | 51 | F51F | A | 1.4e-07 | 2.3e-07 | 1.8e-07 | 1.833E-07 | 4.509E-08 | 1.400E-07 | 2.300E-07 |
| 1015 | 51 | F51P | A | 4.3e-07 | 3.9e-07 | 2.8e-07 | 3.667E-07 | 7.767E-08 | 2.800E-07 | 4.300E-07 |
| 1016 | 51 | F51S | A | 1.8e-07 | 3e-07 | 2.1e-07 | 2.300E-07 | 6.245E-08 | 1.800E-07 | 3.000E-07 |
| 1017 | 51 | F51T | A | 1.7e-07 | 1.6e-07 | 5.1e-07 | 2.800E-07 | 1.992E-07 | 1.600E-07 | 5.100E-07 |
| 1018 | 51 | F51W | A | 2.2e-07 | 2.5e-07 | 2.8e-07 | 2.500E-07 | 3.000E-08 | 2.200E-07 | 2.800E-07 |
| 1019 | 51 | F51Y | A | 5.5e-07 | 2e-07 | 3.4e-07 | 3.633E-07 | 1.762E-07 | 2.000E-07 | 5.500E-07 |

| | | | | | | | | | | |
|---|---|---|---|---|---|---|---|---|---|---|
| 1020 | 51 | F51V | A | 2.9e-07 | 7.4e-08 | 1.4e-07 | 1.680E-07 | 1.107E-07 | 7.400E-08 | 2.900E-07 |
| 1021 | 52 | L52A | A | 2.6e-07 | 2.4e-07 | 1.7e-07 | 2.233E-07 | 4.726E-08 | 1.700E-07 | 2.600E-07 |
| 1022 | 52 | L52R | A | 2.8e-07 | 3.4e-07 | 4.5e-07 | 3.567E-07 | 8.622E-08 | 2.800E-07 | 4.500E-07 |
| 1023 | 52 | L52N | A | 3.4e-07 | 3.6e-07 | 2e-07 | 3.000E-07 | 8.718E-08 | 2.000E-07 | 3.600E-07 |
| 1024 | 52 | L52D | A | 2.3e-07 | 2.3e-07 | 3.4e-07 | 2.667E-07 | 6.351E-08 | 2.300E-07 | 3.400E-07 |
| 1025 | 52 | L52C | A | 1.8e-07 | 1.4e-07 | 2.7e-07 | 1.967E-07 | 6.658E-08 | 1.400E-07 | 2.700E-07 |
| 1026 | 52 | L52E | A | 2.2e-07 | 2.2e-07 | 3.4e-07 | 2.600E-07 | 6.928E-08 | 2.200E-07 | 3.400E-07 |
| 1027 | 52 | L52Q | A | 1.7e-07 | 1.7e-07 | 4.3e-07 | 2.567E-07 | 1.501E-07 | 1.700E-07 | 4.300E-07 |
| 1028 | 52 | L52G | A | 2.4e-07 | 2e-07 | 1.3e-06 | 5.800E-07 | 6.239E-07 | 2.000E-07 | 1.300E-06 |
| 1029 | 52 | L52H | A | 5e-07 | 2.4e-07 | 4.7e-07 | 4.033E-07 | 1.422E-07 | 2.400E-07 | 5.000E-07 |
| 1030 | 52 | L52I | A | 1.9e-07 | 1.8e-07 | 4.4e-07 | 2.700E-07 | 1.473E-07 | 1.800E-07 | 4.400E-07 |
| 1031 | 52 | L52L | A | 4.6e-07 | 1.8e-07 | 4.9e-07 | 3.767E-07 | 1.710E-07 | 1.800E-07 | 4.900E-07 |
| 1032 | 52 | L52K | A | 1.8e-07 | 1.7e-07 | 2.5e-07 | 2.000E-07 | 4.359E-08 | 1.700E-07 | 2.500E-07 |
| 1033 | 52 | L52M | A | 1.6e-07 | 4.5e-07 | 3.5e-07 | 3.200E-07 | 1.473E-07 | 1.600E-07 | 4.500E-07 |
| 1034 | 52 | L52F | A | 4.3e-07 | 4.9e-07 | 3e-07 | 4.067E-07 | 9.713E-08 | 3.000E-07 | 4.900E-07 |
| 1035 | 52 | L52P | A | 1e-07 | 2.4e-07 | 3.1e-07 | 2.167E-07 | 1.069E-07 | 1.000E-07 | 3.100E-07 |
| 1036 | 52 | L52S | A | 2.3e-07 | 7.6e-08 | 1.7e-07 | 1.587E-07 | 7.762E-08 | 7.600E-08 | 2.300E-07 |
| 1037 | 52 | L52T | A | 2.4e-07 | 1.6e-07 | 1.3e-07 | 1.767E-07 | 5.686E-08 | 1.300E-07 | 2.400E-07 |
| 1038 | 52 | L52W | A | 3.2e-07 | 4.1e-07 | 1.7e-07 | 3.000E-07 | 1.212E-07 | 1.700E-07 | 4.100E-07 |
| 1039 | 52 | L52Y | A | 3.2e-07 | 2.2e-07 | 3.1e-07 | 2.833E-07 | 5.508E-08 | 2.200E-07 | 3.200E-07 |
| 1040 | 52 | L52V | A | 1.6e-07 | 3.1e-07 | 1.1e-07 | 1.933E-07 | 1.041E-07 | 1.100E-07 | 3.100E-07 |
| 1041 | 53 | Q53A | A | 4.4e-07 | 4.7e-07 | 4.5e-07 | 4.533E-07 | 1.528E-08 | 4.400E-07 | 4.700E-07 |
| 1042 | 53 | Q53R | A | 2.3e-07 | 2.2e-07 | 4.4e-07 | 2.967E-07 | 1.242E-07 | 2.200E-07 | 4.400E-07 |
| 1043 | 53 | Q53N | A | 2.4e-07 | 1.3e-07 | 1.7e-07 | 1.800E-07 | 5.568E-08 | 1.300E-07 | 2.400E-07 |
| 1044 | 53 | Q53D | A | 3e-07 | 1.3e-07 | 1.6e-07 | 1.967E-07 | 9.074E-08 | 1.300E-07 | 3.000E-07 |
| 1045 | 53 | Q53C | A | 1.8e-07 | 1.7e-07 | 4.8e-07 | 2.767E-07 | 1.762E-07 | 1.700E-07 | 4.800E-07 |
| 1046 | 53 | Q53E | A | 1.8e-07 | 3.4e-07 | 3.6e-07 | 2.933E-07 | 9.866E-08 | 1.800E-07 | 3.600E-07 |
| 1047 | 53 | Q53Q | A | 2.7e-07 | 3e-07 | 1.5e-07 | 2.400E-07 | 7.937E-08 | 1.500E-07 | 3.000E-07 |
| 1048 | 53 | Q53G | A | 1.1e-07 | 1.7e-07 | 1.3e-07 | 1.367E-07 | 3.055E-08 | 1.100E-07 | 1.700E-07 |
| 1049 | 53 | Q53H | A | 1.3e-07 | 1.6e-07 | 2.9e-07 | 1.933E-07 | 8.505E-08 | 1.300E-07 | 2.900E-07 |
| 1050 | 53 | Q53I | A | 1.6e-07 | 2.5e-07 | 4.1e-07 | 2.733E-07 | 1.266E-07 | 1.600E-07 | 4.100E-07 |
| 1051 | 53 | Q53L | A | 2.1e-07 | 7.6e-08 | 2.2e-07 | 1.687E-07 | 8.041E-08 | 7.600E-08 | 2.200E-07 |
| 1052 | 53 | Q53K | A | 2.1e-07 | 1.3e-07 | 3.1e-07 | 2.167E-07 | 9.018E-08 | 1.300E-07 | 3.100E-07 |
| 1053 | 53 | Q53M | A | 2.3e-07 | 2.4e-07 | 3.5e-07 | 2.733E-07 | 6.658E-08 | 2.300E-07 | 3.500E-07 |
| 1054 | 53 | Q53F | A | 2.4e-07 | 4.7e-07 | 1.4e-07 | 2.833E-07 | 1.692E-07 | 1.400E-07 | 4.700E-07 |
| 1055 | 53 | Q53P | A | 1.5e-07 | 1.3e-07 | 5.6e-07 | 2.800E-07 | 2.427E-07 | 1.300E-07 | 5.600E-07 |
| 1056 | 53 | Q53S | A | 4.1e-07 | 5.9e-07 | 2.1e-07 | 4.033E-07 | 1.901E-07 | 2.100E-07 | 5.900E-07 |
| 1057 | 53 | Q53T | A | 2.6e-07 | 2.8e-07 | 1.2e-07 | 2.200E-07 | 8.718E-08 | 1.200E-07 | 2.800E-07 |
| 1058 | 53 | Q53W | A | 3.7e-07 | 1.6e-07 | 4.6e-07 | 3.300E-07 | 1.539E-07 | 1.600E-07 | 4.600E-07 |
| 1059 | 53 | Q53Y | A | 1.9e-07 | 4.9e-07 | 1.5e-07 | 2.767E-07 | 1.858E-07 | 1.500E-07 | 4.900E-07 |
| 1060 | 53 | Q53V | A | 1.9e-07 | 2.5e-07 | 4.2e-07 | 2.867E-07 | 1.193E-07 | 1.900E-07 | 4.200E-07 |
| 1061 | 54 | D54A | A | 2.3e-07 | 1.8e-07 | 2.4e-07 | 2.167E-07 | 3.215E-08 | 1.800E-07 | 2.400E-07 |
| 1062 | 54 | D54R | A | 2.7e-07 | 2.7e-07 | 3.9e-07 | 3.100E-07 | 6.928E-08 | 2.700E-07 | 3.900E-07 |
| 1063 | 54 | D54N | A | 8.5e-07 | 2.1e-07 | 1.5e-07 | 4.033E-07 | 3.880E-07 | 1.500E-07 | 8.500E-07 |
| 1064 | 54 | D54D | A | 4.7e-07 | 4.8e-07 | 1.7e-07 | 3.733E-07 | 1.762E-07 | 1.700E-07 | 4.800E-07 |
| 1065 | 54 | D54C | A | 2.6e-07 | 2.3e-07 | 3.1e-07 | 2.667E-07 | 4.041E-08 | 2.300E-07 | 3.100E-07 |
| 1066 | 54 | D54E | A | 3.8e-07 | 2.8e-07 | 9.8e-07 | 5.467E-07 | 3.786E-07 | 2.800E-07 | 9.800E-07 |
| 1067 | 54 | D54Q | A | 2.4e-07 | 2.5e-07 | 1.6e-07 | 2.167E-07 | 4.933E-08 | 1.600E-07 | 2.500E-07 |
| 1068 | 54 | D54G | A | 3.3e-07 | 1.4e-07 | 2.3e-07 | 2.333E-07 | 9.504E-08 | 1.400E-07 | 3.300E-07 |
| 1069 | 54 | D54H | A | 2.9e-07 | 2.9e-07 | 4.2e-07 | 3.333E-07 | 7.506E-08 | 2.900E-07 | 4.200E-07 |
| 1070 | 54 | D54I | A | 1.6e-07 | 1.4e-07 | 2.9e-07 | 1.967E-07 | 8.145E-08 | 1.400E-07 | 2.900E-07 |
| 1071 | 54 | D54L | A | 4.2e-07 | 1.5e-07 | 4e-07 | 3.233E-07 | 1.504E-07 | 1.500E-07 | 4.200E-07 |
| 1072 | 54 | D54K | A | 1.8e-07 | 4.4e-07 | 1.2e-07 | 2.467E-07 | 1.701E-07 | 1.200E-07 | 4.400E-07 |
| 1073 | 54 | D54M | A | 2.2e-07 | 4.5e-07 | 1.7e-07 | 2.800E-07 | 1.493E-07 | 1.700E-07 | 4.500E-07 |
| 1074 | 54 | D54F | A | 2.4e-07 | 2.8e-07 | 1.7e-07 | 2.300E-07 | 5.568E-08 | 1.700E-07 | 2.800E-07 |
| 1075 | 54 | D54P | A | 1.8e-07 | 3e-07 | 2.9e-07 | 2.567E-07 | 6.658E-08 | 1.800E-07 | 3.000E-07 |
| 1076 | 54 | D54S | A | 1.7e-07 | 1.9e-07 | 2e-07 | 1.867E-07 | 1.528E-08 | 1.700E-07 | 2.000E-07 |
| 1077 | 54 | D54T | A | 2.7e-07 | 3.1e-07 | 3.4e-07 | 3.067E-07 | 3.512E-08 | 2.700E-07 | 3.400E-07 |
| 1078 | 54 | D54W | A | 4e-07 | 2.2e-07 | 2.4e-07 | 2.867E-07 | 9.866E-08 | 2.200E-07 | 4.000E-07 |
| 1079 | 54 | D54Y | A | 1.2e-07 | 1.7e-07 | 3.2e-07 | 2.033E-07 | 1.041E-07 | 1.200E-07 | 3.200E-07 |
| 1080 | 54 | D54V | A | 3.8e-07 | 3e-07 | 2.1e-07 | 2.967E-07 | 8.505E-08 | 2.100E-07 | 3.800E-07 |
| 1081 | 55 | I55A | A | 1.1e-07 | 2.4e-07 | 1.3e-07 | 1.600E-07 | 7.000E-08 | 1.100E-07 | 2.400E-07 |
| 1082 | 55 | I55R | A | 1.7e-07 | 2.4e-07 | 6.8e-07 | 3.633E-07 | 2.765E-07 | 1.700E-07 | 6.800E-07 |
| 1083 | 55 | I55N | A | 4e-07 | 5.1e-07 | 1.7e-07 | 3.600E-07 | 1.735E-07 | 1.700E-07 | 5.100E-07 |
| 1084 | 55 | I55D | A | 1.5e-07 | 2e-07 | 2.2e-07 | 1.900E-07 | 3.606E-08 | 1.500E-07 | 2.200E-07 |
| 1085 | 55 | I55C | A | 2.8e-07 | 2.6e-07 | 3.6e-07 | 3.000E-07 | 5.292E-08 | 2.600E-07 | 3.600E-07 |
| 1086 | 55 | I55E | A | 2.7e-07 | 2.4e-07 | 7.1e-07 | 4.067E-07 | 2.631E-07 | 2.400E-07 | 7.100E-07 |
| 1087 | 55 | I55Q | A | 1.6e-07 | 1.7e-07 | 3.2e-07 | 2.167E-07 | 8.963E-08 | 1.600E-07 | 3.200E-07 |
| 1088 | 55 | I55G | A | 3.1e-07 | 2.3e-07 | 2.7e-07 | 2.700E-07 | 4.000E-08 | 2.300E-07 | 3.100E-07 |
| 1089 | 55 | I55H | A | 3.6e-07 | 2.7e-07 | 3.8e-07 | 3.367E-07 | 5.859E-08 | 2.700E-07 | 3.800E-07 |
| 1090 | 55 | I55I | A | 7.4e-08 | 2.9e-07 | 1.9e-07 | 1.847E-07 | 1.081E-07 | 7.400E-08 | 2.900E-07 |
| 1091 | 55 | I55L | A | 2.4e-07 | 2.5e-07 | 1.9e-07 | 2.267E-07 | 3.215E-08 | 1.900E-07 | 2.500E-07 |
| 1092 | 55 | I55K | A | 1.9e-07 | 3.1e-07 | 2.8e-07 | 2.600E-07 | 6.245E-08 | 1.900E-07 | 3.100E-07 |
| 1093 | 55 | I55M | A | 1.8e-07 | 2.2e-07 | 1e-07 | 1.667E-07 | 6.110E-08 | 1.000E-07 | 2.200E-07 |
| 1094 | 55 | I55F | A | 1.4e-07 | 3.7e-07 | 2.4e-07 | 2.500E-07 | 1.153E-07 | 1.400E-07 | 3.700E-07 |
| 1095 | 55 | I55P | A | 2.6e-07 | 1.7e-07 | 5.8e-07 | 3.367E-07 | 2.155E-07 | 1.700E-07 | 5.800E-07 |
| 1096 | 55 | I55S | A | 1.1e-07 | 2.8e-07 | 2.3e-07 | 2.067E-07 | 8.737E-08 | 1.100E-07 | 2.800E-07 |
| 1097 | 55 | I55T | A | 1.2e-07 | 2.2e-07 | 1.7e-07 | 1.700E-07 | 5.000E-08 | 1.200E-07 | 2.200E-07 |
| 1098 | 55 | I55W | A | 2.7e-07 | 2.9e-07 | 1.2e-07 | 2.267E-07 | 9.292E-08 | 1.200E-07 | 2.900E-07 |
| 1099 | 55 | I55Y | A | 3.1e-07 | 2.2e-07 | 2.3e-07 | 2.533E-07 | 4.933E-08 | 2.200E-07 | 3.100E-07 |
| 1100 | 55 | I55V | A | 2.1e-07 | 1.5e-07 | 2.2e-07 | 1.933E-07 | 3.786E-08 | 1.500E-07 | 2.200E-07 |
| 1101 | 56 | Q56A | A | 4.3e-07 | 2.3e-07 | 3.4e-07 | 3.333E-07 | 1.002E-07 | 2.300E-07 | 4.300E-07 |
| 1102 | 56 | Q56R | A | 3.9e-07 | 2.1e-07 | 2.9e-07 | 2.967E-07 | 9.018E-08 | 2.100E-07 | 3.900E-07 |
| 1103 | 56 | Q56N | A | 3.1e-07 | 3.1e-07 | 2.5e-07 | 2.900E-07 | 3.464E-08 | 2.500E-07 | 3.100E-07 |
| 1104 | 56 | Q56D | A | 2.4e-07 | 3.9e-07 | 4.1e-07 | 3.467E-07 | 9.292E-08 | 2.400E-07 | 4.100E-07 |

| | | | | | | | | | | |
|---|---|---|---|---|---|---|---|---|---|---|
| 1105 | 56 | Q56C | A | 3.4e-07 | 1.1e-07 | 3.1e-07 | 2.533E-07 | 1.250E-07 | 1.100E-07 | 3.400E-07 |
| 1106 | 56 | Q56E | A | 2e-07 | 1.9e-07 | 1.5e-07 | 1.800E-07 | 2.646E-08 | 1.500E-07 | 2.000E-07 |
| 1107 | 56 | Q56Q | A | 1.9e-07 | 2.2e-07 | 3.1e-07 | 2.400E-07 | 6.245E-08 | 1.900E-07 | 3.100E-07 |
| 1108 | 56 | Q56G | A | 1.7e-07 | 1.7e-07 | 3.6e-07 | 2.333E-07 | 1.097E-07 | 1.700E-07 | 3.600E-07 |
| 1109 | 56 | Q56H | A | 2.1e-07 | 1.8e-07 | 3.4e-07 | 2.433E-07 | 8.505E-08 | 1.800E-07 | 3.400E-07 |
| 1110 | 56 | Q56I | A | 2.6e-07 | 2.5e-07 | 2.4e-07 | 2.500E-07 | 1.000E-08 | 2.400E-07 | 2.600E-07 |
| 1111 | 56 | Q56L | A | 2e-07 | 1.4e-07 | 2.4e-07 | 1.933E-07 | 5.033E-08 | 1.400E-07 | 2.400E-07 |
| 1112 | 56 | Q56K | A | 2.9e-07 | 1.9e-07 | 5.1e-08 | 1.770E-07 | 1.200E-07 | 5.100E-08 | 2.900E-07 |
| 1113 | 56 | Q56M | A | 3.2e-07 | 4e-07 | 1.3e-07 | 2.833E-07 | 1.387E-07 | 1.300E-07 | 4.000E-07 |
| 1114 | 56 | Q56F | A | 5e-07 | 4.1e-07 | 2.4e-07 | 3.833E-07 | 1.320E-07 | 2.400E-07 | 5.000E-07 |
| 1115 | 56 | Q56P | A | 4.6e-07 | 2.9e-07 | 2.4e-07 | 3.300E-07 | 1.153E-07 | 2.400E-07 | 4.600E-07 |
| 1116 | 56 | Q56S | A | 1.9e-07 | 2.7e-07 | 3.5e-07 | 2.700E-07 | 8.000E-08 | 1.900E-07 | 3.500E-07 |
| 1117 | 56 | Q56T | A | 8.6e-08 | 1.1e-07 | 1.2e-07 | 1.053E-07 | 1.747E-08 | 8.600E-08 | 1.200E-07 |
| 1118 | 56 | Q56W | A | 1e-07 | 9e-08 | 4e-07 | 1.967E-07 | 1.762E-07 | 9.000E-08 | 4.000E-07 |
| 1119 | 56 | Q56Y | A | 2.4e-07 | 2.6e-07 | 1.1e-07 | 2.033E-07 | 8.145E-08 | 1.100E-07 | 2.600E-07 |
| 1120 | 56 | Q56V | A | 1.4e-07 | 1.5e-07 | 7.8e-08 | 1.227E-07 | 3.900E-08 | 7.800E-08 | 1.500E-07 |
| 1121 | 57 | E57A | A | 2.9e-07 | 1.4e-07 | 2.4e-07 | 2.233E-07 | 7.638E-08 | 1.400E-07 | 2.900E-07 |
| 1122 | 57 | E57R | A | 1.4e-07 | 2e-07 | 2.9e-07 | 2.100E-07 | 7.550E-08 | 1.400E-07 | 2.900E-07 |
| 1123 | 57 | E57N | A | 1.3e-07 | 3.3e-07 | 3.5e-07 | 2.700E-07 | 1.217E-07 | 1.300E-07 | 3.500E-07 |
| 1124 | 57 | E57D | A | 1.1e-07 | 2.5e-07 | 2.3e-07 | 1.967E-07 | 7.572E-08 | 1.100E-07 | 2.500E-07 |
| 1125 | 57 | E57C | A | 1.6e-07 | 1.5e-07 | 1.2e-07 | 1.433E-07 | 2.082E-08 | 1.200E-07 | 1.600E-07 |
| 1126 | 57 | E57E | A | 2.7e-07 | 1.7e-07 | 8.3e-08 | 1.743E-07 | 9.358E-08 | 8.300E-08 | 2.700E-07 |
| 1127 | 57 | E57Q | A | 2e-07 | 1.2e-07 | 1.2e-07 | 1.467E-07 | 4.619E-08 | 1.200E-07 | 2.000E-07 |
| 1128 | 57 | E57G | A | 1.5e-07 | 6e-07 | 1.3e-07 | 2.933E-07 | 2.658E-07 | 1.300E-07 | 6.000E-07 |
| 1129 | 57 | E57H | A | 9.9e-08 | 2.6e-07 | 1.3e-07 | 1.630E-07 | 8.542E-08 | 9.900E-08 | 2.600E-07 |
| 1130 | 57 | E57I | A | 1.7e-07 | 3.3e-07 | 2.2e-07 | 2.400E-07 | 8.185E-08 | 1.700E-07 | 3.300E-07 |
| 1131 | 57 | E57L | A | 3.3e-07 | 1.2e-07 | 1.8e-07 | 2.100E-07 | 1.082E-07 | 1.200E-07 | 3.300E-07 |
| 1132 | 57 | E57K | A | 2.7e-07 | 3.9e-07 | 6e-07 | 4.200E-07 | 1.670E-07 | 2.700E-07 | 6.000E-07 |
| 1133 | 57 | E57M | A | 7.5e-07 | 2.6e-07 | 2.1e-07 | 4.067E-07 | 2.984E-07 | 2.100E-07 | 7.500E-07 |
| 1134 | 57 | E57F | A | 6.6e-07 | 3.3e-07 | 1.9e-07 | 3.933E-07 | 2.413E-07 | 1.900E-07 | 6.600E-07 |
| 1135 | 57 | E57P | A | 7.9e-08 | 3.3e-07 | 2.8e-07 | 2.297E-07 | 1.329E-07 | 7.900E-08 | 3.300E-07 |
| 1136 | 57 | E57S | A | 4.8e-07 | 5.7e-07 | 2.5e-07 | 4.333E-07 | 1.650E-07 | 2.500E-07 | 5.700E-07 |
| 1137 | 57 | E57T | A | 1.9e-07 | 1.9e-07 | 1.5e-07 | 1.767E-07 | 2.309E-08 | 1.500E-07 | 1.900E-07 |
| 1138 | 57 | E57W | A | 2.2e-07 | 3e-07 | 2.5e-07 | 2.567E-07 | 4.041E-08 | 2.200E-07 | 3.000E-07 |
| 1139 | 57 | E57Y | A | 2.1e-07 | 3.5e-07 | 3e-07 | 2.867E-07 | 7.095E-08 | 2.100E-07 | 3.500E-07 |
| 1140 | 57 | E57V | A | 2.7e-07 | 3.7e-07 | 2.9e-07 | 3.100E-07 | 5.292E-08 | 2.700E-07 | 3.700E-07 |
| 1141 | 58 | V58A | A | 1.5e-07 | 1.5e-07 | 5.3e-07 | 2.767E-07 | 2.194E-07 | 1.500E-07 | 5.300E-07 |
| 1142 | 58 | V58R | A | 3.1e-07 | 2.1e-07 | 2e-07 | 2.400E-07 | 6.083E-08 | 2.000E-07 | 3.100E-07 |
| 1143 | 58 | V58N | A | 1.9e-07 | 1.9e-07 | 1.7e-07 | 1.833E-07 | 1.155E-08 | 1.700E-07 | 1.900E-07 |
| 1144 | 58 | V58D | A | 2.4e-07 | 8.5e-07 | 1.9e-07 | 4.267E-07 | 3.675E-07 | 1.900E-07 | 8.500E-07 |
| 1145 | 58 | V58C | A | 3.7e-07 | 2.1e-07 | 3e-07 | 2.933E-07 | 8.021E-08 | 2.100E-07 | 3.700E-07 |
| 1146 | 58 | V58E | A | 2.6e-07 | 1.7e-07 | 4.8e-07 | 3.033E-07 | 1.595E-07 | 1.700E-07 | 4.800E-07 |
| 1147 | 58 | V58Q | A | 1.8e-07 | 2.2e-07 | 4.2e-07 | 2.733E-07 | 1.286E-07 | 1.800E-07 | 4.200E-07 |
| 1148 | 58 | V58G | A | 2.6e-07 | 1.7e-07 | 2.1e-07 | 2.133E-07 | 4.509E-08 | 1.700E-07 | 2.600E-07 |
| 1149 | 58 | V58H | A | 2.3e-07 | 2.6e-07 | 3.3e-07 | 2.733E-07 | 5.132E-08 | 2.300E-07 | 3.300E-07 |
| 1150 | 58 | V58I | A | 4.3e-07 | 1.6e-07 | 1.9e-07 | 2.600E-07 | 1.480E-07 | 1.600E-07 | 4.300E-07 |
| 1151 | 58 | V58L | A | 4.5e-07 | 1.1e-07 | 1.7e-07 | 2.433E-07 | 1.815E-07 | 1.100E-07 | 4.500E-07 |
| 1152 | 58 | V58K | A | 2.2e-07 | 1.6e-07 | 3.7e-07 | 2.500E-07 | 1.082E-07 | 1.600E-07 | 3.700E-07 |
| 1153 | 58 | V58M | A | 1.4e-07 | 2.3e-07 | 3.7e-07 | 2.467E-07 | 1.159E-07 | 1.400E-07 | 3.700E-07 |
| 1154 | 58 | V58F | A | 4.2e-07 | 2.9e-07 | 1.4e-07 | 2.833E-07 | 1.401E-07 | 1.400E-07 | 4.200E-07 |
| 1155 | 58 | V58P | A | 1.8e-07 | 3.5e-07 | 2.5e-07 | 2.600E-07 | 8.544E-08 | 1.800E-07 | 3.500E-07 |
| 1156 | 58 | V58S | A | 3.1e-07 | 2.3e-07 | 2.5e-07 | 2.633E-07 | 4.163E-08 | 2.300E-07 | 3.100E-07 |
| 1157 | 58 | V58T | A | 3.3e-07 | 4.5e-07 | 2.2e-07 | 3.333E-07 | 1.150E-07 | 2.200E-07 | 4.500E-07 |
| 1158 | 58 | V58W | A | 2.5e-07 | 1.4e-07 | 2.7e-07 | 2.200E-07 | 7.000E-08 | 1.400E-07 | 2.700E-07 |
| 1159 | 58 | V58Y | A | 8.6e-08 | 1.9e-07 | 1.9e-07 | 1.553E-07 | 6.004E-08 | 8.600E-08 | 1.900E-07 |
| 1160 | 58 | V58V | A | 2.8e-07 | 2.3e-07 | 4.6e-07 | 3.233E-07 | 1.210E-07 | 2.300E-07 | 4.600E-07 |
| 1161 | 59 | Q59A | A | 2.3e-07 | 2.1e-07 | 2.4e-07 | 2.267E-07 | 1.528E-08 | 2.100E-07 | 2.400E-07 |
| 1162 | 59 | Q59R | A | 2.8e-07 | 1.9e-07 | 1.3e-07 | 2.000E-07 | 7.550E-08 | 1.300E-07 | 2.800E-07 |
| 1163 | 59 | Q59N | A | 3.6e-07 | 1.1e-07 | 2.2e-07 | 2.300E-07 | 1.253E-07 | 1.100E-07 | 3.600E-07 |
| 1164 | 59 | Q59D | A | 1.8e-07 | 1.3e-07 | 2e-07 | 1.700E-07 | 3.606E-08 | 1.300E-07 | 2.000E-07 |
| 1165 | 59 | Q59C | A | 2e-07 | 2.7e-07 | 1.4e-07 | 2.033E-07 | 6.506E-08 | 1.400E-07 | 2.700E-07 |
| 1166 | 59 | Q59E | A | 4.5e-07 | 3e-07 | 2.3e-07 | 3.267E-07 | 1.124E-07 | 2.300E-07 | 4.500E-07 |
| 1167 | 59 | Q59Q | A | 1.8e-07 | 2.9e-07 | 3.4e-07 | 2.700E-07 | 8.185E-08 | 1.800E-07 | 3.400E-07 |
| 1168 | 59 | Q59G | A | 2.4e-07 | 2e-07 | 1.4e-07 | 1.933E-07 | 5.033E-08 | 1.400E-07 | 2.400E-07 |
| 1169 | 59 | Q59H | A | 1.5e-07 | 5e-07 | 2.7e-07 | 3.067E-07 | 1.779E-07 | 1.500E-07 | 5.000E-07 |
| 1170 | 59 | Q59I | A | 3.1e-07 | 1.7e-07 | 2.1e-07 | 2.300E-07 | 7.211E-08 | 1.700E-07 | 3.100E-07 |
| 1171 | 59 | Q59L | A | 1.6e-07 | 1.7e-07 | 2.2e-07 | 1.833E-07 | 3.215E-08 | 1.600E-07 | 2.200E-07 |
| 1172 | 59 | Q59K | A | 2e-07 | 1.4e-07 | 1.8e-07 | 1.733E-07 | 3.055E-08 | 1.400E-07 | 2.000E-07 |
| 1173 | 59 | Q59M | A | 2e-07 | 3.7e-07 | 5.9e-07 | 3.867E-07 | 1.955E-07 | 2.000E-07 | 5.900E-07 |
| 1174 | 59 | Q59F | A | 2.4e-07 | 2.1e-07 | 3e-07 | 2.500E-07 | 4.583E-08 | 2.100E-07 | 3.000E-07 |
| 1175 | 59 | Q59P | A | 1.5e-07 | 5.1e-07 | 1e-07 | 2.533E-07 | 2.237E-07 | 1.000E-07 | 5.100E-07 |
| 1176 | 59 | Q59S | A | 2.6e-07 | 2.7e-07 | 2.6e-07 | 2.633E-07 | 5.774E-09 | 2.600E-07 | 2.700E-07 |
| 1177 | 59 | Q59T | A | 2.6e-07 | 3.3e-07 | 1.6e-07 | 2.500E-07 | 8.544E-08 | 1.600E-07 | 3.300E-07 |
| 1178 | 59 | Q59W | A | 3.7e-07 | 1.3e-07 | 3.3e-07 | 2.767E-07 | 1.286E-07 | 1.300E-07 | 3.700E-07 |
| 1179 | 59 | Q59Y | A | 6.5e-07 | 3e-07 | 6.1e-07 | 5.200E-07 | 1.916E-07 | 3.000E-07 | 6.500E-07 |
| 1180 | 59 | Q59V | A | 1.2e-07 | 5.4e-07 | 4.8e-07 | 3.800E-07 | 2.272E-07 | 1.200E-07 | 5.400E-07 |
| 1181 | 60 | G60A | A | 1.7e-07 | 1.6e-07 | 4.3e-07 | 2.533E-07 | 1.531E-07 | 1.600E-07 | 4.300E-07 |
| 1182 | 60 | G60R | A | 5.1e-07 | 1.1e-07 | 1.4e-07 | 2.533E-07 | 2.228E-07 | 1.100E-07 | 5.100E-07 |
| 1183 | 60 | G60N | A | 1.4e-07 | 5.4e-07 | 1.8e-07 | 2.867E-07 | 2.203E-07 | 1.400E-07 | 5.400E-07 |
| 1184 | 60 | G60D | A | 3.9e-07 | 1.8e-07 | 4.6e-07 | 3.433E-07 | 1.457E-07 | 1.800E-07 | 4.600E-07 |
| 1185 | 60 | G60C | A | 2.2e-07 | 7.2e-07 | 1.7e-07 | 3.700E-07 | 3.041E-07 | 1.700E-07 | 7.200E-07 |
| 1186 | 60 | G60E | A | 9.4e-08 | 2.1e-07 | 4.2e-07 | 2.413E-07 | 1.652E-07 | 9.400E-08 | 4.200E-07 |
| 1187 | 60 | G60Q | A | 2e-07 | 2.8e-07 | 2.2e-07 | 2.333E-07 | 4.163E-08 | 2.000E-07 | 2.800E-07 |
| 1188 | 60 | G60G | A | 2.8e-07 | 1.4e-07 | 4.2e-07 | 2.800E-07 | 1.400E-07 | 1.400E-07 | 4.200E-07 |
| 1189 | 60 | G60H | A | 4.1e-07 | 2e-07 | 2.7e-07 | 2.933E-07 | 1.069E-07 | 2.000E-07 | 4.100E-07 |

| | | | | | | | | | | |
|---|---|---|---|---|---|---|---|---|---|---|
| 1190 | 60 | G60I | A | 1.9e-07 | 5.3e-07 | 4.2e-07 | 3.800E-07 | 1.735E-07 | 1.900E-07 | 5.300E-07 |
| 1191 | 60 | G60L | A | 2.5e-07 | 1.3e-07 | 1.9e-07 | 1.900E-07 | 6.000E-08 | 1.300E-07 | 2.500E-07 |
| 1192 | 60 | G60K | A | 3.3e-07 | 3.7e-07 | 2e-07 | 3.000E-07 | 8.888E-08 | 2.000E-07 | 3.700E-07 |
| 1193 | 60 | G60M | A | 2.6e-07 | 2.8e-07 | 4.8e-07 | 3.400E-07 | 1.217E-07 | 2.600E-07 | 4.800E-07 |
| 1194 | 60 | G60F | A | 2.1e-07 | 2.2e-07 | 1.1e-07 | 1.800E-07 | 6.083E-08 | 1.100E-07 | 2.200E-07 |
| 1195 | 60 | G60P | A | 3.5e-07 | 2.5e-07 | 2.2e-07 | 2.733E-07 | 6.807E-08 | 2.200E-07 | 3.500E-07 |
| 1196 | 60 | G60S | A | 1e-07 | 3.2e-07 | 1.7e-07 | 1.967E-07 | 1.124E-07 | 1.000E-07 | 3.200E-07 |
| 1197 | 60 | G60T | A | 3.4e-07 | 3.4e-07 | 1.1e-07 | 2.633E-07 | 1.328E-07 | 1.100E-07 | 3.400E-07 |
| 1198 | 60 | G60W | A | 1.2e-07 | 3.5e-07 | 3.7e-07 | 2.800E-07 | 1.389E-07 | 1.200E-07 | 3.700E-07 |
| 1199 | 60 | G60Y | A | 2e-07 | 1.7e-07 | 4.7e-07 | 2.800E-07 | 1.652E-07 | 1.700E-07 | 4.700E-07 |
| 1200 | 60 | G60V | A | 3.6e-07 | 2.5e-07 | 3.4e-07 | 3.167E-07 | 5.859E-08 | 2.500E-07 | 3.600E-07 |
| 1201 | 61 | Y61A | A | 3.9e-07 | 2.7e-07 | 2.7e-07 | 3.100E-07 | 6.928E-08 | 2.700E-07 | 3.900E-07 |
| 1202 | 61 | Y61R | A | 3.8e-07 | 4e-07 | 3.1e-07 | 3.633E-07 | 4.726E-08 | 3.100E-07 | 4.000E-07 |
| 1203 | 61 | Y61N | A | 1.5e-07 | 2.8e-07 | 1.9e-07 | 2.067E-07 | 6.658E-08 | 1.500E-07 | 2.800E-07 |
| 1204 | 61 | Y61D | A | 2.5e-07 | 3.2e-07 | 3.2e-07 | 2.967E-07 | 4.041E-08 | 2.500E-07 | 3.200E-07 |
| 1205 | 61 | Y61C | A | 6.1e-07 | 1.3e-07 | 4.2e-07 | 3.867E-07 | 2.417E-07 | 1.300E-07 | 6.100E-07 |
| 1206 | 61 | Y61E | A | 1.7e-07 | 2.8e-07 | 2.6e-07 | 2.367E-07 | 5.859E-08 | 1.700E-07 | 2.800E-07 |
| 1207 | 61 | Y61Q | A | 2.4e-07 | 2.7e-07 | 1.7e-07 | 2.267E-07 | 5.132E-08 | 1.700E-07 | 2.700E-07 |
| 1208 | 61 | Y61G | A | 3.9e-07 | 3.1e-07 | 2.1e-07 | 3.033E-07 | 9.018E-08 | 2.100E-07 | 3.900E-07 |
| 1209 | 61 | Y61H | A | 1.6e-07 | 5e-07 | 2.8e-07 | 3.133E-07 | 1.724E-07 | 1.600E-07 | 5.000E-07 |
| 1210 | 61 | Y61I | A | 1.8e-07 | 4.5e-07 | 3e-07 | 3.100E-07 | 1.353E-07 | 1.800E-07 | 4.500E-07 |
| 1211 | 61 | Y61L | A | 5.5e-07 | 1.5e-07 | 4.5e-07 | 3.833E-07 | 2.082E-07 | 1.500E-07 | 5.500E-07 |
| 1212 | 61 | Y61K | A | 2e-07 | 2.9e-07 | 3.2e-07 | 2.700E-07 | 6.245E-08 | 2.000E-07 | 3.200E-07 |
| 1213 | 61 | Y61M | A | 1.5e-07 | 2.5e-07 | 1.8e-07 | 1.933E-07 | 5.132E-08 | 1.500E-07 | 2.500E-07 |
| 1214 | 61 | Y61F | A | 2.2e-07 | 1.9e-07 | 1.3e-07 | 1.800E-07 | 4.583E-08 | 1.300E-07 | 2.200E-07 |
| 1215 | 61 | Y61P | A | 2.1e-07 | 3e-07 | 2.1e-07 | 2.400E-07 | 5.196E-08 | 2.100E-07 | 3.000E-07 |
| 1216 | 61 | Y61S | A | 7.8e-08 | 2.2e-07 | 2e-07 | 1.660E-07 | 7.686E-08 | 7.800E-08 | 2.200E-07 |
| 1217 | 61 | Y61T | A | 2.5e-07 | 2.6e-07 | 2.7e-07 | 2.600E-07 | 1.000E-08 | 2.500E-07 | 2.700E-07 |
| 1218 | 61 | Y61W | A | 4.4e-07 | 2.7e-07 | 2.4e-07 | 3.167E-07 | 1.079E-07 | 2.400E-07 | 4.400E-07 |
| 1219 | 61 | Y61Y | A | 1.2e-07 | 3.3e-07 | 6.6e-07 | 3.700E-07 | 2.722E-07 | 1.200E-07 | 6.600E-07 |
| 1220 | 61 | Y61V | A | 1.5e-07 | 7.3e-08 | 1.6e-07 | 1.277E-07 | 4.761E-08 | 7.300E-08 | 1.600E-07 |
| 1221 | 62 | V62A | A | 2.4e-07 | 2.7e-07 | 3.1e-07 | 2.733E-07 | 3.512E-08 | 2.400E-07 | 3.100E-07 |
| 1222 | 62 | V62R | A | 1.8e-07 | 3.4e-07 | 1.1e-07 | 2.100E-07 | 1.179E-07 | 1.100E-07 | 3.400E-07 |
| 1223 | 62 | V62N | A | 2.8e-07 | 1.2e-07 | 1.1e-07 | 1.700E-07 | 9.539E-08 | 1.100E-07 | 2.800E-07 |
| 1224 | 62 | V62D | A | 6.1e-07 | 2.9e-07 | 2.3e-07 | 3.767E-07 | 2.043E-07 | 2.300E-07 | 6.100E-07 |
| 1225 | 62 | V62C | A | 1.3e-07 | 2.7e-07 | 3.2e-07 | 2.400E-07 | 9.849E-08 | 1.300E-07 | 3.200E-07 |
| 1226 | 62 | V62E | A | 6.7e-07 | 1.7e-07 | 1.3e-07 | 3.233E-07 | 3.009E-07 | 1.300E-07 | 6.700E-07 |
| 1227 | 62 | V62Q | A | 3.4e-07 | 2.9e-07 | 2.9e-07 | 3.067E-07 | 2.887E-08 | 2.900E-07 | 3.400E-07 |
| 1228 | 62 | V62G | A | 4.6e-07 | 3.4e-07 | 2.5e-07 | 3.500E-07 | 1.054E-07 | 2.500E-07 | 4.600E-07 |
| 1229 | 62 | V62H | A | 1.1e-07 | 2.2e-07 | 3.4e-07 | 2.233E-07 | 1.150E-07 | 1.100E-07 | 3.400E-07 |
| 1230 | 62 | V62I | A | 2.3e-07 | 2.6e-07 | 1.6e-07 | 2.167E-07 | 5.132E-08 | 1.600E-07 | 2.600E-07 |
| 1231 | 62 | V62L | A | 3.9e-07 | 2.1e-07 | 3.1e-07 | 3.033E-07 | 9.018E-08 | 2.100E-07 | 3.900E-07 |
| 1232 | 62 | V62K | A | 1.6e-07 | 1.9e-07 | 1.8e-07 | 1.767E-07 | 1.528E-08 | 1.600E-07 | 1.900E-07 |
| 1233 | 62 | V62M | A | 3.1e-07 | 1.3e-07 | 2.4e-07 | 2.267E-07 | 9.074E-08 | 1.300E-07 | 3.100E-07 |
| 1234 | 62 | V62F | A | 1.9e-07 | 2.3e-07 | 3.5e-07 | 2.567E-07 | 8.327E-08 | 1.900E-07 | 3.500E-07 |
| 1235 | 62 | V62P | A | 2.2e-07 | 2.3e-07 | 2.5e-07 | 2.333E-07 | 1.528E-08 | 2.200E-07 | 2.500E-07 |
| 1236 | 62 | V62S | A | 3.7e-07 | 2.4e-07 | 1.9e-07 | 2.667E-07 | 9.292E-08 | 1.900E-07 | 3.700E-07 |
| 1237 | 62 | V62T | A | 1.9e-07 | 1.8e-07 | 2.5e-07 | 2.067E-07 | 3.786E-08 | 1.800E-07 | 2.500E-07 |
| 1238 | 62 | V62W | A | 1.9e-07 | 2.4e-07 | 1.8e-07 | 2.033E-07 | 3.215E-08 | 1.800E-07 | 2.400E-07 |
| 1239 | 62 | V62Y | A | 2.2e-07 | 1.6e-07 | 2.2e-07 | 2.000E-07 | 3.464E-08 | 1.600E-07 | 2.200E-07 |
| 1240 | 62 | V62V | A | 9.8e-08 | 2.7e-07 | 2.4e-07 | 2.027E-07 | 9.188E-08 | 9.800E-08 | 2.700E-07 |
| 1241 | 63 | L63A | A | 8.4e-07 | 4.6e-07 | 4e-07 | 5.667E-07 | 2.386E-07 | 4.000E-07 | 8.400E-07 |
| 1242 | 63 | L63R | A | 2.2e-07 | 1.2e-07 | 1.8e-07 | 1.733E-07 | 5.033E-08 | 1.200E-07 | 2.200E-07 |
| 1243 | 63 | L63N | A | 1.5e-07 | 5.1e-07 | 3.5e-07 | 3.367E-07 | 1.804E-07 | 1.500E-07 | 5.100E-07 |
| 1244 | 63 | L63D | A | 2.2e-07 | 1.7e-07 | 6.4e-07 | 3.433E-07 | 2.581E-07 | 1.700E-07 | 6.400E-07 |
| 1245 | 63 | L63C | A | 4.2e-07 | 1.9e-07 | 2e-07 | 2.700E-07 | 1.300E-07 | 1.900E-07 | 4.200E-07 |
| 1246 | 63 | L63E | A | 2.7e-07 | 2.6e-07 | 2.6e-07 | 2.633E-07 | 5.774E-09 | 2.600E-07 | 2.700E-07 |
| 1247 | 63 | L63Q | A | 2.9e-07 | 2.2e-07 | 2.1e-07 | 2.400E-07 | 4.359E-08 | 2.100E-07 | 2.900E-07 |
| 1248 | 63 | L63G | A | 4.2e-07 | 1.7e-07 | 2.9e-07 | 2.933E-07 | 1.250E-07 | 1.700E-07 | 4.200E-07 |
| 1249 | 63 | L63H | A | 1.3e-07 | 1.9e-07 | 3.3e-07 | 2.167E-07 | 1.026E-07 | 1.300E-07 | 3.300E-07 |
| 1250 | 63 | L63I | A | 2e-07 | 1.8e-07 | 2.2e-07 | 2.000E-07 | 2.000E-08 | 1.800E-07 | 2.200E-07 |
| 1251 | 63 | L63L | A | 2.7e-07 | 1.4e-07 | 5e-07 | 3.033E-07 | 1.823E-07 | 1.400E-07 | 5.000E-07 |
| 1252 | 63 | L63K | A | 1.4e-07 | 2.4e-07 | 1.7e-07 | 1.833E-07 | 5.132E-08 | 1.400E-07 | 2.400E-07 |
| 1253 | 63 | L63M | A | 3.1e-07 | 3.5e-07 | 2.9e-07 | 3.167E-07 | 3.055E-08 | 2.900E-07 | 3.500E-07 |
| 1254 | 63 | L63F | A | 1.4e-07 | 1.7e-07 | 1.8e-07 | 1.633E-07 | 2.082E-08 | 1.400E-07 | 1.800E-07 |
| 1255 | 63 | L63P | A | 1.7e-07 | 2.6e-07 | 1.6e-07 | 1.967E-07 | 5.508E-08 | 1.600E-07 | 2.600E-07 |
| 1256 | 63 | L63S | A | 2e-07 | 1.7e-07 | 5.3e-07 | 3.000E-07 | 1.997E-07 | 1.700E-07 | 5.300E-07 |
| 1257 | 63 | L63T | A | 3.8e-07 | 1.9e-07 | 2e-07 | 2.567E-07 | 1.069E-07 | 1.900E-07 | 3.800E-07 |
| 1258 | 63 | L63W | A | 2.9e-07 | 4e-07 | 3e-07 | 3.300E-07 | 6.083E-08 | 2.900E-07 | 4.000E-07 |
| 1259 | 63 | L63Y | A | 2.2e-07 | 3.1e-07 | 3e-07 | 2.767E-07 | 4.933E-08 | 2.200E-07 | 3.100E-07 |
| 1260 | 63 | L63V | A | 6.1e-07 | 1e-07 | 2.3e-07 | 3.133E-07 | 2.650E-07 | 1.000E-07 | 6.100E-07 |
| 1261 | 64 | I64A | A | 2.3e-07 | 2.4e-07 | 3.1e-07 | 2.600E-07 | 4.359E-08 | 2.300E-07 | 3.100E-07 |
| 1262 | 64 | I64R | A | 3.1e-07 | 3.9e-07 | 1.8e-07 | 2.933E-07 | 1.060E-07 | 1.800E-07 | 3.900E-07 |
| 1263 | 64 | I64N | A | 3.5e-07 | 1.9e-07 | 2.2e-07 | 2.533E-07 | 8.505E-08 | 1.900E-07 | 3.500E-07 |
| 1264 | 64 | I64D | A | 3.6e-07 | 1.5e-07 | 2e-07 | 2.367E-07 | 1.097E-07 | 1.500E-07 | 3.600E-07 |
| 1265 | 64 | I64C | A | 2.1e-07 | 3e-07 | 3.3e-07 | 2.800E-07 | 6.245E-08 | 2.100E-07 | 3.300E-07 |
| 1266 | 64 | I64E | A | 2.6e-07 | 7.1e-07 | 1.7e-07 | 3.800E-07 | 2.893E-07 | 1.700E-07 | 7.100E-07 |
| 1267 | 64 | I64Q | A | 2.9e-07 | 3.3e-07 | 1.1e-07 | 2.433E-07 | 1.172E-07 | 1.100E-07 | 3.300E-07 |
| 1268 | 64 | I64G | A | 2.4e-07 | 1.5e-07 | 3e-07 | 2.300E-07 | 7.550E-08 | 1.500E-07 | 3.000E-07 |
| 1269 | 64 | I64H | A | 3.4e-07 | 2.8e-07 | 2e-07 | 2.733E-07 | 7.024E-08 | 2.000E-07 | 3.400E-07 |
| 1270 | 64 | I64I | A | 2.9e-07 | 1.7e-07 | 2.1e-07 | 2.233E-07 | 6.110E-08 | 1.700E-07 | 2.900E-07 |
| 1271 | 64 | I64L | A | 3e-07 | 3.7e-07 | 3e-07 | 3.233E-07 | 4.041E-08 | 3.000E-07 | 3.700E-07 |
| 1272 | 64 | I64K | A | 1.9e-07 | 2.5e-07 | 1e-07 | 1.800E-07 | 7.550E-08 | 1.000E-07 | 2.500E-07 |
| 1273 | 64 | I64M | A | 3.7e-07 | 4e-07 | 3e-07 | 3.567E-07 | 5.132E-08 | 3.000E-07 | 4.000E-07 |
| 1274 | 64 | I64F | A | 1.2e-07 | 3e-07 | 1.8e-07 | 2.000E-07 | 9.165E-08 | 1.200E-07 | 3.000E-07 |

| | | | | | | | | | | |
|---|---|---|---|---|---|---|---|---|---|---|
| 1275 | 64 | I64P | A | 2.1e-07 | 3e-07 | 4.7e-07 | 3.267E-07 | 1.320E-07 | 2.100E-07 | 4.700E-07 |
| 1276 | 64 | I64S | A | 2.5e-07 | 5.1e-07 | 1.5e-07 | 3.033E-07 | 1.858E-07 | 1.500E-07 | 5.100E-07 |
| 1277 | 64 | I64T | A | 3.8e-07 | 5e-07 | 2.3e-07 | 3.700E-07 | 1.353E-07 | 2.300E-07 | 5.000E-07 |
| 1278 | 64 | I64W | A | 2.2e-07 | 1.8e-07 | 1.3e-07 | 1.767E-07 | 4.509E-08 | 1.300E-07 | 2.200E-07 |
| 1279 | 64 | I64Y | A | 1.8e-07 | 2e-07 | 1.5e-07 | 1.767E-07 | 2.517E-08 | 1.500E-07 | 2.000E-07 |
| 1280 | 64 | I64V | A | 3e-07 | 2e-07 | 3e-07 | 2.667E-07 | 5.774E-08 | 2.000E-07 | 3.000E-07 |
| 1281 | 65 | A65A | A | 1.3e-07 | 1.6e-07 | 4.5e-07 | 2.467E-07 | 1.767E-07 | 1.300E-07 | 4.500E-07 |
| 1282 | 65 | A65R | A | 3.3e-07 | 3.6e-07 | 3.7e-07 | 3.533E-07 | 2.082E-08 | 3.300E-07 | 3.700E-07 |
| 1283 | 65 | A65N | A | 1.6e-07 | 3.8e-07 | 1.8e-07 | 2.400E-07 | 1.217E-07 | 1.600E-07 | 3.800E-07 |
| 1284 | 65 | A65D | A | 1.8e-07 | 2.6e-07 | 2.6e-07 | 2.333E-07 | 4.619E-08 | 1.800E-07 | 2.600E-07 |
| 1285 | 65 | A65C | A | 3.9e-07 | 3.8e-07 | 3.8e-07 | 3.833E-07 | 5.774E-09 | 3.800E-07 | 3.900E-07 |
| 1286 | 65 | A65E | A | 2e-07 | 1.9e-07 | 2.4e-07 | 2.100E-07 | 2.646E-08 | 1.900E-07 | 2.400E-07 |
| 1287 | 65 | A65Q | A | 3.5e-07 | 3.6e-07 | 2.6e-07 | 3.233E-07 | 5.508E-08 | 2.600E-07 | 3.600E-07 |
| 1288 | 65 | A65G | A | 1.3e-07 | 1.7e-07 | 2.2e-07 | 1.733E-07 | 4.509E-08 | 1.300E-07 | 2.200E-07 |
| 1289 | 65 | A65H | A | 2.4e-07 | 1.9e-07 | 2.9e-07 | 2.400E-07 | 5.000E-08 | 1.900E-07 | 2.900E-07 |
| 1290 | 65 | A65I | A | 2.5e-07 | 1.5e-07 | 1e-07 | 1.667E-07 | 7.638E-08 | 1.000E-07 | 2.500E-07 |
| 1291 | 65 | A65L | A | 2.5e-07 | 3.6e-07 | 3.4e-07 | 3.167E-07 | 5.859E-08 | 2.500E-07 | 3.600E-07 |
| 1292 | 65 | A65K | A | 4e-07 | 2.8e-07 | 2.8e-07 | 3.200E-07 | 6.928E-08 | 2.800E-07 | 4.000E-07 |
| 1293 | 65 | A65M | A | 9.5e-08 | 5.1e-07 | 4.2e-07 | 3.417E-07 | 2.183E-07 | 9.500E-08 | 5.100E-07 |
| 1294 | 65 | A65F | A | 1.7e-07 | 1.7e-07 | 1.7e-07 | 1.700E-07 | 0.000E+00 | 1.700E-07 | 1.700E-07 |
| 1295 | 65 | A65P | A | 1.7e-07 | 1.6e-07 | 3.8e-07 | 2.367E-07 | 1.242E-07 | 1.600E-07 | 3.800E-07 |
| 1296 | 65 | A65S | A | 1.9e-07 | 4.2e-07 | 6.5e-07 | 4.200E-07 | 2.300E-07 | 1.900E-07 | 6.500E-07 |
| 1297 | 65 | A65T | A | 3.7e-07 | 1.4e-07 | 1.5e-07 | 2.200E-07 | 1.300E-07 | 1.400E-07 | 3.700E-07 |
| 1298 | 65 | A65W | A | 4e-07 | 1.8e-07 | 2.8e-07 | 2.867E-07 | 1.102E-07 | 1.800E-07 | 4.000E-07 |
| 1299 | 65 | A65Y | A | 2.1e-07 | 2.7e-07 | 2e-07 | 2.267E-07 | 3.786E-08 | 2.000E-07 | 2.700E-07 |
| 1300 | 65 | A65V | A | 2.2e-07 | 2.1e-07 | 4.3e-07 | 2.867E-07 | 1.242E-07 | 2.100E-07 | 4.300E-07 |
| 1301 | 66 | H66A | A | 1.7e-07 | 2.5e-07 | 2.4e-07 | 2.200E-07 | 4.359E-08 | 1.700E-07 | 2.500E-07 |
| 1302 | 66 | H66R | A | 4.3e-07 | 3.1e-07 | 8.3e-08 | 2.743E-07 | 1.762E-07 | 8.300E-08 | 4.300E-07 |
| 1303 | 66 | H66N | A | 1.2e-07 | 4.6e-07 | 1.8e-07 | 2.533E-07 | 1.815E-07 | 1.200E-07 | 4.600E-07 |
| 1304 | 66 | H66D | A | 1.8e-07 | 2.3e-07 | 2.4e-07 | 2.167E-07 | 3.215E-08 | 1.800E-07 | 2.400E-07 |
| 1305 | 66 | H66C | A | 1.3e-07 | 2.9e-07 | 1.5e-07 | 1.900E-07 | 8.718E-08 | 1.300E-07 | 2.900E-07 |
| 1306 | 66 | H66E | A | 1.1e-07 | 5.3e-07 | 2.8e-07 | 3.067E-07 | 2.113E-07 | 1.100E-07 | 5.300E-07 |
| 1307 | 66 | H66Q | A | 3.4e-07 | 3.6e-07 | 1.2e-07 | 2.733E-07 | 1.332E-07 | 1.200E-07 | 3.600E-07 |
| 1308 | 66 | H66G | A | 2e-07 | 1.7e-07 | 1.5e-07 | 1.733E-07 | 2.517E-08 | 1.500E-07 | 2.000E-07 |
| 1309 | 66 | H66H | A | 3e-07 | 3.9e-07 | 4.3e-07 | 3.733E-07 | 6.658E-08 | 3.000E-07 | 4.300E-07 |
| 1310 | 66 | H66I | A | 3.7e-07 | 2e-07 | 3.8e-07 | 3.167E-07 | 1.012E-07 | 2.000E-07 | 3.800E-07 |
| 1311 | 66 | H66L | A | 5.5e-07 | 9.5e-08 | 1.9e-07 | 2.783E-07 | 2.400E-07 | 9.500E-08 | 5.500E-07 |
| 1312 | 66 | H66K | A | 3.1e-07 | 2.3e-07 | 2.1e-07 | 2.500E-07 | 5.292E-08 | 2.100E-07 | 3.100E-07 |
| 1313 | 66 | H66M | A | 1.2e-07 | 1.7e-07 | 4.1e-07 | 2.333E-07 | 1.550E-07 | 1.200E-07 | 4.100E-07 |
| 1314 | 66 | H66F | A | 1.5e-07 | 3.8e-07 | 3.2e-07 | 2.833E-07 | 1.193E-07 | 1.500E-07 | 3.800E-07 |
| 1315 | 66 | H66P | A | 3.9e-07 | 2.3e-07 | 2.9e-07 | 3.033E-07 | 8.083E-08 | 2.300E-07 | 3.900E-07 |
| 1316 | 66 | H66S | A | 2.1e-07 | 2.7e-07 | 1.8e-07 | 2.200E-07 | 4.583E-08 | 1.800E-07 | 2.700E-07 |
| 1317 | 66 | H66T | A | 3.6e-07 | 5e-07 | 3.1e-07 | 3.900E-07 | 9.849E-08 | 3.100E-07 | 5.000E-07 |
| 1318 | 66 | H66W | A | 2.7e-07 | 2.1e-07 | 4.7e-07 | 3.167E-07 | 1.361E-07 | 2.100E-07 | 4.700E-07 |
| 1319 | 66 | H66Y | A | 7.9e-07 | 4.7e-07 | 2.2e-07 | 4.933E-07 | 2.857E-07 | 2.200E-07 | 7.900E-07 |
| 1320 | 66 | H66V | A | 2.7e-07 | 3.2e-07 | 2.2e-07 | 2.700E-07 | 5.000E-08 | 2.200E-07 | 3.200E-07 |
| 1321 | 67 | N67A | A | 4.7e-07 | 2.1e-07 | 7.3e-07 | 4.700E-07 | 2.600E-07 | 2.100E-07 | 7.300E-07 |
| 1322 | 67 | N67R | A | 7.4e-07 | 1.4e-07 | 1.4e-07 | 3.400E-07 | 3.464E-07 | 1.400E-07 | 7.400E-07 |
| 1323 | 67 | N67N | A | 2.7e-07 | 1.7e-07 | 2.8e-07 | 2.400E-07 | 6.083E-08 | 1.700E-07 | 2.800E-07 |
| 1324 | 67 | N67D | A | 1.2e-07 | 3.7e-07 | 2.1e-07 | 2.333E-07 | 1.266E-07 | 1.200E-07 | 3.700E-07 |
| 1325 | 67 | N67C | A | 2.3e-07 | 1.7e-07 | 3.5e-07 | 2.500E-07 | 9.165E-08 | 1.700E-07 | 3.500E-07 |
| 1326 | 67 | N67E | A | 2e-07 | 2.6e-07 | 2.9e-07 | 2.500E-07 | 4.583E-08 | 2.000E-07 | 2.900E-07 |
| 1327 | 67 | N67Q | A | 2.5e-07 | 1.6e-07 | 3.8e-07 | 2.633E-07 | 1.106E-07 | 1.600E-07 | 3.800E-07 |
| 1328 | 67 | N67G | A | 2.1e-07 | 3.9e-07 | 2.7e-07 | 2.900E-07 | 9.165E-08 | 2.100E-07 | 3.900E-07 |
| 1329 | 67 | N67H | A | 3.8e-07 | 2.3e-07 | 2.9e-07 | 3.000E-07 | 7.550E-08 | 2.300E-07 | 3.800E-07 |
| 1330 | 67 | N67I | A | 1.8e-07 | 2.4e-07 | 9.8e-08 | 1.727E-07 | 7.128E-08 | 9.800E-08 | 2.400E-07 |
| 1331 | 67 | N67L | A | 2e-07 | 1.7e-07 | 1.7e-07 | 1.800E-07 | 1.732E-08 | 1.700E-07 | 2.000E-07 |
| 1332 | 67 | N67K | A | 6.7e-07 | 2.8e-07 | 3.3e-07 | 4.267E-07 | 2.122E-07 | 2.800E-07 | 6.700E-07 |
| 1333 | 67 | N67M | A | 1.9e-07 | 1.7e-07 | 1.9e-07 | 1.833E-07 | 1.155E-08 | 1.700E-07 | 1.900E-07 |
| 1334 | 67 | N67F | A | 4.9e-07 | 1.9e-07 | 2.3e-07 | 3.033E-07 | 1.629E-07 | 1.900E-07 | 4.900E-07 |
| 1335 | 67 | N67P | A | 1.4e-07 | 3e-07 | 4.3e-07 | 2.900E-07 | 1.453E-07 | 1.400E-07 | 4.300E-07 |
| 1336 | 67 | N67S | A | 2.5e-07 | 3.7e-07 | 4.2e-07 | 3.467E-07 | 8.737E-08 | 2.500E-07 | 4.200E-07 |
| 1337 | 67 | N67T | A | 3.6e-07 | 2.1e-07 | 2.1e-07 | 2.600E-07 | 8.660E-08 | 2.100E-07 | 3.600E-07 |
| 1338 | 67 | N67W | A | 1.7e-07 | 2.6e-07 | 2.4e-07 | 2.233E-07 | 4.726E-08 | 1.700E-07 | 2.600E-07 |
| 1339 | 67 | N67Y | A | 3.1e-07 | 2.6e-07 | 2.7e-07 | 2.800E-07 | 2.646E-08 | 2.600E-07 | 3.100E-07 |
| 1340 | 67 | N67V | A | 2.4e-07 | 3.2e-07 | 3.3e-07 | 2.967E-07 | 4.933E-08 | 2.400E-07 | 3.300E-07 |
| 1341 | 68 | Q68A | A | 2.5e-07 | 2.1e-07 | 2.6e-07 | 2.400E-07 | 2.646E-08 | 2.100E-07 | 2.600E-07 |
| 1342 | 68 | Q68R | A | 3.5e-07 | 2.5e-07 | 5.5e-07 | 3.833E-07 | 1.528E-07 | 2.500E-07 | 5.500E-07 |
| 1343 | 68 | Q68N | A | 4.3e-07 | 6.5e-07 | 1.7e-07 | 4.167E-07 | 2.403E-07 | 1.700E-07 | 6.500E-07 |
| 1344 | 68 | Q68D | A | 1.2e-07 | 9.4e-07 | 4.3e-07 | 4.967E-07 | 4.140E-07 | 1.200E-07 | 9.400E-07 |
| 1345 | 68 | Q68C | A | 2.6e-07 | 2.4e-07 | 3e-07 | 2.667E-07 | 3.055E-08 | 2.400E-07 | 3.000E-07 |
| 1346 | 68 | Q68E | A | 5.5e-07 | 1.8e-07 | 1.8e-07 | 3.033E-07 | 2.136E-07 | 1.800E-07 | 5.500E-07 |
| 1347 | 68 | Q68Q | A | 3.1e-07 | 1.6e-07 | 2.9e-07 | 2.533E-07 | 8.145E-08 | 1.600E-07 | 3.100E-07 |
| 1348 | 68 | Q68G | A | 1.8e-07 | 1.2e-07 | 2e-07 | 1.667E-07 | 4.163E-08 | 1.200E-07 | 2.000E-07 |
| 1349 | 68 | Q68H | A | 2.6e-07 | 1.4e-07 | 1.4e-07 | 1.800E-07 | 6.928E-08 | 1.400E-07 | 2.600E-07 |
| 1350 | 68 | Q68I | A | 2.4e-07 | 3.1e-07 | 1.8e-07 | 2.433E-07 | 6.506E-08 | 1.800E-07 | 3.100E-07 |
| 1351 | 68 | Q68L | A | 1.6e-07 | 3.9e-07 | 1.8e-07 | 2.433E-07 | 1.274E-07 | 1.600E-07 | 3.900E-07 |
| 1352 | 68 | Q68K | A | 3.5e-07 | 3.5e-07 | 5e-07 | 4.000E-07 | 8.660E-08 | 3.500E-07 | 5.000E-07 |
| 1353 | 68 | Q68M | A | 6.5e-08 | 5.2e-07 | 2.2e-07 | 2.683E-07 | 2.313E-07 | 6.500E-08 | 5.200E-07 |
| 1354 | 68 | Q68F | A | 2.2e-07 | 1.3e-07 | 5.9e-07 | 3.133E-07 | 2.438E-07 | 1.300E-07 | 5.900E-07 |
| 1355 | 68 | Q68P | A | 1.6e-07 | 4.2e-07 | 1.5e-07 | 2.433E-07 | 1.531E-07 | 1.500E-07 | 4.200E-07 |
| 1356 | 68 | Q68S | A | 1.9e-07 | 2.8e-07 | 3.8e-07 | 2.833E-07 | 9.504E-08 | 1.900E-07 | 3.800E-07 |
| 1357 | 68 | Q68T | A | 2.5e-07 | 2.6e-07 | 1.9e-07 | 2.333E-07 | 3.786E-08 | 1.900E-07 | 2.600E-07 |
| 1358 | 68 | Q68W | A | 2.3e-07 | 1.7e-07 | 2.2e-07 | 2.067E-07 | 3.215E-08 | 1.700E-07 | 2.300E-07 |
| 1359 | 68 | Q68Y | A | 1.7e-07 | 1.9e-07 | 3.4e-07 | 2.333E-07 | 9.292E-08 | 1.700E-07 | 3.400E-07 |

| | | | | | | | | | | |
|---|---|---|---|---|---|---|---|---|---|---|
| 1360 | 68 | Q68V | A | 2.6e-07 | 2.6e-07 | 1.6e-07 | 2.267E-07 | 5.774E-08 | 1.600E-07 | 2.600E-07 |
| 1361 | 69 | V69A | A | 1.6e-07 | 3.4e-07 | 2.4e-07 | 2.467E-07 | 9.018E-08 | 1.600E-07 | 3.400E-07 |
| 1362 | 69 | V69R | A | 1.5e-07 | 2.9e-07 | 2.3e-07 | 2.233E-07 | 7.024E-08 | 1.500E-07 | 2.900E-07 |
| 1363 | 69 | V69N | A | 2.2e-07 | 1.9e-07 | 3.1e-07 | 2.400E-07 | 6.245E-08 | 1.900E-07 | 3.100E-07 |
| 1364 | 69 | V69D | A | 2.3e-07 | 2.2e-07 | 2.3e-07 | 2.267E-07 | 5.774E-09 | 2.200E-07 | 2.300E-07 |
| 1365 | 69 | V69C | A | 2e-07 | 4.3e-07 | 3.7e-07 | 3.333E-07 | 1.193E-07 | 2.000E-07 | 4.300E-07 |
| 1366 | 69 | V69E | A | 1.2e-07 | 3e-07 | 2.8e-07 | 2.333E-07 | 9.866E-08 | 1.200E-07 | 3.000E-07 |
| 1367 | 69 | V69Q | A | 8.3e-08 | 8.1e-08 | 4e-07 | 1.880E-07 | 1.836E-07 | 8.100E-08 | 4.000E-07 |
| 1368 | 69 | V69G | A | 2.1e-07 | 4.9e-07 | 2.1e-07 | 3.033E-07 | 1.617E-07 | 2.100E-07 | 4.900E-07 |
| 1369 | 69 | V69H | A | 1.8e-07 | 5.3e-07 | 2.2e-07 | 3.100E-07 | 1.916E-07 | 1.800E-07 | 5.300E-07 |
| 1370 | 69 | V69I | A | 1.8e-07 | 3.1e-07 | 4.7e-07 | 3.200E-07 | 1.453E-07 | 1.800E-07 | 4.700E-07 |
| 1371 | 69 | V69L | A | 2.3e-07 | 1.6e-07 | 2e-07 | 1.967E-07 | 3.512E-08 | 1.600E-07 | 2.300E-07 |
| 1372 | 69 | V69K | A | 3e-07 | 2.6e-07 | 4.3e-07 | 3.300E-07 | 8.888E-08 | 2.600E-07 | 4.300E-07 |
| 1373 | 69 | V69M | A | 3.8e-07 | 1.8e-07 | 5.6e-07 | 3.733E-07 | 1.901E-07 | 1.800E-07 | 5.600E-07 |
| 1374 | 69 | V69F | A | 1e-07 | 2e-07 | 3.2e-07 | 2.067E-07 | 1.102E-07 | 1.000E-07 | 3.200E-07 |
| 1375 | 69 | V69P | A | 1.7e-07 | 1.6e-07 | 1.4e-07 | 1.567E-07 | 1.528E-08 | 1.400E-07 | 1.700E-07 |
| 1376 | 69 | V69S | A | 1.4e-07 | 3.6e-07 | 2.5e-07 | 2.500E-07 | 1.100E-07 | 1.400E-07 | 3.600E-07 |
| 1377 | 69 | V69T | A | 2.1e-07 | 2.3e-07 | 4.2e-07 | 2.867E-07 | 1.159E-07 | 2.100E-07 | 4.200E-07 |
| 1378 | 69 | V69W | A | 4.7e-07 | 2.3e-07 | 3e-07 | 3.333E-07 | 1.234E-07 | 2.300E-07 | 4.700E-07 |
| 1379 | 69 | V69Y | A | 3.1e-07 | 2.5e-07 | 1.6e-07 | 2.400E-07 | 7.550E-08 | 1.600E-07 | 3.100E-07 |
| 1380 | 69 | V69V | A | 7.5e-07 | 2.8e-07 | 1.6e-07 | 3.967E-07 | 3.118E-07 | 1.600E-07 | 7.500E-07 |
| 1381 | 70 | R70A | A | 3.5e-07 | 2.6e-07 | 2.6e-07 | 2.900E-07 | 5.196E-08 | 2.600E-07 | 3.500E-07 |
| 1382 | 70 | R70R | A | 3.5e-07 | 3.4e-07 | 3e-07 | 3.300E-07 | 2.646E-08 | 3.000E-07 | 3.500E-07 |
| 1383 | 70 | R70N | A | 3.6e-07 | 2.2e-07 | 1.9e-07 | 2.567E-07 | 9.074E-08 | 1.900E-07 | 3.600E-07 |
| 1384 | 70 | R70D | A | 4.6e-07 | 1.1e-07 | 2e-07 | 2.567E-07 | 1.818E-07 | 1.100E-07 | 4.600E-07 |
| 1385 | 70 | R70C | A | 2.4e-07 | 4.5e-07 | 1.7e-07 | 2.867E-07 | 1.457E-07 | 1.700E-07 | 4.500E-07 |
| 1386 | 70 | R70E | A | 2.1e-07 | 3.5e-07 | 3.5e-07 | 3.033E-07 | 8.083E-08 | 2.100E-07 | 3.500E-07 |
| 1387 | 70 | R70Q | A | 3.2e-07 | 6e-07 | 3.6e-07 | 4.267E-07 | 1.514E-07 | 3.200E-07 | 6.000E-07 |
| 1388 | 70 | R70G | A | 2.7e-07 | 3.4e-07 | 2.6e-07 | 2.900E-07 | 4.359E-08 | 2.600E-07 | 3.400E-07 |
| 1389 | 70 | R70H | A | 1.9e-07 | 2.9e-07 | 2.2e-07 | 2.333E-07 | 5.132E-08 | 1.900E-07 | 2.900E-07 |
| 1390 | 70 | R70I | A | 4.5e-07 | 2.3e-07 | 3.9e-07 | 3.567E-07 | 1.137E-07 | 2.300E-07 | 4.500E-07 |
| 1391 | 70 | R70L | A | 2e-07 | 3.1e-07 | 5.9e-08 | 1.897E-07 | 1.258E-07 | 5.900E-08 | 3.100E-07 |
| 1392 | 70 | R70K | A | 4.9e-07 | 1.7e-07 | 2.5e-07 | 3.033E-07 | 1.665E-07 | 1.700E-07 | 4.900E-07 |
| 1393 | 70 | R70M | A | 1.5e-07 | 3.5e-07 | 2.9e-07 | 2.633E-07 | 1.026E-07 | 1.500E-07 | 3.500E-07 |
| 1394 | 70 | R70F | A | 2e-07 | 2.3e-07 | 3.2e-07 | 2.500E-07 | 6.245E-08 | 2.000E-07 | 3.200E-07 |
| 1395 | 70 | R70P | A | 3.2e-07 | 2.3e-07 | 4.4e-07 | 3.300E-07 | 1.054E-07 | 2.300E-07 | 4.400E-07 |
| 1396 | 70 | R70S | A | 2.9e-07 | 3.8e-07 | 1.2e-07 | 2.633E-07 | 1.320E-07 | 1.200E-07 | 3.800E-07 |
| 1397 | 70 | R70T | A | 2.9e-07 | 2.3e-07 | 2.6e-07 | 2.600E-07 | 3.000E-08 | 2.300E-07 | 2.900E-07 |
| 1398 | 70 | R70W | A | 3.2e-07 | 1.8e-07 | 2.1e-07 | 2.367E-07 | 7.371E-08 | 1.800E-07 | 3.200E-07 |
| 1399 | 70 | R70Y | A | 3.3e-07 | 1.2e-07 | 2e-07 | 2.167E-07 | 1.060E-07 | 1.200E-07 | 3.300E-07 |
| 1400 | 70 | R70V | A | 2.8e-07 | 1.7e-07 | 2.9e-07 | 2.467E-07 | 6.658E-08 | 1.700E-07 | 2.900E-07 |
| 1401 | 71 | Q71A | A | 2.5e-07 | 1.6e-07 | 2.1e-07 | 2.067E-07 | 4.509E-08 | 1.600E-07 | 2.500E-07 |
| 1402 | 71 | Q71R | A | 3.4e-07 | 3.7e-07 | 2.8e-07 | 3.300E-07 | 4.583E-08 | 2.800E-07 | 3.700E-07 |
| 1403 | 71 | Q71N | A | 3.9e-07 | 1.4e-07 | 2.4e-07 | 2.567E-07 | 1.258E-07 | 1.400E-07 | 3.900E-07 |
| 1404 | 71 | Q71D | A | 3e-07 | 3.4e-07 | 2.3e-07 | 2.900E-07 | 5.568E-08 | 2.300E-07 | 3.400E-07 |
| 1405 | 71 | Q71C | A | 1.3e-07 | 2.4e-07 | 2.7e-07 | 2.133E-07 | 7.371E-08 | 1.300E-07 | 2.700E-07 |
| 1406 | 71 | Q71E | A | 2.4e-07 | 2e-07 | 1.2e-07 | 1.867E-07 | 6.110E-08 | 1.200E-07 | 2.400E-07 |
| 1407 | 71 | Q71Q | A | 1.2e-07 | 1.9e-07 | 2.4e-07 | 1.833E-07 | 6.028E-08 | 1.200E-07 | 2.400E-07 |
| 1408 | 71 | Q71G | A | 2.1e-07 | 3.3e-07 | 1.6e-07 | 2.333E-07 | 8.737E-08 | 1.600E-07 | 3.300E-07 |
| 1409 | 71 | Q71H | A | 3e-07 | 6.2e-07 | 2.7e-07 | 3.967E-07 | 1.940E-07 | 2.700E-07 | 6.200E-07 |
| 1410 | 71 | Q71I | A | 3.2e-07 | 1.4e-07 | 1.6e-07 | 2.067E-07 | 9.866E-08 | 1.400E-07 | 3.200E-07 |
| 1411 | 71 | Q71L | A | 1.9e-07 | 4.5e-07 | 3.6e-07 | 3.333E-07 | 1.320E-07 | 1.900E-07 | 4.500E-07 |
| 1412 | 71 | Q71K | A | 2.3e-07 | 2.4e-07 | 3.2e-07 | 2.633E-07 | 4.933E-08 | 2.300E-07 | 3.200E-07 |
| 1413 | 71 | Q71M | A | 1.4e-07 | 2.5e-07 | 1.9e-07 | 1.933E-07 | 5.508E-08 | 1.400E-07 | 2.500E-07 |
| 1414 | 71 | Q71F | A | 9.7e-08 | 3.2e-07 | 2.2e-07 | 2.123E-07 | 1.117E-07 | 9.700E-08 | 3.200E-07 |
| 1415 | 71 | Q71P | A | 2.5e-07 | 2.1e-07 | 2.6e-07 | 2.400E-07 | 2.646E-08 | 2.100E-07 | 2.600E-07 |
| 1416 | 71 | Q71S | A | 3.8e-07 | 2.5e-07 | 2e-07 | 2.767E-07 | 9.292E-08 | 2.000E-07 | 3.800E-07 |
| 1417 | 71 | Q71T | A | 2e-07 | 4.1e-07 | 2e-07 | 2.700E-07 | 1.212E-07 | 2.000E-07 | 4.100E-07 |
| 1418 | 71 | Q71W | A | 3.4e-07 | 1.8e-07 | 1.8e-07 | 2.333E-07 | 9.238E-08 | 1.800E-07 | 3.400E-07 |
| 1419 | 71 | Q71Y | A | 1.6e-07 | 4e-07 | 2e-07 | 2.533E-07 | 1.286E-07 | 1.600E-07 | 4.000E-07 |
| 1420 | 71 | Q71V | A | 1.7e-07 | 1.5e-07 | 3e-07 | 2.067E-07 | 8.145E-08 | 1.500E-07 | 3.000E-07 |
| 1421 | 72 | V72A | A | 1.8e-07 | 1.6e-07 | 8.4e-08 | 1.413E-07 | 5.065E-08 | 8.400E-08 | 1.800E-07 |
| 1422 | 72 | V72R | A | 2.7e-07 | 2.7e-07 | 1.9e-07 | 2.433E-07 | 4.619E-08 | 1.900E-07 | 2.700E-07 |
| 1423 | 72 | V72N | A | 3.3e-07 | 3.5e-07 | 2.4e-07 | 3.067E-07 | 5.859E-08 | 2.400E-07 | 3.500E-07 |
| 1424 | 72 | V72D | A | 2.4e-07 | 2.4e-07 | 2.3e-07 | 2.367E-07 | 5.774E-09 | 2.300E-07 | 2.400E-07 |
| 1425 | 72 | V72C | A | 3.3e-07 | 1.8e-07 | 6.5e-07 | 3.867E-07 | 2.401E-07 | 1.800E-07 | 6.500E-07 |
| 1426 | 72 | V72E | A | 1.5e-07 | 2.9e-07 | 4e-07 | 2.800E-07 | 1.253E-07 | 1.500E-07 | 4.000E-07 |
| 1427 | 72 | V72Q | A | 3.2e-07 | 4.1e-07 | 2e-07 | 3.100E-07 | 1.054E-07 | 2.000E-07 | 4.100E-07 |
| 1428 | 72 | V72G | A | 9.6e-07 | 2.6e-07 | 5.2e-07 | 5.800E-07 | 3.538E-07 | 2.600E-07 | 9.600E-07 |
| 1429 | 72 | V72H | A | 4.1e-07 | 2.4e-07 | 2.3e-07 | 2.933E-07 | 1.012E-07 | 2.300E-07 | 4.100E-07 |
| 1430 | 72 | V72I | A | 2.2e-07 | 1.9e-07 | 2.3e-07 | 2.133E-07 | 2.082E-08 | 1.900E-07 | 2.300E-07 |
| 1431 | 72 | V72L | A | 3.3e-07 | 1.7e-07 | 1.7e-07 | 2.233E-07 | 9.238E-08 | 1.700E-07 | 3.300E-07 |
| 1432 | 72 | V72K | A | 1.7e-07 | 3.3e-07 | 2.3e-07 | 2.433E-07 | 8.083E-08 | 1.700E-07 | 3.300E-07 |
| 1433 | 72 | V72M | A | 3.3e-07 | 4.4e-07 | 1.3e-07 | 3.000E-07 | 1.572E-07 | 1.300E-07 | 4.400E-07 |
| 1434 | 72 | V72F | A | 1.4e-07 | 2.2e-07 | 1.8e-07 | 1.800E-07 | 4.000E-08 | 1.400E-07 | 2.200E-07 |
| 1435 | 72 | V72P | A | 2.2e-07 | 4.2e-07 | 1.7e-07 | 2.700E-07 | 1.323E-07 | 1.700E-07 | 4.200E-07 |
| 1436 | 72 | V72S | A | 5.6e-07 | 3e-07 | 1.6e-07 | 3.400E-07 | 2.030E-07 | 1.600E-07 | 5.600E-07 |
| 1437 | 72 | V72T | A | 1.3e-07 | 2.5e-07 | 2.8e-07 | 2.200E-07 | 7.937E-08 | 1.300E-07 | 2.800E-07 |
| 1438 | 72 | V72W | A | 4.4e-07 | 3.1e-07 | 1.5e-07 | 3.000E-07 | 1.453E-07 | 1.500E-07 | 4.400E-07 |
| 1439 | 72 | V72Y | A | 3e-07 | 2.9e-07 | 1.9e-07 | 2.600E-07 | 6.083E-08 | 1.900E-07 | 3.000E-07 |
| 1440 | 72 | V72V | A | 5.4e-07 | 1.8e-07 | 2.8e-07 | 3.333E-07 | 1.858E-07 | 1.800E-07 | 5.400E-07 |
| 1441 | 73 | P73A | A | 2.3e-07 | 2.5e-07 | 2.6e-07 | 2.467E-07 | 1.528E-08 | 2.300E-07 | 2.600E-07 |
| 1442 | 73 | P73R | A | 2.9e-07 | 3e-07 | 1.4e-07 | 2.433E-07 | 8.963E-08 | 1.400E-07 | 3.000E-07 |
| 1443 | 73 | P73N | A | 2.3e-07 | 2.7e-07 | 1.3e-07 | 2.100E-07 | 7.211E-08 | 1.300E-07 | 2.700E-07 |
| 1444 | 73 | P73D | A | 1.9e-07 | 2.8e-07 | 1.7e-07 | 2.133E-07 | 5.859E-08 | 1.700E-07 | 2.800E-07 |

| | | | | | | | | | | |
|---|---|---|---|---|---|---|---|---|---|---|
| 1445 | 73 | P73C | A | 2e-07 | 1.8e-07 | 1.6e-07 | 1.800E-07 | 2.000E-08 | 1.600E-07 | 2.000E-07 |
| 1446 | 73 | P73E | A | 5.3e-07 | 2.3e-07 | 2.2e-07 | 3.267E-07 | 1.762E-07 | 2.200E-07 | 5.300E-07 |
| 1447 | 73 | P73Q | A | 2.1e-07 | 2e-07 | 3.2e-07 | 2.433E-07 | 6.658E-08 | 2.000E-07 | 3.200E-07 |
| 1448 | 73 | P73G | A | 1.5e-07 | 3.7e-07 | 2.5e-07 | 2.567E-07 | 1.102E-07 | 1.500E-07 | 3.700E-07 |
| 1449 | 73 | P73H | A | 8.1e-07 | 3.5e-07 | 3.3e-07 | 4.967E-07 | 2.715E-07 | 3.300E-07 | 8.100E-07 |
| 1450 | 73 | P73I | A | 3.8e-07 | 4.7e-07 | 1.1e-07 | 3.200E-07 | 1.873E-07 | 1.100E-07 | 4.700E-07 |
| 1451 | 73 | P73L | A | 2.3e-07 | 4.4e-07 | 2e-07 | 2.900E-07 | 1.308E-07 | 2.000E-07 | 4.400E-07 |
| 1452 | 73 | P73K | A | 3.9e-07 | 2.7e-07 | 3.9e-07 | 3.500E-07 | 6.928E-08 | 2.700E-07 | 3.900E-07 |
| 1453 | 73 | P73M | A | 8.9e-08 | 3.3e-07 | 3.3e-07 | 2.497E-07 | 1.391E-07 | 8.900E-08 | 3.300E-07 |
| 1454 | 73 | P73F | A | 2.6e-07 | 2.3e-07 | 2.2e-07 | 2.367E-07 | 2.082E-08 | 2.200E-07 | 2.600E-07 |
| 1455 | 73 | P73P | A | 1.7e-07 | 2.8e-07 | 1.6e-07 | 2.033E-07 | 6.658E-08 | 1.600E-07 | 2.800E-07 |
| 1456 | 73 | P73S | A | 1.5e-07 | 3.3e-07 | 4e-07 | 2.933E-07 | 1.290E-07 | 1.500E-07 | 4.000E-07 |
| 1457 | 73 | P73T | A | 2.7e-07 | 2.7e-07 | 3.4e-07 | 2.933E-07 | 4.041E-08 | 2.700E-07 | 3.400E-07 |
| 1458 | 73 | P73W | A | 2.5e-07 | 3.3e-07 | 2e-07 | 2.600E-07 | 6.557E-08 | 2.000E-07 | 3.300E-07 |
| 1459 | 73 | P73Y | A | 1.8e-07 | 2.8e-07 | 2.7e-07 | 2.433E-07 | 5.508E-08 | 1.800E-07 | 2.800E-07 |
| 1460 | 73 | P73V | A | 2.1e-07 | 1.5e-07 | 1.7e-07 | 1.767E-07 | 3.055E-08 | 1.500E-07 | 2.100E-07 |
| 1461 | 74 | L74A | A | 1.4e-07 | 2.2e-07 | 4.2e-07 | 2.600E-07 | 1.442E-07 | 1.400E-07 | 4.200E-07 |
| 1462 | 74 | L74R | A | 2.7e-07 | 1.2e-07 | 1.4e-07 | 1.767E-07 | 8.145E-08 | 1.200E-07 | 2.700E-07 |
| 1463 | 74 | L74N | A | 3.1e-07 | 1.8e-07 | 2e-07 | 2.300E-07 | 7.000E-08 | 1.800E-07 | 3.100E-07 |
| 1464 | 74 | L74D | A | 3.5e-07 | 3.7e-07 | 1.8e-07 | 3.000E-07 | 1.044E-07 | 1.800E-07 | 3.700E-07 |
| 1465 | 74 | L74C | A | 1.7e-07 | 1.6e-07 | 2.9e-07 | 2.067E-07 | 7.234E-08 | 1.600E-07 | 2.900E-07 |
| 1466 | 74 | L74E | A | 4e-07 | 1.7e-07 | 1.1e-07 | 2.267E-07 | 1.531E-07 | 1.100E-07 | 4.000E-07 |
| 1467 | 74 | L74Q | A | 2.2e-07 | 1.6e-07 | 1.8e-07 | 1.867E-07 | 3.055E-08 | 1.600E-07 | 2.200E-07 |
| 1468 | 74 | L74G | A | 2.4e-07 | 1.7e-07 | 1.9e-07 | 2.000E-07 | 3.606E-08 | 1.700E-07 | 2.400E-07 |
| 1469 | 74 | L74H | A | 4e-07 | 4.7e-07 | 3.5e-07 | 4.067E-07 | 6.028E-08 | 3.500E-07 | 4.700E-07 |
| 1470 | 74 | L74I | A | 3.2e-07 | 1.5e-07 | 2.7e-07 | 2.467E-07 | 8.737E-08 | 1.500E-07 | 3.200E-07 |
| 1471 | 74 | L74L | A | 2e-07 | 2.6e-07 | 3.6e-07 | 2.733E-07 | 8.083E-08 | 2.000E-07 | 3.600E-07 |
| 1472 | 74 | L74K | A | 1.6e-07 | 2.7e-07 | 1.4e-07 | 1.900E-07 | 7.000E-08 | 1.400E-07 | 2.700E-07 |
| 1473 | 74 | L74M | A | 3.8e-07 | 3e-07 | 5.4e-07 | 4.067E-07 | 1.222E-07 | 3.000E-07 | 5.400E-07 |
| 1474 | 74 | L74F | A | 4.2e-07 | 2.5e-07 | 2.8e-07 | 3.167E-07 | 9.074E-08 | 2.500E-07 | 4.200E-07 |
| 1475 | 74 | L74P | A | 3.1e-07 | 1.3e-07 | 2.4e-07 | 2.267E-07 | 9.074E-08 | 1.300E-07 | 3.100E-07 |
| 1476 | 74 | L74S | A | 2.1e-07 | 1e-07 | 3.2e-07 | 2.100E-07 | 1.100E-07 | 1.000E-07 | 3.200E-07 |
| 1477 | 74 | L74T | A | 3.6e-07 | 4.2e-07 | 3.4e-07 | 3.733E-07 | 4.163E-08 | 3.400E-07 | 4.200E-07 |
| 1478 | 74 | L74W | A | 2.2e-07 | 3.5e-07 | 1.5e-07 | 2.400E-07 | 1.015E-07 | 1.500E-07 | 3.500E-07 |
| 1479 | 74 | L74Y | A | 3e-07 | 3.1e-07 | 3.3e-07 | 3.133E-07 | 1.528E-08 | 3.000E-07 | 3.300E-07 |
| 1480 | 74 | L74V | A | 3.1e-07 | 2.2e-07 | 2.6e-07 | 2.633E-07 | 4.509E-08 | 2.200E-07 | 3.100E-07 |
| 1481 | 75 | Q75A | A | 2.9e-07 | 2.4e-07 | 2.8e-07 | 2.700E-07 | 2.646E-08 | 2.400E-07 | 2.900E-07 |
| 1482 | 75 | Q75R | A | 2e-07 | 1.5e-07 | 2.9e-07 | 2.133E-07 | 7.095E-08 | 1.500E-07 | 2.900E-07 |
| 1483 | 75 | Q75N | A | 1.8e-07 | 2.5e-07 | 2.4e-07 | 2.233E-07 | 3.786E-08 | 1.800E-07 | 2.500E-07 |
| 1484 | 75 | Q75D | A | 2.6e-07 | 2.1e-07 | 1.7e-07 | 2.133E-07 | 4.509E-08 | 1.700E-07 | 2.600E-07 |
| 1485 | 75 | Q75C | A | 1e-07 | 1.9e-07 | 4.1e-07 | 2.333E-07 | 1.595E-07 | 1.000E-07 | 4.100E-07 |
| 1486 | 75 | Q75E | A | 2.5e-07 | 3.6e-07 | 3.4e-07 | 3.167E-07 | 5.859E-08 | 2.500E-07 | 3.600E-07 |
| 1487 | 75 | Q75Q | A | 3.7e-07 | 3.7e-07 | 3.6e-07 | 3.667E-07 | 5.774E-09 | 3.600E-07 | 3.700E-07 |
| 1488 | 75 | Q75G | A | 2.3e-07 | 3e-07 | 3.3e-07 | 2.867E-07 | 5.132E-08 | 2.300E-07 | 3.300E-07 |
| 1489 | 75 | Q75H | A | 2e-07 | 3.3e-07 | 3e-07 | 2.767E-07 | 6.807E-08 | 2.000E-07 | 3.300E-07 |
| 1490 | 75 | Q75I | A | 1.8e-07 | 3.4e-07 | 1.2e-07 | 2.133E-07 | 1.137E-07 | 1.200E-07 | 3.400E-07 |
| 1491 | 75 | Q75L | A | 3.1e-07 | 2.6e-07 | 3.6e-07 | 3.100E-07 | 5.000E-08 | 2.600E-07 | 3.600E-07 |
| 1492 | 75 | Q75K | A | 5.6e-07 | 1.3e-07 | 2.7e-07 | 3.200E-07 | 2.193E-07 | 1.300E-07 | 5.600E-07 |
| 1493 | 75 | Q75M | A | 5.7e-07 | 1.8e-07 | 1.8e-07 | 3.100E-07 | 2.252E-07 | 1.800E-07 | 5.700E-07 |
| 1494 | 75 | Q75F | A | 4.8e-07 | 1.8e-07 | 3.3e-07 | 3.300E-07 | 1.500E-07 | 1.800E-07 | 4.800E-07 |
| 1495 | 75 | Q75P | A | 1.5e-07 | 4.9e-07 | 2.5e-07 | 2.967E-07 | 1.747E-07 | 1.500E-07 | 4.900E-07 |
| 1496 | 75 | Q75S | A | 9.6e-08 | 1.7e-07 | 1.1e-07 | 1.253E-07 | 3.931E-08 | 9.600E-08 | 1.700E-07 |
| 1497 | 75 | Q75T | A | 3.6e-07 | 3e-07 | 1.8e-07 | 2.800E-07 | 9.165E-08 | 1.800E-07 | 3.600E-07 |
| 1498 | 75 | Q75W | A | 1.5e-07 | 1.8e-07 | 4e-07 | 2.433E-07 | 1.365E-07 | 1.500E-07 | 4.000E-07 |
| 1499 | 75 | Q75Y | A | 2.2e-07 | 4.3e-07 | 1.1e-07 | 2.533E-07 | 1.626E-07 | 1.100E-07 | 4.300E-07 |
| 1500 | 75 | Q75V | A | 1.7e-07 | 2.9e-07 | 1.8e-07 | 2.133E-07 | 6.658E-08 | 1.700E-07 | 2.900E-07 |
| 1501 | 76 | R76A | A | 2.7e-07 | 2.1e-07 | 3.8e-07 | 2.867E-07 | 8.622E-08 | 2.100E-07 | 3.800E-07 |
| 1502 | 76 | R76R | A | 2.7e-07 | 2.3e-07 | 2.8e-07 | 2.600E-07 | 2.646E-08 | 2.300E-07 | 2.800E-07 |
| 1503 | 76 | R76N | A | 1.7e-07 | 1.2e-07 | 1.9e-07 | 1.600E-07 | 3.606E-08 | 1.200E-07 | 1.900E-07 |
| 1504 | 76 | R76D | A | 4.2e-07 | 3.5e-07 | 2.1e-07 | 3.267E-07 | 1.069E-07 | 2.100E-07 | 4.200E-07 |
| 1505 | 76 | R76C | A | 1.5e-07 | 1.5e-07 | 2.2e-07 | 1.733E-07 | 4.041E-08 | 1.500E-07 | 2.200E-07 |
| 1506 | 76 | R76E | A | 6.8e-07 | 3.2e-07 | 1.2e-07 | 3.733E-07 | 2.838E-07 | 1.200E-07 | 6.800E-07 |
| 1507 | 76 | R76Q | A | 3.1e-07 | 1.4e-07 | 1.5e-07 | 2.000E-07 | 9.539E-08 | 1.400E-07 | 3.100E-07 |
| 1508 | 76 | R76G | A | 2.8e-07 | 2.9e-07 | 2e-07 | 2.567E-07 | 4.933E-08 | 2.000E-07 | 2.900E-07 |
| 1509 | 76 | R76H | A | 3.4e-07 | 1.4e-07 | 2.9e-07 | 2.567E-07 | 1.041E-07 | 1.400E-07 | 3.400E-07 |
| 1510 | 76 | R76I | A | 2.1e-07 | 6.5e-07 | 2.5e-07 | 3.700E-07 | 2.433E-07 | 2.100E-07 | 6.500E-07 |
| 1511 | 76 | R76L | A | 2.5e-07 | 3.3e-07 | 1.2e-07 | 2.333E-07 | 1.060E-07 | 1.200E-07 | 3.300E-07 |
| 1512 | 76 | R76K | A | 1.5e-07 | 3.4e-07 | 1.5e-07 | 2.133E-07 | 1.097E-07 | 1.500E-07 | 3.400E-07 |
| 1513 | 76 | R76M | A | 4.4e-07 | 2.3e-07 | 1.4e-07 | 2.700E-07 | 1.539E-07 | 1.400E-07 | 4.400E-07 |
| 1514 | 76 | R76F | A | 4e-07 | 1.8e-07 | 2.4e-07 | 2.733E-07 | 1.137E-07 | 1.800E-07 | 4.000E-07 |
| 1515 | 76 | R76P | A | 3.4e-07 | 3.6e-07 | 1.4e-07 | 2.800E-07 | 1.217E-07 | 1.400E-07 | 3.600E-07 |
| 1516 | 76 | R76S | A | 1.5e-07 | 3.4e-07 | 1.8e-07 | 2.233E-07 | 1.021E-07 | 1.500E-07 | 3.400E-07 |
| 1517 | 76 | R76T | A | 1.6e-07 | 9.6e-08 | 2.9e-07 | 1.820E-07 | 9.885E-08 | 9.600E-08 | 2.900E-07 |
| 1518 | 76 | R76W | A | 9.2e-08 | 3.4e-07 | 2.5e-07 | 2.273E-07 | 1.255E-07 | 9.200E-08 | 3.400E-07 |
| 1519 | 76 | R76Y | A | 2.8e-07 | 3.3e-07 | 1.7e-07 | 2.600E-07 | 8.185E-08 | 1.700E-07 | 3.300E-07 |
| 1520 | 76 | R76V | A | 2.6e-07 | 2.9e-07 | 1.4e-07 | 2.300E-07 | 7.937E-08 | 1.400E-07 | 2.900E-07 |
| 1521 | 77 | L77A | A | 3.5e-07 | 2.2e-07 | 1.4e-07 | 2.367E-07 | 1.060E-07 | 1.400E-07 | 3.500E-07 |
| 1522 | 77 | L77R | A | 2.5e-07 | 1.4e-07 | 2.4e-07 | 2.100E-07 | 6.083E-08 | 1.400E-07 | 2.500E-07 |
| 1523 | 77 | L77N | A | 2.4e-07 | 2.3e-07 | 2.3e-07 | 2.333E-07 | 5.774E-09 | 2.300E-07 | 2.400E-07 |
| 1524 | 77 | L77D | A | 1.7e-07 | 2.4e-07 | 6.3e-07 | 3.467E-07 | 2.479E-07 | 1.700E-07 | 6.300E-07 |
| 1525 | 77 | L77C | A | 1.5e-07 | 1.7e-07 | 2.1e-07 | 1.767E-07 | 3.055E-08 | 1.500E-07 | 2.100E-07 |
| 1526 | 77 | L77E | A | 3.6e-07 | 3.9e-07 | 3.1e-07 | 3.533E-07 | 4.041E-08 | 3.100E-07 | 3.900E-07 |
| 1527 | 77 | L77Q | A | 2.8e-07 | 3.8e-07 | 2.9e-07 | 3.167E-07 | 5.508E-08 | 2.800E-07 | 3.800E-07 |
| 1528 | 77 | L77G | A | 2.4e-07 | 4.2e-07 | 1.7e-07 | 2.767E-07 | 1.290E-07 | 1.700E-07 | 4.200E-07 |
| 1529 | 77 | L77H | A | 1.2e-07 | 2.8e-07 | 1.2e-07 | 1.733E-07 | 9.238E-08 | 1.200E-07 | 2.800E-07 |

| | | | | | | | | | | |
|---|---|---|---|---|---|---|---|---|---|---|
| 1530 | 77 | L77I | A | 6.1e-07 | 4.8e-07 | 2.1e-07 | 4.333E-07 | 2.040E-07 | 2.100E-07 | 6.100E-07 |
| 1531 | 77 | L77L | A | 3.2e-07 | 1.9e-07 | 2e-07 | 2.367E-07 | 7.234E-08 | 1.900E-07 | 3.200E-07 |
| 1532 | 77 | L77K | A | 5.4e-07 | 1.6e-07 | 1.4e-07 | 2.800E-07 | 2.254E-07 | 1.400E-07 | 5.400E-07 |
| 1533 | 77 | L77M | A | 2.2e-07 | 2.5e-07 | 1.5e-07 | 2.067E-07 | 5.132E-08 | 1.500E-07 | 2.500E-07 |
| 1534 | 77 | L77F | A | 2.5e-07 | 3.4e-07 | 2.2e-07 | 2.700E-07 | 6.245E-08 | 2.200E-07 | 3.400E-07 |
| 1535 | 77 | L77P | A | 3e-07 | 5.2e-07 | 3.1e-07 | 3.767E-07 | 1.242E-07 | 3.000E-07 | 5.200E-07 |
| 1536 | 77 | L77S | A | 1.2e-07 | 2.1e-07 | 5.4e-07 | 2.900E-07 | 2.211E-07 | 1.200E-07 | 5.400E-07 |
| 1537 | 77 | L77T | A | 4.1e-07 | 3.1e-07 | 4.6e-07 | 3.933E-07 | 7.638E-08 | 3.100E-07 | 4.600E-07 |
| 1538 | 77 | L77W | A | 9.5e-08 | 2.7e-07 | 1.8e-07 | 1.817E-07 | 8.751E-08 | 9.500E-08 | 2.700E-07 |
| 1539 | 77 | L77Y | A | 3.2e-07 | 2e-07 | 1.5e-07 | 2.233E-07 | 8.737E-08 | 1.500E-07 | 3.200E-07 |
| 1540 | 77 | L77V | A | 1.5e-07 | 3.2e-07 | 4.7e-07 | 3.133E-07 | 1.601E-07 | 1.500E-07 | 4.700E-07 |
| 1541 | 78 | R78A | A | 2.1e-07 | 1.6e-07 | 4.6e-07 | 2.767E-07 | 1.607E-07 | 1.600E-07 | 4.600E-07 |
| 1542 | 78 | R78R | A | 2.2e-07 | 4.4e-07 | 7.9e-08 | 2.463E-07 | 1.819E-07 | 7.900E-08 | 4.400E-07 |
| 1543 | 78 | R78N | A | 3.2e-07 | 1.5e-07 | 1.5e-07 | 2.067E-07 | 9.815E-08 | 1.500E-07 | 3.200E-07 |
| 1544 | 78 | R78D | A | 1.3e-07 | 2e-07 | 2.9e-07 | 2.067E-07 | 8.021E-08 | 1.300E-07 | 2.900E-07 |
| 1545 | 78 | R78C | A | 2e-07 | 2.2e-07 | 3.6e-07 | 2.600E-07 | 8.718E-08 | 2.000E-07 | 3.600E-07 |
| 1546 | 78 | R78E | A | 2.1e-07 | 1.4e-07 | 2.1e-07 | 1.867E-07 | 4.041E-08 | 1.400E-07 | 2.100E-07 |
| 1547 | 78 | R78Q | A | 3.4e-07 | 2.8e-07 | 2e-07 | 2.733E-07 | 7.024E-08 | 2.000E-07 | 3.400E-07 |
| 1548 | 78 | R78G | A | 2.2e-07 | 1.6e-07 | 2.9e-07 | 2.233E-07 | 6.506E-08 | 1.600E-07 | 2.900E-07 |
| 1549 | 78 | R78H | A | 4.2e-07 | 3.5e-07 | 3.2e-07 | 3.633E-07 | 5.132E-08 | 3.200E-07 | 4.200E-07 |
| 1550 | 78 | R78I | A | 1.1e-07 | 3e-07 | 1.7e-07 | 1.933E-07 | 9.713E-08 | 1.100E-07 | 3.000E-07 |
| 1551 | 78 | R78L | A | 1.9e-07 | 1.7e-07 | 1.9e-07 | 1.833E-07 | 1.155E-08 | 1.700E-07 | 1.900E-07 |
| 1552 | 78 | R78K | A | 3.7e-07 | 1.3e-07 | 1.3e-07 | 2.100E-07 | 1.386E-07 | 1.300E-07 | 3.700E-07 |
| 1553 | 78 | R78M | A | 2.8e-07 | 2.3e-07 | 2.2e-07 | 2.433E-07 | 3.215E-08 | 2.200E-07 | 2.800E-07 |
| 1554 | 78 | R78F | A | 4e-07 | 1.9e-07 | 3.3e-07 | 3.067E-07 | 1.069E-07 | 1.900E-07 | 4.000E-07 |
| 1555 | 78 | R78P | A | 1.8e-07 | 3.3e-07 | 2.6e-07 | 2.567E-07 | 7.506E-08 | 1.800E-07 | 3.300E-07 |
| 1556 | 78 | R78S | A | 1.8e-07 | 2.1e-07 | 3.3e-07 | 2.400E-07 | 7.937E-08 | 1.800E-07 | 3.300E-07 |
| 1557 | 78 | R78T | A | 4.8e-07 | 2.8e-07 | 3.8e-07 | 3.800E-07 | 1.000E-07 | 2.800E-07 | 4.800E-07 |
| 1558 | 78 | R78W | A | 1.9e-07 | 3.2e-07 | 1.2e-07 | 2.100E-07 | 1.015E-07 | 1.200E-07 | 3.200E-07 |
| 1559 | 78 | R78Y | A | 3.6e-07 | 3.9e-07 | 1.8e-07 | 3.100E-07 | 1.136E-07 | 1.800E-07 | 3.900E-07 |
| 1560 | 78 | R78V | A | 3e-07 | 2.8e-07 | 4.1e-07 | 3.300E-07 | 7.000E-08 | 2.800E-07 | 4.100E-07 |
| 1561 | 79 | I79A | A | 2.5e-07 | 2e-07 | 6.1e-08 | 1.703E-07 | 9.793E-08 | 6.100E-08 | 2.500E-07 |
| 1562 | 79 | I79R | A | 1.9e-07 | 1.9e-07 | 1.8e-07 | 1.867E-07 | 5.774E-09 | 1.800E-07 | 1.900E-07 |
| 1563 | 79 | I79N | A | 2.6e-07 | 5.3e-07 | 3.3e-07 | 3.733E-07 | 1.401E-07 | 2.600E-07 | 5.300E-07 |
| 1564 | 79 | I79D | A | 1.5e-07 | 1.9e-07 | 3.1e-07 | 2.167E-07 | 8.327E-08 | 1.500E-07 | 3.100E-07 |
| 1565 | 79 | I79C | A | 3.2e-07 | 2.6e-07 | 2.9e-07 | 2.900E-07 | 3.000E-08 | 2.600E-07 | 3.200E-07 |
| 1566 | 79 | I79E | A | 2e-07 | 2.6e-07 | 1.7e-07 | 2.100E-07 | 4.583E-08 | 1.700E-07 | 2.600E-07 |
| 1567 | 79 | I79Q | A | 3.2e-07 | 2.2e-07 | 2.1e-07 | 2.500E-07 | 6.083E-08 | 2.100E-07 | 3.200E-07 |
| 1568 | 79 | I79G | A | 2.5e-07 | 1.6e-07 | 2e-07 | 2.033E-07 | 4.509E-08 | 1.600E-07 | 2.500E-07 |
| 1569 | 79 | I79H | A | 2e-07 | 2e-07 | 1.8e-07 | 1.933E-07 | 1.155E-08 | 1.800E-07 | 2.000E-07 |
| 1570 | 79 | I79I | A | 2.3e-07 | 2.9e-07 | 2.8e-07 | 2.667E-07 | 3.215E-08 | 2.300E-07 | 2.900E-07 |
| 1571 | 79 | I79L | A | 2.6e-07 | 1.9e-07 | 1.2e-07 | 1.900E-07 | 7.000E-08 | 1.200E-07 | 2.600E-07 |
| 1572 | 79 | I79K | A | 2.3e-07 | 2.6e-07 | 1.8e-07 | 2.233E-07 | 4.041E-08 | 1.800E-07 | 2.600E-07 |
| 1573 | 79 | I79M | A | 6.5e-07 | 6e-07 | 1.4e-07 | 4.633E-07 | 2.811E-07 | 1.400E-07 | 6.500E-07 |
| 1574 | 79 | I79F | A | 3.9e-07 | 1.2e-07 | 1.5e-07 | 2.200E-07 | 1.480E-07 | 1.200E-07 | 3.900E-07 |
| 1575 | 79 | I79P | A | 1.6e-07 | 3.3e-07 | 3e-07 | 2.633E-07 | 9.074E-08 | 1.600E-07 | 3.300E-07 |
| 1576 | 79 | I79S | A | 2.4e-07 | 3e-07 | 2.5e-07 | 2.633E-07 | 3.215E-08 | 2.400E-07 | 3.000E-07 |
| 1577 | 79 | I79T | A | 2.5e-07 | 1.4e-07 | 1.3e-07 | 1.733E-07 | 6.658E-08 | 1.300E-07 | 2.500E-07 |
| 1578 | 79 | I79W | A | 4.6e-07 | 2.4e-07 | 1.1e-06 | 6.000E-07 | 4.468E-07 | 2.400E-07 | 1.100E-06 |
| 1579 | 79 | I79Y | A | 3.1e-07 | 2e-07 | 3.3e-07 | 2.800E-07 | 7.000E-08 | 2.000E-07 | 3.300E-07 |
| 1580 | 79 | I79V | A | 2.8e-07 | 2.1e-07 | 3.9e-07 | 2.933E-07 | 9.074E-08 | 2.100E-07 | 3.900E-07 |
| 1581 | 80 | V80A | A | 2.1e-07 | 1.5e-07 | 1.7e-07 | 1.767E-07 | 3.055E-08 | 1.500E-07 | 2.100E-07 |
| 1582 | 80 | V80R | A | 2.5e-07 | 2.3e-07 | 2.3e-07 | 2.367E-07 | 1.155E-08 | 2.300E-07 | 2.500E-07 |
| 1583 | 80 | V80N | A | 8e-07 | 3.8e-07 | 2.1e-07 | 4.633E-07 | 3.037E-07 | 2.100E-07 | 8.000E-07 |
| 1584 | 80 | V80D | A | 1.8e-07 | 3.2e-07 | 5.1e-07 | 3.367E-07 | 1.656E-07 | 1.800E-07 | 5.100E-07 |
| 1585 | 80 | V80C | A | 1.6e-07 | 4.4e-07 | 2.7e-07 | 2.900E-07 | 1.411E-07 | 1.600E-07 | 4.400E-07 |
| 1586 | 80 | V80E | A | 3.4e-07 | 2.8e-07 | 1.8e-07 | 2.667E-07 | 8.083E-08 | 1.800E-07 | 3.400E-07 |
| 1587 | 80 | V80Q | A | 1.4e-07 | 2.8e-07 | 2e-07 | 2.067E-07 | 7.024E-08 | 1.400E-07 | 2.800E-07 |
| 1588 | 80 | V80G | A | 3.4e-07 | 4.5e-07 | 2.3e-07 | 3.400E-07 | 1.100E-07 | 2.300E-07 | 4.500E-07 |
| 1589 | 80 | V80H | A | 1.6e-07 | 3e-07 | 2.4e-07 | 2.333E-07 | 7.024E-08 | 1.600E-07 | 3.000E-07 |
| 1590 | 80 | V80I | A | 2.5e-07 | 3e-07 | 2.5e-07 | 2.667E-07 | 2.887E-08 | 2.500E-07 | 3.000E-07 |
| 1591 | 80 | V80L | A | 1.8e-07 | 2e-07 | 2.1e-07 | 1.967E-07 | 1.528E-08 | 1.800E-07 | 2.100E-07 |
| 1592 | 80 | V80K | A | 2.7e-07 | 3.4e-07 | 1.8e-07 | 2.633E-07 | 8.021E-08 | 1.800E-07 | 3.400E-07 |
| 1593 | 80 | V80M | A | 4.6e-07 | 2e-07 | 2e-07 | 2.867E-07 | 1.501E-07 | 2.000E-07 | 4.600E-07 |
| 1594 | 80 | V80F | A | 5.3e-07 | 2.1e-07 | 2e-07 | 3.133E-07 | 1.877E-07 | 2.000E-07 | 5.300E-07 |
| 1595 | 80 | V80P | A | 5.7e-07 | 3e-07 | 2.5e-07 | 3.733E-07 | 1.721E-07 | 2.500E-07 | 5.700E-07 |
| 1596 | 80 | V80S | A | 1.3e-07 | 2.5e-07 | 4e-07 | 2.600E-07 | 1.353E-07 | 1.300E-07 | 4.000E-07 |
| 1597 | 80 | V80T | A | 1.5e-07 | 2.9e-07 | 9.5e-08 | 1.783E-07 | 1.005E-07 | 9.500E-08 | 2.900E-07 |
| 1598 | 80 | V80W | A | 1.3e-07 | 2.3e-07 | 1.6e-07 | 1.733E-07 | 5.132E-08 | 1.300E-07 | 2.300E-07 |
| 1599 | 80 | V80Y | A | 1.8e-07 | 4e-07 | 1.5e-07 | 2.433E-07 | 1.365E-07 | 1.500E-07 | 4.000E-07 |
| 1600 | 80 | V80V | A | 2.7e-07 | 2.1e-07 | 3.3e-07 | 2.700E-07 | 6.000E-08 | 2.100E-07 | 3.300E-07 |
| 1601 | 81 | R81A | A | 2.4e-07 | 1.9e-07 | 2.5e-07 | 2.267E-07 | 3.215E-08 | 1.900E-07 | 2.500E-07 |
| 1602 | 81 | R81R | A | 2.2e-07 | 1.7e-07 | 1.9e-07 | 1.933E-07 | 2.517E-08 | 1.700E-07 | 2.200E-07 |
| 1603 | 81 | R81N | A | 3.2e-07 | 2.8e-07 | 3.7e-07 | 3.233E-07 | 4.509E-08 | 2.800E-07 | 3.700E-07 |
| 1604 | 81 | R81D | A | 7e-07 | 3e-07 | 3.1e-07 | 4.367E-07 | 2.281E-07 | 3.000E-07 | 7.000E-07 |
| 1605 | 81 | R81C | A | 2.8e-07 | 2.5e-07 | 1.7e-07 | 2.333E-07 | 5.686E-08 | 1.700E-07 | 2.800E-07 |
| 1606 | 81 | R81E | A | 2e-07 | 4.3e-07 | 2e-07 | 2.767E-07 | 1.328E-07 | 2.000E-07 | 4.300E-07 |
| 1607 | 81 | R81Q | A | 2.2e-07 | 1.8e-07 | 1.2e-07 | 1.733E-07 | 5.033E-08 | 1.200E-07 | 2.200E-07 |
| 1608 | 81 | R81G | A | 3e-07 | 6.4e-08 | 5.3e-07 | 2.980E-07 | 2.330E-07 | 6.400E-08 | 5.300E-07 |
| 1609 | 81 | R81H | A | 2.2e-07 | 3.2e-07 | 1.3e-07 | 2.233E-07 | 9.504E-08 | 1.300E-07 | 3.200E-07 |
| 1610 | 81 | R81I | A | 2e-07 | 1.2e-07 | 3.7e-07 | 2.300E-07 | 1.277E-07 | 1.200E-07 | 3.700E-07 |
| 1611 | 81 | R81L | A | 2e-07 | 2.4e-07 | 1.7e-07 | 2.033E-07 | 3.512E-08 | 1.700E-07 | 2.400E-07 |
| 1612 | 81 | R81K | A | 2.2e-07 | 1.3e-07 | 1.8e-07 | 1.767E-07 | 4.509E-08 | 1.300E-07 | 2.200E-07 |
| 1613 | 81 | R81M | A | 2.4e-07 | 1.5e-07 | 3e-07 | 2.300E-07 | 7.550E-08 | 1.500E-07 | 3.000E-07 |
| 1614 | 81 | R81F | A | 4e-07 | 2.2e-07 | 1.6e-07 | 2.600E-07 | 1.249E-07 | 1.600E-07 | 4.000E-07 |

| | | | | | | | | | | |
|---|---|---|---|---|---|---|---|---|---|---|
| 1615 | 81 | R81P | A | 4e-07 | 2.8e-07 | 1.3e-07 | 2.700E-07 | 1.353E-07 | 1.300E-07 | 4.000E-07 |
| 1616 | 81 | R81S | A | 1.7e-07 | 1.6e-07 | 3.9e-07 | 2.400E-07 | 1.300E-07 | 1.600E-07 | 3.900E-07 |
| 1617 | 81 | R81T | A | 3.3e-07 | 2.4e-07 | 1.7e-07 | 2.467E-07 | 8.021E-08 | 1.700E-07 | 3.300E-07 |
| 1618 | 81 | R81W | A | 4.7e-07 | 3e-07 | 2.8e-07 | 3.500E-07 | 1.044E-07 | 2.800E-07 | 4.700E-07 |
| 1619 | 81 | R81Y | A | 2.2e-07 | 1.8e-07 | 3.3e-07 | 2.433E-07 | 7.767E-08 | 1.800E-07 | 3.300E-07 |
| 1620 | 81 | R81V | A | 1.6e-07 | 1.8e-07 | 2.7e-07 | 2.033E-07 | 5.859E-08 | 1.600E-07 | 2.700E-07 |
| 1621 | 82 | G82A | A | 2.7e-07 | 3.1e-07 | 2.4e-07 | 2.733E-07 | 3.512E-08 | 2.400E-07 | 3.100E-07 |
| 1622 | 82 | G82R | A | 2.3e-07 | 1.1e-07 | 1.7e-07 | 1.700E-07 | 6.000E-08 | 1.100E-07 | 2.300E-07 |
| 1623 | 82 | G82N | A | 2.8e-07 | 2.3e-07 | 3.7e-07 | 2.933E-07 | 7.095E-08 | 2.300E-07 | 3.700E-07 |
| 1624 | 82 | G82D | A | 2.3e-07 | 2e-07 | 3.7e-07 | 2.667E-07 | 9.074E-08 | 2.000E-07 | 3.700E-07 |
| 1625 | 82 | G82C | A | 2.9e-07 | 3.1e-07 | 1.9e-07 | 2.633E-07 | 6.429E-08 | 1.900E-07 | 3.100E-07 |
| 1626 | 82 | G82E | A | 4.9e-07 | 4.5e-07 | 6.1e-07 | 5.167E-07 | 8.327E-08 | 4.500E-07 | 6.100E-07 |
| 1627 | 82 | G82Q | A | 4e-07 | 2.9e-07 | 1.6e-07 | 2.833E-07 | 1.201E-07 | 1.600E-07 | 4.000E-07 |
| 1628 | 82 | G82G | A | 4.5e-07 | 3e-07 | 2.6e-07 | 3.367E-07 | 1.002E-07 | 2.600E-07 | 4.500E-07 |
| 1629 | 82 | G82H | A | 2.4e-07 | 1.6e-07 | 2.9e-07 | 2.300E-07 | 6.557E-08 | 1.600E-07 | 2.900E-07 |
| 1630 | 82 | G82I | A | 3.2e-07 | 4.9e-07 | 3.7e-07 | 3.933E-07 | 8.737E-08 | 3.200E-07 | 4.900E-07 |
| 1631 | 82 | G82L | A | 1.6e-07 | 1.4e-07 | 3.8e-07 | 2.267E-07 | 1.332E-07 | 1.400E-07 | 3.800E-07 |
| 1632 | 82 | G82K | A | 1.5e-07 | 3.1e-07 | 1.4e-07 | 2.000E-07 | 9.539E-08 | 1.400E-07 | 3.100E-07 |
| 1633 | 82 | G82M | A | 3.6e-07 | 1.9e-07 | 1.7e-07 | 2.400E-07 | 1.044E-07 | 1.700E-07 | 3.600E-07 |
| 1634 | 82 | G82F | A | 2.1e-07 | 6.7e-07 | 2.1e-07 | 3.633E-07 | 2.656E-07 | 2.100E-07 | 6.700E-07 |
| 1635 | 82 | G82P | A | 1.6e-07 | 1.9e-07 | 3.1e-07 | 2.200E-07 | 7.937E-08 | 1.600E-07 | 3.100E-07 |
| 1636 | 82 | G82S | A | 2.9e-07 | 1.8e-07 | 1.4e-07 | 2.033E-07 | 7.767E-08 | 1.400E-07 | 2.900E-07 |
| 1637 | 82 | G82T | A | 1.9e-07 | 3.6e-07 | 2.9e-07 | 2.800E-07 | 8.544E-08 | 1.900E-07 | 3.600E-07 |
| 1638 | 82 | G82W | A | 2.9e-07 | 2.2e-07 | 3.6e-07 | 2.900E-07 | 7.000E-08 | 2.200E-07 | 3.600E-07 |
| 1639 | 82 | G82Y | A | 5.1e-07 | 4.4e-07 | 1.9e-07 | 3.800E-07 | 1.682E-07 | 1.900E-07 | 5.100E-07 |
| 1640 | 82 | G82V | A | 1.5e-07 | 1.8e-07 | 1.7e-07 | 1.667E-07 | 1.528E-08 | 1.500E-07 | 1.800E-07 |
| 1641 | 83 | T83A | A | 4.8e-07 | 4.1e-07 | 3.5e-07 | 4.133E-07 | 6.506E-08 | 3.500E-07 | 4.800E-07 |
| 1642 | 83 | T83R | A | 3.3e-07 | 1.8e-07 | 2.3e-07 | 2.467E-07 | 7.638E-08 | 1.800E-07 | 3.300E-07 |
| 1643 | 83 | T83N | A | 1.3e-07 | 3e-07 | 2.1e-07 | 2.133E-07 | 8.505E-08 | 1.300E-07 | 3.000E-07 |
| 1644 | 83 | T83D | A | 5.5e-07 | 2.9e-07 | 1.7e-07 | 3.367E-07 | 1.943E-07 | 1.700E-07 | 5.500E-07 |
| 1645 | 83 | T83C | A | 2.3e-07 | 2.8e-07 | 4e-07 | 3.033E-07 | 8.737E-08 | 2.300E-07 | 4.000E-07 |
| 1646 | 83 | T83E | A | 4.6e-07 | 2.6e-07 | 2e-07 | 3.067E-07 | 1.361E-07 | 2.000E-07 | 4.600E-07 |
| 1647 | 83 | T83Q | A | 7.4e-08 | 1.8e-07 | 2.2e-07 | 1.580E-07 | 7.545E-08 | 7.400E-08 | 2.200E-07 |
| 1648 | 83 | T83G | A | 2e-07 | 4.4e-07 | 3e-07 | 3.133E-07 | 1.206E-07 | 2.000E-07 | 4.400E-07 |
| 1649 | 83 | T83H | A | 1.9e-07 | 6.9e-07 | 1.3e-07 | 3.367E-07 | 3.075E-07 | 1.300E-07 | 6.900E-07 |
| 1650 | 83 | T83I | A | 3.4e-07 | 1.5e-07 | 1.8e-07 | 2.233E-07 | 1.021E-07 | 1.500E-07 | 3.400E-07 |
| 1651 | 83 | T83L | A | 1.7e-07 | 2.9e-07 | 4.3e-07 | 2.967E-07 | 1.301E-07 | 1.700E-07 | 4.300E-07 |
| 1652 | 83 | T83K | A | 2.4e-07 | 3e-07 | 2.3e-07 | 2.567E-07 | 3.786E-08 | 2.300E-07 | 3.000E-07 |
| 1653 | 83 | T83M | A | 5.7e-07 | 2.4e-07 | 2.6e-07 | 3.567E-07 | 1.850E-07 | 2.400E-07 | 5.700E-07 |
| 1654 | 83 | T83F | A | 1.5e-07 | 9.5e-08 | 4.1e-07 | 2.183E-07 | 1.683E-07 | 9.500E-08 | 4.100E-07 |
| 1655 | 83 | T83P | A | 3.1e-07 | 3.6e-07 | 3.1e-07 | 3.267E-07 | 2.887E-08 | 3.100E-07 | 3.600E-07 |
| 1656 | 83 | T83S | A | 4.4e-07 | 4.6e-07 | 3.1e-07 | 4.033E-07 | 8.145E-08 | 3.100E-07 | 4.600E-07 |
| 1657 | 83 | T83T | A | 2.4e-07 | 3.2e-07 | 3e-07 | 2.867E-07 | 4.163E-08 | 2.400E-07 | 3.200E-07 |
| 1658 | 83 | T83W | A | 2e-07 | 4.3e-07 | 1.1e-07 | 2.467E-07 | 1.650E-07 | 1.100E-07 | 4.300E-07 |
| 1659 | 83 | T83Y | A | 4.5e-07 | 3.2e-07 | 3.5e-07 | 3.733E-07 | 6.807E-08 | 3.200E-07 | 4.500E-07 |
| 1660 | 83 | T83V | A | 4.9e-07 | 1.6e-07 | 5.3e-07 | 3.933E-07 | 2.031E-07 | 1.600E-07 | 5.300E-07 |
| 1661 | 84 | Q84A | A | 1.8e-07 | 3.4e-07 | 1.9e-07 | 2.367E-07 | 8.963E-08 | 1.800E-07 | 3.400E-07 |
| 1662 | 84 | Q84R | A | 2e-07 | 2.7e-07 | 1.8e-07 | 2.167E-07 | 4.726E-08 | 1.800E-07 | 2.700E-07 |
| 1663 | 84 | Q84N | A | 2.3e-07 | 1.8e-07 | 9e-08 | 1.667E-07 | 7.095E-08 | 9.000E-08 | 2.300E-07 |
| 1664 | 84 | Q84D | A | 2.4e-07 | 3.7e-07 | 2.5e-07 | 2.867E-07 | 7.234E-08 | 2.400E-07 | 3.700E-07 |
| 1665 | 84 | Q84C | A | 2.9e-07 | 3.6e-07 | 3.2e-07 | 3.233E-07 | 3.512E-08 | 2.900E-07 | 3.600E-07 |
| 1666 | 84 | Q84E | A | 1.9e-07 | 4e-07 | 3.8e-07 | 3.233E-07 | 1.159E-07 | 1.900E-07 | 4.000E-07 |
| 1667 | 84 | Q84Q | A | 1.9e-07 | 2.1e-07 | 1e-07 | 1.667E-07 | 5.859E-08 | 1.000E-07 | 2.100E-07 |
| 1668 | 84 | Q84G | A | 4e-07 | 4e-07 | 4.9e-07 | 4.300E-07 | 5.196E-08 | 4.000E-07 | 4.900E-07 |
| 1669 | 84 | Q84H | A | 4.3e-07 | 4.1e-07 | 4.3e-07 | 4.233E-07 | 1.155E-08 | 4.100E-07 | 4.300E-07 |
| 1670 | 84 | Q84I | A | 1.9e-07 | 1.3e-07 | 1e-07 | 1.400E-07 | 4.583E-08 | 1.000E-07 | 1.900E-07 |
| 1671 | 84 | Q84L | A | 1.5e-07 | 1.3e-07 | 2.1e-07 | 1.633E-07 | 4.163E-08 | 1.300E-07 | 2.100E-07 |
| 1672 | 84 | Q84K | A | 2.6e-07 | 1.4e-07 | 1.9e-07 | 1.967E-07 | 6.028E-08 | 1.400E-07 | 2.600E-07 |
| 1673 | 84 | Q84M | A | 2.4e-07 | 1.4e-07 | 1.8e-07 | 1.867E-07 | 5.033E-08 | 1.400E-07 | 2.400E-07 |
| 1674 | 84 | Q84F | A | 2.5e-07 | 3e-07 | 3.6e-07 | 3.033E-07 | 5.508E-08 | 2.500E-07 | 3.600E-07 |
| 1675 | 84 | Q84P | A | 3.5e-07 | 4.6e-07 | 2.3e-07 | 3.467E-07 | 1.150E-07 | 2.300E-07 | 4.600E-07 |
| 1676 | 84 | Q84S | A | 3.9e-07 | 3.1e-07 | 2e-07 | 3.000E-07 | 9.539E-08 | 2.000E-07 | 3.900E-07 |
| 1677 | 84 | Q84T | A | 3.4e-07 | 2.3e-07 | 3.1e-07 | 2.933E-07 | 5.686E-08 | 2.300E-07 | 3.400E-07 |
| 1678 | 84 | Q84W | A | 2e-07 | 4.4e-07 | 1.2e-07 | 2.533E-07 | 1.665E-07 | 1.200E-07 | 4.400E-07 |
| 1679 | 84 | Q84Y | A | 2.2e-07 | 2e-07 | 1.8e-07 | 2.000E-07 | 2.000E-08 | 1.800E-07 | 2.200E-07 |
| 1680 | 84 | Q84V | A | 1.7e-07 | 1.3e-07 | 3.9e-07 | 2.300E-07 | 1.400E-07 | 1.300E-07 | 3.900E-07 |
| 1681 | 85 | L85A | A | 2.6e-07 | 2.1e-07 | 2e-07 | 2.233E-07 | 3.215E-08 | 2.000E-07 | 2.600E-07 |
| 1682 | 85 | L85R | A | 1.4e-07 | 3.5e-07 | 4e-07 | 2.967E-07 | 1.380E-07 | 1.400E-07 | 4.000E-07 |
| 1683 | 85 | L85N | A | 1.9e-07 | 3.9e-07 | 5.6e-07 | 3.800E-07 | 1.852E-07 | 1.900E-07 | 5.600E-07 |
| 1684 | 85 | L85D | A | 2.3e-07 | 5.4e-07 | 6e-07 | 4.567E-07 | 1.986E-07 | 2.300E-07 | 6.000E-07 |
| 1685 | 85 | L85C | A | 1.4e-07 | 2.3e-07 | 4.4e-07 | 2.700E-07 | 1.539E-07 | 1.400E-07 | 4.400E-07 |
| 1686 | 85 | L85E | A | 4.4e-07 | 1.8e-07 | 3e-07 | 3.067E-07 | 1.301E-07 | 1.800E-07 | 4.400E-07 |
| 1687 | 85 | L85Q | A | 3.9e-07 | 3.6e-07 | 3.3e-07 | 3.600E-07 | 3.000E-08 | 3.300E-07 | 3.900E-07 |
| 1688 | 85 | L85G | A | 2.9e-07 | 3.4e-07 | 1.9e-07 | 2.733E-07 | 7.638E-08 | 1.900E-07 | 3.400E-07 |
| 1689 | 85 | L85H | A | 1.5e-07 | 5.2e-07 | 3.6e-07 | 3.433E-07 | 1.856E-07 | 1.500E-07 | 5.200E-07 |
| 1690 | 85 | L85I | A | 3.5e-07 | 2.5e-07 | 1.9e-07 | 2.633E-07 | 8.083E-08 | 1.900E-07 | 3.500E-07 |
| 1691 | 85 | L85L | A | 1.5e-07 | 2.4e-07 | 2.3e-07 | 2.067E-07 | 4.933E-08 | 1.500E-07 | 2.400E-07 |
| 1692 | 85 | L85K | A | 1.5e-07 | 8.3e-08 | 2e-07 | 1.443E-07 | 5.871E-08 | 8.300E-08 | 2.000E-07 |
| 1693 | 85 | L85M | A | 2.3e-07 | 1.7e-07 | 3.6e-07 | 2.533E-07 | 9.713E-08 | 1.700E-07 | 3.600E-07 |
| 1694 | 85 | L85F | A | 4.8e-07 | 1.7e-07 | 1.7e-07 | 2.733E-07 | 1.790E-07 | 1.700E-07 | 4.800E-07 |
| 1695 | 85 | L85P | A | 3e-07 | 2.9e-07 | 2.9e-07 | 2.933E-07 | 5.774E-09 | 2.900E-07 | 3.000E-07 |
| 1696 | 85 | L85S | A | 1.6e-07 | 1.9e-07 | 3.6e-07 | 2.367E-07 | 1.079E-07 | 1.600E-07 | 3.600E-07 |
| 1697 | 85 | L85T | A | 4.4e-07 | 2.2e-07 | 4.7e-07 | 3.767E-07 | 1.365E-07 | 2.200E-07 | 4.700E-07 |
| 1698 | 85 | L85W | A | 2e-07 | 1.9e-07 | 2.8e-07 | 2.233E-07 | 4.933E-08 | 1.900E-07 | 2.800E-07 |
| 1699 | 85 | L85Y | A | 1.5e-07 | 2.1e-07 | 2.5e-07 | 2.033E-07 | 5.033E-08 | 1.500E-07 | 2.500E-07 |

| | | | | | | | | | | |
|---|---|---|---|---|---|---|---|---|---|---|
| 1700 | 85 | L85V | A | 4.8e-07 | 3.7e-07 | 3.7e-07 | 4.067E-07 | 6.351E-08 | 3.700E-07 | 4.800E-07 |
| 1701 | 86 | F86A | A | 3.1e-07 | 2.3e-07 | 3.7e-07 | 3.033E-07 | 7.024E-08 | 2.300E-07 | 3.700E-07 |
| 1702 | 86 | F86R | A | 1.7e-07 | 2.3e-07 | 2.6e-07 | 2.200E-07 | 4.583E-08 | 1.700E-07 | 2.600E-07 |
| 1703 | 86 | F86N | A | 1.3e-07 | 1.8e-07 | 1.8e-07 | 1.633E-07 | 2.887E-08 | 1.300E-07 | 1.800E-07 |
| 1704 | 86 | F86D | A | 2.7e-07 | 2.6e-07 | 1.9e-07 | 2.400E-07 | 4.359E-08 | 1.900E-07 | 2.700E-07 |
| 1705 | 86 | F86C | A | 3.4e-07 | 2.3e-07 | 4.3e-07 | 3.333E-07 | 1.002E-07 | 2.300E-07 | 4.300E-07 |
| 1706 | 86 | F86E | A | 2.4e-07 | 3e-07 | 1.3e-07 | 2.233E-07 | 8.622E-08 | 1.300E-07 | 3.000E-07 |
| 1707 | 86 | F86Q | A | 2.4e-07 | 4.2e-07 | 4.3e-07 | 3.633E-07 | 1.069E-07 | 2.400E-07 | 4.300E-07 |
| 1708 | 86 | F86G | A | 4.5e-07 | 4.7e-07 | 1.6e-07 | 3.600E-07 | 1.735E-07 | 1.600E-07 | 4.700E-07 |
| 1709 | 86 | F86H | A | 3e-07 | 2.4e-07 | 2.9e-07 | 2.767E-07 | 3.215E-08 | 2.400E-07 | 3.000E-07 |
| 1710 | 86 | F86I | A | 2.7e-07 | 5.1e-07 | 3.7e-07 | 3.833E-07 | 1.206E-07 | 2.700E-07 | 5.100E-07 |
| 1711 | 86 | F86L | A | 4.5e-07 | 1.2e-07 | 2.3e-07 | 2.667E-07 | 1.680E-07 | 1.200E-07 | 4.500E-07 |
| 1712 | 86 | F86K | A | 1.6e-07 | 3.5e-07 | 4.3e-07 | 3.133E-07 | 1.387E-07 | 1.600E-07 | 4.300E-07 |
| 1713 | 86 | F86M | A | 1.3e-07 | 2.3e-07 | 1.5e-07 | 1.700E-07 | 5.292E-08 | 1.300E-07 | 2.300E-07 |
| 1714 | 86 | F86F | A | 1.3e-07 | 1.4e-07 | 2.1e-07 | 1.600E-07 | 4.359E-08 | 1.300E-07 | 2.100E-07 |
| 1715 | 86 | F86P | A | 1.9e-07 | 5.6e-07 | 1.5e-07 | 3.000E-07 | 2.261E-07 | 1.500E-07 | 5.600E-07 |
| 1716 | 86 | F86S | A | 2.2e-07 | 1.1e-07 | 2.9e-07 | 2.067E-07 | 9.074E-08 | 1.100E-07 | 2.900E-07 |
| 1717 | 86 | F86T | A | 4.4e-07 | 1.3e-07 | 1.1e-07 | 2.267E-07 | 1.850E-07 | 1.100E-07 | 4.400E-07 |
| 1718 | 86 | F86W | A | 2.1e-07 | 2.7e-07 | 2.6e-07 | 2.467E-07 | 3.215E-08 | 2.100E-07 | 2.700E-07 |
| 1719 | 86 | F86Y | A | 2.8e-07 | 4e-07 | 4.8e-07 | 3.867E-07 | 1.007E-07 | 2.800E-07 | 4.800E-07 |
| 1720 | 86 | F86V | A | 3.4e-07 | 1.2e-07 | 4.3e-07 | 2.967E-07 | 1.595E-07 | 1.200E-07 | 4.300E-07 |
| 1721 | 87 | E87A | A | 3.6e-07 | 2.5e-07 | 2.9e-07 | 3.000E-07 | 5.568E-08 | 2.500E-07 | 3.600E-07 |
| 1722 | 87 | E87R | A | 5.6e-07 | 3.9e-07 | 1.9e-07 | 3.800E-07 | 1.852E-07 | 1.900E-07 | 5.600E-07 |
| 1723 | 87 | E87N | A | 1.7e-07 | 3.6e-07 | 3e-07 | 2.767E-07 | 9.713E-08 | 1.700E-07 | 3.600E-07 |
| 1724 | 87 | E87D | A | 3.5e-07 | 5e-07 | 5.6e-07 | 4.700E-07 | 1.082E-07 | 3.500E-07 | 5.600E-07 |
| 1725 | 87 | E87C | A | 2.9e-07 | 2.3e-07 | 4.8e-07 | 3.333E-07 | 1.305E-07 | 2.300E-07 | 4.800E-07 |
| 1726 | 87 | E87E | A | 3.1e-07 | 2.5e-07 | 1.7e-07 | 2.433E-07 | 7.024E-08 | 1.700E-07 | 3.100E-07 |
| 1727 | 87 | E87Q | A | 1.6e-07 | 1.5e-07 | 1.7e-07 | 1.600E-07 | 1.000E-08 | 1.500E-07 | 1.700E-07 |
| 1728 | 87 | E87G | A | 2.2e-07 | 2.3e-07 | 2e-07 | 2.167E-07 | 1.528E-08 | 2.000E-07 | 2.300E-07 |
| 1729 | 87 | E87H | A | 3e-07 | 2e-07 | 3.5e-07 | 2.833E-07 | 7.638E-08 | 2.000E-07 | 3.500E-07 |
| 1730 | 87 | E87I | A | 5e-07 | 2.3e-07 | 2.6e-07 | 3.300E-07 | 1.480E-07 | 2.300E-07 | 5.000E-07 |
| 1731 | 87 | E87L | A | 1.9e-07 | 3.2e-07 | 1.7e-07 | 2.267E-07 | 8.145E-08 | 1.700E-07 | 3.200E-07 |
| 1732 | 87 | E87K | A | 4.9e-07 | 2.4e-07 | 1.5e-07 | 2.933E-07 | 1.762E-07 | 1.500E-07 | 4.900E-07 |
| 1733 | 87 | E87M | A | 1.3e-07 | 3.8e-07 | 3e-07 | 2.700E-07 | 1.277E-07 | 1.300E-07 | 3.800E-07 |
| 1734 | 87 | E87F | A | 3.4e-07 | 3e-07 | 2.2e-07 | 2.867E-07 | 6.110E-08 | 2.200E-07 | 3.400E-07 |
| 1735 | 87 | E87P | A | 3.7e-07 | 2.5e-07 | 3e-07 | 3.067E-07 | 6.028E-08 | 2.500E-07 | 3.700E-07 |
| 1736 | 87 | E87S | A | 4e-07 | 4e-07 | 1.1e-07 | 3.033E-07 | 1.674E-07 | 1.100E-07 | 4.000E-07 |
| 1737 | 87 | E87T | A | 2.1e-07 | 2.1e-07 | 2.5e-07 | 2.233E-07 | 2.309E-08 | 2.100E-07 | 2.500E-07 |
| 1738 | 87 | E87W | A | 3.7e-07 | 2.5e-07 | 1.9e-07 | 2.700E-07 | 9.165E-08 | 1.900E-07 | 3.700E-07 |
| 1739 | 87 | E87Y | A | 1.9e-07 | 4.5e-07 | 2.9e-07 | 3.100E-07 | 1.311E-07 | 1.900E-07 | 4.500E-07 |
| 1740 | 87 | E87V | A | 2.8e-07 | 1.5e-07 | 3.3e-07 | 2.533E-07 | 9.292E-08 | 1.500E-07 | 3.300E-07 |
| 1741 | 88 | D88A | A | 1.3e-07 | 2.1e-07 | 3.8e-07 | 2.400E-07 | 1.277E-07 | 1.300E-07 | 3.800E-07 |
| 1742 | 88 | D88R | A | 2.3e-07 | 1.1e-07 | 3e-07 | 2.133E-07 | 9.609E-08 | 1.100E-07 | 3.000E-07 |
| 1743 | 88 | D88N | A | 2.1e-07 | 1.2e-07 | 2.8e-07 | 2.033E-07 | 8.021E-08 | 1.200E-07 | 2.800E-07 |
| 1744 | 88 | D88D | A | 2.3e-07 | 1.6e-07 | 1.7e-07 | 1.867E-07 | 3.786E-08 | 1.600E-07 | 2.300E-07 |
| 1745 | 88 | D88C | A | 2.7e-07 | 2.2e-07 | 1.5e-07 | 2.133E-07 | 6.028E-08 | 1.500E-07 | 2.700E-07 |
| 1746 | 88 | D88E | A | 4.5e-07 | 5.8e-07 | 2.9e-07 | 4.400E-07 | 1.453E-07 | 2.900E-07 | 5.800E-07 |
| 1747 | 88 | D88Q | A | 4.2e-07 | 5.9e-07 | 5.3e-07 | 5.133E-07 | 8.622E-08 | 4.200E-07 | 5.900E-07 |
| 1748 | 88 | D88G | A | 2.9e-07 | 3.9e-07 | 2.9e-07 | 3.233E-07 | 5.774E-08 | 2.900E-07 | 3.900E-07 |
| 1749 | 88 | D88H | A | 2.4e-07 | 2.6e-07 | 5.8e-07 | 3.600E-07 | 1.908E-07 | 2.400E-07 | 5.800E-07 |
| 1750 | 88 | D88I | A | 3.3e-07 | 2.4e-07 | 3e-07 | 2.900E-07 | 4.583E-08 | 2.400E-07 | 3.300E-07 |
| 1751 | 88 | D88L | A | 3.8e-07 | 4.6e-07 | 2.1e-07 | 3.500E-07 | 1.277E-07 | 2.100E-07 | 4.600E-07 |
| 1752 | 88 | D88K | A | 4e-07 | 7.3e-07 | 1.1e-07 | 4.133E-07 | 3.102E-07 | 1.100E-07 | 7.300E-07 |
| 1753 | 88 | D88M | A | 6.5e-07 | 1.9e-07 | 2.9e-07 | 3.767E-07 | 2.419E-07 | 1.900E-07 | 6.500E-07 |
| 1754 | 88 | D88F | A | 3.2e-07 | 1.6e-07 | 2.5e-07 | 2.433E-07 | 8.021E-08 | 1.600E-07 | 3.200E-07 |
| 1755 | 88 | D88P | A | 3.6e-07 | 3.9e-07 | 3.7e-07 | 3.733E-07 | 1.528E-08 | 3.600E-07 | 3.900E-07 |
| 1756 | 88 | D88S | A | 3.2e-07 | 1.3e-07 | 2.8e-07 | 2.433E-07 | 1.002E-07 | 1.300E-07 | 3.200E-07 |
| 1757 | 88 | D88T | A | 1.4e-07 | 1.9e-07 | 2.6e-07 | 1.967E-07 | 6.028E-08 | 1.400E-07 | 2.600E-07 |
| 1758 | 88 | D88W | A | 1.5e-07 | 2.3e-07 | 5.6e-07 | 3.133E-07 | 2.173E-07 | 1.500E-07 | 5.600E-07 |
| 1759 | 88 | D88Y | A | 3.7e-07 | 1.7e-07 | 1.6e-07 | 2.333E-07 | 1.185E-07 | 1.600E-07 | 3.700E-07 |
| 1760 | 88 | D88V | A | 2.9e-07 | 9.1e-08 | 1.9e-07 | 1.903E-07 | 9.950E-08 | 9.100E-08 | 2.900E-07 |
| 1761 | 89 | N89A | A | 2.9e-07 | 4e-07 | 1.4e-07 | 2.767E-07 | 1.305E-07 | 1.400E-07 | 4.000E-07 |
| 1762 | 89 | N89R | A | 4.4e-07 | 1.6e-07 | 2.7e-07 | 2.900E-07 | 1.411E-07 | 1.600E-07 | 4.400E-07 |
| 1763 | 89 | N89N | A | 2e-07 | 2.2e-07 | 1.7e-07 | 1.967E-07 | 2.517E-08 | 1.700E-07 | 2.200E-07 |
| 1764 | 89 | N89D | A | 2.6e-07 | 1.9e-07 | 1.7e-07 | 2.067E-07 | 4.726E-08 | 1.700E-07 | 2.600E-07 |
| 1765 | 89 | N89C | A | 3.3e-07 | 1.9e-07 | 1.5e-07 | 2.233E-07 | 9.452E-08 | 1.500E-07 | 3.300E-07 |
| 1766 | 89 | N89E | A | 2.1e-07 | 4.4e-07 | 2.1e-07 | 2.867E-07 | 1.328E-07 | 2.100E-07 | 4.400E-07 |
| 1767 | 89 | N89Q | A | 4e-07 | 2.3e-07 | 2e-07 | 2.767E-07 | 1.079E-07 | 2.000E-07 | 4.000E-07 |
| 1768 | 89 | N89G | A | 2.2e-07 | 3.9e-07 | 2.5e-07 | 2.867E-07 | 9.074E-08 | 2.200E-07 | 3.900E-07 |
| 1769 | 89 | N89H | A | 3e-07 | 5.1e-07 | 2.6e-07 | 3.567E-07 | 1.343E-07 | 2.600E-07 | 5.100E-07 |
| 1770 | 89 | N89I | A | 3.7e-07 | 1.7e-07 | 3.2e-07 | 2.867E-07 | 1.041E-07 | 1.700E-07 | 3.700E-07 |
| 1771 | 89 | N89L | A | 3e-07 | 2.1e-07 | 2.9e-07 | 2.667E-07 | 4.933E-08 | 2.100E-07 | 3.000E-07 |
| 1772 | 89 | N89K | A | 2.9e-07 | 2.1e-07 | 2.2e-07 | 2.400E-07 | 4.359E-08 | 2.100E-07 | 2.900E-07 |
| 1773 | 89 | N89M | A | 4.1e-07 | 9.9e-08 | 1.1e-07 | 2.063E-07 | 1.765E-07 | 9.900E-08 | 4.100E-07 |
| 1774 | 89 | N89F | A | 2.3e-07 | 1.8e-07 | 3e-07 | 2.367E-07 | 6.028E-08 | 1.800E-07 | 3.000E-07 |
| 1775 | 89 | N89P | A | 1.2e-07 | 7.1e-07 | 7e-08 | 3.000E-07 | 3.559E-07 | 7.000E-08 | 7.100E-07 |
| 1776 | 89 | N89S | A | 1e-07 | 3.2e-07 | 1.9e-07 | 2.033E-07 | 1.106E-07 | 1.000E-07 | 3.200E-07 |
| 1777 | 89 | N89T | A | 6.8e-08 | 4.1e-07 | 3e-07 | 2.593E-07 | 1.746E-07 | 6.800E-08 | 4.100E-07 |
| 1778 | 89 | N89W | A | 8.3e-08 | 1.8e-07 | 1.9e-07 | 1.510E-07 | 5.910E-08 | 8.300E-08 | 1.900E-07 |
| 1779 | 89 | N89Y | A | 3.9e-07 | 2.1e-07 | 2.7e-07 | 2.900E-07 | 9.165E-08 | 2.100E-07 | 3.900E-07 |
| 1780 | 89 | N89V | A | 4.6e-07 | 1.6e-07 | 2.2e-07 | 2.800E-07 | 1.587E-07 | 1.600E-07 | 4.600E-07 |
| 1781 | 90 | Y90A | A | 1.3e-07 | 1.5e-07 | 3.1e-07 | 1.967E-07 | 9.866E-08 | 1.300E-07 | 3.100E-07 |
| 1782 | 90 | Y90R | A | 9.4e-08 | 1.6e-07 | 1.9e-07 | 1.480E-07 | 4.911E-08 | 9.400E-08 | 1.900E-07 |
| 1783 | 90 | Y90N | A | 1.5e-07 | 1.5e-07 | 1.8e-07 | 1.600E-07 | 1.732E-08 | 1.500E-07 | 1.800E-07 |
| 1784 | 90 | Y90D | A | 3.4e-07 | 1.5e-07 | 4.2e-07 | 3.033E-07 | 1.387E-07 | 1.500E-07 | 4.200E-07 |

| | | | | | | | | | | |
|---|---|---|---|---|---|---|---|---|---|---|
| 1785 | 90 | Y90C | A | 2.4e-07 | 3.2e-07 | 1.6e-07 | 2.400E-07 | 8.000E-08 | 1.600E-07 | 3.200E-07 |
| 1786 | 90 | Y90E | A | 1.3e-07 | 3e-07 | 2.2e-07 | 2.167E-07 | 8.505E-08 | 1.300E-07 | 3.000E-07 |
| 1787 | 90 | Y90Q | A | 3.3e-07 | 2.2e-07 | 2.5e-07 | 2.667E-07 | 5.686E-08 | 2.200E-07 | 3.300E-07 |
| 1788 | 90 | Y90G | A | 3.1e-07 | 3.3e-07 | 3.9e-07 | 3.433E-07 | 4.163E-08 | 3.100E-07 | 3.900E-07 |
| 1789 | 90 | Y90H | A | 4e-07 | 9.3e-08 | 1.2e-07 | 2.043E-07 | 1.700E-07 | 9.300E-08 | 4.000E-07 |
| 1790 | 90 | Y90I | A | 1.6e-07 | 3.9e-07 | 3.3e-07 | 2.933E-07 | 1.193E-07 | 1.600E-07 | 3.900E-07 |
| 1791 | 90 | Y90L | A | 1.8e-07 | 3.5e-07 | 5.6e-07 | 3.633E-07 | 1.904E-07 | 1.800E-07 | 5.600E-07 |
| 1792 | 90 | Y90K | A | 1.7e-07 | 4.8e-07 | 3.1e-07 | 3.200E-07 | 1.552E-07 | 1.700E-07 | 4.800E-07 |
| 1793 | 90 | Y90M | A | 1.8e-07 | 2.6e-07 | 3.2e-07 | 2.533E-07 | 7.024E-08 | 1.800E-07 | 3.200E-07 |
| 1794 | 90 | Y90F | A | 3.7e-07 | 2e-07 | 4.3e-07 | 3.333E-07 | 1.193E-07 | 2.000E-07 | 4.300E-07 |
| 1795 | 90 | Y90P | A | 2.8e-07 | 2e-07 | 2e-07 | 2.267E-07 | 4.619E-08 | 2.000E-07 | 2.800E-07 |
| 1796 | 90 | Y90S | A | 3.2e-07 | 1.6e-07 | 8.8e-08 | 1.893E-07 | 1.187E-07 | 8.800E-08 | 3.200E-07 |
| 1797 | 90 | Y90T | A | 1.8e-07 | 2.1e-07 | 3.4e-07 | 2.433E-07 | 8.505E-08 | 1.800E-07 | 3.400E-07 |
| 1798 | 90 | Y90W | A | 1.2e-07 | 3.7e-07 | 3.2e-07 | 2.700E-07 | 1.323E-07 | 1.200E-07 | 3.700E-07 |
| 1799 | 90 | Y90Y | A | 2e-07 | 1.5e-07 | 2.5e-07 | 2.000E-07 | 5.000E-08 | 1.500E-07 | 2.500E-07 |
| 1800 | 90 | Y90V | A | 1.1e-07 | 4.2e-07 | 2.7e-07 | 2.667E-07 | 1.550E-07 | 1.100E-07 | 4.200E-07 |
| 1801 | 91 | A91A | A | 2.4e-07 | 4.6e-07 | 1.3e-07 | 2.767E-07 | 1.680E-07 | 1.300E-07 | 4.600E-07 |
| 1802 | 91 | A91R | A | 5.4e-07 | 1.9e-07 | 4.3e-07 | 3.867E-07 | 1.790E-07 | 1.900E-07 | 5.400E-07 |
| 1803 | 91 | A91N | A | 3.7e-07 | 4.9e-07 | 1.5e-07 | 3.367E-07 | 1.724E-07 | 1.500E-07 | 4.900E-07 |
| 1804 | 91 | A91D | A | 2.4e-07 | 1.8e-07 | 1.8e-07 | 2.000E-07 | 3.464E-08 | 1.800E-07 | 2.400E-07 |
| 1805 | 91 | A91C | A | 2.2e-07 | 1.7e-07 | 1.6e-07 | 1.833E-07 | 3.215E-08 | 1.600E-07 | 2.200E-07 |
| 1806 | 91 | A91E | A | 1.5e-07 | 2.1e-07 | 1.4e-07 | 1.667E-07 | 3.786E-08 | 1.400E-07 | 2.100E-07 |
| 1807 | 91 | A91Q | A | 2.8e-07 | 9.9e-08 | 2e-07 | 1.930E-07 | 9.070E-08 | 9.900E-08 | 2.800E-07 |
| 1808 | 91 | A91G | A | 2.7e-07 | 2.2e-07 | 4.6e-07 | 3.167E-07 | 1.266E-07 | 2.200E-07 | 4.600E-07 |
| 1809 | 91 | A91H | A | 4.2e-07 | 2.3e-07 | 2.8e-07 | 3.100E-07 | 9.849E-08 | 2.300E-07 | 4.200E-07 |
| 1810 | 91 | A91I | A | 1.5e-07 | 1.3e-07 | 1.6e-07 | 1.467E-07 | 1.528E-08 | 1.300E-07 | 1.600E-07 |
| 1811 | 91 | A91L | A | 3.6e-07 | 1.4e-07 | 2.8e-07 | 2.600E-07 | 1.114E-07 | 1.400E-07 | 3.600E-07 |
| 1812 | 91 | A91K | A | 4.5e-07 | 2e-07 | 2.6e-07 | 3.033E-07 | 1.305E-07 | 2.000E-07 | 4.500E-07 |
| 1813 | 91 | A91M | A | 1.3e-07 | 1.9e-07 | 1.3e-07 | 1.500E-07 | 3.464E-08 | 1.300E-07 | 1.900E-07 |
| 1814 | 91 | A91F | A | 1.6e-07 | 2.3e-07 | 1.9e-07 | 1.933E-07 | 3.512E-08 | 1.600E-07 | 2.300E-07 |
| 1815 | 91 | A91P | A | 1.5e-07 | 6.6e-07 | 1.5e-07 | 3.200E-07 | 2.944E-07 | 1.500E-07 | 6.600E-07 |
| 1816 | 91 | A91S | A | 1.6e-07 | 2.7e-07 | 4.5e-07 | 2.933E-07 | 1.464E-07 | 1.600E-07 | 4.500E-07 |
| 1817 | 91 | A91T | A | 2.1e-07 | 3.2e-07 | 4.2e-07 | 3.167E-07 | 1.050E-07 | 2.100E-07 | 4.200E-07 |
| 1818 | 91 | A91W | A | 4.9e-07 | 1.6e-07 | 1.7e-07 | 2.733E-07 | 1.877E-07 | 1.600E-07 | 4.900E-07 |
| 1819 | 91 | A91Y | A | 4.7e-07 | 2.5e-07 | 1.9e-07 | 3.033E-07 | 1.474E-07 | 1.900E-07 | 4.700E-07 |
| 1820 | 91 | A91V | A | 2.5e-07 | 3.5e-07 | 3e-07 | 3.000E-07 | 5.000E-08 | 2.500E-07 | 3.500E-07 |
| 1821 | 92 | L92A | A | 2.6e-07 | 2.4e-07 | 1.3e-07 | 2.100E-07 | 7.000E-08 | 1.300E-07 | 2.600E-07 |
| 1822 | 92 | L92R | A | 2e-07 | 1.7e-07 | 2.1e-07 | 1.933E-07 | 2.082E-08 | 1.700E-07 | 2.100E-07 |
| 1823 | 92 | L92N | A | 1.8e-07 | 1.6e-07 | 3.6e-07 | 2.333E-07 | 1.102E-07 | 1.600E-07 | 3.600E-07 |
| 1824 | 92 | L92D | A | 6.2e-07 | 1.7e-07 | 1.2e-07 | 3.033E-07 | 2.754E-07 | 1.200E-07 | 6.200E-07 |
| 1825 | 92 | L92C | A | 2.4e-07 | 2.9e-07 | 4.1e-07 | 3.133E-07 | 8.737E-08 | 2.400E-07 | 4.100E-07 |
| 1826 | 92 | L92E | A | 2.3e-07 | 3.4e-07 | 2.1e-07 | 2.600E-07 | 7.000E-08 | 2.100E-07 | 3.400E-07 |
| 1827 | 92 | L92Q | A | 2.2e-07 | 1.5e-07 | 3.1e-07 | 2.267E-07 | 8.021E-08 | 1.500E-07 | 3.100E-07 |
| 1828 | 92 | L92G | A | 2.9e-07 | 2.4e-07 | 2.8e-07 | 2.700E-07 | 2.646E-08 | 2.400E-07 | 2.900E-07 |
| 1829 | 92 | L92H | A | 4.3e-07 | 1.2e-07 | 1.3e-07 | 2.267E-07 | 1.762E-07 | 1.200E-07 | 4.300E-07 |
| 1830 | 92 | L92I | A | 4.2e-07 | 2.5e-07 | 3.5e-07 | 3.400E-07 | 8.544E-08 | 2.500E-07 | 4.200E-07 |
| 1831 | 92 | L92L | A | 1.9e-07 | 2.1e-07 | 1.7e-07 | 1.900E-07 | 2.000E-08 | 1.700E-07 | 2.100E-07 |
| 1832 | 92 | L92K | A | 3.2e-07 | 2.2e-07 | 5.3e-07 | 3.567E-07 | 1.582E-07 | 2.200E-07 | 5.300E-07 |
| 1833 | 92 | L92M | A | 1.9e-07 | 2.4e-07 | 1.9e-07 | 2.067E-07 | 2.887E-08 | 1.900E-07 | 2.400E-07 |
| 1834 | 92 | L92F | A | 1.2e-07 | 1e-07 | 2.1e-07 | 1.433E-07 | 5.859E-08 | 1.000E-07 | 2.100E-07 |
| 1835 | 92 | L92P | A | 4.5e-07 | 2.6e-07 | 1.8e-07 | 2.967E-07 | 1.387E-07 | 1.800E-07 | 4.500E-07 |
| 1836 | 92 | L92S | A | 3.1e-07 | 1.8e-07 | 3.5e-07 | 2.800E-07 | 8.888E-08 | 1.800E-07 | 3.500E-07 |
| 1837 | 92 | L92T | A | 1.8e-07 | 2.6e-07 | 2.9e-07 | 2.433E-07 | 5.686E-08 | 1.800E-07 | 2.900E-07 |
| 1838 | 92 | L92W | A | 1.3e-07 | 4.1e-07 | 3.2e-07 | 2.867E-07 | 1.429E-07 | 1.300E-07 | 4.100E-07 |
| 1839 | 92 | L92Y | A | 2.2e-07 | 2e-07 | 2.6e-07 | 2.267E-07 | 3.055E-08 | 2.000E-07 | 2.600E-07 |
| 1840 | 92 | L92V | A | 4e-07 | 2.5e-07 | 5.9e-07 | 4.133E-07 | 1.704E-07 | 2.500E-07 | 5.900E-07 |
| 1841 | 93 | A93A | A | 2.9e-07 | 1.6e-07 | 5.4e-07 | 3.300E-07 | 1.931E-07 | 1.600E-07 | 5.400E-07 |
| 1842 | 93 | A93R | A | 1.8e-07 | 2.5e-07 | 1.6e-07 | 1.967E-07 | 4.726E-08 | 1.600E-07 | 2.500E-07 |
| 1843 | 93 | A93N | A | 1.6e-07 | 3.1e-07 | 2.1e-07 | 2.267E-07 | 7.638E-08 | 1.600E-07 | 3.100E-07 |
| 1844 | 93 | A93D | A | 1.8e-07 | 4.8e-07 | 1.8e-07 | 2.800E-07 | 1.732E-07 | 1.800E-07 | 4.800E-07 |
| 1845 | 93 | A93C | A | 2.9e-07 | 3.7e-07 | 3.3e-07 | 3.300E-07 | 4.000E-08 | 2.900E-07 | 3.700E-07 |
| 1846 | 93 | A93E | A | 9.8e-08 | 2.5e-07 | 4e-07 | 2.493E-07 | 1.510E-07 | 9.800E-08 | 4.000E-07 |
| 1847 | 93 | A93Q | A | 1.8e-07 | 9.8e-08 | 3e-07 | 1.927E-07 | 1.016E-07 | 9.800E-08 | 3.000E-07 |
| 1848 | 93 | A93G | A | 3.3e-07 | 1.8e-07 | 1.2e-07 | 2.100E-07 | 1.082E-07 | 1.200E-07 | 3.300E-07 |
| 1849 | 93 | A93H | A | 2.2e-07 | 4.1e-07 | 4.5e-07 | 3.600E-07 | 1.229E-07 | 2.200E-07 | 4.500E-07 |
| 1850 | 93 | A93I | A | 1.3e-07 | 2.7e-07 | 2.3e-07 | 2.100E-07 | 7.211E-08 | 1.300E-07 | 2.700E-07 |
| 1851 | 93 | A93L | A | 1.6e-07 | 3.8e-07 | 1.8e-07 | 2.400E-07 | 1.217E-07 | 1.600E-07 | 3.800E-07 |
| 1852 | 93 | A93K | A | 2.1e-07 | 2e-07 | 2.3e-07 | 2.133E-07 | 1.528E-08 | 2.000E-07 | 2.300E-07 |
| 1853 | 93 | A93M | A | 2.6e-07 | 5.3e-07 | 6.2e-07 | 4.700E-07 | 1.873E-07 | 2.600E-07 | 6.200E-07 |
| 1854 | 93 | A93F | A | 2.3e-07 | 2.3e-07 | 3.7e-07 | 2.767E-07 | 8.083E-08 | 2.300E-07 | 3.700E-07 |
| 1855 | 93 | A93P | A | 5.5e-07 | 2.9e-07 | 2e-07 | 3.467E-07 | 1.818E-07 | 2.000E-07 | 5.500E-07 |
| 1856 | 93 | A93S | A | 1.8e-07 | 1.9e-07 | 1.8e-07 | 1.833E-07 | 5.774E-09 | 1.800E-07 | 1.900E-07 |
| 1857 | 93 | A93T | A | 2.4e-07 | 2.9e-07 | 1.3e-07 | 2.200E-07 | 8.185E-08 | 1.300E-07 | 2.900E-07 |
| 1858 | 93 | A93W | A | 4.2e-07 | 2.6e-07 | 4.7e-07 | 3.833E-07 | 1.097E-07 | 2.600E-07 | 4.700E-07 |
| 1859 | 93 | A93Y | A | 5.4e-07 | 3.1e-07 | 2.2e-07 | 3.567E-07 | 1.650E-07 | 2.200E-07 | 5.400E-07 |
| 1860 | 93 | A93V | A | 2.5e-07 | 2.1e-07 | 2.7e-07 | 2.433E-07 | 3.055E-08 | 2.100E-07 | 2.700E-07 |
| 1861 | 94 | V94A | A | 3.8e-07 | 2.8e-07 | 3.6e-07 | 3.400E-07 | 5.292E-08 | 2.800E-07 | 3.800E-07 |
| 1862 | 94 | V94R | A | 2.3e-07 | 9.2e-08 | 1.7e-07 | 1.640E-07 | 6.920E-08 | 9.200E-08 | 2.300E-07 |
| 1863 | 94 | V94N | A | 3e-07 | 1.5e-07 | 1.7e-07 | 2.067E-07 | 8.145E-08 | 1.500E-07 | 3.000E-07 |
| 1864 | 94 | V94D | A | 2.1e-07 | 3.4e-07 | 5.3e-07 | 3.600E-07 | 1.609E-07 | 2.100E-07 | 5.300E-07 |
| 1865 | 94 | V94C | A | 4.5e-07 | 5.3e-07 | 1.7e-07 | 3.833E-07 | 1.890E-07 | 1.700E-07 | 5.300E-07 |
| 1866 | 94 | V94E | A | 3.1e-07 | 2e-07 | 3.7e-07 | 2.933E-07 | 8.622E-08 | 2.000E-07 | 3.700E-07 |
| 1867 | 94 | V94Q | A | 2.2e-07 | 2.8e-07 | 4.9e-07 | 3.300E-07 | 1.418E-07 | 2.200E-07 | 4.900E-07 |
| 1868 | 94 | V94G | A | 4.5e-07 | 3.8e-07 | 2.2e-07 | 3.500E-07 | 1.179E-07 | 2.200E-07 | 4.500E-07 |
| 1869 | 94 | V94H | A | 2e-07 | 1.7e-07 | 2.1e-07 | 1.933E-07 | 2.082E-08 | 1.700E-07 | 2.100E-07 |

| | | | | | | | | | | |
|---|---|---|---|---|---|---|---|---|---|---|
| 1870 | 94 | V94I | A | 2.4e-07 | 2.2e-07 | 1.5e-07 | 2.033E-07 | 4.726E-08 | 1.500E-07 | 2.400E-07 |
| 1871 | 94 | V94L | A | 2.5e-07 | 1.3e-07 | 4.2e-07 | 2.667E-07 | 1.457E-07 | 1.300E-07 | 4.200E-07 |
| 1872 | 94 | V94K | A | 1.9e-07 | 1.9e-07 | 3e-07 | 2.267E-07 | 6.351E-08 | 1.900E-07 | 3.000E-07 |
| 1873 | 94 | V94M | A | 2.4e-07 | 5.7e-07 | 2.9e-07 | 3.667E-07 | 1.779E-07 | 2.400E-07 | 5.700E-07 |
| 1874 | 94 | V94F | A | 3.5e-07 | 2.5e-07 | 1.8e-07 | 2.600E-07 | 8.544E-08 | 1.800E-07 | 3.500E-07 |
| 1875 | 94 | V94P | A | 1.1e-07 | 2.4e-07 | 2.1e-07 | 1.867E-07 | 6.807E-08 | 1.100E-07 | 2.400E-07 |
| 1876 | 94 | V94S | A | 2.5e-07 | 2.7e-07 | 2.6e-07 | 2.600E-07 | 1.000E-08 | 2.500E-07 | 2.700E-07 |
| 1877 | 94 | V94T | A | 2.8e-07 | 7.2e-07 | 2.3e-07 | 4.100E-07 | 2.696E-07 | 2.300E-07 | 7.200E-07 |
| 1878 | 94 | V94W | A | 2.4e-07 | 2.4e-07 | 2.7e-07 | 2.500E-07 | 1.732E-08 | 2.400E-07 | 2.700E-07 |
| 1879 | 94 | V94Y | A | 1.6e-07 | 3.4e-07 | 3e-07 | 2.667E-07 | 9.452E-08 | 1.600E-07 | 3.400E-07 |
| 1880 | 94 | V94V | A | 1.3e-07 | 1.1e-07 | 2.6e-07 | 1.667E-07 | 8.145E-08 | 1.100E-07 | 2.600E-07 |
| 1881 | 95 | L95A | A | 4e-07 | 1.7e-07 | 4.7e-07 | 3.467E-07 | 1.570E-07 | 1.700E-07 | 4.700E-07 |
| 1882 | 95 | L95R | A | 1.4e-07 | 3.4e-07 | 1.7e-07 | 2.167E-07 | 1.079E-07 | 1.400E-07 | 3.400E-07 |
| 1883 | 95 | L95N | A | 1.9e-07 | 2.2e-07 | 3.3e-07 | 2.467E-07 | 7.371E-08 | 1.900E-07 | 3.300E-07 |
| 1884 | 95 | L95D | A | 3.2e-07 | 2.7e-07 | 1.2e-07 | 2.367E-07 | 1.041E-07 | 1.200E-07 | 3.200E-07 |
| 1885 | 95 | L95C | A | 3.6e-07 | 1.6e-07 | 2.2e-07 | 2.467E-07 | 1.026E-07 | 1.600E-07 | 3.600E-07 |
| 1886 | 95 | L95E | A | 2.5e-07 | 2.8e-07 | 3.4e-07 | 2.900E-07 | 4.583E-08 | 2.500E-07 | 3.400E-07 |
| 1887 | 95 | L95Q | A | 1.9e-07 | 2.1e-07 | 3.5e-07 | 2.500E-07 | 8.718E-08 | 1.900E-07 | 3.500E-07 |
| 1888 | 95 | L95G | A | 3e-07 | 1.9e-07 | 3.4e-07 | 2.767E-07 | 7.767E-08 | 1.900E-07 | 3.400E-07 |
| 1889 | 95 | L95H | A | 1.4e-07 | 3e-07 | 1.7e-07 | 2.033E-07 | 8.505E-08 | 1.400E-07 | 3.000E-07 |
| 1890 | 95 | L95I | A | 4.1e-07 | 2.4e-07 | 2.2e-07 | 2.900E-07 | 1.044E-07 | 2.200E-07 | 4.100E-07 |
| 1891 | 95 | L95L | A | 2.2e-07 | 1.7e-07 | 2.6e-07 | 2.167E-07 | 4.509E-08 | 1.700E-07 | 2.600E-07 |
| 1892 | 95 | L95K | A | 2.5e-07 | 3.1e-07 | 1.8e-07 | 2.467E-07 | 6.506E-08 | 1.800E-07 | 3.100E-07 |
| 1893 | 95 | L95M | A | 3e-07 | 2.2e-07 | 2.1e-07 | 2.433E-07 | 4.933E-08 | 2.100E-07 | 3.000E-07 |
| 1894 | 95 | L95F | A | 2.4e-07 | 2.9e-07 | 3e-07 | 2.767E-07 | 3.215E-08 | 2.400E-07 | 3.000E-07 |
| 1895 | 95 | L95P | A | 1.8e-07 | 4.8e-07 | 2e-07 | 2.867E-07 | 1.677E-07 | 1.800E-07 | 4.800E-07 |
| 1896 | 95 | L95S | A | 5.1e-07 | 1.7e-07 | 5e-07 | 3.933E-07 | 1.935E-07 | 1.700E-07 | 5.100E-07 |
| 1897 | 95 | L95T | A | 2e-07 | 2.4e-07 | 2.6e-07 | 2.333E-07 | 3.055E-08 | 2.000E-07 | 2.600E-07 |
| 1898 | 95 | L95W | A | 9.6e-08 | 1.8e-07 | 2.5e-07 | 1.753E-07 | 7.711E-08 | 9.600E-08 | 2.500E-07 |
| 1899 | 95 | L95Y | A | 4.1e-07 | 1e-07 | 1.4e-07 | 2.167E-07 | 1.686E-07 | 1.000E-07 | 4.100E-07 |
| 1900 | 95 | L95V | A | 1.6e-07 | 4e-07 | 1.9e-07 | 2.500E-07 | 1.308E-07 | 1.600E-07 | 4.000E-07 |
| 1901 | 96 | D96A | A | 3.1e-07 | 2e-07 | 4.5e-07 | 3.200E-07 | 1.253E-07 | 2.000E-07 | 4.500E-07 |
| 1902 | 96 | D96R | A | 1.8e-07 | 2.1e-07 | 2.4e-07 | 2.100E-07 | 3.000E-08 | 1.800E-07 | 2.400E-07 |
| 1903 | 96 | D96N | A | 2.1e-07 | 3.4e-07 | 3.2e-07 | 2.900E-07 | 7.000E-08 | 2.100E-07 | 3.400E-07 |
| 1904 | 96 | D96D | A | 2.4e-07 | 2.9e-07 | 3.9e-07 | 3.067E-07 | 7.638E-08 | 2.400E-07 | 3.900E-07 |
| 1905 | 96 | D96C | A | 3.6e-07 | 3.3e-07 | 1.8e-07 | 2.900E-07 | 9.644E-08 | 1.800E-07 | 3.600E-07 |
| 1906 | 96 | D96E | A | 1.6e-07 | 3.8e-07 | 2.1e-07 | 2.500E-07 | 1.153E-07 | 1.600E-07 | 3.800E-07 |
| 1907 | 96 | D96Q | A | 2e-07 | 2.1e-07 | 2.2e-07 | 2.100E-07 | 1.000E-08 | 2.000E-07 | 2.200E-07 |
| 1908 | 96 | D96G | A | 1.9e-07 | 4.4e-07 | 1.5e-07 | 2.600E-07 | 1.572E-07 | 1.500E-07 | 4.400E-07 |
| 1909 | 96 | D96H | A | 1.4e-07 | 2.4e-07 | 1.6e-07 | 1.800E-07 | 5.292E-08 | 1.400E-07 | 2.400E-07 |
| 1910 | 96 | D96I | A | 2.9e-07 | 3.5e-07 | 3.5e-07 | 3.300E-07 | 3.464E-08 | 2.900E-07 | 3.500E-07 |
| 1911 | 96 | D96L | A | 3.3e-07 | 3.4e-07 | 3.3e-07 | 3.333E-07 | 5.774E-09 | 3.300E-07 | 3.400E-07 |
| 1912 | 96 | D96K | A | 2.6e-07 | 6.6e-07 | 1.9e-07 | 3.700E-07 | 2.536E-07 | 1.900E-07 | 6.600E-07 |
| 1913 | 96 | D96M | A | 2.8e-07 | 1.7e-07 | 1.8e-07 | 2.100E-07 | 6.083E-08 | 1.700E-07 | 2.800E-07 |
| 1914 | 96 | D96F | A | 2.4e-07 | 1.7e-07 | 2.2e-07 | 2.100E-07 | 3.606E-08 | 1.700E-07 | 2.400E-07 |
| 1915 | 96 | D96P | A | 2.3e-07 | 5.8e-07 | 1.9e-07 | 3.333E-07 | 2.146E-07 | 1.900E-07 | 5.800E-07 |
| 1916 | 96 | D96S | A | 2.1e-07 | 2.3e-07 | 2.2e-07 | 2.200E-07 | 1.000E-08 | 2.100E-07 | 2.300E-07 |
| 1917 | 96 | D96T | A | 1.6e-07 | 3e-07 | 2.7e-07 | 2.433E-07 | 7.371E-08 | 1.600E-07 | 3.000E-07 |
| 1918 | 96 | D96W | A | 2.5e-07 | 1.7e-07 | 2.6e-07 | 2.267E-07 | 4.933E-08 | 1.700E-07 | 2.600E-07 |
| 1919 | 96 | D96Y | A | 1.3e-07 | 2.2e-07 | 4.3e-07 | 2.600E-07 | 1.539E-07 | 1.300E-07 | 4.300E-07 |
| 1920 | 96 | D96V | A | 1.7e-07 | 4.6e-07 | 3.7e-07 | 3.333E-07 | 1.484E-07 | 1.700E-07 | 4.600E-07 |
| 1921 | 97 | N97A | A | 2.4e-07 | 1.5e-07 | 3.7e-07 | 2.533E-07 | 1.106E-07 | 1.500E-07 | 3.700E-07 |
| 1922 | 97 | N97R | A | 1.3e-07 | 3.7e-07 | 1.8e-07 | 2.267E-07 | 1.266E-07 | 1.300E-07 | 3.700E-07 |
| 1923 | 97 | N97N | A | 3e-07 | 3.2e-07 | 1.2e-07 | 2.467E-07 | 1.102E-07 | 1.200E-07 | 3.200E-07 |
| 1924 | 97 | N97D | A | 1.6e-07 | 2.8e-07 | 1e-07 | 1.800E-07 | 9.165E-08 | 1.000E-07 | 2.800E-07 |
| 1925 | 97 | N97C | A | 2.6e-07 | 2.7e-07 | 2.8e-07 | 2.700E-07 | 1.000E-08 | 2.600E-07 | 2.800E-07 |
| 1926 | 97 | N97E | A | 3.2e-07 | 3.3e-07 | 3e-07 | 3.167E-07 | 1.528E-08 | 3.000E-07 | 3.300E-07 |
| 1927 | 97 | N97Q | A | 1.3e-07 | 1.7e-07 | 3.5e-07 | 2.167E-07 | 1.172E-07 | 1.300E-07 | 3.500E-07 |
| 1928 | 97 | N97G | A | 1.4e-07 | 3e-07 | 3.3e-07 | 2.567E-07 | 1.021E-07 | 1.400E-07 | 3.300E-07 |
| 1929 | 97 | N97H | A | 1.5e-07 | 2.5e-07 | 1.6e-07 | 1.867E-07 | 5.508E-08 | 1.500E-07 | 2.500E-07 |
| 1930 | 97 | N97I | A | 2.6e-07 | 3.1e-07 | 2.7e-07 | 2.800E-07 | 2.646E-08 | 2.600E-07 | 3.100E-07 |
| 1931 | 97 | N97L | A | 1.9e-07 | 3.8e-07 | 1.3e-07 | 2.333E-07 | 1.305E-07 | 1.300E-07 | 3.800E-07 |
| 1932 | 97 | N97K | A | 5.4e-07 | 3.9e-07 | 4.1e-07 | 4.467E-07 | 8.145E-08 | 3.900E-07 | 5.400E-07 |
| 1933 | 97 | N97M | A | 4.4e-07 | 2.1e-07 | 3.5e-07 | 3.333E-07 | 1.159E-07 | 2.100E-07 | 4.400E-07 |
| 1934 | 97 | N97F | A | 2.7e-07 | 2.2e-07 | 3.6e-07 | 2.833E-07 | 7.095E-08 | 2.200E-07 | 3.600E-07 |
| 1935 | 97 | N97P | A | 2.2e-07 | 1.8e-07 | 3e-07 | 2.333E-07 | 6.110E-08 | 1.800E-07 | 3.000E-07 |
| 1936 | 97 | N97S | A | 2.9e-07 | 3.7e-07 | 4.9e-07 | 3.833E-07 | 1.007E-07 | 2.900E-07 | 4.900E-07 |
| 1937 | 97 | N97T | A | 1.6e-07 | 5.2e-07 | 9.2e-08 | 2.573E-07 | 2.300E-07 | 9.200E-08 | 5.200E-07 |
| 1938 | 97 | N97W | A | 1.2e-07 | 1.7e-07 | 2.6e-07 | 1.833E-07 | 7.095E-08 | 1.200E-07 | 2.600E-07 |
| 1939 | 97 | N97Y | A | 3.6e-07 | 1.9e-07 | 3.1e-07 | 2.867E-07 | 8.737E-08 | 1.900E-07 | 3.600E-07 |
| 1940 | 97 | N97V | A | 2.4e-07 | 1.2e-07 | 1.7e-07 | 1.767E-07 | 6.028E-08 | 1.200E-07 | 2.400E-07 |
| 1941 | 98 | G98A | A | 1.8e-07 | 1.8e-07 | 2.3e-07 | 1.967E-07 | 2.887E-08 | 1.800E-07 | 2.300E-07 |
| 1942 | 98 | G98R | A | 3.3e-07 | 3.5e-07 | 2e-07 | 2.933E-07 | 8.145E-08 | 2.000E-07 | 3.500E-07 |
| 1943 | 98 | G98N | A | 1.6e-07 | 2.8e-07 | 2.2e-07 | 2.200E-07 | 6.000E-08 | 1.600E-07 | 2.800E-07 |
| 1944 | 98 | G98D | A | 4.3e-07 | 4.4e-07 | 2.3e-07 | 3.667E-07 | 1.185E-07 | 2.300E-07 | 4.400E-07 |
| 1945 | 98 | G98C | A | 1.2e-07 | 1.8e-07 | 3e-07 | 2.000E-07 | 9.165E-08 | 1.200E-07 | 3.000E-07 |
| 1946 | 98 | G98E | A | 2.3e-07 | 5.1e-07 | 3.1e-07 | 3.500E-07 | 1.442E-07 | 2.300E-07 | 5.100E-07 |
| 1947 | 98 | G98Q | A | 3.3e-07 | 1.8e-07 | 3.1e-07 | 2.733E-07 | 8.145E-08 | 1.800E-07 | 3.300E-07 |
| 1948 | 98 | G98G | A | 3.7e-07 | 5.4e-07 | 2.8e-07 | 3.967E-07 | 1.320E-07 | 2.800E-07 | 5.400E-07 |
| 1949 | 98 | G98H | A | 1.8e-07 | 2.4e-07 | 9.8e-08 | 1.727E-07 | 7.128E-08 | 9.800E-08 | 2.400E-07 |
| 1950 | 98 | G98I | A | 3.1e-07 | 3.1e-07 | 3e-07 | 3.067E-07 | 5.774E-09 | 3.000E-07 | 3.100E-07 |
| 1951 | 98 | G98L | A | 4e-07 | 1.8e-07 | 2.4e-07 | 2.733E-07 | 1.137E-07 | 1.800E-07 | 4.000E-07 |
| 1952 | 98 | G98K | A | 3e-07 | 1.4e-07 | 2e-07 | 2.133E-07 | 8.083E-08 | 1.400E-07 | 3.000E-07 |
| 1953 | 98 | G98M | A | 1.5e-07 | 1.1e-07 | 8.4e-08 | 1.147E-07 | 3.325E-08 | 8.400E-08 | 1.500E-07 |
| 1954 | 98 | G98F | A | 1e-07 | 1.6e-07 | 2.4e-07 | 1.667E-07 | 7.024E-08 | 1.000E-07 | 2.400E-07 |

| | | | | | | | | | | |
|---|---|---|---|---|---|---|---|---|---|---|
| 1955 | 98 | G98P | A | 1.9e-07 | 1.7e-07 | 4.4e-07 | 2.667E-07 | 1.504E-07 | 1.700E-07 | 4.400E-07 |
| 1956 | 98 | G98S | A | 3.9e-07 | 1.6e-07 | 2.5e-07 | 2.667E-07 | 1.159E-07 | 1.600E-07 | 3.900E-07 |
| 1957 | 98 | G98T | A | 9.1e-08 | 3.6e-07 | 1.3e-07 | 1.937E-07 | 1.454E-07 | 9.100E-08 | 3.600E-07 |
| 1958 | 98 | G98W | A | 2.1e-07 | 8e-07 | 1.3e-07 | 3.800E-07 | 3.659E-07 | 1.300E-07 | 8.000E-07 |
| 1959 | 98 | G98Y | A | 1.9e-07 | 9.9e-08 | 1.8e-07 | 1.563E-07 | 4.990E-08 | 9.900E-08 | 1.900E-07 |
| 1960 | 98 | G98V | A | 1.1e-07 | 2.3e-07 | 3.5e-07 | 2.300E-07 | 1.200E-07 | 1.100E-07 | 3.500E-07 |
| 1961 | 99 | D99A | A | 2.7e-07 | 7.4e-07 | 2.2e-07 | 4.100E-07 | 2.869E-07 | 2.200E-07 | 7.400E-07 |
| 1962 | 99 | D99R | A | 6.5e-07 | 2.1e-07 | 2e-07 | 3.533E-07 | 2.570E-07 | 2.000E-07 | 6.500E-07 |
| 1963 | 99 | D99N | A | 3.1e-07 | 1.9e-07 | 1.4e-07 | 2.133E-07 | 8.737E-08 | 1.400E-07 | 3.100E-07 |
| 1964 | 99 | D99D | A | 2.1e-07 | 1.5e-07 | 1.5e-07 | 1.700E-07 | 3.464E-08 | 1.500E-07 | 2.100E-07 |
| 1965 | 99 | D99C | A | 1.8e-07 | 3.1e-07 | 7.2e-08 | 1.873E-07 | 1.192E-07 | 7.200E-08 | 3.100E-07 |
| 1966 | 99 | D99E | A | 2.2e-07 | 2.6e-07 | 1.9e-07 | 2.233E-07 | 3.512E-08 | 1.900E-07 | 2.600E-07 |
| 1967 | 99 | D99Q | A | 1.5e-07 | 2.2e-07 | 3.8e-07 | 2.500E-07 | 1.179E-07 | 1.500E-07 | 3.800E-07 |
| 1968 | 99 | D99G | A | 5.9e-07 | 4.1e-07 | 2.3e-07 | 4.100E-07 | 1.800E-07 | 2.300E-07 | 5.900E-07 |
| 1969 | 99 | D99H | A | 3.4e-07 | 2.4e-07 | 3.1e-07 | 2.967E-07 | 5.132E-08 | 2.400E-07 | 3.400E-07 |
| 1970 | 99 | D99I | A | 4e-07 | 2e-07 | 3.2e-07 | 3.067E-07 | 1.007E-07 | 2.000E-07 | 4.000E-07 |
| 1971 | 99 | D99L | A | 1.1e-07 | 3.3e-07 | 2e-07 | 2.133E-07 | 1.106E-07 | 1.100E-07 | 3.300E-07 |
| 1972 | 99 | D99K | A | 4.3e-07 | 3.6e-07 | 2e-07 | 3.300E-07 | 1.179E-07 | 2.000E-07 | 4.300E-07 |
| 1973 | 99 | D99M | A | 3.1e-07 | 1.5e-07 | 2.4e-07 | 2.333E-07 | 8.021E-08 | 1.500E-07 | 3.100E-07 |
| 1974 | 99 | D99F | A | 2.7e-07 | 3.3e-07 | 4e-07 | 3.333E-07 | 6.506E-08 | 2.700E-07 | 4.000E-07 |
| 1975 | 99 | D99P | A | 2.1e-07 | 2.2e-07 | 2.2e-07 | 2.167E-07 | 5.774E-09 | 2.100E-07 | 2.200E-07 |
| 1976 | 99 | D99S | A | 2.7e-07 | 2.4e-07 | 1.9e-07 | 2.333E-07 | 4.041E-08 | 1.900E-07 | 2.700E-07 |
| 1977 | 99 | D99T | A | 1.3e-07 | 1.7e-07 | 2e-07 | 1.667E-07 | 3.512E-08 | 1.300E-07 | 2.000E-07 |
| 1978 | 99 | D99W | A | 2.3e-07 | 1.9e-07 | 1.4e-07 | 1.867E-07 | 4.509E-08 | 1.400E-07 | 2.300E-07 |
| 1979 | 99 | D99Y | A | 1.5e-07 | 1.9e-07 | 1.9e-07 | 1.767E-07 | 2.309E-08 | 1.500E-07 | 1.900E-07 |
| 1980 | 99 | D99V | A | 3.3e-07 | 2.2e-07 | 3.7e-07 | 3.067E-07 | 7.767E-08 | 2.200E-07 | 3.700E-07 |
| 1981 | 100 | P100A | A | 2.9e-07 | 2e-07 | 2.4e-07 | 2.433E-07 | 4.509E-08 | 2.000E-07 | 2.900E-07 |
| 1982 | 100 | P100R | A | 4e-07 | 1.9e-07 | 1.4e-07 | 2.433E-07 | 1.380E-07 | 1.400E-07 | 4.000E-07 |
| 1983 | 100 | P100N | A | 3.6e-07 | 3.4e-07 | 1.4e-07 | 2.800E-07 | 1.217E-07 | 1.400E-07 | 3.600E-07 |
| 1984 | 100 | P100D | A | 2.6e-07 | 1.9e-07 | 3.2e-07 | 2.567E-07 | 6.506E-08 | 1.900E-07 | 3.200E-07 |
| 1985 | 100 | P100C | A | 4e-07 | 3.1e-07 | 3.1e-07 | 3.400E-07 | 5.196E-08 | 3.100E-07 | 4.000E-07 |
| 1986 | 100 | P100E | A | 3.4e-07 | 2.8e-07 | 2.2e-07 | 2.800E-07 | 6.000E-08 | 2.200E-07 | 3.400E-07 |
| 1987 | 100 | P100Q | A | 4e-07 | 2e-07 | 5.7e-07 | 3.900E-07 | 1.852E-07 | 2.000E-07 | 5.700E-07 |
| 1988 | 100 | P100G | A | 3.6e-07 | 2.7e-07 | 1.8e-07 | 2.700E-07 | 9.000E-08 | 1.800E-07 | 3.600E-07 |
| 1989 | 100 | P100H | A | 1.6e-07 | 2.9e-07 | 4.5e-07 | 3.000E-07 | 1.453E-07 | 1.600E-07 | 4.500E-07 |
| 1990 | 100 | P100I | A | 3.5e-07 | 2.4e-07 | 1.3e-07 | 2.400E-07 | 1.100E-07 | 1.300E-07 | 3.500E-07 |
| 1991 | 100 | P100L | A | 1.7e-07 | 1.7e-07 | 3e-07 | 2.133E-07 | 7.506E-08 | 1.700E-07 | 3.000E-07 |
| 1992 | 100 | P100K | A | 1.5e-07 | 1.9e-07 | 2.6e-07 | 2.000E-07 | 5.568E-08 | 1.500E-07 | 2.600E-07 |
| 1993 | 100 | P100M | A | 2.8e-07 | 3.9e-07 | 1.7e-07 | 2.800E-07 | 1.100E-07 | 1.700E-07 | 3.900E-07 |
| 1994 | 100 | P100F | A | 1.7e-07 | 2.3e-07 | 1.8e-07 | 1.933E-07 | 3.215E-08 | 1.700E-07 | 2.300E-07 |
| 1995 | 100 | P100P | A | 4.7e-07 | 3.6e-07 | 3.3e-07 | 3.867E-07 | 7.371E-08 | 3.300E-07 | 4.700E-07 |
| 1996 | 100 | P100S | A | 2.9e-07 | 2e-07 | 3e-07 | 2.633E-07 | 5.508E-08 | 2.000E-07 | 3.000E-07 |
| 1997 | 100 | P100T | A | 2.2e-07 | 3.2e-07 | 5.9e-07 | 3.767E-07 | 1.914E-07 | 2.200E-07 | 5.900E-07 |
| 1998 | 100 | P100W | A | 2.9e-07 | 2.4e-07 | 2.4e-07 | 2.567E-07 | 2.887E-08 | 2.400E-07 | 2.900E-07 |
| 1999 | 100 | P100Y | A | 2.4e-07 | 1.4e-07 | 2.3e-07 | 2.033E-07 | 5.508E-08 | 1.400E-07 | 2.400E-07 |
| 2000 | 100 | P100V | A | 3.7e-07 | 2.6e-07 | 3.7e-07 | 3.333E-07 | 6.351E-08 | 2.600E-07 | 3.700E-07 |
| 2001 | 101 | L101A | A | 3.6e-07 | 3e-07 | 3.8e-07 | 3.467E-07 | 4.163E-08 | 3.000E-07 | 3.800E-07 |
| 2002 | 101 | L101R | A | 2.5e-07 | 2.6e-07 | 1.1e-07 | 2.067E-07 | 8.386E-08 | 1.100E-07 | 2.600E-07 |
| 2003 | 101 | L101N | A | 3.4e-07 | 2.8e-07 | 5e-07 | 3.733E-07 | 1.137E-07 | 2.800E-07 | 5.000E-07 |
| 2004 | 101 | L101D | A | 2.9e-07 | 1.6e-07 | 1.1e-07 | 1.867E-07 | 9.292E-08 | 1.100E-07 | 2.900E-07 |
| 2005 | 101 | L101C | A | 1.9e-07 | 3.7e-07 | 2.2e-07 | 2.600E-07 | 9.644E-08 | 1.900E-07 | 3.700E-07 |
| 2006 | 101 | L101E | A | 3e-07 | 2.9e-07 | 2.7e-07 | 2.867E-07 | 1.528E-08 | 2.700E-07 | 3.000E-07 |
| 2007 | 101 | L101Q | A | 2.2e-07 | 4.2e-07 | 3.3e-07 | 3.233E-07 | 1.002E-07 | 2.200E-07 | 4.200E-07 |
| 2008 | 101 | L101G | A | 3.7e-07 | 2.4e-07 | 2.3e-07 | 2.800E-07 | 7.810E-08 | 2.300E-07 | 3.700E-07 |
| 2009 | 101 | L101H | A | 2.6e-07 | 1.1e-07 | 4.7e-07 | 2.800E-07 | 1.808E-07 | 1.100E-07 | 4.700E-07 |
| 2010 | 101 | L101I | A | 2.7e-07 | 3.6e-07 | 3.3e-07 | 3.200E-07 | 4.583E-08 | 2.700E-07 | 3.600E-07 |
| 2011 | 101 | L101L | A | 2e-07 | 1.6e-07 | 2.9e-07 | 2.167E-07 | 6.658E-08 | 1.600E-07 | 2.900E-07 |
| 2012 | 101 | L101K | A | 5.6e-07 | 2.6e-07 | 2e-07 | 3.400E-07 | 1.929E-07 | 2.000E-07 | 5.600E-07 |
| 2013 | 101 | L101M | A | 2.4e-07 | 2.4e-07 | 3e-07 | 2.600E-07 | 3.464E-08 | 2.400E-07 | 3.000E-07 |
| 2014 | 101 | L101F | A | 1.2e-07 | 5e-07 | 1.6e-07 | 2.600E-07 | 2.088E-07 | 1.200E-07 | 5.000E-07 |
| 2015 | 101 | L101P | A | 3e-07 | 3.3e-07 | 5.1e-07 | 3.800E-07 | 1.136E-07 | 3.000E-07 | 5.100E-07 |
| 2016 | 101 | L101S | A | 3e-07 | 1.5e-07 | 4.5e-07 | 3.000E-07 | 1.500E-07 | 1.500E-07 | 4.500E-07 |
| 2017 | 101 | L101T | A | 5.1e-07 | 1.9e-07 | 5.2e-07 | 4.067E-07 | 1.877E-07 | 1.900E-07 | 5.200E-07 |
| 2018 | 101 | L101W | A | 2.3e-07 | 1.3e-07 | 2.2e-07 | 1.933E-07 | 5.508E-08 | 1.300E-07 | 2.300E-07 |
| 2019 | 101 | L101Y | A | 2.4e-07 | 2.1e-07 | 1.5e-07 | 2.000E-07 | 4.583E-08 | 1.500E-07 | 2.400E-07 |
| 2020 | 101 | L101V | A | 1.4e-07 | 5.4e-07 | 3.8e-07 | 3.533E-07 | 2.013E-07 | 1.400E-07 | 5.400E-07 |
| 2021 | 102 | N102A | A | 3.6e-07 | 2.5e-07 | 5.8e-07 | 3.967E-07 | 1.680E-07 | 2.500E-07 | 5.800E-07 |
| 2022 | 102 | N102R | A | 2.7e-07 | 2e-07 | 3.5e-07 | 2.733E-07 | 7.506E-08 | 2.000E-07 | 3.500E-07 |
| 2023 | 102 | N102N | A | 2e-07 | 5.4e-08 | 1.7e-07 | 1.413E-07 | 7.711E-08 | 5.400E-08 | 2.000E-07 |
| 2024 | 102 | N102D | A | 2.4e-07 | 6e-07 | 2.7e-07 | 3.700E-07 | 1.997E-07 | 2.400E-07 | 6.000E-07 |
| 2025 | 102 | N102C | A | 2.9e-07 | 9.3e-08 | 2.4e-07 | 2.077E-07 | 1.024E-07 | 9.300E-08 | 2.900E-07 |
| 2026 | 102 | N102E | A | 2.8e-07 | 1.7e-07 | 4.1e-07 | 2.867E-07 | 1.201E-07 | 1.700E-07 | 4.100E-07 |
| 2027 | 102 | N102Q | A | 3.3e-07 | 2.8e-07 | 6.5e-07 | 4.200E-07 | 2.007E-07 | 2.800E-07 | 6.500E-07 |
| 2028 | 102 | N102G | A | 2.4e-07 | 2.4e-07 | 3.4e-07 | 2.733E-07 | 5.774E-08 | 2.400E-07 | 3.400E-07 |
| 2029 | 102 | N102H | A | 1.7e-07 | 3.4e-07 | 3.2e-07 | 2.767E-07 | 9.292E-08 | 1.700E-07 | 3.400E-07 |
| 2030 | 102 | N102I | A | 2.4e-07 | 2.3e-07 | 2.6e-07 | 2.433E-07 | 1.528E-08 | 2.300E-07 | 2.600E-07 |
| 2031 | 102 | N102L | A | 2.8e-07 | 2.8e-07 | 1.6e-07 | 2.400E-07 | 6.928E-08 | 1.600E-07 | 2.800E-07 |
| 2032 | 102 | N102K | A | 9e-08 | 3.7e-07 | 1.6e-07 | 2.067E-07 | 1.457E-07 | 9.000E-08 | 3.700E-07 |
| 2033 | 102 | N102M | A | 2.3e-07 | 1.9e-07 | 3.2e-07 | 2.467E-07 | 6.658E-08 | 1.900E-07 | 3.200E-07 |
| 2034 | 102 | N102F | A | 1.5e-07 | 4.4e-07 | 3.3e-07 | 3.067E-07 | 1.464E-07 | 1.500E-07 | 4.400E-07 |
| 2035 | 102 | N102P | A | 2.7e-07 | 2.1e-07 | 2.6e-07 | 2.467E-07 | 3.215E-08 | 2.100E-07 | 2.700E-07 |
| 2036 | 102 | N102S | A | 1.8e-07 | 4.4e-07 | 1.1e-07 | 2.433E-07 | 1.739E-07 | 1.100E-07 | 4.400E-07 |
| 2037 | 102 | N102T | A | 1.4e-07 | 5.4e-07 | 2.8e-07 | 3.200E-07 | 2.030E-07 | 1.400E-07 | 5.400E-07 |
| 2038 | 102 | N102W | A | 4.5e-07 | 5.7e-07 | 2.2e-07 | 4.133E-07 | 1.779E-07 | 2.200E-07 | 5.700E-07 |
| 2039 | 102 | N102Y | A | 6.5e-07 | 2.4e-07 | 2.6e-07 | 3.833E-07 | 2.312E-07 | 2.400E-07 | 6.500E-07 |

| | | | | | | | | | | |
|---|---|---|---|---|---|---|---|---|---|---|
| 2040 | 102 | N102V | A | 1.8e-07 | 2.9e-07 | 4.6e-07 | 3.100E-07 | 1.411E-07 | 1.800E-07 | 4.600E-07 |
| 2041 | 103 | N103A | A | 1.7e-07 | 1.3e-07 | 2.6e-07 | 1.867E-07 | 6.658E-08 | 1.300E-07 | 2.600E-07 |
| 2042 | 103 | N103R | A | 2.6e-07 | 6.9e-07 | 2.3e-07 | 3.933E-07 | 2.574E-07 | 2.300E-07 | 6.900E-07 |
| 2043 | 103 | N103N | A | 1.4e-07 | 1.4e-07 | 2e-07 | 1.600E-07 | 3.464E-08 | 1.400E-07 | 2.000E-07 |
| 2044 | 103 | N103D | A | 1.8e-07 | 3.4e-07 | 1.9e-07 | 2.367E-07 | 8.963E-08 | 1.800E-07 | 3.400E-07 |
| 2045 | 103 | N103C | A | 3.2e-07 | 1.3e-07 | 1.7e-07 | 2.067E-07 | 1.002E-07 | 1.300E-07 | 3.200E-07 |
| 2046 | 103 | N103E | A | 2.2e-07 | 2.3e-07 | 1.7e-07 | 2.067E-07 | 3.215E-08 | 1.700E-07 | 2.300E-07 |
| 2047 | 103 | N103Q | A | 4.3e-07 | 2.8e-07 | 3.6e-07 | 3.567E-07 | 7.506E-08 | 2.800E-07 | 4.300E-07 |
| 2048 | 103 | N103G | A | 3e-07 | 1.3e-07 | 1.6e-07 | 1.967E-07 | 9.074E-08 | 1.300E-07 | 3.000E-07 |
| 2049 | 103 | N103H | A | 1.8e-07 | 1.2e-07 | 2.4e-07 | 1.800E-07 | 6.000E-08 | 1.200E-07 | 2.400E-07 |
| 2050 | 103 | N103I | A | 2.3e-07 | 3.5e-07 | 1.8e-07 | 2.533E-07 | 8.737E-08 | 1.800E-07 | 3.500E-07 |
| 2051 | 103 | N103L | A | 3.6e-07 | 1.7e-07 | 1.5e-07 | 2.267E-07 | 1.159E-07 | 1.500E-07 | 3.600E-07 |
| 2052 | 103 | N103K | A | 2.3e-07 | 3.4e-07 | 1.2e-07 | 2.300E-07 | 1.100E-07 | 1.200E-07 | 3.400E-07 |
| 2053 | 103 | N103M | A | 1.4e-07 | 4.4e-07 | 3e-07 | 2.933E-07 | 1.501E-07 | 1.400E-07 | 4.400E-07 |
| 2054 | 103 | N103F | A | 2.4e-07 | 1.1e-07 | 1.6e-07 | 1.700E-07 | 6.557E-08 | 1.100E-07 | 2.400E-07 |
| 2055 | 103 | N103P | A | 1.2e-07 | 4.1e-07 | 1.6e-07 | 2.300E-07 | 1.572E-07 | 1.200E-07 | 4.100E-07 |
| 2056 | 103 | N103S | A | 2.6e-07 | 5e-07 | 2.2e-07 | 3.267E-07 | 1.514E-07 | 2.200E-07 | 5.000E-07 |
| 2057 | 103 | N103T | A | 1.7e-07 | 2.4e-07 | 1.9e-07 | 2.000E-07 | 3.606E-08 | 1.700E-07 | 2.400E-07 |
| 2058 | 103 | N103W | A | 1.8e-07 | 2.8e-07 | 1.7e-07 | 2.100E-07 | 6.083E-08 | 1.700E-07 | 2.800E-07 |
| 2059 | 103 | N103Y | A | 1.9e-07 | 3.9e-07 | 6.8e-07 | 4.200E-07 | 2.464E-07 | 1.900E-07 | 6.800E-07 |
| 2060 | 103 | N103V | A | 7.4e-07 | 2.4e-07 | 2.3e-07 | 4.033E-07 | 2.916E-07 | 2.300E-07 | 7.400E-07 |
| 2061 | 104 | T104A | A | 5.4e-07 | 2.3e-07 | 3.8e-07 | 3.833E-07 | 1.550E-07 | 2.300E-07 | 5.400E-07 |
| 2062 | 104 | T104R | A | 2e-07 | 3.2e-07 | 3.5e-07 | 2.900E-07 | 7.937E-08 | 2.000E-07 | 3.500E-07 |
| 2063 | 104 | T104N | A | 1.1e-07 | 2e-07 | 8.5e-08 | 1.317E-07 | 6.048E-08 | 8.500E-08 | 2.000E-07 |
| 2064 | 104 | T104D | A | 2.2e-07 | 1.4e-07 | 2e-07 | 1.867E-07 | 4.163E-08 | 1.400E-07 | 2.200E-07 |
| 2065 | 104 | T104C | A | 1.2e-07 | 3.6e-07 | 1.7e-07 | 2.167E-07 | 1.266E-07 | 1.200E-07 | 3.600E-07 |
| 2066 | 104 | T104E | A | 3.2e-07 | 9.4e-08 | 3.1e-07 | 2.413E-07 | 1.277E-07 | 9.400E-08 | 3.200E-07 |
| 2067 | 104 | T104Q | A | 3.9e-07 | 2.9e-07 | 1.6e-07 | 2.800E-07 | 1.153E-07 | 1.600E-07 | 3.900E-07 |
| 2068 | 104 | T104G | A | 3.7e-07 | 2.6e-07 | 5.5e-07 | 3.933E-07 | 1.464E-07 | 2.600E-07 | 5.500E-07 |
| 2069 | 104 | T104H | A | 2.1e-07 | 3e-07 | 2.7e-07 | 2.600E-07 | 4.583E-08 | 2.100E-07 | 3.000E-07 |
| 2070 | 104 | T104I | A | 1.2e-07 | 2.3e-07 | 4e-07 | 2.500E-07 | 1.411E-07 | 1.200E-07 | 4.000E-07 |
| 2071 | 104 | T104L | A | 2.4e-07 | 3.2e-07 | 2.1e-07 | 2.567E-07 | 5.686E-08 | 2.100E-07 | 3.200E-07 |
| 2072 | 104 | T104K | A | 2.3e-07 | 4.6e-07 | 2.6e-07 | 3.167E-07 | 1.250E-07 | 2.300E-07 | 4.600E-07 |
| 2073 | 104 | T104M | A | 1.9e-07 | 1.6e-07 | 1.5e-07 | 1.667E-07 | 2.082E-08 | 1.500E-07 | 1.900E-07 |
| 2074 | 104 | T104F | A | 8.7e-08 | 3.5e-07 | 2.2e-07 | 2.190E-07 | 1.315E-07 | 8.700E-08 | 3.500E-07 |
| 2075 | 104 | T104P | A | 1.7e-07 | 2.7e-07 | 2.6e-07 | 2.333E-07 | 5.508E-08 | 1.700E-07 | 2.700E-07 |
| 2076 | 104 | T104S | A | 3.5e-07 | 2.3e-07 | 2.9e-07 | 2.900E-07 | 6.000E-08 | 2.300E-07 | 3.500E-07 |
| 2077 | 104 | T104T | A | 1.5e-07 | 4.1e-07 | 3.9e-07 | 3.167E-07 | 1.447E-07 | 1.500E-07 | 4.100E-07 |
| 2078 | 104 | T104W | A | 5.4e-07 | 1.5e-07 | 4.6e-07 | 3.833E-07 | 2.060E-07 | 1.500E-07 | 5.400E-07 |
| 2079 | 104 | T104Y | A | 2.2e-07 | 3.9e-07 | 1.7e-07 | 2.600E-07 | 1.153E-07 | 1.700E-07 | 3.900E-07 |
| 2080 | 104 | T104V | A | 3.3e-07 | 6.2e-07 | 3e-07 | 4.167E-07 | 1.767E-07 | 3.000E-07 | 6.200E-07 |
| 2081 | 105 | T105A | A | 1e-07 | 3.2e-07 | 1.5e-07 | 1.900E-07 | 1.153E-07 | 1.000E-07 | 3.200E-07 |
| 2082 | 105 | T105R | A | 2.3e-07 | 2.6e-07 | 2.7e-07 | 2.533E-07 | 2.082E-08 | 2.300E-07 | 2.700E-07 |
| 2083 | 105 | T105N | A | 2.1e-07 | 2.7e-07 | 2.6e-07 | 2.467E-07 | 3.215E-08 | 2.100E-07 | 2.700E-07 |
| 2084 | 105 | T105D | A | 2.2e-07 | 3.2e-07 | 2.3e-07 | 2.567E-07 | 5.508E-08 | 2.200E-07 | 3.200E-07 |
| 2085 | 105 | T105C | A | 2e-07 | 4.3e-07 | 2.9e-07 | 3.067E-07 | 1.159E-07 | 2.000E-07 | 4.300E-07 |
| 2086 | 105 | T105E | A | 3.4e-07 | 7e-07 | 3.1e-07 | 4.500E-07 | 2.170E-07 | 3.100E-07 | 7.000E-07 |
| 2087 | 105 | T105Q | A | 4e-07 | 2.6e-07 | 4.2e-07 | 3.600E-07 | 8.718E-08 | 2.600E-07 | 4.200E-07 |
| 2088 | 105 | T105G | A | 3.1e-07 | 1.9e-07 | 3.3e-07 | 2.767E-07 | 7.572E-08 | 1.900E-07 | 3.300E-07 |
| 2089 | 105 | T105H | A | 3.2e-07 | 4e-07 | 1.2e-07 | 2.800E-07 | 1.442E-07 | 1.200E-07 | 4.000E-07 |
| 2090 | 105 | T105I | A | 8.3e-08 | 1.5e-07 | 2.5e-07 | 1.610E-07 | 8.404E-08 | 8.300E-08 | 2.500E-07 |
| 2091 | 105 | T105L | A | 4e-07 | 3.4e-07 | 2.2e-07 | 3.200E-07 | 9.165E-08 | 2.200E-07 | 4.000E-07 |
| 2092 | 105 | T105K | A | 3.4e-07 | 1.8e-07 | 4.4e-07 | 3.200E-07 | 1.311E-07 | 1.800E-07 | 4.400E-07 |
| 2093 | 105 | T105M | A | 1.1e-07 | 2.9e-07 | 2.1e-07 | 2.033E-07 | 9.018E-08 | 1.100E-07 | 2.900E-07 |
| 2094 | 105 | T105F | A | 2e-07 | 3.7e-07 | 5.3e-07 | 3.667E-07 | 1.650E-07 | 2.000E-07 | 5.300E-07 |
| 2095 | 105 | T105P | A | 2.5e-07 | 1.6e-07 | 3e-07 | 2.367E-07 | 7.095E-08 | 1.600E-07 | 3.000E-07 |
| 2096 | 105 | T105S | A | 4e-07 | 2.1e-07 | 3.7e-07 | 3.267E-07 | 1.021E-07 | 2.100E-07 | 4.000E-07 |
| 2097 | 105 | T105T | A | 1.4e-07 | 2.2e-07 | 2.7e-07 | 2.100E-07 | 6.557E-08 | 1.400E-07 | 2.700E-07 |
| 2098 | 105 | T105W | A | 2.6e-07 | 2.9e-07 | 3e-07 | 2.833E-07 | 2.082E-08 | 2.600E-07 | 3.000E-07 |
| 2099 | 105 | T105Y | A | 2e-07 | 6.2e-07 | 1.9e-07 | 3.367E-07 | 2.454E-07 | 1.900E-07 | 6.200E-07 |
| 2100 | 105 | T105V | A | 3.9e-07 | 2e-07 | 1.6e-07 | 2.500E-07 | 1.229E-07 | 1.600E-07 | 3.900E-07 |
| 2101 | 106 | P106A | A | 1.3e-07 | 4e-07 | 2e-07 | 2.433E-07 | 1.401E-07 | 1.300E-07 | 4.000E-07 |
| 2102 | 106 | P106R | A | 3.7e-07 | 9.8e-08 | 4e-07 | 2.893E-07 | 1.664E-07 | 9.800E-08 | 4.000E-07 |
| 2103 | 106 | P106N | A | 2.4e-07 | 3.4e-07 | 1.1e-07 | 2.300E-07 | 1.153E-07 | 1.100E-07 | 3.400E-07 |
| 2104 | 106 | P106D | A | 2.6e-07 | 2.2e-07 | 2.4e-07 | 2.400E-07 | 2.000E-08 | 2.200E-07 | 2.600E-07 |
| 2105 | 106 | P106C | A | 1.5e-07 | 2e-07 | 1.4e-07 | 1.633E-07 | 3.215E-08 | 1.400E-07 | 2.000E-07 |
| 2106 | 106 | P106E | A | 4.1e-07 | 2.6e-07 | 2.8e-07 | 3.167E-07 | 8.145E-08 | 2.600E-07 | 4.100E-07 |
| 2107 | 106 | P106Q | A | 3e-07 | 1.8e-07 | 2.3e-07 | 2.367E-07 | 6.028E-08 | 1.800E-07 | 3.000E-07 |
| 2108 | 106 | P106G | A | 3.1e-07 | 1.8e-07 | 2.4e-07 | 2.433E-07 | 6.506E-08 | 1.800E-07 | 3.100E-07 |
| 2109 | 106 | P106H | A | 2.6e-07 | 3.5e-07 | 1.8e-07 | 2.633E-07 | 8.505E-08 | 1.800E-07 | 3.500E-07 |
| 2110 | 106 | P106I | A | 2.3e-07 | 2.2e-07 | 3.8e-07 | 2.767E-07 | 8.963E-08 | 2.200E-07 | 3.800E-07 |
| 2111 | 106 | P106L | A | 3.8e-07 | 2.8e-07 | 1.6e-07 | 2.733E-07 | 1.102E-07 | 1.600E-07 | 3.800E-07 |
| 2112 | 106 | P106K | A | 1.9e-07 | 5.7e-07 | 1.5e-07 | 3.033E-07 | 2.318E-07 | 1.500E-07 | 5.700E-07 |
| 2113 | 106 | P106M | A | 2.5e-07 | 1.9e-07 | 2.7e-07 | 2.367E-07 | 4.163E-08 | 1.900E-07 | 2.700E-07 |
| 2114 | 106 | P106F | A | 8.4e-07 | 1.5e-07 | 3.8e-07 | 4.567E-07 | 3.513E-07 | 1.500E-07 | 8.400E-07 |
| 2115 | 106 | P106P | A | 2.3e-07 | 2.4e-07 | 2.7e-07 | 2.467E-07 | 2.082E-08 | 2.300E-07 | 2.700E-07 |
| 2116 | 106 | P106S | A | 3.2e-07 | 1.9e-07 | 1.8e-07 | 2.300E-07 | 7.810E-08 | 1.800E-07 | 3.200E-07 |
| 2117 | 106 | P106T | A | 2.6e-07 | 2.2e-07 | 3.7e-07 | 2.833E-07 | 7.767E-08 | 2.200E-07 | 3.700E-07 |
| 2118 | 106 | P106W | A | 6.8e-07 | 2.5e-07 | 2.5e-07 | 3.933E-07 | 2.483E-07 | 2.500E-07 | 6.800E-07 |
| 2119 | 106 | P106Y | A | 3.5e-07 | 3e-07 | 1.5e-07 | 2.667E-07 | 1.041E-07 | 1.500E-07 | 3.500E-07 |
| 2120 | 106 | P106V | A | 4.1e-07 | 2.3e-07 | 3.4e-07 | 3.267E-07 | 9.074E-08 | 2.300E-07 | 4.100E-07 |
| 2121 | 107 | V107A | A | 2.8e-07 | 3.4e-07 | 5.8e-07 | 4.000E-07 | 1.587E-07 | 2.800E-07 | 5.800E-07 |
| 2122 | 107 | V107R | A | 1.4e-07 | 2.1e-07 | 2.9e-07 | 2.133E-07 | 7.506E-08 | 1.400E-07 | 2.900E-07 |
| 2123 | 107 | V107N | A | 3e-07 | 3.3e-07 | 2.7e-07 | 3.000E-07 | 3.000E-08 | 2.700E-07 | 3.300E-07 |
| 2124 | 107 | V107D | A | 5.4e-07 | 2e-07 | 2.1e-07 | 3.167E-07 | 1.935E-07 | 2.000E-07 | 5.400E-07 |

| | | | | | | | | | | |
|---|---|---|---|---|---|---|---|---|---|---|
| 2125 | 107 | V107C | A | 2.9e-07 | 3.1e-07 | 2.3e-07 | 2.767E-07 | 4.163E-08 | 2.300E-07 | 3.100E-07 |
| 2126 | 107 | V107E | A | 4.9e-07 | 6.5e-07 | 1.1e-07 | 4.167E-07 | 2.774E-07 | 1.100E-07 | 6.500E-07 |
| 2127 | 107 | V107Q | A | 2.1e-07 | 5.3e-07 | 3.2e-07 | 3.533E-07 | 1.626E-07 | 2.100E-07 | 5.300E-07 |
| 2128 | 107 | V107G | A | 2.9e-07 | 1.9e-07 | 5.9e-07 | 3.567E-07 | 2.082E-07 | 1.900E-07 | 5.900E-07 |
| 2129 | 107 | V107H | A | 1.5e-07 | 1e-07 | 3.7e-07 | 2.067E-07 | 1.436E-07 | 1.000E-07 | 3.700E-07 |
| 2130 | 107 | V107I | A | 3.4e-07 | 2.6e-07 | 3.5e-07 | 3.167E-07 | 4.933E-08 | 2.600E-07 | 3.500E-07 |
| 2131 | 107 | V107L | A | 2.1e-07 | 2.3e-07 | 1.4e-07 | 1.933E-07 | 4.726E-08 | 1.400E-07 | 2.300E-07 |
| 2132 | 107 | V107K | A | 3.9e-07 | 1.6e-07 | 1.6e-07 | 2.367E-07 | 1.328E-07 | 1.600E-07 | 3.900E-07 |
| 2133 | 107 | V107M | A | 2.6e-07 | 3.7e-07 | 6.3e-07 | 4.200E-07 | 1.900E-07 | 2.600E-07 | 6.300E-07 |
| 2134 | 107 | V107F | A | 2.6e-07 | 2.8e-07 | 4.4e-07 | 3.267E-07 | 9.866E-08 | 2.600E-07 | 4.400E-07 |
| 2135 | 107 | V107P | A | 4e-07 | 6.2e-07 | 4.2e-07 | 4.800E-07 | 1.217E-07 | 4.000E-07 | 6.200E-07 |
| 2136 | 107 | V107S | A | 2.2e-07 | 2.8e-07 | 2.6e-07 | 2.533E-07 | 3.055E-08 | 2.200E-07 | 2.800E-07 |
| 2137 | 107 | V107T | A | 3.7e-07 | 1e-07 | 1.1e-07 | 1.933E-07 | 1.531E-07 | 1.000E-07 | 3.700E-07 |
| 2138 | 107 | V107W | A | 2.1e-07 | 3.9e-07 | 3.5e-07 | 3.167E-07 | 9.452E-08 | 2.100E-07 | 3.900E-07 |
| 2139 | 107 | V107Y | A | 2e-07 | 1.8e-07 | 2e-07 | 1.933E-07 | 1.155E-08 | 1.800E-07 | 2.000E-07 |
| 2140 | 107 | V107V | A | 4.5e-07 | 3.3e-07 | 5.3e-07 | 4.367E-07 | 1.007E-07 | 3.300E-07 | 5.300E-07 |
| 2141 | 108 | T108A | A | 2.7e-07 | 2.5e-07 | 4.6e-07 | 3.267E-07 | 1.159E-07 | 2.500E-07 | 4.600E-07 |
| 2142 | 108 | T108R | A | 3.6e-07 | 4e-07 | 2.1e-07 | 3.233E-07 | 1.002E-07 | 2.100E-07 | 4.000E-07 |
| 2143 | 108 | T108N | A | 1.1e-07 | 2.1e-07 | 3.3e-07 | 2.167E-07 | 1.102E-07 | 1.100E-07 | 3.300E-07 |
| 2144 | 108 | T108D | A | 2.8e-07 | 2e-07 | 2.6e-07 | 2.467E-07 | 4.163E-08 | 2.000E-07 | 2.800E-07 |
| 2145 | 108 | T108C | A | 2.7e-07 | 2.2e-07 | 2.6e-07 | 2.500E-07 | 2.646E-08 | 2.200E-07 | 2.700E-07 |
| 2146 | 108 | T108E | A | 4.5e-07 | 3e-07 | 4e-07 | 3.833E-07 | 7.638E-08 | 3.000E-07 | 4.500E-07 |
| 2147 | 108 | T108Q | A | 2e-07 | 2e-07 | 2.3e-07 | 2.100E-07 | 1.732E-08 | 2.000E-07 | 2.300E-07 |
| 2148 | 108 | T108G | A | 2.4e-07 | 1.4e-07 | 5e-07 | 2.933E-07 | 1.858E-07 | 1.400E-07 | 5.000E-07 |
| 2149 | 108 | T108H | A | 4.8e-07 | 3e-07 | 2.3e-07 | 3.367E-07 | 1.290E-07 | 2.300E-07 | 4.800E-07 |
| 2150 | 108 | T108I | A | 2.8e-07 | 2.3e-07 | 2.8e-07 | 2.633E-07 | 2.887E-08 | 2.300E-07 | 2.800E-07 |
| 2151 | 108 | T108L | A | 2.5e-07 | 3.8e-07 | 2.5e-07 | 2.933E-07 | 7.506E-08 | 2.500E-07 | 3.800E-07 |
| 2152 | 108 | T108K | A | 3.1e-07 | 3.4e-07 | 3.6e-07 | 3.367E-07 | 2.517E-08 | 3.100E-07 | 3.600E-07 |
| 2153 | 108 | T108M | A | 2.6e-07 | 2.4e-07 | 2.5e-07 | 2.500E-07 | 1.000E-08 | 2.400E-07 | 2.600E-07 |
| 2154 | 108 | T108F | A | 1.9e-07 | 1.9e-07 | 3.3e-07 | 2.367E-07 | 8.083E-08 | 1.900E-07 | 3.300E-07 |
| 2155 | 108 | T108P | A | 1.7e-07 | 1.3e-07 | 4.5e-07 | 2.500E-07 | 1.744E-07 | 1.300E-07 | 4.500E-07 |
| 2156 | 108 | T108S | A | 1.3e-07 | 4.5e-07 | 1.4e-07 | 2.400E-07 | 1.819E-07 | 1.300E-07 | 4.500E-07 |
| 2157 | 108 | T108T | A | 1.9e-07 | 3.1e-07 | 3.8e-07 | 2.933E-07 | 9.609E-08 | 1.900E-07 | 3.800E-07 |
| 2158 | 108 | T108W | A | 3.4e-07 | 2.9e-07 | 1.1e-07 | 2.467E-07 | 1.210E-07 | 1.100E-07 | 3.400E-07 |
| 2159 | 108 | T108Y | A | 1.8e-07 | 2.2e-07 | 1.8e-07 | 1.933E-07 | 2.309E-08 | 1.800E-07 | 2.200E-07 |
| 2160 | 108 | T108V | A | 1.3e-07 | 2.1e-07 | 3.1e-07 | 2.167E-07 | 9.018E-08 | 1.300E-07 | 3.100E-07 |
| 2161 | 109 | G109A | A | 2.8e-07 | 3.2e-07 | 4.6e-07 | 3.533E-07 | 9.452E-08 | 2.800E-07 | 4.600E-07 |
| 2162 | 109 | G109R | A | 1.6e-07 | 3.7e-07 | 1.5e-07 | 2.267E-07 | 1.242E-07 | 1.500E-07 | 3.700E-07 |
| 2163 | 109 | G109N | A | 2e-07 | 1.1e-07 | 2.7e-07 | 1.933E-07 | 8.021E-08 | 1.100E-07 | 2.700E-07 |
| 2164 | 109 | G109D | A | 5.7e-07 | 3.5e-07 | 3.9e-07 | 4.367E-07 | 1.172E-07 | 3.500E-07 | 5.700E-07 |
| 2165 | 109 | G109C | A | 2.9e-07 | 3.6e-07 | 3e-07 | 3.167E-07 | 3.786E-08 | 2.900E-07 | 3.600E-07 |
| 2166 | 109 | G109E | A | 1.8e-07 | 2.3e-07 | 1.8e-07 | 1.967E-07 | 2.887E-08 | 1.800E-07 | 2.300E-07 |
| 2167 | 109 | G109Q | A | 3.1e-07 | 2.3e-07 | 1.6e-07 | 2.333E-07 | 7.506E-08 | 1.600E-07 | 3.100E-07 |
| 2168 | 109 | G109G | A | 3.8e-07 | 1.9e-07 | 2.6e-07 | 2.767E-07 | 9.609E-08 | 1.900E-07 | 3.800E-07 |
| 2169 | 109 | G109H | A | 1.8e-07 | 1.5e-07 | 3.2e-07 | 2.167E-07 | 9.074E-08 | 1.500E-07 | 3.200E-07 |
| 2170 | 109 | G109I | A | 3.9e-07 | 2.1e-07 | 1.4e-07 | 2.467E-07 | 1.290E-07 | 1.400E-07 | 3.900E-07 |
| 2171 | 109 | G109L | A | 2.1e-07 | 2.7e-07 | 2.4e-07 | 2.400E-07 | 3.000E-08 | 2.100E-07 | 2.700E-07 |
| 2172 | 109 | G109K | A | 3.8e-07 | 1.8e-07 | 9.3e-08 | 2.177E-07 | 1.472E-07 | 9.300E-08 | 3.800E-07 |
| 2173 | 109 | G109M | A | 2.1e-07 | 1.7e-07 | 3.3e-07 | 2.367E-07 | 8.327E-08 | 1.700E-07 | 3.300E-07 |
| 2174 | 109 | G109F | A | 3.4e-07 | 1.7e-07 | 2.4e-07 | 2.500E-07 | 8.544E-08 | 1.700E-07 | 3.400E-07 |
| 2175 | 109 | G109P | A | 3.2e-07 | 2.2e-07 | 2.6e-07 | 2.667E-07 | 5.033E-08 | 2.200E-07 | 3.200E-07 |
| 2176 | 109 | G109S | A | 3.3e-07 | 2e-07 | 1.6e-07 | 2.300E-07 | 8.888E-08 | 1.600E-07 | 3.300E-07 |
| 2177 | 109 | G109T | A | 2.9e-07 | 2.6e-07 | 2.9e-07 | 2.800E-07 | 1.732E-08 | 2.600E-07 | 2.900E-07 |
| 2178 | 109 | G109W | A | 4.9e-07 | 1.2e-07 | 6.9e-07 | 4.333E-07 | 2.892E-07 | 1.200E-07 | 6.900E-07 |
| 2179 | 109 | G109Y | A | 1.7e-07 | 1.8e-07 | 2.8e-07 | 2.100E-07 | 6.083E-08 | 1.700E-07 | 2.800E-07 |
| 2180 | 109 | G109V | A | 2.4e-07 | 1.2e-07 | 3.6e-07 | 2.400E-07 | 1.200E-07 | 1.200E-07 | 3.600E-07 |
| 2181 | 110 | A110A | A | 2.6e-07 | 2.6e-07 | 2.3e-07 | 2.500E-07 | 1.732E-08 | 2.300E-07 | 2.600E-07 |
| 2182 | 110 | A110R | A | 2.8e-07 | 2.4e-07 | 2.5e-07 | 2.567E-07 | 2.082E-08 | 2.400E-07 | 2.800E-07 |
| 2183 | 110 | A110N | A | 3.1e-07 | 3.9e-07 | 3.3e-07 | 3.433E-07 | 4.163E-08 | 3.100E-07 | 3.900E-07 |
| 2184 | 110 | A110D | A | 3.1e-07 | 1.5e-07 | 1.5e-07 | 2.033E-07 | 9.238E-08 | 1.500E-07 | 3.100E-07 |
| 2185 | 110 | A110C | A | 4.7e-07 | 1.1e-07 | 3.3e-07 | 3.033E-07 | 1.815E-07 | 1.100E-07 | 4.700E-07 |
| 2186 | 110 | A110E | A | 4.3e-07 | 3.1e-07 | 2.2e-07 | 3.200E-07 | 1.054E-07 | 2.200E-07 | 4.300E-07 |
| 2187 | 110 | A110Q | A | 1.5e-07 | 1.5e-07 | 3.7e-07 | 2.233E-07 | 1.270E-07 | 1.500E-07 | 3.700E-07 |
| 2188 | 110 | A110G | A | 4.1e-07 | 3.2e-07 | 1.4e-07 | 2.900E-07 | 1.375E-07 | 1.400E-07 | 4.100E-07 |
| 2189 | 110 | A110H | A | 2.5e-07 | 2.7e-07 | 3.4e-07 | 2.867E-07 | 4.726E-08 | 2.500E-07 | 3.400E-07 |
| 2190 | 110 | A110I | A | 1.7e-07 | 4.7e-07 | 1.8e-07 | 2.733E-07 | 1.704E-07 | 1.700E-07 | 4.700E-07 |
| 2191 | 110 | A110L | A | 2.2e-07 | 4e-07 | 2.8e-07 | 3.000E-07 | 9.165E-08 | 2.200E-07 | 4.000E-07 |
| 2192 | 110 | A110K | A | 1.5e-07 | 1.3e-07 | 1.9e-07 | 1.567E-07 | 3.055E-08 | 1.300E-07 | 1.900E-07 |
| 2193 | 110 | A110M | A | 1.8e-07 | 3e-07 | 1.7e-07 | 2.167E-07 | 7.234E-08 | 1.700E-07 | 3.000E-07 |
| 2194 | 110 | A110F | A | 2.1e-07 | 3.1e-07 | 1.3e-07 | 2.167E-07 | 9.018E-08 | 1.300E-07 | 3.100E-07 |
| 2195 | 110 | A110P | A | 3.9e-07 | 4.1e-07 | 2.6e-07 | 3.533E-07 | 8.145E-08 | 2.600E-07 | 4.100E-07 |
| 2196 | 110 | A110S | A | 4.1e-07 | 1.8e-07 | 4e-07 | 3.300E-07 | 1.300E-07 | 1.800E-07 | 4.100E-07 |
| 2197 | 110 | A110T | A | 2.2e-07 | 3.6e-07 | 2e-07 | 2.600E-07 | 8.718E-08 | 2.000E-07 | 3.600E-07 |
| 2198 | 110 | A110W | A | 3e-07 | 1.5e-07 | 1.2e-07 | 1.900E-07 | 9.644E-08 | 1.200E-07 | 3.000E-07 |
| 2199 | 110 | A110Y | A | 3.3e-07 | 2.3e-07 | 2.5e-07 | 2.700E-07 | 5.292E-08 | 2.300E-07 | 3.300E-07 |
| 2200 | 110 | A110V | A | 3.2e-07 | 1.3e-07 | 3.2e-07 | 2.567E-07 | 1.097E-07 | 1.300E-07 | 3.200E-07 |
| 2201 | 111 | S111A | A | 3.1e-07 | 2.5e-07 | 1.4e-07 | 2.333E-07 | 8.622E-08 | 1.400E-07 | 3.100E-07 |
| 2202 | 111 | S111R | A | 2.3e-07 | 2.4e-07 | 2.8e-07 | 2.500E-07 | 2.646E-08 | 2.300E-07 | 2.800E-07 |
| 2203 | 111 | S111N | A | 4e-07 | 1.8e-07 | 3.9e-07 | 3.233E-07 | 1.242E-07 | 1.800E-07 | 4.000E-07 |
| 2204 | 111 | S111D | A | 1.6e-07 | 2.1e-07 | 1.3e-07 | 1.667E-07 | 4.041E-08 | 1.300E-07 | 2.100E-07 |
| 2205 | 111 | S111C | A | 2.8e-07 | 2e-07 | 2.2e-07 | 2.333E-07 | 4.163E-08 | 2.000E-07 | 2.800E-07 |
| 2206 | 111 | S111E | A | 1.2e-07 | 2e-07 | 2.9e-07 | 2.033E-07 | 8.505E-08 | 1.200E-07 | 2.900E-07 |
| 2207 | 111 | S111Q | A | 4.3e-07 | 2.5e-07 | 2.2e-07 | 3.000E-07 | 1.136E-07 | 2.200E-07 | 4.300E-07 |
| 2208 | 111 | S111G | A | 4.3e-07 | 1.4e-07 | 3.6e-07 | 3.100E-07 | 1.513E-07 | 1.400E-07 | 4.300E-07 |
| 2209 | 111 | S111H | A | 2.5e-07 | 3.5e-07 | 1.7e-07 | 2.567E-07 | 9.018E-08 | 1.700E-07 | 3.500E-07 |

| | | | | | | | | | | |
|---|---|---|---|---|---|---|---|---|---|---|
| 2210 | 111 | S111I | A | 2.6e-07 | 3.2e-07 | 2.6e-07 | 2.800E-07 | 3.464E-08 | 2.600E-07 | 3.200E-07 |
| 2211 | 111 | S111L | A | 4e-07 | 2.3e-07 | 1.3e-07 | 2.533E-07 | 1.365E-07 | 1.300E-07 | 4.000E-07 |
| 2212 | 111 | S111K | A | 1.8e-07 | 1.8e-07 | 1.3e-07 | 1.633E-07 | 2.887E-08 | 1.300E-07 | 1.800E-07 |
| 2213 | 111 | S111M | A | 2.8e-07 | 1.3e-07 | 2.1e-07 | 2.067E-07 | 7.506E-08 | 1.300E-07 | 2.800E-07 |
| 2214 | 111 | S111F | A | 2.2e-07 | 2e-07 | 1.6e-07 | 1.933E-07 | 3.055E-08 | 1.600E-07 | 2.200E-07 |
| 2215 | 111 | S111P | A | 1.7e-07 | 2.3e-07 | 1.7e-07 | 1.900E-07 | 3.464E-08 | 1.700E-07 | 2.300E-07 |
| 2216 | 111 | S111S | A | 2.5e-07 | 3.4e-07 | 3.1e-07 | 3.000E-07 | 4.583E-08 | 2.500E-07 | 3.400E-07 |
| 2217 | 111 | S111T | A | 1.1e-07 | 2.5e-07 | 5.4e-07 | 3.000E-07 | 2.193E-07 | 1.100E-07 | 5.400E-07 |
| 2218 | 111 | S111W | A | 2.9e-07 | 2.2e-07 | 3.2e-07 | 2.767E-07 | 5.132E-08 | 2.200E-07 | 3.200E-07 |
| 2219 | 111 | S111Y | A | 1.4e-07 | 5.3e-07 | 2.5e-07 | 3.067E-07 | 2.011E-07 | 1.400E-07 | 5.300E-07 |
| 2220 | 111 | S111V | A | 3.5e-07 | 2.4e-07 | 3.7e-07 | 3.200E-07 | 7.000E-08 | 2.400E-07 | 3.700E-07 |
| 2221 | 112 | P112A | A | 3.1e-07 | 1.1e-07 | 1.9e-07 | 2.033E-07 | 1.007E-07 | 1.100E-07 | 3.100E-07 |
| 2222 | 112 | P112R | A | 1.8e-07 | 1.8e-07 | 2e-07 | 1.867E-07 | 1.155E-08 | 1.800E-07 | 2.000E-07 |
| 2223 | 112 | P112N | A | 1.9e-07 | 1.7e-07 | 3.4e-07 | 2.333E-07 | 9.292E-08 | 1.700E-07 | 3.400E-07 |
| 2224 | 112 | P112D | A | 1.4e-07 | 2e-07 | 2.2e-07 | 1.867E-07 | 4.163E-08 | 1.400E-07 | 2.200E-07 |
| 2225 | 112 | P112C | A | 3.5e-07 | 1.5e-07 | 4.1e-07 | 3.033E-07 | 1.361E-07 | 1.500E-07 | 4.100E-07 |
| 2226 | 112 | P112E | A | 7e-07 | 3.3e-07 | 1.8e-07 | 4.033E-07 | 2.676E-07 | 1.800E-07 | 7.000E-07 |
| 2227 | 112 | P112Q | A | 3.1e-07 | 1.1e-07 | 1.9e-07 | 2.033E-07 | 1.007E-07 | 1.100E-07 | 3.100E-07 |
| 2228 | 112 | P112G | A | 3.1e-07 | 2.6e-07 | 1.7e-07 | 2.467E-07 | 7.095E-08 | 1.700E-07 | 3.100E-07 |
| 2229 | 112 | P112H | A | 1.1e-07 | 4.5e-07 | 3.3e-07 | 2.967E-07 | 1.724E-07 | 1.100E-07 | 4.500E-07 |
| 2230 | 112 | P112I | A | 2e-07 | 1.6e-07 | 3.2e-07 | 2.267E-07 | 8.327E-08 | 1.600E-07 | 3.200E-07 |
| 2231 | 112 | P112L | A | 2.4e-07 | 1.7e-07 | 2e-07 | 2.033E-07 | 3.512E-08 | 1.700E-07 | 2.400E-07 |
| 2232 | 112 | P112K | A | 5.2e-07 | 3.1e-07 | 3.3e-07 | 3.867E-07 | 1.159E-07 | 3.100E-07 | 5.200E-07 |
| 2233 | 112 | P112M | A | 2.6e-07 | 2.6e-07 | 1.6e-07 | 2.267E-07 | 5.774E-08 | 1.600E-07 | 2.600E-07 |
| 2234 | 112 | P112F | A | 2.4e-07 | 2.7e-07 | 5.7e-07 | 3.600E-07 | 1.825E-07 | 2.400E-07 | 5.700E-07 |
| 2235 | 112 | P112P | A | 2.2e-07 | 1e-07 | 2.4e-07 | 1.867E-07 | 7.572E-08 | 1.000E-07 | 2.400E-07 |
| 2236 | 112 | P112S | A | 2.8e-07 | 1.6e-07 | 1.7e-07 | 2.033E-07 | 6.658E-08 | 1.600E-07 | 2.800E-07 |
| 2237 | 112 | P112T | A | 5e-07 | 4.5e-07 | 2.1e-07 | 3.867E-07 | 1.550E-07 | 2.100E-07 | 5.000E-07 |
| 2238 | 112 | P112W | A | 2.5e-07 | 1.9e-07 | 1.6e-07 | 2.000E-07 | 4.583E-08 | 1.600E-07 | 2.500E-07 |
| 2239 | 112 | P112Y | A | 2.6e-07 | 3.4e-07 | 4.9e-07 | 3.633E-07 | 1.168E-07 | 2.600E-07 | 4.900E-07 |
| 2240 | 112 | P112V | A | 9.8e-08 | 3.1e-07 | 2.7e-07 | 2.260E-07 | 1.126E-07 | 9.800E-08 | 3.100E-07 |
| 2241 | 113 | G113A | A | 2.9e-07 | 1.8e-07 | 2.2e-07 | 2.300E-07 | 5.568E-08 | 1.800E-07 | 2.900E-07 |
| 2242 | 113 | G113R | A | 2.2e-07 | 2.4e-07 | 2.6e-07 | 2.400E-07 | 2.000E-08 | 2.200E-07 | 2.600E-07 |
| 2243 | 113 | G113N | A | 2.1e-07 | 5.3e-07 | 1.4e-07 | 2.933E-07 | 2.079E-07 | 1.400E-07 | 5.300E-07 |
| 2244 | 113 | G113D | A | 4.6e-07 | 8.1e-07 | 1.7e-07 | 4.800E-07 | 3.205E-07 | 1.700E-07 | 8.100E-07 |
| 2245 | 113 | G113C | A | 4.3e-07 | 2.5e-07 | 3.3e-07 | 3.367E-07 | 9.018E-08 | 2.500E-07 | 4.300E-07 |
| 2246 | 113 | G113E | A | 1.5e-07 | 7.7e-08 | 2.4e-07 | 1.557E-07 | 8.165E-08 | 7.700E-08 | 2.400E-07 |
| 2247 | 113 | G113Q | A | 4.6e-07 | 2.8e-07 | 2.7e-07 | 3.367E-07 | 1.069E-07 | 2.700E-07 | 4.600E-07 |
| 2248 | 113 | G113G | A | 2.3e-07 | 2e-07 | 1.1e-07 | 1.800E-07 | 6.245E-08 | 1.100E-07 | 2.300E-07 |
| 2249 | 113 | G113H | A | 1.8e-07 | 1.5e-07 | 3e-07 | 2.100E-07 | 7.937E-08 | 1.500E-07 | 3.000E-07 |
| 2250 | 113 | G113I | A | 2.3e-07 | 2.7e-07 | 5.3e-07 | 3.433E-07 | 1.629E-07 | 2.300E-07 | 5.300E-07 |
| 2251 | 113 | G113L | A | 1.3e-07 | 9.1e-08 | 3.1e-07 | 1.770E-07 | 1.168E-07 | 9.100E-08 | 3.100E-07 |
| 2252 | 113 | G113K | A | 1.7e-07 | 3.3e-07 | 1.4e-07 | 2.133E-07 | 1.021E-07 | 1.400E-07 | 3.300E-07 |
| 2253 | 113 | G113M | A | 3e-07 | 2.2e-07 | 2.2e-07 | 2.467E-07 | 4.619E-08 | 2.200E-07 | 3.000E-07 |
| 2254 | 113 | G113F | A | 2.9e-07 | 4.5e-07 | 1.8e-07 | 3.067E-07 | 1.358E-07 | 1.800E-07 | 4.500E-07 |
| 2255 | 113 | G113P | A | 2.5e-07 | 1.1e-07 | 2.1e-07 | 1.900E-07 | 7.211E-08 | 1.100E-07 | 2.500E-07 |
| 2256 | 113 | G113S | A | 3.2e-07 | 7.9e-07 | 2.6e-07 | 4.567E-07 | 2.902E-07 | 2.600E-07 | 7.900E-07 |
| 2257 | 113 | G113T | A | 4.9e-07 | 2.4e-07 | 3.7e-07 | 3.667E-07 | 1.250E-07 | 2.400E-07 | 4.900E-07 |
| 2258 | 113 | G113W | A | 2.4e-07 | 2.9e-07 | 2.2e-07 | 2.500E-07 | 3.606E-08 | 2.200E-07 | 2.900E-07 |
| 2259 | 113 | G113Y | A | 1.9e-07 | 2e-07 | 2.7e-07 | 2.200E-07 | 4.359E-08 | 1.900E-07 | 2.700E-07 |
| 2260 | 113 | G113V | A | 2.7e-07 | 4.4e-07 | 1.7e-07 | 2.933E-07 | 1.365E-07 | 1.700E-07 | 4.400E-07 |
| 2261 | 114 | G114A | A | 1.1e-07 | 3.2e-07 | 1.1e-07 | 1.800E-07 | 1.212E-07 | 1.100E-07 | 3.200E-07 |
| 2262 | 114 | G114R | A | 3.3e-07 | 1.8e-07 | 2.9e-07 | 2.667E-07 | 7.767E-08 | 1.800E-07 | 3.300E-07 |
| 2263 | 114 | G114N | A | 2.1e-07 | 5e-07 | 6.8e-07 | 4.633E-07 | 2.371E-07 | 2.100E-07 | 6.800E-07 |
| 2264 | 114 | G114D | A | 1.8e-07 | 2.5e-07 | 1.3e-07 | 1.867E-07 | 6.028E-08 | 1.300E-07 | 2.500E-07 |
| 2265 | 114 | G114C | A | 2.6e-07 | 1.7e-07 | 1.4e-07 | 1.900E-07 | 6.245E-08 | 1.400E-07 | 2.600E-07 |
| 2266 | 114 | G114E | A | 2.9e-07 | 2e-07 | 2.1e-07 | 2.333E-07 | 4.933E-08 | 2.000E-07 | 2.900E-07 |
| 2267 | 114 | G114Q | A | 2.7e-07 | 2.5e-07 | 3.3e-07 | 2.833E-07 | 4.163E-08 | 2.500E-07 | 3.300E-07 |
| 2268 | 114 | G114G | A | 2.9e-07 | 4.3e-07 | 2e-07 | 3.067E-07 | 1.159E-07 | 2.000E-07 | 4.300E-07 |
| 2269 | 114 | G114H | A | 5e-07 | 1.8e-07 | 3.6e-07 | 3.467E-07 | 1.604E-07 | 1.800E-07 | 5.000E-07 |
| 2270 | 114 | G114I | A | 1.9e-07 | 2.9e-07 | 1.1e-07 | 1.967E-07 | 9.018E-08 | 1.100E-07 | 2.900E-07 |
| 2271 | 114 | G114L | A | 2.3e-07 | 1.4e-07 | 2.8e-07 | 2.167E-07 | 7.095E-08 | 1.400E-07 | 2.800E-07 |
| 2272 | 114 | G114K | A | 4.6e-07 | 1.1e-07 | 3.4e-07 | 3.033E-07 | 1.779E-07 | 1.100E-07 | 4.600E-07 |
| 2273 | 114 | G114M | A | 5.5e-07 | 2.7e-07 | 1.7e-07 | 3.300E-07 | 1.970E-07 | 1.700E-07 | 5.500E-07 |
| 2274 | 114 | G114F | A | 2.6e-07 | 1.2e-07 | 2e-07 | 1.933E-07 | 7.024E-08 | 1.200E-07 | 2.600E-07 |
| 2275 | 114 | G114P | A | 2e-07 | 1.7e-07 | 1.6e-07 | 1.767E-07 | 2.082E-08 | 1.600E-07 | 2.000E-07 |
| 2276 | 114 | G114S | A | 3.6e-07 | 1.1e-07 | 1.5e-07 | 2.067E-07 | 1.343E-07 | 1.100E-07 | 3.600E-07 |
| 2277 | 114 | G114T | A | 3.5e-07 | 1.6e-07 | 3.4e-07 | 2.833E-07 | 1.069E-07 | 1.600E-07 | 3.500E-07 |
| 2278 | 114 | G114W | A | 2.1e-07 | 4e-07 | 3.3e-07 | 3.133E-07 | 9.609E-08 | 2.100E-07 | 4.000E-07 |
| 2279 | 114 | G114Y | A | 1.5e-07 | 2.8e-07 | 2.5e-07 | 2.267E-07 | 6.807E-08 | 1.500E-07 | 2.800E-07 |
| 2280 | 114 | G114V | A | 1.9e-07 | 5.1e-07 | 1.9e-07 | 2.967E-07 | 1.848E-07 | 1.900E-07 | 5.100E-07 |
| 2281 | 115 | L115A | A | 2.7e-07 | 3.5e-07 | 3e-07 | 3.067E-07 | 4.041E-08 | 2.700E-07 | 3.500E-07 |
| 2282 | 115 | L115R | A | 3.7e-07 | 1.7e-07 | 1.6e-07 | 2.333E-07 | 1.185E-07 | 1.600E-07 | 3.700E-07 |
| 2283 | 115 | L115N | A | 4.4e-07 | 1.4e-07 | 2e-07 | 2.600E-07 | 1.587E-07 | 1.400E-07 | 4.400E-07 |
| 2284 | 115 | L115D | A | 3.6e-07 | 2e-07 | 5e-07 | 3.533E-07 | 1.501E-07 | 2.000E-07 | 5.000E-07 |
| 2285 | 115 | L115C | A | 3.7e-07 | 3.6e-07 | 3.4e-07 | 3.567E-07 | 1.528E-08 | 3.400E-07 | 3.700E-07 |
| 2286 | 115 | L115E | A | 5.3e-07 | 1.9e-07 | 3.1e-07 | 3.433E-07 | 1.724E-07 | 1.900E-07 | 5.300E-07 |
| 2287 | 115 | L115Q | A | 2.1e-07 | 3e-07 | 2.2e-07 | 2.433E-07 | 4.933E-08 | 2.100E-07 | 3.000E-07 |
| 2288 | 115 | L115G | A | 3.8e-07 | 1.7e-07 | 1.3e-07 | 2.267E-07 | 1.343E-07 | 1.300E-07 | 3.800E-07 |
| 2289 | 115 | L115H | A | 1.6e-07 | 3.9e-07 | 2.4e-07 | 2.633E-07 | 1.168E-07 | 1.600E-07 | 3.900E-07 |
| 2290 | 115 | L115I | A | 3e-07 | 1.6e-07 | 3e-07 | 2.533E-07 | 8.083E-08 | 1.600E-07 | 3.000E-07 |
| 2291 | 115 | L115L | A | 4.1e-07 | 1.8e-07 | 1.1e-07 | 2.333E-07 | 1.570E-07 | 1.100E-07 | 4.100E-07 |
| 2292 | 115 | L115K | A | 1.4e-07 | 2.6e-07 | 2.9e-07 | 2.300E-07 | 7.937E-08 | 1.400E-07 | 2.900E-07 |
| 2293 | 115 | L115M | A | 2e-07 | 4.5e-07 | 1.7e-07 | 2.733E-07 | 1.537E-07 | 1.700E-07 | 4.500E-07 |
| 2294 | 115 | L115F | A | 2.9e-07 | 2.3e-07 | 3.7e-07 | 2.967E-07 | 7.024E-08 | 2.300E-07 | 3.700E-07 |

| | | | | | | | | | | |
|---|---|---|---|---|---|---|---|---|---|---|
| 2295 | 115 | L115P | A | 3.8e-07 | 4e-07 | 6.7e-07 | 4.833E-07 | 1.620E-07 | 3.800E-07 | 6.700E-07 |
| 2296 | 115 | L115S | A | 3.4e-07 | 3.7e-07 | 1.4e-07 | 2.833E-07 | 1.250E-07 | 1.400E-07 | 3.700E-07 |
| 2297 | 115 | L115T | A | 4.1e-07 | 2.3e-07 | 2.7e-07 | 3.033E-07 | 9.452E-08 | 2.300E-07 | 4.100E-07 |
| 2298 | 115 | L115W | A | 1.7e-07 | 3.1e-07 | 1.2e-07 | 2.000E-07 | 9.849E-08 | 1.200E-07 | 3.100E-07 |
| 2299 | 115 | L115Y | A | 1.8e-07 | 2.3e-07 | 2.3e-07 | 2.133E-07 | 2.887E-08 | 1.800E-07 | 2.300E-07 |
| 2300 | 115 | L115V | A | 1.9e-07 | 2.2e-07 | 4.2e-07 | 2.767E-07 | 1.250E-07 | 1.900E-07 | 4.200E-07 |
| 2301 | 116 | R116A | A | 1.3e-07 | 3.1e-07 | 4.3e-07 | 2.900E-07 | 1.510E-07 | 1.300E-07 | 4.300E-07 |
| 2302 | 116 | R116R | A | 2.6e-07 | 1.5e-07 | 1.7e-07 | 1.933E-07 | 5.859E-08 | 1.500E-07 | 2.600E-07 |
| 2303 | 116 | R116N | A | 5.2e-07 | 2.1e-07 | 1.3e-07 | 2.867E-07 | 2.060E-07 | 1.300E-07 | 5.200E-07 |
| 2304 | 116 | R116D | A | 4.9e-07 | 2.2e-07 | 2.8e-07 | 3.300E-07 | 1.418E-07 | 2.200E-07 | 4.900E-07 |
| 2305 | 116 | R116C | A | 1.7e-07 | 2.7e-07 | 1.7e-07 | 2.033E-07 | 5.774E-08 | 1.700E-07 | 2.700E-07 |
| 2306 | 116 | R116E | A | 2.6e-07 | 2.2e-07 | 1.9e-07 | 2.233E-07 | 3.512E-08 | 1.900E-07 | 2.600E-07 |
| 2307 | 116 | R116Q | A | 2.6e-07 | 2.4e-07 | 4.4e-07 | 3.133E-07 | 1.102E-07 | 2.400E-07 | 4.400E-07 |
| 2308 | 116 | R116G | A | 1.9e-07 | 3.3e-07 | 3.2e-07 | 2.800E-07 | 7.810E-08 | 1.900E-07 | 3.300E-07 |
| 2309 | 116 | R116H | A | 1.4e-07 | 2.3e-07 | 3.6e-07 | 2.433E-07 | 1.106E-07 | 1.400E-07 | 3.600E-07 |
| 2310 | 116 | R116I | A | 1.2e-07 | 1.1e-07 | 3.1e-07 | 1.800E-07 | 1.127E-07 | 1.100E-07 | 3.100E-07 |
| 2311 | 116 | R116L | A | 1.9e-07 | 3.3e-07 | 1.2e-07 | 2.133E-07 | 1.069E-07 | 1.200E-07 | 3.300E-07 |
| 2312 | 116 | R116K | A | 3.9e-07 | 2.4e-07 | 3e-07 | 3.100E-07 | 7.550E-08 | 2.400E-07 | 3.900E-07 |
| 2313 | 116 | R116M | A | 1.9e-07 | 3.1e-07 | 2.2e-07 | 2.400E-07 | 6.245E-08 | 1.900E-07 | 3.100E-07 |
| 2314 | 116 | R116F | A | 5.3e-07 | 1.9e-07 | 2.3e-07 | 3.167E-07 | 1.858E-07 | 1.900E-07 | 5.300E-07 |
| 2315 | 116 | R116P | A | 2e-07 | 1.7e-07 | 1.6e-07 | 1.767E-07 | 2.082E-08 | 1.600E-07 | 2.000E-07 |
| 2316 | 116 | R116S | A | 2.1e-07 | 2e-07 | 2.1e-07 | 2.067E-07 | 5.774E-09 | 2.000E-07 | 2.100E-07 |
| 2317 | 116 | R116T | A | 2.1e-07 | 2.9e-07 | 3.5e-07 | 2.833E-07 | 7.024E-08 | 2.100E-07 | 3.500E-07 |
| 2318 | 116 | R116W | A | 2.4e-07 | 2.9e-07 | 3.5e-07 | 2.933E-07 | 5.508E-08 | 2.400E-07 | 3.500E-07 |
| 2319 | 116 | R116Y | A | 2.9e-07 | 2.2e-07 | 3.3e-07 | 2.800E-07 | 5.568E-08 | 2.200E-07 | 3.300E-07 |
| 2320 | 116 | R116V | A | 2.2e-07 | 3.3e-07 | 2.6e-07 | 2.700E-07 | 5.568E-08 | 2.200E-07 | 3.300E-07 |
| 2321 | 117 | E117A | A | 2.3e-07 | 2e-07 | 1.8e-07 | 2.033E-07 | 2.517E-08 | 1.800E-07 | 2.300E-07 |
| 2322 | 117 | E117R | A | 2.4e-07 | 3e-07 | 2.8e-07 | 2.733E-07 | 3.055E-08 | 2.400E-07 | 3.000E-07 |
| 2323 | 117 | E117N | A | 4.2e-07 | 3.6e-07 | 3.5e-07 | 3.767E-07 | 3.786E-08 | 3.500E-07 | 4.200E-07 |
| 2324 | 117 | E117D | A | 1.7e-07 | 3e-07 | 1.9e-07 | 2.200E-07 | 7.000E-08 | 1.700E-07 | 3.000E-07 |
| 2325 | 117 | E117C | A | 3.3e-07 | 1.3e-07 | 1.8e-07 | 2.133E-07 | 1.041E-07 | 1.300E-07 | 3.300E-07 |
| 2326 | 117 | E117E | A | 2.3e-07 | 2.2e-07 | 2.2e-07 | 2.233E-07 | 5.774E-09 | 2.200E-07 | 2.300E-07 |
| 2327 | 117 | E117Q | A | 2.7e-07 | 4e-07 | 1.9e-07 | 2.867E-07 | 1.060E-07 | 1.900E-07 | 4.000E-07 |
| 2328 | 117 | E117G | A | 2.3e-07 | 6.6e-07 | 6.8e-07 | 5.233E-07 | 2.542E-07 | 2.300E-07 | 6.800E-07 |
| 2329 | 117 | E117H | A | 2.3e-07 | 2.2e-07 | 3.4e-07 | 2.633E-07 | 6.658E-08 | 2.200E-07 | 3.400E-07 |
| 2330 | 117 | E117I | A | 2e-07 | 4e-07 | 9.7e-08 | 2.323E-07 | 1.541E-07 | 9.700E-08 | 4.000E-07 |
| 2331 | 117 | E117L | A | 2.5e-07 | 3e-07 | 2.8e-07 | 2.767E-07 | 2.517E-08 | 2.500E-07 | 3.000E-07 |
| 2332 | 117 | E117K | A | 1.3e-07 | 3e-07 | 2.6e-07 | 2.300E-07 | 8.888E-08 | 1.300E-07 | 3.000E-07 |
| 2333 | 117 | E117M | A | 2.4e-07 | 1e-07 | 2.1e-07 | 1.833E-07 | 7.371E-08 | 1.000E-07 | 2.400E-07 |
| 2334 | 117 | E117F | A | 5.3e-07 | 4.9e-07 | 3.3e-07 | 4.500E-07 | 1.058E-07 | 3.300E-07 | 5.300E-07 |
| 2335 | 117 | E117P | A | 2.7e-07 | 8e-08 | 2.9e-07 | 2.133E-07 | 1.159E-07 | 8.000E-08 | 2.900E-07 |
| 2336 | 117 | E117S | A | 2.7e-07 | 3.7e-07 | 4.9e-07 | 3.767E-07 | 1.102E-07 | 2.700E-07 | 4.900E-07 |
| 2337 | 117 | E117T | A | 9.2e-08 | 5.1e-07 | 3.8e-07 | 3.273E-07 | 2.139E-07 | 9.200E-08 | 5.100E-07 |
| 2338 | 117 | E117W | A | 3.3e-07 | 2.6e-07 | 3e-07 | 2.967E-07 | 3.512E-08 | 2.600E-07 | 3.300E-07 |
| 2339 | 117 | E117Y | A | 1.8e-07 | 3e-07 | 1.6e-07 | 2.133E-07 | 7.572E-08 | 1.600E-07 | 3.000E-07 |
| 2340 | 117 | E117V | A | 3.2e-07 | 4.5e-07 | 1.5e-07 | 3.067E-07 | 1.504E-07 | 1.500E-07 | 4.500E-07 |
| 2341 | 118 | L118A | A | 3e-07 | 3.6e-07 | 3.6e-07 | 3.400E-07 | 3.464E-08 | 3.000E-07 | 3.600E-07 |
| 2342 | 118 | L118R | A | 1.6e-07 | 3.1e-07 | 1.5e-07 | 2.067E-07 | 8.963E-08 | 1.500E-07 | 3.100E-07 |
| 2343 | 118 | L118N | A | 1.9e-07 | 3.7e-07 | 5.6e-07 | 3.733E-07 | 1.850E-07 | 1.900E-07 | 5.600E-07 |
| 2344 | 118 | L118D | A | 1.9e-07 | 2.8e-07 | 2.3e-07 | 2.333E-07 | 4.509E-08 | 1.900E-07 | 2.800E-07 |
| 2345 | 118 | L118C | A | 2.2e-07 | 3.2e-07 | 4.3e-07 | 3.233E-07 | 1.050E-07 | 2.200E-07 | 4.300E-07 |
| 2346 | 118 | L118E | A | 3.7e-07 | 3e-07 | 2.1e-07 | 2.933E-07 | 8.021E-08 | 2.100E-07 | 3.700E-07 |
| 2347 | 118 | L118Q | A | 3.1e-07 | 2.6e-07 | 1.6e-07 | 2.433E-07 | 7.638E-08 | 1.600E-07 | 3.100E-07 |
| 2348 | 118 | L118G | A | 4.1e-07 | 1.4e-07 | 2.1e-07 | 2.533E-07 | 1.401E-07 | 1.400E-07 | 4.100E-07 |
| 2349 | 118 | L118H | A | 1.8e-07 | 3.3e-07 | 2.1e-07 | 2.400E-07 | 7.937E-08 | 1.800E-07 | 3.300E-07 |
| 2350 | 118 | L118I | A | 4.7e-07 | 2.9e-07 | 2.9e-07 | 3.500E-07 | 1.039E-07 | 2.900E-07 | 4.700E-07 |
| 2351 | 118 | L118L | A | 5.1e-07 | 3.5e-07 | 3.4e-07 | 4.000E-07 | 9.539E-08 | 3.400E-07 | 5.100E-07 |
| 2352 | 118 | L118K | A | 2.7e-07 | 2.8e-07 | 3.6e-07 | 3.033E-07 | 4.933E-08 | 2.700E-07 | 3.600E-07 |
| 2353 | 118 | L118M | A | 2.5e-07 | 1.6e-07 | 3.4e-07 | 2.500E-07 | 9.000E-08 | 1.600E-07 | 3.400E-07 |
| 2354 | 118 | L118F | A | 3.8e-07 | 1.8e-07 | 1.8e-07 | 2.467E-07 | 1.155E-07 | 1.800E-07 | 3.800E-07 |
| 2355 | 118 | L118P | A | 3.1e-07 | 2.7e-07 | 2.6e-07 | 2.800E-07 | 2.646E-08 | 2.600E-07 | 3.100E-07 |
| 2356 | 118 | L118S | A | 3.4e-07 | 1.9e-07 | 2.8e-07 | 2.700E-07 | 7.550E-08 | 1.900E-07 | 3.400E-07 |
| 2357 | 118 | L118T | A | 2.4e-07 | 2.1e-07 | 3e-07 | 2.500E-07 | 4.583E-08 | 2.100E-07 | 3.000E-07 |
| 2358 | 118 | L118W | A | 2.8e-07 | 1.8e-07 | 2.2e-07 | 2.267E-07 | 5.033E-08 | 1.800E-07 | 2.800E-07 |
| 2359 | 118 | L118Y | A | 3.1e-07 | 2.9e-07 | 2.6e-07 | 2.867E-07 | 2.517E-08 | 2.600E-07 | 3.100E-07 |
| 2360 | 118 | L118V | A | 2e-07 | 1.9e-07 | 2.1e-07 | 2.000E-07 | 1.000E-08 | 1.900E-07 | 2.100E-07 |
| 2361 | 119 | Q119A | A | 1.9e-07 | 3.5e-07 | 2.3e-07 | 2.567E-07 | 8.327E-08 | 1.900E-07 | 3.500E-07 |
| 2362 | 119 | Q119R | A | 2.5e-07 | 7.7e-07 | 2.5e-07 | 4.233E-07 | 3.002E-07 | 2.500E-07 | 7.700E-07 |
| 2363 | 119 | Q119N | A | 2.1e-07 | 3.3e-07 | 3.9e-07 | 3.100E-07 | 9.165E-08 | 2.100E-07 | 3.900E-07 |
| 2364 | 119 | Q119D | A | 2.9e-07 | 4.2e-07 | 2e-07 | 3.033E-07 | 1.106E-07 | 2.000E-07 | 4.200E-07 |
| 2365 | 119 | Q119C | A | 3.2e-07 | 3.1e-07 | 2.6e-07 | 2.967E-07 | 3.215E-08 | 2.600E-07 | 3.200E-07 |
| 2366 | 119 | Q119E | A | 3e-07 | 1.6e-07 | 2.5e-07 | 2.367E-07 | 7.095E-08 | 1.600E-07 | 3.000E-07 |
| 2367 | 119 | Q119Q | A | 1.4e-07 | 3.2e-07 | 1.4e-07 | 2.000E-07 | 1.039E-07 | 1.400E-07 | 3.200E-07 |
| 2368 | 119 | Q119G | A | 4.9e-07 | 2.1e-07 | 2.6e-07 | 3.200E-07 | 1.493E-07 | 2.100E-07 | 4.900E-07 |
| 2369 | 119 | Q119H | A | 2.1e-07 | 2.6e-07 | 2.3e-07 | 2.333E-07 | 2.517E-08 | 2.100E-07 | 2.600E-07 |
| 2370 | 119 | Q119I | A | 3.6e-07 | 7.9e-08 | 7.8e-07 | 4.063E-07 | 3.528E-07 | 7.900E-08 | 7.800E-07 |
| 2371 | 119 | Q119L | A | 2.3e-07 | 3e-07 | 2.9e-07 | 2.733E-07 | 3.786E-08 | 2.300E-07 | 3.000E-07 |
| 2372 | 119 | Q119K | A | 3.4e-07 | 2.9e-07 | 2.4e-07 | 2.900E-07 | 5.000E-08 | 2.400E-07 | 3.400E-07 |
| 2373 | 119 | Q119M | A | 1e-07 | 9.3e-08 | 1.6e-07 | 1.177E-07 | 3.683E-08 | 9.300E-08 | 1.600E-07 |
| 2374 | 119 | Q119F | A | 1.2e-07 | 2.2e-07 | 2.5e-07 | 1.967E-07 | 6.807E-08 | 1.200E-07 | 2.500E-07 |
| 2375 | 119 | Q119P | A | 5.3e-07 | 1.3e-07 | 6.8e-07 | 4.467E-07 | 2.843E-07 | 1.300E-07 | 6.800E-07 |
| 2376 | 119 | Q119S | A | 3.2e-07 | 2.8e-07 | 1.7e-07 | 2.567E-07 | 7.767E-08 | 1.700E-07 | 3.200E-07 |
| 2377 | 119 | Q119T | A | 2.1e-07 | 1.6e-07 | 4.2e-07 | 2.633E-07 | 1.380E-07 | 1.600E-07 | 4.200E-07 |
| 2378 | 119 | Q119W | A | 5e-07 | 3.9e-07 | 1.6e-07 | 3.500E-07 | 1.735E-07 | 1.600E-07 | 5.000E-07 |
| 2379 | 119 | Q119Y | A | 1.7e-07 | 3.5e-07 | 1.7e-07 | 2.300E-07 | 1.039E-07 | 1.700E-07 | 3.500E-07 |

| | | | | | | | | | | |
|---|---|---|---|---|---|---|---|---|---|---|
| 2380 | 119 | Q119V | A | 2.5e-07 | 3.4e-07 | 3.1e-07 | 3.000E-07 | 4.583E-08 | 2.500E-07 | 3.400E-07 |
| 2381 | 120 | L120A | A | 1.6e-07 | 1.7e-07 | 1.4e-07 | 1.567E-07 | 1.528E-08 | 1.400E-07 | 1.700E-07 |
| 2382 | 120 | L120R | A | 3.3e-07 | 2.2e-07 | 2.1e-07 | 2.533E-07 | 6.658E-08 | 2.100E-07 | 3.300E-07 |
| 2383 | 120 | L120N | A | 2.3e-07 | 2.3e-07 | 3.8e-07 | 2.800E-07 | 8.660E-08 | 2.300E-07 | 3.800E-07 |
| 2384 | 120 | L120D | A | 1.9e-07 | 2.1e-07 | 2.3e-07 | 2.100E-07 | 2.000E-08 | 1.900E-07 | 2.300E-07 |
| 2385 | 120 | L120C | A | 3.8e-07 | 2.7e-07 | 3.1e-07 | 3.200E-07 | 5.568E-08 | 2.700E-07 | 3.800E-07 |
| 2386 | 120 | L120E | A | 3.4e-07 | 2.2e-07 | 3.9e-07 | 3.167E-07 | 8.737E-08 | 2.200E-07 | 3.900E-07 |
| 2387 | 120 | L120Q | A | 1.7e-07 | 1.5e-07 | 4.1e-07 | 2.433E-07 | 1.447E-07 | 1.500E-07 | 4.100E-07 |
| 2388 | 120 | L120G | A | 2.6e-07 | 1.7e-07 | 1.6e-07 | 1.967E-07 | 5.508E-08 | 1.600E-07 | 2.600E-07 |
| 2389 | 120 | L120H | A | 1.6e-07 | 2.3e-07 | 3.1e-07 | 2.333E-07 | 7.506E-08 | 1.600E-07 | 3.100E-07 |
| 2390 | 120 | L120I | A | 1.2e-07 | 8e-07 | 1.6e-07 | 3.600E-07 | 3.816E-07 | 1.200E-07 | 8.000E-07 |
| 2391 | 120 | L120L | A | 2.2e-07 | 3.7e-07 | 9.1e-07 | 5.000E-07 | 3.629E-07 | 2.200E-07 | 9.100E-07 |
| 2392 | 120 | L120K | A | 1.6e-07 | 2.6e-07 | 3.1e-07 | 2.433E-07 | 7.638E-08 | 1.600E-07 | 3.100E-07 |
| 2393 | 120 | L120M | A | 1.9e-07 | 1.3e-07 | 4.6e-07 | 2.600E-07 | 1.758E-07 | 1.300E-07 | 4.600E-07 |
| 2394 | 120 | L120F | A | 3.1e-07 | 2.4e-07 | 2.6e-07 | 2.700E-07 | 3.606E-08 | 2.400E-07 | 3.100E-07 |
| 2395 | 120 | L120P | A | 1.9e-07 | 3.4e-07 | 2.8e-07 | 2.700E-07 | 7.550E-08 | 1.900E-07 | 3.400E-07 |
| 2396 | 120 | L120S | A | 2.3e-07 | 2.7e-07 | 2.1e-07 | 2.367E-07 | 3.055E-08 | 2.100E-07 | 2.700E-07 |
| 2397 | 120 | L120T | A | 2.3e-07 | 9.9e-08 | 2e-07 | 1.763E-07 | 6.863E-08 | 9.900E-08 | 2.300E-07 |
| 2398 | 120 | L120W | A | 1.6e-07 | 2.6e-07 | 3e-07 | 2.400E-07 | 7.211E-08 | 1.600E-07 | 3.000E-07 |
| 2399 | 120 | L120Y | A | 3.6e-07 | 1.7e-07 | 1.9e-07 | 2.400E-07 | 1.044E-07 | 1.700E-07 | 3.600E-07 |
| 2400 | 120 | L120V | A | 2.4e-07 | 2.9e-07 | 9.6e-08 | 2.087E-07 | 1.007E-07 | 9.600E-08 | 2.900E-07 |
| 2401 | 121 | R121A | A | 2.8e-07 | 1.6e-07 | 1.5e-07 | 1.967E-07 | 7.234E-08 | 1.500E-07 | 2.800E-07 |
| 2402 | 121 | R121R | A | 2.8e-07 | 5.1e-07 | 2.6e-07 | 3.500E-07 | 1.389E-07 | 2.600E-07 | 5.100E-07 |
| 2403 | 121 | R121N | A | 1.1e-07 | 2.3e-07 | 2.4e-07 | 1.933E-07 | 7.234E-08 | 1.100E-07 | 2.400E-07 |
| 2404 | 121 | R121D | A | 1.9e-07 | 2.7e-07 | 2.3e-07 | 2.300E-07 | 4.000E-08 | 1.900E-07 | 2.700E-07 |
| 2405 | 121 | R121C | A | 1.2e-07 | 1.9e-07 | 2.2e-07 | 1.767E-07 | 5.132E-08 | 1.200E-07 | 2.200E-07 |
| 2406 | 121 | R121E | A | 1.5e-07 | 7.3e-08 | 4.2e-07 | 2.143E-07 | 1.822E-07 | 7.300E-08 | 4.200E-07 |
| 2407 | 121 | R121Q | A | 1.9e-07 | 4.4e-07 | 1.9e-07 | 2.733E-07 | 1.443E-07 | 1.900E-07 | 4.400E-07 |
| 2408 | 121 | R121G | A | 2e-07 | 1.5e-07 | 3e-07 | 2.167E-07 | 7.638E-08 | 1.500E-07 | 3.000E-07 |
| 2409 | 121 | R121H | A | 2.6e-07 | 2e-07 | 3.1e-07 | 2.567E-07 | 5.508E-08 | 2.000E-07 | 3.100E-07 |
| 2410 | 121 | R121I | A | 2.2e-07 | 2.1e-07 | 3.8e-07 | 2.700E-07 | 9.539E-08 | 2.100E-07 | 3.800E-07 |
| 2411 | 121 | R121L | A | 2.3e-07 | 1.8e-07 | 2.1e-07 | 2.067E-07 | 2.517E-08 | 1.800E-07 | 2.300E-07 |
| 2412 | 121 | R121K | A | 1.4e-07 | 1.4e-07 | 3.3e-07 | 2.033E-07 | 1.097E-07 | 1.400E-07 | 3.300E-07 |
| 2413 | 121 | R121M | A | 2.1e-07 | 3.3e-07 | 2.4e-07 | 2.600E-07 | 6.245E-08 | 2.100E-07 | 3.300E-07 |
| 2414 | 121 | R121F | A | 2.6e-07 | 2.4e-07 | 2.1e-07 | 2.367E-07 | 2.517E-08 | 2.100E-07 | 2.600E-07 |
| 2415 | 121 | R121P | A | 2.3e-07 | 3.3e-07 | 2.3e-07 | 2.633E-07 | 5.774E-08 | 2.300E-07 | 3.300E-07 |
| 2416 | 121 | R121S | A | 2.8e-07 | 1.9e-07 | 3.6e-07 | 2.767E-07 | 8.505E-08 | 1.900E-07 | 3.600E-07 |
| 2417 | 121 | R121T | A | 2.3e-07 | 1.9e-07 | 3.1e-07 | 2.433E-07 | 6.110E-08 | 1.900E-07 | 3.100E-07 |
| 2418 | 121 | R121W | A | 1.8e-07 | 4.7e-07 | 1.3e-07 | 2.600E-07 | 1.836E-07 | 1.300E-07 | 4.700E-07 |
| 2419 | 121 | R121Y | A | 3.6e-07 | 3.9e-07 | 2.5e-07 | 3.333E-07 | 7.371E-08 | 2.500E-07 | 3.900E-07 |
| 2420 | 121 | R121V | A | 4.5e-07 | 2.9e-07 | 4.2e-07 | 3.867E-07 | 8.505E-08 | 2.900E-07 | 4.500E-07 |
| 2421 | 122 | S122A | A | 1.6e-07 | 3e-07 | 1.1e-07 | 1.900E-07 | 9.849E-08 | 1.100E-07 | 3.000E-07 |
| 2422 | 122 | S122R | A | 1.8e-07 | 1.3e-07 | 3e-07 | 2.033E-07 | 8.737E-08 | 1.300E-07 | 3.000E-07 |
| 2423 | 122 | S122N | A | 5.1e-07 | 1.5e-07 | 1.1e-07 | 2.567E-07 | 2.203E-07 | 1.100E-07 | 5.100E-07 |
| 2424 | 122 | S122D | A | 3.5e-07 | 1.3e-07 | 1.7e-07 | 2.167E-07 | 1.172E-07 | 1.300E-07 | 3.500E-07 |
| 2425 | 122 | S122C | A | 1.1e-07 | 1.1e-07 | 4.6e-07 | 2.267E-07 | 2.021E-07 | 1.100E-07 | 4.600E-07 |
| 2426 | 122 | S122E | A | 1.6e-07 | 1.7e-07 | 2.2e-07 | 1.833E-07 | 3.215E-08 | 1.600E-07 | 2.200E-07 |
| 2427 | 122 | S122Q | A | 2.1e-07 | 2.3e-07 | 1.4e-07 | 1.933E-07 | 4.726E-08 | 1.400E-07 | 2.300E-07 |
| 2428 | 122 | S122G | A | 4.8e-07 | 2.8e-07 | 2.6e-07 | 3.400E-07 | 1.217E-07 | 2.600E-07 | 4.800E-07 |
| 2429 | 122 | S122H | A | 4.1e-07 | 1.4e-07 | 3.1e-07 | 2.867E-07 | 1.365E-07 | 1.400E-07 | 4.100E-07 |
| 2430 | 122 | S122I | A | 2.4e-07 | 2.8e-07 | 2.3e-07 | 2.500E-07 | 2.646E-08 | 2.300E-07 | 2.800E-07 |
| 2431 | 122 | S122L | A | 1.9e-07 | 2.9e-07 | 3.3e-07 | 2.700E-07 | 7.211E-08 | 1.900E-07 | 3.300E-07 |
| 2432 | 122 | S122K | A | 4.6e-07 | 3.2e-07 | 1.6e-07 | 3.133E-07 | 1.501E-07 | 1.600E-07 | 4.600E-07 |
| 2433 | 122 | S122M | A | 2.8e-07 | 2.7e-07 | 2.4e-07 | 2.633E-07 | 2.082E-08 | 2.400E-07 | 2.800E-07 |
| 2434 | 122 | S122F | A | 3.5e-07 | 2.1e-07 | 3e-07 | 2.867E-07 | 7.095E-08 | 2.100E-07 | 3.500E-07 |
| 2435 | 122 | S122P | A | 3.4e-07 | 2.9e-07 | 1.7e-07 | 2.667E-07 | 8.737E-08 | 1.700E-07 | 3.400E-07 |
| 2436 | 122 | S122S | A | 3.2e-07 | 2.7e-07 | 1.1e-07 | 2.333E-07 | 1.097E-07 | 1.100E-07 | 3.200E-07 |
| 2437 | 122 | S122T | A | 1.6e-07 | 2.4e-07 | 2.5e-07 | 2.167E-07 | 4.933E-08 | 1.600E-07 | 2.500E-07 |
| 2438 | 122 | S122W | A | 3.4e-07 | 2.5e-07 | 4.1e-07 | 3.333E-07 | 8.021E-08 | 2.500E-07 | 4.100E-07 |
| 2439 | 122 | S122Y | A | 2.6e-07 | 3.1e-07 | 3.4e-07 | 3.033E-07 | 4.041E-08 | 2.600E-07 | 3.400E-07 |
| 2440 | 122 | S122V | A | 3e-07 | 1.6e-07 | 1.9e-07 | 2.167E-07 | 7.371E-08 | 1.600E-07 | 3.000E-07 |
| 2441 | 123 | L123A | A | 6.7e-07 | 3e-07 | 1.7e-07 | 3.800E-07 | 2.594E-07 | 1.700E-07 | 6.700E-07 |
| 2442 | 123 | L123R | A | 1.4e-07 | 2.5e-07 | 2.5e-07 | 2.133E-07 | 6.351E-08 | 1.400E-07 | 2.500E-07 |
| 2443 | 123 | L123N | A | 9.7e-08 | 2.8e-07 | 2.1e-07 | 1.957E-07 | 9.234E-08 | 9.700E-08 | 2.800E-07 |
| 2444 | 123 | L123D | A | 2.9e-07 | 2.2e-07 | 2.2e-07 | 2.433E-07 | 4.041E-08 | 2.200E-07 | 2.900E-07 |
| 2445 | 123 | L123C | A | 1.8e-07 | 3.8e-07 | 4.1e-07 | 3.233E-07 | 1.250E-07 | 1.800E-07 | 4.100E-07 |
| 2446 | 123 | L123E | A | 3.2e-07 | 2.3e-07 | 1.7e-07 | 2.400E-07 | 7.550E-08 | 1.700E-07 | 3.200E-07 |
| 2447 | 123 | L123Q | A | 3.4e-07 | 4.5e-07 | 1.3e-07 | 3.067E-07 | 1.626E-07 | 1.300E-07 | 4.500E-07 |
| 2448 | 123 | L123G | A | 2.5e-07 | 2.8e-07 | 4.7e-07 | 3.333E-07 | 1.193E-07 | 2.500E-07 | 4.700E-07 |
| 2449 | 123 | L123H | A | 1.5e-07 | 5.7e-07 | 2.2e-07 | 3.133E-07 | 2.250E-07 | 1.500E-07 | 5.700E-07 |
| 2450 | 123 | L123I | A | 3.6e-07 | 3.1e-07 | 3.8e-07 | 3.500E-07 | 3.606E-08 | 3.100E-07 | 3.800E-07 |
| 2451 | 123 | L123L | A | 3.9e-07 | 3e-07 | 2.8e-07 | 3.233E-07 | 5.859E-08 | 2.800E-07 | 3.900E-07 |
| 2452 | 123 | L123K | A | 4.1e-07 | 1.4e-07 | 1.7e-07 | 2.400E-07 | 1.480E-07 | 1.400E-07 | 4.100E-07 |
| 2453 | 123 | L123M | A | 1.8e-07 | 2.4e-07 | 1.1e-07 | 1.767E-07 | 6.506E-08 | 1.100E-07 | 2.400E-07 |
| 2454 | 123 | L123F | A | 1.3e-07 | 1.3e-07 | 2.6e-07 | 1.733E-07 | 7.506E-08 | 1.300E-07 | 2.600E-07 |
| 2455 | 123 | L123P | A | 3.4e-07 | 7.1e-07 | 1.8e-07 | 4.100E-07 | 2.718E-07 | 1.800E-07 | 7.100E-07 |
| 2456 | 123 | L123S | A | 5.4e-07 | 5.2e-07 | 1.1e-07 | 3.900E-07 | 2.427E-07 | 1.100E-07 | 5.400E-07 |
| 2457 | 123 | L123T | A | 1.6e-07 | 4e-07 | 2.3e-07 | 2.633E-07 | 1.234E-07 | 1.600E-07 | 4.000E-07 |
| 2458 | 123 | L123W | A | 1.9e-07 | 2.3e-07 | 2.8e-07 | 2.333E-07 | 4.509E-08 | 1.900E-07 | 2.800E-07 |
| 2459 | 123 | L123Y | A | 6.4e-07 | 2e-07 | 2.3e-07 | 3.567E-07 | 2.458E-07 | 2.000E-07 | 6.400E-07 |
| 2460 | 123 | L123V | A | 1.7e-07 | 2.3e-07 | 2.8e-07 | 2.267E-07 | 5.508E-08 | 1.700E-07 | 2.800E-07 |
| 2461 | 124 | T124A | A | 2.7e-07 | 5.4e-07 | 3.2e-07 | 3.767E-07 | 1.436E-07 | 2.700E-07 | 5.400E-07 |
| 2462 | 124 | T124R | A | 2.9e-07 | 2.8e-07 | 2.4e-07 | 2.700E-07 | 2.646E-08 | 2.400E-07 | 2.900E-07 |
| 2463 | 124 | T124N | A | 2.3e-07 | 4.6e-07 | 2.9e-07 | 3.267E-07 | 1.193E-07 | 2.300E-07 | 4.600E-07 |
| 2464 | 124 | T124D | A | 2.1e-07 | 2.7e-07 | 2.2e-07 | 2.333E-07 | 3.215E-08 | 2.100E-07 | 2.700E-07 |

| | | | | | | | | | | |
|---|---|---|---|---|---|---|---|---|---|---|
| 2465 | 124 | T124C | A | 2.3e-07 | 3.7e-07 | 3e-07 | 3.000E-07 | 7.000E-08 | 2.300E-07 | 3.700E-07 |
| 2466 | 124 | T124E | A | 3.5e-07 | 2.3e-07 | 2.1e-07 | 2.633E-07 | 7.572E-08 | 2.100E-07 | 3.500E-07 |
| 2467 | 124 | T124Q | A | 3.7e-07 | 2.5e-07 | 2.9e-07 | 3.033E-07 | 6.110E-08 | 2.500E-07 | 3.700E-07 |
| 2468 | 124 | T124G | A | 3.5e-07 | 6.7e-07 | 1.9e-07 | 4.033E-07 | 2.444E-07 | 1.900E-07 | 6.700E-07 |
| 2469 | 124 | T124H | A | 6.7e-07 | 3e-07 | 2.1e-07 | 3.933E-07 | 2.438E-07 | 2.100E-07 | 6.700E-07 |
| 2470 | 124 | T124I | A | 1.9e-07 | 1.8e-07 | 2.6e-07 | 2.100E-07 | 4.359E-08 | 1.800E-07 | 2.600E-07 |
| 2471 | 124 | T124L | A | 1.8e-07 | 3.1e-07 | 3.4e-07 | 2.767E-07 | 8.505E-08 | 1.800E-07 | 3.400E-07 |
| 2472 | 124 | T124K | A | 1.6e-07 | 1.6e-07 | 1.3e-07 | 1.500E-07 | 1.732E-08 | 1.300E-07 | 1.600E-07 |
| 2473 | 124 | T124M | A | 3.6e-07 | 1e-07 | 3.5e-07 | 2.700E-07 | 1.473E-07 | 1.000E-07 | 3.600E-07 |
| 2474 | 124 | T124F | A | 3.3e-07 | 1.6e-07 | 2.8e-07 | 2.567E-07 | 8.737E-08 | 1.600E-07 | 3.300E-07 |
| 2475 | 124 | T124P | A | 1.2e-07 | 3.5e-07 | 2.6e-07 | 2.433E-07 | 1.159E-07 | 1.200E-07 | 3.500E-07 |
| 2476 | 124 | T124S | A | 1.7e-07 | 1.9e-07 | 3.5e-07 | 2.367E-07 | 9.866E-08 | 1.700E-07 | 3.500E-07 |
| 2477 | 124 | T124T | A | 1.4e-07 | 2.6e-07 | 3.1e-07 | 2.367E-07 | 8.737E-08 | 1.400E-07 | 3.100E-07 |
| 2478 | 124 | T124W | A | 2.8e-07 | 3.2e-07 | 3.1e-07 | 3.033E-07 | 2.082E-08 | 2.800E-07 | 3.200E-07 |
| 2479 | 124 | T124Y | A | 3.4e-07 | 1.9e-07 | 2.5e-07 | 2.600E-07 | 7.550E-08 | 1.900E-07 | 3.400E-07 |
| 2480 | 124 | T124V | A | 3.7e-07 | 2.1e-07 | 3.2e-07 | 3.000E-07 | 8.185E-08 | 2.100E-07 | 3.700E-07 |
| 2481 | 125 | E125A | A | 4.5e-07 | 1.6e-07 | 3.3e-07 | 3.133E-07 | 1.457E-07 | 1.600E-07 | 4.500E-07 |
| 2482 | 125 | E125R | A | 3e-07 | 1.9e-07 | 2.3e-07 | 2.400E-07 | 5.568E-08 | 1.900E-07 | 3.000E-07 |
| 2483 | 125 | E125N | A | 3.2e-07 | 3.3e-07 | 2.8e-07 | 3.100E-07 | 2.646E-08 | 2.800E-07 | 3.300E-07 |
| 2484 | 125 | E125D | A | 1.1e-07 | 1.9e-07 | 3.9e-07 | 2.300E-07 | 1.442E-07 | 1.100E-07 | 3.900E-07 |
| 2485 | 125 | E125C | A | 3.4e-07 | 2.5e-07 | 2.2e-07 | 2.700E-07 | 6.245E-08 | 2.200E-07 | 3.400E-07 |
| 2486 | 125 | E125E | A | 1.9e-07 | 1.2e-07 | 3.6e-07 | 2.233E-07 | 1.234E-07 | 1.200E-07 | 3.600E-07 |
| 2487 | 125 | E125Q | A | 2.2e-07 | 2.8e-07 | 2.3e-07 | 2.433E-07 | 3.215E-08 | 2.200E-07 | 2.800E-07 |
| 2488 | 125 | E125G | A | 3.5e-07 | 2.9e-07 | 2.2e-07 | 2.867E-07 | 6.506E-08 | 2.200E-07 | 3.500E-07 |
| 2489 | 125 | E125H | A | 3.7e-07 | 3.1e-07 | 2.3e-07 | 3.033E-07 | 7.024E-08 | 2.300E-07 | 3.700E-07 |
| 2490 | 125 | E125I | A | 1.9e-07 | 3.5e-07 | 3.1e-07 | 2.833E-07 | 8.327E-08 | 1.900E-07 | 3.500E-07 |
| 2491 | 125 | E125L | A | 2.1e-07 | 2.2e-07 | 2e-07 | 2.100E-07 | 1.000E-08 | 2.000E-07 | 2.200E-07 |
| 2492 | 125 | E125K | A | 2e-07 | 4e-07 | 3.1e-07 | 3.033E-07 | 1.002E-07 | 2.000E-07 | 4.000E-07 |
| 2493 | 125 | E125M | A | 1.4e-07 | 2.5e-07 | 2.4e-07 | 2.100E-07 | 6.083E-08 | 1.400E-07 | 2.500E-07 |
| 2494 | 125 | E125F | A | 2.5e-07 | 1.9e-07 | 1.8e-07 | 2.067E-07 | 3.786E-08 | 1.800E-07 | 2.500E-07 |
| 2495 | 125 | E125P | A | 1.7e-07 | 6.7e-07 | 5.3e-07 | 4.567E-07 | 2.579E-07 | 1.700E-07 | 6.700E-07 |
| 2496 | 125 | E125S | A | 1e-07 | 2.4e-07 | 3e-07 | 2.133E-07 | 1.026E-07 | 1.000E-07 | 3.000E-07 |
| 2497 | 125 | E125T | A | 2.8e-07 | 1.9e-07 | 2e-07 | 2.233E-07 | 4.933E-08 | 1.900E-07 | 2.800E-07 |
| 2498 | 125 | E125W | A | 1.5e-07 | 1.1e-07 | 2.3e-07 | 1.633E-07 | 6.110E-08 | 1.100E-07 | 2.300E-07 |
| 2499 | 125 | E125Y | A | 2.5e-07 | 1.9e-07 | 2.8e-07 | 2.400E-07 | 4.583E-08 | 1.900E-07 | 2.800E-07 |
| 2500 | 125 | E125V | A | 1.2e-07 | 1.9e-07 | 2.6e-07 | 1.900E-07 | 7.000E-08 | 1.200E-07 | 2.600E-07 |
| 2501 | 126 | I126A | A | 3.4e-07 | 1.5e-07 | 1.3e-07 | 2.067E-07 | 1.159E-07 | 1.300E-07 | 3.400E-07 |
| 2502 | 126 | I126R | A | 1.4e-07 | 2.4e-07 | 1.8e-07 | 1.867E-07 | 5.033E-08 | 1.400E-07 | 2.400E-07 |
| 2503 | 126 | I126N | A | 2.9e-07 | 3.1e-07 | 1.5e-07 | 2.500E-07 | 8.718E-08 | 1.500E-07 | 3.100E-07 |
| 2504 | 126 | I126D | A | 2.7e-07 | 5.4e-07 | 1.7e-07 | 3.267E-07 | 1.914E-07 | 1.700E-07 | 5.400E-07 |
| 2505 | 126 | I126C | A | 1.3e-07 | 4.2e-07 | 1.7e-07 | 2.400E-07 | 1.572E-07 | 1.300E-07 | 4.200E-07 |
| 2506 | 126 | I126E | A | 7e-07 | 3.4e-07 | 1.3e-07 | 3.900E-07 | 2.883E-07 | 1.300E-07 | 7.000E-07 |
| 2507 | 126 | I126Q | A | 2.7e-07 | 9.5e-08 | 4.1e-07 | 2.583E-07 | 1.578E-07 | 9.500E-08 | 4.100E-07 |
| 2508 | 126 | I126G | A | 1.8e-07 | 3.1e-07 | 1.8e-07 | 2.233E-07 | 7.506E-08 | 1.800E-07 | 3.100E-07 |
| 2509 | 126 | I126H | A | 2.4e-07 | 1e-07 | 4.9e-07 | 2.767E-07 | 1.976E-07 | 1.000E-07 | 4.900E-07 |
| 2510 | 126 | I126I | A | 2.7e-07 | 1.6e-07 | 1.6e-07 | 1.967E-07 | 6.351E-08 | 1.600E-07 | 2.700E-07 |
| 2511 | 126 | I126L | A | 2.4e-07 | 2.5e-07 | 5e-07 | 3.300E-07 | 1.473E-07 | 2.400E-07 | 5.000E-07 |
| 2512 | 126 | I126K | A | 2.3e-07 | 2.6e-07 | 1.7e-07 | 2.200E-07 | 4.583E-08 | 1.700E-07 | 2.600E-07 |
| 2513 | 126 | I126M | A | 2.1e-07 | 3.6e-07 | 3.3e-07 | 3.000E-07 | 7.937E-08 | 2.100E-07 | 3.600E-07 |
| 2514 | 126 | I126F | A | 2.5e-07 | 2.4e-07 | 3.1e-07 | 2.667E-07 | 3.786E-08 | 2.400E-07 | 3.100E-07 |
| 2515 | 126 | I126P | A | 4.1e-07 | 1.3e-07 | 1.6e-07 | 2.333E-07 | 1.537E-07 | 1.300E-07 | 4.100E-07 |
| 2516 | 126 | I126S | A | 2.7e-07 | 3.3e-07 | 3.4e-07 | 3.133E-07 | 3.786E-08 | 2.700E-07 | 3.400E-07 |
| 2517 | 126 | I126T | A | 2.7e-07 | 2e-07 | 1.8e-07 | 2.167E-07 | 4.726E-08 | 1.800E-07 | 2.700E-07 |
| 2518 | 126 | I126W | A | 1.8e-07 | 2.2e-07 | 3.7e-07 | 2.567E-07 | 1.002E-07 | 1.800E-07 | 3.700E-07 |
| 2519 | 126 | I126Y | A | 2.2e-07 | 2.6e-07 | 4e-07 | 2.933E-07 | 9.452E-08 | 2.200E-07 | 4.000E-07 |
| 2520 | 126 | I126V | A | 4.2e-07 | 1.1e-07 | 1.3e-07 | 2.200E-07 | 1.735E-07 | 1.100E-07 | 4.200E-07 |
| 2521 | 127 | L127A | A | 2.9e-07 | 1.7e-07 | 6.3e-08 | 1.743E-07 | 1.136E-07 | 6.300E-08 | 2.900E-07 |
| 2522 | 127 | L127R | A | 2.8e-07 | 1.8e-07 | 2.4e-07 | 2.333E-07 | 5.033E-08 | 1.800E-07 | 2.800E-07 |
| 2523 | 127 | L127N | A | 4.7e-07 | 1.7e-07 | 1.9e-07 | 2.767E-07 | 1.677E-07 | 1.700E-07 | 4.700E-07 |
| 2524 | 127 | L127D | A | 2e-07 | 3.1e-07 | 5.2e-07 | 3.433E-07 | 1.626E-07 | 2.000E-07 | 5.200E-07 |
| 2525 | 127 | L127C | A | 1.8e-07 | 4.6e-07 | 6.2e-07 | 4.200E-07 | 2.227E-07 | 1.800E-07 | 6.200E-07 |
| 2526 | 127 | L127E | A | 2.7e-07 | 4.4e-07 | 1.7e-07 | 2.933E-07 | 1.365E-07 | 1.700E-07 | 4.400E-07 |
| 2527 | 127 | L127Q | A | 3.7e-07 | 1.3e-07 | 3.8e-07 | 2.933E-07 | 1.415E-07 | 1.300E-07 | 3.800E-07 |
| 2528 | 127 | L127G | A | 2.8e-07 | 3e-07 | 3e-07 | 2.933E-07 | 1.155E-08 | 2.800E-07 | 3.000E-07 |
| 2529 | 127 | L127H | A | 1.9e-07 | 2.1e-07 | 2.3e-07 | 2.100E-07 | 2.000E-08 | 1.900E-07 | 2.300E-07 |
| 2530 | 127 | L127I | A | 4.8e-07 | 3.5e-07 | 1.5e-07 | 3.267E-07 | 1.662E-07 | 1.500E-07 | 4.800E-07 |
| 2531 | 127 | L127L | A | 1.2e-07 | 1.8e-07 | 2.5e-07 | 1.833E-07 | 6.506E-08 | 1.200E-07 | 2.500E-07 |
| 2532 | 127 | L127K | A | 2.3e-07 | 3e-07 | 2.1e-07 | 2.467E-07 | 4.726E-08 | 2.100E-07 | 3.000E-07 |
| 2533 | 127 | L127M | A | 2.4e-07 | 5.6e-07 | 2.2e-07 | 3.400E-07 | 1.908E-07 | 2.200E-07 | 5.600E-07 |
| 2534 | 127 | L127F | A | 5e-07 | 1.5e-07 | 3.2e-07 | 3.233E-07 | 1.750E-07 | 1.500E-07 | 5.000E-07 |
| 2535 | 127 | L127P | A | 3.8e-07 | 1.3e-07 | 7.6e-07 | 4.233E-07 | 3.172E-07 | 1.300E-07 | 7.600E-07 |
| 2536 | 127 | L127S | A | 1.5e-07 | 3e-07 | 1.3e-07 | 1.933E-07 | 9.292E-08 | 1.300E-07 | 3.000E-07 |
| 2537 | 127 | L127T | A | 4e-07 | 6.7e-07 | 2.7e-07 | 4.467E-07 | 2.040E-07 | 2.700E-07 | 6.700E-07 |
| 2538 | 127 | L127W | A | 3.2e-07 | 3e-07 | 1.5e-07 | 2.567E-07 | 9.292E-08 | 1.500E-07 | 3.200E-07 |
| 2539 | 127 | L127Y | A | 2e-07 | 2.1e-07 | 2.4e-07 | 2.167E-07 | 2.082E-08 | 2.000E-07 | 2.400E-07 |
| 2540 | 127 | L127V | A | 3.9e-07 | 2.3e-07 | 1.4e-07 | 2.533E-07 | 1.266E-07 | 1.400E-07 | 3.900E-07 |
| 2541 | 128 | K128A | A | 3.6e-07 | 1.4e-07 | 7.4e-07 | 4.133E-07 | 3.035E-07 | 1.400E-07 | 7.400E-07 |
| 2542 | 128 | K128R | A | 2.5e-07 | 1.8e-07 | 5.5e-07 | 3.267E-07 | 1.966E-07 | 1.800E-07 | 5.500E-07 |
| 2543 | 128 | K128N | A | 3.1e-07 | 2.6e-07 | 2.8e-07 | 2.833E-07 | 2.517E-08 | 2.600E-07 | 3.100E-07 |
| 2544 | 128 | K128D | A | 3e-07 | 2.5e-07 | 2.6e-07 | 2.700E-07 | 2.646E-08 | 2.500E-07 | 3.000E-07 |
| 2545 | 128 | K128C | A | 1.7e-07 | 2.8e-07 | 3.5e-07 | 2.667E-07 | 9.074E-08 | 1.700E-07 | 3.500E-07 |
| 2546 | 128 | K128E | A | 3.3e-07 | 3e-07 | 2.7e-07 | 3.000E-07 | 3.000E-08 | 2.700E-07 | 3.300E-07 |
| 2547 | 128 | K128Q | A | 2.5e-07 | 2.1e-07 | 2.1e-07 | 2.233E-07 | 2.309E-08 | 2.100E-07 | 2.500E-07 |
| 2548 | 128 | K128G | A | 1.9e-07 | 2.4e-07 | 2.7e-07 | 2.333E-07 | 4.041E-08 | 1.900E-07 | 2.700E-07 |
| 2549 | 128 | K128H | A | 3.4e-07 | 1.4e-07 | 3.6e-07 | 2.800E-07 | 1.217E-07 | 1.400E-07 | 3.600E-07 |

| | | | | | | | | | | |
|---|---|---|---|---|---|---|---|---|---|---|
| 2550 | 128 | K128I | A | 2.2e-07 | 8.4e-07 | 2.2e-07 | 4.267E-07 | 3.580E-07 | 2.200E-07 | 8.400E-07 |
| 2551 | 128 | K128L | A | 3.1e-07 | 2.4e-07 | 3.2e-07 | 2.900E-07 | 4.359E-08 | 2.400E-07 | 3.200E-07 |
| 2552 | 128 | K128K | A | 3.3e-07 | 2.2e-07 | 1.8e-07 | 2.433E-07 | 7.767E-08 | 1.800E-07 | 3.300E-07 |
| 2553 | 128 | K128M | A | 2.6e-07 | 3.6e-07 | 2.7e-07 | 2.967E-07 | 5.508E-08 | 2.600E-07 | 3.600E-07 |
| 2554 | 128 | K128F | A | 1.8e-07 | 1.4e-07 | 3.8e-07 | 2.333E-07 | 1.286E-07 | 1.400E-07 | 3.800E-07 |
| 2555 | 128 | K128P | A | 3.2e-07 | 3.1e-07 | 1.7e-07 | 2.667E-07 | 8.386E-08 | 1.700E-07 | 3.200E-07 |
| 2556 | 128 | K128S | A | 2.4e-07 | 1.4e-07 | 3.2e-07 | 2.333E-07 | 9.018E-08 | 1.400E-07 | 3.200E-07 |
| 2557 | 128 | K128T | A | 3.4e-07 | 4.5e-07 | 3.6e-07 | 3.833E-07 | 5.859E-08 | 3.400E-07 | 4.500E-07 |
| 2558 | 128 | K128W | A | 2.3e-07 | 3.3e-07 | 1.9e-07 | 2.500E-07 | 7.211E-08 | 1.900E-07 | 3.300E-07 |
| 2559 | 128 | K128Y | A | 1.7e-07 | 1.5e-07 | 1.6e-07 | 1.600E-07 | 1.000E-08 | 1.500E-07 | 1.700E-07 |
| 2560 | 128 | K128V | A | 2.8e-07 | 2.6e-07 | 3.5e-07 | 2.967E-07 | 4.726E-08 | 2.600E-07 | 3.500E-07 |
| 2561 | 129 | G129A | A | 1.2e-07 | 1.1e-07 | 2.1e-07 | 1.467E-07 | 5.508E-08 | 1.100E-07 | 2.100E-07 |
| 2562 | 129 | G129R | A | 2.7e-07 | 1.7e-07 | 2.4e-07 | 2.267E-07 | 5.132E-08 | 1.700E-07 | 2.700E-07 |
| 2563 | 129 | G129N | A | 5.2e-07 | 2.7e-07 | 4.6e-07 | 4.167E-07 | 1.305E-07 | 2.700E-07 | 5.200E-07 |
| 2564 | 129 | G129D | A | 2.6e-07 | 2.8e-07 | 3e-07 | 2.800E-07 | 2.000E-08 | 2.600E-07 | 3.000E-07 |
| 2565 | 129 | G129C | A | 2.5e-07 | 5.1e-07 | 2.5e-07 | 3.367E-07 | 1.501E-07 | 2.500E-07 | 5.100E-07 |
| 2566 | 129 | G129E | A | 3.6e-07 | 2.3e-07 | 3.3e-07 | 3.067E-07 | 6.807E-08 | 2.300E-07 | 3.600E-07 |
| 2567 | 129 | G129Q | A | 3e-07 | 3e-07 | 2.4e-07 | 2.800E-07 | 3.464E-08 | 2.400E-07 | 3.000E-07 |
| 2568 | 129 | G129G | A | 2.5e-07 | 3.7e-07 | 4.4e-07 | 3.533E-07 | 9.609E-08 | 2.500E-07 | 4.400E-07 |
| 2569 | 129 | G129H | A | 3.9e-07 | 8.7e-08 | 7.5e-08 | 1.840E-07 | 1.785E-07 | 7.500E-08 | 3.900E-07 |
| 2570 | 129 | G129I | A | 5.5e-07 | 2.8e-07 | 1.8e-07 | 3.367E-07 | 1.914E-07 | 1.800E-07 | 5.500E-07 |
| 2571 | 129 | G129L | A | 3e-07 | 2.9e-07 | 2.4e-07 | 2.767E-07 | 3.215E-08 | 2.400E-07 | 3.000E-07 |
| 2572 | 129 | G129K | A | 1.7e-07 | 2.9e-07 | 4.4e-07 | 3.000E-07 | 1.353E-07 | 1.700E-07 | 4.400E-07 |
| 2573 | 129 | G129M | A | 1.9e-07 | 1.9e-07 | 2.7e-07 | 2.167E-07 | 4.619E-08 | 1.900E-07 | 2.700E-07 |
| 2574 | 129 | G129F | A | 1.8e-07 | 9.2e-07 | 2.4e-07 | 4.467E-07 | 4.110E-07 | 1.800E-07 | 9.200E-07 |
| 2575 | 129 | G129P | A | 3.3e-07 | 2.8e-07 | 1.1e-07 | 2.400E-07 | 1.153E-07 | 1.100E-07 | 3.300E-07 |
| 2576 | 129 | G129S | A | 3.6e-07 | 2.4e-07 | 2.3e-07 | 2.767E-07 | 7.234E-08 | 2.300E-07 | 3.600E-07 |
| 2577 | 129 | G129T | A | 2.6e-07 | 3.2e-07 | 2.3e-07 | 2.700E-07 | 4.583E-08 | 2.300E-07 | 3.200E-07 |
| 2578 | 129 | G129W | A | 2.9e-07 | 5.3e-07 | 3e-07 | 3.733E-07 | 1.358E-07 | 2.900E-07 | 5.300E-07 |
| 2579 | 129 | G129Y | A | 3e-07 | 1.1e-07 | 3.2e-07 | 2.433E-07 | 1.159E-07 | 1.100E-07 | 3.200E-07 |
| 2580 | 129 | G129V | A | 3.2e-07 | 1.8e-07 | 2.2e-07 | 2.400E-07 | 7.211E-08 | 1.800E-07 | 3.200E-07 |
| 2581 | 130 | G130A | A | 2.2e-07 | 4.8e-07 | 1.4e-07 | 2.800E-07 | 1.778E-07 | 1.400E-07 | 4.800E-07 |
| 2582 | 130 | G130R | A | 1.3e-07 | 1.3e-07 | 4.3e-07 | 2.300E-07 | 1.732E-07 | 1.300E-07 | 4.300E-07 |
| 2583 | 130 | G130N | A | 1.9e-07 | 3.3e-07 | 2.7e-07 | 2.633E-07 | 7.024E-08 | 1.900E-07 | 3.300E-07 |
| 2584 | 130 | G130D | A | 6.8e-07 | 3.8e-07 | 3.3e-07 | 4.633E-07 | 1.893E-07 | 3.300E-07 | 6.800E-07 |
| 2585 | 130 | G130C | A | 5.6e-07 | 1.6e-07 | 2.5e-07 | 3.233E-07 | 2.098E-07 | 1.600E-07 | 5.600E-07 |
| 2586 | 130 | G130E | A | 3.1e-07 | 1.8e-07 | 3.3e-07 | 2.733E-07 | 8.145E-08 | 1.800E-07 | 3.300E-07 |
| 2587 | 130 | G130Q | A | 5.9e-07 | 2.2e-07 | 2.8e-07 | 3.633E-07 | 1.986E-07 | 2.200E-07 | 5.900E-07 |
| 2588 | 130 | G130G | A | 4.3e-07 | 2.8e-07 | 2.1e-07 | 3.067E-07 | 1.124E-07 | 2.100E-07 | 4.300E-07 |
| 2589 | 130 | G130H | A | 2.5e-07 | 1.9e-07 | 2.1e-07 | 2.167E-07 | 3.055E-08 | 1.900E-07 | 2.500E-07 |
| 2590 | 130 | G130I | A | 3.6e-07 | 5e-07 | 2.9e-07 | 3.833E-07 | 1.069E-07 | 2.900E-07 | 5.000E-07 |
| 2591 | 130 | G130L | A | 3.1e-07 | 2.5e-07 | 4.4e-07 | 3.333E-07 | 9.713E-08 | 2.500E-07 | 4.400E-07 |
| 2592 | 130 | G130K | A | 3.2e-07 | 1.9e-07 | 1.8e-07 | 2.300E-07 | 7.810E-08 | 1.800E-07 | 3.200E-07 |
| 2593 | 130 | G130M | A | 3.2e-07 | 1.6e-07 | 2.9e-07 | 2.567E-07 | 8.505E-08 | 1.600E-07 | 3.200E-07 |
| 2594 | 130 | G130F | A | 1.5e-07 | 3.5e-07 | 3.4e-07 | 2.800E-07 | 1.127E-07 | 1.500E-07 | 3.500E-07 |
| 2595 | 130 | G130P | A | 1.3e-07 | 2.5e-07 | 3e-07 | 2.267E-07 | 8.737E-08 | 1.300E-07 | 3.000E-07 |
| 2596 | 130 | G130S | A | 1.4e-07 | 3.6e-07 | 3.7e-07 | 2.900E-07 | 1.300E-07 | 1.400E-07 | 3.700E-07 |
| 2597 | 130 | G130T | A | 4.6e-07 | 4e-07 | 4.4e-07 | 4.333E-07 | 3.055E-08 | 4.000E-07 | 4.600E-07 |
| 2598 | 130 | G130W | A | 1.7e-07 | 2.4e-07 | 2.5e-07 | 2.200E-07 | 4.359E-08 | 1.700E-07 | 2.500E-07 |
| 2599 | 130 | G130Y | A | 1.9e-07 | 2.3e-07 | 2.3e-07 | 2.167E-07 | 2.309E-08 | 1.900E-07 | 2.300E-07 |
| 2600 | 130 | G130V | A | 1.8e-07 | 1.4e-07 | 2.5e-07 | 1.900E-07 | 5.568E-08 | 1.400E-07 | 2.500E-07 |
| 2601 | 131 | V131A | A | 2.2e-07 | 1.8e-07 | 4.5e-07 | 2.833E-07 | 1.457E-07 | 1.800E-07 | 4.500E-07 |
| 2602 | 131 | V131R | A | 2.5e-07 | 2.1e-07 | 1.4e-07 | 2.000E-07 | 5.568E-08 | 1.400E-07 | 2.500E-07 |
| 2603 | 131 | V131N | A | 1.5e-07 | 2.2e-07 | 8.9e-07 | 4.200E-07 | 4.085E-07 | 1.500E-07 | 8.900E-07 |
| 2604 | 131 | V131D | A | 2.4e-07 | 1.2e-07 | 2.5e-07 | 2.033E-07 | 7.234E-08 | 1.200E-07 | 2.500E-07 |
| 2605 | 131 | V131C | A | 1.5e-07 | 6.9e-07 | 2.5e-07 | 3.633E-07 | 2.873E-07 | 1.500E-07 | 6.900E-07 |
| 2606 | 131 | V131E | A | 2.6e-07 | 2.6e-07 | 3.2e-07 | 2.800E-07 | 3.464E-08 | 2.600E-07 | 3.200E-07 |
| 2607 | 131 | V131Q | A | 2e-07 | 3.2e-07 | 2.2e-07 | 2.467E-07 | 6.429E-08 | 2.000E-07 | 3.200E-07 |
| 2608 | 131 | V131G | A | 9.8e-08 | 1.9e-07 | 3.3e-07 | 2.060E-07 | 1.168E-07 | 9.800E-08 | 3.300E-07 |
| 2609 | 131 | V131H | A | 1.7e-07 | 2.4e-07 | 2.8e-07 | 2.300E-07 | 5.568E-08 | 1.700E-07 | 2.800E-07 |
| 2610 | 131 | V131I | A | 1.6e-07 | 4.3e-07 | 3e-07 | 2.967E-07 | 1.350E-07 | 1.600E-07 | 4.300E-07 |
| 2611 | 131 | V131L | A | 3.1e-07 | 4.1e-07 | 2.4e-07 | 3.200E-07 | 8.544E-08 | 2.400E-07 | 4.100E-07 |
| 2612 | 131 | V131K | A | 2.6e-07 | 5.2e-07 | 3.3e-07 | 3.700E-07 | 1.345E-07 | 2.600E-07 | 5.200E-07 |
| 2613 | 131 | V131M | A | 1.9e-07 | 3.9e-07 | 2.1e-07 | 2.633E-07 | 1.102E-07 | 1.900E-07 | 3.900E-07 |
| 2614 | 131 | V131F | A | 2.3e-07 | 1.6e-07 | 2.4e-07 | 2.100E-07 | 4.359E-08 | 1.600E-07 | 2.400E-07 |
| 2615 | 131 | V131P | A | 1.5e-07 | 2.5e-07 | 2.1e-07 | 2.033E-07 | 5.033E-08 | 1.500E-07 | 2.500E-07 |
| 2616 | 131 | V131S | A | 2.4e-07 | 1.6e-07 | 1.5e-07 | 1.833E-07 | 4.933E-08 | 1.500E-07 | 2.400E-07 |
| 2617 | 131 | V131T | A | 5.1e-07 | 3.3e-07 | 2.2e-07 | 3.533E-07 | 1.464E-07 | 2.200E-07 | 5.100E-07 |
| 2618 | 131 | V131W | A | 2.6e-07 | 2.2e-07 | 1.5e-07 | 2.100E-07 | 5.568E-08 | 1.500E-07 | 2.600E-07 |
| 2619 | 131 | V131Y | A | 2.4e-07 | 1.3e-07 | 9.8e-08 | 1.560E-07 | 7.448E-08 | 9.800E-08 | 2.400E-07 |
| 2620 | 131 | V131V | A | 3.1e-07 | 3.1e-07 | 2.1e-07 | 2.767E-07 | 5.774E-08 | 2.100E-07 | 3.100E-07 |
| 2621 | 132 | L132A | A | 3.1e-07 | 7e-08 | 5.1e-07 | 2.967E-07 | 2.203E-07 | 7.000E-08 | 5.100E-07 |
| 2622 | 132 | L132R | A | 4.9e-07 | 2e-07 | 2.7e-07 | 3.200E-07 | 1.513E-07 | 2.000E-07 | 4.900E-07 |
| 2623 | 132 | L132N | A | 2.7e-07 | 1.9e-07 | 1.2e-07 | 1.933E-07 | 7.506E-08 | 1.200E-07 | 2.700E-07 |
| 2624 | 132 | L132D | A | 1.4e-07 | 1.3e-07 | 2.3e-07 | 1.667E-07 | 5.508E-08 | 1.300E-07 | 2.300E-07 |
| 2625 | 132 | L132C | A | 3.4e-07 | 3.5e-07 | 2.4e-07 | 3.100E-07 | 6.083E-08 | 2.400E-07 | 3.500E-07 |
| 2626 | 132 | L132E | A | 3.6e-07 | 1.9e-07 | 1.9e-07 | 2.467E-07 | 9.815E-08 | 1.900E-07 | 3.600E-07 |
| 2627 | 132 | L132Q | A | 1.2e-07 | 3e-07 | 2.5e-07 | 2.233E-07 | 9.292E-08 | 1.200E-07 | 3.000E-07 |
| 2628 | 132 | L132G | A | 2.9e-07 | 3.1e-07 | 5e-07 | 3.667E-07 | 1.159E-07 | 2.900E-07 | 5.000E-07 |
| 2629 | 132 | L132H | A | 2e-07 | 3.2e-07 | 4.9e-07 | 3.367E-07 | 1.457E-07 | 2.000E-07 | 4.900E-07 |
| 2630 | 132 | L132I | A | 3.9e-07 | 5.1e-07 | 4.4e-07 | 4.467E-07 | 6.028E-08 | 3.900E-07 | 5.100E-07 |
| 2631 | 132 | L132L | A | 1.1e-07 | 1.6e-07 | 1.4e-07 | 1.367E-07 | 2.517E-08 | 1.100E-07 | 1.600E-07 |
| 2632 | 132 | L132K | A | 4e-07 | 1.8e-07 | 1.9e-07 | 2.567E-07 | 1.242E-07 | 1.800E-07 | 4.000E-07 |
| 2633 | 132 | L132M | A | 3.1e-07 | 2.4e-07 | 3.3e-07 | 2.933E-07 | 4.726E-08 | 2.400E-07 | 3.300E-07 |
| 2634 | 132 | L132F | A | 2.1e-07 | 1.5e-07 | 3.4e-07 | 2.333E-07 | 9.713E-08 | 1.500E-07 | 3.400E-07 |

| | | | | | | | | | | |
|---|---|---|---|---|---|---|---|---|---|---|
| 2635 | 132 | L132P | A | 3.1e-07 | 2.5e-07 | 3.6e-07 | 3.067E-07 | 5.508E-08 | 2.500E-07 | 3.600E-07 |
| 2636 | 132 | L132S | A | 1.8e-07 | 2.2e-07 | 2.2e-07 | 2.067E-07 | 2.309E-08 | 1.800E-07 | 2.200E-07 |
| 2637 | 132 | L132T | A | 9.7e-07 | 3.5e-07 | 1.4e-07 | 4.867E-07 | 4.315E-07 | 1.400E-07 | 9.700E-07 |
| 2638 | 132 | L132W | A | 2.3e-07 | 3.7e-07 | 5.7e-07 | 3.900E-07 | 1.709E-07 | 2.300E-07 | 5.700E-07 |
| 2639 | 132 | L132Y | A | 4.5e-07 | 2.3e-07 | 4.9e-07 | 3.900E-07 | 1.400E-07 | 2.300E-07 | 4.900E-07 |
| 2640 | 132 | L132V | A | 5e-07 | 3.9e-07 | 2.4e-07 | 3.767E-07 | 1.305E-07 | 2.400E-07 | 5.000E-07 |
| 2641 | 133 | I133A | A | 3.8e-07 | 4.3e-07 | 2.5e-07 | 3.533E-07 | 9.292E-08 | 2.500E-07 | 4.300E-07 |
| 2642 | 133 | I133R | A | 2.6e-07 | 3.1e-07 | 1.5e-07 | 2.400E-07 | 8.185E-08 | 1.500E-07 | 3.100E-07 |
| 2643 | 133 | I133N | A | 3.6e-07 | 1.2e-07 | 1.9e-07 | 2.233E-07 | 1.234E-07 | 1.200E-07 | 3.600E-07 |
| 2644 | 133 | I133D | A | 2e-07 | 1.6e-07 | 3.9e-07 | 2.500E-07 | 1.229E-07 | 1.600E-07 | 3.900E-07 |
| 2645 | 133 | I133C | A | 3e-07 | 1.4e-07 | 1.6e-07 | 2.000E-07 | 8.718E-08 | 1.400E-07 | 3.000E-07 |
| 2646 | 133 | I133E | A | 4e-07 | 5.1e-07 | 2.1e-07 | 3.733E-07 | 1.518E-07 | 2.100E-07 | 5.100E-07 |
| 2647 | 133 | I133Q | A | 4.5e-07 | 2.2e-07 | 4.8e-07 | 3.833E-07 | 1.422E-07 | 2.200E-07 | 4.800E-07 |
| 2648 | 133 | I133G | A | 1.3e-07 | 3.7e-07 | 1.7e-07 | 2.233E-07 | 1.286E-07 | 1.300E-07 | 3.700E-07 |
| 2649 | 133 | I133H | A | 2.5e-07 | 1.3e-07 | 1.8e-07 | 1.867E-07 | 6.028E-08 | 1.300E-07 | 2.500E-07 |
| 2650 | 133 | I133I | A | 2.3e-07 | 1.3e-07 | 1.8e-07 | 1.800E-07 | 5.000E-08 | 1.300E-07 | 2.300E-07 |
| 2651 | 133 | I133L | A | 9.7e-08 | 2.2e-07 | 3e-07 | 2.057E-07 | 1.023E-07 | 9.700E-08 | 3.000E-07 |
| 2652 | 133 | I133K | A | 6e-07 | 2.1e-07 | 3.2e-07 | 3.767E-07 | 2.011E-07 | 2.100E-07 | 6.000E-07 |
| 2653 | 133 | I133M | A | 1.5e-07 | 2.6e-07 | 2.6e-07 | 2.233E-07 | 6.351E-08 | 1.500E-07 | 2.600E-07 |
| 2654 | 133 | I133F | A | 3.2e-07 | 2.1e-07 | 2.9e-07 | 2.733E-07 | 5.686E-08 | 2.100E-07 | 3.200E-07 |
| 2655 | 133 | I133P | A | 3.8e-07 | 1.6e-07 | 1.9e-07 | 2.433E-07 | 1.193E-07 | 1.600E-07 | 3.800E-07 |
| 2656 | 133 | I133S | A | 4.1e-07 | 3.5e-07 | 2.1e-07 | 3.233E-07 | 1.026E-07 | 2.100E-07 | 4.100E-07 |
| 2657 | 133 | I133T | A | 1.9e-07 | 1.6e-07 | 1e-07 | 1.500E-07 | 4.583E-08 | 1.000E-07 | 1.900E-07 |
| 2658 | 133 | I133W | A | 3.4e-07 | 1.3e-07 | 3.9e-07 | 2.867E-07 | 1.380E-07 | 1.300E-07 | 3.900E-07 |
| 2659 | 133 | I133Y | A | 2.8e-07 | 1.4e-07 | 8.7e-08 | 1.690E-07 | 9.971E-08 | 8.700E-08 | 2.800E-07 |
| 2660 | 133 | I133V | A | 1.9e-07 | 4.9e-07 | 1.7e-07 | 2.833E-07 | 1.793E-07 | 1.700E-07 | 4.900E-07 |
| 2661 | 134 | Q134A | A | 4.2e-07 | 3e-07 | 2.5e-07 | 3.233E-07 | 8.737E-08 | 2.500E-07 | 4.200E-07 |
| 2662 | 134 | Q134R | A | 1.1e-07 | 6.6e-07 | 3.1e-07 | 3.600E-07 | 2.784E-07 | 1.100E-07 | 6.600E-07 |
| 2663 | 134 | Q134N | A | 2.9e-07 | 3.3e-07 | 2.6e-07 | 2.933E-07 | 3.512E-08 | 2.600E-07 | 3.300E-07 |
| 2664 | 134 | Q134D | A | 3.2e-07 | 5.1e-07 | 2.4e-07 | 3.567E-07 | 1.387E-07 | 2.400E-07 | 5.100E-07 |
| 2665 | 134 | Q134C | A | 2.4e-07 | 2.2e-07 | 2.1e-07 | 2.233E-07 | 1.528E-08 | 2.100E-07 | 2.400E-07 |
| 2666 | 134 | Q134E | A | 1.7e-07 | 2.1e-07 | 1e-07 | 1.600E-07 | 5.568E-08 | 1.000E-07 | 2.100E-07 |
| 2667 | 134 | Q134Q | A | 2.3e-07 | 2.2e-07 | 2.4e-07 | 2.300E-07 | 1.000E-08 | 2.200E-07 | 2.400E-07 |
| 2668 | 134 | Q134G | A | 4.5e-07 | 3.3e-07 | 3.9e-07 | 3.900E-07 | 6.000E-08 | 3.300E-07 | 4.500E-07 |
| 2669 | 134 | Q134H | A | 1.3e-07 | 2.5e-07 | 1.9e-07 | 1.900E-07 | 6.000E-08 | 1.300E-07 | 2.500E-07 |
| 2670 | 134 | Q134I | A | 2.5e-07 | 2.1e-07 | 2.8e-07 | 2.467E-07 | 3.512E-08 | 2.100E-07 | 2.800E-07 |
| 2671 | 134 | Q134L | A | 2.5e-07 | 2.9e-07 | 2.7e-07 | 2.700E-07 | 2.000E-08 | 2.500E-07 | 2.900E-07 |
| 2672 | 134 | Q134K | A | 2.9e-07 | 4e-07 | 2.1e-07 | 3.000E-07 | 9.539E-08 | 2.100E-07 | 4.000E-07 |
| 2673 | 134 | Q134M | A | 3.8e-07 | 1.8e-07 | 2.8e-07 | 2.800E-07 | 1.000E-07 | 1.800E-07 | 3.800E-07 |
| 2674 | 134 | Q134F | A | 2.1e-07 | 2.8e-07 | 4.6e-07 | 3.167E-07 | 1.290E-07 | 2.100E-07 | 4.600E-07 |
| 2675 | 134 | Q134P | A | 4.1e-07 | 2.5e-07 | 1.8e-07 | 2.800E-07 | 1.179E-07 | 1.800E-07 | 4.100E-07 |
| 2676 | 134 | Q134S | A | 2.4e-07 | 2.2e-07 | 2.1e-07 | 2.233E-07 | 1.528E-08 | 2.100E-07 | 2.400E-07 |
| 2677 | 134 | Q134T | A | 4.3e-07 | 2.2e-07 | 1.7e-07 | 2.733E-07 | 1.380E-07 | 1.700E-07 | 4.300E-07 |
| 2678 | 134 | Q134W | A | 1.2e-07 | 2.3e-07 | 5.6e-07 | 3.033E-07 | 2.290E-07 | 1.200E-07 | 5.600E-07 |
| 2679 | 134 | Q134Y | A | 2e-07 | 1.1e-07 | 3.2e-07 | 2.100E-07 | 1.054E-07 | 1.100E-07 | 3.200E-07 |
| 2680 | 134 | Q134V | A | 2.7e-07 | 1.8e-07 | 4.7e-07 | 3.067E-07 | 1.484E-07 | 1.800E-07 | 4.700E-07 |
| 2681 | 135 | R135A | A | 1.6e-07 | 5.6e-07 | 3.4e-07 | 3.533E-07 | 2.003E-07 | 1.600E-07 | 5.600E-07 |
| 2682 | 135 | R135R | A | 4.6e-07 | 3.6e-07 | 4.7e-07 | 4.300E-07 | 6.083E-08 | 3.600E-07 | 4.700E-07 |
| 2683 | 135 | R135N | A | 3e-07 | 1.3e-07 | 1.1e-07 | 1.800E-07 | 1.044E-07 | 1.100E-07 | 3.000E-07 |
| 2684 | 135 | R135D | A | 1.9e-07 | 2.7e-07 | 2.5e-07 | 2.367E-07 | 4.163E-08 | 1.900E-07 | 2.700E-07 |
| 2685 | 135 | R135C | A | 2.4e-07 | 4e-08 | 1.3e-07 | 1.367E-07 | 1.002E-07 | 4.000E-08 | 2.400E-07 |
| 2686 | 135 | R135E | A | 2.4e-07 | 2.2e-07 | 1.5e-07 | 2.033E-07 | 4.726E-08 | 1.500E-07 | 2.400E-07 |
| 2687 | 135 | R135Q | A | 2.1e-07 | 2.6e-07 | 2.9e-07 | 2.533E-07 | 4.041E-08 | 2.100E-07 | 2.900E-07 |
| 2688 | 135 | R135G | A | 1.7e-07 | 3.1e-07 | 1.8e-07 | 2.200E-07 | 7.810E-08 | 1.700E-07 | 3.100E-07 |
| 2689 | 135 | R135H | A | 2.5e-07 | 2.7e-07 | 2.5e-07 | 2.567E-07 | 1.155E-08 | 2.500E-07 | 2.700E-07 |
| 2690 | 135 | R135I | A | 2.4e-07 | 2.2e-07 | 2.2e-07 | 2.267E-07 | 1.155E-08 | 2.200E-07 | 2.400E-07 |
| 2691 | 135 | R135L | A | 6e-07 | 2.9e-07 | 3.8e-07 | 4.233E-07 | 1.595E-07 | 2.900E-07 | 6.000E-07 |
| 2692 | 135 | R135K | A | 2.6e-07 | 1.2e-07 | 2.1e-07 | 1.967E-07 | 7.095E-08 | 1.200E-07 | 2.600E-07 |
| 2693 | 135 | R135M | A | 1.3e-07 | 1.4e-07 | 1.7e-07 | 1.467E-07 | 2.082E-08 | 1.300E-07 | 1.700E-07 |
| 2694 | 135 | R135F | A | 2.4e-07 | 3.3e-07 | 2.6e-07 | 2.767E-07 | 4.726E-08 | 2.400E-07 | 3.300E-07 |
| 2695 | 135 | R135P | A | 1.2e-07 | 2e-07 | 3.3e-07 | 2.167E-07 | 1.060E-07 | 1.200E-07 | 3.300E-07 |
| 2696 | 135 | R135S | A | 1.8e-07 | 4.5e-07 | 2.7e-07 | 3.000E-07 | 1.375E-07 | 1.800E-07 | 4.500E-07 |
| 2697 | 135 | R135T | A | 3.6e-07 | 1.6e-07 | 2.9e-07 | 2.700E-07 | 1.015E-07 | 1.600E-07 | 3.600E-07 |
| 2698 | 135 | R135W | A | 2.5e-07 | 4.2e-07 | 1.5e-07 | 2.733E-07 | 1.365E-07 | 1.500E-07 | 4.200E-07 |
| 2699 | 135 | R135Y | A | 4e-07 | 4e-07 | 2.4e-07 | 3.467E-07 | 9.238E-08 | 2.400E-07 | 4.000E-07 |
| 2700 | 135 | R135V | A | 3.1e-07 | 1.9e-07 | 2.2e-07 | 2.400E-07 | 6.245E-08 | 1.900E-07 | 3.100E-07 |
| 2701 | 136 | N136A | A | 1.5e-07 | 4.6e-07 | 2.1e-07 | 2.733E-07 | 1.644E-07 | 1.500E-07 | 4.600E-07 |
| 2702 | 136 | N136R | A | 3.7e-07 | 2.3e-07 | 1.7e-07 | 2.567E-07 | 1.026E-07 | 1.700E-07 | 3.700E-07 |
| 2703 | 136 | N136N | A | 1.6e-07 | 1.7e-07 | 1.6e-07 | 1.633E-07 | 5.774E-09 | 1.600E-07 | 1.700E-07 |
| 2704 | 136 | N136D | A | 1.6e-07 | 4.8e-07 | 2.9e-07 | 3.100E-07 | 1.609E-07 | 1.600E-07 | 4.800E-07 |
| 2705 | 136 | N136C | A | 2.9e-07 | 3.8e-07 | 2.3e-07 | 3.000E-07 | 7.550E-08 | 2.300E-07 | 3.800E-07 |
| 2706 | 136 | N136E | A | 1.4e-07 | 1.9e-07 | 3.1e-07 | 2.133E-07 | 8.737E-08 | 1.400E-07 | 3.100E-07 |
| 2707 | 136 | N136Q | A | 1.6e-07 | 2.5e-07 | 3.5e-07 | 2.533E-07 | 9.504E-08 | 1.600E-07 | 3.500E-07 |
| 2708 | 136 | N136G | A | 3.2e-07 | 1.8e-07 | 5.2e-07 | 3.400E-07 | 1.709E-07 | 1.800E-07 | 5.200E-07 |
| 2709 | 136 | N136H | A | 1.2e-07 | 1.8e-07 | 7.5e-07 | 3.500E-07 | 3.477E-07 | 1.200E-07 | 7.500E-07 |
| 2710 | 136 | N136I | A | 5.3e-07 | 5.2e-07 | 1.9e-07 | 4.133E-07 | 1.935E-07 | 1.900E-07 | 5.300E-07 |
| 2711 | 136 | N136L | A | 3.1e-07 | 2.8e-07 | 3.1e-07 | 3.000E-07 | 1.732E-08 | 2.800E-07 | 3.100E-07 |
| 2712 | 136 | N136K | A | 2.4e-07 | 3e-07 | 3e-07 | 2.800E-07 | 3.464E-08 | 2.400E-07 | 3.000E-07 |
| 2713 | 136 | N136M | A | 1.1e-07 | 3.3e-07 | 2.2e-07 | 2.200E-07 | 1.100E-07 | 1.100E-07 | 3.300E-07 |
| 2714 | 136 | N136F | A | 1.8e-07 | 2e-07 | 2.3e-07 | 2.033E-07 | 2.517E-08 | 1.800E-07 | 2.300E-07 |
| 2715 | 136 | N136P | A | 3.8e-07 | 9.5e-08 | 2.4e-07 | 2.383E-07 | 1.425E-07 | 9.500E-08 | 3.800E-07 |
| 2716 | 136 | N136S | A | 3.1e-07 | 2.5e-07 | 1.9e-07 | 2.500E-07 | 6.000E-08 | 1.900E-07 | 3.100E-07 |
| 2717 | 136 | N136T | A | 2e-07 | 1.4e-07 | 1.7e-07 | 1.700E-07 | 3.000E-08 | 1.400E-07 | 2.000E-07 |
| 2718 | 136 | N136W | A | 1.8e-07 | 2.3e-07 | 2.3e-07 | 2.133E-07 | 2.887E-08 | 1.800E-07 | 2.300E-07 |
| 2719 | 136 | N136Y | A | 3.3e-07 | 3.2e-07 | 2.2e-07 | 2.900E-07 | 6.083E-08 | 2.200E-07 | 3.300E-07 |

| | | | | | | | | | | |
|---|---|---|---|---|---|---|---|---|---|---|
| 2720 | 136 | N136V | A | 1.3e-07 | 4.6e-07 | 2.6e-07 | 2.833E-07 | 1.662E-07 | 1.300E-07 | 4.600E-07 |
| 2721 | 137 | P137A | A | 2.3e-07 | 6e-07 | 1.6e-07 | 3.300E-07 | 2.364E-07 | 1.600E-07 | 6.000E-07 |
| 2722 | 137 | P137R | A | 2.9e-07 | 2.9e-07 | 1.6e-07 | 2.467E-07 | 7.506E-08 | 1.600E-07 | 2.900E-07 |
| 2723 | 137 | P137N | A | 1.1e-07 | 4.8e-07 | 2e-07 | 2.633E-07 | 1.930E-07 | 1.100E-07 | 4.800E-07 |
| 2724 | 137 | P137D | A | 2.6e-07 | 2.3e-07 | 2.7e-07 | 2.533E-07 | 2.082E-08 | 2.300E-07 | 2.700E-07 |
| 2725 | 137 | P137C | A | 1.5e-07 | 5.2e-07 | 2.4e-07 | 3.033E-07 | 1.930E-07 | 1.500E-07 | 5.200E-07 |
| 2726 | 137 | P137E | A | 3e-07 | 2.3e-07 | 1.9e-07 | 2.400E-07 | 5.568E-08 | 1.900E-07 | 3.000E-07 |
| 2727 | 137 | P137Q | A | 3.1e-07 | 2.8e-07 | 3.6e-07 | 3.167E-07 | 4.041E-08 | 2.800E-07 | 3.600E-07 |
| 2728 | 137 | P137G | A | 3.3e-07 | 1.4e-07 | 2.7e-07 | 2.467E-07 | 9.713E-08 | 1.400E-07 | 3.300E-07 |
| 2729 | 137 | P137H | A | 2.3e-07 | 2.6e-07 | 2e-07 | 2.300E-07 | 3.000E-08 | 2.000E-07 | 2.600E-07 |
| 2730 | 137 | P137I | A | 1.8e-07 | 1.7e-07 | 1.3e-07 | 1.600E-07 | 2.646E-08 | 1.300E-07 | 1.800E-07 |
| 2731 | 137 | P137L | A | 4.3e-07 | 3.2e-07 | 2e-07 | 3.167E-07 | 1.150E-07 | 2.000E-07 | 4.300E-07 |
| 2732 | 137 | P137K | A | 2.1e-07 | 3.2e-07 | 9.9e-08 | 2.097E-07 | 1.105E-07 | 9.900E-08 | 3.200E-07 |
| 2733 | 137 | P137M | A | 2.9e-07 | 4.3e-07 | 1.7e-07 | 2.967E-07 | 1.301E-07 | 1.700E-07 | 4.300E-07 |
| 2734 | 137 | P137F | A | 5.1e-07 | 3.5e-07 | 2.2e-07 | 3.600E-07 | 1.453E-07 | 2.200E-07 | 5.100E-07 |
| 2735 | 137 | P137P | A | 2.4e-07 | 2.8e-07 | 2.4e-07 | 2.533E-07 | 2.309E-08 | 2.400E-07 | 2.800E-07 |
| 2736 | 137 | P137S | A | 2.2e-07 | 1.9e-07 | 2.1e-07 | 2.067E-07 | 1.528E-08 | 1.900E-07 | 2.200E-07 |
| 2737 | 137 | P137T | A | 7e-07 | 1.8e-07 | 1.5e-07 | 3.433E-07 | 3.092E-07 | 1.500E-07 | 7.000E-07 |
| 2738 | 137 | P137W | A | 2.7e-07 | 3.1e-07 | 1.5e-07 | 2.433E-07 | 8.327E-08 | 1.500E-07 | 3.100E-07 |
| 2739 | 137 | P137Y | A | 2.4e-07 | 3.3e-07 | 6.2e-07 | 3.967E-07 | 1.986E-07 | 2.400E-07 | 6.200E-07 |
| 2740 | 137 | P137V | A | 2.2e-07 | 3.1e-07 | 2.7e-07 | 2.667E-07 | 4.509E-08 | 2.200E-07 | 3.100E-07 |
| 2741 | 138 | Q138A | A | 3.3e-07 | 1.6e-07 | 2.8e-07 | 2.567E-07 | 8.737E-08 | 1.600E-07 | 3.300E-07 |
| 2742 | 138 | Q138R | A | 1.1e-07 | 1.1e-07 | 2e-07 | 1.400E-07 | 5.196E-08 | 1.100E-07 | 2.000E-07 |
| 2743 | 138 | Q138N | A | 3.3e-07 | 2.2e-07 | 1.6e-07 | 2.367E-07 | 8.622E-08 | 1.600E-07 | 3.300E-07 |
| 2744 | 138 | Q138D | A | 1.6e-07 | 4.3e-07 | 1.9e-07 | 2.600E-07 | 1.480E-07 | 1.600E-07 | 4.300E-07 |
| 2745 | 138 | Q138C | A | 2.7e-07 | 3.9e-07 | 1.9e-07 | 2.833E-07 | 1.007E-07 | 1.900E-07 | 3.900E-07 |
| 2746 | 138 | Q138E | A | 1.6e-07 | 3.2e-07 | 6.5e-07 | 3.767E-07 | 2.499E-07 | 1.600E-07 | 6.500E-07 |
| 2747 | 138 | Q138Q | A | 5.2e-07 | 3.6e-07 | 4.6e-07 | 4.467E-07 | 8.083E-08 | 3.600E-07 | 5.200E-07 |
| 2748 | 138 | Q138G | A | 2.2e-07 | 2.4e-07 | 1.5e-07 | 2.033E-07 | 4.726E-08 | 1.500E-07 | 2.400E-07 |
| 2749 | 138 | Q138H | A | 2e-07 | 3e-07 | 1.8e-07 | 2.267E-07 | 6.429E-08 | 1.800E-07 | 3.000E-07 |
| 2750 | 138 | Q138I | A | 2.2e-07 | 4.3e-07 | 9.5e-07 | 5.333E-07 | 3.758E-07 | 2.200E-07 | 9.500E-07 |
| 2751 | 138 | Q138L | A | 2.5e-07 | 1.6e-07 | 5.5e-07 | 3.200E-07 | 2.042E-07 | 1.600E-07 | 5.500E-07 |
| 2752 | 138 | Q138K | A | 1.4e-07 | 2.6e-07 | 1.6e-07 | 1.867E-07 | 6.429E-08 | 1.400E-07 | 2.600E-07 |
| 2753 | 138 | Q138M | A | 2.9e-07 | 2.7e-07 | 3.4e-07 | 3.000E-07 | 3.606E-08 | 2.700E-07 | 3.400E-07 |
| 2754 | 138 | Q138F | A | 1.1e-07 | 1.5e-07 | 3.5e-07 | 2.033E-07 | 1.286E-07 | 1.100E-07 | 3.500E-07 |
| 2755 | 138 | Q138P | A | 2.9e-07 | 2.9e-07 | 1.8e-07 | 2.533E-07 | 6.351E-08 | 1.800E-07 | 2.900E-07 |
| 2756 | 138 | Q138S | A | 5.6e-07 | 1.8e-07 | 2.5e-07 | 3.300E-07 | 2.022E-07 | 1.800E-07 | 5.600E-07 |
| 2757 | 138 | Q138T | A | 3.4e-07 | 1.6e-07 | 2.9e-07 | 2.633E-07 | 9.292E-08 | 1.600E-07 | 3.400E-07 |
| 2758 | 138 | Q138W | A | 3.8e-07 | 1.7e-07 | 1.8e-07 | 2.433E-07 | 1.185E-07 | 1.700E-07 | 3.800E-07 |
| 2759 | 138 | Q138Y | A | 1.9e-07 | 4.9e-07 | 2.5e-07 | 3.100E-07 | 1.587E-07 | 1.900E-07 | 4.900E-07 |
| 2760 | 138 | Q138V | A | 2.6e-07 | 2.8e-07 | 3.2e-07 | 2.867E-07 | 3.055E-08 | 2.600E-07 | 3.200E-07 |
| 2761 | 139 | L139A | A | 3.8e-07 | 3.1e-07 | 1.2e-07 | 2.700E-07 | 1.345E-07 | 1.200E-07 | 3.800E-07 |
| 2762 | 139 | L139R | A | 1.6e-07 | 2.3e-07 | 2.2e-07 | 2.033E-07 | 3.786E-08 | 1.600E-07 | 2.300E-07 |
| 2763 | 139 | L139N | A | 1.4e-07 | 2e-07 | 2.4e-07 | 1.933E-07 | 5.033E-08 | 1.400E-07 | 2.400E-07 |
| 2764 | 139 | L139D | A | 4.9e-07 | 2.3e-07 | 3.8e-07 | 3.667E-07 | 1.305E-07 | 2.300E-07 | 4.900E-07 |
| 2765 | 139 | L139C | A | 3.2e-07 | 2.6e-07 | 2.3e-07 | 2.700E-07 | 4.583E-08 | 2.300E-07 | 3.200E-07 |
| 2766 | 139 | L139E | A | 1e-07 | 3.6e-07 | 2.9e-07 | 2.500E-07 | 1.345E-07 | 1.000E-07 | 3.600E-07 |
| 2767 | 139 | L139Q | A | 3e-07 | 1.6e-07 | 1.6e-07 | 2.067E-07 | 8.083E-08 | 1.600E-07 | 3.000E-07 |
| 2768 | 139 | L139G | A | 1.2e-07 | 2e-07 | 2.4e-07 | 1.867E-07 | 6.110E-08 | 1.200E-07 | 2.400E-07 |
| 2769 | 139 | L139H | A | 1.5e-07 | 1.7e-07 | 4e-07 | 2.400E-07 | 1.389E-07 | 1.500E-07 | 4.000E-07 |
| 2770 | 139 | L139I | A | 5.2e-07 | 4.7e-07 | 3.2e-07 | 4.367E-07 | 1.041E-07 | 3.200E-07 | 5.200E-07 |
| 2771 | 139 | L139L | A | 4.4e-07 | 1.9e-07 | 2e-07 | 2.767E-07 | 1.415E-07 | 1.900E-07 | 4.400E-07 |
| 2772 | 139 | L139K | A | 2.5e-07 | 9.7e-08 | 1.3e-07 | 1.590E-07 | 8.052E-08 | 9.700E-08 | 2.500E-07 |
| 2773 | 139 | L139M | A | 1.5e-07 | 3.5e-07 | 8.6e-08 | 1.953E-07 | 1.377E-07 | 8.600E-08 | 3.500E-07 |
| 2774 | 139 | L139F | A | 1.7e-07 | 2e-07 | 3.4e-07 | 2.367E-07 | 9.074E-08 | 1.700E-07 | 3.400E-07 |
| 2775 | 139 | L139P | A | 2.1e-07 | 6e-07 | 3.3e-07 | 3.800E-07 | 1.997E-07 | 2.100E-07 | 6.000E-07 |
| 2776 | 139 | L139S | A | 2.9e-07 | 2e-07 | 2.3e-07 | 2.400E-07 | 4.583E-08 | 2.000E-07 | 2.900E-07 |
| 2777 | 139 | L139T | A | 2.9e-07 | 2.2e-07 | 1.3e-07 | 2.133E-07 | 8.021E-08 | 1.300E-07 | 2.900E-07 |
| 2778 | 139 | L139W | A | 2.4e-07 | 1.8e-07 | 1.4e-07 | 1.867E-07 | 5.033E-08 | 1.400E-07 | 2.400E-07 |
| 2779 | 139 | L139Y | A | 1.4e-07 | 1.7e-07 | 1.2e-07 | 1.433E-07 | 2.517E-08 | 1.200E-07 | 1.700E-07 |
| 2780 | 139 | L139V | A | 2.9e-07 | 3.2e-07 | 1.3e-07 | 2.467E-07 | 1.021E-07 | 1.300E-07 | 3.200E-07 |
| 2781 | 140 | C140A | A | 1.8e-07 | 3.1e-07 | 1.9e-07 | 2.267E-07 | 7.234E-08 | 1.800E-07 | 3.100E-07 |
| 2782 | 140 | C140R | A | 3e-07 | 2.6e-07 | 3.3e-07 | 2.967E-07 | 3.512E-08 | 2.600E-07 | 3.300E-07 |
| 2783 | 140 | C140N | A | 2.8e-07 | 3.4e-07 | 4.1e-07 | 3.433E-07 | 6.506E-08 | 2.800E-07 | 4.100E-07 |
| 2784 | 140 | C140D | A | 2.2e-07 | 2e-07 | 5.1e-07 | 3.100E-07 | 1.735E-07 | 2.000E-07 | 5.100E-07 |
| 2785 | 140 | C140C | A | 3.6e-07 | 1.3e-07 | 4.2e-07 | 3.033E-07 | 1.531E-07 | 1.300E-07 | 4.200E-07 |
| 2786 | 140 | C140E | A | 1.9e-07 | 2.5e-07 | 4.9e-07 | 3.100E-07 | 1.587E-07 | 1.900E-07 | 4.900E-07 |
| 2787 | 140 | C140Q | A | 1.2e-07 | 2.3e-07 | 1.6e-07 | 1.700E-07 | 5.568E-08 | 1.200E-07 | 2.300E-07 |
| 2788 | 140 | C140G | A | 4.7e-07 | 2.3e-07 | 5.4e-07 | 4.133E-07 | 1.626E-07 | 2.300E-07 | 5.400E-07 |
| 2789 | 140 | C140H | A | 1.8e-07 | 2.2e-07 | 2.9e-07 | 2.300E-07 | 5.568E-08 | 1.800E-07 | 2.900E-07 |
| 2790 | 140 | C140I | A | 2.3e-07 | 3.7e-07 | 2.9e-07 | 2.967E-07 | 7.024E-08 | 2.300E-07 | 3.700E-07 |
| 2791 | 140 | C140L | A | 4.4e-07 | 1.8e-07 | 3.1e-07 | 3.100E-07 | 1.300E-07 | 1.800E-07 | 4.400E-07 |
| 2792 | 140 | C140K | A | 1.9e-07 | 1.9e-07 | 2.9e-07 | 2.233E-07 | 5.774E-08 | 1.900E-07 | 2.900E-07 |
| 2793 | 140 | C140M | A | 2.6e-07 | 2.8e-07 | 2.2e-07 | 2.533E-07 | 3.055E-08 | 2.200E-07 | 2.800E-07 |
| 2794 | 140 | C140F | A | 3e-07 | 3.3e-07 | 1.5e-07 | 2.600E-07 | 9.644E-08 | 1.500E-07 | 3.300E-07 |
| 2795 | 140 | C140P | A | 3.5e-07 | 2.1e-07 | 1.4e-07 | 2.333E-07 | 1.069E-07 | 1.400E-07 | 3.500E-07 |
| 2796 | 140 | C140S | A | 2e-07 | 2.7e-07 | 2.8e-07 | 2.500E-07 | 4.359E-08 | 2.000E-07 | 2.800E-07 |
| 2797 | 140 | C140T | A | 2.2e-07 | 1.4e-07 | 7.5e-07 | 3.700E-07 | 3.315E-07 | 1.400E-07 | 7.500E-07 |
| 2798 | 140 | C140W | A | 1.2e-07 | 2.6e-07 | 4.7e-07 | 2.833E-07 | 1.762E-07 | 1.200E-07 | 4.700E-07 |
| 2799 | 140 | C140Y | A | 3.2e-07 | 3e-07 | 2e-07 | 2.733E-07 | 6.429E-08 | 2.000E-07 | 3.200E-07 |
| 2800 | 140 | C140V | A | 2.5e-07 | 2.2e-07 | 2.2e-07 | 2.300E-07 | 1.732E-08 | 2.200E-07 | 2.500E-07 |
| 2801 | 141 | Y141A | A | 1.5e-07 | 3.3e-07 | 4e-07 | 2.933E-07 | 1.290E-07 | 1.500E-07 | 4.000E-07 |
| 2802 | 141 | Y141R | A | 2.2e-07 | 2.3e-07 | 3.2e-07 | 2.567E-07 | 5.508E-08 | 2.200E-07 | 3.200E-07 |
| 2803 | 141 | Y141N | A | 2.9e-07 | 6.1e-07 | 2.8e-07 | 3.933E-07 | 1.877E-07 | 2.800E-07 | 6.100E-07 |
| 2804 | 141 | Y141D | A | 1.9e-07 | 9.9e-08 | 1.2e-07 | 1.363E-07 | 4.765E-08 | 9.900E-08 | 1.900E-07 |

| | | | | | | | | | | |
|---|---|---|---|---|---|---|---|---|---|---|
| 2805 | 141 | Y141C | A | 1.7e-07 | 3.4e-07 | 8.5e-08 | 1.983E-07 | 1.298E-07 | 8.500E-08 | 3.400E-07 |
| 2806 | 141 | Y141E | A | 2.3e-07 | 1.2e-07 | 2.1e-07 | 1.867E-07 | 5.859E-08 | 1.200E-07 | 2.300E-07 |
| 2807 | 141 | Y141Q | A | 1.3e-07 | 2.6e-07 | 2.7e-07 | 2.200E-07 | 7.810E-08 | 1.300E-07 | 2.700E-07 |
| 2808 | 141 | Y141G | A | 2.9e-07 | 3.8e-07 | 2.1e-07 | 2.933E-07 | 8.505E-08 | 2.100E-07 | 3.800E-07 |
| 2809 | 141 | Y141H | A | 4.1e-07 | 3.5e-07 | 3.2e-07 | 3.600E-07 | 4.583E-08 | 3.200E-07 | 4.100E-07 |
| 2810 | 141 | Y141I | A | 3.3e-07 | 2.6e-07 | 4.7e-07 | 3.533E-07 | 1.069E-07 | 2.600E-07 | 4.700E-07 |
| 2811 | 141 | Y141L | A | 4.7e-07 | 3.1e-07 | 3.1e-07 | 3.633E-07 | 9.238E-08 | 3.100E-07 | 4.700E-07 |
| 2812 | 141 | Y141K | A | 1.6e-07 | 1.3e-07 | 2.6e-07 | 1.833E-07 | 6.807E-08 | 1.300E-07 | 2.600E-07 |
| 2813 | 141 | Y141M | A | 5.9e-07 | 2.4e-07 | 2.2e-07 | 3.500E-07 | 2.081E-07 | 2.200E-07 | 5.900E-07 |
| 2814 | 141 | Y141F | A | 4.2e-07 | 1.9e-07 | 1.8e-07 | 2.633E-07 | 1.358E-07 | 1.800E-07 | 4.200E-07 |
| 2815 | 141 | Y141P | A | 3.3e-07 | 3.2e-07 | 2.6e-07 | 3.033E-07 | 3.786E-08 | 2.600E-07 | 3.300E-07 |
| 2816 | 141 | Y141S | A | 3.5e-07 | 4.1e-07 | 1.9e-07 | 3.167E-07 | 1.137E-07 | 1.900E-07 | 4.100E-07 |
| 2817 | 141 | Y141T | A | 3.2e-07 | 2.5e-07 | 1.9e-07 | 2.533E-07 | 6.506E-08 | 1.900E-07 | 3.200E-07 |
| 2818 | 141 | Y141W | A | 4.6e-07 | 1e-07 | 3.8e-07 | 3.133E-07 | 1.890E-07 | 1.000E-07 | 4.600E-07 |
| 2819 | 141 | Y141Y | A | 2.1e-07 | 2.8e-07 | 3.7e-07 | 2.867E-07 | 8.021E-08 | 2.100E-07 | 3.700E-07 |
| 2820 | 141 | Y141V | A | 3.7e-07 | 4.3e-07 | 2.1e-07 | 3.367E-07 | 1.137E-07 | 2.100E-07 | 4.300E-07 |
| 2821 | 142 | Q142A | A | 2.3e-07 | 5.2e-07 | 2.4e-07 | 3.300E-07 | 1.646E-07 | 2.300E-07 | 5.200E-07 |
| 2822 | 142 | Q142R | A | 3.3e-07 | 1.5e-07 | 1.9e-07 | 2.233E-07 | 9.452E-08 | 1.500E-07 | 3.300E-07 |
| 2823 | 142 | Q142N | A | 2.5e-07 | 2.6e-07 | 2.3e-07 | 2.467E-07 | 1.528E-08 | 2.300E-07 | 2.600E-07 |
| 2824 | 142 | Q142D | A | 3.5e-07 | 1.5e-07 | 2.7e-07 | 2.567E-07 | 1.007E-07 | 1.500E-07 | 3.500E-07 |
| 2825 | 142 | Q142C | A | 3.5e-07 | 1.5e-07 | 2.9e-07 | 2.633E-07 | 1.026E-07 | 1.500E-07 | 3.500E-07 |
| 2826 | 142 | Q142E | A | 1.9e-07 | 2e-07 | 2.4e-07 | 2.100E-07 | 2.646E-08 | 1.900E-07 | 2.400E-07 |
| 2827 | 142 | Q142Q | A | 2.2e-07 | 3.1e-07 | 2.1e-07 | 2.467E-07 | 5.508E-08 | 2.100E-07 | 3.100E-07 |
| 2828 | 142 | Q142G | A | 3.1e-07 | 2.5e-07 | 1.3e-07 | 2.300E-07 | 9.165E-08 | 1.300E-07 | 3.100E-07 |
| 2829 | 142 | Q142H | A | 2.3e-07 | 2.7e-07 | 2.1e-07 | 2.367E-07 | 3.055E-08 | 2.100E-07 | 2.700E-07 |
| 2830 | 142 | Q142I | A | 3e-07 | 2.6e-07 | 1.7e-07 | 2.433E-07 | 6.658E-08 | 1.700E-07 | 3.000E-07 |
| 2831 | 142 | Q142L | A | 1.5e-07 | 2.5e-07 | 2.5e-07 | 2.167E-07 | 5.774E-08 | 1.500E-07 | 2.500E-07 |
| 2832 | 142 | Q142K | A | 3.1e-07 | 1.7e-07 | 2e-07 | 2.267E-07 | 7.371E-08 | 1.700E-07 | 3.100E-07 |
| 2833 | 142 | Q142M | A | 4.2e-07 | 2.5e-07 | 1.7e-07 | 2.800E-07 | 1.277E-07 | 1.700E-07 | 4.200E-07 |
| 2834 | 142 | Q142F | A | 2.3e-07 | 1.8e-07 | 3.6e-07 | 2.567E-07 | 9.292E-08 | 1.800E-07 | 3.600E-07 |
| 2835 | 142 | Q142P | A | 1.7e-07 | 3.7e-07 | 2.8e-07 | 2.733E-07 | 1.002E-07 | 1.700E-07 | 3.700E-07 |
| 2836 | 142 | Q142S | A | 1.4e-07 | 2.5e-07 | 5e-07 | 2.967E-07 | 1.845E-07 | 1.400E-07 | 5.000E-07 |
| 2837 | 142 | Q142T | A | 2e-07 | 3.5e-07 | 2.3e-07 | 2.600E-07 | 7.937E-08 | 2.000E-07 | 3.500E-07 |
| 2838 | 142 | Q142W | A | 2.9e-07 | 2.3e-07 | 5.7e-07 | 3.633E-07 | 1.815E-07 | 2.300E-07 | 5.700E-07 |
| 2839 | 142 | Q142Y | A | 2.2e-07 | 2.1e-07 | 2.6e-07 | 2.300E-07 | 2.646E-08 | 2.100E-07 | 2.600E-07 |
| 2840 | 142 | Q142V | A | 3.3e-07 | 3.5e-07 | 2.6e-07 | 3.133E-07 | 4.726E-08 | 2.600E-07 | 3.500E-07 |
| 2841 | 143 | D143A | A | 1.6e-07 | 2.2e-07 | 3.1e-07 | 2.300E-07 | 7.550E-08 | 1.600E-07 | 3.100E-07 |
| 2842 | 143 | D143R | A | 2.9e-07 | 3.3e-07 | 2.4e-07 | 2.867E-07 | 4.509E-08 | 2.400E-07 | 3.300E-07 |
| 2843 | 143 | D143N | A | 2.8e-07 | 1.5e-07 | 1.6e-07 | 1.967E-07 | 7.234E-08 | 1.500E-07 | 2.800E-07 |
| 2844 | 143 | D143D | A | 2.9e-07 | 1.6e-07 | 1.8e-07 | 2.100E-07 | 7.000E-08 | 1.600E-07 | 2.900E-07 |
| 2845 | 143 | D143C | A | 1.3e-07 | 2.8e-07 | 2.4e-07 | 2.167E-07 | 7.767E-08 | 1.300E-07 | 2.800E-07 |
| 2846 | 143 | D143E | A | 2.4e-07 | 2.6e-07 | 3.5e-07 | 2.833E-07 | 5.859E-08 | 2.400E-07 | 3.500E-07 |
| 2847 | 143 | D143Q | A | 2.5e-07 | 1.7e-07 | 3.5e-07 | 2.567E-07 | 9.018E-08 | 1.700E-07 | 3.500E-07 |
| 2848 | 143 | D143G | A | 3.5e-07 | 1.7e-07 | 5.3e-07 | 3.500E-07 | 1.800E-07 | 1.700E-07 | 5.300E-07 |
| 2849 | 143 | D143H | A | 3.4e-07 | 3.3e-07 | 2.2e-07 | 2.967E-07 | 6.658E-08 | 2.200E-07 | 3.400E-07 |
| 2850 | 143 | D143I | A | 2.7e-07 | 2.1e-07 | 3.5e-07 | 2.767E-07 | 7.024E-08 | 2.100E-07 | 3.500E-07 |
| 2851 | 143 | D143L | A | 3.9e-07 | 4.3e-07 | 2.2e-07 | 3.467E-07 | 1.115E-07 | 2.200E-07 | 4.300E-07 |
| 2852 | 143 | D143K | A | 2.8e-07 | 2e-07 | 3e-07 | 2.600E-07 | 5.292E-08 | 2.000E-07 | 3.000E-07 |
| 2853 | 143 | D143M | A | 1.6e-07 | 2.2e-07 | 2.3e-07 | 2.033E-07 | 3.786E-08 | 1.600E-07 | 2.300E-07 |
| 2854 | 143 | D143F | A | 1.1e-07 | 2.3e-07 | 1.3e-07 | 1.567E-07 | 6.429E-08 | 1.100E-07 | 2.300E-07 |
| 2855 | 143 | D143P | A | 1.2e-07 | 4.8e-07 | 2.5e-07 | 2.833E-07 | 1.823E-07 | 1.200E-07 | 4.800E-07 |
| 2856 | 143 | D143S | A | 3.3e-07 | 2.1e-07 | 2.1e-07 | 2.500E-07 | 6.928E-08 | 2.100E-07 | 3.300E-07 |
| 2857 | 143 | D143T | A | 2e-07 | 2.1e-07 | 2.4e-07 | 2.167E-07 | 2.082E-08 | 2.000E-07 | 2.400E-07 |
| 2858 | 143 | D143W | A | 4.3e-07 | 2.1e-07 | 2.4e-07 | 2.933E-07 | 1.193E-07 | 2.100E-07 | 4.300E-07 |
| 2859 | 143 | D143Y | A | 2.5e-07 | 4.2e-07 | 3.1e-07 | 3.267E-07 | 8.622E-08 | 2.500E-07 | 4.200E-07 |
| 2860 | 143 | D143V | A | 1.3e-07 | 3.4e-07 | 2.8e-07 | 2.500E-07 | 1.082E-07 | 1.300E-07 | 3.400E-07 |
| 2861 | 144 | T144A | A | 2.5e-07 | 1.4e-07 | 3.2e-07 | 2.367E-07 | 9.074E-08 | 1.400E-07 | 3.200E-07 |
| 2862 | 144 | T144R | A | 2.6e-07 | 4.3e-07 | 3.2e-07 | 3.367E-07 | 8.622E-08 | 2.600E-07 | 4.300E-07 |
| 2863 | 144 | T144N | A | 2.4e-07 | 3.1e-07 | 2.6e-07 | 2.700E-07 | 3.606E-08 | 2.400E-07 | 3.100E-07 |
| 2864 | 144 | T144D | A | 3.2e-07 | 5.3e-07 | 2.4e-07 | 3.633E-07 | 1.498E-07 | 2.400E-07 | 5.300E-07 |
| 2865 | 144 | T144C | A | 1.9e-07 | 2.4e-07 | 4.3e-07 | 2.867E-07 | 1.266E-07 | 1.900E-07 | 4.300E-07 |
| 2866 | 144 | T144E | A | 3.2e-07 | 3e-07 | 1.9e-07 | 2.700E-07 | 7.000E-08 | 1.900E-07 | 3.200E-07 |
| 2867 | 144 | T144Q | A | 1.8e-07 | 1e-07 | 9.9e-08 | 1.263E-07 | 4.648E-08 | 9.900E-08 | 1.800E-07 |
| 2868 | 144 | T144G | A | 5.4e-07 | 4.5e-07 | 5.1e-07 | 5.000E-07 | 4.583E-08 | 4.500E-07 | 5.400E-07 |
| 2869 | 144 | T144H | A | 7.1e-07 | 2.5e-07 | 2.1e-07 | 3.900E-07 | 2.778E-07 | 2.100E-07 | 7.100E-07 |
| 2870 | 144 | T144I | A | 2.9e-07 | 1.1e-07 | 2.3e-07 | 2.100E-07 | 9.165E-08 | 1.100E-07 | 2.900E-07 |
| 2871 | 144 | T144L | A | 2.4e-07 | 2.5e-07 | 2.3e-07 | 2.400E-07 | 1.000E-08 | 2.300E-07 | 2.500E-07 |
| 2872 | 144 | T144K | A | 1.9e-07 | 2.4e-07 | 4.4e-07 | 2.900E-07 | 1.323E-07 | 1.900E-07 | 4.400E-07 |
| 2873 | 144 | T144M | A | 3.8e-07 | 2.9e-07 | 1.2e-07 | 2.633E-07 | 1.320E-07 | 1.200E-07 | 3.800E-07 |
| 2874 | 144 | T144F | A | 2.3e-07 | 3.1e-07 | 2.1e-07 | 2.500E-07 | 5.292E-08 | 2.100E-07 | 3.100E-07 |
| 2875 | 144 | T144P | A | 4e-07 | 2.9e-07 | 1.4e-07 | 2.767E-07 | 1.305E-07 | 1.400E-07 | 4.000E-07 |
| 2876 | 144 | T144S | A | 2.5e-07 | 2.1e-07 | 4.9e-07 | 3.167E-07 | 1.514E-07 | 2.100E-07 | 4.900E-07 |
| 2877 | 144 | T144T | A | 9.3e-08 | 2.8e-07 | 2e-07 | 1.910E-07 | 9.382E-08 | 9.300E-08 | 2.800E-07 |
| 2878 | 144 | T144W | A | 3.7e-07 | 3e-07 | 3.8e-07 | 3.500E-07 | 4.359E-08 | 3.000E-07 | 3.800E-07 |
| 2879 | 144 | T144Y | A | 3.8e-07 | 5.5e-07 | 1.9e-07 | 3.733E-07 | 1.801E-07 | 1.900E-07 | 5.500E-07 |
| 2880 | 144 | T144V | A | 1.8e-07 | 1.9e-07 | 7e-07 | 3.567E-07 | 2.974E-07 | 1.800E-07 | 7.000E-07 |
| 2881 | 145 | I145A | A | 1.6e-07 | 1.5e-07 | 2.7e-07 | 1.933E-07 | 6.658E-08 | 1.500E-07 | 2.700E-07 |
| 2882 | 145 | I145R | A | 2.4e-07 | 3.5e-07 | 2.1e-07 | 2.667E-07 | 7.371E-08 | 2.100E-07 | 3.500E-07 |
| 2883 | 145 | I145N | A | 1.8e-07 | 6.9e-07 | 1.8e-07 | 3.500E-07 | 2.944E-07 | 1.800E-07 | 6.900E-07 |
| 2884 | 145 | I145D | A | 1e-07 | 1.8e-07 | 1.8e-07 | 1.533E-07 | 4.619E-08 | 1.000E-07 | 1.800E-07 |
| 2885 | 145 | I145C | A | 2.7e-07 | 2.1e-07 | 4.4e-07 | 3.067E-07 | 1.193E-07 | 2.100E-07 | 4.400E-07 |
| 2886 | 145 | I145E | A | 2.8e-07 | 5e-07 | 2.8e-07 | 3.533E-07 | 1.270E-07 | 2.800E-07 | 5.000E-07 |
| 2887 | 145 | I145Q | A | 1.5e-07 | 2.1e-07 | 4.1e-07 | 2.567E-07 | 1.361E-07 | 1.500E-07 | 4.100E-07 |
| 2888 | 145 | I145G | A | 2.3e-07 | 1.3e-07 | 5.9e-07 | 3.167E-07 | 2.419E-07 | 1.300E-07 | 5.900E-07 |
| 2889 | 145 | I145H | A | 1.9e-07 | 4.5e-07 | 4.7e-07 | 3.700E-07 | 1.562E-07 | 1.900E-07 | 4.700E-07 |

| | | | | | | | | | | |
|---|---|---|---|---|---|---|---|---|---|---|
| 2890 | 145 | I145I | A | 1.9e-07 | 2.6e-07 | 3.2e-07 | 2.567E-07 | 6.506E-08 | 1.900E-07 | 3.200E-07 |
| 2891 | 145 | I145L | A | 3.1e-07 | 1.5e-07 | 2.7e-07 | 2.433E-07 | 8.327E-08 | 1.500E-07 | 3.100E-07 |
| 2892 | 145 | I145K | A | 2.1e-07 | 3.7e-07 | 1.5e-07 | 2.433E-07 | 1.137E-07 | 1.500E-07 | 3.700E-07 |
| 2893 | 145 | I145M | A | 3.1e-07 | 9.8e-08 | 2.5e-07 | 2.193E-07 | 1.093E-07 | 9.800E-08 | 3.100E-07 |
| 2894 | 145 | I145F | A | 2.1e-07 | 5e-07 | 2e-07 | 3.033E-07 | 1.704E-07 | 2.000E-07 | 5.000E-07 |
| 2895 | 145 | I145P | A | 3.3e-07 | 2.5e-07 | 2.1e-07 | 2.633E-07 | 6.110E-08 | 2.100E-07 | 3.300E-07 |
| 2896 | 145 | I145S | A | 2.8e-07 | 1.7e-07 | 2.7e-07 | 2.400E-07 | 6.083E-08 | 1.700E-07 | 2.800E-07 |
| 2897 | 145 | I145T | A | 2.5e-07 | 1.5e-07 | 1.2e-07 | 1.733E-07 | 6.807E-08 | 1.200E-07 | 2.500E-07 |
| 2898 | 145 | I145W | A | 3e-07 | 2.9e-07 | 1.9e-07 | 2.600E-07 | 6.083E-08 | 1.900E-07 | 3.000E-07 |
| 2899 | 145 | I145Y | A | 1.2e-07 | 2.6e-07 | 3.8e-07 | 2.533E-07 | 1.301E-07 | 1.200E-07 | 3.800E-07 |
| 2900 | 145 | I145V | A | 2.4e-07 | 1.2e-07 | 2.7e-07 | 2.100E-07 | 7.937E-08 | 1.200E-07 | 2.700E-07 |
| 2901 | 146 | L146A | A | 2.3e-07 | 3.8e-07 | 2.6e-07 | 2.900E-07 | 7.937E-08 | 2.300E-07 | 3.800E-07 |
| 2902 | 146 | L146R | A | 6.4e-07 | 1.6e-07 | 1.5e-07 | 3.167E-07 | 2.801E-07 | 1.500E-07 | 6.400E-07 |
| 2903 | 146 | L146N | A | 2e-07 | 1.3e-07 | 2.1e-07 | 1.800E-07 | 4.359E-08 | 1.300E-07 | 2.100E-07 |
| 2904 | 146 | L146D | A | 3e-07 | 1.7e-07 | 2e-07 | 2.233E-07 | 6.807E-08 | 1.700E-07 | 3.000E-07 |
| 2905 | 146 | L146C | A | 1.6e-07 | 2.7e-07 | 1.3e-07 | 1.867E-07 | 7.371E-08 | 1.300E-07 | 2.700E-07 |
| 2906 | 146 | L146E | A | 3e-07 | 2.4e-07 | 1.5e-07 | 2.300E-07 | 7.550E-08 | 1.500E-07 | 3.000E-07 |
| 2907 | 146 | L146Q | A | 1.2e-07 | 8.5e-08 | 1.1e-07 | 1.050E-07 | 1.803E-08 | 8.500E-08 | 1.200E-07 |
| 2908 | 146 | L146G | A | 3.1e-07 | 2.7e-07 | 4.6e-07 | 3.467E-07 | 1.002E-07 | 2.700E-07 | 4.600E-07 |
| 2909 | 146 | L146H | A | 1.9e-07 | 2e-07 | 1.3e-07 | 1.733E-07 | 3.786E-08 | 1.300E-07 | 2.000E-07 |
| 2910 | 146 | L146I | A | 1.5e-07 | 4.4e-07 | 3.2e-07 | 3.033E-07 | 1.457E-07 | 1.500E-07 | 4.400E-07 |
| 2911 | 146 | L146L | A | 2.7e-07 | 1.7e-07 | 1.8e-07 | 2.067E-07 | 5.508E-08 | 1.700E-07 | 2.700E-07 |
| 2912 | 146 | L146K | A | 3.2e-07 | 2.2e-07 | 1.1e-07 | 2.167E-07 | 1.050E-07 | 1.100E-07 | 3.200E-07 |
| 2913 | 146 | L146M | A | 4e-07 | 1.5e-07 | 3.8e-07 | 3.100E-07 | 1.389E-07 | 1.500E-07 | 4.000E-07 |
| 2914 | 146 | L146F | A | 2.1e-07 | 4.2e-07 | 3.2e-07 | 3.167E-07 | 1.050E-07 | 2.100E-07 | 4.200E-07 |
| 2915 | 146 | L146P | A | 3.4e-07 | 2.8e-07 | 4.1e-07 | 3.433E-07 | 6.506E-08 | 2.800E-07 | 4.100E-07 |
| 2916 | 146 | L146S | A | 4.5e-07 | 1.9e-07 | 1.3e-07 | 2.567E-07 | 1.701E-07 | 1.300E-07 | 4.500E-07 |
| 2917 | 146 | L146T | A | 3.6e-07 | 2.2e-07 | 1.6e-07 | 2.467E-07 | 1.026E-07 | 1.600E-07 | 3.600E-07 |
| 2918 | 146 | L146W | A | 2.3e-07 | 1.8e-07 | 2.7e-07 | 2.267E-07 | 4.509E-08 | 1.800E-07 | 2.700E-07 |
| 2919 | 146 | L146Y | A | 1.5e-07 | 1.5e-07 | 4.2e-07 | 2.400E-07 | 1.559E-07 | 1.500E-07 | 4.200E-07 |
| 2920 | 146 | L146V | A | 4.9e-07 | 1.4e-07 | 3e-07 | 3.100E-07 | 1.752E-07 | 1.400E-07 | 4.900E-07 |
| 2921 | 147 | W147A | A | 2.3e-07 | 2.9e-07 | 4.1e-07 | 3.100E-07 | 9.165E-08 | 2.300E-07 | 4.100E-07 |
| 2922 | 147 | W147R | A | 2.4e-07 | 2e-07 | 3.7e-07 | 2.700E-07 | 8.888E-08 | 2.000E-07 | 3.700E-07 |
| 2923 | 147 | W147N | A | 2.3e-07 | 1.8e-07 | 2.3e-07 | 2.133E-07 | 2.887E-08 | 1.800E-07 | 2.300E-07 |
| 2924 | 147 | W147D | A | 1.9e-07 | 1.5e-07 | 2.6e-07 | 2.000E-07 | 5.568E-08 | 1.500E-07 | 2.600E-07 |
| 2925 | 147 | W147C | A | 1.6e-07 | 2.9e-07 | 2.2e-07 | 2.233E-07 | 6.506E-08 | 1.600E-07 | 2.900E-07 |
| 2926 | 147 | W147E | A | 1.6e-07 | 1.8e-07 | 1e-07 | 1.467E-07 | 4.163E-08 | 1.000E-07 | 1.800E-07 |
| 2927 | 147 | W147Q | A | 3.8e-07 | 1e-07 | 4.9e-07 | 3.233E-07 | 2.011E-07 | 1.000E-07 | 4.900E-07 |
| 2928 | 147 | W147G | A | 4.5e-07 | 2.7e-07 | 3.1e-07 | 3.433E-07 | 9.452E-08 | 2.700E-07 | 4.500E-07 |
| 2929 | 147 | W147H | A | 2.8e-07 | 1.5e-07 | 2.5e-07 | 2.267E-07 | 6.807E-08 | 1.500E-07 | 2.800E-07 |
| 2930 | 147 | W147I | A | 3.6e-07 | 4.1e-07 | 3.7e-07 | 3.800E-07 | 2.646E-08 | 3.600E-07 | 4.100E-07 |
| 2931 | 147 | W147L | A | 2.1e-07 | 3.9e-07 | 6.1e-07 | 4.033E-07 | 2.003E-07 | 2.100E-07 | 6.100E-07 |
| 2932 | 147 | W147K | A | 3.8e-07 | 2.8e-07 | 1.6e-07 | 2.733E-07 | 1.102E-07 | 1.600E-07 | 3.800E-07 |
| 2933 | 147 | W147M | A | 5.1e-07 | 2.6e-07 | 1.8e-07 | 3.167E-07 | 1.721E-07 | 1.800E-07 | 5.100E-07 |
| 2934 | 147 | W147F | A | 2.3e-07 | 3.2e-07 | 2.6e-07 | 2.700E-07 | 4.583E-08 | 2.300E-07 | 3.200E-07 |
| 2935 | 147 | W147P | A | 2.2e-07 | 2.3e-07 | 3.8e-07 | 2.767E-07 | 8.963E-08 | 2.200E-07 | 3.800E-07 |
| 2936 | 147 | W147S | A | 4.7e-07 | 5.2e-07 | 4.7e-07 | 4.867E-07 | 2.887E-08 | 4.700E-07 | 5.200E-07 |
| 2937 | 147 | W147T | A | 1.6e-07 | 8.3e-08 | 1.8e-07 | 1.410E-07 | 5.122E-08 | 8.300E-08 | 1.800E-07 |
| 2938 | 147 | W147W | A | 2.6e-07 | 2e-07 | 4.2e-07 | 2.933E-07 | 1.137E-07 | 2.000E-07 | 4.200E-07 |
| 2939 | 147 | W147Y | A | 1.3e-07 | 2.5e-07 | 3.1e-07 | 2.300E-07 | 9.165E-08 | 1.300E-07 | 3.100E-07 |
| 2940 | 147 | W147V | A | 1.4e-07 | 1.7e-07 | 1.4e-07 | 1.500E-07 | 1.732E-08 | 1.400E-07 | 1.700E-07 |
| 2941 | 148 | K148A | A | 6.6e-07 | 2.4e-07 | 2e-07 | 3.667E-07 | 2.548E-07 | 2.000E-07 | 6.600E-07 |
| 2942 | 148 | K148R | A | 1.3e-07 | 3.3e-07 | 2.6e-07 | 2.400E-07 | 1.015E-07 | 1.300E-07 | 3.300E-07 |
| 2943 | 148 | K148N | A | 2.3e-07 | 3.5e-07 | 2.8e-07 | 2.867E-07 | 6.028E-08 | 2.300E-07 | 3.500E-07 |
| 2944 | 148 | K148D | A | 3.2e-07 | 2.6e-07 | 2.8e-07 | 2.867E-07 | 3.055E-08 | 2.600E-07 | 3.200E-07 |
| 2945 | 148 | K148C | A | 3.6e-07 | 3.8e-07 | 3.3e-07 | 3.567E-07 | 2.517E-08 | 3.300E-07 | 3.800E-07 |
| 2946 | 148 | K148E | A | 2.8e-07 | 1.9e-07 | 5.8e-07 | 3.500E-07 | 2.042E-07 | 1.900E-07 | 5.800E-07 |
| 2947 | 148 | K148Q | A | 2.8e-07 | 2.5e-07 | 2.8e-07 | 2.700E-07 | 1.732E-08 | 2.500E-07 | 2.800E-07 |
| 2948 | 148 | K148G | A | 1.6e-07 | 1.7e-07 | 1.5e-07 | 1.600E-07 | 1.000E-08 | 1.500E-07 | 1.700E-07 |
| 2949 | 148 | K148H | A | 2.6e-07 | 2.8e-07 | 1.1e-07 | 2.167E-07 | 9.292E-08 | 1.100E-07 | 2.800E-07 |
| 2950 | 148 | K148I | A | 1.9e-07 | 2.3e-07 | 6.2e-07 | 3.467E-07 | 2.376E-07 | 1.900E-07 | 6.200E-07 |
| 2951 | 148 | K148L | A | 2.5e-07 | 2.3e-07 | 1.7e-07 | 2.167E-07 | 4.163E-08 | 1.700E-07 | 2.500E-07 |
| 2952 | 148 | K148K | A | 1.7e-07 | 2.8e-07 | 5e-07 | 3.167E-07 | 1.680E-07 | 1.700E-07 | 5.000E-07 |
| 2953 | 148 | K148M | A | 2.9e-07 | 1.3e-07 | 3.7e-07 | 2.633E-07 | 1.222E-07 | 1.300E-07 | 3.700E-07 |
| 2954 | 148 | K148F | A | 3.4e-07 | 2.5e-07 | 2e-07 | 2.633E-07 | 7.095E-08 | 2.000E-07 | 3.400E-07 |
| 2955 | 148 | K148P | A | 2.3e-07 | 4e-07 | 7.9e-07 | 4.733E-07 | 2.871E-07 | 2.300E-07 | 7.900E-07 |
| 2956 | 148 | K148S | A | 3.5e-07 | 5.1e-07 | 4e-07 | 4.200E-07 | 8.185E-08 | 3.500E-07 | 5.100E-07 |
| 2957 | 148 | K148T | A | 1.3e-07 | 3.4e-07 | 2.8e-07 | 2.500E-07 | 1.082E-07 | 1.300E-07 | 3.400E-07 |
| 2958 | 148 | K148W | A | 1.9e-07 | 1.8e-07 | 2e-07 | 1.900E-07 | 1.000E-08 | 1.800E-07 | 2.000E-07 |
| 2959 | 148 | K148Y | A | 3.4e-07 | 3.4e-07 | 1.9e-07 | 2.900E-07 | 8.660E-08 | 1.900E-07 | 3.400E-07 |
| 2960 | 148 | K148V | A | 1.7e-07 | 2.6e-07 | 2.4e-07 | 2.233E-07 | 4.726E-08 | 1.700E-07 | 2.600E-07 |
| 2961 | 149 | D149A | A | 1.9e-07 | 8.5e-08 | 5.7e-07 | 2.817E-07 | 2.552E-07 | 8.500E-08 | 5.700E-07 |
| 2962 | 149 | D149R | A | 3.1e-07 | 1.4e-07 | 3.5e-07 | 2.667E-07 | 1.115E-07 | 1.400E-07 | 3.500E-07 |
| 2963 | 149 | D149N | A | 4.2e-07 | 1.4e-07 | 2.2e-07 | 2.600E-07 | 1.442E-07 | 1.400E-07 | 4.200E-07 |
| 2964 | 149 | D149D | A | 3e-07 | 2.7e-07 | 3.9e-07 | 3.200E-07 | 6.245E-08 | 2.700E-07 | 3.900E-07 |
| 2965 | 149 | D149C | A | 1.7e-07 | 5.6e-07 | 2.1e-07 | 3.133E-07 | 2.146E-07 | 1.700E-07 | 5.600E-07 |
| 2966 | 149 | D149E | A | 1.3e-07 | 3.9e-07 | 1.9e-07 | 2.367E-07 | 1.361E-07 | 1.300E-07 | 3.900E-07 |
| 2967 | 149 | D149Q | A | 2.1e-07 | 2.5e-07 | 2.1e-07 | 2.233E-07 | 2.309E-08 | 2.100E-07 | 2.500E-07 |
| 2968 | 149 | D149G | A | 2.5e-07 | 3e-07 | 5.4e-07 | 3.633E-07 | 1.550E-07 | 2.500E-07 | 5.400E-07 |
| 2969 | 149 | D149H | A | 2e-07 | 1.2e-07 | 5e-07 | 2.733E-07 | 2.003E-07 | 1.200E-07 | 5.000E-07 |
| 2970 | 149 | D149I | A | 2.7e-07 | 2.1e-07 | 3e-07 | 2.600E-07 | 4.583E-08 | 2.100E-07 | 3.000E-07 |
| 2971 | 149 | D149L | A | 1.2e-07 | 3.4e-07 | 3.3e-07 | 2.633E-07 | 1.242E-07 | 1.200E-07 | 3.400E-07 |
| 2972 | 149 | D149K | A | 2e-07 | 2.2e-07 | 4.4e-07 | 2.867E-07 | 1.332E-07 | 2.000E-07 | 4.400E-07 |
| 2973 | 149 | D149M | A | 1.3e-07 | 1.8e-07 | 2.1e-07 | 1.733E-07 | 4.041E-08 | 1.300E-07 | 2.100E-07 |
| 2974 | 149 | D149F | A | 1.2e-07 | 1.3e-07 | 2.3e-07 | 1.600E-07 | 6.083E-08 | 1.200E-07 | 2.300E-07 |

| | | | | | | | | | | |
|---|---|---|---|---|---|---|---|---|---|---|
| 2975 | 149 | D149P | A | 5.1e-07 | 1.7e-07 | 1.9e-07 | 2.900E-07 | 1.908E-07 | 1.700E-07 | 5.100E-07 |
| 2976 | 149 | D149S | A | 2.8e-07 | 1.3e-07 | 4.1e-07 | 2.733E-07 | 1.401E-07 | 1.300E-07 | 4.100E-07 |
| 2977 | 149 | D149T | A | 4.5e-07 | 2.5e-07 | 2.7e-07 | 3.233E-07 | 1.102E-07 | 2.500E-07 | 4.500E-07 |
| 2978 | 149 | D149W | A | 2.4e-07 | 2.4e-07 | 2.5e-07 | 2.433E-07 | 5.774E-09 | 2.400E-07 | 2.500E-07 |
| 2979 | 149 | D149Y | A | 1.7e-07 | 2.6e-07 | 2.1e-07 | 2.133E-07 | 4.509E-08 | 1.700E-07 | 2.600E-07 |
| 2980 | 149 | D149V | A | 1.7e-07 | 3.6e-07 | 1.4e-07 | 2.233E-07 | 1.193E-07 | 1.400E-07 | 3.600E-07 |
| 2981 | 150 | I150A | A | 4.6e-07 | 1.8e-07 | 2.6e-07 | 3.000E-07 | 1.442E-07 | 1.800E-07 | 4.600E-07 |
| 2982 | 150 | I150R | A | 3.2e-07 | 4.8e-07 | 2.6e-07 | 3.533E-07 | 1.137E-07 | 2.600E-07 | 4.800E-07 |
| 2983 | 150 | I150N | A | 2.9e-07 | 1.1e-07 | 1.9e-07 | 1.967E-07 | 9.018E-08 | 1.100E-07 | 2.900E-07 |
| 2984 | 150 | I150D | A | 2.2e-07 | 3.3e-07 | 2.8e-07 | 2.767E-07 | 5.508E-08 | 2.200E-07 | 3.300E-07 |
| 2985 | 150 | I150C | A | 3e-07 | 2.6e-07 | 1.2e-07 | 2.267E-07 | 9.452E-08 | 1.200E-07 | 3.000E-07 |
| 2986 | 150 | I150E | A | 1.5e-07 | 1.6e-07 | 1.2e-07 | 1.433E-07 | 2.082E-08 | 1.200E-07 | 1.600E-07 |
| 2987 | 150 | I150Q | A | 2.8e-07 | 2.9e-07 | 2.7e-07 | 2.800E-07 | 1.000E-08 | 2.700E-07 | 2.900E-07 |
| 2988 | 150 | I150G | A | 2e-07 | 1.5e-07 | 2.2e-07 | 1.900E-07 | 3.606E-08 | 1.500E-07 | 2.200E-07 |
| 2989 | 150 | I150H | A | 3.4e-07 | 1.8e-07 | 3e-07 | 2.733E-07 | 8.327E-08 | 1.800E-07 | 3.400E-07 |
| 2990 | 150 | I150I | A | 2.2e-07 | 2.3e-07 | 2.9e-07 | 2.467E-07 | 3.786E-08 | 2.200E-07 | 2.900E-07 |
| 2991 | 150 | I150L | A | 4.7e-07 | 3e-07 | 3.6e-07 | 3.767E-07 | 8.622E-08 | 3.000E-07 | 4.700E-07 |
| 2992 | 150 | I150K | A | 1.5e-07 | 2.3e-07 | 2.4e-07 | 2.067E-07 | 4.933E-08 | 1.500E-07 | 2.400E-07 |
| 2993 | 150 | I150M | A | 1.7e-07 | 2.9e-07 | 5.7e-07 | 3.433E-07 | 2.053E-07 | 1.700E-07 | 5.700E-07 |
| 2994 | 150 | I150F | A | 1.2e-07 | 3.2e-07 | 2.2e-07 | 2.200E-07 | 1.000E-07 | 1.200E-07 | 3.200E-07 |
| 2995 | 150 | I150P | A | 3.3e-07 | 2.5e-07 | 1.9e-07 | 2.567E-07 | 7.024E-08 | 1.900E-07 | 3.300E-07 |
| 2996 | 150 | I150S | A | 2.7e-07 | 2.2e-07 | 2.3e-07 | 2.400E-07 | 2.646E-08 | 2.200E-07 | 2.700E-07 |
| 2997 | 150 | I150T | A | 2.5e-07 | 2e-07 | 1.3e-07 | 1.933E-07 | 6.028E-08 | 1.300E-07 | 2.500E-07 |
| 2998 | 150 | I150W | A | 3.5e-07 | 1.7e-07 | 3.4e-07 | 2.867E-07 | 1.012E-07 | 1.700E-07 | 3.500E-07 |
| 2999 | 150 | I150Y | A | 3.1e-07 | 3.7e-07 | 2.2e-07 | 3.000E-07 | 7.550E-08 | 2.200E-07 | 3.700E-07 |
| 3000 | 150 | I150V | A | 4.6e-07 | 2.3e-07 | 2.7e-07 | 3.200E-07 | 1.229E-07 | 2.300E-07 | 4.600E-07 |
| 3001 | 151 | F151A | A | 1.1e-07 | 2.3e-07 | 1.3e-07 | 1.567E-07 | 6.429E-08 | 1.100E-07 | 2.300E-07 |
| 3002 | 151 | F151R | A | 2.9e-07 | 2.8e-07 | 9.6e-08 | 2.220E-07 | 1.092E-07 | 9.600E-08 | 2.900E-07 |
| 3003 | 151 | F151N | A | 5.7e-07 | 1.9e-07 | 6.6e-07 | 4.733E-07 | 2.495E-07 | 1.900E-07 | 6.600E-07 |
| 3004 | 151 | F151D | A | 3.4e-07 | 2.6e-07 | 1.4e-07 | 2.467E-07 | 1.007E-07 | 1.400E-07 | 3.400E-07 |
| 3005 | 151 | F151C | A | 2.6e-07 | 1.3e-07 | 1.5e-07 | 1.800E-07 | 7.000E-08 | 1.300E-07 | 2.600E-07 |
| 3006 | 151 | F151E | A | 3.9e-07 | 1.5e-07 | 1.8e-07 | 2.400E-07 | 1.308E-07 | 1.500E-07 | 3.900E-07 |
| 3007 | 151 | F151Q | A | 2.7e-07 | 4e-07 | 4.7e-07 | 3.800E-07 | 1.015E-07 | 2.700E-07 | 4.700E-07 |
| 3008 | 151 | F151G | A | 2.4e-07 | 4.8e-07 | 2e-07 | 3.067E-07 | 1.514E-07 | 2.000E-07 | 4.800E-07 |
| 3009 | 151 | F151H | A | 2.5e-07 | 2.1e-07 | 1.8e-07 | 2.133E-07 | 3.512E-08 | 1.800E-07 | 2.500E-07 |
| 3010 | 151 | F151I | A | 5.2e-07 | 5.9e-07 | 2.4e-07 | 4.500E-07 | 1.852E-07 | 2.400E-07 | 5.900E-07 |
| 3011 | 151 | F151L | A | 4.3e-07 | 3e-07 | 2e-07 | 3.100E-07 | 1.153E-07 | 2.000E-07 | 4.300E-07 |
| 3012 | 151 | F151K | A | 2.2e-07 | 1.8e-07 | 2e-07 | 2.000E-07 | 2.000E-08 | 1.800E-07 | 2.200E-07 |
| 3013 | 151 | F151M | A | 3.7e-07 | 1.8e-07 | 1.7e-07 | 2.400E-07 | 1.127E-07 | 1.700E-07 | 3.700E-07 |
| 3014 | 151 | F151F | A | 3.2e-07 | 2.7e-07 | 2.1e-07 | 2.667E-07 | 5.508E-08 | 2.100E-07 | 3.200E-07 |
| 3015 | 151 | F151P | A | 3.1e-07 | 2.2e-07 | 1.5e-07 | 2.267E-07 | 8.021E-08 | 1.500E-07 | 3.100E-07 |
| 3016 | 151 | F151S | A | 2.3e-07 | 1.8e-07 | 3.4e-07 | 2.500E-07 | 8.185E-08 | 1.800E-07 | 3.400E-07 |
| 3017 | 151 | F151T | A | 4.8e-07 | 1.6e-07 | 1.7e-07 | 2.700E-07 | 1.819E-07 | 1.600E-07 | 4.800E-07 |
| 3018 | 151 | F151W | A | 1.7e-07 | 2.9e-07 | 2.7e-07 | 2.433E-07 | 6.429E-08 | 1.700E-07 | 2.900E-07 |
| 3019 | 151 | F151Y | A | 3.5e-07 | 9.2e-08 | 1.9e-07 | 2.107E-07 | 1.302E-07 | 9.200E-08 | 3.500E-07 |
| 3020 | 151 | F151V | A | 5.1e-07 | 4.8e-07 | 3.3e-07 | 4.400E-07 | 9.644E-08 | 3.300E-07 | 5.100E-07 |
| 3021 | 152 | H152A | A | 2.6e-07 | 2e-07 | 2.5e-07 | 2.367E-07 | 3.215E-08 | 2.000E-07 | 2.600E-07 |
| 3022 | 152 | H152R | A | 3.2e-07 | 3.7e-07 | 4.5e-07 | 3.800E-07 | 6.557E-08 | 3.200E-07 | 4.500E-07 |
| 3023 | 152 | H152N | A | 2.9e-07 | 1.9e-07 | 2.5e-07 | 2.433E-07 | 5.033E-08 | 1.900E-07 | 2.900E-07 |
| 3024 | 152 | H152D | A | 1.4e-07 | 1.2e-07 | 1.1e-07 | 1.233E-07 | 1.528E-08 | 1.100E-07 | 1.400E-07 |
| 3025 | 152 | H152C | A | 2.3e-07 | 1.5e-07 | 1.9e-07 | 1.900E-07 | 4.000E-08 | 1.500E-07 | 2.300E-07 |
| 3026 | 152 | H152E | A | 1.3e-07 | 2.9e-07 | 3.5e-07 | 2.567E-07 | 1.137E-07 | 1.300E-07 | 3.500E-07 |
| 3027 | 152 | H152Q | A | 1.5e-07 | 3.1e-07 | 1.2e-07 | 1.933E-07 | 1.021E-07 | 1.200E-07 | 3.100E-07 |
| 3028 | 152 | H152G | A | 1.8e-07 | 1.8e-07 | 2e-07 | 1.867E-07 | 1.155E-08 | 1.800E-07 | 2.000E-07 |
| 3029 | 152 | H152H | A | 1.3e-07 | 3.9e-07 | 1.4e-07 | 2.200E-07 | 1.473E-07 | 1.300E-07 | 3.900E-07 |
| 3030 | 152 | H152I | A | 1e-07 | 3.4e-07 | 2.2e-07 | 2.200E-07 | 1.200E-07 | 1.000E-07 | 3.400E-07 |
| 3031 | 152 | H152L | A | 3.2e-07 | 1.3e-07 | 2.3e-07 | 2.267E-07 | 9.504E-08 | 1.300E-07 | 3.200E-07 |
| 3032 | 152 | H152K | A | 1.3e-07 | 2.2e-07 | 1.8e-07 | 1.767E-07 | 4.509E-08 | 1.300E-07 | 2.200E-07 |
| 3033 | 152 | H152M | A | 1.3e-07 | 4.1e-07 | 2.4e-07 | 2.600E-07 | 1.411E-07 | 1.300E-07 | 4.100E-07 |
| 3034 | 152 | H152F | A | 2.5e-07 | 5.2e-07 | 1.8e-07 | 3.167E-07 | 1.795E-07 | 1.800E-07 | 5.200E-07 |
| 3035 | 152 | H152P | A | 5.5e-07 | 1.4e-07 | 1.8e-07 | 2.900E-07 | 2.261E-07 | 1.400E-07 | 5.500E-07 |
| 3036 | 152 | H152S | A | 1.5e-07 | 3.7e-07 | 3e-07 | 2.733E-07 | 1.124E-07 | 1.500E-07 | 3.700E-07 |
| 3037 | 152 | H152T | A | 1.6e-07 | 2.8e-07 | 9.7e-08 | 1.790E-07 | 9.297E-08 | 9.700E-08 | 2.800E-07 |
| 3038 | 152 | H152W | A | 4e-07 | 3.3e-07 | 8.1e-08 | 2.703E-07 | 1.677E-07 | 8.100E-08 | 4.000E-07 |
| 3039 | 152 | H152Y | A | 3e-07 | 1.7e-07 | 2.5e-07 | 2.400E-07 | 6.557E-08 | 1.700E-07 | 3.000E-07 |
| 3040 | 152 | H152V | A | 3.6e-07 | 3.2e-07 | 1.5e-07 | 2.767E-07 | 1.115E-07 | 1.500E-07 | 3.600E-07 |
| 3041 | 153 | K153A | A | 2.9e-07 | 4.5e-07 | 2.1e-07 | 3.167E-07 | 1.222E-07 | 2.100E-07 | 4.500E-07 |
| 3042 | 153 | K153R | A | 1.5e-07 | 1.5e-07 | 1.7e-07 | 1.567E-07 | 1.155E-08 | 1.500E-07 | 1.700E-07 |
| 3043 | 153 | K153N | A | 1.4e-07 | 5.9e-07 | 2.9e-07 | 3.400E-07 | 2.291E-07 | 1.400E-07 | 5.900E-07 |
| 3044 | 153 | K153D | A | 2.9e-07 | 1.3e-07 | 1.8e-07 | 2.000E-07 | 8.185E-08 | 1.300E-07 | 2.900E-07 |
| 3045 | 153 | K153C | A | 2.1e-07 | 1.3e-07 | 2.6e-07 | 2.000E-07 | 6.557E-08 | 1.300E-07 | 2.600E-07 |
| 3046 | 153 | K153E | A | 3.2e-07 | 2.2e-07 | 2.8e-07 | 2.733E-07 | 5.033E-08 | 2.200E-07 | 3.200E-07 |
| 3047 | 153 | K153Q | A | 2.1e-07 | 2e-07 | 1.6e-07 | 1.900E-07 | 2.646E-08 | 1.600E-07 | 2.100E-07 |
| 3048 | 153 | K153G | A | 3.4e-07 | 9.4e-08 | 4.4e-07 | 2.913E-07 | 1.781E-07 | 9.400E-08 | 4.400E-07 |
| 3049 | 153 | K153H | A | 2.4e-07 | 3.2e-07 | 1.6e-07 | 2.400E-07 | 8.000E-08 | 1.600E-07 | 3.200E-07 |
| 3050 | 153 | K153I | A | 1.1e-07 | 4.6e-07 | 2.5e-07 | 2.733E-07 | 1.762E-07 | 1.100E-07 | 4.600E-07 |
| 3051 | 153 | K153L | A | 2.2e-07 | 3.7e-07 | 3.3e-07 | 3.067E-07 | 7.767E-08 | 2.200E-07 | 3.700E-07 |
| 3052 | 153 | K153K | A | 1.9e-07 | 1.5e-07 | 2.2e-07 | 1.867E-07 | 3.512E-08 | 1.500E-07 | 2.200E-07 |
| 3053 | 153 | K153M | A | 2.1e-07 | 3e-07 | 2e-07 | 2.367E-07 | 5.508E-08 | 2.000E-07 | 3.000E-07 |
| 3054 | 153 | K153F | A | 9.2e-08 | 3.1e-07 | 3.2e-07 | 2.407E-07 | 1.288E-07 | 9.200E-08 | 3.200E-07 |
| 3055 | 153 | K153P | A | 1.1e-07 | 8.8e-08 | 5.1e-07 | 2.360E-07 | 2.375E-07 | 8.800E-08 | 5.100E-07 |
| 3056 | 153 | K153S | A | 2e-07 | 2.7e-07 | 3.5e-07 | 2.733E-07 | 7.506E-08 | 2.000E-07 | 3.500E-07 |
| 3057 | 153 | K153T | A | 3e-07 | 2e-07 | 4.2e-07 | 3.067E-07 | 1.102E-07 | 2.000E-07 | 4.200E-07 |
| 3058 | 153 | K153W | A | 5.8e-07 | 1.6e-07 | 6.9e-08 | 2.697E-07 | 2.726E-07 | 6.900E-08 | 5.800E-07 |
| 3059 | 153 | K153Y | A | 2.4e-07 | 2.6e-07 | 2.4e-07 | 2.467E-07 | 1.155E-08 | 2.400E-07 | 2.600E-07 |

| | | | | | | | | | | |
|---|---|---|---|---|---|---|---|---|---|---|
| 3060 | 153 | K153V | A | 1.3e-07 | 1.7e-07 | 3.5e-07 | 2.167E-07 | 1.172E-07 | 1.300E-07 | 3.500E-07 |
| 3061 | 154 | N154A | A | 3e-07 | 1.8e-07 | 6.3e-07 | 3.700E-07 | 2.330E-07 | 1.800E-07 | 6.300E-07 |
| 3062 | 154 | N154R | A | 3.6e-07 | 2.4e-07 | 3.2e-07 | 3.067E-07 | 6.110E-08 | 2.400E-07 | 3.600E-07 |
| 3063 | 154 | N154N | A | 4.1e-07 | 2.6e-07 | 7.3e-07 | 4.667E-07 | 2.401E-07 | 2.600E-07 | 7.300E-07 |
| 3064 | 154 | N154D | A | 2.9e-07 | 2.4e-07 | 2.2e-07 | 2.500E-07 | 3.606E-08 | 2.200E-07 | 2.900E-07 |
| 3065 | 154 | N154C | A | 2.4e-07 | 2.2e-07 | 1.7e-07 | 2.100E-07 | 3.606E-08 | 1.700E-07 | 2.400E-07 |
| 3066 | 154 | N154E | A | 2e-07 | 1.4e-07 | 4.6e-07 | 2.667E-07 | 1.701E-07 | 1.400E-07 | 4.600E-07 |
| 3067 | 154 | N154Q | A | 2e-07 | 3.3e-07 | 1.4e-07 | 2.233E-07 | 9.713E-08 | 1.400E-07 | 3.300E-07 |
| 3068 | 154 | N154G | A | 2.8e-07 | 2.8e-07 | 4.5e-07 | 3.367E-07 | 9.815E-08 | 2.800E-07 | 4.500E-07 |
| 3069 | 154 | N154H | A | 5.4e-07 | 3.6e-07 | 3.1e-07 | 4.033E-07 | 1.210E-07 | 3.100E-07 | 5.400E-07 |
| 3070 | 154 | N154I | A | 1.7e-07 | 1.9e-07 | 1.7e-07 | 1.767E-07 | 1.155E-08 | 1.700E-07 | 1.900E-07 |
| 3071 | 154 | N154L | A | 2.8e-07 | 1.7e-07 | 1.4e-07 | 1.967E-07 | 7.371E-08 | 1.400E-07 | 2.800E-07 |
| 3072 | 154 | N154K | A | 1e-07 | 2.6e-07 | 1.9e-07 | 1.833E-07 | 8.021E-08 | 1.000E-07 | 2.600E-07 |
| 3073 | 154 | N154M | A | 4.9e-07 | 1.2e-07 | 2e-07 | 2.700E-07 | 1.947E-07 | 1.200E-07 | 4.900E-07 |
| 3074 | 154 | N154F | A | 1.2e-07 | 2.1e-07 | 9e-08 | 1.400E-07 | 6.245E-08 | 9.000E-08 | 2.100E-07 |
| 3075 | 154 | N154P | A | 2.1e-07 | 5e-07 | 2.3e-07 | 3.133E-07 | 1.620E-07 | 2.100E-07 | 5.000E-07 |
| 3076 | 154 | N154S | A | 5.6e-07 | 1.9e-07 | 2.8e-07 | 3.433E-07 | 1.930E-07 | 1.900E-07 | 5.600E-07 |
| 3077 | 154 | N154T | A | 2.6e-07 | 1.2e-07 | 2.1e-07 | 1.967E-07 | 7.095E-08 | 1.200E-07 | 2.600E-07 |
| 3078 | 154 | N154W | A | 4.3e-07 | 2.7e-07 | 2.6e-07 | 3.200E-07 | 9.539E-08 | 2.600E-07 | 4.300E-07 |
| 3079 | 154 | N154Y | A | 2.6e-07 | 1.8e-07 | 2.3e-07 | 2.233E-07 | 4.041E-08 | 1.800E-07 | 2.600E-07 |
| 3080 | 154 | N154V | A | 2.5e-07 | 1.8e-07 | 2.7e-07 | 2.333E-07 | 4.726E-08 | 1.800E-07 | 2.700E-07 |
| 3081 | 155 | N155A | A | 3.1e-07 | 3.4e-07 | 2.5e-07 | 3.000E-07 | 4.583E-08 | 2.500E-07 | 3.400E-07 |
| 3082 | 155 | N155R | A | 2e-07 | 1.3e-07 | 2.8e-07 | 2.033E-07 | 7.506E-08 | 1.300E-07 | 2.800E-07 |
| 3083 | 155 | N155N | A | 3.3e-07 | 2.1e-07 | 1.4e-07 | 2.267E-07 | 9.609E-08 | 1.400E-07 | 3.300E-07 |
| 3084 | 155 | N155D | A | 1.3e-07 | 3.4e-07 | 1.7e-07 | 2.133E-07 | 1.115E-07 | 1.300E-07 | 3.400E-07 |
| 3085 | 155 | N155C | A | 3.1e-07 | 3.1e-07 | 8.2e-08 | 2.340E-07 | 1.316E-07 | 8.200E-08 | 3.100E-07 |
| 3086 | 155 | N155E | A | 4.9e-07 | 3.3e-07 | 2.8e-07 | 3.667E-07 | 1.097E-07 | 2.800E-07 | 4.900E-07 |
| 3087 | 155 | N155Q | A | 2.3e-07 | 3.3e-07 | 1.6e-07 | 2.400E-07 | 8.544E-08 | 1.600E-07 | 3.300E-07 |
| 3088 | 155 | N155G | A | 2e-07 | 2.6e-07 | 1.2e-07 | 1.933E-07 | 7.024E-08 | 1.200E-07 | 2.600E-07 |
| 3089 | 155 | N155H | A | 2.7e-07 | 2.8e-07 | 3e-07 | 2.833E-07 | 1.528E-08 | 2.700E-07 | 3.000E-07 |
| 3090 | 155 | N155I | A | 4e-07 | 5e-07 | 1.2e-07 | 3.400E-07 | 1.970E-07 | 1.200E-07 | 5.000E-07 |
| 3091 | 155 | N155L | A | 1.6e-07 | 3.9e-07 | 3.1e-07 | 2.867E-07 | 1.168E-07 | 1.600E-07 | 3.900E-07 |
| 3092 | 155 | N155K | A | 1.7e-07 | 2.4e-07 | 2.3e-07 | 2.133E-07 | 3.786E-08 | 1.700E-07 | 2.400E-07 |
| 3093 | 155 | N155M | A | 2.4e-07 | 1.3e-07 | 1.8e-07 | 1.833E-07 | 5.508E-08 | 1.300E-07 | 2.400E-07 |
| 3094 | 155 | N155F | A | 2.2e-07 | 3.5e-07 | 2.4e-07 | 2.700E-07 | 7.000E-08 | 2.200E-07 | 3.500E-07 |
| 3095 | 155 | N155P | A | 2.2e-07 | 2.1e-07 | 3.2e-07 | 2.500E-07 | 6.083E-08 | 2.100E-07 | 3.200E-07 |
| 3096 | 155 | N155S | A | 2.9e-07 | 3.6e-07 | 3.3e-07 | 3.267E-07 | 3.512E-08 | 2.900E-07 | 3.600E-07 |
| 3097 | 155 | N155T | A | 2.8e-07 | 2.1e-07 | 1.7e-07 | 2.200E-07 | 5.568E-08 | 1.700E-07 | 2.800E-07 |
| 3098 | 155 | N155W | A | 2.4e-07 | 2.1e-07 | 2.8e-07 | 2.433E-07 | 3.512E-08 | 2.100E-07 | 2.800E-07 |
| 3099 | 155 | N155Y | A | 2.6e-07 | 1.2e-07 | 1.2e-07 | 1.667E-07 | 8.083E-08 | 1.200E-07 | 2.600E-07 |
| 3100 | 155 | N155V | A | 1.2e-07 | 1.3e-07 | 3.6e-07 | 2.033E-07 | 1.358E-07 | 1.200E-07 | 3.600E-07 |
| 3101 | 156 | Q156A | A | 2.9e-07 | 2.5e-07 | 1.8e-07 | 2.400E-07 | 5.568E-08 | 1.800E-07 | 2.900E-07 |
| 3102 | 156 | Q156R | A | 1.3e-07 | 2.7e-07 | 1.6e-07 | 1.867E-07 | 7.371E-08 | 1.300E-07 | 2.700E-07 |
| 3103 | 156 | Q156N | A | 2.5e-07 | 2.8e-07 | 2.3e-07 | 2.533E-07 | 2.517E-08 | 2.300E-07 | 2.800E-07 |
| 3104 | 156 | Q156D | A | 1.7e-07 | 5e-07 | 2.5e-07 | 3.067E-07 | 1.721E-07 | 1.700E-07 | 5.000E-07 |
| 3105 | 156 | Q156C | A | 2.4e-07 | 2.2e-07 | 1.9e-07 | 2.167E-07 | 2.517E-08 | 1.900E-07 | 2.400E-07 |
| 3106 | 156 | Q156E | A | 1.7e-07 | 1.6e-07 | 2e-07 | 1.767E-07 | 2.082E-08 | 1.600E-07 | 2.000E-07 |
| 3107 | 156 | Q156Q | A | 1.9e-07 | 4.3e-07 | 1.7e-07 | 2.633E-07 | 1.447E-07 | 1.700E-07 | 4.300E-07 |
| 3108 | 156 | Q156G | A | 2.8e-07 | 4.2e-07 | 2.6e-07 | 3.200E-07 | 8.718E-08 | 2.600E-07 | 4.200E-07 |
| 3109 | 156 | Q156H | A | 2e-07 | 5.4e-07 | 2.4e-07 | 3.267E-07 | 1.858E-07 | 2.000E-07 | 5.400E-07 |
| 3110 | 156 | Q156I | A | 1.5e-07 | 3.7e-07 | 3.7e-07 | 2.967E-07 | 1.270E-07 | 1.500E-07 | 3.700E-07 |
| 3111 | 156 | Q156L | A | 2.1e-07 | 2.3e-07 | 1.6e-07 | 2.000E-07 | 3.606E-08 | 1.600E-07 | 2.300E-07 |
| 3112 | 156 | Q156K | A | 2.9e-07 | 2.2e-07 | 2.3e-07 | 2.467E-07 | 3.786E-08 | 2.200E-07 | 2.900E-07 |
| 3113 | 156 | Q156M | A | 2.8e-07 | 4.3e-07 | 3.5e-07 | 3.533E-07 | 7.506E-08 | 2.800E-07 | 4.300E-07 |
| 3114 | 156 | Q156F | A | 2.2e-07 | 3.3e-07 | 8.8e-08 | 2.127E-07 | 1.212E-07 | 8.800E-08 | 3.300E-07 |
| 3115 | 156 | Q156P | A | 2.9e-07 | 1.6e-07 | 2.9e-07 | 2.467E-07 | 7.506E-08 | 1.600E-07 | 2.900E-07 |
| 3116 | 156 | Q156S | A | 2.2e-07 | 3e-07 | 2.3e-07 | 2.500E-07 | 4.359E-08 | 2.200E-07 | 3.000E-07 |
| 3117 | 156 | Q156T | A | 1.9e-07 | 4.4e-07 | 2.6e-07 | 2.967E-07 | 1.290E-07 | 1.900E-07 | 4.400E-07 |
| 3118 | 156 | Q156W | A | 4.7e-07 | 5.7e-07 | 3e-07 | 4.467E-07 | 1.365E-07 | 3.000E-07 | 5.700E-07 |
| 3119 | 156 | Q156Y | A | 1.6e-07 | 4.6e-07 | 2.4e-07 | 2.867E-07 | 1.553E-07 | 1.600E-07 | 4.600E-07 |
| 3120 | 156 | Q156V | A | 7.4e-08 | 1.8e-07 | 4.5e-07 | 2.347E-07 | 1.939E-07 | 7.400E-08 | 4.500E-07 |
| 3121 | 157 | L157A | A | 2.3e-07 | 2.6e-07 | 1.9e-07 | 2.267E-07 | 3.512E-08 | 1.900E-07 | 2.600E-07 |
| 3122 | 157 | L157R | A | 1.6e-07 | 1.5e-07 | 1.8e-07 | 1.633E-07 | 1.528E-08 | 1.500E-07 | 1.800E-07 |
| 3123 | 157 | L157N | A | 1.3e-07 | 2.1e-07 | 2e-07 | 1.800E-07 | 4.359E-08 | 1.300E-07 | 2.100E-07 |
| 3124 | 157 | L157D | A | 3e-07 | 3.1e-07 | 3.8e-07 | 3.300E-07 | 4.359E-08 | 3.000E-07 | 3.800E-07 |
| 3125 | 157 | L157C | A | 1.8e-07 | 7.3e-08 | 2e-07 | 1.510E-07 | 6.829E-08 | 7.300E-08 | 2.000E-07 |
| 3126 | 157 | L157E | A | 5.2e-07 | 6.5e-07 | 4.5e-07 | 5.400E-07 | 1.015E-07 | 4.500E-07 | 6.500E-07 |
| 3127 | 157 | L157Q | A | 1.1e-07 | 2.5e-07 | 2.3e-07 | 1.967E-07 | 7.572E-08 | 1.100E-07 | 2.500E-07 |
| 3128 | 157 | L157G | A | 1.2e-07 | 4.5e-07 | 3e-07 | 2.900E-07 | 1.652E-07 | 1.200E-07 | 4.500E-07 |
| 3129 | 157 | L157H | A | 1.9e-07 | 2.3e-07 | 2.7e-07 | 2.300E-07 | 4.000E-08 | 1.900E-07 | 2.700E-07 |
| 3130 | 157 | L157I | A | 4.9e-07 | 3.3e-07 | 3.2e-07 | 3.800E-07 | 9.539E-08 | 3.200E-07 | 4.900E-07 |
| 3131 | 157 | L157L | A | 3.2e-07 | 1.5e-07 | 2.5e-07 | 2.400E-07 | 8.544E-08 | 1.500E-07 | 3.200E-07 |
| 3132 | 157 | L157K | A | 4e-07 | 1.8e-07 | 1.4e-07 | 2.400E-07 | 1.400E-07 | 1.400E-07 | 4.000E-07 |
| 3133 | 157 | L157M | A | 3.3e-07 | 3.9e-07 | 3.3e-07 | 3.500E-07 | 3.464E-08 | 3.300E-07 | 3.900E-07 |
| 3134 | 157 | L157F | A | 2.6e-07 | 3.5e-07 | 2.5e-07 | 2.867E-07 | 5.508E-08 | 2.500E-07 | 3.500E-07 |
| 3135 | 157 | L157P | A | 1.9e-07 | 1.6e-07 | 3.3e-07 | 2.267E-07 | 9.074E-08 | 1.600E-07 | 3.300E-07 |
| 3136 | 157 | L157S | A | 1.7e-07 | 4.3e-07 | 3.2e-07 | 3.067E-07 | 1.305E-07 | 1.700E-07 | 4.300E-07 |
| 3137 | 157 | L157T | A | 1.7e-07 | 2.9e-07 | 2.6e-07 | 2.400E-07 | 6.245E-08 | 1.700E-07 | 2.900E-07 |
| 3138 | 157 | L157W | A | 3.1e-07 | 1.2e-07 | 1.3e-07 | 1.867E-07 | 1.069E-07 | 1.200E-07 | 3.100E-07 |
| 3139 | 157 | L157Y | A | 1.8e-07 | 2.3e-07 | 2.2e-07 | 2.100E-07 | 2.646E-08 | 1.800E-07 | 2.300E-07 |
| 3140 | 157 | L157V | A | 1.2e-07 | 2.9e-07 | 1.7e-07 | 1.933E-07 | 8.737E-08 | 1.200E-07 | 2.900E-07 |
| 3141 | 158 | A158A | A | 2.3e-07 | 2.1e-07 | 1.5e-07 | 1.967E-07 | 4.163E-08 | 1.500E-07 | 2.300E-07 |
| 3142 | 158 | A158R | A | 8.2e-08 | 1.3e-07 | 1.4e-07 | 1.173E-07 | 3.101E-08 | 8.200E-08 | 1.400E-07 |
| 3143 | 158 | A158N | A | 5.3e-07 | 2.5e-07 | 8.1e-08 | 2.870E-07 | 2.268E-07 | 8.100E-08 | 5.300E-07 |
| 3144 | 158 | A158D | A | 5.3e-07 | 2.2e-07 | 5e-07 | 4.167E-07 | 1.710E-07 | 2.200E-07 | 5.300E-07 |

| | | | | | | | | | | |
|---|---|---|---|---|---|---|---|---|---|---|
| 3145 | 158 | A158C | A | 1.7e-07 | 4e-07 | 1.9e-07 | 2.533E-07 | 1.274E-07 | 1.700E-07 | 4.000E-07 |
| 3146 | 158 | A158E | A | 2.7e-07 | 2e-07 | 4.9e-07 | 3.200E-07 | 1.513E-07 | 2.000E-07 | 4.900E-07 |
| 3147 | 158 | A158Q | A | 1.6e-07 | 4e-07 | 2e-07 | 2.533E-07 | 1.286E-07 | 1.600E-07 | 4.000E-07 |
| 3148 | 158 | A158G | A | 2.8e-07 | 1.3e-07 | 4.2e-07 | 2.767E-07 | 1.450E-07 | 1.300E-07 | 4.200E-07 |
| 3149 | 158 | A158H | A | 1.5e-07 | 1.8e-07 | 1.7e-07 | 1.667E-07 | 1.528E-08 | 1.500E-07 | 1.800E-07 |
| 3150 | 158 | A158I | A | 2.8e-07 | 2.4e-07 | 1.8e-07 | 2.333E-07 | 5.033E-08 | 1.800E-07 | 2.800E-07 |
| 3151 | 158 | A158L | A | 2.9e-07 | 8.9e-07 | 2.7e-07 | 4.833E-07 | 3.523E-07 | 2.700E-07 | 8.900E-07 |
| 3152 | 158 | A158K | A | 1.8e-07 | 2.3e-07 | 3.3e-07 | 2.467E-07 | 7.638E-08 | 1.800E-07 | 3.300E-07 |
| 3153 | 158 | A158M | A | 6.8e-08 | 4.1e-07 | 2.4e-07 | 2.393E-07 | 1.710E-07 | 6.800E-08 | 4.100E-07 |
| 3154 | 158 | A158F | A | 2e-07 | 2.6e-07 | 3e-07 | 2.533E-07 | 5.033E-08 | 2.000E-07 | 3.000E-07 |
| 3155 | 158 | A158P | A | 1.6e-07 | 3.4e-07 | 1.8e-07 | 2.267E-07 | 9.866E-08 | 1.600E-07 | 3.400E-07 |
| 3156 | 158 | A158S | A | 8.2e-07 | 3.1e-07 | 1.2e-07 | 4.167E-07 | 3.620E-07 | 1.200E-07 | 8.200E-07 |
| 3157 | 158 | A158T | A | 3.2e-07 | 3.7e-07 | 5.5e-07 | 4.133E-07 | 1.210E-07 | 3.200E-07 | 5.500E-07 |
| 3158 | 158 | A158W | A | 3.3e-07 | 9.4e-08 | 2.2e-07 | 2.147E-07 | 1.181E-07 | 9.400E-08 | 3.300E-07 |
| 3159 | 158 | A158Y | A | 1.9e-07 | 1.8e-07 | 4.6e-07 | 2.767E-07 | 1.589E-07 | 1.800E-07 | 4.600E-07 |
| 3160 | 158 | A158V | A | 1.2e-07 | 1e-07 | 2.3e-07 | 1.500E-07 | 7.000E-08 | 1.000E-07 | 2.300E-07 |
| 3161 | 159 | L159A | A | 2.3e-07 | 2e-07 | 1.5e-07 | 1.933E-07 | 4.041E-08 | 1.500E-07 | 2.300E-07 |
| 3162 | 159 | L159R | A | 1.9e-07 | 2.1e-07 | 2e-07 | 2.000E-07 | 1.000E-08 | 1.900E-07 | 2.100E-07 |
| 3163 | 159 | L159N | A | 4.3e-07 | 1.9e-07 | 2.9e-07 | 3.033E-07 | 1.206E-07 | 1.900E-07 | 4.300E-07 |
| 3164 | 159 | L159D | A | 1.6e-07 | 1.6e-07 | 1.7e-07 | 1.633E-07 | 5.774E-09 | 1.600E-07 | 1.700E-07 |
| 3165 | 159 | L159C | A | 4e-07 | 1.9e-07 | 2.9e-07 | 2.933E-07 | 1.050E-07 | 1.900E-07 | 4.000E-07 |
| 3166 | 159 | L159E | A | 2.2e-07 | 4.2e-07 | 1.6e-07 | 2.667E-07 | 1.361E-07 | 1.600E-07 | 4.200E-07 |
| 3167 | 159 | L159Q | A | 2.4e-07 | 1.7e-07 | 2.4e-07 | 2.167E-07 | 4.041E-08 | 1.700E-07 | 2.400E-07 |
| 3168 | 159 | L159G | A | 2.8e-07 | 3.3e-07 | 3.1e-07 | 3.067E-07 | 2.517E-08 | 2.800E-07 | 3.300E-07 |
| 3169 | 159 | L159H | A | 2.7e-07 | 7.1e-07 | 5.1e-07 | 4.967E-07 | 2.203E-07 | 2.700E-07 | 7.100E-07 |
| 3170 | 159 | L159I | A | 4.1e-07 | 2.7e-07 | 3e-07 | 3.267E-07 | 7.371E-08 | 2.700E-07 | 4.100E-07 |
| 3171 | 159 | L159L | A | 3e-07 | 2.1e-07 | 1.7e-07 | 2.267E-07 | 6.658E-08 | 1.700E-07 | 3.000E-07 |
| 3172 | 159 | L159K | A | 3e-07 | 4.1e-07 | 1.6e-07 | 2.900E-07 | 1.253E-07 | 1.600E-07 | 4.100E-07 |
| 3173 | 159 | L159M | A | 2.6e-07 | 2.3e-07 | 2.1e-07 | 2.333E-07 | 2.517E-08 | 2.100E-07 | 2.600E-07 |
| 3174 | 159 | L159F | A | 4.3e-07 | 2.3e-07 | 4.9e-07 | 3.833E-07 | 1.361E-07 | 2.300E-07 | 4.900E-07 |
| 3175 | 159 | L159P | A | 2.1e-07 | 4.9e-07 | 1.8e-07 | 2.933E-07 | 1.710E-07 | 1.800E-07 | 4.900E-07 |
| 3176 | 159 | L159S | A | 3.3e-07 | 1.5e-07 | 2e-07 | 2.267E-07 | 9.292E-08 | 1.500E-07 | 3.300E-07 |
| 3177 | 159 | L159T | A | 2.6e-07 | 5.3e-07 | 2.4e-07 | 3.433E-07 | 1.620E-07 | 2.400E-07 | 5.300E-07 |
| 3178 | 159 | L159W | A | 1.7e-07 | 1.1e-07 | 2e-07 | 1.600E-07 | 4.583E-08 | 1.100E-07 | 2.000E-07 |
| 3179 | 159 | L159Y | A | 8.8e-08 | 1.8e-07 | 1.7e-07 | 1.460E-07 | 5.048E-08 | 8.800E-08 | 1.800E-07 |
| 3180 | 159 | L159V | A | 4.3e-07 | 1.4e-07 | 4e-07 | 3.233E-07 | 1.595E-07 | 1.400E-07 | 4.300E-07 |
| 3181 | 160 | T160A | A | 3.1e-07 | 3.7e-07 | 2.8e-07 | 3.200E-07 | 4.583E-08 | 2.800E-07 | 3.700E-07 |
| 3182 | 160 | T160R | A | 2.1e-07 | 2.3e-07 | 5.6e-07 | 3.333E-07 | 1.966E-07 | 2.100E-07 | 5.600E-07 |
| 3183 | 160 | T160N | A | 2.6e-07 | 3.6e-07 | 3.1e-07 | 3.100E-07 | 5.000E-08 | 2.600E-07 | 3.600E-07 |
| 3184 | 160 | T160D | A | 3.9e-07 | 2.8e-07 | 4.1e-07 | 3.600E-07 | 7.000E-08 | 2.800E-07 | 4.100E-07 |
| 3185 | 160 | T160C | A | 1.7e-07 | 3.1e-07 | 1.4e-07 | 2.067E-07 | 9.074E-08 | 1.400E-07 | 3.100E-07 |
| 3186 | 160 | T160E | A | 2.2e-07 | 2.8e-07 | 2.4e-07 | 2.467E-07 | 3.055E-08 | 2.200E-07 | 2.800E-07 |
| 3187 | 160 | T160Q | A | 1.6e-07 | 2.7e-07 | 4.6e-07 | 2.967E-07 | 1.518E-07 | 1.600E-07 | 4.600E-07 |
| 3188 | 160 | T160G | A | 3.6e-07 | 1.2e-07 | 4.6e-07 | 3.133E-07 | 1.747E-07 | 1.200E-07 | 4.600E-07 |
| 3189 | 160 | T160H | A | 1.9e-07 | 2.6e-07 | 2.5e-07 | 2.333E-07 | 3.786E-08 | 1.900E-07 | 2.600E-07 |
| 3190 | 160 | T160I | A | 3.7e-07 | 2.9e-07 | 4e-07 | 3.533E-07 | 5.686E-08 | 2.900E-07 | 4.000E-07 |
| 3191 | 160 | T160L | A | 3.1e-07 | 1e-07 | 2.9e-07 | 2.333E-07 | 1.159E-07 | 1.000E-07 | 3.100E-07 |
| 3192 | 160 | T160K | A | 9e-08 | 3.1e-07 | 2.3e-07 | 2.100E-07 | 1.114E-07 | 9.000E-08 | 3.100E-07 |
| 3193 | 160 | T160M | A | 1.7e-07 | 2.8e-07 | 3.1e-07 | 2.533E-07 | 7.371E-08 | 1.700E-07 | 3.100E-07 |
| 3194 | 160 | T160F | A | 2.1e-07 | 1.9e-07 | 5.5e-07 | 3.167E-07 | 2.023E-07 | 1.900E-07 | 5.500E-07 |
| 3195 | 160 | T160P | A | 2.2e-07 | 2.1e-07 | 4.8e-07 | 3.033E-07 | 1.531E-07 | 2.100E-07 | 4.800E-07 |
| 3196 | 160 | T160S | A | 2.2e-07 | 1.6e-07 | 2.4e-07 | 2.067E-07 | 4.163E-08 | 1.600E-07 | 2.400E-07 |
| 3197 | 160 | T160T | A | 3e-07 | 2.1e-07 | 2.8e-07 | 2.633E-07 | 4.726E-08 | 2.100E-07 | 3.000E-07 |
| 3198 | 160 | T160W | A | 2e-07 | 2.1e-07 | 1.8e-07 | 1.967E-07 | 1.528E-08 | 1.800E-07 | 2.100E-07 |
| 3199 | 160 | T160Y | A | 4.1e-07 | 1.7e-07 | 2.1e-07 | 2.633E-07 | 1.286E-07 | 1.700E-07 | 4.100E-07 |
| 3200 | 160 | T160V | A | 3.3e-07 | 2.1e-07 | 4.1e-07 | 3.167E-07 | 1.007E-07 | 2.100E-07 | 4.100E-07 |
| 3201 | 161 | L161A | A | 5.6e-07 | 3.8e-07 | 1.5e-07 | 3.633E-07 | 2.055E-07 | 1.500E-07 | 5.600E-07 |
| 3202 | 161 | L161R | A | 1.6e-07 | 3.8e-07 | 1.8e-07 | 2.400E-07 | 1.217E-07 | 1.600E-07 | 3.800E-07 |
| 3203 | 161 | L161N | A | 5.6e-07 | 3e-07 | 2e-07 | 3.533E-07 | 1.858E-07 | 2.000E-07 | 5.600E-07 |
| 3204 | 161 | L161D | A | 1.2e-07 | 2.1e-07 | 2.4e-07 | 1.900E-07 | 6.245E-08 | 1.200E-07 | 2.400E-07 |
| 3205 | 161 | L161C | A | 1.9e-07 | 1.7e-07 | 1.7e-07 | 1.767E-07 | 1.155E-08 | 1.700E-07 | 1.900E-07 |
| 3206 | 161 | L161E | A | 3.4e-07 | 1.7e-07 | 2.6e-07 | 2.567E-07 | 8.505E-08 | 1.700E-07 | 3.400E-07 |
| 3207 | 161 | L161Q | A | 2.8e-07 | 5.7e-07 | 3.4e-07 | 3.967E-07 | 1.531E-07 | 2.800E-07 | 5.700E-07 |
| 3208 | 161 | L161G | A | 2.5e-07 | 4.8e-07 | 6.7e-07 | 4.667E-07 | 2.103E-07 | 2.500E-07 | 6.700E-07 |
| 3209 | 161 | L161H | A | 1.3e-07 | 8.5e-07 | 3.3e-07 | 4.367E-07 | 3.717E-07 | 1.300E-07 | 8.500E-07 |
| 3210 | 161 | L161I | A | 1.7e-07 | 2.3e-07 | 4.5e-07 | 2.833E-07 | 1.474E-07 | 1.700E-07 | 4.500E-07 |
| 3211 | 161 | L161L | A | 3.5e-07 | 1.2e-07 | 4e-07 | 2.900E-07 | 1.493E-07 | 1.200E-07 | 4.000E-07 |
| 3212 | 161 | L161K | A | 3.7e-07 | 1.8e-07 | 2.1e-07 | 2.533E-07 | 1.021E-07 | 1.800E-07 | 3.700E-07 |
| 3213 | 161 | L161M | A | 1.5e-07 | 3.2e-07 | 2.3e-07 | 2.333E-07 | 8.505E-08 | 1.500E-07 | 3.200E-07 |
| 3214 | 161 | L161F | A | 2.5e-07 | 3.1e-07 | 4.8e-07 | 3.467E-07 | 1.193E-07 | 2.500E-07 | 4.800E-07 |
| 3215 | 161 | L161P | A | 2.1e-07 | 2.1e-07 | 4.3e-07 | 2.833E-07 | 1.270E-07 | 2.100E-07 | 4.300E-07 |
| 3216 | 161 | L161S | A | 2.8e-07 | 5.3e-07 | 3.1e-07 | 3.733E-07 | 1.365E-07 | 2.800E-07 | 5.300E-07 |
| 3217 | 161 | L161T | A | 2.9e-07 | 1.3e-07 | 2.3e-07 | 2.167E-07 | 8.083E-08 | 1.300E-07 | 2.900E-07 |
| 3218 | 161 | L161W | A | 1.6e-07 | 2.8e-07 | 2.2e-07 | 2.200E-07 | 6.000E-08 | 1.600E-07 | 2.800E-07 |
| 3219 | 161 | L161Y | A | 1.9e-07 | 2.1e-07 | 2.7e-07 | 2.233E-07 | 4.163E-08 | 1.900E-07 | 2.700E-07 |
| 3220 | 161 | L161V | A | 1.4e-07 | 1.7e-07 | 2e-07 | 1.700E-07 | 3.000E-08 | 1.400E-07 | 2.000E-07 |
| 3221 | 162 | I162A | A | 1.4e-07 | 2.5e-07 | 2e-07 | 1.967E-07 | 5.508E-08 | 1.400E-07 | 2.500E-07 |
| 3222 | 162 | I162R | A | 3.5e-07 | 2.2e-07 | 3.6e-07 | 3.100E-07 | 7.810E-08 | 2.200E-07 | 3.600E-07 |
| 3223 | 162 | I162N | A | 4.1e-07 | 1.7e-07 | 2.4e-07 | 2.733E-07 | 1.234E-07 | 1.700E-07 | 4.100E-07 |
| 3224 | 162 | I162D | A | 2.7e-07 | 1.9e-07 | 4.8e-07 | 3.133E-07 | 1.498E-07 | 1.900E-07 | 4.800E-07 |
| 3225 | 162 | I162C | A | 4.8e-07 | 1.7e-07 | 2.7e-07 | 3.067E-07 | 1.582E-07 | 1.700E-07 | 4.800E-07 |
| 3226 | 162 | I162E | A | 6e-07 | 1.7e-07 | 3.4e-07 | 3.700E-07 | 2.166E-07 | 1.700E-07 | 6.000E-07 |
| 3227 | 162 | I162Q | A | 4.1e-07 | 1.7e-07 | 2.9e-07 | 2.900E-07 | 1.200E-07 | 1.700E-07 | 4.100E-07 |
| 3228 | 162 | I162G | A | 2.8e-07 | 3.2e-07 | 2e-07 | 2.667E-07 | 6.110E-08 | 2.000E-07 | 3.200E-07 |
| 3229 | 162 | I162H | A | 3.1e-07 | 1.7e-07 | 1.9e-07 | 2.233E-07 | 7.572E-08 | 1.700E-07 | 3.100E-07 |

| | | | | | | | | | | |
|---|---|---|---|---|---|---|---|---|---|---|
| 3230 | 162 | I162I | A | 2.2e-07 | 2.1e-07 | 2.3e-07 | 2.200E-07 | 1.000E-08 | 2.100E-07 | 2.300E-07 |
| 3231 | 162 | I162L | A | 3e-07 | 4e-07 | 3.6e-07 | 3.533E-07 | 5.033E-08 | 3.000E-07 | 4.000E-07 |
| 3232 | 162 | I162K | A | 1.8e-07 | 1.8e-07 | 4.2e-07 | 2.600E-07 | 1.386E-07 | 1.800E-07 | 4.200E-07 |
| 3233 | 162 | I162M | A | 1.6e-07 | 2.6e-07 | 2.1e-07 | 2.100E-07 | 5.000E-08 | 1.600E-07 | 2.600E-07 |
| 3234 | 162 | I162F | A | 2.5e-07 | 1.9e-07 | 2.8e-07 | 2.400E-07 | 4.583E-08 | 1.900E-07 | 2.800E-07 |
| 3235 | 162 | I162P | A | 3.6e-07 | 1.3e-07 | 2.1e-07 | 2.333E-07 | 1.168E-07 | 1.300E-07 | 3.600E-07 |
| 3236 | 162 | I162S | A | 3.1e-07 | 1.7e-07 | 3.5e-07 | 2.767E-07 | 9.452E-08 | 1.700E-07 | 3.500E-07 |
| 3237 | 162 | I162T | A | 2.5e-07 | 4.2e-07 | 3.6e-07 | 3.433E-07 | 8.622E-08 | 2.500E-07 | 4.200E-07 |
| 3238 | 162 | I162W | A | 1.2e-07 | 8.9e-08 | 2.6e-07 | 1.563E-07 | 9.111E-08 | 8.900E-08 | 2.600E-07 |
| 3239 | 162 | I162Y | A | 1.7e-07 | 2.1e-07 | 2.3e-07 | 2.033E-07 | 3.055E-08 | 1.700E-07 | 2.300E-07 |
| 3240 | 162 | I162V | A | 1.6e-07 | 3.6e-07 | 3.9e-07 | 3.033E-07 | 1.250E-07 | 1.600E-07 | 3.900E-07 |
| 3241 | 163 | D163A | A | 5.5e-07 | 1.1e-07 | 4.3e-07 | 3.633E-07 | 2.274E-07 | 1.100E-07 | 5.500E-07 |
| 3242 | 163 | D163R | A | 2.2e-07 | 3.1e-07 | 2.6e-07 | 2.633E-07 | 4.509E-08 | 2.200E-07 | 3.100E-07 |
| 3243 | 163 | D163N | A | 3.6e-07 | 2.8e-07 | 2.3e-07 | 2.900E-07 | 6.557E-08 | 2.300E-07 | 3.600E-07 |
| 3244 | 163 | D163D | A | 1.2e-07 | 2e-07 | 2.3e-07 | 1.833E-07 | 5.686E-08 | 1.200E-07 | 2.300E-07 |
| 3245 | 163 | D163C | A | 2.9e-07 | 1.3e-07 | 5.2e-07 | 3.133E-07 | 1.960E-07 | 1.300E-07 | 5.200E-07 |
| 3246 | 163 | D163E | A | 2.2e-07 | 3.2e-07 | 2.9e-07 | 2.767E-07 | 5.132E-08 | 2.200E-07 | 3.200E-07 |
| 3247 | 163 | D163Q | A | 2.1e-07 | 2.9e-07 | 2.2e-07 | 2.400E-07 | 4.359E-08 | 2.100E-07 | 2.900E-07 |
| 3248 | 163 | D163G | A | 9.2e-08 | 2.3e-07 | 2.8e-07 | 2.007E-07 | 9.737E-08 | 9.200E-08 | 2.800E-07 |
| 3249 | 163 | D163H | A | 1.3e-07 | 3.3e-07 | 3.7e-07 | 2.767E-07 | 1.286E-07 | 1.300E-07 | 3.700E-07 |
| 3250 | 163 | D163I | A | 4e-07 | 4.5e-07 | 2.9e-07 | 3.800E-07 | 8.185E-08 | 2.900E-07 | 4.500E-07 |
| 3251 | 163 | D163L | A | 1.3e-07 | 1.7e-07 | 2.2e-07 | 1.733E-07 | 4.509E-08 | 1.300E-07 | 2.200E-07 |
| 3252 | 163 | D163K | A | 3.4e-07 | 4e-07 | 4.2e-07 | 3.867E-07 | 4.163E-08 | 3.400E-07 | 4.200E-07 |
| 3253 | 163 | D163M | A | 2.1e-07 | 2.2e-07 | 2.4e-07 | 2.233E-07 | 1.528E-08 | 2.100E-07 | 2.400E-07 |
| 3254 | 163 | D163F | A | 3e-07 | 1.7e-07 | 2e-07 | 2.233E-07 | 6.807E-08 | 1.700E-07 | 3.000E-07 |
| 3255 | 163 | D163P | A | 3e-07 | 2.8e-07 | 2.2e-07 | 2.667E-07 | 4.163E-08 | 2.200E-07 | 3.000E-07 |
| 3256 | 163 | D163S | A | 8.5e-08 | 8.5e-08 | 1.6e-07 | 1.100E-07 | 4.330E-08 | 8.500E-08 | 1.600E-07 |
| 3257 | 163 | D163T | A | 2.3e-07 | 1.4e-07 | 2.5e-07 | 2.067E-07 | 5.859E-08 | 1.400E-07 | 2.500E-07 |
| 3258 | 163 | D163W | A | 4.4e-07 | 1.4e-07 | 3.3e-07 | 3.033E-07 | 1.518E-07 | 1.400E-07 | 4.400E-07 |
| 3259 | 163 | D163Y | A | 1.4e-07 | 1e-07 | 2.3e-07 | 1.567E-07 | 6.658E-08 | 1.000E-07 | 2.300E-07 |
| 3260 | 163 | D163V | A | 2.9e-07 | 2.8e-07 | 2.9e-07 | 2.867E-07 | 5.774E-09 | 2.800E-07 | 2.900E-07 |
| 3261 | 164 | T164A | A | 2.3e-07 | 5.5e-07 | 6.4e-07 | 4.733E-07 | 2.155E-07 | 2.300E-07 | 6.400E-07 |
| 3262 | 164 | T164R | A | 4.9e-07 | 1.9e-07 | 1.7e-07 | 2.833E-07 | 1.793E-07 | 1.700E-07 | 4.900E-07 |
| 3263 | 164 | T164N | A | 1.1e-07 | 4.3e-07 | 2.6e-07 | 2.667E-07 | 1.601E-07 | 1.100E-07 | 4.300E-07 |
| 3264 | 164 | T164D | A | 1.2e-07 | 2.6e-07 | 4.6e-07 | 2.800E-07 | 1.709E-07 | 1.200E-07 | 4.600E-07 |
| 3265 | 164 | T164C | A | 6.2e-07 | 3.3e-07 | 1.8e-07 | 3.767E-07 | 2.237E-07 | 1.800E-07 | 6.200E-07 |
| 3266 | 164 | T164E | A | 1.5e-07 | 1.6e-07 | 4.3e-07 | 2.467E-07 | 1.589E-07 | 1.500E-07 | 4.300E-07 |
| 3267 | 164 | T164Q | A | 2.6e-07 | 2.2e-07 | 1.7e-07 | 2.167E-07 | 4.509E-08 | 1.700E-07 | 2.600E-07 |
| 3268 | 164 | T164G | A | 5.7e-07 | 3e-07 | 4.8e-07 | 4.500E-07 | 1.375E-07 | 3.000E-07 | 5.700E-07 |
| 3269 | 164 | T164H | A | 2.6e-07 | 1.6e-07 | 1.2e-07 | 1.800E-07 | 7.211E-08 | 1.200E-07 | 2.600E-07 |
| 3270 | 164 | T164I | A | 4.7e-07 | 2.5e-07 | 2.6e-07 | 3.267E-07 | 1.242E-07 | 2.500E-07 | 4.700E-07 |
| 3271 | 164 | T164L | A | 2.4e-07 | 4.3e-07 | 2.7e-07 | 3.133E-07 | 1.021E-07 | 2.400E-07 | 4.300E-07 |
| 3272 | 164 | T164K | A | 2.1e-07 | 2.9e-07 | 2.9e-07 | 2.633E-07 | 4.619E-08 | 2.100E-07 | 2.900E-07 |
| 3273 | 164 | T164M | A | 1.9e-07 | 4.4e-07 | 3.3e-07 | 3.200E-07 | 1.253E-07 | 1.900E-07 | 4.400E-07 |
| 3274 | 164 | T164F | A | 2.3e-07 | 3e-07 | 2.1e-07 | 2.467E-07 | 4.726E-08 | 2.100E-07 | 3.000E-07 |
| 3275 | 164 | T164P | A | 3.7e-07 | 1.2e-07 | 2.4e-07 | 2.433E-07 | 1.250E-07 | 1.200E-07 | 3.700E-07 |
| 3276 | 164 | T164S | A | 2.9e-07 | 2.2e-07 | 1.9e-07 | 2.333E-07 | 5.132E-08 | 1.900E-07 | 2.900E-07 |
| 3277 | 164 | T164T | A | 1.4e-07 | 1.7e-07 | 2.9e-07 | 2.000E-07 | 7.937E-08 | 1.400E-07 | 2.900E-07 |
| 3278 | 164 | T164W | A | 4.4e-07 | 3.4e-07 | 2.4e-07 | 3.400E-07 | 1.000E-07 | 2.400E-07 | 4.400E-07 |
| 3279 | 164 | T164Y | A | 3.7e-07 | 3.8e-07 | 2.2e-07 | 3.233E-07 | 8.963E-08 | 2.200E-07 | 3.800E-07 |
| 3280 | 164 | T164V | A | 2.7e-07 | 3.8e-07 | 2.1e-07 | 2.867E-07 | 8.622E-08 | 2.100E-07 | 3.800E-07 |
| 3281 | 165 | N165A | A | 3.1e-07 | 3.7e-07 | 3e-07 | 3.267E-07 | 3.786E-08 | 3.000E-07 | 3.700E-07 |
| 3282 | 165 | N165R | A | 3.4e-07 | 3.8e-07 | 2e-07 | 3.067E-07 | 9.452E-08 | 2.000E-07 | 3.800E-07 |
| 3283 | 165 | N165N | A | 1.4e-07 | 2.2e-07 | 2.4e-07 | 2.000E-07 | 5.292E-08 | 1.400E-07 | 2.400E-07 |
| 3284 | 165 | N165D | A | 6.1e-07 | 2.3e-07 | 4.3e-07 | 4.233E-07 | 1.901E-07 | 2.300E-07 | 6.100E-07 |
| 3285 | 165 | N165C | A | 3.2e-07 | 3.1e-07 | 2.2e-07 | 2.833E-07 | 5.508E-08 | 2.200E-07 | 3.200E-07 |
| 3286 | 165 | N165E | A | 5.7e-07 | 3.3e-07 | 1.8e-07 | 3.600E-07 | 1.967E-07 | 1.800E-07 | 5.700E-07 |
| 3287 | 165 | N165Q | A | 3.1e-07 | 2.9e-07 | 2.2e-07 | 2.733E-07 | 4.726E-08 | 2.200E-07 | 3.100E-07 |
| 3288 | 165 | N165G | A | 1.8e-07 | 2.4e-07 | 1.6e-07 | 1.933E-07 | 4.163E-08 | 1.600E-07 | 2.400E-07 |
| 3289 | 165 | N165H | A | 6.3e-07 | 3.1e-07 | 2e-07 | 3.800E-07 | 2.234E-07 | 2.000E-07 | 6.300E-07 |
| 3290 | 165 | N165I | A | 2.2e-07 | 1.2e-07 | 2.2e-07 | 1.867E-07 | 5.774E-08 | 1.200E-07 | 2.200E-07 |
| 3291 | 165 | N165L | A | 1.9e-07 | 2.7e-07 | 6e-07 | 3.533E-07 | 2.173E-07 | 1.900E-07 | 6.000E-07 |
| 3292 | 165 | N165K | A | 8.7e-08 | 2.8e-07 | 1.1e-07 | 1.590E-07 | 1.054E-07 | 8.700E-08 | 2.800E-07 |
| 3293 | 165 | N165M | A | 3e-07 | 1.3e-07 | 2e-07 | 2.100E-07 | 8.544E-08 | 1.300E-07 | 3.000E-07 |
| 3294 | 165 | N165F | A | 3.5e-07 | 2.2e-07 | 1.5e-07 | 2.400E-07 | 1.015E-07 | 1.500E-07 | 3.500E-07 |
| 3295 | 165 | N165P | A | 4.2e-07 | 1.5e-07 | 2e-07 | 2.567E-07 | 1.436E-07 | 1.500E-07 | 4.200E-07 |
| 3296 | 165 | N165S | A | 2.3e-07 | 4.2e-07 | 3.2e-07 | 3.233E-07 | 9.504E-08 | 2.300E-07 | 4.200E-07 |
| 3297 | 165 | N165T | A | 2.6e-07 | 4.3e-07 | 2.5e-07 | 3.133E-07 | 1.012E-07 | 2.500E-07 | 4.300E-07 |
| 3298 | 165 | N165W | A | 1.5e-07 | 4.5e-07 | 3.9e-07 | 3.300E-07 | 1.587E-07 | 1.500E-07 | 4.500E-07 |
| 3299 | 165 | N165Y | A | 4.5e-07 | 1.5e-07 | 2.1e-07 | 2.700E-07 | 1.587E-07 | 1.500E-07 | 4.500E-07 |
| 3300 | 165 | N165V | A | 1.9e-07 | 1.3e-07 | 1.7e-07 | 1.633E-07 | 3.055E-08 | 1.300E-07 | 1.900E-07 |
| 3301 | 166 | R166A | A | 2.1e-07 | 4.2e-07 | 3.6e-07 | 3.300E-07 | 1.082E-07 | 2.100E-07 | 4.200E-07 |
| 3302 | 166 | R166R | A | 1.4e-07 | 1.4e-07 | 3.1e-07 | 1.967E-07 | 9.815E-08 | 1.400E-07 | 3.100E-07 |
| 3303 | 166 | R166N | A | 2.4e-07 | 3.6e-07 | 1.4e-07 | 2.467E-07 | 1.102E-07 | 1.400E-07 | 3.600E-07 |
| 3304 | 166 | R166D | A | 2.5e-07 | 2.3e-07 | 8.9e-07 | 4.567E-07 | 3.754E-07 | 2.300E-07 | 8.900E-07 |
| 3305 | 166 | R166C | A | 2e-07 | 1.5e-07 | 2.2e-07 | 1.900E-07 | 3.606E-08 | 1.500E-07 | 2.200E-07 |
| 3306 | 166 | R166E | A | 3.3e-07 | 1.3e-07 | 2.2e-07 | 2.267E-07 | 1.002E-07 | 1.300E-07 | 3.300E-07 |
| 3307 | 166 | R166Q | A | 3.8e-07 | 1.3e-07 | 3.7e-07 | 2.933E-07 | 1.415E-07 | 1.300E-07 | 3.800E-07 |
| 3308 | 166 | R166G | A | 4.9e-07 | 6.2e-07 | 3.4e-07 | 4.833E-07 | 1.401E-07 | 3.400E-07 | 6.200E-07 |
| 3309 | 166 | R166H | A | 1.3e-07 | 1.5e-07 | 1.9e-07 | 1.567E-07 | 3.055E-08 | 1.300E-07 | 1.900E-07 |
| 3310 | 166 | R166I | A | 5.6e-07 | 3e-07 | 5.3e-07 | 4.633E-07 | 1.422E-07 | 3.000E-07 | 5.600E-07 |
| 3311 | 166 | R166L | A | 2.4e-07 | 2.5e-07 | 2.6e-07 | 2.500E-07 | 1.000E-08 | 2.400E-07 | 2.600E-07 |
| 3312 | 166 | R166K | A | 3e-07 | 2.2e-07 | 2.3e-07 | 2.500E-07 | 4.359E-08 | 2.200E-07 | 3.000E-07 |
| 3313 | 166 | R166M | A | 2e-07 | 3e-07 | 2.9e-07 | 2.633E-07 | 5.508E-08 | 2.000E-07 | 3.000E-07 |
| 3314 | 166 | R166F | A | 7.8e-08 | 1.4e-07 | 2.8e-07 | 1.660E-07 | 1.035E-07 | 7.800E-08 | 2.800E-07 |

| | | | | | | | | | | |
|---|---|---|---|---|---|---|---|---|---|---|
| 3315 | 166 | R166P | A | 1e-07 | 2.5e-07 | 2.7e-07 | 2.067E-07 | 9.292E-08 | 1.000E-07 | 2.700E-07 |
| 3316 | 166 | R166S | A | 4e-07 | 1.6e-07 | 1.6e-07 | 2.400E-07 | 1.386E-07 | 1.600E-07 | 4.000E-07 |
| 3317 | 166 | R166T | A | 3e-07 | 2.9e-07 | 2e-07 | 2.633E-07 | 5.508E-08 | 2.000E-07 | 3.000E-07 |
| 3318 | 166 | R166W | A | 3.4e-07 | 1.9e-07 | 2.5e-07 | 2.600E-07 | 7.550E-08 | 1.900E-07 | 3.400E-07 |
| 3319 | 166 | R166Y | A | 3.9e-07 | 3.5e-07 | 1.6e-07 | 3.000E-07 | 1.229E-07 | 1.600E-07 | 3.900E-07 |
| 3320 | 166 | R166V | A | 1.6e-07 | 3.1e-07 | 1.6e-07 | 2.100E-07 | 8.660E-08 | 1.600E-07 | 3.100E-07 |
| 3321 | 167 | S167A | A | 1.6e-07 | 3.1e-07 | 1.9e-07 | 2.200E-07 | 7.937E-08 | 1.600E-07 | 3.100E-07 |
| 3322 | 167 | S167R | A | 1.7e-07 | 2.5e-07 | 2.6e-07 | 2.267E-07 | 4.933E-08 | 1.700E-07 | 2.600E-07 |
| 3323 | 167 | S167N | A | 3.2e-07 | 9.2e-08 | 2.6e-07 | 2.240E-07 | 1.182E-07 | 9.200E-08 | 3.200E-07 |
| 3324 | 167 | S167D | A | 2.5e-07 | 5e-07 | 3.6e-07 | 3.700E-07 | 1.253E-07 | 2.500E-07 | 5.000E-07 |
| 3325 | 167 | S167C | A | 4.4e-07 | 5.6e-07 | 1.3e-07 | 3.767E-07 | 2.219E-07 | 1.300E-07 | 5.600E-07 |
| 3326 | 167 | S167E | A | 2.1e-07 | 2.6e-07 | 2.7e-07 | 2.467E-07 | 3.215E-08 | 2.100E-07 | 2.700E-07 |
| 3327 | 167 | S167Q | A | 1.5e-07 | 2e-07 | 2.7e-07 | 2.067E-07 | 6.028E-08 | 1.500E-07 | 2.700E-07 |
| 3328 | 167 | S167G | A | 2.7e-07 | 2e-07 | 5.8e-07 | 3.500E-07 | 2.022E-07 | 2.000E-07 | 5.800E-07 |
| 3329 | 167 | S167H | A | 2.1e-07 | 4.4e-07 | 1.6e-07 | 2.700E-07 | 1.493E-07 | 1.600E-07 | 4.400E-07 |
| 3330 | 167 | S167I | A | 2.9e-07 | 4.7e-07 | 2.7e-07 | 3.433E-07 | 1.102E-07 | 2.700E-07 | 4.700E-07 |
| 3331 | 167 | S167L | A | 1.7e-07 | 1.9e-07 | 1.9e-07 | 1.833E-07 | 1.155E-08 | 1.700E-07 | 1.900E-07 |
| 3332 | 167 | S167K | A | 3.3e-07 | 2e-07 | 2.3e-07 | 2.533E-07 | 6.807E-08 | 2.000E-07 | 3.300E-07 |
| 3333 | 167 | S167M | A | 2.5e-07 | 1.8e-07 | 9.1e-07 | 4.467E-07 | 4.028E-07 | 1.800E-07 | 9.100E-07 |
| 3334 | 167 | S167F | A | 1.9e-07 | 2.5e-07 | 2e-07 | 2.133E-07 | 3.215E-08 | 1.900E-07 | 2.500E-07 |
| 3335 | 167 | S167P | A | 4.3e-07 | 2.6e-07 | 1.9e-07 | 2.933E-07 | 1.234E-07 | 1.900E-07 | 4.300E-07 |
| 3336 | 167 | S167S | A | 3e-07 | 3.7e-07 | 3.9e-07 | 3.533E-07 | 4.726E-08 | 3.000E-07 | 3.900E-07 |
| 3337 | 167 | S167T | A | 1e-07 | 3.4e-07 | 1.7e-07 | 2.033E-07 | 1.234E-07 | 1.000E-07 | 3.400E-07 |
| 3338 | 167 | S167W | A | 5.9e-07 | 2.2e-07 | 1.6e-07 | 3.233E-07 | 2.329E-07 | 1.600E-07 | 5.900E-07 |
| 3339 | 167 | S167Y | A | 2.3e-07 | 1e-07 | 1.7e-07 | 1.667E-07 | 6.506E-08 | 1.000E-07 | 2.300E-07 |
| 3340 | 167 | S167V | A | 4.9e-07 | 2.8e-07 | 1.9e-07 | 3.200E-07 | 1.539E-07 | 1.900E-07 | 4.900E-07 |
| 3341 | 168 | R168A | A | 2.5e-07 | 1.9e-07 | 1.8e-07 | 2.067E-07 | 3.786E-08 | 1.800E-07 | 2.500E-07 |
| 3342 | 168 | R168R | A | 2.4e-07 | 7.7e-08 | 4.7e-07 | 2.623E-07 | 1.974E-07 | 7.700E-08 | 4.700E-07 |
| 3343 | 168 | R168N | A | 1.4e-07 | 1.7e-07 | 2.7e-07 | 1.933E-07 | 6.807E-08 | 1.400E-07 | 2.700E-07 |
| 3344 | 168 | R168D | A | 2.8e-07 | 2e-07 | 2.4e-07 | 2.400E-07 | 4.000E-08 | 2.000E-07 | 2.800E-07 |
| 3345 | 168 | R168C | A | 3e-07 | 3.1e-07 | 2e-07 | 2.700E-07 | 6.083E-08 | 2.000E-07 | 3.100E-07 |
| 3346 | 168 | R168E | A | 2.7e-07 | 3.7e-07 | 1.1e-07 | 2.500E-07 | 1.311E-07 | 1.100E-07 | 3.700E-07 |
| 3347 | 168 | R168Q | A | 2.9e-07 | 3e-07 | 2.3e-07 | 2.733E-07 | 3.786E-08 | 2.300E-07 | 3.000E-07 |
| 3348 | 168 | R168G | A | 6.3e-07 | 1.1e-07 | 2.3e-07 | 3.233E-07 | 2.723E-07 | 1.100E-07 | 6.300E-07 |
| 3349 | 168 | R168H | A | 1.6e-07 | 2.3e-07 | 2e-07 | 1.967E-07 | 3.512E-08 | 1.600E-07 | 2.300E-07 |
| 3350 | 168 | R168I | A | 3.7e-07 | 1.4e-07 | 7.5e-08 | 1.950E-07 | 1.550E-07 | 7.500E-08 | 3.700E-07 |
| 3351 | 168 | R168L | A | 5e-07 | 2.3e-07 | 4.1e-07 | 3.800E-07 | 1.375E-07 | 2.300E-07 | 5.000E-07 |
| 3352 | 168 | R168K | A | 2.4e-07 | 3.1e-07 | 4.5e-08 | 1.983E-07 | 1.373E-07 | 4.500E-08 | 3.100E-07 |
| 3353 | 168 | R168M | A | 2.9e-07 | 1.8e-07 | 3.9e-07 | 2.867E-07 | 1.050E-07 | 1.800E-07 | 3.900E-07 |
| 3354 | 168 | R168F | A | 2.6e-07 | 3e-07 | 5.6e-07 | 3.733E-07 | 1.629E-07 | 2.600E-07 | 5.600E-07 |
| 3355 | 168 | R168P | A | 1.9e-07 | 2.8e-07 | 8.1e-08 | 1.837E-07 | 9.965E-08 | 8.100E-08 | 2.800E-07 |
| 3356 | 168 | R168S | A | 2.4e-07 | 1.6e-07 | 1.2e-07 | 1.733E-07 | 6.110E-08 | 1.200E-07 | 2.400E-07 |
| 3357 | 168 | R168T | A | 1.2e-07 | 2.7e-07 | 1.8e-07 | 1.900E-07 | 7.550E-08 | 1.200E-07 | 2.700E-07 |
| 3358 | 168 | R168W | A | 4.1e-07 | 3.5e-07 | 2.8e-07 | 3.467E-07 | 6.506E-08 | 2.800E-07 | 4.100E-07 |
| 3359 | 168 | R168Y | A | 2.6e-07 | 1.2e-07 | 3.7e-07 | 2.500E-07 | 1.253E-07 | 1.200E-07 | 3.700E-07 |
| 3360 | 168 | R168V | A | 2.4e-07 | 1.9e-07 | 2.2e-07 | 2.167E-07 | 2.517E-08 | 1.900E-07 | 2.400E-07 |
| 3361 | 169 | A169A | A | 2.3e-07 | 1.5e-07 | 2.4e-07 | 2.067E-07 | 4.933E-08 | 1.500E-07 | 2.400E-07 |
| 3362 | 169 | A169R | A | 9.9e-08 | 1.6e-07 | 5.8e-07 | 2.797E-07 | 2.619E-07 | 9.900E-08 | 5.800E-07 |
| 3363 | 169 | A169N | A | 2.3e-07 | 4.3e-07 | 2e-07 | 2.867E-07 | 1.250E-07 | 2.000E-07 | 4.300E-07 |
| 3364 | 169 | A169D | A | 1.3e-07 | 1.5e-07 | 2e-07 | 1.600E-07 | 3.606E-08 | 1.300E-07 | 2.000E-07 |
| 3365 | 169 | A169C | A | 2.7e-07 | 2.2e-07 | 3.6e-07 | 2.833E-07 | 7.095E-08 | 2.200E-07 | 3.600E-07 |
| 3366 | 169 | A169E | A | 1.4e-07 | 4.8e-07 | 1.7e-07 | 2.633E-07 | 1.882E-07 | 1.400E-07 | 4.800E-07 |
| 3367 | 169 | A169Q | A | 2.3e-07 | 2.4e-07 | 2.5e-07 | 2.400E-07 | 1.000E-08 | 2.300E-07 | 2.500E-07 |
| 3368 | 169 | A169G | A | 1.4e-07 | 2.2e-07 | 1.9e-07 | 1.833E-07 | 4.041E-08 | 1.400E-07 | 2.200E-07 |
| 3369 | 169 | A169H | A | 1.5e-07 | 1.2e-07 | 4.1e-07 | 2.267E-07 | 1.595E-07 | 1.200E-07 | 4.100E-07 |
| 3370 | 169 | A169I | A | 2.2e-07 | 2.1e-07 | 1.5e-07 | 1.933E-07 | 3.786E-08 | 1.500E-07 | 2.200E-07 |
| 3371 | 169 | A169L | A | 8.1e-08 | 4.5e-07 | 2e-07 | 2.437E-07 | 1.883E-07 | 8.100E-08 | 4.500E-07 |
| 3372 | 169 | A169K | A | 3e-07 | 2.7e-07 | 2.3e-07 | 2.667E-07 | 3.512E-08 | 2.300E-07 | 3.000E-07 |
| 3373 | 169 | A169M | A | 3.2e-07 | 2.6e-07 | 1.9e-07 | 2.567E-07 | 6.506E-08 | 1.900E-07 | 3.200E-07 |
| 3374 | 169 | A169F | A | 2.2e-07 | 1.5e-07 | 1.3e-07 | 1.667E-07 | 4.726E-08 | 1.300E-07 | 2.200E-07 |
| 3375 | 169 | A169P | A | 1.8e-07 | 1.9e-07 | 2.8e-07 | 2.167E-07 | 5.508E-08 | 1.800E-07 | 2.800E-07 |
| 3376 | 169 | A169S | A | 2.1e-07 | 5.7e-07 | 2.7e-07 | 3.500E-07 | 1.929E-07 | 2.100E-07 | 5.700E-07 |
| 3377 | 169 | A169T | A | 2.9e-07 | 2.7e-07 | 1.6e-07 | 2.400E-07 | 7.000E-08 | 1.600E-07 | 2.900E-07 |
| 3378 | 169 | A169W | A | 9.4e-08 | 3.5e-07 | 1.8e-07 | 2.080E-07 | 1.303E-07 | 9.400E-08 | 3.500E-07 |
| 3379 | 169 | A169Y | A | 2.2e-07 | 1.4e-07 | 4e-07 | 2.533E-07 | 1.332E-07 | 1.400E-07 | 4.000E-07 |
| 3380 | 169 | A169V | A | 3.1e-07 | 1.6e-07 | 2e-07 | 2.233E-07 | 7.767E-08 | 1.600E-07 | 3.100E-07 |
| 3381 | 170 | C170A | A | 2.5e-07 | 3.2e-07 | 2.2e-07 | 2.633E-07 | 5.132E-08 | 2.200E-07 | 3.200E-07 |
| 3382 | 170 | C170R | A | 1.6e-07 | 1.5e-07 | 2.3e-07 | 1.800E-07 | 4.359E-08 | 1.500E-07 | 2.300E-07 |
| 3383 | 170 | C170N | A | 3.4e-07 | 3.2e-07 | 2.6e-07 | 3.067E-07 | 4.163E-08 | 2.600E-07 | 3.400E-07 |
| 3384 | 170 | C170D | A | 3.2e-07 | 3.9e-07 | 3.5e-07 | 3.533E-07 | 3.512E-08 | 3.200E-07 | 3.900E-07 |
| 3385 | 170 | C170C | A | 2.5e-07 | 2.8e-07 | 3.5e-07 | 2.933E-07 | 5.132E-08 | 2.500E-07 | 3.500E-07 |
| 3386 | 170 | C170E | A | 5.4e-07 | 2.3e-07 | 2.8e-07 | 3.500E-07 | 1.664E-07 | 2.300E-07 | 5.400E-07 |
| 3387 | 170 | C170Q | A | 3.5e-07 | 2.4e-07 | 2.7e-07 | 2.867E-07 | 5.686E-08 | 2.400E-07 | 3.500E-07 |
| 3388 | 170 | C170G | A | 2.1e-07 | 1.2e-07 | 1.7e-07 | 1.667E-07 | 4.509E-08 | 1.200E-07 | 2.100E-07 |
| 3389 | 170 | C170H | A | 1.8e-07 | 8.9e-07 | 3.8e-07 | 4.833E-07 | 3.661E-07 | 1.800E-07 | 8.900E-07 |
| 3390 | 170 | C170I | A | 2.7e-07 | 2.8e-07 | 9.4e-08 | 2.147E-07 | 1.046E-07 | 9.400E-08 | 2.800E-07 |
| 3391 | 170 | C170L | A | 1.4e-07 | 2.3e-07 | 2.3e-07 | 2.000E-07 | 5.196E-08 | 1.400E-07 | 2.300E-07 |
| 3392 | 170 | C170K | A | 3e-07 | 4.3e-07 | 6.3e-07 | 4.533E-07 | 1.662E-07 | 3.000E-07 | 6.300E-07 |
| 3393 | 170 | C170M | A | 1.2e-07 | 2.3e-07 | 2.9e-07 | 2.133E-07 | 8.622E-08 | 1.200E-07 | 2.900E-07 |
| 3394 | 170 | C170F | A | 1.5e-07 | 3.8e-07 | 2.1e-07 | 2.467E-07 | 1.193E-07 | 1.500E-07 | 3.800E-07 |
| 3395 | 170 | C170P | A | 2.7e-07 | 3.6e-07 | 4.6e-07 | 3.633E-07 | 9.504E-08 | 2.700E-07 | 4.600E-07 |
| 3396 | 170 | C170S | A | 2.5e-07 | 2.2e-07 | 3.3e-07 | 2.667E-07 | 5.686E-08 | 2.200E-07 | 3.300E-07 |
| 3397 | 170 | C170T | A | 5.8e-07 | 1.6e-07 | 3.3e-07 | 3.567E-07 | 2.113E-07 | 1.600E-07 | 5.800E-07 |
| 3398 | 170 | C170W | A | 1.9e-07 | 2.9e-07 | 3.7e-07 | 2.833E-07 | 9.018E-08 | 1.900E-07 | 3.700E-07 |
| 3399 | 170 | C170Y | A | 2.7e-07 | 1.3e-07 | 2.6e-07 | 2.200E-07 | 7.810E-08 | 1.300E-07 | 2.700E-07 |

| | | | | | | | | | | |
|---|---|---|---|---|---|---|---|---|---|---|
| 3400 | 170 | C170V | A | 3.4e-07 | 2.8e-07 | 3.2e-07 | 3.133E-07 | 3.055E-08 | 2.800E-07 | 3.400E-07 |
| 3401 | 171 | H171A | A | 1.2e-07 | 3.1e-07 | 1.9e-07 | 2.067E-07 | 9.609E-08 | 1.200E-07 | 3.100E-07 |
| 3402 | 171 | H171R | A | 5.7e-07 | 2.9e-07 | 2.5e-07 | 3.700E-07 | 1.744E-07 | 2.500E-07 | 5.700E-07 |
| 3403 | 171 | H171N | A | 3.5e-07 | 1.2e-07 | 5.5e-07 | 3.400E-07 | 2.152E-07 | 1.200E-07 | 5.500E-07 |
| 3404 | 171 | H171D | A | 1.4e-07 | 2.8e-07 | 1.7e-07 | 1.967E-07 | 7.371E-08 | 1.400E-07 | 2.800E-07 |
| 3405 | 171 | H171C | A | 1.4e-07 | 2.8e-07 | 2e-07 | 2.067E-07 | 7.024E-08 | 1.400E-07 | 2.800E-07 |
| 3406 | 171 | H171E | A | 3e-07 | 1e-07 | 6e-07 | 3.333E-07 | 2.517E-07 | 1.000E-07 | 6.000E-07 |
| 3407 | 171 | H171Q | A | 2.1e-07 | 1.8e-07 | 2.5e-07 | 2.133E-07 | 3.512E-08 | 1.800E-07 | 2.500E-07 |
| 3408 | 171 | H171G | A | 1.4e-07 | 5.5e-07 | 2.5e-07 | 3.133E-07 | 2.122E-07 | 1.400E-07 | 5.500E-07 |
| 3409 | 171 | H171H | A | 7.6e-07 | 2e-07 | 3.2e-07 | 4.267E-07 | 2.948E-07 | 2.000E-07 | 7.600E-07 |
| 3410 | 171 | H171I | A | 2e-07 | 3.4e-07 | 2.4e-07 | 2.600E-07 | 7.211E-08 | 2.000E-07 | 3.400E-07 |
| 3411 | 171 | H171L | A | 2.1e-07 | 8.2e-08 | 3.3e-07 | 2.073E-07 | 1.240E-07 | 8.200E-08 | 3.300E-07 |
| 3412 | 171 | H171K | A | 4.1e-07 | 2.1e-07 | 2.2e-07 | 2.800E-07 | 1.127E-07 | 2.100E-07 | 4.100E-07 |
| 3413 | 171 | H171M | A | 1.9e-07 | 1.7e-07 | 2.3e-07 | 1.967E-07 | 3.055E-08 | 1.700E-07 | 2.300E-07 |
| 3414 | 171 | H171F | A | 4.5e-07 | 2e-07 | 3.3e-07 | 3.267E-07 | 1.250E-07 | 2.000E-07 | 4.500E-07 |
| 3415 | 171 | H171P | A | 2.8e-07 | 4.4e-07 | 1.7e-07 | 2.967E-07 | 1.358E-07 | 1.700E-07 | 4.400E-07 |
| 3416 | 171 | H171S | A | 1.5e-07 | 2.7e-07 | 3.2e-07 | 2.467E-07 | 8.737E-08 | 1.500E-07 | 3.200E-07 |
| 3417 | 171 | H171T | A | 2.1e-07 | 3e-07 | 5.5e-07 | 3.533E-07 | 1.762E-07 | 2.100E-07 | 5.500E-07 |
| 3418 | 171 | H171W | A | 3.3e-07 | 2.2e-07 | 1e-07 | 2.167E-07 | 1.150E-07 | 1.000E-07 | 3.300E-07 |
| 3419 | 171 | H171Y | A | 2.3e-07 | 3.1e-07 | 2.6e-07 | 2.667E-07 | 4.041E-08 | 2.300E-07 | 3.100E-07 |
| 3420 | 171 | H171V | A | 1.4e-07 | 2.4e-07 | 1.7e-07 | 1.833E-07 | 5.132E-08 | 1.400E-07 | 2.400E-07 |
| 3421 | 172 | P172A | A | 2.8e-07 | 5.1e-07 | 3.7e-07 | 3.867E-07 | 1.159E-07 | 2.800E-07 | 5.100E-07 |
| 3422 | 172 | P172R | A | 3.7e-07 | 2.7e-07 | 4.5e-07 | 3.633E-07 | 9.018E-08 | 2.700E-07 | 4.500E-07 |
| 3423 | 172 | P172N | A | 7.5e-07 | 1.7e-07 | 5e-07 | 4.733E-07 | 2.909E-07 | 1.700E-07 | 7.500E-07 |
| 3424 | 172 | P172D | A | 8.1e-08 | 1.8e-07 | 5e-07 | 2.537E-07 | 2.190E-07 | 8.100E-08 | 5.000E-07 |
| 3425 | 172 | P172C | A | 1.5e-07 | 7.5e-08 | 9.8e-08 | 1.077E-07 | 3.842E-08 | 7.500E-08 | 1.500E-07 |
| 3426 | 172 | P172E | A | 4.4e-07 | 3.6e-07 | 4.3e-07 | 4.100E-07 | 4.359E-08 | 3.600E-07 | 4.400E-07 |
| 3427 | 172 | P172Q | A | 2.4e-07 | 4.1e-07 | 2.2e-07 | 2.900E-07 | 1.044E-07 | 2.200E-07 | 4.100E-07 |
| 3428 | 172 | P172G | A | 3.7e-07 | 3.4e-07 | 2e-07 | 3.033E-07 | 9.074E-08 | 2.000E-07 | 3.700E-07 |
| 3429 | 172 | P172H | A | 2.6e-07 | 1.7e-07 | 2.5e-07 | 2.267E-07 | 4.933E-08 | 1.700E-07 | 2.600E-07 |
| 3430 | 172 | P172I | A | 1.4e-07 | 3.7e-07 | 3.2e-07 | 2.767E-07 | 1.210E-07 | 1.400E-07 | 3.700E-07 |
| 3431 | 172 | P172L | A | 9e-08 | 2.2e-07 | 3.8e-07 | 2.300E-07 | 1.453E-07 | 9.000E-08 | 3.800E-07 |
| 3432 | 172 | P172K | A | 2.4e-07 | 1.2e-07 | 2.7e-07 | 2.100E-07 | 7.937E-08 | 1.200E-07 | 2.700E-07 |
| 3433 | 172 | P172M | A | 2.4e-07 | 1.4e-07 | 1.6e-07 | 1.800E-07 | 5.292E-08 | 1.400E-07 | 2.400E-07 |
| 3434 | 172 | P172F | A | 4.1e-07 | 9.8e-08 | 3.9e-07 | 2.993E-07 | 1.746E-07 | 9.800E-08 | 4.100E-07 |
| 3435 | 172 | P172P | A | 5.1e-07 | 2.4e-07 | 2.4e-07 | 3.300E-07 | 1.559E-07 | 2.400E-07 | 5.100E-07 |
| 3436 | 172 | P172S | A | 2.9e-07 | 4.8e-07 | 3.8e-07 | 3.833E-07 | 9.504E-08 | 2.900E-07 | 4.800E-07 |
| 3437 | 172 | P172T | A | 1.2e-07 | 3.2e-07 | 1.7e-07 | 2.033E-07 | 1.041E-07 | 1.200E-07 | 3.200E-07 |
| 3438 | 172 | P172W | A | 2.3e-07 | 2.9e-07 | 3e-07 | 2.733E-07 | 3.786E-08 | 2.300E-07 | 3.000E-07 |
| 3439 | 172 | P172Y | A | 1.7e-07 | 1.8e-07 | 4.7e-07 | 2.733E-07 | 1.704E-07 | 1.700E-07 | 4.700E-07 |
| 3440 | 172 | P172V | A | 2.1e-07 | 1.7e-07 | 4.5e-07 | 2.767E-07 | 1.514E-07 | 1.700E-07 | 4.500E-07 |
| 3441 | 173 | C173A | A | 2.2e-07 | 1.7e-07 | 2.1e-07 | 2.000E-07 | 2.646E-08 | 1.700E-07 | 2.200E-07 |
| 3442 | 173 | C173R | A | 2.2e-07 | 2.5e-07 | 9.2e-08 | 1.873E-07 | 8.391E-08 | 9.200E-08 | 2.500E-07 |
| 3443 | 173 | C173N | A | 3.8e-07 | 3.2e-07 | 4.2e-07 | 3.733E-07 | 5.033E-08 | 3.200E-07 | 4.200E-07 |
| 3444 | 173 | C173D | A | 1.7e-07 | 5.4e-07 | 1.2e-07 | 2.767E-07 | 2.294E-07 | 1.200E-07 | 5.400E-07 |
| 3445 | 173 | C173C | A | 1.8e-07 | 3.8e-07 | 2.4e-07 | 2.667E-07 | 1.026E-07 | 1.800E-07 | 3.800E-07 |
| 3446 | 173 | C173E | A | 3e-07 | 3.2e-07 | 3.5e-07 | 3.233E-07 | 2.517E-08 | 3.000E-07 | 3.500E-07 |
| 3447 | 173 | C173Q | A | 2.9e-07 | 4.8e-07 | 1.4e-07 | 3.033E-07 | 1.704E-07 | 1.400E-07 | 4.800E-07 |
| 3448 | 173 | C173G | A | 4.1e-07 | 5.8e-07 | 2.6e-07 | 4.167E-07 | 1.601E-07 | 2.600E-07 | 5.800E-07 |
| 3449 | 173 | C173H | A | 7.6e-07 | 1.9e-07 | 2.7e-07 | 4.067E-07 | 3.086E-07 | 1.900E-07 | 7.600E-07 |
| 3450 | 173 | C173I | A | 3.7e-07 | 3e-07 | 2.9e-07 | 3.200E-07 | 4.359E-08 | 2.900E-07 | 3.700E-07 |
| 3451 | 173 | C173L | A | 2.4e-07 | 1.1e-07 | 4.2e-07 | 2.567E-07 | 1.557E-07 | 1.100E-07 | 4.200E-07 |
| 3452 | 173 | C173K | A | 2.5e-07 | 2.6e-07 | 2.8e-07 | 2.633E-07 | 1.528E-08 | 2.500E-07 | 2.800E-07 |
| 3453 | 173 | C173M | A | 4.8e-07 | 4.9e-07 | 2.5e-07 | 4.067E-07 | 1.358E-07 | 2.500E-07 | 4.900E-07 |
| 3454 | 173 | C173F | A | 3.1e-07 | 3.7e-07 | 5.4e-07 | 4.067E-07 | 1.193E-07 | 3.100E-07 | 5.400E-07 |
| 3455 | 173 | C173P | A | 2.5e-07 | 2.2e-07 | 4.8e-07 | 3.167E-07 | 1.422E-07 | 2.200E-07 | 4.800E-07 |
| 3456 | 173 | C173S | A | 9.8e-08 | 2.4e-07 | 1.3e-07 | 1.560E-07 | 7.448E-08 | 9.800E-08 | 2.400E-07 |
| 3457 | 173 | C173T | A | 4.6e-07 | 1.8e-07 | 1.7e-07 | 2.700E-07 | 1.646E-07 | 1.700E-07 | 4.600E-07 |
| 3458 | 173 | C173W | A | 2.5e-07 | 3.5e-07 | 3.9e-07 | 3.300E-07 | 7.211E-08 | 2.500E-07 | 3.900E-07 |
| 3459 | 173 | C173Y | A | 3.9e-07 | 2.9e-07 | 7.1e-07 | 4.633E-07 | 2.194E-07 | 2.900E-07 | 7.100E-07 |
| 3460 | 173 | C173V | A | 2.3e-07 | 4e-07 | 3.2e-07 | 3.167E-07 | 8.505E-08 | 2.300E-07 | 4.000E-07 |
| 3461 | 174 | S174A | A | 2.7e-07 | 2e-07 | 3.3e-07 | 2.667E-07 | 6.506E-08 | 2.000E-07 | 3.300E-07 |
| 3462 | 174 | S174R | A | 2.5e-07 | 2.6e-07 | 1.1e-07 | 2.067E-07 | 8.386E-08 | 1.100E-07 | 2.600E-07 |
| 3463 | 174 | S174N | A | 1.8e-07 | 3.4e-07 | 3.4e-07 | 2.867E-07 | 9.238E-08 | 1.800E-07 | 3.400E-07 |
| 3464 | 174 | S174D | A | 3.6e-07 | 2.1e-07 | 2.1e-07 | 2.600E-07 | 8.660E-08 | 2.100E-07 | 3.600E-07 |
| 3465 | 174 | S174C | A | 2.1e-07 | 1.9e-07 | 1.5e-07 | 1.833E-07 | 3.055E-08 | 1.500E-07 | 2.100E-07 |
| 3466 | 174 | S174E | A | 3e-07 | 1.7e-07 | 1.6e-07 | 2.100E-07 | 7.810E-08 | 1.600E-07 | 3.000E-07 |
| 3467 | 174 | S174Q | A | 3.7e-07 | 4.5e-07 | 1.3e-07 | 3.167E-07 | 1.665E-07 | 1.300E-07 | 4.500E-07 |
| 3468 | 174 | S174G | A | 7.9e-08 | 1.1e-07 | 5.2e-07 | 2.363E-07 | 2.462E-07 | 7.900E-08 | 5.200E-07 |
| 3469 | 174 | S174H | A | 1.7e-07 | 3.1e-07 | 2.4e-07 | 2.400E-07 | 7.000E-08 | 1.700E-07 | 3.100E-07 |
| 3470 | 174 | S174I | A | 3.6e-07 | 3.8e-07 | 2.5e-07 | 3.300E-07 | 7.000E-08 | 2.500E-07 | 3.800E-07 |
| 3471 | 174 | S174L | A | 2.7e-07 | 2.7e-07 | 2.3e-07 | 2.567E-07 | 2.309E-08 | 2.300E-07 | 2.700E-07 |
| 3472 | 174 | S174K | A | 1.7e-07 | 2.1e-07 | 3e-07 | 2.267E-07 | 6.658E-08 | 1.700E-07 | 3.000E-07 |
| 3473 | 174 | S174M | A | 3.8e-07 | 1.8e-07 | 1e-07 | 2.200E-07 | 1.442E-07 | 1.000E-07 | 3.800E-07 |
| 3474 | 174 | S174F | A | 3.2e-07 | 4.9e-07 | 3.6e-07 | 3.900E-07 | 8.888E-08 | 3.200E-07 | 4.900E-07 |
| 3475 | 174 | S174P | A | 3e-07 | 3.2e-07 | 1.8e-07 | 2.667E-07 | 7.572E-08 | 1.800E-07 | 3.200E-07 |
| 3476 | 174 | S174S | A | 1.7e-07 | 3.8e-07 | 3.9e-07 | 3.133E-07 | 1.242E-07 | 1.700E-07 | 3.900E-07 |
| 3477 | 174 | S174T | A | 1.9e-07 | 1.6e-07 | 5.1e-07 | 2.867E-07 | 1.940E-07 | 1.600E-07 | 5.100E-07 |
| 3478 | 174 | S174W | A | 4.4e-07 | 1.6e-07 | 1.3e-07 | 2.433E-07 | 1.710E-07 | 1.300E-07 | 4.400E-07 |
| 3479 | 174 | S174Y | A | 2.7e-07 | 1.8e-07 | 3.1e-07 | 2.533E-07 | 6.658E-08 | 1.800E-07 | 3.100E-07 |
| 3480 | 174 | S174V | A | 2.5e-07 | 3.2e-07 | 3.2e-07 | 2.967E-07 | 4.041E-08 | 2.500E-07 | 3.200E-07 |
| 3481 | 175 | P175A | A | 3.8e-07 | 8.3e-07 | 2.1e-07 | 4.733E-07 | 3.204E-07 | 2.100E-07 | 8.300E-07 |
| 3482 | 175 | P175R | A | 1.9e-07 | 2e-07 | 3.5e-07 | 2.467E-07 | 8.963E-08 | 1.900E-07 | 3.500E-07 |
| 3483 | 175 | P175N | A | 9.3e-08 | 2e-07 | 1.6e-07 | 1.510E-07 | 5.406E-08 | 9.300E-08 | 2.000E-07 |
| 3484 | 175 | P175D | A | 2.9e-07 | 1.8e-07 | 3e-07 | 2.567E-07 | 6.658E-08 | 1.800E-07 | 3.000E-07 |

| | | | | | | | | | | |
|---|---|---|---|---|---|---|---|---|---|---|
| 3485 | 175 | P175C | A | 4.1e-07 | 3e-07 | 2e-07 | 3.033E-07 | 1.050E-07 | 2.000E-07 | 4.100E-07 |
| 3486 | 175 | P175E | A | 1.8e-07 | 4.4e-07 | 2.8e-07 | 3.000E-07 | 1.311E-07 | 1.800E-07 | 4.400E-07 |
| 3487 | 175 | P175Q | A | 2.6e-07 | 5.5e-07 | 3.2e-07 | 3.767E-07 | 1.531E-07 | 2.600E-07 | 5.500E-07 |
| 3488 | 175 | P175G | A | 1.6e-07 | 2.6e-07 | 1.1e-07 | 1.767E-07 | 7.638E-08 | 1.100E-07 | 2.600E-07 |
| 3489 | 175 | P175H | A | 3.2e-07 | 1.5e-07 | 2.6e-07 | 2.433E-07 | 8.622E-08 | 1.500E-07 | 3.200E-07 |
| 3490 | 175 | P175I | A | 5.2e-07 | 6.5e-07 | 3.1e-07 | 4.933E-07 | 1.716E-07 | 3.100E-07 | 6.500E-07 |
| 3491 | 175 | P175L | A | 1.4e-07 | 1.8e-07 | 2.5e-07 | 1.900E-07 | 5.568E-08 | 1.400E-07 | 2.500E-07 |
| 3492 | 175 | P175K | A | 6.7e-08 | 2.5e-07 | 2.2e-07 | 1.790E-07 | 9.815E-08 | 6.700E-08 | 2.500E-07 |
| 3493 | 175 | P175M | A | 5.2e-07 | 1.2e-07 | 3.3e-07 | 3.233E-07 | 2.001E-07 | 1.200E-07 | 5.200E-07 |
| 3494 | 175 | P175F | A | 2.1e-07 | 2.7e-07 | 1.3e-07 | 2.033E-07 | 7.024E-08 | 1.300E-07 | 2.700E-07 |
| 3495 | 175 | P175P | A | 2e-07 | 1.6e-07 | 2.8e-07 | 2.133E-07 | 6.110E-08 | 1.600E-07 | 2.800E-07 |
| 3496 | 175 | P175S | A | 1.1e-07 | 2.2e-07 | 3.8e-07 | 2.367E-07 | 1.358E-07 | 1.100E-07 | 3.800E-07 |
| 3497 | 175 | P175T | A | 3.4e-07 | 2.3e-07 | 2e-07 | 2.567E-07 | 7.371E-08 | 2.000E-07 | 3.400E-07 |
| 3498 | 175 | P175W | A | 1.3e-07 | 4.1e-07 | 3.1e-07 | 2.833E-07 | 1.419E-07 | 1.300E-07 | 4.100E-07 |
| 3499 | 175 | P175Y | A | 2.1e-07 | 9.1e-08 | 3.3e-07 | 2.103E-07 | 1.195E-07 | 9.100E-08 | 3.300E-07 |
| 3500 | 175 | P175V | A | 2.1e-07 | 2.5e-07 | 1.5e-07 | 2.033E-07 | 5.033E-08 | 1.500E-07 | 2.500E-07 |
| 3501 | 176 | M176A | A | 2.5e-07 | 2.3e-07 | 2.6e-07 | 2.467E-07 | 1.528E-08 | 2.300E-07 | 2.600E-07 |
| 3502 | 176 | M176R | A | 1.9e-07 | 2.2e-07 | 3.9e-07 | 2.667E-07 | 1.079E-07 | 1.900E-07 | 3.900E-07 |
| 3503 | 176 | M176N | A | 1.9e-07 | 3.2e-07 | 1.6e-07 | 2.233E-07 | 8.505E-08 | 1.600E-07 | 3.200E-07 |
| 3504 | 176 | M176D | A | 4.1e-07 | 4e-07 | 2.4e-07 | 3.500E-07 | 9.539E-08 | 2.400E-07 | 4.100E-07 |
| 3505 | 176 | M176C | A | 2.6e-07 | 4.1e-07 | 3.3e-07 | 3.333E-07 | 7.506E-08 | 2.600E-07 | 4.100E-07 |
| 3506 | 176 | M176E | A | 1.9e-07 | 1.3e-07 | 3.5e-07 | 2.233E-07 | 1.137E-07 | 1.300E-07 | 3.500E-07 |
| 3507 | 176 | M176Q | A | 2.9e-07 | 2.4e-07 | 4.8e-07 | 3.367E-07 | 1.266E-07 | 2.400E-07 | 4.800E-07 |
| 3508 | 176 | M176G | A | 1.5e-07 | 2.8e-07 | 2.8e-07 | 2.367E-07 | 7.506E-08 | 1.500E-07 | 2.800E-07 |
| 3509 | 176 | M176H | A | 3.8e-07 | 3.2e-07 | 1.7e-07 | 2.900E-07 | 1.082E-07 | 1.700E-07 | 3.800E-07 |
| 3510 | 176 | M176I | A | 1.4e-07 | 2.4e-07 | 3e-07 | 2.267E-07 | 8.083E-08 | 1.400E-07 | 3.000E-07 |
| 3511 | 176 | M176L | A | 4.3e-07 | 2.1e-07 | 1.1e-07 | 2.500E-07 | 1.637E-07 | 1.100E-07 | 4.300E-07 |
| 3512 | 176 | M176K | A | 1.6e-07 | 2.5e-07 | 2.3e-07 | 2.133E-07 | 4.726E-08 | 1.600E-07 | 2.500E-07 |
| 3513 | 176 | M176M | A | 2.8e-07 | 2.3e-07 | 2.3e-07 | 2.467E-07 | 2.887E-08 | 2.300E-07 | 2.800E-07 |
| 3514 | 176 | M176F | A | 2.1e-07 | 2e-07 | 1.1e-07 | 1.733E-07 | 5.508E-08 | 1.100E-07 | 2.100E-07 |
| 3515 | 176 | M176P | A | 4e-07 | 5.7e-07 | 4.7e-07 | 4.800E-07 | 8.544E-08 | 4.000E-07 | 5.700E-07 |
| 3516 | 176 | M176S | A | 1.5e-07 | 3.2e-07 | 2.2e-07 | 2.300E-07 | 8.544E-08 | 1.500E-07 | 3.200E-07 |
| 3517 | 176 | M176T | A | 5.8e-07 | 2.6e-07 | 2.2e-07 | 3.533E-07 | 1.973E-07 | 2.200E-07 | 5.800E-07 |
| 3518 | 176 | M176W | A | 3.2e-07 | 2.2e-07 | 2e-07 | 2.467E-07 | 6.429E-08 | 2.000E-07 | 3.200E-07 |
| 3519 | 176 | M176Y | A | 2.9e-07 | 1.1e-07 | 1.5e-07 | 1.833E-07 | 9.452E-08 | 1.100E-07 | 2.900E-07 |
| 3520 | 176 | M176V | A | 3.7e-07 | 1.8e-07 | 1.5e-07 | 2.333E-07 | 1.193E-07 | 1.500E-07 | 3.700E-07 |
| 3521 | 177 | C177A | A | 3.2e-07 | 3.9e-07 | 8.4e-07 | 5.167E-07 | 2.822E-07 | 3.200E-07 | 8.400E-07 |
| 3522 | 177 | C177R | A | 4.4e-07 | 2.8e-07 | 5.1e-07 | 4.100E-07 | 1.179E-07 | 2.800E-07 | 5.100E-07 |
| 3523 | 177 | C177N | A | 1.9e-07 | 4.9e-07 | 1.7e-07 | 2.833E-07 | 1.793E-07 | 1.700E-07 | 4.900E-07 |
| 3524 | 177 | C177D | A | 4.4e-07 | 3.2e-07 | 3.3e-07 | 3.633E-07 | 6.658E-08 | 3.200E-07 | 4.400E-07 |
| 3525 | 177 | C177C | A | 1.4e-07 | 6.8e-08 | 3e-07 | 1.693E-07 | 1.187E-07 | 6.800E-08 | 3.000E-07 |
| 3526 | 177 | C177E | A | 4.1e-07 | 4.4e-07 | 2.3e-07 | 3.600E-07 | 1.136E-07 | 2.300E-07 | 4.400E-07 |
| 3527 | 177 | C177Q | A | 3.3e-07 | 3e-07 | 4.1e-07 | 3.467E-07 | 5.686E-08 | 3.000E-07 | 4.100E-07 |
| 3528 | 177 | C177G | A | 4.9e-07 | 4.9e-07 | 3.1e-07 | 4.300E-07 | 1.039E-07 | 3.100E-07 | 4.900E-07 |
| 3529 | 177 | C177H | A | 1.2e-07 | 2.3e-07 | 2.1e-07 | 1.867E-07 | 5.859E-08 | 1.200E-07 | 2.300E-07 |
| 3530 | 177 | C177I | A | 1.5e-07 | 2.4e-07 | 2.2e-07 | 2.033E-07 | 4.726E-08 | 1.500E-07 | 2.400E-07 |
| 3531 | 177 | C177L | A | 1.7e-07 | 4.3e-07 | 3.2e-07 | 3.067E-07 | 1.305E-07 | 1.700E-07 | 4.300E-07 |
| 3532 | 177 | C177K | A | 4.6e-07 | 2.9e-07 | 1.5e-07 | 3.000E-07 | 1.552E-07 | 1.500E-07 | 4.600E-07 |
| 3533 | 177 | C177M | A | 5.9e-07 | 1.8e-07 | 1.8e-07 | 3.167E-07 | 2.367E-07 | 1.800E-07 | 5.900E-07 |
| 3534 | 177 | C177F | A | 6.7e-07 | 2.9e-07 | 3.5e-07 | 4.367E-07 | 2.043E-07 | 2.900E-07 | 6.700E-07 |
| 3535 | 177 | C177P | A | 2.1e-07 | 2.6e-07 | 2.5e-07 | 2.400E-07 | 2.646E-08 | 2.100E-07 | 2.600E-07 |
| 3536 | 177 | C177S | A | 1.1e-07 | 3.4e-07 | 2.8e-07 | 2.433E-07 | 1.193E-07 | 1.100E-07 | 3.400E-07 |
| 3537 | 177 | C177T | A | 2.9e-07 | 2.1e-07 | 1.5e-07 | 2.167E-07 | 7.024E-08 | 1.500E-07 | 2.900E-07 |
| 3538 | 177 | C177W | A | 5.7e-07 | 1.8e-07 | 1.2e-07 | 2.900E-07 | 2.443E-07 | 1.200E-07 | 5.700E-07 |
| 3539 | 177 | C177Y | A | 3.3e-07 | 1.9e-07 | 1.4e-07 | 2.200E-07 | 9.849E-08 | 1.400E-07 | 3.300E-07 |
| 3540 | 177 | C177V | A | 3.9e-07 | 1.4e-07 | 4.5e-07 | 3.267E-07 | 1.644E-07 | 1.400E-07 | 4.500E-07 |
| 3541 | 178 | K178A | A | 2.9e-07 | 1.2e-07 | 1.6e-07 | 1.900E-07 | 8.888E-08 | 1.200E-07 | 2.900E-07 |
| 3542 | 178 | K178R | A | 2.1e-07 | 1.1e-07 | 1.7e-07 | 1.633E-07 | 5.033E-08 | 1.100E-07 | 2.100E-07 |
| 3543 | 178 | K178N | A | 1.7e-07 | 1.5e-07 | 1.8e-07 | 1.667E-07 | 1.528E-08 | 1.500E-07 | 1.800E-07 |
| 3544 | 178 | K178D | A | 1.7e-07 | 2.2e-07 | 3.5e-07 | 2.467E-07 | 9.292E-08 | 1.700E-07 | 3.500E-07 |
| 3545 | 178 | K178C | A | 1.3e-07 | 2.4e-07 | 2.3e-07 | 2.000E-07 | 6.083E-08 | 1.300E-07 | 2.400E-07 |
| 3546 | 178 | K178E | A | 2.7e-07 | 1.3e-07 | 1.7e-07 | 1.900E-07 | 7.211E-08 | 1.300E-07 | 2.700E-07 |
| 3547 | 178 | K178Q | A | 2.5e-07 | 9.4e-08 | 2.3e-07 | 1.913E-07 | 8.488E-08 | 9.400E-08 | 2.500E-07 |
| 3548 | 178 | K178G | A | 3.7e-07 | 4.6e-07 | 2.1e-07 | 3.467E-07 | 1.266E-07 | 2.100E-07 | 4.600E-07 |
| 3549 | 178 | K178H | A | 4e-07 | 2.4e-07 | 8.4e-07 | 4.933E-07 | 3.107E-07 | 2.400E-07 | 8.400E-07 |
| 3550 | 178 | K178I | A | 5.5e-07 | 2.1e-07 | 2.7e-07 | 3.433E-07 | 1.815E-07 | 2.100E-07 | 5.500E-07 |
| 3551 | 178 | K178L | A | 9.9e-07 | 5.1e-07 | 2.4e-07 | 5.800E-07 | 3.799E-07 | 2.400E-07 | 9.900E-07 |
| 3552 | 178 | K178K | A | 1.7e-07 | 3.6e-07 | 2.3e-07 | 2.533E-07 | 9.713E-08 | 1.700E-07 | 3.600E-07 |
| 3553 | 178 | K178M | A | 1.5e-07 | 2.6e-07 | 3.1e-07 | 2.400E-07 | 8.185E-08 | 1.500E-07 | 3.100E-07 |
| 3554 | 178 | K178F | A | 1e-07 | 3.1e-07 | 2.5e-07 | 2.200E-07 | 1.082E-07 | 1.000E-07 | 3.100E-07 |
| 3555 | 178 | K178P | A | 1.4e-07 | 2e-07 | 2.2e-07 | 1.867E-07 | 4.163E-08 | 1.400E-07 | 2.200E-07 |
| 3556 | 178 | K178S | A | 3.3e-07 | 5.5e-07 | 3.6e-07 | 4.133E-07 | 1.193E-07 | 3.300E-07 | 5.500E-07 |
| 3557 | 178 | K178T | A | 3.7e-07 | 2.5e-07 | 2e-07 | 2.733E-07 | 8.737E-08 | 2.000E-07 | 3.700E-07 |
| 3558 | 178 | K178W | A | 2.3e-07 | 1.7e-07 | 2.6e-07 | 2.200E-07 | 4.583E-08 | 1.700E-07 | 2.600E-07 |
| 3559 | 178 | K178Y | A | 2.9e-07 | 2.4e-07 | 2.2e-07 | 2.500E-07 | 3.606E-08 | 2.200E-07 | 2.900E-07 |
| 3560 | 178 | K178V | A | 4.5e-07 | 2.1e-07 | 3.2e-07 | 3.267E-07 | 1.201E-07 | 2.100E-07 | 4.500E-07 |
| 3561 | 179 | G179A | A | 1.2e-07 | 4e-07 | 2.4e-07 | 2.533E-07 | 1.405E-07 | 1.200E-07 | 4.000E-07 |
| 3562 | 179 | G179R | A | 1.9e-07 | 1.6e-07 | 4e-07 | 2.500E-07 | 1.308E-07 | 1.600E-07 | 4.000E-07 |
| 3563 | 179 | G179N | A | 1.2e-07 | 3e-07 | 2.9e-07 | 2.367E-07 | 1.012E-07 | 1.200E-07 | 3.000E-07 |
| 3564 | 179 | G179D | A | 2.6e-07 | 2e-07 | 2.5e-07 | 2.367E-07 | 3.215E-08 | 2.000E-07 | 2.600E-07 |
| 3565 | 179 | G179C | A | 1e-07 | 1.1e-07 | 1.5e-07 | 1.200E-07 | 2.646E-08 | 1.000E-07 | 1.500E-07 |
| 3566 | 179 | G179E | A | 3.7e-07 | 6.7e-07 | 3.4e-07 | 4.600E-07 | 1.825E-07 | 3.400E-07 | 6.700E-07 |
| 3567 | 179 | G179Q | A | 3.2e-07 | 1.5e-07 | 2.7e-07 | 2.467E-07 | 8.737E-08 | 1.500E-07 | 3.200E-07 |
| 3568 | 179 | G179G | A | 1.4e-07 | 1.3e-07 | 1.2e-07 | 1.300E-07 | 1.000E-08 | 1.200E-07 | 1.400E-07 |
| 3569 | 179 | G179H | A | 2.3e-07 | 2.6e-07 | 4.2e-07 | 3.033E-07 | 1.021E-07 | 2.300E-07 | 4.200E-07 |

| | | | | | | | | | | |
|---|---|---|---|---|---|---|---|---|---|---|
| 3570 | 179 | G179I | A | 2.3e-07 | 1.5e-07 | 3.3e-07 | 2.367E-07 | 9.018E-08 | 1.500E-07 | 3.300E-07 |
| 3571 | 179 | G179L | A | 3.1e-07 | 2.6e-07 | 2.8e-07 | 2.833E-07 | 2.517E-08 | 2.600E-07 | 3.100E-07 |
| 3572 | 179 | G179K | A | 2.3e-07 | 2.1e-07 | 3.6e-07 | 2.667E-07 | 8.145E-08 | 2.100E-07 | 3.600E-07 |
| 3573 | 179 | G179M | A | 2.7e-07 | 3.1e-07 | 1.8e-07 | 2.533E-07 | 6.658E-08 | 1.800E-07 | 3.100E-07 |
| 3574 | 179 | G179F | A | 2.9e-07 | 2.7e-07 | 4.4e-07 | 3.333E-07 | 9.292E-08 | 2.700E-07 | 4.400E-07 |
| 3575 | 179 | G179P | A | 2.5e-07 | 4.4e-07 | 3.1e-07 | 3.333E-07 | 9.713E-08 | 2.500E-07 | 4.400E-07 |
| 3576 | 179 | G179S | A | 2e-07 | 2.1e-07 | 5.9e-07 | 3.333E-07 | 2.223E-07 | 2.000E-07 | 5.900E-07 |
| 3577 | 179 | G179T | A | 2.1e-07 | 3.7e-07 | 3.1e-07 | 2.967E-07 | 8.083E-08 | 2.100E-07 | 3.700E-07 |
| 3578 | 179 | G179W | A | 1.8e-07 | 2.7e-07 | 1.2e-07 | 1.900E-07 | 7.550E-08 | 1.200E-07 | 2.700E-07 |
| 3579 | 179 | G179Y | A | 1.3e-07 | 2.2e-07 | 1.9e-07 | 1.800E-07 | 4.583E-08 | 1.300E-07 | 2.200E-07 |
| 3580 | 179 | G179V | A | 1.4e-07 | 3.7e-07 | 3.1e-07 | 2.733E-07 | 1.193E-07 | 1.400E-07 | 3.700E-07 |
| 3581 | 180 | S180A | A | 1.3e-07 | 2.8e-07 | 2e-07 | 2.033E-07 | 7.506E-08 | 1.300E-07 | 2.800E-07 |
| 3582 | 180 | S180R | A | 2.6e-07 | 3.7e-07 | 4.6e-07 | 3.633E-07 | 1.002E-07 | 2.600E-07 | 4.600E-07 |
| 3583 | 180 | S180N | A | 3.3e-07 | 3.9e-07 | 1.5e-07 | 2.900E-07 | 1.249E-07 | 1.500E-07 | 3.900E-07 |
| 3584 | 180 | S180D | A | 3.1e-07 | 1.5e-07 | 4.2e-07 | 2.933E-07 | 1.358E-07 | 1.500E-07 | 4.200E-07 |
| 3585 | 180 | S180C | A | 3.3e-07 | 1.3e-07 | 2.2e-07 | 2.267E-07 | 1.002E-07 | 1.300E-07 | 3.300E-07 |
| 3586 | 180 | S180E | A | 1.9e-07 | 2e-07 | 4.6e-07 | 2.833E-07 | 1.531E-07 | 1.900E-07 | 4.600E-07 |
| 3587 | 180 | S180Q | A | 2.5e-07 | 2.4e-07 | 1.6e-07 | 2.167E-07 | 4.933E-08 | 1.600E-07 | 2.500E-07 |
| 3588 | 180 | S180G | A | 3.3e-07 | 2.7e-07 | 2.6e-07 | 2.867E-07 | 3.786E-08 | 2.600E-07 | 3.300E-07 |
| 3589 | 180 | S180H | A | 3.1e-07 | 2.7e-07 | 3.7e-07 | 3.167E-07 | 5.033E-08 | 2.700E-07 | 3.700E-07 |
| 3590 | 180 | S180I | A | 2.5e-07 | 9.2e-08 | 2e-07 | 1.807E-07 | 8.075E-08 | 9.200E-08 | 2.500E-07 |
| 3591 | 180 | S180L | A | 4.5e-07 | 2.2e-07 | 1.9e-07 | 2.867E-07 | 1.422E-07 | 1.900E-07 | 4.500E-07 |
| 3592 | 180 | S180K | A | 3.1e-07 | 2.4e-07 | 1.4e-07 | 2.300E-07 | 8.544E-08 | 1.400E-07 | 3.100E-07 |
| 3593 | 180 | S180M | A | 2.7e-07 | 3.5e-07 | 4.2e-07 | 3.467E-07 | 7.506E-08 | 2.700E-07 | 4.200E-07 |
| 3594 | 180 | S180F | A | 3e-07 | 3.7e-07 | 3.9e-07 | 3.533E-07 | 4.726E-08 | 3.000E-07 | 3.900E-07 |
| 3595 | 180 | S180P | A | 4.4e-07 | 2.4e-07 | 1.6e-07 | 2.800E-07 | 1.442E-07 | 1.600E-07 | 4.400E-07 |
| 3596 | 180 | S180S | A | 2.3e-07 | 4e-07 | 2.6e-07 | 2.967E-07 | 9.074E-08 | 2.300E-07 | 4.000E-07 |
| 3597 | 180 | S180T | A | 6.7e-07 | 2.1e-07 | 2.7e-07 | 3.833E-07 | 2.501E-07 | 2.100E-07 | 6.700E-07 |
| 3598 | 180 | S180W | A | 4.5e-07 | 3.9e-07 | 2.2e-07 | 3.533E-07 | 1.193E-07 | 2.200E-07 | 4.500E-07 |
| 3599 | 180 | S180Y | A | 6.1e-07 | 4.1e-07 | 2.1e-07 | 4.100E-07 | 2.000E-07 | 2.100E-07 | 6.100E-07 |
| 3600 | 180 | S180V | A | 2.2e-07 | 3e-07 | 2.1e-07 | 2.433E-07 | 4.933E-08 | 2.100E-07 | 3.000E-07 |
| 3601 | 181 | R181A | A | 4.9e-07 | 1.5e-07 | 3.3e-07 | 3.233E-07 | 1.701E-07 | 1.500E-07 | 4.900E-07 |
| 3602 | 181 | R181R | A | 3.4e-07 | 3.8e-07 | 4.3e-07 | 3.833E-07 | 4.509E-08 | 3.400E-07 | 4.300E-07 |
| 3603 | 181 | R181N | A | 2.3e-07 | 2.8e-07 | 3.3e-07 | 2.800E-07 | 5.000E-08 | 2.300E-07 | 3.300E-07 |
| 3604 | 181 | R181D | A | 2.8e-07 | 1.6e-07 | 2.5e-07 | 2.300E-07 | 6.245E-08 | 1.600E-07 | 2.800E-07 |
| 3605 | 181 | R181C | A | 1.8e-07 | 1.5e-07 | 2e-07 | 1.767E-07 | 2.517E-08 | 1.500E-07 | 2.000E-07 |
| 3606 | 181 | R181E | A | 2.9e-07 | 1.1e-07 | 3e-07 | 2.333E-07 | 1.069E-07 | 1.100E-07 | 3.000E-07 |
| 3607 | 181 | R181Q | A | 4.3e-07 | 2.2e-07 | 1.6e-07 | 2.700E-07 | 1.418E-07 | 1.600E-07 | 4.300E-07 |
| 3608 | 181 | R181G | A | 4.1e-07 | 1.1e-06 | 1.4e-07 | 5.500E-07 | 4.951E-07 | 1.400E-07 | 1.100E-06 |
| 3609 | 181 | R181H | A | 5e-07 | 1.9e-07 | 8.5e-08 | 2.583E-07 | 2.158E-07 | 8.500E-08 | 5.000E-07 |
| 3610 | 181 | R181I | A | 2e-07 | 2.1e-07 | 4.6e-07 | 2.900E-07 | 1.473E-07 | 2.000E-07 | 4.600E-07 |
| 3611 | 181 | R181L | A | 1.5e-07 | 1.8e-07 | 4.2e-07 | 2.500E-07 | 1.480E-07 | 1.500E-07 | 4.200E-07 |
| 3612 | 181 | R181K | A | 2.2e-07 | 2.7e-07 | 2.9e-07 | 2.600E-07 | 3.606E-08 | 2.200E-07 | 2.900E-07 |
| 3613 | 181 | R181M | A | 2.3e-07 | 4.7e-07 | 2.2e-07 | 3.067E-07 | 1.415E-07 | 2.200E-07 | 4.700E-07 |
| 3614 | 181 | R181F | A | 7.2e-07 | 2.7e-07 | 2e-07 | 3.967E-07 | 2.822E-07 | 2.000E-07 | 7.200E-07 |
| 3615 | 181 | R181P | A | 1.6e-07 | 8.9e-08 | 2e-07 | 1.497E-07 | 5.622E-08 | 8.900E-08 | 2.000E-07 |
| 3616 | 181 | R181S | A | 3.6e-07 | 3.3e-07 | 1.4e-07 | 2.767E-07 | 1.193E-07 | 1.400E-07 | 3.600E-07 |
| 3617 | 181 | R181T | A | 2.2e-07 | 2.6e-07 | 1.6e-07 | 2.133E-07 | 5.033E-08 | 1.600E-07 | 2.600E-07 |
| 3618 | 181 | R181W | A | 1.6e-07 | 2.8e-07 | 2.7e-07 | 2.367E-07 | 6.658E-08 | 1.600E-07 | 2.800E-07 |
| 3619 | 181 | R181Y | A | 3.4e-07 | 2.7e-07 | 3e-07 | 3.033E-07 | 3.512E-08 | 2.700E-07 | 3.400E-07 |
| 3620 | 181 | R181V | A | 2.9e-07 | 5.9e-07 | 5.6e-07 | 4.800E-07 | 1.652E-07 | 2.900E-07 | 5.900E-07 |
| 3621 | 182 | C182A | A | 2.2e-07 | 4.1e-07 | 1e-07 | 2.433E-07 | 1.563E-07 | 1.000E-07 | 4.100E-07 |
| 3622 | 182 | C182R | A | 3.9e-07 | 3.1e-07 | 2.2e-07 | 3.067E-07 | 8.505E-08 | 2.200E-07 | 3.900E-07 |
| 3623 | 182 | C182N | A | 1.3e-07 | 1.9e-07 | 2.7e-07 | 1.967E-07 | 7.024E-08 | 1.300E-07 | 2.700E-07 |
| 3624 | 182 | C182D | A | 1.7e-07 | 3.8e-07 | 1.4e-07 | 2.300E-07 | 1.308E-07 | 1.400E-07 | 3.800E-07 |
| 3625 | 182 | C182C | A | 2.7e-07 | 4.2e-07 | 3.5e-07 | 3.467E-07 | 7.506E-08 | 2.700E-07 | 4.200E-07 |
| 3626 | 182 | C182E | A | 3.4e-07 | 4.2e-07 | 3.4e-07 | 3.667E-07 | 4.619E-08 | 3.400E-07 | 4.200E-07 |
| 3627 | 182 | C182Q | A | 1.4e-07 | 3.3e-07 | 3.1e-07 | 2.600E-07 | 1.044E-07 | 1.400E-07 | 3.300E-07 |
| 3628 | 182 | C182G | A | 2.4e-07 | 1.7e-07 | 1.4e-07 | 1.833E-07 | 5.132E-08 | 1.400E-07 | 2.400E-07 |
| 3629 | 182 | C182H | A | 5.3e-07 | 2.4e-07 | 2.5e-07 | 3.400E-07 | 1.646E-07 | 2.400E-07 | 5.300E-07 |
| 3630 | 182 | C182I | A | 5e-07 | 2.7e-07 | 3.2e-07 | 3.633E-07 | 1.210E-07 | 2.700E-07 | 5.000E-07 |
| 3631 | 182 | C182L | A | 2e-07 | 3.1e-07 | 3.4e-07 | 2.833E-07 | 7.371E-08 | 2.000E-07 | 3.400E-07 |
| 3632 | 182 | C182K | A | 3.3e-07 | 3.1e-07 | 2.5e-07 | 2.967E-07 | 4.163E-08 | 2.500E-07 | 3.300E-07 |
| 3633 | 182 | C182M | A | 3.3e-07 | 3.2e-07 | 4.7e-07 | 3.733E-07 | 8.386E-08 | 3.200E-07 | 4.700E-07 |
| 3634 | 182 | C182F | A | 2.9e-07 | 2.4e-07 | 1.9e-07 | 2.400E-07 | 5.000E-08 | 1.900E-07 | 2.900E-07 |
| 3635 | 182 | C182P | A | 3.5e-07 | 2.6e-07 | 2.7e-07 | 2.933E-07 | 4.933E-08 | 2.600E-07 | 3.500E-07 |
| 3636 | 182 | C182S | A | 1.9e-07 | 3e-07 | 4.7e-07 | 3.200E-07 | 1.411E-07 | 1.900E-07 | 4.700E-07 |
| 3637 | 182 | C182T | A | 1.8e-07 | 1.2e-07 | 1.7e-07 | 1.567E-07 | 3.215E-08 | 1.200E-07 | 1.800E-07 |
| 3638 | 182 | C182W | A | 1.9e-07 | 1.9e-07 | 3.1e-07 | 2.300E-07 | 6.928E-08 | 1.900E-07 | 3.100E-07 |
| 3639 | 182 | C182Y | A | 2.9e-07 | 2.6e-07 | 2.4e-07 | 2.633E-07 | 2.517E-08 | 2.400E-07 | 2.900E-07 |
| 3640 | 182 | C182V | A | 3.5e-07 | 2.1e-07 | 1.2e-07 | 2.267E-07 | 1.159E-07 | 1.200E-07 | 3.500E-07 |
| 3641 | 183 | W183A | A | 4e-07 | 3.9e-07 | 1.6e-07 | 3.167E-07 | 1.358E-07 | 1.600E-07 | 4.000E-07 |
| 3642 | 183 | W183R | A | 2.3e-07 | 2.8e-07 | 2.1e-07 | 2.400E-07 | 3.606E-08 | 2.100E-07 | 2.800E-07 |
| 3643 | 183 | W183N | A | 2.8e-07 | 2.7e-07 | 3.3e-07 | 2.933E-07 | 3.215E-08 | 2.700E-07 | 3.300E-07 |
| 3644 | 183 | W183D | A | 2.8e-07 | 2.3e-07 | 2.9e-07 | 2.667E-07 | 3.215E-08 | 2.300E-07 | 2.900E-07 |
| 3645 | 183 | W183C | A | 2.9e-07 | 1.8e-07 | 3e-07 | 2.567E-07 | 6.658E-08 | 1.800E-07 | 3.000E-07 |
| 3646 | 183 | W183E | A | 6.7e-07 | 3.1e-07 | 2.2e-07 | 4.000E-07 | 2.381E-07 | 2.200E-07 | 6.700E-07 |
| 3647 | 183 | W183Q | A | 1.9e-07 | 1.6e-07 | 1.6e-07 | 1.700E-07 | 1.732E-08 | 1.600E-07 | 1.900E-07 |
| 3648 | 183 | W183G | A | 2.6e-07 | 3.7e-07 | 4.5e-07 | 3.600E-07 | 9.539E-08 | 2.600E-07 | 4.500E-07 |
| 3649 | 183 | W183H | A | 4.2e-07 | 3.2e-07 | 1.7e-07 | 3.033E-07 | 1.258E-07 | 1.700E-07 | 4.200E-07 |
| 3650 | 183 | W183I | A | 2e-07 | 4.7e-07 | 9e-08 | 2.533E-07 | 1.955E-07 | 9.000E-08 | 4.700E-07 |
| 3651 | 183 | W183L | A | 2.6e-07 | 2.1e-07 | 1.2e-07 | 1.967E-07 | 7.095E-08 | 1.200E-07 | 2.600E-07 |
| 3652 | 183 | W183K | A | 1.9e-07 | 2.6e-07 | 3.4e-07 | 2.633E-07 | 7.506E-08 | 1.900E-07 | 3.400E-07 |
| 3653 | 183 | W183M | A | 1.5e-07 | 5.2e-07 | 2.4e-07 | 3.033E-07 | 1.930E-07 | 1.500E-07 | 5.200E-07 |
| 3654 | 183 | W183F | A | 2.1e-07 | 4e-07 | 5.5e-07 | 3.867E-07 | 1.704E-07 | 2.100E-07 | 5.500E-07 |

| | | | | | | | | | | |
|---|---|---|---|---|---|---|---|---|---|---|
| 3655 | 183 | W183P | A | 2e-07 | 2.9e-07 | 6.2e-08 | 1.840E-07 | 1.148E-07 | 6.200E-08 | 2.900E-07 |
| 3656 | 183 | W183S | A | 2.6e-07 | 2e-07 | 3e-07 | 2.533E-07 | 5.033E-08 | 2.000E-07 | 3.000E-07 |
| 3657 | 183 | W183T | A | 2.5e-07 | 1.9e-07 | 2.3e-07 | 2.233E-07 | 3.055E-08 | 1.900E-07 | 2.500E-07 |
| 3658 | 183 | W183W | A | 1.1e-07 | 3.5e-07 | 2.1e-07 | 2.233E-07 | 1.206E-07 | 1.100E-07 | 3.500E-07 |
| 3659 | 183 | W183Y | A | 1.7e-07 | 4e-07 | 5.2e-07 | 3.633E-07 | 1.779E-07 | 1.700E-07 | 5.200E-07 |
| 3660 | 183 | W183V | A | 3.4e-07 | 2.5e-07 | 4.1e-07 | 3.333E-07 | 8.021E-08 | 2.500E-07 | 4.100E-07 |
| 3661 | 184 | G184A | A | 1.6e-07 | 3.3e-07 | 1.3e-07 | 2.067E-07 | 1.079E-07 | 1.300E-07 | 3.300E-07 |
| 3662 | 184 | G184R | A | 4.6e-07 | 2e-07 | 2.7e-07 | 3.100E-07 | 1.345E-07 | 2.000E-07 | 4.600E-07 |
| 3663 | 184 | G184N | A | 4.1e-07 | 8.8e-07 | 3.6e-07 | 5.500E-07 | 2.869E-07 | 3.600E-07 | 8.800E-07 |
| 3664 | 184 | G184D | A | 3.7e-07 | 2.6e-07 | 5.3e-07 | 3.867E-07 | 1.358E-07 | 2.600E-07 | 5.300E-07 |
| 3665 | 184 | G184C | A | 8.3e-07 | 3.7e-07 | 2.3e-07 | 4.767E-07 | 3.139E-07 | 2.300E-07 | 8.300E-07 |
| 3666 | 184 | G184E | A | 2.4e-07 | 1.6e-07 | 5.5e-07 | 3.167E-07 | 2.060E-07 | 1.600E-07 | 5.500E-07 |
| 3667 | 184 | G184Q | A | 2.3e-07 | 3.1e-07 | 2.2e-07 | 2.533E-07 | 4.933E-08 | 2.200E-07 | 3.100E-07 |
| 3668 | 184 | G184G | A | 1.7e-07 | 4.4e-07 | 4.1e-07 | 3.400E-07 | 1.480E-07 | 1.700E-07 | 4.400E-07 |
| 3669 | 184 | G184H | A | 2.7e-07 | 3e-07 | 4.4e-07 | 3.367E-07 | 9.074E-08 | 2.700E-07 | 4.400E-07 |
| 3670 | 184 | G184I | A | 3.5e-07 | 2.1e-07 | 6.1e-07 | 3.900E-07 | 2.030E-07 | 2.100E-07 | 6.100E-07 |
| 3671 | 184 | G184L | A | 2.2e-07 | 2.2e-07 | 4.1e-07 | 2.833E-07 | 1.097E-07 | 2.200E-07 | 4.100E-07 |
| 3672 | 184 | G184K | A | 2e-07 | 4.9e-07 | 5.9e-07 | 4.267E-07 | 2.026E-07 | 2.000E-07 | 5.900E-07 |
| 3673 | 184 | G184M | A | 4.1e-07 | 3.2e-07 | 3.3e-07 | 3.533E-07 | 4.933E-08 | 3.200E-07 | 4.100E-07 |
| 3674 | 184 | G184F | A | 2.5e-07 | 3.2e-07 | 3.5e-07 | 3.067E-07 | 5.132E-08 | 2.500E-07 | 3.500E-07 |
| 3675 | 184 | G184P | A | 1.1e-07 | 2.4e-07 | 4e-07 | 2.500E-07 | 1.453E-07 | 1.100E-07 | 4.000E-07 |
| 3676 | 184 | G184S | A | 1.9e-07 | 1.9e-07 | 3.7e-07 | 2.500E-07 | 1.039E-07 | 1.900E-07 | 3.700E-07 |
| 3677 | 184 | G184T | A | 2.1e-07 | 3.2e-07 | 2.1e-07 | 2.467E-07 | 6.351E-08 | 2.100E-07 | 3.200E-07 |
| 3678 | 184 | G184W | A | 3e-07 | 2e-07 | 5.1e-07 | 3.367E-07 | 1.582E-07 | 2.000E-07 | 5.100E-07 |
| 3679 | 184 | G184Y | A | 2e-07 | 2.7e-07 | 3.1e-07 | 2.600E-07 | 5.568E-08 | 2.000E-07 | 3.100E-07 |
| 3680 | 184 | G184V | A | 3.2e-07 | 2.5e-07 | 1.7e-07 | 2.467E-07 | 7.506E-08 | 1.700E-07 | 3.200E-07 |
| 3681 | 185 | E185A | A | 5.7e-07 | 3e-07 | 4.4e-07 | 4.367E-07 | 1.350E-07 | 3.000E-07 | 5.700E-07 |
| 3682 | 185 | E185R | A | 1.9e-07 | 3.7e-07 | 3.1e-07 | 2.900E-07 | 9.165E-08 | 1.900E-07 | 3.700E-07 |
| 3683 | 185 | E185N | A | 7.4e-08 | 2.3e-07 | 2.7e-07 | 1.913E-07 | 1.036E-07 | 7.400E-08 | 2.700E-07 |
| 3684 | 185 | E185D | A | 2.4e-07 | 1.2e-07 | 2.8e-07 | 2.133E-07 | 8.327E-08 | 1.200E-07 | 2.800E-07 |
| 3685 | 185 | E185C | A | 2.9e-07 | 2.1e-07 | 1.9e-07 | 2.300E-07 | 5.292E-08 | 1.900E-07 | 2.900E-07 |
| 3686 | 185 | E185E | A | 3.4e-07 | 2.4e-07 | 3.4e-07 | 3.067E-07 | 5.774E-08 | 2.400E-07 | 3.400E-07 |
| 3687 | 185 | E185Q | A | 3.7e-07 | 3.6e-07 | 3.3e-07 | 3.533E-07 | 2.082E-08 | 3.300E-07 | 3.700E-07 |
| 3688 | 185 | E185G | A | 2e-07 | 2.7e-07 | 1e-06 | 4.900E-07 | 4.431E-07 | 2.000E-07 | 1.000E-06 |
| 3689 | 185 | E185H | A | 1.3e-07 | 3.1e-07 | 2.4e-07 | 2.267E-07 | 9.074E-08 | 1.300E-07 | 3.100E-07 |
| 3690 | 185 | E185I | A | 5.1e-07 | 1.1e-07 | 4.2e-07 | 3.467E-07 | 2.098E-07 | 1.100E-07 | 5.100E-07 |
| 3691 | 185 | E185L | A | 2.6e-07 | 3.2e-07 | 2.3e-07 | 2.700E-07 | 4.583E-08 | 2.300E-07 | 3.200E-07 |
| 3692 | 185 | E185K | A | 3.1e-07 | 1.7e-07 | 2.3e-07 | 2.367E-07 | 7.024E-08 | 1.700E-07 | 3.100E-07 |
| 3693 | 185 | E185M | A | 6.2e-07 | 1.4e-07 | 2.9e-07 | 3.500E-07 | 2.456E-07 | 1.400E-07 | 6.200E-07 |
| 3694 | 185 | E185F | A | 2.3e-07 | 3.9e-07 | 3e-07 | 3.067E-07 | 8.021E-08 | 2.300E-07 | 3.900E-07 |
| 3695 | 185 | E185P | A | 3e-07 | 3.2e-07 | 3.3e-07 | 3.167E-07 | 1.528E-08 | 3.000E-07 | 3.300E-07 |
| 3696 | 185 | E185S | A | 3.9e-07 | 1.8e-07 | 3.1e-07 | 2.933E-07 | 1.060E-07 | 1.800E-07 | 3.900E-07 |
| 3697 | 185 | E185T | A | 2.1e-07 | 2.1e-07 | 2.2e-07 | 2.133E-07 | 5.774E-09 | 2.100E-07 | 2.200E-07 |
| 3698 | 185 | E185W | A | 1.8e-07 | 2.4e-07 | 2.2e-07 | 2.133E-07 | 3.055E-08 | 1.800E-07 | 2.400E-07 |
| 3699 | 185 | E185Y | A | 6.7e-07 | 4.3e-07 | 1.4e-07 | 4.133E-07 | 2.654E-07 | 1.400E-07 | 6.700E-07 |
| 3700 | 185 | E185V | A | 2.8e-07 | 4.9e-07 | 2.4e-07 | 3.367E-07 | 1.343E-07 | 2.400E-07 | 4.900E-07 |
| 3701 | 186 | S186A | A | 2.9e-07 | 1.2e-07 | 2.1e-07 | 2.067E-07 | 8.505E-08 | 1.200E-07 | 2.900E-07 |
| 3702 | 186 | S186R | A | 2.7e-07 | 2.3e-07 | 2.3e-07 | 2.433E-07 | 2.309E-08 | 2.300E-07 | 2.700E-07 |
| 3703 | 186 | S186N | A | 3.7e-07 | 5.8e-07 | 2.9e-07 | 4.133E-07 | 1.498E-07 | 2.900E-07 | 5.800E-07 |
| 3704 | 186 | S186D | A | 2e-07 | 2.5e-07 | 4.3e-07 | 2.933E-07 | 1.210E-07 | 2.000E-07 | 4.300E-07 |
| 3705 | 186 | S186C | A | 2.9e-07 | 1.9e-07 | 2e-07 | 2.267E-07 | 5.508E-08 | 1.900E-07 | 2.900E-07 |
| 3706 | 186 | S186E | A | 3e-07 | 2.9e-07 | 2e-07 | 2.633E-07 | 5.508E-08 | 2.000E-07 | 3.000E-07 |
| 3707 | 186 | S186Q | A | 2.8e-07 | 3.9e-07 | 2.3e-07 | 3.000E-07 | 8.185E-08 | 2.300E-07 | 3.900E-07 |
| 3708 | 186 | S186G | A | 6.3e-07 | 1.2e-07 | 2.7e-07 | 3.400E-07 | 2.621E-07 | 1.200E-07 | 6.300E-07 |
| 3709 | 186 | S186H | A | 3.1e-07 | 2e-07 | 3.2e-07 | 2.767E-07 | 6.658E-08 | 2.000E-07 | 3.200E-07 |
| 3710 | 186 | S186I | A | 1.4e-07 | 4.6e-07 | 4.5e-07 | 3.500E-07 | 1.819E-07 | 1.400E-07 | 4.600E-07 |
| 3711 | 186 | S186L | A | 1.9e-07 | 2.5e-07 | 3.1e-07 | 2.500E-07 | 6.000E-08 | 1.900E-07 | 3.100E-07 |
| 3712 | 186 | S186K | A | 3e-07 | 2.3e-07 | 2.9e-07 | 2.733E-07 | 3.786E-08 | 2.300E-07 | 3.000E-07 |
| 3713 | 186 | S186M | A | 3.1e-07 | 1.9e-07 | 4.6e-07 | 3.200E-07 | 1.353E-07 | 1.900E-07 | 4.600E-07 |
| 3714 | 186 | S186F | A | 3.2e-07 | 3e-07 | 2.1e-07 | 2.767E-07 | 5.859E-08 | 2.100E-07 | 3.200E-07 |
| 3715 | 186 | S186P | A | 1.4e-07 | 3.1e-07 | 2.7e-07 | 2.400E-07 | 8.888E-08 | 1.400E-07 | 3.100E-07 |
| 3716 | 186 | S186S | A | 6.8e-07 | 1.4e-07 | 1.7e-07 | 3.300E-07 | 3.035E-07 | 1.400E-07 | 6.800E-07 |
| 3717 | 186 | S186T | A | 2.7e-07 | 2.5e-07 | 1.6e-07 | 2.267E-07 | 5.859E-08 | 1.600E-07 | 2.700E-07 |
| 3718 | 186 | S186W | A | 1.5e-07 | 1.6e-07 | 2.7e-07 | 1.933E-07 | 6.658E-08 | 1.500E-07 | 2.700E-07 |
| 3719 | 186 | S186Y | A | 1.7e-07 | 2.1e-07 | 3.6e-07 | 2.467E-07 | 1.002E-07 | 1.700E-07 | 3.600E-07 |
| 3720 | 186 | S186V | A | 4.1e-07 | 1.8e-07 | 2.3e-07 | 2.733E-07 | 1.210E-07 | 1.800E-07 | 4.100E-07 |
| 3721 | 187 | S187A | A | 1.3e-07 | 1.6e-07 | 4e-07 | 2.300E-07 | 1.480E-07 | 1.300E-07 | 4.000E-07 |
| 3722 | 187 | S187R | A | 2.4e-07 | 3.5e-07 | 1.9e-07 | 2.600E-07 | 8.185E-08 | 1.900E-07 | 3.500E-07 |
| 3723 | 187 | S187N | A | 3.5e-07 | 1.9e-07 | 3.6e-07 | 3.000E-07 | 9.539E-08 | 1.900E-07 | 3.600E-07 |
| 3724 | 187 | S187D | A | 2e-07 | 1.5e-07 | 9.5e-08 | 1.483E-07 | 5.252E-08 | 9.500E-08 | 2.000E-07 |
| 3725 | 187 | S187C | A | 3.2e-07 | 4.1e-07 | 3.2e-07 | 3.500E-07 | 5.196E-08 | 3.200E-07 | 4.100E-07 |
| 3726 | 187 | S187E | A | 2.3e-07 | 1.7e-07 | 1.9e-07 | 1.967E-07 | 3.055E-08 | 1.700E-07 | 2.300E-07 |
| 3727 | 187 | S187Q | A | 2.7e-07 | 1.9e-07 | 2.8e-07 | 2.467E-07 | 4.933E-08 | 1.900E-07 | 2.800E-07 |
| 3728 | 187 | S187G | A | 3.1e-07 | 1.8e-07 | 2.4e-07 | 2.433E-07 | 6.506E-08 | 1.800E-07 | 3.100E-07 |
| 3729 | 187 | S187H | A | 3e-07 | 2.2e-07 | 2.8e-07 | 2.667E-07 | 4.163E-08 | 2.200E-07 | 3.000E-07 |
| 3730 | 187 | S187I | A | 2.3e-07 | 2e-07 | 3.5e-07 | 2.600E-07 | 7.937E-08 | 2.000E-07 | 3.500E-07 |
| 3731 | 187 | S187L | A | 1.7e-07 | 2.8e-07 | 3.6e-07 | 2.700E-07 | 9.539E-08 | 1.700E-07 | 3.600E-07 |
| 3732 | 187 | S187K | A | 1.3e-07 | 2.3e-07 | 3.5e-07 | 2.367E-07 | 1.102E-07 | 1.300E-07 | 3.500E-07 |
| 3733 | 187 | S187M | A | 2.4e-07 | 3.1e-07 | 2.1e-07 | 2.533E-07 | 5.132E-08 | 2.100E-07 | 3.100E-07 |
| 3734 | 187 | S187F | A | 1.4e-07 | 3.6e-07 | 1.7e-07 | 2.233E-07 | 1.193E-07 | 1.400E-07 | 3.600E-07 |
| 3735 | 187 | S187P | A | 2.5e-07 | 3.3e-07 | 4.8e-07 | 3.533E-07 | 1.168E-07 | 2.500E-07 | 4.800E-07 |
| 3736 | 187 | S187S | A | 3.2e-07 | 2.3e-07 | 3.6e-07 | 3.033E-07 | 6.658E-08 | 2.300E-07 | 3.600E-07 |
| 3737 | 187 | S187T | A | 2e-07 | 1.6e-07 | 4.1e-07 | 2.567E-07 | 1.343E-07 | 1.600E-07 | 4.100E-07 |
| 3738 | 187 | S187W | A | 2.5e-07 | 2e-07 | 6.3e-08 | 1.710E-07 | 9.681E-08 | 6.300E-08 | 2.500E-07 |
| 3739 | 187 | S187Y | A | 2.8e-07 | 5.5e-07 | 6.4e-08 | 2.980E-07 | 2.435E-07 | 6.400E-08 | 5.500E-07 |

| | | | | | | | | | | |
|---|---|---|---|---|---|---|---|---|---|---|
| 3740 | 187 | S187V | A | 1.7e-07 | 1.8e-07 | 1.3e-07 | 1.600E-07 | 2.646E-08 | 1.300E-07 | 1.800E-07 |
| 3741 | 188 | E188A | A | 3.5e-07 | 1.7e-07 | 5.2e-07 | 3.467E-07 | 1.750E-07 | 1.700E-07 | 5.200E-07 |
| 3742 | 188 | E188R | A | 3.9e-07 | 2.1e-07 | 9.3e-08 | 2.310E-07 | 1.496E-07 | 9.300E-08 | 3.900E-07 |
| 3743 | 188 | E188N | A | 3.3e-07 | 1.7e-07 | 2.4e-07 | 2.467E-07 | 8.021E-08 | 1.700E-07 | 3.300E-07 |
| 3744 | 188 | E188D | A | 2.4e-07 | 2.5e-07 | 2.6e-07 | 2.500E-07 | 1.000E-08 | 2.400E-07 | 2.600E-07 |
| 3745 | 188 | E188C | A | 2.1e-07 | 1.8e-07 | 2.4e-07 | 2.100E-07 | 3.000E-08 | 1.800E-07 | 2.400E-07 |
| 3746 | 188 | E188E | A | 3.3e-07 | 2.2e-07 | 3e-07 | 2.833E-07 | 5.686E-08 | 2.200E-07 | 3.300E-07 |
| 3747 | 188 | E188Q | A | 1.8e-07 | 4e-07 | 2.5e-07 | 2.767E-07 | 1.124E-07 | 1.800E-07 | 4.000E-07 |
| 3748 | 188 | E188G | A | 1.6e-07 | 4.3e-07 | 3.3e-07 | 3.067E-07 | 1.365E-07 | 1.600E-07 | 4.300E-07 |
| 3749 | 188 | E188H | A | 4.1e-07 | 1.5e-07 | 2e-07 | 2.533E-07 | 1.380E-07 | 1.500E-07 | 4.100E-07 |
| 3750 | 188 | E188I | A | 3.5e-07 | 1.9e-07 | 1.5e-07 | 2.300E-07 | 1.058E-07 | 1.500E-07 | 3.500E-07 |
| 3751 | 188 | E188L | A | 3.1e-07 | 2.4e-07 | 7.5e-07 | 4.333E-07 | 2.765E-07 | 2.400E-07 | 7.500E-07 |
| 3752 | 188 | E188K | A | 2.7e-07 | 1.9e-07 | 1.7e-07 | 2.100E-07 | 5.292E-08 | 1.700E-07 | 2.700E-07 |
| 3753 | 188 | E188M | A | 2.9e-07 | 2.2e-07 | 3.2e-07 | 2.767E-07 | 5.132E-08 | 2.200E-07 | 3.200E-07 |
| 3754 | 188 | E188F | A | 1.8e-07 | 4.7e-07 | 1.2e-07 | 2.567E-07 | 1.872E-07 | 1.200E-07 | 4.700E-07 |
| 3755 | 188 | E188P | A | 2.5e-07 | 2.5e-07 | 4e-07 | 3.000E-07 | 8.660E-08 | 2.500E-07 | 4.000E-07 |
| 3756 | 188 | E188S | A | 1.8e-07 | 3e-07 | 2.5e-07 | 2.433E-07 | 6.028E-08 | 1.800E-07 | 3.000E-07 |
| 3757 | 188 | E188T | A | 2.5e-07 | 1e-07 | 1.6e-07 | 1.700E-07 | 7.550E-08 | 1.000E-07 | 2.500E-07 |
| 3758 | 188 | E188W | A | 2.1e-07 | 5.6e-07 | 1.8e-07 | 3.167E-07 | 2.113E-07 | 1.800E-07 | 5.600E-07 |
| 3759 | 188 | E188Y | A | 1e-07 | 2e-07 | 4e-07 | 2.333E-07 | 1.528E-07 | 1.000E-07 | 4.000E-07 |
| 3760 | 188 | E188V | A | 1.7e-07 | 9.9e-08 | 4e-07 | 2.230E-07 | 1.573E-07 | 9.900E-08 | 4.000E-07 |
| 3761 | 189 | D189A | A | 1e-07 | 2.1e-07 | 2.2e-07 | 1.767E-07 | 6.658E-08 | 1.000E-07 | 2.200E-07 |
| 3762 | 189 | D189R | A | 1.2e-07 | 1.1e-07 | 2e-07 | 1.433E-07 | 4.933E-08 | 1.100E-07 | 2.000E-07 |
| 3763 | 189 | D189N | A | 3.4e-07 | 2.5e-07 | 3.3e-07 | 3.067E-07 | 4.933E-08 | 2.500E-07 | 3.400E-07 |
| 3764 | 189 | D189D | A | 3.6e-07 | 5.2e-07 | 2e-07 | 3.600E-07 | 1.600E-07 | 2.000E-07 | 5.200E-07 |
| 3765 | 189 | D189C | A | 1.8e-07 | 4.3e-07 | 1e-07 | 2.367E-07 | 1.721E-07 | 1.000E-07 | 4.300E-07 |
| 3766 | 189 | D189E | A | 2.7e-07 | 3e-07 | 1.1e-06 | 5.567E-07 | 4.708E-07 | 2.700E-07 | 1.100E-06 |
| 3767 | 189 | D189Q | A | 1.4e-07 | 2.9e-07 | 3.3e-07 | 2.533E-07 | 1.002E-07 | 1.400E-07 | 3.300E-07 |
| 3768 | 189 | D189G | A | 6.2e-07 | 4.6e-07 | 3.3e-07 | 4.700E-07 | 1.453E-07 | 3.300E-07 | 6.200E-07 |
| 3769 | 189 | D189H | A | 2.1e-07 | 6.5e-07 | 1.1e-07 | 3.233E-07 | 2.873E-07 | 1.100E-07 | 6.500E-07 |
| 3770 | 189 | D189I | A | 3.1e-07 | 4.1e-07 | 1.3e-07 | 2.833E-07 | 1.419E-07 | 1.300E-07 | 4.100E-07 |
| 3771 | 189 | D189L | A | 5.7e-07 | 4e-07 | 4.9e-07 | 4.867E-07 | 8.505E-08 | 4.000E-07 | 5.700E-07 |
| 3772 | 189 | D189K | A | 3.2e-07 | 1.7e-07 | 3.4e-07 | 2.767E-07 | 9.292E-08 | 1.700E-07 | 3.400E-07 |
| 3773 | 189 | D189M | A | 1.3e-07 | 2.1e-07 | 2e-07 | 1.800E-07 | 4.359E-08 | 1.300E-07 | 2.100E-07 |
| 3774 | 189 | D189F | A | 4.3e-07 | 3.6e-07 | 2e-07 | 3.300E-07 | 1.179E-07 | 2.000E-07 | 4.300E-07 |
| 3775 | 189 | D189P | A | 1.8e-07 | 3.2e-07 | 1.6e-07 | 2.200E-07 | 8.718E-08 | 1.600E-07 | 3.200E-07 |
| 3776 | 189 | D189S | A | 4.7e-07 | 3.4e-07 | 1.7e-07 | 3.267E-07 | 1.504E-07 | 1.700E-07 | 4.700E-07 |
| 3777 | 189 | D189T | A | 2.1e-07 | 1.7e-07 | 1.1e-07 | 1.633E-07 | 5.033E-08 | 1.100E-07 | 2.100E-07 |
| 3778 | 189 | D189W | A | 1.9e-07 | 3.5e-07 | 3e-07 | 2.800E-07 | 8.185E-08 | 1.900E-07 | 3.500E-07 |
| 3779 | 189 | D189Y | A | 3.2e-07 | 3.8e-07 | 2.8e-07 | 3.267E-07 | 5.033E-08 | 2.800E-07 | 3.800E-07 |
| 3780 | 189 | D189V | A | 2.2e-07 | 2.5e-07 | 4.4e-07 | 3.033E-07 | 1.193E-07 | 2.200E-07 | 4.400E-07 |
| 3781 | 190 | C190A | A | 3.7e-07 | 4.2e-07 | 4.6e-07 | 4.167E-07 | 4.509E-08 | 3.700E-07 | 4.600E-07 |
| 3782 | 190 | C190R | A | 4.2e-07 | 2.1e-07 | 4.6e-07 | 3.633E-07 | 1.343E-07 | 2.100E-07 | 4.600E-07 |
| 3783 | 190 | C190N | A | 2.1e-07 | 5.6e-07 | 3.3e-07 | 3.667E-07 | 1.779E-07 | 2.100E-07 | 5.600E-07 |
| 3784 | 190 | C190D | A | 1.9e-07 | 2.6e-07 | 3.5e-07 | 2.667E-07 | 8.021E-08 | 1.900E-07 | 3.500E-07 |
| 3785 | 190 | C190C | A | 1.6e-07 | 3.1e-07 | 3.3e-07 | 2.667E-07 | 9.292E-08 | 1.600E-07 | 3.300E-07 |
| 3786 | 190 | C190E | A | 2.2e-07 | 2.3e-07 | 3e-07 | 2.500E-07 | 4.359E-08 | 2.200E-07 | 3.000E-07 |
| 3787 | 190 | C190Q | A | 1e-07 | 1.6e-07 | 4.9e-07 | 2.500E-07 | 2.100E-07 | 1.000E-07 | 4.900E-07 |
| 3788 | 190 | C190G | A | 4.1e-07 | 2.1e-07 | 1.3e-07 | 2.500E-07 | 1.442E-07 | 1.300E-07 | 4.100E-07 |
| 3789 | 190 | C190H | A | 2.2e-07 | 2e-07 | 2e-07 | 2.067E-07 | 1.155E-08 | 2.000E-07 | 2.200E-07 |
| 3790 | 190 | C190I | A | 4.1e-07 | 3.2e-07 | 3.5e-07 | 3.600E-07 | 4.583E-08 | 3.200E-07 | 4.100E-07 |
| 3791 | 190 | C190L | A | 1.8e-07 | 3e-07 | 5.2e-07 | 3.333E-07 | 1.724E-07 | 1.800E-07 | 5.200E-07 |
| 3792 | 190 | C190K | A | 3e-07 | 2.7e-07 | 2.8e-07 | 2.833E-07 | 1.528E-08 | 2.700E-07 | 3.000E-07 |
| 3793 | 190 | C190M | A | 3.5e-07 | 2.2e-07 | 3e-07 | 2.900E-07 | 6.557E-08 | 2.200E-07 | 3.500E-07 |
| 3794 | 190 | C190F | A | 2.1e-07 | 7.5e-07 | 3.1e-07 | 4.233E-07 | 2.873E-07 | 2.100E-07 | 7.500E-07 |
| 3795 | 190 | C190P | A | 2.6e-07 | 2.1e-07 | 2.8e-07 | 2.500E-07 | 3.606E-08 | 2.100E-07 | 2.800E-07 |
| 3796 | 190 | C190S | A | 3.2e-07 | 1.9e-07 | 2.1e-07 | 2.400E-07 | 7.000E-08 | 1.900E-07 | 3.200E-07 |
| 3797 | 190 | C190T | A | 2.3e-07 | 1.7e-07 | 1.8e-07 | 1.933E-07 | 3.215E-08 | 1.700E-07 | 2.300E-07 |
| 3798 | 190 | C190W | A | 3.8e-07 | 1.7e-07 | 1.6e-07 | 2.367E-07 | 1.242E-07 | 1.600E-07 | 3.800E-07 |
| 3799 | 190 | C190Y | A | 3e-07 | 3.2e-07 | 2.4e-07 | 2.867E-07 | 4.163E-08 | 2.400E-07 | 3.200E-07 |
| 3800 | 190 | C190V | A | 1.5e-07 | 2e-07 | 2.5e-07 | 2.000E-07 | 5.000E-08 | 1.500E-07 | 2.500E-07 |
| 3801 | 191 | Q191A | A | 2.6e-07 | 3.5e-07 | 1.4e-07 | 2.500E-07 | 1.054E-07 | 1.400E-07 | 3.500E-07 |
| 3802 | 191 | Q191R | A | 1.9e-07 | 3.2e-07 | 2.8e-07 | 2.633E-07 | 6.658E-08 | 1.900E-07 | 3.200E-07 |
| 3803 | 191 | Q191N | A | 2e-07 | 1.4e-07 | 2.9e-07 | 2.100E-07 | 7.550E-08 | 1.400E-07 | 2.900E-07 |
| 3804 | 191 | Q191D | A | 5.6e-07 | 1.9e-07 | 1.4e-07 | 2.967E-07 | 2.294E-07 | 1.400E-07 | 5.600E-07 |
| 3805 | 191 | Q191C | A | 3.7e-07 | 1.9e-07 | 5.1e-07 | 3.567E-07 | 1.604E-07 | 1.900E-07 | 5.100E-07 |
| 3806 | 191 | Q191E | A | 3.1e-07 | 6.6e-07 | 1.6e-07 | 3.767E-07 | 2.566E-07 | 1.600E-07 | 6.600E-07 |
| 3807 | 191 | Q191Q | A | 2.2e-07 | 2e-07 | 2.8e-07 | 2.333E-07 | 4.163E-08 | 2.000E-07 | 2.800E-07 |
| 3808 | 191 | Q191G | A | 2e-07 | 2.1e-07 | 2.1e-07 | 2.067E-07 | 5.774E-09 | 2.000E-07 | 2.100E-07 |
| 3809 | 191 | Q191H | A | 3.3e-07 | 4.8e-07 | 1.6e-07 | 3.233E-07 | 1.601E-07 | 1.600E-07 | 4.800E-07 |
| 3810 | 191 | Q191I | A | 5.5e-07 | 2.9e-07 | 2.7e-07 | 3.700E-07 | 1.562E-07 | 2.700E-07 | 5.500E-07 |
| 3811 | 191 | Q191L | A | 4.5e-07 | 2.6e-07 | 3.2e-07 | 3.433E-07 | 9.713E-08 | 2.600E-07 | 4.500E-07 |
| 3812 | 191 | Q191K | A | 3.2e-07 | 3.7e-07 | 3.2e-07 | 3.367E-07 | 2.887E-08 | 3.200E-07 | 3.700E-07 |
| 3813 | 191 | Q191M | A | 2.2e-07 | 3.6e-07 | 2.4e-07 | 2.733E-07 | 7.572E-08 | 2.200E-07 | 3.600E-07 |
| 3814 | 191 | Q191F | A | 2.7e-07 | 3.2e-07 | 1.8e-07 | 2.567E-07 | 7.095E-08 | 1.800E-07 | 3.200E-07 |
| 3815 | 191 | Q191P | A | 2.1e-07 | 2.4e-07 | 2.6e-07 | 2.367E-07 | 2.517E-08 | 2.100E-07 | 2.600E-07 |
| 3816 | 191 | Q191S | A | 1e-07 | 2.5e-07 | 4e-07 | 2.500E-07 | 1.500E-07 | 1.000E-07 | 4.000E-07 |
| 3817 | 191 | Q191T | A | 3.1e-07 | 1.4e-07 | 1.8e-07 | 2.100E-07 | 8.888E-08 | 1.400E-07 | 3.100E-07 |
| 3818 | 191 | Q191W | A | 2.4e-07 | 1.7e-07 | 1.9e-07 | 2.000E-07 | 3.606E-08 | 1.700E-07 | 2.400E-07 |
| 3819 | 191 | Q191Y | A | 2.7e-07 | 1.3e-07 | 2.1e-07 | 2.033E-07 | 7.024E-08 | 1.300E-07 | 2.700E-07 |
| 3820 | 191 | Q191V | A | 3.4e-07 | 3.1e-07 | 3.7e-07 | 3.400E-07 | 3.000E-08 | 3.100E-07 | 3.700E-07 |
| 3821 | 192 | S192A | A | 5.2e-07 | 2e-07 | 3e-07 | 3.400E-07 | 1.637E-07 | 2.000E-07 | 5.200E-07 |
| 3822 | 192 | S192R | A | 2.3e-07 | 1.1e-07 | 3.6e-07 | 2.333E-07 | 1.250E-07 | 1.100E-07 | 3.600E-07 |
| 3823 | 192 | S192N | A | 2e-07 | 2.7e-07 | 1.8e-07 | 2.167E-07 | 4.726E-08 | 1.800E-07 | 2.700E-07 |
| 3824 | 192 | S192D | A | 2.2e-07 | 1.3e-07 | 1.2e-07 | 1.567E-07 | 5.508E-08 | 1.200E-07 | 2.200E-07 |

| | | | | | | | | | | |
|---|---|---|---|---|---|---|---|---|---|---|
| 3825 | 192 | S192C | A | 5e-07 | 2.1e-07 | 2.5e-07 | 3.200E-07 | 1.572E-07 | 2.100E-07 | 5.000E-07 |
| 3826 | 192 | S192E | A | 2.5e-07 | 3.5e-07 | 3.6e-07 | 3.200E-07 | 6.083E-08 | 2.500E-07 | 3.600E-07 |
| 3827 | 192 | S192Q | A | 2.4e-07 | 1.5e-07 | 1.5e-07 | 1.800E-07 | 5.196E-08 | 1.500E-07 | 2.400E-07 |
| 3828 | 192 | S192G | A | 9.8e-08 | 1.2e-07 | 2.8e-07 | 1.660E-07 | 9.934E-08 | 9.800E-08 | 2.800E-07 |
| 3829 | 192 | S192H | A | 4.8e-07 | 2.1e-07 | 2.6e-07 | 3.167E-07 | 1.436E-07 | 2.100E-07 | 4.800E-07 |
| 3830 | 192 | S192I | A | 3.2e-07 | 4.1e-07 | 3.3e-07 | 3.533E-07 | 4.933E-08 | 3.200E-07 | 4.100E-07 |
| 3831 | 192 | S192L | A | 4.7e-07 | 2.3e-07 | 3.3e-07 | 3.433E-07 | 1.206E-07 | 2.300E-07 | 4.700E-07 |
| 3832 | 192 | S192K | A | 9.1e-08 | 3.8e-07 | 3.7e-07 | 2.803E-07 | 1.640E-07 | 9.100E-08 | 3.800E-07 |
| 3833 | 192 | S192M | A | 1.7e-07 | 3.8e-07 | 2e-07 | 2.500E-07 | 1.136E-07 | 1.700E-07 | 3.800E-07 |
| 3834 | 192 | S192F | A | 3e-07 | 2.6e-07 | 3.1e-07 | 2.900E-07 | 2.646E-08 | 2.600E-07 | 3.100E-07 |
| 3835 | 192 | S192P | A | 3.2e-07 | 1.6e-07 | 1.4e-07 | 2.067E-07 | 9.866E-08 | 1.400E-07 | 3.200E-07 |
| 3836 | 192 | S192S | A | 3.8e-07 | 3.1e-07 | 2.3e-07 | 3.067E-07 | 7.506E-08 | 2.300E-07 | 3.800E-07 |
| 3837 | 192 | S192T | A | 8.9e-08 | 3.8e-07 | 2.4e-07 | 2.363E-07 | 1.455E-07 | 8.900E-08 | 3.800E-07 |
| 3838 | 192 | S192W | A | 3.5e-07 | 1.4e-07 | 2.2e-07 | 2.367E-07 | 1.060E-07 | 1.400E-07 | 3.500E-07 |
| 3839 | 192 | S192Y | A | 1.6e-07 | 2.8e-07 | 2.4e-07 | 2.267E-07 | 6.110E-08 | 1.600E-07 | 2.800E-07 |
| 3840 | 192 | S192V | A | 2.3e-07 | 2.8e-07 | 5.5e-07 | 3.533E-07 | 1.721E-07 | 2.300E-07 | 5.500E-07 |
| 3841 | 193 | L193A | A | 2.8e-07 | 5.3e-07 | 1.5e-07 | 3.200E-07 | 1.931E-07 | 1.500E-07 | 5.300E-07 |
| 3842 | 193 | L193R | A | 2.5e-07 | 2.1e-07 | 2e-07 | 2.200E-07 | 2.646E-08 | 2.000E-07 | 2.500E-07 |
| 3843 | 193 | L193N | A | 1.8e-07 | 4.2e-07 | 7.1e-08 | 2.237E-07 | 1.786E-07 | 7.100E-08 | 4.200E-07 |
| 3844 | 193 | L193D | A | 2.4e-07 | 1.3e-07 | 3.7e-07 | 2.467E-07 | 1.201E-07 | 1.300E-07 | 3.700E-07 |
| 3845 | 193 | L193C | A | 2.3e-07 | 2.7e-07 | 3.8e-07 | 2.933E-07 | 7.767E-08 | 2.300E-07 | 3.800E-07 |
| 3846 | 193 | L193E | A | 1.9e-07 | 3.1e-07 | 2.4e-07 | 2.467E-07 | 6.028E-08 | 1.900E-07 | 3.100E-07 |
| 3847 | 193 | L193Q | A | 2.8e-07 | 2.3e-07 | 3.3e-07 | 2.800E-07 | 5.000E-08 | 2.300E-07 | 3.300E-07 |
| 3848 | 193 | L193G | A | 4.2e-07 | 2.6e-07 | 2.2e-07 | 3.000E-07 | 1.058E-07 | 2.200E-07 | 4.200E-07 |
| 3849 | 193 | L193H | A | 2.9e-07 | 4.9e-07 | 2.4e-07 | 3.400E-07 | 1.323E-07 | 2.400E-07 | 4.900E-07 |
| 3850 | 193 | L193I | A | 1.8e-07 | 2.3e-07 | 2.8e-07 | 2.300E-07 | 5.000E-08 | 1.800E-07 | 2.800E-07 |
| 3851 | 193 | L193L | A | 2e-07 | 3.3e-07 | 4.2e-07 | 3.167E-07 | 1.106E-07 | 2.000E-07 | 4.200E-07 |
| 3852 | 193 | L193K | A | 1.7e-07 | 1.4e-07 | 1.6e-07 | 1.567E-07 | 1.528E-08 | 1.400E-07 | 1.700E-07 |
| 3853 | 193 | L193M | A | 2.1e-07 | 2.9e-07 | 2.1e-07 | 2.367E-07 | 4.619E-08 | 2.100E-07 | 2.900E-07 |
| 3854 | 193 | L193F | A | 2.1e-07 | 2.2e-07 | 2.2e-07 | 2.167E-07 | 5.774E-09 | 2.100E-07 | 2.200E-07 |
| 3855 | 193 | L193P | A | 3.3e-07 | 2.3e-07 | 2.6e-07 | 2.733E-07 | 5.132E-08 | 2.300E-07 | 3.300E-07 |
| 3856 | 193 | L193S | A | 2.6e-07 | 1.6e-07 | 3.4e-07 | 2.533E-07 | 9.018E-08 | 1.600E-07 | 3.400E-07 |
| 3857 | 193 | L193T | A | 2.8e-07 | 1.4e-07 | 6e-07 | 3.400E-07 | 2.358E-07 | 1.400E-07 | 6.000E-07 |
| 3858 | 193 | L193W | A | 3.4e-07 | 2.8e-07 | 2.9e-07 | 3.033E-07 | 3.215E-08 | 2.800E-07 | 3.400E-07 |
| 3859 | 193 | L193Y | A | 2.2e-07 | 1.4e-07 | 1.9e-07 | 1.833E-07 | 4.041E-08 | 1.400E-07 | 2.200E-07 |
| 3860 | 193 | L193V | A | 2.6e-07 | 2.6e-07 | 2.1e-07 | 2.433E-07 | 2.887E-08 | 2.100E-07 | 2.600E-07 |
| 3861 | 194 | T194A | A | 1.4e-07 | 4.4e-07 | 2.7e-07 | 2.833E-07 | 1.504E-07 | 1.400E-07 | 4.400E-07 |
| 3862 | 194 | T194R | A | 3e-07 | 5.1e-07 | 1.6e-07 | 3.233E-07 | 1.762E-07 | 1.600E-07 | 5.100E-07 |
| 3863 | 194 | T194N | A | 2.5e-07 | 3.6e-07 | 3.2e-07 | 3.100E-07 | 5.568E-08 | 2.500E-07 | 3.600E-07 |
| 3864 | 194 | T194D | A | 3.5e-07 | 3.1e-07 | 2.8e-07 | 3.133E-07 | 3.512E-08 | 2.800E-07 | 3.500E-07 |
| 3865 | 194 | T194C | A | 4.3e-07 | 2.9e-07 | 1.4e-07 | 2.867E-07 | 1.450E-07 | 1.400E-07 | 4.300E-07 |
| 3866 | 194 | T194E | A | 3.7e-07 | 1.2e-07 | 4.5e-07 | 3.133E-07 | 1.721E-07 | 1.200E-07 | 4.500E-07 |
| 3867 | 194 | T194Q | A | 1.8e-07 | 4.1e-07 | 1.6e-07 | 2.500E-07 | 1.389E-07 | 1.600E-07 | 4.100E-07 |
| 3868 | 194 | T194G | A | 1.4e-07 | 5.2e-07 | 2.9e-07 | 3.167E-07 | 1.914E-07 | 1.400E-07 | 5.200E-07 |
| 3869 | 194 | T194H | A | 2.5e-07 | 1.4e-07 | 2.6e-07 | 2.167E-07 | 6.658E-08 | 1.400E-07 | 2.600E-07 |
| 3870 | 194 | T194I | A | 2e-07 | 1.3e-07 | 2.8e-07 | 2.033E-07 | 7.506E-08 | 1.300E-07 | 2.800E-07 |
| 3871 | 194 | T194L | A | 1.4e-07 | 1.5e-07 | 2.4e-07 | 1.767E-07 | 5.508E-08 | 1.400E-07 | 2.400E-07 |
| 3872 | 194 | T194K | A | 2.8e-07 | 3.6e-07 | 2.1e-07 | 2.833E-07 | 7.506E-08 | 2.100E-07 | 3.600E-07 |
| 3873 | 194 | T194M | A | 2.8e-07 | 2.7e-07 | 3.2e-07 | 2.900E-07 | 2.646E-08 | 2.700E-07 | 3.200E-07 |
| 3874 | 194 | T194F | A | 2.3e-07 | 2e-07 | 2.2e-07 | 2.167E-07 | 1.528E-08 | 2.000E-07 | 2.300E-07 |
| 3875 | 194 | T194P | A | 4.9e-07 | 1.8e-07 | 2.3e-07 | 3.000E-07 | 1.664E-07 | 1.800E-07 | 4.900E-07 |
| 3876 | 194 | T194S | A | 3e-07 | 2.8e-07 | 1.7e-07 | 2.500E-07 | 7.000E-08 | 1.700E-07 | 3.000E-07 |
| 3877 | 194 | T194T | A | 2.3e-07 | 3.6e-07 | 2.3e-07 | 2.733E-07 | 7.506E-08 | 2.300E-07 | 3.600E-07 |
| 3878 | 194 | T194W | A | 2.3e-07 | 1.4e-07 | 2.5e-07 | 2.067E-07 | 5.859E-08 | 1.400E-07 | 2.500E-07 |
| 3879 | 194 | T194Y | A | 3.3e-07 | 1.9e-07 | 1.6e-07 | 2.267E-07 | 9.074E-08 | 1.600E-07 | 3.300E-07 |
| 3880 | 194 | T194V | A | 1.3e-07 | 1.3e-07 | 2.4e-07 | 1.667E-07 | 6.351E-08 | 1.300E-07 | 2.400E-07 |
| 3881 | 195 | R195A | A | 3.2e-07 | 1.4e-07 | 1.9e-07 | 2.167E-07 | 9.292E-08 | 1.400E-07 | 3.200E-07 |
| 3882 | 195 | R195R | A | 2e-07 | 2.4e-07 | 3e-07 | 2.467E-07 | 5.033E-08 | 2.000E-07 | 3.000E-07 |
| 3883 | 195 | R195N | A | 3.3e-07 | 2.6e-07 | 1.2e-07 | 2.367E-07 | 1.069E-07 | 1.200E-07 | 3.300E-07 |
| 3884 | 195 | R195D | A | 3.4e-07 | 1.3e-07 | 4.9e-07 | 3.200E-07 | 1.808E-07 | 1.300E-07 | 4.900E-07 |
| 3885 | 195 | R195C | A | 1.3e-07 | 1.7e-07 | 2.9e-07 | 1.967E-07 | 8.327E-08 | 1.300E-07 | 2.900E-07 |
| 3886 | 195 | R195E | A | 3.7e-07 | 4.1e-07 | 1.9e-07 | 3.233E-07 | 1.172E-07 | 1.900E-07 | 4.100E-07 |
| 3887 | 195 | R195Q | A | 2.4e-07 | 2.2e-07 | 8e-08 | 1.800E-07 | 8.718E-08 | 8.000E-08 | 2.400E-07 |
| 3888 | 195 | R195G | A | 6.8e-07 | 2e-07 | 2.2e-07 | 3.667E-07 | 2.715E-07 | 2.000E-07 | 6.800E-07 |
| 3889 | 195 | R195H | A | 4.5e-07 | 1.8e-07 | 1.3e-07 | 2.533E-07 | 1.721E-07 | 1.300E-07 | 4.500E-07 |
| 3890 | 195 | R195I | A | 3e-07 | 1.3e-07 | 3.7e-07 | 2.667E-07 | 1.234E-07 | 1.300E-07 | 3.700E-07 |
| 3891 | 195 | R195L | A | 2.3e-07 | 1.5e-07 | 4.1e-07 | 2.633E-07 | 1.332E-07 | 1.500E-07 | 4.100E-07 |
| 3892 | 195 | R195K | A | 1.2e-07 | 2.9e-07 | 2.4e-07 | 2.167E-07 | 8.737E-08 | 1.200E-07 | 2.900E-07 |
| 3893 | 195 | R195M | A | 3.3e-07 | 2.1e-07 | 2.3e-07 | 2.567E-07 | 6.429E-08 | 2.100E-07 | 3.300E-07 |
| 3894 | 195 | R195F | A | 1.9e-07 | 1.9e-07 | 2.4e-07 | 2.067E-07 | 2.887E-08 | 1.900E-07 | 2.400E-07 |
| 3895 | 195 | R195P | A | 8.9e-08 | 3.8e-07 | 4.5e-07 | 3.063E-07 | 1.914E-07 | 8.900E-08 | 4.500E-07 |
| 3896 | 195 | R195S | A | 4.9e-07 | 3.6e-07 | 2.5e-07 | 3.667E-07 | 1.201E-07 | 2.500E-07 | 4.900E-07 |
| 3897 | 195 | R195T | A | 1.7e-07 | 2.9e-07 | 2.1e-07 | 2.233E-07 | 6.110E-08 | 1.700E-07 | 2.900E-07 |
| 3898 | 195 | R195W | A | 2.1e-07 | 1.8e-07 | 3.3e-07 | 2.400E-07 | 7.937E-08 | 1.800E-07 | 3.300E-07 |
| 3899 | 195 | R195Y | A | 3.7e-07 | 1.2e-07 | 6.6e-07 | 3.833E-07 | 2.702E-07 | 1.200E-07 | 6.600E-07 |
| 3900 | 195 | R195V | A | 2.4e-07 | 3.4e-07 | 1.6e-07 | 2.467E-07 | 9.018E-08 | 1.600E-07 | 3.400E-07 |
| 3901 | 196 | T196A | A | 3.4e-07 | 2.3e-07 | 2.2e-07 | 2.633E-07 | 6.658E-08 | 2.200E-07 | 3.400E-07 |
| 3902 | 196 | T196R | A | 2.4e-07 | 3.2e-07 | 2.9e-07 | 2.833E-07 | 4.041E-08 | 2.400E-07 | 3.200E-07 |
| 3903 | 196 | T196N | A | 2.9e-07 | 2e-07 | 4.3e-07 | 3.067E-07 | 1.159E-07 | 2.000E-07 | 4.300E-07 |
| 3904 | 196 | T196D | A | 3.7e-07 | 1.8e-07 | 2.8e-07 | 2.767E-07 | 9.504E-08 | 1.800E-07 | 3.700E-07 |
| 3905 | 196 | T196C | A | 3.5e-07 | 2.7e-07 | 3.2e-07 | 3.133E-07 | 4.041E-08 | 2.700E-07 | 3.500E-07 |
| 3906 | 196 | T196E | A | 2.6e-07 | 2.9e-07 | 4.1e-07 | 3.200E-07 | 7.937E-08 | 2.600E-07 | 4.100E-07 |
| 3907 | 196 | T196Q | A | 2.5e-07 | 6.5e-07 | 3.3e-07 | 4.100E-07 | 2.117E-07 | 2.500E-07 | 6.500E-07 |
| 3908 | 196 | T196G | A | 3.1e-07 | 2.6e-07 | 2.6e-07 | 2.767E-07 | 2.887E-08 | 2.600E-07 | 3.100E-07 |
| 3909 | 196 | T196H | A | 1.4e-07 | 3e-07 | 1.9e-07 | 2.100E-07 | 8.185E-08 | 1.400E-07 | 3.000E-07 |

| | | | | | | | | | | |
|---|---|---|---|---|---|---|---|---|---|---|
| 3910 | 196 | T196I | A | 4.5e-07 | 2.3e-07 | 4.1e-07 | 3.633E-07 | 1.172E-07 | 2.300E-07 | 4.500E-07 |
| 3911 | 196 | T196L | A | 1.4e-07 | 3.1e-07 | 1.9e-07 | 2.133E-07 | 8.737E-08 | 1.400E-07 | 3.100E-07 |
| 3912 | 196 | T196K | A | 1.2e-07 | 1.9e-07 | 2.5e-07 | 1.867E-07 | 6.506E-08 | 1.200E-07 | 2.500E-07 |
| 3913 | 196 | T196M | A | 2.3e-07 | 1.9e-07 | 2e-07 | 2.067E-07 | 2.082E-08 | 1.900E-07 | 2.300E-07 |
| 3914 | 196 | T196F | A | 6.6e-07 | 1.8e-07 | 4.2e-07 | 4.200E-07 | 2.400E-07 | 1.800E-07 | 6.600E-07 |
| 3915 | 196 | T196P | A | 1.9e-07 | 2.8e-07 | 3.1e-07 | 2.600E-07 | 6.245E-08 | 1.900E-07 | 3.100E-07 |
| 3916 | 196 | T196S | A | 3.2e-07 | 2.7e-07 | 5.4e-07 | 3.767E-07 | 1.436E-07 | 2.700E-07 | 5.400E-07 |
| 3917 | 196 | T196T | A | 1.8e-07 | 2e-07 | 9.2e-08 | 1.573E-07 | 5.746E-08 | 9.200E-08 | 2.000E-07 |
| 3918 | 196 | T196W | A | 2.5e-07 | 5.7e-07 | 3.4e-07 | 3.867E-07 | 1.650E-07 | 2.500E-07 | 5.700E-07 |
| 3919 | 196 | T196Y | A | 1.6e-07 | 3.5e-07 | 3.4e-07 | 2.833E-07 | 1.069E-07 | 1.600E-07 | 3.500E-07 |
| 3920 | 196 | T196V | A | 2e-07 | 2.6e-07 | 1.6e-07 | 2.067E-07 | 5.033E-08 | 1.600E-07 | 2.600E-07 |
| 3921 | 197 | V197A | A | 1.6e-07 | 4.5e-07 | 1.6e-07 | 2.567E-07 | 1.674E-07 | 1.600E-07 | 4.500E-07 |
| 3922 | 197 | V197R | A | 7.1e-08 | 9.3e-08 | 2.5e-07 | 1.380E-07 | 9.762E-08 | 7.100E-08 | 2.500E-07 |
| 3923 | 197 | V197N | A | 3.8e-07 | 5e-07 | 4.2e-07 | 4.333E-07 | 6.110E-08 | 3.800E-07 | 5.000E-07 |
| 3924 | 197 | V197D | A | 2e-07 | 5.6e-07 | 2.4e-07 | 3.333E-07 | 1.973E-07 | 2.000E-07 | 5.600E-07 |
| 3925 | 197 | V197C | A | 9.6e-08 | 2.3e-07 | 1.9e-07 | 1.720E-07 | 6.879E-08 | 9.600E-08 | 2.300E-07 |
| 3926 | 197 | V197E | A | 3.5e-07 | 2.8e-07 | 1.5e-07 | 2.600E-07 | 1.015E-07 | 1.500E-07 | 3.500E-07 |
| 3927 | 197 | V197Q | A | 3e-07 | 1.3e-07 | 2e-07 | 2.100E-07 | 8.544E-08 | 1.300E-07 | 3.000E-07 |
| 3928 | 197 | V197G | A | 1e-07 | 2.4e-07 | 3.4e-07 | 2.267E-07 | 1.206E-07 | 1.000E-07 | 3.400E-07 |
| 3929 | 197 | V197H | A | 5.6e-07 | 5.2e-07 | 2.4e-07 | 4.400E-07 | 1.744E-07 | 2.400E-07 | 5.600E-07 |
| 3930 | 197 | V197I | A | 2.8e-07 | 3.3e-07 | 1.3e-07 | 2.467E-07 | 1.041E-07 | 1.300E-07 | 3.300E-07 |
| 3931 | 197 | V197L | A | 2.1e-07 | 2.8e-07 | 2.3e-07 | 2.400E-07 | 3.606E-08 | 2.100E-07 | 2.800E-07 |
| 3932 | 197 | V197K | A | 1.4e-07 | 3.6e-07 | 2.4e-07 | 2.467E-07 | 1.102E-07 | 1.400E-07 | 3.600E-07 |
| 3933 | 197 | V197M | A | 4e-07 | 1.4e-07 | 1.7e-07 | 2.367E-07 | 1.422E-07 | 1.400E-07 | 4.000E-07 |
| 3934 | 197 | V197F | A | 4e-07 | 1.9e-07 | 3.7e-07 | 3.200E-07 | 1.136E-07 | 1.900E-07 | 4.000E-07 |
| 3935 | 197 | V197P | A | 1.4e-07 | 3.2e-07 | 1.7e-07 | 2.100E-07 | 9.644E-08 | 1.400E-07 | 3.200E-07 |
| 3936 | 197 | V197S | A | 2.6e-07 | 1.5e-07 | 1.4e-07 | 1.833E-07 | 6.658E-08 | 1.400E-07 | 2.600E-07 |
| 3937 | 197 | V197T | A | 2e-07 | 2.3e-07 | 2.9e-07 | 2.400E-07 | 4.583E-08 | 2.000E-07 | 2.900E-07 |
| 3938 | 197 | V197W | A | 7.9e-07 | 2.1e-07 | 1.3e-07 | 3.767E-07 | 3.602E-07 | 1.300E-07 | 7.900E-07 |
| 3939 | 197 | V197Y | A | 2.7e-07 | 5.4e-07 | 3.5e-07 | 3.867E-07 | 1.387E-07 | 2.700E-07 | 5.400E-07 |
| 3940 | 197 | V197V | A | 3.5e-07 | 3.5e-07 | 1.3e-07 | 2.767E-07 | 1.270E-07 | 1.300E-07 | 3.500E-07 |
| 3941 | 198 | C198A | A | 1.9e-07 | 6.5e-07 | 1.7e-07 | 3.367E-07 | 2.715E-07 | 1.700E-07 | 6.500E-07 |
| 3942 | 198 | C198R | A | 2.1e-07 | 5.9e-07 | 9.7e-08 | 2.990E-07 | 2.583E-07 | 9.700E-08 | 5.900E-07 |
| 3943 | 198 | C198N | A | 1.9e-07 | 2e-07 | 4.4e-07 | 2.767E-07 | 1.415E-07 | 1.900E-07 | 4.400E-07 |
| 3944 | 198 | C198D | A | 3.1e-07 | 2.2e-07 | 1.5e-07 | 2.267E-07 | 8.021E-08 | 1.500E-07 | 3.100E-07 |
| 3945 | 198 | C198C | A | 1.9e-07 | 1.9e-07 | 2.5e-07 | 2.100E-07 | 3.464E-08 | 1.900E-07 | 2.500E-07 |
| 3946 | 198 | C198E | A | 2.5e-07 | 1.6e-07 | 2.5e-07 | 2.200E-07 | 5.196E-08 | 1.600E-07 | 2.500E-07 |
| 3947 | 198 | C198Q | A | 1.4e-07 | 2.5e-07 | 3e-07 | 2.300E-07 | 8.185E-08 | 1.400E-07 | 3.000E-07 |
| 3948 | 198 | C198G | A | 4.3e-07 | 4.7e-07 | 1.4e-07 | 3.467E-07 | 1.801E-07 | 1.400E-07 | 4.700E-07 |
| 3949 | 198 | C198H | A | 3.2e-07 | 3.6e-07 | 3.6e-07 | 3.467E-07 | 2.309E-08 | 3.200E-07 | 3.600E-07 |
| 3950 | 198 | C198I | A | 1e-07 | 9.5e-08 | 3.2e-07 | 1.717E-07 | 1.285E-07 | 9.500E-08 | 3.200E-07 |
| 3951 | 198 | C198L | A | 2.9e-07 | 2.7e-07 | 2.8e-07 | 2.800E-07 | 1.000E-08 | 2.700E-07 | 2.900E-07 |
| 3952 | 198 | C198K | A | 2.4e-07 | 1.9e-07 | 2.7e-07 | 2.333E-07 | 4.041E-08 | 1.900E-07 | 2.700E-07 |
| 3953 | 198 | C198M | A | 4.7e-07 | 1.4e-07 | 1.5e-07 | 2.533E-07 | 1.877E-07 | 1.400E-07 | 4.700E-07 |
| 3954 | 198 | C198F | A | 2.7e-07 | 5.3e-07 | 3.1e-07 | 3.700E-07 | 1.400E-07 | 2.700E-07 | 5.300E-07 |
| 3955 | 198 | C198P | A | 2e-07 | 2.9e-07 | 2.7e-07 | 2.533E-07 | 4.726E-08 | 2.000E-07 | 2.900E-07 |
| 3956 | 198 | C198S | A | 4e-07 | 2.4e-07 | 3.3e-07 | 3.233E-07 | 8.021E-08 | 2.400E-07 | 4.000E-07 |
| 3957 | 198 | C198T | A | 2.8e-07 | 2e-07 | 1e-07 | 1.933E-07 | 9.018E-08 | 1.000E-07 | 2.800E-07 |
| 3958 | 198 | C198W | A | 2.2e-07 | 1.5e-07 | 3.4e-07 | 2.367E-07 | 9.609E-08 | 1.500E-07 | 3.400E-07 |
| 3959 | 198 | C198Y | A | 1.4e-07 | 1.6e-07 | 3.4e-07 | 2.133E-07 | 1.102E-07 | 1.400E-07 | 3.400E-07 |
| 3960 | 198 | C198V | A | 1.7e-07 | 2.9e-07 | 1e-07 | 1.867E-07 | 9.609E-08 | 1.000E-07 | 2.900E-07 |
| 3961 | 199 | A199A | A | 1.8e-07 | 2.9e-07 | 1.4e-07 | 2.033E-07 | 7.767E-08 | 1.400E-07 | 2.900E-07 |
| 3962 | 199 | A199R | A | 1.6e-07 | 1.7e-07 | 1.6e-07 | 1.633E-07 | 5.774E-09 | 1.600E-07 | 1.700E-07 |
| 3963 | 199 | A199N | A | 3.7e-07 | 1.3e-07 | 2.5e-07 | 2.500E-07 | 1.200E-07 | 1.300E-07 | 3.700E-07 |
| 3964 | 199 | A199D | A | 3.2e-07 | 3.8e-07 | 9.7e-08 | 2.657E-07 | 1.491E-07 | 9.700E-08 | 3.800E-07 |
| 3965 | 199 | A199C | A | 2.5e-07 | 3.7e-07 | 2.4e-07 | 2.867E-07 | 7.234E-08 | 2.400E-07 | 3.700E-07 |
| 3966 | 199 | A199E | A | 4e-07 | 2.6e-07 | 3.3e-07 | 3.300E-07 | 7.000E-08 | 2.600E-07 | 4.000E-07 |
| 3967 | 199 | A199Q | A | 1.7e-07 | 1.1e-07 | 3.2e-07 | 2.000E-07 | 1.082E-07 | 1.100E-07 | 3.200E-07 |
| 3968 | 199 | A199G | A | 2e-07 | 2.1e-07 | 2.6e-07 | 2.233E-07 | 3.215E-08 | 2.000E-07 | 2.600E-07 |
| 3969 | 199 | A199H | A | 1.9e-07 | 4.1e-07 | 2.3e-07 | 2.767E-07 | 1.172E-07 | 1.900E-07 | 4.100E-07 |
| 3970 | 199 | A199I | A | 5.4e-07 | 2.8e-07 | 2.2e-07 | 3.467E-07 | 1.701E-07 | 2.200E-07 | 5.400E-07 |
| 3971 | 199 | A199L | A | 4.7e-07 | 2.5e-07 | 1.7e-07 | 2.967E-07 | 1.553E-07 | 1.700E-07 | 4.700E-07 |
| 3972 | 199 | A199K | A | 3.4e-07 | 2.8e-07 | 3.6e-07 | 3.267E-07 | 4.163E-08 | 2.800E-07 | 3.600E-07 |
| 3973 | 199 | A199M | A | 4e-07 | 9e-08 | 2.6e-07 | 2.500E-07 | 1.552E-07 | 9.000E-08 | 4.000E-07 |
| 3974 | 199 | A199F | A | 1.7e-07 | 1.4e-07 | 3.4e-07 | 2.167E-07 | 1.079E-07 | 1.400E-07 | 3.400E-07 |
| 3975 | 199 | A199P | A | 3.2e-07 | 5.6e-07 | 3.3e-07 | 4.033E-07 | 1.358E-07 | 3.200E-07 | 5.600E-07 |
| 3976 | 199 | A199S | A | 1.7e-07 | 4e-07 | 2.2e-07 | 2.633E-07 | 1.210E-07 | 1.700E-07 | 4.000E-07 |
| 3977 | 199 | A199T | A | 2.2e-07 | 2.5e-07 | 3.6e-07 | 2.767E-07 | 7.371E-08 | 2.200E-07 | 3.600E-07 |
| 3978 | 199 | A199W | A | 2.5e-07 | 2e-07 | 2.6e-07 | 2.367E-07 | 3.215E-08 | 2.000E-07 | 2.600E-07 |
| 3979 | 199 | A199Y | A | 1.4e-07 | 3.9e-07 | 1.6e-07 | 2.300E-07 | 1.389E-07 | 1.400E-07 | 3.900E-07 |
| 3980 | 199 | A199V | A | 3.7e-07 | 2.7e-07 | 4.5e-07 | 3.633E-07 | 9.018E-08 | 2.700E-07 | 4.500E-07 |
| 3981 | 200 | G200A | A | 2.3e-07 | 1.4e-07 | 3.2e-07 | 2.300E-07 | 9.000E-08 | 1.400E-07 | 3.200E-07 |
| 3982 | 200 | G200R | A | 3e-07 | 3.6e-07 | 2.5e-07 | 3.033E-07 | 5.508E-08 | 2.500E-07 | 3.600E-07 |
| 3983 | 200 | G200N | A | 1.4e-07 | 1.5e-07 | 2.3e-07 | 1.733E-07 | 4.933E-08 | 1.400E-07 | 2.300E-07 |
| 3984 | 200 | G200D | A | 3.4e-07 | 1.6e-07 | 2.9e-07 | 2.633E-07 | 9.292E-08 | 1.600E-07 | 3.400E-07 |
| 3985 | 200 | G200C | A | 1.2e-07 | 1.1e-07 | 1.2e-07 | 1.167E-07 | 5.774E-09 | 1.100E-07 | 1.200E-07 |
| 3986 | 200 | G200E | A | 3.1e-07 | 2.2e-07 | 2.4e-07 | 2.567E-07 | 4.726E-08 | 2.200E-07 | 3.100E-07 |
| 3987 | 200 | G200Q | A | 2.7e-07 | 7.7e-08 | 2.5e-07 | 1.990E-07 | 1.061E-07 | 7.700E-08 | 2.700E-07 |
| 3988 | 200 | G200G | A | 7.3e-08 | 3.8e-07 | 2.7e-07 | 2.410E-07 | 1.555E-07 | 7.300E-08 | 3.800E-07 |
| 3989 | 200 | G200H | A | 3e-07 | 3.2e-07 | 1.7e-07 | 2.633E-07 | 8.145E-08 | 1.700E-07 | 3.200E-07 |
| 3990 | 200 | G200I | A | 2.3e-07 | 2.3e-07 | 1.7e-07 | 2.100E-07 | 3.464E-08 | 1.700E-07 | 2.300E-07 |
| 3991 | 200 | G200L | A | 1.9e-07 | 4.5e-07 | 2.5e-07 | 2.967E-07 | 1.361E-07 | 1.900E-07 | 4.500E-07 |
| 3992 | 200 | G200K | A | 3e-07 | 2.3e-07 | 1.1e-07 | 2.133E-07 | 9.609E-08 | 1.100E-07 | 3.000E-07 |
| 3993 | 200 | G200M | A | 2.1e-07 | 5.2e-07 | 4.4e-07 | 3.900E-07 | 1.609E-07 | 2.100E-07 | 5.200E-07 |
| 3994 | 200 | G200F | A | 1e-07 | 4.2e-07 | 2.5e-07 | 2.567E-07 | 1.601E-07 | 1.000E-07 | 4.200E-07 |

| | | | | | | | | | | |
|---|---|---|---|---|---|---|---|---|---|---|
| 3995 | 200 | G200P | A | 2.9e-07 | 2.9e-07 | 3.4e-07 | 3.067E-07 | 2.887E-08 | 2.900E-07 | 3.400E-07 |
| 3996 | 200 | G200S | A | 3e-07 | 1.5e-07 | 1.5e-07 | 2.000E-07 | 8.660E-08 | 1.500E-07 | 3.000E-07 |
| 3997 | 200 | G200T | A | 3.8e-07 | 3.4e-07 | 2.6e-07 | 3.267E-07 | 6.110E-08 | 2.600E-07 | 3.800E-07 |
| 3998 | 200 | G200W | A | 1.1e-07 | 2.7e-07 | 3.5e-07 | 2.433E-07 | 1.222E-07 | 1.100E-07 | 3.500E-07 |
| 3999 | 200 | G200Y | A | 3.2e-07 | 1.8e-07 | 2.1e-07 | 2.367E-07 | 7.371E-08 | 1.800E-07 | 3.200E-07 |
| 4000 | 200 | G200V | A | 3.4e-07 | 1.2e-07 | 1.6e-07 | 2.067E-07 | 1.172E-07 | 1.200E-07 | 3.400E-07 |
| 4001 | 201 | G201A | A | 1.4e-07 | 3.8e-07 | 2.1e-07 | 2.433E-07 | 1.234E-07 | 1.400E-07 | 3.800E-07 |
| 4002 | 201 | G201R | A | 3.6e-07 | 1.7e-07 | 2.1e-07 | 2.467E-07 | 1.002E-07 | 1.700E-07 | 3.600E-07 |
| 4003 | 201 | G201N | A | 1.7e-07 | 3.4e-07 | 2.7e-07 | 2.600E-07 | 8.544E-08 | 1.700E-07 | 3.400E-07 |
| 4004 | 201 | G201D | A | 1.8e-07 | 1.2e-07 | 2.8e-07 | 1.933E-07 | 8.083E-08 | 1.200E-07 | 2.800E-07 |
| 4005 | 201 | G201C | A | 1.9e-07 | 1e-07 | 1.3e-07 | 1.400E-07 | 4.583E-08 | 1.000E-07 | 1.900E-07 |
| 4006 | 201 | G201E | A | 4.9e-07 | 2.6e-07 | 2.4e-07 | 3.300E-07 | 1.389E-07 | 2.400E-07 | 4.900E-07 |
| 4007 | 201 | G201Q | A | 6.3e-07 | 1.5e-07 | 1.3e-07 | 3.033E-07 | 2.831E-07 | 1.300E-07 | 6.300E-07 |
| 4008 | 201 | G201G | A | 3.3e-07 | 3.1e-07 | 2.9e-07 | 3.100E-07 | 2.000E-08 | 2.900E-07 | 3.300E-07 |
| 4009 | 201 | G201H | A | 5e-07 | 3.6e-07 | 1.5e-07 | 3.367E-07 | 1.762E-07 | 1.500E-07 | 5.000E-07 |
| 4010 | 201 | G201I | A | 2.1e-07 | 4.7e-07 | 2.7e-07 | 3.167E-07 | 1.361E-07 | 2.100E-07 | 4.700E-07 |
| 4011 | 201 | G201L | A | 1.8e-07 | 3.2e-07 | 3.3e-07 | 2.767E-07 | 8.386E-08 | 1.800E-07 | 3.300E-07 |
| 4012 | 201 | G201K | A | 4e-07 | 3.8e-07 | 3.3e-07 | 3.700E-07 | 3.606E-08 | 3.300E-07 | 4.000E-07 |
| 4013 | 201 | G201M | A | 1.5e-07 | 4.5e-07 | 4e-07 | 3.333E-07 | 1.607E-07 | 1.500E-07 | 4.500E-07 |
| 4014 | 201 | G201F | A | 4e-07 | 2.9e-07 | 3.6e-07 | 3.500E-07 | 5.568E-08 | 2.900E-07 | 4.000E-07 |
| 4015 | 201 | G201P | A | 3e-07 | 4.2e-07 | 2.5e-07 | 3.233E-07 | 8.737E-08 | 2.500E-07 | 4.200E-07 |
| 4016 | 201 | G201S | A | 2.9e-07 | 2.6e-07 | 2.4e-07 | 2.633E-07 | 2.517E-08 | 2.400E-07 | 2.900E-07 |
| 4017 | 201 | G201T | A | 3.7e-07 | 2.1e-07 | 1.7e-07 | 2.500E-07 | 1.058E-07 | 1.700E-07 | 3.700E-07 |
| 4018 | 201 | G201W | A | 3.4e-07 | 2.3e-07 | 2.4e-07 | 2.700E-07 | 6.083E-08 | 2.300E-07 | 3.400E-07 |
| 4019 | 201 | G201Y | A | 2.1e-07 | 1.7e-07 | 2.3e-07 | 2.033E-07 | 3.055E-08 | 1.700E-07 | 2.300E-07 |
| 4020 | 201 | G201V | A | 3.3e-07 | 3.3e-07 | 1.6e-07 | 2.733E-07 | 9.815E-08 | 1.600E-07 | 3.300E-07 |
| 4021 | 202 | C202A | A | 2.9e-07 | 5.7e-07 | 2.9e-07 | 3.833E-07 | 1.617E-07 | 2.900E-07 | 5.700E-07 |
| 4022 | 202 | C202R | A | 3.2e-07 | 1.8e-07 | 3.2e-07 | 2.733E-07 | 8.083E-08 | 1.800E-07 | 3.200E-07 |
| 4023 | 202 | C202N | A | 2.4e-07 | 2.9e-07 | 2e-07 | 2.433E-07 | 4.509E-08 | 2.000E-07 | 2.900E-07 |
| 4024 | 202 | C202D | A | 3.2e-07 | 2.1e-07 | 2.8e-07 | 2.700E-07 | 5.568E-08 | 2.100E-07 | 3.200E-07 |
| 4025 | 202 | C202C | A | 5.1e-07 | 4e-07 | 9.3e-07 | 6.133E-07 | 2.797E-07 | 4.000E-07 | 9.300E-07 |
| 4026 | 202 | C202E | A | 2.2e-07 | 2.3e-07 | 2e-07 | 2.167E-07 | 1.528E-08 | 2.000E-07 | 2.300E-07 |
| 4027 | 202 | C202Q | A | 2.1e-07 | 2.9e-07 | 2.1e-07 | 2.367E-07 | 4.619E-08 | 2.100E-07 | 2.900E-07 |
| 4028 | 202 | C202G | A | 2.8e-07 | 1.7e-07 | 3.2e-07 | 2.567E-07 | 7.767E-08 | 1.700E-07 | 3.200E-07 |
| 4029 | 202 | C202H | A | 3.2e-07 | 5.5e-07 | 1.3e-07 | 3.333E-07 | 2.103E-07 | 1.300E-07 | 5.500E-07 |
| 4030 | 202 | C202I | A | 3.5e-07 | 2.9e-07 | 6.5e-07 | 4.300E-07 | 1.929E-07 | 2.900E-07 | 6.500E-07 |
| 4031 | 202 | C202L | A | 2.2e-07 | 1.8e-07 | 3.5e-07 | 2.500E-07 | 8.888E-08 | 1.800E-07 | 3.500E-07 |
| 4032 | 202 | C202K | A | 1.6e-07 | 1.6e-07 | 2.8e-07 | 2.000E-07 | 6.928E-08 | 1.600E-07 | 2.800E-07 |
| 4033 | 202 | C202M | A | 3e-07 | 6.2e-08 | 2.1e-07 | 1.907E-07 | 1.202E-07 | 6.200E-08 | 3.000E-07 |
| 4034 | 202 | C202F | A | 2.9e-07 | 2.6e-07 | 1.2e-07 | 2.233E-07 | 9.074E-08 | 1.200E-07 | 2.900E-07 |
| 4035 | 202 | C202P | A | 2.4e-07 | 1.6e-07 | 1.2e-07 | 1.733E-07 | 6.110E-08 | 1.200E-07 | 2.400E-07 |
| 4036 | 202 | C202S | A | 3.1e-07 | 2.7e-07 | 2.9e-07 | 2.900E-07 | 2.000E-08 | 2.700E-07 | 3.100E-07 |
| 4037 | 202 | C202T | A | 2.6e-07 | 5.7e-07 | 1.9e-07 | 3.400E-07 | 2.022E-07 | 1.900E-07 | 5.700E-07 |
| 4038 | 202 | C202W | A | 6.2e-07 | 3.1e-07 | 1.4e-07 | 3.567E-07 | 2.434E-07 | 1.400E-07 | 6.200E-07 |
| 4039 | 202 | C202Y | A | 1.4e-07 | 8.7e-08 | 2.9e-07 | 1.723E-07 | 1.053E-07 | 8.700E-08 | 2.900E-07 |
| 4040 | 202 | C202V | A | 1.5e-07 | 2.8e-07 | 4.7e-07 | 3.000E-07 | 1.609E-07 | 1.500E-07 | 4.700E-07 |
| 4041 | 203 | A203A | A | 1.8e-07 | 2.1e-07 | 2.3e-07 | 2.067E-07 | 2.517E-08 | 1.800E-07 | 2.300E-07 |
| 4042 | 203 | A203R | A | 2e-07 | 1.8e-07 | 1.4e-07 | 1.733E-07 | 3.055E-08 | 1.400E-07 | 2.000E-07 |
| 4043 | 203 | A203N | A | 1.5e-07 | 1.4e-07 | 3.8e-07 | 2.233E-07 | 1.358E-07 | 1.400E-07 | 3.800E-07 |
| 4044 | 203 | A203D | A | 2.9e-07 | 4e-07 | 2.1e-07 | 3.000E-07 | 9.539E-08 | 2.100E-07 | 4.000E-07 |
| 4045 | 203 | A203C | A | 2.3e-07 | 2.1e-07 | 6.8e-08 | 1.693E-07 | 8.833E-08 | 6.800E-08 | 2.300E-07 |
| 4046 | 203 | A203E | A | 3.8e-07 | 1.5e-07 | 2.7e-07 | 2.667E-07 | 1.150E-07 | 1.500E-07 | 3.800E-07 |
| 4047 | 203 | A203Q | A | 6.8e-08 | 2.2e-07 | 1.1e-07 | 1.327E-07 | 7.849E-08 | 6.800E-08 | 2.200E-07 |
| 4048 | 203 | A203G | A | 9.3e-07 | 1.2e-07 | 1.9e-07 | 4.133E-07 | 4.488E-07 | 1.200E-07 | 9.300E-07 |
| 4049 | 203 | A203H | A | 2.6e-07 | 1.2e-07 | 4.5e-07 | 2.767E-07 | 1.656E-07 | 1.200E-07 | 4.500E-07 |
| 4050 | 203 | A203I | A | 3.6e-07 | 2.7e-07 | 2.6e-07 | 2.967E-07 | 5.508E-08 | 2.600E-07 | 3.600E-07 |
| 4051 | 203 | A203L | A | 1e-06 | 2.2e-07 | 3.5e-07 | 5.233E-07 | 4.179E-07 | 2.200E-07 | 1.000E-06 |
| 4052 | 203 | A203K | A | 2e-07 | 2.1e-07 | 4.9e-07 | 3.000E-07 | 1.646E-07 | 2.000E-07 | 4.900E-07 |
| 4053 | 203 | A203M | A | 4.5e-07 | 3.4e-07 | 2.3e-07 | 3.400E-07 | 1.100E-07 | 2.300E-07 | 4.500E-07 |
| 4054 | 203 | A203F | A | 3.4e-07 | 1.8e-07 | 6.6e-07 | 3.933E-07 | 2.444E-07 | 1.800E-07 | 6.600E-07 |
| 4055 | 203 | A203P | A | 3.3e-07 | 2.5e-07 | 2.6e-07 | 2.800E-07 | 4.359E-08 | 2.500E-07 | 3.300E-07 |
| 4056 | 203 | A203S | A | 2.6e-07 | 5e-07 | 1.8e-07 | 3.133E-07 | 1.665E-07 | 1.800E-07 | 5.000E-07 |
| 4057 | 203 | A203T | A | 4.1e-07 | 3.7e-07 | 1.3e-07 | 3.033E-07 | 1.514E-07 | 1.300E-07 | 4.100E-07 |
| 4058 | 203 | A203W | A | 4.1e-07 | 2.4e-07 | 2.4e-07 | 2.967E-07 | 9.815E-08 | 2.400E-07 | 4.100E-07 |
| 4059 | 203 | A203Y | A | 2.3e-07 | 2.8e-07 | 1.6e-07 | 2.233E-07 | 6.028E-08 | 1.600E-07 | 2.800E-07 |
| 4060 | 203 | A203V | A | 3.6e-07 | 3.9e-07 | 1.6e-07 | 3.033E-07 | 1.250E-07 | 1.600E-07 | 3.900E-07 |
| 4061 | 204 | R204A | A | 1.7e-07 | 2e-07 | 1.1e-07 | 1.600E-07 | 4.583E-08 | 1.100E-07 | 2.000E-07 |
| 4062 | 204 | R204R | A | 9.9e-08 | 9.7e-08 | 1.3e-07 | 1.087E-07 | 1.850E-08 | 9.700E-08 | 1.300E-07 |
| 4063 | 204 | R204N | A | 7.8e-07 | 2e-07 | 2.9e-07 | 4.233E-07 | 3.121E-07 | 2.000E-07 | 7.800E-07 |
| 4064 | 204 | R204D | A | 2.6e-07 | 1.7e-07 | 2.7e-07 | 2.333E-07 | 5.508E-08 | 1.700E-07 | 2.700E-07 |
| 4065 | 204 | R204C | A | 2.3e-07 | 2.1e-07 | 4.2e-07 | 2.867E-07 | 1.159E-07 | 2.100E-07 | 4.200E-07 |
| 4066 | 204 | R204E | A | 5e-07 | 3.6e-07 | 4e-07 | 4.200E-07 | 7.211E-08 | 3.600E-07 | 5.000E-07 |
| 4067 | 204 | R204Q | A | 2.2e-07 | 1e-07 | 5.4e-07 | 2.867E-07 | 2.274E-07 | 1.000E-07 | 5.400E-07 |
| 4068 | 204 | R204G | A | 2.3e-07 | 3.9e-07 | 2.3e-07 | 2.833E-07 | 9.238E-08 | 2.300E-07 | 3.900E-07 |
| 4069 | 204 | R204H | A | 2e-07 | 2.5e-07 | 4.3e-07 | 2.933E-07 | 1.210E-07 | 2.000E-07 | 4.300E-07 |
| 4070 | 204 | R204I | A | 1.7e-07 | 2.2e-07 | 3.1e-07 | 2.333E-07 | 7.095E-08 | 1.700E-07 | 3.100E-07 |
| 4071 | 204 | R204L | A | 1.9e-07 | 1.2e-07 | 1.8e-07 | 1.633E-07 | 3.786E-08 | 1.200E-07 | 1.900E-07 |
| 4072 | 204 | R204K | A | 3.3e-07 | 2e-07 | 2.5e-07 | 2.600E-07 | 6.557E-08 | 2.000E-07 | 3.300E-07 |
| 4073 | 204 | R204M | A | 2.2e-07 | 3.3e-07 | 2.4e-07 | 2.633E-07 | 5.859E-08 | 2.200E-07 | 3.300E-07 |
| 4074 | 204 | R204F | A | 3e-07 | 3e-07 | 3.9e-07 | 3.300E-07 | 5.196E-08 | 3.000E-07 | 3.900E-07 |
| 4075 | 204 | R204P | A | 3.3e-07 | 2.7e-07 | 3e-07 | 3.000E-07 | 3.000E-08 | 2.700E-07 | 3.300E-07 |
| 4076 | 204 | R204S | A | 2.4e-07 | 6.5e-07 | 1.4e-07 | 3.433E-07 | 2.702E-07 | 1.400E-07 | 6.500E-07 |
| 4077 | 204 | R204T | A | 3.8e-07 | 3.4e-07 | 1.9e-07 | 3.033E-07 | 1.002E-07 | 1.900E-07 | 3.800E-07 |
| 4078 | 204 | R204W | A | 2.8e-07 | 2.2e-07 | 2.4e-07 | 2.467E-07 | 3.055E-08 | 2.200E-07 | 2.800E-07 |
| 4079 | 204 | R204Y | A | 1.5e-07 | 1.4e-07 | 2.8e-07 | 1.900E-07 | 7.810E-08 | 1.400E-07 | 2.800E-07 |

| | | | | | | | | | | |
|---|---|---|---|---|---|---|---|---|---|---|
| 4080 | 204 | R204V | A | 3.2e-07 | 3.3e-07 | 6.3e-07 | 4.267E-07 | 1.762E-07 | 3.200E-07 | 6.300E-07 |
| 4081 | 205 | C205A | A | 1.2e-07 | 3.6e-07 | 2.9e-07 | 2.567E-07 | 1.234E-07 | 1.200E-07 | 3.600E-07 |
| 4082 | 205 | C205R | A | 3e-07 | 3e-07 | 4.9e-07 | 3.633E-07 | 1.097E-07 | 3.000E-07 | 4.900E-07 |
| 4083 | 205 | C205N | A | 3.9e-07 | 2e-07 | 3.6e-07 | 3.167E-07 | 1.021E-07 | 2.000E-07 | 3.900E-07 |
| 4084 | 205 | C205D | A | 1.9e-07 | 2.2e-07 | 3.3e-07 | 2.467E-07 | 7.371E-08 | 1.900E-07 | 3.300E-07 |
| 4085 | 205 | C205C | A | 5e-07 | 2.4e-07 | 1.5e-07 | 2.967E-07 | 1.818E-07 | 1.500E-07 | 5.000E-07 |
| 4086 | 205 | C205E | A | 3.4e-07 | 4.9e-07 | 5.6e-07 | 4.633E-07 | 1.124E-07 | 3.400E-07 | 5.600E-07 |
| 4087 | 205 | C205Q | A | 3.9e-07 | 1.8e-07 | 2.4e-07 | 2.700E-07 | 1.082E-07 | 1.800E-07 | 3.900E-07 |
| 4088 | 205 | C205G | A | 4.3e-07 | 3.4e-07 | 2.7e-07 | 3.467E-07 | 8.021E-08 | 2.700E-07 | 4.300E-07 |
| 4089 | 205 | C205H | A | 4.7e-07 | 1e-07 | 1.7e-07 | 2.467E-07 | 1.966E-07 | 1.000E-07 | 4.700E-07 |
| 4090 | 205 | C205I | A | 2.6e-07 | 4.9e-07 | 1.5e-07 | 3.000E-07 | 1.735E-07 | 1.500E-07 | 4.900E-07 |
| 4091 | 205 | C205L | A | 1.2e-07 | 3.1e-07 | 1.8e-07 | 2.033E-07 | 9.713E-08 | 1.200E-07 | 3.100E-07 |
| 4092 | 205 | C205K | A | 2.1e-07 | 9.6e-08 | 2.5e-07 | 1.853E-07 | 7.991E-08 | 9.600E-08 | 2.500E-07 |
| 4093 | 205 | C205M | A | 5.1e-07 | 1.8e-07 | 3.2e-07 | 3.367E-07 | 1.656E-07 | 1.800E-07 | 5.100E-07 |
| 4094 | 205 | C205F | A | 2.1e-07 | 2.2e-07 | 1.6e-07 | 1.967E-07 | 3.215E-08 | 1.600E-07 | 2.200E-07 |
| 4095 | 205 | C205P | A | 3e-07 | 1.3e-07 | 2.8e-07 | 2.367E-07 | 9.292E-08 | 1.300E-07 | 3.000E-07 |
| 4096 | 205 | C205S | A | 2.8e-07 | 1.3e-07 | 2.5e-07 | 2.200E-07 | 7.937E-08 | 1.300E-07 | 2.800E-07 |
| 4097 | 205 | C205T | A | 2.8e-07 | 1.4e-07 | 2e-07 | 2.067E-07 | 7.024E-08 | 1.400E-07 | 2.800E-07 |
| 4098 | 205 | C205W | A | 3.6e-07 | 5.3e-07 | 6.8e-07 | 5.233E-07 | 1.601E-07 | 3.600E-07 | 6.800E-07 |
| 4099 | 205 | C205Y | A | 6.7e-07 | 3.2e-07 | 4.7e-07 | 4.867E-07 | 1.756E-07 | 3.200E-07 | 6.700E-07 |
| 4100 | 205 | C205V | A | 5.8e-07 | 2.1e-07 | 1.2e-07 | 3.033E-07 | 2.438E-07 | 1.200E-07 | 5.800E-07 |
| 4101 | 206 | K206A | A | 2.1e-07 | 2e-07 | 1.5e-07 | 1.867E-07 | 3.215E-08 | 1.500E-07 | 2.100E-07 |
| 4102 | 206 | K206R | A | 2.7e-07 | 2.2e-07 | 2.8e-07 | 2.567E-07 | 3.215E-08 | 2.200E-07 | 2.800E-07 |
| 4103 | 206 | K206N | A | 4e-07 | 1.2e-07 | 4e-07 | 3.067E-07 | 1.617E-07 | 1.200E-07 | 4.000E-07 |
| 4104 | 206 | K206D | A | 2.9e-07 | 2.8e-07 | 2e-07 | 2.567E-07 | 4.933E-08 | 2.000E-07 | 2.900E-07 |
| 4105 | 206 | K206C | A | 1.2e-07 | 2.4e-07 | 1.3e-07 | 1.633E-07 | 6.658E-08 | 1.200E-07 | 2.400E-07 |
| 4106 | 206 | K206E | A | 2.8e-07 | 3.2e-07 | 1.5e-07 | 2.500E-07 | 8.888E-08 | 1.500E-07 | 3.200E-07 |
| 4107 | 206 | K206Q | A | 1.2e-07 | 2.5e-07 | 3.2e-07 | 2.300E-07 | 1.015E-07 | 1.200E-07 | 3.200E-07 |
| 4108 | 206 | K206G | A | 4.4e-07 | 2.8e-07 | 2.9e-07 | 3.367E-07 | 8.963E-08 | 2.800E-07 | 4.400E-07 |
| 4109 | 206 | K206H | A | 2.5e-07 | 1.5e-07 | 1e-07 | 1.667E-07 | 7.638E-08 | 1.000E-07 | 2.500E-07 |
| 4110 | 206 | K206I | A | 3.9e-07 | 1.9e-07 | 2.9e-07 | 2.900E-07 | 1.000E-07 | 1.900E-07 | 3.900E-07 |
| 4111 | 206 | K206L | A | 1.8e-07 | 3.4e-07 | 2.2e-07 | 2.467E-07 | 8.327E-08 | 1.800E-07 | 3.400E-07 |
| 4112 | 206 | K206K | A | 1.5e-07 | 2.2e-07 | 2.9e-07 | 2.200E-07 | 7.000E-08 | 1.500E-07 | 2.900E-07 |
| 4113 | 206 | K206M | A | 2e-07 | 3.2e-07 | 1.2e-07 | 2.133E-07 | 1.007E-07 | 1.200E-07 | 3.200E-07 |
| 4114 | 206 | K206F | A | 1.6e-07 | 5.1e-07 | 1.7e-07 | 2.800E-07 | 1.992E-07 | 1.600E-07 | 5.100E-07 |
| 4115 | 206 | K206P | A | 2.6e-07 | 3.1e-07 | 2.5e-07 | 2.733E-07 | 3.215E-08 | 2.500E-07 | 3.100E-07 |
| 4116 | 206 | K206S | A | 2.3e-07 | 3.4e-07 | 2.7e-07 | 2.800E-07 | 5.568E-08 | 2.300E-07 | 3.400E-07 |
| 4117 | 206 | K206T | A | 4.2e-07 | 1.5e-07 | 3.4e-07 | 3.033E-07 | 1.387E-07 | 1.500E-07 | 4.200E-07 |
| 4118 | 206 | K206W | A | 2.5e-07 | 8.6e-07 | 1.4e-07 | 4.167E-07 | 3.879E-07 | 1.400E-07 | 8.600E-07 |
| 4119 | 206 | K206Y | A | 1.2e-07 | 3.3e-07 | 3.2e-07 | 2.567E-07 | 1.185E-07 | 1.200E-07 | 3.300E-07 |
| 4120 | 206 | K206V | A | 1.4e-07 | 2.2e-07 | 1.2e-07 | 1.600E-07 | 5.292E-08 | 1.200E-07 | 2.200E-07 |
| 4121 | 207 | G207A | A | 5.2e-07 | 2.4e-07 | 1.4e-07 | 3.000E-07 | 1.970E-07 | 1.400E-07 | 5.200E-07 |
| 4122 | 207 | G207R | A | 3e-07 | 2.8e-07 | 3.3e-07 | 3.033E-07 | 2.517E-08 | 2.800E-07 | 3.300E-07 |
| 4123 | 207 | G207N | A | 2.7e-07 | 5e-07 | 2.4e-07 | 3.367E-07 | 1.422E-07 | 2.400E-07 | 5.000E-07 |
| 4124 | 207 | G207D | A | 5.4e-07 | 4.1e-07 | 2.7e-07 | 4.067E-07 | 1.350E-07 | 2.700E-07 | 5.400E-07 |
| 4125 | 207 | G207C | A | 2.8e-07 | 2.1e-07 | 1e-07 | 1.967E-07 | 9.074E-08 | 1.000E-07 | 2.800E-07 |
| 4126 | 207 | G207E | A | 2.5e-07 | 2.6e-07 | 1.5e-07 | 2.200E-07 | 6.083E-08 | 1.500E-07 | 2.600E-07 |
| 4127 | 207 | G207Q | A | 2.3e-07 | 3.7e-07 | 1.4e-07 | 2.467E-07 | 1.159E-07 | 1.400E-07 | 3.700E-07 |
| 4128 | 207 | G207G | A | 2.7e-07 | 2.2e-07 | 4.6e-07 | 3.167E-07 | 1.266E-07 | 2.200E-07 | 4.600E-07 |
| 4129 | 207 | G207H | A | 2.5e-07 | 3.7e-07 | 3e-07 | 3.067E-07 | 6.028E-08 | 2.500E-07 | 3.700E-07 |
| 4130 | 207 | G207I | A | 2.1e-07 | 3.8e-07 | 3.3e-07 | 3.067E-07 | 8.737E-08 | 2.100E-07 | 3.800E-07 |
| 4131 | 207 | G207L | A | 2.9e-07 | 2.1e-07 | 2e-07 | 2.333E-07 | 4.933E-08 | 2.000E-07 | 2.900E-07 |
| 4132 | 207 | G207K | A | 2.9e-07 | 3.6e-07 | 2.2e-07 | 2.900E-07 | 7.000E-08 | 2.200E-07 | 3.600E-07 |
| 4133 | 207 | G207M | A | 2.7e-07 | 1.6e-07 | 2.6e-07 | 2.300E-07 | 6.083E-08 | 1.600E-07 | 2.700E-07 |
| 4134 | 207 | G207F | A | 3.3e-07 | 2.1e-07 | 2e-07 | 2.467E-07 | 7.234E-08 | 2.000E-07 | 3.300E-07 |
| 4135 | 207 | G207P | A | 1.6e-07 | 1.9e-07 | 2.9e-07 | 2.133E-07 | 6.807E-08 | 1.600E-07 | 2.900E-07 |
| 4136 | 207 | G207S | A | 2.7e-07 | 9.2e-08 | 4.7e-07 | 2.773E-07 | 1.891E-07 | 9.200E-08 | 4.700E-07 |
| 4137 | 207 | G207T | A | 2.6e-07 | 2.1e-07 | 2.7e-07 | 2.467E-07 | 3.215E-08 | 2.100E-07 | 2.700E-07 |
| 4138 | 207 | G207W | A | 4.8e-07 | 9e-08 | 2.5e-07 | 2.733E-07 | 1.960E-07 | 9.000E-08 | 4.800E-07 |
| 4139 | 207 | G207Y | A | 3.2e-07 | 1.5e-07 | 1.8e-07 | 2.167E-07 | 9.074E-08 | 1.500E-07 | 3.200E-07 |
| 4140 | 207 | G207V | A | 2.2e-07 | 1.6e-07 | 1.2e-07 | 1.667E-07 | 5.033E-08 | 1.200E-07 | 2.200E-07 |
| 4141 | 208 | P208A | A | 3.1e-07 | 1.5e-07 | 3.3e-07 | 2.633E-07 | 9.866E-08 | 1.500E-07 | 3.300E-07 |
| 4142 | 208 | P208R | A | 1.2e-07 | 1.9e-07 | 1.9e-07 | 1.667E-07 | 4.041E-08 | 1.200E-07 | 1.900E-07 |
| 4143 | 208 | P208N | A | 1.7e-07 | 1.8e-07 | 3.8e-07 | 2.433E-07 | 1.185E-07 | 1.700E-07 | 3.800E-07 |
| 4144 | 208 | P208D | A | 4.5e-07 | 2.8e-07 | 2.2e-07 | 3.167E-07 | 1.193E-07 | 2.200E-07 | 4.500E-07 |
| 4145 | 208 | P208C | A | 6.7e-07 | 5.6e-07 | 2e-07 | 4.767E-07 | 2.458E-07 | 2.000E-07 | 6.700E-07 |
| 4146 | 208 | P208E | A | 2.5e-07 | 4.6e-07 | 1.7e-07 | 2.933E-07 | 1.498E-07 | 1.700E-07 | 4.600E-07 |
| 4147 | 208 | P208Q | A | 4.8e-07 | 3.8e-07 | 4.5e-07 | 4.367E-07 | 5.132E-08 | 3.800E-07 | 4.800E-07 |
| 4148 | 208 | P208G | A | 1.8e-07 | 1.7e-07 | 5.8e-07 | 3.100E-07 | 2.339E-07 | 1.700E-07 | 5.800E-07 |
| 4149 | 208 | P208H | A | 2.6e-07 | 2e-07 | 2.3e-07 | 2.300E-07 | 3.000E-08 | 2.000E-07 | 2.600E-07 |
| 4150 | 208 | P208I | A | 2e-07 | 3.4e-07 | 2.5e-07 | 2.633E-07 | 7.095E-08 | 2.000E-07 | 3.400E-07 |
| 4151 | 208 | P208L | A | 1.4e-07 | 8.2e-08 | 1.3e-07 | 1.173E-07 | 3.101E-08 | 8.200E-08 | 1.400E-07 |
| 4152 | 208 | P208K | A | 1.6e-07 | 2.7e-07 | 2.1e-07 | 2.133E-07 | 5.508E-08 | 1.600E-07 | 2.700E-07 |
| 4153 | 208 | P208M | A | 1.8e-07 | 1.6e-07 | 2.1e-07 | 1.833E-07 | 2.517E-08 | 1.600E-07 | 2.100E-07 |
| 4154 | 208 | P208F | A | 2.9e-07 | 1.8e-07 | 2.7e-07 | 2.467E-07 | 5.859E-08 | 1.800E-07 | 2.900E-07 |
| 4155 | 208 | P208P | A | 8.3e-07 | 3.3e-07 | 2.5e-07 | 4.700E-07 | 3.143E-07 | 2.500E-07 | 8.300E-07 |
| 4156 | 208 | P208S | A | 3e-07 | 2.6e-07 | 2.1e-07 | 2.567E-07 | 4.509E-08 | 2.100E-07 | 3.000E-07 |
| 4157 | 208 | P208T | A | 3.2e-07 | 2.5e-07 | 1.6e-07 | 2.433E-07 | 8.021E-08 | 1.600E-07 | 3.200E-07 |
| 4158 | 208 | P208W | A | 4.2e-07 | 2.5e-07 | 1e-07 | 2.567E-07 | 1.601E-07 | 1.000E-07 | 4.200E-07 |
| 4159 | 208 | P208Y | A | 3.3e-07 | 3.8e-07 | 4.8e-07 | 3.967E-07 | 7.638E-08 | 3.300E-07 | 4.800E-07 |
| 4160 | 208 | P208V | A | 2.4e-07 | 2.6e-07 | 1.3e-07 | 2.100E-07 | 7.000E-08 | 1.300E-07 | 2.600E-07 |
| 4161 | 209 | L209A | A | 9.9e-08 | 2.4e-07 | 2.2e-07 | 1.863E-07 | 7.629E-08 | 9.900E-08 | 2.400E-07 |
| 4162 | 209 | L209R | A | 2.2e-07 | 1.3e-07 | 2.5e-07 | 2.000E-07 | 6.245E-08 | 1.300E-07 | 2.500E-07 |
| 4163 | 209 | L209N | A | 3.9e-07 | 3.1e-07 | 2.9e-07 | 3.300E-07 | 5.292E-08 | 2.900E-07 | 3.900E-07 |
| 4164 | 209 | L209D | A | 1.7e-07 | 2.1e-07 | 2.9e-07 | 2.233E-07 | 6.110E-08 | 1.700E-07 | 2.900E-07 |

| | | | | | | | | | | |
|---|---|---|---|---|---|---|---|---|---|---|
| 4165 | 209 | L209C | A | 3.8e-07 | 4.2e-07 | 1.1e-07 | 3.033E-07 | 1.686E-07 | 1.100E-07 | 4.200E-07 |
| 4166 | 209 | L209E | A | 1.7e-07 | 1.7e-07 | 9.7e-08 | 1.457E-07 | 4.215E-08 | 9.700E-08 | 1.700E-07 |
| 4167 | 209 | L209Q | A | 3.3e-07 | 2.2e-07 | 1.9e-07 | 2.467E-07 | 7.371E-08 | 1.900E-07 | 3.300E-07 |
| 4168 | 209 | L209G | A | 2.5e-07 | 2.4e-07 | 1.7e-07 | 2.200E-07 | 4.359E-08 | 1.700E-07 | 2.500E-07 |
| 4169 | 209 | L209H | A | 5.2e-07 | 3.2e-07 | 2.2e-07 | 3.533E-07 | 1.528E-07 | 2.200E-07 | 5.200E-07 |
| 4170 | 209 | L209I | A | 1.1e-06 | 6e-08 | 4e-07 | 5.200E-07 | 5.303E-07 | 6.000E-08 | 1.100E-06 |
| 4171 | 209 | L209L | A | 2.2e-07 | 1.9e-07 | 2e-07 | 2.033E-07 | 1.528E-08 | 1.900E-07 | 2.200E-07 |
| 4172 | 209 | L209K | A | 1.8e-07 | 1.4e-07 | 1.5e-07 | 1.567E-07 | 2.082E-08 | 1.400E-07 | 1.800E-07 |
| 4173 | 209 | L209M | A | 3.7e-07 | 3.7e-07 | 2e-07 | 3.133E-07 | 9.815E-08 | 2.000E-07 | 3.700E-07 |
| 4174 | 209 | L209F | A | 2.4e-07 | 1.7e-07 | 2.9e-07 | 2.333E-07 | 6.028E-08 | 1.700E-07 | 2.900E-07 |
| 4175 | 209 | L209P | A | 2.3e-07 | 4e-07 | 2.6e-07 | 2.967E-07 | 9.074E-08 | 2.300E-07 | 4.000E-07 |
| 4176 | 209 | L209S | A | 1.8e-07 | 2.5e-07 | 1.1e-07 | 1.800E-07 | 7.000E-08 | 1.100E-07 | 2.500E-07 |
| 4177 | 209 | L209T | A | 1.8e-07 | 2e-07 | 4.3e-07 | 2.700E-07 | 1.389E-07 | 1.800E-07 | 4.300E-07 |
| 4178 | 209 | L209W | A | 3.7e-07 | 2.9e-07 | 5.2e-07 | 3.933E-07 | 1.168E-07 | 2.900E-07 | 5.200E-07 |
| 4179 | 209 | L209Y | A | 2.5e-07 | 2e-07 | 2.1e-07 | 2.200E-07 | 2.646E-08 | 2.000E-07 | 2.500E-07 |
| 4180 | 209 | L209V | A | 2.2e-07 | 3.1e-07 | 3.7e-07 | 3.000E-07 | 7.550E-08 | 2.200E-07 | 3.700E-07 |
| 4181 | 210 | P210A | A | 3.4e-07 | 2.6e-07 | 2.6e-07 | 2.867E-07 | 4.619E-08 | 2.600E-07 | 3.400E-07 |
| 4182 | 210 | P210R | A | 9.5e-08 | 1.6e-07 | 3.1e-07 | 1.883E-07 | 1.103E-07 | 9.500E-08 | 3.100E-07 |
| 4183 | 210 | P210N | A | 1.2e-07 | 2e-07 | 3.1e-07 | 2.100E-07 | 9.539E-08 | 1.200E-07 | 3.100E-07 |
| 4184 | 210 | P210D | A | 2.6e-07 | 3.5e-07 | 2.7e-07 | 2.933E-07 | 4.933E-08 | 2.600E-07 | 3.500E-07 |
| 4185 | 210 | P210C | A | 5e-07 | 2.6e-07 | 1.6e-07 | 3.067E-07 | 1.747E-07 | 1.600E-07 | 5.000E-07 |
| 4186 | 210 | P210E | A | 1.3e-07 | 3.9e-07 | 4.7e-07 | 3.300E-07 | 1.778E-07 | 1.300E-07 | 4.700E-07 |
| 4187 | 210 | P210Q | A | 3.5e-07 | 1.1e-07 | 2.1e-07 | 2.233E-07 | 1.206E-07 | 1.100E-07 | 3.500E-07 |
| 4188 | 210 | P210G | A | 2.3e-07 | 1.7e-07 | 2.8e-07 | 2.267E-07 | 5.508E-08 | 1.700E-07 | 2.800E-07 |
| 4189 | 210 | P210H | A | 1.9e-07 | 2.5e-07 | 1.8e-07 | 2.067E-07 | 3.786E-08 | 1.800E-07 | 2.500E-07 |
| 4190 | 210 | P210I | A | 3e-07 | 3.7e-07 | 2.6e-07 | 3.100E-07 | 5.568E-08 | 2.600E-07 | 3.700E-07 |
| 4191 | 210 | P210L | A | 2.3e-07 | 1.8e-07 | 3.3e-07 | 2.467E-07 | 7.638E-08 | 1.800E-07 | 3.300E-07 |
| 4192 | 210 | P210K | A | 4.2e-07 | 1.7e-07 | 3.1e-07 | 3.000E-07 | 1.253E-07 | 1.700E-07 | 4.200E-07 |
| 4193 | 210 | P210M | A | 3e-07 | 2.5e-07 | 4.6e-07 | 3.367E-07 | 1.097E-07 | 2.500E-07 | 4.600E-07 |
| 4194 | 210 | P210F | A | 2.7e-07 | 2.4e-07 | 2.1e-07 | 2.400E-07 | 3.000E-08 | 2.100E-07 | 2.700E-07 |
| 4195 | 210 | P210P | A | 1.7e-07 | 2.2e-07 | 3.5e-07 | 2.467E-07 | 9.292E-08 | 1.700E-07 | 3.500E-07 |
| 4196 | 210 | P210S | A | 2.3e-07 | 2.3e-07 | 1.9e-07 | 2.167E-07 | 2.309E-08 | 1.900E-07 | 2.300E-07 |
| 4197 | 210 | P210T | A | 1.6e-07 | 1.8e-07 | 2.6e-07 | 2.000E-07 | 5.292E-08 | 1.600E-07 | 2.600E-07 |
| 4198 | 210 | P210W | A | 8.7e-07 | 1.8e-07 | 2.5e-07 | 4.333E-07 | 3.798E-07 | 1.800E-07 | 8.700E-07 |
| 4199 | 210 | P210Y | A | 2.1e-07 | 4.1e-07 | 2.3e-07 | 2.833E-07 | 1.102E-07 | 2.100E-07 | 4.100E-07 |
| 4200 | 210 | P210V | A | 1.7e-07 | 3.6e-07 | 3.5e-07 | 2.933E-07 | 1.069E-07 | 1.700E-07 | 3.600E-07 |
| 4201 | 211 | T211A | A | 3.6e-07 | 2.9e-07 | 1.3e-07 | 2.600E-07 | 1.179E-07 | 1.300E-07 | 3.600E-07 |
| 4202 | 211 | T211R | A | 2.9e-07 | 2e-07 | 2.2e-07 | 2.367E-07 | 4.726E-08 | 2.000E-07 | 2.900E-07 |
| 4203 | 211 | T211N | A | 3.1e-07 | 3.3e-07 | 1.6e-07 | 2.667E-07 | 9.292E-08 | 1.600E-07 | 3.300E-07 |
| 4204 | 211 | T211D | A | 1.6e-07 | 4.3e-07 | 4e-07 | 3.300E-07 | 1.480E-07 | 1.600E-07 | 4.300E-07 |
| 4205 | 211 | T211C | A | 1.4e-07 | 4.5e-07 | 2.8e-07 | 2.900E-07 | 1.552E-07 | 1.400E-07 | 4.500E-07 |
| 4206 | 211 | T211E | A | 1.6e-07 | 1.5e-07 | 2e-07 | 1.700E-07 | 2.646E-08 | 1.500E-07 | 2.000E-07 |
| 4207 | 211 | T211Q | A | 2.3e-07 | 2.9e-07 | 1.6e-07 | 2.267E-07 | 6.506E-08 | 1.600E-07 | 2.900E-07 |
| 4208 | 211 | T211G | A | 3.4e-07 | 1.2e-06 | 1.2e-07 | 5.533E-07 | 5.707E-07 | 1.200E-07 | 1.200E-06 |
| 4209 | 211 | T211H | A | 2.9e-07 | 2e-07 | 1e-07 | 1.967E-07 | 9.504E-08 | 1.000E-07 | 2.900E-07 |
| 4210 | 211 | T211I | A | 3.1e-07 | 3.2e-07 | 3.2e-07 | 3.167E-07 | 5.774E-09 | 3.100E-07 | 3.200E-07 |
| 4211 | 211 | T211L | A | 2.7e-07 | 1.6e-07 | 2.8e-07 | 2.367E-07 | 6.658E-08 | 1.600E-07 | 2.800E-07 |
| 4212 | 211 | T211K | A | 1.3e-07 | 1.5e-07 | 1.4e-07 | 1.400E-07 | 1.000E-08 | 1.300E-07 | 1.500E-07 |
| 4213 | 211 | T211M | A | 6.1e-07 | 4.3e-07 | 1.6e-07 | 4.000E-07 | 2.265E-07 | 1.600E-07 | 6.100E-07 |
| 4214 | 211 | T211F | A | 3.2e-07 | 2.6e-07 | 2.3e-07 | 2.700E-07 | 4.583E-08 | 2.300E-07 | 3.200E-07 |
| 4215 | 211 | T211P | A | 1.5e-07 | 1.6e-07 | 4e-07 | 2.367E-07 | 1.415E-07 | 1.500E-07 | 4.000E-07 |
| 4216 | 211 | T211S | A | 3.8e-07 | 1.9e-07 | 2.5e-07 | 2.733E-07 | 9.713E-08 | 1.900E-07 | 3.800E-07 |
| 4217 | 211 | T211T | A | 5.7e-07 | 5.3e-07 | 6.3e-07 | 5.767E-07 | 5.033E-08 | 5.300E-07 | 6.300E-07 |
| 4218 | 211 | T211W | A | 2.7e-07 | 2.4e-07 | 3.1e-07 | 2.733E-07 | 3.512E-08 | 2.400E-07 | 3.100E-07 |
| 4219 | 211 | T211Y | A | 3.3e-07 | 6.1e-07 | 3e-07 | 4.133E-07 | 1.710E-07 | 3.000E-07 | 6.100E-07 |
| 4220 | 211 | T211V | A | 1.4e-07 | 3.2e-07 | 2.4e-07 | 2.333E-07 | 9.018E-08 | 1.400E-07 | 3.200E-07 |
| 4221 | 212 | D212A | A | 2.6e-07 | 2.8e-07 | 2.2e-07 | 2.533E-07 | 3.055E-08 | 2.200E-07 | 2.800E-07 |
| 4222 | 212 | D212R | A | 1.6e-07 | 3.5e-07 | 4.7e-07 | 3.267E-07 | 1.563E-07 | 1.600E-07 | 4.700E-07 |
| 4223 | 212 | D212N | A | 3.2e-07 | 4.7e-07 | 2.4e-07 | 3.433E-07 | 1.168E-07 | 2.400E-07 | 4.700E-07 |
| 4224 | 212 | D212D | A | 1.6e-07 | 3.1e-07 | 3.7e-07 | 2.800E-07 | 1.082E-07 | 1.600E-07 | 3.700E-07 |
| 4225 | 212 | D212C | A | 1.1e-07 | 2.2e-07 | 1.6e-07 | 1.633E-07 | 5.508E-08 | 1.100E-07 | 2.200E-07 |
| 4226 | 212 | D212E | A | 1.7e-07 | 2.8e-07 | 2.6e-07 | 2.367E-07 | 5.859E-08 | 1.700E-07 | 2.800E-07 |
| 4227 | 212 | D212Q | A | 2.8e-07 | 3.4e-07 | 2.7e-07 | 2.967E-07 | 3.786E-08 | 2.700E-07 | 3.400E-07 |
| 4228 | 212 | D212G | A | 1.8e-07 | 3.7e-07 | 2e-07 | 2.500E-07 | 1.044E-07 | 1.800E-07 | 3.700E-07 |
| 4229 | 212 | D212H | A | 2.3e-07 | 2e-07 | 3e-07 | 2.433E-07 | 5.132E-08 | 2.000E-07 | 3.000E-07 |
| 4230 | 212 | D212I | A | 1.3e-07 | 2.3e-07 | 1.3e-07 | 1.633E-07 | 5.774E-08 | 1.300E-07 | 2.300E-07 |
| 4231 | 212 | D212L | A | 1.8e-07 | 1.9e-07 | 3.7e-07 | 2.467E-07 | 1.069E-07 | 1.800E-07 | 3.700E-07 |
| 4232 | 212 | D212K | A | 3.7e-07 | 2e-07 | 1.4e-07 | 2.367E-07 | 1.193E-07 | 1.400E-07 | 3.700E-07 |
| 4233 | 212 | D212M | A | 5e-07 | 9.7e-08 | 1.1e-07 | 2.357E-07 | 2.290E-07 | 9.700E-08 | 5.000E-07 |
| 4234 | 212 | D212F | A | 1.6e-07 | 4.9e-07 | 3.6e-07 | 3.367E-07 | 1.662E-07 | 1.600E-07 | 4.900E-07 |
| 4235 | 212 | D212P | A | 2.3e-07 | 2.8e-07 | 2.7e-07 | 2.600E-07 | 2.646E-08 | 2.300E-07 | 2.800E-07 |
| 4236 | 212 | D212S | A | 1.3e-07 | 1.7e-07 | 1.7e-07 | 1.567E-07 | 2.309E-08 | 1.300E-07 | 1.700E-07 |
| 4237 | 212 | D212T | A | 2.6e-07 | 1.8e-07 | 1.7e-07 | 2.033E-07 | 4.933E-08 | 1.700E-07 | 2.600E-07 |
| 4238 | 212 | D212W | A | 3.3e-07 | 3.8e-07 | 9.9e-07 | 5.667E-07 | 3.675E-07 | 3.300E-07 | 9.900E-07 |
| 4239 | 212 | D212Y | A | 2e-07 | 3.3e-07 | 1.6e-07 | 2.300E-07 | 8.888E-08 | 1.600E-07 | 3.300E-07 |
| 4240 | 212 | D212V | A | 3.7e-07 | 4.5e-07 | 2.1e-07 | 3.433E-07 | 1.222E-07 | 2.100E-07 | 4.500E-07 |
| 4241 | 213 | C213A | A | 2.8e-07 | 3.6e-07 | 1.3e-07 | 2.567E-07 | 1.168E-07 | 1.300E-07 | 3.600E-07 |
| 4242 | 213 | C213R | A | 2.1e-07 | 2.4e-07 | 3.8e-07 | 2.767E-07 | 9.074E-08 | 2.100E-07 | 3.800E-07 |
| 4243 | 213 | C213N | A | 2.4e-07 | 2.6e-07 | 3.1e-07 | 2.700E-07 | 3.606E-08 | 2.400E-07 | 3.100E-07 |
| 4244 | 213 | C213D | A | 3.2e-07 | 2.1e-07 | 3.4e-07 | 2.900E-07 | 7.000E-08 | 2.100E-07 | 3.400E-07 |
| 4245 | 213 | C213C | A | 3.7e-07 | 4.6e-07 | 2.2e-07 | 3.500E-07 | 1.212E-07 | 2.200E-07 | 4.600E-07 |
| 4246 | 213 | C213E | A | 5.1e-07 | 3e-07 | 3.2e-07 | 3.767E-07 | 1.159E-07 | 3.000E-07 | 5.100E-07 |
| 4247 | 213 | C213Q | A | 2e-07 | 3.3e-07 | 1.2e-07 | 2.167E-07 | 1.060E-07 | 1.200E-07 | 3.300E-07 |
| 4248 | 213 | C213G | A | 3.3e-07 | 1.2e-07 | 3.2e-07 | 2.567E-07 | 1.185E-07 | 1.200E-07 | 3.300E-07 |
| 4249 | 213 | C213H | A | 2e-07 | 3.1e-07 | 2.4e-07 | 2.500E-07 | 5.568E-08 | 2.000E-07 | 3.100E-07 |

| | | | | | | | | | | |
|---|---|---|---|---|---|---|---|---|---|---|
| 4250 | 213 | C213I | A | 2.4e-07 | 2.5e-07 | 2.9e-07 | 2.600E-07 | 2.646E-08 | 2.400E-07 | 2.900E-07 |
| 4251 | 213 | C213L | A | 3.5e-07 | 3e-07 | 1.4e-07 | 2.633E-07 | 1.097E-07 | 1.400E-07 | 3.500E-07 |
| 4252 | 213 | C213K | A | 1.1e-07 | 1.6e-07 | 2.4e-07 | 1.700E-07 | 6.557E-08 | 1.100E-07 | 2.400E-07 |
| 4253 | 213 | C213M | A | 2.2e-07 | 1.9e-07 | 2.6e-07 | 2.233E-07 | 3.512E-08 | 1.900E-07 | 2.600E-07 |
| 4254 | 213 | C213F | A | 1.5e-07 | 1.9e-07 | 1.7e-07 | 1.700E-07 | 2.000E-08 | 1.500E-07 | 1.900E-07 |
| 4255 | 213 | C213P | A | 2.3e-07 | 3.4e-07 | 1.6e-07 | 2.433E-07 | 9.074E-08 | 1.600E-07 | 3.400E-07 |
| 4256 | 213 | C213S | A | 2.3e-07 | 2e-07 | 2.9e-07 | 2.400E-07 | 4.583E-08 | 2.000E-07 | 2.900E-07 |
| 4257 | 213 | C213T | A | 3.2e-07 | 2.5e-07 | 1.1e-07 | 2.267E-07 | 1.069E-07 | 1.100E-07 | 3.200E-07 |
| 4258 | 213 | C213W | A | 2e-07 | 1.1e-07 | 8.5e-08 | 1.317E-07 | 6.048E-08 | 8.500E-08 | 2.000E-07 |
| 4259 | 213 | C213Y | A | 1.4e-07 | 2.5e-07 | 1.3e-07 | 1.733E-07 | 6.658E-08 | 1.300E-07 | 2.500E-07 |
| 4260 | 213 | C213V | A | 2.2e-07 | 3.2e-07 | 1.1e-07 | 2.167E-07 | 1.050E-07 | 1.100E-07 | 3.200E-07 |
| 4261 | 214 | C214A | A | 2.8e-07 | 1.7e-07 | 3e-07 | 2.500E-07 | 7.000E-08 | 1.700E-07 | 3.000E-07 |
| 4262 | 214 | C214R | A | 3e-07 | 3.5e-07 | 1.9e-07 | 2.800E-07 | 8.185E-08 | 1.900E-07 | 3.500E-07 |
| 4263 | 214 | C214N | A | 5.1e-07 | 3.1e-07 | 4.9e-07 | 4.367E-07 | 1.102E-07 | 3.100E-07 | 5.100E-07 |
| 4264 | 214 | C214D | A | 2.9e-07 | 5.7e-07 | 4.8e-07 | 4.467E-07 | 1.429E-07 | 2.900E-07 | 5.700E-07 |
| 4265 | 214 | C214C | A | 2.3e-07 | 1.9e-07 | 4.6e-07 | 2.933E-07 | 1.457E-07 | 1.900E-07 | 4.600E-07 |
| 4266 | 214 | C214E | A | 2.3e-07 | 1.7e-07 | 5.8e-07 | 3.267E-07 | 2.214E-07 | 1.700E-07 | 5.800E-07 |
| 4267 | 214 | C214Q | A | 3.3e-07 | 2e-07 | 3.6e-07 | 2.967E-07 | 8.505E-08 | 2.000E-07 | 3.600E-07 |
| 4268 | 214 | C214G | A | 1.8e-07 | 3.7e-07 | 5.1e-07 | 3.533E-07 | 1.656E-07 | 1.800E-07 | 5.100E-07 |
| 4269 | 214 | C214H | A | 1.7e-07 | 2.5e-07 | 2.8e-07 | 2.333E-07 | 5.686E-08 | 1.700E-07 | 2.800E-07 |
| 4270 | 214 | C214I | A | 2.5e-07 | 3.5e-07 | 9.2e-07 | 5.067E-07 | 3.614E-07 | 2.500E-07 | 9.200E-07 |
| 4271 | 214 | C214L | A | 2.3e-07 | 2.7e-07 | 3e-07 | 2.667E-07 | 3.512E-08 | 2.300E-07 | 3.000E-07 |
| 4272 | 214 | C214K | A | 2.5e-07 | 4.3e-07 | 2.3e-07 | 3.033E-07 | 1.102E-07 | 2.300E-07 | 4.300E-07 |
| 4273 | 214 | C214M | A | 2.4e-07 | 2.7e-07 | 2.6e-07 | 2.567E-07 | 1.528E-08 | 2.400E-07 | 2.700E-07 |
| 4274 | 214 | C214F | A | 1.1e-07 | 1.6e-07 | 5e-07 | 2.567E-07 | 2.122E-07 | 1.100E-07 | 5.000E-07 |
| 4275 | 214 | C214P | A | 1.4e-07 | 4.8e-07 | 2e-07 | 2.733E-07 | 1.815E-07 | 1.400E-07 | 4.800E-07 |
| 4276 | 214 | C214S | A | 3.4e-07 | 1.9e-07 | 2.7e-07 | 2.667E-07 | 7.506E-08 | 1.900E-07 | 3.400E-07 |
| 4277 | 214 | C214T | A | 2e-07 | 2.2e-07 | 2.6e-07 | 2.267E-07 | 3.055E-08 | 2.000E-07 | 2.600E-07 |
| 4278 | 214 | C214W | A | 1.6e-07 | 1.7e-07 | 2.2e-07 | 1.833E-07 | 3.215E-08 | 1.600E-07 | 2.200E-07 |
| 4279 | 214 | C214Y | A | 4.1e-07 | 1.7e-07 | 2.8e-07 | 2.867E-07 | 1.201E-07 | 1.700E-07 | 4.100E-07 |
| 4280 | 214 | C214V | A | 3.1e-07 | 3.9e-07 | 2.1e-07 | 3.033E-07 | 9.018E-08 | 2.100E-07 | 3.900E-07 |
| 4281 | 215 | H215A | A | 1.3e-07 | 2.1e-07 | 3.4e-07 | 2.267E-07 | 1.060E-07 | 1.300E-07 | 3.400E-07 |
| 4282 | 215 | H215R | A | 2.2e-07 | 3.3e-07 | 2.5e-07 | 2.667E-07 | 5.686E-08 | 2.200E-07 | 3.300E-07 |
| 4283 | 215 | H215N | A | 3.6e-07 | 2.3e-07 | 3.5e-07 | 3.133E-07 | 7.234E-08 | 2.300E-07 | 3.600E-07 |
| 4284 | 215 | H215D | A | 9.6e-08 | 3.2e-07 | 3.5e-07 | 2.553E-07 | 1.388E-07 | 9.600E-08 | 3.500E-07 |
| 4285 | 215 | H215C | A | 2.7e-07 | 4.4e-07 | 1.5e-07 | 2.867E-07 | 1.457E-07 | 1.500E-07 | 4.400E-07 |
| 4286 | 215 | H215E | A | 4.1e-07 | 3.7e-07 | 4.3e-07 | 4.033E-07 | 3.055E-08 | 3.700E-07 | 4.300E-07 |
| 4287 | 215 | H215Q | A | 1.3e-07 | 1.7e-07 | 2.6e-07 | 1.867E-07 | 6.658E-08 | 1.300E-07 | 2.600E-07 |
| 4288 | 215 | H215G | A | 6.5e-07 | 3.8e-07 | 5.2e-07 | 5.167E-07 | 1.350E-07 | 3.800E-07 | 6.500E-07 |
| 4289 | 215 | H215H | A | 3.5e-07 | 3.4e-07 | 2.4e-07 | 3.100E-07 | 6.083E-08 | 2.400E-07 | 3.500E-07 |
| 4290 | 215 | H215I | A | 2e-07 | 3.6e-07 | 3e-07 | 2.867E-07 | 8.083E-08 | 2.000E-07 | 3.600E-07 |
| 4291 | 215 | H215L | A | 1.1e-07 | 1.9e-07 | 3.2e-07 | 2.067E-07 | 1.060E-07 | 1.100E-07 | 3.200E-07 |
| 4292 | 215 | H215K | A | 1.9e-07 | 2e-07 | 4.8e-07 | 2.900E-07 | 1.646E-07 | 1.900E-07 | 4.800E-07 |
| 4293 | 215 | H215M | A | 3.3e-07 | 1.9e-07 | 1.3e-07 | 2.167E-07 | 1.026E-07 | 1.300E-07 | 3.300E-07 |
| 4294 | 215 | H215F | A | 6.8e-07 | 3.6e-07 | 2.2e-07 | 4.200E-07 | 2.358E-07 | 2.200E-07 | 6.800E-07 |
| 4295 | 215 | H215P | A | 2.5e-07 | 2.8e-07 | 3.6e-07 | 2.967E-07 | 5.686E-08 | 2.500E-07 | 3.600E-07 |
| 4296 | 215 | H215S | A | 4.2e-07 | 3.4e-07 | 2.3e-07 | 3.300E-07 | 9.539E-08 | 2.300E-07 | 4.200E-07 |
| 4297 | 215 | H215T | A | 2.1e-07 | 2.9e-07 | 2.7e-07 | 2.567E-07 | 4.163E-08 | 2.100E-07 | 2.900E-07 |
| 4298 | 215 | H215W | A | 2.6e-07 | 1.3e-07 | 1.5e-07 | 1.800E-07 | 7.000E-08 | 1.300E-07 | 2.600E-07 |
| 4299 | 215 | H215Y | A | 4e-07 | 1.5e-07 | 4.6e-07 | 3.367E-07 | 1.644E-07 | 1.500E-07 | 4.600E-07 |
| 4300 | 215 | H215V | A | 2.8e-07 | 2.9e-07 | 6.3e-07 | 4.000E-07 | 1.992E-07 | 2.800E-07 | 6.300E-07 |
| 4301 | 216 | E216A | A | 4.3e-07 | 2.9e-07 | 2.1e-07 | 3.100E-07 | 1.114E-07 | 2.100E-07 | 4.300E-07 |
| 4302 | 216 | E216R | A | 2.5e-07 | 9.8e-08 | 2.9e-07 | 2.127E-07 | 1.013E-07 | 9.800E-08 | 2.900E-07 |
| 4303 | 216 | E216N | A | 1.7e-07 | 1.4e-07 | 3.8e-07 | 2.300E-07 | 1.308E-07 | 1.400E-07 | 3.800E-07 |
| 4304 | 216 | E216D | A | 2.5e-07 | 3.4e-07 | 2.5e-07 | 2.800E-07 | 5.196E-08 | 2.500E-07 | 3.400E-07 |
| 4305 | 216 | E216C | A | 1.6e-07 | 6.4e-07 | 3.3e-07 | 3.767E-07 | 2.434E-07 | 1.600E-07 | 6.400E-07 |
| 4306 | 216 | E216E | A | 2.7e-07 | 2.3e-07 | 2.7e-07 | 2.567E-07 | 2.309E-08 | 2.300E-07 | 2.700E-07 |
| 4307 | 216 | E216Q | A | 3.5e-07 | 2e-07 | 3.5e-07 | 3.000E-07 | 8.660E-08 | 2.000E-07 | 3.500E-07 |
| 4308 | 216 | E216G | A | 2.9e-07 | 1.8e-07 | 2.2e-07 | 2.300E-07 | 5.568E-08 | 1.800E-07 | 2.900E-07 |
| 4309 | 216 | E216H | A | 2.7e-07 | 4.9e-07 | 1.4e-07 | 3.000E-07 | 1.769E-07 | 1.400E-07 | 4.900E-07 |
| 4310 | 216 | E216I | A | 6.9e-07 | 4.1e-07 | 1.8e-07 | 4.267E-07 | 2.554E-07 | 1.800E-07 | 6.900E-07 |
| 4311 | 216 | E216L | A | 2.2e-07 | 2.2e-07 | 5.2e-07 | 3.200E-07 | 1.732E-07 | 2.200E-07 | 5.200E-07 |
| 4312 | 216 | E216K | A | 2.8e-07 | 3.8e-07 | 2.5e-07 | 3.033E-07 | 6.807E-08 | 2.500E-07 | 3.800E-07 |
| 4313 | 216 | E216M | A | 2.9e-07 | 5.7e-07 | 3.5e-07 | 4.033E-07 | 1.474E-07 | 2.900E-07 | 5.700E-07 |
| 4314 | 216 | E216F | A | 5.9e-07 | 2e-07 | 3.4e-07 | 3.767E-07 | 1.976E-07 | 2.000E-07 | 5.900E-07 |
| 4315 | 216 | E216P | A | 2.7e-07 | 2e-07 | 1.5e-07 | 2.067E-07 | 6.028E-08 | 1.500E-07 | 2.700E-07 |
| 4316 | 216 | E216S | A | 3.5e-07 | 2.1e-07 | 2.4e-07 | 2.667E-07 | 7.371E-08 | 2.100E-07 | 3.500E-07 |
| 4317 | 216 | E216T | A | 1.3e-07 | 3.6e-07 | 6.3e-07 | 3.733E-07 | 2.503E-07 | 1.300E-07 | 6.300E-07 |
| 4318 | 216 | E216W | A | 3.8e-07 | 2.3e-07 | 2.7e-07 | 2.933E-07 | 7.767E-08 | 2.300E-07 | 3.800E-07 |
| 4319 | 216 | E216Y | A | 2.1e-07 | 1.6e-07 | 5.6e-07 | 3.100E-07 | 2.179E-07 | 1.600E-07 | 5.600E-07 |
| 4320 | 216 | E216V | A | 3.2e-07 | 5.6e-07 | 2.4e-07 | 3.733E-07 | 1.665E-07 | 2.400E-07 | 5.600E-07 |
| 4321 | 217 | Q217A | A | 1.3e-07 | 1.2e-07 | 2.6e-07 | 1.700E-07 | 7.810E-08 | 1.200E-07 | 2.600E-07 |
| 4322 | 217 | Q217R | A | 1.3e-07 | 4.3e-07 | 6.1e-07 | 3.900E-07 | 2.425E-07 | 1.300E-07 | 6.100E-07 |
| 4323 | 217 | Q217N | A | 2.8e-07 | 3.5e-07 | 1.4e-07 | 2.567E-07 | 1.069E-07 | 1.400E-07 | 3.500E-07 |
| 4324 | 217 | Q217D | A | 1.8e-07 | 2.8e-07 | 9.8e-08 | 1.860E-07 | 9.115E-08 | 9.800E-08 | 2.800E-07 |
| 4325 | 217 | Q217C | A | 2.5e-07 | 9.5e-08 | 1.1e-07 | 1.517E-07 | 8.549E-08 | 9.500E-08 | 2.500E-07 |
| 4326 | 217 | Q217E | A | 1.9e-07 | 1.8e-07 | 1.9e-07 | 1.867E-07 | 5.774E-09 | 1.800E-07 | 1.900E-07 |
| 4327 | 217 | Q217Q | A | 2.3e-07 | 4e-07 | 3.1e-07 | 3.133E-07 | 8.505E-08 | 2.300E-07 | 4.000E-07 |
| 4328 | 217 | Q217G | A | 5.5e-07 | 2.6e-07 | 1.9e-07 | 3.333E-07 | 1.909E-07 | 1.900E-07 | 5.500E-07 |
| 4329 | 217 | Q217H | A | 2e-07 | 2.4e-07 | 4.2e-07 | 2.867E-07 | 1.172E-07 | 2.000E-07 | 4.200E-07 |
| 4330 | 217 | Q217I | A | 4.2e-07 | 1.9e-07 | 1.7e-07 | 2.600E-07 | 1.389E-07 | 1.700E-07 | 4.200E-07 |
| 4331 | 217 | Q217L | A | 1.7e-07 | 4.4e-07 | 2.6e-07 | 2.900E-07 | 1.375E-07 | 1.700E-07 | 4.400E-07 |
| 4332 | 217 | Q217K | A | 1.1e-07 | 1.2e-07 | 3.6e-07 | 1.967E-07 | 1.415E-07 | 1.100E-07 | 3.600E-07 |
| 4333 | 217 | Q217M | A | 3.8e-07 | 6.9e-07 | 2.7e-07 | 4.467E-07 | 2.178E-07 | 2.700E-07 | 6.900E-07 |
| 4334 | 217 | Q217F | A | 3.3e-07 | 4e-07 | 4.4e-07 | 3.900E-07 | 5.568E-08 | 3.300E-07 | 4.400E-07 |

| | | | | | | | | | | |
|---|---|---|---|---|---|---|---|---|---|---|
| 4335 | 217 | Q217P | A | 1.4e-07 | 2.2e-07 | 3.7e-07 | 2.433E-07 | 1.168E-07 | 1.400E-07 | 3.700E-07 |
| 4336 | 217 | Q217S | A | 2.3e-07 | 1.2e-07 | 1e-07 | 1.500E-07 | 7.000E-08 | 1.000E-07 | 2.300E-07 |
| 4337 | 217 | Q217T | A | 4.1e-07 | 2.3e-07 | 3.3e-07 | 3.233E-07 | 9.018E-08 | 2.300E-07 | 4.100E-07 |
| 4338 | 217 | Q217W | A | 2.4e-07 | 2.5e-07 | 2.3e-07 | 2.400E-07 | 1.000E-08 | 2.300E-07 | 2.500E-07 |
| 4339 | 217 | Q217Y | A | 2.1e-07 | 2.1e-07 | 1.7e-07 | 1.967E-07 | 2.309E-08 | 1.700E-07 | 2.100E-07 |
| 4340 | 217 | Q217V | A | 4.5e-07 | 1.4e-07 | 1.6e-07 | 2.500E-07 | 1.735E-07 | 1.400E-07 | 4.500E-07 |
| 4341 | 218 | C218A | A | 2.5e-07 | 2.6e-07 | 3e-07 | 2.700E-07 | 2.646E-08 | 2.500E-07 | 3.000E-07 |
| 4342 | 218 | C218R | A | 3.4e-07 | 1.5e-07 | 3.2e-07 | 2.700E-07 | 1.044E-07 | 1.500E-07 | 3.400E-07 |
| 4343 | 218 | C218N | A | 4.4e-07 | 2.6e-07 | 1.7e-07 | 2.900E-07 | 1.375E-07 | 1.700E-07 | 4.400E-07 |
| 4344 | 218 | C218D | A | 1.1e-07 | 1.9e-07 | 8.2e-08 | 1.273E-07 | 5.605E-08 | 8.200E-08 | 1.900E-07 |
| 4345 | 218 | C218C | A | 3.9e-07 | 2.7e-07 | 4.1e-07 | 3.567E-07 | 7.572E-08 | 2.700E-07 | 4.100E-07 |
| 4346 | 218 | C218E | A | 2.8e-07 | 1.7e-07 | 3e-07 | 2.500E-07 | 7.000E-08 | 1.700E-07 | 3.000E-07 |
| 4347 | 218 | C218Q | A | 2.1e-07 | 1.5e-07 | 3.8e-07 | 2.467E-07 | 1.193E-07 | 1.500E-07 | 3.800E-07 |
| 4348 | 218 | C218G | A | 2.3e-07 | 6.7e-07 | 1.9e-07 | 3.633E-07 | 2.663E-07 | 1.900E-07 | 6.700E-07 |
| 4349 | 218 | C218H | A | 2.5e-07 | 3.3e-07 | 1.2e-07 | 2.333E-07 | 1.060E-07 | 1.200E-07 | 3.300E-07 |
| 4350 | 218 | C218I | A | 3.6e-07 | 6.5e-07 | 2.9e-07 | 4.333E-07 | 1.909E-07 | 2.900E-07 | 6.500E-07 |
| 4351 | 218 | C218L | A | 1.4e-07 | 2.7e-07 | 3e-07 | 2.367E-07 | 8.505E-08 | 1.400E-07 | 3.000E-07 |
| 4352 | 218 | C218K | A | 3.8e-07 | 2.7e-07 | 3.1e-07 | 3.200E-07 | 5.568E-08 | 2.700E-07 | 3.800E-07 |
| 4353 | 218 | C218M | A | 2.2e-07 | 4e-07 | 1.6e-07 | 2.600E-07 | 1.249E-07 | 1.600E-07 | 4.000E-07 |
| 4354 | 218 | C218F | A | 2.1e-07 | 4.2e-07 | 1.9e-07 | 2.733E-07 | 1.274E-07 | 1.900E-07 | 4.200E-07 |
| 4355 | 218 | C218P | A | 2.2e-07 | 3.1e-07 | 2.4e-07 | 2.567E-07 | 4.726E-08 | 2.200E-07 | 3.100E-07 |
| 4356 | 218 | C218S | A | 3.9e-07 | 3.3e-07 | 2.7e-07 | 3.300E-07 | 6.000E-08 | 2.700E-07 | 3.900E-07 |
| 4357 | 218 | C218T | A | 1.1e-07 | 2e-07 | 3.2e-07 | 2.100E-07 | 1.054E-07 | 1.100E-07 | 3.200E-07 |
| 4358 | 218 | C218W | A | 3.1e-07 | 3.3e-07 | 5.7e-07 | 4.033E-07 | 1.447E-07 | 3.100E-07 | 5.700E-07 |
| 4359 | 218 | C218Y | A | 3.2e-07 | 2.3e-07 | 1.5e-07 | 2.333E-07 | 8.505E-08 | 1.500E-07 | 3.200E-07 |
| 4360 | 218 | C218V | A | 8.6e-08 | 3.1e-07 | 2.2e-07 | 2.053E-07 | 1.127E-07 | 8.600E-08 | 3.100E-07 |
| 4361 | 219 | A219A | A | 2.7e-07 | 3.1e-07 | 2.2e-07 | 2.667E-07 | 4.509E-08 | 2.200E-07 | 3.100E-07 |
| 4362 | 219 | A219R | A | 4.2e-07 | 4e-07 | 4.4e-07 | 4.200E-07 | 2.000E-08 | 4.000E-07 | 4.400E-07 |
| 4363 | 219 | A219N | A | 4.4e-07 | 1.6e-07 | 2.8e-07 | 2.933E-07 | 1.405E-07 | 1.600E-07 | 4.400E-07 |
| 4364 | 219 | A219D | A | 3.3e-07 | 2.8e-07 | 4.2e-07 | 3.433E-07 | 7.095E-08 | 2.800E-07 | 4.200E-07 |
| 4365 | 219 | A219C | A | 7.2e-07 | 4.6e-07 | 1.8e-07 | 4.533E-07 | 2.701E-07 | 1.800E-07 | 7.200E-07 |
| 4366 | 219 | A219E | A | 2.8e-07 | 1.7e-07 | 1.2e-07 | 1.900E-07 | 8.185E-08 | 1.200E-07 | 2.800E-07 |
| 4367 | 219 | A219Q | A | 1.3e-07 | 4.1e-07 | 2.2e-07 | 2.533E-07 | 1.429E-07 | 1.300E-07 | 4.100E-07 |
| 4368 | 219 | A219G | A | 1.9e-07 | 1.3e-07 | 3.5e-07 | 2.233E-07 | 1.137E-07 | 1.300E-07 | 3.500E-07 |
| 4369 | 219 | A219H | A | 1.8e-07 | 3.5e-07 | 3e-07 | 2.767E-07 | 8.737E-08 | 1.800E-07 | 3.500E-07 |
| 4370 | 219 | A219I | A | 4.2e-07 | 1.8e-07 | 7.1e-07 | 4.367E-07 | 2.654E-07 | 1.800E-07 | 7.100E-07 |
| 4371 | 219 | A219L | A | 1.7e-07 | 3.4e-07 | 3.5e-07 | 2.867E-07 | 1.012E-07 | 1.700E-07 | 3.500E-07 |
| 4372 | 219 | A219K | A | 2.5e-07 | 3e-07 | 4e-07 | 3.167E-07 | 7.638E-08 | 2.500E-07 | 4.000E-07 |
| 4373 | 219 | A219M | A | 2.8e-07 | 3.6e-07 | 1.8e-07 | 2.733E-07 | 9.018E-08 | 1.800E-07 | 3.600E-07 |
| 4374 | 219 | A219F | A | 2.4e-07 | 1.2e-07 | 2.2e-07 | 1.933E-07 | 6.429E-08 | 1.200E-07 | 2.400E-07 |
| 4375 | 219 | A219P | A | 1.6e-07 | 3.2e-07 | 6.1e-07 | 3.633E-07 | 2.281E-07 | 1.600E-07 | 6.100E-07 |
| 4376 | 219 | A219S | A | 4.4e-07 | 1.1e-07 | 2.5e-07 | 2.667E-07 | 1.656E-07 | 1.100E-07 | 4.400E-07 |
| 4377 | 219 | A219T | A | 4.4e-07 | 1.9e-07 | 2.9e-07 | 3.067E-07 | 1.258E-07 | 1.900E-07 | 4.400E-07 |
| 4378 | 219 | A219W | A | 2.8e-07 | 1.6e-07 | 1.5e-07 | 1.967E-07 | 7.234E-08 | 1.500E-07 | 2.800E-07 |
| 4379 | 219 | A219Y | A | 3.7e-07 | 3.2e-07 | 2.6e-07 | 3.167E-07 | 5.508E-08 | 2.600E-07 | 3.700E-07 |
| 4380 | 219 | A219V | A | 1.6e-07 | 2.3e-07 | 1.9e-07 | 1.933E-07 | 3.512E-08 | 1.600E-07 | 2.300E-07 |
| 4381 | 220 | A220A | A | 4e-07 | 1.8e-07 | 2.9e-07 | 2.900E-07 | 1.100E-07 | 1.800E-07 | 4.000E-07 |
| 4382 | 220 | A220R | A | 1.5e-07 | 2.2e-07 | 2e-07 | 1.900E-07 | 3.606E-08 | 1.500E-07 | 2.200E-07 |
| 4383 | 220 | A220N | A | 4.1e-07 | 3.6e-07 | 1.4e-07 | 3.033E-07 | 1.436E-07 | 1.400E-07 | 4.100E-07 |
| 4384 | 220 | A220D | A | 1.1e-07 | 3.4e-07 | 2.3e-07 | 2.267E-07 | 1.150E-07 | 1.100E-07 | 3.400E-07 |
| 4385 | 220 | A220C | A | 2e-07 | 1.6e-07 | 2.5e-07 | 2.033E-07 | 4.509E-08 | 1.600E-07 | 2.500E-07 |
| 4386 | 220 | A220E | A | 2.1e-07 | 2.4e-07 | 2.4e-07 | 2.300E-07 | 1.732E-08 | 2.100E-07 | 2.400E-07 |
| 4387 | 220 | A220Q | A | 1.6e-07 | 1.9e-07 | 3.4e-07 | 2.300E-07 | 9.644E-08 | 1.600E-07 | 3.400E-07 |
| 4388 | 220 | A220G | A | 3e-07 | 1.1e-07 | 1.5e-07 | 1.867E-07 | 1.002E-07 | 1.100E-07 | 3.000E-07 |
| 4389 | 220 | A220H | A | 3.4e-07 | 1.8e-07 | 1.7e-07 | 2.300E-07 | 9.539E-08 | 1.700E-07 | 3.400E-07 |
| 4390 | 220 | A220I | A | 2.3e-07 | 8.2e-07 | 2.6e-07 | 4.367E-07 | 3.323E-07 | 2.300E-07 | 8.200E-07 |
| 4391 | 220 | A220L | A | 1.8e-07 | 3e-07 | 1.4e-07 | 2.067E-07 | 8.327E-08 | 1.400E-07 | 3.000E-07 |
| 4392 | 220 | A220K | A | 2.8e-07 | 2.4e-07 | 2.1e-07 | 2.433E-07 | 3.512E-08 | 2.100E-07 | 2.800E-07 |
| 4393 | 220 | A220M | A | 2.8e-07 | 4e-07 | 4.8e-07 | 3.867E-07 | 1.007E-07 | 2.800E-07 | 4.800E-07 |
| 4394 | 220 | A220F | A | 3e-07 | 4.5e-07 | 1.8e-07 | 3.100E-07 | 1.353E-07 | 1.800E-07 | 4.500E-07 |
| 4395 | 220 | A220P | A | 2.4e-07 | 2.3e-07 | 4e-07 | 2.900E-07 | 9.539E-08 | 2.300E-07 | 4.000E-07 |
| 4396 | 220 | A220S | A | 1.9e-07 | 3.4e-07 | 7.2e-07 | 4.167E-07 | 2.732E-07 | 1.900E-07 | 7.200E-07 |
| 4397 | 220 | A220T | A | 5.2e-07 | 4e-07 | 2e-07 | 3.733E-07 | 1.617E-07 | 2.000E-07 | 5.200E-07 |
| 4398 | 220 | A220W | A | 3e-07 | 2.4e-07 | 2.2e-07 | 2.533E-07 | 4.163E-08 | 2.200E-07 | 3.000E-07 |
| 4399 | 220 | A220Y | A | 4.2e-07 | 4e-07 | 6.5e-07 | 4.900E-07 | 1.389E-07 | 4.000E-07 | 6.500E-07 |
| 4400 | 220 | A220V | A | 3.6e-07 | 1.8e-07 | 4.7e-07 | 3.367E-07 | 1.464E-07 | 1.800E-07 | 4.700E-07 |
| 4401 | 221 | G221A | A | 4.4e-07 | 1.8e-07 | 2.1e-07 | 2.767E-07 | 1.422E-07 | 1.800E-07 | 4.400E-07 |
| 4402 | 221 | G221R | A | 2.4e-07 | 2.6e-07 | 1.8e-07 | 2.267E-07 | 4.163E-08 | 1.800E-07 | 2.600E-07 |
| 4403 | 221 | G221N | A | 3e-07 | 1.9e-07 | 3.2e-07 | 2.700E-07 | 7.000E-08 | 1.900E-07 | 3.200E-07 |
| 4404 | 221 | G221D | A | 3.6e-07 | 4.3e-07 | 2.3e-07 | 3.400E-07 | 1.015E-07 | 2.300E-07 | 4.300E-07 |
| 4405 | 221 | G221C | A | 2.8e-07 | 3.8e-07 | 1.7e-07 | 2.767E-07 | 1.050E-07 | 1.700E-07 | 3.800E-07 |
| 4406 | 221 | G221E | A | 2.2e-07 | 2e-07 | 3.5e-07 | 2.567E-07 | 8.145E-08 | 2.000E-07 | 3.500E-07 |
| 4407 | 221 | G221Q | A | 3e-07 | 4.6e-07 | 4.6e-07 | 4.067E-07 | 9.238E-08 | 3.000E-07 | 4.600E-07 |
| 4408 | 221 | G221G | A | 2.7e-07 | 2.7e-07 | 1.3e-07 | 2.233E-07 | 8.083E-08 | 1.300E-07 | 2.700E-07 |
| 4409 | 221 | G221H | A | 2.9e-07 | 1.4e-07 | 2.5e-07 | 2.267E-07 | 7.767E-08 | 1.400E-07 | 2.900E-07 |
| 4410 | 221 | G221I | A | 3.6e-07 | 3.6e-07 | 3.5e-07 | 3.567E-07 | 5.774E-09 | 3.500E-07 | 3.600E-07 |
| 4411 | 221 | G221L | A | 2.1e-07 | 4.3e-07 | 2e-07 | 2.800E-07 | 1.300E-07 | 2.000E-07 | 4.300E-07 |
| 4412 | 221 | G221K | A | 2.6e-07 | 2.5e-07 | 2.3e-07 | 2.467E-07 | 1.528E-08 | 2.300E-07 | 2.600E-07 |
| 4413 | 221 | G221M | A | 2.7e-07 | 2.1e-07 | 1.7e-07 | 2.167E-07 | 5.033E-08 | 1.700E-07 | 2.700E-07 |
| 4414 | 221 | G221F | A | 2.9e-07 | 4e-07 | 2.2e-07 | 3.033E-07 | 9.074E-08 | 2.200E-07 | 4.000E-07 |
| 4415 | 221 | G221P | A | 3.3e-07 | 3.1e-07 | 2.4e-07 | 2.933E-07 | 4.726E-08 | 2.400E-07 | 3.300E-07 |
| 4416 | 221 | G221S | A | 5.3e-07 | 3.8e-07 | 3.2e-07 | 4.100E-07 | 1.082E-07 | 3.200E-07 | 5.300E-07 |
| 4417 | 221 | G221T | A | 2.5e-07 | 1.3e-07 | 1.7e-07 | 1.833E-07 | 6.110E-08 | 1.300E-07 | 2.500E-07 |
| 4418 | 221 | G221W | A | 6.6e-08 | 5.2e-07 | 1.4e-07 | 2.420E-07 | 2.436E-07 | 6.600E-08 | 5.200E-07 |
| 4419 | 221 | G221Y | A | 1.9e-07 | 4.5e-07 | 2.1e-07 | 2.833E-07 | 1.447E-07 | 1.900E-07 | 4.500E-07 |

| | | | | | | | | | | |
|---|---|---|---|---|---|---|---|---|---|---|
| 4420 | 221 | G221V | A | 4.2e-07 | 3.2e-07 | 1.8e-07 | 3.067E-07 | 1.206E-07 | 1.800E-07 | 4.200E-07 |
| 4421 | 222 | C222A | A | 1.7e-07 | 3.7e-07 | 3.1e-07 | 2.833E-07 | 1.026E-07 | 1.700E-07 | 3.700E-07 |
| 4422 | 222 | C222R | A | 2.2e-07 | 3e-07 | 5.4e-07 | 3.533E-07 | 1.665E-07 | 2.200E-07 | 5.400E-07 |
| 4423 | 222 | C222N | A | 5.2e-07 | 8.8e-08 | 3.2e-07 | 3.093E-07 | 2.162E-07 | 8.800E-08 | 5.200E-07 |
| 4424 | 222 | C222D | A | 1.2e-07 | 2.8e-07 | 1.9e-07 | 1.967E-07 | 8.021E-08 | 1.200E-07 | 2.800E-07 |
| 4425 | 222 | C222C | A | 2.5e-07 | 5.9e-08 | 1.5e-07 | 1.530E-07 | 9.554E-08 | 5.900E-08 | 2.500E-07 |
| 4426 | 222 | C222E | A | 4.2e-07 | 2.2e-07 | 1.3e-07 | 2.567E-07 | 1.484E-07 | 1.300E-07 | 4.200E-07 |
| 4427 | 222 | C222Q | A | 1.4e-07 | 3.5e-07 | 6.3e-07 | 3.733E-07 | 2.458E-07 | 1.400E-07 | 6.300E-07 |
| 4428 | 222 | C222G | A | 2.2e-07 | 4.8e-07 | 1.7e-07 | 2.900E-07 | 1.664E-07 | 1.700E-07 | 4.800E-07 |
| 4429 | 222 | C222H | A | 4.7e-07 | 2e-07 | 2.3e-07 | 3.000E-07 | 1.480E-07 | 2.000E-07 | 4.700E-07 |
| 4430 | 222 | C222I | A | 4.5e-07 | 3.3e-07 | 3e-07 | 3.600E-07 | 7.937E-08 | 3.000E-07 | 4.500E-07 |
| 4431 | 222 | C222L | A | 2.1e-07 | 1.8e-07 | 3.7e-07 | 2.533E-07 | 1.021E-07 | 1.800E-07 | 3.700E-07 |
| 4432 | 222 | C222K | A | 7.9e-08 | 5.1e-07 | 3.3e-07 | 3.063E-07 | 2.165E-07 | 7.900E-08 | 5.100E-07 |
| 4433 | 222 | C222M | A | 3.9e-07 | 2e-07 | 8.3e-08 | 2.243E-07 | 1.549E-07 | 8.300E-08 | 3.900E-07 |
| 4434 | 222 | C222F | A | 1.2e-07 | 3.2e-07 | 5.2e-07 | 3.200E-07 | 2.000E-07 | 1.200E-07 | 5.200E-07 |
| 4435 | 222 | C222P | A | 2.2e-07 | 1.5e-07 | 1.9e-07 | 1.867E-07 | 3.512E-08 | 1.500E-07 | 2.200E-07 |
| 4436 | 222 | C222S | A | 1.1e-07 | 2.7e-07 | 2.8e-07 | 2.200E-07 | 9.539E-08 | 1.100E-07 | 2.800E-07 |
| 4437 | 222 | C222T | A | 3.9e-07 | 4.2e-07 | 3.6e-07 | 3.900E-07 | 3.000E-08 | 3.600E-07 | 4.200E-07 |
| 4438 | 222 | C222W | A | 2.8e-07 | 2.7e-07 | 3.3e-07 | 2.933E-07 | 3.215E-08 | 2.700E-07 | 3.300E-07 |
| 4439 | 222 | C222Y | A | 1.3e-07 | 4e-07 | 1.4e-07 | 2.233E-07 | 1.531E-07 | 1.300E-07 | 4.000E-07 |
| 4440 | 222 | C222V | A | 1.1e-07 | 4.5e-07 | 1.6e-07 | 2.400E-07 | 1.836E-07 | 1.100E-07 | 4.500E-07 |
| 4441 | 223 | T223A | A | 1.5e-07 | 4e-07 | 3e-07 | 2.833E-07 | 1.258E-07 | 1.500E-07 | 4.000E-07 |
| 4442 | 223 | T223R | A | 1.8e-07 | 3.7e-07 | 3.2e-07 | 2.900E-07 | 9.849E-08 | 1.800E-07 | 3.700E-07 |
| 4443 | 223 | T223N | A | 2.9e-07 | 3.8e-07 | 2.2e-07 | 2.967E-07 | 8.021E-08 | 2.200E-07 | 3.800E-07 |
| 4444 | 223 | T223D | A | 4.5e-07 | 3e-07 | 3.1e-07 | 3.533E-07 | 8.386E-08 | 3.000E-07 | 4.500E-07 |
| 4445 | 223 | T223C | A | 3.3e-07 | 3.7e-07 | 1.3e-07 | 2.767E-07 | 1.286E-07 | 1.300E-07 | 3.700E-07 |
| 4446 | 223 | T223E | A | 2.8e-07 | 2e-07 | 3.5e-07 | 2.767E-07 | 7.506E-08 | 2.000E-07 | 3.500E-07 |
| 4447 | 223 | T223Q | A | 1.8e-07 | 1.3e-07 | 1.5e-07 | 1.533E-07 | 2.517E-08 | 1.300E-07 | 1.800E-07 |
| 4448 | 223 | T223G | A | 2.2e-07 | 2.7e-07 | 1.6e-07 | 2.167E-07 | 5.508E-08 | 1.600E-07 | 2.700E-07 |
| 4449 | 223 | T223H | A | 2.3e-07 | 2.2e-07 | 2.3e-07 | 2.267E-07 | 5.774E-09 | 2.200E-07 | 2.300E-07 |
| 4450 | 223 | T223I | A | 3.8e-07 | 6.3e-07 | 3.4e-07 | 4.500E-07 | 1.572E-07 | 3.400E-07 | 6.300E-07 |
| 4451 | 223 | T223L | A | 3.4e-07 | 1.7e-07 | 2.1e-07 | 2.400E-07 | 8.888E-08 | 1.700E-07 | 3.400E-07 |
| 4452 | 223 | T223K | A | 2.9e-07 | 2.1e-07 | 1.6e-07 | 2.200E-07 | 6.557E-08 | 1.600E-07 | 2.900E-07 |
| 4453 | 223 | T223M | A | 2.8e-07 | 2.8e-07 | 2e-07 | 2.533E-07 | 4.619E-08 | 2.000E-07 | 2.800E-07 |
| 4454 | 223 | T223F | A | 4.9e-07 | 4.1e-07 | 1.6e-07 | 3.533E-07 | 1.721E-07 | 1.600E-07 | 4.900E-07 |
| 4455 | 223 | T223P | A | 1.4e-07 | 4.9e-07 | 1.7e-07 | 2.667E-07 | 1.940E-07 | 1.400E-07 | 4.900E-07 |
| 4456 | 223 | T223S | A | 1.8e-07 | 1.7e-07 | 2.5e-07 | 2.000E-07 | 4.359E-08 | 1.700E-07 | 2.500E-07 |
| 4457 | 223 | T223T | A | 1.3e-07 | 1.5e-07 | 3.4e-07 | 2.067E-07 | 1.159E-07 | 1.300E-07 | 3.400E-07 |
| 4458 | 223 | T223W | A | 2.2e-07 | 3.3e-07 | 1.8e-07 | 2.433E-07 | 7.767E-08 | 1.800E-07 | 3.300E-07 |
| 4459 | 223 | T223Y | A | 2.1e-07 | 2.8e-07 | 1.9e-07 | 2.267E-07 | 4.726E-08 | 1.900E-07 | 2.800E-07 |
| 4460 | 223 | T223V | A | 3.3e-07 | 3.3e-07 | 3e-07 | 3.200E-07 | 1.732E-08 | 3.000E-07 | 3.300E-07 |
| 4461 | 224 | G224A | A | 1.8e-07 | 3e-07 | 3e-07 | 2.600E-07 | 6.928E-08 | 1.800E-07 | 3.000E-07 |
| 4462 | 224 | G224R | A | 2.3e-07 | 3.7e-07 | 1.2e-07 | 2.400E-07 | 1.253E-07 | 1.200E-07 | 3.700E-07 |
| 4463 | 224 | G224N | A | 2.9e-07 | 1.6e-07 | 2.5e-07 | 2.333E-07 | 6.658E-08 | 1.600E-07 | 2.900E-07 |
| 4464 | 224 | G224D | A | 1.8e-07 | 2.8e-07 | 3.3e-07 | 2.633E-07 | 7.638E-08 | 1.800E-07 | 3.300E-07 |
| 4465 | 224 | G224C | A | 1.4e-07 | 2e-07 | 1.5e-07 | 1.633E-07 | 3.215E-08 | 1.400E-07 | 2.000E-07 |
| 4466 | 224 | G224E | A | 3.2e-07 | 3e-07 | 3.1e-07 | 3.100E-07 | 1.000E-08 | 3.000E-07 | 3.200E-07 |
| 4467 | 224 | G224Q | A | 5.8e-07 | 1.8e-07 | 3.8e-07 | 3.800E-07 | 2.000E-07 | 1.800E-07 | 5.800E-07 |
| 4468 | 224 | G224G | A | 3.8e-07 | 1.2e-07 | 1.7e-07 | 2.233E-07 | 1.380E-07 | 1.200E-07 | 3.800E-07 |
| 4469 | 224 | G224H | A | 9.2e-08 | 1.6e-07 | 3.3e-07 | 1.940E-07 | 1.226E-07 | 9.200E-08 | 3.300E-07 |
| 4470 | 224 | G224I | A | 1.9e-07 | 3e-07 | 1.6e-07 | 2.167E-07 | 7.371E-08 | 1.600E-07 | 3.000E-07 |
| 4471 | 224 | G224L | A | 3.5e-07 | 3.4e-07 | 6.1e-07 | 4.333E-07 | 1.531E-07 | 3.400E-07 | 6.100E-07 |
| 4472 | 224 | G224K | A | 4.8e-07 | 3.2e-07 | 7.1e-07 | 5.033E-07 | 1.960E-07 | 3.200E-07 | 7.100E-07 |
| 4473 | 224 | G224M | A | 2.6e-07 | 2.4e-07 | 3.8e-07 | 2.933E-07 | 7.572E-08 | 2.400E-07 | 3.800E-07 |
| 4474 | 224 | G224F | A | 1.5e-07 | 2.7e-07 | 1.5e-07 | 1.900E-07 | 6.928E-08 | 1.500E-07 | 2.700E-07 |
| 4475 | 224 | G224P | A | 3e-07 | 2.3e-07 | 3.4e-07 | 2.900E-07 | 5.568E-08 | 2.300E-07 | 3.400E-07 |
| 4476 | 224 | G224S | A | 4.4e-07 | 1e-07 | 7e-07 | 4.133E-07 | 3.009E-07 | 1.000E-07 | 7.000E-07 |
| 4477 | 224 | G224T | A | 3e-07 | 4.6e-07 | 6.7e-07 | 4.767E-07 | 1.856E-07 | 3.000E-07 | 6.700E-07 |
| 4478 | 224 | G224W | A | 2.3e-07 | 8.2e-07 | 5.6e-07 | 5.367E-07 | 2.957E-07 | 2.300E-07 | 8.200E-07 |
| 4479 | 224 | G224Y | A | 3.1e-07 | 1.3e-07 | 1.2e-07 | 1.867E-07 | 1.069E-07 | 1.200E-07 | 3.100E-07 |
| 4480 | 224 | G224V | A | 1.8e-07 | 1.4e-07 | 4.7e-07 | 2.633E-07 | 1.801E-07 | 1.400E-07 | 4.700E-07 |
| 4481 | 225 | P225A | A | 2.1e-07 | 3.3e-07 | 3.9e-07 | 3.100E-07 | 9.165E-08 | 2.100E-07 | 3.900E-07 |
| 4482 | 225 | P225R | A | 2.6e-07 | 2e-07 | 1.9e-07 | 2.167E-07 | 3.786E-08 | 1.900E-07 | 2.600E-07 |
| 4483 | 225 | P225N | A | 4.2e-07 | 2.2e-07 | 2.6e-07 | 3.000E-07 | 1.058E-07 | 2.200E-07 | 4.200E-07 |
| 4484 | 225 | P225D | A | 1.9e-07 | 2.7e-07 | 2.8e-07 | 2.467E-07 | 4.933E-08 | 1.900E-07 | 2.800E-07 |
| 4485 | 225 | P225C | A | 3.9e-07 | 5e-07 | 2.8e-07 | 3.900E-07 | 1.100E-07 | 2.800E-07 | 5.000E-07 |
| 4486 | 225 | P225E | A | 2.2e-07 | 2.8e-07 | 2.4e-07 | 2.467E-07 | 3.055E-08 | 2.200E-07 | 2.800E-07 |
| 4487 | 225 | P225Q | A | 2.7e-07 | 1.1e-07 | 2e-07 | 1.933E-07 | 8.021E-08 | 1.100E-07 | 2.700E-07 |
| 4488 | 225 | P225G | A | 1.6e-07 | 1.9e-07 | 2.2e-07 | 1.900E-07 | 3.000E-08 | 1.600E-07 | 2.200E-07 |
| 4489 | 225 | P225H | A | 2e-07 | 2.7e-07 | 3e-07 | 2.567E-07 | 5.132E-08 | 2.000E-07 | 3.000E-07 |
| 4490 | 225 | P225I | A | 3e-07 | 2.8e-07 | 1.2e-07 | 2.333E-07 | 9.866E-08 | 1.200E-07 | 3.000E-07 |
| 4491 | 225 | P225L | A | 2.8e-07 | 1.9e-07 | 2.7e-07 | 2.467E-07 | 4.933E-08 | 1.900E-07 | 2.800E-07 |
| 4492 | 225 | P225K | A | 4.6e-07 | 2.1e-07 | 2e-07 | 2.900E-07 | 1.473E-07 | 2.000E-07 | 4.600E-07 |
| 4493 | 225 | P225M | A | 1.2e-07 | 1.1e-07 | 4e-07 | 2.100E-07 | 1.646E-07 | 1.100E-07 | 4.000E-07 |
| 4494 | 225 | P225F | A | 5.7e-07 | 1.9e-07 | 1.7e-07 | 3.100E-07 | 2.254E-07 | 1.700E-07 | 5.700E-07 |
| 4495 | 225 | P225P | A | 1.7e-07 | 1.6e-07 | 4.1e-07 | 2.467E-07 | 1.415E-07 | 1.600E-07 | 4.100E-07 |
| 4496 | 225 | P225S | A | 2.5e-07 | 4e-07 | 3.8e-07 | 3.433E-07 | 8.145E-08 | 2.500E-07 | 4.000E-07 |
| 4497 | 225 | P225T | A | 3.7e-07 | 1.8e-07 | 2.5e-07 | 2.667E-07 | 9.609E-08 | 1.800E-07 | 3.700E-07 |
| 4498 | 225 | P225W | A | 1.7e-07 | 2.7e-07 | 2.3e-07 | 2.233E-07 | 5.033E-08 | 1.700E-07 | 2.700E-07 |
| 4499 | 225 | P225Y | A | 2e-07 | 3.4e-07 | 1.3e-07 | 2.233E-07 | 1.069E-07 | 1.300E-07 | 3.400E-07 |
| 4500 | 225 | P225V | A | 3.8e-07 | 1.5e-07 | 3.6e-07 | 2.967E-07 | 1.274E-07 | 1.500E-07 | 3.800E-07 |
| 4501 | 226 | K226A | A | 5.8e-07 | 3.2e-07 | 4.1e-07 | 4.367E-07 | 1.320E-07 | 3.200E-07 | 5.800E-07 |
| 4502 | 226 | K226R | A | 2.2e-07 | 2.7e-07 | 3.5e-07 | 2.800E-07 | 6.557E-08 | 2.200E-07 | 3.500E-07 |
| 4503 | 226 | K226N | A | 2.2e-07 | 9.3e-08 | 1.7e-07 | 1.610E-07 | 6.398E-08 | 9.300E-08 | 2.200E-07 |
| 4504 | 226 | K226D | A | 3.4e-07 | 1.6e-07 | 5e-07 | 3.333E-07 | 1.701E-07 | 1.600E-07 | 5.000E-07 |

| | | | | | | | | | | |
|---|---|---|---|---|---|---|---|---|---|---|
| 4505 | 226 | K226C | A | 2e-07 | 2e-07 | 2e-07 | 2.000E-07 | 0.000E+00 | 2.000E-07 | 2.000E-07 |
| 4506 | 226 | K226E | A | 4.9e-07 | 2e-07 | 2.1e-07 | 3.000E-07 | 1.646E-07 | 2.000E-07 | 4.900E-07 |
| 4507 | 226 | K226Q | A | 9.5e-08 | 2.7e-07 | 1.7e-07 | 1.783E-07 | 8.780E-08 | 9.500E-08 | 2.700E-07 |
| 4508 | 226 | K226G | A | 7.6e-07 | 2.4e-07 | 2.1e-07 | 4.033E-07 | 3.092E-07 | 2.100E-07 | 7.600E-07 |
| 4509 | 226 | K226H | A | 1e-07 | 2.8e-07 | 6e-07 | 3.267E-07 | 2.532E-07 | 1.000E-07 | 6.000E-07 |
| 4510 | 226 | K226I | A | 3.5e-07 | 1.8e-07 | 2.2e-07 | 2.500E-07 | 8.888E-08 | 1.800E-07 | 3.500E-07 |
| 4511 | 226 | K226L | A | 2.9e-07 | 2.7e-07 | 2.2e-07 | 2.600E-07 | 3.606E-08 | 2.200E-07 | 2.900E-07 |
| 4512 | 226 | K226K | A | 2e-07 | 1.8e-07 | 2.8e-07 | 2.200E-07 | 5.292E-08 | 1.800E-07 | 2.800E-07 |
| 4513 | 226 | K226M | A | 1.9e-07 | 2.2e-07 | 2e-07 | 2.033E-07 | 1.528E-08 | 1.900E-07 | 2.200E-07 |
| 4514 | 226 | K226F | A | 1.6e-07 | 4.3e-07 | 2e-07 | 2.633E-07 | 1.457E-07 | 1.600E-07 | 4.300E-07 |
| 4515 | 226 | K226P | A | 2e-07 | 8.7e-08 | 2.5e-07 | 1.790E-07 | 8.350E-08 | 8.700E-08 | 2.500E-07 |
| 4516 | 226 | K226S | A | 4.1e-07 | 3e-07 | 1.4e-07 | 2.833E-07 | 1.358E-07 | 1.400E-07 | 4.100E-07 |
| 4517 | 226 | K226T | A | 3.1e-07 | 3.2e-07 | 2.9e-07 | 3.067E-07 | 1.528E-08 | 2.900E-07 | 3.200E-07 |
| 4518 | 226 | K226W | A | 2.8e-07 | 3.6e-07 | 2.7e-07 | 3.033E-07 | 4.933E-08 | 2.700E-07 | 3.600E-07 |
| 4519 | 226 | K226Y | A | 2.9e-07 | 2.1e-07 | 2.1e-07 | 2.367E-07 | 4.619E-08 | 2.100E-07 | 2.900E-07 |
| 4520 | 226 | K226V | A | 3e-07 | 1.4e-07 | 1.7e-07 | 2.033E-07 | 8.505E-08 | 1.400E-07 | 3.000E-07 |
| 4521 | 227 | H227A | A | 3.3e-07 | 2.3e-07 | 2.4e-07 | 2.667E-07 | 5.508E-08 | 2.300E-07 | 3.300E-07 |
| 4522 | 227 | H227R | A | 2.7e-07 | 2.1e-07 | 7.1e-07 | 3.967E-07 | 2.730E-07 | 2.100E-07 | 7.100E-07 |
| 4523 | 227 | H227N | A | 2.3e-07 | 3.1e-07 | 2.7e-07 | 2.700E-07 | 4.000E-08 | 2.300E-07 | 3.100E-07 |
| 4524 | 227 | H227D | A | 2e-07 | 1.7e-07 | 1.6e-07 | 1.767E-07 | 2.082E-08 | 1.600E-07 | 2.000E-07 |
| 4525 | 227 | H227C | A | 4.9e-07 | 1.9e-07 | 4.6e-07 | 3.800E-07 | 1.652E-07 | 1.900E-07 | 4.900E-07 |
| 4526 | 227 | H227E | A | 2.3e-07 | 1.6e-07 | 4.3e-07 | 2.733E-07 | 1.401E-07 | 1.600E-07 | 4.300E-07 |
| 4527 | 227 | H227Q | A | 1.7e-07 | 6.6e-07 | 1.4e-07 | 3.233E-07 | 2.919E-07 | 1.400E-07 | 6.600E-07 |
| 4528 | 227 | H227G | A | 1.4e-07 | 2.6e-07 | 3.5e-07 | 2.500E-07 | 1.054E-07 | 1.400E-07 | 3.500E-07 |
| 4529 | 227 | H227H | A | 2.7e-07 | 1.9e-07 | 1.5e-07 | 2.033E-07 | 6.110E-08 | 1.500E-07 | 2.700E-07 |
| 4530 | 227 | H227I | A | 4.8e-07 | 2.8e-07 | 4.2e-07 | 3.933E-07 | 1.026E-07 | 2.800E-07 | 4.800E-07 |
| 4531 | 227 | H227L | A | 1.3e-07 | 7e-08 | 3.4e-07 | 1.800E-07 | 1.418E-07 | 7.000E-08 | 3.400E-07 |
| 4532 | 227 | H227K | A | 1.9e-07 | 3.4e-07 | 1.6e-07 | 2.300E-07 | 9.644E-08 | 1.600E-07 | 3.400E-07 |
| 4533 | 227 | H227M | A | 1.2e-07 | 4.7e-07 | 3e-07 | 2.967E-07 | 1.750E-07 | 1.200E-07 | 4.700E-07 |
| 4534 | 227 | H227F | A | 1.4e-07 | 2.9e-07 | 2.5e-07 | 2.267E-07 | 7.767E-08 | 1.400E-07 | 2.900E-07 |
| 4535 | 227 | H227P | A | 4.6e-07 | 1.8e-07 | 2.4e-07 | 2.933E-07 | 1.474E-07 | 1.800E-07 | 4.600E-07 |
| 4536 | 227 | H227S | A | 1.4e-07 | 6.1e-07 | 3.2e-07 | 3.567E-07 | 2.371E-07 | 1.400E-07 | 6.100E-07 |
| 4537 | 227 | H227T | A | 2.2e-07 | 1.9e-07 | 2.2e-07 | 2.100E-07 | 1.732E-08 | 1.900E-07 | 2.200E-07 |
| 4538 | 227 | H227W | A | 2.6e-07 | 4.4e-07 | 5.2e-07 | 4.067E-07 | 1.332E-07 | 2.600E-07 | 5.200E-07 |
| 4539 | 227 | H227Y | A | 2.7e-07 | 3.6e-07 | 4.7e-07 | 3.667E-07 | 1.002E-07 | 2.700E-07 | 4.700E-07 |
| 4540 | 227 | H227V | A | 2.8e-07 | 4.1e-07 | 2.1e-07 | 3.000E-07 | 1.015E-07 | 2.100E-07 | 4.100E-07 |
| 4541 | 228 | S228A | A | 2.4e-07 | 8.4e-08 | 1.6e-07 | 1.613E-07 | 7.801E-08 | 8.400E-08 | 2.400E-07 |
| 4542 | 228 | S228R | A | 2.4e-07 | 2.8e-07 | 2.4e-07 | 2.533E-07 | 2.309E-08 | 2.400E-07 | 2.800E-07 |
| 4543 | 228 | S228N | A | 4.7e-07 | 4.8e-07 | 3.7e-07 | 4.400E-07 | 6.083E-08 | 3.700E-07 | 4.800E-07 |
| 4544 | 228 | S228D | A | 4e-07 | 2.9e-07 | 1.6e-07 | 2.833E-07 | 1.201E-07 | 1.600E-07 | 4.000E-07 |
| 4545 | 228 | S228C | A | 1.3e-06 | 4.4e-07 | 2.3e-07 | 6.567E-07 | 5.670E-07 | 2.300E-07 | 1.300E-06 |
| 4546 | 228 | S228E | A | 9.8e-08 | 1.8e-07 | 4.6e-07 | 2.460E-07 | 1.898E-07 | 9.800E-08 | 4.600E-07 |
| 4547 | 228 | S228Q | A | 3.3e-07 | 1.5e-07 | 2e-07 | 2.267E-07 | 9.292E-08 | 1.500E-07 | 3.300E-07 |
| 4548 | 228 | S228G | A | 3.9e-07 | 5e-07 | 1.7e-07 | 3.533E-07 | 1.680E-07 | 1.700E-07 | 5.000E-07 |
| 4549 | 228 | S228H | A | 1.5e-07 | 2.1e-07 | 2.7e-07 | 2.100E-07 | 6.000E-08 | 1.500E-07 | 2.700E-07 |
| 4550 | 228 | S228I | A | 1.9e-07 | 3.8e-07 | 2.7e-07 | 2.800E-07 | 9.539E-08 | 1.900E-07 | 3.800E-07 |
| 4551 | 228 | S228L | A | 2.7e-07 | 3.2e-07 | 1.1e-07 | 2.333E-07 | 1.097E-07 | 1.100E-07 | 3.200E-07 |
| 4552 | 228 | S228K | A | 2.1e-07 | 2.4e-07 | 2.8e-07 | 2.433E-07 | 3.512E-08 | 2.100E-07 | 2.800E-07 |
| 4553 | 228 | S228M | A | 1.9e-07 | 1.4e-07 | 1.4e-07 | 1.567E-07 | 2.887E-08 | 1.400E-07 | 1.900E-07 |
| 4554 | 228 | S228F | A | 2e-07 | 1.1e-07 | 2.9e-07 | 2.000E-07 | 9.000E-08 | 1.100E-07 | 2.900E-07 |
| 4555 | 228 | S228P | A | 2.1e-07 | 2.1e-07 | 1.8e-07 | 2.000E-07 | 1.732E-08 | 1.800E-07 | 2.100E-07 |
| 4556 | 228 | S228S | A | 3.3e-07 | 2.1e-07 | 2.7e-07 | 2.700E-07 | 6.000E-08 | 2.100E-07 | 3.300E-07 |
| 4557 | 228 | S228T | A | 1.6e-07 | 2.1e-07 | 2.2e-07 | 1.967E-07 | 3.215E-08 | 1.600E-07 | 2.200E-07 |
| 4558 | 228 | S228W | A | 2.5e-07 | 1.8e-07 | 3.3e-07 | 2.533E-07 | 7.506E-08 | 1.800E-07 | 3.300E-07 |
| 4559 | 228 | S228Y | A | 4.8e-07 | 2.4e-07 | 1.8e-07 | 3.000E-07 | 1.587E-07 | 1.800E-07 | 4.800E-07 |
| 4560 | 228 | S228V | A | 3.6e-07 | 2.6e-07 | 4.2e-07 | 3.467E-07 | 8.083E-08 | 2.600E-07 | 4.200E-07 |
| 4561 | 229 | D229A | A | 3.1e-07 | 5.2e-07 | 5.2e-07 | 4.500E-07 | 1.212E-07 | 3.100E-07 | 5.200E-07 |
| 4562 | 229 | D229R | A | 3.8e-07 | 1.4e-07 | 1.1e-07 | 2.100E-07 | 1.480E-07 | 1.100E-07 | 3.800E-07 |
| 4563 | 229 | D229N | A | 5.7e-07 | 2.2e-07 | 3.5e-07 | 3.800E-07 | 1.769E-07 | 2.200E-07 | 5.700E-07 |
| 4564 | 229 | D229D | A | 2.8e-07 | 3.4e-07 | 2.3e-07 | 2.833E-07 | 5.508E-08 | 2.300E-07 | 3.400E-07 |
| 4565 | 229 | D229C | A | 5.6e-07 | 1.4e-07 | 1.9e-07 | 2.967E-07 | 2.294E-07 | 1.400E-07 | 5.600E-07 |
| 4566 | 229 | D229E | A | 2.4e-07 | 3.8e-07 | 4.9e-07 | 3.700E-07 | 1.253E-07 | 2.400E-07 | 4.900E-07 |
| 4567 | 229 | D229Q | A | 1.5e-07 | 2.6e-07 | 2.1e-07 | 2.067E-07 | 5.508E-08 | 1.500E-07 | 2.600E-07 |
| 4568 | 229 | D229G | A | 3.1e-07 | 3e-07 | 2.8e-07 | 2.967E-07 | 1.528E-08 | 2.800E-07 | 3.100E-07 |
| 4569 | 229 | D229H | A | 1.3e-07 | 1.1e-07 | 2.4e-07 | 1.600E-07 | 7.000E-08 | 1.100E-07 | 2.400E-07 |
| 4570 | 229 | D229I | A | 2.6e-07 | 2.1e-07 | 3e-07 | 2.567E-07 | 4.509E-08 | 2.100E-07 | 3.000E-07 |
| 4571 | 229 | D229L | A | 4.6e-07 | 3e-07 | 2.7e-07 | 3.433E-07 | 1.021E-07 | 2.700E-07 | 4.600E-07 |
| 4572 | 229 | D229K | A | 3.3e-07 | 3.4e-07 | 9.9e-08 | 2.563E-07 | 1.363E-07 | 9.900E-08 | 3.400E-07 |
| 4573 | 229 | D229M | A | 1.5e-07 | 2e-07 | 1.3e-07 | 1.600E-07 | 3.606E-08 | 1.300E-07 | 2.000E-07 |
| 4574 | 229 | D229F | A | 2.3e-07 | 1.9e-07 | 1.5e-07 | 1.900E-07 | 4.000E-08 | 1.500E-07 | 2.300E-07 |
| 4575 | 229 | D229P | A | 5.6e-07 | 3.3e-07 | 3e-07 | 3.967E-07 | 1.422E-07 | 3.000E-07 | 5.600E-07 |
| 4576 | 229 | D229S | A | 3e-07 | 2.3e-07 | 4.3e-07 | 3.200E-07 | 1.015E-07 | 2.300E-07 | 4.300E-07 |
| 4577 | 229 | D229T | A | 2.8e-07 | 1.8e-07 | 3.8e-07 | 2.800E-07 | 1.000E-07 | 1.800E-07 | 3.800E-07 |
| 4578 | 229 | D229W | A | 3.1e-07 | 2.9e-07 | 1.5e-07 | 2.500E-07 | 8.718E-08 | 1.500E-07 | 3.100E-07 |
| 4579 | 229 | D229Y | A | 1.5e-07 | 3.2e-07 | 2.5e-07 | 2.400E-07 | 8.544E-08 | 1.500E-07 | 3.200E-07 |
| 4580 | 229 | D229V | A | 3.3e-07 | 3e-07 | 1.6e-07 | 2.633E-07 | 9.074E-08 | 1.600E-07 | 3.300E-07 |
| 4581 | 230 | C230A | A | 1.1e-07 | 1.4e-07 | 2.6e-07 | 1.700E-07 | 7.937E-08 | 1.100E-07 | 2.600E-07 |
| 4582 | 230 | C230R | A | 4.8e-07 | 1.4e-07 | 5.1e-07 | 3.767E-07 | 2.055E-07 | 1.400E-07 | 5.100E-07 |
| 4583 | 230 | C230N | A | 1.3e-07 | 1.4e-07 | 3.5e-07 | 2.067E-07 | 1.242E-07 | 1.300E-07 | 3.500E-07 |
| 4584 | 230 | C230D | A | 2.2e-07 | 3.8e-07 | 2.4e-07 | 2.800E-07 | 8.718E-08 | 2.200E-07 | 3.800E-07 |
| 4585 | 230 | C230C | A | 3.4e-07 | 2.3e-07 | 1.8e-07 | 2.500E-07 | 8.185E-08 | 1.800E-07 | 3.400E-07 |
| 4586 | 230 | C230E | A | 4.3e-07 | 1.8e-07 | 3.7e-07 | 3.267E-07 | 1.305E-07 | 1.800E-07 | 4.300E-07 |
| 4587 | 230 | C230Q | A | 2.3e-07 | 3.9e-07 | 3.6e-07 | 3.267E-07 | 8.505E-08 | 2.300E-07 | 3.900E-07 |
| 4588 | 230 | C230G | A | 1.9e-07 | 4.9e-07 | 1.2e-07 | 2.667E-07 | 1.966E-07 | 1.200E-07 | 4.900E-07 |
| 4589 | 230 | C230H | A | 1.3e-07 | 1.5e-07 | 1.3e-07 | 1.367E-07 | 1.155E-08 | 1.300E-07 | 1.500E-07 |

| | | | | | | | | | | |
|---|---|---|---|---|---|---|---|---|---|---|
| 4590 | 230 | C230I | A | 2.6e-07 | 2.6e-07 | 2.4e-07 | 2.533E-07 | 1.155E-08 | 2.400E-07 | 2.600E-07 |
| 4591 | 230 | C230L | A | 3.7e-07 | 4.1e-07 | 3.6e-07 | 3.800E-07 | 2.646E-08 | 3.600E-07 | 4.100E-07 |
| 4592 | 230 | C230K | A | 2.9e-07 | 5.1e-07 | 2.3e-07 | 3.433E-07 | 1.474E-07 | 2.300E-07 | 5.100E-07 |
| 4593 | 230 | C230M | A | 1.5e-07 | 3.2e-07 | 2e-07 | 2.233E-07 | 8.737E-08 | 1.500E-07 | 3.200E-07 |
| 4594 | 230 | C230F | A | 5.7e-07 | 3.2e-07 | 2.6e-07 | 3.833E-07 | 1.644E-07 | 2.600E-07 | 5.700E-07 |
| 4595 | 230 | C230P | A | 4e-07 | 3.3e-07 | 2.6e-07 | 3.300E-07 | 7.000E-08 | 2.600E-07 | 4.000E-07 |
| 4596 | 230 | C230S | A | 2.6e-07 | 2.2e-07 | 2.7e-07 | 2.500E-07 | 2.646E-08 | 2.200E-07 | 2.700E-07 |
| 4597 | 230 | C230T | A | 3.8e-07 | 2.4e-07 | 1.8e-07 | 2.667E-07 | 1.026E-07 | 1.800E-07 | 3.800E-07 |
| 4598 | 230 | C230W | A | 3.7e-07 | 4.7e-07 | 2.3e-07 | 3.567E-07 | 1.206E-07 | 2.300E-07 | 4.700E-07 |
| 4599 | 230 | C230Y | A | 2.7e-07 | 1.4e-07 | 2e-07 | 2.033E-07 | 6.506E-08 | 1.400E-07 | 2.700E-07 |
| 4600 | 230 | C230V | A | 3.6e-07 | 2.4e-07 | 1.1e-07 | 2.367E-07 | 1.250E-07 | 1.100E-07 | 3.600E-07 |
| 4601 | 231 | L231A | A | 2.2e-07 | 1.2e-07 | 1.6e-07 | 1.667E-07 | 5.033E-08 | 1.200E-07 | 2.200E-07 |
| 4602 | 231 | L231R | A | 2.5e-07 | 2.5e-07 | 3.1e-07 | 2.700E-07 | 3.464E-08 | 2.500E-07 | 3.100E-07 |
| 4603 | 231 | L231N | A | 2.1e-07 | 1.9e-07 | 1.8e-07 | 1.933E-07 | 1.528E-08 | 1.800E-07 | 2.100E-07 |
| 4604 | 231 | L231D | A | 3.5e-07 | 2e-07 | 1.7e-07 | 2.400E-07 | 9.644E-08 | 1.700E-07 | 3.500E-07 |
| 4605 | 231 | L231C | A | 2.5e-07 | 2e-07 | 2.8e-07 | 2.433E-07 | 4.041E-08 | 2.000E-07 | 2.800E-07 |
| 4606 | 231 | L231E | A | 2.4e-07 | 2.1e-07 | 2.8e-07 | 2.433E-07 | 3.512E-08 | 2.100E-07 | 2.800E-07 |
| 4607 | 231 | L231Q | A | 2.4e-07 | 5.4e-07 | 1.8e-07 | 3.200E-07 | 1.929E-07 | 1.800E-07 | 5.400E-07 |
| 4608 | 231 | L231G | A | 1.4e-07 | 2.8e-07 | 2e-07 | 2.067E-07 | 7.024E-08 | 1.400E-07 | 2.800E-07 |
| 4609 | 231 | L231H | A | 2.7e-07 | 4.1e-07 | 3.2e-07 | 3.333E-07 | 7.095E-08 | 2.700E-07 | 4.100E-07 |
| 4610 | 231 | L231I | A | 2.6e-07 | 2.5e-07 | 2.2e-07 | 2.433E-07 | 2.082E-08 | 2.200E-07 | 2.600E-07 |
| 4611 | 231 | L231L | A | 3.8e-07 | 3.1e-07 | 4.2e-07 | 3.700E-07 | 5.568E-08 | 3.100E-07 | 4.200E-07 |
| 4612 | 231 | L231K | A | 3e-07 | 1.3e-07 | 1.7e-07 | 2.000E-07 | 8.888E-08 | 1.300E-07 | 3.000E-07 |
| 4613 | 231 | L231M | A | 1.6e-07 | 4e-07 | 3.3e-07 | 2.967E-07 | 1.234E-07 | 1.600E-07 | 4.000E-07 |
| 4614 | 231 | L231F | A | 3.6e-07 | 3e-07 | 2.6e-07 | 3.067E-07 | 5.033E-08 | 2.600E-07 | 3.600E-07 |
| 4615 | 231 | L231P | A | 2.3e-07 | 1.1e-07 | 3.2e-07 | 2.200E-07 | 1.054E-07 | 1.100E-07 | 3.200E-07 |
| 4616 | 231 | L231S | A | 3.4e-07 | 3.7e-07 | 2.4e-07 | 3.167E-07 | 6.807E-08 | 2.400E-07 | 3.700E-07 |
| 4617 | 231 | L231T | A | 2e-07 | 1e-07 | 2.9e-07 | 1.967E-07 | 9.504E-08 | 1.000E-07 | 2.900E-07 |
| 4618 | 231 | L231W | A | 1.7e-07 | 1.4e-07 | 5.4e-07 | 2.833E-07 | 2.228E-07 | 1.400E-07 | 5.400E-07 |
| 4619 | 231 | L231Y | A | 1.6e-07 | 3.9e-07 | 2.4e-07 | 2.633E-07 | 1.168E-07 | 1.600E-07 | 3.900E-07 |
| 4620 | 231 | L231V | A | 2.3e-07 | 4.8e-07 | 1.9e-07 | 3.000E-07 | 1.572E-07 | 1.900E-07 | 4.800E-07 |
| 4621 | 232 | A232A | A | 1.8e-07 | 3.3e-07 | 1.7e-07 | 2.267E-07 | 8.963E-08 | 1.700E-07 | 3.300E-07 |
| 4622 | 232 | A232R | A | 2.4e-07 | 3.9e-07 | 3.5e-07 | 3.267E-07 | 7.767E-08 | 2.400E-07 | 3.900E-07 |
| 4623 | 232 | A232N | A | 1.8e-07 | 3.8e-07 | 2.9e-07 | 2.833E-07 | 1.002E-07 | 1.800E-07 | 3.800E-07 |
| 4624 | 232 | A232D | A | 3.6e-07 | 1.3e-07 | 2.2e-07 | 2.367E-07 | 1.159E-07 | 1.300E-07 | 3.600E-07 |
| 4625 | 232 | A232C | A | 5.8e-07 | 2.3e-07 | 2.3e-07 | 3.467E-07 | 2.021E-07 | 2.300E-07 | 5.800E-07 |
| 4626 | 232 | A232E | A | 2.7e-07 | 1.8e-07 | 2.5e-07 | 2.333E-07 | 4.726E-08 | 1.800E-07 | 2.700E-07 |
| 4627 | 232 | A232Q | A | 2.5e-07 | 5.1e-07 | 1.7e-07 | 3.100E-07 | 1.778E-07 | 1.700E-07 | 5.100E-07 |
| 4628 | 232 | A232G | A | 2.5e-07 | 2.8e-07 | 1.3e-07 | 2.200E-07 | 7.937E-08 | 1.300E-07 | 2.800E-07 |
| 4629 | 232 | A232H | A | 4.8e-07 | 1.4e-07 | 3.3e-07 | 3.167E-07 | 1.704E-07 | 1.400E-07 | 4.800E-07 |
| 4630 | 232 | A232I | A | 5.7e-07 | 3.9e-07 | 2.2e-07 | 3.933E-07 | 1.750E-07 | 2.200E-07 | 5.700E-07 |
| 4631 | 232 | A232L | A | 2.3e-07 | 4.1e-07 | 1.8e-07 | 2.733E-07 | 1.210E-07 | 1.800E-07 | 4.100E-07 |
| 4632 | 232 | A232K | A | 2e-07 | 3.2e-07 | 5.3e-07 | 3.500E-07 | 1.670E-07 | 2.000E-07 | 5.300E-07 |
| 4633 | 232 | A232M | A | 2.9e-07 | 2.7e-07 | 2.8e-07 | 2.800E-07 | 1.000E-08 | 2.700E-07 | 2.900E-07 |
| 4634 | 232 | A232F | A | 1.6e-07 | 1.5e-07 | 3.6e-07 | 2.233E-07 | 1.185E-07 | 1.500E-07 | 3.600E-07 |
| 4635 | 232 | A232P | A | 6.7e-07 | 2.5e-07 | 2.8e-07 | 4.000E-07 | 2.343E-07 | 2.500E-07 | 6.700E-07 |
| 4636 | 232 | A232S | A | 6.8e-07 | 1.8e-07 | 2e-07 | 3.533E-07 | 2.831E-07 | 1.800E-07 | 6.800E-07 |
| 4637 | 232 | A232T | A | 2e-07 | 3.2e-07 | 4.5e-07 | 3.233E-07 | 1.250E-07 | 2.000E-07 | 4.500E-07 |
| 4638 | 232 | A232W | A | 7.3e-07 | 3.2e-07 | 2.9e-07 | 4.467E-07 | 2.458E-07 | 2.900E-07 | 7.300E-07 |
| 4639 | 232 | A232Y | A | 3.8e-07 | 5.2e-07 | 1.9e-07 | 3.633E-07 | 1.656E-07 | 1.900E-07 | 5.200E-07 |
| 4640 | 232 | A232V | A | 1.7e-07 | 7.4e-08 | 2.2e-07 | 1.547E-07 | 7.420E-08 | 7.400E-08 | 2.200E-07 |
| 4641 | 233 | C233A | A | 2.7e-07 | 1.7e-07 | 2.6e-07 | 2.333E-07 | 5.508E-08 | 1.700E-07 | 2.700E-07 |
| 4642 | 233 | C233R | A | 1.2e-07 | 1.5e-07 | 2.6e-07 | 1.767E-07 | 7.371E-08 | 1.200E-07 | 2.600E-07 |
| 4643 | 233 | C233N | A | 2.3e-07 | 1.8e-07 | 1.9e-07 | 2.000E-07 | 2.646E-08 | 1.800E-07 | 2.300E-07 |
| 4644 | 233 | C233D | A | 3.5e-07 | 1.4e-07 | 2.7e-07 | 2.533E-07 | 1.060E-07 | 1.400E-07 | 3.500E-07 |
| 4645 | 233 | C233C | A | 2.3e-07 | 2e-07 | 3.7e-07 | 2.667E-07 | 9.074E-08 | 2.000E-07 | 3.700E-07 |
| 4646 | 233 | C233E | A | 4.8e-07 | 3e-07 | 3.9e-07 | 3.900E-07 | 9.000E-08 | 3.000E-07 | 4.800E-07 |
| 4647 | 233 | C233Q | A | 3.2e-07 | 2.8e-07 | 2.4e-07 | 2.800E-07 | 4.000E-08 | 2.400E-07 | 3.200E-07 |
| 4648 | 233 | C233G | A | 3.2e-07 | 1.7e-07 | 5.5e-07 | 3.467E-07 | 1.914E-07 | 1.700E-07 | 5.500E-07 |
| 4649 | 233 | C233H | A | 2.3e-07 | 2.4e-07 | 2.3e-07 | 2.333E-07 | 5.774E-09 | 2.300E-07 | 2.400E-07 |
| 4650 | 233 | C233I | A | 2.6e-07 | 2.3e-07 | 6.1e-07 | 3.667E-07 | 2.113E-07 | 2.300E-07 | 6.100E-07 |
| 4651 | 233 | C233L | A | 1.4e-07 | 3.4e-07 | 2.6e-07 | 2.467E-07 | 1.007E-07 | 1.400E-07 | 3.400E-07 |
| 4652 | 233 | C233K | A | 1.1e-06 | 1.9e-07 | 2.6e-07 | 5.167E-07 | 5.064E-07 | 1.900E-07 | 1.100E-06 |
| 4653 | 233 | C233M | A | 4e-07 | 2.6e-07 | 2.2e-07 | 2.933E-07 | 9.452E-08 | 2.200E-07 | 4.000E-07 |
| 4654 | 233 | C233F | A | 1.4e-07 | 3.7e-07 | 5.5e-07 | 3.533E-07 | 2.055E-07 | 1.400E-07 | 5.500E-07 |
| 4655 | 233 | C233P | A | 3.9e-07 | 6.3e-07 | 2.1e-07 | 4.100E-07 | 2.107E-07 | 2.100E-07 | 6.300E-07 |
| 4656 | 233 | C233S | A | 2.6e-07 | 2.9e-07 | 2.5e-07 | 2.667E-07 | 2.082E-08 | 2.500E-07 | 2.900E-07 |
| 4657 | 233 | C233T | A | 4.9e-07 | 2.4e-07 | 6.5e-07 | 4.600E-07 | 2.066E-07 | 2.400E-07 | 6.500E-07 |
| 4658 | 233 | C233W | A | 4e-07 | 3e-07 | 4.3e-07 | 3.767E-07 | 6.807E-08 | 3.000E-07 | 4.300E-07 |
| 4659 | 233 | C233Y | A | 7e-07 | 2.7e-07 | 1.4e-07 | 3.700E-07 | 2.931E-07 | 1.400E-07 | 7.000E-07 |
| 4660 | 233 | C233V | A | 3e-07 | 2.4e-07 | 3.7e-07 | 3.033E-07 | 6.506E-08 | 2.400E-07 | 3.700E-07 |
| 4661 | 234 | L234A | A | 2.9e-07 | 1.1e-07 | 3.3e-07 | 2.433E-07 | 1.172E-07 | 1.100E-07 | 3.300E-07 |
| 4662 | 234 | L234R | A | 2.7e-07 | 1.4e-07 | 2.3e-07 | 2.133E-07 | 6.658E-08 | 1.400E-07 | 2.700E-07 |
| 4663 | 234 | L234N | A | 3e-07 | 6.1e-07 | 2.7e-07 | 3.933E-07 | 1.882E-07 | 2.700E-07 | 6.100E-07 |
| 4664 | 234 | L234D | A | 1.1e-06 | 5e-07 | 2.7e-07 | 6.233E-07 | 4.285E-07 | 2.700E-07 | 1.100E-06 |
| 4665 | 234 | L234C | A | 3.1e-07 | 4.3e-07 | 1.6e-07 | 3.000E-07 | 1.353E-07 | 1.600E-07 | 4.300E-07 |
| 4666 | 234 | L234E | A | 2.2e-07 | 2.6e-07 | 2.7e-07 | 2.500E-07 | 2.646E-08 | 2.200E-07 | 2.700E-07 |
| 4667 | 234 | L234Q | A | 3.5e-07 | 3e-07 | 2.9e-07 | 3.133E-07 | 3.215E-08 | 2.900E-07 | 3.500E-07 |
| 4668 | 234 | L234G | A | 2.4e-07 | 4.8e-07 | 9e-08 | 2.700E-07 | 1.967E-07 | 9.000E-08 | 4.800E-07 |
| 4669 | 234 | L234H | A | 3.4e-07 | 3e-07 | 3.1e-07 | 3.167E-07 | 2.082E-08 | 3.000E-07 | 3.400E-07 |
| 4670 | 234 | L234I | A | 1.8e-07 | 2.2e-07 | 2.1e-07 | 2.033E-07 | 2.082E-08 | 1.800E-07 | 2.200E-07 |
| 4671 | 234 | L234L | A | 3.5e-07 | 3.2e-07 | 4.9e-07 | 3.867E-07 | 9.074E-08 | 3.200E-07 | 4.900E-07 |
| 4672 | 234 | L234K | A | 1.9e-07 | 1.7e-07 | 2.3e-07 | 1.967E-07 | 3.055E-08 | 1.700E-07 | 2.300E-07 |
| 4673 | 234 | L234M | A | 1.6e-07 | 4.3e-07 | 2.4e-07 | 2.767E-07 | 1.387E-07 | 1.600E-07 | 4.300E-07 |
| 4674 | 234 | L234F | A | 2.1e-07 | 3.1e-07 | 1.4e-07 | 2.200E-07 | 8.544E-08 | 1.400E-07 | 3.100E-07 |

| | | | | | | | | | | |
|---|---|---|---|---|---|---|---|---|---|---|
| 4675 | 234 | L234P | A | 5.1e-07 | 4.9e-07 | 3.1e-07 | 4.367E-07 | 1.102E-07 | 3.100E-07 | 5.100E-07 |
| 4676 | 234 | L234S | A | 3.5e-07 | 2.6e-07 | 3e-07 | 3.033E-07 | 4.509E-08 | 2.600E-07 | 3.500E-07 |
| 4677 | 234 | L234T | A | 1.1e-07 | 2.8e-07 | 2.1e-07 | 2.000E-07 | 8.544E-08 | 1.100E-07 | 2.800E-07 |
| 4678 | 234 | L234W | A | 1.3e-07 | 4.2e-07 | 3e-07 | 2.833E-07 | 1.457E-07 | 1.300E-07 | 4.200E-07 |
| 4679 | 234 | L234Y | A | 2e-07 | 3.4e-07 | 2.6e-07 | 2.667E-07 | 7.024E-08 | 2.000E-07 | 3.400E-07 |
| 4680 | 234 | L234V | A | 3.4e-07 | 2.3e-07 | 3.2e-07 | 2.967E-07 | 5.859E-08 | 2.300E-07 | 3.400E-07 |
| 4681 | 235 | H235A | A | 2.3e-07 | 2.4e-07 | 2.2e-07 | 2.300E-07 | 1.000E-08 | 2.200E-07 | 2.400E-07 |
| 4682 | 235 | H235R | A | 3e-07 | 2.5e-07 | 1.2e-07 | 2.233E-07 | 9.292E-08 | 1.200E-07 | 3.000E-07 |
| 4683 | 235 | H235N | A | 2.3e-07 | 2.9e-07 | 1.6e-07 | 2.267E-07 | 6.506E-08 | 1.600E-07 | 2.900E-07 |
| 4684 | 235 | H235D | A | 3.3e-07 | 1.1e-07 | 2.4e-07 | 2.267E-07 | 1.106E-07 | 1.100E-07 | 3.300E-07 |
| 4685 | 235 | H235C | A | 4.9e-07 | 5.2e-07 | 1.5e-07 | 3.867E-07 | 2.055E-07 | 1.500E-07 | 5.200E-07 |
| 4686 | 235 | H235E | A | 2.5e-07 | 2.4e-07 | 2.3e-07 | 2.400E-07 | 1.000E-08 | 2.300E-07 | 2.500E-07 |
| 4687 | 235 | H235Q | A | 1.8e-07 | 1.4e-07 | 2.2e-07 | 1.800E-07 | 4.000E-08 | 1.400E-07 | 2.200E-07 |
| 4688 | 235 | H235G | A | 4.5e-07 | 2.9e-07 | 4.1e-07 | 3.833E-07 | 8.327E-08 | 2.900E-07 | 4.500E-07 |
| 4689 | 235 | H235H | A | 4.8e-07 | 2.7e-07 | 3.6e-07 | 3.700E-07 | 1.054E-07 | 2.700E-07 | 4.800E-07 |
| 4690 | 235 | H235I | A | 3.4e-07 | 1.7e-07 | 2.8e-07 | 2.633E-07 | 8.622E-08 | 1.700E-07 | 3.400E-07 |
| 4691 | 235 | H235L | A | 1.7e-07 | 2.4e-07 | 2.8e-07 | 2.300E-07 | 5.568E-08 | 1.700E-07 | 2.800E-07 |
| 4692 | 235 | H235K | A | 1.8e-07 | 1.7e-07 | 1.9e-07 | 1.800E-07 | 1.000E-08 | 1.700E-07 | 1.900E-07 |
| 4693 | 235 | H235M | A | 3.7e-07 | 1.9e-07 | 3.7e-07 | 3.100E-07 | 1.039E-07 | 1.900E-07 | 3.700E-07 |
| 4694 | 235 | H235F | A | 3.5e-07 | 2.3e-07 | 2.7e-07 | 2.833E-07 | 6.110E-08 | 2.300E-07 | 3.500E-07 |
| 4695 | 235 | H235P | A | 1.3e-07 | 2.3e-07 | 9.2e-08 | 1.507E-07 | 7.128E-08 | 9.200E-08 | 2.300E-07 |
| 4696 | 235 | H235S | A | 1.6e-07 | 1.1e-07 | 3.1e-07 | 1.933E-07 | 1.041E-07 | 1.100E-07 | 3.100E-07 |
| 4697 | 235 | H235T | A | 3.2e-07 | 2.1e-07 | 3.6e-07 | 2.967E-07 | 7.767E-08 | 2.100E-07 | 3.600E-07 |
| 4698 | 235 | H235W | A | 3.2e-07 | 5e-07 | 4.4e-07 | 4.200E-07 | 9.165E-08 | 3.200E-07 | 5.000E-07 |
| 4699 | 235 | H235Y | A | 3.2e-07 | 4.6e-07 | 2.5e-07 | 3.433E-07 | 1.069E-07 | 2.500E-07 | 4.600E-07 |
| 4700 | 235 | H235V | A | 2.7e-07 | 2e-07 | 2.2e-07 | 2.300E-07 | 3.606E-08 | 2.000E-07 | 2.700E-07 |
| 4701 | 236 | F236A | A | 4.9e-07 | 2.1e-07 | 3.3e-07 | 3.433E-07 | 1.405E-07 | 2.100E-07 | 4.900E-07 |
| 4702 | 236 | F236R | A | 2.3e-07 | 1.8e-07 | 1.7e-07 | 1.933E-07 | 3.215E-08 | 1.700E-07 | 2.300E-07 |
| 4703 | 236 | F236N | A | 3.1e-07 | 3.8e-07 | 3e-07 | 3.300E-07 | 4.359E-08 | 3.000E-07 | 3.800E-07 |
| 4704 | 236 | F236D | A | 4.4e-07 | 3.2e-07 | 3.3e-07 | 3.633E-07 | 6.658E-08 | 3.200E-07 | 4.400E-07 |
| 4705 | 236 | F236C | A | 3.2e-07 | 2.2e-07 | 1.4e-07 | 2.267E-07 | 9.018E-08 | 1.400E-07 | 3.200E-07 |
| 4706 | 236 | F236E | A | 2e-07 | 2e-07 | 1.2e-07 | 1.733E-07 | 4.619E-08 | 1.200E-07 | 2.000E-07 |
| 4707 | 236 | F236Q | A | 2.4e-07 | 2.4e-07 | 1.1e-07 | 1.967E-07 | 7.506E-08 | 1.100E-07 | 2.400E-07 |
| 4708 | 236 | F236G | A | 4.9e-07 | 3e-07 | 2.5e-07 | 3.467E-07 | 1.266E-07 | 2.500E-07 | 4.900E-07 |
| 4709 | 236 | F236H | A | 1.6e-07 | 4.5e-07 | 4.2e-07 | 3.433E-07 | 1.595E-07 | 1.600E-07 | 4.500E-07 |
| 4710 | 236 | F236I | A | 3.8e-07 | 1.8e-07 | 1e-07 | 2.200E-07 | 1.442E-07 | 1.000E-07 | 3.800E-07 |
| 4711 | 236 | F236L | A | 1.9e-07 | 1.2e-07 | 1.9e-07 | 1.667E-07 | 4.041E-08 | 1.200E-07 | 1.900E-07 |
| 4712 | 236 | F236K | A | 3.1e-07 | 3.5e-07 | 3.7e-07 | 3.433E-07 | 3.055E-08 | 3.100E-07 | 3.700E-07 |
| 4713 | 236 | F236M | A | 7.4e-07 | 4.2e-07 | 1.9e-07 | 4.500E-07 | 2.762E-07 | 1.900E-07 | 7.400E-07 |
| 4714 | 236 | F236F | A | 2.9e-07 | 1.7e-07 | 2.4e-07 | 2.333E-07 | 6.028E-08 | 1.700E-07 | 2.900E-07 |
| 4715 | 236 | F236P | A | 3.1e-07 | 1.3e-07 | 2.1e-07 | 2.167E-07 | 9.018E-08 | 1.300E-07 | 3.100E-07 |
| 4716 | 236 | F236S | A | 2.4e-07 | 1.3e-07 | 2.7e-07 | 2.133E-07 | 7.371E-08 | 1.300E-07 | 2.700E-07 |
| 4717 | 236 | F236T | A | 1.7e-07 | 2.5e-07 | 2.5e-07 | 2.233E-07 | 4.619E-08 | 1.700E-07 | 2.500E-07 |
| 4718 | 236 | F236W | A | 3.8e-07 | 3.7e-07 | 2.5e-07 | 3.333E-07 | 7.234E-08 | 2.500E-07 | 3.800E-07 |
| 4719 | 236 | F236Y | A | 1.6e-07 | 2.3e-07 | 2.7e-07 | 2.200E-07 | 5.568E-08 | 1.600E-07 | 2.700E-07 |
| 4720 | 236 | F236V | A | 2.2e-07 | 1.7e-07 | 1.3e-07 | 1.733E-07 | 4.509E-08 | 1.300E-07 | 2.200E-07 |
| 4721 | 237 | N237A | A | 1.9e-07 | 4.3e-07 | 2e-07 | 2.733E-07 | 1.358E-07 | 1.900E-07 | 4.300E-07 |
| 4722 | 237 | N237R | A | 2.5e-07 | 1.2e-07 | 1.8e-07 | 1.833E-07 | 6.506E-08 | 1.200E-07 | 2.500E-07 |
| 4723 | 237 | N237N | A | 1.9e-07 | 2.6e-07 | 1.7e-07 | 2.067E-07 | 4.726E-08 | 1.700E-07 | 2.600E-07 |
| 4724 | 237 | N237D | A | 2.4e-07 | 2.6e-07 | 9.2e-08 | 1.973E-07 | 9.177E-08 | 9.200E-08 | 2.600E-07 |
| 4725 | 237 | N237C | A | 2.5e-07 | 4.2e-07 | 1e-07 | 2.567E-07 | 1.601E-07 | 1.000E-07 | 4.200E-07 |
| 4726 | 237 | N237E | A | 2.4e-07 | 2.6e-07 | 1.1e-07 | 2.033E-07 | 8.145E-08 | 1.100E-07 | 2.600E-07 |
| 4727 | 237 | N237Q | A | 3.1e-07 | 1.1e-07 | 4.6e-07 | 2.933E-07 | 1.756E-07 | 1.100E-07 | 4.600E-07 |
| 4728 | 237 | N237G | A | 2.6e-07 | 4.3e-07 | 2.1e-07 | 3.000E-07 | 1.153E-07 | 2.100E-07 | 4.300E-07 |
| 4729 | 237 | N237H | A | 2.5e-07 | 2.6e-07 | 1.9e-07 | 2.333E-07 | 3.786E-08 | 1.900E-07 | 2.600E-07 |
| 4730 | 237 | N237I | A | 2e-07 | 2e-07 | 3.4e-07 | 2.467E-07 | 8.083E-08 | 2.000E-07 | 3.400E-07 |
| 4731 | 237 | N237L | A | 3.2e-07 | 7.6e-07 | 2.4e-07 | 4.400E-07 | 2.800E-07 | 2.400E-07 | 7.600E-07 |
| 4732 | 237 | N237K | A | 3.4e-07 | 4.1e-07 | 2.1e-07 | 3.200E-07 | 1.015E-07 | 2.100E-07 | 4.100E-07 |
| 4733 | 237 | N237M | A | 1.3e-07 | 1.8e-07 | 2.4e-07 | 1.833E-07 | 5.508E-08 | 1.300E-07 | 2.400E-07 |
| 4734 | 237 | N237F | A | 5.7e-08 | 2e-07 | 1.8e-07 | 1.457E-07 | 7.744E-08 | 5.700E-08 | 2.000E-07 |
| 4735 | 237 | N237P | A | 2.1e-07 | 1.1e-07 | 2.4e-07 | 1.867E-07 | 6.807E-08 | 1.100E-07 | 2.400E-07 |
| 4736 | 237 | N237S | A | 1.8e-07 | 3e-07 | 4.9e-07 | 3.233E-07 | 1.563E-07 | 1.800E-07 | 4.900E-07 |
| 4737 | 237 | N237T | A | 3.9e-07 | 1.7e-07 | 4.3e-07 | 3.300E-07 | 1.400E-07 | 1.700E-07 | 4.300E-07 |
| 4738 | 237 | N237W | A | 3.8e-07 | 2.7e-07 | 4e-07 | 3.500E-07 | 7.000E-08 | 2.700E-07 | 4.000E-07 |
| 4739 | 237 | N237Y | A | 6.3e-07 | 3.4e-07 | 2.7e-07 | 4.133E-07 | 1.909E-07 | 2.700E-07 | 6.300E-07 |
| 4740 | 237 | N237V | A | 1.9e-07 | 5.2e-07 | 2.9e-07 | 3.333E-07 | 1.692E-07 | 1.900E-07 | 5.200E-07 |
| 4741 | 238 | H238A | A | 5.3e-07 | 1.7e-07 | 7.4e-07 | 4.800E-07 | 2.883E-07 | 1.700E-07 | 7.400E-07 |
| 4742 | 238 | H238R | A | 4.5e-07 | 2e-07 | 2.1e-07 | 2.867E-07 | 1.415E-07 | 2.000E-07 | 4.500E-07 |
| 4743 | 238 | H238N | A | 2.7e-07 | 2.7e-07 | 3.8e-07 | 3.067E-07 | 6.351E-08 | 2.700E-07 | 3.800E-07 |
| 4744 | 238 | H238D | A | 2.8e-07 | 2.1e-07 | 2.7e-07 | 2.533E-07 | 3.786E-08 | 2.100E-07 | 2.800E-07 |
| 4745 | 238 | H238C | A | 2e-07 | 2.3e-07 | 2.7e-07 | 2.333E-07 | 3.512E-08 | 2.000E-07 | 2.700E-07 |
| 4746 | 238 | H238E | A | 1.7e-07 | 3.7e-07 | 1.4e-07 | 2.267E-07 | 1.250E-07 | 1.400E-07 | 3.700E-07 |
| 4747 | 238 | H238Q | A | 5.7e-07 | 1.4e-07 | 2.1e-07 | 3.067E-07 | 2.307E-07 | 1.400E-07 | 5.700E-07 |
| 4748 | 238 | H238G | A | 2.2e-07 | 3e-07 | 2.7e-07 | 2.633E-07 | 4.041E-08 | 2.200E-07 | 3.000E-07 |
| 4749 | 238 | H238H | A | 3.8e-07 | 2.2e-07 | 1.9e-07 | 2.633E-07 | 1.021E-07 | 1.900E-07 | 3.800E-07 |
| 4750 | 238 | H238I | A | 1.7e-07 | 2.5e-07 | 3.2e-07 | 2.467E-07 | 7.506E-08 | 1.700E-07 | 3.200E-07 |
| 4751 | 238 | H238L | A | 3.6e-07 | 5.4e-07 | 2.8e-07 | 3.933E-07 | 1.332E-07 | 2.800E-07 | 5.400E-07 |
| 4752 | 238 | H238K | A | 1.5e-07 | 1.5e-07 | 4.8e-07 | 2.600E-07 | 1.905E-07 | 1.500E-07 | 4.800E-07 |
| 4753 | 238 | H238M | A | 2.2e-07 | 2.3e-07 | 5.4e-07 | 3.300E-07 | 1.819E-07 | 2.200E-07 | 5.400E-07 |
| 4754 | 238 | H238F | A | 1.4e-07 | 3.8e-07 | 3.3e-07 | 2.833E-07 | 1.266E-07 | 1.400E-07 | 3.800E-07 |
| 4755 | 238 | H238P | A | 2.9e-07 | 2e-07 | 1.8e-07 | 2.233E-07 | 5.859E-08 | 1.800E-07 | 2.900E-07 |
| 4756 | 238 | H238S | A | 1.7e-07 | 2e-07 | 2.5e-07 | 2.067E-07 | 4.041E-08 | 1.700E-07 | 2.500E-07 |
| 4757 | 238 | H238T | A | 1.3e-07 | 2.5e-07 | 2.9e-07 | 2.233E-07 | 8.327E-08 | 1.300E-07 | 2.900E-07 |
| 4758 | 238 | H238W | A | 2.3e-07 | 6.8e-08 | 3e-07 | 1.993E-07 | 1.190E-07 | 6.800E-08 | 3.000E-07 |
| 4759 | 238 | H238Y | A | 3.3e-07 | 5.1e-07 | 2.6e-07 | 3.667E-07 | 1.290E-07 | 2.600E-07 | 5.100E-07 |

| | | | | | | | | | | |
|---|---|---|---|---|---|---|---|---|---|---|
| 4760 | 238 | H238V | A | 4.1e-07 | 2.6e-07 | 1.8e-07 | 2.833E-07 | 1.168E-07 | 1.800E-07 | 4.100E-07 |
| 4761 | 239 | S239A | A | 1.9e-07 | 2.2e-07 | 3.3e-07 | 2.467E-07 | 7.371E-08 | 1.900E-07 | 3.300E-07 |
| 4762 | 239 | S239R | A | 5.3e-07 | 2.4e-07 | 2.8e-07 | 3.500E-07 | 1.572E-07 | 2.400E-07 | 5.300E-07 |
| 4763 | 239 | S239N | A | 2.4e-07 | 4.8e-07 | 2.3e-07 | 3.167E-07 | 1.415E-07 | 2.300E-07 | 4.800E-07 |
| 4764 | 239 | S239D | A | 2.4e-07 | 3.6e-07 | 2.7e-07 | 2.900E-07 | 6.245E-08 | 2.400E-07 | 3.600E-07 |
| 4765 | 239 | S239C | A | 5e-07 | 4.7e-07 | 4.7e-07 | 4.800E-07 | 1.732E-08 | 4.700E-07 | 5.000E-07 |
| 4766 | 239 | S239E | A | 3.2e-07 | 3.2e-07 | 3.2e-07 | 3.200E-07 | 6.484E-23 | 3.200E-07 | 3.200E-07 |
| 4767 | 239 | S239Q | A | 4.9e-07 | 1.6e-07 | 2e-07 | 2.833E-07 | 1.801E-07 | 1.600E-07 | 4.900E-07 |
| 4768 | 239 | S239G | A | 2.7e-07 | 1.8e-07 | 3.9e-07 | 2.800E-07 | 1.054E-07 | 1.800E-07 | 3.900E-07 |
| 4769 | 239 | S239H | A | 2.9e-07 | 4.3e-07 | 3.5e-07 | 3.567E-07 | 7.024E-08 | 2.900E-07 | 4.300E-07 |
| 4770 | 239 | S239I | A | 1.3e-07 | 2.3e-07 | 2e-07 | 1.867E-07 | 5.132E-08 | 1.300E-07 | 2.300E-07 |
| 4771 | 239 | S239L | A | 1.4e-07 | 2.9e-07 | 2.1e-07 | 2.133E-07 | 7.506E-08 | 1.400E-07 | 2.900E-07 |
| 4772 | 239 | S239K | A | 3.8e-07 | 2.4e-07 | 1.6e-07 | 2.600E-07 | 1.114E-07 | 1.600E-07 | 3.800E-07 |
| 4773 | 239 | S239M | A | 2.3e-07 | 1.5e-07 | 3.6e-07 | 2.467E-07 | 1.060E-07 | 1.500E-07 | 3.600E-07 |
| 4774 | 239 | S239F | A | 3e-07 | 2.6e-07 | 1.3e-07 | 2.300E-07 | 8.888E-08 | 1.300E-07 | 3.000E-07 |
| 4775 | 239 | S239P | A | 3.5e-07 | 5.6e-07 | 4.9e-07 | 4.667E-07 | 1.069E-07 | 3.500E-07 | 5.600E-07 |
| 4776 | 239 | S239S | A | 4.3e-07 | 2.2e-07 | 4e-07 | 3.500E-07 | 1.136E-07 | 2.200E-07 | 4.300E-07 |
| 4777 | 239 | S239T | A | 1.9e-07 | 2.8e-07 | 3.9e-07 | 2.867E-07 | 1.002E-07 | 1.900E-07 | 3.900E-07 |
| 4778 | 239 | S239W | A | 2.6e-07 | 2.4e-07 | 3.4e-07 | 2.800E-07 | 5.292E-08 | 2.400E-07 | 3.400E-07 |
| 4779 | 239 | S239Y | A | 1.9e-07 | 2.8e-07 | 1.7e-07 | 2.133E-07 | 5.859E-08 | 1.700E-07 | 2.800E-07 |
| 4780 | 239 | S239V | A | 5.9e-07 | 4.2e-07 | 2e-07 | 4.033E-07 | 1.955E-07 | 2.000E-07 | 5.900E-07 |
| 4781 | 240 | G240A | A | 1.5e-07 | 1.8e-07 | 3.1e-07 | 2.133E-07 | 8.505E-08 | 1.500E-07 | 3.100E-07 |
| 4782 | 240 | G240R | A | 1.8e-07 | 1.8e-07 | 1.8e-07 | 1.800E-07 | 0.000E+00 | 1.800E-07 | 1.800E-07 |
| 4783 | 240 | G240N | A | 2.4e-07 | 2.1e-07 | 1.2e-07 | 1.900E-07 | 6.245E-08 | 1.200E-07 | 2.400E-07 |
| 4784 | 240 | G240D | A | 1.5e-07 | 1.9e-07 | 3.5e-07 | 2.300E-07 | 1.058E-07 | 1.500E-07 | 3.500E-07 |
| 4785 | 240 | G240C | A | 1.8e-07 | 1.6e-07 | 2.2e-07 | 1.867E-07 | 3.055E-08 | 1.600E-07 | 2.200E-07 |
| 4786 | 240 | G240E | A | 2.3e-07 | 2.6e-07 | 2.4e-07 | 2.433E-07 | 1.528E-08 | 2.300E-07 | 2.600E-07 |
| 4787 | 240 | G240Q | A | 3e-07 | 3.8e-07 | 3.1e-07 | 3.300E-07 | 4.359E-08 | 3.000E-07 | 3.800E-07 |
| 4788 | 240 | G240G | A | 3.6e-07 | 1.2e-07 | 6e-07 | 3.600E-07 | 2.400E-07 | 1.200E-07 | 6.000E-07 |
| 4789 | 240 | G240H | A | 3.9e-07 | 1.7e-07 | 3.6e-07 | 3.067E-07 | 1.193E-07 | 1.700E-07 | 3.900E-07 |
| 4790 | 240 | G240I | A | 2.1e-07 | 9.4e-08 | 2.1e-07 | 1.713E-07 | 6.697E-08 | 9.400E-08 | 2.100E-07 |
| 4791 | 240 | G240L | A | 2.2e-07 | 2.7e-07 | 1.6e-07 | 2.167E-07 | 5.508E-08 | 1.600E-07 | 2.700E-07 |
| 4792 | 240 | G240K | A | 1.2e-07 | 7.3e-08 | 1.9e-07 | 1.277E-07 | 5.888E-08 | 7.300E-08 | 1.900E-07 |
| 4793 | 240 | G240M | A | 3.3e-07 | 4.6e-07 | 4e-07 | 3.967E-07 | 6.506E-08 | 3.300E-07 | 4.600E-07 |
| 4794 | 240 | G240F | A | 1.4e-07 | 1.8e-07 | 8.1e-07 | 3.767E-07 | 3.758E-07 | 1.400E-07 | 8.100E-07 |
| 4795 | 240 | G240P | A | 5.7e-07 | 1.2e-07 | 3e-07 | 3.300E-07 | 2.265E-07 | 1.200E-07 | 5.700E-07 |
| 4796 | 240 | G240S | A | 3.2e-07 | 1.9e-07 | 8.7e-08 | 1.990E-07 | 1.168E-07 | 8.700E-08 | 3.200E-07 |
| 4797 | 240 | G240T | A | 2.2e-07 | 3.4e-07 | 2.5e-07 | 2.700E-07 | 6.245E-08 | 2.200E-07 | 3.400E-07 |
| 4798 | 240 | G240W | A | 3.4e-07 | 3.3e-07 | 1.3e-07 | 2.667E-07 | 1.185E-07 | 1.300E-07 | 3.400E-07 |
| 4799 | 240 | G240Y | A | 1e-07 | 2.3e-07 | 2.5e-07 | 1.933E-07 | 8.145E-08 | 1.000E-07 | 2.500E-07 |
| 4800 | 240 | G240V | A | 1.8e-07 | 2.6e-07 | 2.5e-07 | 2.300E-07 | 4.359E-08 | 1.800E-07 | 2.600E-07 |
| 4801 | 241 | I241A | A | 2.7e-07 | 1.8e-07 | 4.3e-07 | 2.933E-07 | 1.266E-07 | 1.800E-07 | 4.300E-07 |
| 4802 | 241 | I241R | A | 2.5e-07 | 2.6e-07 | 3.8e-07 | 2.967E-07 | 7.234E-08 | 2.500E-07 | 3.800E-07 |
| 4803 | 241 | I241N | A | 2.3e-07 | 2.4e-07 | 3.2e-07 | 2.633E-07 | 4.933E-08 | 2.300E-07 | 3.200E-07 |
| 4804 | 241 | I241D | A | 2e-07 | 3.8e-07 | 2.9e-07 | 2.900E-07 | 9.000E-08 | 2.000E-07 | 3.800E-07 |
| 4805 | 241 | I241C | A | 3.1e-07 | 1.9e-07 | 3.1e-07 | 2.700E-07 | 6.928E-08 | 1.900E-07 | 3.100E-07 |
| 4806 | 241 | I241E | A | 8e-08 | 1.2e-07 | 4.7e-07 | 2.233E-07 | 2.146E-07 | 8.000E-08 | 4.700E-07 |
| 4807 | 241 | I241Q | A | 6.3e-07 | 2.6e-07 | 1.6e-07 | 3.500E-07 | 2.476E-07 | 1.600E-07 | 6.300E-07 |
| 4808 | 241 | I241G | A | 2.3e-07 | 8.3e-07 | 3.8e-07 | 4.800E-07 | 3.122E-07 | 2.300E-07 | 8.300E-07 |
| 4809 | 241 | I241H | A | 2e-07 | 1.6e-07 | 1.4e-07 | 1.667E-07 | 3.055E-08 | 1.400E-07 | 2.000E-07 |
| 4810 | 241 | I241I | A | 2.2e-07 | 3.2e-07 | 2.9e-07 | 2.767E-07 | 5.132E-08 | 2.200E-07 | 3.200E-07 |
| 4811 | 241 | I241L | A | 3.1e-07 | 1.4e-07 | 1.4e-07 | 1.967E-07 | 9.815E-08 | 1.400E-07 | 3.100E-07 |
| 4812 | 241 | I241K | A | 3e-07 | 3.3e-07 | 2.3e-07 | 2.867E-07 | 5.132E-08 | 2.300E-07 | 3.300E-07 |
| 4813 | 241 | I241M | A | 2.1e-07 | 4.7e-07 | 4.7e-07 | 3.833E-07 | 1.501E-07 | 2.100E-07 | 4.700E-07 |
| 4814 | 241 | I241F | A | 1.9e-07 | 2.6e-07 | 3.4e-07 | 2.633E-07 | 7.506E-08 | 1.900E-07 | 3.400E-07 |
| 4815 | 241 | I241P | A | 1.8e-07 | 2e-07 | 2e-07 | 1.933E-07 | 1.155E-08 | 1.800E-07 | 2.000E-07 |
| 4816 | 241 | I241S | A | 1.5e-07 | 3.3e-07 | 1.5e-07 | 2.100E-07 | 1.039E-07 | 1.500E-07 | 3.300E-07 |
| 4817 | 241 | I241T | A | 3.8e-07 | 3.2e-07 | 2.6e-07 | 3.200E-07 | 6.000E-08 | 2.600E-07 | 3.800E-07 |
| 4818 | 241 | I241W | A | 2.8e-07 | 1.6e-07 | 1.5e-07 | 1.967E-07 | 7.234E-08 | 1.500E-07 | 2.800E-07 |
| 4819 | 241 | I241Y | A | 3.5e-07 | 2.5e-07 | 1.9e-07 | 2.633E-07 | 8.083E-08 | 1.900E-07 | 3.500E-07 |
| 4820 | 241 | I241V | A | 2.9e-07 | 1.2e-07 | 6.4e-07 | 3.500E-07 | 2.651E-07 | 1.200E-07 | 6.400E-07 |
| 4821 | 242 | C242A | A | 4.1e-07 | 9.6e-08 | 1.1e-07 | 2.053E-07 | 1.774E-07 | 9.600E-08 | 4.100E-07 |
| 4822 | 242 | C242R | A | 1.4e-07 | 1.4e-07 | 1.5e-07 | 1.433E-07 | 5.774E-09 | 1.400E-07 | 1.500E-07 |
| 4823 | 242 | C242N | A | 1.7e-07 | 3e-07 | 1.3e-07 | 2.000E-07 | 8.888E-08 | 1.300E-07 | 3.000E-07 |
| 4824 | 242 | C242D | A | 4e-07 | 5.6e-07 | 1.5e-07 | 3.700E-07 | 2.066E-07 | 1.500E-07 | 5.600E-07 |
| 4825 | 242 | C242C | A | 8.2e-08 | 2.6e-07 | 2.6e-07 | 2.007E-07 | 1.028E-07 | 8.200E-08 | 2.600E-07 |
| 4826 | 242 | C242E | A | 3e-07 | 3.9e-07 | 2.5e-07 | 3.133E-07 | 7.095E-08 | 2.500E-07 | 3.900E-07 |
| 4827 | 242 | C242Q | A | 3.5e-07 | 1.2e-07 | 2.1e-07 | 2.267E-07 | 1.159E-07 | 1.200E-07 | 3.500E-07 |
| 4828 | 242 | C242G | A | 2.8e-07 | 2.5e-07 | 1.9e-07 | 2.400E-07 | 4.583E-08 | 1.900E-07 | 2.800E-07 |
| 4829 | 242 | C242H | A | 1.5e-07 | 3.2e-07 | 3.3e-07 | 2.667E-07 | 1.012E-07 | 1.500E-07 | 3.300E-07 |
| 4830 | 242 | C242I | A | 4.8e-07 | 1.6e-07 | 1.1e-07 | 2.500E-07 | 2.007E-07 | 1.100E-07 | 4.800E-07 |
| 4831 | 242 | C242L | A | 2.7e-07 | 1.9e-07 | 1.8e-07 | 2.133E-07 | 4.933E-08 | 1.800E-07 | 2.700E-07 |
| 4832 | 242 | C242K | A | 4.4e-07 | 2.9e-07 | 1.4e-07 | 2.900E-07 | 1.500E-07 | 1.400E-07 | 4.400E-07 |
| 4833 | 242 | C242M | A | 2.7e-07 | 1.6e-07 | 3.2e-07 | 2.500E-07 | 8.185E-08 | 1.600E-07 | 3.200E-07 |
| 4834 | 242 | C242F | A | 2.1e-07 | 1e-07 | 2.1e-07 | 1.733E-07 | 6.351E-08 | 1.000E-07 | 2.100E-07 |
| 4835 | 242 | C242P | A | 2e-07 | 1.1e-07 | 2.2e-07 | 1.767E-07 | 5.859E-08 | 1.100E-07 | 2.200E-07 |
| 4836 | 242 | C242S | A | 4.3e-07 | 3.3e-07 | 2.8e-07 | 3.467E-07 | 7.638E-08 | 2.800E-07 | 4.300E-07 |
| 4837 | 242 | C242T | A | 2.1e-07 | 2.2e-07 | 2.3e-07 | 2.200E-07 | 1.000E-08 | 2.100E-07 | 2.300E-07 |
| 4838 | 242 | C242W | A | 3.8e-07 | 1.6e-07 | 1.1e-07 | 2.167E-07 | 1.436E-07 | 1.100E-07 | 3.800E-07 |
| 4839 | 242 | C242Y | A | 3.5e-07 | 2e-07 | 2.1e-07 | 2.533E-07 | 8.386E-08 | 2.000E-07 | 3.500E-07 |
| 4840 | 242 | C242V | A | 1.6e-07 | 1.9e-07 | 3.5e-07 | 2.333E-07 | 1.021E-07 | 1.600E-07 | 3.500E-07 |
| 4841 | 243 | E243A | A | 1.6e-07 | 6e-07 | 3.9e-07 | 3.833E-07 | 2.201E-07 | 1.600E-07 | 6.000E-07 |
| 4842 | 243 | E243R | A | 2.7e-07 | 2.7e-07 | 1.3e-07 | 2.233E-07 | 8.083E-08 | 1.300E-07 | 2.700E-07 |
| 4843 | 243 | E243N | A | 1e-07 | 2.2e-07 | 4.2e-07 | 2.467E-07 | 1.617E-07 | 1.000E-07 | 4.200E-07 |
| 4844 | 243 | E243D | A | 2.5e-07 | 1.6e-07 | 2.9e-07 | 2.333E-07 | 6.658E-08 | 1.600E-07 | 2.900E-07 |

| | | | | | | | | | | |
|---|---|---|---|---|---|---|---|---|---|---|
| 4845 | 243 | E243C | A | 4.6e-07 | 1.7e-07 | 4e-07 | 3.433E-07 | 1.531E-07 | 1.700E-07 | 4.600E-07 |
| 4846 | 243 | E243E | A | 2.6e-07 | 2.6e-07 | 1.6e-07 | 2.267E-07 | 5.774E-08 | 1.600E-07 | 2.600E-07 |
| 4847 | 243 | E243Q | A | 1.5e-07 | 1.6e-07 | 2.9e-07 | 2.000E-07 | 7.810E-08 | 1.500E-07 | 2.900E-07 |
| 4848 | 243 | E243G | A | 3.9e-07 | 4.9e-07 | 1.7e-07 | 3.500E-07 | 1.637E-07 | 1.700E-07 | 4.900E-07 |
| 4849 | 243 | E243H | A | 2.3e-07 | 1.3e-07 | 3.4e-07 | 2.333E-07 | 1.050E-07 | 1.300E-07 | 3.400E-07 |
| 4850 | 243 | E243I | A | 2.6e-07 | 3.7e-07 | 1.4e-07 | 2.567E-07 | 1.150E-07 | 1.400E-07 | 3.700E-07 |
| 4851 | 243 | E243L | A | 6e-07 | 2.2e-07 | 2.4e-07 | 3.533E-07 | 2.139E-07 | 2.200E-07 | 6.000E-07 |
| 4852 | 243 | E243K | A | 3.1e-07 | 2.7e-07 | 4.1e-07 | 3.300E-07 | 7.211E-08 | 2.700E-07 | 4.100E-07 |
| 4853 | 243 | E243M | A | 2.2e-07 | 1.6e-07 | 3.4e-07 | 2.400E-07 | 9.165E-08 | 1.600E-07 | 3.400E-07 |
| 4854 | 243 | E243F | A | 5.3e-07 | 2.6e-07 | 2.4e-07 | 3.433E-07 | 1.620E-07 | 2.400E-07 | 5.300E-07 |
| 4855 | 243 | E243P | A | 3.4e-07 | 3e-07 | 3e-07 | 3.133E-07 | 2.309E-08 | 3.000E-07 | 3.400E-07 |
| 4856 | 243 | E243S | A | 2.9e-07 | 2.3e-07 | 1.9e-07 | 2.367E-07 | 5.033E-08 | 1.900E-07 | 2.900E-07 |
| 4857 | 243 | E243T | A | 2.4e-07 | 2.6e-07 | 6.5e-07 | 3.833E-07 | 2.312E-07 | 2.400E-07 | 6.500E-07 |
| 4858 | 243 | E243W | A | 3.5e-07 | 2.5e-07 | 3e-07 | 3.000E-07 | 5.000E-08 | 2.500E-07 | 3.500E-07 |
| 4859 | 243 | E243Y | A | 3.7e-07 | 2.1e-07 | 2.3e-07 | 2.700E-07 | 8.718E-08 | 2.100E-07 | 3.700E-07 |
| 4860 | 243 | E243V | A | 3.4e-07 | 9.2e-07 | 2.2e-07 | 4.933E-07 | 3.743E-07 | 2.200E-07 | 9.200E-07 |
| 4861 | 244 | L244A | A | 2.5e-07 | 3e-07 | 2.3e-07 | 2.600E-07 | 3.606E-08 | 2.300E-07 | 3.000E-07 |
| 4862 | 244 | L244R | A | 1.6e-07 | 2.2e-07 | 1.6e-07 | 1.800E-07 | 3.464E-08 | 1.600E-07 | 2.200E-07 |
| 4863 | 244 | L244N | A | 1.7e-07 | 1.9e-07 | 1.6e-07 | 1.733E-07 | 1.528E-08 | 1.600E-07 | 1.900E-07 |
| 4864 | 244 | L244D | A | 2.7e-07 | 1.3e-07 | 2.5e-07 | 2.167E-07 | 7.572E-08 | 1.300E-07 | 2.700E-07 |
| 4865 | 244 | L244C | A | 2e-07 | 4.5e-07 | 2.2e-07 | 2.900E-07 | 1.389E-07 | 2.000E-07 | 4.500E-07 |
| 4866 | 244 | L244E | A | 1.6e-07 | 2.7e-07 | 1.3e-07 | 1.867E-07 | 7.371E-08 | 1.300E-07 | 2.700E-07 |
| 4867 | 244 | L244Q | A | 4.9e-07 | 3.2e-07 | 2.2e-07 | 3.433E-07 | 1.365E-07 | 2.200E-07 | 4.900E-07 |
| 4868 | 244 | L244G | A | 2.9e-07 | 6.6e-08 | 3.8e-07 | 2.453E-07 | 1.617E-07 | 6.600E-08 | 3.800E-07 |
| 4869 | 244 | L244H | A | 6.3e-07 | 2.2e-07 | 3.1e-07 | 3.867E-07 | 2.155E-07 | 2.200E-07 | 6.300E-07 |
| 4870 | 244 | L244I | A | 6.6e-07 | 2.7e-07 | 2.8e-07 | 4.033E-07 | 2.223E-07 | 2.700E-07 | 6.600E-07 |
| 4871 | 244 | L244L | A | 4.2e-07 | 3.2e-07 | 2.9e-07 | 3.433E-07 | 6.807E-08 | 2.900E-07 | 4.200E-07 |
| 4872 | 244 | L244K | A | 1.8e-07 | 3.3e-07 | 2.3e-07 | 2.467E-07 | 7.638E-08 | 1.800E-07 | 3.300E-07 |
| 4873 | 244 | L244M | A | 3.6e-07 | 2.4e-07 | 1.6e-07 | 2.533E-07 | 1.007E-07 | 1.600E-07 | 3.600E-07 |
| 4874 | 244 | L244F | A | 3.2e-07 | 1.6e-07 | 1.6e-07 | 2.133E-07 | 9.238E-08 | 1.600E-07 | 3.200E-07 |
| 4875 | 244 | L244P | A | 2.9e-07 | 4.3e-07 | 3.5e-07 | 3.567E-07 | 7.024E-08 | 2.900E-07 | 4.300E-07 |
| 4876 | 244 | L244S | A | 1.4e-07 | 1.2e-07 | 2e-07 | 1.533E-07 | 4.163E-08 | 1.200E-07 | 2.000E-07 |
| 4877 | 244 | L244T | A | 2.7e-07 | 1.2e-07 | 2.1e-07 | 2.000E-07 | 7.550E-08 | 1.200E-07 | 2.700E-07 |
| 4878 | 244 | L244W | A | 2.5e-07 | 2.9e-07 | 2.7e-07 | 2.700E-07 | 2.000E-08 | 2.500E-07 | 2.900E-07 |
| 4879 | 244 | L244Y | A | 1.9e-07 | 3e-07 | 4.3e-07 | 3.067E-07 | 1.201E-07 | 1.900E-07 | 4.300E-07 |
| 4880 | 244 | L244V | A | 5.1e-07 | 2.5e-07 | 3.2e-07 | 3.600E-07 | 1.345E-07 | 2.500E-07 | 5.100E-07 |
| 4881 | 245 | H245A | A | 3.8e-07 | 2.1e-07 | 1.8e-07 | 2.567E-07 | 1.079E-07 | 1.800E-07 | 3.800E-07 |
| 4882 | 245 | H245R | A | 2.8e-07 | 1.7e-07 | 2.3e-07 | 2.267E-07 | 5.508E-08 | 1.700E-07 | 2.800E-07 |
| 4883 | 245 | H245N | A | 4.2e-07 | 2e-07 | 3e-07 | 3.067E-07 | 1.102E-07 | 2.000E-07 | 4.200E-07 |
| 4884 | 245 | H245D | A | 3.1e-07 | 1.9e-07 | 1.5e-07 | 2.167E-07 | 8.327E-08 | 1.500E-07 | 3.100E-07 |
| 4885 | 245 | H245C | A | 1.7e-07 | 2.6e-07 | 4.6e-07 | 2.967E-07 | 1.484E-07 | 1.700E-07 | 4.600E-07 |
| 4886 | 245 | H245E | A | 1.6e-07 | 2.6e-07 | 1.2e-07 | 1.800E-07 | 7.211E-08 | 1.200E-07 | 2.600E-07 |
| 4887 | 245 | H245Q | A | 1.9e-07 | 4.3e-07 | 1.6e-07 | 2.600E-07 | 1.480E-07 | 1.600E-07 | 4.300E-07 |
| 4888 | 245 | H245G | A | 2.5e-07 | 2e-07 | 2.1e-07 | 2.200E-07 | 2.646E-08 | 2.000E-07 | 2.500E-07 |
| 4889 | 245 | H245H | A | 2.6e-07 | 2.2e-07 | 2e-07 | 2.267E-07 | 3.055E-08 | 2.000E-07 | 2.600E-07 |
| 4890 | 245 | H245I | A | 8.6e-08 | 1.3e-07 | 4.1e-07 | 2.087E-07 | 1.757E-07 | 8.600E-08 | 4.100E-07 |
| 4891 | 245 | H245L | A | 3.6e-07 | 1.3e-07 | 3.1e-07 | 2.667E-07 | 1.210E-07 | 1.300E-07 | 3.600E-07 |
| 4892 | 245 | H245K | A | 2.8e-07 | 3.5e-07 | 3.1e-07 | 3.133E-07 | 3.512E-08 | 2.800E-07 | 3.500E-07 |
| 4893 | 245 | H245M | A | 1.7e-07 | 5e-07 | 3.3e-07 | 3.333E-07 | 1.650E-07 | 1.700E-07 | 5.000E-07 |
| 4894 | 245 | H245F | A | 1.4e-07 | 1e-07 | 2.4e-07 | 1.600E-07 | 7.211E-08 | 1.000E-07 | 2.400E-07 |
| 4895 | 245 | H245P | A | 2.3e-07 | 1.9e-07 | 1.5e-07 | 1.900E-07 | 4.000E-08 | 1.500E-07 | 2.300E-07 |
| 4896 | 245 | H245S | A | 1.6e-07 | 2.8e-07 | 1.4e-07 | 1.933E-07 | 7.572E-08 | 1.400E-07 | 2.800E-07 |
| 4897 | 245 | H245T | A | 4e-07 | 2.1e-07 | 3.3e-07 | 3.133E-07 | 9.609E-08 | 2.100E-07 | 4.000E-07 |
| 4898 | 245 | H245W | A | 4e-07 | 2.5e-07 | 3.6e-07 | 3.367E-07 | 7.767E-08 | 2.500E-07 | 4.000E-07 |
| 4899 | 245 | H245Y | A | 4.4e-07 | 3.5e-07 | 2.1e-07 | 3.333E-07 | 1.159E-07 | 2.100E-07 | 4.400E-07 |
| 4900 | 245 | H245V | A | 4.5e-07 | 1.9e-07 | 4.1e-07 | 3.500E-07 | 1.400E-07 | 1.900E-07 | 4.500E-07 |
| 4901 | 246 | C246A | A | 3.1e-07 | 1.8e-07 | 2.4e-07 | 2.433E-07 | 6.506E-08 | 1.800E-07 | 3.100E-07 |
| 4902 | 246 | C246R | A | 3.3e-07 | 2.2e-07 | 1.9e-07 | 2.467E-07 | 7.371E-08 | 1.900E-07 | 3.300E-07 |
| 4903 | 246 | C246N | A | 1.4e-07 | 1.6e-07 | 3.8e-07 | 2.267E-07 | 1.332E-07 | 1.400E-07 | 3.800E-07 |
| 4904 | 246 | C246D | A | 3.4e-07 | 5.3e-07 | 2.1e-07 | 3.600E-07 | 1.609E-07 | 2.100E-07 | 5.300E-07 |
| 4905 | 246 | C246C | A | 1.7e-07 | 4e-07 | 3.5e-07 | 3.067E-07 | 1.210E-07 | 1.700E-07 | 4.000E-07 |
| 4906 | 246 | C246E | A | 1.7e-07 | 4e-07 | 1.6e-07 | 2.433E-07 | 1.358E-07 | 1.600E-07 | 4.000E-07 |
| 4907 | 246 | C246Q | A | 1.6e-07 | 7.3e-07 | 5.6e-07 | 4.833E-07 | 2.926E-07 | 1.600E-07 | 7.300E-07 |
| 4908 | 246 | C246G | A | 1.6e-07 | 2.2e-07 | 2.3e-07 | 2.033E-07 | 3.786E-08 | 1.600E-07 | 2.300E-07 |
| 4909 | 246 | C246H | A | 2.1e-07 | 2.2e-07 | 3.1e-07 | 2.467E-07 | 5.508E-08 | 2.100E-07 | 3.100E-07 |
| 4910 | 246 | C246I | A | 2.4e-07 | 2.8e-07 | 2.4e-07 | 2.533E-07 | 2.309E-08 | 2.400E-07 | 2.800E-07 |
| 4911 | 246 | C246L | A | 2e-07 | 2.6e-07 | 3.9e-07 | 2.833E-07 | 9.713E-08 | 2.000E-07 | 3.900E-07 |
| 4912 | 246 | C246K | A | 1.3e-07 | 2.6e-07 | 3.1e-07 | 2.333E-07 | 9.292E-08 | 1.300E-07 | 3.100E-07 |
| 4913 | 246 | C246M | A | 4e-07 | 3.7e-07 | 3.9e-07 | 3.867E-07 | 1.528E-08 | 3.700E-07 | 4.000E-07 |
| 4914 | 246 | C246F | A | 1.6e-07 | 3.8e-07 | 2.7e-07 | 2.700E-07 | 1.100E-07 | 1.600E-07 | 3.800E-07 |
| 4915 | 246 | C246P | A | 3.2e-07 | 2.4e-07 | 1.7e-07 | 2.433E-07 | 7.506E-08 | 1.700E-07 | 3.200E-07 |
| 4916 | 246 | C246S | A | 2.6e-07 | 3.5e-07 | 3.3e-07 | 3.133E-07 | 4.726E-08 | 2.600E-07 | 3.500E-07 |
| 4917 | 246 | C246T | A | 2.1e-07 | 5.1e-07 | 1.8e-07 | 3.000E-07 | 1.825E-07 | 1.800E-07 | 5.100E-07 |
| 4918 | 246 | C246W | A | 4.1e-07 | 4.9e-07 | 3.4e-07 | 4.133E-07 | 7.506E-08 | 3.400E-07 | 4.900E-07 |
| 4919 | 246 | C246Y | A | 1.7e-07 | 3.5e-07 | 3e-07 | 2.733E-07 | 9.292E-08 | 1.700E-07 | 3.500E-07 |
| 4920 | 246 | C246V | A | 9.1e-08 | 2.1e-07 | 2.9e-07 | 1.970E-07 | 1.001E-07 | 9.100E-08 | 2.900E-07 |
| 4921 | 247 | P247A | A | 2.4e-07 | 3.5e-07 | 2.3e-07 | 2.733E-07 | 6.658E-08 | 2.300E-07 | 3.500E-07 |
| 4922 | 247 | P247R | A | 1.6e-07 | 2.9e-07 | 3.1e-07 | 2.533E-07 | 8.145E-08 | 1.600E-07 | 3.100E-07 |
| 4923 | 247 | P247N | A | 2.4e-07 | 2.9e-07 | 1e-07 | 2.100E-07 | 9.849E-08 | 1.000E-07 | 2.900E-07 |
| 4924 | 247 | P247D | A | 3.8e-07 | 2.1e-07 | 1.8e-07 | 2.567E-07 | 1.079E-07 | 1.800E-07 | 3.800E-07 |
| 4925 | 247 | P247C | A | 2e-07 | 3.5e-07 | 1.6e-07 | 2.367E-07 | 1.002E-07 | 1.600E-07 | 3.500E-07 |
| 4926 | 247 | P247E | A | 3.1e-07 | 1.1e-07 | 2.6e-07 | 2.267E-07 | 1.041E-07 | 1.100E-07 | 3.100E-07 |
| 4927 | 247 | P247Q | A | 3.7e-07 | 2.3e-07 | 1.9e-07 | 2.633E-07 | 9.452E-08 | 1.900E-07 | 3.700E-07 |
| 4928 | 247 | P247G | A | 2e-07 | 1.4e-07 | 1.5e-07 | 1.633E-07 | 3.215E-08 | 1.400E-07 | 2.000E-07 |
| 4929 | 247 | P247H | A | 4.4e-07 | 2e-07 | 3e-07 | 3.133E-07 | 1.206E-07 | 2.000E-07 | 4.400E-07 |

| | | | | | | | | | | |
|---|---|---|---|---|---|---|---|---|---|---|
| 4930 | 247 | P247I | A | 2.5e-07 | 4.3e-07 | 1.5e-07 | 2.767E-07 | 1.419E-07 | 1.500E-07 | 4.300E-07 |
| 4931 | 247 | P247L | A | 2.9e-07 | 1.9e-07 | 7e-07 | 3.933E-07 | 2.702E-07 | 1.900E-07 | 7.000E-07 |
| 4932 | 247 | P247K | A | 1.7e-07 | 3.6e-07 | 2.5e-07 | 2.600E-07 | 9.539E-08 | 1.700E-07 | 3.600E-07 |
| 4933 | 247 | P247M | A | 3.8e-07 | 3.7e-07 | 1.7e-07 | 3.067E-07 | 1.185E-07 | 1.700E-07 | 3.800E-07 |
| 4934 | 247 | P247F | A | 2.6e-07 | 1.5e-07 | 1.7e-07 | 1.933E-07 | 5.859E-08 | 1.500E-07 | 2.600E-07 |
| 4935 | 247 | P247P | A | 2e-07 | 3.3e-07 | 1.1e-07 | 2.133E-07 | 1.106E-07 | 1.100E-07 | 3.300E-07 |
| 4936 | 247 | P247S | A | 1.4e-07 | 2.8e-07 | 4.4e-07 | 2.867E-07 | 1.501E-07 | 1.400E-07 | 4.400E-07 |
| 4937 | 247 | P247T | A | 1.9e-07 | 2.9e-07 | 1e-07 | 1.933E-07 | 9.504E-08 | 1.000E-07 | 2.900E-07 |
| 4938 | 247 | P247W | A | 1.4e-07 | 6.2e-07 | 1.4e-07 | 3.000E-07 | 2.771E-07 | 1.400E-07 | 6.200E-07 |
| 4939 | 247 | P247Y | A | 2.8e-07 | 4.2e-07 | 1.9e-07 | 2.967E-07 | 1.159E-07 | 1.900E-07 | 4.200E-07 |
| 4940 | 247 | P247V | A | 3.1e-07 | 4.8e-07 | 3e-07 | 3.633E-07 | 1.012E-07 | 3.000E-07 | 4.800E-07 |
| 4941 | 248 | A248A | A | 3.7e-07 | 4.3e-07 | 1.4e-07 | 3.133E-07 | 1.531E-07 | 1.400E-07 | 4.300E-07 |
| 4942 | 248 | A248R | A | 5.3e-07 | 2e-07 | 1.9e-07 | 3.067E-07 | 1.935E-07 | 1.900E-07 | 5.300E-07 |
| 4943 | 248 | A248N | A | 1.8e-07 | 1.7e-07 | 1.7e-07 | 1.733E-07 | 5.774E-09 | 1.700E-07 | 1.800E-07 |
| 4944 | 248 | A248D | A | 1.9e-07 | 2.9e-07 | 1.7e-07 | 2.167E-07 | 6.429E-08 | 1.700E-07 | 2.900E-07 |
| 4945 | 248 | A248C | A | 1.6e-07 | 2.2e-07 | 6.2e-07 | 3.333E-07 | 2.501E-07 | 1.600E-07 | 6.200E-07 |
| 4946 | 248 | A248E | A | 1.9e-07 | 3.1e-07 | 2.8e-07 | 2.600E-07 | 6.245E-08 | 1.900E-07 | 3.100E-07 |
| 4947 | 248 | A248Q | A | 4.1e-07 | 8.6e-08 | 2.1e-07 | 2.353E-07 | 1.635E-07 | 8.600E-08 | 4.100E-07 |
| 4948 | 248 | A248G | A | 3.8e-07 | 4.2e-07 | 3e-07 | 3.667E-07 | 6.110E-08 | 3.000E-07 | 4.200E-07 |
| 4949 | 248 | A248H | A | 2.9e-07 | 3.1e-07 | 3.3e-07 | 3.100E-07 | 2.000E-08 | 2.900E-07 | 3.300E-07 |
| 4950 | 248 | A248I | A | 8e-07 | 3.2e-07 | 1.4e-07 | 4.200E-07 | 3.412E-07 | 1.400E-07 | 8.000E-07 |
| 4951 | 248 | A248L | A | 9.9e-08 | 2.2e-07 | 2.1e-07 | 1.763E-07 | 6.716E-08 | 9.900E-08 | 2.200E-07 |
| 4952 | 248 | A248K | A | 2.5e-07 | 3.9e-07 | 2.2e-07 | 2.867E-07 | 9.074E-08 | 2.200E-07 | 3.900E-07 |
| 4953 | 248 | A248M | A | 4.1e-07 | 3.5e-07 | 4.2e-07 | 3.933E-07 | 3.786E-08 | 3.500E-07 | 4.200E-07 |
| 4954 | 248 | A248F | A | 4.9e-07 | 2.4e-07 | 4.3e-07 | 3.867E-07 | 1.305E-07 | 2.400E-07 | 4.900E-07 |
| 4955 | 248 | A248P | A | 3.4e-07 | 3.1e-07 | 1.1e-07 | 2.533E-07 | 1.250E-07 | 1.100E-07 | 3.400E-07 |
| 4956 | 248 | A248S | A | 1.8e-07 | 3.2e-07 | 5.7e-08 | 1.857E-07 | 1.316E-07 | 5.700E-08 | 3.200E-07 |
| 4957 | 248 | A248T | A | 4.6e-07 | 2e-07 | 4.4e-07 | 3.667E-07 | 1.447E-07 | 2.000E-07 | 4.600E-07 |
| 4958 | 248 | A248W | A | 3.2e-07 | 2.3e-07 | 3.2e-07 | 2.900E-07 | 5.196E-08 | 2.300E-07 | 3.200E-07 |
| 4959 | 248 | A248Y | A | 3.2e-07 | 2.8e-07 | 2.5e-07 | 2.833E-07 | 3.512E-08 | 2.500E-07 | 3.200E-07 |
| 4960 | 248 | A248V | A | 3.3e-07 | 2.3e-07 | 6.3e-07 | 3.967E-07 | 2.082E-07 | 2.300E-07 | 6.300E-07 |
| 4961 | 249 | L249A | A | 2.2e-07 | 4e-07 | 3.6e-07 | 3.267E-07 | 9.452E-08 | 2.200E-07 | 4.000E-07 |
| 4962 | 249 | L249R | A | 3.1e-07 | 2.1e-07 | 4.1e-07 | 3.100E-07 | 1.000E-07 | 2.100E-07 | 4.100E-07 |
| 4963 | 249 | L249N | A | 1.3e-07 | 3.6e-07 | 3.9e-07 | 2.933E-07 | 1.422E-07 | 1.300E-07 | 3.900E-07 |
| 4964 | 249 | L249D | A | 3.2e-07 | 2.2e-07 | 2.7e-07 | 2.700E-07 | 5.000E-08 | 2.200E-07 | 3.200E-07 |
| 4965 | 249 | L249C | A | 2.4e-07 | 1.2e-07 | 5.2e-07 | 2.933E-07 | 2.053E-07 | 1.200E-07 | 5.200E-07 |
| 4966 | 249 | L249E | A | 3.3e-07 | 6.4e-07 | 6.2e-07 | 5.300E-07 | 1.735E-07 | 3.300E-07 | 6.400E-07 |
| 4967 | 249 | L249Q | A | 1.2e-07 | 2.4e-07 | 1.9e-07 | 1.833E-07 | 6.028E-08 | 1.200E-07 | 2.400E-07 |
| 4968 | 249 | L249G | A | 3.2e-07 | 1.9e-07 | 3.9e-07 | 3.000E-07 | 1.015E-07 | 1.900E-07 | 3.900E-07 |
| 4969 | 249 | L249H | A | 2.5e-07 | 4e-07 | 2.5e-07 | 3.000E-07 | 8.660E-08 | 2.500E-07 | 4.000E-07 |
| 4970 | 249 | L249I | A | 2.3e-07 | 3.6e-07 | 2.7e-07 | 2.867E-07 | 6.658E-08 | 2.300E-07 | 3.600E-07 |
| 4971 | 249 | L249L | A | 8.9e-07 | 2.3e-07 | 1.1e-07 | 4.100E-07 | 4.200E-07 | 1.100E-07 | 8.900E-07 |
| 4972 | 249 | L249K | A | 2.6e-07 | 1.8e-07 | 2.6e-07 | 2.333E-07 | 4.619E-08 | 1.800E-07 | 2.600E-07 |
| 4973 | 249 | L249M | A | 3.9e-07 | 2.7e-07 | 2.2e-07 | 2.933E-07 | 8.737E-08 | 2.200E-07 | 3.900E-07 |
| 4974 | 249 | L249F | A | 4.4e-07 | 2.6e-07 | 2.9e-07 | 3.300E-07 | 9.644E-08 | 2.600E-07 | 4.400E-07 |
| 4975 | 249 | L249P | A | 2.6e-07 | 3e-07 | 1.8e-07 | 2.467E-07 | 6.110E-08 | 1.800E-07 | 3.000E-07 |
| 4976 | 249 | L249S | A | 1.3e-07 | 1.3e-07 | 2.5e-07 | 1.700E-07 | 6.928E-08 | 1.300E-07 | 2.500E-07 |
| 4977 | 249 | L249T | A | 2.2e-07 | 1.1e-07 | 2.7e-07 | 2.000E-07 | 8.185E-08 | 1.100E-07 | 2.700E-07 |
| 4978 | 249 | L249W | A | 1.1e-07 | 3.4e-07 | 2.6e-07 | 2.367E-07 | 1.168E-07 | 1.100E-07 | 3.400E-07 |
| 4979 | 249 | L249Y | A | 1e-07 | 2.8e-07 | 2.3e-07 | 2.033E-07 | 9.292E-08 | 1.000E-07 | 2.800E-07 |
| 4980 | 249 | L249V | A | 4.6e-07 | 3.5e-07 | 1.2e-07 | 3.100E-07 | 1.735E-07 | 1.200E-07 | 4.600E-07 |
| 4981 | 250 | V250A | A | 3.7e-07 | 1.4e-07 | 1.6e-07 | 2.233E-07 | 1.274E-07 | 1.400E-07 | 3.700E-07 |
| 4982 | 250 | V250R | A | 1.5e-07 | 2e-07 | 2.2e-07 | 1.900E-07 | 3.606E-08 | 1.500E-07 | 2.200E-07 |
| 4983 | 250 | V250N | A | 3.7e-07 | 1.7e-07 | 2.4e-07 | 2.600E-07 | 1.015E-07 | 1.700E-07 | 3.700E-07 |
| 4984 | 250 | V250D | A | 1.9e-07 | 2.3e-07 | 3.7e-07 | 2.633E-07 | 9.452E-08 | 1.900E-07 | 3.700E-07 |
| 4985 | 250 | V250C | A | 2.2e-07 | 2.4e-07 | 1.3e-07 | 1.967E-07 | 5.859E-08 | 1.300E-07 | 2.400E-07 |
| 4986 | 250 | V250E | A | 3.7e-07 | 5.8e-07 | 1.6e-07 | 3.700E-07 | 2.100E-07 | 1.600E-07 | 5.800E-07 |
| 4987 | 250 | V250Q | A | 2.5e-07 | 3.1e-07 | 8.8e-08 | 2.160E-07 | 1.148E-07 | 8.800E-08 | 3.100E-07 |
| 4988 | 250 | V250G | A | 1e-07 | 2.6e-07 | 2e-07 | 1.867E-07 | 8.083E-08 | 1.000E-07 | 2.600E-07 |
| 4989 | 250 | V250H | A | 3.1e-07 | 1.1e-07 | 1.9e-07 | 2.033E-07 | 1.007E-07 | 1.100E-07 | 3.100E-07 |
| 4990 | 250 | V250I | A | 2.9e-07 | 1.7e-07 | 8.8e-07 | 4.467E-07 | 3.800E-07 | 1.700E-07 | 8.800E-07 |
| 4991 | 250 | V250L | A | 5.4e-07 | 2.8e-07 | 3.5e-07 | 3.900E-07 | 1.345E-07 | 2.800E-07 | 5.400E-07 |
| 4992 | 250 | V250K | A | 3.2e-07 | 3.5e-07 | 4.1e-07 | 3.600E-07 | 4.583E-08 | 3.200E-07 | 4.100E-07 |
| 4993 | 250 | V250M | A | 1.8e-07 | 3.2e-07 | 2.3e-07 | 2.433E-07 | 7.095E-08 | 1.800E-07 | 3.200E-07 |
| 4994 | 250 | V250F | A | 3e-07 | 1.8e-07 | 1.8e-07 | 2.200E-07 | 6.928E-08 | 1.800E-07 | 3.000E-07 |
| 4995 | 250 | V250P | A | 3.9e-07 | 2.3e-07 | 2.1e-07 | 2.767E-07 | 9.866E-08 | 2.100E-07 | 3.900E-07 |
| 4996 | 250 | V250S | A | 1.3e-07 | 2.6e-07 | 2.9e-07 | 2.267E-07 | 8.505E-08 | 1.300E-07 | 2.900E-07 |
| 4997 | 250 | V250T | A | 1.5e-07 | 3.5e-07 | 1.1e-07 | 2.033E-07 | 1.286E-07 | 1.100E-07 | 3.500E-07 |
| 4998 | 250 | V250W | A | 8.9e-08 | 2.5e-07 | 3.4e-07 | 2.263E-07 | 1.272E-07 | 8.900E-08 | 3.400E-07 |
| 4999 | 250 | V250Y | A | 2.1e-07 | 3.9e-07 | 3.6e-07 | 3.200E-07 | 9.644E-08 | 2.100E-07 | 3.900E-07 |
| 5000 | 250 | V250V | A | 2.5e-07 | 6.2e-07 | 2.9e-07 | 3.867E-07 | 2.031E-07 | 2.500E-07 | 6.200E-07 |
| 5001 | 251 | T251A | A | 3.1e-07 | 3.1e-07 | 1.7e-07 | 2.633E-07 | 8.083E-08 | 1.700E-07 | 3.100E-07 |
| 5002 | 251 | T251R | A | 4.1e-07 | 2.8e-07 | 1.9e-07 | 2.933E-07 | 1.106E-07 | 1.900E-07 | 4.100E-07 |
| 5003 | 251 | T251N | A | 5e-07 | 2.6e-07 | 3.5e-07 | 3.700E-07 | 1.212E-07 | 2.600E-07 | 5.000E-07 |
| 5004 | 251 | T251D | A | 2.4e-07 | 2.3e-07 | 4.2e-07 | 2.967E-07 | 1.069E-07 | 2.300E-07 | 4.200E-07 |
| 5005 | 251 | T251C | A | 4.6e-07 | 2.7e-07 | 3.4e-07 | 3.567E-07 | 9.609E-08 | 2.700E-07 | 4.600E-07 |
| 5006 | 251 | T251E | A | 1.3e-07 | 2.5e-07 | 2.3e-07 | 2.033E-07 | 6.429E-08 | 1.300E-07 | 2.500E-07 |
| 5007 | 251 | T251Q | A | 2.1e-07 | 3.2e-07 | 2.9e-07 | 2.733E-07 | 5.686E-08 | 2.100E-07 | 3.200E-07 |
| 5008 | 251 | T251G | A | 3.3e-07 | 4.1e-07 | 9.2e-08 | 2.773E-07 | 1.654E-07 | 9.200E-08 | 4.100E-07 |
| 5009 | 251 | T251H | A | 2e-07 | 2.4e-07 | 2.1e-07 | 2.167E-07 | 2.082E-08 | 2.000E-07 | 2.400E-07 |
| 5010 | 251 | T251I | A | 2e-07 | 7.2e-08 | 4.2e-07 | 2.307E-07 | 1.760E-07 | 7.200E-08 | 4.200E-07 |
| 5011 | 251 | T251L | A | 2.8e-07 | 4.1e-07 | 2.9e-07 | 3.267E-07 | 7.234E-08 | 2.800E-07 | 4.100E-07 |
| 5012 | 251 | T251K | A | 3.1e-07 | 2.6e-07 | 3.6e-07 | 3.100E-07 | 5.000E-08 | 2.600E-07 | 3.600E-07 |
| 5013 | 251 | T251M | A | 3.3e-07 | 1.2e-07 | 2e-07 | 2.167E-07 | 1.060E-07 | 1.200E-07 | 3.300E-07 |
| 5014 | 251 | T251F | A | 2.2e-07 | 3.9e-07 | 2.7e-07 | 2.933E-07 | 8.737E-08 | 2.200E-07 | 3.900E-07 |

| | | | | | | | | | | |
|---|---|---|---|---|---|---|---|---|---|---|
| 5015 | 251 | T251P | A | 2.8e-07 | 3.4e-07 | 1.8e-07 | 2.667E-07 | 8.083E-08 | 1.800E-07 | 3.400E-07 |
| 5016 | 251 | T251S | A | 9.4e-08 | 2e-07 | 1.7e-07 | 1.547E-07 | 5.464E-08 | 9.400E-08 | 2.000E-07 |
| 5017 | 251 | T251T | A | 1.9e-07 | 2.4e-07 | 1.8e-07 | 2.033E-07 | 3.215E-08 | 1.800E-07 | 2.400E-07 |
| 5018 | 251 | T251W | A | 5.1e-07 | 4.6e-07 | 2.3e-07 | 4.000E-07 | 1.493E-07 | 2.300E-07 | 5.100E-07 |
| 5019 | 251 | T251Y | A | 1.9e-07 | 2.4e-07 | 1.4e-07 | 1.900E-07 | 5.000E-08 | 1.400E-07 | 2.400E-07 |
| 5020 | 251 | T251V | A | 1.5e-07 | 2.1e-07 | 4.1e-07 | 2.567E-07 | 1.361E-07 | 1.500E-07 | 4.100E-07 |
| 5021 | 252 | Y252A | A | 1.8e-07 | 9.3e-08 | 1.6e-07 | 1.443E-07 | 4.557E-08 | 9.300E-08 | 1.800E-07 |
| 5022 | 252 | Y252R | A | 2.5e-07 | 3.2e-07 | 1.9e-07 | 2.533E-07 | 6.506E-08 | 1.900E-07 | 3.200E-07 |
| 5023 | 252 | Y252N | A | 2.7e-07 | 1.9e-07 | 2.3e-07 | 2.300E-07 | 4.000E-08 | 1.900E-07 | 2.700E-07 |
| 5024 | 252 | Y252D | A | 2.1e-07 | 5e-07 | 1.3e-07 | 2.800E-07 | 1.947E-07 | 1.300E-07 | 5.000E-07 |
| 5025 | 252 | Y252C | A | 1.4e-07 | 2.9e-07 | 1e-07 | 1.767E-07 | 1.002E-07 | 1.000E-07 | 2.900E-07 |
| 5026 | 252 | Y252E | A | 2.4e-07 | 2.2e-07 | 3.3e-07 | 2.633E-07 | 5.859E-08 | 2.200E-07 | 3.300E-07 |
| 5027 | 252 | Y252Q | A | 1.2e-07 | 1.1e-07 | 1.3e-07 | 1.200E-07 | 1.000E-08 | 1.100E-07 | 1.300E-07 |
| 5028 | 252 | Y252G | A | 1.3e-07 | 2.4e-07 | 2.5e-07 | 2.067E-07 | 6.658E-08 | 1.300E-07 | 2.500E-07 |
| 5029 | 252 | Y252H | A | 1.7e-07 | 2.6e-07 | 3.3e-07 | 2.533E-07 | 8.021E-08 | 1.700E-07 | 3.300E-07 |
| 5030 | 252 | Y252I | A | 5.1e-07 | 1.5e-07 | 3.1e-07 | 3.233E-07 | 1.804E-07 | 1.500E-07 | 5.100E-07 |
| 5031 | 252 | Y252L | A | 2.9e-07 | 3.5e-07 | 4.2e-07 | 3.533E-07 | 6.506E-08 | 2.900E-07 | 4.200E-07 |
| 5032 | 252 | Y252K | A | 2.2e-07 | 2.4e-07 | 1.1e-07 | 1.900E-07 | 7.000E-08 | 1.100E-07 | 2.400E-07 |
| 5033 | 252 | Y252M | A | 2.1e-07 | 1.1e-07 | 1.7e-07 | 1.633E-07 | 5.033E-08 | 1.100E-07 | 2.100E-07 |
| 5034 | 252 | Y252F | A | 2e-07 | 2.7e-07 | 2.4e-07 | 2.367E-07 | 3.512E-08 | 2.000E-07 | 2.700E-07 |
| 5035 | 252 | Y252P | A | 8.3e-07 | 1.8e-07 | 6.8e-07 | 5.633E-07 | 3.403E-07 | 1.800E-07 | 8.300E-07 |
| 5036 | 252 | Y252S | A | 2.2e-07 | 7.7e-07 | 3.4e-07 | 4.433E-07 | 2.892E-07 | 2.200E-07 | 7.700E-07 |
| 5037 | 252 | Y252T | A | 1.4e-07 | 5.7e-07 | 1.8e-07 | 2.967E-07 | 2.376E-07 | 1.400E-07 | 5.700E-07 |
| 5038 | 252 | Y252W | A | 2.7e-07 | 5.4e-07 | 3.1e-07 | 3.733E-07 | 1.457E-07 | 2.700E-07 | 5.400E-07 |
| 5039 | 252 | Y252Y | A | 1.9e-07 | 3.3e-07 | 1.9e-07 | 2.367E-07 | 8.083E-08 | 1.900E-07 | 3.300E-07 |
| 5040 | 252 | Y252V | A | 1.4e-07 | 4.4e-07 | 2.2e-07 | 2.667E-07 | 1.553E-07 | 1.400E-07 | 4.400E-07 |
| 5041 | 253 | N253A | A | 2.2e-07 | 2.8e-07 | 3e-07 | 2.667E-07 | 4.163E-08 | 2.200E-07 | 3.000E-07 |
| 5042 | 253 | N253R | A | 3.3e-07 | 4.7e-07 | 2.2e-07 | 3.400E-07 | 1.253E-07 | 2.200E-07 | 4.700E-07 |
| 5043 | 253 | N253N | A | 3e-07 | 3e-07 | 4.7e-07 | 3.567E-07 | 9.815E-08 | 3.000E-07 | 4.700E-07 |
| 5044 | 253 | N253D | A | 3.6e-07 | 2.3e-07 | 1.9e-07 | 2.600E-07 | 8.888E-08 | 1.900E-07 | 3.600E-07 |
| 5045 | 253 | N253C | A | 9e-08 | 1.8e-07 | 4.6e-07 | 2.433E-07 | 1.930E-07 | 9.000E-08 | 4.600E-07 |
| 5046 | 253 | N253E | A | 2.4e-07 | 2.7e-07 | 4.1e-07 | 3.067E-07 | 9.074E-08 | 2.400E-07 | 4.100E-07 |
| 5047 | 253 | N253Q | A | 5.2e-07 | 2.3e-07 | 5.6e-07 | 4.367E-07 | 1.801E-07 | 2.300E-07 | 5.600E-07 |
| 5048 | 253 | N253G | A | 2e-07 | 3.3e-07 | 2e-07 | 2.433E-07 | 7.506E-08 | 2.000E-07 | 3.300E-07 |
| 5049 | 253 | N253H | A | 1.8e-07 | 4.8e-07 | 3.6e-07 | 3.400E-07 | 1.510E-07 | 1.800E-07 | 4.800E-07 |
| 5050 | 253 | N253I | A | 2.1e-07 | 2.6e-07 | 3.1e-07 | 2.600E-07 | 5.000E-08 | 2.100E-07 | 3.100E-07 |
| 5051 | 253 | N253L | A | 4e-07 | 3.5e-07 | 3.3e-07 | 3.600E-07 | 3.606E-08 | 3.300E-07 | 4.000E-07 |
| 5052 | 253 | N253K | A | 4.1e-07 | 7.2e-07 | 1.6e-07 | 4.300E-07 | 2.805E-07 | 1.600E-07 | 7.200E-07 |
| 5053 | 253 | N253M | A | 3.9e-07 | 5.6e-07 | 3.5e-07 | 4.333E-07 | 1.115E-07 | 3.500E-07 | 5.600E-07 |
| 5054 | 253 | N253F | A | 4.8e-07 | 3.4e-07 | 1.4e-07 | 3.200E-07 | 1.709E-07 | 1.400E-07 | 4.800E-07 |
| 5055 | 253 | N253P | A | 3e-07 | 3.3e-07 | 3e-07 | 3.100E-07 | 1.732E-08 | 3.000E-07 | 3.300E-07 |
| 5056 | 253 | N253S | A | 2.8e-07 | 5.1e-07 | 9.1e-07 | 5.667E-07 | 3.188E-07 | 2.800E-07 | 9.100E-07 |
| 5057 | 253 | N253T | A | 4.6e-07 | 2e-07 | 2.5e-07 | 3.033E-07 | 1.380E-07 | 2.000E-07 | 4.600E-07 |
| 5058 | 253 | N253W | A | 9.6e-08 | 2.5e-07 | 1.6e-07 | 1.687E-07 | 7.736E-08 | 9.600E-08 | 2.500E-07 |
| 5059 | 253 | N253Y | A | 2e-07 | 1.1e-07 | 1.3e-07 | 1.467E-07 | 4.726E-08 | 1.100E-07 | 2.000E-07 |
| 5060 | 253 | N253V | A | 4.2e-07 | 2.7e-07 | 3e-07 | 3.300E-07 | 7.937E-08 | 2.700E-07 | 4.200E-07 |
| 5061 | 254 | T254A | A | 2.5e-07 | 6.1e-07 | 6.3e-07 | 4.967E-07 | 2.139E-07 | 2.500E-07 | 6.300E-07 |
| 5062 | 254 | T254R | A | 2.9e-07 | 2.1e-07 | 4.7e-07 | 3.233E-07 | 1.332E-07 | 2.100E-07 | 4.700E-07 |
| 5063 | 254 | T254N | A | 2.9e-07 | 6.6e-07 | 3.3e-07 | 4.267E-07 | 2.031E-07 | 2.900E-07 | 6.600E-07 |
| 5064 | 254 | T254D | A | 1.9e-07 | 4.8e-07 | 2.4e-07 | 3.033E-07 | 1.550E-07 | 1.900E-07 | 4.800E-07 |
| 5065 | 254 | T254C | A | 2.8e-07 | 2e-07 | 2.6e-07 | 2.467E-07 | 4.163E-08 | 2.000E-07 | 2.800E-07 |
| 5066 | 254 | T254E | A | 3.9e-07 | 4.4e-07 | 2.3e-07 | 3.533E-07 | 1.097E-07 | 2.300E-07 | 4.400E-07 |
| 5067 | 254 | T254Q | A | 2.2e-07 | 1.8e-07 | 2.1e-07 | 2.033E-07 | 2.082E-08 | 1.800E-07 | 2.200E-07 |
| 5068 | 254 | T254G | A | 7e-07 | 1.6e-07 | 3.5e-07 | 4.033E-07 | 2.739E-07 | 1.600E-07 | 7.000E-07 |
| 5069 | 254 | T254H | A | 2.5e-07 | 3e-07 | 2e-07 | 2.500E-07 | 5.000E-08 | 2.000E-07 | 3.000E-07 |
| 5070 | 254 | T254I | A | 2.1e-07 | 6.8e-07 | 4.4e-07 | 4.433E-07 | 2.350E-07 | 2.100E-07 | 6.800E-07 |
| 5071 | 254 | T254L | A | 3.3e-07 | 1e-07 | 4.3e-07 | 2.867E-07 | 1.692E-07 | 1.000E-07 | 4.300E-07 |
| 5072 | 254 | T254K | A | 2.4e-07 | 5.1e-07 | 3.8e-07 | 3.767E-07 | 1.350E-07 | 2.400E-07 | 5.100E-07 |
| 5073 | 254 | T254M | A | 2.8e-07 | 4.6e-07 | 2.3e-07 | 3.233E-07 | 1.210E-07 | 2.300E-07 | 4.600E-07 |
| 5074 | 254 | T254F | A | 4.8e-07 | 1.7e-07 | 4.1e-07 | 3.533E-07 | 1.626E-07 | 1.700E-07 | 4.800E-07 |
| 5075 | 254 | T254P | A | 1.7e-07 | 4.8e-07 | 2.7e-07 | 3.067E-07 | 1.582E-07 | 1.700E-07 | 4.800E-07 |
| 5076 | 254 | T254S | A | 2.5e-07 | 3.2e-07 | 4.3e-07 | 3.333E-07 | 9.074E-08 | 2.500E-07 | 4.300E-07 |
| 5077 | 254 | T254T | A | 1.6e-07 | 4.6e-07 | 1.3e-07 | 2.500E-07 | 1.825E-07 | 1.300E-07 | 4.600E-07 |
| 5078 | 254 | T254W | A | 2.9e-07 | 4.3e-07 | 2.3e-07 | 3.167E-07 | 1.026E-07 | 2.300E-07 | 4.300E-07 |
| 5079 | 254 | T254Y | A | 2.7e-07 | 1.5e-07 | 4.5e-07 | 2.900E-07 | 1.510E-07 | 1.500E-07 | 4.500E-07 |
| 5080 | 254 | T254V | A | 2.7e-07 | 2.7e-07 | 2e-07 | 2.467E-07 | 4.041E-08 | 2.000E-07 | 2.700E-07 |
| 5081 | 255 | D255A | A | 1.9e-07 | 4.4e-07 | 2.6e-07 | 2.967E-07 | 1.290E-07 | 1.900E-07 | 4.400E-07 |
| 5082 | 255 | D255R | A | 1.6e-07 | 1.9e-07 | 2.5e-07 | 2.000E-07 | 4.583E-08 | 1.600E-07 | 2.500E-07 |
| 5083 | 255 | D255N | A | 1.6e-07 | 4.4e-07 | 1.7e-07 | 2.567E-07 | 1.589E-07 | 1.600E-07 | 4.400E-07 |
| 5084 | 255 | D255D | A | 2.3e-07 | 3.2e-07 | 2.7e-07 | 2.733E-07 | 4.509E-08 | 2.300E-07 | 3.200E-07 |
| 5085 | 255 | D255C | A | 3.3e-07 | 2.8e-07 | 2.4e-07 | 2.833E-07 | 4.509E-08 | 2.400E-07 | 3.300E-07 |
| 5086 | 255 | D255E | A | 2.3e-07 | 2.7e-07 | 1.6e-07 | 2.200E-07 | 5.568E-08 | 1.600E-07 | 2.700E-07 |
| 5087 | 255 | D255Q | A | 2.7e-07 | 1.6e-07 | 3.4e-07 | 2.567E-07 | 9.074E-08 | 1.600E-07 | 3.400E-07 |
| 5088 | 255 | D255G | A | 3.2e-07 | 2.7e-07 | 3.2e-07 | 3.033E-07 | 2.887E-08 | 2.700E-07 | 3.200E-07 |
| 5089 | 255 | D255H | A | 2.4e-07 | 5.3e-07 | 1.4e-07 | 3.033E-07 | 2.026E-07 | 1.400E-07 | 5.300E-07 |
| 5090 | 255 | D255I | A | 4.7e-07 | 2.4e-07 | 3.4e-07 | 3.500E-07 | 1.153E-07 | 2.400E-07 | 4.700E-07 |
| 5091 | 255 | D255L | A | 1.7e-07 | 2e-07 | 1.4e-07 | 1.700E-07 | 3.000E-08 | 1.400E-07 | 2.000E-07 |
| 5092 | 255 | D255K | A | 1.4e-07 | 2.2e-07 | 3e-07 | 2.200E-07 | 8.000E-08 | 1.400E-07 | 3.000E-07 |
| 5093 | 255 | D255M | A | 1.8e-07 | 9.9e-08 | 2.6e-07 | 1.797E-07 | 8.050E-08 | 9.900E-08 | 2.600E-07 |
| 5094 | 255 | D255F | A | 2.9e-07 | 2.6e-07 | 1.4e-07 | 2.300E-07 | 7.937E-08 | 1.400E-07 | 2.900E-07 |
| 5095 | 255 | D255P | A | 1.5e-07 | 3.7e-07 | 1.1e-07 | 2.100E-07 | 1.400E-07 | 1.100E-07 | 3.700E-07 |
| 5096 | 255 | D255S | A | 2.1e-07 | 1.9e-07 | 2.8e-07 | 2.267E-07 | 4.726E-08 | 1.900E-07 | 2.800E-07 |
| 5097 | 255 | D255T | A | 1e-07 | 4.3e-07 | 2.7e-07 | 2.667E-07 | 1.650E-07 | 1.000E-07 | 4.300E-07 |
| 5098 | 255 | D255W | A | 1.8e-07 | 2.3e-07 | 4.1e-07 | 2.733E-07 | 1.210E-07 | 1.800E-07 | 4.100E-07 |
| 5099 | 255 | D255Y | A | 9.2e-07 | 1.5e-07 | 2.2e-07 | 4.300E-07 | 4.258E-07 | 1.500E-07 | 9.200E-07 |

| | | | | | | | | | | |
|---|---|---|---|---|---|---|---|---|---|---|
| 5100 | 255 | D255V | A | 2.9e-07 | 2.4e-07 | 2.4e-07 | 2.567E-07 | 2.887E-08 | 2.400E-07 | 2.900E-07 |
| 5101 | 256 | T256A | A | 1.8e-07 | 3.6e-07 | 3e-07 | 2.800E-07 | 9.165E-08 | 1.800E-07 | 3.600E-07 |
| 5102 | 256 | T256R | A | 1.7e-07 | 2.4e-07 | 2.2e-07 | 2.100E-07 | 3.606E-08 | 1.700E-07 | 2.400E-07 |
| 5103 | 256 | T256N | A | 1.6e-07 | 2.8e-07 | 1.7e-07 | 2.033E-07 | 6.658E-08 | 1.600E-07 | 2.800E-07 |
| 5104 | 256 | T256D | A | 1.6e-07 | 4.4e-07 | 3.4e-07 | 3.133E-07 | 1.419E-07 | 1.600E-07 | 4.400E-07 |
| 5105 | 256 | T256C | A | 2.3e-07 | 1.3e-07 | 2.5e-07 | 2.033E-07 | 6.429E-08 | 1.300E-07 | 2.500E-07 |
| 5106 | 256 | T256E | A | 1.7e-07 | 3.4e-07 | 6.9e-07 | 4.000E-07 | 2.651E-07 | 1.700E-07 | 6.900E-07 |
| 5107 | 256 | T256Q | A | 2.9e-07 | 1.8e-07 | 5e-07 | 3.233E-07 | 1.626E-07 | 1.800E-07 | 5.000E-07 |
| 5108 | 256 | T256G | A | 2.3e-07 | 5.8e-08 | 3.4e-07 | 2.093E-07 | 1.421E-07 | 5.800E-08 | 3.400E-07 |
| 5109 | 256 | T256H | A | 1.9e-07 | 2.4e-07 | 3.1e-07 | 2.467E-07 | 6.028E-08 | 1.900E-07 | 3.100E-07 |
| 5110 | 256 | T256I | A | 3.1e-07 | 4.5e-07 | 9e-08 | 2.833E-07 | 1.815E-07 | 9.000E-08 | 4.500E-07 |
| 5111 | 256 | T256L | A | 3.2e-07 | 2.3e-07 | 2e-07 | 2.500E-07 | 6.245E-08 | 2.000E-07 | 3.200E-07 |
| 5112 | 256 | T256K | A | 3.2e-07 | 1.7e-07 | 2.3e-07 | 2.400E-07 | 7.550E-08 | 1.700E-07 | 3.200E-07 |
| 5113 | 256 | T256M | A | 2.3e-07 | 1.7e-07 | 2.2e-07 | 2.067E-07 | 3.215E-08 | 1.700E-07 | 2.300E-07 |
| 5114 | 256 | T256F | A | 3.3e-07 | 1.8e-07 | 3.5e-07 | 2.867E-07 | 9.292E-08 | 1.800E-07 | 3.500E-07 |
| 5115 | 256 | T256P | A | 2.4e-07 | 2.2e-07 | 3e-07 | 2.533E-07 | 4.163E-08 | 2.200E-07 | 3.000E-07 |
| 5116 | 256 | T256S | A | 2.2e-07 | 2.4e-07 | 3e-07 | 2.533E-07 | 4.163E-08 | 2.200E-07 | 3.000E-07 |
| 5117 | 256 | T256T | A | 2.6e-07 | 2.3e-07 | 2.2e-07 | 2.367E-07 | 2.082E-08 | 2.200E-07 | 2.600E-07 |
| 5118 | 256 | T256W | A | 1.6e-07 | 2.6e-07 | 5.5e-07 | 3.233E-07 | 2.026E-07 | 1.600E-07 | 5.500E-07 |
| 5119 | 256 | T256Y | A | 2e-07 | 5.1e-07 | 4.1e-07 | 3.733E-07 | 1.582E-07 | 2.000E-07 | 5.100E-07 |
| 5120 | 256 | T256V | A | 2.5e-07 | 1.6e-07 | 4e-07 | 2.700E-07 | 1.212E-07 | 1.600E-07 | 4.000E-07 |
| 5121 | 257 | F257A | A | 3.6e-07 | 2.4e-07 | 3.2e-07 | 3.067E-07 | 6.110E-08 | 2.400E-07 | 3.600E-07 |
| 5122 | 257 | F257R | A | 1.7e-07 | 3.8e-07 | 2e-07 | 2.500E-07 | 1.136E-07 | 1.700E-07 | 3.800E-07 |
| 5123 | 257 | F257N | A | 2e-07 | 1.7e-07 | 2.4e-07 | 2.033E-07 | 3.512E-08 | 1.700E-07 | 2.400E-07 |
| 5124 | 257 | F257D | A | 2.7e-07 | 4.3e-07 | 3e-07 | 3.333E-07 | 8.505E-08 | 2.700E-07 | 4.300E-07 |
| 5125 | 257 | F257C | A | 3.4e-07 | 3e-07 | 1.3e-07 | 2.567E-07 | 1.115E-07 | 1.300E-07 | 3.400E-07 |
| 5126 | 257 | F257E | A | 8.8e-08 | 5.6e-07 | 1.6e-07 | 2.693E-07 | 2.543E-07 | 8.800E-08 | 5.600E-07 |
| 5127 | 257 | F257Q | A | 5.6e-08 | 4.4e-07 | 2e-07 | 2.320E-07 | 1.940E-07 | 5.600E-08 | 4.400E-07 |
| 5128 | 257 | F257G | A | 2.1e-07 | 1.6e-07 | 2.1e-07 | 1.933E-07 | 2.887E-08 | 1.600E-07 | 2.100E-07 |
| 5129 | 257 | F257H | A | 2.1e-07 | 2.9e-07 | 3e-07 | 2.667E-07 | 4.933E-08 | 2.100E-07 | 3.000E-07 |
| 5130 | 257 | F257I | A | 3.8e-07 | 2.2e-07 | 2.5e-07 | 2.833E-07 | 8.505E-08 | 2.200E-07 | 3.800E-07 |
| 5131 | 257 | F257L | A | 1.8e-07 | 2.3e-07 | 9.5e-07 | 4.533E-07 | 4.309E-07 | 1.800E-07 | 9.500E-07 |
| 5132 | 257 | F257K | A | 3.4e-07 | 1.5e-07 | 2.5e-07 | 2.467E-07 | 9.504E-08 | 1.500E-07 | 3.400E-07 |
| 5133 | 257 | F257M | A | 2e-07 | 1.9e-07 | 3.3e-07 | 2.400E-07 | 7.810E-08 | 1.900E-07 | 3.300E-07 |
| 5134 | 257 | F257F | A | 2e-07 | 2.8e-07 | 4.2e-07 | 3.000E-07 | 1.114E-07 | 2.000E-07 | 4.200E-07 |
| 5135 | 257 | F257P | A | 1.2e-07 | 3e-07 | 2.8e-07 | 2.333E-07 | 9.866E-08 | 1.200E-07 | 3.000E-07 |
| 5136 | 257 | F257S | A | 3.8e-07 | 3.5e-07 | 6.2e-08 | 2.640E-07 | 1.756E-07 | 6.200E-08 | 3.800E-07 |
| 5137 | 257 | F257T | A | 2e-07 | 1.6e-07 | 1.8e-07 | 1.800E-07 | 2.000E-08 | 1.600E-07 | 2.000E-07 |
| 5138 | 257 | F257W | A | 2.8e-07 | 1.4e-07 | 1.4e-07 | 1.867E-07 | 8.083E-08 | 1.400E-07 | 2.800E-07 |
| 5139 | 257 | F257Y | A | 1.8e-07 | 1.7e-07 | 1.7e-07 | 1.733E-07 | 5.774E-09 | 1.700E-07 | 1.800E-07 |
| 5140 | 257 | F257V | A | 2e-07 | 5.8e-07 | 2.8e-07 | 3.533E-07 | 2.003E-07 | 2.000E-07 | 5.800E-07 |
| 5141 | 258 | E258A | A | 6.8e-07 | 4.8e-07 | 1.3e-07 | 4.300E-07 | 2.784E-07 | 1.300E-07 | 6.800E-07 |
| 5142 | 258 | E258R | A | 1.8e-07 | 2.7e-07 | 6.6e-07 | 3.700E-07 | 2.551E-07 | 1.800E-07 | 6.600E-07 |
| 5143 | 258 | E258N | A | 2.1e-07 | 1.5e-07 | 2.5e-07 | 2.033E-07 | 5.033E-08 | 1.500E-07 | 2.500E-07 |
| 5144 | 258 | E258D | A | 2e-07 | 4e-07 | 3.9e-07 | 3.300E-07 | 1.127E-07 | 2.000E-07 | 4.000E-07 |
| 5145 | 258 | E258C | A | 1.9e-07 | 3.1e-07 | 4.4e-07 | 3.133E-07 | 1.250E-07 | 1.900E-07 | 4.400E-07 |
| 5146 | 258 | E258E | A | 1.7e-07 | 1.4e-07 | 1.7e-07 | 1.600E-07 | 1.732E-08 | 1.400E-07 | 1.700E-07 |
| 5147 | 258 | E258Q | A | 2.5e-07 | 1.2e-07 | 4.6e-07 | 2.767E-07 | 1.716E-07 | 1.200E-07 | 4.600E-07 |
| 5148 | 258 | E258G | A | 3.3e-07 | 2.8e-07 | 3.8e-07 | 3.300E-07 | 5.000E-08 | 2.800E-07 | 3.800E-07 |
| 5149 | 258 | E258H | A | 3.4e-07 | 3.5e-07 | 1.2e-07 | 2.700E-07 | 1.300E-07 | 1.200E-07 | 3.500E-07 |
| 5150 | 258 | E258I | A | 1.8e-07 | 3.7e-07 | 1.7e-07 | 2.400E-07 | 1.127E-07 | 1.700E-07 | 3.700E-07 |
| 5151 | 258 | E258L | A | 2.3e-07 | 3.1e-07 | 2.7e-07 | 2.700E-07 | 4.000E-08 | 2.300E-07 | 3.100E-07 |
| 5152 | 258 | E258K | A | 3.7e-07 | 2.5e-07 | 1.6e-07 | 2.600E-07 | 1.054E-07 | 1.600E-07 | 3.700E-07 |
| 5153 | 258 | E258M | A | 2e-07 | 3e-07 | 4.6e-07 | 3.200E-07 | 1.311E-07 | 2.000E-07 | 4.600E-07 |
| 5154 | 258 | E258F | A | 2.7e-07 | 3.5e-07 | 3.7e-07 | 3.300E-07 | 5.292E-08 | 2.700E-07 | 3.700E-07 |
| 5155 | 258 | E258P | A | 2.5e-07 | 3.4e-07 | 3.1e-07 | 3.000E-07 | 4.583E-08 | 2.500E-07 | 3.400E-07 |
| 5156 | 258 | E258S | A | 2.1e-07 | 1.6e-07 | 2.2e-07 | 1.967E-07 | 3.215E-08 | 1.600E-07 | 2.200E-07 |
| 5157 | 258 | E258T | A | 3.1e-07 | 2.7e-07 | 3.4e-07 | 3.067E-07 | 3.512E-08 | 2.700E-07 | 3.400E-07 |
| 5158 | 258 | E258W | A | 3.5e-07 | 1.5e-07 | 1.8e-07 | 2.267E-07 | 1.079E-07 | 1.500E-07 | 3.500E-07 |
| 5159 | 258 | E258Y | A | 2.6e-07 | 6.1e-07 | 1.8e-07 | 3.500E-07 | 2.287E-07 | 1.800E-07 | 6.100E-07 |
| 5160 | 258 | E258V | A | 2e-07 | 2.4e-07 | 3.1e-07 | 2.500E-07 | 5.568E-08 | 2.000E-07 | 3.100E-07 |
| 5161 | 259 | S259A | A | 2.1e-07 | 3.2e-07 | 1.4e-07 | 2.233E-07 | 9.074E-08 | 1.400E-07 | 3.200E-07 |
| 5162 | 259 | S259R | A | 5.3e-07 | 3.6e-07 | 1.3e-07 | 3.400E-07 | 2.007E-07 | 1.300E-07 | 5.300E-07 |
| 5163 | 259 | S259N | A | 3.2e-07 | 3.3e-07 | 2.2e-07 | 2.900E-07 | 6.083E-08 | 2.200E-07 | 3.300E-07 |
| 5164 | 259 | S259D | A | 3.1e-07 | 6.6e-07 | 3.2e-07 | 4.300E-07 | 1.992E-07 | 3.100E-07 | 6.600E-07 |
| 5165 | 259 | S259C | A | 2.2e-07 | 1.3e-07 | 2.7e-07 | 2.067E-07 | 7.095E-08 | 1.300E-07 | 2.700E-07 |
| 5166 | 259 | S259E | A | 2.5e-07 | 1.8e-07 | 1.8e-07 | 2.033E-07 | 4.041E-08 | 1.800E-07 | 2.500E-07 |
| 5167 | 259 | S259Q | A | 2.3e-07 | 4.5e-07 | 1.6e-07 | 2.800E-07 | 1.513E-07 | 1.600E-07 | 4.500E-07 |
| 5168 | 259 | S259G | A | 5.7e-07 | 1.8e-07 | 2.7e-07 | 3.400E-07 | 2.042E-07 | 1.800E-07 | 5.700E-07 |
| 5169 | 259 | S259H | A | 1.3e-07 | 2e-07 | 1.5e-07 | 1.600E-07 | 3.606E-08 | 1.300E-07 | 2.000E-07 |
| 5170 | 259 | S259I | A | 3.4e-07 | 1e-07 | 4.5e-07 | 2.967E-07 | 1.790E-07 | 1.000E-07 | 4.500E-07 |
| 5171 | 259 | S259L | A | 2.8e-07 | 2.8e-07 | 1.9e-07 | 2.500E-07 | 5.196E-08 | 1.900E-07 | 2.800E-07 |
| 5172 | 259 | S259K | A | 2.8e-07 | 1.3e-07 | 1.9e-07 | 2.000E-07 | 7.550E-08 | 1.300E-07 | 2.800E-07 |
| 5173 | 259 | S259M | A | 3.2e-07 | 2.8e-07 | 2.8e-07 | 2.933E-07 | 2.309E-08 | 2.800E-07 | 3.200E-07 |
| 5174 | 259 | S259F | A | 1.4e-07 | 2e-07 | 2.3e-07 | 1.900E-07 | 4.583E-08 | 1.400E-07 | 2.300E-07 |
| 5175 | 259 | S259P | A | 4.1e-07 | 3.6e-07 | 2.9e-07 | 3.533E-07 | 6.028E-08 | 2.900E-07 | 4.100E-07 |
| 5176 | 259 | S259S | A | 2e-07 | 2.4e-07 | 3.5e-07 | 2.633E-07 | 7.767E-08 | 2.000E-07 | 3.500E-07 |
| 5177 | 259 | S259T | A | 1.2e-07 | 3.2e-07 | 3.8e-07 | 2.733E-07 | 1.361E-07 | 1.200E-07 | 3.800E-07 |
| 5178 | 259 | S259W | A | 2e-07 | 3.9e-07 | 2.3e-07 | 2.733E-07 | 1.021E-07 | 2.000E-07 | 3.900E-07 |
| 5179 | 259 | S259Y | A | 3e-07 | 2.4e-07 | 2.7e-07 | 2.700E-07 | 3.000E-08 | 2.400E-07 | 3.000E-07 |
| 5180 | 259 | S259V | A | 3.6e-07 | 2.9e-07 | 1.5e-07 | 2.667E-07 | 1.069E-07 | 1.500E-07 | 3.600E-07 |
| 5181 | 260 | M260A | A | 1.8e-07 | 2.5e-07 | 6e-07 | 3.433E-07 | 2.250E-07 | 1.800E-07 | 6.000E-07 |
| 5182 | 260 | M260R | A | 4.5e-07 | 2.2e-07 | 1.4e-07 | 2.700E-07 | 1.609E-07 | 1.400E-07 | 4.500E-07 |
| 5183 | 260 | M260N | A | 3.1e-07 | 3e-07 | 1.8e-07 | 2.633E-07 | 7.234E-08 | 1.800E-07 | 3.100E-07 |
| 5184 | 260 | M260D | A | 2e-07 | 2.3e-07 | 1.2e-07 | 1.833E-07 | 5.686E-08 | 1.200E-07 | 2.300E-07 |

| | | | | | | | | | | |
|---|---|---|---|---|---|---|---|---|---|---|
| 5185 | 260 | M260C | A | 1.7e-07 | 3.4e-07 | 1.5e-07 | 2.200E-07 | 1.044E-07 | 1.500E-07 | 3.400E-07 |
| 5186 | 260 | M260E | A | 3.1e-07 | 3e-07 | 2.3e-07 | 2.800E-07 | 4.359E-08 | 2.300E-07 | 3.100E-07 |
| 5187 | 260 | M260Q | A | 2.3e-07 | 1.3e-07 | 2.8e-07 | 2.133E-07 | 7.638E-08 | 1.300E-07 | 2.800E-07 |
| 5188 | 260 | M260G | A | 1.7e-07 | 1.4e-07 | 2.7e-07 | 1.933E-07 | 6.807E-08 | 1.400E-07 | 2.700E-07 |
| 5189 | 260 | M260H | A | 4.2e-07 | 3.1e-07 | 3.5e-07 | 3.600E-07 | 5.568E-08 | 3.100E-07 | 4.200E-07 |
| 5190 | 260 | M260I | A | 2.1e-07 | 3.4e-07 | 1.6e-07 | 2.367E-07 | 9.292E-08 | 1.600E-07 | 3.400E-07 |
| 5191 | 260 | M260L | A | 3e-07 | 8.9e-08 | 2.5e-07 | 2.130E-07 | 1.103E-07 | 8.900E-08 | 3.000E-07 |
| 5192 | 260 | M260K | A | 2.1e-07 | 2.4e-07 | 2e-07 | 2.167E-07 | 2.082E-08 | 2.000E-07 | 2.400E-07 |
| 5193 | 260 | M260M | A | 3.5e-07 | 6.3e-07 | 2.3e-07 | 4.033E-07 | 2.053E-07 | 2.300E-07 | 6.300E-07 |
| 5194 | 260 | M260F | A | 1.1e-07 | 2.6e-07 | 2.3e-07 | 2.000E-07 | 7.937E-08 | 1.100E-07 | 2.600E-07 |
| 5195 | 260 | M260P | A | 2.8e-07 | 4.9e-07 | 1.9e-07 | 3.200E-07 | 1.539E-07 | 1.900E-07 | 4.900E-07 |
| 5196 | 260 | M260S | A | 1.4e-07 | 2.1e-07 | 4.4e-07 | 2.633E-07 | 1.570E-07 | 1.400E-07 | 4.400E-07 |
| 5197 | 260 | M260T | A | 4.1e-07 | 1.5e-07 | 3.9e-07 | 3.167E-07 | 1.447E-07 | 1.500E-07 | 4.100E-07 |
| 5198 | 260 | M260W | A | 1.1e-07 | 3.2e-07 | 2.6e-07 | 2.300E-07 | 1.082E-07 | 1.100E-07 | 3.200E-07 |
| 5199 | 260 | M260Y | A | 4.1e-07 | 2.5e-07 | 1.4e-07 | 2.667E-07 | 1.358E-07 | 1.400E-07 | 4.100E-07 |
| 5200 | 260 | M260V | A | 1e-07 | 4e-07 | 7e-07 | 4.000E-07 | 3.000E-07 | 1.000E-07 | 7.000E-07 |
| 5201 | 261 | P261A | A | 3e-07 | 3.1e-07 | 1.9e-07 | 2.667E-07 | 6.658E-08 | 1.900E-07 | 3.100E-07 |
| 5202 | 261 | P261R | A | 8e-08 | 3e-07 | 2.2e-07 | 2.000E-07 | 1.114E-07 | 8.000E-08 | 3.000E-07 |
| 5203 | 261 | P261N | A | 1.8e-07 | 1.5e-07 | 1.2e-07 | 1.500E-07 | 3.000E-08 | 1.200E-07 | 1.800E-07 |
| 5204 | 261 | P261D | A | 3.5e-07 | 2.5e-07 | 1.5e-07 | 2.500E-07 | 1.000E-07 | 1.500E-07 | 3.500E-07 |
| 5205 | 261 | P261C | A | 3.1e-07 | 2.5e-07 | 4e-07 | 3.200E-07 | 7.550E-08 | 2.500E-07 | 4.000E-07 |
| 5206 | 261 | P261E | A | 4.1e-07 | 3.1e-07 | 6.4e-07 | 4.533E-07 | 1.692E-07 | 3.100E-07 | 6.400E-07 |
| 5207 | 261 | P261Q | A | 3.1e-07 | 1.7e-07 | 5.7e-07 | 3.500E-07 | 2.030E-07 | 1.700E-07 | 5.700E-07 |
| 5208 | 261 | P261G | A | 2.4e-07 | 2.5e-07 | 2.9e-07 | 2.600E-07 | 2.646E-08 | 2.400E-07 | 2.900E-07 |
| 5209 | 261 | P261H | A | 4.1e-07 | 2.7e-07 | 8.6e-08 | 2.553E-07 | 1.625E-07 | 8.600E-08 | 4.100E-07 |
| 5210 | 261 | P261I | A | 2.5e-07 | 3.5e-07 | 2.5e-07 | 2.833E-07 | 5.774E-08 | 2.500E-07 | 3.500E-07 |
| 5211 | 261 | P261L | A | 3e-07 | 3.2e-07 | 4.6e-07 | 3.600E-07 | 8.718E-08 | 3.000E-07 | 4.600E-07 |
| 5212 | 261 | P261K | A | 2.3e-07 | 2.5e-07 | 3.9e-07 | 2.900E-07 | 8.718E-08 | 2.300E-07 | 3.900E-07 |
| 5213 | 261 | P261M | A | 2.2e-07 | 3.4e-07 | 1.2e-07 | 2.267E-07 | 1.102E-07 | 1.200E-07 | 3.400E-07 |
| 5214 | 261 | P261F | A | 3.1e-07 | 1.6e-07 | 4.4e-07 | 3.033E-07 | 1.401E-07 | 1.600E-07 | 4.400E-07 |
| 5215 | 261 | P261P | A | 4.6e-07 | 2.4e-07 | 1.7e-07 | 2.900E-07 | 1.513E-07 | 1.700E-07 | 4.600E-07 |
| 5216 | 261 | P261S | A | 1.8e-07 | 3.2e-07 | 1.5e-07 | 2.167E-07 | 9.074E-08 | 1.500E-07 | 3.200E-07 |
| 5217 | 261 | P261T | A | 1.4e-07 | 2e-07 | 3.6e-07 | 2.333E-07 | 1.137E-07 | 1.400E-07 | 3.600E-07 |
| 5218 | 261 | P261W | A | 8.4e-08 | 1.7e-07 | 3.4e-07 | 1.980E-07 | 1.303E-07 | 8.400E-08 | 3.400E-07 |
| 5219 | 261 | P261Y | A | 1.6e-07 | 1.5e-07 | 3.4e-07 | 2.167E-07 | 1.069E-07 | 1.500E-07 | 3.400E-07 |
| 5220 | 261 | P261V | A | 4.4e-07 | 2.8e-07 | 9.3e-08 | 2.710E-07 | 1.737E-07 | 9.300E-08 | 4.400E-07 |
| 5221 | 262 | N262A | A | 3.3e-07 | 3.5e-07 | 1e-07 | 2.600E-07 | 1.389E-07 | 1.000E-07 | 3.500E-07 |
| 5222 | 262 | N262R | A | 2.7e-07 | 2.7e-07 | 1.9e-07 | 2.433E-07 | 4.619E-08 | 1.900E-07 | 2.700E-07 |
| 5223 | 262 | N262N | A | 2.1e-07 | 3.9e-07 | 2.4e-07 | 2.800E-07 | 9.644E-08 | 2.100E-07 | 3.900E-07 |
| 5224 | 262 | N262D | A | 2.5e-07 | 2.6e-07 | 2.5e-07 | 2.533E-07 | 5.774E-09 | 2.500E-07 | 2.600E-07 |
| 5225 | 262 | N262C | A | 5.1e-07 | 4.1e-07 | 1.8e-07 | 3.667E-07 | 1.692E-07 | 1.800E-07 | 5.100E-07 |
| 5226 | 262 | N262E | A | 2e-07 | 2.5e-07 | 2.4e-07 | 2.300E-07 | 2.646E-08 | 2.000E-07 | 2.500E-07 |
| 5227 | 262 | N262Q | A | 1.5e-07 | 1.7e-07 | 3e-07 | 2.067E-07 | 8.145E-08 | 1.500E-07 | 3.000E-07 |
| 5228 | 262 | N262G | A | 1.4e-07 | 1.3e-07 | 2.4e-07 | 1.700E-07 | 6.083E-08 | 1.300E-07 | 2.400E-07 |
| 5229 | 262 | N262H | A | 3.9e-07 | 2.5e-07 | 1.9e-07 | 2.767E-07 | 1.026E-07 | 1.900E-07 | 3.900E-07 |
| 5230 | 262 | N262I | A | 2.5e-07 | 3.4e-07 | 2.9e-07 | 2.933E-07 | 4.509E-08 | 2.500E-07 | 3.400E-07 |
| 5231 | 262 | N262L | A | 1.8e-07 | 3.8e-07 | 2.3e-07 | 2.633E-07 | 1.041E-07 | 1.800E-07 | 3.800E-07 |
| 5232 | 262 | N262K | A | 2.4e-07 | 2e-07 | 2.7e-07 | 2.367E-07 | 3.512E-08 | 2.000E-07 | 2.700E-07 |
| 5233 | 262 | N262M | A | 1.2e-07 | 3e-07 | 1.2e-07 | 1.800E-07 | 1.039E-07 | 1.200E-07 | 3.000E-07 |
| 5234 | 262 | N262F | A | 1.9e-07 | 5e-07 | 1.7e-07 | 2.867E-07 | 1.850E-07 | 1.700E-07 | 5.000E-07 |
| 5235 | 262 | N262P | A | 3.9e-07 | 1.4e-07 | 2.2e-07 | 2.500E-07 | 1.277E-07 | 1.400E-07 | 3.900E-07 |
| 5236 | 262 | N262S | A | 2.1e-07 | 3.2e-07 | 5.9e-07 | 3.733E-07 | 1.955E-07 | 2.100E-07 | 5.900E-07 |
| 5237 | 262 | N262T | A | 2.1e-07 | 1e-06 | 1.5e-07 | 4.533E-07 | 4.744E-07 | 1.500E-07 | 1.000E-06 |
| 5238 | 262 | N262W | A | 2.3e-07 | 1.4e-07 | 5.9e-07 | 3.200E-07 | 2.381E-07 | 1.400E-07 | 5.900E-07 |
| 5239 | 262 | N262Y | A | 1.8e-07 | 1.9e-07 | 3.2e-07 | 2.300E-07 | 7.810E-08 | 1.800E-07 | 3.200E-07 |
| 5240 | 262 | N262V | A | 2.8e-07 | 4e-07 | 4.2e-07 | 3.667E-07 | 7.572E-08 | 2.800E-07 | 4.200E-07 |
| 5241 | 263 | P263A | A | 5.1e-07 | 2.9e-07 | 1.4e-07 | 3.133E-07 | 1.861E-07 | 1.400E-07 | 5.100E-07 |
| 5242 | 263 | P263R | A | 1.5e-07 | 2.3e-07 | 1.4e-07 | 1.733E-07 | 4.933E-08 | 1.400E-07 | 2.300E-07 |
| 5243 | 263 | P263N | A | 1.9e-07 | 3.6e-07 | 3.4e-07 | 2.967E-07 | 9.292E-08 | 1.900E-07 | 3.600E-07 |
| 5244 | 263 | P263D | A | 3.4e-07 | 1.6e-07 | 2.9e-07 | 2.633E-07 | 9.292E-08 | 1.600E-07 | 3.400E-07 |
| 5245 | 263 | P263C | A | 4.2e-07 | 1.8e-07 | 2.8e-07 | 2.933E-07 | 1.206E-07 | 1.800E-07 | 4.200E-07 |
| 5246 | 263 | P263E | A | 2.2e-07 | 2.3e-07 | 2.4e-07 | 2.300E-07 | 1.000E-08 | 2.200E-07 | 2.400E-07 |
| 5247 | 263 | P263Q | A | 2.3e-07 | 1.7e-07 | 1.5e-07 | 1.833E-07 | 4.163E-08 | 1.500E-07 | 2.300E-07 |
| 5248 | 263 | P263G | A | 9.7e-08 | 2.6e-07 | 8e-08 | 1.457E-07 | 9.938E-08 | 8.000E-08 | 2.600E-07 |
| 5249 | 263 | P263H | A | 3.3e-07 | 2.5e-07 | 1.1e-07 | 2.300E-07 | 1.114E-07 | 1.100E-07 | 3.300E-07 |
| 5250 | 263 | P263I | A | 1.9e-07 | 4.5e-07 | 3.9e-07 | 3.433E-07 | 1.361E-07 | 1.900E-07 | 4.500E-07 |
| 5251 | 263 | P263L | A | 3.7e-07 | 2.9e-07 | 9.2e-07 | 5.267E-07 | 3.430E-07 | 2.900E-07 | 9.200E-07 |
| 5252 | 263 | P263K | A | 2.2e-07 | 1.9e-07 | 6.2e-07 | 3.433E-07 | 2.401E-07 | 1.900E-07 | 6.200E-07 |
| 5253 | 263 | P263M | A | 3.6e-07 | 2.1e-07 | 4.4e-07 | 3.367E-07 | 1.168E-07 | 2.100E-07 | 4.400E-07 |
| 5254 | 263 | P263F | A | 2.8e-07 | 1.5e-07 | 1.9e-07 | 2.067E-07 | 6.658E-08 | 1.500E-07 | 2.800E-07 |
| 5255 | 263 | P263P | A | 1.1e-07 | 2.8e-07 | 1.5e-07 | 1.800E-07 | 8.888E-08 | 1.100E-07 | 2.800E-07 |
| 5256 | 263 | P263S | A | 5.9e-07 | 2.6e-07 | 3.8e-07 | 4.100E-07 | 1.670E-07 | 2.600E-07 | 5.900E-07 |
| 5257 | 263 | P263T | A | 8.6e-08 | 3e-07 | 3e-07 | 2.287E-07 | 1.236E-07 | 8.600E-08 | 3.000E-07 |
| 5258 | 263 | P263W | A | 1.1e-07 | 2e-07 | 3.7e-07 | 2.267E-07 | 1.320E-07 | 1.100E-07 | 3.700E-07 |
| 5259 | 263 | P263Y | A | 4.3e-07 | 3.4e-07 | 4e-07 | 3.900E-07 | 4.583E-08 | 3.400E-07 | 4.300E-07 |
| 5260 | 263 | P263V | A | 4e-07 | 6e-07 | 2e-07 | 4.000E-07 | 2.000E-07 | 2.000E-07 | 6.000E-07 |
| 5261 | 264 | E264A | A | 4.3e-07 | 3e-07 | 2.7e-07 | 3.333E-07 | 8.505E-08 | 2.700E-07 | 4.300E-07 |
| 5262 | 264 | E264R | A | 4.2e-07 | 2.4e-07 | 2.8e-07 | 3.133E-07 | 9.452E-08 | 2.400E-07 | 4.200E-07 |
| 5263 | 264 | E264N | A | 4.8e-07 | 5.1e-07 | 1.9e-07 | 3.933E-07 | 1.767E-07 | 1.900E-07 | 5.100E-07 |
| 5264 | 264 | E264D | A | 2.1e-07 | 3.9e-07 | 2.8e-07 | 2.933E-07 | 9.074E-08 | 2.100E-07 | 3.900E-07 |
| 5265 | 264 | E264C | A | 3e-07 | 2.1e-07 | 2.5e-07 | 2.533E-07 | 4.509E-08 | 2.100E-07 | 3.000E-07 |
| 5266 | 264 | E264E | A | 4.4e-07 | 2.3e-07 | 1.5e-07 | 2.733E-07 | 1.498E-07 | 1.500E-07 | 4.400E-07 |
| 5267 | 264 | E264Q | A | 2.5e-07 | 1.5e-07 | 2e-07 | 2.000E-07 | 5.000E-08 | 1.500E-07 | 2.500E-07 |
| 5268 | 264 | E264G | A | 1.8e-07 | 2.2e-07 | 3.6e-07 | 2.533E-07 | 9.452E-08 | 1.800E-07 | 3.600E-07 |
| 5269 | 264 | E264H | A | 2.1e-07 | 3.1e-07 | 2.7e-07 | 2.633E-07 | 5.033E-08 | 2.100E-07 | 3.100E-07 |

| | | | | | | | | | | |
|---|---|---|---|---|---|---|---|---|---|---|
| 5270 | 264 | E264I | A | 4.6e-07 | 2.7e-07 | 1.4e-07 | 2.900E-07 | 1.609E-07 | 1.400E-07 | 4.600E-07 |
| 5271 | 264 | E264L | A | 2.7e-07 | 1.2e-07 | 1.6e-07 | 1.833E-07 | 7.767E-08 | 1.200E-07 | 2.700E-07 |
| 5272 | 264 | E264K | A | 2.3e-07 | 3.1e-07 | 2.8e-07 | 2.733E-07 | 4.041E-08 | 2.300E-07 | 3.100E-07 |
| 5273 | 264 | E264M | A | 3.4e-07 | 2.5e-07 | 3.2e-07 | 3.033E-07 | 4.726E-08 | 2.500E-07 | 3.400E-07 |
| 5274 | 264 | E264F | A | 1.4e-07 | 1.5e-07 | 1.6e-07 | 1.500E-07 | 1.000E-08 | 1.400E-07 | 1.600E-07 |
| 5275 | 264 | E264P | A | 4.1e-07 | 4.1e-07 | 7.6e-07 | 5.267E-07 | 2.021E-07 | 4.100E-07 | 7.600E-07 |
| 5276 | 264 | E264S | A | 3.1e-07 | 2.6e-07 | 1.9e-07 | 2.533E-07 | 6.028E-08 | 1.900E-07 | 3.100E-07 |
| 5277 | 264 | E264T | A | 2.5e-07 | 3.5e-07 | 3.2e-07 | 3.067E-07 | 5.132E-08 | 2.500E-07 | 3.500E-07 |
| 5278 | 264 | E264W | A | 1.4e-07 | 1.8e-07 | 4.4e-07 | 2.533E-07 | 1.629E-07 | 1.400E-07 | 4.400E-07 |
| 5279 | 264 | E264Y | A | 3.4e-07 | 1.4e-07 | 1.3e-07 | 2.033E-07 | 1.185E-07 | 1.300E-07 | 3.400E-07 |
| 5280 | 264 | E264V | A | 5.6e-07 | 3.4e-07 | 1.3e-07 | 3.433E-07 | 2.150E-07 | 1.300E-07 | 5.600E-07 |
| 5281 | 265 | G265A | A | 2.8e-07 | 2e-07 | 3.6e-07 | 2.800E-07 | 8.000E-08 | 2.000E-07 | 3.600E-07 |
| 5282 | 265 | G265R | A | 3.2e-07 | 1.8e-07 | 1.7e-07 | 2.233E-07 | 8.386E-08 | 1.700E-07 | 3.200E-07 |
| 5283 | 265 | G265N | A | 2.7e-07 | 2.3e-07 | 5.1e-07 | 3.367E-07 | 1.514E-07 | 2.300E-07 | 5.100E-07 |
| 5284 | 265 | G265D | A | 2.5e-07 | 9.6e-08 | 2.5e-07 | 1.987E-07 | 8.891E-08 | 9.600E-08 | 2.500E-07 |
| 5285 | 265 | G265C | A | 2.5e-07 | 9e-07 | 1.4e-07 | 4.300E-07 | 4.107E-07 | 1.400E-07 | 9.000E-07 |
| 5286 | 265 | G265E | A | 4.9e-07 | 3.7e-07 | 1.9e-07 | 3.500E-07 | 1.510E-07 | 1.900E-07 | 4.900E-07 |
| 5287 | 265 | G265Q | A | 8.4e-08 | 2e-07 | 4.6e-07 | 2.480E-07 | 1.925E-07 | 8.400E-08 | 4.600E-07 |
| 5288 | 265 | G265G | A | 2.5e-07 | 3.5e-07 | 4.9e-07 | 3.633E-07 | 1.206E-07 | 2.500E-07 | 4.900E-07 |
| 5289 | 265 | G265H | A | 3.1e-07 | 2.4e-07 | 1.4e-07 | 2.300E-07 | 8.544E-08 | 1.400E-07 | 3.100E-07 |
| 5290 | 265 | G265I | A | 3.2e-07 | 2.3e-07 | 3.4e-07 | 2.967E-07 | 5.859E-08 | 2.300E-07 | 3.400E-07 |
| 5291 | 265 | G265L | A | 2.1e-07 | 3e-07 | 5.1e-07 | 3.400E-07 | 1.539E-07 | 2.100E-07 | 5.100E-07 |
| 5292 | 265 | G265K | A | 1.5e-07 | 5.9e-07 | 1.3e-07 | 2.900E-07 | 2.600E-07 | 1.300E-07 | 5.900E-07 |
| 5293 | 265 | G265M | A | 3e-07 | 1.4e-07 | 3.1e-07 | 2.500E-07 | 9.539E-08 | 1.400E-07 | 3.100E-07 |
| 5294 | 265 | G265F | A | 2.6e-07 | 1.4e-07 | 1.7e-07 | 1.900E-07 | 6.245E-08 | 1.400E-07 | 2.600E-07 |
| 5295 | 265 | G265P | A | 1.7e-07 | 3.4e-07 | 2e-07 | 2.367E-07 | 9.074E-08 | 1.700E-07 | 3.400E-07 |
| 5296 | 265 | G265S | A | 5e-07 | 2.4e-07 | 3e-07 | 3.467E-07 | 1.361E-07 | 2.400E-07 | 5.000E-07 |
| 5297 | 265 | G265T | A | 4e-07 | 2.6e-07 | 3.4e-07 | 3.333E-07 | 7.024E-08 | 2.600E-07 | 4.000E-07 |
| 5298 | 265 | G265W | A | 2.7e-07 | 3.9e-07 | 1.6e-07 | 2.733E-07 | 1.150E-07 | 1.600E-07 | 3.900E-07 |
| 5299 | 265 | G265Y | A | 2.2e-07 | 2.4e-07 | 3.1e-07 | 2.567E-07 | 4.726E-08 | 2.200E-07 | 3.100E-07 |
| 5300 | 265 | G265V | A | 2.8e-07 | 2.6e-07 | 2.6e-07 | 2.667E-07 | 1.155E-08 | 2.600E-07 | 2.800E-07 |
| 5301 | 266 | R266A | A | 2.1e-07 | 2.3e-07 | 1.3e-07 | 1.900E-07 | 5.292E-08 | 1.300E-07 | 2.300E-07 |
| 5302 | 266 | R266R | A | 6.4e-07 | 3.8e-07 | 3.3e-07 | 4.500E-07 | 1.664E-07 | 3.300E-07 | 6.400E-07 |
| 5303 | 266 | R266N | A | 4e-07 | 4.7e-07 | 5.1e-07 | 4.600E-07 | 5.568E-08 | 4.000E-07 | 5.100E-07 |
| 5304 | 266 | R266D | A | 3.6e-07 | 5.7e-07 | 2.2e-07 | 3.833E-07 | 1.762E-07 | 2.200E-07 | 5.700E-07 |
| 5305 | 266 | R266C | A | 1.5e-07 | 1.7e-07 | 2.6e-07 | 1.933E-07 | 5.859E-08 | 1.500E-07 | 2.600E-07 |
| 5306 | 266 | R266E | A | 2e-07 | 2.1e-07 | 4.5e-07 | 2.867E-07 | 1.415E-07 | 2.000E-07 | 4.500E-07 |
| 5307 | 266 | R266Q | A | 1.3e-07 | 2.6e-07 | 2.8e-07 | 2.233E-07 | 8.145E-08 | 1.300E-07 | 2.800E-07 |
| 5308 | 266 | R266G | A | 2.4e-07 | 2.1e-07 | 2.1e-07 | 2.200E-07 | 1.732E-08 | 2.100E-07 | 2.400E-07 |
| 5309 | 266 | R266H | A | 3.5e-07 | 2.8e-07 | 2.8e-07 | 3.033E-07 | 4.041E-08 | 2.800E-07 | 3.500E-07 |
| 5310 | 266 | R266I | A | 4.8e-07 | 1.4e-07 | 2.6e-07 | 2.933E-07 | 1.724E-07 | 1.400E-07 | 4.800E-07 |
| 5311 | 266 | R266L | A | 2.8e-07 | 1.5e-07 | 4.7e-07 | 3.000E-07 | 1.609E-07 | 1.500E-07 | 4.700E-07 |
| 5312 | 266 | R266K | A | 3.2e-07 | 2e-07 | 2e-07 | 2.400E-07 | 6.928E-08 | 2.000E-07 | 3.200E-07 |
| 5313 | 266 | R266M | A | 3.1e-07 | 1.9e-07 | 1.4e-07 | 2.133E-07 | 8.737E-08 | 1.400E-07 | 3.100E-07 |
| 5314 | 266 | R266F | A | 2e-07 | 8.1e-07 | 1.3e-07 | 3.800E-07 | 3.740E-07 | 1.300E-07 | 8.100E-07 |
| 5315 | 266 | R266P | A | 2e-07 | 2.1e-07 | 2.3e-07 | 2.133E-07 | 1.528E-08 | 2.000E-07 | 2.300E-07 |
| 5316 | 266 | R266S | A | 2e-07 | 2e-07 | 1.9e-07 | 1.967E-07 | 5.774E-09 | 1.900E-07 | 2.000E-07 |
| 5317 | 266 | R266T | A | 3e-07 | 1.9e-07 | 2.7e-07 | 2.533E-07 | 5.686E-08 | 1.900E-07 | 3.000E-07 |
| 5318 | 266 | R266W | A | 2.1e-07 | 2.6e-07 | 1.4e-07 | 2.033E-07 | 6.028E-08 | 1.400E-07 | 2.600E-07 |
| 5319 | 266 | R266Y | A | 2e-07 | 1.7e-07 | 4e-07 | 2.567E-07 | 1.250E-07 | 1.700E-07 | 4.000E-07 |
| 5320 | 266 | R266V | A | 3.9e-07 | 1.7e-07 | 3.6e-07 | 3.067E-07 | 1.193E-07 | 1.700E-07 | 3.900E-07 |
| 5321 | 267 | Y267A | A | 4.1e-07 | 1.5e-07 | 3e-07 | 2.867E-07 | 1.305E-07 | 1.500E-07 | 4.100E-07 |
| 5322 | 267 | Y267R | A | 1.9e-07 | 4.2e-07 | 2.5e-07 | 2.867E-07 | 1.193E-07 | 1.900E-07 | 4.200E-07 |
| 5323 | 267 | Y267N | A | 1.7e-07 | 3.6e-07 | 2.6e-07 | 2.633E-07 | 9.504E-08 | 1.700E-07 | 3.600E-07 |
| 5324 | 267 | Y267D | A | 2.3e-07 | 1.7e-07 | 7.4e-07 | 3.800E-07 | 3.132E-07 | 1.700E-07 | 7.400E-07 |
| 5325 | 267 | Y267C | A | 1.2e-07 | 5.1e-07 | 3e-07 | 3.100E-07 | 1.952E-07 | 1.200E-07 | 5.100E-07 |
| 5326 | 267 | Y267E | A | 2.4e-07 | 3.3e-07 | 2.9e-07 | 2.867E-07 | 4.509E-08 | 2.400E-07 | 3.300E-07 |
| 5327 | 267 | Y267Q | A | 1.9e-07 | 3.8e-07 | 3.7e-07 | 3.133E-07 | 1.069E-07 | 1.900E-07 | 3.800E-07 |
| 5328 | 267 | Y267G | A | 1.8e-07 | 1.8e-07 | 1.8e-07 | 1.800E-07 | 0.000E+00 | 1.800E-07 | 1.800E-07 |
| 5329 | 267 | Y267H | A | 1.9e-07 | 2.4e-07 | 3.4e-07 | 2.567E-07 | 7.638E-08 | 1.900E-07 | 3.400E-07 |
| 5330 | 267 | Y267I | A | 3.7e-07 | 9.8e-08 | 5e-07 | 3.227E-07 | 2.051E-07 | 9.800E-08 | 5.000E-07 |
| 5331 | 267 | Y267L | A | 2.4e-07 | 2.6e-07 | 1.7e-07 | 2.233E-07 | 4.726E-08 | 1.700E-07 | 2.600E-07 |
| 5332 | 267 | Y267K | A | 1.7e-07 | 2.1e-07 | 1.2e-07 | 1.667E-07 | 4.509E-08 | 1.200E-07 | 2.100E-07 |
| 5333 | 267 | Y267M | A | 1.3e-07 | 2.2e-07 | 5.8e-07 | 3.100E-07 | 2.381E-07 | 1.300E-07 | 5.800E-07 |
| 5334 | 267 | Y267F | A | 9.4e-08 | 3.3e-07 | 2.5e-07 | 2.247E-07 | 1.200E-07 | 9.400E-08 | 3.300E-07 |
| 5335 | 267 | Y267P | A | 3.2e-07 | 3.9e-07 | 4.7e-07 | 3.933E-07 | 7.506E-08 | 3.200E-07 | 4.700E-07 |
| 5336 | 267 | Y267S | A | 2.1e-07 | 1.4e-07 | 3.4e-07 | 2.300E-07 | 1.015E-07 | 1.400E-07 | 3.400E-07 |
| 5337 | 267 | Y267T | A | 1.9e-07 | 1.6e-07 | 2.6e-07 | 2.033E-07 | 5.132E-08 | 1.600E-07 | 2.600E-07 |
| 5338 | 267 | Y267W | A | 2.2e-07 | 2.2e-07 | 1.9e-07 | 2.100E-07 | 1.732E-08 | 1.900E-07 | 2.200E-07 |
| 5339 | 267 | Y267Y | A | 2.1e-07 | 6e-08 | 3.6e-08 | 1.020E-07 | 9.430E-08 | 3.600E-08 | 2.100E-07 |
| 5340 | 267 | Y267V | A | 1.8e-07 | 5.5e-07 | 4e-07 | 3.767E-07 | 1.861E-07 | 1.800E-07 | 5.500E-07 |
| 5341 | 268 | T268A | A | 3e-07 | 2e-07 | 1.6e-07 | 2.200E-07 | 7.211E-08 | 1.600E-07 | 3.000E-07 |
| 5342 | 268 | T268R | A | 2.3e-07 | 3.4e-07 | 1.3e-07 | 2.333E-07 | 1.050E-07 | 1.300E-07 | 3.400E-07 |
| 5343 | 268 | T268N | A | 1.7e-07 | 4.8e-07 | 2.3e-07 | 2.933E-07 | 1.644E-07 | 1.700E-07 | 4.800E-07 |
| 5344 | 268 | T268D | A | 1.8e-07 | 2.9e-07 | 3.3e-07 | 2.667E-07 | 7.767E-08 | 1.800E-07 | 3.300E-07 |
| 5345 | 268 | T268C | A | 3.5e-07 | 3e-07 | 3.3e-07 | 3.267E-07 | 2.517E-08 | 3.000E-07 | 3.500E-07 |
| 5346 | 268 | T268E | A | 5.9e-07 | 4.5e-07 | 1.9e-07 | 4.100E-07 | 2.030E-07 | 1.900E-07 | 5.900E-07 |
| 5347 | 268 | T268Q | A | 2.6e-07 | 3.9e-07 | 2.6e-07 | 3.033E-07 | 7.506E-08 | 2.600E-07 | 3.900E-07 |
| 5348 | 268 | T268G | A | 1.5e-07 | 1.9e-07 | 3.5e-07 | 2.300E-07 | 1.058E-07 | 1.500E-07 | 3.500E-07 |
| 5349 | 268 | T268H | A | 3.8e-07 | 1.5e-07 | 2e-07 | 2.433E-07 | 1.210E-07 | 1.500E-07 | 3.800E-07 |
| 5350 | 268 | T268I | A | 1.5e-07 | 2.4e-07 | 2.4e-07 | 2.100E-07 | 5.196E-08 | 1.500E-07 | 2.400E-07 |
| 5351 | 268 | T268L | A | 1.2e-07 | 1.4e-07 | 1.6e-07 | 1.400E-07 | 2.000E-08 | 1.200E-07 | 1.600E-07 |
| 5352 | 268 | T268K | A | 2.7e-07 | 4e-07 | 2.9e-07 | 3.200E-07 | 7.000E-08 | 2.700E-07 | 4.000E-07 |
| 5353 | 268 | T268M | A | 1.8e-07 | 2.6e-07 | 1.2e-07 | 1.867E-07 | 7.024E-08 | 1.200E-07 | 2.600E-07 |
| 5354 | 268 | T268F | A | 3.3e-07 | 1.3e-07 | 2.8e-07 | 2.467E-07 | 1.041E-07 | 1.300E-07 | 3.300E-07 |

| | | | | | | | | | | |
|---|---|---|---|---|---|---|---|---|---|---|
| 5355 | 268 | T268P | A | 1.3e-07 | 3.6e-07 | 3.3e-07 | 2.733E-07 | 1.250E-07 | 1.300E-07 | 3.600E-07 |
| 5356 | 268 | T268S | A | 1.9e-07 | 2.7e-07 | 2.2e-07 | 2.267E-07 | 4.041E-08 | 1.900E-07 | 2.700E-07 |
| 5357 | 268 | T268T | A | 3.3e-07 | 1.5e-07 | 5.6e-07 | 3.467E-07 | 2.055E-07 | 1.500E-07 | 5.600E-07 |
| 5358 | 268 | T268W | A | 2.4e-07 | 1.4e-07 | 1.3e-07 | 1.700E-07 | 6.083E-08 | 1.300E-07 | 2.400E-07 |
| 5359 | 268 | T268Y | A | 2.5e-07 | 2.4e-07 | 1.8e-07 | 2.233E-07 | 3.786E-08 | 1.800E-07 | 2.500E-07 |
| 5360 | 268 | T268V | A | 2.9e-07 | 2.5e-07 | 6.7e-07 | 4.033E-07 | 2.318E-07 | 2.500E-07 | 6.700E-07 |
| 5361 | 269 | F269A | A | 3e-07 | 2e-07 | 1.7e-07 | 2.233E-07 | 6.807E-08 | 1.700E-07 | 3.000E-07 |
| 5362 | 269 | F269R | A | 1.9e-07 | 2.9e-07 | 4.7e-07 | 3.167E-07 | 1.419E-07 | 1.900E-07 | 4.700E-07 |
| 5363 | 269 | F269N | A | 2.3e-07 | 2.3e-07 | 2.5e-07 | 2.367E-07 | 1.155E-08 | 2.300E-07 | 2.500E-07 |
| 5364 | 269 | F269D | A | 2.8e-07 | 1.1e-07 | 1.3e-07 | 1.733E-07 | 9.292E-08 | 1.100E-07 | 2.800E-07 |
| 5365 | 269 | F269C | A | 2.6e-07 | 3.2e-07 | 2.4e-07 | 2.733E-07 | 4.163E-08 | 2.400E-07 | 3.200E-07 |
| 5366 | 269 | F269E | A | 2.9e-07 | 1.4e-07 | 4.3e-07 | 2.867E-07 | 1.450E-07 | 1.400E-07 | 4.300E-07 |
| 5367 | 269 | F269Q | A | 4.4e-07 | 1.3e-07 | 2.3e-07 | 2.667E-07 | 1.582E-07 | 1.300E-07 | 4.400E-07 |
| 5368 | 269 | F269G | A | 3.3e-07 | 3.8e-07 | 2e-07 | 3.033E-07 | 9.292E-08 | 2.000E-07 | 3.800E-07 |
| 5369 | 269 | F269H | A | 1.7e-07 | 4.5e-07 | 1.7e-07 | 2.633E-07 | 1.617E-07 | 1.700E-07 | 4.500E-07 |
| 5370 | 269 | F269I | A | 4e-07 | 3.2e-07 | 2.4e-07 | 3.200E-07 | 8.000E-08 | 2.400E-07 | 4.000E-07 |
| 5371 | 269 | F269L | A | 1.8e-07 | 2.6e-07 | 2.2e-07 | 2.200E-07 | 4.000E-08 | 1.800E-07 | 2.600E-07 |
| 5372 | 269 | F269K | A | 3.1e-07 | 2.5e-07 | 2.5e-07 | 2.700E-07 | 3.464E-08 | 2.500E-07 | 3.100E-07 |
| 5373 | 269 | F269M | A | 1.7e-07 | 4.3e-07 | 2.4e-07 | 2.800E-07 | 1.345E-07 | 1.700E-07 | 4.300E-07 |
| 5374 | 269 | F269F | A | 1.3e-07 | 1.7e-07 | 3.4e-07 | 2.133E-07 | 1.115E-07 | 1.300E-07 | 3.400E-07 |
| 5375 | 269 | F269P | A | 2.5e-07 | 2.1e-07 | 3.5e-07 | 2.700E-07 | 7.211E-08 | 2.100E-07 | 3.500E-07 |
| 5376 | 269 | F269S | A | 1.1e-07 | 1.7e-07 | 1.8e-07 | 1.533E-07 | 3.786E-08 | 1.100E-07 | 1.800E-07 |
| 5377 | 269 | F269T | A | 2.7e-07 | 2.8e-07 | 1.7e-07 | 2.400E-07 | 6.083E-08 | 1.700E-07 | 2.800E-07 |
| 5378 | 269 | F269W | A | 1.3e-07 | 2.4e-07 | 3.9e-07 | 2.533E-07 | 1.305E-07 | 1.300E-07 | 3.900E-07 |
| 5379 | 269 | F269Y | A | 2.6e-07 | 2.5e-07 | 1.2e-07 | 2.100E-07 | 7.810E-08 | 1.200E-07 | 2.600E-07 |
| 5380 | 269 | F269V | A | 2.8e-07 | 1.4e-07 | 3.8e-07 | 2.667E-07 | 1.206E-07 | 1.400E-07 | 3.800E-07 |
| 5381 | 270 | G270A | A | 1.5e-07 | 7.1e-07 | 2.7e-07 | 3.767E-07 | 2.948E-07 | 1.500E-07 | 7.100E-07 |
| 5382 | 270 | G270R | A | 2.2e-07 | 3.2e-07 | 1.2e-07 | 2.200E-07 | 1.000E-07 | 1.200E-07 | 3.200E-07 |
| 5383 | 270 | G270N | A | 2.3e-07 | 4.2e-07 | 2.4e-07 | 2.967E-07 | 1.069E-07 | 2.300E-07 | 4.200E-07 |
| 5384 | 270 | G270D | A | 3.2e-07 | 2.8e-07 | 3.7e-07 | 3.233E-07 | 4.509E-08 | 2.800E-07 | 3.700E-07 |
| 5385 | 270 | G270C | A | 3.1e-07 | 3.9e-07 | 2.3e-07 | 3.100E-07 | 8.000E-08 | 2.300E-07 | 3.900E-07 |
| 5386 | 270 | G270E | A | 2.7e-07 | 1.3e-07 | 8.2e-07 | 4.067E-07 | 3.647E-07 | 1.300E-07 | 8.200E-07 |
| 5387 | 270 | G270Q | A | 3e-07 | 2.8e-07 | 3.3e-07 | 3.033E-07 | 2.517E-08 | 2.800E-07 | 3.300E-07 |
| 5388 | 270 | G270G | A | 5.1e-07 | 3.3e-07 | 2.3e-07 | 3.567E-07 | 1.419E-07 | 2.300E-07 | 5.100E-07 |
| 5389 | 270 | G270H | A | 1.1e-07 | 1.1e-07 | 1.6e-07 | 1.267E-07 | 2.887E-08 | 1.100E-07 | 1.600E-07 |
| 5390 | 270 | G270I | A | 3.7e-07 | 2.2e-07 | 2.7e-07 | 2.867E-07 | 7.638E-08 | 2.200E-07 | 3.700E-07 |
| 5391 | 270 | G270L | A | 1.7e-07 | 2.2e-07 | 2.5e-07 | 2.133E-07 | 4.041E-08 | 1.700E-07 | 2.500E-07 |
| 5392 | 270 | G270K | A | 1.2e-07 | 2.5e-07 | 7.6e-08 | 1.487E-07 | 9.047E-08 | 7.600E-08 | 2.500E-07 |
| 5393 | 270 | G270M | A | 3.6e-07 | 2.4e-07 | 2.5e-07 | 2.833E-07 | 6.658E-08 | 2.400E-07 | 3.600E-07 |
| 5394 | 270 | G270F | A | 2.2e-07 | 2.9e-07 | 3.2e-07 | 2.767E-07 | 5.132E-08 | 2.200E-07 | 3.200E-07 |
| 5395 | 270 | G270P | A | 1.2e-07 | 4e-07 | 1.4e-07 | 2.200E-07 | 1.562E-07 | 1.200E-07 | 4.000E-07 |
| 5396 | 270 | G270S | A | 6.4e-08 | 3.2e-07 | 2.6e-07 | 2.147E-07 | 1.339E-07 | 6.400E-08 | 3.200E-07 |
| 5397 | 270 | G270T | A | 2.1e-07 | 1.9e-07 | 3.6e-07 | 2.533E-07 | 9.292E-08 | 1.900E-07 | 3.600E-07 |
| 5398 | 270 | G270W | A | 4.6e-07 | 2.5e-07 | 1.9e-07 | 3.000E-07 | 1.418E-07 | 1.900E-07 | 4.600E-07 |
| 5399 | 270 | G270Y | A | 3.1e-07 | 1.7e-07 | 1.7e-07 | 2.167E-07 | 8.083E-08 | 1.700E-07 | 3.100E-07 |
| 5400 | 270 | G270V | A | 2.9e-07 | 1.6e-07 | 6.7e-07 | 3.733E-07 | 2.650E-07 | 1.600E-07 | 6.700E-07 |
| 5401 | 271 | A271A | A | 1.6e-07 | 2.8e-07 | 2.7e-07 | 2.367E-07 | 6.658E-08 | 1.600E-07 | 2.800E-07 |
| 5402 | 271 | A271R | A | 8e-07 | 3.8e-07 | 3.1e-07 | 4.967E-07 | 2.650E-07 | 3.100E-07 | 8.000E-07 |
| 5403 | 271 | A271N | A | 2.7e-07 | 1.8e-07 | 2e-07 | 2.167E-07 | 4.726E-08 | 1.800E-07 | 2.700E-07 |
| 5404 | 271 | A271D | A | 2.9e-07 | 2.7e-07 | 1.5e-07 | 2.367E-07 | 7.572E-08 | 1.500E-07 | 2.900E-07 |
| 5405 | 271 | A271C | A | 2.4e-07 | 2.6e-07 | 2.3e-07 | 2.433E-07 | 1.528E-08 | 2.300E-07 | 2.600E-07 |
| 5406 | 271 | A271E | A | 3.2e-07 | 1.7e-07 | 3.9e-07 | 2.933E-07 | 1.124E-07 | 1.700E-07 | 3.900E-07 |
| 5407 | 271 | A271Q | A | 2.6e-07 | 2.1e-07 | 2.5e-07 | 2.400E-07 | 2.646E-08 | 2.100E-07 | 2.600E-07 |
| 5408 | 271 | A271G | A | 2.5e-07 | 2.7e-07 | 3.4e-07 | 2.867E-07 | 4.726E-08 | 2.500E-07 | 3.400E-07 |
| 5409 | 271 | A271H | A | 3.5e-07 | 3.1e-07 | 3.7e-07 | 3.433E-07 | 3.055E-08 | 3.100E-07 | 3.700E-07 |
| 5410 | 271 | A271I | A | 3.9e-07 | 1.7e-07 | 3.1e-07 | 2.900E-07 | 1.114E-07 | 1.700E-07 | 3.900E-07 |
| 5411 | 271 | A271L | A | 2.5e-07 | 2.8e-07 | 2.8e-07 | 2.700E-07 | 1.732E-08 | 2.500E-07 | 2.800E-07 |
| 5412 | 271 | A271K | A | 2e-07 | 2.4e-07 | 2.5e-07 | 2.300E-07 | 2.646E-08 | 2.000E-07 | 2.500E-07 |
| 5413 | 271 | A271M | A | 1.6e-07 | 3.7e-07 | 2.2e-07 | 2.500E-07 | 1.082E-07 | 1.600E-07 | 3.700E-07 |
| 5414 | 271 | A271F | A | 2.3e-07 | 2.3e-07 | 3.7e-07 | 2.767E-07 | 8.083E-08 | 2.300E-07 | 3.700E-07 |
| 5415 | 271 | A271P | A | 2.6e-07 | 3.2e-07 | 1.9e-07 | 2.567E-07 | 6.506E-08 | 1.900E-07 | 3.200E-07 |
| 5416 | 271 | A271S | A | 9.7e-08 | 5.5e-07 | 2.6e-07 | 3.023E-07 | 2.294E-07 | 9.700E-08 | 5.500E-07 |
| 5417 | 271 | A271T | A | 1.7e-07 | 3.4e-07 | 1.6e-07 | 2.233E-07 | 1.012E-07 | 1.600E-07 | 3.400E-07 |
| 5418 | 271 | A271W | A | 2.9e-07 | 1.8e-07 | 2.4e-07 | 2.367E-07 | 5.508E-08 | 1.800E-07 | 2.900E-07 |
| 5419 | 271 | A271Y | A | 5e-07 | 1.6e-07 | 1.3e-07 | 2.633E-07 | 2.055E-07 | 1.300E-07 | 5.000E-07 |
| 5420 | 271 | A271V | A | 3.1e-07 | 1.2e-07 | 2.7e-07 | 2.333E-07 | 1.002E-07 | 1.200E-07 | 3.100E-07 |
| 5421 | 272 | S272A | A | 6e-07 | 3.5e-07 | 1.6e-07 | 3.700E-07 | 2.207E-07 | 1.600E-07 | 6.000E-07 |
| 5422 | 272 | S272R | A | 1.5e-07 | 2.5e-07 | 2.8e-07 | 2.267E-07 | 6.807E-08 | 1.500E-07 | 2.800E-07 |
| 5423 | 272 | S272N | A | 9.8e-08 | 2.1e-07 | 5.4e-07 | 2.827E-07 | 2.298E-07 | 9.800E-08 | 5.400E-07 |
| 5424 | 272 | S272D | A | 4.8e-07 | 2.4e-07 | 4.1e-07 | 3.767E-07 | 1.234E-07 | 2.400E-07 | 4.800E-07 |
| 5425 | 272 | S272C | A | 3e-07 | 1.7e-07 | 5.7e-07 | 3.467E-07 | 2.040E-07 | 1.700E-07 | 5.700E-07 |
| 5426 | 272 | S272E | A | 2.7e-07 | 3.1e-07 | 4.9e-07 | 3.567E-07 | 1.172E-07 | 2.700E-07 | 4.900E-07 |
| 5427 | 272 | S272Q | A | 2e-07 | 1.9e-07 | 5.9e-07 | 3.267E-07 | 2.281E-07 | 1.900E-07 | 5.900E-07 |
| 5428 | 272 | S272G | A | 3.4e-07 | 2.6e-07 | 2.4e-07 | 2.800E-07 | 5.292E-08 | 2.400E-07 | 3.400E-07 |
| 5429 | 272 | S272H | A | 2.5e-07 | 2.4e-07 | 2.9e-07 | 2.600E-07 | 2.646E-08 | 2.400E-07 | 2.900E-07 |
| 5430 | 272 | S272I | A | 4.5e-07 | 2.5e-07 | 2.9e-07 | 3.300E-07 | 1.058E-07 | 2.500E-07 | 4.500E-07 |
| 5431 | 272 | S272L | A | 1.6e-07 | 3.5e-07 | 2.1e-07 | 2.400E-07 | 9.849E-08 | 1.600E-07 | 3.500E-07 |
| 5432 | 272 | S272K | A | 8.5e-07 | 3.9e-07 | 4.5e-07 | 5.633E-07 | 2.501E-07 | 3.900E-07 | 8.500E-07 |
| 5433 | 272 | S272M | A | 1.7e-07 | 5.4e-07 | 1.3e-07 | 2.800E-07 | 2.261E-07 | 1.300E-07 | 5.400E-07 |
| 5434 | 272 | S272F | A | 4.6e-07 | 3.6e-07 | 3e-07 | 3.733E-07 | 8.083E-08 | 3.000E-07 | 4.600E-07 |
| 5435 | 272 | S272P | A | 3.2e-07 | 3e-07 | 2.4e-07 | 2.867E-07 | 4.163E-08 | 2.400E-07 | 3.200E-07 |
| 5436 | 272 | S272S | A | 1.1e-07 | 3.4e-07 | 1.2e-07 | 1.900E-07 | 1.300E-07 | 1.100E-07 | 3.400E-07 |
| 5437 | 272 | S272T | A | 1.6e-07 | 4.9e-07 | 2.4e-07 | 2.967E-07 | 1.721E-07 | 1.600E-07 | 4.900E-07 |
| 5438 | 272 | S272W | A | 2.1e-07 | 1.7e-07 | 1.1e-06 | 4.933E-07 | 5.258E-07 | 1.700E-07 | 1.100E-06 |
| 5439 | 272 | S272Y | A | 1.6e-07 | 2e-07 | 3.6e-07 | 2.400E-07 | 1.058E-07 | 1.600E-07 | 3.600E-07 |

| | | | | | | | | | | |
|---|---|---|---|---|---|---|---|---|---|---|
| 5440 | 272 | S272V | A | 6.4e-07 | 3.3e-07 | 2.6e-07 | 4.100E-07 | 2.022E-07 | 2.600E-07 | 6.400E-07 |
| 5441 | 273 | C273A | A | 2.7e-07 | 4.8e-07 | 2.6e-07 | 3.367E-07 | 1.242E-07 | 2.600E-07 | 4.800E-07 |
| 5442 | 273 | C273R | A | 1e-07 | 1.3e-07 | 2.2e-07 | 1.500E-07 | 6.245E-08 | 1.000E-07 | 2.200E-07 |
| 5443 | 273 | C273N | A | 1.3e-07 | 3.2e-07 | 2e-07 | 2.167E-07 | 9.609E-08 | 1.300E-07 | 3.200E-07 |
| 5444 | 273 | C273D | A | 2.7e-07 | 2.5e-07 | 2.1e-07 | 2.433E-07 | 3.055E-08 | 2.100E-07 | 2.700E-07 |
| 5445 | 273 | C273C | A | 2e-07 | 3.4e-07 | 1.8e-07 | 2.400E-07 | 8.718E-08 | 1.800E-07 | 3.400E-07 |
| 5446 | 273 | C273E | A | 2.3e-07 | 4.4e-07 | 3.2e-07 | 3.300E-07 | 1.054E-07 | 2.300E-07 | 4.400E-07 |
| 5447 | 273 | C273Q | A | 4e-07 | 1.7e-07 | 9.8e-08 | 2.227E-07 | 1.577E-07 | 9.800E-08 | 4.000E-07 |
| 5448 | 273 | C273G | A | 2e-07 | 1.1e-07 | 2.8e-07 | 1.967E-07 | 8.505E-08 | 1.100E-07 | 2.800E-07 |
| 5449 | 273 | C273H | A | 2.1e-07 | 2e-07 | 2.8e-07 | 2.300E-07 | 4.359E-08 | 2.000E-07 | 2.800E-07 |
| 5450 | 273 | C273I | A | 2.4e-07 | 2.5e-07 | 4.2e-07 | 3.033E-07 | 1.012E-07 | 2.400E-07 | 4.200E-07 |
| 5451 | 273 | C273L | A | 4.4e-07 | 2.7e-07 | 3.7e-07 | 3.600E-07 | 8.544E-08 | 2.700E-07 | 4.400E-07 |
| 5452 | 273 | C273K | A | 5.4e-07 | 1.4e-07 | 3.1e-07 | 3.300E-07 | 2.007E-07 | 1.400E-07 | 5.400E-07 |
| 5453 | 273 | C273M | A | 2.2e-07 | 3.2e-07 | 4.7e-07 | 3.367E-07 | 1.258E-07 | 2.200E-07 | 4.700E-07 |
| 5454 | 273 | C273F | A | 2.1e-07 | 3.2e-07 | 2e-07 | 2.433E-07 | 6.658E-08 | 2.000E-07 | 3.200E-07 |
| 5455 | 273 | C273P | A | 4.7e-07 | 4.3e-07 | 1.9e-07 | 3.633E-07 | 1.514E-07 | 1.900E-07 | 4.700E-07 |
| 5456 | 273 | C273S | A | 1.8e-07 | 1.5e-07 | 4.2e-07 | 2.500E-07 | 1.480E-07 | 1.500E-07 | 4.200E-07 |
| 5457 | 273 | C273T | A | 1.1e-07 | 7.2e-08 | 2.2e-07 | 1.340E-07 | 7.686E-08 | 7.200E-08 | 2.200E-07 |
| 5458 | 273 | C273W | A | 3.3e-07 | 3.4e-07 | 3e-07 | 3.233E-07 | 2.082E-08 | 3.000E-07 | 3.400E-07 |
| 5459 | 273 | C273Y | A | 2e-07 | 3.9e-07 | 6.1e-07 | 4.000E-07 | 2.052E-07 | 2.000E-07 | 6.100E-07 |
| 5460 | 273 | C273V | A | 4.6e-07 | 2.3e-07 | 2.2e-07 | 3.033E-07 | 1.358E-07 | 2.200E-07 | 4.600E-07 |
| 5461 | 274 | V274A | A | 2.8e-07 | 2.4e-07 | 2e-07 | 2.400E-07 | 4.000E-08 | 2.000E-07 | 2.800E-07 |
| 5462 | 274 | V274R | A | 3.4e-07 | 3.5e-07 | 3.6e-07 | 3.500E-07 | 1.000E-08 | 3.400E-07 | 3.600E-07 |
| 5463 | 274 | V274N | A | 2.4e-07 | 9.1e-08 | 1.7e-07 | 1.670E-07 | 7.455E-08 | 9.100E-08 | 2.400E-07 |
| 5464 | 274 | V274D | A | 2.2e-07 | 1.9e-07 | 3.4e-07 | 2.500E-07 | 7.937E-08 | 1.900E-07 | 3.400E-07 |
| 5465 | 274 | V274C | A | 1.5e-07 | 1.1e-07 | 1.6e-07 | 1.400E-07 | 2.646E-08 | 1.100E-07 | 1.600E-07 |
| 5466 | 274 | V274E | A | 4.3e-07 | 4.4e-07 | 2.6e-07 | 3.767E-07 | 1.012E-07 | 2.600E-07 | 4.400E-07 |
| 5467 | 274 | V274Q | A | 1.8e-07 | 1e-07 | 4e-07 | 2.267E-07 | 1.553E-07 | 1.000E-07 | 4.000E-07 |
| 5468 | 274 | V274G | A | 1.5e-07 | 2e-07 | 8.3e-08 | 1.443E-07 | 5.871E-08 | 8.300E-08 | 2.000E-07 |
| 5469 | 274 | V274H | A | 2.2e-07 | 1.2e-07 | 1.9e-07 | 1.767E-07 | 5.132E-08 | 1.200E-07 | 2.200E-07 |
| 5470 | 274 | V274I | A | 4.2e-07 | 2.9e-07 | 2.4e-07 | 3.167E-07 | 9.292E-08 | 2.400E-07 | 4.200E-07 |
| 5471 | 274 | V274L | A | 4.8e-07 | 2.1e-07 | 1.6e-07 | 2.833E-07 | 1.721E-07 | 1.600E-07 | 4.800E-07 |
| 5472 | 274 | V274K | A | 1.5e-07 | 2.8e-07 | 4.1e-07 | 2.800E-07 | 1.300E-07 | 1.500E-07 | 4.100E-07 |
| 5473 | 274 | V274M | A | 2e-07 | 3.1e-07 | 4.2e-07 | 3.100E-07 | 1.100E-07 | 2.000E-07 | 4.200E-07 |
| 5474 | 274 | V274F | A | 1.5e-07 | 2.9e-07 | 2.4e-07 | 2.267E-07 | 7.095E-08 | 1.500E-07 | 2.900E-07 |
| 5475 | 274 | V274P | A | 1.5e-07 | 3e-07 | 4.7e-07 | 3.067E-07 | 1.601E-07 | 1.500E-07 | 4.700E-07 |
| 5476 | 274 | V274S | A | 1.1e-07 | 3.7e-07 | 3.7e-07 | 2.833E-07 | 1.501E-07 | 1.100E-07 | 3.700E-07 |
| 5477 | 274 | V274T | A | 3.8e-07 | 2.4e-07 | 3.1e-07 | 3.100E-07 | 7.000E-08 | 2.400E-07 | 3.800E-07 |
| 5478 | 274 | V274W | A | 1.7e-07 | 3.4e-07 | 3.6e-07 | 2.900E-07 | 1.044E-07 | 1.700E-07 | 3.600E-07 |
| 5479 | 274 | V274Y | A | 3.8e-07 | 1.9e-07 | 2.2e-07 | 2.633E-07 | 1.021E-07 | 1.900E-07 | 3.800E-07 |
| 5480 | 274 | V274V | A | 2.5e-07 | 3.1e-07 | 1.9e-07 | 2.500E-07 | 6.000E-08 | 1.900E-07 | 3.100E-07 |
| 5481 | 275 | T275A | A | 4.5e-07 | 5e-07 | 9.1e-08 | 3.470E-07 | 2.231E-07 | 9.100E-08 | 5.000E-07 |
| 5482 | 275 | T275R | A | 2.6e-07 | 2.5e-07 | 2.6e-07 | 2.567E-07 | 5.774E-09 | 2.500E-07 | 2.600E-07 |
| 5483 | 275 | T275N | A | 1.7e-07 | 2e-07 | 2.1e-07 | 1.933E-07 | 2.082E-08 | 1.700E-07 | 2.100E-07 |
| 5484 | 275 | T275D | A | 1.9e-07 | 2.9e-07 | 3.7e-07 | 2.833E-07 | 9.018E-08 | 1.900E-07 | 3.700E-07 |
| 5485 | 275 | T275C | A | 2.1e-07 | 3.3e-07 | 4e-07 | 3.133E-07 | 9.609E-08 | 2.100E-07 | 4.000E-07 |
| 5486 | 275 | T275E | A | 3.4e-07 | 3.3e-07 | 3.3e-07 | 3.333E-07 | 5.774E-09 | 3.300E-07 | 3.400E-07 |
| 5487 | 275 | T275Q | A | 2.2e-07 | 1.6e-07 | 3.3e-07 | 2.367E-07 | 8.622E-08 | 1.600E-07 | 3.300E-07 |
| 5488 | 275 | T275G | A | 1.7e-07 | 2.7e-07 | 4.8e-07 | 3.067E-07 | 1.582E-07 | 1.700E-07 | 4.800E-07 |
| 5489 | 275 | T275H | A | 1.4e-07 | 3.5e-07 | 4e-07 | 2.967E-07 | 1.380E-07 | 1.400E-07 | 4.000E-07 |
| 5490 | 275 | T275I | A | 7.5e-07 | 2.5e-07 | 1.9e-07 | 3.967E-07 | 3.075E-07 | 1.900E-07 | 7.500E-07 |
| 5491 | 275 | T275L | A | 2.1e-07 | 2e-07 | 2.5e-07 | 2.200E-07 | 2.646E-08 | 2.000E-07 | 2.500E-07 |
| 5492 | 275 | T275K | A | 3.6e-07 | 3.2e-07 | 1.5e-07 | 2.767E-07 | 1.115E-07 | 1.500E-07 | 3.600E-07 |
| 5493 | 275 | T275M | A | 1.4e-07 | 2.9e-07 | 2.1e-07 | 2.133E-07 | 7.506E-08 | 1.400E-07 | 2.900E-07 |
| 5494 | 275 | T275F | A | 2.2e-07 | 3.7e-07 | 2.1e-07 | 2.667E-07 | 8.963E-08 | 2.100E-07 | 3.700E-07 |
| 5495 | 275 | T275P | A | 1.2e-07 | 1.7e-07 | 3.7e-07 | 2.200E-07 | 1.323E-07 | 1.200E-07 | 3.700E-07 |
| 5496 | 275 | T275S | A | 4.1e-07 | 8.7e-08 | 4.3e-07 | 3.090E-07 | 1.925E-07 | 8.700E-08 | 4.300E-07 |
| 5497 | 275 | T275T | A | 1.4e-07 | 1.8e-07 | 1.2e-07 | 1.467E-07 | 3.055E-08 | 1.200E-07 | 1.800E-07 |
| 5498 | 275 | T275W | A | 1.2e-07 | 2.9e-07 | 3e-07 | 2.367E-07 | 1.012E-07 | 1.200E-07 | 3.000E-07 |
| 5499 | 275 | T275Y | A | 3e-07 | 1.5e-07 | 2.8e-07 | 2.433E-07 | 8.145E-08 | 1.500E-07 | 3.000E-07 |
| 5500 | 275 | T275V | A | 2.9e-07 | 1.7e-07 | 2.9e-07 | 2.500E-07 | 6.928E-08 | 1.700E-07 | 2.900E-07 |
| 5501 | 276 | A276A | A | 1.5e-07 | 3e-07 | 2.9e-07 | 2.467E-07 | 8.386E-08 | 1.500E-07 | 3.000E-07 |
| 5502 | 276 | A276R | A | 2e-07 | 4.8e-07 | 3.1e-07 | 3.300E-07 | 1.411E-07 | 2.000E-07 | 4.800E-07 |
| 5503 | 276 | A276N | A | 1.7e-07 | 4.8e-07 | 2.9e-07 | 3.133E-07 | 1.563E-07 | 1.700E-07 | 4.800E-07 |
| 5504 | 276 | A276D | A | 2.6e-07 | 3.6e-07 | 2.2e-07 | 2.800E-07 | 7.211E-08 | 2.200E-07 | 3.600E-07 |
| 5505 | 276 | A276C | A | 2.8e-07 | 3.5e-07 | 1.7e-07 | 2.667E-07 | 9.074E-08 | 1.700E-07 | 3.500E-07 |
| 5506 | 276 | A276E | A | 4.6e-07 | 3.1e-07 | 6.1e-07 | 4.600E-07 | 1.500E-07 | 3.100E-07 | 6.100E-07 |
| 5507 | 276 | A276Q | A | 1.2e-07 | 2.9e-07 | 2.6e-07 | 2.233E-07 | 9.074E-08 | 1.200E-07 | 2.900E-07 |
| 5508 | 276 | A276G | A | 3.6e-07 | 1.9e-07 | 2.2e-07 | 2.567E-07 | 9.074E-08 | 1.900E-07 | 3.600E-07 |
| 5509 | 276 | A276H | A | 2.5e-07 | 2.2e-07 | 3.1e-07 | 2.600E-07 | 4.583E-08 | 2.200E-07 | 3.100E-07 |
| 5510 | 276 | A276I | A | 5.5e-07 | 3.4e-07 | 3.1e-07 | 4.000E-07 | 1.308E-07 | 3.100E-07 | 5.500E-07 |
| 5511 | 276 | A276L | A | 4.1e-07 | 1.7e-07 | 1.2e-07 | 2.333E-07 | 1.550E-07 | 1.200E-07 | 4.100E-07 |
| 5512 | 276 | A276K | A | 2.6e-07 | 3.3e-07 | 1.7e-07 | 2.533E-07 | 8.021E-08 | 1.700E-07 | 3.300E-07 |
| 5513 | 276 | A276M | A | 3.1e-07 | 1.2e-07 | 3.9e-07 | 2.733E-07 | 1.387E-07 | 1.200E-07 | 3.900E-07 |
| 5514 | 276 | A276F | A | 1.7e-07 | 4.3e-07 | 2.4e-07 | 2.800E-07 | 1.345E-07 | 1.700E-07 | 4.300E-07 |
| 5515 | 276 | A276P | A | 3.1e-07 | 3.1e-07 | 2.4e-07 | 2.867E-07 | 4.041E-08 | 2.400E-07 | 3.100E-07 |
| 5516 | 276 | A276S | A | 1.7e-07 | 1.9e-07 | 4.2e-07 | 2.600E-07 | 1.389E-07 | 1.700E-07 | 4.200E-07 |
| 5517 | 276 | A276T | A | 1.7e-07 | 2.2e-07 | 1.8e-07 | 1.900E-07 | 2.646E-08 | 1.700E-07 | 2.200E-07 |
| 5518 | 276 | A276W | A | 1.8e-07 | 2.3e-07 | 6.1e-07 | 3.400E-07 | 2.352E-07 | 1.800E-07 | 6.100E-07 |
| 5519 | 276 | A276Y | A | 2.3e-07 | 2.8e-07 | 2.8e-07 | 2.633E-07 | 2.887E-08 | 2.300E-07 | 2.800E-07 |
| 5520 | 276 | A276V | A | 1.4e-07 | 4.1e-07 | 3.2e-07 | 2.900E-07 | 1.375E-07 | 1.400E-07 | 4.100E-07 |
| 5521 | 277 | C277A | A | 1.5e-07 | 6e-07 | 1.9e-07 | 3.133E-07 | 2.491E-07 | 1.500E-07 | 6.000E-07 |
| 5522 | 277 | C277R | A | 1.3e-07 | 4e-07 | 2.3e-07 | 2.533E-07 | 1.365E-07 | 1.300E-07 | 4.000E-07 |
| 5523 | 277 | C277N | A | 3e-07 | 2.3e-07 | 2.2e-07 | 2.500E-07 | 4.359E-08 | 2.200E-07 | 3.000E-07 |
| 5524 | 277 | C277D | A | 3.2e-07 | 3.6e-07 | 1.7e-07 | 2.833E-07 | 1.002E-07 | 1.700E-07 | 3.600E-07 |

| | | | | | | | | | | |
|---|---|---|---|---|---|---|---|---|---|---|
| 5525 | 277 | C277C | A | 3.3e-07 | 3.2e-07 | 3.3e-07 | 3.267E-07 | 5.774E-09 | 3.200E-07 | 3.300E-07 |
| 5526 | 277 | C277E | A | 1.9e-07 | 2.9e-07 | 1.6e-07 | 2.133E-07 | 6.807E-08 | 1.600E-07 | 2.900E-07 |
| 5527 | 277 | C277Q | A | 9.5e-08 | 3.4e-07 | 1.1e-07 | 1.817E-07 | 1.373E-07 | 9.500E-08 | 3.400E-07 |
| 5528 | 277 | C277G | A | 3.4e-07 | 1.4e-07 | 4e-07 | 2.933E-07 | 1.361E-07 | 1.400E-07 | 4.000E-07 |
| 5529 | 277 | C277H | A | 2.2e-07 | 2.1e-07 | 2.8e-07 | 2.367E-07 | 3.786E-08 | 2.100E-07 | 2.800E-07 |
| 5530 | 277 | C277I | A | 2.3e-07 | 3e-07 | 2.1e-07 | 2.467E-07 | 4.726E-08 | 2.100E-07 | 3.000E-07 |
| 5531 | 277 | C277L | A | 3.5e-07 | 1.9e-07 | 2.3e-07 | 2.567E-07 | 8.327E-08 | 1.900E-07 | 3.500E-07 |
| 5532 | 277 | C277K | A | 2.5e-07 | 3.2e-07 | 2.6e-07 | 2.767E-07 | 3.786E-08 | 2.500E-07 | 3.200E-07 |
| 5533 | 277 | C277M | A | 4.1e-07 | 2e-07 | 1.5e-07 | 2.533E-07 | 1.380E-07 | 1.500E-07 | 4.100E-07 |
| 5534 | 277 | C277F | A | 1.8e-07 | 1.8e-07 | 2.8e-07 | 2.133E-07 | 5.774E-08 | 1.800E-07 | 2.800E-07 |
| 5535 | 277 | C277P | A | 2.5e-07 | 5.3e-07 | 1.8e-07 | 3.200E-07 | 1.852E-07 | 1.800E-07 | 5.300E-07 |
| 5536 | 277 | C277S | A | 2.2e-07 | 1.6e-07 | 4.6e-07 | 2.800E-07 | 1.587E-07 | 1.600E-07 | 4.600E-07 |
| 5537 | 277 | C277T | A | 3.9e-07 | 1.7e-07 | 2.4e-07 | 2.667E-07 | 1.124E-07 | 1.700E-07 | 3.900E-07 |
| 5538 | 277 | C277W | A | 2.8e-07 | 1.2e-07 | 2.3e-07 | 2.100E-07 | 8.185E-08 | 1.200E-07 | 2.800E-07 |
| 5539 | 277 | C277Y | A | 4.7e-07 | 5.7e-07 | 3.4e-07 | 4.600E-07 | 1.153E-07 | 3.400E-07 | 5.700E-07 |
| 5540 | 277 | C277V | A | 1.7e-07 | 3.5e-07 | 5e-07 | 3.400E-07 | 1.652E-07 | 1.700E-07 | 5.000E-07 |
| 5541 | 278 | P278A | A | 2.6e-07 | 2.4e-07 | 1.6e-07 | 2.200E-07 | 5.292E-08 | 1.600E-07 | 2.600E-07 |
| 5542 | 278 | P278R | A | 1.5e-07 | 1.4e-07 | 7.9e-07 | 3.600E-07 | 3.724E-07 | 1.400E-07 | 7.900E-07 |
| 5543 | 278 | P278N | A | 2.2e-07 | 1.5e-07 | 2.3e-07 | 2.000E-07 | 4.359E-08 | 1.500E-07 | 2.300E-07 |
| 5544 | 278 | P278D | A | 1.3e-07 | 2.4e-07 | 3.5e-07 | 2.400E-07 | 1.100E-07 | 1.300E-07 | 3.500E-07 |
| 5545 | 278 | P278C | A | 2.7e-07 | 3.3e-07 | 3.7e-07 | 3.233E-07 | 5.033E-08 | 2.700E-07 | 3.700E-07 |
| 5546 | 278 | P278E | A | 4.2e-07 | 3.5e-07 | 3.5e-07 | 3.733E-07 | 4.041E-08 | 3.500E-07 | 4.200E-07 |
| 5547 | 278 | P278Q | A | 2.6e-07 | 1.6e-07 | 3.5e-07 | 2.567E-07 | 9.504E-08 | 1.600E-07 | 3.500E-07 |
| 5548 | 278 | P278G | A | 3.5e-07 | 6.2e-07 | 1.4e-07 | 3.700E-07 | 2.406E-07 | 1.400E-07 | 6.200E-07 |
| 5549 | 278 | P278H | A | 1.5e-07 | 2.4e-07 | 2.7e-07 | 2.200E-07 | 6.245E-08 | 1.500E-07 | 2.700E-07 |
| 5550 | 278 | P278I | A | 2.1e-07 | 2.7e-07 | 3.6e-07 | 2.800E-07 | 7.550E-08 | 2.100E-07 | 3.600E-07 |
| 5551 | 278 | P278L | A | 2.2e-07 | 5.1e-07 | 1.9e-07 | 3.067E-07 | 1.767E-07 | 1.900E-07 | 5.100E-07 |
| 5552 | 278 | P278K | A | 2.8e-07 | 3.6e-07 | 3.1e-07 | 3.167E-07 | 4.041E-08 | 2.800E-07 | 3.600E-07 |
| 5553 | 278 | P278M | A | 1.7e-07 | 3.9e-07 | 2e-07 | 2.533E-07 | 1.193E-07 | 1.700E-07 | 3.900E-07 |
| 5554 | 278 | P278F | A | 2.2e-07 | 2.5e-07 | 2.5e-07 | 2.400E-07 | 1.732E-08 | 2.200E-07 | 2.500E-07 |
| 5555 | 278 | P278P | A | 4.6e-07 | 3e-07 | 1.4e-07 | 3.000E-07 | 1.600E-07 | 1.400E-07 | 4.600E-07 |
| 5556 | 278 | P278S | A | 3.4e-07 | 1.7e-07 | 2e-07 | 2.367E-07 | 9.074E-08 | 1.700E-07 | 3.400E-07 |
| 5557 | 278 | P278T | A | 1.9e-07 | 1.4e-07 | 1.5e-07 | 1.600E-07 | 2.646E-08 | 1.400E-07 | 1.900E-07 |
| 5558 | 278 | P278W | A | 2.9e-07 | 1.4e-07 | 3.7e-07 | 2.667E-07 | 1.168E-07 | 1.400E-07 | 3.700E-07 |
| 5559 | 278 | P278Y | A | 1.4e-07 | 2.1e-07 | 3.4e-07 | 2.300E-07 | 1.015E-07 | 1.400E-07 | 3.400E-07 |
| 5560 | 278 | P278V | A | 1.8e-07 | 2.4e-07 | 2.6e-07 | 2.267E-07 | 4.163E-08 | 1.800E-07 | 2.600E-07 |
| 5561 | 279 | Y279A | A | 2.3e-07 | 3.5e-07 | 2.4e-07 | 2.733E-07 | 6.658E-08 | 2.300E-07 | 3.500E-07 |
| 5562 | 279 | Y279R | A | 1.9e-07 | 1.2e-07 | 1.4e-07 | 1.500E-07 | 3.606E-08 | 1.200E-07 | 1.900E-07 |
| 5563 | 279 | Y279N | A | 9.8e-08 | 2.8e-07 | 4.5e-07 | 2.760E-07 | 1.760E-07 | 9.800E-08 | 4.500E-07 |
| 5564 | 279 | Y279D | A | 2.2e-07 | 3.4e-07 | 4.4e-07 | 3.333E-07 | 1.102E-07 | 2.200E-07 | 4.400E-07 |
| 5565 | 279 | Y279C | A | 2.9e-07 | 3.6e-07 | 2.3e-07 | 2.933E-07 | 6.506E-08 | 2.300E-07 | 3.600E-07 |
| 5566 | 279 | Y279E | A | 2.7e-07 | 3.6e-07 | 4.4e-07 | 3.567E-07 | 8.505E-08 | 2.700E-07 | 4.400E-07 |
| 5567 | 279 | Y279Q | A | 2.3e-07 | 2.2e-07 | 2.5e-07 | 2.333E-07 | 1.528E-08 | 2.200E-07 | 2.500E-07 |
| 5568 | 279 | Y279G | A | 1.9e-07 | 1.9e-07 | 1.2e-07 | 1.667E-07 | 4.041E-08 | 1.200E-07 | 1.900E-07 |
| 5569 | 279 | Y279H | A | 2.5e-07 | 1.3e-07 | 3.6e-07 | 2.467E-07 | 1.150E-07 | 1.300E-07 | 3.600E-07 |
| 5570 | 279 | Y279I | A | 2.2e-07 | 2.1e-07 | 2.3e-07 | 2.200E-07 | 1.000E-08 | 2.100E-07 | 2.300E-07 |
| 5571 | 279 | Y279L | A | 2.7e-07 | 1.8e-07 | 1.3e-07 | 1.933E-07 | 7.095E-08 | 1.300E-07 | 2.700E-07 |
| 5572 | 279 | Y279K | A | 3e-07 | 2.3e-07 | 2.1e-07 | 2.467E-07 | 4.726E-08 | 2.100E-07 | 3.000E-07 |
| 5573 | 279 | Y279M | A | 3.2e-07 | 2.8e-07 | 2.2e-07 | 2.733E-07 | 5.033E-08 | 2.200E-07 | 3.200E-07 |
| 5574 | 279 | Y279F | A | 4.5e-07 | 1.9e-07 | 3.4e-07 | 3.267E-07 | 1.305E-07 | 1.900E-07 | 4.500E-07 |
| 5575 | 279 | Y279P | A | 1e-07 | 2.9e-07 | 4.4e-07 | 2.767E-07 | 1.704E-07 | 1.000E-07 | 4.400E-07 |
| 5576 | 279 | Y279S | A | 5.1e-07 | 2.4e-07 | 1.7e-07 | 3.067E-07 | 1.795E-07 | 1.700E-07 | 5.100E-07 |
| 5577 | 279 | Y279T | A | 4.8e-07 | 1.7e-07 | 2e-07 | 2.833E-07 | 1.710E-07 | 1.700E-07 | 4.800E-07 |
| 5578 | 279 | Y279W | A | 4.6e-07 | 1.3e-06 | 1.7e-07 | 6.433E-07 | 5.869E-07 | 1.700E-07 | 1.300E-06 |
| 5579 | 279 | Y279Y | A | 3.4e-07 | 1.8e-07 | 3.4e-07 | 2.867E-07 | 9.238E-08 | 1.800E-07 | 3.400E-07 |
| 5580 | 279 | Y279V | A | 2.7e-07 | 1.4e-07 | 1.9e-07 | 2.000E-07 | 6.557E-08 | 1.400E-07 | 2.700E-07 |
| 5581 | 280 | N280A | A | 6.3e-07 | 2.4e-07 | 2.1e-07 | 3.600E-07 | 2.343E-07 | 2.100E-07 | 6.300E-07 |
| 5582 | 280 | N280R | A | 2.7e-07 | 1.5e-07 | 2.2e-07 | 2.133E-07 | 6.028E-08 | 1.500E-07 | 2.700E-07 |
| 5583 | 280 | N280N | A | 4.3e-07 | 2.2e-07 | 5.5e-07 | 4.000E-07 | 1.670E-07 | 2.200E-07 | 5.500E-07 |
| 5584 | 280 | N280D | A | 2.5e-07 | 2.3e-07 | 8.9e-08 | 1.897E-07 | 8.775E-08 | 8.900E-08 | 2.500E-07 |
| 5585 | 280 | N280C | A | 2.9e-07 | 5.2e-07 | 4.2e-07 | 4.100E-07 | 1.153E-07 | 2.900E-07 | 5.200E-07 |
| 5586 | 280 | N280E | A | 3e-07 | 2.4e-07 | 1.9e-07 | 2.433E-07 | 5.508E-08 | 1.900E-07 | 3.000E-07 |
| 5587 | 280 | N280Q | A | 2.7e-07 | 3.1e-07 | 2.6e-07 | 2.800E-07 | 2.646E-08 | 2.600E-07 | 3.100E-07 |
| 5588 | 280 | N280G | A | 2.1e-07 | 1.3e-07 | 2.4e-07 | 1.933E-07 | 5.686E-08 | 1.300E-07 | 2.400E-07 |
| 5589 | 280 | N280H | A | 3.1e-07 | 2.8e-07 | 1.4e-07 | 2.433E-07 | 9.074E-08 | 1.400E-07 | 3.100E-07 |
| 5590 | 280 | N280I | A | 2.1e-07 | 1.9e-07 | 1.4e-07 | 1.800E-07 | 3.606E-08 | 1.400E-07 | 2.100E-07 |
| 5591 | 280 | N280L | A | 2.8e-07 | 2.3e-07 | 1.9e-07 | 2.333E-07 | 4.509E-08 | 1.900E-07 | 2.800E-07 |
| 5592 | 280 | N280K | A | 3.9e-07 | 1.7e-07 | 3e-07 | 2.867E-07 | 1.106E-07 | 1.700E-07 | 3.900E-07 |
| 5593 | 280 | N280M | A | 4.7e-07 | 1.7e-07 | 1.6e-07 | 2.667E-07 | 1.762E-07 | 1.600E-07 | 4.700E-07 |
| 5594 | 280 | N280F | A | 2e-07 | 4e-07 | 3e-07 | 3.000E-07 | 1.000E-07 | 2.000E-07 | 4.000E-07 |
| 5595 | 280 | N280P | A | 3.1e-07 | 3.9e-07 | 3.1e-07 | 3.367E-07 | 4.619E-08 | 3.100E-07 | 3.900E-07 |
| 5596 | 280 | N280S | A | 1.1e-07 | 1.5e-07 | 1.6e-07 | 1.400E-07 | 2.646E-08 | 1.100E-07 | 1.600E-07 |
| 5597 | 280 | N280T | A | 4.9e-07 | 1.7e-07 | 2.8e-07 | 3.133E-07 | 1.626E-07 | 1.700E-07 | 4.900E-07 |
| 5598 | 280 | N280W | A | 1.5e-07 | 2.4e-07 | 2.8e-07 | 2.233E-07 | 6.658E-08 | 1.500E-07 | 2.800E-07 |
| 5599 | 280 | N280Y | A | 2.1e-07 | 3.6e-07 | 9.2e-07 | 4.967E-07 | 3.742E-07 | 2.100E-07 | 9.200E-07 |
| 5600 | 280 | N280V | A | 2.3e-07 | 2.5e-07 | 2.2e-07 | 2.333E-07 | 1.528E-08 | 2.200E-07 | 2.500E-07 |
| 5601 | 281 | Y281A | A | 2.1e-07 | 1.2e-07 | 1.4e-07 | 1.567E-07 | 4.726E-08 | 1.200E-07 | 2.100E-07 |
| 5602 | 281 | Y281R | A | 7.3e-07 | 2e-07 | 2.5e-07 | 3.933E-07 | 2.926E-07 | 2.000E-07 | 7.300E-07 |
| 5603 | 281 | Y281N | A | 2.5e-07 | 1.7e-07 | 2.1e-07 | 2.100E-07 | 4.000E-08 | 1.700E-07 | 2.500E-07 |
| 5604 | 281 | Y281D | A | 2.7e-07 | 2.2e-07 | 6.8e-07 | 3.900E-07 | 2.524E-07 | 2.200E-07 | 6.800E-07 |
| 5605 | 281 | Y281C | A | 1.3e-07 | 1.7e-07 | 2.5e-07 | 1.833E-07 | 6.110E-08 | 1.300E-07 | 2.500E-07 |
| 5606 | 281 | Y281E | A | 4.7e-07 | 1.2e-07 | 2.4e-07 | 2.767E-07 | 1.779E-07 | 1.200E-07 | 4.700E-07 |
| 5607 | 281 | Y281Q | A | 2.6e-07 | 1.1e-07 | 3.6e-07 | 2.433E-07 | 1.258E-07 | 1.100E-07 | 3.600E-07 |
| 5608 | 281 | Y281G | A | 1.7e-07 | 1.6e-07 | 1.1e-07 | 1.467E-07 | 3.215E-08 | 1.100E-07 | 1.700E-07 |
| 5609 | 281 | Y281H | A | 1.6e-07 | 4.1e-07 | 1.7e-07 | 2.467E-07 | 1.415E-07 | 1.600E-07 | 4.100E-07 |

| | | | | | | | | | | |
|---|---|---|---|---|---|---|---|---|---|---|
| 5610 | 281 | Y281I | A | 2e-07 | 4e-07 | 5.3e-07 | 3.767E-07 | 1.662E-07 | 2.000E-07 | 5.300E-07 |
| 5611 | 281 | Y281L | A | 1.5e-07 | 1.1e-07 | 2.4e-07 | 1.667E-07 | 6.658E-08 | 1.100E-07 | 2.400E-07 |
| 5612 | 281 | Y281K | A | 2.7e-07 | 3.2e-07 | 2.5e-07 | 2.800E-07 | 3.606E-08 | 2.500E-07 | 3.200E-07 |
| 5613 | 281 | Y281M | A | 2.2e-07 | 3.2e-07 | 2.9e-07 | 2.767E-07 | 5.132E-08 | 2.200E-07 | 3.200E-07 |
| 5614 | 281 | Y281F | A | 3.1e-07 | 5.1e-07 | 2.5e-07 | 3.567E-07 | 1.361E-07 | 2.500E-07 | 5.100E-07 |
| 5615 | 281 | Y281P | A | 1.9e-07 | 1.6e-07 | 2.9e-07 | 2.133E-07 | 6.807E-08 | 1.600E-07 | 2.900E-07 |
| 5616 | 281 | Y281S | A | 1.8e-07 | 2e-07 | 1.3e-07 | 1.700E-07 | 3.606E-08 | 1.300E-07 | 2.000E-07 |
| 5617 | 281 | Y281T | A | 2e-07 | 4e-07 | 2.3e-07 | 2.767E-07 | 1.079E-07 | 2.000E-07 | 4.000E-07 |
| 5618 | 281 | Y281W | A | 5.4e-07 | 2.6e-07 | 1.9e-07 | 3.300E-07 | 1.852E-07 | 1.900E-07 | 5.400E-07 |
| 5619 | 281 | Y281Y | A | 2.6e-07 | 1.6e-07 | 3.8e-07 | 2.667E-07 | 1.102E-07 | 1.600E-07 | 3.800E-07 |
| 5620 | 281 | Y281V | A | 1.6e-07 | 1.4e-07 | 2.9e-07 | 1.967E-07 | 8.145E-08 | 1.400E-07 | 2.900E-07 |
| 5621 | 282 | L282A | A | 3.3e-07 | 1.4e-07 | 2.1e-07 | 2.267E-07 | 9.609E-08 | 1.400E-07 | 3.300E-07 |
| 5622 | 282 | L282R | A | 1.1e-07 | 4.4e-07 | 4.4e-07 | 3.300E-07 | 1.905E-07 | 1.100E-07 | 4.400E-07 |
| 5623 | 282 | L282N | A | 1.9e-07 | 1.6e-07 | 2.2e-07 | 1.900E-07 | 3.000E-08 | 1.600E-07 | 2.200E-07 |
| 5624 | 282 | L282D | A | 1.1e-07 | 3.9e-07 | 1.5e-07 | 2.167E-07 | 1.514E-07 | 1.100E-07 | 3.900E-07 |
| 5625 | 282 | L282C | A | 1.8e-07 | 1.2e-07 | 2.4e-07 | 1.800E-07 | 6.000E-08 | 1.200E-07 | 2.400E-07 |
| 5626 | 282 | L282E | A | 2.2e-07 | 3e-07 | 1.8e-07 | 2.333E-07 | 6.110E-08 | 1.800E-07 | 3.000E-07 |
| 5627 | 282 | L282Q | A | 2e-07 | 2.1e-07 | 1.8e-07 | 1.967E-07 | 1.528E-08 | 1.800E-07 | 2.100E-07 |
| 5628 | 282 | L282G | A | 3.1e-07 | 2e-07 | 3.1e-07 | 2.733E-07 | 6.351E-08 | 2.000E-07 | 3.100E-07 |
| 5629 | 282 | L282H | A | 2.3e-07 | 3.2e-07 | 1.4e-07 | 2.300E-07 | 9.000E-08 | 1.400E-07 | 3.200E-07 |
| 5630 | 282 | L282I | A | 1.9e-07 | 1.7e-07 | 2.5e-07 | 2.033E-07 | 4.163E-08 | 1.700E-07 | 2.500E-07 |
| 5631 | 282 | L282L | A | 5.7e-07 | 4.4e-07 | 4.9e-07 | 5.000E-07 | 6.557E-08 | 4.400E-07 | 5.700E-07 |
| 5632 | 282 | L282K | A | 3e-07 | 2.6e-07 | 1.7e-07 | 2.433E-07 | 6.658E-08 | 1.700E-07 | 3.000E-07 |
| 5633 | 282 | L282M | A | 1.2e-07 | 1.9e-07 | 3.5e-07 | 2.200E-07 | 1.179E-07 | 1.200E-07 | 3.500E-07 |
| 5634 | 282 | L282F | A | 4.2e-07 | 4.9e-07 | 3.5e-07 | 4.200E-07 | 7.000E-08 | 3.500E-07 | 4.900E-07 |
| 5635 | 282 | L282P | A | 3.2e-07 | 2.5e-07 | 1.8e-07 | 2.500E-07 | 7.000E-08 | 1.800E-07 | 3.200E-07 |
| 5636 | 282 | L282S | A | 1.2e-07 | 3.5e-07 | 5e-07 | 3.233E-07 | 1.914E-07 | 1.200E-07 | 5.000E-07 |
| 5637 | 282 | L282T | A | 2.4e-07 | 4e-07 | 3e-07 | 3.133E-07 | 8.083E-08 | 2.400E-07 | 4.000E-07 |
| 5638 | 282 | L282W | A | 1.9e-07 | 2.4e-07 | 3.4e-07 | 2.567E-07 | 7.638E-08 | 1.900E-07 | 3.400E-07 |
| 5639 | 282 | L282Y | A | 2.1e-07 | 2.1e-07 | 2.7e-07 | 2.300E-07 | 3.464E-08 | 2.100E-07 | 2.700E-07 |
| 5640 | 282 | L282V | A | 4.1e-07 | 3e-07 | 3e-07 | 3.367E-07 | 6.351E-08 | 3.000E-07 | 4.100E-07 |
| 5641 | 283 | S283A | A | 3.5e-07 | 2.3e-07 | 2e-07 | 2.600E-07 | 7.937E-08 | 2.000E-07 | 3.500E-07 |
| 5642 | 283 | S283R | A | 5.6e-07 | 1.3e-07 | 6.5e-07 | 4.467E-07 | 2.779E-07 | 1.300E-07 | 6.500E-07 |
| 5643 | 283 | S283N | A | 2.6e-07 | 3.7e-07 | 2.5e-07 | 2.933E-07 | 6.658E-08 | 2.500E-07 | 3.700E-07 |
| 5644 | 283 | S283D | A | 1.5e-07 | 1.6e-07 | 2.8e-07 | 1.967E-07 | 7.234E-08 | 1.500E-07 | 2.800E-07 |
| 5645 | 283 | S283C | A | 4.1e-07 | 1.1e-07 | 8.2e-07 | 4.467E-07 | 3.564E-07 | 1.100E-07 | 8.200E-07 |
| 5646 | 283 | S283E | A | 3.7e-07 | 1.7e-07 | 4.7e-07 | 3.367E-07 | 1.528E-07 | 1.700E-07 | 4.700E-07 |
| 5647 | 283 | S283Q | A | 1.9e-07 | 4.2e-07 | 2.7e-07 | 2.933E-07 | 1.168E-07 | 1.900E-07 | 4.200E-07 |
| 5648 | 283 | S283G | A | 2.6e-07 | 1.5e-07 | 1.6e-07 | 1.900E-07 | 6.083E-08 | 1.500E-07 | 2.600E-07 |
| 5649 | 283 | S283H | A | 9e-07 | 1.9e-07 | 2.4e-07 | 4.433E-07 | 3.963E-07 | 1.900E-07 | 9.000E-07 |
| 5650 | 283 | S283I | A | 2.5e-07 | 1.7e-07 | 2.6e-07 | 2.267E-07 | 4.933E-08 | 1.700E-07 | 2.600E-07 |
| 5651 | 283 | S283L | A | 1.3e-07 | 2e-07 | 2.4e-07 | 1.900E-07 | 5.568E-08 | 1.300E-07 | 2.400E-07 |
| 5652 | 283 | S283K | A | 2.4e-07 | 1.7e-07 | 3.2e-07 | 2.433E-07 | 7.506E-08 | 1.700E-07 | 3.200E-07 |
| 5653 | 283 | S283M | A | 4.9e-07 | 1.3e-07 | 1.7e-07 | 2.633E-07 | 1.973E-07 | 1.300E-07 | 4.900E-07 |
| 5654 | 283 | S283F | A | 5e-07 | 2.2e-07 | 2.8e-07 | 3.333E-07 | 1.474E-07 | 2.200E-07 | 5.000E-07 |
| 5655 | 283 | S283P | A | 1.4e-07 | 3.9e-07 | 1.7e-07 | 2.333E-07 | 1.365E-07 | 1.400E-07 | 3.900E-07 |
| 5656 | 283 | S283S | A | 1.4e-07 | 1e-07 | 1.7e-07 | 1.367E-07 | 3.512E-08 | 1.000E-07 | 1.700E-07 |
| 5657 | 283 | S283T | A | 2.1e-07 | 1.7e-07 | 2.6e-07 | 2.133E-07 | 4.509E-08 | 1.700E-07 | 2.600E-07 |
| 5658 | 283 | S283W | A | 2.4e-07 | 3e-07 | 2.8e-07 | 2.733E-07 | 3.055E-08 | 2.400E-07 | 3.000E-07 |
| 5659 | 283 | S283Y | A | 2.3e-07 | 3.4e-07 | 3e-07 | 2.900E-07 | 5.568E-08 | 2.300E-07 | 3.400E-07 |
| 5660 | 283 | S283V | A | 5.6e-07 | 1.5e-07 | 2.5e-07 | 3.200E-07 | 2.138E-07 | 1.500E-07 | 5.600E-07 |
| 5661 | 284 | T284A | A | 2.5e-07 | 5.3e-07 | 2.6e-07 | 3.467E-07 | 1.589E-07 | 2.500E-07 | 5.300E-07 |
| 5662 | 284 | T284R | A | 2.6e-07 | 4.2e-07 | 1.7e-07 | 2.833E-07 | 1.266E-07 | 1.700E-07 | 4.200E-07 |
| 5663 | 284 | T284N | A | 2.2e-07 | 4e-07 | 2.2e-07 | 2.800E-07 | 1.039E-07 | 2.200E-07 | 4.000E-07 |
| 5664 | 284 | T284D | A | 1.4e-07 | 4.6e-07 | 3.5e-07 | 3.167E-07 | 1.626E-07 | 1.400E-07 | 4.600E-07 |
| 5665 | 284 | T284C | A | 3.4e-07 | 2.2e-07 | 2e-07 | 2.533E-07 | 7.572E-08 | 2.000E-07 | 3.400E-07 |
| 5666 | 284 | T284E | A | 3.8e-07 | 8.8e-07 | 3.8e-07 | 5.467E-07 | 2.887E-07 | 3.800E-07 | 8.800E-07 |
| 5667 | 284 | T284Q | A | 2.1e-07 | 1.8e-07 | 1.2e-07 | 1.700E-07 | 4.583E-08 | 1.200E-07 | 2.100E-07 |
| 5668 | 284 | T284G | A | 2.3e-07 | 3e-07 | 1.1e-07 | 2.133E-07 | 9.609E-08 | 1.100E-07 | 3.000E-07 |
| 5669 | 284 | T284H | A | 2.3e-07 | 2.3e-07 | 3.5e-07 | 2.700E-07 | 6.928E-08 | 2.300E-07 | 3.500E-07 |
| 5670 | 284 | T284I | A | 4.8e-07 | 3.9e-07 | 2.8e-07 | 3.833E-07 | 1.002E-07 | 2.800E-07 | 4.800E-07 |
| 5671 | 284 | T284L | A | 3.8e-07 | 3.7e-07 | 2e-07 | 3.167E-07 | 1.012E-07 | 2.000E-07 | 3.800E-07 |
| 5672 | 284 | T284K | A | 2.8e-07 | 1.9e-07 | 2.7e-07 | 2.467E-07 | 4.933E-08 | 1.900E-07 | 2.800E-07 |
| 5673 | 284 | T284M | A | 2.5e-07 | 3e-07 | 3.7e-07 | 3.067E-07 | 6.028E-08 | 2.500E-07 | 3.700E-07 |
| 5674 | 284 | T284F | A | 2.4e-07 | 3.5e-07 | 1.9e-07 | 2.600E-07 | 8.185E-08 | 1.900E-07 | 3.500E-07 |
| 5675 | 284 | T284P | A | 2.4e-07 | 2e-07 | 2.4e-07 | 2.267E-07 | 2.309E-08 | 2.000E-07 | 2.400E-07 |
| 5676 | 284 | T284S | A | 1.5e-07 | 1.4e-07 | 1.2e-07 | 1.367E-07 | 1.528E-08 | 1.200E-07 | 1.500E-07 |
| 5677 | 284 | T284T | A | 4.4e-07 | 3e-07 | 1.9e-07 | 3.100E-07 | 1.253E-07 | 1.900E-07 | 4.400E-07 |
| 5678 | 284 | T284W | A | 3e-07 | 2e-07 | 2.2e-07 | 2.400E-07 | 5.292E-08 | 2.000E-07 | 3.000E-07 |
| 5679 | 284 | T284Y | A | 2.6e-07 | 5e-07 | 2.6e-07 | 3.400E-07 | 1.386E-07 | 2.600E-07 | 5.000E-07 |
| 5680 | 284 | T284V | A | 1.7e-07 | 2e-07 | 1.8e-07 | 1.833E-07 | 1.528E-08 | 1.700E-07 | 2.000E-07 |
| 5681 | 285 | D285A | A | 6.2e-07 | 3e-07 | 1.7e-07 | 3.633E-07 | 2.316E-07 | 1.700E-07 | 6.200E-07 |
| 5682 | 285 | D285R | A | 3.5e-07 | 4.5e-07 | 2.5e-07 | 3.500E-07 | 1.000E-07 | 2.500E-07 | 4.500E-07 |
| 5683 | 285 | D285N | A | 1.4e-07 | 1.7e-07 | 3.1e-07 | 2.067E-07 | 9.074E-08 | 1.400E-07 | 3.100E-07 |
| 5684 | 285 | D285D | A | 3e-07 | 1.6e-07 | 3.5e-07 | 2.700E-07 | 9.849E-08 | 1.600E-07 | 3.500E-07 |
| 5685 | 285 | D285C | A | 1.4e-07 | 5.9e-07 | 2.8e-07 | 3.367E-07 | 2.303E-07 | 1.400E-07 | 5.900E-07 |
| 5686 | 285 | D285E | A | 2.5e-07 | 3.7e-07 | 2.1e-07 | 2.767E-07 | 8.327E-08 | 2.100E-07 | 3.700E-07 |
| 5687 | 285 | D285Q | A | 2.6e-07 | 2.4e-07 | 3e-07 | 2.667E-07 | 3.055E-08 | 2.400E-07 | 3.000E-07 |
| 5688 | 285 | D285G | A | 2.7e-07 | 1.3e-07 | 1.3e-07 | 1.767E-07 | 8.083E-08 | 1.300E-07 | 2.700E-07 |
| 5689 | 285 | D285H | A | 3.2e-07 | 1.8e-07 | 4.6e-07 | 3.200E-07 | 1.400E-07 | 1.800E-07 | 4.600E-07 |
| 5690 | 285 | D285I | A | 1.2e-07 | 1.2e-07 | 7.2e-07 | 3.200E-07 | 3.464E-07 | 1.200E-07 | 7.200E-07 |
| 5691 | 285 | D285L | A | 3.1e-07 | 2.6e-07 | 4.5e-07 | 3.400E-07 | 9.849E-08 | 2.600E-07 | 4.500E-07 |
| 5692 | 285 | D285K | A | 2.4e-07 | 2.7e-07 | 3.1e-07 | 2.733E-07 | 3.512E-08 | 2.400E-07 | 3.100E-07 |
| 5693 | 285 | D285M | A | 3e-07 | 2.5e-07 | 2.4e-07 | 2.633E-07 | 3.215E-08 | 2.400E-07 | 3.000E-07 |
| 5694 | 285 | D285F | A | 5e-07 | 1.6e-07 | 3.6e-07 | 3.400E-07 | 1.709E-07 | 1.600E-07 | 5.000E-07 |

| | | | | | | | | | | |
|---|---|---|---|---|---|---|---|---|---|---|
| 5695 | 285 | D285P | A | 1.8e-07 | 2.2e-07 | 1.3e-07 | 1.767E-07 | 4.509E-08 | 1.300E-07 | 2.200E-07 |
| 5696 | 285 | D285S | A | 2.5e-07 | 7e-07 | 1.9e-07 | 3.800E-07 | 2.787E-07 | 1.900E-07 | 7.000E-07 |
| 5697 | 285 | D285T | A | 3.3e-07 | 2.5e-07 | 5.5e-07 | 3.767E-07 | 1.553E-07 | 2.500E-07 | 5.500E-07 |
| 5698 | 285 | D285W | A | 1.5e-07 | 2.3e-07 | 1.5e-07 | 1.767E-07 | 4.619E-08 | 1.500E-07 | 2.300E-07 |
| 5699 | 285 | D285Y | A | 2.1e-07 | 2.2e-07 | 2.3e-07 | 2.200E-07 | 1.000E-08 | 2.100E-07 | 2.300E-07 |
| 5700 | 285 | D285V | A | 1.1e-07 | 2.8e-07 | 3e-07 | 2.300E-07 | 1.044E-07 | 1.100E-07 | 3.000E-07 |
| 5701 | 286 | V286A | A | 2.3e-07 | 3.6e-07 | 2.2e-07 | 2.700E-07 | 7.810E-08 | 2.200E-07 | 3.600E-07 |
| 5702 | 286 | V286R | A | 2.8e-07 | 9e-08 | 1.5e-07 | 1.733E-07 | 9.713E-08 | 9.000E-08 | 2.800E-07 |
| 5703 | 286 | V286N | A | 2.1e-07 | 2.3e-07 | 2.8e-07 | 2.400E-07 | 3.606E-08 | 2.100E-07 | 2.800E-07 |
| 5704 | 286 | V286D | A | 3.2e-07 | 2.5e-07 | 3.3e-07 | 3.000E-07 | 4.359E-08 | 2.500E-07 | 3.300E-07 |
| 5705 | 286 | V286C | A | 3.2e-07 | 1.4e-07 | 2.7e-07 | 2.433E-07 | 9.292E-08 | 1.400E-07 | 3.200E-07 |
| 5706 | 286 | V286E | A | 3e-07 | 5.5e-07 | 1.7e-07 | 3.400E-07 | 1.931E-07 | 1.700E-07 | 5.500E-07 |
| 5707 | 286 | V286Q | A | 2.9e-07 | 2.5e-07 | 3.3e-07 | 2.900E-07 | 4.000E-08 | 2.500E-07 | 3.300E-07 |
| 5708 | 286 | V286G | A | 2.5e-07 | 1.1e-07 | 4e-07 | 2.533E-07 | 1.450E-07 | 1.100E-07 | 4.000E-07 |
| 5709 | 286 | V286H | A | 1.8e-07 | 3e-07 | 2e-07 | 2.267E-07 | 6.429E-08 | 1.800E-07 | 3.000E-07 |
| 5710 | 286 | V286I | A | 2.5e-07 | 1.8e-07 | 1.4e-07 | 1.900E-07 | 5.568E-08 | 1.400E-07 | 2.500E-07 |
| 5711 | 286 | V286L | A | 2.4e-07 | 3e-07 | 1.7e-07 | 2.367E-07 | 6.506E-08 | 1.700E-07 | 3.000E-07 |
| 5712 | 286 | V286K | A | 1.3e-07 | 2.2e-07 | 2.3e-07 | 1.933E-07 | 5.508E-08 | 1.300E-07 | 2.300E-07 |
| 5713 | 286 | V286M | A | 1.6e-07 | 1.3e-07 | 1.1e-07 | 1.333E-07 | 2.517E-08 | 1.100E-07 | 1.600E-07 |
| 5714 | 286 | V286F | A | 2.9e-07 | 2.9e-07 | 1.7e-07 | 2.500E-07 | 6.928E-08 | 1.700E-07 | 2.900E-07 |
| 5715 | 286 | V286P | A | 2.5e-07 | 3e-07 | 1.9e-07 | 2.467E-07 | 5.508E-08 | 1.900E-07 | 3.000E-07 |
| 5716 | 286 | V286S | A | 2.3e-07 | 3.5e-07 | 2.8e-07 | 2.867E-07 | 6.028E-08 | 2.300E-07 | 3.500E-07 |
| 5717 | 286 | V286T | A | 1.7e-07 | 2.4e-07 | 2.8e-07 | 2.300E-07 | 5.568E-08 | 1.700E-07 | 2.800E-07 |
| 5718 | 286 | V286W | A | 3.9e-07 | 2.3e-07 | 3.4e-07 | 3.200E-07 | 8.185E-08 | 2.300E-07 | 3.900E-07 |
| 5719 | 286 | V286Y | A | 5.1e-07 | 2.3e-07 | 3.2e-07 | 3.533E-07 | 1.429E-07 | 2.300E-07 | 5.100E-07 |
| 5720 | 286 | V286V | A | 3e-07 | 1.4e-07 | 2.4e-07 | 2.267E-07 | 8.083E-08 | 1.400E-07 | 3.000E-07 |
| 5721 | 287 | G287A | A | 2.9e-07 | 4.7e-07 | 4.4e-07 | 4.000E-07 | 9.644E-08 | 2.900E-07 | 4.700E-07 |
| 5722 | 287 | G287R | A | 1.3e-07 | 1.6e-07 | 1.7e-07 | 1.533E-07 | 2.082E-08 | 1.300E-07 | 1.700E-07 |
| 5723 | 287 | G287N | A | 2.9e-07 | 2.3e-07 | 3.6e-07 | 2.933E-07 | 6.506E-08 | 2.300E-07 | 3.600E-07 |
| 5724 | 287 | G287D | A | 2.3e-07 | 2.4e-07 | 5.6e-07 | 3.433E-07 | 1.877E-07 | 2.300E-07 | 5.600E-07 |
| 5725 | 287 | G287C | A | 2.1e-07 | 5.9e-07 | 1.7e-07 | 3.233E-07 | 2.318E-07 | 1.700E-07 | 5.900E-07 |
| 5726 | 287 | G287E | A | 2.8e-07 | 3.5e-07 | 1.6e-07 | 2.633E-07 | 9.609E-08 | 1.600E-07 | 3.500E-07 |
| 5727 | 287 | G287Q | A | 1.1e-07 | 1.4e-07 | 5.4e-07 | 2.633E-07 | 2.401E-07 | 1.100E-07 | 5.400E-07 |
| 5728 | 287 | G287G | A | 2.1e-07 | 4.7e-07 | 1.8e-07 | 2.867E-07 | 1.595E-07 | 1.800E-07 | 4.700E-07 |
| 5729 | 287 | G287H | A | 2.3e-07 | 6.4e-08 | 2.8e-07 | 1.913E-07 | 1.131E-07 | 6.400E-08 | 2.800E-07 |
| 5730 | 287 | G287I | A | 1.3e-07 | 3.2e-07 | 4.1e-07 | 2.867E-07 | 1.429E-07 | 1.300E-07 | 4.100E-07 |
| 5731 | 287 | G287L | A | 5.3e-07 | 1.4e-07 | 2.4e-07 | 3.033E-07 | 2.026E-07 | 1.400E-07 | 5.300E-07 |
| 5732 | 287 | G287K | A | 2.5e-07 | 1.6e-07 | 3.1e-07 | 2.400E-07 | 7.550E-08 | 1.600E-07 | 3.100E-07 |
| 5733 | 287 | G287M | A | 2.5e-07 | 3e-07 | 1.4e-07 | 2.300E-07 | 8.185E-08 | 1.400E-07 | 3.000E-07 |
| 5734 | 287 | G287F | A | 2.6e-07 | 3.2e-07 | 1.8e-07 | 2.533E-07 | 7.024E-08 | 1.800E-07 | 3.200E-07 |
| 5735 | 287 | G287P | A | 1.1e-07 | 1.7e-07 | 2.3e-07 | 1.700E-07 | 6.000E-08 | 1.100E-07 | 2.300E-07 |
| 5736 | 287 | G287S | A | 2.5e-07 | 1.5e-07 | 1.7e-07 | 1.900E-07 | 5.292E-08 | 1.500E-07 | 2.500E-07 |
| 5737 | 287 | G287T | A | 5.2e-07 | 1.3e-07 | 2.6e-07 | 3.033E-07 | 1.986E-07 | 1.300E-07 | 5.200E-07 |
| 5738 | 287 | G287W | A | 2.7e-07 | 4e-07 | 3.4e-07 | 3.367E-07 | 6.506E-08 | 2.700E-07 | 4.000E-07 |
| 5739 | 287 | G287Y | A | 9.4e-08 | 5.6e-07 | 2.7e-07 | 3.080E-07 | 2.353E-07 | 9.400E-08 | 5.600E-07 |
| 5740 | 287 | G287V | A | 1.4e-07 | 3e-07 | 3.8e-07 | 2.733E-07 | 1.222E-07 | 1.400E-07 | 3.800E-07 |
| 5741 | 288 | S288A | A | 3e-07 | 4.6e-07 | 3.9e-07 | 3.833E-07 | 8.021E-08 | 3.000E-07 | 4.600E-07 |
| 5742 | 288 | S288R | A | 1.1e-07 | 8.8e-08 | 2.6e-07 | 1.527E-07 | 9.360E-08 | 8.800E-08 | 2.600E-07 |
| 5743 | 288 | S288N | A | 4.9e-07 | 2.3e-07 | 9.4e-08 | 2.713E-07 | 2.012E-07 | 9.400E-08 | 4.900E-07 |
| 5744 | 288 | S288D | A | 1.3e-07 | 1.9e-07 | 2.9e-07 | 2.033E-07 | 8.083E-08 | 1.300E-07 | 2.900E-07 |
| 5745 | 288 | S288C | A | 2.5e-07 | 5.3e-07 | 4.5e-07 | 4.100E-07 | 1.442E-07 | 2.500E-07 | 5.300E-07 |
| 5746 | 288 | S288E | A | 2e-07 | 1.3e-07 | 2.9e-07 | 2.067E-07 | 8.021E-08 | 1.300E-07 | 2.900E-07 |
| 5747 | 288 | S288Q | A | 2.4e-07 | 3.9e-07 | 1.8e-07 | 2.700E-07 | 1.082E-07 | 1.800E-07 | 3.900E-07 |
| 5748 | 288 | S288G | A | 3.7e-07 | 4.5e-07 | 2.5e-07 | 3.567E-07 | 1.007E-07 | 2.500E-07 | 4.500E-07 |
| 5749 | 288 | S288H | A | 1.9e-07 | 1.2e-07 | 2.8e-07 | 1.967E-07 | 8.021E-08 | 1.200E-07 | 2.800E-07 |
| 5750 | 288 | S288I | A | 1.8e-07 | 5.7e-07 | 1.3e-07 | 2.933E-07 | 2.409E-07 | 1.300E-07 | 5.700E-07 |
| 5751 | 288 | S288L | A | 4.2e-07 | 5e-07 | 4.3e-07 | 4.500E-07 | 4.359E-08 | 4.200E-07 | 5.000E-07 |
| 5752 | 288 | S288K | A | 2.4e-07 | 5.9e-07 | 4.6e-07 | 4.300E-07 | 1.769E-07 | 2.400E-07 | 5.900E-07 |
| 5753 | 288 | S288M | A | 4.4e-07 | 1.9e-07 | 4.3e-07 | 3.533E-07 | 1.415E-07 | 1.900E-07 | 4.400E-07 |
| 5754 | 288 | S288F | A | 2.7e-07 | 1.9e-07 | 3e-07 | 2.533E-07 | 5.686E-08 | 1.900E-07 | 3.000E-07 |
| 5755 | 288 | S288P | A | 7.4e-07 | 1.5e-07 | 4.5e-07 | 4.467E-07 | 2.950E-07 | 1.500E-07 | 7.400E-07 |
| 5756 | 288 | S288S | A | 6.6e-07 | 3.3e-07 | 3.7e-07 | 4.533E-07 | 1.801E-07 | 3.300E-07 | 6.600E-07 |
| 5757 | 288 | S288T | A | 2.7e-07 | 2.6e-07 | 2.7e-07 | 2.667E-07 | 5.774E-09 | 2.600E-07 | 2.700E-07 |
| 5758 | 288 | S288W | A | 2.2e-07 | 4.5e-07 | 2.9e-07 | 3.200E-07 | 1.179E-07 | 2.200E-07 | 4.500E-07 |
| 5759 | 288 | S288Y | A | 2.7e-07 | 1.6e-07 | 1.7e-07 | 2.000E-07 | 6.083E-08 | 1.600E-07 | 2.700E-07 |
| 5760 | 288 | S288V | A | 3.4e-07 | 3e-07 | 2.8e-07 | 3.067E-07 | 3.055E-08 | 2.800E-07 | 3.400E-07 |
| 5761 | 289 | C289A | A | 2.5e-07 | 2.2e-07 | 3.8e-07 | 2.833E-07 | 8.505E-08 | 2.200E-07 | 3.800E-07 |
| 5762 | 289 | C289R | A | 6.2e-07 | 6.3e-08 | 2.7e-07 | 3.177E-07 | 2.815E-07 | 6.300E-08 | 6.200E-07 |
| 5763 | 289 | C289N | A | 1.2e-07 | 4.3e-07 | 1.6e-07 | 2.367E-07 | 1.686E-07 | 1.200E-07 | 4.300E-07 |
| 5764 | 289 | C289D | A | 3.1e-07 | 2.3e-07 | 2.7e-07 | 2.700E-07 | 4.000E-08 | 2.300E-07 | 3.100E-07 |
| 5765 | 289 | C289C | A | 1.2e-07 | 2.4e-07 | 2e-07 | 1.867E-07 | 6.110E-08 | 1.200E-07 | 2.400E-07 |
| 5766 | 289 | C289E | A | 2.3e-07 | 1.6e-07 | 2.8e-07 | 2.233E-07 | 6.028E-08 | 1.600E-07 | 2.800E-07 |
| 5767 | 289 | C289Q | A | 4.9e-07 | 1.8e-07 | 2.5e-07 | 3.067E-07 | 1.626E-07 | 1.800E-07 | 4.900E-07 |
| 5768 | 289 | C289G | A | 5.5e-07 | 2.3e-07 | 3.1e-07 | 3.633E-07 | 1.665E-07 | 2.300E-07 | 5.500E-07 |
| 5769 | 289 | C289H | A | 2.8e-07 | 1.9e-07 | 2e-07 | 2.233E-07 | 4.933E-08 | 1.900E-07 | 2.800E-07 |
| 5770 | 289 | C289I | A | 2.2e-07 | 2.6e-07 | 3.4e-07 | 2.733E-07 | 6.110E-08 | 2.200E-07 | 3.400E-07 |
| 5771 | 289 | C289L | A | 4.7e-07 | 2.8e-07 | 2.6e-07 | 3.367E-07 | 1.159E-07 | 2.600E-07 | 4.700E-07 |
| 5772 | 289 | C289K | A | 2.5e-07 | 1.3e-07 | 1.8e-07 | 1.867E-07 | 6.028E-08 | 1.300E-07 | 2.500E-07 |
| 5773 | 289 | C289M | A | 4.9e-07 | 2.7e-07 | 2.4e-07 | 3.333E-07 | 1.365E-07 | 2.400E-07 | 4.900E-07 |
| 5774 | 289 | C289F | A | 2.7e-07 | 1.3e-06 | 2.9e-07 | 6.200E-07 | 5.890E-07 | 2.700E-07 | 1.300E-06 |
| 5775 | 289 | C289P | A | 2.2e-07 | 2.7e-07 | 3.7e-07 | 2.867E-07 | 7.638E-08 | 2.200E-07 | 3.700E-07 |
| 5776 | 289 | C289S | A | 2.1e-07 | 6.1e-07 | 3.2e-07 | 3.800E-07 | 2.066E-07 | 2.100E-07 | 6.100E-07 |
| 5777 | 289 | C289T | A | 6.9e-07 | 1.7e-07 | 2.3e-07 | 3.633E-07 | 2.845E-07 | 1.700E-07 | 6.900E-07 |
| 5778 | 289 | C289W | A | 1.6e-07 | 2.2e-07 | 2.4e-07 | 2.067E-07 | 4.163E-08 | 1.600E-07 | 2.400E-07 |
| 5779 | 289 | C289Y | A | 2.4e-07 | 5.2e-07 | 2.5e-07 | 3.367E-07 | 1.589E-07 | 2.400E-07 | 5.200E-07 |

| | | | | | | | | | | |
|---|---|---|---|---|---|---|---|---|---|---|
| 5780 | 289 | C289V | A | 2.2e-07 | 1.7e-07 | 1.4e-07 | 1.767E-07 | 4.041E-08 | 1.400E-07 | 2.200E-07 |
| 5781 | 290 | T290A | A | 2.3e-07 | 2.1e-07 | 2.1e-07 | 2.167E-07 | 1.155E-08 | 2.100E-07 | 2.300E-07 |
| 5782 | 290 | T290R | A | 3.4e-07 | 2.6e-07 | 2.4e-07 | 2.800E-07 | 5.292E-08 | 2.400E-07 | 3.400E-07 |
| 5783 | 290 | T290N | A | 4.1e-07 | 1.7e-07 | 3.8e-07 | 3.200E-07 | 1.308E-07 | 1.700E-07 | 4.100E-07 |
| 5784 | 290 | T290D | A | 2.8e-07 | 2.2e-07 | 2.9e-07 | 2.633E-07 | 3.786E-08 | 2.200E-07 | 2.900E-07 |
| 5785 | 290 | T290C | A | 4e-07 | 4.4e-07 | 2.2e-07 | 3.533E-07 | 1.172E-07 | 2.200E-07 | 4.400E-07 |
| 5786 | 290 | T290E | A | 2.1e-07 | 1.2e-07 | 4.1e-07 | 2.467E-07 | 1.484E-07 | 1.200E-07 | 4.100E-07 |
| 5787 | 290 | T290Q | A | 3.3e-07 | 5.5e-07 | 1.7e-07 | 3.500E-07 | 1.908E-07 | 1.700E-07 | 5.500E-07 |
| 5788 | 290 | T290G | A | 5.3e-07 | 1.9e-07 | 3.6e-07 | 3.600E-07 | 1.700E-07 | 1.900E-07 | 5.300E-07 |
| 5789 | 290 | T290H | A | 1.5e-07 | 1.1e-07 | 2.4e-07 | 1.667E-07 | 6.658E-08 | 1.100E-07 | 2.400E-07 |
| 5790 | 290 | T290I | A | 1.1e-07 | 2e-07 | 2.6e-07 | 1.900E-07 | 7.550E-08 | 1.100E-07 | 2.600E-07 |
| 5791 | 290 | T290L | A | 3.3e-07 | 2.4e-07 | 2.3e-07 | 2.667E-07 | 5.508E-08 | 2.300E-07 | 3.300E-07 |
| 5792 | 290 | T290K | A | 1.4e-07 | 3.8e-07 | 4.1e-07 | 3.100E-07 | 1.480E-07 | 1.400E-07 | 4.100E-07 |
| 5793 | 290 | T290M | A | 1.4e-07 | 2.9e-07 | 1.3e-07 | 1.867E-07 | 8.963E-08 | 1.300E-07 | 2.900E-07 |
| 5794 | 290 | T290F | A | 3.8e-07 | 1.8e-07 | 2.4e-07 | 2.667E-07 | 1.026E-07 | 1.800E-07 | 3.800E-07 |
| 5795 | 290 | T290P | A | 4.3e-07 | 3.1e-07 | 4.5e-07 | 3.967E-07 | 7.572E-08 | 3.100E-07 | 4.500E-07 |
| 5796 | 290 | T290S | A | 7.3e-08 | 1.5e-07 | 2.8e-07 | 1.677E-07 | 1.046E-07 | 7.300E-08 | 2.800E-07 |
| 5797 | 290 | T290T | A | 3.3e-07 | 2.5e-07 | 3e-07 | 2.933E-07 | 4.041E-08 | 2.500E-07 | 3.300E-07 |
| 5798 | 290 | T290W | A | 1.7e-07 | 3.1e-07 | 3.1e-07 | 2.633E-07 | 8.083E-08 | 1.700E-07 | 3.100E-07 |
| 5799 | 290 | T290Y | A | 1.9e-07 | 1.7e-07 | 1.6e-07 | 1.733E-07 | 1.528E-08 | 1.600E-07 | 1.900E-07 |
| 5800 | 290 | T290V | A | 1.3e-07 | 3.7e-07 | 1.6e-07 | 2.200E-07 | 1.308E-07 | 1.300E-07 | 3.700E-07 |
| 5801 | 291 | L291A | A | 2.2e-07 | 4.3e-07 | 2e-07 | 2.833E-07 | 1.274E-07 | 2.000E-07 | 4.300E-07 |
| 5802 | 291 | L291R | A | 1.4e-07 | 2e-07 | 2.4e-07 | 1.933E-07 | 5.033E-08 | 1.400E-07 | 2.400E-07 |
| 5803 | 291 | L291N | A | 1.4e-07 | 2.9e-07 | 1.7e-07 | 2.000E-07 | 7.937E-08 | 1.400E-07 | 2.900E-07 |
| 5804 | 291 | L291D | A | 2.9e-07 | 2.7e-07 | 2.3e-07 | 2.633E-07 | 3.055E-08 | 2.300E-07 | 2.900E-07 |
| 5805 | 291 | L291C | A | 2.4e-07 | 3.3e-07 | 1.6e-07 | 2.433E-07 | 8.505E-08 | 1.600E-07 | 3.300E-07 |
| 5806 | 291 | L291E | A | 2e-07 | 2e-07 | 2.3e-07 | 2.100E-07 | 1.732E-08 | 2.000E-07 | 2.300E-07 |
| 5807 | 291 | L291Q | A | 2e-07 | 3.1e-07 | 2e-07 | 2.367E-07 | 6.351E-08 | 2.000E-07 | 3.100E-07 |
| 5808 | 291 | L291G | A | 2.5e-07 | 1.9e-07 | 1.2e-07 | 1.867E-07 | 6.506E-08 | 1.200E-07 | 2.500E-07 |
| 5809 | 291 | L291H | A | 4.8e-07 | 4.6e-07 | 4.8e-07 | 4.733E-07 | 1.155E-08 | 4.600E-07 | 4.800E-07 |
| 5810 | 291 | L291I | A | 3.7e-07 | 2.5e-07 | 2.7e-07 | 2.967E-07 | 6.429E-08 | 2.500E-07 | 3.700E-07 |
| 5811 | 291 | L291L | A | 1.4e-07 | 2.7e-07 | 4.9e-07 | 3.000E-07 | 1.769E-07 | 1.400E-07 | 4.900E-07 |
| 5812 | 291 | L291K | A | 3.3e-07 | 1.6e-07 | 2.9e-07 | 2.600E-07 | 8.888E-08 | 1.600E-07 | 3.300E-07 |
| 5813 | 291 | L291M | A | 2.4e-07 | 2e-07 | 2.3e-07 | 2.233E-07 | 2.082E-08 | 2.000E-07 | 2.400E-07 |
| 5814 | 291 | L291F | A | 2.3e-07 | 1e-07 | 2.3e-07 | 1.867E-07 | 7.506E-08 | 1.000E-07 | 2.300E-07 |
| 5815 | 291 | L291P | A | 3.8e-07 | 3.2e-07 | 1.5e-07 | 2.833E-07 | 1.193E-07 | 1.500E-07 | 3.800E-07 |
| 5816 | 291 | L291S | A | 2.7e-07 | 3.2e-07 | 6.6e-07 | 4.167E-07 | 2.122E-07 | 2.700E-07 | 6.600E-07 |
| 5817 | 291 | L291T | A | 1.6e-07 | 2.3e-07 | 1e-07 | 1.633E-07 | 6.506E-08 | 1.000E-07 | 2.300E-07 |
| 5818 | 291 | L291W | A | 1.2e-07 | 2.7e-07 | 4.7e-07 | 2.867E-07 | 1.756E-07 | 1.200E-07 | 4.700E-07 |
| 5819 | 291 | L291Y | A | 1.5e-07 | 1.7e-07 | 3.4e-07 | 2.200E-07 | 1.044E-07 | 1.500E-07 | 3.400E-07 |
| 5820 | 291 | L291V | A | 1.3e-07 | 2.5e-07 | 2.4e-07 | 2.067E-07 | 6.658E-08 | 1.300E-07 | 2.500E-07 |
| 5821 | 292 | V292A | A | 1.8e-07 | 1.1e-07 | 2e-07 | 1.633E-07 | 4.726E-08 | 1.100E-07 | 2.000E-07 |
| 5822 | 292 | V292R | A | 3.4e-07 | 4e-07 | 2.9e-07 | 3.433E-07 | 5.508E-08 | 2.900E-07 | 4.000E-07 |
| 5823 | 292 | V292N | A | 1.2e-07 | 1.8e-07 | 3.7e-07 | 2.233E-07 | 1.305E-07 | 1.200E-07 | 3.700E-07 |
| 5824 | 292 | V292D | A | 1.9e-07 | 1.7e-07 | 1.2e-07 | 1.600E-07 | 3.606E-08 | 1.200E-07 | 1.900E-07 |
| 5825 | 292 | V292C | A | 7e-07 | 3.4e-07 | 2.3e-07 | 4.233E-07 | 2.458E-07 | 2.300E-07 | 7.000E-07 |
| 5826 | 292 | V292E | A | 2.2e-07 | 3.2e-07 | 1.8e-07 | 2.400E-07 | 7.211E-08 | 1.800E-07 | 3.200E-07 |
| 5827 | 292 | V292Q | A | 1.6e-07 | 1.2e-07 | 4.5e-07 | 2.433E-07 | 1.801E-07 | 1.200E-07 | 4.500E-07 |
| 5828 | 292 | V292G | A | 1.2e-07 | 3.5e-07 | 2e-07 | 2.233E-07 | 1.168E-07 | 1.200E-07 | 3.500E-07 |
| 5829 | 292 | V292H | A | 2.2e-07 | 2.1e-07 | 2.5e-07 | 2.267E-07 | 2.082E-08 | 2.100E-07 | 2.500E-07 |
| 5830 | 292 | V292I | A | 3.6e-07 | 3.5e-07 | 4.6e-07 | 3.900E-07 | 6.083E-08 | 3.500E-07 | 4.600E-07 |
| 5831 | 292 | V292L | A | 1.5e-07 | 2.2e-07 | 4.6e-07 | 2.767E-07 | 1.626E-07 | 1.500E-07 | 4.600E-07 |
| 5832 | 292 | V292K | A | 1.3e-07 | 2.2e-07 | 2.5e-07 | 2.000E-07 | 6.245E-08 | 1.300E-07 | 2.500E-07 |
| 5833 | 292 | V292M | A | 2.1e-07 | 4.2e-07 | 4.7e-07 | 3.667E-07 | 1.380E-07 | 2.100E-07 | 4.700E-07 |
| 5834 | 292 | V292F | A | 2e-07 | 2.2e-07 | 2.3e-07 | 2.167E-07 | 1.528E-08 | 2.000E-07 | 2.300E-07 |
| 5835 | 292 | V292P | A | 4.3e-07 | 1.3e-07 | 2.8e-07 | 2.800E-07 | 1.500E-07 | 1.300E-07 | 4.300E-07 |
| 5836 | 292 | V292S | A | 1.8e-07 | 1.5e-07 | 8.9e-08 | 1.397E-07 | 4.637E-08 | 8.900E-08 | 1.800E-07 |
| 5837 | 292 | V292T | A | 2.5e-07 | 1.5e-07 | 3e-07 | 2.333E-07 | 7.638E-08 | 1.500E-07 | 3.000E-07 |
| 5838 | 292 | V292W | A | 2.5e-07 | 2.1e-07 | 4.2e-07 | 2.933E-07 | 1.115E-07 | 2.100E-07 | 4.200E-07 |
| 5839 | 292 | V292Y | A | 5e-07 | 2.4e-07 | 2e-07 | 3.133E-07 | 1.629E-07 | 2.000E-07 | 5.000E-07 |
| 5840 | 292 | V292V | A | 1.6e-07 | 2.4e-07 | 2.8e-07 | 2.267E-07 | 6.110E-08 | 1.600E-07 | 2.800E-07 |
| 5841 | 293 | C293A | A | 4.1e-07 | 2.7e-07 | 3.1e-07 | 3.300E-07 | 7.211E-08 | 2.700E-07 | 4.100E-07 |
| 5842 | 293 | C293R | A | 5.5e-07 | 3.7e-07 | 2e-07 | 3.733E-07 | 1.750E-07 | 2.000E-07 | 5.500E-07 |
| 5843 | 293 | C293N | A | 1.6e-07 | 2.5e-07 | 3.1e-07 | 2.400E-07 | 7.550E-08 | 1.600E-07 | 3.100E-07 |
| 5844 | 293 | C293D | A | 2.9e-07 | 1.7e-07 | 1.2e-07 | 1.933E-07 | 8.737E-08 | 1.200E-07 | 2.900E-07 |
| 5845 | 293 | C293C | A | 1.3e-07 | 2.6e-07 | 1.9e-07 | 1.933E-07 | 6.506E-08 | 1.300E-07 | 2.600E-07 |
| 5846 | 293 | C293E | A | 1.7e-07 | 5.9e-07 | 4.9e-07 | 4.167E-07 | 2.194E-07 | 1.700E-07 | 5.900E-07 |
| 5847 | 293 | C293Q | A | 2.2e-07 | 1.8e-07 | 2.1e-07 | 2.033E-07 | 2.082E-08 | 1.800E-07 | 2.200E-07 |
| 5848 | 293 | C293G | A | 2.4e-07 | 3.4e-07 | 1.4e-07 | 2.400E-07 | 1.000E-07 | 1.400E-07 | 3.400E-07 |
| 5849 | 293 | C293H | A | 2.7e-07 | 3.4e-07 | 1.1e-07 | 2.400E-07 | 1.179E-07 | 1.100E-07 | 3.400E-07 |
| 5850 | 293 | C293I | A | 2.1e-07 | 2.7e-07 | 2e-07 | 2.267E-07 | 3.786E-08 | 2.000E-07 | 2.700E-07 |
| 5851 | 293 | C293L | A | 5.5e-07 | 1.3e-07 | 4.9e-07 | 3.900E-07 | 2.272E-07 | 1.300E-07 | 5.500E-07 |
| 5852 | 293 | C293K | A | 5.1e-07 | 2.3e-07 | 1.7e-07 | 3.033E-07 | 1.815E-07 | 1.700E-07 | 5.100E-07 |
| 5853 | 293 | C293M | A | 1.5e-07 | 2.1e-07 | 3.8e-07 | 2.467E-07 | 1.193E-07 | 1.500E-07 | 3.800E-07 |
| 5854 | 293 | C293F | A | 3.5e-07 | 2.2e-07 | 1.9e-07 | 2.533E-07 | 8.505E-08 | 1.900E-07 | 3.500E-07 |
| 5855 | 293 | C293P | A | 2.1e-07 | 3.3e-07 | 5.6e-07 | 3.667E-07 | 1.779E-07 | 2.100E-07 | 5.600E-07 |
| 5856 | 293 | C293S | A | 5.5e-08 | 2.5e-07 | 1.2e-07 | 1.417E-07 | 9.929E-08 | 5.500E-08 | 2.500E-07 |
| 5857 | 293 | C293T | A | 1.8e-07 | 1.7e-07 | 1.5e-07 | 1.667E-07 | 1.528E-08 | 1.500E-07 | 1.800E-07 |
| 5858 | 293 | C293W | A | 2.5e-07 | 2.9e-07 | 2e-07 | 2.467E-07 | 4.509E-08 | 2.000E-07 | 2.900E-07 |
| 5859 | 293 | C293Y | A | 3.8e-07 | 2.5e-07 | 5.6e-07 | 3.967E-07 | 1.557E-07 | 2.500E-07 | 5.600E-07 |
| 5860 | 293 | C293V | A | 2.2e-07 | 2.2e-07 | 1.5e-07 | 1.967E-07 | 4.041E-08 | 1.500E-07 | 2.200E-07 |
| 5861 | 294 | P294A | A | 2.4e-07 | 2.6e-07 | 1.5e-07 | 2.167E-07 | 5.859E-08 | 1.500E-07 | 2.600E-07 |
| 5862 | 294 | P294R | A | 1.7e-07 | 2.6e-07 | 2.6e-07 | 2.300E-07 | 5.196E-08 | 1.700E-07 | 2.600E-07 |
| 5863 | 294 | P294N | A | 5.2e-07 | 2.3e-07 | 1.5e-07 | 3.000E-07 | 1.947E-07 | 1.500E-07 | 5.200E-07 |
| 5864 | 294 | P294D | A | 1.9e-07 | 3.2e-07 | 2.7e-07 | 2.600E-07 | 6.557E-08 | 1.900E-07 | 3.200E-07 |

| | | | | | | | | | | |
|---|---|---|---|---|---|---|---|---|---|---|
| 5865 | 294 | P294C | A | 3.6e-07 | 4.9e-07 | 2.1e-07 | 3.533E-07 | 1.401E-07 | 2.100E-07 | 4.900E-07 |
| 5866 | 294 | P294E | A | 2.8e-07 | 3.2e-07 | 3e-07 | 3.000E-07 | 2.000E-08 | 2.800E-07 | 3.200E-07 |
| 5867 | 294 | P294Q | A | 2.1e-07 | 1.2e-07 | 3.5e-07 | 2.267E-07 | 1.159E-07 | 1.200E-07 | 3.500E-07 |
| 5868 | 294 | P294G | A | 3e-07 | 2.8e-07 | 1e-07 | 2.267E-07 | 1.102E-07 | 1.000E-07 | 3.000E-07 |
| 5869 | 294 | P294H | A | 2.4e-07 | 1.4e-07 | 3.2e-07 | 2.333E-07 | 9.018E-08 | 1.400E-07 | 3.200E-07 |
| 5870 | 294 | P294I | A | 3.5e-07 | 1.2e-07 | 4.7e-07 | 3.133E-07 | 1.779E-07 | 1.200E-07 | 4.700E-07 |
| 5871 | 294 | P294L | A | 3.1e-07 | 2.4e-07 | 2.3e-07 | 2.600E-07 | 4.359E-08 | 2.300E-07 | 3.100E-07 |
| 5872 | 294 | P294K | A | 2.6e-07 | 2.8e-07 | 2.2e-07 | 2.533E-07 | 3.055E-08 | 2.200E-07 | 2.800E-07 |
| 5873 | 294 | P294M | A | 2.6e-07 | 1.9e-07 | 1.9e-07 | 2.133E-07 | 4.041E-08 | 1.900E-07 | 2.600E-07 |
| 5874 | 294 | P294F | A | 3.1e-07 | 1.4e-07 | 1.7e-07 | 2.067E-07 | 9.074E-08 | 1.400E-07 | 3.100E-07 |
| 5875 | 294 | P294P | A | 2.8e-07 | 1.1e-07 | 5.7e-07 | 3.200E-07 | 2.326E-07 | 1.100E-07 | 5.700E-07 |
| 5876 | 294 | P294S | A | 2.5e-07 | 1.9e-07 | 1.6e-07 | 2.000E-07 | 4.583E-08 | 1.600E-07 | 2.500E-07 |
| 5877 | 294 | P294T | A | 2.5e-07 | 4.7e-07 | 3.9e-07 | 3.700E-07 | 1.114E-07 | 2.500E-07 | 4.700E-07 |
| 5878 | 294 | P294W | A | 3e-07 | 2.3e-07 | 1.8e-07 | 2.367E-07 | 6.028E-08 | 1.800E-07 | 3.000E-07 |
| 5879 | 294 | P294Y | A | 2.3e-07 | 2.6e-07 | 2.7e-07 | 2.533E-07 | 2.082E-08 | 2.300E-07 | 2.700E-07 |
| 5880 | 294 | P294V | A | 2.6e-07 | 4.6e-07 | 3.5e-07 | 3.567E-07 | 1.002E-07 | 2.600E-07 | 4.600E-07 |
| 5881 | 295 | L295A | A | 3.1e-07 | 5e-07 | 3.5e-07 | 3.867E-07 | 1.002E-07 | 3.100E-07 | 5.000E-07 |
| 5882 | 295 | L295R | A | 2.4e-07 | 2.1e-07 | 1.1e-07 | 1.867E-07 | 6.807E-08 | 1.100E-07 | 2.400E-07 |
| 5883 | 295 | L295N | A | 1.2e-07 | 4e-07 | 1.5e-07 | 2.233E-07 | 1.537E-07 | 1.200E-07 | 4.000E-07 |
| 5884 | 295 | L295D | A | 2e-07 | 2.8e-07 | 4.6e-07 | 3.133E-07 | 1.332E-07 | 2.000E-07 | 4.600E-07 |
| 5885 | 295 | L295C | A | 1.8e-07 | 2.6e-07 | 1.8e-07 | 2.067E-07 | 4.619E-08 | 1.800E-07 | 2.600E-07 |
| 5886 | 295 | L295E | A | 2e-07 | 5.2e-07 | 4.8e-07 | 4.000E-07 | 1.744E-07 | 2.000E-07 | 5.200E-07 |
| 5887 | 295 | L295Q | A | 2.6e-07 | 4.3e-07 | 2.5e-07 | 3.133E-07 | 1.012E-07 | 2.500E-07 | 4.300E-07 |
| 5888 | 295 | L295G | A | 2.6e-07 | 8.6e-08 | 3.3e-07 | 2.253E-07 | 1.256E-07 | 8.600E-08 | 3.300E-07 |
| 5889 | 295 | L295H | A | 2.7e-07 | 1.4e-07 | 3.9e-07 | 2.667E-07 | 1.250E-07 | 1.400E-07 | 3.900E-07 |
| 5890 | 295 | L295I | A | 1.2e-07 | 1.9e-07 | 3.7e-07 | 2.267E-07 | 1.290E-07 | 1.200E-07 | 3.700E-07 |
| 5891 | 295 | L295L | A | 1.3e-07 | 2.6e-07 | 1.9e-07 | 1.933E-07 | 6.506E-08 | 1.300E-07 | 2.600E-07 |
| 5892 | 295 | L295K | A | 2.8e-07 | 1.6e-07 | 1.9e-07 | 2.100E-07 | 6.245E-08 | 1.600E-07 | 2.800E-07 |
| 5893 | 295 | L295M | A | 3.4e-07 | 3e-07 | 2.7e-07 | 3.033E-07 | 3.512E-08 | 2.700E-07 | 3.400E-07 |
| 5894 | 295 | L295F | A | 2.1e-07 | 2.8e-07 | 2.3e-07 | 2.400E-07 | 3.606E-08 | 2.100E-07 | 2.800E-07 |
| 5895 | 295 | L295P | A | 1.2e-07 | 1.6e-07 | 3.5e-07 | 2.100E-07 | 1.229E-07 | 1.200E-07 | 3.500E-07 |
| 5896 | 295 | L295S | A | 2.2e-07 | 3.4e-07 | 4.4e-07 | 3.333E-07 | 1.102E-07 | 2.200E-07 | 4.400E-07 |
| 5897 | 295 | L295T | A | 3.1e-07 | 2.3e-07 | 1.9e-07 | 2.433E-07 | 6.110E-08 | 1.900E-07 | 3.100E-07 |
| 5898 | 295 | L295W | A | 2.5e-07 | 2.2e-07 | 3.9e-07 | 2.867E-07 | 9.074E-08 | 2.200E-07 | 3.900E-07 |
| 5899 | 295 | L295Y | A | 1.7e-07 | 3.8e-07 | 1.3e-07 | 2.267E-07 | 1.343E-07 | 1.300E-07 | 3.800E-07 |
| 5900 | 295 | L295V | A | 2.9e-07 | 3e-07 | 3.4e-07 | 3.100E-07 | 2.646E-08 | 2.900E-07 | 3.400E-07 |
| 5901 | 296 | H296A | A | 3e-07 | 1.4e-07 | 2e-07 | 2.133E-07 | 8.083E-08 | 1.400E-07 | 3.000E-07 |
| 5902 | 296 | H296R | A | 3.2e-07 | 3e-07 | 5.8e-07 | 4.000E-07 | 1.562E-07 | 3.000E-07 | 5.800E-07 |
| 5903 | 296 | H296N | A | 2.4e-07 | 3.4e-07 | 1.7e-07 | 2.500E-07 | 8.544E-08 | 1.700E-07 | 3.400E-07 |
| 5904 | 296 | H296D | A | 2.2e-07 | 3e-07 | 2.2e-07 | 2.467E-07 | 4.619E-08 | 2.200E-07 | 3.000E-07 |
| 5905 | 296 | H296C | A | 3.3e-07 | 2.1e-07 | 1.1e-07 | 2.167E-07 | 1.102E-07 | 1.100E-07 | 3.300E-07 |
| 5906 | 296 | H296E | A | 3.2e-07 | 3.8e-07 | 4.6e-07 | 3.867E-07 | 7.024E-08 | 3.200E-07 | 4.600E-07 |
| 5907 | 296 | H296Q | A | 4e-07 | 3e-07 | 9.9e-08 | 2.663E-07 | 1.533E-07 | 9.900E-08 | 4.000E-07 |
| 5908 | 296 | H296G | A | 1.8e-07 | 6.6e-07 | 2e-07 | 3.467E-07 | 2.715E-07 | 1.800E-07 | 6.600E-07 |
| 5909 | 296 | H296H | A | 2.3e-07 | 1e-07 | 2e-07 | 1.767E-07 | 6.807E-08 | 1.000E-07 | 2.300E-07 |
| 5910 | 296 | H296I | A | 2.9e-07 | 3.5e-07 | 1.3e-07 | 2.567E-07 | 1.137E-07 | 1.300E-07 | 3.500E-07 |
| 5911 | 296 | H296L | A | 2.7e-07 | 3.1e-07 | 1.9e-07 | 2.567E-07 | 6.110E-08 | 1.900E-07 | 3.100E-07 |
| 5912 | 296 | H296K | A | 2.1e-07 | 2.5e-07 | 2.3e-07 | 2.300E-07 | 2.000E-08 | 2.100E-07 | 2.500E-07 |
| 5913 | 296 | H296M | A | 2.8e-07 | 6.1e-07 | 1.7e-07 | 3.533E-07 | 2.290E-07 | 1.700E-07 | 6.100E-07 |
| 5914 | 296 | H296F | A | 3.5e-07 | 3.5e-07 | 1.6e-07 | 2.867E-07 | 1.097E-07 | 1.600E-07 | 3.500E-07 |
| 5915 | 296 | H296P | A | 1.6e-07 | 2.6e-07 | 3.2e-07 | 2.467E-07 | 8.083E-08 | 1.600E-07 | 3.200E-07 |
| 5916 | 296 | H296S | A | 2e-07 | 4.7e-07 | 3.4e-07 | 3.367E-07 | 1.350E-07 | 2.000E-07 | 4.700E-07 |
| 5917 | 296 | H296T | A | 1.9e-07 | 1.2e-07 | 2e-07 | 1.700E-07 | 4.359E-08 | 1.200E-07 | 2.000E-07 |
| 5918 | 296 | H296W | A | 3.6e-07 | 2.4e-07 | 4.1e-07 | 3.367E-07 | 8.737E-08 | 2.400E-07 | 4.100E-07 |
| 5919 | 296 | H296Y | A | 2e-07 | 3.5e-07 | 3.2e-07 | 2.900E-07 | 7.937E-08 | 2.000E-07 | 3.500E-07 |
| 5920 | 296 | H296V | A | 3.6e-07 | 3.1e-07 | 3.2e-07 | 3.300E-07 | 2.646E-08 | 3.100E-07 | 3.600E-07 |
| 5921 | 297 | N297A | A | 7.9e-07 | 2.1e-07 | 1.8e-07 | 3.933E-07 | 3.439E-07 | 1.800E-07 | 7.900E-07 |
| 5922 | 297 | N297R | A | 2.8e-07 | 2.4e-07 | 3.7e-07 | 2.967E-07 | 6.658E-08 | 2.400E-07 | 3.700E-07 |
| 5923 | 297 | N297N | A | 2.5e-07 | 6.9e-07 | 2.9e-07 | 4.100E-07 | 2.433E-07 | 2.500E-07 | 6.900E-07 |
| 5924 | 297 | N297D | A | 3.1e-07 | 2.2e-07 | 2.8e-07 | 2.700E-07 | 4.583E-08 | 2.200E-07 | 3.100E-07 |
| 5925 | 297 | N297C | A | 4.3e-07 | 3.2e-07 | 1e-07 | 2.833E-07 | 1.680E-07 | 1.000E-07 | 4.300E-07 |
| 5926 | 297 | N297E | A | 1.5e-07 | 4.1e-07 | 1e-07 | 2.200E-07 | 1.664E-07 | 1.000E-07 | 4.100E-07 |
| 5927 | 297 | N297Q | A | 1.5e-07 | 4.3e-07 | 2.2e-07 | 2.667E-07 | 1.457E-07 | 1.500E-07 | 4.300E-07 |
| 5928 | 297 | N297G | A | 3.6e-07 | 2.9e-07 | 3e-07 | 3.167E-07 | 3.786E-08 | 2.900E-07 | 3.600E-07 |
| 5929 | 297 | N297H | A | 1.1e-07 | 2e-07 | 4.4e-07 | 2.500E-07 | 1.706E-07 | 1.100E-07 | 4.400E-07 |
| 5930 | 297 | N297I | A | 2.3e-07 | 4.7e-07 | 4.2e-07 | 3.733E-07 | 1.266E-07 | 2.300E-07 | 4.700E-07 |
| 5931 | 297 | N297L | A | 1.4e-07 | 4.6e-07 | 2.6e-07 | 2.867E-07 | 1.617E-07 | 1.400E-07 | 4.600E-07 |
| 5932 | 297 | N297K | A | 3.3e-07 | 3.1e-07 | 3e-07 | 3.133E-07 | 1.528E-08 | 3.000E-07 | 3.300E-07 |
| 5933 | 297 | N297M | A | 2.2e-07 | 3.8e-07 | 2e-07 | 2.667E-07 | 9.866E-08 | 2.000E-07 | 3.800E-07 |
| 5934 | 297 | N297F | A | 2.3e-07 | 3e-07 | 3.5e-07 | 2.933E-07 | 6.028E-08 | 2.300E-07 | 3.500E-07 |
| 5935 | 297 | N297P | A | 1.8e-07 | 5.2e-07 | 2.6e-07 | 3.200E-07 | 1.778E-07 | 1.800E-07 | 5.200E-07 |
| 5936 | 297 | N297S | A | 5.5e-07 | 4.6e-07 | 1.4e-07 | 3.833E-07 | 2.155E-07 | 1.400E-07 | 5.500E-07 |
| 5937 | 297 | N297T | A | 3.6e-07 | 3.5e-07 | 3e-07 | 3.367E-07 | 3.215E-08 | 3.000E-07 | 3.600E-07 |
| 5938 | 297 | N297W | A | 3.9e-07 | 2.9e-07 | 7.4e-07 | 4.733E-07 | 2.363E-07 | 2.900E-07 | 7.400E-07 |
| 5939 | 297 | N297Y | A | 4.1e-07 | 2.8e-07 | 4.6e-07 | 3.833E-07 | 9.292E-08 | 2.800E-07 | 4.600E-07 |
| 5940 | 297 | N297V | A | 4.3e-07 | 3e-07 | 2.1e-07 | 3.133E-07 | 1.106E-07 | 2.100E-07 | 4.300E-07 |
| 5941 | 298 | Q298A | A | 4.2e-07 | 2.4e-07 | 1.5e-07 | 2.700E-07 | 1.375E-07 | 1.500E-07 | 4.200E-07 |
| 5942 | 298 | Q298R | A | 2e-07 | 5.9e-07 | 1.8e-07 | 3.233E-07 | 2.312E-07 | 1.800E-07 | 5.900E-07 |
| 5943 | 298 | Q298N | A | 3e-07 | 1.4e-07 | 1.1e-07 | 1.833E-07 | 1.021E-07 | 1.100E-07 | 3.000E-07 |
| 5944 | 298 | Q298D | A | 2.9e-07 | 2.5e-07 | 1.5e-07 | 2.300E-07 | 7.211E-08 | 1.500E-07 | 2.900E-07 |
| 5945 | 298 | Q298C | A | 2.4e-07 | 3.8e-07 | 3.4e-07 | 3.200E-07 | 7.211E-08 | 2.400E-07 | 3.800E-07 |
| 5946 | 298 | Q298E | A | 3.2e-07 | 1.8e-07 | 1.5e-07 | 2.167E-07 | 9.074E-08 | 1.500E-07 | 3.200E-07 |
| 5947 | 298 | Q298Q | A | 4.2e-07 | 5.1e-07 | 7.3e-07 | 5.533E-07 | 1.595E-07 | 4.200E-07 | 7.300E-07 |
| 5948 | 298 | Q298G | A | 4.6e-07 | 3.4e-07 | 4e-07 | 4.000E-07 | 6.000E-08 | 3.400E-07 | 4.600E-07 |
| 5949 | 298 | Q298H | A | 9.7e-08 | 1.8e-07 | 2.6e-07 | 1.790E-07 | 8.150E-08 | 9.700E-08 | 2.600E-07 |

| | | | | | | | | | | |
|---|---|---|---|---|---|---|---|---|---|---|
| 5950 | 298 | Q298I | A | 2.2e-07 | 3.1e-07 | 2.9e-07 | 2.733E-07 | 4.726E-08 | 2.200E-07 | 3.100E-07 |
| 5951 | 298 | Q298L | A | 3e-07 | 3.2e-07 | 2.9e-07 | 3.033E-07 | 1.528E-08 | 2.900E-07 | 3.200E-07 |
| 5952 | 298 | Q298K | A | 1.4e-07 | 2.4e-07 | 3.7e-07 | 2.500E-07 | 1.153E-07 | 1.400E-07 | 3.700E-07 |
| 5953 | 298 | Q298M | A | 2.3e-07 | 1.6e-07 | 3e-07 | 2.300E-07 | 7.000E-08 | 1.600E-07 | 3.000E-07 |
| 5954 | 298 | Q298F | A | 4.6e-07 | 1.9e-07 | 4.3e-07 | 3.600E-07 | 1.480E-07 | 1.900E-07 | 4.600E-07 |
| 5955 | 298 | Q298P | A | 2.9e-07 | 8.2e-07 | 4.5e-07 | 5.200E-07 | 2.718E-07 | 2.900E-07 | 8.200E-07 |
| 5956 | 298 | Q298S | A | 4e-07 | 2.4e-07 | 2.7e-07 | 3.033E-07 | 8.505E-08 | 2.400E-07 | 4.000E-07 |
| 5957 | 298 | Q298T | A | 2.9e-07 | 2.8e-07 | 1.7e-07 | 2.467E-07 | 6.658E-08 | 1.700E-07 | 2.900E-07 |
| 5958 | 298 | Q298W | A | 3.2e-07 | 1.7e-07 | 2.9e-07 | 2.600E-07 | 7.937E-08 | 1.700E-07 | 3.200E-07 |
| 5959 | 298 | Q298Y | A | 2.3e-07 | 1.7e-07 | 4.4e-07 | 2.800E-07 | 1.418E-07 | 1.700E-07 | 4.400E-07 |
| 5960 | 298 | Q298V | A | 2.4e-07 | 2.2e-07 | 1.6e-07 | 2.067E-07 | 4.163E-08 | 1.600E-07 | 2.400E-07 |
| 5961 | 299 | E299A | A | 1.7e-07 | 1.1e-07 | 4e-07 | 2.267E-07 | 1.531E-07 | 1.100E-07 | 4.000E-07 |
| 5962 | 299 | E299R | A | 2.8e-07 | 5.9e-07 | 1.6e-07 | 3.433E-07 | 2.219E-07 | 1.600E-07 | 5.900E-07 |
| 5963 | 299 | E299N | A | 3e-07 | 4.9e-07 | 8.5e-08 | 2.917E-07 | 2.026E-07 | 8.500E-08 | 4.900E-07 |
| 5964 | 299 | E299D | A | 2e-07 | 1.9e-07 | 1.6e-07 | 1.833E-07 | 2.082E-08 | 1.600E-07 | 2.000E-07 |
| 5965 | 299 | E299C | A | 3.6e-07 | 2.3e-07 | 2.2e-07 | 2.700E-07 | 7.810E-08 | 2.200E-07 | 3.600E-07 |
| 5966 | 299 | E299E | A | 3.9e-07 | 1.8e-07 | 4e-07 | 3.233E-07 | 1.242E-07 | 1.800E-07 | 4.000E-07 |
| 5967 | 299 | E299Q | A | 2.1e-07 | 3.2e-07 | 1.6e-07 | 2.300E-07 | 8.185E-08 | 1.600E-07 | 3.200E-07 |
| 5968 | 299 | E299G | A | 2.2e-07 | 3.4e-07 | 2.8e-07 | 2.800E-07 | 6.000E-08 | 2.200E-07 | 3.400E-07 |
| 5969 | 299 | E299H | A | 4.2e-07 | 4.2e-07 | 3e-07 | 3.800E-07 | 6.928E-08 | 3.000E-07 | 4.200E-07 |
| 5970 | 299 | E299I | A | 3e-07 | 2.8e-07 | 2.5e-07 | 2.767E-07 | 2.517E-08 | 2.500E-07 | 3.000E-07 |
| 5971 | 299 | E299L | A | 1.3e-07 | 1.7e-07 | 1.8e-07 | 1.600E-07 | 2.646E-08 | 1.300E-07 | 1.800E-07 |
| 5972 | 299 | E299K | A | 2.2e-07 | 4.8e-07 | 2.9e-07 | 3.300E-07 | 1.345E-07 | 2.200E-07 | 4.800E-07 |
| 5973 | 299 | E299M | A | 1.4e-07 | 3.5e-07 | 3e-07 | 2.633E-07 | 1.097E-07 | 1.400E-07 | 3.500E-07 |
| 5974 | 299 | E299F | A | 3.5e-07 | 3.3e-07 | 1.5e-07 | 2.767E-07 | 1.102E-07 | 1.500E-07 | 3.500E-07 |
| 5975 | 299 | E299P | A | 3.3e-07 | 2.2e-07 | 1.7e-07 | 2.400E-07 | 8.185E-08 | 1.700E-07 | 3.300E-07 |
| 5976 | 299 | E299S | A | 3.3e-07 | 2.2e-07 | 1.1e-07 | 2.200E-07 | 1.100E-07 | 1.100E-07 | 3.300E-07 |
| 5977 | 299 | E299T | A | 2.8e-07 | 5e-07 | 2.8e-07 | 3.533E-07 | 1.270E-07 | 2.800E-07 | 5.000E-07 |
| 5978 | 299 | E299W | A | 1.1e-07 | 4e-07 | 2.2e-07 | 2.433E-07 | 1.464E-07 | 1.100E-07 | 4.000E-07 |
| 5979 | 299 | E299Y | A | 2.2e-07 | 4e-07 | 5.3e-07 | 3.833E-07 | 1.557E-07 | 2.200E-07 | 5.300E-07 |
| 5980 | 299 | E299V | A | 1.8e-07 | 2.3e-07 | 2.4e-07 | 2.167E-07 | 3.215E-08 | 1.800E-07 | 2.400E-07 |
| 5981 | 300 | V300A | A | 7.2e-07 | 3.2e-07 | 3e-07 | 4.467E-07 | 2.369E-07 | 3.000E-07 | 7.200E-07 |
| 5982 | 300 | V300R | A | 9.3e-08 | 4.8e-07 | 3.1e-07 | 2.943E-07 | 1.940E-07 | 9.300E-08 | 4.800E-07 |
| 5983 | 300 | V300N | A | 3.7e-07 | 2.9e-07 | 2.3e-07 | 2.967E-07 | 7.024E-08 | 2.300E-07 | 3.700E-07 |
| 5984 | 300 | V300D | A | 2.3e-07 | 2.6e-07 | 3.3e-07 | 2.733E-07 | 5.132E-08 | 2.300E-07 | 3.300E-07 |
| 5985 | 300 | V300C | A | 1.7e-07 | 1.3e-07 | 3.3e-07 | 2.100E-07 | 1.058E-07 | 1.300E-07 | 3.300E-07 |
| 5986 | 300 | V300E | A | 2e-07 | 2.3e-07 | 3.3e-07 | 2.533E-07 | 6.807E-08 | 2.000E-07 | 3.300E-07 |
| 5987 | 300 | V300Q | A | 3e-07 | 6.3e-07 | 1.9e-07 | 3.733E-07 | 2.290E-07 | 1.900E-07 | 6.300E-07 |
| 5988 | 300 | V300G | A | 2.3e-07 | 8.4e-08 | 3.2e-07 | 2.113E-07 | 1.191E-07 | 8.400E-08 | 3.200E-07 |
| 5989 | 300 | V300H | A | 1.8e-07 | 1.8e-07 | 3.1e-07 | 2.233E-07 | 7.506E-08 | 1.800E-07 | 3.100E-07 |
| 5990 | 300 | V300I | A | 1.9e-07 | 5.2e-07 | 1.3e-07 | 2.800E-07 | 2.100E-07 | 1.300E-07 | 5.200E-07 |
| 5991 | 300 | V300L | A | 2.5e-07 | 2.2e-07 | 3.2e-07 | 2.633E-07 | 5.132E-08 | 2.200E-07 | 3.200E-07 |
| 5992 | 300 | V300K | A | 1.6e-07 | 4e-07 | 1.5e-07 | 2.367E-07 | 1.415E-07 | 1.500E-07 | 4.000E-07 |
| 5993 | 300 | V300M | A | 1.2e-07 | 2.3e-07 | 2.2e-07 | 1.900E-07 | 6.083E-08 | 1.200E-07 | 2.300E-07 |
| 5994 | 300 | V300F | A | 4e-07 | 2.3e-07 | 1.7e-07 | 2.667E-07 | 1.193E-07 | 1.700E-07 | 4.000E-07 |
| 5995 | 300 | V300P | A | 1.5e-07 | 2e-07 | 3.5e-07 | 2.333E-07 | 1.041E-07 | 1.500E-07 | 3.500E-07 |
| 5996 | 300 | V300S | A | 1.7e-07 | 2.1e-07 | 2.4e-07 | 2.067E-07 | 3.512E-08 | 1.700E-07 | 2.400E-07 |
| 5997 | 300 | V300T | A | 4.6e-07 | 3e-07 | 2e-07 | 3.200E-07 | 1.311E-07 | 2.000E-07 | 4.600E-07 |
| 5998 | 300 | V300W | A | 4.5e-07 | 2.1e-07 | 1.2e-07 | 2.600E-07 | 1.706E-07 | 1.200E-07 | 4.500E-07 |
| 5999 | 300 | V300Y | A | 3e-07 | 2.5e-07 | 2.9e-07 | 2.800E-07 | 2.646E-08 | 2.500E-07 | 3.000E-07 |
| 6000 | 300 | V300V | A | 3.4e-07 | 2.5e-07 | 2.9e-07 | 2.933E-07 | 4.509E-08 | 2.500E-07 | 3.400E-07 |
| 6001 | 301 | T301A | A | 3.2e-07 | 3.4e-07 | 3.1e-07 | 3.233E-07 | 1.528E-08 | 3.100E-07 | 3.400E-07 |
| 6002 | 301 | T301R | A | 4.1e-07 | 2.2e-07 | 4.7e-07 | 3.667E-07 | 1.305E-07 | 2.200E-07 | 4.700E-07 |
| 6003 | 301 | T301N | A | 2.4e-07 | 1.5e-07 | 1.1e-07 | 1.667E-07 | 6.658E-08 | 1.100E-07 | 2.400E-07 |
| 6004 | 301 | T301D | A | 7.6e-07 | 2.7e-07 | 2.3e-07 | 4.200E-07 | 2.951E-07 | 2.300E-07 | 7.600E-07 |
| 6005 | 301 | T301C | A | 1.5e-07 | 2e-07 | 2.1e-07 | 1.867E-07 | 3.215E-08 | 1.500E-07 | 2.100E-07 |
| 6006 | 301 | T301E | A | 1.8e-07 | 2.8e-07 | 2.8e-07 | 2.467E-07 | 5.774E-08 | 1.800E-07 | 2.800E-07 |
| 6007 | 301 | T301Q | A | 3.9e-07 | 3.5e-07 | 2e-07 | 3.133E-07 | 1.002E-07 | 2.000E-07 | 3.900E-07 |
| 6008 | 301 | T301G | A | 2.5e-07 | 2.2e-07 | 3.8e-07 | 2.833E-07 | 8.505E-08 | 2.200E-07 | 3.800E-07 |
| 6009 | 301 | T301H | A | 1.1e-07 | 2.1e-07 | 1.3e-07 | 1.500E-07 | 5.292E-08 | 1.100E-07 | 2.100E-07 |
| 6010 | 301 | T301I | A | 1.4e-07 | 2.4e-07 | 3e-07 | 2.267E-07 | 8.083E-08 | 1.400E-07 | 3.000E-07 |
| 6011 | 301 | T301L | A | 2.1e-07 | 3.4e-07 | 2.2e-07 | 2.567E-07 | 7.234E-08 | 2.100E-07 | 3.400E-07 |
| 6012 | 301 | T301K | A | 6.4e-07 | 2.1e-07 | 2.1e-07 | 3.533E-07 | 2.483E-07 | 2.100E-07 | 6.400E-07 |
| 6013 | 301 | T301M | A | 1.5e-07 | 4.7e-07 | 3.3e-07 | 3.167E-07 | 1.604E-07 | 1.500E-07 | 4.700E-07 |
| 6014 | 301 | T301F | A | 1.8e-07 | 4.6e-07 | 2.2e-07 | 2.867E-07 | 1.514E-07 | 1.800E-07 | 4.600E-07 |
| 6015 | 301 | T301P | A | 8.5e-08 | 2.2e-07 | 2.8e-07 | 1.950E-07 | 9.987E-08 | 8.500E-08 | 2.800E-07 |
| 6016 | 301 | T301S | A | 3.2e-07 | 3e-07 | 6e-07 | 4.067E-07 | 1.677E-07 | 3.000E-07 | 6.000E-07 |
| 6017 | 301 | T301T | A | 2.3e-07 | 1.4e-07 | 2.4e-07 | 2.033E-07 | 5.508E-08 | 1.400E-07 | 2.400E-07 |
| 6018 | 301 | T301W | A | 4.5e-07 | 1.2e-07 | 2.8e-07 | 2.833E-07 | 1.650E-07 | 1.200E-07 | 4.500E-07 |
| 6019 | 301 | T301Y | A | 1.4e-07 | 3.2e-07 | 1.7e-07 | 2.100E-07 | 9.644E-08 | 1.400E-07 | 3.200E-07 |
| 6020 | 301 | T301V | A | 2.2e-07 | 2.4e-07 | 3.5e-07 | 2.700E-07 | 7.000E-08 | 2.200E-07 | 3.500E-07 |
| 6021 | 302 | A302A | A | 3e-07 | 1.3e-07 | 3.2e-07 | 2.500E-07 | 1.044E-07 | 1.300E-07 | 3.200E-07 |
| 6022 | 302 | A302R | A | 2.6e-07 | 3.1e-07 | 2.2e-07 | 2.633E-07 | 4.509E-08 | 2.200E-07 | 3.100E-07 |
| 6023 | 302 | A302N | A | 9.8e-08 | 3.6e-07 | 4.2e-07 | 2.927E-07 | 1.712E-07 | 9.800E-08 | 4.200E-07 |
| 6024 | 302 | A302D | A | 3.3e-07 | 2.4e-07 | 2.4e-07 | 2.700E-07 | 5.196E-08 | 2.400E-07 | 3.300E-07 |
| 6025 | 302 | A302C | A | 2.1e-07 | 2.3e-07 | 1.7e-07 | 2.033E-07 | 3.055E-08 | 1.700E-07 | 2.300E-07 |
| 6026 | 302 | A302E | A | 2.5e-07 | 3.8e-07 | 1.2e-07 | 2.500E-07 | 1.300E-07 | 1.200E-07 | 3.800E-07 |
| 6027 | 302 | A302Q | A | 1.6e-07 | 4.1e-07 | 5.1e-07 | 3.600E-07 | 1.803E-07 | 1.600E-07 | 5.100E-07 |
| 6028 | 302 | A302G | A | 1.2e-07 | 3.4e-07 | 4.9e-07 | 3.167E-07 | 1.861E-07 | 1.200E-07 | 4.900E-07 |
| 6029 | 302 | A302H | A | 6e-07 | 3.6e-07 | 1.7e-07 | 3.767E-07 | 2.155E-07 | 1.700E-07 | 6.000E-07 |
| 6030 | 302 | A302I | A | 2.6e-07 | 2.4e-07 | 2.5e-07 | 2.500E-07 | 1.000E-08 | 2.400E-07 | 2.600E-07 |
| 6031 | 302 | A302L | A | 3.9e-07 | 6.5e-07 | 4.6e-07 | 5.000E-07 | 1.345E-07 | 3.900E-07 | 6.500E-07 |
| 6032 | 302 | A302K | A | 1.9e-07 | 5.4e-07 | 2.5e-07 | 3.267E-07 | 1.872E-07 | 1.900E-07 | 5.400E-07 |
| 6033 | 302 | A302M | A | 3.5e-07 | 2.5e-07 | 2e-07 | 2.667E-07 | 7.638E-08 | 2.000E-07 | 3.500E-07 |
| 6034 | 302 | A302F | A | 3.9e-07 | 4.2e-07 | 3.7e-07 | 3.933E-07 | 2.517E-08 | 3.700E-07 | 4.200E-07 |

| | | | | | | | | | | |
|---|---|---|---|---|---|---|---|---|---|---|
| 6035 | 302 | A302P | A | 7.5e-07 | 2.2e-07 | 4.6e-07 | 4.767E-07 | 2.654E-07 | 2.200E-07 | 7.500E-07 |
| 6036 | 302 | A302S | A | 2.5e-07 | 3.2e-07 | 1.8e-07 | 2.500E-07 | 7.000E-08 | 1.800E-07 | 3.200E-07 |
| 6037 | 302 | A302T | A | 2.1e-07 | 1.9e-07 | 3.8e-07 | 2.600E-07 | 1.044E-07 | 1.900E-07 | 3.800E-07 |
| 6038 | 302 | A302W | A | 9.9e-08 | 2e-07 | 1.8e-07 | 1.597E-07 | 5.348E-08 | 9.900E-08 | 2.000E-07 |
| 6039 | 302 | A302Y | A | 1.3e-07 | 3e-07 | 4e-07 | 2.767E-07 | 1.365E-07 | 1.300E-07 | 4.000E-07 |
| 6040 | 302 | A302V | A | 2.8e-07 | 5.5e-07 | 2.6e-07 | 3.633E-07 | 1.620E-07 | 2.600E-07 | 5.500E-07 |
| 6041 | 303 | E303A | A | 2.8e-07 | 1.8e-07 | 4.3e-08 | 1.677E-07 | 1.190E-07 | 4.300E-08 | 2.800E-07 |
| 6042 | 303 | E303R | A | 5.1e-07 | 1.5e-07 | 2.2e-07 | 2.933E-07 | 1.909E-07 | 1.500E-07 | 5.100E-07 |
| 6043 | 303 | E303N | A | 2.7e-07 | 2.9e-07 | 4.1e-07 | 3.233E-07 | 7.572E-08 | 2.700E-07 | 4.100E-07 |
| 6044 | 303 | E303D | A | 2.2e-07 | 9.7e-08 | 2.7e-07 | 1.957E-07 | 8.903E-08 | 9.700E-08 | 2.700E-07 |
| 6045 | 303 | E303C | A | 2.7e-07 | 2.1e-07 | 8.4e-07 | 4.400E-07 | 3.477E-07 | 2.100E-07 | 8.400E-07 |
| 6046 | 303 | E303E | A | 3.3e-07 | 1.4e-07 | 1.7e-07 | 2.133E-07 | 1.021E-07 | 1.400E-07 | 3.300E-07 |
| 6047 | 303 | E303Q | A | 1.3e-07 | 2.3e-07 | 3.6e-07 | 2.400E-07 | 1.153E-07 | 1.300E-07 | 3.600E-07 |
| 6048 | 303 | E303G | A | 4.1e-07 | 1.9e-07 | 2.7e-07 | 2.900E-07 | 1.114E-07 | 1.900E-07 | 4.100E-07 |
| 6049 | 303 | E303H | A | 4.5e-07 | 1.8e-07 | 1.6e-07 | 2.633E-07 | 1.620E-07 | 1.600E-07 | 4.500E-07 |
| 6050 | 303 | E303I | A | 2.8e-07 | 2.1e-07 | 1.8e-07 | 2.233E-07 | 5.132E-08 | 1.800E-07 | 2.800E-07 |
| 6051 | 303 | E303L | A | 3.4e-07 | 2.4e-07 | 2.3e-07 | 2.700E-07 | 6.083E-08 | 2.300E-07 | 3.400E-07 |
| 6052 | 303 | E303K | A | 3e-07 | 2.1e-07 | 1.6e-07 | 2.233E-07 | 7.095E-08 | 1.600E-07 | 3.000E-07 |
| 6053 | 303 | E303M | A | 1.7e-07 | 3.4e-07 | 2.2e-07 | 2.433E-07 | 8.737E-08 | 1.700E-07 | 3.400E-07 |
| 6054 | 303 | E303F | A | 1.6e-07 | 1.5e-07 | 2.1e-07 | 1.733E-07 | 3.215E-08 | 1.500E-07 | 2.100E-07 |
| 6055 | 303 | E303P | A | 1e-07 | 1.4e-07 | 1.2e-07 | 1.200E-07 | 2.000E-08 | 1.000E-07 | 1.400E-07 |
| 6056 | 303 | E303S | A | 2.5e-07 | 2.6e-07 | 2.4e-07 | 2.500E-07 | 1.000E-08 | 2.400E-07 | 2.600E-07 |
| 6057 | 303 | E303T | A | 3e-07 | 2.8e-07 | 2e-07 | 2.600E-07 | 5.292E-08 | 2.000E-07 | 3.000E-07 |
| 6058 | 303 | E303W | A | 4e-07 | 1.9e-07 | 3.2e-07 | 3.033E-07 | 1.060E-07 | 1.900E-07 | 4.000E-07 |
| 6059 | 303 | E303Y | A | 2.2e-07 | 8.6e-08 | 1.6e-07 | 1.553E-07 | 6.712E-08 | 8.600E-08 | 2.200E-07 |
| 6060 | 303 | E303V | A | 3e-07 | 2.1e-07 | 4.1e-07 | 3.067E-07 | 1.002E-07 | 2.100E-07 | 4.100E-07 |
| 6061 | 304 | D304A | A | 3.6e-07 | 1.2e-07 | 3.1e-07 | 2.633E-07 | 1.266E-07 | 1.200E-07 | 3.600E-07 |
| 6062 | 304 | D304R | A | 2.9e-07 | 1.5e-07 | 1.5e-07 | 1.967E-07 | 8.083E-08 | 1.500E-07 | 2.900E-07 |
| 6063 | 304 | D304N | A | 2.7e-07 | 8.6e-07 | 6.5e-07 | 5.933E-07 | 2.991E-07 | 2.700E-07 | 8.600E-07 |
| 6064 | 304 | D304D | A | 1.6e-07 | 1.9e-07 | 1.1e-07 | 1.533E-07 | 4.041E-08 | 1.100E-07 | 1.900E-07 |
| 6065 | 304 | D304C | A | 1.6e-07 | 1.5e-07 | 2.4e-07 | 1.833E-07 | 4.933E-08 | 1.500E-07 | 2.400E-07 |
| 6066 | 304 | D304E | A | 2.3e-07 | 1.4e-07 | 1.6e-07 | 1.767E-07 | 4.726E-08 | 1.400E-07 | 2.300E-07 |
| 6067 | 304 | D304Q | A | 2.9e-07 | 2e-07 | 4.6e-07 | 3.167E-07 | 1.320E-07 | 2.000E-07 | 4.600E-07 |
| 6068 | 304 | D304G | A | 2.8e-07 | 2.8e-07 | 1.9e-07 | 2.500E-07 | 5.196E-08 | 1.900E-07 | 2.800E-07 |
| 6069 | 304 | D304H | A | 2.5e-07 | 2.1e-07 | 7.7e-07 | 4.100E-07 | 3.124E-07 | 2.100E-07 | 7.700E-07 |
| 6070 | 304 | D304I | A | 3.2e-07 | 3.3e-07 | 1.9e-07 | 2.800E-07 | 7.810E-08 | 1.900E-07 | 3.300E-07 |
| 6071 | 304 | D304L | A | 9e-08 | 1.3e-07 | 3.7e-07 | 1.967E-07 | 1.514E-07 | 9.000E-08 | 3.700E-07 |
| 6072 | 304 | D304K | A | 2.4e-07 | 3e-07 | 1.8e-07 | 2.400E-07 | 6.000E-08 | 1.800E-07 | 3.000E-07 |
| 6073 | 304 | D304M | A | 5.5e-07 | 3.9e-07 | 2.3e-07 | 3.900E-07 | 1.600E-07 | 2.300E-07 | 5.500E-07 |
| 6074 | 304 | D304F | A | 9.2e-08 | 9.8e-08 | 2.5e-07 | 1.467E-07 | 8.954E-08 | 9.200E-08 | 2.500E-07 |
| 6075 | 304 | D304P | A | 2.3e-07 | 3e-07 | 4e-07 | 3.100E-07 | 8.544E-08 | 2.300E-07 | 4.000E-07 |
| 6076 | 304 | D304S | A | 5.6e-07 | 1.7e-07 | 2.3e-07 | 3.200E-07 | 2.100E-07 | 1.700E-07 | 5.600E-07 |
| 6077 | 304 | D304T | A | 2.5e-07 | 3e-07 | 1.6e-07 | 2.367E-07 | 7.095E-08 | 1.600E-07 | 3.000E-07 |
| 6078 | 304 | D304W | A | 2.1e-07 | 2.7e-07 | 1.4e-07 | 2.067E-07 | 6.506E-08 | 1.400E-07 | 2.700E-07 |
| 6079 | 304 | D304Y | A | 8.6e-08 | 4.4e-07 | 2.7e-07 | 2.653E-07 | 1.770E-07 | 8.600E-08 | 4.400E-07 |
| 6080 | 304 | D304V | A | 3.2e-07 | 1.7e-07 | 1.7e-07 | 2.200E-07 | 8.660E-08 | 1.700E-07 | 3.200E-07 |
| 6081 | 305 | G305A | A | 2e-07 | 1.8e-07 | 1.7e-07 | 1.833E-07 | 1.528E-08 | 1.700E-07 | 2.000E-07 |
| 6082 | 305 | G305R | A | 2.1e-07 | 3.7e-07 | 1.9e-07 | 2.567E-07 | 9.866E-08 | 1.900E-07 | 3.700E-07 |
| 6083 | 305 | G305N | A | 2e-07 | 2.7e-07 | 2.3e-07 | 2.333E-07 | 3.512E-08 | 2.000E-07 | 2.700E-07 |
| 6084 | 305 | G305D | A | 2.2e-07 | 2.2e-07 | 4.3e-07 | 2.900E-07 | 1.212E-07 | 2.200E-07 | 4.300E-07 |
| 6085 | 305 | G305C | A | 1.6e-07 | 2.6e-07 | 2e-07 | 2.067E-07 | 5.033E-08 | 1.600E-07 | 2.600E-07 |
| 6086 | 305 | G305E | A | 1.7e-07 | 2.7e-07 | 4.4e-07 | 2.933E-07 | 1.365E-07 | 1.700E-07 | 4.400E-07 |
| 6087 | 305 | G305Q | A | 2.3e-07 | 3.3e-07 | 3e-07 | 2.867E-07 | 5.132E-08 | 2.300E-07 | 3.300E-07 |
| 6088 | 305 | G305G | A | 5.3e-07 | 3.1e-07 | 8.1e-08 | 3.070E-07 | 2.245E-07 | 8.100E-08 | 5.300E-07 |
| 6089 | 305 | G305H | A | 6.7e-07 | 1.3e-07 | 2.7e-07 | 3.567E-07 | 2.802E-07 | 1.300E-07 | 6.700E-07 |
| 6090 | 305 | G305I | A | 1.4e-07 | 2.9e-07 | 1.4e-07 | 1.900E-07 | 8.660E-08 | 1.400E-07 | 2.900E-07 |
| 6091 | 305 | G305L | A | 3.6e-07 | 4.9e-07 | 1.7e-07 | 3.400E-07 | 1.609E-07 | 1.700E-07 | 4.900E-07 |
| 6092 | 305 | G305K | A | 2.6e-07 | 4.2e-07 | 2.3e-07 | 3.033E-07 | 1.021E-07 | 2.300E-07 | 4.200E-07 |
| 6093 | 305 | G305M | A | 1.3e-07 | 1.8e-07 | 4.8e-07 | 2.633E-07 | 1.893E-07 | 1.300E-07 | 4.800E-07 |
| 6094 | 305 | G305F | A | 1.3e-07 | 3.5e-07 | 2e-07 | 2.267E-07 | 1.124E-07 | 1.300E-07 | 3.500E-07 |
| 6095 | 305 | G305P | A | 2.5e-07 | 1.8e-07 | 1.8e-07 | 2.033E-07 | 4.041E-08 | 1.800E-07 | 2.500E-07 |
| 6096 | 305 | G305S | A | 2.7e-07 | 1.7e-07 | 2.7e-07 | 2.367E-07 | 5.774E-08 | 1.700E-07 | 2.700E-07 |
| 6097 | 305 | G305T | A | 3.1e-07 | 3.4e-07 | 2.3e-07 | 2.933E-07 | 5.686E-08 | 2.300E-07 | 3.400E-07 |
| 6098 | 305 | G305W | A | 2.5e-07 | 2.3e-07 | 2.7e-07 | 2.500E-07 | 2.000E-08 | 2.300E-07 | 2.700E-07 |
| 6099 | 305 | G305Y | A | 3.3e-07 | 3.2e-07 | 1.7e-07 | 2.733E-07 | 8.963E-08 | 1.700E-07 | 3.300E-07 |
| 6100 | 305 | G305V | A | 3.1e-07 | 1.4e-07 | 1.9e-07 | 2.133E-07 | 8.737E-08 | 1.400E-07 | 3.100E-07 |
| 6101 | 306 | T306A | A | 2.4e-07 | 3.4e-07 | 2.2e-07 | 2.667E-07 | 6.429E-08 | 2.200E-07 | 3.400E-07 |
| 6102 | 306 | T306R | A | 4.5e-07 | 1.4e-07 | 2.5e-07 | 2.800E-07 | 1.572E-07 | 1.400E-07 | 4.500E-07 |
| 6103 | 306 | T306N | A | 2.6e-07 | 4.1e-07 | 1.9e-07 | 2.867E-07 | 1.124E-07 | 1.900E-07 | 4.100E-07 |
| 6104 | 306 | T306D | A | 9.7e-08 | 2.2e-07 | 1.4e-07 | 1.523E-07 | 6.242E-08 | 9.700E-08 | 2.200E-07 |
| 6105 | 306 | T306C | A | 1.1e-07 | 1.6e-07 | 2.6e-07 | 1.767E-07 | 7.638E-08 | 1.100E-07 | 2.600E-07 |
| 6106 | 306 | T306E | A | 4.1e-07 | 2.5e-07 | 3.6e-07 | 3.400E-07 | 8.185E-08 | 2.500E-07 | 4.100E-07 |
| 6107 | 306 | T306Q | A | 1.6e-07 | 1.2e-07 | 5.2e-07 | 2.667E-07 | 2.203E-07 | 1.200E-07 | 5.200E-07 |
| 6108 | 306 | T306G | A | 2.7e-07 | 3.7e-07 | 3e-07 | 3.133E-07 | 5.132E-08 | 2.700E-07 | 3.700E-07 |
| 6109 | 306 | T306H | A | 3.2e-07 | 2.5e-07 | 3.4e-07 | 3.033E-07 | 4.726E-08 | 2.500E-07 | 3.400E-07 |
| 6110 | 306 | T306I | A | 2.1e-07 | 3.2e-07 | 1.5e-07 | 2.267E-07 | 8.622E-08 | 1.500E-07 | 3.200E-07 |
| 6111 | 306 | T306L | A | 4.7e-07 | 2.2e-07 | 1.4e-07 | 2.767E-07 | 1.721E-07 | 1.400E-07 | 4.700E-07 |
| 6112 | 306 | T306K | A | 3.7e-07 | 2e-07 | 1.8e-07 | 2.500E-07 | 1.044E-07 | 1.800E-07 | 3.700E-07 |
| 6113 | 306 | T306M | A | 2.2e-07 | 1.6e-07 | 2.7e-07 | 2.167E-07 | 5.508E-08 | 1.600E-07 | 2.700E-07 |
| 6114 | 306 | T306F | A | 3.9e-07 | 1.2e-07 | 2.5e-07 | 2.533E-07 | 1.350E-07 | 1.200E-07 | 3.900E-07 |
| 6115 | 306 | T306P | A | 3.8e-07 | 3e-07 | 2.5e-07 | 3.100E-07 | 6.557E-08 | 2.500E-07 | 3.800E-07 |
| 6116 | 306 | T306S | A | 2.3e-07 | 2.3e-07 | 2.3e-07 | 2.300E-07 | 0.000E+00 | 2.300E-07 | 2.300E-07 |
| 6117 | 306 | T306T | A | 2.1e-07 | 3.3e-07 | 7.3e-08 | 2.043E-07 | 1.286E-07 | 7.300E-08 | 3.300E-07 |
| 6118 | 306 | T306W | A | 1.8e-07 | 3.5e-07 | 2.3e-07 | 2.533E-07 | 8.737E-08 | 1.800E-07 | 3.500E-07 |
| 6119 | 306 | T306Y | A | 2.3e-07 | 1.4e-07 | 4.6e-07 | 2.767E-07 | 1.650E-07 | 1.400E-07 | 4.600E-07 |

| | | | | | | | | | | |
|---|---|---|---|---|---|---|---|---|---|---|
| 6120 | 306 | T306V | A | 2.8e-07 | 2.5e-07 | 3e-07 | 2.767E-07 | 2.517E-08 | 2.500E-07 | 3.000E-07 |
| 6121 | 307 | Q307A | A | 1.7e-07 | 1.7e-07 | 3.2e-07 | 2.200E-07 | 8.660E-08 | 1.700E-07 | 3.200E-07 |
| 6122 | 307 | Q307R | A | 3.3e-07 | 2e-07 | 2.8e-07 | 2.700E-07 | 6.557E-08 | 2.000E-07 | 3.300E-07 |
| 6123 | 307 | Q307N | A | 3.9e-07 | 3e-07 | 2.9e-07 | 3.267E-07 | 5.508E-08 | 2.900E-07 | 3.900E-07 |
| 6124 | 307 | Q307D | A | 9.1e-08 | 5.6e-07 | 4e-07 | 3.503E-07 | 2.384E-07 | 9.100E-08 | 5.600E-07 |
| 6125 | 307 | Q307C | A | 1.9e-07 | 1.3e-07 | 2.1e-07 | 1.767E-07 | 4.163E-08 | 1.300E-07 | 2.100E-07 |
| 6126 | 307 | Q307E | A | 2.2e-07 | 2.6e-07 | 2.3e-07 | 2.367E-07 | 2.082E-08 | 2.200E-07 | 2.600E-07 |
| 6127 | 307 | Q307Q | A | 4e-07 | 1.7e-07 | 1.7e-07 | 2.467E-07 | 1.328E-07 | 1.700E-07 | 4.000E-07 |
| 6128 | 307 | Q307G | A | 2e-07 | 3.9e-07 | 3.5e-07 | 3.133E-07 | 1.002E-07 | 2.000E-07 | 3.900E-07 |
| 6129 | 307 | Q307H | A | 1.8e-07 | 1.1e-07 | 2.6e-07 | 1.833E-07 | 7.506E-08 | 1.100E-07 | 2.600E-07 |
| 6130 | 307 | Q307I | A | 1.8e-07 | 6e-07 | 2.4e-07 | 3.400E-07 | 2.272E-07 | 1.800E-07 | 6.000E-07 |
| 6131 | 307 | Q307L | A | 2e-07 | 8.1e-08 | 2.6e-07 | 1.803E-07 | 9.111E-08 | 8.100E-08 | 2.600E-07 |
| 6132 | 307 | Q307K | A | 1.8e-07 | 1.7e-07 | 3.9e-07 | 2.467E-07 | 1.242E-07 | 1.700E-07 | 3.900E-07 |
| 6133 | 307 | Q307M | A | 1.3e-07 | 2.7e-07 | 1.5e-07 | 1.833E-07 | 7.572E-08 | 1.300E-07 | 2.700E-07 |
| 6134 | 307 | Q307F | A | 4.1e-07 | 2e-07 | 1.9e-07 | 2.667E-07 | 1.242E-07 | 1.900E-07 | 4.100E-07 |
| 6135 | 307 | Q307P | A | 3.1e-07 | 2e-07 | 3.2e-07 | 2.767E-07 | 6.658E-08 | 2.000E-07 | 3.200E-07 |
| 6136 | 307 | Q307S | A | 2.5e-07 | 2.6e-07 | 3e-07 | 2.700E-07 | 2.646E-08 | 2.500E-07 | 3.000E-07 |
| 6137 | 307 | Q307T | A | 2.6e-07 | 3.3e-07 | 2.6e-07 | 2.833E-07 | 4.041E-08 | 2.600E-07 | 3.300E-07 |
| 6138 | 307 | Q307W | A | 3.2e-07 | 3.2e-07 | 1.6e-07 | 2.667E-07 | 9.238E-08 | 1.600E-07 | 3.200E-07 |
| 6139 | 307 | Q307Y | A | 1.6e-07 | 2.2e-07 | 3.4e-07 | 2.400E-07 | 9.165E-08 | 1.600E-07 | 3.400E-07 |
| 6140 | 307 | Q307V | A | 1.5e-07 | 1.5e-07 | 1.8e-07 | 1.600E-07 | 1.732E-08 | 1.500E-07 | 1.800E-07 |
| 6141 | 308 | R308A | A | 2.9e-07 | 2.5e-07 | 4.2e-07 | 3.200E-07 | 8.888E-08 | 2.500E-07 | 4.200E-07 |
| 6142 | 308 | R308R | A | 1.1e-07 | 3.4e-07 | 2e-07 | 2.167E-07 | 1.159E-07 | 1.100E-07 | 3.400E-07 |
| 6143 | 308 | R308N | A | 1.2e-07 | 3.9e-07 | 2.5e-07 | 2.533E-07 | 1.350E-07 | 1.200E-07 | 3.900E-07 |
| 6144 | 308 | R308D | A | 3.4e-07 | 9.3e-08 | 2.5e-07 | 2.277E-07 | 1.250E-07 | 9.300E-08 | 3.400E-07 |
| 6145 | 308 | R308C | A | 1.5e-07 | 2.8e-07 | 2e-07 | 2.100E-07 | 6.557E-08 | 1.500E-07 | 2.800E-07 |
| 6146 | 308 | R308E | A | 1.6e-07 | 5.3e-07 | 2.2e-07 | 3.033E-07 | 1.986E-07 | 1.600E-07 | 5.300E-07 |
| 6147 | 308 | R308Q | A | 2e-07 | 3e-07 | 3.3e-07 | 2.767E-07 | 6.807E-08 | 2.000E-07 | 3.300E-07 |
| 6148 | 308 | R308G | A | 1.6e-07 | 4e-07 | 5.5e-07 | 3.700E-07 | 1.967E-07 | 1.600E-07 | 5.500E-07 |
| 6149 | 308 | R308H | A | 3.3e-07 | 2.8e-07 | 2.2e-07 | 2.767E-07 | 5.508E-08 | 2.200E-07 | 3.300E-07 |
| 6150 | 308 | R308I | A | 1.9e-07 | 2e-07 | 3.2e-07 | 2.367E-07 | 7.234E-08 | 1.900E-07 | 3.200E-07 |
| 6151 | 308 | R308L | A | 5.9e-07 | 2.1e-07 | 1.6e-07 | 3.200E-07 | 2.352E-07 | 1.600E-07 | 5.900E-07 |
| 6152 | 308 | R308K | A | 2.7e-07 | 3.2e-07 | 2.5e-07 | 2.800E-07 | 3.606E-08 | 2.500E-07 | 3.200E-07 |
| 6153 | 308 | R308M | A | 2.2e-07 | 2.8e-07 | 2.6e-07 | 2.533E-07 | 3.055E-08 | 2.200E-07 | 2.800E-07 |
| 6154 | 308 | R308F | A | 2.4e-07 | 1.4e-07 | 2.2e-07 | 2.000E-07 | 5.292E-08 | 1.400E-07 | 2.400E-07 |
| 6155 | 308 | R308P | A | 8.4e-07 | 2.9e-07 | 3.4e-07 | 4.900E-07 | 3.041E-07 | 2.900E-07 | 8.400E-07 |
| 6156 | 308 | R308S | A | 5.6e-07 | 2.2e-07 | 3e-07 | 3.600E-07 | 1.778E-07 | 2.200E-07 | 5.600E-07 |
| 6157 | 308 | R308T | A | 2.3e-07 | 1.7e-07 | 9.6e-08 | 1.653E-07 | 6.712E-08 | 9.600E-08 | 2.300E-07 |
| 6158 | 308 | R308W | A | 5.3e-07 | 2.5e-07 | 3.5e-07 | 3.767E-07 | 1.419E-07 | 2.500E-07 | 5.300E-07 |
| 6159 | 308 | R308Y | A | 1.7e-07 | 2.9e-07 | 3e-07 | 2.533E-07 | 7.234E-08 | 1.700E-07 | 3.000E-07 |
| 6160 | 308 | R308V | A | 1.6e-07 | 3.4e-07 | 3.3e-07 | 2.767E-07 | 1.012E-07 | 1.600E-07 | 3.400E-07 |
| 6161 | 309 | C309A | A | 2.5e-07 | 2.2e-07 | 3.6e-07 | 2.767E-07 | 7.371E-08 | 2.200E-07 | 3.600E-07 |
| 6162 | 309 | C309R | A | 2.4e-07 | 2.1e-07 | 2.6e-07 | 2.367E-07 | 2.517E-08 | 2.100E-07 | 2.600E-07 |
| 6163 | 309 | C309N | A | 3.2e-07 | 4.2e-07 | 5.2e-07 | 4.200E-07 | 1.000E-07 | 3.200E-07 | 5.200E-07 |
| 6164 | 309 | C309D | A | 2.6e-07 | 3e-07 | 1.9e-07 | 2.500E-07 | 5.568E-08 | 1.900E-07 | 3.000E-07 |
| 6165 | 309 | C309C | A | 2.2e-07 | 2.8e-07 | 2.7e-07 | 2.567E-07 | 3.215E-08 | 2.200E-07 | 2.800E-07 |
| 6166 | 309 | C309E | A | 3.5e-07 | 1.6e-07 | 2e-07 | 2.367E-07 | 1.002E-07 | 1.600E-07 | 3.500E-07 |
| 6167 | 309 | C309Q | A | 2e-07 | 3.8e-07 | 1.9e-07 | 2.567E-07 | 1.069E-07 | 1.900E-07 | 3.800E-07 |
| 6168 | 309 | C309G | A | 3.3e-07 | 2.2e-07 | 7.6e-07 | 4.367E-07 | 2.854E-07 | 2.200E-07 | 7.600E-07 |
| 6169 | 309 | C309H | A | 2.2e-07 | 2e-07 | 4.5e-07 | 2.900E-07 | 1.389E-07 | 2.000E-07 | 4.500E-07 |
| 6170 | 309 | C309I | A | 1.6e-07 | 6.1e-07 | 4.4e-07 | 4.033E-07 | 2.272E-07 | 1.600E-07 | 6.100E-07 |
| 6171 | 309 | C309L | A | 4.3e-07 | 3.1e-07 | 2.5e-07 | 3.300E-07 | 9.165E-08 | 2.500E-07 | 4.300E-07 |
| 6172 | 309 | C309K | A | 1.4e-07 | 2.2e-07 | 1.2e-07 | 1.600E-07 | 5.292E-08 | 1.200E-07 | 2.200E-07 |
| 6173 | 309 | C309M | A | 3.5e-07 | 3.7e-07 | 3.5e-07 | 3.567E-07 | 1.155E-08 | 3.500E-07 | 3.700E-07 |
| 6174 | 309 | C309F | A | 3.2e-07 | 2.3e-07 | 4.6e-07 | 3.367E-07 | 1.159E-07 | 2.300E-07 | 4.600E-07 |
| 6175 | 309 | C309P | A | 2.3e-07 | 2.4e-07 | 2.8e-07 | 2.500E-07 | 2.646E-08 | 2.300E-07 | 2.800E-07 |
| 6176 | 309 | C309S | A | 1.8e-07 | 1.6e-07 | 2.8e-07 | 2.067E-07 | 6.429E-08 | 1.600E-07 | 2.800E-07 |
| 6177 | 309 | C309T | A | 1.9e-07 | 1.3e-07 | 3.8e-07 | 2.333E-07 | 1.305E-07 | 1.300E-07 | 3.800E-07 |
| 6178 | 309 | C309W | A | 3.8e-07 | 2.1e-07 | 3.1e-07 | 3.000E-07 | 8.544E-08 | 2.100E-07 | 3.800E-07 |
| 6179 | 309 | C309Y | A | 3.8e-07 | 1.2e-07 | 2e-07 | 2.333E-07 | 1.332E-07 | 1.200E-07 | 3.800E-07 |
| 6180 | 309 | C309V | A | 3.6e-07 | 2e-07 | 1e-07 | 2.200E-07 | 1.311E-07 | 1.000E-07 | 3.600E-07 |
| 6181 | 310 | E310A | A | 2.7e-07 | 4.7e-07 | 2.1e-07 | 3.167E-07 | 1.361E-07 | 2.100E-07 | 4.700E-07 |
| 6182 | 310 | E310R | A | 3.4e-07 | 3.1e-07 | 1.3e-07 | 2.600E-07 | 1.136E-07 | 1.300E-07 | 3.400E-07 |
| 6183 | 310 | E310N | A | 1.5e-07 | 3.4e-07 | 2e-07 | 2.300E-07 | 9.849E-08 | 1.500E-07 | 3.400E-07 |
| 6184 | 310 | E310D | A | 1e-07 | 1.8e-07 | 2e-07 | 1.600E-07 | 5.292E-08 | 1.000E-07 | 2.000E-07 |
| 6185 | 310 | E310C | A | 3.6e-07 | 3e-07 | 3.6e-07 | 3.400E-07 | 3.464E-08 | 3.000E-07 | 3.600E-07 |
| 6186 | 310 | E310E | A | 2.7e-07 | 1e-07 | 2.4e-07 | 2.033E-07 | 9.074E-08 | 1.000E-07 | 2.700E-07 |
| 6187 | 310 | E310Q | A | 2.2e-07 | 2.2e-07 | 2.6e-07 | 2.333E-07 | 2.309E-08 | 2.200E-07 | 2.600E-07 |
| 6188 | 310 | E310G | A | 1.8e-07 | 7e-07 | 1.7e-07 | 3.500E-07 | 3.032E-07 | 1.700E-07 | 7.000E-07 |
| 6189 | 310 | E310H | A | 1.9e-07 | 2.3e-07 | 1.4e-07 | 1.867E-07 | 4.509E-08 | 1.400E-07 | 2.300E-07 |
| 6190 | 310 | E310I | A | 1.7e-07 | 3.9e-07 | 1.7e-07 | 2.433E-07 | 1.270E-07 | 1.700E-07 | 3.900E-07 |
| 6191 | 310 | E310L | A | 1.4e-07 | 3.4e-07 | 3e-07 | 2.600E-07 | 1.058E-07 | 1.400E-07 | 3.400E-07 |
| 6192 | 310 | E310K | A | 1.8e-07 | 2.1e-07 | 1.7e-07 | 1.867E-07 | 2.082E-08 | 1.700E-07 | 2.100E-07 |
| 6193 | 310 | E310M | A | 2.2e-07 | 1.9e-07 | 3.9e-07 | 2.667E-07 | 1.079E-07 | 1.900E-07 | 3.900E-07 |
| 6194 | 310 | E310F | A | 3e-07 | 2.5e-07 | 1.8e-07 | 2.433E-07 | 6.028E-08 | 1.800E-07 | 3.000E-07 |
| 6195 | 310 | E310P | A | 1.1e-07 | 3.2e-07 | 1.2e-07 | 1.833E-07 | 1.185E-07 | 1.100E-07 | 3.200E-07 |
| 6196 | 310 | E310S | A | 8.4e-08 | 2.3e-07 | 3.8e-07 | 2.313E-07 | 1.480E-07 | 8.400E-08 | 3.800E-07 |
| 6197 | 310 | E310T | A | 2.9e-07 | 2e-07 | 3.2e-07 | 2.700E-07 | 6.245E-08 | 2.000E-07 | 3.200E-07 |
| 6198 | 310 | E310W | A | 2.3e-07 | 2.2e-07 | 1.8e-07 | 2.100E-07 | 2.646E-08 | 1.800E-07 | 2.300E-07 |
| 6199 | 310 | E310Y | A | 1.4e-07 | 3.1e-07 | 2.1e-07 | 2.200E-07 | 8.544E-08 | 1.400E-07 | 3.100E-07 |
| 6200 | 310 | E310V | A | 2.3e-07 | 2.3e-07 | 2.5e-07 | 2.367E-07 | 1.155E-08 | 2.300E-07 | 2.500E-07 |
| 6201 | 311 | K311A | A | 9.9e-08 | 2.2e-07 | 1.4e-07 | 1.530E-07 | 6.154E-08 | 9.900E-08 | 2.200E-07 |
| 6202 | 311 | K311R | A | 4.1e-07 | 2.3e-07 | 1.5e-07 | 2.633E-07 | 1.332E-07 | 1.500E-07 | 4.100E-07 |
| 6203 | 311 | K311N | A | 1.3e-07 | 2.7e-07 | 1.4e-07 | 1.800E-07 | 7.810E-08 | 1.300E-07 | 2.700E-07 |
| 6204 | 311 | K311D | A | 2.7e-07 | 2.7e-07 | 1.6e-07 | 2.333E-07 | 6.351E-08 | 1.600E-07 | 2.700E-07 |

| | | | | | | | | | | |
|---|---|---|---|---|---|---|---|---|---|---|
| 6205 | 311 | K311C | A | 1.1e-07 | 1.6e-07 | 2.7e-07 | 1.800E-07 | 8.185E-08 | 1.100E-07 | 2.700E-07 |
| 6206 | 311 | K311E | A | 2.2e-07 | 3.8e-07 | 3.5e-07 | 3.167E-07 | 8.505E-08 | 2.200E-07 | 3.800E-07 |
| 6207 | 311 | K311Q | A | 4.5e-07 | 2.6e-07 | 2.7e-07 | 3.267E-07 | 1.069E-07 | 2.600E-07 | 4.500E-07 |
| 6208 | 311 | K311G | A | 4e-07 | 2.9e-07 | 2.5e-07 | 3.133E-07 | 7.767E-08 | 2.500E-07 | 4.000E-07 |
| 6209 | 311 | K311H | A | 1.9e-07 | 1.8e-07 | 1.3e-07 | 1.667E-07 | 3.215E-08 | 1.300E-07 | 1.900E-07 |
| 6210 | 311 | K311I | A | 1.1e-07 | 3e-07 | 2.7e-07 | 2.267E-07 | 1.021E-07 | 1.100E-07 | 3.000E-07 |
| 6211 | 311 | K311L | A | 2.6e-07 | 3.3e-07 | 1.3e-07 | 2.400E-07 | 1.015E-07 | 1.300E-07 | 3.300E-07 |
| 6212 | 311 | K311K | A | 2.6e-07 | 1.1e-07 | 2.7e-07 | 2.133E-07 | 8.963E-08 | 1.100E-07 | 2.700E-07 |
| 6213 | 311 | K311M | A | 2.7e-07 | 2.1e-07 | 3.4e-07 | 2.733E-07 | 6.506E-08 | 2.100E-07 | 3.400E-07 |
| 6214 | 311 | K311F | A | 1.7e-07 | 4.3e-07 | 1.7e-07 | 2.567E-07 | 1.501E-07 | 1.700E-07 | 4.300E-07 |
| 6215 | 311 | K311P | A | 3.6e-07 | 3.1e-07 | 3e-07 | 3.233E-07 | 3.215E-08 | 3.000E-07 | 3.600E-07 |
| 6216 | 311 | K311S | A | 1.7e-07 | 2.4e-07 | 4.3e-07 | 2.800E-07 | 1.345E-07 | 1.700E-07 | 4.300E-07 |
| 6217 | 311 | K311T | A | 2.6e-07 | 2.4e-07 | 2.6e-07 | 2.533E-07 | 1.155E-08 | 2.400E-07 | 2.600E-07 |
| 6218 | 311 | K311W | A | 2.8e-07 | 2.1e-07 | 5.9e-07 | 3.600E-07 | 2.022E-07 | 2.100E-07 | 5.900E-07 |
| 6219 | 311 | K311Y | A | 2.7e-07 | 3.9e-07 | 3.6e-07 | 3.400E-07 | 6.245E-08 | 2.700E-07 | 3.900E-07 |
| 6220 | 311 | K311V | A | 3.2e-07 | 2.1e-07 | 2.5e-07 | 2.600E-07 | 5.568E-08 | 2.100E-07 | 3.200E-07 |
| 6221 | 312 | C312A | A | 3.9e-07 | 1.8e-07 | 2.4e-07 | 2.700E-07 | 1.082E-07 | 1.800E-07 | 3.900E-07 |
| 6222 | 312 | C312R | A | 2.7e-07 | 2.3e-07 | 2.7e-07 | 2.567E-07 | 2.309E-08 | 2.300E-07 | 2.700E-07 |
| 6223 | 312 | C312N | A | 2.3e-07 | 5e-07 | 2e-07 | 3.100E-07 | 1.652E-07 | 2.000E-07 | 5.000E-07 |
| 6224 | 312 | C312D | A | 5.7e-07 | 1.1e-07 | 1.8e-07 | 2.867E-07 | 2.479E-07 | 1.100E-07 | 5.700E-07 |
| 6225 | 312 | C312C | A | 3.1e-07 | 2.2e-07 | 3.9e-07 | 3.067E-07 | 8.505E-08 | 2.200E-07 | 3.900E-07 |
| 6226 | 312 | C312E | A | 2.6e-07 | 3.6e-07 | 3.7e-07 | 3.300E-07 | 6.083E-08 | 2.600E-07 | 3.700E-07 |
| 6227 | 312 | C312Q | A | 2.9e-07 | 3.4e-07 | 1.9e-07 | 2.733E-07 | 7.638E-08 | 1.900E-07 | 3.400E-07 |
| 6228 | 312 | C312G | A | 1.3e-07 | 4.2e-07 | 1.8e-07 | 2.433E-07 | 1.550E-07 | 1.300E-07 | 4.200E-07 |
| 6229 | 312 | C312H | A | 2.6e-07 | 3.4e-07 | 3e-07 | 3.000E-07 | 4.000E-08 | 2.600E-07 | 3.400E-07 |
| 6230 | 312 | C312I | A | 2e-07 | 3.2e-07 | 2.4e-07 | 2.533E-07 | 6.110E-08 | 2.000E-07 | 3.200E-07 |
| 6231 | 312 | C312L | A | 3.2e-07 | 1.7e-07 | 1.7e-07 | 2.200E-07 | 8.660E-08 | 1.700E-07 | 3.200E-07 |
| 6232 | 312 | C312K | A | 2.1e-07 | 1.4e-07 | 2e-07 | 1.833E-07 | 3.786E-08 | 1.400E-07 | 2.100E-07 |
| 6233 | 312 | C312M | A | 2.7e-07 | 7.5e-08 | 3.4e-07 | 2.283E-07 | 1.373E-07 | 7.500E-08 | 3.400E-07 |
| 6234 | 312 | C312F | A | 3.7e-07 | 2.1e-07 | 3.1e-07 | 2.967E-07 | 8.083E-08 | 2.100E-07 | 3.700E-07 |
| 6235 | 312 | C312P | A | 4.8e-07 | 1.7e-07 | 3.8e-07 | 3.433E-07 | 1.582E-07 | 1.700E-07 | 4.800E-07 |
| 6236 | 312 | C312S | A | 5.7e-07 | 1.7e-07 | 2e-07 | 3.133E-07 | 2.228E-07 | 1.700E-07 | 5.700E-07 |
| 6237 | 312 | C312T | A | 2e-07 | 3.6e-07 | 3.3e-07 | 2.967E-07 | 8.505E-08 | 2.000E-07 | 3.600E-07 |
| 6238 | 312 | C312W | A | 2.1e-07 | 2e-07 | 1.5e-07 | 1.867E-07 | 3.215E-08 | 1.500E-07 | 2.100E-07 |
| 6239 | 312 | C312Y | A | 1.7e-07 | 1.6e-07 | 2.8e-07 | 2.033E-07 | 6.658E-08 | 1.600E-07 | 2.800E-07 |
| 6240 | 312 | C312V | A | 1.8e-07 | 4.5e-07 | 2.3e-07 | 2.867E-07 | 1.436E-07 | 1.800E-07 | 4.500E-07 |
| 6241 | 313 | S313A | A | 1.1e-07 | 2e-07 | 3.5e-07 | 2.200E-07 | 1.212E-07 | 1.100E-07 | 3.500E-07 |
| 6242 | 313 | S313R | A | 8.3e-08 | 3.7e-07 | 3e-07 | 2.510E-07 | 1.496E-07 | 8.300E-08 | 3.700E-07 |
| 6243 | 313 | S313N | A | 1.6e-07 | 1.4e-07 | 3.5e-07 | 2.167E-07 | 1.159E-07 | 1.400E-07 | 3.500E-07 |
| 6244 | 313 | S313D | A | 2.7e-07 | 1.6e-07 | 2.1e-07 | 2.133E-07 | 5.508E-08 | 1.600E-07 | 2.700E-07 |
| 6245 | 313 | S313C | A | 3e-07 | 3e-07 | 3.4e-07 | 3.133E-07 | 2.309E-08 | 3.000E-07 | 3.400E-07 |
| 6246 | 313 | S313E | A | 1.9e-07 | 5.6e-08 | 4.4e-07 | 2.287E-07 | 1.949E-07 | 5.600E-08 | 4.400E-07 |
| 6247 | 313 | S313Q | A | 3.2e-07 | 2.1e-07 | 3.4e-07 | 2.900E-07 | 7.000E-08 | 2.100E-07 | 3.400E-07 |
| 6248 | 313 | S313G | A | 6.9e-07 | 1.6e-07 | 6.1e-07 | 4.867E-07 | 2.857E-07 | 1.600E-07 | 6.900E-07 |
| 6249 | 313 | S313H | A | 3.1e-07 | 2.5e-07 | 1.7e-07 | 2.433E-07 | 7.024E-08 | 1.700E-07 | 3.100E-07 |
| 6250 | 313 | S313I | A | 1.2e-07 | 1.3e-07 | 1.6e-07 | 1.367E-07 | 2.082E-08 | 1.200E-07 | 1.600E-07 |
| 6251 | 313 | S313L | A | 6.1e-07 | 2.3e-07 | 3.6e-07 | 4.000E-07 | 1.931E-07 | 2.300E-07 | 6.100E-07 |
| 6252 | 313 | S313K | A | 7.7e-08 | 2.4e-07 | 5.1e-07 | 2.757E-07 | 2.187E-07 | 7.700E-08 | 5.100E-07 |
| 6253 | 313 | S313M | A | 2.8e-07 | 2.2e-07 | 5.7e-07 | 3.567E-07 | 1.872E-07 | 2.200E-07 | 5.700E-07 |
| 6254 | 313 | S313F | A | 1.3e-07 | 5.1e-07 | 3.2e-07 | 3.200E-07 | 1.900E-07 | 1.300E-07 | 5.100E-07 |
| 6255 | 313 | S313P | A | 8.6e-08 | 9.7e-08 | 1.4e-07 | 1.077E-07 | 2.854E-08 | 8.600E-08 | 1.400E-07 |
| 6256 | 313 | S313S | A | 3.5e-07 | 1.6e-07 | 1.3e-07 | 2.133E-07 | 1.193E-07 | 1.300E-07 | 3.500E-07 |
| 6257 | 313 | S313T | A | 1.5e-07 | 1.9e-07 | 3e-07 | 2.133E-07 | 7.767E-08 | 1.500E-07 | 3.000E-07 |
| 6258 | 313 | S313W | A | 2.4e-07 | 9.4e-07 | 4e-07 | 5.267E-07 | 3.668E-07 | 2.400E-07 | 9.400E-07 |
| 6259 | 313 | S313Y | A | 3.8e-07 | 3.2e-07 | 2.3e-07 | 3.100E-07 | 7.550E-08 | 2.300E-07 | 3.800E-07 |
| 6260 | 313 | S313V | A | 6.2e-07 | 3e-07 | 4.9e-07 | 4.700E-07 | 1.609E-07 | 3.000E-07 | 6.200E-07 |
| 6261 | 314 | K314A | A | 1e-07 | 3.4e-07 | 2.4e-07 | 2.267E-07 | 1.206E-07 | 1.000E-07 | 3.400E-07 |
| 6262 | 314 | K314R | A | 1.1e-07 | 1.7e-07 | 1.4e-07 | 1.400E-07 | 3.000E-08 | 1.100E-07 | 1.700E-07 |
| 6263 | 314 | K314N | A | 6.4e-07 | 2.2e-07 | 5.6e-07 | 4.733E-07 | 2.230E-07 | 2.200E-07 | 6.400E-07 |
| 6264 | 314 | K314D | A | 2.4e-07 | 9.8e-08 | 9.5e-08 | 1.443E-07 | 8.286E-08 | 9.500E-08 | 2.400E-07 |
| 6265 | 314 | K314C | A | 4.2e-07 | 3.2e-07 | 4e-07 | 3.800E-07 | 5.292E-08 | 3.200E-07 | 4.200E-07 |
| 6266 | 314 | K314E | A | 2e-07 | 2.4e-07 | 3.4e-07 | 2.600E-07 | 7.211E-08 | 2.000E-07 | 3.400E-07 |
| 6267 | 314 | K314Q | A | 2.1e-07 | 2.1e-07 | 2.9e-07 | 2.367E-07 | 4.619E-08 | 2.100E-07 | 2.900E-07 |
| 6268 | 314 | K314G | A | 3.2e-07 | 8.5e-08 | 2.3e-07 | 2.117E-07 | 1.186E-07 | 8.500E-08 | 3.200E-07 |
| 6269 | 314 | K314H | A | 5e-07 | 2.6e-07 | 2.3e-07 | 3.300E-07 | 1.480E-07 | 2.300E-07 | 5.000E-07 |
| 6270 | 314 | K314I | A | 1.9e-07 | 2.2e-07 | 3.8e-07 | 2.633E-07 | 1.021E-07 | 1.900E-07 | 3.800E-07 |
| 6271 | 314 | K314L | A | 3.6e-07 | 1.7e-07 | 2.2e-07 | 2.500E-07 | 9.849E-08 | 1.700E-07 | 3.600E-07 |
| 6272 | 314 | K314K | A | 2.4e-07 | 2e-07 | 1.3e-07 | 1.900E-07 | 5.568E-08 | 1.300E-07 | 2.400E-07 |
| 6273 | 314 | K314M | A | 3.7e-07 | 2.4e-07 | 2.6e-07 | 2.900E-07 | 7.000E-08 | 2.400E-07 | 3.700E-07 |
| 6274 | 314 | K314F | A | 9.2e-08 | 1.8e-07 | 2.3e-07 | 1.673E-07 | 6.987E-08 | 9.200E-08 | 2.300E-07 |
| 6275 | 314 | K314P | A | 7.1e-08 | 8.3e-08 | 2e-07 | 1.180E-07 | 7.127E-08 | 7.100E-08 | 2.000E-07 |
| 6276 | 314 | K314S | A | 2.3e-07 | 4.5e-07 | 2.5e-07 | 3.100E-07 | 1.217E-07 | 2.300E-07 | 4.500E-07 |
| 6277 | 314 | K314T | A | 3.5e-07 | 1.8e-07 | 3.1e-07 | 2.800E-07 | 8.888E-08 | 1.800E-07 | 3.500E-07 |
| 6278 | 314 | K314W | A | 2.5e-07 | 5.7e-07 | 1.4e-07 | 3.200E-07 | 2.234E-07 | 1.400E-07 | 5.700E-07 |
| 6279 | 314 | K314Y | A | 1.3e-07 | 2.3e-07 | 4.7e-07 | 2.767E-07 | 1.747E-07 | 1.300E-07 | 4.700E-07 |
| 6280 | 314 | K314V | A | 2.2e-07 | 4.1e-07 | 4.1e-07 | 3.467E-07 | 1.097E-07 | 2.200E-07 | 4.100E-07 |
| 6281 | 315 | P315A | A | 3.3e-07 | 3.6e-07 | 3.7e-07 | 3.533E-07 | 2.082E-08 | 3.300E-07 | 3.700E-07 |
| 6282 | 315 | P315R | A | 1.1e-07 | 2.1e-07 | 1.1e-07 | 1.433E-07 | 5.774E-08 | 1.100E-07 | 2.100E-07 |
| 6283 | 315 | P315N | A | 1e-07 | 1.3e-07 | 3e-07 | 1.767E-07 | 1.079E-07 | 1.000E-07 | 3.000E-07 |
| 6284 | 315 | P315D | A | 2.3e-07 | 1.3e-07 | 2.2e-07 | 1.933E-07 | 5.508E-08 | 1.300E-07 | 2.300E-07 |
| 6285 | 315 | P315C | A | 6.8e-07 | 2.1e-07 | 3.6e-07 | 4.167E-07 | 2.401E-07 | 2.100E-07 | 6.800E-07 |
| 6286 | 315 | P315E | A | 3.1e-07 | 2.9e-07 | 3e-07 | 3.000E-07 | 1.000E-08 | 2.900E-07 | 3.100E-07 |
| 6287 | 315 | P315Q | A | 8.4e-08 | 1.4e-07 | 1.8e-07 | 1.347E-07 | 4.822E-08 | 8.400E-08 | 1.800E-07 |
| 6288 | 315 | P315G | A | 2.6e-07 | 1.2e-07 | 3.2e-07 | 2.333E-07 | 1.026E-07 | 1.200E-07 | 3.200E-07 |
| 6289 | 315 | P315H | A | 3.3e-07 | 2.9e-07 | 6.1e-08 | 2.270E-07 | 1.451E-07 | 6.100E-08 | 3.300E-07 |

| | | | | | | | | | | |
|---|---|---|---|---|---|---|---|---|---|---|
| 6290 | 315 | P315I | A | 1.6e-07 | 1.9e-07 | 2.2e-07 | 1.900E-07 | 3.000E-08 | 1.600E-07 | 2.200E-07 |
| 6291 | 315 | P315L | A | 5e-07 | 2.6e-07 | 4.5e-07 | 4.033E-07 | 1.266E-07 | 2.600E-07 | 5.000E-07 |
| 6292 | 315 | P315K | A | 2.2e-07 | 4e-07 | 2.4e-07 | 2.867E-07 | 9.866E-08 | 2.200E-07 | 4.000E-07 |
| 6293 | 315 | P315M | A | 1.8e-07 | 2.9e-07 | 2e-07 | 2.233E-07 | 5.859E-08 | 1.800E-07 | 2.900E-07 |
| 6294 | 315 | P315F | A | 2.3e-07 | 3e-07 | 3.1e-07 | 2.800E-07 | 4.359E-08 | 2.300E-07 | 3.100E-07 |
| 6295 | 315 | P315P | A | 1.7e-07 | 1.7e-07 | 1.7e-07 | 1.700E-07 | 0.000E+00 | 1.700E-07 | 1.700E-07 |
| 6296 | 315 | P315S | A | 1.1e-07 | 3.4e-07 | 3e-07 | 2.500E-07 | 1.229E-07 | 1.100E-07 | 3.400E-07 |
| 6297 | 315 | P315T | A | 2.3e-07 | 3.9e-07 | 1.5e-07 | 2.567E-07 | 1.222E-07 | 1.500E-07 | 3.900E-07 |
| 6298 | 315 | P315W | A | 3.9e-07 | 3.3e-07 | 2.4e-07 | 3.200E-07 | 7.550E-08 | 2.400E-07 | 3.900E-07 |
| 6299 | 315 | P315Y | A | 2.3e-07 | 3e-07 | 1.7e-07 | 2.333E-07 | 6.506E-08 | 1.700E-07 | 3.000E-07 |
| 6300 | 315 | P315V | A | 1.4e-07 | 2.7e-07 | 4.7e-07 | 2.933E-07 | 1.662E-07 | 1.400E-07 | 4.700E-07 |
| 6301 | 316 | C316A | A | 3.7e-07 | 3.8e-07 | 1e-07 | 2.833E-07 | 1.589E-07 | 1.000E-07 | 3.800E-07 |
| 6302 | 316 | C316R | A | 4e-07 | 2e-07 | 3.4e-07 | 3.133E-07 | 1.026E-07 | 2.000E-07 | 4.000E-07 |
| 6303 | 316 | C316N | A | 2.9e-07 | 2.8e-07 | 3.5e-07 | 3.067E-07 | 3.786E-08 | 2.800E-07 | 3.500E-07 |
| 6304 | 316 | C316D | A | 1.5e-07 | 1.8e-07 | 2.4e-07 | 1.900E-07 | 4.583E-08 | 1.500E-07 | 2.400E-07 |
| 6305 | 316 | C316C | A | 3.7e-07 | 1.4e-07 | 2.2e-07 | 2.433E-07 | 1.168E-07 | 1.400E-07 | 3.700E-07 |
| 6306 | 316 | C316E | A | 1.6e-07 | 1.7e-07 | 4e-07 | 2.433E-07 | 1.358E-07 | 1.600E-07 | 4.000E-07 |
| 6307 | 316 | C316Q | A | 3e-07 | 3.1e-07 | 5.1e-07 | 3.733E-07 | 1.185E-07 | 3.000E-07 | 5.100E-07 |
| 6308 | 316 | C316G | A | 1.7e-07 | 2.7e-07 | 3.2e-07 | 2.533E-07 | 7.638E-08 | 1.700E-07 | 3.200E-07 |
| 6309 | 316 | C316H | A | 3.1e-07 | 2e-07 | 2.7e-07 | 2.600E-07 | 5.568E-08 | 2.000E-07 | 3.100E-07 |
| 6310 | 316 | C316I | A | 3.6e-07 | 1.1e-07 | 2.6e-07 | 2.433E-07 | 1.258E-07 | 1.100E-07 | 3.600E-07 |
| 6311 | 316 | C316L | A | 4.7e-07 | 2.9e-07 | 2.4e-07 | 3.333E-07 | 1.210E-07 | 2.400E-07 | 4.700E-07 |
| 6312 | 316 | C316K | A | 2.9e-07 | 2.8e-07 | 2.6e-07 | 2.767E-07 | 1.528E-08 | 2.600E-07 | 2.900E-07 |
| 6313 | 316 | C316M | A | 1.5e-07 | 5.4e-07 | 1.1e-07 | 2.667E-07 | 2.376E-07 | 1.100E-07 | 5.400E-07 |
| 6314 | 316 | C316F | A | 2.4e-07 | 7.5e-07 | 3.2e-07 | 4.367E-07 | 2.743E-07 | 2.400E-07 | 7.500E-07 |
| 6315 | 316 | C316P | A | 7.4e-08 | 4.7e-07 | 2.2e-07 | 2.547E-07 | 2.003E-07 | 7.400E-08 | 4.700E-07 |
| 6316 | 316 | C316S | A | 2.4e-07 | 1.3e-07 | 1.9e-07 | 1.867E-07 | 5.508E-08 | 1.300E-07 | 2.400E-07 |
| 6317 | 316 | C316T | A | 2.7e-07 | 2.5e-07 | 1.1e-07 | 2.100E-07 | 8.718E-08 | 1.100E-07 | 2.700E-07 |
| 6318 | 316 | C316W | A | 1.7e-07 | 2.5e-07 | 3.5e-07 | 2.567E-07 | 9.018E-08 | 1.700E-07 | 3.500E-07 |
| 6319 | 316 | C316Y | A | 1.5e-07 | 5.4e-07 | 2.2e-07 | 3.033E-07 | 2.079E-07 | 1.500E-07 | 5.400E-07 |
| 6320 | 316 | C316V | A | 4.4e-07 | 2.5e-07 | 2.7e-07 | 3.200E-07 | 1.044E-07 | 2.500E-07 | 4.400E-07 |
| 6321 | 317 | A317A | A | 2.8e-07 | 1.1e-07 | 1.7e-07 | 1.867E-07 | 8.622E-08 | 1.100E-07 | 2.800E-07 |
| 6322 | 317 | A317R | A | 2.9e-07 | 7.6e-07 | 1.9e-07 | 4.133E-07 | 3.044E-07 | 1.900E-07 | 7.600E-07 |
| 6323 | 317 | A317N | A | 1.7e-07 | 5.2e-07 | 2.5e-07 | 3.133E-07 | 1.834E-07 | 1.700E-07 | 5.200E-07 |
| 6324 | 317 | A317D | A | 6.5e-07 | 3.5e-07 | 2.4e-07 | 4.133E-07 | 2.122E-07 | 2.400E-07 | 6.500E-07 |
| 6325 | 317 | A317C | A | 2.4e-07 | 3.1e-07 | 3.4e-07 | 2.967E-07 | 5.132E-08 | 2.400E-07 | 3.400E-07 |
| 6326 | 317 | A317E | A | 6.2e-07 | 3.5e-07 | 2.2e-07 | 3.967E-07 | 2.040E-07 | 2.200E-07 | 6.200E-07 |
| 6327 | 317 | A317Q | A | 4.1e-07 | 2e-07 | 2.6e-07 | 2.900E-07 | 1.082E-07 | 2.000E-07 | 4.100E-07 |
| 6328 | 317 | A317G | A | 2e-07 | 1.5e-07 | 4.3e-07 | 2.600E-07 | 1.493E-07 | 1.500E-07 | 4.300E-07 |
| 6329 | 317 | A317H | A | 1.2e-06 | 3e-07 | 2.4e-07 | 5.800E-07 | 5.378E-07 | 2.400E-07 | 1.200E-06 |
| 6330 | 317 | A317I | A | 8.4e-07 | 2.3e-07 | 1.2e-07 | 3.967E-07 | 3.879E-07 | 1.200E-07 | 8.400E-07 |
| 6331 | 317 | A317L | A | 3.6e-07 | 3.6e-07 | 2.3e-07 | 3.167E-07 | 7.506E-08 | 2.300E-07 | 3.600E-07 |
| 6332 | 317 | A317K | A | 2.5e-07 | 1.9e-07 | 2.3e-07 | 2.233E-07 | 3.055E-08 | 1.900E-07 | 2.500E-07 |
| 6333 | 317 | A317M | A | 4.1e-07 | 3.1e-07 | 2.6e-07 | 3.267E-07 | 7.638E-08 | 2.600E-07 | 4.100E-07 |
| 6334 | 317 | A317F | A | 2.9e-07 | 1.8e-07 | 2.1e-07 | 2.267E-07 | 5.686E-08 | 1.800E-07 | 2.900E-07 |
| 6335 | 317 | A317P | A | 2.6e-07 | 3.2e-07 | 2.6e-07 | 2.800E-07 | 3.464E-08 | 2.600E-07 | 3.200E-07 |
| 6336 | 317 | A317S | A | 3e-07 | 3.4e-07 | 9.3e-08 | 2.443E-07 | 1.326E-07 | 9.300E-08 | 3.400E-07 |
| 6337 | 317 | A317T | A | 3e-07 | 2.4e-07 | 2.4e-07 | 2.600E-07 | 3.464E-08 | 2.400E-07 | 3.000E-07 |
| 6338 | 317 | A317W | A | 3.1e-07 | 1.1e-07 | 2.9e-07 | 2.367E-07 | 1.102E-07 | 1.100E-07 | 3.100E-07 |
| 6339 | 317 | A317Y | A | 4.1e-07 | 1.6e-07 | 7.5e-08 | 2.150E-07 | 1.741E-07 | 7.500E-08 | 4.100E-07 |
| 6340 | 317 | A317V | A | 3.2e-07 | 2.3e-07 | 4.2e-07 | 3.233E-07 | 9.504E-08 | 2.300E-07 | 4.200E-07 |
| 6341 | 318 | R318A | A | 3e-07 | 5.1e-07 | 6.7e-07 | 4.933E-07 | 1.856E-07 | 3.000E-07 | 6.700E-07 |
| 6342 | 318 | R318R | A | 1.6e-07 | 3.7e-07 | 3.4e-07 | 2.900E-07 | 1.136E-07 | 1.600E-07 | 3.700E-07 |
| 6343 | 318 | R318N | A | 1.3e-07 | 2.7e-07 | 4e-07 | 2.667E-07 | 1.350E-07 | 1.300E-07 | 4.000E-07 |
| 6344 | 318 | R318D | A | 2.8e-07 | 2.9e-07 | 2.3e-07 | 2.667E-07 | 3.215E-08 | 2.300E-07 | 2.900E-07 |
| 6345 | 318 | R318C | A | 1.9e-07 | 4.4e-07 | 4.1e-07 | 3.467E-07 | 1.365E-07 | 1.900E-07 | 4.400E-07 |
| 6346 | 318 | R318E | A | 2.6e-07 | 1.3e-07 | 2.1e-07 | 2.000E-07 | 6.557E-08 | 1.300E-07 | 2.600E-07 |
| 6347 | 318 | R318Q | A | 3e-07 | 2.6e-07 | 2.5e-07 | 2.700E-07 | 2.646E-08 | 2.500E-07 | 3.000E-07 |
| 6348 | 318 | R318G | A | 2.2e-07 | 2.6e-07 | 3.5e-07 | 2.767E-07 | 6.658E-08 | 2.200E-07 | 3.500E-07 |
| 6349 | 318 | R318H | A | 1.9e-07 | 3.6e-07 | 2.7e-07 | 2.733E-07 | 8.505E-08 | 1.900E-07 | 3.600E-07 |
| 6350 | 318 | R318I | A | 2.2e-07 | 1.4e-07 | 2.9e-07 | 2.167E-07 | 7.506E-08 | 1.400E-07 | 2.900E-07 |
| 6351 | 318 | R318L | A | 3e-07 | 2.1e-07 | 2.2e-07 | 2.433E-07 | 4.933E-08 | 2.100E-07 | 3.000E-07 |
| 6352 | 318 | R318K | A | 1.6e-07 | 3.4e-07 | 2.2e-07 | 2.400E-07 | 9.165E-08 | 1.600E-07 | 3.400E-07 |
| 6353 | 318 | R318M | A | 3.4e-07 | 9.4e-08 | 1.2e-07 | 1.847E-07 | 1.351E-07 | 9.400E-08 | 3.400E-07 |
| 6354 | 318 | R318F | A | 2.2e-07 | 2.5e-07 | 2.8e-07 | 2.500E-07 | 3.000E-08 | 2.200E-07 | 2.800E-07 |
| 6355 | 318 | R318P | A | 2.7e-07 | 3e-07 | 2.5e-07 | 2.733E-07 | 2.517E-08 | 2.500E-07 | 3.000E-07 |
| 6356 | 318 | R318S | A | 3.1e-07 | 3.6e-07 | 2.1e-07 | 2.933E-07 | 7.638E-08 | 2.100E-07 | 3.600E-07 |
| 6357 | 318 | R318T | A | 2.8e-07 | 4e-07 | 1.6e-07 | 2.800E-07 | 1.200E-07 | 1.600E-07 | 4.000E-07 |
| 6358 | 318 | R318W | A | 2.1e-07 | 3.3e-07 | 2.7e-07 | 2.700E-07 | 6.000E-08 | 2.100E-07 | 3.300E-07 |
| 6359 | 318 | R318Y | A | 2.1e-07 | 1.9e-07 | 2.1e-07 | 2.033E-07 | 1.155E-08 | 1.900E-07 | 2.100E-07 |
| 6360 | 318 | R318V | A | 5.1e-07 | 1.7e-07 | 2.7e-07 | 3.167E-07 | 1.747E-07 | 1.700E-07 | 5.100E-07 |
| 6361 | 319 | V319A | A | 1.4e-07 | 1.3e-07 | 2e-07 | 1.567E-07 | 3.786E-08 | 1.300E-07 | 2.000E-07 |
| 6362 | 319 | V319R | A | 3.9e-07 | 4.6e-07 | 3.5e-07 | 4.000E-07 | 5.568E-08 | 3.500E-07 | 4.600E-07 |
| 6363 | 319 | V319N | A | 3.8e-07 | 3.6e-07 | 2.3e-07 | 3.233E-07 | 8.145E-08 | 2.300E-07 | 3.800E-07 |
| 6364 | 319 | V319D | A | 2.4e-07 | 3.8e-07 | 2.7e-07 | 2.967E-07 | 7.371E-08 | 2.400E-07 | 3.800E-07 |
| 6365 | 319 | V319C | A | 4.4e-07 | 4.4e-07 | 4.1e-07 | 4.300E-07 | 1.732E-08 | 4.100E-07 | 4.400E-07 |
| 6366 | 319 | V319E | A | 4.2e-07 | 2.5e-07 | 9.6e-08 | 2.553E-07 | 1.621E-07 | 9.600E-08 | 4.200E-07 |
| 6367 | 319 | V319Q | A | 2.6e-07 | 2.1e-07 | 1.5e-07 | 2.067E-07 | 5.508E-08 | 1.500E-07 | 2.600E-07 |
| 6368 | 319 | V319G | A | 1.8e-07 | 7.3e-08 | 3.9e-07 | 2.143E-07 | 1.613E-07 | 7.300E-08 | 3.900E-07 |
| 6369 | 319 | V319H | A | 1.3e-07 | 3.1e-07 | 2e-07 | 2.133E-07 | 9.074E-08 | 1.300E-07 | 3.100E-07 |
| 6370 | 319 | V319I | A | 1.7e-07 | 2.1e-07 | 5.5e-07 | 3.100E-07 | 2.088E-07 | 1.700E-07 | 5.500E-07 |
| 6371 | 319 | V319L | A | 2.4e-07 | 4.3e-07 | 2.4e-07 | 3.033E-07 | 1.097E-07 | 2.400E-07 | 4.300E-07 |
| 6372 | 319 | V319K | A | 3.4e-07 | 1.9e-07 | 1.7e-07 | 2.333E-07 | 9.292E-08 | 1.700E-07 | 3.400E-07 |
| 6373 | 319 | V319M | A | 1.3e-07 | 2.4e-07 | 3.7e-07 | 2.467E-07 | 1.201E-07 | 1.300E-07 | 3.700E-07 |
| 6374 | 319 | V319F | A | 2.6e-07 | 4e-07 | 3.1e-07 | 3.233E-07 | 7.095E-08 | 2.600E-07 | 4.000E-07 |

| | | | | | | | | | | |
|---|---|---|---|---|---|---|---|---|---|---|
| 6375 | 319 | V319P | A | 2.5e-07 | 3.4e-07 | 2.6e-07 | 2.833E-07 | 4.933E-08 | 2.500E-07 | 3.400E-07 |
| 6376 | 319 | V319S | A | 6.5e-08 | 2.4e-07 | 2.8e-07 | 1.950E-07 | 1.143E-07 | 6.500E-08 | 2.800E-07 |
| 6377 | 319 | V319T | A | 3.6e-07 | 2.2e-07 | 1.1e-07 | 2.300E-07 | 1.253E-07 | 1.100E-07 | 3.600E-07 |
| 6378 | 319 | V319W | A | 2.9e-07 | 8.8e-08 | 1.7e-07 | 1.827E-07 | 1.016E-07 | 8.800E-08 | 2.900E-07 |
| 6379 | 319 | V319Y | A | 2.8e-07 | 2.3e-07 | 1.3e-07 | 2.133E-07 | 7.638E-08 | 1.300E-07 | 2.800E-07 |
| 6380 | 319 | V319V | A | 3.2e-07 | 2.7e-07 | 1.1e-07 | 2.333E-07 | 1.097E-07 | 1.100E-07 | 3.200E-07 |
| 6381 | 320 | C320A | A | 2.2e-07 | 2e-07 | 2.8e-07 | 2.333E-07 | 4.163E-08 | 2.000E-07 | 2.800E-07 |
| 6382 | 320 | C320R | A | 2.3e-07 | 3.7e-07 | 2.4e-07 | 2.800E-07 | 7.810E-08 | 2.300E-07 | 3.700E-07 |
| 6383 | 320 | C320N | A | 2.7e-07 | 9.3e-08 | 3.6e-07 | 2.410E-07 | 1.358E-07 | 9.300E-08 | 3.600E-07 |
| 6384 | 320 | C320D | A | 1.9e-07 | 2.9e-07 | 2.8e-07 | 2.533E-07 | 5.508E-08 | 1.900E-07 | 2.900E-07 |
| 6385 | 320 | C320C | A | 2.4e-07 | 2e-07 | 2.3e-07 | 2.233E-07 | 2.082E-08 | 2.000E-07 | 2.400E-07 |
| 6386 | 320 | C320E | A | 2.5e-07 | 3.7e-07 | 2.5e-07 | 2.900E-07 | 6.928E-08 | 2.500E-07 | 3.700E-07 |
| 6387 | 320 | C320Q | A | 2.8e-07 | 2.8e-07 | 2.3e-07 | 2.633E-07 | 2.887E-08 | 2.300E-07 | 2.800E-07 |
| 6388 | 320 | C320G | A | 1.8e-07 | 2.4e-07 | 1e-07 | 1.733E-07 | 7.024E-08 | 1.000E-07 | 2.400E-07 |
| 6389 | 320 | C320H | A | 3.8e-07 | 2.4e-07 | 3e-07 | 3.067E-07 | 7.024E-08 | 2.400E-07 | 3.800E-07 |
| 6390 | 320 | C320I | A | 2.6e-07 | 1.6e-07 | 2.2e-07 | 2.133E-07 | 5.033E-08 | 1.600E-07 | 2.600E-07 |
| 6391 | 320 | C320L | A | 1.8e-07 | 4.8e-07 | 2.7e-07 | 3.100E-07 | 1.539E-07 | 1.800E-07 | 4.800E-07 |
| 6392 | 320 | C320K | A | 1.4e-07 | 2.3e-07 | 2.1e-07 | 1.933E-07 | 4.726E-08 | 1.400E-07 | 2.300E-07 |
| 6393 | 320 | C320M | A | 4.2e-07 | 1.3e-07 | 4e-07 | 3.167E-07 | 1.620E-07 | 1.300E-07 | 4.200E-07 |
| 6394 | 320 | C320F | A | 1.7e-07 | 3.2e-07 | 3.7e-07 | 2.867E-07 | 1.041E-07 | 1.700E-07 | 3.700E-07 |
| 6395 | 320 | C320P | A | 2.2e-07 | 2.4e-07 | 1.1e-07 | 1.900E-07 | 7.000E-08 | 1.100E-07 | 2.400E-07 |
| 6396 | 320 | C320S | A | 1.2e-07 | 1.6e-07 | 2e-07 | 1.600E-07 | 4.000E-08 | 1.200E-07 | 2.000E-07 |
| 6397 | 320 | C320T | A | 3.3e-07 | 1.6e-07 | 1.3e-07 | 2.067E-07 | 1.079E-07 | 1.300E-07 | 3.300E-07 |
| 6398 | 320 | C320W | A | 2.4e-07 | 2.6e-07 | 2.8e-07 | 2.600E-07 | 2.000E-08 | 2.400E-07 | 2.800E-07 |
| 6399 | 320 | C320Y | A | 3.5e-07 | 1.9e-07 | 1.8e-07 | 2.400E-07 | 9.539E-08 | 1.800E-07 | 3.500E-07 |
| 6400 | 320 | C320V | A | 1.4e-07 | 1.6e-07 | 2.2e-07 | 1.733E-07 | 4.163E-08 | 1.400E-07 | 2.200E-07 |
| 6401 | 321 | Y321A | A | 1.9e-07 | 4.7e-07 | 1.9e-07 | 2.833E-07 | 1.617E-07 | 1.900E-07 | 4.700E-07 |
| 6402 | 321 | Y321R | A | 3.6e-07 | 9.6e-08 | 3.5e-07 | 2.687E-07 | 1.496E-07 | 9.600E-08 | 3.600E-07 |
| 6403 | 321 | Y321N | A | 2e-07 | 2.2e-07 | 5e-07 | 3.067E-07 | 1.677E-07 | 2.000E-07 | 5.000E-07 |
| 6404 | 321 | Y321D | A | 2.7e-07 | 3.5e-07 | 3.6e-07 | 3.267E-07 | 4.933E-08 | 2.700E-07 | 3.600E-07 |
| 6405 | 321 | Y321C | A | 3.1e-07 | 1.6e-07 | 3.4e-07 | 2.700E-07 | 9.644E-08 | 1.600E-07 | 3.400E-07 |
| 6406 | 321 | Y321E | A | 2e-07 | 2.2e-07 | 1.9e-07 | 2.033E-07 | 1.528E-08 | 1.900E-07 | 2.200E-07 |
| 6407 | 321 | Y321Q | A | 2.3e-07 | 3.2e-07 | 2.5e-07 | 2.667E-07 | 4.726E-08 | 2.300E-07 | 3.200E-07 |
| 6408 | 321 | Y321G | A | 3.8e-07 | 3.8e-07 | 1.9e-07 | 3.167E-07 | 1.097E-07 | 1.900E-07 | 3.800E-07 |
| 6409 | 321 | Y321H | A | 1.7e-07 | 3.6e-07 | 2.5e-07 | 2.600E-07 | 9.539E-08 | 1.700E-07 | 3.600E-07 |
| 6410 | 321 | Y321I | A | 2.9e-07 | 1.7e-07 | 1.6e-07 | 2.067E-07 | 7.234E-08 | 1.600E-07 | 2.900E-07 |
| 6411 | 321 | Y321L | A | 1.5e-07 | 3.2e-07 | 3.5e-07 | 2.733E-07 | 1.079E-07 | 1.500E-07 | 3.500E-07 |
| 6412 | 321 | Y321K | A | 5.2e-07 | 1.9e-07 | 2.5e-07 | 3.200E-07 | 1.758E-07 | 1.900E-07 | 5.200E-07 |
| 6413 | 321 | Y321M | A | 1.8e-07 | 2.8e-07 | 1.9e-07 | 2.167E-07 | 5.508E-08 | 1.800E-07 | 2.800E-07 |
| 6414 | 321 | Y321F | A | 2.8e-07 | 2.9e-07 | 1.4e-07 | 2.367E-07 | 8.386E-08 | 1.400E-07 | 2.900E-07 |
| 6415 | 321 | Y321P | A | 3.5e-07 | 2.6e-07 | 1.5e-07 | 2.533E-07 | 1.002E-07 | 1.500E-07 | 3.500E-07 |
| 6416 | 321 | Y321S | A | 3.3e-07 | 2.2e-07 | 2.2e-07 | 2.567E-07 | 6.351E-08 | 2.200E-07 | 3.300E-07 |
| 6417 | 321 | Y321T | A | 1.9e-07 | 2.6e-07 | 2.1e-07 | 2.200E-07 | 3.606E-08 | 1.900E-07 | 2.600E-07 |
| 6418 | 321 | Y321W | A | 3.7e-07 | 2e-07 | 3.7e-07 | 3.133E-07 | 9.815E-08 | 2.000E-07 | 3.700E-07 |
| 6419 | 321 | Y321Y | A | 3.3e-07 | 2.7e-07 | 4.1e-07 | 3.367E-07 | 7.024E-08 | 2.700E-07 | 4.100E-07 |
| 6420 | 321 | Y321V | A | 5.2e-07 | 3e-07 | 1.8e-07 | 3.333E-07 | 1.724E-07 | 1.800E-07 | 5.200E-07 |
| 6421 | 322 | G322A | A | 1.1e-07 | 1.6e-07 | 2.8e-07 | 1.833E-07 | 8.737E-08 | 1.100E-07 | 2.800E-07 |
| 6422 | 322 | G322R | A | 3.9e-07 | 3.3e-07 | 1.1e-07 | 2.767E-07 | 1.474E-07 | 1.100E-07 | 3.900E-07 |
| 6423 | 322 | G322N | A | 2.2e-07 | 4.1e-07 | 2.4e-07 | 2.900E-07 | 1.044E-07 | 2.200E-07 | 4.100E-07 |
| 6424 | 322 | G322D | A | 2.3e-07 | 4.2e-07 | 2e-07 | 2.833E-07 | 1.193E-07 | 2.000E-07 | 4.200E-07 |
| 6425 | 322 | G322C | A | 2e-07 | 2e-07 | 9.4e-08 | 1.647E-07 | 6.120E-08 | 9.400E-08 | 2.000E-07 |
| 6426 | 322 | G322E | A | 1.9e-07 | 3.7e-07 | 1.8e-07 | 2.467E-07 | 1.069E-07 | 1.800E-07 | 3.700E-07 |
| 6427 | 322 | G322Q | A | 2.3e-07 | 1.3e-07 | 2.3e-07 | 1.967E-07 | 5.774E-08 | 1.300E-07 | 2.300E-07 |
| 6428 | 322 | G322G | A | 2.3e-07 | 2.8e-07 | 1.4e-07 | 2.167E-07 | 7.095E-08 | 1.400E-07 | 2.800E-07 |
| 6429 | 322 | G322H | A | 4.7e-07 | 1.7e-07 | 3.3e-07 | 3.233E-07 | 1.501E-07 | 1.700E-07 | 4.700E-07 |
| 6430 | 322 | G322I | A | 2.4e-07 | 3.9e-07 | 1.4e-07 | 2.567E-07 | 1.258E-07 | 1.400E-07 | 3.900E-07 |
| 6431 | 322 | G322L | A | 3.3e-07 | 2.3e-07 | 3e-07 | 2.867E-07 | 5.132E-08 | 2.300E-07 | 3.300E-07 |
| 6432 | 322 | G322K | A | 2.4e-07 | 2.2e-07 | 2.3e-07 | 2.300E-07 | 1.000E-08 | 2.200E-07 | 2.400E-07 |
| 6433 | 322 | G322M | A | 2.6e-07 | 1.4e-07 | 3e-07 | 2.333E-07 | 8.327E-08 | 1.400E-07 | 3.000E-07 |
| 6434 | 322 | G322F | A | 2.4e-07 | 2.8e-07 | 2.7e-07 | 2.633E-07 | 2.082E-08 | 2.400E-07 | 2.800E-07 |
| 6435 | 322 | G322P | A | 9.3e-08 | 1.6e-07 | 1.5e-07 | 1.343E-07 | 3.614E-08 | 9.300E-08 | 1.600E-07 |
| 6436 | 322 | G322S | A | 1.5e-07 | 3.1e-07 | 3.6e-07 | 2.733E-07 | 1.097E-07 | 1.500E-07 | 3.600E-07 |
| 6437 | 322 | G322T | A | 2.1e-07 | 2e-07 | 8.8e-08 | 1.660E-07 | 6.773E-08 | 8.800E-08 | 2.100E-07 |
| 6438 | 322 | G322W | A | 3.5e-07 | 1.5e-07 | 2.1e-07 | 2.367E-07 | 1.026E-07 | 1.500E-07 | 3.500E-07 |
| 6439 | 322 | G322Y | A | 3.1e-07 | 4.6e-07 | 1e-07 | 2.900E-07 | 1.808E-07 | 1.000E-07 | 4.600E-07 |
| 6440 | 322 | G322V | A | 4.2e-07 | 1.6e-07 | 3.3e-07 | 3.033E-07 | 1.320E-07 | 1.600E-07 | 4.200E-07 |
| 6441 | 323 | L323A | A | 2e-07 | 5e-07 | 1.8e-07 | 2.933E-07 | 1.793E-07 | 1.800E-07 | 5.000E-07 |
| 6442 | 323 | L323R | A | 2.4e-07 | 2.1e-07 | 3.1e-07 | 2.533E-07 | 5.132E-08 | 2.100E-07 | 3.100E-07 |
| 6443 | 323 | L323N | A | 2.2e-07 | 2.8e-07 | 1.3e-07 | 2.100E-07 | 7.550E-08 | 1.300E-07 | 2.800E-07 |
| 6444 | 323 | L323D | A | 7.4e-07 | 2.1e-07 | 3.8e-07 | 4.433E-07 | 2.706E-07 | 2.100E-07 | 7.400E-07 |
| 6445 | 323 | L323C | A | 1.4e-07 | 4.1e-07 | 3.1e-07 | 2.867E-07 | 1.365E-07 | 1.400E-07 | 4.100E-07 |
| 6446 | 323 | L323E | A | 2.1e-07 | 1.7e-07 | 7.1e-08 | 1.503E-07 | 7.156E-08 | 7.100E-08 | 2.100E-07 |
| 6447 | 323 | L323Q | A | 3e-07 | 1.1e-07 | 2.9e-07 | 2.333E-07 | 1.069E-07 | 1.100E-07 | 3.000E-07 |
| 6448 | 323 | L323G | A | 1.2e-07 | 3.2e-07 | 1.6e-07 | 2.000E-07 | 1.058E-07 | 1.200E-07 | 3.200E-07 |
| 6449 | 323 | L323H | A | 2.2e-07 | 2.2e-07 | 1.5e-07 | 1.967E-07 | 4.041E-08 | 1.500E-07 | 2.200E-07 |
| 6450 | 323 | L323I | A | 1.7e-07 | 3.7e-07 | 3.2e-07 | 2.867E-07 | 1.041E-07 | 1.700E-07 | 3.700E-07 |
| 6451 | 323 | L323L | A | 4e-07 | 1.6e-07 | 1.5e-07 | 2.367E-07 | 1.415E-07 | 1.500E-07 | 4.000E-07 |
| 6452 | 323 | L323K | A | 1.8e-07 | 3.8e-07 | 2e-07 | 2.533E-07 | 1.102E-07 | 1.800E-07 | 3.800E-07 |
| 6453 | 323 | L323M | A | 2.3e-07 | 2.2e-07 | 1.1e-07 | 1.867E-07 | 6.658E-08 | 1.100E-07 | 2.300E-07 |
| 6454 | 323 | L323F | A | 3e-07 | 1.6e-07 | 2.1e-07 | 2.233E-07 | 7.095E-08 | 1.600E-07 | 3.000E-07 |
| 6455 | 323 | L323P | A | 1.9e-07 | 2.8e-07 | 1.9e-07 | 2.200E-07 | 5.196E-08 | 1.900E-07 | 2.800E-07 |
| 6456 | 323 | L323S | A | 1.4e-07 | 1.6e-07 | 2.2e-07 | 1.733E-07 | 4.163E-08 | 1.400E-07 | 2.200E-07 |
| 6457 | 323 | L323T | A | 2.7e-07 | 2.4e-07 | 1.8e-07 | 2.300E-07 | 4.583E-08 | 1.800E-07 | 2.700E-07 |
| 6458 | 323 | L323W | A | 2.2e-07 | 2.3e-07 | 1.7e-07 | 2.067E-07 | 3.215E-08 | 1.700E-07 | 2.300E-07 |
| 6459 | 323 | L323Y | A | 2.4e-07 | 4.1e-07 | 3.4e-07 | 3.300E-07 | 8.544E-08 | 2.400E-07 | 4.100E-07 |

| | | | | | | | | | | |
|---|---|---|---|---|---|---|---|---|---|---|
| 6460 | 323 | L323V | A | 2.3e-07 | 1.1e-07 | 2e-07 | 1.800E-07 | 6.245E-08 | 1.100E-07 | 2.300E-07 |
| 6461 | 324 | G324A | A | 1e-07 | 1.2e-07 | 3.4e-07 | 1.867E-07 | 1.332E-07 | 1.000E-07 | 3.400E-07 |
| 6462 | 324 | G324R | A | 4e-07 | 6.6e-07 | 2.8e-07 | 4.467E-07 | 1.943E-07 | 2.800E-07 | 6.600E-07 |
| 6463 | 324 | G324N | A | 1.1e-07 | 1.5e-07 | 1.8e-07 | 1.467E-07 | 3.512E-08 | 1.100E-07 | 1.800E-07 |
| 6464 | 324 | G324D | A | 2.1e-07 | 4.3e-07 | 2.3e-07 | 2.900E-07 | 1.217E-07 | 2.100E-07 | 4.300E-07 |
| 6465 | 324 | G324C | A | 1.6e-07 | 2.4e-07 | 2.9e-07 | 2.300E-07 | 6.557E-08 | 1.600E-07 | 2.900E-07 |
| 6466 | 324 | G324E | A | 1.2e-07 | 1.5e-07 | 2.5e-07 | 1.733E-07 | 6.807E-08 | 1.200E-07 | 2.500E-07 |
| 6467 | 324 | G324Q | A | 4.6e-07 | 2.1e-07 | 1.3e-07 | 2.667E-07 | 1.721E-07 | 1.300E-07 | 4.600E-07 |
| 6468 | 324 | G324G | A | 4e-07 | 2.2e-07 | 1.8e-07 | 2.667E-07 | 1.172E-07 | 1.800E-07 | 4.000E-07 |
| 6469 | 324 | G324H | A | 1.4e-07 | 1.9e-07 | 4.7e-07 | 2.667E-07 | 1.779E-07 | 1.400E-07 | 4.700E-07 |
| 6470 | 324 | G324I | A | 1.7e-07 | 3.5e-07 | 1.8e-07 | 2.333E-07 | 1.012E-07 | 1.700E-07 | 3.500E-07 |
| 6471 | 324 | G324L | A | 2.5e-07 | 2e-07 | 3.7e-07 | 2.733E-07 | 8.737E-08 | 2.000E-07 | 3.700E-07 |
| 6472 | 324 | G324K | A | 1.5e-07 | 2.1e-07 | 3.5e-07 | 2.367E-07 | 1.026E-07 | 1.500E-07 | 3.500E-07 |
| 6473 | 324 | G324M | A | 3.2e-07 | 1.9e-07 | 2.6e-07 | 2.567E-07 | 6.506E-08 | 1.900E-07 | 3.200E-07 |
| 6474 | 324 | G324F | A | 2.8e-07 | 2e-07 | 3e-07 | 2.600E-07 | 5.292E-08 | 2.000E-07 | 3.000E-07 |
| 6475 | 324 | G324P | A | 2e-07 | 1.3e-07 | 1.7e-07 | 1.667E-07 | 3.512E-08 | 1.300E-07 | 2.000E-07 |
| 6476 | 324 | G324S | A | 1.5e-07 | 1.7e-07 | 1.5e-07 | 1.567E-07 | 1.155E-08 | 1.500E-07 | 1.700E-07 |
| 6477 | 324 | G324T | A | 2.6e-07 | 4.5e-07 | 5.3e-08 | 2.543E-07 | 1.986E-07 | 5.300E-08 | 4.500E-07 |
| 6478 | 324 | G324W | A | 1.8e-07 | 3.5e-07 | 1.5e-07 | 2.267E-07 | 1.079E-07 | 1.500E-07 | 3.500E-07 |
| 6479 | 324 | G324Y | A | 2.9e-07 | 5e-07 | 1.8e-07 | 3.233E-07 | 1.626E-07 | 1.800E-07 | 5.000E-07 |
| 6480 | 324 | G324V | A | 2.1e-07 | 3.1e-07 | 8.8e-08 | 2.027E-07 | 1.112E-07 | 8.800E-08 | 3.100E-07 |
| 6481 | 325 | M325A | A | 2.1e-07 | 2.1e-07 | 1.7e-07 | 1.967E-07 | 2.309E-08 | 1.700E-07 | 2.100E-07 |
| 6482 | 325 | M325R | A | 2.1e-07 | 3.3e-07 | 1.4e-07 | 2.267E-07 | 9.609E-08 | 1.400E-07 | 3.300E-07 |
| 6483 | 325 | M325N | A | 4e-07 | 4e-07 | 9.6e-08 | 2.987E-07 | 1.755E-07 | 9.600E-08 | 4.000E-07 |
| 6484 | 325 | M325D | A | 1.8e-07 | 9.8e-08 | 6.3e-08 | 1.137E-07 | 6.005E-08 | 6.300E-08 | 1.800E-07 |
| 6485 | 325 | M325C | A | 1.3e-07 | 2.2e-07 | 2.9e-07 | 2.133E-07 | 8.021E-08 | 1.300E-07 | 2.900E-07 |
| 6486 | 325 | M325E | A | 2.4e-07 | 1.9e-07 | 2.1e-07 | 2.133E-07 | 2.517E-08 | 1.900E-07 | 2.400E-07 |
| 6487 | 325 | M325Q | A | 1.6e-07 | 1.4e-07 | 2e-07 | 1.667E-07 | 3.055E-08 | 1.400E-07 | 2.000E-07 |
| 6488 | 325 | M325G | A | 3.4e-07 | 6.7e-07 | 9e-08 | 3.667E-07 | 2.909E-07 | 9.000E-08 | 6.700E-07 |
| 6489 | 325 | M325H | A | 3e-07 | 3.7e-07 | 1.4e-07 | 2.700E-07 | 1.179E-07 | 1.400E-07 | 3.700E-07 |
| 6490 | 325 | M325I | A | 4e-07 | 1.7e-07 | 3.4e-07 | 3.033E-07 | 1.193E-07 | 1.700E-07 | 4.000E-07 |
| 6491 | 325 | M325L | A | 3.8e-07 | 2.7e-07 | 8.2e-08 | 2.440E-07 | 1.507E-07 | 8.200E-08 | 3.800E-07 |
| 6492 | 325 | M325K | A | 2.1e-07 | 2.7e-07 | 2.5e-07 | 2.433E-07 | 3.055E-08 | 2.100E-07 | 2.700E-07 |
| 6493 | 325 | M325M | A | 2.5e-07 | 5.8e-07 | 3.4e-07 | 3.900E-07 | 1.706E-07 | 2.500E-07 | 5.800E-07 |
| 6494 | 325 | M325F | A | 2.8e-07 | 3.2e-07 | 2.4e-07 | 2.800E-07 | 4.000E-08 | 2.400E-07 | 3.200E-07 |
| 6495 | 325 | M325P | A | 1.1e-07 | 2.2e-07 | 1.9e-07 | 1.733E-07 | 5.686E-08 | 1.100E-07 | 2.200E-07 |
| 6496 | 325 | M325S | A | 1.6e-07 | 2.2e-07 | 2.5e-07 | 2.100E-07 | 4.583E-08 | 1.600E-07 | 2.500E-07 |
| 6497 | 325 | M325T | A | 2e-07 | 2.4e-07 | 1.5e-07 | 1.967E-07 | 4.509E-08 | 1.500E-07 | 2.400E-07 |
| 6498 | 325 | M325W | A | 3.3e-07 | 4.6e-07 | 2.9e-07 | 3.600E-07 | 8.888E-08 | 2.900E-07 | 4.600E-07 |
| 6499 | 325 | M325Y | A | 1.7e-07 | 2.7e-07 | 1.1e-07 | 1.833E-07 | 8.083E-08 | 1.100E-07 | 2.700E-07 |
| 6500 | 325 | M325V | A | 1.5e-07 | 4.6e-07 | 1.8e-07 | 2.633E-07 | 1.710E-07 | 1.500E-07 | 4.600E-07 |
| 6501 | 326 | E326A | A | 2.6e-07 | 1.9e-07 | 1.2e-07 | 1.900E-07 | 7.000E-08 | 1.200E-07 | 2.600E-07 |
| 6502 | 326 | E326R | A | 1.7e-07 | 2.7e-07 | 4.5e-07 | 2.967E-07 | 1.419E-07 | 1.700E-07 | 4.500E-07 |
| 6503 | 326 | E326N | A | 5.4e-07 | 1.4e-07 | 1.8e-07 | 2.867E-07 | 2.203E-07 | 1.400E-07 | 5.400E-07 |
| 6504 | 326 | E326D | A | 3.2e-07 | 3.1e-07 | 4.9e-07 | 3.733E-07 | 1.012E-07 | 3.100E-07 | 4.900E-07 |
| 6505 | 326 | E326C | A | 1.5e-07 | 3.1e-07 | 2.9e-07 | 2.500E-07 | 8.718E-08 | 1.500E-07 | 3.100E-07 |
| 6506 | 326 | E326E | A | 4.1e-07 | 1.5e-07 | 2.4e-07 | 2.667E-07 | 1.320E-07 | 1.500E-07 | 4.100E-07 |
| 6507 | 326 | E326Q | A | 1.7e-07 | 2.1e-07 | 3.8e-07 | 2.533E-07 | 1.115E-07 | 1.700E-07 | 3.800E-07 |
| 6508 | 326 | E326G | A | 2.4e-07 | 3.2e-07 | 2.6e-07 | 2.733E-07 | 4.163E-08 | 2.400E-07 | 3.200E-07 |
| 6509 | 326 | E326H | A | 2.9e-07 | 1.6e-07 | 1.3e-07 | 1.933E-07 | 8.505E-08 | 1.300E-07 | 2.900E-07 |
| 6510 | 326 | E326I | A | 2.7e-07 | 1.5e-07 | 4.5e-07 | 2.900E-07 | 1.510E-07 | 1.500E-07 | 4.500E-07 |
| 6511 | 326 | E326L | A | 2.2e-07 | 8e-08 | 2.2e-07 | 1.733E-07 | 8.083E-08 | 8.000E-08 | 2.200E-07 |
| 6512 | 326 | E326K | A | 1.8e-07 | 3e-07 | 5.4e-07 | 3.400E-07 | 1.833E-07 | 1.800E-07 | 5.400E-07 |
| 6513 | 326 | E326M | A | 4.5e-07 | 4.1e-07 | 2.4e-07 | 3.667E-07 | 1.115E-07 | 2.400E-07 | 4.500E-07 |
| 6514 | 326 | E326F | A | 2.3e-07 | 1.7e-07 | 1.9e-07 | 1.967E-07 | 3.055E-08 | 1.700E-07 | 2.300E-07 |
| 6515 | 326 | E326P | A | 3.2e-07 | 2.9e-07 | 2.6e-07 | 2.900E-07 | 3.000E-08 | 2.600E-07 | 3.200E-07 |
| 6516 | 326 | E326S | A | 1.5e-07 | 1e-07 | 2e-07 | 1.500E-07 | 5.000E-08 | 1.000E-07 | 2.000E-07 |
| 6517 | 326 | E326T | A | 2.3e-07 | 4e-07 | 4.2e-07 | 3.500E-07 | 1.044E-07 | 2.300E-07 | 4.200E-07 |
| 6518 | 326 | E326W | A | 1.2e-07 | 2.4e-07 | 2.4e-07 | 2.000E-07 | 6.928E-08 | 1.200E-07 | 2.400E-07 |
| 6519 | 326 | E326Y | A | 1.9e-07 | 3e-07 | 2.5e-07 | 2.467E-07 | 5.508E-08 | 1.900E-07 | 3.000E-07 |
| 6520 | 326 | E326V | A | 2.7e-07 | 2e-07 | 3.8e-07 | 2.833E-07 | 9.074E-08 | 2.000E-07 | 3.800E-07 |
| 6521 | 327 | H327A | A | 2.7e-07 | 2.5e-07 | 3.7e-07 | 2.967E-07 | 6.429E-08 | 2.500E-07 | 3.700E-07 |
| 6522 | 327 | H327R | A | 1.8e-07 | 4.9e-07 | 5.4e-07 | 4.033E-07 | 1.950E-07 | 1.800E-07 | 5.400E-07 |
| 6523 | 327 | H327N | A | 1.8e-07 | 2.9e-07 | 2.5e-07 | 2.400E-07 | 5.568E-08 | 1.800E-07 | 2.900E-07 |
| 6524 | 327 | H327D | A | 1.2e-07 | 2.3e-07 | 2.8e-07 | 2.100E-07 | 8.185E-08 | 1.200E-07 | 2.800E-07 |
| 6525 | 327 | H327C | A | 1.7e-07 | 3.4e-07 | 4.2e-07 | 3.100E-07 | 1.277E-07 | 1.700E-07 | 4.200E-07 |
| 6526 | 327 | H327E | A | 1.7e-07 | 2.9e-07 | 2.2e-07 | 2.267E-07 | 6.028E-08 | 1.700E-07 | 2.900E-07 |
| 6527 | 327 | H327Q | A | 1.1e-07 | 2.3e-07 | 2.2e-07 | 1.867E-07 | 6.658E-08 | 1.100E-07 | 2.300E-07 |
| 6528 | 327 | H327G | A | 3.6e-07 | 4.5e-07 | 3.3e-07 | 3.800E-07 | 6.245E-08 | 3.300E-07 | 4.500E-07 |
| 6529 | 327 | H327H | A | 2.2e-07 | 9.7e-08 | 2.4e-07 | 1.857E-07 | 7.744E-08 | 9.700E-08 | 2.400E-07 |
| 6530 | 327 | H327I | A | 3.5e-07 | 3.1e-07 | 2.9e-07 | 3.167E-07 | 3.055E-08 | 2.900E-07 | 3.500E-07 |
| 6531 | 327 | H327L | A | 3.2e-07 | 2.1e-07 | 5e-07 | 3.433E-07 | 1.464E-07 | 2.100E-07 | 5.000E-07 |
| 6532 | 327 | H327K | A | 1.8e-07 | 2.3e-07 | 1.5e-07 | 1.867E-07 | 4.041E-08 | 1.500E-07 | 2.300E-07 |
| 6533 | 327 | H327M | A | 2.9e-07 | 1.5e-07 | 3.7e-07 | 2.700E-07 | 1.114E-07 | 1.500E-07 | 3.700E-07 |
| 6534 | 327 | H327F | A | 2e-07 | 2.7e-07 | 2e-07 | 2.233E-07 | 4.041E-08 | 2.000E-07 | 2.700E-07 |
| 6535 | 327 | H327P | A | 1.3e-07 | 2e-07 | 1.9e-07 | 1.733E-07 | 3.786E-08 | 1.300E-07 | 2.000E-07 |
| 6536 | 327 | H327S | A | 2.2e-07 | 2.4e-07 | 3.8e-07 | 2.800E-07 | 8.718E-08 | 2.200E-07 | 3.800E-07 |
| 6537 | 327 | H327T | A | 2.5e-07 | 2.3e-07 | 3.7e-07 | 2.833E-07 | 7.572E-08 | 2.300E-07 | 3.700E-07 |
| 6538 | 327 | H327W | A | 2.5e-07 | 3.2e-07 | 1.5e-07 | 2.400E-07 | 8.544E-08 | 1.500E-07 | 3.200E-07 |
| 6539 | 327 | H327Y | A | 1.3e-07 | 4.1e-07 | 1.6e-07 | 2.333E-07 | 1.537E-07 | 1.300E-07 | 4.100E-07 |
| 6540 | 327 | H327V | A | 1.8e-07 | 2.6e-07 | 5.2e-07 | 3.200E-07 | 1.778E-07 | 1.800E-07 | 5.200E-07 |
| 6541 | 328 | L328A | A | 9.7e-08 | 1.1e-07 | 2.4e-07 | 1.490E-07 | 7.908E-08 | 9.700E-08 | 2.400E-07 |
| 6542 | 328 | L328R | A | 3.2e-07 | 1e-07 | 3.1e-07 | 2.433E-07 | 1.242E-07 | 1.000E-07 | 3.200E-07 |
| 6543 | 328 | L328N | A | 2.2e-07 | 1.9e-07 | 2.9e-07 | 2.333E-07 | 5.132E-08 | 1.900E-07 | 2.900E-07 |
| 6544 | 328 | L328D | A | 2.2e-07 | 1.7e-07 | 2.9e-07 | 2.267E-07 | 6.028E-08 | 1.700E-07 | 2.900E-07 |

| | | | | | | | | | | |
|---|---|---|---|---|---|---|---|---|---|---|
| 6545 | 328 | L328C | A | 1.9e-07 | 4.2e-07 | 2.5e-07 | 2.867E-07 | 1.193E-07 | 1.900E-07 | 4.200E-07 |
| 6546 | 328 | L328E | A | 4.2e-07 | 3.1e-07 | 2.7e-07 | 3.333E-07 | 7.767E-08 | 2.700E-07 | 4.200E-07 |
| 6547 | 328 | L328Q | A | 3e-07 | 2.5e-07 | 1.1e-07 | 2.200E-07 | 9.849E-08 | 1.100E-07 | 3.000E-07 |
| 6548 | 328 | L328G | A | 2.1e-07 | 2.5e-07 | 3.3e-07 | 2.633E-07 | 6.110E-08 | 2.100E-07 | 3.300E-07 |
| 6549 | 328 | L328H | A | 3.1e-07 | 4.2e-07 | 2.2e-07 | 3.167E-07 | 1.002E-07 | 2.200E-07 | 4.200E-07 |
| 6550 | 328 | L328I | A | 3.2e-07 | 2.4e-07 | 1.8e-07 | 2.467E-07 | 7.024E-08 | 1.800E-07 | 3.200E-07 |
| 6551 | 328 | L328L | A | 1.3e-07 | 1.8e-07 | 2.7e-07 | 1.933E-07 | 7.095E-08 | 1.300E-07 | 2.700E-07 |
| 6552 | 328 | L328K | A | 4.2e-07 | 2.5e-07 | 2.3e-07 | 3.000E-07 | 1.044E-07 | 2.300E-07 | 4.200E-07 |
| 6553 | 328 | L328M | A | 4.5e-07 | 5.8e-07 | 2.8e-07 | 4.367E-07 | 1.504E-07 | 2.800E-07 | 5.800E-07 |
| 6554 | 328 | L328F | A | 9.3e-08 | 2.8e-07 | 2e-07 | 1.910E-07 | 9.382E-08 | 9.300E-08 | 2.800E-07 |
| 6555 | 328 | L328P | A | 9.2e-08 | 1.8e-07 | 1.1e-07 | 1.273E-07 | 4.649E-08 | 9.200E-08 | 1.800E-07 |
| 6556 | 328 | L328S | A | 3.7e-07 | 3.8e-07 | 3.5e-07 | 3.667E-07 | 1.528E-08 | 3.500E-07 | 3.800E-07 |
| 6557 | 328 | L328T | A | 3.2e-07 | 2.7e-07 | 2e-07 | 2.633E-07 | 6.028E-08 | 2.000E-07 | 3.200E-07 |
| 6558 | 328 | L328W | A | 2.9e-07 | 6.8e-08 | 1.5e-07 | 1.693E-07 | 1.123E-07 | 6.800E-08 | 2.900E-07 |
| 6559 | 328 | L328Y | A | 3.4e-07 | 2.7e-07 | 1.3e-07 | 2.467E-07 | 1.069E-07 | 1.300E-07 | 3.400E-07 |
| 6560 | 328 | L328V | A | 2e-07 | 3e-07 | 1.6e-07 | 2.200E-07 | 7.211E-08 | 1.600E-07 | 3.000E-07 |
| 6561 | 329 | R329A | A | 2.2e-07 | 1.7e-07 | 2.7e-07 | 2.200E-07 | 5.000E-08 | 1.700E-07 | 2.700E-07 |
| 6562 | 329 | R329R | A | 4.5e-07 | 2.5e-07 | 1.6e-07 | 2.867E-07 | 1.484E-07 | 1.600E-07 | 4.500E-07 |
| 6563 | 329 | R329N | A | 2.7e-07 | 3.4e-07 | 7.7e-07 | 4.600E-07 | 2.707E-07 | 2.700E-07 | 7.700E-07 |
| 6564 | 329 | R329D | A | 1.5e-07 | 1.5e-07 | 1.8e-07 | 1.600E-07 | 1.732E-08 | 1.500E-07 | 1.800E-07 |
| 6565 | 329 | R329C | A | 2.3e-07 | 3.3e-07 | 1.8e-07 | 2.467E-07 | 7.638E-08 | 1.800E-07 | 3.300E-07 |
| 6566 | 329 | R329E | A | 4.7e-07 | 3.1e-07 | 4.1e-07 | 3.967E-07 | 8.083E-08 | 3.100E-07 | 4.700E-07 |
| 6567 | 329 | R329Q | A | 3.4e-07 | 1.8e-07 | 3.4e-07 | 2.867E-07 | 9.238E-08 | 1.800E-07 | 3.400E-07 |
| 6568 | 329 | R329G | A | 2e-07 | 3.2e-07 | 2.9e-07 | 2.700E-07 | 6.245E-08 | 2.000E-07 | 3.200E-07 |
| 6569 | 329 | R329H | A | 3.8e-07 | 3.9e-07 | 2.2e-07 | 3.300E-07 | 9.539E-08 | 2.200E-07 | 3.900E-07 |
| 6570 | 329 | R329I | A | 1.8e-07 | 3.3e-07 | 5.6e-07 | 3.567E-07 | 1.914E-07 | 1.800E-07 | 5.600E-07 |
| 6571 | 329 | R329L | A | 4.2e-07 | 2.2e-07 | 4.6e-07 | 3.667E-07 | 1.286E-07 | 2.200E-07 | 4.600E-07 |
| 6572 | 329 | R329K | A | 3.2e-07 | 1.7e-07 | 1.8e-07 | 2.233E-07 | 8.386E-08 | 1.700E-07 | 3.200E-07 |
| 6573 | 329 | R329M | A | 2.2e-07 | 1.5e-07 | 1.1e-07 | 1.600E-07 | 5.568E-08 | 1.100E-07 | 2.200E-07 |
| 6574 | 329 | R329F | A | 2.3e-07 | 2.9e-07 | 2.8e-07 | 2.667E-07 | 3.215E-08 | 2.300E-07 | 2.900E-07 |
| 6575 | 329 | R329P | A | 1.7e-07 | 3.4e-07 | 4.2e-07 | 3.100E-07 | 1.277E-07 | 1.700E-07 | 4.200E-07 |
| 6576 | 329 | R329S | A | 1.5e-07 | 1.8e-07 | 3.7e-07 | 2.333E-07 | 1.193E-07 | 1.500E-07 | 3.700E-07 |
| 6577 | 329 | R329T | A | 3.2e-07 | 4e-07 | 3.3e-07 | 3.500E-07 | 4.359E-08 | 3.200E-07 | 4.000E-07 |
| 6578 | 329 | R329W | A | 2.2e-07 | 7.7e-07 | 1.9e-07 | 3.933E-07 | 3.265E-07 | 1.900E-07 | 7.700E-07 |
| 6579 | 329 | R329Y | A | 1.8e-07 | 1.4e-07 | 1.6e-07 | 1.600E-07 | 2.000E-08 | 1.400E-07 | 1.800E-07 |
| 6580 | 329 | R329V | A | 2.5e-07 | 3.1e-07 | 1.9e-07 | 2.500E-07 | 6.000E-08 | 1.900E-07 | 3.100E-07 |
| 6581 | 330 | E330A | A | 3.9e-07 | 2.8e-07 | 2.8e-07 | 3.167E-07 | 6.351E-08 | 2.800E-07 | 3.900E-07 |
| 6582 | 330 | E330R | A | 1.4e-07 | 4e-07 | 1.3e-07 | 2.233E-07 | 1.531E-07 | 1.300E-07 | 4.000E-07 |
| 6583 | 330 | E330N | A | 2e-07 | 1.8e-07 | 2.5e-07 | 2.100E-07 | 3.606E-08 | 1.800E-07 | 2.500E-07 |
| 6584 | 330 | E330D | A | 5.1e-07 | 2.3e-07 | 2.7e-07 | 3.367E-07 | 1.514E-07 | 2.300E-07 | 5.100E-07 |
| 6585 | 330 | E330C | A | 2.2e-07 | 4.7e-07 | 1.4e-07 | 2.767E-07 | 1.721E-07 | 1.400E-07 | 4.700E-07 |
| 6586 | 330 | E330E | A | 1.8e-07 | 1.9e-07 | 2.8e-07 | 2.167E-07 | 5.508E-08 | 1.800E-07 | 2.800E-07 |
| 6587 | 330 | E330Q | A | 3e-07 | 8.4e-08 | 1.7e-07 | 1.847E-07 | 1.087E-07 | 8.400E-08 | 3.000E-07 |
| 6588 | 330 | E330G | A | 1.4e-07 | 5.9e-07 | 2.3e-07 | 3.200E-07 | 2.381E-07 | 1.400E-07 | 5.900E-07 |
| 6589 | 330 | E330H | A | 2.9e-07 | 3.9e-07 | 2e-07 | 2.933E-07 | 9.504E-08 | 2.000E-07 | 3.900E-07 |
| 6590 | 330 | E330I | A | 2.7e-07 | 3.2e-07 | 1.1e-07 | 2.333E-07 | 1.097E-07 | 1.100E-07 | 3.200E-07 |
| 6591 | 330 | E330L | A | 3.5e-07 | 2.7e-07 | 2.3e-07 | 2.833E-07 | 6.110E-08 | 2.300E-07 | 3.500E-07 |
| 6592 | 330 | E330K | A | 1.8e-07 | 3.8e-07 | 1.9e-07 | 2.500E-07 | 1.127E-07 | 1.800E-07 | 3.800E-07 |
| 6593 | 330 | E330M | A | 3.4e-07 | 2.9e-07 | 1.9e-07 | 2.733E-07 | 7.638E-08 | 1.900E-07 | 3.400E-07 |
| 6594 | 330 | E330F | A | 1.3e-07 | 1.8e-07 | 2.3e-07 | 1.800E-07 | 5.000E-08 | 1.300E-07 | 2.300E-07 |
| 6595 | 330 | E330P | A | 2.4e-07 | 2.5e-07 | 6.5e-08 | 1.850E-07 | 1.040E-07 | 6.500E-08 | 2.500E-07 |
| 6596 | 330 | E330S | A | 3.2e-07 | 2.6e-07 | 1.4e-07 | 2.400E-07 | 9.165E-08 | 1.400E-07 | 3.200E-07 |
| 6597 | 330 | E330T | A | 1.8e-07 | 1.7e-07 | 1.6e-07 | 1.700E-07 | 1.000E-08 | 1.600E-07 | 1.800E-07 |
| 6598 | 330 | E330W | A | 3.6e-07 | 3.8e-07 | 1.7e-07 | 3.033E-07 | 1.159E-07 | 1.700E-07 | 3.800E-07 |
| 6599 | 330 | E330Y | A | 1.7e-07 | 3.5e-07 | 2e-07 | 2.400E-07 | 9.644E-08 | 1.700E-07 | 3.500E-07 |
| 6600 | 330 | E330V | A | 4e-07 | 2.9e-07 | 5.9e-07 | 4.267E-07 | 1.518E-07 | 2.900E-07 | 5.900E-07 |
| 6601 | 331 | V331A | A | 2.5e-07 | 2.4e-07 | 3e-07 | 2.633E-07 | 3.215E-08 | 2.400E-07 | 3.000E-07 |
| 6602 | 331 | V331R | A | 3e-07 | 2.2e-07 | 3.7e-07 | 2.967E-07 | 7.506E-08 | 2.200E-07 | 3.700E-07 |
| 6603 | 331 | V331N | A | 2e-07 | 2.3e-07 | 3.3e-07 | 2.533E-07 | 6.807E-08 | 2.000E-07 | 3.300E-07 |
| 6604 | 331 | V331D | A | 2.1e-07 | 9.2e-08 | 1.6e-07 | 1.540E-07 | 5.923E-08 | 9.200E-08 | 2.100E-07 |
| 6605 | 331 | V331C | A | 2.9e-07 | 2.6e-07 | 3.5e-07 | 3.000E-07 | 4.583E-08 | 2.600E-07 | 3.500E-07 |
| 6606 | 331 | V331E | A | 2.3e-07 | 1.4e-07 | 4.4e-07 | 2.700E-07 | 1.539E-07 | 1.400E-07 | 4.400E-07 |
| 6607 | 331 | V331Q | A | 4e-07 | 2.2e-07 | 2e-07 | 2.733E-07 | 1.102E-07 | 2.000E-07 | 4.000E-07 |
| 6608 | 331 | V331G | A | 1.9e-07 | 1.6e-07 | 1.6e-07 | 1.700E-07 | 1.732E-08 | 1.600E-07 | 1.900E-07 |
| 6609 | 331 | V331H | A | 3.2e-07 | 2.7e-07 | 6.1e-07 | 4.000E-07 | 1.836E-07 | 2.700E-07 | 6.100E-07 |
| 6610 | 331 | V331I | A | 3.4e-07 | 4.1e-07 | 2.5e-07 | 3.333E-07 | 8.021E-08 | 2.500E-07 | 4.100E-07 |
| 6611 | 331 | V331L | A | 2.2e-07 | 2.9e-07 | 2.5e-07 | 2.533E-07 | 3.512E-08 | 2.200E-07 | 2.900E-07 |
| 6612 | 331 | V331K | A | 3.5e-07 | 2e-07 | 2e-07 | 2.500E-07 | 8.660E-08 | 2.000E-07 | 3.500E-07 |
| 6613 | 331 | V331M | A | 6.4e-07 | 2.7e-07 | 3.2e-07 | 4.100E-07 | 2.007E-07 | 2.700E-07 | 6.400E-07 |
| 6614 | 331 | V331F | A | 1.8e-07 | 4.1e-07 | 1e-07 | 2.300E-07 | 1.609E-07 | 1.000E-07 | 4.100E-07 |
| 6615 | 331 | V331P | A | 3e-07 | 2.2e-07 | 3.1e-07 | 2.767E-07 | 4.933E-08 | 2.200E-07 | 3.100E-07 |
| 6616 | 331 | V331S | A | 1.5e-07 | 4.8e-07 | 4.7e-07 | 3.667E-07 | 1.877E-07 | 1.500E-07 | 4.800E-07 |
| 6617 | 331 | V331T | A | 4.1e-07 | 4.2e-07 | 3.9e-07 | 4.067E-07 | 1.528E-08 | 3.900E-07 | 4.200E-07 |
| 6618 | 331 | V331W | A | 8.4e-08 | 2.4e-07 | 3.4e-07 | 2.213E-07 | 1.290E-07 | 8.400E-08 | 3.400E-07 |
| 6619 | 331 | V331Y | A | 1.6e-07 | 1.7e-07 | 3e-07 | 2.100E-07 | 7.810E-08 | 1.600E-07 | 3.000E-07 |
| 6620 | 331 | V331V | A | 1.6e-07 | 2.9e-07 | 2e-07 | 2.167E-07 | 6.658E-08 | 1.600E-07 | 2.900E-07 |
| 6621 | 332 | R332A | A | 5.1e-07 | 3.3e-07 | 2e-07 | 3.467E-07 | 1.557E-07 | 2.000E-07 | 5.100E-07 |
| 6622 | 332 | R332R | A | 3.8e-07 | 2.8e-07 | 8.4e-08 | 2.480E-07 | 1.506E-07 | 8.400E-08 | 3.800E-07 |
| 6623 | 332 | R332N | A | 2.4e-07 | 3.2e-07 | 1.7e-07 | 2.433E-07 | 7.506E-08 | 1.700E-07 | 3.200E-07 |
| 6624 | 332 | R332D | A | 4.8e-07 | 3.9e-07 | 4.6e-08 | 3.053E-07 | 2.291E-07 | 4.600E-08 | 4.800E-07 |
| 6625 | 332 | R332C | A | 1.6e-07 | 3.2e-07 | 4.4e-07 | 3.067E-07 | 1.405E-07 | 1.600E-07 | 4.400E-07 |
| 6626 | 332 | R332E | A | 2.3e-07 | 4.4e-07 | 1.5e-07 | 2.733E-07 | 1.498E-07 | 1.500E-07 | 4.400E-07 |
| 6627 | 332 | R332Q | A | 1.7e-07 | 2.6e-07 | 1.8e-07 | 2.033E-07 | 4.933E-08 | 1.700E-07 | 2.600E-07 |
| 6628 | 332 | R332G | A | 3.3e-07 | 1.9e-07 | 3.9e-07 | 3.033E-07 | 1.026E-07 | 1.900E-07 | 3.900E-07 |
| 6629 | 332 | R332H | A | 1.6e-07 | 2.5e-07 | 3.9e-07 | 2.667E-07 | 1.159E-07 | 1.600E-07 | 3.900E-07 |

| | | | | | | | | | | |
|---|---|---|---|---|---|---|---|---|---|---|
| 6630 | 332 | R332I | A | 4.2e-07 | 2.4e-07 | 1.3e-07 | 2.633E-07 | 1.464E-07 | 1.300E-07 | 4.200E-07 |
| 6631 | 332 | R332L | A | 1.8e-07 | 2.2e-07 | 1.5e-07 | 1.833E-07 | 3.512E-08 | 1.500E-07 | 2.200E-07 |
| 6632 | 332 | R332K | A | 9.6e-08 | 2.8e-07 | 2e-07 | 1.920E-07 | 9.226E-08 | 9.600E-08 | 2.800E-07 |
| 6633 | 332 | R332M | A | 1.5e-07 | 1.5e-07 | 2.7e-07 | 1.900E-07 | 6.928E-08 | 1.500E-07 | 2.700E-07 |
| 6634 | 332 | R332F | A | 3.3e-07 | 3.2e-07 | 2.6e-07 | 3.033E-07 | 3.786E-08 | 2.600E-07 | 3.300E-07 |
| 6635 | 332 | R332P | A | 2.1e-07 | 2.4e-07 | 1.6e-07 | 2.033E-07 | 4.041E-08 | 1.600E-07 | 2.400E-07 |
| 6636 | 332 | R332S | A | 3.1e-07 | 2.8e-07 | 4.2e-07 | 3.367E-07 | 7.371E-08 | 2.800E-07 | 4.200E-07 |
| 6637 | 332 | R332T | A | 8.1e-07 | 2.6e-07 | 1.2e-07 | 3.967E-07 | 3.647E-07 | 1.200E-07 | 8.100E-07 |
| 6638 | 332 | R332W | A | 1.8e-07 | 2.6e-07 | 2.1e-07 | 2.167E-07 | 4.041E-08 | 1.800E-07 | 2.600E-07 |
| 6639 | 332 | R332Y | A | 1.9e-07 | 5.2e-07 | 2.8e-07 | 3.300E-07 | 1.706E-07 | 1.900E-07 | 5.200E-07 |
| 6640 | 332 | R332V | A | 3.8e-07 | 2.6e-07 | 1.7e-07 | 2.700E-07 | 1.054E-07 | 1.700E-07 | 3.800E-07 |
| 6641 | 333 | A333A | A | 2.1e-07 | 6.1e-07 | 1.5e-07 | 3.233E-07 | 2.501E-07 | 1.500E-07 | 6.100E-07 |
| 6642 | 333 | A333R | A | 2e-07 | 2.2e-07 | 1.7e-07 | 1.967E-07 | 2.517E-08 | 1.700E-07 | 2.200E-07 |
| 6643 | 333 | A333N | A | 1.5e-07 | 2.7e-07 | 1e-07 | 1.733E-07 | 8.737E-08 | 1.000E-07 | 2.700E-07 |
| 6644 | 333 | A333D | A | 3.7e-07 | 3.6e-07 | 3e-07 | 3.433E-07 | 3.786E-08 | 3.000E-07 | 3.700E-07 |
| 6645 | 333 | A333C | A | 3.4e-07 | 1.8e-07 | 4.1e-07 | 3.100E-07 | 1.179E-07 | 1.800E-07 | 4.100E-07 |
| 6646 | 333 | A333E | A | 2.2e-07 | 3e-07 | 1.6e-07 | 2.267E-07 | 7.024E-08 | 1.600E-07 | 3.000E-07 |
| 6647 | 333 | A333Q | A | 1.4e-07 | 2.5e-07 | 3.7e-07 | 2.533E-07 | 1.150E-07 | 1.400E-07 | 3.700E-07 |
| 6648 | 333 | A333G | A | 2.3e-07 | 5.8e-07 | 2.2e-07 | 3.433E-07 | 2.050E-07 | 2.200E-07 | 5.800E-07 |
| 6649 | 333 | A333H | A | 7.6e-08 | 3.4e-07 | 3.7e-07 | 2.620E-07 | 1.618E-07 | 7.600E-08 | 3.700E-07 |
| 6650 | 333 | A333I | A | 1.1e-07 | 1.6e-07 | 1.7e-07 | 1.467E-07 | 3.215E-08 | 1.100E-07 | 1.700E-07 |
| 6651 | 333 | A333L | A | 1.7e-07 | 1.8e-07 | 2.5e-07 | 2.000E-07 | 4.359E-08 | 1.700E-07 | 2.500E-07 |
| 6652 | 333 | A333K | A | 1.8e-07 | 3.1e-07 | 6.5e-07 | 3.800E-07 | 2.427E-07 | 1.800E-07 | 6.500E-07 |
| 6653 | 333 | A333M | A | 2.2e-07 | 2.4e-07 | 2e-07 | 2.200E-07 | 2.000E-08 | 2.000E-07 | 2.400E-07 |
| 6654 | 333 | A333F | A | 4.2e-07 | 2.2e-07 | 2.1e-07 | 2.833E-07 | 1.185E-07 | 2.100E-07 | 4.200E-07 |
| 6655 | 333 | A333P | A | 2.1e-07 | 3e-07 | 2.1e-07 | 2.400E-07 | 5.196E-08 | 2.100E-07 | 3.000E-07 |
| 6656 | 333 | A333S | A | 2.9e-07 | 2.9e-07 | 2e-07 | 2.600E-07 | 5.196E-08 | 2.000E-07 | 2.900E-07 |
| 6657 | 333 | A333T | A | 1.5e-07 | 2.1e-07 | 1.6e-07 | 1.733E-07 | 3.215E-08 | 1.500E-07 | 2.100E-07 |
| 6658 | 333 | A333W | A | 2.3e-07 | 2.7e-07 | 3.1e-07 | 2.700E-07 | 4.000E-08 | 2.300E-07 | 3.100E-07 |
| 6659 | 333 | A333Y | A | 1.7e-07 | 2.2e-07 | 3.4e-07 | 2.433E-07 | 8.737E-08 | 1.700E-07 | 3.400E-07 |
| 6660 | 333 | A333V | A | 1.7e-07 | 5.4e-07 | 2e-07 | 3.033E-07 | 2.055E-07 | 1.700E-07 | 5.400E-07 |
| 6661 | 334 | V334A | A | 2.9e-07 | 2e-07 | 2.8e-07 | 2.567E-07 | 4.933E-08 | 2.000E-07 | 2.900E-07 |
| 6662 | 334 | V334R | A | 5.8e-07 | 9.9e-08 | 1.6e-07 | 2.797E-07 | 2.619E-07 | 9.900E-08 | 5.800E-07 |
| 6663 | 334 | V334N | A | 1.3e-07 | 1.9e-07 | 4.5e-07 | 2.567E-07 | 1.701E-07 | 1.300E-07 | 4.500E-07 |
| 6664 | 334 | V334D | A | 3.4e-07 | 3.3e-07 | 2.3e-07 | 3.000E-07 | 6.083E-08 | 2.300E-07 | 3.400E-07 |
| 6665 | 334 | V334C | A | 3e-07 | 2.3e-07 | 2e-07 | 2.433E-07 | 5.132E-08 | 2.000E-07 | 3.000E-07 |
| 6666 | 334 | V334E | A | 1.6e-07 | 3.8e-07 | 1.1e-07 | 2.167E-07 | 1.436E-07 | 1.100E-07 | 3.800E-07 |
| 6667 | 334 | V334Q | A | 1.7e-07 | 1.5e-07 | 2.3e-07 | 1.833E-07 | 4.163E-08 | 1.500E-07 | 2.300E-07 |
| 6668 | 334 | V334G | A | 2.3e-07 | 4.5e-07 | 4.1e-07 | 3.633E-07 | 1.172E-07 | 2.300E-07 | 4.500E-07 |
| 6669 | 334 | V334H | A | 1.7e-07 | 1.6e-07 | 1.7e-07 | 1.667E-07 | 5.774E-09 | 1.600E-07 | 1.700E-07 |
| 6670 | 334 | V334I | A | 2.4e-07 | 3.4e-07 | 3.4e-07 | 3.067E-07 | 5.774E-08 | 2.400E-07 | 3.400E-07 |
| 6671 | 334 | V334L | A | 3.4e-07 | 3.5e-07 | 2e-07 | 2.967E-07 | 8.386E-08 | 2.000E-07 | 3.500E-07 |
| 6672 | 334 | V334K | A | 3.6e-07 | 3.4e-07 | 2.6e-07 | 3.200E-07 | 5.292E-08 | 2.600E-07 | 3.600E-07 |
| 6673 | 334 | V334M | A | 3.9e-07 | 1.7e-07 | 4.2e-07 | 3.267E-07 | 1.365E-07 | 1.700E-07 | 4.200E-07 |
| 6674 | 334 | V334F | A | 2.7e-07 | 1.2e-07 | 3.2e-07 | 2.367E-07 | 1.041E-07 | 1.200E-07 | 3.200E-07 |
| 6675 | 334 | V334P | A | 1.6e-07 | 1.5e-07 | 3.1e-07 | 2.067E-07 | 8.963E-08 | 1.500E-07 | 3.100E-07 |
| 6676 | 334 | V334S | A | 3.2e-07 | 1.9e-07 | 1.6e-07 | 2.233E-07 | 8.505E-08 | 1.600E-07 | 3.200E-07 |
| 6677 | 334 | V334T | A | 2.8e-07 | 3.6e-07 | 2.3e-07 | 2.900E-07 | 6.557E-08 | 2.300E-07 | 3.600E-07 |
| 6678 | 334 | V334W | A | 2.6e-07 | 3.8e-07 | 2.6e-07 | 3.000E-07 | 6.928E-08 | 2.600E-07 | 3.800E-07 |
| 6679 | 334 | V334Y | A | 3.8e-07 | 3e-07 | 1.2e-07 | 2.667E-07 | 1.332E-07 | 1.200E-07 | 3.800E-07 |
| 6680 | 334 | V334V | A | 3.3e-07 | 3.5e-07 | 1.6e-07 | 2.800E-07 | 1.044E-07 | 1.600E-07 | 3.500E-07 |
| 6681 | 335 | T335A | A | 1e-07 | 2.7e-07 | 1.6e-07 | 1.767E-07 | 8.622E-08 | 1.000E-07 | 2.700E-07 |
| 6682 | 335 | T335R | A | 4.4e-07 | 9.2e-08 | 1.1e-07 | 2.140E-07 | 1.959E-07 | 9.200E-08 | 4.400E-07 |
| 6683 | 335 | T335N | A | 2.8e-07 | 4e-07 | 2.5e-07 | 3.100E-07 | 7.937E-08 | 2.500E-07 | 4.000E-07 |
| 6684 | 335 | T335D | A | 1.2e-07 | 3.3e-07 | 1.9e-07 | 2.133E-07 | 1.069E-07 | 1.200E-07 | 3.300E-07 |
| 6685 | 335 | T335C | A | 1.7e-07 | 1.9e-07 | 1.6e-07 | 1.733E-07 | 1.528E-08 | 1.600E-07 | 1.900E-07 |
| 6686 | 335 | T335E | A | 2.6e-07 | 2.1e-07 | 1.7e-07 | 2.133E-07 | 4.509E-08 | 1.700E-07 | 2.600E-07 |
| 6687 | 335 | T335Q | A | 2.5e-07 | 1.4e-07 | 2.2e-07 | 2.033E-07 | 5.686E-08 | 1.400E-07 | 2.500E-07 |
| 6688 | 335 | T335G | A | 2.1e-07 | 6.4e-07 | 2.2e-07 | 3.567E-07 | 2.454E-07 | 2.100E-07 | 6.400E-07 |
| 6689 | 335 | T335H | A | 6.3e-07 | 3.3e-07 | 2.6e-07 | 4.067E-07 | 1.966E-07 | 2.600E-07 | 6.300E-07 |
| 6690 | 335 | T335I | A | 3.4e-07 | 1.7e-07 | 1.7e-07 | 2.267E-07 | 9.815E-08 | 1.700E-07 | 3.400E-07 |
| 6691 | 335 | T335L | A | 5.6e-07 | 3.1e-07 | 3.2e-07 | 3.967E-07 | 1.415E-07 | 3.100E-07 | 5.600E-07 |
| 6692 | 335 | T335K | A | 6.1e-07 | 2.8e-07 | 4.5e-07 | 4.467E-07 | 1.650E-07 | 2.800E-07 | 6.100E-07 |
| 6693 | 335 | T335M | A | 4.1e-07 | 3.3e-07 | 5.7e-07 | 4.367E-07 | 1.222E-07 | 3.300E-07 | 5.700E-07 |
| 6694 | 335 | T335F | A | 2e-07 | 1.7e-07 | 1.6e-07 | 1.767E-07 | 2.082E-08 | 1.600E-07 | 2.000E-07 |
| 6695 | 335 | T335P | A | 1.5e-07 | 2.7e-07 | 1.4e-07 | 1.867E-07 | 7.234E-08 | 1.400E-07 | 2.700E-07 |
| 6696 | 335 | T335S | A | 4.4e-07 | 3.3e-07 | 2.3e-07 | 3.333E-07 | 1.050E-07 | 2.300E-07 | 4.400E-07 |
| 6697 | 335 | T335T | A | 1.8e-07 | 1.6e-07 | 7.6e-07 | 3.667E-07 | 3.408E-07 | 1.600E-07 | 7.600E-07 |
| 6698 | 335 | T335W | A | 1.1e-07 | 2.8e-07 | 4e-07 | 2.633E-07 | 1.457E-07 | 1.100E-07 | 4.000E-07 |
| 6699 | 335 | T335Y | A | 3.3e-07 | 8.1e-08 | 3.2e-07 | 2.437E-07 | 1.410E-07 | 8.100E-08 | 3.300E-07 |
| 6700 | 335 | T335V | A | 3.1e-07 | 1.9e-07 | 3.4e-07 | 2.800E-07 | 7.937E-08 | 1.900E-07 | 3.400E-07 |
| 6701 | 336 | S336A | A | 1.8e-07 | 1.3e-07 | 1.1e-07 | 1.400E-07 | 3.606E-08 | 1.100E-07 | 1.800E-07 |
| 6702 | 336 | S336R | A | 1.3e-07 | 3.8e-07 | 2.7e-07 | 2.600E-07 | 1.253E-07 | 1.300E-07 | 3.800E-07 |
| 6703 | 336 | S336N | A | 2.1e-07 | 5.3e-07 | 2.7e-07 | 3.367E-07 | 1.701E-07 | 2.100E-07 | 5.300E-07 |
| 6704 | 336 | S336D | A | 2.7e-07 | 2.1e-07 | 2.2e-07 | 2.333E-07 | 3.215E-08 | 2.100E-07 | 2.700E-07 |
| 6705 | 336 | S336C | A | 2.4e-07 | 3e-07 | 2.6e-07 | 2.667E-07 | 3.055E-08 | 2.400E-07 | 3.000E-07 |
| 6706 | 336 | S336E | A | 7.5e-07 | 3.1e-07 | 2.4e-07 | 4.333E-07 | 2.765E-07 | 2.400E-07 | 7.500E-07 |
| 6707 | 336 | S336Q | A | 1.2e-07 | 2.9e-07 | 4.7e-07 | 2.933E-07 | 1.750E-07 | 1.200E-07 | 4.700E-07 |
| 6708 | 336 | S336G | A | 3.6e-07 | 3.2e-07 | 2.7e-07 | 3.167E-07 | 4.509E-08 | 2.700E-07 | 3.600E-07 |
| 6709 | 336 | S336H | A | 2.7e-07 | 2.7e-07 | 6.6e-08 | 2.020E-07 | 1.178E-07 | 6.600E-08 | 2.700E-07 |
| 6710 | 336 | S336I | A | 2.8e-07 | 2.8e-07 | 1.4e-07 | 2.333E-07 | 8.083E-08 | 1.400E-07 | 2.800E-07 |
| 6711 | 336 | S336L | A | 2.2e-07 | 2.8e-07 | 2.6e-07 | 2.533E-07 | 3.055E-08 | 2.200E-07 | 2.800E-07 |
| 6712 | 336 | S336K | A | 1.8e-07 | 2.5e-07 | 3.1e-07 | 2.467E-07 | 6.506E-08 | 1.800E-07 | 3.100E-07 |
| 6713 | 336 | S336M | A | 3.3e-07 | 4e-07 | 1.5e-07 | 2.933E-07 | 1.290E-07 | 1.500E-07 | 4.000E-07 |
| 6714 | 336 | S336F | A | 4e-07 | 2.5e-07 | 2.5e-07 | 3.000E-07 | 8.660E-08 | 2.500E-07 | 4.000E-07 |

| | | | | | | | | | | |
|---|---|---|---|---|---|---|---|---|---|---|
| 6715 | 336 | S336P | A | 1.8e-07 | 3.5e-07 | 2.4e-07 | 2.567E-07 | 8.622E-08 | 1.800E-07 | 3.500E-07 |
| 6716 | 336 | S336S | A | 2.6e-07 | 3.4e-07 | 1.1e-07 | 2.367E-07 | 1.168E-07 | 1.100E-07 | 3.400E-07 |
| 6717 | 336 | S336T | A | 6.9e-07 | 3e-07 | 1.9e-07 | 3.933E-07 | 2.627E-07 | 1.900E-07 | 6.900E-07 |
| 6718 | 336 | S336W | A | 4.4e-07 | 2.1e-07 | 1.7e-07 | 2.733E-07 | 1.457E-07 | 1.700E-07 | 4.400E-07 |
| 6719 | 336 | S336Y | A | 2e-07 | 4.6e-07 | 1.5e-07 | 2.700E-07 | 1.664E-07 | 1.500E-07 | 4.600E-07 |
| 6720 | 336 | S336V | A | 5.6e-07 | 6.6e-07 | 5.3e-07 | 5.833E-07 | 6.807E-08 | 5.300E-07 | 6.600E-07 |
| 6721 | 337 | A337A | A | 2e-07 | 7.2e-07 | 1.4e-07 | 3.533E-07 | 3.190E-07 | 1.400E-07 | 7.200E-07 |
| 6722 | 337 | A337R | A | 3e-07 | 1.3e-07 | 4.8e-07 | 3.033E-07 | 1.750E-07 | 1.300E-07 | 4.800E-07 |
| 6723 | 337 | A337N | A | 2.1e-07 | 2.2e-07 | 3e-07 | 2.433E-07 | 4.933E-08 | 2.100E-07 | 3.000E-07 |
| 6724 | 337 | A337D | A | 3.2e-07 | 1.7e-07 | 4.6e-07 | 3.167E-07 | 1.450E-07 | 1.700E-07 | 4.600E-07 |
| 6725 | 337 | A337C | A | 2.3e-07 | 2.9e-07 | 2e-07 | 2.400E-07 | 4.583E-08 | 2.000E-07 | 2.900E-07 |
| 6726 | 337 | A337E | A | 3.2e-07 | 3.8e-07 | 3.2e-07 | 3.400E-07 | 3.464E-08 | 3.200E-07 | 3.800E-07 |
| 6727 | 337 | A337Q | A | 4e-07 | 1.6e-07 | 1.4e-07 | 2.333E-07 | 1.447E-07 | 1.400E-07 | 4.000E-07 |
| 6728 | 337 | A337G | A | 1.8e-07 | 1.2e-07 | 9.3e-08 | 1.310E-07 | 4.453E-08 | 9.300E-08 | 1.800E-07 |
| 6729 | 337 | A337H | A | 2.9e-07 | 1.5e-07 | 4.6e-07 | 3.000E-07 | 1.552E-07 | 1.500E-07 | 4.600E-07 |
| 6730 | 337 | A337I | A | 1.9e-07 | 2.4e-07 | 2.5e-07 | 2.267E-07 | 3.215E-08 | 1.900E-07 | 2.500E-07 |
| 6731 | 337 | A337L | A | 4.1e-07 | 3e-07 | 3.5e-07 | 3.533E-07 | 5.508E-08 | 3.000E-07 | 4.100E-07 |
| 6732 | 337 | A337K | A | 3.5e-07 | 1.5e-07 | 1.9e-07 | 2.300E-07 | 1.058E-07 | 1.500E-07 | 3.500E-07 |
| 6733 | 337 | A337M | A | 2.5e-07 | 2.9e-07 | 2.4e-07 | 2.600E-07 | 2.646E-08 | 2.400E-07 | 2.900E-07 |
| 6734 | 337 | A337F | A | 4.1e-07 | 1.8e-07 | 5.1e-07 | 3.667E-07 | 1.692E-07 | 1.800E-07 | 5.100E-07 |
| 6735 | 337 | A337P | A | 3.1e-07 | 2.8e-07 | 2e-07 | 2.633E-07 | 5.686E-08 | 2.000E-07 | 3.100E-07 |
| 6736 | 337 | A337S | A | 4e-07 | 4.4e-07 | 3.8e-07 | 4.067E-07 | 3.055E-08 | 3.800E-07 | 4.400E-07 |
| 6737 | 337 | A337T | A | 1.3e-07 | 1.5e-07 | 2.9e-07 | 1.900E-07 | 8.718E-08 | 1.300E-07 | 2.900E-07 |
| 6738 | 337 | A337W | A | 2.2e-07 | 2.8e-07 | 2e-07 | 2.333E-07 | 4.163E-08 | 2.000E-07 | 2.800E-07 |
| 6739 | 337 | A337Y | A | 1.7e-07 | 1.2e-07 | 4.1e-07 | 2.333E-07 | 1.550E-07 | 1.200E-07 | 4.100E-07 |
| 6740 | 337 | A337V | A | 5.3e-07 | 4.3e-07 | 2.7e-07 | 4.100E-07 | 1.311E-07 | 2.700E-07 | 5.300E-07 |
| 6741 | 338 | N338A | A | 2e-07 | 2.4e-07 | 4e-07 | 2.800E-07 | 1.058E-07 | 2.000E-07 | 4.000E-07 |
| 6742 | 338 | N338R | A | 2.6e-07 | 2.1e-07 | 1.2e-07 | 1.967E-07 | 7.095E-08 | 1.200E-07 | 2.600E-07 |
| 6743 | 338 | N338N | A | 2.9e-07 | 3.4e-07 | 1.5e-07 | 2.600E-07 | 9.849E-08 | 1.500E-07 | 3.400E-07 |
| 6744 | 338 | N338D | A | 1.4e-07 | 3.3e-07 | 1.6e-07 | 2.100E-07 | 1.044E-07 | 1.400E-07 | 3.300E-07 |
| 6745 | 338 | N338C | A | 3.4e-07 | 2.2e-07 | 1.9e-07 | 2.500E-07 | 7.937E-08 | 1.900E-07 | 3.400E-07 |
| 6746 | 338 | N338E | A | 1.7e-07 | 2.3e-07 | 2.2e-07 | 2.067E-07 | 3.215E-08 | 1.700E-07 | 2.300E-07 |
| 6747 | 338 | N338Q | A | 2.5e-07 | 1.3e-07 | 1.4e-07 | 1.733E-07 | 6.658E-08 | 1.300E-07 | 2.500E-07 |
| 6748 | 338 | N338G | A | 2.5e-07 | 4.3e-07 | 2.3e-07 | 3.033E-07 | 1.102E-07 | 2.300E-07 | 4.300E-07 |
| 6749 | 338 | N338H | A | 2.4e-07 | 3.4e-07 | 2.9e-07 | 2.900E-07 | 5.000E-08 | 2.400E-07 | 3.400E-07 |
| 6750 | 338 | N338I | A | 2.1e-07 | 3.3e-07 | 2.4e-07 | 2.600E-07 | 6.245E-08 | 2.100E-07 | 3.300E-07 |
| 6751 | 338 | N338L | A | 3e-07 | 2.7e-07 | 2.3e-07 | 2.667E-07 | 3.512E-08 | 2.300E-07 | 3.000E-07 |
| 6752 | 338 | N338K | A | 2.4e-07 | 8.8e-08 | 4e-07 | 2.427E-07 | 1.560E-07 | 8.800E-08 | 4.000E-07 |
| 6753 | 338 | N338M | A | 1.8e-07 | 1.9e-07 | 2.5e-07 | 2.067E-07 | 3.786E-08 | 1.800E-07 | 2.500E-07 |
| 6754 | 338 | N338F | A | 6.2e-07 | 2.8e-07 | 1.7e-07 | 3.567E-07 | 2.346E-07 | 1.700E-07 | 6.200E-07 |
| 6755 | 338 | N338P | A | 7.3e-08 | 4.8e-07 | 2.8e-07 | 2.777E-07 | 2.035E-07 | 7.300E-08 | 4.800E-07 |
| 6756 | 338 | N338S | A | 2.3e-07 | 2e-07 | 2.1e-07 | 2.133E-07 | 1.528E-08 | 2.000E-07 | 2.300E-07 |
| 6757 | 338 | N338T | A | 1.5e-07 | 3e-07 | 3e-07 | 2.500E-07 | 8.660E-08 | 1.500E-07 | 3.000E-07 |
| 6758 | 338 | N338W | A | 1.8e-07 | 1.8e-07 | 2.2e-07 | 1.933E-07 | 2.309E-08 | 1.800E-07 | 2.200E-07 |
| 6759 | 338 | N338Y | A | 5.9e-07 | 2.7e-07 | 1.8e-07 | 3.467E-07 | 2.155E-07 | 1.800E-07 | 5.900E-07 |
| 6760 | 338 | N338V | A | 2.7e-07 | 2.8e-07 | 2.6e-07 | 2.700E-07 | 1.000E-08 | 2.600E-07 | 2.800E-07 |
| 6761 | 339 | I339A | A | 5.1e-07 | 4e-07 | 4.1e-07 | 4.400E-07 | 6.083E-08 | 4.000E-07 | 5.100E-07 |
| 6762 | 339 | I339R | A | 2.4e-07 | 2.9e-07 | 3.5e-07 | 2.933E-07 | 5.508E-08 | 2.400E-07 | 3.500E-07 |
| 6763 | 339 | I339N | A | 3.5e-07 | 2.1e-07 | 3e-07 | 2.867E-07 | 7.095E-08 | 2.100E-07 | 3.500E-07 |
| 6764 | 339 | I339D | A | 3.2e-07 | 1.2e-07 | 1.8e-07 | 2.067E-07 | 1.026E-07 | 1.200E-07 | 3.200E-07 |
| 6765 | 339 | I339C | A | 3.8e-07 | 2.2e-07 | 1.9e-07 | 2.633E-07 | 1.021E-07 | 1.900E-07 | 3.800E-07 |
| 6766 | 339 | I339E | A | 4.2e-07 | 2e-07 | 2.2e-07 | 2.800E-07 | 1.217E-07 | 2.000E-07 | 4.200E-07 |
| 6767 | 339 | I339Q | A | 2.9e-07 | 3.2e-07 | 2.6e-07 | 2.900E-07 | 3.000E-08 | 2.600E-07 | 3.200E-07 |
| 6768 | 339 | I339G | A | 2.1e-07 | 3.2e-07 | 2.5e-07 | 2.600E-07 | 5.568E-08 | 2.100E-07 | 3.200E-07 |
| 6769 | 339 | I339H | A | 3.4e-07 | 3.9e-07 | 2.7e-07 | 3.333E-07 | 6.028E-08 | 2.700E-07 | 3.900E-07 |
| 6770 | 339 | I339I | A | 2e-07 | 1.5e-07 | 4e-07 | 2.500E-07 | 1.323E-07 | 1.500E-07 | 4.000E-07 |
| 6771 | 339 | I339L | A | 1.6e-07 | 9.7e-08 | 1.6e-07 | 1.390E-07 | 3.637E-08 | 9.700E-08 | 1.600E-07 |
| 6772 | 339 | I339K | A | 3.1e-07 | 2.4e-07 | 2.1e-07 | 2.533E-07 | 5.132E-08 | 2.100E-07 | 3.100E-07 |
| 6773 | 339 | I339M | A | 1.6e-07 | 1.9e-07 | 3.2e-07 | 2.233E-07 | 8.505E-08 | 1.600E-07 | 3.200E-07 |
| 6774 | 339 | I339F | A | 4e-07 | 2.2e-07 | 4.5e-07 | 3.567E-07 | 1.210E-07 | 2.200E-07 | 4.500E-07 |
| 6775 | 339 | I339P | A | 1.2e-07 | 2.2e-07 | 2.2e-07 | 1.867E-07 | 5.774E-08 | 1.200E-07 | 2.200E-07 |
| 6776 | 339 | I339S | A | 2.7e-07 | 2.1e-07 | 1.7e-07 | 2.167E-07 | 5.033E-08 | 1.700E-07 | 2.700E-07 |
| 6777 | 339 | I339T | A | 2.1e-07 | 2.1e-07 | 3.9e-07 | 2.700E-07 | 1.039E-07 | 2.100E-07 | 3.900E-07 |
| 6778 | 339 | I339W | A | 3e-07 | 2.9e-07 | 2.2e-07 | 2.700E-07 | 4.359E-08 | 2.200E-07 | 3.000E-07 |
| 6779 | 339 | I339Y | A | 2.3e-07 | 1.3e-07 | 1.8e-07 | 1.800E-07 | 5.000E-08 | 1.300E-07 | 2.300E-07 |
| 6780 | 339 | I339V | A | 9e-08 | 1.9e-07 | 3.5e-07 | 2.100E-07 | 1.311E-07 | 9.000E-08 | 3.500E-07 |
| 6781 | 340 | Q340A | A | 5.4e-07 | 1.1e-07 | 2.9e-07 | 3.133E-07 | 2.159E-07 | 1.100E-07 | 5.400E-07 |
| 6782 | 340 | Q340R | A | 1.5e-07 | 4.5e-07 | 6.1e-07 | 4.033E-07 | 2.335E-07 | 1.500E-07 | 6.100E-07 |
| 6783 | 340 | Q340N | A | 3.2e-07 | 3.6e-07 | 3.5e-07 | 3.433E-07 | 2.082E-08 | 3.200E-07 | 3.600E-07 |
| 6784 | 340 | Q340D | A | 3.4e-07 | 1.9e-07 | 2e-07 | 2.433E-07 | 8.386E-08 | 1.900E-07 | 3.400E-07 |
| 6785 | 340 | Q340C | A | 1.9e-07 | 4.3e-07 | 2.3e-07 | 2.833E-07 | 1.286E-07 | 1.900E-07 | 4.300E-07 |
| 6786 | 340 | Q340E | A | 4.6e-07 | 3.6e-07 | 3.2e-07 | 3.800E-07 | 7.211E-08 | 3.200E-07 | 4.600E-07 |
| 6787 | 340 | Q340Q | A | 3.2e-07 | 3.2e-07 | 2.2e-07 | 2.867E-07 | 5.774E-08 | 2.200E-07 | 3.200E-07 |
| 6788 | 340 | Q340G | A | 2.5e-07 | 4.7e-07 | 2.7e-07 | 3.300E-07 | 1.217E-07 | 2.500E-07 | 4.700E-07 |
| 6789 | 340 | Q340H | A | 3.2e-07 | 3.1e-07 | 1.6e-07 | 2.633E-07 | 8.963E-08 | 1.600E-07 | 3.200E-07 |
| 6790 | 340 | Q340I | A | 3.9e-07 | 1.7e-07 | 4.3e-07 | 3.300E-07 | 1.400E-07 | 1.700E-07 | 4.300E-07 |
| 6791 | 340 | Q340L | A | 4.3e-07 | 3.1e-07 | 1.5e-07 | 2.967E-07 | 1.405E-07 | 1.500E-07 | 4.300E-07 |
| 6792 | 340 | Q340K | A | 4.1e-07 | 3e-07 | 1.9e-07 | 3.000E-07 | 1.100E-07 | 1.900E-07 | 4.100E-07 |
| 6793 | 340 | Q340M | A | 2.1e-07 | 1.6e-07 | 3.3e-07 | 2.333E-07 | 8.737E-08 | 1.600E-07 | 3.300E-07 |
| 6794 | 340 | Q340F | A | 1.5e-07 | 1.7e-07 | 1.2e-07 | 1.467E-07 | 2.517E-08 | 1.200E-07 | 1.700E-07 |
| 6795 | 340 | Q340P | A | 2.1e-07 | 4.6e-07 | 2.7e-07 | 3.133E-07 | 1.305E-07 | 2.100E-07 | 4.600E-07 |
| 6796 | 340 | Q340S | A | 3.9e-07 | 1.2e-07 | 3.2e-07 | 2.767E-07 | 1.401E-07 | 1.200E-07 | 3.900E-07 |
| 6797 | 340 | Q340T | A | 2.2e-07 | 2.6e-07 | 3.1e-07 | 2.633E-07 | 4.509E-08 | 2.200E-07 | 3.100E-07 |
| 6798 | 340 | Q340W | A | 3.6e-07 | 2.5e-07 | 2.5e-07 | 2.867E-07 | 6.351E-08 | 2.500E-07 | 3.600E-07 |
| 6799 | 340 | Q340Y | A | 2e-07 | 3.1e-07 | 1.9e-07 | 2.333E-07 | 6.658E-08 | 1.900E-07 | 3.100E-07 |

| | | | | | | | | | | |
|---|---|---|---|---|---|---|---|---|---|---|
| 6800 | 340 | Q340V | A | 1.5e-07 | 2e-07 | 1.8e-07 | 1.767E-07 | 2.517E-08 | 1.500E-07 | 2.000E-07 |
| 6801 | 341 | E341A | A | 2.2e-07 | 1.2e-07 | 8e-07 | 3.800E-07 | 3.672E-07 | 1.200E-07 | 8.000E-07 |
| 6802 | 341 | E341R | A | 3.9e-07 | 2e-07 | 3.4e-07 | 3.100E-07 | 9.849E-08 | 2.000E-07 | 3.900E-07 |
| 6803 | 341 | E341N | A | 2.9e-07 | 1.5e-07 | 3.7e-07 | 2.700E-07 | 1.114E-07 | 1.500E-07 | 3.700E-07 |
| 6804 | 341 | E341D | A | 2.3e-07 | 3.3e-07 | 2.7e-07 | 2.767E-07 | 5.033E-08 | 2.300E-07 | 3.300E-07 |
| 6805 | 341 | E341C | A | 2.8e-07 | 4.7e-07 | 2.2e-07 | 3.233E-07 | 1.305E-07 | 2.200E-07 | 4.700E-07 |
| 6806 | 341 | E341E | A | 1.5e-07 | 3.3e-07 | 4e-07 | 2.933E-07 | 1.290E-07 | 1.500E-07 | 4.000E-07 |
| 6807 | 341 | E341Q | A | 1.6e-07 | 2.9e-07 | 9.9e-08 | 1.830E-07 | 9.756E-08 | 9.900E-08 | 2.900E-07 |
| 6808 | 341 | E341G | A | 1.4e-07 | 3.4e-07 | 2.7e-07 | 2.500E-07 | 1.015E-07 | 1.400E-07 | 3.400E-07 |
| 6809 | 341 | E341H | A | 6.1e-07 | 2.3e-07 | 5.3e-07 | 4.567E-07 | 2.003E-07 | 2.300E-07 | 6.100E-07 |
| 6810 | 341 | E341I | A | 2e-07 | 2.5e-07 | 7.4e-07 | 3.967E-07 | 2.984E-07 | 2.000E-07 | 7.400E-07 |
| 6811 | 341 | E341L | A | 1.7e-07 | 3.9e-07 | 2.5e-07 | 2.700E-07 | 1.114E-07 | 1.700E-07 | 3.900E-07 |
| 6812 | 341 | E341K | A | 1.6e-07 | 1.8e-07 | 3.5e-07 | 2.300E-07 | 1.044E-07 | 1.600E-07 | 3.500E-07 |
| 6813 | 341 | E341M | A | 2.8e-07 | 3.4e-07 | 2.2e-07 | 2.800E-07 | 6.000E-08 | 2.200E-07 | 3.400E-07 |
| 6814 | 341 | E341F | A | 2.3e-07 | 2.2e-07 | 2.5e-07 | 2.333E-07 | 1.528E-08 | 2.200E-07 | 2.500E-07 |
| 6815 | 341 | E341P | A | 2e-07 | 1.8e-07 | 2.2e-07 | 2.000E-07 | 2.000E-08 | 1.800E-07 | 2.200E-07 |
| 6816 | 341 | E341S | A | 2.6e-07 | 1.7e-07 | 2.2e-07 | 2.167E-07 | 4.509E-08 | 1.700E-07 | 2.600E-07 |
| 6817 | 341 | E341T | A | 3.3e-07 | 3.5e-07 | 3.9e-07 | 3.567E-07 | 3.055E-08 | 3.300E-07 | 3.900E-07 |
| 6818 | 341 | E341W | A | 9.6e-08 | 1.1e-07 | 2.2e-07 | 1.420E-07 | 6.791E-08 | 9.600E-08 | 2.200E-07 |
| 6819 | 341 | E341Y | A | 2.6e-07 | 4.7e-07 | 2.5e-07 | 3.267E-07 | 1.242E-07 | 2.500E-07 | 4.700E-07 |
| 6820 | 341 | E341V | A | 3.5e-07 | 1.9e-07 | 3.6e-07 | 3.000E-07 | 9.539E-08 | 1.900E-07 | 3.600E-07 |
| 6821 | 342 | F342A | A | 2.4e-07 | 3.3e-07 | 3.3e-07 | 3.000E-07 | 5.196E-08 | 2.400E-07 | 3.300E-07 |
| 6822 | 342 | F342R | A | 2.5e-07 | 1.8e-07 | 2.4e-07 | 2.233E-07 | 3.786E-08 | 1.800E-07 | 2.500E-07 |
| 6823 | 342 | F342N | A | 3.8e-07 | 5e-07 | 3.2e-07 | 4.000E-07 | 9.165E-08 | 3.200E-07 | 5.000E-07 |
| 6824 | 342 | F342D | A | 2.5e-07 | 5.3e-07 | 2.2e-07 | 3.333E-07 | 1.710E-07 | 2.200E-07 | 5.300E-07 |
| 6825 | 342 | F342C | A | 4.2e-07 | 1.1e-07 | 3.2e-07 | 2.833E-07 | 1.582E-07 | 1.100E-07 | 4.200E-07 |
| 6826 | 342 | F342E | A | 1.4e-07 | 9.8e-08 | 1.4e-07 | 1.260E-07 | 2.425E-08 | 9.800E-08 | 1.400E-07 |
| 6827 | 342 | F342Q | A | 1.5e-07 | 4.2e-07 | 1.3e-07 | 2.333E-07 | 1.620E-07 | 1.300E-07 | 4.200E-07 |
| 6828 | 342 | F342G | A | 1.5e-07 | 3e-07 | 7.7e-08 | 1.757E-07 | 1.137E-07 | 7.700E-08 | 3.000E-07 |
| 6829 | 342 | F342H | A | 2.4e-07 | 3e-07 | 4.8e-07 | 3.400E-07 | 1.249E-07 | 2.400E-07 | 4.800E-07 |
| 6830 | 342 | F342I | A | 1.9e-07 | 4.7e-07 | 2.9e-07 | 3.167E-07 | 1.419E-07 | 1.900E-07 | 4.700E-07 |
| 6831 | 342 | F342L | A | 4.9e-07 | 2.7e-07 | 2.6e-07 | 3.400E-07 | 1.300E-07 | 2.600E-07 | 4.900E-07 |
| 6832 | 342 | F342K | A | 1.2e-07 | 3e-07 | 3.1e-07 | 2.433E-07 | 1.069E-07 | 1.200E-07 | 3.100E-07 |
| 6833 | 342 | F342M | A | 1.6e-07 | 3.1e-07 | 2.9e-07 | 2.533E-07 | 8.145E-08 | 1.600E-07 | 3.100E-07 |
| 6834 | 342 | F342F | A | 3.5e-07 | 1.2e-07 | 1e-07 | 1.900E-07 | 1.389E-07 | 1.000E-07 | 3.500E-07 |
| 6835 | 342 | F342P | A | 5e-07 | 2e-07 | 1.8e-07 | 2.933E-07 | 1.793E-07 | 1.800E-07 | 5.000E-07 |
| 6836 | 342 | F342S | A | 3.3e-07 | 2.1e-07 | 3.3e-07 | 2.900E-07 | 6.928E-08 | 2.100E-07 | 3.300E-07 |
| 6837 | 342 | F342T | A | 3.9e-07 | 2.4e-07 | 2.2e-07 | 2.833E-07 | 9.292E-08 | 2.200E-07 | 3.900E-07 |
| 6838 | 342 | F342W | A | 1.2e-07 | 3.2e-07 | 1.6e-07 | 2.000E-07 | 1.058E-07 | 1.200E-07 | 3.200E-07 |
| 6839 | 342 | F342Y | A | 2e-07 | 1.8e-07 | 2.6e-07 | 2.133E-07 | 4.163E-08 | 1.800E-07 | 2.600E-07 |
| 6840 | 342 | F342V | A | 2e-07 | 1.9e-07 | 3.7e-07 | 2.533E-07 | 1.012E-07 | 1.900E-07 | 3.700E-07 |
| 6841 | 343 | A343A | A | 3.3e-07 | 2.3e-07 | 2.9e-07 | 2.833E-07 | 5.033E-08 | 2.300E-07 | 3.300E-07 |
| 6842 | 343 | A343R | A | 2.8e-07 | 1.8e-07 | 2.6e-07 | 2.400E-07 | 5.292E-08 | 1.800E-07 | 2.800E-07 |
| 6843 | 343 | A343N | A | 1.2e-07 | 2.3e-07 | 1.9e-07 | 1.800E-07 | 5.568E-08 | 1.200E-07 | 2.300E-07 |
| 6844 | 343 | A343D | A | 2.5e-07 | 2.6e-07 | 2.3e-07 | 2.467E-07 | 1.528E-08 | 2.300E-07 | 2.600E-07 |
| 6845 | 343 | A343C | A | 5e-07 | 4e-07 | 1.8e-07 | 3.600E-07 | 1.637E-07 | 1.800E-07 | 5.000E-07 |
| 6846 | 343 | A343E | A | 4.9e-07 | 1.4e-07 | 1.5e-07 | 2.600E-07 | 1.992E-07 | 1.400E-07 | 4.900E-07 |
| 6847 | 343 | A343Q | A | 2.7e-07 | 4.2e-07 | 8.7e-08 | 2.590E-07 | 1.668E-07 | 8.700E-08 | 4.200E-07 |
| 6848 | 343 | A343G | A | 3.4e-07 | 2.7e-07 | 2.7e-07 | 2.933E-07 | 4.041E-08 | 2.700E-07 | 3.400E-07 |
| 6849 | 343 | A343H | A | 3e-07 | 3e-07 | 1.3e-07 | 2.433E-07 | 9.815E-08 | 1.300E-07 | 3.000E-07 |
| 6850 | 343 | A343I | A | 1.4e-07 | 2.6e-07 | 8.4e-08 | 1.613E-07 | 8.992E-08 | 8.400E-08 | 2.600E-07 |
| 6851 | 343 | A343L | A | 3.6e-07 | 1.4e-07 | 2.1e-07 | 2.367E-07 | 1.124E-07 | 1.400E-07 | 3.600E-07 |
| 6852 | 343 | A343K | A | 3e-07 | 2.2e-07 | 2.9e-07 | 2.700E-07 | 4.359E-08 | 2.200E-07 | 3.000E-07 |
| 6853 | 343 | A343M | A | 2.5e-07 | 2.5e-07 | 2e-07 | 2.333E-07 | 2.887E-08 | 2.000E-07 | 2.500E-07 |
| 6854 | 343 | A343F | A | 2.9e-07 | 3.3e-07 | 1.3e-07 | 2.500E-07 | 1.058E-07 | 1.300E-07 | 3.300E-07 |
| 6855 | 343 | A343P | A | 1.4e-07 | 4e-07 | 2.3e-07 | 2.567E-07 | 1.320E-07 | 1.400E-07 | 4.000E-07 |
| 6856 | 343 | A343S | A | 2.5e-07 | 1.9e-07 | 1.2e-07 | 1.867E-07 | 6.506E-08 | 1.200E-07 | 2.500E-07 |
| 6857 | 343 | A343T | A | 2.6e-07 | 2.5e-07 | 3.3e-07 | 2.800E-07 | 4.359E-08 | 2.500E-07 | 3.300E-07 |
| 6858 | 343 | A343W | A | 1.4e-07 | 1.8e-07 | 4.8e-07 | 2.667E-07 | 1.858E-07 | 1.400E-07 | 4.800E-07 |
| 6859 | 343 | A343Y | A | 2.2e-07 | 3.9e-07 | 3.9e-07 | 3.333E-07 | 9.815E-08 | 2.200E-07 | 3.900E-07 |
| 6860 | 343 | A343V | A | 1.8e-07 | 3.2e-07 | 1.8e-07 | 2.267E-07 | 8.083E-08 | 1.800E-07 | 3.200E-07 |
| 6861 | 344 | G344A | A | 6.6e-07 | 6.7e-07 | 2.8e-07 | 5.367E-07 | 2.223E-07 | 2.800E-07 | 6.700E-07 |
| 6862 | 344 | G344R | A | 2.9e-07 | 1.4e-07 | 4.1e-07 | 2.800E-07 | 1.353E-07 | 1.400E-07 | 4.100E-07 |
| 6863 | 344 | G344N | A | 3e-07 | 2.9e-07 | 2.6e-07 | 2.833E-07 | 2.082E-08 | 2.600E-07 | 3.000E-07 |
| 6864 | 344 | G344D | A | 1.5e-07 | 3.1e-07 | 2.4e-07 | 2.333E-07 | 8.021E-08 | 1.500E-07 | 3.100E-07 |
| 6865 | 344 | G344C | A | 2.4e-07 | 2.5e-07 | 1.4e-07 | 2.100E-07 | 6.083E-08 | 1.400E-07 | 2.500E-07 |
| 6866 | 344 | G344E | A | 1.9e-07 | 3.1e-07 | 3e-07 | 2.667E-07 | 6.658E-08 | 1.900E-07 | 3.100E-07 |
| 6867 | 344 | G344Q | A | 3e-07 | 1.6e-07 | 1.6e-07 | 2.067E-07 | 8.083E-08 | 1.600E-07 | 3.000E-07 |
| 6868 | 344 | G344G | A | 1.5e-07 | 3.6e-07 | 4.1e-07 | 3.067E-07 | 1.380E-07 | 1.500E-07 | 4.100E-07 |
| 6869 | 344 | G344H | A | 2.6e-07 | 1.9e-07 | 4.7e-07 | 3.067E-07 | 1.457E-07 | 1.900E-07 | 4.700E-07 |
| 6870 | 344 | G344I | A | 2.5e-07 | 2.3e-07 | 4.3e-07 | 3.033E-07 | 1.102E-07 | 2.300E-07 | 4.300E-07 |
| 6871 | 344 | G344L | A | 4.5e-07 | 1.9e-07 | 1.8e-07 | 2.733E-07 | 1.531E-07 | 1.800E-07 | 4.500E-07 |
| 6872 | 344 | G344K | A | 2.2e-07 | 3.3e-07 | 4.3e-07 | 3.267E-07 | 1.050E-07 | 2.200E-07 | 4.300E-07 |
| 6873 | 344 | G344M | A | 2.1e-07 | 3.3e-07 | 2.7e-07 | 2.700E-07 | 6.000E-08 | 2.100E-07 | 3.300E-07 |
| 6874 | 344 | G344F | A | 3.7e-07 | 2.8e-07 | 1e-07 | 2.500E-07 | 1.375E-07 | 1.000E-07 | 3.700E-07 |
| 6875 | 344 | G344P | A | 3.4e-07 | 1.6e-07 | 3.7e-07 | 2.900E-07 | 1.136E-07 | 1.600E-07 | 3.700E-07 |
| 6876 | 344 | G344S | A | 2.4e-07 | 1.5e-07 | 1.9e-07 | 1.933E-07 | 4.509E-08 | 1.500E-07 | 2.400E-07 |
| 6877 | 344 | G344T | A | 1.8e-07 | 1.8e-07 | 3.4e-07 | 2.333E-07 | 9.238E-08 | 1.800E-07 | 3.400E-07 |
| 6878 | 344 | G344W | A | 3.1e-07 | 2.2e-07 | 3.3e-07 | 2.867E-07 | 5.859E-08 | 2.200E-07 | 3.300E-07 |
| 6879 | 344 | G344Y | A | 1.7e-07 | 2.1e-07 | 2.8e-07 | 2.200E-07 | 5.568E-08 | 1.700E-07 | 2.800E-07 |
| 6880 | 344 | G344V | A | 2.9e-07 | 2.1e-07 | 2e-07 | 2.333E-07 | 4.933E-08 | 2.000E-07 | 2.900E-07 |
| 6881 | 345 | C345A | A | 2.9e-07 | 3.3e-07 | 4.4e-07 | 3.533E-07 | 7.767E-08 | 2.900E-07 | 4.400E-07 |
| 6882 | 345 | C345R | A | 9.7e-08 | 3.4e-07 | 1.7e-07 | 2.023E-07 | 1.247E-07 | 9.700E-08 | 3.400E-07 |
| 6883 | 345 | C345N | A | 3e-07 | 1.7e-07 | 2.9e-07 | 2.533E-07 | 7.234E-08 | 1.700E-07 | 3.000E-07 |
| 6884 | 345 | C345D | A | 1.9e-07 | 3.7e-07 | 1.9e-07 | 2.500E-07 | 1.039E-07 | 1.900E-07 | 3.700E-07 |

| | | | | | | | | | | |
|---|---|---|---|---|---|---|---|---|---|---|
| 6885 | 345 | C345C | A | 2.9e-07 | 2.6e-07 | 5.8e-07 | 3.767E-07 | 1.767E-07 | 2.600E-07 | 5.800E-07 |
| 6886 | 345 | C345E | A | 1.3e-07 | 1.2e-07 | 2.7e-07 | 1.733E-07 | 8.386E-08 | 1.200E-07 | 2.700E-07 |
| 6887 | 345 | C345Q | A | 2e-07 | 2.5e-07 | 2.2e-07 | 2.233E-07 | 2.517E-08 | 2.000E-07 | 2.500E-07 |
| 6888 | 345 | C345G | A | 1.5e-07 | 3e-07 | 1.9e-07 | 2.133E-07 | 7.767E-08 | 1.500E-07 | 3.000E-07 |
| 6889 | 345 | C345H | A | 2.1e-07 | 2.3e-07 | 2.4e-07 | 2.267E-07 | 1.528E-08 | 2.100E-07 | 2.400E-07 |
| 6890 | 345 | C345I | A | 2.2e-07 | 2.8e-07 | 3.5e-07 | 2.833E-07 | 6.506E-08 | 2.200E-07 | 3.500E-07 |
| 6891 | 345 | C345L | A | 4.8e-07 | 4.2e-07 | 2.8e-07 | 3.933E-07 | 1.026E-07 | 2.800E-07 | 4.800E-07 |
| 6892 | 345 | C345K | A | 2.4e-07 | 1.4e-07 | 1.4e-07 | 1.733E-07 | 5.774E-08 | 1.400E-07 | 2.400E-07 |
| 6893 | 345 | C345M | A | 2.4e-07 | 2.1e-07 | 3.1e-07 | 2.533E-07 | 5.132E-08 | 2.100E-07 | 3.100E-07 |
| 6894 | 345 | C345F | A | 2e-07 | 2.8e-07 | 4.4e-07 | 3.067E-07 | 1.222E-07 | 2.000E-07 | 4.400E-07 |
| 6895 | 345 | C345P | A | 8.4e-08 | 1.4e-07 | 4.2e-07 | 2.147E-07 | 1.800E-07 | 8.400E-08 | 4.200E-07 |
| 6896 | 345 | C345S | A | 1.9e-07 | 2.6e-07 | 1.8e-07 | 2.100E-07 | 4.359E-08 | 1.800E-07 | 2.600E-07 |
| 6897 | 345 | C345T | A | 1.6e-07 | 2e-07 | 3.6e-07 | 2.400E-07 | 1.058E-07 | 1.600E-07 | 3.600E-07 |
| 6898 | 345 | C345W | A | 3.2e-07 | 1e-07 | 6.6e-07 | 3.600E-07 | 2.821E-07 | 1.000E-07 | 6.600E-07 |
| 6899 | 345 | C345Y | A | 9.7e-08 | 3.1e-07 | 2.4e-07 | 2.157E-07 | 1.086E-07 | 9.700E-08 | 3.100E-07 |
| 6900 | 345 | C345V | A | 5.2e-07 | 5.1e-07 | 1.5e-07 | 3.933E-07 | 2.108E-07 | 1.500E-07 | 5.200E-07 |
| 6901 | 346 | K346A | A | 2.1e-07 | 2.3e-07 | 4e-07 | 2.800E-07 | 1.044E-07 | 2.100E-07 | 4.000E-07 |
| 6902 | 346 | K346R | A | 2e-07 | 1.8e-07 | 1.5e-07 | 1.767E-07 | 2.517E-08 | 1.500E-07 | 2.000E-07 |
| 6903 | 346 | K346N | A | 1.1e-07 | 1.6e-07 | 1.6e-07 | 1.433E-07 | 2.887E-08 | 1.100E-07 | 1.600E-07 |
| 6904 | 346 | K346D | A | 5.3e-07 | 2.6e-07 | 2.6e-07 | 3.500E-07 | 1.559E-07 | 2.600E-07 | 5.300E-07 |
| 6905 | 346 | K346C | A | 5.7e-07 | 2.4e-07 | 2.3e-07 | 3.467E-07 | 1.935E-07 | 2.300E-07 | 5.700E-07 |
| 6906 | 346 | K346E | A | 3.7e-07 | 3.5e-07 | 1.7e-07 | 2.967E-07 | 1.102E-07 | 1.700E-07 | 3.700E-07 |
| 6907 | 346 | K346Q | A | 3.5e-07 | 2.2e-07 | 2.3e-07 | 2.667E-07 | 7.234E-08 | 2.200E-07 | 3.500E-07 |
| 6908 | 346 | K346G | A | 2.2e-07 | 2.2e-07 | 2.6e-07 | 2.333E-07 | 2.309E-08 | 2.200E-07 | 2.600E-07 |
| 6909 | 346 | K346H | A | 4.3e-07 | 2.3e-07 | 1.9e-07 | 2.833E-07 | 1.286E-07 | 1.900E-07 | 4.300E-07 |
| 6910 | 346 | K346I | A | 2.9e-07 | 3.8e-07 | 3.2e-07 | 3.300E-07 | 4.583E-08 | 2.900E-07 | 3.800E-07 |
| 6911 | 346 | K346L | A | 1.4e-07 | 4e-07 | 1.7e-07 | 2.367E-07 | 1.422E-07 | 1.400E-07 | 4.000E-07 |
| 6912 | 346 | K346K | A | 1.9e-07 | 4.5e-07 | 4.3e-07 | 3.567E-07 | 1.447E-07 | 1.900E-07 | 4.500E-07 |
| 6913 | 346 | K346M | A | 3.6e-07 | 2.1e-07 | 1.3e-07 | 2.333E-07 | 1.168E-07 | 1.300E-07 | 3.600E-07 |
| 6914 | 346 | K346F | A | 2.9e-07 | 1.8e-07 | 4.2e-07 | 2.967E-07 | 1.201E-07 | 1.800E-07 | 4.200E-07 |
| 6915 | 346 | K346P | A | 2.4e-07 | 3.4e-07 | 1.7e-07 | 2.500E-07 | 8.544E-08 | 1.700E-07 | 3.400E-07 |
| 6916 | 346 | K346S | A | 1.6e-07 | 2.8e-07 | 2.3e-07 | 2.233E-07 | 6.028E-08 | 1.600E-07 | 2.800E-07 |
| 6917 | 346 | K346T | A | 2.2e-07 | 3.4e-07 | 4.5e-07 | 3.367E-07 | 1.150E-07 | 2.200E-07 | 4.500E-07 |
| 6918 | 346 | K346W | A | 6.5e-08 | 3.1e-07 | 2.4e-07 | 2.050E-07 | 1.262E-07 | 6.500E-08 | 3.100E-07 |
| 6919 | 346 | K346Y | A | 3.5e-07 | 1.9e-07 | 1.3e-07 | 2.233E-07 | 1.137E-07 | 1.300E-07 | 3.500E-07 |
| 6920 | 346 | K346V | A | 3.8e-07 | 4.3e-07 | 3.5e-07 | 3.867E-07 | 4.041E-08 | 3.500E-07 | 4.300E-07 |
| 6921 | 347 | K347A | A | 2.8e-07 | 2.9e-07 | 1.9e-07 | 2.533E-07 | 5.508E-08 | 1.900E-07 | 2.900E-07 |
| 6922 | 347 | K347R | A | 1.8e-07 | 2.5e-07 | 3.6e-07 | 2.633E-07 | 9.074E-08 | 1.800E-07 | 3.600E-07 |
| 6923 | 347 | K347N | A | 3.4e-07 | 4.5e-07 | 3e-07 | 3.633E-07 | 7.767E-08 | 3.000E-07 | 4.500E-07 |
| 6924 | 347 | K347D | A | 1.6e-07 | 1.8e-07 | 1.6e-07 | 1.667E-07 | 1.155E-08 | 1.600E-07 | 1.800E-07 |
| 6925 | 347 | K347C | A | 1.5e-07 | 2.9e-07 | 6.4e-07 | 3.600E-07 | 2.524E-07 | 1.500E-07 | 6.400E-07 |
| 6926 | 347 | K347E | A | 2.5e-07 | 1.7e-07 | 1.8e-07 | 2.000E-07 | 4.359E-08 | 1.700E-07 | 2.500E-07 |
| 6927 | 347 | K347Q | A | 1.8e-07 | 4.3e-07 | 2.2e-07 | 2.767E-07 | 1.343E-07 | 1.800E-07 | 4.300E-07 |
| 6928 | 347 | K347G | A | 3.3e-07 | 2.7e-07 | 1.4e-07 | 2.467E-07 | 9.713E-08 | 1.400E-07 | 3.300E-07 |
| 6929 | 347 | K347H | A | 2.1e-07 | 1.4e-07 | 4e-07 | 2.500E-07 | 1.345E-07 | 1.400E-07 | 4.000E-07 |
| 6930 | 347 | K347I | A | 2.8e-07 | 1.9e-07 | 2e-07 | 2.233E-07 | 4.933E-08 | 1.900E-07 | 2.800E-07 |
| 6931 | 347 | K347L | A | 3.3e-07 | 1.8e-07 | 2.9e-07 | 2.667E-07 | 7.767E-08 | 1.800E-07 | 3.300E-07 |
| 6932 | 347 | K347K | A | 3.3e-07 | 1.2e-07 | 2.1e-07 | 2.200E-07 | 1.054E-07 | 1.200E-07 | 3.300E-07 |
| 6933 | 347 | K347M | A | 6.9e-08 | 1.5e-07 | 1.8e-07 | 1.330E-07 | 5.742E-08 | 6.900E-08 | 1.800E-07 |
| 6934 | 347 | K347F | A | 2.4e-07 | 2.2e-07 | 3.6e-07 | 2.733E-07 | 7.572E-08 | 2.200E-07 | 3.600E-07 |
| 6935 | 347 | K347P | A | 2.3e-07 | 2.8e-07 | 2.6e-07 | 2.567E-07 | 2.517E-08 | 2.300E-07 | 2.800E-07 |
| 6936 | 347 | K347S | A | 1.4e-07 | 3.7e-07 | 1.8e-07 | 2.300E-07 | 1.229E-07 | 1.400E-07 | 3.700E-07 |
| 6937 | 347 | K347T | A | 3.6e-07 | 2.6e-07 | 1.5e-07 | 2.567E-07 | 1.050E-07 | 1.500E-07 | 3.600E-07 |
| 6938 | 347 | K347W | A | 2.4e-07 | 2.4e-07 | 2.2e-07 | 2.333E-07 | 1.155E-08 | 2.200E-07 | 2.400E-07 |
| 6939 | 347 | K347Y | A | 1.3e-07 | 2.2e-07 | 2.2e-07 | 1.900E-07 | 5.196E-08 | 1.300E-07 | 2.200E-07 |
| 6940 | 347 | K347V | A | 2.6e-07 | 2.7e-07 | 1.8e-07 | 2.367E-07 | 4.933E-08 | 1.800E-07 | 2.700E-07 |
| 6941 | 348 | I348A | A | 1.9e-07 | 2e-07 | 1.8e-07 | 1.900E-07 | 1.000E-08 | 1.800E-07 | 2.000E-07 |
| 6942 | 348 | I348R | A | 1.8e-07 | 3e-07 | 1.9e-07 | 2.233E-07 | 6.658E-08 | 1.800E-07 | 3.000E-07 |
| 6943 | 348 | I348N | A | 3e-07 | 3.4e-07 | 6.5e-07 | 4.300E-07 | 1.916E-07 | 3.000E-07 | 6.500E-07 |
| 6944 | 348 | I348D | A | 2.4e-07 | 1.9e-07 | 3.3e-07 | 2.533E-07 | 7.095E-08 | 1.900E-07 | 3.300E-07 |
| 6945 | 348 | I348C | A | 2.6e-07 | 3.2e-07 | 5.1e-07 | 3.633E-07 | 1.305E-07 | 2.600E-07 | 5.100E-07 |
| 6946 | 348 | I348E | A | 1.9e-07 | 4.4e-07 | 3.7e-07 | 3.333E-07 | 1.290E-07 | 1.900E-07 | 4.400E-07 |
| 6947 | 348 | I348Q | A | 3.9e-07 | 2.4e-07 | 3.1e-07 | 3.133E-07 | 7.506E-08 | 2.400E-07 | 3.900E-07 |
| 6948 | 348 | I348G | A | 1.3e-07 | 4e-07 | 3.4e-07 | 2.900E-07 | 1.418E-07 | 1.300E-07 | 4.000E-07 |
| 6949 | 348 | I348H | A | 2.8e-07 | 5.6e-07 | 3.1e-07 | 3.833E-07 | 1.537E-07 | 2.800E-07 | 5.600E-07 |
| 6950 | 348 | I348I | A | 3.7e-07 | 2.8e-07 | 4.6e-07 | 3.700E-07 | 9.000E-08 | 2.800E-07 | 4.600E-07 |
| 6951 | 348 | I348L | A | 5.6e-07 | 3.7e-07 | 1.9e-07 | 3.733E-07 | 1.850E-07 | 1.900E-07 | 5.600E-07 |
| 6952 | 348 | I348K | A | 2.1e-07 | 1.6e-07 | 2.4e-07 | 2.033E-07 | 4.041E-08 | 1.600E-07 | 2.400E-07 |
| 6953 | 348 | I348M | A | 5e-07 | 1.9e-07 | 3.4e-07 | 3.433E-07 | 1.550E-07 | 1.900E-07 | 5.000E-07 |
| 6954 | 348 | I348F | A | 2.1e-07 | 2.9e-07 | 2.7e-07 | 2.567E-07 | 4.163E-08 | 2.100E-07 | 2.900E-07 |
| 6955 | 348 | I348P | A | 4e-07 | 1.3e-07 | 2e-07 | 2.433E-07 | 1.401E-07 | 1.300E-07 | 4.000E-07 |
| 6956 | 348 | I348S | A | 1.1e-06 | 2.8e-07 | 2.5e-07 | 5.433E-07 | 4.823E-07 | 2.500E-07 | 1.100E-06 |
| 6957 | 348 | I348T | A | 2.9e-07 | 1.2e-07 | 2.4e-07 | 2.167E-07 | 8.737E-08 | 1.200E-07 | 2.900E-07 |
| 6958 | 348 | I348W | A | 4.4e-07 | 2.1e-07 | 1.5e-07 | 2.667E-07 | 1.531E-07 | 1.500E-07 | 4.400E-07 |
| 6959 | 348 | I348Y | A | 2.2e-07 | 2.5e-07 | 2.7e-07 | 2.467E-07 | 2.517E-08 | 2.200E-07 | 2.700E-07 |
| 6960 | 348 | I348V | A | 3.2e-07 | 1.9e-07 | 3.4e-07 | 2.833E-07 | 8.145E-08 | 1.900E-07 | 3.400E-07 |
| 6961 | 349 | F349A | A | 1.6e-07 | 3e-07 | 3.9e-07 | 2.833E-07 | 1.159E-07 | 1.600E-07 | 3.900E-07 |
| 6962 | 349 | F349R | A | 4.4e-07 | 3.4e-07 | 1.5e-07 | 3.100E-07 | 1.473E-07 | 1.500E-07 | 4.400E-07 |
| 6963 | 349 | F349N | A | 2.3e-07 | 9.7e-08 | 1.6e-07 | 1.623E-07 | 6.653E-08 | 9.700E-08 | 2.300E-07 |
| 6964 | 349 | F349D | A | 2.3e-07 | 2.3e-07 | 3.4e-07 | 2.667E-07 | 6.351E-08 | 2.300E-07 | 3.400E-07 |
| 6965 | 349 | F349C | A | 3.5e-07 | 5.5e-07 | 6.7e-07 | 5.233E-07 | 1.617E-07 | 3.500E-07 | 6.700E-07 |
| 6966 | 349 | F349E | A | 5.3e-07 | 2.9e-07 | 3.1e-07 | 3.767E-07 | 1.332E-07 | 2.900E-07 | 5.300E-07 |
| 6967 | 349 | F349Q | A | 4.2e-07 | 1.6e-07 | 2.3e-07 | 2.700E-07 | 1.345E-07 | 1.600E-07 | 4.200E-07 |
| 6968 | 349 | F349G | A | 5.3e-07 | 1.9e-07 | 2.6e-07 | 3.267E-07 | 1.795E-07 | 1.900E-07 | 5.300E-07 |
| 6969 | 349 | F349H | A | 5.9e-07 | 2.8e-07 | 1.6e-07 | 3.433E-07 | 2.219E-07 | 1.600E-07 | 5.900E-07 |

| | | | | | | | | | | |
|---|---|---|---|---|---|---|---|---|---|---|
| 6970 | 349 | F349I | A | 3.6e-07 | 1.7e-07 | 2.2e-07 | 2.500E-07 | 9.849E-08 | 1.700E-07 | 3.600E-07 |
| 6971 | 349 | F349L | A | 3.7e-07 | 1.7e-07 | 1.8e-07 | 2.400E-07 | 1.127E-07 | 1.700E-07 | 3.700E-07 |
| 6972 | 349 | F349K | A | 2.6e-07 | 2.1e-07 | 3.9e-07 | 2.867E-07 | 9.292E-08 | 2.100E-07 | 3.900E-07 |
| 6973 | 349 | F349M | A | 2.6e-07 | 2.8e-07 | 4.5e-07 | 3.300E-07 | 1.044E-07 | 2.600E-07 | 4.500E-07 |
| 6974 | 349 | F349F | A | 2.3e-07 | 3.1e-07 | 2.6e-07 | 2.667E-07 | 4.041E-08 | 2.300E-07 | 3.100E-07 |
| 6975 | 349 | F349P | A | 6.7e-07 | 5.1e-07 | 3e-07 | 4.933E-07 | 1.856E-07 | 3.000E-07 | 6.700E-07 |
| 6976 | 349 | F349S | A | 2.4e-07 | 2.5e-07 | 2.6e-07 | 2.500E-07 | 1.000E-08 | 2.400E-07 | 2.600E-07 |
| 6977 | 349 | F349T | A | 4e-07 | 2.2e-07 | 2.7e-07 | 2.967E-07 | 9.292E-08 | 2.200E-07 | 4.000E-07 |
| 6978 | 349 | F349W | A | 2.7e-07 | 2.2e-07 | 1.4e-07 | 2.100E-07 | 6.557E-08 | 1.400E-07 | 2.700E-07 |
| 6979 | 349 | F349Y | A | 1.9e-07 | 3.5e-07 | 1.8e-07 | 2.400E-07 | 9.539E-08 | 1.800E-07 | 3.500E-07 |
| 6980 | 349 | F349V | A | 1.4e-07 | 2.4e-07 | 5.2e-07 | 3.000E-07 | 1.970E-07 | 1.400E-07 | 5.200E-07 |
| 6981 | 350 | G350A | A | 4.8e-07 | 1.6e-07 | 1.7e-07 | 2.700E-07 | 1.819E-07 | 1.600E-07 | 4.800E-07 |
| 6982 | 350 | G350R | A | 2.4e-07 | 3.1e-07 | 2e-07 | 2.500E-07 | 5.568E-08 | 2.000E-07 | 3.100E-07 |
| 6983 | 350 | G350N | A | 1.7e-07 | 5.5e-07 | 1.8e-07 | 3.000E-07 | 2.166E-07 | 1.700E-07 | 5.500E-07 |
| 6984 | 350 | G350D | A | 1.1e-07 | 1.9e-07 | 3.1e-07 | 2.033E-07 | 1.007E-07 | 1.100E-07 | 3.100E-07 |
| 6985 | 350 | G350C | A | 2.5e-07 | 2.3e-07 | 4.9e-07 | 3.233E-07 | 1.447E-07 | 2.300E-07 | 4.900E-07 |
| 6986 | 350 | G350E | A | 1.3e-07 | 2.2e-07 | 2.1e-07 | 1.867E-07 | 4.933E-08 | 1.300E-07 | 2.200E-07 |
| 6987 | 350 | G350Q | A | 3e-07 | 1.5e-07 | 2.5e-07 | 2.333E-07 | 7.638E-08 | 1.500E-07 | 3.000E-07 |
| 6988 | 350 | G350G | A | 2.6e-07 | 1.7e-07 | 1.6e-07 | 1.967E-07 | 5.508E-08 | 1.600E-07 | 2.600E-07 |
| 6989 | 350 | G350H | A | 1.3e-06 | 2.5e-07 | 1.5e-07 | 5.667E-07 | 6.371E-07 | 1.500E-07 | 1.300E-06 |
| 6990 | 350 | G350I | A | 9.8e-08 | 2.2e-07 | 2.2e-07 | 1.793E-07 | 7.044E-08 | 9.800E-08 | 2.200E-07 |
| 6991 | 350 | G350L | A | 4.3e-07 | 1.6e-07 | 3.6e-07 | 3.167E-07 | 1.401E-07 | 1.600E-07 | 4.300E-07 |
| 6992 | 350 | G350K | A | 2.8e-07 | 1.6e-07 | 2.1e-07 | 2.167E-07 | 6.028E-08 | 1.600E-07 | 2.800E-07 |
| 6993 | 350 | G350M | A | 1.7e-07 | 1.4e-07 | 3.4e-07 | 2.167E-07 | 1.079E-07 | 1.400E-07 | 3.400E-07 |
| 6994 | 350 | G350F | A | 1.6e-07 | 3.7e-07 | 2.5e-07 | 2.600E-07 | 1.054E-07 | 1.600E-07 | 3.700E-07 |
| 6995 | 350 | G350P | A | 2.4e-07 | 1.5e-07 | 2.5e-07 | 2.133E-07 | 5.508E-08 | 1.500E-07 | 2.500E-07 |
| 6996 | 350 | G350S | A | 1.7e-07 | 1.4e-07 | 5.1e-07 | 2.733E-07 | 2.055E-07 | 1.400E-07 | 5.100E-07 |
| 6997 | 350 | G350T | A | 3.4e-07 | 3e-07 | 8.1e-08 | 2.403E-07 | 1.394E-07 | 8.100E-08 | 3.400E-07 |
| 6998 | 350 | G350W | A | 1.3e-07 | 2.8e-07 | 2.8e-07 | 2.300E-07 | 8.660E-08 | 1.300E-07 | 2.800E-07 |
| 6999 | 350 | G350Y | A | 2.9e-07 | 2.2e-07 | 3.8e-07 | 2.967E-07 | 8.021E-08 | 2.200E-07 | 3.800E-07 |
| 7000 | 350 | G350V | A | 2.7e-07 | 4.4e-07 | 2.6e-07 | 3.233E-07 | 1.012E-07 | 2.600E-07 | 4.400E-07 |
| 7001 | 351 | S351A | A | 4.1e-07 | 2.5e-07 | 1.2e-07 | 2.600E-07 | 1.453E-07 | 1.200E-07 | 4.100E-07 |
| 7002 | 351 | S351R | A | 3.7e-07 | 2.5e-07 | 2.5e-07 | 2.900E-07 | 6.928E-08 | 2.500E-07 | 3.700E-07 |
| 7003 | 351 | S351N | A | 4.3e-07 | 1.5e-07 | 1.8e-07 | 2.533E-07 | 1.537E-07 | 1.500E-07 | 4.300E-07 |
| 7004 | 351 | S351D | A | 2.4e-07 | 2.7e-07 | 2.1e-07 | 2.400E-07 | 3.000E-08 | 2.100E-07 | 2.700E-07 |
| 7005 | 351 | S351C | A | 1.8e-07 | 3.2e-07 | 1.4e-07 | 2.133E-07 | 9.452E-08 | 1.400E-07 | 3.200E-07 |
| 7006 | 351 | S351E | A | 2.5e-07 | 3.9e-07 | 5.6e-07 | 4.000E-07 | 1.552E-07 | 2.500E-07 | 5.600E-07 |
| 7007 | 351 | S351Q | A | 3.7e-07 | 1.6e-07 | 1.8e-07 | 2.367E-07 | 1.159E-07 | 1.600E-07 | 3.700E-07 |
| 7008 | 351 | S351G | A | 2e-07 | 3.4e-07 | 3e-07 | 2.800E-07 | 7.211E-08 | 2.000E-07 | 3.400E-07 |
| 7009 | 351 | S351H | A | 1.5e-07 | 2.1e-07 | 8.1e-07 | 3.900E-07 | 3.650E-07 | 1.500E-07 | 8.100E-07 |
| 7010 | 351 | S351I | A | 1.4e-07 | 2.1e-07 | 3.2e-07 | 2.233E-07 | 9.074E-08 | 1.400E-07 | 3.200E-07 |
| 7011 | 351 | S351L | A | 1.3e-07 | 3.1e-07 | 3.2e-07 | 2.533E-07 | 1.069E-07 | 1.300E-07 | 3.200E-07 |
| 7012 | 351 | S351K | A | 2.2e-07 | 2.7e-07 | 2e-07 | 2.300E-07 | 3.606E-08 | 2.000E-07 | 2.700E-07 |
| 7013 | 351 | S351M | A | 3.8e-07 | 3.1e-07 | 2.1e-07 | 3.000E-07 | 8.544E-08 | 2.100E-07 | 3.800E-07 |
| 7014 | 351 | S351F | A | 4.8e-07 | 1.7e-07 | 2e-07 | 2.833E-07 | 1.710E-07 | 1.700E-07 | 4.800E-07 |
| 7015 | 351 | S351P | A | 2.4e-07 | 1.8e-07 | 3.8e-07 | 2.667E-07 | 1.026E-07 | 1.800E-07 | 3.800E-07 |
| 7016 | 351 | S351S | A | 2.6e-07 | 1.5e-07 | 1.6e-07 | 1.900E-07 | 6.083E-08 | 1.500E-07 | 2.600E-07 |
| 7017 | 351 | S351T | A | 3.1e-07 | 3.1e-07 | 9.4e-08 | 2.380E-07 | 1.247E-07 | 9.400E-08 | 3.100E-07 |
| 7018 | 351 | S351W | A | 4.9e-07 | 4.8e-07 | 2.8e-07 | 4.167E-07 | 1.185E-07 | 2.800E-07 | 4.900E-07 |
| 7019 | 351 | S351Y | A | 4.5e-07 | 2.7e-07 | 2e-07 | 3.067E-07 | 1.290E-07 | 2.000E-07 | 4.500E-07 |
| 7020 | 351 | S351V | A | 2.7e-07 | 1.3e-07 | 2.2e-07 | 2.067E-07 | 7.095E-08 | 1.300E-07 | 2.700E-07 |
| 7021 | 352 | L352A | A | 3.9e-07 | 2.3e-07 | 3.7e-07 | 3.300E-07 | 8.718E-08 | 2.300E-07 | 3.900E-07 |
| 7022 | 352 | L352R | A | 2.8e-07 | 2.1e-07 | 1e-07 | 1.967E-07 | 9.074E-08 | 1.000E-07 | 2.800E-07 |
| 7023 | 352 | L352N | A | 2.6e-07 | 1.6e-07 | 3.3e-07 | 2.500E-07 | 8.544E-08 | 1.600E-07 | 3.300E-07 |
| 7024 | 352 | L352D | A | 2.6e-07 | 6.3e-07 | 2.3e-07 | 3.733E-07 | 2.228E-07 | 2.300E-07 | 6.300E-07 |
| 7025 | 352 | L352C | A | 2.7e-07 | 1.4e-07 | 3.3e-07 | 2.467E-07 | 9.713E-08 | 1.400E-07 | 3.300E-07 |
| 7026 | 352 | L352E | A | 2.9e-07 | 2.5e-07 | 3.5e-07 | 2.967E-07 | 5.033E-08 | 2.500E-07 | 3.500E-07 |
| 7027 | 352 | L352Q | A | 1.2e-07 | 3.7e-07 | 1.8e-07 | 2.233E-07 | 1.305E-07 | 1.200E-07 | 3.700E-07 |
| 7028 | 352 | L352G | A | 1.5e-07 | 3.9e-07 | 2.6e-07 | 2.667E-07 | 1.201E-07 | 1.500E-07 | 3.900E-07 |
| 7029 | 352 | L352H | A | 1.5e-07 | 2.8e-07 | 1.9e-07 | 2.067E-07 | 6.658E-08 | 1.500E-07 | 2.800E-07 |
| 7030 | 352 | L352I | A | 2.5e-07 | 2.8e-07 | 2.5e-07 | 2.600E-07 | 1.732E-08 | 2.500E-07 | 2.800E-07 |
| 7031 | 352 | L352L | A | 1.8e-07 | 3e-07 | 2.8e-07 | 2.533E-07 | 6.429E-08 | 1.800E-07 | 3.000E-07 |
| 7032 | 352 | L352K | A | 1.4e-07 | 4.2e-07 | 1.7e-07 | 2.433E-07 | 1.537E-07 | 1.400E-07 | 4.200E-07 |
| 7033 | 352 | L352M | A | 1.4e-07 | 3.3e-07 | 7e-07 | 3.900E-07 | 2.848E-07 | 1.400E-07 | 7.000E-07 |
| 7034 | 352 | L352F | A | 2e-07 | 4.3e-07 | 4.9e-07 | 3.733E-07 | 1.531E-07 | 2.000E-07 | 4.900E-07 |
| 7035 | 352 | L352P | A | 2.7e-07 | 2.2e-07 | 2.3e-07 | 2.400E-07 | 2.646E-08 | 2.200E-07 | 2.700E-07 |
| 7036 | 352 | L352S | A | 2.1e-07 | 4.1e-07 | 3.5e-07 | 3.233E-07 | 1.026E-07 | 2.100E-07 | 4.100E-07 |
| 7037 | 352 | L352T | A | 2.3e-07 | 1.9e-07 | 3.2e-07 | 2.467E-07 | 6.658E-08 | 1.900E-07 | 3.200E-07 |
| 7038 | 352 | L352W | A | 1.5e-07 | 2.7e-07 | 1.6e-07 | 1.933E-07 | 6.658E-08 | 1.500E-07 | 2.700E-07 |
| 7039 | 352 | L352Y | A | 7.2e-08 | 3.6e-07 | 2.4e-07 | 2.240E-07 | 1.447E-07 | 7.200E-08 | 3.600E-07 |
| 7040 | 352 | L352V | A | 3.7e-07 | 2.6e-07 | 2.9e-07 | 3.067E-07 | 5.686E-08 | 2.600E-07 | 3.700E-07 |
| 7041 | 353 | A353A | A | 2.1e-07 | 3.3e-07 | 2.4e-07 | 2.600E-07 | 6.245E-08 | 2.100E-07 | 3.300E-07 |
| 7042 | 353 | A353R | A | 1.7e-07 | 3.4e-07 | 2.9e-07 | 2.667E-07 | 8.737E-08 | 1.700E-07 | 3.400E-07 |
| 7043 | 353 | A353N | A | 2.9e-07 | 1.1e-07 | 3.9e-07 | 2.633E-07 | 1.419E-07 | 1.100E-07 | 3.900E-07 |
| 7044 | 353 | A353D | A | 2.8e-07 | 2e-07 | 4.7e-07 | 3.167E-07 | 1.387E-07 | 2.000E-07 | 4.700E-07 |
| 7045 | 353 | A353C | A | 3e-07 | 3.8e-07 | 7.6e-08 | 2.520E-07 | 1.576E-07 | 7.600E-08 | 3.800E-07 |
| 7046 | 353 | A353E | A | 1.9e-07 | 4.1e-07 | 2.1e-07 | 2.700E-07 | 1.217E-07 | 1.900E-07 | 4.100E-07 |
| 7047 | 353 | A353Q | A | 1.8e-07 | 2.8e-07 | 2.9e-07 | 2.500E-07 | 6.083E-08 | 1.800E-07 | 2.900E-07 |
| 7048 | 353 | A353G | A | 2.1e-07 | 3.7e-07 | 1.9e-07 | 2.567E-07 | 9.866E-08 | 1.900E-07 | 3.700E-07 |
| 7049 | 353 | A353H | A | 3.7e-07 | 3.2e-07 | 1.9e-07 | 2.933E-07 | 9.292E-08 | 1.900E-07 | 3.700E-07 |
| 7050 | 353 | A353I | A | 5.6e-07 | 2e-07 | 1.9e-07 | 3.167E-07 | 2.108E-07 | 1.900E-07 | 5.600E-07 |
| 7051 | 353 | A353L | A | 2.9e-07 | 3e-07 | 2.8e-07 | 2.900E-07 | 1.000E-08 | 2.800E-07 | 3.000E-07 |
| 7052 | 353 | A353K | A | 3.6e-07 | 5.2e-07 | 1.8e-07 | 3.533E-07 | 1.701E-07 | 1.800E-07 | 5.200E-07 |
| 7053 | 353 | A353M | A | 3.5e-07 | 2.2e-07 | 1.9e-07 | 2.533E-07 | 8.505E-08 | 1.900E-07 | 3.500E-07 |
| 7054 | 353 | A353F | A | 2.7e-07 | 4e-07 | 3.9e-07 | 3.533E-07 | 7.234E-08 | 2.700E-07 | 4.000E-07 |

| | | | | | | | | | | |
|---|---|---|---|---|---|---|---|---|---|---|
| 7055 | 353 | A353P | A | 1.5e-07 | 6.8e-07 | 4.3e-07 | 4.200E-07 | 2.651E-07 | 1.500E-07 | 6.800E-07 |
| 7056 | 353 | A353S | A | 9.7e-08 | 6.2e-07 | 1.4e-07 | 2.857E-07 | 2.903E-07 | 9.700E-08 | 6.200E-07 |
| 7057 | 353 | A353T | A | 1.6e-07 | 2e-07 | 3.5e-07 | 2.367E-07 | 1.002E-07 | 1.600E-07 | 3.500E-07 |
| 7058 | 353 | A353W | A | 2e-07 | 2e-07 | 2.3e-07 | 2.100E-07 | 1.732E-08 | 2.000E-07 | 2.300E-07 |
| 7059 | 353 | A353Y | A | 2e-07 | 2.9e-07 | 1.1e-07 | 2.000E-07 | 9.000E-08 | 1.100E-07 | 2.900E-07 |
| 7060 | 353 | A353V | A | 1.7e-07 | 1.9e-07 | 2.5e-07 | 2.033E-07 | 4.163E-08 | 1.700E-07 | 2.500E-07 |
| 7061 | 354 | F354A | A | 1.9e-07 | 2.8e-07 | 3.6e-07 | 2.767E-07 | 8.505E-08 | 1.900E-07 | 3.600E-07 |
| 7062 | 354 | F354R | A | 6.3e-07 | 3.6e-07 | 2.4e-07 | 4.100E-07 | 1.997E-07 | 2.400E-07 | 6.300E-07 |
| 7063 | 354 | F354N | A | 1.4e-07 | 9.2e-08 | 4.2e-07 | 2.173E-07 | 1.771E-07 | 9.200E-08 | 4.200E-07 |
| 7064 | 354 | F354D | A | 3.4e-07 | 7.3e-07 | 2.1e-07 | 4.267E-07 | 2.706E-07 | 2.100E-07 | 7.300E-07 |
| 7065 | 354 | F354C | A | 2e-07 | 6.5e-07 | 3.3e-07 | 3.933E-07 | 2.316E-07 | 2.000E-07 | 6.500E-07 |
| 7066 | 354 | F354E | A | 5.5e-07 | 1e-07 | 2.4e-07 | 2.967E-07 | 2.303E-07 | 1.000E-07 | 5.500E-07 |
| 7067 | 354 | F354Q | A | 2.7e-07 | 4.9e-07 | 2.8e-07 | 3.467E-07 | 1.242E-07 | 2.700E-07 | 4.900E-07 |
| 7068 | 354 | F354G | A | 3.5e-07 | 2.4e-07 | 3.7e-07 | 3.200E-07 | 7.000E-08 | 2.400E-07 | 3.700E-07 |
| 7069 | 354 | F354H | A | 4.8e-07 | 2.6e-07 | 2.9e-07 | 3.433E-07 | 1.193E-07 | 2.600E-07 | 4.800E-07 |
| 7070 | 354 | F354I | A | 2.9e-07 | 2.4e-07 | 1.6e-07 | 2.300E-07 | 6.557E-08 | 1.600E-07 | 2.900E-07 |
| 7071 | 354 | F354L | A | 3e-07 | 2.6e-07 | 2.7e-07 | 2.767E-07 | 2.082E-08 | 2.600E-07 | 3.000E-07 |
| 7072 | 354 | F354K | A | 2.3e-07 | 1.9e-07 | 2.4e-07 | 2.200E-07 | 2.646E-08 | 1.900E-07 | 2.400E-07 |
| 7073 | 354 | F354M | A | 2.1e-07 | 2.6e-07 | 3.1e-07 | 2.600E-07 | 5.000E-08 | 2.100E-07 | 3.100E-07 |
| 7074 | 354 | F354F | A | 1.2e-07 | 1.2e-07 | 3e-07 | 1.800E-07 | 1.039E-07 | 1.200E-07 | 3.000E-07 |
| 7075 | 354 | F354P | A | 3.2e-07 | 4.1e-07 | 2.7e-07 | 3.333E-07 | 7.095E-08 | 2.700E-07 | 4.100E-07 |
| 7076 | 354 | F354S | A | 2.9e-07 | 1.1e-07 | 2.6e-07 | 2.200E-07 | 9.644E-08 | 1.100E-07 | 2.900E-07 |
| 7077 | 354 | F354T | A | 1.2e-07 | 1.9e-07 | 3.3e-07 | 2.133E-07 | 1.069E-07 | 1.200E-07 | 3.300E-07 |
| 7078 | 354 | F354W | A | 3.9e-07 | 3.6e-07 | 2.2e-07 | 3.233E-07 | 9.074E-08 | 2.200E-07 | 3.900E-07 |
| 7079 | 354 | F354Y | A | 3.2e-07 | 2.4e-07 | 1.4e-07 | 2.333E-07 | 9.018E-08 | 1.400E-07 | 3.200E-07 |
| 7080 | 354 | F354V | A | 1.4e-07 | 5.6e-07 | 3.2e-07 | 3.400E-07 | 2.107E-07 | 1.400E-07 | 5.600E-07 |
| 7081 | 355 | L355A | A | 1.1e-07 | 1.2e-07 | 1.6e-07 | 1.300E-07 | 2.646E-08 | 1.100E-07 | 1.600E-07 |
| 7082 | 355 | L355R | A | 2.1e-07 | 1.2e-07 | 2.7e-07 | 2.000E-07 | 7.550E-08 | 1.200E-07 | 2.700E-07 |
| 7083 | 355 | L355N | A | 2.5e-07 | 1.1e-07 | 6e-07 | 3.200E-07 | 2.524E-07 | 1.100E-07 | 6.000E-07 |
| 7084 | 355 | L355D | A | 2.9e-07 | 3.5e-07 | 2.3e-07 | 2.900E-07 | 6.000E-08 | 2.300E-07 | 3.500E-07 |
| 7085 | 355 | L355C | A | 2.2e-07 | 1.4e-07 | 3.1e-07 | 2.233E-07 | 8.505E-08 | 1.400E-07 | 3.100E-07 |
| 7086 | 355 | L355E | A | 3.2e-07 | 4.3e-07 | 1.4e-07 | 2.967E-07 | 1.464E-07 | 1.400E-07 | 4.300E-07 |
| 7087 | 355 | L355Q | A | 2.5e-07 | 3.6e-07 | 8.1e-08 | 2.303E-07 | 1.405E-07 | 8.100E-08 | 3.600E-07 |
| 7088 | 355 | L355G | A | 5.5e-07 | 1.4e-07 | 1.5e-07 | 2.800E-07 | 2.339E-07 | 1.400E-07 | 5.500E-07 |
| 7089 | 355 | L355H | A | 2.1e-07 | 2.2e-07 | 2.1e-07 | 2.133E-07 | 5.774E-09 | 2.100E-07 | 2.200E-07 |
| 7090 | 355 | L355I | A | 2.2e-07 | 1.3e-07 | 1.3e-07 | 1.600E-07 | 5.196E-08 | 1.300E-07 | 2.200E-07 |
| 7091 | 355 | L355L | A | 1.6e-07 | 3.1e-07 | 4e-07 | 2.900E-07 | 1.212E-07 | 1.600E-07 | 4.000E-07 |
| 7092 | 355 | L355K | A | 2.3e-07 | 1.6e-07 | 2.5e-07 | 2.133E-07 | 4.726E-08 | 1.600E-07 | 2.500E-07 |
| 7093 | 355 | L355M | A | 2.1e-07 | 1.5e-07 | 2e-07 | 1.867E-07 | 3.215E-08 | 1.500E-07 | 2.100E-07 |
| 7094 | 355 | L355F | A | 2.2e-07 | 3.5e-07 | 2e-07 | 2.567E-07 | 8.145E-08 | 2.000E-07 | 3.500E-07 |
| 7095 | 355 | L355P | A | 5.6e-07 | 2.4e-07 | 1.9e-07 | 3.300E-07 | 2.007E-07 | 1.900E-07 | 5.600E-07 |
| 7096 | 355 | L355S | A | 1.7e-07 | 2.9e-07 | 1.3e-07 | 1.967E-07 | 8.327E-08 | 1.300E-07 | 2.900E-07 |
| 7097 | 355 | L355T | A | 3e-07 | 1.5e-07 | 3.6e-07 | 2.700E-07 | 1.082E-07 | 1.500E-07 | 3.600E-07 |
| 7098 | 355 | L355W | A | 2e-07 | 2.6e-07 | 1.5e-07 | 2.033E-07 | 5.508E-08 | 1.500E-07 | 2.600E-07 |
| 7099 | 355 | L355Y | A | 3e-07 | 1.1e-07 | 2.1e-07 | 2.067E-07 | 9.504E-08 | 1.100E-07 | 3.000E-07 |
| 7100 | 355 | L355V | A | 3.5e-07 | 1.7e-07 | 3.9e-07 | 3.033E-07 | 1.172E-07 | 1.700E-07 | 3.900E-07 |
| 7101 | 356 | P356A | A | 2e-07 | 4.6e-07 | 3.9e-07 | 3.500E-07 | 1.345E-07 | 2.000E-07 | 4.600E-07 |
| 7102 | 356 | P356R | A | 3.9e-07 | 3.1e-07 | 4.1e-07 | 3.700E-07 | 5.292E-08 | 3.100E-07 | 4.100E-07 |
| 7103 | 356 | P356N | A | 2.5e-07 | 3.4e-07 | 1.2e-07 | 2.367E-07 | 1.106E-07 | 1.200E-07 | 3.400E-07 |
| 7104 | 356 | P356D | A | 2.7e-07 | 3e-07 | 3.1e-07 | 2.933E-07 | 2.082E-08 | 2.700E-07 | 3.100E-07 |
| 7105 | 356 | P356C | A | 4.4e-07 | 1.6e-07 | 1.6e-07 | 2.533E-07 | 1.617E-07 | 1.600E-07 | 4.400E-07 |
| 7106 | 356 | P356E | A | 2.3e-07 | 3.4e-07 | 2.8e-07 | 2.833E-07 | 5.508E-08 | 2.300E-07 | 3.400E-07 |
| 7107 | 356 | P356Q | A | 3.1e-07 | 2.6e-07 | 3e-07 | 2.900E-07 | 2.646E-08 | 2.600E-07 | 3.100E-07 |
| 7108 | 356 | P356G | A | 2.5e-07 | 1.8e-07 | 1.5e-07 | 1.933E-07 | 5.132E-08 | 1.500E-07 | 2.500E-07 |
| 7109 | 356 | P356H | A | 2.4e-07 | 2e-07 | 1e-07 | 1.800E-07 | 7.211E-08 | 1.000E-07 | 2.400E-07 |
| 7110 | 356 | P356I | A | 2.8e-07 | 3.2e-07 | 2.4e-07 | 2.800E-07 | 4.000E-08 | 2.400E-07 | 3.200E-07 |
| 7111 | 356 | P356L | A | 3e-07 | 2.1e-07 | 1.9e-07 | 2.333E-07 | 5.859E-08 | 1.900E-07 | 3.000E-07 |
| 7112 | 356 | P356K | A | 2.4e-07 | 2.4e-07 | 2.8e-07 | 2.533E-07 | 2.309E-08 | 2.400E-07 | 2.800E-07 |
| 7113 | 356 | P356M | A | 1.5e-07 | 2.5e-07 | 3e-07 | 2.333E-07 | 7.638E-08 | 1.500E-07 | 3.000E-07 |
| 7114 | 356 | P356F | A | 1.6e-07 | 5.6e-07 | 2.1e-07 | 3.100E-07 | 2.179E-07 | 1.600E-07 | 5.600E-07 |
| 7115 | 356 | P356P | A | 1.2e-07 | 2.1e-07 | 1.8e-07 | 1.700E-07 | 4.583E-08 | 1.200E-07 | 2.100E-07 |
| 7116 | 356 | P356S | A | 1.3e-07 | 2.7e-07 | 2.2e-07 | 2.067E-07 | 7.095E-08 | 1.300E-07 | 2.700E-07 |
| 7117 | 356 | P356T | A | 2.8e-07 | 1e-07 | 1.8e-07 | 1.867E-07 | 9.018E-08 | 1.000E-07 | 2.800E-07 |
| 7118 | 356 | P356W | A | 2.7e-07 | 2.1e-07 | 1.4e-07 | 2.067E-07 | 6.506E-08 | 1.400E-07 | 2.700E-07 |
| 7119 | 356 | P356Y | A | 2.5e-07 | 2.3e-07 | 3.5e-07 | 2.767E-07 | 6.429E-08 | 2.300E-07 | 3.500E-07 |
| 7120 | 356 | P356V | A | 2.1e-07 | 1.7e-07 | 2.4e-07 | 2.067E-07 | 3.512E-08 | 1.700E-07 | 2.400E-07 |
| 7121 | 357 | E357A | A | 1.6e-07 | 2.2e-07 | 2.6e-07 | 2.133E-07 | 5.033E-08 | 1.600E-07 | 2.600E-07 |
| 7122 | 357 | E357R | A | 3.6e-07 | 2.1e-07 | 5.2e-07 | 3.633E-07 | 1.550E-07 | 2.100E-07 | 5.200E-07 |
| 7123 | 357 | E357N | A | 2.4e-07 | 3e-07 | 2.7e-07 | 2.700E-07 | 3.000E-08 | 2.400E-07 | 3.000E-07 |
| 7124 | 357 | E357D | A | 1.8e-07 | 2.4e-07 | 2.1e-07 | 2.100E-07 | 3.000E-08 | 1.800E-07 | 2.400E-07 |
| 7125 | 357 | E357C | A | 1.9e-07 | 2.7e-07 | 1.9e-07 | 2.167E-07 | 4.619E-08 | 1.900E-07 | 2.700E-07 |
| 7126 | 357 | E357E | A | 3.1e-07 | 3.7e-07 | 2.5e-07 | 3.100E-07 | 6.000E-08 | 2.500E-07 | 3.700E-07 |
| 7127 | 357 | E357Q | A | 2.3e-07 | 2.2e-07 | 1.8e-07 | 2.100E-07 | 2.646E-08 | 1.800E-07 | 2.300E-07 |
| 7128 | 357 | E357G | A | 3.6e-07 | 2.9e-07 | 1.7e-07 | 2.733E-07 | 9.609E-08 | 1.700E-07 | 3.600E-07 |
| 7129 | 357 | E357H | A | 1.8e-07 | 2.4e-07 | 1.8e-07 | 2.000E-07 | 3.464E-08 | 1.800E-07 | 2.400E-07 |
| 7130 | 357 | E357I | A | 1.6e-07 | 1.9e-07 | 2.8e-07 | 2.100E-07 | 6.245E-08 | 1.600E-07 | 2.800E-07 |
| 7131 | 357 | E357L | A | 2.7e-07 | 1.9e-07 | 3.1e-07 | 2.567E-07 | 6.110E-08 | 1.900E-07 | 3.100E-07 |
| 7132 | 357 | E357K | A | 3.1e-07 | 2.8e-07 | 2.3e-07 | 2.733E-07 | 4.041E-08 | 2.300E-07 | 3.100E-07 |
| 7133 | 357 | E357M | A | 1e-07 | 2.8e-07 | 1.7e-07 | 1.833E-07 | 9.074E-08 | 1.000E-07 | 2.800E-07 |
| 7134 | 357 | E357F | A | 6.2e-07 | 3.5e-07 | 3.5e-07 | 4.400E-07 | 1.559E-07 | 3.500E-07 | 6.200E-07 |
| 7135 | 357 | E357P | A | 2.6e-07 | 6.9e-08 | 1.3e-07 | 1.530E-07 | 9.756E-08 | 6.900E-08 | 2.600E-07 |
| 7136 | 357 | E357S | A | 2.2e-07 | 3.3e-07 | 1.6e-07 | 2.367E-07 | 8.622E-08 | 1.600E-07 | 3.300E-07 |
| 7137 | 357 | E357T | A | 8.2e-07 | 4.3e-07 | 5.7e-07 | 6.067E-07 | 1.976E-07 | 4.300E-07 | 8.200E-07 |
| 7138 | 357 | E357W | A | 1.3e-07 | 2.1e-07 | 1e-07 | 1.467E-07 | 5.686E-08 | 1.000E-07 | 2.100E-07 |
| 7139 | 357 | E357Y | A | 3.4e-07 | 3.5e-07 | 5e-07 | 3.967E-07 | 8.963E-08 | 3.400E-07 | 5.000E-07 |

| | | | | | | | | | | |
|---|---|---|---|---|---|---|---|---|---|---|
| 7140 | 357 | E357V | A | 3.4e-07 | 2.7e-07 | 3e-07 | 3.033E-07 | 3.512E-08 | 2.700E-07 | 3.400E-07 |
| 7141 | 358 | S358A | A | 5.5e-07 | 5.1e-07 | 1.5e-07 | 4.033E-07 | 2.203E-07 | 1.500E-07 | 5.500E-07 |
| 7142 | 358 | S358R | A | 2.4e-07 | 5.1e-07 | 2.2e-07 | 3.233E-07 | 1.620E-07 | 2.200E-07 | 5.100E-07 |
| 7143 | 358 | S358N | A | 2.2e-07 | 2.5e-07 | 2.4e-07 | 2.367E-07 | 1.528E-08 | 2.200E-07 | 2.500E-07 |
| 7144 | 358 | S358D | A | 1.1e-07 | 2.5e-07 | 2e-07 | 1.867E-07 | 7.095E-08 | 1.100E-07 | 2.500E-07 |
| 7145 | 358 | S358C | A | 9.7e-08 | 6.7e-07 | 2.2e-07 | 3.290E-07 | 3.017E-07 | 9.700E-08 | 6.700E-07 |
| 7146 | 358 | S358E | A | 3.3e-07 | 2.7e-07 | 1.8e-07 | 2.600E-07 | 7.550E-08 | 1.800E-07 | 3.300E-07 |
| 7147 | 358 | S358Q | A | 2.6e-07 | 2.1e-07 | 2.4e-07 | 2.367E-07 | 2.517E-08 | 2.100E-07 | 2.600E-07 |
| 7148 | 358 | S358G | A | 3.1e-07 | 3.2e-07 | 2.9e-07 | 3.067E-07 | 1.528E-08 | 2.900E-07 | 3.200E-07 |
| 7149 | 358 | S358H | A | 5.9e-07 | 3.2e-07 | 1.5e-07 | 3.533E-07 | 2.219E-07 | 1.500E-07 | 5.900E-07 |
| 7150 | 358 | S358I | A | 2.5e-07 | 1.8e-07 | 2.1e-07 | 2.133E-07 | 3.512E-08 | 1.800E-07 | 2.500E-07 |
| 7151 | 358 | S358L | A | 1.2e-07 | 1.8e-07 | 2.7e-07 | 1.900E-07 | 7.550E-08 | 1.200E-07 | 2.700E-07 |
| 7152 | 358 | S358K | A | 3e-07 | 1.1e-07 | 1.4e-07 | 1.833E-07 | 1.021E-07 | 1.100E-07 | 3.000E-07 |
| 7153 | 358 | S358M | A | 3.2e-07 | 2.3e-07 | 4.3e-07 | 3.267E-07 | 1.002E-07 | 2.300E-07 | 4.300E-07 |
| 7154 | 358 | S358F | A | 3.8e-07 | 2.2e-07 | 2.1e-07 | 2.700E-07 | 9.539E-08 | 2.100E-07 | 3.800E-07 |
| 7155 | 358 | S358P | A | 4.1e-07 | 2e-07 | 4.6e-07 | 3.567E-07 | 1.380E-07 | 2.000E-07 | 4.600E-07 |
| 7156 | 358 | S358S | A | 2.7e-07 | 1.1e-07 | 2.9e-07 | 2.233E-07 | 9.866E-08 | 1.100E-07 | 2.900E-07 |
| 7157 | 358 | S358T | A | 1.1e-07 | 1.3e-07 | 2.9e-07 | 1.767E-07 | 9.866E-08 | 1.100E-07 | 2.900E-07 |
| 7158 | 358 | S358W | A | 3.3e-07 | 2.1e-07 | 2.8e-07 | 2.733E-07 | 6.028E-08 | 2.100E-07 | 3.300E-07 |
| 7159 | 358 | S358Y | A | 1.2e-07 | 1.7e-07 | 2.1e-07 | 1.667E-07 | 4.509E-08 | 1.200E-07 | 2.100E-07 |
| 7160 | 358 | S358V | A | 5.2e-07 | 2.1e-07 | 2.9e-07 | 3.400E-07 | 1.609E-07 | 2.100E-07 | 5.200E-07 |
| 7161 | 359 | F359A | A | 3.8e-07 | 2e-07 | 2.4e-07 | 2.733E-07 | 9.452E-08 | 2.000E-07 | 3.800E-07 |
| 7162 | 359 | F359R | A | 2.1e-07 | 2.9e-07 | 1.9e-07 | 2.300E-07 | 5.292E-08 | 1.900E-07 | 2.900E-07 |
| 7163 | 359 | F359N | A | 2.7e-07 | 3.2e-07 | 3.5e-07 | 3.133E-07 | 4.041E-08 | 2.700E-07 | 3.500E-07 |
| 7164 | 359 | F359D | A | 3.3e-07 | 3.5e-07 | 2.3e-07 | 3.033E-07 | 6.429E-08 | 2.300E-07 | 3.500E-07 |
| 7165 | 359 | F359C | A | 3.2e-07 | 2.1e-07 | 1.2e-07 | 2.167E-07 | 1.002E-07 | 1.200E-07 | 3.200E-07 |
| 7166 | 359 | F359E | A | 1.8e-07 | 1.2e-07 | 3.2e-07 | 2.067E-07 | 1.026E-07 | 1.200E-07 | 3.200E-07 |
| 7167 | 359 | F359Q | A | 1.2e-07 | 3e-07 | 1.3e-07 | 1.833E-07 | 1.012E-07 | 1.200E-07 | 3.000E-07 |
| 7168 | 359 | F359G | A | 3.1e-07 | 2.7e-07 | 3.9e-07 | 3.233E-07 | 6.110E-08 | 2.700E-07 | 3.900E-07 |
| 7169 | 359 | F359H | A | 3e-07 | 4.6e-07 | 1.6e-07 | 3.067E-07 | 1.501E-07 | 1.600E-07 | 4.600E-07 |
| 7170 | 359 | F359I | A | 3.5e-07 | 5.5e-07 | 1.4e-07 | 3.467E-07 | 2.050E-07 | 1.400E-07 | 5.500E-07 |
| 7171 | 359 | F359L | A | 4.3e-07 | 3.8e-07 | 2.1e-07 | 3.400E-07 | 1.153E-07 | 2.100E-07 | 4.300E-07 |
| 7172 | 359 | F359K | A | 1.4e-07 | 3.4e-07 | 2.8e-07 | 2.533E-07 | 1.026E-07 | 1.400E-07 | 3.400E-07 |
| 7173 | 359 | F359M | A | 2e-07 | 1.8e-07 | 1.4e-07 | 1.733E-07 | 3.055E-08 | 1.400E-07 | 2.000E-07 |
| 7174 | 359 | F359F | A | 3.4e-07 | 2.2e-07 | 2.6e-07 | 2.733E-07 | 6.110E-08 | 2.200E-07 | 3.400E-07 |
| 7175 | 359 | F359P | A | 1.6e-07 | 8.7e-08 | 5.6e-07 | 2.690E-07 | 2.546E-07 | 8.700E-08 | 5.600E-07 |
| 7176 | 359 | F359S | A | 3.9e-07 | 2.5e-07 | 2.5e-07 | 2.967E-07 | 8.083E-08 | 2.500E-07 | 3.900E-07 |
| 7177 | 359 | F359T | A | 2.9e-07 | 7.4e-08 | 1.6e-07 | 1.747E-07 | 1.087E-07 | 7.400E-08 | 2.900E-07 |
| 7178 | 359 | F359W | A | 2e-07 | 2.2e-07 | 2.2e-07 | 2.133E-07 | 1.155E-08 | 2.000E-07 | 2.200E-07 |
| 7179 | 359 | F359Y | A | 1.8e-07 | 2e-07 | 4.2e-07 | 2.667E-07 | 1.332E-07 | 1.800E-07 | 4.200E-07 |
| 7180 | 359 | F359V | A | 1.5e-07 | 1.7e-07 | 1.9e-07 | 1.700E-07 | 2.000E-08 | 1.500E-07 | 1.900E-07 |
| 7181 | 360 | D360A | A | 2.1e-07 | 2.9e-07 | 3e-07 | 2.667E-07 | 4.933E-08 | 2.100E-07 | 3.000E-07 |
| 7182 | 360 | D360R | A | 2e-07 | 2e-07 | 2.4e-07 | 2.133E-07 | 2.309E-08 | 2.000E-07 | 2.400E-07 |
| 7183 | 360 | D360N | A | 3.8e-07 | 2.1e-07 | 4.6e-07 | 3.500E-07 | 1.277E-07 | 2.100E-07 | 4.600E-07 |
| 7184 | 360 | D360D | A | 2.5e-07 | 2.5e-07 | 2.1e-07 | 2.367E-07 | 2.309E-08 | 2.100E-07 | 2.500E-07 |
| 7185 | 360 | D360C | A | 1.6e-07 | 3.8e-07 | 1.5e-07 | 2.300E-07 | 1.300E-07 | 1.500E-07 | 3.800E-07 |
| 7186 | 360 | D360E | A | 2e-07 | 3.2e-07 | 1.9e-07 | 2.367E-07 | 7.234E-08 | 1.900E-07 | 3.200E-07 |
| 7187 | 360 | D360Q | A | 1.3e-07 | 4.6e-07 | 2.9e-07 | 2.933E-07 | 1.650E-07 | 1.300E-07 | 4.600E-07 |
| 7188 | 360 | D360G | A | 2.5e-07 | 2.2e-07 | 3.2e-07 | 2.633E-07 | 5.132E-08 | 2.200E-07 | 3.200E-07 |
| 7189 | 360 | D360H | A | 1.9e-07 | 2.2e-07 | 2.2e-07 | 2.100E-07 | 1.732E-08 | 1.900E-07 | 2.200E-07 |
| 7190 | 360 | D360I | A | 1.7e-07 | 1.8e-07 | 2.8e-07 | 2.100E-07 | 6.083E-08 | 1.700E-07 | 2.800E-07 |
| 7191 | 360 | D360L | A | 9.4e-08 | 2.4e-07 | 3.1e-07 | 2.147E-07 | 1.102E-07 | 9.400E-08 | 3.100E-07 |
| 7192 | 360 | D360K | A | 1.7e-07 | 3.2e-07 | 3.1e-07 | 2.667E-07 | 8.386E-08 | 1.700E-07 | 3.200E-07 |
| 7193 | 360 | D360M | A | 2.1e-07 | 3.1e-07 | 1.7e-07 | 2.300E-07 | 7.211E-08 | 1.700E-07 | 3.100E-07 |
| 7194 | 360 | D360F | A | 1.5e-07 | 8.4e-08 | 4.2e-07 | 2.180E-07 | 1.780E-07 | 8.400E-08 | 4.200E-07 |
| 7195 | 360 | D360P | A | 4.2e-07 | 2.6e-07 | 3.3e-07 | 3.367E-07 | 8.021E-08 | 2.600E-07 | 4.200E-07 |
| 7196 | 360 | D360S | A | 2.4e-07 | 1.8e-07 | 1.8e-07 | 2.000E-07 | 3.464E-08 | 1.800E-07 | 2.400E-07 |
| 7197 | 360 | D360T | A | 2.3e-07 | 4.5e-07 | 3.4e-07 | 3.400E-07 | 1.100E-07 | 2.300E-07 | 4.500E-07 |
| 7198 | 360 | D360W | A | 1.4e-07 | 1.2e-07 | 1.7e-07 | 1.433E-07 | 2.517E-08 | 1.200E-07 | 1.700E-07 |
| 7199 | 360 | D360Y | A | 3.7e-07 | 2e-07 | 4.8e-07 | 3.500E-07 | 1.411E-07 | 2.000E-07 | 4.800E-07 |
| 7200 | 360 | D360V | A | 2e-07 | 5e-07 | 5.7e-07 | 4.233E-07 | 1.966E-07 | 2.000E-07 | 5.700E-07 |
| 7201 | 361 | G361A | A | 4.7e-07 | 3.5e-07 | 3.4e-07 | 3.867E-07 | 7.234E-08 | 3.400E-07 | 4.700E-07 |
| 7202 | 361 | G361R | A | 2.8e-07 | 1.5e-07 | 2.3e-07 | 2.200E-07 | 6.557E-08 | 1.500E-07 | 2.800E-07 |
| 7203 | 361 | G361N | A | 3.7e-07 | 2e-07 | 1.1e-07 | 2.267E-07 | 1.320E-07 | 1.100E-07 | 3.700E-07 |
| 7204 | 361 | G361D | A | 1.9e-07 | 2.2e-07 | 1.6e-07 | 1.900E-07 | 3.000E-08 | 1.600E-07 | 2.200E-07 |
| 7205 | 361 | G361C | A | 1.2e-07 | 4.7e-07 | 3.1e-07 | 3.000E-07 | 1.752E-07 | 1.200E-07 | 4.700E-07 |
| 7206 | 361 | G361E | A | 2.1e-07 | 4.8e-07 | 1.4e-07 | 2.767E-07 | 1.795E-07 | 1.400E-07 | 4.800E-07 |
| 7207 | 361 | G361Q | A | 2.6e-07 | 2.9e-07 | 1.9e-07 | 2.467E-07 | 5.132E-08 | 1.900E-07 | 2.900E-07 |
| 7208 | 361 | G361G | A | 3.1e-07 | 3.5e-07 | 2.3e-07 | 2.967E-07 | 6.110E-08 | 2.300E-07 | 3.500E-07 |
| 7209 | 361 | G361H | A | 5e-07 | 3.8e-07 | 1.7e-07 | 3.500E-07 | 1.670E-07 | 1.700E-07 | 5.000E-07 |
| 7210 | 361 | G361I | A | 2.8e-07 | 2.7e-07 | 2.2e-07 | 2.567E-07 | 3.215E-08 | 2.200E-07 | 2.800E-07 |
| 7211 | 361 | G361L | A | 3.9e-07 | 2.5e-07 | 3.9e-07 | 3.433E-07 | 8.083E-08 | 2.500E-07 | 3.900E-07 |
| 7212 | 361 | G361K | A | 2.4e-07 | 4.2e-07 | 2.7e-07 | 3.100E-07 | 9.644E-08 | 2.400E-07 | 4.200E-07 |
| 7213 | 361 | G361M | A | 2.5e-07 | 2.6e-07 | 1.7e-07 | 2.267E-07 | 4.933E-08 | 1.700E-07 | 2.600E-07 |
| 7214 | 361 | G361F | A | 4.7e-07 | 4e-07 | 6.5e-07 | 5.067E-07 | 1.290E-07 | 4.000E-07 | 6.500E-07 |
| 7215 | 361 | G361P | A | 1.4e-07 | 3.8e-07 | 3.6e-07 | 2.933E-07 | 1.332E-07 | 1.400E-07 | 3.800E-07 |
| 7216 | 361 | G361S | A | 5.4e-07 | 3.8e-07 | 1.4e-07 | 3.533E-07 | 2.013E-07 | 1.400E-07 | 5.400E-07 |
| 7217 | 361 | G361T | A | 3.2e-07 | 3.1e-07 | 1.8e-07 | 2.700E-07 | 7.810E-08 | 1.800E-07 | 3.200E-07 |
| 7218 | 361 | G361W | A | 1.2e-07 | 2.4e-07 | 4.1e-07 | 2.567E-07 | 1.457E-07 | 1.200E-07 | 4.100E-07 |
| 7219 | 361 | G361Y | A | 1.7e-07 | 2.3e-07 | 2.4e-07 | 2.133E-07 | 3.786E-08 | 1.700E-07 | 2.400E-07 |
| 7220 | 361 | G361V | A | 1.9e-07 | 2.4e-07 | 1.6e-07 | 1.967E-07 | 4.041E-08 | 1.600E-07 | 2.400E-07 |
| 7221 | 362 | D362A | A | 2.8e-07 | 1e-07 | 2.4e-07 | 2.067E-07 | 9.452E-08 | 1.000E-07 | 2.800E-07 |
| 7222 | 362 | D362R | A | 2.6e-07 | 4.7e-07 | 1.6e-07 | 2.967E-07 | 1.582E-07 | 1.600E-07 | 4.700E-07 |
| 7223 | 362 | D362N | A | 1.1e-07 | 1.6e-07 | 3.8e-07 | 2.167E-07 | 1.436E-07 | 1.100E-07 | 3.800E-07 |
| 7224 | 362 | D362D | A | 2.2e-07 | 2.6e-07 | 4.1e-07 | 2.967E-07 | 1.002E-07 | 2.200E-07 | 4.100E-07 |

| | | | | | | | | | | |
|---|---|---|---|---|---|---|---|---|---|---|
| 7225 | 362 | D362C | A | 3.1e-07 | 2e-07 | 2e-07 | 2.367E-07 | 6.351E-08 | 2.000E-07 | 3.100E-07 |
| 7226 | 362 | D362E | A | 4.2e-07 | 2.5e-07 | 2.6e-07 | 3.100E-07 | 9.539E-08 | 2.500E-07 | 4.200E-07 |
| 7227 | 362 | D362Q | A | 1.3e-07 | 9.2e-08 | 2.1e-07 | 1.440E-07 | 6.023E-08 | 9.200E-08 | 2.100E-07 |
| 7228 | 362 | D362G | A | 5.8e-08 | 3.2e-07 | 3.9e-07 | 2.560E-07 | 1.750E-07 | 5.800E-08 | 3.900E-07 |
| 7229 | 362 | D362H | A | 3.1e-07 | 1.5e-07 | 9.7e-08 | 1.857E-07 | 1.109E-07 | 9.700E-08 | 3.100E-07 |
| 7230 | 362 | D362I | A | 2.9e-07 | 2e-07 | 2.1e-07 | 2.333E-07 | 4.933E-08 | 2.000E-07 | 2.900E-07 |
| 7231 | 362 | D362L | A | 5.5e-07 | 2.3e-07 | 3.6e-07 | 3.800E-07 | 1.609E-07 | 2.300E-07 | 5.500E-07 |
| 7232 | 362 | D362K | A | 3.2e-07 | 3.5e-07 | 1.7e-07 | 2.800E-07 | 9.644E-08 | 1.700E-07 | 3.500E-07 |
| 7233 | 362 | D362M | A | 1.5e-07 | 1.7e-07 | 3.4e-07 | 2.200E-07 | 1.044E-07 | 1.500E-07 | 3.400E-07 |
| 7234 | 362 | D362F | A | 1.4e-07 | 2e-07 | 2.5e-07 | 1.967E-07 | 5.508E-08 | 1.400E-07 | 2.500E-07 |
| 7235 | 362 | D362P | A | 1.7e-07 | 2.3e-07 | 1.8e-07 | 1.933E-07 | 3.215E-08 | 1.700E-07 | 2.300E-07 |
| 7236 | 362 | D362S | A | 1.1e-07 | 3.8e-07 | 2.8e-07 | 2.567E-07 | 1.365E-07 | 1.100E-07 | 3.800E-07 |
| 7237 | 362 | D362T | A | 2.1e-07 | 1.1e-07 | 3e-07 | 2.067E-07 | 9.504E-08 | 1.100E-07 | 3.000E-07 |
| 7238 | 362 | D362W | A | 2.5e-07 | 3.5e-07 | 2.5e-07 | 2.833E-07 | 5.774E-08 | 2.500E-07 | 3.500E-07 |
| 7239 | 362 | D362Y | A | 3.2e-07 | 1.7e-07 | 1.6e-07 | 2.167E-07 | 8.963E-08 | 1.600E-07 | 3.200E-07 |
| 7240 | 362 | D362V | A | 3.2e-07 | 5.5e-07 | 1.4e-07 | 3.367E-07 | 2.055E-07 | 1.400E-07 | 5.500E-07 |
| 7241 | 363 | P363A | A | 2.3e-07 | 2.1e-07 | 3.9e-07 | 2.767E-07 | 9.866E-08 | 2.100E-07 | 3.900E-07 |
| 7242 | 363 | P363R | A | 4.7e-07 | 2e-07 | 2.9e-07 | 3.200E-07 | 1.375E-07 | 2.000E-07 | 4.700E-07 |
| 7243 | 363 | P363N | A | 4.2e-07 | 1.7e-07 | 2.3e-07 | 2.733E-07 | 1.305E-07 | 1.700E-07 | 4.200E-07 |
| 7244 | 363 | P363D | A | 1.9e-07 | 3.6e-07 | 3.3e-07 | 2.933E-07 | 9.074E-08 | 1.900E-07 | 3.600E-07 |
| 7245 | 363 | P363C | A | 4.5e-07 | 9.7e-08 | 2.1e-07 | 2.523E-07 | 1.803E-07 | 9.700E-08 | 4.500E-07 |
| 7246 | 363 | P363E | A | 2.5e-07 | 2.5e-07 | 1.6e-07 | 2.200E-07 | 5.196E-08 | 1.600E-07 | 2.500E-07 |
| 7247 | 363 | P363Q | A | 2.2e-07 | 9.3e-08 | 4.5e-07 | 2.543E-07 | 1.810E-07 | 9.300E-08 | 4.500E-07 |
| 7248 | 363 | P363G | A | 1.8e-07 | 1.5e-07 | 3.4e-07 | 2.233E-07 | 1.021E-07 | 1.500E-07 | 3.400E-07 |
| 7249 | 363 | P363H | A | 2.8e-07 | 2e-07 | 1.5e-07 | 2.100E-07 | 6.557E-08 | 1.500E-07 | 2.800E-07 |
| 7250 | 363 | P363I | A | 1.8e-07 | 3.9e-07 | 3.8e-07 | 3.167E-07 | 1.185E-07 | 1.800E-07 | 3.900E-07 |
| 7251 | 363 | P363L | A | 2.4e-07 | 1.2e-07 | 3.1e-07 | 2.233E-07 | 9.609E-08 | 1.200E-07 | 3.100E-07 |
| 7252 | 363 | P363K | A | 3.5e-07 | 2.3e-07 | 2.3e-07 | 2.700E-07 | 6.928E-08 | 2.300E-07 | 3.500E-07 |
| 7253 | 363 | P363M | A | 4.1e-07 | 1.5e-07 | 1.5e-07 | 2.367E-07 | 1.501E-07 | 1.500E-07 | 4.100E-07 |
| 7254 | 363 | P363F | A | 7.4e-08 | 3.4e-07 | 3.4e-07 | 2.513E-07 | 1.536E-07 | 7.400E-08 | 3.400E-07 |
| 7255 | 363 | P363P | A | 6.1e-08 | 1.8e-07 | 2.9e-07 | 1.770E-07 | 1.145E-07 | 6.100E-08 | 2.900E-07 |
| 7256 | 363 | P363S | A | 2.9e-07 | 3e-07 | 2.3e-07 | 2.733E-07 | 3.786E-08 | 2.300E-07 | 3.000E-07 |
| 7257 | 363 | P363T | A | 3.5e-07 | 5.6e-07 | 4.5e-07 | 4.533E-07 | 1.050E-07 | 3.500E-07 | 5.600E-07 |
| 7258 | 363 | P363W | A | 2e-07 | 8.5e-08 | 2.7e-07 | 1.850E-07 | 9.341E-08 | 8.500E-08 | 2.700E-07 |
| 7259 | 363 | P363Y | A | 8.9e-08 | 9.1e-07 | 2.7e-07 | 4.230E-07 | 4.314E-07 | 8.900E-08 | 9.100E-07 |
| 7260 | 363 | P363V | A | 2.2e-07 | 4.1e-07 | 2.4e-07 | 2.900E-07 | 1.044E-07 | 2.200E-07 | 4.100E-07 |
| 7261 | 364 | A364A | A | 1.3e-07 | 2e-07 | 1.1e-07 | 1.467E-07 | 4.726E-08 | 1.100E-07 | 2.000E-07 |
| 7262 | 364 | A364R | A | 3e-07 | 2.6e-07 | 3.4e-07 | 3.000E-07 | 4.000E-08 | 2.600E-07 | 3.400E-07 |
| 7263 | 364 | A364N | A | 1.5e-07 | 4.2e-07 | 2.2e-07 | 2.633E-07 | 1.401E-07 | 1.500E-07 | 4.200E-07 |
| 7264 | 364 | A364D | A | 3.5e-07 | 1.6e-07 | 2.2e-07 | 2.433E-07 | 9.713E-08 | 1.600E-07 | 3.500E-07 |
| 7265 | 364 | A364C | A | 1.7e-07 | 2.6e-07 | 2.9e-07 | 2.400E-07 | 6.245E-08 | 1.700E-07 | 2.900E-07 |
| 7266 | 364 | A364E | A | 1.9e-07 | 5.7e-07 | 2.1e-07 | 3.233E-07 | 2.139E-07 | 1.900E-07 | 5.700E-07 |
| 7267 | 364 | A364Q | A | 1.7e-07 | 3.1e-07 | 1.9e-07 | 2.233E-07 | 7.572E-08 | 1.700E-07 | 3.100E-07 |
| 7268 | 364 | A364G | A | 1.7e-07 | 4e-07 | 1.8e-07 | 2.500E-07 | 1.300E-07 | 1.700E-07 | 4.000E-07 |
| 7269 | 364 | A364H | A | 2.7e-07 | 2.8e-07 | 4.1e-07 | 3.200E-07 | 7.810E-08 | 2.700E-07 | 4.100E-07 |
| 7270 | 364 | A364I | A | 3.5e-07 | 2.1e-07 | 4.7e-07 | 3.433E-07 | 1.301E-07 | 2.100E-07 | 4.700E-07 |
| 7271 | 364 | A364L | A | 5.4e-07 | 2.2e-07 | 1.4e-07 | 3.000E-07 | 2.117E-07 | 1.400E-07 | 5.400E-07 |
| 7272 | 364 | A364K | A | 2.1e-07 | 2e-07 | 4.3e-07 | 2.800E-07 | 1.300E-07 | 2.000E-07 | 4.300E-07 |
| 7273 | 364 | A364M | A | 3.4e-07 | 3.8e-07 | 2.8e-07 | 3.333E-07 | 5.033E-08 | 2.800E-07 | 3.800E-07 |
| 7274 | 364 | A364F | A | 4e-07 | 4e-07 | 3.3e-07 | 3.767E-07 | 4.041E-08 | 3.300E-07 | 4.000E-07 |
| 7275 | 364 | A364P | A | 1.6e-07 | 2.4e-07 | 1.7e-07 | 1.900E-07 | 4.359E-08 | 1.600E-07 | 2.400E-07 |
| 7276 | 364 | A364S | A | 3e-07 | 3.5e-07 | 1.9e-07 | 2.800E-07 | 8.185E-08 | 1.900E-07 | 3.500E-07 |
| 7277 | 364 | A364T | A | 2.2e-07 | 7.8e-08 | 2.4e-07 | 1.793E-07 | 8.833E-08 | 7.800E-08 | 2.400E-07 |
| 7278 | 364 | A364W | A | 2.1e-07 | 2e-07 | 2.1e-07 | 2.067E-07 | 5.774E-09 | 2.000E-07 | 2.100E-07 |
| 7279 | 364 | A364Y | A | 3.3e-07 | 5.7e-07 | 3.3e-07 | 4.100E-07 | 1.386E-07 | 3.300E-07 | 5.700E-07 |
| 7280 | 364 | A364V | A | 1.8e-07 | 2.9e-07 | 2.6e-07 | 2.433E-07 | 5.686E-08 | 1.800E-07 | 2.900E-07 |
| 7281 | 365 | S365A | A | 3.3e-07 | 1.1e-07 | 1.1e-07 | 1.833E-07 | 1.270E-07 | 1.100E-07 | 3.300E-07 |
| 7282 | 365 | S365R | A | 3e-07 | 3.3e-07 | 1.8e-07 | 2.700E-07 | 7.937E-08 | 1.800E-07 | 3.300E-07 |
| 7283 | 365 | S365N | A | 3.8e-07 | 2.5e-07 | 4.2e-07 | 3.500E-07 | 8.888E-08 | 2.500E-07 | 4.200E-07 |
| 7284 | 365 | S365D | A | 4.1e-07 | 1.4e-07 | 3.1e-07 | 2.867E-07 | 1.365E-07 | 1.400E-07 | 4.100E-07 |
| 7285 | 365 | S365C | A | 5.4e-07 | 1.8e-07 | 4.4e-07 | 3.867E-07 | 1.858E-07 | 1.800E-07 | 5.400E-07 |
| 7286 | 365 | S365E | A | 1.9e-07 | 1.4e-07 | 2.1e-07 | 1.800E-07 | 3.606E-08 | 1.400E-07 | 2.100E-07 |
| 7287 | 365 | S365Q | A | 2.4e-07 | 3.9e-07 | 1.8e-07 | 2.700E-07 | 1.082E-07 | 1.800E-07 | 3.900E-07 |
| 7288 | 365 | S365G | A | 6.4e-07 | 3.6e-07 | 3.9e-07 | 4.633E-07 | 1.537E-07 | 3.600E-07 | 6.400E-07 |
| 7289 | 365 | S365H | A | 3.8e-07 | 2.1e-07 | 2.8e-07 | 2.900E-07 | 8.544E-08 | 2.100E-07 | 3.800E-07 |
| 7290 | 365 | S365I | A | 3.7e-07 | 5.9e-07 | 4.3e-07 | 4.633E-07 | 1.137E-07 | 3.700E-07 | 5.900E-07 |
| 7291 | 365 | S365L | A | 3.7e-07 | 3.6e-07 | 2.3e-07 | 3.200E-07 | 7.810E-08 | 2.300E-07 | 3.700E-07 |
| 7292 | 365 | S365K | A | 1.6e-07 | 1.6e-07 | 1.2e-07 | 1.467E-07 | 2.309E-08 | 1.200E-07 | 1.600E-07 |
| 7293 | 365 | S365M | A | 3.3e-07 | 3.1e-07 | 5.3e-07 | 3.900E-07 | 1.217E-07 | 3.100E-07 | 5.300E-07 |
| 7294 | 365 | S365F | A | 1.2e-07 | 3.6e-07 | 1.8e-07 | 2.200E-07 | 1.249E-07 | 1.200E-07 | 3.600E-07 |
| 7295 | 365 | S365P | A | 1.2e-07 | 2.4e-07 | 2.3e-07 | 1.967E-07 | 6.658E-08 | 1.200E-07 | 2.400E-07 |
| 7296 | 365 | S365S | A | 4.5e-07 | 5.7e-07 | 2.3e-07 | 4.167E-07 | 1.724E-07 | 2.300E-07 | 5.700E-07 |
| 7297 | 365 | S365T | A | 3.5e-07 | 2.2e-07 | 2.8e-07 | 2.833E-07 | 6.506E-08 | 2.200E-07 | 3.500E-07 |
| 7298 | 365 | S365W | A | 2.7e-07 | 2.2e-07 | 3.6e-07 | 2.833E-07 | 7.095E-08 | 2.200E-07 | 3.600E-07 |
| 7299 | 365 | S365Y | A | 3.4e-07 | 2.7e-07 | 1.6e-07 | 2.567E-07 | 9.074E-08 | 1.600E-07 | 3.400E-07 |
| 7300 | 365 | S365V | A | 3.6e-07 | 1.9e-07 | 2.7e-07 | 2.733E-07 | 8.505E-08 | 1.900E-07 | 3.600E-07 |
| 7301 | 366 | N366A | A | 3.1e-07 | 2.3e-07 | 1.9e-07 | 2.433E-07 | 6.110E-08 | 1.900E-07 | 3.100E-07 |
| 7302 | 366 | N366R | A | 2.5e-07 | 3.5e-07 | 2.8e-07 | 2.933E-07 | 5.132E-08 | 2.500E-07 | 3.500E-07 |
| 7303 | 366 | N366N | A | 1.2e-07 | 4.8e-07 | 3e-07 | 3.000E-07 | 1.800E-07 | 1.200E-07 | 4.800E-07 |
| 7304 | 366 | N366D | A | 2.1e-07 | 1.2e-07 | 1.2e-07 | 1.500E-07 | 5.196E-08 | 1.200E-07 | 2.100E-07 |
| 7305 | 366 | N366C | A | 1.7e-07 | 4.5e-07 | 2.3e-07 | 2.833E-07 | 1.474E-07 | 1.700E-07 | 4.500E-07 |
| 7306 | 366 | N366E | A | 1.9e-07 | 2.6e-07 | 4.3e-07 | 2.933E-07 | 1.234E-07 | 1.900E-07 | 4.300E-07 |
| 7307 | 366 | N366Q | A | 2.5e-07 | 1.1e-07 | 3e-07 | 2.200E-07 | 9.849E-08 | 1.100E-07 | 3.000E-07 |
| 7308 | 366 | N366G | A | 2.7e-07 | 5.1e-07 | 3.4e-07 | 3.733E-07 | 1.234E-07 | 2.700E-07 | 5.100E-07 |
| 7309 | 366 | N366H | A | 2e-07 | 2.5e-07 | 2.6e-07 | 2.367E-07 | 3.215E-08 | 2.000E-07 | 2.600E-07 |

| | | | | | | | | | | |
|---|---|---|---|---|---|---|---|---|---|---|
| 7310 | 366 | N366I | A | 1.6e-07 | 3.8e-07 | 3.2e-07 | 2.867E-07 | 1.137E-07 | 1.600E-07 | 3.800E-07 |
| 7311 | 366 | N366L | A | 2.3e-07 | 2.3e-07 | 1.6e-07 | 2.067E-07 | 4.041E-08 | 1.600E-07 | 2.300E-07 |
| 7312 | 366 | N366K | A | 3.4e-07 | 1.3e-07 | 1.7e-07 | 2.133E-07 | 1.115E-07 | 1.300E-07 | 3.400E-07 |
| 7313 | 366 | N366M | A | 2.9e-07 | 1.7e-07 | 3.1e-07 | 2.567E-07 | 7.572E-08 | 1.700E-07 | 3.100E-07 |
| 7314 | 366 | N366F | A | 5.2e-07 | 3.6e-07 | 1.4e-07 | 3.400E-07 | 1.908E-07 | 1.400E-07 | 5.200E-07 |
| 7315 | 366 | N366P | A | 2.4e-07 | 3.5e-07 | 2.3e-07 | 2.733E-07 | 6.658E-08 | 2.300E-07 | 3.500E-07 |
| 7316 | 366 | N366S | A | 3.1e-07 | 2.6e-07 | 2.4e-07 | 2.700E-07 | 3.606E-08 | 2.400E-07 | 3.100E-07 |
| 7317 | 366 | N366T | A | 5.2e-07 | 1.7e-07 | 3e-07 | 3.300E-07 | 1.769E-07 | 1.700E-07 | 5.200E-07 |
| 7318 | 366 | N366W | A | 3.1e-07 | 2.2e-07 | 2.3e-07 | 2.533E-07 | 4.933E-08 | 2.200E-07 | 3.100E-07 |
| 7319 | 366 | N366Y | A | 2.7e-07 | 2.4e-07 | 2.1e-07 | 2.400E-07 | 3.000E-08 | 2.100E-07 | 2.700E-07 |
| 7320 | 366 | N366V | A | 4.2e-07 | 2.5e-07 | 2.5e-07 | 3.067E-07 | 9.815E-08 | 2.500E-07 | 4.200E-07 |
| 7321 | 367 | T367A | A | 3.9e-07 | 2.4e-07 | 3.3e-07 | 3.200E-07 | 7.550E-08 | 2.400E-07 | 3.900E-07 |
| 7322 | 367 | T367R | A | 1.8e-07 | 2.5e-07 | 1.3e-07 | 1.867E-07 | 6.028E-08 | 1.300E-07 | 2.500E-07 |
| 7323 | 367 | T367N | A | 2.1e-07 | 1.7e-07 | 3.5e-07 | 2.433E-07 | 9.452E-08 | 1.700E-07 | 3.500E-07 |
| 7324 | 367 | T367D | A | 3.2e-07 | 2.6e-07 | 3.4e-07 | 3.067E-07 | 4.163E-08 | 2.600E-07 | 3.400E-07 |
| 7325 | 367 | T367C | A | 1.7e-07 | 1.4e-07 | 3.1e-07 | 2.067E-07 | 9.074E-08 | 1.400E-07 | 3.100E-07 |
| 7326 | 367 | T367E | A | 3e-07 | 4e-07 | 2.4e-07 | 3.133E-07 | 8.083E-08 | 2.400E-07 | 4.000E-07 |
| 7327 | 367 | T367Q | A | 2.7e-07 | 1.6e-07 | 2e-07 | 2.100E-07 | 5.568E-08 | 1.600E-07 | 2.700E-07 |
| 7328 | 367 | T367G | A | 2.5e-07 | 3e-07 | 2.4e-07 | 2.633E-07 | 3.215E-08 | 2.400E-07 | 3.000E-07 |
| 7329 | 367 | T367H | A | 3.3e-07 | 1.8e-07 | 1.2e-07 | 2.100E-07 | 1.082E-07 | 1.200E-07 | 3.300E-07 |
| 7330 | 367 | T367I | A | 2.9e-07 | 1.7e-07 | 1.2e-07 | 1.933E-07 | 8.737E-08 | 1.200E-07 | 2.900E-07 |
| 7331 | 367 | T367L | A | 2.5e-07 | 1.8e-07 | 2.4e-07 | 2.233E-07 | 3.786E-08 | 1.800E-07 | 2.500E-07 |
| 7332 | 367 | T367K | A | 3.2e-07 | 3.3e-07 | 1e-07 | 2.500E-07 | 1.300E-07 | 1.000E-07 | 3.300E-07 |
| 7333 | 367 | T367M | A | 2.3e-07 | 6.2e-08 | 3.4e-07 | 2.107E-07 | 1.400E-07 | 6.200E-08 | 3.400E-07 |
| 7334 | 367 | T367F | A | 2.5e-07 | 3e-07 | 1.4e-07 | 2.300E-07 | 8.185E-08 | 1.400E-07 | 3.000E-07 |
| 7335 | 367 | T367P | A | 3.9e-07 | 1.9e-07 | 2.2e-07 | 2.667E-07 | 1.079E-07 | 1.900E-07 | 3.900E-07 |
| 7336 | 367 | T367S | A | 5.9e-07 | 3.3e-07 | 4.5e-07 | 4.567E-07 | 1.301E-07 | 3.300E-07 | 5.900E-07 |
| 7337 | 367 | T367T | A | 4.5e-07 | 6e-07 | 2.4e-07 | 4.300E-07 | 1.808E-07 | 2.400E-07 | 6.000E-07 |
| 7338 | 367 | T367W | A | 2.4e-07 | 2.4e-07 | 1.4e-07 | 2.067E-07 | 5.774E-08 | 1.400E-07 | 2.400E-07 |
| 7339 | 367 | T367Y | A | 1.8e-07 | 5.2e-07 | 1e-07 | 2.667E-07 | 2.230E-07 | 1.000E-07 | 5.200E-07 |
| 7340 | 367 | T367V | A | 6.6e-07 | 4.3e-07 | 1.2e-07 | 4.033E-07 | 2.710E-07 | 1.200E-07 | 6.600E-07 |
| 7341 | 368 | A368A | A | 3.5e-07 | 4.1e-07 | 2.5e-07 | 3.367E-07 | 8.083E-08 | 2.500E-07 | 4.100E-07 |
| 7342 | 368 | A368R | A | 4.1e-07 | 3e-07 | 5e-07 | 4.033E-07 | 1.002E-07 | 3.000E-07 | 5.000E-07 |
| 7343 | 368 | A368N | A | 9.4e-08 | 3.8e-07 | 2.1e-07 | 2.280E-07 | 1.438E-07 | 9.400E-08 | 3.800E-07 |
| 7344 | 368 | A368D | A | 2.5e-07 | 3.8e-07 | 2.6e-07 | 2.967E-07 | 7.234E-08 | 2.500E-07 | 3.800E-07 |
| 7345 | 368 | A368C | A | 2.4e-07 | 1.3e-07 | 2.1e-07 | 1.933E-07 | 5.686E-08 | 1.300E-07 | 2.400E-07 |
| 7346 | 368 | A368E | A | 2.9e-07 | 4.5e-07 | 1.5e-07 | 2.967E-07 | 1.501E-07 | 1.500E-07 | 4.500E-07 |
| 7347 | 368 | A368Q | A | 3.4e-07 | 3.4e-07 | 3.3e-07 | 3.367E-07 | 5.774E-09 | 3.300E-07 | 3.400E-07 |
| 7348 | 368 | A368G | A | 3.6e-07 | 2.4e-07 | 3.1e-07 | 3.033E-07 | 6.028E-08 | 2.400E-07 | 3.600E-07 |
| 7349 | 368 | A368H | A | 3e-07 | 1.5e-07 | 1e-07 | 1.833E-07 | 1.041E-07 | 1.000E-07 | 3.000E-07 |
| 7350 | 368 | A368I | A | 2.5e-07 | 2.3e-07 | 1.7e-07 | 2.167E-07 | 4.163E-08 | 1.700E-07 | 2.500E-07 |
| 7351 | 368 | A368L | A | 2.7e-07 | 3.2e-07 | 5.3e-07 | 3.733E-07 | 1.380E-07 | 2.700E-07 | 5.300E-07 |
| 7352 | 368 | A368K | A | 1.1e-07 | 3.9e-07 | 3.3e-07 | 2.767E-07 | 1.474E-07 | 1.100E-07 | 3.900E-07 |
| 7353 | 368 | A368M | A | 2.6e-07 | 1.2e-07 | 2.6e-07 | 2.133E-07 | 8.083E-08 | 1.200E-07 | 2.600E-07 |
| 7354 | 368 | A368F | A | 1.4e-07 | 1.5e-07 | 2.7e-07 | 1.867E-07 | 7.234E-08 | 1.400E-07 | 2.700E-07 |
| 7355 | 368 | A368P | A | 2.8e-07 | 7.6e-08 | 2e-07 | 1.853E-07 | 1.028E-07 | 7.600E-08 | 2.800E-07 |
| 7356 | 368 | A368S | A | 3.2e-07 | 1.5e-07 | 2.8e-07 | 2.500E-07 | 8.888E-08 | 1.500E-07 | 3.200E-07 |
| 7357 | 368 | A368T | A | 2.7e-07 | 2.8e-07 | 1.9e-07 | 2.467E-07 | 4.933E-08 | 1.900E-07 | 2.800E-07 |
| 7358 | 368 | A368W | A | 2.1e-07 | 2e-07 | 2.3e-07 | 2.133E-07 | 1.528E-08 | 2.000E-07 | 2.300E-07 |
| 7359 | 368 | A368Y | A | 1.4e-07 | 3.1e-07 | 1.6e-07 | 2.033E-07 | 9.292E-08 | 1.400E-07 | 3.100E-07 |
| 7360 | 368 | A368V | A | 2.8e-07 | 2e-07 | 1.9e-07 | 2.233E-07 | 4.933E-08 | 1.900E-07 | 2.800E-07 |
| 7361 | 369 | P369A | A | 1.9e-07 | 3.3e-07 | 1.7e-07 | 2.300E-07 | 8.718E-08 | 1.700E-07 | 3.300E-07 |
| 7362 | 369 | P369R | A | 1.5e-07 | 2.2e-07 | 2.2e-07 | 1.967E-07 | 4.041E-08 | 1.500E-07 | 2.200E-07 |
| 7363 | 369 | P369N | A | 3.6e-07 | 1.8e-07 | 1.5e-07 | 2.300E-07 | 1.136E-07 | 1.500E-07 | 3.600E-07 |
| 7364 | 369 | P369D | A | 1.3e-07 | 1.8e-07 | 5.2e-07 | 2.767E-07 | 2.122E-07 | 1.300E-07 | 5.200E-07 |
| 7365 | 369 | P369C | A | 2.1e-07 | 1.5e-07 | 2.3e-07 | 1.967E-07 | 4.163E-08 | 1.500E-07 | 2.300E-07 |
| 7366 | 369 | P369E | A | 1.5e-07 | 3.9e-07 | 2.5e-07 | 2.633E-07 | 1.206E-07 | 1.500E-07 | 3.900E-07 |
| 7367 | 369 | P369Q | A | 3.9e-07 | 8.5e-08 | 2.5e-07 | 2.417E-07 | 1.527E-07 | 8.500E-08 | 3.900E-07 |
| 7368 | 369 | P369G | A | 4.3e-07 | 2.3e-07 | 1.6e-07 | 2.733E-07 | 1.401E-07 | 1.600E-07 | 4.300E-07 |
| 7369 | 369 | P369H | A | 2.3e-07 | 2.1e-07 | 5.3e-07 | 3.233E-07 | 1.793E-07 | 2.100E-07 | 5.300E-07 |
| 7370 | 369 | P369I | A | 5.1e-07 | 2.4e-07 | 3.1e-07 | 3.533E-07 | 1.401E-07 | 2.400E-07 | 5.100E-07 |
| 7371 | 369 | P369L | A | 1.7e-07 | 2.4e-07 | 1.7e-07 | 1.933E-07 | 4.041E-08 | 1.700E-07 | 2.400E-07 |
| 7372 | 369 | P369K | A | 1.4e-07 | 1.7e-07 | 3e-07 | 2.033E-07 | 8.505E-08 | 1.400E-07 | 3.000E-07 |
| 7373 | 369 | P369M | A | 2.8e-07 | 3.8e-07 | 9.3e-08 | 2.510E-07 | 1.457E-07 | 9.300E-08 | 3.800E-07 |
| 7374 | 369 | P369F | A | 4e-07 | 3e-07 | 2e-07 | 3.000E-07 | 1.000E-07 | 2.000E-07 | 4.000E-07 |
| 7375 | 369 | P369P | A | 2.4e-07 | 3.7e-07 | 1.5e-07 | 2.533E-07 | 1.106E-07 | 1.500E-07 | 3.700E-07 |
| 7376 | 369 | P369S | A | 1.6e-07 | 3.4e-07 | 4.6e-07 | 3.200E-07 | 1.510E-07 | 1.600E-07 | 4.600E-07 |
| 7377 | 369 | P369T | A | 3.4e-07 | 3.7e-07 | 1e-07 | 2.700E-07 | 1.480E-07 | 1.000E-07 | 3.700E-07 |
| 7378 | 369 | P369W | A | 2.2e-07 | 2.2e-07 | 2.9e-07 | 2.433E-07 | 4.041E-08 | 2.200E-07 | 2.900E-07 |
| 7379 | 369 | P369Y | A | 3.5e-07 | 5.5e-07 | 5e-07 | 4.667E-07 | 1.041E-07 | 3.500E-07 | 5.500E-07 |
| 7380 | 369 | P369V | A | 2.2e-07 | 1.6e-07 | 3e-07 | 2.267E-07 | 7.024E-08 | 1.600E-07 | 3.000E-07 |
| 7381 | 370 | L370A | A | 7.9e-08 | 2.8e-07 | 5e-07 | 2.863E-07 | 2.106E-07 | 7.900E-08 | 5.000E-07 |
| 7382 | 370 | L370R | A | 1.7e-07 | 2.9e-07 | 4.6e-07 | 3.067E-07 | 1.457E-07 | 1.700E-07 | 4.600E-07 |
| 7383 | 370 | L370N | A | 6.9e-07 | 1.6e-07 | 1.3e-07 | 3.267E-07 | 3.150E-07 | 1.300E-07 | 6.900E-07 |
| 7384 | 370 | L370D | A | 3.6e-07 | 7.6e-08 | 5.4e-07 | 3.253E-07 | 2.339E-07 | 7.600E-08 | 5.400E-07 |
| 7385 | 370 | L370C | A | 1.9e-07 | 3.3e-07 | 1.1e-07 | 2.100E-07 | 1.114E-07 | 1.100E-07 | 3.300E-07 |
| 7386 | 370 | L370E | A | 3.1e-07 | 2.9e-07 | 1.5e-07 | 2.500E-07 | 8.718E-08 | 1.500E-07 | 3.100E-07 |
| 7387 | 370 | L370Q | A | 3e-07 | 1.6e-07 | 1.9e-07 | 2.167E-07 | 7.371E-08 | 1.600E-07 | 3.000E-07 |
| 7388 | 370 | L370G | A | 3.8e-07 | 4.5e-07 | 2.3e-07 | 3.533E-07 | 1.124E-07 | 2.300E-07 | 4.500E-07 |
| 7389 | 370 | L370H | A | 2e-07 | 1.8e-07 | 3.3e-07 | 2.367E-07 | 8.145E-08 | 1.800E-07 | 3.300E-07 |
| 7390 | 370 | L370I | A | 1.6e-07 | 1.1e-07 | 2e-07 | 1.567E-07 | 4.509E-08 | 1.100E-07 | 2.000E-07 |
| 7391 | 370 | L370L | A | 3.4e-07 | 1.9e-07 | 4.6e-07 | 3.300E-07 | 1.353E-07 | 1.900E-07 | 4.600E-07 |
| 7392 | 370 | L370K | A | 6.6e-07 | 2.4e-07 | 2.2e-07 | 3.733E-07 | 2.485E-07 | 2.200E-07 | 6.600E-07 |
| 7393 | 370 | L370M | A | 8.7e-07 | 1.6e-07 | 2.4e-07 | 4.233E-07 | 3.889E-07 | 1.600E-07 | 8.700E-07 |
| 7394 | 370 | L370F | A | 1.5e-07 | 2.9e-07 | 1.4e-07 | 1.933E-07 | 8.386E-08 | 1.400E-07 | 2.900E-07 |

| | | | | | | | | | | |
|---|---|---|---|---|---|---|---|---|---|---|
| 7395 | 370 | L370P | A | 2.7e-07 | 2.1e-07 | 4.9e-07 | 3.233E-07 | 1.474E-07 | 2.100E-07 | 4.900E-07 |
| 7396 | 370 | L370S | A | 3e-07 | 3.1e-07 | 3.3e-07 | 3.133E-07 | 1.528E-08 | 3.000E-07 | 3.300E-07 |
| 7397 | 370 | L370T | A | 2.5e-07 | 3.4e-07 | 5.9e-07 | 3.933E-07 | 1.762E-07 | 2.500E-07 | 5.900E-07 |
| 7398 | 370 | L370W | A | 1.2e-07 | 5e-07 | 2.7e-07 | 2.967E-07 | 1.914E-07 | 1.200E-07 | 5.000E-07 |
| 7399 | 370 | L370Y | A | 2.3e-07 | 1.2e-07 | 3.7e-07 | 2.400E-07 | 1.253E-07 | 1.200E-07 | 3.700E-07 |
| 7400 | 370 | L370V | A | 4.1e-07 | 1.6e-07 | 2.7e-07 | 2.800E-07 | 1.253E-07 | 1.600E-07 | 4.100E-07 |
| 7401 | 371 | Q371A | A | 1.6e-07 | 2.2e-07 | 2.1e-07 | 1.967E-07 | 3.215E-08 | 1.600E-07 | 2.200E-07 |
| 7402 | 371 | Q371R | A | 5.2e-07 | 2.8e-07 | 2.2e-07 | 3.400E-07 | 1.587E-07 | 2.200E-07 | 5.200E-07 |
| 7403 | 371 | Q371N | A | 2.2e-07 | 1.3e-07 | 2.9e-07 | 2.133E-07 | 8.021E-08 | 1.300E-07 | 2.900E-07 |
| 7404 | 371 | Q371D | A | 2.7e-07 | 1.8e-07 | 2.3e-07 | 2.267E-07 | 4.509E-08 | 1.800E-07 | 2.700E-07 |
| 7405 | 371 | Q371C | A | 2.4e-07 | 4.3e-07 | 2.7e-07 | 3.133E-07 | 1.021E-07 | 2.400E-07 | 4.300E-07 |
| 7406 | 371 | Q371E | A | 2.4e-07 | 3.8e-07 | 5e-07 | 3.733E-07 | 1.301E-07 | 2.400E-07 | 5.000E-07 |
| 7407 | 371 | Q371Q | A | 4.3e-07 | 1.6e-07 | 4.6e-07 | 3.500E-07 | 1.652E-07 | 1.600E-07 | 4.600E-07 |
| 7408 | 371 | Q371G | A | 2.1e-07 | 2e-07 | 2.5e-07 | 2.200E-07 | 2.646E-08 | 2.000E-07 | 2.500E-07 |
| 7409 | 371 | Q371H | A | 1.1e-07 | 2.2e-07 | 2.9e-07 | 2.067E-07 | 9.074E-08 | 1.100E-07 | 2.900E-07 |
| 7410 | 371 | Q371I | A | 7.3e-08 | 2.3e-07 | 2.4e-07 | 1.810E-07 | 9.366E-08 | 7.300E-08 | 2.400E-07 |
| 7411 | 371 | Q371L | A | 2.3e-07 | 3.4e-07 | 2.9e-07 | 2.867E-07 | 5.508E-08 | 2.300E-07 | 3.400E-07 |
| 7412 | 371 | Q371K | A | 3.4e-07 | 1.3e-07 | 3.3e-07 | 2.667E-07 | 1.185E-07 | 1.300E-07 | 3.400E-07 |
| 7413 | 371 | Q371M | A | 2.9e-07 | 2.1e-07 | 2.7e-07 | 2.567E-07 | 4.163E-08 | 2.100E-07 | 2.900E-07 |
| 7414 | 371 | Q371F | A | 2.2e-07 | 2.7e-07 | 4.3e-07 | 3.067E-07 | 1.097E-07 | 2.200E-07 | 4.300E-07 |
| 7415 | 371 | Q371P | A | 2e-07 | 2.7e-07 | 1.3e-07 | 2.000E-07 | 7.000E-08 | 1.300E-07 | 2.700E-07 |
| 7416 | 371 | Q371S | A | 3.7e-07 | 1.8e-07 | 3e-07 | 2.833E-07 | 9.609E-08 | 1.800E-07 | 3.700E-07 |
| 7417 | 371 | Q371T | A | 4.2e-07 | 3.1e-07 | 1.2e-07 | 2.833E-07 | 1.518E-07 | 1.200E-07 | 4.200E-07 |
| 7418 | 371 | Q371W | A | 2.9e-07 | 1.7e-07 | 1.8e-07 | 2.133E-07 | 6.658E-08 | 1.700E-07 | 2.900E-07 |
| 7419 | 371 | Q371Y | A | 4.1e-07 | 2e-07 | 1.7e-07 | 2.600E-07 | 1.308E-07 | 1.700E-07 | 4.100E-07 |
| 7420 | 371 | Q371V | A | 1.6e-07 | 6.4e-07 | 4.1e-07 | 4.033E-07 | 2.401E-07 | 1.600E-07 | 6.400E-07 |
| 7421 | 372 | P372A | A | 3.2e-07 | 4.7e-07 | 2e-07 | 3.300E-07 | 1.353E-07 | 2.000E-07 | 4.700E-07 |
| 7422 | 372 | P372R | A | 3e-07 | 2.8e-07 | 1.3e-07 | 2.367E-07 | 9.292E-08 | 1.300E-07 | 3.000E-07 |
| 7423 | 372 | P372N | A | 1.3e-07 | 4e-07 | 1.6e-07 | 2.300E-07 | 1.480E-07 | 1.300E-07 | 4.000E-07 |
| 7424 | 372 | P372D | A | 1.9e-07 | 4.7e-07 | 2.2e-07 | 2.933E-07 | 1.537E-07 | 1.900E-07 | 4.700E-07 |
| 7425 | 372 | P372C | A | 1.4e-07 | 2.2e-07 | 3e-07 | 2.200E-07 | 8.000E-08 | 1.400E-07 | 3.000E-07 |
| 7426 | 372 | P372E | A | 2.3e-07 | 9.3e-08 | 2e-07 | 1.743E-07 | 7.202E-08 | 9.300E-08 | 2.300E-07 |
| 7427 | 372 | P372Q | A | 3.7e-07 | 1.2e-07 | 1.9e-07 | 2.267E-07 | 1.290E-07 | 1.200E-07 | 3.700E-07 |
| 7428 | 372 | P372G | A | 3.3e-07 | 3.7e-07 | 5.4e-07 | 4.133E-07 | 1.115E-07 | 3.300E-07 | 5.400E-07 |
| 7429 | 372 | P372H | A | 2.9e-07 | 3e-07 | 2.6e-07 | 2.833E-07 | 2.082E-08 | 2.600E-07 | 3.000E-07 |
| 7430 | 372 | P372I | A | 3.2e-07 | 2.3e-07 | 2.2e-07 | 2.567E-07 | 5.508E-08 | 2.200E-07 | 3.200E-07 |
| 7431 | 372 | P372L | A | 3.1e-07 | 3.1e-07 | 2.9e-07 | 3.033E-07 | 1.155E-08 | 2.900E-07 | 3.100E-07 |
| 7432 | 372 | P372K | A | 2.9e-07 | 2.9e-07 | 2.6e-07 | 2.800E-07 | 1.732E-08 | 2.600E-07 | 2.900E-07 |
| 7433 | 372 | P372M | A | 3.9e-07 | 2.2e-07 | 4.4e-07 | 3.500E-07 | 1.153E-07 | 2.200E-07 | 4.400E-07 |
| 7434 | 372 | P372F | A | 2.9e-07 | 2.2e-07 | 3.2e-07 | 2.767E-07 | 5.132E-08 | 2.200E-07 | 3.200E-07 |
| 7435 | 372 | P372P | A | 1.8e-07 | 4.9e-07 | 2.1e-07 | 2.933E-07 | 1.710E-07 | 1.800E-07 | 4.900E-07 |
| 7436 | 372 | P372S | A | 3.4e-07 | 2.3e-07 | 1.9e-07 | 2.533E-07 | 7.767E-08 | 1.900E-07 | 3.400E-07 |
| 7437 | 372 | P372T | A | 4.2e-07 | 2.7e-07 | 1.2e-07 | 2.700E-07 | 1.500E-07 | 1.200E-07 | 4.200E-07 |
| 7438 | 372 | P372W | A | 1.5e-07 | 2.1e-07 | 2.1e-07 | 1.900E-07 | 3.464E-08 | 1.500E-07 | 2.100E-07 |
| 7439 | 372 | P372Y | A | 2.5e-07 | 2.6e-07 | 8.3e-08 | 1.977E-07 | 9.943E-08 | 8.300E-08 | 2.600E-07 |
| 7440 | 372 | P372V | A | 1.7e-07 | 2.6e-07 | 4e-07 | 2.767E-07 | 1.159E-07 | 1.700E-07 | 4.000E-07 |
| 7441 | 373 | E373A | A | 2.9e-07 | 3e-07 | 1.9e-07 | 2.600E-07 | 6.083E-08 | 1.900E-07 | 3.000E-07 |
| 7442 | 373 | E373R | A | 3.1e-07 | 2.6e-07 | 2e-07 | 2.567E-07 | 5.508E-08 | 2.000E-07 | 3.100E-07 |
| 7443 | 373 | E373N | A | 3.4e-07 | 2.1e-07 | 2.3e-07 | 2.600E-07 | 7.000E-08 | 2.100E-07 | 3.400E-07 |
| 7444 | 373 | E373D | A | 3.7e-07 | 2.8e-07 | 3.3e-07 | 3.267E-07 | 4.509E-08 | 2.800E-07 | 3.700E-07 |
| 7445 | 373 | E373C | A | 2.2e-07 | 3.7e-07 | 3.3e-07 | 3.067E-07 | 7.767E-08 | 2.200E-07 | 3.700E-07 |
| 7446 | 373 | E373E | A | 2.2e-07 | 2.9e-07 | 2.6e-07 | 2.567E-07 | 3.512E-08 | 2.200E-07 | 2.900E-07 |
| 7447 | 373 | E373Q | A | 3.7e-07 | 3.2e-07 | 2.5e-07 | 3.133E-07 | 6.028E-08 | 2.500E-07 | 3.700E-07 |
| 7448 | 373 | E373G | A | 2.2e-07 | 2.5e-07 | 1.7e-07 | 2.133E-07 | 4.041E-08 | 1.700E-07 | 2.500E-07 |
| 7449 | 373 | E373H | A | 2.6e-07 | 3.7e-07 | 1.9e-07 | 2.733E-07 | 9.074E-08 | 1.900E-07 | 3.700E-07 |
| 7450 | 373 | E373I | A | 2.3e-07 | 1.9e-07 | 5.3e-07 | 3.167E-07 | 1.858E-07 | 1.900E-07 | 5.300E-07 |
| 7451 | 373 | E373L | A | 3.7e-07 | 3.7e-07 | 2.7e-07 | 3.367E-07 | 5.774E-08 | 2.700E-07 | 3.700E-07 |
| 7452 | 373 | E373K | A | 2.1e-07 | 3e-07 | 3.3e-07 | 2.800E-07 | 6.245E-08 | 2.100E-07 | 3.300E-07 |
| 7453 | 373 | E373M | A | 1.8e-07 | 4e-07 | 2.1e-07 | 2.633E-07 | 1.193E-07 | 1.800E-07 | 4.000E-07 |
| 7454 | 373 | E373F | A | 2.6e-07 | 2.7e-07 | 1.8e-07 | 2.367E-07 | 4.933E-08 | 1.800E-07 | 2.700E-07 |
| 7455 | 373 | E373P | A | 2e-07 | 2.3e-07 | 1.5e-07 | 1.933E-07 | 4.041E-08 | 1.500E-07 | 2.300E-07 |
| 7456 | 373 | E373S | A | 3.7e-07 | 3.6e-07 | 2.4e-07 | 3.233E-07 | 7.234E-08 | 2.400E-07 | 3.700E-07 |
| 7457 | 373 | E373T | A | 1.5e-07 | 3.5e-07 | 3.6e-07 | 2.867E-07 | 1.185E-07 | 1.500E-07 | 3.600E-07 |
| 7458 | 373 | E373W | A | 2.6e-07 | 4.1e-07 | 2.6e-07 | 3.100E-07 | 8.660E-08 | 2.600E-07 | 4.100E-07 |
| 7459 | 373 | E373Y | A | 2.2e-07 | 3.7e-07 | 4.1e-07 | 3.333E-07 | 1.002E-07 | 2.200E-07 | 4.100E-07 |
| 7460 | 373 | E373V | A | 2.3e-07 | 1.5e-07 | 1.9e-07 | 1.900E-07 | 4.000E-08 | 1.500E-07 | 2.300E-07 |
| 7461 | 374 | Q374A | A | 2.7e-07 | 2.2e-07 | 4.5e-07 | 3.133E-07 | 1.210E-07 | 2.200E-07 | 4.500E-07 |
| 7462 | 374 | Q374R | A | 7.1e-07 | 2.9e-07 | 3.2e-07 | 4.400E-07 | 2.343E-07 | 2.900E-07 | 7.100E-07 |
| 7463 | 374 | Q374N | A | 2.4e-07 | 2e-07 | 2.3e-07 | 2.233E-07 | 2.082E-08 | 2.000E-07 | 2.400E-07 |
| 7464 | 374 | Q374D | A | 2.4e-07 | 2.6e-07 | 3.1e-07 | 2.700E-07 | 3.606E-08 | 2.400E-07 | 3.100E-07 |
| 7465 | 374 | Q374C | A | 2.3e-07 | 2.8e-07 | 2.3e-07 | 2.467E-07 | 2.887E-08 | 2.300E-07 | 2.800E-07 |
| 7466 | 374 | Q374E | A | 1.6e-07 | 2.1e-07 | 2.4e-07 | 2.033E-07 | 4.041E-08 | 1.600E-07 | 2.400E-07 |
| 7467 | 374 | Q374Q | A | 4e-07 | 7.4e-08 | 2.5e-07 | 2.413E-07 | 1.632E-07 | 7.400E-08 | 4.000E-07 |
| 7468 | 374 | Q374G | A | 2.8e-07 | 5.2e-07 | 3.2e-07 | 3.733E-07 | 1.286E-07 | 2.800E-07 | 5.200E-07 |
| 7469 | 374 | Q374H | A | 3.9e-07 | 1.3e-07 | 2.4e-07 | 2.533E-07 | 1.305E-07 | 1.300E-07 | 3.900E-07 |
| 7470 | 374 | Q374I | A | 6.1e-07 | 3e-07 | 3.9e-07 | 4.333E-07 | 1.595E-07 | 3.000E-07 | 6.100E-07 |
| 7471 | 374 | Q374L | A | 3.1e-07 | 2.1e-07 | 7.4e-08 | 1.980E-07 | 1.185E-07 | 7.400E-08 | 3.100E-07 |
| 7472 | 374 | Q374K | A | 2.9e-07 | 3.2e-07 | 4.3e-07 | 3.467E-07 | 7.371E-08 | 2.900E-07 | 4.300E-07 |
| 7473 | 374 | Q374M | A | 2.5e-07 | 1.6e-07 | 2.3e-07 | 2.133E-07 | 4.726E-08 | 1.600E-07 | 2.500E-07 |
| 7474 | 374 | Q374F | A | 1.6e-07 | 1.6e-07 | 3.6e-07 | 2.267E-07 | 1.155E-07 | 1.600E-07 | 3.600E-07 |
| 7475 | 374 | Q374P | A | 2.4e-07 | 2.5e-07 | 4.4e-07 | 3.100E-07 | 1.127E-07 | 2.400E-07 | 4.400E-07 |
| 7476 | 374 | Q374S | A | 2.4e-07 | 3.1e-07 | 3.1e-07 | 2.867E-07 | 4.041E-08 | 2.400E-07 | 3.100E-07 |
| 7477 | 374 | Q374T | A | 2.9e-07 | 3.4e-07 | 2.3e-07 | 2.867E-07 | 5.508E-08 | 2.300E-07 | 3.400E-07 |
| 7478 | 374 | Q374W | A | 9.4e-08 | 9.3e-08 | 2.2e-07 | 1.357E-07 | 7.304E-08 | 9.300E-08 | 2.200E-07 |
| 7479 | 374 | Q374Y | A | 2.1e-07 | 3.5e-07 | 2.2e-07 | 2.600E-07 | 7.810E-08 | 2.100E-07 | 3.500E-07 |

| | | | | | | | | | | |
|---|---|---|---|---|---|---|---|---|---|---|
| 7480 | 374 | Q374V | A | 6e-07 | 2.2e-07 | 2.1e-07 | 3.433E-07 | 2.223E-07 | 2.100E-07 | 6.000E-07 |
| 7481 | 375 | L375A | A | 5.8e-07 | 1e-07 | 3e-07 | 3.267E-07 | 2.411E-07 | 1.000E-07 | 5.800E-07 |
| 7482 | 375 | L375R | A | 7.4e-07 | 6.9e-08 | 2.1e-07 | 3.397E-07 | 3.538E-07 | 6.900E-08 | 7.400E-07 |
| 7483 | 375 | L375N | A | 3.7e-07 | 2e-07 | 2.3e-07 | 2.667E-07 | 9.074E-08 | 2.000E-07 | 3.700E-07 |
| 7484 | 375 | L375D | A | 4.2e-07 | 1.4e-07 | 3.2e-07 | 2.933E-07 | 1.419E-07 | 1.400E-07 | 4.200E-07 |
| 7485 | 375 | L375C | A | 2.3e-07 | 2.4e-07 | 2.3e-07 | 2.333E-07 | 5.774E-09 | 2.300E-07 | 2.400E-07 |
| 7486 | 375 | L375E | A | 1.6e-07 | 3.7e-07 | 3.3e-07 | 2.867E-07 | 1.115E-07 | 1.600E-07 | 3.700E-07 |
| 7487 | 375 | L375Q | A | 2.7e-07 | 1.1e-07 | 2.2e-07 | 2.000E-07 | 8.185E-08 | 1.100E-07 | 2.700E-07 |
| 7488 | 375 | L375G | A | 1.1e-07 | 1.8e-07 | 2e-07 | 1.633E-07 | 4.726E-08 | 1.100E-07 | 2.000E-07 |
| 7489 | 375 | L375H | A | 2.2e-07 | 3.1e-07 | 4.4e-07 | 3.233E-07 | 1.106E-07 | 2.200E-07 | 4.400E-07 |
| 7490 | 375 | L375I | A | 1.2e-07 | 1.5e-07 | 3.6e-07 | 2.100E-07 | 1.308E-07 | 1.200E-07 | 3.600E-07 |
| 7491 | 375 | L375L | A | 2.8e-07 | 1.5e-07 | 2.9e-07 | 2.400E-07 | 7.810E-08 | 1.500E-07 | 2.900E-07 |
| 7492 | 375 | L375K | A | 2.1e-07 | 2.1e-07 | 3.6e-07 | 2.600E-07 | 8.660E-08 | 2.100E-07 | 3.600E-07 |
| 7493 | 375 | L375M | A | 2.8e-07 | 3.6e-07 | 1.5e-07 | 2.633E-07 | 1.060E-07 | 1.500E-07 | 3.600E-07 |
| 7494 | 375 | L375F | A | 1.1e-07 | 3.1e-07 | 3.7e-07 | 2.633E-07 | 1.361E-07 | 1.100E-07 | 3.700E-07 |
| 7495 | 375 | L375P | A | 1.5e-07 | 1.7e-07 | 4e-07 | 2.400E-07 | 1.389E-07 | 1.500E-07 | 4.000E-07 |
| 7496 | 375 | L375S | A | 2.6e-07 | 3.3e-07 | 5.7e-07 | 3.867E-07 | 1.626E-07 | 2.600E-07 | 5.700E-07 |
| 7497 | 375 | L375T | A | 1.9e-07 | 5.6e-07 | 2.8e-07 | 3.433E-07 | 1.930E-07 | 1.900E-07 | 5.600E-07 |
| 7498 | 375 | L375W | A | 2.7e-07 | 5e-07 | 3.7e-07 | 3.800E-07 | 1.153E-07 | 2.700E-07 | 5.000E-07 |
| 7499 | 375 | L375Y | A | 3.2e-07 | 2.4e-07 | 2.8e-07 | 2.800E-07 | 4.000E-08 | 2.400E-07 | 3.200E-07 |
| 7500 | 375 | L375V | A | 3.1e-07 | 1e-07 | 4.8e-07 | 2.967E-07 | 1.904E-07 | 1.000E-07 | 4.800E-07 |
| 7501 | 376 | Q376A | A | 1.6e-07 | 2.1e-07 | 2.9e-07 | 2.200E-07 | 6.557E-08 | 1.600E-07 | 2.900E-07 |
| 7502 | 376 | Q376R | A | 1.5e-07 | 2e-07 | 4.8e-07 | 2.767E-07 | 1.779E-07 | 1.500E-07 | 4.800E-07 |
| 7503 | 376 | Q376N | A | 2.5e-07 | 2.8e-07 | 3.6e-07 | 2.967E-07 | 5.686E-08 | 2.500E-07 | 3.600E-07 |
| 7504 | 376 | Q376D | A | 2.1e-07 | 3.1e-07 | 2.5e-07 | 2.567E-07 | 5.033E-08 | 2.100E-07 | 3.100E-07 |
| 7505 | 376 | Q376C | A | 2.8e-07 | 3.9e-07 | 2.9e-07 | 3.200E-07 | 6.083E-08 | 2.800E-07 | 3.900E-07 |
| 7506 | 376 | Q376E | A | 1.4e-07 | 3.4e-07 | 4.2e-07 | 3.000E-07 | 1.442E-07 | 1.400E-07 | 4.200E-07 |
| 7507 | 376 | Q376Q | A | 2.5e-07 | 6.8e-07 | 1.9e-07 | 3.733E-07 | 2.673E-07 | 1.900E-07 | 6.800E-07 |
| 7508 | 376 | Q376G | A | 3.1e-07 | 1.7e-07 | 2e-07 | 2.267E-07 | 7.371E-08 | 1.700E-07 | 3.100E-07 |
| 7509 | 376 | Q376H | A | 1.6e-07 | 8.7e-07 | 2.9e-07 | 4.400E-07 | 3.780E-07 | 1.600E-07 | 8.700E-07 |
| 7510 | 376 | Q376I | A | 4.1e-07 | 1.9e-07 | 4e-07 | 3.333E-07 | 1.242E-07 | 1.900E-07 | 4.100E-07 |
| 7511 | 376 | Q376L | A | 5.4e-07 | 2.6e-07 | 1.7e-07 | 3.233E-07 | 1.930E-07 | 1.700E-07 | 5.400E-07 |
| 7512 | 376 | Q376K | A | 2.3e-07 | 2.5e-07 | 3.6e-07 | 2.800E-07 | 7.000E-08 | 2.300E-07 | 3.600E-07 |
| 7513 | 376 | Q376M | A | 4.5e-07 | 3e-07 | 2.5e-07 | 3.333E-07 | 1.041E-07 | 2.500E-07 | 4.500E-07 |
| 7514 | 376 | Q376F | A | 1.6e-07 | 3.5e-07 | 1.2e-07 | 2.100E-07 | 1.229E-07 | 1.200E-07 | 3.500E-07 |
| 7515 | 376 | Q376P | A | 2.7e-07 | 2e-07 | 2.3e-07 | 2.333E-07 | 3.512E-08 | 2.000E-07 | 2.700E-07 |
| 7516 | 376 | Q376S | A | 2.1e-07 | 1.4e-07 | 3.3e-07 | 2.267E-07 | 9.609E-08 | 1.400E-07 | 3.300E-07 |
| 7517 | 376 | Q376T | A | 3.5e-07 | 3.7e-07 | 3.1e-07 | 3.433E-07 | 3.055E-08 | 3.100E-07 | 3.700E-07 |
| 7518 | 376 | Q376W | A | 2.6e-07 | 1.2e-07 | 2e-07 | 1.933E-07 | 7.024E-08 | 1.200E-07 | 2.600E-07 |
| 7519 | 376 | Q376Y | A | 3.2e-07 | 2.2e-07 | 2.6e-07 | 2.667E-07 | 5.033E-08 | 2.200E-07 | 3.200E-07 |
| 7520 | 376 | Q376V | A | 1.7e-07 | 2e-07 | 4e-07 | 2.567E-07 | 1.250E-07 | 1.700E-07 | 4.000E-07 |
| 7521 | 377 | V377A | A | 7.9e-07 | 1.6e-07 | 3.7e-07 | 4.400E-07 | 3.208E-07 | 1.600E-07 | 7.900E-07 |
| 7522 | 377 | V377R | A | 2.8e-07 | 3.1e-07 | 3.4e-07 | 3.100E-07 | 3.000E-08 | 2.800E-07 | 3.400E-07 |
| 7523 | 377 | V377N | A | 2.1e-07 | 2.1e-07 | 1.5e-07 | 1.900E-07 | 3.464E-08 | 1.500E-07 | 2.100E-07 |
| 7524 | 377 | V377D | A | 3.8e-07 | 2.8e-07 | 2.6e-07 | 3.067E-07 | 6.429E-08 | 2.600E-07 | 3.800E-07 |
| 7525 | 377 | V377C | A | 1.8e-07 | 1.7e-07 | 2.2e-07 | 1.900E-07 | 2.646E-08 | 1.700E-07 | 2.200E-07 |
| 7526 | 377 | V377E | A | 1.1e-07 | 3.7e-07 | 4.4e-07 | 3.067E-07 | 1.739E-07 | 1.100E-07 | 4.400E-07 |
| 7527 | 377 | V377Q | A | 3.8e-07 | 7.3e-07 | 2.5e-07 | 4.533E-07 | 2.483E-07 | 2.500E-07 | 7.300E-07 |
| 7528 | 377 | V377G | A | 2e-07 | 5e-07 | 2.2e-07 | 3.067E-07 | 1.677E-07 | 2.000E-07 | 5.000E-07 |
| 7529 | 377 | V377H | A | 3.9e-07 | 3.9e-07 | 2.6e-07 | 3.467E-07 | 7.506E-08 | 2.600E-07 | 3.900E-07 |
| 7530 | 377 | V377I | A | 2.1e-07 | 2.5e-07 | 1.9e-07 | 2.167E-07 | 3.055E-08 | 1.900E-07 | 2.500E-07 |
| 7531 | 377 | V377L | A | 2.2e-07 | 1.9e-07 | 1.2e-07 | 1.767E-07 | 5.132E-08 | 1.200E-07 | 2.200E-07 |
| 7532 | 377 | V377K | A | 2.9e-07 | 2e-07 | 1.5e-07 | 2.133E-07 | 7.095E-08 | 1.500E-07 | 2.900E-07 |
| 7533 | 377 | V377M | A | 2.7e-07 | 4.4e-07 | 3.6e-07 | 3.567E-07 | 8.505E-08 | 2.700E-07 | 4.400E-07 |
| 7534 | 377 | V377F | A | 8.5e-08 | 3.6e-07 | 3.1e-07 | 2.517E-07 | 1.465E-07 | 8.500E-08 | 3.600E-07 |
| 7535 | 377 | V377P | A | 2e-07 | 1.9e-07 | 3.1e-07 | 2.333E-07 | 6.658E-08 | 1.900E-07 | 3.100E-07 |
| 7536 | 377 | V377S | A | 1.5e-07 | 2.6e-07 | 1.9e-07 | 2.000E-07 | 5.568E-08 | 1.500E-07 | 2.600E-07 |
| 7537 | 377 | V377T | A | 2.6e-07 | 2.6e-07 | 5.1e-07 | 3.433E-07 | 1.443E-07 | 2.600E-07 | 5.100E-07 |
| 7538 | 377 | V377W | A | 2.3e-07 | 2.5e-07 | 1.7e-07 | 2.167E-07 | 4.163E-08 | 1.700E-07 | 2.500E-07 |
| 7539 | 377 | V377Y | A | 2.6e-07 | 2.7e-07 | 2.7e-07 | 2.667E-07 | 5.774E-09 | 2.600E-07 | 2.700E-07 |
| 7540 | 377 | V377V | A | 2.1e-07 | 3.1e-07 | 1.5e-07 | 2.233E-07 | 8.083E-08 | 1.500E-07 | 3.100E-07 |
| 7541 | 378 | F378A | A | 4.7e-07 | 4.3e-07 | 2.6e-07 | 3.867E-07 | 1.115E-07 | 2.600E-07 | 4.700E-07 |
| 7542 | 378 | F378R | A | 1e-06 | 9.6e-08 | 4.4e-07 | 5.120E-07 | 4.563E-07 | 9.600E-08 | 1.000E-06 |
| 7543 | 378 | F378N | A | 4.1e-07 | 1.7e-07 | 2.5e-07 | 2.767E-07 | 1.222E-07 | 1.700E-07 | 4.100E-07 |
| 7544 | 378 | F378D | A | 1.7e-07 | 4.4e-07 | 3e-07 | 3.033E-07 | 1.350E-07 | 1.700E-07 | 4.400E-07 |
| 7545 | 378 | F378C | A | 2.2e-07 | 4.2e-07 | 2e-07 | 2.800E-07 | 1.217E-07 | 2.000E-07 | 4.200E-07 |
| 7546 | 378 | F378E | A | 1.9e-07 | 1.9e-07 | 4.2e-07 | 2.667E-07 | 1.328E-07 | 1.900E-07 | 4.200E-07 |
| 7547 | 378 | F378Q | A | 1.5e-07 | 2.5e-07 | 1.4e-07 | 1.800E-07 | 6.083E-08 | 1.400E-07 | 2.500E-07 |
| 7548 | 378 | F378G | A | 1.8e-07 | 3.1e-07 | 1.2e-07 | 2.033E-07 | 9.713E-08 | 1.200E-07 | 3.100E-07 |
| 7549 | 378 | F378H | A | 2.1e-07 | 1.6e-07 | 2e-07 | 1.900E-07 | 2.646E-08 | 1.600E-07 | 2.100E-07 |
| 7550 | 378 | F378I | A | 1.9e-07 | 2.3e-07 | 3.1e-07 | 2.433E-07 | 6.110E-08 | 1.900E-07 | 3.100E-07 |
| 7551 | 378 | F378L | A | 1.5e-07 | 8.7e-08 | 3.3e-07 | 1.890E-07 | 1.261E-07 | 8.700E-08 | 3.300E-07 |
| 7552 | 378 | F378K | A | 5.3e-07 | 3e-07 | 3.3e-07 | 3.867E-07 | 1.250E-07 | 3.000E-07 | 5.300E-07 |
| 7553 | 378 | F378M | A | 2.3e-07 | 1.8e-07 | 2.5e-07 | 2.200E-07 | 3.606E-08 | 1.800E-07 | 2.500E-07 |
| 7554 | 378 | F378F | A | 5.4e-07 | 1.7e-07 | 1.5e-07 | 2.867E-07 | 2.196E-07 | 1.500E-07 | 5.400E-07 |
| 7555 | 378 | F378P | A | 2.3e-07 | 1.4e-07 | 1e-07 | 1.567E-07 | 6.658E-08 | 1.000E-07 | 2.300E-07 |
| 7556 | 378 | F378S | A | 3.7e-07 | 2e-07 | 3.8e-07 | 3.167E-07 | 1.012E-07 | 2.000E-07 | 3.800E-07 |
| 7557 | 378 | F378T | A | 1.6e-07 | 2.2e-07 | 2.5e-07 | 2.100E-07 | 4.583E-08 | 1.600E-07 | 2.500E-07 |
| 7558 | 378 | F378W | A | 2e-07 | 1.6e-07 | 5.2e-07 | 2.933E-07 | 1.973E-07 | 1.600E-07 | 5.200E-07 |
| 7559 | 378 | F378Y | A | 1.5e-07 | 1.8e-07 | 1.9e-07 | 1.733E-07 | 2.082E-08 | 1.500E-07 | 1.900E-07 |
| 7560 | 378 | F378V | A | 2.7e-07 | 2.9e-07 | 2.3e-07 | 2.633E-07 | 3.055E-08 | 2.300E-07 | 2.900E-07 |
| 7561 | 379 | E379A | A | 1.8e-07 | 2.3e-07 | 6.1e-07 | 3.400E-07 | 2.352E-07 | 1.800E-07 | 6.100E-07 |
| 7562 | 379 | E379R | A | 1e-07 | 4e-07 | 1.3e-07 | 2.100E-07 | 1.652E-07 | 1.000E-07 | 4.000E-07 |
| 7563 | 379 | E379N | A | 1.7e-07 | 4.6e-07 | 9.8e-07 | 5.367E-07 | 4.104E-07 | 1.700E-07 | 9.800E-07 |
| 7564 | 379 | E379D | A | 2.1e-07 | 2.5e-07 | 2.5e-07 | 2.367E-07 | 2.309E-08 | 2.100E-07 | 2.500E-07 |

| | | | | | | | | | | |
|---|---|---|---|---|---|---|---|---|---|---|
| 7565 | 379 | E379C | A | 3.2e-07 | 1.9e-07 | 3.7e-07 | 2.933E-07 | 9.292E-08 | 1.900E-07 | 3.700E-07 |
| 7566 | 379 | E379E | A | 2.8e-07 | 2.4e-07 | 1.8e-07 | 2.333E-07 | 5.033E-08 | 1.800E-07 | 2.800E-07 |
| 7567 | 379 | E379Q | A | 4.1e-07 | 1.3e-07 | 1.6e-07 | 2.333E-07 | 1.537E-07 | 1.300E-07 | 4.100E-07 |
| 7568 | 379 | E379G | A | 3.5e-07 | 2.6e-07 | 3e-07 | 3.033E-07 | 4.509E-08 | 2.600E-07 | 3.500E-07 |
| 7569 | 379 | E379H | A | 1.6e-07 | 1.8e-07 | 2.6e-07 | 2.000E-07 | 5.292E-08 | 1.600E-07 | 2.600E-07 |
| 7570 | 379 | E379I | A | 3.2e-07 | 3.1e-07 | 1.9e-07 | 2.733E-07 | 7.234E-08 | 1.900E-07 | 3.200E-07 |
| 7571 | 379 | E379L | A | 2.5e-07 | 1.5e-07 | 1.8e-07 | 1.933E-07 | 5.132E-08 | 1.500E-07 | 2.500E-07 |
| 7572 | 379 | E379K | A | 2.8e-07 | 2.1e-07 | 5e-07 | 3.300E-07 | 1.513E-07 | 2.100E-07 | 5.000E-07 |
| 7573 | 379 | E379M | A | 2.4e-07 | 1.9e-07 | 5.1e-07 | 3.133E-07 | 1.721E-07 | 1.900E-07 | 5.100E-07 |
| 7574 | 379 | E379F | A | 3e-07 | 2.5e-07 | 2.7e-07 | 2.733E-07 | 2.517E-08 | 2.500E-07 | 3.000E-07 |
| 7575 | 379 | E379P | A | 3.2e-07 | 3.1e-07 | 1.8e-07 | 2.700E-07 | 7.810E-08 | 1.800E-07 | 3.200E-07 |
| 7576 | 379 | E379S | A | 3.4e-07 | 3.1e-07 | 2.7e-07 | 3.067E-07 | 3.512E-08 | 2.700E-07 | 3.400E-07 |
| 7577 | 379 | E379T | A | 2.3e-07 | 7.5e-07 | 1.8e-07 | 3.867E-07 | 3.156E-07 | 1.800E-07 | 7.500E-07 |
| 7578 | 379 | E379W | A | 3.4e-07 | 3.7e-07 | 4e-07 | 3.700E-07 | 3.000E-08 | 3.400E-07 | 4.000E-07 |
| 7579 | 379 | E379Y | A | 6.7e-07 | 3.7e-07 | 2e-07 | 4.133E-07 | 2.380E-07 | 2.000E-07 | 6.700E-07 |
| 7580 | 379 | E379V | A | 1.9e-07 | 4.2e-07 | 3.3e-07 | 3.133E-07 | 1.159E-07 | 1.900E-07 | 4.200E-07 |
| 7581 | 380 | T380A | A | 8e-07 | 2e-07 | 2.8e-07 | 4.267E-07 | 3.258E-07 | 2.000E-07 | 8.000E-07 |
| 7582 | 380 | T380R | A | 1.9e-07 | 2.2e-07 | 2.2e-07 | 2.100E-07 | 1.732E-08 | 1.900E-07 | 2.200E-07 |
| 7583 | 380 | T380N | A | 4e-07 | 1.2e-07 | 2.7e-07 | 2.633E-07 | 1.401E-07 | 1.200E-07 | 4.000E-07 |
| 7584 | 380 | T380D | A | 3.9e-07 | 2.1e-07 | 3.4e-07 | 3.133E-07 | 9.292E-08 | 2.100E-07 | 3.900E-07 |
| 7585 | 380 | T380C | A | 4.7e-07 | 3.6e-07 | 5.8e-07 | 4.700E-07 | 1.100E-07 | 3.600E-07 | 5.800E-07 |
| 7586 | 380 | T380E | A | 4e-07 | 4.9e-07 | 2e-07 | 3.633E-07 | 1.484E-07 | 2.000E-07 | 4.900E-07 |
| 7587 | 380 | T380Q | A | 1e-07 | 2.6e-07 | 1.5e-07 | 1.700E-07 | 8.185E-08 | 1.000E-07 | 2.600E-07 |
| 7588 | 380 | T380G | A | 2.8e-07 | 1.9e-07 | 1.8e-07 | 2.167E-07 | 5.508E-08 | 1.800E-07 | 2.800E-07 |
| 7589 | 380 | T380H | A | 1.3e-07 | 2e-07 | 2.8e-07 | 2.033E-07 | 7.506E-08 | 1.300E-07 | 2.800E-07 |
| 7590 | 380 | T380I | A | 2.2e-07 | 5.4e-07 | 2.2e-07 | 3.267E-07 | 1.848E-07 | 2.200E-07 | 5.400E-07 |
| 7591 | 380 | T380L | A | 2e-07 | 8.5e-08 | 3.1e-07 | 1.983E-07 | 1.125E-07 | 8.500E-08 | 3.100E-07 |
| 7592 | 380 | T380K | A | 2.5e-07 | 2.3e-07 | 1.3e-07 | 2.033E-07 | 6.429E-08 | 1.300E-07 | 2.500E-07 |
| 7593 | 380 | T380M | A | 3.2e-07 | 3.3e-07 | 3e-07 | 3.167E-07 | 1.528E-08 | 3.000E-07 | 3.300E-07 |
| 7594 | 380 | T380F | A | 1.7e-07 | 5.3e-07 | 1.2e-07 | 2.733E-07 | 2.237E-07 | 1.200E-07 | 5.300E-07 |
| 7595 | 380 | T380P | A | 2.4e-07 | 5.8e-07 | 3.7e-07 | 3.967E-07 | 1.716E-07 | 2.400E-07 | 5.800E-07 |
| 7596 | 380 | T380S | A | 3.2e-07 | 1.9e-07 | 5e-07 | 3.367E-07 | 1.557E-07 | 1.900E-07 | 5.000E-07 |
| 7597 | 380 | T380T | A | 2.8e-07 | 1.4e-07 | 1.5e-07 | 1.900E-07 | 7.810E-08 | 1.400E-07 | 2.800E-07 |
| 7598 | 380 | T380W | A | 3.6e-07 | 3.2e-07 | 4.1e-07 | 3.633E-07 | 4.509E-08 | 3.200E-07 | 4.100E-07 |
| 7599 | 380 | T380Y | A | 3.9e-07 | 1.7e-07 | 2.3e-07 | 2.633E-07 | 1.137E-07 | 1.700E-07 | 3.900E-07 |
| 7600 | 380 | T380V | A | 4.5e-07 | 2.1e-07 | 2.5e-07 | 3.033E-07 | 1.286E-07 | 2.100E-07 | 4.500E-07 |
| 7601 | 381 | L381A | A | 2.8e-07 | 3.3e-07 | 5.6e-07 | 3.900E-07 | 1.493E-07 | 2.800E-07 | 5.600E-07 |
| 7602 | 381 | L381R | A | 3.9e-07 | 2.8e-07 | 3.2e-07 | 3.300E-07 | 5.568E-08 | 2.800E-07 | 3.900E-07 |
| 7603 | 381 | L381N | A | 1.6e-07 | 3e-07 | 2.3e-07 | 2.300E-07 | 7.000E-08 | 1.600E-07 | 3.000E-07 |
| 7604 | 381 | L381D | A | 4.9e-07 | 3.3e-07 | 1.8e-07 | 3.333E-07 | 1.550E-07 | 1.800E-07 | 4.900E-07 |
| 7605 | 381 | L381C | A | 3.8e-07 | 2.9e-07 | 7e-07 | 4.567E-07 | 2.155E-07 | 2.900E-07 | 7.000E-07 |
| 7606 | 381 | L381E | A | 2.4e-07 | 4.9e-07 | 1e-07 | 2.767E-07 | 1.976E-07 | 1.000E-07 | 4.900E-07 |
| 7607 | 381 | L381Q | A | 8.9e-08 | 2.8e-07 | 1.7e-07 | 1.797E-07 | 9.587E-08 | 8.900E-08 | 2.800E-07 |
| 7608 | 381 | L381G | A | 3.8e-07 | 2.4e-07 | 3.4e-07 | 3.200E-07 | 7.211E-08 | 2.400E-07 | 3.800E-07 |
| 7609 | 381 | L381H | A | 1.6e-07 | 3.8e-07 | 2.2e-07 | 2.533E-07 | 1.137E-07 | 1.600E-07 | 3.800E-07 |
| 7610 | 381 | L381I | A | 3e-07 | 2e-07 | 3.9e-07 | 2.967E-07 | 9.504E-08 | 2.000E-07 | 3.900E-07 |
| 7611 | 381 | L381L | A | 3e-07 | 2.4e-07 | 1.8e-07 | 2.400E-07 | 6.000E-08 | 1.800E-07 | 3.000E-07 |
| 7612 | 381 | L381K | A | 2.2e-07 | 3.8e-07 | 2.2e-07 | 2.733E-07 | 9.238E-08 | 2.200E-07 | 3.800E-07 |
| 7613 | 381 | L381M | A | 2.8e-07 | 2.1e-07 | 1.5e-07 | 2.133E-07 | 6.506E-08 | 1.500E-07 | 2.800E-07 |
| 7614 | 381 | L381F | A | 1.9e-07 | 2.1e-07 | 2.4e-07 | 2.133E-07 | 2.517E-08 | 1.900E-07 | 2.400E-07 |
| 7615 | 381 | L381P | A | 1.3e-07 | 1.5e-07 | 2.9e-07 | 1.900E-07 | 8.718E-08 | 1.300E-07 | 2.900E-07 |
| 7616 | 381 | L381S | A | 2.5e-07 | 2.5e-07 | 1.5e-07 | 2.167E-07 | 5.774E-08 | 1.500E-07 | 2.500E-07 |
| 7617 | 381 | L381T | A | 5e-07 | 2.3e-07 | 4.1e-07 | 3.800E-07 | 1.375E-07 | 2.300E-07 | 5.000E-07 |
| 7618 | 381 | L381W | A | 3e-07 | 3.6e-07 | 1.8e-07 | 2.800E-07 | 9.165E-08 | 1.800E-07 | 3.600E-07 |
| 7619 | 381 | L381Y | A | 4.2e-07 | 3.7e-07 | 1.5e-07 | 3.133E-07 | 1.436E-07 | 1.500E-07 | 4.200E-07 |
| 7620 | 381 | L381V | A | 3e-07 | 1.2e-07 | 4.3e-07 | 2.833E-07 | 1.557E-07 | 1.200E-07 | 4.300E-07 |
| 7621 | 382 | E382A | A | 4e-07 | 1.7e-07 | 3.4e-07 | 3.033E-07 | 1.193E-07 | 1.700E-07 | 4.000E-07 |
| 7622 | 382 | E382R | A | 3.1e-07 | 3.9e-07 | 2e-07 | 3.000E-07 | 9.539E-08 | 2.000E-07 | 3.900E-07 |
| 7623 | 382 | E382N | A | 3.4e-07 | 1.8e-07 | 2.6e-07 | 2.600E-07 | 8.000E-08 | 1.800E-07 | 3.400E-07 |
| 7624 | 382 | E382D | A | 1.9e-07 | 1.2e-07 | 1.4e-07 | 1.500E-07 | 3.606E-08 | 1.200E-07 | 1.900E-07 |
| 7625 | 382 | E382C | A | 1.3e-07 | 5.7e-07 | 3.2e-07 | 3.400E-07 | 2.207E-07 | 1.300E-07 | 5.700E-07 |
| 7626 | 382 | E382E | A | 2.9e-07 | 3e-07 | 8.6e-08 | 2.253E-07 | 1.208E-07 | 8.600E-08 | 3.000E-07 |
| 7627 | 382 | E382Q | A | 5.3e-07 | 4.1e-07 | 2.5e-07 | 3.967E-07 | 1.405E-07 | 2.500E-07 | 5.300E-07 |
| 7628 | 382 | E382G | A | 3.2e-07 | 2.8e-07 | 3.3e-07 | 3.100E-07 | 2.646E-08 | 2.800E-07 | 3.300E-07 |
| 7629 | 382 | E382H | A | 1.6e-06 | 1.3e-07 | 9.6e-08 | 6.087E-07 | 8.587E-07 | 9.600E-08 | 1.600E-06 |
| 7630 | 382 | E382I | A | 3.4e-07 | 1.1e-07 | 2.4e-07 | 2.300E-07 | 1.153E-07 | 1.100E-07 | 3.400E-07 |
| 7631 | 382 | E382L | A | 2.6e-07 | 4.4e-07 | 1.8e-07 | 2.933E-07 | 1.332E-07 | 1.800E-07 | 4.400E-07 |
| 7632 | 382 | E382K | A | 2.2e-07 | 3e-07 | 1.9e-07 | 2.367E-07 | 5.686E-08 | 1.900E-07 | 3.000E-07 |
| 7633 | 382 | E382M | A | 1.7e-07 | 2.5e-07 | 2.9e-07 | 2.367E-07 | 6.110E-08 | 1.700E-07 | 2.900E-07 |
| 7634 | 382 | E382F | A | 3.8e-07 | 4.5e-07 | 2.9e-07 | 3.733E-07 | 8.021E-08 | 2.900E-07 | 4.500E-07 |
| 7635 | 382 | E382P | A | 2.2e-07 | 6.4e-07 | 2.9e-07 | 3.833E-07 | 2.250E-07 | 2.200E-07 | 6.400E-07 |
| 7636 | 382 | E382S | A | 1.3e-07 | 4.7e-07 | 1.1e-07 | 2.367E-07 | 2.023E-07 | 1.100E-07 | 4.700E-07 |
| 7637 | 382 | E382T | A | 1.8e-07 | 3.2e-07 | 1.4e-07 | 2.133E-07 | 9.452E-08 | 1.400E-07 | 3.200E-07 |
| 7638 | 382 | E382W | A | 1.7e-07 | 3.1e-07 | 2.5e-07 | 2.433E-07 | 7.024E-08 | 1.700E-07 | 3.100E-07 |
| 7639 | 382 | E382Y | A | 2e-07 | 2.1e-07 | 1.5e-07 | 1.867E-07 | 3.215E-08 | 1.500E-07 | 2.100E-07 |
| 7640 | 382 | E382V | A | 3.5e-07 | 2.5e-07 | 3.1e-07 | 3.033E-07 | 5.033E-08 | 2.500E-07 | 3.500E-07 |
| 7641 | 383 | E383A | A | 4.2e-07 | 1.2e-07 | 2.8e-07 | 2.733E-07 | 1.501E-07 | 1.200E-07 | 4.200E-07 |
| 7642 | 383 | E383R | A | 1.6e-07 | 1.2e-07 | 1.7e-07 | 1.500E-07 | 2.646E-08 | 1.200E-07 | 1.700E-07 |
| 7643 | 383 | E383N | A | 2.8e-07 | 1.9e-07 | 9.7e-08 | 1.890E-07 | 9.150E-08 | 9.700E-08 | 2.800E-07 |
| 7644 | 383 | E383D | A | 1.9e-07 | 3.6e-07 | 2.2e-07 | 2.567E-07 | 9.074E-08 | 1.900E-07 | 3.600E-07 |
| 7645 | 383 | E383C | A | 2e-07 | 2.6e-07 | 1.8e-07 | 2.133E-07 | 4.163E-08 | 1.800E-07 | 2.600E-07 |
| 7646 | 383 | E383E | A | 2.2e-07 | 4.1e-07 | 2.6e-07 | 2.967E-07 | 1.002E-07 | 2.200E-07 | 4.100E-07 |
| 7647 | 383 | E383Q | A | 1.5e-07 | 2.1e-07 | 7.2e-07 | 3.600E-07 | 3.132E-07 | 1.500E-07 | 7.200E-07 |
| 7648 | 383 | E383G | A | 1.8e-07 | 3e-07 | 2.2e-07 | 2.333E-07 | 6.110E-08 | 1.800E-07 | 3.000E-07 |
| 7649 | 383 | E383H | A | 6.6e-07 | 2.6e-07 | 2.4e-07 | 3.867E-07 | 2.369E-07 | 2.400E-07 | 6.600E-07 |

| | | | | | | | | | | |
|---|---|---|---|---|---|---|---|---|---|---|
| 7650 | 383 | E383I | A | 1.3e-07 | 2.1e-07 | 3.4e-07 | 2.267E-07 | 1.060E-07 | 1.300E-07 | 3.400E-07 |
| 7651 | 383 | E383L | A | 3.8e-07 | 3.6e-07 | 4e-07 | 3.800E-07 | 2.000E-08 | 3.600E-07 | 4.000E-07 |
| 7652 | 383 | E383K | A | 2.6e-07 | 3.5e-07 | 3.7e-07 | 3.267E-07 | 5.859E-08 | 2.600E-07 | 3.700E-07 |
| 7653 | 383 | E383M | A | 2.1e-07 | 2.2e-07 | 2e-07 | 2.100E-07 | 1.000E-08 | 2.000E-07 | 2.200E-07 |
| 7654 | 383 | E383F | A | 2.9e-07 | 1.7e-07 | 1.6e-07 | 2.067E-07 | 7.234E-08 | 1.600E-07 | 2.900E-07 |
| 7655 | 383 | E383P | A | 2.9e-07 | 6.1e-07 | 4.5e-07 | 4.500E-07 | 1.600E-07 | 2.900E-07 | 6.100E-07 |
| 7656 | 383 | E383S | A | 6.7e-07 | 2.4e-07 | 2.5e-07 | 3.867E-07 | 2.454E-07 | 2.400E-07 | 6.700E-07 |
| 7657 | 383 | E383T | A | 2.4e-07 | 7.5e-07 | 3e-07 | 4.300E-07 | 2.787E-07 | 2.400E-07 | 7.500E-07 |
| 7658 | 383 | E383W | A | 3.7e-07 | 2.5e-07 | 2.9e-07 | 3.033E-07 | 6.110E-08 | 2.500E-07 | 3.700E-07 |
| 7659 | 383 | E383Y | A | 1.8e-07 | 3.6e-07 | 2.6e-07 | 2.667E-07 | 9.018E-08 | 1.800E-07 | 3.600E-07 |
| 7660 | 383 | E383V | A | 2.4e-07 | 2.6e-07 | 4.4e-07 | 3.133E-07 | 1.102E-07 | 2.400E-07 | 4.400E-07 |
| 7661 | 384 | I384A | A | 7.6e-07 | 3.6e-07 | 2.6e-07 | 4.600E-07 | 2.646E-07 | 2.600E-07 | 7.600E-07 |
| 7662 | 384 | I384R | A | 2.3e-07 | 1.2e-07 | 1.3e-07 | 1.600E-07 | 6.083E-08 | 1.200E-07 | 2.300E-07 |
| 7663 | 384 | I384N | A | 2.9e-07 | 3.9e-07 | 3.3e-07 | 3.367E-07 | 5.033E-08 | 2.900E-07 | 3.900E-07 |
| 7664 | 384 | I384D | A | 1.6e-07 | 1.2e-07 | 5.7e-07 | 2.833E-07 | 2.491E-07 | 1.200E-07 | 5.700E-07 |
| 7665 | 384 | I384C | A | 5e-07 | 3.3e-07 | 2.2e-07 | 3.500E-07 | 1.411E-07 | 2.200E-07 | 5.000E-07 |
| 7666 | 384 | I384E | A | 3.7e-07 | 2.6e-07 | 2.6e-07 | 2.967E-07 | 6.351E-08 | 2.600E-07 | 3.700E-07 |
| 7667 | 384 | I384Q | A | 1.9e-07 | 2.4e-07 | 2.5e-07 | 2.267E-07 | 3.215E-08 | 1.900E-07 | 2.500E-07 |
| 7668 | 384 | I384G | A | 2.9e-07 | 2e-07 | 1.9e-07 | 2.267E-07 | 5.508E-08 | 1.900E-07 | 2.900E-07 |
| 7669 | 384 | I384H | A | 2.2e-07 | 4.2e-07 | 2.8e-07 | 3.067E-07 | 1.026E-07 | 2.200E-07 | 4.200E-07 |
| 7670 | 384 | I384I | A | 2.9e-07 | 5.7e-07 | 2.4e-07 | 3.667E-07 | 1.779E-07 | 2.400E-07 | 5.700E-07 |
| 7671 | 384 | I384L | A | 2.5e-07 | 2.6e-07 | 6.5e-07 | 3.867E-07 | 2.281E-07 | 2.500E-07 | 6.500E-07 |
| 7672 | 384 | I384K | A | 3.9e-07 | 1.7e-07 | 1.8e-07 | 2.467E-07 | 1.242E-07 | 1.700E-07 | 3.900E-07 |
| 7673 | 384 | I384M | A | 2.3e-07 | 3e-07 | 3.7e-07 | 3.000E-07 | 7.000E-08 | 2.300E-07 | 3.700E-07 |
| 7674 | 384 | I384F | A | 4e-07 | 2.2e-07 | 1.9e-07 | 2.700E-07 | 1.136E-07 | 1.900E-07 | 4.000E-07 |
| 7675 | 384 | I384P | A | 2.6e-07 | 2.7e-07 | 2.3e-07 | 2.533E-07 | 2.082E-08 | 2.300E-07 | 2.700E-07 |
| 7676 | 384 | I384S | A | 1.9e-07 | 2.3e-07 | 1.6e-07 | 1.933E-07 | 3.512E-08 | 1.600E-07 | 2.300E-07 |
| 7677 | 384 | I384T | A | 2.5e-07 | 4.6e-07 | 3.9e-07 | 3.667E-07 | 1.069E-07 | 2.500E-07 | 4.600E-07 |
| 7678 | 384 | I384W | A | 3.7e-07 | 4.2e-07 | 3.1e-07 | 3.667E-07 | 5.508E-08 | 3.100E-07 | 4.200E-07 |
| 7679 | 384 | I384Y | A | 4.7e-07 | 1.2e-07 | 3.4e-07 | 3.100E-07 | 1.769E-07 | 1.200E-07 | 4.700E-07 |
| 7680 | 384 | I384V | A | 2.5e-07 | 2.3e-07 | 4e-07 | 2.933E-07 | 9.292E-08 | 2.300E-07 | 4.000E-07 |
| 7681 | 385 | T385A | A | 1.8e-07 | 2.5e-07 | 2.7e-07 | 2.333E-07 | 4.726E-08 | 1.800E-07 | 2.700E-07 |
| 7682 | 385 | T385R | A | 1.7e-07 | 1.9e-07 | 3.3e-07 | 2.300E-07 | 8.718E-08 | 1.700E-07 | 3.300E-07 |
| 7683 | 385 | T385N | A | 1.1e-07 | 3.8e-07 | 2.1e-07 | 2.333E-07 | 1.365E-07 | 1.100E-07 | 3.800E-07 |
| 7684 | 385 | T385D | A | 2.5e-07 | 4.9e-07 | 1.6e-07 | 3.000E-07 | 1.706E-07 | 1.600E-07 | 4.900E-07 |
| 7685 | 385 | T385C | A | 3e-07 | 1.1e-07 | 2.3e-07 | 2.133E-07 | 9.609E-08 | 1.100E-07 | 3.000E-07 |
| 7686 | 385 | T385E | A | 2.4e-07 | 2.2e-07 | 2.5e-07 | 2.367E-07 | 1.528E-08 | 2.200E-07 | 2.500E-07 |
| 7687 | 385 | T385Q | A | 2.5e-07 | 1.2e-07 | 3.1e-07 | 2.267E-07 | 9.713E-08 | 1.200E-07 | 3.100E-07 |
| 7688 | 385 | T385G | A | 4.3e-07 | 2.1e-07 | 1.3e-07 | 2.567E-07 | 1.553E-07 | 1.300E-07 | 4.300E-07 |
| 7689 | 385 | T385H | A | 3.1e-07 | 1.3e-07 | 2.9e-07 | 2.433E-07 | 9.866E-08 | 1.300E-07 | 3.100E-07 |
| 7690 | 385 | T385I | A | 1.1e-06 | 2.2e-07 | 2.6e-07 | 5.267E-07 | 4.969E-07 | 2.200E-07 | 1.100E-06 |
| 7691 | 385 | T385L | A | 4.7e-07 | 2.8e-07 | 1.2e-07 | 2.900E-07 | 1.752E-07 | 1.200E-07 | 4.700E-07 |
| 7692 | 385 | T385K | A | 2.7e-07 | 3.8e-07 | 1.5e-07 | 2.667E-07 | 1.150E-07 | 1.500E-07 | 3.800E-07 |
| 7693 | 385 | T385M | A | 3e-07 | 3.2e-07 | 3e-07 | 3.067E-07 | 1.155E-08 | 3.000E-07 | 3.200E-07 |
| 7694 | 385 | T385F | A | 1.6e-07 | 4.4e-07 | 2.5e-07 | 2.833E-07 | 1.429E-07 | 1.600E-07 | 4.400E-07 |
| 7695 | 385 | T385P | A | 3e-07 | 2.9e-07 | 2.7e-07 | 2.867E-07 | 1.528E-08 | 2.700E-07 | 3.000E-07 |
| 7696 | 385 | T385S | A | 1.9e-07 | 2.4e-07 | 2e-07 | 2.100E-07 | 2.646E-08 | 1.900E-07 | 2.400E-07 |
| 7697 | 385 | T385T | A | 2.6e-07 | 3.3e-07 | 2.1e-07 | 2.667E-07 | 6.028E-08 | 2.100E-07 | 3.300E-07 |
| 7698 | 385 | T385W | A | 2e-07 | 5.2e-07 | 1.3e-07 | 2.833E-07 | 2.079E-07 | 1.300E-07 | 5.200E-07 |
| 7699 | 385 | T385Y | A | 2.2e-07 | 3.9e-07 | 2.3e-07 | 2.800E-07 | 9.539E-08 | 2.200E-07 | 3.900E-07 |
| 7700 | 385 | T385V | A | 3.8e-07 | 1.8e-07 | 3.3e-07 | 2.967E-07 | 1.041E-07 | 1.800E-07 | 3.800E-07 |
| 7701 | 386 | G386A | A | 3.5e-07 | 1.8e-07 | 9.5e-07 | 4.933E-07 | 4.045E-07 | 1.800E-07 | 9.500E-07 |
| 7702 | 386 | G386R | A | 1.4e-07 | 2.3e-07 | 2.7e-07 | 2.133E-07 | 6.658E-08 | 1.400E-07 | 2.700E-07 |
| 7703 | 386 | G386N | A | 1.6e-07 | 1.8e-07 | 3.4e-07 | 2.267E-07 | 9.866E-08 | 1.600E-07 | 3.400E-07 |
| 7704 | 386 | G386D | A | 1.7e-07 | 2.2e-07 | 3.5e-07 | 2.467E-07 | 9.292E-08 | 1.700E-07 | 3.500E-07 |
| 7705 | 386 | G386C | A | 1.6e-07 | 1.8e-07 | 3.1e-07 | 2.167E-07 | 8.145E-08 | 1.600E-07 | 3.100E-07 |
| 7706 | 386 | G386E | A | 3.6e-07 | 2.3e-07 | 3.3e-07 | 3.067E-07 | 6.807E-08 | 2.300E-07 | 3.600E-07 |
| 7707 | 386 | G386Q | A | 2.8e-07 | 3.2e-07 | 1.7e-07 | 2.567E-07 | 7.767E-08 | 1.700E-07 | 3.200E-07 |
| 7708 | 386 | G386G | A | 1.7e-07 | 1.8e-07 | 5e-07 | 2.833E-07 | 1.877E-07 | 1.700E-07 | 5.000E-07 |
| 7709 | 386 | G386H | A | 1.6e-07 | 1.1e-07 | 3.1e-07 | 1.933E-07 | 1.041E-07 | 1.100E-07 | 3.100E-07 |
| 7710 | 386 | G386I | A | 1.3e-07 | 2.2e-07 | 1.7e-07 | 1.733E-07 | 4.509E-08 | 1.300E-07 | 2.200E-07 |
| 7711 | 386 | G386L | A | 2.1e-07 | 1.6e-07 | 2.2e-07 | 1.967E-07 | 3.215E-08 | 1.600E-07 | 2.200E-07 |
| 7712 | 386 | G386K | A | 3.2e-07 | 1.9e-07 | 1.6e-07 | 2.233E-07 | 8.505E-08 | 1.600E-07 | 3.200E-07 |
| 7713 | 386 | G386M | A | 1.4e-07 | 3.5e-07 | 3.4e-07 | 2.767E-07 | 1.185E-07 | 1.400E-07 | 3.500E-07 |
| 7714 | 386 | G386F | A | 1.3e-07 | 2.6e-07 | 8.9e-08 | 1.597E-07 | 8.928E-08 | 8.900E-08 | 2.600E-07 |
| 7715 | 386 | G386P | A | 2.4e-07 | 4.2e-07 | 1.9e-07 | 2.833E-07 | 1.210E-07 | 1.900E-07 | 4.200E-07 |
| 7716 | 386 | G386S | A | 6.7e-07 | 2e-07 | 2.2e-07 | 3.633E-07 | 2.658E-07 | 2.000E-07 | 6.700E-07 |
| 7717 | 386 | G386T | A | 2.2e-07 | 3.8e-07 | 1.3e-07 | 2.433E-07 | 1.266E-07 | 1.300E-07 | 3.800E-07 |
| 7718 | 386 | G386W | A | 1.6e-07 | 4.9e-07 | 2.3e-07 | 2.933E-07 | 1.739E-07 | 1.600E-07 | 4.900E-07 |
| 7719 | 386 | G386Y | A | 2.6e-07 | 1.9e-07 | 1.5e-07 | 2.000E-07 | 5.568E-08 | 1.500E-07 | 2.600E-07 |
| 7720 | 386 | G386V | A | 3.7e-07 | 2.1e-07 | 8.9e-08 | 2.230E-07 | 1.410E-07 | 8.900E-08 | 3.700E-07 |
| 7721 | 387 | Y387A | A | 4.6e-07 | 2.5e-07 | 2.5e-07 | 3.200E-07 | 1.212E-07 | 2.500E-07 | 4.600E-07 |
| 7722 | 387 | Y387R | A | 3.4e-07 | 3.1e-07 | 1.9e-07 | 2.800E-07 | 7.937E-08 | 1.900E-07 | 3.400E-07 |
| 7723 | 387 | Y387N | A | 5.3e-08 | 3.2e-07 | 2.3e-07 | 2.010E-07 | 1.358E-07 | 5.300E-08 | 3.200E-07 |
| 7724 | 387 | Y387D | A | 1.7e-07 | 1.4e-07 | 2.9e-07 | 2.000E-07 | 7.937E-08 | 1.400E-07 | 2.900E-07 |
| 7725 | 387 | Y387C | A | 2e-07 | 4.8e-07 | 1.6e-07 | 2.800E-07 | 1.744E-07 | 1.600E-07 | 4.800E-07 |
| 7726 | 387 | Y387E | A | 1.9e-07 | 1.8e-07 | 3.7e-07 | 2.467E-07 | 1.069E-07 | 1.800E-07 | 3.700E-07 |
| 7727 | 387 | Y387Q | A | 5.2e-07 | 2.7e-07 | 2.4e-07 | 3.433E-07 | 1.537E-07 | 2.400E-07 | 5.200E-07 |
| 7728 | 387 | Y387G | A | 3.2e-07 | 1.9e-07 | 2.8e-07 | 2.633E-07 | 6.658E-08 | 1.900E-07 | 3.200E-07 |
| 7729 | 387 | Y387H | A | 4.7e-07 | 4e-07 | 2.2e-07 | 3.633E-07 | 1.290E-07 | 2.200E-07 | 4.700E-07 |
| 7730 | 387 | Y387I | A | 2.1e-07 | 2.3e-07 | 3.1e-07 | 2.500E-07 | 5.292E-08 | 2.100E-07 | 3.100E-07 |
| 7731 | 387 | Y387L | A | 3.1e-07 | 1.9e-07 | 1.8e-07 | 2.267E-07 | 7.234E-08 | 1.800E-07 | 3.100E-07 |
| 7732 | 387 | Y387K | A | 2.3e-07 | 2.8e-07 | 7.1e-07 | 4.067E-07 | 2.639E-07 | 2.300E-07 | 7.100E-07 |
| 7733 | 387 | Y387M | A | 2.8e-07 | 2.1e-07 | 3e-07 | 2.633E-07 | 4.726E-08 | 2.100E-07 | 3.000E-07 |
| 7734 | 387 | Y387F | A | 7.1e-07 | 3.8e-07 | 3.5e-07 | 4.800E-07 | 1.997E-07 | 3.500E-07 | 7.100E-07 |

| | | | | | | | | | | |
|---|---|---|---|---|---|---|---|---|---|---|
| 7735 | 387 | Y387P | A | 2.3e-07 | 4.1e-07 | 5.9e-07 | 4.100E-07 | 1.800E-07 | 2.300E-07 | 5.900E-07 |
| 7736 | 387 | Y387S | A | 1.3e-07 | 2.4e-07 | 2.1e-07 | 1.933E-07 | 5.686E-08 | 1.300E-07 | 2.400E-07 |
| 7737 | 387 | Y387T | A | 3.3e-07 | 5e-07 | 4e-07 | 4.100E-07 | 8.544E-08 | 3.300E-07 | 5.000E-07 |
| 7738 | 387 | Y387W | A | 2.3e-07 | 2.2e-07 | 2.5e-07 | 2.333E-07 | 1.528E-08 | 2.200E-07 | 2.500E-07 |
| 7739 | 387 | Y387Y | A | 8.8e-08 | 1.6e-07 | 2.1e-07 | 1.527E-07 | 6.133E-08 | 8.800E-08 | 2.100E-07 |
| 7740 | 387 | Y387V | A | 3.2e-07 | 2.5e-07 | 1.8e-07 | 2.500E-07 | 7.000E-08 | 1.800E-07 | 3.200E-07 |
| 7741 | 388 | L388A | A | 1.5e-07 | 3e-07 | 2.6e-07 | 2.367E-07 | 7.767E-08 | 1.500E-07 | 3.000E-07 |
| 7742 | 388 | L388R | A | 3.4e-07 | 2.3e-07 | 2.3e-07 | 2.667E-07 | 6.351E-08 | 2.300E-07 | 3.400E-07 |
| 7743 | 388 | L388N | A | 2.6e-07 | 2.9e-07 | 1.4e-07 | 2.300E-07 | 7.937E-08 | 1.400E-07 | 2.900E-07 |
| 7744 | 388 | L388D | A | 3.2e-07 | 4.7e-07 | 1.8e-07 | 3.233E-07 | 1.450E-07 | 1.800E-07 | 4.700E-07 |
| 7745 | 388 | L388C | A | 2.8e-07 | 2.4e-07 | 2.8e-07 | 2.667E-07 | 2.309E-08 | 2.400E-07 | 2.800E-07 |
| 7746 | 388 | L388E | A | 2.3e-07 | 2.2e-07 | 2.6e-07 | 2.367E-07 | 2.082E-08 | 2.200E-07 | 2.600E-07 |
| 7747 | 388 | L388Q | A | 1.4e-07 | 3e-07 | 1.9e-07 | 2.100E-07 | 8.185E-08 | 1.400E-07 | 3.000E-07 |
| 7748 | 388 | L388G | A | 5.5e-07 | 2.4e-07 | 3.7e-07 | 3.867E-07 | 1.557E-07 | 2.400E-07 | 5.500E-07 |
| 7749 | 388 | L388H | A | 2.1e-07 | 3.2e-07 | 1e-07 | 2.100E-07 | 1.100E-07 | 1.000E-07 | 3.200E-07 |
| 7750 | 388 | L388I | A | 2.2e-07 | 1.3e-07 | 6e-07 | 3.167E-07 | 2.495E-07 | 1.300E-07 | 6.000E-07 |
| 7751 | 388 | L388L | A | 2e-07 | 6.5e-07 | 3.4e-07 | 3.967E-07 | 2.303E-07 | 2.000E-07 | 6.500E-07 |
| 7752 | 388 | L388K | A | 2.1e-07 | 3e-07 | 1.9e-07 | 2.333E-07 | 5.859E-08 | 1.900E-07 | 3.000E-07 |
| 7753 | 388 | L388M | A | 3e-07 | 3.3e-07 | 3.8e-07 | 3.367E-07 | 4.041E-08 | 3.000E-07 | 3.800E-07 |
| 7754 | 388 | L388F | A | 2.5e-07 | 2.2e-07 | 4e-07 | 2.900E-07 | 9.644E-08 | 2.200E-07 | 4.000E-07 |
| 7755 | 388 | L388P | A | 1.8e-07 | 3.2e-07 | 2.9e-07 | 2.633E-07 | 7.371E-08 | 1.800E-07 | 3.200E-07 |
| 7756 | 388 | L388S | A | 1.1e-07 | 1.4e-07 | 1.9e-07 | 1.467E-07 | 4.041E-08 | 1.100E-07 | 1.900E-07 |
| 7757 | 388 | L388T | A | 2.9e-07 | 2.1e-07 | 3.3e-07 | 2.767E-07 | 6.110E-08 | 2.100E-07 | 3.300E-07 |
| 7758 | 388 | L388W | A | 2.6e-07 | 1.6e-07 | 2.6e-07 | 2.267E-07 | 5.774E-08 | 1.600E-07 | 2.600E-07 |
| 7759 | 388 | L388Y | A | 2.5e-07 | 1.6e-07 | 2.4e-07 | 2.167E-07 | 4.933E-08 | 1.600E-07 | 2.500E-07 |
| 7760 | 388 | L388V | A | 2.2e-07 | 1.4e-07 | 2.7e-07 | 2.100E-07 | 6.557E-08 | 1.400E-07 | 2.700E-07 |
| 7761 | 389 | Y389A | A | 2.3e-07 | 3.4e-07 | 3.1e-07 | 2.933E-07 | 5.686E-08 | 2.300E-07 | 3.400E-07 |
| 7762 | 389 | Y389R | A | 2.2e-07 | 3.2e-07 | 3.2e-07 | 2.867E-07 | 5.774E-08 | 2.200E-07 | 3.200E-07 |
| 7763 | 389 | Y389N | A | 3e-07 | 2.2e-07 | 2.1e-07 | 2.433E-07 | 4.933E-08 | 2.100E-07 | 3.000E-07 |
| 7764 | 389 | Y389D | A | 5.4e-07 | 1.9e-07 | 1.8e-07 | 3.033E-07 | 2.050E-07 | 1.800E-07 | 5.400E-07 |
| 7765 | 389 | Y389C | A | 4.9e-07 | 3.1e-07 | 1.9e-07 | 3.300E-07 | 1.510E-07 | 1.900E-07 | 4.900E-07 |
| 7766 | 389 | Y389E | A | 2.7e-07 | 1.9e-07 | 5.3e-07 | 3.300E-07 | 1.778E-07 | 1.900E-07 | 5.300E-07 |
| 7767 | 389 | Y389Q | A | 3.8e-07 | 2.6e-07 | 4.2e-07 | 3.533E-07 | 8.327E-08 | 2.600E-07 | 4.200E-07 |
| 7768 | 389 | Y389G | A | 2e-07 | 3.7e-07 | 1.9e-07 | 2.533E-07 | 1.012E-07 | 1.900E-07 | 3.700E-07 |
| 7769 | 389 | Y389H | A | 1.9e-07 | 2e-07 | 2.7e-07 | 2.200E-07 | 4.359E-08 | 1.900E-07 | 2.700E-07 |
| 7770 | 389 | Y389I | A | 8.6e-08 | 3.6e-07 | 3.7e-07 | 2.720E-07 | 1.612E-07 | 8.600E-08 | 3.700E-07 |
| 7771 | 389 | Y389L | A | 2.7e-07 | 5.6e-07 | 2.4e-07 | 3.567E-07 | 1.767E-07 | 2.400E-07 | 5.600E-07 |
| 7772 | 389 | Y389K | A | 2.7e-07 | 2.7e-07 | 2.9e-07 | 2.767E-07 | 1.155E-08 | 2.700E-07 | 2.900E-07 |
| 7773 | 389 | Y389M | A | 6.5e-07 | 2.8e-07 | 2.1e-07 | 3.800E-07 | 2.364E-07 | 2.100E-07 | 6.500E-07 |
| 7774 | 389 | Y389F | A | 4.6e-07 | 1.7e-07 | 1.1e-07 | 2.467E-07 | 1.872E-07 | 1.100E-07 | 4.600E-07 |
| 7775 | 389 | Y389P | A | 4.3e-07 | 3.9e-07 | 2.8e-07 | 3.667E-07 | 7.767E-08 | 2.800E-07 | 4.300E-07 |
| 7776 | 389 | Y389S | A | 3.7e-07 | 3.4e-07 | 2.6e-07 | 3.233E-07 | 5.686E-08 | 2.600E-07 | 3.700E-07 |
| 7777 | 389 | Y389T | A | 3.1e-07 | 2.1e-07 | 1.2e-07 | 2.133E-07 | 9.504E-08 | 1.200E-07 | 3.100E-07 |
| 7778 | 389 | Y389W | A | 4.4e-07 | 1.9e-07 | 2.8e-07 | 3.033E-07 | 1.266E-07 | 1.900E-07 | 4.400E-07 |
| 7779 | 389 | Y389Y | A | 2.8e-07 | 2e-07 | 3.1e-07 | 2.633E-07 | 5.686E-08 | 2.000E-07 | 3.100E-07 |
| 7780 | 389 | Y389V | A | 2.3e-07 | 3e-07 | 5.3e-07 | 3.533E-07 | 1.570E-07 | 2.300E-07 | 5.300E-07 |
| 7781 | 390 | I390A | A | 3.6e-07 | 4.8e-07 | 3.8e-07 | 4.067E-07 | 6.429E-08 | 3.600E-07 | 4.800E-07 |
| 7782 | 390 | I390R | A | 2.2e-07 | 3.9e-07 | 3.8e-07 | 3.300E-07 | 9.539E-08 | 2.200E-07 | 3.900E-07 |
| 7783 | 390 | I390N | A | 3.8e-07 | 1.6e-07 | 1.3e-07 | 2.233E-07 | 1.365E-07 | 1.300E-07 | 3.800E-07 |
| 7784 | 390 | I390D | A | 2.1e-07 | 3.2e-07 | 2.6e-07 | 2.633E-07 | 5.508E-08 | 2.100E-07 | 3.200E-07 |
| 7785 | 390 | I390C | A | 2.4e-07 | 2.8e-07 | 4.5e-07 | 3.233E-07 | 1.115E-07 | 2.400E-07 | 4.500E-07 |
| 7786 | 390 | I390E | A | 2.5e-07 | 1.6e-07 | 3.5e-07 | 2.533E-07 | 9.504E-08 | 1.600E-07 | 3.500E-07 |
| 7787 | 390 | I390Q | A | 3e-07 | 1.5e-07 | 3.4e-07 | 2.633E-07 | 1.002E-07 | 1.500E-07 | 3.400E-07 |
| 7788 | 390 | I390G | A | 3.1e-07 | 2.4e-07 | 2e-07 | 2.500E-07 | 5.568E-08 | 2.000E-07 | 3.100E-07 |
| 7789 | 390 | I390H | A | 2.8e-07 | 4.2e-07 | 3.7e-07 | 3.567E-07 | 7.095E-08 | 2.800E-07 | 4.200E-07 |
| 7790 | 390 | I390I | A | 4.3e-07 | 2.4e-07 | 2e-07 | 2.900E-07 | 1.229E-07 | 2.000E-07 | 4.300E-07 |
| 7791 | 390 | I390L | A | 5.8e-07 | 1.6e-07 | 4.3e-07 | 3.900E-07 | 2.128E-07 | 1.600E-07 | 5.800E-07 |
| 7792 | 390 | I390K | A | 2e-07 | 3.3e-07 | 1.4e-07 | 2.233E-07 | 9.713E-08 | 1.400E-07 | 3.300E-07 |
| 7793 | 390 | I390M | A | 2e-07 | 1.5e-07 | 2.9e-07 | 2.133E-07 | 7.095E-08 | 1.500E-07 | 2.900E-07 |
| 7794 | 390 | I390F | A | 1.5e-07 | 1.6e-07 | 4.5e-07 | 2.533E-07 | 1.704E-07 | 1.500E-07 | 4.500E-07 |
| 7795 | 390 | I390P | A | 3.2e-07 | 2.8e-07 | 1.2e-07 | 2.400E-07 | 1.058E-07 | 1.200E-07 | 3.200E-07 |
| 7796 | 390 | I390S | A | 2.1e-07 | 1.8e-07 | 1.9e-07 | 1.933E-07 | 1.528E-08 | 1.800E-07 | 2.100E-07 |
| 7797 | 390 | I390T | A | 3.3e-07 | 2.5e-07 | 2.9e-07 | 2.900E-07 | 4.000E-08 | 2.500E-07 | 3.300E-07 |
| 7798 | 390 | I390W | A | 2.6e-07 | 3e-07 | 2.8e-07 | 2.800E-07 | 2.000E-08 | 2.600E-07 | 3.000E-07 |
| 7799 | 390 | I390Y | A | 2.3e-07 | 5.3e-07 | 1.4e-07 | 3.000E-07 | 2.042E-07 | 1.400E-07 | 5.300E-07 |
| 7800 | 390 | I390V | A | 1.8e-07 | 2.5e-07 | 1.3e-07 | 1.867E-07 | 6.028E-08 | 1.300E-07 | 2.500E-07 |
| 7801 | 391 | S391A | A | 2.2e-07 | 2.4e-07 | 5.1e-07 | 3.233E-07 | 1.620E-07 | 2.200E-07 | 5.100E-07 |
| 7802 | 391 | S391R | A | 2.2e-07 | 2.6e-07 | 2e-07 | 2.267E-07 | 3.055E-08 | 2.000E-07 | 2.600E-07 |
| 7803 | 391 | S391N | A | 2e-07 | 2e-07 | 2.3e-07 | 2.100E-07 | 1.732E-08 | 2.000E-07 | 2.300E-07 |
| 7804 | 391 | S391D | A | 3e-07 | 2.3e-07 | 1.7e-07 | 2.333E-07 | 6.506E-08 | 1.700E-07 | 3.000E-07 |
| 7805 | 391 | S391C | A | 2e-07 | 2.8e-07 | 3e-07 | 2.600E-07 | 5.292E-08 | 2.000E-07 | 3.000E-07 |
| 7806 | 391 | S391E | A | 1.9e-07 | 1.1e-07 | 5.1e-07 | 2.700E-07 | 2.117E-07 | 1.100E-07 | 5.100E-07 |
| 7807 | 391 | S391Q | A | 6.5e-08 | 1.7e-07 | 1.8e-07 | 1.383E-07 | 6.371E-08 | 6.500E-08 | 1.800E-07 |
| 7808 | 391 | S391G | A | 2.4e-07 | 2.9e-07 | 2.4e-07 | 2.567E-07 | 2.887E-08 | 2.400E-07 | 2.900E-07 |
| 7809 | 391 | S391H | A | 5.1e-07 | 2.8e-07 | 2.5e-07 | 3.467E-07 | 1.422E-07 | 2.500E-07 | 5.100E-07 |
| 7810 | 391 | S391I | A | 1.4e-07 | 2.4e-07 | 3.8e-07 | 2.533E-07 | 1.206E-07 | 1.400E-07 | 3.800E-07 |
| 7811 | 391 | S391L | A | 5.1e-07 | 2.3e-07 | 1.7e-07 | 3.033E-07 | 1.815E-07 | 1.700E-07 | 5.100E-07 |
| 7812 | 391 | S391K | A | 2.4e-07 | 3.5e-07 | 2e-07 | 2.633E-07 | 7.767E-08 | 2.000E-07 | 3.500E-07 |
| 7813 | 391 | S391M | A | 2.6e-07 | 2.8e-07 | 1.6e-07 | 2.333E-07 | 6.429E-08 | 1.600E-07 | 2.800E-07 |
| 7814 | 391 | S391F | A | 7e-08 | 1.5e-07 | 1.5e-07 | 1.233E-07 | 4.619E-08 | 7.000E-08 | 1.500E-07 |
| 7815 | 391 | S391P | A | 2.4e-07 | 2e-07 | 3.6e-07 | 2.667E-07 | 8.327E-08 | 2.000E-07 | 3.600E-07 |
| 7816 | 391 | S391S | A | 4.2e-07 | 3.1e-07 | 4.1e-07 | 3.800E-07 | 6.083E-08 | 3.100E-07 | 4.200E-07 |
| 7817 | 391 | S391T | A | 2.1e-07 | 3.1e-07 | 1.5e-07 | 2.233E-07 | 8.083E-08 | 1.500E-07 | 3.100E-07 |
| 7818 | 391 | S391W | A | 4e-07 | 2.8e-07 | 1.7e-07 | 2.833E-07 | 1.150E-07 | 1.700E-07 | 4.000E-07 |
| 7819 | 391 | S391Y | A | 2.1e-07 | 2.3e-07 | 1.4e-07 | 1.933E-07 | 4.726E-08 | 1.400E-07 | 2.300E-07 |

| | | | | | | | | | | |
|---|---|---|---|---|---|---|---|---|---|---|
| 7820 | 391 | S391V | A | 2.2e-07 | 2.8e-07 | 3.8e-07 | 2.933E-07 | 8.083E-08 | 2.200E-07 | 3.800E-07 |
| 7821 | 392 | A392A | A | 3.3e-07 | 3.2e-07 | 3.5e-07 | 3.333E-07 | 1.528E-08 | 3.200E-07 | 3.500E-07 |
| 7822 | 392 | A392R | A | 8.8e-08 | 2.7e-07 | 2.7e-07 | 2.093E-07 | 1.051E-07 | 8.800E-08 | 2.700E-07 |
| 7823 | 392 | A392N | A | 2.1e-07 | 1.3e-07 | 3.8e-07 | 2.400E-07 | 1.277E-07 | 1.300E-07 | 3.800E-07 |
| 7824 | 392 | A392D | A | 1.5e-07 | 2.9e-07 | 8.9e-08 | 1.763E-07 | 1.031E-07 | 8.900E-08 | 2.900E-07 |
| 7825 | 392 | A392C | A | 3.3e-07 | 2.9e-07 | 2.8e-07 | 3.000E-07 | 2.646E-08 | 2.800E-07 | 3.300E-07 |
| 7826 | 392 | A392E | A | 3.4e-07 | 7.5e-07 | 2.8e-07 | 4.567E-07 | 2.558E-07 | 2.800E-07 | 7.500E-07 |
| 7827 | 392 | A392Q | A | 2.9e-07 | 5.6e-07 | 9.9e-08 | 3.163E-07 | 2.316E-07 | 9.900E-08 | 5.600E-07 |
| 7828 | 392 | A392G | A | 1.5e-07 | 1.6e-07 | 2.5e-07 | 1.867E-07 | 5.508E-08 | 1.500E-07 | 2.500E-07 |
| 7829 | 392 | A392H | A | 1.5e-07 | 1.7e-07 | 3.8e-07 | 2.333E-07 | 1.274E-07 | 1.500E-07 | 3.800E-07 |
| 7830 | 392 | A392I | A | 1.7e-07 | 2.6e-07 | 2.6e-07 | 2.300E-07 | 5.196E-08 | 1.700E-07 | 2.600E-07 |
| 7831 | 392 | A392L | A | 1.1e-07 | 2.1e-07 | 4.1e-07 | 2.433E-07 | 1.528E-07 | 1.100E-07 | 4.100E-07 |
| 7832 | 392 | A392K | A | 2.6e-07 | 1.5e-07 | 2.9e-07 | 2.333E-07 | 7.371E-08 | 1.500E-07 | 2.900E-07 |
| 7833 | 392 | A392M | A | 1.3e-07 | 2.2e-07 | 3.3e-07 | 2.267E-07 | 1.002E-07 | 1.300E-07 | 3.300E-07 |
| 7834 | 392 | A392F | A | 1.2e-07 | 2.4e-07 | 3.4e-07 | 2.333E-07 | 1.102E-07 | 1.200E-07 | 3.400E-07 |
| 7835 | 392 | A392P | A | 2.6e-07 | 2.4e-07 | 3.9e-07 | 2.967E-07 | 8.145E-08 | 2.400E-07 | 3.900E-07 |
| 7836 | 392 | A392S | A | 1.7e-07 | 1.7e-07 | 7.2e-07 | 3.533E-07 | 3.175E-07 | 1.700E-07 | 7.200E-07 |
| 7837 | 392 | A392T | A | 3.7e-07 | 2.5e-07 | 1.6e-07 | 2.600E-07 | 1.054E-07 | 1.600E-07 | 3.700E-07 |
| 7838 | 392 | A392W | A | 3.8e-07 | 1e-07 | 3e-07 | 2.600E-07 | 1.442E-07 | 1.000E-07 | 3.800E-07 |
| 7839 | 392 | A392Y | A | 1.7e-07 | 2.8e-07 | 3.5e-07 | 2.667E-07 | 9.074E-08 | 1.700E-07 | 3.500E-07 |
| 7840 | 392 | A392V | A | 1.4e-07 | 1.5e-07 | 2.6e-07 | 1.833E-07 | 6.658E-08 | 1.400E-07 | 2.600E-07 |
| 7841 | 393 | W393A | A | 1.8e-07 | 2.9e-07 | 3.4e-07 | 2.700E-07 | 8.185E-08 | 1.800E-07 | 3.400E-07 |
| 7842 | 393 | W393R | A | 3.3e-07 | 2.9e-07 | 4.1e-07 | 3.433E-07 | 6.110E-08 | 2.900E-07 | 4.100E-07 |
| 7843 | 393 | W393N | A | 2.8e-07 | 1.5e-07 | 5.2e-07 | 3.167E-07 | 1.877E-07 | 1.500E-07 | 5.200E-07 |
| 7844 | 393 | W393D | A | 2.3e-07 | 2.2e-07 | 1.3e-07 | 1.933E-07 | 5.508E-08 | 1.300E-07 | 2.300E-07 |
| 7845 | 393 | W393C | A | 3.3e-07 | 3e-07 | 1.6e-07 | 2.633E-07 | 9.074E-08 | 1.600E-07 | 3.300E-07 |
| 7846 | 393 | W393E | A | 2.9e-07 | 2.8e-07 | 1.9e-07 | 2.533E-07 | 5.508E-08 | 1.900E-07 | 2.900E-07 |
| 7847 | 393 | W393Q | A | 1.9e-07 | 5.2e-07 | 1.7e-07 | 2.933E-07 | 1.966E-07 | 1.700E-07 | 5.200E-07 |
| 7848 | 393 | W393G | A | 3.5e-07 | 2.2e-07 | 2.7e-07 | 2.800E-07 | 6.557E-08 | 2.200E-07 | 3.500E-07 |
| 7849 | 393 | W393H | A | 6.2e-07 | 2.4e-07 | 2.7e-07 | 3.767E-07 | 2.113E-07 | 2.400E-07 | 6.200E-07 |
| 7850 | 393 | W393I | A | 1.6e-07 | 4.3e-07 | 3.5e-07 | 3.133E-07 | 1.387E-07 | 1.600E-07 | 4.300E-07 |
| 7851 | 393 | W393L | A | 3.2e-07 | 2.2e-07 | 2.8e-07 | 2.733E-07 | 5.033E-08 | 2.200E-07 | 3.200E-07 |
| 7852 | 393 | W393K | A | 2.1e-07 | 7.9e-07 | 1.8e-07 | 3.933E-07 | 3.439E-07 | 1.800E-07 | 7.900E-07 |
| 7853 | 393 | W393M | A | 3.8e-07 | 1.4e-07 | 2.9e-07 | 2.700E-07 | 1.212E-07 | 1.400E-07 | 3.800E-07 |
| 7854 | 393 | W393F | A | 8.6e-08 | 2e-07 | 3e-07 | 1.953E-07 | 1.071E-07 | 8.600E-08 | 3.000E-07 |
| 7855 | 393 | W393P | A | 3.7e-07 | 2.3e-07 | 3.9e-07 | 3.300E-07 | 8.718E-08 | 2.300E-07 | 3.900E-07 |
| 7856 | 393 | W393S | A | 5.1e-07 | 3.2e-07 | 1.4e-07 | 3.233E-07 | 1.850E-07 | 1.400E-07 | 5.100E-07 |
| 7857 | 393 | W393T | A | 2.6e-07 | 5.2e-07 | 2.1e-07 | 3.300E-07 | 1.664E-07 | 2.100E-07 | 5.200E-07 |
| 7858 | 393 | W393W | A | 2.4e-07 | 5.9e-07 | 1.5e-07 | 3.267E-07 | 2.325E-07 | 1.500E-07 | 5.900E-07 |
| 7859 | 393 | W393Y | A | 9.2e-08 | 2.4e-07 | 3e-07 | 2.107E-07 | 1.071E-07 | 9.200E-08 | 3.000E-07 |
| 7860 | 393 | W393V | A | 3.4e-07 | 1.4e-07 | 2.2e-07 | 2.333E-07 | 1.007E-07 | 1.400E-07 | 3.400E-07 |
| 7861 | 394 | P394A | A | 4.1e-07 | 2.7e-07 | 3.9e-07 | 3.567E-07 | 7.572E-08 | 2.700E-07 | 4.100E-07 |
| 7862 | 394 | P394R | A | 1.1e-07 | 8.8e-07 | 2.5e-07 | 4.133E-07 | 4.102E-07 | 1.100E-07 | 8.800E-07 |
| 7863 | 394 | P394N | A | 3e-07 | 2.6e-07 | 1.6e-07 | 2.400E-07 | 7.211E-08 | 1.600E-07 | 3.000E-07 |
| 7864 | 394 | P394D | A | 3.4e-07 | 1.5e-07 | 3.2e-07 | 2.700E-07 | 1.044E-07 | 1.500E-07 | 3.400E-07 |
| 7865 | 394 | P394C | A | 5.5e-07 | 2.4e-07 | 2.5e-07 | 3.467E-07 | 1.762E-07 | 2.400E-07 | 5.500E-07 |
| 7866 | 394 | P394E | A | 3.9e-07 | 1.6e-07 | 2.2e-07 | 2.567E-07 | 1.193E-07 | 1.600E-07 | 3.900E-07 |
| 7867 | 394 | P394Q | A | 2.5e-07 | 2.7e-07 | 3.9e-07 | 3.033E-07 | 7.572E-08 | 2.500E-07 | 3.900E-07 |
| 7868 | 394 | P394G | A | 6.2e-07 | 3e-07 | 2.6e-07 | 3.933E-07 | 1.973E-07 | 2.600E-07 | 6.200E-07 |
| 7869 | 394 | P394H | A | 3.3e-07 | 3.3e-07 | 3.9e-07 | 3.500E-07 | 3.464E-08 | 3.300E-07 | 3.900E-07 |
| 7870 | 394 | P394I | A | 2.8e-07 | 1.6e-07 | 4.5e-07 | 2.967E-07 | 1.457E-07 | 1.600E-07 | 4.500E-07 |
| 7871 | 394 | P394L | A | 1.6e-07 | 2.3e-07 | 2e-07 | 1.967E-07 | 3.512E-08 | 1.600E-07 | 2.300E-07 |
| 7872 | 394 | P394K | A | 1.6e-07 | 1.5e-07 | 1.8e-07 | 1.633E-07 | 1.528E-08 | 1.500E-07 | 1.800E-07 |
| 7873 | 394 | P394M | A | 2.2e-07 | 5e-07 | 1.5e-07 | 2.900E-07 | 1.852E-07 | 1.500E-07 | 5.000E-07 |
| 7874 | 394 | P394F | A | 3e-07 | 2.9e-07 | 1.3e-07 | 2.400E-07 | 9.539E-08 | 1.300E-07 | 3.000E-07 |
| 7875 | 394 | P394P | A | 1.4e-07 | 1.9e-07 | 1.6e-07 | 1.633E-07 | 2.517E-08 | 1.400E-07 | 1.900E-07 |
| 7876 | 394 | P394S | A | 1.6e-07 | 2.6e-07 | 2.4e-07 | 2.200E-07 | 5.292E-08 | 1.600E-07 | 2.600E-07 |
| 7877 | 394 | P394T | A | 3.9e-07 | 3.5e-07 | 1.4e-07 | 2.933E-07 | 1.343E-07 | 1.400E-07 | 3.900E-07 |
| 7878 | 394 | P394W | A | 1.5e-07 | 2.3e-07 | 2.7e-07 | 2.167E-07 | 6.110E-08 | 1.500E-07 | 2.700E-07 |
| 7879 | 394 | P394Y | A | 9.8e-08 | 2.4e-07 | 2.6e-07 | 1.993E-07 | 8.833E-08 | 9.800E-08 | 2.600E-07 |
| 7880 | 394 | P394V | A | 7.3e-07 | 1.9e-07 | 2.7e-07 | 3.967E-07 | 2.914E-07 | 1.900E-07 | 7.300E-07 |
| 7881 | 395 | D395A | A | 1.4e-07 | 1.6e-07 | 1.3e-07 | 1.433E-07 | 1.528E-08 | 1.300E-07 | 1.600E-07 |
| 7882 | 395 | D395R | A | 2.8e-07 | 2.5e-07 | 1.5e-07 | 2.267E-07 | 6.807E-08 | 1.500E-07 | 2.800E-07 |
| 7883 | 395 | D395N | A | 1.2e-07 | 3.9e-07 | 1.9e-07 | 2.333E-07 | 1.401E-07 | 1.200E-07 | 3.900E-07 |
| 7884 | 395 | D395D | A | 2e-07 | 4.9e-07 | 2.4e-07 | 3.100E-07 | 1.572E-07 | 2.000E-07 | 4.900E-07 |
| 7885 | 395 | D395C | A | 2.5e-07 | 2.2e-07 | 4.3e-07 | 3.000E-07 | 1.136E-07 | 2.200E-07 | 4.300E-07 |
| 7886 | 395 | D395E | A | 1.1e-07 | 3.1e-07 | 2.7e-07 | 2.300E-07 | 1.058E-07 | 1.100E-07 | 3.100E-07 |
| 7887 | 395 | D395Q | A | 3.1e-07 | 4.6e-07 | 4.6e-07 | 4.100E-07 | 8.660E-08 | 3.100E-07 | 4.600E-07 |
| 7888 | 395 | D395G | A | 1.6e-07 | 3.1e-07 | 3.4e-07 | 2.700E-07 | 9.644E-08 | 1.600E-07 | 3.400E-07 |
| 7889 | 395 | D395H | A | 1.5e-07 | 3.7e-07 | 1.7e-07 | 2.300E-07 | 1.217E-07 | 1.500E-07 | 3.700E-07 |
| 7890 | 395 | D395I | A | 1.5e-07 | 3.7e-07 | 2.6e-07 | 2.600E-07 | 1.100E-07 | 1.500E-07 | 3.700E-07 |
| 7891 | 395 | D395L | A | 1.2e-07 | 2e-07 | 3.1e-07 | 2.100E-07 | 9.539E-08 | 1.200E-07 | 3.100E-07 |
| 7892 | 395 | D395K | A | 2.2e-07 | 2.1e-07 | 2.8e-07 | 2.367E-07 | 3.786E-08 | 2.100E-07 | 2.800E-07 |
| 7893 | 395 | D395M | A | 4.1e-07 | 3.9e-07 | 2.3e-07 | 3.433E-07 | 9.866E-08 | 2.300E-07 | 4.100E-07 |
| 7894 | 395 | D395F | A | 3.8e-07 | 3.4e-07 | 2.5e-07 | 3.233E-07 | 6.658E-08 | 2.500E-07 | 3.800E-07 |
| 7895 | 395 | D395P | A | 3.7e-07 | 2.3e-07 | 3.1e-07 | 3.033E-07 | 7.024E-08 | 2.300E-07 | 3.700E-07 |
| 7896 | 395 | D395S | A | 2.4e-07 | 3.1e-07 | 9.7e-08 | 2.157E-07 | 1.086E-07 | 9.700E-08 | 3.100E-07 |
| 7897 | 395 | D395T | A | 2.6e-07 | 2.8e-07 | 1.9e-07 | 2.433E-07 | 4.726E-08 | 1.900E-07 | 2.800E-07 |
| 7898 | 395 | D395W | A | 2.3e-07 | 1.6e-07 | 1.7e-07 | 1.867E-07 | 3.786E-08 | 1.600E-07 | 2.300E-07 |
| 7899 | 395 | D395Y | A | 5.1e-07 | 3e-07 | 2.2e-07 | 3.433E-07 | 1.498E-07 | 2.200E-07 | 5.100E-07 |
| 7900 | 395 | D395V | A | 3.6e-07 | 2.2e-07 | 4.3e-07 | 3.367E-07 | 1.069E-07 | 2.200E-07 | 4.300E-07 |
| 7901 | 396 | S396A | A | 2.5e-07 | 3.9e-07 | 2.3e-07 | 2.900E-07 | 8.718E-08 | 2.300E-07 | 3.900E-07 |
| 7902 | 396 | S396R | A | 3.8e-07 | 2.2e-07 | 2.8e-07 | 2.933E-07 | 8.083E-08 | 2.200E-07 | 3.800E-07 |
| 7903 | 396 | S396N | A | 2e-07 | 3.7e-07 | 4.8e-07 | 3.500E-07 | 1.411E-07 | 2.000E-07 | 4.800E-07 |
| 7904 | 396 | S396D | A | 3.2e-07 | 2.3e-07 | 2e-07 | 2.500E-07 | 6.245E-08 | 2.000E-07 | 3.200E-07 |

| | | | | | | | | | | |
|---|---|---|---|---|---|---|---|---|---|---|
| 7905 | 396 | S396C | A | 2.4e-07 | 1e-07 | 3.3e-07 | 2.233E-07 | 1.159E-07 | 1.000E-07 | 3.300E-07 |
| 7906 | 396 | S396E | A | 1.1e-06 | 1.7e-07 | 1.1e-07 | 4.600E-07 | 5.551E-07 | 1.100E-07 | 1.100E-06 |
| 7907 | 396 | S396Q | A | 1.9e-07 | 1.8e-07 | 3.1e-07 | 2.267E-07 | 7.234E-08 | 1.800E-07 | 3.100E-07 |
| 7908 | 396 | S396G | A | 2.6e-07 | 2.4e-07 | 2.1e-07 | 2.367E-07 | 2.517E-08 | 2.100E-07 | 2.600E-07 |
| 7909 | 396 | S396H | A | 3.2e-07 | 1.7e-07 | 2.1e-07 | 2.333E-07 | 7.767E-08 | 1.700E-07 | 3.200E-07 |
| 7910 | 396 | S396I | A | 3.3e-07 | 3.7e-07 | 3.3e-07 | 3.433E-07 | 2.309E-08 | 3.300E-07 | 3.700E-07 |
| 7911 | 396 | S396L | A | 1.6e-07 | 2.2e-07 | 1.6e-07 | 1.800E-07 | 3.464E-08 | 1.600E-07 | 2.200E-07 |
| 7912 | 396 | S396K | A | 2.5e-07 | 3.6e-07 | 2e-07 | 2.700E-07 | 8.185E-08 | 2.000E-07 | 3.600E-07 |
| 7913 | 396 | S396M | A | 3.7e-07 | 2.8e-07 | 3e-07 | 3.167E-07 | 4.726E-08 | 2.800E-07 | 3.700E-07 |
| 7914 | 396 | S396F | A | 2.2e-07 | 2.3e-07 | 2e-07 | 2.167E-07 | 1.528E-08 | 2.000E-07 | 2.300E-07 |
| 7915 | 396 | S396P | A | 6.3e-07 | 1.8e-07 | 1.2e-07 | 3.100E-07 | 2.787E-07 | 1.200E-07 | 6.300E-07 |
| 7916 | 396 | S396S | A | 4.7e-07 | 3.8e-07 | 3.2e-07 | 3.900E-07 | 7.550E-08 | 3.200E-07 | 4.700E-07 |
| 7917 | 396 | S396T | A | 3.1e-07 | 2.7e-07 | 2e-07 | 2.600E-07 | 5.568E-08 | 2.000E-07 | 3.100E-07 |
| 7918 | 396 | S396W | A | 1.9e-07 | 2.9e-07 | 1.9e-07 | 2.233E-07 | 5.774E-08 | 1.900E-07 | 2.900E-07 |
| 7919 | 396 | S396Y | A | 1.7e-07 | 2.2e-07 | 2.5e-07 | 2.133E-07 | 4.041E-08 | 1.700E-07 | 2.500E-07 |
| 7920 | 396 | S396V | A | 2.4e-07 | 2.8e-07 | 3.5e-07 | 2.900E-07 | 5.568E-08 | 2.400E-07 | 3.500E-07 |
| 7921 | 397 | L397A | A | 2.2e-07 | 2e-07 | 2.6e-07 | 2.267E-07 | 3.055E-08 | 2.000E-07 | 2.600E-07 |
| 7922 | 397 | L397R | A | 2.2e-07 | 3.2e-07 | 3.4e-07 | 2.933E-07 | 6.429E-08 | 2.200E-07 | 3.400E-07 |
| 7923 | 397 | L397N | A | 5.4e-07 | 4.7e-07 | 1.3e-07 | 3.800E-07 | 2.193E-07 | 1.300E-07 | 5.400E-07 |
| 7924 | 397 | L397D | A | 4.4e-07 | 5.4e-07 | 1.1e-07 | 3.633E-07 | 2.250E-07 | 1.100E-07 | 5.400E-07 |
| 7925 | 397 | L397C | A | 1.2e-07 | 1.6e-07 | 2.1e-07 | 1.633E-07 | 4.509E-08 | 1.200E-07 | 2.100E-07 |
| 7926 | 397 | L397E | A | 1.9e-07 | 1.4e-07 | 1.7e-07 | 1.667E-07 | 2.517E-08 | 1.400E-07 | 1.900E-07 |
| 7927 | 397 | L397Q | A | 2.5e-07 | 1.4e-07 | 2.5e-07 | 2.133E-07 | 6.351E-08 | 1.400E-07 | 2.500E-07 |
| 7928 | 397 | L397G | A | 2.9e-07 | 5.9e-07 | 1.7e-07 | 3.500E-07 | 2.163E-07 | 1.700E-07 | 5.900E-07 |
| 7929 | 397 | L397H | A | 2.3e-07 | 4.9e-08 | 1.9e-07 | 1.563E-07 | 9.508E-08 | 4.900E-08 | 2.300E-07 |
| 7930 | 397 | L397I | A | 1.9e-07 | 1.8e-07 | 1.9e-07 | 1.867E-07 | 5.774E-09 | 1.800E-07 | 1.900E-07 |
| 7931 | 397 | L397L | A | 3.2e-07 | 2.4e-07 | 2.8e-07 | 2.800E-07 | 4.000E-08 | 2.400E-07 | 3.200E-07 |
| 7932 | 397 | L397K | A | 2.6e-07 | 1.8e-07 | 3.3e-07 | 2.567E-07 | 7.506E-08 | 1.800E-07 | 3.300E-07 |
| 7933 | 397 | L397M | A | 3.8e-07 | 1.3e-07 | 2.6e-07 | 2.567E-07 | 1.250E-07 | 1.300E-07 | 3.800E-07 |
| 7934 | 397 | L397F | A | 2.4e-07 | 1.2e-07 | 1.7e-07 | 1.767E-07 | 6.028E-08 | 1.200E-07 | 2.400E-07 |
| 7935 | 397 | L397P | A | 3.2e-07 | 2.3e-07 | 4.2e-07 | 3.233E-07 | 9.504E-08 | 2.300E-07 | 4.200E-07 |
| 7936 | 397 | L397S | A | 1.6e-07 | 3.8e-07 | 2.6e-07 | 2.667E-07 | 1.102E-07 | 1.600E-07 | 3.800E-07 |
| 7937 | 397 | L397T | A | 2e-07 | 6e-07 | 2.3e-07 | 3.433E-07 | 2.228E-07 | 2.000E-07 | 6.000E-07 |
| 7938 | 397 | L397W | A | 4e-07 | 2.2e-07 | 2.2e-07 | 2.800E-07 | 1.039E-07 | 2.200E-07 | 4.000E-07 |
| 7939 | 397 | L397Y | A | 2.3e-07 | 2.7e-07 | 3e-07 | 2.667E-07 | 3.512E-08 | 2.300E-07 | 3.000E-07 |
| 7940 | 397 | L397V | A | 1.4e-07 | 1.9e-07 | 3.3e-07 | 2.200E-07 | 9.849E-08 | 1.400E-07 | 3.300E-07 |
| 7941 | 398 | P398A | A | 1.9e-07 | 1e-07 | 4.3e-07 | 2.400E-07 | 1.706E-07 | 1.000E-07 | 4.300E-07 |
| 7942 | 398 | P398R | A | 2.9e-07 | 2.7e-07 | 3.9e-07 | 3.167E-07 | 6.429E-08 | 2.700E-07 | 3.900E-07 |
| 7943 | 398 | P398N | A | 1.2e-07 | 1.6e-07 | 2.1e-07 | 1.633E-07 | 4.509E-08 | 1.200E-07 | 2.100E-07 |
| 7944 | 398 | P398D | A | 2.7e-07 | 2.1e-07 | 4e-07 | 2.933E-07 | 9.713E-08 | 2.100E-07 | 4.000E-07 |
| 7945 | 398 | P398C | A | 1e-06 | 2.7e-07 | 5.3e-07 | 6.000E-07 | 3.700E-07 | 2.700E-07 | 1.000E-06 |
| 7946 | 398 | P398E | A | 3.9e-07 | 2.1e-07 | 1.2e-07 | 2.400E-07 | 1.375E-07 | 1.200E-07 | 3.900E-07 |
| 7947 | 398 | P398Q | A | 2.2e-07 | 1.2e-07 | 1.9e-07 | 1.767E-07 | 5.132E-08 | 1.200E-07 | 2.200E-07 |
| 7948 | 398 | P398G | A | 4.2e-07 | 1.8e-07 | 1.8e-07 | 2.600E-07 | 1.386E-07 | 1.800E-07 | 4.200E-07 |
| 7949 | 398 | P398H | A | 1.7e-07 | 1.5e-07 | 2.7e-07 | 1.967E-07 | 6.429E-08 | 1.500E-07 | 2.700E-07 |
| 7950 | 398 | P398I | A | 8.5e-08 | 2.4e-07 | 1.2e-07 | 1.483E-07 | 8.129E-08 | 8.500E-08 | 2.400E-07 |
| 7951 | 398 | P398L | A | 2.9e-07 | 1.6e-07 | 2.6e-07 | 2.367E-07 | 6.807E-08 | 1.600E-07 | 2.900E-07 |
| 7952 | 398 | P398K | A | 1.8e-07 | 2.2e-07 | 1.8e-07 | 1.933E-07 | 2.309E-08 | 1.800E-07 | 2.200E-07 |
| 7953 | 398 | P398M | A | 2.5e-07 | 2.3e-07 | 2.1e-07 | 2.300E-07 | 2.000E-08 | 2.100E-07 | 2.500E-07 |
| 7954 | 398 | P398F | A | 1.8e-07 | 2e-07 | 2.1e-07 | 1.967E-07 | 1.528E-08 | 1.800E-07 | 2.100E-07 |
| 7955 | 398 | P398P | A | 2.3e-07 | 3.2e-07 | 5.6e-07 | 3.700E-07 | 1.706E-07 | 2.300E-07 | 5.600E-07 |
| 7956 | 398 | P398S | A | 4.6e-07 | 2.3e-07 | 1.6e-07 | 2.833E-07 | 1.570E-07 | 1.600E-07 | 4.600E-07 |
| 7957 | 398 | P398T | A | 1.8e-07 | 5.8e-07 | 1.2e-07 | 2.933E-07 | 2.501E-07 | 1.200E-07 | 5.800E-07 |
| 7958 | 398 | P398W | A | 3.6e-07 | 1.6e-07 | 5.2e-07 | 3.467E-07 | 1.804E-07 | 1.600E-07 | 5.200E-07 |
| 7959 | 398 | P398Y | A | 2e-07 | 2.4e-07 | 2.8e-07 | 2.400E-07 | 4.000E-08 | 2.000E-07 | 2.800E-07 |
| 7960 | 398 | P398V | A | 1.9e-07 | 2.2e-07 | 2.6e-07 | 2.233E-07 | 3.512E-08 | 1.900E-07 | 2.600E-07 |
| 7961 | 399 | D399A | A | 2.1e-07 | 3.6e-07 | 1.7e-07 | 2.467E-07 | 1.002E-07 | 1.700E-07 | 3.600E-07 |
| 7962 | 399 | D399R | A | 1.9e-07 | 1.4e-07 | 1.3e-07 | 1.533E-07 | 3.215E-08 | 1.300E-07 | 1.900E-07 |
| 7963 | 399 | D399N | A | 1.8e-07 | 3.9e-07 | 1.8e-07 | 2.500E-07 | 1.212E-07 | 1.800E-07 | 3.900E-07 |
| 7964 | 399 | D399D | A | 4.3e-07 | 2.6e-07 | 2.6e-07 | 3.167E-07 | 9.815E-08 | 2.600E-07 | 4.300E-07 |
| 7965 | 399 | D399C | A | 1.5e-07 | 9.8e-08 | 1.7e-07 | 1.393E-07 | 3.717E-08 | 9.800E-08 | 1.700E-07 |
| 7966 | 399 | D399E | A | 5.8e-07 | 3.4e-07 | 1.2e-07 | 3.467E-07 | 2.301E-07 | 1.200E-07 | 5.800E-07 |
| 7967 | 399 | D399Q | A | 2e-07 | 4.6e-07 | 2.1e-07 | 2.900E-07 | 1.473E-07 | 2.000E-07 | 4.600E-07 |
| 7968 | 399 | D399G | A | 8.9e-08 | 4.4e-07 | 2.5e-07 | 2.597E-07 | 1.757E-07 | 8.900E-08 | 4.400E-07 |
| 7969 | 399 | D399H | A | 3.1e-07 | 1.2e-07 | 4.4e-07 | 2.900E-07 | 1.609E-07 | 1.200E-07 | 4.400E-07 |
| 7970 | 399 | D399I | A | 3.5e-07 | 2.7e-07 | 1.7e-07 | 2.633E-07 | 9.018E-08 | 1.700E-07 | 3.500E-07 |
| 7971 | 399 | D399L | A | 2.5e-07 | 2.8e-07 | 4.6e-07 | 3.300E-07 | 1.136E-07 | 2.500E-07 | 4.600E-07 |
| 7972 | 399 | D399K | A | 4.7e-07 | 1.8e-07 | 2.1e-07 | 2.867E-07 | 1.595E-07 | 1.800E-07 | 4.700E-07 |
| 7973 | 399 | D399M | A | 3.8e-07 | 2.7e-07 | 2.4e-07 | 2.967E-07 | 7.371E-08 | 2.400E-07 | 3.800E-07 |
| 7974 | 399 | D399F | A | 2.1e-07 | 2e-07 | 9e-08 | 1.667E-07 | 6.658E-08 | 9.000E-08 | 2.100E-07 |
| 7975 | 399 | D399P | A | 2.4e-07 | 1.1e-07 | 2.5e-07 | 2.000E-07 | 7.810E-08 | 1.100E-07 | 2.500E-07 |
| 7976 | 399 | D399S | A | 3.3e-07 | 2.8e-07 | 3e-07 | 3.033E-07 | 2.517E-08 | 2.800E-07 | 3.300E-07 |
| 7977 | 399 | D399T | A | 3e-07 | 2.1e-07 | 2.5e-07 | 2.533E-07 | 4.509E-08 | 2.100E-07 | 3.000E-07 |
| 7978 | 399 | D399W | A | 9.6e-08 | 1.5e-07 | 2.4e-07 | 1.620E-07 | 7.275E-08 | 9.600E-08 | 2.400E-07 |
| 7979 | 399 | D399Y | A | 5.4e-07 | 2.2e-07 | 2.2e-07 | 3.267E-07 | 1.848E-07 | 2.200E-07 | 5.400E-07 |
| 7980 | 399 | D399V | A | 2.6e-07 | 4.9e-07 | 6.2e-07 | 4.567E-07 | 1.823E-07 | 2.600E-07 | 6.200E-07 |
| 7981 | 400 | L400A | A | 3.6e-07 | 2.3e-07 | 2e-07 | 2.633E-07 | 8.505E-08 | 2.000E-07 | 3.600E-07 |
| 7982 | 400 | L400R | A | 1.2e-07 | 1.3e-07 | 1.9e-07 | 1.467E-07 | 3.786E-08 | 1.200E-07 | 1.900E-07 |
| 7983 | 400 | L400N | A | 2.3e-07 | 3e-07 | 3e-07 | 2.767E-07 | 4.041E-08 | 2.300E-07 | 3.000E-07 |
| 7984 | 400 | L400D | A | 3.1e-07 | 5.6e-07 | 2.4e-07 | 3.700E-07 | 1.682E-07 | 2.400E-07 | 5.600E-07 |
| 7985 | 400 | L400C | A | 1.1e-07 | 3.2e-07 | 3e-07 | 2.433E-07 | 1.159E-07 | 1.100E-07 | 3.200E-07 |
| 7986 | 400 | L400E | A | 2.6e-07 | 2.8e-07 | 2.7e-07 | 2.700E-07 | 1.000E-08 | 2.600E-07 | 2.800E-07 |
| 7987 | 400 | L400Q | A | 2.1e-07 | 1.9e-07 | 2.3e-07 | 2.100E-07 | 2.000E-08 | 1.900E-07 | 2.300E-07 |
| 7988 | 400 | L400G | A | 4.7e-07 | 2.3e-07 | 6.7e-07 | 4.567E-07 | 2.203E-07 | 2.300E-07 | 6.700E-07 |
| 7989 | 400 | L400H | A | 3.3e-07 | 1.9e-07 | 4.4e-07 | 3.200E-07 | 1.253E-07 | 1.900E-07 | 4.400E-07 |

| | | | | | | | | | | |
|---|---|---|---|---|---|---|---|---|---|---|
| 7990 | 400 | L400I | A | 7.1e-07 | 2.2e-07 | 2.5e-07 | 3.933E-07 | 2.747E-07 | 2.200E-07 | 7.100E-07 |
| 7991 | 400 | L400L | A | 1.1e-07 | 2.6e-07 | 2.6e-07 | 2.100E-07 | 8.660E-08 | 1.100E-07 | 2.600E-07 |
| 7992 | 400 | L400K | A | 1e-07 | 1.6e-07 | 2e-07 | 1.533E-07 | 5.033E-08 | 1.000E-07 | 2.000E-07 |
| 7993 | 400 | L400M | A | 2.8e-07 | 2.7e-07 | 2.7e-07 | 2.733E-07 | 5.774E-09 | 2.700E-07 | 2.800E-07 |
| 7994 | 400 | L400F | A | 2.8e-07 | 2e-07 | 1.6e-07 | 2.133E-07 | 6.110E-08 | 1.600E-07 | 2.800E-07 |
| 7995 | 400 | L400P | A | 2.8e-07 | 3.7e-07 | 2.4e-07 | 2.967E-07 | 6.658E-08 | 2.400E-07 | 3.700E-07 |
| 7996 | 400 | L400S | A | 2.7e-07 | 3.9e-07 | 1.7e-07 | 2.767E-07 | 1.102E-07 | 1.700E-07 | 3.900E-07 |
| 7997 | 400 | L400T | A | 2.5e-07 | 7.8e-07 | 2.2e-07 | 4.167E-07 | 3.150E-07 | 2.200E-07 | 7.800E-07 |
| 7998 | 400 | L400W | A | 3e-07 | 1.5e-07 | 1.1e-07 | 1.867E-07 | 1.002E-07 | 1.100E-07 | 3.000E-07 |
| 7999 | 400 | L400Y | A | 2.9e-07 | 1.7e-07 | 3.8e-07 | 2.800E-07 | 1.054E-07 | 1.700E-07 | 3.800E-07 |
| 8000 | 400 | L400V | A | 1.9e-07 | 8.9e-07 | 1e-07 | 3.933E-07 | 4.325E-07 | 1.000E-07 | 8.900E-07 |
| 8001 | 401 | S401A | A | 3.8e-07 | 3.5e-07 | 1.5e-07 | 2.933E-07 | 1.250E-07 | 1.500E-07 | 3.800E-07 |
| 8002 | 401 | S401R | A | 5.5e-07 | 3.2e-07 | 4e-07 | 4.233E-07 | 1.168E-07 | 3.200E-07 | 5.500E-07 |
| 8003 | 401 | S401N | A | 1.8e-07 | 3e-07 | 1.8e-07 | 2.200E-07 | 6.928E-08 | 1.800E-07 | 3.000E-07 |
| 8004 | 401 | S401D | A | 5.1e-07 | 1.6e-07 | 3.5e-07 | 3.400E-07 | 1.752E-07 | 1.600E-07 | 5.100E-07 |
| 8005 | 401 | S401C | A | 3.4e-07 | 2.4e-07 | 1.7e-07 | 2.500E-07 | 8.544E-08 | 1.700E-07 | 3.400E-07 |
| 8006 | 401 | S401E | A | 3.8e-07 | 2e-07 | 2.9e-07 | 2.900E-07 | 9.000E-08 | 2.000E-07 | 3.800E-07 |
| 8007 | 401 | S401Q | A | 2.1e-07 | 2.1e-07 | 1.8e-07 | 2.000E-07 | 1.732E-08 | 1.800E-07 | 2.100E-07 |
| 8008 | 401 | S401G | A | 2.7e-07 | 2.8e-07 | 7.7e-07 | 4.400E-07 | 2.858E-07 | 2.700E-07 | 7.700E-07 |
| 8009 | 401 | S401H | A | 2e-07 | 2.9e-07 | 3.5e-07 | 2.800E-07 | 7.550E-08 | 2.000E-07 | 3.500E-07 |
| 8010 | 401 | S401I | A | 3.9e-07 | 2.3e-07 | 1.3e-07 | 2.500E-07 | 1.311E-07 | 1.300E-07 | 3.900E-07 |
| 8011 | 401 | S401L | A | 2.1e-07 | 2.4e-07 | 2.5e-07 | 2.333E-07 | 2.082E-08 | 2.100E-07 | 2.500E-07 |
| 8012 | 401 | S401K | A | 1.4e-07 | 2.5e-07 | 5.3e-07 | 3.067E-07 | 2.011E-07 | 1.400E-07 | 5.300E-07 |
| 8013 | 401 | S401M | A | 2.1e-07 | 1.2e-07 | 2.7e-07 | 2.000E-07 | 7.550E-08 | 1.200E-07 | 2.700E-07 |
| 8014 | 401 | S401F | A | 1.8e-07 | 1.4e-07 | 1.7e-07 | 1.633E-07 | 2.082E-08 | 1.400E-07 | 1.800E-07 |
| 8015 | 401 | S401P | A | 3.5e-07 | 2.3e-07 | 1.3e-07 | 2.367E-07 | 1.102E-07 | 1.300E-07 | 3.500E-07 |
| 8016 | 401 | S401S | A | 3.9e-07 | 4e-07 | 3.2e-07 | 3.700E-07 | 4.359E-08 | 3.200E-07 | 4.000E-07 |
| 8017 | 401 | S401T | A | 2.2e-07 | 1.5e-07 | 3.5e-07 | 2.400E-07 | 1.015E-07 | 1.500E-07 | 3.500E-07 |
| 8018 | 401 | S401W | A | 1.8e-07 | 1.7e-07 | 1.7e-07 | 1.733E-07 | 5.774E-09 | 1.700E-07 | 1.800E-07 |
| 8019 | 401 | S401Y | A | 1.7e-07 | 2.4e-07 | 2.2e-07 | 2.100E-07 | 3.606E-08 | 1.700E-07 | 2.400E-07 |
| 8020 | 401 | S401V | A | 1.7e-07 | 1.4e-07 | 4.1e-07 | 2.400E-07 | 1.480E-07 | 1.400E-07 | 4.100E-07 |
| 8021 | 402 | V402A | A | 4.8e-07 | 3.6e-07 | 1.3e-07 | 3.233E-07 | 1.779E-07 | 1.300E-07 | 4.800E-07 |
| 8022 | 402 | V402R | A | 3.6e-07 | 1.7e-07 | 5.8e-07 | 3.700E-07 | 2.052E-07 | 1.700E-07 | 5.800E-07 |
| 8023 | 402 | V402N | A | 1.5e-07 | 2.4e-07 | 8.2e-07 | 4.033E-07 | 3.636E-07 | 1.500E-07 | 8.200E-07 |
| 8024 | 402 | V402D | A | 3e-07 | 3e-07 | 1.4e-07 | 2.467E-07 | 9.238E-08 | 1.400E-07 | 3.000E-07 |
| 8025 | 402 | V402C | A | 3.5e-07 | 1e-07 | 1.7e-07 | 2.067E-07 | 1.290E-07 | 1.000E-07 | 3.500E-07 |
| 8026 | 402 | V402E | A | 3.1e-07 | 3.2e-07 | 3.7e-07 | 3.333E-07 | 3.215E-08 | 3.100E-07 | 3.700E-07 |
| 8027 | 402 | V402Q | A | 3e-07 | 6.6e-07 | 2.9e-07 | 4.167E-07 | 2.108E-07 | 2.900E-07 | 6.600E-07 |
| 8028 | 402 | V402G | A | 4.2e-07 | 4e-07 | 6.6e-07 | 4.933E-07 | 1.447E-07 | 4.000E-07 | 6.600E-07 |
| 8029 | 402 | V402H | A | 4.8e-07 | 2.7e-07 | 4e-07 | 3.833E-07 | 1.060E-07 | 2.700E-07 | 4.800E-07 |
| 8030 | 402 | V402I | A | 6.6e-07 | 3e-07 | 1.8e-07 | 3.800E-07 | 2.498E-07 | 1.800E-07 | 6.600E-07 |
| 8031 | 402 | V402L | A | 3.4e-07 | 3.4e-07 | 1.2e-07 | 2.667E-07 | 1.270E-07 | 1.200E-07 | 3.400E-07 |
| 8032 | 402 | V402K | A | 1.5e-07 | 3.1e-07 | 1.6e-07 | 2.067E-07 | 8.963E-08 | 1.500E-07 | 3.100E-07 |
| 8033 | 402 | V402M | A | 2.1e-07 | 4.5e-07 | 2e-07 | 2.867E-07 | 1.415E-07 | 2.000E-07 | 4.500E-07 |
| 8034 | 402 | V402F | A | 2.1e-07 | 4.4e-07 | 3.4e-07 | 3.300E-07 | 1.153E-07 | 2.100E-07 | 4.400E-07 |
| 8035 | 402 | V402P | A | 3.1e-07 | 3.2e-07 | 1.8e-07 | 2.700E-07 | 7.810E-08 | 1.800E-07 | 3.200E-07 |
| 8036 | 402 | V402S | A | 1.1e-07 | 1.3e-07 | 3.2e-07 | 1.867E-07 | 1.159E-07 | 1.100E-07 | 3.200E-07 |
| 8037 | 402 | V402T | A | 2.5e-07 | 2.7e-07 | 2.6e-07 | 2.600E-07 | 1.000E-08 | 2.500E-07 | 2.700E-07 |
| 8038 | 402 | V402W | A | 4.2e-07 | 1.1e-07 | 4.2e-07 | 3.167E-07 | 1.790E-07 | 1.100E-07 | 4.200E-07 |
| 8039 | 402 | V402Y | A | 2.3e-07 | 3.4e-07 | 2.5e-07 | 2.733E-07 | 5.859E-08 | 2.300E-07 | 3.400E-07 |
| 8040 | 402 | V402V | A | 2.9e-07 | 2.9e-07 | 1.3e-07 | 2.367E-07 | 9.238E-08 | 1.300E-07 | 2.900E-07 |
| 8041 | 403 | F403A | A | 5.4e-07 | 2e-07 | 4e-07 | 3.800E-07 | 1.709E-07 | 2.000E-07 | 5.400E-07 |
| 8042 | 403 | F403R | A | 2e-07 | 2.2e-07 | 1.6e-07 | 1.933E-07 | 3.055E-08 | 1.600E-07 | 2.200E-07 |
| 8043 | 403 | F403N | A | 3e-07 | 2.5e-07 | 2.9e-07 | 2.800E-07 | 2.646E-08 | 2.500E-07 | 3.000E-07 |
| 8044 | 403 | F403D | A | 2.6e-07 | 1.8e-07 | 2.4e-07 | 2.267E-07 | 4.163E-08 | 1.800E-07 | 2.600E-07 |
| 8045 | 403 | F403C | A | 4.8e-07 | 1.2e-07 | 2.5e-07 | 2.833E-07 | 1.823E-07 | 1.200E-07 | 4.800E-07 |
| 8046 | 403 | F403E | A | 4.8e-07 | 3.8e-07 | 2.6e-07 | 3.733E-07 | 1.102E-07 | 2.600E-07 | 4.800E-07 |
| 8047 | 403 | F403Q | A | 3.2e-07 | 8.5e-08 | 3e-07 | 2.350E-07 | 1.303E-07 | 8.500E-08 | 3.200E-07 |
| 8048 | 403 | F403G | A | 2.6e-07 | 1.9e-07 | 4.4e-07 | 2.967E-07 | 1.290E-07 | 1.900E-07 | 4.400E-07 |
| 8049 | 403 | F403H | A | 1.6e-07 | 2.4e-07 | 1.3e-07 | 1.767E-07 | 5.686E-08 | 1.300E-07 | 2.400E-07 |
| 8050 | 403 | F403I | A | 2.6e-07 | 1.7e-07 | 3.2e-07 | 2.500E-07 | 7.550E-08 | 1.700E-07 | 3.200E-07 |
| 8051 | 403 | F403L | A | 5.3e-07 | 1.7e-07 | 3.5e-07 | 3.500E-07 | 1.800E-07 | 1.700E-07 | 5.300E-07 |
| 8052 | 403 | F403K | A | 1.9e-07 | 3.1e-07 | 2.8e-07 | 2.600E-07 | 6.245E-08 | 1.900E-07 | 3.100E-07 |
| 8053 | 403 | F403M | A | 1.9e-07 | 2.5e-07 | 2e-07 | 2.133E-07 | 3.215E-08 | 1.900E-07 | 2.500E-07 |
| 8054 | 403 | F403F | A | 2.5e-07 | 2e-07 | 2.2e-07 | 2.233E-07 | 2.517E-08 | 2.000E-07 | 2.500E-07 |
| 8055 | 403 | F403P | A | 4.5e-07 | 2.1e-07 | 3.6e-07 | 3.400E-07 | 1.212E-07 | 2.100E-07 | 4.500E-07 |
| 8056 | 403 | F403S | A | 3.3e-07 | 5.9e-07 | 2.5e-07 | 3.900E-07 | 1.778E-07 | 2.500E-07 | 5.900E-07 |
| 8057 | 403 | F403T | A | 2.5e-07 | 2.1e-07 | 3.3e-07 | 2.633E-07 | 6.110E-08 | 2.100E-07 | 3.300E-07 |
| 8058 | 403 | F403W | A | 3.2e-07 | 1.2e-07 | 3.7e-07 | 2.700E-07 | 1.323E-07 | 1.200E-07 | 3.700E-07 |
| 8059 | 403 | F403Y | A | 2.2e-07 | 3.4e-07 | 5.6e-07 | 3.733E-07 | 1.724E-07 | 2.200E-07 | 5.600E-07 |
| 8060 | 403 | F403V | A | 9.7e-08 | 3.1e-07 | 3.4e-07 | 2.490E-07 | 1.325E-07 | 9.700E-08 | 3.400E-07 |
| 8061 | 404 | Q404A | A | 3.8e-07 | 4.3e-07 | 4.2e-07 | 4.100E-07 | 2.646E-08 | 3.800E-07 | 4.300E-07 |
| 8062 | 404 | Q404R | A | 4.1e-07 | 3e-07 | 3.2e-07 | 3.433E-07 | 5.859E-08 | 3.000E-07 | 4.100E-07 |
| 8063 | 404 | Q404N | A | 1.6e-07 | 4.8e-07 | 1.3e-07 | 2.567E-07 | 1.940E-07 | 1.300E-07 | 4.800E-07 |
| 8064 | 404 | Q404D | A | 1.2e-07 | 3.9e-07 | 3.1e-07 | 2.733E-07 | 1.387E-07 | 1.200E-07 | 3.900E-07 |
| 8065 | 404 | Q404C | A | 1.6e-07 | 1.7e-07 | 2.7e-07 | 2.000E-07 | 6.083E-08 | 1.600E-07 | 2.700E-07 |
| 8066 | 404 | Q404E | A | 2e-07 | 3.1e-07 | 4e-07 | 3.033E-07 | 1.002E-07 | 2.000E-07 | 4.000E-07 |
| 8067 | 404 | Q404Q | A | 1.4e-07 | 3.6e-07 | 3e-07 | 2.667E-07 | 1.137E-07 | 1.400E-07 | 3.600E-07 |
| 8068 | 404 | Q404G | A | 2.8e-07 | 2.4e-07 | 4.8e-07 | 3.333E-07 | 1.286E-07 | 2.400E-07 | 4.800E-07 |
| 8069 | 404 | Q404H | A | 3e-07 | 2.4e-07 | 2.8e-07 | 2.733E-07 | 3.055E-08 | 2.400E-07 | 3.000E-07 |
| 8070 | 404 | Q404I | A | 3.5e-07 | 1.6e-07 | 1.3e-07 | 2.133E-07 | 1.193E-07 | 1.300E-07 | 3.500E-07 |
| 8071 | 404 | Q404L | A | 3.2e-07 | 2.5e-07 | 3.1e-07 | 2.933E-07 | 3.786E-08 | 2.500E-07 | 3.200E-07 |
| 8072 | 404 | Q404K | A | 3.6e-07 | 2.5e-07 | 1.7e-07 | 2.600E-07 | 9.539E-08 | 1.700E-07 | 3.600E-07 |
| 8073 | 404 | Q404M | A | 4.4e-07 | 3.2e-07 | 1.3e-07 | 2.967E-07 | 1.563E-07 | 1.300E-07 | 4.400E-07 |
| 8074 | 404 | Q404F | A | 4.5e-07 | 4.7e-07 | 3.4e-07 | 4.200E-07 | 7.000E-08 | 3.400E-07 | 4.700E-07 |

| | | | | | | | | | | |
|---|---|---|---|---|---|---|---|---|---|---|
| 8075 | 404 | Q404P | A | 2.9e-07 | 2.5e-07 | 1.6e-07 | 2.333E-07 | 6.658E-08 | 1.600E-07 | 2.900E-07 |
| 8076 | 404 | Q404S | A | 3.5e-07 | 3e-07 | 2.5e-07 | 3.000E-07 | 5.000E-08 | 2.500E-07 | 3.500E-07 |
| 8077 | 404 | Q404T | A | 9.9e-08 | 2.1e-07 | 2.1e-07 | 1.730E-07 | 6.409E-08 | 9.900E-08 | 2.100E-07 |
| 8078 | 404 | Q404W | A | 2.2e-07 | 1.3e-07 | 1e-07 | 1.500E-07 | 6.245E-08 | 1.000E-07 | 2.200E-07 |
| 8079 | 404 | Q404Y | A | 1.9e-07 | 2.6e-07 | 1.3e-07 | 1.933E-07 | 6.506E-08 | 1.300E-07 | 2.600E-07 |
| 8080 | 404 | Q404V | A | 1.2e-07 | 2.3e-07 | 1.6e-07 | 1.700E-07 | 5.568E-08 | 1.200E-07 | 2.300E-07 |
| 8081 | 405 | N405A | A | 1.8e-07 | 2.1e-07 | 1.3e-07 | 1.733E-07 | 4.041E-08 | 1.300E-07 | 2.100E-07 |
| 8082 | 405 | N405R | A | 1.4e-07 | 1.7e-07 | 2.7e-07 | 1.933E-07 | 6.807E-08 | 1.400E-07 | 2.700E-07 |
| 8083 | 405 | N405N | A | 2e-07 | 3.2e-07 | 6.1e-07 | 3.767E-07 | 2.108E-07 | 2.000E-07 | 6.100E-07 |
| 8084 | 405 | N405D | A | 2.7e-07 | 2.3e-07 | 3.6e-07 | 2.867E-07 | 6.658E-08 | 2.300E-07 | 3.600E-07 |
| 8085 | 405 | N405C | A | 5.6e-07 | 2.2e-07 | 4e-07 | 3.933E-07 | 1.701E-07 | 2.200E-07 | 5.600E-07 |
| 8086 | 405 | N405E | A | 2.3e-07 | 6.3e-08 | 2.4e-07 | 1.777E-07 | 9.943E-08 | 6.300E-08 | 2.400E-07 |
| 8087 | 405 | N405Q | A | 2.5e-07 | 1.2e-07 | 5.1e-07 | 2.933E-07 | 1.986E-07 | 1.200E-07 | 5.100E-07 |
| 8088 | 405 | N405G | A | 1.5e-07 | 2.6e-07 | 1.8e-07 | 1.967E-07 | 5.686E-08 | 1.500E-07 | 2.600E-07 |
| 8089 | 405 | N405H | A | 2.2e-07 | 5.4e-07 | 6.7e-07 | 4.767E-07 | 2.316E-07 | 2.200E-07 | 6.700E-07 |
| 8090 | 405 | N405I | A | 4.5e-07 | 4.1e-07 | 1.2e-07 | 3.267E-07 | 1.801E-07 | 1.200E-07 | 4.500E-07 |
| 8091 | 405 | N405L | A | 2.5e-07 | 3.4e-07 | 3.1e-07 | 3.000E-07 | 4.583E-08 | 2.500E-07 | 3.400E-07 |
| 8092 | 405 | N405K | A | 1.6e-07 | 1.5e-07 | 2.3e-07 | 1.800E-07 | 4.359E-08 | 1.500E-07 | 2.300E-07 |
| 8093 | 405 | N405M | A | 2.5e-07 | 1.6e-07 | 2.7e-07 | 2.267E-07 | 5.859E-08 | 1.600E-07 | 2.700E-07 |
| 8094 | 405 | N405F | A | 2.2e-07 | 3.8e-07 | 1.5e-07 | 2.500E-07 | 1.179E-07 | 1.500E-07 | 3.800E-07 |
| 8095 | 405 | N405P | A | 1.5e-07 | 6.5e-07 | 2.5e-07 | 3.500E-07 | 2.646E-07 | 1.500E-07 | 6.500E-07 |
| 8096 | 405 | N405S | A | 1.1e-07 | 2.9e-07 | 2.4e-07 | 2.133E-07 | 9.292E-08 | 1.100E-07 | 2.900E-07 |
| 8097 | 405 | N405T | A | 2.2e-07 | 1.8e-07 | 3.4e-07 | 2.467E-07 | 8.327E-08 | 1.800E-07 | 3.400E-07 |
| 8098 | 405 | N405W | A | 3.9e-07 | 2.9e-07 | 2.8e-07 | 3.200E-07 | 6.083E-08 | 2.800E-07 | 3.900E-07 |
| 8099 | 405 | N405Y | A | 2.3e-07 | 2.7e-07 | 2.7e-07 | 2.567E-07 | 2.309E-08 | 2.300E-07 | 2.700E-07 |
| 8100 | 405 | N405V | A | 8.4e-07 | 4.7e-07 | 3e-07 | 5.367E-07 | 2.761E-07 | 3.000E-07 | 8.400E-07 |
| 8101 | 406 | L406A | A | 3.4e-07 | 1.9e-07 | 2.3e-07 | 2.533E-07 | 7.767E-08 | 1.900E-07 | 3.400E-07 |
| 8102 | 406 | L406R | A | 3.6e-07 | 1.2e-07 | 4.5e-07 | 3.100E-07 | 1.706E-07 | 1.200E-07 | 4.500E-07 |
| 8103 | 406 | L406N | A | 2.5e-07 | 2.1e-07 | 2.3e-07 | 2.300E-07 | 2.000E-08 | 2.100E-07 | 2.500E-07 |
| 8104 | 406 | L406D | A | 2.7e-07 | 5e-07 | 2.6e-07 | 3.433E-07 | 1.358E-07 | 2.600E-07 | 5.000E-07 |
| 8105 | 406 | L406C | A | 3.1e-07 | 9.3e-08 | 4.5e-07 | 2.843E-07 | 1.799E-07 | 9.300E-08 | 4.500E-07 |
| 8106 | 406 | L406E | A | 2.1e-07 | 1.7e-07 | 1.2e-07 | 1.667E-07 | 4.509E-08 | 1.200E-07 | 2.100E-07 |
| 8107 | 406 | L406Q | A | 2.5e-07 | 4.8e-07 | 2e-07 | 3.100E-07 | 1.493E-07 | 2.000E-07 | 4.800E-07 |
| 8108 | 406 | L406G | A | 1.4e-07 | 2.8e-07 | 1.8e-07 | 2.000E-07 | 7.211E-08 | 1.400E-07 | 2.800E-07 |
| 8109 | 406 | L406H | A | 4.3e-07 | 2.5e-07 | 4.1e-07 | 3.633E-07 | 9.866E-08 | 2.500E-07 | 4.300E-07 |
| 8110 | 406 | L406I | A | 2.7e-07 | 3.5e-07 | 4.9e-07 | 3.700E-07 | 1.114E-07 | 2.700E-07 | 4.900E-07 |
| 8111 | 406 | L406L | A | 1.9e-07 | 4.5e-07 | 9.3e-08 | 2.443E-07 | 1.846E-07 | 9.300E-08 | 4.500E-07 |
| 8112 | 406 | L406K | A | 3.6e-07 | 1.8e-07 | 3e-07 | 2.800E-07 | 9.165E-08 | 1.800E-07 | 3.600E-07 |
| 8113 | 406 | L406M | A | 2.9e-07 | 3e-07 | 3.4e-07 | 3.100E-07 | 2.646E-08 | 2.900E-07 | 3.400E-07 |
| 8114 | 406 | L406F | A | 1.5e-07 | 2.3e-07 | 2.9e-07 | 2.233E-07 | 7.024E-08 | 1.500E-07 | 2.900E-07 |
| 8115 | 406 | L406P | A | 3.8e-07 | 1.6e-07 | 2.2e-07 | 2.533E-07 | 1.137E-07 | 1.600E-07 | 3.800E-07 |
| 8116 | 406 | L406S | A | 2.1e-07 | 2.6e-07 | 1.7e-07 | 2.133E-07 | 4.509E-08 | 1.700E-07 | 2.600E-07 |
| 8117 | 406 | L406T | A | 2.6e-07 | 3.3e-07 | 2.6e-07 | 2.833E-07 | 4.041E-08 | 2.600E-07 | 3.300E-07 |
| 8118 | 406 | L406W | A | 3.2e-07 | 2.9e-07 | 1.2e-07 | 2.433E-07 | 1.079E-07 | 1.200E-07 | 3.200E-07 |
| 8119 | 406 | L406Y | A | 6.6e-08 | 1.2e-07 | 1.7e-07 | 1.187E-07 | 5.201E-08 | 6.600E-08 | 1.700E-07 |
| 8120 | 406 | L406V | A | 1.3e-07 | 2.5e-07 | 2e-07 | 1.933E-07 | 6.028E-08 | 1.300E-07 | 2.500E-07 |
| 8121 | 407 | Q407A | A | 3.4e-07 | 3.7e-07 | 3.8e-07 | 3.633E-07 | 2.082E-08 | 3.400E-07 | 3.800E-07 |
| 8122 | 407 | Q407R | A | 4.5e-07 | 2.3e-07 | 3.6e-07 | 3.467E-07 | 1.106E-07 | 2.300E-07 | 4.500E-07 |
| 8123 | 407 | Q407N | A | 4.5e-07 | 2.3e-07 | 1.3e-07 | 2.700E-07 | 1.637E-07 | 1.300E-07 | 4.500E-07 |
| 8124 | 407 | Q407D | A | 3.5e-07 | 4.1e-07 | 1.3e-07 | 2.967E-07 | 1.474E-07 | 1.300E-07 | 4.100E-07 |
| 8125 | 407 | Q407C | A | 2.2e-07 | 3.3e-07 | 1.4e-07 | 2.300E-07 | 9.539E-08 | 1.400E-07 | 3.300E-07 |
| 8126 | 407 | Q407E | A | 2.6e-07 | 1.6e-07 | 1.8e-07 | 2.000E-07 | 5.292E-08 | 1.600E-07 | 2.600E-07 |
| 8127 | 407 | Q407Q | A | 5.9e-07 | 4.6e-07 | 1.9e-07 | 4.133E-07 | 2.040E-07 | 1.900E-07 | 5.900E-07 |
| 8128 | 407 | Q407G | A | 2.7e-07 | 2.2e-07 | 2.9e-07 | 2.600E-07 | 3.606E-08 | 2.200E-07 | 2.900E-07 |
| 8129 | 407 | Q407H | A | 8.7e-08 | 1.9e-07 | 2.3e-07 | 1.690E-07 | 7.378E-08 | 8.700E-08 | 2.300E-07 |
| 8130 | 407 | Q407I | A | 1.7e-07 | 3e-07 | 4.7e-07 | 3.133E-07 | 1.504E-07 | 1.700E-07 | 4.700E-07 |
| 8131 | 407 | Q407L | A | 4.3e-07 | 2e-07 | 4.7e-07 | 3.667E-07 | 1.457E-07 | 2.000E-07 | 4.700E-07 |
| 8132 | 407 | Q407K | A | 2.2e-07 | 1.9e-07 | 2.6e-07 | 2.233E-07 | 3.512E-08 | 1.900E-07 | 2.600E-07 |
| 8133 | 407 | Q407M | A | 2.9e-07 | 1.8e-07 | 4e-07 | 2.900E-07 | 1.100E-07 | 1.800E-07 | 4.000E-07 |
| 8134 | 407 | Q407F | A | 1.3e-07 | 2.7e-07 | 2.2e-07 | 2.067E-07 | 7.095E-08 | 1.300E-07 | 2.700E-07 |
| 8135 | 407 | Q407P | A | 2.6e-07 | 2.4e-07 | 1.6e-07 | 2.200E-07 | 5.292E-08 | 1.600E-07 | 2.600E-07 |
| 8136 | 407 | Q407S | A | 3e-07 | 1.9e-07 | 4.5e-07 | 3.133E-07 | 1.305E-07 | 1.900E-07 | 4.500E-07 |
| 8137 | 407 | Q407T | A | 2.8e-07 | 1.5e-07 | 2.2e-07 | 2.167E-07 | 6.506E-08 | 1.500E-07 | 2.800E-07 |
| 8138 | 407 | Q407W | A | 4.2e-07 | 1.9e-07 | 2.6e-07 | 2.900E-07 | 1.179E-07 | 1.900E-07 | 4.200E-07 |
| 8139 | 407 | Q407Y | A | 1.7e-07 | 4.3e-07 | 2e-07 | 2.667E-07 | 1.422E-07 | 1.700E-07 | 4.300E-07 |
| 8140 | 407 | Q407V | A | 4.4e-07 | 6.4e-07 | 6.6e-08 | 3.820E-07 | 2.914E-07 | 6.600E-08 | 6.400E-07 |
| 8141 | 408 | V408A | A | 1.3e-07 | 3.5e-07 | 1.3e-07 | 2.033E-07 | 1.270E-07 | 1.300E-07 | 3.500E-07 |
| 8142 | 408 | V408R | A | 1.9e-07 | 5e-07 | 2.3e-07 | 3.067E-07 | 1.686E-07 | 1.900E-07 | 5.000E-07 |
| 8143 | 408 | V408N | A | 1.7e-07 | 1.8e-07 | 7e-08 | 1.400E-07 | 6.083E-08 | 7.000E-08 | 1.800E-07 |
| 8144 | 408 | V408D | A | 1.8e-07 | 2.2e-07 | 1.1e-07 | 1.700E-07 | 5.568E-08 | 1.100E-07 | 2.200E-07 |
| 8145 | 408 | V408C | A | 3.6e-07 | 1.7e-07 | 2.4e-07 | 2.567E-07 | 9.609E-08 | 1.700E-07 | 3.600E-07 |
| 8146 | 408 | V408E | A | 1.9e-07 | 2e-07 | 3.2e-07 | 2.367E-07 | 7.234E-08 | 1.900E-07 | 3.200E-07 |
| 8147 | 408 | V408Q | A | 1.5e-07 | 6.5e-07 | 4.7e-07 | 4.233E-07 | 2.532E-07 | 1.500E-07 | 6.500E-07 |
| 8148 | 408 | V408G | A | 2.6e-07 | 1.5e-07 | 4e-07 | 2.700E-07 | 1.253E-07 | 1.500E-07 | 4.000E-07 |
| 8149 | 408 | V408H | A | 2.6e-07 | 3.2e-07 | 2e-07 | 2.600E-07 | 6.000E-08 | 2.000E-07 | 3.200E-07 |
| 8150 | 408 | V408I | A | 2e-07 | 4e-07 | 4.7e-07 | 3.567E-07 | 1.401E-07 | 2.000E-07 | 4.700E-07 |
| 8151 | 408 | V408L | A | 4.3e-07 | 5.3e-07 | 2.6e-07 | 4.067E-07 | 1.365E-07 | 2.600E-07 | 5.300E-07 |
| 8152 | 408 | V408K | A | 2.3e-07 | 4.2e-07 | 1.5e-07 | 2.667E-07 | 1.387E-07 | 1.500E-07 | 4.200E-07 |
| 8153 | 408 | V408M | A | 1.3e-07 | 2.8e-07 | 2.7e-07 | 2.267E-07 | 8.386E-08 | 1.300E-07 | 2.800E-07 |
| 8154 | 408 | V408F | A | 1.8e-07 | 1.4e-07 | 1.2e-07 | 1.467E-07 | 3.055E-08 | 1.200E-07 | 1.800E-07 |
| 8155 | 408 | V408P | A | 1.3e-07 | 4.2e-07 | 5.4e-07 | 3.633E-07 | 2.108E-07 | 1.300E-07 | 5.400E-07 |
| 8156 | 408 | V408S | A | 3.2e-07 | 2.8e-07 | 1.7e-07 | 2.567E-07 | 7.767E-08 | 1.700E-07 | 3.200E-07 |
| 8157 | 408 | V408T | A | 2.7e-07 | 2.1e-07 | 2.5e-07 | 2.433E-07 | 3.055E-08 | 2.100E-07 | 2.700E-07 |
| 8158 | 408 | V408W | A | 4.5e-07 | 2.9e-07 | 2.1e-07 | 3.167E-07 | 1.222E-07 | 2.100E-07 | 4.500E-07 |
| 8159 | 408 | V408Y | A | 2.6e-07 | 1.8e-07 | 1.8e-07 | 2.067E-07 | 4.619E-08 | 1.800E-07 | 2.600E-07 |

| | | | | | | | | | | |
|---|---|---|---|---|---|---|---|---|---|---|
| 8160 | 408 | V408V | A | 6.9e-07 | 2e-07 | 2.2e-07 | 3.700E-07 | 2.773E-07 | 2.000E-07 | 6.900E-07 |
| 8161 | 409 | I409A | A | 6.8e-07 | 3.9e-07 | 3.4e-07 | 4.700E-07 | 1.836E-07 | 3.400E-07 | 6.800E-07 |
| 8162 | 409 | I409R | A | 3e-07 | 1.3e-07 | 2.9e-07 | 2.400E-07 | 9.539E-08 | 1.300E-07 | 3.000E-07 |
| 8163 | 409 | I409N | A | 2.8e-07 | 3.8e-07 | 9.2e-08 | 2.507E-07 | 1.462E-07 | 9.200E-08 | 3.800E-07 |
| 8164 | 409 | I409D | A | 2.8e-07 | 1.7e-07 | 3.5e-07 | 2.667E-07 | 9.074E-08 | 1.700E-07 | 3.500E-07 |
| 8165 | 409 | I409C | A | 1.8e-07 | 1.5e-07 | 2.6e-07 | 1.967E-07 | 5.686E-08 | 1.500E-07 | 2.600E-07 |
| 8166 | 409 | I409E | A | 2.2e-07 | 1.9e-07 | 1.2e-07 | 1.767E-07 | 5.132E-08 | 1.200E-07 | 2.200E-07 |
| 8167 | 409 | I409Q | A | 4.5e-07 | 2.9e-07 | 3.4e-07 | 3.600E-07 | 8.185E-08 | 2.900E-07 | 4.500E-07 |
| 8168 | 409 | I409G | A | 4.6e-07 | 2.2e-07 | 1.5e-07 | 2.767E-07 | 1.626E-07 | 1.500E-07 | 4.600E-07 |
| 8169 | 409 | I409H | A | 1.8e-07 | 3.1e-07 | 2.3e-07 | 2.400E-07 | 6.557E-08 | 1.800E-07 | 3.100E-07 |
| 8170 | 409 | I409I | A | 1.9e-07 | 2.1e-07 | 1.5e-07 | 1.833E-07 | 3.055E-08 | 1.500E-07 | 2.100E-07 |
| 8171 | 409 | I409L | A | 2.8e-07 | 2.1e-07 | 1.7e-07 | 2.200E-07 | 5.568E-08 | 1.700E-07 | 2.800E-07 |
| 8172 | 409 | I409K | A | 3.2e-07 | 2.1e-07 | 3e-07 | 2.767E-07 | 5.859E-08 | 2.100E-07 | 3.200E-07 |
| 8173 | 409 | I409M | A | 5.6e-07 | 2.5e-07 | 3.6e-07 | 3.900E-07 | 1.572E-07 | 2.500E-07 | 5.600E-07 |
| 8174 | 409 | I409F | A | 2.2e-07 | 8.9e-08 | 4.3e-07 | 2.463E-07 | 1.720E-07 | 8.900E-08 | 4.300E-07 |
| 8175 | 409 | I409P | A | 2.2e-07 | 6.5e-08 | 2.1e-07 | 1.650E-07 | 8.675E-08 | 6.500E-08 | 2.200E-07 |
| 8176 | 409 | I409S | A | 2.3e-07 | 4.9e-07 | 2.6e-07 | 3.267E-07 | 1.422E-07 | 2.300E-07 | 4.900E-07 |
| 8177 | 409 | I409T | A | 8.6e-07 | 3.2e-07 | 4e-07 | 5.267E-07 | 2.914E-07 | 3.200E-07 | 8.600E-07 |
| 8178 | 409 | I409W | A | 1.3e-07 | 4.3e-07 | 2.4e-07 | 2.667E-07 | 1.518E-07 | 1.300E-07 | 4.300E-07 |
| 8179 | 409 | I409Y | A | 1.1e-07 | 1.2e-07 | 4.3e-07 | 2.200E-07 | 1.819E-07 | 1.100E-07 | 4.300E-07 |
| 8180 | 409 | I409V | A | 4.5e-07 | 2.1e-07 | 2.8e-07 | 3.133E-07 | 1.234E-07 | 2.100E-07 | 4.500E-07 |
| 8181 | 410 | R410A | A | 2.5e-07 | 1.6e-07 | 2.5e-07 | 2.200E-07 | 5.196E-08 | 1.600E-07 | 2.500E-07 |
| 8182 | 410 | R410R | A | 4.8e-07 | 2.7e-07 | 2.6e-07 | 3.367E-07 | 1.242E-07 | 2.600E-07 | 4.800E-07 |
| 8183 | 410 | R410N | A | 2e-07 | 2.1e-07 | 1.8e-07 | 1.967E-07 | 1.528E-08 | 1.800E-07 | 2.100E-07 |
| 8184 | 410 | R410D | A | 3.5e-07 | 2.2e-07 | 6.7e-07 | 4.133E-07 | 2.316E-07 | 2.200E-07 | 6.700E-07 |
| 8185 | 410 | R410C | A | 1.9e-07 | 2.7e-07 | 2e-07 | 2.200E-07 | 4.359E-08 | 1.900E-07 | 2.700E-07 |
| 8186 | 410 | R410E | A | 1.6e-07 | 4.4e-07 | 3.3e-07 | 3.100E-07 | 1.411E-07 | 1.600E-07 | 4.400E-07 |
| 8187 | 410 | R410Q | A | 1.8e-07 | 4e-07 | 2e-07 | 2.600E-07 | 1.217E-07 | 1.800E-07 | 4.000E-07 |
| 8188 | 410 | R410G | A | 2.1e-07 | 3.9e-07 | 3.3e-07 | 3.100E-07 | 9.165E-08 | 2.100E-07 | 3.900E-07 |
| 8189 | 410 | R410H | A | 2.8e-07 | 3.4e-07 | 2.2e-07 | 2.800E-07 | 6.000E-08 | 2.200E-07 | 3.400E-07 |
| 8190 | 410 | R410I | A | 1.9e-07 | 4.2e-07 | 2.1e-07 | 2.733E-07 | 1.274E-07 | 1.900E-07 | 4.200E-07 |
| 8191 | 410 | R410L | A | 2.9e-07 | 1.9e-07 | 2.3e-07 | 2.367E-07 | 5.033E-08 | 1.900E-07 | 2.900E-07 |
| 8192 | 410 | R410K | A | 1.9e-07 | 2.3e-07 | 3.5e-07 | 2.567E-07 | 8.327E-08 | 1.900E-07 | 3.500E-07 |
| 8193 | 410 | R410M | A | 3.1e-07 | 2e-07 | 1.1e-07 | 2.067E-07 | 1.002E-07 | 1.100E-07 | 3.100E-07 |
| 8194 | 410 | R410F | A | 2.5e-07 | 3.8e-07 | 4e-07 | 3.433E-07 | 8.145E-08 | 2.500E-07 | 4.000E-07 |
| 8195 | 410 | R410P | A | 1.4e-07 | 2.7e-07 | 2.6e-07 | 2.233E-07 | 7.234E-08 | 1.400E-07 | 2.700E-07 |
| 8196 | 410 | R410S | A | 2.8e-07 | 2.8e-07 | 3e-07 | 2.867E-07 | 1.155E-08 | 2.800E-07 | 3.000E-07 |
| 8197 | 410 | R410T | A | 2.3e-07 | 2.7e-07 | 2.1e-07 | 2.367E-07 | 3.055E-08 | 2.100E-07 | 2.700E-07 |
| 8198 | 410 | R410W | A | 4.3e-07 | 2.3e-07 | 8.4e-08 | 2.480E-07 | 1.737E-07 | 8.400E-08 | 4.300E-07 |
| 8199 | 410 | R410Y | A | 5.2e-07 | 4.5e-07 | 2.3e-07 | 4.000E-07 | 1.513E-07 | 2.300E-07 | 5.200E-07 |
| 8200 | 410 | R410V | A | 2.8e-07 | 2.5e-07 | 1.9e-07 | 2.400E-07 | 4.583E-08 | 1.900E-07 | 2.800E-07 |
| 8201 | 411 | G411A | A | 2.1e-07 | 4.1e-07 | 5.8e-07 | 4.000E-07 | 1.852E-07 | 2.100E-07 | 5.800E-07 |
| 8202 | 411 | G411R | A | 7.1e-07 | 1.4e-07 | 1.2e-07 | 3.233E-07 | 3.350E-07 | 1.200E-07 | 7.100E-07 |
| 8203 | 411 | G411N | A | 2.1e-07 | 3e-07 | 1.2e-07 | 2.100E-07 | 9.000E-08 | 1.200E-07 | 3.000E-07 |
| 8204 | 411 | G411D | A | 2.4e-07 | 3.2e-07 | 1.2e-07 | 2.267E-07 | 1.007E-07 | 1.200E-07 | 3.200E-07 |
| 8205 | 411 | G411C | A | 3e-07 | 3.1e-07 | 2.2e-07 | 2.767E-07 | 4.933E-08 | 2.200E-07 | 3.100E-07 |
| 8206 | 411 | G411E | A | 4.5e-07 | 6.5e-07 | 2.6e-07 | 4.533E-07 | 1.950E-07 | 2.600E-07 | 6.500E-07 |
| 8207 | 411 | G411Q | A | 1.9e-07 | 4.2e-07 | 1.8e-07 | 2.633E-07 | 1.358E-07 | 1.800E-07 | 4.200E-07 |
| 8208 | 411 | G411G | A | 3.5e-07 | 2.1e-07 | 4e-07 | 3.200E-07 | 9.849E-08 | 2.100E-07 | 4.000E-07 |
| 8209 | 411 | G411H | A | 2.2e-07 | 2.4e-07 | 4.2e-07 | 2.933E-07 | 1.102E-07 | 2.200E-07 | 4.200E-07 |
| 8210 | 411 | G411I | A | 1.7e-07 | 1.2e-07 | 1.6e-07 | 1.500E-07 | 2.646E-08 | 1.200E-07 | 1.700E-07 |
| 8211 | 411 | G411L | A | 7.1e-07 | 2.7e-07 | 3.7e-07 | 4.500E-07 | 2.307E-07 | 2.700E-07 | 7.100E-07 |
| 8212 | 411 | G411K | A | 1.8e-07 | 2.5e-07 | 3.5e-07 | 2.600E-07 | 8.544E-08 | 1.800E-07 | 3.500E-07 |
| 8213 | 411 | G411M | A | 2.4e-07 | 7.8e-07 | 4.1e-07 | 4.767E-07 | 2.761E-07 | 2.400E-07 | 7.800E-07 |
| 8214 | 411 | G411F | A | 2.1e-07 | 2.3e-07 | 1.3e-07 | 1.900E-07 | 5.292E-08 | 1.300E-07 | 2.300E-07 |
| 8215 | 411 | G411P | A | 4.3e-07 | 2.5e-07 | 5.5e-07 | 4.100E-07 | 1.510E-07 | 2.500E-07 | 5.500E-07 |
| 8216 | 411 | G411S | A | 4.1e-07 | 2e-07 | 1.3e-07 | 2.467E-07 | 1.457E-07 | 1.300E-07 | 4.100E-07 |
| 8217 | 411 | G411T | A | 4.4e-07 | 1.8e-07 | 4.1e-07 | 3.433E-07 | 1.422E-07 | 1.800E-07 | 4.400E-07 |
| 8218 | 411 | G411W | A | 2.5e-07 | 5.8e-07 | 5.9e-07 | 4.733E-07 | 1.935E-07 | 2.500E-07 | 5.900E-07 |
| 8219 | 411 | G411Y | A | 2.5e-07 | 1.9e-07 | 2.1e-07 | 2.167E-07 | 3.055E-08 | 1.900E-07 | 2.500E-07 |
| 8220 | 411 | G411V | A | 3.2e-07 | 4.5e-07 | 1.1e-07 | 2.933E-07 | 1.716E-07 | 1.100E-07 | 4.500E-07 |
| 8221 | 412 | R412A | A | 2.3e-07 | 4.1e-07 | 1.6e-07 | 2.667E-07 | 1.290E-07 | 1.600E-07 | 4.100E-07 |
| 8222 | 412 | R412R | A | 3.8e-07 | 3.6e-07 | 1.4e-07 | 2.933E-07 | 1.332E-07 | 1.400E-07 | 3.800E-07 |
| 8223 | 412 | R412N | A | 1.2e-07 | 2.7e-07 | 3.5e-07 | 2.467E-07 | 1.168E-07 | 1.200E-07 | 3.500E-07 |
| 8224 | 412 | R412D | A | 4.1e-07 | 3.1e-07 | 2e-07 | 3.067E-07 | 1.050E-07 | 2.000E-07 | 4.100E-07 |
| 8225 | 412 | R412C | A | 4.9e-07 | 2e-07 | 1.3e-07 | 2.733E-07 | 1.909E-07 | 1.300E-07 | 4.900E-07 |
| 8226 | 412 | R412E | A | 2.4e-07 | 2e-07 | 1.7e-07 | 2.033E-07 | 3.512E-08 | 1.700E-07 | 2.400E-07 |
| 8227 | 412 | R412Q | A | 2.1e-07 | 2.6e-07 | 3.3e-07 | 2.667E-07 | 6.028E-08 | 2.100E-07 | 3.300E-07 |
| 8228 | 412 | R412G | A | 2.3e-07 | 2.8e-07 | 2.2e-07 | 2.433E-07 | 3.215E-08 | 2.200E-07 | 2.800E-07 |
| 8229 | 412 | R412H | A | 3.1e-07 | 1.3e-07 | 5.9e-07 | 3.433E-07 | 2.318E-07 | 1.300E-07 | 5.900E-07 |
| 8230 | 412 | R412I | A | 2.3e-07 | 2.7e-07 | 2.3e-07 | 2.433E-07 | 2.309E-08 | 2.300E-07 | 2.700E-07 |
| 8231 | 412 | R412L | A | 2.9e-07 | 1.4e-07 | 2.7e-07 | 2.333E-07 | 8.145E-08 | 1.400E-07 | 2.900E-07 |
| 8232 | 412 | R412K | A | 2e-07 | 1.2e-07 | 2e-07 | 1.733E-07 | 4.619E-08 | 1.200E-07 | 2.000E-07 |
| 8233 | 412 | R412M | A | 2e-07 | 4e-07 | 2e-07 | 2.667E-07 | 1.155E-07 | 2.000E-07 | 4.000E-07 |
| 8234 | 412 | R412F | A | 3.3e-07 | 3.6e-07 | 1.4e-07 | 2.767E-07 | 1.193E-07 | 1.400E-07 | 3.600E-07 |
| 8235 | 412 | R412P | A | 2.8e-07 | 4.6e-07 | 1.3e-07 | 2.900E-07 | 1.652E-07 | 1.300E-07 | 4.600E-07 |
| 8236 | 412 | R412S | A | 1.7e-07 | 2.3e-07 | 2.9e-07 | 2.300E-07 | 6.000E-08 | 1.700E-07 | 2.900E-07 |
| 8237 | 412 | R412T | A | 2.7e-07 | 2.9e-07 | 2.7e-07 | 2.767E-07 | 1.155E-08 | 2.700E-07 | 2.900E-07 |
| 8238 | 412 | R412W | A | 1.3e-07 | 2.4e-07 | 3.2e-07 | 2.300E-07 | 9.539E-08 | 1.300E-07 | 3.200E-07 |
| 8239 | 412 | R412Y | A | 2.4e-07 | 2.3e-07 | 3e-07 | 2.567E-07 | 3.786E-08 | 2.300E-07 | 3.000E-07 |
| 8240 | 412 | R412V | A | 2.1e-07 | 8.7e-07 | 6.1e-07 | 5.633E-07 | 3.325E-07 | 2.100E-07 | 8.700E-07 |
| 8241 | 413 | I413A | A | 5.1e-07 | 1.5e-07 | 3.9e-07 | 3.500E-07 | 1.833E-07 | 1.500E-07 | 5.100E-07 |
| 8242 | 413 | I413R | A | 3e-07 | 2.7e-07 | 3e-07 | 2.900E-07 | 1.732E-08 | 2.700E-07 | 3.000E-07 |
| 8243 | 413 | I413N | A | 2e-07 | 1.8e-07 | 5.2e-07 | 3.000E-07 | 1.908E-07 | 1.800E-07 | 5.200E-07 |
| 8244 | 413 | I413D | A | 1.4e-07 | 3.6e-07 | 2.6e-07 | 2.533E-07 | 1.102E-07 | 1.400E-07 | 3.600E-07 |

| | | | | | | | | | | |
|---|---|---|---|---|---|---|---|---|---|---|
| 8245 | 413 | I413C | A | 2.7e-07 | 3.1e-07 | 4.6e-07 | 3.467E-07 | 1.002E-07 | 2.700E-07 | 4.600E-07 |
| 8246 | 413 | I413E | A | 2.4e-07 | 3.8e-07 | 6.7e-07 | 4.300E-07 | 2.193E-07 | 2.400E-07 | 6.700E-07 |
| 8247 | 413 | I413Q | A | 2.6e-07 | 4.4e-07 | 4.9e-07 | 3.967E-07 | 1.210E-07 | 2.600E-07 | 4.900E-07 |
| 8248 | 413 | I413G | A | 2.4e-07 | 1.9e-07 | 2.2e-07 | 2.167E-07 | 2.517E-08 | 1.900E-07 | 2.400E-07 |
| 8249 | 413 | I413H | A | 2.3e-07 | 2.6e-07 | 1.4e-07 | 2.100E-07 | 6.245E-08 | 1.400E-07 | 2.600E-07 |
| 8250 | 413 | I413I | A | 3.2e-07 | 1.4e-07 | 1.7e-07 | 2.100E-07 | 9.644E-08 | 1.400E-07 | 3.200E-07 |
| 8251 | 413 | I413L | A | 1.8e-07 | 3.4e-07 | 9.8e-08 | 2.060E-07 | 1.231E-07 | 9.800E-08 | 3.400E-07 |
| 8252 | 413 | I413K | A | 1.9e-07 | 2.2e-07 | 5.5e-07 | 3.200E-07 | 1.997E-07 | 1.900E-07 | 5.500E-07 |
| 8253 | 413 | I413M | A | 3.7e-07 | 4.1e-07 | 4e-07 | 3.933E-07 | 2.082E-08 | 3.700E-07 | 4.100E-07 |
| 8254 | 413 | I413F | A | 2.9e-07 | 2.5e-07 | 1.1e-07 | 2.167E-07 | 9.452E-08 | 1.100E-07 | 2.900E-07 |
| 8255 | 413 | I413P | A | 3.6e-07 | 1.6e-07 | 4.9e-07 | 3.367E-07 | 1.662E-07 | 1.600E-07 | 4.900E-07 |
| 8256 | 413 | I413S | A | 4e-07 | 2.6e-07 | 5.1e-07 | 3.900E-07 | 1.253E-07 | 2.600E-07 | 5.100E-07 |
| 8257 | 413 | I413T | A | 1.6e-07 | 1.8e-07 | 2.3e-07 | 1.900E-07 | 3.606E-08 | 1.600E-07 | 2.300E-07 |
| 8258 | 413 | I413W | A | 3.6e-07 | 2e-07 | 3.7e-07 | 3.100E-07 | 9.539E-08 | 2.000E-07 | 3.700E-07 |
| 8259 | 413 | I413Y | A | 4e-07 | 5e-07 | 1.7e-07 | 3.567E-07 | 1.692E-07 | 1.700E-07 | 5.000E-07 |
| 8260 | 413 | I413V | A | 1.4e-07 | 2.1e-07 | 2.2e-07 | 1.900E-07 | 4.359E-08 | 1.400E-07 | 2.200E-07 |
| 8261 | 414 | L414A | A | 1e-06 | 2.7e-07 | 1.9e-07 | 4.867E-07 | 4.464E-07 | 1.900E-07 | 1.000E-06 |
| 8262 | 414 | L414R | A | 1.5e-07 | 2.7e-07 | 2.7e-07 | 2.300E-07 | 6.928E-08 | 1.500E-07 | 2.700E-07 |
| 8263 | 414 | L414N | A | 1.7e-07 | 9.1e-08 | 2.3e-07 | 1.637E-07 | 6.972E-08 | 9.100E-08 | 2.300E-07 |
| 8264 | 414 | L414D | A | 3.7e-07 | 2.1e-07 | 6.4e-07 | 4.067E-07 | 2.173E-07 | 2.100E-07 | 6.400E-07 |
| 8265 | 414 | L414C | A | 4.2e-07 | 2.8e-07 | 4.2e-07 | 3.733E-07 | 8.083E-08 | 2.800E-07 | 4.200E-07 |
| 8266 | 414 | L414E | A | 2.4e-07 | 2.5e-07 | 6.2e-07 | 3.700E-07 | 2.166E-07 | 2.400E-07 | 6.200E-07 |
| 8267 | 414 | L414Q | A | 1.8e-07 | 1.9e-07 | 1.5e-07 | 1.733E-07 | 2.082E-08 | 1.500E-07 | 1.900E-07 |
| 8268 | 414 | L414G | A | 1.7e-07 | 1.9e-07 | 2.1e-07 | 1.900E-07 | 2.000E-08 | 1.700E-07 | 2.100E-07 |
| 8269 | 414 | L414H | A | 2e-07 | 3.7e-07 | 2.9e-07 | 2.867E-07 | 8.505E-08 | 2.000E-07 | 3.700E-07 |
| 8270 | 414 | L414I | A | 2.8e-08 | 2e-07 | 3.2e-07 | 1.827E-07 | 1.468E-07 | 2.800E-08 | 3.200E-07 |
| 8271 | 414 | L414L | A | 1.2e-07 | 3.1e-07 | 2e-07 | 2.100E-07 | 9.539E-08 | 1.200E-07 | 3.100E-07 |
| 8272 | 414 | L414K | A | 2.1e-07 | 1.5e-07 | 3.5e-07 | 2.367E-07 | 1.026E-07 | 1.500E-07 | 3.500E-07 |
| 8273 | 414 | L414M | A | 4.9e-07 | 3.3e-07 | 1.8e-07 | 3.333E-07 | 1.550E-07 | 1.800E-07 | 4.900E-07 |
| 8274 | 414 | L414F | A | 1.9e-07 | 1.8e-07 | 2.6e-07 | 2.100E-07 | 4.359E-08 | 1.800E-07 | 2.600E-07 |
| 8275 | 414 | L414P | A | 2.1e-07 | 2.1e-07 | 1.1e-07 | 1.767E-07 | 5.774E-08 | 1.100E-07 | 2.100E-07 |
| 8276 | 414 | L414S | A | 3.4e-07 | 3.5e-07 | 2.8e-07 | 3.233E-07 | 3.786E-08 | 2.800E-07 | 3.500E-07 |
| 8277 | 414 | L414T | A | 1.3e-07 | 6.5e-08 | 2e-07 | 1.317E-07 | 6.752E-08 | 6.500E-08 | 2.000E-07 |
| 8278 | 414 | L414W | A | 2.2e-07 | 1.3e-07 | 3.1e-07 | 2.200E-07 | 9.000E-08 | 1.300E-07 | 3.100E-07 |
| 8279 | 414 | L414Y | A | 4.2e-07 | 8.3e-07 | 3.1e-07 | 5.200E-07 | 2.740E-07 | 3.100E-07 | 8.300E-07 |
| 8280 | 414 | L414V | A | 2.8e-07 | 4.8e-07 | 5.9e-07 | 4.500E-07 | 1.572E-07 | 2.800E-07 | 5.900E-07 |
| 8281 | 415 | H415A | A | 2e-07 | 1.8e-07 | 2.1e-07 | 1.967E-07 | 1.528E-08 | 1.800E-07 | 2.100E-07 |
| 8282 | 415 | H415R | A | 3.6e-07 | 2.6e-07 | 2e-07 | 2.733E-07 | 8.083E-08 | 2.000E-07 | 3.600E-07 |
| 8283 | 415 | H415N | A | 1.7e-07 | 3.3e-07 | 1.7e-07 | 2.233E-07 | 9.238E-08 | 1.700E-07 | 3.300E-07 |
| 8284 | 415 | H415D | A | 1.5e-07 | 4.4e-07 | 2e-07 | 2.633E-07 | 1.550E-07 | 1.500E-07 | 4.400E-07 |
| 8285 | 415 | H415C | A | 1.9e-07 | 2.8e-07 | 2.7e-07 | 2.467E-07 | 4.933E-08 | 1.900E-07 | 2.800E-07 |
| 8286 | 415 | H415E | A | 2.4e-07 | 2.1e-07 | 3.3e-07 | 2.600E-07 | 6.245E-08 | 2.100E-07 | 3.300E-07 |
| 8287 | 415 | H415Q | A | 2e-07 | 7.8e-08 | 1.3e-07 | 1.360E-07 | 6.122E-08 | 7.800E-08 | 2.000E-07 |
| 8288 | 415 | H415G | A | 4.1e-07 | 2.2e-07 | 2e-07 | 2.767E-07 | 1.159E-07 | 2.000E-07 | 4.100E-07 |
| 8289 | 415 | H415H | A | 2.2e-07 | 1.7e-07 | 4.5e-07 | 2.800E-07 | 1.493E-07 | 1.700E-07 | 4.500E-07 |
| 8290 | 415 | H415I | A | 3.7e-07 | 1.5e-07 | 2.6e-07 | 2.600E-07 | 1.100E-07 | 1.500E-07 | 3.700E-07 |
| 8291 | 415 | H415L | A | 4.2e-07 | 2.1e-07 | 1.8e-07 | 2.700E-07 | 1.308E-07 | 1.800E-07 | 4.200E-07 |
| 8292 | 415 | H415K | A | 5.6e-07 | 3.8e-07 | 1.3e-07 | 3.567E-07 | 2.159E-07 | 1.300E-07 | 5.600E-07 |
| 8293 | 415 | H415M | A | 3.5e-07 | 2.1e-07 | 1.8e-07 | 2.467E-07 | 9.074E-08 | 1.800E-07 | 3.500E-07 |
| 8294 | 415 | H415F | A | 6.1e-07 | 3.2e-07 | 3.3e-07 | 4.200E-07 | 1.646E-07 | 3.200E-07 | 6.100E-07 |
| 8295 | 415 | H415P | A | 2e-07 | 2.1e-07 | 2.9e-07 | 2.333E-07 | 4.933E-08 | 2.000E-07 | 2.900E-07 |
| 8296 | 415 | H415S | A | 5.8e-07 | 1.9e-07 | 1.6e-07 | 3.100E-07 | 2.343E-07 | 1.600E-07 | 5.800E-07 |
| 8297 | 415 | H415T | A | 4.8e-07 | 1.2e-07 | 2.5e-07 | 2.833E-07 | 1.823E-07 | 1.200E-07 | 4.800E-07 |
| 8298 | 415 | H415W | A | 2.4e-07 | 7e-07 | 3.3e-07 | 4.233E-07 | 2.438E-07 | 2.400E-07 | 7.000E-07 |
| 8299 | 415 | H415Y | A | 3.2e-07 | 2.1e-07 | 1.4e-07 | 2.233E-07 | 9.074E-08 | 1.400E-07 | 3.200E-07 |
| 8300 | 415 | H415V | A | 4e-07 | 1.6e-07 | 4.3e-07 | 3.300E-07 | 1.480E-07 | 1.600E-07 | 4.300E-07 |
| 8301 | 416 | N416A | A | 3.6e-07 | 1.9e-07 | 8e-08 | 2.100E-07 | 1.411E-07 | 8.000E-08 | 3.600E-07 |
| 8302 | 416 | N416R | A | 1.3e-07 | 7.6e-07 | 3.1e-07 | 4.000E-07 | 3.245E-07 | 1.300E-07 | 7.600E-07 |
| 8303 | 416 | N416N | A | 6.6e-07 | 3.4e-07 | 2.7e-07 | 4.233E-07 | 2.079E-07 | 2.700E-07 | 6.600E-07 |
| 8304 | 416 | N416D | A | 4.7e-07 | 1.4e-07 | 3.8e-07 | 3.300E-07 | 1.706E-07 | 1.400E-07 | 4.700E-07 |
| 8305 | 416 | N416C | A | 1.3e-07 | 3e-07 | 1.3e-07 | 1.867E-07 | 9.815E-08 | 1.300E-07 | 3.000E-07 |
| 8306 | 416 | N416E | A | 3e-07 | 7.3e-07 | 1.1e-07 | 3.800E-07 | 3.176E-07 | 1.100E-07 | 7.300E-07 |
| 8307 | 416 | N416Q | A | 6.2e-07 | 3.6e-07 | 2.2e-07 | 4.000E-07 | 2.030E-07 | 2.200E-07 | 6.200E-07 |
| 8308 | 416 | N416G | A | 8.9e-08 | 3.2e-07 | 1.7e-07 | 1.930E-07 | 1.172E-07 | 8.900E-08 | 3.200E-07 |
| 8309 | 416 | N416H | A | 2.2e-07 | 1.6e-07 | 2.1e-07 | 1.967E-07 | 3.215E-08 | 1.600E-07 | 2.200E-07 |
| 8310 | 416 | N416I | A | 1.6e-07 | 2.3e-07 | 2.8e-07 | 2.233E-07 | 6.028E-08 | 1.600E-07 | 2.800E-07 |
| 8311 | 416 | N416L | A | 2.1e-07 | 4.5e-07 | 2.6e-07 | 3.067E-07 | 1.266E-07 | 2.100E-07 | 4.500E-07 |
| 8312 | 416 | N416K | A | 4.9e-07 | 5.9e-07 | 1.1e-07 | 3.967E-07 | 2.532E-07 | 1.100E-07 | 5.900E-07 |
| 8313 | 416 | N416M | A | 3.5e-07 | 1.3e-07 | 1.6e-07 | 2.133E-07 | 1.193E-07 | 1.300E-07 | 3.500E-07 |
| 8314 | 416 | N416F | A | 1.7e-07 | 1.2e-07 | 1.1e-07 | 1.333E-07 | 3.215E-08 | 1.100E-07 | 1.700E-07 |
| 8315 | 416 | N416P | A | 2.2e-07 | 2.4e-07 | 3.3e-07 | 2.633E-07 | 5.859E-08 | 2.200E-07 | 3.300E-07 |
| 8316 | 416 | N416S | A | 1.3e-07 | 4.5e-07 | 3.1e-07 | 2.967E-07 | 1.604E-07 | 1.300E-07 | 4.500E-07 |
| 8317 | 416 | N416T | A | 2.3e-07 | 1.6e-07 | 2.3e-07 | 2.067E-07 | 4.041E-08 | 1.600E-07 | 2.300E-07 |
| 8318 | 416 | N416W | A | 4e-07 | 4.4e-07 | 3.2e-07 | 3.867E-07 | 6.110E-08 | 3.200E-07 | 4.400E-07 |
| 8319 | 416 | N416Y | A | 2.7e-07 | 2.3e-07 | 1.1e-07 | 2.033E-07 | 8.327E-08 | 1.100E-07 | 2.700E-07 |
| 8320 | 416 | N416V | A | 2.1e-07 | 2.6e-07 | 2e-07 | 2.233E-07 | 3.215E-08 | 2.000E-07 | 2.600E-07 |
| 8321 | 417 | G417A | A | 2e-07 | 2.4e-07 | 2e-07 | 2.133E-07 | 2.309E-08 | 2.000E-07 | 2.400E-07 |
| 8322 | 417 | G417R | A | 2.4e-07 | 3.8e-07 | 1.1e-07 | 2.433E-07 | 1.350E-07 | 1.100E-07 | 3.800E-07 |
| 8323 | 417 | G417N | A | 2.7e-07 | 1.4e-07 | 2.3e-07 | 2.133E-07 | 6.658E-08 | 1.400E-07 | 2.700E-07 |
| 8324 | 417 | G417D | A | 2.1e-07 | 4.3e-07 | 1.9e-07 | 2.767E-07 | 1.332E-07 | 1.900E-07 | 4.300E-07 |
| 8325 | 417 | G417C | A | 2.8e-07 | 3.7e-07 | 1.8e-07 | 2.767E-07 | 9.504E-08 | 1.800E-07 | 3.700E-07 |
| 8326 | 417 | G417E | A | 1.1e-07 | 4.6e-07 | 2.9e-07 | 2.867E-07 | 1.750E-07 | 1.100E-07 | 4.600E-07 |
| 8327 | 417 | G417Q | A | 6.5e-08 | 4.5e-07 | 4.9e-07 | 3.350E-07 | 2.347E-07 | 6.500E-08 | 4.900E-07 |
| 8328 | 417 | G417G | A | 3.5e-07 | 1.3e-07 | 4e-07 | 2.933E-07 | 1.436E-07 | 1.300E-07 | 4.000E-07 |
| 8329 | 417 | G417H | A | 2.8e-07 | 2.6e-07 | 2.1e-07 | 2.500E-07 | 3.606E-08 | 2.100E-07 | 2.800E-07 |

| | | | | | | | | | | |
|---|---|---|---|---|---|---|---|---|---|---|
| 8330 | 417 | G417I | A | 2.2e-07 | 1.4e-07 | 4e-07 | 2.533E-07 | 1.332E-07 | 1.400E-07 | 4.000E-07 |
| 8331 | 417 | G417L | A | 2.5e-07 | 1.5e-07 | 2.5e-07 | 2.167E-07 | 5.774E-08 | 1.500E-07 | 2.500E-07 |
| 8332 | 417 | G417K | A | 1.1e-07 | 3e-07 | 2.1e-07 | 2.067E-07 | 9.504E-08 | 1.100E-07 | 3.000E-07 |
| 8333 | 417 | G417M | A | 1.5e-07 | 2.4e-07 | 8.5e-08 | 1.583E-07 | 7.784E-08 | 8.500E-08 | 2.400E-07 |
| 8334 | 417 | G417F | A | 1.5e-07 | 2.8e-07 | 2.7e-07 | 2.333E-07 | 7.234E-08 | 1.500E-07 | 2.800E-07 |
| 8335 | 417 | G417P | A | 2.5e-07 | 1.3e-07 | 3.1e-07 | 2.300E-07 | 9.165E-08 | 1.300E-07 | 3.100E-07 |
| 8336 | 417 | G417S | A | 3e-07 | 1.5e-07 | 2.5e-07 | 2.333E-07 | 7.638E-08 | 1.500E-07 | 3.000E-07 |
| 8337 | 417 | G417T | A | 2.5e-07 | 1.9e-07 | 3.4e-07 | 2.600E-07 | 7.550E-08 | 1.900E-07 | 3.400E-07 |
| 8338 | 417 | G417W | A | 1.3e-07 | 1.3e-07 | 1.9e-07 | 1.500E-07 | 3.464E-08 | 1.300E-07 | 1.900E-07 |
| 8339 | 417 | G417Y | A | 6.2e-07 | 3.2e-07 | 3.3e-07 | 4.233E-07 | 1.704E-07 | 3.200E-07 | 6.200E-07 |
| 8340 | 417 | G417V | A | 2.7e-07 | 2.2e-07 | 6.3e-07 | 3.733E-07 | 2.237E-07 | 2.200E-07 | 6.300E-07 |
| 8341 | 418 | A418A | A | 3.4e-07 | 1e-07 | 2.1e-07 | 2.167E-07 | 1.201E-07 | 1.000E-07 | 3.400E-07 |
| 8342 | 418 | A418R | A | 2e-07 | 5.8e-07 | 1.6e-07 | 3.133E-07 | 2.318E-07 | 1.600E-07 | 5.800E-07 |
| 8343 | 418 | A418N | A | 1.7e-07 | 3e-07 | 3e-07 | 2.567E-07 | 7.506E-08 | 1.700E-07 | 3.000E-07 |
| 8344 | 418 | A418D | A | 2.4e-07 | 1.4e-07 | 3.2e-07 | 2.333E-07 | 9.018E-08 | 1.400E-07 | 3.200E-07 |
| 8345 | 418 | A418C | A | 1.3e-07 | 3.5e-07 | 6.5e-07 | 3.767E-07 | 2.610E-07 | 1.300E-07 | 6.500E-07 |
| 8346 | 418 | A418E | A | 2.2e-07 | 4e-07 | 4.6e-07 | 3.600E-07 | 1.249E-07 | 2.200E-07 | 4.600E-07 |
| 8347 | 418 | A418Q | A | 1.9e-07 | 4.7e-07 | 3.4e-07 | 3.333E-07 | 1.401E-07 | 1.900E-07 | 4.700E-07 |
| 8348 | 418 | A418G | A | 2.1e-07 | 8.9e-07 | 3.9e-07 | 4.967E-07 | 3.523E-07 | 2.100E-07 | 8.900E-07 |
| 8349 | 418 | A418H | A | 1.3e-07 | 1.9e-07 | 2.5e-07 | 1.900E-07 | 6.000E-08 | 1.300E-07 | 2.500E-07 |
| 8350 | 418 | A418I | A | 1.9e-07 | 1.9e-07 | 4e-07 | 2.600E-07 | 1.212E-07 | 1.900E-07 | 4.000E-07 |
| 8351 | 418 | A418L | A | 2e-07 | 1.2e-07 | 2.4e-07 | 1.867E-07 | 6.110E-08 | 1.200E-07 | 2.400E-07 |
| 8352 | 418 | A418K | A | 1.9e-07 | 5.9e-07 | 1e-07 | 2.933E-07 | 2.608E-07 | 1.000E-07 | 5.900E-07 |
| 8353 | 418 | A418M | A | 1.8e-07 | 2.5e-07 | 9.4e-08 | 1.747E-07 | 7.814E-08 | 9.400E-08 | 2.500E-07 |
| 8354 | 418 | A418F | A | 2.2e-07 | 2.1e-07 | 1e-07 | 1.767E-07 | 6.658E-08 | 1.000E-07 | 2.200E-07 |
| 8355 | 418 | A418P | A | 3.2e-07 | 1.6e-07 | 1.6e-07 | 2.133E-07 | 9.238E-08 | 1.600E-07 | 3.200E-07 |
| 8356 | 418 | A418S | A | 3.4e-07 | 2.6e-07 | 3.3e-07 | 3.100E-07 | 4.359E-08 | 2.600E-07 | 3.400E-07 |
| 8357 | 418 | A418T | A | 4.1e-07 | 2.8e-07 | 1.8e-07 | 2.900E-07 | 1.153E-07 | 1.800E-07 | 4.100E-07 |
| 8358 | 418 | A418W | A | 2.6e-07 | 3.5e-07 | 2.5e-07 | 2.867E-07 | 5.508E-08 | 2.500E-07 | 3.500E-07 |
| 8359 | 418 | A418Y | A | 3.3e-07 | 3.6e-07 | 2.1e-07 | 3.000E-07 | 7.937E-08 | 2.100E-07 | 3.600E-07 |
| 8360 | 418 | A418V | A | 1.3e-07 | 1.8e-07 | 5.4e-07 | 2.833E-07 | 2.237E-07 | 1.300E-07 | 5.400E-07 |
| 8361 | 419 | Y419A | A | 2.1e-07 | 3.4e-07 | 1.1e-07 | 2.200E-07 | 1.153E-07 | 1.100E-07 | 3.400E-07 |
| 8362 | 419 | Y419R | A | 4.8e-07 | 2.7e-07 | 5.6e-07 | 4.367E-07 | 1.498E-07 | 2.700E-07 | 5.600E-07 |
| 8363 | 419 | Y419N | A | 3.9e-07 | 2e-07 | 3.7e-07 | 3.200E-07 | 1.044E-07 | 2.000E-07 | 3.900E-07 |
| 8364 | 419 | Y419D | A | 1.3e-07 | 5.7e-07 | 4.2e-07 | 3.733E-07 | 2.237E-07 | 1.300E-07 | 5.700E-07 |
| 8365 | 419 | Y419C | A | 7e-07 | 3.5e-07 | 1.6e-07 | 4.033E-07 | 2.739E-07 | 1.600E-07 | 7.000E-07 |
| 8366 | 419 | Y419E | A | 4.1e-07 | 2.1e-07 | 3.1e-07 | 3.100E-07 | 1.000E-07 | 2.100E-07 | 4.100E-07 |
| 8367 | 419 | Y419Q | A | 3e-07 | 1.4e-07 | 2.2e-07 | 2.200E-07 | 8.000E-08 | 1.400E-07 | 3.000E-07 |
| 8368 | 419 | Y419G | A | 2.2e-07 | 2.3e-07 | 2.6e-07 | 2.367E-07 | 2.082E-08 | 2.200E-07 | 2.600E-07 |
| 8369 | 419 | Y419H | A | 2.1e-07 | 2.1e-07 | 2e-07 | 2.067E-07 | 5.774E-09 | 2.000E-07 | 2.100E-07 |
| 8370 | 419 | Y419I | A | 1.1e-07 | 2.8e-07 | 3e-07 | 2.300E-07 | 1.044E-07 | 1.100E-07 | 3.000E-07 |
| 8371 | 419 | Y419L | A | 6.2e-08 | 1.9e-07 | 5.9e-07 | 2.807E-07 | 2.754E-07 | 6.200E-08 | 5.900E-07 |
| 8372 | 419 | Y419K | A | 3.9e-07 | 4.6e-07 | 1.5e-07 | 3.333E-07 | 1.626E-07 | 1.500E-07 | 4.600E-07 |
| 8373 | 419 | Y419M | A | 2.9e-07 | 3e-07 | 1.4e-07 | 2.433E-07 | 8.963E-08 | 1.400E-07 | 3.000E-07 |
| 8374 | 419 | Y419F | A | 3.7e-07 | 3e-07 | 1.6e-07 | 2.767E-07 | 1.069E-07 | 1.600E-07 | 3.700E-07 |
| 8375 | 419 | Y419P | A | 1.7e-07 | 5.8e-07 | 2e-07 | 3.167E-07 | 2.285E-07 | 1.700E-07 | 5.800E-07 |
| 8376 | 419 | Y419S | A | 2.5e-07 | 2.4e-07 | 1.7e-07 | 2.200E-07 | 4.359E-08 | 1.700E-07 | 2.500E-07 |
| 8377 | 419 | Y419T | A | 5.7e-07 | 3.1e-07 | 1.4e-07 | 3.400E-07 | 2.166E-07 | 1.400E-07 | 5.700E-07 |
| 8378 | 419 | Y419W | A | 2.4e-07 | 2.5e-07 | 3.7e-07 | 2.867E-07 | 7.234E-08 | 2.400E-07 | 3.700E-07 |
| 8379 | 419 | Y419Y | A | 3.8e-07 | 2.6e-07 | 2.5e-07 | 2.967E-07 | 7.234E-08 | 2.500E-07 | 3.800E-07 |
| 8380 | 419 | Y419V | A | 2.9e-07 | 1.1e-07 | 1.2e-07 | 1.733E-07 | 1.012E-07 | 1.100E-07 | 2.900E-07 |
| 8381 | 420 | S420A | A | 2.6e-07 | 4.1e-07 | 3.7e-07 | 3.467E-07 | 7.767E-08 | 2.600E-07 | 4.100E-07 |
| 8382 | 420 | S420R | A | 3.3e-07 | 2.5e-07 | 2e-07 | 2.600E-07 | 6.557E-08 | 2.000E-07 | 3.300E-07 |
| 8383 | 420 | S420N | A | 2.2e-07 | 2.5e-07 | 1.7e-07 | 2.133E-07 | 4.041E-08 | 1.700E-07 | 2.500E-07 |
| 8384 | 420 | S420D | A | 1.8e-07 | 2.2e-07 | 3.5e-07 | 2.500E-07 | 8.888E-08 | 1.800E-07 | 3.500E-07 |
| 8385 | 420 | S420C | A | 3.5e-07 | 1.5e-07 | 2.9e-07 | 2.633E-07 | 1.026E-07 | 1.500E-07 | 3.500E-07 |
| 8386 | 420 | S420E | A | 1.3e-07 | 2.6e-07 | 4.3e-07 | 2.733E-07 | 1.504E-07 | 1.300E-07 | 4.300E-07 |
| 8387 | 420 | S420Q | A | 2.3e-07 | 2e-07 | 3.5e-07 | 2.600E-07 | 7.937E-08 | 2.000E-07 | 3.500E-07 |
| 8388 | 420 | S420G | A | 1.8e-07 | 2.5e-07 | 1.5e-07 | 1.933E-07 | 5.132E-08 | 1.500E-07 | 2.500E-07 |
| 8389 | 420 | S420H | A | 3.4e-07 | 2e-07 | 2.1e-07 | 2.500E-07 | 7.810E-08 | 2.000E-07 | 3.400E-07 |
| 8390 | 420 | S420I | A | 1e-07 | 2.3e-07 | 3.4e-07 | 2.233E-07 | 1.201E-07 | 1.000E-07 | 3.400E-07 |
| 8391 | 420 | S420L | A | 5.5e-07 | 2.8e-07 | 1.8e-07 | 3.367E-07 | 1.914E-07 | 1.800E-07 | 5.500E-07 |
| 8392 | 420 | S420K | A | 5.9e-07 | 3.9e-07 | 2.7e-07 | 4.167E-07 | 1.617E-07 | 2.700E-07 | 5.900E-07 |
| 8393 | 420 | S420M | A | 1.9e-07 | 1.8e-07 | 2.7e-07 | 2.133E-07 | 4.933E-08 | 1.800E-07 | 2.700E-07 |
| 8394 | 420 | S420F | A | 5.5e-07 | 3.5e-07 | 3.2e-07 | 4.067E-07 | 1.250E-07 | 3.200E-07 | 5.500E-07 |
| 8395 | 420 | S420P | A | 2e-07 | 2e-07 | 2.5e-07 | 2.167E-07 | 2.887E-08 | 2.000E-07 | 2.500E-07 |
| 8396 | 420 | S420S | A | 3.4e-07 | 2.1e-07 | 1.1e-07 | 2.200E-07 | 1.153E-07 | 1.100E-07 | 3.400E-07 |
| 8397 | 420 | S420T | A | 1.6e-07 | 2.3e-07 | 2.2e-07 | 2.033E-07 | 3.786E-08 | 1.600E-07 | 2.300E-07 |
| 8398 | 420 | S420W | A | 1.8e-07 | 2.2e-07 | 2.7e-07 | 2.233E-07 | 4.509E-08 | 1.800E-07 | 2.700E-07 |
| 8399 | 420 | S420Y | A | 3.1e-07 | 2.4e-07 | 2e-07 | 2.500E-07 | 5.568E-08 | 2.000E-07 | 3.100E-07 |
| 8400 | 420 | S420V | A | 2.7e-07 | 3.6e-07 | 1.9e-07 | 2.733E-07 | 8.505E-08 | 1.900E-07 | 3.600E-07 |
| 8401 | 421 | L421A | A | 1.1e-07 | 1.4e-07 | 4.5e-07 | 2.333E-07 | 1.882E-07 | 1.100E-07 | 4.500E-07 |
| 8402 | 421 | L421R | A | 3.9e-07 | 1.1e-07 | 3.5e-07 | 2.833E-07 | 1.514E-07 | 1.100E-07 | 3.900E-07 |
| 8403 | 421 | L421N | A | 2.4e-07 | 2.9e-07 | 4.1e-07 | 3.133E-07 | 8.737E-08 | 2.400E-07 | 4.100E-07 |
| 8404 | 421 | L421D | A | 3.1e-07 | 2.4e-07 | 3.9e-07 | 3.133E-07 | 7.506E-08 | 2.400E-07 | 3.900E-07 |
| 8405 | 421 | L421C | A | 2.8e-07 | 1.6e-07 | 3.8e-07 | 2.733E-07 | 1.102E-07 | 1.600E-07 | 3.800E-07 |
| 8406 | 421 | L421E | A | 3.1e-07 | 6.3e-08 | 8.5e-08 | 1.527E-07 | 1.367E-07 | 6.300E-08 | 3.100E-07 |
| 8407 | 421 | L421Q | A | 3.2e-07 | 2.4e-07 | 2e-07 | 2.533E-07 | 6.110E-08 | 2.000E-07 | 3.200E-07 |
| 8408 | 421 | L421G | A | 1.5e-07 | 3e-07 | 4.6e-07 | 3.033E-07 | 1.550E-07 | 1.500E-07 | 4.600E-07 |
| 8409 | 421 | L421H | A | 5.1e-07 | 3.6e-07 | 1.8e-07 | 3.500E-07 | 1.652E-07 | 1.800E-07 | 5.100E-07 |
| 8410 | 421 | L421I | A | 4.3e-07 | 8.9e-08 | 2.2e-07 | 2.463E-07 | 1.720E-07 | 8.900E-08 | 4.300E-07 |
| 8411 | 421 | L421L | A | 4e-07 | 3.7e-07 | 2e-07 | 3.233E-07 | 1.079E-07 | 2.000E-07 | 4.000E-07 |
| 8412 | 421 | L421K | A | 5.8e-07 | 2.5e-07 | 4.1e-07 | 4.133E-07 | 1.650E-07 | 2.500E-07 | 5.800E-07 |
| 8413 | 421 | L421M | A | 1.1e-07 | 3.1e-07 | 1.4e-07 | 1.867E-07 | 1.079E-07 | 1.100E-07 | 3.100E-07 |
| 8414 | 421 | L421F | A | 2.8e-07 | 1.4e-07 | 7.2e-08 | 1.640E-07 | 1.061E-07 | 7.200E-08 | 2.800E-07 |

| | | | | | | | | | | |
|---|---|---|---|---|---|---|---|---|---|---|
| 8415 | 421 | L421P | A | 4.1e-07 | 3.3e-07 | 6.6e-07 | 4.667E-07 | 1.721E-07 | 3.300E-07 | 6.600E-07 |
| 8416 | 421 | L421S | A | 1.5e-07 | 3.9e-07 | 2.1e-07 | 2.500E-07 | 1.249E-07 | 1.500E-07 | 3.900E-07 |
| 8417 | 421 | L421T | A | 3.5e-07 | 4.4e-07 | 3.3e-07 | 3.733E-07 | 5.859E-08 | 3.300E-07 | 4.400E-07 |
| 8418 | 421 | L421W | A | 1.8e-07 | 3.7e-07 | 2.6e-07 | 2.700E-07 | 9.539E-08 | 1.800E-07 | 3.700E-07 |
| 8419 | 421 | L421Y | A | 1.1e-07 | 4.7e-07 | 2.7e-07 | 2.833E-07 | 1.804E-07 | 1.100E-07 | 4.700E-07 |
| 8420 | 421 | L421V | A | 2.6e-07 | 2.6e-07 | 2.3e-07 | 2.500E-07 | 1.732E-08 | 2.300E-07 | 2.600E-07 |
| 8421 | 422 | T422A | A | 2.8e-07 | 3.2e-07 | 3.1e-07 | 3.033E-07 | 2.082E-08 | 2.800E-07 | 3.200E-07 |
| 8422 | 422 | T422R | A | 1.7e-07 | 2.4e-07 | 4.2e-07 | 2.767E-07 | 1.290E-07 | 1.700E-07 | 4.200E-07 |
| 8423 | 422 | T422N | A | 2.6e-07 | 1.6e-07 | 3.2e-07 | 2.467E-07 | 8.083E-08 | 1.600E-07 | 3.200E-07 |
| 8424 | 422 | T422D | A | 2.7e-07 | 1.3e-07 | 2e-07 | 2.000E-07 | 7.000E-08 | 1.300E-07 | 2.700E-07 |
| 8425 | 422 | T422C | A | 3.3e-07 | 1.2e-07 | 1.7e-07 | 2.067E-07 | 1.097E-07 | 1.200E-07 | 3.300E-07 |
| 8426 | 422 | T422E | A | 1.7e-07 | 4.1e-07 | 1.2e-07 | 2.333E-07 | 1.550E-07 | 1.200E-07 | 4.100E-07 |
| 8427 | 422 | T422Q | A | 1.9e-07 | 4e-07 | 1.7e-07 | 2.533E-07 | 1.274E-07 | 1.700E-07 | 4.000E-07 |
| 8428 | 422 | T422G | A | 1.7e-07 | 1.4e-07 | 2.1e-07 | 1.733E-07 | 3.512E-08 | 1.400E-07 | 2.100E-07 |
| 8429 | 422 | T422H | A | 1.7e-07 | 2.8e-07 | 4.8e-07 | 3.100E-07 | 1.572E-07 | 1.700E-07 | 4.800E-07 |
| 8430 | 422 | T422I | A | 3.9e-07 | 1.7e-07 | 2.1e-07 | 2.567E-07 | 1.172E-07 | 1.700E-07 | 3.900E-07 |
| 8431 | 422 | T422L | A | 3.1e-07 | 2e-07 | 4.1e-07 | 3.067E-07 | 1.050E-07 | 2.000E-07 | 4.100E-07 |
| 8432 | 422 | T422K | A | 1.8e-07 | 2.6e-07 | 1.9e-07 | 2.100E-07 | 4.359E-08 | 1.800E-07 | 2.600E-07 |
| 8433 | 422 | T422M | A | 5.3e-07 | 2.4e-07 | 1.4e-07 | 3.033E-07 | 2.026E-07 | 1.400E-07 | 5.300E-07 |
| 8434 | 422 | T422F | A | 1.3e-07 | 1.8e-07 | 7.8e-07 | 3.633E-07 | 3.617E-07 | 1.300E-07 | 7.800E-07 |
| 8435 | 422 | T422P | A | 2.7e-07 | 2.4e-07 | 1.7e-07 | 2.267E-07 | 5.132E-08 | 1.700E-07 | 2.700E-07 |
| 8436 | 422 | T422S | A | 2.9e-07 | 2.6e-07 | 1.3e-07 | 2.267E-07 | 8.505E-08 | 1.300E-07 | 2.900E-07 |
| 8437 | 422 | T422T | A | 2.6e-07 | 1.9e-07 | 2.8e-07 | 2.433E-07 | 4.726E-08 | 1.900E-07 | 2.800E-07 |
| 8438 | 422 | T422W | A | 2.5e-07 | 4e-07 | 3.6e-07 | 3.367E-07 | 7.767E-08 | 2.500E-07 | 4.000E-07 |
| 8439 | 422 | T422Y | A | 2.6e-07 | 3.4e-07 | 2.2e-07 | 2.733E-07 | 6.110E-08 | 2.200E-07 | 3.400E-07 |
| 8440 | 422 | T422V | A | 6.5e-07 | 3.1e-07 | 2.1e-07 | 3.900E-07 | 2.307E-07 | 2.100E-07 | 6.500E-07 |
| 8441 | 423 | L423A | A | 3.4e-07 | 1.2e-07 | 1.7e-07 | 2.100E-07 | 1.153E-07 | 1.200E-07 | 3.400E-07 |
| 8442 | 423 | L423R | A | 1.6e-07 | 3.6e-07 | 2.7e-07 | 2.633E-07 | 1.002E-07 | 1.600E-07 | 3.600E-07 |
| 8443 | 423 | L423N | A | 3.1e-07 | 1.8e-07 | 4.9e-07 | 3.267E-07 | 1.557E-07 | 1.800E-07 | 4.900E-07 |
| 8444 | 423 | L423D | A | 2.8e-07 | 1.1e-07 | 5.5e-07 | 3.133E-07 | 2.219E-07 | 1.100E-07 | 5.500E-07 |
| 8445 | 423 | L423C | A | 3.2e-07 | 1.2e-07 | 5.1e-07 | 3.167E-07 | 1.950E-07 | 1.200E-07 | 5.100E-07 |
| 8446 | 423 | L423E | A | 3.2e-07 | 1.9e-07 | 2.3e-07 | 2.467E-07 | 6.658E-08 | 1.900E-07 | 3.200E-07 |
| 8447 | 423 | L423Q | A | 3.3e-07 | 3.3e-07 | 2.4e-07 | 3.000E-07 | 5.196E-08 | 2.400E-07 | 3.300E-07 |
| 8448 | 423 | L423G | A | 3.9e-07 | 4e-07 | 1.4e-07 | 3.100E-07 | 1.473E-07 | 1.400E-07 | 4.000E-07 |
| 8449 | 423 | L423H | A | 1.7e-07 | 1.9e-07 | 2e-07 | 1.867E-07 | 1.528E-08 | 1.700E-07 | 2.000E-07 |
| 8450 | 423 | L423I | A | 4.1e-07 | 1.6e-07 | 2.5e-07 | 2.733E-07 | 1.266E-07 | 1.600E-07 | 4.100E-07 |
| 8451 | 423 | L423L | A | 1.6e-07 | 5.6e-07 | 2.9e-07 | 3.367E-07 | 2.040E-07 | 1.600E-07 | 5.600E-07 |
| 8452 | 423 | L423K | A | 4.7e-07 | 3.2e-07 | 1.7e-07 | 3.200E-07 | 1.500E-07 | 1.700E-07 | 4.700E-07 |
| 8453 | 423 | L423M | A | 4.4e-07 | 4.3e-07 | 2.6e-07 | 3.767E-07 | 1.012E-07 | 2.600E-07 | 4.400E-07 |
| 8454 | 423 | L423F | A | 3.2e-07 | 1.1e-07 | 3.3e-07 | 2.533E-07 | 1.242E-07 | 1.100E-07 | 3.300E-07 |
| 8455 | 423 | L423P | A | 2.6e-07 | 2.4e-07 | 1.1e-07 | 2.033E-07 | 8.145E-08 | 1.100E-07 | 2.600E-07 |
| 8456 | 423 | L423S | A | 2.6e-07 | 1.8e-07 | 2.4e-07 | 2.267E-07 | 4.163E-08 | 1.800E-07 | 2.600E-07 |
| 8457 | 423 | L423T | A | 5.4e-07 | 1.6e-07 | 2.8e-07 | 3.267E-07 | 1.943E-07 | 1.600E-07 | 5.400E-07 |
| 8458 | 423 | L423W | A | 3.9e-07 | 1.9e-07 | 1.9e-07 | 2.567E-07 | 1.155E-07 | 1.900E-07 | 3.900E-07 |
| 8459 | 423 | L423Y | A | 1.5e-07 | 3.7e-07 | 8.5e-08 | 2.017E-07 | 1.494E-07 | 8.500E-08 | 3.700E-07 |
| 8460 | 423 | L423V | A | 6.3e-07 | 1.5e-07 | 1.8e-07 | 3.200E-07 | 2.689E-07 | 1.500E-07 | 6.300E-07 |
| 8461 | 424 | Q424A | A | 2.3e-07 | 2.1e-07 | 3.9e-07 | 2.767E-07 | 9.866E-08 | 2.100E-07 | 3.900E-07 |
| 8462 | 424 | Q424R | A | 1.9e-07 | 1.3e-07 | 4.9e-07 | 2.700E-07 | 1.929E-07 | 1.300E-07 | 4.900E-07 |
| 8463 | 424 | Q424N | A | 2.6e-07 | 1.7e-07 | 2.3e-07 | 2.200E-07 | 4.583E-08 | 1.700E-07 | 2.600E-07 |
| 8464 | 424 | Q424D | A | 3.4e-07 | 3.2e-07 | 2.7e-07 | 3.100E-07 | 3.606E-08 | 2.700E-07 | 3.400E-07 |
| 8465 | 424 | Q424C | A | 3.1e-07 | 3.5e-07 | 3.3e-07 | 3.300E-07 | 2.000E-08 | 3.100E-07 | 3.500E-07 |
| 8466 | 424 | Q424E | A | 1.3e-07 | 4.2e-07 | 1e-07 | 2.167E-07 | 1.767E-07 | 1.000E-07 | 4.200E-07 |
| 8467 | 424 | Q424Q | A | 3.3e-07 | 1.6e-07 | 1.9e-07 | 2.267E-07 | 9.074E-08 | 1.600E-07 | 3.300E-07 |
| 8468 | 424 | Q424G | A | 2.8e-07 | 2.2e-07 | 1.8e-07 | 2.267E-07 | 5.033E-08 | 1.800E-07 | 2.800E-07 |
| 8469 | 424 | Q424H | A | 2.1e-07 | 2.6e-07 | 2.6e-07 | 2.433E-07 | 2.887E-08 | 2.100E-07 | 2.600E-07 |
| 8470 | 424 | Q424I | A | 2e-07 | 6.2e-08 | 9.8e-08 | 1.200E-07 | 7.158E-08 | 6.200E-08 | 2.000E-07 |
| 8471 | 424 | Q424L | A | 1.6e-07 | 2.1e-07 | 3.2e-07 | 2.300E-07 | 8.185E-08 | 1.600E-07 | 3.200E-07 |
| 8472 | 424 | Q424K | A | 3.6e-07 | 3.3e-07 | 3.6e-07 | 3.500E-07 | 1.732E-08 | 3.300E-07 | 3.600E-07 |
| 8473 | 424 | Q424M | A | 2.2e-07 | 1.4e-07 | 6.5e-07 | 3.367E-07 | 2.743E-07 | 1.400E-07 | 6.500E-07 |
| 8474 | 424 | Q424F | A | 1.4e-07 | 4.1e-07 | 2.2e-07 | 2.567E-07 | 1.387E-07 | 1.400E-07 | 4.100E-07 |
| 8475 | 424 | Q424P | A | 2.4e-07 | 2.4e-07 | 5.9e-07 | 3.567E-07 | 2.021E-07 | 2.400E-07 | 5.900E-07 |
| 8476 | 424 | Q424S | A | 3.1e-07 | 1.8e-07 | 2.6e-07 | 2.500E-07 | 6.557E-08 | 1.800E-07 | 3.100E-07 |
| 8477 | 424 | Q424T | A | 1.9e-07 | 1.2e-07 | 1.3e-07 | 1.467E-07 | 3.786E-08 | 1.200E-07 | 1.900E-07 |
| 8478 | 424 | Q424W | A | 4.1e-07 | 3.7e-07 | 6.2e-07 | 4.667E-07 | 1.343E-07 | 3.700E-07 | 6.200E-07 |
| 8479 | 424 | Q424Y | A | 2.2e-07 | 1.3e-07 | 3e-07 | 2.167E-07 | 8.505E-08 | 1.300E-07 | 3.000E-07 |
| 8480 | 424 | Q424V | A | 1.5e-07 | 3e-07 | 2.8e-07 | 2.433E-07 | 8.145E-08 | 1.500E-07 | 3.000E-07 |
| 8481 | 425 | G425A | A | 2.6e-07 | 1.3e-07 | 2.5e-07 | 2.133E-07 | 7.234E-08 | 1.300E-07 | 2.600E-07 |
| 8482 | 425 | G425R | A | 1.3e-07 | 2.6e-07 | 3e-07 | 2.300E-07 | 8.888E-08 | 1.300E-07 | 3.000E-07 |
| 8483 | 425 | G425N | A | 1.7e-07 | 2.2e-07 | 3e-07 | 2.300E-07 | 6.557E-08 | 1.700E-07 | 3.000E-07 |
| 8484 | 425 | G425D | A | 9.5e-08 | 4.3e-07 | 2.6e-07 | 2.617E-07 | 1.675E-07 | 9.500E-08 | 4.300E-07 |
| 8485 | 425 | G425C | A | 9.7e-08 | 1.5e-07 | 3.2e-07 | 1.890E-07 | 1.165E-07 | 9.700E-08 | 3.200E-07 |
| 8486 | 425 | G425E | A | 3.9e-07 | 1.5e-07 | 3.9e-07 | 3.100E-07 | 1.386E-07 | 1.500E-07 | 3.900E-07 |
| 8487 | 425 | G425Q | A | 2.1e-07 | 1.3e-07 | 1.3e-07 | 1.567E-07 | 4.619E-08 | 1.300E-07 | 2.100E-07 |
| 8488 | 425 | G425G | A | 2.9e-07 | 2.1e-07 | 3e-07 | 2.667E-07 | 4.933E-08 | 2.100E-07 | 3.000E-07 |
| 8489 | 425 | G425H | A | 2.2e-07 | 1.4e-07 | 8.4e-08 | 1.480E-07 | 6.835E-08 | 8.400E-08 | 2.200E-07 |
| 8490 | 425 | G425I | A | 2.8e-07 | 2.9e-07 | 2e-07 | 2.567E-07 | 4.933E-08 | 2.000E-07 | 2.900E-07 |
| 8491 | 425 | G425L | A | 3.5e-07 | 1.9e-07 | 3.1e-07 | 2.833E-07 | 8.327E-08 | 1.900E-07 | 3.500E-07 |
| 8492 | 425 | G425K | A | 2.4e-07 | 2.4e-07 | 1.3e-07 | 2.033E-07 | 6.351E-08 | 1.300E-07 | 2.400E-07 |
| 8493 | 425 | G425M | A | 2.5e-07 | 2.6e-07 | 2.5e-07 | 2.533E-07 | 5.774E-09 | 2.500E-07 | 2.600E-07 |
| 8494 | 425 | G425F | A | 2.5e-07 | 1.8e-07 | 2.7e-07 | 2.333E-07 | 4.726E-08 | 1.800E-07 | 2.700E-07 |
| 8495 | 425 | G425P | A | 1e-07 | 5.1e-07 | 1.4e-07 | 2.500E-07 | 2.261E-07 | 1.000E-07 | 5.100E-07 |
| 8496 | 425 | G425S | A | 3.4e-07 | 1.7e-07 | 2.6e-07 | 2.567E-07 | 8.505E-08 | 1.700E-07 | 3.400E-07 |
| 8497 | 425 | G425T | A | 2.2e-07 | 1.5e-07 | 5.3e-08 | 1.410E-07 | 8.386E-08 | 5.300E-08 | 2.200E-07 |
| 8498 | 425 | G425W | A | 2.1e-07 | 1.9e-07 | 1.2e-07 | 1.733E-07 | 4.726E-08 | 1.200E-07 | 2.100E-07 |
| 8499 | 425 | G425Y | A | 3e-07 | 1.7e-07 | 5.2e-07 | 3.300E-07 | 1.769E-07 | 1.700E-07 | 5.200E-07 |

| | | | | | | | | | | |
|---|---|---|---|---|---|---|---|---|---|---|
| 8500 | 425 | G425V | A | 1.9e-07 | 2.3e-07 | 1.5e-07 | 1.900E-07 | 4.000E-08 | 1.500E-07 | 2.300E-07 |
| 8501 | 426 | L426A | A | 1.3e-07 | 1.2e-07 | 2e-07 | 1.500E-07 | 4.359E-08 | 1.200E-07 | 2.000E-07 |
| 8502 | 426 | L426R | A | 2e-07 | 1.8e-07 | 2.8e-07 | 2.200E-07 | 5.292E-08 | 1.800E-07 | 2.800E-07 |
| 8503 | 426 | L426N | A | 1.5e-07 | 2e-07 | 2e-07 | 1.833E-07 | 2.887E-08 | 1.500E-07 | 2.000E-07 |
| 8504 | 426 | L426D | A | 1.8e-07 | 3.4e-07 | 3e-07 | 2.733E-07 | 8.327E-08 | 1.800E-07 | 3.400E-07 |
| 8505 | 426 | L426C | A | 1.5e-07 | 6.4e-07 | 5.7e-07 | 4.533E-07 | 2.650E-07 | 1.500E-07 | 6.400E-07 |
| 8506 | 426 | L426E | A | 2.2e-07 | 1.4e-07 | 2e-07 | 1.867E-07 | 4.163E-08 | 1.400E-07 | 2.200E-07 |
| 8507 | 426 | L426Q | A | 2.3e-07 | 1e-07 | 2.4e-07 | 1.900E-07 | 7.810E-08 | 1.000E-07 | 2.400E-07 |
| 8508 | 426 | L426G | A | 1.4e-07 | 2.4e-07 | 3.3e-07 | 2.367E-07 | 9.504E-08 | 1.400E-07 | 3.300E-07 |
| 8509 | 426 | L426H | A | 2.3e-07 | 1.3e-07 | 2.5e-07 | 2.033E-07 | 6.429E-08 | 1.300E-07 | 2.500E-07 |
| 8510 | 426 | L426I | A | 2.5e-07 | 3e-07 | 2.6e-07 | 2.700E-07 | 2.646E-08 | 2.500E-07 | 3.000E-07 |
| 8511 | 426 | L426L | A | 1.3e-07 | 2.1e-07 | 9.1e-08 | 1.437E-07 | 6.067E-08 | 9.100E-08 | 2.100E-07 |
| 8512 | 426 | L426K | A | 1.7e-07 | 1.8e-07 | 8.8e-08 | 1.460E-07 | 5.048E-08 | 8.800E-08 | 1.800E-07 |
| 8513 | 426 | L426M | A | 2.5e-07 | 5.9e-07 | 1.5e-07 | 3.300E-07 | 2.307E-07 | 1.500E-07 | 5.900E-07 |
| 8514 | 426 | L426F | A | 1.1e-07 | 2.9e-07 | 2.7e-07 | 2.233E-07 | 9.866E-08 | 1.100E-07 | 2.900E-07 |
| 8515 | 426 | L426P | A | 3.9e-07 | 2.4e-07 | 2.3e-07 | 2.867E-07 | 8.963E-08 | 2.300E-07 | 3.900E-07 |
| 8516 | 426 | L426S | A | 1.6e-07 | 5.1e-07 | 8.4e-08 | 2.513E-07 | 2.272E-07 | 8.400E-08 | 5.100E-07 |
| 8517 | 426 | L426T | A | 5.9e-07 | 2.7e-07 | 1.6e-07 | 3.400E-07 | 2.234E-07 | 1.600E-07 | 5.900E-07 |
| 8518 | 426 | L426W | A | 2.9e-07 | 3e-07 | 3.6e-07 | 3.167E-07 | 3.786E-08 | 2.900E-07 | 3.600E-07 |
| 8519 | 426 | L426Y | A | 6.2e-07 | 1.2e-07 | 3.7e-07 | 3.700E-07 | 2.500E-07 | 1.200E-07 | 6.200E-07 |
| 8520 | 426 | L426V | A | 1.7e-07 | 1.3e-07 | 3.8e-07 | 2.267E-07 | 1.343E-07 | 1.300E-07 | 3.800E-07 |
| 8521 | 427 | G427A | A | 3.9e-07 | 3.9e-07 | 1.6e-07 | 3.133E-07 | 1.328E-07 | 1.600E-07 | 3.900E-07 |
| 8522 | 427 | G427R | A | 2.1e-07 | 2.9e-07 | 1.8e-07 | 2.267E-07 | 5.686E-08 | 1.800E-07 | 2.900E-07 |
| 8523 | 427 | G427N | A | 2.7e-07 | 1.7e-07 | 2.2e-07 | 2.200E-07 | 5.000E-08 | 1.700E-07 | 2.700E-07 |
| 8524 | 427 | G427D | A | 2.2e-07 | 2e-07 | 1.7e-07 | 1.967E-07 | 2.517E-08 | 1.700E-07 | 2.200E-07 |
| 8525 | 427 | G427C | A | 2.9e-07 | 1.1e-07 | 1.9e-07 | 1.967E-07 | 9.018E-08 | 1.100E-07 | 2.900E-07 |
| 8526 | 427 | G427E | A | 4.3e-07 | 1.7e-07 | 4.5e-07 | 3.500E-07 | 1.562E-07 | 1.700E-07 | 4.500E-07 |
| 8527 | 427 | G427Q | A | 1.4e-07 | 2.6e-07 | 2.1e-07 | 2.033E-07 | 6.028E-08 | 1.400E-07 | 2.600E-07 |
| 8528 | 427 | G427G | A | 2.7e-07 | 2.2e-07 | 2.3e-07 | 2.400E-07 | 2.646E-08 | 2.200E-07 | 2.700E-07 |
| 8529 | 427 | G427H | A | 1.8e-07 | 3.2e-07 | 1.7e-07 | 2.233E-07 | 8.386E-08 | 1.700E-07 | 3.200E-07 |
| 8530 | 427 | G427I | A | 1.8e-07 | 3.3e-07 | 2.6e-07 | 2.567E-07 | 7.506E-08 | 1.800E-07 | 3.300E-07 |
| 8531 | 427 | G427L | A | 2.3e-07 | 3.3e-07 | 1.2e-07 | 2.267E-07 | 1.050E-07 | 1.200E-07 | 3.300E-07 |
| 8532 | 427 | G427K | A | 1.6e-07 | 3.4e-07 | 2.8e-07 | 2.600E-07 | 9.165E-08 | 1.600E-07 | 3.400E-07 |
| 8533 | 427 | G427M | A | 4.7e-07 | 1.4e-07 | 2.4e-07 | 2.833E-07 | 1.692E-07 | 1.400E-07 | 4.700E-07 |
| 8534 | 427 | G427F | A | 1.6e-07 | 1.5e-07 | 3.9e-07 | 2.333E-07 | 1.358E-07 | 1.500E-07 | 3.900E-07 |
| 8535 | 427 | G427P | A | 1.8e-07 | 3.3e-07 | 3.6e-07 | 2.900E-07 | 9.644E-08 | 1.800E-07 | 3.600E-07 |
| 8536 | 427 | G427S | A | 1.9e-07 | 3.4e-07 | 1.1e-07 | 2.133E-07 | 1.168E-07 | 1.100E-07 | 3.400E-07 |
| 8537 | 427 | G427T | A | 3.6e-07 | 1.1e-07 | 2.7e-07 | 2.467E-07 | 1.266E-07 | 1.100E-07 | 3.600E-07 |
| 8538 | 427 | G427W | A | 3.8e-07 | 2.9e-07 | 2.7e-07 | 3.133E-07 | 5.859E-08 | 2.700E-07 | 3.800E-07 |
| 8539 | 427 | G427Y | A | 1.9e-07 | 2.2e-07 | 3.4e-07 | 2.500E-07 | 7.937E-08 | 1.900E-07 | 3.400E-07 |
| 8540 | 427 | G427V | A | 1.4e-07 | 2.5e-07 | 2.2e-07 | 2.033E-07 | 5.686E-08 | 1.400E-07 | 2.500E-07 |
| 8541 | 428 | I428A | A | 4.7e-07 | 4e-07 | 1.9e-07 | 3.533E-07 | 1.457E-07 | 1.900E-07 | 4.700E-07 |
| 8542 | 428 | I428R | A | 2.2e-07 | 1.1e-07 | 3.3e-07 | 2.200E-07 | 1.100E-07 | 1.100E-07 | 3.300E-07 |
| 8543 | 428 | I428N | A | 2.5e-07 | 4.2e-07 | 2.8e-07 | 3.167E-07 | 9.074E-08 | 2.500E-07 | 4.200E-07 |
| 8544 | 428 | I428D | A | 1e-07 | 3e-07 | 5e-07 | 3.000E-07 | 2.000E-07 | 1.000E-07 | 5.000E-07 |
| 8545 | 428 | I428C | A | 2.1e-07 | 1.6e-07 | 4.7e-07 | 2.800E-07 | 1.664E-07 | 1.600E-07 | 4.700E-07 |
| 8546 | 428 | I428E | A | 3.7e-07 | 2.6e-07 | 3.3e-07 | 3.200E-07 | 5.568E-08 | 2.600E-07 | 3.700E-07 |
| 8547 | 428 | I428Q | A | 1.7e-07 | 2.4e-07 | 2.4e-07 | 2.167E-07 | 4.041E-08 | 1.700E-07 | 2.400E-07 |
| 8548 | 428 | I428G | A | 4e-07 | 1.3e-07 | 2e-07 | 2.433E-07 | 1.401E-07 | 1.300E-07 | 4.000E-07 |
| 8549 | 428 | I428H | A | 1.2e-07 | 1.1e-07 | 1.1e-07 | 1.133E-07 | 5.774E-09 | 1.100E-07 | 1.200E-07 |
| 8550 | 428 | I428I | A | 1.7e-07 | 2.8e-07 | 3e-07 | 2.500E-07 | 7.000E-08 | 1.700E-07 | 3.000E-07 |
| 8551 | 428 | I428L | A | 2.4e-07 | 2e-07 | 3.5e-07 | 2.633E-07 | 7.767E-08 | 2.000E-07 | 3.500E-07 |
| 8552 | 428 | I428K | A | 2.8e-07 | 2.8e-07 | 1.4e-07 | 2.333E-07 | 8.083E-08 | 1.400E-07 | 2.800E-07 |
| 8553 | 428 | I428M | A | 3.7e-07 | 3.2e-07 | 4.9e-07 | 3.933E-07 | 8.737E-08 | 3.200E-07 | 4.900E-07 |
| 8554 | 428 | I428F | A | 2.7e-07 | 4e-07 | 2.7e-07 | 3.133E-07 | 7.506E-08 | 2.700E-07 | 4.000E-07 |
| 8555 | 428 | I428P | A | 4.7e-07 | 2.2e-07 | 2.6e-07 | 3.167E-07 | 1.343E-07 | 2.200E-07 | 4.700E-07 |
| 8556 | 428 | I428S | A | 3.1e-07 | 1.6e-07 | 3.4e-07 | 2.700E-07 | 9.644E-08 | 1.600E-07 | 3.400E-07 |
| 8557 | 428 | I428T | A | 1.3e-07 | 5.6e-07 | 1.8e-07 | 2.900E-07 | 2.352E-07 | 1.300E-07 | 5.600E-07 |
| 8558 | 428 | I428W | A | 2.6e-07 | 4.1e-07 | 1.5e-07 | 2.733E-07 | 1.305E-07 | 1.500E-07 | 4.100E-07 |
| 8559 | 428 | I428Y | A | 2.6e-07 | 1.9e-07 | 3.6e-07 | 2.700E-07 | 8.544E-08 | 1.900E-07 | 3.600E-07 |
| 8560 | 428 | I428V | A | 1.5e-07 | 1.7e-07 | 2.5e-07 | 1.900E-07 | 5.292E-08 | 1.500E-07 | 2.500E-07 |
| 8561 | 429 | S429A | A | 3.4e-07 | 5e-07 | 2.6e-07 | 3.667E-07 | 1.222E-07 | 2.600E-07 | 5.000E-07 |
| 8562 | 429 | S429R | A | 5e-07 | 1.9e-07 | 1.9e-07 | 2.933E-07 | 1.790E-07 | 1.900E-07 | 5.000E-07 |
| 8563 | 429 | S429N | A | 2.1e-07 | 2.3e-07 | 1.9e-07 | 2.100E-07 | 2.000E-08 | 1.900E-07 | 2.300E-07 |
| 8564 | 429 | S429D | A | 3.6e-07 | 1.8e-07 | 1.3e-07 | 2.233E-07 | 1.210E-07 | 1.300E-07 | 3.600E-07 |
| 8565 | 429 | S429C | A | 9.8e-08 | 1.1e-07 | 3.2e-07 | 1.760E-07 | 1.249E-07 | 9.800E-08 | 3.200E-07 |
| 8566 | 429 | S429E | A | 4e-07 | 1.9e-07 | 1.5e-07 | 2.467E-07 | 1.343E-07 | 1.500E-07 | 4.000E-07 |
| 8567 | 429 | S429Q | A | 5.9e-07 | 2.8e-07 | 2e-07 | 3.567E-07 | 2.060E-07 | 2.000E-07 | 5.900E-07 |
| 8568 | 429 | S429G | A | 2.7e-07 | 2.3e-07 | 3e-07 | 2.667E-07 | 3.512E-08 | 2.300E-07 | 3.000E-07 |
| 8569 | 429 | S429H | A | 3.6e-07 | 4e-07 | 7.1e-07 | 4.900E-07 | 1.916E-07 | 3.600E-07 | 7.100E-07 |
| 8570 | 429 | S429I | A | 2.3e-07 | 2.5e-07 | 2.9e-07 | 2.567E-07 | 3.055E-08 | 2.300E-07 | 2.900E-07 |
| 8571 | 429 | S429L | A | 5.8e-07 | 2e-07 | 3.3e-07 | 3.700E-07 | 1.931E-07 | 2.000E-07 | 5.800E-07 |
| 8572 | 429 | S429K | A | 2.1e-07 | 2.8e-07 | 1.1e-07 | 2.000E-07 | 8.544E-08 | 1.100E-07 | 2.800E-07 |
| 8573 | 429 | S429M | A | 1.8e-07 | 2.7e-07 | 3.7e-07 | 2.733E-07 | 9.504E-08 | 1.800E-07 | 3.700E-07 |
| 8574 | 429 | S429F | A | 2.5e-07 | 2.3e-07 | 5.5e-07 | 3.433E-07 | 1.793E-07 | 2.300E-07 | 5.500E-07 |
| 8575 | 429 | S429P | A | 2.7e-07 | 1.6e-07 | 2.9e-07 | 2.400E-07 | 7.000E-08 | 1.600E-07 | 2.900E-07 |
| 8576 | 429 | S429S | A | 2.2e-07 | 5.4e-07 | 1.9e-07 | 3.167E-07 | 1.940E-07 | 1.900E-07 | 5.400E-07 |
| 8577 | 429 | S429T | A | 1.9e-07 | 8.1e-08 | 2.2e-07 | 1.637E-07 | 7.315E-08 | 8.100E-08 | 2.200E-07 |
| 8578 | 429 | S429W | A | 2.5e-07 | 3e-07 | 3.3e-07 | 2.933E-07 | 4.041E-08 | 2.500E-07 | 3.300E-07 |
| 8579 | 429 | S429Y | A | 3.8e-07 | 2.4e-07 | 3.5e-07 | 3.233E-07 | 7.371E-08 | 2.400E-07 | 3.800E-07 |
| 8580 | 429 | S429V | A | 2e-07 | 9e-08 | 2.3e-07 | 1.733E-07 | 7.371E-08 | 9.000E-08 | 2.300E-07 |
| 8581 | 430 | W430A | A | 2.9e-07 | 2.9e-07 | 4e-07 | 3.267E-07 | 6.351E-08 | 2.900E-07 | 4.000E-07 |
| 8582 | 430 | W430R | A | 8.9e-08 | 2e-07 | 2.3e-07 | 1.730E-07 | 7.428E-08 | 8.900E-08 | 2.300E-07 |
| 8583 | 430 | W430N | A | 1.4e-07 | 2.3e-07 | 2.8e-07 | 2.167E-07 | 7.095E-08 | 1.400E-07 | 2.800E-07 |
| 8584 | 430 | W430D | A | 3.6e-07 | 2.4e-07 | 5.5e-07 | 3.833E-07 | 1.563E-07 | 2.400E-07 | 5.500E-07 |

| | | | | | | | | | | |
|---|---|---|---|---|---|---|---|---|---|---|
| 8585 | 430 | W430C | A | 4.5e-07 | 5e-07 | 3.5e-07 | 4.333E-07 | 7.638E-08 | 3.500E-07 | 5.000E-07 |
| 8586 | 430 | W430E | A | 5.1e-07 | 7.5e-07 | 1.8e-07 | 4.800E-07 | 2.862E-07 | 1.800E-07 | 7.500E-07 |
| 8587 | 430 | W430Q | A | 1.3e-07 | 3.9e-07 | 2.2e-07 | 2.467E-07 | 1.320E-07 | 1.300E-07 | 3.900E-07 |
| 8588 | 430 | W430G | A | 4e-07 | 4.7e-07 | 2.4e-07 | 3.700E-07 | 1.179E-07 | 2.400E-07 | 4.700E-07 |
| 8589 | 430 | W430H | A | 2.3e-07 | 4.1e-07 | 1.2e-07 | 2.533E-07 | 1.464E-07 | 1.200E-07 | 4.100E-07 |
| 8590 | 430 | W430I | A | 3.2e-07 | 2.9e-07 | 2.4e-07 | 2.833E-07 | 4.041E-08 | 2.400E-07 | 3.200E-07 |
| 8591 | 430 | W430L | A | 4.6e-07 | 8.5e-08 | 1.4e-07 | 2.283E-07 | 2.025E-07 | 8.500E-08 | 4.600E-07 |
| 8592 | 430 | W430K | A | 2.2e-07 | 2.2e-07 | 1.6e-07 | 2.000E-07 | 3.464E-08 | 1.600E-07 | 2.200E-07 |
| 8593 | 430 | W430M | A | 1.5e-07 | 1.9e-07 | 1.3e-07 | 1.567E-07 | 3.055E-08 | 1.300E-07 | 1.900E-07 |
| 8594 | 430 | W430F | A | 1.6e-07 | 8e-08 | 1.4e-07 | 1.267E-07 | 4.163E-08 | 8.000E-08 | 1.600E-07 |
| 8595 | 430 | W430P | A | 2.9e-07 | 1.9e-07 | 1.3e-07 | 2.033E-07 | 8.083E-08 | 1.300E-07 | 2.900E-07 |
| 8596 | 430 | W430S | A | 1.8e-07 | 3.7e-07 | 1.9e-07 | 2.467E-07 | 1.069E-07 | 1.800E-07 | 3.700E-07 |
| 8597 | 430 | W430T | A | 2.4e-07 | 2.8e-07 | 2.2e-07 | 2.467E-07 | 3.055E-08 | 2.200E-07 | 2.800E-07 |
| 8598 | 430 | W430W | A | 2.6e-07 | 3.2e-07 | 3.2e-07 | 3.000E-07 | 3.464E-08 | 2.600E-07 | 3.200E-07 |
| 8599 | 430 | W430Y | A | 4.2e-07 | 3.2e-07 | 9.6e-08 | 2.787E-07 | 1.659E-07 | 9.600E-08 | 4.200E-07 |
| 8600 | 430 | W430V | A | 2.6e-07 | 3e-07 | 2.2e-07 | 2.600E-07 | 4.000E-08 | 2.200E-07 | 3.000E-07 |
| 8601 | 431 | L431A | A | 5.6e-07 | 4.6e-07 | 5.7e-07 | 5.300E-07 | 6.083E-08 | 4.600E-07 | 5.700E-07 |
| 8602 | 431 | L431R | A | 2.8e-07 | 1.4e-07 | 2.6e-07 | 2.267E-07 | 7.572E-08 | 1.400E-07 | 2.800E-07 |
| 8603 | 431 | L431N | A | 3.4e-07 | 1.5e-07 | 5e-07 | 3.300E-07 | 1.752E-07 | 1.500E-07 | 5.000E-07 |
| 8604 | 431 | L431D | A | 2.7e-07 | 2.7e-07 | 2.3e-07 | 2.567E-07 | 2.309E-08 | 2.300E-07 | 2.700E-07 |
| 8605 | 431 | L431C | A | 5.9e-07 | 1.7e-07 | 1.9e-07 | 3.167E-07 | 2.369E-07 | 1.700E-07 | 5.900E-07 |
| 8606 | 431 | L431E | A | 1.3e-07 | 2.2e-07 | 1.3e-07 | 1.600E-07 | 5.196E-08 | 1.300E-07 | 2.200E-07 |
| 8607 | 431 | L431Q | A | 3.9e-07 | 1.6e-07 | 1.5e-07 | 2.333E-07 | 1.358E-07 | 1.500E-07 | 3.900E-07 |
| 8608 | 431 | L431G | A | 2.7e-07 | 3.6e-07 | 3e-07 | 3.100E-07 | 4.583E-08 | 2.700E-07 | 3.600E-07 |
| 8609 | 431 | L431H | A | 2.1e-07 | 1.9e-07 | 3.6e-07 | 2.533E-07 | 9.292E-08 | 1.900E-07 | 3.600E-07 |
| 8610 | 431 | L431I | A | 3.5e-07 | 3.1e-07 | 2.5e-07 | 3.033E-07 | 5.033E-08 | 2.500E-07 | 3.500E-07 |
| 8611 | 431 | L431L | A | 2.4e-07 | 3.4e-07 | 3e-07 | 2.933E-07 | 5.033E-08 | 2.400E-07 | 3.400E-07 |
| 8612 | 431 | L431K | A | 3.6e-07 | 2.4e-07 | 2.4e-07 | 2.800E-07 | 6.928E-08 | 2.400E-07 | 3.600E-07 |
| 8613 | 431 | L431M | A | 3.3e-07 | 2.7e-07 | 4.2e-07 | 3.400E-07 | 7.550E-08 | 2.700E-07 | 4.200E-07 |
| 8614 | 431 | L431F | A | 2.9e-07 | 3.9e-07 | 3.2e-07 | 3.333E-07 | 5.132E-08 | 2.900E-07 | 3.900E-07 |
| 8615 | 431 | L431P | A | 4.5e-07 | 1.5e-07 | 2.9e-07 | 2.967E-07 | 1.501E-07 | 1.500E-07 | 4.500E-07 |
| 8616 | 431 | L431S | A | 2e-07 | 3.6e-07 | 3.7e-07 | 3.100E-07 | 9.539E-08 | 2.000E-07 | 3.700E-07 |
| 8617 | 431 | L431T | A | 1.2e-07 | 2.5e-07 | 1.7e-07 | 1.800E-07 | 6.557E-08 | 1.200E-07 | 2.500E-07 |
| 8618 | 431 | L431W | A | 3.2e-07 | 2.8e-07 | 2.7e-07 | 2.900E-07 | 2.646E-08 | 2.700E-07 | 3.200E-07 |
| 8619 | 431 | L431Y | A | 3.2e-07 | 1.6e-07 | 1.3e-07 | 2.033E-07 | 1.021E-07 | 1.300E-07 | 3.200E-07 |
| 8620 | 431 | L431V | A | 3.1e-07 | 3.2e-07 | 1.8e-07 | 2.700E-07 | 7.810E-08 | 1.800E-07 | 3.200E-07 |
| 8621 | 432 | G432A | A | 3.2e-07 | 2.8e-07 | 1.9e-07 | 2.633E-07 | 6.658E-08 | 1.900E-07 | 3.200E-07 |
| 8622 | 432 | G432R | A | 3e-07 | 2.9e-07 | 3.6e-07 | 3.167E-07 | 3.786E-08 | 2.900E-07 | 3.600E-07 |
| 8623 | 432 | G432N | A | 1.7e-07 | 1.6e-07 | 3e-07 | 2.100E-07 | 7.810E-08 | 1.600E-07 | 3.000E-07 |
| 8624 | 432 | G432D | A | 6.7e-08 | 2.4e-07 | 2.5e-07 | 1.857E-07 | 1.029E-07 | 6.700E-08 | 2.500E-07 |
| 8625 | 432 | G432C | A | 3e-07 | 2.2e-07 | 3.3e-07 | 2.833E-07 | 5.686E-08 | 2.200E-07 | 3.300E-07 |
| 8626 | 432 | G432E | A | 1.5e-07 | 1.9e-07 | 1.5e-07 | 1.633E-07 | 2.309E-08 | 1.500E-07 | 1.900E-07 |
| 8627 | 432 | G432Q | A | 3.4e-07 | 2.9e-07 | 1.7e-07 | 2.667E-07 | 8.737E-08 | 1.700E-07 | 3.400E-07 |
| 8628 | 432 | G432G | A | 2.6e-07 | 2.4e-07 | 3.9e-07 | 2.967E-07 | 8.145E-08 | 2.400E-07 | 3.900E-07 |
| 8629 | 432 | G432H | A | 3e-07 | 3.7e-07 | 2e-07 | 2.900E-07 | 8.544E-08 | 2.000E-07 | 3.700E-07 |
| 8630 | 432 | G432I | A | 1.7e-07 | 6e-07 | 1.5e-07 | 3.067E-07 | 2.542E-07 | 1.500E-07 | 6.000E-07 |
| 8631 | 432 | G432L | A | 1.4e-07 | 2e-07 | 2.1e-07 | 1.833E-07 | 3.786E-08 | 1.400E-07 | 2.100E-07 |
| 8632 | 432 | G432K | A | 2.3e-07 | 3.9e-07 | 4.1e-07 | 3.433E-07 | 9.866E-08 | 2.300E-07 | 4.100E-07 |
| 8633 | 432 | G432M | A | 3.6e-07 | 3.4e-07 | 1.7e-07 | 2.900E-07 | 1.044E-07 | 1.700E-07 | 3.600E-07 |
| 8634 | 432 | G432F | A | 1.7e-07 | 5.1e-07 | 2.1e-07 | 2.967E-07 | 1.858E-07 | 1.700E-07 | 5.100E-07 |
| 8635 | 432 | G432P | A | 1.8e-07 | 1.9e-07 | 3.5e-07 | 2.400E-07 | 9.539E-08 | 1.800E-07 | 3.500E-07 |
| 8636 | 432 | G432S | A | 3.8e-07 | 4.3e-07 | 1.2e-07 | 3.100E-07 | 1.664E-07 | 1.200E-07 | 4.300E-07 |
| 8637 | 432 | G432T | A | 9.9e-08 | 2.1e-07 | 1.7e-07 | 1.597E-07 | 5.622E-08 | 9.900E-08 | 2.100E-07 |
| 8638 | 432 | G432W | A | 1.4e-07 | 1.9e-07 | 2.8e-07 | 2.033E-07 | 7.095E-08 | 1.400E-07 | 2.800E-07 |
| 8639 | 432 | G432Y | A | 4.3e-07 | 3.6e-07 | 1.9e-07 | 3.267E-07 | 1.234E-07 | 1.900E-07 | 4.300E-07 |
| 8640 | 432 | G432V | A | 4.1e-07 | 3.5e-07 | 2.7e-07 | 3.433E-07 | 7.024E-08 | 2.700E-07 | 4.100E-07 |
| 8641 | 433 | L433A | A | 3.7e-07 | 8.8e-07 | 1.8e-07 | 4.767E-07 | 3.620E-07 | 1.800E-07 | 8.800E-07 |
| 8642 | 433 | L433R | A | 3.5e-07 | 2.8e-07 | 4.2e-07 | 3.500E-07 | 7.000E-08 | 2.800E-07 | 4.200E-07 |
| 8643 | 433 | L433N | A | 1.9e-07 | 2.3e-07 | 2.2e-07 | 2.133E-07 | 2.082E-08 | 1.900E-07 | 2.300E-07 |
| 8644 | 433 | L433D | A | 4.1e-07 | 2.6e-07 | 2.9e-07 | 3.200E-07 | 7.937E-08 | 2.600E-07 | 4.100E-07 |
| 8645 | 433 | L433C | A | 4.2e-07 | 3.3e-07 | 3e-07 | 3.500E-07 | 6.245E-08 | 3.000E-07 | 4.200E-07 |
| 8646 | 433 | L433E | A | 2.4e-07 | 1.9e-07 | 2.1e-07 | 2.133E-07 | 2.517E-08 | 1.900E-07 | 2.400E-07 |
| 8647 | 433 | L433Q | A | 3.7e-07 | 1.5e-07 | 1.2e-07 | 2.133E-07 | 1.365E-07 | 1.200E-07 | 3.700E-07 |
| 8648 | 433 | L433G | A | 2.5e-07 | 1e-07 | 2.6e-07 | 2.033E-07 | 8.963E-08 | 1.000E-07 | 2.600E-07 |
| 8649 | 433 | L433H | A | 2.4e-07 | 2.2e-07 | 6.8e-07 | 3.800E-07 | 2.600E-07 | 2.200E-07 | 6.800E-07 |
| 8650 | 433 | L433I | A | 2.9e-07 | 2.8e-07 | 2.8e-07 | 2.833E-07 | 5.774E-09 | 2.800E-07 | 2.900E-07 |
| 8651 | 433 | L433L | A | 1.7e-07 | 2.7e-07 | 2.1e-07 | 2.167E-07 | 5.033E-08 | 1.700E-07 | 2.700E-07 |
| 8652 | 433 | L433K | A | 4.9e-07 | 2.4e-07 | 2.3e-07 | 3.200E-07 | 1.473E-07 | 2.300E-07 | 4.900E-07 |
| 8653 | 433 | L433M | A | 1.5e-07 | 2.6e-07 | 1.8e-07 | 1.967E-07 | 5.686E-08 | 1.500E-07 | 2.600E-07 |
| 8654 | 433 | L433F | A | 2e-07 | 2.7e-07 | 2.6e-07 | 2.433E-07 | 3.786E-08 | 2.000E-07 | 2.700E-07 |
| 8655 | 433 | L433P | A | 6.3e-07 | 2e-07 | 2e-07 | 3.433E-07 | 2.483E-07 | 2.000E-07 | 6.300E-07 |
| 8656 | 433 | L433S | A | 3.3e-07 | 1.5e-07 | 1.9e-07 | 2.233E-07 | 9.452E-08 | 1.500E-07 | 3.300E-07 |
| 8657 | 433 | L433T | A | 1.2e-07 | 2e-07 | 2.8e-07 | 2.000E-07 | 8.000E-08 | 1.200E-07 | 2.800E-07 |
| 8658 | 433 | L433W | A | 1.6e-07 | 2e-07 | 5.3e-07 | 2.967E-07 | 2.031E-07 | 1.600E-07 | 5.300E-07 |
| 8659 | 433 | L433Y | A | 3.4e-07 | 1.8e-07 | 4.2e-07 | 3.133E-07 | 1.222E-07 | 1.800E-07 | 4.200E-07 |
| 8660 | 433 | L433V | A | 2.3e-07 | 1.5e-07 | 2.3e-07 | 2.033E-07 | 4.619E-08 | 1.500E-07 | 2.300E-07 |
| 8661 | 434 | R434A | A | 1.5e-07 | 3.6e-07 | 1.3e-07 | 2.133E-07 | 1.274E-07 | 1.300E-07 | 3.600E-07 |
| 8662 | 434 | R434R | A | 4.3e-07 | 2.4e-07 | 2e-07 | 2.900E-07 | 1.229E-07 | 2.000E-07 | 4.300E-07 |
| 8663 | 434 | R434N | A | 3.2e-07 | 2.3e-07 | 3e-07 | 2.833E-07 | 4.726E-08 | 2.300E-07 | 3.200E-07 |
| 8664 | 434 | R434D | A | 3.3e-07 | 2.4e-07 | 2.9e-07 | 2.867E-07 | 4.509E-08 | 2.400E-07 | 3.300E-07 |
| 8665 | 434 | R434C | A | 5.1e-07 | 1.1e-07 | 3e-07 | 3.067E-07 | 2.001E-07 | 1.100E-07 | 5.100E-07 |
| 8666 | 434 | R434E | A | 3.2e-07 | 2.1e-07 | 4.7e-07 | 3.333E-07 | 1.305E-07 | 2.100E-07 | 4.700E-07 |
| 8667 | 434 | R434Q | A | 3.1e-07 | 1.5e-07 | 4.4e-07 | 3.000E-07 | 1.453E-07 | 1.500E-07 | 4.400E-07 |
| 8668 | 434 | R434G | A | 1.3e-07 | 4.7e-07 | 4e-07 | 3.333E-07 | 1.795E-07 | 1.300E-07 | 4.700E-07 |
| 8669 | 434 | R434H | A | 1.6e-07 | 4.5e-07 | 3.7e-07 | 3.267E-07 | 1.498E-07 | 1.600E-07 | 4.500E-07 |

| | | | | | | | | | | |
|---|---|---|---|---|---|---|---|---|---|---|
| 8670 | 434 | R434I | A | 2.8e-07 | 1.3e-07 | 1.2e-07 | 1.767E-07 | 8.963E-08 | 1.200E-07 | 2.800E-07 |
| 8671 | 434 | R434L | A | 1.1e-07 | 2.5e-07 | 2.2e-07 | 1.933E-07 | 7.371E-08 | 1.100E-07 | 2.500E-07 |
| 8672 | 434 | R434K | A | 5e-07 | 2.7e-07 | 2.8e-07 | 3.500E-07 | 1.300E-07 | 2.700E-07 | 5.000E-07 |
| 8673 | 434 | R434M | A | 3.2e-07 | 1.7e-07 | 1.4e-07 | 2.100E-07 | 9.644E-08 | 1.400E-07 | 3.200E-07 |
| 8674 | 434 | R434F | A | 4.4e-07 | 4.9e-07 | 1.5e-07 | 3.600E-07 | 1.836E-07 | 1.500E-07 | 4.900E-07 |
| 8675 | 434 | R434P | A | 1.5e-07 | 1.8e-07 | 1.6e-07 | 1.633E-07 | 1.528E-08 | 1.500E-07 | 1.800E-07 |
| 8676 | 434 | R434S | A | 1.7e-07 | 1.1e-07 | 4.8e-07 | 2.533E-07 | 1.986E-07 | 1.100E-07 | 4.800E-07 |
| 8677 | 434 | R434T | A | 1.4e-07 | 3.9e-07 | 1.2e-07 | 2.167E-07 | 1.504E-07 | 1.200E-07 | 3.900E-07 |
| 8678 | 434 | R434W | A | 2.3e-07 | 2.6e-07 | 4.5e-07 | 3.133E-07 | 1.193E-07 | 2.300E-07 | 4.500E-07 |
| 8679 | 434 | R434Y | A | 1.2e-07 | 1.7e-07 | 4.9e-07 | 2.600E-07 | 2.007E-07 | 1.200E-07 | 4.900E-07 |
| 8680 | 434 | R434V | A | 1.6e-07 | 3.6e-07 | 6e-07 | 3.733E-07 | 2.203E-07 | 1.600E-07 | 6.000E-07 |
| 8681 | 435 | S435A | A | 2.1e-07 | 2.9e-07 | 2.3e-07 | 2.433E-07 | 4.163E-08 | 2.100E-07 | 2.900E-07 |
| 8682 | 435 | S435R | A | 9.1e-08 | 2e-07 | 3.3e-07 | 2.070E-07 | 1.197E-07 | 9.100E-08 | 3.300E-07 |
| 8683 | 435 | S435N | A | 3.5e-07 | 1.8e-07 | 4.6e-07 | 3.300E-07 | 1.411E-07 | 1.800E-07 | 4.600E-07 |
| 8684 | 435 | S435D | A | 1.4e-07 | 3.6e-07 | 3.1e-07 | 2.700E-07 | 1.153E-07 | 1.400E-07 | 3.600E-07 |
| 8685 | 435 | S435C | A | 2.9e-07 | 2.4e-07 | 1.4e-07 | 2.233E-07 | 7.638E-08 | 1.400E-07 | 2.900E-07 |
| 8686 | 435 | S435E | A | 1.6e-07 | 3.9e-07 | 1.4e-07 | 2.300E-07 | 1.389E-07 | 1.400E-07 | 3.900E-07 |
| 8687 | 435 | S435Q | A | 5.3e-07 | 2.3e-07 | 2.6e-07 | 3.400E-07 | 1.652E-07 | 2.300E-07 | 5.300E-07 |
| 8688 | 435 | S435G | A | 9.7e-08 | 2.5e-07 | 1.7e-07 | 1.723E-07 | 7.653E-08 | 9.700E-08 | 2.500E-07 |
| 8689 | 435 | S435H | A | 1.8e-07 | 2.7e-07 | 2.3e-07 | 2.267E-07 | 4.509E-08 | 1.800E-07 | 2.700E-07 |
| 8690 | 435 | S435I | A | 1.5e-07 | 1.5e-07 | 1.8e-07 | 1.600E-07 | 1.732E-08 | 1.500E-07 | 1.800E-07 |
| 8691 | 435 | S435L | A | 2.5e-07 | 2.1e-07 | 3.6e-07 | 2.733E-07 | 7.767E-08 | 2.100E-07 | 3.600E-07 |
| 8692 | 435 | S435K | A | 2e-07 | 2.7e-07 | 9.7e-08 | 1.890E-07 | 8.702E-08 | 9.700E-08 | 2.700E-07 |
| 8693 | 435 | S435M | A | 2.3e-07 | 3.7e-07 | 2e-07 | 2.667E-07 | 9.074E-08 | 2.000E-07 | 3.700E-07 |
| 8694 | 435 | S435F | A | 2.4e-07 | 3e-07 | 2.9e-07 | 2.767E-07 | 3.215E-08 | 2.400E-07 | 3.000E-07 |
| 8695 | 435 | S435P | A | 1.4e-07 | 2.9e-07 | 1.4e-07 | 1.900E-07 | 8.660E-08 | 1.400E-07 | 2.900E-07 |
| 8696 | 435 | S435S | A | 1.9e-07 | 2.3e-07 | 8.2e-08 | 1.673E-07 | 7.656E-08 | 8.200E-08 | 2.300E-07 |
| 8697 | 435 | S435T | A | 5.5e-07 | 1.3e-07 | 2.2e-07 | 3.000E-07 | 2.211E-07 | 1.300E-07 | 5.500E-07 |
| 8698 | 435 | S435W | A | 1.6e-07 | 3.9e-07 | 1.7e-07 | 2.400E-07 | 1.300E-07 | 1.600E-07 | 3.900E-07 |
| 8699 | 435 | S435Y | A | 3.7e-07 | 2.9e-07 | 2.6e-07 | 3.067E-07 | 5.686E-08 | 2.600E-07 | 3.700E-07 |
| 8700 | 435 | S435V | A | 2.3e-07 | 2.5e-07 | 4.6e-07 | 3.133E-07 | 1.274E-07 | 2.300E-07 | 4.600E-07 |
| 8701 | 436 | L436A | A | 2.6e-07 | 6.7e-07 | 2.3e-07 | 3.867E-07 | 2.458E-07 | 2.300E-07 | 6.700E-07 |
| 8702 | 436 | L436R | A | 1.6e-07 | 2.8e-07 | 2.3e-07 | 2.233E-07 | 6.028E-08 | 1.600E-07 | 2.800E-07 |
| 8703 | 436 | L436N | A | 1.5e-07 | 3.3e-07 | 3.1e-07 | 2.633E-07 | 9.866E-08 | 1.500E-07 | 3.300E-07 |
| 8704 | 436 | L436D | A | 3.6e-07 | 4e-07 | 4.5e-07 | 4.033E-07 | 4.509E-08 | 3.600E-07 | 4.500E-07 |
| 8705 | 436 | L436C | A | 1.7e-07 | 1.7e-07 | 2.9e-07 | 2.100E-07 | 6.928E-08 | 1.700E-07 | 2.900E-07 |
| 8706 | 436 | L436E | A | 1.6e-07 | 3.9e-07 | 2.3e-07 | 2.600E-07 | 1.179E-07 | 1.600E-07 | 3.900E-07 |
| 8707 | 436 | L436Q | A | 5.4e-07 | 2e-07 | 3.4e-07 | 3.600E-07 | 1.709E-07 | 2.000E-07 | 5.400E-07 |
| 8708 | 436 | L436G | A | 1.9e-07 | 3.2e-07 | 2.5e-07 | 2.533E-07 | 6.506E-08 | 1.900E-07 | 3.200E-07 |
| 8709 | 436 | L436H | A | 3e-07 | 5.9e-07 | 1.1e-07 | 3.333E-07 | 2.417E-07 | 1.100E-07 | 5.900E-07 |
| 8710 | 436 | L436I | A | 4.4e-07 | 1.8e-07 | 2.6e-07 | 2.933E-07 | 1.332E-07 | 1.800E-07 | 4.400E-07 |
| 8711 | 436 | L436L | A | 4.4e-07 | 2.1e-07 | 3.7e-07 | 3.400E-07 | 1.179E-07 | 2.100E-07 | 4.400E-07 |
| 8712 | 436 | L436K | A | 1.3e-07 | 1.9e-07 | 3.4e-07 | 2.200E-07 | 1.082E-07 | 1.300E-07 | 3.400E-07 |
| 8713 | 436 | L436M | A | 4.5e-07 | 1.7e-07 | 1.4e-07 | 2.533E-07 | 1.710E-07 | 1.400E-07 | 4.500E-07 |
| 8714 | 436 | L436F | A | 1.4e-07 | 2.7e-07 | 2.6e-07 | 2.233E-07 | 7.234E-08 | 1.400E-07 | 2.700E-07 |
| 8715 | 436 | L436P | A | 3.2e-07 | 4e-07 | 1.7e-07 | 2.967E-07 | 1.168E-07 | 1.700E-07 | 4.000E-07 |
| 8716 | 436 | L436S | A | 2.2e-07 | 4e-07 | 5.9e-07 | 4.033E-07 | 1.850E-07 | 2.200E-07 | 5.900E-07 |
| 8717 | 436 | L436T | A | 2.8e-07 | 1.5e-07 | 1.3e-07 | 1.867E-07 | 8.145E-08 | 1.300E-07 | 2.800E-07 |
| 8718 | 436 | L436W | A | 3.4e-07 | 2.8e-07 | 3.1e-07 | 3.100E-07 | 3.000E-08 | 2.800E-07 | 3.400E-07 |
| 8719 | 436 | L436Y | A | 3.1e-07 | 1.4e-07 | 1.6e-07 | 2.033E-07 | 9.292E-08 | 1.400E-07 | 3.100E-07 |
| 8720 | 436 | L436V | A | 9e-08 | 2.8e-07 | 2.6e-07 | 2.100E-07 | 1.044E-07 | 9.000E-08 | 2.800E-07 |
| 8721 | 437 | R437A | A | 2.3e-07 | 4.7e-07 | 1.3e-07 | 2.767E-07 | 1.747E-07 | 1.300E-07 | 4.700E-07 |
| 8722 | 437 | R437R | A | 2.5e-07 | 1.2e-07 | 3.3e-07 | 2.333E-07 | 1.060E-07 | 1.200E-07 | 3.300E-07 |
| 8723 | 437 | R437N | A | 4.1e-07 | 1.2e-07 | 2.3e-07 | 2.533E-07 | 1.464E-07 | 1.200E-07 | 4.100E-07 |
| 8724 | 437 | R437D | A | 2.4e-07 | 7.3e-07 | 1.5e-07 | 3.733E-07 | 3.121E-07 | 1.500E-07 | 7.300E-07 |
| 8725 | 437 | R437C | A | 2.5e-07 | 2.6e-07 | 1.8e-07 | 2.300E-07 | 4.359E-08 | 1.800E-07 | 2.600E-07 |
| 8726 | 437 | R437E | A | 3.1e-07 | 2e-07 | 3e-07 | 2.700E-07 | 6.083E-08 | 2.000E-07 | 3.100E-07 |
| 8727 | 437 | R437Q | A | 2.5e-07 | 1.9e-07 | 2.6e-07 | 2.333E-07 | 3.786E-08 | 1.900E-07 | 2.600E-07 |
| 8728 | 437 | R437G | A | 1.5e-07 | 2.8e-07 | 2.4e-07 | 2.233E-07 | 6.658E-08 | 1.500E-07 | 2.800E-07 |
| 8729 | 437 | R437H | A | 5.5e-07 | 1.6e-07 | 1.4e-07 | 2.833E-07 | 2.312E-07 | 1.400E-07 | 5.500E-07 |
| 8730 | 437 | R437I | A | 2.4e-07 | 3.2e-07 | 1.5e-07 | 2.367E-07 | 8.505E-08 | 1.500E-07 | 3.200E-07 |
| 8731 | 437 | R437L | A | 1.7e-07 | 2.4e-07 | 3.8e-07 | 2.633E-07 | 1.069E-07 | 1.700E-07 | 3.800E-07 |
| 8732 | 437 | R437K | A | 2.6e-07 | 7.3e-08 | 1.2e-07 | 1.510E-07 | 9.728E-08 | 7.300E-08 | 2.600E-07 |
| 8733 | 437 | R437M | A | 1.6e-07 | 1.2e-07 | 2.7e-07 | 1.833E-07 | 7.767E-08 | 1.200E-07 | 2.700E-07 |
| 8734 | 437 | R437F | A | 2e-07 | 2.2e-07 | 1.5e-07 | 1.900E-07 | 3.606E-08 | 1.500E-07 | 2.200E-07 |
| 8735 | 437 | R437P | A | 1.2e-07 | 2.7e-07 | 2.7e-07 | 2.200E-07 | 8.660E-08 | 1.200E-07 | 2.700E-07 |
| 8736 | 437 | R437S | A | 2.6e-07 | 2.3e-07 | 1.4e-07 | 2.100E-07 | 6.245E-08 | 1.400E-07 | 2.600E-07 |
| 8737 | 437 | R437T | A | 2.9e-07 | 1.8e-07 | 2.2e-07 | 2.300E-07 | 5.568E-08 | 1.800E-07 | 2.900E-07 |
| 8738 | 437 | R437W | A | 1.3e-07 | 3.5e-07 | 3.9e-07 | 2.900E-07 | 1.400E-07 | 1.300E-07 | 3.900E-07 |
| 8739 | 437 | R437Y | A | 2.3e-07 | 2.1e-07 | 1.7e-07 | 2.033E-07 | 3.055E-08 | 1.700E-07 | 2.300E-07 |
| 8740 | 437 | R437V | A | 3.5e-07 | 2.5e-07 | 2.4e-07 | 2.800E-07 | 6.083E-08 | 2.400E-07 | 3.500E-07 |
| 8741 | 438 | E438A | A | 4.4e-07 | 3.8e-07 | 1.6e-07 | 3.267E-07 | 1.474E-07 | 1.600E-07 | 4.400E-07 |
| 8742 | 438 | E438R | A | 2e-07 | 1.7e-07 | 3.8e-07 | 2.500E-07 | 1.136E-07 | 1.700E-07 | 3.800E-07 |
| 8743 | 438 | E438N | A | 4.2e-07 | 3.1e-07 | 4.6e-07 | 3.967E-07 | 7.767E-08 | 3.100E-07 | 4.600E-07 |
| 8744 | 438 | E438D | A | 7.2e-07 | 3.4e-07 | 4.6e-07 | 5.067E-07 | 1.943E-07 | 3.400E-07 | 7.200E-07 |
| 8745 | 438 | E438C | A | 5e-07 | 1.8e-07 | 2.1e-07 | 2.967E-07 | 1.767E-07 | 1.800E-07 | 5.000E-07 |
| 8746 | 438 | E438E | A | 3.5e-07 | 4.1e-07 | 4e-07 | 3.867E-07 | 3.215E-08 | 3.500E-07 | 4.100E-07 |
| 8747 | 438 | E438Q | A | 2.1e-07 | 4.9e-07 | 2.3e-07 | 3.100E-07 | 1.562E-07 | 2.100E-07 | 4.900E-07 |
| 8748 | 438 | E438G | A | 2.5e-07 | 2.6e-07 | 3.2e-07 | 2.767E-07 | 3.786E-08 | 2.500E-07 | 3.200E-07 |
| 8749 | 438 | E438H | A | 1.3e-07 | 1.1e-07 | 2.1e-07 | 1.500E-07 | 5.292E-08 | 1.100E-07 | 2.100E-07 |
| 8750 | 438 | E438I | A | 2.8e-07 | 2.6e-07 | 4.7e-07 | 3.367E-07 | 1.159E-07 | 2.600E-07 | 4.700E-07 |
| 8751 | 438 | E438L | A | 2e-07 | 2.3e-07 | 4.7e-07 | 3.000E-07 | 1.480E-07 | 2.000E-07 | 4.700E-07 |
| 8752 | 438 | E438K | A | 1.7e-07 | 4.7e-07 | 3.9e-07 | 3.433E-07 | 1.553E-07 | 1.700E-07 | 4.700E-07 |
| 8753 | 438 | E438M | A | 1.6e-07 | 1.2e-07 | 2.4e-07 | 1.733E-07 | 6.110E-08 | 1.200E-07 | 2.400E-07 |
| 8754 | 438 | E438F | A | 3.9e-07 | 1.8e-07 | 1.8e-07 | 2.500E-07 | 1.212E-07 | 1.800E-07 | 3.900E-07 |

| | | | | | | | | | | |
|---|---|---|---|---|---|---|---|---|---|---|
| 8755 | 438 | E438P | A | 3.8e-07 | 2.1e-07 | 4.5e-07 | 3.467E-07 | 1.234E-07 | 2.100E-07 | 4.500E-07 |
| 8756 | 438 | E438S | A | 1.8e-07 | 8.6e-08 | 2.8e-07 | 1.820E-07 | 9.702E-08 | 8.600E-08 | 2.800E-07 |
| 8757 | 438 | E438T | A | 2.2e-07 | 2.5e-07 | 1.3e-07 | 2.000E-07 | 6.245E-08 | 1.300E-07 | 2.500E-07 |
| 8758 | 438 | E438W | A | 2.4e-07 | 1e-07 | 2.9e-07 | 2.100E-07 | 9.849E-08 | 1.000E-07 | 2.900E-07 |
| 8759 | 438 | E438Y | A | 1.7e-07 | 3.1e-07 | 3e-07 | 2.600E-07 | 7.810E-08 | 1.700E-07 | 3.100E-07 |
| 8760 | 438 | E438V | A | 4.2e-07 | 2.2e-07 | 4.7e-07 | 3.700E-07 | 1.323E-07 | 2.200E-07 | 4.700E-07 |
| 8761 | 439 | L439A | A | 5.6e-07 | 2.5e-07 | 4.4e-07 | 4.167E-07 | 1.563E-07 | 2.500E-07 | 5.600E-07 |
| 8762 | 439 | L439R | A | 1.6e-07 | 1.4e-07 | 2.2e-07 | 1.733E-07 | 4.163E-08 | 1.400E-07 | 2.200E-07 |
| 8763 | 439 | L439N | A | 3.5e-07 | 2.6e-07 | 3.5e-07 | 3.200E-07 | 5.196E-08 | 2.600E-07 | 3.500E-07 |
| 8764 | 439 | L439D | A | 2e-07 | 5.7e-07 | 2.5e-07 | 3.400E-07 | 2.007E-07 | 2.000E-07 | 5.700E-07 |
| 8765 | 439 | L439C | A | 4.4e-07 | 2.7e-07 | 2.3e-07 | 3.133E-07 | 1.115E-07 | 2.300E-07 | 4.400E-07 |
| 8766 | 439 | L439E | A | 1.9e-07 | 1.3e-07 | 1.8e-07 | 1.667E-07 | 3.215E-08 | 1.300E-07 | 1.900E-07 |
| 8767 | 439 | L439Q | A | 1.6e-07 | 1.2e-07 | 2.3e-07 | 1.700E-07 | 5.568E-08 | 1.200E-07 | 2.300E-07 |
| 8768 | 439 | L439G | A | 1.7e-07 | 1.6e-07 | 4.5e-07 | 2.600E-07 | 1.646E-07 | 1.600E-07 | 4.500E-07 |
| 8769 | 439 | L439H | A | 2.1e-07 | 2.6e-07 | 1.1e-07 | 1.933E-07 | 7.638E-08 | 1.100E-07 | 2.600E-07 |
| 8770 | 439 | L439I | A | 2.9e-07 | 2.6e-07 | 2.4e-07 | 2.633E-07 | 2.517E-08 | 2.400E-07 | 2.900E-07 |
| 8771 | 439 | L439L | A | 1.3e-07 | 2.2e-07 | 1.6e-07 | 1.700E-07 | 4.583E-08 | 1.300E-07 | 2.200E-07 |
| 8772 | 439 | L439K | A | 1.1e-07 | 1.7e-07 | 2.2e-07 | 1.667E-07 | 5.508E-08 | 1.100E-07 | 2.200E-07 |
| 8773 | 439 | L439M | A | 3.4e-07 | 2.9e-07 | 2.6e-07 | 2.967E-07 | 4.041E-08 | 2.600E-07 | 3.400E-07 |
| 8774 | 439 | L439F | A | 2.7e-07 | 2.5e-07 | 3.5e-07 | 2.900E-07 | 5.292E-08 | 2.500E-07 | 3.500E-07 |
| 8775 | 439 | L439P | A | 2.7e-07 | 8.9e-08 | 1.5e-07 | 1.697E-07 | 9.209E-08 | 8.900E-08 | 2.700E-07 |
| 8776 | 439 | L439S | A | 2.1e-07 | 2.9e-07 | 2.4e-07 | 2.467E-07 | 4.041E-08 | 2.100E-07 | 2.900E-07 |
| 8777 | 439 | L439T | A | 2.6e-07 | 2e-07 | 1.8e-07 | 2.133E-07 | 4.163E-08 | 1.800E-07 | 2.600E-07 |
| 8778 | 439 | L439W | A | 2.2e-07 | 3e-07 | 2.1e-07 | 2.433E-07 | 4.933E-08 | 2.100E-07 | 3.000E-07 |
| 8779 | 439 | L439Y | A | 1.4e-07 | 3.9e-07 | 2.7e-07 | 2.667E-07 | 1.250E-07 | 1.400E-07 | 3.900E-07 |
| 8780 | 439 | L439V | A | 1.6e-07 | 1.5e-07 | 2.9e-07 | 2.000E-07 | 7.810E-08 | 1.500E-07 | 2.900E-07 |
| 8781 | 440 | G440A | A | 8.9e-08 | 2.2e-07 | 3.9e-07 | 2.330E-07 | 1.509E-07 | 8.900E-08 | 3.900E-07 |
| 8782 | 440 | G440R | A | 1.5e-07 | 4e-07 | 1.3e-07 | 2.267E-07 | 1.504E-07 | 1.300E-07 | 4.000E-07 |
| 8783 | 440 | G440N | A | 1.7e-07 | 2.3e-07 | 2.9e-07 | 2.300E-07 | 6.000E-08 | 1.700E-07 | 2.900E-07 |
| 8784 | 440 | G440D | A | 2.6e-07 | 4.8e-07 | 3e-07 | 3.467E-07 | 1.172E-07 | 2.600E-07 | 4.800E-07 |
| 8785 | 440 | G440C | A | 2.4e-07 | 2e-07 | 2e-07 | 2.133E-07 | 2.309E-08 | 2.000E-07 | 2.400E-07 |
| 8786 | 440 | G440E | A | 6.3e-07 | 2.1e-07 | 1.3e-07 | 3.233E-07 | 2.686E-07 | 1.300E-07 | 6.300E-07 |
| 8787 | 440 | G440Q | A | 3.9e-07 | 2e-07 | 2.1e-07 | 2.667E-07 | 1.069E-07 | 2.000E-07 | 3.900E-07 |
| 8788 | 440 | G440G | A | 1.7e-07 | 4.2e-07 | 3.9e-07 | 3.267E-07 | 1.365E-07 | 1.700E-07 | 4.200E-07 |
| 8789 | 440 | G440H | A | 1.9e-07 | 4.4e-07 | 3.8e-07 | 3.367E-07 | 1.305E-07 | 1.900E-07 | 4.400E-07 |
| 8790 | 440 | G440I | A | 1.6e-07 | 3.2e-07 | 1.8e-07 | 2.200E-07 | 8.718E-08 | 1.600E-07 | 3.200E-07 |
| 8791 | 440 | G440L | A | 3.4e-07 | 8.4e-08 | 6.4e-07 | 3.547E-07 | 2.783E-07 | 8.400E-08 | 6.400E-07 |
| 8792 | 440 | G440K | A | 4.6e-07 | 3.5e-07 | 2e-07 | 3.367E-07 | 1.305E-07 | 2.000E-07 | 4.600E-07 |
| 8793 | 440 | G440M | A | 1.7e-07 | 2.9e-07 | 3.3e-07 | 2.633E-07 | 8.327E-08 | 1.700E-07 | 3.300E-07 |
| 8794 | 440 | G440F | A | 1.7e-07 | 1.9e-07 | 1.5e-07 | 1.700E-07 | 2.000E-08 | 1.500E-07 | 1.900E-07 |
| 8795 | 440 | G440P | A | 2.1e-07 | 1.6e-07 | 1.7e-07 | 1.800E-07 | 2.646E-08 | 1.600E-07 | 2.100E-07 |
| 8796 | 440 | G440S | A | 3e-07 | 3e-07 | 2.9e-07 | 2.967E-07 | 5.774E-09 | 2.900E-07 | 3.000E-07 |
| 8797 | 440 | G440T | A | 1.2e-07 | 2e-07 | 1.5e-07 | 1.567E-07 | 4.041E-08 | 1.200E-07 | 2.000E-07 |
| 8798 | 440 | G440W | A | 2.1e-07 | 6.6e-07 | 2.4e-07 | 3.700E-07 | 2.516E-07 | 2.100E-07 | 6.600E-07 |
| 8799 | 440 | G440Y | A | 2.6e-07 | 2.5e-07 | 3.2e-07 | 2.767E-07 | 3.786E-08 | 2.500E-07 | 3.200E-07 |
| 8800 | 440 | G440V | A | 1.9e-07 | 2.3e-07 | 2.3e-07 | 2.167E-07 | 2.309E-08 | 1.900E-07 | 2.300E-07 |
| 8801 | 441 | S441A | A | 3.8e-07 | 2.1e-07 | 2.4e-07 | 2.767E-07 | 9.074E-08 | 2.100E-07 | 3.800E-07 |
| 8802 | 441 | S441R | A | 6.8e-07 | 5.4e-07 | 3e-07 | 5.067E-07 | 1.922E-07 | 3.000E-07 | 6.800E-07 |
| 8803 | 441 | S441N | A | 4.4e-07 | 1.8e-07 | 2.7e-07 | 2.967E-07 | 1.320E-07 | 1.800E-07 | 4.400E-07 |
| 8804 | 441 | S441D | A | 1.3e-07 | 4.6e-07 | 1.5e-07 | 2.467E-07 | 1.850E-07 | 1.300E-07 | 4.600E-07 |
| 8805 | 441 | S441C | A | 2.7e-07 | 3.9e-07 | 1.7e-07 | 2.767E-07 | 1.102E-07 | 1.700E-07 | 3.900E-07 |
| 8806 | 441 | S441E | A | 2.1e-07 | 4.1e-07 | 3.6e-07 | 3.267E-07 | 1.041E-07 | 2.100E-07 | 4.100E-07 |
| 8807 | 441 | S441Q | A | 6.9e-07 | 4.1e-07 | 2.4e-07 | 4.467E-07 | 2.272E-07 | 2.400E-07 | 6.900E-07 |
| 8808 | 441 | S441G | A | 4.5e-07 | 1.4e-07 | 1.3e-07 | 2.400E-07 | 1.819E-07 | 1.300E-07 | 4.500E-07 |
| 8809 | 441 | S441H | A | 1.2e-07 | 3e-07 | 3.2e-07 | 2.467E-07 | 1.102E-07 | 1.200E-07 | 3.200E-07 |
| 8810 | 441 | S441I | A | 2.5e-07 | 1.7e-07 | 4.4e-07 | 2.867E-07 | 1.387E-07 | 1.700E-07 | 4.400E-07 |
| 8811 | 441 | S441L | A | 2.8e-07 | 4.3e-07 | 2.4e-07 | 3.167E-07 | 1.002E-07 | 2.400E-07 | 4.300E-07 |
| 8812 | 441 | S441K | A | 1.8e-07 | 2.4e-07 | 3.2e-07 | 2.467E-07 | 7.024E-08 | 1.800E-07 | 3.200E-07 |
| 8813 | 441 | S441M | A | 2.7e-07 | 1.8e-07 | 1.9e-07 | 2.133E-07 | 4.933E-08 | 1.800E-07 | 2.700E-07 |
| 8814 | 441 | S441F | A | 2.7e-07 | 3.1e-07 | 2.2e-07 | 2.667E-07 | 4.509E-08 | 2.200E-07 | 3.100E-07 |
| 8815 | 441 | S441P | A | 1.3e-07 | 2.8e-07 | 2.9e-07 | 2.333E-07 | 8.963E-08 | 1.300E-07 | 2.900E-07 |
| 8816 | 441 | S441S | A | 1.7e-07 | 4.7e-07 | 2e-07 | 2.800E-07 | 1.652E-07 | 1.700E-07 | 4.700E-07 |
| 8817 | 441 | S441T | A | 2.4e-07 | 4.1e-07 | 2.1e-07 | 2.867E-07 | 1.079E-07 | 2.100E-07 | 4.100E-07 |
| 8818 | 441 | S441W | A | 2.4e-07 | 2.2e-07 | 2e-07 | 2.200E-07 | 2.000E-08 | 2.000E-07 | 2.400E-07 |
| 8819 | 441 | S441Y | A | 2.4e-07 | 4.8e-07 | 3.2e-07 | 3.467E-07 | 1.222E-07 | 2.400E-07 | 4.800E-07 |
| 8820 | 441 | S441V | A | 5.6e-07 | 1.9e-07 | 5.2e-07 | 4.233E-07 | 2.031E-07 | 1.900E-07 | 5.600E-07 |
| 8821 | 442 | G442A | A | 1.4e-07 | 1.6e-07 | 1.9e-07 | 1.633E-07 | 2.517E-08 | 1.400E-07 | 1.900E-07 |
| 8822 | 442 | G442R | A | 2.8e-07 | 2.7e-07 | 3.3e-07 | 2.933E-07 | 3.215E-08 | 2.700E-07 | 3.300E-07 |
| 8823 | 442 | G442N | A | 2e-07 | 3.8e-07 | 1.3e-07 | 2.367E-07 | 1.290E-07 | 1.300E-07 | 3.800E-07 |
| 8824 | 442 | G442D | A | 2.5e-07 | 1.8e-07 | 2.5e-07 | 2.267E-07 | 4.041E-08 | 1.800E-07 | 2.500E-07 |
| 8825 | 442 | G442C | A | 4.7e-07 | 2.9e-07 | 2.3e-07 | 3.300E-07 | 1.249E-07 | 2.300E-07 | 4.700E-07 |
| 8826 | 442 | G442E | A | 2.4e-07 | 1.1e-07 | 2e-07 | 1.833E-07 | 6.658E-08 | 1.100E-07 | 2.400E-07 |
| 8827 | 442 | G442Q | A | 4.8e-07 | 2.5e-07 | 8.5e-07 | 5.267E-07 | 3.027E-07 | 2.500E-07 | 8.500E-07 |
| 8828 | 442 | G442G | A | 1.3e-07 | 3e-07 | 2.9e-07 | 2.400E-07 | 9.539E-08 | 1.300E-07 | 3.000E-07 |
| 8829 | 442 | G442H | A | 1.7e-07 | 5e-07 | 2e-07 | 2.900E-07 | 1.825E-07 | 1.700E-07 | 5.000E-07 |
| 8830 | 442 | G442I | A | 7.8e-08 | 1.8e-07 | 2.8e-07 | 1.793E-07 | 1.010E-07 | 7.800E-08 | 2.800E-07 |
| 8831 | 442 | G442L | A | 2.9e-07 | 2.3e-07 | 3.1e-07 | 2.767E-07 | 4.163E-08 | 2.300E-07 | 3.100E-07 |
| 8832 | 442 | G442K | A | 2e-07 | 3.2e-07 | 2.8e-07 | 2.667E-07 | 6.110E-08 | 2.000E-07 | 3.200E-07 |
| 8833 | 442 | G442M | A | 1.2e-07 | 6.8e-08 | 3.4e-07 | 1.760E-07 | 1.444E-07 | 6.800E-08 | 3.400E-07 |
| 8834 | 442 | G442F | A | 2.2e-07 | 3e-07 | 1.9e-07 | 2.367E-07 | 5.686E-08 | 1.900E-07 | 3.000E-07 |
| 8835 | 442 | G442P | A | 3.6e-07 | 2.3e-07 | 3.6e-07 | 3.167E-07 | 7.506E-08 | 2.300E-07 | 3.600E-07 |
| 8836 | 442 | G442S | A | 1.2e-07 | 3.3e-07 | 4.9e-07 | 3.133E-07 | 1.856E-07 | 1.200E-07 | 4.900E-07 |
| 8837 | 442 | G442T | A | 3.7e-07 | 4.3e-07 | 2.3e-07 | 3.433E-07 | 1.026E-07 | 2.300E-07 | 4.300E-07 |
| 8838 | 442 | G442W | A | 1.4e-07 | 1.2e-07 | 2.1e-07 | 1.567E-07 | 4.726E-08 | 1.200E-07 | 2.100E-07 |
| 8839 | 442 | G442Y | A | 4.6e-07 | 1.1e-07 | 2.6e-07 | 2.767E-07 | 1.756E-07 | 1.100E-07 | 4.600E-07 |

| | | | | | | | | | | |
|---|---|---|---|---|---|---|---|---|---|---|
| 8840 | 442 | G442V | A | 2e-07 | 1.9e-07 | 2.7e-07 | 2.200E-07 | 4.359E-08 | 1.900E-07 | 2.700E-07 |
| 8841 | 443 | L443A | A | 4e-07 | 2.1e-07 | 1.8e-07 | 2.633E-07 | 1.193E-07 | 1.800E-07 | 4.000E-07 |
| 8842 | 443 | L443R | A | 2.7e-07 | 1.9e-07 | 2.4e-07 | 2.333E-07 | 4.041E-08 | 1.900E-07 | 2.700E-07 |
| 8843 | 443 | L443N | A | 2.1e-07 | 2.4e-07 | 2.1e-07 | 2.200E-07 | 1.732E-08 | 2.100E-07 | 2.400E-07 |
| 8844 | 443 | L443D | A | 6.2e-07 | 3.4e-07 | 1.3e-07 | 3.633E-07 | 2.458E-07 | 1.300E-07 | 6.200E-07 |
| 8845 | 443 | L443C | A | 3.8e-07 | 2.1e-07 | 2e-07 | 2.633E-07 | 1.012E-07 | 2.000E-07 | 3.800E-07 |
| 8846 | 443 | L443E | A | 1.1e-07 | 1.3e-07 | 3.7e-07 | 2.033E-07 | 1.447E-07 | 1.100E-07 | 3.700E-07 |
| 8847 | 443 | L443Q | A | 1.9e-07 | 3.3e-07 | 2.1e-07 | 2.433E-07 | 7.572E-08 | 1.900E-07 | 3.300E-07 |
| 8848 | 443 | L443G | A | 3.4e-07 | 1.9e-07 | 3.8e-07 | 3.033E-07 | 1.002E-07 | 1.900E-07 | 3.800E-07 |
| 8849 | 443 | L443H | A | 3.6e-07 | 3.5e-07 | 3.4e-07 | 3.500E-07 | 1.000E-08 | 3.400E-07 | 3.600E-07 |
| 8850 | 443 | L443I | A | 3.5e-07 | 2e-07 | 2.4e-07 | 2.633E-07 | 7.767E-08 | 2.000E-07 | 3.500E-07 |
| 8851 | 443 | L443L | A | 2.7e-07 | 3.4e-07 | 2.8e-07 | 2.967E-07 | 3.786E-08 | 2.700E-07 | 3.400E-07 |
| 8852 | 443 | L443K | A | 3.4e-07 | 3.9e-07 | 2.8e-07 | 3.367E-07 | 5.508E-08 | 2.800E-07 | 3.900E-07 |
| 8853 | 443 | L443M | A | 2e-07 | 2.6e-07 | 2.5e-07 | 2.367E-07 | 3.215E-08 | 2.000E-07 | 2.600E-07 |
| 8854 | 443 | L443F | A | 2.3e-07 | 1.8e-07 | 9.5e-07 | 4.533E-07 | 4.309E-07 | 1.800E-07 | 9.500E-07 |
| 8855 | 443 | L443P | A | 2.4e-07 | 1.4e-07 | 3.2e-07 | 2.333E-07 | 9.018E-08 | 1.400E-07 | 3.200E-07 |
| 8856 | 443 | L443S | A | 1.1e-07 | 4.3e-07 | 4.5e-07 | 3.300E-07 | 1.908E-07 | 1.100E-07 | 4.500E-07 |
| 8857 | 443 | L443T | A | 4.9e-07 | 3.4e-07 | 5e-07 | 4.433E-07 | 8.963E-08 | 3.400E-07 | 5.000E-07 |
| 8858 | 443 | L443W | A | 3e-07 | 2.7e-07 | 1.9e-07 | 2.533E-07 | 5.686E-08 | 1.900E-07 | 3.000E-07 |
| 8859 | 443 | L443Y | A | 1.9e-07 | 2.6e-07 | 1e-07 | 1.833E-07 | 8.021E-08 | 1.000E-07 | 2.600E-07 |
| 8860 | 443 | L443V | A | 2.9e-07 | 2.4e-07 | 2.2e-07 | 2.500E-07 | 3.606E-08 | 2.200E-07 | 2.900E-07 |
| 8861 | 444 | A444A | A | 3.2e-07 | 3.7e-07 | 1.9e-07 | 2.933E-07 | 9.292E-08 | 1.900E-07 | 3.700E-07 |
| 8862 | 444 | A444R | A | 1.7e-07 | 4e-07 | 3.3e-07 | 3.000E-07 | 1.179E-07 | 1.700E-07 | 4.000E-07 |
| 8863 | 444 | A444N | A | 2.7e-07 | 5.2e-07 | 2.5e-07 | 3.467E-07 | 1.504E-07 | 2.500E-07 | 5.200E-07 |
| 8864 | 444 | A444D | A | 2.8e-07 | 3.9e-07 | 3e-07 | 3.233E-07 | 5.859E-08 | 2.800E-07 | 3.900E-07 |
| 8865 | 444 | A444C | A | 2.6e-07 | 1.3e-07 | 2.1e-07 | 2.000E-07 | 6.557E-08 | 1.300E-07 | 2.600E-07 |
| 8866 | 444 | A444E | A | 2e-07 | 1.4e-07 | 2.5e-07 | 1.967E-07 | 5.508E-08 | 1.400E-07 | 2.500E-07 |
| 8867 | 444 | A444Q | A | 2.3e-07 | 2.5e-07 | 1.3e-07 | 2.033E-07 | 6.429E-08 | 1.300E-07 | 2.500E-07 |
| 8868 | 444 | A444G | A | 2.6e-07 | 1.6e-07 | 2.6e-07 | 2.267E-07 | 5.774E-08 | 1.600E-07 | 2.600E-07 |
| 8869 | 444 | A444H | A | 2.2e-07 | 2.1e-07 | 3.5e-07 | 2.600E-07 | 7.810E-08 | 2.100E-07 | 3.500E-07 |
| 8870 | 444 | A444I | A | 1.4e-07 | 1e-07 | 1.5e-07 | 1.300E-07 | 2.646E-08 | 1.000E-07 | 1.500E-07 |
| 8871 | 444 | A444L | A | 2.1e-07 | 2.6e-07 | 9.8e-08 | 1.893E-07 | 8.295E-08 | 9.800E-08 | 2.600E-07 |
| 8872 | 444 | A444K | A | 2.3e-07 | 2e-07 | 1e-07 | 1.767E-07 | 6.807E-08 | 1.000E-07 | 2.300E-07 |
| 8873 | 444 | A444M | A | 4e-07 | 7.1e-07 | 2.8e-07 | 4.633E-07 | 2.219E-07 | 2.800E-07 | 7.100E-07 |
| 8874 | 444 | A444F | A | 2.2e-07 | 2.3e-07 | 1.8e-07 | 2.100E-07 | 2.646E-08 | 1.800E-07 | 2.300E-07 |
| 8875 | 444 | A444P | A | 5.2e-07 | 2.8e-07 | 3e-07 | 3.667E-07 | 1.332E-07 | 2.800E-07 | 5.200E-07 |
| 8876 | 444 | A444S | A | 1.5e-07 | 2e-07 | 3.5e-07 | 2.333E-07 | 1.041E-07 | 1.500E-07 | 3.500E-07 |
| 8877 | 444 | A444T | A | 3.4e-07 | 2.8e-07 | 6e-07 | 4.067E-07 | 1.701E-07 | 2.800E-07 | 6.000E-07 |
| 8878 | 444 | A444W | A | 4.8e-07 | 4.4e-07 | 1.3e-07 | 3.500E-07 | 1.916E-07 | 1.300E-07 | 4.800E-07 |
| 8879 | 444 | A444Y | A | 2.5e-07 | 3.7e-07 | 3.4e-07 | 3.200E-07 | 6.245E-08 | 2.500E-07 | 3.700E-07 |
| 8880 | 444 | A444V | A | 2.9e-07 | 1.4e-07 | 5.9e-07 | 3.400E-07 | 2.291E-07 | 1.400E-07 | 5.900E-07 |
| 8881 | 445 | L445A | A | 2.2e-07 | 2.3e-07 | 3e-07 | 2.500E-07 | 4.359E-08 | 2.200E-07 | 3.000E-07 |
| 8882 | 445 | L445R | A | 3.5e-07 | 1.2e-07 | 1.9e-07 | 2.200E-07 | 1.179E-07 | 1.200E-07 | 3.500E-07 |
| 8883 | 445 | L445N | A | 1.9e-07 | 4e-07 | 4.7e-07 | 3.533E-07 | 1.457E-07 | 1.900E-07 | 4.700E-07 |
| 8884 | 445 | L445D | A | 2e-07 | 3.2e-07 | 9.8e-08 | 2.060E-07 | 1.111E-07 | 9.800E-08 | 3.200E-07 |
| 8885 | 445 | L445C | A | 6.9e-07 | 2.8e-07 | 2.4e-07 | 4.033E-07 | 2.491E-07 | 2.400E-07 | 6.900E-07 |
| 8886 | 445 | L445E | A | 4.5e-07 | 2.5e-07 | 1.7e-07 | 2.900E-07 | 1.442E-07 | 1.700E-07 | 4.500E-07 |
| 8887 | 445 | L445Q | A | 3.8e-07 | 1.8e-07 | 4e-07 | 3.200E-07 | 1.217E-07 | 1.800E-07 | 4.000E-07 |
| 8888 | 445 | L445G | A | 1.9e-07 | 2.2e-07 | 2.4e-07 | 2.167E-07 | 2.517E-08 | 1.900E-07 | 2.400E-07 |
| 8889 | 445 | L445H | A | 4.4e-07 | 3.8e-07 | 3.2e-07 | 3.800E-07 | 6.000E-08 | 3.200E-07 | 4.400E-07 |
| 8890 | 445 | L445I | A | 1.5e-07 | 2.4e-07 | 2.6e-07 | 2.167E-07 | 5.859E-08 | 1.500E-07 | 2.600E-07 |
| 8891 | 445 | L445L | A | 2.5e-07 | 1.9e-07 | 3.7e-07 | 2.700E-07 | 9.165E-08 | 1.900E-07 | 3.700E-07 |
| 8892 | 445 | L445K | A | 3.9e-07 | 2.7e-07 | 2e-07 | 2.867E-07 | 9.609E-08 | 2.000E-07 | 3.900E-07 |
| 8893 | 445 | L445M | A | 3.6e-07 | 1.4e-07 | 1.7e-07 | 2.233E-07 | 1.193E-07 | 1.400E-07 | 3.600E-07 |
| 8894 | 445 | L445F | A | 5.8e-07 | 2.6e-07 | 4e-07 | 4.133E-07 | 1.604E-07 | 2.600E-07 | 5.800E-07 |
| 8895 | 445 | L445P | A | 1.1e-07 | 1.6e-07 | 1.5e-07 | 1.400E-07 | 2.646E-08 | 1.100E-07 | 1.600E-07 |
| 8896 | 445 | L445S | A | 1.5e-07 | 3.1e-07 | 4.6e-07 | 3.067E-07 | 1.550E-07 | 1.500E-07 | 4.600E-07 |
| 8897 | 445 | L445T | A | 1.9e-07 | 9.9e-08 | 4.7e-07 | 2.530E-07 | 1.934E-07 | 9.900E-08 | 4.700E-07 |
| 8898 | 445 | L445W | A | 2.4e-07 | 2.8e-07 | 2.2e-07 | 2.467E-07 | 3.055E-08 | 2.200E-07 | 2.800E-07 |
| 8899 | 445 | L445Y | A | 2.6e-07 | 2.5e-07 | 2.4e-07 | 2.500E-07 | 1.000E-08 | 2.400E-07 | 2.600E-07 |
| 8900 | 445 | L445V | A | 3.2e-07 | 2e-07 | 2.5e-07 | 2.567E-07 | 6.028E-08 | 2.000E-07 | 3.200E-07 |
| 8901 | 446 | I446A | A | 3.2e-07 | 2.1e-07 | 1.8e-07 | 2.367E-07 | 7.371E-08 | 1.800E-07 | 3.200E-07 |
| 8902 | 446 | I446R | A | 2.2e-07 | 9.6e-08 | 2.9e-07 | 2.020E-07 | 9.824E-08 | 9.600E-08 | 2.900E-07 |
| 8903 | 446 | I446N | A | 2.1e-07 | 1.9e-07 | 2.4e-07 | 2.133E-07 | 2.517E-08 | 1.900E-07 | 2.400E-07 |
| 8904 | 446 | I446D | A | 4e-07 | 3.2e-07 | 4.2e-07 | 3.800E-07 | 5.292E-08 | 3.200E-07 | 4.200E-07 |
| 8905 | 446 | I446C | A | 1.7e-07 | 1.8e-07 | 2.7e-07 | 2.067E-07 | 5.508E-08 | 1.700E-07 | 2.700E-07 |
| 8906 | 446 | I446E | A | 1.2e-07 | 1.5e-07 | 2e-07 | 1.567E-07 | 4.041E-08 | 1.200E-07 | 2.000E-07 |
| 8907 | 446 | I446Q | A | 2.6e-07 | 3.3e-07 | 2.4e-07 | 2.767E-07 | 4.726E-08 | 2.400E-07 | 3.300E-07 |
| 8908 | 446 | I446G | A | 2.3e-07 | 2e-07 | 2.5e-07 | 2.267E-07 | 2.517E-08 | 2.000E-07 | 2.500E-07 |
| 8909 | 446 | I446H | A | 2.8e-07 | 2.4e-07 | 2.4e-07 | 2.533E-07 | 2.309E-08 | 2.400E-07 | 2.800E-07 |
| 8910 | 446 | I446I | A | 5.7e-07 | 2.8e-07 | 6.3e-07 | 4.933E-07 | 1.872E-07 | 2.800E-07 | 6.300E-07 |
| 8911 | 446 | I446L | A | 1.4e-07 | 3.7e-07 | 2.9e-07 | 2.667E-07 | 1.168E-07 | 1.400E-07 | 3.700E-07 |
| 8912 | 446 | I446K | A | 9.5e-08 | 1.9e-07 | 1.8e-07 | 1.550E-07 | 5.220E-08 | 9.500E-08 | 1.900E-07 |
| 8913 | 446 | I446M | A | 3.8e-07 | 3.5e-07 | 2.3e-07 | 3.200E-07 | 7.937E-08 | 2.300E-07 | 3.800E-07 |
| 8914 | 446 | I446F | A | 1.1e-07 | 4e-07 | 2.8e-07 | 2.633E-07 | 1.457E-07 | 1.100E-07 | 4.000E-07 |
| 8915 | 446 | I446P | A | 2.2e-07 | 2.5e-07 | 1.3e-07 | 2.000E-07 | 6.245E-08 | 1.300E-07 | 2.500E-07 |
| 8916 | 446 | I446S | A | 3.5e-07 | 3.1e-07 | 4.3e-07 | 3.633E-07 | 6.110E-08 | 3.100E-07 | 4.300E-07 |
| 8917 | 446 | I446T | A | 4.7e-07 | 3.8e-07 | 3e-07 | 3.833E-07 | 8.505E-08 | 3.000E-07 | 4.700E-07 |
| 8918 | 446 | I446W | A | 3.2e-07 | 1.8e-07 | 1.9e-07 | 2.300E-07 | 7.810E-08 | 1.800E-07 | 3.200E-07 |
| 8919 | 446 | I446Y | A | 2e-07 | 3.2e-07 | 4.6e-07 | 3.267E-07 | 1.301E-07 | 2.000E-07 | 4.600E-07 |
| 8920 | 446 | I446V | A | 3.3e-07 | 2e-07 | 1.5e-07 | 2.267E-07 | 9.292E-08 | 1.500E-07 | 3.300E-07 |
| 8921 | 447 | H447A | A | 4.4e-07 | 2.9e-07 | 3e-07 | 3.433E-07 | 8.386E-08 | 2.900E-07 | 4.400E-07 |
| 8922 | 447 | H447R | A | 8.8e-08 | 8e-08 | 1.7e-07 | 1.127E-07 | 4.981E-08 | 8.000E-08 | 1.700E-07 |
| 8923 | 447 | H447N | A | 3.3e-07 | 2.5e-07 | 2.9e-07 | 2.900E-07 | 4.000E-08 | 2.500E-07 | 3.300E-07 |
| 8924 | 447 | H447D | A | 5.8e-07 | 1.7e-07 | 2.6e-07 | 3.367E-07 | 2.155E-07 | 1.700E-07 | 5.800E-07 |

| | | | | | | | | | | |
|---|---|---|---|---|---|---|---|---|---|---|
| 8925 | 447 | H447C | A | 2.1e-07 | 2.2e-07 | 2.5e-07 | 2.267E-07 | 2.082E-08 | 2.100E-07 | 2.500E-07 |
| 8926 | 447 | H447E | A | 3.1e-07 | 2.3e-07 | 1.7e-07 | 2.367E-07 | 7.024E-08 | 1.700E-07 | 3.100E-07 |
| 8927 | 447 | H447Q | A | 1.4e-07 | 1.8e-07 | 3.4e-07 | 2.200E-07 | 1.058E-07 | 1.400E-07 | 3.400E-07 |
| 8928 | 447 | H447G | A | 2.7e-07 | 2e-07 | 2.5e-07 | 2.400E-07 | 3.606E-08 | 2.000E-07 | 2.700E-07 |
| 8929 | 447 | H447H | A | 3.8e-07 | 2.6e-07 | 2.9e-07 | 3.100E-07 | 6.245E-08 | 2.600E-07 | 3.800E-07 |
| 8930 | 447 | H447I | A | 2.2e-07 | 2.4e-07 | 2.7e-07 | 2.433E-07 | 2.517E-08 | 2.200E-07 | 2.700E-07 |
| 8931 | 447 | H447L | A | 2.5e-07 | 2.6e-07 | 3.3e-07 | 2.800E-07 | 4.359E-08 | 2.500E-07 | 3.300E-07 |
| 8932 | 447 | H447K | A | 2.6e-07 | 2.2e-07 | 2.2e-07 | 2.333E-07 | 2.309E-08 | 2.200E-07 | 2.600E-07 |
| 8933 | 447 | H447M | A | 3.7e-07 | 2.6e-07 | 2.8e-07 | 3.033E-07 | 5.859E-08 | 2.600E-07 | 3.700E-07 |
| 8934 | 447 | H447F | A | 1.6e-07 | 4.4e-07 | 3.2e-07 | 3.067E-07 | 1.405E-07 | 1.600E-07 | 4.400E-07 |
| 8935 | 447 | H447P | A | 2.2e-07 | 3.6e-07 | 1.2e-07 | 2.333E-07 | 1.206E-07 | 1.200E-07 | 3.600E-07 |
| 8936 | 447 | H447S | A | 4.8e-07 | 3e-07 | 4.3e-07 | 4.033E-07 | 9.292E-08 | 3.000E-07 | 4.800E-07 |
| 8937 | 447 | H447T | A | 3e-07 | 2e-07 | 3.9e-07 | 2.967E-07 | 9.504E-08 | 2.000E-07 | 3.900E-07 |
| 8938 | 447 | H447W | A | 2.8e-07 | 1.3e-07 | 2.8e-07 | 2.300E-07 | 8.660E-08 | 1.300E-07 | 2.800E-07 |
| 8939 | 447 | H447Y | A | 1.4e-07 | 1.4e-07 | 1.3e-07 | 1.367E-07 | 5.774E-09 | 1.300E-07 | 1.400E-07 |
| 8940 | 447 | H447V | A | 3.1e-07 | 4.3e-07 | 3.6e-07 | 3.667E-07 | 6.028E-08 | 3.100E-07 | 4.300E-07 |
| 8941 | 448 | H448A | A | 1.9e-07 | 1.7e-07 | 1.7e-07 | 1.767E-07 | 1.155E-08 | 1.700E-07 | 1.900E-07 |
| 8942 | 448 | H448R | A | 1.5e-07 | 3.8e-07 | 3.1e-07 | 2.800E-07 | 1.179E-07 | 1.500E-07 | 3.800E-07 |
| 8943 | 448 | H448N | A | 2.1e-07 | 2.2e-07 | 2.2e-07 | 2.167E-07 | 5.774E-09 | 2.100E-07 | 2.200E-07 |
| 8944 | 448 | H448D | A | 1.6e-07 | 4.3e-07 | 4.3e-07 | 3.400E-07 | 1.559E-07 | 1.600E-07 | 4.300E-07 |
| 8945 | 448 | H448C | A | 2.3e-07 | 1.1e-07 | 1.6e-07 | 1.667E-07 | 6.028E-08 | 1.100E-07 | 2.300E-07 |
| 8946 | 448 | H448E | A | 6.4e-07 | 6.2e-07 | 3.3e-07 | 5.300E-07 | 1.735E-07 | 3.300E-07 | 6.400E-07 |
| 8947 | 448 | H448Q | A | 1.7e-07 | 1.7e-07 | 4.6e-07 | 2.667E-07 | 1.674E-07 | 1.700E-07 | 4.600E-07 |
| 8948 | 448 | H448G | A | 1.8e-07 | 3.7e-07 | 5.3e-07 | 3.600E-07 | 1.752E-07 | 1.800E-07 | 5.300E-07 |
| 8949 | 448 | H448H | A | 9.9e-08 | 2.8e-07 | 2.4e-07 | 2.063E-07 | 9.508E-08 | 9.900E-08 | 2.800E-07 |
| 8950 | 448 | H448I | A | 2.2e-07 | 2.1e-07 | 2.5e-07 | 2.267E-07 | 2.082E-08 | 2.100E-07 | 2.500E-07 |
| 8951 | 448 | H448L | A | 4.3e-07 | 3.9e-07 | 1.3e-07 | 3.167E-07 | 1.629E-07 | 1.300E-07 | 4.300E-07 |
| 8952 | 448 | H448K | A | 1.9e-07 | 3.1e-07 | 1.2e-07 | 2.067E-07 | 9.609E-08 | 1.200E-07 | 3.100E-07 |
| 8953 | 448 | H448M | A | 3.2e-07 | 3.1e-07 | 3.6e-07 | 3.300E-07 | 2.646E-08 | 3.100E-07 | 3.600E-07 |
| 8954 | 448 | H448F | A | 3.5e-07 | 2.3e-07 | 2.8e-07 | 2.867E-07 | 6.028E-08 | 2.300E-07 | 3.500E-07 |
| 8955 | 448 | H448P | A | 2.4e-07 | 2.6e-07 | 4.1e-07 | 3.033E-07 | 9.292E-08 | 2.400E-07 | 4.100E-07 |
| 8956 | 448 | H448S | A | 4.9e-07 | 2.5e-07 | 2.3e-07 | 3.233E-07 | 1.447E-07 | 2.300E-07 | 4.900E-07 |
| 8957 | 448 | H448T | A | 2.3e-07 | 2.7e-07 | 1.1e-07 | 2.033E-07 | 8.327E-08 | 1.100E-07 | 2.700E-07 |
| 8958 | 448 | H448W | A | 3.3e-07 | 4.1e-07 | 2.7e-07 | 3.367E-07 | 7.024E-08 | 2.700E-07 | 4.100E-07 |
| 8959 | 448 | H448Y | A | 2.9e-07 | 2.9e-07 | 2.2e-07 | 2.667E-07 | 4.041E-08 | 2.200E-07 | 2.900E-07 |
| 8960 | 448 | H448V | A | 2.4e-07 | 3.3e-07 | 3.4e-07 | 3.033E-07 | 5.508E-08 | 2.400E-07 | 3.400E-07 |
| 8961 | 449 | N449A | A | 3.1e-07 | 2.4e-07 | 2.3e-07 | 2.600E-07 | 4.359E-08 | 2.300E-07 | 3.100E-07 |
| 8962 | 449 | N449R | A | 2.1e-07 | 1.6e-07 | 2.1e-07 | 1.933E-07 | 2.887E-08 | 1.600E-07 | 2.100E-07 |
| 8963 | 449 | N449N | A | 2.9e-07 | 2.7e-07 | 2.7e-07 | 2.767E-07 | 1.155E-08 | 2.700E-07 | 2.900E-07 |
| 8964 | 449 | N449D | A | 2.7e-07 | 2.3e-07 | 1e-07 | 2.000E-07 | 8.888E-08 | 1.000E-07 | 2.700E-07 |
| 8965 | 449 | N449C | A | 2.2e-07 | 3.9e-07 | 1.9e-07 | 2.667E-07 | 1.079E-07 | 1.900E-07 | 3.900E-07 |
| 8966 | 449 | N449E | A | 1.1e-07 | 1.7e-07 | 3.8e-07 | 2.200E-07 | 1.418E-07 | 1.100E-07 | 3.800E-07 |
| 8967 | 449 | N449Q | A | 2.4e-07 | 3.4e-07 | 2e-07 | 2.600E-07 | 7.211E-08 | 2.000E-07 | 3.400E-07 |
| 8968 | 449 | N449G | A | 6e-07 | 1.7e-07 | 1.1e-07 | 2.933E-07 | 2.673E-07 | 1.100E-07 | 6.000E-07 |
| 8969 | 449 | N449H | A | 4e-07 | 1.5e-07 | 4.4e-07 | 3.300E-07 | 1.572E-07 | 1.500E-07 | 4.400E-07 |
| 8970 | 449 | N449I | A | 2.2e-07 | 1.2e-07 | 1.5e-07 | 1.633E-07 | 5.132E-08 | 1.200E-07 | 2.200E-07 |
| 8971 | 449 | N449L | A | 1.7e-07 | 1.6e-07 | 3.3e-07 | 2.200E-07 | 9.539E-08 | 1.600E-07 | 3.300E-07 |
| 8972 | 449 | N449K | A | 2.3e-07 | 6.3e-07 | 1.3e-07 | 3.300E-07 | 2.646E-07 | 1.300E-07 | 6.300E-07 |
| 8973 | 449 | N449M | A | 1.1e-07 | 1.4e-07 | 1.6e-07 | 1.367E-07 | 2.517E-08 | 1.100E-07 | 1.600E-07 |
| 8974 | 449 | N449F | A | 3.4e-07 | 1.7e-07 | 1.4e-07 | 2.167E-07 | 1.079E-07 | 1.400E-07 | 3.400E-07 |
| 8975 | 449 | N449P | A | 1.8e-07 | 1e-07 | 2.8e-07 | 1.867E-07 | 9.018E-08 | 1.000E-07 | 2.800E-07 |
| 8976 | 449 | N449S | A | 2.7e-07 | 1.6e-07 | 4.3e-07 | 2.867E-07 | 1.358E-07 | 1.600E-07 | 4.300E-07 |
| 8977 | 449 | N449T | A | 6.5e-07 | 3.7e-07 | 3.5e-07 | 4.567E-07 | 1.677E-07 | 3.500E-07 | 6.500E-07 |
| 8978 | 449 | N449W | A | 1.8e-07 | 3.3e-07 | 1.7e-07 | 2.267E-07 | 8.963E-08 | 1.700E-07 | 3.300E-07 |
| 8979 | 449 | N449Y | A | 2.2e-07 | 3.7e-07 | 1.4e-07 | 2.433E-07 | 1.168E-07 | 1.400E-07 | 3.700E-07 |
| 8980 | 449 | N449V | A | 3.3e-07 | 1.8e-07 | 1.6e-07 | 2.233E-07 | 9.292E-08 | 1.600E-07 | 3.300E-07 |
| 8981 | 450 | T450A | A | 2.4e-07 | 2.4e-07 | 3.2e-07 | 2.667E-07 | 4.619E-08 | 2.400E-07 | 3.200E-07 |
| 8982 | 450 | T450R | A | 2.8e-07 | 2.5e-07 | 2.6e-07 | 2.633E-07 | 1.528E-08 | 2.500E-07 | 2.800E-07 |
| 8983 | 450 | T450N | A | 2.8e-07 | 2.9e-07 | 1.9e-07 | 2.533E-07 | 5.508E-08 | 1.900E-07 | 2.900E-07 |
| 8984 | 450 | T450D | A | 1.5e-07 | 3e-07 | 1.5e-07 | 2.000E-07 | 8.660E-08 | 1.500E-07 | 3.000E-07 |
| 8985 | 450 | T450C | A | 2.4e-07 | 1.7e-07 | 1.9e-07 | 2.000E-07 | 3.606E-08 | 1.700E-07 | 2.400E-07 |
| 8986 | 450 | T450E | A | 2.3e-07 | 2.9e-07 | 2e-07 | 2.400E-07 | 4.583E-08 | 2.000E-07 | 2.900E-07 |
| 8987 | 450 | T450Q | A | 2.7e-07 | 2.3e-07 | 3.5e-07 | 2.833E-07 | 6.110E-08 | 2.300E-07 | 3.500E-07 |
| 8988 | 450 | T450G | A | 5e-07 | 1.9e-07 | 1.1e-07 | 2.667E-07 | 2.060E-07 | 1.100E-07 | 5.000E-07 |
| 8989 | 450 | T450H | A | 1.7e-07 | 3.3e-07 | 1.6e-07 | 2.200E-07 | 9.539E-08 | 1.600E-07 | 3.300E-07 |
| 8990 | 450 | T450I | A | 2e-07 | 2.5e-07 | 2.6e-07 | 2.367E-07 | 3.215E-08 | 2.000E-07 | 2.600E-07 |
| 8991 | 450 | T450L | A | 5.2e-07 | 5.2e-07 | 2e-07 | 4.133E-07 | 1.848E-07 | 2.000E-07 | 5.200E-07 |
| 8992 | 450 | T450K | A | 2.9e-07 | 9.1e-08 | 3.2e-07 | 2.337E-07 | 1.245E-07 | 9.100E-08 | 3.200E-07 |
| 8993 | 450 | T450M | A | 1.3e-07 | 2.5e-07 | 6.4e-07 | 3.400E-07 | 2.666E-07 | 1.300E-07 | 6.400E-07 |
| 8994 | 450 | T450F | A | 2.1e-07 | 2.2e-07 | 2.1e-07 | 2.133E-07 | 5.774E-09 | 2.100E-07 | 2.200E-07 |
| 8995 | 450 | T450P | A | 3.1e-07 | 3.3e-07 | 4.6e-07 | 3.667E-07 | 8.145E-08 | 3.100E-07 | 4.600E-07 |
| 8996 | 450 | T450S | A | 3.6e-07 | 5e-07 | 2.5e-07 | 3.700E-07 | 1.253E-07 | 2.500E-07 | 5.000E-07 |
| 8997 | 450 | T450T | A | 1.5e-07 | 2.3e-07 | 3.3e-07 | 2.367E-07 | 9.018E-08 | 1.500E-07 | 3.300E-07 |
| 8998 | 450 | T450W | A | 1.5e-07 | 2.5e-07 | 1.7e-07 | 1.900E-07 | 5.292E-08 | 1.500E-07 | 2.500E-07 |
| 8999 | 450 | T450Y | A | 2.4e-07 | 4.1e-07 | 1.4e-07 | 2.633E-07 | 1.365E-07 | 1.400E-07 | 4.100E-07 |
| 9000 | 450 | T450V | A | 1.6e-07 | 2.7e-07 | 2.8e-07 | 2.367E-07 | 6.658E-08 | 1.600E-07 | 2.800E-07 |
| 9001 | 451 | H451A | A | 2.7e-07 | 3.9e-07 | 2e-07 | 2.867E-07 | 9.609E-08 | 2.000E-07 | 3.900E-07 |
| 9002 | 451 | H451R | A | 1.8e-07 | 1.5e-07 | 5.7e-07 | 3.000E-07 | 2.343E-07 | 1.500E-07 | 5.700E-07 |
| 9003 | 451 | H451N | A | 2.3e-07 | 5.8e-07 | 3.4e-07 | 3.833E-07 | 1.790E-07 | 2.300E-07 | 5.800E-07 |
| 9004 | 451 | H451D | A | 3.5e-07 | 1.4e-07 | 2.7e-07 | 2.533E-07 | 1.060E-07 | 1.400E-07 | 3.500E-07 |
| 9005 | 451 | H451C | A | 2.1e-07 | 8.1e-08 | 2.7e-07 | 1.870E-07 | 9.658E-08 | 8.100E-08 | 2.700E-07 |
| 9006 | 451 | H451E | A | 3.8e-07 | 3.8e-07 | 3e-07 | 3.533E-07 | 4.619E-08 | 3.000E-07 | 3.800E-07 |
| 9007 | 451 | H451Q | A | 3.8e-07 | 2.7e-07 | 3.6e-07 | 3.367E-07 | 5.859E-08 | 2.700E-07 | 3.800E-07 |
| 9008 | 451 | H451G | A | 1.9e-07 | 2.2e-07 | 2.2e-07 | 2.100E-07 | 1.732E-08 | 1.900E-07 | 2.200E-07 |
| 9009 | 451 | H451H | A | 1.4e-07 | 5.1e-07 | 3.3e-07 | 3.267E-07 | 1.850E-07 | 1.400E-07 | 5.100E-07 |

| | | | | | | | | | | |
|---|---|---|---|---|---|---|---|---|---|---|
| 9010 | 451 | H451I | A | 3.4e-07 | 1.6e-07 | 3.3e-07 | 2.767E-07 | 1.012E-07 | 1.600E-07 | 3.400E-07 |
| 9011 | 451 | H451L | A | 4.1e-07 | 1.6e-07 | 2.4e-07 | 2.700E-07 | 1.277E-07 | 1.600E-07 | 4.100E-07 |
| 9012 | 451 | H451K | A | 9.8e-08 | 3.2e-07 | 1.5e-07 | 1.893E-07 | 1.161E-07 | 9.800E-08 | 3.200E-07 |
| 9013 | 451 | H451M | A | 3.5e-07 | 2.8e-07 | 1.5e-07 | 2.600E-07 | 1.015E-07 | 1.500E-07 | 3.500E-07 |
| 9014 | 451 | H451F | A | 3.3e-07 | 2.1e-07 | 2.7e-07 | 2.700E-07 | 6.000E-08 | 2.100E-07 | 3.300E-07 |
| 9015 | 451 | H451P | A | 1.6e-07 | 4e-07 | 3.1e-07 | 2.900E-07 | 1.212E-07 | 1.600E-07 | 4.000E-07 |
| 9016 | 451 | H451S | A | 3e-07 | 3.2e-07 | 2.5e-07 | 2.900E-07 | 3.606E-08 | 2.500E-07 | 3.200E-07 |
| 9017 | 451 | H451T | A | 1.6e-07 | 1.9e-07 | 2.4e-07 | 1.967E-07 | 4.041E-08 | 1.600E-07 | 2.400E-07 |
| 9018 | 451 | H451W | A | 1.8e-07 | 2.9e-07 | 1.9e-07 | 2.200E-07 | 6.083E-08 | 1.800E-07 | 2.900E-07 |
| 9019 | 451 | H451Y | A | 2.8e-07 | 8.9e-07 | 1.8e-07 | 4.500E-07 | 3.843E-07 | 1.800E-07 | 8.900E-07 |
| 9020 | 451 | H451V | A | 2e-07 | 3.5e-07 | 4.8e-07 | 3.433E-07 | 1.401E-07 | 2.000E-07 | 4.800E-07 |
| 9021 | 452 | L452A | A | 1.5e-07 | 4.2e-07 | 3.3e-07 | 3.000E-07 | 1.375E-07 | 1.500E-07 | 4.200E-07 |
| 9022 | 452 | L452R | A | 2.3e-07 | 3e-07 | 2.4e-07 | 2.567E-07 | 3.786E-08 | 2.300E-07 | 3.000E-07 |
| 9023 | 452 | L452N | A | 2.3e-07 | 2.4e-07 | 2.3e-07 | 2.333E-07 | 5.774E-09 | 2.300E-07 | 2.400E-07 |
| 9024 | 452 | L452D | A | 2.7e-07 | 4.4e-07 | 4.8e-07 | 3.967E-07 | 1.115E-07 | 2.700E-07 | 4.800E-07 |
| 9025 | 452 | L452C | A | 3.2e-07 | 2.6e-07 | 3.3e-07 | 3.033E-07 | 3.786E-08 | 2.600E-07 | 3.300E-07 |
| 9026 | 452 | L452E | A | 3.1e-07 | 4.8e-07 | 1.8e-07 | 3.233E-07 | 1.504E-07 | 1.800E-07 | 4.800E-07 |
| 9027 | 452 | L452Q | A | 1.5e-07 | 2.7e-07 | 2.2e-07 | 2.133E-07 | 6.028E-08 | 1.500E-07 | 2.700E-07 |
| 9028 | 452 | L452G | A | 2.3e-07 | 2.9e-07 | 1.9e-07 | 2.367E-07 | 5.033E-08 | 1.900E-07 | 2.900E-07 |
| 9029 | 452 | L452H | A | 4.4e-07 | 2.3e-07 | 3.8e-07 | 3.500E-07 | 1.082E-07 | 2.300E-07 | 4.400E-07 |
| 9030 | 452 | L452I | A | 2.5e-07 | 1.5e-07 | 2.7e-07 | 2.233E-07 | 6.429E-08 | 1.500E-07 | 2.700E-07 |
| 9031 | 452 | L452L | A | 3.1e-07 | 2.8e-07 | 2.9e-07 | 2.933E-07 | 1.528E-08 | 2.800E-07 | 3.100E-07 |
| 9032 | 452 | L452K | A | 1e-07 | 1.3e-07 | 3.2e-07 | 1.833E-07 | 1.193E-07 | 1.000E-07 | 3.200E-07 |
| 9033 | 452 | L452M | A | 3.4e-07 | 2.4e-07 | 3.8e-07 | 3.200E-07 | 7.211E-08 | 2.400E-07 | 3.800E-07 |
| 9034 | 452 | L452F | A | 4.8e-07 | 1.7e-07 | 2.3e-07 | 2.933E-07 | 1.644E-07 | 1.700E-07 | 4.800E-07 |
| 9035 | 452 | L452P | A | 1.7e-07 | 3e-07 | 3.8e-07 | 2.833E-07 | 1.060E-07 | 1.700E-07 | 3.800E-07 |
| 9036 | 452 | L452S | A | 2.8e-07 | 3.7e-07 | 2.3e-07 | 2.933E-07 | 7.095E-08 | 2.300E-07 | 3.700E-07 |
| 9037 | 452 | L452T | A | 2.7e-07 | 4.1e-07 | 3.6e-07 | 3.467E-07 | 7.095E-08 | 2.700E-07 | 4.100E-07 |
| 9038 | 452 | L452W | A | 2.4e-07 | 3.7e-07 | 2.8e-07 | 2.967E-07 | 6.658E-08 | 2.400E-07 | 3.700E-07 |
| 9039 | 452 | L452Y | A | 2.6e-07 | 2.5e-07 | 2.3e-07 | 2.467E-07 | 1.528E-08 | 2.300E-07 | 2.600E-07 |
| 9040 | 452 | L452V | A | 1.1e-07 | 3.9e-07 | 4e-07 | 3.000E-07 | 1.646E-07 | 1.100E-07 | 4.000E-07 |
| 9041 | 453 | C453A | A | 3.7e-07 | 1.1e-07 | 4.3e-07 | 3.033E-07 | 1.701E-07 | 1.100E-07 | 4.300E-07 |
| 9042 | 453 | C453R | A | 1.6e-07 | 1.7e-07 | 1.6e-07 | 1.633E-07 | 5.774E-09 | 1.600E-07 | 1.700E-07 |
| 9043 | 453 | C453N | A | 3.3e-07 | 1.7e-07 | 1.7e-07 | 2.233E-07 | 9.238E-08 | 1.700E-07 | 3.300E-07 |
| 9044 | 453 | C453D | A | 4.1e-07 | 2.8e-07 | 2.2e-07 | 3.033E-07 | 9.713E-08 | 2.200E-07 | 4.100E-07 |
| 9045 | 453 | C453C | A | 2.3e-07 | 9.4e-08 | 2e-07 | 1.747E-07 | 7.145E-08 | 9.400E-08 | 2.300E-07 |
| 9046 | 453 | C453E | A | 3.5e-07 | 2.9e-07 | 3.4e-07 | 3.267E-07 | 3.215E-08 | 2.900E-07 | 3.500E-07 |
| 9047 | 453 | C453Q | A | 5.7e-07 | 1.9e-07 | 2.4e-07 | 3.333E-07 | 2.065E-07 | 1.900E-07 | 5.700E-07 |
| 9048 | 453 | C453G | A | 2.9e-07 | 2.9e-07 | 3.2e-07 | 3.000E-07 | 1.732E-08 | 2.900E-07 | 3.200E-07 |
| 9049 | 453 | C453H | A | 1.7e-07 | 1.5e-07 | 2.9e-07 | 2.033E-07 | 7.572E-08 | 1.500E-07 | 2.900E-07 |
| 9050 | 453 | C453I | A | 2.4e-07 | 1.9e-07 | 6.4e-07 | 3.567E-07 | 2.466E-07 | 1.900E-07 | 6.400E-07 |
| 9051 | 453 | C453L | A | 1e-07 | 3e-07 | 1.8e-07 | 1.933E-07 | 1.007E-07 | 1.000E-07 | 3.000E-07 |
| 9052 | 453 | C453K | A | 2.5e-07 | 3.8e-07 | 1.9e-07 | 2.733E-07 | 9.713E-08 | 1.900E-07 | 3.800E-07 |
| 9053 | 453 | C453M | A | 3.8e-07 | 2.4e-07 | 2.7e-07 | 2.967E-07 | 7.371E-08 | 2.400E-07 | 3.800E-07 |
| 9054 | 453 | C453F | A | 2.5e-07 | 6.3e-07 | 2.9e-07 | 3.900E-07 | 2.088E-07 | 2.500E-07 | 6.300E-07 |
| 9055 | 453 | C453P | A | 4.5e-07 | 2.9e-07 | 1.7e-07 | 3.033E-07 | 1.405E-07 | 1.700E-07 | 4.500E-07 |
| 9056 | 453 | C453S | A | 3.1e-07 | 3.6e-07 | 2.6e-07 | 3.100E-07 | 5.000E-08 | 2.600E-07 | 3.600E-07 |
| 9057 | 453 | C453T | A | 2.8e-07 | 3.2e-07 | 2.2e-07 | 2.733E-07 | 5.033E-08 | 2.200E-07 | 3.200E-07 |
| 9058 | 453 | C453W | A | 1.5e-07 | 2.1e-07 | 3.4e-07 | 2.333E-07 | 9.713E-08 | 1.500E-07 | 3.400E-07 |
| 9059 | 453 | C453Y | A | 2.1e-07 | 1.4e-07 | 2.6e-07 | 2.033E-07 | 6.028E-08 | 1.400E-07 | 2.600E-07 |
| 9060 | 453 | C453V | A | 4e-07 | 1.7e-07 | 3.8e-07 | 3.167E-07 | 1.274E-07 | 1.700E-07 | 4.000E-07 |
| 9061 | 454 | F454A | A | 1.6e-07 | 2.2e-07 | 2.1e-07 | 1.967E-07 | 3.215E-08 | 1.600E-07 | 2.200E-07 |
| 9062 | 454 | F454R | A | 4.5e-07 | 2.8e-07 | 3.1e-07 | 3.467E-07 | 9.074E-08 | 2.800E-07 | 4.500E-07 |
| 9063 | 454 | F454N | A | 1.9e-07 | 4.1e-07 | 4e-07 | 3.333E-07 | 1.242E-07 | 1.900E-07 | 4.100E-07 |
| 9064 | 454 | F454D | A | 3.9e-07 | 2e-07 | 2e-07 | 2.633E-07 | 1.097E-07 | 2.000E-07 | 3.900E-07 |
| 9065 | 454 | F454C | A | 4e-07 | 1.9e-07 | 4.4e-07 | 3.433E-07 | 1.343E-07 | 1.900E-07 | 4.400E-07 |
| 9066 | 454 | F454E | A | 3e-07 | 2.4e-07 | 3.3e-07 | 2.900E-07 | 4.583E-08 | 2.400E-07 | 3.300E-07 |
| 9067 | 454 | F454Q | A | 1.6e-07 | 1.9e-07 | 2.1e-07 | 1.867E-07 | 2.517E-08 | 1.600E-07 | 2.100E-07 |
| 9068 | 454 | F454G | A | 1.7e-07 | 1.5e-07 | 1.4e-07 | 1.533E-07 | 1.528E-08 | 1.400E-07 | 1.700E-07 |
| 9069 | 454 | F454H | A | 2.2e-07 | 2.2e-07 | 2.5e-07 | 2.300E-07 | 1.732E-08 | 2.200E-07 | 2.500E-07 |
| 9070 | 454 | F454I | A | 2.1e-07 | 3.1e-07 | 2.2e-07 | 2.467E-07 | 5.508E-08 | 2.100E-07 | 3.100E-07 |
| 9071 | 454 | F454L | A | 3.5e-07 | 2.9e-07 | 5.5e-07 | 3.967E-07 | 1.361E-07 | 2.900E-07 | 5.500E-07 |
| 9072 | 454 | F454K | A | 1.8e-07 | 2.1e-07 | 1.6e-07 | 1.833E-07 | 2.517E-08 | 1.600E-07 | 2.100E-07 |
| 9073 | 454 | F454M | A | 3.2e-07 | 3.9e-07 | 7.6e-08 | 2.620E-07 | 1.648E-07 | 7.600E-08 | 3.900E-07 |
| 9074 | 454 | F454F | A | 2.5e-07 | 2.5e-07 | 1.4e-07 | 2.133E-07 | 6.351E-08 | 1.400E-07 | 2.500E-07 |
| 9075 | 454 | F454P | A | 1.8e-07 | 1.9e-07 | 2.8e-07 | 2.167E-07 | 5.508E-08 | 1.800E-07 | 2.800E-07 |
| 9076 | 454 | F454S | A | 2e-07 | 3.2e-07 | 7.5e-07 | 4.233E-07 | 2.892E-07 | 2.000E-07 | 7.500E-07 |
| 9077 | 454 | F454T | A | 1.5e-07 | 2.4e-07 | 2.6e-07 | 2.167E-07 | 5.859E-08 | 1.500E-07 | 2.600E-07 |
| 9078 | 454 | F454W | A | 3.2e-07 | 1.6e-07 | 2e-07 | 2.267E-07 | 8.327E-08 | 1.600E-07 | 3.200E-07 |
| 9079 | 454 | F454Y | A | 3.9e-07 | 2.2e-07 | 1.5e-07 | 2.533E-07 | 1.234E-07 | 1.500E-07 | 3.900E-07 |
| 9080 | 454 | F454V | A | 6.7e-07 | 3e-07 | 3e-07 | 4.233E-07 | 2.136E-07 | 3.000E-07 | 6.700E-07 |
| 9081 | 455 | V455A | A | 2.2e-07 | 2.3e-07 | 1.5e-07 | 2.000E-07 | 4.359E-08 | 1.500E-07 | 2.300E-07 |
| 9082 | 455 | V455R | A | 4.7e-07 | 2.3e-07 | 2.7e-07 | 3.233E-07 | 1.286E-07 | 2.300E-07 | 4.700E-07 |
| 9083 | 455 | V455N | A | 3e-07 | 2.9e-07 | 1.6e-07 | 2.500E-07 | 7.810E-08 | 1.600E-07 | 3.000E-07 |
| 9084 | 455 | V455D | A | 1.6e-07 | 2.1e-07 | 1.6e-07 | 1.767E-07 | 2.887E-08 | 1.600E-07 | 2.100E-07 |
| 9085 | 455 | V455C | A | 3.4e-07 | 1.1e-07 | 3.3e-07 | 2.600E-07 | 1.300E-07 | 1.100E-07 | 3.400E-07 |
| 9086 | 455 | V455E | A | 1e-07 | 3e-07 | 1.7e-07 | 1.900E-07 | 1.015E-07 | 1.000E-07 | 3.000E-07 |
| 9087 | 455 | V455Q | A | 4.3e-07 | 3.5e-07 | 3.4e-07 | 3.733E-07 | 4.933E-08 | 3.400E-07 | 4.300E-07 |
| 9088 | 455 | V455G | A | 2e-07 | 1.6e-07 | 3.8e-07 | 2.467E-07 | 1.172E-07 | 1.600E-07 | 3.800E-07 |
| 9089 | 455 | V455H | A | 1.2e-07 | 2.5e-07 | 2.5e-07 | 2.067E-07 | 7.506E-08 | 1.200E-07 | 2.500E-07 |
| 9090 | 455 | V455I | A | 2.9e-07 | 6.2e-07 | 1.8e-07 | 3.633E-07 | 2.290E-07 | 1.800E-07 | 6.200E-07 |
| 9091 | 455 | V455L | A | 2.8e-07 | 2.8e-07 | 2e-07 | 2.533E-07 | 4.619E-08 | 2.000E-07 | 2.800E-07 |
| 9092 | 455 | V455K | A | 2.4e-07 | 4.1e-07 | 1.8e-07 | 2.767E-07 | 1.193E-07 | 1.800E-07 | 4.100E-07 |
| 9093 | 455 | V455M | A | 6.3e-07 | 2.2e-07 | 1.9e-07 | 3.467E-07 | 2.458E-07 | 1.900E-07 | 6.300E-07 |
| 9094 | 455 | V455F | A | 4.4e-07 | 1.6e-07 | 4e-07 | 3.333E-07 | 1.514E-07 | 1.600E-07 | 4.400E-07 |

| | | | | | | | | | | |
|---|---|---|---|---|---|---|---|---|---|---|
| 9095 | 455 | V455P | A | 2e-07 | 3.4e-07 | 2.3e-07 | 2.567E-07 | 7.371E-08 | 2.000E-07 | 3.400E-07 |
| 9096 | 455 | V455S | A | 1.9e-07 | 2.2e-07 | 5.4e-07 | 3.167E-07 | 1.940E-07 | 1.900E-07 | 5.400E-07 |
| 9097 | 455 | V455T | A | 2.8e-07 | 4.5e-07 | 1.5e-07 | 2.933E-07 | 1.504E-07 | 1.500E-07 | 4.500E-07 |
| 9098 | 455 | V455W | A | 2.4e-07 | 1.8e-07 | 2.3e-07 | 2.167E-07 | 3.215E-08 | 1.800E-07 | 2.400E-07 |
| 9099 | 455 | V455Y | A | 1.5e-07 | 1.3e-07 | 4.9e-07 | 2.567E-07 | 2.023E-07 | 1.300E-07 | 4.900E-07 |
| 9100 | 455 | V455V | A | 1.9e-07 | 5e-07 | 2.7e-07 | 3.200E-07 | 1.609E-07 | 1.900E-07 | 5.000E-07 |
| 9101 | 456 | H456A | A | 2.8e-07 | 1.7e-07 | 3.4e-07 | 2.633E-07 | 8.622E-08 | 1.700E-07 | 3.400E-07 |
| 9102 | 456 | H456R | A | 3.5e-07 | 1.8e-07 | 6.4e-08 | 1.980E-07 | 1.438E-07 | 6.400E-08 | 3.500E-07 |
| 9103 | 456 | H456N | A | 2.9e-07 | 2e-07 | 3.1e-07 | 2.667E-07 | 5.859E-08 | 2.000E-07 | 3.100E-07 |
| 9104 | 456 | H456D | A | 2.8e-07 | 1.9e-07 | 2e-07 | 2.233E-07 | 4.933E-08 | 1.900E-07 | 2.800E-07 |
| 9105 | 456 | H456C | A | 2.2e-07 | 3.8e-07 | 1.9e-07 | 2.633E-07 | 1.021E-07 | 1.900E-07 | 3.800E-07 |
| 9106 | 456 | H456E | A | 2.5e-07 | 1.3e-07 | 2.2e-07 | 2.000E-07 | 6.245E-08 | 1.300E-07 | 2.500E-07 |
| 9107 | 456 | H456Q | A | 3.4e-07 | 1.2e-07 | 3.3e-07 | 2.633E-07 | 1.242E-07 | 1.200E-07 | 3.400E-07 |
| 9108 | 456 | H456G | A | 3.3e-07 | 4.3e-07 | 2.3e-07 | 3.300E-07 | 1.000E-07 | 2.300E-07 | 4.300E-07 |
| 9109 | 456 | H456H | A | 1.2e-07 | 2.7e-07 | 9.8e-08 | 1.627E-07 | 9.360E-08 | 9.800E-08 | 2.700E-07 |
| 9110 | 456 | H456I | A | 4.1e-07 | 3e-07 | 4.8e-07 | 3.967E-07 | 9.074E-08 | 3.000E-07 | 4.800E-07 |
| 9111 | 456 | H456L | A | 1.8e-07 | 2.5e-07 | 2.4e-07 | 2.233E-07 | 3.786E-08 | 1.800E-07 | 2.500E-07 |
| 9112 | 456 | H456K | A | 4e-07 | 3e-07 | 3.5e-07 | 3.500E-07 | 5.000E-08 | 3.000E-07 | 4.000E-07 |
| 9113 | 456 | H456M | A | 2.9e-07 | 3.2e-07 | 1.9e-07 | 2.667E-07 | 6.807E-08 | 1.900E-07 | 3.200E-07 |
| 9114 | 456 | H456F | A | 8.5e-07 | 1.9e-07 | 1.9e-07 | 4.100E-07 | 3.811E-07 | 1.900E-07 | 8.500E-07 |
| 9115 | 456 | H456P | A | 1.6e-07 | 5.7e-07 | 3.1e-07 | 3.467E-07 | 2.074E-07 | 1.600E-07 | 5.700E-07 |
| 9116 | 456 | H456S | A | 1.9e-07 | 2.1e-07 | 1.8e-07 | 1.933E-07 | 1.528E-08 | 1.800E-07 | 2.100E-07 |
| 9117 | 456 | H456T | A | 1.4e-07 | 3.6e-07 | 3.5e-07 | 2.833E-07 | 1.242E-07 | 1.400E-07 | 3.600E-07 |
| 9118 | 456 | H456W | A | 3.5e-07 | 1.9e-07 | 7.6e-08 | 2.053E-07 | 1.376E-07 | 7.600E-08 | 3.500E-07 |
| 9119 | 456 | H456Y | A | 2.1e-07 | 4.3e-07 | 2.4e-07 | 2.933E-07 | 1.193E-07 | 2.100E-07 | 4.300E-07 |
| 9120 | 456 | H456V | A | 1.6e-07 | 2.8e-07 | 4.2e-07 | 2.867E-07 | 1.301E-07 | 1.600E-07 | 4.200E-07 |
| 9121 | 457 | T457A | A | 6.8e-08 | 3.9e-07 | 2.2e-07 | 2.260E-07 | 1.611E-07 | 6.800E-08 | 3.900E-07 |
| 9122 | 457 | T457R | A | 2.3e-07 | 4.1e-07 | 2.2e-07 | 2.867E-07 | 1.069E-07 | 2.200E-07 | 4.100E-07 |
| 9123 | 457 | T457N | A | 1.5e-07 | 2e-07 | 4.1e-07 | 2.533E-07 | 1.380E-07 | 1.500E-07 | 4.100E-07 |
| 9124 | 457 | T457D | A | 1.2e-07 | 3.4e-07 | 1.3e-07 | 1.967E-07 | 1.242E-07 | 1.200E-07 | 3.400E-07 |
| 9125 | 457 | T457C | A | 3.1e-07 | 3.2e-07 | 2.6e-07 | 2.967E-07 | 3.215E-08 | 2.600E-07 | 3.200E-07 |
| 9126 | 457 | T457E | A | 5.5e-07 | 4e-07 | 2.8e-07 | 4.100E-07 | 1.353E-07 | 2.800E-07 | 5.500E-07 |
| 9127 | 457 | T457Q | A | 2.5e-07 | 3.3e-07 | 2.1e-07 | 2.633E-07 | 6.110E-08 | 2.100E-07 | 3.300E-07 |
| 9128 | 457 | T457G | A | 3.8e-07 | 2.9e-07 | 2.4e-07 | 3.033E-07 | 7.095E-08 | 2.400E-07 | 3.800E-07 |
| 9129 | 457 | T457H | A | 2.4e-07 | 2.4e-07 | 2.5e-07 | 2.433E-07 | 5.774E-09 | 2.400E-07 | 2.500E-07 |
| 9130 | 457 | T457I | A | 2.7e-07 | 3.5e-07 | 2.1e-07 | 2.767E-07 | 7.024E-08 | 2.100E-07 | 3.500E-07 |
| 9131 | 457 | T457L | A | 2.1e-07 | 2.1e-07 | 2.4e-07 | 2.200E-07 | 1.732E-08 | 2.100E-07 | 2.400E-07 |
| 9132 | 457 | T457K | A | 1.9e-07 | 2.6e-07 | 1.4e-07 | 1.967E-07 | 6.028E-08 | 1.400E-07 | 2.600E-07 |
| 9133 | 457 | T457M | A | 4.4e-07 | 2.3e-07 | 1.2e-07 | 2.633E-07 | 1.626E-07 | 1.200E-07 | 4.400E-07 |
| 9134 | 457 | T457F | A | 2.2e-07 | 3.2e-07 | 1.9e-07 | 2.433E-07 | 6.807E-08 | 1.900E-07 | 3.200E-07 |
| 9135 | 457 | T457P | A | 2e-07 | 3.7e-07 | 1.6e-07 | 2.433E-07 | 1.115E-07 | 1.600E-07 | 3.700E-07 |
| 9136 | 457 | T457S | A | 2.4e-07 | 1.4e-07 | 2.6e-07 | 2.133E-07 | 6.429E-08 | 1.400E-07 | 2.600E-07 |
| 9137 | 457 | T457T | A | 2.5e-07 | 1.5e-07 | 3.2e-07 | 2.400E-07 | 8.544E-08 | 1.500E-07 | 3.200E-07 |
| 9138 | 457 | T457W | A | 3.5e-07 | 2.2e-07 | 1.3e-07 | 2.333E-07 | 1.106E-07 | 1.300E-07 | 3.500E-07 |
| 9139 | 457 | T457Y | A | 2.4e-07 | 1.7e-07 | 1.9e-07 | 2.000E-07 | 3.606E-08 | 1.700E-07 | 2.400E-07 |
| 9140 | 457 | T457V | A | 2.7e-07 | 2.5e-07 | 2.4e-07 | 2.533E-07 | 1.528E-08 | 2.400E-07 | 2.700E-07 |
| 9141 | 458 | V458A | A | 1.9e-07 | 2.1e-07 | 3.6e-07 | 2.533E-07 | 9.292E-08 | 1.900E-07 | 3.600E-07 |
| 9142 | 458 | V458R | A | 2.9e-07 | 2.1e-07 | 2.3e-07 | 2.433E-07 | 4.163E-08 | 2.100E-07 | 2.900E-07 |
| 9143 | 458 | V458N | A | 3.3e-07 | 3.5e-07 | 2.1e-07 | 2.967E-07 | 7.572E-08 | 2.100E-07 | 3.500E-07 |
| 9144 | 458 | V458D | A | 2.4e-07 | 3.4e-07 | 1.7e-07 | 2.500E-07 | 8.544E-08 | 1.700E-07 | 3.400E-07 |
| 9145 | 458 | V458C | A | 1.7e-07 | 4.6e-07 | 2.7e-07 | 3.000E-07 | 1.473E-07 | 1.700E-07 | 4.600E-07 |
| 9146 | 458 | V458E | A | 5.7e-07 | 3.4e-07 | 1.9e-07 | 3.667E-07 | 1.914E-07 | 1.900E-07 | 5.700E-07 |
| 9147 | 458 | V458Q | A | 2.8e-07 | 2e-07 | 2.6e-07 | 2.467E-07 | 4.163E-08 | 2.000E-07 | 2.800E-07 |
| 9148 | 458 | V458G | A | 3.9e-07 | 2.5e-07 | 3.9e-07 | 3.433E-07 | 8.083E-08 | 2.500E-07 | 3.900E-07 |
| 9149 | 458 | V458H | A | 2.3e-07 | 2.4e-07 | 1.1e-07 | 1.933E-07 | 7.234E-08 | 1.100E-07 | 2.400E-07 |
| 9150 | 458 | V458I | A | 2.3e-07 | 3.3e-07 | 3.4e-07 | 3.000E-07 | 6.083E-08 | 2.300E-07 | 3.400E-07 |
| 9151 | 458 | V458L | A | 4.7e-07 | 1.3e-07 | 2.5e-07 | 2.833E-07 | 1.724E-07 | 1.300E-07 | 4.700E-07 |
| 9152 | 458 | V458K | A | 4.2e-07 | 2.4e-07 | 2.7e-07 | 3.100E-07 | 9.644E-08 | 2.400E-07 | 4.200E-07 |
| 9153 | 458 | V458M | A | 4.2e-07 | 1.3e-07 | 3.4e-07 | 2.967E-07 | 1.498E-07 | 1.300E-07 | 4.200E-07 |
| 9154 | 458 | V458F | A | 2.4e-07 | 1.4e-07 | 1.9e-07 | 1.900E-07 | 5.000E-08 | 1.400E-07 | 2.400E-07 |
| 9155 | 458 | V458P | A | 5.1e-07 | 3.6e-07 | 2.3e-07 | 3.667E-07 | 1.401E-07 | 2.300E-07 | 5.100E-07 |
| 9156 | 458 | V458S | A | 3.2e-07 | 3.5e-07 | 3.3e-07 | 3.333E-07 | 1.528E-08 | 3.200E-07 | 3.500E-07 |
| 9157 | 458 | V458T | A | 1.9e-07 | 1.4e-07 | 2.8e-07 | 2.033E-07 | 7.095E-08 | 1.400E-07 | 2.800E-07 |
| 9158 | 458 | V458W | A | 3e-07 | 1.4e-07 | 4.7e-07 | 3.033E-07 | 1.650E-07 | 1.400E-07 | 4.700E-07 |
| 9159 | 458 | V458Y | A | 1.8e-07 | 3.6e-07 | 2.4e-07 | 2.600E-07 | 9.165E-08 | 1.800E-07 | 3.600E-07 |
| 9160 | 458 | V458V | A | 2.4e-07 | 2.3e-07 | 2.8e-07 | 2.500E-07 | 2.646E-08 | 2.300E-07 | 2.800E-07 |
| 9161 | 459 | P459A | A | 2.1e-07 | 1.4e-07 | 9.8e-08 | 1.493E-07 | 5.658E-08 | 9.800E-08 | 2.100E-07 |
| 9162 | 459 | P459R | A | 2.5e-07 | 1.8e-07 | 2.3e-07 | 2.200E-07 | 3.606E-08 | 1.800E-07 | 2.500E-07 |
| 9163 | 459 | P459N | A | 2.5e-07 | 9.7e-08 | 2.7e-07 | 2.057E-07 | 9.464E-08 | 9.700E-08 | 2.700E-07 |
| 9164 | 459 | P459D | A | 3.4e-07 | 2.7e-07 | 1.8e-07 | 2.633E-07 | 8.021E-08 | 1.800E-07 | 3.400E-07 |
| 9165 | 459 | P459C | A | 3.6e-07 | 7.9e-08 | 1.7e-07 | 2.030E-07 | 1.434E-07 | 7.900E-08 | 3.600E-07 |
| 9166 | 459 | P459E | A | 1.7e-07 | 1.7e-07 | 2.7e-07 | 2.033E-07 | 5.774E-08 | 1.700E-07 | 2.700E-07 |
| 9167 | 459 | P459Q | A | 2.7e-07 | 2.5e-07 | 3.3e-07 | 2.833E-07 | 4.163E-08 | 2.500E-07 | 3.300E-07 |
| 9168 | 459 | P459G | A | 4.7e-07 | 2e-07 | 2.4e-07 | 3.033E-07 | 1.457E-07 | 2.000E-07 | 4.700E-07 |
| 9169 | 459 | P459H | A | 1.9e-07 | 3e-07 | 2.3e-07 | 2.400E-07 | 5.568E-08 | 1.900E-07 | 3.000E-07 |
| 9170 | 459 | P459I | A | 1.3e-07 | 3.3e-07 | 1.8e-07 | 2.133E-07 | 1.041E-07 | 1.300E-07 | 3.300E-07 |
| 9171 | 459 | P459L | A | 2.7e-07 | 1.8e-07 | 3.3e-07 | 2.600E-07 | 7.550E-08 | 1.800E-07 | 3.300E-07 |
| 9172 | 459 | P459K | A | 4.6e-07 | 1.1e-07 | 4.6e-07 | 3.433E-07 | 2.021E-07 | 1.100E-07 | 4.600E-07 |
| 9173 | 459 | P459M | A | 4.4e-07 | 1.6e-07 | 2.6e-07 | 2.867E-07 | 1.419E-07 | 1.600E-07 | 4.400E-07 |
| 9174 | 459 | P459F | A | 3.7e-07 | 1e-07 | 9.3e-08 | 1.877E-07 | 1.579E-07 | 9.300E-08 | 3.700E-07 |
| 9175 | 459 | P459P | A | 1.4e-07 | 1.9e-07 | 2.5e-07 | 1.933E-07 | 5.508E-08 | 1.400E-07 | 2.500E-07 |
| 9176 | 459 | P459S | A | 5.3e-07 | 1e-07 | 2.3e-07 | 2.867E-07 | 2.205E-07 | 1.000E-07 | 5.300E-07 |
| 9177 | 459 | P459T | A | 5.5e-07 | 2.5e-07 | 3.4e-07 | 3.800E-07 | 1.539E-07 | 2.500E-07 | 5.500E-07 |
| 9178 | 459 | P459W | A | 3.7e-07 | 1.7e-07 | 3.2e-07 | 2.867E-07 | 1.041E-07 | 1.700E-07 | 3.700E-07 |
| 9179 | 459 | P459Y | A | 8.8e-08 | 1.4e-07 | 4.3e-07 | 2.193E-07 | 1.843E-07 | 8.800E-08 | 4.300E-07 |

| | | | | | | | | | | |
|---|---|---|---|---|---|---|---|---|---|---|
| 9180 | 459 | P459V | A | 2.2e-07 | 3.2e-07 | 1.9e-07 | 2.433E-07 | 6.807E-08 | 1.900E-07 | 3.200E-07 |
| 9181 | 460 | W460A | A | 2.2e-07 | 2.4e-07 | 2.1e-07 | 2.233E-07 | 1.528E-08 | 2.100E-07 | 2.400E-07 |
| 9182 | 460 | W460R | A | 6e-07 | 2.9e-07 | 1.2e-07 | 3.367E-07 | 2.434E-07 | 1.200E-07 | 6.000E-07 |
| 9183 | 460 | W460N | A | 2.7e-07 | 3.3e-07 | 4.1e-07 | 3.367E-07 | 7.024E-08 | 2.700E-07 | 4.100E-07 |
| 9184 | 460 | W460D | A | 5.2e-07 | 3.1e-07 | 1.2e-07 | 3.167E-07 | 2.001E-07 | 1.200E-07 | 5.200E-07 |
| 9185 | 460 | W460C | A | 3.4e-07 | 3.1e-07 | 4.6e-07 | 3.700E-07 | 7.937E-08 | 3.100E-07 | 4.600E-07 |
| 9186 | 460 | W460E | A | 4.1e-07 | 1.7e-07 | 1.3e-07 | 2.367E-07 | 1.514E-07 | 1.300E-07 | 4.100E-07 |
| 9187 | 460 | W460Q | A | 4.5e-07 | 2.2e-07 | 2.2e-07 | 2.967E-07 | 1.328E-07 | 2.200E-07 | 4.500E-07 |
| 9188 | 460 | W460G | A | 1.9e-07 | 1.2e-07 | 1.9e-07 | 1.667E-07 | 4.041E-08 | 1.200E-07 | 1.900E-07 |
| 9189 | 460 | W460H | A | 1.7e-07 | 2.1e-07 | 2.2e-07 | 2.000E-07 | 2.646E-08 | 1.700E-07 | 2.200E-07 |
| 9190 | 460 | W460I | A | 1.1e-07 | 2.2e-07 | 1.7e-07 | 1.667E-07 | 5.508E-08 | 1.100E-07 | 2.200E-07 |
| 9191 | 460 | W460L | A | 1.9e-07 | 3.6e-07 | 1.1e-07 | 2.200E-07 | 1.277E-07 | 1.100E-07 | 3.600E-07 |
| 9192 | 460 | W460K | A | 4.1e-07 | 3.6e-07 | 3.5e-07 | 3.733E-07 | 3.215E-08 | 3.500E-07 | 4.100E-07 |
| 9193 | 460 | W460M | A | 3.9e-07 | 1.2e-07 | 2.7e-07 | 2.600E-07 | 1.353E-07 | 1.200E-07 | 3.900E-07 |
| 9194 | 460 | W460F | A | 4e-07 | 1.3e-07 | 1.6e-07 | 2.300E-07 | 1.480E-07 | 1.300E-07 | 4.000E-07 |
| 9195 | 460 | W460P | A | 2.5e-07 | 5.9e-07 | 2.8e-07 | 3.733E-07 | 1.882E-07 | 2.500E-07 | 5.900E-07 |
| 9196 | 460 | W460S | A | 3.6e-07 | 5.4e-07 | 4.2e-07 | 4.400E-07 | 9.165E-08 | 3.600E-07 | 5.400E-07 |
| 9197 | 460 | W460T | A | 2.2e-07 | 2.6e-07 | 4e-07 | 2.933E-07 | 9.452E-08 | 2.200E-07 | 4.000E-07 |
| 9198 | 460 | W460W | A | 1.6e-07 | 1.2e-07 | 1.2e-07 | 1.333E-07 | 2.309E-08 | 1.200E-07 | 1.600E-07 |
| 9199 | 460 | W460Y | A | 3.8e-07 | 1.9e-07 | 3.2e-07 | 2.967E-07 | 9.713E-08 | 1.900E-07 | 3.800E-07 |
| 9200 | 460 | W460V | A | 1.6e-07 | 3.1e-07 | 2.4e-07 | 2.367E-07 | 7.506E-08 | 1.600E-07 | 3.100E-07 |
| 9201 | 461 | D461A | A | 2.7e-07 | 1.9e-07 | 3.1e-07 | 2.567E-07 | 6.110E-08 | 1.900E-07 | 3.100E-07 |
| 9202 | 461 | D461R | A | 2.8e-07 | 3.5e-07 | 2.2e-07 | 2.833E-07 | 6.506E-08 | 2.200E-07 | 3.500E-07 |
| 9203 | 461 | D461N | A | 3.2e-07 | 1.5e-07 | 2.2e-07 | 2.300E-07 | 8.544E-08 | 1.500E-07 | 3.200E-07 |
| 9204 | 461 | D461D | A | 2e-07 | 2.7e-07 | 1.8e-07 | 2.167E-07 | 4.726E-08 | 1.800E-07 | 2.700E-07 |
| 9205 | 461 | D461C | A | 1.3e-07 | 2e-07 | 3.4e-07 | 2.233E-07 | 1.069E-07 | 1.300E-07 | 3.400E-07 |
| 9206 | 461 | D461E | A | 1.4e-07 | 3.7e-07 | 2.7e-07 | 2.600E-07 | 1.153E-07 | 1.400E-07 | 3.700E-07 |
| 9207 | 461 | D461Q | A | 2.8e-07 | 4.9e-07 | 2.2e-07 | 3.300E-07 | 1.418E-07 | 2.200E-07 | 4.900E-07 |
| 9208 | 461 | D461G | A | 1.4e-07 | 3.3e-07 | 3.7e-07 | 2.800E-07 | 1.229E-07 | 1.400E-07 | 3.700E-07 |
| 9209 | 461 | D461H | A | 3.2e-07 | 1.9e-07 | 2.8e-07 | 2.633E-07 | 6.658E-08 | 1.900E-07 | 3.200E-07 |
| 9210 | 461 | D461I | A | 3.4e-07 | 1.3e-07 | 3.8e-07 | 2.833E-07 | 1.343E-07 | 1.300E-07 | 3.800E-07 |
| 9211 | 461 | D461L | A | 1.2e-07 | 4e-07 | 3.3e-07 | 2.833E-07 | 1.457E-07 | 1.200E-07 | 4.000E-07 |
| 9212 | 461 | D461K | A | 2.7e-07 | 2.6e-07 | 5.2e-07 | 3.500E-07 | 1.473E-07 | 2.600E-07 | 5.200E-07 |
| 9213 | 461 | D461M | A | 2.4e-07 | 2.8e-07 | 2.4e-07 | 2.533E-07 | 2.309E-08 | 2.400E-07 | 2.800E-07 |
| 9214 | 461 | D461F | A | 1.1e-07 | 3.3e-07 | 2.5e-07 | 2.300E-07 | 1.114E-07 | 1.100E-07 | 3.300E-07 |
| 9215 | 461 | D461P | A | 3.3e-07 | 3.3e-07 | 4.9e-07 | 3.833E-07 | 9.238E-08 | 3.300E-07 | 4.900E-07 |
| 9216 | 461 | D461S | A | 2.5e-07 | 2.9e-07 | 3.6e-07 | 3.000E-07 | 5.568E-08 | 2.500E-07 | 3.600E-07 |
| 9217 | 461 | D461T | A | 1.9e-07 | 2.6e-07 | 3e-07 | 2.500E-07 | 5.568E-08 | 1.900E-07 | 3.000E-07 |
| 9218 | 461 | D461W | A | 2e-07 | 1.6e-07 | 4.2e-07 | 2.600E-07 | 1.400E-07 | 1.600E-07 | 4.200E-07 |
| 9219 | 461 | D461Y | A | 2.4e-07 | 3.2e-07 | 2e-07 | 2.533E-07 | 6.110E-08 | 2.000E-07 | 3.200E-07 |
| 9220 | 461 | D461V | A | 1.9e-07 | 1.3e-07 | 1.6e-07 | 1.600E-07 | 3.000E-08 | 1.300E-07 | 1.900E-07 |
| 9221 | 462 | Q462A | A | 2.2e-07 | 1.7e-07 | 2.4e-07 | 2.100E-07 | 3.606E-08 | 1.700E-07 | 2.400E-07 |
| 9222 | 462 | Q462R | A | 2.6e-07 | 2.7e-07 | 3e-07 | 2.767E-07 | 2.082E-08 | 2.600E-07 | 3.000E-07 |
| 9223 | 462 | Q462N | A | 2.4e-07 | 1.4e-07 | 2.8e-07 | 2.200E-07 | 7.211E-08 | 1.400E-07 | 2.800E-07 |
| 9224 | 462 | Q462D | A | 2.7e-07 | 2.2e-07 | 2.9e-07 | 2.600E-07 | 3.606E-08 | 2.200E-07 | 2.900E-07 |
| 9225 | 462 | Q462C | A | 3.2e-07 | 2.4e-07 | 3.5e-07 | 3.033E-07 | 5.686E-08 | 2.400E-07 | 3.500E-07 |
| 9226 | 462 | Q462E | A | 3.4e-07 | 5.2e-07 | 4.2e-07 | 4.267E-07 | 9.018E-08 | 3.400E-07 | 5.200E-07 |
| 9227 | 462 | Q462Q | A | 4.4e-07 | 4.7e-07 | 2.8e-07 | 3.967E-07 | 1.021E-07 | 2.800E-07 | 4.700E-07 |
| 9228 | 462 | Q462G | A | 3.8e-07 | 1.4e-07 | 4e-07 | 3.067E-07 | 1.447E-07 | 1.400E-07 | 4.000E-07 |
| 9229 | 462 | Q462H | A | 2.8e-07 | 1.5e-07 | 2.8e-07 | 2.367E-07 | 7.506E-08 | 1.500E-07 | 2.800E-07 |
| 9230 | 462 | Q462I | A | 2.4e-07 | 2.1e-07 | 5.1e-07 | 3.200E-07 | 1.652E-07 | 2.100E-07 | 5.100E-07 |
| 9231 | 462 | Q462L | A | 1.5e-07 | 2.8e-07 | 2.7e-07 | 2.333E-07 | 7.234E-08 | 1.500E-07 | 2.800E-07 |
| 9232 | 462 | Q462K | A | 3.7e-07 | 1.2e-07 | 2e-07 | 2.300E-07 | 1.277E-07 | 1.200E-07 | 3.700E-07 |
| 9233 | 462 | Q462M | A | 8.2e-08 | 1.3e-07 | 4.3e-07 | 2.140E-07 | 1.886E-07 | 8.200E-08 | 4.300E-07 |
| 9234 | 462 | Q462F | A | 1.7e-07 | 1.5e-07 | 4.3e-07 | 2.500E-07 | 1.562E-07 | 1.500E-07 | 4.300E-07 |
| 9235 | 462 | Q462P | A | 3.4e-07 | 6.1e-07 | 1.8e-07 | 3.767E-07 | 2.173E-07 | 1.800E-07 | 6.100E-07 |
| 9236 | 462 | Q462S | A | 1.6e-07 | 4.4e-07 | 2.5e-07 | 2.833E-07 | 1.429E-07 | 1.600E-07 | 4.400E-07 |
| 9237 | 462 | Q462T | A | 2.5e-07 | 2.5e-07 | 1.5e-07 | 2.167E-07 | 5.774E-08 | 1.500E-07 | 2.500E-07 |
| 9238 | 462 | Q462W | A | 2.9e-07 | 9.8e-08 | 4.8e-07 | 2.893E-07 | 1.910E-07 | 9.800E-08 | 4.800E-07 |
| 9239 | 462 | Q462Y | A | 1.6e-07 | 9.9e-07 | 1.6e-07 | 4.367E-07 | 4.792E-07 | 1.600E-07 | 9.900E-07 |
| 9240 | 462 | Q462V | A | 3.8e-07 | 1.7e-07 | 1.9e-07 | 2.467E-07 | 1.159E-07 | 1.700E-07 | 3.800E-07 |
| 9241 | 463 | L463A | A | 3.6e-07 | 3e-07 | 4.5e-07 | 3.700E-07 | 7.550E-08 | 3.000E-07 | 4.500E-07 |
| 9242 | 463 | L463R | A | 1.6e-07 | 3e-07 | 2.6e-07 | 2.400E-07 | 7.211E-08 | 1.600E-07 | 3.000E-07 |
| 9243 | 463 | L463N | A | 4.6e-07 | 2.7e-07 | 1.9e-07 | 3.067E-07 | 1.387E-07 | 1.900E-07 | 4.600E-07 |
| 9244 | 463 | L463D | A | 2.8e-07 | 1.2e-07 | 2.9e-07 | 2.300E-07 | 9.539E-08 | 1.200E-07 | 2.900E-07 |
| 9245 | 463 | L463C | A | 4.2e-07 | 2.2e-07 | 1.4e-07 | 2.600E-07 | 1.442E-07 | 1.400E-07 | 4.200E-07 |
| 9246 | 463 | L463E | A | 3.6e-07 | 6.9e-08 | 2.7e-07 | 2.330E-07 | 1.490E-07 | 6.900E-08 | 3.600E-07 |
| 9247 | 463 | L463Q | A | 3.3e-07 | 1.5e-07 | 2.4e-07 | 2.400E-07 | 9.000E-08 | 1.500E-07 | 3.300E-07 |
| 9248 | 463 | L463G | A | 2.8e-07 | 1.3e-07 | 2.9e-07 | 2.333E-07 | 8.963E-08 | 1.300E-07 | 2.900E-07 |
| 9249 | 463 | L463H | A | 2.8e-07 | 1.5e-07 | 1.5e-07 | 1.933E-07 | 7.506E-08 | 1.500E-07 | 2.800E-07 |
| 9250 | 463 | L463I | A | 1.9e-07 | 2.1e-07 | 4.1e-07 | 2.700E-07 | 1.217E-07 | 1.900E-07 | 4.100E-07 |
| 9251 | 463 | L463L | A | 3.4e-07 | 4.8e-07 | 1.5e-07 | 3.233E-07 | 1.656E-07 | 1.500E-07 | 4.800E-07 |
| 9252 | 463 | L463K | A | 1.4e-07 | 1.4e-07 | 1.6e-07 | 1.467E-07 | 1.155E-08 | 1.400E-07 | 1.600E-07 |
| 9253 | 463 | L463M | A | 4e-07 | 1.5e-07 | 1.4e-07 | 2.300E-07 | 1.473E-07 | 1.400E-07 | 4.000E-07 |
| 9254 | 463 | L463F | A | 3.6e-07 | 3.2e-07 | 3.6e-07 | 3.467E-07 | 2.309E-08 | 3.200E-07 | 3.600E-07 |
| 9255 | 463 | L463P | A | 2.7e-07 | 2.4e-07 | 5.4e-07 | 3.500E-07 | 1.652E-07 | 2.400E-07 | 5.400E-07 |
| 9256 | 463 | L463S | A | 1.8e-07 | 1.5e-07 | 1.7e-07 | 1.667E-07 | 1.528E-08 | 1.500E-07 | 1.800E-07 |
| 9257 | 463 | L463T | A | 1.8e-07 | 1.4e-07 | 3.5e-07 | 2.233E-07 | 1.115E-07 | 1.400E-07 | 3.500E-07 |
| 9258 | 463 | L463W | A | 3e-07 | 2.1e-07 | 2.7e-07 | 2.600E-07 | 4.583E-08 | 2.100E-07 | 3.000E-07 |
| 9259 | 463 | L463Y | A | 1.8e-07 | 3e-07 | 1.7e-07 | 2.167E-07 | 7.234E-08 | 1.700E-07 | 3.000E-07 |
| 9260 | 463 | L463V | A | 7.3e-07 | 2.5e-07 | 2.4e-07 | 4.067E-07 | 2.801E-07 | 2.400E-07 | 7.300E-07 |
| 9261 | 464 | F464A | A | 3.6e-07 | 2.6e-07 | 4e-07 | 3.400E-07 | 7.211E-08 | 2.600E-07 | 4.000E-07 |
| 9262 | 464 | F464R | A | 2.2e-07 | 4e-07 | 2e-07 | 2.733E-07 | 1.102E-07 | 2.000E-07 | 4.000E-07 |
| 9263 | 464 | F464N | A | 6.6e-07 | 2.8e-07 | 2.8e-07 | 4.067E-07 | 2.194E-07 | 2.800E-07 | 6.600E-07 |
| 9264 | 464 | F464D | A | 2.3e-07 | 4.1e-07 | 2.7e-07 | 3.033E-07 | 9.452E-08 | 2.300E-07 | 4.100E-07 |

| | | | | | | | | | | |
|---|---|---|---|---|---|---|---|---|---|---|
| 9265 | 464 | F464C | A | 4.2e-07 | 2.2e-07 | 2.5e-07 | 2.967E-07 | 1.079E-07 | 2.200E-07 | 4.200E-07 |
| 9266 | 464 | F464E | A | 1.5e-07 | 4.3e-07 | 2.1e-07 | 2.633E-07 | 1.474E-07 | 1.500E-07 | 4.300E-07 |
| 9267 | 464 | F464Q | A | 2.2e-07 | 1.3e-07 | 3e-07 | 2.167E-07 | 8.505E-08 | 1.300E-07 | 3.000E-07 |
| 9268 | 464 | F464G | A | 1.8e-07 | 2.7e-07 | 2.6e-07 | 2.367E-07 | 4.933E-08 | 1.800E-07 | 2.700E-07 |
| 9269 | 464 | F464H | A | 6.6e-07 | 2.2e-07 | 2.4e-07 | 3.733E-07 | 2.485E-07 | 2.200E-07 | 6.600E-07 |
| 9270 | 464 | F464I | A | 1.8e-07 | 2.4e-07 | 2.9e-07 | 2.367E-07 | 5.508E-08 | 1.800E-07 | 2.900E-07 |
| 9271 | 464 | F464L | A | 2e-07 | 3.9e-07 | 1.5e-07 | 2.467E-07 | 1.266E-07 | 1.500E-07 | 3.900E-07 |
| 9272 | 464 | F464K | A | 2e-07 | 8.6e-08 | 3.5e-07 | 2.120E-07 | 1.324E-07 | 8.600E-08 | 3.500E-07 |
| 9273 | 464 | F464M | A | 2.2e-07 | 3e-07 | 2.4e-07 | 2.533E-07 | 4.163E-08 | 2.200E-07 | 3.000E-07 |
| 9274 | 464 | F464F | A | 2e-07 | 3.1e-07 | 2.4e-07 | 2.500E-07 | 5.568E-08 | 2.000E-07 | 3.100E-07 |
| 9275 | 464 | F464P | A | 3.1e-07 | 1.6e-07 | 2.4e-07 | 2.367E-07 | 7.506E-08 | 1.600E-07 | 3.100E-07 |
| 9276 | 464 | F464S | A | 2.8e-07 | 1.7e-07 | 4.2e-07 | 2.900E-07 | 1.253E-07 | 1.700E-07 | 4.200E-07 |
| 9277 | 464 | F464T | A | 4.2e-07 | 2.6e-07 | 3.2e-07 | 3.333E-07 | 8.083E-08 | 2.600E-07 | 4.200E-07 |
| 9278 | 464 | F464W | A | 2.1e-07 | 1.9e-07 | 1.5e-07 | 1.833E-07 | 3.055E-08 | 1.500E-07 | 2.100E-07 |
| 9279 | 464 | F464Y | A | 1.5e-07 | 2.7e-07 | 3.3e-07 | 2.500E-07 | 9.165E-08 | 1.500E-07 | 3.300E-07 |
| 9280 | 464 | F464V | A | 2.4e-07 | 3.3e-07 | 2.3e-07 | 2.667E-07 | 5.508E-08 | 2.300E-07 | 3.300E-07 |
| 9281 | 465 | R465A | A | 4.5e-07 | 1.7e-07 | 5e-07 | 3.733E-07 | 1.779E-07 | 1.700E-07 | 5.000E-07 |
| 9282 | 465 | R465R | A | 1.1e-07 | 1.4e-07 | 4.8e-07 | 2.433E-07 | 2.055E-07 | 1.100E-07 | 4.800E-07 |
| 9283 | 465 | R465N | A | 3.2e-07 | 1.7e-07 | 2.5e-07 | 2.467E-07 | 7.506E-08 | 1.700E-07 | 3.200E-07 |
| 9284 | 465 | R465D | A | 3e-07 | 3.9e-07 | 2.6e-07 | 3.167E-07 | 6.658E-08 | 2.600E-07 | 3.900E-07 |
| 9285 | 465 | R465C | A | 2.6e-07 | 2.4e-07 | 3.3e-07 | 2.767E-07 | 4.726E-08 | 2.400E-07 | 3.300E-07 |
| 9286 | 465 | R465E | A | 2.4e-07 | 2.6e-07 | 3.2e-07 | 2.733E-07 | 4.163E-08 | 2.400E-07 | 3.200E-07 |
| 9287 | 465 | R465Q | A | 2.6e-07 | 1.1e-07 | 1.6e-07 | 1.767E-07 | 7.638E-08 | 1.100E-07 | 2.600E-07 |
| 9288 | 465 | R465G | A | 5e-07 | 3.3e-07 | 1.8e-07 | 3.367E-07 | 1.601E-07 | 1.800E-07 | 5.000E-07 |
| 9289 | 465 | R465H | A | 2.4e-07 | 7.9e-08 | 3.6e-07 | 2.263E-07 | 1.410E-07 | 7.900E-08 | 3.600E-07 |
| 9290 | 465 | R465I | A | 2.2e-07 | 1.7e-07 | 3.5e-07 | 2.467E-07 | 9.292E-08 | 1.700E-07 | 3.500E-07 |
| 9291 | 465 | R465L | A | 2.8e-07 | 2e-07 | 3.7e-07 | 2.833E-07 | 8.505E-08 | 2.000E-07 | 3.700E-07 |
| 9292 | 465 | R465K | A | 5.7e-07 | 2.8e-07 | 1.4e-07 | 3.300E-07 | 2.193E-07 | 1.400E-07 | 5.700E-07 |
| 9293 | 465 | R465M | A | 2.6e-07 | 2.8e-07 | 3.3e-07 | 2.900E-07 | 3.606E-08 | 2.600E-07 | 3.300E-07 |
| 9294 | 465 | R465F | A | 2.1e-07 | 1.6e-07 | 2.6e-07 | 2.100E-07 | 5.000E-08 | 1.600E-07 | 2.600E-07 |
| 9295 | 465 | R465P | A | 6.5e-07 | 3.5e-07 | 2.4e-07 | 4.133E-07 | 2.122E-07 | 2.400E-07 | 6.500E-07 |
| 9296 | 465 | R465S | A | 1.9e-07 | 3.3e-07 | 4.4e-07 | 3.200E-07 | 1.253E-07 | 1.900E-07 | 4.400E-07 |
| 9297 | 465 | R465T | A | 2.8e-07 | 2.2e-07 | 5.7e-07 | 3.567E-07 | 1.872E-07 | 2.200E-07 | 5.700E-07 |
| 9298 | 465 | R465W | A | 3.2e-07 | 2.6e-07 | 2.3e-07 | 2.700E-07 | 4.583E-08 | 2.300E-07 | 3.200E-07 |
| 9299 | 465 | R465Y | A | 2.5e-07 | 4.6e-07 | 1.2e-07 | 2.767E-07 | 1.716E-07 | 1.200E-07 | 4.600E-07 |
| 9300 | 465 | R465V | A | 4.6e-07 | 6.6e-07 | 1.5e-07 | 4.233E-07 | 2.570E-07 | 1.500E-07 | 6.600E-07 |
| 9301 | 466 | N466A | A | 2.3e-07 | 1.8e-07 | 4.2e-07 | 2.767E-07 | 1.266E-07 | 1.800E-07 | 4.200E-07 |
| 9302 | 466 | N466R | A | 3.1e-07 | 3.9e-07 | 2.1e-07 | 3.033E-07 | 9.018E-08 | 2.100E-07 | 3.900E-07 |
| 9303 | 466 | N466N | A | 1.9e-07 | 4.9e-07 | 1.7e-07 | 2.833E-07 | 1.793E-07 | 1.700E-07 | 4.900E-07 |
| 9304 | 466 | N466D | A | 1.5e-07 | 2.1e-07 | 1.1e-07 | 1.567E-07 | 5.033E-08 | 1.100E-07 | 2.100E-07 |
| 9305 | 466 | N466C | A | 2.2e-07 | 2.3e-07 | 1.5e-07 | 2.000E-07 | 4.359E-08 | 1.500E-07 | 2.300E-07 |
| 9306 | 466 | N466E | A | 3.2e-07 | 3.4e-07 | 4.4e-07 | 3.667E-07 | 6.429E-08 | 3.200E-07 | 4.400E-07 |
| 9307 | 466 | N466Q | A | 5.3e-07 | 3.2e-07 | 2.7e-07 | 3.733E-07 | 1.380E-07 | 2.700E-07 | 5.300E-07 |
| 9308 | 466 | N466G | A | 5.6e-07 | 3e-07 | 1.8e-07 | 3.467E-07 | 1.943E-07 | 1.800E-07 | 5.600E-07 |
| 9309 | 466 | N466H | A | 3.8e-07 | 1.5e-07 | 2.7e-07 | 2.667E-07 | 1.150E-07 | 1.500E-07 | 3.800E-07 |
| 9310 | 466 | N466I | A | 2.5e-07 | 2e-07 | 3.4e-07 | 2.633E-07 | 7.095E-08 | 2.000E-07 | 3.400E-07 |
| 9311 | 466 | N466L | A | 2.4e-07 | 5.4e-07 | 1.5e-07 | 3.100E-07 | 2.042E-07 | 1.500E-07 | 5.400E-07 |
| 9312 | 466 | N466K | A | 3.6e-07 | 1.6e-07 | 2.3e-07 | 2.500E-07 | 1.015E-07 | 1.600E-07 | 3.600E-07 |
| 9313 | 466 | N466M | A | 1.8e-07 | 2.9e-07 | 3.2e-07 | 2.633E-07 | 7.371E-08 | 1.800E-07 | 3.200E-07 |
| 9314 | 466 | N466F | A | 2.3e-07 | 3.9e-07 | 2.5e-07 | 2.900E-07 | 8.718E-08 | 2.300E-07 | 3.900E-07 |
| 9315 | 466 | N466P | A | 1.2e-07 | 3e-07 | 2e-07 | 2.067E-07 | 9.018E-08 | 1.200E-07 | 3.000E-07 |
| 9316 | 466 | N466S | A | 1.4e-07 | 3.8e-07 | 2.1e-07 | 2.433E-07 | 1.234E-07 | 1.400E-07 | 3.800E-07 |
| 9317 | 466 | N466T | A | 1.3e-07 | 3e-07 | 2.7e-07 | 2.333E-07 | 9.074E-08 | 1.300E-07 | 3.000E-07 |
| 9318 | 466 | N466W | A | 2.8e-07 | 2e-07 | 1.9e-07 | 2.233E-07 | 4.933E-08 | 1.900E-07 | 2.800E-07 |
| 9319 | 466 | N466Y | A | 1.7e-07 | 6.9e-08 | 2.1e-07 | 1.497E-07 | 7.267E-08 | 6.900E-08 | 2.100E-07 |
| 9320 | 466 | N466V | A | 3e-07 | 1.7e-07 | 2.6e-07 | 2.433E-07 | 6.658E-08 | 1.700E-07 | 3.000E-07 |
| 9321 | 467 | P467A | A | 1.9e-07 | 2.2e-07 | 1.4e-07 | 1.833E-07 | 4.041E-08 | 1.400E-07 | 2.200E-07 |
| 9322 | 467 | P467R | A | 2.4e-07 | 5.5e-07 | 1.8e-07 | 3.233E-07 | 1.986E-07 | 1.800E-07 | 5.500E-07 |
| 9323 | 467 | P467N | A | 1.7e-07 | 4.4e-07 | 7.6e-07 | 4.567E-07 | 2.954E-07 | 1.700E-07 | 7.600E-07 |
| 9324 | 467 | P467D | A | 1.9e-07 | 2.1e-07 | 5.7e-07 | 3.233E-07 | 2.139E-07 | 1.900E-07 | 5.700E-07 |
| 9325 | 467 | P467C | A | 1.7e-07 | 2.9e-07 | 3e-07 | 2.533E-07 | 7.234E-08 | 1.700E-07 | 3.000E-07 |
| 9326 | 467 | P467E | A | 2.7e-07 | 1.6e-07 | 5.8e-07 | 3.367E-07 | 2.178E-07 | 1.600E-07 | 5.800E-07 |
| 9327 | 467 | P467Q | A | 9.3e-08 | 1.1e-07 | 2.2e-07 | 1.410E-07 | 6.894E-08 | 9.300E-08 | 2.200E-07 |
| 9328 | 467 | P467G | A | 2.3e-07 | 4.8e-07 | 2.9e-07 | 3.333E-07 | 1.305E-07 | 2.300E-07 | 4.800E-07 |
| 9329 | 467 | P467H | A | 3.3e-07 | 1.6e-07 | 2.6e-07 | 2.500E-07 | 8.544E-08 | 1.600E-07 | 3.300E-07 |
| 9330 | 467 | P467I | A | 2.6e-07 | 2.7e-07 | 2.4e-07 | 2.567E-07 | 1.528E-08 | 2.400E-07 | 2.700E-07 |
| 9331 | 467 | P467L | A | 3.2e-07 | 2.8e-07 | 8.4e-08 | 2.280E-07 | 1.263E-07 | 8.400E-08 | 3.200E-07 |
| 9332 | 467 | P467K | A | 1.4e-07 | 1.9e-07 | 6e-07 | 3.100E-07 | 2.524E-07 | 1.400E-07 | 6.000E-07 |
| 9333 | 467 | P467M | A | 2.3e-07 | 1.8e-07 | 2.8e-07 | 2.300E-07 | 5.000E-08 | 1.800E-07 | 2.800E-07 |
| 9334 | 467 | P467F | A | 3.2e-07 | 3.6e-07 | 1.9e-07 | 2.900E-07 | 8.888E-08 | 1.900E-07 | 3.600E-07 |
| 9335 | 467 | P467P | A | 2e-07 | 2.1e-07 | 1.2e-07 | 1.767E-07 | 4.933E-08 | 1.200E-07 | 2.100E-07 |
| 9336 | 467 | P467S | A | 1.3e-07 | 1.8e-07 | 1.1e-07 | 1.400E-07 | 3.606E-08 | 1.100E-07 | 1.800E-07 |
| 9337 | 467 | P467T | A | 2.1e-07 | 2.5e-07 | 2e-07 | 2.200E-07 | 2.646E-08 | 2.000E-07 | 2.500E-07 |
| 9338 | 467 | P467W | A | 3.8e-07 | 3.1e-07 | 2.4e-07 | 3.100E-07 | 7.000E-08 | 2.400E-07 | 3.800E-07 |
| 9339 | 467 | P467Y | A | 2e-07 | 3.4e-07 | 3e-07 | 2.800E-07 | 7.211E-08 | 2.000E-07 | 3.400E-07 |
| 9340 | 467 | P467V | A | 3.6e-07 | 2.8e-07 | 2.2e-07 | 2.867E-07 | 7.024E-08 | 2.200E-07 | 3.600E-07 |
| 9341 | 468 | H468A | A | 2.4e-07 | 1.8e-07 | 3.9e-07 | 2.700E-07 | 1.082E-07 | 1.800E-07 | 3.900E-07 |
| 9342 | 468 | H468R | A | 2.3e-07 | 1.9e-07 | 2.8e-07 | 2.333E-07 | 4.509E-08 | 1.900E-07 | 2.800E-07 |
| 9343 | 468 | H468N | A | 2.2e-07 | 2.5e-07 | 2.7e-07 | 2.467E-07 | 2.517E-08 | 2.200E-07 | 2.700E-07 |
| 9344 | 468 | H468D | A | 6e-07 | 1.5e-07 | 2e-07 | 3.167E-07 | 2.466E-07 | 1.500E-07 | 6.000E-07 |
| 9345 | 468 | H468C | A | 2e-07 | 9.7e-07 | 4.4e-07 | 5.367E-07 | 3.940E-07 | 2.000E-07 | 9.700E-07 |
| 9346 | 468 | H468E | A | 2.7e-07 | 3.5e-07 | 1.6e-07 | 2.600E-07 | 9.539E-08 | 1.600E-07 | 3.500E-07 |
| 9347 | 468 | H468Q | A | 1.5e-07 | 3e-07 | 1.7e-07 | 2.067E-07 | 8.145E-08 | 1.500E-07 | 3.000E-07 |
| 9348 | 468 | H468G | A | 2.4e-07 | 2.6e-07 | 4.6e-07 | 3.200E-07 | 1.217E-07 | 2.400E-07 | 4.600E-07 |
| 9349 | 468 | H468H | A | 2.6e-07 | 3.3e-07 | 3.4e-07 | 3.100E-07 | 4.359E-08 | 2.600E-07 | 3.400E-07 |

| | | | | | | | | | | |
|---|---|---|---|---|---|---|---|---|---|---|
| 9350 | 468 | H468I | A | 3.5e-07 | 3e-07 | 3.3e-07 | 3.267E-07 | 2.517E-08 | 3.000E-07 | 3.500E-07 |
| 9351 | 468 | H468L | A | 2.4e-07 | 2.7e-07 | 1.8e-07 | 2.300E-07 | 4.583E-08 | 1.800E-07 | 2.700E-07 |
| 9352 | 468 | H468K | A | 3.3e-07 | 3.9e-07 | 1.5e-07 | 2.900E-07 | 1.249E-07 | 1.500E-07 | 3.900E-07 |
| 9353 | 468 | H468M | A | 5.8e-07 | 1.7e-07 | 3.4e-07 | 3.633E-07 | 2.060E-07 | 1.700E-07 | 5.800E-07 |
| 9354 | 468 | H468F | A | 1.7e-07 | 1.3e-07 | 2.9e-07 | 1.967E-07 | 8.327E-08 | 1.300E-07 | 2.900E-07 |
| 9355 | 468 | H468P | A | 3.3e-07 | 2.4e-07 | 8.3e-08 | 2.177E-07 | 1.250E-07 | 8.300E-08 | 3.300E-07 |
| 9356 | 468 | H468S | A | 3.6e-07 | 2.3e-07 | 9.1e-08 | 2.270E-07 | 1.345E-07 | 9.100E-08 | 3.600E-07 |
| 9357 | 468 | H468T | A | 3.2e-07 | 2e-07 | 1.9e-07 | 2.367E-07 | 7.234E-08 | 1.900E-07 | 3.200E-07 |
| 9358 | 468 | H468W | A | 1.6e-07 | 2.4e-07 | 2.1e-07 | 2.033E-07 | 4.041E-08 | 1.600E-07 | 2.400E-07 |
| 9359 | 468 | H468Y | A | 3e-07 | 2e-07 | 3.2e-07 | 2.733E-07 | 6.429E-08 | 2.000E-07 | 3.200E-07 |
| 9360 | 468 | H468V | A | 2.4e-07 | 2.1e-07 | 1.3e-07 | 1.933E-07 | 5.686E-08 | 1.300E-07 | 2.400E-07 |
| 9361 | 469 | Q469A | A | 1.6e-07 | 2.3e-07 | 1.3e-07 | 1.733E-07 | 5.132E-08 | 1.300E-07 | 2.300E-07 |
| 9362 | 469 | Q469R | A | 4.1e-07 | 4.5e-07 | 1.9e-07 | 3.500E-07 | 1.400E-07 | 1.900E-07 | 4.500E-07 |
| 9363 | 469 | Q469N | A | 5.9e-07 | 2.2e-07 | 1.9e-07 | 3.333E-07 | 2.228E-07 | 1.900E-07 | 5.900E-07 |
| 9364 | 469 | Q469D | A | 3e-07 | 4.6e-07 | 3.2e-07 | 3.600E-07 | 8.718E-08 | 3.000E-07 | 4.600E-07 |
| 9365 | 469 | Q469C | A | 2.5e-07 | 1.2e-07 | 5.1e-07 | 2.933E-07 | 1.986E-07 | 1.200E-07 | 5.100E-07 |
| 9366 | 469 | Q469E | A | 1.3e-07 | 1.7e-07 | 4.4e-07 | 2.467E-07 | 1.686E-07 | 1.300E-07 | 4.400E-07 |
| 9367 | 469 | Q469Q | A | 3.1e-07 | 2.2e-07 | 2.6e-07 | 2.633E-07 | 4.509E-08 | 2.200E-07 | 3.100E-07 |
| 9368 | 469 | Q469G | A | 1.7e-07 | 2e-07 | 2e-07 | 1.900E-07 | 1.732E-08 | 1.700E-07 | 2.000E-07 |
| 9369 | 469 | Q469H | A | 2.3e-07 | 4.4e-07 | 3.1e-07 | 3.267E-07 | 1.060E-07 | 2.300E-07 | 4.400E-07 |
| 9370 | 469 | Q469I | A | 2.8e-07 | 2.6e-07 | 2.5e-07 | 2.633E-07 | 1.528E-08 | 2.500E-07 | 2.800E-07 |
| 9371 | 469 | Q469L | A | 1.9e-07 | 3.3e-07 | 2.8e-07 | 2.667E-07 | 7.095E-08 | 1.900E-07 | 3.300E-07 |
| 9372 | 469 | Q469K | A | 3.5e-07 | 2.5e-07 | 1.4e-07 | 2.467E-07 | 1.050E-07 | 1.400E-07 | 3.500E-07 |
| 9373 | 469 | Q469M | A | 2.2e-07 | 9.3e-07 | 2.2e-07 | 4.567E-07 | 4.099E-07 | 2.200E-07 | 9.300E-07 |
| 9374 | 469 | Q469F | A | 2.4e-07 | 3.2e-07 | 1.4e-07 | 2.333E-07 | 9.018E-08 | 1.400E-07 | 3.200E-07 |
| 9375 | 469 | Q469P | A | 1.3e-07 | 6.2e-07 | 7.2e-08 | 2.740E-07 | 3.010E-07 | 7.200E-08 | 6.200E-07 |
| 9376 | 469 | Q469S | A | 5.5e-07 | 1.9e-07 | 3.5e-07 | 3.633E-07 | 1.804E-07 | 1.900E-07 | 5.500E-07 |
| 9377 | 469 | Q469T | A | 1.8e-07 | 3.4e-07 | 2.5e-07 | 2.567E-07 | 8.021E-08 | 1.800E-07 | 3.400E-07 |
| 9378 | 469 | Q469W | A | 5.1e-07 | 2.1e-07 | 4.4e-07 | 3.867E-07 | 1.570E-07 | 2.100E-07 | 5.100E-07 |
| 9379 | 469 | Q469Y | A | 3.1e-07 | 1.6e-07 | 2.4e-07 | 2.367E-07 | 7.506E-08 | 1.600E-07 | 3.100E-07 |
| 9380 | 469 | Q469V | A | 3.3e-07 | 2.6e-07 | 3.1e-07 | 3.000E-07 | 3.606E-08 | 2.600E-07 | 3.300E-07 |
| 9381 | 470 | A470A | A | 1.3e-07 | 1.7e-07 | 2.4e-07 | 1.800E-07 | 5.568E-08 | 1.300E-07 | 2.400E-07 |
| 9382 | 470 | A470R | A | 1.7e-07 | 1.4e-07 | 3.8e-07 | 2.300E-07 | 1.308E-07 | 1.400E-07 | 3.800E-07 |
| 9383 | 470 | A470N | A | 1.9e-07 | 1.9e-07 | 1.8e-07 | 1.867E-07 | 5.774E-09 | 1.800E-07 | 1.900E-07 |
| 9384 | 470 | A470D | A | 2.7e-07 | 2.1e-07 | 4.1e-07 | 2.967E-07 | 1.026E-07 | 2.100E-07 | 4.100E-07 |
| 9385 | 470 | A470C | A | 1.4e-07 | 3.6e-07 | 1.6e-07 | 2.200E-07 | 1.217E-07 | 1.400E-07 | 3.600E-07 |
| 9386 | 470 | A470E | A | 4.8e-07 | 1.5e-07 | 3e-07 | 3.100E-07 | 1.652E-07 | 1.500E-07 | 4.800E-07 |
| 9387 | 470 | A470Q | A | 3.1e-07 | 2.9e-07 | 1.5e-07 | 2.500E-07 | 8.718E-08 | 1.500E-07 | 3.100E-07 |
| 9388 | 470 | A470G | A | 2.6e-07 | 2.3e-07 | 4.3e-07 | 3.067E-07 | 1.079E-07 | 2.300E-07 | 4.300E-07 |
| 9389 | 470 | A470H | A | 1.1e-07 | 2.4e-07 | 3.2e-07 | 2.233E-07 | 1.060E-07 | 1.100E-07 | 3.200E-07 |
| 9390 | 470 | A470I | A | 3.7e-07 | 1.7e-07 | 2.7e-07 | 2.700E-07 | 1.000E-07 | 1.700E-07 | 3.700E-07 |
| 9391 | 470 | A470L | A | 1.6e-07 | 1.4e-07 | 1.8e-07 | 1.600E-07 | 2.000E-08 | 1.400E-07 | 1.800E-07 |
| 9392 | 470 | A470K | A | 1.8e-07 | 7.6e-08 | 2.7e-07 | 1.753E-07 | 9.708E-08 | 7.600E-08 | 2.700E-07 |
| 9393 | 470 | A470M | A | 1.7e-07 | 3.6e-07 | 4.2e-07 | 3.167E-07 | 1.305E-07 | 1.700E-07 | 4.200E-07 |
| 9394 | 470 | A470F | A | 3.3e-07 | 2.8e-07 | 2.3e-07 | 2.800E-07 | 5.000E-08 | 2.300E-07 | 3.300E-07 |
| 9395 | 470 | A470P | A | 4.7e-07 | 2.2e-07 | 2.2e-07 | 3.033E-07 | 1.443E-07 | 2.200E-07 | 4.700E-07 |
| 9396 | 470 | A470S | A | 2.1e-07 | 2.1e-07 | 4.5e-07 | 2.900E-07 | 1.386E-07 | 2.100E-07 | 4.500E-07 |
| 9397 | 470 | A470T | A | 3.3e-07 | 2.2e-07 | 1.3e-07 | 2.267E-07 | 1.002E-07 | 1.300E-07 | 3.300E-07 |
| 9398 | 470 | A470W | A | 3.5e-07 | 1.7e-07 | 1.5e-07 | 2.233E-07 | 1.102E-07 | 1.500E-07 | 3.500E-07 |
| 9399 | 470 | A470Y | A | 1.2e-07 | 2.2e-07 | 1.9e-07 | 1.767E-07 | 5.132E-08 | 1.200E-07 | 2.200E-07 |
| 9400 | 470 | A470V | A | 5.4e-07 | 4.3e-07 | 9.8e-08 | 3.560E-07 | 2.301E-07 | 9.800E-08 | 5.400E-07 |
| 9401 | 471 | L471A | A | 2.2e-07 | 2.6e-07 | 1.2e-07 | 2.000E-07 | 7.211E-08 | 1.200E-07 | 2.600E-07 |
| 9402 | 471 | L471R | A | 2.7e-07 | 1.3e-07 | 3.4e-07 | 2.467E-07 | 1.069E-07 | 1.300E-07 | 3.400E-07 |
| 9403 | 471 | L471N | A | 1.9e-07 | 1.9e-07 | 3e-07 | 2.267E-07 | 6.351E-08 | 1.900E-07 | 3.000E-07 |
| 9404 | 471 | L471D | A | 1.9e-07 | 2.2e-07 | 9.8e-07 | 4.633E-07 | 4.477E-07 | 1.900E-07 | 9.800E-07 |
| 9405 | 471 | L471C | A | 2.5e-07 | 1.7e-07 | 3.3e-07 | 2.500E-07 | 8.000E-08 | 1.700E-07 | 3.300E-07 |
| 9406 | 471 | L471E | A | 1.9e-07 | 2.3e-07 | 3.9e-07 | 2.700E-07 | 1.058E-07 | 1.900E-07 | 3.900E-07 |
| 9407 | 471 | L471Q | A | 2.2e-07 | 2.2e-07 | 2.7e-07 | 2.367E-07 | 2.887E-08 | 2.200E-07 | 2.700E-07 |
| 9408 | 471 | L471G | A | 1.7e-07 | 1.2e-07 | 3.4e-07 | 2.100E-07 | 1.153E-07 | 1.200E-07 | 3.400E-07 |
| 9409 | 471 | L471H | A | 1.8e-07 | 2e-07 | 1.2e-07 | 1.667E-07 | 4.163E-08 | 1.200E-07 | 2.000E-07 |
| 9410 | 471 | L471I | A | 2.5e-07 | 3.3e-07 | 2.7e-07 | 2.833E-07 | 4.163E-08 | 2.500E-07 | 3.300E-07 |
| 9411 | 471 | L471L | A | 2e-07 | 2.9e-07 | 4.2e-07 | 3.033E-07 | 1.106E-07 | 2.000E-07 | 4.200E-07 |
| 9412 | 471 | L471K | A | 2.2e-07 | 4.1e-07 | 6.4e-07 | 4.233E-07 | 2.103E-07 | 2.200E-07 | 6.400E-07 |
| 9413 | 471 | L471M | A | 2.1e-07 | 4.8e-07 | 1.6e-07 | 2.833E-07 | 1.721E-07 | 1.600E-07 | 4.800E-07 |
| 9414 | 471 | L471F | A | 3.6e-07 | 2.7e-07 | 4.2e-07 | 3.500E-07 | 7.550E-08 | 2.700E-07 | 4.200E-07 |
| 9415 | 471 | L471P | A | 2.7e-07 | 2.4e-07 | 3e-07 | 2.700E-07 | 3.000E-08 | 2.400E-07 | 3.000E-07 |
| 9416 | 471 | L471S | A | 3.8e-07 | 2e-07 | 2.5e-07 | 2.767E-07 | 9.292E-08 | 2.000E-07 | 3.800E-07 |
| 9417 | 471 | L471T | A | 1.1e-07 | 1.4e-07 | 2.5e-07 | 1.667E-07 | 7.371E-08 | 1.100E-07 | 2.500E-07 |
| 9418 | 471 | L471W | A | 6.6e-07 | 2.3e-07 | 3.4e-07 | 4.100E-07 | 2.234E-07 | 2.300E-07 | 6.600E-07 |
| 9419 | 471 | L471Y | A | 7.1e-07 | 2.8e-07 | 2.7e-07 | 4.200E-07 | 2.512E-07 | 2.700E-07 | 7.100E-07 |
| 9420 | 471 | L471V | A | 2.8e-07 | 1.6e-07 | 2.4e-07 | 2.267E-07 | 6.110E-08 | 1.600E-07 | 2.800E-07 |
| 9421 | 472 | L472A | A | 1.7e-07 | 2.4e-07 | 2.9e-07 | 2.333E-07 | 6.028E-08 | 1.700E-07 | 2.900E-07 |
| 9422 | 472 | L472R | A | 4e-07 | 2.9e-07 | 1.6e-07 | 2.833E-07 | 1.201E-07 | 1.600E-07 | 4.000E-07 |
| 9423 | 472 | L472N | A | 2e-07 | 3.1e-07 | 1.1e-07 | 2.067E-07 | 1.002E-07 | 1.100E-07 | 3.100E-07 |
| 9424 | 472 | L472D | A | 1.8e-07 | 1.9e-07 | 1.3e-07 | 1.667E-07 | 3.215E-08 | 1.300E-07 | 1.900E-07 |
| 9425 | 472 | L472C | A | 1.9e-07 | 3.8e-07 | 2e-07 | 2.567E-07 | 1.069E-07 | 1.900E-07 | 3.800E-07 |
| 9426 | 472 | L472E | A | 2.7e-07 | 1.8e-07 | 2.9e-07 | 2.467E-07 | 5.859E-08 | 1.800E-07 | 2.900E-07 |
| 9427 | 472 | L472Q | A | 1e-07 | 1.4e-07 | 3.7e-07 | 2.033E-07 | 1.457E-07 | 1.000E-07 | 3.700E-07 |
| 9428 | 472 | L472G | A | 2.3e-07 | 1.9e-07 | 8.9e-08 | 1.697E-07 | 7.267E-08 | 8.900E-08 | 2.300E-07 |
| 9429 | 472 | L472H | A | 2.3e-07 | 2.7e-07 | 7.5e-08 | 1.917E-07 | 1.030E-07 | 7.500E-08 | 2.700E-07 |
| 9430 | 472 | L472I | A | 3.3e-07 | 2.2e-07 | 1.8e-07 | 2.433E-07 | 7.767E-08 | 1.800E-07 | 3.300E-07 |
| 9431 | 472 | L472L | A | 2.4e-07 | 1.8e-07 | 1.3e-07 | 1.833E-07 | 5.508E-08 | 1.300E-07 | 2.400E-07 |
| 9432 | 472 | L472K | A | 1.8e-07 | 1.6e-07 | 3.3e-07 | 2.233E-07 | 9.292E-08 | 1.600E-07 | 3.300E-07 |
| 9433 | 472 | L472M | A | 2e-07 | 2.2e-07 | 2.3e-07 | 2.167E-07 | 1.528E-08 | 2.000E-07 | 2.300E-07 |
| 9434 | 472 | L472F | A | 1.9e-07 | 1.7e-07 | 2.6e-07 | 2.067E-07 | 4.726E-08 | 1.700E-07 | 2.600E-07 |

| | | | | | | | | | | |
|---|---|---|---|---|---|---|---|---|---|---|
| 9435 | 472 | L472P | A | 2.4e-07 | 3.5e-07 | 4.9e-07 | 3.600E-07 | 1.253E-07 | 2.400E-07 | 4.900E-07 |
| 9436 | 472 | L472S | A | 3.2e-07 | 1.3e-07 | 1.7e-07 | 2.067E-07 | 1.002E-07 | 1.300E-07 | 3.200E-07 |
| 9437 | 472 | L472T | A | 1.7e-07 | 2.4e-07 | 4.8e-07 | 2.967E-07 | 1.626E-07 | 1.700E-07 | 4.800E-07 |
| 9438 | 472 | L472W | A | 3.3e-07 | 4.1e-07 | 2.3e-07 | 3.233E-07 | 9.018E-08 | 2.300E-07 | 4.100E-07 |
| 9439 | 472 | L472Y | A | 3.5e-07 | 3.7e-07 | 5.1e-08 | 2.570E-07 | 1.787E-07 | 5.100E-08 | 3.700E-07 |
| 9440 | 472 | L472V | A | 3.1e-07 | 5e-07 | 1.2e-07 | 3.100E-07 | 1.900E-07 | 1.200E-07 | 5.000E-07 |
| 9441 | 473 | H473A | A | 2.4e-07 | 1.1e-07 | 1.9e-07 | 1.800E-07 | 6.557E-08 | 1.100E-07 | 2.400E-07 |
| 9442 | 473 | H473R | A | 3.7e-07 | 3.4e-07 | 2.2e-07 | 3.100E-07 | 7.937E-08 | 2.200E-07 | 3.700E-07 |
| 9443 | 473 | H473N | A | 1.5e-07 | 2.1e-07 | 1.6e-07 | 1.733E-07 | 3.215E-08 | 1.500E-07 | 2.100E-07 |
| 9444 | 473 | H473D | A | 2.7e-07 | 4.1e-07 | 1.9e-07 | 2.900E-07 | 1.114E-07 | 1.900E-07 | 4.100E-07 |
| 9445 | 473 | H473C | A | 3.3e-07 | 1.8e-07 | 2.3e-07 | 2.467E-07 | 7.638E-08 | 1.800E-07 | 3.300E-07 |
| 9446 | 473 | H473E | A | 3.9e-07 | 1.6e-07 | 2.9e-07 | 2.800E-07 | 1.153E-07 | 1.600E-07 | 3.900E-07 |
| 9447 | 473 | H473Q | A | 3.3e-07 | 2.3e-07 | 1.3e-07 | 2.300E-07 | 1.000E-07 | 1.300E-07 | 3.300E-07 |
| 9448 | 473 | H473G | A | 4e-07 | 1.9e-07 | 1.5e-07 | 2.467E-07 | 1.343E-07 | 1.500E-07 | 4.000E-07 |
| 9449 | 473 | H473H | A | 2.9e-07 | 4.2e-07 | 2e-07 | 3.033E-07 | 1.106E-07 | 2.000E-07 | 4.200E-07 |
| 9450 | 473 | H473I | A | 4.1e-07 | 3e-07 | 2.2e-07 | 3.100E-07 | 9.539E-08 | 2.200E-07 | 4.100E-07 |
| 9451 | 473 | H473L | A | 2.3e-07 | 3.8e-07 | 2.7e-07 | 2.933E-07 | 7.767E-08 | 2.300E-07 | 3.800E-07 |
| 9452 | 473 | H473K | A | 1.8e-07 | 1.7e-07 | 2.6e-07 | 2.033E-07 | 4.933E-08 | 1.700E-07 | 2.600E-07 |
| 9453 | 473 | H473M | A | 3.6e-07 | 3.6e-07 | 2e-07 | 3.067E-07 | 9.238E-08 | 2.000E-07 | 3.600E-07 |
| 9454 | 473 | H473F | A | 3.1e-07 | 1.6e-07 | 1.6e-07 | 2.100E-07 | 8.660E-08 | 1.600E-07 | 3.100E-07 |
| 9455 | 473 | H473P | A | 2.9e-07 | 2e-07 | 1.4e-07 | 2.100E-07 | 7.550E-08 | 1.400E-07 | 2.900E-07 |
| 9456 | 473 | H473S | A | 2.1e-07 | 4e-07 | 2.9e-07 | 3.000E-07 | 9.539E-08 | 2.100E-07 | 4.000E-07 |
| 9457 | 473 | H473T | A | 1.6e-07 | 4.2e-07 | 1.9e-07 | 2.567E-07 | 1.422E-07 | 1.600E-07 | 4.200E-07 |
| 9458 | 473 | H473W | A | 4.1e-07 | 1.9e-07 | 2.1e-07 | 2.700E-07 | 1.217E-07 | 1.900E-07 | 4.100E-07 |
| 9459 | 473 | H473Y | A | 4.2e-07 | 3.2e-07 | 1.9e-07 | 3.100E-07 | 1.153E-07 | 1.900E-07 | 4.200E-07 |
| 9460 | 473 | H473V | A | 1.7e-07 | 3e-07 | 2.4e-07 | 2.367E-07 | 6.506E-08 | 1.700E-07 | 3.000E-07 |
| 9461 | 474 | T474A | A | 2.4e-07 | 6.9e-07 | 2.9e-07 | 4.067E-07 | 2.466E-07 | 2.400E-07 | 6.900E-07 |
| 9462 | 474 | T474R | A | 2e-07 | 2e-07 | 1.8e-07 | 1.933E-07 | 1.155E-08 | 1.800E-07 | 2.000E-07 |
| 9463 | 474 | T474N | A | 4.4e-07 | 1.6e-07 | 3.3e-07 | 3.100E-07 | 1.411E-07 | 1.600E-07 | 4.400E-07 |
| 9464 | 474 | T474D | A | 2.4e-07 | 6e-07 | 3.4e-07 | 3.933E-07 | 1.858E-07 | 2.400E-07 | 6.000E-07 |
| 9465 | 474 | T474C | A | 2.9e-07 | 1.7e-07 | 2.3e-07 | 2.300E-07 | 6.000E-08 | 1.700E-07 | 2.900E-07 |
| 9466 | 474 | T474E | A | 4.4e-07 | 3.4e-07 | 7.4e-07 | 5.067E-07 | 2.082E-07 | 3.400E-07 | 7.400E-07 |
| 9467 | 474 | T474Q | A | 2.9e-07 | 3e-07 | 2.5e-07 | 2.800E-07 | 2.646E-08 | 2.500E-07 | 3.000E-07 |
| 9468 | 474 | T474G | A | 3.9e-07 | 3e-07 | 1e-06 | 5.633E-07 | 3.808E-07 | 3.000E-07 | 1.000E-06 |
| 9469 | 474 | T474H | A | 3.2e-07 | 2.5e-07 | 8e-07 | 4.567E-07 | 2.994E-07 | 2.500E-07 | 8.000E-07 |
| 9470 | 474 | T474I | A | 1.4e-07 | 3.8e-07 | 2.9e-07 | 2.700E-07 | 1.212E-07 | 1.400E-07 | 3.800E-07 |
| 9471 | 474 | T474L | A | 4.7e-07 | 4.8e-07 | 2.8e-07 | 4.100E-07 | 1.127E-07 | 2.800E-07 | 4.800E-07 |
| 9472 | 474 | T474K | A | 4.2e-07 | 3.6e-07 | 1.8e-07 | 3.200E-07 | 1.249E-07 | 1.800E-07 | 4.200E-07 |
| 9473 | 474 | T474M | A | 3.6e-07 | 5.4e-07 | 7.8e-08 | 3.260E-07 | 2.329E-07 | 7.800E-08 | 5.400E-07 |
| 9474 | 474 | T474F | A | 4.3e-07 | 3.6e-07 | 1.9e-07 | 3.267E-07 | 1.234E-07 | 1.900E-07 | 4.300E-07 |
| 9475 | 474 | T474P | A | 2.4e-07 | 4.9e-07 | 2.7e-07 | 3.333E-07 | 1.365E-07 | 2.400E-07 | 4.900E-07 |
| 9476 | 474 | T474S | A | 1.1e-07 | 2.6e-07 | 1.2e-07 | 1.633E-07 | 8.386E-08 | 1.100E-07 | 2.600E-07 |
| 9477 | 474 | T474T | A | 3.6e-07 | 2.6e-07 | 7.4e-07 | 4.533E-07 | 2.532E-07 | 2.600E-07 | 7.400E-07 |
| 9478 | 474 | T474W | A | 2.8e-07 | 2.2e-07 | 1.4e-07 | 2.133E-07 | 7.024E-08 | 1.400E-07 | 2.800E-07 |
| 9479 | 474 | T474Y | A | 2.3e-07 | 3.8e-07 | 3.8e-07 | 3.300E-07 | 8.660E-08 | 2.300E-07 | 3.800E-07 |
| 9480 | 474 | T474V | A | 3.1e-07 | 1.7e-07 | 3.9e-07 | 2.900E-07 | 1.114E-07 | 1.700E-07 | 3.900E-07 |
| 9481 | 475 | A475A | A | 4e-07 | 2.1e-07 | 3.2e-07 | 3.100E-07 | 9.539E-08 | 2.100E-07 | 4.000E-07 |
| 9482 | 475 | A475R | A | 1.5e-07 | 3.4e-07 | 1.5e-07 | 2.133E-07 | 1.097E-07 | 1.500E-07 | 3.400E-07 |
| 9483 | 475 | A475N | A | 3.6e-07 | 1.6e-07 | 4.6e-07 | 3.267E-07 | 1.528E-07 | 1.600E-07 | 4.600E-07 |
| 9484 | 475 | A475D | A | 1.2e-07 | 2.5e-07 | 4.6e-07 | 2.767E-07 | 1.716E-07 | 1.200E-07 | 4.600E-07 |
| 9485 | 475 | A475C | A | 2.7e-07 | 3.3e-07 | 2.5e-07 | 2.833E-07 | 4.163E-08 | 2.500E-07 | 3.300E-07 |
| 9486 | 475 | A475E | A | 3e-07 | 2.9e-07 | 3.9e-07 | 3.267E-07 | 5.508E-08 | 2.900E-07 | 3.900E-07 |
| 9487 | 475 | A475Q | A | 2.3e-07 | 3.4e-07 | 2.1e-07 | 2.600E-07 | 7.000E-08 | 2.100E-07 | 3.400E-07 |
| 9488 | 475 | A475G | A | 3e-07 | 4.1e-07 | 2.1e-07 | 3.067E-07 | 1.002E-07 | 2.100E-07 | 4.100E-07 |
| 9489 | 475 | A475H | A | 1.9e-07 | 1.3e-07 | 2.7e-07 | 1.967E-07 | 7.024E-08 | 1.300E-07 | 2.700E-07 |
| 9490 | 475 | A475I | A | 2.6e-07 | 2.6e-07 | 3.5e-07 | 2.900E-07 | 5.196E-08 | 2.600E-07 | 3.500E-07 |
| 9491 | 475 | A475L | A | 1.3e-07 | 1.7e-07 | 3.2e-07 | 2.067E-07 | 1.002E-07 | 1.300E-07 | 3.200E-07 |
| 9492 | 475 | A475K | A | 4.7e-07 | 2e-07 | 1.9e-07 | 2.867E-07 | 1.589E-07 | 1.900E-07 | 4.700E-07 |
| 9493 | 475 | A475M | A | 2.3e-07 | 2.3e-07 | 1.6e-07 | 2.067E-07 | 4.041E-08 | 1.600E-07 | 2.300E-07 |
| 9494 | 475 | A475F | A | 5.6e-07 | 1.4e-07 | 2.1e-07 | 3.033E-07 | 2.250E-07 | 1.400E-07 | 5.600E-07 |
| 9495 | 475 | A475P | A | 5.4e-07 | 2.3e-07 | 1.2e-07 | 2.967E-07 | 2.178E-07 | 1.200E-07 | 5.400E-07 |
| 9496 | 475 | A475S | A | 2e-07 | 2.9e-07 | 2.3e-07 | 2.400E-07 | 4.583E-08 | 2.000E-07 | 2.900E-07 |
| 9497 | 475 | A475T | A | 5.3e-07 | 1.4e-07 | 4.1e-07 | 3.600E-07 | 1.997E-07 | 1.400E-07 | 5.300E-07 |
| 9498 | 475 | A475W | A | 1.3e-07 | 2.8e-07 | 1.3e-07 | 1.800E-07 | 8.660E-08 | 1.300E-07 | 2.800E-07 |
| 9499 | 475 | A475Y | A | 3.3e-07 | 3.5e-07 | 8.9e-08 | 2.563E-07 | 1.453E-07 | 8.900E-08 | 3.500E-07 |
| 9500 | 475 | A475V | A | 2.3e-07 | 4.6e-07 | 3.7e-07 | 3.533E-07 | 1.159E-07 | 2.300E-07 | 4.600E-07 |
| 9501 | 476 | N476A | A | 1.4e-07 | 2.1e-07 | 3.3e-07 | 2.267E-07 | 9.609E-08 | 1.400E-07 | 3.300E-07 |
| 9502 | 476 | N476R | A | 1.2e-07 | 3.9e-07 | 2.2e-07 | 2.433E-07 | 1.365E-07 | 1.200E-07 | 3.900E-07 |
| 9503 | 476 | N476N | A | 4e-07 | 9.3e-08 | 2.2e-07 | 2.377E-07 | 1.543E-07 | 9.300E-08 | 4.000E-07 |
| 9504 | 476 | N476D | A | 5.1e-07 | 1.4e-07 | 3.8e-07 | 3.433E-07 | 1.877E-07 | 1.400E-07 | 5.100E-07 |
| 9505 | 476 | N476C | A | 2.3e-07 | 1.6e-07 | 2.4e-07 | 2.100E-07 | 4.359E-08 | 1.600E-07 | 2.400E-07 |
| 9506 | 476 | N476E | A | 3e-07 | 2.9e-07 | 2.5e-07 | 2.800E-07 | 2.646E-08 | 2.500E-07 | 3.000E-07 |
| 9507 | 476 | N476Q | A | 2.7e-07 | 2.2e-07 | 1.2e-07 | 2.033E-07 | 7.638E-08 | 1.200E-07 | 2.700E-07 |
| 9508 | 476 | N476G | A | 3.4e-07 | 4.1e-07 | 1.4e-07 | 2.967E-07 | 1.401E-07 | 1.400E-07 | 4.100E-07 |
| 9509 | 476 | N476H | A | 2.2e-07 | 1.8e-07 | 1.9e-07 | 1.967E-07 | 2.082E-08 | 1.800E-07 | 2.200E-07 |
| 9510 | 476 | N476I | A | 2.2e-07 | 2.5e-07 | 5.1e-07 | 3.267E-07 | 1.595E-07 | 2.200E-07 | 5.100E-07 |
| 9511 | 476 | N476L | A | 1.2e-07 | 6.3e-07 | 2e-07 | 3.167E-07 | 2.743E-07 | 1.200E-07 | 6.300E-07 |
| 9512 | 476 | N476K | A | 1.8e-07 | 3.5e-07 | 4.4e-07 | 3.233E-07 | 1.320E-07 | 1.800E-07 | 4.400E-07 |
| 9513 | 476 | N476M | A | 2.3e-07 | 3.1e-07 | 3.2e-07 | 2.867E-07 | 4.933E-08 | 2.300E-07 | 3.200E-07 |
| 9514 | 476 | N476F | A | 3.8e-07 | 2.2e-07 | 2.6e-07 | 2.867E-07 | 8.327E-08 | 2.200E-07 | 3.800E-07 |
| 9515 | 476 | N476P | A | 2.2e-07 | 1.4e-07 | 2.9e-07 | 2.167E-07 | 7.506E-08 | 1.400E-07 | 2.900E-07 |
| 9516 | 476 | N476S | A | 4.4e-07 | 2e-07 | 2.1e-07 | 2.833E-07 | 1.358E-07 | 2.000E-07 | 4.400E-07 |
| 9517 | 476 | N476T | A | 4.6e-07 | 2.6e-07 | 2.6e-07 | 3.267E-07 | 1.155E-07 | 2.600E-07 | 4.600E-07 |
| 9518 | 476 | N476W | A | 7.6e-07 | 2.4e-07 | 1.6e-07 | 3.867E-07 | 3.258E-07 | 1.600E-07 | 7.600E-07 |
| 9519 | 476 | N476Y | A | 4.4e-07 | 2.1e-07 | 9.3e-08 | 2.477E-07 | 1.765E-07 | 9.300E-08 | 4.400E-07 |

| | | | | | | | | | | |
|---|---|---|---|---|---|---|---|---|---|---|
| 9520 | 476 | N476V | A | 1.9e-07 | 2.7e-07 | 1e-07 | 1.867E-07 | 8.505E-08 | 1.000E-07 | 2.700E-07 |
| 9521 | 477 | R477A | A | 3.9e-07 | 5.3e-07 | 1.4e-07 | 3.533E-07 | 1.976E-07 | 1.400E-07 | 5.300E-07 |
| 9522 | 477 | R477R | A | 1.1e-07 | 1.3e-07 | 1.8e-07 | 1.400E-07 | 3.606E-08 | 1.100E-07 | 1.800E-07 |
| 9523 | 477 | R477N | A | 1.4e-07 | 1.6e-07 | 2.8e-07 | 1.933E-07 | 7.572E-08 | 1.400E-07 | 2.800E-07 |
| 9524 | 477 | R477D | A | 1.6e-07 | 1.6e-07 | 3.2e-07 | 2.133E-07 | 9.238E-08 | 1.600E-07 | 3.200E-07 |
| 9525 | 477 | R477C | A | 1.3e-07 | 5.6e-07 | 2.1e-07 | 3.000E-07 | 2.287E-07 | 1.300E-07 | 5.600E-07 |
| 9526 | 477 | R477E | A | 1.6e-07 | 4.1e-07 | 1.8e-07 | 2.500E-07 | 1.389E-07 | 1.600E-07 | 4.100E-07 |
| 9527 | 477 | R477Q | A | 2.8e-07 | 3.1e-07 | 5.5e-07 | 3.800E-07 | 1.480E-07 | 2.800E-07 | 5.500E-07 |
| 9528 | 477 | R477G | A | 1.8e-07 | 2.7e-07 | 1.6e-07 | 2.033E-07 | 5.859E-08 | 1.600E-07 | 2.700E-07 |
| 9529 | 477 | R477H | A | 3.1e-07 | 5.2e-07 | 2.9e-07 | 3.733E-07 | 1.274E-07 | 2.900E-07 | 5.200E-07 |
| 9530 | 477 | R477I | A | 2.7e-07 | 2.7e-07 | 1.5e-07 | 2.300E-07 | 6.928E-08 | 1.500E-07 | 2.700E-07 |
| 9531 | 477 | R477L | A | 1.9e-07 | 1.9e-07 | 3.4e-07 | 2.400E-07 | 8.660E-08 | 1.900E-07 | 3.400E-07 |
| 9532 | 477 | R477K | A | 1.5e-07 | 1.4e-07 | 3.2e-07 | 2.033E-07 | 1.012E-07 | 1.400E-07 | 3.200E-07 |
| 9533 | 477 | R477M | A | 3.3e-07 | 1.7e-07 | 1e-07 | 2.000E-07 | 1.179E-07 | 1.000E-07 | 3.300E-07 |
| 9534 | 477 | R477F | A | 5.7e-07 | 3.4e-07 | 2.3e-07 | 3.800E-07 | 1.735E-07 | 2.300E-07 | 5.700E-07 |
| 9535 | 477 | R477P | A | 3.4e-07 | 2.4e-07 | 2.1e-07 | 2.633E-07 | 6.807E-08 | 2.100E-07 | 3.400E-07 |
| 9536 | 477 | R477S | A | 1.2e-07 | 3.5e-07 | 1.9e-07 | 2.200E-07 | 1.179E-07 | 1.200E-07 | 3.500E-07 |
| 9537 | 477 | R477T | A | 3.7e-07 | 3.1e-07 | 3.1e-07 | 3.300E-07 | 3.464E-08 | 3.100E-07 | 3.700E-07 |
| 9538 | 477 | R477W | A | 2.4e-07 | 1.9e-07 | 3.5e-07 | 2.600E-07 | 8.185E-08 | 1.900E-07 | 3.500E-07 |
| 9539 | 477 | R477Y | A | 3.5e-07 | 2.3e-07 | 2.9e-07 | 2.900E-07 | 6.000E-08 | 2.300E-07 | 3.500E-07 |
| 9540 | 477 | R477V | A | 1.7e-07 | 2.1e-07 | 5.4e-07 | 3.067E-07 | 2.031E-07 | 1.700E-07 | 5.400E-07 |
| 9541 | 478 | P478A | A | 1.7e-07 | 2.3e-07 | 2.8e-07 | 2.267E-07 | 5.508E-08 | 1.700E-07 | 2.800E-07 |
| 9542 | 478 | P478R | A | 1.5e-07 | 3.3e-07 | 2e-07 | 2.267E-07 | 9.292E-08 | 1.500E-07 | 3.300E-07 |
| 9543 | 478 | P478N | A | 2.4e-07 | 3.1e-07 | 3.5e-07 | 3.000E-07 | 5.568E-08 | 2.400E-07 | 3.500E-07 |
| 9544 | 478 | P478D | A | 1.6e-07 | 2.3e-07 | 3e-07 | 2.300E-07 | 7.000E-08 | 1.600E-07 | 3.000E-07 |
| 9545 | 478 | P478C | A | 2.7e-07 | 4.2e-07 | 2.5e-07 | 3.133E-07 | 9.292E-08 | 2.500E-07 | 4.200E-07 |
| 9546 | 478 | P478E | A | 1.7e-07 | 3.3e-07 | 4.8e-07 | 3.267E-07 | 1.550E-07 | 1.700E-07 | 4.800E-07 |
| 9547 | 478 | P478Q | A | 2.4e-07 | 2.2e-07 | 3.6e-07 | 2.733E-07 | 7.572E-08 | 2.200E-07 | 3.600E-07 |
| 9548 | 478 | P478G | A | 3.3e-07 | 1.5e-07 | 5e-07 | 3.267E-07 | 1.750E-07 | 1.500E-07 | 5.000E-07 |
| 9549 | 478 | P478H | A | 1.9e-07 | 1.2e-07 | 3.1e-07 | 2.067E-07 | 9.609E-08 | 1.200E-07 | 3.100E-07 |
| 9550 | 478 | P478I | A | 1.7e-07 | 3.1e-07 | 3.6e-07 | 2.800E-07 | 9.849E-08 | 1.700E-07 | 3.600E-07 |
| 9551 | 478 | P478L | A | 3.7e-07 | 2.3e-07 | 1.9e-07 | 2.633E-07 | 9.452E-08 | 1.900E-07 | 3.700E-07 |
| 9552 | 478 | P478K | A | 3.3e-07 | 3.7e-07 | 4.3e-07 | 3.767E-07 | 5.033E-08 | 3.300E-07 | 4.300E-07 |
| 9553 | 478 | P478M | A | 2.1e-07 | 3.1e-07 | 1.4e-07 | 2.200E-07 | 8.544E-08 | 1.400E-07 | 3.100E-07 |
| 9554 | 478 | P478F | A | 3e-07 | 4.6e-07 | 5e-07 | 4.200E-07 | 1.058E-07 | 3.000E-07 | 5.000E-07 |
| 9555 | 478 | P478P | A | 3.6e-07 | 5.9e-07 | 1.1e-07 | 3.533E-07 | 2.401E-07 | 1.100E-07 | 5.900E-07 |
| 9556 | 478 | P478S | A | 2.6e-07 | 1.7e-07 | 1.9e-07 | 2.067E-07 | 4.726E-08 | 1.700E-07 | 2.600E-07 |
| 9557 | 478 | P478T | A | 3.2e-07 | 2e-07 | 1.2e-07 | 2.133E-07 | 1.007E-07 | 1.200E-07 | 3.200E-07 |
| 9558 | 478 | P478W | A | 3.1e-07 | 5.7e-08 | 5.8e-07 | 3.157E-07 | 2.615E-07 | 5.700E-08 | 5.800E-07 |
| 9559 | 478 | P478Y | A | 1.3e-07 | 5.2e-07 | 2.9e-07 | 3.133E-07 | 1.960E-07 | 1.300E-07 | 5.200E-07 |
| 9560 | 478 | P478V | A | 2e-07 | 3.3e-07 | 4.4e-07 | 3.233E-07 | 1.201E-07 | 2.000E-07 | 4.400E-07 |
| 9561 | 479 | E479A | A | 1.7e-07 | 1.5e-07 | 2.3e-07 | 1.833E-07 | 4.163E-08 | 1.500E-07 | 2.300E-07 |
| 9562 | 479 | E479R | A | 2.5e-07 | 2.2e-07 | 1.6e-07 | 2.100E-07 | 4.583E-08 | 1.600E-07 | 2.500E-07 |
| 9563 | 479 | E479N | A | 3.3e-07 | 3.3e-07 | 3e-07 | 3.200E-07 | 1.732E-08 | 3.000E-07 | 3.300E-07 |
| 9564 | 479 | E479D | A | 1.1e-07 | 1.9e-07 | 3.4e-07 | 2.133E-07 | 1.168E-07 | 1.100E-07 | 3.400E-07 |
| 9565 | 479 | E479C | A | 3.6e-07 | 2.1e-07 | 1.9e-07 | 2.533E-07 | 9.292E-08 | 1.900E-07 | 3.600E-07 |
| 9566 | 479 | E479E | A | 5.3e-08 | 2.1e-07 | 1.5e-07 | 1.377E-07 | 7.922E-08 | 5.300E-08 | 2.100E-07 |
| 9567 | 479 | E479Q | A | 1.6e-07 | 2e-07 | 2.8e-07 | 2.133E-07 | 6.110E-08 | 1.600E-07 | 2.800E-07 |
| 9568 | 479 | E479G | A | 1.5e-07 | 5e-07 | 2.5e-07 | 3.000E-07 | 1.803E-07 | 1.500E-07 | 5.000E-07 |
| 9569 | 479 | E479H | A | 4.1e-07 | 3.5e-07 | 1.7e-07 | 3.100E-07 | 1.249E-07 | 1.700E-07 | 4.100E-07 |
| 9570 | 479 | E479I | A | 1.6e-07 | 1.8e-07 | 3.3e-07 | 2.233E-07 | 9.292E-08 | 1.600E-07 | 3.300E-07 |
| 9571 | 479 | E479L | A | 2.4e-07 | 2.3e-07 | 2.2e-07 | 2.300E-07 | 1.000E-08 | 2.200E-07 | 2.400E-07 |
| 9572 | 479 | E479K | A | 2.3e-07 | 3.6e-07 | 2.1e-07 | 2.667E-07 | 8.145E-08 | 2.100E-07 | 3.600E-07 |
| 9573 | 479 | E479M | A | 4.6e-07 | 2.3e-07 | 2e-07 | 2.967E-07 | 1.422E-07 | 2.000E-07 | 4.600E-07 |
| 9574 | 479 | E479F | A | 2.8e-07 | 2.8e-07 | 1.9e-07 | 2.500E-07 | 5.196E-08 | 1.900E-07 | 2.800E-07 |
| 9575 | 479 | E479P | A | 4.3e-07 | 1.7e-07 | 5.6e-07 | 3.867E-07 | 1.986E-07 | 1.700E-07 | 5.600E-07 |
| 9576 | 479 | E479S | A | 3.5e-07 | 2.7e-07 | 5.2e-07 | 3.800E-07 | 1.277E-07 | 2.700E-07 | 5.200E-07 |
| 9577 | 479 | E479T | A | 2.2e-07 | 9.2e-08 | 2.2e-07 | 1.773E-07 | 7.390E-08 | 9.200E-08 | 2.200E-07 |
| 9578 | 479 | E479W | A | 3.6e-07 | 4.8e-07 | 1.7e-07 | 3.367E-07 | 1.563E-07 | 1.700E-07 | 4.800E-07 |
| 9579 | 479 | E479Y | A | 3.4e-07 | 1.4e-07 | 1.4e-07 | 2.067E-07 | 1.155E-07 | 1.400E-07 | 3.400E-07 |
| 9580 | 479 | E479V | A | 4.3e-07 | 2.4e-07 | 4.2e-07 | 3.633E-07 | 1.069E-07 | 2.400E-07 | 4.300E-07 |
| 9581 | 480 | D480A | A | 3.5e-07 | 2.8e-07 | 4.9e-07 | 3.733E-07 | 1.069E-07 | 2.800E-07 | 4.900E-07 |
| 9582 | 480 | D480R | A | 4.7e-07 | 4.3e-07 | 1.6e-07 | 3.533E-07 | 1.686E-07 | 1.600E-07 | 4.700E-07 |
| 9583 | 480 | D480N | A | 2.2e-07 | 2.9e-07 | 4.9e-07 | 3.333E-07 | 1.401E-07 | 2.200E-07 | 4.900E-07 |
| 9584 | 480 | D480D | A | 1.3e-07 | 2.4e-07 | 3.6e-07 | 2.433E-07 | 1.150E-07 | 1.300E-07 | 3.600E-07 |
| 9585 | 480 | D480C | A | 2.6e-07 | 3e-07 | 1.1e-07 | 2.233E-07 | 1.002E-07 | 1.100E-07 | 3.000E-07 |
| 9586 | 480 | D480E | A | 3.4e-07 | 5.3e-07 | 1.9e-07 | 3.533E-07 | 1.704E-07 | 1.900E-07 | 5.300E-07 |
| 9587 | 480 | D480Q | A | 3.5e-07 | 3.8e-07 | 2e-07 | 3.100E-07 | 9.644E-08 | 2.000E-07 | 3.800E-07 |
| 9588 | 480 | D480G | A | 2.4e-07 | 4e-07 | 1.8e-07 | 2.733E-07 | 1.137E-07 | 1.800E-07 | 4.000E-07 |
| 9589 | 480 | D480H | A | 4.5e-07 | 3.5e-07 | 3e-07 | 3.667E-07 | 7.638E-08 | 3.000E-07 | 4.500E-07 |
| 9590 | 480 | D480I | A | 2.9e-07 | 4.9e-07 | 1.5e-07 | 3.100E-07 | 1.709E-07 | 1.500E-07 | 4.900E-07 |
| 9591 | 480 | D480L | A | 2.3e-07 | 1.1e-07 | 1.7e-07 | 1.700E-07 | 6.000E-08 | 1.100E-07 | 2.300E-07 |
| 9592 | 480 | D480K | A | 1.6e-07 | 2.8e-07 | 3.6e-07 | 2.667E-07 | 1.007E-07 | 1.600E-07 | 3.600E-07 |
| 9593 | 480 | D480M | A | 9.2e-08 | 3.5e-07 | 2.2e-07 | 2.207E-07 | 1.290E-07 | 9.200E-08 | 3.500E-07 |
| 9594 | 480 | D480F | A | 2.4e-07 | 5.3e-07 | 4.7e-07 | 4.133E-07 | 1.531E-07 | 2.400E-07 | 5.300E-07 |
| 9595 | 480 | D480P | A | 4.5e-07 | 5.2e-07 | 1.6e-07 | 3.767E-07 | 1.909E-07 | 1.600E-07 | 5.200E-07 |
| 9596 | 480 | D480S | A | 3.1e-07 | 2.2e-07 | 2.1e-07 | 2.467E-07 | 5.508E-08 | 2.100E-07 | 3.100E-07 |
| 9597 | 480 | D480T | A | 2.8e-07 | 1.6e-07 | 2.8e-07 | 2.400E-07 | 6.928E-08 | 1.600E-07 | 2.800E-07 |
| 9598 | 480 | D480W | A | 2.3e-07 | 2.5e-07 | 1.5e-07 | 2.100E-07 | 5.292E-08 | 1.500E-07 | 2.500E-07 |
| 9599 | 480 | D480Y | A | 2.3e-07 | 2.6e-07 | 1.8e-07 | 2.233E-07 | 4.041E-08 | 1.800E-07 | 2.600E-07 |
| 9600 | 480 | D480V | A | 4.3e-07 | 3.4e-07 | 2.5e-07 | 3.400E-07 | 9.000E-08 | 2.500E-07 | 4.300E-07 |
| 9601 | 481 | E481A | A | 3.8e-07 | 1.9e-07 | 3.8e-07 | 3.167E-07 | 1.097E-07 | 1.900E-07 | 3.800E-07 |
| 9602 | 481 | E481R | A | 1.5e-07 | 2.5e-07 | 2.7e-07 | 2.233E-07 | 6.429E-08 | 1.500E-07 | 2.700E-07 |
| 9603 | 481 | E481N | A | 7e-07 | 3.8e-07 | 3.4e-07 | 4.733E-07 | 1.973E-07 | 3.400E-07 | 7.000E-07 |
| 9604 | 481 | E481D | A | 4.2e-07 | 2.2e-07 | 4.4e-07 | 3.600E-07 | 1.217E-07 | 2.200E-07 | 4.400E-07 |

| | | | | | | | | | | |
|---|---|---|---|---|---|---|---|---|---|---|
| 9605 | 481 | E481C | A | 2.2e-07 | 4.2e-07 | 5.6e-07 | 4.000E-07 | 1.709E-07 | 2.200E-07 | 5.600E-07 |
| 9606 | 481 | E481E | A | 1.7e-07 | 1.6e-07 | 8e-07 | 3.767E-07 | 3.667E-07 | 1.600E-07 | 8.000E-07 |
| 9607 | 481 | E481Q | A | 1.8e-07 | 1.4e-07 | 1.4e-07 | 1.533E-07 | 2.309E-08 | 1.400E-07 | 1.800E-07 |
| 9608 | 481 | E481G | A | 2.1e-07 | 3.9e-07 | 2.3e-07 | 2.767E-07 | 9.866E-08 | 2.100E-07 | 3.900E-07 |
| 9609 | 481 | E481H | A | 2.9e-07 | 2.5e-07 | 2.6e-07 | 2.667E-07 | 2.082E-08 | 2.500E-07 | 2.900E-07 |
| 9610 | 481 | E481I | A | 2.5e-07 | 2.1e-07 | 1.6e-07 | 2.067E-07 | 4.509E-08 | 1.600E-07 | 2.500E-07 |
| 9611 | 481 | E481L | A | 3e-07 | 2e-07 | 3.1e-07 | 2.700E-07 | 6.083E-08 | 2.000E-07 | 3.100E-07 |
| 9612 | 481 | E481K | A | 6.8e-08 | 2.6e-07 | 2.8e-07 | 2.027E-07 | 1.171E-07 | 6.800E-08 | 2.800E-07 |
| 9613 | 481 | E481M | A | 2.5e-07 | 3.1e-07 | 3.8e-07 | 3.133E-07 | 6.506E-08 | 2.500E-07 | 3.800E-07 |
| 9614 | 481 | E481F | A | 1.9e-07 | 3.8e-07 | 3.2e-07 | 2.967E-07 | 9.713E-08 | 1.900E-07 | 3.800E-07 |
| 9615 | 481 | E481P | A | 2.1e-07 | 4e-07 | 2.4e-07 | 2.833E-07 | 1.021E-07 | 2.100E-07 | 4.000E-07 |
| 9616 | 481 | E481S | A | 3.7e-07 | 2.3e-07 | 1.1e-07 | 2.367E-07 | 1.301E-07 | 1.100E-07 | 3.700E-07 |
| 9617 | 481 | E481T | A | 2.6e-07 | 1.1e-07 | 4.2e-07 | 2.633E-07 | 1.550E-07 | 1.100E-07 | 4.200E-07 |
| 9618 | 481 | E481W | A | 1.6e-07 | 2.6e-07 | 4e-07 | 2.733E-07 | 1.206E-07 | 1.600E-07 | 4.000E-07 |
| 9619 | 481 | E481Y | A | 3.9e-07 | 3.4e-07 | 2.3e-07 | 3.200E-07 | 8.185E-08 | 2.300E-07 | 3.900E-07 |
| 9620 | 481 | E481V | A | 1.6e-07 | 2.3e-07 | 3.7e-07 | 2.533E-07 | 1.069E-07 | 1.600E-07 | 3.700E-07 |
| 9621 | 482 | C482A | A | 3.3e-07 | 1.8e-07 | 3.2e-07 | 2.767E-07 | 8.386E-08 | 1.800E-07 | 3.300E-07 |
| 9622 | 482 | C482R | A | 1.8e-07 | 9.4e-08 | 5.8e-07 | 2.847E-07 | 2.594E-07 | 9.400E-08 | 5.800E-07 |
| 9623 | 482 | C482N | A | 3.8e-07 | 3.5e-07 | 1.1e-07 | 2.800E-07 | 1.480E-07 | 1.100E-07 | 3.800E-07 |
| 9624 | 482 | C482D | A | 1.8e-07 | 2e-07 | 2.5e-07 | 2.100E-07 | 3.606E-08 | 1.800E-07 | 2.500E-07 |
| 9625 | 482 | C482C | A | 4.6e-07 | 2.2e-07 | 1.7e-07 | 2.833E-07 | 1.550E-07 | 1.700E-07 | 4.600E-07 |
| 9626 | 482 | C482E | A | 2.5e-07 | 3.2e-07 | 1.5e-07 | 2.400E-07 | 8.544E-08 | 1.500E-07 | 3.200E-07 |
| 9627 | 482 | C482Q | A | 3.5e-07 | 4.1e-07 | 2.1e-07 | 3.233E-07 | 1.026E-07 | 2.100E-07 | 4.100E-07 |
| 9628 | 482 | C482G | A | 2.1e-07 | 5.6e-07 | 3.8e-07 | 3.833E-07 | 1.750E-07 | 2.100E-07 | 5.600E-07 |
| 9629 | 482 | C482H | A | 2.1e-07 | 1.9e-07 | 3.5e-07 | 2.500E-07 | 8.718E-08 | 1.900E-07 | 3.500E-07 |
| 9630 | 482 | C482I | A | 3.3e-07 | 3.1e-07 | 5.3e-07 | 3.900E-07 | 1.217E-07 | 3.100E-07 | 5.300E-07 |
| 9631 | 482 | C482L | A | 3.4e-07 | 2.8e-07 | 2.1e-07 | 2.767E-07 | 6.506E-08 | 2.100E-07 | 3.400E-07 |
| 9632 | 482 | C482K | A | 3.2e-07 | 2.4e-07 | 2.9e-07 | 2.833E-07 | 4.041E-08 | 2.400E-07 | 3.200E-07 |
| 9633 | 482 | C482M | A | 2.6e-07 | 3.1e-07 | 1.7e-07 | 2.467E-07 | 7.095E-08 | 1.700E-07 | 3.100E-07 |
| 9634 | 482 | C482F | A | 2.9e-07 | 2e-07 | 4.8e-07 | 3.233E-07 | 1.429E-07 | 2.000E-07 | 4.800E-07 |
| 9635 | 482 | C482P | A | 3.4e-07 | 3.1e-07 | 5.4e-07 | 3.967E-07 | 1.250E-07 | 3.100E-07 | 5.400E-07 |
| 9636 | 482 | C482S | A | 6.3e-07 | 3.7e-07 | 2.8e-07 | 4.267E-07 | 1.818E-07 | 2.800E-07 | 6.300E-07 |
| 9637 | 482 | C482T | A | 1.8e-07 | 2.1e-07 | 2.1e-07 | 2.000E-07 | 1.732E-08 | 1.800E-07 | 2.100E-07 |
| 9638 | 482 | C482W | A | 4.4e-07 | 1.3e-07 | 7.5e-08 | 2.150E-07 | 1.968E-07 | 7.500E-08 | 4.400E-07 |
| 9639 | 482 | C482Y | A | 1.5e-07 | 1.8e-07 | 3.7e-07 | 2.333E-07 | 1.193E-07 | 1.500E-07 | 3.700E-07 |
| 9640 | 482 | C482V | A | 3.7e-07 | 1.8e-07 | 1.9e-07 | 2.467E-07 | 1.069E-07 | 1.800E-07 | 3.700E-07 |
| 9641 | 483 | V483A | A | 1.6e-07 | 2.5e-07 | 2.4e-07 | 2.167E-07 | 4.933E-08 | 1.600E-07 | 2.500E-07 |
| 9642 | 483 | V483R | A | 5e-07 | 2.9e-07 | 2.2e-07 | 3.367E-07 | 1.457E-07 | 2.200E-07 | 5.000E-07 |
| 9643 | 483 | V483N | A | 1.8e-07 | 2.1e-07 | 5.9e-07 | 3.267E-07 | 2.285E-07 | 1.800E-07 | 5.900E-07 |
| 9644 | 483 | V483D | A | 1.4e-07 | 1e-07 | 1.3e-07 | 1.233E-07 | 2.082E-08 | 1.000E-07 | 1.400E-07 |
| 9645 | 483 | V483C | A | 4.8e-07 | 1.7e-07 | 2.9e-07 | 3.133E-07 | 1.563E-07 | 1.700E-07 | 4.800E-07 |
| 9646 | 483 | V483E | A | 5.3e-07 | 3.9e-07 | 2.7e-07 | 3.967E-07 | 1.301E-07 | 2.700E-07 | 5.300E-07 |
| 9647 | 483 | V483Q | A | 2e-07 | 2.2e-07 | 2.4e-07 | 2.200E-07 | 2.000E-08 | 2.000E-07 | 2.400E-07 |
| 9648 | 483 | V483G | A | 3.8e-07 | 3.4e-07 | 1.7e-07 | 2.967E-07 | 1.115E-07 | 1.700E-07 | 3.800E-07 |
| 9649 | 483 | V483H | A | 5.7e-07 | 5e-07 | 2.8e-07 | 4.500E-07 | 1.513E-07 | 2.800E-07 | 5.700E-07 |
| 9650 | 483 | V483I | A | 4.1e-07 | 2.5e-07 | 4.8e-07 | 3.800E-07 | 1.179E-07 | 2.500E-07 | 4.800E-07 |
| 9651 | 483 | V483L | A | 2.9e-07 | 2.1e-07 | 6.1e-07 | 3.700E-07 | 2.117E-07 | 2.100E-07 | 6.100E-07 |
| 9652 | 483 | V483K | A | 4.3e-07 | 3.2e-07 | 4e-07 | 3.833E-07 | 5.686E-08 | 3.200E-07 | 4.300E-07 |
| 9653 | 483 | V483M | A | 2.5e-07 | 4.7e-07 | 4.8e-07 | 4.000E-07 | 1.300E-07 | 2.500E-07 | 4.800E-07 |
| 9654 | 483 | V483F | A | 1.8e-07 | 4e-07 | 1.2e-07 | 2.333E-07 | 1.474E-07 | 1.200E-07 | 4.000E-07 |
| 9655 | 483 | V483P | A | 1.4e-07 | 3.1e-07 | 2.2e-07 | 2.233E-07 | 8.505E-08 | 1.400E-07 | 3.100E-07 |
| 9656 | 483 | V483S | A | 2.4e-07 | 2.1e-07 | 3.1e-07 | 2.533E-07 | 5.132E-08 | 2.100E-07 | 3.100E-07 |
| 9657 | 483 | V483T | A | 2.9e-07 | 3.4e-07 | 5.2e-07 | 3.833E-07 | 1.210E-07 | 2.900E-07 | 5.200E-07 |
| 9658 | 483 | V483W | A | 2.4e-07 | 1.2e-07 | 4e-07 | 2.533E-07 | 1.405E-07 | 1.200E-07 | 4.000E-07 |
| 9659 | 483 | V483Y | A | 2.9e-07 | 1.7e-07 | 3.3e-07 | 2.633E-07 | 8.327E-08 | 1.700E-07 | 3.300E-07 |
| 9660 | 483 | V483V | A | 1.3e-07 | 3.1e-07 | 1.8e-07 | 2.067E-07 | 9.292E-08 | 1.300E-07 | 3.100E-07 |
| 9661 | 484 | G484A | A | 5.1e-07 | 1.6e-07 | 2.6e-07 | 3.100E-07 | 1.803E-07 | 1.600E-07 | 5.100E-07 |
| 9662 | 484 | G484R | A | 2.4e-07 | 3.1e-07 | 4.5e-07 | 3.333E-07 | 1.069E-07 | 2.400E-07 | 4.500E-07 |
| 9663 | 484 | G484N | A | 1.6e-07 | 1.5e-07 | 2.6e-07 | 1.900E-07 | 6.083E-08 | 1.500E-07 | 2.600E-07 |
| 9664 | 484 | G484D | A | 2.9e-07 | 2.2e-07 | 2.3e-07 | 2.467E-07 | 3.786E-08 | 2.200E-07 | 2.900E-07 |
| 9665 | 484 | G484C | A | 1.9e-07 | 1.7e-07 | 2.6e-07 | 2.067E-07 | 4.726E-08 | 1.700E-07 | 2.600E-07 |
| 9666 | 484 | G484E | A | 4.2e-07 | 4.2e-07 | 2.2e-07 | 3.533E-07 | 1.155E-07 | 2.200E-07 | 4.200E-07 |
| 9667 | 484 | G484Q | A | 2.2e-07 | 1.3e-07 | 2e-07 | 1.833E-07 | 4.726E-08 | 1.300E-07 | 2.200E-07 |
| 9668 | 484 | G484G | A | 4.1e-07 | 2.3e-07 | 1.2e-07 | 2.533E-07 | 1.464E-07 | 1.200E-07 | 4.100E-07 |
| 9669 | 484 | G484H | A | 5.2e-07 | 2e-07 | 2.1e-07 | 3.100E-07 | 1.819E-07 | 2.000E-07 | 5.200E-07 |
| 9670 | 484 | G484I | A | 2.1e-07 | 2.5e-07 | 5.2e-07 | 3.267E-07 | 1.686E-07 | 2.100E-07 | 5.200E-07 |
| 9671 | 484 | G484L | A | 1.2e-07 | 3.7e-07 | 1.9e-07 | 2.267E-07 | 1.290E-07 | 1.200E-07 | 3.700E-07 |
| 9672 | 484 | G484K | A | 4.2e-07 | 3.2e-07 | 2.1e-07 | 3.167E-07 | 1.050E-07 | 2.100E-07 | 4.200E-07 |
| 9673 | 484 | G484M | A | 1.4e-07 | 4.6e-07 | 1.5e-07 | 2.500E-07 | 1.819E-07 | 1.400E-07 | 4.600E-07 |
| 9674 | 484 | G484F | A | 4.1e-07 | 2.8e-07 | 2.9e-07 | 3.267E-07 | 7.234E-08 | 2.800E-07 | 4.100E-07 |
| 9675 | 484 | G484P | A | 4e-07 | 2.1e-07 | 2.3e-07 | 2.800E-07 | 1.044E-07 | 2.100E-07 | 4.000E-07 |
| 9676 | 484 | G484S | A | 5.5e-07 | 1e-07 | 1.8e-07 | 2.767E-07 | 2.401E-07 | 1.000E-07 | 5.500E-07 |
| 9677 | 484 | G484T | A | 1.7e-07 | 2.3e-07 | 5.9e-07 | 3.300E-07 | 2.272E-07 | 1.700E-07 | 5.900E-07 |
| 9678 | 484 | G484W | A | 2.1e-07 | 4.9e-07 | 3e-07 | 3.333E-07 | 1.429E-07 | 2.100E-07 | 4.900E-07 |
| 9679 | 484 | G484Y | A | 5.1e-07 | 7.1e-07 | 2.7e-07 | 4.967E-07 | 2.203E-07 | 2.700E-07 | 7.100E-07 |
| 9680 | 484 | G484V | A | 1.2e-07 | 9.9e-08 | 2.8e-07 | 1.663E-07 | 9.900E-08 | 9.900E-08 | 2.800E-07 |
| 9681 | 485 | E485A | A | 2.2e-07 | 3e-07 | 1.1e-07 | 2.100E-07 | 9.539E-08 | 1.100E-07 | 3.000E-07 |
| 9682 | 485 | E485R | A | 3.1e-07 | 4e-07 | 2.3e-07 | 3.133E-07 | 8.505E-08 | 2.300E-07 | 4.000E-07 |
| 9683 | 485 | E485N | A | 1.4e-07 | 4.4e-07 | 4.5e-07 | 3.433E-07 | 1.762E-07 | 1.400E-07 | 4.500E-07 |
| 9684 | 485 | E485D | A | 3.8e-07 | 3.7e-07 | 4e-07 | 3.833E-07 | 1.528E-08 | 3.700E-07 | 4.000E-07 |
| 9685 | 485 | E485C | A | 6.6e-07 | 5.2e-07 | 2.4e-07 | 4.733E-07 | 2.139E-07 | 2.400E-07 | 6.600E-07 |
| 9686 | 485 | E485E | A | 2.6e-07 | 8.8e-08 | 2.8e-07 | 2.093E-07 | 1.056E-07 | 8.800E-08 | 2.800E-07 |
| 9687 | 485 | E485Q | A | 3.4e-07 | 1.3e-07 | 1.5e-07 | 2.067E-07 | 1.159E-07 | 1.300E-07 | 3.400E-07 |
| 9688 | 485 | E485G | A | 2.4e-07 | 2.5e-07 | 2e-07 | 2.300E-07 | 2.646E-08 | 2.000E-07 | 2.500E-07 |
| 9689 | 485 | E485H | A | 3.8e-07 | 3.1e-07 | 1.9e-07 | 2.933E-07 | 9.609E-08 | 1.900E-07 | 3.800E-07 |

| | | | | | | | | | | |
|---|---|---|---|---|---|---|---|---|---|---|
| 9690 | 485 | E485I | A | 6.5e-07 | 5.7e-07 | 3.3e-07 | 5.167E-07 | 1.665E-07 | 3.300E-07 | 6.500E-07 |
| 9691 | 485 | E485L | A | 1.4e-07 | 7.7e-07 | 3.2e-07 | 4.100E-07 | 3.245E-07 | 1.400E-07 | 7.700E-07 |
| 9692 | 485 | E485K | A | 2.5e-07 | 9.8e-08 | 2.6e-07 | 2.027E-07 | 9.078E-08 | 9.800E-08 | 2.600E-07 |
| 9693 | 485 | E485M | A | 1.6e-07 | 2.6e-07 | 2.5e-07 | 2.233E-07 | 5.508E-08 | 1.600E-07 | 2.600E-07 |
| 9694 | 485 | E485F | A | 3e-07 | 1.9e-07 | 2.5e-07 | 2.467E-07 | 5.508E-08 | 1.900E-07 | 3.000E-07 |
| 9695 | 485 | E485P | A | 2.9e-07 | 5.1e-07 | 3.4e-07 | 3.800E-07 | 1.153E-07 | 2.900E-07 | 5.100E-07 |
| 9696 | 485 | E485S | A | 2e-07 | 2.3e-07 | 2.5e-07 | 2.267E-07 | 2.517E-08 | 2.000E-07 | 2.500E-07 |
| 9697 | 485 | E485T | A | 1.6e-07 | 3.5e-07 | 3.3e-07 | 2.800E-07 | 1.044E-07 | 1.600E-07 | 3.500E-07 |
| 9698 | 485 | E485W | A | 7.6e-08 | 1.1e-07 | 2.9e-07 | 1.587E-07 | 1.150E-07 | 7.600E-08 | 2.900E-07 |
| 9699 | 485 | E485Y | A | 2e-07 | 3e-07 | 2.7e-07 | 2.567E-07 | 5.132E-08 | 2.000E-07 | 3.000E-07 |
| 9700 | 485 | E485V | A | 4.3e-07 | 4.2e-07 | 2.7e-07 | 3.733E-07 | 8.963E-08 | 2.700E-07 | 4.300E-07 |
| 9701 | 486 | G486A | A | 1.5e-07 | 3.8e-07 | 2.9e-07 | 2.733E-07 | 1.159E-07 | 1.500E-07 | 3.800E-07 |
| 9702 | 486 | G486R | A | 2.3e-07 | 2.7e-07 | 3.8e-07 | 2.933E-07 | 7.767E-08 | 2.300E-07 | 3.800E-07 |
| 9703 | 486 | G486N | A | 3.5e-07 | 2e-07 | 1.9e-07 | 2.467E-07 | 8.963E-08 | 1.900E-07 | 3.500E-07 |
| 9704 | 486 | G486D | A | 5.5e-07 | 1e-07 | 5.3e-07 | 3.933E-07 | 2.542E-07 | 1.000E-07 | 5.500E-07 |
| 9705 | 486 | G486C | A | 3.7e-07 | 2.2e-07 | 1.4e-07 | 2.433E-07 | 1.168E-07 | 1.400E-07 | 3.700E-07 |
| 9706 | 486 | G486E | A | 1.6e-07 | 2.3e-07 | 3e-07 | 2.300E-07 | 7.000E-08 | 1.600E-07 | 3.000E-07 |
| 9707 | 486 | G486Q | A | 1.6e-07 | 2.2e-07 | 4.5e-07 | 2.767E-07 | 1.531E-07 | 1.600E-07 | 4.500E-07 |
| 9708 | 486 | G486G | A | 4e-07 | 1.2e-07 | 3.9e-07 | 3.033E-07 | 1.589E-07 | 1.200E-07 | 4.000E-07 |
| 9709 | 486 | G486H | A | 1.9e-07 | 2.8e-07 | 1.5e-07 | 2.067E-07 | 6.658E-08 | 1.500E-07 | 2.800E-07 |
| 9710 | 486 | G486I | A | 2.8e-07 | 3.2e-07 | 7.7e-07 | 4.567E-07 | 2.721E-07 | 2.800E-07 | 7.700E-07 |
| 9711 | 486 | G486L | A | 2.8e-07 | 1.3e-07 | 2.2e-07 | 2.100E-07 | 7.550E-08 | 1.300E-07 | 2.800E-07 |
| 9712 | 486 | G486K | A | 2e-07 | 9.8e-08 | 3.7e-07 | 2.227E-07 | 1.374E-07 | 9.800E-08 | 3.700E-07 |
| 9713 | 486 | G486M | A | 3.9e-07 | 2.9e-07 | 6.1e-07 | 4.300E-07 | 1.637E-07 | 2.900E-07 | 6.100E-07 |
| 9714 | 486 | G486F | A | 1.6e-07 | 1.8e-07 | 3.9e-07 | 2.433E-07 | 1.274E-07 | 1.600E-07 | 3.900E-07 |
| 9715 | 486 | G486P | A | 1.3e-07 | 1.7e-07 | 5.5e-07 | 2.833E-07 | 2.318E-07 | 1.300E-07 | 5.500E-07 |
| 9716 | 486 | G486S | A | 2.3e-07 | 2.5e-07 | 2.2e-07 | 2.333E-07 | 1.528E-08 | 2.200E-07 | 2.500E-07 |
| 9717 | 486 | G486T | A | 2.6e-07 | 2.3e-07 | 2.7e-07 | 2.533E-07 | 2.082E-08 | 2.300E-07 | 2.700E-07 |
| 9718 | 486 | G486W | A | 2.1e-07 | 2.7e-07 | 2.1e-07 | 2.300E-07 | 3.464E-08 | 2.100E-07 | 2.700E-07 |
| 9719 | 486 | G486Y | A | 2.2e-07 | 5.6e-07 | 2.7e-07 | 3.500E-07 | 1.836E-07 | 2.200E-07 | 5.600E-07 |
| 9720 | 486 | G486V | A | 2.3e-07 | 3e-07 | 1.8e-07 | 2.367E-07 | 6.028E-08 | 1.800E-07 | 3.000E-07 |
| 9721 | 487 | L487A | A | 4.5e-07 | 2.6e-07 | 5e-08 | 2.533E-07 | 2.001E-07 | 5.000E-08 | 4.500E-07 |
| 9722 | 487 | L487R | A | 2.1e-07 | 1.5e-07 | 1.3e-07 | 1.633E-07 | 4.163E-08 | 1.300E-07 | 2.100E-07 |
| 9723 | 487 | L487N | A | 1.3e-07 | 3e-07 | 3.3e-07 | 2.533E-07 | 1.079E-07 | 1.300E-07 | 3.300E-07 |
| 9724 | 487 | L487D | A | 2.4e-07 | 1.4e-07 | 1.6e-07 | 1.800E-07 | 5.292E-08 | 1.400E-07 | 2.400E-07 |
| 9725 | 487 | L487C | A | 1.8e-07 | 5.9e-07 | 1.6e-07 | 3.100E-07 | 2.427E-07 | 1.600E-07 | 5.900E-07 |
| 9726 | 487 | L487E | A | 2e-07 | 2.2e-07 | 3.5e-07 | 2.567E-07 | 8.145E-08 | 2.000E-07 | 3.500E-07 |
| 9727 | 487 | L487Q | A | 4.2e-07 | 2.9e-07 | 2e-07 | 3.033E-07 | 1.106E-07 | 2.000E-07 | 4.200E-07 |
| 9728 | 487 | L487G | A | 2.9e-07 | 1.9e-07 | 4.3e-07 | 3.033E-07 | 1.206E-07 | 1.900E-07 | 4.300E-07 |
| 9729 | 487 | L487H | A | 9.3e-08 | 3.8e-07 | 2.9e-07 | 2.543E-07 | 1.468E-07 | 9.300E-08 | 3.800E-07 |
| 9730 | 487 | L487I | A | 2.6e-07 | 2.8e-07 | 2.5e-07 | 2.633E-07 | 1.528E-08 | 2.500E-07 | 2.800E-07 |
| 9731 | 487 | L487L | A | 1.6e-07 | 3.9e-07 | 2.1e-07 | 2.533E-07 | 1.210E-07 | 1.600E-07 | 3.900E-07 |
| 9732 | 487 | L487K | A | 1.6e-07 | 4.5e-07 | 5e-07 | 3.700E-07 | 1.836E-07 | 1.600E-07 | 5.000E-07 |
| 9733 | 487 | L487M | A | 1.7e-07 | 2.9e-07 | 2.5e-07 | 2.367E-07 | 6.110E-08 | 1.700E-07 | 2.900E-07 |
| 9734 | 487 | L487F | A | 1.9e-07 | 1.9e-07 | 1.8e-07 | 1.867E-07 | 5.774E-09 | 1.800E-07 | 1.900E-07 |
| 9735 | 487 | L487P | A | 2.7e-07 | 2.9e-07 | 2.8e-07 | 2.800E-07 | 1.000E-08 | 2.700E-07 | 2.900E-07 |
| 9736 | 487 | L487S | A | 4.7e-07 | 5.3e-07 | 1.7e-07 | 3.900E-07 | 1.929E-07 | 1.700E-07 | 5.300E-07 |
| 9737 | 487 | L487T | A | 3.6e-07 | 3.9e-07 | 3.3e-07 | 3.600E-07 | 3.000E-08 | 3.300E-07 | 3.900E-07 |
| 9738 | 487 | L487W | A | 3.8e-07 | 2.1e-07 | 1.9e-07 | 2.600E-07 | 1.044E-07 | 1.900E-07 | 3.800E-07 |
| 9739 | 487 | L487Y | A | 9.2e-08 | 2.1e-07 | 3.5e-07 | 2.173E-07 | 1.292E-07 | 9.200E-08 | 3.500E-07 |
| 9740 | 487 | L487V | A | 2.4e-07 | 3.7e-07 | 4.2e-07 | 3.433E-07 | 9.292E-08 | 2.400E-07 | 4.200E-07 |
| 9741 | 488 | A488A | A | 3.3e-07 | 5.6e-07 | 4.3e-07 | 4.400E-07 | 1.153E-07 | 3.300E-07 | 5.600E-07 |
| 9742 | 488 | A488R | A | 2.6e-07 | 3.6e-07 | 1.9e-07 | 2.700E-07 | 8.544E-08 | 1.900E-07 | 3.600E-07 |
| 9743 | 488 | A488N | A | 2e-07 | 2.4e-07 | 3.2e-07 | 2.533E-07 | 6.110E-08 | 2.000E-07 | 3.200E-07 |
| 9744 | 488 | A488D | A | 2.3e-07 | 2.1e-07 | 2e-07 | 2.133E-07 | 1.528E-08 | 2.000E-07 | 2.300E-07 |
| 9745 | 488 | A488C | A | 2e-07 | 1.6e-07 | 1.6e-07 | 1.733E-07 | 2.309E-08 | 1.600E-07 | 2.000E-07 |
| 9746 | 488 | A488E | A | 1.5e-07 | 7.6e-07 | 3.7e-07 | 4.267E-07 | 3.089E-07 | 1.500E-07 | 7.600E-07 |
| 9747 | 488 | A488Q | A | 1.7e-07 | 3.6e-07 | 4e-07 | 3.100E-07 | 1.229E-07 | 1.700E-07 | 4.000E-07 |
| 9748 | 488 | A488G | A | 3.8e-07 | 2.8e-07 | 1.4e-07 | 2.667E-07 | 1.206E-07 | 1.400E-07 | 3.800E-07 |
| 9749 | 488 | A488H | A | 3.1e-07 | 1.5e-07 | 4.7e-07 | 3.100E-07 | 1.600E-07 | 1.500E-07 | 4.700E-07 |
| 9750 | 488 | A488I | A | 3.7e-07 | 2.8e-07 | 1.8e-07 | 2.767E-07 | 9.504E-08 | 1.800E-07 | 3.700E-07 |
| 9751 | 488 | A488L | A | 4.3e-07 | 2.5e-07 | 4.9e-07 | 3.900E-07 | 1.249E-07 | 2.500E-07 | 4.900E-07 |
| 9752 | 488 | A488K | A | 1.5e-07 | 2.2e-07 | 2.1e-07 | 1.933E-07 | 3.786E-08 | 1.500E-07 | 2.200E-07 |
| 9753 | 488 | A488M | A | 3.3e-07 | 1.4e-07 | 2e-07 | 2.233E-07 | 9.713E-08 | 1.400E-07 | 3.300E-07 |
| 9754 | 488 | A488F | A | 1.2e-07 | 4.1e-07 | 3.1e-07 | 2.800E-07 | 1.473E-07 | 1.200E-07 | 4.100E-07 |
| 9755 | 488 | A488P | A | 1e-07 | 1.9e-07 | 1.6e-07 | 1.500E-07 | 4.583E-08 | 1.000E-07 | 1.900E-07 |
| 9756 | 488 | A488S | A | 4.3e-07 | 2.5e-07 | 1.1e-07 | 2.633E-07 | 1.604E-07 | 1.100E-07 | 4.300E-07 |
| 9757 | 488 | A488T | A | 2.1e-07 | 2.5e-07 | 4.9e-07 | 3.167E-07 | 1.514E-07 | 2.100E-07 | 4.900E-07 |
| 9758 | 488 | A488W | A | 1.6e-07 | 3.8e-07 | 1.3e-07 | 2.233E-07 | 1.365E-07 | 1.300E-07 | 3.800E-07 |
| 9759 | 488 | A488Y | A | 2.4e-07 | 2.5e-07 | 2.5e-07 | 2.467E-07 | 5.774E-09 | 2.400E-07 | 2.500E-07 |
| 9760 | 488 | A488V | A | 1.9e-07 | 1.8e-07 | 1.5e-07 | 1.733E-07 | 2.082E-08 | 1.500E-07 | 1.900E-07 |
| 9761 | 489 | C489A | A | 2.8e-07 | 1.8e-07 | 1.9e-07 | 2.167E-07 | 5.508E-08 | 1.800E-07 | 2.800E-07 |
| 9762 | 489 | C489R | A | 2.7e-07 | 1.6e-07 | 1.9e-07 | 2.067E-07 | 5.686E-08 | 1.600E-07 | 2.700E-07 |
| 9763 | 489 | C489N | A | 1.8e-07 | 2.4e-07 | 2.2e-07 | 2.133E-07 | 3.055E-08 | 1.800E-07 | 2.400E-07 |
| 9764 | 489 | C489D | A | 3.2e-07 | 3.4e-07 | 3.4e-07 | 3.333E-07 | 1.155E-08 | 3.200E-07 | 3.400E-07 |
| 9765 | 489 | C489C | A | 5.7e-07 | 2.1e-07 | 1.4e-07 | 3.067E-07 | 2.307E-07 | 1.400E-07 | 5.700E-07 |
| 9766 | 489 | C489E | A | 1.8e-07 | 3.1e-07 | 2.1e-07 | 2.333E-07 | 6.807E-08 | 1.800E-07 | 3.100E-07 |
| 9767 | 489 | C489Q | A | 2.9e-07 | 3.6e-07 | 5.4e-07 | 3.967E-07 | 1.290E-07 | 2.900E-07 | 5.400E-07 |
| 9768 | 489 | C489G | A | 3e-07 | 3.3e-07 | 5.5e-07 | 3.933E-07 | 1.365E-07 | 3.000E-07 | 5.500E-07 |
| 9769 | 489 | C489H | A | 4e-07 | 2.4e-07 | 2.2e-07 | 2.867E-07 | 9.866E-08 | 2.200E-07 | 4.000E-07 |
| 9770 | 489 | C489I | A | 4.2e-07 | 3e-07 | 2.2e-07 | 3.133E-07 | 1.007E-07 | 2.200E-07 | 4.200E-07 |
| 9771 | 489 | C489L | A | 3.4e-07 | 3.3e-07 | 5.1e-07 | 3.933E-07 | 1.012E-07 | 3.300E-07 | 5.100E-07 |
| 9772 | 489 | C489K | A | 2.3e-07 | 2.2e-07 | 1.3e-07 | 1.933E-07 | 5.508E-08 | 1.300E-07 | 2.300E-07 |
| 9773 | 489 | C489M | A | 2.3e-07 | 9.2e-08 | 2.1e-07 | 1.773E-07 | 7.457E-08 | 9.200E-08 | 2.300E-07 |
| 9774 | 489 | C489F | A | 5.8e-07 | 2.6e-07 | 1.9e-07 | 3.433E-07 | 2.079E-07 | 1.900E-07 | 5.800E-07 |

| | | | | | | | | | | |
|---|---|---|---|---|---|---|---|---|---|---|
| 9775 | 489 | C489P | A | 2.3e-07 | 9.8e-08 | 1.5e-07 | 1.593E-07 | 6.649E-08 | 9.800E-08 | 2.300E-07 |
| 9776 | 489 | C489S | A | 2e-07 | 2.3e-07 | 3.5e-07 | 2.600E-07 | 7.937E-08 | 2.000E-07 | 3.500E-07 |
| 9777 | 489 | C489T | A | 2.6e-07 | 3e-07 | 2.9e-07 | 2.833E-07 | 2.082E-08 | 2.600E-07 | 3.000E-07 |
| 9778 | 489 | C489W | A | 1.8e-07 | 3.7e-07 | 5.3e-07 | 3.600E-07 | 1.752E-07 | 1.800E-07 | 5.300E-07 |
| 9779 | 489 | C489Y | A | 2.9e-07 | 2.3e-07 | 2e-07 | 2.400E-07 | 4.583E-08 | 2.000E-07 | 2.900E-07 |
| 9780 | 489 | C489V | A | 3.3e-07 | 2.6e-07 | 4.7e-07 | 3.533E-07 | 1.069E-07 | 2.600E-07 | 4.700E-07 |
| 9781 | 490 | H490A | A | 2.5e-07 | 5.1e-07 | 3.9e-07 | 3.833E-07 | 1.301E-07 | 2.500E-07 | 5.100E-07 |
| 9782 | 490 | H490R | A | 1.9e-07 | 1.5e-07 | 1.9e-07 | 1.767E-07 | 2.309E-08 | 1.500E-07 | 1.900E-07 |
| 9783 | 490 | H490N | A | 3.8e-07 | 1.2e-07 | 2.6e-07 | 2.533E-07 | 1.301E-07 | 1.200E-07 | 3.800E-07 |
| 9784 | 490 | H490D | A | 4.1e-07 | 2.2e-07 | 2.5e-07 | 2.933E-07 | 1.021E-07 | 2.200E-07 | 4.100E-07 |
| 9785 | 490 | H490C | A | 2.9e-07 | 5.5e-07 | 3.9e-07 | 4.100E-07 | 1.311E-07 | 2.900E-07 | 5.500E-07 |
| 9786 | 490 | H490E | A | 3.2e-07 | 2.1e-07 | 2.5e-07 | 2.600E-07 | 5.568E-08 | 2.100E-07 | 3.200E-07 |
| 9787 | 490 | H490Q | A | 1.7e-07 | 1.6e-07 | 2.8e-07 | 2.033E-07 | 6.658E-08 | 1.600E-07 | 2.800E-07 |
| 9788 | 490 | H490G | A | 1.6e-07 | 3.4e-07 | 2.1e-07 | 2.367E-07 | 9.292E-08 | 1.600E-07 | 3.400E-07 |
| 9789 | 490 | H490H | A | 3.2e-07 | 4.4e-07 | 2.7e-07 | 3.433E-07 | 8.737E-08 | 2.700E-07 | 4.400E-07 |
| 9790 | 490 | H490I | A | 3.4e-07 | 2.6e-07 | 1.8e-07 | 2.600E-07 | 8.000E-08 | 1.800E-07 | 3.400E-07 |
| 9791 | 490 | H490L | A | 1.3e-07 | 4.5e-07 | 5.8e-07 | 3.867E-07 | 2.316E-07 | 1.300E-07 | 5.800E-07 |
| 9792 | 490 | H490K | A | 5.2e-07 | 2.4e-07 | 2.9e-07 | 3.500E-07 | 1.493E-07 | 2.400E-07 | 5.200E-07 |
| 9793 | 490 | H490M | A | 3.1e-07 | 3.5e-07 | 2.3e-07 | 2.967E-07 | 6.110E-08 | 2.300E-07 | 3.500E-07 |
| 9794 | 490 | H490F | A | 1.4e-07 | 4.6e-07 | 5e-07 | 3.667E-07 | 1.973E-07 | 1.400E-07 | 5.000E-07 |
| 9795 | 490 | H490P | A | 1.5e-07 | 2.9e-07 | 3.7e-07 | 2.700E-07 | 1.114E-07 | 1.500E-07 | 3.700E-07 |
| 9796 | 490 | H490S | A | 2.7e-07 | 1.6e-07 | 1.7e-07 | 2.000E-07 | 6.083E-08 | 1.600E-07 | 2.700E-07 |
| 9797 | 490 | H490T | A | 2.7e-07 | 2.3e-07 | 2.5e-07 | 2.500E-07 | 2.000E-08 | 2.300E-07 | 2.700E-07 |
| 9798 | 490 | H490W | A | 4.1e-07 | 2.8e-07 | 2.8e-07 | 3.233E-07 | 7.506E-08 | 2.800E-07 | 4.100E-07 |
| 9799 | 490 | H490Y | A | 2e-07 | 1.9e-07 | 9.6e-08 | 1.620E-07 | 5.738E-08 | 9.600E-08 | 2.000E-07 |
| 9800 | 490 | H490V | A | 1.7e-07 | 1.4e-07 | 3.5e-07 | 2.200E-07 | 1.136E-07 | 1.400E-07 | 3.500E-07 |
| 9801 | 491 | Q491A | A | 3.1e-07 | 3.2e-07 | 3.9e-07 | 3.400E-07 | 4.359E-08 | 3.100E-07 | 3.900E-07 |
| 9802 | 491 | Q491R | A | 1.6e-07 | 4e-07 | 2.5e-07 | 2.700E-07 | 1.212E-07 | 1.600E-07 | 4.000E-07 |
| 9803 | 491 | Q491N | A | 5.3e-07 | 1.9e-07 | 2e-07 | 3.067E-07 | 1.935E-07 | 1.900E-07 | 5.300E-07 |
| 9804 | 491 | Q491D | A | 3.5e-07 | 2.1e-07 | 2.2e-07 | 2.600E-07 | 7.810E-08 | 2.100E-07 | 3.500E-07 |
| 9805 | 491 | Q491C | A | 4.5e-07 | 2.7e-07 | 1.9e-07 | 3.033E-07 | 1.332E-07 | 1.900E-07 | 4.500E-07 |
| 9806 | 491 | Q491E | A | 4.1e-07 | 1.5e-07 | 3.2e-07 | 2.933E-07 | 1.320E-07 | 1.500E-07 | 4.100E-07 |
| 9807 | 491 | Q491Q | A | 5.5e-07 | 2.8e-07 | 2.8e-07 | 3.700E-07 | 1.559E-07 | 2.800E-07 | 5.500E-07 |
| 9808 | 491 | Q491G | A | 1.6e-07 | 2e-07 | 5.9e-07 | 3.167E-07 | 2.376E-07 | 1.600E-07 | 5.900E-07 |
| 9809 | 491 | Q491H | A | 5.5e-07 | 2.1e-07 | 3.5e-07 | 3.700E-07 | 1.709E-07 | 2.100E-07 | 5.500E-07 |
| 9810 | 491 | Q491I | A | 2.1e-07 | 9.5e-08 | 4.8e-07 | 2.617E-07 | 1.976E-07 | 9.500E-08 | 4.800E-07 |
| 9811 | 491 | Q491L | A | 4.3e-07 | 5.6e-07 | 5.7e-07 | 5.200E-07 | 7.810E-08 | 4.300E-07 | 5.700E-07 |
| 9812 | 491 | Q491K | A | 2.2e-07 | 2.8e-07 | 2e-07 | 2.333E-07 | 4.163E-08 | 2.000E-07 | 2.800E-07 |
| 9813 | 491 | Q491M | A | 2.1e-07 | 1.7e-07 | 3.3e-07 | 2.367E-07 | 8.327E-08 | 1.700E-07 | 3.300E-07 |
| 9814 | 491 | Q491F | A | 3.8e-07 | 4.6e-07 | 1.3e-07 | 3.233E-07 | 1.721E-07 | 1.300E-07 | 4.600E-07 |
| 9815 | 491 | Q491P | A | 4.4e-07 | 4.5e-07 | 2.1e-07 | 3.667E-07 | 1.358E-07 | 2.100E-07 | 4.500E-07 |
| 9816 | 491 | Q491S | A | 3.4e-07 | 1.6e-07 | 2.5e-07 | 2.500E-07 | 9.000E-08 | 1.600E-07 | 3.400E-07 |
| 9817 | 491 | Q491T | A | 2.2e-07 | 2.2e-07 | 5e-07 | 3.133E-07 | 1.617E-07 | 2.200E-07 | 5.000E-07 |
| 9818 | 491 | Q491W | A | 1.7e-07 | 2.7e-07 | 1.9e-07 | 2.100E-07 | 5.292E-08 | 1.700E-07 | 2.700E-07 |
| 9819 | 491 | Q491Y | A | 4e-07 | 8.4e-08 | 2.1e-07 | 2.313E-07 | 1.591E-07 | 8.400E-08 | 4.000E-07 |
| 9820 | 491 | Q491V | A | 4e-07 | 1.4e-07 | 3.9e-07 | 3.100E-07 | 1.473E-07 | 1.400E-07 | 4.000E-07 |
| 9821 | 492 | L492A | A | 1.5e-07 | 1.3e-07 | 1.9e-07 | 1.567E-07 | 3.055E-08 | 1.300E-07 | 1.900E-07 |
| 9822 | 492 | L492R | A | 7.2e-07 | 1.7e-07 | 1.2e-07 | 3.367E-07 | 3.329E-07 | 1.200E-07 | 7.200E-07 |
| 9823 | 492 | L492N | A | 5.7e-07 | 3e-07 | 2.5e-07 | 3.733E-07 | 1.721E-07 | 2.500E-07 | 5.700E-07 |
| 9824 | 492 | L492D | A | 1.6e-07 | 2.5e-07 | 1.2e-07 | 1.767E-07 | 6.658E-08 | 1.200E-07 | 2.500E-07 |
| 9825 | 492 | L492C | A | 1.6e-07 | 4.1e-07 | 2.4e-07 | 2.700E-07 | 1.277E-07 | 1.600E-07 | 4.100E-07 |
| 9826 | 492 | L492E | A | 1.7e-07 | 2.4e-07 | 4.7e-07 | 2.933E-07 | 1.570E-07 | 1.700E-07 | 4.700E-07 |
| 9827 | 492 | L492Q | A | 4.4e-07 | 3.9e-07 | 1.1e-07 | 3.133E-07 | 1.779E-07 | 1.100E-07 | 4.400E-07 |
| 9828 | 492 | L492G | A | 2.4e-07 | 3.8e-07 | 4.5e-07 | 3.567E-07 | 1.069E-07 | 2.400E-07 | 4.500E-07 |
| 9829 | 492 | L492H | A | 2.8e-07 | 2.7e-07 | 1.4e-07 | 2.300E-07 | 7.810E-08 | 1.400E-07 | 2.800E-07 |
| 9830 | 492 | L492I | A | 2.9e-07 | 3.4e-07 | 3.1e-07 | 3.133E-07 | 2.517E-08 | 2.900E-07 | 3.400E-07 |
| 9831 | 492 | L492L | A | 1.6e-07 | 3.5e-07 | 4.5e-07 | 3.200E-07 | 1.473E-07 | 1.600E-07 | 4.500E-07 |
| 9832 | 492 | L492K | A | 3.2e-07 | 1.6e-07 | 1.6e-07 | 2.133E-07 | 9.238E-08 | 1.600E-07 | 3.200E-07 |
| 9833 | 492 | L492M | A | 2.2e-07 | 2.6e-07 | 4e-07 | 2.933E-07 | 9.452E-08 | 2.200E-07 | 4.000E-07 |
| 9834 | 492 | L492F | A | 2.9e-07 | 1.7e-07 | 2.4e-07 | 2.333E-07 | 6.028E-08 | 1.700E-07 | 2.900E-07 |
| 9835 | 492 | L492P | A | 1.4e-07 | 1.5e-07 | 1.9e-07 | 1.600E-07 | 2.646E-08 | 1.400E-07 | 1.900E-07 |
| 9836 | 492 | L492S | A | 2.2e-07 | 2.6e-07 | 2.2e-07 | 2.333E-07 | 2.309E-08 | 2.200E-07 | 2.600E-07 |
| 9837 | 492 | L492T | A | 2.8e-07 | 6.3e-07 | 1.8e-07 | 3.633E-07 | 2.363E-07 | 1.800E-07 | 6.300E-07 |
| 9838 | 492 | L492W | A | 1.4e-07 | 1.3e-07 | 2.9e-07 | 1.867E-07 | 8.963E-08 | 1.300E-07 | 2.900E-07 |
| 9839 | 492 | L492Y | A | 3.2e-07 | 2.7e-07 | 3.4e-07 | 3.100E-07 | 3.606E-08 | 2.700E-07 | 3.400E-07 |
| 9840 | 492 | L492V | A | 5.5e-07 | 2e-07 | 3e-07 | 3.500E-07 | 1.803E-07 | 2.000E-07 | 5.500E-07 |
| 9841 | 493 | C493A | A | 3e-07 | 2.3e-07 | 2.4e-07 | 2.567E-07 | 3.786E-08 | 2.300E-07 | 3.000E-07 |
| 9842 | 493 | C493R | A | 2e-07 | 1.5e-07 | 1.4e-07 | 1.633E-07 | 3.215E-08 | 1.400E-07 | 2.000E-07 |
| 9843 | 493 | C493N | A | 5.2e-07 | 3.5e-07 | 2.8e-07 | 3.833E-07 | 1.234E-07 | 2.800E-07 | 5.200E-07 |
| 9844 | 493 | C493D | A | 5.4e-07 | 2.5e-07 | 2.8e-07 | 3.567E-07 | 1.595E-07 | 2.500E-07 | 5.400E-07 |
| 9845 | 493 | C493C | A | 6.3e-07 | 2.2e-07 | 2.3e-07 | 3.600E-07 | 2.339E-07 | 2.200E-07 | 6.300E-07 |
| 9846 | 493 | C493E | A | 5.6e-07 | 2e-07 | 2e-07 | 3.200E-07 | 2.078E-07 | 2.000E-07 | 5.600E-07 |
| 9847 | 493 | C493Q | A | 2.1e-07 | 3.6e-07 | 3.9e-07 | 3.200E-07 | 9.644E-08 | 2.100E-07 | 3.900E-07 |
| 9848 | 493 | C493G | A | 2.9e-07 | 5.6e-07 | 3.4e-07 | 3.967E-07 | 1.436E-07 | 2.900E-07 | 5.600E-07 |
| 9849 | 493 | C493H | A | 1.9e-07 | 4.6e-07 | 3.7e-07 | 3.400E-07 | 1.375E-07 | 1.900E-07 | 4.600E-07 |
| 9850 | 493 | C493I | A | 4.7e-07 | 7.9e-08 | 3.9e-07 | 3.130E-07 | 2.066E-07 | 7.900E-08 | 4.700E-07 |
| 9851 | 493 | C493L | A | 2e-07 | 3.9e-07 | 3.1e-07 | 3.000E-07 | 9.539E-08 | 2.000E-07 | 3.900E-07 |
| 9852 | 493 | C493K | A | 2.6e-07 | 1.8e-07 | 2e-07 | 2.133E-07 | 4.163E-08 | 1.800E-07 | 2.600E-07 |
| 9853 | 493 | C493M | A | 3.9e-07 | 2.3e-07 | 1.4e-07 | 2.533E-07 | 1.266E-07 | 1.400E-07 | 3.900E-07 |
| 9854 | 493 | C493F | A | 2.9e-07 | 1.9e-07 | 3.3e-07 | 2.700E-07 | 7.211E-08 | 1.900E-07 | 3.300E-07 |
| 9855 | 493 | C493P | A | 4.8e-07 | 1.4e-07 | 3.2e-07 | 3.133E-07 | 1.701E-07 | 1.400E-07 | 4.800E-07 |
| 9856 | 493 | C493S | A | 6.5e-07 | 5.7e-07 | 1.8e-07 | 4.667E-07 | 2.515E-07 | 1.800E-07 | 6.500E-07 |
| 9857 | 493 | C493T | A | 3.4e-07 | 2.8e-07 | 1.5e-07 | 2.567E-07 | 9.713E-08 | 1.500E-07 | 3.400E-07 |
| 9858 | 493 | C493W | A | 1.9e-07 | 1.6e-07 | 1.4e-07 | 1.633E-07 | 2.517E-08 | 1.400E-07 | 1.900E-07 |
| 9859 | 493 | C493Y | A | 3.7e-07 | 3.1e-07 | 2.7e-07 | 3.167E-07 | 5.033E-08 | 2.700E-07 | 3.700E-07 |

| | | | | | | | | | | |
|---|---|---|---|---|---|---|---|---|---|---|
| 9860 | 493 | C493V | A | 3.5e-07 | 1.8e-07 | 4.5e-07 | 3.267E-07 | 1.365E-07 | 1.800E-07 | 4.500E-07 |
| 9861 | 494 | A494A | A | 4e-07 | 4.3e-07 | 2.4e-07 | 3.567E-07 | 1.021E-07 | 2.400E-07 | 4.300E-07 |
| 9862 | 494 | A494R | A | 1.6e-07 | 3.3e-07 | 2.1e-07 | 2.333E-07 | 8.737E-08 | 1.600E-07 | 3.300E-07 |
| 9863 | 494 | A494N | A | 2.1e-07 | 6.4e-07 | 2.8e-07 | 3.767E-07 | 2.307E-07 | 2.100E-07 | 6.400E-07 |
| 9864 | 494 | A494D | A | 1.7e-07 | 4.2e-07 | 2.1e-07 | 2.667E-07 | 1.343E-07 | 1.700E-07 | 4.200E-07 |
| 9865 | 494 | A494C | A | 1.4e-07 | 4.5e-07 | 1.8e-07 | 2.567E-07 | 1.686E-07 | 1.400E-07 | 4.500E-07 |
| 9866 | 494 | A494E | A | 2.2e-07 | 2.6e-07 | 5.4e-07 | 3.400E-07 | 1.744E-07 | 2.200E-07 | 5.400E-07 |
| 9867 | 494 | A494Q | A | 2.2e-07 | 2.1e-07 | 1.9e-07 | 2.067E-07 | 1.528E-08 | 1.900E-07 | 2.200E-07 |
| 9868 | 494 | A494G | A | 6.6e-07 | 8.1e-07 | 2.4e-07 | 5.700E-07 | 2.955E-07 | 2.400E-07 | 8.100E-07 |
| 9869 | 494 | A494H | A | 2.6e-07 | 2.1e-07 | 1.3e-07 | 2.000E-07 | 6.557E-08 | 1.300E-07 | 2.600E-07 |
| 9870 | 494 | A494I | A | 5.6e-07 | 3.9e-07 | 3.1e-07 | 4.200E-07 | 1.277E-07 | 3.100E-07 | 5.600E-07 |
| 9871 | 494 | A494L | A | 1.5e-07 | 4.4e-07 | 2.5e-07 | 2.800E-07 | 1.473E-07 | 1.500E-07 | 4.400E-07 |
| 9872 | 494 | A494K | A | 2e-07 | 2.7e-07 | 4.5e-07 | 3.067E-07 | 1.290E-07 | 2.000E-07 | 4.500E-07 |
| 9873 | 494 | A494M | A | 2.6e-07 | 4e-07 | 2.5e-07 | 3.033E-07 | 8.386E-08 | 2.500E-07 | 4.000E-07 |
| 9874 | 494 | A494F | A | 4.4e-07 | 2.9e-07 | 1.4e-07 | 2.900E-07 | 1.500E-07 | 1.400E-07 | 4.400E-07 |
| 9875 | 494 | A494P | A | 2.6e-07 | 3.3e-07 | 3.2e-07 | 3.033E-07 | 3.786E-08 | 2.600E-07 | 3.300E-07 |
| 9876 | 494 | A494S | A | 3.6e-07 | 3.4e-07 | 2.9e-07 | 3.300E-07 | 3.606E-08 | 2.900E-07 | 3.600E-07 |
| 9877 | 494 | A494T | A | 3.1e-07 | 2.6e-07 | 4.7e-07 | 3.467E-07 | 1.097E-07 | 2.600E-07 | 4.700E-07 |
| 9878 | 494 | A494W | A | 4.8e-07 | 2.5e-07 | 3.5e-07 | 3.600E-07 | 1.153E-07 | 2.500E-07 | 4.800E-07 |
| 9879 | 494 | A494Y | A | 2.2e-07 | 2.6e-07 | 2.7e-07 | 2.500E-07 | 2.646E-08 | 2.200E-07 | 2.700E-07 |
| 9880 | 494 | A494V | A | 1.3e-07 | 2.3e-07 | 1.5e-07 | 1.700E-07 | 5.292E-08 | 1.300E-07 | 2.300E-07 |
| 9881 | 495 | R495A | A | 1.2e-07 | 3.4e-07 | 5.2e-07 | 3.267E-07 | 2.003E-07 | 1.200E-07 | 5.200E-07 |
| 9882 | 495 | R495R | A | 2.4e-07 | 1.5e-07 | 2.1e-07 | 2.000E-07 | 4.583E-08 | 1.500E-07 | 2.400E-07 |
| 9883 | 495 | R495N | A | 3e-07 | 5e-07 | 5.5e-07 | 4.500E-07 | 1.323E-07 | 3.000E-07 | 5.500E-07 |
| 9884 | 495 | R495D | A | 3.9e-07 | 2e-07 | 2.1e-07 | 2.667E-07 | 1.069E-07 | 2.000E-07 | 3.900E-07 |
| 9885 | 495 | R495C | A | 1.8e-07 | 1.5e-07 | 3.4e-07 | 2.233E-07 | 1.021E-07 | 1.500E-07 | 3.400E-07 |
| 9886 | 495 | R495E | A | 2.1e-07 | 6.2e-07 | 3.2e-07 | 3.833E-07 | 2.122E-07 | 2.100E-07 | 6.200E-07 |
| 9887 | 495 | R495Q | A | 3.5e-07 | 2.3e-07 | 1.5e-07 | 2.433E-07 | 1.007E-07 | 1.500E-07 | 3.500E-07 |
| 9888 | 495 | R495G | A | 3.3e-07 | 1.3e-07 | 2.9e-07 | 2.500E-07 | 1.058E-07 | 1.300E-07 | 3.300E-07 |
| 9889 | 495 | R495H | A | 2.4e-07 | 3.9e-07 | 2.7e-07 | 3.000E-07 | 7.937E-08 | 2.400E-07 | 3.900E-07 |
| 9890 | 495 | R495I | A | 3.3e-07 | 2.6e-07 | 2e-07 | 2.633E-07 | 6.506E-08 | 2.000E-07 | 3.300E-07 |
| 9891 | 495 | R495L | A | 3.3e-07 | 5.4e-07 | 2.2e-07 | 3.633E-07 | 1.626E-07 | 2.200E-07 | 5.400E-07 |
| 9892 | 495 | R495K | A | 2.3e-07 | 1.8e-07 | 2.5e-07 | 2.200E-07 | 3.606E-08 | 1.800E-07 | 2.500E-07 |
| 9893 | 495 | R495M | A | 2.7e-07 | 4e-07 | 3.8e-07 | 3.500E-07 | 7.000E-08 | 2.700E-07 | 4.000E-07 |
| 9894 | 495 | R495F | A | 1.9e-07 | 4.1e-07 | 3.2e-07 | 3.067E-07 | 1.106E-07 | 1.900E-07 | 4.100E-07 |
| 9895 | 495 | R495P | A | 6.7e-08 | 3e-07 | 1.7e-07 | 1.790E-07 | 1.168E-07 | 6.700E-08 | 3.000E-07 |
| 9896 | 495 | R495S | A | 9.3e-08 | 2.2e-07 | 1.8e-07 | 1.643E-07 | 6.493E-08 | 9.300E-08 | 2.200E-07 |
| 9897 | 495 | R495T | A | 1.7e-07 | 2.1e-07 | 3.9e-07 | 2.567E-07 | 1.172E-07 | 1.700E-07 | 3.900E-07 |
| 9898 | 495 | R495W | A | 4.1e-07 | 2.5e-07 | 2.5e-07 | 3.033E-07 | 9.238E-08 | 2.500E-07 | 4.100E-07 |
| 9899 | 495 | R495Y | A | 1.8e-07 | 2e-07 | 4.3e-07 | 2.700E-07 | 1.389E-07 | 1.800E-07 | 4.300E-07 |
| 9900 | 495 | R495V | A | 1.8e-07 | 2.4e-07 | 1.6e-07 | 1.933E-07 | 4.163E-08 | 1.600E-07 | 2.400E-07 |
| 9901 | 496 | G496A | A | 2e-07 | 3.3e-07 | 1.7e-07 | 2.333E-07 | 8.505E-08 | 1.700E-07 | 3.300E-07 |
| 9902 | 496 | G496R | A | 1.5e-07 | 2.1e-07 | 2.3e-07 | 1.967E-07 | 4.163E-08 | 1.500E-07 | 2.300E-07 |
| 9903 | 496 | G496N | A | 1.7e-07 | 5.1e-07 | 2.1e-07 | 2.967E-07 | 1.858E-07 | 1.700E-07 | 5.100E-07 |
| 9904 | 496 | G496D | A | 3.6e-07 | 1.3e-07 | 3e-07 | 2.633E-07 | 1.193E-07 | 1.300E-07 | 3.600E-07 |
| 9905 | 496 | G496C | A | 5.5e-07 | 2e-07 | 1.5e-07 | 3.000E-07 | 2.179E-07 | 1.500E-07 | 5.500E-07 |
| 9906 | 496 | G496E | A | 1.9e-07 | 1.6e-07 | 3e-07 | 2.167E-07 | 7.371E-08 | 1.600E-07 | 3.000E-07 |
| 9907 | 496 | G496Q | A | 2.5e-07 | 1.1e-07 | 3.7e-07 | 2.433E-07 | 1.301E-07 | 1.100E-07 | 3.700E-07 |
| 9908 | 496 | G496G | A | 1.4e-07 | 2.8e-07 | 5.6e-07 | 3.267E-07 | 2.139E-07 | 1.400E-07 | 5.600E-07 |
| 9909 | 496 | G496H | A | 2.8e-07 | 2.1e-07 | 1.5e-07 | 2.133E-07 | 6.506E-08 | 1.500E-07 | 2.800E-07 |
| 9910 | 496 | G496I | A | 3.7e-07 | 2e-07 | 2e-07 | 2.567E-07 | 9.815E-08 | 2.000E-07 | 3.700E-07 |
| 9911 | 496 | G496L | A | 2.7e-07 | 3.2e-07 | 2.2e-07 | 2.700E-07 | 5.000E-08 | 2.200E-07 | 3.200E-07 |
| 9912 | 496 | G496K | A | 3.6e-07 | 1.3e-07 | 3.1e-07 | 2.667E-07 | 1.210E-07 | 1.300E-07 | 3.600E-07 |
| 9913 | 496 | G496M | A | 3.3e-07 | 3.5e-07 | 2.5e-07 | 3.100E-07 | 5.292E-08 | 2.500E-07 | 3.500E-07 |
| 9914 | 496 | G496F | A | 4.8e-07 | 5.1e-07 | 3e-07 | 4.300E-07 | 1.136E-07 | 3.000E-07 | 5.100E-07 |
| 9915 | 496 | G496P | A | 1.5e-07 | 3.7e-07 | 4.8e-07 | 3.333E-07 | 1.680E-07 | 1.500E-07 | 4.800E-07 |
| 9916 | 496 | G496S | A | 3.2e-07 | 5e-07 | 2.3e-07 | 3.500E-07 | 1.375E-07 | 2.300E-07 | 5.000E-07 |
| 9917 | 496 | G496T | A | 3e-07 | 1.4e-07 | 3.3e-07 | 2.567E-07 | 1.021E-07 | 1.400E-07 | 3.300E-07 |
| 9918 | 496 | G496W | A | 3.7e-07 | 4.2e-07 | 2.1e-07 | 3.333E-07 | 1.097E-07 | 2.100E-07 | 4.200E-07 |
| 9919 | 496 | G496Y | A | 3.5e-07 | 1.7e-07 | 2.8e-07 | 2.667E-07 | 9.074E-08 | 1.700E-07 | 3.500E-07 |
| 9920 | 496 | G496V | A | 3.9e-07 | 6.2e-07 | 2.2e-07 | 4.100E-07 | 2.007E-07 | 2.200E-07 | 6.200E-07 |
| 9921 | 497 | H497A | A | 5e-07 | 2.6e-07 | 1e-07 | 2.867E-07 | 2.013E-07 | 1.000E-07 | 5.000E-07 |
| 9922 | 497 | H497R | A | 2.7e-07 | 2e-07 | 8.1e-07 | 4.267E-07 | 3.338E-07 | 2.000E-07 | 8.100E-07 |
| 9923 | 497 | H497N | A | 1.5e-07 | 3.1e-07 | 2.5e-07 | 2.367E-07 | 8.083E-08 | 1.500E-07 | 3.100E-07 |
| 9924 | 497 | H497D | A | 2.4e-07 | 1.2e-07 | 2.5e-07 | 2.033E-07 | 7.234E-08 | 1.200E-07 | 2.500E-07 |
| 9925 | 497 | H497C | A | 4.5e-07 | 3.3e-07 | 1.5e-07 | 3.100E-07 | 1.510E-07 | 1.500E-07 | 4.500E-07 |
| 9926 | 497 | H497E | A | 1.9e-07 | 1.9e-07 | 4.1e-07 | 2.633E-07 | 1.270E-07 | 1.900E-07 | 4.100E-07 |
| 9927 | 497 | H497Q | A | 2.9e-07 | 3.3e-07 | 2.6e-07 | 2.933E-07 | 3.512E-08 | 2.600E-07 | 3.300E-07 |
| 9928 | 497 | H497G | A | 2.2e-07 | 2.2e-07 | 5.5e-07 | 3.300E-07 | 1.905E-07 | 2.200E-07 | 5.500E-07 |
| 9929 | 497 | H497H | A | 2.6e-07 | 3.3e-07 | 3.1e-07 | 3.000E-07 | 3.606E-08 | 2.600E-07 | 3.300E-07 |
| 9930 | 497 | H497I | A | 3e-07 | 1.5e-07 | 2.3e-07 | 2.267E-07 | 7.506E-08 | 1.500E-07 | 3.000E-07 |
| 9931 | 497 | H497L | A | 1.5e-07 | 2.9e-07 | 1.9e-07 | 2.100E-07 | 7.211E-08 | 1.500E-07 | 2.900E-07 |
| 9932 | 497 | H497K | A | 3.7e-07 | 2.7e-07 | 4.2e-07 | 3.533E-07 | 7.638E-08 | 2.700E-07 | 4.200E-07 |
| 9933 | 497 | H497M | A | 3.3e-07 | 1.6e-07 | 2.9e-07 | 2.600E-07 | 8.888E-08 | 1.600E-07 | 3.300E-07 |
| 9934 | 497 | H497F | A | 2.8e-07 | 1.7e-07 | 3.1e-07 | 2.533E-07 | 7.371E-08 | 1.700E-07 | 3.100E-07 |
| 9935 | 497 | H497P | A | 2.4e-07 | 2.6e-07 | 2.2e-07 | 2.400E-07 | 2.000E-08 | 2.200E-07 | 2.600E-07 |
| 9936 | 497 | H497S | A | 7.3e-08 | 3e-07 | 2.6e-07 | 2.110E-07 | 1.212E-07 | 7.300E-08 | 3.000E-07 |
| 9937 | 497 | H497T | A | 1.6e-07 | 1.4e-07 | 3e-07 | 2.000E-07 | 8.718E-08 | 1.400E-07 | 3.000E-07 |
| 9938 | 497 | H497W | A | 3.2e-07 | 5.3e-07 | 3.8e-07 | 4.100E-07 | 1.082E-07 | 3.200E-07 | 5.300E-07 |
| 9939 | 497 | H497Y | A | 2.7e-07 | 2.7e-07 | 1.7e-07 | 2.367E-07 | 5.774E-08 | 1.700E-07 | 2.700E-07 |
| 9940 | 497 | H497V | A | 3.1e-07 | 2.3e-07 | 2e-07 | 2.467E-07 | 5.686E-08 | 2.000E-07 | 3.100E-07 |
| 9941 | 498 | C498A | A | 3.8e-07 | 3.9e-07 | 2.1e-07 | 3.267E-07 | 1.012E-07 | 2.100E-07 | 3.900E-07 |
| 9942 | 498 | C498R | A | 1.2e-07 | 1.7e-07 | 3e-07 | 1.967E-07 | 9.292E-08 | 1.200E-07 | 3.000E-07 |
| 9943 | 498 | C498N | A | 2.4e-07 | 1.9e-07 | 2.8e-07 | 2.367E-07 | 4.509E-08 | 1.900E-07 | 2.800E-07 |
| 9944 | 498 | C498D | A | 1.4e-07 | 3.5e-07 | 1.9e-07 | 2.267E-07 | 1.097E-07 | 1.400E-07 | 3.500E-07 |

| | | | | | | | | | | |
|---|---|---|---|---|---|---|---|---|---|---|
| 9945 | 498 | C498C | A | 4.2e-07 | 3.5e-07 | 2.1e-07 | 3.267E-07 | 1.069E-07 | 2.100E-07 | 4.200E-07 |
| 9946 | 498 | C498E | A | 1.2e-07 | 2.3e-07 | 2.4e-07 | 1.967E-07 | 6.658E-08 | 1.200E-07 | 2.400E-07 |
| 9947 | 498 | C498Q | A | 2.2e-07 | 2.2e-07 | 2e-07 | 2.133E-07 | 1.155E-08 | 2.000E-07 | 2.200E-07 |
| 9948 | 498 | C498G | A | 3.1e-07 | 2.8e-07 | 3.6e-07 | 3.167E-07 | 4.041E-08 | 2.800E-07 | 3.600E-07 |
| 9949 | 498 | C498H | A | 3.2e-07 | 2.2e-07 | 2.5e-07 | 2.633E-07 | 5.132E-08 | 2.200E-07 | 3.200E-07 |
| 9950 | 498 | C498I | A | 1e-07 | 3.9e-07 | 3.8e-07 | 2.900E-07 | 1.646E-07 | 1.000E-07 | 3.900E-07 |
| 9951 | 498 | C498L | A | 1.6e-07 | 1.3e-07 | 1e-07 | 1.300E-07 | 3.000E-08 | 1.000E-07 | 1.600E-07 |
| 9952 | 498 | C498K | A | 2e-07 | 3.8e-07 | 1.2e-07 | 2.333E-07 | 1.332E-07 | 1.200E-07 | 3.800E-07 |
| 9953 | 498 | C498M | A | 2.3e-07 | 2.8e-07 | 1.7e-07 | 2.267E-07 | 5.508E-08 | 1.700E-07 | 2.800E-07 |
| 9954 | 498 | C498F | A | 8e-07 | 1.1e-07 | 1.8e-07 | 3.633E-07 | 3.798E-07 | 1.100E-07 | 8.000E-07 |
| 9955 | 498 | C498P | A | 2.7e-07 | 2.2e-07 | 2.2e-07 | 2.367E-07 | 2.887E-08 | 2.200E-07 | 2.700E-07 |
| 9956 | 498 | C498S | A | 2.3e-07 | 4.3e-07 | 2.3e-07 | 2.967E-07 | 1.155E-07 | 2.300E-07 | 4.300E-07 |
| 9957 | 498 | C498T | A | 4.4e-07 | 2.3e-07 | 3.4e-07 | 3.367E-07 | 1.050E-07 | 2.300E-07 | 4.400E-07 |
| 9958 | 498 | C498W | A | 3.5e-07 | 1.1e-07 | 2.2e-07 | 2.267E-07 | 1.201E-07 | 1.100E-07 | 3.500E-07 |
| 9959 | 498 | C498Y | A | 3.3e-07 | 2e-07 | 1.2e-07 | 2.167E-07 | 1.060E-07 | 1.200E-07 | 3.300E-07 |
| 9960 | 498 | C498V | A | 2e-07 | 2.1e-07 | 2.5e-07 | 2.200E-07 | 2.646E-08 | 2.000E-07 | 2.500E-07 |
| 9961 | 499 | W499A | A | 4e-07 | 1.2e-07 | 3e-07 | 2.733E-07 | 1.419E-07 | 1.200E-07 | 4.000E-07 |
| 9962 | 499 | W499R | A | 1.6e-07 | 2.8e-07 | 1.5e-07 | 1.967E-07 | 7.234E-08 | 1.500E-07 | 2.800E-07 |
| 9963 | 499 | W499N | A | 2.1e-07 | 3.1e-07 | 1.5e-07 | 2.233E-07 | 8.083E-08 | 1.500E-07 | 3.100E-07 |
| 9964 | 499 | W499D | A | 3.8e-07 | 2.1e-07 | 1.3e-07 | 2.400E-07 | 1.277E-07 | 1.300E-07 | 3.800E-07 |
| 9965 | 499 | W499C | A | 2.9e-07 | 2e-07 | 2.5e-07 | 2.467E-07 | 4.509E-08 | 2.000E-07 | 2.900E-07 |
| 9966 | 499 | W499E | A | 2.6e-07 | 5.7e-07 | 1.3e-07 | 3.200E-07 | 2.261E-07 | 1.300E-07 | 5.700E-07 |
| 9967 | 499 | W499Q | A | 2e-07 | 3.1e-07 | 2.1e-07 | 2.400E-07 | 6.083E-08 | 2.000E-07 | 3.100E-07 |
| 9968 | 499 | W499G | A | 3.9e-07 | 1.9e-07 | 2.5e-07 | 2.767E-07 | 1.026E-07 | 1.900E-07 | 3.900E-07 |
| 9969 | 499 | W499H | A | 2e-07 | 2.7e-07 | 5.1e-07 | 3.267E-07 | 1.626E-07 | 2.000E-07 | 5.100E-07 |
| 9970 | 499 | W499I | A | 3.5e-07 | 4.1e-07 | 3.3e-07 | 3.633E-07 | 4.163E-08 | 3.300E-07 | 4.100E-07 |
| 9971 | 499 | W499L | A | 1.9e-07 | 3.8e-07 | 2.4e-07 | 2.700E-07 | 9.849E-08 | 1.900E-07 | 3.800E-07 |
| 9972 | 499 | W499K | A | 2.3e-07 | 2.5e-07 | 4.1e-07 | 2.967E-07 | 9.866E-08 | 2.300E-07 | 4.100E-07 |
| 9973 | 499 | W499M | A | 2e-07 | 9.9e-08 | 2.9e-07 | 1.963E-07 | 9.555E-08 | 9.900E-08 | 2.900E-07 |
| 9974 | 499 | W499F | A | 1.9e-07 | 1.3e-07 | 4.2e-07 | 2.467E-07 | 1.531E-07 | 1.300E-07 | 4.200E-07 |
| 9975 | 499 | W499P | A | 4.3e-07 | 6e-07 | 2.7e-07 | 4.333E-07 | 1.650E-07 | 2.700E-07 | 6.000E-07 |
| 9976 | 499 | W499S | A | 2.9e-07 | 1.7e-07 | 4.7e-07 | 3.100E-07 | 1.510E-07 | 1.700E-07 | 4.700E-07 |
| 9977 | 499 | W499T | A | 2.1e-07 | 3.3e-07 | 3.2e-07 | 2.867E-07 | 6.658E-08 | 2.100E-07 | 3.300E-07 |
| 9978 | 499 | W499W | A | 4.7e-07 | 1.5e-07 | 3.2e-07 | 3.133E-07 | 1.601E-07 | 1.500E-07 | 4.700E-07 |
| 9979 | 499 | W499Y | A | 2.3e-07 | 1.9e-07 | 2.2e-07 | 2.133E-07 | 2.082E-08 | 1.900E-07 | 2.300E-07 |
| 9980 | 499 | W499V | A | 2e-07 | 2.4e-07 | 5.5e-07 | 3.300E-07 | 1.916E-07 | 2.000E-07 | 5.500E-07 |
| 9981 | 500 | G500A | A | 1.6e-07 | 1.4e-07 | 3e-07 | 2.000E-07 | 8.718E-08 | 1.400E-07 | 3.000E-07 |
| 9982 | 500 | G500R | A | 2.3e-07 | 1.6e-07 | 2.8e-07 | 2.233E-07 | 6.028E-08 | 1.600E-07 | 2.800E-07 |
| 9983 | 500 | G500N | A | 2e-07 | 2.9e-07 | 1.9e-07 | 2.267E-07 | 5.508E-08 | 1.900E-07 | 2.900E-07 |
| 9984 | 500 | G500D | A | 3.6e-07 | 3.1e-07 | 1.2e-07 | 2.633E-07 | 1.266E-07 | 1.200E-07 | 3.600E-07 |
| 9985 | 500 | G500C | A | 2.1e-07 | 3.6e-07 | 3.4e-07 | 3.033E-07 | 8.145E-08 | 2.100E-07 | 3.600E-07 |
| 9986 | 500 | G500E | A | 3e-07 | 2.7e-07 | 2.7e-07 | 2.800E-07 | 1.732E-08 | 2.700E-07 | 3.000E-07 |
| 9987 | 500 | G500Q | A | 1.5e-07 | 1.3e-07 | 3.6e-07 | 2.133E-07 | 1.274E-07 | 1.300E-07 | 3.600E-07 |
| 9988 | 500 | G500G | A | 2.8e-07 | 2.5e-07 | 2.7e-07 | 2.667E-07 | 1.528E-08 | 2.500E-07 | 2.800E-07 |
| 9989 | 500 | G500H | A | 1.9e-07 | 4.5e-07 | 1.4e-07 | 2.600E-07 | 1.664E-07 | 1.400E-07 | 4.500E-07 |
| 9990 | 500 | G500I | A | 2.3e-07 | 2e-07 | 1.8e-07 | 2.033E-07 | 2.517E-08 | 1.800E-07 | 2.300E-07 |
| 9991 | 500 | G500L | A | 1.8e-07 | 2.7e-07 | 4.2e-07 | 2.900E-07 | 1.212E-07 | 1.800E-07 | 4.200E-07 |
| 9992 | 500 | G500K | A | 2.5e-07 | 2e-07 | 3.5e-07 | 2.667E-07 | 7.638E-08 | 2.000E-07 | 3.500E-07 |
| 9993 | 500 | G500M | A | 1.3e-07 | 3e-07 | 2e-07 | 2.100E-07 | 8.544E-08 | 1.300E-07 | 3.000E-07 |
| 9994 | 500 | G500F | A | 3.9e-07 | 3.4e-07 | 2.9e-07 | 3.400E-07 | 5.000E-08 | 2.900E-07 | 3.900E-07 |
| 9995 | 500 | G500P | A | 3.9e-07 | 3.3e-07 | 2.4e-07 | 3.200E-07 | 7.550E-08 | 2.400E-07 | 3.900E-07 |
| 9996 | 500 | G500S | A | 3e-07 | 3e-07 | 1.9e-07 | 2.633E-07 | 6.351E-08 | 1.900E-07 | 3.000E-07 |
| 9997 | 500 | G500T | A | 2.5e-07 | 2.1e-07 | 1.3e-07 | 1.967E-07 | 6.110E-08 | 1.300E-07 | 2.500E-07 |
| 9998 | 500 | G500W | A | 7.4e-07 | 3.5e-07 | 1.1e-07 | 4.000E-07 | 3.180E-07 | 1.100E-07 | 7.400E-07 |
| 9999 | 500 | G500Y | A | 1.6e-07 | 2e-07 | 2.5e-07 | 2.033E-07 | 4.509E-08 | 1.600E-07 | 2.500E-07 |
| 10000 | 500 | G500V | A | 1.3e-07 | 2.5e-07 | 1.6e-07 | 1.800E-07 | 6.245E-08 | 1.300E-07 | 2.500E-07 |
| 10001 | 501 | P501A | A | 3.9e-07 | 2.7e-07 | 1.9e-07 | 2.833E-07 | 1.007E-07 | 1.900E-07 | 3.900E-07 |
| 10002 | 501 | P501R | A | 4.8e-07 | 2.4e-07 | 2.7e-07 | 3.300E-07 | 1.308E-07 | 2.400E-07 | 4.800E-07 |
| 10003 | 501 | P501N | A | 2.2e-07 | 1.8e-07 | 4.7e-07 | 2.900E-07 | 1.572E-07 | 1.800E-07 | 4.700E-07 |
| 10004 | 501 | P501D | A | 1.5e-07 | 3.5e-07 | 3.9e-07 | 2.967E-07 | 1.286E-07 | 1.500E-07 | 3.900E-07 |
| 10005 | 501 | P501C | A | 5.3e-07 | 2.6e-07 | 1.4e-07 | 3.100E-07 | 1.997E-07 | 1.400E-07 | 5.300E-07 |
| 10006 | 501 | P501E | A | 3e-07 | 2.3e-07 | 1.6e-07 | 2.300E-07 | 7.000E-08 | 1.600E-07 | 3.000E-07 |
| 10007 | 501 | P501Q | A | 4.1e-07 | 4.4e-07 | 1.5e-07 | 3.333E-07 | 1.595E-07 | 1.500E-07 | 4.400E-07 |
| 10008 | 501 | P501G | A | 2.2e-07 | 3.3e-07 | 2.9e-07 | 2.800E-07 | 5.568E-08 | 2.200E-07 | 3.300E-07 |
| 10009 | 501 | P501H | A | 7.5e-08 | 2.6e-07 | 1.2e-07 | 1.517E-07 | 9.648E-08 | 7.500E-08 | 2.600E-07 |
| 10010 | 501 | P501I | A | 1.2e-07 | 2.9e-07 | 2.5e-07 | 2.200E-07 | 8.888E-08 | 1.200E-07 | 2.900E-07 |
| 10011 | 501 | P501L | A | 2.3e-07 | 2.1e-07 | 2.8e-07 | 2.400E-07 | 3.606E-08 | 2.100E-07 | 2.800E-07 |
| 10012 | 501 | P501K | A | 2.4e-07 | 1.7e-07 | 2.7e-07 | 2.267E-07 | 5.132E-08 | 1.700E-07 | 2.700E-07 |
| 10013 | 501 | P501M | A | 1.4e-07 | 3e-07 | 3.6e-07 | 2.667E-07 | 1.137E-07 | 1.400E-07 | 3.600E-07 |
| 10014 | 501 | P501F | A | 1.6e-07 | 3e-07 | 2.6e-07 | 2.400E-07 | 7.211E-08 | 1.600E-07 | 3.000E-07 |
| 10015 | 501 | P501P | A | 3.8e-07 | 2.9e-07 | 2.6e-07 | 3.100E-07 | 6.245E-08 | 2.600E-07 | 3.800E-07 |
| 10016 | 501 | P501S | A | 3.4e-07 | 4.3e-07 | 1.5e-07 | 3.067E-07 | 1.429E-07 | 1.500E-07 | 4.300E-07 |
| 10017 | 501 | P501T | A | 1.2e-07 | 3.3e-07 | 2.5e-07 | 2.333E-07 | 1.060E-07 | 1.200E-07 | 3.300E-07 |
| 10018 | 501 | P501W | A | 3.2e-07 | 2.7e-07 | 1.8e-07 | 2.567E-07 | 7.095E-08 | 1.800E-07 | 3.200E-07 |
| 10019 | 501 | P501Y | A | 1.5e-07 | 1.2e-07 | 3.9e-07 | 2.200E-07 | 1.480E-07 | 1.200E-07 | 3.900E-07 |
| 10020 | 501 | P501V | A | 2.5e-07 | 2.5e-07 | 1.5e-07 | 2.167E-07 | 5.774E-08 | 1.500E-07 | 2.500E-07 |
| 10021 | 502 | G502A | A | 1.6e-07 | 1.3e-07 | 3.3e-07 | 2.067E-07 | 1.079E-07 | 1.300E-07 | 3.300E-07 |
| 10022 | 502 | G502R | A | 3.4e-07 | 6.9e-08 | 1.9e-07 | 1.997E-07 | 1.358E-07 | 6.900E-08 | 3.400E-07 |
| 10023 | 502 | G502N | A | 3.6e-07 | 4.3e-07 | 5.6e-07 | 4.500E-07 | 1.015E-07 | 3.600E-07 | 5.600E-07 |
| 10024 | 502 | G502D | A | 1.4e-07 | 1.7e-07 | 5.5e-07 | 2.867E-07 | 2.285E-07 | 1.400E-07 | 5.500E-07 |
| 10025 | 502 | G502C | A | 4.2e-07 | 1.6e-07 | 1.8e-07 | 2.533E-07 | 1.447E-07 | 1.600E-07 | 4.200E-07 |
| 10026 | 502 | G502E | A | 3.8e-07 | 1.9e-07 | 1.3e-07 | 2.333E-07 | 1.305E-07 | 1.300E-07 | 3.800E-07 |
| 10027 | 502 | G502Q | A | 1.6e-07 | 3e-07 | 2.7e-07 | 2.433E-07 | 7.371E-08 | 1.600E-07 | 3.000E-07 |
| 10028 | 502 | G502G | A | 3e-07 | 1.4e-07 | 1.4e-07 | 1.933E-07 | 9.238E-08 | 1.400E-07 | 3.000E-07 |
| 10029 | 502 | G502H | A | 1.6e-07 | 3.2e-07 | 2.3e-07 | 2.367E-07 | 8.021E-08 | 1.600E-07 | 3.200E-07 |

| | | | | | | | | | | |
|---|---|---|---|---|---|---|---|---|---|---|
| 10030 | 502 | G502I | A | 3.6e-07 | 2.3e-07 | 1.9e-07 | 2.600E-07 | 8.888E-08 | 1.900E-07 | 3.600E-07 |
| 10031 | 502 | G502L | A | 3.1e-07 | 5.9e-07 | 4.6e-07 | 4.533E-07 | 1.401E-07 | 3.100E-07 | 5.900E-07 |
| 10032 | 502 | G502K | A | 1.1e-07 | 4.2e-07 | 1.9e-07 | 2.400E-07 | 1.609E-07 | 1.100E-07 | 4.200E-07 |
| 10033 | 502 | G502M | A | 4.2e-07 | 4.7e-07 | 2.4e-07 | 3.767E-07 | 1.210E-07 | 2.400E-07 | 4.700E-07 |
| 10034 | 502 | G502F | A | 4.8e-07 | 1.9e-07 | 1.8e-07 | 2.833E-07 | 1.704E-07 | 1.800E-07 | 4.800E-07 |
| 10035 | 502 | G502P | A | 2.2e-07 | 1.6e-07 | 2.2e-07 | 2.000E-07 | 3.464E-08 | 1.600E-07 | 2.200E-07 |
| 10036 | 502 | G502S | A | 2e-07 | 1.6e-07 | 2.4e-07 | 2.000E-07 | 4.000E-08 | 1.600E-07 | 2.400E-07 |
| 10037 | 502 | G502T | A | 5.8e-07 | 4.8e-07 | 1.9e-07 | 4.167E-07 | 2.026E-07 | 1.900E-07 | 5.800E-07 |
| 10038 | 502 | G502W | A | 4.3e-07 | 2.4e-07 | 1.4e-07 | 2.700E-07 | 1.473E-07 | 1.400E-07 | 4.300E-07 |
| 10039 | 502 | G502Y | A | 4.9e-07 | 2.4e-07 | 3.2e-07 | 3.500E-07 | 1.277E-07 | 2.400E-07 | 4.900E-07 |
| 10040 | 502 | G502V | A | 2e-07 | 2.7e-07 | 2e-07 | 2.233E-07 | 4.041E-08 | 2.000E-07 | 2.700E-07 |
| 10041 | 503 | P503A | A | 7.1e-07 | 3e-07 | 2e-07 | 4.033E-07 | 2.702E-07 | 2.000E-07 | 7.100E-07 |
| 10042 | 503 | P503R | A | 1e-07 | 4.5e-07 | 7.8e-07 | 4.433E-07 | 3.400E-07 | 1.000E-07 | 7.800E-07 |
| 10043 | 503 | P503N | A | 6.9e-08 | 5.6e-07 | 2.2e-07 | 2.830E-07 | 2.515E-07 | 6.900E-08 | 5.600E-07 |
| 10044 | 503 | P503D | A | 1.2e-07 | 1.5e-07 | 2.9e-07 | 1.867E-07 | 9.074E-08 | 1.200E-07 | 2.900E-07 |
| 10045 | 503 | P503C | A | 1.8e-07 | 1.8e-07 | 4.1e-07 | 2.567E-07 | 1.328E-07 | 1.800E-07 | 4.100E-07 |
| 10046 | 503 | P503E | A | 2.2e-07 | 2.8e-07 | 2.4e-07 | 2.467E-07 | 3.055E-08 | 2.200E-07 | 2.800E-07 |
| 10047 | 503 | P503Q | A | 1.6e-07 | 2.1e-07 | 3.1e-07 | 2.267E-07 | 7.638E-08 | 1.600E-07 | 3.100E-07 |
| 10048 | 503 | P503G | A | 2.9e-07 | 5e-07 | 2.3e-07 | 3.400E-07 | 1.418E-07 | 2.300E-07 | 5.000E-07 |
| 10049 | 503 | P503H | A | 3.7e-07 | 2.6e-07 | 4.8e-07 | 3.700E-07 | 1.100E-07 | 2.600E-07 | 4.800E-07 |
| 10050 | 503 | P503I | A | 2.1e-07 | 1.4e-07 | 3.1e-07 | 2.200E-07 | 8.544E-08 | 1.400E-07 | 3.100E-07 |
| 10051 | 503 | P503L | A | 2.7e-07 | 2.5e-07 | 5e-07 | 3.400E-07 | 1.389E-07 | 2.500E-07 | 5.000E-07 |
| 10052 | 503 | P503K | A | 3.3e-07 | 3.5e-07 | 3.4e-07 | 3.400E-07 | 1.000E-08 | 3.300E-07 | 3.500E-07 |
| 10053 | 503 | P503M | A | 3.9e-07 | 2.1e-07 | 2.6e-07 | 2.867E-07 | 9.292E-08 | 2.100E-07 | 3.900E-07 |
| 10054 | 503 | P503F | A | 9.9e-08 | 1.8e-07 | 4.3e-07 | 2.363E-07 | 1.725E-07 | 9.900E-08 | 4.300E-07 |
| 10055 | 503 | P503P | A | 3e-07 | 2.3e-07 | 2.3e-07 | 2.533E-07 | 4.041E-08 | 2.300E-07 | 3.000E-07 |
| 10056 | 503 | P503S | A | 1.5e-07 | 2.5e-07 | 1.8e-07 | 1.933E-07 | 5.132E-08 | 1.500E-07 | 2.500E-07 |
| 10057 | 503 | P503T | A | 2.2e-07 | 2e-07 | 3e-07 | 2.400E-07 | 5.292E-08 | 2.000E-07 | 3.000E-07 |
| 10058 | 503 | P503W | A | 3.8e-07 | 2.8e-07 | 2.2e-07 | 2.933E-07 | 8.083E-08 | 2.200E-07 | 3.800E-07 |
| 10059 | 503 | P503Y | A | 3.1e-07 | 1.6e-07 | 3.1e-07 | 2.600E-07 | 8.660E-08 | 1.600E-07 | 3.100E-07 |
| 10060 | 503 | P503V | A | 3.1e-07 | 2.1e-07 | 3.4e-07 | 2.867E-07 | 6.807E-08 | 2.100E-07 | 3.400E-07 |
| 10061 | 504 | T504A | A | 1.4e-07 | 3.1e-07 | 2.2e-07 | 2.233E-07 | 8.505E-08 | 1.400E-07 | 3.100E-07 |
| 10062 | 504 | T504R | A | 3.2e-07 | 1.7e-07 | 2.4e-07 | 2.433E-07 | 7.506E-08 | 1.700E-07 | 3.200E-07 |
| 10063 | 504 | T504N | A | 1.1e-07 | 2e-07 | 2.6e-07 | 1.900E-07 | 7.550E-08 | 1.100E-07 | 2.600E-07 |
| 10064 | 504 | T504D | A | 1.9e-07 | 2.3e-07 | 2.7e-07 | 2.300E-07 | 4.000E-08 | 1.900E-07 | 2.700E-07 |
| 10065 | 504 | T504C | A | 2.1e-07 | 3.3e-07 | 2e-07 | 2.467E-07 | 7.234E-08 | 2.000E-07 | 3.300E-07 |
| 10066 | 504 | T504E | A | 1.4e-07 | 2.4e-07 | 3.5e-07 | 2.433E-07 | 1.050E-07 | 1.400E-07 | 3.500E-07 |
| 10067 | 504 | T504Q | A | 2.3e-07 | 2.4e-07 | 2.5e-07 | 2.400E-07 | 1.000E-08 | 2.300E-07 | 2.500E-07 |
| 10068 | 504 | T504G | A | 1.2e-07 | 3.1e-07 | 2.1e-07 | 2.133E-07 | 9.504E-08 | 1.200E-07 | 3.100E-07 |
| 10069 | 504 | T504H | A | 2.4e-07 | 2.5e-07 | 1.3e-07 | 2.067E-07 | 6.658E-08 | 1.300E-07 | 2.500E-07 |
| 10070 | 504 | T504I | A | 1.7e-07 | 5.5e-07 | 3.1e-07 | 3.433E-07 | 1.922E-07 | 1.700E-07 | 5.500E-07 |
| 10071 | 504 | T504L | A | 1.5e-07 | 5.4e-07 | 1.5e-07 | 2.800E-07 | 2.252E-07 | 1.500E-07 | 5.400E-07 |
| 10072 | 504 | T504K | A | 2.5e-07 | 3.2e-07 | 1.2e-07 | 2.300E-07 | 1.015E-07 | 1.200E-07 | 3.200E-07 |
| 10073 | 504 | T504M | A | 2.4e-07 | 3.5e-07 | 3.3e-07 | 3.067E-07 | 5.859E-08 | 2.400E-07 | 3.500E-07 |
| 10074 | 504 | T504F | A | 3.5e-07 | 2.3e-07 | 1.7e-07 | 2.500E-07 | 9.165E-08 | 1.700E-07 | 3.500E-07 |
| 10075 | 504 | T504P | A | 3e-07 | 2.2e-07 | 2.4e-07 | 2.533E-07 | 4.163E-08 | 2.200E-07 | 3.000E-07 |
| 10076 | 504 | T504S | A | 1.6e-07 | 3.4e-07 | 2.2e-07 | 2.400E-07 | 9.165E-08 | 1.600E-07 | 3.400E-07 |
| 10077 | 504 | T504T | A | 1.7e-07 | 3.1e-07 | 3.6e-07 | 2.800E-07 | 9.849E-08 | 1.700E-07 | 3.600E-07 |
| 10078 | 504 | T504W | A | 2.1e-07 | 3e-07 | 2.2e-07 | 2.433E-07 | 4.933E-08 | 2.100E-07 | 3.000E-07 |
| 10079 | 504 | T504Y | A | 2.8e-07 | 1.6e-07 | 3.2e-07 | 2.533E-07 | 8.327E-08 | 1.600E-07 | 3.200E-07 |
| 10080 | 504 | T504V | A | 1.4e-07 | 2.6e-07 | 2.4e-07 | 2.133E-07 | 6.429E-08 | 1.400E-07 | 2.600E-07 |
| 10081 | 505 | Q505A | A | 3.3e-07 | 1.3e-07 | 1.8e-07 | 2.133E-07 | 1.041E-07 | 1.300E-07 | 3.300E-07 |
| 10082 | 505 | Q505R | A | 3.2e-07 | 2.2e-07 | 2.5e-07 | 2.633E-07 | 5.132E-08 | 2.200E-07 | 3.200E-07 |
| 10083 | 505 | Q505N | A | 4e-07 | 3.2e-07 | 2e-07 | 3.067E-07 | 1.007E-07 | 2.000E-07 | 4.000E-07 |
| 10084 | 505 | Q505D | A | 2.6e-07 | 1.3e-07 | 2.1e-07 | 2.000E-07 | 6.557E-08 | 1.300E-07 | 2.600E-07 |
| 10085 | 505 | Q505C | A | 2.2e-07 | 1.3e-07 | 3.9e-07 | 2.467E-07 | 1.320E-07 | 1.300E-07 | 3.900E-07 |
| 10086 | 505 | Q505E | A | 4.3e-07 | 2.8e-07 | 7.2e-07 | 4.767E-07 | 2.237E-07 | 2.800E-07 | 7.200E-07 |
| 10087 | 505 | Q505Q | A | 2.9e-07 | 2.2e-07 | 2.5e-07 | 2.533E-07 | 3.512E-08 | 2.200E-07 | 2.900E-07 |
| 10088 | 505 | Q505G | A | 2e-07 | 2.7e-07 | 7.2e-08 | 1.807E-07 | 1.004E-07 | 7.200E-08 | 2.700E-07 |
| 10089 | 505 | Q505H | A | 9.1e-08 | 2e-07 | 1.7e-07 | 1.537E-07 | 5.631E-08 | 9.100E-08 | 2.000E-07 |
| 10090 | 505 | Q505I | A | 1.9e-07 | 1.6e-07 | 2.2e-07 | 1.900E-07 | 3.000E-08 | 1.600E-07 | 2.200E-07 |
| 10091 | 505 | Q505L | A | 2.5e-07 | 1.5e-07 | 4.9e-07 | 2.967E-07 | 1.747E-07 | 1.500E-07 | 4.900E-07 |
| 10092 | 505 | Q505K | A | 2e-07 | 1.6e-07 | 1.8e-07 | 1.800E-07 | 2.000E-08 | 1.600E-07 | 2.000E-07 |
| 10093 | 505 | Q505M | A | 2.4e-07 | 2.2e-07 | 5e-07 | 3.200E-07 | 1.562E-07 | 2.200E-07 | 5.000E-07 |
| 10094 | 505 | Q505F | A | 3e-07 | 2.1e-07 | 2.2e-07 | 2.433E-07 | 4.933E-08 | 2.100E-07 | 3.000E-07 |
| 10095 | 505 | Q505P | A | 2.8e-07 | 2.7e-07 | 5.9e-08 | 2.030E-07 | 1.248E-07 | 5.900E-08 | 2.800E-07 |
| 10096 | 505 | Q505S | A | 1.6e-07 | 3.2e-07 | 2e-07 | 2.267E-07 | 8.327E-08 | 1.600E-07 | 3.200E-07 |
| 10097 | 505 | Q505T | A | 1.9e-07 | 2e-07 | 1.7e-07 | 1.867E-07 | 1.528E-08 | 1.700E-07 | 2.000E-07 |
| 10098 | 505 | Q505W | A | 2.9e-07 | 6.4e-08 | 5.4e-07 | 2.980E-07 | 2.381E-07 | 6.400E-08 | 5.400E-07 |
| 10099 | 505 | Q505Y | A | 1.7e-07 | 2e-07 | 2e-07 | 1.900E-07 | 1.732E-08 | 1.700E-07 | 2.000E-07 |
| 10100 | 505 | Q505V | A | 3.5e-07 | 1.6e-07 | 2.7e-07 | 2.600E-07 | 9.539E-08 | 1.600E-07 | 3.500E-07 |
| 10101 | 506 | C506A | A | 2.5e-07 | 1.8e-07 | 3.5e-07 | 2.600E-07 | 8.544E-08 | 1.800E-07 | 3.500E-07 |
| 10102 | 506 | C506R | A | 1.3e-07 | 4.4e-07 | 2.8e-07 | 2.833E-07 | 1.550E-07 | 1.300E-07 | 4.400E-07 |
| 10103 | 506 | C506N | A | 2.4e-07 | 2.5e-07 | 2.3e-07 | 2.400E-07 | 1.000E-08 | 2.300E-07 | 2.500E-07 |
| 10104 | 506 | C506D | A | 1.7e-07 | 3.5e-07 | 6.9e-07 | 4.033E-07 | 2.641E-07 | 1.700E-07 | 6.900E-07 |
| 10105 | 506 | C506C | A | 5e-07 | 1.8e-07 | 2.1e-07 | 2.967E-07 | 1.767E-07 | 1.800E-07 | 5.000E-07 |
| 10106 | 506 | C506E | A | 9.6e-07 | 3.3e-07 | 2.2e-07 | 5.033E-07 | 3.993E-07 | 2.200E-07 | 9.600E-07 |
| 10107 | 506 | C506Q | A | 1.5e-07 | 2.7e-07 | 3.4e-07 | 2.533E-07 | 9.609E-08 | 1.500E-07 | 3.400E-07 |
| 10108 | 506 | C506G | A | 1.2e-07 | 3.2e-07 | 2.7e-07 | 2.367E-07 | 1.041E-07 | 1.200E-07 | 3.200E-07 |
| 10109 | 506 | C506H | A | 1.5e-07 | 1.7e-07 | 2.5e-07 | 1.900E-07 | 5.292E-08 | 1.500E-07 | 2.500E-07 |
| 10110 | 506 | C506I | A | 2.9e-07 | 2.2e-07 | 1.7e-07 | 2.267E-07 | 6.028E-08 | 1.700E-07 | 2.900E-07 |
| 10111 | 506 | C506L | A | 4.8e-07 | 1.9e-07 | 3.2e-07 | 3.300E-07 | 1.453E-07 | 1.900E-07 | 4.800E-07 |
| 10112 | 506 | C506K | A | 1.2e-07 | 6.9e-07 | 4.1e-07 | 4.067E-07 | 2.850E-07 | 1.200E-07 | 6.900E-07 |
| 10113 | 506 | C506M | A | 3e-07 | 3.6e-07 | 5.5e-07 | 4.033E-07 | 1.305E-07 | 3.000E-07 | 5.500E-07 |
| 10114 | 506 | C506F | A | 3.1e-07 | 3e-07 | 3.8e-07 | 3.300E-07 | 4.359E-08 | 3.000E-07 | 3.800E-07 |

| | | | | | | | | | | |
|---|---|---|---|---|---|---|---|---|---|---|
| 10115 | 506 | C506P | A | 3.4e-07 | 1.2e-07 | 1.9e-07 | 2.167E-07 | 1.124E-07 | 1.200E-07 | 3.400E-07 |
| 10116 | 506 | C506S | A | 4.7e-07 | 3e-07 | 3.3e-07 | 3.667E-07 | 9.074E-08 | 3.000E-07 | 4.700E-07 |
| 10117 | 506 | C506T | A | 1.9e-07 | 1.8e-07 | 1.6e-07 | 1.767E-07 | 1.528E-08 | 1.600E-07 | 1.900E-07 |
| 10118 | 506 | C506W | A | 3e-07 | 1.4e-07 | 1.9e-07 | 2.100E-07 | 8.185E-08 | 1.400E-07 | 3.000E-07 |
| 10119 | 506 | C506Y | A | 2.8e-07 | 4.4e-07 | 3.4e-07 | 3.533E-07 | 8.083E-08 | 2.800E-07 | 4.400E-07 |
| 10120 | 506 | C506V | A | 2.6e-07 | 4.7e-07 | 1.8e-07 | 3.033E-07 | 1.498E-07 | 1.800E-07 | 4.700E-07 |
| 10121 | 507 | V507A | A | 2e-07 | 1.1e-07 | 1.9e-07 | 1.667E-07 | 4.933E-08 | 1.100E-07 | 2.000E-07 |
| 10122 | 507 | V507R | A | 1e-07 | 1.9e-07 | 2.5e-07 | 1.800E-07 | 7.550E-08 | 1.000E-07 | 2.500E-07 |
| 10123 | 507 | V507N | A | 2.8e-07 | 4.5e-07 | 1.8e-07 | 3.033E-07 | 1.365E-07 | 1.800E-07 | 4.500E-07 |
| 10124 | 507 | V507D | A | 2.9e-07 | 2.2e-07 | 3.1e-07 | 2.733E-07 | 4.726E-08 | 2.200E-07 | 3.100E-07 |
| 10125 | 507 | V507C | A | 1.7e-07 | 2.6e-07 | 3e-07 | 2.433E-07 | 6.658E-08 | 1.700E-07 | 3.000E-07 |
| 10126 | 507 | V507E | A | 1.5e-07 | 2.2e-07 | 1.5e-07 | 1.733E-07 | 4.041E-08 | 1.500E-07 | 2.200E-07 |
| 10127 | 507 | V507Q | A | 2.6e-07 | 3e-07 | 1.9e-07 | 2.500E-07 | 5.568E-08 | 1.900E-07 | 3.000E-07 |
| 10128 | 507 | V507G | A | 2.1e-07 | 2.3e-07 | 4e-07 | 2.800E-07 | 1.044E-07 | 2.100E-07 | 4.000E-07 |
| 10129 | 507 | V507H | A | 2.7e-07 | 3.2e-07 | 3.4e-07 | 3.100E-07 | 3.606E-08 | 2.700E-07 | 3.400E-07 |
| 10130 | 507 | V507I | A | 3.5e-07 | 1.9e-07 | 2.2e-07 | 2.533E-07 | 8.505E-08 | 1.900E-07 | 3.500E-07 |
| 10131 | 507 | V507L | A | 2.1e-07 | 4.1e-07 | 2.6e-07 | 2.933E-07 | 1.041E-07 | 2.100E-07 | 4.100E-07 |
| 10132 | 507 | V507K | A | 5e-07 | 2.8e-07 | 7.5e-07 | 5.100E-07 | 2.352E-07 | 2.800E-07 | 7.500E-07 |
| 10133 | 507 | V507M | A | 2e-07 | 1.3e-07 | 2.8e-07 | 2.033E-07 | 7.506E-08 | 1.300E-07 | 2.800E-07 |
| 10134 | 507 | V507F | A | 3.7e-07 | 6.3e-07 | 2.6e-07 | 4.200E-07 | 1.900E-07 | 2.600E-07 | 6.300E-07 |
| 10135 | 507 | V507P | A | 6.3e-07 | 2.9e-07 | 2.7e-07 | 3.967E-07 | 2.023E-07 | 2.700E-07 | 6.300E-07 |
| 10136 | 507 | V507S | A | 2.9e-07 | 3.6e-07 | 1.8e-07 | 2.767E-07 | 9.074E-08 | 1.800E-07 | 3.600E-07 |
| 10137 | 507 | V507T | A | 2.8e-07 | 2.2e-07 | 3e-07 | 2.667E-07 | 4.163E-08 | 2.200E-07 | 3.000E-07 |
| 10138 | 507 | V507W | A | 4.1e-07 | 4e-07 | 2e-07 | 3.367E-07 | 1.185E-07 | 2.000E-07 | 4.100E-07 |
| 10139 | 507 | V507Y | A | 2.2e-07 | 1.9e-07 | 1.5e-07 | 1.867E-07 | 3.512E-08 | 1.500E-07 | 2.200E-07 |
| 10140 | 507 | V507V | A | 2.3e-07 | 4e-07 | 2.5e-07 | 2.933E-07 | 9.292E-08 | 2.300E-07 | 4.000E-07 |
| 10141 | 508 | N508A | A | 3.5e-07 | 3.6e-07 | 4e-07 | 3.700E-07 | 2.646E-08 | 3.500E-07 | 4.000E-07 |
| 10142 | 508 | N508R | A | 3.8e-07 | 1.6e-07 | 1.4e-07 | 2.267E-07 | 1.332E-07 | 1.400E-07 | 3.800E-07 |
| 10143 | 508 | N508N | A | 3.1e-07 | 2.8e-07 | 4.7e-07 | 3.533E-07 | 1.021E-07 | 2.800E-07 | 4.700E-07 |
| 10144 | 508 | N508D | A | 1.7e-07 | 4.5e-07 | 1.5e-07 | 2.567E-07 | 1.677E-07 | 1.500E-07 | 4.500E-07 |
| 10145 | 508 | N508C | A | 1.6e-07 | 1.1e-07 | 1.2e-07 | 1.300E-07 | 2.646E-08 | 1.100E-07 | 1.600E-07 |
| 10146 | 508 | N508E | A | 8.3e-08 | 2.5e-07 | 1.3e-07 | 1.543E-07 | 8.612E-08 | 8.300E-08 | 2.500E-07 |
| 10147 | 508 | N508Q | A | 1.5e-07 | 1.7e-07 | 1.3e-07 | 1.500E-07 | 2.000E-08 | 1.300E-07 | 1.700E-07 |
| 10148 | 508 | N508G | A | 5.6e-07 | 3.1e-07 | 4.8e-07 | 4.500E-07 | 1.277E-07 | 3.100E-07 | 5.600E-07 |
| 10149 | 508 | N508H | A | 2.9e-07 | 1.6e-07 | 3.8e-07 | 2.767E-07 | 1.106E-07 | 1.600E-07 | 3.800E-07 |
| 10150 | 508 | N508I | A | 3e-07 | 2.4e-07 | 2.3e-07 | 2.567E-07 | 3.786E-08 | 2.300E-07 | 3.000E-07 |
| 10151 | 508 | N508L | A | 1.7e-07 | 1.2e-07 | 7.6e-08 | 1.220E-07 | 4.703E-08 | 7.600E-08 | 1.700E-07 |
| 10152 | 508 | N508K | A | 1.4e-07 | 6e-07 | 3e-07 | 3.467E-07 | 2.335E-07 | 1.400E-07 | 6.000E-07 |
| 10153 | 508 | N508M | A | 1.1e-07 | 2.2e-07 | 3.2e-07 | 2.167E-07 | 1.050E-07 | 1.100E-07 | 3.200E-07 |
| 10154 | 508 | N508F | A | 1.7e-07 | 7.4e-07 | 2.6e-07 | 3.900E-07 | 3.064E-07 | 1.700E-07 | 7.400E-07 |
| 10155 | 508 | N508P | A | 2e-07 | 2.8e-07 | 2.2e-07 | 2.333E-07 | 4.163E-08 | 2.000E-07 | 2.800E-07 |
| 10156 | 508 | N508S | A | 3.1e-07 | 1.3e-07 | 3.4e-07 | 2.600E-07 | 1.136E-07 | 1.300E-07 | 3.400E-07 |
| 10157 | 508 | N508T | A | 1.8e-07 | 7.7e-07 | 2.8e-07 | 4.100E-07 | 3.158E-07 | 1.800E-07 | 7.700E-07 |
| 10158 | 508 | N508W | A | 2.2e-07 | 2.9e-07 | 9.6e-08 | 2.020E-07 | 9.824E-08 | 9.600E-08 | 2.900E-07 |
| 10159 | 508 | N508Y | A | 2.4e-07 | 3e-07 | 6e-07 | 3.800E-07 | 1.929E-07 | 2.400E-07 | 6.000E-07 |
| 10160 | 508 | N508V | A | 4.5e-07 | 7.4e-07 | 4.5e-07 | 5.467E-07 | 1.674E-07 | 4.500E-07 | 7.400E-07 |
| 10161 | 509 | C509A | A | 3.4e-07 | 5.6e-07 | 2.5e-07 | 3.833E-07 | 1.595E-07 | 2.500E-07 | 5.600E-07 |
| 10162 | 509 | C509R | A | 4.1e-07 | 1.4e-07 | 2.4e-07 | 2.633E-07 | 1.365E-07 | 1.400E-07 | 4.100E-07 |
| 10163 | 509 | C509N | A | 1.2e-07 | 1.3e-07 | 6.1e-07 | 2.867E-07 | 2.801E-07 | 1.200E-07 | 6.100E-07 |
| 10164 | 509 | C509D | A | 2.8e-07 | 3.3e-07 | 1.5e-07 | 2.533E-07 | 9.292E-08 | 1.500E-07 | 3.300E-07 |
| 10165 | 509 | C509C | A | 2e-07 | 2.2e-07 | 1.1e-07 | 1.767E-07 | 5.859E-08 | 1.100E-07 | 2.200E-07 |
| 10166 | 509 | C509E | A | 4.3e-07 | 1.8e-07 | 5.7e-07 | 3.933E-07 | 1.976E-07 | 1.800E-07 | 5.700E-07 |
| 10167 | 509 | C509Q | A | 2.5e-07 | 2.4e-07 | 3.5e-07 | 2.800E-07 | 6.083E-08 | 2.400E-07 | 3.500E-07 |
| 10168 | 509 | C509G | A | 3.7e-07 | 2.6e-07 | 5.7e-07 | 4.000E-07 | 1.572E-07 | 2.600E-07 | 5.700E-07 |
| 10169 | 509 | C509H | A | 4.7e-07 | 1e-07 | 1.9e-07 | 2.533E-07 | 1.930E-07 | 1.000E-07 | 4.700E-07 |
| 10170 | 509 | C509I | A | 3.2e-07 | 1.2e-07 | 2.6e-07 | 2.333E-07 | 1.026E-07 | 1.200E-07 | 3.200E-07 |
| 10171 | 509 | C509L | A | 1.9e-07 | 4.9e-07 | 3e-07 | 3.267E-07 | 1.518E-07 | 1.900E-07 | 4.900E-07 |
| 10172 | 509 | C509K | A | 2.4e-07 | 2.9e-07 | 3.5e-07 | 2.933E-07 | 5.508E-08 | 2.400E-07 | 3.500E-07 |
| 10173 | 509 | C509M | A | 2.1e-07 | 1.7e-07 | 7.1e-07 | 3.633E-07 | 3.009E-07 | 1.700E-07 | 7.100E-07 |
| 10174 | 509 | C509F | A | 6.2e-07 | 1.2e-07 | 2.3e-07 | 3.233E-07 | 2.627E-07 | 1.200E-07 | 6.200E-07 |
| 10175 | 509 | C509P | A | 3.6e-07 | 1.7e-07 | 3e-07 | 2.767E-07 | 9.713E-08 | 1.700E-07 | 3.600E-07 |
| 10176 | 509 | C509S | A | 2.3e-07 | 2.1e-07 | 2.3e-07 | 2.233E-07 | 1.155E-08 | 2.100E-07 | 2.300E-07 |
| 10177 | 509 | C509T | A | 2.1e-07 | 1.4e-07 | 1e-07 | 1.500E-07 | 5.568E-08 | 1.000E-07 | 2.100E-07 |
| 10178 | 509 | C509W | A | 1.9e-07 | 2.5e-07 | 3e-07 | 2.467E-07 | 5.508E-08 | 1.900E-07 | 3.000E-07 |
| 10179 | 509 | C509Y | A | 2.2e-07 | 1.1e-07 | 2.3e-07 | 1.867E-07 | 6.658E-08 | 1.100E-07 | 2.300E-07 |
| 10180 | 509 | C509V | A | 2.5e-07 | 4.1e-07 | 2.6e-07 | 3.067E-07 | 8.963E-08 | 2.500E-07 | 4.100E-07 |
| 10181 | 510 | S510A | A | 1.8e-07 | 1.2e-07 | 3.5e-07 | 2.167E-07 | 1.193E-07 | 1.200E-07 | 3.500E-07 |
| 10182 | 510 | S510R | A | 2.1e-07 | 4e-07 | 6.3e-07 | 4.133E-07 | 2.103E-07 | 2.100E-07 | 6.300E-07 |
| 10183 | 510 | S510N | A | 2.8e-07 | 2.4e-07 | 4.9e-07 | 3.367E-07 | 1.343E-07 | 2.400E-07 | 4.900E-07 |
| 10184 | 510 | S510D | A | 2.7e-07 | 2.4e-07 | 8.6e-07 | 4.567E-07 | 3.496E-07 | 2.400E-07 | 8.600E-07 |
| 10185 | 510 | S510C | A | 2e-07 | 1.8e-07 | 2.2e-07 | 2.000E-07 | 2.000E-08 | 1.800E-07 | 2.200E-07 |
| 10186 | 510 | S510E | A | 1.8e-07 | 2.3e-07 | 2.1e-07 | 2.067E-07 | 2.517E-08 | 1.800E-07 | 2.300E-07 |
| 10187 | 510 | S510Q | A | 1.4e-07 | 1.4e-07 | 3.5e-07 | 2.100E-07 | 1.212E-07 | 1.400E-07 | 3.500E-07 |
| 10188 | 510 | S510G | A | 6.2e-07 | 4.8e-07 | 2.4e-07 | 4.467E-07 | 1.922E-07 | 2.400E-07 | 6.200E-07 |
| 10189 | 510 | S510H | A | 3.9e-07 | 1.4e-07 | 8.2e-08 | 2.040E-07 | 1.637E-07 | 8.200E-08 | 3.900E-07 |
| 10190 | 510 | S510I | A | 4.1e-07 | 2.5e-07 | 1.5e-07 | 2.700E-07 | 1.311E-07 | 1.500E-07 | 4.100E-07 |
| 10191 | 510 | S510L | A | 5.8e-07 | 3.8e-07 | 1.4e-07 | 3.667E-07 | 2.203E-07 | 1.400E-07 | 5.800E-07 |
| 10192 | 510 | S510K | A | 1.7e-07 | 2.9e-07 | 4.2e-07 | 2.933E-07 | 1.250E-07 | 1.700E-07 | 4.200E-07 |
| 10193 | 510 | S510M | A | 1.5e-07 | 2.3e-07 | 2.2e-07 | 2.000E-07 | 4.359E-08 | 1.500E-07 | 2.300E-07 |
| 10194 | 510 | S510F | A | 2.9e-07 | 2e-07 | 4e-07 | 2.967E-07 | 1.002E-07 | 2.000E-07 | 4.000E-07 |
| 10195 | 510 | S510P | A | 2.1e-07 | 2.7e-07 | 5.2e-07 | 3.333E-07 | 1.644E-07 | 2.100E-07 | 5.200E-07 |
| 10196 | 510 | S510S | A | 2.1e-07 | 3e-07 | 2.9e-07 | 2.667E-07 | 4.933E-08 | 2.100E-07 | 3.000E-07 |
| 10197 | 510 | S510T | A | 2e-07 | 1.6e-07 | 2.7e-07 | 2.100E-07 | 5.568E-08 | 1.600E-07 | 2.700E-07 |
| 10198 | 510 | S510W | A | 2.3e-07 | 2.1e-07 | 2.6e-07 | 2.333E-07 | 2.517E-08 | 2.100E-07 | 2.600E-07 |
| 10199 | 510 | S510Y | A | 9.5e-08 | 1.6e-07 | 2.1e-07 | 1.550E-07 | 5.766E-08 | 9.500E-08 | 2.100E-07 |

| | | | | | | | | | | |
|---|---|---|---|---|---|---|---|---|---|---|
| 10200 | 510 | S510V | A | 1.6e-07 | 2e-07 | 2.4e-07 | 2.000E-07 | 4.000E-08 | 1.600E-07 | 2.400E-07 |
| 10201 | 511 | Q511A | A | 1.9e-07 | 3.7e-07 | 3.1e-07 | 2.900E-07 | 9.165E-08 | 1.900E-07 | 3.700E-07 |
| 10202 | 511 | Q511R | A | 1.7e-07 | 1.2e-07 | 2.5e-07 | 1.800E-07 | 6.557E-08 | 1.200E-07 | 2.500E-07 |
| 10203 | 511 | Q511N | A | 2.2e-07 | 1.7e-07 | 1.7e-07 | 1.867E-07 | 2.887E-08 | 1.700E-07 | 2.200E-07 |
| 10204 | 511 | Q511D | A | 5.7e-07 | 3.5e-07 | 2.8e-07 | 4.000E-07 | 1.513E-07 | 2.800E-07 | 5.700E-07 |
| 10205 | 511 | Q511C | A | 1.8e-07 | 2.6e-07 | 2.6e-07 | 2.333E-07 | 4.619E-08 | 1.800E-07 | 2.600E-07 |
| 10206 | 511 | Q511E | A | 1.6e-07 | 3.2e-07 | 3.3e-07 | 2.700E-07 | 9.539E-08 | 1.600E-07 | 3.300E-07 |
| 10207 | 511 | Q511Q | A | 3.2e-07 | 3.9e-07 | 2.9e-07 | 3.333E-07 | 5.132E-08 | 2.900E-07 | 3.900E-07 |
| 10208 | 511 | Q511G | A | 1.1e-07 | 2.9e-07 | 3.3e-07 | 2.433E-07 | 1.172E-07 | 1.100E-07 | 3.300E-07 |
| 10209 | 511 | Q511H | A | 2.3e-07 | 3.7e-07 | 3.6e-07 | 3.200E-07 | 7.810E-08 | 2.300E-07 | 3.700E-07 |
| 10210 | 511 | Q511I | A | 1.9e-07 | 2.3e-07 | 5e-07 | 3.067E-07 | 1.686E-07 | 1.900E-07 | 5.000E-07 |
| 10211 | 511 | Q511L | A | 2.5e-07 | 2.5e-07 | 1.3e-07 | 2.100E-07 | 6.928E-08 | 1.300E-07 | 2.500E-07 |
| 10212 | 511 | Q511K | A | 2.9e-07 | 4.1e-07 | 4.8e-07 | 3.933E-07 | 9.609E-08 | 2.900E-07 | 4.800E-07 |
| 10213 | 511 | Q511M | A | 6e-07 | 5.1e-07 | 9.2e-07 | 6.767E-07 | 2.155E-07 | 5.100E-07 | 9.200E-07 |
| 10214 | 511 | Q511F | A | 2e-07 | 2.7e-07 | 1.9e-07 | 2.200E-07 | 4.359E-08 | 1.900E-07 | 2.700E-07 |
| 10215 | 511 | Q511P | A | 2.6e-07 | 1.6e-07 | 2.4e-07 | 2.200E-07 | 5.292E-08 | 1.600E-07 | 2.600E-07 |
| 10216 | 511 | Q511S | A | 1.6e-07 | 4.6e-07 | 4.3e-07 | 3.500E-07 | 1.652E-07 | 1.600E-07 | 4.600E-07 |
| 10217 | 511 | Q511T | A | 2.8e-07 | 4.1e-07 | 1.1e-07 | 2.667E-07 | 1.504E-07 | 1.100E-07 | 4.100E-07 |
| 10218 | 511 | Q511W | A | 1.7e-07 | 1.7e-07 | 3.3e-07 | 2.233E-07 | 9.238E-08 | 1.700E-07 | 3.300E-07 |
| 10219 | 511 | Q511Y | A | 2.6e-07 | 2.6e-07 | 3.7e-07 | 2.967E-07 | 6.351E-08 | 2.600E-07 | 3.700E-07 |
| 10220 | 511 | Q511V | A | 6.4e-07 | 2.2e-07 | 1.5e-07 | 3.367E-07 | 2.650E-07 | 1.500E-07 | 6.400E-07 |
| 10221 | 512 | F512A | A | 1.8e-07 | 2.3e-07 | 1.7e-07 | 1.933E-07 | 3.215E-08 | 1.700E-07 | 2.300E-07 |
| 10222 | 512 | F512R | A | 2e-07 | 1.2e-07 | 6.4e-07 | 3.200E-07 | 2.800E-07 | 1.200E-07 | 6.400E-07 |
| 10223 | 512 | F512N | A | 2.2e-07 | 1e-07 | 1.4e-07 | 1.533E-07 | 6.110E-08 | 1.000E-07 | 2.200E-07 |
| 10224 | 512 | F512D | A | 2.6e-07 | 3e-07 | 2.4e-07 | 2.667E-07 | 3.055E-08 | 2.400E-07 | 3.000E-07 |
| 10225 | 512 | F512C | A | 2.3e-07 | 6.2e-07 | 2.3e-07 | 3.600E-07 | 2.252E-07 | 2.300E-07 | 6.200E-07 |
| 10226 | 512 | F512E | A | 3.6e-07 | 3.1e-07 | 2.6e-07 | 3.100E-07 | 5.000E-08 | 2.600E-07 | 3.600E-07 |
| 10227 | 512 | F512Q | A | 2.2e-07 | 3.1e-07 | 9.1e-08 | 2.070E-07 | 1.101E-07 | 9.100E-08 | 3.100E-07 |
| 10228 | 512 | F512G | A | 2.8e-07 | 1.5e-07 | 3.4e-07 | 2.567E-07 | 9.713E-08 | 1.500E-07 | 3.400E-07 |
| 10229 | 512 | F512H | A | 1.8e-07 | 1.6e-07 | 2.9e-07 | 2.100E-07 | 7.000E-08 | 1.600E-07 | 2.900E-07 |
| 10230 | 512 | F512I | A | 3.8e-07 | 2e-07 | 1.2e-07 | 2.333E-07 | 1.332E-07 | 1.200E-07 | 3.800E-07 |
| 10231 | 512 | F512L | A | 4.8e-07 | 1.7e-07 | 6.4e-07 | 4.300E-07 | 2.390E-07 | 1.700E-07 | 6.400E-07 |
| 10232 | 512 | F512K | A | 2.5e-07 | 4.4e-07 | 4.1e-07 | 3.667E-07 | 1.021E-07 | 2.500E-07 | 4.400E-07 |
| 10233 | 512 | F512M | A | 3.4e-07 | 2.4e-07 | 3e-07 | 2.933E-07 | 5.033E-08 | 2.400E-07 | 3.400E-07 |
| 10234 | 512 | F512F | A | 3.1e-07 | 1.3e-07 | 3.6e-07 | 2.667E-07 | 1.210E-07 | 1.300E-07 | 3.600E-07 |
| 10235 | 512 | F512P | A | 3.5e-07 | 3.2e-07 | 8.3e-08 | 2.510E-07 | 1.463E-07 | 8.300E-08 | 3.500E-07 |
| 10236 | 512 | F512S | A | 1.8e-07 | 3.1e-07 | 2.4e-07 | 2.433E-07 | 6.506E-08 | 1.800E-07 | 3.100E-07 |
| 10237 | 512 | F512T | A | 3e-07 | 2.4e-07 | 1.7e-07 | 2.367E-07 | 6.506E-08 | 1.700E-07 | 3.000E-07 |
| 10238 | 512 | F512W | A | 2.4e-07 | 1.6e-07 | 1.8e-07 | 1.933E-07 | 4.163E-08 | 1.600E-07 | 2.400E-07 |
| 10239 | 512 | F512Y | A | 5.4e-07 | 3.7e-07 | 3.3e-07 | 4.133E-07 | 1.115E-07 | 3.300E-07 | 5.400E-07 |
| 10240 | 512 | F512V | A | 4.8e-07 | 2.3e-07 | 1.7e-07 | 2.933E-07 | 1.644E-07 | 1.700E-07 | 4.800E-07 |
| 10241 | 513 | L513A | A | 1.6e-07 | 1.9e-07 | 3.5e-07 | 2.333E-07 | 1.021E-07 | 1.600E-07 | 3.500E-07 |
| 10242 | 513 | L513R | A | 2.8e-07 | 2.3e-07 | 1.8e-07 | 2.300E-07 | 5.000E-08 | 1.800E-07 | 2.800E-07 |
| 10243 | 513 | L513N | A | 2.2e-07 | 3.2e-07 | 4.4e-07 | 3.267E-07 | 1.102E-07 | 2.200E-07 | 4.400E-07 |
| 10244 | 513 | L513D | A | 2.6e-07 | 3.4e-07 | 3.1e-07 | 3.033E-07 | 4.041E-08 | 2.600E-07 | 3.400E-07 |
| 10245 | 513 | L513C | A | 2.2e-07 | 1.8e-07 | 4.4e-07 | 2.800E-07 | 1.400E-07 | 1.800E-07 | 4.400E-07 |
| 10246 | 513 | L513E | A | 3.3e-07 | 1.6e-07 | 2e-07 | 2.300E-07 | 8.888E-08 | 1.600E-07 | 3.300E-07 |
| 10247 | 513 | L513Q | A | 3.6e-07 | 1.9e-07 | 1.3e-07 | 2.267E-07 | 1.193E-07 | 1.300E-07 | 3.600E-07 |
| 10248 | 513 | L513G | A | 2.1e-07 | 3.6e-07 | 2.1e-07 | 2.600E-07 | 8.660E-08 | 2.100E-07 | 3.600E-07 |
| 10249 | 513 | L513H | A | 2.6e-07 | 1.7e-07 | 4.9e-07 | 3.067E-07 | 1.650E-07 | 1.700E-07 | 4.900E-07 |
| 10250 | 513 | L513I | A | 3.4e-07 | 2.2e-07 | 2.5e-07 | 2.700E-07 | 6.245E-08 | 2.200E-07 | 3.400E-07 |
| 10251 | 513 | L513L | A | 1.6e-07 | 3.8e-08 | 2.3e-07 | 1.427E-07 | 9.717E-08 | 3.800E-08 | 2.300E-07 |
| 10252 | 513 | L513K | A | 9.5e-08 | 7.6e-08 | 2.1e-07 | 1.270E-07 | 7.251E-08 | 7.600E-08 | 2.100E-07 |
| 10253 | 513 | L513M | A | 2.8e-07 | 2.3e-07 | 1.8e-07 | 2.300E-07 | 5.000E-08 | 1.800E-07 | 2.800E-07 |
| 10254 | 513 | L513F | A | 3.2e-07 | 1.4e-07 | 2.2e-07 | 2.267E-07 | 9.018E-08 | 1.400E-07 | 3.200E-07 |
| 10255 | 513 | L513P | A | 3.1e-07 | 2.8e-07 | 3.3e-07 | 3.067E-07 | 2.517E-08 | 2.800E-07 | 3.300E-07 |
| 10256 | 513 | L513S | A | 2.3e-07 | 1.1e-07 | 2.1e-07 | 1.833E-07 | 6.429E-08 | 1.100E-07 | 2.300E-07 |
| 10257 | 513 | L513T | A | 3.6e-07 | 3.2e-07 | 1.7e-07 | 2.833E-07 | 1.002E-07 | 1.700E-07 | 3.600E-07 |
| 10258 | 513 | L513W | A | 1.6e-07 | 2.4e-07 | 5.3e-07 | 3.100E-07 | 1.947E-07 | 1.600E-07 | 5.300E-07 |
| 10259 | 513 | L513Y | A | 4.2e-07 | 5.2e-07 | 3e-07 | 4.133E-07 | 1.102E-07 | 3.000E-07 | 5.200E-07 |
| 10260 | 513 | L513V | A | 2.6e-07 | 2.4e-07 | 3.1e-07 | 2.700E-07 | 3.606E-08 | 2.400E-07 | 3.100E-07 |
| 10261 | 514 | R514A | A | 1.7e-07 | 2e-07 | 2.4e-07 | 2.033E-07 | 3.512E-08 | 1.700E-07 | 2.400E-07 |
| 10262 | 514 | R514R | A | 2.5e-07 | 3.3e-07 | 2.1e-07 | 2.633E-07 | 6.110E-08 | 2.100E-07 | 3.300E-07 |
| 10263 | 514 | R514N | A | 1.7e-07 | 1.3e-07 | 3.1e-07 | 2.033E-07 | 9.452E-08 | 1.300E-07 | 3.100E-07 |
| 10264 | 514 | R514D | A | 3e-07 | 3.6e-07 | 2.2e-07 | 2.933E-07 | 7.024E-08 | 2.200E-07 | 3.600E-07 |
| 10265 | 514 | R514C | A | 1.4e-07 | 2.4e-07 | 2.6e-07 | 2.133E-07 | 6.429E-08 | 1.400E-07 | 2.600E-07 |
| 10266 | 514 | R514E | A | 4e-07 | 1.6e-07 | 1.9e-07 | 2.500E-07 | 1.308E-07 | 1.600E-07 | 4.000E-07 |
| 10267 | 514 | R514Q | A | 2.9e-07 | 2.7e-07 | 2e-07 | 2.533E-07 | 4.726E-08 | 2.000E-07 | 2.900E-07 |
| 10268 | 514 | R514G | A | 2.3e-07 | 3.3e-07 | 3.9e-07 | 3.167E-07 | 8.083E-08 | 2.300E-07 | 3.900E-07 |
| 10269 | 514 | R514H | A | 4.6e-07 | 2.2e-07 | 9.5e-08 | 2.583E-07 | 1.855E-07 | 9.500E-08 | 4.600E-07 |
| 10270 | 514 | R514I | A | 2.4e-07 | 2.3e-07 | 7.3e-07 | 4.000E-07 | 2.858E-07 | 2.300E-07 | 7.300E-07 |
| 10271 | 514 | R514L | A | 6.2e-07 | 1.6e-07 | 2e-07 | 3.267E-07 | 2.548E-07 | 1.600E-07 | 6.200E-07 |
| 10272 | 514 | R514K | A | 3.9e-07 | 1.7e-07 | 1.4e-07 | 2.333E-07 | 1.365E-07 | 1.400E-07 | 3.900E-07 |
| 10273 | 514 | R514M | A | 1.7e-07 | 2.2e-07 | 1.7e-07 | 1.867E-07 | 2.887E-08 | 1.700E-07 | 2.200E-07 |
| 10274 | 514 | R514F | A | 2.3e-07 | 4e-07 | 1.2e-07 | 2.500E-07 | 1.411E-07 | 1.200E-07 | 4.000E-07 |
| 10275 | 514 | R514P | A | 4.6e-07 | 3.9e-07 | 2.9e-07 | 3.800E-07 | 8.544E-08 | 2.900E-07 | 4.600E-07 |
| 10276 | 514 | R514S | A | 3.3e-07 | 1.9e-07 | 2.8e-07 | 2.667E-07 | 7.095E-08 | 1.900E-07 | 3.300E-07 |
| 10277 | 514 | R514T | A | 3.5e-07 | 2.7e-07 | 2e-07 | 2.733E-07 | 7.506E-08 | 2.000E-07 | 3.500E-07 |
| 10278 | 514 | R514W | A | 3.3e-07 | 2.4e-07 | 1.2e-07 | 2.300E-07 | 1.054E-07 | 1.200E-07 | 3.300E-07 |
| 10279 | 514 | R514Y | A | 1.3e-07 | 2.5e-07 | 2.6e-07 | 2.133E-07 | 7.234E-08 | 1.300E-07 | 2.600E-07 |
| 10280 | 514 | R514V | A | 7.7e-08 | 2.5e-07 | 4.1e-07 | 2.457E-07 | 1.665E-07 | 7.700E-08 | 4.100E-07 |
| 10281 | 515 | G515A | A | 1.7e-07 | 2.8e-07 | 3.3e-07 | 2.600E-07 | 8.185E-08 | 1.700E-07 | 3.300E-07 |
| 10282 | 515 | G515R | A | 3.9e-07 | 2.8e-07 | 3.4e-07 | 3.367E-07 | 5.508E-08 | 2.800E-07 | 3.900E-07 |
| 10283 | 515 | G515N | A | 1.7e-07 | 2e-07 | 9.6e-08 | 1.553E-07 | 5.353E-08 | 9.600E-08 | 2.000E-07 |
| 10284 | 515 | G515D | A | 3e-07 | 2.5e-07 | 2.7e-07 | 2.733E-07 | 2.517E-08 | 2.500E-07 | 3.000E-07 |

| | | | | | | | | | | |
|---|---|---|---|---|---|---|---|---|---|---|
| 10285 | 515 | G515C | A | 1.3e-07 | 2.9e-07 | 8e-08 | 1.667E-07 | 1.097E-07 | 8.000E-08 | 2.900E-07 |
| 10286 | 515 | G515E | A | 2.7e-07 | 1.9e-07 | 2e-07 | 2.200E-07 | 4.359E-08 | 1.900E-07 | 2.700E-07 |
| 10287 | 515 | G515Q | A | 2.6e-07 | 2.3e-07 | 2e-07 | 2.300E-07 | 3.000E-08 | 2.000E-07 | 2.600E-07 |
| 10288 | 515 | G515G | A | 4.7e-07 | 4e-07 | 1.5e-07 | 3.400E-07 | 1.682E-07 | 1.500E-07 | 4.700E-07 |
| 10289 | 515 | G515H | A | 2.6e-07 | 2.5e-07 | 1.9e-07 | 2.333E-07 | 3.786E-08 | 1.900E-07 | 2.600E-07 |
| 10290 | 515 | G515I | A | 1.5e-07 | 2.6e-07 | 1.7e-07 | 1.933E-07 | 5.859E-08 | 1.500E-07 | 2.600E-07 |
| 10291 | 515 | G515L | A | 3.1e-07 | 2.1e-07 | 2.3e-07 | 2.500E-07 | 5.292E-08 | 2.100E-07 | 3.100E-07 |
| 10292 | 515 | G515K | A | 4.2e-07 | 4.1e-07 | 4e-07 | 4.100E-07 | 1.000E-08 | 4.000E-07 | 4.200E-07 |
| 10293 | 515 | G515M | A | 3.1e-07 | 2.5e-07 | 6.1e-07 | 3.900E-07 | 1.929E-07 | 2.500E-07 | 6.100E-07 |
| 10294 | 515 | G515F | A | 3e-07 | 4.3e-07 | 6.3e-07 | 4.533E-07 | 1.662E-07 | 3.000E-07 | 6.300E-07 |
| 10295 | 515 | G515P | A | 2.1e-07 | 3.4e-07 | 1.9e-07 | 2.467E-07 | 8.145E-08 | 1.900E-07 | 3.400E-07 |
| 10296 | 515 | G515S | A | 2.7e-07 | 2e-07 | 1.6e-07 | 2.100E-07 | 5.568E-08 | 1.600E-07 | 2.700E-07 |
| 10297 | 515 | G515T | A | 3e-07 | 2.3e-07 | 1.8e-07 | 2.367E-07 | 6.028E-08 | 1.800E-07 | 3.000E-07 |
| 10298 | 515 | G515W | A | 1.8e-07 | 4e-07 | 3.3e-07 | 3.033E-07 | 1.124E-07 | 1.800E-07 | 4.000E-07 |
| 10299 | 515 | G515Y | A | 4.7e-07 | 2e-07 | 3.2e-07 | 3.300E-07 | 1.353E-07 | 2.000E-07 | 4.700E-07 |
| 10300 | 515 | G515V | A | 1.1e-07 | 1.3e-07 | 4.9e-07 | 2.433E-07 | 2.139E-07 | 1.100E-07 | 4.900E-07 |
| 10301 | 516 | Q516A | A | 2.4e-07 | 4.8e-07 | 1.5e-07 | 2.900E-07 | 1.706E-07 | 1.500E-07 | 4.800E-07 |
| 10302 | 516 | Q516R | A | 4.4e-07 | 3e-07 | 1.5e-07 | 2.967E-07 | 1.450E-07 | 1.500E-07 | 4.400E-07 |
| 10303 | 516 | Q516N | A | 4.9e-07 | 2e-07 | 4e-07 | 3.633E-07 | 1.484E-07 | 2.000E-07 | 4.900E-07 |
| 10304 | 516 | Q516D | A | 2.6e-07 | 1.9e-07 | 2.5e-07 | 2.333E-07 | 3.786E-08 | 1.900E-07 | 2.600E-07 |
| 10305 | 516 | Q516C | A | 2.5e-07 | 3.4e-07 | 1.4e-07 | 2.433E-07 | 1.002E-07 | 1.400E-07 | 3.400E-07 |
| 10306 | 516 | Q516E | A | 2.5e-07 | 4e-07 | 2.9e-07 | 3.133E-07 | 7.767E-08 | 2.500E-07 | 4.000E-07 |
| 10307 | 516 | Q516Q | A | 2.3e-07 | 3.8e-07 | 3e-07 | 3.033E-07 | 7.506E-08 | 2.300E-07 | 3.800E-07 |
| 10308 | 516 | Q516G | A | 3.3e-07 | 2.9e-07 | 1.4e-07 | 2.533E-07 | 1.002E-07 | 1.400E-07 | 3.300E-07 |
| 10309 | 516 | Q516H | A | 2.1e-07 | 7.7e-07 | 2.2e-07 | 4.000E-07 | 3.205E-07 | 2.100E-07 | 7.700E-07 |
| 10310 | 516 | Q516I | A | 1.2e-07 | 2.8e-07 | 2.5e-07 | 2.167E-07 | 8.505E-08 | 1.200E-07 | 2.800E-07 |
| 10311 | 516 | Q516L | A | 2.5e-07 | 2.6e-07 | 1.4e-07 | 2.167E-07 | 6.658E-08 | 1.400E-07 | 2.600E-07 |
| 10312 | 516 | Q516K | A | 2e-07 | 1e-07 | 2.4e-07 | 1.800E-07 | 7.211E-08 | 1.000E-07 | 2.400E-07 |
| 10313 | 516 | Q516M | A | 2.5e-07 | 1.7e-07 | 2e-07 | 2.067E-07 | 4.041E-08 | 1.700E-07 | 2.500E-07 |
| 10314 | 516 | Q516F | A | 1.7e-07 | 3e-07 | 1.9e-07 | 2.200E-07 | 7.000E-08 | 1.700E-07 | 3.000E-07 |
| 10315 | 516 | Q516P | A | 1.7e-07 | 3e-07 | 2.6e-07 | 2.433E-07 | 6.658E-08 | 1.700E-07 | 3.000E-07 |
| 10316 | 516 | Q516S | A | 2.2e-07 | 1.6e-07 | 7.5e-07 | 3.767E-07 | 3.247E-07 | 1.600E-07 | 7.500E-07 |
| 10317 | 516 | Q516T | A | 3.2e-07 | 1.2e-07 | 2.3e-07 | 2.233E-07 | 1.002E-07 | 1.200E-07 | 3.200E-07 |
| 10318 | 516 | Q516W | A | 2.4e-07 | 2.6e-07 | 3.6e-07 | 2.867E-07 | 6.429E-08 | 2.400E-07 | 3.600E-07 |
| 10319 | 516 | Q516Y | A | 2.6e-07 | 2.2e-07 | 3.5e-07 | 2.767E-07 | 6.658E-08 | 2.200E-07 | 3.500E-07 |
| 10320 | 516 | Q516V | A | 2.1e-07 | 3.1e-07 | 2.5e-07 | 2.567E-07 | 5.033E-08 | 2.100E-07 | 3.100E-07 |
| 10321 | 517 | E517A | A | 3.6e-07 | 3.2e-07 | 3.7e-07 | 3.500E-07 | 2.646E-08 | 3.200E-07 | 3.700E-07 |
| 10322 | 517 | E517R | A | 3.9e-07 | 2.4e-07 | 1.5e-07 | 2.600E-07 | 1.212E-07 | 1.500E-07 | 3.900E-07 |
| 10323 | 517 | E517N | A | 4.2e-07 | 2.1e-07 | 6.2e-07 | 4.167E-07 | 2.050E-07 | 2.100E-07 | 6.200E-07 |
| 10324 | 517 | E517D | A | 2.4e-07 | 2.4e-07 | 1e-07 | 1.933E-07 | 8.083E-08 | 1.000E-07 | 2.400E-07 |
| 10325 | 517 | E517C | A | 2.5e-07 | 2.6e-07 | 2e-07 | 2.367E-07 | 3.215E-08 | 2.000E-07 | 2.600E-07 |
| 10326 | 517 | E517E | A | 2.8e-07 | 3.8e-07 | 2.6e-07 | 3.067E-07 | 6.429E-08 | 2.600E-07 | 3.800E-07 |
| 10327 | 517 | E517Q | A | 1.5e-07 | 3.3e-07 | 6.2e-07 | 3.667E-07 | 2.371E-07 | 1.500E-07 | 6.200E-07 |
| 10328 | 517 | E517G | A | 2.5e-07 | 1.1e-07 | 2e-07 | 1.867E-07 | 7.095E-08 | 1.100E-07 | 2.500E-07 |
| 10329 | 517 | E517H | A | 3.4e-07 | 2.4e-07 | 2.2e-07 | 2.667E-07 | 6.429E-08 | 2.200E-07 | 3.400E-07 |
| 10330 | 517 | E517I | A | 3.5e-07 | 4e-07 | 2.6e-07 | 3.367E-07 | 7.095E-08 | 2.600E-07 | 4.000E-07 |
| 10331 | 517 | E517L | A | 1.2e-06 | 3.3e-07 | 5.4e-07 | 6.900E-07 | 4.540E-07 | 3.300E-07 | 1.200E-06 |
| 10332 | 517 | E517K | A | 2.4e-07 | 1.6e-07 | 2.5e-07 | 2.167E-07 | 4.933E-08 | 1.600E-07 | 2.500E-07 |
| 10333 | 517 | E517M | A | 3.2e-07 | 1.1e-07 | 2e-07 | 2.100E-07 | 1.054E-07 | 1.100E-07 | 3.200E-07 |
| 10334 | 517 | E517F | A | 1.7e-07 | 2.7e-07 | 1.6e-07 | 2.000E-07 | 6.083E-08 | 1.600E-07 | 2.700E-07 |
| 10335 | 517 | E517P | A | 1.6e-07 | 1.1e-07 | 4e-07 | 2.233E-07 | 1.550E-07 | 1.100E-07 | 4.000E-07 |
| 10336 | 517 | E517S | A | 3.5e-07 | 2.4e-07 | 3e-07 | 2.967E-07 | 5.508E-08 | 2.400E-07 | 3.500E-07 |
| 10337 | 517 | E517T | A | 2.3e-07 | 3.2e-07 | 2.5e-07 | 2.667E-07 | 4.726E-08 | 2.300E-07 | 3.200E-07 |
| 10338 | 517 | E517W | A | 2.2e-07 | 2.2e-07 | 2.9e-07 | 2.433E-07 | 4.041E-08 | 2.200E-07 | 2.900E-07 |
| 10339 | 517 | E517Y | A | 1.4e-07 | 3.3e-07 | 5.5e-07 | 3.400E-07 | 2.052E-07 | 1.400E-07 | 5.500E-07 |
| 10340 | 517 | E517V | A | 8.8e-08 | 3.1e-07 | 2.9e-07 | 2.293E-07 | 1.228E-07 | 8.800E-08 | 3.100E-07 |
| 10341 | 518 | C518A | A | 2.9e-07 | 7.9e-07 | 1.7e-07 | 4.167E-07 | 3.288E-07 | 1.700E-07 | 7.900E-07 |
| 10342 | 518 | C518R | A | 3.3e-07 | 2.2e-07 | 4.3e-07 | 3.267E-07 | 1.050E-07 | 2.200E-07 | 4.300E-07 |
| 10343 | 518 | C518N | A | 1.6e-07 | 1.7e-07 | 2e-07 | 1.767E-07 | 2.082E-08 | 1.600E-07 | 2.000E-07 |
| 10344 | 518 | C518D | A | 3.4e-07 | 3.3e-07 | 1.4e-07 | 2.700E-07 | 1.127E-07 | 1.400E-07 | 3.400E-07 |
| 10345 | 518 | C518C | A | 2.5e-07 | 1.8e-07 | 3e-07 | 2.433E-07 | 6.028E-08 | 1.800E-07 | 3.000E-07 |
| 10346 | 518 | C518E | A | 2.4e-07 | 3.3e-07 | 2.7e-07 | 2.800E-07 | 4.583E-08 | 2.400E-07 | 3.300E-07 |
| 10347 | 518 | C518Q | A | 1.6e-07 | 2.2e-07 | 3.2e-07 | 2.333E-07 | 8.083E-08 | 1.600E-07 | 3.200E-07 |
| 10348 | 518 | C518G | A | 2.2e-07 | 1.5e-07 | 1.9e-07 | 1.867E-07 | 3.512E-08 | 1.500E-07 | 2.200E-07 |
| 10349 | 518 | C518H | A | 3.4e-07 | 2.2e-07 | 1.2e-07 | 2.267E-07 | 1.102E-07 | 1.200E-07 | 3.400E-07 |
| 10350 | 518 | C518I | A | 1.7e-07 | 4.9e-07 | 2.2e-07 | 2.933E-07 | 1.721E-07 | 1.700E-07 | 4.900E-07 |
| 10351 | 518 | C518L | A | 1.6e-07 | 3.4e-07 | 3.2e-07 | 2.733E-07 | 9.866E-08 | 1.600E-07 | 3.400E-07 |
| 10352 | 518 | C518K | A | 2.3e-07 | 6.6e-07 | 2e-07 | 3.633E-07 | 2.574E-07 | 2.000E-07 | 6.600E-07 |
| 10353 | 518 | C518M | A | 2.4e-07 | 3.5e-07 | 4e-07 | 3.300E-07 | 8.185E-08 | 2.400E-07 | 4.000E-07 |
| 10354 | 518 | C518F | A | 2.6e-07 | 2.1e-07 | 2.6e-07 | 2.433E-07 | 2.887E-08 | 2.100E-07 | 2.600E-07 |
| 10355 | 518 | C518P | A | 3.2e-07 | 3.1e-07 | 1.7e-07 | 2.667E-07 | 8.386E-08 | 1.700E-07 | 3.200E-07 |
| 10356 | 518 | C518S | A | 1.6e-07 | 1.9e-07 | 2.1e-07 | 1.867E-07 | 2.517E-08 | 1.600E-07 | 2.100E-07 |
| 10357 | 518 | C518T | A | 2.4e-07 | 9e-08 | 2.5e-07 | 1.933E-07 | 8.963E-08 | 9.000E-08 | 2.500E-07 |
| 10358 | 518 | C518W | A | 2.6e-07 | 3.8e-07 | 2.2e-07 | 2.867E-07 | 8.327E-08 | 2.200E-07 | 3.800E-07 |
| 10359 | 518 | C518Y | A | 2.8e-07 | 3.7e-07 | 6.5e-07 | 4.333E-07 | 1.930E-07 | 2.800E-07 | 6.500E-07 |
| 10360 | 518 | C518V | A | 1.9e-07 | 1.2e-07 | 4.9e-07 | 2.667E-07 | 1.966E-07 | 1.200E-07 | 4.900E-07 |
| 10361 | 519 | V519A | A | 2.1e-07 | 1.1e-07 | 1.8e-07 | 1.667E-07 | 5.132E-08 | 1.100E-07 | 2.100E-07 |
| 10362 | 519 | V519R | A | 1e-06 | 2e-07 | 1.5e-07 | 4.500E-07 | 4.770E-07 | 1.500E-07 | 1.000E-06 |
| 10363 | 519 | V519N | A | 1.7e-07 | 1.6e-07 | 1.5e-07 | 1.600E-07 | 1.000E-08 | 1.500E-07 | 1.700E-07 |
| 10364 | 519 | V519D | A | 2.6e-07 | 1.9e-07 | 3.6e-07 | 2.700E-07 | 8.544E-08 | 1.900E-07 | 3.600E-07 |
| 10365 | 519 | V519C | A | 2.9e-07 | 2.7e-07 | 2.5e-07 | 2.700E-07 | 2.000E-08 | 2.500E-07 | 2.900E-07 |
| 10366 | 519 | V519E | A | 3.1e-07 | 1.7e-07 | 4.6e-07 | 3.133E-07 | 1.450E-07 | 1.700E-07 | 4.600E-07 |
| 10367 | 519 | V519Q | A | 2.5e-07 | 2.1e-07 | 2.4e-07 | 2.333E-07 | 2.082E-08 | 2.100E-07 | 2.500E-07 |
| 10368 | 519 | V519G | A | 1.2e-07 | 2.4e-07 | 2e-07 | 1.867E-07 | 6.110E-08 | 1.200E-07 | 2.400E-07 |
| 10369 | 519 | V519H | A | 5.5e-07 | 1.3e-07 | 1.3e-07 | 2.700E-07 | 2.425E-07 | 1.300E-07 | 5.500E-07 |

| | | | | | | | | | | |
|---|---|---|---|---|---|---|---|---|---|---|
| 10370 | 519 | V519I | A | 3.1e-07 | 8.2e-08 | 6.6e-07 | 3.507E-07 | 2.911E-07 | 8.200E-08 | 6.600E-07 |
| 10371 | 519 | V519L | A | 2.1e-07 | 1.2e-07 | 2.7e-07 | 2.000E-07 | 7.550E-08 | 1.200E-07 | 2.700E-07 |
| 10372 | 519 | V519K | A | 1.8e-07 | 1.2e-07 | 3.3e-07 | 2.100E-07 | 1.082E-07 | 1.200E-07 | 3.300E-07 |
| 10373 | 519 | V519M | A | 2.1e-07 | 2.5e-07 | 6e-07 | 3.533E-07 | 2.146E-07 | 2.100E-07 | 6.000E-07 |
| 10374 | 519 | V519F | A | 2.4e-07 | 2.2e-07 | 1.6e-07 | 2.067E-07 | 4.163E-08 | 1.600E-07 | 2.400E-07 |
| 10375 | 519 | V519P | A | 1.8e-07 | 8e-08 | 2.9e-07 | 1.833E-07 | 1.050E-07 | 8.000E-08 | 2.900E-07 |
| 10376 | 519 | V519S | A | 2.2e-07 | 1.8e-07 | 2.9e-07 | 2.300E-07 | 5.568E-08 | 1.800E-07 | 2.900E-07 |
| 10377 | 519 | V519T | A | 4.5e-07 | 2.8e-07 | 1.9e-07 | 3.067E-07 | 1.320E-07 | 1.900E-07 | 4.500E-07 |
| 10378 | 519 | V519W | A | 4.1e-07 | 5.2e-07 | 2.6e-07 | 3.967E-07 | 1.305E-07 | 2.600E-07 | 5.200E-07 |
| 10379 | 519 | V519Y | A | 1.4e-07 | 1.8e-07 | 4.1e-07 | 2.433E-07 | 1.457E-07 | 1.400E-07 | 4.100E-07 |
| 10380 | 519 | V519V | A | 2.3e-07 | 4.4e-07 | 1.8e-07 | 2.833E-07 | 1.380E-07 | 1.800E-07 | 4.400E-07 |
| 10381 | 520 | E520A | A | 1.8e-07 | 2e-07 | 6.9e-07 | 3.567E-07 | 2.888E-07 | 1.800E-07 | 6.900E-07 |
| 10382 | 520 | E520R | A | 2.7e-07 | 3.7e-07 | 1.1e-07 | 2.500E-07 | 1.311E-07 | 1.100E-07 | 3.700E-07 |
| 10383 | 520 | E520N | A | 2.1e-07 | 4e-07 | 2e-07 | 2.700E-07 | 1.127E-07 | 2.000E-07 | 4.000E-07 |
| 10384 | 520 | E520D | A | 4.3e-07 | 2.6e-07 | 2.7e-07 | 3.200E-07 | 9.539E-08 | 2.600E-07 | 4.300E-07 |
| 10385 | 520 | E520C | A | 2.3e-07 | 3e-07 | 4.9e-07 | 3.400E-07 | 1.345E-07 | 2.300E-07 | 4.900E-07 |
| 10386 | 520 | E520E | A | 1.3e-07 | 1.1e-07 | 2.5e-07 | 1.633E-07 | 7.572E-08 | 1.100E-07 | 2.500E-07 |
| 10387 | 520 | E520Q | A | 1.5e-07 | 1.5e-07 | 2.1e-07 | 1.700E-07 | 3.464E-08 | 1.500E-07 | 2.100E-07 |
| 10388 | 520 | E520G | A | 2.6e-07 | 3.5e-07 | 4.1e-07 | 3.400E-07 | 7.550E-08 | 2.600E-07 | 4.100E-07 |
| 10389 | 520 | E520H | A | 4e-07 | 3.8e-07 | 1.8e-07 | 3.200E-07 | 1.217E-07 | 1.800E-07 | 4.000E-07 |
| 10390 | 520 | E520I | A | 2.9e-07 | 3.8e-07 | 4.9e-07 | 3.867E-07 | 1.002E-07 | 2.900E-07 | 4.900E-07 |
| 10391 | 520 | E520L | A | 2.6e-07 | 3e-07 | 3.4e-07 | 3.000E-07 | 4.000E-08 | 2.600E-07 | 3.400E-07 |
| 10392 | 520 | E520K | A | 8.3e-08 | 1.9e-07 | 2.5e-07 | 1.743E-07 | 8.460E-08 | 8.300E-08 | 2.500E-07 |
| 10393 | 520 | E520M | A | 3e-07 | 2e-07 | 2.6e-07 | 2.533E-07 | 5.033E-08 | 2.000E-07 | 3.000E-07 |
| 10394 | 520 | E520F | A | 5e-07 | 2.3e-07 | 2.3e-07 | 3.200E-07 | 1.559E-07 | 2.300E-07 | 5.000E-07 |
| 10395 | 520 | E520P | A | 3.4e-07 | 1.7e-07 | 2.8e-07 | 2.633E-07 | 8.622E-08 | 1.700E-07 | 3.400E-07 |
| 10396 | 520 | E520S | A | 5.3e-07 | 4.1e-07 | 2.7e-07 | 4.033E-07 | 1.301E-07 | 2.700E-07 | 5.300E-07 |
| 10397 | 520 | E520T | A | 2.8e-07 | 1e-07 | 3.5e-07 | 2.433E-07 | 1.290E-07 | 1.000E-07 | 3.500E-07 |
| 10398 | 520 | E520W | A | 5.8e-07 | 3e-07 | 2.5e-07 | 3.767E-07 | 1.779E-07 | 2.500E-07 | 5.800E-07 |
| 10399 | 520 | E520Y | A | 3.2e-07 | 1.5e-07 | 2.7e-07 | 2.467E-07 | 8.737E-08 | 1.500E-07 | 3.200E-07 |
| 10400 | 520 | E520V | A | 2.1e-07 | 1.6e-07 | 2.9e-07 | 2.200E-07 | 6.557E-08 | 1.600E-07 | 2.900E-07 |
| 10401 | 521 | E521A | A | 3.4e-07 | 6.8e-07 | 3.4e-07 | 4.533E-07 | 1.963E-07 | 3.400E-07 | 6.800E-07 |
| 10402 | 521 | E521R | A | 1.8e-07 | 1.7e-07 | 3e-07 | 2.167E-07 | 7.234E-08 | 1.700E-07 | 3.000E-07 |
| 10403 | 521 | E521N | A | 2.8e-07 | 6.6e-07 | 2.2e-07 | 3.867E-07 | 2.386E-07 | 2.200E-07 | 6.600E-07 |
| 10404 | 521 | E521D | A | 3.8e-07 | 4.8e-07 | 2.7e-07 | 3.767E-07 | 1.050E-07 | 2.700E-07 | 4.800E-07 |
| 10405 | 521 | E521C | A | 2.9e-07 | 3.2e-07 | 3.9e-07 | 3.333E-07 | 5.132E-08 | 2.900E-07 | 3.900E-07 |
| 10406 | 521 | E521E | A | 2.4e-07 | 1.7e-07 | 1.9e-07 | 2.000E-07 | 3.606E-08 | 1.700E-07 | 2.400E-07 |
| 10407 | 521 | E521Q | A | 1.9e-07 | 1.7e-07 | 1.4e-07 | 1.667E-07 | 2.517E-08 | 1.400E-07 | 1.900E-07 |
| 10408 | 521 | E521G | A | 1.6e-07 | 5.8e-07 | 1.2e-07 | 2.867E-07 | 2.548E-07 | 1.200E-07 | 5.800E-07 |
| 10409 | 521 | E521H | A | 7.6e-07 | 3.5e-07 | 2.8e-07 | 4.633E-07 | 2.593E-07 | 2.800E-07 | 7.600E-07 |
| 10410 | 521 | E521I | A | 1.8e-07 | 3.5e-07 | 6.2e-07 | 3.833E-07 | 2.219E-07 | 1.800E-07 | 6.200E-07 |
| 10411 | 521 | E521L | A | 6.8e-07 | 1.8e-07 | 2.4e-07 | 3.667E-07 | 2.730E-07 | 1.800E-07 | 6.800E-07 |
| 10412 | 521 | E521K | A | 3.3e-07 | 1.9e-07 | 1.4e-07 | 2.200E-07 | 9.849E-08 | 1.400E-07 | 3.300E-07 |
| 10413 | 521 | E521M | A | 3.3e-07 | 2.2e-07 | 4.9e-07 | 3.467E-07 | 1.358E-07 | 2.200E-07 | 4.900E-07 |
| 10414 | 521 | E521F | A | 3.8e-07 | 2.8e-07 | 7.2e-08 | 2.440E-07 | 1.571E-07 | 7.200E-08 | 3.800E-07 |
| 10415 | 521 | E521P | A | 3.1e-07 | 1.8e-07 | 2.1e-07 | 2.333E-07 | 6.807E-08 | 1.800E-07 | 3.100E-07 |
| 10416 | 521 | E521S | A | 2.1e-07 | 3.3e-07 | 3.2e-07 | 2.867E-07 | 6.658E-08 | 2.100E-07 | 3.300E-07 |
| 10417 | 521 | E521T | A | 3.6e-07 | 2.6e-07 | 3.7e-07 | 3.300E-07 | 6.083E-08 | 2.600E-07 | 3.700E-07 |
| 10418 | 521 | E521W | A | 2.9e-07 | 4.6e-07 | 2.5e-07 | 3.333E-07 | 1.115E-07 | 2.500E-07 | 4.600E-07 |
| 10419 | 521 | E521Y | A | 3e-07 | 5.2e-07 | 3e-07 | 3.733E-07 | 1.270E-07 | 3.000E-07 | 5.200E-07 |
| 10420 | 521 | E521V | A | 3.4e-07 | 2.8e-07 | 3.2e-07 | 3.133E-07 | 3.055E-08 | 2.800E-07 | 3.400E-07 |
| 10421 | 522 | C522A | A | 4.2e-07 | 3.4e-07 | 3.1e-07 | 3.567E-07 | 5.686E-08 | 3.100E-07 | 4.200E-07 |
| 10422 | 522 | C522R | A | 2.8e-07 | 3.7e-07 | 3.7e-07 | 3.400E-07 | 5.196E-08 | 2.800E-07 | 3.700E-07 |
| 10423 | 522 | C522N | A | 7e-08 | 1.4e-07 | 2.4e-07 | 1.500E-07 | 8.544E-08 | 7.000E-08 | 2.400E-07 |
| 10424 | 522 | C522D | A | 4.2e-07 | 3.6e-07 | 1.2e-07 | 3.000E-07 | 1.587E-07 | 1.200E-07 | 4.200E-07 |
| 10425 | 522 | C522C | A | 1.3e-07 | 4.7e-07 | 2.1e-07 | 2.700E-07 | 1.778E-07 | 1.300E-07 | 4.700E-07 |
| 10426 | 522 | C522E | A | 1.8e-07 | 2.4e-07 | 2.8e-07 | 2.333E-07 | 5.033E-08 | 1.800E-07 | 2.800E-07 |
| 10427 | 522 | C522Q | A | 1.3e-07 | 6.4e-07 | 3.1e-07 | 3.600E-07 | 2.587E-07 | 1.300E-07 | 6.400E-07 |
| 10428 | 522 | C522G | A | 4.1e-07 | 1.5e-07 | 5.4e-07 | 3.667E-07 | 1.986E-07 | 1.500E-07 | 5.400E-07 |
| 10429 | 522 | C522H | A | 2.2e-07 | 1.8e-07 | 2.4e-07 | 2.133E-07 | 3.055E-08 | 1.800E-07 | 2.400E-07 |
| 10430 | 522 | C522I | A | 3.6e-07 | 3.3e-07 | 4.9e-07 | 3.933E-07 | 8.505E-08 | 3.300E-07 | 4.900E-07 |
| 10431 | 522 | C522L | A | 2.2e-07 | 2.5e-07 | 2.7e-07 | 2.467E-07 | 2.517E-08 | 2.200E-07 | 2.700E-07 |
| 10432 | 522 | C522K | A | 2.9e-07 | 1.9e-07 | 2e-07 | 2.267E-07 | 5.508E-08 | 1.900E-07 | 2.900E-07 |
| 10433 | 522 | C522M | A | 3.4e-07 | 6.4e-07 | 2.5e-07 | 4.100E-07 | 2.042E-07 | 2.500E-07 | 6.400E-07 |
| 10434 | 522 | C522F | A | 2.1e-07 | 3.3e-07 | 3e-07 | 2.800E-07 | 6.245E-08 | 2.100E-07 | 3.300E-07 |
| 10435 | 522 | C522P | A | 2e-07 | 3.3e-07 | 2.9e-07 | 2.733E-07 | 6.658E-08 | 2.000E-07 | 3.300E-07 |
| 10436 | 522 | C522S | A | 2e-07 | 2.5e-07 | 2.7e-07 | 2.400E-07 | 3.606E-08 | 2.000E-07 | 2.700E-07 |
| 10437 | 522 | C522T | A | 3.9e-07 | 1.7e-07 | 2e-07 | 2.533E-07 | 1.193E-07 | 1.700E-07 | 3.900E-07 |
| 10438 | 522 | C522W | A | 2.6e-07 | 2.9e-07 | 2.2e-07 | 2.567E-07 | 3.512E-08 | 2.200E-07 | 2.900E-07 |
| 10439 | 522 | C522Y | A | 4.3e-07 | 1.9e-07 | 2e-07 | 2.733E-07 | 1.358E-07 | 1.900E-07 | 4.300E-07 |
| 10440 | 522 | C522V | A | 4.7e-07 | 2.2e-07 | 2.8e-07 | 3.233E-07 | 1.305E-07 | 2.200E-07 | 4.700E-07 |
| 10441 | 523 | R523A | A | 1.2e-07 | 2.2e-07 | 4e-07 | 2.467E-07 | 1.419E-07 | 1.200E-07 | 4.000E-07 |
| 10442 | 523 | R523R | A | 1.7e-07 | 7.9e-07 | 1.4e-07 | 3.667E-07 | 3.669E-07 | 1.400E-07 | 7.900E-07 |
| 10443 | 523 | R523N | A | 1.7e-07 | 3e-07 | 1.4e-07 | 2.033E-07 | 8.505E-08 | 1.400E-07 | 3.000E-07 |
| 10444 | 523 | R523D | A | 1.9e-07 | 1.6e-07 | 4.8e-07 | 2.767E-07 | 1.767E-07 | 1.600E-07 | 4.800E-07 |
| 10445 | 523 | R523C | A | 3.8e-07 | 2.1e-07 | 2.3e-07 | 2.733E-07 | 9.292E-08 | 2.100E-07 | 3.800E-07 |
| 10446 | 523 | R523E | A | 2.1e-07 | 3.1e-07 | 2.4e-07 | 2.533E-07 | 5.132E-08 | 2.100E-07 | 3.100E-07 |
| 10447 | 523 | R523Q | A | 2e-07 | 3e-07 | 1.8e-07 | 2.267E-07 | 6.429E-08 | 1.800E-07 | 3.000E-07 |
| 10448 | 523 | R523G | A | 2.9e-07 | 3e-07 | 2.5e-07 | 2.800E-07 | 2.646E-08 | 2.500E-07 | 3.000E-07 |
| 10449 | 523 | R523H | A | 1.7e-07 | 2.5e-07 | 1.7e-07 | 1.967E-07 | 4.619E-08 | 1.700E-07 | 2.500E-07 |
| 10450 | 523 | R523I | A | 2e-07 | 1.9e-07 | 4.2e-07 | 2.700E-07 | 1.300E-07 | 1.900E-07 | 4.200E-07 |
| 10451 | 523 | R523L | A | 3.3e-07 | 2.8e-07 | 2.1e-07 | 2.733E-07 | 6.028E-08 | 2.100E-07 | 3.300E-07 |
| 10452 | 523 | R523K | A | 2.6e-07 | 4.4e-07 | 1.9e-07 | 2.967E-07 | 1.290E-07 | 1.900E-07 | 4.400E-07 |
| 10453 | 523 | R523M | A | 2.5e-07 | 1.2e-07 | 1.6e-07 | 1.767E-07 | 6.658E-08 | 1.200E-07 | 2.500E-07 |
| 10454 | 523 | R523F | A | 1.2e-07 | 1.4e-07 | 3e-07 | 1.867E-07 | 9.866E-08 | 1.200E-07 | 3.000E-07 |

| | | | | | | | | | | |
|---|---|---|---|---|---|---|---|---|---|---|
| 10455 | 523 | R523P | A | 1.8e-07 | 1.8e-07 | 2.4e-07 | 2.000E-07 | 3.464E-08 | 1.800E-07 | 2.400E-07 |
| 10456 | 523 | R523S | A | 2.2e-07 | 5.3e-07 | 4.5e-07 | 4.000E-07 | 1.609E-07 | 2.200E-07 | 5.300E-07 |
| 10457 | 523 | R523T | A | 1.6e-07 | 2.1e-07 | 3.3e-07 | 2.333E-07 | 8.737E-08 | 1.600E-07 | 3.300E-07 |
| 10458 | 523 | R523W | A | 4e-07 | 1.6e-07 | 2.6e-07 | 2.733E-07 | 1.206E-07 | 1.600E-07 | 4.000E-07 |
| 10459 | 523 | R523Y | A | 2.7e-07 | 1.8e-07 | 2.4e-07 | 2.300E-07 | 4.583E-08 | 1.800E-07 | 2.700E-07 |
| 10460 | 523 | R523V | A | 3.5e-07 | 3.3e-07 | 2.9e-07 | 3.233E-07 | 3.055E-08 | 2.900E-07 | 3.500E-07 |
| 10461 | 524 | V524A | A | 1.6e-07 | 1.4e-07 | 1.5e-07 | 1.500E-07 | 1.000E-08 | 1.400E-07 | 1.600E-07 |
| 10462 | 524 | V524R | A | 1.7e-07 | 7.1e-08 | 1.5e-07 | 1.303E-07 | 5.235E-08 | 7.100E-08 | 1.700E-07 |
| 10463 | 524 | V524N | A | 2.2e-07 | 1.3e-07 | 1.9e-07 | 1.800E-07 | 4.583E-08 | 1.300E-07 | 2.200E-07 |
| 10464 | 524 | V524D | A | 1.9e-07 | 1.7e-07 | 2.9e-07 | 2.167E-07 | 6.429E-08 | 1.700E-07 | 2.900E-07 |
| 10465 | 524 | V524C | A | 1.4e-07 | 5e-07 | 2.5e-07 | 2.967E-07 | 1.845E-07 | 1.400E-07 | 5.000E-07 |
| 10466 | 524 | V524E | A | 2.1e-07 | 2.8e-07 | 1.9e-07 | 2.267E-07 | 4.726E-08 | 1.900E-07 | 2.800E-07 |
| 10467 | 524 | V524Q | A | 3.7e-07 | 4.4e-07 | 2e-07 | 3.367E-07 | 1.234E-07 | 2.000E-07 | 4.400E-07 |
| 10468 | 524 | V524G | A | 2e-07 | 3.4e-07 | 2.5e-07 | 2.633E-07 | 7.095E-08 | 2.000E-07 | 3.400E-07 |
| 10469 | 524 | V524H | A | 2.2e-07 | 2.4e-07 | 1.8e-07 | 2.133E-07 | 3.055E-08 | 1.800E-07 | 2.400E-07 |
| 10470 | 524 | V524I | A | 3.2e-07 | 4e-07 | 4.8e-07 | 4.000E-07 | 8.000E-08 | 3.200E-07 | 4.800E-07 |
| 10471 | 524 | V524L | A | 2.9e-07 | 2.2e-07 | 3.8e-07 | 2.967E-07 | 8.021E-08 | 2.200E-07 | 3.800E-07 |
| 10472 | 524 | V524K | A | 3e-07 | 1.9e-07 | 1.9e-07 | 2.267E-07 | 6.351E-08 | 1.900E-07 | 3.000E-07 |
| 10473 | 524 | V524M | A | 3.3e-07 | 3.4e-07 | 3.2e-07 | 3.300E-07 | 1.000E-08 | 3.200E-07 | 3.400E-07 |
| 10474 | 524 | V524F | A | 9.7e-08 | 2.8e-07 | 4.6e-07 | 2.790E-07 | 1.815E-07 | 9.700E-08 | 4.600E-07 |
| 10475 | 524 | V524P | A | 3.4e-07 | 3.5e-07 | 1.1e-07 | 2.667E-07 | 1.358E-07 | 1.100E-07 | 3.500E-07 |
| 10476 | 524 | V524S | A | 3.6e-07 | 3.7e-07 | 2e-07 | 3.100E-07 | 9.539E-08 | 2.000E-07 | 3.700E-07 |
| 10477 | 524 | V524T | A | 2.4e-07 | 3e-07 | 2.8e-07 | 2.733E-07 | 3.055E-08 | 2.400E-07 | 3.000E-07 |
| 10478 | 524 | V524W | A | 2.5e-07 | 2.1e-07 | 2.2e-07 | 2.267E-07 | 2.082E-08 | 2.100E-07 | 2.500E-07 |
| 10479 | 524 | V524Y | A | 3.3e-07 | 1.3e-07 | 1.3e-07 | 1.967E-07 | 1.155E-07 | 1.300E-07 | 3.300E-07 |
| 10480 | 524 | V524V | A | 2.2e-07 | 3.9e-07 | 3.1e-07 | 3.067E-07 | 8.505E-08 | 2.200E-07 | 3.900E-07 |
| 10481 | 525 | L525A | A | 2.5e-07 | 1.4e-07 | 3.2e-07 | 2.367E-07 | 9.074E-08 | 1.400E-07 | 3.200E-07 |
| 10482 | 525 | L525R | A | 2e-07 | 3.8e-07 | 2.2e-07 | 2.667E-07 | 9.866E-08 | 2.000E-07 | 3.800E-07 |
| 10483 | 525 | L525N | A | 6.7e-08 | 1.8e-07 | 2.4e-07 | 1.623E-07 | 8.784E-08 | 6.700E-08 | 2.400E-07 |
| 10484 | 525 | L525D | A | 1.4e-07 | 2.3e-07 | 2.9e-07 | 2.200E-07 | 7.550E-08 | 1.400E-07 | 2.900E-07 |
| 10485 | 525 | L525C | A | 2.9e-07 | 2.2e-07 | 6.3e-07 | 3.800E-07 | 2.193E-07 | 2.200E-07 | 6.300E-07 |
| 10486 | 525 | L525E | A | 1.6e-07 | 4.8e-07 | 2.1e-07 | 2.833E-07 | 1.721E-07 | 1.600E-07 | 4.800E-07 |
| 10487 | 525 | L525Q | A | 1.2e-07 | 3.3e-07 | 4.6e-07 | 3.033E-07 | 1.716E-07 | 1.200E-07 | 4.600E-07 |
| 10488 | 525 | L525G | A | 2.8e-07 | 1.3e-07 | 5.7e-08 | 1.557E-07 | 1.137E-07 | 5.700E-08 | 2.800E-07 |
| 10489 | 525 | L525H | A | 1.1e-07 | 2e-07 | 2.1e-07 | 1.733E-07 | 5.508E-08 | 1.100E-07 | 2.100E-07 |
| 10490 | 525 | L525I | A | 3.9e-07 | 3.5e-07 | 3.2e-07 | 3.533E-07 | 3.512E-08 | 3.200E-07 | 3.900E-07 |
| 10491 | 525 | L525L | A | 1.7e-07 | 1.7e-07 | 3.6e-07 | 2.333E-07 | 1.097E-07 | 1.700E-07 | 3.600E-07 |
| 10492 | 525 | L525K | A | 1.1e-07 | 5.3e-07 | 4.5e-07 | 3.633E-07 | 2.230E-07 | 1.100E-07 | 5.300E-07 |
| 10493 | 525 | L525M | A | 2.3e-07 | 1.3e-07 | 2.3e-07 | 1.967E-07 | 5.774E-08 | 1.300E-07 | 2.300E-07 |
| 10494 | 525 | L525F | A | 1.8e-07 | 2.8e-07 | 1.9e-07 | 2.167E-07 | 5.508E-08 | 1.800E-07 | 2.800E-07 |
| 10495 | 525 | L525P | A | 1.7e-07 | 2.4e-07 | 2.3e-07 | 2.133E-07 | 3.786E-08 | 1.700E-07 | 2.400E-07 |
| 10496 | 525 | L525S | A | 1.3e-07 | 4e-07 | 4.4e-07 | 3.233E-07 | 1.686E-07 | 1.300E-07 | 4.400E-07 |
| 10497 | 525 | L525T | A | 1e-07 | 2.6e-07 | 1.5e-07 | 1.700E-07 | 8.185E-08 | 1.000E-07 | 2.600E-07 |
| 10498 | 525 | L525W | A | 4.6e-07 | 5.2e-07 | 6.1e-07 | 5.300E-07 | 7.550E-08 | 4.600E-07 | 6.100E-07 |
| 10499 | 525 | L525Y | A | 3.1e-07 | 2e-07 | 2.8e-07 | 2.633E-07 | 5.686E-08 | 2.000E-07 | 3.100E-07 |
| 10500 | 525 | L525V | A | 2.1e-07 | 2.4e-07 | 1.9e-07 | 2.133E-07 | 2.517E-08 | 1.900E-07 | 2.400E-07 |
| 10501 | 526 | Q526A | A | 1.7e-07 | 3.2e-07 | 4.5e-07 | 3.133E-07 | 1.401E-07 | 1.700E-07 | 4.500E-07 |
| 10502 | 526 | Q526R | A | 3.3e-07 | 2e-07 | 3.3e-07 | 2.867E-07 | 7.506E-08 | 2.000E-07 | 3.300E-07 |
| 10503 | 526 | Q526N | A | 2.6e-07 | 1.9e-07 | 5.9e-07 | 3.467E-07 | 2.136E-07 | 1.900E-07 | 5.900E-07 |
| 10504 | 526 | Q526D | A | 2.4e-07 | 4.2e-07 | 2.9e-07 | 3.167E-07 | 9.292E-08 | 2.400E-07 | 4.200E-07 |
| 10505 | 526 | Q526C | A | 3.4e-07 | 3.7e-07 | 3.3e-07 | 3.467E-07 | 2.082E-08 | 3.300E-07 | 3.700E-07 |
| 10506 | 526 | Q526E | A | 1.2e-07 | 2.2e-07 | 2.1e-07 | 1.833E-07 | 5.508E-08 | 1.200E-07 | 2.200E-07 |
| 10507 | 526 | Q526Q | A | 2.7e-07 | 2.9e-07 | 2.4e-07 | 2.667E-07 | 2.517E-08 | 2.400E-07 | 2.900E-07 |
| 10508 | 526 | Q526G | A | 2.8e-07 | 4.7e-07 | 6e-07 | 4.500E-07 | 1.609E-07 | 2.800E-07 | 6.000E-07 |
| 10509 | 526 | Q526H | A | 1.9e-07 | 2.2e-07 | 1.5e-07 | 1.867E-07 | 3.512E-08 | 1.500E-07 | 2.200E-07 |
| 10510 | 526 | Q526I | A | 5.7e-07 | 3.2e-07 | 3.6e-07 | 4.167E-07 | 1.343E-07 | 3.200E-07 | 5.700E-07 |
| 10511 | 526 | Q526L | A | 1.9e-07 | 1.9e-07 | 2.3e-07 | 2.033E-07 | 2.309E-08 | 1.900E-07 | 2.300E-07 |
| 10512 | 526 | Q526K | A | 1.7e-07 | 2.9e-07 | 2e-07 | 2.200E-07 | 6.245E-08 | 1.700E-07 | 2.900E-07 |
| 10513 | 526 | Q526M | A | 2.8e-07 | 1.5e-07 | 1.6e-07 | 1.967E-07 | 7.234E-08 | 1.500E-07 | 2.800E-07 |
| 10514 | 526 | Q526F | A | 2.4e-07 | 3.2e-07 | 6.2e-08 | 2.073E-07 | 1.321E-07 | 6.200E-08 | 3.200E-07 |
| 10515 | 526 | Q526P | A | 3.8e-07 | 2.5e-07 | 1.3e-07 | 2.533E-07 | 1.250E-07 | 1.300E-07 | 3.800E-07 |
| 10516 | 526 | Q526S | A | 1.6e-07 | 4.8e-07 | 8.9e-08 | 2.430E-07 | 2.083E-07 | 8.900E-08 | 4.800E-07 |
| 10517 | 526 | Q526T | A | 1.7e-07 | 1.7e-07 | 3.9e-07 | 2.433E-07 | 1.270E-07 | 1.700E-07 | 3.900E-07 |
| 10518 | 526 | Q526W | A | 1.7e-07 | 2e-07 | 1.1e-07 | 1.600E-07 | 4.583E-08 | 1.100E-07 | 2.000E-07 |
| 10519 | 526 | Q526Y | A | 2.8e-07 | 1.2e-07 | 2e-07 | 2.000E-07 | 8.000E-08 | 1.200E-07 | 2.800E-07 |
| 10520 | 526 | Q526V | A | 2.5e-07 | 3.8e-07 | 6.1e-07 | 4.133E-07 | 1.823E-07 | 2.500E-07 | 6.100E-07 |
| 10521 | 527 | G527A | A | 4.2e-07 | 3.3e-07 | 2e-07 | 3.167E-07 | 1.106E-07 | 2.000E-07 | 4.200E-07 |
| 10522 | 527 | G527R | A | 1.9e-07 | 3.1e-07 | 2.7e-07 | 2.567E-07 | 6.110E-08 | 1.900E-07 | 3.100E-07 |
| 10523 | 527 | G527N | A | 3.4e-07 | 3.7e-07 | 1.5e-07 | 2.867E-07 | 1.193E-07 | 1.500E-07 | 3.700E-07 |
| 10524 | 527 | G527D | A | 2.5e-07 | 3.7e-07 | 1.6e-07 | 2.600E-07 | 1.054E-07 | 1.600E-07 | 3.700E-07 |
| 10525 | 527 | G527C | A | 6.8e-07 | 2e-07 | 4e-07 | 4.267E-07 | 2.411E-07 | 2.000E-07 | 6.800E-07 |
| 10526 | 527 | G527E | A | 3.5e-07 | 4.7e-07 | 2.8e-07 | 3.667E-07 | 9.609E-08 | 2.800E-07 | 4.700E-07 |
| 10527 | 527 | G527Q | A | 1.9e-07 | 1.7e-07 | 2.5e-07 | 2.033E-07 | 4.163E-08 | 1.700E-07 | 2.500E-07 |
| 10528 | 527 | G527G | A | 3e-07 | 4.7e-07 | 1.5e-07 | 3.067E-07 | 1.601E-07 | 1.500E-07 | 4.700E-07 |
| 10529 | 527 | G527H | A | 3.8e-07 | 3.2e-07 | 2.3e-07 | 3.100E-07 | 7.550E-08 | 2.300E-07 | 3.800E-07 |
| 10530 | 527 | G527I | A | 2.3e-07 | 2.2e-07 | 1.8e-07 | 2.100E-07 | 2.646E-08 | 1.800E-07 | 2.300E-07 |
| 10531 | 527 | G527L | A | 3.3e-07 | 2.6e-07 | 3e-07 | 2.967E-07 | 3.512E-08 | 2.600E-07 | 3.300E-07 |
| 10532 | 527 | G527K | A | 4.7e-07 | 3.4e-07 | 1.5e-07 | 3.200E-07 | 1.609E-07 | 1.500E-07 | 4.700E-07 |
| 10533 | 527 | G527M | A | 3.8e-07 | 2.1e-07 | 3e-07 | 2.967E-07 | 8.505E-08 | 2.100E-07 | 3.800E-07 |
| 10534 | 527 | G527F | A | 2.3e-07 | 5.6e-07 | 2.8e-07 | 3.567E-07 | 1.779E-07 | 2.300E-07 | 5.600E-07 |
| 10535 | 527 | G527P | A | 2.9e-07 | 1.7e-07 | 2e-07 | 2.200E-07 | 6.245E-08 | 1.700E-07 | 2.900E-07 |
| 10536 | 527 | G527S | A | 2.6e-07 | 2.6e-07 | 3.7e-07 | 2.967E-07 | 6.351E-08 | 2.600E-07 | 3.700E-07 |
| 10537 | 527 | G527T | A | 4.2e-07 | 2.5e-07 | 3.8e-07 | 3.500E-07 | 8.888E-08 | 2.500E-07 | 4.200E-07 |
| 10538 | 527 | G527W | A | 2.9e-07 | 1.3e-07 | 2.7e-07 | 2.300E-07 | 8.718E-08 | 1.300E-07 | 2.900E-07 |
| 10539 | 527 | G527Y | A | 2.7e-07 | 8.2e-08 | 2.7e-07 | 2.073E-07 | 1.085E-07 | 8.200E-08 | 2.700E-07 |

| | | | | | | | | | | |
|---|---|---|---|---|---|---|---|---|---|---|
| 10540 | 527 | G527V | A | 2.3e-07 | 1.6e-07 | 2.5e-07 | 2.133E-07 | 4.726E-08 | 1.600E-07 | 2.500E-07 |
| 10541 | 528 | L528A | A | 1.3e-07 | 1.9e-07 | 2.3e-07 | 1.833E-07 | 5.033E-08 | 1.300E-07 | 2.300E-07 |
| 10542 | 528 | L528R | A | 3.5e-07 | 1.7e-07 | 1.1e-07 | 2.100E-07 | 1.249E-07 | 1.100E-07 | 3.500E-07 |
| 10543 | 528 | L528N | A | 2.8e-07 | 2.8e-07 | 2e-07 | 2.533E-07 | 4.619E-08 | 2.000E-07 | 2.800E-07 |
| 10544 | 528 | L528D | A | 6.4e-07 | 6e-07 | 2.7e-07 | 5.033E-07 | 2.031E-07 | 2.700E-07 | 6.400E-07 |
| 10545 | 528 | L528C | A | 2.6e-07 | 1.2e-07 | 1.4e-07 | 1.733E-07 | 7.572E-08 | 1.200E-07 | 2.600E-07 |
| 10546 | 528 | L528E | A | 3.8e-07 | 2.4e-07 | 1.2e-07 | 2.467E-07 | 1.301E-07 | 1.200E-07 | 3.800E-07 |
| 10547 | 528 | L528Q | A | 3e-07 | 4.5e-07 | 3.7e-07 | 3.733E-07 | 7.506E-08 | 3.000E-07 | 4.500E-07 |
| 10548 | 528 | L528G | A | 2.5e-07 | 1e-07 | 1.5e-07 | 1.667E-07 | 7.638E-08 | 1.000E-07 | 2.500E-07 |
| 10549 | 528 | L528H | A | 2.9e-07 | 3.1e-07 | 1.9e-07 | 2.633E-07 | 6.429E-08 | 1.900E-07 | 3.100E-07 |
| 10550 | 528 | L528I | A | 2.9e-07 | 3.1e-07 | 3.7e-07 | 3.233E-07 | 4.163E-08 | 2.900E-07 | 3.700E-07 |
| 10551 | 528 | L528L | A | 4.3e-07 | 3.8e-07 | 3.8e-07 | 3.967E-07 | 2.887E-08 | 3.800E-07 | 4.300E-07 |
| 10552 | 528 | L528K | A | 3.3e-07 | 3e-07 | 5.1e-07 | 3.800E-07 | 1.136E-07 | 3.000E-07 | 5.100E-07 |
| 10553 | 528 | L528M | A | 5.3e-07 | 1.7e-07 | 3e-07 | 3.333E-07 | 1.823E-07 | 1.700E-07 | 5.300E-07 |
| 10554 | 528 | L528F | A | 3e-07 | 3.7e-07 | 4.4e-07 | 3.700E-07 | 7.000E-08 | 3.000E-07 | 4.400E-07 |
| 10555 | 528 | L528P | A | 3.9e-07 | 2.9e-07 | 2e-07 | 2.933E-07 | 9.504E-08 | 2.000E-07 | 3.900E-07 |
| 10556 | 528 | L528S | A | 2.4e-07 | 2.4e-07 | 1.3e-07 | 2.033E-07 | 6.351E-08 | 1.300E-07 | 2.400E-07 |
| 10557 | 528 | L528T | A | 1.1e-07 | 2.6e-07 | 2.5e-07 | 2.067E-07 | 8.386E-08 | 1.100E-07 | 2.600E-07 |
| 10558 | 528 | L528W | A | 2.7e-07 | 3e-07 | 6.1e-07 | 3.933E-07 | 1.882E-07 | 2.700E-07 | 6.100E-07 |
| 10559 | 528 | L528Y | A | 2.3e-07 | 1.6e-07 | 2.1e-07 | 2.000E-07 | 3.606E-08 | 1.600E-07 | 2.300E-07 |
| 10560 | 528 | L528V | A | 2.3e-07 | 3.8e-07 | 1.7e-07 | 2.600E-07 | 1.082E-07 | 1.700E-07 | 3.800E-07 |
| 10561 | 529 | P529A | A | 1.7e-07 | 1.5e-07 | 1.8e-07 | 1.667E-07 | 1.528E-08 | 1.500E-07 | 1.800E-07 |
| 10562 | 529 | P529R | A | 2.3e-07 | 3.4e-07 | 3.6e-07 | 3.100E-07 | 7.000E-08 | 2.300E-07 | 3.600E-07 |
| 10563 | 529 | P529N | A | 2.1e-07 | 5e-07 | 1.3e-07 | 2.800E-07 | 1.947E-07 | 1.300E-07 | 5.000E-07 |
| 10564 | 529 | P529D | A | 3.6e-07 | 3e-07 | 1.4e-07 | 2.667E-07 | 1.137E-07 | 1.400E-07 | 3.600E-07 |
| 10565 | 529 | P529C | A | 2.2e-07 | 1.5e-07 | 3.4e-07 | 2.367E-07 | 9.609E-08 | 1.500E-07 | 3.400E-07 |
| 10566 | 529 | P529E | A | 2.7e-07 | 1.5e-07 | 3.1e-07 | 2.433E-07 | 8.327E-08 | 1.500E-07 | 3.100E-07 |
| 10567 | 529 | P529Q | A | 3.3e-07 | 3e-07 | 3.3e-07 | 3.200E-07 | 1.732E-08 | 3.000E-07 | 3.300E-07 |
| 10568 | 529 | P529G | A | 1.9e-07 | 2.4e-07 | 3.4e-07 | 2.567E-07 | 7.638E-08 | 1.900E-07 | 3.400E-07 |
| 10569 | 529 | P529H | A | 3.3e-07 | 3e-07 | 1.7e-07 | 2.667E-07 | 8.505E-08 | 1.700E-07 | 3.300E-07 |
| 10570 | 529 | P529I | A | 4.8e-07 | 2.2e-07 | 2.1e-07 | 3.033E-07 | 1.531E-07 | 2.100E-07 | 4.800E-07 |
| 10571 | 529 | P529L | A | 3.6e-07 | 1.4e-07 | 3e-07 | 2.667E-07 | 1.137E-07 | 1.400E-07 | 3.600E-07 |
| 10572 | 529 | P529K | A | 1.9e-07 | 2.1e-07 | 1.3e-07 | 1.767E-07 | 4.163E-08 | 1.300E-07 | 2.100E-07 |
| 10573 | 529 | P529M | A | 4.2e-07 | 1.3e-07 | 3.5e-07 | 3.000E-07 | 1.513E-07 | 1.300E-07 | 4.200E-07 |
| 10574 | 529 | P529F | A | 2.7e-07 | 1.6e-07 | 2.1e-07 | 2.133E-07 | 5.508E-08 | 1.600E-07 | 2.700E-07 |
| 10575 | 529 | P529P | A | 3.8e-07 | 3.4e-07 | 2.3e-07 | 3.167E-07 | 7.767E-08 | 2.300E-07 | 3.800E-07 |
| 10576 | 529 | P529S | A | 1.8e-07 | 1.9e-07 | 3e-07 | 2.233E-07 | 6.658E-08 | 1.800E-07 | 3.000E-07 |
| 10577 | 529 | P529T | A | 1.7e-07 | 2.7e-07 | 1.8e-07 | 2.067E-07 | 5.508E-08 | 1.700E-07 | 2.700E-07 |
| 10578 | 529 | P529W | A | 1.3e-07 | 4.4e-07 | 2.3e-07 | 2.667E-07 | 1.582E-07 | 1.300E-07 | 4.400E-07 |
| 10579 | 529 | P529Y | A | 1.5e-07 | 1.7e-07 | 1.7e-07 | 1.633E-07 | 1.155E-08 | 1.500E-07 | 1.700E-07 |
| 10580 | 529 | P529V | A | 2e-07 | 2.4e-07 | 1.7e-07 | 2.033E-07 | 3.512E-08 | 1.700E-07 | 2.400E-07 |
| 10581 | 530 | R530A | A | 1.7e-07 | 5e-08 | 4.5e-07 | 2.233E-07 | 2.053E-07 | 5.000E-08 | 4.500E-07 |
| 10582 | 530 | R530R | A | 4.2e-07 | 4.4e-07 | 1.4e-07 | 3.333E-07 | 1.677E-07 | 1.400E-07 | 4.400E-07 |
| 10583 | 530 | R530N | A | 4.4e-07 | 1.5e-07 | 4e-07 | 3.300E-07 | 1.572E-07 | 1.500E-07 | 4.400E-07 |
| 10584 | 530 | R530D | A | 4e-07 | 2.7e-07 | 2.6e-07 | 3.100E-07 | 7.810E-08 | 2.600E-07 | 4.000E-07 |
| 10585 | 530 | R530C | A | 1.8e-07 | 1.9e-07 | 1.7e-07 | 1.800E-07 | 1.000E-08 | 1.700E-07 | 1.900E-07 |
| 10586 | 530 | R530E | A | 1.5e-07 | 3.2e-07 | 2.3e-07 | 2.333E-07 | 8.505E-08 | 1.500E-07 | 3.200E-07 |
| 10587 | 530 | R530Q | A | 2.5e-07 | 1.8e-07 | 3.3e-07 | 2.533E-07 | 7.506E-08 | 1.800E-07 | 3.300E-07 |
| 10588 | 530 | R530G | A | 2.6e-07 | 2.7e-07 | 4.8e-07 | 3.367E-07 | 1.242E-07 | 2.600E-07 | 4.800E-07 |
| 10589 | 530 | R530H | A | 3e-07 | 1.7e-07 | 3.1e-07 | 2.600E-07 | 7.810E-08 | 1.700E-07 | 3.100E-07 |
| 10590 | 530 | R530I | A | 1.9e-07 | 2.7e-07 | 2.9e-07 | 2.500E-07 | 5.292E-08 | 1.900E-07 | 2.900E-07 |
| 10591 | 530 | R530L | A | 2.4e-07 | 9.7e-08 | 1.6e-07 | 1.657E-07 | 7.167E-08 | 9.700E-08 | 2.400E-07 |
| 10592 | 530 | R530K | A | 2.3e-07 | 3.3e-07 | 3.7e-07 | 3.100E-07 | 7.211E-08 | 2.300E-07 | 3.700E-07 |
| 10593 | 530 | R530M | A | 3.5e-07 | 3.9e-07 | 5.2e-07 | 4.200E-07 | 8.888E-08 | 3.500E-07 | 5.200E-07 |
| 10594 | 530 | R530F | A | 2.4e-07 | 3.4e-07 | 2e-07 | 2.600E-07 | 7.211E-08 | 2.000E-07 | 3.400E-07 |
| 10595 | 530 | R530P | A | 6.2e-07 | 4.2e-07 | 3e-07 | 4.467E-07 | 1.617E-07 | 3.000E-07 | 6.200E-07 |
| 10596 | 530 | R530S | A | 2.2e-07 | 1.1e-07 | 2.7e-07 | 2.000E-07 | 8.185E-08 | 1.100E-07 | 2.700E-07 |
| 10597 | 530 | R530T | A | 3.2e-07 | 3.3e-07 | 2.1e-07 | 2.867E-07 | 6.658E-08 | 2.100E-07 | 3.300E-07 |
| 10598 | 530 | R530W | A | 2.1e-07 | 2.7e-07 | 3.2e-07 | 2.667E-07 | 5.508E-08 | 2.100E-07 | 3.200E-07 |
| 10599 | 530 | R530Y | A | 2.5e-07 | 2.5e-07 | 6.3e-07 | 3.767E-07 | 2.194E-07 | 2.500E-07 | 6.300E-07 |
| 10600 | 530 | R530V | A | 5.2e-07 | 9.1e-08 | 2e-07 | 2.703E-07 | 2.230E-07 | 9.100E-08 | 5.200E-07 |
| 10601 | 531 | E531A | A | 4.4e-07 | 2.8e-07 | 2e-07 | 3.067E-07 | 1.222E-07 | 2.000E-07 | 4.400E-07 |
| 10602 | 531 | E531R | A | 2.4e-07 | 2.7e-07 | 1.7e-07 | 2.267E-07 | 5.132E-08 | 1.700E-07 | 2.700E-07 |
| 10603 | 531 | E531N | A | 5.6e-07 | 3.3e-07 | 3.2e-07 | 4.033E-07 | 1.358E-07 | 3.200E-07 | 5.600E-07 |
| 10604 | 531 | E531D | A | 4.5e-07 | 3.1e-07 | 3.1e-07 | 3.567E-07 | 8.083E-08 | 3.100E-07 | 4.500E-07 |
| 10605 | 531 | E531C | A | 3.6e-07 | 5.6e-07 | 3.8e-07 | 4.333E-07 | 1.102E-07 | 3.600E-07 | 5.600E-07 |
| 10606 | 531 | E531E | A | 1.4e-07 | 2e-07 | 2.1e-07 | 1.833E-07 | 3.786E-08 | 1.400E-07 | 2.100E-07 |
| 10607 | 531 | E531Q | A | 2.1e-07 | 2.3e-07 | 2.1e-07 | 2.167E-07 | 1.155E-08 | 2.100E-07 | 2.300E-07 |
| 10608 | 531 | E531G | A | 2.5e-07 | 4.1e-07 | 3.2e-07 | 3.267E-07 | 8.021E-08 | 2.500E-07 | 4.100E-07 |
| 10609 | 531 | E531H | A | 3.9e-07 | 2e-07 | 1.8e-07 | 2.567E-07 | 1.159E-07 | 1.800E-07 | 3.900E-07 |
| 10610 | 531 | E531I | A | 5.7e-07 | 4.2e-07 | 2.3e-07 | 4.067E-07 | 1.704E-07 | 2.300E-07 | 5.700E-07 |
| 10611 | 531 | E531L | A | 3.7e-07 | 2.2e-07 | 2.4e-07 | 2.767E-07 | 8.145E-08 | 2.200E-07 | 3.700E-07 |
| 10612 | 531 | E531K | A | 4.8e-07 | 3e-07 | 2.3e-07 | 3.367E-07 | 1.290E-07 | 2.300E-07 | 4.800E-07 |
| 10613 | 531 | E531M | A | 1.8e-07 | 1.8e-07 | 3.9e-07 | 2.500E-07 | 1.212E-07 | 1.800E-07 | 3.900E-07 |
| 10614 | 531 | E531F | A | 1.9e-07 | 4.4e-07 | 1.1e-07 | 2.467E-07 | 1.721E-07 | 1.100E-07 | 4.400E-07 |
| 10615 | 531 | E531P | A | 6.1e-07 | 2.8e-07 | 1.7e-07 | 3.533E-07 | 2.290E-07 | 1.700E-07 | 6.100E-07 |
| 10616 | 531 | E531S | A | 1.9e-07 | 1.6e-07 | 5e-07 | 2.833E-07 | 1.882E-07 | 1.600E-07 | 5.000E-07 |
| 10617 | 531 | E531T | A | 2.1e-07 | 2.1e-07 | 2.9e-07 | 2.367E-07 | 4.619E-08 | 2.100E-07 | 2.900E-07 |
| 10618 | 531 | E531W | A | 2.6e-07 | 1.9e-07 | 1.5e-07 | 2.000E-07 | 5.568E-08 | 1.500E-07 | 2.600E-07 |
| 10619 | 531 | E531Y | A | 2.1e-07 | 2.5e-07 | 4.2e-07 | 2.933E-07 | 1.115E-07 | 2.100E-07 | 4.200E-07 |
| 10620 | 531 | E531V | A | 2.8e-07 | 6.6e-07 | 1.6e-07 | 3.667E-07 | 2.610E-07 | 1.600E-07 | 6.600E-07 |
| 10621 | 532 | Y532A | A | 2.5e-07 | 2.8e-07 | 5.2e-07 | 3.500E-07 | 1.480E-07 | 2.500E-07 | 5.200E-07 |
| 10622 | 532 | Y532R | A | 2.1e-07 | 8.1e-08 | 2.8e-07 | 1.903E-07 | 1.009E-07 | 8.100E-08 | 2.800E-07 |
| 10623 | 532 | Y532N | A | 2.1e-07 | 1.7e-07 | 2.9e-07 | 2.233E-07 | 6.110E-08 | 1.700E-07 | 2.900E-07 |
| 10624 | 532 | Y532D | A | 1.8e-07 | 4.9e-07 | 2.3e-07 | 3.000E-07 | 1.664E-07 | 1.800E-07 | 4.900E-07 |

| | | | | | | | | | | |
|---|---|---|---|---|---|---|---|---|---|---|
| 10625 | 532 | Y532C | A | 1.8e-07 | 2.2e-07 | 2.3e-07 | 2.100E-07 | 2.646E-08 | 1.800E-07 | 2.300E-07 |
| 10626 | 532 | Y532E | A | 2.1e-07 | 3.8e-07 | 2.1e-07 | 2.667E-07 | 9.815E-08 | 2.100E-07 | 3.800E-07 |
| 10627 | 532 | Y532Q | A | 2.9e-07 | 2.3e-07 | 2.6e-07 | 2.600E-07 | 3.000E-08 | 2.300E-07 | 2.900E-07 |
| 10628 | 532 | Y532G | A | 2.9e-07 | 2.1e-07 | 2.1e-07 | 2.367E-07 | 4.619E-08 | 2.100E-07 | 2.900E-07 |
| 10629 | 532 | Y532H | A | 2e-07 | 1.4e-07 | 2.2e-07 | 1.867E-07 | 4.163E-08 | 1.400E-07 | 2.200E-07 |
| 10630 | 532 | Y532I | A | 2e-07 | 2.4e-07 | 4.4e-07 | 2.933E-07 | 1.286E-07 | 2.000E-07 | 4.400E-07 |
| 10631 | 532 | Y532L | A | 2.3e-07 | 2.6e-07 | 1.2e-07 | 2.033E-07 | 7.371E-08 | 1.200E-07 | 2.600E-07 |
| 10632 | 532 | Y532K | A | 1.8e-07 | 2.8e-07 | 2.2e-07 | 2.267E-07 | 5.033E-08 | 1.800E-07 | 2.800E-07 |
| 10633 | 532 | Y532M | A | 2.8e-07 | 2.8e-07 | 3.6e-07 | 3.067E-07 | 4.619E-08 | 2.800E-07 | 3.600E-07 |
| 10634 | 532 | Y532F | A | 4.8e-07 | 4.4e-07 | 2.1e-07 | 3.767E-07 | 1.457E-07 | 2.100E-07 | 4.800E-07 |
| 10635 | 532 | Y532P | A | 2.6e-07 | 3.1e-07 | 4.4e-07 | 3.367E-07 | 9.292E-08 | 2.600E-07 | 4.400E-07 |
| 10636 | 532 | Y532S | A | 3.5e-07 | 6.6e-07 | 3.2e-07 | 4.433E-07 | 1.882E-07 | 3.200E-07 | 6.600E-07 |
| 10637 | 532 | Y532T | A | 2e-07 | 4.1e-07 | 1.8e-07 | 2.633E-07 | 1.274E-07 | 1.800E-07 | 4.100E-07 |
| 10638 | 532 | Y532W | A | 3.4e-07 | 1.7e-07 | 1.7e-07 | 2.267E-07 | 9.815E-08 | 1.700E-07 | 3.400E-07 |
| 10639 | 532 | Y532Y | A | 2.3e-07 | 1.7e-07 | 2.4e-07 | 2.133E-07 | 3.786E-08 | 1.700E-07 | 2.400E-07 |
| 10640 | 532 | Y532V | A | 3.1e-07 | 4.2e-07 | 2.4e-07 | 3.233E-07 | 9.074E-08 | 2.400E-07 | 4.200E-07 |
| 10641 | 533 | V533A | A | 2.1e-07 | 2.2e-07 | 1.7e-07 | 2.000E-07 | 2.646E-08 | 1.700E-07 | 2.200E-07 |
| 10642 | 533 | V533R | A | 3.5e-07 | 2.9e-07 | 2.8e-07 | 3.067E-07 | 3.786E-08 | 2.800E-07 | 3.500E-07 |
| 10643 | 533 | V533N | A | 3.2e-07 | 2.1e-07 | 9.4e-08 | 2.080E-07 | 1.130E-07 | 9.400E-08 | 3.200E-07 |
| 10644 | 533 | V533D | A | 1.4e-07 | 2.8e-07 | 1.6e-07 | 1.933E-07 | 7.572E-08 | 1.400E-07 | 2.800E-07 |
| 10645 | 533 | V533C | A | 2.3e-07 | 4.1e-07 | 2.5e-07 | 2.967E-07 | 9.866E-08 | 2.300E-07 | 4.100E-07 |
| 10646 | 533 | V533E | A | 2e-07 | 2.9e-07 | 3.5e-07 | 2.800E-07 | 7.550E-08 | 2.000E-07 | 3.500E-07 |
| 10647 | 533 | V533Q | A | 1.7e-07 | 2.5e-07 | 2.1e-07 | 2.100E-07 | 4.000E-08 | 1.700E-07 | 2.500E-07 |
| 10648 | 533 | V533G | A | 7.6e-08 | 4.9e-07 | 2.2e-07 | 2.620E-07 | 2.102E-07 | 7.600E-08 | 4.900E-07 |
| 10649 | 533 | V533H | A | 2.2e-07 | 2.4e-07 | 2e-07 | 2.200E-07 | 2.000E-08 | 2.000E-07 | 2.400E-07 |
| 10650 | 533 | V533I | A | 1.8e-07 | 6.4e-07 | 3.7e-07 | 3.967E-07 | 2.312E-07 | 1.800E-07 | 6.400E-07 |
| 10651 | 533 | V533L | A | 3.4e-07 | 4.3e-07 | 3.1e-07 | 3.600E-07 | 6.245E-08 | 3.100E-07 | 4.300E-07 |
| 10652 | 533 | V533K | A | 2.2e-07 | 2.1e-07 | 1.7e-07 | 2.000E-07 | 2.646E-08 | 1.700E-07 | 2.200E-07 |
| 10653 | 533 | V533M | A | 2.7e-07 | 2.3e-07 | 2.1e-07 | 2.367E-07 | 3.055E-08 | 2.100E-07 | 2.700E-07 |
| 10654 | 533 | V533F | A | 2e-07 | 8e-07 | 4.3e-07 | 4.767E-07 | 3.027E-07 | 2.000E-07 | 8.000E-07 |
| 10655 | 533 | V533P | A | 3.4e-07 | 2.3e-07 | 4.7e-07 | 3.467E-07 | 1.201E-07 | 2.300E-07 | 4.700E-07 |
| 10656 | 533 | V533S | A | 2.1e-07 | 1.1e-07 | 1.8e-07 | 1.667E-07 | 5.132E-08 | 1.100E-07 | 2.100E-07 |
| 10657 | 533 | V533T | A | 2.9e-07 | 2e-07 | 2.4e-07 | 2.433E-07 | 4.509E-08 | 2.000E-07 | 2.900E-07 |
| 10658 | 533 | V533W | A | 2.7e-07 | 4.1e-07 | 3.2e-07 | 3.333E-07 | 7.095E-08 | 2.700E-07 | 4.100E-07 |
| 10659 | 533 | V533Y | A | 1.6e-07 | 2e-07 | 1.6e-07 | 1.733E-07 | 2.309E-08 | 1.600E-07 | 2.000E-07 |
| 10660 | 533 | V533V | A | 2.3e-07 | 1.6e-07 | 2e-07 | 1.967E-07 | 3.512E-08 | 1.600E-07 | 2.300E-07 |
| 10661 | 534 | N534A | A | 1.5e-07 | 2.3e-07 | 2.7e-07 | 2.167E-07 | 6.110E-08 | 1.500E-07 | 2.700E-07 |
| 10662 | 534 | N534R | A | 3.5e-07 | 1.9e-07 | 3.6e-07 | 3.000E-07 | 9.539E-08 | 1.900E-07 | 3.600E-07 |
| 10663 | 534 | N534N | A | 1.9e-07 | 4.9e-07 | 1.1e-07 | 2.633E-07 | 2.003E-07 | 1.100E-07 | 4.900E-07 |
| 10664 | 534 | N534D | A | 3e-07 | 3.4e-07 | 1.7e-07 | 2.700E-07 | 8.888E-08 | 1.700E-07 | 3.400E-07 |
| 10665 | 534 | N534C | A | 2.4e-07 | 2.9e-07 | 2.1e-07 | 2.467E-07 | 4.041E-08 | 2.100E-07 | 2.900E-07 |
| 10666 | 534 | N534E | A | 2.5e-07 | 2e-07 | 1.8e-07 | 2.100E-07 | 3.606E-08 | 1.800E-07 | 2.500E-07 |
| 10667 | 534 | N534Q | A | 2.7e-07 | 1.5e-07 | 1.7e-07 | 1.967E-07 | 6.429E-08 | 1.500E-07 | 2.700E-07 |
| 10668 | 534 | N534G | A | 2e-07 | 3.5e-07 | 2.2e-07 | 2.567E-07 | 8.145E-08 | 2.000E-07 | 3.500E-07 |
| 10669 | 534 | N534H | A | 1.5e-07 | 1.6e-07 | 1.9e-07 | 1.667E-07 | 2.082E-08 | 1.500E-07 | 1.900E-07 |
| 10670 | 534 | N534I | A | 2.4e-07 | 2.3e-07 | 3e-07 | 2.567E-07 | 3.786E-08 | 2.300E-07 | 3.000E-07 |
| 10671 | 534 | N534L | A | 1.1e-07 | 4.3e-07 | 3.1e-07 | 2.833E-07 | 1.617E-07 | 1.100E-07 | 4.300E-07 |
| 10672 | 534 | N534K | A | 2.1e-07 | 5.3e-07 | 2.7e-07 | 3.367E-07 | 1.701E-07 | 2.100E-07 | 5.300E-07 |
| 10673 | 534 | N534M | A | 2.5e-07 | 1.2e-07 | 3.9e-07 | 2.533E-07 | 1.350E-07 | 1.200E-07 | 3.900E-07 |
| 10674 | 534 | N534F | A | 2.3e-07 | 2.7e-07 | 7.4e-07 | 4.133E-07 | 2.836E-07 | 2.300E-07 | 7.400E-07 |
| 10675 | 534 | N534P | A | 2.3e-07 | 3.6e-07 | 1.5e-07 | 2.467E-07 | 1.060E-07 | 1.500E-07 | 3.600E-07 |
| 10676 | 534 | N534S | A | 2e-07 | 1.8e-07 | 1e-06 | 4.600E-07 | 4.678E-07 | 1.800E-07 | 1.000E-06 |
| 10677 | 534 | N534T | A | 3e-07 | 6.7e-07 | 4.4e-07 | 4.700E-07 | 1.868E-07 | 3.000E-07 | 6.700E-07 |
| 10678 | 534 | N534W | A | 3e-07 | 2.4e-07 | 2.9e-07 | 2.767E-07 | 3.215E-08 | 2.400E-07 | 3.000E-07 |
| 10679 | 534 | N534Y | A | 3.7e-07 | 2.2e-07 | 1.8e-07 | 2.567E-07 | 1.002E-07 | 1.800E-07 | 3.700E-07 |
| 10680 | 534 | N534V | A | 1.2e-07 | 3.6e-07 | 3.5e-07 | 2.767E-07 | 1.358E-07 | 1.200E-07 | 3.600E-07 |
| 10681 | 535 | A535A | A | 5.1e-07 | 2.5e-07 | 1.4e-07 | 3.000E-07 | 1.900E-07 | 1.400E-07 | 5.100E-07 |
| 10682 | 535 | A535R | A | 1.3e-07 | 3.9e-07 | 2.5e-07 | 2.567E-07 | 1.301E-07 | 1.300E-07 | 3.900E-07 |
| 10683 | 535 | A535N | A | 2.3e-07 | 4.1e-07 | 3.7e-07 | 3.367E-07 | 9.452E-08 | 2.300E-07 | 4.100E-07 |
| 10684 | 535 | A535D | A | 2.7e-07 | 2e-07 | 1.6e-07 | 2.100E-07 | 5.568E-08 | 1.600E-07 | 2.700E-07 |
| 10685 | 535 | A535C | A | 2.4e-07 | 1.6e-07 | 3.6e-07 | 2.533E-07 | 1.007E-07 | 1.600E-07 | 3.600E-07 |
| 10686 | 535 | A535E | A | 4.5e-07 | 2.1e-07 | 3.5e-07 | 3.367E-07 | 1.206E-07 | 2.100E-07 | 4.500E-07 |
| 10687 | 535 | A535Q | A | 1.8e-07 | 3.6e-07 | 3e-07 | 2.800E-07 | 9.165E-08 | 1.800E-07 | 3.600E-07 |
| 10688 | 535 | A535G | A | 3.8e-07 | 2.9e-07 | 1.8e-07 | 2.833E-07 | 1.002E-07 | 1.800E-07 | 3.800E-07 |
| 10689 | 535 | A535H | A | 3.2e-07 | 2.1e-07 | 8.2e-08 | 2.040E-07 | 1.191E-07 | 8.200E-08 | 3.200E-07 |
| 10690 | 535 | A535I | A | 3.2e-07 | 2.5e-07 | 4e-07 | 3.233E-07 | 7.506E-08 | 2.500E-07 | 4.000E-07 |
| 10691 | 535 | A535L | A | 2.5e-07 | 8.1e-08 | 3.7e-07 | 2.337E-07 | 1.452E-07 | 8.100E-08 | 3.700E-07 |
| 10692 | 535 | A535K | A | 1.6e-07 | 3.1e-07 | 4.3e-07 | 3.000E-07 | 1.353E-07 | 1.600E-07 | 4.300E-07 |
| 10693 | 535 | A535M | A | 4.2e-07 | 2.9e-07 | 1.7e-07 | 2.933E-07 | 1.250E-07 | 1.700E-07 | 4.200E-07 |
| 10694 | 535 | A535F | A | 1.5e-07 | 1e-07 | 2.1e-07 | 1.533E-07 | 5.508E-08 | 1.000E-07 | 2.100E-07 |
| 10695 | 535 | A535P | A | 1.5e-07 | 1.4e-07 | 3.4e-07 | 2.100E-07 | 1.127E-07 | 1.400E-07 | 3.400E-07 |
| 10696 | 535 | A535S | A | 2e-07 | 2.9e-07 | 1.6e-07 | 2.167E-07 | 6.658E-08 | 1.600E-07 | 2.900E-07 |
| 10697 | 535 | A535T | A | 1.7e-07 | 4e-07 | 1.5e-07 | 2.400E-07 | 1.389E-07 | 1.500E-07 | 4.000E-07 |
| 10698 | 535 | A535W | A | 1e-07 | 2.3e-07 | 2e-07 | 1.767E-07 | 6.807E-08 | 1.000E-07 | 2.300E-07 |
| 10699 | 535 | A535Y | A | 3e-07 | 3.4e-07 | 9.6e-08 | 2.453E-07 | 1.309E-07 | 9.600E-08 | 3.400E-07 |
| 10700 | 535 | A535V | A | 3.4e-07 | 3.3e-07 | 1.9e-07 | 2.867E-07 | 8.386E-08 | 1.900E-07 | 3.400E-07 |
| 10701 | 536 | R536A | A | 3.9e-07 | 2.7e-07 | 4.8e-07 | 3.800E-07 | 1.054E-07 | 2.700E-07 | 4.800E-07 |
| 10702 | 536 | R536R | A | 2.8e-07 | 7e-07 | 1.5e-07 | 3.767E-07 | 2.875E-07 | 1.500E-07 | 7.000E-07 |
| 10703 | 536 | R536N | A | 2.1e-07 | 8.7e-07 | 2.2e-07 | 4.333E-07 | 3.782E-07 | 2.100E-07 | 8.700E-07 |
| 10704 | 536 | R536D | A | 1.6e-07 | 2.9e-07 | 2.3e-07 | 2.267E-07 | 6.506E-08 | 1.600E-07 | 2.900E-07 |
| 10705 | 536 | R536C | A | 2.8e-07 | 1.5e-07 | 2.8e-07 | 2.367E-07 | 7.506E-08 | 1.500E-07 | 2.800E-07 |
| 10706 | 536 | R536E | A | 5e-07 | 1.6e-07 | 5.3e-07 | 3.967E-07 | 2.055E-07 | 1.600E-07 | 5.300E-07 |
| 10707 | 536 | R536Q | A | 2.3e-07 | 2.4e-07 | 2.9e-07 | 2.533E-07 | 3.215E-08 | 2.300E-07 | 2.900E-07 |
| 10708 | 536 | R536G | A | 4.5e-07 | 4.1e-07 | 2.4e-07 | 3.667E-07 | 1.115E-07 | 2.400E-07 | 4.500E-07 |
| 10709 | 536 | R536H | A | 1.1e-07 | 1.2e-07 | 1.4e-07 | 1.233E-07 | 1.528E-08 | 1.100E-07 | 1.400E-07 |

| | | | | | | | | | | |
|---|---|---|---|---|---|---|---|---|---|---|
| 10710 | 536 | R536I | A | 3.7e-07 | 2.3e-07 | 4.4e-07 | 3.467E-07 | 1.069E-07 | 2.300E-07 | 4.400E-07 |
| 10711 | 536 | R536L | A | 3.1e-07 | 1.3e-07 | 2.1e-07 | 2.167E-07 | 9.018E-08 | 1.300E-07 | 3.100E-07 |
| 10712 | 536 | R536K | A | 2e-07 | 3.6e-07 | 2e-07 | 2.533E-07 | 9.238E-08 | 2.000E-07 | 3.600E-07 |
| 10713 | 536 | R536M | A | 4e-07 | 2.2e-07 | 3.3e-07 | 3.167E-07 | 9.074E-08 | 2.200E-07 | 4.000E-07 |
| 10714 | 536 | R536F | A | 2e-07 | 1e-07 | 2.1e-07 | 1.700E-07 | 6.083E-08 | 1.000E-07 | 2.100E-07 |
| 10715 | 536 | R536P | A | 1.3e-07 | 1.5e-07 | 2.2e-07 | 1.667E-07 | 4.726E-08 | 1.300E-07 | 2.200E-07 |
| 10716 | 536 | R536S | A | 2.1e-07 | 4.1e-07 | 1.6e-07 | 2.600E-07 | 1.323E-07 | 1.600E-07 | 4.100E-07 |
| 10717 | 536 | R536T | A | 2.5e-07 | 3.8e-07 | 2.3e-07 | 2.867E-07 | 8.145E-08 | 2.300E-07 | 3.800E-07 |
| 10718 | 536 | R536W | A | 1.7e-07 | 1.9e-07 | 2.3e-07 | 1.967E-07 | 3.055E-08 | 1.700E-07 | 2.300E-07 |
| 10719 | 536 | R536Y | A | 2.6e-07 | 8.9e-08 | 2e-07 | 1.830E-07 | 8.676E-08 | 8.900E-08 | 2.600E-07 |
| 10720 | 536 | R536V | A | 2.4e-07 | 3.5e-07 | 3.6e-07 | 3.167E-07 | 6.658E-08 | 2.400E-07 | 3.600E-07 |
| 10721 | 537 | H537A | A | 3.8e-07 | 2.3e-07 | 2.4e-07 | 2.833E-07 | 8.386E-08 | 2.300E-07 | 3.800E-07 |
| 10722 | 537 | H537R | A | 2.6e-07 | 2.6e-07 | 1e-07 | 2.067E-07 | 9.238E-08 | 1.000E-07 | 2.600E-07 |
| 10723 | 537 | H537N | A | 1.4e-07 | 1.7e-07 | 2.7e-07 | 1.933E-07 | 6.807E-08 | 1.400E-07 | 2.700E-07 |
| 10724 | 537 | H537D | A | 3e-07 | 3.7e-07 | 3.6e-07 | 3.433E-07 | 3.786E-08 | 3.000E-07 | 3.700E-07 |
| 10725 | 537 | H537C | A | 2.4e-07 | 6.3e-07 | 1.5e-07 | 3.400E-07 | 2.551E-07 | 1.500E-07 | 6.300E-07 |
| 10726 | 537 | H537E | A | 3.5e-07 | 1.3e-07 | 1.5e-07 | 2.100E-07 | 1.217E-07 | 1.300E-07 | 3.500E-07 |
| 10727 | 537 | H537Q | A | 5.5e-07 | 2.6e-07 | 3.6e-07 | 3.900E-07 | 1.473E-07 | 2.600E-07 | 5.500E-07 |
| 10728 | 537 | H537G | A | 1.4e-07 | 2.5e-07 | 2.4e-07 | 2.100E-07 | 6.083E-08 | 1.400E-07 | 2.500E-07 |
| 10729 | 537 | H537H | A | 3.8e-07 | 2.8e-07 | 3.5e-07 | 3.367E-07 | 5.132E-08 | 2.800E-07 | 3.800E-07 |
| 10730 | 537 | H537I | A | 3.1e-07 | 1.4e-07 | 3.1e-07 | 2.533E-07 | 9.815E-08 | 1.400E-07 | 3.100E-07 |
| 10731 | 537 | H537L | A | 4.5e-07 | 2.1e-07 | 5.9e-07 | 4.167E-07 | 1.922E-07 | 2.100E-07 | 5.900E-07 |
| 10732 | 537 | H537K | A | 3.1e-07 | 3.9e-07 | 2.4e-07 | 3.133E-07 | 7.506E-08 | 2.400E-07 | 3.900E-07 |
| 10733 | 537 | H537M | A | 1.7e-07 | 1.4e-07 | 1.6e-07 | 1.567E-07 | 1.528E-08 | 1.400E-07 | 1.700E-07 |
| 10734 | 537 | H537F | A | 2.6e-07 | 2.5e-07 | 2.7e-07 | 2.600E-07 | 1.000E-08 | 2.500E-07 | 2.700E-07 |
| 10735 | 537 | H537P | A | 2.4e-07 | 4.3e-07 | 2.3e-07 | 3.000E-07 | 1.127E-07 | 2.300E-07 | 4.300E-07 |
| 10736 | 537 | H537S | A | 1.6e-07 | 5.6e-07 | 2.2e-07 | 3.133E-07 | 2.157E-07 | 1.600E-07 | 5.600E-07 |
| 10737 | 537 | H537T | A | 3.2e-07 | 2.7e-07 | 3.7e-07 | 3.200E-07 | 5.000E-08 | 2.700E-07 | 3.700E-07 |
| 10738 | 537 | H537W | A | 2e-07 | 4.8e-07 | 1.3e-07 | 2.700E-07 | 1.852E-07 | 1.300E-07 | 4.800E-07 |
| 10739 | 537 | H537Y | A | 2.8e-07 | 1.6e-07 | 1.6e-07 | 2.000E-07 | 6.928E-08 | 1.600E-07 | 2.800E-07 |
| 10740 | 537 | H537V | A | 4.3e-07 | 2.9e-07 | 2e-07 | 3.067E-07 | 1.159E-07 | 2.000E-07 | 4.300E-07 |
| 10741 | 538 | C538A | A | 1.7e-07 | 2.7e-07 | 2e-07 | 2.133E-07 | 5.132E-08 | 1.700E-07 | 2.700E-07 |
| 10742 | 538 | C538R | A | 2.3e-07 | 3.2e-07 | 3.3e-07 | 2.933E-07 | 5.508E-08 | 2.300E-07 | 3.300E-07 |
| 10743 | 538 | C538N | A | 8.7e-08 | 1.2e-07 | 1.2e-07 | 1.090E-07 | 1.905E-08 | 8.700E-08 | 1.200E-07 |
| 10744 | 538 | C538D | A | 3.4e-07 | 3e-07 | 2.4e-07 | 2.933E-07 | 5.033E-08 | 2.400E-07 | 3.400E-07 |
| 10745 | 538 | C538C | A | 4.3e-07 | 1.9e-07 | 1.2e-07 | 2.467E-07 | 1.626E-07 | 1.200E-07 | 4.300E-07 |
| 10746 | 538 | C538E | A | 3.2e-07 | 3.2e-07 | 5.8e-07 | 4.067E-07 | 1.501E-07 | 3.200E-07 | 5.800E-07 |
| 10747 | 538 | C538Q | A | 3.5e-07 | 3.9e-07 | 1.1e-07 | 2.833E-07 | 1.514E-07 | 1.100E-07 | 3.900E-07 |
| 10748 | 538 | C538G | A | 3.5e-07 | 1.2e-07 | 2.6e-07 | 2.433E-07 | 1.159E-07 | 1.200E-07 | 3.500E-07 |
| 10749 | 538 | C538H | A | 1.1e-07 | 1.7e-07 | 3.2e-07 | 2.000E-07 | 1.082E-07 | 1.100E-07 | 3.200E-07 |
| 10750 | 538 | C538I | A | 3e-07 | 2.3e-07 | 1.7e-07 | 2.333E-07 | 6.506E-08 | 1.700E-07 | 3.000E-07 |
| 10751 | 538 | C538L | A | 1.2e-07 | 1.2e-07 | 2.1e-07 | 1.500E-07 | 5.196E-08 | 1.200E-07 | 2.100E-07 |
| 10752 | 538 | C538K | A | 2.1e-07 | 3.4e-07 | 2.5e-07 | 2.667E-07 | 6.658E-08 | 2.100E-07 | 3.400E-07 |
| 10753 | 538 | C538M | A | 3.4e-07 | 3e-07 | 3.1e-07 | 3.167E-07 | 2.082E-08 | 3.000E-07 | 3.400E-07 |
| 10754 | 538 | C538F | A | 3.6e-07 | 1.8e-07 | 7.3e-07 | 4.233E-07 | 2.804E-07 | 1.800E-07 | 7.300E-07 |
| 10755 | 538 | C538P | A | 3.4e-07 | 2.3e-07 | 1.9e-07 | 2.533E-07 | 7.767E-08 | 1.900E-07 | 3.400E-07 |
| 10756 | 538 | C538S | A | 2.8e-07 | 3e-07 | 1.8e-07 | 2.533E-07 | 6.429E-08 | 1.800E-07 | 3.000E-07 |
| 10757 | 538 | C538T | A | 3.8e-07 | 1.7e-07 | 2.1e-07 | 2.533E-07 | 1.115E-07 | 1.700E-07 | 3.800E-07 |
| 10758 | 538 | C538W | A | 3.9e-07 | 1.6e-07 | 4.9e-07 | 3.467E-07 | 1.692E-07 | 1.600E-07 | 4.900E-07 |
| 10759 | 538 | C538Y | A | 4.2e-07 | 2.6e-07 | 1.9e-07 | 2.900E-07 | 1.179E-07 | 1.900E-07 | 4.200E-07 |
| 10760 | 538 | C538V | A | 2.8e-07 | 1.5e-07 | 2.5e-07 | 2.267E-07 | 6.807E-08 | 1.500E-07 | 2.800E-07 |
| 10761 | 539 | L539A | A | 2.1e-07 | 1.7e-07 | 1.3e-07 | 1.700E-07 | 4.000E-08 | 1.300E-07 | 2.100E-07 |
| 10762 | 539 | L539R | A | 2.1e-07 | 1.2e-07 | 1.6e-07 | 1.633E-07 | 4.509E-08 | 1.200E-07 | 2.100E-07 |
| 10763 | 539 | L539N | A | 2.5e-07 | 7.7e-07 | 2.5e-07 | 4.233E-07 | 3.002E-07 | 2.500E-07 | 7.700E-07 |
| 10764 | 539 | L539D | A | 2.4e-07 | 2.8e-07 | 2.2e-07 | 2.467E-07 | 3.055E-08 | 2.200E-07 | 2.800E-07 |
| 10765 | 539 | L539C | A | 1.1e-07 | 2.2e-07 | 1.9e-07 | 1.733E-07 | 5.686E-08 | 1.100E-07 | 2.200E-07 |
| 10766 | 539 | L539E | A | 2.1e-07 | 7.5e-08 | 2.8e-07 | 1.883E-07 | 1.042E-07 | 7.500E-08 | 2.800E-07 |
| 10767 | 539 | L539Q | A | 4.3e-07 | 1.7e-07 | 1.5e-07 | 2.500E-07 | 1.562E-07 | 1.500E-07 | 4.300E-07 |
| 10768 | 539 | L539G | A | 8.7e-07 | 3.8e-07 | 1.7e-07 | 4.733E-07 | 3.592E-07 | 1.700E-07 | 8.700E-07 |
| 10769 | 539 | L539H | A | 2.7e-07 | 3.8e-07 | 1.1e-07 | 2.533E-07 | 1.358E-07 | 1.100E-07 | 3.800E-07 |
| 10770 | 539 | L539I | A | 1.8e-07 | 1.8e-07 | 2.7e-07 | 2.100E-07 | 5.196E-08 | 1.800E-07 | 2.700E-07 |
| 10771 | 539 | L539L | A | 9.4e-08 | 2.1e-07 | 3e-07 | 2.013E-07 | 1.033E-07 | 9.400E-08 | 3.000E-07 |
| 10772 | 539 | L539K | A | 1.4e-07 | 1.7e-07 | 1.4e-07 | 1.500E-07 | 1.732E-08 | 1.400E-07 | 1.700E-07 |
| 10773 | 539 | L539M | A | 2.6e-07 | 5.2e-07 | 2.7e-07 | 3.500E-07 | 1.473E-07 | 2.600E-07 | 5.200E-07 |
| 10774 | 539 | L539F | A | 6.2e-07 | 4.5e-07 | 2.6e-07 | 4.433E-07 | 1.801E-07 | 2.600E-07 | 6.200E-07 |
| 10775 | 539 | L539P | A | 3.2e-07 | 4.6e-07 | 2.2e-07 | 3.333E-07 | 1.206E-07 | 2.200E-07 | 4.600E-07 |
| 10776 | 539 | L539S | A | 1.6e-07 | 1.5e-07 | 1.8e-07 | 1.633E-07 | 1.528E-08 | 1.500E-07 | 1.800E-07 |
| 10777 | 539 | L539T | A | 1.5e-07 | 2.4e-07 | 3.2e-07 | 2.367E-07 | 8.505E-08 | 1.500E-07 | 3.200E-07 |
| 10778 | 539 | L539W | A | 3.2e-07 | 1.3e-07 | 1.5e-07 | 2.000E-07 | 1.044E-07 | 1.300E-07 | 3.200E-07 |
| 10779 | 539 | L539Y | A | 2.4e-07 | 1.6e-07 | 1.6e-07 | 1.867E-07 | 4.619E-08 | 1.600E-07 | 2.400E-07 |
| 10780 | 539 | L539V | A | 1.2e-07 | 5.2e-07 | 1.6e-07 | 2.667E-07 | 2.203E-07 | 1.200E-07 | 5.200E-07 |
| 10781 | 540 | P540A | A | 2.5e-07 | 2.4e-07 | 3.6e-07 | 2.833E-07 | 6.658E-08 | 2.400E-07 | 3.600E-07 |
| 10782 | 540 | P540R | A | 2.3e-07 | 4.6e-07 | 2.6e-07 | 3.167E-07 | 1.250E-07 | 2.300E-07 | 4.600E-07 |
| 10783 | 540 | P540N | A | 8.2e-08 | 1.7e-07 | 4.5e-07 | 2.340E-07 | 1.922E-07 | 8.200E-08 | 4.500E-07 |
| 10784 | 540 | P540D | A | 4e-07 | 1.8e-07 | 3.1e-07 | 2.967E-07 | 1.106E-07 | 1.800E-07 | 4.000E-07 |
| 10785 | 540 | P540C | A | 2.4e-07 | 2.5e-07 | 9.3e-08 | 1.943E-07 | 8.790E-08 | 9.300E-08 | 2.500E-07 |
| 10786 | 540 | P540E | A | 3.7e-07 | 2e-07 | 1.2e-07 | 2.300E-07 | 1.277E-07 | 1.200E-07 | 3.700E-07 |
| 10787 | 540 | P540Q | A | 3.5e-07 | 1.7e-07 | 1.4e-07 | 2.200E-07 | 1.136E-07 | 1.400E-07 | 3.500E-07 |
| 10788 | 540 | P540G | A | 2.9e-07 | 6.4e-07 | 4.5e-07 | 4.600E-07 | 1.752E-07 | 2.900E-07 | 6.400E-07 |
| 10789 | 540 | P540H | A | 2.9e-07 | 3.3e-07 | 1.4e-07 | 2.533E-07 | 1.002E-07 | 1.400E-07 | 3.300E-07 |
| 10790 | 540 | P540I | A | 3e-07 | 3.5e-07 | 3.9e-07 | 3.467E-07 | 4.509E-08 | 3.000E-07 | 3.900E-07 |
| 10791 | 540 | P540L | A | 2.7e-07 | 3.2e-07 | 4.6e-07 | 3.500E-07 | 9.849E-08 | 2.700E-07 | 4.600E-07 |
| 10792 | 540 | P540K | A | 5.5e-07 | 1.9e-07 | 1.1e-07 | 2.833E-07 | 2.344E-07 | 1.100E-07 | 5.500E-07 |
| 10793 | 540 | P540M | A | 3.9e-07 | 2.6e-07 | 1.4e-07 | 2.633E-07 | 1.250E-07 | 1.400E-07 | 3.900E-07 |
| 10794 | 540 | P540F | A | 8.7e-07 | 5.4e-07 | 1.6e-07 | 5.233E-07 | 3.553E-07 | 1.600E-07 | 8.700E-07 |

| | | | | | | | | | | |
|---|---|---|---|---|---|---|---|---|---|---|
| 10795 | 540 | P540P | A | 2.4e-07 | 6.6e-07 | 4.3e-07 | 4.433E-07 | 2.103E-07 | 2.400E-07 | 6.600E-07 |
| 10796 | 540 | P540S | A | 3.2e-07 | 2.7e-07 | 5.7e-07 | 3.867E-07 | 1.607E-07 | 2.700E-07 | 5.700E-07 |
| 10797 | 540 | P540T | A | 1.3e-07 | 1.1e-07 | 3e-07 | 1.800E-07 | 1.044E-07 | 1.100E-07 | 3.000E-07 |
| 10798 | 540 | P540W | A | 4e-07 | 2.4e-07 | 3.9e-07 | 3.433E-07 | 8.963E-08 | 2.400E-07 | 4.000E-07 |
| 10799 | 540 | P540Y | A | 1.4e-07 | 2.8e-07 | 1.8e-07 | 2.000E-07 | 7.211E-08 | 1.400E-07 | 2.800E-07 |
| 10800 | 540 | P540V | A | 2.1e-07 | 3.1e-07 | 1.6e-07 | 2.267E-07 | 7.638E-08 | 1.600E-07 | 3.100E-07 |
| 10801 | 541 | C541A | A | 1.5e-07 | 2e-07 | 2.3e-07 | 1.933E-07 | 4.041E-08 | 1.500E-07 | 2.300E-07 |
| 10802 | 541 | C541R | A | 2.3e-07 | 2.8e-07 | 1.6e-07 | 2.233E-07 | 6.028E-08 | 1.600E-07 | 2.800E-07 |
| 10803 | 541 | C541N | A | 2.1e-07 | 3.6e-07 | 1.6e-07 | 2.433E-07 | 1.041E-07 | 1.600E-07 | 3.600E-07 |
| 10804 | 541 | C541D | A | 2.3e-07 | 1.1e-07 | 3.2e-07 | 2.200E-07 | 1.054E-07 | 1.100E-07 | 3.200E-07 |
| 10805 | 541 | C541C | A | 1.9e-07 | 2.1e-07 | 4.5e-07 | 2.833E-07 | 1.447E-07 | 1.900E-07 | 4.500E-07 |
| 10806 | 541 | C541E | A | 2.3e-07 | 1.7e-07 | 1.7e-07 | 1.900E-07 | 3.464E-08 | 1.700E-07 | 2.300E-07 |
| 10807 | 541 | C541Q | A | 2e-07 | 2.1e-07 | 3.4e-07 | 2.500E-07 | 7.810E-08 | 2.000E-07 | 3.400E-07 |
| 10808 | 541 | C541G | A | 3.4e-07 | 3.4e-07 | 3.4e-07 | 3.400E-07 | 0.000E+00 | 3.400E-07 | 3.400E-07 |
| 10809 | 541 | C541H | A | 1.7e-07 | 1.6e-07 | 1.8e-07 | 1.700E-07 | 1.000E-08 | 1.600E-07 | 1.800E-07 |
| 10810 | 541 | C541I | A | 3.2e-07 | 2.7e-07 | 2.3e-07 | 2.733E-07 | 4.509E-08 | 2.300E-07 | 3.200E-07 |
| 10811 | 541 | C541L | A | 2.9e-07 | 1.9e-07 | 3e-07 | 2.600E-07 | 6.083E-08 | 1.900E-07 | 3.000E-07 |
| 10812 | 541 | C541K | A | 1.3e-07 | 1.7e-07 | 1.5e-07 | 1.500E-07 | 2.000E-08 | 1.300E-07 | 1.700E-07 |
| 10813 | 541 | C541M | A | 6.6e-07 | 2.8e-07 | 2.3e-07 | 3.900E-07 | 2.352E-07 | 2.300E-07 | 6.600E-07 |
| 10814 | 541 | C541F | A | 2.1e-07 | 3.5e-07 | 3.4e-07 | 3.000E-07 | 7.810E-08 | 2.100E-07 | 3.500E-07 |
| 10815 | 541 | C541P | A | 4.4e-07 | 1.9e-07 | 2.4e-07 | 2.900E-07 | 1.323E-07 | 1.900E-07 | 4.400E-07 |
| 10816 | 541 | C541S | A | 2.2e-07 | 1.6e-07 | 2.9e-07 | 2.233E-07 | 6.506E-08 | 1.600E-07 | 2.900E-07 |
| 10817 | 541 | C541T | A | 3.3e-07 | 3.1e-07 | 1.5e-07 | 2.633E-07 | 9.866E-08 | 1.500E-07 | 3.300E-07 |
| 10818 | 541 | C541W | A | 3e-07 | 1.9e-07 | 2.7e-07 | 2.533E-07 | 5.686E-08 | 1.900E-07 | 3.000E-07 |
| 10819 | 541 | C541Y | A | 4.5e-07 | 2.9e-07 | 2.7e-07 | 3.367E-07 | 9.866E-08 | 2.700E-07 | 4.500E-07 |
| 10820 | 541 | C541V | A | 8.2e-08 | 3.5e-07 | 2.8e-07 | 2.373E-07 | 1.390E-07 | 8.200E-08 | 3.500E-07 |
| 10821 | 542 | H542A | A | 1.5e-07 | 2.1e-07 | 2.2e-07 | 1.933E-07 | 3.786E-08 | 1.500E-07 | 2.200E-07 |
| 10822 | 542 | H542R | A | 1.5e-07 | 2.5e-07 | 1.9e-07 | 1.967E-07 | 5.033E-08 | 1.500E-07 | 2.500E-07 |
| 10823 | 542 | H542N | A | 2.9e-07 | 3.1e-07 | 2.9e-07 | 2.967E-07 | 1.155E-08 | 2.900E-07 | 3.100E-07 |
| 10824 | 542 | H542D | A | 4.9e-07 | 4.9e-07 | 3.9e-07 | 4.567E-07 | 5.774E-08 | 3.900E-07 | 4.900E-07 |
| 10825 | 542 | H542C | A | 1.1e-07 | 2e-07 | 2e-07 | 1.700E-07 | 5.196E-08 | 1.100E-07 | 2.000E-07 |
| 10826 | 542 | H542E | A | 1.4e-07 | 1.4e-07 | 1.5e-07 | 1.433E-07 | 5.774E-09 | 1.400E-07 | 1.500E-07 |
| 10827 | 542 | H542Q | A | 2.9e-07 | 1.8e-07 | 3.6e-07 | 2.767E-07 | 9.074E-08 | 1.800E-07 | 3.600E-07 |
| 10828 | 542 | H542G | A | 1.9e-07 | 2.2e-07 | 3.7e-07 | 2.600E-07 | 9.644E-08 | 1.900E-07 | 3.700E-07 |
| 10829 | 542 | H542H | A | 2.6e-07 | 5.5e-07 | 2.5e-07 | 3.533E-07 | 1.704E-07 | 2.500E-07 | 5.500E-07 |
| 10830 | 542 | H542I | A | 2.7e-07 | 2.9e-07 | 1.6e-07 | 2.400E-07 | 7.000E-08 | 1.600E-07 | 2.900E-07 |
| 10831 | 542 | H542L | A | 2.2e-07 | 7.6e-07 | 1.7e-07 | 3.833E-07 | 3.272E-07 | 1.700E-07 | 7.600E-07 |
| 10832 | 542 | H542K | A | 1.4e-07 | 2.5e-07 | 1.5e-07 | 1.800E-07 | 6.083E-08 | 1.400E-07 | 2.500E-07 |
| 10833 | 542 | H542M | A | 4.2e-07 | 2.7e-07 | 1e-06 | 5.633E-07 | 3.855E-07 | 2.700E-07 | 1.000E-06 |
| 10834 | 542 | H542F | A | 2.4e-07 | 2.6e-07 | 1.6e-07 | 2.200E-07 | 5.292E-08 | 1.600E-07 | 2.600E-07 |
| 10835 | 542 | H542P | A | 1.7e-07 | 1.5e-07 | 4.8e-07 | 2.667E-07 | 1.850E-07 | 1.500E-07 | 4.800E-07 |
| 10836 | 542 | H542S | A | 2.1e-07 | 3e-07 | 4.1e-07 | 3.067E-07 | 1.002E-07 | 2.100E-07 | 4.100E-07 |
| 10837 | 542 | H542T | A | 2.4e-07 | 3.5e-07 | 3e-07 | 2.967E-07 | 5.508E-08 | 2.400E-07 | 3.500E-07 |
| 10838 | 542 | H542W | A | 2.4e-07 | 4e-07 | 1.7e-07 | 2.700E-07 | 1.179E-07 | 1.700E-07 | 4.000E-07 |
| 10839 | 542 | H542Y | A | 2.5e-07 | 2.6e-07 | 3.4e-07 | 2.833E-07 | 4.933E-08 | 2.500E-07 | 3.400E-07 |
| 10840 | 542 | H542V | A | 2.4e-07 | 3.7e-07 | 1.7e-07 | 2.600E-07 | 1.015E-07 | 1.700E-07 | 3.700E-07 |
| 10841 | 543 | P543A | A | 1.3e-07 | 2.2e-07 | 1.6e-07 | 1.700E-07 | 4.583E-08 | 1.300E-07 | 2.200E-07 |
| 10842 | 543 | P543R | A | 4.8e-07 | 8.8e-07 | 2.6e-07 | 5.400E-07 | 3.143E-07 | 2.600E-07 | 8.800E-07 |
| 10843 | 543 | P543N | A | 5.3e-07 | 4.2e-07 | 2.2e-07 | 3.900E-07 | 1.572E-07 | 2.200E-07 | 5.300E-07 |
| 10844 | 543 | P543D | A | 1.1e-07 | 1.7e-07 | 3.3e-07 | 2.033E-07 | 1.137E-07 | 1.100E-07 | 3.300E-07 |
| 10845 | 543 | P543C | A | 6.1e-07 | 1.2e-07 | 1.1e-07 | 2.800E-07 | 2.858E-07 | 1.100E-07 | 6.100E-07 |
| 10846 | 543 | P543E | A | 1.3e-07 | 1.6e-07 | 1.6e-07 | 1.500E-07 | 1.732E-08 | 1.300E-07 | 1.600E-07 |
| 10847 | 543 | P543Q | A | 1.9e-07 | 4.4e-07 | 2.1e-07 | 2.800E-07 | 1.389E-07 | 1.900E-07 | 4.400E-07 |
| 10848 | 543 | P543G | A | 2.4e-07 | 1.3e-07 | 4.1e-07 | 2.600E-07 | 1.411E-07 | 1.300E-07 | 4.100E-07 |
| 10849 | 543 | P543H | A | 3.2e-07 | 4.6e-07 | 3.4e-07 | 3.733E-07 | 7.572E-08 | 3.200E-07 | 4.600E-07 |
| 10850 | 543 | P543I | A | 3.4e-07 | 2.6e-07 | 4.8e-07 | 3.600E-07 | 1.114E-07 | 2.600E-07 | 4.800E-07 |
| 10851 | 543 | P543L | A | 2.9e-07 | 2.3e-07 | 2e-07 | 2.400E-07 | 4.583E-08 | 2.000E-07 | 2.900E-07 |
| 10852 | 543 | P543K | A | 2.4e-07 | 2.1e-07 | 3.8e-07 | 2.767E-07 | 9.074E-08 | 2.100E-07 | 3.800E-07 |
| 10853 | 543 | P543M | A | 2.1e-07 | 1.5e-07 | 5.7e-07 | 3.100E-07 | 2.272E-07 | 1.500E-07 | 5.700E-07 |
| 10854 | 543 | P543F | A | 2.1e-07 | 5.4e-07 | 2.3e-07 | 3.267E-07 | 1.850E-07 | 2.100E-07 | 5.400E-07 |
| 10855 | 543 | P543P | A | 2.5e-07 | 3.5e-07 | 2e-07 | 2.667E-07 | 7.638E-08 | 2.000E-07 | 3.500E-07 |
| 10856 | 543 | P543S | A | 2.8e-07 | 2.4e-07 | 2.4e-07 | 2.533E-07 | 2.309E-08 | 2.400E-07 | 2.800E-07 |
| 10857 | 543 | P543T | A | 1.2e-07 | 1.5e-07 | 2.1e-07 | 1.600E-07 | 4.583E-08 | 1.200E-07 | 2.100E-07 |
| 10858 | 543 | P543W | A | 2.5e-07 | 1e-07 | 2.6e-07 | 2.033E-07 | 8.963E-08 | 1.000E-07 | 2.600E-07 |
| 10859 | 543 | P543Y | A | 3e-07 | 2e-07 | 1.3e-07 | 2.100E-07 | 8.544E-08 | 1.300E-07 | 3.000E-07 |
| 10860 | 543 | P543V | A | 3.5e-07 | 1.8e-07 | 2.4e-07 | 2.567E-07 | 8.622E-08 | 1.800E-07 | 3.500E-07 |
| 10861 | 544 | E544A | A | 4e-07 | 3.2e-07 | 3.8e-07 | 3.667E-07 | 4.163E-08 | 3.200E-07 | 4.000E-07 |
| 10862 | 544 | E544R | A | 1.4e-07 | 1.9e-07 | 1.1e-07 | 1.467E-07 | 4.041E-08 | 1.100E-07 | 1.900E-07 |
| 10863 | 544 | E544N | A | 3.2e-07 | 1.3e-07 | 2e-07 | 2.167E-07 | 9.609E-08 | 1.300E-07 | 3.200E-07 |
| 10864 | 544 | E544D | A | 3e-07 | 1.7e-07 | 3.5e-07 | 2.733E-07 | 9.292E-08 | 1.700E-07 | 3.500E-07 |
| 10865 | 544 | E544C | A | 2.9e-07 | 6.2e-07 | 5.6e-07 | 4.900E-07 | 1.758E-07 | 2.900E-07 | 6.200E-07 |
| 10866 | 544 | E544E | A | 1.7e-07 | 2.3e-07 | 1.3e-07 | 1.767E-07 | 5.033E-08 | 1.300E-07 | 2.300E-07 |
| 10867 | 544 | E544Q | A | 1.9e-07 | 1.1e-07 | 2.5e-07 | 1.833E-07 | 7.024E-08 | 1.100E-07 | 2.500E-07 |
| 10868 | 544 | E544G | A | 2e-07 | 3.4e-07 | 1.4e-07 | 2.267E-07 | 1.026E-07 | 1.400E-07 | 3.400E-07 |
| 10869 | 544 | E544H | A | 2.3e-07 | 2.5e-07 | 2.3e-07 | 2.367E-07 | 1.155E-08 | 2.300E-07 | 2.500E-07 |
| 10870 | 544 | E544I | A | 5e-07 | 1.8e-07 | 3.2e-07 | 3.333E-07 | 1.604E-07 | 1.800E-07 | 5.000E-07 |
| 10871 | 544 | E544L | A | 2.4e-07 | 2.4e-07 | 2.6e-07 | 2.467E-07 | 1.155E-08 | 2.400E-07 | 2.600E-07 |
| 10872 | 544 | E544K | A | 2.7e-07 | 1.9e-07 | 5.8e-07 | 3.467E-07 | 2.060E-07 | 1.900E-07 | 5.800E-07 |
| 10873 | 544 | E544M | A | 2.6e-07 | 3e-07 | 1.7e-07 | 2.433E-07 | 6.658E-08 | 1.700E-07 | 3.000E-07 |
| 10874 | 544 | E544F | A | 3.6e-07 | 1.8e-07 | 2.7e-07 | 2.700E-07 | 9.000E-08 | 1.800E-07 | 3.600E-07 |
| 10875 | 544 | E544P | A | 4.8e-07 | 1.6e-07 | 5.6e-07 | 4.000E-07 | 2.117E-07 | 1.600E-07 | 5.600E-07 |
| 10876 | 544 | E544S | A | 2.6e-07 | 2.2e-07 | 1.5e-07 | 2.100E-07 | 5.568E-08 | 1.500E-07 | 2.600E-07 |
| 10877 | 544 | E544T | A | 2e-07 | 3.1e-07 | 2.4e-07 | 2.500E-07 | 5.568E-08 | 2.000E-07 | 3.100E-07 |
| 10878 | 544 | E544W | A | 3.5e-07 | 3.4e-07 | 1.9e-07 | 2.933E-07 | 8.963E-08 | 1.900E-07 | 3.500E-07 |
| 10879 | 544 | E544Y | A | 2e-07 | 4.6e-07 | 4.6e-07 | 3.733E-07 | 1.501E-07 | 2.000E-07 | 4.600E-07 |

| | | | | | | | | | | |
|---|---|---|---|---|---|---|---|---|---|---|
| 10880 | 544 | E544V | A | 4e-07 | 2.1e-07 | 4.4e-07 | 3.500E-07 | 1.229E-07 | 2.100E-07 | 4.400E-07 |
| 10881 | 545 | C545A | A | 1.8e-07 | 2.5e-07 | 3.2e-07 | 2.500E-07 | 7.000E-08 | 1.800E-07 | 3.200E-07 |
| 10882 | 545 | C545R | A | 1.9e-07 | 4e-07 | 6e-07 | 3.967E-07 | 2.050E-07 | 1.900E-07 | 6.000E-07 |
| 10883 | 545 | C545N | A | 2.1e-07 | 2.3e-07 | 2.8e-07 | 2.400E-07 | 3.606E-08 | 2.100E-07 | 2.800E-07 |
| 10884 | 545 | C545D | A | 2.9e-07 | 2.9e-07 | 2e-07 | 2.600E-07 | 5.196E-08 | 2.000E-07 | 2.900E-07 |
| 10885 | 545 | C545C | A | 3.3e-07 | 4.1e-07 | 1.4e-07 | 2.933E-07 | 1.387E-07 | 1.400E-07 | 4.100E-07 |
| 10886 | 545 | C545E | A | 2.8e-07 | 1.1e-07 | 1.9e-07 | 1.933E-07 | 8.505E-08 | 1.100E-07 | 2.800E-07 |
| 10887 | 545 | C545Q | A | 3.2e-07 | 2.1e-07 | 2.2e-07 | 2.500E-07 | 6.083E-08 | 2.100E-07 | 3.200E-07 |
| 10888 | 545 | C545G | A | 9.6e-08 | 3e-07 | 4.6e-08 | 1.473E-07 | 1.346E-07 | 4.600E-08 | 3.000E-07 |
| 10889 | 545 | C545H | A | 3.2e-07 | 1.3e-07 | 4.5e-07 | 3.000E-07 | 1.609E-07 | 1.300E-07 | 4.500E-07 |
| 10890 | 545 | C545I | A | 2.7e-07 | 2.6e-07 | 2.4e-07 | 2.567E-07 | 1.528E-08 | 2.400E-07 | 2.700E-07 |
| 10891 | 545 | C545L | A | 1.3e-07 | 3.6e-07 | 4.7e-07 | 3.200E-07 | 1.735E-07 | 1.300E-07 | 4.700E-07 |
| 10892 | 545 | C545K | A | 1.4e-07 | 2.7e-07 | 2.7e-07 | 2.267E-07 | 7.506E-08 | 1.400E-07 | 2.700E-07 |
| 10893 | 545 | C545M | A | 1.3e-07 | 2.2e-07 | 4.5e-07 | 2.667E-07 | 1.650E-07 | 1.300E-07 | 4.500E-07 |
| 10894 | 545 | C545F | A | 1e-06 | 1.9e-07 | 1.9e-07 | 4.600E-07 | 4.677E-07 | 1.900E-07 | 1.000E-06 |
| 10895 | 545 | C545P | A | 3e-07 | 2.6e-07 | 1.9e-07 | 2.500E-07 | 5.568E-08 | 1.900E-07 | 3.000E-07 |
| 10896 | 545 | C545S | A | 1.2e-07 | 2.1e-07 | 5.4e-07 | 2.900E-07 | 2.211E-07 | 1.200E-07 | 5.400E-07 |
| 10897 | 545 | C545T | A | 3.7e-07 | 3.8e-07 | 1.7e-07 | 3.067E-07 | 1.185E-07 | 1.700E-07 | 3.800E-07 |
| 10898 | 545 | C545W | A | 2.8e-07 | 2.6e-07 | 1.4e-07 | 2.267E-07 | 7.572E-08 | 1.400E-07 | 2.800E-07 |
| 10899 | 545 | C545Y | A | 2.9e-07 | 2.2e-07 | 1.9e-07 | 2.333E-07 | 5.132E-08 | 1.900E-07 | 2.900E-07 |
| 10900 | 545 | C545V | A | 3.5e-07 | 1.9e-07 | 3.8e-07 | 3.067E-07 | 1.021E-07 | 1.900E-07 | 3.800E-07 |
| 10901 | 546 | Q546A | A | 2.5e-07 | 3.4e-07 | 1.5e-07 | 2.467E-07 | 9.504E-08 | 1.500E-07 | 3.400E-07 |
| 10902 | 546 | Q546R | A | 1.8e-07 | 1.8e-07 | 2e-07 | 1.867E-07 | 1.155E-08 | 1.800E-07 | 2.000E-07 |
| 10903 | 546 | Q546N | A | 1.2e-07 | 1.9e-07 | 4.4e-07 | 2.500E-07 | 1.682E-07 | 1.200E-07 | 4.400E-07 |
| 10904 | 546 | Q546D | A | 2.3e-07 | 1.8e-07 | 2.1e-07 | 2.067E-07 | 2.517E-08 | 1.800E-07 | 2.300E-07 |
| 10905 | 546 | Q546C | A | 1.7e-07 | 5.8e-08 | 8.8e-08 | 1.053E-07 | 5.798E-08 | 5.800E-08 | 1.700E-07 |
| 10906 | 546 | Q546E | A | 3.8e-07 | 4e-07 | 2.9e-07 | 3.567E-07 | 5.859E-08 | 2.900E-07 | 4.000E-07 |
| 10907 | 546 | Q546Q | A | 1.9e-07 | 3.3e-07 | 1.5e-07 | 2.233E-07 | 9.452E-08 | 1.500E-07 | 3.300E-07 |
| 10908 | 546 | Q546G | A | 1.1e-07 | 2.2e-07 | 4.1e-07 | 2.467E-07 | 1.518E-07 | 1.100E-07 | 4.100E-07 |
| 10909 | 546 | Q546H | A | 4.8e-07 | 4.1e-07 | 2.3e-07 | 3.733E-07 | 1.290E-07 | 2.300E-07 | 4.800E-07 |
| 10910 | 546 | Q546I | A | 2.9e-07 | 1e-07 | 2.7e-07 | 2.200E-07 | 1.044E-07 | 1.000E-07 | 2.900E-07 |
| 10911 | 546 | Q546L | A | 1.8e-07 | 5.2e-07 | 4.1e-07 | 3.700E-07 | 1.735E-07 | 1.800E-07 | 5.200E-07 |
| 10912 | 546 | Q546K | A | 1.8e-07 | 1.4e-07 | 2.8e-07 | 2.000E-07 | 7.211E-08 | 1.400E-07 | 2.800E-07 |
| 10913 | 546 | Q546M | A | 2.7e-07 | 2.4e-07 | 1.9e-07 | 2.333E-07 | 4.041E-08 | 1.900E-07 | 2.700E-07 |
| 10914 | 546 | Q546F | A | 2.3e-07 | 2e-07 | 1.4e-07 | 1.900E-07 | 4.583E-08 | 1.400E-07 | 2.300E-07 |
| 10915 | 546 | Q546P | A | 2e-07 | 4.1e-07 | 2e-07 | 2.700E-07 | 1.212E-07 | 2.000E-07 | 4.100E-07 |
| 10916 | 546 | Q546S | A | 2e-07 | 2.6e-07 | 3.4e-07 | 2.667E-07 | 7.024E-08 | 2.000E-07 | 3.400E-07 |
| 10917 | 546 | Q546T | A | 2e-07 | 1.6e-07 | 5.6e-07 | 3.067E-07 | 2.203E-07 | 1.600E-07 | 5.600E-07 |
| 10918 | 546 | Q546W | A | 2.2e-07 | 1.4e-07 | 3.1e-07 | 2.233E-07 | 8.505E-08 | 1.400E-07 | 3.100E-07 |
| 10919 | 546 | Q546Y | A | 8.5e-07 | 2.4e-07 | 3e-07 | 4.633E-07 | 3.362E-07 | 2.400E-07 | 8.500E-07 |
| 10920 | 546 | Q546V | A | 1.8e-07 | 2.1e-07 | 3.5e-07 | 2.467E-07 | 9.074E-08 | 1.800E-07 | 3.500E-07 |
| 10921 | 547 | P547A | A | 1.7e-07 | 2.7e-07 | 2.1e-07 | 2.167E-07 | 5.033E-08 | 1.700E-07 | 2.700E-07 |
| 10922 | 547 | P547R | A | 5.8e-07 | 1.2e-07 | 2.9e-07 | 3.300E-07 | 2.326E-07 | 1.200E-07 | 5.800E-07 |
| 10923 | 547 | P547N | A | 2.1e-07 | 1.4e-07 | 1.2e-07 | 1.567E-07 | 4.726E-08 | 1.200E-07 | 2.100E-07 |
| 10924 | 547 | P547D | A | 1.4e-07 | 1.4e-07 | 1.7e-07 | 1.500E-07 | 1.732E-08 | 1.400E-07 | 1.700E-07 |
| 10925 | 547 | P547C | A | 2.4e-07 | 2.9e-07 | 2.1e-07 | 2.467E-07 | 4.041E-08 | 2.100E-07 | 2.900E-07 |
| 10926 | 547 | P547E | A | 1.2e-07 | 1.5e-07 | 2.4e-07 | 1.700E-07 | 6.245E-08 | 1.200E-07 | 2.400E-07 |
| 10927 | 547 | P547Q | A | 3.3e-07 | 4.7e-07 | 3.4e-07 | 3.800E-07 | 7.810E-08 | 3.300E-07 | 4.700E-07 |
| 10928 | 547 | P547G | A | 1.8e-07 | 7.7e-08 | 1.5e-07 | 1.357E-07 | 5.297E-08 | 7.700E-08 | 1.800E-07 |
| 10929 | 547 | P547H | A | 1.2e-07 | 3.7e-07 | 1.5e-07 | 2.133E-07 | 1.365E-07 | 1.200E-07 | 3.700E-07 |
| 10930 | 547 | P547I | A | 1.9e-07 | 4.3e-07 | 2.6e-07 | 2.933E-07 | 1.234E-07 | 1.900E-07 | 4.300E-07 |
| 10931 | 547 | P547L | A | 2.4e-07 | 3.1e-07 | 1.7e-07 | 2.400E-07 | 7.000E-08 | 1.700E-07 | 3.100E-07 |
| 10932 | 547 | P547K | A | 2.1e-07 | 1.1e-07 | 3.6e-07 | 2.267E-07 | 1.258E-07 | 1.100E-07 | 3.600E-07 |
| 10933 | 547 | P547M | A | 4.1e-07 | 2.6e-07 | 2.6e-07 | 3.100E-07 | 8.660E-08 | 2.600E-07 | 4.100E-07 |
| 10934 | 547 | P547F | A | 5.8e-07 | 3.8e-07 | 1.3e-07 | 3.633E-07 | 2.255E-07 | 1.300E-07 | 5.800E-07 |
| 10935 | 547 | P547P | A | 3.7e-07 | 1.4e-07 | 3.7e-07 | 2.933E-07 | 1.328E-07 | 1.400E-07 | 3.700E-07 |
| 10936 | 547 | P547S | A | 3.2e-07 | 2.9e-07 | 1.8e-07 | 2.633E-07 | 7.371E-08 | 1.800E-07 | 3.200E-07 |
| 10937 | 547 | P547T | A | 4.2e-07 | 1.3e-07 | 1.1e-07 | 2.200E-07 | 1.735E-07 | 1.100E-07 | 4.200E-07 |
| 10938 | 547 | P547W | A | 2.1e-07 | 2.3e-07 | 2e-07 | 2.133E-07 | 1.528E-08 | 2.000E-07 | 2.300E-07 |
| 10939 | 547 | P547Y | A | 2.1e-07 | 4.7e-07 | 8.1e-07 | 4.967E-07 | 3.009E-07 | 2.100E-07 | 8.100E-07 |
| 10940 | 547 | P547V | A | 3.7e-07 | 5.5e-07 | 3.2e-07 | 4.133E-07 | 1.210E-07 | 3.200E-07 | 5.500E-07 |
| 10941 | 548 | Q548A | A | 2.5e-07 | 3.8e-07 | 4.6e-07 | 3.633E-07 | 1.060E-07 | 2.500E-07 | 4.600E-07 |
| 10942 | 548 | Q548R | A | 3.4e-07 | 2.7e-07 | 2.1e-07 | 2.733E-07 | 6.506E-08 | 2.100E-07 | 3.400E-07 |
| 10943 | 548 | Q548N | A | 1.8e-07 | 2.5e-07 | 3.1e-07 | 2.467E-07 | 6.506E-08 | 1.800E-07 | 3.100E-07 |
| 10944 | 548 | Q548D | A | 2.8e-07 | 1.7e-07 | 1.3e-07 | 1.933E-07 | 7.767E-08 | 1.300E-07 | 2.800E-07 |
| 10945 | 548 | Q548C | A | 1.6e-07 | 3.7e-07 | 2.9e-07 | 2.733E-07 | 1.060E-07 | 1.600E-07 | 3.700E-07 |
| 10946 | 548 | Q548E | A | 3.5e-07 | 3e-07 | 1.7e-07 | 2.733E-07 | 9.292E-08 | 1.700E-07 | 3.500E-07 |
| 10947 | 548 | Q548Q | A | 1.7e-07 | 1.8e-07 | 2.7e-07 | 2.067E-07 | 5.508E-08 | 1.700E-07 | 2.700E-07 |
| 10948 | 548 | Q548G | A | 4.1e-07 | 2.7e-07 | 2.2e-07 | 3.000E-07 | 9.849E-08 | 2.200E-07 | 4.100E-07 |
| 10949 | 548 | Q548H | A | 1.9e-07 | 2.3e-07 | 3.7e-07 | 2.633E-07 | 9.452E-08 | 1.900E-07 | 3.700E-07 |
| 10950 | 548 | Q548I | A | 2.2e-07 | 2e-07 | 2.1e-07 | 2.100E-07 | 1.000E-08 | 2.000E-07 | 2.200E-07 |
| 10951 | 548 | Q548L | A | 9.3e-08 | 1.9e-07 | 4.5e-07 | 2.443E-07 | 1.846E-07 | 9.300E-08 | 4.500E-07 |
| 10952 | 548 | Q548K | A | 2.4e-07 | 1e-07 | 4.7e-07 | 2.700E-07 | 1.868E-07 | 1.000E-07 | 4.700E-07 |
| 10953 | 548 | Q548M | A | 5.1e-07 | 3.1e-07 | 1.2e-07 | 3.133E-07 | 1.950E-07 | 1.200E-07 | 5.100E-07 |
| 10954 | 548 | Q548F | A | 3e-07 | 2e-07 | 1.9e-07 | 2.300E-07 | 6.083E-08 | 1.900E-07 | 3.000E-07 |
| 10955 | 548 | Q548P | A | 1.2e-07 | 3.4e-07 | 3.9e-07 | 2.833E-07 | 1.436E-07 | 1.200E-07 | 3.900E-07 |
| 10956 | 548 | Q548S | A | 2e-07 | 1.9e-07 | 3.2e-07 | 2.367E-07 | 7.234E-08 | 1.900E-07 | 3.200E-07 |
| 10957 | 548 | Q548T | A | 1.9e-07 | 3.5e-07 | 3.8e-07 | 3.067E-07 | 1.021E-07 | 1.900E-07 | 3.800E-07 |
| 10958 | 548 | Q548W | A | 4.1e-07 | 5.5e-07 | 2.2e-07 | 3.933E-07 | 1.656E-07 | 2.200E-07 | 5.500E-07 |
| 10959 | 548 | Q548Y | A | 1.9e-07 | 1.5e-07 | 3.9e-07 | 2.433E-07 | 1.286E-07 | 1.500E-07 | 3.900E-07 |
| 10960 | 548 | Q548V | A | 4.5e-07 | 5.2e-08 | 3.5e-07 | 2.840E-07 | 2.070E-07 | 5.200E-08 | 4.500E-07 |
| 10961 | 549 | N549A | A | 2e-07 | 1.5e-07 | 2.1e-07 | 1.867E-07 | 3.215E-08 | 1.500E-07 | 2.100E-07 |
| 10962 | 549 | N549R | A | 1.9e-07 | 2e-07 | 3.6e-07 | 2.500E-07 | 9.539E-08 | 1.900E-07 | 3.600E-07 |
| 10963 | 549 | N549N | A | 1.7e-07 | 1e-07 | 3.2e-07 | 1.967E-07 | 1.124E-07 | 1.000E-07 | 3.200E-07 |
| 10964 | 549 | N549D | A | 5.3e-07 | 1.3e-07 | 3.7e-07 | 3.433E-07 | 2.013E-07 | 1.300E-07 | 5.300E-07 |

| | | | | | | | | | | |
|---|---|---|---|---|---|---|---|---|---|---|
| 10965 | 549 | N549C | A | 5.3e-07 | 2.5e-07 | 3e-07 | 3.600E-07 | 1.493E-07 | 2.500E-07 | 5.300E-07 |
| 10966 | 549 | N549E | A | 4.7e-07 | 1.4e-07 | 3.4e-07 | 3.167E-07 | 1.662E-07 | 1.400E-07 | 4.700E-07 |
| 10967 | 549 | N549Q | A | 2e-07 | 1.4e-07 | 2.1e-07 | 1.833E-07 | 3.786E-08 | 1.400E-07 | 2.100E-07 |
| 10968 | 549 | N549G | A | 1.8e-07 | 3.3e-07 | 4.5e-07 | 3.200E-07 | 1.353E-07 | 1.800E-07 | 4.500E-07 |
| 10969 | 549 | N549H | A | 2.3e-07 | 3.7e-07 | 1.7e-07 | 2.567E-07 | 1.026E-07 | 1.700E-07 | 3.700E-07 |
| 10970 | 549 | N549I | A | 7e-07 | 1.4e-07 | 3.6e-07 | 4.000E-07 | 2.821E-07 | 1.400E-07 | 7.000E-07 |
| 10971 | 549 | N549L | A | 2e-07 | 2.8e-07 | 6.3e-07 | 3.700E-07 | 2.287E-07 | 2.000E-07 | 6.300E-07 |
| 10972 | 549 | N549K | A | 1.4e-07 | 2.7e-07 | 5.3e-07 | 3.133E-07 | 1.986E-07 | 1.400E-07 | 5.300E-07 |
| 10973 | 549 | N549M | A | 1.4e-07 | 2.8e-07 | 3.1e-07 | 2.433E-07 | 9.074E-08 | 1.400E-07 | 3.100E-07 |
| 10974 | 549 | N549F | A | 4.2e-07 | 5.1e-07 | 2.9e-07 | 4.067E-07 | 1.106E-07 | 2.900E-07 | 5.100E-07 |
| 10975 | 549 | N549P | A | 3.2e-07 | 3.3e-07 | 1.4e-07 | 2.633E-07 | 1.069E-07 | 1.400E-07 | 3.300E-07 |
| 10976 | 549 | N549S | A | 2.3e-07 | 1.6e-07 | 1.6e-07 | 1.833E-07 | 4.041E-08 | 1.600E-07 | 2.300E-07 |
| 10977 | 549 | N549T | A | 3.1e-07 | 1.4e-07 | 2.7e-07 | 2.400E-07 | 8.888E-08 | 1.400E-07 | 3.100E-07 |
| 10978 | 549 | N549W | A | 1e-07 | 1.9e-07 | 2e-07 | 1.633E-07 | 5.508E-08 | 1.000E-07 | 2.000E-07 |
| 10979 | 549 | N549Y | A | 2.5e-07 | 1.3e-07 | 4.3e-07 | 2.700E-07 | 1.510E-07 | 1.300E-07 | 4.300E-07 |
| 10980 | 549 | N549V | A | 2e-07 | 3e-07 | 2.8e-07 | 2.600E-07 | 5.292E-08 | 2.000E-07 | 3.000E-07 |
| 10981 | 550 | G550A | A | 2.2e-07 | 1.6e-07 | 4.4e-07 | 2.733E-07 | 1.474E-07 | 1.600E-07 | 4.400E-07 |
| 10982 | 550 | G550R | A | 4.2e-07 | 3.8e-07 | 2.5e-07 | 3.500E-07 | 8.888E-08 | 2.500E-07 | 4.200E-07 |
| 10983 | 550 | G550N | A | 2.9e-07 | 2.1e-07 | 2.7e-07 | 2.567E-07 | 4.163E-08 | 2.100E-07 | 2.900E-07 |
| 10984 | 550 | G550D | A | 2.6e-07 | 3.6e-07 | 2e-07 | 2.733E-07 | 8.083E-08 | 2.000E-07 | 3.600E-07 |
| 10985 | 550 | G550C | A | 4e-07 | 1.7e-07 | 2.5e-07 | 2.733E-07 | 1.168E-07 | 1.700E-07 | 4.000E-07 |
| 10986 | 550 | G550E | A | 2.4e-07 | 1.9e-07 | 2.7e-07 | 2.333E-07 | 4.041E-08 | 1.900E-07 | 2.700E-07 |
| 10987 | 550 | G550Q | A | 3.8e-07 | 3.3e-07 | 1.9e-07 | 3.000E-07 | 9.849E-08 | 1.900E-07 | 3.800E-07 |
| 10988 | 550 | G550G | A | 2.5e-07 | 1.7e-07 | 9.5e-08 | 1.717E-07 | 7.751E-08 | 9.500E-08 | 2.500E-07 |
| 10989 | 550 | G550H | A | 3e-07 | 1.1e-07 | 2.9e-07 | 2.333E-07 | 1.069E-07 | 1.100E-07 | 3.000E-07 |
| 10990 | 550 | G550I | A | 2.6e-07 | 1.5e-07 | 3.2e-07 | 2.433E-07 | 8.622E-08 | 1.500E-07 | 3.200E-07 |
| 10991 | 550 | G550L | A | 1e-07 | 2.7e-07 | 2e-07 | 1.900E-07 | 8.544E-08 | 1.000E-07 | 2.700E-07 |
| 10992 | 550 | G550K | A | 7.4e-07 | 3.6e-07 | 3.9e-07 | 4.967E-07 | 2.113E-07 | 3.600E-07 | 7.400E-07 |
| 10993 | 550 | G550M | A | 2.2e-07 | 3.5e-07 | 1.7e-07 | 2.467E-07 | 9.292E-08 | 1.700E-07 | 3.500E-07 |
| 10994 | 550 | G550F | A | 1.2e-07 | 2.1e-07 | 1.6e-07 | 1.633E-07 | 4.509E-08 | 1.200E-07 | 2.100E-07 |
| 10995 | 550 | G550P | A | 1.5e-07 | 2.7e-07 | 3.2e-07 | 2.467E-07 | 8.737E-08 | 1.500E-07 | 3.200E-07 |
| 10996 | 550 | G550S | A | 2.1e-07 | 2e-07 | 2.8e-07 | 2.300E-07 | 4.359E-08 | 2.000E-07 | 2.800E-07 |
| 10997 | 550 | G550T | A | 1.7e-07 | 2.7e-07 | 3.1e-07 | 2.500E-07 | 7.211E-08 | 1.700E-07 | 3.100E-07 |
| 10998 | 550 | G550W | A | 1.1e-07 | 2.9e-07 | 1.1e-07 | 1.700E-07 | 1.039E-07 | 1.100E-07 | 2.900E-07 |
| 10999 | 550 | G550Y | A | 4.5e-07 | 1e-07 | 2e-07 | 2.500E-07 | 1.803E-07 | 1.000E-07 | 4.500E-07 |
| 11000 | 550 | G550V | A | 2.3e-07 | 3.1e-07 | 2.7e-07 | 2.700E-07 | 4.000E-08 | 2.300E-07 | 3.100E-07 |
| 11001 | 551 | S551A | A | 1.9e-07 | 4.5e-07 | 1.6e-07 | 2.667E-07 | 1.595E-07 | 1.600E-07 | 4.500E-07 |
| 11002 | 551 | S551R | A | 2.3e-07 | 2.6e-07 | 5.2e-07 | 3.367E-07 | 1.595E-07 | 2.300E-07 | 5.200E-07 |
| 11003 | 551 | S551N | A | 2.4e-07 | 1.9e-07 | 2.4e-07 | 2.233E-07 | 2.887E-08 | 1.900E-07 | 2.400E-07 |
| 11004 | 551 | S551D | A | 1.4e-07 | 3.9e-07 | 2.4e-07 | 2.567E-07 | 1.258E-07 | 1.400E-07 | 3.900E-07 |
| 11005 | 551 | S551C | A | 2e-07 | 2.4e-07 | 1.8e-07 | 2.067E-07 | 3.055E-08 | 1.800E-07 | 2.400E-07 |
| 11006 | 551 | S551E | A | 1.7e-07 | 6.7e-07 | 1.2e-07 | 3.200E-07 | 3.041E-07 | 1.200E-07 | 6.700E-07 |
| 11007 | 551 | S551Q | A | 5.5e-07 | 2.8e-07 | 1.5e-07 | 3.267E-07 | 2.040E-07 | 1.500E-07 | 5.500E-07 |
| 11008 | 551 | S551G | A | 3.1e-07 | 2.2e-07 | 3e-07 | 2.767E-07 | 4.933E-08 | 2.200E-07 | 3.100E-07 |
| 11009 | 551 | S551H | A | 6.8e-07 | 1.6e-07 | 2.6e-07 | 3.667E-07 | 2.759E-07 | 1.600E-07 | 6.800E-07 |
| 11010 | 551 | S551I | A | 2.4e-07 | 1.1e-07 | 1.8e-07 | 1.767E-07 | 6.506E-08 | 1.100E-07 | 2.400E-07 |
| 11011 | 551 | S551L | A | 2.1e-07 | 4.5e-07 | 3.7e-07 | 3.433E-07 | 1.222E-07 | 2.100E-07 | 4.500E-07 |
| 11012 | 551 | S551K | A | 4.7e-07 | 2.9e-07 | 3.1e-07 | 3.567E-07 | 9.866E-08 | 2.900E-07 | 4.700E-07 |
| 11013 | 551 | S551M | A | 4.1e-07 | 4.2e-07 | 2e-07 | 3.433E-07 | 1.242E-07 | 2.000E-07 | 4.200E-07 |
| 11014 | 551 | S551F | A | 2.9e-07 | 4.4e-07 | 2.7e-07 | 3.333E-07 | 9.292E-08 | 2.700E-07 | 4.400E-07 |
| 11015 | 551 | S551P | A | 4.5e-07 | 2.4e-07 | 1.7e-07 | 2.867E-07 | 1.457E-07 | 1.700E-07 | 4.500E-07 |
| 11016 | 551 | S551S | A | 2.6e-07 | 3.4e-07 | 2.7e-07 | 2.900E-07 | 4.359E-08 | 2.600E-07 | 3.400E-07 |
| 11017 | 551 | S551T | A | 4.2e-07 | 3.8e-07 | 2.6e-07 | 3.533E-07 | 8.327E-08 | 2.600E-07 | 4.200E-07 |
| 11018 | 551 | S551W | A | 2.2e-07 | 2.5e-07 | 1.6e-07 | 2.100E-07 | 4.583E-08 | 1.600E-07 | 2.500E-07 |
| 11019 | 551 | S551Y | A | 2.3e-07 | 2.6e-07 | 3.7e-07 | 2.867E-07 | 7.371E-08 | 2.300E-07 | 3.700E-07 |
| 11020 | 551 | S551V | A | 2.5e-07 | 2.3e-07 | 8e-07 | 4.267E-07 | 3.235E-07 | 2.300E-07 | 8.000E-07 |
| 11021 | 552 | V552A | A | 3.1e-07 | 1.4e-07 | 2.4e-07 | 2.300E-07 | 8.544E-08 | 1.400E-07 | 3.100E-07 |
| 11022 | 552 | V552R | A | 1.8e-07 | 3.4e-07 | 2.1e-07 | 2.433E-07 | 8.505E-08 | 1.800E-07 | 3.400E-07 |
| 11023 | 552 | V552N | A | 2.3e-07 | 3.2e-07 | 2.3e-07 | 2.600E-07 | 5.196E-08 | 2.300E-07 | 3.200E-07 |
| 11024 | 552 | V552D | A | 2.5e-07 | 1.2e-07 | 3.8e-07 | 2.500E-07 | 1.300E-07 | 1.200E-07 | 3.800E-07 |
| 11025 | 552 | V552C | A | 2e-07 | 2.7e-07 | 2.7e-07 | 2.467E-07 | 4.041E-08 | 2.000E-07 | 2.700E-07 |
| 11026 | 552 | V552E | A | 2.8e-07 | 2e-07 | 1.4e-07 | 2.067E-07 | 7.024E-08 | 1.400E-07 | 2.800E-07 |
| 11027 | 552 | V552Q | A | 1.8e-07 | 3.5e-07 | 1.7e-07 | 2.333E-07 | 1.012E-07 | 1.700E-07 | 3.500E-07 |
| 11028 | 552 | V552G | A | 2.1e-07 | 2.1e-07 | 2.8e-07 | 2.333E-07 | 4.041E-08 | 2.100E-07 | 2.800E-07 |
| 11029 | 552 | V552H | A | 2.3e-07 | 2.4e-07 | 2e-07 | 2.233E-07 | 2.082E-08 | 2.000E-07 | 2.400E-07 |
| 11030 | 552 | V552I | A | 2.4e-07 | 2.7e-07 | 3.1e-07 | 2.733E-07 | 3.512E-08 | 2.400E-07 | 3.100E-07 |
| 11031 | 552 | V552L | A | 2.1e-07 | 3.6e-07 | 4.9e-07 | 3.533E-07 | 1.401E-07 | 2.100E-07 | 4.900E-07 |
| 11032 | 552 | V552K | A | 3.9e-07 | 1.3e-07 | 1.7e-07 | 2.300E-07 | 1.400E-07 | 1.300E-07 | 3.900E-07 |
| 11033 | 552 | V552M | A | 1.1e-07 | 3e-07 | 3.4e-07 | 2.500E-07 | 1.229E-07 | 1.100E-07 | 3.400E-07 |
| 11034 | 552 | V552F | A | 2.8e-07 | 1.4e-07 | 2.3e-07 | 2.167E-07 | 7.095E-08 | 1.400E-07 | 2.800E-07 |
| 11035 | 552 | V552P | A | 1.3e-07 | 2.6e-07 | 3.1e-07 | 2.333E-07 | 9.292E-08 | 1.300E-07 | 3.100E-07 |
| 11036 | 552 | V552S | A | 2.2e-07 | 6.9e-07 | 2.8e-07 | 3.967E-07 | 2.558E-07 | 2.200E-07 | 6.900E-07 |
| 11037 | 552 | V552T | A | 5.7e-07 | 2.7e-07 | 1.5e-07 | 3.300E-07 | 2.163E-07 | 1.500E-07 | 5.700E-07 |
| 11038 | 552 | V552W | A | 2.4e-07 | 3e-07 | 1.1e-07 | 2.167E-07 | 9.713E-08 | 1.100E-07 | 3.000E-07 |
| 11039 | 552 | V552Y | A | 5.1e-07 | 4.2e-07 | 3.6e-07 | 4.300E-07 | 7.550E-08 | 3.600E-07 | 5.100E-07 |
| 11040 | 552 | V552V | A | 1.7e-07 | 3e-07 | 4.2e-07 | 2.967E-07 | 1.250E-07 | 1.700E-07 | 4.200E-07 |
| 11041 | 553 | T553A | A | 2e-07 | 1.8e-07 | 2.2e-07 | 2.000E-07 | 2.000E-08 | 1.800E-07 | 2.200E-07 |
| 11042 | 553 | T553R | A | 2.4e-07 | 1.8e-07 | 2.5e-07 | 2.233E-07 | 3.786E-08 | 1.800E-07 | 2.500E-07 |
| 11043 | 553 | T553N | A | 3.4e-07 | 2.1e-07 | 2.9e-07 | 2.800E-07 | 6.557E-08 | 2.100E-07 | 3.400E-07 |
| 11044 | 553 | T553D | A | 2e-07 | 1.8e-07 | 3.3e-07 | 2.367E-07 | 8.145E-08 | 1.800E-07 | 3.300E-07 |
| 11045 | 553 | T553C | A | 1.5e-07 | 2.2e-07 | 2.5e-07 | 2.067E-07 | 5.132E-08 | 1.500E-07 | 2.500E-07 |
| 11046 | 553 | T553E | A | 2.6e-07 | 2.3e-07 | 2.2e-07 | 2.367E-07 | 2.082E-08 | 2.200E-07 | 2.600E-07 |
| 11047 | 553 | T553Q | A | 2.6e-07 | 4.3e-07 | 2.3e-07 | 3.067E-07 | 1.079E-07 | 2.300E-07 | 4.300E-07 |
| 11048 | 553 | T553G | A | 5e-07 | 1.6e-07 | 2.1e-07 | 2.900E-07 | 1.836E-07 | 1.600E-07 | 5.000E-07 |
| 11049 | 553 | T553H | A | 2.1e-07 | 1.7e-07 | 2.1e-07 | 1.967E-07 | 2.309E-08 | 1.700E-07 | 2.100E-07 |

| | | | | | | | | | | |
|---|---|---|---|---|---|---|---|---|---|---|
| 11050 | 553 | T553I | A | 4.8e-07 | 2e-07 | 3.3e-07 | 3.367E-07 | 1.401E-07 | 2.000E-07 | 4.800E-07 |
| 11051 | 553 | T553L | A | 1.4e-07 | 2.2e-07 | 1.6e-07 | 1.733E-07 | 4.163E-08 | 1.400E-07 | 2.200E-07 |
| 11052 | 553 | T553K | A | 2.6e-07 | 6.1e-07 | 2.6e-07 | 3.767E-07 | 2.021E-07 | 2.600E-07 | 6.100E-07 |
| 11053 | 553 | T553M | A | 2.4e-07 | 7e-08 | 3e-07 | 2.033E-07 | 1.193E-07 | 7.000E-08 | 3.000E-07 |
| 11054 | 553 | T553F | A | 2.8e-07 | 2.1e-07 | 1.6e-07 | 2.167E-07 | 6.028E-08 | 1.600E-07 | 2.800E-07 |
| 11055 | 553 | T553P | A | 1.7e-07 | 2.4e-07 | 2.2e-07 | 2.100E-07 | 3.606E-08 | 1.700E-07 | 2.400E-07 |
| 11056 | 553 | T553S | A | 3.6e-07 | 1.2e-07 | 3.8e-07 | 2.867E-07 | 1.447E-07 | 1.200E-07 | 3.800E-07 |
| 11057 | 553 | T553T | A | 3.7e-07 | 2.1e-07 | 3.7e-07 | 3.167E-07 | 9.238E-08 | 2.100E-07 | 3.700E-07 |
| 11058 | 553 | T553W | A | 4.1e-07 | 1.8e-07 | 4.8e-07 | 3.567E-07 | 1.570E-07 | 1.800E-07 | 4.800E-07 |
| 11059 | 553 | T553Y | A | 3e-07 | 3.2e-07 | 2.8e-07 | 3.000E-07 | 2.000E-08 | 2.800E-07 | 3.200E-07 |
| 11060 | 553 | T553V | A | 2.6e-07 | 1.7e-07 | 2.5e-07 | 2.267E-07 | 4.933E-08 | 1.700E-07 | 2.600E-07 |
| 11061 | 554 | C554A | A | 2.6e-07 | 1.1e-07 | 2.6e-07 | 2.100E-07 | 8.660E-08 | 1.100E-07 | 2.600E-07 |
| 11062 | 554 | C554R | A | 3.1e-07 | 4.7e-07 | 1.6e-07 | 3.133E-07 | 1.550E-07 | 1.600E-07 | 4.700E-07 |
| 11063 | 554 | C554N | A | 2e-07 | 1.6e-07 | 7.2e-07 | 3.600E-07 | 3.124E-07 | 1.600E-07 | 7.200E-07 |
| 11064 | 554 | C554D | A | 4.2e-07 | 1.8e-07 | 2.1e-07 | 2.700E-07 | 1.308E-07 | 1.800E-07 | 4.200E-07 |
| 11065 | 554 | C554C | A | 2.2e-07 | 2.7e-07 | 1.7e-07 | 2.200E-07 | 5.000E-08 | 1.700E-07 | 2.700E-07 |
| 11066 | 554 | C554E | A | 1.2e-07 | 2e-07 | 3.5e-07 | 2.233E-07 | 1.168E-07 | 1.200E-07 | 3.500E-07 |
| 11067 | 554 | C554Q | A | 2.5e-07 | 3.1e-07 | 1.2e-07 | 2.267E-07 | 9.713E-08 | 1.200E-07 | 3.100E-07 |
| 11068 | 554 | C554G | A | 2.4e-07 | 2.1e-07 | 2.2e-07 | 2.233E-07 | 1.528E-08 | 2.100E-07 | 2.400E-07 |
| 11069 | 554 | C554H | A | 1.3e-07 | 3.2e-07 | 4e-07 | 2.833E-07 | 1.387E-07 | 1.300E-07 | 4.000E-07 |
| 11070 | 554 | C554I | A | 2.6e-07 | 1.8e-07 | 3.2e-07 | 2.533E-07 | 7.024E-08 | 1.800E-07 | 3.200E-07 |
| 11071 | 554 | C554L | A | 2.2e-07 | 3.8e-07 | 4.6e-07 | 3.533E-07 | 1.222E-07 | 2.200E-07 | 4.600E-07 |
| 11072 | 554 | C554K | A | 2.7e-07 | 2.4e-07 | 4.2e-07 | 3.100E-07 | 9.644E-08 | 2.400E-07 | 4.200E-07 |
| 11073 | 554 | C554M | A | 1.7e-07 | 2.7e-07 | 2.8e-07 | 2.400E-07 | 6.083E-08 | 1.700E-07 | 2.800E-07 |
| 11074 | 554 | C554F | A | 2.5e-07 | 1.8e-07 | 3.2e-07 | 2.500E-07 | 7.000E-08 | 1.800E-07 | 3.200E-07 |
| 11075 | 554 | C554P | A | 1.3e-07 | 2.4e-07 | 4.5e-07 | 2.733E-07 | 1.626E-07 | 1.300E-07 | 4.500E-07 |
| 11076 | 554 | C554S | A | 2.5e-07 | 4.6e-07 | 2.8e-07 | 3.300E-07 | 1.136E-07 | 2.500E-07 | 4.600E-07 |
| 11077 | 554 | C554T | A | 8.1e-08 | 3.2e-07 | 1.3e-07 | 1.770E-07 | 1.262E-07 | 8.100E-08 | 3.200E-07 |
| 11078 | 554 | C554W | A | 8.8e-08 | 2e-07 | 1.7e-07 | 1.527E-07 | 5.798E-08 | 8.800E-08 | 2.000E-07 |
| 11079 | 554 | C554Y | A | 2.6e-07 | 1.1e-07 | 2.1e-07 | 1.933E-07 | 7.638E-08 | 1.100E-07 | 2.600E-07 |
| 11080 | 554 | C554V | A | 2.3e-07 | 2.6e-07 | 2.8e-07 | 2.567E-07 | 2.517E-08 | 2.300E-07 | 2.800E-07 |
| 11081 | 555 | F555A | A | 2.6e-07 | 1.7e-07 | 1.4e-07 | 1.900E-07 | 6.245E-08 | 1.400E-07 | 2.600E-07 |
| 11082 | 555 | F555R | A | 1.7e-07 | 1.9e-07 | 3.9e-07 | 2.500E-07 | 1.217E-07 | 1.700E-07 | 3.900E-07 |
| 11083 | 555 | F555N | A | 2.3e-07 | 1.5e-07 | 1.7e-07 | 1.833E-07 | 4.163E-08 | 1.500E-07 | 2.300E-07 |
| 11084 | 555 | F555D | A | 6.2e-07 | 4.7e-07 | 2.4e-07 | 4.433E-07 | 1.914E-07 | 2.400E-07 | 6.200E-07 |
| 11085 | 555 | F555C | A | 2.4e-07 | 1.6e-07 | 6.2e-07 | 3.400E-07 | 2.458E-07 | 1.600E-07 | 6.200E-07 |
| 11086 | 555 | F555E | A | 3.5e-07 | 1.7e-07 | 4.2e-07 | 3.133E-07 | 1.290E-07 | 1.700E-07 | 4.200E-07 |
| 11087 | 555 | F555Q | A | 1.2e-07 | 2.9e-07 | 1.7e-07 | 1.933E-07 | 8.737E-08 | 1.200E-07 | 2.900E-07 |
| 11088 | 555 | F555G | A | 3e-07 | 2.9e-07 | 2.4e-07 | 2.767E-07 | 3.215E-08 | 2.400E-07 | 3.000E-07 |
| 11089 | 555 | F555H | A | 1.5e-07 | 4.6e-07 | 2e-07 | 2.700E-07 | 1.664E-07 | 1.500E-07 | 4.600E-07 |
| 11090 | 555 | F555I | A | 1.6e-07 | 2.4e-07 | 2.6e-07 | 2.200E-07 | 5.292E-08 | 1.600E-07 | 2.600E-07 |
| 11091 | 555 | F555L | A | 2.6e-07 | 2.5e-07 | 1.7e-07 | 2.267E-07 | 4.933E-08 | 1.700E-07 | 2.600E-07 |
| 11092 | 555 | F555K | A | 2.9e-07 | 1.9e-07 | 2.9e-07 | 2.567E-07 | 5.774E-08 | 1.900E-07 | 2.900E-07 |
| 11093 | 555 | F555M | A | 2.4e-07 | 3.6e-07 | 1.9e-07 | 2.633E-07 | 8.737E-08 | 1.900E-07 | 3.600E-07 |
| 11094 | 555 | F555F | A | 1.4e-07 | 2.4e-07 | 1.2e-07 | 1.667E-07 | 6.429E-08 | 1.200E-07 | 2.400E-07 |
| 11095 | 555 | F555P | A | 1.3e-07 | 3e-07 | 1.4e-07 | 1.900E-07 | 9.539E-08 | 1.300E-07 | 3.000E-07 |
| 11096 | 555 | F555S | A | 2.7e-07 | 2e-07 | 1.9e-07 | 2.200E-07 | 4.359E-08 | 1.900E-07 | 2.700E-07 |
| 11097 | 555 | F555T | A | 1.4e-07 | 4.6e-07 | 9.7e-08 | 2.323E-07 | 1.983E-07 | 9.700E-08 | 4.600E-07 |
| 11098 | 555 | F555W | A | 1.2e-07 | 2.5e-07 | 2.5e-07 | 2.067E-07 | 7.506E-08 | 1.200E-07 | 2.500E-07 |
| 11099 | 555 | F555Y | A | 1.9e-07 | 2.5e-07 | 1.7e-07 | 2.033E-07 | 4.163E-08 | 1.700E-07 | 2.500E-07 |
| 11100 | 555 | F555V | A | 1.2e-07 | 4.2e-07 | 3.1e-07 | 2.833E-07 | 1.518E-07 | 1.200E-07 | 4.200E-07 |
| 11101 | 556 | G556A | A | 1.7e-07 | 1.1e-07 | 1.4e-07 | 1.400E-07 | 3.000E-08 | 1.100E-07 | 1.700E-07 |
| 11102 | 556 | G556R | A | 2.2e-07 | 2.1e-07 | 2.5e-07 | 2.267E-07 | 2.082E-08 | 2.100E-07 | 2.500E-07 |
| 11103 | 556 | G556N | A | 5.5e-07 | 2.1e-07 | 3.4e-07 | 3.667E-07 | 1.716E-07 | 2.100E-07 | 5.500E-07 |
| 11104 | 556 | G556D | A | 8.9e-08 | 9.1e-08 | 2.5e-07 | 1.433E-07 | 9.238E-08 | 8.900E-08 | 2.500E-07 |
| 11105 | 556 | G556C | A | 2.1e-07 | 1.4e-07 | 1.8e-07 | 1.767E-07 | 3.512E-08 | 1.400E-07 | 2.100E-07 |
| 11106 | 556 | G556E | A | 1.5e-07 | 2.1e-07 | 1.6e-07 | 1.733E-07 | 3.215E-08 | 1.500E-07 | 2.100E-07 |
| 11107 | 556 | G556Q | A | 3.4e-07 | 9.3e-08 | 2.5e-07 | 2.277E-07 | 1.250E-07 | 9.300E-08 | 3.400E-07 |
| 11108 | 556 | G556G | A | 1.7e-07 | 1.1e-07 | 4.6e-07 | 2.467E-07 | 1.872E-07 | 1.100E-07 | 4.600E-07 |
| 11109 | 556 | G556H | A | 1.5e-07 | 2.8e-07 | 2.7e-07 | 2.333E-07 | 7.234E-08 | 1.500E-07 | 2.800E-07 |
| 11110 | 556 | G556I | A | 2.9e-07 | 1.7e-07 | 1.1e-07 | 1.900E-07 | 9.165E-08 | 1.100E-07 | 2.900E-07 |
| 11111 | 556 | G556L | A | 1.2e-07 | 1.2e-07 | 1.2e-07 | 1.200E-07 | 0.000E+00 | 1.200E-07 | 1.200E-07 |
| 11112 | 556 | G556K | A | 2.5e-07 | 1.8e-07 | 2.8e-07 | 2.367E-07 | 5.132E-08 | 1.800E-07 | 2.800E-07 |
| 11113 | 556 | G556M | A | 2.9e-07 | 1.1e-07 | 6.7e-08 | 1.557E-07 | 1.183E-07 | 6.700E-08 | 2.900E-07 |
| 11114 | 556 | G556F | A | 1.5e-07 | 3.3e-07 | 3.3e-07 | 2.700E-07 | 1.039E-07 | 1.500E-07 | 3.300E-07 |
| 11115 | 556 | G556P | A | 5.4e-07 | 7.8e-07 | 2.7e-07 | 5.300E-07 | 2.551E-07 | 2.700E-07 | 7.800E-07 |
| 11116 | 556 | G556S | A | 1.9e-07 | 2.8e-07 | 5.3e-07 | 3.333E-07 | 1.762E-07 | 1.900E-07 | 5.300E-07 |
| 11117 | 556 | G556T | A | 2.2e-07 | 2.2e-07 | 1.7e-07 | 2.033E-07 | 2.887E-08 | 1.700E-07 | 2.200E-07 |
| 11118 | 556 | G556W | A | 1.1e-07 | 2e-07 | 9.4e-08 | 1.347E-07 | 5.714E-08 | 9.400E-08 | 2.000E-07 |
| 11119 | 556 | G556Y | A | 4.8e-08 | 1.8e-07 | 1.8e-07 | 1.360E-07 | 7.621E-08 | 4.800E-08 | 1.800E-07 |
| 11120 | 556 | G556V | A | 2.3e-07 | 2.4e-07 | 1.2e-07 | 1.967E-07 | 6.658E-08 | 1.200E-07 | 2.400E-07 |
| 11121 | 557 | P557A | A | 1.8e-07 | 1.4e-07 | 3.1e-07 | 2.100E-07 | 8.888E-08 | 1.400E-07 | 3.100E-07 |
| 11122 | 557 | P557R | A | 1.7e-07 | 2.9e-07 | 8.5e-08 | 1.817E-07 | 1.030E-07 | 8.500E-08 | 2.900E-07 |
| 11123 | 557 | P557N | A | 1.6e-07 | 3.6e-07 | 3.2e-07 | 2.800E-07 | 1.058E-07 | 1.600E-07 | 3.600E-07 |
| 11124 | 557 | P557D | A | 1.9e-07 | 2.1e-07 | 1.3e-07 | 1.767E-07 | 4.163E-08 | 1.300E-07 | 2.100E-07 |
| 11125 | 557 | P557C | A | 2.6e-07 | 3.1e-07 | 2.6e-07 | 2.767E-07 | 2.887E-08 | 2.600E-07 | 3.100E-07 |
| 11126 | 557 | P557E | A | 2.1e-07 | 3e-07 | 2.1e-07 | 2.400E-07 | 5.196E-08 | 2.100E-07 | 3.000E-07 |
| 11127 | 557 | P557Q | A | 2.2e-07 | 1.5e-07 | 1.5e-07 | 1.733E-07 | 4.041E-08 | 1.500E-07 | 2.200E-07 |
| 11128 | 557 | P557G | A | 2.3e-07 | 1.1e-07 | 9.8e-08 | 1.460E-07 | 7.299E-08 | 9.800E-08 | 2.300E-07 |
| 11129 | 557 | P557H | A | 3.3e-07 | 2.3e-07 | 2.3e-07 | 2.633E-07 | 5.774E-08 | 2.300E-07 | 3.300E-07 |
| 11130 | 557 | P557I | A | 1.5e-07 | 1.4e-07 | 1.8e-07 | 1.567E-07 | 2.082E-08 | 1.400E-07 | 1.800E-07 |
| 11131 | 557 | P557L | A | 2.3e-07 | 1.8e-07 | 3.5e-07 | 2.533E-07 | 8.737E-08 | 1.800E-07 | 3.500E-07 |
| 11132 | 557 | P557K | A | 5.8e-07 | 3.4e-07 | 1.3e-07 | 3.500E-07 | 2.252E-07 | 1.300E-07 | 5.800E-07 |
| 11133 | 557 | P557M | A | 2.7e-07 | 1.8e-07 | 2.2e-07 | 2.233E-07 | 4.509E-08 | 1.800E-07 | 2.700E-07 |
| 11134 | 557 | P557F | A | 1.2e-07 | 1.7e-07 | 2.6e-07 | 1.833E-07 | 7.095E-08 | 1.200E-07 | 2.600E-07 |

| | | | | | | | | | | |
|---|---|---|---|---|---|---|---|---|---|---|
| 11135 | 557 | P557P | A | 1.6e-07 | 1.7e-07 | 2.4e-07 | 1.900E-07 | 4.359E-08 | 1.600E-07 | 2.400E-07 |
| 11136 | 557 | P557S | A | 2.5e-07 | 1.1e-07 | 2.9e-07 | 2.167E-07 | 9.452E-08 | 1.100E-07 | 2.900E-07 |
| 11137 | 557 | P557T | A | 7.6e-08 | 3.1e-07 | 1.7e-07 | 1.853E-07 | 1.178E-07 | 7.600E-08 | 3.100E-07 |
| 11138 | 557 | P557W | A | 2.3e-07 | 1.5e-07 | 1.2e-07 | 1.667E-07 | 5.686E-08 | 1.200E-07 | 2.300E-07 |
| 11139 | 557 | P557Y | A | 2.9e-07 | 2.1e-07 | 3.4e-07 | 2.800E-07 | 6.557E-08 | 2.100E-07 | 3.400E-07 |
| 11140 | 557 | P557V | A | 3.2e-07 | 2e-07 | 2.3e-07 | 2.500E-07 | 6.245E-08 | 2.000E-07 | 3.200E-07 |
| 11141 | 558 | E558A | A | 3.5e-07 | 3.3e-07 | 3.2e-07 | 3.333E-07 | 1.528E-08 | 3.200E-07 | 3.500E-07 |
| 11142 | 558 | E558R | A | 2.9e-07 | 2.8e-07 | 3.4e-07 | 3.033E-07 | 3.215E-08 | 2.800E-07 | 3.400E-07 |
| 11143 | 558 | E558N | A | 3e-07 | 1.8e-07 | 3.2e-07 | 2.667E-07 | 7.572E-08 | 1.800E-07 | 3.200E-07 |
| 11144 | 558 | E558D | A | 3e-07 | 5.8e-07 | 3.5e-07 | 4.100E-07 | 1.493E-07 | 3.000E-07 | 5.800E-07 |
| 11145 | 558 | E558C | A | 2.2e-07 | 2.1e-07 | 5.3e-07 | 3.200E-07 | 1.819E-07 | 2.100E-07 | 5.300E-07 |
| 11146 | 558 | E558E | A | 2.2e-07 | 1.6e-07 | 4.4e-07 | 2.733E-07 | 1.474E-07 | 1.600E-07 | 4.400E-07 |
| 11147 | 558 | E558Q | A | 2.7e-07 | 1.1e-07 | 7.3e-07 | 3.700E-07 | 3.219E-07 | 1.100E-07 | 7.300E-07 |
| 11148 | 558 | E558G | A | 4.2e-07 | 6.2e-07 | 7.2e-07 | 5.867E-07 | 1.528E-07 | 4.200E-07 | 7.200E-07 |
| 11149 | 558 | E558H | A | 8.3e-08 | 1.2e-07 | 1.3e-07 | 1.110E-07 | 2.476E-08 | 8.300E-08 | 1.300E-07 |
| 11150 | 558 | E558I | A | 3.6e-07 | 2.6e-07 | 1e-06 | 5.400E-07 | 4.015E-07 | 2.600E-07 | 1.000E-06 |
| 11151 | 558 | E558L | A | 2e-07 | 3e-07 | 2.9e-07 | 2.633E-07 | 5.508E-08 | 2.000E-07 | 3.000E-07 |
| 11152 | 558 | E558K | A | 2.8e-07 | 1.8e-07 | 1.4e-07 | 2.000E-07 | 7.211E-08 | 1.400E-07 | 2.800E-07 |
| 11153 | 558 | E558M | A | 1.4e-07 | 3.1e-07 | 3.8e-07 | 2.767E-07 | 1.234E-07 | 1.400E-07 | 3.800E-07 |
| 11154 | 558 | E558F | A | 2.8e-07 | 1.4e-07 | 2e-07 | 2.067E-07 | 7.024E-08 | 1.400E-07 | 2.800E-07 |
| 11155 | 558 | E558P | A | 3.6e-07 | 1.9e-07 | 5.5e-07 | 3.667E-07 | 1.801E-07 | 1.900E-07 | 5.500E-07 |
| 11156 | 558 | E558S | A | 6e-07 | 4.5e-07 | 7.9e-07 | 6.133E-07 | 1.704E-07 | 4.500E-07 | 7.900E-07 |
| 11157 | 558 | E558T | A | 1.7e-07 | 1.6e-07 | 2.5e-07 | 1.933E-07 | 4.933E-08 | 1.600E-07 | 2.500E-07 |
| 11158 | 558 | E558W | A | 3.6e-07 | 2.9e-07 | 3.9e-07 | 3.467E-07 | 5.132E-08 | 2.900E-07 | 3.900E-07 |
| 11159 | 558 | E558Y | A | 1.8e-07 | 3e-07 | 1.6e-07 | 2.133E-07 | 7.572E-08 | 1.600E-07 | 3.000E-07 |
| 11160 | 558 | E558V | A | 4.1e-07 | 8.4e-07 | 3.1e-07 | 5.200E-07 | 2.816E-07 | 3.100E-07 | 8.400E-07 |
| 11161 | 559 | A559A | A | 5.4e-07 | 8.3e-08 | 4.2e-07 | 3.477E-07 | 2.369E-07 | 8.300E-08 | 5.400E-07 |
| 11162 | 559 | A559R | A | 2.4e-07 | 2.8e-07 | 3.6e-07 | 2.933E-07 | 6.110E-08 | 2.400E-07 | 3.600E-07 |
| 11163 | 559 | A559N | A | 5.5e-07 | 2.4e-07 | 1.9e-07 | 3.267E-07 | 1.950E-07 | 1.900E-07 | 5.500E-07 |
| 11164 | 559 | A559D | A | 2.7e-07 | 1.8e-07 | 2.4e-07 | 2.300E-07 | 4.583E-08 | 1.800E-07 | 2.700E-07 |
| 11165 | 559 | A559C | A | 2.5e-07 | 3.2e-07 | 3.7e-07 | 3.133E-07 | 6.028E-08 | 2.500E-07 | 3.700E-07 |
| 11166 | 559 | A559E | A | 2.2e-07 | 4.8e-07 | 2.5e-07 | 3.167E-07 | 1.422E-07 | 2.200E-07 | 4.800E-07 |
| 11167 | 559 | A559Q | A | 1.9e-07 | 2e-07 | 4e-07 | 2.633E-07 | 1.185E-07 | 1.900E-07 | 4.000E-07 |
| 11168 | 559 | A559G | A | 1.4e-07 | 3.1e-07 | 1.2e-07 | 1.900E-07 | 1.044E-07 | 1.200E-07 | 3.100E-07 |
| 11169 | 559 | A559H | A | 1.8e-07 | 3e-07 | 8.2e-07 | 4.333E-07 | 3.402E-07 | 1.800E-07 | 8.200E-07 |
| 11170 | 559 | A559I | A | 3.3e-07 | 1.6e-07 | 3.8e-07 | 2.900E-07 | 1.153E-07 | 1.600E-07 | 3.800E-07 |
| 11171 | 559 | A559L | A | 1.7e-07 | 2.4e-07 | 2.4e-07 | 2.167E-07 | 4.041E-08 | 1.700E-07 | 2.400E-07 |
| 11172 | 559 | A559K | A | 4.2e-07 | 4e-07 | 2.1e-07 | 3.433E-07 | 1.159E-07 | 2.100E-07 | 4.200E-07 |
| 11173 | 559 | A559M | A | 3.3e-07 | 3.7e-07 | 2e-07 | 3.000E-07 | 8.888E-08 | 2.000E-07 | 3.700E-07 |
| 11174 | 559 | A559F | A | 2.6e-07 | 2.4e-07 | 2.9e-07 | 2.633E-07 | 2.517E-08 | 2.400E-07 | 2.900E-07 |
| 11175 | 559 | A559P | A | 2.8e-07 | 4e-07 | 2.6e-07 | 3.133E-07 | 7.572E-08 | 2.600E-07 | 4.000E-07 |
| 11176 | 559 | A559S | A | 4.3e-07 | 2.2e-07 | 1.8e-07 | 2.767E-07 | 1.343E-07 | 1.800E-07 | 4.300E-07 |
| 11177 | 559 | A559T | A | 1.9e-07 | 3.2e-07 | 3.4e-07 | 2.833E-07 | 8.145E-08 | 1.900E-07 | 3.400E-07 |
| 11178 | 559 | A559W | A | 3e-07 | 3.4e-07 | 3.1e-07 | 3.167E-07 | 2.082E-08 | 3.000E-07 | 3.400E-07 |
| 11179 | 559 | A559Y | A | 3.9e-07 | 2.8e-07 | 2.5e-07 | 3.067E-07 | 7.371E-08 | 2.500E-07 | 3.900E-07 |
| 11180 | 559 | A559V | A | 2.6e-07 | 3.3e-07 | 1.6e-07 | 2.500E-07 | 8.544E-08 | 1.600E-07 | 3.300E-07 |
| 11181 | 560 | D560A | A | 9.8e-08 | 2.4e-07 | 2.4e-07 | 1.927E-07 | 8.198E-08 | 9.800E-08 | 2.400E-07 |
| 11182 | 560 | D560R | A | 1.9e-07 | 2e-07 | 1.9e-07 | 1.933E-07 | 5.774E-09 | 1.900E-07 | 2.000E-07 |
| 11183 | 560 | D560N | A | 4.1e-07 | 2.8e-07 | 1e-06 | 5.633E-07 | 3.837E-07 | 2.800E-07 | 1.000E-06 |
| 11184 | 560 | D560D | A | 2e-07 | 2e-07 | 2e-07 | 2.000E-07 | 0.000E+00 | 2.000E-07 | 2.000E-07 |
| 11185 | 560 | D560C | A | 1.2e-07 | 1.7e-07 | 2.1e-07 | 1.667E-07 | 4.509E-08 | 1.200E-07 | 2.100E-07 |
| 11186 | 560 | D560E | A | 1.7e-07 | 3.3e-07 | 1.5e-07 | 2.167E-07 | 9.866E-08 | 1.500E-07 | 3.300E-07 |
| 11187 | 560 | D560Q | A | 4.9e-07 | 6.8e-07 | 7.7e-07 | 6.467E-07 | 1.429E-07 | 4.900E-07 | 7.700E-07 |
| 11188 | 560 | D560G | A | 2.1e-07 | 2e-07 | 7.1e-07 | 3.733E-07 | 2.916E-07 | 2.000E-07 | 7.100E-07 |
| 11189 | 560 | D560H | A | 2.5e-07 | 2.8e-07 | 3.3e-07 | 2.867E-07 | 4.041E-08 | 2.500E-07 | 3.300E-07 |
| 11190 | 560 | D560I | A | 2.4e-07 | 1.3e-07 | 1.2e-07 | 1.633E-07 | 6.658E-08 | 1.200E-07 | 2.400E-07 |
| 11191 | 560 | D560L | A | 7.2e-08 | 9.1e-08 | 6.1e-08 | 7.467E-08 | 1.518E-08 | 6.100E-08 | 9.100E-08 |
| 11192 | 560 | D560K | A | 3.4e-07 | 3.7e-07 | 4.2e-07 | 3.767E-07 | 4.041E-08 | 3.400E-07 | 4.200E-07 |
| 11193 | 560 | D560M | A | 1e-07 | 1.5e-07 | 1.1e-07 | 1.200E-07 | 2.646E-08 | 1.000E-07 | 1.500E-07 |
| 11194 | 560 | D560F | A | 6.9e-08 | 9.7e-08 | 1.9e-07 | 1.187E-07 | 6.334E-08 | 6.900E-08 | 1.900E-07 |
| 11195 | 560 | D560P | A | 2.5e-07 | 3.8e-07 | 2.4e-07 | 2.900E-07 | 7.810E-08 | 2.400E-07 | 3.800E-07 |
| 11196 | 560 | D560S | A | 5.1e-07 | 3.7e-07 | 7.1e-07 | 5.300E-07 | 1.709E-07 | 3.700E-07 | 7.100E-07 |
| 11197 | 560 | D560T | A | 5.9e-07 | 5.4e-07 | 3.5e-07 | 4.933E-07 | 1.266E-07 | 3.500E-07 | 5.900E-07 |
| 11198 | 560 | D560W | A | 1e-07 | 1.1e-07 | 1.1e-07 | 1.067E-07 | 5.774E-09 | 1.000E-07 | 1.100E-07 |
| 11199 | 560 | D560Y | A | 7.8e-08 | 1.7e-07 | 6.2e-08 | 1.033E-07 | 5.829E-08 | 6.200E-08 | 1.700E-07 |
| 11200 | 560 | D560V | A | 2.1e-07 | 1.9e-07 | 1.8e-07 | 1.933E-07 | 1.528E-08 | 1.800E-07 | 2.100E-07 |
| 11201 | 561 | Q561A | A | 2.8e-07 | 1.8e-07 | 2.6e-07 | 2.400E-07 | 5.292E-08 | 1.800E-07 | 2.800E-07 |
| 11202 | 561 | Q561R | A | 2.3e-07 | 1.1e-07 | 1.5e-07 | 1.633E-07 | 6.110E-08 | 1.100E-07 | 2.300E-07 |
| 11203 | 561 | Q561N | A | 2.2e-07 | 2.2e-07 | 4.3e-07 | 2.900E-07 | 1.212E-07 | 2.200E-07 | 4.300E-07 |
| 11204 | 561 | Q561D | A | 1.9e-07 | 1.2e-07 | 2.5e-07 | 1.867E-07 | 6.506E-08 | 1.200E-07 | 2.500E-07 |
| 11205 | 561 | Q561C | A | 8.7e-08 | 2.9e-07 | 2.6e-07 | 2.123E-07 | 1.096E-07 | 8.700E-08 | 2.900E-07 |
| 11206 | 561 | Q561E | A | 1.4e-07 | 1.1e-07 | 2.4e-07 | 1.633E-07 | 6.807E-08 | 1.100E-07 | 2.400E-07 |
| 11207 | 561 | Q561Q | A | 2.3e-07 | 4e-07 | 3.9e-07 | 3.400E-07 | 9.539E-08 | 2.300E-07 | 4.000E-07 |
| 11208 | 561 | Q561G | A | 1.1e-07 | 4.3e-07 | 2.8e-07 | 2.733E-07 | 1.601E-07 | 1.100E-07 | 4.300E-07 |
| 11209 | 561 | Q561H | A | 7.9e-08 | 4.2e-07 | 1.3e-07 | 2.097E-07 | 1.839E-07 | 7.900E-08 | 4.200E-07 |
| 11210 | 561 | Q561I | A | 1.9e-07 | 2.5e-07 | 9.9e-08 | 1.797E-07 | 7.603E-08 | 9.900E-08 | 2.500E-07 |
| 11211 | 561 | Q561L | A | 1.2e-07 | 6e-08 | 5.7e-08 | 7.900E-08 | 3.554E-08 | 5.700E-08 | 1.200E-07 |
| 11212 | 561 | Q561K | A | 4e-07 | 1.8e-07 | 2.2e-07 | 2.667E-07 | 1.172E-07 | 1.800E-07 | 4.000E-07 |
| 11213 | 561 | Q561M | A | 1.1e-07 | 9.2e-08 | 7.4e-08 | 9.200E-08 | 1.800E-08 | 7.400E-08 | 1.100E-07 |
| 11214 | 561 | Q561F | A | 9.3e-08 | 2.3e-07 | 1.7e-07 | 1.643E-07 | 6.868E-08 | 9.300E-08 | 2.300E-07 |
| 11215 | 561 | Q561P | A | 1.6e-07 | 2.8e-07 | 1.9e-07 | 2.100E-07 | 6.245E-08 | 1.600E-07 | 2.800E-07 |
| 11216 | 561 | Q561S | A | 2e-07 | 2.2e-07 | 2.2e-07 | 2.133E-07 | 1.155E-08 | 2.000E-07 | 2.200E-07 |
| 11217 | 561 | Q561T | A | 2.5e-07 | 5e-07 | 2.1e-07 | 3.200E-07 | 1.572E-07 | 2.100E-07 | 5.000E-07 |
| 11218 | 561 | Q561W | A | 7.5e-08 | 7.5e-08 | 1.1e-07 | 8.667E-08 | 2.021E-08 | 7.500E-08 | 1.100E-07 |
| 11219 | 561 | Q561Y | A | 1.3e-07 | 1.2e-07 | 1.4e-07 | 1.300E-07 | 1.000E-08 | 1.200E-07 | 1.400E-07 |

| | | | | | | | | | | |
|---|---|---|---|---|---|---|---|---|---|---|
| 11220 | 561 | Q561V | A | 1.6e-07 | 1.3e-07 | 2.5e-07 | 1.800E-07 | 6.245E-08 | 1.300E-07 | 2.500E-07 |
| 11221 | 562 | C562A | A | 3.1e-07 | 1.6e-07 | 3.2e-07 | 2.633E-07 | 8.963E-08 | 1.600E-07 | 3.200E-07 |
| 11222 | 562 | C562R | A | 2.5e-07 | 3.7e-07 | 1.3e-07 | 2.500E-07 | 1.200E-07 | 1.300E-07 | 3.700E-07 |
| 11223 | 562 | C562N | A | 2.6e-07 | 4.8e-07 | 2.5e-07 | 3.300E-07 | 1.300E-07 | 2.500E-07 | 4.800E-07 |
| 11224 | 562 | C562D | A | 1.8e-07 | 2.5e-07 | 2.1e-07 | 2.133E-07 | 3.512E-08 | 1.800E-07 | 2.500E-07 |
| 11225 | 562 | C562C | A | 3.1e-07 | 1.9e-07 | 3.4e-07 | 2.800E-07 | 7.937E-08 | 1.900E-07 | 3.400E-07 |
| 11226 | 562 | C562E | A | 2.9e-07 | 4.5e-07 | 3.2e-07 | 3.533E-07 | 8.505E-08 | 2.900E-07 | 4.500E-07 |
| 11227 | 562 | C562Q | A | 5.9e-08 | 3.5e-07 | 3.1e-07 | 2.397E-07 | 1.577E-07 | 5.900E-08 | 3.500E-07 |
| 11228 | 562 | C562G | A | 3.6e-07 | 1.8e-07 | 5.2e-07 | 3.533E-07 | 1.701E-07 | 1.800E-07 | 5.200E-07 |
| 11229 | 562 | C562H | A | 1.6e-07 | 3.7e-07 | 2.6e-07 | 2.633E-07 | 1.050E-07 | 1.600E-07 | 3.700E-07 |
| 11230 | 562 | C562I | A | 2.7e-07 | 2.1e-07 | 6.4e-07 | 3.733E-07 | 2.329E-07 | 2.100E-07 | 6.400E-07 |
| 11231 | 562 | C562L | A | 3.4e-07 | 2.6e-07 | 2.3e-07 | 2.767E-07 | 5.686E-08 | 2.300E-07 | 3.400E-07 |
| 11232 | 562 | C562K | A | 1.1e-07 | 2.5e-07 | 2.7e-07 | 2.100E-07 | 8.718E-08 | 1.100E-07 | 2.700E-07 |
| 11233 | 562 | C562M | A | 4.1e-07 | 3.2e-07 | 2.7e-07 | 3.333E-07 | 7.095E-08 | 2.700E-07 | 4.100E-07 |
| 11234 | 562 | C562F | A | 1.1e-07 | 2.6e-07 | 3.8e-07 | 2.500E-07 | 1.353E-07 | 1.100E-07 | 3.800E-07 |
| 11235 | 562 | C562P | A | 1.5e-07 | 2e-07 | 2.1e-07 | 1.867E-07 | 3.215E-08 | 1.500E-07 | 2.100E-07 |
| 11236 | 562 | C562S | A | 2.3e-07 | 1.7e-07 | 7.7e-07 | 3.900E-07 | 3.305E-07 | 1.700E-07 | 7.700E-07 |
| 11237 | 562 | C562T | A | 1e-07 | 3.2e-07 | 3.1e-07 | 2.433E-07 | 1.242E-07 | 1.000E-07 | 3.200E-07 |
| 11238 | 562 | C562W | A | 1.5e-07 | 3.8e-07 | 1.4e-07 | 2.233E-07 | 1.358E-07 | 1.400E-07 | 3.800E-07 |
| 11239 | 562 | C562Y | A | 8.7e-08 | 4.1e-07 | 1.4e-07 | 2.123E-07 | 1.732E-07 | 8.700E-08 | 4.100E-07 |
| 11240 | 562 | C562V | A | 1.5e-07 | 4.2e-07 | 2.4e-07 | 2.700E-07 | 1.375E-07 | 1.500E-07 | 4.200E-07 |
| 11241 | 563 | V563A | A | 1.6e-07 | 2.8e-07 | 3.7e-07 | 2.700E-07 | 1.054E-07 | 1.600E-07 | 3.700E-07 |
| 11242 | 563 | V563R | A | 3.3e-07 | 1e-07 | 8.3e-08 | 1.710E-07 | 1.380E-07 | 8.300E-08 | 3.300E-07 |
| 11243 | 563 | V563N | A | 4e-07 | 2.4e-07 | 3e-07 | 3.133E-07 | 8.083E-08 | 2.400E-07 | 4.000E-07 |
| 11244 | 563 | V563D | A | 2.6e-07 | 1.7e-07 | 4.5e-07 | 2.933E-07 | 1.429E-07 | 1.700E-07 | 4.500E-07 |
| 11245 | 563 | V563C | A | 3.4e-07 | 1.4e-07 | 2.1e-07 | 2.300E-07 | 1.015E-07 | 1.400E-07 | 3.400E-07 |
| 11246 | 563 | V563E | A | 1.6e-07 | 2.8e-07 | 4.9e-07 | 3.100E-07 | 1.670E-07 | 1.600E-07 | 4.900E-07 |
| 11247 | 563 | V563Q | A | 3.1e-07 | 1.8e-07 | 5.8e-07 | 3.567E-07 | 2.040E-07 | 1.800E-07 | 5.800E-07 |
| 11248 | 563 | V563G | A | 2.7e-07 | 6.5e-07 | 3.7e-07 | 4.300E-07 | 1.970E-07 | 2.700E-07 | 6.500E-07 |
| 11249 | 563 | V563H | A | 2.3e-07 | 2.5e-07 | 2.6e-07 | 2.467E-07 | 1.528E-08 | 2.300E-07 | 2.600E-07 |
| 11250 | 563 | V563I | A | 3.9e-07 | 4.7e-07 | 3e-07 | 3.867E-07 | 8.505E-08 | 3.000E-07 | 4.700E-07 |
| 11251 | 563 | V563L | A | 3.3e-07 | 1.3e-07 | 3e-07 | 2.533E-07 | 1.079E-07 | 1.300E-07 | 3.300E-07 |
| 11252 | 563 | V563K | A | 3.6e-07 | 2e-07 | 2e-07 | 2.533E-07 | 9.238E-08 | 2.000E-07 | 3.600E-07 |
| 11253 | 563 | V563M | A | 1.4e-07 | 4.5e-07 | 3e-07 | 2.967E-07 | 1.550E-07 | 1.400E-07 | 4.500E-07 |
| 11254 | 563 | V563F | A | 6.2e-07 | 2.7e-07 | 3.2e-07 | 4.033E-07 | 1.893E-07 | 2.700E-07 | 6.200E-07 |
| 11255 | 563 | V563P | A | 4.3e-07 | 1.8e-07 | 4.4e-07 | 3.500E-07 | 1.473E-07 | 1.800E-07 | 4.400E-07 |
| 11256 | 563 | V563S | A | 2.6e-07 | 5.6e-07 | 3e-07 | 3.733E-07 | 1.629E-07 | 2.600E-07 | 5.600E-07 |
| 11257 | 563 | V563T | A | 2e-07 | 2.7e-07 | 2.4e-07 | 2.367E-07 | 3.512E-08 | 2.000E-07 | 2.700E-07 |
| 11258 | 563 | V563W | A | 2e-07 | 2e-07 | 2.1e-07 | 2.033E-07 | 5.774E-09 | 2.000E-07 | 2.100E-07 |
| 11259 | 563 | V563Y | A | 2.3e-07 | 1.6e-07 | 2e-07 | 1.967E-07 | 3.512E-08 | 1.600E-07 | 2.300E-07 |
| 11260 | 563 | V563V | A | 1.7e-07 | 2.5e-07 | 3.1e-07 | 2.433E-07 | 7.024E-08 | 1.700E-07 | 3.100E-07 |
| 11261 | 564 | A564A | A | 3.3e-07 | 1.9e-07 | 3.2e-07 | 2.800E-07 | 7.810E-08 | 1.900E-07 | 3.300E-07 |
| 11262 | 564 | A564R | A | 1.3e-07 | 2e-07 | 2.1e-07 | 1.800E-07 | 4.359E-08 | 1.300E-07 | 2.100E-07 |
| 11263 | 564 | A564N | A | 8e-08 | 1.8e-07 | 1.7e-07 | 1.433E-07 | 5.508E-08 | 8.000E-08 | 1.800E-07 |
| 11264 | 564 | A564D | A | 2e-07 | 1.1e-07 | 1.4e-07 | 1.500E-07 | 4.583E-08 | 1.100E-07 | 2.000E-07 |
| 11265 | 564 | A564C | A | 1.8e-07 | 2.9e-07 | 9.7e-08 | 1.890E-07 | 9.681E-08 | 9.700E-08 | 2.900E-07 |
| 11266 | 564 | A564E | A | 2.5e-07 | 2.1e-07 | 2.1e-07 | 2.233E-07 | 2.309E-08 | 2.100E-07 | 2.500E-07 |
| 11267 | 564 | A564Q | A | 2e-07 | 2.4e-07 | 2.3e-07 | 2.233E-07 | 2.082E-08 | 2.000E-07 | 2.400E-07 |
| 11268 | 564 | A564G | A | 2.6e-07 | 1.4e-07 | 1.3e-07 | 1.767E-07 | 7.234E-08 | 1.300E-07 | 2.600E-07 |
| 11269 | 564 | A564H | A | 1.3e-07 | 1.2e-07 | 2.4e-07 | 1.633E-07 | 6.658E-08 | 1.200E-07 | 2.400E-07 |
| 11270 | 564 | A564I | A | 2.8e-07 | 1.7e-07 | 2.1e-07 | 2.200E-07 | 5.568E-08 | 1.700E-07 | 2.800E-07 |
| 11271 | 564 | A564L | A | 4.8e-07 | 2.5e-07 | 2.1e-07 | 3.133E-07 | 1.457E-07 | 2.100E-07 | 4.800E-07 |
| 11272 | 564 | A564K | A | 1.2e-07 | 1.6e-07 | 1.5e-07 | 1.433E-07 | 2.082E-08 | 1.200E-07 | 1.600E-07 |
| 11273 | 564 | A564M | A | 2.8e-07 | 1.4e-07 | 2e-07 | 2.067E-07 | 7.024E-08 | 1.400E-07 | 2.800E-07 |
| 11274 | 564 | A564F | A | 1.8e-07 | 1.8e-07 | 1.2e-07 | 1.600E-07 | 3.464E-08 | 1.200E-07 | 1.800E-07 |
| 11275 | 564 | A564P | A | 2.8e-07 | 1.1e-07 | 3.1e-07 | 2.333E-07 | 1.079E-07 | 1.100E-07 | 3.100E-07 |
| 11276 | 564 | A564S | A | 4.1e-07 | 1.1e-07 | 3.5e-07 | 2.900E-07 | 1.587E-07 | 1.100E-07 | 4.100E-07 |
| 11277 | 564 | A564T | A | 2.4e-07 | 1.6e-07 | 1.5e-07 | 1.833E-07 | 4.933E-08 | 1.500E-07 | 2.400E-07 |
| 11278 | 564 | A564W | A | 3.1e-07 | 2.4e-07 | 3.2e-07 | 2.900E-07 | 4.359E-08 | 2.400E-07 | 3.200E-07 |
| 11279 | 564 | A564Y | A | 1e-07 | 4e-07 | 1.9e-07 | 2.300E-07 | 1.539E-07 | 1.000E-07 | 4.000E-07 |
| 11280 | 564 | A564V | A | 2.3e-07 | 1.9e-07 | 2.7e-07 | 2.300E-07 | 4.000E-08 | 1.900E-07 | 2.700E-07 |
| 11281 | 565 | C565A | A | 6.9e-07 | 2.2e-07 | 3.3e-07 | 4.133E-07 | 2.458E-07 | 2.200E-07 | 6.900E-07 |
| 11282 | 565 | C565R | A | 3.2e-07 | 3.4e-07 | 1.9e-07 | 2.833E-07 | 8.145E-08 | 1.900E-07 | 3.400E-07 |
| 11283 | 565 | C565N | A | 1.2e-06 | 1.1e-07 | 1.1e-07 | 4.733E-07 | 6.293E-07 | 1.100E-07 | 1.200E-06 |
| 11284 | 565 | C565D | A | 4.3e-07 | 3e-07 | 1.5e-07 | 2.933E-07 | 1.401E-07 | 1.500E-07 | 4.300E-07 |
| 11285 | 565 | C565C | A | 1.9e-07 | 2.3e-07 | 2.3e-07 | 2.167E-07 | 2.309E-08 | 1.900E-07 | 2.300E-07 |
| 11286 | 565 | C565E | A | 2.3e-07 | 5.2e-07 | 3.1e-07 | 3.533E-07 | 1.498E-07 | 2.300E-07 | 5.200E-07 |
| 11287 | 565 | C565Q | A | 4.1e-07 | 3.2e-07 | 4.5e-07 | 3.933E-07 | 6.658E-08 | 3.200E-07 | 4.500E-07 |
| 11288 | 565 | C565G | A | 2.1e-07 | 1e-07 | 2.6e-07 | 1.900E-07 | 8.185E-08 | 1.000E-07 | 2.600E-07 |
| 11289 | 565 | C565H | A | 2.2e-07 | 2.4e-07 | 3.9e-07 | 2.833E-07 | 9.292E-08 | 2.200E-07 | 3.900E-07 |
| 11290 | 565 | C565I | A | 4.1e-07 | 3.9e-07 | 3.3e-07 | 3.767E-07 | 4.163E-08 | 3.300E-07 | 4.100E-07 |
| 11291 | 565 | C565L | A | 3.6e-07 | 1.2e-07 | 1.8e-07 | 2.200E-07 | 1.249E-07 | 1.200E-07 | 3.600E-07 |
| 11292 | 565 | C565K | A | 2.7e-07 | 2.1e-07 | 1.6e-07 | 2.133E-07 | 5.508E-08 | 1.600E-07 | 2.700E-07 |
| 11293 | 565 | C565M | A | 2.9e-07 | 1.3e-07 | 1.6e-07 | 1.933E-07 | 8.505E-08 | 1.300E-07 | 2.900E-07 |
| 11294 | 565 | C565F | A | 2.9e-07 | 2e-07 | 1.6e-07 | 2.167E-07 | 6.658E-08 | 1.600E-07 | 2.900E-07 |
| 11295 | 565 | C565P | A | 1.8e-07 | 3.7e-07 | 3e-07 | 2.833E-07 | 9.609E-08 | 1.800E-07 | 3.700E-07 |
| 11296 | 565 | C565S | A | 2.2e-07 | 1.1e-07 | 7.7e-07 | 3.667E-07 | 3.536E-07 | 1.100E-07 | 7.700E-07 |
| 11297 | 565 | C565T | A | 2.8e-07 | 5e-07 | 1.4e-07 | 3.067E-07 | 1.815E-07 | 1.400E-07 | 5.000E-07 |
| 11298 | 565 | C565W | A | 3.6e-07 | 3e-07 | 2.6e-07 | 3.067E-07 | 5.033E-08 | 2.600E-07 | 3.600E-07 |
| 11299 | 565 | C565Y | A | 1.9e-07 | 2.6e-07 | 4.9e-07 | 3.133E-07 | 1.570E-07 | 1.900E-07 | 4.900E-07 |
| 11300 | 565 | C565V | A | 4e-07 | 1.6e-07 | 1.7e-07 | 2.433E-07 | 1.358E-07 | 1.600E-07 | 4.000E-07 |
| 11301 | 566 | A566A | A | 5e-07 | 2e-07 | 1.5e-07 | 2.833E-07 | 1.893E-07 | 1.500E-07 | 5.000E-07 |
| 11302 | 566 | A566R | A | 1.4e-07 | 3.2e-07 | 1.3e-07 | 1.967E-07 | 1.069E-07 | 1.300E-07 | 3.200E-07 |
| 11303 | 566 | A566N | A | 2.9e-07 | 2e-07 | 1.9e-07 | 2.267E-07 | 5.508E-08 | 1.900E-07 | 2.900E-07 |
| 11304 | 566 | A566D | A | 2.1e-07 | 2.1e-07 | 2.7e-07 | 2.300E-07 | 3.464E-08 | 2.100E-07 | 2.700E-07 |

| | | | | | | | | | | |
|---|---|---|---|---|---|---|---|---|---|---|
| 11305 | 566 | A566C | A | 9e-08 | 1.8e-07 | 2.8e-07 | 1.833E-07 | 9.504E-08 | 9.000E-08 | 2.800E-07 |
| 11306 | 566 | A566E | A | 2.8e-07 | 3.1e-07 | 2.1e-07 | 2.667E-07 | 5.132E-08 | 2.100E-07 | 3.100E-07 |
| 11307 | 566 | A566Q | A | 3.7e-07 | 1.7e-07 | 3.5e-07 | 2.967E-07 | 1.102E-07 | 1.700E-07 | 3.700E-07 |
| 11308 | 566 | A566G | A | 2.6e-07 | 5.7e-07 | 4.3e-07 | 4.200E-07 | 1.552E-07 | 2.600E-07 | 5.700E-07 |
| 11309 | 566 | A566H | A | 2.4e-07 | 3.1e-07 | 3.4e-07 | 2.967E-07 | 5.132E-08 | 2.400E-07 | 3.400E-07 |
| 11310 | 566 | A566I | A | 1.8e-07 | 3.4e-07 | 3.7e-07 | 2.967E-07 | 1.021E-07 | 1.800E-07 | 3.700E-07 |
| 11311 | 566 | A566L | A | 1.4e-07 | 2e-07 | 7.7e-08 | 1.390E-07 | 6.151E-08 | 7.700E-08 | 2.000E-07 |
| 11312 | 566 | A566K | A | 2.5e-07 | 4.9e-07 | 4.6e-07 | 4.000E-07 | 1.308E-07 | 2.500E-07 | 4.900E-07 |
| 11313 | 566 | A566M | A | 2.6e-07 | 1.1e-07 | 2e-07 | 1.900E-07 | 7.550E-08 | 1.100E-07 | 2.600E-07 |
| 11314 | 566 | A566F | A | 3.1e-07 | 3.1e-07 | 2.8e-07 | 3.000E-07 | 1.732E-08 | 2.800E-07 | 3.100E-07 |
| 11315 | 566 | A566P | A | 1.8e-07 | 1.4e-07 | 2.7e-07 | 1.967E-07 | 6.658E-08 | 1.400E-07 | 2.700E-07 |
| 11316 | 566 | A566S | A | 2.5e-07 | 2.4e-07 | 2.6e-07 | 2.500E-07 | 1.000E-08 | 2.400E-07 | 2.600E-07 |
| 11317 | 566 | A566T | A | 1.8e-07 | 1.1e-07 | 2.4e-07 | 1.767E-07 | 6.506E-08 | 1.100E-07 | 2.400E-07 |
| 11318 | 566 | A566W | A | 1.7e-07 | 2.5e-07 | 1.7e-07 | 1.967E-07 | 4.619E-08 | 1.700E-07 | 2.500E-07 |
| 11319 | 566 | A566Y | A | 1.1e-07 | 2.8e-07 | 3.3e-07 | 2.400E-07 | 1.153E-07 | 1.100E-07 | 3.300E-07 |
| 11320 | 566 | A566V | A | 2.1e-07 | 1.9e-07 | 8.6e-08 | 1.620E-07 | 6.657E-08 | 8.600E-08 | 2.100E-07 |
| 11321 | 567 | H567A | A | 2.9e-07 | 2e-07 | 4e-07 | 2.967E-07 | 1.002E-07 | 2.000E-07 | 4.000E-07 |
| 11322 | 567 | H567R | A | 2.4e-07 | 1.3e-07 | 2.7e-07 | 2.133E-07 | 7.371E-08 | 1.300E-07 | 2.700E-07 |
| 11323 | 567 | H567N | A | 1.7e-07 | 2.2e-07 | 1.9e-07 | 1.933E-07 | 2.517E-08 | 1.700E-07 | 2.200E-07 |
| 11324 | 567 | H567D | A | 1.5e-07 | 1.5e-07 | 6.6e-07 | 3.200E-07 | 2.944E-07 | 1.500E-07 | 6.600E-07 |
| 11325 | 567 | H567C | A | 3.1e-07 | 2.6e-07 | 6.3e-07 | 4.000E-07 | 2.007E-07 | 2.600E-07 | 6.300E-07 |
| 11326 | 567 | H567E | A | 3.8e-07 | 7.4e-07 | 2.2e-07 | 4.467E-07 | 2.663E-07 | 2.200E-07 | 7.400E-07 |
| 11327 | 567 | H567Q | A | 2.5e-07 | 3.1e-07 | 2.2e-07 | 2.600E-07 | 4.583E-08 | 2.200E-07 | 3.100E-07 |
| 11328 | 567 | H567G | A | 2.2e-07 | 2.1e-07 | 4.3e-07 | 2.867E-07 | 1.242E-07 | 2.100E-07 | 4.300E-07 |
| 11329 | 567 | H567H | A | 5e-07 | 7.9e-07 | 2.7e-07 | 5.200E-07 | 2.606E-07 | 2.700E-07 | 7.900E-07 |
| 11330 | 567 | H567I | A | 2.9e-07 | 1.7e-07 | 2.9e-07 | 2.500E-07 | 6.928E-08 | 1.700E-07 | 2.900E-07 |
| 11331 | 567 | H567L | A | 3.4e-07 | 1e-07 | 7e-07 | 3.800E-07 | 3.020E-07 | 1.000E-07 | 7.000E-07 |
| 11332 | 567 | H567K | A | 3.2e-07 | 2.3e-07 | 2.4e-07 | 2.633E-07 | 4.933E-08 | 2.300E-07 | 3.200E-07 |
| 11333 | 567 | H567M | A | 3.1e-07 | 4.6e-07 | 1.6e-07 | 3.100E-07 | 1.500E-07 | 1.600E-07 | 4.600E-07 |
| 11334 | 567 | H567F | A | 1.6e-07 | 5.5e-07 | 3.1e-07 | 3.400E-07 | 1.967E-07 | 1.600E-07 | 5.500E-07 |
| 11335 | 567 | H567P | A | 2.7e-07 | 3.1e-07 | 4.2e-07 | 3.333E-07 | 7.767E-08 | 2.700E-07 | 4.200E-07 |
| 11336 | 567 | H567S | A | 1.7e-07 | 3e-07 | 4.7e-07 | 3.133E-07 | 1.504E-07 | 1.700E-07 | 4.700E-07 |
| 11337 | 567 | H567T | A | 3e-07 | 3.1e-07 | 9.5e-08 | 2.350E-07 | 1.213E-07 | 9.500E-08 | 3.100E-07 |
| 11338 | 567 | H567W | A | 2.7e-07 | 2.7e-07 | 4.2e-07 | 3.200E-07 | 8.660E-08 | 2.700E-07 | 4.200E-07 |
| 11339 | 567 | H567Y | A | 1.9e-07 | 1.8e-07 | 2.6e-07 | 2.100E-07 | 4.359E-08 | 1.800E-07 | 2.600E-07 |
| 11340 | 567 | H567V | A | 3.2e-07 | 2.4e-07 | 3e-07 | 2.867E-07 | 4.163E-08 | 2.400E-07 | 3.200E-07 |
| 11341 | 568 | Y568A | A | 1.9e-07 | 3.1e-07 | 2.3e-07 | 2.433E-07 | 6.110E-08 | 1.900E-07 | 3.100E-07 |
| 11342 | 568 | Y568R | A | 2.6e-07 | 1.5e-07 | 2.6e-07 | 2.233E-07 | 6.351E-08 | 1.500E-07 | 2.600E-07 |
| 11343 | 568 | Y568N | A | 3.8e-07 | 2.1e-07 | 1.2e-07 | 2.367E-07 | 1.320E-07 | 1.200E-07 | 3.800E-07 |
| 11344 | 568 | Y568D | A | 1.8e-07 | 1.6e-07 | 2.4e-07 | 1.933E-07 | 4.163E-08 | 1.600E-07 | 2.400E-07 |
| 11345 | 568 | Y568C | A | 1.8e-07 | 2.5e-07 | 2.7e-07 | 2.333E-07 | 4.726E-08 | 1.800E-07 | 2.700E-07 |
| 11346 | 568 | Y568E | A | 2.3e-07 | 2.8e-07 | 2.4e-07 | 2.500E-07 | 2.646E-08 | 2.300E-07 | 2.800E-07 |
| 11347 | 568 | Y568Q | A | 9.5e-08 | 2e-07 | 3.7e-07 | 2.217E-07 | 1.388E-07 | 9.500E-08 | 3.700E-07 |
| 11348 | 568 | Y568G | A | 1.3e-07 | 1.5e-07 | 2.2e-07 | 1.667E-07 | 4.726E-08 | 1.300E-07 | 2.200E-07 |
| 11349 | 568 | Y568H | A | 1.6e-07 | 1.3e-07 | 2.7e-07 | 1.867E-07 | 7.371E-08 | 1.300E-07 | 2.700E-07 |
| 11350 | 568 | Y568I | A | 2.1e-07 | 2.8e-07 | 2.2e-07 | 2.367E-07 | 3.786E-08 | 2.100E-07 | 2.800E-07 |
| 11351 | 568 | Y568L | A | 5.3e-07 | 1.9e-07 | 1.6e-07 | 2.933E-07 | 2.055E-07 | 1.600E-07 | 5.300E-07 |
| 11352 | 568 | Y568K | A | 1.2e-07 | 1.7e-07 | 3.4e-07 | 2.100E-07 | 1.153E-07 | 1.200E-07 | 3.400E-07 |
| 11353 | 568 | Y568M | A | 2.9e-07 | 1.6e-07 | 2.1e-07 | 2.200E-07 | 6.557E-08 | 1.600E-07 | 2.900E-07 |
| 11354 | 568 | Y568F | A | 2.5e-07 | 2.2e-07 | 4.3e-07 | 3.000E-07 | 1.136E-07 | 2.200E-07 | 4.300E-07 |
| 11355 | 568 | Y568P | A | 1.5e-07 | 3e-07 | 2.4e-07 | 2.300E-07 | 7.550E-08 | 1.500E-07 | 3.000E-07 |
| 11356 | 568 | Y568S | A | 1.1e-07 | 1.8e-07 | 5.5e-07 | 2.800E-07 | 2.364E-07 | 1.100E-07 | 5.500E-07 |
| 11357 | 568 | Y568T | A | 1.5e-07 | 1.8e-07 | 1.9e-07 | 1.733E-07 | 2.082E-08 | 1.500E-07 | 1.900E-07 |
| 11358 | 568 | Y568W | A | 5.2e-07 | 1.6e-07 | 2.9e-07 | 3.233E-07 | 1.823E-07 | 1.600E-07 | 5.200E-07 |
| 11359 | 568 | Y568Y | A | 1.8e-07 | 2.5e-07 | 2.4e-07 | 2.233E-07 | 3.786E-08 | 1.800E-07 | 2.500E-07 |
| 11360 | 568 | Y568V | A | 2.5e-07 | 3e-07 | 2.7e-07 | 2.733E-07 | 2.517E-08 | 2.500E-07 | 3.000E-07 |
| 11361 | 569 | K569A | A | 2.4e-07 | 3.4e-07 | 2.2e-07 | 2.667E-07 | 6.429E-08 | 2.200E-07 | 3.400E-07 |
| 11362 | 569 | K569R | A | 4.6e-07 | 1.3e-07 | 2.2e-07 | 2.700E-07 | 1.706E-07 | 1.300E-07 | 4.600E-07 |
| 11363 | 569 | K569N | A | 4.4e-07 | 1.5e-07 | 3.6e-07 | 3.167E-07 | 1.498E-07 | 1.500E-07 | 4.400E-07 |
| 11364 | 569 | K569D | A | 1.8e-07 | 3e-07 | 3.6e-07 | 2.800E-07 | 9.165E-08 | 1.800E-07 | 3.600E-07 |
| 11365 | 569 | K569C | A | 2.3e-07 | 1.7e-07 | 1.6e-07 | 1.867E-07 | 3.786E-08 | 1.600E-07 | 2.300E-07 |
| 11366 | 569 | K569E | A | 1.5e-07 | 2.3e-07 | 6.2e-07 | 3.333E-07 | 2.515E-07 | 1.500E-07 | 6.200E-07 |
| 11367 | 569 | K569Q | A | 2.1e-07 | 1.8e-07 | 8e-08 | 1.567E-07 | 6.807E-08 | 8.000E-08 | 2.100E-07 |
| 11368 | 569 | K569G | A | 2.9e-07 | 2.6e-07 | 5.1e-07 | 3.533E-07 | 1.365E-07 | 2.600E-07 | 5.100E-07 |
| 11369 | 569 | K569H | A | 3.3e-07 | 3.5e-07 | 3e-07 | 3.267E-07 | 2.517E-08 | 3.000E-07 | 3.500E-07 |
| 11370 | 569 | K569I | A | 2.1e-07 | 2.3e-07 | 5.1e-07 | 3.167E-07 | 1.677E-07 | 2.100E-07 | 5.100E-07 |
| 11371 | 569 | K569L | A | 2.2e-07 | 2.3e-07 | 4.9e-07 | 3.133E-07 | 1.531E-07 | 2.200E-07 | 4.900E-07 |
| 11372 | 569 | K569K | A | 1.7e-07 | 3.7e-07 | 4.9e-07 | 3.433E-07 | 1.617E-07 | 1.700E-07 | 4.900E-07 |
| 11373 | 569 | K569M | A | 2.7e-07 | 2e-07 | 2.3e-07 | 2.333E-07 | 3.512E-08 | 2.000E-07 | 2.700E-07 |
| 11374 | 569 | K569F | A | 2.9e-07 | 9.5e-08 | 3.8e-07 | 2.550E-07 | 1.457E-07 | 9.500E-08 | 3.800E-07 |
| 11375 | 569 | K569P | A | 2.2e-07 | 1.7e-07 | 5.4e-07 | 3.100E-07 | 2.007E-07 | 1.700E-07 | 5.400E-07 |
| 11376 | 569 | K569S | A | 2.2e-07 | 2.1e-07 | 6e-07 | 3.433E-07 | 2.223E-07 | 2.100E-07 | 6.000E-07 |
| 11377 | 569 | K569T | A | 2.1e-07 | 1.9e-07 | 1e-07 | 1.667E-07 | 5.859E-08 | 1.000E-07 | 2.100E-07 |
| 11378 | 569 | K569W | A | 1.8e-07 | 4.6e-07 | 1.6e-07 | 2.667E-07 | 1.677E-07 | 1.600E-07 | 4.600E-07 |
| 11379 | 569 | K569Y | A | 1.6e-07 | 1.5e-07 | 2.8e-07 | 1.967E-07 | 7.234E-08 | 1.500E-07 | 2.800E-07 |
| 11380 | 569 | K569V | A | 2.9e-07 | 3.2e-07 | 1.4e-07 | 2.500E-07 | 9.644E-08 | 1.400E-07 | 3.200E-07 |
| 11381 | 570 | D570A | A | 2.2e-07 | 3.2e-07 | 3.8e-07 | 3.067E-07 | 8.083E-08 | 2.200E-07 | 3.800E-07 |
| 11382 | 570 | D570R | A | 5.5e-07 | 6.2e-07 | 1.7e-07 | 4.467E-07 | 2.421E-07 | 1.700E-07 | 6.200E-07 |
| 11383 | 570 | D570N | A | 2.7e-07 | 1.5e-07 | 3.7e-07 | 2.633E-07 | 1.102E-07 | 1.500E-07 | 3.700E-07 |
| 11384 | 570 | D570D | A | 2.1e-07 | 2e-07 | 1.3e-07 | 1.800E-07 | 4.359E-08 | 1.300E-07 | 2.100E-07 |
| 11385 | 570 | D570C | A | 1.3e-07 | 7e-08 | 3.5e-07 | 1.833E-07 | 1.474E-07 | 7.000E-08 | 3.500E-07 |
| 11386 | 570 | D570E | A | 2.2e-07 | 1.6e-07 | 9.8e-08 | 1.593E-07 | 6.100E-08 | 9.800E-08 | 2.200E-07 |
| 11387 | 570 | D570Q | A | 2.9e-07 | 2.7e-07 | 2.1e-07 | 2.567E-07 | 4.163E-08 | 2.100E-07 | 2.900E-07 |
| 11388 | 570 | D570G | A | 2.7e-07 | 1.3e-07 | 1.6e-06 | 6.667E-07 | 8.113E-07 | 1.300E-07 | 1.600E-06 |
| 11389 | 570 | D570H | A | 3.9e-07 | 7.9e-08 | 2.4e-07 | 2.363E-07 | 1.555E-07 | 7.900E-08 | 3.900E-07 |

| | | | | | | | | | | |
|---|---|---|---|---|---|---|---|---|---|---|
| 11390 | 570 | D570I | A | 4.5e-07 | 3.5e-07 | 4.9e-07 | 4.300E-07 | 7.211E-08 | 3.500E-07 | 4.900E-07 |
| 11391 | 570 | D570L | A | 6.1e-07 | 3.1e-07 | 3.9e-07 | 4.367E-07 | 1.553E-07 | 3.100E-07 | 6.100E-07 |
| 11392 | 570 | D570K | A | 2.7e-07 | 4.4e-07 | 2.9e-07 | 3.333E-07 | 9.292E-08 | 2.700E-07 | 4.400E-07 |
| 11393 | 570 | D570M | A | 2.3e-07 | 4.3e-07 | 5.2e-07 | 3.933E-07 | 1.484E-07 | 2.300E-07 | 5.200E-07 |
| 11394 | 570 | D570F | A | 3.6e-07 | 1.8e-07 | 3.1e-07 | 2.833E-07 | 9.292E-08 | 1.800E-07 | 3.600E-07 |
| 11395 | 570 | D570P | A | 6.6e-07 | 5.2e-07 | 1.9e-07 | 4.567E-07 | 2.413E-07 | 1.900E-07 | 6.600E-07 |
| 11396 | 570 | D570S | A | 1.6e-07 | 1.7e-07 | 1.5e-07 | 1.600E-07 | 1.000E-08 | 1.500E-07 | 1.700E-07 |
| 11397 | 570 | D570T | A | 2.1e-07 | 1.3e-07 | 1.2e-07 | 1.533E-07 | 4.933E-08 | 1.200E-07 | 2.100E-07 |
| 11398 | 570 | D570W | A | 7.3e-07 | 1.5e-07 | 2.4e-07 | 3.733E-07 | 3.121E-07 | 1.500E-07 | 7.300E-07 |
| 11399 | 570 | D570Y | A | 2e-07 | 2e-07 | 7.9e-07 | 3.967E-07 | 3.406E-07 | 2.000E-07 | 7.900E-07 |
| 11400 | 570 | D570V | A | 2.4e-07 | 2.6e-07 | 4.1e-07 | 3.033E-07 | 9.292E-08 | 2.400E-07 | 4.100E-07 |
| 11401 | 571 | P571A | A | 2.7e-07 | 3.1e-07 | 3.7e-07 | 3.167E-07 | 5.033E-08 | 2.700E-07 | 3.700E-07 |
| 11402 | 571 | P571R | A | 2.2e-07 | 1.1e-07 | 1.1e-07 | 1.467E-07 | 6.351E-08 | 1.100E-07 | 2.200E-07 |
| 11403 | 571 | P571N | A | 2.3e-07 | 3.2e-07 | 2.4e-07 | 2.633E-07 | 4.933E-08 | 2.300E-07 | 3.200E-07 |
| 11404 | 571 | P571D | A | 1.4e-07 | 3.3e-07 | 1.9e-07 | 2.200E-07 | 9.849E-08 | 1.400E-07 | 3.300E-07 |
| 11405 | 571 | P571C | A | 2.6e-07 | 1e-07 | 1.8e-07 | 1.800E-07 | 8.000E-08 | 1.000E-07 | 2.600E-07 |
| 11406 | 571 | P571E | A | 1.8e-07 | 2e-07 | 2.6e-07 | 2.133E-07 | 4.163E-08 | 1.800E-07 | 2.600E-07 |
| 11407 | 571 | P571Q | A | 2.6e-07 | 2.1e-07 | 2.2e-07 | 2.300E-07 | 2.646E-08 | 2.100E-07 | 2.600E-07 |
| 11408 | 571 | P571G | A | 4.5e-07 | 3e-07 | 2.5e-07 | 3.333E-07 | 1.041E-07 | 2.500E-07 | 4.500E-07 |
| 11409 | 571 | P571H | A | 2.2e-07 | 2e-07 | 1.2e-07 | 1.800E-07 | 5.292E-08 | 1.200E-07 | 2.200E-07 |
| 11410 | 571 | P571I | A | 2.5e-07 | 2.4e-07 | 9.1e-08 | 1.937E-07 | 8.905E-08 | 9.100E-08 | 2.500E-07 |
| 11411 | 571 | P571L | A | 8.8e-08 | 1.5e-07 | 7.7e-08 | 1.050E-07 | 3.936E-08 | 7.700E-08 | 1.500E-07 |
| 11412 | 571 | P571K | A | 3.3e-07 | 2e-07 | 2.6e-07 | 2.633E-07 | 6.506E-08 | 2.000E-07 | 3.300E-07 |
| 11413 | 571 | P571M | A | 1.8e-07 | 7.9e-08 | 1.2e-07 | 1.263E-07 | 5.080E-08 | 7.900E-08 | 1.800E-07 |
| 11414 | 571 | P571F | A | 1.1e-07 | 2.4e-07 | 9.1e-08 | 1.470E-07 | 8.110E-08 | 9.100E-08 | 2.400E-07 |
| 11415 | 571 | P571P | A | 2.1e-07 | 6.1e-07 | 1.5e-07 | 3.233E-07 | 2.501E-07 | 1.500E-07 | 6.100E-07 |
| 11416 | 571 | P571S | A | 1.5e-07 | 1.4e-07 | 1.7e-07 | 1.533E-07 | 1.528E-08 | 1.400E-07 | 1.700E-07 |
| 11417 | 571 | P571T | A | 1.6e-07 | 1.2e-07 | 1.6e-07 | 1.467E-07 | 2.309E-08 | 1.200E-07 | 1.600E-07 |
| 11418 | 571 | P571W | A | 5.5e-08 | 2.7e-07 | 9.6e-08 | 1.403E-07 | 1.142E-07 | 5.500E-08 | 2.700E-07 |
| 11419 | 571 | P571Y | A | 1.4e-07 | 1.8e-07 | 2.2e-07 | 1.800E-07 | 4.000E-08 | 1.400E-07 | 2.200E-07 |
| 11420 | 571 | P571V | A | 9.8e-08 | 7.6e-08 | 1.9e-07 | 1.213E-07 | 6.048E-08 | 7.600E-08 | 1.900E-07 |
| 11421 | 572 | P572A | A | 6.1e-07 | 8.6e-08 | 3e-07 | 3.320E-07 | 2.635E-07 | 8.600E-08 | 6.100E-07 |
| 11422 | 572 | P572R | A | 2.2e-07 | 4.7e-07 | 1.4e-07 | 2.767E-07 | 1.721E-07 | 1.400E-07 | 4.700E-07 |
| 11423 | 572 | P572N | A | 5e-07 | 6.9e-07 | 3.7e-07 | 5.200E-07 | 1.609E-07 | 3.700E-07 | 6.900E-07 |
| 11424 | 572 | P572D | A | 2.1e-07 | 4.3e-07 | 3.3e-07 | 3.233E-07 | 1.102E-07 | 2.100E-07 | 4.300E-07 |
| 11425 | 572 | P572C | A | 2.1e-07 | 3.1e-07 | 3.5e-07 | 2.900E-07 | 7.211E-08 | 2.100E-07 | 3.500E-07 |
| 11426 | 572 | P572E | A | 5.9e-07 | 1e-07 | 1.5e-07 | 2.800E-07 | 2.696E-07 | 1.000E-07 | 5.900E-07 |
| 11427 | 572 | P572Q | A | 5.7e-07 | 9.6e-08 | 6.7e-07 | 4.453E-07 | 3.066E-07 | 9.600E-08 | 6.700E-07 |
| 11428 | 572 | P572G | A | 2e-07 | 4.6e-07 | 2.3e-07 | 2.967E-07 | 1.422E-07 | 2.000E-07 | 4.600E-07 |
| 11429 | 572 | P572H | A | 4.3e-07 | 3.6e-07 | 1.7e-07 | 3.200E-07 | 1.345E-07 | 1.700E-07 | 4.300E-07 |
| 11430 | 572 | P572I | A | 1.7e-07 | 2.1e-07 | 2.2e-07 | 2.000E-07 | 2.646E-08 | 1.700E-07 | 2.200E-07 |
| 11431 | 572 | P572L | A | 8.9e-08 | 4.2e-07 | 2.1e-07 | 2.397E-07 | 1.675E-07 | 8.900E-08 | 4.200E-07 |
| 11432 | 572 | P572K | A | 2e-07 | 4.2e-07 | 1.6e-07 | 2.600E-07 | 1.400E-07 | 1.600E-07 | 4.200E-07 |
| 11433 | 572 | P572M | A | 5.5e-07 | 3.3e-07 | 1.8e-07 | 3.533E-07 | 1.861E-07 | 1.800E-07 | 5.500E-07 |
| 11434 | 572 | P572F | A | 1.3e-07 | 1.8e-07 | 2.5e-07 | 1.867E-07 | 6.028E-08 | 1.300E-07 | 2.500E-07 |
| 11435 | 572 | P572P | A | 6.5e-07 | 2e-07 | 2.5e-07 | 3.667E-07 | 2.466E-07 | 2.000E-07 | 6.500E-07 |
| 11436 | 572 | P572S | A | 9.8e-07 | 5.5e-07 | 5.7e-07 | 7.000E-07 | 2.427E-07 | 5.500E-07 | 9.800E-07 |
| 11437 | 572 | P572T | A | 3.7e-07 | 8.8e-07 | 6.2e-07 | 6.233E-07 | 2.550E-07 | 3.700E-07 | 8.800E-07 |
| 11438 | 572 | P572W | A | 1.1e-07 | 2.2e-07 | 1.3e-07 | 1.533E-07 | 5.859E-08 | 1.100E-07 | 2.200E-07 |
| 11439 | 572 | P572Y | A | 1.8e-07 | 2.3e-07 | 1.7e-07 | 1.933E-07 | 3.215E-08 | 1.700E-07 | 2.300E-07 |
| 11440 | 572 | P572V | A | 1.5e-07 | 2.8e-07 | 1.6e-07 | 1.967E-07 | 7.234E-08 | 1.500E-07 | 2.800E-07 |
| 11441 | 573 | F573A | A | 3.2e-07 | 3.9e-07 | 2e-07 | 3.033E-07 | 9.609E-08 | 2.000E-07 | 3.900E-07 |
| 11442 | 573 | F573R | A | 1.2e-07 | 1.5e-07 | 1.7e-07 | 1.467E-07 | 2.517E-08 | 1.200E-07 | 1.700E-07 |
| 11443 | 573 | F573N | A | 3.9e-07 | 1.9e-07 | 1.2e-07 | 2.333E-07 | 1.401E-07 | 1.200E-07 | 3.900E-07 |
| 11444 | 573 | F573D | A | 2e-07 | 1.8e-07 | 1.4e-07 | 1.733E-07 | 3.055E-08 | 1.400E-07 | 2.000E-07 |
| 11445 | 573 | F573C | A | 2e-07 | 3e-07 | 4.5e-07 | 3.167E-07 | 1.258E-07 | 2.000E-07 | 4.500E-07 |
| 11446 | 573 | F573E | A | 1.4e-07 | 1.4e-07 | 2.8e-07 | 1.867E-07 | 8.083E-08 | 1.400E-07 | 2.800E-07 |
| 11447 | 573 | F573Q | A | 1.3e-07 | 1.7e-07 | 7.9e-08 | 1.263E-07 | 4.561E-08 | 7.900E-08 | 1.700E-07 |
| 11448 | 573 | F573G | A | 2.3e-07 | 3.8e-07 | 2.1e-07 | 2.733E-07 | 9.292E-08 | 2.100E-07 | 3.800E-07 |
| 11449 | 573 | F573H | A | 1.7e-07 | 1.1e-07 | 2.4e-07 | 1.733E-07 | 6.506E-08 | 1.100E-07 | 2.400E-07 |
| 11450 | 573 | F573I | A | 5.1e-07 | 4e-07 | 2.7e-07 | 3.933E-07 | 1.201E-07 | 2.700E-07 | 5.100E-07 |
| 11451 | 573 | F573L | A | 2.3e-07 | 2.4e-07 | 3.4e-07 | 2.700E-07 | 6.083E-08 | 2.300E-07 | 3.400E-07 |
| 11452 | 573 | F573K | A | 2e-07 | 2.7e-07 | 1.9e-07 | 2.200E-07 | 4.359E-08 | 1.900E-07 | 2.700E-07 |
| 11453 | 573 | F573M | A | 2.1e-07 | 2.1e-07 | 2.5e-07 | 2.233E-07 | 2.309E-08 | 2.100E-07 | 2.500E-07 |
| 11454 | 573 | F573F | A | 4.9e-07 | 2.4e-07 | 2.4e-07 | 3.233E-07 | 1.443E-07 | 2.400E-07 | 4.900E-07 |
| 11455 | 573 | F573P | A | 9.9e-07 | 4.6e-07 | 3.6e-07 | 6.033E-07 | 3.386E-07 | 3.600E-07 | 9.900E-07 |
| 11456 | 573 | F573S | A | 2.5e-07 | 2.3e-07 | 4.7e-07 | 3.167E-07 | 1.332E-07 | 2.300E-07 | 4.700E-07 |
| 11457 | 573 | F573T | A | 1.9e-07 | 1.5e-07 | 1.3e-07 | 1.567E-07 | 3.055E-08 | 1.300E-07 | 1.900E-07 |
| 11458 | 573 | F573W | A | 5.3e-07 | 1.6e-07 | 1.5e-07 | 2.800E-07 | 2.166E-07 | 1.500E-07 | 5.300E-07 |
| 11459 | 573 | F573Y | A | 1.7e-07 | 6.9e-07 | 2.5e-07 | 3.700E-07 | 2.800E-07 | 1.700E-07 | 6.900E-07 |
| 11460 | 573 | F573V | A | 3e-07 | 5.1e-07 | 1.5e-07 | 3.200E-07 | 1.808E-07 | 1.500E-07 | 5.100E-07 |
| 11461 | 574 | C574A | A | 2.9e-07 | 4.2e-07 | 7.8e-07 | 4.967E-07 | 2.538E-07 | 2.900E-07 | 7.800E-07 |
| 11462 | 574 | C574R | A | 2.3e-07 | 3.3e-07 | 7.7e-07 | 4.433E-07 | 2.873E-07 | 2.300E-07 | 7.700E-07 |
| 11463 | 574 | C574N | A | 1.2e-07 | 1.8e-07 | 1.9e-07 | 1.633E-07 | 3.786E-08 | 1.200E-07 | 1.900E-07 |
| 11464 | 574 | C574D | A | 2.5e-07 | 2.1e-07 | 2.1e-07 | 2.233E-07 | 2.309E-08 | 2.100E-07 | 2.500E-07 |
| 11465 | 574 | C574C | A | 2.1e-07 | 3.5e-07 | 2.3e-07 | 2.633E-07 | 7.572E-08 | 2.100E-07 | 3.500E-07 |
| 11466 | 574 | C574E | A | 2.2e-07 | 2.9e-07 | 2.4e-07 | 2.500E-07 | 3.606E-08 | 2.200E-07 | 2.900E-07 |
| 11467 | 574 | C574Q | A | 3.3e-07 | 3.3e-07 | 9.4e-08 | 2.513E-07 | 1.363E-07 | 9.400E-08 | 3.300E-07 |
| 11468 | 574 | C574G | A | 1.2e-07 | 1.4e-07 | 3.7e-07 | 2.100E-07 | 1.389E-07 | 1.200E-07 | 3.700E-07 |
| 11469 | 574 | C574H | A | 3.1e-07 | 1.7e-07 | 3.1e-07 | 2.633E-07 | 8.083E-08 | 1.700E-07 | 3.100E-07 |
| 11470 | 574 | C574I | A | 1.8e-07 | 1.5e-07 | 1.1e-07 | 1.467E-07 | 3.512E-08 | 1.100E-07 | 1.800E-07 |
| 11471 | 574 | C574L | A | 2.4e-07 | 1.9e-07 | 1.5e-07 | 1.933E-07 | 4.509E-08 | 1.500E-07 | 2.400E-07 |
| 11472 | 574 | C574K | A | 2.9e-07 | 2.2e-07 | 3.6e-07 | 2.900E-07 | 7.000E-08 | 2.200E-07 | 3.600E-07 |
| 11473 | 574 | C574M | A | 2.8e-07 | 2.3e-07 | 2.5e-07 | 2.533E-07 | 2.517E-08 | 2.300E-07 | 2.800E-07 |
| 11474 | 574 | C574F | A | 2.1e-07 | 3.2e-07 | 3.2e-07 | 2.833E-07 | 6.351E-08 | 2.100E-07 | 3.200E-07 |

| | | | | | | | | | | |
|---|---|---|---|---|---|---|---|---|---|---|
| 11475 | 574 | C574P | A | 5.8e-07 | 1.5e-07 | 3.7e-07 | 3.667E-07 | 2.150E-07 | 1.500E-07 | 5.800E-07 |
| 11476 | 574 | C574S | A | 4e-07 | 2.1e-07 | 9e-08 | 2.333E-07 | 1.563E-07 | 9.000E-08 | 4.000E-07 |
| 11477 | 574 | C574T | A | 2.6e-07 | 1.1e-07 | 3.6e-07 | 2.433E-07 | 1.258E-07 | 1.100E-07 | 3.600E-07 |
| 11478 | 574 | C574W | A | 1.3e-07 | 4.5e-07 | 1.5e-07 | 2.433E-07 | 1.793E-07 | 1.300E-07 | 4.500E-07 |
| 11479 | 574 | C574Y | A | 2.8e-07 | 2.7e-07 | 2.1e-07 | 2.533E-07 | 3.786E-08 | 2.100E-07 | 2.800E-07 |
| 11480 | 574 | C574V | A | 2.3e-07 | 3.3e-07 | 1.9e-07 | 2.500E-07 | 7.211E-08 | 1.900E-07 | 3.300E-07 |
| 11481 | 575 | V575A | A | 4.8e-07 | 1.7e-07 | 2.6e-07 | 3.033E-07 | 1.595E-07 | 1.700E-07 | 4.800E-07 |
| 11482 | 575 | V575R | A | 1.7e-07 | 1.7e-07 | 1.5e-07 | 1.633E-07 | 1.155E-08 | 1.500E-07 | 1.700E-07 |
| 11483 | 575 | V575N | A | 2.1e-07 | 1.2e-07 | 1.2e-07 | 1.500E-07 | 5.196E-08 | 1.200E-07 | 2.100E-07 |
| 11484 | 575 | V575D | A | 4.5e-07 | 3.4e-07 | 1.9e-07 | 3.267E-07 | 1.305E-07 | 1.900E-07 | 4.500E-07 |
| 11485 | 575 | V575C | A | 1.5e-07 | 1.7e-07 | 1.4e-07 | 1.533E-07 | 1.528E-08 | 1.400E-07 | 1.700E-07 |
| 11486 | 575 | V575E | A | 1.1e-07 | 1.8e-07 | 8.6e-08 | 1.253E-07 | 4.884E-08 | 8.600E-08 | 1.800E-07 |
| 11487 | 575 | V575Q | A | 2.6e-07 | 2.7e-07 | 2.1e-07 | 2.467E-07 | 3.215E-08 | 2.100E-07 | 2.700E-07 |
| 11488 | 575 | V575G | A | 3.4e-07 | 1.9e-07 | 1.3e-07 | 2.200E-07 | 1.082E-07 | 1.300E-07 | 3.400E-07 |
| 11489 | 575 | V575H | A | 1.7e-07 | 1.6e-07 | 1.7e-07 | 1.667E-07 | 5.774E-09 | 1.600E-07 | 1.700E-07 |
| 11490 | 575 | V575I | A | 1.5e-07 | 3e-07 | 1e-07 | 1.833E-07 | 1.041E-07 | 1.000E-07 | 3.000E-07 |
| 11491 | 575 | V575L | A | 3.2e-07 | 5.8e-07 | 2.2e-07 | 3.733E-07 | 1.858E-07 | 2.200E-07 | 5.800E-07 |
| 11492 | 575 | V575K | A | 3e-07 | 1.9e-07 | 2.1e-07 | 2.333E-07 | 5.859E-08 | 1.900E-07 | 3.000E-07 |
| 11493 | 575 | V575M | A | 4.2e-07 | 2e-07 | 2e-07 | 2.733E-07 | 1.270E-07 | 2.000E-07 | 4.200E-07 |
| 11494 | 575 | V575F | A | 3.4e-07 | 3.3e-07 | 2.3e-07 | 3.000E-07 | 6.083E-08 | 2.300E-07 | 3.400E-07 |
| 11495 | 575 | V575P | A | 1.5e-07 | 5.3e-07 | 2.5e-07 | 3.100E-07 | 1.970E-07 | 1.500E-07 | 5.300E-07 |
| 11496 | 575 | V575S | A | 1.8e-07 | 3.4e-07 | 3.3e-07 | 2.833E-07 | 8.963E-08 | 1.800E-07 | 3.400E-07 |
| 11497 | 575 | V575T | A | 1.4e-07 | 2.8e-07 | 1.4e-07 | 1.867E-07 | 8.083E-08 | 1.400E-07 | 2.800E-07 |
| 11498 | 575 | V575W | A | 6.6e-08 | 1.9e-07 | 1.5e-07 | 1.353E-07 | 6.329E-08 | 6.600E-08 | 1.900E-07 |
| 11499 | 575 | V575Y | A | 3.1e-07 | 2.7e-07 | 8.9e-08 | 2.230E-07 | 1.178E-07 | 8.900E-08 | 3.100E-07 |
| 11500 | 575 | V575V | A | 1.2e-07 | 5.9e-07 | 3e-07 | 3.367E-07 | 2.371E-07 | 1.200E-07 | 5.900E-07 |
| 11501 | 576 | A576A | A | 1.6e-07 | 1.7e-07 | 1.2e-07 | 1.500E-07 | 2.646E-08 | 1.200E-07 | 1.700E-07 |
| 11502 | 576 | A576R | A | 2.3e-07 | 5.2e-07 | 2.4e-07 | 3.300E-07 | 1.646E-07 | 2.300E-07 | 5.200E-07 |
| 11503 | 576 | A576N | A | 2.6e-07 | 2.8e-07 | 2.2e-07 | 2.533E-07 | 3.055E-08 | 2.200E-07 | 2.800E-07 |
| 11504 | 576 | A576D | A | 2.2e-07 | 2.7e-07 | 5.3e-07 | 3.400E-07 | 1.664E-07 | 2.200E-07 | 5.300E-07 |
| 11505 | 576 | A576C | A | 3.3e-07 | 4.3e-07 | 1.3e-07 | 2.967E-07 | 1.528E-07 | 1.300E-07 | 4.300E-07 |
| 11506 | 576 | A576E | A | 2e-07 | 2.1e-07 | 1.7e-07 | 1.933E-07 | 2.082E-08 | 1.700E-07 | 2.100E-07 |
| 11507 | 576 | A576Q | A | 3.9e-07 | 2e-07 | 3.2e-07 | 3.033E-07 | 9.609E-08 | 2.000E-07 | 3.900E-07 |
| 11508 | 576 | A576G | A | 2.8e-07 | 1.9e-07 | 1.9e-07 | 2.200E-07 | 5.196E-08 | 1.900E-07 | 2.800E-07 |
| 11509 | 576 | A576H | A | 1.6e-07 | 2.4e-07 | 1.9e-07 | 1.967E-07 | 4.041E-08 | 1.600E-07 | 2.400E-07 |
| 11510 | 576 | A576I | A | 2.1e-07 | 1.2e-07 | 3.7e-07 | 2.333E-07 | 1.266E-07 | 1.200E-07 | 3.700E-07 |
| 11511 | 576 | A576L | A | 2e-07 | 2.5e-07 | 4e-07 | 2.833E-07 | 1.041E-07 | 2.000E-07 | 4.000E-07 |
| 11512 | 576 | A576K | A | 2.2e-07 | 1.9e-07 | 1.4e-07 | 1.833E-07 | 4.041E-08 | 1.400E-07 | 2.200E-07 |
| 11513 | 576 | A576M | A | 7.3e-08 | 1.5e-07 | 2.4e-07 | 1.543E-07 | 8.358E-08 | 7.300E-08 | 2.400E-07 |
| 11514 | 576 | A576F | A | 1.9e-07 | 3.1e-07 | 4.2e-07 | 3.067E-07 | 1.150E-07 | 1.900E-07 | 4.200E-07 |
| 11515 | 576 | A576P | A | 2.5e-07 | 2.2e-07 | 2.8e-07 | 2.500E-07 | 3.000E-08 | 2.200E-07 | 2.800E-07 |
| 11516 | 576 | A576S | A | 3.8e-07 | 3.1e-07 | 7.3e-08 | 2.543E-07 | 1.609E-07 | 7.300E-08 | 3.800E-07 |
| 11517 | 576 | A576T | A | 3.9e-07 | 2.4e-07 | 3.6e-07 | 3.300E-07 | 7.937E-08 | 2.400E-07 | 3.900E-07 |
| 11518 | 576 | A576W | A | 1.9e-07 | 2.8e-07 | 1.6e-07 | 2.100E-07 | 6.245E-08 | 1.600E-07 | 2.800E-07 |
| 11519 | 576 | A576Y | A | 2.8e-07 | 3.4e-07 | 4.1e-07 | 3.433E-07 | 6.506E-08 | 2.800E-07 | 4.100E-07 |
| 11520 | 576 | A576V | A | 2.6e-07 | 1.3e-07 | 5.5e-07 | 3.133E-07 | 2.150E-07 | 1.300E-07 | 5.500E-07 |
| 11521 | 577 | R577A | A | 5.3e-07 | 2.2e-07 | 2.9e-07 | 3.467E-07 | 1.626E-07 | 2.200E-07 | 5.300E-07 |
| 11522 | 577 | R577R | A | 5.3e-07 | 1.4e-07 | 1.2e-07 | 2.633E-07 | 2.312E-07 | 1.200E-07 | 5.300E-07 |
| 11523 | 577 | R577N | A | 2.4e-07 | 1.3e-07 | 2.6e-07 | 2.100E-07 | 7.000E-08 | 1.300E-07 | 2.600E-07 |
| 11524 | 577 | R577D | A | 2.3e-07 | 6.6e-07 | 3.5e-07 | 4.133E-07 | 2.219E-07 | 2.300E-07 | 6.600E-07 |
| 11525 | 577 | R577C | A | 2.6e-07 | 2.1e-07 | 2.2e-07 | 2.300E-07 | 2.646E-08 | 2.100E-07 | 2.600E-07 |
| 11526 | 577 | R577E | A | 3.5e-07 | 2.8e-07 | 1.6e-07 | 2.633E-07 | 9.609E-08 | 1.600E-07 | 3.500E-07 |
| 11527 | 577 | R577Q | A | 1.6e-07 | 2.2e-07 | 2.4e-07 | 2.067E-07 | 4.163E-08 | 1.600E-07 | 2.400E-07 |
| 11528 | 577 | R577G | A | 1.6e-07 | 1.4e-07 | 1.8e-07 | 1.600E-07 | 2.000E-08 | 1.400E-07 | 1.800E-07 |
| 11529 | 577 | R577H | A | 1.8e-07 | 6.7e-07 | 5.6e-07 | 4.700E-07 | 2.571E-07 | 1.800E-07 | 6.700E-07 |
| 11530 | 577 | R577I | A | 1.6e-07 | 3.6e-07 | 2e-07 | 2.400E-07 | 1.058E-07 | 1.600E-07 | 3.600E-07 |
| 11531 | 577 | R577L | A | 4.9e-07 | 2.5e-07 | 2.6e-07 | 3.333E-07 | 1.358E-07 | 2.500E-07 | 4.900E-07 |
| 11532 | 577 | R577K | A | 1.9e-07 | 2.6e-07 | 2.2e-07 | 2.233E-07 | 3.512E-08 | 1.900E-07 | 2.600E-07 |
| 11533 | 577 | R577M | A | 3.1e-07 | 1.1e-07 | 2e-07 | 2.067E-07 | 1.002E-07 | 1.100E-07 | 3.100E-07 |
| 11534 | 577 | R577F | A | 2.2e-07 | 1.9e-07 | 3.8e-07 | 2.633E-07 | 1.021E-07 | 1.900E-07 | 3.800E-07 |
| 11535 | 577 | R577P | A | 3.2e-07 | 2e-07 | 1.7e-07 | 2.300E-07 | 7.937E-08 | 1.700E-07 | 3.200E-07 |
| 11536 | 577 | R577S | A | 2.9e-07 | 7.2e-07 | 1.2e-07 | 3.767E-07 | 3.092E-07 | 1.200E-07 | 7.200E-07 |
| 11537 | 577 | R577T | A | 2e-07 | 2.9e-07 | 1.3e-07 | 2.067E-07 | 8.021E-08 | 1.300E-07 | 2.900E-07 |
| 11538 | 577 | R577W | A | 2.7e-07 | 1.6e-07 | 1.8e-07 | 2.033E-07 | 5.859E-08 | 1.600E-07 | 2.700E-07 |
| 11539 | 577 | R577Y | A | 1.9e-07 | 1.6e-07 | 2.2e-07 | 1.900E-07 | 3.000E-08 | 1.600E-07 | 2.200E-07 |
| 11540 | 577 | R577V | A | 2.1e-07 | 3.1e-07 | 2.9e-07 | 2.700E-07 | 5.292E-08 | 2.100E-07 | 3.100E-07 |
| 11541 | 578 | C578A | A | 6.7e-07 | 4.3e-07 | 4.4e-07 | 5.133E-07 | 1.358E-07 | 4.300E-07 | 6.700E-07 |
| 11542 | 578 | C578R | A | 2.9e-07 | 4e-07 | 2.7e-07 | 3.200E-07 | 7.000E-08 | 2.700E-07 | 4.000E-07 |
| 11543 | 578 | C578N | A | 1.6e-07 | 1.5e-07 | 3e-07 | 2.033E-07 | 8.386E-08 | 1.500E-07 | 3.000E-07 |
| 11544 | 578 | C578D | A | 1.7e-07 | 1.7e-07 | 2.3e-07 | 1.900E-07 | 3.464E-08 | 1.700E-07 | 2.300E-07 |
| 11545 | 578 | C578C | A | 2.6e-07 | 1.8e-07 | 2.7e-07 | 2.367E-07 | 4.933E-08 | 1.800E-07 | 2.700E-07 |
| 11546 | 578 | C578E | A | 1.1e-07 | 1.7e-07 | 2.1e-07 | 1.633E-07 | 5.033E-08 | 1.100E-07 | 2.100E-07 |
| 11547 | 578 | C578Q | A | 4.7e-07 | 3e-07 | 4.4e-07 | 4.033E-07 | 9.074E-08 | 3.000E-07 | 4.700E-07 |
| 11548 | 578 | C578G | A | 3e-07 | 2.5e-07 | 1.3e-07 | 2.267E-07 | 8.737E-08 | 1.300E-07 | 3.000E-07 |
| 11549 | 578 | C578H | A | 1.3e-07 | 4.3e-07 | 1.7e-07 | 2.433E-07 | 1.629E-07 | 1.300E-07 | 4.300E-07 |
| 11550 | 578 | C578I | A | 3.8e-07 | 4.3e-07 | 3.1e-07 | 3.733E-07 | 6.028E-08 | 3.100E-07 | 4.300E-07 |
| 11551 | 578 | C578L | A | 2.9e-07 | 2.4e-07 | 1.8e-07 | 2.367E-07 | 5.508E-08 | 1.800E-07 | 2.900E-07 |
| 11552 | 578 | C578K | A | 3.3e-07 | 2.6e-07 | 1.3e-07 | 2.400E-07 | 1.015E-07 | 1.300E-07 | 3.300E-07 |
| 11553 | 578 | C578M | A | 1.3e-07 | 1.7e-07 | 2.3e-07 | 1.767E-07 | 5.033E-08 | 1.300E-07 | 2.300E-07 |
| 11554 | 578 | C578F | A | 5.7e-07 | 1.2e-07 | 3.6e-07 | 3.500E-07 | 2.252E-07 | 1.200E-07 | 5.700E-07 |
| 11555 | 578 | C578P | A | 2.9e-07 | 1.9e-07 | 1.2e-06 | 5.600E-07 | 5.565E-07 | 1.900E-07 | 1.200E-06 |
| 11556 | 578 | C578S | A | 1.3e-07 | 1.1e-07 | 7.3e-08 | 1.043E-07 | 2.892E-08 | 7.300E-08 | 1.300E-07 |
| 11557 | 578 | C578T | A | 4.1e-07 | 4.1e-07 | 2.8e-07 | 3.667E-07 | 7.506E-08 | 2.800E-07 | 4.100E-07 |
| 11558 | 578 | C578W | A | 1.2e-07 | 9.8e-08 | 1.2e-07 | 1.127E-07 | 1.270E-08 | 9.800E-08 | 1.200E-07 |
| 11559 | 578 | C578Y | A | 1.5e-07 | 2.2e-07 | 1.8e-07 | 1.833E-07 | 3.512E-08 | 1.500E-07 | 2.200E-07 |

| | | | | | | | | | | |
|---|---|---|---|---|---|---|---|---|---|---|
| 11560 | 578 | C578V | A | 3.7e-07 | 2e-07 | 2.1e-07 | 2.600E-07 | 9.539E-08 | 2.000E-07 | 3.700E-07 |
| 11561 | 579 | P579A | A | 2.9e-07 | 6.3e-07 | 4.5e-07 | 4.567E-07 | 1.701E-07 | 2.900E-07 | 6.300E-07 |
| 11562 | 579 | P579R | A | 2.1e-07 | 1.7e-07 | 2.3e-07 | 2.033E-07 | 3.055E-08 | 1.700E-07 | 2.300E-07 |
| 11563 | 579 | P579N | A | 1.7e-07 | 2.5e-07 | 1.1e-06 | 5.067E-07 | 5.154E-07 | 1.700E-07 | 1.100E-06 |
| 11564 | 579 | P579D | A | 2.2e-07 | 2.6e-07 | 1.3e-07 | 2.033E-07 | 6.658E-08 | 1.300E-07 | 2.600E-07 |
| 11565 | 579 | P579C | A | 2e-07 | 2.3e-07 | 1.2e-07 | 1.833E-07 | 5.686E-08 | 1.200E-07 | 2.300E-07 |
| 11566 | 579 | P579E | A | 6e-07 | 3.1e-07 | 3.1e-07 | 4.067E-07 | 1.674E-07 | 3.100E-07 | 6.000E-07 |
| 11567 | 579 | P579Q | A | 2.3e-07 | 1.3e-07 | 2.9e-07 | 2.167E-07 | 8.083E-08 | 1.300E-07 | 2.900E-07 |
| 11568 | 579 | P579G | A | 2.8e-07 | 3.7e-07 | 4.1e-07 | 3.533E-07 | 6.658E-08 | 2.800E-07 | 4.100E-07 |
| 11569 | 579 | P579H | A | 3.3e-07 | 2.8e-07 | 1.3e-07 | 2.467E-07 | 1.041E-07 | 1.300E-07 | 3.300E-07 |
| 11570 | 579 | P579I | A | 1.9e-07 | 1.6e-07 | 2.2e-07 | 1.900E-07 | 3.000E-08 | 1.600E-07 | 2.200E-07 |
| 11571 | 579 | P579L | A | 9.7e-08 | 1.2e-07 | 1.6e-07 | 1.257E-07 | 3.188E-08 | 9.700E-08 | 1.600E-07 |
| 11572 | 579 | P579K | A | 2.8e-07 | 3.7e-07 | 3.4e-07 | 3.300E-07 | 4.583E-08 | 2.800E-07 | 3.700E-07 |
| 11573 | 579 | P579M | A | 5.1e-07 | 5.2e-07 | 2.1e-07 | 4.133E-07 | 1.762E-07 | 2.100E-07 | 5.200E-07 |
| 11574 | 579 | P579F | A | 1.9e-07 | 2.8e-07 | 2.6e-07 | 2.433E-07 | 4.726E-08 | 1.900E-07 | 2.800E-07 |
| 11575 | 579 | P579P | A | 4.5e-07 | 1.6e-07 | 3.3e-07 | 3.133E-07 | 1.457E-07 | 1.600E-07 | 4.500E-07 |
| 11576 | 579 | P579S | A | 3.9e-07 | 7.8e-08 | 2.6e-07 | 2.427E-07 | 1.567E-07 | 7.800E-08 | 3.900E-07 |
| 11577 | 579 | P579T | A | 1.3e-07 | 2.3e-07 | 1.3e-07 | 1.633E-07 | 5.774E-08 | 1.300E-07 | 2.300E-07 |
| 11578 | 579 | P579W | A | 3.1e-07 | 1.7e-07 | 5.1e-07 | 3.300E-07 | 1.709E-07 | 1.700E-07 | 5.100E-07 |
| 11579 | 579 | P579Y | A | 1.7e-07 | 1.2e-07 | 1.1e-07 | 1.333E-07 | 3.215E-08 | 1.100E-07 | 1.700E-07 |
| 11580 | 579 | P579V | A | 2.4e-07 | 2.5e-07 | 1.7e-07 | 2.200E-07 | 4.359E-08 | 1.700E-07 | 2.500E-07 |
| 11581 | 580 | S580A | A | 1.9e-07 | 1.9e-07 | 1.6e-07 | 1.800E-07 | 1.732E-08 | 1.600E-07 | 1.900E-07 |
| 11582 | 580 | S580R | A | 1.4e-07 | 1.2e-07 | 2.5e-07 | 1.700E-07 | 7.000E-08 | 1.200E-07 | 2.500E-07 |
| 11583 | 580 | S580N | A | 7.2e-07 | 2.2e-07 | 2.4e-07 | 3.933E-07 | 2.831E-07 | 2.200E-07 | 7.200E-07 |
| 11584 | 580 | S580D | A | 1.6e-07 | 2.1e-07 | 3.5e-07 | 2.400E-07 | 9.849E-08 | 1.600E-07 | 3.500E-07 |
| 11585 | 580 | S580C | A | 1.9e-07 | 1.3e-07 | 3e-07 | 2.067E-07 | 8.622E-08 | 1.300E-07 | 3.000E-07 |
| 11586 | 580 | S580E | A | 3.9e-07 | 2.5e-07 | 1.6e-07 | 2.667E-07 | 1.159E-07 | 1.600E-07 | 3.900E-07 |
| 11587 | 580 | S580Q | A | 2.1e-07 | 1.7e-07 | 2.4e-07 | 2.067E-07 | 3.512E-08 | 1.700E-07 | 2.400E-07 |
| 11588 | 580 | S580G | A | 2.2e-07 | 2.8e-07 | 3.6e-07 | 2.867E-07 | 7.024E-08 | 2.200E-07 | 3.600E-07 |
| 11589 | 580 | S580H | A | 2.7e-07 | 3.9e-07 | 2.7e-07 | 3.100E-07 | 6.928E-08 | 2.700E-07 | 3.900E-07 |
| 11590 | 580 | S580I | A | 1.7e-07 | 2.9e-07 | 1.6e-07 | 2.067E-07 | 7.234E-08 | 1.600E-07 | 2.900E-07 |
| 11591 | 580 | S580L | A | 4.8e-07 | 2.9e-07 | 1.3e-07 | 3.000E-07 | 1.752E-07 | 1.300E-07 | 4.800E-07 |
| 11592 | 580 | S580K | A | 1.9e-07 | 1.9e-07 | 2.5e-07 | 2.100E-07 | 3.464E-08 | 1.900E-07 | 2.500E-07 |
| 11593 | 580 | S580M | A | 2.4e-07 | 4.2e-07 | 2.5e-07 | 3.033E-07 | 1.012E-07 | 2.400E-07 | 4.200E-07 |
| 11594 | 580 | S580F | A | 2.4e-07 | 3.3e-07 | 3.3e-07 | 3.000E-07 | 5.196E-08 | 2.400E-07 | 3.300E-07 |
| 11595 | 580 | S580P | A | 3.2e-07 | 3.1e-07 | 3.3e-07 | 3.200E-07 | 1.000E-08 | 3.100E-07 | 3.300E-07 |
| 11596 | 580 | S580S | A | 2.4e-07 | 1.5e-07 | 2.5e-07 | 2.133E-07 | 5.508E-08 | 1.500E-07 | 2.500E-07 |
| 11597 | 580 | S580T | A | 2.5e-07 | 2.8e-07 | 2.1e-07 | 2.467E-07 | 3.512E-08 | 2.100E-07 | 2.800E-07 |
| 11598 | 580 | S580W | A | 8.4e-08 | 4.2e-07 | 1.4e-07 | 2.147E-07 | 1.800E-07 | 8.400E-08 | 4.200E-07 |
| 11599 | 580 | S580Y | A | 2.3e-07 | 2.6e-07 | 3.4e-07 | 2.767E-07 | 5.686E-08 | 2.300E-07 | 3.400E-07 |
| 11600 | 580 | S580V | A | 5.6e-07 | 2.6e-07 | 2.1e-07 | 3.433E-07 | 1.893E-07 | 2.100E-07 | 5.600E-07 |
| 11601 | 581 | G581A | A | 2.9e-07 | 1.9e-07 | 5.7e-07 | 3.500E-07 | 1.970E-07 | 1.900E-07 | 5.700E-07 |
| 11602 | 581 | G581R | A | 2.4e-07 | 1.6e-07 | 1.1e-07 | 1.700E-07 | 6.557E-08 | 1.100E-07 | 2.400E-07 |
| 11603 | 581 | G581N | A | 1.4e-07 | 4.4e-07 | 1.4e-07 | 2.400E-07 | 1.732E-07 | 1.400E-07 | 4.400E-07 |
| 11604 | 581 | G581D | A | 3.9e-07 | 1.4e-07 | 4.5e-07 | 3.267E-07 | 1.644E-07 | 1.400E-07 | 4.500E-07 |
| 11605 | 581 | G581C | A | 1.3e-07 | 1.9e-07 | 1.7e-07 | 1.633E-07 | 3.055E-08 | 1.300E-07 | 1.900E-07 |
| 11606 | 581 | G581E | A | 1.4e-07 | 1.8e-07 | 1.2e-07 | 1.467E-07 | 3.055E-08 | 1.200E-07 | 1.800E-07 |
| 11607 | 581 | G581Q | A | 2.3e-07 | 4.2e-07 | 3.6e-07 | 3.367E-07 | 9.713E-08 | 2.300E-07 | 4.200E-07 |
| 11608 | 581 | G581G | A | 2.5e-07 | 1.7e-07 | 2.6e-07 | 2.267E-07 | 4.933E-08 | 1.700E-07 | 2.600E-07 |
| 11609 | 581 | G581H | A | 1.9e-07 | 1.9e-07 | 9.7e-08 | 1.590E-07 | 5.369E-08 | 9.700E-08 | 1.900E-07 |
| 11610 | 581 | G581I | A | 2.2e-07 | 2e-07 | 1.9e-07 | 2.033E-07 | 1.528E-08 | 1.900E-07 | 2.200E-07 |
| 11611 | 581 | G581L | A | 1.4e-07 | 1.8e-07 | 2.3e-07 | 1.833E-07 | 4.509E-08 | 1.400E-07 | 2.300E-07 |
| 11612 | 581 | G581K | A | 1.2e-07 | 1.6e-07 | 3.1e-07 | 1.967E-07 | 1.002E-07 | 1.200E-07 | 3.100E-07 |
| 11613 | 581 | G581M | A | 1.9e-07 | 1.8e-07 | 2.2e-07 | 1.967E-07 | 2.082E-08 | 1.800E-07 | 2.200E-07 |
| 11614 | 581 | G581F | A | 2e-07 | 4e-07 | 2e-07 | 2.667E-07 | 1.155E-07 | 2.000E-07 | 4.000E-07 |
| 11615 | 581 | G581P | A | 2.8e-07 | 2.3e-07 | 1.6e-07 | 2.233E-07 | 6.028E-08 | 1.600E-07 | 2.800E-07 |
| 11616 | 581 | G581S | A | 2.6e-07 | 6.1e-07 | 1.5e-07 | 3.400E-07 | 2.402E-07 | 1.500E-07 | 6.100E-07 |
| 11617 | 581 | G581T | A | 9.9e-08 | 2.2e-07 | 1.8e-07 | 1.663E-07 | 6.165E-08 | 9.900E-08 | 2.200E-07 |
| 11618 | 581 | G581W | A | 2.2e-07 | 7.7e-08 | 1.6e-07 | 1.523E-07 | 7.181E-08 | 7.700E-08 | 2.200E-07 |
| 11619 | 581 | G581Y | A | 7.8e-08 | 4.2e-07 | 1.5e-07 | 2.160E-07 | 1.803E-07 | 7.800E-08 | 4.200E-07 |
| 11620 | 581 | G581V | A | 2.4e-07 | 1.9e-07 | 2.6e-07 | 2.300E-07 | 3.606E-08 | 1.900E-07 | 2.600E-07 |
| 11621 | 582 | V582A | A | 6.1e-07 | 1.9e-07 | 1.8e-07 | 3.267E-07 | 2.454E-07 | 1.800E-07 | 6.100E-07 |
| 11622 | 582 | V582R | A | 2.3e-07 | 2.4e-07 | 2e-07 | 2.233E-07 | 2.082E-08 | 2.000E-07 | 2.400E-07 |
| 11623 | 582 | V582N | A | 2.2e-07 | 2e-07 | 1.8e-07 | 2.000E-07 | 2.000E-08 | 1.800E-07 | 2.200E-07 |
| 11624 | 582 | V582D | A | 2.9e-07 | 2.4e-07 | 1.9e-07 | 2.400E-07 | 5.000E-08 | 1.900E-07 | 2.900E-07 |
| 11625 | 582 | V582C | A | 2.9e-07 | 2.4e-07 | 2.2e-07 | 2.500E-07 | 3.606E-08 | 2.200E-07 | 2.900E-07 |
| 11626 | 582 | V582E | A | 2.6e-07 | 3.7e-07 | 1.8e-07 | 2.700E-07 | 9.539E-08 | 1.800E-07 | 3.700E-07 |
| 11627 | 582 | V582Q | A | 2.6e-07 | 1.5e-07 | 7.7e-08 | 1.623E-07 | 9.212E-08 | 7.700E-08 | 2.600E-07 |
| 11628 | 582 | V582G | A | 1.8e-07 | 2.1e-07 | 3.2e-07 | 2.367E-07 | 7.371E-08 | 1.800E-07 | 3.200E-07 |
| 11629 | 582 | V582H | A | 1.7e-07 | 3.4e-07 | 1.3e-07 | 2.133E-07 | 1.115E-07 | 1.300E-07 | 3.400E-07 |
| 11630 | 582 | V582I | A | 1e-07 | 1.2e-07 | 3.4e-07 | 1.867E-07 | 1.332E-07 | 1.000E-07 | 3.400E-07 |
| 11631 | 582 | V582L | A | 2.3e-07 | 3.3e-07 | 1.8e-07 | 2.467E-07 | 7.638E-08 | 1.800E-07 | 3.300E-07 |
| 11632 | 582 | V582K | A | 1.9e-07 | 1.6e-07 | 1.5e-07 | 1.667E-07 | 2.082E-08 | 1.500E-07 | 1.900E-07 |
| 11633 | 582 | V582M | A | 4.8e-07 | 3.4e-07 | 2.9e-07 | 3.700E-07 | 9.849E-08 | 2.900E-07 | 4.800E-07 |
| 11634 | 582 | V582F | A | 2.5e-07 | 2.7e-07 | 1.8e-07 | 2.333E-07 | 4.726E-08 | 1.800E-07 | 2.700E-07 |
| 11635 | 582 | V582P | A | 4.8e-08 | 2.8e-07 | 1e-07 | 1.427E-07 | 1.217E-07 | 4.800E-08 | 2.800E-07 |
| 11636 | 582 | V582S | A | 2.3e-07 | 2.7e-07 | 2.8e-07 | 2.600E-07 | 2.646E-08 | 2.300E-07 | 2.800E-07 |
| 11637 | 582 | V582T | A | 2e-07 | 1.5e-07 | 1.9e-07 | 1.800E-07 | 2.646E-08 | 1.500E-07 | 2.000E-07 |
| 11638 | 582 | V582W | A | 1.7e-07 | 3.1e-07 | 3.9e-07 | 2.900E-07 | 1.114E-07 | 1.700E-07 | 3.900E-07 |
| 11639 | 582 | V582Y | A | 2.1e-07 | 3.4e-07 | 3.4e-07 | 2.967E-07 | 7.506E-08 | 2.100E-07 | 3.400E-07 |
| 11640 | 582 | V582V | A | 2.2e-07 | 3.4e-07 | 3e-07 | 2.867E-07 | 6.110E-08 | 2.200E-07 | 3.400E-07 |
| 11641 | 583 | K583A | A | 1.7e-07 | 3.4e-07 | 2.8e-07 | 2.633E-07 | 8.622E-08 | 1.700E-07 | 3.400E-07 |
| 11642 | 583 | K583R | A | 2.1e-07 | 3.2e-07 | 1.5e-07 | 2.267E-07 | 8.622E-08 | 1.500E-07 | 3.200E-07 |
| 11643 | 583 | K583N | A | 2.7e-07 | 4e-07 | 1.6e-07 | 2.767E-07 | 1.201E-07 | 1.600E-07 | 4.000E-07 |
| 11644 | 583 | K583D | A | 5.1e-07 | 4e-07 | 2.9e-07 | 4.000E-07 | 1.100E-07 | 2.900E-07 | 5.100E-07 |

| | | | | | | | | | | |
|---|---|---|---|---|---|---|---|---|---|---|
| 11645 | 583 | K583C | A | 2.1e-07 | 3.2e-07 | 2.4e-07 | 2.567E-07 | 5.686E-08 | 2.100E-07 | 3.200E-07 |
| 11646 | 583 | K583E | A | 4.6e-07 | 3e-07 | 2.8e-07 | 3.467E-07 | 9.866E-08 | 2.800E-07 | 4.600E-07 |
| 11647 | 583 | K583Q | A | 7.2e-07 | 7.2e-07 | 1.7e-07 | 5.367E-07 | 3.175E-07 | 1.700E-07 | 7.200E-07 |
| 11648 | 583 | K583G | A | 1.9e-07 | 2e-07 | 1.7e-07 | 1.867E-07 | 1.528E-08 | 1.700E-07 | 2.000E-07 |
| 11649 | 583 | K583H | A | 4.8e-07 | 1.5e-07 | 2.6e-07 | 2.967E-07 | 1.680E-07 | 1.500E-07 | 4.800E-07 |
| 11650 | 583 | K583I | A | 2.3e-07 | 2.4e-07 | 3.5e-07 | 2.733E-07 | 6.658E-08 | 2.300E-07 | 3.500E-07 |
| 11651 | 583 | K583L | A | 2.3e-07 | 3.4e-07 | 4.5e-07 | 3.400E-07 | 1.100E-07 | 2.300E-07 | 4.500E-07 |
| 11652 | 583 | K583K | A | 2.5e-07 | 2.8e-07 | 2e-07 | 2.433E-07 | 4.041E-08 | 2.000E-07 | 2.800E-07 |
| 11653 | 583 | K583M | A | 4e-07 | 1e-07 | 4e-07 | 3.000E-07 | 1.732E-07 | 1.000E-07 | 4.000E-07 |
| 11654 | 583 | K583F | A | 2.1e-07 | 2.3e-07 | 4.1e-07 | 2.833E-07 | 1.102E-07 | 2.100E-07 | 4.100E-07 |
| 11655 | 583 | K583P | A | 4e-07 | 9.9e-08 | 1.8e-07 | 2.263E-07 | 1.558E-07 | 9.900E-08 | 4.000E-07 |
| 11656 | 583 | K583S | A | 1.8e-07 | 1.8e-07 | 2e-07 | 1.867E-07 | 1.155E-08 | 1.800E-07 | 2.000E-07 |
| 11657 | 583 | K583T | A | 2.9e-07 | 3.3e-07 | 3.5e-07 | 3.233E-07 | 3.055E-08 | 2.900E-07 | 3.500E-07 |
| 11658 | 583 | K583W | A | 4.5e-07 | 3.2e-07 | 2.9e-07 | 3.533E-07 | 8.505E-08 | 2.900E-07 | 4.500E-07 |
| 11659 | 583 | K583Y | A | 2.5e-07 | 1.6e-07 | 2.3e-07 | 2.133E-07 | 4.726E-08 | 1.600E-07 | 2.500E-07 |
| 11660 | 583 | K583V | A | 1.2e-07 | 4e-07 | 1.5e-07 | 2.233E-07 | 1.537E-07 | 1.200E-07 | 4.000E-07 |
| 11661 | 584 | P584A | A | 2.2e-07 | 3.3e-07 | 1.4e-07 | 2.300E-07 | 9.539E-08 | 1.400E-07 | 3.300E-07 |
| 11662 | 584 | P584R | A | 1.9e-07 | 1.5e-07 | 3.1e-07 | 2.167E-07 | 8.327E-08 | 1.500E-07 | 3.100E-07 |
| 11663 | 584 | P584N | A | 5.2e-07 | 2.9e-07 | 1.5e-07 | 3.200E-07 | 1.868E-07 | 1.500E-07 | 5.200E-07 |
| 11664 | 584 | P584D | A | 1.7e-07 | 4.5e-07 | 1.6e-07 | 2.600E-07 | 1.646E-07 | 1.600E-07 | 4.500E-07 |
| 11665 | 584 | P584C | A | 2.1e-07 | 3.1e-07 | 3.6e-07 | 2.933E-07 | 7.638E-08 | 2.100E-07 | 3.600E-07 |
| 11666 | 584 | P584E | A | 2.1e-07 | 2.8e-07 | 2.8e-07 | 2.567E-07 | 4.041E-08 | 2.100E-07 | 2.800E-07 |
| 11667 | 584 | P584Q | A | 3e-07 | 1.9e-07 | 3.4e-07 | 2.767E-07 | 7.767E-08 | 1.900E-07 | 3.400E-07 |
| 11668 | 584 | P584G | A | 1.6e-07 | 2e-07 | 2.6e-07 | 2.067E-07 | 5.033E-08 | 1.600E-07 | 2.600E-07 |
| 11669 | 584 | P584H | A | 2.8e-07 | 1.7e-07 | 2.1e-07 | 2.200E-07 | 5.568E-08 | 1.700E-07 | 2.800E-07 |
| 11670 | 584 | P584I | A | 4.4e-07 | 5.3e-07 | 1.6e-07 | 3.767E-07 | 1.930E-07 | 1.600E-07 | 5.300E-07 |
| 11671 | 584 | P584L | A | 1.6e-07 | 5.8e-07 | 1.5e-07 | 2.967E-07 | 2.454E-07 | 1.500E-07 | 5.800E-07 |
| 11672 | 584 | P584K | A | 4.5e-07 | 1.8e-07 | 2.3e-07 | 2.867E-07 | 1.436E-07 | 1.800E-07 | 4.500E-07 |
| 11673 | 584 | P584M | A | 3.1e-07 | 1.5e-07 | 1.8e-07 | 2.133E-07 | 8.505E-08 | 1.500E-07 | 3.100E-07 |
| 11674 | 584 | P584F | A | 7.1e-08 | 4.4e-07 | 3.3e-07 | 2.803E-07 | 1.894E-07 | 7.100E-08 | 4.400E-07 |
| 11675 | 584 | P584P | A | 1.2e-07 | 2.6e-07 | 2.1e-07 | 1.967E-07 | 7.095E-08 | 1.200E-07 | 2.600E-07 |
| 11676 | 584 | P584S | A | 2.2e-07 | 2.3e-07 | 1.1e-07 | 1.867E-07 | 6.658E-08 | 1.100E-07 | 2.300E-07 |
| 11677 | 584 | P584T | A | 4.5e-07 | 1e-07 | 5.3e-07 | 3.600E-07 | 2.287E-07 | 1.000E-07 | 5.300E-07 |
| 11678 | 584 | P584W | A | 1.7e-07 | 7.4e-08 | 1.6e-07 | 1.347E-07 | 5.278E-08 | 7.400E-08 | 1.700E-07 |
| 11679 | 584 | P584Y | A | 4.9e-07 | 3.2e-07 | 1.1e-07 | 3.067E-07 | 1.904E-07 | 1.100E-07 | 4.900E-07 |
| 11680 | 584 | P584V | A | 3e-07 | 3.1e-07 | 3.1e-07 | 3.067E-07 | 5.774E-09 | 3.000E-07 | 3.100E-07 |
| 11681 | 585 | D585A | A | 1.9e-07 | 2.3e-07 | 3.2e-07 | 2.467E-07 | 6.658E-08 | 1.900E-07 | 3.200E-07 |
| 11682 | 585 | D585R | A | 5.2e-08 | 3.2e-07 | 1.2e-07 | 1.640E-07 | 1.393E-07 | 5.200E-08 | 3.200E-07 |
| 11683 | 585 | D585N | A | 7.1e-07 | 3.1e-07 | 2.5e-07 | 4.233E-07 | 2.501E-07 | 2.500E-07 | 7.100E-07 |
| 11684 | 585 | D585D | A | 2.1e-07 | 2.2e-07 | 3.5e-07 | 2.600E-07 | 7.810E-08 | 2.100E-07 | 3.500E-07 |
| 11685 | 585 | D585C | A | 7.5e-07 | 1.9e-07 | 1.3e-07 | 3.567E-07 | 3.420E-07 | 1.300E-07 | 7.500E-07 |
| 11686 | 585 | D585E | A | 1.6e-07 | 1.5e-07 | 1.5e-07 | 1.533E-07 | 5.774E-09 | 1.500E-07 | 1.600E-07 |
| 11687 | 585 | D585Q | A | 1.8e-07 | 1.2e-07 | 2.8e-07 | 1.933E-07 | 8.083E-08 | 1.200E-07 | 2.800E-07 |
| 11688 | 585 | D585G | A | 2.7e-07 | 2.9e-07 | 3.6e-07 | 3.067E-07 | 4.726E-08 | 2.700E-07 | 3.600E-07 |
| 11689 | 585 | D585H | A | 1.5e-07 | 2.2e-07 | 2e-07 | 1.900E-07 | 3.606E-08 | 1.500E-07 | 2.200E-07 |
| 11690 | 585 | D585I | A | 1.8e-07 | 3.9e-07 | 9.2e-08 | 2.207E-07 | 1.531E-07 | 9.200E-08 | 3.900E-07 |
| 11691 | 585 | D585L | A | 1.2e-07 | 4.1e-07 | 4e-07 | 3.100E-07 | 1.646E-07 | 1.200E-07 | 4.100E-07 |
| 11692 | 585 | D585K | A | 2.3e-07 | 4.3e-07 | 4.8e-07 | 3.800E-07 | 1.323E-07 | 2.300E-07 | 4.800E-07 |
| 11693 | 585 | D585M | A | 2.3e-07 | 2.7e-07 | 2.2e-07 | 2.400E-07 | 2.646E-08 | 2.200E-07 | 2.700E-07 |
| 11694 | 585 | D585F | A | 2.1e-07 | 9.9e-08 | 1.3e-07 | 1.463E-07 | 5.727E-08 | 9.900E-08 | 2.100E-07 |
| 11695 | 585 | D585P | A | 4.4e-07 | 3.4e-07 | 4.9e-07 | 4.233E-07 | 7.638E-08 | 3.400E-07 | 4.900E-07 |
| 11696 | 585 | D585S | A | 3.9e-07 | 1.4e-07 | 3.8e-07 | 3.033E-07 | 1.415E-07 | 1.400E-07 | 3.900E-07 |
| 11697 | 585 | D585T | A | 1.8e-07 | 1.3e-07 | 1.8e-07 | 1.633E-07 | 2.887E-08 | 1.300E-07 | 1.800E-07 |
| 11698 | 585 | D585W | A | 2.1e-07 | 1.7e-07 | 2.4e-07 | 2.067E-07 | 3.512E-08 | 1.700E-07 | 2.400E-07 |
| 11699 | 585 | D585Y | A | 1.4e-07 | 3.1e-07 | 1.4e-07 | 1.967E-07 | 9.815E-08 | 1.400E-07 | 3.100E-07 |
| 11700 | 585 | D585V | A | 2.3e-07 | 3.2e-07 | 2.1e-07 | 2.533E-07 | 5.859E-08 | 2.100E-07 | 3.200E-07 |
| 11701 | 586 | L586A | A | 1.9e-07 | 6.2e-07 | 2.5e-07 | 3.533E-07 | 2.329E-07 | 1.900E-07 | 6.200E-07 |
| 11702 | 586 | L586R | A | 6.8e-08 | 1.4e-07 | 1.5e-07 | 1.193E-07 | 4.474E-08 | 6.800E-08 | 1.500E-07 |
| 11703 | 586 | L586N | A | 1.2e-07 | 1.9e-07 | 7e-08 | 1.267E-07 | 6.028E-08 | 7.000E-08 | 1.900E-07 |
| 11704 | 586 | L586D | A | 1e-07 | 1.7e-07 | 1.7e-07 | 1.467E-07 | 4.041E-08 | 1.000E-07 | 1.700E-07 |
| 11705 | 586 | L586C | A | 3.1e-07 | 3e-07 | 2.4e-07 | 2.833E-07 | 3.786E-08 | 2.400E-07 | 3.100E-07 |
| 11706 | 586 | L586E | A | 2.7e-07 | 1.9e-07 | 1.4e-07 | 2.000E-07 | 6.557E-08 | 1.400E-07 | 2.700E-07 |
| 11707 | 586 | L586Q | A | 9.2e-08 | 7.6e-08 | 1.4e-07 | 1.027E-07 | 3.331E-08 | 7.600E-08 | 1.400E-07 |
| 11708 | 586 | L586G | A | 6.2e-07 | 2.5e-07 | 2.2e-07 | 3.633E-07 | 2.228E-07 | 2.200E-07 | 6.200E-07 |
| 11709 | 586 | L586H | A | 2.7e-07 | 1.2e-07 | 2e-07 | 1.967E-07 | 7.506E-08 | 1.200E-07 | 2.700E-07 |
| 11710 | 586 | L586I | A | 2.3e-07 | 1.9e-07 | 1.8e-07 | 2.000E-07 | 2.646E-08 | 1.800E-07 | 2.300E-07 |
| 11711 | 586 | L586L | A | 2.4e-07 | 4.2e-07 | 4.9e-07 | 3.833E-07 | 1.290E-07 | 2.400E-07 | 4.900E-07 |
| 11712 | 586 | L586K | A | 1.7e-07 | 1.3e-07 | 1e-07 | 1.333E-07 | 3.512E-08 | 1.000E-07 | 1.700E-07 |
| 11713 | 586 | L586M | A | 1.2e-07 | 4.1e-07 | 2.5e-07 | 2.600E-07 | 1.453E-07 | 1.200E-07 | 4.100E-07 |
| 11714 | 586 | L586F | A | 4e-07 | 1.9e-07 | 2.2e-07 | 2.700E-07 | 1.136E-07 | 1.900E-07 | 4.000E-07 |
| 11715 | 586 | L586P | A | 2.9e-07 | 3e-07 | 4.3e-07 | 3.400E-07 | 7.810E-08 | 2.900E-07 | 4.300E-07 |
| 11716 | 586 | L586S | A | 3.7e-07 | 1.9e-07 | 1.5e-07 | 2.367E-07 | 1.172E-07 | 1.500E-07 | 3.700E-07 |
| 11717 | 586 | L586T | A | 9e-08 | 1.7e-07 | 1.6e-07 | 1.400E-07 | 4.359E-08 | 9.000E-08 | 1.700E-07 |
| 11718 | 586 | L586W | A | 1.5e-07 | 1.3e-07 | 2.9e-07 | 1.900E-07 | 8.718E-08 | 1.300E-07 | 2.900E-07 |
| 11719 | 586 | L586Y | A | 1e-07 | 2.3e-07 | 1.8e-07 | 1.700E-07 | 6.557E-08 | 1.000E-07 | 2.300E-07 |
| 11720 | 586 | L586V | A | 2.3e-07 | 2.7e-07 | 3.4e-07 | 2.800E-07 | 5.568E-08 | 2.300E-07 | 3.400E-07 |
| 11721 | 587 | S587A | A | 3.7e-07 | 3.5e-07 | 2e-07 | 3.067E-07 | 9.292E-08 | 2.000E-07 | 3.700E-07 |
| 11722 | 587 | S587R | A | 2.1e-07 | 3.9e-07 | 1.4e-07 | 2.467E-07 | 1.290E-07 | 1.400E-07 | 3.900E-07 |
| 11723 | 587 | S587N | A | 2.8e-07 | 4e-07 | 1.3e-07 | 2.700E-07 | 1.353E-07 | 1.300E-07 | 4.000E-07 |
| 11724 | 587 | S587D | A | 2.9e-07 | 4.1e-07 | 2e-07 | 3.000E-07 | 1.054E-07 | 2.000E-07 | 4.100E-07 |
| 11725 | 587 | S587C | A | 3.6e-07 | 1.8e-07 | 1.3e-07 | 2.233E-07 | 1.210E-07 | 1.300E-07 | 3.600E-07 |
| 11726 | 587 | S587E | A | 2e-07 | 2.6e-07 | 2.4e-07 | 2.333E-07 | 3.055E-08 | 2.000E-07 | 2.600E-07 |
| 11727 | 587 | S587Q | A | 2.8e-07 | 3.4e-07 | 2.5e-07 | 2.900E-07 | 4.583E-08 | 2.500E-07 | 3.400E-07 |
| 11728 | 587 | S587G | A | 2.9e-07 | 2.6e-07 | 5e-07 | 3.500E-07 | 1.308E-07 | 2.600E-07 | 5.000E-07 |
| 11729 | 587 | S587H | A | 1.9e-07 | 3.2e-07 | 5.3e-07 | 3.467E-07 | 1.716E-07 | 1.900E-07 | 5.300E-07 |

| | | | | | | | | | | |
|---|---|---|---|---|---|---|---|---|---|---|
| 11730 | 587 | S587I | A | 1.8e-07 | 1.7e-07 | 3.7e-07 | 2.400E-07 | 1.127E-07 | 1.700E-07 | 3.700E-07 |
| 11731 | 587 | S587L | A | 1.4e-07 | 1.5e-07 | 2.2e-07 | 1.700E-07 | 4.359E-08 | 1.400E-07 | 2.200E-07 |
| 11732 | 587 | S587K | A | 3e-07 | 2.1e-07 | 1.6e-07 | 2.233E-07 | 7.095E-08 | 1.600E-07 | 3.000E-07 |
| 11733 | 587 | S587M | A | 3.3e-07 | 2.9e-07 | 2.6e-07 | 2.933E-07 | 3.512E-08 | 2.600E-07 | 3.300E-07 |
| 11734 | 587 | S587F | A | 1.8e-07 | 1.9e-07 | 3.7e-07 | 2.467E-07 | 1.069E-07 | 1.800E-07 | 3.700E-07 |
| 11735 | 587 | S587P | A | 3.3e-07 | 2.5e-07 | 5.8e-07 | 3.867E-07 | 1.721E-07 | 2.500E-07 | 5.800E-07 |
| 11736 | 587 | S587S | A | 1.4e-07 | 1.9e-07 | 1.5e-07 | 1.600E-07 | 2.646E-08 | 1.400E-07 | 1.900E-07 |
| 11737 | 587 | S587T | A | 2e-07 | 3.1e-07 | 2.4e-07 | 2.500E-07 | 5.568E-08 | 2.000E-07 | 3.100E-07 |
| 11738 | 587 | S587W | A | 3.6e-07 | 1.4e-07 | 2.7e-07 | 2.567E-07 | 1.106E-07 | 1.400E-07 | 3.600E-07 |
| 11739 | 587 | S587Y | A | 1.6e-07 | 4.3e-07 | 2e-07 | 2.633E-07 | 1.457E-07 | 1.600E-07 | 4.300E-07 |
| 11740 | 587 | S587V | A | 2e-07 | 2.4e-07 | 2.1e-07 | 2.167E-07 | 2.082E-08 | 2.000E-07 | 2.400E-07 |
| 11741 | 588 | Y588A | A | 4.9e-07 | 4e-07 | 3.7e-07 | 4.200E-07 | 6.245E-08 | 3.700E-07 | 4.900E-07 |
| 11742 | 588 | Y588R | A | 2.7e-07 | 1.6e-07 | 2.7e-07 | 2.333E-07 | 6.351E-08 | 1.600E-07 | 2.700E-07 |
| 11743 | 588 | Y588N | A | 7.5e-07 | 1.3e-07 | 5e-07 | 4.600E-07 | 3.119E-07 | 1.300E-07 | 7.500E-07 |
| 11744 | 588 | Y588D | A | 2.4e-07 | 2.9e-07 | 3.2e-07 | 2.833E-07 | 4.041E-08 | 2.400E-07 | 3.200E-07 |
| 11745 | 588 | Y588C | A | 3.9e-07 | 3e-07 | 3.3e-07 | 3.400E-07 | 4.583E-08 | 3.000E-07 | 3.900E-07 |
| 11746 | 588 | Y588E | A | 1.4e-07 | 2.1e-07 | 5.7e-07 | 3.067E-07 | 2.307E-07 | 1.400E-07 | 5.700E-07 |
| 11747 | 588 | Y588Q | A | 2.5e-07 | 9.4e-08 | 1e-07 | 1.480E-07 | 8.839E-08 | 9.400E-08 | 2.500E-07 |
| 11748 | 588 | Y588G | A | 7.9e-07 | 2.6e-07 | 3.3e-07 | 4.600E-07 | 2.879E-07 | 2.600E-07 | 7.900E-07 |
| 11749 | 588 | Y588H | A | 5.1e-07 | 2.9e-07 | 3.4e-07 | 3.800E-07 | 1.153E-07 | 2.900E-07 | 5.100E-07 |
| 11750 | 588 | Y588I | A | 5.2e-07 | 6.6e-07 | 6.7e-07 | 6.167E-07 | 8.386E-08 | 5.200E-07 | 6.700E-07 |
| 11751 | 588 | Y588L | A | 4e-07 | 4.4e-07 | 2.1e-07 | 3.500E-07 | 1.229E-07 | 2.100E-07 | 4.400E-07 |
| 11752 | 588 | Y588K | A | 1.6e-07 | 3e-07 | 1.4e-07 | 2.000E-07 | 8.718E-08 | 1.400E-07 | 3.000E-07 |
| 11753 | 588 | Y588M | A | 4.5e-07 | 5.4e-07 | 2e-07 | 3.967E-07 | 1.762E-07 | 2.000E-07 | 5.400E-07 |
| 11754 | 588 | Y588F | A | 2.9e-07 | 9.9e-08 | 4.3e-07 | 2.730E-07 | 1.662E-07 | 9.900E-08 | 4.300E-07 |
| 11755 | 588 | Y588P | A | 4.1e-07 | 2.3e-07 | 3.4e-07 | 3.267E-07 | 9.074E-08 | 2.300E-07 | 4.100E-07 |
| 11756 | 588 | Y588S | A | 3.9e-07 | 1.1e-07 | 2.2e-07 | 2.400E-07 | 1.411E-07 | 1.100E-07 | 3.900E-07 |
| 11757 | 588 | Y588T | A | 3.3e-07 | 8.3e-07 | 8.6e-07 | 6.733E-07 | 2.977E-07 | 3.300E-07 | 8.600E-07 |
| 11758 | 588 | Y588W | A | 3.6e-07 | 9.4e-07 | 4.7e-07 | 5.900E-07 | 3.081E-07 | 3.600E-07 | 9.400E-07 |
| 11759 | 588 | Y588Y | A | 2.2e-07 | 1.8e-07 | 3.8e-07 | 2.600E-07 | 1.058E-07 | 1.800E-07 | 3.800E-07 |
| 11760 | 588 | Y588V | A | 2.1e-07 | 6e-07 | 3.3e-07 | 3.800E-07 | 1.997E-07 | 2.100E-07 | 6.000E-07 |
| 11761 | 589 | M589A | A | 2.9e-07 | 1.2e-07 | 2.3e-07 | 2.133E-07 | 8.622E-08 | 1.200E-07 | 2.900E-07 |
| 11762 | 589 | M589R | A | 3.3e-07 | 1.5e-07 | 2.5e-07 | 2.433E-07 | 9.018E-08 | 1.500E-07 | 3.300E-07 |
| 11763 | 589 | M589N | A | 1.6e-07 | 3.5e-07 | 1.7e-07 | 2.267E-07 | 1.069E-07 | 1.600E-07 | 3.500E-07 |
| 11764 | 589 | M589D | A | 3.6e-07 | 2.4e-07 | 2.4e-07 | 2.800E-07 | 6.928E-08 | 2.400E-07 | 3.600E-07 |
| 11765 | 589 | M589C | A | 7.4e-07 | 3.5e-07 | 3.1e-07 | 4.667E-07 | 2.376E-07 | 3.100E-07 | 7.400E-07 |
| 11766 | 589 | M589E | A | 2.8e-07 | 3.4e-07 | 1.2e-07 | 2.467E-07 | 1.137E-07 | 1.200E-07 | 3.400E-07 |
| 11767 | 589 | M589Q | A | 2.3e-07 | 1.3e-07 | 3.2e-07 | 2.267E-07 | 9.504E-08 | 1.300E-07 | 3.200E-07 |
| 11768 | 589 | M589G | A | 2.6e-07 | 3.8e-07 | 1.6e-07 | 2.667E-07 | 1.102E-07 | 1.600E-07 | 3.800E-07 |
| 11769 | 589 | M589H | A | 2.6e-07 | 2.2e-07 | 2.1e-07 | 2.300E-07 | 2.646E-08 | 2.100E-07 | 2.600E-07 |
| 11770 | 589 | M589I | A | 1.3e-07 | 1.9e-07 | 1.1e-07 | 1.433E-07 | 4.163E-08 | 1.100E-07 | 1.900E-07 |
| 11771 | 589 | M589L | A | 1.7e-07 | 3.1e-07 | 1.2e-07 | 2.000E-07 | 9.849E-08 | 1.200E-07 | 3.100E-07 |
| 11772 | 589 | M589K | A | 4.3e-07 | 3e-07 | 1.8e-07 | 3.033E-07 | 1.250E-07 | 1.800E-07 | 4.300E-07 |
| 11773 | 589 | M589M | A | 2.3e-07 | 4.5e-07 | 3.1e-07 | 3.300E-07 | 1.114E-07 | 2.300E-07 | 4.500E-07 |
| 11774 | 589 | M589F | A | 1.4e-07 | 2.3e-07 | 2.2e-07 | 1.967E-07 | 4.933E-08 | 1.400E-07 | 2.300E-07 |
| 11775 | 589 | M589P | A | 1.7e-07 | 1.2e-07 | 2.7e-07 | 1.867E-07 | 7.638E-08 | 1.200E-07 | 2.700E-07 |
| 11776 | 589 | M589S | A | 2.3e-07 | 3.3e-07 | 2.2e-07 | 2.600E-07 | 6.083E-08 | 2.200E-07 | 3.300E-07 |
| 11777 | 589 | M589T | A | 2.3e-07 | 3.1e-07 | 3.2e-07 | 2.867E-07 | 4.933E-08 | 2.300E-07 | 3.200E-07 |
| 11778 | 589 | M589W | A | 4.1e-07 | 1.5e-07 | 2.8e-07 | 2.800E-07 | 1.300E-07 | 1.500E-07 | 4.100E-07 |
| 11779 | 589 | M589Y | A | 3.1e-07 | 1.8e-07 | 1.9e-07 | 2.267E-07 | 7.234E-08 | 1.800E-07 | 3.100E-07 |
| 11780 | 589 | M589V | A | 1.9e-07 | 5e-07 | 3.2e-07 | 3.367E-07 | 1.557E-07 | 1.900E-07 | 5.000E-07 |
| 11781 | 590 | P590A | A | 2.7e-07 | 4.5e-07 | 1.9e-07 | 3.033E-07 | 1.332E-07 | 1.900E-07 | 4.500E-07 |
| 11782 | 590 | P590R | A | 2.4e-07 | 1e-07 | 2e-07 | 1.800E-07 | 7.211E-08 | 1.000E-07 | 2.400E-07 |
| 11783 | 590 | P590N | A | 2.7e-07 | 2.2e-07 | 8.1e-08 | 1.903E-07 | 9.793E-08 | 8.100E-08 | 2.700E-07 |
| 11784 | 590 | P590D | A | 1.4e-07 | 6.1e-08 | 2e-07 | 1.337E-07 | 6.972E-08 | 6.100E-08 | 2.000E-07 |
| 11785 | 590 | P590C | A | 1.3e-07 | 2.3e-07 | 1.5e-07 | 1.700E-07 | 5.292E-08 | 1.300E-07 | 2.300E-07 |
| 11786 | 590 | P590E | A | 9.1e-08 | 1.7e-07 | 2.5e-07 | 1.703E-07 | 7.950E-08 | 9.100E-08 | 2.500E-07 |
| 11787 | 590 | P590Q | A | 2.2e-07 | 1.9e-07 | 2.7e-07 | 2.267E-07 | 4.041E-08 | 1.900E-07 | 2.700E-07 |
| 11788 | 590 | P590G | A | 5.2e-07 | 2e-07 | 1.6e-07 | 2.933E-07 | 1.973E-07 | 1.600E-07 | 5.200E-07 |
| 11789 | 590 | P590H | A | 1.2e-07 | 1e-07 | 1.3e-07 | 1.167E-07 | 1.528E-08 | 1.000E-07 | 1.300E-07 |
| 11790 | 590 | P590I | A | 4.3e-07 | 3.1e-07 | 2.5e-07 | 3.300E-07 | 9.165E-08 | 2.500E-07 | 4.300E-07 |
| 11791 | 590 | P590L | A | 3e-07 | 3.1e-07 | 1.1e-07 | 2.400E-07 | 1.127E-07 | 1.100E-07 | 3.100E-07 |
| 11792 | 590 | P590K | A | 1.9e-07 | 1.5e-07 | 2.3e-07 | 1.900E-07 | 4.000E-08 | 1.500E-07 | 2.300E-07 |
| 11793 | 590 | P590M | A | 3.3e-07 | 2.7e-07 | 1.4e-07 | 2.467E-07 | 9.713E-08 | 1.400E-07 | 3.300E-07 |
| 11794 | 590 | P590F | A | 3.9e-07 | 1.5e-07 | 1.5e-07 | 2.300E-07 | 1.386E-07 | 1.500E-07 | 3.900E-07 |
| 11795 | 590 | P590P | A | 2.2e-07 | 3e-07 | 4e-07 | 3.067E-07 | 9.018E-08 | 2.200E-07 | 4.000E-07 |
| 11796 | 590 | P590S | A | 1.6e-07 | 9.1e-08 | 2.1e-07 | 1.537E-07 | 5.975E-08 | 9.100E-08 | 2.100E-07 |
| 11797 | 590 | P590T | A | 8.2e-08 | 1.9e-07 | 2.1e-07 | 1.607E-07 | 6.886E-08 | 8.200E-08 | 2.100E-07 |
| 11798 | 590 | P590W | A | 1.9e-07 | 1.5e-07 | 2.8e-07 | 2.067E-07 | 6.658E-08 | 1.500E-07 | 2.800E-07 |
| 11799 | 590 | P590Y | A | 3.2e-07 | 1.4e-07 | 2.6e-07 | 2.400E-07 | 9.165E-08 | 1.400E-07 | 3.200E-07 |
| 11800 | 590 | P590V | A | 2.2e-07 | 2.8e-07 | 2.1e-07 | 2.367E-07 | 3.786E-08 | 2.100E-07 | 2.800E-07 |
| 11801 | 591 | I591A | A | 2.2e-07 | 2.2e-07 | 2.9e-07 | 2.433E-07 | 4.041E-08 | 2.200E-07 | 2.900E-07 |
| 11802 | 591 | I591R | A | 3.2e-07 | 2.2e-07 | 2.4e-07 | 2.600E-07 | 5.292E-08 | 2.200E-07 | 3.200E-07 |
| 11803 | 591 | I591N | A | 3.3e-07 | 4.9e-07 | 3.5e-07 | 3.900E-07 | 8.718E-08 | 3.300E-07 | 4.900E-07 |
| 11804 | 591 | I591D | A | 2.9e-07 | 1.4e-07 | 2.2e-07 | 2.167E-07 | 7.506E-08 | 1.400E-07 | 2.900E-07 |
| 11805 | 591 | I591C | A | 2.2e-07 | 3.9e-07 | 2.1e-07 | 2.733E-07 | 1.012E-07 | 2.100E-07 | 3.900E-07 |
| 11806 | 591 | I591E | A | 2.5e-07 | 5.6e-07 | 2.5e-07 | 3.533E-07 | 1.790E-07 | 2.500E-07 | 5.600E-07 |
| 11807 | 591 | I591Q | A | 2.3e-07 | 3.6e-07 | 2.8e-07 | 2.900E-07 | 6.557E-08 | 2.300E-07 | 3.600E-07 |
| 11808 | 591 | I591G | A | 9.2e-08 | 2.6e-07 | 3.6e-07 | 2.373E-07 | 1.354E-07 | 9.200E-08 | 3.600E-07 |
| 11809 | 591 | I591H | A | 2.3e-07 | 2.1e-07 | 3.5e-07 | 2.633E-07 | 7.572E-08 | 2.100E-07 | 3.500E-07 |
| 11810 | 591 | I591I | A | 4.3e-07 | 3.8e-07 | 3.1e-07 | 3.733E-07 | 6.028E-08 | 3.100E-07 | 4.300E-07 |
| 11811 | 591 | I591L | A | 1.9e-07 | 2.2e-07 | 2.4e-07 | 2.167E-07 | 2.517E-08 | 1.900E-07 | 2.400E-07 |
| 11812 | 591 | I591K | A | 3.5e-07 | 2e-07 | 2.7e-07 | 2.733E-07 | 7.506E-08 | 2.000E-07 | 3.500E-07 |
| 11813 | 591 | I591M | A | 2.2e-07 | 2.3e-07 | 1.7e-07 | 2.067E-07 | 3.215E-08 | 1.700E-07 | 2.300E-07 |
| 11814 | 591 | I591F | A | 3.2e-07 | 2.4e-07 | 4.3e-07 | 3.300E-07 | 9.539E-08 | 2.400E-07 | 4.300E-07 |

| | | | | | | | | | | |
|---|---|---|---|---|---|---|---|---|---|---|
| 11815 | 591 | I591P | A | 3.8e-07 | 5e-07 | 1.1e-07 | 3.300E-07 | 1.997E-07 | 1.100E-07 | 5.000E-07 |
| 11816 | 591 | I591S | A | 2.7e-07 | 1.4e-07 | 1.8e-07 | 1.967E-07 | 6.658E-08 | 1.400E-07 | 2.700E-07 |
| 11817 | 591 | I591T | A | 1.3e-07 | 1.9e-07 | 1.9e-07 | 1.700E-07 | 3.464E-08 | 1.300E-07 | 1.900E-07 |
| 11818 | 591 | I591W | A | 2.9e-07 | 3.3e-07 | 2.3e-07 | 2.833E-07 | 5.033E-08 | 2.300E-07 | 3.300E-07 |
| 11819 | 591 | I591Y | A | 7.2e-07 | 2.2e-07 | 2.3e-07 | 3.900E-07 | 2.858E-07 | 2.200E-07 | 7.200E-07 |
| 11820 | 591 | I591V | A | 9.9e-08 | 2.3e-07 | 2.1e-07 | 1.797E-07 | 7.057E-08 | 9.900E-08 | 2.300E-07 |
| 11821 | 592 | W592A | A | 1.5e-07 | 2e-07 | 4.7e-07 | 2.733E-07 | 1.721E-07 | 1.500E-07 | 4.700E-07 |
| 11822 | 592 | W592R | A | 3.1e-07 | 2.6e-07 | 3.3e-07 | 3.000E-07 | 3.606E-08 | 2.600E-07 | 3.300E-07 |
| 11823 | 592 | W592N | A | 3.6e-07 | 2.3e-07 | 5.5e-07 | 3.800E-07 | 1.609E-07 | 2.300E-07 | 5.500E-07 |
| 11824 | 592 | W592D | A | 2e-07 | 2e-07 | 9.4e-08 | 1.647E-07 | 6.120E-08 | 9.400E-08 | 2.000E-07 |
| 11825 | 592 | W592C | A | 4.1e-07 | 1.4e-07 | 6.3e-07 | 3.933E-07 | 2.454E-07 | 1.400E-07 | 6.300E-07 |
| 11826 | 592 | W592E | A | 2.8e-07 | 2e-07 | 2.6e-07 | 2.467E-07 | 4.163E-08 | 2.000E-07 | 2.800E-07 |
| 11827 | 592 | W592Q | A | 3.5e-07 | 2.7e-07 | 3.5e-07 | 3.233E-07 | 4.619E-08 | 2.700E-07 | 3.500E-07 |
| 11828 | 592 | W592G | A | 4.2e-07 | 3.9e-07 | 5.8e-07 | 4.633E-07 | 1.021E-07 | 3.900E-07 | 5.800E-07 |
| 11829 | 592 | W592H | A | 3.1e-07 | 3e-07 | 4.1e-07 | 3.400E-07 | 6.083E-08 | 3.000E-07 | 4.100E-07 |
| 11830 | 592 | W592I | A | 2.6e-07 | 7.7e-07 | 3.5e-07 | 4.600E-07 | 2.722E-07 | 2.600E-07 | 7.700E-07 |
| 11831 | 592 | W592L | A | 3.4e-07 | 1.6e-07 | 3.3e-07 | 2.767E-07 | 1.012E-07 | 1.600E-07 | 3.400E-07 |
| 11832 | 592 | W592K | A | 1.6e-07 | 1.8e-07 | 1.9e-07 | 1.767E-07 | 1.528E-08 | 1.600E-07 | 1.900E-07 |
| 11833 | 592 | W592M | A | 1.5e-07 | 1.9e-07 | 4.3e-07 | 2.567E-07 | 1.514E-07 | 1.500E-07 | 4.300E-07 |
| 11834 | 592 | W592F | A | 3.1e-07 | 4.4e-07 | 5.5e-07 | 4.333E-07 | 1.201E-07 | 3.100E-07 | 5.500E-07 |
| 11835 | 592 | W592P | A | 2.2e-07 | 2.2e-07 | 2e-07 | 2.133E-07 | 1.155E-08 | 2.000E-07 | 2.200E-07 |
| 11836 | 592 | W592S | A | 2.8e-07 | 1.6e-07 | 2.1e-07 | 2.167E-07 | 6.028E-08 | 1.600E-07 | 2.800E-07 |
| 11837 | 592 | W592T | A | 4.2e-07 | 2.6e-07 | 9.3e-07 | 5.367E-07 | 3.499E-07 | 2.600E-07 | 9.300E-07 |
| 11838 | 592 | W592W | A | 9e-08 | 1.1e-07 | 3.5e-07 | 1.833E-07 | 1.447E-07 | 9.000E-08 | 3.500E-07 |
| 11839 | 592 | W592Y | A | 4.7e-07 | 4e-07 | 5.3e-07 | 4.667E-07 | 6.506E-08 | 4.000E-07 | 5.300E-07 |
| 11840 | 592 | W592V | A | 1.6e-07 | 2e-07 | 1.6e-07 | 1.733E-07 | 2.309E-08 | 1.600E-07 | 2.000E-07 |
| 11841 | 593 | K593A | A | 5.9e-07 | 4e-07 | 5.2e-07 | 5.033E-07 | 9.609E-08 | 4.000E-07 | 5.900E-07 |
| 11842 | 593 | K593R | A | 5.6e-07 | 2.5e-07 | 2e-07 | 3.367E-07 | 1.950E-07 | 2.000E-07 | 5.600E-07 |
| 11843 | 593 | K593N | A | 1.4e-07 | 4.8e-07 | 3.2e-07 | 3.133E-07 | 1.701E-07 | 1.400E-07 | 4.800E-07 |
| 11844 | 593 | K593D | A | 2.8e-07 | 1.5e-07 | 3.3e-07 | 2.533E-07 | 9.292E-08 | 1.500E-07 | 3.300E-07 |
| 11845 | 593 | K593C | A | 2.3e-07 | 3.2e-07 | 5.7e-07 | 3.733E-07 | 1.762E-07 | 2.300E-07 | 5.700E-07 |
| 11846 | 593 | K593E | A | 5.6e-07 | 2.5e-07 | 2.5e-07 | 3.533E-07 | 1.790E-07 | 2.500E-07 | 5.600E-07 |
| 11847 | 593 | K593Q | A | 2.9e-07 | 1.8e-07 | 7.1e-07 | 3.933E-07 | 2.797E-07 | 1.800E-07 | 7.100E-07 |
| 11848 | 593 | K593G | A | 6.6e-07 | 4.1e-07 | 7.7e-07 | 6.133E-07 | 1.845E-07 | 4.100E-07 | 7.700E-07 |
| 11849 | 593 | K593H | A | 2.9e-07 | 4e-07 | 2e-07 | 2.967E-07 | 1.002E-07 | 2.000E-07 | 4.000E-07 |
| 11850 | 593 | K593I | A | 8e-07 | 5.7e-07 | 4.7e-07 | 6.133E-07 | 1.692E-07 | 4.700E-07 | 8.000E-07 |
| 11851 | 593 | K593L | A | 2.2e-07 | 5.6e-07 | 5e-07 | 4.267E-07 | 1.815E-07 | 2.200E-07 | 5.600E-07 |
| 11852 | 593 | K593K | A | 1.6e-07 | 5.6e-07 | 2.7e-07 | 3.300E-07 | 2.066E-07 | 1.600E-07 | 5.600E-07 |
| 11853 | 593 | K593M | A | 1.8e-07 | 4.3e-07 | 3.1e-07 | 3.067E-07 | 1.250E-07 | 1.800E-07 | 4.300E-07 |
| 11854 | 593 | K593F | A | 3.6e-07 | 2.8e-07 | 5.2e-07 | 3.867E-07 | 1.222E-07 | 2.800E-07 | 5.200E-07 |
| 11855 | 593 | K593P | A | 3.8e-07 | 6.1e-07 | 3.9e-07 | 4.600E-07 | 1.300E-07 | 3.800E-07 | 6.100E-07 |
| 11856 | 593 | K593S | A | 3.2e-07 | 4.2e-07 | 3.7e-07 | 3.700E-07 | 5.000E-08 | 3.200E-07 | 4.200E-07 |
| 11857 | 593 | K593T | A | 6.8e-07 | 5.6e-07 | 5.6e-07 | 6.000E-07 | 6.928E-08 | 5.600E-07 | 6.800E-07 |
| 11858 | 593 | K593W | A | 3.7e-07 | 3.3e-07 | 2.2e-07 | 3.067E-07 | 7.767E-08 | 2.200E-07 | 3.700E-07 |
| 11859 | 593 | K593Y | A | 7.7e-07 | 4.3e-07 | 3.2e-07 | 5.067E-07 | 2.346E-07 | 3.200E-07 | 7.700E-07 |
| 11860 | 593 | K593V | A | 6.5e-07 | 1.5e-07 | 2.9e-07 | 3.633E-07 | 2.579E-07 | 1.500E-07 | 6.500E-07 |
| 11861 | 594 | F594A | A | 2.7e-07 | 4.6e-07 | 2.4e-07 | 3.233E-07 | 1.193E-07 | 2.400E-07 | 4.600E-07 |
| 11862 | 594 | F594R | A | 2.5e-07 | 2.2e-07 | 2.7e-07 | 2.467E-07 | 2.517E-08 | 2.200E-07 | 2.700E-07 |
| 11863 | 594 | F594N | A | 1.3e-07 | 5e-07 | 2.5e-07 | 2.933E-07 | 1.888E-07 | 1.300E-07 | 5.000E-07 |
| 11864 | 594 | F594D | A | 1.7e-07 | 4.7e-07 | 7.3e-07 | 4.567E-07 | 2.802E-07 | 1.700E-07 | 7.300E-07 |
| 11865 | 594 | F594C | A | 5.1e-07 | 1.5e-07 | 4.2e-07 | 3.600E-07 | 1.873E-07 | 1.500E-07 | 5.100E-07 |
| 11866 | 594 | F594E | A | 1.4e-07 | 2.4e-07 | 4.3e-07 | 2.700E-07 | 1.473E-07 | 1.400E-07 | 4.300E-07 |
| 11867 | 594 | F594Q | A | 1.3e-07 | 2.3e-07 | 2.1e-07 | 1.900E-07 | 5.292E-08 | 1.300E-07 | 2.300E-07 |
| 11868 | 594 | F594G | A | 1.6e-07 | 2.2e-07 | 2.3e-07 | 2.033E-07 | 3.786E-08 | 1.600E-07 | 2.300E-07 |
| 11869 | 594 | F594H | A | 2.8e-07 | 4e-07 | 2.5e-07 | 3.100E-07 | 7.937E-08 | 2.500E-07 | 4.000E-07 |
| 11870 | 594 | F594I | A | 2.1e-07 | 1.5e-07 | 2.2e-07 | 1.933E-07 | 3.786E-08 | 1.500E-07 | 2.200E-07 |
| 11871 | 594 | F594L | A | 2.5e-07 | 1.7e-07 | 2.4e-07 | 2.200E-07 | 4.359E-08 | 1.700E-07 | 2.500E-07 |
| 11872 | 594 | F594K | A | 3.9e-07 | 2.6e-07 | 2.6e-07 | 3.033E-07 | 7.506E-08 | 2.600E-07 | 3.900E-07 |
| 11873 | 594 | F594M | A | 2.5e-07 | 2.1e-07 | 1.9e-07 | 2.167E-07 | 3.055E-08 | 1.900E-07 | 2.500E-07 |
| 11874 | 594 | F594F | A | 3.9e-07 | 1.9e-07 | 2.7e-07 | 2.833E-07 | 1.007E-07 | 1.900E-07 | 3.900E-07 |
| 11875 | 594 | F594P | A | 2.6e-07 | 1.7e-07 | 2.4e-07 | 2.233E-07 | 4.726E-08 | 1.700E-07 | 2.600E-07 |
| 11876 | 594 | F594S | A | 1.3e-07 | 1.4e-07 | 1e-07 | 1.233E-07 | 2.082E-08 | 1.000E-07 | 1.400E-07 |
| 11877 | 594 | F594T | A | 2.7e-07 | 1.3e-07 | 2.8e-07 | 2.267E-07 | 8.386E-08 | 1.300E-07 | 2.800E-07 |
| 11878 | 594 | F594W | A | 3e-07 | 2.1e-07 | 2.5e-07 | 2.533E-07 | 4.509E-08 | 2.100E-07 | 3.000E-07 |
| 11879 | 594 | F594Y | A | 9e-08 | 3.6e-07 | 2.5e-07 | 2.333E-07 | 1.358E-07 | 9.000E-08 | 3.600E-07 |
| 11880 | 594 | F594V | A | 3e-07 | 3.5e-07 | 1.9e-07 | 2.800E-07 | 8.185E-08 | 1.900E-07 | 3.500E-07 |
| 11881 | 595 | P595A | A | 3e-07 | 3.3e-07 | 3.5e-07 | 3.267E-07 | 2.517E-08 | 3.000E-07 | 3.500E-07 |
| 11882 | 595 | P595R | A | 3.6e-07 | 2.2e-07 | 2e-07 | 2.600E-07 | 8.718E-08 | 2.000E-07 | 3.600E-07 |
| 11883 | 595 | P595N | A | 5.6e-07 | 2.1e-07 | 9e-08 | 2.867E-07 | 2.442E-07 | 9.000E-08 | 5.600E-07 |
| 11884 | 595 | P595D | A | 3.8e-07 | 3e-07 | 2e-07 | 2.933E-07 | 9.018E-08 | 2.000E-07 | 3.800E-07 |
| 11885 | 595 | P595C | A | 2.7e-07 | 2.4e-07 | 1.1e-07 | 2.067E-07 | 8.505E-08 | 1.100E-07 | 2.700E-07 |
| 11886 | 595 | P595E | A | 1.5e-07 | 2.3e-07 | 3.6e-07 | 2.467E-07 | 1.060E-07 | 1.500E-07 | 3.600E-07 |
| 11887 | 595 | P595Q | A | 1.8e-07 | 2e-07 | 2.6e-07 | 2.133E-07 | 4.163E-08 | 1.800E-07 | 2.600E-07 |
| 11888 | 595 | P595G | A | 5.3e-07 | 1.7e-07 | 4.5e-07 | 3.833E-07 | 1.890E-07 | 1.700E-07 | 5.300E-07 |
| 11889 | 595 | P595H | A | 4.8e-07 | 2e-07 | 1.9e-07 | 2.900E-07 | 1.646E-07 | 1.900E-07 | 4.800E-07 |
| 11890 | 595 | P595I | A | 2.5e-07 | 2e-07 | 3.1e-07 | 2.533E-07 | 5.508E-08 | 2.000E-07 | 3.100E-07 |
| 11891 | 595 | P595L | A | 1.3e-07 | 9.8e-08 | 1.6e-07 | 1.293E-07 | 3.101E-08 | 9.800E-08 | 1.600E-07 |
| 11892 | 595 | P595K | A | 1.6e-07 | 2.3e-07 | 2.6e-07 | 2.167E-07 | 5.132E-08 | 1.600E-07 | 2.600E-07 |
| 11893 | 595 | P595M | A | 2.7e-07 | 3.7e-07 | 3.8e-07 | 3.400E-07 | 6.083E-08 | 2.700E-07 | 3.800E-07 |
| 11894 | 595 | P595F | A | 2.2e-07 | 2.9e-07 | 3.3e-07 | 2.800E-07 | 5.568E-08 | 2.200E-07 | 3.300E-07 |
| 11895 | 595 | P595P | A | 1.4e-07 | 2.2e-07 | 2.5e-07 | 2.033E-07 | 5.686E-08 | 1.400E-07 | 2.500E-07 |
| 11896 | 595 | P595S | A | 4.2e-07 | 2.7e-07 | 4e-07 | 3.633E-07 | 8.145E-08 | 2.700E-07 | 4.200E-07 |
| 11897 | 595 | P595T | A | 1.2e-07 | 4.8e-07 | 2e-07 | 2.667E-07 | 1.890E-07 | 1.200E-07 | 4.800E-07 |
| 11898 | 595 | P595W | A | 2.4e-07 | 1.5e-07 | 1.7e-07 | 1.867E-07 | 4.726E-08 | 1.500E-07 | 2.400E-07 |
| 11899 | 595 | P595Y | A | 1.5e-07 | 5.9e-07 | 1.9e-07 | 3.100E-07 | 2.433E-07 | 1.500E-07 | 5.900E-07 |

| | | | | | | | | | | |
|---|---|---|---|---|---|---|---|---|---|---|
| 11900 | 595 | P595V | A | 1.7e-07 | 2.3e-07 | 2.3e-07 | 2.100E-07 | 3.464E-08 | 1.700E-07 | 2.300E-07 |
| 11901 | 596 | D596A | A | 1.4e-07 | 1.7e-07 | 2.1e-07 | 1.733E-07 | 3.512E-08 | 1.400E-07 | 2.100E-07 |
| 11902 | 596 | D596R | A | 1.3e-07 | 2.1e-07 | 2.7e-07 | 2.033E-07 | 7.024E-08 | 1.300E-07 | 2.700E-07 |
| 11903 | 596 | D596N | A | 1.2e-07 | 3.3e-07 | 7.8e-07 | 4.100E-07 | 3.372E-07 | 1.200E-07 | 7.800E-07 |
| 11904 | 596 | D596D | A | 1.6e-07 | 4.1e-07 | 5.2e-07 | 3.633E-07 | 1.845E-07 | 1.600E-07 | 5.200E-07 |
| 11905 | 596 | D596C | A | 5.5e-08 | 1.3e-07 | 3.9e-07 | 1.917E-07 | 1.758E-07 | 5.500E-08 | 3.900E-07 |
| 11906 | 596 | D596E | A | 3.5e-07 | 2.3e-07 | 2.5e-07 | 2.767E-07 | 6.429E-08 | 2.300E-07 | 3.500E-07 |
| 11907 | 596 | D596Q | A | 7.8e-07 | 3.8e-07 | 5.2e-07 | 5.600E-07 | 2.030E-07 | 3.800E-07 | 7.800E-07 |
| 11908 | 596 | D596G | A | 5.9e-07 | 2.8e-07 | 1.9e-07 | 3.533E-07 | 2.098E-07 | 1.900E-07 | 5.900E-07 |
| 11909 | 596 | D596H | A | 3.2e-07 | 1e-06 | 3.5e-07 | 5.567E-07 | 3.842E-07 | 3.200E-07 | 1.000E-06 |
| 11910 | 596 | D596I | A | 2.7e-07 | 2.4e-07 | 1.1e-07 | 2.067E-07 | 8.505E-08 | 1.100E-07 | 2.700E-07 |
| 11911 | 596 | D596L | A | 1.2e-07 | 1.2e-07 | 1.4e-07 | 1.267E-07 | 1.155E-08 | 1.200E-07 | 1.400E-07 |
| 11912 | 596 | D596K | A | 2.7e-07 | 1.3e-07 | 4.5e-07 | 2.833E-07 | 1.604E-07 | 1.300E-07 | 4.500E-07 |
| 11913 | 596 | D596M | A | 1.9e-07 | 6.6e-08 | 2e-07 | 1.520E-07 | 7.465E-08 | 6.600E-08 | 2.000E-07 |
| 11914 | 596 | D596F | A | 2e-07 | 1.3e-07 | 6.6e-08 | 1.320E-07 | 6.702E-08 | 6.600E-08 | 2.000E-07 |
| 11915 | 596 | D596P | A | 6.3e-07 | 6.8e-07 | 2e-07 | 5.033E-07 | 2.639E-07 | 2.000E-07 | 6.800E-07 |
| 11916 | 596 | D596S | A | 3.4e-07 | 2.8e-07 | 2.1e-07 | 2.767E-07 | 6.506E-08 | 2.100E-07 | 3.400E-07 |
| 11917 | 596 | D596T | A | 3.7e-07 | 3.9e-07 | 2e-07 | 3.200E-07 | 1.044E-07 | 2.000E-07 | 3.900E-07 |
| 11918 | 596 | D596W | A | 1.9e-07 | 2.4e-07 | 2.3e-07 | 2.200E-07 | 2.646E-08 | 1.900E-07 | 2.400E-07 |
| 11919 | 596 | D596Y | A | 1.2e-07 | 1.9e-07 | 1.3e-07 | 1.467E-07 | 3.786E-08 | 1.200E-07 | 1.900E-07 |
| 11920 | 596 | D596V | A | 3.2e-07 | 8.5e-07 | 1.6e-07 | 4.433E-07 | 3.612E-07 | 1.600E-07 | 8.500E-07 |
| 11921 | 597 | E597A | A | 3.1e-07 | 3.2e-07 | 2.5e-07 | 2.933E-07 | 3.786E-08 | 2.500E-07 | 3.200E-07 |
| 11922 | 597 | E597R | A | 1.4e-07 | 1.9e-07 | 3.6e-07 | 2.300E-07 | 1.153E-07 | 1.400E-07 | 3.600E-07 |
| 11923 | 597 | E597N | A | 3.8e-07 | 2.8e-07 | 5.4e-07 | 4.000E-07 | 1.311E-07 | 2.800E-07 | 5.400E-07 |
| 11924 | 597 | E597D | A | 3.4e-07 | 2.1e-07 | 2.1e-07 | 2.533E-07 | 7.506E-08 | 2.100E-07 | 3.400E-07 |
| 11925 | 597 | E597C | A | 3e-07 | 1.6e-07 | 6.5e-07 | 3.700E-07 | 2.524E-07 | 1.600E-07 | 6.500E-07 |
| 11926 | 597 | E597E | A | 2.1e-07 | 2.3e-07 | 1.2e-07 | 1.867E-07 | 5.859E-08 | 1.200E-07 | 2.300E-07 |
| 11927 | 597 | E597Q | A | 1.7e-07 | 4.1e-07 | 1.8e-07 | 2.533E-07 | 1.358E-07 | 1.700E-07 | 4.100E-07 |
| 11928 | 597 | E597G | A | 5.8e-07 | 2.3e-07 | 4e-07 | 4.033E-07 | 1.750E-07 | 2.300E-07 | 5.800E-07 |
| 11929 | 597 | E597H | A | 3.9e-07 | 2.3e-07 | 3.4e-07 | 3.200E-07 | 8.185E-08 | 2.300E-07 | 3.900E-07 |
| 11930 | 597 | E597I | A | 1.8e-07 | 1.4e-07 | 3.4e-07 | 2.200E-07 | 1.058E-07 | 1.400E-07 | 3.400E-07 |
| 11931 | 597 | E597L | A | 2e-07 | 1e-07 | 2.4e-07 | 1.800E-07 | 7.211E-08 | 1.000E-07 | 2.400E-07 |
| 11932 | 597 | E597K | A | 2e-07 | 2.7e-07 | 2.3e-07 | 2.333E-07 | 3.512E-08 | 2.000E-07 | 2.700E-07 |
| 11933 | 597 | E597M | A | 3e-07 | 3.4e-07 | 1.5e-07 | 2.633E-07 | 1.002E-07 | 1.500E-07 | 3.400E-07 |
| 11934 | 597 | E597F | A | 1.9e-07 | 2e-07 | 1.8e-07 | 1.900E-07 | 1.000E-08 | 1.800E-07 | 2.000E-07 |
| 11935 | 597 | E597P | A | 2.9e-07 | 6.9e-07 | 1.2e-07 | 3.667E-07 | 2.926E-07 | 1.200E-07 | 6.900E-07 |
| 11936 | 597 | E597S | A | 2e-07 | 2.5e-07 | 5e-07 | 3.167E-07 | 1.607E-07 | 2.000E-07 | 5.000E-07 |
| 11937 | 597 | E597T | A | 3.6e-07 | 1.5e-07 | 1.4e-07 | 2.167E-07 | 1.242E-07 | 1.400E-07 | 3.600E-07 |
| 11938 | 597 | E597W | A | 2.5e-07 | 1.5e-07 | 3.3e-07 | 2.433E-07 | 9.018E-08 | 1.500E-07 | 3.300E-07 |
| 11939 | 597 | E597Y | A | 1.3e-07 | 1.7e-07 | 2.3e-07 | 1.767E-07 | 5.033E-08 | 1.300E-07 | 2.300E-07 |
| 11940 | 597 | E597V | A | 1.8e-07 | 4.1e-07 | 3.2e-07 | 3.033E-07 | 1.159E-07 | 1.800E-07 | 4.100E-07 |
| 11941 | 598 | E598A | A | 2.4e-07 | 7.6e-08 | 1.4e-07 | 1.520E-07 | 8.266E-08 | 7.600E-08 | 2.400E-07 |
| 11942 | 598 | E598R | A | 2.4e-07 | 1e-07 | 1.9e-07 | 1.767E-07 | 7.095E-08 | 1.000E-07 | 2.400E-07 |
| 11943 | 598 | E598N | A | 3.9e-07 | 1.7e-07 | 2.5e-07 | 2.700E-07 | 1.114E-07 | 1.700E-07 | 3.900E-07 |
| 11944 | 598 | E598D | A | 3.4e-07 | 1.4e-07 | 1.8e-07 | 2.200E-07 | 1.058E-07 | 1.400E-07 | 3.400E-07 |
| 11945 | 598 | E598C | A | 1.1e-07 | 2.4e-07 | 1.5e-07 | 1.667E-07 | 6.658E-08 | 1.100E-07 | 2.400E-07 |
| 11946 | 598 | E598E | A | 2.3e-07 | 2.1e-07 | 3e-07 | 2.467E-07 | 4.726E-08 | 2.100E-07 | 3.000E-07 |
| 11947 | 598 | E598Q | A | 3.1e-07 | 2.7e-07 | 2.1e-07 | 2.633E-07 | 5.033E-08 | 2.100E-07 | 3.100E-07 |
| 11948 | 598 | E598G | A | 3e-07 | 1.9e-07 | 2e-07 | 2.300E-07 | 6.083E-08 | 1.900E-07 | 3.000E-07 |
| 11949 | 598 | E598H | A | 3.1e-07 | 2.9e-07 | 6.5e-07 | 4.167E-07 | 2.023E-07 | 2.900E-07 | 6.500E-07 |
| 11950 | 598 | E598I | A | 9.9e-08 | 3e-07 | 1.2e-07 | 1.730E-07 | 1.105E-07 | 9.900E-08 | 3.000E-07 |
| 11951 | 598 | E598L | A | 3.4e-07 | 1.9e-07 | 9.5e-08 | 2.083E-07 | 1.235E-07 | 9.500E-08 | 3.400E-07 |
| 11952 | 598 | E598K | A | 2.3e-07 | 2.9e-07 | 6.4e-07 | 3.867E-07 | 2.214E-07 | 2.300E-07 | 6.400E-07 |
| 11953 | 598 | E598M | A | 2.3e-07 | 3.3e-07 | 3.5e-07 | 3.033E-07 | 6.429E-08 | 2.300E-07 | 3.500E-07 |
| 11954 | 598 | E598F | A | 2.1e-07 | 1.6e-07 | 2e-07 | 1.900E-07 | 2.646E-08 | 1.600E-07 | 2.100E-07 |
| 11955 | 598 | E598P | A | 4.2e-07 | 2.3e-07 | 1.1e-07 | 2.533E-07 | 1.563E-07 | 1.100E-07 | 4.200E-07 |
| 11956 | 598 | E598S | A | 3.2e-07 | 4.3e-07 | 7e-07 | 4.833E-07 | 1.955E-07 | 3.200E-07 | 7.000E-07 |
| 11957 | 598 | E598T | A | 3e-07 | 7e-07 | 4.4e-07 | 4.800E-07 | 2.030E-07 | 3.000E-07 | 7.000E-07 |
| 11958 | 598 | E598W | A | 2.7e-07 | 1.6e-07 | 1.1e-07 | 1.800E-07 | 8.185E-08 | 1.100E-07 | 2.700E-07 |
| 11959 | 598 | E598Y | A | 3.3e-07 | 2.2e-07 | 2.1e-07 | 2.533E-07 | 6.658E-08 | 2.100E-07 | 3.300E-07 |
| 11960 | 598 | E598V | A | 2.7e-07 | 2.6e-07 | 2.8e-07 | 2.700E-07 | 1.000E-08 | 2.600E-07 | 2.800E-07 |
| 11961 | 599 | G599A | A | 1.8e-07 | 3.7e-07 | 1.7e-07 | 2.400E-07 | 1.127E-07 | 1.700E-07 | 3.700E-07 |
| 11962 | 599 | G599R | A | 2.1e-07 | 2.3e-07 | 2.6e-07 | 2.333E-07 | 2.517E-08 | 2.100E-07 | 2.600E-07 |
| 11963 | 599 | G599N | A | 1.9e-07 | 2.2e-07 | 1.9e-07 | 2.000E-07 | 1.732E-08 | 1.900E-07 | 2.200E-07 |
| 11964 | 599 | G599D | A | 2.4e-07 | 2.6e-07 | 1.5e-07 | 2.167E-07 | 5.859E-08 | 1.500E-07 | 2.600E-07 |
| 11965 | 599 | G599C | A | 3.8e-07 | 3.1e-07 | 1.7e-07 | 2.867E-07 | 1.069E-07 | 1.700E-07 | 3.800E-07 |
| 11966 | 599 | G599E | A | 2e-07 | 1.4e-07 | 1.9e-07 | 1.767E-07 | 3.215E-08 | 1.400E-07 | 2.000E-07 |
| 11967 | 599 | G599Q | A | 1.6e-07 | 5.6e-07 | 1.8e-07 | 3.000E-07 | 2.254E-07 | 1.600E-07 | 5.600E-07 |
| 11968 | 599 | G599G | A | 2.1e-07 | 1.8e-07 | 2.7e-07 | 2.200E-07 | 4.583E-08 | 1.800E-07 | 2.700E-07 |
| 11969 | 599 | G599H | A | 2.1e-07 | 2.5e-07 | 3.1e-07 | 2.567E-07 | 5.033E-08 | 2.100E-07 | 3.100E-07 |
| 11970 | 599 | G599I | A | 1.3e-07 | 2.4e-07 | 2.1e-07 | 1.933E-07 | 5.686E-08 | 1.300E-07 | 2.400E-07 |
| 11971 | 599 | G599L | A | 1.7e-07 | 2.8e-07 | 2.8e-07 | 2.433E-07 | 6.351E-08 | 1.700E-07 | 2.800E-07 |
| 11972 | 599 | G599K | A | 1.8e-07 | 1.4e-07 | 1.1e-07 | 1.433E-07 | 3.512E-08 | 1.100E-07 | 1.800E-07 |
| 11973 | 599 | G599M | A | 1.2e-07 | 1.5e-07 | 2.7e-07 | 1.800E-07 | 7.937E-08 | 1.200E-07 | 2.700E-07 |
| 11974 | 599 | G599F | A | 2.6e-07 | 2.1e-07 | 1.9e-07 | 2.200E-07 | 3.606E-08 | 1.900E-07 | 2.600E-07 |
| 11975 | 599 | G599P | A | 1.1e-07 | 1.1e-07 | 2e-07 | 1.400E-07 | 5.196E-08 | 1.100E-07 | 2.000E-07 |
| 11976 | 599 | G599S | A | 2.6e-07 | 2.3e-07 | 2.8e-07 | 2.567E-07 | 2.517E-08 | 2.300E-07 | 2.800E-07 |
| 11977 | 599 | G599T | A | 2.5e-07 | 5.9e-07 | 5.4e-07 | 4.600E-07 | 1.836E-07 | 2.500E-07 | 5.900E-07 |
| 11978 | 599 | G599W | A | 6.5e-07 | 1.4e-07 | 3.3e-07 | 3.733E-07 | 2.577E-07 | 1.400E-07 | 6.500E-07 |
| 11979 | 599 | G599Y | A | 3.4e-07 | 2.4e-07 | 3.6e-07 | 3.133E-07 | 6.429E-08 | 2.400E-07 | 3.600E-07 |
| 11980 | 599 | G599V | A | 2.5e-07 | 6.4e-07 | 2.4e-07 | 3.767E-07 | 2.281E-07 | 2.400E-07 | 6.400E-07 |
| 11981 | 600 | A600A | A | 2.4e-07 | 4.3e-07 | 2.8e-07 | 3.167E-07 | 1.002E-07 | 2.400E-07 | 4.300E-07 |
| 11982 | 600 | A600R | A | 3.6e-07 | 2.1e-07 | 3.4e-07 | 3.033E-07 | 8.145E-08 | 2.100E-07 | 3.600E-07 |
| 11983 | 600 | A600N | A | 8.1e-07 | 3.9e-07 | 3.5e-07 | 5.167E-07 | 2.548E-07 | 3.500E-07 | 8.100E-07 |
| 11984 | 600 | A600D | A | 4.5e-07 | 6e-07 | 2.5e-07 | 4.333E-07 | 1.756E-07 | 2.500E-07 | 6.000E-07 |

| | | | | | | | | | | |
|---|---|---|---|---|---|---|---|---|---|---|
| 11985 | 600 | A600C | A | 2.5e-07 | 2.5e-07 | 4.4e-07 | 3.133E-07 | 1.097E-07 | 2.500E-07 | 4.400E-07 |
| 11986 | 600 | A600E | A | 5.7e-07 | 3.3e-07 | 4.5e-07 | 4.500E-07 | 1.200E-07 | 3.300E-07 | 5.700E-07 |
| 11987 | 600 | A600Q | A | 1.2e-06 | 4.2e-07 | 8.5e-07 | 8.233E-07 | 3.907E-07 | 4.200E-07 | 1.200E-06 |
| 11988 | 600 | A600G | A | 5e-07 | 6.3e-07 | 2.8e-07 | 4.700E-07 | 1.769E-07 | 2.800E-07 | 6.300E-07 |
| 11989 | 600 | A600H | A | 2.7e-07 | 3.8e-07 | 6e-07 | 4.167E-07 | 1.680E-07 | 2.700E-07 | 6.000E-07 |
| 11990 | 600 | A600I | A | 2.9e-07 | 1.7e-07 | 1.7e-07 | 2.100E-07 | 6.928E-08 | 1.700E-07 | 2.900E-07 |
| 11991 | 600 | A600L | A | 1.7e-07 | 1.6e-07 | 1.6e-07 | 1.633E-07 | 5.774E-09 | 1.600E-07 | 1.700E-07 |
| 11992 | 600 | A600K | A | 7.6e-07 | 2e-07 | 4.3e-07 | 4.633E-07 | 2.815E-07 | 2.000E-07 | 7.600E-07 |
| 11993 | 600 | A600M | A | 4e-07 | 3.8e-07 | 4.8e-07 | 4.200E-07 | 5.292E-08 | 3.800E-07 | 4.800E-07 |
| 11994 | 600 | A600F | A | 2e-07 | 2.3e-07 | 4.1e-07 | 2.800E-07 | 1.136E-07 | 2.000E-07 | 4.100E-07 |
| 11995 | 600 | A600P | A | 1.7e-07 | 1.1e-07 | 1.1e-07 | 1.300E-07 | 3.464E-08 | 1.100E-07 | 1.700E-07 |
| 11996 | 600 | A600S | A | 1.7e-06 | 4.3e-07 | 2.5e-06 | 1.543E-06 | 1.044E-06 | 4.300E-07 | 2.500E-06 |
| 11997 | 600 | A600T | A | 1.2e-06 | 7.7e-07 | 4.9e-07 | 8.200E-07 | 3.576E-07 | 4.900E-07 | 1.200E-06 |
| 11998 | 600 | A600W | A | 2.7e-07 | 4.7e-07 | 1.4e-07 | 2.933E-07 | 1.662E-07 | 1.400E-07 | 4.700E-07 |
| 11999 | 600 | A600Y | A | 3.2e-07 | 2.9e-07 | 4.9e-07 | 3.667E-07 | 1.079E-07 | 2.900E-07 | 4.900E-07 |
| 12000 | 600 | A600V | A | 2.1e-07 | 1.7e-07 | 2.1e-07 | 1.967E-07 | 2.309E-08 | 1.700E-07 | 2.100E-07 |
| 12001 | 601 | C601A | A | 1.8e-07 | 1.5e-07 | 3.1e-07 | 2.133E-07 | 8.505E-08 | 1.500E-07 | 3.100E-07 |
| 12002 | 601 | C601R | A | 1e-07 | 1e-07 | 1.1e-07 | 1.033E-07 | 5.774E-09 | 1.000E-07 | 1.100E-07 |
| 12003 | 601 | C601N | A | 2e-07 | 1.3e-07 | 3.9e-07 | 2.400E-07 | 1.345E-07 | 1.300E-07 | 3.900E-07 |
| 12004 | 601 | C601D | A | 2e-07 | 4.5e-07 | 2.6e-07 | 3.033E-07 | 1.305E-07 | 2.000E-07 | 4.500E-07 |
| 12005 | 601 | C601C | A | 1.8e-07 | 3e-07 | 3.1e-07 | 2.633E-07 | 7.234E-08 | 1.800E-07 | 3.100E-07 |
| 12006 | 601 | C601E | A | 1.9e-07 | 1.1e-07 | 2.7e-07 | 1.900E-07 | 8.000E-08 | 1.100E-07 | 2.700E-07 |
| 12007 | 601 | C601Q | A | 3.3e-07 | 1.3e-07 | 8.8e-08 | 1.827E-07 | 1.293E-07 | 8.800E-08 | 3.300E-07 |
| 12008 | 601 | C601G | A | 3.1e-07 | 3e-07 | 1.3e-07 | 2.467E-07 | 1.012E-07 | 1.300E-07 | 3.100E-07 |
| 12009 | 601 | C601H | A | 1.3e-07 | 1.4e-07 | 1.6e-07 | 1.433E-07 | 1.528E-08 | 1.300E-07 | 1.600E-07 |
| 12010 | 601 | C601I | A | 1.6e-07 | 4e-07 | 3.4e-07 | 3.000E-07 | 1.249E-07 | 1.600E-07 | 4.000E-07 |
| 12011 | 601 | C601L | A | 3.1e-07 | 4e-07 | 2.9e-07 | 3.333E-07 | 5.859E-08 | 2.900E-07 | 4.000E-07 |
| 12012 | 601 | C601K | A | 2.9e-07 | 2.4e-07 | 2.2e-07 | 2.500E-07 | 3.606E-08 | 2.200E-07 | 2.900E-07 |
| 12013 | 601 | C601M | A | 2.1e-07 | 2.7e-07 | 3.1e-07 | 2.633E-07 | 5.033E-08 | 2.100E-07 | 3.100E-07 |
| 12014 | 601 | C601F | A | 1.6e-07 | 2.3e-07 | 4.4e-07 | 2.767E-07 | 1.457E-07 | 1.600E-07 | 4.400E-07 |
| 12015 | 601 | C601P | A | 2.5e-07 | 4.6e-07 | 2.5e-07 | 3.200E-07 | 1.212E-07 | 2.500E-07 | 4.600E-07 |
| 12016 | 601 | C601S | A | 2.9e-07 | 1.3e-07 | 3.5e-07 | 2.567E-07 | 1.137E-07 | 1.300E-07 | 3.500E-07 |
| 12017 | 601 | C601T | A | 3.5e-07 | 3.6e-07 | 2.8e-07 | 3.300E-07 | 4.359E-08 | 2.800E-07 | 3.600E-07 |
| 12018 | 601 | C601W | A | 1.1e-07 | 5.4e-07 | 1.7e-07 | 2.733E-07 | 2.329E-07 | 1.100E-07 | 5.400E-07 |
| 12019 | 601 | C601Y | A | 1.3e-07 | 2.3e-07 | 1.3e-07 | 1.633E-07 | 5.774E-08 | 1.300E-07 | 2.300E-07 |
| 12020 | 601 | C601V | A | 2.8e-07 | 2.4e-07 | 1.7e-07 | 2.300E-07 | 5.568E-08 | 1.700E-07 | 2.800E-07 |
| 12021 | 602 | Q602A | A | 2.4e-07 | 2.1e-07 | 7.2e-08 | 1.740E-07 | 8.960E-08 | 7.200E-08 | 2.400E-07 |
| 12022 | 602 | Q602R | A | 6.9e-08 | 1.8e-07 | 1e-07 | 1.163E-07 | 5.727E-08 | 6.900E-08 | 1.800E-07 |
| 12023 | 602 | Q602N | A | 2.8e-07 | 2.5e-07 | 3.9e-07 | 3.067E-07 | 7.371E-08 | 2.500E-07 | 3.900E-07 |
| 12024 | 602 | Q602D | A | 1.4e-07 | 9.9e-08 | 1.2e-07 | 1.197E-07 | 2.050E-08 | 9.900E-08 | 1.400E-07 |
| 12025 | 602 | Q602C | A | 5.6e-08 | 8.8e-08 | 1.6e-07 | 1.013E-07 | 5.327E-08 | 5.600E-08 | 1.600E-07 |
| 12026 | 602 | Q602E | A | 1.8e-07 | 1.3e-07 | 1.6e-07 | 1.567E-07 | 2.517E-08 | 1.300E-07 | 1.800E-07 |
| 12027 | 602 | Q602Q | A | 7.8e-07 | 1.6e-07 | 5.2e-07 | 4.867E-07 | 3.113E-07 | 1.600E-07 | 7.800E-07 |
| 12028 | 602 | Q602G | A | 1e-07 | 6.7e-08 | 1.6e-07 | 1.090E-07 | 4.715E-08 | 6.700E-08 | 1.600E-07 |
| 12029 | 602 | Q602H | A | 1.4e-07 | 1.1e-07 | 2.2e-07 | 1.567E-07 | 5.686E-08 | 1.100E-07 | 2.200E-07 |
| 12030 | 602 | Q602I | A | 7.4e-08 | 6.2e-08 | 9.3e-08 | 7.633E-08 | 1.563E-08 | 6.200E-08 | 9.300E-08 |
| 12031 | 602 | Q602L | A | 1.2e-07 | 1.1e-07 | 1.1e-07 | 1.133E-07 | 5.774E-09 | 1.100E-07 | 1.200E-07 |
| 12032 | 602 | Q602K | A | 6.8e-08 | 9.8e-08 | 2e-07 | 1.220E-07 | 6.920E-08 | 6.800E-08 | 2.000E-07 |
| 12033 | 602 | Q602M | A | 2.1e-07 | 7.1e-08 | 1.6e-07 | 1.470E-07 | 7.041E-08 | 7.100E-08 | 2.100E-07 |
| 12034 | 602 | Q602F | A | 3.4e-08 | 1.7e-08 | 5e-08 | 3.367E-08 | 1.650E-08 | 1.700E-08 | 5.000E-08 |
| 12035 | 602 | Q602P | A | 1.7e-07 | 1.1e-07 | 8.2e-08 | 1.207E-07 | 4.496E-08 | 8.200E-08 | 1.700E-07 |
| 12036 | 602 | Q602S | A | 1.7e-07 | 1.4e-07 | 1.1e-07 | 1.400E-07 | 3.000E-08 | 1.100E-07 | 1.700E-07 |
| 12037 | 602 | Q602T | A | 9.6e-08 | 1.1e-07 | 9.2e-08 | 9.933E-08 | 9.452E-09 | 9.200E-08 | 1.100E-07 |
| 12038 | 602 | Q602W | A | 9.3e-08 | 4.1e-08 | 2.3e-08 | 5.233E-08 | 3.635E-08 | 2.300E-08 | 9.300E-08 |
| 12039 | 602 | Q602Y | A | 3.8e-08 | 8.1e-08 | 4.9e-08 | 5.600E-08 | 2.234E-08 | 3.800E-08 | 8.100E-08 |
| 12040 | 602 | Q602V | A | 9.6e-08 | 1.4e-07 | 1.2e-07 | 1.187E-07 | 2.203E-08 | 9.600E-08 | 1.400E-07 |
| 12041 | 603 | P603A | A | 4.6e-07 | 3.4e-07 | 1.9e-07 | 3.300E-07 | 1.353E-07 | 1.900E-07 | 4.600E-07 |
| 12042 | 603 | P603R | A | 2.8e-07 | 2.1e-07 | 9.1e-08 | 1.937E-07 | 9.555E-08 | 9.100E-08 | 2.800E-07 |
| 12043 | 603 | P603N | A | 2.4e-07 | 1e-07 | 2.9e-07 | 2.100E-07 | 9.849E-08 | 1.000E-07 | 2.900E-07 |
| 12044 | 603 | P603D | A | 2.4e-07 | 3.7e-07 | 3.7e-07 | 3.267E-07 | 7.506E-08 | 2.400E-07 | 3.700E-07 |
| 12045 | 603 | P603C | A | 1.6e-07 | 1.5e-07 | 2.2e-07 | 1.767E-07 | 3.786E-08 | 1.500E-07 | 2.200E-07 |
| 12046 | 603 | P603E | A | 1.2e-07 | 2.9e-07 | 2.1e-07 | 2.067E-07 | 8.505E-08 | 1.200E-07 | 2.900E-07 |
| 12047 | 603 | P603Q | A | 4.1e-07 | 1.6e-07 | 1.7e-07 | 2.467E-07 | 1.415E-07 | 1.600E-07 | 4.100E-07 |
| 12048 | 603 | P603G | A | 3.2e-07 | 4.6e-07 | 5.7e-07 | 4.500E-07 | 1.253E-07 | 3.200E-07 | 5.700E-07 |
| 12049 | 603 | P603H | A | 4e-07 | 3.6e-07 | 6.8e-07 | 4.800E-07 | 1.744E-07 | 3.600E-07 | 6.800E-07 |
| 12050 | 603 | P603I | A | 2.1e-07 | 1.3e-07 | 3.4e-07 | 2.267E-07 | 1.060E-07 | 1.300E-07 | 3.400E-07 |
| 12051 | 603 | P603L | A | 2.9e-07 | 2.3e-07 | 4.5e-07 | 3.233E-07 | 1.137E-07 | 2.300E-07 | 4.500E-07 |
| 12052 | 603 | P603K | A | 3e-07 | 2e-07 | 2.3e-07 | 2.433E-07 | 5.132E-08 | 2.000E-07 | 3.000E-07 |
| 12053 | 603 | P603M | A | 1e-07 | 4.2e-07 | 1.3e-07 | 2.167E-07 | 1.767E-07 | 1.000E-07 | 4.200E-07 |
| 12054 | 603 | P603F | A | 1.9e-07 | 2.9e-07 | 5.7e-07 | 3.500E-07 | 1.970E-07 | 1.900E-07 | 5.700E-07 |
| 12055 | 603 | P603P | A | 8.1e-08 | 1.9e-07 | 1.3e-07 | 1.337E-07 | 5.459E-08 | 8.100E-08 | 1.900E-07 |
| 12056 | 603 | P603S | A | 3e-07 | 3.8e-07 | 3.4e-07 | 3.400E-07 | 4.000E-08 | 3.000E-07 | 3.800E-07 |
| 12057 | 603 | P603T | A | 1.5e-07 | 2.8e-07 | 2.2e-07 | 2.167E-07 | 6.506E-08 | 1.500E-07 | 2.800E-07 |
| 12058 | 603 | P603W | A | 2.7e-07 | 1.9e-07 | 2.5e-07 | 2.367E-07 | 4.163E-08 | 1.900E-07 | 2.700E-07 |
| 12059 | 603 | P603Y | A | 1.3e-07 | 1.3e-07 | 2.2e-07 | 1.600E-07 | 5.196E-08 | 1.300E-07 | 2.200E-07 |
| 12060 | 603 | P603V | A | 2.2e-07 | 2.9e-07 | 2.8e-07 | 2.633E-07 | 3.786E-08 | 2.200E-07 | 2.900E-07 |
| 12061 | 604 | C604A | A | 3.2e-07 | 1.9e-07 | 2.3e-07 | 2.467E-07 | 6.658E-08 | 1.900E-07 | 3.200E-07 |
| 12062 | 604 | C604R | A | 2.4e-07 | 4.1e-07 | 2.2e-07 | 2.900E-07 | 1.044E-07 | 2.200E-07 | 4.100E-07 |
| 12063 | 604 | C604N | A | 2.1e-07 | 1.9e-07 | 2.5e-07 | 2.167E-07 | 3.055E-08 | 1.900E-07 | 2.500E-07 |
| 12064 | 604 | C604D | A | 2.2e-07 | 2.2e-07 | 2.5e-07 | 2.300E-07 | 1.732E-08 | 2.200E-07 | 2.500E-07 |
| 12065 | 604 | C604C | A | 2.5e-07 | 1.7e-07 | 3.1e-07 | 2.433E-07 | 7.024E-08 | 1.700E-07 | 3.100E-07 |
| 12066 | 604 | C604E | A | 2.9e-07 | 4e-07 | 1.2e-07 | 2.700E-07 | 1.411E-07 | 1.200E-07 | 4.000E-07 |
| 12067 | 604 | C604Q | A | 1.5e-07 | 1.7e-07 | 1.8e-07 | 1.667E-07 | 1.528E-08 | 1.500E-07 | 1.800E-07 |
| 12068 | 604 | C604G | A | 4e-07 | 2.4e-07 | 2.1e-07 | 2.833E-07 | 1.021E-07 | 2.100E-07 | 4.000E-07 |
| 12069 | 604 | C604H | A | 1.7e-07 | 3.9e-07 | 2.1e-07 | 2.567E-07 | 1.172E-07 | 1.700E-07 | 3.900E-07 |

| | | | | | | | | | | |
|---|---|---|---|---|---|---|---|---|---|---|
| 12070 | 604 | C604I | A | 2.7e-07 | 2.4e-07 | 4.6e-07 | 3.233E-07 | 1.193E-07 | 2.400E-07 | 4.600E-07 |
| 12071 | 604 | C604L | A | 4.9e-07 | 4.4e-07 | 2.2e-07 | 3.833E-07 | 1.436E-07 | 2.200E-07 | 4.900E-07 |
| 12072 | 604 | C604K | A | 1.7e-07 | 4.1e-07 | 4.3e-07 | 3.367E-07 | 1.447E-07 | 1.700E-07 | 4.300E-07 |
| 12073 | 604 | C604M | A | 1.2e-07 | 2.5e-07 | 1.6e-07 | 1.767E-07 | 6.658E-08 | 1.200E-07 | 2.500E-07 |
| 12074 | 604 | C604F | A | 4.1e-07 | 7e-08 | 3.1e-07 | 2.633E-07 | 1.747E-07 | 7.000E-08 | 4.100E-07 |
| 12075 | 604 | C604P | A | 4.1e-07 | 2.7e-07 | 2.5e-07 | 3.100E-07 | 8.718E-08 | 2.500E-07 | 4.100E-07 |
| 12076 | 604 | C604S | A | 1.6e-07 | 2e-07 | 4.7e-07 | 2.767E-07 | 1.686E-07 | 1.600E-07 | 4.700E-07 |
| 12077 | 604 | C604T | A | 2.4e-07 | 1.5e-07 | 2.3e-07 | 2.067E-07 | 4.933E-08 | 1.500E-07 | 2.400E-07 |
| 12078 | 604 | C604W | A | 4.2e-07 | 3.1e-07 | 2.8e-07 | 3.367E-07 | 7.371E-08 | 2.800E-07 | 4.200E-07 |
| 12079 | 604 | C604Y | A | 3.1e-07 | 1.5e-07 | 3e-07 | 2.533E-07 | 8.963E-08 | 1.500E-07 | 3.100E-07 |
| 12080 | 604 | C604V | A | 3.5e-07 | 1.1e-07 | 2.8e-07 | 2.467E-07 | 1.234E-07 | 1.100E-07 | 3.500E-07 |
| 12081 | 605 | P605A | A | 2e-07 | 2.2e-07 | 1.1e-07 | 1.767E-07 | 5.859E-08 | 1.100E-07 | 2.200E-07 |
| 12082 | 605 | P605R | A | 3.3e-07 | 2e-07 | 2.3e-07 | 2.533E-07 | 6.807E-08 | 2.000E-07 | 3.300E-07 |
| 12083 | 605 | P605N | A | 1.5e-07 | 2.5e-07 | 2.3e-07 | 2.100E-07 | 5.292E-08 | 1.500E-07 | 2.500E-07 |
| 12084 | 605 | P605D | A | 2.4e-07 | 2.6e-07 | 1.6e-07 | 2.200E-07 | 5.292E-08 | 1.600E-07 | 2.600E-07 |
| 12085 | 605 | P605C | A | 1.8e-07 | 3.9e-07 | 2.6e-07 | 2.767E-07 | 1.060E-07 | 1.800E-07 | 3.900E-07 |
| 12086 | 605 | P605E | A | 2.8e-07 | 7.4e-07 | 3.5e-07 | 4.567E-07 | 2.479E-07 | 2.800E-07 | 7.400E-07 |
| 12087 | 605 | P605Q | A | 2e-07 | 1.7e-07 | 2.3e-07 | 2.000E-07 | 3.000E-08 | 1.700E-07 | 2.300E-07 |
| 12088 | 605 | P605G | A | 1.8e-07 | 1.8e-07 | 4e-07 | 2.533E-07 | 1.270E-07 | 1.800E-07 | 4.000E-07 |
| 12089 | 605 | P605H | A | 2.2e-07 | 2.8e-07 | 8.6e-08 | 1.953E-07 | 9.932E-08 | 8.600E-08 | 2.800E-07 |
| 12090 | 605 | P605I | A | 9e-08 | 3.8e-07 | 2e-07 | 2.233E-07 | 1.464E-07 | 9.000E-08 | 3.800E-07 |
| 12091 | 605 | P605L | A | 3.4e-07 | 5.2e-07 | 1.9e-07 | 3.500E-07 | 1.652E-07 | 1.900E-07 | 5.200E-07 |
| 12092 | 605 | P605K | A | 2.5e-07 | 5.5e-07 | 2.1e-07 | 3.367E-07 | 1.858E-07 | 2.100E-07 | 5.500E-07 |
| 12093 | 605 | P605M | A | 2.4e-07 | 2.5e-07 | 1.4e-07 | 2.100E-07 | 6.083E-08 | 1.400E-07 | 2.500E-07 |
| 12094 | 605 | P605F | A | 2.1e-07 | 1.2e-07 | 3.5e-07 | 2.267E-07 | 1.159E-07 | 1.200E-07 | 3.500E-07 |
| 12095 | 605 | P605P | A | 2.3e-07 | 2.2e-07 | 3e-07 | 2.500E-07 | 4.359E-08 | 2.200E-07 | 3.000E-07 |
| 12096 | 605 | P605S | A | 1.8e-07 | 2.3e-07 | 1.6e-07 | 1.900E-07 | 3.606E-08 | 1.600E-07 | 2.300E-07 |
| 12097 | 605 | P605T | A | 1.5e-07 | 3.8e-07 | 9e-08 | 2.067E-07 | 1.531E-07 | 9.000E-08 | 3.800E-07 |
| 12098 | 605 | P605W | A | 2.7e-07 | 1.8e-07 | 5.1e-07 | 3.200E-07 | 1.706E-07 | 1.800E-07 | 5.100E-07 |
| 12099 | 605 | P605Y | A | 1.9e-07 | 2.7e-07 | 1.4e-07 | 2.000E-07 | 6.557E-08 | 1.400E-07 | 2.700E-07 |
| 12100 | 605 | P605V | A | 1.8e-07 | 2e-07 | 2.6e-07 | 2.133E-07 | 4.163E-08 | 1.800E-07 | 2.600E-07 |
| 12101 | 606 | I606A | A | 8.6e-08 | 1.6e-07 | 1.7e-07 | 1.387E-07 | 4.588E-08 | 8.600E-08 | 1.700E-07 |
| 12102 | 606 | I606R | A | 2.2e-07 | 3.5e-07 | 1.8e-07 | 2.500E-07 | 8.888E-08 | 1.800E-07 | 3.500E-07 |
| 12103 | 606 | I606N | A | 1.6e-07 | 1.2e-07 | 1.3e-07 | 1.367E-07 | 2.082E-08 | 1.200E-07 | 1.600E-07 |
| 12104 | 606 | I606D | A | 1.7e-07 | 2.1e-07 | 2.6e-07 | 2.133E-07 | 4.509E-08 | 1.700E-07 | 2.600E-07 |
| 12105 | 606 | I606C | A | 3.2e-07 | 1.4e-07 | 2.8e-07 | 2.467E-07 | 9.452E-08 | 1.400E-07 | 3.200E-07 |
| 12106 | 606 | I606E | A | 2.3e-07 | 1.4e-07 | 1.8e-07 | 1.833E-07 | 4.509E-08 | 1.400E-07 | 2.300E-07 |
| 12107 | 606 | I606Q | A | 3.8e-07 | 1.5e-07 | 1.7e-07 | 2.333E-07 | 1.274E-07 | 1.500E-07 | 3.800E-07 |
| 12108 | 606 | I606G | A | 2.2e-07 | 2.8e-07 | 4e-07 | 3.000E-07 | 9.165E-08 | 2.200E-07 | 4.000E-07 |
| 12109 | 606 | I606H | A | 2.5e-07 | 1.4e-07 | 1.5e-07 | 1.800E-07 | 6.083E-08 | 1.400E-07 | 2.500E-07 |
| 12110 | 606 | I606I | A | 1.7e-07 | 7.4e-07 | 2.5e-07 | 3.867E-07 | 3.086E-07 | 1.700E-07 | 7.400E-07 |
| 12111 | 606 | I606L | A | 2.5e-07 | 3.8e-07 | 1.7e-07 | 2.667E-07 | 1.060E-07 | 1.700E-07 | 3.800E-07 |
| 12112 | 606 | I606K | A | 4.7e-07 | 2.9e-07 | 1.6e-07 | 3.067E-07 | 1.557E-07 | 1.600E-07 | 4.700E-07 |
| 12113 | 606 | I606M | A | 1.9e-07 | 4.4e-07 | 3e-07 | 3.100E-07 | 1.253E-07 | 1.900E-07 | 4.400E-07 |
| 12114 | 606 | I606F | A | 2.6e-07 | 4.1e-07 | 2e-07 | 2.900E-07 | 1.082E-07 | 2.000E-07 | 4.100E-07 |
| 12115 | 606 | I606P | A | 3e-07 | 2.3e-07 | 1.9e-07 | 2.400E-07 | 5.568E-08 | 1.900E-07 | 3.000E-07 |
| 12116 | 606 | I606S | A | 1.1e-07 | 3.9e-07 | 3.5e-07 | 2.833E-07 | 1.514E-07 | 1.100E-07 | 3.900E-07 |
| 12117 | 606 | I606T | A | 2.5e-07 | 1.6e-07 | 1.5e-07 | 1.867E-07 | 5.508E-08 | 1.500E-07 | 2.500E-07 |
| 12118 | 606 | I606W | A | 2.5e-07 | 2.5e-07 | 2.2e-07 | 2.400E-07 | 1.732E-08 | 2.200E-07 | 2.500E-07 |
| 12119 | 606 | I606Y | A | 1.7e-07 | 1.3e-07 | 5.4e-07 | 2.800E-07 | 2.261E-07 | 1.300E-07 | 5.400E-07 |
| 12120 | 606 | I606V | A | 2.3e-07 | 1.4e-07 | 1.3e-07 | 1.667E-07 | 5.508E-08 | 1.300E-07 | 2.300E-07 |
| 12121 | 607 | N607A | A | 2.8e-07 | 2.7e-07 | 3.1e-07 | 2.867E-07 | 2.082E-08 | 2.700E-07 | 3.100E-07 |
| 12122 | 607 | N607R | A | 2.5e-07 | 1.7e-07 | 2.9e-07 | 2.367E-07 | 6.110E-08 | 1.700E-07 | 2.900E-07 |
| 12123 | 607 | N607N | A | 3.2e-07 | 1e-07 | 3.5e-07 | 2.567E-07 | 1.365E-07 | 1.000E-07 | 3.500E-07 |
| 12124 | 607 | N607D | A | 3.5e-07 | 2.6e-07 | 2.3e-07 | 2.800E-07 | 6.245E-08 | 2.300E-07 | 3.500E-07 |
| 12125 | 607 | N607C | A | 6.9e-07 | 1.8e-07 | 3e-07 | 3.900E-07 | 2.666E-07 | 1.800E-07 | 6.900E-07 |
| 12126 | 607 | N607E | A | 1.6e-07 | 1.6e-07 | 2.5e-07 | 1.900E-07 | 5.196E-08 | 1.600E-07 | 2.500E-07 |
| 12127 | 607 | N607Q | A | 5.9e-07 | 2.4e-07 | 2.5e-07 | 3.600E-07 | 1.992E-07 | 2.400E-07 | 5.900E-07 |
| 12128 | 607 | N607G | A | 1.8e-07 | 2.2e-07 | 4.5e-07 | 2.833E-07 | 1.457E-07 | 1.800E-07 | 4.500E-07 |
| 12129 | 607 | N607H | A | 1.4e-07 | 1.3e-07 | 3.2e-07 | 1.967E-07 | 1.069E-07 | 1.300E-07 | 3.200E-07 |
| 12130 | 607 | N607I | A | 7.6e-07 | 2.1e-07 | 2.2e-07 | 3.967E-07 | 3.147E-07 | 2.100E-07 | 7.600E-07 |
| 12131 | 607 | N607L | A | 1e-07 | 1.7e-07 | 2.4e-07 | 1.700E-07 | 7.000E-08 | 1.000E-07 | 2.400E-07 |
| 12132 | 607 | N607K | A | 1.9e-07 | 3.2e-07 | 4e-07 | 3.033E-07 | 1.060E-07 | 1.900E-07 | 4.000E-07 |
| 12133 | 607 | N607M | A | 3.2e-07 | 1.9e-07 | 2.3e-07 | 2.467E-07 | 6.658E-08 | 1.900E-07 | 3.200E-07 |
| 12134 | 607 | N607F | A | 2.7e-07 | 1.8e-07 | 2.2e-07 | 2.233E-07 | 4.509E-08 | 1.800E-07 | 2.700E-07 |
| 12135 | 607 | N607P | A | 2.1e-07 | 1.1e-07 | 2.9e-07 | 2.033E-07 | 9.018E-08 | 1.100E-07 | 2.900E-07 |
| 12136 | 607 | N607S | A | 2.3e-07 | 4.7e-07 | 2.5e-07 | 3.167E-07 | 1.332E-07 | 2.300E-07 | 4.700E-07 |
| 12137 | 607 | N607T | A | 2.5e-07 | 4.8e-07 | 2.1e-07 | 3.133E-07 | 1.457E-07 | 2.100E-07 | 4.800E-07 |
| 12138 | 607 | N607W | A | 1.6e-07 | 2.3e-07 | 2.2e-07 | 2.033E-07 | 3.786E-08 | 1.600E-07 | 2.300E-07 |
| 12139 | 607 | N607Y | A | 2.5e-07 | 2.4e-07 | 2.4e-07 | 2.433E-07 | 5.774E-09 | 2.400E-07 | 2.500E-07 |
| 12140 | 607 | N607V | A | 3.5e-07 | 1.6e-07 | 5.1e-07 | 3.400E-07 | 1.752E-07 | 1.600E-07 | 5.100E-07 |
| 12141 | 608 | C608A | A | 4.7e-07 | 1.7e-07 | 2.6e-07 | 3.000E-07 | 1.539E-07 | 1.700E-07 | 4.700E-07 |
| 12142 | 608 | C608R | A | 1.2e-07 | 3e-07 | 2.1e-07 | 2.100E-07 | 9.000E-08 | 1.200E-07 | 3.000E-07 |
| 12143 | 608 | C608N | A | 1.7e-07 | 2.5e-07 | 2.2e-07 | 2.133E-07 | 4.041E-08 | 1.700E-07 | 2.500E-07 |
| 12144 | 608 | C608D | A | 1.8e-07 | 1.4e-07 | 3.7e-07 | 2.300E-07 | 1.229E-07 | 1.400E-07 | 3.700E-07 |
| 12145 | 608 | C608C | A | 1.9e-07 | 1.7e-07 | 3.1e-07 | 2.233E-07 | 7.572E-08 | 1.700E-07 | 3.100E-07 |
| 12146 | 608 | C608E | A | 3.2e-07 | 3e-07 | 2.8e-07 | 3.000E-07 | 2.000E-08 | 2.800E-07 | 3.200E-07 |
| 12147 | 608 | C608Q | A | 5.2e-07 | 4.8e-07 | 5.3e-07 | 5.100E-07 | 2.646E-08 | 4.800E-07 | 5.300E-07 |
| 12148 | 608 | C608G | A | 2.5e-07 | 3e-07 | 2.4e-07 | 2.633E-07 | 3.215E-08 | 2.400E-07 | 3.000E-07 |
| 12149 | 608 | C608H | A | 1.4e-07 | 9.5e-08 | 2.1e-07 | 1.483E-07 | 5.795E-08 | 9.500E-08 | 2.100E-07 |
| 12150 | 608 | C608I | A | 4e-07 | 4.2e-07 | 2.7e-07 | 3.633E-07 | 8.145E-08 | 2.700E-07 | 4.200E-07 |
| 12151 | 608 | C608L | A | 1.9e-07 | 2.3e-07 | 4.4e-07 | 2.867E-07 | 1.343E-07 | 1.900E-07 | 4.400E-07 |
| 12152 | 608 | C608K | A | 2.8e-07 | 3.5e-07 | 3.6e-07 | 3.300E-07 | 4.359E-08 | 2.800E-07 | 3.600E-07 |
| 12153 | 608 | C608M | A | 2.2e-07 | 2.1e-07 | 1.8e-07 | 2.033E-07 | 2.082E-08 | 1.800E-07 | 2.200E-07 |
| 12154 | 608 | C608F | A | 1.9e-07 | 3.1e-07 | 3.4e-07 | 2.800E-07 | 7.937E-08 | 1.900E-07 | 3.400E-07 |

| | | | | | | | | | | |
|---|---|---|---|---|---|---|---|---|---|---|
| 12155 | 608 | C608P | A | 1.1e-07 | 2.6e-07 | 2e-07 | 1.900E-07 | 7.550E-08 | 1.100E-07 | 2.600E-07 |
| 12156 | 608 | C608S | A | 2.7e-07 | 2.9e-07 | 1.7e-07 | 2.433E-07 | 6.429E-08 | 1.700E-07 | 2.900E-07 |
| 12157 | 608 | C608T | A | 2e-07 | 3.7e-07 | 8.9e-07 | 4.867E-07 | 3.595E-07 | 2.000E-07 | 8.900E-07 |
| 12158 | 608 | C608W | A | 2.7e-07 | 1.5e-07 | 3.5e-07 | 2.567E-07 | 1.007E-07 | 1.500E-07 | 3.500E-07 |
| 12159 | 608 | C608Y | A | 4.7e-07 | 1.4e-07 | 3e-07 | 3.033E-07 | 1.650E-07 | 1.400E-07 | 4.700E-07 |
| 12160 | 608 | C608V | A | 3.1e-07 | 1.5e-07 | 2.4e-07 | 2.333E-07 | 8.021E-08 | 1.500E-07 | 3.100E-07 |
| 12161 | 609 | T609A | A | 1.8e-07 | 1.3e-07 | 1.8e-07 | 1.633E-07 | 2.887E-08 | 1.300E-07 | 1.800E-07 |
| 12162 | 609 | T609R | A | 2.8e-07 | 2.3e-07 | 2.5e-07 | 2.533E-07 | 2.517E-08 | 2.300E-07 | 2.800E-07 |
| 12163 | 609 | T609N | A | 3.3e-07 | 4.6e-07 | 4.1e-07 | 4.000E-07 | 6.557E-08 | 3.300E-07 | 4.600E-07 |
| 12164 | 609 | T609D | A | 1.5e-07 | 1.3e-07 | 1.6e-07 | 1.467E-07 | 1.528E-08 | 1.300E-07 | 1.600E-07 |
| 12165 | 609 | T609C | A | 4e-07 | 4.1e-07 | 1.5e-07 | 3.200E-07 | 1.473E-07 | 1.500E-07 | 4.100E-07 |
| 12166 | 609 | T609E | A | 6.9e-07 | 1.9e-07 | 2.7e-07 | 3.833E-07 | 2.686E-07 | 1.900E-07 | 6.900E-07 |
| 12167 | 609 | T609Q | A | 3.7e-07 | 7.1e-07 | 3.2e-07 | 4.667E-07 | 2.122E-07 | 3.200E-07 | 7.100E-07 |
| 12168 | 609 | T609G | A | 4.2e-07 | 4.1e-07 | 3.7e-07 | 4.000E-07 | 2.646E-08 | 3.700E-07 | 4.200E-07 |
| 12169 | 609 | T609H | A | 3.7e-07 | 3.2e-07 | 2.3e-07 | 3.067E-07 | 7.095E-08 | 2.300E-07 | 3.700E-07 |
| 12170 | 609 | T609I | A | 7.5e-07 | 3.1e-07 | 2.1e-07 | 4.233E-07 | 2.873E-07 | 2.100E-07 | 7.500E-07 |
| 12171 | 609 | T609L | A | 1.4e-07 | 1.2e-07 | 2.3e-07 | 1.633E-07 | 5.859E-08 | 1.200E-07 | 2.300E-07 |
| 12172 | 609 | T609K | A | 3.1e-07 | 2.4e-07 | 2.4e-07 | 2.633E-07 | 4.041E-08 | 2.400E-07 | 3.100E-07 |
| 12173 | 609 | T609M | A | 2e-07 | 1.7e-07 | 2.5e-07 | 2.067E-07 | 4.041E-08 | 1.700E-07 | 2.500E-07 |
| 12174 | 609 | T609F | A | 1.1e-07 | 2.6e-07 | 1.5e-07 | 1.733E-07 | 7.767E-08 | 1.100E-07 | 2.600E-07 |
| 12175 | 609 | T609P | A | 3.3e-07 | 1.9e-07 | 4e-07 | 3.067E-07 | 1.069E-07 | 1.900E-07 | 4.000E-07 |
| 12176 | 609 | T609S | A | 3.4e-07 | 1.9e-07 | 2.7e-07 | 2.667E-07 | 7.506E-08 | 1.900E-07 | 3.400E-07 |
| 12177 | 609 | T609T | A | 2.4e-07 | 2.4e-07 | 3.6e-07 | 2.800E-07 | 6.928E-08 | 2.400E-07 | 3.600E-07 |
| 12178 | 609 | T609W | A | 3.3e-07 | 1.4e-07 | 3.1e-07 | 2.600E-07 | 1.044E-07 | 1.400E-07 | 3.300E-07 |
| 12179 | 609 | T609Y | A | 2.9e-07 | 3.4e-07 | 2.7e-07 | 3.000E-07 | 3.606E-08 | 2.700E-07 | 3.400E-07 |
| 12180 | 609 | T609V | A | 1.9e-07 | 2.6e-07 | 1.2e-07 | 1.900E-07 | 7.000E-08 | 1.200E-07 | 2.600E-07 |
| 12181 | 610 | H610A | A | 1.7e-07 | 4.9e-07 | 2e-07 | 2.867E-07 | 1.767E-07 | 1.700E-07 | 4.900E-07 |
| 12182 | 610 | H610R | A | 1.6e-07 | 2.6e-07 | 3.2e-07 | 2.467E-07 | 8.083E-08 | 1.600E-07 | 3.200E-07 |
| 12183 | 610 | H610N | A | 2.4e-07 | 3.7e-07 | 2.2e-07 | 2.767E-07 | 8.145E-08 | 2.200E-07 | 3.700E-07 |
| 12184 | 610 | H610D | A | 4.2e-07 | 1.2e-07 | 1.8e-07 | 2.400E-07 | 1.587E-07 | 1.200E-07 | 4.200E-07 |
| 12185 | 610 | H610C | A | 3e-07 | 3.5e-07 | 2.1e-07 | 2.867E-07 | 7.095E-08 | 2.100E-07 | 3.500E-07 |
| 12186 | 610 | H610E | A | 1.4e-07 | 2.7e-07 | 5.4e-07 | 3.167E-07 | 2.040E-07 | 1.400E-07 | 5.400E-07 |
| 12187 | 610 | H610Q | A | 1.9e-07 | 3.2e-07 | 2e-07 | 2.367E-07 | 7.234E-08 | 1.900E-07 | 3.200E-07 |
| 12188 | 610 | H610G | A | 2.6e-07 | 3e-07 | 2.4e-07 | 2.667E-07 | 3.055E-08 | 2.400E-07 | 3.000E-07 |
| 12189 | 610 | H610H | A | 3e-07 | 1.8e-07 | 1.4e-07 | 2.067E-07 | 8.327E-08 | 1.400E-07 | 3.000E-07 |
| 12190 | 610 | H610I | A | 4.6e-07 | 3.2e-07 | 5.1e-07 | 4.300E-07 | 9.849E-08 | 3.200E-07 | 5.100E-07 |
| 12191 | 610 | H610L | A | 6e-07 | 1.6e-07 | 3.2e-07 | 3.600E-07 | 2.227E-07 | 1.600E-07 | 6.000E-07 |
| 12192 | 610 | H610K | A | 2.2e-07 | 3.1e-07 | 1.8e-07 | 2.367E-07 | 6.658E-08 | 1.800E-07 | 3.100E-07 |
| 12193 | 610 | H610M | A | 1.6e-07 | 4.7e-07 | 1.6e-07 | 2.633E-07 | 1.790E-07 | 1.600E-07 | 4.700E-07 |
| 12194 | 610 | H610F | A | 2.6e-07 | 3.7e-07 | 2.6e-07 | 2.967E-07 | 6.351E-08 | 2.600E-07 | 3.700E-07 |
| 12195 | 610 | H610P | A | 2.6e-07 | 1.9e-07 | 3e-07 | 2.500E-07 | 5.568E-08 | 1.900E-07 | 3.000E-07 |
| 12196 | 610 | H610S | A | 2.3e-07 | 2.4e-07 | 5.8e-07 | 3.500E-07 | 1.992E-07 | 2.300E-07 | 5.800E-07 |
| 12197 | 610 | H610T | A | 4.5e-07 | 3.4e-07 | 2.9e-07 | 3.600E-07 | 8.185E-08 | 2.900E-07 | 4.500E-07 |
| 12198 | 610 | H610W | A | 3.2e-07 | 2.1e-07 | 1.6e-07 | 2.300E-07 | 8.185E-08 | 1.600E-07 | 3.200E-07 |
| 12199 | 610 | H610Y | A | 4.8e-07 | 1.3e-07 | 1.4e-07 | 2.500E-07 | 1.992E-07 | 1.300E-07 | 4.800E-07 |
| 12200 | 610 | H610V | A | 3.3e-07 | 2e-07 | 2e-07 | 2.433E-07 | 7.506E-08 | 2.000E-07 | 3.300E-07 |
| 12201 | 611 | S611A | A | 2.1e-07 | 2.8e-07 | 4.8e-07 | 3.233E-07 | 1.401E-07 | 2.100E-07 | 4.800E-07 |
| 12202 | 611 | S611R | A | 2e-07 | 1.5e-07 | 4.6e-07 | 2.700E-07 | 1.664E-07 | 1.500E-07 | 4.600E-07 |
| 12203 | 611 | S611N | A | 1.7e-07 | 1.8e-07 | 3.3e-07 | 2.267E-07 | 8.963E-08 | 1.700E-07 | 3.300E-07 |
| 12204 | 611 | S611D | A | 2e-07 | 2e-07 | 2.5e-07 | 2.167E-07 | 2.887E-08 | 2.000E-07 | 2.500E-07 |
| 12205 | 611 | S611C | A | 5.2e-07 | 1.2e-07 | 3.6e-07 | 3.333E-07 | 2.013E-07 | 1.200E-07 | 5.200E-07 |
| 12206 | 611 | S611E | A | 3.1e-07 | 1.5e-07 | 2e-07 | 2.200E-07 | 8.185E-08 | 1.500E-07 | 3.100E-07 |
| 12207 | 611 | S611Q | A | 1.4e-07 | 1.6e-07 | 3.5e-07 | 2.167E-07 | 1.159E-07 | 1.400E-07 | 3.500E-07 |
| 12208 | 611 | S611G | A | 3.9e-07 | 5.4e-07 | 3.7e-07 | 4.333E-07 | 9.292E-08 | 3.700E-07 | 5.400E-07 |
| 12209 | 611 | S611H | A | 3.3e-07 | 1.3e-07 | 2.9e-07 | 2.500E-07 | 1.058E-07 | 1.300E-07 | 3.300E-07 |
| 12210 | 611 | S611I | A | 3.8e-07 | 5.8e-07 | 2e-07 | 3.867E-07 | 1.901E-07 | 2.000E-07 | 5.800E-07 |
| 12211 | 611 | S611L | A | 3.2e-07 | 4.6e-07 | 3.8e-07 | 3.867E-07 | 7.024E-08 | 3.200E-07 | 4.600E-07 |
| 12212 | 611 | S611K | A | 2.3e-07 | 4e-07 | 1.3e-07 | 2.533E-07 | 1.365E-07 | 1.300E-07 | 4.000E-07 |
| 12213 | 611 | S611M | A | 1.8e-07 | 2.5e-07 | 2.6e-07 | 2.300E-07 | 4.359E-08 | 1.800E-07 | 2.600E-07 |
| 12214 | 611 | S611F | A | 2.8e-07 | 1.3e-07 | 3.4e-07 | 2.500E-07 | 1.082E-07 | 1.300E-07 | 3.400E-07 |
| 12215 | 611 | S611P | A | 2.7e-07 | 3.3e-07 | 3.6e-07 | 3.200E-07 | 4.583E-08 | 2.700E-07 | 3.600E-07 |
| 12216 | 611 | S611S | A | 1.2e-07 | 1.7e-07 | 3.9e-07 | 2.267E-07 | 1.436E-07 | 1.200E-07 | 3.900E-07 |
| 12217 | 611 | S611T | A | 1.2e-07 | 4.4e-07 | 2.4e-07 | 2.667E-07 | 1.617E-07 | 1.200E-07 | 4.400E-07 |
| 12218 | 611 | S611W | A | 1.2e-07 | 1.1e-07 | 2.1e-07 | 1.467E-07 | 5.508E-08 | 1.100E-07 | 2.100E-07 |
| 12219 | 611 | S611Y | A | 2.2e-07 | 4.9e-07 | 6e-07 | 4.367E-07 | 1.955E-07 | 2.200E-07 | 6.000E-07 |
| 12220 | 611 | S611V | A | 1.6e-07 | 2.4e-07 | 1.9e-07 | 1.967E-07 | 4.041E-08 | 1.600E-07 | 2.400E-07 |
| 12221 | 612 | C612A | A | 1.5e-07 | 4.3e-07 | 2.6e-07 | 2.800E-07 | 1.411E-07 | 1.500E-07 | 4.300E-07 |
| 12222 | 612 | C612R | A | 3.1e-07 | 2.4e-07 | 2.5e-07 | 2.667E-07 | 3.786E-08 | 2.400E-07 | 3.100E-07 |
| 12223 | 612 | C612N | A | 1.9e-07 | 8e-07 | 2.8e-07 | 4.233E-07 | 3.293E-07 | 1.900E-07 | 8.000E-07 |
| 12224 | 612 | C612D | A | 2.3e-07 | 5.2e-07 | 1.9e-07 | 3.133E-07 | 1.801E-07 | 1.900E-07 | 5.200E-07 |
| 12225 | 612 | C612C | A | 2.6e-07 | 4e-07 | 3.5e-07 | 3.367E-07 | 7.095E-08 | 2.600E-07 | 4.000E-07 |
| 12226 | 612 | C612E | A | 3.2e-07 | 4.4e-07 | 2e-07 | 3.200E-07 | 1.200E-07 | 2.000E-07 | 4.400E-07 |
| 12227 | 612 | C612Q | A | 2e-07 | 3.2e-07 | 1.7e-07 | 2.300E-07 | 7.937E-08 | 1.700E-07 | 3.200E-07 |
| 12228 | 612 | C612G | A | 1.5e-07 | 3.8e-07 | 3.1e-07 | 2.800E-07 | 1.179E-07 | 1.500E-07 | 3.800E-07 |
| 12229 | 612 | C612H | A | 2.9e-07 | 3.6e-07 | 2.8e-07 | 3.100E-07 | 4.359E-08 | 2.800E-07 | 3.600E-07 |
| 12230 | 612 | C612I | A | 1.8e-07 | 2.1e-07 | 2.2e-07 | 2.033E-07 | 2.082E-08 | 1.800E-07 | 2.200E-07 |
| 12231 | 612 | C612L | A | 1.6e-07 | 2.6e-07 | 1.9e-07 | 2.033E-07 | 5.132E-08 | 1.600E-07 | 2.600E-07 |
| 12232 | 612 | C612K | A | 3.9e-07 | 2.8e-07 | 5.2e-07 | 3.967E-07 | 1.201E-07 | 2.800E-07 | 5.200E-07 |
| 12233 | 612 | C612M | A | 1.9e-07 | 1.9e-07 | 4.1e-07 | 2.633E-07 | 1.270E-07 | 1.900E-07 | 4.100E-07 |
| 12234 | 612 | C612F | A | 1.5e-07 | 2e-07 | 1.7e-07 | 1.733E-07 | 2.517E-08 | 1.500E-07 | 2.000E-07 |
| 12235 | 612 | C612P | A | 2.1e-07 | 2.7e-07 | 3.1e-07 | 2.633E-07 | 5.033E-08 | 2.100E-07 | 3.100E-07 |
| 12236 | 612 | C612S | A | 1.2e-07 | 1.7e-07 | 4.9e-07 | 2.600E-07 | 2.007E-07 | 1.200E-07 | 4.900E-07 |
| 12237 | 612 | C612T | A | 2.9e-07 | 1.6e-07 | 1.2e-07 | 1.900E-07 | 8.888E-08 | 1.200E-07 | 2.900E-07 |
| 12238 | 612 | C612W | A | 1.9e-07 | 2.3e-07 | 9.6e-08 | 1.720E-07 | 6.879E-08 | 9.600E-08 | 2.300E-07 |
| 12239 | 612 | C612Y | A | 2.4e-07 | 1.4e-07 | 2.8e-07 | 2.200E-07 | 7.211E-08 | 1.400E-07 | 2.800E-07 |

| | | | | | | | | | | |
|---|---|---|---|---|---|---|---|---|---|---|
| 12240 | 612 | C612V | A | 1.5e-07 | 1.5e-07 | 4.5e-07 | 2.500E-07 | 1.732E-07 | 1.500E-07 | 4.500E-07 |
| 12241 | 613 | V613A | A | 2.6e-07 | 2.9e-07 | 6.2e-07 | 3.900E-07 | 1.997E-07 | 2.600E-07 | 6.200E-07 |
| 12242 | 613 | V613R | A | 1.5e-07 | 2e-07 | 3.6e-07 | 2.367E-07 | 1.097E-07 | 1.500E-07 | 3.600E-07 |
| 12243 | 613 | V613N | A | 2.4e-07 | 1.7e-07 | 2.2e-07 | 2.100E-07 | 3.606E-08 | 1.700E-07 | 2.400E-07 |
| 12244 | 613 | V613D | A | 3.8e-07 | 2.5e-07 | 1.3e-07 | 2.533E-07 | 1.250E-07 | 1.300E-07 | 3.800E-07 |
| 12245 | 613 | V613C | A | 3.3e-07 | 2e-07 | 1.8e-07 | 2.367E-07 | 8.145E-08 | 1.800E-07 | 3.300E-07 |
| 12246 | 613 | V613E | A | 3.1e-07 | 2.7e-07 | 1.6e-07 | 2.467E-07 | 7.767E-08 | 1.600E-07 | 3.100E-07 |
| 12247 | 613 | V613Q | A | 2.2e-07 | 3e-07 | 3.4e-07 | 2.867E-07 | 6.110E-08 | 2.200E-07 | 3.400E-07 |
| 12248 | 613 | V613G | A | 1.1e-07 | 1.1e-07 | 2.5e-07 | 1.567E-07 | 8.083E-08 | 1.100E-07 | 2.500E-07 |
| 12249 | 613 | V613H | A | 3.8e-07 | 2.1e-07 | 2e-07 | 2.633E-07 | 1.012E-07 | 2.000E-07 | 3.800E-07 |
| 12250 | 613 | V613I | A | 2.2e-07 | 3.1e-07 | 6.8e-07 | 4.033E-07 | 2.438E-07 | 2.200E-07 | 6.800E-07 |
| 12251 | 613 | V613L | A | 1.6e-07 | 2.7e-07 | 4e-07 | 2.767E-07 | 1.201E-07 | 1.600E-07 | 4.000E-07 |
| 12252 | 613 | V613K | A | 1.9e-07 | 3.9e-07 | 1.6e-07 | 2.467E-07 | 1.250E-07 | 1.600E-07 | 3.900E-07 |
| 12253 | 613 | V613M | A | 6.5e-07 | 2.1e-07 | 4.2e-07 | 4.267E-07 | 2.201E-07 | 2.100E-07 | 6.500E-07 |
| 12254 | 613 | V613F | A | 3.1e-07 | 2.3e-07 | 5.9e-07 | 3.767E-07 | 1.890E-07 | 2.300E-07 | 5.900E-07 |
| 12255 | 613 | V613P | A | 4.7e-07 | 3.9e-07 | 1.4e-07 | 3.333E-07 | 1.721E-07 | 1.400E-07 | 4.700E-07 |
| 12256 | 613 | V613S | A | 3.1e-07 | 3.4e-07 | 3.4e-07 | 3.300E-07 | 1.732E-08 | 3.100E-07 | 3.400E-07 |
| 12257 | 613 | V613T | A | 3.5e-07 | 1.9e-07 | 2.3e-07 | 2.567E-07 | 8.327E-08 | 1.900E-07 | 3.500E-07 |
| 12258 | 613 | V613W | A | 5.2e-07 | 5.2e-07 | 4.3e-07 | 4.900E-07 | 5.196E-08 | 4.300E-07 | 5.200E-07 |
| 12259 | 613 | V613Y | A | 2.7e-07 | 4.8e-07 | 3.6e-07 | 3.700E-07 | 1.054E-07 | 2.700E-07 | 4.800E-07 |
| 12260 | 613 | V613V | A | 2.1e-07 | 1.5e-07 | 2.6e-07 | 2.067E-07 | 5.508E-08 | 1.500E-07 | 2.600E-07 |
| 12261 | 614 | D614A | A | 9.6e-08 | 2.8e-07 | 3.5e-07 | 2.420E-07 | 1.312E-07 | 9.600E-08 | 3.500E-07 |
| 12262 | 614 | D614R | A | 3.5e-07 | 5.3e-07 | 1.7e-07 | 3.500E-07 | 1.800E-07 | 1.700E-07 | 5.300E-07 |
| 12263 | 614 | D614N | A | 2e-07 | 1.9e-07 | 2e-07 | 1.967E-07 | 5.774E-09 | 1.900E-07 | 2.000E-07 |
| 12264 | 614 | D614D | A | 1.5e-07 | 3e-07 | 1.2e-07 | 1.900E-07 | 9.644E-08 | 1.200E-07 | 3.000E-07 |
| 12265 | 614 | D614C | A | 1.6e-07 | 2.9e-07 | 2.9e-07 | 2.467E-07 | 7.506E-08 | 1.600E-07 | 2.900E-07 |
| 12266 | 614 | D614E | A | 2.1e-07 | 5e-07 | 2.9e-07 | 3.333E-07 | 1.498E-07 | 2.100E-07 | 5.000E-07 |
| 12267 | 614 | D614Q | A | 3.1e-07 | 7.5e-07 | 3.2e-07 | 4.600E-07 | 2.512E-07 | 3.100E-07 | 7.500E-07 |
| 12268 | 614 | D614G | A | 3.7e-07 | 3e-07 | 1.5e-07 | 2.733E-07 | 1.124E-07 | 1.500E-07 | 3.700E-07 |
| 12269 | 614 | D614H | A | 1.5e-07 | 3.8e-07 | 2.8e-07 | 2.700E-07 | 1.153E-07 | 1.500E-07 | 3.800E-07 |
| 12270 | 614 | D614I | A | 5.8e-07 | 3.3e-07 | 2.1e-07 | 3.733E-07 | 1.888E-07 | 2.100E-07 | 5.800E-07 |
| 12271 | 614 | D614L | A | 2.4e-07 | 2.9e-07 | 3.1e-07 | 2.800E-07 | 3.606E-08 | 2.400E-07 | 3.100E-07 |
| 12272 | 614 | D614K | A | 1.8e-07 | 2.4e-07 | 1.2e-07 | 1.800E-07 | 6.000E-08 | 1.200E-07 | 2.400E-07 |
| 12273 | 614 | D614M | A | 2e-07 | 1.9e-07 | 1.7e-07 | 1.867E-07 | 1.528E-08 | 1.700E-07 | 2.000E-07 |
| 12274 | 614 | D614F | A | 1.7e-07 | 1.4e-07 | 2.7e-07 | 1.933E-07 | 6.807E-08 | 1.400E-07 | 2.700E-07 |
| 12275 | 614 | D614P | A | 3.4e-07 | 2.8e-07 | 2.4e-07 | 2.867E-07 | 5.033E-08 | 2.400E-07 | 3.400E-07 |
| 12276 | 614 | D614S | A | 5e-07 | 2.7e-07 | 2.9e-07 | 3.533E-07 | 1.274E-07 | 2.700E-07 | 5.000E-07 |
| 12277 | 614 | D614T | A | 2e-07 | 3.3e-07 | 1.8e-07 | 2.367E-07 | 8.145E-08 | 1.800E-07 | 3.300E-07 |
| 12278 | 614 | D614W | A | 3.5e-07 | 2.4e-07 | 1.3e-07 | 2.400E-07 | 1.100E-07 | 1.300E-07 | 3.500E-07 |
| 12279 | 614 | D614Y | A | 2.5e-07 | 2.3e-07 | 3.8e-07 | 2.867E-07 | 8.145E-08 | 2.300E-07 | 3.800E-07 |
| 12280 | 614 | D614V | A | 2.5e-07 | 1.6e-07 | 1.4e-07 | 1.833E-07 | 5.859E-08 | 1.400E-07 | 2.500E-07 |
| 12281 | 615 | L615A | A | 3.9e-07 | 2.8e-07 | 1.1e-07 | 2.600E-07 | 1.411E-07 | 1.100E-07 | 3.900E-07 |
| 12282 | 615 | L615R | A | 1.8e-07 | 8e-08 | 2.6e-07 | 1.733E-07 | 9.018E-08 | 8.000E-08 | 2.600E-07 |
| 12283 | 615 | L615N | A | 6.1e-07 | 1.8e-07 | 1.6e-07 | 3.167E-07 | 2.542E-07 | 1.600E-07 | 6.100E-07 |
| 12284 | 615 | L615D | A | 8.7e-07 | 1.8e-07 | 3e-07 | 4.500E-07 | 3.686E-07 | 1.800E-07 | 8.700E-07 |
| 12285 | 615 | L615C | A | 2.8e-07 | 3.1e-07 | 1.8e-07 | 2.567E-07 | 6.807E-08 | 1.800E-07 | 3.100E-07 |
| 12286 | 615 | L615E | A | 4.4e-07 | 3.7e-07 | 4.5e-07 | 4.200E-07 | 4.359E-08 | 3.700E-07 | 4.500E-07 |
| 12287 | 615 | L615Q | A | 2.9e-07 | 2.5e-07 | 1.6e-07 | 2.333E-07 | 6.658E-08 | 1.600E-07 | 2.900E-07 |
| 12288 | 615 | L615G | A | 7.7e-07 | 4e-07 | 1.4e-07 | 4.367E-07 | 3.166E-07 | 1.400E-07 | 7.700E-07 |
| 12289 | 615 | L615H | A | 2e-07 | 2.7e-07 | 1.9e-07 | 2.200E-07 | 4.359E-08 | 1.900E-07 | 2.700E-07 |
| 12290 | 615 | L615I | A | 1.9e-07 | 2.8e-07 | 2.2e-07 | 2.300E-07 | 4.583E-08 | 1.900E-07 | 2.800E-07 |
| 12291 | 615 | L615L | A | 1.7e-07 | 1.8e-07 | 1.1e-07 | 1.533E-07 | 3.786E-08 | 1.100E-07 | 1.800E-07 |
| 12292 | 615 | L615K | A | 1.7e-07 | 1.5e-07 | 2.8e-07 | 2.000E-07 | 7.000E-08 | 1.500E-07 | 2.800E-07 |
| 12293 | 615 | L615M | A | 2.3e-07 | 1.2e-07 | 2.1e-07 | 1.867E-07 | 5.859E-08 | 1.200E-07 | 2.300E-07 |
| 12294 | 615 | L615F | A | 1.4e-07 | 3.2e-07 | 1.1e-07 | 1.900E-07 | 1.136E-07 | 1.100E-07 | 3.200E-07 |
| 12295 | 615 | L615P | A | 4.1e-07 | 1.5e-07 | 2.3e-07 | 2.633E-07 | 1.332E-07 | 1.500E-07 | 4.100E-07 |
| 12296 | 615 | L615S | A | 4.7e-07 | 1.4e-07 | 2.6e-07 | 2.900E-07 | 1.670E-07 | 1.400E-07 | 4.700E-07 |
| 12297 | 615 | L615T | A | 4e-07 | 2.3e-07 | 1.9e-07 | 2.733E-07 | 1.115E-07 | 1.900E-07 | 4.000E-07 |
| 12298 | 615 | L615W | A | 2.3e-07 | 3.4e-07 | 4.2e-07 | 3.300E-07 | 9.539E-08 | 2.300E-07 | 4.200E-07 |
| 12299 | 615 | L615Y | A | 4.8e-07 | 2.5e-07 | 1.6e-07 | 2.967E-07 | 1.650E-07 | 1.600E-07 | 4.800E-07 |
| 12300 | 615 | L615V | A | 4.2e-07 | 1.3e-07 | 4.4e-07 | 3.300E-07 | 1.735E-07 | 1.300E-07 | 4.400E-07 |
| 12301 | 616 | D616A | A | 1.5e-07 | 2.8e-07 | 2.3e-07 | 2.200E-07 | 6.557E-08 | 1.500E-07 | 2.800E-07 |
| 12302 | 616 | D616R | A | 4e-07 | 4.3e-07 | 3.4e-07 | 3.900E-07 | 4.583E-08 | 3.400E-07 | 4.300E-07 |
| 12303 | 616 | D616N | A | 6.1e-07 | 2.8e-07 | 1.8e-07 | 3.567E-07 | 2.250E-07 | 1.800E-07 | 6.100E-07 |
| 12304 | 616 | D616D | A | 8.8e-08 | 3.7e-07 | 3.8e-07 | 2.793E-07 | 1.658E-07 | 8.800E-08 | 3.800E-07 |
| 12305 | 616 | D616C | A | 1.7e-07 | 3.5e-07 | 2.2e-07 | 2.467E-07 | 9.292E-08 | 1.700E-07 | 3.500E-07 |
| 12306 | 616 | D616E | A | 2.8e-07 | 1.9e-07 | 1e-07 | 1.900E-07 | 9.000E-08 | 1.000E-07 | 2.800E-07 |
| 12307 | 616 | D616Q | A | 9.8e-08 | 2.4e-07 | 1.6e-07 | 1.660E-07 | 7.119E-08 | 9.800E-08 | 2.400E-07 |
| 12308 | 616 | D616G | A | 3.5e-07 | 1.9e-07 | 1.9e-07 | 2.433E-07 | 9.238E-08 | 1.900E-07 | 3.500E-07 |
| 12309 | 616 | D616H | A | 3.7e-07 | 2.6e-07 | 1.3e-07 | 2.533E-07 | 1.201E-07 | 1.300E-07 | 3.700E-07 |
| 12310 | 616 | D616I | A | 3.2e-07 | 2e-07 | 1.8e-07 | 2.333E-07 | 7.572E-08 | 1.800E-07 | 3.200E-07 |
| 12311 | 616 | D616L | A | 3.9e-07 | 4.1e-07 | 2.7e-07 | 3.567E-07 | 7.572E-08 | 2.700E-07 | 4.100E-07 |
| 12312 | 616 | D616K | A | 1.9e-07 | 2.4e-07 | 1.5e-07 | 1.933E-07 | 4.509E-08 | 1.500E-07 | 2.400E-07 |
| 12313 | 616 | D616M | A | 3.8e-07 | 1.8e-07 | 2.9e-07 | 2.833E-07 | 1.002E-07 | 1.800E-07 | 3.800E-07 |
| 12314 | 616 | D616F | A | 3.1e-07 | 1.7e-07 | 1.4e-07 | 2.067E-07 | 9.074E-08 | 1.400E-07 | 3.100E-07 |
| 12315 | 616 | D616P | A | 2.6e-07 | 2e-07 | 1.5e-07 | 2.033E-07 | 5.508E-08 | 1.500E-07 | 2.600E-07 |
| 12316 | 616 | D616S | A | 1.5e-07 | 1e-07 | 3.1e-07 | 1.867E-07 | 1.097E-07 | 1.000E-07 | 3.100E-07 |
| 12317 | 616 | D616T | A | 3.1e-07 | 2.8e-07 | 2.9e-07 | 2.933E-07 | 1.528E-08 | 2.800E-07 | 3.100E-07 |
| 12318 | 616 | D616W | A | 1.8e-07 | 3.3e-07 | 1.1e-07 | 2.067E-07 | 1.124E-07 | 1.100E-07 | 3.300E-07 |
| 12319 | 616 | D616Y | A | 2.3e-07 | 9e-08 | 3.1e-07 | 2.100E-07 | 1.114E-07 | 9.000E-08 | 3.100E-07 |
| 12320 | 616 | D616V | A | 2.2e-07 | 1.6e-07 | 1.8e-07 | 1.867E-07 | 3.055E-08 | 1.600E-07 | 2.200E-07 |
| 12321 | 617 | D617A | A | 2e-07 | 2.2e-07 | 2.2e-07 | 2.133E-07 | 1.155E-08 | 2.000E-07 | 2.200E-07 |
| 12322 | 617 | D617R | A | 2.2e-07 | 1.2e-07 | 4.7e-07 | 2.700E-07 | 1.803E-07 | 1.200E-07 | 4.700E-07 |
| 12323 | 617 | D617N | A | 1.8e-07 | 1.6e-07 | 1.7e-07 | 1.700E-07 | 1.000E-08 | 1.600E-07 | 1.800E-07 |
| 12324 | 617 | D617D | A | 3.2e-07 | 2.7e-07 | 2.3e-07 | 2.733E-07 | 4.509E-08 | 2.300E-07 | 3.200E-07 |

| | | | | | | | | | | |
|---|---|---|---|---|---|---|---|---|---|---|
| 12325 | 617 | D617C | A | 3.8e-07 | 1.4e-07 | 2.4e-07 | 2.533E-07 | 1.206E-07 | 1.400E-07 | 3.800E-07 |
| 12326 | 617 | D617E | A | 2.6e-07 | 1.5e-07 | 1.8e-07 | 1.967E-07 | 5.686E-08 | 1.500E-07 | 2.600E-07 |
| 12327 | 617 | D617Q | A | 3.1e-07 | 1.7e-07 | 3.3e-07 | 2.700E-07 | 8.718E-08 | 1.700E-07 | 3.300E-07 |
| 12328 | 617 | D617G | A | 3e-07 | 2.8e-07 | 2.6e-07 | 2.800E-07 | 2.000E-08 | 2.600E-07 | 3.000E-07 |
| 12329 | 617 | D617H | A | 3.6e-07 | 2.5e-07 | 1.5e-07 | 2.533E-07 | 1.050E-07 | 1.500E-07 | 3.600E-07 |
| 12330 | 617 | D617I | A | 1.8e-07 | 2.6e-07 | 3.6e-07 | 2.667E-07 | 9.018E-08 | 1.800E-07 | 3.600E-07 |
| 12331 | 617 | D617L | A | 3.1e-07 | 7.2e-07 | 1.6e-07 | 3.967E-07 | 2.899E-07 | 1.600E-07 | 7.200E-07 |
| 12332 | 617 | D617K | A | 2.2e-07 | 1.7e-07 | 3.4e-07 | 2.433E-07 | 8.737E-08 | 1.700E-07 | 3.400E-07 |
| 12333 | 617 | D617M | A | 3.7e-07 | 1.2e-07 | 3.3e-07 | 2.733E-07 | 1.343E-07 | 1.200E-07 | 3.700E-07 |
| 12334 | 617 | D617F | A | 2e-07 | 5.7e-07 | 2e-07 | 3.233E-07 | 2.136E-07 | 2.000E-07 | 5.700E-07 |
| 12335 | 617 | D617P | A | 5.3e-07 | 2.5e-07 | 1.4e-07 | 3.067E-07 | 2.011E-07 | 1.400E-07 | 5.300E-07 |
| 12336 | 617 | D617S | A | 2.8e-07 | 2.7e-07 | 2.2e-07 | 2.567E-07 | 3.215E-08 | 2.200E-07 | 2.800E-07 |
| 12337 | 617 | D617T | A | 3.2e-07 | 1.6e-07 | 3.9e-07 | 2.900E-07 | 1.179E-07 | 1.600E-07 | 3.900E-07 |
| 12338 | 617 | D617W | A | 2.5e-07 | 1.9e-07 | 1.9e-07 | 2.100E-07 | 3.464E-08 | 1.900E-07 | 2.500E-07 |
| 12339 | 617 | D617Y | A | 2.6e-07 | 3.9e-07 | 2.9e-07 | 3.133E-07 | 6.807E-08 | 2.600E-07 | 3.900E-07 |
| 12340 | 617 | D617V | A | 3.5e-07 | 4.5e-07 | 5e-07 | 4.333E-07 | 7.638E-08 | 3.500E-07 | 5.000E-07 |
| 12341 | 618 | K618A | A | 1.6e-07 | 3.6e-07 | 4e-07 | 3.067E-07 | 1.286E-07 | 1.600E-07 | 4.000E-07 |
| 12342 | 618 | K618R | A | 3.7e-07 | 1.4e-07 | 2.5e-07 | 2.533E-07 | 1.150E-07 | 1.400E-07 | 3.700E-07 |
| 12343 | 618 | K618N | A | 2.3e-07 | 2.8e-07 | 2.3e-07 | 2.467E-07 | 2.887E-08 | 2.300E-07 | 2.800E-07 |
| 12344 | 618 | K618D | A | 2e-07 | 2.3e-07 | 1.8e-07 | 2.033E-07 | 2.517E-08 | 1.800E-07 | 2.300E-07 |
| 12345 | 618 | K618C | A | 4.1e-07 | 1.4e-07 | 1.9e-07 | 2.467E-07 | 1.436E-07 | 1.400E-07 | 4.100E-07 |
| 12346 | 618 | K618E | A | 2.5e-07 | 3.2e-07 | 1.9e-07 | 2.533E-07 | 6.506E-08 | 1.900E-07 | 3.200E-07 |
| 12347 | 618 | K618Q | A | 3e-07 | 2.6e-07 | 2.1e-07 | 2.567E-07 | 4.509E-08 | 2.100E-07 | 3.000E-07 |
| 12348 | 618 | K618G | A | 1.5e-07 | 2.7e-07 | 2.9e-07 | 2.367E-07 | 7.572E-08 | 1.500E-07 | 2.900E-07 |
| 12349 | 618 | K618H | A | 3.6e-07 | 2.4e-07 | 2.3e-07 | 2.767E-07 | 7.234E-08 | 2.300E-07 | 3.600E-07 |
| 12350 | 618 | K618I | A | 2.7e-07 | 2.7e-07 | 3.3e-07 | 2.900E-07 | 3.464E-08 | 2.700E-07 | 3.300E-07 |
| 12351 | 618 | K618L | A | 1.9e-07 | 1.9e-07 | 2.6e-07 | 2.133E-07 | 4.041E-08 | 1.900E-07 | 2.600E-07 |
| 12352 | 618 | K618K | A | 3.4e-07 | 4.4e-07 | 2.8e-07 | 3.533E-07 | 8.083E-08 | 2.800E-07 | 4.400E-07 |
| 12353 | 618 | K618M | A | 2.2e-07 | 2.6e-07 | 3.3e-07 | 2.700E-07 | 5.568E-08 | 2.200E-07 | 3.300E-07 |
| 12354 | 618 | K618F | A | 2e-07 | 1.9e-07 | 1.9e-07 | 1.933E-07 | 5.774E-09 | 1.900E-07 | 2.000E-07 |
| 12355 | 618 | K618P | A | 2.1e-07 | 4.7e-07 | 5.2e-07 | 4.000E-07 | 1.664E-07 | 2.100E-07 | 5.200E-07 |
| 12356 | 618 | K618S | A | 1.7e-07 | 1.2e-07 | 2.4e-07 | 1.767E-07 | 6.028E-08 | 1.200E-07 | 2.400E-07 |
| 12357 | 618 | K618T | A | 3.2e-07 | 1.1e-07 | 2.7e-07 | 2.333E-07 | 1.097E-07 | 1.100E-07 | 3.200E-07 |
| 12358 | 618 | K618W | A | 1.6e-07 | 2e-07 | 2.4e-07 | 2.000E-07 | 4.000E-08 | 1.600E-07 | 2.400E-07 |
| 12359 | 618 | K618Y | A | 4.3e-07 | 2.6e-07 | 2.8e-07 | 3.233E-07 | 9.292E-08 | 2.600E-07 | 4.300E-07 |
| 12360 | 618 | K618V | A | 1.4e-07 | 1.8e-07 | 3.5e-07 | 2.233E-07 | 1.115E-07 | 1.400E-07 | 3.500E-07 |
| 12361 | 619 | G619A | A | 2.3e-07 | 2.1e-07 | 3.2e-07 | 2.533E-07 | 5.859E-08 | 2.100E-07 | 3.200E-07 |
| 12362 | 619 | G619R | A | 2.1e-07 | 2.2e-07 | 1.4e-07 | 1.900E-07 | 4.359E-08 | 1.400E-07 | 2.200E-07 |
| 12363 | 619 | G619N | A | 2.6e-07 | 2.9e-07 | 1.9e-07 | 2.467E-07 | 5.132E-08 | 1.900E-07 | 2.900E-07 |
| 12364 | 619 | G619D | A | 5.4e-07 | 1.7e-07 | 2.7e-07 | 3.267E-07 | 1.914E-07 | 1.700E-07 | 5.400E-07 |
| 12365 | 619 | G619C | A | 1.8e-07 | 2.9e-07 | 2.1e-07 | 2.267E-07 | 5.686E-08 | 1.800E-07 | 2.900E-07 |
| 12366 | 619 | G619E | A | 3.9e-07 | 4.1e-07 | 1.7e-07 | 3.233E-07 | 1.332E-07 | 1.700E-07 | 4.100E-07 |
| 12367 | 619 | G619Q | A | 2.1e-07 | 1.2e-07 | 2.1e-07 | 1.800E-07 | 5.196E-08 | 1.200E-07 | 2.100E-07 |
| 12368 | 619 | G619G | A | 1.1e-07 | 3.2e-07 | 2.9e-07 | 2.400E-07 | 1.136E-07 | 1.100E-07 | 3.200E-07 |
| 12369 | 619 | G619H | A | 1.4e-07 | 2.2e-07 | 3.2e-07 | 2.267E-07 | 9.018E-08 | 1.400E-07 | 3.200E-07 |
| 12370 | 619 | G619I | A | 2.3e-07 | 2.1e-07 | 5e-07 | 3.133E-07 | 1.620E-07 | 2.100E-07 | 5.000E-07 |
| 12371 | 619 | G619L | A | 2.8e-07 | 4.2e-07 | 3.7e-07 | 3.567E-07 | 7.095E-08 | 2.800E-07 | 4.200E-07 |
| 12372 | 619 | G619K | A | 3e-07 | 6e-08 | 1.2e-07 | 1.600E-07 | 1.249E-07 | 6.000E-08 | 3.000E-07 |
| 12373 | 619 | G619M | A | 2.2e-07 | 4.4e-07 | 1.5e-07 | 2.700E-07 | 1.513E-07 | 1.500E-07 | 4.400E-07 |
| 12374 | 619 | G619F | A | 2.6e-07 | 1.6e-07 | 4.1e-07 | 2.767E-07 | 1.258E-07 | 1.600E-07 | 4.100E-07 |
| 12375 | 619 | G619P | A | 2.1e-07 | 2.3e-07 | 8.4e-08 | 1.747E-07 | 7.915E-08 | 8.400E-08 | 2.300E-07 |
| 12376 | 619 | G619S | A | 3.8e-07 | 1.4e-07 | 3.8e-07 | 3.000E-07 | 1.386E-07 | 1.400E-07 | 3.800E-07 |
| 12377 | 619 | G619T | A | 1.5e-07 | 1.5e-07 | 2.9e-07 | 1.967E-07 | 8.083E-08 | 1.500E-07 | 2.900E-07 |
| 12378 | 619 | G619W | A | 2e-07 | 1.5e-07 | 2.9e-07 | 2.133E-07 | 7.095E-08 | 1.500E-07 | 2.900E-07 |
| 12379 | 619 | G619Y | A | 9.5e-08 | 2.2e-07 | 2.2e-07 | 1.783E-07 | 7.217E-08 | 9.500E-08 | 2.200E-07 |
| 12380 | 619 | G619V | A | 1.6e-07 | 3.5e-07 | 1.7e-07 | 2.267E-07 | 1.069E-07 | 1.600E-07 | 3.500E-07 |
| 12381 | 620 | C620A | A | 2.3e-07 | 2.4e-07 | 1.5e-07 | 2.067E-07 | 4.933E-08 | 1.500E-07 | 2.400E-07 |
| 12382 | 620 | C620R | A | 2.1e-07 | 2.6e-07 | 1.2e-07 | 1.967E-07 | 7.095E-08 | 1.200E-07 | 2.600E-07 |
| 12383 | 620 | C620N | A | 4.1e-07 | 5.7e-07 | 2e-07 | 3.933E-07 | 1.856E-07 | 2.000E-07 | 5.700E-07 |
| 12384 | 620 | C620D | A | 2.1e-07 | 2e-07 | 3.3e-07 | 2.467E-07 | 7.234E-08 | 2.000E-07 | 3.300E-07 |
| 12385 | 620 | C620C | A | 1.7e-07 | 2.3e-07 | 3.1e-07 | 2.367E-07 | 7.024E-08 | 1.700E-07 | 3.100E-07 |
| 12386 | 620 | C620E | A | 2.4e-07 | 2e-07 | 2.7e-07 | 2.367E-07 | 3.512E-08 | 2.000E-07 | 2.700E-07 |
| 12387 | 620 | C620Q | A | 3.6e-07 | 2.4e-07 | 2.3e-07 | 2.767E-07 | 7.234E-08 | 2.300E-07 | 3.600E-07 |
| 12388 | 620 | C620G | A | 2.9e-07 | 3.1e-07 | 1.7e-07 | 2.567E-07 | 7.572E-08 | 1.700E-07 | 3.100E-07 |
| 12389 | 620 | C620H | A | 1.7e-07 | 2.8e-07 | 2.4e-07 | 2.300E-07 | 5.568E-08 | 1.700E-07 | 2.800E-07 |
| 12390 | 620 | C620I | A | 2.2e-07 | 2.6e-07 | 3.6e-07 | 2.800E-07 | 7.211E-08 | 2.200E-07 | 3.600E-07 |
| 12391 | 620 | C620L | A | 4.2e-07 | 4.6e-07 | 2.8e-07 | 3.867E-07 | 9.452E-08 | 2.800E-07 | 4.600E-07 |
| 12392 | 620 | C620K | A | 3.3e-07 | 3.4e-07 | 2.8e-07 | 3.167E-07 | 3.215E-08 | 2.800E-07 | 3.400E-07 |
| 12393 | 620 | C620M | A | 1.5e-07 | 5.7e-07 | 1.4e-07 | 2.867E-07 | 2.454E-07 | 1.400E-07 | 5.700E-07 |
| 12394 | 620 | C620F | A | 2.2e-07 | 4.5e-07 | 2.3e-07 | 3.000E-07 | 1.300E-07 | 2.200E-07 | 4.500E-07 |
| 12395 | 620 | C620P | A | 4.3e-07 | 2.5e-07 | 4.2e-07 | 3.667E-07 | 1.012E-07 | 2.500E-07 | 4.300E-07 |
| 12396 | 620 | C620S | A | 1.7e-07 | 1.4e-07 | 5.7e-07 | 2.933E-07 | 2.401E-07 | 1.400E-07 | 5.700E-07 |
| 12397 | 620 | C620T | A | 2.8e-07 | 3.6e-07 | 3.6e-07 | 3.333E-07 | 4.619E-08 | 2.800E-07 | 3.600E-07 |
| 12398 | 620 | C620W | A | 3.6e-07 | 2.7e-07 | 3.4e-07 | 3.233E-07 | 4.726E-08 | 2.700E-07 | 3.600E-07 |
| 12399 | 620 | C620Y | A | 5.7e-07 | 2.1e-07 | 2.4e-07 | 3.400E-07 | 1.997E-07 | 2.100E-07 | 5.700E-07 |
| 12400 | 620 | C620V | A | 3.6e-07 | 1.9e-07 | 2e-07 | 2.500E-07 | 9.539E-08 | 1.900E-07 | 3.600E-07 |
| 12401 | 621 | P621A | A | 4e-07 | 2.1e-07 | 3.4e-07 | 3.167E-07 | 9.713E-08 | 2.100E-07 | 4.000E-07 |
| 12402 | 621 | P621R | A | 1.3e-07 | 1.6e-07 | 5.5e-07 | 2.800E-07 | 2.343E-07 | 1.300E-07 | 5.500E-07 |
| 12403 | 621 | P621N | A | 5e-07 | 2.1e-07 | 4.5e-07 | 3.867E-07 | 1.550E-07 | 2.100E-07 | 5.000E-07 |
| 12404 | 621 | P621D | A | 4.1e-07 | 2.4e-07 | 4.9e-07 | 3.800E-07 | 1.277E-07 | 2.400E-07 | 4.900E-07 |
| 12405 | 621 | P621C | A | 1.9e-07 | 2e-07 | 2.1e-07 | 2.000E-07 | 1.000E-08 | 1.900E-07 | 2.100E-07 |
| 12406 | 621 | P621E | A | 1.4e-07 | 4.3e-07 | 1.5e-07 | 2.400E-07 | 1.646E-07 | 1.400E-07 | 4.300E-07 |
| 12407 | 621 | P621Q | A | 3.5e-07 | 1.5e-07 | 9.3e-08 | 1.977E-07 | 1.350E-07 | 9.300E-08 | 3.500E-07 |
| 12408 | 621 | P621G | A | 1.5e-07 | 2.5e-07 | 2.6e-07 | 2.200E-07 | 6.083E-08 | 1.500E-07 | 2.600E-07 |
| 12409 | 621 | P621H | A | 1.5e-07 | 3.9e-07 | 2.8e-07 | 2.733E-07 | 1.201E-07 | 1.500E-07 | 3.900E-07 |

| | | | | | | | | | | |
|---|---|---|---|---|---|---|---|---|---|---|
| 12410 | 621 | P621I | A | 2.9e-07 | 1.1e-07 | 1.2e-07 | 1.733E-07 | 1.012E-07 | 1.100E-07 | 2.900E-07 |
| 12411 | 621 | P621L | A | 3.3e-07 | 3.1e-07 | 1.9e-07 | 2.767E-07 | 7.572E-08 | 1.900E-07 | 3.300E-07 |
| 12412 | 621 | P621K | A | 1.8e-07 | 1.6e-07 | 7.5e-07 | 3.633E-07 | 3.350E-07 | 1.600E-07 | 7.500E-07 |
| 12413 | 621 | P621M | A | 2e-07 | 1.2e-07 | 2e-07 | 1.733E-07 | 4.619E-08 | 1.200E-07 | 2.000E-07 |
| 12414 | 621 | P621F | A | 2.4e-07 | 3.3e-07 | 3.7e-07 | 3.133E-07 | 6.658E-08 | 2.400E-07 | 3.700E-07 |
| 12415 | 621 | P621P | A | 1.1e-07 | 2.1e-07 | 2.7e-07 | 1.967E-07 | 8.083E-08 | 1.100E-07 | 2.700E-07 |
| 12416 | 621 | P621S | A | 1.9e-07 | 4.2e-07 | 2.3e-07 | 2.800E-07 | 1.229E-07 | 1.900E-07 | 4.200E-07 |
| 12417 | 621 | P621T | A | 2.1e-07 | 2.5e-07 | 1.4e-07 | 2.000E-07 | 5.568E-08 | 1.400E-07 | 2.500E-07 |
| 12418 | 621 | P621W | A | 4.3e-07 | 1.4e-07 | 4.2e-07 | 3.300E-07 | 1.646E-07 | 1.400E-07 | 4.300E-07 |
| 12419 | 621 | P621Y | A | 2e-07 | 4.7e-07 | 1.4e-07 | 2.700E-07 | 1.758E-07 | 1.400E-07 | 4.700E-07 |
| 12420 | 621 | P621V | A | 4e-07 | 1.7e-07 | 6.3e-07 | 4.000E-07 | 2.300E-07 | 1.700E-07 | 6.300E-07 |
| 12421 | 622 | A622A | A | 2.9e-07 | 2.4e-07 | 1.9e-07 | 2.400E-07 | 5.000E-08 | 1.900E-07 | 2.900E-07 |
| 12422 | 622 | A622R | A | 1.6e-07 | 1.6e-07 | 2.1e-07 | 1.767E-07 | 2.887E-08 | 1.600E-07 | 2.100E-07 |
| 12423 | 622 | A622N | A | 3.6e-07 | 1.6e-07 | 1.6e-07 | 2.267E-07 | 1.155E-07 | 1.600E-07 | 3.600E-07 |
| 12424 | 622 | A622D | A | 3.4e-07 | 2.7e-07 | 1.8e-07 | 2.633E-07 | 8.021E-08 | 1.800E-07 | 3.400E-07 |
| 12425 | 622 | A622C | A | 2.8e-07 | 2.7e-07 | 9.7e-08 | 2.157E-07 | 1.029E-07 | 9.700E-08 | 2.800E-07 |
| 12426 | 622 | A622E | A | 9.6e-08 | 3.9e-07 | 2.3e-07 | 2.387E-07 | 1.472E-07 | 9.600E-08 | 3.900E-07 |
| 12427 | 622 | A622Q | A | 2.1e-07 | 3.8e-07 | 3.1e-07 | 3.000E-07 | 8.544E-08 | 2.100E-07 | 3.800E-07 |
| 12428 | 622 | A622G | A | 1.8e-07 | 1.4e-07 | 3.1e-07 | 2.100E-07 | 8.888E-08 | 1.400E-07 | 3.100E-07 |
| 12429 | 622 | A622H | A | 2.1e-07 | 2.6e-07 | 1.6e-07 | 2.100E-07 | 5.000E-08 | 1.600E-07 | 2.600E-07 |
| 12430 | 622 | A622I | A | 2.1e-07 | 1e-07 | 2.3e-07 | 1.800E-07 | 7.000E-08 | 1.000E-07 | 2.300E-07 |
| 12431 | 622 | A622L | A | 3.7e-07 | 2.2e-07 | 4.7e-07 | 3.533E-07 | 1.258E-07 | 2.200E-07 | 4.700E-07 |
| 12432 | 622 | A622K | A | 2.4e-07 | 2.2e-07 | 5.2e-07 | 3.267E-07 | 1.677E-07 | 2.200E-07 | 5.200E-07 |
| 12433 | 622 | A622M | A | 1.4e-07 | 1.9e-07 | 3e-07 | 2.100E-07 | 8.185E-08 | 1.400E-07 | 3.000E-07 |
| 12434 | 622 | A622F | A | 3.2e-07 | 9.7e-08 | 3.9e-07 | 2.690E-07 | 1.530E-07 | 9.700E-08 | 3.900E-07 |
| 12435 | 622 | A622P | A | 2.1e-07 | 1.6e-07 | 3.8e-07 | 2.500E-07 | 1.153E-07 | 1.600E-07 | 3.800E-07 |
| 12436 | 622 | A622S | A | 2.6e-07 | 3.8e-07 | 4.5e-07 | 3.633E-07 | 9.609E-08 | 2.600E-07 | 4.500E-07 |
| 12437 | 622 | A622T | A | 3e-07 | 4.6e-07 | 2.5e-07 | 3.367E-07 | 1.097E-07 | 2.500E-07 | 4.600E-07 |
| 12438 | 622 | A622W | A | 2.4e-07 | 1.8e-07 | 1.5e-07 | 1.900E-07 | 4.583E-08 | 1.500E-07 | 2.400E-07 |
| 12439 | 622 | A622Y | A | 3.8e-07 | 1.9e-07 | 2.3e-07 | 2.667E-07 | 1.002E-07 | 1.900E-07 | 3.800E-07 |
| 12440 | 622 | A622V | A | 1.7e-07 | 1.7e-07 | 2.3e-07 | 1.900E-07 | 3.464E-08 | 1.700E-07 | 2.300E-07 |
| 12441 | 623 | D623A | B | 4.3e-07 | 2.5e-07 | 2.2e-07 | 3.000E-07 | 1.136E-07 | 2.200E-07 | 4.300E-07 |
| 12442 | 623 | D623R | B | 4.1e-07 | 2.6e-07 | 3.8e-07 | 3.500E-07 | 7.937E-08 | 2.600E-07 | 4.100E-07 |
| 12443 | 623 | D623N | B | 2e-07 | 3.2e-07 | 4.7e-07 | 3.300E-07 | 1.353E-07 | 2.000E-07 | 4.700E-07 |
| 12444 | 623 | D623D | B | 2.4e-07 | 2e-07 | 1.8e-07 | 2.067E-07 | 3.055E-08 | 1.800E-07 | 2.400E-07 |
| 12445 | 623 | D623C | B | 4.6e-07 | 3.3e-07 | 2.5e-07 | 3.467E-07 | 1.060E-07 | 2.500E-07 | 4.600E-07 |
| 12446 | 623 | D623E | B | 1.7e-07 | 3.3e-07 | 2.7e-07 | 2.567E-07 | 8.083E-08 | 1.700E-07 | 3.300E-07 |
| 12447 | 623 | D623Q | B | 1.8e-07 | 4.5e-07 | 2.8e-07 | 3.033E-07 | 1.365E-07 | 1.800E-07 | 4.500E-07 |
| 12448 | 623 | D623G | B | 3.3e-07 | 4.7e-07 | 2e-07 | 3.333E-07 | 1.350E-07 | 2.000E-07 | 4.700E-07 |
| 12449 | 623 | D623H | B | 2.3e-07 | 2e-07 | 2e-07 | 2.100E-07 | 1.732E-08 | 2.000E-07 | 2.300E-07 |
| 12450 | 623 | D623I | B | 2.8e-07 | 7.9e-08 | 1.4e-07 | 1.663E-07 | 1.031E-07 | 7.900E-08 | 2.800E-07 |
| 12451 | 623 | D623L | B | 2.6e-07 | 1.1e-07 | 2.1e-07 | 1.933E-07 | 7.638E-08 | 1.100E-07 | 2.600E-07 |
| 12452 | 623 | D623K | B | 3e-07 | 4.4e-07 | 3e-07 | 3.467E-07 | 8.083E-08 | 3.000E-07 | 4.400E-07 |
| 12453 | 623 | D623M | B | 2e-07 | 1.4e-07 | 2.9e-07 | 2.100E-07 | 7.550E-08 | 1.400E-07 | 2.900E-07 |
| 12454 | 623 | D623F | B | 4.3e-07 | 3.4e-07 | 2.3e-07 | 3.333E-07 | 1.002E-07 | 2.300E-07 | 4.300E-07 |
| 12455 | 623 | D623P | B | 4.4e-07 | 2.4e-07 | 1.9e-07 | 2.900E-07 | 1.323E-07 | 1.900E-07 | 4.400E-07 |
| 12456 | 623 | D623S | B | 7.3e-07 | 3.1e-07 | 2.7e-07 | 4.367E-07 | 2.548E-07 | 2.700E-07 | 7.300E-07 |
| 12457 | 623 | D623T | B | 3.4e-07 | 2.3e-07 | 2.1e-07 | 2.600E-07 | 7.000E-08 | 2.100E-07 | 3.400E-07 |
| 12458 | 623 | D623W | B | 3.5e-07 | 5.2e-07 | 3.3e-07 | 4.000E-07 | 1.044E-07 | 3.300E-07 | 5.200E-07 |
| 12459 | 623 | D623Y | B | 3.7e-07 | 6.3e-07 | 3.8e-07 | 4.600E-07 | 1.473E-07 | 3.700E-07 | 6.300E-07 |
| 12460 | 623 | D623V | B | 4.6e-07 | 4.3e-07 | 3e-07 | 3.967E-07 | 8.505E-08 | 3.000E-07 | 4.600E-07 |
| 12461 | 624 | I624A | B | 1.1e-07 | 3.2e-07 | 5.6e-07 | 3.300E-07 | 2.252E-07 | 1.100E-07 | 5.600E-07 |
| 12462 | 624 | I624R | B | 3.5e-07 | 2.6e-07 | 4e-07 | 3.367E-07 | 7.095E-08 | 2.600E-07 | 4.000E-07 |
| 12463 | 624 | I624N | B | 2.1e-07 | 3.1e-07 | 1.8e-07 | 2.333E-07 | 6.807E-08 | 1.800E-07 | 3.100E-07 |
| 12464 | 624 | I624D | B | 2.6e-07 | 4.1e-07 | 3.2e-07 | 3.300E-07 | 7.550E-08 | 2.600E-07 | 4.100E-07 |
| 12465 | 624 | I624C | B | 2.3e-07 | 3.2e-07 | 3.4e-07 | 2.967E-07 | 5.859E-08 | 2.300E-07 | 3.400E-07 |
| 12466 | 624 | I624E | B | 2.5e-07 | 2.6e-07 | 3.7e-07 | 2.933E-07 | 6.658E-08 | 2.500E-07 | 3.700E-07 |
| 12467 | 624 | I624Q | B | 1.8e-07 | 2.8e-07 | 3.3e-07 | 2.633E-07 | 7.638E-08 | 1.800E-07 | 3.300E-07 |
| 12468 | 624 | I624G | B | 4.7e-07 | 1.2e-07 | 1.6e-07 | 2.500E-07 | 1.916E-07 | 1.200E-07 | 4.700E-07 |
| 12469 | 624 | I624H | B | 3.4e-07 | 2.5e-07 | 3.2e-07 | 3.033E-07 | 4.726E-08 | 2.500E-07 | 3.400E-07 |
| 12470 | 624 | I624I | B | 1.8e-07 | 3.1e-07 | 1.5e-07 | 2.133E-07 | 8.505E-08 | 1.500E-07 | 3.100E-07 |
| 12471 | 624 | I624L | B | 1.6e-07 | 2.6e-07 | 1.7e-07 | 1.967E-07 | 5.508E-08 | 1.600E-07 | 2.600E-07 |
| 12472 | 624 | I624K | B | 2.7e-07 | 2.3e-07 | 3.1e-07 | 2.700E-07 | 4.000E-08 | 2.300E-07 | 3.100E-07 |
| 12473 | 624 | I624M | B | 3.7e-07 | 2.7e-07 | 3.9e-07 | 3.433E-07 | 6.429E-08 | 2.700E-07 | 3.900E-07 |
| 12474 | 624 | I624F | B | 9.4e-08 | 1.9e-07 | 1.3e-07 | 1.380E-07 | 4.850E-08 | 9.400E-08 | 1.900E-07 |
| 12475 | 624 | I624P | B | 2.8e-07 | 4.6e-07 | 1.8e-07 | 3.067E-07 | 1.419E-07 | 1.800E-07 | 4.600E-07 |
| 12476 | 624 | I624S | B | 3.9e-07 | 3.4e-07 | 2.1e-07 | 3.133E-07 | 9.292E-08 | 2.100E-07 | 3.900E-07 |
| 12477 | 624 | I624T | B | 2.3e-07 | 1.8e-07 | 3.4e-07 | 2.500E-07 | 8.185E-08 | 1.800E-07 | 3.400E-07 |
| 12478 | 624 | I624W | B | 1.4e-07 | 1.2e-07 | 1.5e-07 | 1.367E-07 | 1.528E-08 | 1.200E-07 | 1.500E-07 |
| 12479 | 624 | I624Y | B | 2.6e-07 | 2.1e-07 | 1.4e-07 | 2.033E-07 | 6.028E-08 | 1.400E-07 | 2.600E-07 |
| 12480 | 624 | I624V | B | 1.7e-07 | 1.6e-07 | 3.3e-07 | 2.200E-07 | 9.539E-08 | 1.600E-07 | 3.300E-07 |
| 12481 | 625 | Q625A | B | 1.8e-07 | 2.7e-07 | 3.1e-07 | 2.533E-07 | 6.658E-08 | 1.800E-07 | 3.100E-07 |
| 12482 | 625 | Q625R | B | 2.5e-07 | 2.9e-07 | 3.8e-07 | 3.067E-07 | 6.658E-08 | 2.500E-07 | 3.800E-07 |
| 12483 | 625 | Q625N | B | 1.3e-07 | 2.1e-07 | 3.2e-07 | 2.200E-07 | 9.539E-08 | 1.300E-07 | 3.200E-07 |
| 12484 | 625 | Q625D | B | 3e-07 | 3.2e-07 | 1.6e-07 | 2.600E-07 | 8.718E-08 | 1.600E-07 | 3.200E-07 |
| 12485 | 625 | Q625C | B | 1.5e-07 | 2e-07 | 2.2e-07 | 1.900E-07 | 3.606E-08 | 1.500E-07 | 2.200E-07 |
| 12486 | 625 | Q625E | B | 7.1e-07 | 4.1e-07 | 5.3e-07 | 5.500E-07 | 1.510E-07 | 4.100E-07 | 7.100E-07 |
| 12487 | 625 | Q625Q | B | 3.2e-07 | 2.1e-07 | 2.2e-07 | 2.500E-07 | 6.083E-08 | 2.100E-07 | 3.200E-07 |
| 12488 | 625 | Q625G | B | 3.7e-07 | 2.4e-07 | 2.3e-07 | 2.800E-07 | 7.810E-08 | 2.300E-07 | 3.700E-07 |
| 12489 | 625 | Q625H | B | 3.7e-07 | 4.6e-07 | 4.1e-07 | 4.133E-07 | 4.509E-08 | 3.700E-07 | 4.600E-07 |
| 12490 | 625 | Q625I | B | 3.7e-07 | 2.4e-07 | 1.2e-07 | 2.433E-07 | 1.250E-07 | 1.200E-07 | 3.700E-07 |
| 12491 | 625 | Q625L | B | 2.5e-07 | 2.7e-07 | 3.2e-07 | 2.800E-07 | 3.606E-08 | 2.500E-07 | 3.200E-07 |
| 12492 | 625 | Q625K | B | 2.6e-07 | 3.4e-07 | 3.1e-07 | 3.033E-07 | 4.041E-08 | 2.600E-07 | 3.400E-07 |
| 12493 | 625 | Q625M | B | 1.7e-07 | 1.5e-07 | 1.8e-07 | 1.667E-07 | 1.528E-08 | 1.500E-07 | 1.800E-07 |
| 12494 | 625 | Q625F | B | 1.9e-07 | 3.4e-07 | 1.2e-07 | 2.167E-07 | 1.124E-07 | 1.200E-07 | 3.400E-07 |

| | | | | | | | | | | |
|---|---|---|---|---|---|---|---|---|---|---|
| 12495 | 625 | Q625P | B | 2.6e-07 | 2.6e-07 | 1.2e-07 | 2.133E-07 | 8.083E-08 | 1.200E-07 | 2.600E-07 |
| 12496 | 625 | Q625S | B | 2.1e-07 | 4.1e-07 | 1.8e-07 | 2.667E-07 | 1.250E-07 | 1.800E-07 | 4.100E-07 |
| 12497 | 625 | Q625T | B | 2.8e-07 | 4.4e-07 | 2.2e-07 | 3.133E-07 | 1.137E-07 | 2.200E-07 | 4.400E-07 |
| 12498 | 625 | Q625W | B | 1.9e-07 | 2e-07 | 3.3e-07 | 2.400E-07 | 7.810E-08 | 1.900E-07 | 3.300E-07 |
| 12499 | 625 | Q625Y | B | 6.5e-07 | 6.4e-07 | 4.2e-07 | 5.700E-07 | 1.300E-07 | 4.200E-07 | 6.500E-07 |
| 12500 | 625 | Q625V | B | 1.7e-07 | 1.2e-07 | 2.8e-07 | 1.900E-07 | 8.185E-08 | 1.200E-07 | 2.800E-07 |
| 12501 | 626 | M626A | B | 4.1e-07 | 4.6e-07 | 2.7e-07 | 3.800E-07 | 9.849E-08 | 2.700E-07 | 4.600E-07 |
| 12502 | 626 | M626R | B | 2.7e-07 | 2.8e-07 | 1.8e-07 | 2.433E-07 | 5.508E-08 | 1.800E-07 | 2.800E-07 |
| 12503 | 626 | M626N | B | 3e-07 | 2e-07 | 2.3e-07 | 2.433E-07 | 5.132E-08 | 2.000E-07 | 3.000E-07 |
| 12504 | 626 | M626D | B | 3.5e-07 | 2.2e-07 | 4.5e-07 | 3.400E-07 | 1.153E-07 | 2.200E-07 | 4.500E-07 |
| 12505 | 626 | M626C | B | 2.2e-07 | 2.9e-07 | 6.2e-07 | 3.767E-07 | 2.136E-07 | 2.200E-07 | 6.200E-07 |
| 12506 | 626 | M626E | B | 5e-07 | 3.7e-07 | 2.3e-07 | 3.667E-07 | 1.350E-07 | 2.300E-07 | 5.000E-07 |
| 12507 | 626 | M626Q | B | 3.2e-07 | 3e-07 | 2.9e-07 | 3.033E-07 | 1.528E-08 | 2.900E-07 | 3.200E-07 |
| 12508 | 626 | M626G | B | 2.8e-07 | 3.4e-07 | 3.3e-07 | 3.167E-07 | 3.215E-08 | 2.800E-07 | 3.400E-07 |
| 12509 | 626 | M626H | B | 2.8e-07 | 1.9e-07 | 1.5e-07 | 2.067E-07 | 6.658E-08 | 1.500E-07 | 2.800E-07 |
| 12510 | 626 | M626I | B | 3.4e-07 | 1.8e-07 | 3.8e-07 | 3.000E-07 | 1.058E-07 | 1.800E-07 | 3.800E-07 |
| 12511 | 626 | M626L | B | 1.2e-07 | 2.6e-07 | 1.5e-07 | 1.767E-07 | 7.371E-08 | 1.200E-07 | 2.600E-07 |
| 12512 | 626 | M626K | B | 1.7e-07 | 1.7e-07 | 1.9e-07 | 1.767E-07 | 1.155E-08 | 1.700E-07 | 1.900E-07 |
| 12513 | 626 | M626M | B | 3.8e-07 | 1.1e-07 | 1.2e-07 | 2.033E-07 | 1.531E-07 | 1.100E-07 | 3.800E-07 |
| 12514 | 626 | M626F | B | 1e-07 | 2.2e-07 | 1.5e-07 | 1.567E-07 | 6.028E-08 | 1.000E-07 | 2.200E-07 |
| 12515 | 626 | M626P | B | 5e-07 | 1.7e-07 | 1.9e-07 | 2.867E-07 | 1.850E-07 | 1.700E-07 | 5.000E-07 |
| 12516 | 626 | M626S | B | 3.1e-07 | 3.6e-07 | 3.8e-07 | 3.500E-07 | 3.606E-08 | 3.100E-07 | 3.800E-07 |
| 12517 | 626 | M626T | B | 3.4e-07 | 2.9e-07 | 1.6e-07 | 2.633E-07 | 9.292E-08 | 1.600E-07 | 3.400E-07 |
| 12518 | 626 | M626W | B | 2e-07 | 2.2e-07 | 2.2e-07 | 2.133E-07 | 1.155E-08 | 2.000E-07 | 2.200E-07 |
| 12519 | 626 | M626Y | B | 6.3e-08 | 1.5e-07 | 3.7e-07 | 1.943E-07 | 1.582E-07 | 6.300E-08 | 3.700E-07 |
| 12520 | 626 | M626V | B | 3.1e-07 | 2.6e-07 | 2.4e-07 | 2.700E-07 | 3.606E-08 | 2.400E-07 | 3.100E-07 |
| 12521 | 627 | T627A | B | 2.2e-07 | 5.6e-07 | 4e-07 | 3.933E-07 | 1.701E-07 | 2.200E-07 | 5.600E-07 |
| 12522 | 627 | T627R | B | 2.3e-07 | 2e-07 | 3.4e-07 | 2.567E-07 | 7.371E-08 | 2.000E-07 | 3.400E-07 |
| 12523 | 627 | T627N | B | 2.8e-07 | 3e-07 | 3e-07 | 2.933E-07 | 1.155E-08 | 2.800E-07 | 3.000E-07 |
| 12524 | 627 | T627D | B | 2.4e-07 | 2.7e-07 | 1.6e-07 | 2.233E-07 | 5.686E-08 | 1.600E-07 | 2.700E-07 |
| 12525 | 627 | T627C | B | 5.5e-07 | 2e-07 | 6.1e-07 | 4.533E-07 | 2.214E-07 | 2.000E-07 | 6.100E-07 |
| 12526 | 627 | T627E | B | 2.5e-07 | 2.1e-07 | 4e-07 | 2.867E-07 | 1.002E-07 | 2.100E-07 | 4.000E-07 |
| 12527 | 627 | T627Q | B | 1.6e-07 | 2.6e-07 | 2.5e-07 | 2.233E-07 | 5.508E-08 | 1.600E-07 | 2.600E-07 |
| 12528 | 627 | T627G | B | 3.6e-07 | 1.6e-07 | 3.2e-07 | 2.800E-07 | 1.058E-07 | 1.600E-07 | 3.600E-07 |
| 12529 | 627 | T627H | B | 3.3e-07 | 2.3e-07 | 2.4e-07 | 2.667E-07 | 5.508E-08 | 2.300E-07 | 3.300E-07 |
| 12530 | 627 | T627I | B | 2.8e-07 | 3.7e-07 | 1.8e-07 | 2.767E-07 | 9.504E-08 | 1.800E-07 | 3.700E-07 |
| 12531 | 627 | T627L | B | 1.1e-07 | 2.3e-07 | 1.9e-07 | 1.767E-07 | 6.110E-08 | 1.100E-07 | 2.300E-07 |
| 12532 | 627 | T627K | B | 5.1e-07 | 2.2e-07 | 2.3e-07 | 3.200E-07 | 1.646E-07 | 2.200E-07 | 5.100E-07 |
| 12533 | 627 | T627M | B | 3.2e-07 | 3.6e-07 | 2.7e-07 | 3.167E-07 | 4.509E-08 | 2.700E-07 | 3.600E-07 |
| 12534 | 627 | T627F | B | 2.2e-07 | 2.8e-07 | 1.7e-07 | 2.233E-07 | 5.508E-08 | 1.700E-07 | 2.800E-07 |
| 12535 | 627 | T627P | B | 1.1e-07 | 5.9e-07 | 3.6e-07 | 3.533E-07 | 2.401E-07 | 1.100E-07 | 5.900E-07 |
| 12536 | 627 | T627S | B | 1.6e-07 | 2.3e-07 | 1.8e-07 | 1.900E-07 | 3.606E-08 | 1.600E-07 | 2.300E-07 |
| 12537 | 627 | T627T | B | 1.7e-07 | 1.4e-07 | 2.9e-07 | 2.000E-07 | 7.937E-08 | 1.400E-07 | 2.900E-07 |
| 12538 | 627 | T627W | B | 3.3e-07 | 2.5e-07 | 1.8e-07 | 2.533E-07 | 7.506E-08 | 1.800E-07 | 3.300E-07 |
| 12539 | 627 | T627Y | B | 3.5e-07 | 1.1e-07 | 3e-07 | 2.533E-07 | 1.266E-07 | 1.100E-07 | 3.500E-07 |
| 12540 | 627 | T627V | B | 1.7e-07 | 2.8e-07 | 2.5e-07 | 2.333E-07 | 5.686E-08 | 1.700E-07 | 2.800E-07 |
| 12541 | 628 | Q628A | B | 3.9e-07 | 2.7e-07 | 1.5e-07 | 2.700E-07 | 1.200E-07 | 1.500E-07 | 3.900E-07 |
| 12542 | 628 | Q628R | B | 2e-07 | 9.9e-08 | 1.3e-07 | 1.430E-07 | 5.174E-08 | 9.900E-08 | 2.000E-07 |
| 12543 | 628 | Q628N | B | 2.7e-07 | 1.5e-07 | 3.2e-07 | 2.467E-07 | 8.737E-08 | 1.500E-07 | 3.200E-07 |
| 12544 | 628 | Q628D | B | 1.4e-07 | 2.4e-07 | 2.2e-07 | 2.000E-07 | 5.292E-08 | 1.400E-07 | 2.400E-07 |
| 12545 | 628 | Q628C | B | 5.5e-07 | 1.6e-07 | 1.3e-07 | 2.800E-07 | 2.343E-07 | 1.300E-07 | 5.500E-07 |
| 12546 | 628 | Q628E | B | 1.4e-07 | 2.7e-07 | 1.5e-07 | 1.867E-07 | 7.234E-08 | 1.400E-07 | 2.700E-07 |
| 12547 | 628 | Q628Q | B | 9.5e-07 | 6.9e-07 | 1.4e-07 | 5.933E-07 | 4.136E-07 | 1.400E-07 | 9.500E-07 |
| 12548 | 628 | Q628G | B | 1.9e-07 | 3.7e-07 | 1.4e-07 | 2.333E-07 | 1.210E-07 | 1.400E-07 | 3.700E-07 |
| 12549 | 628 | Q628H | B | 1.2e-07 | 4.2e-07 | 3.4e-07 | 2.933E-07 | 1.553E-07 | 1.200E-07 | 4.200E-07 |
| 12550 | 628 | Q628I | B | 3.3e-07 | 4.8e-07 | 2.1e-07 | 3.400E-07 | 1.353E-07 | 2.100E-07 | 4.800E-07 |
| 12551 | 628 | Q628L | B | 4e-07 | 2.2e-07 | 3.6e-07 | 3.267E-07 | 9.452E-08 | 2.200E-07 | 4.000E-07 |
| 12552 | 628 | Q628K | B | 1.4e-07 | 3.1e-07 | 1.6e-07 | 2.033E-07 | 9.292E-08 | 1.400E-07 | 3.100E-07 |
| 12553 | 628 | Q628M | B | 1.9e-07 | 2.2e-07 | 2.9e-07 | 2.333E-07 | 5.132E-08 | 1.900E-07 | 2.900E-07 |
| 12554 | 628 | Q628F | B | 2e-07 | 2.2e-07 | 5e-07 | 3.067E-07 | 1.677E-07 | 2.000E-07 | 5.000E-07 |
| 12555 | 628 | Q628P | B | 1.4e-07 | 2.8e-07 | 7e-07 | 3.733E-07 | 2.914E-07 | 1.400E-07 | 7.000E-07 |
| 12556 | 628 | Q628S | B | 3.1e-07 | 2.9e-07 | 2.2e-07 | 2.733E-07 | 4.726E-08 | 2.200E-07 | 3.100E-07 |
| 12557 | 628 | Q628T | B | 2.1e-07 | 2.9e-07 | 3.4e-07 | 2.800E-07 | 6.557E-08 | 2.100E-07 | 3.400E-07 |
| 12558 | 628 | Q628W | B | 2.2e-07 | 2.1e-07 | 1.9e-07 | 2.067E-07 | 1.528E-08 | 1.900E-07 | 2.200E-07 |
| 12559 | 628 | Q628Y | B | 2.4e-07 | 3.2e-07 | 2.8e-07 | 2.800E-07 | 4.000E-08 | 2.400E-07 | 3.200E-07 |
| 12560 | 628 | Q628V | B | 2.6e-07 | 2.5e-07 | 5e-07 | 3.367E-07 | 1.415E-07 | 2.500E-07 | 5.000E-07 |
| 12561 | 629 | S629A | B | 3.4e-07 | 1.8e-07 | 5.4e-07 | 3.533E-07 | 1.804E-07 | 1.800E-07 | 5.400E-07 |
| 12562 | 629 | S629R | B | 2.8e-07 | 2.7e-07 | 3.8e-07 | 3.100E-07 | 6.083E-08 | 2.700E-07 | 3.800E-07 |
| 12563 | 629 | S629N | B | 5.7e-07 | 2.9e-07 | 1.4e-07 | 3.333E-07 | 2.183E-07 | 1.400E-07 | 5.700E-07 |
| 12564 | 629 | S629D | B | 2.2e-07 | 3e-07 | 3.1e-07 | 2.767E-07 | 4.933E-08 | 2.200E-07 | 3.100E-07 |
| 12565 | 629 | S629C | B | 1.8e-07 | 2.3e-07 | 4.9e-07 | 3.000E-07 | 1.664E-07 | 1.800E-07 | 4.900E-07 |
| 12566 | 629 | S629E | B | 1.6e-07 | 2e-07 | 2.3e-07 | 1.967E-07 | 3.512E-08 | 1.600E-07 | 2.300E-07 |
| 12567 | 629 | S629Q | B | 1.9e-07 | 3.6e-07 | 2.7e-07 | 2.733E-07 | 8.505E-08 | 1.900E-07 | 3.600E-07 |
| 12568 | 629 | S629G | B | 3.7e-07 | 2.9e-07 | 2.3e-07 | 2.967E-07 | 7.024E-08 | 2.300E-07 | 3.700E-07 |
| 12569 | 629 | S629H | B | 1.3e-07 | 1.4e-07 | 1.9e-07 | 1.533E-07 | 3.215E-08 | 1.300E-07 | 1.900E-07 |
| 12570 | 629 | S629I | B | 2.3e-07 | 3.7e-07 | 2.5e-07 | 2.833E-07 | 7.572E-08 | 2.300E-07 | 3.700E-07 |
| 12571 | 629 | S629L | B | 1.7e-07 | 2.4e-07 | 3.7e-07 | 2.600E-07 | 1.015E-07 | 1.700E-07 | 3.700E-07 |
| 12572 | 629 | S629K | B | 4.5e-07 | 2.3e-07 | 2.6e-07 | 3.133E-07 | 1.193E-07 | 2.300E-07 | 4.500E-07 |
| 12573 | 629 | S629M | B | 3.4e-07 | 1.9e-07 | 2.4e-07 | 2.567E-07 | 7.638E-08 | 1.900E-07 | 3.400E-07 |
| 12574 | 629 | S629F | B | 3e-07 | 5.2e-07 | 5e-07 | 4.400E-07 | 1.217E-07 | 3.000E-07 | 5.200E-07 |
| 12575 | 629 | S629P | B | 1.7e-07 | 3.4e-07 | 3.5e-07 | 2.867E-07 | 1.012E-07 | 1.700E-07 | 3.500E-07 |
| 12576 | 629 | S629S | B | 3.4e-07 | 3.4e-07 | 1.9e-07 | 2.900E-07 | 8.660E-08 | 1.900E-07 | 3.400E-07 |
| 12577 | 629 | S629T | B | 2.6e-07 | 4.7e-07 | 2.8e-07 | 3.367E-07 | 1.159E-07 | 2.600E-07 | 4.700E-07 |
| 12578 | 629 | S629W | B | 2.3e-07 | 1.4e-07 | 2.3e-07 | 2.000E-07 | 5.196E-08 | 1.400E-07 | 2.300E-07 |
| 12579 | 629 | S629Y | B | 3.9e-07 | 2.5e-07 | 3.6e-07 | 3.333E-07 | 7.371E-08 | 2.500E-07 | 3.900E-07 |

| | | | | | | | | | | |
|---|---|---|---|---|---|---|---|---|---|---|
| 12580 | 629 | S629V | B | 5.3e-07 | 3.2e-07 | 2.6e-07 | 3.700E-07 | 1.418E-07 | 2.600E-07 | 5.300E-07 |
| 12581 | 630 | P630A | B | 2.3e-07 | 3.5e-07 | 2e-07 | 2.600E-07 | 7.937E-08 | 2.000E-07 | 3.500E-07 |
| 12582 | 630 | P630R | B | 3.5e-07 | 2e-07 | 2.1e-07 | 2.533E-07 | 8.386E-08 | 2.000E-07 | 3.500E-07 |
| 12583 | 630 | P630N | B | 3.7e-07 | 2.6e-07 | 2.1e-07 | 2.800E-07 | 8.185E-08 | 2.100E-07 | 3.700E-07 |
| 12584 | 630 | P630D | B | 1.4e-07 | 1.6e-07 | 1.9e-07 | 1.633E-07 | 2.517E-08 | 1.400E-07 | 1.900E-07 |
| 12585 | 630 | P630C | B | 1.8e-07 | 3.2e-07 | 3.9e-07 | 2.967E-07 | 1.069E-07 | 1.800E-07 | 3.900E-07 |
| 12586 | 630 | P630E | B | 4e-07 | 2.6e-07 | 2.2e-07 | 2.933E-07 | 9.452E-08 | 2.200E-07 | 4.000E-07 |
| 12587 | 630 | P630Q | B | 1.9e-07 | 3.9e-07 | 2.5e-07 | 2.767E-07 | 1.026E-07 | 1.900E-07 | 3.900E-07 |
| 12588 | 630 | P630G | B | 4.2e-07 | 1.7e-07 | 4.4e-07 | 3.433E-07 | 1.504E-07 | 1.700E-07 | 4.400E-07 |
| 12589 | 630 | P630H | B | 7.7e-07 | 3.1e-07 | 4.1e-07 | 4.967E-07 | 2.419E-07 | 3.100E-07 | 7.700E-07 |
| 12590 | 630 | P630I | B | 1.8e-07 | 3.4e-07 | 2.9e-07 | 2.700E-07 | 8.185E-08 | 1.800E-07 | 3.400E-07 |
| 12591 | 630 | P630L | B | 2.1e-07 | 2.5e-07 | 3.9e-07 | 2.833E-07 | 9.452E-08 | 2.100E-07 | 3.900E-07 |
| 12592 | 630 | P630K | B | 2.8e-07 | 7.7e-07 | 3.2e-07 | 4.567E-07 | 2.721E-07 | 2.800E-07 | 7.700E-07 |
| 12593 | 630 | P630M | B | 4e-07 | 1.8e-07 | 2.2e-07 | 2.667E-07 | 1.172E-07 | 1.800E-07 | 4.000E-07 |
| 12594 | 630 | P630F | B | 2.8e-07 | 2.8e-07 | 1.1e-07 | 2.233E-07 | 9.815E-08 | 1.100E-07 | 2.800E-07 |
| 12595 | 630 | P630P | B | 4.9e-07 | 3e-07 | 3.4e-07 | 3.767E-07 | 1.002E-07 | 3.000E-07 | 4.900E-07 |
| 12596 | 630 | P630S | B | 1.8e-07 | 3.9e-07 | 2.7e-07 | 2.800E-07 | 1.054E-07 | 1.800E-07 | 3.900E-07 |
| 12597 | 630 | P630T | B | 3.5e-07 | 1.8e-07 | 3.9e-07 | 3.067E-07 | 1.115E-07 | 1.800E-07 | 3.900E-07 |
| 12598 | 630 | P630W | B | 5.3e-07 | 2.3e-07 | 3.6e-07 | 3.733E-07 | 1.504E-07 | 2.300E-07 | 5.300E-07 |
| 12599 | 630 | P630Y | B | 1.8e-07 | 2.1e-07 | 3.3e-07 | 2.400E-07 | 7.937E-08 | 1.800E-07 | 3.300E-07 |
| 12600 | 630 | P630V | B | 2.4e-07 | 1.9e-07 | 1.8e-07 | 2.033E-07 | 3.215E-08 | 1.800E-07 | 2.400E-07 |
| 12601 | 631 | S631A | B | 4.4e-07 | 1.3e-07 | 5.3e-07 | 3.667E-07 | 2.098E-07 | 1.300E-07 | 5.300E-07 |
| 12602 | 631 | S631R | B | 3.2e-07 | 4.1e-07 | 1.8e-07 | 3.033E-07 | 1.159E-07 | 1.800E-07 | 4.100E-07 |
| 12603 | 631 | S631N | B | 2.9e-07 | 3.4e-07 | 1.5e-07 | 2.600E-07 | 9.849E-08 | 1.500E-07 | 3.400E-07 |
| 12604 | 631 | S631D | B | 1.4e-07 | 2.9e-07 | 3.9e-07 | 2.733E-07 | 1.258E-07 | 1.400E-07 | 3.900E-07 |
| 12605 | 631 | S631C | B | 1.7e-07 | 5.3e-07 | 2.5e-07 | 3.167E-07 | 1.890E-07 | 1.700E-07 | 5.300E-07 |
| 12606 | 631 | S631E | B | 1.7e-07 | 1.9e-07 | 1.5e-07 | 1.700E-07 | 2.000E-08 | 1.500E-07 | 1.900E-07 |
| 12607 | 631 | S631Q | B | 2.3e-07 | 2.9e-07 | 2.4e-07 | 2.533E-07 | 3.215E-08 | 2.300E-07 | 2.900E-07 |
| 12608 | 631 | S631G | B | 3.4e-07 | 2.5e-07 | 4.7e-07 | 3.533E-07 | 1.106E-07 | 2.500E-07 | 4.700E-07 |
| 12609 | 631 | S631H | B | 1.7e-07 | 2.1e-07 | 4.3e-07 | 2.700E-07 | 1.400E-07 | 1.700E-07 | 4.300E-07 |
| 12610 | 631 | S631I | B | 1.4e-07 | 4.5e-07 | 2e-07 | 2.633E-07 | 1.644E-07 | 1.400E-07 | 4.500E-07 |
| 12611 | 631 | S631L | B | 1.4e-07 | 1.5e-07 | 5.5e-07 | 2.800E-07 | 2.339E-07 | 1.400E-07 | 5.500E-07 |
| 12612 | 631 | S631K | B | 1.5e-07 | 1.9e-07 | 1.7e-07 | 1.700E-07 | 2.000E-08 | 1.500E-07 | 1.900E-07 |
| 12613 | 631 | S631M | B | 1.7e-07 | 3.2e-07 | 2.3e-07 | 2.400E-07 | 7.550E-08 | 1.700E-07 | 3.200E-07 |
| 12614 | 631 | S631F | B | 3.5e-07 | 3.4e-07 | 3.3e-07 | 3.400E-07 | 1.000E-08 | 3.300E-07 | 3.500E-07 |
| 12615 | 631 | S631P | B | 2e-07 | 2.5e-07 | 2.3e-07 | 2.267E-07 | 2.517E-08 | 2.000E-07 | 2.500E-07 |
| 12616 | 631 | S631S | B | 3.4e-07 | 2.2e-07 | 1.8e-07 | 2.467E-07 | 8.327E-08 | 1.800E-07 | 3.400E-07 |
| 12617 | 631 | S631T | B | 2.1e-07 | 2.4e-07 | 3.3e-07 | 2.600E-07 | 6.245E-08 | 2.100E-07 | 3.300E-07 |
| 12618 | 631 | S631W | B | 5.5e-08 | 1.3e-07 | 2.4e-07 | 1.417E-07 | 9.305E-08 | 5.500E-08 | 2.400E-07 |
| 12619 | 631 | S631Y | B | 1.5e-07 | 3.2e-07 | 2e-07 | 2.233E-07 | 8.737E-08 | 1.500E-07 | 3.200E-07 |
| 12620 | 631 | S631V | B | 4.2e-07 | 3e-07 | 5.1e-07 | 4.100E-07 | 1.054E-07 | 3.000E-07 | 5.100E-07 |
| 12621 | 632 | S632A | B | 3.9e-07 | 1.4e-07 | 2.2e-07 | 2.500E-07 | 1.277E-07 | 1.400E-07 | 3.900E-07 |
| 12622 | 632 | S632R | B | 1.6e-07 | 2.6e-07 | 2.7e-07 | 2.300E-07 | 6.083E-08 | 1.600E-07 | 2.700E-07 |
| 12623 | 632 | S632N | B | 3.3e-07 | 2e-07 | 4.6e-07 | 3.300E-07 | 1.300E-07 | 2.000E-07 | 4.600E-07 |
| 12624 | 632 | S632D | B | 4.3e-07 | 4.1e-07 | 4e-07 | 4.133E-07 | 1.528E-08 | 4.000E-07 | 4.300E-07 |
| 12625 | 632 | S632C | B | 2.1e-07 | 2.9e-07 | 1.8e-07 | 2.267E-07 | 5.686E-08 | 1.800E-07 | 2.900E-07 |
| 12626 | 632 | S632E | B | 3.4e-07 | 2.6e-07 | 1.5e-07 | 2.500E-07 | 9.539E-08 | 1.500E-07 | 3.400E-07 |
| 12627 | 632 | S632Q | B | 1.4e-07 | 2.5e-07 | 4.4e-07 | 2.767E-07 | 1.518E-07 | 1.400E-07 | 4.400E-07 |
| 12628 | 632 | S632G | B | 2.2e-07 | 3.9e-07 | 2e-07 | 2.700E-07 | 1.044E-07 | 2.000E-07 | 3.900E-07 |
| 12629 | 632 | S632H | B | 2.7e-07 | 3.3e-07 | 2.5e-07 | 2.833E-07 | 4.163E-08 | 2.500E-07 | 3.300E-07 |
| 12630 | 632 | S632I | B | 4.9e-07 | 2e-07 | 2e-07 | 2.967E-07 | 1.674E-07 | 2.000E-07 | 4.900E-07 |
| 12631 | 632 | S632L | B | 4.7e-07 | 3.1e-07 | 1.4e-07 | 3.067E-07 | 1.650E-07 | 1.400E-07 | 4.700E-07 |
| 12632 | 632 | S632K | B | 4.7e-07 | 1e-07 | 3.2e-07 | 2.967E-07 | 1.861E-07 | 1.000E-07 | 4.700E-07 |
| 12633 | 632 | S632M | B | 1.9e-07 | 3.2e-07 | 2.1e-07 | 2.400E-07 | 7.000E-08 | 1.900E-07 | 3.200E-07 |
| 12634 | 632 | S632F | B | 1.2e-07 | 1.7e-07 | 2.5e-07 | 1.800E-07 | 6.557E-08 | 1.200E-07 | 2.500E-07 |
| 12635 | 632 | S632P | B | 2.9e-07 | 4.2e-07 | 1.6e-07 | 2.900E-07 | 1.300E-07 | 1.600E-07 | 4.200E-07 |
| 12636 | 632 | S632S | B | 4.8e-07 | 4.4e-07 | 1.5e-07 | 3.567E-07 | 1.801E-07 | 1.500E-07 | 4.800E-07 |
| 12637 | 632 | S632T | B | 2.9e-07 | 2.3e-07 | 2.1e-07 | 2.433E-07 | 4.163E-08 | 2.100E-07 | 2.900E-07 |
| 12638 | 632 | S632W | B | 1.2e-07 | 2.2e-07 | 3.9e-07 | 2.433E-07 | 1.365E-07 | 1.200E-07 | 3.900E-07 |
| 12639 | 632 | S632Y | B | 2.4e-07 | 2.3e-07 | 1.4e-07 | 2.033E-07 | 5.508E-08 | 1.400E-07 | 2.400E-07 |
| 12640 | 632 | S632V | B | 2.6e-07 | 2.2e-07 | 4.5e-07 | 3.100E-07 | 1.229E-07 | 2.200E-07 | 4.500E-07 |
| 12641 | 633 | L633A | B | 2.3e-07 | 4.6e-07 | 1.7e-07 | 2.867E-07 | 1.531E-07 | 1.700E-07 | 4.600E-07 |
| 12642 | 633 | L633R | B | 1.3e-07 | 3.5e-07 | 1.8e-07 | 2.200E-07 | 1.153E-07 | 1.300E-07 | 3.500E-07 |
| 12643 | 633 | L633N | B | 3.3e-07 | 2.5e-07 | 1.7e-07 | 2.500E-07 | 8.000E-08 | 1.700E-07 | 3.300E-07 |
| 12644 | 633 | L633D | B | 2.5e-07 | 3.1e-07 | 1.3e-07 | 2.300E-07 | 9.165E-08 | 1.300E-07 | 3.100E-07 |
| 12645 | 633 | L633C | B | 2.2e-07 | 4.3e-07 | 2e-07 | 2.833E-07 | 1.274E-07 | 2.000E-07 | 4.300E-07 |
| 12646 | 633 | L633E | B | 2.2e-07 | 2.8e-07 | 3.8e-07 | 2.933E-07 | 8.083E-08 | 2.200E-07 | 3.800E-07 |
| 12647 | 633 | L633Q | B | 2.2e-07 | 4e-07 | 2.7e-07 | 2.967E-07 | 9.292E-08 | 2.200E-07 | 4.000E-07 |
| 12648 | 633 | L633G | B | 1.6e-07 | 2.7e-07 | 2.7e-07 | 2.333E-07 | 6.351E-08 | 1.600E-07 | 2.700E-07 |
| 12649 | 633 | L633H | B | 2.6e-07 | 2e-07 | 2.6e-07 | 2.400E-07 | 3.464E-08 | 2.000E-07 | 2.600E-07 |
| 12650 | 633 | L633I | B | 3.2e-07 | 1e-07 | 2.9e-07 | 2.367E-07 | 1.193E-07 | 1.000E-07 | 3.200E-07 |
| 12651 | 633 | L633L | B | 2.6e-07 | 3.2e-07 | 6.5e-07 | 4.100E-07 | 2.100E-07 | 2.600E-07 | 6.500E-07 |
| 12652 | 633 | L633K | B | 3.8e-07 | 2.3e-07 | 3.2e-07 | 3.100E-07 | 7.550E-08 | 2.300E-07 | 3.800E-07 |
| 12653 | 633 | L633M | B | 3.1e-07 | 5.3e-07 | 3.2e-07 | 3.867E-07 | 1.242E-07 | 3.100E-07 | 5.300E-07 |
| 12654 | 633 | L633F | B | 2e-07 | 4.9e-07 | 1.4e-07 | 2.767E-07 | 1.872E-07 | 1.400E-07 | 4.900E-07 |
| 12655 | 633 | L633P | B | 5.2e-07 | 8.2e-08 | 7.5e-07 | 4.507E-07 | 3.394E-07 | 8.200E-08 | 7.500E-07 |
| 12656 | 633 | L633S | B | 2.5e-07 | 5.3e-07 | 3.1e-07 | 3.633E-07 | 1.474E-07 | 2.500E-07 | 5.300E-07 |
| 12657 | 633 | L633T | B | 2.4e-07 | 3.5e-07 | 2.1e-07 | 2.667E-07 | 7.371E-08 | 2.100E-07 | 3.500E-07 |
| 12658 | 633 | L633W | B | 2.8e-07 | 3.6e-07 | 2e-07 | 2.800E-07 | 8.000E-08 | 2.000E-07 | 3.600E-07 |
| 12659 | 633 | L633Y | B | 3.8e-07 | 3.1e-07 | 2.5e-07 | 3.133E-07 | 6.506E-08 | 2.500E-07 | 3.800E-07 |
| 12660 | 633 | L633V | B | 2.5e-07 | 2.1e-07 | 1.5e-07 | 2.033E-07 | 5.033E-08 | 1.500E-07 | 2.500E-07 |
| 12661 | 634 | S634A | B | 2e-07 | 1.2e-07 | 3.8e-07 | 2.333E-07 | 1.332E-07 | 1.200E-07 | 3.800E-07 |
| 12662 | 634 | S634R | B | 3.5e-07 | 1.7e-07 | 2.1e-07 | 2.433E-07 | 9.452E-08 | 1.700E-07 | 3.500E-07 |
| 12663 | 634 | S634N | B | 2.1e-07 | 3.8e-07 | 1.7e-07 | 2.533E-07 | 1.115E-07 | 1.700E-07 | 3.800E-07 |
| 12664 | 634 | S634D | B | 1.9e-07 | 3.4e-07 | 3.6e-07 | 2.967E-07 | 9.292E-08 | 1.900E-07 | 3.600E-07 |

| | | | | | | | | | | |
|---|---|---|---|---|---|---|---|---|---|---|
| 12665 | 634 | S634C | B | 1.8e-07 | 2.4e-07 | 2.8e-07 | 2.333E-07 | 5.033E-08 | 1.800E-07 | 2.800E-07 |
| 12666 | 634 | S634E | B | 2.9e-07 | 2.5e-07 | 2.1e-07 | 2.500E-07 | 4.000E-08 | 2.100E-07 | 2.900E-07 |
| 12667 | 634 | S634Q | B | 1.8e-07 | 2.2e-07 | 3.3e-07 | 2.433E-07 | 7.767E-08 | 1.800E-07 | 3.300E-07 |
| 12668 | 634 | S634G | B | 2.5e-07 | 3.6e-07 | 3.3e-07 | 3.133E-07 | 5.686E-08 | 2.500E-07 | 3.600E-07 |
| 12669 | 634 | S634H | B | 1.4e-07 | 2.6e-07 | 3.3e-07 | 2.433E-07 | 9.609E-08 | 1.400E-07 | 3.300E-07 |
| 12670 | 634 | S634I | B | 1.7e-07 | 2.3e-07 | 6.6e-07 | 3.533E-07 | 2.673E-07 | 1.700E-07 | 6.600E-07 |
| 12671 | 634 | S634L | B | 4.7e-07 | 3.2e-07 | 2.8e-07 | 3.567E-07 | 1.002E-07 | 2.800E-07 | 4.700E-07 |
| 12672 | 634 | S634K | B | 4.6e-07 | 3.7e-07 | 4e-07 | 4.100E-07 | 4.583E-08 | 3.700E-07 | 4.600E-07 |
| 12673 | 634 | S634M | B | 2.3e-07 | 1.9e-07 | 1.4e-07 | 1.867E-07 | 4.509E-08 | 1.400E-07 | 2.300E-07 |
| 12674 | 634 | S634F | B | 2.8e-07 | 2.7e-07 | 2.3e-07 | 2.600E-07 | 2.646E-08 | 2.300E-07 | 2.800E-07 |
| 12675 | 634 | S634P | B | 6.6e-08 | 1.7e-07 | 2.4e-07 | 1.587E-07 | 8.755E-08 | 6.600E-08 | 2.400E-07 |
| 12676 | 634 | S634S | B | 2.6e-07 | 3.1e-07 | 7e-07 | 4.233E-07 | 2.409E-07 | 2.600E-07 | 7.000E-07 |
| 12677 | 634 | S634T | B | 3e-07 | 2.9e-07 | 2.4e-07 | 2.767E-07 | 3.215E-08 | 2.400E-07 | 3.000E-07 |
| 12678 | 634 | S634W | B | 2.3e-07 | 4.1e-07 | 2.3e-07 | 2.900E-07 | 1.039E-07 | 2.300E-07 | 4.100E-07 |
| 12679 | 634 | S634Y | B | 3.4e-07 | 1.8e-07 | 2.6e-07 | 2.600E-07 | 8.000E-08 | 1.800E-07 | 3.400E-07 |
| 12680 | 634 | S634V | B | 2.5e-07 | 1.2e-07 | 4.3e-07 | 2.667E-07 | 1.557E-07 | 1.200E-07 | 4.300E-07 |
| 12681 | 635 | A635A | B | 3.3e-07 | 1.7e-07 | 1.8e-07 | 2.267E-07 | 8.963E-08 | 1.700E-07 | 3.300E-07 |
| 12682 | 635 | A635R | B | 4.8e-07 | 3.4e-07 | 2.1e-07 | 3.433E-07 | 1.350E-07 | 2.100E-07 | 4.800E-07 |
| 12683 | 635 | A635N | B | 2.4e-07 | 3.1e-07 | 3.2e-07 | 2.900E-07 | 4.359E-08 | 2.400E-07 | 3.200E-07 |
| 12684 | 635 | A635D | B | 3.9e-07 | 2e-07 | 1.1e-07 | 2.333E-07 | 1.429E-07 | 1.100E-07 | 3.900E-07 |
| 12685 | 635 | A635C | B | 4.5e-07 | 1.6e-07 | 4.3e-07 | 3.467E-07 | 1.620E-07 | 1.600E-07 | 4.500E-07 |
| 12686 | 635 | A635E | B | 2.6e-07 | 2.9e-07 | 2.3e-07 | 2.600E-07 | 3.000E-08 | 2.300E-07 | 2.900E-07 |
| 12687 | 635 | A635Q | B | 4e-07 | 1.6e-07 | 1.9e-07 | 2.500E-07 | 1.308E-07 | 1.600E-07 | 4.000E-07 |
| 12688 | 635 | A635G | B | 1.9e-07 | 2.2e-07 | 2.2e-07 | 2.100E-07 | 1.732E-08 | 1.900E-07 | 2.200E-07 |
| 12689 | 635 | A635H | B | 1.2e-07 | 3.4e-07 | 2.1e-07 | 2.233E-07 | 1.106E-07 | 1.200E-07 | 3.400E-07 |
| 12690 | 635 | A635I | B | 1.3e-07 | 1.2e-07 | 4.6e-07 | 2.367E-07 | 1.935E-07 | 1.200E-07 | 4.600E-07 |
| 12691 | 635 | A635L | B | 1.8e-07 | 1.6e-07 | 3.2e-07 | 2.200E-07 | 8.718E-08 | 1.600E-07 | 3.200E-07 |
| 12692 | 635 | A635K | B | 6.2e-07 | 6.9e-07 | 2.1e-07 | 5.067E-07 | 2.593E-07 | 2.100E-07 | 6.900E-07 |
| 12693 | 635 | A635M | B | 3.1e-07 | 2.6e-07 | 4.7e-07 | 3.467E-07 | 1.097E-07 | 2.600E-07 | 4.700E-07 |
| 12694 | 635 | A635F | B | 2.7e-07 | 3.4e-07 | 3.1e-07 | 3.067E-07 | 3.512E-08 | 2.700E-07 | 3.400E-07 |
| 12695 | 635 | A635P | B | 1.7e-07 | 5.7e-07 | 1.4e-07 | 2.933E-07 | 2.401E-07 | 1.400E-07 | 5.700E-07 |
| 12696 | 635 | A635S | B | 1.5e-07 | 2.9e-07 | 3e-07 | 2.467E-07 | 8.386E-08 | 1.500E-07 | 3.000E-07 |
| 12697 | 635 | A635T | B | 3.3e-07 | 3.3e-07 | 2.7e-07 | 3.100E-07 | 3.464E-08 | 2.700E-07 | 3.300E-07 |
| 12698 | 635 | A635W | B | 1.8e-07 | 2.2e-07 | 7.1e-07 | 3.700E-07 | 2.951E-07 | 1.800E-07 | 7.100E-07 |
| 12699 | 635 | A635Y | B | 1.5e-07 | 1.4e-07 | 2e-07 | 1.633E-07 | 3.215E-08 | 1.400E-07 | 2.000E-07 |
| 12700 | 635 | A635V | B | 2e-07 | 4.7e-07 | 1.7e-07 | 2.800E-07 | 1.652E-07 | 1.700E-07 | 4.700E-07 |
| 12701 | 636 | S636A | B | 1.3e-07 | 2.9e-07 | 1.8e-07 | 2.000E-07 | 8.185E-08 | 1.300E-07 | 2.900E-07 |
| 12702 | 636 | S636R | B | 4.2e-07 | 3e-07 | 3.8e-07 | 3.667E-07 | 6.110E-08 | 3.000E-07 | 4.200E-07 |
| 12703 | 636 | S636N | B | 3.1e-07 | 1.7e-07 | 1.2e-07 | 2.000E-07 | 9.849E-08 | 1.200E-07 | 3.100E-07 |
| 12704 | 636 | S636D | B | 1.4e-07 | 2.6e-07 | 4.1e-07 | 2.700E-07 | 1.353E-07 | 1.400E-07 | 4.100E-07 |
| 12705 | 636 | S636C | B | 2e-07 | 2.5e-07 | 3.2e-07 | 2.567E-07 | 6.028E-08 | 2.000E-07 | 3.200E-07 |
| 12706 | 636 | S636E | B | 2.1e-07 | 3.2e-07 | 3.1e-07 | 2.800E-07 | 6.083E-08 | 2.100E-07 | 3.200E-07 |
| 12707 | 636 | S636Q | B | 2.3e-07 | 2.6e-07 | 2.7e-07 | 2.533E-07 | 2.082E-08 | 2.300E-07 | 2.700E-07 |
| 12708 | 636 | S636G | B | 5.2e-07 | 2.5e-07 | 2.8e-07 | 3.500E-07 | 1.480E-07 | 2.500E-07 | 5.200E-07 |
| 12709 | 636 | S636H | B | 1.6e-07 | 2.1e-07 | 3.7e-07 | 2.467E-07 | 1.097E-07 | 1.600E-07 | 3.700E-07 |
| 12710 | 636 | S636I | B | 2.3e-07 | 3.6e-07 | 2.7e-07 | 2.867E-07 | 6.658E-08 | 2.300E-07 | 3.600E-07 |
| 12711 | 636 | S636L | B | 3.6e-07 | 2.4e-07 | 2.7e-07 | 2.900E-07 | 6.245E-08 | 2.400E-07 | 3.600E-07 |
| 12712 | 636 | S636K | B | 4.3e-07 | 1.8e-07 | 3.6e-07 | 3.233E-07 | 1.290E-07 | 1.800E-07 | 4.300E-07 |
| 12713 | 636 | S636M | B | 4e-07 | 1.9e-07 | 4.7e-07 | 3.533E-07 | 1.457E-07 | 1.900E-07 | 4.700E-07 |
| 12714 | 636 | S636F | B | 3.9e-07 | 2.5e-07 | 3e-07 | 3.133E-07 | 7.095E-08 | 2.500E-07 | 3.900E-07 |
| 12715 | 636 | S636P | B | 4.7e-07 | 2.9e-07 | 3.2e-07 | 3.600E-07 | 9.644E-08 | 2.900E-07 | 4.700E-07 |
| 12716 | 636 | S636S | B | 3.2e-07 | 1.7e-07 | 1.5e-07 | 2.133E-07 | 9.292E-08 | 1.500E-07 | 3.200E-07 |
| 12717 | 636 | S636T | B | 2.2e-07 | 3.3e-07 | 2.5e-07 | 2.667E-07 | 5.686E-08 | 2.200E-07 | 3.300E-07 |
| 12718 | 636 | S636W | B | 1.7e-07 | 2.4e-07 | 3.9e-07 | 2.667E-07 | 1.124E-07 | 1.700E-07 | 3.900E-07 |
| 12719 | 636 | S636Y | B | 2.1e-07 | 5.3e-07 | 3e-07 | 3.467E-07 | 1.650E-07 | 2.100E-07 | 5.300E-07 |
| 12720 | 636 | S636V | B | 2.2e-07 | 1.4e-07 | 1.6e-07 | 1.733E-07 | 4.163E-08 | 1.400E-07 | 2.200E-07 |
| 12721 | 637 | V637A | B | 2.6e-07 | 3.1e-07 | 3e-07 | 2.900E-07 | 2.646E-08 | 2.600E-07 | 3.100E-07 |
| 12722 | 637 | V637R | B | 3.7e-07 | 2.1e-07 | 4.1e-07 | 3.300E-07 | 1.058E-07 | 2.100E-07 | 4.100E-07 |
| 12723 | 637 | V637N | B | 1.8e-07 | 3.3e-07 | 3.8e-07 | 2.967E-07 | 1.041E-07 | 1.800E-07 | 3.800E-07 |
| 12724 | 637 | V637D | B | 5e-07 | 1.7e-07 | 2.8e-07 | 3.167E-07 | 1.680E-07 | 1.700E-07 | 5.000E-07 |
| 12725 | 637 | V637C | B | 3.1e-07 | 3.2e-07 | 2.7e-07 | 3.000E-07 | 2.646E-08 | 2.700E-07 | 3.200E-07 |
| 12726 | 637 | V637E | B | 1.5e-07 | 2.4e-07 | 1.8e-07 | 1.900E-07 | 4.583E-08 | 1.500E-07 | 2.400E-07 |
| 12727 | 637 | V637Q | B | 3.1e-07 | 9.4e-08 | 3.5e-07 | 2.513E-07 | 1.377E-07 | 9.400E-08 | 3.500E-07 |
| 12728 | 637 | V637G | B | 2.7e-07 | 2.6e-07 | 4.6e-07 | 3.300E-07 | 1.127E-07 | 2.600E-07 | 4.600E-07 |
| 12729 | 637 | V637H | B | 4.7e-07 | 2.4e-07 | 2.1e-07 | 3.067E-07 | 1.422E-07 | 2.100E-07 | 4.700E-07 |
| 12730 | 637 | V637I | B | 2.4e-07 | 3.2e-07 | 2.1e-07 | 2.567E-07 | 5.686E-08 | 2.100E-07 | 3.200E-07 |
| 12731 | 637 | V637L | B | 2.4e-07 | 2.7e-07 | 1.8e-07 | 2.300E-07 | 4.583E-08 | 1.800E-07 | 2.700E-07 |
| 12732 | 637 | V637K | B | 2.9e-07 | 1.1e-07 | 2.2e-07 | 2.067E-07 | 9.074E-08 | 1.100E-07 | 2.900E-07 |
| 12733 | 637 | V637M | B | 1.5e-07 | 2.4e-07 | 1.8e-07 | 1.900E-07 | 4.583E-08 | 1.500E-07 | 2.400E-07 |
| 12734 | 637 | V637F | B | 1.8e-07 | 3.6e-07 | 3.2e-07 | 2.867E-07 | 9.452E-08 | 1.800E-07 | 3.600E-07 |
| 12735 | 637 | V637P | B | 3e-07 | 2.5e-07 | 2.4e-07 | 2.633E-07 | 3.215E-08 | 2.400E-07 | 3.000E-07 |
| 12736 | 637 | V637S | B | 1.7e-07 | 8.1e-08 | 1.8e-07 | 1.437E-07 | 5.450E-08 | 8.100E-08 | 1.800E-07 |
| 12737 | 637 | V637T | B | 2.7e-07 | 2e-07 | 2.7e-07 | 2.467E-07 | 4.041E-08 | 2.000E-07 | 2.700E-07 |
| 12738 | 637 | V637W | B | 2e-07 | 1.3e-07 | 2.9e-07 | 2.067E-07 | 8.021E-08 | 1.300E-07 | 2.900E-07 |
| 12739 | 637 | V637Y | B | 1.7e-07 | 2.8e-07 | 2.3e-07 | 2.267E-07 | 5.508E-08 | 1.700E-07 | 2.800E-07 |
| 12740 | 637 | V637V | B | 3.8e-07 | 2.5e-07 | 3.2e-07 | 3.167E-07 | 6.506E-08 | 2.500E-07 | 3.800E-07 |
| 12741 | 638 | G638A | B | 1.5e-07 | 2.4e-07 | 5.2e-07 | 3.033E-07 | 1.930E-07 | 1.500E-07 | 5.200E-07 |
| 12742 | 638 | G638R | B | 2.6e-07 | 1.2e-07 | 2.8e-07 | 2.200E-07 | 8.718E-08 | 1.200E-07 | 2.800E-07 |
| 12743 | 638 | G638N | B | 2e-07 | 3.9e-07 | 1.4e-07 | 2.433E-07 | 1.305E-07 | 1.400E-07 | 3.900E-07 |
| 12744 | 638 | G638D | B | 1.6e-07 | 2.2e-07 | 4.4e-07 | 2.733E-07 | 1.474E-07 | 1.600E-07 | 4.400E-07 |
| 12745 | 638 | G638C | B | 2.1e-07 | 1.3e-07 | 3.3e-07 | 2.233E-07 | 1.007E-07 | 1.300E-07 | 3.300E-07 |
| 12746 | 638 | G638E | B | 1.9e-07 | 2.2e-07 | 2.8e-07 | 2.300E-07 | 4.583E-08 | 1.900E-07 | 2.800E-07 |
| 12747 | 638 | G638Q | B | 3e-07 | 3.6e-07 | 2.6e-07 | 3.067E-07 | 5.033E-08 | 2.600E-07 | 3.600E-07 |
| 12748 | 638 | G638G | B | 3.7e-07 | 4.8e-07 | 4.2e-07 | 4.233E-07 | 5.508E-08 | 3.700E-07 | 4.800E-07 |
| 12749 | 638 | G638H | B | 2.1e-07 | 1.8e-07 | 2.8e-07 | 2.233E-07 | 5.132E-08 | 1.800E-07 | 2.800E-07 |

| | | | | | | | | | | |
|---|---|---|---|---|---|---|---|---|---|---|
| 12750 | 638 | G638I | B | 2.1e-07 | 9.6e-08 | 2.5e-07 | 1.853E-07 | 7.991E-08 | 9.600E-08 | 2.500E-07 |
| 12751 | 638 | G638L | B | 3.1e-07 | 2e-07 | 2.5e-07 | 2.533E-07 | 5.508E-08 | 2.000E-07 | 3.100E-07 |
| 12752 | 638 | G638K | B | 1.9e-07 | 1.5e-07 | 3.3e-07 | 2.233E-07 | 9.452E-08 | 1.500E-07 | 3.300E-07 |
| 12753 | 638 | G638M | B | 1.7e-07 | 3e-07 | 3e-07 | 2.567E-07 | 7.506E-08 | 1.700E-07 | 3.000E-07 |
| 12754 | 638 | G638F | B | 4.6e-07 | 2.8e-07 | 2.6e-07 | 3.333E-07 | 1.102E-07 | 2.600E-07 | 4.600E-07 |
| 12755 | 638 | G638P | B | 5.8e-07 | 1.9e-07 | 2.8e-07 | 3.500E-07 | 2.042E-07 | 1.900E-07 | 5.800E-07 |
| 12756 | 638 | G638S | B | 8.2e-08 | 2.6e-07 | 7.6e-08 | 1.393E-07 | 1.045E-07 | 7.600E-08 | 2.600E-07 |
| 12757 | 638 | G638T | B | 3.5e-07 | 2.4e-07 | 1.1e-07 | 2.333E-07 | 1.201E-07 | 1.100E-07 | 3.500E-07 |
| 12758 | 638 | G638W | B | 2.9e-07 | 7.9e-08 | 1.5e-07 | 1.730E-07 | 1.074E-07 | 7.900E-08 | 2.900E-07 |
| 12759 | 638 | G638Y | B | 1.7e-07 | 2.9e-07 | 2.2e-07 | 2.267E-07 | 6.028E-08 | 1.700E-07 | 2.900E-07 |
| 12760 | 638 | G638V | B | 1.6e-07 | 3.3e-07 | 3.3e-07 | 2.733E-07 | 9.815E-08 | 1.600E-07 | 3.300E-07 |
| 12761 | 639 | D639A | B | 1.8e-07 | 4.9e-07 | 5.7e-07 | 4.133E-07 | 2.060E-07 | 1.800E-07 | 5.700E-07 |
| 12762 | 639 | D639R | B | 1.5e-07 | 2.3e-07 | 2.6e-07 | 2.133E-07 | 5.686E-08 | 1.500E-07 | 2.600E-07 |
| 12763 | 639 | D639N | B | 1.4e-07 | 1.8e-07 | 2e-07 | 1.733E-07 | 3.055E-08 | 1.400E-07 | 2.000E-07 |
| 12764 | 639 | D639D | B | 2.8e-07 | 2.2e-07 | 2.4e-07 | 2.467E-07 | 3.055E-08 | 2.200E-07 | 2.800E-07 |
| 12765 | 639 | D639C | B | 2.4e-07 | 2.4e-07 | 2.4e-07 | 2.400E-07 | 0.000E+00 | 2.400E-07 | 2.400E-07 |
| 12766 | 639 | D639E | B | 2.1e-07 | 1.9e-07 | 2e-07 | 2.000E-07 | 1.000E-08 | 1.900E-07 | 2.100E-07 |
| 12767 | 639 | D639Q | B | 4.5e-07 | 1.6e-07 | 2.5e-07 | 2.867E-07 | 1.484E-07 | 1.600E-07 | 4.500E-07 |
| 12768 | 639 | D639G | B | 1.9e-07 | 2e-07 | 2.2e-07 | 2.033E-07 | 1.528E-08 | 1.900E-07 | 2.200E-07 |
| 12769 | 639 | D639H | B | 2.3e-07 | 3.8e-07 | 2.1e-07 | 2.733E-07 | 9.292E-08 | 2.100E-07 | 3.800E-07 |
| 12770 | 639 | D639I | B | 1.1e-07 | 2.4e-07 | 1.9e-07 | 1.800E-07 | 6.557E-08 | 1.100E-07 | 2.400E-07 |
| 12771 | 639 | D639L | B | 2.4e-07 | 2.1e-07 | 2.6e-07 | 2.367E-07 | 2.517E-08 | 2.100E-07 | 2.600E-07 |
| 12772 | 639 | D639K | B | 2.4e-07 | 4.4e-07 | 2.1e-07 | 2.967E-07 | 1.250E-07 | 2.100E-07 | 4.400E-07 |
| 12773 | 639 | D639M | B | 3.7e-07 | 2e-07 | 2e-07 | 2.567E-07 | 9.815E-08 | 2.000E-07 | 3.700E-07 |
| 12774 | 639 | D639F | B | 1.6e-07 | 3.8e-07 | 1.6e-07 | 2.333E-07 | 1.270E-07 | 1.600E-07 | 3.800E-07 |
| 12775 | 639 | D639P | B | 2.4e-07 | 2.6e-07 | 2.5e-07 | 2.500E-07 | 1.000E-08 | 2.400E-07 | 2.600E-07 |
| 12776 | 639 | D639S | B | 3e-07 | 1.8e-07 | 4.8e-07 | 3.200E-07 | 1.510E-07 | 1.800E-07 | 4.800E-07 |
| 12777 | 639 | D639T | B | 3.1e-07 | 2.7e-07 | 3.9e-07 | 3.233E-07 | 6.110E-08 | 2.700E-07 | 3.900E-07 |
| 12778 | 639 | D639W | B | 2.8e-07 | 2e-07 | 1.8e-07 | 2.200E-07 | 5.292E-08 | 1.800E-07 | 2.800E-07 |
| 12779 | 639 | D639Y | B | 4.7e-07 | 2.8e-07 | 3.3e-07 | 3.600E-07 | 9.849E-08 | 2.800E-07 | 4.700E-07 |
| 12780 | 639 | D639V | B | 1.4e-07 | 4.4e-07 | 3.9e-07 | 3.233E-07 | 1.607E-07 | 1.400E-07 | 4.400E-07 |
| 12781 | 640 | R640A | B | 4.1e-07 | 2.4e-07 | 3.2e-07 | 3.233E-07 | 8.505E-08 | 2.400E-07 | 4.100E-07 |
| 12782 | 640 | R640R | B | 3.4e-07 | 4.6e-07 | 3.3e-07 | 3.767E-07 | 7.234E-08 | 3.300E-07 | 4.600E-07 |
| 12783 | 640 | R640N | B | 3e-07 | 2.7e-07 | 4e-07 | 3.233E-07 | 6.807E-08 | 2.700E-07 | 4.000E-07 |
| 12784 | 640 | R640D | B | 2.9e-07 | 9.4e-08 | 2.1e-07 | 1.980E-07 | 9.855E-08 | 9.400E-08 | 2.900E-07 |
| 12785 | 640 | R640C | B | 2.8e-07 | 6.5e-07 | 2.5e-07 | 3.933E-07 | 2.228E-07 | 2.500E-07 | 6.500E-07 |
| 12786 | 640 | R640E | B | 1.9e-07 | 4.2e-07 | 2.6e-07 | 2.900E-07 | 1.179E-07 | 1.900E-07 | 4.200E-07 |
| 12787 | 640 | R640Q | B | 6.3e-07 | 1.5e-07 | 1.9e-07 | 3.233E-07 | 2.663E-07 | 1.500E-07 | 6.300E-07 |
| 12788 | 640 | R640G | B | 2.3e-07 | 1.9e-07 | 1.5e-07 | 1.900E-07 | 4.000E-08 | 1.500E-07 | 2.300E-07 |
| 12789 | 640 | R640H | B | 2.4e-07 | 3.3e-07 | 3e-07 | 2.900E-07 | 4.583E-08 | 2.400E-07 | 3.300E-07 |
| 12790 | 640 | R640I | B | 1.8e-07 | 1.2e-07 | 1.6e-07 | 1.533E-07 | 3.055E-08 | 1.200E-07 | 1.800E-07 |
| 12791 | 640 | R640L | B | 2.6e-07 | 3.3e-07 | 2.9e-07 | 2.933E-07 | 3.512E-08 | 2.600E-07 | 3.300E-07 |
| 12792 | 640 | R640K | B | 2.6e-07 | 2.6e-07 | 3.4e-07 | 2.867E-07 | 4.619E-08 | 2.600E-07 | 3.400E-07 |
| 12793 | 640 | R640M | B | 3e-07 | 2.4e-07 | 2e-07 | 2.467E-07 | 5.033E-08 | 2.000E-07 | 3.000E-07 |
| 12794 | 640 | R640F | B | 1.4e-07 | 2.9e-07 | 4.2e-07 | 2.833E-07 | 1.401E-07 | 1.400E-07 | 4.200E-07 |
| 12795 | 640 | R640P | B | 3.1e-07 | 1.8e-07 | 4.4e-07 | 3.100E-07 | 1.300E-07 | 1.800E-07 | 4.400E-07 |
| 12796 | 640 | R640S | B | 2.7e-07 | 3.2e-07 | 2.2e-07 | 2.700E-07 | 5.000E-08 | 2.200E-07 | 3.200E-07 |
| 12797 | 640 | R640T | B | 3.5e-07 | 1.1e-07 | 2.4e-07 | 2.333E-07 | 1.201E-07 | 1.100E-07 | 3.500E-07 |
| 12798 | 640 | R640W | B | 2e-07 | 4.4e-07 | 3.2e-07 | 3.200E-07 | 1.200E-07 | 2.000E-07 | 4.400E-07 |
| 12799 | 640 | R640Y | B | 2.6e-07 | 2.9e-07 | 1.1e-07 | 2.200E-07 | 9.644E-08 | 1.100E-07 | 2.900E-07 |
| 12800 | 640 | R640V | B | 2.7e-07 | 2.3e-07 | 4.3e-07 | 3.100E-07 | 1.058E-07 | 2.300E-07 | 4.300E-07 |
| 12801 | 641 | V641A | B | 2.1e-07 | 1.1e-07 | 2.5e-07 | 1.900E-07 | 7.211E-08 | 1.100E-07 | 2.500E-07 |
| 12802 | 641 | V641R | B | 4.1e-07 | 2.4e-07 | 3.5e-07 | 3.333E-07 | 8.622E-08 | 2.400E-07 | 4.100E-07 |
| 12803 | 641 | V641N | B | 3.7e-07 | 4.1e-07 | 3.1e-07 | 3.633E-07 | 5.033E-08 | 3.100E-07 | 4.100E-07 |
| 12804 | 641 | V641D | B | 2.7e-07 | 1.4e-07 | 1.6e-07 | 1.900E-07 | 7.000E-08 | 1.400E-07 | 2.700E-07 |
| 12805 | 641 | V641C | B | 2.4e-07 | 3.6e-07 | 4.9e-07 | 3.633E-07 | 1.250E-07 | 2.400E-07 | 4.900E-07 |
| 12806 | 641 | V641E | B | 2.1e-07 | 1.5e-07 | 2.4e-07 | 2.000E-07 | 4.583E-08 | 1.500E-07 | 2.400E-07 |
| 12807 | 641 | V641Q | B | 1.9e-07 | 5.4e-07 | 1.9e-07 | 3.067E-07 | 2.021E-07 | 1.900E-07 | 5.400E-07 |
| 12808 | 641 | V641G | B | 3.7e-07 | 2e-07 | 3e-07 | 2.900E-07 | 8.544E-08 | 2.000E-07 | 3.700E-07 |
| 12809 | 641 | V641H | B | 3.2e-07 | 3.5e-07 | 5e-07 | 3.900E-07 | 9.644E-08 | 3.200E-07 | 5.000E-07 |
| 12810 | 641 | V641I | B | 2.7e-07 | 1.4e-07 | 3e-07 | 2.367E-07 | 8.505E-08 | 1.400E-07 | 3.000E-07 |
| 12811 | 641 | V641L | B | 2.4e-07 | 4.3e-07 | 2.3e-07 | 3.000E-07 | 1.127E-07 | 2.300E-07 | 4.300E-07 |
| 12812 | 641 | V641K | B | 1.9e-07 | 1.9e-07 | 1.1e-07 | 1.633E-07 | 4.619E-08 | 1.100E-07 | 1.900E-07 |
| 12813 | 641 | V641M | B | 9.5e-08 | 2.6e-07 | 5.7e-07 | 3.083E-07 | 2.412E-07 | 9.500E-08 | 5.700E-07 |
| 12814 | 641 | V641F | B | 1.4e-07 | 1.7e-07 | 1.8e-07 | 1.633E-07 | 2.082E-08 | 1.400E-07 | 1.800E-07 |
| 12815 | 641 | V641P | B | 4.7e-07 | 4e-07 | 7.1e-07 | 5.267E-07 | 1.626E-07 | 4.000E-07 | 7.100E-07 |
| 12816 | 641 | V641S | B | 1.8e-07 | 1.9e-07 | 2.2e-07 | 1.967E-07 | 2.082E-08 | 1.800E-07 | 2.200E-07 |
| 12817 | 641 | V641T | B | 1.8e-07 | 6.7e-08 | 5.8e-07 | 2.757E-07 | 2.695E-07 | 6.700E-08 | 5.800E-07 |
| 12818 | 641 | V641W | B | 2.9e-07 | 1.1e-07 | 3.4e-07 | 2.467E-07 | 1.210E-07 | 1.100E-07 | 3.400E-07 |
| 12819 | 641 | V641Y | B | 1.5e-07 | 2.4e-07 | 1e-07 | 1.633E-07 | 7.095E-08 | 1.000E-07 | 2.400E-07 |
| 12820 | 641 | V641V | B | 1.8e-07 | 4.4e-07 | 2.6e-07 | 2.933E-07 | 1.332E-07 | 1.800E-07 | 4.400E-07 |
| 12821 | 642 | T642A | B | 1.7e-07 | 2.1e-07 | 2.4e-07 | 2.067E-07 | 3.512E-08 | 1.700E-07 | 2.400E-07 |
| 12822 | 642 | T642R | B | 4.4e-07 | 2.3e-07 | 2.4e-07 | 3.033E-07 | 1.185E-07 | 2.300E-07 | 4.400E-07 |
| 12823 | 642 | T642N | B | 2.4e-07 | 3.7e-07 | 2.4e-07 | 2.833E-07 | 7.506E-08 | 2.400E-07 | 3.700E-07 |
| 12824 | 642 | T642D | B | 1.7e-07 | 1.8e-07 | 2.9e-07 | 2.133E-07 | 6.658E-08 | 1.700E-07 | 2.900E-07 |
| 12825 | 642 | T642C | B | 3.3e-07 | 3.5e-07 | 2.4e-07 | 3.067E-07 | 5.859E-08 | 2.400E-07 | 3.500E-07 |
| 12826 | 642 | T642E | B | 1.9e-07 | 2.6e-07 | 3.6e-07 | 2.700E-07 | 8.544E-08 | 1.900E-07 | 3.600E-07 |
| 12827 | 642 | T642Q | B | 2.9e-07 | 2.7e-07 | 2.8e-07 | 2.800E-07 | 1.000E-08 | 2.700E-07 | 2.900E-07 |
| 12828 | 642 | T642G | B | 3.5e-07 | 2.1e-07 | 1.5e-07 | 2.367E-07 | 1.026E-07 | 1.500E-07 | 3.500E-07 |
| 12829 | 642 | T642H | B | 3.8e-07 | 2.2e-07 | 2e-07 | 2.667E-07 | 9.866E-08 | 2.000E-07 | 3.800E-07 |
| 12830 | 642 | T642I | B | 3.3e-07 | 4.2e-07 | 3.7e-07 | 3.733E-07 | 4.509E-08 | 3.300E-07 | 4.200E-07 |
| 12831 | 642 | T642L | B | 5e-07 | 2.2e-07 | 1.4e-07 | 2.867E-07 | 1.890E-07 | 1.400E-07 | 5.000E-07 |
| 12832 | 642 | T642K | B | 4.6e-07 | 2.1e-07 | 2.4e-07 | 3.033E-07 | 1.365E-07 | 2.100E-07 | 4.600E-07 |
| 12833 | 642 | T642M | B | 2.9e-07 | 1.5e-07 | 3.2e-07 | 2.533E-07 | 9.074E-08 | 1.500E-07 | 3.200E-07 |
| 12834 | 642 | T642F | B | 1.8e-07 | 3.5e-07 | 2.3e-07 | 2.533E-07 | 8.737E-08 | 1.800E-07 | 3.500E-07 |

| | | | | | | | | | | |
|---|---|---|---|---|---|---|---|---|---|---|
| 12835 | 642 | T642P | B | 5.1e-07 | 4.4e-07 | 1.4e-07 | 3.633E-07 | 1.966E-07 | 1.400E-07 | 5.100E-07 |
| 12836 | 642 | T642S | B | 2e-07 | 2.9e-07 | 2.3e-07 | 2.400E-07 | 4.583E-08 | 2.000E-07 | 2.900E-07 |
| 12837 | 642 | T642T | B | 1.9e-07 | 2.3e-07 | 2.5e-07 | 2.233E-07 | 3.055E-08 | 1.900E-07 | 2.500E-07 |
| 12838 | 642 | T642W | B | 6.6e-08 | 3.2e-07 | 2.8e-07 | 2.220E-07 | 1.366E-07 | 6.600E-08 | 3.200E-07 |
| 12839 | 642 | T642Y | B | 1.2e-07 | 3.3e-07 | 1.9e-07 | 2.133E-07 | 1.069E-07 | 1.200E-07 | 3.300E-07 |
| 12840 | 642 | T642V | B | 1.5e-07 | 1.6e-07 | 1.6e-07 | 1.567E-07 | 5.774E-09 | 1.500E-07 | 1.600E-07 |
| 12841 | 643 | I643A | B | 2.2e-07 | 1.9e-07 | 1e-07 | 1.700E-07 | 6.245E-08 | 1.000E-07 | 2.200E-07 |
| 12842 | 643 | I643R | B | 1.3e-07 | 2.3e-07 | 4.2e-07 | 2.600E-07 | 1.473E-07 | 1.300E-07 | 4.200E-07 |
| 12843 | 643 | I643N | B | 1.4e-07 | 3.2e-07 | 1.5e-07 | 2.033E-07 | 1.012E-07 | 1.400E-07 | 3.200E-07 |
| 12844 | 643 | I643D | B | 4.6e-07 | 1.9e-07 | 2.5e-07 | 3.000E-07 | 1.418E-07 | 1.900E-07 | 4.600E-07 |
| 12845 | 643 | I643C | B | 2.9e-07 | 8.3e-07 | 1.4e-07 | 4.200E-07 | 3.629E-07 | 1.400E-07 | 8.300E-07 |
| 12846 | 643 | I643E | B | 2.6e-07 | 2.6e-07 | 1.7e-07 | 2.300E-07 | 5.196E-08 | 1.700E-07 | 2.600E-07 |
| 12847 | 643 | I643Q | B | 1.8e-07 | 2.8e-07 | 1.4e-07 | 2.000E-07 | 7.211E-08 | 1.400E-07 | 2.800E-07 |
| 12848 | 643 | I643G | B | 1.6e-07 | 7.3e-07 | 3.2e-07 | 4.033E-07 | 2.940E-07 | 1.600E-07 | 7.300E-07 |
| 12849 | 643 | I643H | B | 3.7e-07 | 2.5e-07 | 1.3e-07 | 2.500E-07 | 1.200E-07 | 1.300E-07 | 3.700E-07 |
| 12850 | 643 | I643I | B | 3.2e-07 | 1.8e-07 | 1.7e-07 | 2.233E-07 | 8.386E-08 | 1.700E-07 | 3.200E-07 |
| 12851 | 643 | I643L | B | 1.2e-07 | 2e-07 | 2.2e-07 | 1.800E-07 | 5.292E-08 | 1.200E-07 | 2.200E-07 |
| 12852 | 643 | I643K | B | 2.5e-07 | 3.4e-07 | 1.4e-07 | 2.433E-07 | 1.002E-07 | 1.400E-07 | 3.400E-07 |
| 12853 | 643 | I643M | B | 2.1e-07 | 8.2e-08 | 9.4e-08 | 1.287E-07 | 7.069E-08 | 8.200E-08 | 2.100E-07 |
| 12854 | 643 | I643F | B | 2.4e-07 | 1.3e-07 | 2.4e-07 | 2.033E-07 | 6.351E-08 | 1.300E-07 | 2.400E-07 |
| 12855 | 643 | I643P | B | 5.5e-07 | 1.7e-07 | 3.6e-07 | 3.600E-07 | 1.900E-07 | 1.700E-07 | 5.500E-07 |
| 12856 | 643 | I643S | B | 3.9e-07 | 1.4e-07 | 2.1e-07 | 2.467E-07 | 1.290E-07 | 1.400E-07 | 3.900E-07 |
| 12857 | 643 | I643T | B | 3.2e-07 | 3.3e-07 | 2.1e-07 | 2.867E-07 | 6.658E-08 | 2.100E-07 | 3.300E-07 |
| 12858 | 643 | I643W | B | 3.3e-07 | 1.6e-07 | 3.7e-07 | 2.867E-07 | 1.115E-07 | 1.600E-07 | 3.700E-07 |
| 12859 | 643 | I643Y | B | 2.8e-07 | 4e-07 | 2.3e-07 | 3.033E-07 | 8.737E-08 | 2.300E-07 | 4.000E-07 |
| 12860 | 643 | I643V | B | 1.1e-07 | 4.8e-07 | 1.2e-07 | 2.367E-07 | 2.108E-07 | 1.100E-07 | 4.800E-07 |
| 12861 | 644 | T644A | B | 3.5e-07 | 2.2e-07 | 3.7e-07 | 3.133E-07 | 8.145E-08 | 2.200E-07 | 3.700E-07 |
| 12862 | 644 | T644R | B | 2e-07 | 3.2e-07 | 3.1e-07 | 2.767E-07 | 6.658E-08 | 2.000E-07 | 3.200E-07 |
| 12863 | 644 | T644N | B | 2e-07 | 1.6e-07 | 2.5e-07 | 2.033E-07 | 4.509E-08 | 1.600E-07 | 2.500E-07 |
| 12864 | 644 | T644D | B | 2.9e-07 | 1.4e-07 | 3.4e-07 | 2.567E-07 | 1.041E-07 | 1.400E-07 | 3.400E-07 |
| 12865 | 644 | T644C | B | 2.1e-07 | 1.9e-07 | 1.4e-07 | 1.800E-07 | 3.606E-08 | 1.400E-07 | 2.100E-07 |
| 12866 | 644 | T644E | B | 2.5e-07 | 4e-07 | 3.7e-07 | 3.400E-07 | 7.937E-08 | 2.500E-07 | 4.000E-07 |
| 12867 | 644 | T644Q | B | 5.4e-07 | 2.1e-07 | 1e-06 | 5.833E-07 | 3.968E-07 | 2.100E-07 | 1.000E-06 |
| 12868 | 644 | T644G | B | 2.2e-07 | 2e-07 | 4.8e-07 | 3.000E-07 | 1.562E-07 | 2.000E-07 | 4.800E-07 |
| 12869 | 644 | T644H | B | 1.9e-07 | 1.5e-07 | 1.4e-07 | 1.600E-07 | 2.646E-08 | 1.400E-07 | 1.900E-07 |
| 12870 | 644 | T644I | B | 4.5e-07 | 3.4e-07 | 1.2e-07 | 3.033E-07 | 1.680E-07 | 1.200E-07 | 4.500E-07 |
| 12871 | 644 | T644L | B | 3.3e-07 | 1.9e-07 | 1.8e-07 | 2.333E-07 | 8.386E-08 | 1.800E-07 | 3.300E-07 |
| 12872 | 644 | T644K | B | 2.5e-07 | 1.9e-07 | 4.4e-07 | 2.933E-07 | 1.305E-07 | 1.900E-07 | 4.400E-07 |
| 12873 | 644 | T644M | B | 2.3e-07 | 1.6e-07 | 3.3e-07 | 2.400E-07 | 8.544E-08 | 1.600E-07 | 3.300E-07 |
| 12874 | 644 | T644F | B | 1.7e-07 | 2.5e-07 | 1.2e-07 | 1.800E-07 | 6.557E-08 | 1.200E-07 | 2.500E-07 |
| 12875 | 644 | T644P | B | 4.8e-07 | 1.8e-07 | 5.6e-07 | 4.067E-07 | 2.003E-07 | 1.800E-07 | 5.600E-07 |
| 12876 | 644 | T644S | B | 1.4e-07 | 1.1e-07 | 2e-07 | 1.500E-07 | 4.583E-08 | 1.100E-07 | 2.000E-07 |
| 12877 | 644 | T644T | B | 4e-07 | 3.4e-07 | 3.3e-07 | 3.567E-07 | 3.786E-08 | 3.300E-07 | 4.000E-07 |
| 12878 | 644 | T644W | B | 2.5e-07 | 2.9e-07 | 8.8e-07 | 4.733E-07 | 3.528E-07 | 2.500E-07 | 8.800E-07 |
| 12879 | 644 | T644Y | B | 2.2e-07 | 3.7e-07 | 2.2e-07 | 2.700E-07 | 8.660E-08 | 2.200E-07 | 3.700E-07 |
| 12880 | 644 | T644V | B | 2.3e-07 | 2.2e-07 | 3.4e-07 | 2.633E-07 | 6.658E-08 | 2.200E-07 | 3.400E-07 |
| 12881 | 645 | C645A | B | 2.8e-07 | 2.2e-07 | 3.3e-07 | 2.767E-07 | 5.508E-08 | 2.200E-07 | 3.300E-07 |
| 12882 | 645 | C645R | B | 3.7e-07 | 1.7e-07 | 4.4e-07 | 3.267E-07 | 1.401E-07 | 1.700E-07 | 4.400E-07 |
| 12883 | 645 | C645N | B | 2.5e-07 | 2.2e-07 | 2.9e-07 | 2.533E-07 | 3.512E-08 | 2.200E-07 | 2.900E-07 |
| 12884 | 645 | C645D | B | 3.1e-07 | 3.3e-07 | 2.2e-07 | 2.867E-07 | 5.859E-08 | 2.200E-07 | 3.300E-07 |
| 12885 | 645 | C645C | B | 2.2e-07 | 2.3e-07 | 1.6e-07 | 2.033E-07 | 3.786E-08 | 1.600E-07 | 2.300E-07 |
| 12886 | 645 | C645E | B | 3.8e-07 | 4.9e-07 | 2.9e-07 | 3.867E-07 | 1.002E-07 | 2.900E-07 | 4.900E-07 |
| 12887 | 645 | C645Q | B | 2.9e-07 | 2.1e-07 | 3.8e-07 | 2.933E-07 | 8.505E-08 | 2.100E-07 | 3.800E-07 |
| 12888 | 645 | C645G | B | 1e-07 | 3.6e-07 | 2.6e-07 | 2.400E-07 | 1.311E-07 | 1.000E-07 | 3.600E-07 |
| 12889 | 645 | C645H | B | 1.3e-07 | 5.1e-07 | 1.9e-07 | 2.767E-07 | 2.043E-07 | 1.300E-07 | 5.100E-07 |
| 12890 | 645 | C645I | B | 4.5e-07 | 2e-07 | 2.2e-07 | 2.900E-07 | 1.389E-07 | 2.000E-07 | 4.500E-07 |
| 12891 | 645 | C645L | B | 1.9e-07 | 2.7e-07 | 3e-07 | 2.533E-07 | 5.686E-08 | 1.900E-07 | 3.000E-07 |
| 12892 | 645 | C645K | B | 4.5e-07 | 3.4e-07 | 2.8e-07 | 3.567E-07 | 8.622E-08 | 2.800E-07 | 4.500E-07 |
| 12893 | 645 | C645M | B | 2.3e-07 | 1.6e-07 | 3.6e-07 | 2.500E-07 | 1.015E-07 | 1.600E-07 | 3.600E-07 |
| 12894 | 645 | C645F | B | 3.1e-07 | 3e-07 | 2.6e-07 | 2.900E-07 | 2.646E-08 | 2.600E-07 | 3.100E-07 |
| 12895 | 645 | C645P | B | 3.4e-07 | 2.1e-07 | 3.8e-07 | 3.100E-07 | 8.888E-08 | 2.100E-07 | 3.800E-07 |
| 12896 | 645 | C645S | B | 2.5e-07 | 2.4e-07 | 1.5e-07 | 2.133E-07 | 5.508E-08 | 1.500E-07 | 2.500E-07 |
| 12897 | 645 | C645T | B | 1.7e-07 | 3.3e-07 | 1.4e-07 | 2.133E-07 | 1.021E-07 | 1.400E-07 | 3.300E-07 |
| 12898 | 645 | C645W | B | 1.6e-07 | 2.2e-07 | 1.8e-07 | 1.867E-07 | 3.055E-08 | 1.600E-07 | 2.200E-07 |
| 12899 | 645 | C645Y | B | 1.4e-07 | 1.7e-07 | 2.1e-07 | 1.733E-07 | 3.512E-08 | 1.400E-07 | 2.100E-07 |
| 12900 | 645 | C645V | B | 2.5e-07 | 3.3e-07 | 1.6e-07 | 2.467E-07 | 8.505E-08 | 1.600E-07 | 3.300E-07 |
| 12901 | 646 | K646A | B | 1.5e-07 | 1.6e-07 | 3.9e-07 | 2.333E-07 | 1.358E-07 | 1.500E-07 | 3.900E-07 |
| 12902 | 646 | K646R | B | 6.4e-07 | 1.5e-07 | 1.7e-07 | 3.200E-07 | 2.773E-07 | 1.500E-07 | 6.400E-07 |
| 12903 | 646 | K646N | B | 4.6e-07 | 2.4e-07 | 3.1e-07 | 3.367E-07 | 1.124E-07 | 2.400E-07 | 4.600E-07 |
| 12904 | 646 | K646D | B | 2.5e-07 | 1.3e-07 | 1.4e-07 | 1.733E-07 | 6.658E-08 | 1.300E-07 | 2.500E-07 |
| 12905 | 646 | K646C | B | 1.9e-07 | 3.2e-07 | 3.1e-07 | 2.733E-07 | 7.234E-08 | 1.900E-07 | 3.200E-07 |
| 12906 | 646 | K646E | B | 2.3e-07 | 3.2e-07 | 1.6e-07 | 2.367E-07 | 8.021E-08 | 1.600E-07 | 3.200E-07 |
| 12907 | 646 | K646Q | B | 1.6e-07 | 2.2e-07 | 3.2e-07 | 2.333E-07 | 8.083E-08 | 1.600E-07 | 3.200E-07 |
| 12908 | 646 | K646G | B | 2.7e-07 | 4.5e-07 | 2e-07 | 3.067E-07 | 1.290E-07 | 2.000E-07 | 4.500E-07 |
| 12909 | 646 | K646H | B | 2e-07 | 2.7e-07 | 5.5e-07 | 3.400E-07 | 1.852E-07 | 2.000E-07 | 5.500E-07 |
| 12910 | 646 | K646I | B | 1e-07 | 2.4e-07 | 4.3e-07 | 2.567E-07 | 1.656E-07 | 1.000E-07 | 4.300E-07 |
| 12911 | 646 | K646L | B | 2.9e-07 | 1.2e-07 | 1.9e-07 | 2.000E-07 | 8.544E-08 | 1.200E-07 | 2.900E-07 |
| 12912 | 646 | K646K | B | 2.3e-07 | 3.3e-07 | 5.6e-07 | 3.733E-07 | 1.692E-07 | 2.300E-07 | 5.600E-07 |
| 12913 | 646 | K646M | B | 3.1e-07 | 3.7e-07 | 3.5e-07 | 3.433E-07 | 3.055E-08 | 3.100E-07 | 3.700E-07 |
| 12914 | 646 | K646F | B | 1.6e-07 | 2.3e-07 | 2e-07 | 1.967E-07 | 3.512E-08 | 1.600E-07 | 2.300E-07 |
| 12915 | 646 | K646P | B | 3.2e-07 | 3.1e-07 | 2e-07 | 2.767E-07 | 6.658E-08 | 2.000E-07 | 3.200E-07 |
| 12916 | 646 | K646S | B | 2.3e-07 | 3.4e-07 | 1.6e-07 | 2.433E-07 | 9.074E-08 | 1.600E-07 | 3.400E-07 |
| 12917 | 646 | K646T | B | 1.7e-07 | 3.4e-07 | 2.4e-07 | 2.500E-07 | 8.544E-08 | 1.700E-07 | 3.400E-07 |
| 12918 | 646 | K646W | B | 1.2e-07 | 1.8e-07 | 3.1e-07 | 2.033E-07 | 9.713E-08 | 1.200E-07 | 3.100E-07 |
| 12919 | 646 | K646Y | B | 1.3e-07 | 3.4e-07 | 2.7e-07 | 2.467E-07 | 1.069E-07 | 1.300E-07 | 3.400E-07 |

| | | | | | | | | | | |
|---|---|---|---|---|---|---|---|---|---|---|
| 12920 | 646 | K646V | B | 2.9e-07 | 2.1e-07 | 4.6e-07 | 3.200E-07 | 1.277E-07 | 2.100E-07 | 4.600E-07 |
| 12921 | 647 | A647A | B | 2.3e-07 | 4.4e-07 | 3.8e-07 | 3.500E-07 | 1.082E-07 | 2.300E-07 | 4.400E-07 |
| 12922 | 647 | A647R | B | 6.1e-07 | 4.2e-07 | 1.7e-07 | 4.000E-07 | 2.207E-07 | 1.700E-07 | 6.100E-07 |
| 12923 | 647 | A647N | B | 3e-07 | 3.3e-07 | 2e-07 | 2.767E-07 | 6.807E-08 | 2.000E-07 | 3.300E-07 |
| 12924 | 647 | A647D | B | 2e-07 | 3.1e-07 | 1.6e-07 | 2.233E-07 | 7.767E-08 | 1.600E-07 | 3.100E-07 |
| 12925 | 647 | A647C | B | 1.9e-07 | 2.1e-07 | 3e-07 | 2.333E-07 | 5.859E-08 | 1.900E-07 | 3.000E-07 |
| 12926 | 647 | A647E | B | 1.7e-07 | 2.1e-07 | 3.1e-07 | 2.300E-07 | 7.211E-08 | 1.700E-07 | 3.100E-07 |
| 12927 | 647 | A647Q | B | 1.4e-07 | 3.7e-07 | 3.9e-07 | 3.000E-07 | 1.389E-07 | 1.400E-07 | 3.900E-07 |
| 12928 | 647 | A647G | B | 1.3e-07 | 2.2e-07 | 3.2e-07 | 2.233E-07 | 9.504E-08 | 1.300E-07 | 3.200E-07 |
| 12929 | 647 | A647H | B | 1.4e-07 | 9.7e-08 | 3.1e-07 | 1.823E-07 | 1.126E-07 | 9.700E-08 | 3.100E-07 |
| 12930 | 647 | A647I | B | 4.4e-07 | 4.7e-07 | 1.1e-07 | 3.400E-07 | 1.997E-07 | 1.100E-07 | 4.700E-07 |
| 12931 | 647 | A647L | B | 1.4e-07 | 2.7e-07 | 4.8e-07 | 2.967E-07 | 1.716E-07 | 1.400E-07 | 4.800E-07 |
| 12932 | 647 | A647K | B | 1.6e-07 | 1.1e-07 | 1.5e-07 | 1.400E-07 | 2.646E-08 | 1.100E-07 | 1.600E-07 |
| 12933 | 647 | A647M | B | 3.5e-07 | 2.8e-07 | 2.6e-07 | 2.967E-07 | 4.726E-08 | 2.600E-07 | 3.500E-07 |
| 12934 | 647 | A647F | B | 1.8e-07 | 1.3e-07 | 1e-07 | 1.367E-07 | 4.041E-08 | 1.000E-07 | 1.800E-07 |
| 12935 | 647 | A647P | B | 1.4e-07 | 1.7e-07 | 3.1e-07 | 2.067E-07 | 9.074E-08 | 1.400E-07 | 3.100E-07 |
| 12936 | 647 | A647S | B | 5.3e-07 | 3.1e-07 | 1.9e-07 | 3.433E-07 | 1.724E-07 | 1.900E-07 | 5.300E-07 |
| 12937 | 647 | A647T | B | 9.7e-08 | 3.8e-07 | 1.1e-07 | 1.957E-07 | 1.598E-07 | 9.700E-08 | 3.800E-07 |
| 12938 | 647 | A647W | B | 3.8e-07 | 4.1e-07 | 1.1e-07 | 3.000E-07 | 1.652E-07 | 1.100E-07 | 4.100E-07 |
| 12939 | 647 | A647Y | B | 1.9e-07 | 3.5e-07 | 1.8e-07 | 2.400E-07 | 9.539E-08 | 1.800E-07 | 3.500E-07 |
| 12940 | 647 | A647V | B | 1.8e-07 | 2.2e-07 | 4.1e-07 | 2.700E-07 | 1.229E-07 | 1.800E-07 | 4.100E-07 |
| 12941 | 648 | S648A | B | 1.8e-07 | 4e-07 | 2.5e-07 | 2.767E-07 | 1.124E-07 | 1.800E-07 | 4.000E-07 |
| 12942 | 648 | S648R | B | 2.2e-07 | 2.4e-07 | 1.5e-07 | 2.033E-07 | 4.726E-08 | 1.500E-07 | 2.400E-07 |
| 12943 | 648 | S648N | B | 4.1e-07 | 4.8e-07 | 1.3e-07 | 3.400E-07 | 1.852E-07 | 1.300E-07 | 4.800E-07 |
| 12944 | 648 | S648D | B | 1.9e-07 | 4.4e-07 | 1e-07 | 2.433E-07 | 1.762E-07 | 1.000E-07 | 4.400E-07 |
| 12945 | 648 | S648C | B | 2.6e-07 | 1.8e-07 | 6.7e-07 | 3.700E-07 | 2.629E-07 | 1.800E-07 | 6.700E-07 |
| 12946 | 648 | S648E | B | 1.6e-07 | 1.9e-07 | 2.4e-07 | 1.967E-07 | 4.041E-08 | 1.600E-07 | 2.400E-07 |
| 12947 | 648 | S648Q | B | 2.3e-07 | 9.5e-08 | 5.4e-07 | 2.883E-07 | 2.282E-07 | 9.500E-08 | 5.400E-07 |
| 12948 | 648 | S648G | B | 1.9e-07 | 2.9e-07 | 2.1e-07 | 2.300E-07 | 5.292E-08 | 1.900E-07 | 2.900E-07 |
| 12949 | 648 | S648H | B | 2.9e-07 | 2.9e-07 | 2e-07 | 2.600E-07 | 5.196E-08 | 2.000E-07 | 2.900E-07 |
| 12950 | 648 | S648I | B | 1.4e-07 | 1.6e-07 | 2.6e-07 | 1.867E-07 | 6.429E-08 | 1.400E-07 | 2.600E-07 |
| 12951 | 648 | S648L | B | 3.9e-07 | 9.8e-08 | 3.5e-07 | 2.793E-07 | 1.583E-07 | 9.800E-08 | 3.900E-07 |
| 12952 | 648 | S648K | B | 4e-07 | 1.6e-07 | 1.8e-07 | 2.467E-07 | 1.332E-07 | 1.600E-07 | 4.000E-07 |
| 12953 | 648 | S648M | B | 1.8e-07 | 2.9e-07 | 2.2e-07 | 2.300E-07 | 5.568E-08 | 1.800E-07 | 2.900E-07 |
| 12954 | 648 | S648F | B | 3e-07 | 3e-07 | 8.1e-08 | 2.270E-07 | 1.264E-07 | 8.100E-08 | 3.000E-07 |
| 12955 | 648 | S648P | B | 2.9e-07 | 3.4e-07 | 2.4e-07 | 2.900E-07 | 5.000E-08 | 2.400E-07 | 3.400E-07 |
| 12956 | 648 | S648S | B | 3.6e-07 | 3.6e-07 | 2.1e-07 | 3.100E-07 | 8.660E-08 | 2.100E-07 | 3.600E-07 |
| 12957 | 648 | S648T | B | 2.7e-07 | 3e-07 | 1.4e-07 | 2.367E-07 | 8.505E-08 | 1.400E-07 | 3.000E-07 |
| 12958 | 648 | S648W | B | 2.4e-07 | 2.6e-07 | 2.4e-07 | 2.467E-07 | 1.155E-08 | 2.400E-07 | 2.600E-07 |
| 12959 | 648 | S648Y | B | 2.5e-07 | 2.5e-07 | 3.5e-07 | 2.833E-07 | 5.774E-08 | 2.500E-07 | 3.500E-07 |
| 12960 | 648 | S648V | B | 2.1e-07 | 3.3e-07 | 4.2e-07 | 3.200E-07 | 1.054E-07 | 2.100E-07 | 4.200E-07 |
| 12961 | 649 | Q649A | B | 1.8e-07 | 3.2e-07 | 2.2e-07 | 2.400E-07 | 7.211E-08 | 1.800E-07 | 3.200E-07 |
| 12962 | 649 | Q649R | B | 3.1e-07 | 2.2e-07 | 7.2e-07 | 4.167E-07 | 2.665E-07 | 2.200E-07 | 7.200E-07 |
| 12963 | 649 | Q649N | B | 5.5e-07 | 2.3e-07 | 3.2e-07 | 3.667E-07 | 1.650E-07 | 2.300E-07 | 5.500E-07 |
| 12964 | 649 | Q649D | B | 2.2e-07 | 1.8e-07 | 3.1e-07 | 2.367E-07 | 6.658E-08 | 1.800E-07 | 3.100E-07 |
| 12965 | 649 | Q649C | B | 1.5e-07 | 2.7e-07 | 1.8e-07 | 2.000E-07 | 6.245E-08 | 1.500E-07 | 2.700E-07 |
| 12966 | 649 | Q649E | B | 1.9e-07 | 2.9e-07 | 4.1e-07 | 2.967E-07 | 1.102E-07 | 1.900E-07 | 4.100E-07 |
| 12967 | 649 | Q649Q | B | 5.8e-07 | 3.2e-07 | 2.3e-07 | 3.767E-07 | 1.818E-07 | 2.300E-07 | 5.800E-07 |
| 12968 | 649 | Q649G | B | 2.7e-07 | 4.1e-07 | 1.9e-07 | 2.900E-07 | 1.114E-07 | 1.900E-07 | 4.100E-07 |
| 12969 | 649 | Q649H | B | 1.7e-07 | 2.1e-07 | 3.3e-07 | 2.367E-07 | 8.327E-08 | 1.700E-07 | 3.300E-07 |
| 12970 | 649 | Q649I | B | 2.1e-07 | 1.6e-07 | 3.4e-07 | 2.367E-07 | 9.292E-08 | 1.600E-07 | 3.400E-07 |
| 12971 | 649 | Q649L | B | 2.1e-07 | 4e-07 | 4.1e-07 | 3.400E-07 | 1.127E-07 | 2.100E-07 | 4.100E-07 |
| 12972 | 649 | Q649K | B | 3.2e-07 | 2.1e-07 | 2.1e-07 | 2.467E-07 | 6.351E-08 | 2.100E-07 | 3.200E-07 |
| 12973 | 649 | Q649M | B | 1.4e-07 | 2.8e-07 | 2.2e-07 | 2.133E-07 | 7.024E-08 | 1.400E-07 | 2.800E-07 |
| 12974 | 649 | Q649F | B | 2.6e-07 | 2e-07 | 2.1e-07 | 2.233E-07 | 3.215E-08 | 2.000E-07 | 2.600E-07 |
| 12975 | 649 | Q649P | B | 4e-07 | 1.8e-07 | 3.9e-07 | 3.233E-07 | 1.242E-07 | 1.800E-07 | 4.000E-07 |
| 12976 | 649 | Q649S | B | 2.4e-07 | 2.9e-07 | 3.1e-07 | 2.800E-07 | 3.606E-08 | 2.400E-07 | 3.100E-07 |
| 12977 | 649 | Q649T | B | 3e-07 | 2.7e-07 | 6.9e-07 | 4.200E-07 | 2.343E-07 | 2.700E-07 | 6.900E-07 |
| 12978 | 649 | Q649W | B | 2.2e-07 | 2.8e-07 | 1.8e-07 | 2.267E-07 | 5.033E-08 | 1.800E-07 | 2.800E-07 |
| 12979 | 649 | Q649Y | B | 3.2e-07 | 4.9e-07 | 4.2e-07 | 4.100E-07 | 8.544E-08 | 3.200E-07 | 4.900E-07 |
| 12980 | 649 | Q649V | B | 3.8e-07 | 2e-07 | 1.2e-07 | 2.333E-07 | 1.332E-07 | 1.200E-07 | 3.800E-07 |
| 12981 | 650 | D650A | B | 2.7e-07 | 2.9e-07 | 1.9e-07 | 2.500E-07 | 5.292E-08 | 1.900E-07 | 2.900E-07 |
| 12982 | 650 | D650R | B | 1.9e-07 | 2.5e-07 | 3.6e-07 | 2.667E-07 | 8.622E-08 | 1.900E-07 | 3.600E-07 |
| 12983 | 650 | D650N | B | 1.3e-07 | 1.2e-07 | 4.3e-07 | 2.267E-07 | 1.762E-07 | 1.200E-07 | 4.300E-07 |
| 12984 | 650 | D650D | B | 3e-07 | 2.1e-07 | 1.3e-07 | 2.133E-07 | 8.505E-08 | 1.300E-07 | 3.000E-07 |
| 12985 | 650 | D650C | B | 1.4e-07 | 1.9e-07 | 2.4e-07 | 1.900E-07 | 5.000E-08 | 1.400E-07 | 2.400E-07 |
| 12986 | 650 | D650E | B | 2.6e-07 | 4.5e-07 | 2.4e-07 | 3.167E-07 | 1.159E-07 | 2.400E-07 | 4.500E-07 |
| 12987 | 650 | D650Q | B | 2.7e-07 | 1.4e-07 | 2.6e-07 | 2.233E-07 | 7.234E-08 | 1.400E-07 | 2.700E-07 |
| 12988 | 650 | D650G | B | 6.2e-07 | 1.5e-07 | 3.4e-07 | 3.700E-07 | 2.364E-07 | 1.500E-07 | 6.200E-07 |
| 12989 | 650 | D650H | B | 1.4e-07 | 2.5e-07 | 2.8e-07 | 2.233E-07 | 7.371E-08 | 1.400E-07 | 2.800E-07 |
| 12990 | 650 | D650I | B | 2.2e-07 | 9.8e-08 | 2.5e-07 | 1.893E-07 | 8.051E-08 | 9.800E-08 | 2.500E-07 |
| 12991 | 650 | D650L | B | 1.1e-07 | 6.1e-07 | 1.4e-07 | 2.867E-07 | 2.804E-07 | 1.100E-07 | 6.100E-07 |
| 12992 | 650 | D650K | B | 2.5e-07 | 2.3e-07 | 1.7e-07 | 2.167E-07 | 4.163E-08 | 1.700E-07 | 2.500E-07 |
| 12993 | 650 | D650M | B | 4.3e-07 | 3.3e-07 | 1.9e-07 | 3.167E-07 | 1.206E-07 | 1.900E-07 | 4.300E-07 |
| 12994 | 650 | D650F | B | 2.8e-07 | 1.7e-07 | 1.3e-07 | 1.933E-07 | 7.767E-08 | 1.300E-07 | 2.800E-07 |
| 12995 | 650 | D650P | B | 3.9e-07 | 2.3e-07 | 1.3e-07 | 2.500E-07 | 1.311E-07 | 1.300E-07 | 3.900E-07 |
| 12996 | 650 | D650S | B | 2.7e-07 | 1.3e-07 | 1.7e-07 | 1.900E-07 | 7.211E-08 | 1.300E-07 | 2.700E-07 |
| 12997 | 650 | D650T | B | 7.7e-08 | 4.6e-07 | 4.7e-07 | 3.357E-07 | 2.241E-07 | 7.700E-08 | 4.700E-07 |
| 12998 | 650 | D650W | B | 7.2e-08 | 3.3e-07 | 8.5e-07 | 4.173E-07 | 3.963E-07 | 7.200E-08 | 8.500E-07 |
| 12999 | 650 | D650Y | B | 1.5e-07 | 2.8e-07 | 2.4e-07 | 2.233E-07 | 6.658E-08 | 1.500E-07 | 2.800E-07 |
| 13000 | 650 | D650V | B | 1.2e-07 | 5.4e-07 | 1.6e-07 | 2.733E-07 | 2.318E-07 | 1.200E-07 | 5.400E-07 |
| 13001 | 651 | V651A | B | 3.9e-07 | 9.4e-08 | 4.7e-07 | 3.180E-07 | 1.981E-07 | 9.400E-08 | 4.700E-07 |
| 13002 | 651 | V651R | B | 1.6e-07 | 1.7e-07 | 1.4e-07 | 1.567E-07 | 1.528E-08 | 1.400E-07 | 1.700E-07 |
| 13003 | 651 | V651N | B | 1.2e-07 | 3.3e-07 | 3.1e-07 | 2.533E-07 | 1.159E-07 | 1.200E-07 | 3.300E-07 |
| 13004 | 651 | V651D | B | 1.7e-07 | 3.5e-07 | 3.7e-07 | 2.967E-07 | 1.102E-07 | 1.700E-07 | 3.700E-07 |

| | | | | | | | | | | |
|---|---|---|---|---|---|---|---|---|---|---|
| 13005 | 651 | V651C | B | 3.7e-07 | 2.6e-07 | 3.6e-07 | 3.300E-07 | 6.083E-08 | 2.600E-07 | 3.700E-07 |
| 13006 | 651 | V651E | B | 2.5e-07 | 1.7e-07 | 2.9e-07 | 2.367E-07 | 6.110E-08 | 1.700E-07 | 2.900E-07 |
| 13007 | 651 | V651Q | B | 1.4e-07 | 6.1e-07 | 1.7e-07 | 3.067E-07 | 2.631E-07 | 1.400E-07 | 6.100E-07 |
| 13008 | 651 | V651G | B | 1.4e-07 | 2.7e-07 | 5.3e-07 | 3.133E-07 | 1.986E-07 | 1.400E-07 | 5.300E-07 |
| 13009 | 651 | V651H | B | 1.5e-07 | 2.1e-07 | 3.1e-07 | 2.233E-07 | 8.083E-08 | 1.500E-07 | 3.100E-07 |
| 13010 | 651 | V651I | B | 2.9e-07 | 2.1e-07 | 3e-07 | 2.667E-07 | 4.933E-08 | 2.100E-07 | 3.000E-07 |
| 13011 | 651 | V651L | B | 3.3e-07 | 5e-07 | 3e-07 | 3.767E-07 | 1.079E-07 | 3.000E-07 | 5.000E-07 |
| 13012 | 651 | V651K | B | 2.5e-07 | 2.2e-07 | 1.7e-07 | 2.133E-07 | 4.041E-08 | 1.700E-07 | 2.500E-07 |
| 13013 | 651 | V651M | B | 9.7e-08 | 3.7e-07 | 2.2e-07 | 2.290E-07 | 1.367E-07 | 9.700E-08 | 3.700E-07 |
| 13014 | 651 | V651F | B | 2.5e-07 | 3e-07 | 2.7e-07 | 2.733E-07 | 2.517E-08 | 2.500E-07 | 3.000E-07 |
| 13015 | 651 | V651P | B | 3.1e-07 | 2.7e-07 | 1.2e-07 | 2.333E-07 | 1.002E-07 | 1.200E-07 | 3.100E-07 |
| 13016 | 651 | V651S | B | 2.8e-07 | 3.2e-07 | 1.7e-07 | 2.567E-07 | 7.767E-08 | 1.700E-07 | 3.200E-07 |
| 13017 | 651 | V651T | B | 2.8e-07 | 2.8e-07 | 2e-07 | 2.533E-07 | 4.619E-08 | 2.000E-07 | 2.800E-07 |
| 13018 | 651 | V651W | B | 2.8e-07 | 3.3e-07 | 1.2e-07 | 2.433E-07 | 1.097E-07 | 1.200E-07 | 3.300E-07 |
| 13019 | 651 | V651Y | B | 2.5e-07 | 1.1e-07 | 2e-07 | 1.867E-07 | 7.095E-08 | 1.100E-07 | 2.500E-07 |
| 13020 | 651 | V651V | B | 2.4e-07 | 4.1e-07 | 1.8e-07 | 2.767E-07 | 1.193E-07 | 1.800E-07 | 4.100E-07 |
| 13021 | 652 | S652A | B | 2.4e-07 | 5.6e-07 | 3.6e-07 | 3.867E-07 | 1.617E-07 | 2.400E-07 | 5.600E-07 |
| 13022 | 652 | S652R | B | 2.4e-07 | 3.9e-07 | 5.4e-07 | 3.900E-07 | 1.500E-07 | 2.400E-07 | 5.400E-07 |
| 13023 | 652 | S652N | B | 3.9e-07 | 2.2e-07 | 2.8e-07 | 2.967E-07 | 8.622E-08 | 2.200E-07 | 3.900E-07 |
| 13024 | 652 | S652D | B | 2.4e-07 | 2.5e-07 | 2.5e-07 | 2.467E-07 | 5.774E-09 | 2.400E-07 | 2.500E-07 |
| 13025 | 652 | S652C | B | 2.6e-07 | 2.7e-07 | 1.6e-07 | 2.300E-07 | 6.083E-08 | 1.600E-07 | 2.700E-07 |
| 13026 | 652 | S652E | B | 1.7e-07 | 2.1e-07 | 1.6e-07 | 1.800E-07 | 2.646E-08 | 1.600E-07 | 2.100E-07 |
| 13027 | 652 | S652Q | B | 2.2e-07 | 2.1e-07 | 2.1e-07 | 2.133E-07 | 5.774E-09 | 2.100E-07 | 2.200E-07 |
| 13028 | 652 | S652G | B | 2.7e-07 | 3.2e-07 | 3.3e-07 | 3.067E-07 | 3.215E-08 | 2.700E-07 | 3.300E-07 |
| 13029 | 652 | S652H | B | 1.9e-07 | 3.2e-07 | 1.2e-07 | 2.100E-07 | 1.015E-07 | 1.200E-07 | 3.200E-07 |
| 13030 | 652 | S652I | B | 2e-07 | 2.3e-07 | 3.8e-07 | 2.700E-07 | 9.644E-08 | 2.000E-07 | 3.800E-07 |
| 13031 | 652 | S652L | B | 7.4e-08 | 2.2e-07 | 4.2e-07 | 2.380E-07 | 1.737E-07 | 7.400E-08 | 4.200E-07 |
| 13032 | 652 | S652K | B | 3.9e-07 | 3e-07 | 4.2e-07 | 3.700E-07 | 6.245E-08 | 3.000E-07 | 4.200E-07 |
| 13033 | 652 | S652M | B | 1.7e-07 | 3.2e-07 | 5.5e-07 | 3.467E-07 | 1.914E-07 | 1.700E-07 | 5.500E-07 |
| 13034 | 652 | S652F | B | 4.3e-07 | 1.6e-07 | 2.1e-07 | 2.667E-07 | 1.436E-07 | 1.600E-07 | 4.300E-07 |
| 13035 | 652 | S652P | B | 3.1e-07 | 2.4e-07 | 3.8e-07 | 3.100E-07 | 7.000E-08 | 2.400E-07 | 3.800E-07 |
| 13036 | 652 | S652S | B | 1.5e-07 | 5.3e-07 | 2.1e-07 | 2.967E-07 | 2.043E-07 | 1.500E-07 | 5.300E-07 |
| 13037 | 652 | S652T | B | 4.3e-07 | 2.6e-07 | 1.2e-07 | 2.700E-07 | 1.552E-07 | 1.200E-07 | 4.300E-07 |
| 13038 | 652 | S652W | B | 1.2e-07 | 2e-07 | 2.6e-07 | 1.933E-07 | 7.024E-08 | 1.200E-07 | 2.600E-07 |
| 13039 | 652 | S652Y | B | 2.7e-07 | 1.5e-07 | 2.4e-07 | 2.200E-07 | 6.245E-08 | 1.500E-07 | 2.700E-07 |
| 13040 | 652 | S652V | B | 1.3e-07 | 3.7e-07 | 1.9e-07 | 2.300E-07 | 1.249E-07 | 1.300E-07 | 3.700E-07 |
| 13041 | 653 | I653A | B | 3.3e-07 | 6e-07 | 9.7e-08 | 3.423E-07 | 2.517E-07 | 9.700E-08 | 6.000E-07 |
| 13042 | 653 | I653R | B | 1.6e-07 | 3.1e-07 | 5.6e-07 | 3.433E-07 | 2.021E-07 | 1.600E-07 | 5.600E-07 |
| 13043 | 653 | I653N | B | 2.6e-07 | 6.8e-08 | 4e-07 | 2.427E-07 | 1.667E-07 | 6.800E-08 | 4.000E-07 |
| 13044 | 653 | I653D | B | 3.3e-07 | 5.2e-07 | 2.2e-07 | 3.567E-07 | 1.518E-07 | 2.200E-07 | 5.200E-07 |
| 13045 | 653 | I653C | B | 4.7e-07 | 9.7e-08 | 3.9e-07 | 3.190E-07 | 1.964E-07 | 9.700E-08 | 4.700E-07 |
| 13046 | 653 | I653E | B | 3.3e-07 | 2e-07 | 2.9e-07 | 2.733E-07 | 6.658E-08 | 2.000E-07 | 3.300E-07 |
| 13047 | 653 | I653Q | B | 2.2e-07 | 2.8e-07 | 2.4e-07 | 2.467E-07 | 3.055E-08 | 2.200E-07 | 2.800E-07 |
| 13048 | 653 | I653G | B | 3.3e-07 | 2.7e-07 | 1.8e-07 | 2.600E-07 | 7.550E-08 | 1.800E-07 | 3.300E-07 |
| 13049 | 653 | I653H | B | 1.7e-07 | 2.2e-07 | 2.6e-07 | 2.167E-07 | 4.509E-08 | 1.700E-07 | 2.600E-07 |
| 13050 | 653 | I653I | B | 2.3e-07 | 1.6e-07 | 1.3e-07 | 1.733E-07 | 5.132E-08 | 1.300E-07 | 2.300E-07 |
| 13051 | 653 | I653L | B | 3.1e-07 | 3.2e-07 | 1.3e-07 | 2.533E-07 | 1.069E-07 | 1.300E-07 | 3.200E-07 |
| 13052 | 653 | I653K | B | 2.4e-07 | 1.5e-07 | 1.7e-07 | 1.867E-07 | 4.726E-08 | 1.500E-07 | 2.400E-07 |
| 13053 | 653 | I653M | B | 2.9e-07 | 3.3e-07 | 4.5e-07 | 3.567E-07 | 8.327E-08 | 2.900E-07 | 4.500E-07 |
| 13054 | 653 | I653F | B | 1.4e-07 | 1.5e-07 | 2.7e-07 | 1.867E-07 | 7.234E-08 | 1.400E-07 | 2.700E-07 |
| 13055 | 653 | I653P | B | 3.6e-07 | 2e-07 | 3.5e-07 | 3.033E-07 | 8.963E-08 | 2.000E-07 | 3.600E-07 |
| 13056 | 653 | I653S | B | 1.5e-07 | 3.6e-07 | 2e-07 | 2.367E-07 | 1.097E-07 | 1.500E-07 | 3.600E-07 |
| 13057 | 653 | I653T | B | 2.3e-07 | 2.4e-07 | 4.2e-07 | 2.967E-07 | 1.069E-07 | 2.300E-07 | 4.200E-07 |
| 13058 | 653 | I653W | B | 2.3e-07 | 1.3e-07 | 3.8e-07 | 2.467E-07 | 1.258E-07 | 1.300E-07 | 3.800E-07 |
| 13059 | 653 | I653Y | B | 4.5e-07 | 1.1e-07 | 3.2e-07 | 2.933E-07 | 1.716E-07 | 1.100E-07 | 4.500E-07 |
| 13060 | 653 | I653V | B | 2e-07 | 2.6e-07 | 2.4e-07 | 2.333E-07 | 3.055E-08 | 2.000E-07 | 2.600E-07 |
| 13061 | 654 | G654A | B | 3.3e-07 | 2.7e-07 | 3.9e-07 | 3.300E-07 | 6.000E-08 | 2.700E-07 | 3.900E-07 |
| 13062 | 654 | G654R | B | 2.2e-07 | 2.7e-07 | 2.7e-07 | 2.533E-07 | 2.887E-08 | 2.200E-07 | 2.700E-07 |
| 13063 | 654 | G654N | B | 4.3e-07 | 9.3e-08 | 2.3e-07 | 2.510E-07 | 1.695E-07 | 9.300E-08 | 4.300E-07 |
| 13064 | 654 | G654D | B | 3.4e-07 | 1.7e-07 | 1.9e-07 | 2.333E-07 | 9.292E-08 | 1.700E-07 | 3.400E-07 |
| 13065 | 654 | G654C | B | 4.2e-07 | 1.2e-07 | 1.6e-07 | 2.333E-07 | 1.629E-07 | 1.200E-07 | 4.200E-07 |
| 13066 | 654 | G654E | B | 1.9e-07 | 2.9e-07 | 2e-07 | 2.267E-07 | 5.508E-08 | 1.900E-07 | 2.900E-07 |
| 13067 | 654 | G654Q | B | 2.8e-07 | 3.4e-07 | 4.6e-07 | 3.600E-07 | 9.165E-08 | 2.800E-07 | 4.600E-07 |
| 13068 | 654 | G654G | B | 1.3e-07 | 2.7e-07 | 1.9e-07 | 1.967E-07 | 7.024E-08 | 1.300E-07 | 2.700E-07 |
| 13069 | 654 | G654H | B | 1.2e-07 | 1.6e-07 | 2.5e-07 | 1.767E-07 | 6.658E-08 | 1.200E-07 | 2.500E-07 |
| 13070 | 654 | G654I | B | 2.9e-07 | 2.2e-07 | 7.1e-07 | 4.067E-07 | 2.650E-07 | 2.200E-07 | 7.100E-07 |
| 13071 | 654 | G654L | B | 3.6e-07 | 2.2e-07 | 4e-07 | 3.267E-07 | 9.452E-08 | 2.200E-07 | 4.000E-07 |
| 13072 | 654 | G654K | B | 4.1e-07 | 1.8e-07 | 2.4e-07 | 2.767E-07 | 1.193E-07 | 1.800E-07 | 4.100E-07 |
| 13073 | 654 | G654M | B | 4.8e-07 | 2.9e-07 | 2.1e-07 | 3.267E-07 | 1.387E-07 | 2.100E-07 | 4.800E-07 |
| 13074 | 654 | G654F | B | 4.2e-07 | 1.5e-07 | 2.6e-07 | 2.767E-07 | 1.358E-07 | 1.500E-07 | 4.200E-07 |
| 13075 | 654 | G654P | B | 3e-07 | 3.8e-07 | 2.4e-07 | 3.067E-07 | 7.024E-08 | 2.400E-07 | 3.800E-07 |
| 13076 | 654 | G654S | B | 2e-07 | 3.4e-07 | 1.7e-07 | 2.367E-07 | 9.074E-08 | 1.700E-07 | 3.400E-07 |
| 13077 | 654 | G654T | B | 2.4e-07 | 2.5e-07 | 2e-07 | 2.300E-07 | 2.646E-08 | 2.000E-07 | 2.500E-07 |
| 13078 | 654 | G654W | B | 2.8e-07 | 8.1e-07 | 2.4e-07 | 4.433E-07 | 3.182E-07 | 2.400E-07 | 8.100E-07 |
| 13079 | 654 | G654Y | B | 4.3e-07 | 3.5e-07 | 4.4e-07 | 4.067E-07 | 4.933E-08 | 3.500E-07 | 4.400E-07 |
| 13080 | 654 | G654V | B | 2.3e-07 | 1.3e-07 | 2.1e-07 | 1.900E-07 | 5.292E-08 | 1.300E-07 | 2.300E-07 |
| 13081 | 655 | V655A | B | 1.7e-07 | 2e-07 | 3.1e-07 | 2.267E-07 | 7.371E-08 | 1.700E-07 | 3.100E-07 |
| 13082 | 655 | V655R | B | 3.5e-07 | 1.9e-07 | 2.4e-07 | 2.600E-07 | 8.185E-08 | 1.900E-07 | 3.500E-07 |
| 13083 | 655 | V655N | B | 4e-07 | 3.6e-07 | 2.1e-07 | 3.233E-07 | 1.002E-07 | 2.100E-07 | 4.000E-07 |
| 13084 | 655 | V655D | B | 3.4e-07 | 2.9e-07 | 9.4e-08 | 2.413E-07 | 1.300E-07 | 9.400E-08 | 3.400E-07 |
| 13085 | 655 | V655C | B | 1.7e-07 | 7.1e-07 | 2.2e-07 | 3.667E-07 | 2.984E-07 | 1.700E-07 | 7.100E-07 |
| 13086 | 655 | V655E | B | 3.9e-07 | 4.4e-07 | 5.8e-07 | 4.700E-07 | 9.849E-08 | 3.900E-07 | 5.800E-07 |
| 13087 | 655 | V655Q | B | 2.2e-07 | 3.8e-07 | 1.6e-07 | 2.533E-07 | 1.137E-07 | 1.600E-07 | 3.800E-07 |
| 13088 | 655 | V655G | B | 3.1e-07 | 2e-07 | 2.9e-07 | 2.667E-07 | 5.859E-08 | 2.000E-07 | 3.100E-07 |
| 13089 | 655 | V655H | B | 2.4e-07 | 2.2e-07 | 3.1e-07 | 2.567E-07 | 4.726E-08 | 2.200E-07 | 3.100E-07 |

| | | | | | | | | | | |
|---|---|---|---|---|---|---|---|---|---|---|
| 13090 | 655 | V655I | B | 1.6e-07 | 1.4e-07 | 2.5e-07 | 1.833E-07 | 5.859E-08 | 1.400E-07 | 2.500E-07 |
| 13091 | 655 | V655L | B | 2.5e-07 | 4e-07 | 2.6e-07 | 3.033E-07 | 8.386E-08 | 2.500E-07 | 4.000E-07 |
| 13092 | 655 | V655K | B | 3e-07 | 4.2e-07 | 6.7e-07 | 4.633E-07 | 1.888E-07 | 3.000E-07 | 6.700E-07 |
| 13093 | 655 | V655M | B | 3.9e-07 | 2.1e-07 | 2.1e-07 | 2.700E-07 | 1.039E-07 | 2.100E-07 | 3.900E-07 |
| 13094 | 655 | V655F | B | 3.7e-07 | 3.3e-07 | 3.7e-07 | 3.567E-07 | 2.309E-08 | 3.300E-07 | 3.700E-07 |
| 13095 | 655 | V655P | B | 1.3e-07 | 3.3e-07 | 4.5e-07 | 3.033E-07 | 1.617E-07 | 1.300E-07 | 4.500E-07 |
| 13096 | 655 | V655S | B | 3.8e-07 | 4.1e-07 | 1e-07 | 2.967E-07 | 1.710E-07 | 1.000E-07 | 4.100E-07 |
| 13097 | 655 | V655T | B | 7.7e-08 | 1.4e-07 | 5.8e-07 | 2.657E-07 | 2.740E-07 | 7.700E-08 | 5.800E-07 |
| 13098 | 655 | V655W | B | 2.8e-07 | 5.7e-07 | 5.4e-07 | 4.633E-07 | 1.595E-07 | 2.800E-07 | 5.700E-07 |
| 13099 | 655 | V655Y | B | 4.2e-07 | 3.6e-07 | 1.8e-07 | 3.200E-07 | 1.249E-07 | 1.800E-07 | 4.200E-07 |
| 13100 | 655 | V655V | B | 3.8e-07 | 2.6e-07 | 8.5e-07 | 4.967E-07 | 3.118E-07 | 2.600E-07 | 8.500E-07 |
| 13101 | 656 | A656A | B | 4.8e-07 | 4.9e-07 | 3.1e-07 | 4.267E-07 | 1.012E-07 | 3.100E-07 | 4.900E-07 |
| 13102 | 656 | A656R | B | 1.3e-07 | 1.4e-07 | 1.2e-07 | 1.300E-07 | 1.000E-08 | 1.200E-07 | 1.400E-07 |
| 13103 | 656 | A656N | B | 3e-07 | 3.1e-07 | 2.9e-07 | 3.000E-07 | 1.000E-08 | 2.900E-07 | 3.100E-07 |
| 13104 | 656 | A656D | B | 1.5e-07 | 2e-07 | 4.7e-07 | 2.733E-07 | 1.721E-07 | 1.500E-07 | 4.700E-07 |
| 13105 | 656 | A656C | B | 2.9e-07 | 2.2e-07 | 2.1e-07 | 2.400E-07 | 4.359E-08 | 2.100E-07 | 2.900E-07 |
| 13106 | 656 | A656E | B | 3.4e-07 | 4.2e-07 | 1.8e-07 | 3.133E-07 | 1.222E-07 | 1.800E-07 | 4.200E-07 |
| 13107 | 656 | A656Q | B | 1.8e-07 | 2.8e-07 | 3.5e-07 | 2.700E-07 | 8.544E-08 | 1.800E-07 | 3.500E-07 |
| 13108 | 656 | A656G | B | 9.4e-07 | 1.7e-07 | 2e-07 | 4.367E-07 | 4.362E-07 | 1.700E-07 | 9.400E-07 |
| 13109 | 656 | A656H | B | 1.5e-07 | 5.9e-07 | 2.9e-07 | 3.433E-07 | 2.248E-07 | 1.500E-07 | 5.900E-07 |
| 13110 | 656 | A656I | B | 3e-07 | 8.3e-08 | 4.5e-07 | 2.777E-07 | 1.845E-07 | 8.300E-08 | 4.500E-07 |
| 13111 | 656 | A656L | B | 4.7e-07 | 1.3e-07 | 1.5e-07 | 2.500E-07 | 1.908E-07 | 1.300E-07 | 4.700E-07 |
| 13112 | 656 | A656K | B | 1.4e-07 | 2.2e-07 | 2.9e-07 | 2.167E-07 | 7.506E-08 | 1.400E-07 | 2.900E-07 |
| 13113 | 656 | A656M | B | 2.2e-07 | 1.9e-07 | 3.2e-07 | 2.433E-07 | 6.807E-08 | 1.900E-07 | 3.200E-07 |
| 13114 | 656 | A656F | B | 4.5e-07 | 3e-07 | 2.3e-07 | 3.267E-07 | 1.124E-07 | 2.300E-07 | 4.500E-07 |
| 13115 | 656 | A656P | B | 6.5e-07 | 4.4e-07 | 1.9e-07 | 4.267E-07 | 2.303E-07 | 1.900E-07 | 6.500E-07 |
| 13116 | 656 | A656S | B | 3.9e-07 | 3e-07 | 1.8e-07 | 2.900E-07 | 1.054E-07 | 1.800E-07 | 3.900E-07 |
| 13117 | 656 | A656T | B | 6.1e-08 | 6.7e-07 | 1.3e-07 | 2.870E-07 | 3.335E-07 | 6.100E-08 | 6.700E-07 |
| 13118 | 656 | A656W | B | 2.3e-07 | 2.3e-07 | 1.7e-07 | 2.100E-07 | 3.464E-08 | 1.700E-07 | 2.300E-07 |
| 13119 | 656 | A656Y | B | 1.3e-07 | 1.4e-07 | 5.7e-07 | 2.800E-07 | 2.512E-07 | 1.300E-07 | 5.700E-07 |
| 13120 | 656 | A656V | B | 2.3e-07 | 1.3e-07 | 3.7e-07 | 2.433E-07 | 1.206E-07 | 1.300E-07 | 3.700E-07 |
| 13121 | 657 | W657A | B | 5e-07 | 1.8e-07 | 3.1e-07 | 3.300E-07 | 1.609E-07 | 1.800E-07 | 5.000E-07 |
| 13122 | 657 | W657R | B | 2.8e-07 | 1.7e-07 | 3.8e-07 | 2.767E-07 | 1.050E-07 | 1.700E-07 | 3.800E-07 |
| 13123 | 657 | W657N | B | 1.5e-07 | 2.2e-07 | 1.7e-07 | 1.800E-07 | 3.606E-08 | 1.500E-07 | 2.200E-07 |
| 13124 | 657 | W657D | B | 1.4e-07 | 2.4e-07 | 2.6e-07 | 2.133E-07 | 6.429E-08 | 1.400E-07 | 2.600E-07 |
| 13125 | 657 | W657C | B | 1.2e-07 | 3.9e-07 | 1.7e-07 | 2.267E-07 | 1.436E-07 | 1.200E-07 | 3.900E-07 |
| 13126 | 657 | W657E | B | 2.4e-07 | 3.3e-07 | 2.3e-07 | 2.667E-07 | 5.508E-08 | 2.300E-07 | 3.300E-07 |
| 13127 | 657 | W657Q | B | 2.7e-07 | 5.5e-07 | 4.4e-07 | 4.200E-07 | 1.411E-07 | 2.700E-07 | 5.500E-07 |
| 13128 | 657 | W657G | B | 2.9e-07 | 2e-07 | 2.7e-07 | 2.533E-07 | 4.726E-08 | 2.000E-07 | 2.900E-07 |
| 13129 | 657 | W657H | B | 5.3e-07 | 2.8e-07 | 2.6e-07 | 3.567E-07 | 1.504E-07 | 2.600E-07 | 5.300E-07 |
| 13130 | 657 | W657I | B | 3.5e-07 | 2.2e-07 | 2.6e-07 | 2.767E-07 | 6.658E-08 | 2.200E-07 | 3.500E-07 |
| 13131 | 657 | W657L | B | 3e-07 | 1.5e-07 | 2.6e-07 | 2.367E-07 | 7.767E-08 | 1.500E-07 | 3.000E-07 |
| 13132 | 657 | W657K | B | 2.9e-07 | 3.2e-07 | 2e-07 | 2.700E-07 | 6.245E-08 | 2.000E-07 | 3.200E-07 |
| 13133 | 657 | W657M | B | 1.8e-07 | 4.3e-07 | 2.7e-07 | 2.933E-07 | 1.266E-07 | 1.800E-07 | 4.300E-07 |
| 13134 | 657 | W657F | B | 3e-07 | 3.2e-07 | 1.6e-07 | 2.600E-07 | 8.718E-08 | 1.600E-07 | 3.200E-07 |
| 13135 | 657 | W657P | B | 2.2e-07 | 4e-07 | 3.7e-07 | 3.300E-07 | 9.644E-08 | 2.200E-07 | 4.000E-07 |
| 13136 | 657 | W657S | B | 1.6e-07 | 4.3e-07 | 4.7e-07 | 3.533E-07 | 1.686E-07 | 1.600E-07 | 4.700E-07 |
| 13137 | 657 | W657T | B | 2.8e-07 | 2.8e-07 | 5.4e-07 | 3.667E-07 | 1.501E-07 | 2.800E-07 | 5.400E-07 |
| 13138 | 657 | W657W | B | 1.6e-07 | 1.9e-07 | 3.2e-07 | 2.233E-07 | 8.505E-08 | 1.600E-07 | 3.200E-07 |
| 13139 | 657 | W657Y | B | 2.6e-07 | 5.5e-07 | 1.9e-07 | 3.333E-07 | 1.909E-07 | 1.900E-07 | 5.500E-07 |
| 13140 | 657 | W657V | B | 3.1e-07 | 1.8e-07 | 2.9e-07 | 2.600E-07 | 7.000E-08 | 1.800E-07 | 3.100E-07 |
| 13141 | 658 | Y658A | B | 2.2e-07 | 1.6e-07 | 1.5e-07 | 1.767E-07 | 3.786E-08 | 1.500E-07 | 2.200E-07 |
| 13142 | 658 | Y658R | B | 3.3e-07 | 2.1e-07 | 3.7e-07 | 3.033E-07 | 8.327E-08 | 2.100E-07 | 3.700E-07 |
| 13143 | 658 | Y658N | B | 3.1e-07 | 1.7e-07 | 2.7e-07 | 2.500E-07 | 7.211E-08 | 1.700E-07 | 3.100E-07 |
| 13144 | 658 | Y658D | B | 1.6e-07 | 2.4e-07 | 3.3e-07 | 2.433E-07 | 8.505E-08 | 1.600E-07 | 3.300E-07 |
| 13145 | 658 | Y658C | B | 3e-07 | 1.6e-07 | 2.2e-07 | 2.267E-07 | 7.024E-08 | 1.600E-07 | 3.000E-07 |
| 13146 | 658 | Y658E | B | 3.2e-07 | 2.7e-07 | 1.9e-07 | 2.600E-07 | 6.557E-08 | 1.900E-07 | 3.200E-07 |
| 13147 | 658 | Y658Q | B | 2.2e-07 | 3.6e-07 | 2.5e-07 | 2.767E-07 | 7.371E-08 | 2.200E-07 | 3.600E-07 |
| 13148 | 658 | Y658G | B | 1.5e-07 | 3.6e-07 | 2.8e-07 | 2.633E-07 | 1.060E-07 | 1.500E-07 | 3.600E-07 |
| 13149 | 658 | Y658H | B | 1.5e-07 | 2.3e-07 | 1.3e-07 | 1.700E-07 | 5.292E-08 | 1.300E-07 | 2.300E-07 |
| 13150 | 658 | Y658I | B | 2.9e-07 | 3e-07 | 3.2e-07 | 3.033E-07 | 1.528E-08 | 2.900E-07 | 3.200E-07 |
| 13151 | 658 | Y658L | B | 2.3e-07 | 2.4e-07 | 1.9e-07 | 2.200E-07 | 2.646E-08 | 1.900E-07 | 2.400E-07 |
| 13152 | 658 | Y658K | B | 2.8e-07 | 1.6e-07 | 4e-07 | 2.800E-07 | 1.200E-07 | 1.600E-07 | 4.000E-07 |
| 13153 | 658 | Y658M | B | 1.1e-07 | 2.5e-07 | 1.8e-07 | 1.800E-07 | 7.000E-08 | 1.100E-07 | 2.500E-07 |
| 13154 | 658 | Y658F | B | 2.5e-07 | 4.4e-07 | 5e-07 | 3.967E-07 | 1.305E-07 | 2.500E-07 | 5.000E-07 |
| 13155 | 658 | Y658P | B | 3.5e-07 | 3.4e-07 | 5.2e-07 | 4.033E-07 | 1.012E-07 | 3.400E-07 | 5.200E-07 |
| 13156 | 658 | Y658S | B | 2.4e-07 | 4.7e-07 | 2.5e-07 | 3.200E-07 | 1.300E-07 | 2.400E-07 | 4.700E-07 |
| 13157 | 658 | Y658T | B | 3.6e-07 | 2.3e-07 | 1.9e-07 | 2.600E-07 | 8.888E-08 | 1.900E-07 | 3.600E-07 |
| 13158 | 658 | Y658W | B | 2e-07 | 2.1e-07 | 1.9e-07 | 2.000E-07 | 1.000E-08 | 1.900E-07 | 2.100E-07 |
| 13159 | 658 | Y658Y | B | 2.5e-07 | 4.8e-07 | 4.2e-07 | 3.833E-07 | 1.193E-07 | 2.500E-07 | 4.800E-07 |
| 13160 | 658 | Y658V | B | 2.6e-07 | 2.1e-07 | 2.3e-07 | 2.333E-07 | 2.517E-08 | 2.100E-07 | 2.600E-07 |
| 13161 | 659 | Q659A | B | 2.8e-07 | 2e-07 | 3.2e-07 | 2.667E-07 | 6.110E-08 | 2.000E-07 | 3.200E-07 |
| 13162 | 659 | Q659R | B | 2.3e-07 | 2.4e-07 | 1.4e-07 | 2.033E-07 | 5.508E-08 | 1.400E-07 | 2.400E-07 |
| 13163 | 659 | Q659N | B | 1.5e-07 | 1.1e-07 | 2.4e-07 | 1.667E-07 | 6.658E-08 | 1.100E-07 | 2.400E-07 |
| 13164 | 659 | Q659D | B | 5.3e-07 | 2.6e-07 | 1.5e-07 | 3.133E-07 | 1.955E-07 | 1.500E-07 | 5.300E-07 |
| 13165 | 659 | Q659C | B | 3.3e-07 | 1.9e-07 | 4.4e-07 | 3.200E-07 | 1.253E-07 | 1.900E-07 | 4.400E-07 |
| 13166 | 659 | Q659E | B | 1.7e-07 | 4.3e-07 | 2e-07 | 2.667E-07 | 1.422E-07 | 1.700E-07 | 4.300E-07 |
| 13167 | 659 | Q659Q | B | 5.1e-07 | 1.8e-07 | 2.6e-07 | 3.167E-07 | 1.721E-07 | 1.800E-07 | 5.100E-07 |
| 13168 | 659 | Q659G | B | 1.9e-07 | 3.7e-07 | 3.5e-07 | 3.033E-07 | 9.866E-08 | 1.900E-07 | 3.700E-07 |
| 13169 | 659 | Q659H | B | 1.8e-07 | 4.3e-07 | 1.3e-07 | 2.467E-07 | 1.607E-07 | 1.300E-07 | 4.300E-07 |
| 13170 | 659 | Q659I | B | 3.4e-07 | 5.8e-07 | 2.7e-07 | 3.967E-07 | 1.626E-07 | 2.700E-07 | 5.800E-07 |
| 13171 | 659 | Q659L | B | 4.5e-07 | 2.4e-07 | 3.3e-07 | 3.400E-07 | 1.054E-07 | 2.400E-07 | 4.500E-07 |
| 13172 | 659 | Q659K | B | 1.8e-07 | 2.7e-07 | 4e-07 | 2.833E-07 | 1.106E-07 | 1.800E-07 | 4.000E-07 |
| 13173 | 659 | Q659M | B | 3.2e-07 | 2e-07 | 1.4e-07 | 2.200E-07 | 9.165E-08 | 1.400E-07 | 3.200E-07 |
| 13174 | 659 | Q659F | B | 2.4e-07 | 4.2e-07 | 3.2e-07 | 3.267E-07 | 9.018E-08 | 2.400E-07 | 4.200E-07 |

| | | | | | | | | | | |
|---|---|---|---|---|---|---|---|---|---|---|
| 13175 | 659 | Q659P | B | 1.9e-07 | 3.4e-07 | 1e-07 | 2.100E-07 | 1.212E-07 | 1.000E-07 | 3.400E-07 |
| 13176 | 659 | Q659S | B | 2.1e-07 | 2.4e-07 | 7e-07 | 3.833E-07 | 2.747E-07 | 2.100E-07 | 7.000E-07 |
| 13177 | 659 | Q659T | B | 1.1e-07 | 2.7e-07 | 1.2e-07 | 1.667E-07 | 8.963E-08 | 1.100E-07 | 2.700E-07 |
| 13178 | 659 | Q659W | B | 2.3e-07 | 2.8e-07 | 4.1e-07 | 3.067E-07 | 9.292E-08 | 2.300E-07 | 4.100E-07 |
| 13179 | 659 | Q659Y | B | 1.2e-07 | 3.6e-07 | 1e-07 | 1.933E-07 | 1.447E-07 | 1.000E-07 | 3.600E-07 |
| 13180 | 659 | Q659V | B | 1.6e-07 | 2.4e-07 | 1.9e-07 | 1.967E-07 | 4.041E-08 | 1.600E-07 | 2.400E-07 |
| 13181 | 660 | Q660A | B | 2.5e-07 | 3.6e-07 | 1.7e-07 | 2.600E-07 | 9.539E-08 | 1.700E-07 | 3.600E-07 |
| 13182 | 660 | Q660R | B | 3.2e-07 | 2.8e-07 | 2.6e-07 | 2.867E-07 | 3.055E-08 | 2.600E-07 | 3.200E-07 |
| 13183 | 660 | Q660N | B | 1.8e-07 | 4.1e-07 | 2e-07 | 2.633E-07 | 1.274E-07 | 1.800E-07 | 4.100E-07 |
| 13184 | 660 | Q660D | B | 4e-07 | 2.6e-07 | 2e-07 | 2.867E-07 | 1.026E-07 | 2.000E-07 | 4.000E-07 |
| 13185 | 660 | Q660C | B | 4e-07 | 2.4e-07 | 2.6e-07 | 3.000E-07 | 8.718E-08 | 2.400E-07 | 4.000E-07 |
| 13186 | 660 | Q660E | B | 4e-07 | 2.3e-07 | 1.7e-07 | 2.667E-07 | 1.193E-07 | 1.700E-07 | 4.000E-07 |
| 13187 | 660 | Q660Q | B | 6.1e-07 | 2.7e-07 | 2.7e-07 | 3.833E-07 | 1.963E-07 | 2.700E-07 | 6.100E-07 |
| 13188 | 660 | Q660G | B | 2.5e-07 | 3.1e-07 | 4.2e-07 | 3.267E-07 | 8.622E-08 | 2.500E-07 | 4.200E-07 |
| 13189 | 660 | Q660H | B | 2.3e-07 | 1e-07 | 2.5e-07 | 1.933E-07 | 8.145E-08 | 1.000E-07 | 2.500E-07 |
| 13190 | 660 | Q660I | B | 2.2e-07 | 3.1e-07 | 3.2e-07 | 2.833E-07 | 5.508E-08 | 2.200E-07 | 3.200E-07 |
| 13191 | 660 | Q660L | B | 2.9e-07 | 5.4e-07 | 2.3e-07 | 3.533E-07 | 1.644E-07 | 2.300E-07 | 5.400E-07 |
| 13192 | 660 | Q660K | B | 1.7e-07 | 2.1e-07 | 2e-07 | 1.933E-07 | 2.082E-08 | 1.700E-07 | 2.100E-07 |
| 13193 | 660 | Q660M | B | 1.6e-07 | 1.8e-07 | 1.8e-07 | 1.733E-07 | 1.155E-08 | 1.600E-07 | 1.800E-07 |
| 13194 | 660 | Q660F | B | 2e-07 | 4.7e-07 | 3.8e-07 | 3.500E-07 | 1.375E-07 | 2.000E-07 | 4.700E-07 |
| 13195 | 660 | Q660P | B | 1.5e-07 | 4.6e-07 | 2.1e-07 | 2.733E-07 | 1.644E-07 | 1.500E-07 | 4.600E-07 |
| 13196 | 660 | Q660S | B | 2.7e-07 | 3.7e-07 | 3.2e-07 | 3.200E-07 | 5.000E-08 | 2.700E-07 | 3.700E-07 |
| 13197 | 660 | Q660T | B | 1.4e-07 | 1.3e-07 | 2.6e-07 | 1.767E-07 | 7.234E-08 | 1.300E-07 | 2.600E-07 |
| 13198 | 660 | Q660W | B | 3e-07 | 3.9e-07 | 1.3e-07 | 2.733E-07 | 1.320E-07 | 1.300E-07 | 3.900E-07 |
| 13199 | 660 | Q660Y | B | 1.9e-07 | 2.3e-07 | 3e-07 | 2.400E-07 | 5.568E-08 | 1.900E-07 | 3.000E-07 |
| 13200 | 660 | Q660V | B | 2.8e-07 | 4.7e-07 | 3.2e-07 | 3.567E-07 | 1.002E-07 | 2.800E-07 | 4.700E-07 |
| 13201 | 661 | K661A | B | 2.8e-07 | 1.2e-07 | 1.1e-07 | 1.700E-07 | 9.539E-08 | 1.100E-07 | 2.800E-07 |
| 13202 | 661 | K661R | B | 2.1e-07 | 2.9e-07 | 1.2e-07 | 2.067E-07 | 8.505E-08 | 1.200E-07 | 2.900E-07 |
| 13203 | 661 | K661N | B | 2.7e-07 | 2.9e-07 | 1.7e-07 | 2.433E-07 | 6.429E-08 | 1.700E-07 | 2.900E-07 |
| 13204 | 661 | K661D | B | 6.2e-07 | 2e-07 | 1.2e-07 | 3.133E-07 | 2.686E-07 | 1.200E-07 | 6.200E-07 |
| 13205 | 661 | K661C | B | 1.5e-07 | 2.8e-07 | 3.8e-07 | 2.700E-07 | 1.153E-07 | 1.500E-07 | 3.800E-07 |
| 13206 | 661 | K661E | B | 4.5e-07 | 1.8e-07 | 2.1e-07 | 2.800E-07 | 1.480E-07 | 1.800E-07 | 4.500E-07 |
| 13207 | 661 | K661Q | B | 3.3e-07 | 5.7e-07 | 2.9e-07 | 3.967E-07 | 1.514E-07 | 2.900E-07 | 5.700E-07 |
| 13208 | 661 | K661G | B | 3.8e-07 | 2.9e-07 | 3.4e-07 | 3.367E-07 | 4.509E-08 | 2.900E-07 | 3.800E-07 |
| 13209 | 661 | K661H | B | 1.6e-07 | 2.6e-07 | 7.2e-07 | 3.800E-07 | 2.987E-07 | 1.600E-07 | 7.200E-07 |
| 13210 | 661 | K661I | B | 2.1e-07 | 1e-07 | 1.6e-07 | 1.567E-07 | 5.508E-08 | 1.000E-07 | 2.100E-07 |
| 13211 | 661 | K661L | B | 3.5e-07 | 5.1e-07 | 2.1e-07 | 3.567E-07 | 1.501E-07 | 2.100E-07 | 5.100E-07 |
| 13212 | 661 | K661K | B | 3.4e-07 | 2e-07 | 2.8e-07 | 2.733E-07 | 7.024E-08 | 2.000E-07 | 3.400E-07 |
| 13213 | 661 | K661M | B | 3.2e-07 | 5.6e-07 | 5.5e-07 | 4.767E-07 | 1.358E-07 | 3.200E-07 | 5.600E-07 |
| 13214 | 661 | K661F | B | 1.8e-07 | 1.5e-07 | 3.8e-07 | 2.367E-07 | 1.250E-07 | 1.500E-07 | 3.800E-07 |
| 13215 | 661 | K661P | B | 4.7e-07 | 1.1e-07 | 1.6e-07 | 2.467E-07 | 1.950E-07 | 1.100E-07 | 4.700E-07 |
| 13216 | 661 | K661S | B | 2.1e-07 | 3.7e-07 | 1.1e-07 | 2.300E-07 | 1.311E-07 | 1.100E-07 | 3.700E-07 |
| 13217 | 661 | K661T | B | 3.3e-07 | 2e-07 | 1.8e-07 | 2.367E-07 | 8.145E-08 | 1.800E-07 | 3.300E-07 |
| 13218 | 661 | K661W | B | 3.9e-07 | 3.4e-07 | 4.3e-07 | 3.867E-07 | 4.509E-08 | 3.400E-07 | 4.300E-07 |
| 13219 | 661 | K661Y | B | 1.9e-07 | 2.2e-07 | 2.1e-07 | 2.067E-07 | 1.528E-08 | 1.900E-07 | 2.200E-07 |
| 13220 | 661 | K661V | B | 2.5e-07 | 3.8e-07 | 3.8e-07 | 3.367E-07 | 7.506E-08 | 2.500E-07 | 3.800E-07 |
| 13221 | 662 | P662A | B | 3.5e-07 | 3.1e-07 | 4.3e-07 | 3.633E-07 | 6.110E-08 | 3.100E-07 | 4.300E-07 |
| 13222 | 662 | P662R | B | 2.2e-07 | 3.4e-07 | 3.1e-07 | 2.900E-07 | 6.245E-08 | 2.200E-07 | 3.400E-07 |
| 13223 | 662 | P662N | B | 4e-07 | 2.7e-07 | 1.7e-07 | 2.800E-07 | 1.153E-07 | 1.700E-07 | 4.000E-07 |
| 13224 | 662 | P662D | B | 4.8e-07 | 2.2e-07 | 2.1e-07 | 3.033E-07 | 1.531E-07 | 2.100E-07 | 4.800E-07 |
| 13225 | 662 | P662C | B | 3e-07 | 2.7e-07 | 3.6e-07 | 3.100E-07 | 4.583E-08 | 2.700E-07 | 3.600E-07 |
| 13226 | 662 | P662E | B | 1.6e-07 | 5.2e-07 | 2e-07 | 2.933E-07 | 1.973E-07 | 1.600E-07 | 5.200E-07 |
| 13227 | 662 | P662Q | B | 3.5e-07 | 1.8e-07 | 2.4e-07 | 2.567E-07 | 8.622E-08 | 1.800E-07 | 3.500E-07 |
| 13228 | 662 | P662G | B | 4e-07 | 1.9e-07 | 4.6e-07 | 3.500E-07 | 1.418E-07 | 1.900E-07 | 4.600E-07 |
| 13229 | 662 | P662H | B | 3e-07 | 2.9e-07 | 2.2e-07 | 2.700E-07 | 4.359E-08 | 2.200E-07 | 3.000E-07 |
| 13230 | 662 | P662I | B | 1.1e-07 | 1.7e-07 | 3.1e-07 | 1.967E-07 | 1.026E-07 | 1.100E-07 | 3.100E-07 |
| 13231 | 662 | P662L | B | 5.4e-07 | 2.8e-07 | 1.1e-07 | 3.100E-07 | 2.166E-07 | 1.100E-07 | 5.400E-07 |
| 13232 | 662 | P662K | B | 2.6e-07 | 1.8e-07 | 2.3e-07 | 2.233E-07 | 4.041E-08 | 1.800E-07 | 2.600E-07 |
| 13233 | 662 | P662M | B | 1.8e-07 | 3.3e-07 | 4.3e-07 | 3.133E-07 | 1.258E-07 | 1.800E-07 | 4.300E-07 |
| 13234 | 662 | P662F | B | 2.5e-07 | 4.3e-07 | 2.9e-07 | 3.233E-07 | 9.452E-08 | 2.500E-07 | 4.300E-07 |
| 13235 | 662 | P662P | B | 1.7e-07 | 4.2e-07 | 4.9e-07 | 3.600E-07 | 1.682E-07 | 1.700E-07 | 4.900E-07 |
| 13236 | 662 | P662S | B | 3.3e-07 | 2.3e-07 | 1.7e-07 | 2.433E-07 | 8.083E-08 | 1.700E-07 | 3.300E-07 |
| 13237 | 662 | P662T | B | 2.3e-07 | 2.7e-07 | 2.3e-07 | 2.433E-07 | 2.309E-08 | 2.300E-07 | 2.700E-07 |
| 13238 | 662 | P662W | B | 7.2e-07 | 2.9e-07 | 2.1e-07 | 4.067E-07 | 2.743E-07 | 2.100E-07 | 7.200E-07 |
| 13239 | 662 | P662Y | B | 2.4e-07 | 4.3e-07 | 2.6e-07 | 3.100E-07 | 1.044E-07 | 2.400E-07 | 4.300E-07 |
| 13240 | 662 | P662V | B | 2.2e-07 | 3.1e-07 | 2.4e-07 | 2.567E-07 | 4.726E-08 | 2.200E-07 | 3.100E-07 |
| 13241 | 663 | G663A | B | 2e-07 | 3.4e-07 | 1.6e-07 | 2.333E-07 | 9.452E-08 | 1.600E-07 | 3.400E-07 |
| 13242 | 663 | G663R | B | 2.1e-07 | 3.6e-07 | 2e-07 | 2.567E-07 | 8.963E-08 | 2.000E-07 | 3.600E-07 |
| 13243 | 663 | G663N | B | 2.1e-07 | 4.1e-07 | 4.4e-07 | 3.533E-07 | 1.250E-07 | 2.100E-07 | 4.400E-07 |
| 13244 | 663 | G663D | B | 1.4e-07 | 2.9e-07 | 2e-07 | 2.100E-07 | 7.550E-08 | 1.400E-07 | 2.900E-07 |
| 13245 | 663 | G663C | B | 1.3e-07 | 2.1e-07 | 3.2e-07 | 2.200E-07 | 9.539E-08 | 1.300E-07 | 3.200E-07 |
| 13246 | 663 | G663E | B | 3.7e-07 | 5.3e-07 | 3.8e-07 | 4.267E-07 | 8.963E-08 | 3.700E-07 | 5.300E-07 |
| 13247 | 663 | G663Q | B | 1.4e-07 | 2.1e-07 | 1.8e-07 | 1.767E-07 | 3.512E-08 | 1.400E-07 | 2.100E-07 |
| 13248 | 663 | G663G | B | 2.8e-07 | 2.1e-07 | 2.5e-07 | 2.467E-07 | 3.512E-08 | 2.100E-07 | 2.800E-07 |
| 13249 | 663 | G663H | B | 1.9e-07 | 2.8e-07 | 1.9e-07 | 2.200E-07 | 5.196E-08 | 1.900E-07 | 2.800E-07 |
| 13250 | 663 | G663I | B | 1.6e-07 | 3.3e-07 | 1.8e-07 | 2.233E-07 | 9.292E-08 | 1.600E-07 | 3.300E-07 |
| 13251 | 663 | G663L | B | 1.8e-07 | 3.2e-07 | 3.3e-07 | 2.767E-07 | 8.386E-08 | 1.800E-07 | 3.300E-07 |
| 13252 | 663 | G663K | B | 2.5e-07 | 2.9e-07 | 3.7e-07 | 3.033E-07 | 6.110E-08 | 2.500E-07 | 3.700E-07 |
| 13253 | 663 | G663M | B | 2.6e-07 | 2.6e-07 | 3.7e-07 | 2.967E-07 | 6.351E-08 | 2.600E-07 | 3.700E-07 |
| 13254 | 663 | G663F | B | 3.3e-07 | 1.5e-07 | 1.3e-07 | 2.033E-07 | 1.102E-07 | 1.300E-07 | 3.300E-07 |
| 13255 | 663 | G663P | B | 1.9e-07 | 3.9e-07 | 2.6e-07 | 2.800E-07 | 1.015E-07 | 1.900E-07 | 3.900E-07 |
| 13256 | 663 | G663S | B | 4.8e-07 | 2.5e-07 | 1.2e-07 | 2.833E-07 | 1.823E-07 | 1.200E-07 | 4.800E-07 |
| 13257 | 663 | G663T | B | 2.6e-07 | 3.3e-07 | 3.3e-07 | 3.067E-07 | 4.041E-08 | 2.600E-07 | 3.300E-07 |
| 13258 | 663 | G663W | B | 3.2e-07 | 1.8e-07 | 2.6e-07 | 2.533E-07 | 7.024E-08 | 1.800E-07 | 3.200E-07 |
| 13259 | 663 | G663Y | B | 1.6e-07 | 2.9e-07 | 2.7e-07 | 2.400E-07 | 7.000E-08 | 1.600E-07 | 2.900E-07 |

| | | | | | | | | | | |
|---|---|---|---|---|---|---|---|---|---|---|
| 13260 | 663 | G663V | B | 2e-07 | 5.5e-07 | 2.5e-07 | 3.333E-07 | 1.893E-07 | 2.000E-07 | 5.500E-07 |
| 13261 | 664 | K664A | B | 2.2e-07 | 2.1e-07 | 1.3e-07 | 1.867E-07 | 4.933E-08 | 1.300E-07 | 2.200E-07 |
| 13262 | 664 | K664R | B | 2.8e-07 | 1.7e-07 | 9.8e-08 | 1.827E-07 | 9.166E-08 | 9.800E-08 | 2.800E-07 |
| 13263 | 664 | K664N | B | 2.3e-07 | 2.1e-07 | 2.6e-07 | 2.333E-07 | 2.517E-08 | 2.100E-07 | 2.600E-07 |
| 13264 | 664 | K664D | B | 1.9e-07 | 2.5e-07 | 1.7e-07 | 2.033E-07 | 4.163E-08 | 1.700E-07 | 2.500E-07 |
| 13265 | 664 | K664C | B | 1.5e-07 | 3.3e-07 | 1.4e-07 | 2.067E-07 | 1.069E-07 | 1.400E-07 | 3.300E-07 |
| 13266 | 664 | K664E | B | 5.1e-07 | 4.1e-07 | 1.4e-07 | 3.533E-07 | 1.914E-07 | 1.400E-07 | 5.100E-07 |
| 13267 | 664 | K664Q | B | 2.7e-07 | 2.8e-07 | 2.9e-07 | 2.800E-07 | 1.000E-08 | 2.700E-07 | 2.900E-07 |
| 13268 | 664 | K664G | B | 1.7e-07 | 2.3e-07 | 1.2e-07 | 1.733E-07 | 5.508E-08 | 1.200E-07 | 2.300E-07 |
| 13269 | 664 | K664H | B | 3.7e-07 | 2.4e-07 | 1.5e-07 | 2.533E-07 | 1.106E-07 | 1.500E-07 | 3.700E-07 |
| 13270 | 664 | K664I | B | 3.2e-07 | 1.3e-07 | 1.9e-07 | 2.133E-07 | 9.713E-08 | 1.300E-07 | 3.200E-07 |
| 13271 | 664 | K664L | B | 1.5e-07 | 4.4e-07 | 2.4e-07 | 2.767E-07 | 1.484E-07 | 1.500E-07 | 4.400E-07 |
| 13272 | 664 | K664K | B | 3.5e-07 | 6e-07 | 1.4e-07 | 3.633E-07 | 2.303E-07 | 1.400E-07 | 6.000E-07 |
| 13273 | 664 | K664M | B | 3.3e-07 | 1.7e-07 | 3.2e-07 | 2.733E-07 | 8.963E-08 | 1.700E-07 | 3.300E-07 |
| 13274 | 664 | K664F | B | 2.5e-07 | 2.8e-07 | 1.4e-07 | 2.233E-07 | 7.371E-08 | 1.400E-07 | 2.800E-07 |
| 13275 | 664 | K664P | B | 2.8e-07 | 1.6e-07 | 2.4e-07 | 2.267E-07 | 6.110E-08 | 1.600E-07 | 2.800E-07 |
| 13276 | 664 | K664S | B | 1.6e-07 | 2.1e-07 | 4.8e-07 | 2.833E-07 | 1.721E-07 | 1.600E-07 | 4.800E-07 |
| 13277 | 664 | K664T | B | 2.3e-07 | 4e-07 | 1.4e-07 | 2.567E-07 | 1.320E-07 | 1.400E-07 | 4.000E-07 |
| 13278 | 664 | K664W | B | 8.8e-08 | 1.3e-07 | 1.6e-07 | 1.260E-07 | 3.617E-08 | 8.800E-08 | 1.600E-07 |
| 13279 | 664 | K664Y | B | 2.7e-07 | 6.4e-07 | 2.8e-07 | 3.967E-07 | 2.108E-07 | 2.700E-07 | 6.400E-07 |
| 13280 | 664 | K664V | B | 3.8e-07 | 2.1e-07 | 4.5e-07 | 3.467E-07 | 1.234E-07 | 2.100E-07 | 4.500E-07 |
| 13281 | 665 | A665A | B | 1.4e-07 | 2.9e-07 | 3.1e-07 | 2.467E-07 | 9.292E-08 | 1.400E-07 | 3.100E-07 |
| 13282 | 665 | A665R | B | 2.6e-07 | 3e-07 | 7.4e-08 | 2.113E-07 | 1.206E-07 | 7.400E-08 | 3.000E-07 |
| 13283 | 665 | A665N | B | 3.1e-07 | 3.7e-07 | 2.3e-07 | 3.033E-07 | 7.024E-08 | 2.300E-07 | 3.700E-07 |
| 13284 | 665 | A665D | B | 1.7e-07 | 2.7e-07 | 4.5e-07 | 2.967E-07 | 1.419E-07 | 1.700E-07 | 4.500E-07 |
| 13285 | 665 | A665C | B | 1.5e-07 | 3.4e-07 | 2.7e-07 | 2.533E-07 | 9.609E-08 | 1.500E-07 | 3.400E-07 |
| 13286 | 665 | A665E | B | 2e-07 | 1.5e-07 | 4.4e-07 | 2.633E-07 | 1.550E-07 | 1.500E-07 | 4.400E-07 |
| 13287 | 665 | A665Q | B | 4e-07 | 1.7e-07 | 4.5e-07 | 3.400E-07 | 1.493E-07 | 1.700E-07 | 4.500E-07 |
| 13288 | 665 | A665G | B | 1.8e-07 | 1.1e-06 | 5.2e-07 | 6.000E-07 | 4.652E-07 | 1.800E-07 | 1.100E-06 |
| 13289 | 665 | A665H | B | 2.2e-07 | 2.3e-07 | 3.1e-07 | 2.533E-07 | 4.933E-08 | 2.200E-07 | 3.100E-07 |
| 13290 | 665 | A665I | B | 2.3e-07 | 2.2e-07 | 1.6e-07 | 2.033E-07 | 3.786E-08 | 1.600E-07 | 2.300E-07 |
| 13291 | 665 | A665L | B | 3.9e-07 | 2.4e-07 | 1.1e-07 | 2.467E-07 | 1.401E-07 | 1.100E-07 | 3.900E-07 |
| 13292 | 665 | A665K | B | 2e-07 | 3.6e-07 | 1.3e-07 | 2.300E-07 | 1.179E-07 | 1.300E-07 | 3.600E-07 |
| 13293 | 665 | A665M | B | 7.9e-07 | 1.5e-07 | 2.4e-07 | 3.933E-07 | 3.465E-07 | 1.500E-07 | 7.900E-07 |
| 13294 | 665 | A665F | B | 3e-07 | 2.4e-07 | 2.2e-07 | 2.533E-07 | 4.163E-08 | 2.200E-07 | 3.000E-07 |
| 13295 | 665 | A665P | B | 3e-07 | 1.6e-07 | 2.6e-07 | 2.400E-07 | 7.211E-08 | 1.600E-07 | 3.000E-07 |
| 13296 | 665 | A665S | B | 2.2e-07 | 1.7e-07 | 2.5e-07 | 2.133E-07 | 4.041E-08 | 1.700E-07 | 2.500E-07 |
| 13297 | 665 | A665T | B | 1.5e-07 | 2.7e-07 | 2.7e-07 | 2.300E-07 | 6.928E-08 | 1.500E-07 | 2.700E-07 |
| 13298 | 665 | A665W | B | 3.2e-07 | 1.1e-07 | 2.5e-07 | 2.267E-07 | 1.069E-07 | 1.100E-07 | 3.200E-07 |
| 13299 | 665 | A665Y | B | 4e-07 | 2.6e-07 | 2e-07 | 2.867E-07 | 1.026E-07 | 2.000E-07 | 4.000E-07 |
| 13300 | 665 | A665V | B | 2.7e-07 | 2.5e-07 | 2.5e-07 | 2.567E-07 | 1.155E-08 | 2.500E-07 | 2.700E-07 |
| 13301 | 666 | P666A | B | 2.9e-07 | 1.9e-07 | 3.5e-07 | 2.767E-07 | 8.083E-08 | 1.900E-07 | 3.500E-07 |
| 13302 | 666 | P666R | B | 4e-07 | 2.3e-07 | 2.6e-07 | 2.967E-07 | 9.074E-08 | 2.300E-07 | 4.000E-07 |
| 13303 | 666 | P666N | B | 4.4e-07 | 1.6e-07 | 1.9e-07 | 2.633E-07 | 1.537E-07 | 1.600E-07 | 4.400E-07 |
| 13304 | 666 | P666D | B | 5e-07 | 2.3e-07 | 2.2e-07 | 3.167E-07 | 1.589E-07 | 2.200E-07 | 5.000E-07 |
| 13305 | 666 | P666C | B | 4.8e-07 | 3.9e-07 | 2e-07 | 3.567E-07 | 1.429E-07 | 2.000E-07 | 4.800E-07 |
| 13306 | 666 | P666E | B | 2.7e-07 | 2.8e-07 | 2.3e-07 | 2.600E-07 | 2.646E-08 | 2.300E-07 | 2.800E-07 |
| 13307 | 666 | P666Q | B | 1.2e-07 | 2.6e-07 | 2.5e-07 | 2.100E-07 | 7.810E-08 | 1.200E-07 | 2.600E-07 |
| 13308 | 666 | P666G | B | 1.6e-07 | 1.6e-07 | 2e-07 | 1.733E-07 | 2.309E-08 | 1.600E-07 | 2.000E-07 |
| 13309 | 666 | P666H | B | 1.5e-07 | 4.2e-07 | 9.1e-08 | 2.203E-07 | 1.754E-07 | 9.100E-08 | 4.200E-07 |
| 13310 | 666 | P666I | B | 1.9e-07 | 6e-07 | 2e-07 | 3.300E-07 | 2.339E-07 | 1.900E-07 | 6.000E-07 |
| 13311 | 666 | P666L | B | 3.4e-07 | 5.6e-08 | 1.4e-07 | 1.787E-07 | 1.459E-07 | 5.600E-08 | 3.400E-07 |
| 13312 | 666 | P666K | B | 2.6e-07 | 1.9e-07 | 2.1e-07 | 2.200E-07 | 3.606E-08 | 1.900E-07 | 2.600E-07 |
| 13313 | 666 | P666M | B | 2.2e-07 | 1.7e-07 | 2e-07 | 1.967E-07 | 2.517E-08 | 1.700E-07 | 2.200E-07 |
| 13314 | 666 | P666F | B | 1.5e-07 | 2.1e-07 | 3.1e-07 | 2.233E-07 | 8.083E-08 | 1.500E-07 | 3.100E-07 |
| 13315 | 666 | P666P | B | 1.7e-07 | 1.7e-07 | 1.8e-07 | 1.733E-07 | 5.774E-09 | 1.700E-07 | 1.800E-07 |
| 13316 | 666 | P666S | B | 5e-07 | 1.4e-07 | 1.7e-07 | 2.700E-07 | 1.997E-07 | 1.400E-07 | 5.000E-07 |
| 13317 | 666 | P666T | B | 1.4e-07 | 3.7e-07 | 1.8e-07 | 2.300E-07 | 1.229E-07 | 1.400E-07 | 3.700E-07 |
| 13318 | 666 | P666W | B | 3.4e-07 | 2.7e-07 | 3e-07 | 3.033E-07 | 3.512E-08 | 2.700E-07 | 3.400E-07 |
| 13319 | 666 | P666Y | B | 1.4e-07 | 1.3e-07 | 4.8e-07 | 2.500E-07 | 1.992E-07 | 1.300E-07 | 4.800E-07 |
| 13320 | 666 | P666V | B | 2.3e-07 | 2.5e-07 | 1.1e-07 | 1.967E-07 | 7.572E-08 | 1.100E-07 | 2.500E-07 |
| 13321 | 667 | K667A | B | 2.9e-07 | 2.4e-07 | 2.4e-07 | 2.567E-07 | 2.887E-08 | 2.400E-07 | 2.900E-07 |
| 13322 | 667 | K667R | B | 1.4e-07 | 1.6e-07 | 1.7e-07 | 1.567E-07 | 1.528E-08 | 1.400E-07 | 1.700E-07 |
| 13323 | 667 | K667N | B | 1.6e-07 | 2.6e-07 | 3.7e-07 | 2.633E-07 | 1.050E-07 | 1.600E-07 | 3.700E-07 |
| 13324 | 667 | K667D | B | 1.2e-07 | 4.2e-07 | 2.6e-07 | 2.667E-07 | 1.501E-07 | 1.200E-07 | 4.200E-07 |
| 13325 | 667 | K667C | B | 6.8e-08 | 1.3e-07 | 3.6e-07 | 1.860E-07 | 1.538E-07 | 6.800E-08 | 3.600E-07 |
| 13326 | 667 | K667E | B | 2.6e-07 | 3.6e-07 | 2.7e-07 | 2.967E-07 | 5.508E-08 | 2.600E-07 | 3.600E-07 |
| 13327 | 667 | K667Q | B | 2.1e-07 | 2.1e-07 | 2.6e-07 | 2.267E-07 | 2.887E-08 | 2.100E-07 | 2.600E-07 |
| 13328 | 667 | K667G | B | 1.8e-07 | 1.8e-07 | 2.4e-07 | 2.000E-07 | 3.464E-08 | 1.800E-07 | 2.400E-07 |
| 13329 | 667 | K667H | B | 5.7e-07 | 2.3e-07 | 4.4e-07 | 4.133E-07 | 1.716E-07 | 2.300E-07 | 5.700E-07 |
| 13330 | 667 | K667I | B | 3.3e-07 | 1.1e-07 | 3.6e-07 | 2.667E-07 | 1.365E-07 | 1.100E-07 | 3.600E-07 |
| 13331 | 667 | K667L | B | 1.5e-07 | 2.4e-07 | 1.3e-07 | 1.733E-07 | 5.859E-08 | 1.300E-07 | 2.400E-07 |
| 13332 | 667 | K667K | B | 2.7e-07 | 2.3e-07 | 2.1e-07 | 2.367E-07 | 3.055E-08 | 2.100E-07 | 2.700E-07 |
| 13333 | 667 | K667M | B | 3.1e-07 | 2.7e-07 | 2e-07 | 2.600E-07 | 5.568E-08 | 2.000E-07 | 3.100E-07 |
| 13334 | 667 | K667F | B | 6.5e-07 | 3e-07 | 2.7e-07 | 4.067E-07 | 2.113E-07 | 2.700E-07 | 6.500E-07 |
| 13335 | 667 | K667P | B | 4.7e-07 | 3e-07 | 5.4e-07 | 4.367E-07 | 1.234E-07 | 3.000E-07 | 5.400E-07 |
| 13336 | 667 | K667S | B | 2.8e-07 | 2.4e-07 | 1.7e-07 | 2.300E-07 | 5.568E-08 | 1.700E-07 | 2.800E-07 |
| 13337 | 667 | K667T | B | 1.6e-07 | 2.7e-07 | 2.3e-07 | 2.200E-07 | 5.568E-08 | 1.600E-07 | 2.700E-07 |
| 13338 | 667 | K667W | B | 2.6e-07 | 3e-07 | 3.3e-07 | 2.967E-07 | 3.512E-08 | 2.600E-07 | 3.300E-07 |
| 13339 | 667 | K667Y | B | 5.8e-07 | 3.9e-07 | 1.6e-07 | 3.767E-07 | 2.103E-07 | 1.600E-07 | 5.800E-07 |
| 13340 | 667 | K667V | B | 1.7e-07 | 2.6e-07 | 3.9e-07 | 2.733E-07 | 1.106E-07 | 1.700E-07 | 3.900E-07 |
| 13341 | 668 | L668A | B | 3.1e-07 | 3.2e-07 | 2e-07 | 2.767E-07 | 6.658E-08 | 2.000E-07 | 3.200E-07 |
| 13342 | 668 | L668R | B | 1.9e-07 | 1.9e-07 | 3.8e-07 | 2.533E-07 | 1.097E-07 | 1.900E-07 | 3.800E-07 |
| 13343 | 668 | L668N | B | 1.1e-07 | 1.7e-07 | 5.6e-08 | 1.120E-07 | 5.703E-08 | 5.600E-08 | 1.700E-07 |
| 13344 | 668 | L668D | B | 3.5e-07 | 3.5e-07 | 2.6e-07 | 3.200E-07 | 5.196E-08 | 2.600E-07 | 3.500E-07 |

| | | | | | | | | | | |
|---|---|---|---|---|---|---|---|---|---|---|
| 13345 | 668 | L668C | B | 3e-07 | 2.3e-07 | 2.5e-07 | 2.600E-07 | 3.606E-08 | 2.300E-07 | 3.000E-07 |
| 13346 | 668 | L668E | B | 3.8e-07 | 1.6e-07 | 4.2e-07 | 3.200E-07 | 1.400E-07 | 1.600E-07 | 4.200E-07 |
| 13347 | 668 | L668Q | B | 2e-07 | 2.8e-07 | 2.4e-07 | 2.400E-07 | 4.000E-08 | 2.000E-07 | 2.800E-07 |
| 13348 | 668 | L668G | B | 2.9e-07 | 1.7e-07 | 1.5e-07 | 2.033E-07 | 7.572E-08 | 1.500E-07 | 2.900E-07 |
| 13349 | 668 | L668H | B | 3.2e-07 | 3.6e-07 | 4.8e-07 | 3.867E-07 | 8.327E-08 | 3.200E-07 | 4.800E-07 |
| 13350 | 668 | L668I | B | 2.3e-07 | 1.6e-07 | 1.9e-07 | 1.933E-07 | 3.512E-08 | 1.600E-07 | 2.300E-07 |
| 13351 | 668 | L668L | B | 2.9e-07 | 4.7e-07 | 2.3e-07 | 3.300E-07 | 1.249E-07 | 2.300E-07 | 4.700E-07 |
| 13352 | 668 | L668K | B | 1e-07 | 5.9e-07 | 3.5e-07 | 3.467E-07 | 2.450E-07 | 1.000E-07 | 5.900E-07 |
| 13353 | 668 | L668M | B | 4e-07 | 1.7e-07 | 7.4e-07 | 4.367E-07 | 2.868E-07 | 1.700E-07 | 7.400E-07 |
| 13354 | 668 | L668F | B | 1.8e-07 | 2.7e-07 | 5.3e-07 | 3.267E-07 | 1.818E-07 | 1.800E-07 | 5.300E-07 |
| 13355 | 668 | L668P | B | 2e-07 | 6.8e-07 | 3.1e-07 | 3.967E-07 | 2.515E-07 | 2.000E-07 | 6.800E-07 |
| 13356 | 668 | L668S | B | 2.7e-07 | 3.7e-07 | 1.7e-07 | 2.700E-07 | 1.000E-07 | 1.700E-07 | 3.700E-07 |
| 13357 | 668 | L668T | B | 3.3e-07 | 2.9e-07 | 2.4e-07 | 2.867E-07 | 4.509E-08 | 2.400E-07 | 3.300E-07 |
| 13358 | 668 | L668W | B | 2.1e-07 | 1.8e-07 | 7.2e-07 | 3.700E-07 | 3.035E-07 | 1.800E-07 | 7.200E-07 |
| 13359 | 668 | L668Y | B | 2.1e-07 | 1.2e-07 | 2.7e-07 | 2.000E-07 | 7.550E-08 | 1.200E-07 | 2.700E-07 |
| 13360 | 668 | L668V | B | 2.9e-07 | 3.3e-07 | 3.5e-07 | 3.233E-07 | 3.055E-08 | 2.900E-07 | 3.500E-07 |
| 13361 | 669 | L669A | B | 1.7e-07 | 2.1e-07 | 1.6e-07 | 1.800E-07 | 2.646E-08 | 1.600E-07 | 2.100E-07 |
| 13362 | 669 | L669R | B | 1.1e-07 | 2.2e-07 | 4e-07 | 2.433E-07 | 1.464E-07 | 1.100E-07 | 4.000E-07 |
| 13363 | 669 | L669N | B | 3.4e-07 | 2.1e-07 | 3.3e-07 | 2.933E-07 | 7.234E-08 | 2.100E-07 | 3.400E-07 |
| 13364 | 669 | L669D | B | 1.7e-07 | 2.5e-07 | 2.1e-07 | 2.100E-07 | 4.000E-08 | 1.700E-07 | 2.500E-07 |
| 13365 | 669 | L669C | B | 2.5e-07 | 1.6e-07 | 4.1e-07 | 2.733E-07 | 1.266E-07 | 1.600E-07 | 4.100E-07 |
| 13366 | 669 | L669E | B | 1.2e-07 | 2e-07 | 1.5e-07 | 1.567E-07 | 4.041E-08 | 1.200E-07 | 2.000E-07 |
| 13367 | 669 | L669Q | B | 2.3e-07 | 4.3e-07 | 1.7e-07 | 2.767E-07 | 1.361E-07 | 1.700E-07 | 4.300E-07 |
| 13368 | 669 | L669G | B | 2.2e-07 | 2e-07 | 8.1e-07 | 4.100E-07 | 3.466E-07 | 2.000E-07 | 8.100E-07 |
| 13369 | 669 | L669H | B | 2.9e-07 | 2e-07 | 3.1e-07 | 2.667E-07 | 5.859E-08 | 2.000E-07 | 3.100E-07 |
| 13370 | 669 | L669I | B | 2.5e-07 | 2.9e-07 | 2.3e-07 | 2.567E-07 | 3.055E-08 | 2.300E-07 | 2.900E-07 |
| 13371 | 669 | L669L | B | 4.2e-07 | 4.4e-07 | 2e-07 | 3.533E-07 | 1.332E-07 | 2.000E-07 | 4.400E-07 |
| 13372 | 669 | L669K | B | 2e-07 | 1.7e-07 | 2.9e-07 | 2.200E-07 | 6.245E-08 | 1.700E-07 | 2.900E-07 |
| 13373 | 669 | L669M | B | 4.4e-07 | 1.9e-07 | 1.8e-07 | 2.700E-07 | 1.473E-07 | 1.800E-07 | 4.400E-07 |
| 13374 | 669 | L669F | B | 2.2e-07 | 1.4e-07 | 1.6e-07 | 1.733E-07 | 4.163E-08 | 1.400E-07 | 2.200E-07 |
| 13375 | 669 | L669P | B | 3.7e-07 | 1.5e-07 | 1.7e-07 | 2.300E-07 | 1.217E-07 | 1.500E-07 | 3.700E-07 |
| 13376 | 669 | L669S | B | 2e-07 | 6e-07 | 1.5e-07 | 3.167E-07 | 2.466E-07 | 1.500E-07 | 6.000E-07 |
| 13377 | 669 | L669T | B | 2.4e-07 | 1.8e-07 | 2.9e-07 | 2.367E-07 | 5.508E-08 | 1.800E-07 | 2.900E-07 |
| 13378 | 669 | L669W | B | 4.1e-07 | 3.4e-07 | 2.3e-07 | 3.267E-07 | 9.074E-08 | 2.300E-07 | 4.100E-07 |
| 13379 | 669 | L669Y | B | 3.3e-07 | 2.3e-07 | 3.7e-07 | 3.100E-07 | 7.211E-08 | 2.300E-07 | 3.700E-07 |
| 13380 | 669 | L669V | B | 1.1e-07 | 6.8e-08 | 1.1e-07 | 9.600E-08 | 2.425E-08 | 6.800E-08 | 1.100E-07 |
| 13381 | 670 | I670A | B | 5.2e-07 | 4e-07 | 1.3e-07 | 3.500E-07 | 1.997E-07 | 1.300E-07 | 5.200E-07 |
| 13382 | 670 | I670R | B | 4e-07 | 2.1e-07 | 1.6e-07 | 2.567E-07 | 1.266E-07 | 1.600E-07 | 4.000E-07 |
| 13383 | 670 | I670N | B | 2.6e-07 | 2e-07 | 1.9e-07 | 2.167E-07 | 3.786E-08 | 1.900E-07 | 2.600E-07 |
| 13384 | 670 | I670D | B | 4.4e-07 | 2e-07 | 2.7e-07 | 3.033E-07 | 1.234E-07 | 2.000E-07 | 4.400E-07 |
| 13385 | 670 | I670C | B | 1.1e-07 | 4.2e-07 | 4.2e-07 | 3.167E-07 | 1.790E-07 | 1.100E-07 | 4.200E-07 |
| 13386 | 670 | I670E | B | 3.6e-07 | 4.3e-07 | 1.4e-07 | 3.100E-07 | 1.513E-07 | 1.400E-07 | 4.300E-07 |
| 13387 | 670 | I670Q | B | 3.2e-07 | 2.1e-07 | 2.1e-07 | 2.467E-07 | 6.351E-08 | 2.100E-07 | 3.200E-07 |
| 13388 | 670 | I670G | B | 2e-07 | 5.7e-07 | 2.1e-07 | 3.267E-07 | 2.108E-07 | 2.000E-07 | 5.700E-07 |
| 13389 | 670 | I670H | B | 2.9e-07 | 6.2e-07 | 3.5e-07 | 4.200E-07 | 1.758E-07 | 2.900E-07 | 6.200E-07 |
| 13390 | 670 | I670I | B | 2.2e-07 | 4.1e-07 | 2e-07 | 2.767E-07 | 1.159E-07 | 2.000E-07 | 4.100E-07 |
| 13391 | 670 | I670L | B | 3e-07 | 1.7e-07 | 2.3e-07 | 2.333E-07 | 6.506E-08 | 1.700E-07 | 3.000E-07 |
| 13392 | 670 | I670K | B | 2e-07 | 2.9e-07 | 3.7e-07 | 2.867E-07 | 8.505E-08 | 2.000E-07 | 3.700E-07 |
| 13393 | 670 | I670M | B | 1.9e-07 | 1.5e-07 | 2.8e-07 | 2.067E-07 | 6.658E-08 | 1.500E-07 | 2.800E-07 |
| 13394 | 670 | I670F | B | 2.3e-07 | 2.4e-07 | 3.4e-07 | 2.700E-07 | 6.083E-08 | 2.300E-07 | 3.400E-07 |
| 13395 | 670 | I670P | B | 1.6e-07 | 4e-07 | 6e-07 | 3.867E-07 | 2.203E-07 | 1.600E-07 | 6.000E-07 |
| 13396 | 670 | I670S | B | 3.5e-07 | 3.4e-07 | 2.7e-07 | 3.200E-07 | 4.359E-08 | 2.700E-07 | 3.500E-07 |
| 13397 | 670 | I670T | B | 1.9e-07 | 2.6e-07 | 1.6e-07 | 2.033E-07 | 5.132E-08 | 1.600E-07 | 2.600E-07 |
| 13398 | 670 | I670W | B | 1.5e-07 | 1.6e-07 | 2.7e-07 | 1.933E-07 | 6.658E-08 | 1.500E-07 | 2.700E-07 |
| 13399 | 670 | I670Y | B | 2.1e-07 | 2.7e-07 | 2.1e-07 | 2.300E-07 | 3.464E-08 | 2.100E-07 | 2.700E-07 |
| 13400 | 670 | I670V | B | 2e-07 | 1.8e-07 | 1.7e-07 | 1.833E-07 | 1.528E-08 | 1.700E-07 | 2.000E-07 |
| 13401 | 671 | Y671A | B | 1.5e-07 | 3.3e-07 | 1.3e-07 | 2.033E-07 | 1.102E-07 | 1.300E-07 | 3.300E-07 |
| 13402 | 671 | Y671R | B | 3.1e-07 | 1.6e-07 | 3.3e-07 | 2.667E-07 | 9.292E-08 | 1.600E-07 | 3.300E-07 |
| 13403 | 671 | Y671N | B | 4.6e-07 | 3e-07 | 3.1e-07 | 3.567E-07 | 8.963E-08 | 3.000E-07 | 4.600E-07 |
| 13404 | 671 | Y671D | B | 2.5e-07 | 1.9e-07 | 3.7e-07 | 2.700E-07 | 9.165E-08 | 1.900E-07 | 3.700E-07 |
| 13405 | 671 | Y671C | B | 1.5e-07 | 1.4e-07 | 2.7e-07 | 1.867E-07 | 7.234E-08 | 1.400E-07 | 2.700E-07 |
| 13406 | 671 | Y671E | B | 4e-07 | 2e-07 | 6.3e-07 | 4.100E-07 | 2.152E-07 | 2.000E-07 | 6.300E-07 |
| 13407 | 671 | Y671Q | B | 1.4e-07 | 1.9e-07 | 2e-07 | 1.767E-07 | 3.215E-08 | 1.400E-07 | 2.000E-07 |
| 13408 | 671 | Y671G | B | 2.1e-07 | 2.4e-07 | 2e-07 | 2.167E-07 | 2.082E-08 | 2.000E-07 | 2.400E-07 |
| 13409 | 671 | Y671H | B | 4.6e-07 | 2.1e-07 | 4.4e-07 | 3.700E-07 | 1.389E-07 | 2.100E-07 | 4.600E-07 |
| 13410 | 671 | Y671I | B | 1.9e-07 | 3.6e-07 | 1.9e-07 | 2.467E-07 | 9.815E-08 | 1.900E-07 | 3.600E-07 |
| 13411 | 671 | Y671L | B | 3.9e-07 | 3.9e-07 | 1.7e-07 | 3.167E-07 | 1.270E-07 | 1.700E-07 | 3.900E-07 |
| 13412 | 671 | Y671K | B | 4e-07 | 1.8e-07 | 1.3e-07 | 2.367E-07 | 1.436E-07 | 1.300E-07 | 4.000E-07 |
| 13413 | 671 | Y671M | B | 1.7e-07 | 3.1e-07 | 3.3e-07 | 2.700E-07 | 8.718E-08 | 1.700E-07 | 3.300E-07 |
| 13414 | 671 | Y671F | B | 3.2e-07 | 3.3e-07 | 1.2e-07 | 2.567E-07 | 1.185E-07 | 1.200E-07 | 3.300E-07 |
| 13415 | 671 | Y671P | B | 1.8e-07 | 5.1e-07 | 2e-07 | 2.967E-07 | 1.850E-07 | 1.800E-07 | 5.100E-07 |
| 13416 | 671 | Y671S | B | 4e-07 | 2.6e-07 | 1.2e-07 | 2.600E-07 | 1.400E-07 | 1.200E-07 | 4.000E-07 |
| 13417 | 671 | Y671T | B | 1.6e-07 | 1.9e-07 | 1.3e-07 | 1.600E-07 | 3.000E-08 | 1.300E-07 | 1.900E-07 |
| 13418 | 671 | Y671W | B | 2.2e-07 | 1.3e-07 | 2e-07 | 1.833E-07 | 4.726E-08 | 1.300E-07 | 2.200E-07 |
| 13419 | 671 | Y671Y | B | 3.6e-07 | 7.4e-07 | 2.1e-07 | 4.367E-07 | 2.732E-07 | 2.100E-07 | 7.400E-07 |
| 13420 | 671 | Y671V | B | 4.5e-07 | 2.3e-07 | 2e-07 | 2.933E-07 | 1.365E-07 | 2.000E-07 | 4.500E-07 |
| 13421 | 672 | S672A | B | 1.4e-07 | 2e-07 | 1.7e-07 | 1.700E-07 | 3.000E-08 | 1.400E-07 | 2.000E-07 |
| 13422 | 672 | S672R | B | 3.8e-07 | 3.3e-07 | 1.3e-07 | 2.800E-07 | 1.323E-07 | 1.300E-07 | 3.800E-07 |
| 13423 | 672 | S672N | B | 4.3e-07 | 3.7e-07 | 4.4e-07 | 4.133E-07 | 3.786E-08 | 3.700E-07 | 4.400E-07 |
| 13424 | 672 | S672D | B | 2.1e-07 | 1.7e-07 | 3.4e-07 | 2.400E-07 | 8.888E-08 | 1.700E-07 | 3.400E-07 |
| 13425 | 672 | S672C | B | 2.3e-07 | 3e-07 | 2.4e-07 | 2.567E-07 | 3.786E-08 | 2.300E-07 | 3.000E-07 |
| 13426 | 672 | S672E | B | 2.9e-07 | 1.7e-07 | 2.6e-07 | 2.400E-07 | 6.245E-08 | 1.700E-07 | 2.900E-07 |
| 13427 | 672 | S672Q | B | 3.4e-07 | 2e-07 | 1.6e-07 | 2.333E-07 | 9.452E-08 | 1.600E-07 | 3.400E-07 |
| 13428 | 672 | S672G | B | 2.4e-07 | 1.5e-07 | 2.5e-07 | 2.133E-07 | 5.508E-08 | 1.500E-07 | 2.500E-07 |
| 13429 | 672 | S672H | B | 1.2e-07 | 3.5e-07 | 1.9e-07 | 2.200E-07 | 1.179E-07 | 1.200E-07 | 3.500E-07 |

| | | | | | | | | | | |
|---|---|---|---|---|---|---|---|---|---|---|
| 13430 | 672 | S672I | B | 2.4e-07 | 2.3e-07 | 2.4e-07 | 2.367E-07 | 5.774E-09 | 2.300E-07 | 2.400E-07 |
| 13431 | 672 | S672L | B | 1.4e-07 | 3.9e-07 | 5.4e-07 | 3.567E-07 | 2.021E-07 | 1.400E-07 | 5.400E-07 |
| 13432 | 672 | S672K | B | 1.8e-07 | 3.6e-07 | 2.1e-07 | 2.500E-07 | 9.644E-08 | 1.800E-07 | 3.600E-07 |
| 13433 | 672 | S672M | B | 1.7e-07 | 1.8e-07 | 2.4e-07 | 1.967E-07 | 3.786E-08 | 1.700E-07 | 2.400E-07 |
| 13434 | 672 | S672F | B | 2.5e-07 | 3.7e-07 | 2e-07 | 2.733E-07 | 8.737E-08 | 2.000E-07 | 3.700E-07 |
| 13435 | 672 | S672P | B | 1.9e-07 | 2e-07 | 1.6e-07 | 1.833E-07 | 2.082E-08 | 1.600E-07 | 2.000E-07 |
| 13436 | 672 | S672S | B | 1.1e-07 | 2.7e-07 | 1.6e-07 | 1.800E-07 | 8.185E-08 | 1.100E-07 | 2.700E-07 |
| 13437 | 672 | S672T | B | 3e-07 | 1.3e-07 | 1.6e-07 | 1.967E-07 | 9.074E-08 | 1.300E-07 | 3.000E-07 |
| 13438 | 672 | S672W | B | 9.5e-08 | 4.7e-07 | 2e-07 | 2.550E-07 | 1.935E-07 | 9.500E-08 | 4.700E-07 |
| 13439 | 672 | S672Y | B | 3.2e-07 | 5.2e-07 | 3.5e-07 | 3.967E-07 | 1.079E-07 | 3.200E-07 | 5.200E-07 |
| 13440 | 672 | S672V | B | 1.9e-07 | 3e-07 | 2e-07 | 2.300E-07 | 6.083E-08 | 1.900E-07 | 3.000E-07 |
| 13441 | 673 | A673A | B | 4.2e-07 | 1.6e-07 | 2.2e-07 | 2.667E-07 | 1.361E-07 | 1.600E-07 | 4.200E-07 |
| 13442 | 673 | A673R | B | 1.4e-07 | 2.7e-07 | 5.3e-07 | 3.133E-07 | 1.986E-07 | 1.400E-07 | 5.300E-07 |
| 13443 | 673 | A673N | B | 1.8e-07 | 3.4e-07 | 2.7e-07 | 2.633E-07 | 8.021E-08 | 1.800E-07 | 3.400E-07 |
| 13444 | 673 | A673D | B | 2.4e-07 | 1.9e-07 | 1.4e-07 | 1.900E-07 | 5.000E-08 | 1.400E-07 | 2.400E-07 |
| 13445 | 673 | A673C | B | 2.1e-07 | 3.2e-07 | 2e-07 | 2.433E-07 | 6.658E-08 | 2.000E-07 | 3.200E-07 |
| 13446 | 673 | A673E | B | 2.7e-07 | 6.3e-07 | 4e-07 | 4.333E-07 | 1.823E-07 | 2.700E-07 | 6.300E-07 |
| 13447 | 673 | A673Q | B | 2.4e-07 | 1.7e-07 | 1.7e-07 | 1.933E-07 | 4.041E-08 | 1.700E-07 | 2.400E-07 |
| 13448 | 673 | A673G | B | 2.6e-07 | 4.8e-07 | 3.9e-07 | 3.767E-07 | 1.106E-07 | 2.600E-07 | 4.800E-07 |
| 13449 | 673 | A673H | B | 2e-07 | 2.4e-07 | 4.5e-07 | 2.967E-07 | 1.343E-07 | 2.000E-07 | 4.500E-07 |
| 13450 | 673 | A673I | B | 7.2e-07 | 1.2e-07 | 2e-07 | 3.467E-07 | 3.258E-07 | 1.200E-07 | 7.200E-07 |
| 13451 | 673 | A673L | B | 2.6e-07 | 3.7e-07 | 3.3e-07 | 3.200E-07 | 5.568E-08 | 2.600E-07 | 3.700E-07 |
| 13452 | 673 | A673K | B | 3.5e-07 | 2.4e-07 | 2.6e-07 | 2.833E-07 | 5.859E-08 | 2.400E-07 | 3.500E-07 |
| 13453 | 673 | A673M | B | 3.3e-07 | 1.4e-07 | 1.9e-07 | 2.200E-07 | 9.849E-08 | 1.400E-07 | 3.300E-07 |
| 13454 | 673 | A673F | B | 2.1e-07 | 3.7e-07 | 2.6e-07 | 2.800E-07 | 8.185E-08 | 2.100E-07 | 3.700E-07 |
| 13455 | 673 | A673P | B | 1.3e-07 | 1.8e-07 | 2.3e-07 | 1.800E-07 | 5.000E-08 | 1.300E-07 | 2.300E-07 |
| 13456 | 673 | A673S | B | 2e-07 | 1.2e-07 | 2e-07 | 1.733E-07 | 4.619E-08 | 1.200E-07 | 2.000E-07 |
| 13457 | 673 | A673T | B | 2e-07 | 1.8e-07 | 2.6e-07 | 2.133E-07 | 4.163E-08 | 1.800E-07 | 2.600E-07 |
| 13458 | 673 | A673W | B | 2.3e-07 | 4e-07 | 2e-07 | 2.767E-07 | 1.079E-07 | 2.000E-07 | 4.000E-07 |
| 13459 | 673 | A673Y | B | 2.7e-07 | 9.8e-08 | 3.2e-07 | 2.293E-07 | 1.165E-07 | 9.800E-08 | 3.200E-07 |
| 13460 | 673 | A673V | B | 2.7e-07 | 3.1e-07 | 2.5e-07 | 2.767E-07 | 3.055E-08 | 2.500E-07 | 3.100E-07 |
| 13461 | 674 | S674A | B | 2.8e-07 | 3.5e-07 | 1.9e-07 | 2.733E-07 | 8.021E-08 | 1.900E-07 | 3.500E-07 |
| 13462 | 674 | S674R | B | 1.8e-07 | 4.2e-07 | 3.1e-07 | 3.033E-07 | 1.201E-07 | 1.800E-07 | 4.200E-07 |
| 13463 | 674 | S674N | B | 2.4e-07 | 2.3e-07 | 2.8e-07 | 2.500E-07 | 2.646E-08 | 2.300E-07 | 2.800E-07 |
| 13464 | 674 | S674D | B | 9.7e-07 | 2.8e-07 | 1.8e-07 | 4.767E-07 | 4.302E-07 | 1.800E-07 | 9.700E-07 |
| 13465 | 674 | S674C | B | 1.9e-07 | 1.3e-07 | 2.7e-07 | 1.967E-07 | 7.024E-08 | 1.300E-07 | 2.700E-07 |
| 13466 | 674 | S674E | B | 3.7e-07 | 1.1e-07 | 1.4e-07 | 2.067E-07 | 1.422E-07 | 1.100E-07 | 3.700E-07 |
| 13467 | 674 | S674Q | B | 1.8e-07 | 1.4e-07 | 2.6e-07 | 1.933E-07 | 6.110E-08 | 1.400E-07 | 2.600E-07 |
| 13468 | 674 | S674G | B | 1.6e-07 | 1.6e-07 | 1.5e-07 | 1.567E-07 | 5.774E-09 | 1.500E-07 | 1.600E-07 |
| 13469 | 674 | S674H | B | 1.4e-07 | 1.7e-07 | 2.9e-07 | 2.000E-07 | 7.937E-08 | 1.400E-07 | 2.900E-07 |
| 13470 | 674 | S674I | B | 5.7e-07 | 3.3e-07 | 2.7e-07 | 3.900E-07 | 1.587E-07 | 2.700E-07 | 5.700E-07 |
| 13471 | 674 | S674L | B | 2.7e-07 | 3.2e-07 | 3.2e-07 | 3.033E-07 | 2.887E-08 | 2.700E-07 | 3.200E-07 |
| 13472 | 674 | S674K | B | 2.4e-07 | 2.5e-07 | 4.7e-07 | 3.200E-07 | 1.300E-07 | 2.400E-07 | 4.700E-07 |
| 13473 | 674 | S674M | B | 3.1e-07 | 1.8e-07 | 1.4e-07 | 2.100E-07 | 8.888E-08 | 1.400E-07 | 3.100E-07 |
| 13474 | 674 | S674F | B | 4.1e-07 | 2.8e-07 | 4.6e-07 | 3.833E-07 | 9.292E-08 | 2.800E-07 | 4.600E-07 |
| 13475 | 674 | S674P | B | 2.3e-07 | 3.2e-07 | 3.7e-07 | 3.067E-07 | 7.095E-08 | 2.300E-07 | 3.700E-07 |
| 13476 | 674 | S674S | B | 2.1e-07 | 2.7e-07 | 2.2e-07 | 2.333E-07 | 3.215E-08 | 2.100E-07 | 2.700E-07 |
| 13477 | 674 | S674T | B | 3.1e-07 | 2.5e-07 | 1.5e-07 | 2.367E-07 | 8.083E-08 | 1.500E-07 | 3.100E-07 |
| 13478 | 674 | S674W | B | 6.2e-07 | 4.1e-07 | 2.1e-07 | 4.133E-07 | 2.050E-07 | 2.100E-07 | 6.200E-07 |
| 13479 | 674 | S674Y | B | 8.5e-07 | 2.7e-07 | 2e-07 | 4.400E-07 | 3.568E-07 | 2.000E-07 | 8.500E-07 |
| 13480 | 674 | S674V | B | 2.5e-07 | 1.9e-07 | 1.9e-07 | 2.100E-07 | 3.464E-08 | 1.900E-07 | 2.500E-07 |
| 13481 | 675 | Y675A | B | 2.8e-07 | 6.8e-08 | 3.3e-07 | 2.260E-07 | 1.391E-07 | 6.800E-08 | 3.300E-07 |
| 13482 | 675 | Y675R | B | 4.8e-07 | 3e-07 | 2.3e-07 | 3.367E-07 | 1.290E-07 | 2.300E-07 | 4.800E-07 |
| 13483 | 675 | Y675N | B | 1.6e-07 | 2e-07 | 1.7e-07 | 1.767E-07 | 2.082E-08 | 1.600E-07 | 2.000E-07 |
| 13484 | 675 | Y675D | B | 2.4e-07 | 1.5e-07 | 3.3e-07 | 2.400E-07 | 9.000E-08 | 1.500E-07 | 3.300E-07 |
| 13485 | 675 | Y675C | B | 2.6e-07 | 1.5e-07 | 3.2e-07 | 2.433E-07 | 8.622E-08 | 1.500E-07 | 3.200E-07 |
| 13486 | 675 | Y675E | B | 1.5e-07 | 2.9e-07 | 3e-07 | 2.467E-07 | 8.386E-08 | 1.500E-07 | 3.000E-07 |
| 13487 | 675 | Y675Q | B | 3.8e-07 | 2.5e-07 | 2.7e-07 | 3.000E-07 | 7.000E-08 | 2.500E-07 | 3.800E-07 |
| 13488 | 675 | Y675G | B | 3e-07 | 2.4e-07 | 2e-07 | 2.467E-07 | 5.033E-08 | 2.000E-07 | 3.000E-07 |
| 13489 | 675 | Y675H | B | 3.8e-07 | 3e-07 | 3.4e-07 | 3.400E-07 | 4.000E-08 | 3.000E-07 | 3.800E-07 |
| 13490 | 675 | Y675I | B | 3e-07 | 4.6e-07 | 2.3e-07 | 3.300E-07 | 1.179E-07 | 2.300E-07 | 4.600E-07 |
| 13491 | 675 | Y675L | B | 4.4e-07 | 3.9e-07 | 3.7e-07 | 4.000E-07 | 3.606E-08 | 3.700E-07 | 4.400E-07 |
| 13492 | 675 | Y675K | B | 2e-07 | 2.3e-07 | 2.7e-07 | 2.333E-07 | 3.512E-08 | 2.000E-07 | 2.700E-07 |
| 13493 | 675 | Y675M | B | 2.4e-07 | 3.5e-07 | 1.6e-07 | 2.500E-07 | 9.539E-08 | 1.600E-07 | 3.500E-07 |
| 13494 | 675 | Y675F | B | 2.5e-07 | 2e-07 | 4.1e-07 | 2.867E-07 | 1.097E-07 | 2.000E-07 | 4.100E-07 |
| 13495 | 675 | Y675P | B | 2e-07 | 1.5e-07 | 6.1e-07 | 3.200E-07 | 2.524E-07 | 1.500E-07 | 6.100E-07 |
| 13496 | 675 | Y675S | B | 2.9e-07 | 2.6e-07 | 1.7e-07 | 2.400E-07 | 6.245E-08 | 1.700E-07 | 2.900E-07 |
| 13497 | 675 | Y675T | B | 2.7e-07 | 4.7e-07 | 1.1e-07 | 2.833E-07 | 1.804E-07 | 1.100E-07 | 4.700E-07 |
| 13498 | 675 | Y675W | B | 4e-07 | 2.6e-07 | 3.2e-07 | 3.267E-07 | 7.024E-08 | 2.600E-07 | 4.000E-07 |
| 13499 | 675 | Y675Y | B | 1.4e-07 | 2.9e-07 | 2.8e-07 | 2.367E-07 | 8.386E-08 | 1.400E-07 | 2.900E-07 |
| 13500 | 675 | Y675V | B | 3.1e-07 | 3e-07 | 2e-07 | 2.700E-07 | 6.083E-08 | 2.000E-07 | 3.100E-07 |
| 13501 | 676 | R676A | B | 2.9e-07 | 1.7e-07 | 1.6e-07 | 2.067E-07 | 7.234E-08 | 1.600E-07 | 2.900E-07 |
| 13502 | 676 | R676R | B | 1.9e-07 | 3.9e-07 | 3.2e-07 | 3.000E-07 | 1.015E-07 | 1.900E-07 | 3.900E-07 |
| 13503 | 676 | R676N | B | 2.7e-07 | 3.4e-07 | 2.3e-07 | 2.800E-07 | 5.568E-08 | 2.300E-07 | 3.400E-07 |
| 13504 | 676 | R676D | B | 3.1e-07 | 2.4e-07 | 2.8e-07 | 2.767E-07 | 3.512E-08 | 2.400E-07 | 3.100E-07 |
| 13505 | 676 | R676C | B | 2e-07 | 1.8e-07 | 1.3e-07 | 1.700E-07 | 3.606E-08 | 1.300E-07 | 2.000E-07 |
| 13506 | 676 | R676E | B | 1e-07 | 3.2e-07 | 1.8e-07 | 2.000E-07 | 1.114E-07 | 1.000E-07 | 3.200E-07 |
| 13507 | 676 | R676Q | B | 2.8e-07 | 2.3e-07 | 2.7e-07 | 2.600E-07 | 2.646E-08 | 2.300E-07 | 2.800E-07 |
| 13508 | 676 | R676G | B | 5e-07 | 2.8e-07 | 1.3e-07 | 3.033E-07 | 1.861E-07 | 1.300E-07 | 5.000E-07 |
| 13509 | 676 | R676H | B | 3.5e-07 | 2.5e-07 | 2.4e-07 | 2.800E-07 | 6.083E-08 | 2.400E-07 | 3.500E-07 |
| 13510 | 676 | R676I | B | 2.9e-07 | 2.4e-07 | 1.8e-07 | 2.367E-07 | 5.508E-08 | 1.800E-07 | 2.900E-07 |
| 13511 | 676 | R676L | B | 2.3e-07 | 2.8e-07 | 3.7e-07 | 2.933E-07 | 7.095E-08 | 2.300E-07 | 3.700E-07 |
| 13512 | 676 | R676K | B | 1.3e-07 | 5.1e-07 | 1.8e-07 | 2.733E-07 | 2.065E-07 | 1.300E-07 | 5.100E-07 |
| 13513 | 676 | R676M | B | 3.3e-07 | 2.7e-07 | 2.7e-07 | 2.900E-07 | 3.464E-08 | 2.700E-07 | 3.300E-07 |
| 13514 | 676 | R676F | B | 2.6e-07 | 6e-07 | 2.4e-07 | 3.667E-07 | 2.023E-07 | 2.400E-07 | 6.000E-07 |

| | | | | | | | | | | |
|---|---|---|---|---|---|---|---|---|---|---|
| 13515 | 676 | R676P | B | 2.3e-07 | 1.8e-07 | 3.8e-07 | 2.633E-07 | 1.041E-07 | 1.800E-07 | 3.800E-07 |
| 13516 | 676 | R676S | B | 5.2e-07 | 4.4e-07 | 4.6e-07 | 4.733E-07 | 4.163E-08 | 4.400E-07 | 5.200E-07 |
| 13517 | 676 | R676T | B | 2.7e-07 | 2.8e-07 | 2e-07 | 2.500E-07 | 4.359E-08 | 2.000E-07 | 2.800E-07 |
| 13518 | 676 | R676W | B | 3e-07 | 2.4e-07 | 3.2e-07 | 2.867E-07 | 4.163E-08 | 2.400E-07 | 3.200E-07 |
| 13519 | 676 | R676Y | B | 2.1e-07 | 2.1e-07 | 3e-07 | 2.400E-07 | 5.196E-08 | 2.100E-07 | 3.000E-07 |
| 13520 | 676 | R676V | B | 2.6e-07 | 9.2e-08 | 5.1e-07 | 2.873E-07 | 2.103E-07 | 9.200E-08 | 5.100E-07 |
| 13521 | 677 | Y677A | B | 4.1e-07 | 1e-07 | 2.4e-07 | 2.500E-07 | 1.552E-07 | 1.000E-07 | 4.100E-07 |
| 13522 | 677 | Y677R | B | 3e-07 | 4.1e-07 | 3.8e-07 | 3.633E-07 | 5.686E-08 | 3.000E-07 | 4.100E-07 |
| 13523 | 677 | Y677N | B | 2.6e-07 | 3.7e-07 | 2.2e-07 | 2.833E-07 | 7.767E-08 | 2.200E-07 | 3.700E-07 |
| 13524 | 677 | Y677D | B | 2.9e-07 | 3.1e-07 | 1.6e-07 | 2.533E-07 | 8.145E-08 | 1.600E-07 | 3.100E-07 |
| 13525 | 677 | Y677C | B | 2.7e-07 | 2.9e-07 | 5.5e-07 | 3.700E-07 | 1.562E-07 | 2.700E-07 | 5.500E-07 |
| 13526 | 677 | Y677E | B | 2.9e-07 | 2.5e-07 | 9e-08 | 2.100E-07 | 1.058E-07 | 9.000E-08 | 2.900E-07 |
| 13527 | 677 | Y677Q | B | 1.4e-07 | 2.2e-07 | 2.2e-07 | 1.933E-07 | 4.619E-08 | 1.400E-07 | 2.200E-07 |
| 13528 | 677 | Y677G | B | 2.1e-07 | 2.1e-07 | 3.7e-07 | 2.633E-07 | 9.238E-08 | 2.100E-07 | 3.700E-07 |
| 13529 | 677 | Y677H | B | 2.5e-07 | 1.7e-07 | 4.1e-07 | 2.767E-07 | 1.222E-07 | 1.700E-07 | 4.100E-07 |
| 13530 | 677 | Y677I | B | 2.2e-07 | 3.8e-07 | 1.1e-07 | 2.367E-07 | 1.358E-07 | 1.100E-07 | 3.800E-07 |
| 13531 | 677 | Y677L | B | 2.1e-07 | 1.5e-07 | 2.7e-07 | 2.100E-07 | 6.000E-08 | 1.500E-07 | 2.700E-07 |
| 13532 | 677 | Y677K | B | 2e-07 | 2.9e-07 | 2.5e-07 | 2.467E-07 | 4.509E-08 | 2.000E-07 | 2.900E-07 |
| 13533 | 677 | Y677M | B | 1.8e-07 | 6.5e-07 | 5.2e-07 | 4.500E-07 | 2.427E-07 | 1.800E-07 | 6.500E-07 |
| 13534 | 677 | Y677F | B | 1.2e-07 | 1.8e-07 | 2.3e-07 | 1.767E-07 | 5.508E-08 | 1.200E-07 | 2.300E-07 |
| 13535 | 677 | Y677P | B | 4e-07 | 2.8e-07 | 1.7e-07 | 2.833E-07 | 1.150E-07 | 1.700E-07 | 4.000E-07 |
| 13536 | 677 | Y677S | B | 1.2e-07 | 4.4e-07 | 3.4e-07 | 3.000E-07 | 1.637E-07 | 1.200E-07 | 4.400E-07 |
| 13537 | 677 | Y677T | B | 1.9e-07 | 2.3e-07 | 3.1e-07 | 2.433E-07 | 6.110E-08 | 1.900E-07 | 3.100E-07 |
| 13538 | 677 | Y677W | B | 3.3e-07 | 4.3e-07 | 2.7e-07 | 3.433E-07 | 8.083E-08 | 2.700E-07 | 4.300E-07 |
| 13539 | 677 | Y677Y | B | 2.7e-07 | 3.2e-07 | 1.6e-07 | 2.500E-07 | 8.185E-08 | 1.600E-07 | 3.200E-07 |
| 13540 | 677 | Y677V | B | 3.7e-07 | 1.7e-07 | 2.5e-07 | 2.633E-07 | 1.007E-07 | 1.700E-07 | 3.700E-07 |
| 13541 | 678 | T678A | B | 2.6e-07 | 1.6e-07 | 3e-07 | 2.400E-07 | 7.211E-08 | 1.600E-07 | 3.000E-07 |
| 13542 | 678 | T678R | B | 3.6e-07 | 2.3e-07 | 1.7e-07 | 2.533E-07 | 9.713E-08 | 1.700E-07 | 3.600E-07 |
| 13543 | 678 | T678N | B | 4.8e-07 | 1.3e-07 | 1.9e-07 | 2.667E-07 | 1.872E-07 | 1.300E-07 | 4.800E-07 |
| 13544 | 678 | T678D | B | 1.5e-07 | 4.3e-07 | 4.5e-07 | 3.433E-07 | 1.677E-07 | 1.500E-07 | 4.500E-07 |
| 13545 | 678 | T678C | B | 3.2e-07 | 5.2e-07 | 2.5e-07 | 3.633E-07 | 1.401E-07 | 2.500E-07 | 5.200E-07 |
| 13546 | 678 | T678E | B | 4.4e-07 | 2.3e-07 | 4.2e-07 | 3.633E-07 | 1.159E-07 | 2.300E-07 | 4.400E-07 |
| 13547 | 678 | T678Q | B | 2.9e-07 | 2e-07 | 5e-07 | 3.300E-07 | 1.539E-07 | 2.000E-07 | 5.000E-07 |
| 13548 | 678 | T678G | B | 7.9e-07 | 6e-07 | 3.2e-07 | 5.700E-07 | 2.364E-07 | 3.200E-07 | 7.900E-07 |
| 13549 | 678 | T678H | B | 3.4e-07 | 3.2e-07 | 3.1e-07 | 3.233E-07 | 1.528E-08 | 3.100E-07 | 3.400E-07 |
| 13550 | 678 | T678I | B | 4.2e-07 | 2.8e-07 | 2e-07 | 3.000E-07 | 1.114E-07 | 2.000E-07 | 4.200E-07 |
| 13551 | 678 | T678L | B | 2.1e-07 | 2.5e-07 | 1.4e-07 | 2.000E-07 | 5.568E-08 | 1.400E-07 | 2.500E-07 |
| 13552 | 678 | T678K | B | 4e-07 | 2.9e-07 | 1.7e-07 | 2.867E-07 | 1.150E-07 | 1.700E-07 | 4.000E-07 |
| 13553 | 678 | T678M | B | 3.1e-07 | 2.2e-07 | 4e-07 | 3.100E-07 | 9.000E-08 | 2.200E-07 | 4.000E-07 |
| 13554 | 678 | T678F | B | 4.6e-07 | 3e-07 | 1.4e-07 | 3.000E-07 | 1.600E-07 | 1.400E-07 | 4.600E-07 |
| 13555 | 678 | T678P | B | 2e-07 | 1.7e-07 | 2.2e-07 | 1.967E-07 | 2.517E-08 | 1.700E-07 | 2.200E-07 |
| 13556 | 678 | T678S | B | 1.1e-07 | 6e-07 | 4.7e-07 | 3.933E-07 | 2.538E-07 | 1.100E-07 | 6.000E-07 |
| 13557 | 678 | T678T | B | 3.1e-07 | 2.7e-07 | 1.6e-07 | 2.467E-07 | 7.767E-08 | 1.600E-07 | 3.100E-07 |
| 13558 | 678 | T678W | B | 3.1e-07 | 3.6e-07 | 3.1e-07 | 3.267E-07 | 2.887E-08 | 3.100E-07 | 3.600E-07 |
| 13559 | 678 | T678Y | B | 2.7e-07 | 9.3e-08 | 2.3e-07 | 1.977E-07 | 9.282E-08 | 9.300E-08 | 2.700E-07 |
| 13560 | 678 | T678V | B | 2.9e-07 | 2.3e-07 | 1.2e-07 | 2.133E-07 | 8.622E-08 | 1.200E-07 | 2.900E-07 |
| 13561 | 679 | G679A | B | 4.7e-07 | 3.6e-07 | 1.9e-07 | 3.400E-07 | 1.411E-07 | 1.900E-07 | 4.700E-07 |
| 13562 | 679 | G679R | B | 3.4e-07 | 1.9e-07 | 3.4e-07 | 2.900E-07 | 8.660E-08 | 1.900E-07 | 3.400E-07 |
| 13563 | 679 | G679N | B | 5.6e-07 | 2.1e-07 | 1.2e-07 | 2.967E-07 | 2.325E-07 | 1.200E-07 | 5.600E-07 |
| 13564 | 679 | G679D | B | 3.2e-07 | 1.9e-07 | 1.9e-07 | 2.333E-07 | 7.506E-08 | 1.900E-07 | 3.200E-07 |
| 13565 | 679 | G679C | B | 3.6e-07 | 3.4e-07 | 3.2e-07 | 3.400E-07 | 2.000E-08 | 3.200E-07 | 3.600E-07 |
| 13566 | 679 | G679E | B | 9.2e-07 | 1.3e-07 | 1.5e-07 | 4.000E-07 | 4.504E-07 | 1.300E-07 | 9.200E-07 |
| 13567 | 679 | G679Q | B | 3.4e-07 | 2.5e-07 | 1.9e-07 | 2.600E-07 | 7.550E-08 | 1.900E-07 | 3.400E-07 |
| 13568 | 679 | G679G | B | 2e-07 | 2.4e-07 | 1.8e-07 | 2.067E-07 | 3.055E-08 | 1.800E-07 | 2.400E-07 |
| 13569 | 679 | G679H | B | 3e-07 | 2.5e-07 | 1.4e-07 | 2.300E-07 | 8.185E-08 | 1.400E-07 | 3.000E-07 |
| 13570 | 679 | G679I | B | 1.7e-07 | 2e-07 | 2.1e-07 | 1.933E-07 | 2.082E-08 | 1.700E-07 | 2.100E-07 |
| 13571 | 679 | G679L | B | 4.6e-07 | 1.1e-07 | 8.9e-08 | 2.197E-07 | 2.084E-07 | 8.900E-08 | 4.600E-07 |
| 13572 | 679 | G679K | B | 5.6e-07 | 3.1e-07 | 2e-07 | 3.567E-07 | 1.845E-07 | 2.000E-07 | 5.600E-07 |
| 13573 | 679 | G679M | B | 1.6e-07 | 2e-07 | 1.4e-07 | 1.667E-07 | 3.055E-08 | 1.400E-07 | 2.000E-07 |
| 13574 | 679 | G679F | B | 9.9e-08 | 4.7e-07 | 3.4e-07 | 3.030E-07 | 1.882E-07 | 9.900E-08 | 4.700E-07 |
| 13575 | 679 | G679P | B | 1.2e-07 | 1.7e-07 | 2.5e-07 | 1.800E-07 | 6.557E-08 | 1.200E-07 | 2.500E-07 |
| 13576 | 679 | G679S | B | 2.8e-07 | 2.1e-07 | 2.8e-07 | 2.567E-07 | 4.041E-08 | 2.100E-07 | 2.800E-07 |
| 13577 | 679 | G679T | B | 4.1e-07 | 2.1e-07 | 1.4e-07 | 2.533E-07 | 1.401E-07 | 1.400E-07 | 4.100E-07 |
| 13578 | 679 | G679W | B | 1.3e-07 | 2.7e-07 | 2.2e-07 | 2.067E-07 | 7.095E-08 | 1.300E-07 | 2.700E-07 |
| 13579 | 679 | G679Y | B | 3.3e-07 | 1.8e-07 | 2.4e-07 | 2.500E-07 | 7.550E-08 | 1.800E-07 | 3.300E-07 |
| 13580 | 679 | G679V | B | 1.6e-07 | 1.9e-07 | 9.1e-08 | 1.470E-07 | 5.076E-08 | 9.100E-08 | 1.900E-07 |
| 13581 | 680 | V680A | B | 1.4e-07 | 3e-07 | 1.2e-07 | 1.867E-07 | 9.866E-08 | 1.200E-07 | 3.000E-07 |
| 13582 | 680 | V680R | B | 9.6e-08 | 3e-07 | 1.1e-07 | 1.687E-07 | 1.140E-07 | 9.600E-08 | 3.000E-07 |
| 13583 | 680 | V680N | B | 1.4e-07 | 1.5e-07 | 3.8e-07 | 2.233E-07 | 1.358E-07 | 1.400E-07 | 3.800E-07 |
| 13584 | 680 | V680D | B | 2.7e-07 | 3.2e-07 | 2e-07 | 2.633E-07 | 6.028E-08 | 2.000E-07 | 3.200E-07 |
| 13585 | 680 | V680C | B | 1.5e-07 | 3.5e-07 | 2.3e-07 | 2.433E-07 | 1.007E-07 | 1.500E-07 | 3.500E-07 |
| 13586 | 680 | V680E | B | 3.8e-07 | 2.5e-07 | 1.5e-07 | 2.600E-07 | 1.153E-07 | 1.500E-07 | 3.800E-07 |
| 13587 | 680 | V680Q | B | 1.1e-07 | 2.8e-07 | 2e-07 | 1.967E-07 | 8.505E-08 | 1.100E-07 | 2.800E-07 |
| 13588 | 680 | V680G | B | 2.3e-07 | 4e-07 | 2.5e-07 | 2.933E-07 | 9.292E-08 | 2.300E-07 | 4.000E-07 |
| 13589 | 680 | V680H | B | 2e-07 | 4.4e-07 | 2e-07 | 2.800E-07 | 1.386E-07 | 2.000E-07 | 4.400E-07 |
| 13590 | 680 | V680I | B | 3.9e-07 | 2.9e-07 | 2.3e-07 | 3.033E-07 | 8.083E-08 | 2.300E-07 | 3.900E-07 |
| 13591 | 680 | V680L | B | 3.8e-07 | 2.8e-07 | 3e-07 | 3.200E-07 | 5.292E-08 | 2.800E-07 | 3.800E-07 |
| 13592 | 680 | V680K | B | 2.7e-07 | 2.4e-07 | 2.6e-07 | 2.567E-07 | 1.528E-08 | 2.400E-07 | 2.700E-07 |
| 13593 | 680 | V680M | B | 2.8e-07 | 1.6e-07 | 1.1e-07 | 1.833E-07 | 8.737E-08 | 1.100E-07 | 2.800E-07 |
| 13594 | 680 | V680F | B | 1.8e-07 | 4.7e-07 | 2.2e-07 | 2.900E-07 | 1.572E-07 | 1.800E-07 | 4.700E-07 |
| 13595 | 680 | V680P | B | 1.2e-07 | 2.1e-07 | 3.8e-07 | 2.367E-07 | 1.320E-07 | 1.200E-07 | 3.800E-07 |
| 13596 | 680 | V680S | B | 1.6e-07 | 6.9e-08 | 4e-07 | 2.097E-07 | 1.710E-07 | 6.900E-08 | 4.000E-07 |
| 13597 | 680 | V680T | B | 8.9e-08 | 1.9e-07 | 2.7e-07 | 1.830E-07 | 9.070E-08 | 8.900E-08 | 2.700E-07 |
| 13598 | 680 | V680W | B | 4.1e-07 | 1.4e-07 | 2.7e-07 | 2.733E-07 | 1.350E-07 | 1.400E-07 | 4.100E-07 |
| 13599 | 680 | V680Y | B | 3.3e-07 | 2.5e-07 | 1.8e-07 | 2.533E-07 | 7.506E-08 | 1.800E-07 | 3.300E-07 |

| | | | | | | | | | | |
|---|---|---|---|---|---|---|---|---|---|---|
| 13600 | 680 | V680V | B | 4.1e-07 | 3.1e-07 | 3.3e-07 | 3.500E-07 | 5.292E-08 | 3.100E-07 | 4.100E-07 |
| 13601 | 681 | P681A | B | 3.3e-07 | 3.2e-07 | 3.5e-07 | 3.333E-07 | 1.528E-08 | 3.200E-07 | 3.500E-07 |
| 13602 | 681 | P681R | B | 2e-07 | 3.5e-07 | 2.5e-07 | 2.667E-07 | 7.638E-08 | 2.000E-07 | 3.500E-07 |
| 13603 | 681 | P681N | B | 1.3e-07 | 2.8e-07 | 2e-07 | 2.033E-07 | 7.506E-08 | 1.300E-07 | 2.800E-07 |
| 13604 | 681 | P681D | B | 1.9e-07 | 2e-07 | 4e-07 | 2.633E-07 | 1.185E-07 | 1.900E-07 | 4.000E-07 |
| 13605 | 681 | P681C | B | 3.3e-07 | 2.7e-07 | 1.2e-07 | 2.400E-07 | 1.082E-07 | 1.200E-07 | 3.300E-07 |
| 13606 | 681 | P681E | B | 2.9e-07 | 1.9e-07 | 1.9e-07 | 2.233E-07 | 5.774E-08 | 1.900E-07 | 2.900E-07 |
| 13607 | 681 | P681Q | B | 2.5e-07 | 3.5e-07 | 2.8e-07 | 2.933E-07 | 5.132E-08 | 2.500E-07 | 3.500E-07 |
| 13608 | 681 | P681G | B | 1.7e-07 | 5.3e-07 | 4.5e-07 | 3.833E-07 | 1.890E-07 | 1.700E-07 | 5.300E-07 |
| 13609 | 681 | P681H | B | 1.9e-07 | 3.4e-07 | 2.6e-07 | 2.633E-07 | 7.506E-08 | 1.900E-07 | 3.400E-07 |
| 13610 | 681 | P681I | B | 2.2e-07 | 9.7e-08 | 6.2e-07 | 3.123E-07 | 2.735E-07 | 9.700E-08 | 6.200E-07 |
| 13611 | 681 | P681L | B | 8.4e-08 | 3.2e-07 | 2.7e-07 | 2.247E-07 | 1.244E-07 | 8.400E-08 | 3.200E-07 |
| 13612 | 681 | P681K | B | 4.7e-07 | 3.6e-07 | 2e-07 | 3.433E-07 | 1.358E-07 | 2.000E-07 | 4.700E-07 |
| 13613 | 681 | P681M | B | 2.3e-07 | 2.3e-07 | 1.3e-07 | 1.967E-07 | 5.774E-08 | 1.300E-07 | 2.300E-07 |
| 13614 | 681 | P681F | B | 2.9e-07 | 2.4e-07 | 3.1e-07 | 2.800E-07 | 3.606E-08 | 2.400E-07 | 3.100E-07 |
| 13615 | 681 | P681P | B | 4.4e-07 | 4.6e-07 | 2.6e-07 | 3.867E-07 | 1.102E-07 | 2.600E-07 | 4.600E-07 |
| 13616 | 681 | P681S | B | 3.1e-07 | 1.5e-07 | 2.4e-07 | 2.333E-07 | 8.021E-08 | 1.500E-07 | 3.100E-07 |
| 13617 | 681 | P681T | B | 8.7e-07 | 1.7e-07 | 4.8e-07 | 5.067E-07 | 3.508E-07 | 1.700E-07 | 8.700E-07 |
| 13618 | 681 | P681W | B | 4.6e-07 | 2.6e-07 | 2.4e-07 | 3.200E-07 | 1.217E-07 | 2.400E-07 | 4.600E-07 |
| 13619 | 681 | P681Y | B | 2.8e-07 | 2e-07 | 4.4e-07 | 3.067E-07 | 1.222E-07 | 2.000E-07 | 4.400E-07 |
| 13620 | 681 | P681V | B | 3.5e-07 | 2.5e-07 | 5.7e-07 | 3.900E-07 | 1.637E-07 | 2.500E-07 | 5.700E-07 |
| 13621 | 682 | S682A | B | 1.3e-07 | 1.9e-07 | 2.8e-07 | 2.000E-07 | 7.550E-08 | 1.300E-07 | 2.800E-07 |
| 13622 | 682 | S682R | B | 2.7e-07 | 1.6e-07 | 1.5e-07 | 1.933E-07 | 6.658E-08 | 1.500E-07 | 2.700E-07 |
| 13623 | 682 | S682N | B | 2.2e-07 | 2.5e-07 | 3.1e-07 | 2.600E-07 | 4.583E-08 | 2.200E-07 | 3.100E-07 |
| 13624 | 682 | S682D | B | 3.2e-07 | 1.8e-07 | 3.2e-07 | 2.733E-07 | 8.083E-08 | 1.800E-07 | 3.200E-07 |
| 13625 | 682 | S682C | B | 2.6e-07 | 3.2e-07 | 6.1e-07 | 3.967E-07 | 1.872E-07 | 2.600E-07 | 6.100E-07 |
| 13626 | 682 | S682E | B | 1.8e-07 | 2.8e-07 | 1.2e-07 | 1.933E-07 | 8.083E-08 | 1.200E-07 | 2.800E-07 |
| 13627 | 682 | S682Q | B | 3.7e-07 | 2.7e-07 | 1.6e-07 | 2.667E-07 | 1.050E-07 | 1.600E-07 | 3.700E-07 |
| 13628 | 682 | S682G | B | 3.4e-07 | 4e-07 | 3.6e-07 | 3.667E-07 | 3.055E-08 | 3.400E-07 | 4.000E-07 |
| 13629 | 682 | S682H | B | 4.4e-07 | 2e-07 | 2.6e-07 | 3.000E-07 | 1.249E-07 | 2.000E-07 | 4.400E-07 |
| 13630 | 682 | S682I | B | 2.2e-07 | 5.7e-07 | 1.2e-07 | 3.033E-07 | 2.363E-07 | 1.200E-07 | 5.700E-07 |
| 13631 | 682 | S682L | B | 2.6e-07 | 1.5e-07 | 1.7e-07 | 1.933E-07 | 5.859E-08 | 1.500E-07 | 2.600E-07 |
| 13632 | 682 | S682K | B | 2.1e-07 | 3e-07 | 2.1e-07 | 2.400E-07 | 5.196E-08 | 2.100E-07 | 3.000E-07 |
| 13633 | 682 | S682M | B | 2.3e-07 | 3.8e-07 | 2.1e-07 | 2.733E-07 | 9.292E-08 | 2.100E-07 | 3.800E-07 |
| 13634 | 682 | S682F | B | 3.3e-07 | 2.9e-07 | 1.8e-07 | 2.667E-07 | 7.767E-08 | 1.800E-07 | 3.300E-07 |
| 13635 | 682 | S682P | B | 1.3e-07 | 1.8e-07 | 4.2e-07 | 2.433E-07 | 1.550E-07 | 1.300E-07 | 4.200E-07 |
| 13636 | 682 | S682S | B | 1.4e-07 | 3.8e-07 | 4.6e-07 | 3.267E-07 | 1.665E-07 | 1.400E-07 | 4.600E-07 |
| 13637 | 682 | S682T | B | 4e-07 | 1.9e-07 | 1.9e-07 | 2.600E-07 | 1.212E-07 | 1.900E-07 | 4.000E-07 |
| 13638 | 682 | S682W | B | 1.6e-07 | 2.1e-07 | 5.9e-07 | 3.200E-07 | 2.352E-07 | 1.600E-07 | 5.900E-07 |
| 13639 | 682 | S682Y | B | 2.1e-07 | 2.7e-07 | 2.2e-07 | 2.333E-07 | 3.215E-08 | 2.100E-07 | 2.700E-07 |
| 13640 | 682 | S682V | B | 2e-07 | 1.5e-07 | 2e-07 | 1.833E-07 | 2.887E-08 | 1.500E-07 | 2.000E-07 |
| 13641 | 683 | R683A | B | 2.7e-07 | 2.9e-07 | 2.8e-07 | 2.800E-07 | 1.000E-08 | 2.700E-07 | 2.900E-07 |
| 13642 | 683 | R683R | B | 2e-07 | 1.5e-07 | 2.4e-07 | 1.967E-07 | 4.509E-08 | 1.500E-07 | 2.400E-07 |
| 13643 | 683 | R683N | B | 2.8e-07 | 1.4e-07 | 1.2e-07 | 1.800E-07 | 8.718E-08 | 1.200E-07 | 2.800E-07 |
| 13644 | 683 | R683D | B | 3.2e-07 | 4.8e-07 | 3.3e-07 | 3.767E-07 | 8.963E-08 | 3.200E-07 | 4.800E-07 |
| 13645 | 683 | R683C | B | 2.5e-07 | 1.3e-07 | 2e-07 | 1.933E-07 | 6.028E-08 | 1.300E-07 | 2.500E-07 |
| 13646 | 683 | R683E | B | 3.1e-07 | 2.5e-07 | 2.8e-07 | 2.800E-07 | 3.000E-08 | 2.500E-07 | 3.100E-07 |
| 13647 | 683 | R683Q | B | 2.2e-07 | 1.6e-07 | 1.9e-07 | 1.900E-07 | 3.000E-08 | 1.600E-07 | 2.200E-07 |
| 13648 | 683 | R683G | B | 4.1e-07 | 2.9e-07 | 2.8e-07 | 3.267E-07 | 7.234E-08 | 2.800E-07 | 4.100E-07 |
| 13649 | 683 | R683H | B | 4.1e-07 | 4.8e-07 | 1.7e-07 | 3.533E-07 | 1.626E-07 | 1.700E-07 | 4.800E-07 |
| 13650 | 683 | R683I | B | 3.8e-07 | 4.2e-07 | 1.8e-07 | 3.267E-07 | 1.286E-07 | 1.800E-07 | 4.200E-07 |
| 13651 | 683 | R683L | B | 1.9e-07 | 2.1e-07 | 3.4e-07 | 2.467E-07 | 8.145E-08 | 1.900E-07 | 3.400E-07 |
| 13652 | 683 | R683K | B | 1.2e-07 | 1.7e-07 | 4e-07 | 2.300E-07 | 1.493E-07 | 1.200E-07 | 4.000E-07 |
| 13653 | 683 | R683M | B | 3.9e-07 | 2.3e-07 | 4e-07 | 3.400E-07 | 9.539E-08 | 2.300E-07 | 4.000E-07 |
| 13654 | 683 | R683F | B | 3.3e-07 | 2.3e-07 | 1.3e-07 | 2.300E-07 | 1.000E-07 | 1.300E-07 | 3.300E-07 |
| 13655 | 683 | R683P | B | 2.4e-07 | 9.5e-08 | 1.5e-07 | 1.617E-07 | 7.320E-08 | 9.500E-08 | 2.400E-07 |
| 13656 | 683 | R683S | B | 1.5e-07 | 1.8e-07 | 3.2e-07 | 2.167E-07 | 9.074E-08 | 1.500E-07 | 3.200E-07 |
| 13657 | 683 | R683T | B | 2.4e-07 | 3.1e-07 | 4.1e-07 | 3.200E-07 | 8.544E-08 | 2.400E-07 | 4.100E-07 |
| 13658 | 683 | R683W | B | 1.3e-07 | 4.2e-07 | 2.8e-07 | 2.767E-07 | 1.450E-07 | 1.300E-07 | 4.200E-07 |
| 13659 | 683 | R683Y | B | 3.4e-07 | 1.9e-07 | 5.5e-07 | 3.600E-07 | 1.808E-07 | 1.900E-07 | 5.500E-07 |
| 13660 | 683 | R683V | B | 3e-07 | 2.3e-07 | 2.9e-07 | 2.733E-07 | 3.786E-08 | 2.300E-07 | 3.000E-07 |
| 13661 | 684 | F684A | B | 3.6e-07 | 3e-07 | 1.6e-07 | 2.733E-07 | 1.026E-07 | 1.600E-07 | 3.600E-07 |
| 13662 | 684 | F684R | B | 3.2e-07 | 5.2e-07 | 4.8e-07 | 4.400E-07 | 1.058E-07 | 3.200E-07 | 5.200E-07 |
| 13663 | 684 | F684N | B | 4.6e-07 | 2.3e-07 | 3.1e-07 | 3.333E-07 | 1.168E-07 | 2.300E-07 | 4.600E-07 |
| 13664 | 684 | F684D | B | 2.2e-07 | 3.1e-07 | 2.6e-07 | 2.633E-07 | 4.509E-08 | 2.200E-07 | 3.100E-07 |
| 13665 | 684 | F684C | B | 2.1e-07 | 1.3e-07 | 2e-07 | 1.800E-07 | 4.359E-08 | 1.300E-07 | 2.100E-07 |
| 13666 | 684 | F684E | B | 3.3e-07 | 3.3e-07 | 2.3e-07 | 2.967E-07 | 5.774E-08 | 2.300E-07 | 3.300E-07 |
| 13667 | 684 | F684Q | B | 3e-07 | 1.2e-07 | 3.4e-07 | 2.533E-07 | 1.172E-07 | 1.200E-07 | 3.400E-07 |
| 13668 | 684 | F684G | B | 1.5e-07 | 2.7e-07 | 1.7e-07 | 1.967E-07 | 6.429E-08 | 1.500E-07 | 2.700E-07 |
| 13669 | 684 | F684H | B | 3.3e-07 | 3.6e-07 | 3.1e-07 | 3.333E-07 | 2.517E-08 | 3.100E-07 | 3.600E-07 |
| 13670 | 684 | F684I | B | 1.5e-07 | 1.8e-07 | 4.1e-07 | 2.467E-07 | 1.422E-07 | 1.500E-07 | 4.100E-07 |
| 13671 | 684 | F684L | B | 1.7e-07 | 2.6e-07 | 2.1e-07 | 2.133E-07 | 4.509E-08 | 1.700E-07 | 2.600E-07 |
| 13672 | 684 | F684K | B | 4e-07 | 1.9e-07 | 2.4e-07 | 2.767E-07 | 1.097E-07 | 1.900E-07 | 4.000E-07 |
| 13673 | 684 | F684M | B | 1.7e-07 | 1.6e-07 | 2.6e-07 | 1.967E-07 | 5.508E-08 | 1.600E-07 | 2.600E-07 |
| 13674 | 684 | F684F | B | 3.9e-07 | 3.8e-07 | 1.9e-07 | 3.200E-07 | 1.127E-07 | 1.900E-07 | 3.900E-07 |
| 13675 | 684 | F684P | B | 1.5e-07 | 4.3e-07 | 2.5e-07 | 2.767E-07 | 1.419E-07 | 1.500E-07 | 4.300E-07 |
| 13676 | 684 | F684S | B | 1.6e-07 | 2e-07 | 1.8e-07 | 1.800E-07 | 2.000E-08 | 1.600E-07 | 2.000E-07 |
| 13677 | 684 | F684T | B | 1.7e-07 | 3.5e-07 | 3.3e-07 | 2.833E-07 | 9.866E-08 | 1.700E-07 | 3.500E-07 |
| 13678 | 684 | F684W | B | 2.2e-07 | 1.4e-07 | 1.3e-07 | 1.633E-07 | 4.933E-08 | 1.300E-07 | 2.200E-07 |
| 13679 | 684 | F684Y | B | 5.1e-08 | 3e-07 | 1e-07 | 1.503E-07 | 1.319E-07 | 5.100E-08 | 3.000E-07 |
| 13680 | 684 | F684V | B | 3.1e-07 | 2.8e-07 | 1.4e-07 | 2.433E-07 | 9.074E-08 | 1.400E-07 | 3.100E-07 |
| 13681 | 685 | S685A | B | 5.8e-07 | 2.1e-07 | 2.1e-07 | 3.333E-07 | 2.136E-07 | 2.100E-07 | 5.800E-07 |
| 13682 | 685 | S685R | B | 1.3e-07 | 1.2e-07 | 2.1e-07 | 1.533E-07 | 4.933E-08 | 1.200E-07 | 2.100E-07 |
| 13683 | 685 | S685N | B | 2e-07 | 3.6e-07 | 2.3e-07 | 2.633E-07 | 8.505E-08 | 2.000E-07 | 3.600E-07 |
| 13684 | 685 | S685D | B | 2.2e-07 | 3.9e-07 | 3.3e-07 | 3.133E-07 | 8.622E-08 | 2.200E-07 | 3.900E-07 |

| | | | | | | | | | | |
|---|---|---|---|---|---|---|---|---|---|---|
| 13685 | 685 | S685C | B | 2.2e-07 | 2.3e-07 | 1.8e-07 | 2.100E-07 | 2.646E-08 | 1.800E-07 | 2.300E-07 |
| 13686 | 685 | S685E | B | 4.2e-07 | 4e-07 | 3.4e-07 | 3.867E-07 | 4.163E-08 | 3.400E-07 | 4.200E-07 |
| 13687 | 685 | S685Q | B | 2.5e-07 | 4.2e-07 | 1.3e-07 | 2.667E-07 | 1.457E-07 | 1.300E-07 | 4.200E-07 |
| 13688 | 685 | S685G | B | 5e-07 | 3.2e-07 | 3.6e-07 | 3.933E-07 | 9.452E-08 | 3.200E-07 | 5.000E-07 |
| 13689 | 685 | S685H | B | 3.1e-07 | 2.7e-07 | 2.1e-07 | 2.633E-07 | 5.033E-08 | 2.100E-07 | 3.100E-07 |
| 13690 | 685 | S685I | B | 3.4e-07 | 1.7e-07 | 3e-07 | 2.700E-07 | 8.888E-08 | 1.700E-07 | 3.400E-07 |
| 13691 | 685 | S685L | B | 2.7e-07 | 1.7e-07 | 3.8e-07 | 2.733E-07 | 1.050E-07 | 1.700E-07 | 3.800E-07 |
| 13692 | 685 | S685K | B | 4.4e-07 | 4e-07 | 3.4e-07 | 3.933E-07 | 5.033E-08 | 3.400E-07 | 4.400E-07 |
| 13693 | 685 | S685M | B | 2.3e-07 | 2.6e-07 | 2.7e-07 | 2.533E-07 | 2.082E-08 | 2.300E-07 | 2.700E-07 |
| 13694 | 685 | S685F | B | 3.5e-07 | 2.8e-07 | 3.4e-07 | 3.233E-07 | 3.786E-08 | 2.800E-07 | 3.500E-07 |
| 13695 | 685 | S685P | B | 1.5e-07 | 2.8e-07 | 2.6e-07 | 2.300E-07 | 7.000E-08 | 1.500E-07 | 2.800E-07 |
| 13696 | 685 | S685S | B | 4.6e-07 | 2.4e-07 | 3e-07 | 3.333E-07 | 1.137E-07 | 2.400E-07 | 4.600E-07 |
| 13697 | 685 | S685T | B | 2.8e-07 | 2.1e-07 | 1.6e-07 | 2.167E-07 | 6.028E-08 | 1.600E-07 | 2.800E-07 |
| 13698 | 685 | S685W | B | 1.4e-07 | 1.4e-07 | 3.6e-07 | 2.133E-07 | 1.270E-07 | 1.400E-07 | 3.600E-07 |
| 13699 | 685 | S685Y | B | 4.4e-07 | 5.6e-07 | 3.5e-07 | 4.500E-07 | 1.054E-07 | 3.500E-07 | 5.600E-07 |
| 13700 | 685 | S685V | B | 1.8e-07 | 3.5e-07 | 3.8e-07 | 3.033E-07 | 1.079E-07 | 1.800E-07 | 3.800E-07 |
| 13701 | 686 | G686A | B | 4.2e-07 | 2.5e-07 | 3e-07 | 3.233E-07 | 8.737E-08 | 2.500E-07 | 4.200E-07 |
| 13702 | 686 | G686R | B | 2.8e-07 | 2.1e-07 | 3.5e-07 | 2.800E-07 | 7.000E-08 | 2.100E-07 | 3.500E-07 |
| 13703 | 686 | G686N | B | 2.8e-07 | 8.8e-08 | 3.8e-07 | 2.493E-07 | 1.484E-07 | 8.800E-08 | 3.800E-07 |
| 13704 | 686 | G686D | B | 3.1e-07 | 1.8e-07 | 2.5e-07 | 2.467E-07 | 6.506E-08 | 1.800E-07 | 3.100E-07 |
| 13705 | 686 | G686C | B | 2e-07 | 1.7e-07 | 1.8e-07 | 1.833E-07 | 1.528E-08 | 1.700E-07 | 2.000E-07 |
| 13706 | 686 | G686E | B | 1.6e-07 | 2.9e-07 | 2.9e-07 | 2.467E-07 | 7.506E-08 | 1.600E-07 | 2.900E-07 |
| 13707 | 686 | G686Q | B | 2e-07 | 2.9e-07 | 2.1e-07 | 2.333E-07 | 4.933E-08 | 2.000E-07 | 2.900E-07 |
| 13708 | 686 | G686G | B | 3.4e-07 | 2.7e-07 | 2.2e-07 | 2.767E-07 | 6.028E-08 | 2.200E-07 | 3.400E-07 |
| 13709 | 686 | G686H | B | 1.9e-07 | 1.1e-07 | 2.8e-07 | 1.933E-07 | 8.505E-08 | 1.100E-07 | 2.800E-07 |
| 13710 | 686 | G686I | B | 2.9e-07 | 3e-07 | 2.5e-07 | 2.800E-07 | 2.646E-08 | 2.500E-07 | 3.000E-07 |
| 13711 | 686 | G686L | B | 1.9e-07 | 2.8e-07 | 3.6e-07 | 2.767E-07 | 8.505E-08 | 1.900E-07 | 3.600E-07 |
| 13712 | 686 | G686K | B | 2.8e-07 | 3e-07 | 2.7e-07 | 2.833E-07 | 1.528E-08 | 2.700E-07 | 3.000E-07 |
| 13713 | 686 | G686M | B | 4.1e-07 | 3.7e-07 | 1.3e-07 | 3.033E-07 | 1.514E-07 | 1.300E-07 | 4.100E-07 |
| 13714 | 686 | G686F | B | 3.2e-07 | 1.7e-07 | 2.1e-07 | 2.333E-07 | 7.767E-08 | 1.700E-07 | 3.200E-07 |
| 13715 | 686 | G686P | B | 1.9e-07 | 3.6e-07 | 1.3e-07 | 2.267E-07 | 1.193E-07 | 1.300E-07 | 3.600E-07 |
| 13716 | 686 | G686S | B | 2.5e-07 | 1.3e-07 | 2.7e-07 | 2.167E-07 | 7.572E-08 | 1.300E-07 | 2.700E-07 |
| 13717 | 686 | G686T | B | 4.3e-07 | 1.3e-07 | 2.6e-07 | 2.733E-07 | 1.504E-07 | 1.300E-07 | 4.300E-07 |
| 13718 | 686 | G686W | B | 3.2e-07 | 3.7e-07 | 1.4e-07 | 2.767E-07 | 1.210E-07 | 1.400E-07 | 3.700E-07 |
| 13719 | 686 | G686Y | B | 2.7e-07 | 1.5e-07 | 3e-07 | 2.400E-07 | 7.937E-08 | 1.500E-07 | 3.000E-07 |
| 13720 | 686 | G686V | B | 1.2e-07 | 2.8e-07 | 4e-07 | 2.667E-07 | 1.405E-07 | 1.200E-07 | 4.000E-07 |
| 13721 | 687 | S687A | B | 2.5e-07 | 1.2e-07 | 2e-07 | 1.900E-07 | 6.557E-08 | 1.200E-07 | 2.500E-07 |
| 13722 | 687 | S687R | B | 2e-07 | 2.7e-07 | 1.1e-07 | 1.933E-07 | 8.021E-08 | 1.100E-07 | 2.700E-07 |
| 13723 | 687 | S687N | B | 2.2e-07 | 3e-07 | 2.5e-07 | 2.567E-07 | 4.041E-08 | 2.200E-07 | 3.000E-07 |
| 13724 | 687 | S687D | B | 3.2e-07 | 1.9e-07 | 3.8e-07 | 2.967E-07 | 9.713E-08 | 1.900E-07 | 3.800E-07 |
| 13725 | 687 | S687C | B | 2.6e-07 | 2e-07 | 2.8e-07 | 2.467E-07 | 4.163E-08 | 2.000E-07 | 2.800E-07 |
| 13726 | 687 | S687E | B | 6.7e-07 | 3.7e-07 | 9.9e-08 | 3.797E-07 | 2.856E-07 | 9.900E-08 | 6.700E-07 |
| 13727 | 687 | S687Q | B | 1.2e-07 | 1.7e-07 | 2.5e-07 | 1.800E-07 | 6.557E-08 | 1.200E-07 | 2.500E-07 |
| 13728 | 687 | S687G | B | 2.8e-07 | 4.1e-07 | 5.1e-07 | 4.000E-07 | 1.153E-07 | 2.800E-07 | 5.100E-07 |
| 13729 | 687 | S687H | B | 1.9e-07 | 1.5e-07 | 3e-07 | 2.133E-07 | 7.767E-08 | 1.500E-07 | 3.000E-07 |
| 13730 | 687 | S687I | B | 2.4e-07 | 1.2e-07 | 3.7e-07 | 2.433E-07 | 1.250E-07 | 1.200E-07 | 3.700E-07 |
| 13731 | 687 | S687L | B | 1.3e-07 | 2e-07 | 3e-07 | 2.100E-07 | 8.544E-08 | 1.300E-07 | 3.000E-07 |
| 13732 | 687 | S687K | B | 2.4e-07 | 1.5e-07 | 2e-07 | 1.967E-07 | 4.509E-08 | 1.500E-07 | 2.400E-07 |
| 13733 | 687 | S687M | B | 3.8e-07 | 3.7e-07 | 2.5e-07 | 3.333E-07 | 7.234E-08 | 2.500E-07 | 3.800E-07 |
| 13734 | 687 | S687F | B | 2.7e-07 | 2.8e-07 | 1.6e-07 | 2.367E-07 | 6.658E-08 | 1.600E-07 | 2.800E-07 |
| 13735 | 687 | S687P | B | 5.5e-07 | 2.4e-07 | 1.5e-07 | 3.133E-07 | 2.098E-07 | 1.500E-07 | 5.500E-07 |
| 13736 | 687 | S687S | B | 3.9e-07 | 3.2e-07 | 2.4e-07 | 3.167E-07 | 7.506E-08 | 2.400E-07 | 3.900E-07 |
| 13737 | 687 | S687T | B | 2.6e-07 | 6e-07 | 2.1e-07 | 3.567E-07 | 2.122E-07 | 2.100E-07 | 6.000E-07 |
| 13738 | 687 | S687W | B | 2.6e-07 | 3.2e-07 | 1.4e-07 | 2.400E-07 | 9.165E-08 | 1.400E-07 | 3.200E-07 |
| 13739 | 687 | S687Y | B | 3.6e-07 | 1.3e-07 | 1.8e-07 | 2.233E-07 | 1.210E-07 | 1.300E-07 | 3.600E-07 |
| 13740 | 687 | S687V | B | 2.9e-07 | 3.4e-07 | 3.9e-07 | 3.400E-07 | 5.000E-08 | 2.900E-07 | 3.900E-07 |
| 13741 | 688 | G688A | B | 3e-07 | 1.8e-07 | 2.8e-07 | 2.533E-07 | 6.429E-08 | 1.800E-07 | 3.000E-07 |
| 13742 | 688 | G688R | B | 1.5e-07 | 1.8e-07 | 2.6e-07 | 1.967E-07 | 5.686E-08 | 1.500E-07 | 2.600E-07 |
| 13743 | 688 | G688N | B | 1.7e-07 | 5.8e-07 | 2.5e-07 | 3.333E-07 | 2.173E-07 | 1.700E-07 | 5.800E-07 |
| 13744 | 688 | G688D | B | 2.3e-07 | 3.1e-07 | 2.2e-07 | 2.533E-07 | 4.933E-08 | 2.200E-07 | 3.100E-07 |
| 13745 | 688 | G688C | B | 1.9e-07 | 1.9e-07 | 1.9e-07 | 1.900E-07 | 0.000E+00 | 1.900E-07 | 1.900E-07 |
| 13746 | 688 | G688E | B | 3.7e-07 | 1.4e-07 | 2.3e-07 | 2.467E-07 | 1.159E-07 | 1.400E-07 | 3.700E-07 |
| 13747 | 688 | G688Q | B | 6.2e-07 | 1.5e-07 | 3.9e-07 | 3.867E-07 | 2.350E-07 | 1.500E-07 | 6.200E-07 |
| 13748 | 688 | G688G | B | 5.3e-07 | 7.3e-07 | 3.3e-07 | 5.300E-07 | 2.000E-07 | 3.300E-07 | 7.300E-07 |
| 13749 | 688 | G688H | B | 3.6e-07 | 1e-07 | 2e-07 | 2.200E-07 | 1.311E-07 | 1.000E-07 | 3.600E-07 |
| 13750 | 688 | G688I | B | 2.3e-07 | 2.8e-07 | 5.4e-07 | 3.500E-07 | 1.664E-07 | 2.300E-07 | 5.400E-07 |
| 13751 | 688 | G688L | B | 1.4e-07 | 2.3e-07 | 4.1e-07 | 2.600E-07 | 1.375E-07 | 1.400E-07 | 4.100E-07 |
| 13752 | 688 | G688K | B | 3.5e-07 | 2.9e-07 | 3.4e-07 | 3.267E-07 | 3.215E-08 | 2.900E-07 | 3.500E-07 |
| 13753 | 688 | G688M | B | 2.9e-07 | 2e-07 | 2.8e-07 | 2.567E-07 | 4.933E-08 | 2.000E-07 | 2.900E-07 |
| 13754 | 688 | G688F | B | 6e-07 | 1.2e-07 | 5.3e-07 | 4.167E-07 | 2.593E-07 | 1.200E-07 | 6.000E-07 |
| 13755 | 688 | G688P | B | 3.5e-07 | 1.3e-07 | 2.8e-07 | 2.533E-07 | 1.124E-07 | 1.300E-07 | 3.500E-07 |
| 13756 | 688 | G688S | B | 1.1e-07 | 2.1e-07 | 2.6e-07 | 1.933E-07 | 7.638E-08 | 1.100E-07 | 2.600E-07 |
| 13757 | 688 | G688T | B | 5.9e-07 | 4.1e-07 | 2.1e-07 | 4.033E-07 | 1.901E-07 | 2.100E-07 | 5.900E-07 |
| 13758 | 688 | G688W | B | 2.6e-07 | 4.9e-07 | 3.4e-07 | 3.633E-07 | 1.168E-07 | 2.600E-07 | 4.900E-07 |
| 13759 | 688 | G688Y | B | 4.2e-07 | 2.6e-07 | 2e-07 | 2.933E-07 | 1.137E-07 | 2.000E-07 | 4.200E-07 |
| 13760 | 688 | G688V | B | 4.2e-07 | 2.7e-07 | 2.2e-07 | 3.033E-07 | 1.041E-07 | 2.200E-07 | 4.200E-07 |
| 13761 | 689 | S689A | B | 2e-07 | 2.9e-07 | 2.5e-07 | 2.467E-07 | 4.509E-08 | 2.000E-07 | 2.900E-07 |
| 13762 | 689 | S689R | B | 2.1e-07 | 1.1e-07 | 3.1e-07 | 2.100E-07 | 1.000E-07 | 1.100E-07 | 3.100E-07 |
| 13763 | 689 | S689N | B | 4.3e-07 | 2.6e-07 | 1.7e-07 | 2.867E-07 | 1.320E-07 | 1.700E-07 | 4.300E-07 |
| 13764 | 689 | S689D | B | 1.5e-07 | 1.5e-07 | 1.8e-07 | 1.600E-07 | 1.732E-08 | 1.500E-07 | 1.800E-07 |
| 13765 | 689 | S689C | B | 3.4e-07 | 3.9e-07 | 3e-07 | 3.433E-07 | 4.509E-08 | 3.000E-07 | 3.900E-07 |
| 13766 | 689 | S689E | B | 2.7e-07 | 3.3e-07 | 3.7e-07 | 3.233E-07 | 5.033E-08 | 2.700E-07 | 3.700E-07 |
| 13767 | 689 | S689Q | B | 4e-07 | 2.9e-07 | 2.1e-07 | 3.000E-07 | 9.539E-08 | 2.100E-07 | 4.000E-07 |
| 13768 | 689 | S689G | B | 3.8e-07 | 4.4e-07 | 1.6e-07 | 3.267E-07 | 1.474E-07 | 1.600E-07 | 4.400E-07 |
| 13769 | 689 | S689H | B | 2.6e-07 | 1.4e-07 | 2.7e-07 | 2.233E-07 | 7.234E-08 | 1.400E-07 | 2.700E-07 |

| | | | | | | | | | | |
|---|---|---|---|---|---|---|---|---|---|---|
| 13770 | 689 | S689I | B | 2.4e-07 | 2.7e-07 | 2e-07 | 2.367E-07 | 3.512E-08 | 2.000E-07 | 2.700E-07 |
| 13771 | 689 | S689L | B | 3.6e-07 | 1.7e-07 | 3.2e-07 | 2.833E-07 | 1.002E-07 | 1.700E-07 | 3.600E-07 |
| 13772 | 689 | S689K | B | 1.7e-07 | 1.1e-07 | 1.6e-07 | 1.467E-07 | 3.215E-08 | 1.100E-07 | 1.700E-07 |
| 13773 | 689 | S689M | B | 5.1e-07 | 3.8e-07 | 1.3e-07 | 3.400E-07 | 1.931E-07 | 1.300E-07 | 5.100E-07 |
| 13774 | 689 | S689F | B | 1.9e-07 | 5.4e-07 | 9.4e-08 | 2.747E-07 | 2.347E-07 | 9.400E-08 | 5.400E-07 |
| 13775 | 689 | S689P | B | 1.3e-07 | 2.1e-07 | 1.9e-07 | 1.767E-07 | 4.163E-08 | 1.300E-07 | 2.100E-07 |
| 13776 | 689 | S689S | B | 2.5e-07 | 2.9e-07 | 2.8e-07 | 2.733E-07 | 2.082E-08 | 2.500E-07 | 2.900E-07 |
| 13777 | 689 | S689T | B | 4.1e-07 | 2.5e-07 | 1.3e-07 | 2.633E-07 | 1.405E-07 | 1.300E-07 | 4.100E-07 |
| 13778 | 689 | S689W | B | 5.8e-07 | 1.8e-07 | 3.2e-07 | 3.600E-07 | 2.030E-07 | 1.800E-07 | 5.800E-07 |
| 13779 | 689 | S689Y | B | 3.6e-07 | 1.7e-07 | 1.9e-07 | 2.400E-07 | 1.044E-07 | 1.700E-07 | 3.600E-07 |
| 13780 | 689 | S689V | B | 5e-07 | 4e-07 | 1e-07 | 3.333E-07 | 2.082E-07 | 1.000E-07 | 5.000E-07 |
| 13781 | 690 | G690A | B | 3.8e-07 | 3.9e-07 | 3.8e-07 | 3.833E-07 | 5.774E-09 | 3.800E-07 | 3.900E-07 |
| 13782 | 690 | G690R | B | 3.1e-07 | 2.6e-07 | 3.9e-07 | 3.200E-07 | 6.557E-08 | 2.600E-07 | 3.900E-07 |
| 13783 | 690 | G690N | B | 4.4e-07 | 3e-07 | 4.2e-07 | 3.867E-07 | 7.572E-08 | 3.000E-07 | 4.400E-07 |
| 13784 | 690 | G690D | B | 5.8e-07 | 5.9e-07 | 1e-07 | 4.233E-07 | 2.801E-07 | 1.000E-07 | 5.900E-07 |
| 13785 | 690 | G690C | B | 1.2e-07 | 1.3e-07 | 2.7e-07 | 1.733E-07 | 8.386E-08 | 1.200E-07 | 2.700E-07 |
| 13786 | 690 | G690E | B | 3.3e-07 | 2.7e-07 | 2.8e-07 | 2.933E-07 | 3.215E-08 | 2.700E-07 | 3.300E-07 |
| 13787 | 690 | G690Q | B | 3e-07 | 2.4e-07 | 2.3e-07 | 2.567E-07 | 3.786E-08 | 2.300E-07 | 3.000E-07 |
| 13788 | 690 | G690G | B | 5.6e-07 | 1.6e-07 | 4e-07 | 3.733E-07 | 2.013E-07 | 1.600E-07 | 5.600E-07 |
| 13789 | 690 | G690H | B | 1.3e-07 | 1.6e-07 | 3.3e-07 | 2.067E-07 | 1.079E-07 | 1.300E-07 | 3.300E-07 |
| 13790 | 690 | G690I | B | 3e-07 | 2.2e-07 | 3.3e-07 | 2.833E-07 | 5.686E-08 | 2.200E-07 | 3.300E-07 |
| 13791 | 690 | G690L | B | 3.4e-07 | 2.4e-07 | 4.7e-07 | 3.500E-07 | 1.153E-07 | 2.400E-07 | 4.700E-07 |
| 13792 | 690 | G690K | B | 1.6e-07 | 6.8e-07 | 1.8e-07 | 3.400E-07 | 2.946E-07 | 1.600E-07 | 6.800E-07 |
| 13793 | 690 | G690M | B | 2.1e-07 | 3.5e-07 | 2.3e-07 | 2.633E-07 | 7.572E-08 | 2.100E-07 | 3.500E-07 |
| 13794 | 690 | G690F | B | 2.9e-07 | 1.9e-07 | 2.6e-07 | 2.467E-07 | 5.132E-08 | 1.900E-07 | 2.900E-07 |
| 13795 | 690 | G690P | B | 1.2e-07 | 3.6e-07 | 1.9e-07 | 2.233E-07 | 1.234E-07 | 1.200E-07 | 3.600E-07 |
| 13796 | 690 | G690S | B | 1.9e-07 | 3e-07 | 4.3e-08 | 1.777E-07 | 1.289E-07 | 4.300E-08 | 3.000E-07 |
| 13797 | 690 | G690T | B | 2e-07 | 1.7e-07 | 4e-07 | 2.567E-07 | 1.250E-07 | 1.700E-07 | 4.000E-07 |
| 13798 | 690 | G690W | B | 2.5e-07 | 3.7e-07 | 2.2e-07 | 2.800E-07 | 7.937E-08 | 2.200E-07 | 3.700E-07 |
| 13799 | 690 | G690Y | B | 2.6e-07 | 2.1e-07 | 2.3e-07 | 2.333E-07 | 2.517E-08 | 2.100E-07 | 2.600E-07 |
| 13800 | 690 | G690V | B | 4e-07 | 2.1e-07 | 1.7e-07 | 2.600E-07 | 1.229E-07 | 1.700E-07 | 4.000E-07 |
| 13801 | 691 | T691A | B | 3.4e-07 | 3e-07 | 3.8e-07 | 3.400E-07 | 4.000E-08 | 3.000E-07 | 3.800E-07 |
| 13802 | 691 | T691R | B | 4.4e-07 | 1.9e-07 | 6.1e-07 | 4.133E-07 | 2.113E-07 | 1.900E-07 | 6.100E-07 |
| 13803 | 691 | T691N | B | 3.9e-07 | 2.3e-07 | 2.4e-07 | 2.867E-07 | 8.963E-08 | 2.300E-07 | 3.900E-07 |
| 13804 | 691 | T691D | B | 2.8e-07 | 7.5e-07 | 3.1e-07 | 4.467E-07 | 2.631E-07 | 2.800E-07 | 7.500E-07 |
| 13805 | 691 | T691C | B | 3.6e-07 | 1.9e-07 | 1.7e-07 | 2.400E-07 | 1.044E-07 | 1.700E-07 | 3.600E-07 |
| 13806 | 691 | T691E | B | 1.3e-07 | 1.7e-07 | 1.5e-07 | 1.500E-07 | 2.000E-08 | 1.300E-07 | 1.700E-07 |
| 13807 | 691 | T691Q | B | 2.2e-07 | 2.2e-07 | 1.8e-07 | 2.067E-07 | 2.309E-08 | 1.800E-07 | 2.200E-07 |
| 13808 | 691 | T691G | B | 3.6e-07 | 2.9e-07 | 2.5e-07 | 3.000E-07 | 5.568E-08 | 2.500E-07 | 3.600E-07 |
| 13809 | 691 | T691H | B | 4.7e-07 | 2.3e-07 | 4.4e-07 | 3.800E-07 | 1.308E-07 | 2.300E-07 | 4.700E-07 |
| 13810 | 691 | T691I | B | 1.2e-07 | 2.9e-07 | 2.5e-07 | 2.200E-07 | 8.888E-08 | 1.200E-07 | 2.900E-07 |
| 13811 | 691 | T691L | B | 1.4e-07 | 2.4e-07 | 3e-07 | 2.267E-07 | 8.083E-08 | 1.400E-07 | 3.000E-07 |
| 13812 | 691 | T691K | B | 3.6e-07 | 1.7e-07 | 1.5e-07 | 2.267E-07 | 1.159E-07 | 1.500E-07 | 3.600E-07 |
| 13813 | 691 | T691M | B | 2.7e-07 | 4.5e-07 | 1.7e-07 | 2.967E-07 | 1.419E-07 | 1.700E-07 | 4.500E-07 |
| 13814 | 691 | T691F | B | 3.7e-07 | 3.4e-07 | 1.9e-07 | 3.000E-07 | 9.644E-08 | 1.900E-07 | 3.700E-07 |
| 13815 | 691 | T691P | B | 3e-07 | 3e-07 | 1.9e-07 | 2.633E-07 | 6.351E-08 | 1.900E-07 | 3.000E-07 |
| 13816 | 691 | T691S | B | 2.3e-07 | 3e-07 | 1.6e-07 | 2.300E-07 | 7.000E-08 | 1.600E-07 | 3.000E-07 |
| 13817 | 691 | T691T | B | 1.8e-07 | 2.5e-07 | 1.5e-07 | 1.933E-07 | 5.132E-08 | 1.500E-07 | 2.500E-07 |
| 13818 | 691 | T691W | B | 1.8e-07 | 2.9e-07 | 2.2e-07 | 2.300E-07 | 5.568E-08 | 1.800E-07 | 2.900E-07 |
| 13819 | 691 | T691Y | B | 2.1e-07 | 4.2e-07 | 4.2e-07 | 3.500E-07 | 1.212E-07 | 2.100E-07 | 4.200E-07 |
| 13820 | 691 | T691V | B | 4.2e-07 | 4.1e-07 | 3.3e-07 | 3.867E-07 | 4.933E-08 | 3.300E-07 | 4.200E-07 |
| 13821 | 692 | D692A | B | 6.4e-07 | 1.6e-07 | 1.4e-07 | 3.133E-07 | 2.831E-07 | 1.400E-07 | 6.400E-07 |
| 13822 | 692 | D692R | B | 4e-07 | 2.4e-07 | 1.6e-07 | 2.667E-07 | 1.222E-07 | 1.600E-07 | 4.000E-07 |
| 13823 | 692 | D692N | B | 2.4e-07 | 2.2e-07 | 1.2e-07 | 1.933E-07 | 6.429E-08 | 1.200E-07 | 2.400E-07 |
| 13824 | 692 | D692D | B | 3.2e-07 | 1.7e-07 | 3e-07 | 2.633E-07 | 8.145E-08 | 1.700E-07 | 3.200E-07 |
| 13825 | 692 | D692C | B | 3e-07 | 1.4e-07 | 3.7e-07 | 2.700E-07 | 1.179E-07 | 1.400E-07 | 3.700E-07 |
| 13826 | 692 | D692E | B | 4.6e-07 | 1.8e-07 | 5e-07 | 3.800E-07 | 1.744E-07 | 1.800E-07 | 5.000E-07 |
| 13827 | 692 | D692Q | B | 2.2e-07 | 2.4e-07 | 5.3e-07 | 3.300E-07 | 1.735E-07 | 2.200E-07 | 5.300E-07 |
| 13828 | 692 | D692G | B | 3.3e-07 | 3.2e-07 | 3.8e-07 | 3.433E-07 | 3.215E-08 | 3.200E-07 | 3.800E-07 |
| 13829 | 692 | D692H | B | 1.7e-07 | 2.8e-07 | 2.1e-07 | 2.200E-07 | 5.568E-08 | 1.700E-07 | 2.800E-07 |
| 13830 | 692 | D692I | B | 4.5e-07 | 2e-07 | 2.9e-07 | 3.133E-07 | 1.266E-07 | 2.000E-07 | 4.500E-07 |
| 13831 | 692 | D692L | B | 2.7e-07 | 1.8e-07 | 1.3e-07 | 1.933E-07 | 7.095E-08 | 1.300E-07 | 2.700E-07 |
| 13832 | 692 | D692K | B | 2.2e-07 | 3.6e-07 | 3.5e-07 | 3.100E-07 | 7.810E-08 | 2.200E-07 | 3.600E-07 |
| 13833 | 692 | D692M | B | 4.6e-07 | 1.9e-07 | 1.3e-07 | 2.600E-07 | 1.758E-07 | 1.300E-07 | 4.600E-07 |
| 13834 | 692 | D692F | B | 1.2e-07 | 1.7e-07 | 2.4e-07 | 1.767E-07 | 6.028E-08 | 1.200E-07 | 2.400E-07 |
| 13835 | 692 | D692P | B | 1.7e-07 | 2.2e-07 | 2.5e-07 | 2.133E-07 | 4.041E-08 | 1.700E-07 | 2.500E-07 |
| 13836 | 692 | D692S | B | 2.1e-07 | 4.6e-07 | 1.6e-07 | 2.767E-07 | 1.607E-07 | 1.600E-07 | 4.600E-07 |
| 13837 | 692 | D692T | B | 4.1e-07 | 4e-07 | 2.7e-07 | 3.600E-07 | 7.810E-08 | 2.700E-07 | 4.100E-07 |
| 13838 | 692 | D692W | B | 2.9e-07 | 2.8e-07 | 2.5e-07 | 2.733E-07 | 2.082E-08 | 2.500E-07 | 2.900E-07 |
| 13839 | 692 | D692Y | B | 1.7e-07 | 2.3e-07 | 2.3e-07 | 2.100E-07 | 3.464E-08 | 1.700E-07 | 2.300E-07 |
| 13840 | 692 | D692V | B | 2e-07 | 2.6e-07 | 6.8e-07 | 3.800E-07 | 2.615E-07 | 2.000E-07 | 6.800E-07 |
| 13841 | 693 | F693A | B | 2.8e-07 | 2.7e-07 | 2.4e-07 | 2.633E-07 | 2.082E-08 | 2.400E-07 | 2.800E-07 |
| 13842 | 693 | F693R | B | 1.5e-07 | 7.9e-08 | 2.1e-07 | 1.463E-07 | 6.558E-08 | 7.900E-08 | 2.100E-07 |
| 13843 | 693 | F693N | B | 2.1e-07 | 2.6e-07 | 4.1e-07 | 2.933E-07 | 1.041E-07 | 2.100E-07 | 4.100E-07 |
| 13844 | 693 | F693D | B | 2.5e-07 | 2e-07 | 2.9e-07 | 2.467E-07 | 4.509E-08 | 2.000E-07 | 2.900E-07 |
| 13845 | 693 | F693C | B | 3.7e-07 | 1.5e-07 | 3.4e-07 | 2.867E-07 | 1.193E-07 | 1.500E-07 | 3.700E-07 |
| 13846 | 693 | F693E | B | 3.4e-07 | 3.7e-07 | 3.9e-07 | 3.667E-07 | 2.517E-08 | 3.400E-07 | 3.900E-07 |
| 13847 | 693 | F693Q | B | 2.2e-07 | 3.8e-07 | 1.5e-07 | 2.500E-07 | 1.179E-07 | 1.500E-07 | 3.800E-07 |
| 13848 | 693 | F693G | B | 3.2e-07 | 3.4e-07 | 3.1e-07 | 3.233E-07 | 1.528E-08 | 3.100E-07 | 3.400E-07 |
| 13849 | 693 | F693H | B | 1.6e-07 | 7.1e-08 | 1.1e-07 | 1.137E-07 | 4.461E-08 | 7.100E-08 | 1.600E-07 |
| 13850 | 693 | F693I | B | 1.8e-07 | 2.3e-07 | 2.9e-07 | 2.333E-07 | 5.508E-08 | 1.800E-07 | 2.900E-07 |
| 13851 | 693 | F693L | B | 7.1e-07 | 1.4e-07 | 3e-07 | 3.833E-07 | 2.940E-07 | 1.400E-07 | 7.100E-07 |
| 13852 | 693 | F693K | B | 2.8e-07 | 2e-07 | 3.5e-07 | 2.767E-07 | 7.506E-08 | 2.000E-07 | 3.500E-07 |
| 13853 | 693 | F693M | B | 1.3e-07 | 3.6e-07 | 4.7e-07 | 3.200E-07 | 1.735E-07 | 1.300E-07 | 4.700E-07 |
| 13854 | 693 | F693F | B | 3.2e-07 | 2.8e-07 | 2.7e-07 | 2.900E-07 | 2.646E-08 | 2.700E-07 | 3.200E-07 |

| | | | | | | | | | | |
|---|---|---|---|---|---|---|---|---|---|---|
| 13855 | 693 | F693P | B | 2.6e-07 | 2.9e-07 | 2.4e-07 | 2.633E-07 | 2.517E-08 | 2.400E-07 | 2.900E-07 |
| 13856 | 693 | F693S | B | 3e-07 | 1.4e-07 | 1.8e-07 | 2.067E-07 | 8.327E-08 | 1.400E-07 | 3.000E-07 |
| 13857 | 693 | F693T | B | 3e-07 | 4.4e-07 | 3.4e-07 | 3.600E-07 | 7.211E-08 | 3.000E-07 | 4.400E-07 |
| 13858 | 693 | F693W | B | 3.2e-07 | 2.5e-07 | 2.1e-07 | 2.600E-07 | 5.568E-08 | 2.100E-07 | 3.200E-07 |
| 13859 | 693 | F693Y | B | 2.7e-07 | 1.6e-07 | 1.4e-07 | 1.900E-07 | 7.000E-08 | 1.400E-07 | 2.700E-07 |
| 13860 | 693 | F693V | B | 3.2e-07 | 1.5e-07 | 2.7e-07 | 2.467E-07 | 8.737E-08 | 1.500E-07 | 3.200E-07 |
| 13861 | 694 | T694A | B | 5.8e-07 | 5.8e-07 | 5.1e-07 | 5.567E-07 | 4.041E-08 | 5.100E-07 | 5.800E-07 |
| 13862 | 694 | T694R | B | 1.8e-07 | 1.1e-07 | 4.8e-07 | 2.567E-07 | 1.966E-07 | 1.100E-07 | 4.800E-07 |
| 13863 | 694 | T694N | B | 5.2e-07 | 1.1e-07 | 2.8e-07 | 3.033E-07 | 2.060E-07 | 1.100E-07 | 5.200E-07 |
| 13864 | 694 | T694D | B | 1.1e-07 | 2.8e-07 | 4.9e-07 | 2.933E-07 | 1.904E-07 | 1.100E-07 | 4.900E-07 |
| 13865 | 694 | T694C | B | 1.4e-07 | 2.3e-07 | 6.9e-08 | 1.463E-07 | 8.069E-08 | 6.900E-08 | 2.300E-07 |
| 13866 | 694 | T694E | B | 2.8e-07 | 5.2e-07 | 1.3e-07 | 3.100E-07 | 1.967E-07 | 1.300E-07 | 5.200E-07 |
| 13867 | 694 | T694Q | B | 1.5e-07 | 4.7e-07 | 2.3e-07 | 2.833E-07 | 1.665E-07 | 1.500E-07 | 4.700E-07 |
| 13868 | 694 | T694G | B | 1.5e-07 | 1.4e-07 | 4.7e-07 | 2.533E-07 | 1.877E-07 | 1.400E-07 | 4.700E-07 |
| 13869 | 694 | T694H | B | 1.7e-07 | 3.1e-07 | 1.5e-07 | 2.100E-07 | 8.718E-08 | 1.500E-07 | 3.100E-07 |
| 13870 | 694 | T694I | B | 2e-07 | 4.7e-07 | 1.8e-07 | 2.833E-07 | 1.620E-07 | 1.800E-07 | 4.700E-07 |
| 13871 | 694 | T694L | B | 1.4e-07 | 2.9e-07 | 3.9e-07 | 2.733E-07 | 1.258E-07 | 1.400E-07 | 3.900E-07 |
| 13872 | 694 | T694K | B | 2.3e-07 | 1.5e-07 | 2.3e-07 | 2.033E-07 | 4.619E-08 | 1.500E-07 | 2.300E-07 |
| 13873 | 694 | T694M | B | 3.3e-07 | 2.8e-07 | 1.6e-07 | 2.567E-07 | 8.737E-08 | 1.600E-07 | 3.300E-07 |
| 13874 | 694 | T694F | B | 1.2e-07 | 2e-07 | 3.5e-07 | 2.233E-07 | 1.168E-07 | 1.200E-07 | 3.500E-07 |
| 13875 | 694 | T694P | B | 1.1e-07 | 1.3e-07 | 3.2e-07 | 1.867E-07 | 1.159E-07 | 1.100E-07 | 3.200E-07 |
| 13876 | 694 | T694S | B | 3.2e-07 | 4.4e-07 | 2.8e-07 | 3.467E-07 | 8.327E-08 | 2.800E-07 | 4.400E-07 |
| 13877 | 694 | T694T | B | 2.6e-07 | 2.3e-07 | 4.8e-07 | 3.233E-07 | 1.365E-07 | 2.300E-07 | 4.800E-07 |
| 13878 | 694 | T694W | B | 1.4e-07 | 1.3e-07 | 3.4e-07 | 2.033E-07 | 1.185E-07 | 1.300E-07 | 3.400E-07 |
| 13879 | 694 | T694Y | B | 2.2e-07 | 4.7e-07 | 3e-07 | 3.300E-07 | 1.277E-07 | 2.200E-07 | 4.700E-07 |
| 13880 | 694 | T694V | B | 2.2e-07 | 4.9e-07 | 1.9e-07 | 3.000E-07 | 1.652E-07 | 1.900E-07 | 4.900E-07 |
| 13881 | 695 | L695A | B | 5.2e-07 | 4.4e-07 | 2.7e-07 | 4.100E-07 | 1.277E-07 | 2.700E-07 | 5.200E-07 |
| 13882 | 695 | L695R | B | 1.3e-07 | 1.4e-07 | 2.3e-07 | 1.667E-07 | 5.508E-08 | 1.300E-07 | 2.300E-07 |
| 13883 | 695 | L695N | B | 1.7e-07 | 3.3e-07 | 2e-07 | 2.333E-07 | 8.505E-08 | 1.700E-07 | 3.300E-07 |
| 13884 | 695 | L695D | B | 2e-07 | 2.5e-07 | 2.9e-07 | 2.467E-07 | 4.509E-08 | 2.000E-07 | 2.900E-07 |
| 13885 | 695 | L695C | B | 1.5e-07 | 2e-07 | 3.3e-07 | 2.267E-07 | 9.292E-08 | 1.500E-07 | 3.300E-07 |
| 13886 | 695 | L695E | B | 2.5e-07 | 1.7e-07 | 2e-07 | 2.067E-07 | 4.041E-08 | 1.700E-07 | 2.500E-07 |
| 13887 | 695 | L695Q | B | 3.1e-07 | 2.3e-07 | 1.9e-07 | 2.433E-07 | 6.110E-08 | 1.900E-07 | 3.100E-07 |
| 13888 | 695 | L695G | B | 1.5e-07 | 8.6e-08 | 1.8e-07 | 1.387E-07 | 4.801E-08 | 8.600E-08 | 1.800E-07 |
| 13889 | 695 | L695H | B | 2.2e-07 | 4.1e-07 | 2e-07 | 2.767E-07 | 1.159E-07 | 2.000E-07 | 4.100E-07 |
| 13890 | 695 | L695I | B | 1.5e-07 | 4.5e-07 | 2.6e-07 | 2.867E-07 | 1.518E-07 | 1.500E-07 | 4.500E-07 |
| 13891 | 695 | L695L | B | 2.6e-07 | 1.8e-07 | 1.9e-07 | 2.100E-07 | 4.359E-08 | 1.800E-07 | 2.600E-07 |
| 13892 | 695 | L695K | B | 4.1e-07 | 1.5e-07 | 1.1e-07 | 2.233E-07 | 1.629E-07 | 1.100E-07 | 4.100E-07 |
| 13893 | 695 | L695M | B | 1.6e-07 | 2.5e-07 | 3.1e-07 | 2.400E-07 | 7.550E-08 | 1.600E-07 | 3.100E-07 |
| 13894 | 695 | L695F | B | 6.1e-07 | 1.8e-07 | 3.2e-07 | 3.700E-07 | 2.193E-07 | 1.800E-07 | 6.100E-07 |
| 13895 | 695 | L695P | B | 2.3e-07 | 5.3e-07 | 4.1e-07 | 3.900E-07 | 1.510E-07 | 2.300E-07 | 5.300E-07 |
| 13896 | 695 | L695S | B | 3.1e-07 | 3.4e-07 | 2.2e-07 | 2.900E-07 | 6.245E-08 | 2.200E-07 | 3.400E-07 |
| 13897 | 695 | L695T | B | 5e-07 | 2.8e-07 | 2.1e-07 | 3.300E-07 | 1.513E-07 | 2.100E-07 | 5.000E-07 |
| 13898 | 695 | L695W | B | 2.6e-07 | 1.8e-07 | 2.3e-07 | 2.233E-07 | 4.041E-08 | 1.800E-07 | 2.600E-07 |
| 13899 | 695 | L695Y | B | 4.6e-07 | 4.7e-07 | 3.1e-07 | 4.133E-07 | 8.963E-08 | 3.100E-07 | 4.700E-07 |
| 13900 | 695 | L695V | B | 6.4e-07 | 1.9e-07 | 4.4e-07 | 4.233E-07 | 2.255E-07 | 1.900E-07 | 6.400E-07 |
| 13901 | 696 | T696A | B | 3.2e-07 | 6.5e-08 | 5.6e-07 | 3.150E-07 | 2.475E-07 | 6.500E-08 | 5.600E-07 |
| 13902 | 696 | T696R | B | 1.2e-07 | 1.4e-07 | 1.9e-07 | 1.500E-07 | 3.606E-08 | 1.200E-07 | 1.900E-07 |
| 13903 | 696 | T696N | B | 4.1e-07 | 3.9e-07 | 1.5e-07 | 3.167E-07 | 1.447E-07 | 1.500E-07 | 4.100E-07 |
| 13904 | 696 | T696D | B | 2.9e-07 | 5e-07 | 3e-07 | 3.633E-07 | 1.185E-07 | 2.900E-07 | 5.000E-07 |
| 13905 | 696 | T696C | B | 4.3e-07 | 1.7e-07 | 2.7e-07 | 2.900E-07 | 1.311E-07 | 1.700E-07 | 4.300E-07 |
| 13906 | 696 | T696E | B | 2.5e-07 | 8.3e-08 | 4e-07 | 2.443E-07 | 1.586E-07 | 8.300E-08 | 4.000E-07 |
| 13907 | 696 | T696Q | B | 2.4e-07 | 2e-07 | 2.1e-07 | 2.167E-07 | 2.082E-08 | 2.000E-07 | 2.400E-07 |
| 13908 | 696 | T696G | B | 1.3e-07 | 1.3e-07 | 2.4e-07 | 1.667E-07 | 6.351E-08 | 1.300E-07 | 2.400E-07 |
| 13909 | 696 | T696H | B | 3e-07 | 1.8e-07 | 3.7e-07 | 2.833E-07 | 9.609E-08 | 1.800E-07 | 3.700E-07 |
| 13910 | 696 | T696I | B | 2.3e-07 | 4.6e-07 | 5.2e-07 | 4.033E-07 | 1.531E-07 | 2.300E-07 | 5.200E-07 |
| 13911 | 696 | T696L | B | 7e-07 | 3.1e-07 | 1.2e-07 | 3.767E-07 | 2.957E-07 | 1.200E-07 | 7.000E-07 |
| 13912 | 696 | T696K | B | 4.1e-07 | 4.3e-07 | 1.6e-07 | 3.333E-07 | 1.504E-07 | 1.600E-07 | 4.300E-07 |
| 13913 | 696 | T696M | B | 6.4e-07 | 2e-07 | 2e-07 | 3.467E-07 | 2.540E-07 | 2.000E-07 | 6.400E-07 |
| 13914 | 696 | T696F | B | 2.8e-07 | 1.7e-07 | 2.3e-07 | 2.267E-07 | 5.508E-08 | 1.700E-07 | 2.800E-07 |
| 13915 | 696 | T696P | B | 3.8e-07 | 3e-07 | 1.3e-07 | 2.700E-07 | 1.277E-07 | 1.300E-07 | 3.800E-07 |
| 13916 | 696 | T696S | B | 3.1e-07 | 2.8e-07 | 1e-07 | 2.300E-07 | 1.136E-07 | 1.000E-07 | 3.100E-07 |
| 13917 | 696 | T696T | B | 3.3e-07 | 2.2e-07 | 2.1e-07 | 2.533E-07 | 6.658E-08 | 2.100E-07 | 3.300E-07 |
| 13918 | 696 | T696W | B | 1.3e-07 | 5.2e-07 | 1.9e-07 | 2.800E-07 | 2.100E-07 | 1.300E-07 | 5.200E-07 |
| 13919 | 696 | T696Y | B | 3.2e-07 | 1.2e-07 | 5.2e-07 | 3.200E-07 | 2.000E-07 | 1.200E-07 | 5.200E-07 |
| 13920 | 696 | T696V | B | 2.5e-07 | 2.1e-07 | 2.9e-07 | 2.500E-07 | 4.000E-08 | 2.100E-07 | 2.900E-07 |
| 13921 | 697 | I697A | B | 1.8e-07 | 3e-07 | 2.2e-07 | 2.333E-07 | 6.110E-08 | 1.800E-07 | 3.000E-07 |
| 13922 | 697 | I697R | B | 2.9e-07 | 3.3e-07 | 2.8e-07 | 3.000E-07 | 2.646E-08 | 2.800E-07 | 3.300E-07 |
| 13923 | 697 | I697N | B | 5.5e-07 | 3.4e-07 | 4.7e-07 | 4.533E-07 | 1.060E-07 | 3.400E-07 | 5.500E-07 |
| 13924 | 697 | I697D | B | 3.9e-07 | 2.2e-07 | 2.4e-07 | 2.833E-07 | 9.292E-08 | 2.200E-07 | 3.900E-07 |
| 13925 | 697 | I697C | B | 1.3e-07 | 4.2e-07 | 2e-07 | 2.500E-07 | 1.513E-07 | 1.300E-07 | 4.200E-07 |
| 13926 | 697 | I697E | B | 2.3e-07 | 5.5e-07 | 3.3e-07 | 3.700E-07 | 1.637E-07 | 2.300E-07 | 5.500E-07 |
| 13927 | 697 | I697Q | B | 3e-07 | 2.3e-07 | 2.2e-07 | 2.500E-07 | 4.359E-08 | 2.200E-07 | 3.000E-07 |
| 13928 | 697 | I697G | B | 1.9e-07 | 1.6e-07 | 4.9e-07 | 2.800E-07 | 1.825E-07 | 1.600E-07 | 4.900E-07 |
| 13929 | 697 | I697H | B | 1.1e-07 | 3.8e-07 | 1.9e-07 | 2.267E-07 | 1.387E-07 | 1.100E-07 | 3.800E-07 |
| 13930 | 697 | I697I | B | 8.9e-08 | 3.2e-07 | 1.8e-07 | 1.963E-07 | 1.164E-07 | 8.900E-08 | 3.200E-07 |
| 13931 | 697 | I697L | B | 1.6e-07 | 1.8e-07 | 4.1e-07 | 2.500E-07 | 1.389E-07 | 1.600E-07 | 4.100E-07 |
| 13932 | 697 | I697K | B | 2.4e-07 | 2.6e-07 | 4.2e-07 | 3.067E-07 | 9.866E-08 | 2.400E-07 | 4.200E-07 |
| 13933 | 697 | I697M | B | 2.4e-07 | 2.4e-07 | 1.7e-07 | 2.167E-07 | 4.041E-08 | 1.700E-07 | 2.400E-07 |
| 13934 | 697 | I697F | B | 5.2e-07 | 1.8e-07 | 2.1e-07 | 3.033E-07 | 1.882E-07 | 1.800E-07 | 5.200E-07 |
| 13935 | 697 | I697P | B | 5e-07 | 2.9e-07 | 2.2e-07 | 3.367E-07 | 1.457E-07 | 2.200E-07 | 5.000E-07 |
| 13936 | 697 | I697S | B | 1.2e-07 | 2.7e-07 | 1.6e-07 | 1.833E-07 | 7.767E-08 | 1.200E-07 | 2.700E-07 |
| 13937 | 697 | I697T | B | 3.9e-07 | 2.4e-07 | 2.9e-07 | 3.067E-07 | 7.638E-08 | 2.400E-07 | 3.900E-07 |
| 13938 | 697 | I697W | B | 1.8e-07 | 3.2e-07 | 2e-07 | 2.333E-07 | 7.572E-08 | 1.800E-07 | 3.200E-07 |
| 13939 | 697 | I697Y | B | 2.7e-07 | 1.6e-07 | 1.1e-07 | 1.800E-07 | 8.185E-08 | 1.100E-07 | 2.700E-07 |

| | | | | | | | | | | |
|---|---|---|---|---|---|---|---|---|---|---|
| 13940 | 697 | I697V | B | 2.3e-07 | 1.8e-07 | 1.6e-07 | 1.900E-07 | 3.606E-08 | 1.600E-07 | 2.300E-07 |
| 13941 | 698 | S698A | B | 1.7e-07 | 2.6e-07 | 1.1e-07 | 1.800E-07 | 7.550E-08 | 1.100E-07 | 2.600E-07 |
| 13942 | 698 | S698R | B | 5e-07 | 3.9e-07 | 2.3e-07 | 3.733E-07 | 1.358E-07 | 2.300E-07 | 5.000E-07 |
| 13943 | 698 | S698N | B | 2.2e-07 | 2.8e-07 | 2.1e-07 | 2.367E-07 | 3.786E-08 | 2.100E-07 | 2.800E-07 |
| 13944 | 698 | S698D | B | 2.8e-07 | 2.9e-07 | 3.1e-07 | 2.933E-07 | 1.528E-08 | 2.800E-07 | 3.100E-07 |
| 13945 | 698 | S698C | B | 3.4e-07 | 2.8e-07 | 1.4e-07 | 2.533E-07 | 1.026E-07 | 1.400E-07 | 3.400E-07 |
| 13946 | 698 | S698E | B | 1.9e-07 | 3.1e-07 | 1.9e-07 | 2.300E-07 | 6.928E-08 | 1.900E-07 | 3.100E-07 |
| 13947 | 698 | S698Q | B | 2e-07 | 2.7e-07 | 3.2e-07 | 2.633E-07 | 6.028E-08 | 2.000E-07 | 3.200E-07 |
| 13948 | 698 | S698G | B | 3e-07 | 3.7e-07 | 2.3e-07 | 3.000E-07 | 7.000E-08 | 2.300E-07 | 3.700E-07 |
| 13949 | 698 | S698H | B | 1.5e-07 | 1.4e-07 | 2.9e-07 | 1.933E-07 | 8.386E-08 | 1.400E-07 | 2.900E-07 |
| 13950 | 698 | S698I | B | 5.4e-07 | 4.4e-07 | 5.3e-07 | 5.033E-07 | 5.508E-08 | 4.400E-07 | 5.400E-07 |
| 13951 | 698 | S698L | B | 1.5e-07 | 4e-07 | 3.2e-07 | 2.900E-07 | 1.277E-07 | 1.500E-07 | 4.000E-07 |
| 13952 | 698 | S698K | B | 2.7e-07 | 2.1e-07 | 2.4e-07 | 2.400E-07 | 3.000E-08 | 2.100E-07 | 2.700E-07 |
| 13953 | 698 | S698M | B | 2.5e-07 | 3e-07 | 4.7e-07 | 3.400E-07 | 1.153E-07 | 2.500E-07 | 4.700E-07 |
| 13954 | 698 | S698F | B | 1.4e-07 | 2.9e-07 | 2.4e-07 | 2.233E-07 | 7.638E-08 | 1.400E-07 | 2.900E-07 |
| 13955 | 698 | S698P | B | 1.6e-07 | 2.5e-07 | 4.4e-07 | 2.833E-07 | 1.429E-07 | 1.600E-07 | 4.400E-07 |
| 13956 | 698 | S698S | B | 1.7e-07 | 1.6e-07 | 3.1e-07 | 2.133E-07 | 8.386E-08 | 1.600E-07 | 3.100E-07 |
| 13957 | 698 | S698T | B | 4.7e-07 | 5.4e-07 | 2e-07 | 4.033E-07 | 1.795E-07 | 2.000E-07 | 5.400E-07 |
| 13958 | 698 | S698W | B | 1.5e-07 | 1.8e-07 | 2.5e-07 | 1.933E-07 | 5.132E-08 | 1.500E-07 | 2.500E-07 |
| 13959 | 698 | S698Y | B | 2.3e-07 | 4.4e-07 | 3.4e-07 | 3.367E-07 | 1.050E-07 | 2.300E-07 | 4.400E-07 |
| 13960 | 698 | S698V | B | 2.3e-07 | 3.1e-07 | 5.5e-07 | 3.633E-07 | 1.665E-07 | 2.300E-07 | 5.500E-07 |
| 13961 | 699 | S699A | B | 1.6e-07 | 1.9e-07 | 1.5e-07 | 1.667E-07 | 2.082E-08 | 1.500E-07 | 1.900E-07 |
| 13962 | 699 | S699R | B | 2.4e-07 | 3.2e-07 | 3.9e-07 | 3.167E-07 | 7.506E-08 | 2.400E-07 | 3.900E-07 |
| 13963 | 699 | S699N | B | 6.4e-08 | 3.9e-07 | 2.8e-07 | 2.447E-07 | 1.658E-07 | 6.400E-08 | 3.900E-07 |
| 13964 | 699 | S699D | B | 4.5e-07 | 2.2e-07 | 6.1e-07 | 4.267E-07 | 1.960E-07 | 2.200E-07 | 6.100E-07 |
| 13965 | 699 | S699C | B | 1.6e-07 | 2.1e-07 | 2.4e-07 | 2.033E-07 | 4.041E-08 | 1.600E-07 | 2.400E-07 |
| 13966 | 699 | S699E | B | 1.6e-07 | 5e-07 | 4.4e-07 | 3.667E-07 | 1.815E-07 | 1.600E-07 | 5.000E-07 |
| 13967 | 699 | S699Q | B | 2.7e-07 | 1.9e-07 | 1.4e-07 | 2.000E-07 | 6.557E-08 | 1.400E-07 | 2.700E-07 |
| 13968 | 699 | S699G | B | 2.6e-07 | 1.6e-07 | 2.8e-07 | 2.333E-07 | 6.429E-08 | 1.600E-07 | 2.800E-07 |
| 13969 | 699 | S699H | B | 1e-07 | 1.8e-07 | 2.1e-07 | 1.633E-07 | 5.686E-08 | 1.000E-07 | 2.100E-07 |
| 13970 | 699 | S699I | B | 3.6e-07 | 6.8e-07 | 1.2e-07 | 3.867E-07 | 2.810E-07 | 1.200E-07 | 6.800E-07 |
| 13971 | 699 | S699L | B | 4.2e-07 | 2.9e-07 | 2.4e-07 | 3.167E-07 | 9.292E-08 | 2.400E-07 | 4.200E-07 |
| 13972 | 699 | S699K | B | 1.2e-07 | 2.1e-07 | 2.8e-07 | 2.033E-07 | 8.021E-08 | 1.200E-07 | 2.800E-07 |
| 13973 | 699 | S699M | B | 3.3e-07 | 2.2e-07 | 1.4e-07 | 2.300E-07 | 9.539E-08 | 1.400E-07 | 3.300E-07 |
| 13974 | 699 | S699F | B | 2.7e-07 | 5.6e-07 | 2.7e-07 | 3.667E-07 | 1.674E-07 | 2.700E-07 | 5.600E-07 |
| 13975 | 699 | S699P | B | 2.1e-07 | 5.2e-07 | 1.3e-07 | 2.867E-07 | 2.060E-07 | 1.300E-07 | 5.200E-07 |
| 13976 | 699 | S699S | B | 1.7e-07 | 3.1e-07 | 1.8e-07 | 2.200E-07 | 7.810E-08 | 1.700E-07 | 3.100E-07 |
| 13977 | 699 | S699T | B | 3.6e-07 | 1.3e-07 | 2.9e-07 | 2.600E-07 | 1.179E-07 | 1.300E-07 | 3.600E-07 |
| 13978 | 699 | S699W | B | 3.9e-07 | 1.5e-07 | 2.1e-07 | 2.500E-07 | 1.249E-07 | 1.500E-07 | 3.900E-07 |
| 13979 | 699 | S699Y | B | 2.9e-07 | 2.6e-07 | 2.3e-07 | 2.600E-07 | 3.000E-08 | 2.300E-07 | 2.900E-07 |
| 13980 | 699 | S699V | B | 1.7e-07 | 2.7e-07 | 2.5e-07 | 2.300E-07 | 5.292E-08 | 1.700E-07 | 2.700E-07 |
| 13981 | 700 | L700A | B | 3.1e-07 | 2.7e-07 | 1.3e-07 | 2.367E-07 | 9.452E-08 | 1.300E-07 | 3.100E-07 |
| 13982 | 700 | L700R | B | 4.9e-07 | 2.4e-07 | 1.8e-07 | 3.033E-07 | 1.644E-07 | 1.800E-07 | 4.900E-07 |
| 13983 | 700 | L700N | B | 2.6e-07 | 1.3e-07 | 2.3e-07 | 2.067E-07 | 6.807E-08 | 1.300E-07 | 2.600E-07 |
| 13984 | 700 | L700D | B | 1.6e-07 | 2.6e-07 | 1e-07 | 1.733E-07 | 8.083E-08 | 1.000E-07 | 2.600E-07 |
| 13985 | 700 | L700C | B | 2.5e-07 | 4.5e-07 | 1.2e-07 | 2.733E-07 | 1.662E-07 | 1.200E-07 | 4.500E-07 |
| 13986 | 700 | L700E | B | 3.1e-07 | 4e-07 | 1.8e-07 | 2.967E-07 | 1.106E-07 | 1.800E-07 | 4.000E-07 |
| 13987 | 700 | L700Q | B | 3.2e-07 | 1.7e-07 | 2.4e-07 | 2.433E-07 | 7.506E-08 | 1.700E-07 | 3.200E-07 |
| 13988 | 700 | L700G | B | 7.8e-07 | 1.8e-07 | 1.5e-07 | 3.700E-07 | 3.554E-07 | 1.500E-07 | 7.800E-07 |
| 13989 | 700 | L700H | B | 3.2e-07 | 4.7e-07 | 1.8e-07 | 3.233E-07 | 1.450E-07 | 1.800E-07 | 4.700E-07 |
| 13990 | 700 | L700I | B | 2.7e-07 | 1.7e-07 | 2e-07 | 2.133E-07 | 5.132E-08 | 1.700E-07 | 2.700E-07 |
| 13991 | 700 | L700L | B | 1.4e-07 | 2.2e-07 | 1.2e-07 | 1.600E-07 | 5.292E-08 | 1.200E-07 | 2.200E-07 |
| 13992 | 700 | L700K | B | 2.9e-07 | 1.9e-07 | 3.2e-07 | 2.667E-07 | 6.807E-08 | 1.900E-07 | 3.200E-07 |
| 13993 | 700 | L700M | B | 2.5e-07 | 3.4e-07 | 3.5e-07 | 3.133E-07 | 5.508E-08 | 2.500E-07 | 3.500E-07 |
| 13994 | 700 | L700F | B | 3.1e-07 | 3.3e-07 | 3.8e-07 | 3.400E-07 | 3.606E-08 | 3.100E-07 | 3.800E-07 |
| 13995 | 700 | L700P | B | 9.4e-08 | 2.1e-07 | 1.5e-07 | 1.513E-07 | 5.801E-08 | 9.400E-08 | 2.100E-07 |
| 13996 | 700 | L700S | B | 3.4e-07 | 1.8e-07 | 1.2e-07 | 2.133E-07 | 1.137E-07 | 1.200E-07 | 3.400E-07 |
| 13997 | 700 | L700T | B | 1.4e-07 | 2e-07 | 1.6e-07 | 1.667E-07 | 3.055E-08 | 1.400E-07 | 2.000E-07 |
| 13998 | 700 | L700W | B | 2.9e-07 | 2.1e-07 | 5e-07 | 3.333E-07 | 1.498E-07 | 2.100E-07 | 5.000E-07 |
| 13999 | 700 | L700Y | B | 2.1e-07 | 1.3e-07 | 2.6e-07 | 2.000E-07 | 6.557E-08 | 1.300E-07 | 2.600E-07 |
| 14000 | 700 | L700V | B | 3.7e-07 | 4.4e-07 | 2.2e-07 | 3.433E-07 | 1.124E-07 | 2.200E-07 | 4.400E-07 |
| 14001 | 701 | Q701A | B | 3.3e-07 | 2.8e-07 | 2.6e-07 | 2.900E-07 | 3.606E-08 | 2.600E-07 | 3.300E-07 |
| 14002 | 701 | Q701R | B | 2.7e-07 | 2.1e-07 | 5.1e-07 | 3.300E-07 | 1.587E-07 | 2.100E-07 | 5.100E-07 |
| 14003 | 701 | Q701N | B | 2.5e-07 | 2.2e-07 | 2e-07 | 2.233E-07 | 2.517E-08 | 2.000E-07 | 2.500E-07 |
| 14004 | 701 | Q701D | B | 3.3e-07 | 2.6e-07 | 5.1e-07 | 3.667E-07 | 1.290E-07 | 2.600E-07 | 5.100E-07 |
| 14005 | 701 | Q701C | B | 2.5e-07 | 2.4e-07 | 2.3e-07 | 2.400E-07 | 1.000E-08 | 2.300E-07 | 2.500E-07 |
| 14006 | 701 | Q701E | B | 2e-07 | 1.3e-07 | 6.2e-07 | 3.167E-07 | 2.650E-07 | 1.300E-07 | 6.200E-07 |
| 14007 | 701 | Q701Q | B | 2.3e-07 | 2.6e-07 | 1.4e-07 | 2.100E-07 | 6.245E-08 | 1.400E-07 | 2.600E-07 |
| 14008 | 701 | Q701G | B | 3.4e-07 | 2.3e-07 | 2.7e-07 | 2.800E-07 | 5.568E-08 | 2.300E-07 | 3.400E-07 |
| 14009 | 701 | Q701H | B | 2.1e-07 | 4e-07 | 2.4e-07 | 2.833E-07 | 1.021E-07 | 2.100E-07 | 4.000E-07 |
| 14010 | 701 | Q701I | B | 2e-07 | 2.8e-07 | 2.8e-07 | 2.533E-07 | 4.619E-08 | 2.000E-07 | 2.800E-07 |
| 14011 | 701 | Q701L | B | 1.1e-07 | 5.1e-07 | 1.6e-07 | 2.600E-07 | 2.179E-07 | 1.100E-07 | 5.100E-07 |
| 14012 | 701 | Q701K | B | 1.8e-07 | 4.5e-07 | 2.3e-07 | 2.867E-07 | 1.436E-07 | 1.800E-07 | 4.500E-07 |
| 14013 | 701 | Q701M | B | 2.3e-07 | 3.6e-07 | 2.2e-07 | 2.700E-07 | 7.810E-08 | 2.200E-07 | 3.600E-07 |
| 14014 | 701 | Q701F | B | 1.5e-07 | 5.1e-07 | 1.9e-07 | 2.833E-07 | 1.973E-07 | 1.500E-07 | 5.100E-07 |
| 14015 | 701 | Q701P | B | 2.2e-07 | 2.1e-07 | 1.7e-07 | 2.000E-07 | 2.646E-08 | 1.700E-07 | 2.200E-07 |
| 14016 | 701 | Q701S | B | 1.3e-07 | 1.4e-07 | 1.5e-07 | 1.400E-07 | 1.000E-08 | 1.300E-07 | 1.500E-07 |
| 14017 | 701 | Q701T | B | 1.3e-07 | 3.8e-07 | 3.3e-07 | 2.800E-07 | 1.323E-07 | 1.300E-07 | 3.800E-07 |
| 14018 | 701 | Q701W | B | 2.7e-07 | 2.3e-07 | 2.2e-07 | 2.400E-07 | 2.646E-08 | 2.200E-07 | 2.700E-07 |
| 14019 | 701 | Q701Y | B | 1.4e-07 | 3.6e-07 | 2e-07 | 2.333E-07 | 1.137E-07 | 1.400E-07 | 3.600E-07 |
| 14020 | 701 | Q701V | B | 2.9e-07 | 1.6e-07 | 2.1e-07 | 2.200E-07 | 6.557E-08 | 1.600E-07 | 2.900E-07 |
| 14021 | 702 | P702A | B | 2.2e-07 | 2.5e-07 | 1.9e-07 | 2.200E-07 | 3.000E-08 | 1.900E-07 | 2.500E-07 |
| 14022 | 702 | P702R | B | 2.4e-07 | 2.3e-07 | 1.4e-07 | 2.033E-07 | 5.508E-08 | 1.400E-07 | 2.400E-07 |
| 14023 | 702 | P702N | B | 2e-07 | 3.7e-07 | 3e-07 | 2.900E-07 | 8.544E-08 | 2.000E-07 | 3.700E-07 |
| 14024 | 702 | P702D | B | 2.9e-07 | 2.4e-07 | 8.3e-08 | 2.043E-07 | 1.080E-07 | 8.300E-08 | 2.900E-07 |

| | | | | | | | | | | |
|---|---|---|---|---|---|---|---|---|---|---|
| 14025 | 702 | P702C | B | 1.2e-07 | 2.4e-07 | 3.3e-07 | 2.300E-07 | 1.054E-07 | 1.200E-07 | 3.300E-07 |
| 14026 | 702 | P702E | B | 2.3e-07 | 2.3e-07 | 1.4e-07 | 2.000E-07 | 5.196E-08 | 1.400E-07 | 2.300E-07 |
| 14027 | 702 | P702Q | B | 1.7e-07 | 4.7e-07 | 1.7e-07 | 2.700E-07 | 1.732E-07 | 1.700E-07 | 4.700E-07 |
| 14028 | 702 | P702G | B | 2.7e-07 | 1.5e-07 | 1.6e-07 | 1.933E-07 | 6.658E-08 | 1.500E-07 | 2.700E-07 |
| 14029 | 702 | P702H | B | 1.9e-07 | 2.2e-07 | 2.4e-07 | 2.167E-07 | 2.517E-08 | 1.900E-07 | 2.400E-07 |
| 14030 | 702 | P702I | B | 9.3e-08 | 3.2e-07 | 2.6e-07 | 2.243E-07 | 1.176E-07 | 9.300E-08 | 3.200E-07 |
| 14031 | 702 | P702L | B | 2.8e-07 | 1.8e-07 | 1.1e-07 | 1.900E-07 | 8.544E-08 | 1.100E-07 | 2.800E-07 |
| 14032 | 702 | P702K | B | 3.5e-07 | 2.1e-07 | 3.4e-07 | 3.000E-07 | 7.810E-08 | 2.100E-07 | 3.500E-07 |
| 14033 | 702 | P702M | B | 4.9e-07 | 1e-07 | 2.5e-07 | 2.800E-07 | 1.967E-07 | 1.000E-07 | 4.900E-07 |
| 14034 | 702 | P702F | B | 3.9e-07 | 3.9e-07 | 1.3e-07 | 3.033E-07 | 1.501E-07 | 1.300E-07 | 3.900E-07 |
| 14035 | 702 | P702P | B | 2.5e-07 | 2.3e-07 | 1.1e-07 | 1.967E-07 | 7.572E-08 | 1.100E-07 | 2.500E-07 |
| 14036 | 702 | P702S | B | 1.7e-07 | 2.6e-07 | 1.8e-07 | 2.033E-07 | 4.933E-08 | 1.700E-07 | 2.600E-07 |
| 14037 | 702 | P702T | B | 1.6e-07 | 2.2e-07 | 1.5e-07 | 1.767E-07 | 3.786E-08 | 1.500E-07 | 2.200E-07 |
| 14038 | 702 | P702W | B | 3.5e-07 | 2.9e-07 | 4.8e-07 | 3.733E-07 | 9.713E-08 | 2.900E-07 | 4.800E-07 |
| 14039 | 702 | P702Y | B | 3.3e-07 | 2.2e-07 | 1.6e-07 | 2.367E-07 | 8.622E-08 | 1.600E-07 | 3.300E-07 |
| 14040 | 702 | P702V | B | 2.1e-07 | 2.2e-07 | 2.3e-07 | 2.200E-07 | 1.000E-08 | 2.100E-07 | 2.300E-07 |
| 14041 | 703 | E703A | B | 2.4e-07 | 3.6e-07 | 2.7e-07 | 2.900E-07 | 6.245E-08 | 2.400E-07 | 3.600E-07 |
| 14042 | 703 | E703R | B | 3.4e-07 | 2.4e-07 | 2e-07 | 2.600E-07 | 7.211E-08 | 2.000E-07 | 3.400E-07 |
| 14043 | 703 | E703N | B | 1.5e-07 | 2.3e-07 | 1.9e-07 | 1.900E-07 | 4.000E-08 | 1.500E-07 | 2.300E-07 |
| 14044 | 703 | E703D | B | 2.6e-07 | 2.1e-07 | 1.9e-07 | 2.200E-07 | 3.606E-08 | 1.900E-07 | 2.600E-07 |
| 14045 | 703 | E703C | B | 5e-07 | 4.2e-07 | 4.1e-07 | 4.433E-07 | 4.933E-08 | 4.100E-07 | 5.000E-07 |
| 14046 | 703 | E703E | B | 3.2e-07 | 3.2e-07 | 4.3e-07 | 3.567E-07 | 6.351E-08 | 3.200E-07 | 4.300E-07 |
| 14047 | 703 | E703Q | B | 2.2e-07 | 1.4e-07 | 1.9e-07 | 1.833E-07 | 4.041E-08 | 1.400E-07 | 2.200E-07 |
| 14048 | 703 | E703G | B | 3.9e-07 | 1.7e-07 | 1.2e-07 | 2.267E-07 | 1.436E-07 | 1.200E-07 | 3.900E-07 |
| 14049 | 703 | E703H | B | 2.1e-07 | 1.1e-07 | 9.8e-08 | 1.393E-07 | 6.149E-08 | 9.800E-08 | 2.100E-07 |
| 14050 | 703 | E703I | B | 2.2e-07 | 1.8e-07 | 2.3e-07 | 2.100E-07 | 2.646E-08 | 1.800E-07 | 2.300E-07 |
| 14051 | 703 | E703L | B | 2.7e-07 | 1.4e-07 | 3.7e-07 | 2.600E-07 | 1.153E-07 | 1.400E-07 | 3.700E-07 |
| 14052 | 703 | E703K | B | 2.7e-07 | 3.2e-07 | 2.8e-07 | 2.900E-07 | 2.646E-08 | 2.700E-07 | 3.200E-07 |
| 14053 | 703 | E703M | B | 2e-07 | 2.8e-07 | 3.2e-07 | 2.667E-07 | 6.110E-08 | 2.000E-07 | 3.200E-07 |
| 14054 | 703 | E703F | B | 3.7e-07 | 1.8e-07 | 2e-07 | 2.500E-07 | 1.044E-07 | 1.800E-07 | 3.700E-07 |
| 14055 | 703 | E703P | B | 2.4e-07 | 1.5e-07 | 2.4e-07 | 2.100E-07 | 5.196E-08 | 1.500E-07 | 2.400E-07 |
| 14056 | 703 | E703S | B | 5.8e-07 | 4.5e-07 | 3.5e-07 | 4.600E-07 | 1.153E-07 | 3.500E-07 | 5.800E-07 |
| 14057 | 703 | E703T | B | 1.7e-07 | 2.5e-07 | 1.2e-07 | 1.800E-07 | 6.557E-08 | 1.200E-07 | 2.500E-07 |
| 14058 | 703 | E703W | B | 1.8e-07 | 1.7e-07 | 5.4e-07 | 2.967E-07 | 2.108E-07 | 1.700E-07 | 5.400E-07 |
| 14059 | 703 | E703Y | B | 3.4e-07 | 2.7e-07 | 3.5e-07 | 3.200E-07 | 4.359E-08 | 2.700E-07 | 3.500E-07 |
| 14060 | 703 | E703V | B | 2.7e-07 | 3.4e-07 | 3.6e-07 | 3.233E-07 | 4.726E-08 | 2.700E-07 | 3.600E-07 |
| 14061 | 704 | D704A | B | 2e-07 | 3.2e-07 | 2.2e-07 | 2.467E-07 | 6.429E-08 | 2.000E-07 | 3.200E-07 |
| 14062 | 704 | D704R | B | 2.2e-07 | 1.2e-07 | 3.7e-07 | 2.367E-07 | 1.258E-07 | 1.200E-07 | 3.700E-07 |
| 14063 | 704 | D704N | B | 1.1e-07 | 1.5e-07 | 4.4e-07 | 2.333E-07 | 1.801E-07 | 1.100E-07 | 4.400E-07 |
| 14064 | 704 | D704D | B | 2.9e-07 | 1.9e-07 | 1.6e-07 | 2.133E-07 | 6.807E-08 | 1.600E-07 | 2.900E-07 |
| 14065 | 704 | D704C | B | 4.3e-07 | 1.8e-07 | 1.3e-07 | 2.467E-07 | 1.607E-07 | 1.300E-07 | 4.300E-07 |
| 14066 | 704 | D704E | B | 2.6e-07 | 2.7e-07 | 6.2e-07 | 3.833E-07 | 2.050E-07 | 2.600E-07 | 6.200E-07 |
| 14067 | 704 | D704Q | B | 2e-07 | 2.3e-07 | 2e-07 | 2.100E-07 | 1.732E-08 | 2.000E-07 | 2.300E-07 |
| 14068 | 704 | D704G | B | 2.6e-07 | 3.5e-07 | 3.6e-07 | 3.233E-07 | 5.508E-08 | 2.600E-07 | 3.600E-07 |
| 14069 | 704 | D704H | B | 2.9e-07 | 4.7e-07 | 2.9e-07 | 3.500E-07 | 1.039E-07 | 2.900E-07 | 4.700E-07 |
| 14070 | 704 | D704I | B | 1.8e-07 | 2.4e-07 | 1.4e-07 | 1.867E-07 | 5.033E-08 | 1.400E-07 | 2.400E-07 |
| 14071 | 704 | D704L | B | 3.7e-07 | 2.6e-07 | 2e-07 | 2.767E-07 | 8.622E-08 | 2.000E-07 | 3.700E-07 |
| 14072 | 704 | D704K | B | 3.4e-07 | 1.6e-07 | 4.8e-07 | 3.267E-07 | 1.604E-07 | 1.600E-07 | 4.800E-07 |
| 14073 | 704 | D704M | B | 1.8e-07 | 2.6e-07 | 2.7e-07 | 2.367E-07 | 4.933E-08 | 1.800E-07 | 2.700E-07 |
| 14074 | 704 | D704F | B | 3.2e-07 | 2.6e-07 | 2.9e-07 | 2.900E-07 | 3.000E-08 | 2.600E-07 | 3.200E-07 |
| 14075 | 704 | D704P | B | 2.3e-07 | 4e-07 | 3e-07 | 3.100E-07 | 8.544E-08 | 2.300E-07 | 4.000E-07 |
| 14076 | 704 | D704S | B | 3.4e-07 | 1.8e-07 | 1.6e-07 | 2.267E-07 | 9.866E-08 | 1.600E-07 | 3.400E-07 |
| 14077 | 704 | D704T | B | 3.8e-07 | 3.1e-07 | 2.6e-07 | 3.167E-07 | 6.028E-08 | 2.600E-07 | 3.800E-07 |
| 14078 | 704 | D704W | B | 4.5e-07 | 1.9e-07 | 1.8e-07 | 2.733E-07 | 1.531E-07 | 1.800E-07 | 4.500E-07 |
| 14079 | 704 | D704Y | B | 1.5e-07 | 2.4e-07 | 4.6e-07 | 2.833E-07 | 1.595E-07 | 1.500E-07 | 4.600E-07 |
| 14080 | 704 | D704V | B | 2.3e-07 | 1.4e-07 | 1.7e-07 | 1.800E-07 | 4.583E-08 | 1.400E-07 | 2.300E-07 |
| 14081 | 705 | F705A | B | 4.1e-07 | 2.8e-07 | 2.2e-07 | 3.033E-07 | 9.713E-08 | 2.200E-07 | 4.100E-07 |
| 14082 | 705 | F705R | B | 2e-07 | 1.4e-07 | 2.8e-07 | 2.067E-07 | 7.024E-08 | 1.400E-07 | 2.800E-07 |
| 14083 | 705 | F705N | B | 2e-07 | 3.3e-07 | 4e-07 | 3.100E-07 | 1.015E-07 | 2.000E-07 | 4.000E-07 |
| 14084 | 705 | F705D | B | 2.4e-07 | 5e-07 | 3.7e-07 | 3.700E-07 | 1.300E-07 | 2.400E-07 | 5.000E-07 |
| 14085 | 705 | F705C | B | 2.5e-07 | 1.5e-07 | 4.6e-07 | 2.867E-07 | 1.582E-07 | 1.500E-07 | 4.600E-07 |
| 14086 | 705 | F705E | B | 1.3e-07 | 3.5e-07 | 1.2e-07 | 2.000E-07 | 1.300E-07 | 1.200E-07 | 3.500E-07 |
| 14087 | 705 | F705Q | B | 3.2e-07 | 2.3e-07 | 1.6e-07 | 2.367E-07 | 8.021E-08 | 1.600E-07 | 3.200E-07 |
| 14088 | 705 | F705G | B | 3.1e-07 | 3.8e-07 | 3.6e-07 | 3.500E-07 | 3.606E-08 | 3.100E-07 | 3.800E-07 |
| 14089 | 705 | F705H | B | 2.7e-07 | 2.1e-07 | 3.2e-07 | 2.667E-07 | 5.508E-08 | 2.100E-07 | 3.200E-07 |
| 14090 | 705 | F705I | B | 2.4e-07 | 5.1e-07 | 1.6e-07 | 3.033E-07 | 1.834E-07 | 1.600E-07 | 5.100E-07 |
| 14091 | 705 | F705L | B | 2.2e-07 | 2.9e-07 | 9.1e-08 | 2.003E-07 | 1.009E-07 | 9.100E-08 | 2.900E-07 |
| 14092 | 705 | F705K | B | 2e-07 | 1e-07 | 4.2e-07 | 2.400E-07 | 1.637E-07 | 1.000E-07 | 4.200E-07 |
| 14093 | 705 | F705M | B | 4e-07 | 3.3e-07 | 2.3e-07 | 3.200E-07 | 8.544E-08 | 2.300E-07 | 4.000E-07 |
| 14094 | 705 | F705F | B | 2.5e-07 | 2.1e-07 | 3.1e-07 | 2.567E-07 | 5.033E-08 | 2.100E-07 | 3.100E-07 |
| 14095 | 705 | F705P | B | 2.3e-07 | 2.2e-07 | 4.3e-07 | 2.933E-07 | 1.185E-07 | 2.200E-07 | 4.300E-07 |
| 14096 | 705 | F705S | B | 3.4e-07 | 2.3e-07 | 2.1e-07 | 2.600E-07 | 7.000E-08 | 2.100E-07 | 3.400E-07 |
| 14097 | 705 | F705T | B | 2.4e-07 | 2.1e-07 | 2.2e-07 | 2.233E-07 | 1.528E-08 | 2.100E-07 | 2.400E-07 |
| 14098 | 705 | F705W | B | 2e-07 | 3.2e-07 | 4.7e-07 | 3.300E-07 | 1.353E-07 | 2.000E-07 | 4.700E-07 |
| 14099 | 705 | F705Y | B | 4.4e-07 | 1.4e-07 | 2.6e-07 | 2.800E-07 | 1.510E-07 | 1.400E-07 | 4.400E-07 |
| 14100 | 705 | F705V | B | 3e-07 | 1.8e-07 | 2.7e-07 | 2.500E-07 | 6.245E-08 | 1.800E-07 | 3.000E-07 |
| 14101 | 706 | A706A | B | 1.9e-07 | 2.2e-07 | 2.3e-07 | 2.133E-07 | 2.082E-08 | 1.900E-07 | 2.300E-07 |
| 14102 | 706 | A706R | B | 2.1e-07 | 2.5e-07 | 2e-07 | 2.200E-07 | 2.646E-08 | 2.000E-07 | 2.500E-07 |
| 14103 | 706 | A706N | B | 7.9e-08 | 1.3e-07 | 1.7e-07 | 1.263E-07 | 4.561E-08 | 7.900E-08 | 1.700E-07 |
| 14104 | 706 | A706D | B | 2.5e-07 | 3.6e-07 | 2.2e-07 | 2.767E-07 | 7.371E-08 | 2.200E-07 | 3.600E-07 |
| 14105 | 706 | A706C | B | 2.4e-07 | 2.4e-07 | 1.2e-07 | 2.000E-07 | 6.928E-08 | 1.200E-07 | 2.400E-07 |
| 14106 | 706 | A706E | B | 3.2e-07 | 1.7e-07 | 2.7e-07 | 2.533E-07 | 7.638E-08 | 1.700E-07 | 3.200E-07 |
| 14107 | 706 | A706Q | B | 2.7e-07 | 2.4e-07 | 1.8e-07 | 2.300E-07 | 4.583E-08 | 1.800E-07 | 2.700E-07 |
| 14108 | 706 | A706G | B | 1.4e-07 | 1.6e-07 | 1.4e-07 | 1.467E-07 | 1.155E-08 | 1.400E-07 | 1.600E-07 |
| 14109 | 706 | A706H | B | 2.6e-07 | 2e-07 | 1.5e-07 | 2.033E-07 | 5.508E-08 | 1.500E-07 | 2.600E-07 |

| | | | | | | | | | | |
|---|---|---|---|---|---|---|---|---|---|---|
| 14110 | 706 | A706I | B | 3.3e-07 | 3.7e-07 | 1e-07 | 2.667E-07 | 1.457E-07 | 1.000E-07 | 3.700E-07 |
| 14111 | 706 | A706L | B | 1.8e-07 | 3.6e-07 | 1.7e-07 | 2.367E-07 | 1.069E-07 | 1.700E-07 | 3.600E-07 |
| 14112 | 706 | A706K | B | 3.5e-07 | 1.8e-07 | 1.5e-07 | 2.267E-07 | 1.079E-07 | 1.500E-07 | 3.500E-07 |
| 14113 | 706 | A706M | B | 4.7e-07 | 2.4e-07 | 7.2e-07 | 4.767E-07 | 2.401E-07 | 2.400E-07 | 7.200E-07 |
| 14114 | 706 | A706F | B | 5.6e-07 | 9e-08 | 1.2e-07 | 2.567E-07 | 2.631E-07 | 9.000E-08 | 5.600E-07 |
| 14115 | 706 | A706P | B | 3e-07 | 5.1e-08 | 2.2e-07 | 1.903E-07 | 1.271E-07 | 5.100E-08 | 3.000E-07 |
| 14116 | 706 | A706S | B | 6.5e-07 | 2.5e-07 | 2.5e-07 | 3.833E-07 | 2.309E-07 | 2.500E-07 | 6.500E-07 |
| 14117 | 706 | A706T | B | 2.9e-07 | 2.8e-07 | 1e-07 | 2.233E-07 | 1.069E-07 | 1.000E-07 | 2.900E-07 |
| 14118 | 706 | A706W | B | 2.5e-07 | 1.1e-07 | 1.6e-07 | 1.733E-07 | 7.095E-08 | 1.100E-07 | 2.500E-07 |
| 14119 | 706 | A706Y | B | 2.3e-07 | 1.9e-07 | 5.3e-07 | 3.167E-07 | 1.858E-07 | 1.900E-07 | 5.300E-07 |
| 14120 | 706 | A706V | B | 1.8e-07 | 2e-07 | 5.6e-07 | 3.133E-07 | 2.139E-07 | 1.800E-07 | 5.600E-07 |
| 14121 | 707 | T707A | B | 1.5e-07 | 1.4e-07 | 2e-07 | 1.633E-07 | 3.215E-08 | 1.400E-07 | 2.000E-07 |
| 14122 | 707 | T707R | B | 2.9e-07 | 2.7e-07 | 1.8e-07 | 2.467E-07 | 5.859E-08 | 1.800E-07 | 2.900E-07 |
| 14123 | 707 | T707N | B | 4.4e-07 | 2.5e-07 | 3e-07 | 3.300E-07 | 9.849E-08 | 2.500E-07 | 4.400E-07 |
| 14124 | 707 | T707D | B | 2.3e-07 | 2.5e-07 | 1.4e-07 | 2.067E-07 | 5.859E-08 | 1.400E-07 | 2.500E-07 |
| 14125 | 707 | T707C | B | 3.8e-07 | 3.8e-07 | 1.2e-07 | 2.933E-07 | 1.501E-07 | 1.200E-07 | 3.800E-07 |
| 14126 | 707 | T707E | B | 1.2e-07 | 2.7e-07 | 3.9e-07 | 2.600E-07 | 1.353E-07 | 1.200E-07 | 3.900E-07 |
| 14127 | 707 | T707Q | B | 4.5e-07 | 2.1e-07 | 4.5e-07 | 3.700E-07 | 1.386E-07 | 2.100E-07 | 4.500E-07 |
| 14128 | 707 | T707G | B | 2.3e-07 | 2.7e-07 | 2.2e-07 | 2.400E-07 | 2.646E-08 | 2.200E-07 | 2.700E-07 |
| 14129 | 707 | T707H | B | 3.4e-07 | 4e-07 | 2.8e-07 | 3.400E-07 | 6.000E-08 | 2.800E-07 | 4.000E-07 |
| 14130 | 707 | T707I | B | 2.9e-07 | 2.4e-07 | 1.4e-07 | 2.233E-07 | 7.638E-08 | 1.400E-07 | 2.900E-07 |
| 14131 | 707 | T707L | B | 2.9e-07 | 2.8e-07 | 2.2e-07 | 2.633E-07 | 3.786E-08 | 2.200E-07 | 2.900E-07 |
| 14132 | 707 | T707K | B | 4.2e-07 | 2.8e-07 | 2.6e-07 | 3.200E-07 | 8.718E-08 | 2.600E-07 | 4.200E-07 |
| 14133 | 707 | T707M | B | 2.6e-07 | 1.9e-07 | 2.7e-07 | 2.400E-07 | 4.359E-08 | 1.900E-07 | 2.700E-07 |
| 14134 | 707 | T707F | B | 4.4e-07 | 2.7e-07 | 2.5e-07 | 3.200E-07 | 1.044E-07 | 2.500E-07 | 4.400E-07 |
| 14135 | 707 | T707P | B | 3.4e-07 | 1.9e-07 | 2e-07 | 2.433E-07 | 8.386E-08 | 1.900E-07 | 3.400E-07 |
| 14136 | 707 | T707S | B | 2.4e-07 | 2.8e-07 | 3.1e-07 | 2.767E-07 | 3.512E-08 | 2.400E-07 | 3.100E-07 |
| 14137 | 707 | T707T | B | 2.2e-07 | 2.2e-07 | 4.1e-07 | 2.833E-07 | 1.097E-07 | 2.200E-07 | 4.100E-07 |
| 14138 | 707 | T707W | B | 2.1e-07 | 3.2e-07 | 3.6e-07 | 2.967E-07 | 7.767E-08 | 2.100E-07 | 3.600E-07 |
| 14139 | 707 | T707Y | B | 3.5e-07 | 4e-07 | 1.6e-07 | 3.033E-07 | 1.266E-07 | 1.600E-07 | 4.000E-07 |
| 14140 | 707 | T707V | B | 1.5e-07 | 5.9e-07 | 2.7e-07 | 3.367E-07 | 2.274E-07 | 1.500E-07 | 5.900E-07 |
| 14141 | 708 | Y708A | B | 3.6e-07 | 1.4e-07 | 2.5e-07 | 2.500E-07 | 1.100E-07 | 1.400E-07 | 3.600E-07 |
| 14142 | 708 | Y708R | B | 1.6e-07 | 4e-07 | 3.4e-07 | 3.000E-07 | 1.249E-07 | 1.600E-07 | 4.000E-07 |
| 14143 | 708 | Y708N | B | 5e-07 | 3.1e-07 | 2.4e-07 | 3.500E-07 | 1.345E-07 | 2.400E-07 | 5.000E-07 |
| 14144 | 708 | Y708D | B | 2.5e-07 | 1.8e-07 | 5.2e-07 | 3.167E-07 | 1.795E-07 | 1.800E-07 | 5.200E-07 |
| 14145 | 708 | Y708C | B | 1.7e-07 | 2.3e-07 | 1.7e-07 | 1.900E-07 | 3.464E-08 | 1.700E-07 | 2.300E-07 |
| 14146 | 708 | Y708E | B | 2.7e-07 | 2.6e-07 | 2.3e-07 | 2.533E-07 | 2.082E-08 | 2.300E-07 | 2.700E-07 |
| 14147 | 708 | Y708Q | B | 2.5e-07 | 5.2e-07 | 3.3e-07 | 3.667E-07 | 1.387E-07 | 2.500E-07 | 5.200E-07 |
| 14148 | 708 | Y708G | B | 2.7e-07 | 2.7e-07 | 2e-07 | 2.467E-07 | 4.041E-08 | 2.000E-07 | 2.700E-07 |
| 14149 | 708 | Y708H | B | 1.7e-07 | 2.3e-07 | 2.1e-07 | 2.033E-07 | 3.055E-08 | 1.700E-07 | 2.300E-07 |
| 14150 | 708 | Y708I | B | 1.6e-07 | 2.1e-07 | 1.8e-07 | 1.833E-07 | 2.517E-08 | 1.600E-07 | 2.100E-07 |
| 14151 | 708 | Y708L | B | 1.2e-07 | 4.8e-07 | 4e-07 | 3.333E-07 | 1.890E-07 | 1.200E-07 | 4.800E-07 |
| 14152 | 708 | Y708K | B | 1.4e-07 | 2e-07 | 2.9e-07 | 2.100E-07 | 7.550E-08 | 1.400E-07 | 2.900E-07 |
| 14153 | 708 | Y708M | B | 1.6e-07 | 1.4e-07 | 2.5e-07 | 1.833E-07 | 5.859E-08 | 1.400E-07 | 2.500E-07 |
| 14154 | 708 | Y708F | B | 3.9e-07 | 4.9e-07 | 4.1e-07 | 4.300E-07 | 5.292E-08 | 3.900E-07 | 4.900E-07 |
| 14155 | 708 | Y708P | B | 1.7e-07 | 1.5e-07 | 1.2e-07 | 1.467E-07 | 2.517E-08 | 1.200E-07 | 1.700E-07 |
| 14156 | 708 | Y708S | B | 1.3e-06 | 3.1e-07 | 4.9e-07 | 7.000E-07 | 5.274E-07 | 3.100E-07 | 1.300E-06 |
| 14157 | 708 | Y708T | B | 4.4e-07 | 2.4e-07 | 2.2e-07 | 3.000E-07 | 1.217E-07 | 2.200E-07 | 4.400E-07 |
| 14158 | 708 | Y708W | B | 4.1e-07 | 1e-07 | 5.7e-07 | 3.600E-07 | 2.390E-07 | 1.000E-07 | 5.700E-07 |
| 14159 | 708 | Y708Y | B | 3.4e-07 | 2.4e-07 | 1.7e-07 | 2.500E-07 | 8.544E-08 | 1.700E-07 | 3.400E-07 |
| 14160 | 708 | Y708V | B | 3.6e-07 | 1.8e-07 | 9.9e-08 | 2.130E-07 | 1.336E-07 | 9.900E-08 | 3.600E-07 |
| 14161 | 709 | Y709A | B | 4.9e-07 | 3.9e-07 | 1.7e-07 | 3.500E-07 | 1.637E-07 | 1.700E-07 | 4.900E-07 |
| 14162 | 709 | Y709R | B | 2.6e-07 | 3.1e-07 | 4e-07 | 3.233E-07 | 7.095E-08 | 2.600E-07 | 4.000E-07 |
| 14163 | 709 | Y709N | B | 2e-07 | 2.3e-07 | 2.9e-07 | 2.400E-07 | 4.583E-08 | 2.000E-07 | 2.900E-07 |
| 14164 | 709 | Y709D | B | 6.7e-07 | 2.3e-07 | 2.7e-07 | 3.900E-07 | 2.433E-07 | 2.300E-07 | 6.700E-07 |
| 14165 | 709 | Y709C | B | 1.2e-07 | 1.5e-07 | 3.3e-07 | 2.000E-07 | 1.136E-07 | 1.200E-07 | 3.300E-07 |
| 14166 | 709 | Y709E | B | 1.2e-07 | 1.5e-07 | 1.9e-07 | 1.533E-07 | 3.512E-08 | 1.200E-07 | 1.900E-07 |
| 14167 | 709 | Y709Q | B | 1.9e-07 | 6.6e-07 | 2.4e-07 | 3.633E-07 | 2.581E-07 | 1.900E-07 | 6.600E-07 |
| 14168 | 709 | Y709G | B | 4.3e-07 | 1.9e-07 | 3.4e-07 | 3.200E-07 | 1.212E-07 | 1.900E-07 | 4.300E-07 |
| 14169 | 709 | Y709H | B | 4.9e-07 | 2.3e-07 | 3.7e-07 | 3.633E-07 | 1.301E-07 | 2.300E-07 | 4.900E-07 |
| 14170 | 709 | Y709I | B | 2.7e-07 | 2.4e-07 | 1.7e-07 | 2.267E-07 | 5.132E-08 | 1.700E-07 | 2.700E-07 |
| 14171 | 709 | Y709L | B | 5e-07 | 2e-07 | 1.4e-07 | 2.800E-07 | 1.929E-07 | 1.400E-07 | 5.000E-07 |
| 14172 | 709 | Y709K | B | 5e-07 | 2.6e-07 | 2e-07 | 3.200E-07 | 1.587E-07 | 2.000E-07 | 5.000E-07 |
| 14173 | 709 | Y709M | B | 1.5e-07 | 1.8e-07 | 1.3e-07 | 1.533E-07 | 2.517E-08 | 1.300E-07 | 1.800E-07 |
| 14174 | 709 | Y709F | B | 2.3e-07 | 1.4e-07 | 3.1e-07 | 2.267E-07 | 8.505E-08 | 1.400E-07 | 3.100E-07 |
| 14175 | 709 | Y709P | B | 6.6e-07 | 2.1e-07 | 6.6e-07 | 5.100E-07 | 2.598E-07 | 2.100E-07 | 6.600E-07 |
| 14176 | 709 | Y709S | B | 4.1e-07 | 2.1e-07 | 2e-07 | 2.733E-07 | 1.185E-07 | 2.000E-07 | 4.100E-07 |
| 14177 | 709 | Y709T | B | 3.2e-07 | 2.4e-07 | 2.1e-07 | 2.567E-07 | 5.686E-08 | 2.100E-07 | 3.200E-07 |
| 14178 | 709 | Y709W | B | 3.2e-07 | 2.1e-07 | 2.2e-07 | 2.500E-07 | 6.083E-08 | 2.100E-07 | 3.200E-07 |
| 14179 | 709 | Y709Y | B | 1.6e-07 | 3.6e-07 | 1.7e-07 | 2.300E-07 | 1.127E-07 | 1.600E-07 | 3.600E-07 |
| 14180 | 709 | Y709V | B | 3.1e-07 | 2.6e-07 | 1.4e-07 | 2.367E-07 | 8.737E-08 | 1.400E-07 | 3.100E-07 |
| 14181 | 710 | C710A | B | 2.7e-07 | 1.5e-07 | 4.4e-07 | 2.867E-07 | 1.457E-07 | 1.500E-07 | 4.400E-07 |
| 14182 | 710 | C710R | B | 3.1e-07 | 2.6e-07 | 5.5e-07 | 3.733E-07 | 1.550E-07 | 2.600E-07 | 5.500E-07 |
| 14183 | 710 | C710N | B | 2.5e-07 | 2.7e-07 | 2.6e-07 | 2.600E-07 | 1.000E-08 | 2.500E-07 | 2.700E-07 |
| 14184 | 710 | C710D | B | 2.3e-07 | 1e-07 | 4.8e-07 | 2.700E-07 | 1.931E-07 | 1.000E-07 | 4.800E-07 |
| 14185 | 710 | C710C | B | 2.5e-07 | 2.2e-07 | 4.6e-07 | 3.100E-07 | 1.308E-07 | 2.200E-07 | 4.600E-07 |
| 14186 | 710 | C710E | B | 1.3e-07 | 1.6e-07 | 5.2e-07 | 2.700E-07 | 2.170E-07 | 1.300E-07 | 5.200E-07 |
| 14187 | 710 | C710Q | B | 2.4e-07 | 1.9e-07 | 2.2e-07 | 2.167E-07 | 2.517E-08 | 1.900E-07 | 2.400E-07 |
| 14188 | 710 | C710G | B | 2e-07 | 1.9e-07 | 2.5e-07 | 2.133E-07 | 3.215E-08 | 1.900E-07 | 2.500E-07 |
| 14189 | 710 | C710H | B | 1.8e-07 | 2.5e-07 | 1.8e-07 | 2.033E-07 | 4.041E-08 | 1.800E-07 | 2.500E-07 |
| 14190 | 710 | C710I | B | 2.4e-07 | 1.8e-07 | 2e-07 | 2.067E-07 | 3.055E-08 | 1.800E-07 | 2.400E-07 |
| 14191 | 710 | C710L | B | 2.7e-07 | 1.8e-07 | 4.1e-07 | 2.867E-07 | 1.159E-07 | 1.800E-07 | 4.100E-07 |
| 14192 | 710 | C710K | B | 2.3e-07 | 3.1e-07 | 1.7e-07 | 2.367E-07 | 7.024E-08 | 1.700E-07 | 3.100E-07 |
| 14193 | 710 | C710M | B | 2e-07 | 3.3e-07 | 4.2e-07 | 3.167E-07 | 1.106E-07 | 2.000E-07 | 4.200E-07 |
| 14194 | 710 | C710F | B | 2.2e-07 | 1.2e-07 | 4.3e-07 | 2.567E-07 | 1.582E-07 | 1.200E-07 | 4.300E-07 |

| | | | | | | | | | | |
|---|---|---|---|---|---|---|---|---|---|---|
| 14195 | 710 | C710P | B | 2.4e-07 | 5.3e-07 | 3.4e-07 | 3.700E-07 | 1.473E-07 | 2.400E-07 | 5.300E-07 |
| 14196 | 710 | C710S | B | 1.8e-07 | 2.6e-07 | 3.3e-07 | 2.567E-07 | 7.506E-08 | 1.800E-07 | 3.300E-07 |
| 14197 | 710 | C710T | B | 1.7e-07 | 2.8e-07 | 1.4e-07 | 1.967E-07 | 7.371E-08 | 1.400E-07 | 2.800E-07 |
| 14198 | 710 | C710W | B | 1.7e-07 | 1.8e-07 | 2.8e-07 | 2.100E-07 | 6.083E-08 | 1.700E-07 | 2.800E-07 |
| 14199 | 710 | C710Y | B | 2.1e-07 | 3e-07 | 2.9e-07 | 2.667E-07 | 4.933E-08 | 2.100E-07 | 3.000E-07 |
| 14200 | 710 | C710V | B | 2.9e-07 | 3e-07 | 3.3e-07 | 3.067E-07 | 2.082E-08 | 2.900E-07 | 3.300E-07 |
| 14201 | 711 | Q711A | B | 3.5e-07 | 2.6e-07 | 1.8e-07 | 2.633E-07 | 8.505E-08 | 1.800E-07 | 3.500E-07 |
| 14202 | 711 | Q711R | B | 2e-07 | 2.7e-07 | 2.6e-07 | 2.433E-07 | 3.786E-08 | 2.000E-07 | 2.700E-07 |
| 14203 | 711 | Q711N | B | 5.6e-07 | 4.1e-07 | 2.3e-07 | 4.000E-07 | 1.652E-07 | 2.300E-07 | 5.600E-07 |
| 14204 | 711 | Q711D | B | 2.9e-07 | 1.8e-07 | 3e-07 | 2.567E-07 | 6.658E-08 | 1.800E-07 | 3.000E-07 |
| 14205 | 711 | Q711C | B | 1.5e-07 | 1e-07 | 1.9e-07 | 1.467E-07 | 4.509E-08 | 1.000E-07 | 1.900E-07 |
| 14206 | 711 | Q711E | B | 3.9e-07 | 1.7e-07 | 2e-07 | 2.533E-07 | 1.193E-07 | 1.700E-07 | 3.900E-07 |
| 14207 | 711 | Q711Q | B | 2.9e-07 | 1.3e-07 | 3.5e-07 | 2.567E-07 | 1.137E-07 | 1.300E-07 | 3.500E-07 |
| 14208 | 711 | Q711G | B | 2.3e-07 | 2.7e-07 | 2.6e-07 | 2.533E-07 | 2.082E-08 | 2.300E-07 | 2.700E-07 |
| 14209 | 711 | Q711H | B | 3.4e-07 | 3.8e-07 | 7.9e-08 | 2.663E-07 | 1.635E-07 | 7.900E-08 | 3.800E-07 |
| 14210 | 711 | Q711I | B | 1.8e-07 | 1.5e-07 | 1.7e-07 | 1.667E-07 | 1.528E-08 | 1.500E-07 | 1.800E-07 |
| 14211 | 711 | Q711L | B | 1.7e-07 | 5.2e-07 | 1.5e-07 | 2.800E-07 | 2.081E-07 | 1.500E-07 | 5.200E-07 |
| 14212 | 711 | Q711K | B | 3.9e-07 | 2e-07 | 3.4e-07 | 3.100E-07 | 9.849E-08 | 2.000E-07 | 3.900E-07 |
| 14213 | 711 | Q711M | B | 1.6e-07 | 5.7e-07 | 6.2e-07 | 4.500E-07 | 2.524E-07 | 1.600E-07 | 6.200E-07 |
| 14214 | 711 | Q711F | B | 1.9e-07 | 3.8e-07 | 2e-07 | 2.567E-07 | 1.069E-07 | 1.900E-07 | 3.800E-07 |
| 14215 | 711 | Q711P | B | 5.7e-07 | 3.1e-07 | 1.9e-07 | 3.567E-07 | 1.943E-07 | 1.900E-07 | 5.700E-07 |
| 14216 | 711 | Q711S | B | 3.8e-07 | 1.4e-07 | 3.3e-07 | 2.833E-07 | 1.266E-07 | 1.400E-07 | 3.800E-07 |
| 14217 | 711 | Q711T | B | 2.8e-07 | 1.9e-07 | 1.5e-07 | 2.067E-07 | 6.658E-08 | 1.500E-07 | 2.800E-07 |
| 14218 | 711 | Q711W | B | 2.5e-07 | 2.7e-07 | 2.4e-07 | 2.533E-07 | 1.528E-08 | 2.400E-07 | 2.700E-07 |
| 14219 | 711 | Q711Y | B | 1e-07 | 2.7e-07 | 2.4e-07 | 2.033E-07 | 9.074E-08 | 1.000E-07 | 2.700E-07 |
| 14220 | 711 | Q711V | B | 2.3e-07 | 4.1e-07 | 3.6e-07 | 3.333E-07 | 9.292E-08 | 2.300E-07 | 4.100E-07 |
| 14221 | 712 | Q712A | B | 6.2e-07 | 6.4e-08 | 4e-07 | 3.613E-07 | 2.800E-07 | 6.400E-08 | 6.200E-07 |
| 14222 | 712 | Q712R | B | 1.6e-07 | 1.7e-07 | 3.7e-07 | 2.333E-07 | 1.185E-07 | 1.600E-07 | 3.700E-07 |
| 14223 | 712 | Q712N | B | 4e-07 | 2.5e-07 | 2.5e-07 | 3.000E-07 | 8.660E-08 | 2.500E-07 | 4.000E-07 |
| 14224 | 712 | Q712D | B | 4.3e-07 | 2.1e-07 | 1.1e-07 | 2.500E-07 | 1.637E-07 | 1.100E-07 | 4.300E-07 |
| 14225 | 712 | Q712C | B | 1.5e-07 | 1.6e-07 | 1.3e-07 | 1.467E-07 | 1.528E-08 | 1.300E-07 | 1.600E-07 |
| 14226 | 712 | Q712E | B | 2.3e-07 | 1.2e-07 | 2e-07 | 1.833E-07 | 5.686E-08 | 1.200E-07 | 2.300E-07 |
| 14227 | 712 | Q712Q | B | 1.7e-07 | 3.4e-07 | 2.7e-07 | 2.600E-07 | 8.544E-08 | 1.700E-07 | 3.400E-07 |
| 14228 | 712 | Q712G | B | 1.8e-07 | 1.9e-07 | 1.8e-07 | 1.833E-07 | 5.774E-09 | 1.800E-07 | 1.900E-07 |
| 14229 | 712 | Q712H | B | 1.5e-07 | 2.7e-07 | 3.9e-07 | 2.700E-07 | 1.200E-07 | 1.500E-07 | 3.900E-07 |
| 14230 | 712 | Q712I | B | 4.9e-07 | 6.1e-07 | 1e-07 | 4.000E-07 | 2.666E-07 | 1.000E-07 | 6.100E-07 |
| 14231 | 712 | Q712L | B | 4.2e-07 | 3e-07 | 1.4e-07 | 2.867E-07 | 1.405E-07 | 1.400E-07 | 4.200E-07 |
| 14232 | 712 | Q712K | B | 6.1e-07 | 2.2e-07 | 2.5e-07 | 3.600E-07 | 2.170E-07 | 2.200E-07 | 6.100E-07 |
| 14233 | 712 | Q712M | B | 1.4e-07 | 1.7e-07 | 3e-07 | 2.033E-07 | 8.505E-08 | 1.400E-07 | 3.000E-07 |
| 14234 | 712 | Q712F | B | 4.2e-07 | 2.1e-07 | 3e-07 | 3.100E-07 | 1.054E-07 | 2.100E-07 | 4.200E-07 |
| 14235 | 712 | Q712P | B | 1.7e-07 | 3.2e-07 | 2e-07 | 2.300E-07 | 7.937E-08 | 1.700E-07 | 3.200E-07 |
| 14236 | 712 | Q712S | B | 2.5e-07 | 5.5e-07 | 1.9e-07 | 3.300E-07 | 1.929E-07 | 1.900E-07 | 5.500E-07 |
| 14237 | 712 | Q712T | B | 3e-07 | 2.6e-07 | 2.4e-07 | 2.667E-07 | 3.055E-08 | 2.400E-07 | 3.000E-07 |
| 14238 | 712 | Q712W | B | 1.6e-07 | 3.8e-07 | 2.2e-07 | 2.533E-07 | 1.137E-07 | 1.600E-07 | 3.800E-07 |
| 14239 | 712 | Q712Y | B | 2e-07 | 2.2e-07 | 3.1e-07 | 2.433E-07 | 5.859E-08 | 2.000E-07 | 3.100E-07 |
| 14240 | 712 | Q712V | B | 2e-07 | 3.7e-07 | 2.3e-07 | 2.667E-07 | 9.074E-08 | 2.000E-07 | 3.700E-07 |
| 14241 | 713 | Y713A | B | 4.8e-07 | 2e-07 | 3.7e-07 | 3.500E-07 | 1.411E-07 | 2.000E-07 | 4.800E-07 |
| 14242 | 713 | Y713R | B | 5.7e-07 | 3.7e-07 | 2.9e-07 | 4.100E-07 | 1.442E-07 | 2.900E-07 | 5.700E-07 |
| 14243 | 713 | Y713N | B | 5.4e-07 | 9.9e-07 | 2.3e-07 | 5.867E-07 | 3.821E-07 | 2.300E-07 | 9.900E-07 |
| 14244 | 713 | Y713D | B | 3.4e-07 | 4.2e-07 | 3.6e-07 | 3.733E-07 | 4.163E-08 | 3.400E-07 | 4.200E-07 |
| 14245 | 713 | Y713C | B | 2.2e-07 | 2.2e-07 | 3.4e-07 | 2.600E-07 | 6.928E-08 | 2.200E-07 | 3.400E-07 |
| 14246 | 713 | Y713E | B | 2e-07 | 2.2e-07 | 3e-07 | 2.400E-07 | 5.292E-08 | 2.000E-07 | 3.000E-07 |
| 14247 | 713 | Y713Q | B | 1.2e-07 | 2.7e-07 | 3.2e-07 | 2.367E-07 | 1.041E-07 | 1.200E-07 | 3.200E-07 |
| 14248 | 713 | Y713G | B | 3.4e-07 | 5.8e-07 | 8.1e-08 | 3.337E-07 | 2.496E-07 | 8.100E-08 | 5.800E-07 |
| 14249 | 713 | Y713H | B | 1.5e-07 | 1.9e-07 | 5.4e-07 | 2.933E-07 | 2.146E-07 | 1.500E-07 | 5.400E-07 |
| 14250 | 713 | Y713I | B | 3.5e-07 | 3.6e-07 | 1.9e-07 | 3.000E-07 | 9.539E-08 | 1.900E-07 | 3.600E-07 |
| 14251 | 713 | Y713L | B | 2.1e-07 | 1e-07 | 2.4e-07 | 1.833E-07 | 7.371E-08 | 1.000E-07 | 2.400E-07 |
| 14252 | 713 | Y713K | B | 4e-07 | 2.3e-07 | 2.7e-07 | 3.000E-07 | 8.888E-08 | 2.300E-07 | 4.000E-07 |
| 14253 | 713 | Y713M | B | 2.5e-07 | 2.6e-07 | 2.5e-07 | 2.533E-07 | 5.774E-09 | 2.500E-07 | 2.600E-07 |
| 14254 | 713 | Y713F | B | 2.9e-07 | 3.4e-07 | 2.6e-07 | 2.967E-07 | 4.041E-08 | 2.600E-07 | 3.400E-07 |
| 14255 | 713 | Y713P | B | 3.2e-07 | 2.2e-07 | 3e-07 | 2.800E-07 | 5.292E-08 | 2.200E-07 | 3.200E-07 |
| 14256 | 713 | Y713S | B | 3.8e-07 | 1.4e-07 | 1.9e-07 | 2.367E-07 | 1.266E-07 | 1.400E-07 | 3.800E-07 |
| 14257 | 713 | Y713T | B | 1e-07 | 2.3e-07 | 2.5e-07 | 1.933E-07 | 8.145E-08 | 1.000E-07 | 2.500E-07 |
| 14258 | 713 | Y713W | B | 4.9e-07 | 2.8e-07 | 2.4e-07 | 3.367E-07 | 1.343E-07 | 2.400E-07 | 4.900E-07 |
| 14259 | 713 | Y713Y | B | 3e-07 | 3.4e-07 | 1.6e-07 | 2.667E-07 | 9.452E-08 | 1.600E-07 | 3.400E-07 |
| 14260 | 713 | Y713V | B | 2.2e-07 | 1.8e-07 | 2.7e-07 | 2.233E-07 | 4.509E-08 | 1.800E-07 | 2.700E-07 |
| 14261 | 714 | Y714A | B | 1.4e-07 | 4.1e-07 | 3e-07 | 2.833E-07 | 1.358E-07 | 1.400E-07 | 4.100E-07 |
| 14262 | 714 | Y714R | B | 3.8e-07 | 4.1e-07 | 3e-07 | 3.633E-07 | 5.686E-08 | 3.000E-07 | 4.100E-07 |
| 14263 | 714 | Y714N | B | 2.2e-07 | 5.2e-07 | 1.4e-07 | 2.933E-07 | 2.003E-07 | 1.400E-07 | 5.200E-07 |
| 14264 | 714 | Y714D | B | 3.8e-07 | 2.8e-07 | 8.8e-08 | 2.493E-07 | 1.484E-07 | 8.800E-08 | 3.800E-07 |
| 14265 | 714 | Y714C | B | 1.3e-07 | 3.4e-07 | 3.1e-07 | 2.600E-07 | 1.136E-07 | 1.300E-07 | 3.400E-07 |
| 14266 | 714 | Y714E | B | 6.4e-07 | 2.8e-07 | 1.5e-07 | 3.567E-07 | 2.538E-07 | 1.500E-07 | 6.400E-07 |
| 14267 | 714 | Y714Q | B | 5.4e-07 | 3.3e-07 | 2.3e-07 | 3.667E-07 | 1.582E-07 | 2.300E-07 | 5.400E-07 |
| 14268 | 714 | Y714G | B | 1.6e-07 | 3.6e-07 | 1.7e-07 | 2.300E-07 | 1.127E-07 | 1.600E-07 | 3.600E-07 |
| 14269 | 714 | Y714H | B | 1.9e-07 | 3.2e-07 | 4e-07 | 3.033E-07 | 1.060E-07 | 1.900E-07 | 4.000E-07 |
| 14270 | 714 | Y714I | B | 4.7e-07 | 2.3e-07 | 2.1e-07 | 3.033E-07 | 1.447E-07 | 2.100E-07 | 4.700E-07 |
| 14271 | 714 | Y714L | B | 1.6e-07 | 2e-07 | 2.4e-07 | 2.000E-07 | 4.000E-08 | 1.600E-07 | 2.400E-07 |
| 14272 | 714 | Y714K | B | 2.8e-07 | 2.7e-07 | 1.7e-07 | 2.400E-07 | 6.083E-08 | 1.700E-07 | 2.800E-07 |
| 14273 | 714 | Y714M | B | 2.8e-07 | 2.7e-07 | 3.7e-07 | 3.067E-07 | 5.508E-08 | 2.700E-07 | 3.700E-07 |
| 14274 | 714 | Y714F | B | 3.6e-07 | 2.6e-07 | 1.6e-07 | 2.600E-07 | 1.000E-07 | 1.600E-07 | 3.600E-07 |
| 14275 | 714 | Y714P | B | 2.9e-07 | 2.8e-07 | 3e-07 | 2.900E-07 | 1.000E-08 | 2.800E-07 | 3.000E-07 |
| 14276 | 714 | Y714S | B | 1.3e-07 | 2.2e-07 | 2.8e-07 | 2.100E-07 | 7.550E-08 | 1.300E-07 | 2.800E-07 |
| 14277 | 714 | Y714T | B | 3.1e-07 | 1.1e-07 | 2.1e-07 | 2.100E-07 | 1.000E-07 | 1.100E-07 | 3.100E-07 |
| 14278 | 714 | Y714W | B | 1.9e-07 | 2.3e-07 | 2.6e-07 | 2.267E-07 | 3.512E-08 | 1.900E-07 | 2.600E-07 |
| 14279 | 714 | Y714Y | B | 1.4e-07 | 2.2e-07 | 1.7e-07 | 1.767E-07 | 4.041E-08 | 1.400E-07 | 2.200E-07 |

| | | | | | | | | | | |
|---|---|---|---|---|---|---|---|---|---|---|
| 14280 | 714 | Y714V | B | 1.2e-07 | 2.3e-07 | 2.4e-07 | 1.967E-07 | 6.658E-08 | 1.200E-07 | 2.400E-07 |
| 14281 | 715 | I715A | B | 4.3e-07 | 1.7e-07 | 2.4e-07 | 2.800E-07 | 1.345E-07 | 1.700E-07 | 4.300E-07 |
| 14282 | 715 | I715R | B | 2.3e-07 | 2.8e-07 | 1.6e-07 | 2.233E-07 | 6.028E-08 | 1.600E-07 | 2.800E-07 |
| 14283 | 715 | I715N | B | 1.7e-07 | 2.8e-07 | 2.9e-07 | 2.467E-07 | 6.658E-08 | 1.700E-07 | 2.900E-07 |
| 14284 | 715 | I715D | B | 2.4e-07 | 1.5e-07 | 3.4e-07 | 2.433E-07 | 9.504E-08 | 1.500E-07 | 3.400E-07 |
| 14285 | 715 | I715C | B | 4.3e-07 | 3.1e-07 | 2.9e-07 | 3.433E-07 | 7.572E-08 | 2.900E-07 | 4.300E-07 |
| 14286 | 715 | I715E | B | 2.9e-07 | 1.5e-07 | 1.6e-07 | 2.000E-07 | 7.810E-08 | 1.500E-07 | 2.900E-07 |
| 14287 | 715 | I715Q | B | 2.7e-07 | 1.4e-07 | 3.5e-07 | 2.533E-07 | 1.060E-07 | 1.400E-07 | 3.500E-07 |
| 14288 | 715 | I715G | B | 2.7e-07 | 4.9e-07 | 4e-07 | 3.867E-07 | 1.106E-07 | 2.700E-07 | 4.900E-07 |
| 14289 | 715 | I715H | B | 2.3e-07 | 3e-07 | 3e-07 | 2.767E-07 | 4.041E-08 | 2.300E-07 | 3.000E-07 |
| 14290 | 715 | I715I | B | 2e-07 | 1.6e-07 | 3.6e-07 | 2.400E-07 | 1.058E-07 | 1.600E-07 | 3.600E-07 |
| 14291 | 715 | I715L | B | 1.6e-07 | 2.8e-07 | 3e-07 | 2.467E-07 | 7.572E-08 | 1.600E-07 | 3.000E-07 |
| 14292 | 715 | I715K | B | 6.8e-07 | 2.2e-07 | 3.8e-07 | 4.267E-07 | 2.335E-07 | 2.200E-07 | 6.800E-07 |
| 14293 | 715 | I715M | B | 2.6e-07 | 2.6e-07 | 2.9e-07 | 2.700E-07 | 1.732E-08 | 2.600E-07 | 2.900E-07 |
| 14294 | 715 | I715F | B | 2.6e-07 | 3.2e-07 | 2.4e-07 | 2.733E-07 | 4.163E-08 | 2.400E-07 | 3.200E-07 |
| 14295 | 715 | I715P | B | 1.9e-07 | 3.7e-07 | 3e-07 | 2.867E-07 | 9.074E-08 | 1.900E-07 | 3.700E-07 |
| 14296 | 715 | I715S | B | 3.1e-07 | 1.1e-07 | 2.3e-07 | 2.167E-07 | 1.007E-07 | 1.100E-07 | 3.100E-07 |
| 14297 | 715 | I715T | B | 3.2e-07 | 2.3e-07 | 1.9e-07 | 2.467E-07 | 6.658E-08 | 1.900E-07 | 3.200E-07 |
| 14298 | 715 | I715W | B | 2.8e-07 | 2.2e-07 | 1.9e-07 | 2.300E-07 | 4.583E-08 | 1.900E-07 | 2.800E-07 |
| 14299 | 715 | I715Y | B | 2.3e-07 | 2e-07 | 2.7e-07 | 2.333E-07 | 3.512E-08 | 2.000E-07 | 2.700E-07 |
| 14300 | 715 | I715V | B | 1.4e-07 | 1.9e-07 | 1.5e-07 | 1.600E-07 | 2.646E-08 | 1.400E-07 | 1.900E-07 |
| 14301 | 716 | Y716A | B | 2.3e-07 | 3.6e-07 | 2.1e-07 | 2.667E-07 | 8.145E-08 | 2.100E-07 | 3.600E-07 |
| 14302 | 716 | Y716R | B | 3.8e-07 | 1.4e-07 | 3e-07 | 2.733E-07 | 1.222E-07 | 1.400E-07 | 3.800E-07 |
| 14303 | 716 | Y716N | B | 1.5e-07 | 5.9e-07 | 3.9e-07 | 3.767E-07 | 2.203E-07 | 1.500E-07 | 5.900E-07 |
| 14304 | 716 | Y716D | B | 1.5e-07 | 2.9e-07 | 2.7e-07 | 2.367E-07 | 7.572E-08 | 1.500E-07 | 2.900E-07 |
| 14305 | 716 | Y716C | B | 2.6e-07 | 3.3e-07 | 5.2e-07 | 3.700E-07 | 1.345E-07 | 2.600E-07 | 5.200E-07 |
| 14306 | 716 | Y716E | B | 2.3e-07 | 2.1e-07 | 3.6e-07 | 2.667E-07 | 8.145E-08 | 2.100E-07 | 3.600E-07 |
| 14307 | 716 | Y716Q | B | 2.7e-07 | 2.4e-07 | 2.8e-07 | 2.633E-07 | 2.082E-08 | 2.400E-07 | 2.800E-07 |
| 14308 | 716 | Y716G | B | 2.7e-07 | 3.7e-07 | 2.8e-07 | 3.067E-07 | 5.508E-08 | 2.700E-07 | 3.700E-07 |
| 14309 | 716 | Y716H | B | 2e-07 | 2.4e-07 | 5.2e-07 | 3.200E-07 | 1.744E-07 | 2.000E-07 | 5.200E-07 |
| 14310 | 716 | Y716I | B | 2.8e-07 | 1.3e-07 | 4.8e-07 | 2.967E-07 | 1.756E-07 | 1.300E-07 | 4.800E-07 |
| 14311 | 716 | Y716L | B | 2.3e-07 | 1.8e-07 | 2.8e-07 | 2.300E-07 | 5.000E-08 | 1.800E-07 | 2.800E-07 |
| 14312 | 716 | Y716K | B | 3.5e-07 | 4.5e-07 | 3.2e-07 | 3.733E-07 | 6.807E-08 | 3.200E-07 | 4.500E-07 |
| 14313 | 716 | Y716M | B | 2.4e-07 | 3.2e-07 | 1e-07 | 2.200E-07 | 1.114E-07 | 1.000E-07 | 3.200E-07 |
| 14314 | 716 | Y716F | B | 1.8e-07 | 9.5e-08 | 1.7e-07 | 1.483E-07 | 4.646E-08 | 9.500E-08 | 1.800E-07 |
| 14315 | 716 | Y716P | B | 6.5e-08 | 1.3e-07 | 1.5e-07 | 1.150E-07 | 4.444E-08 | 6.500E-08 | 1.500E-07 |
| 14316 | 716 | Y716S | B | 2.7e-07 | 2.8e-07 | 2e-07 | 2.500E-07 | 4.359E-08 | 2.000E-07 | 2.800E-07 |
| 14317 | 716 | Y716T | B | 3.5e-07 | 2.5e-07 | 9.8e-08 | 2.327E-07 | 1.269E-07 | 9.800E-08 | 3.500E-07 |
| 14318 | 716 | Y716W | B | 2.1e-07 | 5.6e-07 | 2.2e-07 | 3.300E-07 | 1.992E-07 | 2.100E-07 | 5.600E-07 |
| 14319 | 716 | Y716Y | B | 8.9e-08 | 1.9e-07 | 3.4e-07 | 2.063E-07 | 1.263E-07 | 8.900E-08 | 3.400E-07 |
| 14320 | 716 | Y716V | B | 2.3e-07 | 1.2e-07 | 3.5e-07 | 2.333E-07 | 1.150E-07 | 1.200E-07 | 3.500E-07 |
| 14321 | 717 | P717A | B | 2.2e-07 | 1.8e-07 | 6.3e-07 | 3.433E-07 | 2.491E-07 | 1.800E-07 | 6.300E-07 |
| 14322 | 717 | P717R | B | 3.6e-07 | 1.4e-07 | 8.2e-08 | 1.940E-07 | 1.467E-07 | 8.200E-08 | 3.600E-07 |
| 14323 | 717 | P717N | B | 1.5e-07 | 1.9e-07 | 2.6e-07 | 2.000E-07 | 5.568E-08 | 1.500E-07 | 2.600E-07 |
| 14324 | 717 | P717D | B | 3.1e-07 | 5e-07 | 2.3e-07 | 3.467E-07 | 1.387E-07 | 2.300E-07 | 5.000E-07 |
| 14325 | 717 | P717C | B | 2.2e-07 | 1.9e-07 | 2.1e-07 | 2.067E-07 | 1.528E-08 | 1.900E-07 | 2.200E-07 |
| 14326 | 717 | P717E | B | 2.4e-07 | 2.3e-07 | 1.8e-07 | 2.167E-07 | 3.215E-08 | 1.800E-07 | 2.400E-07 |
| 14327 | 717 | P717Q | B | 2.2e-07 | 3.4e-07 | 2.9e-07 | 2.833E-07 | 6.028E-08 | 2.200E-07 | 3.400E-07 |
| 14328 | 717 | P717G | B | 3e-07 | 1.9e-07 | 1.7e-07 | 2.200E-07 | 7.000E-08 | 1.700E-07 | 3.000E-07 |
| 14329 | 717 | P717H | B | 1.2e-07 | 4.2e-07 | 1.2e-07 | 2.200E-07 | 1.732E-07 | 1.200E-07 | 4.200E-07 |
| 14330 | 717 | P717I | B | 5.1e-07 | 2.3e-07 | 3.9e-07 | 3.767E-07 | 1.405E-07 | 2.300E-07 | 5.100E-07 |
| 14331 | 717 | P717L | B | 4.6e-07 | 2.3e-07 | 1.9e-07 | 2.933E-07 | 1.457E-07 | 1.900E-07 | 4.600E-07 |
| 14332 | 717 | P717K | B | 2.2e-07 | 3.1e-07 | 1.9e-07 | 2.400E-07 | 6.245E-08 | 1.900E-07 | 3.100E-07 |
| 14333 | 717 | P717M | B | 1.8e-07 | 5.7e-07 | 2.5e-07 | 3.333E-07 | 2.079E-07 | 1.800E-07 | 5.700E-07 |
| 14334 | 717 | P717F | B | 1.5e-07 | 2.3e-07 | 2.2e-07 | 2.000E-07 | 4.359E-08 | 1.500E-07 | 2.300E-07 |
| 14335 | 717 | P717P | B | 2.2e-07 | 2.3e-07 | 1.5e-07 | 2.000E-07 | 4.359E-08 | 1.500E-07 | 2.300E-07 |
| 14336 | 717 | P717S | B | 1.9e-07 | 3e-07 | 4.6e-07 | 3.167E-07 | 1.358E-07 | 1.900E-07 | 4.600E-07 |
| 14337 | 717 | P717T | B | 1.5e-07 | 2.4e-07 | 3.1e-07 | 2.333E-07 | 8.021E-08 | 1.500E-07 | 3.100E-07 |
| 14338 | 717 | P717W | B | 3.4e-07 | 1.7e-07 | 3e-07 | 2.700E-07 | 8.888E-08 | 1.700E-07 | 3.400E-07 |
| 14339 | 717 | P717Y | B | 1.2e-07 | 1.8e-07 | 4.1e-07 | 2.367E-07 | 1.531E-07 | 1.200E-07 | 4.100E-07 |
| 14340 | 717 | P717V | B | 1.5e-07 | 1.1e-07 | 2.7e-07 | 1.767E-07 | 8.327E-08 | 1.100E-07 | 2.700E-07 |
| 14341 | 718 | Y718A | B | 2.9e-07 | 4e-07 | 1.7e-07 | 2.867E-07 | 1.150E-07 | 1.700E-07 | 4.000E-07 |
| 14342 | 718 | Y718R | B | 2.6e-07 | 2.3e-07 | 2.3e-07 | 2.400E-07 | 1.732E-08 | 2.300E-07 | 2.600E-07 |
| 14343 | 718 | Y718N | B | 1.6e-07 | 2.2e-07 | 1.9e-07 | 1.900E-07 | 3.000E-08 | 1.600E-07 | 2.200E-07 |
| 14344 | 718 | Y718D | B | 8.4e-07 | 2.5e-07 | 1.8e-07 | 4.233E-07 | 3.625E-07 | 1.800E-07 | 8.400E-07 |
| 14345 | 718 | Y718C | B | 3.8e-07 | 6.2e-07 | 1.4e-07 | 3.800E-07 | 2.400E-07 | 1.400E-07 | 6.200E-07 |
| 14346 | 718 | Y718E | B | 1.8e-07 | 5e-07 | 6.4e-07 | 4.400E-07 | 2.358E-07 | 1.800E-07 | 6.400E-07 |
| 14347 | 718 | Y718Q | B | 4e-07 | 3e-07 | 1.6e-07 | 2.867E-07 | 1.206E-07 | 1.600E-07 | 4.000E-07 |
| 14348 | 718 | Y718G | B | 2.3e-07 | 1.9e-07 | 4.4e-07 | 2.867E-07 | 1.343E-07 | 1.900E-07 | 4.400E-07 |
| 14349 | 718 | Y718H | B | 1.1e-07 | 3.4e-07 | 1.9e-07 | 2.133E-07 | 1.168E-07 | 1.100E-07 | 3.400E-07 |
| 14350 | 718 | Y718I | B | 2.1e-07 | 2.3e-07 | 2.5e-07 | 2.300E-07 | 2.000E-08 | 2.100E-07 | 2.500E-07 |
| 14351 | 718 | Y718L | B | 3.7e-07 | 1.5e-07 | 2.9e-07 | 2.700E-07 | 1.114E-07 | 1.500E-07 | 3.700E-07 |
| 14352 | 718 | Y718K | B | 3.4e-07 | 1.7e-07 | 1.9e-07 | 2.333E-07 | 9.292E-08 | 1.700E-07 | 3.400E-07 |
| 14353 | 718 | Y718M | B | 1.4e-07 | 4.2e-07 | 2e-07 | 2.533E-07 | 1.474E-07 | 1.400E-07 | 4.200E-07 |
| 14354 | 718 | Y718F | B | 3.6e-07 | 2.7e-07 | 6e-07 | 4.100E-07 | 1.706E-07 | 2.700E-07 | 6.000E-07 |
| 14355 | 718 | Y718P | B | 2.2e-07 | 1.6e-07 | 3.8e-07 | 2.533E-07 | 1.137E-07 | 1.600E-07 | 3.800E-07 |
| 14356 | 718 | Y718S | B | 2e-07 | 5e-07 | 2.6e-07 | 3.200E-07 | 1.587E-07 | 2.000E-07 | 5.000E-07 |
| 14357 | 718 | Y718T | B | 1.5e-07 | 3.6e-07 | 1.1e-07 | 2.067E-07 | 1.343E-07 | 1.100E-07 | 3.600E-07 |
| 14358 | 718 | Y718W | B | 8.1e-08 | 2.9e-07 | 5.4e-07 | 3.037E-07 | 2.298E-07 | 8.100E-08 | 5.400E-07 |
| 14359 | 718 | Y718Y | B | 7e-08 | 1.9e-07 | 5.2e-07 | 2.600E-07 | 2.330E-07 | 7.000E-08 | 5.200E-07 |
| 14360 | 718 | Y718V | B | 4.5e-07 | 8.8e-08 | 2.2e-07 | 2.527E-07 | 1.832E-07 | 8.800E-08 | 4.500E-07 |
| 14361 | 719 | T719A | B | 2.5e-07 | 4e-07 | 4.4e-07 | 3.633E-07 | 1.002E-07 | 2.500E-07 | 4.400E-07 |
| 14362 | 719 | T719R | B | 1.8e-07 | 2.7e-07 | 3.3e-07 | 2.600E-07 | 7.550E-08 | 1.800E-07 | 3.300E-07 |
| 14363 | 719 | T719N | B | 3.9e-07 | 1e-07 | 3e-07 | 2.633E-07 | 1.484E-07 | 1.000E-07 | 3.900E-07 |
| 14364 | 719 | T719D | B | 2.3e-07 | 4.3e-07 | 3.6e-07 | 3.400E-07 | 1.015E-07 | 2.300E-07 | 4.300E-07 |

| | | | | | | | | | | |
|---|---|---|---|---|---|---|---|---|---|---|
| 14365 | 719 | T719C | B | 3e-07 | 4e-07 | 3.1e-07 | 3.367E-07 | 5.508E-08 | 3.000E-07 | 4.000E-07 |
| 14366 | 719 | T719E | B | 5.1e-07 | 3.7e-07 | 4e-07 | 4.267E-07 | 7.371E-08 | 3.700E-07 | 5.100E-07 |
| 14367 | 719 | T719Q | B | 2.4e-07 | 8.1e-08 | 3.4e-07 | 2.203E-07 | 1.306E-07 | 8.100E-08 | 3.400E-07 |
| 14368 | 719 | T719G | B | 2.5e-07 | 2.4e-07 | 1.8e-07 | 2.233E-07 | 3.786E-08 | 1.800E-07 | 2.500E-07 |
| 14369 | 719 | T719H | B | 2.4e-07 | 3e-07 | 1.4e-07 | 2.267E-07 | 8.083E-08 | 1.400E-07 | 3.000E-07 |
| 14370 | 719 | T719I | B | 3.6e-07 | 7.3e-08 | 2.1e-07 | 2.143E-07 | 1.435E-07 | 7.300E-08 | 3.600E-07 |
| 14371 | 719 | T719L | B | 2.7e-07 | 7.8e-08 | 2.2e-07 | 1.893E-07 | 9.961E-08 | 7.800E-08 | 2.700E-07 |
| 14372 | 719 | T719K | B | 1.8e-07 | 3.8e-07 | 3e-07 | 2.867E-07 | 1.007E-07 | 1.800E-07 | 3.800E-07 |
| 14373 | 719 | T719M | B | 4.9e-07 | 1.7e-07 | 1.6e-07 | 2.733E-07 | 1.877E-07 | 1.600E-07 | 4.900E-07 |
| 14374 | 719 | T719F | B | 4.2e-07 | 1.1e-07 | 1.9e-07 | 2.400E-07 | 1.609E-07 | 1.100E-07 | 4.200E-07 |
| 14375 | 719 | T719P | B | 2.6e-07 | 6.7e-07 | 2.1e-07 | 3.800E-07 | 2.524E-07 | 2.100E-07 | 6.700E-07 |
| 14376 | 719 | T719S | B | 2e-07 | 2.7e-07 | 2.3e-07 | 2.333E-07 | 3.512E-08 | 2.000E-07 | 2.700E-07 |
| 14377 | 719 | T719T | B | 1.1e-07 | 1.8e-07 | 3.3e-07 | 2.067E-07 | 1.124E-07 | 1.100E-07 | 3.300E-07 |
| 14378 | 719 | T719W | B | 1.2e-07 | 4.4e-07 | 1.6e-07 | 2.400E-07 | 1.744E-07 | 1.200E-07 | 4.400E-07 |
| 14379 | 719 | T719Y | B | 1.8e-07 | 2e-07 | 6.8e-07 | 3.533E-07 | 2.831E-07 | 1.800E-07 | 6.800E-07 |
| 14380 | 719 | T719V | B | 4.8e-07 | 2e-07 | 4.5e-07 | 3.767E-07 | 1.537E-07 | 2.000E-07 | 4.800E-07 |
| 14381 | 720 | F720A | B | 3.8e-07 | 2.1e-07 | 1.2e-07 | 2.367E-07 | 1.320E-07 | 1.200E-07 | 3.800E-07 |
| 14382 | 720 | F720R | B | 3e-07 | 3.2e-07 | 1.9e-07 | 2.700E-07 | 7.000E-08 | 1.900E-07 | 3.200E-07 |
| 14383 | 720 | F720N | B | 1.1e-07 | 1.8e-07 | 6.1e-07 | 3.000E-07 | 2.707E-07 | 1.100E-07 | 6.100E-07 |
| 14384 | 720 | F720D | B | 2.9e-07 | 2.1e-07 | 2.6e-07 | 2.533E-07 | 4.041E-08 | 2.100E-07 | 2.900E-07 |
| 14385 | 720 | F720C | B | 7.9e-08 | 2.2e-07 | 2.6e-07 | 1.863E-07 | 9.508E-08 | 7.900E-08 | 2.600E-07 |
| 14386 | 720 | F720E | B | 1.8e-07 | 2.4e-07 | 3.2e-07 | 2.467E-07 | 7.024E-08 | 1.800E-07 | 3.200E-07 |
| 14387 | 720 | F720Q | B | 3.4e-07 | 2.9e-07 | 1.3e-07 | 2.533E-07 | 1.097E-07 | 1.300E-07 | 3.400E-07 |
| 14388 | 720 | F720G | B | 2.8e-07 | 5.1e-07 | 3e-07 | 3.633E-07 | 1.274E-07 | 2.800E-07 | 5.100E-07 |
| 14389 | 720 | F720H | B | 3.5e-07 | 2.3e-07 | 1.5e-07 | 2.433E-07 | 1.007E-07 | 1.500E-07 | 3.500E-07 |
| 14390 | 720 | F720I | B | 2.3e-07 | 2.6e-07 | 1.6e-07 | 2.167E-07 | 5.132E-08 | 1.600E-07 | 2.600E-07 |
| 14391 | 720 | F720L | B | 3e-07 | 3.1e-07 | 3.1e-07 | 3.067E-07 | 5.774E-09 | 3.000E-07 | 3.100E-07 |
| 14392 | 720 | F720K | B | 2.9e-07 | 2e-07 | 9.4e-08 | 1.947E-07 | 9.811E-08 | 9.400E-08 | 2.900E-07 |
| 14393 | 720 | F720M | B | 1.4e-07 | 2e-07 | 2.9e-07 | 2.100E-07 | 7.550E-08 | 1.400E-07 | 2.900E-07 |
| 14394 | 720 | F720F | B | 3.8e-07 | 1.5e-07 | 1.9e-07 | 2.400E-07 | 1.229E-07 | 1.500E-07 | 3.800E-07 |
| 14395 | 720 | F720P | B | 3.1e-07 | 1.7e-07 | 2e-07 | 2.267E-07 | 7.371E-08 | 1.700E-07 | 3.100E-07 |
| 14396 | 720 | F720S | B | 1.7e-07 | 2.5e-07 | 1.9e-07 | 2.033E-07 | 4.163E-08 | 1.700E-07 | 2.500E-07 |
| 14397 | 720 | F720T | B | 3.4e-07 | 2.8e-07 | 2.7e-07 | 2.967E-07 | 3.786E-08 | 2.700E-07 | 3.400E-07 |
| 14398 | 720 | F720W | B | 6.1e-07 | 1.9e-07 | 3.1e-07 | 3.700E-07 | 2.163E-07 | 1.900E-07 | 6.100E-07 |
| 14399 | 720 | F720Y | B | 2.5e-07 | 1.4e-07 | 1.9e-07 | 1.933E-07 | 5.508E-08 | 1.400E-07 | 2.500E-07 |
| 14400 | 720 | F720V | B | 1.7e-07 | 2e-07 | 1.5e-07 | 1.733E-07 | 2.517E-08 | 1.500E-07 | 2.000E-07 |
| 14401 | 721 | G721A | B | 2.7e-07 | 2.3e-07 | 3.3e-07 | 2.767E-07 | 5.033E-08 | 2.300E-07 | 3.300E-07 |
| 14402 | 721 | G721R | B | 1.3e-07 | 2.5e-07 | 2.1e-07 | 1.967E-07 | 6.110E-08 | 1.300E-07 | 2.500E-07 |
| 14403 | 721 | G721N | B | 2.5e-07 | 1.1e-07 | 1.3e-07 | 1.633E-07 | 7.572E-08 | 1.100E-07 | 2.500E-07 |
| 14404 | 721 | G721D | B | 3.4e-07 | 1.2e-07 | 1.9e-07 | 2.167E-07 | 1.124E-07 | 1.200E-07 | 3.400E-07 |
| 14405 | 721 | G721C | B | 2e-07 | 2.9e-07 | 3.4e-07 | 2.767E-07 | 7.095E-08 | 2.000E-07 | 3.400E-07 |
| 14406 | 721 | G721E | B | 2.6e-07 | 1.9e-07 | 1.5e-07 | 2.000E-07 | 5.568E-08 | 1.500E-07 | 2.600E-07 |
| 14407 | 721 | G721Q | B | 3.5e-07 | 2.9e-07 | 2.5e-07 | 2.967E-07 | 5.033E-08 | 2.500E-07 | 3.500E-07 |
| 14408 | 721 | G721G | B | 2.9e-07 | 3.4e-07 | 3.7e-07 | 3.333E-07 | 4.041E-08 | 2.900E-07 | 3.700E-07 |
| 14409 | 721 | G721H | B | 3.9e-07 | 3.2e-07 | 1.9e-07 | 3.000E-07 | 1.015E-07 | 1.900E-07 | 3.900E-07 |
| 14410 | 721 | G721I | B | 2.2e-07 | 3.1e-07 | 1.4e-07 | 2.233E-07 | 8.505E-08 | 1.400E-07 | 3.100E-07 |
| 14411 | 721 | G721L | B | 3.3e-07 | 1.9e-07 | 2.2e-07 | 2.467E-07 | 7.371E-08 | 1.900E-07 | 3.300E-07 |
| 14412 | 721 | G721K | B | 1.8e-07 | 2.1e-07 | 2.8e-07 | 2.233E-07 | 5.132E-08 | 1.800E-07 | 2.800E-07 |
| 14413 | 721 | G721M | B | 1.6e-07 | 2.3e-07 | 2.8e-07 | 2.233E-07 | 6.028E-08 | 1.600E-07 | 2.800E-07 |
| 14414 | 721 | G721F | B | 2.5e-07 | 3.1e-07 | 1.7e-07 | 2.433E-07 | 7.024E-08 | 1.700E-07 | 3.100E-07 |
| 14415 | 721 | G721P | B | 3.6e-07 | 1.9e-07 | 2.3e-07 | 2.600E-07 | 8.888E-08 | 1.900E-07 | 3.600E-07 |
| 14416 | 721 | G721S | B | 8.9e-07 | 2.2e-07 | 2.1e-07 | 4.400E-07 | 3.897E-07 | 2.100E-07 | 8.900E-07 |
| 14417 | 721 | G721T | B | 1.2e-07 | 3.2e-07 | 9.1e-08 | 1.770E-07 | 1.247E-07 | 9.100E-08 | 3.200E-07 |
| 14418 | 721 | G721W | B | 1.8e-07 | 2.7e-07 | 1.6e-07 | 2.033E-07 | 5.859E-08 | 1.600E-07 | 2.700E-07 |
| 14419 | 721 | G721Y | B | 3.2e-07 | 3e-07 | 2.8e-07 | 3.000E-07 | 2.000E-08 | 2.800E-07 | 3.200E-07 |
| 14420 | 721 | G721V | B | 2.2e-07 | 3.7e-07 | 2.9e-07 | 2.933E-07 | 7.506E-08 | 2.200E-07 | 3.700E-07 |
| 14421 | 722 | Q722A | B | 3.9e-07 | 2.8e-07 | 4.2e-07 | 3.633E-07 | 7.371E-08 | 2.800E-07 | 4.200E-07 |
| 14422 | 722 | Q722R | B | 2.7e-07 | 3.4e-07 | 2.2e-07 | 2.767E-07 | 6.028E-08 | 2.200E-07 | 3.400E-07 |
| 14423 | 722 | Q722N | B | 3.7e-07 | 3.7e-07 | 2.4e-07 | 3.267E-07 | 7.506E-08 | 2.400E-07 | 3.700E-07 |
| 14424 | 722 | Q722D | B | 3e-07 | 1.9e-07 | 1.9e-07 | 2.267E-07 | 6.351E-08 | 1.900E-07 | 3.000E-07 |
| 14425 | 722 | Q722C | B | 2.8e-07 | 2.6e-07 | 2.3e-07 | 2.567E-07 | 2.517E-08 | 2.300E-07 | 2.800E-07 |
| 14426 | 722 | Q722E | B | 2.8e-07 | 2.6e-07 | 2.5e-07 | 2.633E-07 | 1.528E-08 | 2.500E-07 | 2.800E-07 |
| 14427 | 722 | Q722Q | B | 2.7e-07 | 2.5e-07 | 1.9e-07 | 2.367E-07 | 4.163E-08 | 1.900E-07 | 2.700E-07 |
| 14428 | 722 | Q722G | B | 3.3e-07 | 2.5e-07 | 3.5e-07 | 3.100E-07 | 5.292E-08 | 2.500E-07 | 3.500E-07 |
| 14429 | 722 | Q722H | B | 4.1e-07 | 2.6e-07 | 5.1e-07 | 3.933E-07 | 1.258E-07 | 2.600E-07 | 5.100E-07 |
| 14430 | 722 | Q722I | B | 3.7e-07 | 3e-07 | 1.6e-07 | 2.767E-07 | 1.069E-07 | 1.600E-07 | 3.700E-07 |
| 14431 | 722 | Q722L | B | 2.8e-07 | 2.9e-07 | 2.2e-07 | 2.633E-07 | 3.786E-08 | 2.200E-07 | 2.900E-07 |
| 14432 | 722 | Q722K | B | 2.1e-07 | 2.3e-07 | 2.9e-07 | 2.433E-07 | 4.163E-08 | 2.100E-07 | 2.900E-07 |
| 14433 | 722 | Q722M | B | 1.4e-07 | 3.9e-07 | 2.5e-07 | 2.600E-07 | 1.253E-07 | 1.400E-07 | 3.900E-07 |
| 14434 | 722 | Q722F | B | 1.3e-07 | 1.9e-07 | 4e-07 | 2.400E-07 | 1.418E-07 | 1.300E-07 | 4.000E-07 |
| 14435 | 722 | Q722P | B | 2.5e-07 | 3.8e-07 | 1.9e-07 | 2.733E-07 | 9.713E-08 | 1.900E-07 | 3.800E-07 |
| 14436 | 722 | Q722S | B | 3.7e-07 | 2.9e-07 | 1.4e-07 | 2.667E-07 | 1.168E-07 | 1.400E-07 | 3.700E-07 |
| 14437 | 722 | Q722T | B | 2.2e-07 | 1.5e-07 | 2.2e-07 | 1.967E-07 | 4.041E-08 | 1.500E-07 | 2.200E-07 |
| 14438 | 722 | Q722W | B | 2.7e-07 | 3.2e-07 | 5.9e-07 | 3.933E-07 | 1.721E-07 | 2.700E-07 | 5.900E-07 |
| 14439 | 722 | Q722Y | B | 3.5e-07 | 2.2e-07 | 6.3e-07 | 4.000E-07 | 2.095E-07 | 2.200E-07 | 6.300E-07 |
| 14440 | 722 | Q722V | B | 5.5e-07 | 3.5e-07 | 3.1e-07 | 4.033E-07 | 1.286E-07 | 3.100E-07 | 5.500E-07 |
| 14441 | 723 | G723A | B | 1.5e-07 | 2.1e-07 | 3.1e-07 | 2.233E-07 | 8.083E-08 | 1.500E-07 | 3.100E-07 |
| 14442 | 723 | G723R | B | 6.6e-08 | 2.2e-07 | 1.6e-07 | 1.487E-07 | 7.762E-08 | 6.600E-08 | 2.200E-07 |
| 14443 | 723 | G723N | B | 1.9e-07 | 3.4e-07 | 1.5e-07 | 2.267E-07 | 1.002E-07 | 1.500E-07 | 3.400E-07 |
| 14444 | 723 | G723D | B | 2e-07 | 1.8e-07 | 5.6e-07 | 3.133E-07 | 2.139E-07 | 1.800E-07 | 5.600E-07 |
| 14445 | 723 | G723C | B | 2e-07 | 2.6e-07 | 2.2e-07 | 2.267E-07 | 3.055E-08 | 2.000E-07 | 2.600E-07 |
| 14446 | 723 | G723E | B | 2.5e-07 | 2.4e-07 | 1.7e-07 | 2.200E-07 | 4.359E-08 | 1.700E-07 | 2.500E-07 |
| 14447 | 723 | G723Q | B | 3.1e-07 | 4.9e-07 | 2e-07 | 3.333E-07 | 1.464E-07 | 2.000E-07 | 4.900E-07 |
| 14448 | 723 | G723G | B | 1.5e-07 | 3.2e-07 | 3.5e-07 | 2.733E-07 | 1.079E-07 | 1.500E-07 | 3.500E-07 |
| 14449 | 723 | G723H | B | 4.4e-07 | 3.3e-07 | 2e-07 | 3.233E-07 | 1.201E-07 | 2.000E-07 | 4.400E-07 |

| | | | | | | | | | | |
|---|---|---|---|---|---|---|---|---|---|---|
| 14450 | 723 | G723I | B | 1.6e-07 | 2.5e-07 | 2.4e-07 | 2.167E-07 | 4.933E-08 | 1.600E-07 | 2.500E-07 |
| 14451 | 723 | G723L | B | 2.2e-07 | 1.5e-07 | 1.6e-07 | 1.767E-07 | 3.786E-08 | 1.500E-07 | 2.200E-07 |
| 14452 | 723 | G723K | B | 2.9e-07 | 4e-07 | 2.1e-07 | 3.000E-07 | 9.539E-08 | 2.100E-07 | 4.000E-07 |
| 14453 | 723 | G723M | B | 2.4e-07 | 1.3e-07 | 1.8e-07 | 1.833E-07 | 5.508E-08 | 1.300E-07 | 2.400E-07 |
| 14454 | 723 | G723F | B | 1.8e-07 | 3e-07 | 3.5e-07 | 2.767E-07 | 8.737E-08 | 1.800E-07 | 3.500E-07 |
| 14455 | 723 | G723P | B | 2.9e-07 | 2.7e-07 | 2e-07 | 2.533E-07 | 4.726E-08 | 2.000E-07 | 2.900E-07 |
| 14456 | 723 | G723S | B | 3.4e-07 | 1.8e-07 | 1.9e-07 | 2.367E-07 | 8.963E-08 | 1.800E-07 | 3.400E-07 |
| 14457 | 723 | G723T | B | 3.3e-07 | 2.4e-07 | 2.7e-07 | 2.800E-07 | 4.583E-08 | 2.400E-07 | 3.300E-07 |
| 14458 | 723 | G723W | B | 2.7e-07 | 2.5e-07 | 2.5e-07 | 2.567E-07 | 1.155E-08 | 2.500E-07 | 2.700E-07 |
| 14459 | 723 | G723Y | B | 2.5e-07 | 2e-07 | 3.4e-07 | 2.633E-07 | 7.095E-08 | 2.000E-07 | 3.400E-07 |
| 14460 | 723 | G723V | B | 3e-07 | 1.6e-07 | 2.1e-07 | 2.233E-07 | 7.095E-08 | 1.600E-07 | 3.000E-07 |
| 14461 | 724 | T724A | B | 4.6e-07 | 1.1e-07 | 3.1e-07 | 2.933E-07 | 1.756E-07 | 1.100E-07 | 4.600E-07 |
| 14462 | 724 | T724R | B | 1.7e-07 | 1.6e-07 | 3.7e-07 | 2.333E-07 | 1.185E-07 | 1.600E-07 | 3.700E-07 |
| 14463 | 724 | T724N | B | 1.6e-07 | 1.8e-07 | 2.4e-07 | 1.933E-07 | 4.163E-08 | 1.600E-07 | 2.400E-07 |
| 14464 | 724 | T724D | B | 1.7e-07 | 5.9e-07 | 3.5e-07 | 3.700E-07 | 2.107E-07 | 1.700E-07 | 5.900E-07 |
| 14465 | 724 | T724C | B | 1.2e-07 | 2.4e-07 | 2.4e-07 | 2.000E-07 | 6.928E-08 | 1.200E-07 | 2.400E-07 |
| 14466 | 724 | T724E | B | 3.4e-07 | 1.5e-07 | 1e-07 | 1.967E-07 | 1.266E-07 | 1.000E-07 | 3.400E-07 |
| 14467 | 724 | T724Q | B | 1.2e-07 | 3e-07 | 1.6e-07 | 1.933E-07 | 9.452E-08 | 1.200E-07 | 3.000E-07 |
| 14468 | 724 | T724G | B | 1.8e-07 | 5e-07 | 8.1e-08 | 2.537E-07 | 2.190E-07 | 8.100E-08 | 5.000E-07 |
| 14469 | 724 | T724H | B | 1.1e-07 | 2.2e-07 | 4.1e-07 | 2.467E-07 | 1.518E-07 | 1.100E-07 | 4.100E-07 |
| 14470 | 724 | T724I | B | 2.3e-07 | 2.3e-07 | 1.6e-07 | 2.067E-07 | 4.041E-08 | 1.600E-07 | 2.300E-07 |
| 14471 | 724 | T724L | B | 2.3e-07 | 1.7e-07 | 2e-07 | 2.000E-07 | 3.000E-08 | 1.700E-07 | 2.300E-07 |
| 14472 | 724 | T724K | B | 1.3e-07 | 1.6e-07 | 2.8e-07 | 1.900E-07 | 7.937E-08 | 1.300E-07 | 2.800E-07 |
| 14473 | 724 | T724M | B | 3.1e-07 | 1.2e-07 | 2.3e-07 | 2.200E-07 | 9.539E-08 | 1.200E-07 | 3.100E-07 |
| 14474 | 724 | T724F | B | 1.6e-07 | 2.2e-07 | 1.3e-07 | 1.700E-07 | 4.583E-08 | 1.300E-07 | 2.200E-07 |
| 14475 | 724 | T724P | B | 1.6e-07 | 3.5e-07 | 9.1e-08 | 2.003E-07 | 1.341E-07 | 9.100E-08 | 3.500E-07 |
| 14476 | 724 | T724S | B | 2.4e-07 | 2.1e-07 | 3.6e-07 | 2.700E-07 | 7.937E-08 | 2.100E-07 | 3.600E-07 |
| 14477 | 724 | T724T | B | 2.6e-07 | 1.8e-07 | 3.8e-07 | 2.733E-07 | 1.007E-07 | 1.800E-07 | 3.800E-07 |
| 14478 | 724 | T724W | B | 3.7e-07 | 4.6e-07 | 7.4e-07 | 5.233E-07 | 1.930E-07 | 3.700E-07 | 7.400E-07 |
| 14479 | 724 | T724Y | B | 5.8e-07 | 4.2e-07 | 3.4e-07 | 4.467E-07 | 1.222E-07 | 3.400E-07 | 5.800E-07 |
| 14480 | 724 | T724V | B | 3.6e-07 | 1.7e-07 | 3.4e-07 | 2.900E-07 | 1.044E-07 | 1.700E-07 | 3.600E-07 |
| 14481 | 725 | K725A | B | 3e-07 | 2.4e-07 | 2.8e-07 | 2.733E-07 | 3.055E-08 | 2.400E-07 | 3.000E-07 |
| 14482 | 725 | K725R | B | 1.8e-07 | 1.9e-07 | 2.4e-07 | 2.033E-07 | 3.215E-08 | 1.800E-07 | 2.400E-07 |
| 14483 | 725 | K725N | B | 2e-07 | 2.1e-07 | 2.7e-07 | 2.267E-07 | 3.786E-08 | 2.000E-07 | 2.700E-07 |
| 14484 | 725 | K725D | B | 7.9e-08 | 4.1e-07 | 1.6e-07 | 2.163E-07 | 1.725E-07 | 7.900E-08 | 4.100E-07 |
| 14485 | 725 | K725C | B | 1.7e-07 | 2.4e-07 | 2.4e-07 | 2.167E-07 | 4.041E-08 | 1.700E-07 | 2.400E-07 |
| 14486 | 725 | K725E | B | 3e-07 | 1.4e-07 | 2.7e-07 | 2.367E-07 | 8.505E-08 | 1.400E-07 | 3.000E-07 |
| 14487 | 725 | K725Q | B | 1.2e-07 | 9.7e-08 | 5.7e-07 | 2.623E-07 | 2.667E-07 | 9.700E-08 | 5.700E-07 |
| 14488 | 725 | K725G | B | 3.3e-07 | 2.2e-07 | 1.6e-07 | 2.367E-07 | 8.622E-08 | 1.600E-07 | 3.300E-07 |
| 14489 | 725 | K725H | B | 1.6e-07 | 1.9e-07 | 5.5e-07 | 3.000E-07 | 2.170E-07 | 1.600E-07 | 5.500E-07 |
| 14490 | 725 | K725I | B | 2.7e-07 | 2.7e-07 | 3.6e-07 | 3.000E-07 | 5.196E-08 | 2.700E-07 | 3.600E-07 |
| 14491 | 725 | K725L | B | 1.4e-07 | 1.5e-07 | 1.7e-07 | 1.533E-07 | 1.528E-08 | 1.400E-07 | 1.700E-07 |
| 14492 | 725 | K725K | B | 1.3e-07 | 1.9e-07 | 1.9e-07 | 1.700E-07 | 3.464E-08 | 1.300E-07 | 1.900E-07 |
| 14493 | 725 | K725M | B | 2.3e-07 | 3.7e-07 | 4.2e-07 | 3.400E-07 | 9.849E-08 | 2.300E-07 | 4.200E-07 |
| 14494 | 725 | K725F | B | 2e-07 | 1.5e-07 | 2.5e-07 | 2.000E-07 | 5.000E-08 | 1.500E-07 | 2.500E-07 |
| 14495 | 725 | K725P | B | 1.8e-07 | 1.6e-07 | 2.7e-07 | 2.033E-07 | 5.859E-08 | 1.600E-07 | 2.700E-07 |
| 14496 | 725 | K725S | B | 3e-07 | 2.6e-07 | 2.5e-07 | 2.700E-07 | 2.646E-08 | 2.500E-07 | 3.000E-07 |
| 14497 | 725 | K725T | B | 2.2e-07 | 1.9e-07 | 3.3e-07 | 2.467E-07 | 7.371E-08 | 1.900E-07 | 3.300E-07 |
| 14498 | 725 | K725W | B | 2.6e-07 | 1.6e-07 | 3.5e-07 | 2.567E-07 | 9.504E-08 | 1.600E-07 | 3.500E-07 |
| 14499 | 725 | K725Y | B | 2.4e-07 | 2.7e-07 | 4.7e-07 | 3.267E-07 | 1.250E-07 | 2.400E-07 | 4.700E-07 |
| 14500 | 725 | K725V | B | 1.4e-07 | 3.1e-07 | 2.8e-07 | 2.433E-07 | 9.074E-08 | 1.400E-07 | 3.100E-07 |
| 14501 | 726 | V726A | B | 2.5e-07 | 2.2e-07 | 1.7e-07 | 2.133E-07 | 4.041E-08 | 1.700E-07 | 2.500E-07 |
| 14502 | 726 | V726R | B | 2.2e-07 | 1.8e-07 | 1.2e-07 | 1.733E-07 | 5.033E-08 | 1.200E-07 | 2.200E-07 |
| 14503 | 726 | V726N | B | 4.2e-07 | 2.9e-07 | 1e-07 | 2.700E-07 | 1.609E-07 | 1.000E-07 | 4.200E-07 |
| 14504 | 726 | V726D | B | 3.4e-07 | 7.5e-07 | 2e-07 | 4.300E-07 | 2.858E-07 | 2.000E-07 | 7.500E-07 |
| 14505 | 726 | V726C | B | 2.4e-07 | 1.4e-07 | 2.6e-07 | 2.133E-07 | 6.429E-08 | 1.400E-07 | 2.600E-07 |
| 14506 | 726 | V726E | B | 2.7e-07 | 2.1e-07 | 2.1e-07 | 2.300E-07 | 3.464E-08 | 2.100E-07 | 2.700E-07 |
| 14507 | 726 | V726Q | B | 2.6e-07 | 1.6e-07 | 2e-07 | 2.067E-07 | 5.033E-08 | 1.600E-07 | 2.600E-07 |
| 14508 | 726 | V726G | B | 3.2e-07 | 1.8e-07 | 2.1e-07 | 2.367E-07 | 7.371E-08 | 1.800E-07 | 3.200E-07 |
| 14509 | 726 | V726H | B | 3e-07 | 1.9e-07 | 9.7e-08 | 1.957E-07 | 1.016E-07 | 9.700E-08 | 3.000E-07 |
| 14510 | 726 | V726I | B | 3.1e-07 | 6.5e-07 | 2.5e-07 | 4.033E-07 | 2.157E-07 | 2.500E-07 | 6.500E-07 |
| 14511 | 726 | V726L | B | 1.8e-07 | 2.2e-07 | 2.7e-07 | 2.233E-07 | 4.509E-08 | 1.800E-07 | 2.700E-07 |
| 14512 | 726 | V726K | B | 2.3e-07 | 3.9e-07 | 4.8e-07 | 3.667E-07 | 1.266E-07 | 2.300E-07 | 4.800E-07 |
| 14513 | 726 | V726M | B | 4.9e-07 | 2.3e-07 | 2.1e-07 | 3.100E-07 | 1.562E-07 | 2.100E-07 | 4.900E-07 |
| 14514 | 726 | V726F | B | 2.4e-07 | 1.5e-07 | 2.7e-07 | 2.200E-07 | 6.245E-08 | 1.500E-07 | 2.700E-07 |
| 14515 | 726 | V726P | B | 2.8e-07 | 1.2e-07 | 2e-07 | 2.000E-07 | 8.000E-08 | 1.200E-07 | 2.800E-07 |
| 14516 | 726 | V726S | B | 5.7e-07 | 2.6e-07 | 2.2e-07 | 3.500E-07 | 1.916E-07 | 2.200E-07 | 5.700E-07 |
| 14517 | 726 | V726T | B | 3.9e-07 | 1.6e-07 | 1.9e-07 | 2.467E-07 | 1.250E-07 | 1.600E-07 | 3.900E-07 |
| 14518 | 726 | V726W | B | 1.8e-07 | 5.6e-07 | 8.8e-08 | 2.760E-07 | 2.502E-07 | 8.800E-08 | 5.600E-07 |
| 14519 | 726 | V726Y | B | 1.5e-07 | 1.2e-07 | 1.9e-07 | 1.533E-07 | 3.512E-08 | 1.200E-07 | 1.900E-07 |
| 14520 | 726 | V726V | B | 1e-07 | 1.4e-07 | 2.7e-07 | 1.700E-07 | 8.888E-08 | 1.000E-07 | 2.700E-07 |
| 14521 | 727 | E727A | B | 1.2e-07 | 2.3e-07 | 1.9e-07 | 1.800E-07 | 5.568E-08 | 1.200E-07 | 2.300E-07 |
| 14522 | 727 | E727R | B | 3.7e-07 | 1.2e-07 | 3.2e-07 | 2.700E-07 | 1.323E-07 | 1.200E-07 | 3.700E-07 |
| 14523 | 727 | E727N | B | 1.8e-07 | 2.3e-07 | 5.4e-07 | 3.167E-07 | 1.950E-07 | 1.800E-07 | 5.400E-07 |
| 14524 | 727 | E727D | B | 2.3e-07 | 3.8e-07 | 3.2e-07 | 3.100E-07 | 7.550E-08 | 2.300E-07 | 3.800E-07 |
| 14525 | 727 | E727C | B | 1.4e-07 | 3e-07 | 2.2e-07 | 2.200E-07 | 8.000E-08 | 1.400E-07 | 3.000E-07 |
| 14526 | 727 | E727E | B | 1.5e-07 | 1.7e-07 | 3.3e-07 | 2.167E-07 | 9.866E-08 | 1.500E-07 | 3.300E-07 |
| 14527 | 727 | E727Q | B | 3.6e-07 | 5e-07 | 2.4e-07 | 3.667E-07 | 1.301E-07 | 2.400E-07 | 5.000E-07 |
| 14528 | 727 | E727G | B | 3.3e-07 | 2.9e-07 | 2.2e-07 | 2.800E-07 | 5.568E-08 | 2.200E-07 | 3.300E-07 |
| 14529 | 727 | E727H | B | 2.5e-07 | 1.7e-07 | 1.4e-07 | 1.867E-07 | 5.686E-08 | 1.400E-07 | 2.500E-07 |
| 14530 | 727 | E727I | B | 4e-07 | 2.5e-07 | 3.4e-07 | 3.300E-07 | 7.550E-08 | 2.500E-07 | 4.000E-07 |
| 14531 | 727 | E727L | B | 2.7e-07 | 2.7e-07 | 2.9e-07 | 2.767E-07 | 1.155E-08 | 2.700E-07 | 2.900E-07 |
| 14532 | 727 | E727K | B | 3.3e-07 | 1.8e-07 | 3.5e-07 | 2.867E-07 | 9.292E-08 | 1.800E-07 | 3.500E-07 |
| 14533 | 727 | E727M | B | 2.3e-07 | 2.6e-07 | 2e-07 | 2.300E-07 | 3.000E-08 | 2.000E-07 | 2.600E-07 |
| 14534 | 727 | E727F | B | 2.5e-07 | 1.2e-07 | 2.2e-07 | 1.967E-07 | 6.807E-08 | 1.200E-07 | 2.500E-07 |

| | | | | | | | | | | |
|---|---|---|---|---|---|---|---|---|---|---|
| 14535 | 727 | E727P | B | 1.7e-07 | 2.7e-07 | 2.3e-07 | 2.233E-07 | 5.033E-08 | 1.700E-07 | 2.700E-07 |
| 14536 | 727 | E727S | B | 1.3e-07 | 3.3e-07 | 2e-07 | 2.200E-07 | 1.015E-07 | 1.300E-07 | 3.300E-07 |
| 14537 | 727 | E727T | B | 2.7e-07 | 9.8e-08 | 2.7e-07 | 2.127E-07 | 9.930E-08 | 9.800E-08 | 2.700E-07 |
| 14538 | 727 | E727W | B | 5.4e-07 | 3.3e-07 | 2.6e-07 | 3.767E-07 | 1.457E-07 | 2.600E-07 | 5.400E-07 |
| 14539 | 727 | E727Y | B | 3.2e-07 | 1.7e-07 | 3e-07 | 2.633E-07 | 8.145E-08 | 1.700E-07 | 3.200E-07 |
| 14540 | 727 | E727V | B | 5.3e-07 | 1.9e-07 | 3.8e-07 | 3.667E-07 | 1.704E-07 | 1.900E-07 | 5.300E-07 |
| 14541 | 728 | I728A | B | 1.6e-07 | 3.3e-07 | 8.5e-08 | 1.917E-07 | 1.255E-07 | 8.500E-08 | 3.300E-07 |
| 14542 | 728 | I728R | B | 1.7e-07 | 2.7e-07 | 3.4e-07 | 2.600E-07 | 8.544E-08 | 1.700E-07 | 3.400E-07 |
| 14543 | 728 | I728N | B | 2.1e-07 | 1.7e-07 | 4.3e-07 | 2.700E-07 | 1.400E-07 | 1.700E-07 | 4.300E-07 |
| 14544 | 728 | I728D | B | 3.4e-07 | 3e-07 | 4e-07 | 3.467E-07 | 5.033E-08 | 3.000E-07 | 4.000E-07 |
| 14545 | 728 | I728C | B | 2.8e-07 | 2.8e-07 | 1.7e-07 | 2.433E-07 | 6.351E-08 | 1.700E-07 | 2.800E-07 |
| 14546 | 728 | I728E | B | 1.7e-07 | 5.5e-07 | 9.1e-08 | 2.703E-07 | 2.454E-07 | 9.100E-08 | 5.500E-07 |
| 14547 | 728 | I728Q | B | 2.4e-07 | 1.5e-07 | 3.6e-07 | 2.500E-07 | 1.054E-07 | 1.500E-07 | 3.600E-07 |
| 14548 | 728 | I728G | B | 3.8e-07 | 3.1e-07 | 2.2e-07 | 3.033E-07 | 8.021E-08 | 2.200E-07 | 3.800E-07 |
| 14549 | 728 | I728H | B | 2.8e-07 | 3e-07 | 8e-08 | 2.200E-07 | 1.217E-07 | 8.000E-08 | 3.000E-07 |
| 14550 | 728 | I728I | B | 2.1e-07 | 2.3e-07 | 2.1e-07 | 2.167E-07 | 1.155E-08 | 2.100E-07 | 2.300E-07 |
| 14551 | 728 | I728L | B | 1.8e-07 | 3.8e-07 | 1.8e-07 | 2.467E-07 | 1.155E-07 | 1.800E-07 | 3.800E-07 |
| 14552 | 728 | I728K | B | 2.2e-07 | 2e-07 | 2.2e-07 | 2.133E-07 | 1.155E-08 | 2.000E-07 | 2.200E-07 |
| 14553 | 728 | I728M | B | 4.4e-07 | 1.5e-07 | 4.6e-07 | 3.500E-07 | 1.735E-07 | 1.500E-07 | 4.600E-07 |
| 14554 | 728 | I728F | B | 3.8e-07 | 2.7e-07 | 3.3e-07 | 3.267E-07 | 5.508E-08 | 2.700E-07 | 3.800E-07 |
| 14555 | 728 | I728P | B | 2.5e-07 | 2.1e-07 | 2.4e-07 | 2.333E-07 | 2.082E-08 | 2.100E-07 | 2.500E-07 |
| 14556 | 728 | I728S | B | 3.5e-07 | 1.3e-07 | 1.5e-07 | 2.100E-07 | 1.217E-07 | 1.300E-07 | 3.500E-07 |
| 14557 | 728 | I728T | B | 1.5e-07 | 1.6e-07 | 3.2e-07 | 2.100E-07 | 9.539E-08 | 1.500E-07 | 3.200E-07 |
| 14558 | 728 | I728W | B | 1.3e-07 | 3e-07 | 3.7e-07 | 2.667E-07 | 1.234E-07 | 1.300E-07 | 3.700E-07 |
| 14559 | 728 | I728Y | B | 2.8e-07 | 4.2e-07 | 3.6e-07 | 3.533E-07 | 7.024E-08 | 2.800E-07 | 4.200E-07 |
| 14560 | 728 | I728V | B | 9.9e-08 | 3.6e-07 | 1.8e-07 | 2.130E-07 | 1.336E-07 | 9.900E-08 | 3.600E-07 |
| 14561 | 729 | K729A | B | 2.4e-07 | 1.9e-07 | 1.8e-07 | 2.033E-07 | 3.215E-08 | 1.800E-07 | 2.400E-07 |
| 14562 | 729 | K729R | B | 1.6e-07 | 2.9e-07 | 1.7e-07 | 2.067E-07 | 7.234E-08 | 1.600E-07 | 2.900E-07 |
| 14563 | 729 | K729N | B | 2.6e-07 | 4.9e-07 | 5.4e-07 | 4.300E-07 | 1.493E-07 | 2.600E-07 | 5.400E-07 |
| 14564 | 729 | K729D | B | 4e-07 | 1.6e-07 | 2e-07 | 2.533E-07 | 1.286E-07 | 1.600E-07 | 4.000E-07 |
| 14565 | 729 | K729C | B | 4.5e-07 | 2.4e-07 | 8e-08 | 2.567E-07 | 1.856E-07 | 8.000E-08 | 4.500E-07 |
| 14566 | 729 | K729E | B | 9e-08 | 2.6e-07 | 2.5e-07 | 2.000E-07 | 9.539E-08 | 9.000E-08 | 2.600E-07 |
| 14567 | 729 | K729Q | B | 2.2e-07 | 3.2e-07 | 5.2e-07 | 3.533E-07 | 1.528E-07 | 2.200E-07 | 5.200E-07 |
| 14568 | 729 | K729G | B | 1e-07 | 1.9e-07 | 1.4e-07 | 1.433E-07 | 4.509E-08 | 1.000E-07 | 1.900E-07 |
| 14569 | 729 | K729H | B | 5.1e-07 | 2.2e-07 | 2.2e-07 | 3.167E-07 | 1.674E-07 | 2.200E-07 | 5.100E-07 |
| 14570 | 729 | K729I | B | 2.8e-07 | 7.3e-07 | 2.9e-07 | 4.333E-07 | 2.570E-07 | 2.800E-07 | 7.300E-07 |
| 14571 | 729 | K729L | B | 2.8e-07 | 1.3e-07 | 3.2e-07 | 2.433E-07 | 1.002E-07 | 1.300E-07 | 3.200E-07 |
| 14572 | 729 | K729K | B | 2.2e-07 | 1.6e-07 | 3.5e-07 | 2.433E-07 | 9.713E-08 | 1.600E-07 | 3.500E-07 |
| 14573 | 729 | K729M | B | 3.3e-07 | 2.2e-07 | 1.9e-07 | 2.467E-07 | 7.371E-08 | 1.900E-07 | 3.300E-07 |
| 14574 | 729 | K729F | B | 1.4e-07 | 2.3e-07 | 4.2e-07 | 2.633E-07 | 1.429E-07 | 1.400E-07 | 4.200E-07 |
| 14575 | 729 | K729P | B | 2.7e-07 | 2e-07 | 1.3e-07 | 2.000E-07 | 7.000E-08 | 1.300E-07 | 2.700E-07 |
| 14576 | 729 | K729S | B | 4.8e-07 | 1e-07 | 3.2e-07 | 3.000E-07 | 1.908E-07 | 1.000E-07 | 4.800E-07 |
| 14577 | 729 | K729T | B | 1.6e-07 | 2.9e-07 | 3.3e-07 | 2.600E-07 | 8.888E-08 | 1.600E-07 | 3.300E-07 |
| 14578 | 729 | K729W | B | 3.7e-07 | 2.6e-07 | 1.4e-07 | 2.567E-07 | 1.150E-07 | 1.400E-07 | 3.700E-07 |
| 14579 | 729 | K729Y | B | 2.8e-07 | 1.8e-07 | 1.6e-07 | 2.067E-07 | 6.429E-08 | 1.600E-07 | 2.800E-07 |
| 14580 | 729 | K729V | B | 2.6e-07 | 1.6e-07 | 1.8e-07 | 2.000E-07 | 5.292E-08 | 1.600E-07 | 2.600E-07 |
| 14581 | 730 | R730A | B | 2.2e-07 | 2.5e-07 | 4.3e-07 | 3.000E-07 | 1.136E-07 | 2.200E-07 | 4.300E-07 |
| 14582 | 730 | R730R | B | 1.7e-07 | 9.2e-08 | 2e-07 | 1.540E-07 | 5.575E-08 | 9.200E-08 | 2.000E-07 |
| 14583 | 730 | R730N | B | 3.4e-07 | 1.3e-07 | 3.6e-07 | 2.767E-07 | 1.274E-07 | 1.300E-07 | 3.600E-07 |
| 14584 | 730 | R730D | B | 2.4e-07 | 2.1e-07 | 2.7e-07 | 2.400E-07 | 3.000E-08 | 2.100E-07 | 2.700E-07 |
| 14585 | 730 | R730C | B | 2.3e-07 | 4.1e-07 | 1.8e-07 | 2.733E-07 | 1.210E-07 | 1.800E-07 | 4.100E-07 |
| 14586 | 730 | R730E | B | 2e-07 | 1.2e-07 | 1.3e-07 | 1.500E-07 | 4.359E-08 | 1.200E-07 | 2.000E-07 |
| 14587 | 730 | R730Q | B | 4.8e-07 | 1.7e-07 | 1.5e-07 | 2.667E-07 | 1.850E-07 | 1.500E-07 | 4.800E-07 |
| 14588 | 730 | R730G | B | 3.5e-07 | 1.7e-07 | 1.7e-07 | 2.300E-07 | 1.039E-07 | 1.700E-07 | 3.500E-07 |
| 14589 | 730 | R730H | B | 3.8e-07 | 4.3e-07 | 9.8e-08 | 3.027E-07 | 1.790E-07 | 9.800E-08 | 4.300E-07 |
| 14590 | 730 | R730I | B | 3.6e-07 | 1.5e-07 | 4.1e-07 | 3.067E-07 | 1.380E-07 | 1.500E-07 | 4.100E-07 |
| 14591 | 730 | R730L | B | 3.9e-07 | 2.4e-07 | 2.3e-07 | 2.867E-07 | 8.963E-08 | 2.300E-07 | 3.900E-07 |
| 14592 | 730 | R730K | B | 1.4e-07 | 3.6e-07 | 5e-07 | 3.333E-07 | 1.815E-07 | 1.400E-07 | 5.000E-07 |
| 14593 | 730 | R730M | B | 1.5e-07 | 3e-07 | 2.3e-07 | 2.267E-07 | 7.506E-08 | 1.500E-07 | 3.000E-07 |
| 14594 | 730 | R730F | B | 1.9e-07 | 1.4e-07 | 3.4e-07 | 2.233E-07 | 1.041E-07 | 1.400E-07 | 3.400E-07 |
| 14595 | 730 | R730P | B | 1.5e-07 | 1.8e-07 | 2.6e-07 | 1.967E-07 | 5.686E-08 | 1.500E-07 | 2.600E-07 |
| 14596 | 730 | R730S | B | 3e-07 | 3.3e-07 | 3.4e-07 | 3.233E-07 | 2.082E-08 | 3.000E-07 | 3.400E-07 |
| 14597 | 730 | R730T | B | 5.1e-07 | 3.7e-07 | 2.9e-07 | 3.900E-07 | 1.114E-07 | 2.900E-07 | 5.100E-07 |
| 14598 | 730 | R730W | B | 2.4e-07 | 3e-07 | 1.9e-07 | 2.433E-07 | 5.508E-08 | 1.900E-07 | 3.000E-07 |
| 14599 | 730 | R730Y | B | 7.3e-07 | 4.7e-07 | 2.4e-07 | 4.800E-07 | 2.452E-07 | 2.400E-07 | 7.300E-07 |
| 14600 | 730 | R730V | B | 3.5e-07 | 2.6e-07 | 6.1e-07 | 4.067E-07 | 1.818E-07 | 2.600E-07 | 6.100E-07 |
| 14601 | 731 | T731A | B | 5.4e-07 | 2.3e-07 | 4.1e-07 | 3.933E-07 | 1.557E-07 | 2.300E-07 | 5.400E-07 |
| 14602 | 731 | T731R | B | 3.1e-07 | 2.8e-07 | 2.4e-07 | 2.767E-07 | 3.512E-08 | 2.400E-07 | 3.100E-07 |
| 14603 | 731 | T731N | B | 9.3e-08 | 3.9e-07 | 1.7e-07 | 2.177E-07 | 1.541E-07 | 9.300E-08 | 3.900E-07 |
| 14604 | 731 | T731D | B | 3.9e-07 | 2.5e-07 | 2e-07 | 2.800E-07 | 9.849E-08 | 2.000E-07 | 3.900E-07 |
| 14605 | 731 | T731C | B | 8.6e-08 | 3e-07 | 1.9e-07 | 1.920E-07 | 1.070E-07 | 8.600E-08 | 3.000E-07 |
| 14606 | 731 | T731E | B | 3.9e-07 | 3.3e-07 | 2.2e-07 | 3.133E-07 | 8.622E-08 | 2.200E-07 | 3.900E-07 |
| 14607 | 731 | T731Q | B | 1.2e-07 | 1.7e-07 | 2.1e-07 | 1.667E-07 | 4.509E-08 | 1.200E-07 | 2.100E-07 |
| 14608 | 731 | T731G | B | 9.9e-08 | 1.9e-07 | 2.4e-07 | 1.763E-07 | 7.149E-08 | 9.900E-08 | 2.400E-07 |
| 14609 | 731 | T731H | B | 1.4e-07 | 1.8e-07 | 1.1e-07 | 1.433E-07 | 3.512E-08 | 1.100E-07 | 1.800E-07 |
| 14610 | 731 | T731I | B | 3.4e-07 | 1.8e-07 | 3.2e-07 | 2.800E-07 | 8.718E-08 | 1.800E-07 | 3.400E-07 |
| 14611 | 731 | T731L | B | 5.2e-07 | 3.1e-07 | 1.6e-07 | 3.300E-07 | 1.808E-07 | 1.600E-07 | 5.200E-07 |
| 14612 | 731 | T731K | B | 3.1e-07 | 3.8e-07 | 3.4e-07 | 3.433E-07 | 3.512E-08 | 3.100E-07 | 3.800E-07 |
| 14613 | 731 | T731M | B | 2.3e-07 | 1.5e-07 | 1.4e-07 | 1.733E-07 | 4.933E-08 | 1.400E-07 | 2.300E-07 |
| 14614 | 731 | T731F | B | 9.4e-08 | 2.5e-07 | 5.7e-07 | 3.047E-07 | 2.427E-07 | 9.400E-08 | 5.700E-07 |
| 14615 | 731 | T731P | B | 5.2e-07 | 3.8e-07 | 2.2e-07 | 3.733E-07 | 1.501E-07 | 2.200E-07 | 5.200E-07 |
| 14616 | 731 | T731S | B | 1.3e-07 | 1.2e-07 | 2e-07 | 1.500E-07 | 4.359E-08 | 1.200E-07 | 2.000E-07 |
| 14617 | 731 | T731T | B | 2.2e-07 | 2.1e-07 | 3.7e-07 | 2.667E-07 | 8.963E-08 | 2.100E-07 | 3.700E-07 |
| 14618 | 731 | T731W | B | 3.9e-07 | 2e-07 | 2.7e-07 | 2.867E-07 | 9.609E-08 | 2.000E-07 | 3.900E-07 |
| 14619 | 731 | T731Y | B | 2.7e-07 | 4.8e-07 | 3.1e-07 | 3.533E-07 | 1.115E-07 | 2.700E-07 | 4.800E-07 |

| | | | | | | | | | | |
|---|---|---|---|---|---|---|---|---|---|---|
| 14620 | 731 | T731V | B | 1e-07 | 2.3e-07 | 3.3e-07 | 2.200E-07 | 1.153E-07 | 1.000E-07 | 3.300E-07 |
| 14621 | 732 | V732A | B | 1.2e-07 | 3e-07 | 3.4e-07 | 2.533E-07 | 1.172E-07 | 1.200E-07 | 3.400E-07 |
| 14622 | 732 | V732R | B | 3.8e-07 | 3.4e-07 | 2.4e-07 | 3.200E-07 | 7.211E-08 | 2.400E-07 | 3.800E-07 |
| 14623 | 732 | V732N | B | 2e-07 | 2.2e-07 | 3.6e-07 | 2.600E-07 | 8.718E-08 | 2.000E-07 | 3.600E-07 |
| 14624 | 732 | V732D | B | 3.6e-07 | 1e-07 | 6.3e-07 | 3.633E-07 | 2.650E-07 | 1.000E-07 | 6.300E-07 |
| 14625 | 732 | V732C | B | 2.9e-07 | 3.3e-07 | 3.4e-07 | 3.200E-07 | 2.646E-08 | 2.900E-07 | 3.400E-07 |
| 14626 | 732 | V732E | B | 3e-07 | 2e-07 | 2.3e-07 | 2.433E-07 | 5.132E-08 | 2.000E-07 | 3.000E-07 |
| 14627 | 732 | V732Q | B | 2e-07 | 3.8e-07 | 2e-07 | 2.600E-07 | 1.039E-07 | 2.000E-07 | 3.800E-07 |
| 14628 | 732 | V732G | B | 1.2e-07 | 1e-07 | 2e-07 | 1.400E-07 | 5.292E-08 | 1.000E-07 | 2.000E-07 |
| 14629 | 732 | V732H | B | 1.4e-07 | 2.8e-07 | 3.2e-07 | 2.467E-07 | 9.452E-08 | 1.400E-07 | 3.200E-07 |
| 14630 | 732 | V732I | B | 1.7e-07 | 5.7e-07 | 3.6e-07 | 3.667E-07 | 2.001E-07 | 1.700E-07 | 5.700E-07 |
| 14631 | 732 | V732L | B | 2.1e-07 | 3.2e-07 | 2.2e-07 | 2.500E-07 | 6.083E-08 | 2.100E-07 | 3.200E-07 |
| 14632 | 732 | V732K | B | 2.2e-07 | 3.7e-07 | 2e-07 | 2.633E-07 | 9.292E-08 | 2.000E-07 | 3.700E-07 |
| 14633 | 732 | V732M | B | 4.3e-07 | 2.7e-07 | 2.8e-07 | 3.267E-07 | 8.963E-08 | 2.700E-07 | 4.300E-07 |
| 14634 | 732 | V732F | B | 1.4e-07 | 3.3e-07 | 1.3e-07 | 2.000E-07 | 1.127E-07 | 1.300E-07 | 3.300E-07 |
| 14635 | 732 | V732P | B | 2.7e-07 | 3.3e-07 | 4.8e-07 | 3.600E-07 | 1.082E-07 | 2.700E-07 | 4.800E-07 |
| 14636 | 732 | V732S | B | 1.8e-07 | 2.1e-07 | 2.6e-07 | 2.167E-07 | 4.041E-08 | 1.800E-07 | 2.600E-07 |
| 14637 | 732 | V732T | B | 1.8e-07 | 2.7e-07 | 2e-07 | 2.167E-07 | 4.726E-08 | 1.800E-07 | 2.700E-07 |
| 14638 | 732 | V732W | B | 1.3e-07 | 2.5e-07 | 1.2e-07 | 1.667E-07 | 7.234E-08 | 1.200E-07 | 2.500E-07 |
| 14639 | 732 | V732Y | B | 3.6e-07 | 2.7e-07 | 2.2e-07 | 2.833E-07 | 7.095E-08 | 2.200E-07 | 3.600E-07 |
| 14640 | 732 | V732V | B | 3.8e-07 | 9.5e-08 | 5e-07 | 3.250E-07 | 2.080E-07 | 9.500E-08 | 5.000E-07 |
| 14641 | 733 | A733A | B | 3e-07 | 2.3e-07 | 4e-07 | 3.100E-07 | 8.544E-08 | 2.300E-07 | 4.000E-07 |
| 14642 | 733 | A733R | B | 3e-07 | 1.7e-07 | 1.5e-07 | 2.067E-07 | 8.145E-08 | 1.500E-07 | 3.000E-07 |
| 14643 | 733 | A733N | B | 2.8e-07 | 1.4e-07 | 3.1e-07 | 2.433E-07 | 9.074E-08 | 1.400E-07 | 3.100E-07 |
| 14644 | 733 | A733D | B | 2.9e-07 | 6.7e-07 | 2.7e-07 | 4.100E-07 | 2.254E-07 | 2.700E-07 | 6.700E-07 |
| 14645 | 733 | A733C | B | 1e-07 | 2.6e-07 | 1.9e-07 | 1.833E-07 | 8.021E-08 | 1.000E-07 | 2.600E-07 |
| 14646 | 733 | A733E | B | 2.8e-07 | 1.4e-07 | 3.1e-07 | 2.433E-07 | 9.074E-08 | 1.400E-07 | 3.100E-07 |
| 14647 | 733 | A733Q | B | 1.2e-07 | 2.3e-07 | 4.3e-07 | 2.600E-07 | 1.572E-07 | 1.200E-07 | 4.300E-07 |
| 14648 | 733 | A733G | B | 3.2e-07 | 4.3e-07 | 2.6e-07 | 3.367E-07 | 8.622E-08 | 2.600E-07 | 4.300E-07 |
| 14649 | 733 | A733H | B | 1.7e-07 | 3.1e-07 | 1.1e-07 | 1.967E-07 | 1.026E-07 | 1.100E-07 | 3.100E-07 |
| 14650 | 733 | A733I | B | 2.4e-07 | 1.9e-07 | 6.6e-08 | 1.653E-07 | 8.958E-08 | 6.600E-08 | 2.400E-07 |
| 14651 | 733 | A733L | B | 2.4e-07 | 6.1e-07 | 2.4e-07 | 3.633E-07 | 2.136E-07 | 2.400E-07 | 6.100E-07 |
| 14652 | 733 | A733K | B | 3.5e-07 | 2e-07 | 3.9e-07 | 3.133E-07 | 1.002E-07 | 2.000E-07 | 3.900E-07 |
| 14653 | 733 | A733M | B | 2.4e-07 | 1.6e-07 | 2.5e-07 | 2.167E-07 | 4.933E-08 | 1.600E-07 | 2.500E-07 |
| 14654 | 733 | A733F | B | 3.9e-07 | 2e-07 | 5.5e-07 | 3.800E-07 | 1.752E-07 | 2.000E-07 | 5.500E-07 |
| 14655 | 733 | A733P | B | 3.8e-07 | 3.8e-07 | 1.6e-07 | 3.067E-07 | 1.270E-07 | 1.600E-07 | 3.800E-07 |
| 14656 | 733 | A733S | B | 1.9e-07 | 5.6e-07 | 2.4e-07 | 3.300E-07 | 2.007E-07 | 1.900E-07 | 5.600E-07 |
| 14657 | 733 | A733T | B | 2.8e-07 | 1.8e-07 | 1.2e-07 | 1.933E-07 | 8.083E-08 | 1.200E-07 | 2.800E-07 |
| 14658 | 733 | A733W | B | 3.1e-07 | 2.9e-07 | 3.4e-07 | 3.133E-07 | 2.517E-08 | 2.900E-07 | 3.400E-07 |
| 14659 | 733 | A733Y | B | 3.9e-07 | 2e-07 | 3.9e-07 | 3.267E-07 | 1.097E-07 | 2.000E-07 | 3.900E-07 |
| 14660 | 733 | A733V | B | 5.5e-07 | 3.6e-07 | 2.3e-07 | 3.800E-07 | 1.609E-07 | 2.300E-07 | 5.500E-07 |
| 14661 | 734 | A734A | B | 2e-07 | 1.9e-07 | 4.9e-07 | 2.933E-07 | 1.704E-07 | 1.900E-07 | 4.900E-07 |
| 14662 | 734 | A734R | B | 2.4e-07 | 1.9e-07 | 1.6e-07 | 1.967E-07 | 4.041E-08 | 1.600E-07 | 2.400E-07 |
| 14663 | 734 | A734N | B | 3.7e-07 | 3.4e-07 | 3.5e-07 | 3.533E-07 | 1.528E-08 | 3.400E-07 | 3.700E-07 |
| 14664 | 734 | A734D | B | 2.1e-07 | 3.2e-07 | 1.9e-07 | 2.400E-07 | 7.000E-08 | 1.900E-07 | 3.200E-07 |
| 14665 | 734 | A734C | B | 2.1e-07 | 4.1e-07 | 2e-07 | 2.733E-07 | 1.185E-07 | 2.000E-07 | 4.100E-07 |
| 14666 | 734 | A734E | B | 3.6e-07 | 4.6e-07 | 1.6e-07 | 3.267E-07 | 1.528E-07 | 1.600E-07 | 4.600E-07 |
| 14667 | 734 | A734Q | B | 4.9e-07 | 2e-07 | 1.6e-07 | 2.833E-07 | 1.801E-07 | 1.600E-07 | 4.900E-07 |
| 14668 | 734 | A734G | B | 4.2e-07 | 2.6e-07 | 3e-07 | 3.267E-07 | 8.327E-08 | 2.600E-07 | 4.200E-07 |
| 14669 | 734 | A734H | B | 1.8e-07 | 2e-07 | 2.2e-07 | 2.000E-07 | 2.000E-08 | 1.800E-07 | 2.200E-07 |
| 14670 | 734 | A734I | B | 2.2e-07 | 2.5e-07 | 1.5e-07 | 2.067E-07 | 5.132E-08 | 1.500E-07 | 2.500E-07 |
| 14671 | 734 | A734L | B | 1.8e-07 | 1.7e-07 | 2.6e-07 | 2.033E-07 | 4.933E-08 | 1.700E-07 | 2.600E-07 |
| 14672 | 734 | A734K | B | 1.7e-07 | 3.4e-07 | 4.5e-07 | 3.200E-07 | 1.411E-07 | 1.700E-07 | 4.500E-07 |
| 14673 | 734 | A734M | B | 6.1e-08 | 2.5e-07 | 3.1e-07 | 2.070E-07 | 1.299E-07 | 6.100E-08 | 3.100E-07 |
| 14674 | 734 | A734F | B | 2.5e-07 | 2.2e-07 | 9e-08 | 1.867E-07 | 8.505E-08 | 9.000E-08 | 2.500E-07 |
| 14675 | 734 | A734P | B | 2.3e-07 | 9.2e-08 | 1.2e-07 | 1.473E-07 | 7.295E-08 | 9.200E-08 | 2.300E-07 |
| 14676 | 734 | A734S | B | 3.7e-07 | 3.3e-07 | 1.6e-07 | 2.867E-07 | 1.115E-07 | 1.600E-07 | 3.700E-07 |
| 14677 | 734 | A734T | B | 3.2e-07 | 4.6e-07 | 2.2e-07 | 3.333E-07 | 1.206E-07 | 2.200E-07 | 4.600E-07 |
| 14678 | 734 | A734W | B | 1.9e-07 | 2.5e-07 | 3.9e-07 | 2.767E-07 | 1.026E-07 | 1.900E-07 | 3.900E-07 |
| 14679 | 734 | A734Y | B | 1.8e-07 | 4.7e-07 | 2e-07 | 2.833E-07 | 1.620E-07 | 1.800E-07 | 4.700E-07 |
| 14680 | 734 | A734V | B | 3.9e-07 | 2.6e-07 | 2e-07 | 2.833E-07 | 9.713E-08 | 2.000E-07 | 3.900E-07 |
| 14681 | 735 | P735A | B | 9.6e-07 | 4.1e-07 | 3.6e-07 | 5.767E-07 | 3.329E-07 | 3.600E-07 | 9.600E-07 |
| 14682 | 735 | P735R | B | 2.6e-07 | 3.6e-07 | 1.8e-07 | 2.667E-07 | 9.018E-08 | 1.800E-07 | 3.600E-07 |
| 14683 | 735 | P735N | B | 2e-07 | 3.9e-07 | 1.5e-07 | 2.467E-07 | 1.266E-07 | 1.500E-07 | 3.900E-07 |
| 14684 | 735 | P735D | B | 3.8e-07 | 8.2e-08 | 3.4e-07 | 2.673E-07 | 1.617E-07 | 8.200E-08 | 3.800E-07 |
| 14685 | 735 | P735C | B | 5.1e-07 | 4.6e-07 | 1.9e-07 | 3.867E-07 | 1.721E-07 | 1.900E-07 | 5.100E-07 |
| 14686 | 735 | P735E | B | 4e-07 | 1.6e-07 | 3.3e-07 | 2.967E-07 | 1.234E-07 | 1.600E-07 | 4.000E-07 |
| 14687 | 735 | P735Q | B | 1.4e-07 | 2.2e-07 | 4.1e-07 | 2.567E-07 | 1.387E-07 | 1.400E-07 | 4.100E-07 |
| 14688 | 735 | P735G | B | 2.1e-07 | 2.6e-07 | 3.6e-07 | 2.767E-07 | 7.638E-08 | 2.100E-07 | 3.600E-07 |
| 14689 | 735 | P735H | B | 2.1e-07 | 4.3e-07 | 2.5e-07 | 2.967E-07 | 1.172E-07 | 2.100E-07 | 4.300E-07 |
| 14690 | 735 | P735I | B | 2.6e-07 | 5.4e-07 | 2.3e-07 | 3.433E-07 | 1.710E-07 | 2.300E-07 | 5.400E-07 |
| 14691 | 735 | P735L | B | 1.4e-07 | 3.5e-07 | 1.5e-07 | 2.133E-07 | 1.185E-07 | 1.400E-07 | 3.500E-07 |
| 14692 | 735 | P735K | B | 1.7e-07 | 4.6e-07 | 3.8e-07 | 3.367E-07 | 1.498E-07 | 1.700E-07 | 4.600E-07 |
| 14693 | 735 | P735M | B | 1.2e-07 | 3e-07 | 1.6e-07 | 1.933E-07 | 9.452E-08 | 1.200E-07 | 3.000E-07 |
| 14694 | 735 | P735F | B | 2.6e-07 | 4.6e-07 | 1.1e-07 | 2.767E-07 | 1.756E-07 | 1.100E-07 | 4.600E-07 |
| 14695 | 735 | P735P | B | 1.7e-07 | 2.7e-07 | 2.3e-07 | 2.233E-07 | 5.033E-08 | 1.700E-07 | 2.700E-07 |
| 14696 | 735 | P735S | B | 5.1e-07 | 1.6e-07 | 2.7e-07 | 3.133E-07 | 1.790E-07 | 1.600E-07 | 5.100E-07 |
| 14697 | 735 | P735T | B | 4.4e-07 | 3.3e-07 | 1.2e-07 | 2.967E-07 | 1.626E-07 | 1.200E-07 | 4.400E-07 |
| 14698 | 735 | P735W | B | 2.2e-07 | 1.7e-07 | 2.8e-07 | 2.233E-07 | 5.508E-08 | 1.700E-07 | 2.800E-07 |
| 14699 | 735 | P735Y | B | 3e-07 | 4.2e-07 | 2.5e-07 | 3.233E-07 | 8.737E-08 | 2.500E-07 | 4.200E-07 |
| 14700 | 735 | P735V | B | 1.7e-07 | 4.4e-07 | 3e-07 | 3.033E-07 | 1.350E-07 | 1.700E-07 | 4.400E-07 |
| 14701 | 736 | S736A | B | 4.6e-07 | 4e-07 | 3.3e-07 | 3.967E-07 | 6.506E-08 | 3.300E-07 | 4.600E-07 |
| 14702 | 736 | S736R | B | 2.2e-07 | 2e-07 | 2.9e-07 | 2.367E-07 | 4.726E-08 | 2.000E-07 | 2.900E-07 |
| 14703 | 736 | S736N | B | 4.2e-07 | 2e-07 | 2.9e-07 | 3.033E-07 | 1.106E-07 | 2.000E-07 | 4.200E-07 |
| 14704 | 736 | S736D | B | 2.3e-07 | 3.6e-07 | 2.3e-07 | 2.733E-07 | 7.506E-08 | 2.300E-07 | 3.600E-07 |

| | | | | | | | | | | |
|---|---|---|---|---|---|---|---|---|---|---|
| 14705 | 736 | S736C | B | 2.1e-07 | 1.7e-07 | 1.7e-07 | 1.833E-07 | 2.309E-08 | 1.700E-07 | 2.100E-07 |
| 14706 | 736 | S736E | B | 2.1e-07 | 1.8e-07 | 6.8e-07 | 3.567E-07 | 2.804E-07 | 1.800E-07 | 6.800E-07 |
| 14707 | 736 | S736Q | B | 1.4e-07 | 2.5e-07 | 2.7e-07 | 2.200E-07 | 7.000E-08 | 1.400E-07 | 2.700E-07 |
| 14708 | 736 | S736G | B | 4.7e-07 | 2.7e-07 | 5.1e-07 | 4.167E-07 | 1.286E-07 | 2.700E-07 | 5.100E-07 |
| 14709 | 736 | S736H | B | 5.7e-07 | 2.4e-07 | 4.2e-07 | 4.100E-07 | 1.652E-07 | 2.400E-07 | 5.700E-07 |
| 14710 | 736 | S736I | B | 3.1e-07 | 2.5e-07 | 2e-07 | 2.533E-07 | 5.508E-08 | 2.000E-07 | 3.100E-07 |
| 14711 | 736 | S736L | B | 2.2e-07 | 2.6e-07 | 2.3e-07 | 2.367E-07 | 2.082E-08 | 2.200E-07 | 2.600E-07 |
| 14712 | 736 | S736K | B | 1.3e-07 | 4e-07 | 2.3e-07 | 2.533E-07 | 1.365E-07 | 1.300E-07 | 4.000E-07 |
| 14713 | 736 | S736M | B | 1.7e-07 | 4.2e-07 | 1.4e-07 | 2.433E-07 | 1.537E-07 | 1.400E-07 | 4.200E-07 |
| 14714 | 736 | S736F | B | 4.1e-07 | 1.2e-07 | 1.7e-07 | 2.333E-07 | 1.550E-07 | 1.200E-07 | 4.100E-07 |
| 14715 | 736 | S736P | B | 3e-07 | 2.5e-07 | 3.2e-07 | 2.900E-07 | 3.606E-08 | 2.500E-07 | 3.200E-07 |
| 14716 | 736 | S736S | B | 1.3e-07 | 4.9e-07 | 2.1e-07 | 2.767E-07 | 1.890E-07 | 1.300E-07 | 4.900E-07 |
| 14717 | 736 | S736T | B | 4.2e-07 | 1.5e-07 | 9.9e-08 | 2.230E-07 | 1.725E-07 | 9.900E-08 | 4.200E-07 |
| 14718 | 736 | S736W | B | 1.9e-07 | 2.8e-07 | 2.5e-07 | 2.400E-07 | 4.583E-08 | 1.900E-07 | 2.800E-07 |
| 14719 | 736 | S736Y | B | 4.1e-07 | 2.7e-07 | 1.4e-07 | 2.733E-07 | 1.350E-07 | 1.400E-07 | 4.100E-07 |
| 14720 | 736 | S736V | B | 2.4e-07 | 3.7e-07 | 3.5e-07 | 3.200E-07 | 7.000E-08 | 2.400E-07 | 3.700E-07 |
| 14721 | 737 | V737A | B | 2.8e-07 | 3e-07 | 2.8e-07 | 2.867E-07 | 1.155E-08 | 2.800E-07 | 3.000E-07 |
| 14722 | 737 | V737R | B | 1.5e-07 | 1.3e-07 | 4e-07 | 2.267E-07 | 1.504E-07 | 1.300E-07 | 4.000E-07 |
| 14723 | 737 | V737N | B | 3.3e-07 | 3.8e-07 | 2.7e-07 | 3.267E-07 | 5.508E-08 | 2.700E-07 | 3.800E-07 |
| 14724 | 737 | V737D | B | 2.2e-07 | 2.6e-07 | 1.3e-07 | 2.033E-07 | 6.658E-08 | 1.300E-07 | 2.600E-07 |
| 14725 | 737 | V737C | B | 2.1e-07 | 2.5e-07 | 1.4e-07 | 2.000E-07 | 5.568E-08 | 1.400E-07 | 2.500E-07 |
| 14726 | 737 | V737E | B | 1.1e-07 | 1.9e-07 | 2.4e-07 | 1.800E-07 | 6.557E-08 | 1.100E-07 | 2.400E-07 |
| 14727 | 737 | V737Q | B | 4.1e-07 | 4.6e-07 | 2.7e-07 | 3.800E-07 | 9.849E-08 | 2.700E-07 | 4.600E-07 |
| 14728 | 737 | V737G | B | 4.5e-07 | 1.5e-07 | 3.9e-07 | 3.300E-07 | 1.587E-07 | 1.500E-07 | 4.500E-07 |
| 14729 | 737 | V737H | B | 1.4e-07 | 2.6e-07 | 2.2e-07 | 2.067E-07 | 6.110E-08 | 1.400E-07 | 2.600E-07 |
| 14730 | 737 | V737I | B | 4.3e-07 | 3.2e-07 | 9.9e-08 | 2.830E-07 | 1.686E-07 | 9.900E-08 | 4.300E-07 |
| 14731 | 737 | V737L | B | 2.6e-07 | 3.9e-07 | 2.1e-07 | 2.867E-07 | 9.292E-08 | 2.100E-07 | 3.900E-07 |
| 14732 | 737 | V737K | B | 3.4e-07 | 2.8e-07 | 3e-07 | 3.067E-07 | 3.055E-08 | 2.800E-07 | 3.400E-07 |
| 14733 | 737 | V737M | B | 2.3e-07 | 3.1e-07 | 2.8e-07 | 2.733E-07 | 4.041E-08 | 2.300E-07 | 3.100E-07 |
| 14734 | 737 | V737F | B | 3e-07 | 2.9e-07 | 2.2e-07 | 2.700E-07 | 4.359E-08 | 2.200E-07 | 3.000E-07 |
| 14735 | 737 | V737P | B | 4.3e-07 | 3.8e-07 | 3.1e-07 | 3.733E-07 | 6.028E-08 | 3.100E-07 | 4.300E-07 |
| 14736 | 737 | V737S | B | 1.6e-07 | 2.4e-07 | 3.2e-07 | 2.400E-07 | 8.000E-08 | 1.600E-07 | 3.200E-07 |
| 14737 | 737 | V737T | B | 2.4e-07 | 2.7e-07 | 2.9e-07 | 2.667E-07 | 2.517E-08 | 2.400E-07 | 2.900E-07 |
| 14738 | 737 | V737W | B | 1.9e-07 | 1.8e-07 | 1.3e-07 | 1.667E-07 | 3.215E-08 | 1.300E-07 | 1.900E-07 |
| 14739 | 737 | V737Y | B | 2.3e-07 | 2e-07 | 1.4e-07 | 1.900E-07 | 4.583E-08 | 1.400E-07 | 2.300E-07 |
| 14740 | 737 | V737V | B | 3.8e-07 | 2.4e-07 | 4.6e-07 | 3.600E-07 | 1.114E-07 | 2.400E-07 | 4.600E-07 |
| 14741 | 738 | F738A | B | 1.7e-07 | 2.2e-07 | 2.1e-07 | 2.000E-07 | 2.646E-08 | 1.700E-07 | 2.200E-07 |
| 14742 | 738 | F738R | B | 3.3e-07 | 1.8e-07 | 4.2e-07 | 3.100E-07 | 1.212E-07 | 1.800E-07 | 4.200E-07 |
| 14743 | 738 | F738N | B | 2e-07 | 2.7e-07 | 3.8e-07 | 2.833E-07 | 9.074E-08 | 2.000E-07 | 3.800E-07 |
| 14744 | 738 | F738D | B | 4.4e-07 | 1.7e-07 | 5.3e-07 | 3.800E-07 | 1.873E-07 | 1.700E-07 | 5.300E-07 |
| 14745 | 738 | F738C | B | 3e-07 | 1.3e-07 | 2e-07 | 2.100E-07 | 8.544E-08 | 1.300E-07 | 3.000E-07 |
| 14746 | 738 | F738E | B | 2.9e-07 | 2.5e-07 | 2.5e-07 | 2.633E-07 | 2.309E-08 | 2.500E-07 | 2.900E-07 |
| 14747 | 738 | F738Q | B | 1.5e-07 | 3.2e-07 | 2.5e-07 | 2.400E-07 | 8.544E-08 | 1.500E-07 | 3.200E-07 |
| 14748 | 738 | F738G | B | 1.9e-07 | 2.2e-07 | 2.1e-07 | 2.067E-07 | 1.528E-08 | 1.900E-07 | 2.200E-07 |
| 14749 | 738 | F738H | B | 2.3e-07 | 4.1e-07 | 1.6e-07 | 2.667E-07 | 1.290E-07 | 1.600E-07 | 4.100E-07 |
| 14750 | 738 | F738I | B | 3.3e-07 | 3.2e-07 | 1.6e-07 | 2.700E-07 | 9.539E-08 | 1.600E-07 | 3.300E-07 |
| 14751 | 738 | F738L | B | 1.4e-07 | 7e-08 | 4e-07 | 2.033E-07 | 1.739E-07 | 7.000E-08 | 4.000E-07 |
| 14752 | 738 | F738K | B | 2.2e-07 | 3.3e-07 | 3.5e-07 | 3.000E-07 | 7.000E-08 | 2.200E-07 | 3.500E-07 |
| 14753 | 738 | F738M | B | 8.4e-08 | 2.1e-07 | 3.2e-07 | 2.047E-07 | 1.181E-07 | 8.400E-08 | 3.200E-07 |
| 14754 | 738 | F738F | B | 1.5e-07 | 2.4e-07 | 4.1e-07 | 2.667E-07 | 1.320E-07 | 1.500E-07 | 4.100E-07 |
| 14755 | 738 | F738P | B | 5.1e-07 | 1.7e-07 | 2e-07 | 2.933E-07 | 1.882E-07 | 1.700E-07 | 5.100E-07 |
| 14756 | 738 | F738S | B | 3.6e-07 | 1.6e-07 | 1.5e-07 | 2.233E-07 | 1.185E-07 | 1.500E-07 | 3.600E-07 |
| 14757 | 738 | F738T | B | 1.2e-07 | 4.6e-07 | 1.4e-07 | 2.400E-07 | 1.908E-07 | 1.200E-07 | 4.600E-07 |
| 14758 | 738 | F738W | B | 4.1e-07 | 1.6e-07 | 5.3e-07 | 3.667E-07 | 1.888E-07 | 1.600E-07 | 5.300E-07 |
| 14759 | 738 | F738Y | B | 1.7e-07 | 3.3e-07 | 8.7e-08 | 1.957E-07 | 1.235E-07 | 8.700E-08 | 3.300E-07 |
| 14760 | 738 | F738V | B | 2.1e-07 | 2.3e-07 | 2.1e-07 | 2.167E-07 | 1.155E-08 | 2.100E-07 | 2.300E-07 |
| 14761 | 739 | I739A | B | 1.2e-06 | 2.1e-07 | 1.2e-07 | 5.100E-07 | 5.992E-07 | 1.200E-07 | 1.200E-06 |
| 14762 | 739 | I739R | B | 3.7e-07 | 4.3e-07 | 3.3e-07 | 3.767E-07 | 5.033E-08 | 3.300E-07 | 4.300E-07 |
| 14763 | 739 | I739N | B | 3e-07 | 1.3e-07 | 3.4e-07 | 2.567E-07 | 1.115E-07 | 1.300E-07 | 3.400E-07 |
| 14764 | 739 | I739D | B | 3e-07 | 2.4e-07 | 3e-07 | 2.800E-07 | 3.464E-08 | 2.400E-07 | 3.000E-07 |
| 14765 | 739 | I739C | B | 2.9e-07 | 2e-07 | 2.6e-07 | 2.500E-07 | 4.583E-08 | 2.000E-07 | 2.900E-07 |
| 14766 | 739 | I739E | B | 3.2e-07 | 2.8e-07 | 1.8e-07 | 2.600E-07 | 7.211E-08 | 1.800E-07 | 3.200E-07 |
| 14767 | 739 | I739Q | B | 3.5e-07 | 1.9e-07 | 2.3e-07 | 2.567E-07 | 8.327E-08 | 1.900E-07 | 3.500E-07 |
| 14768 | 739 | I739G | B | 4.1e-07 | 2e-07 | 2.9e-07 | 3.000E-07 | 1.054E-07 | 2.000E-07 | 4.100E-07 |
| 14769 | 739 | I739H | B | 3.4e-07 | 2.6e-07 | 3.1e-07 | 3.033E-07 | 4.041E-08 | 2.600E-07 | 3.400E-07 |
| 14770 | 739 | I739I | B | 2.7e-07 | 1.5e-07 | 2.4e-07 | 2.200E-07 | 6.245E-08 | 1.500E-07 | 2.700E-07 |
| 14771 | 739 | I739L | B | 1.6e-07 | 1.8e-07 | 2e-07 | 1.800E-07 | 2.000E-08 | 1.600E-07 | 2.000E-07 |
| 14772 | 739 | I739K | B | 4.9e-07 | 5.3e-07 | 2e-07 | 4.067E-07 | 1.801E-07 | 2.000E-07 | 5.300E-07 |
| 14773 | 739 | I739M | B | 2.1e-07 | 4.2e-07 | 5.5e-07 | 3.933E-07 | 1.716E-07 | 2.100E-07 | 5.500E-07 |
| 14774 | 739 | I739F | B | 2.1e-07 | 2.2e-07 | 1.1e-07 | 1.800E-07 | 6.083E-08 | 1.100E-07 | 2.200E-07 |
| 14775 | 739 | I739P | B | 6.1e-07 | 2.7e-07 | 2.4e-07 | 3.733E-07 | 2.055E-07 | 2.400E-07 | 6.100E-07 |
| 14776 | 739 | I739S | B | 2.1e-07 | 3.9e-07 | 5.7e-07 | 3.900E-07 | 1.800E-07 | 2.100E-07 | 5.700E-07 |
| 14777 | 739 | I739T | B | 2.1e-07 | 2e-07 | 2.1e-07 | 2.067E-07 | 5.774E-09 | 2.000E-07 | 2.100E-07 |
| 14778 | 739 | I739W | B | 1.2e-07 | 1.6e-07 | 5e-07 | 2.600E-07 | 2.088E-07 | 1.200E-07 | 5.000E-07 |
| 14779 | 739 | I739Y | B | 2.3e-07 | 1.3e-07 | 3.6e-07 | 2.400E-07 | 1.153E-07 | 1.300E-07 | 3.600E-07 |
| 14780 | 739 | I739V | B | 3.4e-07 | 6.5e-07 | 2.8e-07 | 4.233E-07 | 1.986E-07 | 2.800E-07 | 6.500E-07 |
| 14781 | 740 | F740A | B | 1.7e-07 | 2.5e-07 | 3.1e-07 | 2.433E-07 | 7.024E-08 | 1.700E-07 | 3.100E-07 |
| 14782 | 740 | F740R | B | 1.4e-07 | 1.7e-07 | 5.6e-07 | 2.900E-07 | 2.343E-07 | 1.400E-07 | 5.600E-07 |
| 14783 | 740 | F740N | B | 3.2e-07 | 1.9e-07 | 1.8e-07 | 2.300E-07 | 7.810E-08 | 1.800E-07 | 3.200E-07 |
| 14784 | 740 | F740D | B | 3e-07 | 2e-07 | 1.5e-07 | 2.167E-07 | 7.638E-08 | 1.500E-07 | 3.000E-07 |
| 14785 | 740 | F740C | B | 8.2e-08 | 3.8e-07 | 2.6e-07 | 2.407E-07 | 1.499E-07 | 8.200E-08 | 3.800E-07 |
| 14786 | 740 | F740E | B | 2e-07 | 1.4e-07 | 6e-07 | 3.133E-07 | 2.501E-07 | 1.400E-07 | 6.000E-07 |
| 14787 | 740 | F740Q | B | 1.7e-07 | 5.2e-07 | 3.4e-07 | 3.433E-07 | 1.750E-07 | 1.700E-07 | 5.200E-07 |
| 14788 | 740 | F740G | B | 6.8e-07 | 3.3e-07 | 2.4e-07 | 4.167E-07 | 2.325E-07 | 2.400E-07 | 6.800E-07 |
| 14789 | 740 | F740H | B | 2.7e-07 | 2.9e-07 | 2.7e-07 | 2.767E-07 | 1.155E-08 | 2.700E-07 | 2.900E-07 |

| | | | | | | | | | | |
|---|---|---|---|---|---|---|---|---|---|---|
| 14790 | 740 | F740I | B | 3.6e-07 | 2.2e-07 | 1.4e-07 | 2.400E-07 | 1.114E-07 | 1.400E-07 | 3.600E-07 |
| 14791 | 740 | F740L | B | 1.6e-07 | 1.4e-07 | 3.3e-07 | 2.100E-07 | 1.044E-07 | 1.400E-07 | 3.300E-07 |
| 14792 | 740 | F740K | B | 1.6e-07 | 1.7e-07 | 2.6e-07 | 1.967E-07 | 5.508E-08 | 1.600E-07 | 2.600E-07 |
| 14793 | 740 | F740M | B | 2.3e-07 | 1.1e-07 | 1.9e-07 | 1.767E-07 | 6.110E-08 | 1.100E-07 | 2.300E-07 |
| 14794 | 740 | F740F | B | 2.4e-07 | 3.1e-07 | 1.9e-07 | 2.467E-07 | 6.028E-08 | 1.900E-07 | 3.100E-07 |
| 14795 | 740 | F740P | B | 1.6e-07 | 1.9e-07 | 2.4e-07 | 1.967E-07 | 4.041E-08 | 1.600E-07 | 2.400E-07 |
| 14796 | 740 | F740S | B | 4.3e-07 | 2.8e-07 | 1.8e-07 | 2.967E-07 | 1.258E-07 | 1.800E-07 | 4.300E-07 |
| 14797 | 740 | F740T | B | 2.8e-07 | 2.5e-07 | 2.9e-07 | 2.733E-07 | 2.082E-08 | 2.500E-07 | 2.900E-07 |
| 14798 | 740 | F740W | B | 1.4e-07 | 1.6e-07 | 2.8e-07 | 1.933E-07 | 7.572E-08 | 1.400E-07 | 2.800E-07 |
| 14799 | 740 | F740Y | B | 1.4e-07 | 2.8e-07 | 1.1e-07 | 1.767E-07 | 9.074E-08 | 1.100E-07 | 2.800E-07 |
| 14800 | 740 | F740V | B | 3.1e-07 | 1.7e-07 | 1.4e-07 | 2.067E-07 | 9.074E-08 | 1.400E-07 | 3.100E-07 |
| 14801 | 741 | P741A | B | 5.5e-07 | 1.3e-07 | 1.6e-07 | 2.800E-07 | 2.343E-07 | 1.300E-07 | 5.500E-07 |
| 14802 | 741 | P741R | B | 2.9e-07 | 1.5e-07 | 1.8e-07 | 2.067E-07 | 7.371E-08 | 1.500E-07 | 2.900E-07 |
| 14803 | 741 | P741N | B | 2.8e-07 | 3e-07 | 2.9e-07 | 2.900E-07 | 1.000E-08 | 2.800E-07 | 3.000E-07 |
| 14804 | 741 | P741D | B | 2e-07 | 1.8e-07 | 4.2e-07 | 2.667E-07 | 1.332E-07 | 1.800E-07 | 4.200E-07 |
| 14805 | 741 | P741C | B | 2.2e-07 | 1.6e-07 | 1.6e-07 | 1.800E-07 | 3.464E-08 | 1.600E-07 | 2.200E-07 |
| 14806 | 741 | P741E | B | 3.5e-07 | 2.6e-07 | 2.4e-07 | 2.833E-07 | 5.859E-08 | 2.400E-07 | 3.500E-07 |
| 14807 | 741 | P741Q | B | 1.8e-07 | 1e-06 | 1.8e-07 | 4.533E-07 | 4.734E-07 | 1.800E-07 | 1.000E-06 |
| 14808 | 741 | P741G | B | 3e-07 | 1.8e-07 | 1.1e-07 | 1.967E-07 | 9.609E-08 | 1.100E-07 | 3.000E-07 |
| 14809 | 741 | P741H | B | 2.5e-07 | 3e-07 | 2.3e-07 | 2.600E-07 | 3.606E-08 | 2.300E-07 | 3.000E-07 |
| 14810 | 741 | P741I | B | 2.1e-07 | 2e-07 | 2.3e-07 | 2.133E-07 | 1.528E-08 | 2.000E-07 | 2.300E-07 |
| 14811 | 741 | P741L | B | 1.9e-07 | 1.8e-07 | 1.1e-07 | 1.600E-07 | 4.359E-08 | 1.100E-07 | 1.900E-07 |
| 14812 | 741 | P741K | B | 2e-07 | 2.5e-07 | 3e-07 | 2.500E-07 | 5.000E-08 | 2.000E-07 | 3.000E-07 |
| 14813 | 741 | P741M | B | 1.6e-07 | 3e-07 | 2.7e-07 | 2.433E-07 | 7.371E-08 | 1.600E-07 | 3.000E-07 |
| 14814 | 741 | P741F | B | 2.1e-07 | 6.6e-07 | 1.9e-07 | 3.533E-07 | 2.658E-07 | 1.900E-07 | 6.600E-07 |
| 14815 | 741 | P741P | B | 3.7e-07 | 1.5e-07 | 2.4e-07 | 2.533E-07 | 1.106E-07 | 1.500E-07 | 3.700E-07 |
| 14816 | 741 | P741S | B | 2e-07 | 2.2e-07 | 6.6e-07 | 3.600E-07 | 2.600E-07 | 2.000E-07 | 6.600E-07 |
| 14817 | 741 | P741T | B | 2.4e-07 | 1.6e-07 | 1.6e-07 | 1.867E-07 | 4.619E-08 | 1.600E-07 | 2.400E-07 |
| 14818 | 741 | P741W | B | 2.6e-07 | 3.3e-07 | 2e-07 | 2.633E-07 | 6.506E-08 | 2.000E-07 | 3.300E-07 |
| 14819 | 741 | P741Y | B | 1.6e-07 | 4.2e-07 | 2.8e-07 | 2.867E-07 | 1.301E-07 | 1.600E-07 | 4.200E-07 |
| 14820 | 741 | P741V | B | 2.3e-07 | 2.2e-07 | 2.7e-07 | 2.400E-07 | 2.646E-08 | 2.200E-07 | 2.700E-07 |
| 14821 | 742 | P742A | B | 1.9e-07 | 5.7e-07 | 1.8e-07 | 3.133E-07 | 2.223E-07 | 1.800E-07 | 5.700E-07 |
| 14822 | 742 | P742R | B | 4e-07 | 2.6e-07 | 4.5e-07 | 3.700E-07 | 9.849E-08 | 2.600E-07 | 4.500E-07 |
| 14823 | 742 | P742N | B | 1.5e-07 | 3.7e-07 | 1.6e-07 | 2.267E-07 | 1.242E-07 | 1.500E-07 | 3.700E-07 |
| 14824 | 742 | P742D | B | 2.6e-07 | 2.4e-07 | 1.6e-07 | 2.200E-07 | 5.292E-08 | 1.600E-07 | 2.600E-07 |
| 14825 | 742 | P742C | B | 1.7e-07 | 2.1e-07 | 1.8e-07 | 1.867E-07 | 2.082E-08 | 1.700E-07 | 2.100E-07 |
| 14826 | 742 | P742E | B | 2.8e-07 | 2e-07 | 2.9e-07 | 2.567E-07 | 4.933E-08 | 2.000E-07 | 2.900E-07 |
| 14827 | 742 | P742Q | B | 2.1e-07 | 2.3e-07 | 3.7e-07 | 2.700E-07 | 8.718E-08 | 2.100E-07 | 3.700E-07 |
| 14828 | 742 | P742G | B | 3.8e-07 | 2e-07 | 5.6e-07 | 3.800E-07 | 1.800E-07 | 2.000E-07 | 5.600E-07 |
| 14829 | 742 | P742H | B | 1.3e-07 | 2.2e-07 | 2.2e-07 | 1.900E-07 | 5.196E-08 | 1.300E-07 | 2.200E-07 |
| 14830 | 742 | P742I | B | 3.5e-07 | 1.9e-07 | 2.6e-07 | 2.667E-07 | 8.021E-08 | 1.900E-07 | 3.500E-07 |
| 14831 | 742 | P742L | B | 3e-07 | 2.7e-07 | 2.2e-07 | 2.633E-07 | 4.041E-08 | 2.200E-07 | 3.000E-07 |
| 14832 | 742 | P742K | B | 2.2e-07 | 1.8e-07 | 2.9e-07 | 2.300E-07 | 5.568E-08 | 1.800E-07 | 2.900E-07 |
| 14833 | 742 | P742M | B | 2.7e-07 | 1.9e-07 | 2e-07 | 2.200E-07 | 4.359E-08 | 1.900E-07 | 2.700E-07 |
| 14834 | 742 | P742F | B | 1.9e-07 | 4.5e-07 | 1.6e-07 | 2.667E-07 | 1.595E-07 | 1.600E-07 | 4.500E-07 |
| 14835 | 742 | P742P | B | 3e-07 | 1.5e-07 | 4.4e-07 | 2.967E-07 | 1.450E-07 | 1.500E-07 | 4.400E-07 |
| 14836 | 742 | P742S | B | 1.6e-07 | 1.7e-07 | 1.4e-07 | 1.567E-07 | 1.528E-08 | 1.400E-07 | 1.700E-07 |
| 14837 | 742 | P742T | B | 1.9e-07 | 2.8e-07 | 2.2e-07 | 2.300E-07 | 4.583E-08 | 1.900E-07 | 2.800E-07 |
| 14838 | 742 | P742W | B | 2.6e-07 | 2.3e-07 | 2e-07 | 2.300E-07 | 3.000E-08 | 2.000E-07 | 2.600E-07 |
| 14839 | 742 | P742Y | B | 2e-07 | 5.9e-07 | 3.9e-07 | 3.933E-07 | 1.950E-07 | 2.000E-07 | 5.900E-07 |
| 14840 | 742 | P742V | B | 4.1e-07 | 1.8e-07 | 2e-07 | 2.633E-07 | 1.274E-07 | 1.800E-07 | 4.100E-07 |
| 14841 | 743 | S743A | B | 2.9e-07 | 1.4e-07 | 3.6e-07 | 2.633E-07 | 1.124E-07 | 1.400E-07 | 3.600E-07 |
| 14842 | 743 | S743R | B | 1.6e-07 | 2e-07 | 2.1e-07 | 1.900E-07 | 2.646E-08 | 1.600E-07 | 2.100E-07 |
| 14843 | 743 | S743N | B | 2.3e-07 | 3e-07 | 1.5e-07 | 2.267E-07 | 7.506E-08 | 1.500E-07 | 3.000E-07 |
| 14844 | 743 | S743D | B | 2e-07 | 9.5e-08 | 3.7e-07 | 2.217E-07 | 1.388E-07 | 9.500E-08 | 3.700E-07 |
| 14845 | 743 | S743C | B | 1.9e-07 | 3.8e-07 | 3.9e-07 | 3.200E-07 | 1.127E-07 | 1.900E-07 | 3.900E-07 |
| 14846 | 743 | S743E | B | 4e-07 | 3.3e-07 | 5e-07 | 4.100E-07 | 8.544E-08 | 3.300E-07 | 5.000E-07 |
| 14847 | 743 | S743Q | B | 3.4e-07 | 2.6e-07 | 1.1e-07 | 2.367E-07 | 1.168E-07 | 1.100E-07 | 3.400E-07 |
| 14848 | 743 | S743G | B | 2.1e-07 | 3e-07 | 4.8e-07 | 3.300E-07 | 1.375E-07 | 2.100E-07 | 4.800E-07 |
| 14849 | 743 | S743H | B | 1.2e-07 | 6.9e-08 | 2e-07 | 1.297E-07 | 6.603E-08 | 6.900E-08 | 2.000E-07 |
| 14850 | 743 | S743I | B | 3.3e-07 | 2.7e-07 | 3e-07 | 3.000E-07 | 3.000E-08 | 2.700E-07 | 3.300E-07 |
| 14851 | 743 | S743L | B | 8.6e-08 | 3.1e-07 | 1.4e-07 | 1.787E-07 | 1.169E-07 | 8.600E-08 | 3.100E-07 |
| 14852 | 743 | S743K | B | 2.7e-07 | 2.6e-07 | 1.1e-07 | 2.133E-07 | 8.963E-08 | 1.100E-07 | 2.700E-07 |
| 14853 | 743 | S743M | B | 1.6e-07 | 3e-07 | 2e-07 | 2.200E-07 | 7.211E-08 | 1.600E-07 | 3.000E-07 |
| 14854 | 743 | S743F | B | 1.9e-07 | 1.5e-07 | 4.4e-07 | 2.600E-07 | 1.572E-07 | 1.500E-07 | 4.400E-07 |
| 14855 | 743 | S743P | B | 3.4e-07 | 1.3e-07 | 2.8e-07 | 2.500E-07 | 1.082E-07 | 1.300E-07 | 3.400E-07 |
| 14856 | 743 | S743S | B | 2.2e-07 | 1.8e-07 | 2e-07 | 2.000E-07 | 2.000E-08 | 1.800E-07 | 2.200E-07 |
| 14857 | 743 | S743T | B | 3.7e-07 | 9.9e-08 | 3.2e-07 | 2.630E-07 | 1.442E-07 | 9.900E-08 | 3.700E-07 |
| 14858 | 743 | S743W | B | 1.8e-07 | 1.5e-07 | 2e-07 | 1.767E-07 | 2.517E-08 | 1.500E-07 | 2.000E-07 |
| 14859 | 743 | S743Y | B | 2.8e-07 | 2.4e-07 | 3.2e-07 | 2.800E-07 | 4.000E-08 | 2.400E-07 | 3.200E-07 |
| 14860 | 743 | S743V | B | 3e-07 | 3.3e-07 | 2.3e-07 | 2.867E-07 | 5.132E-08 | 2.300E-07 | 3.300E-07 |
| 14861 | 744 | D744A | B | 1.2e-07 | 2.6e-07 | 4.9e-07 | 2.900E-07 | 1.868E-07 | 1.200E-07 | 4.900E-07 |
| 14862 | 744 | D744R | B | 3.1e-07 | 2.9e-07 | 2.2e-07 | 2.733E-07 | 4.726E-08 | 2.200E-07 | 3.100E-07 |
| 14863 | 744 | D744N | B | 1.8e-07 | 4.2e-07 | 1.6e-07 | 2.533E-07 | 1.447E-07 | 1.600E-07 | 4.200E-07 |
| 14864 | 744 | D744D | B | 2e-07 | 2.7e-07 | 1e-07 | 1.900E-07 | 8.544E-08 | 1.000E-07 | 2.700E-07 |
| 14865 | 744 | D744C | B | 3.2e-07 | 1.8e-07 | 2.3e-07 | 2.433E-07 | 7.095E-08 | 1.800E-07 | 3.200E-07 |
| 14866 | 744 | D744E | B | 2.1e-07 | 1.9e-07 | 1e-07 | 1.667E-07 | 5.859E-08 | 1.000E-07 | 2.100E-07 |
| 14867 | 744 | D744Q | B | 1.4e-07 | 1.9e-07 | 3e-07 | 2.100E-07 | 8.185E-08 | 1.400E-07 | 3.000E-07 |
| 14868 | 744 | D744G | B | 2.2e-07 | 1.9e-07 | 2.5e-07 | 2.200E-07 | 3.000E-08 | 1.900E-07 | 2.500E-07 |
| 14869 | 744 | D744H | B | 2.5e-07 | 3.3e-07 | 2e-07 | 2.600E-07 | 6.557E-08 | 2.000E-07 | 3.300E-07 |
| 14870 | 744 | D744I | B | 3.3e-07 | 2.5e-07 | 3.2e-07 | 3.000E-07 | 4.359E-08 | 2.500E-07 | 3.300E-07 |
| 14871 | 744 | D744L | B | 4.7e-07 | 1.4e-07 | 1.4e-07 | 2.500E-07 | 1.905E-07 | 1.400E-07 | 4.700E-07 |
| 14872 | 744 | D744K | B | 2.6e-07 | 4.6e-07 | 4.7e-07 | 3.967E-07 | 1.185E-07 | 2.600E-07 | 4.700E-07 |
| 14873 | 744 | D744M | B | 2.4e-07 | 2.3e-07 | 1.7e-07 | 2.133E-07 | 3.786E-08 | 1.700E-07 | 2.400E-07 |
| 14874 | 744 | D744F | B | 5.4e-07 | 2e-07 | 2.2e-07 | 3.200E-07 | 1.908E-07 | 2.000E-07 | 5.400E-07 |

| | | | | | | | | | | |
|---|---|---|---|---|---|---|---|---|---|---|
| 14875 | 744 | D744P | B | 1.4e-07 | 4.1e-07 | 3.4e-07 | 2.967E-07 | 1.401E-07 | 1.400E-07 | 4.100E-07 |
| 14876 | 744 | D744S | B | 3.5e-07 | 1.5e-07 | 1.9e-07 | 2.300E-07 | 1.058E-07 | 1.500E-07 | 3.500E-07 |
| 14877 | 744 | D744T | B | 2.2e-07 | 2.9e-07 | 3e-07 | 2.700E-07 | 4.359E-08 | 2.200E-07 | 3.000E-07 |
| 14878 | 744 | D744W | B | 3.1e-07 | 3e-07 | 3.7e-07 | 3.267E-07 | 3.786E-08 | 3.000E-07 | 3.700E-07 |
| 14879 | 744 | D744Y | B | 2.3e-07 | 2.1e-07 | 3.1e-07 | 2.500E-07 | 5.292E-08 | 2.100E-07 | 3.100E-07 |
| 14880 | 744 | D744V | B | 2.5e-07 | 3.4e-07 | 2.3e-07 | 2.733E-07 | 5.859E-08 | 2.300E-07 | 3.400E-07 |
| 14881 | 745 | E745A | B | 1.4e-07 | 1.7e-07 | 2.7e-07 | 1.933E-07 | 6.807E-08 | 1.400E-07 | 2.700E-07 |
| 14882 | 745 | E745R | B | 1.9e-07 | 4.6e-07 | 2.9e-07 | 3.133E-07 | 1.365E-07 | 1.900E-07 | 4.600E-07 |
| 14883 | 745 | E745N | B | 3.9e-07 | 2.8e-07 | 6.4e-07 | 4.367E-07 | 1.845E-07 | 2.800E-07 | 6.400E-07 |
| 14884 | 745 | E745D | B | 3.1e-07 | 2e-07 | 3.5e-07 | 2.867E-07 | 7.767E-08 | 2.000E-07 | 3.500E-07 |
| 14885 | 745 | E745C | B | 2.1e-07 | 2.8e-07 | 1.3e-07 | 2.067E-07 | 7.506E-08 | 1.300E-07 | 2.800E-07 |
| 14886 | 745 | E745E | B | 1.9e-07 | 2.6e-07 | 1.4e-07 | 1.967E-07 | 6.028E-08 | 1.400E-07 | 2.600E-07 |
| 14887 | 745 | E745Q | B | 2.9e-07 | 2.4e-07 | 1.3e-07 | 2.200E-07 | 8.185E-08 | 1.300E-07 | 2.900E-07 |
| 14888 | 745 | E745G | B | 1.3e-07 | 4e-07 | 3.1e-07 | 2.800E-07 | 1.375E-07 | 1.300E-07 | 4.000E-07 |
| 14889 | 745 | E745H | B | 3.8e-07 | 2.2e-07 | 1.8e-07 | 2.600E-07 | 1.058E-07 | 1.800E-07 | 3.800E-07 |
| 14890 | 745 | E745I | B | 4.1e-07 | 3.1e-07 | 1.5e-07 | 2.900E-07 | 1.311E-07 | 1.500E-07 | 4.100E-07 |
| 14891 | 745 | E745L | B | 3.2e-07 | 3e-07 | 4e-07 | 3.400E-07 | 5.292E-08 | 3.000E-07 | 4.000E-07 |
| 14892 | 745 | E745K | B | 1.4e-07 | 2.7e-07 | 1.5e-07 | 1.867E-07 | 7.234E-08 | 1.400E-07 | 2.700E-07 |
| 14893 | 745 | E745M | B | 1.7e-07 | 1.9e-07 | 4.2e-07 | 2.600E-07 | 1.389E-07 | 1.700E-07 | 4.200E-07 |
| 14894 | 745 | E745F | B | 4.5e-07 | 4.6e-07 | 4.3e-07 | 4.467E-07 | 1.528E-08 | 4.300E-07 | 4.600E-07 |
| 14895 | 745 | E745P | B | 3.4e-07 | 4.1e-07 | 3.9e-07 | 3.800E-07 | 3.606E-08 | 3.400E-07 | 4.100E-07 |
| 14896 | 745 | E745S | B | 2.9e-07 | 2.3e-07 | 5.6e-07 | 3.600E-07 | 1.758E-07 | 2.300E-07 | 5.600E-07 |
| 14897 | 745 | E745T | B | 1.9e-07 | 6.1e-07 | 2.1e-07 | 3.367E-07 | 2.369E-07 | 1.900E-07 | 6.100E-07 |
| 14898 | 745 | E745W | B | 1.8e-07 | 4.2e-07 | 2.1e-07 | 2.700E-07 | 1.308E-07 | 1.800E-07 | 4.200E-07 |
| 14899 | 745 | E745Y | B | 6e-07 | 2.5e-07 | 4.3e-07 | 4.267E-07 | 1.750E-07 | 2.500E-07 | 6.000E-07 |
| 14900 | 745 | E745V | B | 3.9e-07 | 2.9e-07 | 2.4e-07 | 3.067E-07 | 7.638E-08 | 2.400E-07 | 3.900E-07 |
| 14901 | 746 | Q746A | B | 4.1e-07 | 1.2e-07 | 8.2e-08 | 2.040E-07 | 1.794E-07 | 8.200E-08 | 4.100E-07 |
| 14902 | 746 | Q746R | B | 2.3e-07 | 2.3e-07 | 2.8e-07 | 2.467E-07 | 2.887E-08 | 2.300E-07 | 2.800E-07 |
| 14903 | 746 | Q746N | B | 1.8e-07 | 1.7e-07 | 3.8e-07 | 2.433E-07 | 1.185E-07 | 1.700E-07 | 3.800E-07 |
| 14904 | 746 | Q746D | B | 1.4e-07 | 1.8e-07 | 3.3e-07 | 2.167E-07 | 1.002E-07 | 1.400E-07 | 3.300E-07 |
| 14905 | 746 | Q746C | B | 2.1e-07 | 1.7e-07 | 3.6e-07 | 2.467E-07 | 1.002E-07 | 1.700E-07 | 3.600E-07 |
| 14906 | 746 | Q746E | B | 1.8e-07 | 3.5e-07 | 3.5e-07 | 2.933E-07 | 9.815E-08 | 1.800E-07 | 3.500E-07 |
| 14907 | 746 | Q746Q | B | 3.4e-07 | 3.6e-07 | 4.3e-07 | 3.767E-07 | 4.726E-08 | 3.400E-07 | 4.300E-07 |
| 14908 | 746 | Q746G | B | 1.5e-07 | 3.2e-07 | 3.8e-07 | 2.833E-07 | 1.193E-07 | 1.500E-07 | 3.800E-07 |
| 14909 | 746 | Q746H | B | 3.1e-07 | 1.7e-07 | 1.2e-07 | 2.000E-07 | 9.849E-08 | 1.200E-07 | 3.100E-07 |
| 14910 | 746 | Q746I | B | 3.7e-07 | 2.4e-07 | 3.3e-07 | 3.133E-07 | 6.658E-08 | 2.400E-07 | 3.700E-07 |
| 14911 | 746 | Q746L | B | 3.4e-07 | 2.5e-07 | 1.2e-07 | 2.367E-07 | 1.106E-07 | 1.200E-07 | 3.400E-07 |
| 14912 | 746 | Q746K | B | 3.3e-07 | 5.1e-07 | 4.8e-07 | 4.400E-07 | 9.644E-08 | 3.300E-07 | 5.100E-07 |
| 14913 | 746 | Q746M | B | 2.5e-07 | 2.2e-07 | 1.7e-07 | 2.133E-07 | 4.041E-08 | 1.700E-07 | 2.500E-07 |
| 14914 | 746 | Q746F | B | 6.4e-07 | 6.3e-07 | 1.8e-07 | 4.833E-07 | 2.627E-07 | 1.800E-07 | 6.400E-07 |
| 14915 | 746 | Q746P | B | 1.9e-07 | 3e-07 | 3.9e-07 | 2.933E-07 | 1.002E-07 | 1.900E-07 | 3.900E-07 |
| 14916 | 746 | Q746S | B | 1.6e-07 | 1.9e-07 | 1e-07 | 1.500E-07 | 4.583E-08 | 1.000E-07 | 1.900E-07 |
| 14917 | 746 | Q746T | B | 1.9e-07 | 1.9e-07 | 5.3e-07 | 3.033E-07 | 1.963E-07 | 1.900E-07 | 5.300E-07 |
| 14918 | 746 | Q746W | B | 2.5e-07 | 1.8e-07 | 4.2e-07 | 2.833E-07 | 1.234E-07 | 1.800E-07 | 4.200E-07 |
| 14919 | 746 | Q746Y | B | 3.1e-07 | 6e-07 | 3.5e-07 | 4.200E-07 | 1.572E-07 | 3.100E-07 | 6.000E-07 |
| 14920 | 746 | Q746V | B | 1.4e-07 | 1.4e-07 | 1.3e-07 | 1.367E-07 | 5.774E-09 | 1.300E-07 | 1.400E-07 |
| 14921 | 747 | L747A | B | 1.9e-07 | 8.4e-08 | 1.9e-07 | 1.547E-07 | 6.120E-08 | 8.400E-08 | 1.900E-07 |
| 14922 | 747 | L747R | B | 2.5e-07 | 1.3e-07 | 1.8e-07 | 1.867E-07 | 6.028E-08 | 1.300E-07 | 2.500E-07 |
| 14923 | 747 | L747N | B | 3.6e-07 | 3e-07 | 5.4e-07 | 4.000E-07 | 1.249E-07 | 3.000E-07 | 5.400E-07 |
| 14924 | 747 | L747D | B | 3.2e-07 | 1.4e-07 | 3.1e-07 | 2.567E-07 | 1.012E-07 | 1.400E-07 | 3.200E-07 |
| 14925 | 747 | L747C | B | 2.1e-07 | 2e-07 | 2e-07 | 2.033E-07 | 5.774E-09 | 2.000E-07 | 2.100E-07 |
| 14926 | 747 | L747E | B | 1.4e-07 | 2.2e-07 | 4.9e-07 | 2.833E-07 | 1.834E-07 | 1.400E-07 | 4.900E-07 |
| 14927 | 747 | L747Q | B | 1.2e-07 | 2.2e-07 | 2.1e-07 | 1.833E-07 | 5.508E-08 | 1.200E-07 | 2.200E-07 |
| 14928 | 747 | L747G | B | 5.9e-07 | 4.3e-07 | 6.1e-07 | 5.433E-07 | 9.866E-08 | 4.300E-07 | 6.100E-07 |
| 14929 | 747 | L747H | B | 2.3e-07 | 1.5e-07 | 1.3e-07 | 1.700E-07 | 5.292E-08 | 1.300E-07 | 2.300E-07 |
| 14930 | 747 | L747I | B | 1.8e-07 | 1.7e-07 | 4.3e-07 | 2.600E-07 | 1.473E-07 | 1.700E-07 | 4.300E-07 |
| 14931 | 747 | L747L | B | 1.8e-07 | 2.7e-07 | 3.2e-07 | 2.567E-07 | 7.095E-08 | 1.800E-07 | 3.200E-07 |
| 14932 | 747 | L747K | B | 1.4e-07 | 2.1e-07 | 2.1e-07 | 1.867E-07 | 4.041E-08 | 1.400E-07 | 2.100E-07 |
| 14933 | 747 | L747M | B | 3.5e-07 | 2.3e-07 | 1.2e-07 | 2.333E-07 | 1.150E-07 | 1.200E-07 | 3.500E-07 |
| 14934 | 747 | L747F | B | 2.4e-07 | 1.8e-07 | 2.7e-07 | 2.300E-07 | 4.583E-08 | 1.800E-07 | 2.700E-07 |
| 14935 | 747 | L747P | B | 2.2e-07 | 3.8e-07 | 1.9e-07 | 2.633E-07 | 1.021E-07 | 1.900E-07 | 3.800E-07 |
| 14936 | 747 | L747S | B | 1.7e-07 | 3.6e-07 | 1.9e-07 | 2.400E-07 | 1.044E-07 | 1.700E-07 | 3.600E-07 |
| 14937 | 747 | L747T | B | 2.2e-07 | 3.3e-07 | 1.3e-07 | 2.267E-07 | 1.002E-07 | 1.300E-07 | 3.300E-07 |
| 14938 | 747 | L747W | B | 1.9e-07 | 3.2e-07 | 2.1e-07 | 2.400E-07 | 7.000E-08 | 1.900E-07 | 3.200E-07 |
| 14939 | 747 | L747Y | B | 1.7e-07 | 3.3e-07 | 2.1e-07 | 2.367E-07 | 8.327E-08 | 1.700E-07 | 3.300E-07 |
| 14940 | 747 | L747V | B | 1.9e-07 | 1.6e-07 | 1.1e-07 | 1.533E-07 | 4.041E-08 | 1.100E-07 | 1.900E-07 |
| 14941 | 748 | K748A | B | 8.7e-08 | 3.5e-07 | 2.5e-07 | 2.290E-07 | 1.328E-07 | 8.700E-08 | 3.500E-07 |
| 14942 | 748 | K748R | B | 3.5e-07 | 9.5e-08 | 2.3e-07 | 2.250E-07 | 1.276E-07 | 9.500E-08 | 3.500E-07 |
| 14943 | 748 | K748N | B | 3.3e-07 | 7.4e-08 | 3.1e-07 | 2.380E-07 | 1.424E-07 | 7.400E-08 | 3.300E-07 |
| 14944 | 748 | K748D | B | 2e-07 | 1.9e-07 | 2.3e-07 | 2.067E-07 | 2.082E-08 | 1.900E-07 | 2.300E-07 |
| 14945 | 748 | K748C | B | 1.6e-07 | 1.7e-07 | 1.8e-07 | 1.700E-07 | 1.000E-08 | 1.600E-07 | 1.800E-07 |
| 14946 | 748 | K748E | B | 2.3e-07 | 1.2e-07 | 2.4e-07 | 1.967E-07 | 6.658E-08 | 1.200E-07 | 2.400E-07 |
| 14947 | 748 | K748Q | B | 5e-07 | 9.6e-08 | 2.5e-07 | 2.820E-07 | 2.039E-07 | 9.600E-08 | 5.000E-07 |
| 14948 | 748 | K748G | B | 2.7e-07 | 2.9e-07 | 2.5e-07 | 2.700E-07 | 2.000E-08 | 2.500E-07 | 2.900E-07 |
| 14949 | 748 | K748H | B | 2.9e-07 | 1.5e-07 | 2.6e-07 | 2.333E-07 | 7.371E-08 | 1.500E-07 | 2.900E-07 |
| 14950 | 748 | K748I | B | 2.4e-07 | 1.3e-07 | 1.9e-07 | 1.867E-07 | 5.508E-08 | 1.300E-07 | 2.400E-07 |
| 14951 | 748 | K748L | B | 1.7e-07 | 2.2e-07 | 3.2e-07 | 2.367E-07 | 7.638E-08 | 1.700E-07 | 3.200E-07 |
| 14952 | 748 | K748K | B | 1.7e-07 | 3.3e-07 | 1.9e-07 | 2.300E-07 | 8.718E-08 | 1.700E-07 | 3.300E-07 |
| 14953 | 748 | K748M | B | 4.1e-07 | 3.2e-07 | 2.9e-07 | 3.400E-07 | 6.245E-08 | 2.900E-07 | 4.100E-07 |
| 14954 | 748 | K748F | B | 1.1e-06 | 2.5e-07 | 6.1e-07 | 6.533E-07 | 4.267E-07 | 2.500E-07 | 1.100E-06 |
| 14955 | 748 | K748P | B | 3.7e-07 | 1.5e-07 | 1.5e-07 | 2.233E-07 | 1.270E-07 | 1.500E-07 | 3.700E-07 |
| 14956 | 748 | K748S | B | 3.1e-07 | 2.2e-07 | 2.7e-07 | 2.667E-07 | 4.509E-08 | 2.200E-07 | 3.100E-07 |
| 14957 | 748 | K748T | B | 2.2e-07 | 2e-07 | 2e-07 | 2.067E-07 | 1.155E-08 | 2.000E-07 | 2.200E-07 |
| 14958 | 748 | K748W | B | 3e-07 | 2.4e-07 | 2.1e-07 | 2.500E-07 | 4.583E-08 | 2.100E-07 | 3.000E-07 |
| 14959 | 748 | K748Y | B | 1e-07 | 3.3e-07 | 3.3e-07 | 2.533E-07 | 1.328E-07 | 1.000E-07 | 3.300E-07 |

| | | | | | | | | | | |
|---|---|---|---|---|---|---|---|---|---|---|
| 14960 | 748 | K748V | B | 1.1e-07 | 3.2e-07 | 3e-07 | 2.433E-07 | 1.159E-07 | 1.100E-07 | 3.200E-07 |
| 14961 | 749 | S749A | B | 1.7e-07 | 2.3e-07 | 2.9e-07 | 2.300E-07 | 6.000E-08 | 1.700E-07 | 2.900E-07 |
| 14962 | 749 | S749R | B | 2.1e-07 | 1.3e-07 | 2.1e-07 | 1.833E-07 | 4.619E-08 | 1.300E-07 | 2.100E-07 |
| 14963 | 749 | S749N | B | 2e-07 | 2.4e-07 | 1.2e-07 | 1.867E-07 | 6.110E-08 | 1.200E-07 | 2.400E-07 |
| 14964 | 749 | S749D | B | 3.1e-07 | 4.8e-07 | 1.8e-07 | 3.233E-07 | 1.504E-07 | 1.800E-07 | 4.800E-07 |
| 14965 | 749 | S749C | B | 2.3e-07 | 3.3e-07 | 4.3e-07 | 3.300E-07 | 1.000E-07 | 2.300E-07 | 4.300E-07 |
| 14966 | 749 | S749E | B | 9.1e-08 | 2.1e-07 | 1.7e-07 | 1.570E-07 | 6.056E-08 | 9.100E-08 | 2.100E-07 |
| 14967 | 749 | S749Q | B | 2.2e-07 | 2.6e-07 | 2.6e-07 | 2.467E-07 | 2.309E-08 | 2.200E-07 | 2.600E-07 |
| 14968 | 749 | S749G | B | 3.3e-07 | 3.4e-07 | 1e-07 | 2.567E-07 | 1.358E-07 | 1.000E-07 | 3.400E-07 |
| 14969 | 749 | S749H | B | 2.6e-07 | 4.2e-07 | 3.7e-07 | 3.500E-07 | 8.185E-08 | 2.600E-07 | 4.200E-07 |
| 14970 | 749 | S749I | B | 1.9e-07 | 2.7e-07 | 4.6e-07 | 3.067E-07 | 1.387E-07 | 1.900E-07 | 4.600E-07 |
| 14971 | 749 | S749L | B | 2.9e-07 | 1.5e-07 | 2.6e-07 | 2.333E-07 | 7.371E-08 | 1.500E-07 | 2.900E-07 |
| 14972 | 749 | S749K | B | 2.4e-07 | 2.6e-07 | 1.9e-07 | 2.300E-07 | 3.606E-08 | 1.900E-07 | 2.600E-07 |
| 14973 | 749 | S749M | B | 3.2e-07 | 2.8e-07 | 2e-07 | 2.667E-07 | 6.110E-08 | 2.000E-07 | 3.200E-07 |
| 14974 | 749 | S749F | B | 2.4e-07 | 6.9e-07 | 4.9e-07 | 4.733E-07 | 2.255E-07 | 2.400E-07 | 6.900E-07 |
| 14975 | 749 | S749P | B | 3.6e-07 | 5.7e-07 | 1.8e-07 | 3.700E-07 | 1.952E-07 | 1.800E-07 | 5.700E-07 |
| 14976 | 749 | S749S | B | 2.1e-07 | 3.4e-07 | 1.8e-07 | 2.433E-07 | 8.505E-08 | 1.800E-07 | 3.400E-07 |
| 14977 | 749 | S749T | B | 9.5e-08 | 2e-07 | 1.7e-07 | 1.550E-07 | 5.408E-08 | 9.500E-08 | 2.000E-07 |
| 14978 | 749 | S749W | B | 3.7e-07 | 1.7e-07 | 2e-07 | 2.467E-07 | 1.079E-07 | 1.700E-07 | 3.700E-07 |
| 14979 | 749 | S749Y | B | 4.9e-07 | 2e-07 | 6.6e-07 | 4.500E-07 | 2.326E-07 | 2.000E-07 | 6.600E-07 |
| 14980 | 749 | S749V | B | 2.6e-07 | 4e-07 | 2.2e-07 | 2.933E-07 | 9.452E-08 | 2.200E-07 | 4.000E-07 |
| 14981 | 750 | G750A | B | 2e-07 | 1.6e-07 | 4.6e-07 | 2.733E-07 | 1.629E-07 | 1.600E-07 | 4.600E-07 |
| 14982 | 750 | G750R | B | 1.7e-07 | 2.3e-07 | 1.6e-07 | 1.867E-07 | 3.786E-08 | 1.600E-07 | 2.300E-07 |
| 14983 | 750 | G750N | B | 2.7e-07 | 2.8e-07 | 4.2e-07 | 3.233E-07 | 8.386E-08 | 2.700E-07 | 4.200E-07 |
| 14984 | 750 | G750D | B | 2e-07 | 4.7e-07 | 3.1e-07 | 3.267E-07 | 1.358E-07 | 2.000E-07 | 4.700E-07 |
| 14985 | 750 | G750C | B | 2.1e-07 | 2.7e-07 | 2e-07 | 2.267E-07 | 3.786E-08 | 2.000E-07 | 2.700E-07 |
| 14986 | 750 | G750E | B | 2.5e-07 | 3.1e-07 | 3.5e-07 | 3.033E-07 | 5.033E-08 | 2.500E-07 | 3.500E-07 |
| 14987 | 750 | G750Q | B | 3.5e-07 | 3.9e-07 | 1.8e-07 | 3.067E-07 | 1.115E-07 | 1.800E-07 | 3.900E-07 |
| 14988 | 750 | G750G | B | 1.7e-07 | 1.2e-07 | 2.7e-07 | 1.867E-07 | 7.638E-08 | 1.200E-07 | 2.700E-07 |
| 14989 | 750 | G750H | B | 1.5e-07 | 5.5e-07 | 1.8e-07 | 2.933E-07 | 2.228E-07 | 1.500E-07 | 5.500E-07 |
| 14990 | 750 | G750I | B | 3.5e-07 | 2.6e-07 | 5e-07 | 3.700E-07 | 1.212E-07 | 2.600E-07 | 5.000E-07 |
| 14991 | 750 | G750L | B | 1.3e-07 | 1.2e-07 | 3.3e-07 | 1.933E-07 | 1.185E-07 | 1.200E-07 | 3.300E-07 |
| 14992 | 750 | G750K | B | 4.6e-07 | 3.7e-07 | 1.9e-07 | 3.400E-07 | 1.375E-07 | 1.900E-07 | 4.600E-07 |
| 14993 | 750 | G750M | B | 1.1e-07 | 3.2e-07 | 1.6e-07 | 1.967E-07 | 1.097E-07 | 1.100E-07 | 3.200E-07 |
| 14994 | 750 | G750F | B | 1.9e-07 | 1.1e-07 | 4.2e-07 | 2.400E-07 | 1.609E-07 | 1.100E-07 | 4.200E-07 |
| 14995 | 750 | G750P | B | 2e-07 | 2.9e-07 | 1.5e-07 | 2.133E-07 | 7.095E-08 | 1.500E-07 | 2.900E-07 |
| 14996 | 750 | G750S | B | 1.4e-07 | 4.2e-07 | 2.2e-07 | 2.600E-07 | 1.442E-07 | 1.400E-07 | 4.200E-07 |
| 14997 | 750 | G750T | B | 2.5e-07 | 1.4e-07 | 2.7e-07 | 2.200E-07 | 7.000E-08 | 1.400E-07 | 2.700E-07 |
| 14998 | 750 | G750W | B | 2.5e-07 | 5.1e-08 | 5e-07 | 2.670E-07 | 2.250E-07 | 5.100E-08 | 5.000E-07 |
| 14999 | 750 | G750Y | B | 2.1e-07 | 3.4e-07 | 2.1e-07 | 2.533E-07 | 7.506E-08 | 2.100E-07 | 3.400E-07 |
| 15000 | 750 | G750V | B | 1.9e-07 | 2.3e-07 | 2.1e-07 | 2.100E-07 | 2.000E-08 | 1.900E-07 | 2.300E-07 |
| 15001 | 751 | T751A | B | 2.3e-07 | 2.6e-07 | 2e-07 | 2.300E-07 | 3.000E-08 | 2.000E-07 | 2.600E-07 |
| 15002 | 751 | T751R | B | 5.8e-07 | 2.5e-07 | 9.7e-08 | 3.090E-07 | 2.468E-07 | 9.700E-08 | 5.800E-07 |
| 15003 | 751 | T751N | B | 2.5e-07 | 2.2e-07 | 3e-07 | 2.567E-07 | 4.041E-08 | 2.200E-07 | 3.000E-07 |
| 15004 | 751 | T751D | B | 1.3e-07 | 3.4e-07 | 1.8e-07 | 2.167E-07 | 1.097E-07 | 1.300E-07 | 3.400E-07 |
| 15005 | 751 | T751C | B | 2.9e-07 | 1.7e-07 | 2.6e-07 | 2.400E-07 | 6.245E-08 | 1.700E-07 | 2.900E-07 |
| 15006 | 751 | T751E | B | 2.1e-07 | 1.9e-07 | 2.8e-07 | 2.267E-07 | 4.726E-08 | 1.900E-07 | 2.800E-07 |
| 15007 | 751 | T751Q | B | 2.8e-07 | 5.2e-07 | 3.7e-07 | 3.900E-07 | 1.212E-07 | 2.800E-07 | 5.200E-07 |
| 15008 | 751 | T751G | B | 2.2e-07 | 3.1e-07 | 2.7e-07 | 2.667E-07 | 4.509E-08 | 2.200E-07 | 3.100E-07 |
| 15009 | 751 | T751H | B | 2.6e-07 | 3.6e-07 | 1.9e-07 | 2.700E-07 | 8.544E-08 | 1.900E-07 | 3.600E-07 |
| 15010 | 751 | T751I | B | 2.2e-07 | 2.9e-07 | 3.8e-07 | 2.967E-07 | 8.021E-08 | 2.200E-07 | 3.800E-07 |
| 15011 | 751 | T751L | B | 2.2e-07 | 3.8e-07 | 1e-07 | 2.333E-07 | 1.405E-07 | 1.000E-07 | 3.800E-07 |
| 15012 | 751 | T751K | B | 1.8e-07 | 1.2e-07 | 3.9e-07 | 2.300E-07 | 1.418E-07 | 1.200E-07 | 3.900E-07 |
| 15013 | 751 | T751M | B | 2e-07 | 2.1e-07 | 2.6e-07 | 2.233E-07 | 3.215E-08 | 2.000E-07 | 2.600E-07 |
| 15014 | 751 | T751F | B | 1.4e-07 | 3.7e-07 | 2.8e-07 | 2.633E-07 | 1.159E-07 | 1.400E-07 | 3.700E-07 |
| 15015 | 751 | T751P | B | 2.7e-07 | 1.9e-07 | 2.5e-07 | 2.367E-07 | 4.163E-08 | 1.900E-07 | 2.700E-07 |
| 15016 | 751 | T751S | B | 1.6e-07 | 2.7e-07 | 2.6e-07 | 2.300E-07 | 6.083E-08 | 1.600E-07 | 2.700E-07 |
| 15017 | 751 | T751T | B | 2.4e-07 | 2e-07 | 3.2e-07 | 2.533E-07 | 6.110E-08 | 2.000E-07 | 3.200E-07 |
| 15018 | 751 | T751W | B | 1.8e-07 | 2.2e-07 | 3.3e-07 | 2.433E-07 | 7.767E-08 | 1.800E-07 | 3.300E-07 |
| 15019 | 751 | T751Y | B | 1.5e-07 | 6.8e-07 | 1.8e-07 | 3.367E-07 | 2.977E-07 | 1.500E-07 | 6.800E-07 |
| 15020 | 751 | T751V | B | 1.3e-07 | 2.1e-07 | 3.2e-07 | 2.200E-07 | 9.539E-08 | 1.300E-07 | 3.200E-07 |
| 15021 | 752 | A752A | B | 1.1e-07 | 3e-07 | 1.1e-07 | 1.733E-07 | 1.097E-07 | 1.100E-07 | 3.000E-07 |
| 15022 | 752 | A752R | B | 3.1e-07 | 1.7e-07 | 2.3e-07 | 2.367E-07 | 7.024E-08 | 1.700E-07 | 3.100E-07 |
| 15023 | 752 | A752N | B | 2.4e-07 | 2.4e-07 | 4.1e-07 | 2.967E-07 | 9.815E-08 | 2.400E-07 | 4.100E-07 |
| 15024 | 752 | A752D | B | 4.4e-07 | 2.4e-07 | 9.8e-08 | 2.593E-07 | 1.718E-07 | 9.800E-08 | 4.400E-07 |
| 15025 | 752 | A752C | B | 1.8e-07 | 5e-07 | 1.5e-07 | 2.767E-07 | 1.940E-07 | 1.500E-07 | 5.000E-07 |
| 15026 | 752 | A752E | B | 1.4e-07 | 9.2e-07 | 2.9e-07 | 4.500E-07 | 4.139E-07 | 1.400E-07 | 9.200E-07 |
| 15027 | 752 | A752Q | B | 4.3e-07 | 3e-07 | 5.3e-07 | 4.200E-07 | 1.153E-07 | 3.000E-07 | 5.300E-07 |
| 15028 | 752 | A752G | B | 2.9e-07 | 3.8e-07 | 2.3e-07 | 3.000E-07 | 7.550E-08 | 2.300E-07 | 3.800E-07 |
| 15029 | 752 | A752H | B | 2.1e-07 | 2.7e-07 | 5.8e-07 | 3.533E-07 | 1.986E-07 | 2.100E-07 | 5.800E-07 |
| 15030 | 752 | A752I | B | 1.8e-07 | 2.8e-07 | 4.5e-07 | 3.033E-07 | 1.365E-07 | 1.800E-07 | 4.500E-07 |
| 15031 | 752 | A752L | B | 6.8e-08 | 2e-07 | 2.4e-07 | 1.693E-07 | 9.001E-08 | 6.800E-08 | 2.400E-07 |
| 15032 | 752 | A752K | B | 1.8e-07 | 1.8e-07 | 1.9e-07 | 1.833E-07 | 5.774E-09 | 1.800E-07 | 1.900E-07 |
| 15033 | 752 | A752M | B | 2e-07 | 2.2e-07 | 4.7e-07 | 2.967E-07 | 1.504E-07 | 2.000E-07 | 4.700E-07 |
| 15034 | 752 | A752F | B | 2.2e-07 | 2.9e-07 | 1.8e-07 | 2.300E-07 | 5.568E-08 | 1.800E-07 | 2.900E-07 |
| 15035 | 752 | A752P | B | 2.4e-07 | 3e-07 | 1.7e-07 | 2.367E-07 | 6.506E-08 | 1.700E-07 | 3.000E-07 |
| 15036 | 752 | A752S | B | 2.4e-07 | 2.5e-07 | 1.7e-07 | 2.200E-07 | 4.359E-08 | 1.700E-07 | 2.500E-07 |
| 15037 | 752 | A752T | B | 2.5e-07 | 2.9e-07 | 3.6e-07 | 3.000E-07 | 5.568E-08 | 2.500E-07 | 3.600E-07 |
| 15038 | 752 | A752W | B | 1.2e-07 | 2.7e-07 | 5.3e-07 | 3.067E-07 | 2.074E-07 | 1.200E-07 | 5.300E-07 |
| 15039 | 752 | A752Y | B | 6e-07 | 1.8e-07 | 1.7e-07 | 3.167E-07 | 2.454E-07 | 1.700E-07 | 6.000E-07 |
| 15040 | 752 | A752V | B | 9.5e-08 | 2.3e-07 | 2.2e-07 | 1.817E-07 | 7.522E-08 | 9.500E-08 | 2.300E-07 |
| 15041 | 753 | S753A | B | 3.1e-07 | 3.1e-07 | 1.6e-07 | 2.600E-07 | 8.660E-08 | 1.600E-07 | 3.100E-07 |
| 15042 | 753 | S753R | B | 1.2e-07 | 1.2e-07 | 3.2e-07 | 1.867E-07 | 1.155E-07 | 1.200E-07 | 3.200E-07 |
| 15043 | 753 | S753N | B | 1.4e-07 | 7.2e-07 | 1.7e-07 | 3.433E-07 | 3.265E-07 | 1.400E-07 | 7.200E-07 |
| 15044 | 753 | S753D | B | 2.7e-07 | 1.1e-07 | 2.1e-07 | 1.967E-07 | 8.083E-08 | 1.100E-07 | 2.700E-07 |

| | | | | | | | | | | |
|---|---|---|---|---|---|---|---|---|---|---|
| 15045 | 753 | S753C | B | 3.6e-07 | 2.2e-07 | 1.5e-07 | 2.433E-07 | 1.069E-07 | 1.500E-07 | 3.600E-07 |
| 15046 | 753 | S753E | B | 2.7e-07 | 2.9e-07 | 2.1e-07 | 2.567E-07 | 4.163E-08 | 2.100E-07 | 2.900E-07 |
| 15047 | 753 | S753Q | B | 2.1e-07 | 2e-07 | 4.2e-07 | 2.767E-07 | 1.242E-07 | 2.000E-07 | 4.200E-07 |
| 15048 | 753 | S753G | B | 2e-07 | 3.2e-07 | 2.4e-07 | 2.533E-07 | 6.110E-08 | 2.000E-07 | 3.200E-07 |
| 15049 | 753 | S753H | B | 2e-07 | 2.1e-07 | 2.1e-07 | 2.067E-07 | 5.774E-09 | 2.000E-07 | 2.100E-07 |
| 15050 | 753 | S753I | B | 2.4e-07 | 2.5e-07 | 4e-07 | 2.967E-07 | 8.963E-08 | 2.400E-07 | 4.000E-07 |
| 15051 | 753 | S753L | B | 1.9e-07 | 2.5e-07 | 1.7e-07 | 2.033E-07 | 4.163E-08 | 1.700E-07 | 2.500E-07 |
| 15052 | 753 | S753K | B | 3.6e-07 | 4.2e-07 | 4.2e-07 | 4.000E-07 | 3.464E-08 | 3.600E-07 | 4.200E-07 |
| 15053 | 753 | S753M | B | 9.4e-08 | 1.4e-07 | 3.1e-07 | 1.813E-07 | 1.138E-07 | 9.400E-08 | 3.100E-07 |
| 15054 | 753 | S753F | B | 2.7e-07 | 1.6e-07 | 1.7e-07 | 2.000E-07 | 6.083E-08 | 1.600E-07 | 2.700E-07 |
| 15055 | 753 | S753P | B | 2e-07 | 8.3e-08 | 1.2e-07 | 1.343E-07 | 5.980E-08 | 8.300E-08 | 2.000E-07 |
| 15056 | 753 | S753S | B | 3.4e-07 | 1.7e-07 | 1.1e-07 | 2.067E-07 | 1.193E-07 | 1.100E-07 | 3.400E-07 |
| 15057 | 753 | S753T | B | 2.1e-07 | 1.8e-07 | 4.8e-07 | 2.900E-07 | 1.652E-07 | 1.800E-07 | 4.800E-07 |
| 15058 | 753 | S753W | B | 2.5e-07 | 2.1e-07 | 4.9e-07 | 3.167E-07 | 1.514E-07 | 2.100E-07 | 4.900E-07 |
| 15059 | 753 | S753Y | B | 1e-07 | 3e-07 | 1.7e-07 | 1.900E-07 | 1.015E-07 | 1.000E-07 | 3.000E-07 |
| 15060 | 753 | S753V | B | 2.6e-07 | 1.8e-07 | 6.8e-07 | 3.733E-07 | 2.686E-07 | 1.800E-07 | 6.800E-07 |
| 15061 | 754 | V754A | B | 6e-07 | 6.4e-07 | 6.7e-07 | 6.367E-07 | 3.512E-08 | 6.000E-07 | 6.700E-07 |
| 15062 | 754 | V754R | B | 1.2e-07 | 2.4e-07 | 3.9e-07 | 2.500E-07 | 1.353E-07 | 1.200E-07 | 3.900E-07 |
| 15063 | 754 | V754N | B | 2e-07 | 9.1e-08 | 1.8e-07 | 1.570E-07 | 5.803E-08 | 9.100E-08 | 2.000E-07 |
| 15064 | 754 | V754D | B | 2.5e-07 | 2.2e-07 | 1.1e-07 | 1.933E-07 | 7.371E-08 | 1.100E-07 | 2.500E-07 |
| 15065 | 754 | V754C | B | 2.2e-07 | 1.4e-07 | 2.7e-07 | 2.100E-07 | 6.557E-08 | 1.400E-07 | 2.700E-07 |
| 15066 | 754 | V754E | B | 2.1e-07 | 1.8e-07 | 1.4e-07 | 1.767E-07 | 3.512E-08 | 1.400E-07 | 2.100E-07 |
| 15067 | 754 | V754Q | B | 2.9e-07 | 2.3e-07 | 1.9e-07 | 2.367E-07 | 5.033E-08 | 1.900E-07 | 2.900E-07 |
| 15068 | 754 | V754G | B | 3.6e-07 | 3.6e-07 | 9.8e-08 | 2.727E-07 | 1.513E-07 | 9.800E-08 | 3.600E-07 |
| 15069 | 754 | V754H | B | 2.1e-07 | 2e-07 | 2.8e-07 | 2.300E-07 | 4.359E-08 | 2.000E-07 | 2.800E-07 |
| 15070 | 754 | V754I | B | 2.3e-07 | 3.5e-07 | 2.3e-07 | 2.700E-07 | 6.928E-08 | 2.300E-07 | 3.500E-07 |
| 15071 | 754 | V754L | B | 1.9e-07 | 1.2e-07 | 1.4e-07 | 1.500E-07 | 3.606E-08 | 1.200E-07 | 1.900E-07 |
| 15072 | 754 | V754K | B | 8e-08 | 3.6e-07 | 1.8e-07 | 2.067E-07 | 1.419E-07 | 8.000E-08 | 3.600E-07 |
| 15073 | 754 | V754M | B | 9.2e-08 | 3.6e-07 | 3.4e-07 | 2.640E-07 | 1.493E-07 | 9.200E-08 | 3.600E-07 |
| 15074 | 754 | V754F | B | 2.4e-07 | 2.5e-07 | 2.9e-07 | 2.600E-07 | 2.646E-08 | 2.400E-07 | 2.900E-07 |
| 15075 | 754 | V754P | B | 4.5e-07 | 2.5e-07 | 2.9e-07 | 3.300E-07 | 1.058E-07 | 2.500E-07 | 4.500E-07 |
| 15076 | 754 | V754S | B | 3.4e-07 | 2.7e-07 | 2.3e-07 | 2.800E-07 | 5.568E-08 | 2.300E-07 | 3.400E-07 |
| 15077 | 754 | V754T | B | 2.1e-07 | 2.4e-07 | 1e-07 | 1.833E-07 | 7.371E-08 | 1.000E-07 | 2.400E-07 |
| 15078 | 754 | V754W | B | 1.8e-07 | 1.4e-07 | 2e-07 | 1.733E-07 | 3.055E-08 | 1.400E-07 | 2.000E-07 |
| 15079 | 754 | V754Y | B | 2.1e-07 | 3.5e-07 | 2e-07 | 2.533E-07 | 8.386E-08 | 2.000E-07 | 3.500E-07 |
| 15080 | 754 | V754V | B | 1.2e-07 | 3e-07 | 2.2e-07 | 2.133E-07 | 9.018E-08 | 1.200E-07 | 3.000E-07 |
| 15081 | 755 | V755A | B | 3.3e-07 | 7.7e-08 | 3.5e-07 | 2.523E-07 | 1.522E-07 | 7.700E-08 | 3.500E-07 |
| 15082 | 755 | V755R | B | 1.3e-07 | 2.4e-07 | 4.5e-07 | 2.733E-07 | 1.626E-07 | 1.300E-07 | 4.500E-07 |
| 15083 | 755 | V755N | B | 1.4e-07 | 3.6e-07 | 1.8e-07 | 2.267E-07 | 1.172E-07 | 1.400E-07 | 3.600E-07 |
| 15084 | 755 | V755D | B | 4e-07 | 2.4e-07 | 2.6e-07 | 3.000E-07 | 8.718E-08 | 2.400E-07 | 4.000E-07 |
| 15085 | 755 | V755C | B | 2.2e-07 | 1.8e-07 | 3e-07 | 2.333E-07 | 6.110E-08 | 1.800E-07 | 3.000E-07 |
| 15086 | 755 | V755E | B | 1.6e-07 | 1.8e-07 | 2.9e-07 | 2.100E-07 | 7.000E-08 | 1.600E-07 | 2.900E-07 |
| 15087 | 755 | V755Q | B | 1.2e-07 | 3.3e-07 | 6.4e-07 | 3.633E-07 | 2.616E-07 | 1.200E-07 | 6.400E-07 |
| 15088 | 755 | V755G | B | 3.5e-07 | 1.8e-07 | 3.4e-07 | 2.900E-07 | 9.539E-08 | 1.800E-07 | 3.500E-07 |
| 15089 | 755 | V755H | B | 2.1e-07 | 1.9e-07 | 2.7e-07 | 2.233E-07 | 4.163E-08 | 1.900E-07 | 2.700E-07 |
| 15090 | 755 | V755I | B | 1.7e-07 | 2.3e-07 | 1.4e-07 | 1.800E-07 | 4.583E-08 | 1.400E-07 | 2.300E-07 |
| 15091 | 755 | V755L | B | 2.1e-07 | 2.9e-07 | 3.2e-07 | 2.733E-07 | 5.686E-08 | 2.100E-07 | 3.200E-07 |
| 15092 | 755 | V755K | B | 2.5e-07 | 4.1e-07 | 2.6e-07 | 3.067E-07 | 8.963E-08 | 2.500E-07 | 4.100E-07 |
| 15093 | 755 | V755M | B | 2.4e-07 | 5.2e-07 | 2e-07 | 3.200E-07 | 1.744E-07 | 2.000E-07 | 5.200E-07 |
| 15094 | 755 | V755F | B | 3.3e-07 | 2.9e-07 | 2e-07 | 2.733E-07 | 6.658E-08 | 2.000E-07 | 3.300E-07 |
| 15095 | 755 | V755P | B | 2.6e-07 | 1.9e-07 | 5.4e-07 | 3.300E-07 | 1.852E-07 | 1.900E-07 | 5.400E-07 |
| 15096 | 755 | V755S | B | 7.2e-08 | 2e-07 | 3.8e-07 | 2.173E-07 | 1.547E-07 | 7.200E-08 | 3.800E-07 |
| 15097 | 755 | V755T | B | 2.3e-07 | 1.9e-07 | 1.8e-07 | 2.000E-07 | 2.646E-08 | 1.800E-07 | 2.300E-07 |
| 15098 | 755 | V755W | B | 2.6e-07 | 2.1e-07 | 1.5e-07 | 2.067E-07 | 5.508E-08 | 1.500E-07 | 2.600E-07 |
| 15099 | 755 | V755Y | B | 1.8e-07 | 1e-07 | 3.6e-07 | 2.133E-07 | 1.332E-07 | 1.000E-07 | 3.600E-07 |
| 15100 | 755 | V755V | B | 3.8e-07 | 4.6e-07 | 2.1e-07 | 3.500E-07 | 1.277E-07 | 2.100E-07 | 4.600E-07 |
| 15101 | 756 | C756A | B | 1.4e-07 | 2.5e-07 | 2.2e-07 | 2.033E-07 | 5.686E-08 | 1.400E-07 | 2.500E-07 |
| 15102 | 756 | C756R | B | 2.7e-07 | 1.5e-07 | 2.2e-07 | 2.133E-07 | 6.028E-08 | 1.500E-07 | 2.700E-07 |
| 15103 | 756 | C756N | B | 1.9e-07 | 3.5e-07 | 1.4e-07 | 2.267E-07 | 1.097E-07 | 1.400E-07 | 3.500E-07 |
| 15104 | 756 | C756D | B | 2.5e-07 | 2.1e-07 | 4e-07 | 2.867E-07 | 1.002E-07 | 2.100E-07 | 4.000E-07 |
| 15105 | 756 | C756C | B | 2.9e-07 | 6.5e-07 | 4.5e-07 | 4.633E-07 | 1.804E-07 | 2.900E-07 | 6.500E-07 |
| 15106 | 756 | C756E | B | 2.5e-07 | 2.3e-07 | 1.6e-07 | 2.133E-07 | 4.726E-08 | 1.600E-07 | 2.500E-07 |
| 15107 | 756 | C756Q | B | 2.8e-07 | 9.1e-08 | 2.3e-07 | 2.003E-07 | 9.793E-08 | 9.100E-08 | 2.800E-07 |
| 15108 | 756 | C756G | B | 4.2e-07 | 1.9e-07 | 2.4e-07 | 2.833E-07 | 1.210E-07 | 1.900E-07 | 4.200E-07 |
| 15109 | 756 | C756H | B | 3.4e-07 | 2e-07 | 1.6e-07 | 2.333E-07 | 9.452E-08 | 1.600E-07 | 3.400E-07 |
| 15110 | 756 | C756I | B | 3.7e-07 | 3.4e-07 | 2.3e-07 | 3.133E-07 | 7.371E-08 | 2.300E-07 | 3.700E-07 |
| 15111 | 756 | C756L | B | 1.8e-07 | 2.2e-07 | 1.8e-07 | 1.933E-07 | 2.309E-08 | 1.800E-07 | 2.200E-07 |
| 15112 | 756 | C756K | B | 2.8e-07 | 4.6e-07 | 2.7e-07 | 3.367E-07 | 1.069E-07 | 2.700E-07 | 4.600E-07 |
| 15113 | 756 | C756M | B | 2.6e-07 | 2.1e-07 | 2.8e-07 | 2.500E-07 | 3.606E-08 | 2.100E-07 | 2.800E-07 |
| 15114 | 756 | C756F | B | 3.3e-07 | 3.8e-07 | 5.1e-07 | 4.067E-07 | 9.292E-08 | 3.300E-07 | 5.100E-07 |
| 15115 | 756 | C756P | B | 2e-07 | 2.1e-07 | 2.9e-07 | 2.333E-07 | 4.933E-08 | 2.000E-07 | 2.900E-07 |
| 15116 | 756 | C756S | B | 2.9e-07 | 1.9e-07 | 2.2e-07 | 2.333E-07 | 5.132E-08 | 1.900E-07 | 2.900E-07 |
| 15117 | 756 | C756T | B | 2e-07 | 2e-07 | 3.8e-07 | 2.600E-07 | 1.039E-07 | 2.000E-07 | 3.800E-07 |
| 15118 | 756 | C756W | B | 3.2e-07 | 1.7e-07 | 2.8e-07 | 2.567E-07 | 7.767E-08 | 1.700E-07 | 3.200E-07 |
| 15119 | 756 | C756Y | B | 1.5e-07 | 6.3e-07 | 3.4e-07 | 3.733E-07 | 2.417E-07 | 1.500E-07 | 6.300E-07 |
| 15120 | 756 | C756V | B | 3.8e-07 | 4e-07 | 1.7e-07 | 3.167E-07 | 1.274E-07 | 1.700E-07 | 4.000E-07 |
| 15121 | 757 | L757A | B | 1.9e-07 | 1.6e-07 | 4.4e-07 | 2.633E-07 | 1.537E-07 | 1.600E-07 | 4.400E-07 |
| 15122 | 757 | L757R | B | 4.1e-07 | 1.7e-07 | 1.4e-07 | 2.400E-07 | 1.480E-07 | 1.400E-07 | 4.100E-07 |
| 15123 | 757 | L757N | B | 2.3e-07 | 2.2e-07 | 2.3e-07 | 2.267E-07 | 5.774E-09 | 2.200E-07 | 2.300E-07 |
| 15124 | 757 | L757D | B | 2.5e-07 | 1.4e-07 | 2.5e-07 | 2.133E-07 | 6.351E-08 | 1.400E-07 | 2.500E-07 |
| 15125 | 757 | L757C | B | 3.5e-07 | 1.3e-07 | 3.5e-07 | 2.767E-07 | 1.270E-07 | 1.300E-07 | 3.500E-07 |
| 15126 | 757 | L757E | B | 2.4e-07 | 4.2e-07 | 2.3e-07 | 2.967E-07 | 1.069E-07 | 2.300E-07 | 4.200E-07 |
| 15127 | 757 | L757Q | B | 1.6e-07 | 7.5e-07 | 2.5e-07 | 3.867E-07 | 3.179E-07 | 1.600E-07 | 7.500E-07 |
| 15128 | 757 | L757G | B | 2.7e-07 | 2.2e-07 | 2.3e-07 | 2.400E-07 | 2.646E-08 | 2.200E-07 | 2.700E-07 |
| 15129 | 757 | L757H | B | 1.7e-07 | 1.5e-07 | 2.3e-07 | 1.833E-07 | 4.163E-08 | 1.500E-07 | 2.300E-07 |

| | | | | | | | | | | |
|---|---|---|---|---|---|---|---|---|---|---|
| 15130 | 757 | L757I | B | 1.7e-07 | 2e-07 | 1.1e-07 | 1.600E-07 | 4.583E-08 | 1.100E-07 | 2.000E-07 |
| 15131 | 757 | L757L | B | 2.1e-07 | 4.1e-07 | 2.9e-07 | 3.033E-07 | 1.007E-07 | 2.100E-07 | 4.100E-07 |
| 15132 | 757 | L757K | B | 1.7e-07 | 2.2e-07 | 1.6e-07 | 1.833E-07 | 3.215E-08 | 1.600E-07 | 2.200E-07 |
| 15133 | 757 | L757M | B | 2.7e-07 | 3.5e-07 | 2.1e-07 | 2.767E-07 | 7.024E-08 | 2.100E-07 | 3.500E-07 |
| 15134 | 757 | L757F | B | 1.5e-07 | 2.9e-07 | 1.4e-07 | 1.933E-07 | 8.386E-08 | 1.400E-07 | 2.900E-07 |
| 15135 | 757 | L757P | B | 3.9e-07 | 2e-07 | 4.1e-07 | 3.333E-07 | 1.159E-07 | 2.000E-07 | 4.100E-07 |
| 15136 | 757 | L757S | B | 3.6e-07 | 3.9e-07 | 1.7e-07 | 3.067E-07 | 1.193E-07 | 1.700E-07 | 3.900E-07 |
| 15137 | 757 | L757T | B | 2.4e-07 | 1.5e-07 | 4.4e-07 | 2.767E-07 | 1.484E-07 | 1.500E-07 | 4.400E-07 |
| 15138 | 757 | L757W | B | 1.1e-07 | 2.3e-07 | 3.6e-07 | 2.333E-07 | 1.250E-07 | 1.100E-07 | 3.600E-07 |
| 15139 | 757 | L757Y | B | 2.1e-07 | 2.6e-07 | 1.9e-07 | 2.200E-07 | 3.606E-08 | 1.900E-07 | 2.600E-07 |
| 15140 | 757 | L757V | B | 3e-07 | 2.1e-07 | 2.9e-07 | 2.667E-07 | 4.933E-08 | 2.100E-07 | 3.000E-07 |
| 15141 | 758 | L758A | B | 2.2e-07 | 1.3e-07 | 2.9e-07 | 2.133E-07 | 8.021E-08 | 1.300E-07 | 2.900E-07 |
| 15142 | 758 | L758R | B | 1.5e-07 | 2.4e-07 | 2.6e-07 | 2.167E-07 | 5.859E-08 | 1.500E-07 | 2.600E-07 |
| 15143 | 758 | L758N | B | 1.8e-07 | 2.9e-07 | 5.8e-07 | 3.500E-07 | 2.066E-07 | 1.800E-07 | 5.800E-07 |
| 15144 | 758 | L758D | B | 6.2e-07 | 6e-07 | 3.6e-07 | 5.267E-07 | 1.447E-07 | 3.600E-07 | 6.200E-07 |
| 15145 | 758 | L758C | B | 2.6e-07 | 2e-07 | 2.3e-07 | 2.300E-07 | 3.000E-08 | 2.000E-07 | 2.600E-07 |
| 15146 | 758 | L758E | B | 2.7e-07 | 1.6e-07 | 3.7e-07 | 2.667E-07 | 1.050E-07 | 1.600E-07 | 3.700E-07 |
| 15147 | 758 | L758Q | B | 3.8e-07 | 1.6e-07 | 1.6e-07 | 2.333E-07 | 1.270E-07 | 1.600E-07 | 3.800E-07 |
| 15148 | 758 | L758G | B | 2.1e-07 | 1.6e-07 | 3.1e-07 | 2.267E-07 | 7.638E-08 | 1.600E-07 | 3.100E-07 |
| 15149 | 758 | L758H | B | 3e-07 | 5.3e-07 | 1.7e-07 | 3.333E-07 | 1.823E-07 | 1.700E-07 | 5.300E-07 |
| 15150 | 758 | L758I | B | 2.1e-07 | 3e-07 | 3.5e-07 | 2.867E-07 | 7.095E-08 | 2.100E-07 | 3.500E-07 |
| 15151 | 758 | L758L | B | 1.6e-07 | 2.6e-07 | 7.2e-08 | 1.640E-07 | 9.406E-08 | 7.200E-08 | 2.600E-07 |
| 15152 | 758 | L758K | B | 1.4e-07 | 3.9e-07 | 3.2e-07 | 2.833E-07 | 1.290E-07 | 1.400E-07 | 3.900E-07 |
| 15153 | 758 | L758M | B | 1.9e-07 | 2.9e-07 | 6.4e-07 | 3.733E-07 | 2.363E-07 | 1.900E-07 | 6.400E-07 |
| 15154 | 758 | L758F | B | 3.1e-07 | 2.5e-07 | 2.8e-07 | 2.800E-07 | 3.000E-08 | 2.500E-07 | 3.100E-07 |
| 15155 | 758 | L758P | B | 4.4e-07 | 5.3e-07 | 4.9e-07 | 4.867E-07 | 4.509E-08 | 4.400E-07 | 5.300E-07 |
| 15156 | 758 | L758S | B | 3e-07 | 3.7e-07 | 4.8e-07 | 3.833E-07 | 9.074E-08 | 3.000E-07 | 4.800E-07 |
| 15157 | 758 | L758T | B | 2.5e-07 | 4.1e-07 | 1.4e-07 | 2.667E-07 | 1.358E-07 | 1.400E-07 | 4.100E-07 |
| 15158 | 758 | L758W | B | 2.1e-07 | 5.8e-07 | 1.6e-07 | 3.167E-07 | 2.294E-07 | 1.600E-07 | 5.800E-07 |
| 15159 | 758 | L758Y | B | 1.4e-07 | 2.3e-07 | 1.4e-07 | 1.700E-07 | 5.196E-08 | 1.400E-07 | 2.300E-07 |
| 15160 | 758 | L758V | B | 3e-07 | 2.9e-07 | 9.1e-08 | 2.270E-07 | 1.179E-07 | 9.100E-08 | 3.000E-07 |
| 15161 | 759 | N759A | B | 2.7e-07 | 2e-07 | 3.2e-07 | 2.633E-07 | 6.028E-08 | 2.000E-07 | 3.200E-07 |
| 15162 | 759 | N759R | B | 1.3e-07 | 2.3e-07 | 3.2e-07 | 2.267E-07 | 9.504E-08 | 1.300E-07 | 3.200E-07 |
| 15163 | 759 | N759N | B | 4.5e-07 | 2.5e-07 | 1.4e-07 | 2.800E-07 | 1.572E-07 | 1.400E-07 | 4.500E-07 |
| 15164 | 759 | N759D | B | 3.9e-07 | 2.3e-07 | 3.4e-07 | 3.200E-07 | 8.185E-08 | 2.300E-07 | 3.900E-07 |
| 15165 | 759 | N759C | B | 2.7e-07 | 2.3e-07 | 2.5e-07 | 2.500E-07 | 2.000E-08 | 2.300E-07 | 2.700E-07 |
| 15166 | 759 | N759E | B | 3.5e-07 | 4e-07 | 2.5e-07 | 3.333E-07 | 7.638E-08 | 2.500E-07 | 4.000E-07 |
| 15167 | 759 | N759Q | B | 1.1e-07 | 2.6e-07 | 2.5e-07 | 2.067E-07 | 8.386E-08 | 1.100E-07 | 2.600E-07 |
| 15168 | 759 | N759G | B | 1.9e-07 | 3.2e-07 | 4.5e-07 | 3.200E-07 | 1.300E-07 | 1.900E-07 | 4.500E-07 |
| 15169 | 759 | N759H | B | 2.4e-07 | 4.1e-07 | 1.6e-07 | 2.700E-07 | 1.277E-07 | 1.600E-07 | 4.100E-07 |
| 15170 | 759 | N759I | B | 1.8e-07 | 2.8e-07 | 2e-07 | 2.200E-07 | 5.292E-08 | 1.800E-07 | 2.800E-07 |
| 15171 | 759 | N759L | B | 3.2e-07 | 2.9e-07 | 4e-07 | 3.367E-07 | 5.686E-08 | 2.900E-07 | 4.000E-07 |
| 15172 | 759 | N759K | B | 1.9e-07 | 1e-07 | 2e-07 | 1.633E-07 | 5.508E-08 | 1.000E-07 | 2.000E-07 |
| 15173 | 759 | N759M | B | 2.9e-07 | 2.3e-07 | 9e-08 | 2.033E-07 | 1.026E-07 | 9.000E-08 | 2.900E-07 |
| 15174 | 759 | N759F | B | 3.4e-07 | 4.3e-07 | 3e-07 | 3.567E-07 | 6.658E-08 | 3.000E-07 | 4.300E-07 |
| 15175 | 759 | N759P | B | 5e-07 | 3.3e-07 | 2.3e-07 | 3.533E-07 | 1.365E-07 | 2.300E-07 | 5.000E-07 |
| 15176 | 759 | N759S | B | 4.2e-07 | 1.3e-07 | 3.9e-07 | 3.133E-07 | 1.595E-07 | 1.300E-07 | 4.200E-07 |
| 15177 | 759 | N759T | B | 1.4e-07 | 3.5e-07 | 2.6e-07 | 2.500E-07 | 1.054E-07 | 1.400E-07 | 3.500E-07 |
| 15178 | 759 | N759W | B | 2.3e-07 | 2.1e-07 | 1.8e-07 | 2.067E-07 | 2.517E-08 | 1.800E-07 | 2.300E-07 |
| 15179 | 759 | N759Y | B | 2.5e-07 | 1.8e-07 | 1.3e-07 | 1.867E-07 | 6.028E-08 | 1.300E-07 | 2.500E-07 |
| 15180 | 759 | N759V | B | 2.2e-07 | 1.7e-07 | 1.5e-07 | 1.800E-07 | 3.606E-08 | 1.500E-07 | 2.200E-07 |
| 15181 | 760 | N760A | B | 3.3e-07 | 2.1e-07 | 5.9e-07 | 3.767E-07 | 1.943E-07 | 2.100E-07 | 5.900E-07 |
| 15182 | 760 | N760R | B | 2.5e-07 | 6.3e-07 | 2.1e-07 | 3.633E-07 | 2.318E-07 | 2.100E-07 | 6.300E-07 |
| 15183 | 760 | N760N | B | 2.3e-07 | 1.9e-07 | 1.7e-07 | 1.967E-07 | 3.055E-08 | 1.700E-07 | 2.300E-07 |
| 15184 | 760 | N760D | B | 2e-07 | 6.6e-07 | 2.2e-07 | 3.600E-07 | 2.600E-07 | 2.000E-07 | 6.600E-07 |
| 15185 | 760 | N760C | B | 3.2e-07 | 2.6e-07 | 3.3e-07 | 3.033E-07 | 3.786E-08 | 2.600E-07 | 3.300E-07 |
| 15186 | 760 | N760E | B | 1.7e-07 | 2.2e-07 | 1.3e-07 | 1.733E-07 | 4.509E-08 | 1.300E-07 | 2.200E-07 |
| 15187 | 760 | N760Q | B | 2.8e-07 | 1.5e-07 | 1.4e-07 | 1.900E-07 | 7.810E-08 | 1.400E-07 | 2.800E-07 |
| 15188 | 760 | N760G | B | 1.4e-07 | 2.6e-07 | 2.4e-07 | 2.133E-07 | 6.429E-08 | 1.400E-07 | 2.600E-07 |
| 15189 | 760 | N760H | B | 1.4e-07 | 2.9e-07 | 2.6e-07 | 2.300E-07 | 7.937E-08 | 1.400E-07 | 2.900E-07 |
| 15190 | 760 | N760I | B | 4.8e-07 | 2.7e-07 | 5.2e-07 | 4.233E-07 | 1.343E-07 | 2.700E-07 | 5.200E-07 |
| 15191 | 760 | N760L | B | 4.7e-07 | 2.6e-07 | 2e-07 | 3.100E-07 | 1.418E-07 | 2.000E-07 | 4.700E-07 |
| 15192 | 760 | N760K | B | 1.2e-07 | 3.1e-07 | 4.2e-07 | 2.833E-07 | 1.518E-07 | 1.200E-07 | 4.200E-07 |
| 15193 | 760 | N760M | B | 2.1e-07 | 4.7e-07 | 2e-07 | 2.933E-07 | 1.531E-07 | 2.000E-07 | 4.700E-07 |
| 15194 | 760 | N760F | B | 2.6e-07 | 3.1e-07 | 2.1e-07 | 2.600E-07 | 5.000E-08 | 2.100E-07 | 3.100E-07 |
| 15195 | 760 | N760P | B | 1.7e-07 | 1.9e-07 | 2.2e-07 | 1.933E-07 | 2.517E-08 | 1.700E-07 | 2.200E-07 |
| 15196 | 760 | N760S | B | 2.4e-07 | 2.8e-07 | 2.1e-07 | 2.433E-07 | 3.512E-08 | 2.100E-07 | 2.800E-07 |
| 15197 | 760 | N760T | B | 1.8e-07 | 3e-07 | 2.7e-07 | 2.500E-07 | 6.245E-08 | 1.800E-07 | 3.000E-07 |
| 15198 | 760 | N760W | B | 2.4e-07 | 3.3e-07 | 1.5e-07 | 2.400E-07 | 9.000E-08 | 1.500E-07 | 3.300E-07 |
| 15199 | 760 | N760Y | B | 4.1e-07 | 6e-07 | 3.3e-07 | 4.467E-07 | 1.387E-07 | 3.300E-07 | 6.000E-07 |
| 15200 | 760 | N760V | B | 2.2e-07 | 5.3e-07 | 2.7e-07 | 3.400E-07 | 1.664E-07 | 2.200E-07 | 5.300E-07 |
| 15201 | 761 | F761A | B | 2e-07 | 1.8e-07 | 4e-07 | 2.600E-07 | 1.217E-07 | 1.800E-07 | 4.000E-07 |
| 15202 | 761 | F761R | B | 2.7e-07 | 3.1e-07 | 2.7e-07 | 2.833E-07 | 2.309E-08 | 2.700E-07 | 3.100E-07 |
| 15203 | 761 | F761N | B | 4.4e-07 | 3.5e-07 | 2.3e-07 | 3.400E-07 | 1.054E-07 | 2.300E-07 | 4.400E-07 |
| 15204 | 761 | F761D | B | 1.1e-07 | 2.1e-07 | 2.6e-07 | 1.933E-07 | 7.638E-08 | 1.100E-07 | 2.600E-07 |
| 15205 | 761 | F761C | B | 1.4e-07 | 3.2e-07 | 3.1e-07 | 2.567E-07 | 1.012E-07 | 1.400E-07 | 3.200E-07 |
| 15206 | 761 | F761E | B | 1.3e-07 | 1.6e-07 | 1.5e-07 | 1.467E-07 | 1.528E-08 | 1.300E-07 | 1.600E-07 |
| 15207 | 761 | F761Q | B | 1.2e-07 | 2.6e-07 | 3.8e-07 | 2.533E-07 | 1.301E-07 | 1.200E-07 | 3.800E-07 |
| 15208 | 761 | F761G | B | 2.9e-07 | 1.5e-07 | 2.3e-07 | 2.233E-07 | 7.024E-08 | 1.500E-07 | 2.900E-07 |
| 15209 | 761 | F761H | B | 2.5e-07 | 2.6e-07 | 3e-07 | 2.700E-07 | 2.646E-08 | 2.500E-07 | 3.000E-07 |
| 15210 | 761 | F761I | B | 5e-07 | 3e-07 | 4.6e-07 | 4.200E-07 | 1.058E-07 | 3.000E-07 | 5.000E-07 |
| 15211 | 761 | F761L | B | 2.6e-07 | 8.7e-08 | 7.2e-08 | 1.397E-07 | 1.045E-07 | 7.200E-08 | 2.600E-07 |
| 15212 | 761 | F761K | B | 2e-07 | 2.9e-07 | 6.9e-07 | 3.933E-07 | 2.608E-07 | 2.000E-07 | 6.900E-07 |
| 15213 | 761 | F761M | B | 4.5e-07 | 1.7e-07 | 1.2e-07 | 2.467E-07 | 1.779E-07 | 1.200E-07 | 4.500E-07 |
| 15214 | 761 | F761F | B | 3.9e-07 | 2e-07 | 2.6e-07 | 2.833E-07 | 9.713E-08 | 2.000E-07 | 3.900E-07 |

| | | | | | | | | | | |
|---|---|---|---|---|---|---|---|---|---|---|
| 15215 | 761 | F761P | B | 5.1e-07 | 2.5e-07 | 2.4e-07 | 3.333E-07 | 1.531E-07 | 2.400E-07 | 5.100E-07 |
| 15216 | 761 | F761S | B | 2.1e-07 | 6.2e-08 | 1.6e-07 | 1.440E-07 | 7.529E-08 | 6.200E-08 | 2.100E-07 |
| 15217 | 761 | F761T | B | 1.3e-07 | 1.6e-07 | 2.1e-07 | 1.667E-07 | 4.041E-08 | 1.300E-07 | 2.100E-07 |
| 15218 | 761 | F761W | B | 3.8e-07 | 1e-07 | 3.5e-07 | 2.767E-07 | 1.537E-07 | 1.000E-07 | 3.800E-07 |
| 15219 | 761 | F761Y | B | 3e-07 | 4.6e-07 | 4.5e-07 | 4.033E-07 | 8.963E-08 | 3.000E-07 | 4.600E-07 |
| 15220 | 761 | F761V | B | 2.3e-07 | 1.5e-07 | 2e-07 | 1.933E-07 | 4.041E-08 | 1.500E-07 | 2.300E-07 |
| 15221 | 762 | Y762A | B | 1.7e-07 | 2.5e-07 | 1.9e-07 | 2.033E-07 | 4.163E-08 | 1.700E-07 | 2.500E-07 |
| 15222 | 762 | Y762R | B | 3.5e-07 | 2.5e-07 | 2.3e-07 | 2.767E-07 | 6.429E-08 | 2.300E-07 | 3.500E-07 |
| 15223 | 762 | Y762N | B | 1e-07 | 2.8e-07 | 4.3e-07 | 2.700E-07 | 1.652E-07 | 1.000E-07 | 4.300E-07 |
| 15224 | 762 | Y762D | B | 2.2e-07 | 6.9e-07 | 5.3e-07 | 4.800E-07 | 2.390E-07 | 2.200E-07 | 6.900E-07 |
| 15225 | 762 | Y762C | B | 2.9e-07 | 1.8e-07 | 2e-07 | 2.233E-07 | 5.859E-08 | 1.800E-07 | 2.900E-07 |
| 15226 | 762 | Y762E | B | 2.7e-07 | 5.6e-07 | 3.1e-07 | 3.800E-07 | 1.572E-07 | 2.700E-07 | 5.600E-07 |
| 15227 | 762 | Y762Q | B | 3.4e-07 | 2.5e-07 | 1.6e-07 | 2.500E-07 | 9.000E-08 | 1.600E-07 | 3.400E-07 |
| 15228 | 762 | Y762G | B | 2.8e-07 | 3.9e-07 | 1.4e-07 | 2.700E-07 | 1.253E-07 | 1.400E-07 | 3.900E-07 |
| 15229 | 762 | Y762H | B | 2.9e-07 | 1.6e-07 | 2.7e-07 | 2.400E-07 | 7.000E-08 | 1.600E-07 | 2.900E-07 |
| 15230 | 762 | Y762I | B | 2.7e-07 | 2.5e-07 | 2e-07 | 2.400E-07 | 3.606E-08 | 2.000E-07 | 2.700E-07 |
| 15231 | 762 | Y762L | B | 1.5e-07 | 2.9e-07 | 4e-07 | 2.800E-07 | 1.253E-07 | 1.500E-07 | 4.000E-07 |
| 15232 | 762 | Y762K | B | 3.2e-07 | 5.5e-07 | 5.6e-07 | 4.767E-07 | 1.358E-07 | 3.200E-07 | 5.600E-07 |
| 15233 | 762 | Y762M | B | 2.8e-07 | 3e-07 | 2e-07 | 2.600E-07 | 5.292E-08 | 2.000E-07 | 3.000E-07 |
| 15234 | 762 | Y762F | B | 4.3e-07 | 1.8e-07 | 2.8e-07 | 2.967E-07 | 1.258E-07 | 1.800E-07 | 4.300E-07 |
| 15235 | 762 | Y762P | B | 1.7e-07 | 2.8e-07 | 2e-07 | 2.167E-07 | 5.686E-08 | 1.700E-07 | 2.800E-07 |
| 15236 | 762 | Y762S | B | 2.4e-07 | 4.5e-07 | 2e-07 | 2.967E-07 | 1.343E-07 | 2.000E-07 | 4.500E-07 |
| 15237 | 762 | Y762T | B | 1.8e-07 | 5.5e-07 | 4.1e-07 | 3.800E-07 | 1.868E-07 | 1.800E-07 | 5.500E-07 |
| 15238 | 762 | Y762W | B | 3.5e-07 | 1.9e-07 | 2.1e-07 | 2.500E-07 | 8.718E-08 | 1.900E-07 | 3.500E-07 |
| 15239 | 762 | Y762Y | B | 3.9e-07 | 1.7e-07 | 2.1e-07 | 2.567E-07 | 1.172E-07 | 1.700E-07 | 3.900E-07 |
| 15240 | 762 | Y762V | B | 3.4e-07 | 2.1e-07 | 2.1e-07 | 2.533E-07 | 7.506E-08 | 2.100E-07 | 3.400E-07 |
| 15241 | 763 | P763A | B | 2.3e-07 | 2.5e-07 | 2.7e-07 | 2.500E-07 | 2.000E-08 | 2.300E-07 | 2.700E-07 |
| 15242 | 763 | P763R | B | 3e-07 | 4.2e-07 | 1.3e-07 | 2.833E-07 | 1.457E-07 | 1.300E-07 | 4.200E-07 |
| 15243 | 763 | P763N | B | 2e-07 | 4.9e-07 | 6.6e-07 | 4.500E-07 | 2.326E-07 | 2.000E-07 | 6.600E-07 |
| 15244 | 763 | P763D | B | 3.2e-07 | 1.3e-07 | 3.7e-07 | 2.733E-07 | 1.266E-07 | 1.300E-07 | 3.700E-07 |
| 15245 | 763 | P763C | B | 1.6e-07 | 2.9e-07 | 2.7e-07 | 2.400E-07 | 7.000E-08 | 1.600E-07 | 2.900E-07 |
| 15246 | 763 | P763E | B | 3.8e-07 | 2.2e-07 | 3e-07 | 3.000E-07 | 8.000E-08 | 2.200E-07 | 3.800E-07 |
| 15247 | 763 | P763Q | B | 2.5e-07 | 2e-07 | 3.4e-07 | 2.633E-07 | 7.095E-08 | 2.000E-07 | 3.400E-07 |
| 15248 | 763 | P763G | B | 1.8e-07 | 5.8e-07 | 1.4e-07 | 3.000E-07 | 2.433E-07 | 1.400E-07 | 5.800E-07 |
| 15249 | 763 | P763H | B | 1.5e-07 | 3.2e-07 | 7.3e-07 | 4.000E-07 | 2.982E-07 | 1.500E-07 | 7.300E-07 |
| 15250 | 763 | P763I | B | 1.8e-07 | 2.3e-07 | 1.8e-07 | 1.967E-07 | 2.887E-08 | 1.800E-07 | 2.300E-07 |
| 15251 | 763 | P763L | B | 3.2e-07 | 2.3e-07 | 3.4e-07 | 2.967E-07 | 5.859E-08 | 2.300E-07 | 3.400E-07 |
| 15252 | 763 | P763K | B | 3.6e-07 | 1.9e-07 | 1.5e-07 | 2.333E-07 | 1.115E-07 | 1.500E-07 | 3.600E-07 |
| 15253 | 763 | P763M | B | 2.2e-07 | 1.8e-07 | 3.8e-07 | 2.600E-07 | 1.058E-07 | 1.800E-07 | 3.800E-07 |
| 15254 | 763 | P763F | B | 1.7e-07 | 2.9e-07 | 2.4e-07 | 2.333E-07 | 6.028E-08 | 1.700E-07 | 2.900E-07 |
| 15255 | 763 | P763P | B | 2.2e-07 | 1.6e-07 | 2.4e-07 | 2.067E-07 | 4.163E-08 | 1.600E-07 | 2.400E-07 |
| 15256 | 763 | P763S | B | 2.7e-07 | 1.5e-07 | 2.6e-07 | 2.267E-07 | 6.658E-08 | 1.500E-07 | 2.700E-07 |
| 15257 | 763 | P763T | B | 1.4e-07 | 5.4e-07 | 3.9e-07 | 3.567E-07 | 2.021E-07 | 1.400E-07 | 5.400E-07 |
| 15258 | 763 | P763W | B | 1.8e-07 | 2.6e-07 | 3.2e-07 | 2.533E-07 | 7.024E-08 | 1.800E-07 | 3.200E-07 |
| 15259 | 763 | P763Y | B | 2.4e-07 | 2.3e-07 | 2.7e-07 | 2.467E-07 | 2.082E-08 | 2.300E-07 | 2.700E-07 |
| 15260 | 763 | P763V | B | 3.1e-07 | 1e-07 | 3.1e-07 | 2.400E-07 | 1.212E-07 | 1.000E-07 | 3.100E-07 |
| 15261 | 764 | R764A | B | 3.8e-07 | 1.1e-07 | 4.6e-07 | 3.167E-07 | 1.834E-07 | 1.100E-07 | 4.600E-07 |
| 15262 | 764 | R764R | B | 3.7e-07 | 1.7e-07 | 2.1e-07 | 2.500E-07 | 1.058E-07 | 1.700E-07 | 3.700E-07 |
| 15263 | 764 | R764N | B | 1.4e-07 | 3e-07 | 1.7e-07 | 2.033E-07 | 8.505E-08 | 1.400E-07 | 3.000E-07 |
| 15264 | 764 | R764D | B | 1e-07 | 2.2e-07 | 2e-07 | 1.733E-07 | 6.429E-08 | 1.000E-07 | 2.200E-07 |
| 15265 | 764 | R764C | B | 3.8e-07 | 7.3e-08 | 2e-07 | 2.177E-07 | 1.543E-07 | 7.300E-08 | 3.800E-07 |
| 15266 | 764 | R764E | B | 3e-07 | 2e-07 | 2.7e-07 | 2.567E-07 | 5.132E-08 | 2.000E-07 | 3.000E-07 |
| 15267 | 764 | R764Q | B | 1.4e-07 | 3.9e-07 | 1.6e-07 | 2.300E-07 | 1.389E-07 | 1.400E-07 | 3.900E-07 |
| 15268 | 764 | R764G | B | 1.7e-07 | 4e-07 | 2.3e-07 | 2.667E-07 | 1.193E-07 | 1.700E-07 | 4.000E-07 |
| 15269 | 764 | R764H | B | 1.9e-07 | 4.5e-07 | 2.3e-07 | 2.900E-07 | 1.400E-07 | 1.900E-07 | 4.500E-07 |
| 15270 | 764 | R764I | B | 2.4e-07 | 3.3e-07 | 3e-07 | 2.900E-07 | 4.583E-08 | 2.400E-07 | 3.300E-07 |
| 15271 | 764 | R764L | B | 4.8e-07 | 2.9e-07 | 4.1e-07 | 3.933E-07 | 9.609E-08 | 2.900E-07 | 4.800E-07 |
| 15272 | 764 | R764K | B | 2.6e-07 | 2.6e-07 | 3.6e-07 | 2.933E-07 | 5.774E-08 | 2.600E-07 | 3.600E-07 |
| 15273 | 764 | R764M | B | 3.2e-07 | 1.5e-07 | 1.9e-07 | 2.200E-07 | 8.888E-08 | 1.500E-07 | 3.200E-07 |
| 15274 | 764 | R764F | B | 2.8e-07 | 2.8e-07 | 1.7e-07 | 2.433E-07 | 6.351E-08 | 1.700E-07 | 2.800E-07 |
| 15275 | 764 | R764P | B | 3.9e-07 | 3.4e-07 | 2.1e-07 | 3.133E-07 | 9.292E-08 | 2.100E-07 | 3.900E-07 |
| 15276 | 764 | R764S | B | 2.2e-07 | 3.1e-07 | 2.5e-07 | 2.600E-07 | 4.583E-08 | 2.200E-07 | 3.100E-07 |
| 15277 | 764 | R764T | B | 1.6e-07 | 1.5e-07 | 3.5e-07 | 2.200E-07 | 1.127E-07 | 1.500E-07 | 3.500E-07 |
| 15278 | 764 | R764W | B | 2e-07 | 2.9e-07 | 3.1e-07 | 2.667E-07 | 5.859E-08 | 2.000E-07 | 3.100E-07 |
| 15279 | 764 | R764Y | B | 3e-07 | 3.5e-07 | 4.2e-07 | 3.567E-07 | 6.028E-08 | 3.000E-07 | 4.200E-07 |
| 15280 | 764 | R764V | B | 3.5e-07 | 2.4e-07 | 2.9e-07 | 2.933E-07 | 5.508E-08 | 2.400E-07 | 3.500E-07 |
| 15281 | 765 | E765A | B | 1.7e-07 | 1.2e-07 | 1.3e-07 | 1.400E-07 | 2.646E-08 | 1.200E-07 | 1.700E-07 |
| 15282 | 765 | E765R | B | 1.9e-07 | 3.9e-07 | 2.5e-07 | 2.767E-07 | 1.026E-07 | 1.900E-07 | 3.900E-07 |
| 15283 | 765 | E765N | B | 1.7e-07 | 2.5e-07 | 2e-07 | 2.067E-07 | 4.041E-08 | 1.700E-07 | 2.500E-07 |
| 15284 | 765 | E765D | B | 2.8e-07 | 3.8e-07 | 5.3e-07 | 3.967E-07 | 1.258E-07 | 2.800E-07 | 5.300E-07 |
| 15285 | 765 | E765C | B | 2.4e-07 | 1.1e-07 | 2.3e-07 | 1.933E-07 | 7.234E-08 | 1.100E-07 | 2.400E-07 |
| 15286 | 765 | E765E | B | 2.4e-07 | 2.7e-07 | 1.4e-07 | 2.167E-07 | 6.807E-08 | 1.400E-07 | 2.700E-07 |
| 15287 | 765 | E765Q | B | 1.1e-07 | 2.1e-07 | 5e-07 | 2.733E-07 | 2.026E-07 | 1.100E-07 | 5.000E-07 |
| 15288 | 765 | E765G | B | 3.4e-07 | 2.5e-07 | 3.1e-07 | 3.000E-07 | 4.583E-08 | 2.500E-07 | 3.400E-07 |
| 15289 | 765 | E765H | B | 1.8e-07 | 1.1e-07 | 1e-07 | 1.300E-07 | 4.359E-08 | 1.000E-07 | 1.800E-07 |
| 15290 | 765 | E765I | B | 3.4e-07 | 3.4e-07 | 1.1e-07 | 2.633E-07 | 1.328E-07 | 1.100E-07 | 3.400E-07 |
| 15291 | 765 | E765L | B | 4.4e-07 | 4.5e-07 | 3.2e-07 | 4.033E-07 | 7.234E-08 | 3.200E-07 | 4.500E-07 |
| 15292 | 765 | E765K | B | 2.4e-07 | 4.2e-07 | 5.2e-07 | 3.933E-07 | 1.419E-07 | 2.400E-07 | 5.200E-07 |
| 15293 | 765 | E765M | B | 2.3e-07 | 4.3e-07 | 1.6e-07 | 2.733E-07 | 1.401E-07 | 1.600E-07 | 4.300E-07 |
| 15294 | 765 | E765F | B | 1.7e-07 | 2.7e-07 | 5.2e-07 | 3.200E-07 | 1.803E-07 | 1.700E-07 | 5.200E-07 |
| 15295 | 765 | E765P | B | 2.6e-07 | 2.1e-07 | 2.3e-07 | 2.333E-07 | 2.517E-08 | 2.100E-07 | 2.600E-07 |
| 15296 | 765 | E765S | B | 1.8e-07 | 1.8e-07 | 2.4e-07 | 2.000E-07 | 3.464E-08 | 1.800E-07 | 2.400E-07 |
| 15297 | 765 | E765T | B | 2.6e-07 | 2.2e-07 | 1.9e-07 | 2.233E-07 | 3.512E-08 | 1.900E-07 | 2.600E-07 |
| 15298 | 765 | E765W | B | 2.3e-07 | 3.8e-07 | 1.2e-07 | 2.433E-07 | 1.305E-07 | 1.200E-07 | 3.800E-07 |
| 15299 | 765 | E765Y | B | 2.7e-07 | 2.7e-07 | 2.8e-07 | 2.733E-07 | 5.774E-09 | 2.700E-07 | 2.800E-07 |

| | | | | | | | | | | |
|---|---|---|---|---|---|---|---|---|---|---|
| 15300 | 765 | E765V | B | 4.7e-07 | 4.6e-07 | 2.7e-07 | 4.000E-07 | 1.127E-07 | 2.700E-07 | 4.700E-07 |
| 15301 | 766 | A766A | B | 5.4e-07 | 1.6e-07 | 3.7e-07 | 3.567E-07 | 1.904E-07 | 1.600E-07 | 5.400E-07 |
| 15302 | 766 | A766R | B | 3.7e-07 | 1.5e-07 | 3.4e-07 | 2.867E-07 | 1.193E-07 | 1.500E-07 | 3.700E-07 |
| 15303 | 766 | A766N | B | 2.4e-07 | 1.7e-07 | 5.1e-07 | 3.067E-07 | 1.795E-07 | 1.700E-07 | 5.100E-07 |
| 15304 | 766 | A766D | B | 3.3e-07 | 4.2e-07 | 4.3e-07 | 3.933E-07 | 5.508E-08 | 3.300E-07 | 4.300E-07 |
| 15305 | 766 | A766C | B | 1.6e-07 | 1.8e-07 | 1.7e-07 | 1.700E-07 | 1.000E-08 | 1.600E-07 | 1.800E-07 |
| 15306 | 766 | A766E | B | 2.4e-07 | 2.8e-07 | 2.5e-07 | 2.567E-07 | 2.082E-08 | 2.400E-07 | 2.800E-07 |
| 15307 | 766 | A766Q | B | 2.1e-07 | 3.2e-07 | 4e-07 | 3.100E-07 | 9.539E-08 | 2.100E-07 | 4.000E-07 |
| 15308 | 766 | A766G | B | 2.7e-07 | 3.3e-07 | 3e-07 | 3.000E-07 | 3.000E-08 | 2.700E-07 | 3.300E-07 |
| 15309 | 766 | A766H | B | 1.9e-07 | 4.4e-07 | 1.7e-07 | 2.667E-07 | 1.504E-07 | 1.700E-07 | 4.400E-07 |
| 15310 | 766 | A766I | B | 4.3e-07 | 3.5e-07 | 1.1e-07 | 2.967E-07 | 1.665E-07 | 1.100E-07 | 4.300E-07 |
| 15311 | 766 | A766L | B | 3e-07 | 3.7e-07 | 2.1e-07 | 2.933E-07 | 8.021E-08 | 2.100E-07 | 3.700E-07 |
| 15312 | 766 | A766K | B | 2.3e-07 | 2e-07 | 2.1e-07 | 2.133E-07 | 1.528E-08 | 2.000E-07 | 2.300E-07 |
| 15313 | 766 | A766M | B | 2.4e-07 | 2.9e-07 | 5.7e-07 | 3.667E-07 | 1.779E-07 | 2.400E-07 | 5.700E-07 |
| 15314 | 766 | A766F | B | 4.9e-07 | 2.1e-07 | 4.8e-07 | 3.933E-07 | 1.589E-07 | 2.100E-07 | 4.900E-07 |
| 15315 | 766 | A766P | B | 2.4e-07 | 1.9e-07 | 5.5e-07 | 3.267E-07 | 1.950E-07 | 1.900E-07 | 5.500E-07 |
| 15316 | 766 | A766S | B | 1.3e-07 | 1.6e-07 | 3.1e-07 | 2.000E-07 | 9.644E-08 | 1.300E-07 | 3.100E-07 |
| 15317 | 766 | A766T | B | 3.7e-07 | 2.3e-07 | 1.9e-07 | 2.633E-07 | 9.452E-08 | 1.900E-07 | 3.700E-07 |
| 15318 | 766 | A766W | B | 2.7e-07 | 8.1e-08 | 2.8e-07 | 2.103E-07 | 1.121E-07 | 8.100E-08 | 2.800E-07 |
| 15319 | 766 | A766Y | B | 1e-07 | 2.3e-07 | 2e-07 | 1.767E-07 | 6.807E-08 | 1.000E-07 | 2.300E-07 |
| 15320 | 766 | A766V | B | 2.1e-07 | 2.5e-07 | 1.8e-07 | 2.133E-07 | 3.512E-08 | 1.800E-07 | 2.500E-07 |
| 15321 | 767 | K767A | B | 5.2e-07 | 1.7e-07 | 3.8e-07 | 3.567E-07 | 1.762E-07 | 1.700E-07 | 5.200E-07 |
| 15322 | 767 | K767R | B | 1.3e-07 | 3.9e-07 | 2.3e-07 | 2.500E-07 | 1.311E-07 | 1.300E-07 | 3.900E-07 |
| 15323 | 767 | K767N | B | 3.7e-07 | 3.4e-07 | 2.4e-07 | 3.167E-07 | 6.807E-08 | 2.400E-07 | 3.700E-07 |
| 15324 | 767 | K767D | B | 6e-07 | 2e-07 | 2.7e-07 | 3.567E-07 | 2.136E-07 | 2.000E-07 | 6.000E-07 |
| 15325 | 767 | K767C | B | 4.1e-07 | 2.5e-07 | 1.6e-07 | 2.733E-07 | 1.266E-07 | 1.600E-07 | 4.100E-07 |
| 15326 | 767 | K767E | B | 2e-07 | 2.6e-07 | 3e-07 | 2.533E-07 | 5.033E-08 | 2.000E-07 | 3.000E-07 |
| 15327 | 767 | K767Q | B | 1.5e-07 | 2.8e-07 | 2.7e-07 | 2.333E-07 | 7.234E-08 | 1.500E-07 | 2.800E-07 |
| 15328 | 767 | K767G | B | 1.8e-07 | 1e-07 | 2e-07 | 1.600E-07 | 5.292E-08 | 1.000E-07 | 2.000E-07 |
| 15329 | 767 | K767H | B | 3.6e-07 | 7.3e-07 | 3.1e-07 | 4.667E-07 | 2.294E-07 | 3.100E-07 | 7.300E-07 |
| 15330 | 767 | K767I | B | 3.5e-07 | 3.3e-07 | 3.1e-07 | 3.300E-07 | 2.000E-08 | 3.100E-07 | 3.500E-07 |
| 15331 | 767 | K767L | B | 2.6e-07 | 4.1e-07 | 1.7e-07 | 2.800E-07 | 1.212E-07 | 1.700E-07 | 4.100E-07 |
| 15332 | 767 | K767K | B | 2.7e-07 | 2.5e-07 | 2.1e-07 | 2.433E-07 | 3.055E-08 | 2.100E-07 | 2.700E-07 |
| 15333 | 767 | K767M | B | 1.7e-07 | 2.6e-07 | 2.1e-07 | 2.133E-07 | 4.509E-08 | 1.700E-07 | 2.600E-07 |
| 15334 | 767 | K767F | B | 1.6e-07 | 2.5e-07 | 2.7e-07 | 2.267E-07 | 5.859E-08 | 1.600E-07 | 2.700E-07 |
| 15335 | 767 | K767P | B | 3.3e-07 | 2.7e-07 | 1.6e-07 | 2.533E-07 | 8.622E-08 | 1.600E-07 | 3.300E-07 |
| 15336 | 767 | K767S | B | 1.5e-07 | 3.1e-07 | 2.1e-07 | 2.233E-07 | 8.083E-08 | 1.500E-07 | 3.100E-07 |
| 15337 | 767 | K767T | B | 3.6e-07 | 1.1e-07 | 2.1e-07 | 2.267E-07 | 1.258E-07 | 1.100E-07 | 3.600E-07 |
| 15338 | 767 | K767W | B | 2.7e-07 | 4e-07 | 2.3e-07 | 3.000E-07 | 8.888E-08 | 2.300E-07 | 4.000E-07 |
| 15339 | 767 | K767Y | B | 8.5e-08 | 3.9e-07 | 1.2e-07 | 1.983E-07 | 1.669E-07 | 8.500E-08 | 3.900E-07 |
| 15340 | 767 | K767V | B | 5.5e-07 | 1.6e-07 | 2.5e-07 | 3.200E-07 | 2.042E-07 | 1.600E-07 | 5.500E-07 |
| 15341 | 768 | V768A | B | 1.8e-07 | 4.4e-07 | 3.4e-07 | 3.200E-07 | 1.311E-07 | 1.800E-07 | 4.400E-07 |
| 15342 | 768 | V768R | B | 3.1e-07 | 1.5e-07 | 2.4e-07 | 2.333E-07 | 8.021E-08 | 1.500E-07 | 3.100E-07 |
| 15343 | 768 | V768N | B | 1.8e-07 | 4.5e-07 | 3.5e-07 | 3.267E-07 | 1.365E-07 | 1.800E-07 | 4.500E-07 |
| 15344 | 768 | V768D | B | 1.5e-07 | 2.1e-07 | 2e-07 | 1.867E-07 | 3.215E-08 | 1.500E-07 | 2.100E-07 |
| 15345 | 768 | V768C | B | 2.8e-07 | 3.7e-07 | 2.2e-07 | 2.900E-07 | 7.550E-08 | 2.200E-07 | 3.700E-07 |
| 15346 | 768 | V768E | B | 6.1e-07 | 2.6e-07 | 2.7e-07 | 3.800E-07 | 1.992E-07 | 2.600E-07 | 6.100E-07 |
| 15347 | 768 | V768Q | B | 4.8e-07 | 1.9e-07 | 4.3e-07 | 3.667E-07 | 1.550E-07 | 1.900E-07 | 4.800E-07 |
| 15348 | 768 | V768G | B | 1.9e-07 | 2.9e-07 | 3e-07 | 2.600E-07 | 6.083E-08 | 1.900E-07 | 3.000E-07 |
| 15349 | 768 | V768H | B | 1.3e-07 | 4.2e-07 | 2.1e-07 | 2.533E-07 | 1.498E-07 | 1.300E-07 | 4.200E-07 |
| 15350 | 768 | V768I | B | 2.3e-07 | 2.3e-07 | 3e-07 | 2.533E-07 | 4.041E-08 | 2.300E-07 | 3.000E-07 |
| 15351 | 768 | V768L | B | 3.3e-07 | 2.5e-07 | 6.2e-07 | 4.000E-07 | 1.947E-07 | 2.500E-07 | 6.200E-07 |
| 15352 | 768 | V768K | B | 1.8e-07 | 8.5e-08 | 3.1e-07 | 1.917E-07 | 1.130E-07 | 8.500E-08 | 3.100E-07 |
| 15353 | 768 | V768M | B | 4e-07 | 2.4e-07 | 2.4e-07 | 2.933E-07 | 9.238E-08 | 2.400E-07 | 4.000E-07 |
| 15354 | 768 | V768F | B | 2.9e-07 | 2.6e-07 | 3.3e-07 | 2.933E-07 | 3.512E-08 | 2.600E-07 | 3.300E-07 |
| 15355 | 768 | V768P | B | 5e-07 | 2.5e-07 | 4.5e-07 | 4.000E-07 | 1.323E-07 | 2.500E-07 | 5.000E-07 |
| 15356 | 768 | V768S | B | 3e-07 | 3e-07 | 2.5e-07 | 2.833E-07 | 2.887E-08 | 2.500E-07 | 3.000E-07 |
| 15357 | 768 | V768T | B | 3.6e-07 | 1.9e-07 | 9.2e-08 | 2.140E-07 | 1.356E-07 | 9.200E-08 | 3.600E-07 |
| 15358 | 768 | V768W | B | 1.2e-07 | 2e-07 | 1.5e-07 | 1.567E-07 | 4.041E-08 | 1.200E-07 | 2.000E-07 |
| 15359 | 768 | V768Y | B | 2e-07 | 1.9e-07 | 1.2e-07 | 1.700E-07 | 4.359E-08 | 1.200E-07 | 2.000E-07 |
| 15360 | 768 | V768V | B | 1.6e-07 | 3.7e-07 | 1.8e-07 | 2.367E-07 | 1.159E-07 | 1.600E-07 | 3.700E-07 |
| 15361 | 769 | Q769A | B | 1.7e-07 | 2e-07 | 2.2e-07 | 1.967E-07 | 2.517E-08 | 1.700E-07 | 2.200E-07 |
| 15362 | 769 | Q769R | B | 1.4e-07 | 1.4e-07 | 2.6e-07 | 1.800E-07 | 6.928E-08 | 1.400E-07 | 2.600E-07 |
| 15363 | 769 | Q769N | B | 3e-07 | 2.1e-07 | 3.6e-07 | 2.900E-07 | 7.550E-08 | 2.100E-07 | 3.600E-07 |
| 15364 | 769 | Q769D | B | 2.2e-07 | 3.2e-07 | 2.1e-07 | 2.500E-07 | 6.083E-08 | 2.100E-07 | 3.200E-07 |
| 15365 | 769 | Q769C | B | 2.1e-07 | 1.7e-07 | 2.4e-07 | 2.067E-07 | 3.512E-08 | 1.700E-07 | 2.400E-07 |
| 15366 | 769 | Q769E | B | 1.4e-07 | 1.5e-07 | 1.6e-07 | 1.500E-07 | 1.000E-08 | 1.400E-07 | 1.600E-07 |
| 15367 | 769 | Q769Q | B | 2.3e-07 | 2.1e-07 | 1.4e-07 | 1.933E-07 | 4.726E-08 | 1.400E-07 | 2.300E-07 |
| 15368 | 769 | Q769G | B | 1.7e-07 | 2.7e-07 | 2.4e-07 | 2.267E-07 | 5.132E-08 | 1.700E-07 | 2.700E-07 |
| 15369 | 769 | Q769H | B | 6.7e-07 | 8.5e-08 | 4.9e-07 | 4.150E-07 | 2.996E-07 | 8.500E-08 | 6.700E-07 |
| 15370 | 769 | Q769I | B | 2e-07 | 3.1e-07 | 4.2e-07 | 3.100E-07 | 1.100E-07 | 2.000E-07 | 4.200E-07 |
| 15371 | 769 | Q769L | B | 2e-07 | 3.2e-07 | 2.8e-07 | 2.667E-07 | 6.110E-08 | 2.000E-07 | 3.200E-07 |
| 15372 | 769 | Q769K | B | 1.8e-07 | 1.5e-07 | 1.5e-07 | 1.600E-07 | 1.732E-08 | 1.500E-07 | 1.800E-07 |
| 15373 | 769 | Q769M | B | 3.7e-07 | 2.9e-07 | 1.7e-07 | 2.767E-07 | 1.007E-07 | 1.700E-07 | 3.700E-07 |
| 15374 | 769 | Q769F | B | 3.2e-07 | 7.9e-07 | 2.5e-07 | 4.533E-07 | 2.937E-07 | 2.500E-07 | 7.900E-07 |
| 15375 | 769 | Q769P | B | 3e-07 | 2.9e-07 | 1.5e-07 | 2.467E-07 | 8.386E-08 | 1.500E-07 | 3.000E-07 |
| 15376 | 769 | Q769S | B | 3.8e-07 | 1.9e-07 | 4.1e-07 | 3.267E-07 | 1.193E-07 | 1.900E-07 | 4.100E-07 |
| 15377 | 769 | Q769T | B | 1.4e-07 | 3.8e-07 | 3.8e-07 | 3.000E-07 | 1.386E-07 | 1.400E-07 | 3.800E-07 |
| 15378 | 769 | Q769W | B | 2.1e-07 | 1.6e-07 | 2.8e-07 | 2.167E-07 | 6.028E-08 | 1.600E-07 | 2.800E-07 |
| 15379 | 769 | Q769Y | B | 2.3e-07 | 2.6e-07 | 1.3e-07 | 2.067E-07 | 6.807E-08 | 1.300E-07 | 2.600E-07 |
| 15380 | 769 | Q769V | B | 2.7e-07 | 3.6e-07 | 4.8e-07 | 3.700E-07 | 1.054E-07 | 2.700E-07 | 4.800E-07 |
| 15381 | 770 | W770A | B | 6.3e-07 | 1.4e-07 | 3.3e-07 | 3.667E-07 | 2.470E-07 | 1.400E-07 | 6.300E-07 |
| 15382 | 770 | W770R | B | 3.3e-07 | 4.5e-07 | 2.4e-07 | 3.400E-07 | 1.054E-07 | 2.400E-07 | 4.500E-07 |
| 15383 | 770 | W770N | B | 3.4e-07 | 2e-07 | 2.4e-07 | 2.600E-07 | 7.211E-08 | 2.000E-07 | 3.400E-07 |
| 15384 | 770 | W770D | B | 3.3e-07 | 1.9e-07 | 2.2e-07 | 2.467E-07 | 7.371E-08 | 1.900E-07 | 3.300E-07 |

| | | | | | | | | | | |
|---|---|---|---|---|---|---|---|---|---|---|
| 15385 | 770 | W770C | B | 3.3e-07 | 3.2e-07 | 1.7e-07 | 2.733E-07 | 8.963E-08 | 1.700E-07 | 3.300E-07 |
| 15386 | 770 | W770E | B | 9.8e-08 | 5.9e-07 | 1.2e-07 | 2.693E-07 | 2.779E-07 | 9.800E-08 | 5.900E-07 |
| 15387 | 770 | W770Q | B | 3.4e-07 | 1.7e-07 | 2.7e-07 | 2.600E-07 | 8.544E-08 | 1.700E-07 | 3.400E-07 |
| 15388 | 770 | W770G | B | 2.8e-07 | 2.2e-07 | 2.5e-07 | 2.500E-07 | 3.000E-08 | 2.200E-07 | 2.800E-07 |
| 15389 | 770 | W770H | B | 2.6e-07 | 4.1e-07 | 9.7e-08 | 2.557E-07 | 1.565E-07 | 9.700E-08 | 4.100E-07 |
| 15390 | 770 | W770I | B | 2.8e-07 | 3e-07 | 1.5e-07 | 2.433E-07 | 8.145E-08 | 1.500E-07 | 3.000E-07 |
| 15391 | 770 | W770L | B | 4e-07 | 3e-07 | 3.1e-07 | 3.367E-07 | 5.508E-08 | 3.000E-07 | 4.000E-07 |
| 15392 | 770 | W770K | B | 1.3e-07 | 2.8e-07 | 4e-07 | 2.700E-07 | 1.353E-07 | 1.300E-07 | 4.000E-07 |
| 15393 | 770 | W770M | B | 3.1e-07 | 2.7e-07 | 4.5e-07 | 3.433E-07 | 9.452E-08 | 2.700E-07 | 4.500E-07 |
| 15394 | 770 | W770F | B | 3.5e-07 | 2.8e-07 | 2.8e-07 | 3.033E-07 | 4.041E-08 | 2.800E-07 | 3.500E-07 |
| 15395 | 770 | W770P | B | 2e-07 | 3.3e-07 | 1.5e-07 | 2.267E-07 | 9.292E-08 | 1.500E-07 | 3.300E-07 |
| 15396 | 770 | W770S | B | 2.2e-07 | 3.1e-07 | 3.7e-07 | 3.000E-07 | 7.550E-08 | 2.200E-07 | 3.700E-07 |
| 15397 | 770 | W770T | B | 2.6e-07 | 2.4e-07 | 2.5e-07 | 2.500E-07 | 1.000E-08 | 2.400E-07 | 2.600E-07 |
| 15398 | 770 | W770W | B | 3.8e-07 | 2e-07 | 2.2e-07 | 2.667E-07 | 9.866E-08 | 2.000E-07 | 3.800E-07 |
| 15399 | 770 | W770Y | B | 3.2e-07 | 1.7e-07 | 1.5e-07 | 2.133E-07 | 9.292E-08 | 1.500E-07 | 3.200E-07 |
| 15400 | 770 | W770V | B | 1.8e-07 | 3.8e-07 | 2.2e-07 | 2.600E-07 | 1.058E-07 | 1.800E-07 | 3.800E-07 |
| 15401 | 771 | K771A | B | 1.3e-07 | 4e-07 | 2.4e-07 | 2.567E-07 | 1.358E-07 | 1.300E-07 | 4.000E-07 |
| 15402 | 771 | K771R | B | 4.2e-07 | 2.9e-07 | 2.8e-07 | 3.300E-07 | 7.810E-08 | 2.800E-07 | 4.200E-07 |
| 15403 | 771 | K771N | B | 1.7e-07 | 2e-07 | 1.6e-07 | 1.767E-07 | 2.082E-08 | 1.600E-07 | 2.000E-07 |
| 15404 | 771 | K771D | B | 6.6e-07 | 1.6e-07 | 2.5e-07 | 3.567E-07 | 2.665E-07 | 1.600E-07 | 6.600E-07 |
| 15405 | 771 | K771C | B | 1.8e-07 | 2.9e-07 | 3.8e-07 | 2.833E-07 | 1.002E-07 | 1.800E-07 | 3.800E-07 |
| 15406 | 771 | K771E | B | 2.5e-07 | 4.1e-07 | 4.4e-07 | 3.667E-07 | 1.021E-07 | 2.500E-07 | 4.400E-07 |
| 15407 | 771 | K771Q | B | 3.6e-07 | 3e-07 | 1.9e-07 | 2.833E-07 | 8.622E-08 | 1.900E-07 | 3.600E-07 |
| 15408 | 771 | K771G | B | 5.8e-07 | 1.3e-07 | 2.6e-07 | 3.233E-07 | 2.316E-07 | 1.300E-07 | 5.800E-07 |
| 15409 | 771 | K771H | B | 2e-07 | 2.7e-07 | 5.7e-07 | 3.467E-07 | 1.966E-07 | 2.000E-07 | 5.700E-07 |
| 15410 | 771 | K771I | B | 1.3e-07 | 2.6e-07 | 1.5e-07 | 1.800E-07 | 7.000E-08 | 1.300E-07 | 2.600E-07 |
| 15411 | 771 | K771L | B | 2.4e-07 | 2.5e-07 | 1.9e-07 | 2.267E-07 | 3.215E-08 | 1.900E-07 | 2.500E-07 |
| 15412 | 771 | K771K | B | 2.6e-07 | 3.3e-07 | 3.4e-07 | 3.100E-07 | 4.359E-08 | 2.600E-07 | 3.400E-07 |
| 15413 | 771 | K771M | B | 3.1e-07 | 2.4e-07 | 2.5e-07 | 2.667E-07 | 3.786E-08 | 2.400E-07 | 3.100E-07 |
| 15414 | 771 | K771F | B | 1.9e-07 | 1.4e-07 | 1.9e-07 | 1.733E-07 | 2.887E-08 | 1.400E-07 | 1.900E-07 |
| 15415 | 771 | K771P | B | 3.1e-07 | 2.2e-07 | 2.7e-07 | 2.667E-07 | 4.509E-08 | 2.200E-07 | 3.100E-07 |
| 15416 | 771 | K771S | B | 1.7e-07 | 1.7e-07 | 2.2e-07 | 1.867E-07 | 2.887E-08 | 1.700E-07 | 2.200E-07 |
| 15417 | 771 | K771T | B | 2.6e-07 | 3.4e-07 | 2.7e-07 | 2.900E-07 | 4.359E-08 | 2.600E-07 | 3.400E-07 |
| 15418 | 771 | K771W | B | 3.1e-07 | 2.3e-07 | 2e-07 | 2.467E-07 | 5.686E-08 | 2.000E-07 | 3.100E-07 |
| 15419 | 771 | K771Y | B | 3.3e-07 | 2.2e-07 | 2.2e-07 | 2.567E-07 | 6.351E-08 | 2.200E-07 | 3.300E-07 |
| 15420 | 771 | K771V | B | 2.3e-07 | 2.9e-07 | 1.2e-07 | 2.133E-07 | 8.622E-08 | 1.200E-07 | 2.900E-07 |
| 15421 | 772 | V772A | B | 1.8e-07 | 9.1e-08 | 2.4e-07 | 1.703E-07 | 7.497E-08 | 9.100E-08 | 2.400E-07 |
| 15422 | 772 | V772R | B | 2.6e-07 | 2.7e-07 | 2e-07 | 2.433E-07 | 3.786E-08 | 2.000E-07 | 2.700E-07 |
| 15423 | 772 | V772N | B | 2.1e-07 | 4.1e-07 | 3.6e-07 | 3.267E-07 | 1.041E-07 | 2.100E-07 | 4.100E-07 |
| 15424 | 772 | V772D | B | 1.2e-07 | 1.2e-07 | 2.3e-07 | 1.567E-07 | 6.351E-08 | 1.200E-07 | 2.300E-07 |
| 15425 | 772 | V772C | B | 2.1e-07 | 2.4e-07 | 2.9e-07 | 2.467E-07 | 4.041E-08 | 2.100E-07 | 2.900E-07 |
| 15426 | 772 | V772E | B | 3.9e-07 | 1.5e-07 | 2.5e-07 | 2.633E-07 | 1.206E-07 | 1.500E-07 | 3.900E-07 |
| 15427 | 772 | V772Q | B | 3.7e-07 | 2.3e-07 | 2e-07 | 2.667E-07 | 9.074E-08 | 2.000E-07 | 3.700E-07 |
| 15428 | 772 | V772G | B | 2.7e-07 | 1.8e-07 | 2.4e-07 | 2.300E-07 | 4.583E-08 | 1.800E-07 | 2.700E-07 |
| 15429 | 772 | V772H | B | 2.5e-07 | 1.8e-07 | 2.7e-07 | 2.333E-07 | 4.726E-08 | 1.800E-07 | 2.700E-07 |
| 15430 | 772 | V772I | B | 2.4e-07 | 3.6e-07 | 2e-07 | 2.667E-07 | 8.327E-08 | 2.000E-07 | 3.600E-07 |
| 15431 | 772 | V772L | B | 2.4e-07 | 3e-07 | 3.1e-07 | 2.833E-07 | 3.786E-08 | 2.400E-07 | 3.100E-07 |
| 15432 | 772 | V772K | B | 1.6e-07 | 2.3e-07 | 9.3e-08 | 1.610E-07 | 6.851E-08 | 9.300E-08 | 2.300E-07 |
| 15433 | 772 | V772M | B | 1.9e-07 | 3.7e-07 | 2.3e-07 | 2.633E-07 | 9.452E-08 | 1.900E-07 | 3.700E-07 |
| 15434 | 772 | V772F | B | 2.1e-07 | 1.7e-07 | 2.9e-07 | 2.233E-07 | 6.110E-08 | 1.700E-07 | 2.900E-07 |
| 15435 | 772 | V772P | B | 2e-07 | 3.6e-07 | 2.8e-07 | 2.800E-07 | 8.000E-08 | 2.000E-07 | 3.600E-07 |
| 15436 | 772 | V772S | B | 5.2e-07 | 1.5e-07 | 2.7e-07 | 3.133E-07 | 1.888E-07 | 1.500E-07 | 5.200E-07 |
| 15437 | 772 | V772T | B | 2.5e-07 | 2.5e-07 | 2.6e-07 | 2.533E-07 | 5.774E-09 | 2.500E-07 | 2.600E-07 |
| 15438 | 772 | V772W | B | 2e-07 | 4.1e-07 | 2.2e-07 | 2.767E-07 | 1.159E-07 | 2.000E-07 | 4.100E-07 |
| 15439 | 772 | V772Y | B | 2.3e-07 | 2.2e-07 | 1.1e-07 | 1.867E-07 | 6.658E-08 | 1.100E-07 | 2.300E-07 |
| 15440 | 772 | V772V | B | 1.7e-07 | 1.4e-07 | 2e-07 | 1.700E-07 | 3.000E-08 | 1.400E-07 | 2.000E-07 |
| 15441 | 773 | D773A | B | 1e-07 | 4.1e-07 | 2.4e-07 | 2.500E-07 | 1.552E-07 | 1.000E-07 | 4.100E-07 |
| 15442 | 773 | D773R | B | 1.9e-07 | 3.4e-07 | 3e-07 | 2.767E-07 | 7.767E-08 | 1.900E-07 | 3.400E-07 |
| 15443 | 773 | D773N | B | 2.7e-07 | 8.6e-07 | 1.5e-07 | 4.267E-07 | 3.800E-07 | 1.500E-07 | 8.600E-07 |
| 15444 | 773 | D773D | B | 2.9e-07 | 2.1e-07 | 2.1e-07 | 2.367E-07 | 4.619E-08 | 2.100E-07 | 2.900E-07 |
| 15445 | 773 | D773C | B | 3.6e-07 | 5e-07 | 2.8e-07 | 3.800E-07 | 1.114E-07 | 2.800E-07 | 5.000E-07 |
| 15446 | 773 | D773E | B | 1.4e-07 | 2.4e-07 | 2.5e-07 | 2.100E-07 | 6.083E-08 | 1.400E-07 | 2.500E-07 |
| 15447 | 773 | D773Q | B | 2.2e-07 | 4.7e-07 | 1.9e-07 | 2.933E-07 | 1.537E-07 | 1.900E-07 | 4.700E-07 |
| 15448 | 773 | D773G | B | 2.6e-07 | 3.4e-07 | 2.1e-07 | 2.700E-07 | 6.557E-08 | 2.100E-07 | 3.400E-07 |
| 15449 | 773 | D773H | B | 2.5e-07 | 2.7e-07 | 1.9e-07 | 2.367E-07 | 4.163E-08 | 1.900E-07 | 2.700E-07 |
| 15450 | 773 | D773I | B | 3.6e-07 | 2.8e-07 | 2.3e-07 | 2.900E-07 | 6.557E-08 | 2.300E-07 | 3.600E-07 |
| 15451 | 773 | D773L | B | 2.3e-07 | 1.1e-07 | 2.1e-07 | 1.833E-07 | 6.429E-08 | 1.100E-07 | 2.300E-07 |
| 15452 | 773 | D773K | B | 2.4e-07 | 3.7e-07 | 3.4e-07 | 3.167E-07 | 6.807E-08 | 2.400E-07 | 3.700E-07 |
| 15453 | 773 | D773M | B | 2.9e-07 | 3.3e-07 | 2.8e-07 | 3.000E-07 | 2.646E-08 | 2.800E-07 | 3.300E-07 |
| 15454 | 773 | D773F | B | 2.2e-07 | 2e-07 | 1.8e-07 | 2.000E-07 | 2.000E-08 | 1.800E-07 | 2.200E-07 |
| 15455 | 773 | D773P | B | 4.8e-07 | 2.7e-07 | 2.1e-07 | 3.200E-07 | 1.418E-07 | 2.100E-07 | 4.800E-07 |
| 15456 | 773 | D773S | B | 2.2e-07 | 1.9e-07 | 1.9e-07 | 2.000E-07 | 1.732E-08 | 1.900E-07 | 2.200E-07 |
| 15457 | 773 | D773T | B | 1.9e-07 | 1.6e-07 | 3.6e-07 | 2.367E-07 | 1.079E-07 | 1.600E-07 | 3.600E-07 |
| 15458 | 773 | D773W | B | 4.6e-07 | 2e-07 | 1.3e-07 | 2.633E-07 | 1.739E-07 | 1.300E-07 | 4.600E-07 |
| 15459 | 773 | D773Y | B | 2.1e-07 | 2.1e-07 | 2.6e-07 | 2.267E-07 | 2.887E-08 | 2.100E-07 | 2.600E-07 |
| 15460 | 773 | D773V | B | 2.9e-07 | 2.3e-07 | 3.5e-07 | 2.900E-07 | 6.000E-08 | 2.300E-07 | 3.500E-07 |
| 15461 | 774 | N774A | B | 1.8e-07 | 2e-07 | 1.7e-07 | 1.833E-07 | 1.528E-08 | 1.700E-07 | 2.000E-07 |
| 15462 | 774 | N774R | B | 3.7e-07 | 4.1e-07 | 1.9e-07 | 3.233E-07 | 1.172E-07 | 1.900E-07 | 4.100E-07 |
| 15463 | 774 | N774N | B | 2.6e-07 | 1.5e-07 | 2e-07 | 2.033E-07 | 5.508E-08 | 1.500E-07 | 2.600E-07 |
| 15464 | 774 | N774D | B | 1.9e-07 | 2.3e-07 | 2.9e-07 | 2.367E-07 | 5.033E-08 | 1.900E-07 | 2.900E-07 |
| 15465 | 774 | N774C | B | 1.9e-07 | 4e-07 | 1.2e-07 | 2.367E-07 | 1.457E-07 | 1.200E-07 | 4.000E-07 |
| 15466 | 774 | N774E | B | 3.9e-07 | 2.4e-07 | 2.5e-07 | 2.933E-07 | 8.386E-08 | 2.400E-07 | 3.900E-07 |
| 15467 | 774 | N774Q | B | 2.2e-07 | 1.7e-07 | 3.1e-07 | 2.333E-07 | 7.095E-08 | 1.700E-07 | 3.100E-07 |
| 15468 | 774 | N774G | B | 4.9e-07 | 2.5e-07 | 1.4e-07 | 2.933E-07 | 1.790E-07 | 1.400E-07 | 4.900E-07 |
| 15469 | 774 | N774H | B | 2.7e-07 | 2.3e-07 | 3.1e-07 | 2.700E-07 | 4.000E-08 | 2.300E-07 | 3.100E-07 |

| | | | | | | | | | | |
|---|---|---|---|---|---|---|---|---|---|---|
| 15470 | 774 | N774I | B | 1.3e-07 | 1.3e-07 | 1.6e-07 | 1.400E-07 | 1.732E-08 | 1.300E-07 | 1.600E-07 |
| 15471 | 774 | N774L | B | 3.4e-07 | 3e-07 | 2.7e-07 | 3.033E-07 | 3.512E-08 | 2.700E-07 | 3.400E-07 |
| 15472 | 774 | N774K | B | 3.2e-07 | 1.7e-07 | 2.8e-07 | 2.567E-07 | 7.767E-08 | 1.700E-07 | 3.200E-07 |
| 15473 | 774 | N774M | B | 4.4e-07 | 2.3e-07 | 2.2e-07 | 2.967E-07 | 1.242E-07 | 2.200E-07 | 4.400E-07 |
| 15474 | 774 | N774F | B | 2e-07 | 2.9e-07 | 2.5e-07 | 2.467E-07 | 4.509E-08 | 2.000E-07 | 2.900E-07 |
| 15475 | 774 | N774P | B | 3.3e-07 | 1.2e-07 | 3.4e-07 | 2.633E-07 | 1.242E-07 | 1.200E-07 | 3.400E-07 |
| 15476 | 774 | N774S | B | 1.2e-07 | 1.3e-07 | 2.3e-07 | 1.600E-07 | 6.083E-08 | 1.200E-07 | 2.300E-07 |
| 15477 | 774 | N774T | B | 2.5e-07 | 2e-07 | 3.9e-07 | 2.800E-07 | 9.849E-08 | 2.000E-07 | 3.900E-07 |
| 15478 | 774 | N774W | B | 3.1e-07 | 2e-07 | 3e-07 | 2.700E-07 | 6.083E-08 | 2.000E-07 | 3.100E-07 |
| 15479 | 774 | N774Y | B | 3.3e-07 | 4e-07 | 2.2e-07 | 3.167E-07 | 9.074E-08 | 2.200E-07 | 4.000E-07 |
| 15480 | 774 | N774V | B | 2.6e-07 | 1.4e-07 | 1.5e-07 | 1.833E-07 | 6.658E-08 | 1.400E-07 | 2.600E-07 |
| 15481 | 775 | A775A | B | 1.3e-07 | 2.2e-07 | 4.7e-07 | 2.733E-07 | 1.762E-07 | 1.300E-07 | 4.700E-07 |
| 15482 | 775 | A775R | B | 3.5e-07 | 8.6e-08 | 2.5e-07 | 2.287E-07 | 1.333E-07 | 8.600E-08 | 3.500E-07 |
| 15483 | 775 | A775N | B | 6.7e-08 | 2.1e-07 | 2e-07 | 1.590E-07 | 7.983E-08 | 6.700E-08 | 2.100E-07 |
| 15484 | 775 | A775D | B | 3.2e-07 | 2.7e-07 | 2.9e-07 | 2.933E-07 | 2.517E-08 | 2.700E-07 | 3.200E-07 |
| 15485 | 775 | A775C | B | 3e-07 | 2.2e-07 | 4.8e-07 | 3.333E-07 | 1.332E-07 | 2.200E-07 | 4.800E-07 |
| 15486 | 775 | A775E | B | 2.1e-07 | 2.8e-07 | 3.2e-07 | 2.700E-07 | 5.568E-08 | 2.100E-07 | 3.200E-07 |
| 15487 | 775 | A775Q | B | 1.8e-07 | 2.3e-07 | 3e-07 | 2.367E-07 | 6.028E-08 | 1.800E-07 | 3.000E-07 |
| 15488 | 775 | A775G | B | 1.6e-07 | 4.8e-07 | 2.1e-07 | 2.833E-07 | 1.721E-07 | 1.600E-07 | 4.800E-07 |
| 15489 | 775 | A775H | B | 1.4e-07 | 2.6e-07 | 2.1e-07 | 2.033E-07 | 6.028E-08 | 1.400E-07 | 2.600E-07 |
| 15490 | 775 | A775I | B | 1.6e-07 | 4.5e-07 | 3e-07 | 3.033E-07 | 1.450E-07 | 1.600E-07 | 4.500E-07 |
| 15491 | 775 | A775L | B | 4.2e-07 | 1.4e-07 | 3.5e-07 | 3.033E-07 | 1.457E-07 | 1.400E-07 | 4.200E-07 |
| 15492 | 775 | A775K | B | 1.1e-07 | 2.2e-07 | 2.4e-07 | 1.900E-07 | 7.000E-08 | 1.100E-07 | 2.400E-07 |
| 15493 | 775 | A775M | B | 2.7e-07 | 3.6e-07 | 1.2e-07 | 2.500E-07 | 1.212E-07 | 1.200E-07 | 3.600E-07 |
| 15494 | 775 | A775F | B | 2.4e-07 | 2.4e-07 | 3.7e-07 | 2.833E-07 | 7.506E-08 | 2.400E-07 | 3.700E-07 |
| 15495 | 775 | A775P | B | 5.2e-07 | 2.2e-07 | 3.3e-07 | 3.567E-07 | 1.518E-07 | 2.200E-07 | 5.200E-07 |
| 15496 | 775 | A775S | B | 2.3e-07 | 1.3e-07 | 4e-07 | 2.533E-07 | 1.365E-07 | 1.300E-07 | 4.000E-07 |
| 15497 | 775 | A775T | B | 5.7e-07 | 1.1e-07 | 1.5e-07 | 2.767E-07 | 2.548E-07 | 1.100E-07 | 5.700E-07 |
| 15498 | 775 | A775W | B | 2.8e-07 | 3.5e-07 | 2.4e-07 | 2.900E-07 | 5.568E-08 | 2.400E-07 | 3.500E-07 |
| 15499 | 775 | A775Y | B | 2.7e-07 | 2e-07 | 2.8e-07 | 2.500E-07 | 4.359E-08 | 2.000E-07 | 2.800E-07 |
| 15500 | 775 | A775V | B | 2.1e-07 | 4.8e-07 | 2.5e-07 | 3.133E-07 | 1.457E-07 | 2.100E-07 | 4.800E-07 |
| 15501 | 776 | L776A | B | 2.7e-07 | 2.6e-07 | 2.6e-07 | 2.633E-07 | 5.774E-09 | 2.600E-07 | 2.700E-07 |
| 15502 | 776 | L776R | B | 5.2e-07 | 2.6e-07 | 3.3e-07 | 3.700E-07 | 1.345E-07 | 2.600E-07 | 5.200E-07 |
| 15503 | 776 | L776N | B | 2.8e-07 | 3e-07 | 1.9e-07 | 2.567E-07 | 5.859E-08 | 1.900E-07 | 3.000E-07 |
| 15504 | 776 | L776D | B | 2.1e-07 | 2.7e-07 | 1.1e-07 | 1.967E-07 | 8.083E-08 | 1.100E-07 | 2.700E-07 |
| 15505 | 776 | L776C | B | 2.8e-07 | 1.5e-07 | 2.4e-07 | 2.233E-07 | 6.658E-08 | 1.500E-07 | 2.800E-07 |
| 15506 | 776 | L776E | B | 2.3e-07 | 2.3e-07 | 2.5e-07 | 2.367E-07 | 1.155E-08 | 2.300E-07 | 2.500E-07 |
| 15507 | 776 | L776Q | B | 4e-07 | 2.9e-07 | 2.4e-07 | 3.100E-07 | 8.185E-08 | 2.400E-07 | 4.000E-07 |
| 15508 | 776 | L776G | B | 1.5e-07 | 3e-07 | 1.6e-07 | 2.033E-07 | 8.386E-08 | 1.500E-07 | 3.000E-07 |
| 15509 | 776 | L776H | B | 3e-07 | 5.8e-07 | 3.1e-07 | 3.967E-07 | 1.589E-07 | 3.000E-07 | 5.800E-07 |
| 15510 | 776 | L776I | B | 2.8e-07 | 3e-07 | 1.3e-07 | 2.367E-07 | 9.292E-08 | 1.300E-07 | 3.000E-07 |
| 15511 | 776 | L776L | B | 2.8e-07 | 2.9e-07 | 5.9e-07 | 3.867E-07 | 1.762E-07 | 2.800E-07 | 5.900E-07 |
| 15512 | 776 | L776K | B | 3.6e-07 | 3e-07 | 1.6e-07 | 2.733E-07 | 1.026E-07 | 1.600E-07 | 3.600E-07 |
| 15513 | 776 | L776M | B | 2.4e-07 | 2.5e-07 | 1.9e-07 | 2.267E-07 | 3.215E-08 | 1.900E-07 | 2.500E-07 |
| 15514 | 776 | L776F | B | 3e-07 | 5.4e-07 | 4.1e-07 | 4.167E-07 | 1.201E-07 | 3.000E-07 | 5.400E-07 |
| 15515 | 776 | L776P | B | 2e-07 | 2.7e-07 | 2.5e-07 | 2.400E-07 | 3.606E-08 | 2.000E-07 | 2.700E-07 |
| 15516 | 776 | L776S | B | 3.2e-07 | 2.2e-07 | 2.1e-07 | 2.500E-07 | 6.083E-08 | 2.100E-07 | 3.200E-07 |
| 15517 | 776 | L776T | B | 2.2e-07 | 2.1e-07 | 3.8e-07 | 2.700E-07 | 9.539E-08 | 2.100E-07 | 3.800E-07 |
| 15518 | 776 | L776W | B | 2.9e-07 | 3.3e-07 | 1.6e-07 | 2.600E-07 | 8.888E-08 | 1.600E-07 | 3.300E-07 |
| 15519 | 776 | L776Y | B | 3.4e-07 | 3.7e-07 | 3.5e-07 | 3.533E-07 | 1.528E-08 | 3.400E-07 | 3.700E-07 |
| 15520 | 776 | L776V | B | 3.6e-07 | 2.8e-07 | 3.8e-07 | 3.400E-07 | 5.292E-08 | 2.800E-07 | 3.800E-07 |
| 15521 | 777 | Q777A | B | 2.2e-07 | 3.8e-07 | 2.4e-07 | 2.800E-07 | 8.718E-08 | 2.200E-07 | 3.800E-07 |
| 15522 | 777 | Q777R | B | 2.2e-07 | 1.3e-07 | 1.7e-07 | 1.733E-07 | 4.509E-08 | 1.300E-07 | 2.200E-07 |
| 15523 | 777 | Q777N | B | 1.5e-07 | 2e-07 | 2.7e-07 | 2.067E-07 | 6.028E-08 | 1.500E-07 | 2.700E-07 |
| 15524 | 777 | Q777D | B | 8.5e-08 | 2.7e-07 | 1.7e-07 | 1.750E-07 | 9.260E-08 | 8.500E-08 | 2.700E-07 |
| 15525 | 777 | Q777C | B | 1.1e-07 | 3e-07 | 5.2e-07 | 3.100E-07 | 2.052E-07 | 1.100E-07 | 5.200E-07 |
| 15526 | 777 | Q777E | B | 2e-07 | 5.5e-07 | 3.7e-07 | 3.733E-07 | 1.750E-07 | 2.000E-07 | 5.500E-07 |
| 15527 | 777 | Q777Q | B | 2.6e-07 | 3.5e-07 | 6e-07 | 4.033E-07 | 1.762E-07 | 2.600E-07 | 6.000E-07 |
| 15528 | 777 | Q777G | B | 2.3e-07 | 4.9e-07 | 2.1e-07 | 3.100E-07 | 1.562E-07 | 2.100E-07 | 4.900E-07 |
| 15529 | 777 | Q777H | B | 1.9e-07 | 1.8e-07 | 3e-07 | 2.233E-07 | 6.658E-08 | 1.800E-07 | 3.000E-07 |
| 15530 | 777 | Q777I | B | 2.6e-07 | 2.3e-07 | 3.6e-07 | 2.833E-07 | 6.807E-08 | 2.300E-07 | 3.600E-07 |
| 15531 | 777 | Q777L | B | 1.9e-07 | 1.4e-07 | 1.4e-07 | 1.567E-07 | 2.887E-08 | 1.400E-07 | 1.900E-07 |
| 15532 | 777 | Q777K | B | 4.5e-07 | 3.2e-07 | 3.6e-07 | 3.767E-07 | 6.658E-08 | 3.200E-07 | 4.500E-07 |
| 15533 | 777 | Q777M | B | 2.5e-07 | 2.1e-07 | 2.4e-07 | 2.333E-07 | 2.082E-08 | 2.100E-07 | 2.500E-07 |
| 15534 | 777 | Q777F | B | 5e-07 | 2.2e-07 | 1.5e-07 | 2.900E-07 | 1.852E-07 | 1.500E-07 | 5.000E-07 |
| 15535 | 777 | Q777P | B | 1.5e-07 | 2.7e-07 | 2.5e-07 | 2.233E-07 | 6.429E-08 | 1.500E-07 | 2.700E-07 |
| 15536 | 777 | Q777S | B | 2.4e-07 | 2.1e-07 | 2e-07 | 2.167E-07 | 2.082E-08 | 2.000E-07 | 2.400E-07 |
| 15537 | 777 | Q777T | B | 3.5e-07 | 2.8e-07 | 2.6e-07 | 2.967E-07 | 4.726E-08 | 2.600E-07 | 3.500E-07 |
| 15538 | 777 | Q777W | B | 2.6e-07 | 1.8e-07 | 2e-07 | 2.133E-07 | 4.163E-08 | 1.800E-07 | 2.600E-07 |
| 15539 | 777 | Q777Y | B | 3.3e-07 | 1.3e-07 | 2.1e-07 | 2.233E-07 | 1.007E-07 | 1.300E-07 | 3.300E-07 |
| 15540 | 777 | Q777V | B | 4.3e-07 | 3e-07 | 2.8e-07 | 3.367E-07 | 8.145E-08 | 2.800E-07 | 4.300E-07 |
| 15541 | 778 | S778A | B | 2.6e-07 | 2.3e-07 | 3.6e-07 | 2.833E-07 | 6.807E-08 | 2.300E-07 | 3.600E-07 |
| 15542 | 778 | S778R | B | 2.7e-07 | 4.3e-07 | 3.4e-07 | 3.467E-07 | 8.021E-08 | 2.700E-07 | 4.300E-07 |
| 15543 | 778 | S778N | B | 3.3e-07 | 2e-07 | 2.5e-07 | 2.600E-07 | 6.557E-08 | 2.000E-07 | 3.300E-07 |
| 15544 | 778 | S778D | B | 3e-07 | 1.9e-07 | 2.3e-07 | 2.400E-07 | 5.568E-08 | 1.900E-07 | 3.000E-07 |
| 15545 | 778 | S778C | B | 2.7e-07 | 1.2e-07 | 2.4e-07 | 2.100E-07 | 7.937E-08 | 1.200E-07 | 2.700E-07 |
| 15546 | 778 | S778E | B | 3.1e-07 | 3e-07 | 3.5e-07 | 3.200E-07 | 2.646E-08 | 3.000E-07 | 3.500E-07 |
| 15547 | 778 | S778Q | B | 1.6e-07 | 1.5e-07 | 2.8e-07 | 1.967E-07 | 7.234E-08 | 1.500E-07 | 2.800E-07 |
| 15548 | 778 | S778G | B | 3.1e-07 | 1.8e-07 | 9e-08 | 1.933E-07 | 1.106E-07 | 9.000E-08 | 3.100E-07 |
| 15549 | 778 | S778H | B | 7.7e-07 | 3.8e-07 | 2.9e-07 | 4.800E-07 | 2.551E-07 | 2.900E-07 | 7.700E-07 |
| 15550 | 778 | S778I | B | 2.3e-07 | 1.6e-07 | 4.2e-07 | 2.700E-07 | 1.345E-07 | 1.600E-07 | 4.200E-07 |
| 15551 | 778 | S778L | B | 1.7e-07 | 2.1e-07 | 2.4e-07 | 2.067E-07 | 3.512E-08 | 1.700E-07 | 2.400E-07 |
| 15552 | 778 | S778K | B | 3.5e-07 | 2.9e-07 | 1.8e-07 | 2.733E-07 | 8.622E-08 | 1.800E-07 | 3.500E-07 |
| 15553 | 778 | S778M | B | 1.8e-07 | 1.9e-07 | 2.5e-07 | 2.067E-07 | 3.786E-08 | 1.800E-07 | 2.500E-07 |
| 15554 | 778 | S778F | B | 3.9e-07 | 2e-07 | 4.4e-07 | 3.433E-07 | 1.266E-07 | 2.000E-07 | 4.400E-07 |

| | | | | | | | | | | |
|---|---|---|---|---|---|---|---|---|---|---|
| 15555 | 778 | S778P | B | 1.9e-07 | 2.8e-07 | 2.2e-07 | 2.300E-07 | 4.583E-08 | 1.900E-07 | 2.800E-07 |
| 15556 | 778 | S778S | B | 1.4e-07 | 1.4e-07 | 3.8e-07 | 2.200E-07 | 1.386E-07 | 1.400E-07 | 3.800E-07 |
| 15557 | 778 | S778T | B | 2.5e-07 | 3e-07 | 1.7e-07 | 2.400E-07 | 6.557E-08 | 1.700E-07 | 3.000E-07 |
| 15558 | 778 | S778W | B | 2.4e-07 | 3.4e-07 | 2.6e-07 | 2.800E-07 | 5.292E-08 | 2.400E-07 | 3.400E-07 |
| 15559 | 778 | S778Y | B | 2.4e-07 | 2e-07 | 8.5e-08 | 1.750E-07 | 8.047E-08 | 8.500E-08 | 2.400E-07 |
| 15560 | 778 | S778V | B | 6.4e-07 | 1.4e-07 | 1.9e-07 | 3.233E-07 | 2.754E-07 | 1.400E-07 | 6.400E-07 |
| 15561 | 779 | G779A | B | 3.3e-07 | 1.6e-07 | 3.3e-07 | 2.733E-07 | 9.815E-08 | 1.600E-07 | 3.300E-07 |
| 15562 | 779 | G779R | B | 5.5e-07 | 1.4e-07 | 3.3e-07 | 3.400E-07 | 2.052E-07 | 1.400E-07 | 5.500E-07 |
| 15563 | 779 | G779N | B | 4.5e-07 | 2e-07 | 3.4e-07 | 3.300E-07 | 1.253E-07 | 2.000E-07 | 4.500E-07 |
| 15564 | 779 | G779D | B | 2.5e-07 | 2.2e-07 | 2.3e-07 | 2.333E-07 | 1.528E-08 | 2.200E-07 | 2.500E-07 |
| 15565 | 779 | G779C | B | 5.5e-07 | 2.5e-07 | 4.1e-07 | 4.033E-07 | 1.501E-07 | 2.500E-07 | 5.500E-07 |
| 15566 | 779 | G779E | B | 1.7e-07 | 2.8e-07 | 2.1e-07 | 2.200E-07 | 5.568E-08 | 1.700E-07 | 2.800E-07 |
| 15567 | 779 | G779Q | B | 2.4e-07 | 2.2e-07 | 3.1e-07 | 2.567E-07 | 4.726E-08 | 2.200E-07 | 3.100E-07 |
| 15568 | 779 | G779G | B | 9e-08 | 3e-07 | 2.1e-07 | 2.000E-07 | 1.054E-07 | 9.000E-08 | 3.000E-07 |
| 15569 | 779 | G779H | B | 2.7e-07 | 3.1e-07 | 4.4e-07 | 3.400E-07 | 8.888E-08 | 2.700E-07 | 4.400E-07 |
| 15570 | 779 | G779I | B | 2.2e-07 | 2.9e-07 | 5.1e-07 | 3.400E-07 | 1.513E-07 | 2.200E-07 | 5.100E-07 |
| 15571 | 779 | G779L | B | 3.1e-07 | 2.3e-07 | 1.6e-07 | 2.333E-07 | 7.506E-08 | 1.600E-07 | 3.100E-07 |
| 15572 | 779 | G779K | B | 4.2e-07 | 3.3e-07 | 2e-07 | 3.167E-07 | 1.106E-07 | 2.000E-07 | 4.200E-07 |
| 15573 | 779 | G779M | B | 9.8e-08 | 3.5e-07 | 3.1e-07 | 2.527E-07 | 1.354E-07 | 9.800E-08 | 3.500E-07 |
| 15574 | 779 | G779F | B | 3.5e-07 | 2.9e-07 | 2.8e-07 | 3.067E-07 | 3.786E-08 | 2.800E-07 | 3.500E-07 |
| 15575 | 779 | G779P | B | 3.3e-07 | 4.3e-07 | 1.2e-07 | 2.933E-07 | 1.582E-07 | 1.200E-07 | 4.300E-07 |
| 15576 | 779 | G779S | B | 2.3e-07 | 2.3e-07 | 3.7e-07 | 2.767E-07 | 8.083E-08 | 2.300E-07 | 3.700E-07 |
| 15577 | 779 | G779T | B | 1.9e-07 | 3.8e-07 | 1.7e-07 | 2.467E-07 | 1.159E-07 | 1.700E-07 | 3.800E-07 |
| 15578 | 779 | G779W | B | 2.6e-07 | 2e-07 | 2.7e-07 | 2.433E-07 | 3.786E-08 | 2.000E-07 | 2.700E-07 |
| 15579 | 779 | G779Y | B | 4.4e-07 | 2e-07 | 1.6e-07 | 2.667E-07 | 1.514E-07 | 1.600E-07 | 4.400E-07 |
| 15580 | 779 | G779V | B | 1.6e-07 | 2.4e-07 | 1.4e-07 | 1.800E-07 | 5.292E-08 | 1.400E-07 | 2.400E-07 |
| 15581 | 780 | N780A | B | 3e-07 | 1.6e-07 | 5e-07 | 3.200E-07 | 1.709E-07 | 1.600E-07 | 5.000E-07 |
| 15582 | 780 | N780R | B | 3.3e-07 | 2.7e-07 | 2.3e-07 | 2.767E-07 | 5.033E-08 | 2.300E-07 | 3.300E-07 |
| 15583 | 780 | N780N | B | 2.6e-07 | 1.3e-07 | 2e-07 | 1.967E-07 | 6.506E-08 | 1.300E-07 | 2.600E-07 |
| 15584 | 780 | N780D | B | 5.7e-07 | 2.2e-07 | 3.2e-07 | 3.700E-07 | 1.803E-07 | 2.200E-07 | 5.700E-07 |
| 15585 | 780 | N780C | B | 1.6e-07 | 9.2e-08 | 1.6e-07 | 1.373E-07 | 3.926E-08 | 9.200E-08 | 1.600E-07 |
| 15586 | 780 | N780E | B | 1.4e-07 | 2.4e-07 | 1.6e-07 | 1.800E-07 | 5.292E-08 | 1.400E-07 | 2.400E-07 |
| 15587 | 780 | N780Q | B | 1.6e-07 | 1.3e-07 | 7.2e-07 | 3.367E-07 | 3.323E-07 | 1.300E-07 | 7.200E-07 |
| 15588 | 780 | N780G | B | 2.8e-07 | 3.5e-07 | 2.2e-07 | 2.833E-07 | 6.506E-08 | 2.200E-07 | 3.500E-07 |
| 15589 | 780 | N780H | B | 3.6e-07 | 2.4e-07 | 1.7e-07 | 2.567E-07 | 9.609E-08 | 1.700E-07 | 3.600E-07 |
| 15590 | 780 | N780I | B | 2.8e-07 | 4.8e-07 | 3.7e-07 | 3.767E-07 | 1.002E-07 | 2.800E-07 | 4.800E-07 |
| 15591 | 780 | N780L | B | 5.3e-07 | 3.8e-07 | 1.9e-07 | 3.667E-07 | 1.704E-07 | 1.900E-07 | 5.300E-07 |
| 15592 | 780 | N780K | B | 3.9e-07 | 1.3e-07 | 2.5e-07 | 2.567E-07 | 1.301E-07 | 1.300E-07 | 3.900E-07 |
| 15593 | 780 | N780M | B | 6.4e-07 | 2.3e-07 | 6.4e-07 | 5.033E-07 | 2.367E-07 | 2.300E-07 | 6.400E-07 |
| 15594 | 780 | N780F | B | 3.7e-07 | 2.7e-07 | 3.3e-07 | 3.233E-07 | 5.033E-08 | 2.700E-07 | 3.700E-07 |
| 15595 | 780 | N780P | B | 2.7e-07 | 1.4e-07 | 2.2e-07 | 2.100E-07 | 6.557E-08 | 1.400E-07 | 2.700E-07 |
| 15596 | 780 | N780S | B | 2.3e-07 | 2.8e-07 | 2.4e-07 | 2.500E-07 | 2.646E-08 | 2.300E-07 | 2.800E-07 |
| 15597 | 780 | N780T | B | 3.8e-07 | 2.1e-07 | 2.5e-07 | 2.800E-07 | 8.888E-08 | 2.100E-07 | 3.800E-07 |
| 15598 | 780 | N780W | B | 1.5e-07 | 3.9e-07 | 1.4e-07 | 2.267E-07 | 1.415E-07 | 1.400E-07 | 3.900E-07 |
| 15599 | 780 | N780Y | B | 2e-07 | 2.4e-07 | 6.6e-08 | 1.687E-07 | 9.113E-08 | 6.600E-08 | 2.400E-07 |
| 15600 | 780 | N780V | B | 3.2e-07 | 4.6e-07 | 4.6e-07 | 4.133E-07 | 8.083E-08 | 3.200E-07 | 4.600E-07 |
| 15601 | 781 | S781A | B | 1.5e-07 | 2e-07 | 1.8e-07 | 1.767E-07 | 2.517E-08 | 1.500E-07 | 2.000E-07 |
| 15602 | 781 | S781R | B | 4.3e-07 | 3.8e-07 | 2.3e-07 | 3.467E-07 | 1.041E-07 | 2.300E-07 | 4.300E-07 |
| 15603 | 781 | S781N | B | 2e-07 | 2.2e-07 | 2.4e-07 | 2.200E-07 | 2.000E-08 | 2.000E-07 | 2.400E-07 |
| 15604 | 781 | S781D | B | 2.7e-07 | 2.7e-07 | 4.7e-07 | 3.367E-07 | 1.155E-07 | 2.700E-07 | 4.700E-07 |
| 15605 | 781 | S781C | B | 2.7e-07 | 4.3e-07 | 1.1e-07 | 2.700E-07 | 1.600E-07 | 1.100E-07 | 4.300E-07 |
| 15606 | 781 | S781E | B | 3.4e-07 | 2.2e-07 | 4e-07 | 3.200E-07 | 9.165E-08 | 2.200E-07 | 4.000E-07 |
| 15607 | 781 | S781Q | B | 1.3e-07 | 1.3e-07 | 2.3e-07 | 1.633E-07 | 5.774E-08 | 1.300E-07 | 2.300E-07 |
| 15608 | 781 | S781G | B | 4.6e-07 | 1.4e-07 | 1.4e-07 | 2.467E-07 | 1.848E-07 | 1.400E-07 | 4.600E-07 |
| 15609 | 781 | S781H | B | 3.1e-07 | 2.2e-07 | 2.5e-07 | 2.600E-07 | 4.583E-08 | 2.200E-07 | 3.100E-07 |
| 15610 | 781 | S781I | B | 6.5e-07 | 2.3e-07 | 1.1e-07 | 3.300E-07 | 2.835E-07 | 1.100E-07 | 6.500E-07 |
| 15611 | 781 | S781L | B | 4.7e-07 | 2.6e-07 | 1.1e-07 | 2.800E-07 | 1.808E-07 | 1.100E-07 | 4.700E-07 |
| 15612 | 781 | S781K | B | 3.8e-07 | 1.7e-07 | 2.9e-07 | 2.800E-07 | 1.054E-07 | 1.700E-07 | 3.800E-07 |
| 15613 | 781 | S781M | B | 3e-07 | 3.7e-07 | 2.2e-07 | 2.967E-07 | 7.506E-08 | 2.200E-07 | 3.700E-07 |
| 15614 | 781 | S781F | B | 2.7e-07 | 4.9e-07 | 3.2e-07 | 3.600E-07 | 1.153E-07 | 2.700E-07 | 4.900E-07 |
| 15615 | 781 | S781P | B | 4.5e-07 | 2.1e-07 | 3.1e-07 | 3.233E-07 | 1.206E-07 | 2.100E-07 | 4.500E-07 |
| 15616 | 781 | S781S | B | 1.4e-07 | 1.6e-07 | 2.8e-07 | 1.933E-07 | 7.572E-08 | 1.400E-07 | 2.800E-07 |
| 15617 | 781 | S781T | B | 1.6e-07 | 3.5e-07 | 2.5e-07 | 2.533E-07 | 9.504E-08 | 1.600E-07 | 3.500E-07 |
| 15618 | 781 | S781W | B | 2.3e-07 | 2.9e-07 | 9.2e-08 | 2.040E-07 | 1.015E-07 | 9.200E-08 | 2.900E-07 |
| 15619 | 781 | S781Y | B | 3.1e-07 | 2.7e-07 | 6.3e-07 | 4.033E-07 | 1.973E-07 | 2.700E-07 | 6.300E-07 |
| 15620 | 781 | S781V | B | 1.5e-07 | 3.4e-07 | 2.2e-07 | 2.367E-07 | 9.609E-08 | 1.500E-07 | 3.400E-07 |
| 15621 | 782 | Q782A | B | 3.2e-07 | 8.2e-08 | 1.9e-07 | 1.973E-07 | 1.192E-07 | 8.200E-08 | 3.200E-07 |
| 15622 | 782 | Q782R | B | 3.4e-07 | 1.1e-07 | 2.5e-07 | 2.333E-07 | 1.159E-07 | 1.100E-07 | 3.400E-07 |
| 15623 | 782 | Q782N | B | 1.5e-07 | 4.1e-07 | 2e-07 | 2.533E-07 | 1.380E-07 | 1.500E-07 | 4.100E-07 |
| 15624 | 782 | Q782D | B | 1.8e-07 | 2.9e-07 | 2.3e-07 | 2.333E-07 | 5.508E-08 | 1.800E-07 | 2.900E-07 |
| 15625 | 782 | Q782C | B | 1.8e-07 | 3.4e-07 | 2.4e-07 | 2.533E-07 | 8.083E-08 | 1.800E-07 | 3.400E-07 |
| 15626 | 782 | Q782E | B | 1.8e-07 | 2.8e-07 | 3e-07 | 2.533E-07 | 6.429E-08 | 1.800E-07 | 3.000E-07 |
| 15627 | 782 | Q782Q | B | 2.5e-07 | 3.8e-07 | 3.5e-07 | 3.267E-07 | 6.807E-08 | 2.500E-07 | 3.800E-07 |
| 15628 | 782 | Q782G | B | 1.9e-07 | 1.6e-07 | 2.3e-07 | 1.933E-07 | 3.512E-08 | 1.600E-07 | 2.300E-07 |
| 15629 | 782 | Q782H | B | 1e-07 | 1.9e-07 | 2.4e-07 | 1.767E-07 | 7.095E-08 | 1.000E-07 | 2.400E-07 |
| 15630 | 782 | Q782I | B | 4.9e-07 | 3.3e-07 | 6.6e-07 | 4.933E-07 | 1.650E-07 | 3.300E-07 | 6.600E-07 |
| 15631 | 782 | Q782L | B | 2.6e-07 | 1.8e-07 | 1.9e-07 | 2.100E-07 | 4.359E-08 | 1.800E-07 | 2.600E-07 |
| 15632 | 782 | Q782K | B | 1.1e-07 | 2.8e-07 | 3.3e-07 | 2.400E-07 | 1.153E-07 | 1.100E-07 | 3.300E-07 |
| 15633 | 782 | Q782M | B | 3.7e-07 | 2.1e-07 | 3.1e-07 | 2.967E-07 | 8.083E-08 | 2.100E-07 | 3.700E-07 |
| 15634 | 782 | Q782F | B | 3.2e-07 | 2.2e-07 | 3.9e-07 | 3.100E-07 | 8.544E-08 | 2.200E-07 | 3.900E-07 |
| 15635 | 782 | Q782P | B | 3.3e-07 | 2.9e-07 | 2.3e-07 | 2.833E-07 | 5.033E-08 | 2.300E-07 | 3.300E-07 |
| 15636 | 782 | Q782S | B | 1.9e-07 | 1.4e-07 | 3.1e-07 | 2.133E-07 | 8.737E-08 | 1.400E-07 | 3.100E-07 |
| 15637 | 782 | Q782T | B | 3.5e-07 | 1.7e-07 | 2e-07 | 2.400E-07 | 9.644E-08 | 1.700E-07 | 3.500E-07 |
| 15638 | 782 | Q782W | B | 3e-07 | 2e-07 | 2.3e-07 | 2.433E-07 | 5.132E-08 | 2.000E-07 | 3.000E-07 |
| 15639 | 782 | Q782Y | B | 4.1e-07 | 2.3e-07 | 2.8e-07 | 3.067E-07 | 9.292E-08 | 2.300E-07 | 4.100E-07 |

| | | | | | | | | | | |
|---|---|---|---|---|---|---|---|---|---|---|
| 15640 | 782 | Q782V | B | 1.9e-07 | 4.7e-07 | 2.4e-07 | 3.000E-07 | 1.493E-07 | 1.900E-07 | 4.700E-07 |
| 15641 | 783 | E783A | B | 3.2e-07 | 1.1e-07 | 2.3e-07 | 2.200E-07 | 1.054E-07 | 1.100E-07 | 3.200E-07 |
| 15642 | 783 | E783R | B | 2.1e-07 | 5e-07 | 3.7e-07 | 3.600E-07 | 1.453E-07 | 2.100E-07 | 5.000E-07 |
| 15643 | 783 | E783N | B | 3.9e-07 | 2.9e-07 | 3.6e-07 | 3.467E-07 | 5.132E-08 | 2.900E-07 | 3.900E-07 |
| 15644 | 783 | E783D | B | 7.9e-07 | 2.1e-07 | 1.9e-07 | 3.967E-07 | 3.408E-07 | 1.900E-07 | 7.900E-07 |
| 15645 | 783 | E783C | B | 1.9e-07 | 6.6e-08 | 1.7e-07 | 1.420E-07 | 6.657E-08 | 6.600E-08 | 1.900E-07 |
| 15646 | 783 | E783E | B | 9.7e-08 | 1.8e-07 | 1.3e-07 | 1.357E-07 | 4.179E-08 | 9.700E-08 | 1.800E-07 |
| 15647 | 783 | E783Q | B | 3.4e-07 | 3.1e-07 | 2.4e-07 | 2.967E-07 | 5.132E-08 | 2.400E-07 | 3.400E-07 |
| 15648 | 783 | E783G | B | 1.8e-07 | 1.8e-07 | 7.1e-07 | 3.567E-07 | 3.060E-07 | 1.800E-07 | 7.100E-07 |
| 15649 | 783 | E783H | B | 3.3e-07 | 5.3e-07 | 2.3e-07 | 3.633E-07 | 1.528E-07 | 2.300E-07 | 5.300E-07 |
| 15650 | 783 | E783I | B | 3.9e-07 | 2.7e-07 | 5.6e-07 | 4.067E-07 | 1.457E-07 | 2.700E-07 | 5.600E-07 |
| 15651 | 783 | E783L | B | 2.3e-07 | 2.5e-07 | 2.1e-07 | 2.300E-07 | 2.000E-08 | 2.100E-07 | 2.500E-07 |
| 15652 | 783 | E783K | B | 1.3e-07 | 2.9e-07 | 2.3e-07 | 2.167E-07 | 8.083E-08 | 1.300E-07 | 2.900E-07 |
| 15653 | 783 | E783M | B | 5.1e-07 | 1.4e-07 | 2.4e-07 | 2.967E-07 | 1.914E-07 | 1.400E-07 | 5.100E-07 |
| 15654 | 783 | E783F | B | 3e-07 | 2.3e-07 | 1.3e-07 | 2.200E-07 | 8.544E-08 | 1.300E-07 | 3.000E-07 |
| 15655 | 783 | E783P | B | 2.1e-07 | 1.7e-07 | 1.6e-07 | 1.800E-07 | 2.646E-08 | 1.600E-07 | 2.100E-07 |
| 15656 | 783 | E783S | B | 2e-07 | 1.9e-07 | 3.9e-07 | 2.600E-07 | 1.127E-07 | 1.900E-07 | 3.900E-07 |
| 15657 | 783 | E783T | B | 2.4e-07 | 1.3e-07 | 1.7e-07 | 1.800E-07 | 5.568E-08 | 1.300E-07 | 2.400E-07 |
| 15658 | 783 | E783W | B | 4.4e-07 | 1.1e-07 | 3.3e-07 | 2.933E-07 | 1.680E-07 | 1.100E-07 | 4.400E-07 |
| 15659 | 783 | E783Y | B | 1.5e-07 | 1.2e-07 | 2.4e-07 | 1.700E-07 | 6.245E-08 | 1.200E-07 | 2.400E-07 |
| 15660 | 783 | E783V | B | 1.9e-07 | 4e-07 | 2.7e-07 | 2.867E-07 | 1.060E-07 | 1.900E-07 | 4.000E-07 |
| 15661 | 784 | S784A | B | 2e-07 | 1.8e-07 | 2.7e-07 | 2.167E-07 | 4.726E-08 | 1.800E-07 | 2.700E-07 |
| 15662 | 784 | S784R | B | 4.3e-07 | 2.1e-07 | 3.4e-07 | 3.267E-07 | 1.106E-07 | 2.100E-07 | 4.300E-07 |
| 15663 | 784 | S784N | B | 4.9e-07 | 6.4e-07 | 1.5e-07 | 4.267E-07 | 2.511E-07 | 1.500E-07 | 6.400E-07 |
| 15664 | 784 | S784D | B | 2.2e-07 | 3.7e-07 | 7.7e-07 | 4.533E-07 | 2.843E-07 | 2.200E-07 | 7.700E-07 |
| 15665 | 784 | S784C | B | 2.2e-07 | 1.7e-07 | 3.2e-07 | 2.367E-07 | 7.638E-08 | 1.700E-07 | 3.200E-07 |
| 15666 | 784 | S784E | B | 3.8e-07 | 1.4e-07 | 1.9e-07 | 2.367E-07 | 1.266E-07 | 1.400E-07 | 3.800E-07 |
| 15667 | 784 | S784Q | B | 5.1e-07 | 2.3e-07 | 3.4e-07 | 3.600E-07 | 1.411E-07 | 2.300E-07 | 5.100E-07 |
| 15668 | 784 | S784G | B | 2.3e-07 | 2.9e-07 | 2.1e-07 | 2.433E-07 | 4.163E-08 | 2.100E-07 | 2.900E-07 |
| 15669 | 784 | S784H | B | 2.1e-07 | 2.1e-07 | 2.3e-07 | 2.167E-07 | 1.155E-08 | 2.100E-07 | 2.300E-07 |
| 15670 | 784 | S784I | B | 3.8e-07 | 2.9e-07 | 1.4e-07 | 2.700E-07 | 1.212E-07 | 1.400E-07 | 3.800E-07 |
| 15671 | 784 | S784L | B | 5.2e-07 | 3.2e-07 | 4.3e-07 | 4.233E-07 | 1.002E-07 | 3.200E-07 | 5.200E-07 |
| 15672 | 784 | S784K | B | 2.6e-07 | 1.9e-07 | 1e-07 | 1.833E-07 | 8.021E-08 | 1.000E-07 | 2.600E-07 |
| 15673 | 784 | S784M | B | 1.6e-07 | 3e-07 | 2.5e-07 | 2.367E-07 | 7.095E-08 | 1.600E-07 | 3.000E-07 |
| 15674 | 784 | S784F | B | 5.5e-07 | 2.2e-07 | 2.2e-07 | 3.300E-07 | 1.905E-07 | 2.200E-07 | 5.500E-07 |
| 15675 | 784 | S784P | B | 1.3e-07 | 4.2e-07 | 4.6e-07 | 3.367E-07 | 1.801E-07 | 1.300E-07 | 4.600E-07 |
| 15676 | 784 | S784S | B | 2.2e-07 | 5e-07 | 3.8e-07 | 3.667E-07 | 1.405E-07 | 2.200E-07 | 5.000E-07 |
| 15677 | 784 | S784T | B | 1.2e-07 | 6.1e-07 | 2.4e-07 | 3.233E-07 | 2.554E-07 | 1.200E-07 | 6.100E-07 |
| 15678 | 784 | S784W | B | 3.2e-07 | 3.6e-07 | 5.2e-07 | 4.000E-07 | 1.058E-07 | 3.200E-07 | 5.200E-07 |
| 15679 | 784 | S784Y | B | 9e-08 | 3.3e-07 | 9e-08 | 1.700E-07 | 1.386E-07 | 9.000E-08 | 3.300E-07 |
| 15680 | 784 | S784V | B | 3.1e-07 | 4.3e-07 | 3.6e-07 | 3.667E-07 | 6.028E-08 | 3.100E-07 | 4.300E-07 |
| 15681 | 785 | V785A | B | 1.3e-07 | 5.7e-07 | 1.7e-07 | 2.900E-07 | 2.433E-07 | 1.300E-07 | 5.700E-07 |
| 15682 | 785 | V785R | B | 4.2e-07 | 1.5e-07 | 4.6e-07 | 3.433E-07 | 1.686E-07 | 1.500E-07 | 4.600E-07 |
| 15683 | 785 | V785N | B | 1.6e-07 | 1.7e-07 | 2.2e-07 | 1.833E-07 | 3.215E-08 | 1.600E-07 | 2.200E-07 |
| 15684 | 785 | V785D | B | 5.1e-07 | 3.2e-07 | 2.1e-07 | 3.467E-07 | 1.518E-07 | 2.100E-07 | 5.100E-07 |
| 15685 | 785 | V785C | B | 3.8e-07 | 1.5e-07 | 2.1e-07 | 2.467E-07 | 1.193E-07 | 1.500E-07 | 3.800E-07 |
| 15686 | 785 | V785E | B | 2.3e-07 | 1.4e-07 | 2.4e-07 | 2.033E-07 | 5.508E-08 | 1.400E-07 | 2.400E-07 |
| 15687 | 785 | V785Q | B | 1.5e-07 | 1.2e-07 | 2.2e-07 | 1.633E-07 | 5.132E-08 | 1.200E-07 | 2.200E-07 |
| 15688 | 785 | V785G | B | 1.7e-07 | 2.6e-07 | 3.5e-07 | 2.600E-07 | 9.000E-08 | 1.700E-07 | 3.500E-07 |
| 15689 | 785 | V785H | B | 2.2e-07 | 1.6e-07 | 2.5e-07 | 2.100E-07 | 4.583E-08 | 1.600E-07 | 2.500E-07 |
| 15690 | 785 | V785I | B | 2.4e-07 | 1.8e-07 | 2.9e-07 | 2.367E-07 | 5.508E-08 | 1.800E-07 | 2.900E-07 |
| 15691 | 785 | V785L | B | 1.7e-07 | 1.5e-07 | 2.3e-07 | 1.833E-07 | 4.163E-08 | 1.500E-07 | 2.300E-07 |
| 15692 | 785 | V785K | B | 3.2e-07 | 2.7e-07 | 1.7e-07 | 2.533E-07 | 7.638E-08 | 1.700E-07 | 3.200E-07 |
| 15693 | 785 | V785M | B | 3.1e-07 | 3.9e-07 | 2e-07 | 3.000E-07 | 9.539E-08 | 2.000E-07 | 3.900E-07 |
| 15694 | 785 | V785F | B | 1.9e-07 | 2.1e-07 | 1.5e-07 | 1.833E-07 | 3.055E-08 | 1.500E-07 | 2.100E-07 |
| 15695 | 785 | V785P | B | 4.2e-07 | 3.3e-07 | 1.1e-07 | 2.867E-07 | 1.595E-07 | 1.100E-07 | 4.200E-07 |
| 15696 | 785 | V785S | B | 1.7e-07 | 4.1e-07 | 1.3e-07 | 2.367E-07 | 1.514E-07 | 1.300E-07 | 4.100E-07 |
| 15697 | 785 | V785T | B | 4.4e-07 | 4.9e-07 | 1.1e-07 | 3.467E-07 | 2.065E-07 | 1.100E-07 | 4.900E-07 |
| 15698 | 785 | V785W | B | 3.5e-07 | 1.3e-07 | 2.8e-07 | 2.533E-07 | 1.124E-07 | 1.300E-07 | 3.500E-07 |
| 15699 | 785 | V785Y | B | 3e-07 | 6.8e-08 | 2.1e-07 | 1.927E-07 | 1.170E-07 | 6.800E-08 | 3.000E-07 |
| 15700 | 785 | V785V | B | 1.1e-07 | 3.6e-07 | 1.7e-07 | 2.133E-07 | 1.305E-07 | 1.100E-07 | 3.600E-07 |
| 15701 | 786 | T786A | B | 3.1e-07 | 4.5e-07 | 3.8e-07 | 3.800E-07 | 7.000E-08 | 3.100E-07 | 4.500E-07 |
| 15702 | 786 | T786R | B | 1.3e-07 | 2.2e-07 | 4.5e-07 | 2.667E-07 | 1.650E-07 | 1.300E-07 | 4.500E-07 |
| 15703 | 786 | T786N | B | 2e-07 | 2.3e-07 | 2.4e-07 | 2.233E-07 | 2.082E-08 | 2.000E-07 | 2.400E-07 |
| 15704 | 786 | T786D | B | 1.8e-07 | 3.1e-07 | 2.3e-07 | 2.400E-07 | 6.557E-08 | 1.800E-07 | 3.100E-07 |
| 15705 | 786 | T786C | B | 1.7e-07 | 1.4e-07 | 3.4e-07 | 2.167E-07 | 1.079E-07 | 1.400E-07 | 3.400E-07 |
| 15706 | 786 | T786E | B | 1.1e-07 | 1.3e-07 | 2.4e-07 | 1.600E-07 | 7.000E-08 | 1.100E-07 | 2.400E-07 |
| 15707 | 786 | T786Q | B | 3.3e-07 | 2.8e-07 | 2e-07 | 2.700E-07 | 6.557E-08 | 2.000E-07 | 3.300E-07 |
| 15708 | 786 | T786G | B | 1.9e-07 | 2.5e-07 | 2.2e-07 | 2.200E-07 | 3.000E-08 | 1.900E-07 | 2.500E-07 |
| 15709 | 786 | T786H | B | 2.6e-07 | 2.4e-07 | 1.9e-07 | 2.300E-07 | 3.606E-08 | 1.900E-07 | 2.600E-07 |
| 15710 | 786 | T786I | B | 7.3e-07 | 9.7e-08 | 3.8e-07 | 4.023E-07 | 3.171E-07 | 9.700E-08 | 7.300E-07 |
| 15711 | 786 | T786L | B | 2.2e-07 | 1.3e-06 | 2e-07 | 5.733E-07 | 6.294E-07 | 2.000E-07 | 1.300E-06 |
| 15712 | 786 | T786K | B | 2.2e-07 | 3.8e-07 | 2.8e-07 | 2.933E-07 | 8.083E-08 | 2.200E-07 | 3.800E-07 |
| 15713 | 786 | T786M | B | 2.5e-07 | 2.7e-07 | 3.2e-07 | 2.800E-07 | 3.606E-08 | 2.500E-07 | 3.200E-07 |
| 15714 | 786 | T786F | B | 6.5e-07 | 1.8e-07 | 3.3e-07 | 3.867E-07 | 2.401E-07 | 1.800E-07 | 6.500E-07 |
| 15715 | 786 | T786P | B | 4e-07 | 1.8e-07 | 3.6e-07 | 3.133E-07 | 1.172E-07 | 1.800E-07 | 4.000E-07 |
| 15716 | 786 | T786S | B | 1.4e-07 | 2e-07 | 1.7e-07 | 1.700E-07 | 3.000E-08 | 1.400E-07 | 2.000E-07 |
| 15717 | 786 | T786T | B | 2e-07 | 1.7e-07 | 2.3e-07 | 2.000E-07 | 3.000E-08 | 1.700E-07 | 2.300E-07 |
| 15718 | 786 | T786W | B | 1.9e-07 | 4.8e-07 | 3.3e-07 | 3.333E-07 | 1.450E-07 | 1.900E-07 | 4.800E-07 |
| 15719 | 786 | T786Y | B | 1.2e-07 | 3.7e-07 | 9.3e-08 | 1.943E-07 | 1.527E-07 | 9.300E-08 | 3.700E-07 |
| 15720 | 786 | T786V | B | 3.3e-07 | 2.1e-07 | 2.3e-07 | 2.567E-07 | 6.429E-08 | 2.100E-07 | 3.300E-07 |
| 15721 | 787 | E787A | B | 1.5e-07 | 1.5e-07 | 2.1e-07 | 1.700E-07 | 3.464E-08 | 1.500E-07 | 2.100E-07 |
| 15722 | 787 | E787R | B | 3.2e-07 | 1.9e-07 | 1.1e-07 | 2.067E-07 | 1.060E-07 | 1.100E-07 | 3.200E-07 |
| 15723 | 787 | E787N | B | 4.8e-07 | 3.9e-07 | 1.3e-07 | 3.333E-07 | 1.818E-07 | 1.300E-07 | 4.800E-07 |
| 15724 | 787 | E787D | B | 4.2e-07 | 3.2e-07 | 3.4e-07 | 3.600E-07 | 5.292E-08 | 3.200E-07 | 4.200E-07 |

| | | | | | | | | | | |
|---|---|---|---|---|---|---|---|---|---|---|
| 15725 | 787 | E787C | B | 3.1e-07 | 2.5e-07 | 2.9e-07 | 2.833E-07 | 3.055E-08 | 2.500E-07 | 3.100E-07 |
| 15726 | 787 | E787E | B | 1.6e-07 | 3e-07 | 3.1e-07 | 2.567E-07 | 8.386E-08 | 1.600E-07 | 3.100E-07 |
| 15727 | 787 | E787Q | B | 2.8e-07 | 2.3e-07 | 1.6e-07 | 2.233E-07 | 6.028E-08 | 1.600E-07 | 2.800E-07 |
| 15728 | 787 | E787G | B | 2.4e-07 | 2.4e-07 | 3.9e-07 | 2.900E-07 | 8.660E-08 | 2.400E-07 | 3.900E-07 |
| 15729 | 787 | E787H | B | 2e-07 | 2.5e-07 | 2.5e-07 | 2.333E-07 | 2.887E-08 | 2.000E-07 | 2.500E-07 |
| 15730 | 787 | E787I | B | 1.6e-07 | 2e-07 | 1.4e-07 | 1.667E-07 | 3.055E-08 | 1.400E-07 | 2.000E-07 |
| 15731 | 787 | E787L | B | 2.8e-07 | 3.8e-07 | 2.9e-07 | 3.167E-07 | 5.508E-08 | 2.800E-07 | 3.800E-07 |
| 15732 | 787 | E787K | B | 2e-07 | 9.2e-08 | 1.6e-07 | 1.507E-07 | 5.460E-08 | 9.200E-08 | 2.000E-07 |
| 15733 | 787 | E787M | B | 1.2e-07 | 1.9e-07 | 2.5e-07 | 1.867E-07 | 6.506E-08 | 1.200E-07 | 2.500E-07 |
| 15734 | 787 | E787F | B | 2.4e-07 | 2.4e-07 | 2.2e-07 | 2.333E-07 | 1.155E-08 | 2.200E-07 | 2.400E-07 |
| 15735 | 787 | E787P | B | 2.1e-07 | 1.9e-07 | 1.3e-07 | 1.767E-07 | 4.163E-08 | 1.300E-07 | 2.100E-07 |
| 15736 | 787 | E787S | B | 1.4e-07 | 2.2e-07 | 1.7e-07 | 1.767E-07 | 4.041E-08 | 1.400E-07 | 2.200E-07 |
| 15737 | 787 | E787T | B | 1.3e-07 | 2.7e-07 | 3.4e-07 | 2.467E-07 | 1.069E-07 | 1.300E-07 | 3.400E-07 |
| 15738 | 787 | E787W | B | 4.5e-07 | 1.7e-07 | 1.5e-07 | 2.567E-07 | 1.677E-07 | 1.500E-07 | 4.500E-07 |
| 15739 | 787 | E787Y | B | 1.3e-07 | 3.7e-07 | 3.1e-07 | 2.700E-07 | 1.249E-07 | 1.300E-07 | 3.700E-07 |
| 15740 | 787 | E787V | B | 4.9e-07 | 2.1e-07 | 2e-07 | 3.000E-07 | 1.646E-07 | 2.000E-07 | 4.900E-07 |
| 15741 | 788 | Q788A | B | 2.3e-07 | 3.6e-07 | 1.4e-07 | 2.433E-07 | 1.106E-07 | 1.400E-07 | 3.600E-07 |
| 15742 | 788 | Q788R | B | 2.9e-07 | 4.2e-07 | 1.4e-07 | 2.833E-07 | 1.401E-07 | 1.400E-07 | 4.200E-07 |
| 15743 | 788 | Q788N | B | 2e-07 | 5.2e-08 | 1.7e-07 | 1.407E-07 | 7.824E-08 | 5.200E-08 | 2.000E-07 |
| 15744 | 788 | Q788D | B | 2.3e-07 | 4.6e-07 | 2.7e-07 | 3.200E-07 | 1.229E-07 | 2.300E-07 | 4.600E-07 |
| 15745 | 788 | Q788C | B | 2.4e-07 | 2.8e-07 | 1.8e-07 | 2.333E-07 | 5.033E-08 | 1.800E-07 | 2.800E-07 |
| 15746 | 788 | Q788E | B | 2.9e-07 | 4.1e-07 | 1.3e-07 | 2.767E-07 | 1.405E-07 | 1.300E-07 | 4.100E-07 |
| 15747 | 788 | Q788Q | B | 2.1e-07 | 2.3e-07 | 4.8e-07 | 3.067E-07 | 1.504E-07 | 2.100E-07 | 4.800E-07 |
| 15748 | 788 | Q788G | B | 1.5e-07 | 2.3e-07 | 2e-07 | 1.933E-07 | 4.041E-08 | 1.500E-07 | 2.300E-07 |
| 15749 | 788 | Q788H | B | 3.5e-07 | 2e-07 | 2.4e-07 | 2.633E-07 | 7.767E-08 | 2.000E-07 | 3.500E-07 |
| 15750 | 788 | Q788I | B | 4.9e-07 | 4.8e-07 | 9.3e-08 | 3.543E-07 | 2.264E-07 | 9.300E-08 | 4.900E-07 |
| 15751 | 788 | Q788L | B | 1.3e-07 | 4.5e-07 | 1.3e-07 | 2.367E-07 | 1.848E-07 | 1.300E-07 | 4.500E-07 |
| 15752 | 788 | Q788K | B | 4.9e-07 | 2e-07 | 1.9e-07 | 2.933E-07 | 1.704E-07 | 1.900E-07 | 4.900E-07 |
| 15753 | 788 | Q788M | B | 1.8e-07 | 2.9e-07 | 2.1e-07 | 2.267E-07 | 5.686E-08 | 1.800E-07 | 2.900E-07 |
| 15754 | 788 | Q788F | B | 3.5e-07 | 1.3e-07 | 1.5e-07 | 2.100E-07 | 1.217E-07 | 1.300E-07 | 3.500E-07 |
| 15755 | 788 | Q788P | B | 3e-07 | 3.6e-07 | 2.1e-07 | 2.900E-07 | 7.550E-08 | 2.100E-07 | 3.600E-07 |
| 15756 | 788 | Q788S | B | 5.3e-07 | 1.6e-07 | 3.5e-07 | 3.467E-07 | 1.850E-07 | 1.600E-07 | 5.300E-07 |
| 15757 | 788 | Q788T | B | 2.8e-07 | 1.5e-07 | 2.6e-07 | 2.300E-07 | 7.000E-08 | 1.500E-07 | 2.800E-07 |
| 15758 | 788 | Q788W | B | 2.3e-07 | 1.9e-07 | 2.6e-07 | 2.267E-07 | 3.512E-08 | 1.900E-07 | 2.600E-07 |
| 15759 | 788 | Q788Y | B | 4.3e-07 | 3.3e-07 | 3.9e-07 | 3.833E-07 | 5.033E-08 | 3.300E-07 | 4.300E-07 |
| 15760 | 788 | Q788V | B | 1.9e-07 | 1.7e-07 | 2.5e-07 | 2.033E-07 | 4.163E-08 | 1.700E-07 | 2.500E-07 |
| 15761 | 789 | D789A | B | 1.6e-07 | 1.6e-07 | 2.7e-07 | 1.967E-07 | 6.351E-08 | 1.600E-07 | 2.700E-07 |
| 15762 | 789 | D789R | B | 1.6e-07 | 2.9e-07 | 2.7e-07 | 2.400E-07 | 7.000E-08 | 1.600E-07 | 2.900E-07 |
| 15763 | 789 | D789N | B | 7.1e-07 | 4e-07 | 2.2e-07 | 4.433E-07 | 2.479E-07 | 2.200E-07 | 7.100E-07 |
| 15764 | 789 | D789D | B | 1.8e-07 | 1.8e-07 | 1.5e-07 | 1.700E-07 | 1.732E-08 | 1.500E-07 | 1.800E-07 |
| 15765 | 789 | D789C | B | 5.6e-07 | 3.7e-07 | 1.7e-07 | 3.667E-07 | 1.950E-07 | 1.700E-07 | 5.600E-07 |
| 15766 | 789 | D789E | B | 3.5e-07 | 1.5e-07 | 2.4e-07 | 2.467E-07 | 1.002E-07 | 1.500E-07 | 3.500E-07 |
| 15767 | 789 | D789Q | B | 2.3e-07 | 2.2e-07 | 4.2e-07 | 2.900E-07 | 1.127E-07 | 2.200E-07 | 4.200E-07 |
| 15768 | 789 | D789G | B | 2.1e-07 | 3.1e-07 | 2.6e-07 | 2.600E-07 | 5.000E-08 | 2.100E-07 | 3.100E-07 |
| 15769 | 789 | D789H | B | 3.4e-07 | 4.5e-07 | 2.4e-07 | 3.433E-07 | 1.050E-07 | 2.400E-07 | 4.500E-07 |
| 15770 | 789 | D789I | B | 2.2e-07 | 4.4e-07 | 1.5e-07 | 2.700E-07 | 1.513E-07 | 1.500E-07 | 4.400E-07 |
| 15771 | 789 | D789L | B | 3.7e-07 | 2.1e-07 | 4.9e-07 | 3.567E-07 | 1.405E-07 | 2.100E-07 | 4.900E-07 |
| 15772 | 789 | D789K | B | 2.2e-07 | 3e-07 | 3.2e-07 | 2.800E-07 | 5.292E-08 | 2.200E-07 | 3.200E-07 |
| 15773 | 789 | D789M | B | 1.7e-07 | 2.5e-07 | 8.2e-08 | 1.673E-07 | 8.403E-08 | 8.200E-08 | 2.500E-07 |
| 15774 | 789 | D789F | B | 3.5e-07 | 5.3e-07 | 2.7e-07 | 3.833E-07 | 1.332E-07 | 2.700E-07 | 5.300E-07 |
| 15775 | 789 | D789P | B | 3.1e-07 | 2.8e-07 | 4.2e-07 | 3.367E-07 | 7.371E-08 | 2.800E-07 | 4.200E-07 |
| 15776 | 789 | D789S | B | 2.6e-07 | 3e-07 | 3.3e-07 | 2.967E-07 | 3.512E-08 | 2.600E-07 | 3.300E-07 |
| 15777 | 789 | D789T | B | 2.3e-07 | 2.1e-07 | 2.2e-07 | 2.200E-07 | 1.000E-08 | 2.100E-07 | 2.300E-07 |
| 15778 | 789 | D789W | B | 1.4e-07 | 4e-07 | 2.7e-07 | 2.700E-07 | 1.300E-07 | 1.400E-07 | 4.000E-07 |
| 15779 | 789 | D789Y | B | 2.3e-07 | 1.9e-07 | 2.2e-07 | 2.133E-07 | 2.082E-08 | 1.900E-07 | 2.300E-07 |
| 15780 | 789 | D789V | B | 4.8e-07 | 2.7e-07 | 3.8e-07 | 3.767E-07 | 1.050E-07 | 2.700E-07 | 4.800E-07 |
| 15781 | 790 | S790A | B | 4.2e-07 | 3.8e-07 | 1.9e-07 | 3.300E-07 | 1.229E-07 | 1.900E-07 | 4.200E-07 |
| 15782 | 790 | S790R | B | 5.5e-07 | 2.2e-07 | 1.8e-07 | 3.167E-07 | 2.031E-07 | 1.800E-07 | 5.500E-07 |
| 15783 | 790 | S790N | B | 1.1e-07 | 1.7e-07 | 2e-07 | 1.600E-07 | 4.583E-08 | 1.100E-07 | 2.000E-07 |
| 15784 | 790 | S790D | B | 1.3e-07 | 3.3e-07 | 3.2e-07 | 2.600E-07 | 1.127E-07 | 1.300E-07 | 3.300E-07 |
| 15785 | 790 | S790C | B | 4e-07 | 2e-07 | 1.1e-07 | 2.367E-07 | 1.484E-07 | 1.100E-07 | 4.000E-07 |
| 15786 | 790 | S790E | B | 3.7e-07 | 7.2e-07 | 1.4e-07 | 4.100E-07 | 2.921E-07 | 1.400E-07 | 7.200E-07 |
| 15787 | 790 | S790Q | B | 1.5e-07 | 5.1e-07 | 2.3e-07 | 2.967E-07 | 1.890E-07 | 1.500E-07 | 5.100E-07 |
| 15788 | 790 | S790G | B | 2.4e-07 | 2.9e-07 | 3e-07 | 2.767E-07 | 3.215E-08 | 2.400E-07 | 3.000E-07 |
| 15789 | 790 | S790H | B | 4e-07 | 1e-07 | 1e-07 | 2.000E-07 | 1.732E-07 | 1.000E-07 | 4.000E-07 |
| 15790 | 790 | S790I | B | 4.8e-07 | 2.1e-07 | 2.3e-07 | 3.067E-07 | 1.504E-07 | 2.100E-07 | 4.800E-07 |
| 15791 | 790 | S790L | B | 3.9e-07 | 2.5e-07 | 1.7e-07 | 2.700E-07 | 1.114E-07 | 1.700E-07 | 3.900E-07 |
| 15792 | 790 | S790K | B | 3.8e-07 | 1.9e-07 | 2.2e-07 | 2.633E-07 | 1.021E-07 | 1.900E-07 | 3.800E-07 |
| 15793 | 790 | S790M | B | 2.2e-07 | 4.5e-07 | 2.7e-07 | 3.133E-07 | 1.210E-07 | 2.200E-07 | 4.500E-07 |
| 15794 | 790 | S790F | B | 7.2e-07 | 5.1e-07 | 1e-07 | 4.433E-07 | 3.153E-07 | 1.000E-07 | 7.200E-07 |
| 15795 | 790 | S790P | B | 2.8e-07 | 2e-07 | 4e-07 | 2.933E-07 | 1.007E-07 | 2.000E-07 | 4.000E-07 |
| 15796 | 790 | S790S | B | 6.3e-07 | 4e-07 | 3.2e-07 | 4.500E-07 | 1.609E-07 | 3.200E-07 | 6.300E-07 |
| 15797 | 790 | S790T | B | 1.4e-07 | 2.1e-07 | 1.6e-07 | 1.700E-07 | 3.606E-08 | 1.400E-07 | 2.100E-07 |
| 15798 | 790 | S790W | B | 2.7e-07 | 1.5e-07 | 2.4e-07 | 2.200E-07 | 6.245E-08 | 1.500E-07 | 2.700E-07 |
| 15799 | 790 | S790Y | B | 2.3e-07 | 1e-07 | 2.7e-07 | 2.000E-07 | 8.888E-08 | 1.000E-07 | 2.700E-07 |
| 15800 | 790 | S790V | B | 1.5e-07 | 1.1e-07 | 3.2e-07 | 1.933E-07 | 1.115E-07 | 1.100E-07 | 3.200E-07 |
| 15801 | 791 | K791A | B | 5.1e-07 | 4.3e-07 | 2.1e-07 | 3.833E-07 | 1.553E-07 | 2.100E-07 | 5.100E-07 |
| 15802 | 791 | K791R | B | 2.4e-07 | 2.1e-07 | 5.4e-07 | 3.300E-07 | 1.825E-07 | 2.100E-07 | 5.400E-07 |
| 15803 | 791 | K791N | B | 2.7e-07 | 2.5e-07 | 6.3e-07 | 3.833E-07 | 2.139E-07 | 2.500E-07 | 6.300E-07 |
| 15804 | 791 | K791D | B | 2e-07 | 2.2e-07 | 2e-07 | 2.067E-07 | 1.155E-08 | 2.000E-07 | 2.200E-07 |
| 15805 | 791 | K791C | B | 1.2e-07 | 2.6e-07 | 1.9e-07 | 1.900E-07 | 7.000E-08 | 1.200E-07 | 2.600E-07 |
| 15806 | 791 | K791E | B | 1.7e-07 | 3e-07 | 5.4e-07 | 3.367E-07 | 1.877E-07 | 1.700E-07 | 5.400E-07 |
| 15807 | 791 | K791Q | B | 2.2e-07 | 1.2e-07 | 3.6e-07 | 2.333E-07 | 1.206E-07 | 1.200E-07 | 3.600E-07 |
| 15808 | 791 | K791G | B | 5.8e-07 | 2.4e-07 | 4e-07 | 4.067E-07 | 1.701E-07 | 2.400E-07 | 5.800E-07 |
| 15809 | 791 | K791H | B | 3.3e-07 | 1.6e-07 | 3.7e-07 | 2.867E-07 | 1.115E-07 | 1.600E-07 | 3.700E-07 |

| | | | | | | | | | | |
|---|---|---|---|---|---|---|---|---|---|---|
| 15810 | 791 | K791I | B | 5e-07 | 1.8e-07 | 3e-07 | 3.267E-07 | 1.617E-07 | 1.800E-07 | 5.000E-07 |
| 15811 | 791 | K791L | B | 1.9e-07 | 7.2e-07 | 2.3e-07 | 3.800E-07 | 2.951E-07 | 1.900E-07 | 7.200E-07 |
| 15812 | 791 | K791K | B | 1.1e-07 | 2.4e-07 | 3.1e-07 | 2.200E-07 | 1.015E-07 | 1.100E-07 | 3.100E-07 |
| 15813 | 791 | K791M | B | 1.8e-07 | 3.2e-07 | 3.2e-07 | 2.733E-07 | 8.083E-08 | 1.800E-07 | 3.200E-07 |
| 15814 | 791 | K791F | B | 1.4e-07 | 2.3e-07 | 2e-07 | 1.900E-07 | 4.583E-08 | 1.400E-07 | 2.300E-07 |
| 15815 | 791 | K791P | B | 1.7e-07 | 2.2e-07 | 2e-07 | 1.967E-07 | 2.517E-08 | 1.700E-07 | 2.200E-07 |
| 15816 | 791 | K791S | B | 2.9e-07 | 3.9e-07 | 1.7e-07 | 2.833E-07 | 1.102E-07 | 1.700E-07 | 3.900E-07 |
| 15817 | 791 | K791T | B | 1.9e-07 | 3.2e-07 | 2.2e-07 | 2.433E-07 | 6.807E-08 | 1.900E-07 | 3.200E-07 |
| 15818 | 791 | K791W | B | 3.4e-07 | 1.9e-07 | 3.3e-07 | 2.867E-07 | 8.386E-08 | 1.900E-07 | 3.400E-07 |
| 15819 | 791 | K791Y | B | 2.4e-07 | 2.8e-07 | 4e-07 | 3.067E-07 | 8.327E-08 | 2.400E-07 | 4.000E-07 |
| 15820 | 791 | K791V | B | 1.6e-07 | 3.2e-07 | 4.4e-07 | 3.067E-07 | 1.405E-07 | 1.600E-07 | 4.400E-07 |
| 15821 | 792 | D792A | B | 2.6e-07 | 2.4e-07 | 1.9e-07 | 2.300E-07 | 3.606E-08 | 1.900E-07 | 2.600E-07 |
| 15822 | 792 | D792R | B | 1.2e-07 | 5.9e-07 | 2.3e-07 | 3.133E-07 | 2.458E-07 | 1.200E-07 | 5.900E-07 |
| 15823 | 792 | D792N | B | 3.3e-07 | 2.1e-07 | 2.4e-07 | 2.600E-07 | 6.245E-08 | 2.100E-07 | 3.300E-07 |
| 15824 | 792 | D792D | B | 7.9e-07 | 5.8e-07 | 2.3e-07 | 5.333E-07 | 2.829E-07 | 2.300E-07 | 7.900E-07 |
| 15825 | 792 | D792C | B | 2e-07 | 2.6e-07 | 1.5e-07 | 2.033E-07 | 5.508E-08 | 1.500E-07 | 2.600E-07 |
| 15826 | 792 | D792E | B | 2.3e-07 | 2.9e-07 | 2.3e-07 | 2.500E-07 | 3.464E-08 | 2.300E-07 | 2.900E-07 |
| 15827 | 792 | D792Q | B | 4.6e-07 | 2.3e-07 | 3.1e-07 | 3.333E-07 | 1.168E-07 | 2.300E-07 | 4.600E-07 |
| 15828 | 792 | D792G | B | 2.4e-07 | 1.8e-07 | 1.4e-07 | 1.867E-07 | 5.033E-08 | 1.400E-07 | 2.400E-07 |
| 15829 | 792 | D792H | B | 2.2e-07 | 3.3e-07 | 2.7e-07 | 2.733E-07 | 5.508E-08 | 2.200E-07 | 3.300E-07 |
| 15830 | 792 | D792I | B | 1.7e-07 | 2.3e-07 | 2.5e-07 | 2.167E-07 | 4.163E-08 | 1.700E-07 | 2.500E-07 |
| 15831 | 792 | D792L | B | 2.2e-07 | 1.9e-07 | 3.2e-07 | 2.433E-07 | 6.807E-08 | 1.900E-07 | 3.200E-07 |
| 15832 | 792 | D792K | B | 2.4e-07 | 2.5e-07 | 3.2e-07 | 2.700E-07 | 4.359E-08 | 2.400E-07 | 3.200E-07 |
| 15833 | 792 | D792M | B | 2.4e-07 | 1.2e-07 | 6.8e-07 | 3.467E-07 | 2.948E-07 | 1.200E-07 | 6.800E-07 |
| 15834 | 792 | D792F | B | 2.5e-07 | 9.1e-08 | 2.6e-07 | 2.003E-07 | 9.482E-08 | 9.100E-08 | 2.600E-07 |
| 15835 | 792 | D792P | B | 3.1e-07 | 6.2e-07 | 3.7e-07 | 4.333E-07 | 1.644E-07 | 3.100E-07 | 6.200E-07 |
| 15836 | 792 | D792S | B | 1.6e-07 | 3.5e-07 | 3e-07 | 2.700E-07 | 9.849E-08 | 1.600E-07 | 3.500E-07 |
| 15837 | 792 | D792T | B | 3.1e-07 | 1.7e-07 | 1.9e-07 | 2.233E-07 | 7.572E-08 | 1.700E-07 | 3.100E-07 |
| 15838 | 792 | D792W | B | 4.1e-07 | 2.3e-07 | 3e-07 | 3.133E-07 | 9.074E-08 | 2.300E-07 | 4.100E-07 |
| 15839 | 792 | D792Y | B | 5e-07 | 2.9e-07 | 3.3e-07 | 3.733E-07 | 1.115E-07 | 2.900E-07 | 5.000E-07 |
| 15840 | 792 | D792V | B | 2.3e-07 | 2.1e-07 | 2.4e-07 | 2.267E-07 | 1.528E-08 | 2.100E-07 | 2.400E-07 |
| 15841 | 793 | S793A | B | 1.4e-07 | 2.5e-07 | 3e-07 | 2.300E-07 | 8.185E-08 | 1.400E-07 | 3.000E-07 |
| 15842 | 793 | S793R | B | 2e-07 | 1.3e-07 | 1.4e-07 | 1.567E-07 | 3.786E-08 | 1.300E-07 | 2.000E-07 |
| 15843 | 793 | S793N | B | 3.5e-07 | 2e-07 | 1.5e-07 | 2.333E-07 | 1.041E-07 | 1.500E-07 | 3.500E-07 |
| 15844 | 793 | S793D | B | 1.3e-07 | 2e-07 | 1.6e-07 | 1.633E-07 | 3.512E-08 | 1.300E-07 | 2.000E-07 |
| 15845 | 793 | S793C | B | 3.9e-07 | 1.3e-07 | 1.7e-07 | 2.300E-07 | 1.400E-07 | 1.300E-07 | 3.900E-07 |
| 15846 | 793 | S793E | B | 1.7e-07 | 3.2e-07 | 2.2e-07 | 2.367E-07 | 7.638E-08 | 1.700E-07 | 3.200E-07 |
| 15847 | 793 | S793Q | B | 4.9e-07 | 1.3e-07 | 4.6e-07 | 3.600E-07 | 1.997E-07 | 1.300E-07 | 4.900E-07 |
| 15848 | 793 | S793G | B | 2.5e-07 | 2.6e-07 | 1.1e-07 | 2.067E-07 | 8.386E-08 | 1.100E-07 | 2.600E-07 |
| 15849 | 793 | S793H | B | 4.3e-07 | 2.3e-07 | 3.1e-07 | 3.233E-07 | 1.007E-07 | 2.300E-07 | 4.300E-07 |
| 15850 | 793 | S793I | B | 2.4e-07 | 3e-07 | 1.7e-07 | 2.367E-07 | 6.506E-08 | 1.700E-07 | 3.000E-07 |
| 15851 | 793 | S793L | B | 2.1e-07 | 2.2e-07 | 2.5e-07 | 2.267E-07 | 2.082E-08 | 2.100E-07 | 2.500E-07 |
| 15852 | 793 | S793K | B | 1.5e-07 | 1.6e-07 | 1.9e-07 | 1.667E-07 | 2.082E-08 | 1.500E-07 | 1.900E-07 |
| 15853 | 793 | S793M | B | 1.6e-07 | 2e-07 | 4.4e-07 | 2.667E-07 | 1.514E-07 | 1.600E-07 | 4.400E-07 |
| 15854 | 793 | S793F | B | 6.9e-07 | 2.7e-07 | 2.9e-07 | 4.167E-07 | 2.369E-07 | 2.700E-07 | 6.900E-07 |
| 15855 | 793 | S793P | B | 3.5e-07 | 5.4e-07 | 2.4e-07 | 3.767E-07 | 1.518E-07 | 2.400E-07 | 5.400E-07 |
| 15856 | 793 | S793S | B | 2e-07 | 1.3e-07 | 3.5e-07 | 2.267E-07 | 1.124E-07 | 1.300E-07 | 3.500E-07 |
| 15857 | 793 | S793T | B | 1.5e-07 | 1.5e-07 | 3.5e-07 | 2.167E-07 | 1.155E-07 | 1.500E-07 | 3.500E-07 |
| 15858 | 793 | S793W | B | 1.9e-07 | 1.6e-07 | 4e-07 | 2.500E-07 | 1.308E-07 | 1.600E-07 | 4.000E-07 |
| 15859 | 793 | S793Y | B | 1.5e-07 | 3.1e-07 | 2.7e-07 | 2.433E-07 | 8.327E-08 | 1.500E-07 | 3.100E-07 |
| 15860 | 793 | S793V | B | 1.9e-07 | 2.8e-07 | 2.4e-07 | 2.367E-07 | 4.509E-08 | 1.900E-07 | 2.800E-07 |
| 15861 | 794 | T794A | B | 1.3e-07 | 2.2e-07 | 3.7e-07 | 2.400E-07 | 1.212E-07 | 1.300E-07 | 3.700E-07 |
| 15862 | 794 | T794R | B | 3e-07 | 4.2e-07 | 4.1e-07 | 3.767E-07 | 6.658E-08 | 3.000E-07 | 4.200E-07 |
| 15863 | 794 | T794N | B | 3e-07 | 3.6e-07 | 2.3e-07 | 2.967E-07 | 6.506E-08 | 2.300E-07 | 3.600E-07 |
| 15864 | 794 | T794D | B | 3.3e-07 | 2.4e-07 | 1.5e-07 | 2.400E-07 | 9.000E-08 | 1.500E-07 | 3.300E-07 |
| 15865 | 794 | T794C | B | 3.1e-07 | 3.2e-07 | 1.4e-07 | 2.567E-07 | 1.012E-07 | 1.400E-07 | 3.200E-07 |
| 15866 | 794 | T794E | B | 2.2e-07 | 3.5e-07 | 1.4e-07 | 2.367E-07 | 1.060E-07 | 1.400E-07 | 3.500E-07 |
| 15867 | 794 | T794Q | B | 3.9e-07 | 3.2e-07 | 2.3e-07 | 3.133E-07 | 8.021E-08 | 2.300E-07 | 3.900E-07 |
| 15868 | 794 | T794G | B | 3.1e-07 | 2.7e-07 | 1.7e-07 | 2.500E-07 | 7.211E-08 | 1.700E-07 | 3.100E-07 |
| 15869 | 794 | T794H | B | 6.6e-07 | 1.8e-07 | 4.4e-07 | 4.267E-07 | 2.403E-07 | 1.800E-07 | 6.600E-07 |
| 15870 | 794 | T794I | B | 3e-07 | 3.4e-07 | 2.6e-07 | 3.000E-07 | 4.000E-08 | 2.600E-07 | 3.400E-07 |
| 15871 | 794 | T794L | B | 4.5e-07 | 2.2e-07 | 1.8e-07 | 2.833E-07 | 1.457E-07 | 1.800E-07 | 4.500E-07 |
| 15872 | 794 | T794K | B | 2.5e-07 | 4.4e-07 | 2.3e-07 | 3.067E-07 | 1.159E-07 | 2.300E-07 | 4.400E-07 |
| 15873 | 794 | T794M | B | 2.4e-07 | 2.4e-07 | 2.1e-07 | 2.300E-07 | 1.732E-08 | 2.100E-07 | 2.400E-07 |
| 15874 | 794 | T794F | B | 2.4e-07 | 3e-07 | 4.1e-07 | 3.167E-07 | 8.622E-08 | 2.400E-07 | 4.100E-07 |
| 15875 | 794 | T794P | B | 2e-07 | 1.9e-07 | 2.1e-07 | 2.000E-07 | 1.000E-08 | 1.900E-07 | 2.100E-07 |
| 15876 | 794 | T794S | B | 4.5e-07 | 3.9e-07 | 4.9e-07 | 4.433E-07 | 5.033E-08 | 3.900E-07 | 4.900E-07 |
| 15877 | 794 | T794T | B | 2.1e-07 | 1.2e-07 | 1.8e-07 | 1.700E-07 | 4.583E-08 | 1.200E-07 | 2.100E-07 |
| 15878 | 794 | T794W | B | 2.1e-07 | 1.7e-07 | 1.8e-07 | 1.867E-07 | 2.082E-08 | 1.700E-07 | 2.100E-07 |
| 15879 | 794 | T794Y | B | 1.1e-07 | 4.8e-07 | 1.6e-07 | 2.500E-07 | 2.007E-07 | 1.100E-07 | 4.800E-07 |
| 15880 | 794 | T794V | B | 2.4e-07 | 2.3e-07 | 6.6e-07 | 3.767E-07 | 2.454E-07 | 2.300E-07 | 6.600E-07 |
| 15881 | 795 | Y795A | B | 2.6e-07 | 1.7e-07 | 3e-07 | 2.433E-07 | 6.658E-08 | 1.700E-07 | 3.000E-07 |
| 15882 | 795 | Y795R | B | 5.7e-07 | 5.6e-07 | 1.2e-07 | 4.167E-07 | 2.570E-07 | 1.200E-07 | 5.700E-07 |
| 15883 | 795 | Y795N | B | 1.4e-07 | 2.2e-07 | 2e-07 | 1.867E-07 | 4.163E-08 | 1.400E-07 | 2.200E-07 |
| 15884 | 795 | Y795D | B | 2.7e-07 | 2.8e-07 | 1.5e-07 | 2.333E-07 | 7.234E-08 | 1.500E-07 | 2.800E-07 |
| 15885 | 795 | Y795C | B | 2.4e-07 | 2.6e-07 | 7e-07 | 4.000E-07 | 2.600E-07 | 2.400E-07 | 7.000E-07 |
| 15886 | 795 | Y795E | B | 1.8e-07 | 3e-07 | 1.1e-07 | 1.967E-07 | 9.609E-08 | 1.100E-07 | 3.000E-07 |
| 15887 | 795 | Y795Q | B | 3.4e-07 | 2.5e-07 | 2e-07 | 2.633E-07 | 7.095E-08 | 2.000E-07 | 3.400E-07 |
| 15888 | 795 | Y795G | B | 1.9e-07 | 1.3e-07 | 1.9e-07 | 1.700E-07 | 3.464E-08 | 1.300E-07 | 1.900E-07 |
| 15889 | 795 | Y795H | B | 1.9e-07 | 3.2e-07 | 1.3e-07 | 2.133E-07 | 9.713E-08 | 1.300E-07 | 3.200E-07 |
| 15890 | 795 | Y795I | B | 2.1e-07 | 1.4e-07 | 3e-07 | 2.167E-07 | 8.021E-08 | 1.400E-07 | 3.000E-07 |
| 15891 | 795 | Y795L | B | 2.6e-07 | 1.2e-07 | 2.8e-07 | 2.200E-07 | 8.718E-08 | 1.200E-07 | 2.800E-07 |
| 15892 | 795 | Y795K | B | 2.2e-07 | 1.7e-07 | 4e-07 | 2.633E-07 | 1.210E-07 | 1.700E-07 | 4.000E-07 |
| 15893 | 795 | Y795M | B | 4.9e-07 | 1.7e-07 | 2.7e-07 | 3.100E-07 | 1.637E-07 | 1.700E-07 | 4.900E-07 |
| 15894 | 795 | Y795F | B | 3.1e-07 | 3.6e-07 | 2e-07 | 2.900E-07 | 8.185E-08 | 2.000E-07 | 3.600E-07 |

| | | | | | | | | | | |
|---|---|---|---|---|---|---|---|---|---|---|
| 15895 | 795 | Y795P | B | 2.4e-07 | 7.1e-07 | 2.8e-07 | 4.100E-07 | 2.606E-07 | 2.400E-07 | 7.100E-07 |
| 15896 | 795 | Y795S | B | 2.4e-07 | 2.9e-07 | 2.3e-07 | 2.533E-07 | 3.215E-08 | 2.300E-07 | 2.900E-07 |
| 15897 | 795 | Y795T | B | 1.8e-07 | 2.1e-07 | 2.1e-07 | 2.000E-07 | 1.732E-08 | 1.800E-07 | 2.100E-07 |
| 15898 | 795 | Y795W | B | 3.6e-07 | 9.7e-08 | 2.9e-07 | 2.490E-07 | 1.362E-07 | 9.700E-08 | 3.600E-07 |
| 15899 | 795 | Y795Y | B | 2.5e-07 | 2.1e-07 | 1.6e-07 | 2.067E-07 | 4.509E-08 | 1.600E-07 | 2.500E-07 |
| 15900 | 795 | Y795V | B | 3.5e-07 | 4e-07 | 2.9e-07 | 3.467E-07 | 5.508E-08 | 2.900E-07 | 4.000E-07 |
| 15901 | 796 | S796A | B | 1e-07 | 2.3e-07 | 1.7e-07 | 1.667E-07 | 6.506E-08 | 1.000E-07 | 2.300E-07 |
| 15902 | 796 | S796R | B | 2.3e-07 | 2.3e-07 | 3.2e-07 | 2.600E-07 | 5.196E-08 | 2.300E-07 | 3.200E-07 |
| 15903 | 796 | S796N | B | 3.1e-07 | 1.1e-07 | 8.7e-08 | 1.690E-07 | 1.226E-07 | 8.700E-08 | 3.100E-07 |
| 15904 | 796 | S796D | B | 1.9e-07 | 2e-07 | 9.2e-07 | 4.367E-07 | 4.186E-07 | 1.900E-07 | 9.200E-07 |
| 15905 | 796 | S796C | B | 3.1e-07 | 1.3e-07 | 1.2e-07 | 1.867E-07 | 1.069E-07 | 1.200E-07 | 3.100E-07 |
| 15906 | 796 | S796E | B | 2.1e-07 | 2.8e-07 | 1.9e-07 | 2.267E-07 | 4.726E-08 | 1.900E-07 | 2.800E-07 |
| 15907 | 796 | S796Q | B | 1.7e-07 | 5e-07 | 2.2e-07 | 2.967E-07 | 1.779E-07 | 1.700E-07 | 5.000E-07 |
| 15908 | 796 | S796G | B | 2.7e-07 | 4.3e-07 | 3.1e-07 | 3.367E-07 | 8.327E-08 | 2.700E-07 | 4.300E-07 |
| 15909 | 796 | S796H | B | 3e-07 | 3.5e-07 | 3.6e-07 | 3.367E-07 | 3.215E-08 | 3.000E-07 | 3.600E-07 |
| 15910 | 796 | S796I | B | 1.4e-07 | 4.5e-07 | 3.7e-07 | 3.200E-07 | 1.609E-07 | 1.400E-07 | 4.500E-07 |
| 15911 | 796 | S796L | B | 3.9e-07 | 5.1e-07 | 2.8e-07 | 3.933E-07 | 1.150E-07 | 2.800E-07 | 5.100E-07 |
| 15912 | 796 | S796K | B | 8.1e-08 | 1.6e-07 | 2.3e-07 | 1.570E-07 | 7.455E-08 | 8.100E-08 | 2.300E-07 |
| 15913 | 796 | S796M | B | 3e-07 | 4e-07 | 1.5e-07 | 2.833E-07 | 1.258E-07 | 1.500E-07 | 4.000E-07 |
| 15914 | 796 | S796F | B | 1.9e-07 | 2.2e-07 | 1.6e-07 | 1.900E-07 | 3.000E-08 | 1.600E-07 | 2.200E-07 |
| 15915 | 796 | S796P | B | 1.8e-07 | 1.9e-07 | 3.2e-07 | 2.300E-07 | 7.810E-08 | 1.800E-07 | 3.200E-07 |
| 15916 | 796 | S796S | B | 2.1e-07 | 2.1e-07 | 1.5e-07 | 1.900E-07 | 3.464E-08 | 1.500E-07 | 2.100E-07 |
| 15917 | 796 | S796T | B | 1.5e-07 | 1.7e-07 | 2.3e-07 | 1.833E-07 | 4.163E-08 | 1.500E-07 | 2.300E-07 |
| 15918 | 796 | S796W | B | 3.3e-07 | 4.3e-07 | 2.4e-07 | 3.333E-07 | 9.504E-08 | 2.400E-07 | 4.300E-07 |
| 15919 | 796 | S796Y | B | 2.2e-07 | 7.7e-07 | 1.2e-07 | 3.700E-07 | 3.500E-07 | 1.200E-07 | 7.700E-07 |
| 15920 | 796 | S796V | B | 2.7e-07 | 3.2e-07 | 3.6e-07 | 3.167E-07 | 4.509E-08 | 2.700E-07 | 3.600E-07 |
| 15921 | 797 | L797A | B | 2.2e-07 | 2.5e-07 | 2.6e-07 | 2.433E-07 | 2.082E-08 | 2.200E-07 | 2.600E-07 |
| 15922 | 797 | L797R | B | 1.4e-07 | 2.3e-07 | 3.1e-07 | 2.267E-07 | 8.505E-08 | 1.400E-07 | 3.100E-07 |
| 15923 | 797 | L797N | B | 5.5e-07 | 1.5e-07 | 2.3e-07 | 3.100E-07 | 2.117E-07 | 1.500E-07 | 5.500E-07 |
| 15924 | 797 | L797D | B | 2.1e-07 | 2.3e-07 | 4.8e-07 | 3.067E-07 | 1.504E-07 | 2.100E-07 | 4.800E-07 |
| 15925 | 797 | L797C | B | 1.4e-07 | 2.3e-07 | 3.7e-07 | 2.467E-07 | 1.159E-07 | 1.400E-07 | 3.700E-07 |
| 15926 | 797 | L797E | B | 3.9e-08 | 2.4e-07 | 1.3e-07 | 1.363E-07 | 1.006E-07 | 3.900E-08 | 2.400E-07 |
| 15927 | 797 | L797Q | B | 2.1e-07 | 3.7e-07 | 9.8e-08 | 2.260E-07 | 1.367E-07 | 9.800E-08 | 3.700E-07 |
| 15928 | 797 | L797G | B | 1.4e-07 | 2.5e-07 | 2.6e-07 | 2.167E-07 | 6.658E-08 | 1.400E-07 | 2.600E-07 |
| 15929 | 797 | L797H | B | 1.2e-07 | 8.9e-08 | 2.6e-07 | 1.563E-07 | 9.111E-08 | 8.900E-08 | 2.600E-07 |
| 15930 | 797 | L797I | B | 4.5e-07 | 2.6e-07 | 2.8e-07 | 3.300E-07 | 1.044E-07 | 2.600E-07 | 4.500E-07 |
| 15931 | 797 | L797L | B | 2.1e-07 | 2.3e-07 | 3.9e-07 | 2.767E-07 | 9.866E-08 | 2.100E-07 | 3.900E-07 |
| 15932 | 797 | L797K | B | 2.7e-07 | 4e-07 | 6.5e-08 | 2.450E-07 | 1.689E-07 | 6.500E-08 | 4.000E-07 |
| 15933 | 797 | L797M | B | 2.6e-07 | 1.1e-07 | 2.5e-07 | 2.067E-07 | 8.386E-08 | 1.100E-07 | 2.600E-07 |
| 15934 | 797 | L797F | B | 1.8e-07 | 1.5e-07 | 2.6e-07 | 1.967E-07 | 5.686E-08 | 1.500E-07 | 2.600E-07 |
| 15935 | 797 | L797P | B | 1.6e-07 | 3e-07 | 3.8e-07 | 2.800E-07 | 1.114E-07 | 1.600E-07 | 3.800E-07 |
| 15936 | 797 | L797S | B | 2.7e-07 | 2.1e-07 | 1.8e-07 | 2.200E-07 | 4.583E-08 | 1.800E-07 | 2.700E-07 |
| 15937 | 797 | L797T | B | 3e-07 | 2.2e-07 | 2.4e-07 | 2.533E-07 | 4.163E-08 | 2.200E-07 | 3.000E-07 |
| 15938 | 797 | L797W | B | 1.2e-07 | 1.4e-07 | 2.7e-07 | 1.767E-07 | 8.145E-08 | 1.200E-07 | 2.700E-07 |
| 15939 | 797 | L797Y | B | 1.9e-07 | 2.3e-07 | 1.4e-07 | 1.867E-07 | 4.509E-08 | 1.400E-07 | 2.300E-07 |
| 15940 | 797 | L797V | B | 2.1e-07 | 1.7e-07 | 3.4e-07 | 2.400E-07 | 8.888E-08 | 1.700E-07 | 3.400E-07 |
| 15941 | 798 | S798A | B | 1.9e-07 | 2.8e-07 | 3.9e-07 | 2.867E-07 | 1.002E-07 | 1.900E-07 | 3.900E-07 |
| 15942 | 798 | S798R | B | 3.3e-07 | 1.8e-07 | 1.7e-07 | 2.267E-07 | 8.963E-08 | 1.700E-07 | 3.300E-07 |
| 15943 | 798 | S798N | B | 2.2e-07 | 2.4e-07 | 3.4e-07 | 2.667E-07 | 6.429E-08 | 2.200E-07 | 3.400E-07 |
| 15944 | 798 | S798D | B | 1.9e-07 | 3.2e-07 | 1.2e-07 | 2.100E-07 | 1.015E-07 | 1.200E-07 | 3.200E-07 |
| 15945 | 798 | S798C | B | 3.1e-07 | 3.3e-07 | 5.9e-07 | 4.100E-07 | 1.562E-07 | 3.100E-07 | 5.900E-07 |
| 15946 | 798 | S798E | B | 4.1e-07 | 2.7e-07 | 3.1e-07 | 3.300E-07 | 7.211E-08 | 2.700E-07 | 4.100E-07 |
| 15947 | 798 | S798Q | B | 4.9e-07 | 2e-07 | 1.3e-07 | 2.733E-07 | 1.909E-07 | 1.300E-07 | 4.900E-07 |
| 15948 | 798 | S798G | B | 2e-07 | 3e-07 | 2e-07 | 2.333E-07 | 5.774E-08 | 2.000E-07 | 3.000E-07 |
| 15949 | 798 | S798H | B | 1.4e-07 | 3.9e-07 | 2.7e-07 | 2.667E-07 | 1.250E-07 | 1.400E-07 | 3.900E-07 |
| 15950 | 798 | S798I | B | 2.3e-07 | 1.7e-07 | 1.3e-07 | 1.767E-07 | 5.033E-08 | 1.300E-07 | 2.300E-07 |
| 15951 | 798 | S798L | B | 2.9e-07 | 1.6e-07 | 3.4e-07 | 2.633E-07 | 9.292E-08 | 1.600E-07 | 3.400E-07 |
| 15952 | 798 | S798K | B | 1.8e-07 | 1e-07 | 1.6e-07 | 1.467E-07 | 4.163E-08 | 1.000E-07 | 1.800E-07 |
| 15953 | 798 | S798M | B | 1.6e-07 | 3.9e-07 | 2.4e-07 | 2.633E-07 | 1.168E-07 | 1.600E-07 | 3.900E-07 |
| 15954 | 798 | S798F | B | 1.3e-07 | 4.4e-07 | 3.9e-07 | 3.200E-07 | 1.664E-07 | 1.300E-07 | 4.400E-07 |
| 15955 | 798 | S798P | B | 2.8e-07 | 3e-07 | 3.5e-07 | 3.100E-07 | 3.606E-08 | 2.800E-07 | 3.500E-07 |
| 15956 | 798 | S798S | B | 2.4e-07 | 5e-07 | 2e-07 | 3.133E-07 | 1.629E-07 | 2.000E-07 | 5.000E-07 |
| 15957 | 798 | S798T | B | 1.2e-07 | 1.5e-07 | 2e-07 | 1.567E-07 | 4.041E-08 | 1.200E-07 | 2.000E-07 |
| 15958 | 798 | S798W | B | 3.3e-07 | 1.7e-07 | 2.7e-07 | 2.567E-07 | 8.083E-08 | 1.700E-07 | 3.300E-07 |
| 15959 | 798 | S798Y | B | 1.4e-07 | 3.1e-07 | 2e-07 | 2.167E-07 | 8.622E-08 | 1.400E-07 | 3.100E-07 |
| 15960 | 798 | S798V | B | 1.7e-07 | 2.9e-07 | 4.8e-07 | 3.133E-07 | 1.563E-07 | 1.700E-07 | 4.800E-07 |
| 15961 | 799 | S799A | B | 3.6e-07 | 2.4e-07 | 1.9e-07 | 2.633E-07 | 8.737E-08 | 1.900E-07 | 3.600E-07 |
| 15962 | 799 | S799R | B | 2.9e-07 | 3.3e-07 | 3.2e-07 | 3.133E-07 | 2.082E-08 | 2.900E-07 | 3.300E-07 |
| 15963 | 799 | S799N | B | 2.1e-07 | 3.3e-07 | 3e-07 | 2.800E-07 | 6.245E-08 | 2.100E-07 | 3.300E-07 |
| 15964 | 799 | S799D | B | 4.2e-07 | 1.1e-07 | 3.1e-07 | 2.800E-07 | 1.572E-07 | 1.100E-07 | 4.200E-07 |
| 15965 | 799 | S799C | B | 1.4e-07 | 3.3e-07 | 4.9e-07 | 3.200E-07 | 1.752E-07 | 1.400E-07 | 4.900E-07 |
| 15966 | 799 | S799E | B | 1.4e-07 | 3e-07 | 3.7e-07 | 2.700E-07 | 1.179E-07 | 1.400E-07 | 3.700E-07 |
| 15967 | 799 | S799Q | B | 3e-07 | 1.8e-07 | 1.2e-07 | 2.000E-07 | 9.165E-08 | 1.200E-07 | 3.000E-07 |
| 15968 | 799 | S799G | B | 1.1e-07 | 2e-07 | 2.5e-07 | 1.867E-07 | 7.095E-08 | 1.100E-07 | 2.500E-07 |
| 15969 | 799 | S799H | B | 9.9e-08 | 2.9e-07 | 2.8e-07 | 2.230E-07 | 1.075E-07 | 9.900E-08 | 2.900E-07 |
| 15970 | 799 | S799I | B | 4.3e-07 | 4e-07 | 3.7e-07 | 4.000E-07 | 3.000E-08 | 3.700E-07 | 4.300E-07 |
| 15971 | 799 | S799L | B | 1.8e-07 | 2.2e-07 | 3.5e-07 | 2.500E-07 | 8.888E-08 | 1.800E-07 | 3.500E-07 |
| 15972 | 799 | S799K | B | 9.3e-07 | 2.5e-07 | 2.4e-07 | 4.733E-07 | 3.955E-07 | 2.400E-07 | 9.300E-07 |
| 15973 | 799 | S799M | B | 2.5e-07 | 2.8e-07 | 5.5e-07 | 3.600E-07 | 1.652E-07 | 2.500E-07 | 5.500E-07 |
| 15974 | 799 | S799F | B | 2.8e-07 | 2.6e-07 | 2.4e-07 | 2.600E-07 | 2.000E-08 | 2.400E-07 | 2.800E-07 |
| 15975 | 799 | S799P | B | 3.3e-07 | 3e-07 | 2.4e-07 | 2.900E-07 | 4.583E-08 | 2.400E-07 | 3.300E-07 |
| 15976 | 799 | S799S | B | 2e-07 | 7.4e-07 | 1.7e-07 | 3.700E-07 | 3.208E-07 | 1.700E-07 | 7.400E-07 |
| 15977 | 799 | S799T | B | 2.8e-07 | 4.3e-07 | 2.1e-07 | 3.067E-07 | 1.124E-07 | 2.100E-07 | 4.300E-07 |
| 15978 | 799 | S799W | B | 2.7e-07 | 3.1e-07 | 3.9e-07 | 3.233E-07 | 6.110E-08 | 2.700E-07 | 3.900E-07 |
| 15979 | 799 | S799Y | B | 2.2e-07 | 2.9e-07 | 3.8e-07 | 2.967E-07 | 8.021E-08 | 2.200E-07 | 3.800E-07 |

| | | | | | | | | | | |
|---|---|---|---|---|---|---|---|---|---|---|
| 15980 | 799 | S799V | B | 3.8e-07 | 3.4e-07 | 1.6e-07 | 2.933E-07 | 1.172E-07 | 1.600E-07 | 3.800E-07 |
| 15981 | 800 | T800A | B | 1.7e-07 | 1.7e-07 | 2.5e-07 | 1.967E-07 | 4.619E-08 | 1.700E-07 | 2.500E-07 |
| 15982 | 800 | T800R | B | 2.4e-07 | 2.7e-07 | 1.9e-07 | 2.333E-07 | 4.041E-08 | 1.900E-07 | 2.700E-07 |
| 15983 | 800 | T800N | B | 1.6e-07 | 3.5e-07 | 2.6e-07 | 2.567E-07 | 9.504E-08 | 1.600E-07 | 3.500E-07 |
| 15984 | 800 | T800D | B | 2.4e-07 | 2.9e-07 | 2.9e-07 | 2.733E-07 | 2.887E-08 | 2.400E-07 | 2.900E-07 |
| 15985 | 800 | T800C | B | 2.8e-07 | 2.5e-07 | 1.4e-07 | 2.233E-07 | 7.371E-08 | 1.400E-07 | 2.800E-07 |
| 15986 | 800 | T800E | B | 1.6e-07 | 3.2e-07 | 3.7e-07 | 2.833E-07 | 1.097E-07 | 1.600E-07 | 3.700E-07 |
| 15987 | 800 | T800Q | B | 3.3e-07 | 2.8e-07 | 1e-07 | 2.367E-07 | 1.210E-07 | 1.000E-07 | 3.300E-07 |
| 15988 | 800 | T800G | B | 4.4e-07 | 3e-07 | 4.6e-07 | 4.000E-07 | 8.718E-08 | 3.000E-07 | 4.600E-07 |
| 15989 | 800 | T800H | B | 4e-07 | 4.1e-07 | 3.4e-07 | 3.833E-07 | 3.786E-08 | 3.400E-07 | 4.100E-07 |
| 15990 | 800 | T800I | B | 1.5e-07 | 2.6e-07 | 9.7e-07 | 4.600E-07 | 4.451E-07 | 1.500E-07 | 9.700E-07 |
| 15991 | 800 | T800L | B | 3.5e-07 | 1e-07 | 3.2e-07 | 2.567E-07 | 1.365E-07 | 1.000E-07 | 3.500E-07 |
| 15992 | 800 | T800K | B | 3.2e-07 | 3.3e-07 | 3.3e-07 | 3.267E-07 | 5.774E-09 | 3.200E-07 | 3.300E-07 |
| 15993 | 800 | T800M | B | 2.5e-07 | 2.5e-07 | 3.2e-07 | 2.733E-07 | 4.041E-08 | 2.500E-07 | 3.200E-07 |
| 15994 | 800 | T800F | B | 2.8e-07 | 3.7e-07 | 3.5e-07 | 3.333E-07 | 4.726E-08 | 2.800E-07 | 3.700E-07 |
| 15995 | 800 | T800P | B | 3.1e-07 | 1e-07 | 2.7e-07 | 2.267E-07 | 1.115E-07 | 1.000E-07 | 3.100E-07 |
| 15996 | 800 | T800S | B | 2.7e-07 | 4e-07 | 3.4e-07 | 3.367E-07 | 6.506E-08 | 2.700E-07 | 4.000E-07 |
| 15997 | 800 | T800T | B | 3.4e-07 | 2.7e-07 | 1e-07 | 2.367E-07 | 1.234E-07 | 1.000E-07 | 3.400E-07 |
| 15998 | 800 | T800W | B | 2.6e-07 | 2e-07 | 3.7e-07 | 2.767E-07 | 8.622E-08 | 2.000E-07 | 3.700E-07 |
| 15999 | 800 | T800Y | B | 1.5e-07 | 3.8e-07 | 3.1e-07 | 2.800E-07 | 1.179E-07 | 1.500E-07 | 3.800E-07 |
| 16000 | 800 | T800V | B | 3.5e-07 | 4.4e-07 | 1.5e-07 | 3.133E-07 | 1.484E-07 | 1.500E-07 | 4.400E-07 |
| 16001 | 801 | L801A | B | 1.6e-07 | 2.5e-07 | 2.2e-07 | 2.100E-07 | 4.583E-08 | 1.600E-07 | 2.500E-07 |
| 16002 | 801 | L801R | B | 1e-07 | 2.1e-07 | 1.5e-07 | 1.533E-07 | 5.508E-08 | 1.000E-07 | 2.100E-07 |
| 16003 | 801 | L801N | B | 2.4e-07 | 3.6e-07 | 2.1e-07 | 2.700E-07 | 7.937E-08 | 2.100E-07 | 3.600E-07 |
| 16004 | 801 | L801D | B | 1.1e-07 | 3.3e-07 | 2.7e-07 | 2.367E-07 | 1.137E-07 | 1.100E-07 | 3.300E-07 |
| 16005 | 801 | L801C | B | 2.9e-07 | 2.1e-07 | 2.3e-07 | 2.433E-07 | 4.163E-08 | 2.100E-07 | 2.900E-07 |
| 16006 | 801 | L801E | B | 6e-07 | 2e-07 | 2.4e-07 | 3.467E-07 | 2.203E-07 | 2.000E-07 | 6.000E-07 |
| 16007 | 801 | L801Q | B | 2.3e-07 | 8.7e-08 | 3.6e-07 | 2.257E-07 | 1.366E-07 | 8.700E-08 | 3.600E-07 |
| 16008 | 801 | L801G | B | 2.1e-07 | 1.5e-07 | 1.7e-07 | 1.767E-07 | 3.055E-08 | 1.500E-07 | 2.100E-07 |
| 16009 | 801 | L801H | B | 2.5e-07 | 3.5e-07 | 3.6e-07 | 3.200E-07 | 6.083E-08 | 2.500E-07 | 3.600E-07 |
| 16010 | 801 | L801I | B | 5.4e-07 | 4e-07 | 2.6e-07 | 4.000E-07 | 1.400E-07 | 2.600E-07 | 5.400E-07 |
| 16011 | 801 | L801L | B | 8.1e-07 | 4e-07 | 1.3e-07 | 4.467E-07 | 3.424E-07 | 1.300E-07 | 8.100E-07 |
| 16012 | 801 | L801K | B | 4e-07 | 2.2e-07 | 3.3e-07 | 3.167E-07 | 9.074E-08 | 2.200E-07 | 4.000E-07 |
| 16013 | 801 | L801M | B | 3.6e-07 | 1.3e-07 | 2.6e-07 | 2.500E-07 | 1.153E-07 | 1.300E-07 | 3.600E-07 |
| 16014 | 801 | L801F | B | 3.8e-07 | 3.2e-07 | 2.5e-07 | 3.167E-07 | 6.506E-08 | 2.500E-07 | 3.800E-07 |
| 16015 | 801 | L801P | B | 4e-07 | 2.3e-07 | 2.5e-07 | 2.933E-07 | 9.292E-08 | 2.300E-07 | 4.000E-07 |
| 16016 | 801 | L801S | B | 4.1e-07 | 4.2e-07 | 2.8e-07 | 3.700E-07 | 7.810E-08 | 2.800E-07 | 4.200E-07 |
| 16017 | 801 | L801T | B | 6.7e-07 | 3.1e-07 | 1.5e-07 | 3.767E-07 | 2.663E-07 | 1.500E-07 | 6.700E-07 |
| 16018 | 801 | L801W | B | 3.2e-07 | 4.2e-07 | 7.7e-07 | 5.033E-07 | 2.363E-07 | 3.200E-07 | 7.700E-07 |
| 16019 | 801 | L801Y | B | 3e-07 | 1.2e-07 | 2.8e-07 | 2.333E-07 | 9.866E-08 | 1.200E-07 | 3.000E-07 |
| 16020 | 801 | L801V | B | 1.7e-07 | 3.1e-07 | 1.4e-07 | 2.067E-07 | 9.074E-08 | 1.400E-07 | 3.100E-07 |
| 16021 | 802 | T802A | B | 4.1e-07 | 2.4e-07 | 2.1e-07 | 2.867E-07 | 1.079E-07 | 2.100E-07 | 4.100E-07 |
| 16022 | 802 | T802R | B | 4.5e-07 | 2.6e-07 | 3.7e-07 | 3.600E-07 | 9.539E-08 | 2.600E-07 | 4.500E-07 |
| 16023 | 802 | T802N | B | 1.7e-07 | 2.4e-07 | 1.2e-07 | 1.767E-07 | 6.028E-08 | 1.200E-07 | 2.400E-07 |
| 16024 | 802 | T802D | B | 2.2e-07 | 3e-07 | 1.5e-07 | 2.233E-07 | 7.506E-08 | 1.500E-07 | 3.000E-07 |
| 16025 | 802 | T802C | B | 2.5e-07 | 2.7e-07 | 2.4e-07 | 2.533E-07 | 1.528E-08 | 2.400E-07 | 2.700E-07 |
| 16026 | 802 | T802E | B | 1.1e-07 | 3.3e-07 | 1.5e-07 | 1.967E-07 | 1.172E-07 | 1.100E-07 | 3.300E-07 |
| 16027 | 802 | T802Q | B | 3.6e-07 | 1.7e-07 | 3.3e-07 | 2.867E-07 | 1.021E-07 | 1.700E-07 | 3.600E-07 |
| 16028 | 802 | T802G | B | 1.6e-07 | 4.3e-07 | 3.1e-07 | 3.000E-07 | 1.353E-07 | 1.600E-07 | 4.300E-07 |
| 16029 | 802 | T802H | B | 2.9e-07 | 2e-07 | 2e-07 | 2.300E-07 | 5.196E-08 | 2.000E-07 | 2.900E-07 |
| 16030 | 802 | T802I | B | 1.5e-07 | 2.4e-07 | 3.3e-07 | 2.400E-07 | 9.000E-08 | 1.500E-07 | 3.300E-07 |
| 16031 | 802 | T802L | B | 2.6e-07 | 3.5e-07 | 3e-07 | 3.033E-07 | 4.509E-08 | 2.600E-07 | 3.500E-07 |
| 16032 | 802 | T802K | B | 2.1e-07 | 1.7e-07 | 2.7e-07 | 2.167E-07 | 5.033E-08 | 1.700E-07 | 2.700E-07 |
| 16033 | 802 | T802M | B | 3.5e-07 | 8.2e-07 | 4.8e-07 | 5.500E-07 | 2.427E-07 | 3.500E-07 | 8.200E-07 |
| 16034 | 802 | T802F | B | 4e-07 | 2.5e-07 | 2.4e-07 | 2.967E-07 | 8.963E-08 | 2.400E-07 | 4.000E-07 |
| 16035 | 802 | T802P | B | 2.2e-07 | 4e-07 | 3.5e-07 | 3.233E-07 | 9.292E-08 | 2.200E-07 | 4.000E-07 |
| 16036 | 802 | T802S | B | 2e-07 | 2.7e-07 | 2.2e-07 | 2.300E-07 | 3.606E-08 | 2.000E-07 | 2.700E-07 |
| 16037 | 802 | T802T | B | 3.6e-07 | 1.4e-07 | 9.3e-08 | 1.977E-07 | 1.425E-07 | 9.300E-08 | 3.600E-07 |
| 16038 | 802 | T802W | B | 7.5e-07 | 3.8e-07 | 2e-07 | 4.433E-07 | 2.804E-07 | 2.000E-07 | 7.500E-07 |
| 16039 | 802 | T802Y | B | 1.7e-07 | 2e-07 | 2.6e-07 | 2.100E-07 | 4.583E-08 | 1.700E-07 | 2.600E-07 |
| 16040 | 802 | T802V | B | 1.8e-07 | 2e-07 | 4.8e-07 | 2.867E-07 | 1.677E-07 | 1.800E-07 | 4.800E-07 |
| 16041 | 803 | L803A | B | 3.5e-07 | 2.2e-07 | 1.6e-07 | 2.433E-07 | 9.713E-08 | 1.600E-07 | 3.500E-07 |
| 16042 | 803 | L803R | B | 2.6e-07 | 2e-07 | 5.5e-07 | 3.367E-07 | 1.872E-07 | 2.000E-07 | 5.500E-07 |
| 16043 | 803 | L803N | B | 2.7e-07 | 1.8e-07 | 3e-07 | 2.500E-07 | 6.245E-08 | 1.800E-07 | 3.000E-07 |
| 16044 | 803 | L803D | B | 1.1e-07 | 3e-07 | 1.5e-07 | 1.867E-07 | 1.002E-07 | 1.100E-07 | 3.000E-07 |
| 16045 | 803 | L803C | B | 3.4e-07 | 1.6e-07 | 4.8e-07 | 3.267E-07 | 1.604E-07 | 1.600E-07 | 4.800E-07 |
| 16046 | 803 | L803E | B | 3.6e-07 | 3.3e-07 | 4.3e-07 | 3.733E-07 | 5.132E-08 | 3.300E-07 | 4.300E-07 |
| 16047 | 803 | L803Q | B | 2.6e-07 | 1.2e-07 | 1.7e-07 | 1.833E-07 | 7.095E-08 | 1.200E-07 | 2.600E-07 |
| 16048 | 803 | L803G | B | 1.5e-07 | 3.9e-07 | 4.9e-07 | 3.433E-07 | 1.747E-07 | 1.500E-07 | 4.900E-07 |
| 16049 | 803 | L803H | B | 2.5e-07 | 1.5e-07 | 2.4e-07 | 2.133E-07 | 5.508E-08 | 1.500E-07 | 2.500E-07 |
| 16050 | 803 | L803I | B | 3e-07 | 2.6e-07 | 2.2e-07 | 2.600E-07 | 4.000E-08 | 2.200E-07 | 3.000E-07 |
| 16051 | 803 | L803L | B | 1.7e-07 | 1.7e-07 | 1.5e-07 | 1.633E-07 | 1.155E-08 | 1.500E-07 | 1.700E-07 |
| 16052 | 803 | L803K | B | 2e-07 | 5.7e-07 | 1.7e-07 | 3.133E-07 | 2.228E-07 | 1.700E-07 | 5.700E-07 |
| 16053 | 803 | L803M | B | 1.9e-07 | 3e-07 | 1.9e-07 | 2.267E-07 | 6.351E-08 | 1.900E-07 | 3.000E-07 |
| 16054 | 803 | L803F | B | 1.1e-07 | 4.8e-07 | 2e-07 | 2.633E-07 | 1.930E-07 | 1.100E-07 | 4.800E-07 |
| 16055 | 803 | L803P | B | 4.5e-07 | 1.7e-07 | 4.1e-07 | 3.433E-07 | 1.514E-07 | 1.700E-07 | 4.500E-07 |
| 16056 | 803 | L803S | B | 4e-07 | 1.8e-07 | 4e-07 | 3.267E-07 | 1.270E-07 | 1.800E-07 | 4.000E-07 |
| 16057 | 803 | L803T | B | 2.4e-07 | 4.2e-07 | 3.6e-07 | 3.400E-07 | 9.165E-08 | 2.400E-07 | 4.200E-07 |
| 16058 | 803 | L803W | B | 3.3e-07 | 2.8e-07 | 1.1e-07 | 2.400E-07 | 1.153E-07 | 1.100E-07 | 3.300E-07 |
| 16059 | 803 | L803Y | B | 1.5e-07 | 8.3e-08 | 1.5e-07 | 1.277E-07 | 3.868E-08 | 8.300E-08 | 1.500E-07 |
| 16060 | 803 | L803V | B | 3.6e-07 | 2.5e-07 | 1.6e-07 | 2.567E-07 | 1.002E-07 | 1.600E-07 | 3.600E-07 |
| 16061 | 804 | S804A | B | 1.2e-07 | 1.7e-07 | 2.2e-07 | 1.700E-07 | 5.000E-08 | 1.200E-07 | 2.200E-07 |
| 16062 | 804 | S804R | B | 1.5e-07 | 4.4e-07 | 3.1e-07 | 3.000E-07 | 1.453E-07 | 1.500E-07 | 4.400E-07 |
| 16063 | 804 | S804N | B | 1.8e-07 | 3e-07 | 1.8e-07 | 2.200E-07 | 6.928E-08 | 1.800E-07 | 3.000E-07 |
| 16064 | 804 | S804D | B | 1.7e-07 | 6e-07 | 3.4e-07 | 3.700E-07 | 2.166E-07 | 1.700E-07 | 6.000E-07 |

| | | | | | | | | | | |
|---|---|---|---|---|---|---|---|---|---|---|
| 16065 | 804 | S804C | B | 2.3e-07 | 7.8e-07 | 4.5e-07 | 4.867E-07 | 2.768E-07 | 2.300E-07 | 7.800E-07 |
| 16066 | 804 | S804E | B | 1.7e-07 | 2.1e-07 | 1.2e-07 | 1.667E-07 | 4.509E-08 | 1.200E-07 | 2.100E-07 |
| 16067 | 804 | S804Q | B | 3.8e-07 | 1.3e-07 | 1.8e-07 | 2.300E-07 | 1.323E-07 | 1.300E-07 | 3.800E-07 |
| 16068 | 804 | S804G | B | 2.6e-07 | 3.6e-07 | 1.4e-07 | 2.533E-07 | 1.102E-07 | 1.400E-07 | 3.600E-07 |
| 16069 | 804 | S804H | B | 3.2e-07 | 4.7e-07 | 2.7e-07 | 3.533E-07 | 1.041E-07 | 2.700E-07 | 4.700E-07 |
| 16070 | 804 | S804I | B | 3e-07 | 1.4e-07 | 3.6e-07 | 2.667E-07 | 1.137E-07 | 1.400E-07 | 3.600E-07 |
| 16071 | 804 | S804L | B | 1.4e-07 | 3.5e-07 | 2.3e-07 | 2.400E-07 | 1.054E-07 | 1.400E-07 | 3.500E-07 |
| 16072 | 804 | S804K | B | 4e-07 | 2.1e-07 | 4.9e-07 | 3.667E-07 | 1.429E-07 | 2.100E-07 | 4.900E-07 |
| 16073 | 804 | S804M | B | 1.5e-07 | 2.7e-07 | 2.3e-07 | 2.167E-07 | 6.110E-08 | 1.500E-07 | 2.700E-07 |
| 16074 | 804 | S804F | B | 2.9e-07 | 1.7e-07 | 3.4e-07 | 2.667E-07 | 8.737E-08 | 1.700E-07 | 3.400E-07 |
| 16075 | 804 | S804P | B | 1.5e-07 | 2.9e-07 | 1.3e-07 | 1.900E-07 | 8.718E-08 | 1.300E-07 | 2.900E-07 |
| 16076 | 804 | S804S | B | 3.9e-07 | 2.1e-07 | 3.3e-07 | 3.100E-07 | 9.165E-08 | 2.100E-07 | 3.900E-07 |
| 16077 | 804 | S804T | B | 2.9e-07 | 6.6e-07 | 4.2e-07 | 4.567E-07 | 1.877E-07 | 2.900E-07 | 6.600E-07 |
| 16078 | 804 | S804W | B | 3.2e-07 | 1.6e-07 | 2.1e-07 | 2.300E-07 | 8.185E-08 | 1.600E-07 | 3.200E-07 |
| 16079 | 804 | S804Y | B | 1.4e-07 | 2.5e-07 | 3.9e-07 | 2.600E-07 | 1.253E-07 | 1.400E-07 | 3.900E-07 |
| 16080 | 804 | S804V | B | 2.1e-07 | 1.6e-07 | 2.2e-07 | 1.967E-07 | 3.215E-08 | 1.600E-07 | 2.200E-07 |
| 16081 | 805 | K805A | B | 2.3e-07 | 5e-07 | 3e-07 | 3.433E-07 | 1.401E-07 | 2.300E-07 | 5.000E-07 |
| 16082 | 805 | K805R | B | 1.8e-07 | 2e-07 | 3.2e-07 | 2.333E-07 | 7.572E-08 | 1.800E-07 | 3.200E-07 |
| 16083 | 805 | K805N | B | 3.5e-07 | 1.6e-07 | 2.1e-07 | 2.400E-07 | 9.849E-08 | 1.600E-07 | 3.500E-07 |
| 16084 | 805 | K805D | B | 1.9e-07 | 2.6e-07 | 1.8e-07 | 2.100E-07 | 4.359E-08 | 1.800E-07 | 2.600E-07 |
| 16085 | 805 | K805C | B | 2e-07 | 5.4e-07 | 1.6e-07 | 3.000E-07 | 2.088E-07 | 1.600E-07 | 5.400E-07 |
| 16086 | 805 | K805E | B | 3e-07 | 3.8e-07 | 3.3e-07 | 3.367E-07 | 4.041E-08 | 3.000E-07 | 3.800E-07 |
| 16087 | 805 | K805Q | B | 2.3e-07 | 3.2e-07 | 2.7e-07 | 2.733E-07 | 4.509E-08 | 2.300E-07 | 3.200E-07 |
| 16088 | 805 | K805G | B | 1.8e-07 | 2.2e-07 | 1.5e-07 | 1.833E-07 | 3.512E-08 | 1.500E-07 | 2.200E-07 |
| 16089 | 805 | K805H | B | 5.3e-07 | 3.2e-07 | 1.7e-07 | 3.400E-07 | 1.808E-07 | 1.700E-07 | 5.300E-07 |
| 16090 | 805 | K805I | B | 1.7e-07 | 2.4e-07 | 4.3e-07 | 2.800E-07 | 1.345E-07 | 1.700E-07 | 4.300E-07 |
| 16091 | 805 | K805L | B | 1.2e-07 | 2.3e-07 | 9.3e-08 | 1.477E-07 | 7.257E-08 | 9.300E-08 | 2.300E-07 |
| 16092 | 805 | K805K | B | 6.8e-08 | 2.9e-07 | 3.7e-07 | 2.427E-07 | 1.565E-07 | 6.800E-08 | 3.700E-07 |
| 16093 | 805 | K805M | B | 2.4e-07 | 1.2e-07 | 3.3e-07 | 2.300E-07 | 1.054E-07 | 1.200E-07 | 3.300E-07 |
| 16094 | 805 | K805F | B | 2.4e-07 | 2.3e-07 | 1.6e-07 | 2.100E-07 | 4.359E-08 | 1.600E-07 | 2.400E-07 |
| 16095 | 805 | K805P | B | 4.3e-07 | 3.2e-07 | 2e-07 | 3.167E-07 | 1.150E-07 | 2.000E-07 | 4.300E-07 |
| 16096 | 805 | K805S | B | 1.5e-07 | 3.3e-07 | 2.4e-07 | 2.400E-07 | 9.000E-08 | 1.500E-07 | 3.300E-07 |
| 16097 | 805 | K805T | B | 1.3e-07 | 3e-07 | 2.6e-07 | 2.300E-07 | 8.888E-08 | 1.300E-07 | 3.000E-07 |
| 16098 | 805 | K805W | B | 2e-07 | 2.7e-07 | 6.2e-07 | 3.633E-07 | 2.250E-07 | 2.000E-07 | 6.200E-07 |
| 16099 | 805 | K805Y | B | 2.9e-07 | 5.1e-07 | 3.9e-07 | 3.967E-07 | 1.102E-07 | 2.900E-07 | 5.100E-07 |
| 16100 | 805 | K805V | B | 1.5e-07 | 2.2e-07 | 9.8e-08 | 1.560E-07 | 6.122E-08 | 9.800E-08 | 2.200E-07 |
| 16101 | 806 | A806A | B | 1.9e-07 | 1.9e-07 | 1.9e-07 | 1.900E-07 | 0.000E+00 | 1.900E-07 | 1.900E-07 |
| 16102 | 806 | A806R | B | 1.9e-07 | 2.3e-07 | 2.5e-07 | 2.233E-07 | 3.055E-08 | 1.900E-07 | 2.500E-07 |
| 16103 | 806 | A806N | B | 3.7e-07 | 1.6e-07 | 1.6e-07 | 2.300E-07 | 1.212E-07 | 1.600E-07 | 3.700E-07 |
| 16104 | 806 | A806D | B | 2.2e-07 | 3.1e-07 | 2e-07 | 2.433E-07 | 5.859E-08 | 2.000E-07 | 3.100E-07 |
| 16105 | 806 | A806C | B | 4e-07 | 1.6e-07 | 1.1e-07 | 2.233E-07 | 1.550E-07 | 1.100E-07 | 4.000E-07 |
| 16106 | 806 | A806E | B | 2.4e-07 | 2.4e-07 | 1.9e-07 | 2.233E-07 | 2.887E-08 | 1.900E-07 | 2.400E-07 |
| 16107 | 806 | A806Q | B | 2.2e-07 | 1.6e-07 | 2.6e-07 | 2.133E-07 | 5.033E-08 | 1.600E-07 | 2.600E-07 |
| 16108 | 806 | A806G | B | 2.8e-07 | 2e-07 | 3.2e-07 | 2.667E-07 | 6.110E-08 | 2.000E-07 | 3.200E-07 |
| 16109 | 806 | A806H | B | 1.7e-07 | 2.7e-07 | 1.7e-07 | 2.033E-07 | 5.774E-08 | 1.700E-07 | 2.700E-07 |
| 16110 | 806 | A806I | B | 3.5e-07 | 2.3e-07 | 1.4e-07 | 2.400E-07 | 1.054E-07 | 1.400E-07 | 3.500E-07 |
| 16111 | 806 | A806L | B | 3.2e-07 | 2.9e-07 | 2.3e-07 | 2.800E-07 | 4.583E-08 | 2.300E-07 | 3.200E-07 |
| 16112 | 806 | A806K | B | 4.7e-07 | 4.1e-07 | 3.7e-07 | 4.167E-07 | 5.033E-08 | 3.700E-07 | 4.700E-07 |
| 16113 | 806 | A806M | B | 1.5e-07 | 3e-07 | 1.7e-07 | 2.067E-07 | 8.145E-08 | 1.500E-07 | 3.000E-07 |
| 16114 | 806 | A806F | B | 3.6e-07 | 3.1e-07 | 1.5e-07 | 2.733E-07 | 1.097E-07 | 1.500E-07 | 3.600E-07 |
| 16115 | 806 | A806P | B | 1.7e-07 | 5e-07 | 5.1e-07 | 3.933E-07 | 1.935E-07 | 1.700E-07 | 5.100E-07 |
| 16116 | 806 | A806S | B | 3.6e-07 | 3e-07 | 1e-07 | 2.533E-07 | 1.361E-07 | 1.000E-07 | 3.600E-07 |
| 16117 | 806 | A806T | B | 3e-07 | 1.8e-07 | 1.6e-07 | 2.133E-07 | 7.572E-08 | 1.600E-07 | 3.000E-07 |
| 16118 | 806 | A806W | B | 2e-07 | 2.2e-07 | 6.8e-08 | 1.627E-07 | 8.259E-08 | 6.800E-08 | 2.200E-07 |
| 16119 | 806 | A806Y | B | 1.7e-07 | 1.3e-07 | 3.6e-07 | 2.200E-07 | 1.229E-07 | 1.300E-07 | 3.600E-07 |
| 16120 | 806 | A806V | B | 3.9e-07 | 1.4e-07 | 1.9e-07 | 2.400E-07 | 1.323E-07 | 1.400E-07 | 3.900E-07 |
| 16121 | 807 | D807A | B | 1.6e-07 | 3.6e-07 | 1.7e-07 | 2.300E-07 | 1.127E-07 | 1.600E-07 | 3.600E-07 |
| 16122 | 807 | D807R | B | 4.3e-07 | 3.6e-07 | 1.3e-07 | 3.067E-07 | 1.570E-07 | 1.300E-07 | 4.300E-07 |
| 16123 | 807 | D807N | B | 3.2e-07 | 3.6e-07 | 9.2e-07 | 5.333E-07 | 3.355E-07 | 3.200E-07 | 9.200E-07 |
| 16124 | 807 | D807D | B | 3.7e-07 | 1.6e-07 | 2e-07 | 2.433E-07 | 1.115E-07 | 1.600E-07 | 3.700E-07 |
| 16125 | 807 | D807C | B | 3e-07 | 2.4e-07 | 1.9e-07 | 2.433E-07 | 5.508E-08 | 1.900E-07 | 3.000E-07 |
| 16126 | 807 | D807E | B | 1.1e-07 | 2.9e-07 | 2e-07 | 2.000E-07 | 9.000E-08 | 1.100E-07 | 2.900E-07 |
| 16127 | 807 | D807Q | B | 2.3e-07 | 2.9e-07 | 1.6e-07 | 2.267E-07 | 6.506E-08 | 1.600E-07 | 2.900E-07 |
| 16128 | 807 | D807G | B | 3.9e-07 | 1.2e-07 | 1.5e-07 | 2.200E-07 | 1.480E-07 | 1.200E-07 | 3.900E-07 |
| 16129 | 807 | D807H | B | 5.3e-07 | 3.4e-07 | 1.7e-07 | 3.467E-07 | 1.801E-07 | 1.700E-07 | 5.300E-07 |
| 16130 | 807 | D807I | B | 2.6e-07 | 2.4e-07 | 2.3e-07 | 2.433E-07 | 1.528E-08 | 2.300E-07 | 2.600E-07 |
| 16131 | 807 | D807L | B | 1.5e-07 | 1.9e-07 | 2.2e-07 | 1.867E-07 | 3.512E-08 | 1.500E-07 | 2.200E-07 |
| 16132 | 807 | D807K | B | 3.3e-07 | 1.8e-07 | 1.8e-07 | 2.300E-07 | 8.660E-08 | 1.800E-07 | 3.300E-07 |
| 16133 | 807 | D807M | B | 1.5e-07 | 2.2e-07 | 1.8e-07 | 1.833E-07 | 3.512E-08 | 1.500E-07 | 2.200E-07 |
| 16134 | 807 | D807F | B | 3.3e-07 | 2e-07 | 2.5e-07 | 2.600E-07 | 6.557E-08 | 2.000E-07 | 3.300E-07 |
| 16135 | 807 | D807P | B | 2.2e-07 | 2.2e-07 | 2.5e-07 | 2.300E-07 | 1.732E-08 | 2.200E-07 | 2.500E-07 |
| 16136 | 807 | D807S | B | 1.8e-07 | 1e-07 | 3.4e-07 | 2.067E-07 | 1.222E-07 | 1.000E-07 | 3.400E-07 |
| 16137 | 807 | D807T | B | 4.5e-07 | 1.5e-07 | 2e-07 | 2.667E-07 | 1.607E-07 | 1.500E-07 | 4.500E-07 |
| 16138 | 807 | D807W | B | 1.9e-07 | 1.4e-07 | 1.9e-07 | 1.733E-07 | 2.887E-08 | 1.400E-07 | 1.900E-07 |
| 16139 | 807 | D807Y | B | 1.9e-07 | 3.2e-07 | 2.7e-07 | 2.600E-07 | 6.557E-08 | 1.900E-07 | 3.200E-07 |
| 16140 | 807 | D807V | B | 2.9e-07 | 2.2e-07 | 1.5e-07 | 2.200E-07 | 7.000E-08 | 1.500E-07 | 2.900E-07 |
| 16141 | 808 | Y808A | B | 2.9e-07 | 1.8e-07 | 2e-07 | 2.233E-07 | 5.859E-08 | 1.800E-07 | 2.900E-07 |
| 16142 | 808 | Y808R | B | 1.4e-07 | 2.2e-07 | 3e-07 | 2.200E-07 | 8.000E-08 | 1.400E-07 | 3.000E-07 |
| 16143 | 808 | Y808N | B | 1.3e-07 | 4e-07 | 1.3e-07 | 2.200E-07 | 1.559E-07 | 1.300E-07 | 4.000E-07 |
| 16144 | 808 | Y808D | B | 7.5e-08 | 5.5e-07 | 2e-07 | 2.750E-07 | 2.462E-07 | 7.500E-08 | 5.500E-07 |
| 16145 | 808 | Y808C | B | 3.5e-07 | 1.8e-07 | 2.3e-07 | 2.533E-07 | 8.737E-08 | 1.800E-07 | 3.500E-07 |
| 16146 | 808 | Y808E | B | 3.9e-07 | 1.6e-07 | 2.6e-07 | 2.700E-07 | 1.153E-07 | 1.600E-07 | 3.900E-07 |
| 16147 | 808 | Y808Q | B | 1.6e-07 | 3.8e-07 | 2.1e-07 | 2.500E-07 | 1.153E-07 | 1.600E-07 | 3.800E-07 |
| 16148 | 808 | Y808G | B | 4.6e-07 | 1.1e-07 | 4.7e-07 | 3.467E-07 | 2.050E-07 | 1.100E-07 | 4.700E-07 |
| 16149 | 808 | Y808H | B | 2e-07 | 2.2e-07 | 3.3e-07 | 2.500E-07 | 7.000E-08 | 2.000E-07 | 3.300E-07 |

| | | | | | | | | | | |
|---|---|---|---|---|---|---|---|---|---|---|
| 16150 | 808 | Y808I | B | 5.2e-07 | 2.4e-07 | 3.7e-07 | 3.767E-07 | 1.401E-07 | 2.400E-07 | 5.200E-07 |
| 16151 | 808 | Y808L | B | 4.5e-07 | 4.6e-07 | 3e-07 | 4.033E-07 | 8.963E-08 | 3.000E-07 | 4.600E-07 |
| 16152 | 808 | Y808K | B | 4.7e-07 | 2.2e-07 | 2.7e-07 | 3.200E-07 | 1.323E-07 | 2.200E-07 | 4.700E-07 |
| 16153 | 808 | Y808M | B | 4.4e-07 | 2.5e-07 | 2.8e-07 | 3.233E-07 | 1.021E-07 | 2.500E-07 | 4.400E-07 |
| 16154 | 808 | Y808F | B | 8.1e-08 | 1.8e-07 | 3e-07 | 1.870E-07 | 1.097E-07 | 8.100E-08 | 3.000E-07 |
| 16155 | 808 | Y808P | B | 3.5e-07 | 4.3e-07 | 2.9e-07 | 3.567E-07 | 7.024E-08 | 2.900E-07 | 4.300E-07 |
| 16156 | 808 | Y808S | B | 1.5e-07 | 2e-07 | 2e-07 | 1.833E-07 | 2.887E-08 | 1.500E-07 | 2.000E-07 |
| 16157 | 808 | Y808T | B | 2.5e-07 | 3.2e-07 | 2.8e-07 | 2.833E-07 | 3.512E-08 | 2.500E-07 | 3.200E-07 |
| 16158 | 808 | Y808W | B | 1.5e-07 | 3.2e-07 | 3e-07 | 2.567E-07 | 9.292E-08 | 1.500E-07 | 3.200E-07 |
| 16159 | 808 | Y808Y | B | 2.6e-07 | 2.4e-07 | 2.2e-07 | 2.400E-07 | 2.000E-08 | 2.200E-07 | 2.600E-07 |
| 16160 | 808 | Y808V | B | 2.7e-07 | 1.9e-07 | 1.9e-07 | 2.167E-07 | 4.619E-08 | 1.900E-07 | 2.700E-07 |
| 16161 | 809 | E809A | B | 1.6e-07 | 3.4e-07 | 4.9e-07 | 3.300E-07 | 1.652E-07 | 1.600E-07 | 4.900E-07 |
| 16162 | 809 | E809R | B | 4.9e-08 | 2.3e-07 | 1.3e-07 | 1.363E-07 | 9.067E-08 | 4.900E-08 | 2.300E-07 |
| 16163 | 809 | E809N | B | 2.5e-07 | 5.1e-07 | 2.1e-07 | 3.233E-07 | 1.629E-07 | 2.100E-07 | 5.100E-07 |
| 16164 | 809 | E809D | B | 2.3e-07 | 4.7e-07 | 2.6e-07 | 3.200E-07 | 1.308E-07 | 2.300E-07 | 4.700E-07 |
| 16165 | 809 | E809C | B | 2.2e-07 | 1e-07 | 3.5e-07 | 2.233E-07 | 1.250E-07 | 1.000E-07 | 3.500E-07 |
| 16166 | 809 | E809E | B | 7.4e-07 | 5e-07 | 2.8e-07 | 5.067E-07 | 2.301E-07 | 2.800E-07 | 7.400E-07 |
| 16167 | 809 | E809Q | B | 2e-07 | 1.2e-07 | 1.6e-07 | 1.600E-07 | 4.000E-08 | 1.200E-07 | 2.000E-07 |
| 16168 | 809 | E809G | B | 1.8e-07 | 3.5e-07 | 1.7e-07 | 2.333E-07 | 1.012E-07 | 1.700E-07 | 3.500E-07 |
| 16169 | 809 | E809H | B | 2.3e-07 | 2.9e-07 | 1.8e-07 | 2.333E-07 | 5.508E-08 | 1.800E-07 | 2.900E-07 |
| 16170 | 809 | E809I | B | 1.2e-07 | 2.4e-07 | 4e-07 | 2.533E-07 | 1.405E-07 | 1.200E-07 | 4.000E-07 |
| 16171 | 809 | E809L | B | 3.9e-07 | 4.1e-07 | 2.8e-07 | 3.600E-07 | 7.000E-08 | 2.800E-07 | 4.100E-07 |
| 16172 | 809 | E809K | B | 2.4e-07 | 2.9e-07 | 5.8e-07 | 3.700E-07 | 1.836E-07 | 2.400E-07 | 5.800E-07 |
| 16173 | 809 | E809M | B | 3.7e-07 | 1.2e-07 | 4e-07 | 2.967E-07 | 1.537E-07 | 1.200E-07 | 4.000E-07 |
| 16174 | 809 | E809F | B | 2.3e-07 | 2.2e-07 | 3e-07 | 2.500E-07 | 4.359E-08 | 2.200E-07 | 3.000E-07 |
| 16175 | 809 | E809P | B | 3e-07 | 6.3e-07 | 1.4e-07 | 3.567E-07 | 2.499E-07 | 1.400E-07 | 6.300E-07 |
| 16176 | 809 | E809S | B | 4.7e-07 | 3.3e-07 | 3.4e-07 | 3.800E-07 | 7.810E-08 | 3.300E-07 | 4.700E-07 |
| 16177 | 809 | E809T | B | 2.6e-07 | 3.7e-07 | 1.9e-07 | 2.733E-07 | 9.074E-08 | 1.900E-07 | 3.700E-07 |
| 16178 | 809 | E809W | B | 1.3e-07 | 3.4e-07 | 1.9e-07 | 2.200E-07 | 1.082E-07 | 1.300E-07 | 3.400E-07 |
| 16179 | 809 | E809Y | B | 1.5e-07 | 3e-07 | 1.8e-07 | 2.100E-07 | 7.937E-08 | 1.500E-07 | 3.000E-07 |
| 16180 | 809 | E809V | B | 4e-07 | 3.5e-07 | 1.3e-07 | 2.933E-07 | 1.436E-07 | 1.300E-07 | 4.000E-07 |
| 16181 | 810 | K810A | B | 3.2e-07 | 2.3e-07 | 2.5e-07 | 2.667E-07 | 4.726E-08 | 2.300E-07 | 3.200E-07 |
| 16182 | 810 | K810R | B | 3.1e-07 | 2.5e-07 | 1.7e-07 | 2.433E-07 | 7.024E-08 | 1.700E-07 | 3.100E-07 |
| 16183 | 810 | K810N | B | 1.6e-07 | 2.1e-07 | 3.4e-07 | 2.367E-07 | 9.292E-08 | 1.600E-07 | 3.400E-07 |
| 16184 | 810 | K810D | B | 2e-07 | 3e-07 | 2e-07 | 2.333E-07 | 5.774E-08 | 2.000E-07 | 3.000E-07 |
| 16185 | 810 | K810C | B | 2e-07 | 2.4e-07 | 2.7e-07 | 2.367E-07 | 3.512E-08 | 2.000E-07 | 2.700E-07 |
| 16186 | 810 | K810E | B | 2.6e-07 | 2.2e-07 | 3.8e-07 | 2.867E-07 | 8.327E-08 | 2.200E-07 | 3.800E-07 |
| 16187 | 810 | K810Q | B | 4.5e-07 | 2.3e-07 | 4.6e-07 | 3.800E-07 | 1.300E-07 | 2.300E-07 | 4.600E-07 |
| 16188 | 810 | K810G | B | 3.2e-07 | 2.6e-07 | 4.7e-07 | 3.500E-07 | 1.082E-07 | 2.600E-07 | 4.700E-07 |
| 16189 | 810 | K810H | B | 4.1e-07 | 1.1e-07 | 5e-07 | 3.400E-07 | 2.042E-07 | 1.100E-07 | 5.000E-07 |
| 16190 | 810 | K810I | B | 2.8e-07 | 1.7e-07 | 4.1e-07 | 2.867E-07 | 1.201E-07 | 1.700E-07 | 4.100E-07 |
| 16191 | 810 | K810L | B | 4.2e-07 | 2.8e-07 | 1.5e-07 | 2.833E-07 | 1.350E-07 | 1.500E-07 | 4.200E-07 |
| 16192 | 810 | K810K | B | 1.7e-07 | 2.7e-07 | 3.4e-07 | 2.600E-07 | 8.544E-08 | 1.700E-07 | 3.400E-07 |
| 16193 | 810 | K810M | B | 1.7e-07 | 1.9e-07 | 2.9e-07 | 2.167E-07 | 6.429E-08 | 1.700E-07 | 2.900E-07 |
| 16194 | 810 | K810F | B | 7.2e-07 | 1.8e-07 | 1.7e-07 | 3.567E-07 | 3.147E-07 | 1.700E-07 | 7.200E-07 |
| 16195 | 810 | K810P | B | 2.2e-07 | 1.8e-07 | 1.6e-07 | 1.867E-07 | 3.055E-08 | 1.600E-07 | 2.200E-07 |
| 16196 | 810 | K810S | B | 4.2e-07 | 3.4e-07 | 2.2e-07 | 3.267E-07 | 1.007E-07 | 2.200E-07 | 4.200E-07 |
| 16197 | 810 | K810T | B | 3.1e-07 | 2.1e-07 | 3.3e-07 | 2.833E-07 | 6.429E-08 | 2.100E-07 | 3.300E-07 |
| 16198 | 810 | K810W | B | 1.5e-07 | 5.9e-07 | 2.3e-07 | 3.233E-07 | 2.344E-07 | 1.500E-07 | 5.900E-07 |
| 16199 | 810 | K810Y | B | 1.8e-07 | 2.5e-07 | 2.1e-07 | 2.133E-07 | 3.512E-08 | 1.800E-07 | 2.500E-07 |
| 16200 | 810 | K810V | B | 3.4e-07 | 1.3e-07 | 1.6e-07 | 2.100E-07 | 1.136E-07 | 1.300E-07 | 3.400E-07 |
| 16201 | 811 | H811A | B | 2.6e-07 | 2.7e-07 | 1.9e-07 | 2.400E-07 | 4.359E-08 | 1.900E-07 | 2.700E-07 |
| 16202 | 811 | H811R | B | 9.7e-08 | 2.6e-07 | 5.5e-07 | 3.023E-07 | 2.294E-07 | 9.700E-08 | 5.500E-07 |
| 16203 | 811 | H811N | B | 7.8e-07 | 2.1e-07 | 1.5e-07 | 3.800E-07 | 3.477E-07 | 1.500E-07 | 7.800E-07 |
| 16204 | 811 | H811D | B | 3.6e-07 | 1.7e-07 | 2.9e-07 | 2.733E-07 | 9.609E-08 | 1.700E-07 | 3.600E-07 |
| 16205 | 811 | H811C | B | 1.4e-07 | 4.6e-07 | 2.6e-07 | 2.867E-07 | 1.617E-07 | 1.400E-07 | 4.600E-07 |
| 16206 | 811 | H811E | B | 1.2e-07 | 2.1e-07 | 2.6e-07 | 1.967E-07 | 7.095E-08 | 1.200E-07 | 2.600E-07 |
| 16207 | 811 | H811Q | B | 1.2e-06 | 2.7e-07 | 1.9e-07 | 5.533E-07 | 5.615E-07 | 1.900E-07 | 1.200E-06 |
| 16208 | 811 | H811G | B | 2.6e-07 | 2.2e-07 | 2.1e-07 | 2.300E-07 | 2.646E-08 | 2.100E-07 | 2.600E-07 |
| 16209 | 811 | H811H | B | 2.3e-07 | 3.9e-07 | 4.4e-07 | 3.533E-07 | 1.097E-07 | 2.300E-07 | 4.400E-07 |
| 16210 | 811 | H811I | B | 2.8e-07 | 2.8e-07 | 7.1e-07 | 4.233E-07 | 2.483E-07 | 2.800E-07 | 7.100E-07 |
| 16211 | 811 | H811L | B | 3e-07 | 1e-07 | 1.9e-07 | 1.967E-07 | 1.002E-07 | 1.000E-07 | 3.000E-07 |
| 16212 | 811 | H811K | B | 4.3e-07 | 3.1e-07 | 8.3e-08 | 2.743E-07 | 1.762E-07 | 8.300E-08 | 4.300E-07 |
| 16213 | 811 | H811M | B | 3.9e-07 | 1.6e-07 | 1.2e-07 | 2.233E-07 | 1.457E-07 | 1.200E-07 | 3.900E-07 |
| 16214 | 811 | H811F | B | 3.8e-07 | 3.7e-07 | 2.4e-07 | 3.300E-07 | 7.810E-08 | 2.400E-07 | 3.800E-07 |
| 16215 | 811 | H811P | B | 3.9e-07 | 2.1e-07 | 2.2e-07 | 2.733E-07 | 1.012E-07 | 2.100E-07 | 3.900E-07 |
| 16216 | 811 | H811S | B | 3.1e-07 | 5.8e-07 | 3.3e-07 | 4.067E-07 | 1.504E-07 | 3.100E-07 | 5.800E-07 |
| 16217 | 811 | H811T | B | 3e-07 | 2.8e-07 | 2.1e-07 | 2.633E-07 | 4.726E-08 | 2.100E-07 | 3.000E-07 |
| 16218 | 811 | H811W | B | 4.2e-07 | 2.3e-07 | 2.1e-07 | 2.867E-07 | 1.159E-07 | 2.100E-07 | 4.200E-07 |
| 16219 | 811 | H811Y | B | 3.5e-07 | 3.7e-07 | 2.1e-07 | 3.100E-07 | 8.718E-08 | 2.100E-07 | 3.700E-07 |
| 16220 | 811 | H811V | B | 2.1e-07 | 2.5e-07 | 4.4e-07 | 3.000E-07 | 1.229E-07 | 2.100E-07 | 4.400E-07 |
| 16221 | 812 | K812A | B | 1.3e-07 | 4.1e-07 | 2.2e-07 | 2.533E-07 | 1.429E-07 | 1.300E-07 | 4.100E-07 |
| 16222 | 812 | K812R | B | 2.7e-07 | 2.5e-07 | 3.3e-07 | 2.833E-07 | 4.163E-08 | 2.500E-07 | 3.300E-07 |
| 16223 | 812 | K812N | B | 4e-07 | 3.7e-07 | 2.2e-07 | 3.300E-07 | 9.644E-08 | 2.200E-07 | 4.000E-07 |
| 16224 | 812 | K812D | B | 2.9e-07 | 1.8e-07 | 1.3e-07 | 2.000E-07 | 8.185E-08 | 1.300E-07 | 2.900E-07 |
| 16225 | 812 | K812C | B | 2.7e-07 | 2.8e-07 | 4.3e-07 | 3.267E-07 | 8.963E-08 | 2.700E-07 | 4.300E-07 |
| 16226 | 812 | K812E | B | 9.6e-08 | 3.9e-07 | 3.6e-07 | 2.820E-07 | 1.618E-07 | 9.600E-08 | 3.900E-07 |
| 16227 | 812 | K812Q | B | 1.5e-07 | 3.8e-07 | 7.8e-08 | 2.027E-07 | 1.577E-07 | 7.800E-08 | 3.800E-07 |
| 16228 | 812 | K812G | B | 3.7e-07 | 3.9e-07 | 1.7e-07 | 3.100E-07 | 1.217E-07 | 1.700E-07 | 3.900E-07 |
| 16229 | 812 | K812H | B | 3.1e-07 | 2e-07 | 2e-07 | 2.367E-07 | 6.351E-08 | 2.000E-07 | 3.100E-07 |
| 16230 | 812 | K812I | B | 9.1e-07 | 2.3e-07 | 4.3e-07 | 5.233E-07 | 3.495E-07 | 2.300E-07 | 9.100E-07 |
| 16231 | 812 | K812L | B | 2.8e-07 | 1.9e-07 | 1.2e-07 | 1.967E-07 | 8.021E-08 | 1.200E-07 | 2.800E-07 |
| 16232 | 812 | K812K | B | 1.6e-07 | 3e-07 | 2.2e-07 | 2.267E-07 | 7.024E-08 | 1.600E-07 | 3.000E-07 |
| 16233 | 812 | K812M | B | 2.9e-07 | 2.3e-07 | 2e-07 | 2.400E-07 | 4.583E-08 | 2.000E-07 | 2.900E-07 |
| 16234 | 812 | K812F | B | 5.4e-07 | 3.2e-07 | 1.5e-07 | 3.367E-07 | 1.955E-07 | 1.500E-07 | 5.400E-07 |

| | | | | | | | | | | |
|---|---|---|---|---|---|---|---|---|---|---|
| 16235 | 812 | K812P | B | 2.7e-07 | 1.8e-07 | 2e-07 | 2.167E-07 | 4.726E-08 | 1.800E-07 | 2.700E-07 |
| 16236 | 812 | K812S | B | 3e-07 | 4.5e-07 | 1.5e-07 | 3.000E-07 | 1.500E-07 | 1.500E-07 | 4.500E-07 |
| 16237 | 812 | K812T | B | 3e-07 | 3.2e-07 | 1.6e-07 | 2.600E-07 | 8.718E-08 | 1.600E-07 | 3.200E-07 |
| 16238 | 812 | K812W | B | 5.7e-07 | 1.7e-07 | 2.6e-07 | 3.333E-07 | 2.098E-07 | 1.700E-07 | 5.700E-07 |
| 16239 | 812 | K812Y | B | 5.5e-07 | 3e-07 | 2.9e-07 | 3.800E-07 | 1.473E-07 | 2.900E-07 | 5.500E-07 |
| 16240 | 812 | K812V | B | 3.2e-07 | 3.3e-07 | 1.5e-07 | 2.667E-07 | 1.012E-07 | 1.500E-07 | 3.300E-07 |
| 16241 | 813 | V813A | B | 2.5e-07 | 2.6e-07 | 2.1e-07 | 2.400E-07 | 2.646E-08 | 2.100E-07 | 2.600E-07 |
| 16242 | 813 | V813R | B | 1.3e-07 | 1.8e-07 | 2.1e-07 | 1.733E-07 | 4.041E-08 | 1.300E-07 | 2.100E-07 |
| 16243 | 813 | V813N | B | 2.7e-07 | 2.6e-07 | 2e-07 | 2.433E-07 | 3.786E-08 | 2.000E-07 | 2.700E-07 |
| 16244 | 813 | V813D | B | 2.4e-07 | 1.4e-07 | 2e-07 | 1.933E-07 | 5.033E-08 | 1.400E-07 | 2.400E-07 |
| 16245 | 813 | V813C | B | 3.8e-07 | 2.8e-07 | 4.2e-07 | 3.600E-07 | 7.211E-08 | 2.800E-07 | 4.200E-07 |
| 16246 | 813 | V813E | B | 3.6e-07 | 1.7e-07 | 4.2e-07 | 3.167E-07 | 1.305E-07 | 1.700E-07 | 4.200E-07 |
| 16247 | 813 | V813Q | B | 7.6e-07 | 4.5e-07 | 4.9e-07 | 5.667E-07 | 1.686E-07 | 4.500E-07 | 7.600E-07 |
| 16248 | 813 | V813G | B | 2.2e-07 | 2.4e-07 | 2.3e-07 | 2.300E-07 | 1.000E-08 | 2.200E-07 | 2.400E-07 |
| 16249 | 813 | V813H | B | 1.3e-07 | 1.4e-07 | 2.8e-07 | 1.833E-07 | 8.386E-08 | 1.300E-07 | 2.800E-07 |
| 16250 | 813 | V813I | B | 2.4e-07 | 7.9e-07 | 1.6e-07 | 3.967E-07 | 3.430E-07 | 1.600E-07 | 7.900E-07 |
| 16251 | 813 | V813L | B | 1.4e-07 | 3.1e-07 | 2.4e-07 | 2.300E-07 | 8.544E-08 | 1.400E-07 | 3.100E-07 |
| 16252 | 813 | V813K | B | 1.6e-07 | 3.1e-07 | 4.5e-07 | 3.067E-07 | 1.450E-07 | 1.600E-07 | 4.500E-07 |
| 16253 | 813 | V813M | B | 1.9e-07 | 4.3e-07 | 4.2e-07 | 3.467E-07 | 1.358E-07 | 1.900E-07 | 4.300E-07 |
| 16254 | 813 | V813F | B | 6.6e-07 | 2.6e-07 | 3.3e-07 | 4.167E-07 | 2.136E-07 | 2.600E-07 | 6.600E-07 |
| 16255 | 813 | V813P | B | 2.3e-07 | 2.2e-07 | 2.9e-07 | 2.467E-07 | 3.786E-08 | 2.200E-07 | 2.900E-07 |
| 16256 | 813 | V813S | B | 3.1e-07 | 2.6e-07 | 2.7e-07 | 2.800E-07 | 2.646E-08 | 2.600E-07 | 3.100E-07 |
| 16257 | 813 | V813T | B | 3e-07 | 1.9e-07 | 4.5e-07 | 3.133E-07 | 1.305E-07 | 1.900E-07 | 4.500E-07 |
| 16258 | 813 | V813W | B | 3.7e-07 | 2.2e-07 | 1.9e-07 | 2.600E-07 | 9.644E-08 | 1.900E-07 | 3.700E-07 |
| 16259 | 813 | V813Y | B | 2.4e-07 | 3.2e-07 | 2.9e-07 | 2.833E-07 | 4.041E-08 | 2.400E-07 | 3.200E-07 |
| 16260 | 813 | V813V | B | 6.2e-08 | 1.4e-07 | 2.8e-07 | 1.607E-07 | 1.105E-07 | 6.200E-08 | 2.800E-07 |
| 16261 | 814 | Y814A | B | 4e-07 | 2.7e-07 | 2.7e-07 | 3.133E-07 | 7.506E-08 | 2.700E-07 | 4.000E-07 |
| 16262 | 814 | Y814R | B | 3.7e-07 | 2.9e-07 | 2.4e-07 | 3.000E-07 | 6.557E-08 | 2.400E-07 | 3.700E-07 |
| 16263 | 814 | Y814N | B | 3e-07 | 6.9e-08 | 4.9e-07 | 2.863E-07 | 2.108E-07 | 6.900E-08 | 4.900E-07 |
| 16264 | 814 | Y814D | B | 3.9e-07 | 2.3e-07 | 1.6e-07 | 2.600E-07 | 1.179E-07 | 1.600E-07 | 3.900E-07 |
| 16265 | 814 | Y814C | B | 1.5e-07 | 2.6e-07 | 8.1e-08 | 1.637E-07 | 9.028E-08 | 8.100E-08 | 2.600E-07 |
| 16266 | 814 | Y814E | B | 1.6e-07 | 5.2e-07 | 1.4e-07 | 2.733E-07 | 2.139E-07 | 1.400E-07 | 5.200E-07 |
| 16267 | 814 | Y814Q | B | 3e-07 | 1.3e-07 | 1.1e-07 | 1.800E-07 | 1.044E-07 | 1.100E-07 | 3.000E-07 |
| 16268 | 814 | Y814G | B | 3.4e-07 | 2.9e-07 | 1e-07 | 2.433E-07 | 1.266E-07 | 1.000E-07 | 3.400E-07 |
| 16269 | 814 | Y814H | B | 2.1e-07 | 1.4e-07 | 1.8e-07 | 1.767E-07 | 3.512E-08 | 1.400E-07 | 2.100E-07 |
| 16270 | 814 | Y814I | B | 3.4e-07 | 1.8e-07 | 1.6e-07 | 2.267E-07 | 9.866E-08 | 1.600E-07 | 3.400E-07 |
| 16271 | 814 | Y814L | B | 2e-07 | 5e-07 | 3.3e-07 | 3.433E-07 | 1.504E-07 | 2.000E-07 | 5.000E-07 |
| 16272 | 814 | Y814K | B | 2e-07 | 3.9e-07 | 1.4e-07 | 2.433E-07 | 1.305E-07 | 1.400E-07 | 3.900E-07 |
| 16273 | 814 | Y814M | B | 9.6e-08 | 3e-07 | 1.8e-07 | 1.920E-07 | 1.025E-07 | 9.600E-08 | 3.000E-07 |
| 16274 | 814 | Y814F | B | 3.7e-07 | 1.5e-07 | 3.2e-07 | 2.800E-07 | 1.153E-07 | 1.500E-07 | 3.700E-07 |
| 16275 | 814 | Y814P | B | 2.9e-07 | 8.1e-07 | 2.1e-07 | 4.367E-07 | 3.258E-07 | 2.100E-07 | 8.100E-07 |
| 16276 | 814 | Y814S | B | 1.5e-07 | 1.7e-07 | 4.1e-07 | 2.433E-07 | 1.447E-07 | 1.500E-07 | 4.100E-07 |
| 16277 | 814 | Y814T | B | 1.8e-07 | 1.4e-07 | 1.6e-07 | 1.600E-07 | 2.000E-08 | 1.400E-07 | 1.800E-07 |
| 16278 | 814 | Y814W | B | 1.8e-07 | 6.2e-07 | 3.6e-07 | 3.867E-07 | 2.212E-07 | 1.800E-07 | 6.200E-07 |
| 16279 | 814 | Y814Y | B | 1.5e-07 | 1.4e-07 | 2.8e-07 | 1.900E-07 | 7.810E-08 | 1.400E-07 | 2.800E-07 |
| 16280 | 814 | Y814V | B | 1.8e-07 | 3.9e-07 | 2.3e-07 | 2.667E-07 | 1.097E-07 | 1.800E-07 | 3.900E-07 |
| 16281 | 815 | A815A | B | 3.8e-07 | 1.2e-07 | 1.7e-07 | 2.233E-07 | 1.380E-07 | 1.200E-07 | 3.800E-07 |
| 16282 | 815 | A815R | B | 8.6e-07 | 2.2e-07 | 2.6e-07 | 4.467E-07 | 3.585E-07 | 2.200E-07 | 8.600E-07 |
| 16283 | 815 | A815N | B | 1.9e-07 | 1.5e-07 | 2.4e-07 | 1.933E-07 | 4.509E-08 | 1.500E-07 | 2.400E-07 |
| 16284 | 815 | A815D | B | 4.5e-07 | 1.7e-07 | 3.1e-07 | 3.100E-07 | 1.400E-07 | 1.700E-07 | 4.500E-07 |
| 16285 | 815 | A815C | B | 2.5e-07 | 8.3e-07 | 1.6e-07 | 4.133E-07 | 3.636E-07 | 1.600E-07 | 8.300E-07 |
| 16286 | 815 | A815E | B | 4e-07 | 1.4e-07 | 2e-07 | 2.467E-07 | 1.361E-07 | 1.400E-07 | 4.000E-07 |
| 16287 | 815 | A815Q | B | 2.8e-07 | 3.9e-07 | 4e-07 | 3.567E-07 | 6.658E-08 | 2.800E-07 | 4.000E-07 |
| 16288 | 815 | A815G | B | 2.4e-07 | 1.2e-07 | 2.6e-07 | 2.067E-07 | 7.572E-08 | 1.200E-07 | 2.600E-07 |
| 16289 | 815 | A815H | B | 4e-07 | 2.6e-07 | 2.3e-07 | 2.967E-07 | 9.074E-08 | 2.300E-07 | 4.000E-07 |
| 16290 | 815 | A815I | B | 2.3e-07 | 1.9e-07 | 4.7e-07 | 2.967E-07 | 1.514E-07 | 1.900E-07 | 4.700E-07 |
| 16291 | 815 | A815L | B | 1.7e-07 | 9.2e-08 | 1.7e-07 | 1.440E-07 | 4.503E-08 | 9.200E-08 | 1.700E-07 |
| 16292 | 815 | A815K | B | 3.8e-07 | 2.3e-07 | 1.6e-07 | 2.567E-07 | 1.124E-07 | 1.600E-07 | 3.800E-07 |
| 16293 | 815 | A815M | B | 3.3e-07 | 3.9e-07 | 1.7e-07 | 2.967E-07 | 1.137E-07 | 1.700E-07 | 3.900E-07 |
| 16294 | 815 | A815F | B | 2.9e-07 | 1.8e-07 | 1.6e-07 | 2.100E-07 | 7.000E-08 | 1.600E-07 | 2.900E-07 |
| 16295 | 815 | A815P | B | 3.9e-07 | 4.9e-07 | 1.7e-07 | 3.500E-07 | 1.637E-07 | 1.700E-07 | 4.900E-07 |
| 16296 | 815 | A815S | B | 2.1e-07 | 2.7e-07 | 2.6e-07 | 2.467E-07 | 3.215E-08 | 2.100E-07 | 2.700E-07 |
| 16297 | 815 | A815T | B | 2.5e-07 | 1.8e-07 | 3.5e-07 | 2.600E-07 | 8.544E-08 | 1.800E-07 | 3.500E-07 |
| 16298 | 815 | A815W | B | 3e-07 | 1.7e-07 | 2.6e-07 | 2.433E-07 | 6.658E-08 | 1.700E-07 | 3.000E-07 |
| 16299 | 815 | A815Y | B | 2.6e-07 | 1.5e-07 | 2.9e-07 | 2.333E-07 | 7.371E-08 | 1.500E-07 | 2.900E-07 |
| 16300 | 815 | A815V | B | 5.8e-07 | 2e-07 | 4.9e-07 | 4.233E-07 | 1.986E-07 | 2.000E-07 | 5.800E-07 |
| 16301 | 816 | C816A | B | 4.2e-07 | 1.7e-07 | 1.7e-07 | 2.533E-07 | 1.443E-07 | 1.700E-07 | 4.200E-07 |
| 16302 | 816 | C816R | B | 3.9e-07 | 3e-07 | 9.7e-08 | 2.623E-07 | 1.501E-07 | 9.700E-08 | 3.900E-07 |
| 16303 | 816 | C816N | B | 4.6e-07 | 2.3e-07 | 3.9e-07 | 3.600E-07 | 1.179E-07 | 2.300E-07 | 4.600E-07 |
| 16304 | 816 | C816D | B | 8.2e-08 | 1.9e-07 | 2.3e-07 | 1.673E-07 | 7.656E-08 | 8.200E-08 | 2.300E-07 |
| 16305 | 816 | C816C | B | 1.9e-07 | 3e-07 | 9.8e-08 | 1.960E-07 | 1.011E-07 | 9.800E-08 | 3.000E-07 |
| 16306 | 816 | C816E | B | 2e-07 | 2.7e-07 | 8.7e-08 | 1.857E-07 | 9.234E-08 | 8.700E-08 | 2.700E-07 |
| 16307 | 816 | C816Q | B | 3.1e-07 | 4.3e-07 | 5.4e-07 | 4.267E-07 | 1.150E-07 | 3.100E-07 | 5.400E-07 |
| 16308 | 816 | C816G | B | 1.2e-07 | 4.4e-07 | 2.5e-07 | 2.700E-07 | 1.609E-07 | 1.200E-07 | 4.400E-07 |
| 16309 | 816 | C816H | B | 1.5e-07 | 2.6e-07 | 2.7e-07 | 2.267E-07 | 6.658E-08 | 1.500E-07 | 2.700E-07 |
| 16310 | 816 | C816I | B | 3.9e-07 | 3.1e-07 | 2.1e-07 | 3.033E-07 | 9.018E-08 | 2.100E-07 | 3.900E-07 |
| 16311 | 816 | C816L | B | 2.4e-07 | 2.3e-07 | 4.1e-07 | 2.933E-07 | 1.012E-07 | 2.300E-07 | 4.100E-07 |
| 16312 | 816 | C816K | B | 1.7e-07 | 2.5e-07 | 1.7e-07 | 1.967E-07 | 4.619E-08 | 1.700E-07 | 2.500E-07 |
| 16313 | 816 | C816M | B | 1.8e-07 | 3.5e-07 | 1.3e-07 | 2.200E-07 | 1.153E-07 | 1.300E-07 | 3.500E-07 |
| 16314 | 816 | C816F | B | 3.1e-07 | 3.4e-07 | 2e-07 | 2.833E-07 | 7.371E-08 | 2.000E-07 | 3.400E-07 |
| 16315 | 816 | C816P | B | 3e-07 | 4.2e-07 | 3.1e-07 | 3.433E-07 | 6.658E-08 | 3.000E-07 | 4.200E-07 |
| 16316 | 816 | C816S | B | 1.6e-07 | 1.6e-07 | 2.8e-07 | 2.000E-07 | 6.928E-08 | 1.600E-07 | 2.800E-07 |
| 16317 | 816 | C816T | B | 1.8e-07 | 2.3e-07 | 6.5e-07 | 3.533E-07 | 2.581E-07 | 1.800E-07 | 6.500E-07 |
| 16318 | 816 | C816W | B | 3e-07 | 1.8e-07 | 1.3e-07 | 2.033E-07 | 8.737E-08 | 1.300E-07 | 3.000E-07 |
| 16319 | 816 | C816Y | B | 2.3e-07 | 2.9e-07 | 2.4e-07 | 2.533E-07 | 3.215E-08 | 2.300E-07 | 2.900E-07 |

| | | | | | | | | | | |
|---|---|---|---|---|---|---|---|---|---|---|
| 16320 | 816 | C816V | B | 2.2e-07 | 2.2e-07 | 2e-07 | 2.133E-07 | 1.155E-08 | 2.000E-07 | 2.200E-07 |
| 16321 | 817 | E817A | B | 2.5e-07 | 3.1e-07 | 3.7e-07 | 3.100E-07 | 6.000E-08 | 2.500E-07 | 3.700E-07 |
| 16322 | 817 | E817R | B | 4.5e-07 | 2e-07 | 2.6e-07 | 3.033E-07 | 1.305E-07 | 2.000E-07 | 4.500E-07 |
| 16323 | 817 | E817N | B | 1.4e-07 | 3.4e-07 | 3.3e-07 | 2.700E-07 | 1.127E-07 | 1.400E-07 | 3.400E-07 |
| 16324 | 817 | E817D | B | 4.5e-07 | 5.5e-07 | 2.4e-07 | 4.133E-07 | 1.582E-07 | 2.400E-07 | 5.500E-07 |
| 16325 | 817 | E817C | B | 1e-07 | 3.1e-07 | 1.4e-07 | 1.833E-07 | 1.115E-07 | 1.000E-07 | 3.100E-07 |
| 16326 | 817 | E817E | B | 3e-07 | 2.3e-07 | 9.9e-08 | 2.097E-07 | 1.020E-07 | 9.900E-08 | 3.000E-07 |
| 16327 | 817 | E817Q | B | 2.8e-07 | 3.3e-07 | 4.4e-07 | 3.500E-07 | 8.185E-08 | 2.800E-07 | 4.400E-07 |
| 16328 | 817 | E817G | B | 1.1e-07 | 2.1e-07 | 1.7e-07 | 1.633E-07 | 5.033E-08 | 1.100E-07 | 2.100E-07 |
| 16329 | 817 | E817H | B | 5.9e-07 | 2.6e-07 | 2.3e-07 | 3.600E-07 | 1.997E-07 | 2.300E-07 | 5.900E-07 |
| 16330 | 817 | E817I | B | 2.7e-07 | 3e-07 | 2.1e-07 | 2.600E-07 | 4.583E-08 | 2.100E-07 | 3.000E-07 |
| 16331 | 817 | E817L | B | 1.9e-07 | 2.6e-07 | 3.4e-07 | 2.633E-07 | 7.506E-08 | 1.900E-07 | 3.400E-07 |
| 16332 | 817 | E817K | B | 3.6e-07 | 4.4e-07 | 2e-07 | 3.333E-07 | 1.222E-07 | 2.000E-07 | 4.400E-07 |
| 16333 | 817 | E817M | B | 3.1e-07 | 1.6e-07 | 1.9e-07 | 2.200E-07 | 7.937E-08 | 1.600E-07 | 3.100E-07 |
| 16334 | 817 | E817F | B | 2.1e-07 | 4.9e-07 | 1.5e-07 | 2.833E-07 | 1.815E-07 | 1.500E-07 | 4.900E-07 |
| 16335 | 817 | E817P | B | 9.9e-07 | 2.4e-07 | 2.3e-07 | 4.867E-07 | 4.359E-07 | 2.300E-07 | 9.900E-07 |
| 16336 | 817 | E817S | B | 2e-07 | 2.3e-07 | 4.2e-07 | 2.833E-07 | 1.193E-07 | 2.000E-07 | 4.200E-07 |
| 16337 | 817 | E817T | B | 2.5e-07 | 2.5e-07 | 1.5e-07 | 2.167E-07 | 5.774E-08 | 1.500E-07 | 2.500E-07 |
| 16338 | 817 | E817W | B | 1.9e-07 | 1.9e-07 | 2e-07 | 1.933E-07 | 5.774E-09 | 1.900E-07 | 2.000E-07 |
| 16339 | 817 | E817Y | B | 2.3e-07 | 2.7e-07 | 2.3e-07 | 2.433E-07 | 2.309E-08 | 2.300E-07 | 2.700E-07 |
| 16340 | 817 | E817V | B | 2.9e-07 | 3.1e-07 | 2.4e-07 | 2.800E-07 | 3.606E-08 | 2.400E-07 | 3.100E-07 |
| 16341 | 818 | V818A | B | 2.1e-07 | 3.8e-07 | 6.4e-07 | 4.100E-07 | 2.166E-07 | 2.100E-07 | 6.400E-07 |
| 16342 | 818 | V818R | B | 3e-07 | 3.1e-07 | 5.5e-07 | 3.867E-07 | 1.415E-07 | 3.000E-07 | 5.500E-07 |
| 16343 | 818 | V818N | B | 1.7e-07 | 3e-07 | 2.9e-07 | 2.533E-07 | 7.234E-08 | 1.700E-07 | 3.000E-07 |
| 16344 | 818 | V818D | B | 1.9e-07 | 4.3e-07 | 1.5e-07 | 2.567E-07 | 1.514E-07 | 1.500E-07 | 4.300E-07 |
| 16345 | 818 | V818C | B | 2.9e-07 | 2.6e-07 | 1.8e-07 | 2.433E-07 | 5.686E-08 | 1.800E-07 | 2.900E-07 |
| 16346 | 818 | V818E | B | 2.6e-07 | 1.5e-07 | 2.3e-07 | 2.133E-07 | 5.686E-08 | 1.500E-07 | 2.600E-07 |
| 16347 | 818 | V818Q | B | 4.5e-07 | 2.2e-07 | 2.4e-07 | 3.033E-07 | 1.274E-07 | 2.200E-07 | 4.500E-07 |
| 16348 | 818 | V818G | B | 1.8e-07 | 3.7e-07 | 1.5e-07 | 2.333E-07 | 1.193E-07 | 1.500E-07 | 3.700E-07 |
| 16349 | 818 | V818H | B | 2.2e-07 | 2.2e-07 | 1.7e-07 | 2.033E-07 | 2.887E-08 | 1.700E-07 | 2.200E-07 |
| 16350 | 818 | V818I | B | 1.8e-07 | 3.4e-07 | 2.5e-07 | 2.567E-07 | 8.021E-08 | 1.800E-07 | 3.400E-07 |
| 16351 | 818 | V818L | B | 2.7e-07 | 2.3e-07 | 1.9e-07 | 2.300E-07 | 4.000E-08 | 1.900E-07 | 2.700E-07 |
| 16352 | 818 | V818K | B | 2.7e-07 | 1.2e-07 | 1.9e-07 | 1.933E-07 | 7.506E-08 | 1.200E-07 | 2.700E-07 |
| 16353 | 818 | V818M | B | 7.2e-07 | 2.9e-07 | 1.8e-07 | 3.967E-07 | 2.854E-07 | 1.800E-07 | 7.200E-07 |
| 16354 | 818 | V818F | B | 9.3e-08 | 2.4e-07 | 1.6e-07 | 1.643E-07 | 7.360E-08 | 9.300E-08 | 2.400E-07 |
| 16355 | 818 | V818P | B | 2.9e-07 | 3e-07 | 1.7e-07 | 2.533E-07 | 7.234E-08 | 1.700E-07 | 3.000E-07 |
| 16356 | 818 | V818S | B | 1.5e-07 | 2.9e-07 | 7.5e-08 | 1.717E-07 | 1.091E-07 | 7.500E-08 | 2.900E-07 |
| 16357 | 818 | V818T | B | 3.1e-07 | 2.5e-07 | 3.5e-07 | 3.033E-07 | 5.033E-08 | 2.500E-07 | 3.500E-07 |
| 16358 | 818 | V818W | B | 5.1e-07 | 1.9e-07 | 3.4e-07 | 3.467E-07 | 1.601E-07 | 1.900E-07 | 5.100E-07 |
| 16359 | 818 | V818Y | B | 1.5e-07 | 3e-07 | 2.8e-07 | 2.433E-07 | 8.145E-08 | 1.500E-07 | 3.000E-07 |
| 16360 | 818 | V818V | B | 1.8e-07 | 2e-07 | 4.7e-07 | 2.833E-07 | 1.620E-07 | 1.800E-07 | 4.700E-07 |
| 16361 | 819 | T819A | B | 3e-07 | 2.7e-07 | 2.6e-07 | 2.767E-07 | 2.082E-08 | 2.600E-07 | 3.000E-07 |
| 16362 | 819 | T819R | B | 9.8e-08 | 4.5e-07 | 2e-07 | 2.493E-07 | 1.811E-07 | 9.800E-08 | 4.500E-07 |
| 16363 | 819 | T819N | B | 3.9e-07 | 1.7e-07 | 2.8e-07 | 2.800E-07 | 1.100E-07 | 1.700E-07 | 3.900E-07 |
| 16364 | 819 | T819D | B | 3.6e-07 | 3.7e-07 | 1.7e-07 | 3.000E-07 | 1.127E-07 | 1.700E-07 | 3.700E-07 |
| 16365 | 819 | T819C | B | 2.6e-07 | 2.5e-07 | 2.9e-07 | 2.667E-07 | 2.082E-08 | 2.500E-07 | 2.900E-07 |
| 16366 | 819 | T819E | B | 2.7e-07 | 1.8e-07 | 1.8e-07 | 2.100E-07 | 5.196E-08 | 1.800E-07 | 2.700E-07 |
| 16367 | 819 | T819Q | B | 1.7e-07 | 2.2e-07 | 1.6e-07 | 1.833E-07 | 3.215E-08 | 1.600E-07 | 2.200E-07 |
| 16368 | 819 | T819G | B | 1.7e-07 | 2.5e-07 | 2.1e-07 | 2.100E-07 | 4.000E-08 | 1.700E-07 | 2.500E-07 |
| 16369 | 819 | T819H | B | 2.1e-07 | 2.6e-07 | 2.7e-07 | 2.467E-07 | 3.215E-08 | 2.100E-07 | 2.700E-07 |
| 16370 | 819 | T819I | B | 2.6e-07 | 1.1e-07 | 3.7e-07 | 2.467E-07 | 1.305E-07 | 1.100E-07 | 3.700E-07 |
| 16371 | 819 | T819L | B | 2.9e-07 | 1.5e-07 | 2.9e-07 | 2.433E-07 | 8.083E-08 | 1.500E-07 | 2.900E-07 |
| 16372 | 819 | T819K | B | 3.2e-07 | 2.7e-07 | 3.1e-07 | 3.000E-07 | 2.646E-08 | 2.700E-07 | 3.200E-07 |
| 16373 | 819 | T819M | B | 2.6e-07 | 2.3e-07 | 2.8e-07 | 2.567E-07 | 2.517E-08 | 2.300E-07 | 2.800E-07 |
| 16374 | 819 | T819F | B | 3.6e-07 | 1.6e-07 | 1.2e-07 | 2.133E-07 | 1.286E-07 | 1.200E-07 | 3.600E-07 |
| 16375 | 819 | T819P | B | 1.9e-07 | 2e-07 | 3.2e-07 | 2.367E-07 | 7.234E-08 | 1.900E-07 | 3.200E-07 |
| 16376 | 819 | T819S | B | 1.6e-07 | 1.9e-07 | 3.7e-07 | 2.400E-07 | 1.136E-07 | 1.600E-07 | 3.700E-07 |
| 16377 | 819 | T819T | B | 2.3e-07 | 1.4e-07 | 3.1e-07 | 2.267E-07 | 8.505E-08 | 1.400E-07 | 3.100E-07 |
| 16378 | 819 | T819W | B | 1.9e-07 | 2e-07 | 2.2e-07 | 2.033E-07 | 1.528E-08 | 1.900E-07 | 2.200E-07 |
| 16379 | 819 | T819Y | B | 2e-07 | 2.2e-07 | 2.3e-07 | 2.167E-07 | 1.528E-08 | 2.000E-07 | 2.300E-07 |
| 16380 | 819 | T819V | B | 1.9e-07 | 5.8e-07 | 9.8e-07 | 5.833E-07 | 3.950E-07 | 1.900E-07 | 9.800E-07 |
| 16381 | 820 | H820A | B | 5.6e-07 | 2.3e-07 | 1.8e-07 | 3.233E-07 | 2.065E-07 | 1.800E-07 | 5.600E-07 |
| 16382 | 820 | H820R | B | 6.1e-07 | 9.8e-08 | 2e-07 | 3.027E-07 | 2.710E-07 | 9.800E-08 | 6.100E-07 |
| 16383 | 820 | H820N | B | 1.7e-07 | 2.8e-07 | 2.2e-07 | 2.233E-07 | 5.508E-08 | 1.700E-07 | 2.800E-07 |
| 16384 | 820 | H820D | B | 2.6e-07 | 2.9e-07 | 1.7e-07 | 2.400E-07 | 6.245E-08 | 1.700E-07 | 2.900E-07 |
| 16385 | 820 | H820C | B | 2e-07 | 3.7e-07 | 3.5e-07 | 3.067E-07 | 9.292E-08 | 2.000E-07 | 3.700E-07 |
| 16386 | 820 | H820E | B | 1.2e-07 | 1.3e-07 | 8.7e-08 | 1.123E-07 | 2.250E-08 | 8.700E-08 | 1.300E-07 |
| 16387 | 820 | H820Q | B | 2.5e-07 | 1.6e-07 | 9.5e-07 | 4.533E-07 | 4.325E-07 | 1.600E-07 | 9.500E-07 |
| 16388 | 820 | H820G | B | 1.9e-07 | 2.5e-07 | 1.5e-07 | 1.967E-07 | 5.033E-08 | 1.500E-07 | 2.500E-07 |
| 16389 | 820 | H820H | B | 2.2e-07 | 2.1e-07 | 4.2e-07 | 2.833E-07 | 1.185E-07 | 2.100E-07 | 4.200E-07 |
| 16390 | 820 | H820I | B | 1.2e-07 | 2e-07 | 1.7e-07 | 1.633E-07 | 4.041E-08 | 1.200E-07 | 2.000E-07 |
| 16391 | 820 | H820L | B | 2.2e-07 | 3.3e-07 | 4.5e-07 | 3.333E-07 | 1.150E-07 | 2.200E-07 | 4.500E-07 |
| 16392 | 820 | H820K | B | 1.5e-07 | 1.9e-07 | 2.3e-07 | 1.900E-07 | 4.000E-08 | 1.500E-07 | 2.300E-07 |
| 16393 | 820 | H820M | B | 4.8e-07 | 3.1e-07 | 1.8e-07 | 3.233E-07 | 1.504E-07 | 1.800E-07 | 4.800E-07 |
| 16394 | 820 | H820F | B | 3.7e-07 | 3.5e-07 | 3.5e-07 | 3.567E-07 | 1.155E-08 | 3.500E-07 | 3.700E-07 |
| 16395 | 820 | H820P | B | 1.3e-07 | 2.8e-07 | 1.4e-07 | 1.833E-07 | 8.386E-08 | 1.300E-07 | 2.800E-07 |
| 16396 | 820 | H820S | B | 4.6e-07 | 3.3e-07 | 2.8e-07 | 3.567E-07 | 9.292E-08 | 2.800E-07 | 4.600E-07 |
| 16397 | 820 | H820T | B | 3.5e-07 | 2.1e-07 | 1.6e-07 | 2.400E-07 | 9.849E-08 | 1.600E-07 | 3.500E-07 |
| 16398 | 820 | H820W | B | 3.3e-07 | 2.6e-07 | 5.3e-07 | 3.733E-07 | 1.401E-07 | 2.600E-07 | 5.300E-07 |
| 16399 | 820 | H820Y | B | 2.4e-07 | 2.2e-07 | 8.8e-08 | 1.827E-07 | 8.259E-08 | 8.800E-08 | 2.400E-07 |
| 16400 | 820 | H820V | B | 2.3e-07 | 3.2e-07 | 5.7e-07 | 3.733E-07 | 1.762E-07 | 2.300E-07 | 5.700E-07 |
| 16401 | 821 | Q821A | B | 1.9e-07 | 3.3e-07 | 3.5e-07 | 2.900E-07 | 8.718E-08 | 1.900E-07 | 3.500E-07 |
| 16402 | 821 | Q821R | B | 1.9e-07 | 3e-07 | 1.9e-07 | 2.267E-07 | 6.351E-08 | 1.900E-07 | 3.000E-07 |
| 16403 | 821 | Q821N | B | 1.7e-07 | 4.2e-07 | 4.7e-07 | 3.533E-07 | 1.607E-07 | 1.700E-07 | 4.700E-07 |
| 16404 | 821 | Q821D | B | 1.6e-07 | 8.1e-08 | 2.8e-07 | 1.737E-07 | 1.002E-07 | 8.100E-08 | 2.800E-07 |

| | | | | | | | | | | |
|---|---|---|---|---|---|---|---|---|---|---|
| 16405 | 821 | Q821C | B | 2.7e-07 | 1.5e-07 | 2e-07 | 2.067E-07 | 6.028E-08 | 1.500E-07 | 2.700E-07 |
| 16406 | 821 | Q821E | B | 2.5e-07 | 1.8e-07 | 2e-07 | 2.100E-07 | 3.606E-08 | 1.800E-07 | 2.500E-07 |
| 16407 | 821 | Q821Q | B | 1.6e-07 | 2.3e-07 | 2e-07 | 1.967E-07 | 3.512E-08 | 1.600E-07 | 2.300E-07 |
| 16408 | 821 | Q821G | B | 2.4e-07 | 3.6e-07 | 3.4e-07 | 3.133E-07 | 6.429E-08 | 2.400E-07 | 3.600E-07 |
| 16409 | 821 | Q821H | B | 2.1e-07 | 1.8e-07 | 1.2e-07 | 1.700E-07 | 4.583E-08 | 1.200E-07 | 2.100E-07 |
| 16410 | 821 | Q821I | B | 1.8e-07 | 3.7e-07 | 1.3e-07 | 2.267E-07 | 1.266E-07 | 1.300E-07 | 3.700E-07 |
| 16411 | 821 | Q821L | B | 3e-07 | 2.5e-07 | 2.8e-07 | 2.767E-07 | 2.517E-08 | 2.500E-07 | 3.000E-07 |
| 16412 | 821 | Q821K | B | 1.2e-07 | 2.8e-07 | 4.4e-07 | 2.800E-07 | 1.600E-07 | 1.200E-07 | 4.400E-07 |
| 16413 | 821 | Q821M | B | 2.2e-07 | 5.8e-07 | 2.2e-07 | 3.400E-07 | 2.078E-07 | 2.200E-07 | 5.800E-07 |
| 16414 | 821 | Q821F | B | 2.8e-07 | 1.6e-07 | 2e-07 | 2.133E-07 | 6.110E-08 | 1.600E-07 | 2.800E-07 |
| 16415 | 821 | Q821P | B | 5e-07 | 2.2e-07 | 3.6e-07 | 3.600E-07 | 1.400E-07 | 2.200E-07 | 5.000E-07 |
| 16416 | 821 | Q821S | B | 2e-07 | 8.5e-08 | 3.1e-07 | 1.983E-07 | 1.125E-07 | 8.500E-08 | 3.100E-07 |
| 16417 | 821 | Q821T | B | 5.6e-07 | 1.9e-07 | 2.4e-07 | 3.300E-07 | 2.007E-07 | 1.900E-07 | 5.600E-07 |
| 16418 | 821 | Q821W | B | 2.7e-07 | 3.8e-07 | 2e-07 | 2.833E-07 | 9.074E-08 | 2.000E-07 | 3.800E-07 |
| 16419 | 821 | Q821Y | B | 5.6e-07 | 3e-07 | 3.9e-07 | 4.167E-07 | 1.320E-07 | 3.000E-07 | 5.600E-07 |
| 16420 | 821 | Q821V | B | 1.7e-07 | 2.3e-07 | 2.4e-07 | 2.133E-07 | 3.786E-08 | 1.700E-07 | 2.400E-07 |
| 16421 | 822 | G822A | B | 1.4e-07 | 2.6e-07 | 4.4e-07 | 2.800E-07 | 1.510E-07 | 1.400E-07 | 4.400E-07 |
| 16422 | 822 | G822R | B | 1.4e-07 | 3e-07 | 2.6e-07 | 2.333E-07 | 8.327E-08 | 1.400E-07 | 3.000E-07 |
| 16423 | 822 | G822N | B | 2e-07 | 7e-07 | 1.1e-07 | 3.367E-07 | 3.179E-07 | 1.100E-07 | 7.000E-07 |
| 16424 | 822 | G822D | B | 1.4e-07 | 2.6e-07 | 3e-07 | 2.333E-07 | 8.327E-08 | 1.400E-07 | 3.000E-07 |
| 16425 | 822 | G822C | B | 3.9e-07 | 3.6e-07 | 3e-07 | 3.500E-07 | 4.583E-08 | 3.000E-07 | 3.900E-07 |
| 16426 | 822 | G822E | B | 1.9e-07 | 1.7e-07 | 5.6e-07 | 3.067E-07 | 2.196E-07 | 1.700E-07 | 5.600E-07 |
| 16427 | 822 | G822Q | B | 4.3e-07 | 3.8e-07 | 1.1e-07 | 3.067E-07 | 1.721E-07 | 1.100E-07 | 4.300E-07 |
| 16428 | 822 | G822G | B | 1e-06 | 2.1e-07 | 3.8e-07 | 5.300E-07 | 4.158E-07 | 2.100E-07 | 1.000E-06 |
| 16429 | 822 | G822H | B | 2.1e-07 | 2.5e-07 | 2.9e-07 | 2.500E-07 | 4.000E-08 | 2.100E-07 | 2.900E-07 |
| 16430 | 822 | G822I | B | 5.6e-07 | 3.5e-07 | 2.4e-07 | 3.833E-07 | 1.626E-07 | 2.400E-07 | 5.600E-07 |
| 16431 | 822 | G822L | B | 2.7e-07 | 4.8e-07 | 1.6e-07 | 3.033E-07 | 1.626E-07 | 1.600E-07 | 4.800E-07 |
| 16432 | 822 | G822K | B | 3e-07 | 2.1e-07 | 2.5e-07 | 2.533E-07 | 4.509E-08 | 2.100E-07 | 3.000E-07 |
| 16433 | 822 | G822M | B | 5.9e-07 | 2.3e-07 | 2.7e-07 | 3.633E-07 | 1.973E-07 | 2.300E-07 | 5.900E-07 |
| 16434 | 822 | G822F | B | 4.6e-07 | 2.7e-07 | 1.8e-07 | 3.033E-07 | 1.429E-07 | 1.800E-07 | 4.600E-07 |
| 16435 | 822 | G822P | B | 2.4e-07 | 2.4e-07 | 2.1e-07 | 2.300E-07 | 1.732E-08 | 2.100E-07 | 2.400E-07 |
| 16436 | 822 | G822S | B | 3.2e-07 | 2.8e-07 | 1.3e-07 | 2.433E-07 | 1.002E-07 | 1.300E-07 | 3.200E-07 |
| 16437 | 822 | G822T | B | 2.5e-07 | 2.5e-07 | 2.4e-07 | 2.467E-07 | 5.774E-09 | 2.400E-07 | 2.500E-07 |
| 16438 | 822 | G822W | B | 3.6e-07 | 1.4e-07 | 3.1e-07 | 2.700E-07 | 1.153E-07 | 1.400E-07 | 3.600E-07 |
| 16439 | 822 | G822Y | B | 2e-07 | 2.9e-07 | 2.3e-07 | 2.400E-07 | 4.583E-08 | 2.000E-07 | 2.900E-07 |
| 16440 | 822 | G822V | B | 1.3e-07 | 3.4e-07 | 2.6e-07 | 2.433E-07 | 1.060E-07 | 1.300E-07 | 3.400E-07 |
| 16441 | 823 | L823A | B | 1.8e-07 | 5.3e-07 | 4.4e-07 | 3.833E-07 | 1.818E-07 | 1.800E-07 | 5.300E-07 |
| 16442 | 823 | L823R | B | 1.9e-07 | 2e-07 | 1.9e-07 | 1.933E-07 | 5.774E-09 | 1.900E-07 | 2.000E-07 |
| 16443 | 823 | L823N | B | 3.1e-07 | 4.2e-07 | 2.6e-07 | 3.300E-07 | 8.185E-08 | 2.600E-07 | 4.200E-07 |
| 16444 | 823 | L823D | B | 1.7e-07 | 4.2e-07 | 3.8e-07 | 3.233E-07 | 1.343E-07 | 1.700E-07 | 4.200E-07 |
| 16445 | 823 | L823C | B | 9.3e-08 | 3.9e-07 | 4.5e-07 | 3.110E-07 | 1.912E-07 | 9.300E-08 | 4.500E-07 |
| 16446 | 823 | L823E | B | 2.6e-07 | 2.5e-07 | 2e-07 | 2.367E-07 | 3.215E-08 | 2.000E-07 | 2.600E-07 |
| 16447 | 823 | L823Q | B | 1.5e-07 | 2.1e-07 | 1.9e-07 | 1.833E-07 | 3.055E-08 | 1.500E-07 | 2.100E-07 |
| 16448 | 823 | L823G | B | 4.7e-07 | 2.1e-07 | 1.7e-07 | 2.833E-07 | 1.629E-07 | 1.700E-07 | 4.700E-07 |
| 16449 | 823 | L823H | B | 3.9e-07 | 2.6e-07 | 4.3e-07 | 3.600E-07 | 8.888E-08 | 2.600E-07 | 4.300E-07 |
| 16450 | 823 | L823I | B | 2.8e-07 | 2.6e-07 | 2.9e-07 | 2.767E-07 | 1.528E-08 | 2.600E-07 | 2.900E-07 |
| 16451 | 823 | L823L | B | 3.3e-07 | 5.1e-07 | 1.4e-07 | 3.267E-07 | 1.850E-07 | 1.400E-07 | 5.100E-07 |
| 16452 | 823 | L823K | B | 2.9e-07 | 1.5e-07 | 3.3e-07 | 2.567E-07 | 9.452E-08 | 1.500E-07 | 3.300E-07 |
| 16453 | 823 | L823M | B | 4.8e-07 | 3.1e-07 | 1.6e-07 | 3.167E-07 | 1.601E-07 | 1.600E-07 | 4.800E-07 |
| 16454 | 823 | L823F | B | 1.9e-07 | 2.6e-07 | 9.1e-08 | 1.803E-07 | 8.491E-08 | 9.100E-08 | 2.600E-07 |
| 16455 | 823 | L823P | B | 7.1e-07 | 2.2e-07 | 1.8e-07 | 3.700E-07 | 2.951E-07 | 1.800E-07 | 7.100E-07 |
| 16456 | 823 | L823S | B | 2.8e-07 | 2.3e-07 | 1.9e-07 | 2.333E-07 | 4.509E-08 | 1.900E-07 | 2.800E-07 |
| 16457 | 823 | L823T | B | 2.4e-07 | 1.4e-07 | 2e-07 | 1.933E-07 | 5.033E-08 | 1.400E-07 | 2.400E-07 |
| 16458 | 823 | L823W | B | 1.8e-07 | 1.3e-07 | 2.5e-07 | 1.867E-07 | 6.028E-08 | 1.300E-07 | 2.500E-07 |
| 16459 | 823 | L823Y | B | 2.7e-07 | 2.9e-07 | 3.1e-07 | 2.900E-07 | 2.000E-08 | 2.700E-07 | 3.100E-07 |
| 16460 | 823 | L823V | B | 4.4e-07 | 4.9e-07 | 3.2e-07 | 4.167E-07 | 8.737E-08 | 3.200E-07 | 4.900E-07 |
| 16461 | 824 | S824A | B | 2.6e-07 | 2.9e-07 | 2.4e-07 | 2.633E-07 | 2.517E-08 | 2.400E-07 | 2.900E-07 |
| 16462 | 824 | S824R | B | 3.3e-07 | 8.4e-08 | 1.8e-07 | 1.980E-07 | 1.240E-07 | 8.400E-08 | 3.300E-07 |
| 16463 | 824 | S824N | B | 2.5e-07 | 3.5e-07 | 1.6e-07 | 2.533E-07 | 9.504E-08 | 1.600E-07 | 3.500E-07 |
| 16464 | 824 | S824D | B | 1.3e-07 | 1.5e-07 | 3.2e-07 | 2.000E-07 | 1.044E-07 | 1.300E-07 | 3.200E-07 |
| 16465 | 824 | S824C | B | 2.8e-07 | 1.9e-07 | 4e-07 | 2.900E-07 | 1.054E-07 | 1.900E-07 | 4.000E-07 |
| 16466 | 824 | S824E | B | 1.8e-07 | 4.5e-07 | 2.4e-07 | 2.900E-07 | 1.418E-07 | 1.800E-07 | 4.500E-07 |
| 16467 | 824 | S824Q | B | 1.6e-07 | 3e-07 | 2.3e-07 | 2.300E-07 | 7.000E-08 | 1.600E-07 | 3.000E-07 |
| 16468 | 824 | S824G | B | 1.5e-07 | 1.9e-07 | 1.9e-07 | 1.767E-07 | 2.309E-08 | 1.500E-07 | 1.900E-07 |
| 16469 | 824 | S824H | B | 5e-07 | 3.7e-07 | 5.1e-07 | 4.600E-07 | 7.810E-08 | 3.700E-07 | 5.100E-07 |
| 16470 | 824 | S824I | B | 2.5e-07 | 1.6e-07 | 9.5e-08 | 1.683E-07 | 7.784E-08 | 9.500E-08 | 2.500E-07 |
| 16471 | 824 | S824L | B | 5.9e-07 | 4e-07 | 5.2e-07 | 5.033E-07 | 9.609E-08 | 4.000E-07 | 5.900E-07 |
| 16472 | 824 | S824K | B | 2.4e-07 | 2e-07 | 1.7e-07 | 2.033E-07 | 3.512E-08 | 1.700E-07 | 2.400E-07 |
| 16473 | 824 | S824M | B | 1.4e-07 | 2.7e-07 | 4.7e-07 | 2.933E-07 | 1.662E-07 | 1.400E-07 | 4.700E-07 |
| 16474 | 824 | S824F | B | 2.1e-07 | 9.8e-08 | 2.3e-07 | 1.793E-07 | 7.114E-08 | 9.800E-08 | 2.300E-07 |
| 16475 | 824 | S824P | B | 1.7e-07 | 8.2e-07 | 2.3e-07 | 4.067E-07 | 3.592E-07 | 1.700E-07 | 8.200E-07 |
| 16476 | 824 | S824S | B | 1.1e-07 | 2.6e-07 | 2.9e-07 | 2.200E-07 | 9.644E-08 | 1.100E-07 | 2.900E-07 |
| 16477 | 824 | S824T | B | 2.4e-07 | 1.1e-07 | 1.7e-07 | 1.733E-07 | 6.506E-08 | 1.100E-07 | 2.400E-07 |
| 16478 | 824 | S824W | B | 2.1e-07 | 4.3e-07 | 1.8e-07 | 2.733E-07 | 1.365E-07 | 1.800E-07 | 4.300E-07 |
| 16479 | 824 | S824Y | B | 4e-07 | 2.7e-07 | 1.9e-07 | 2.867E-07 | 1.060E-07 | 1.900E-07 | 4.000E-07 |
| 16480 | 824 | S824V | B | 3.8e-07 | 1.6e-07 | 3.3e-07 | 2.900E-07 | 1.153E-07 | 1.600E-07 | 3.800E-07 |
| 16481 | 825 | S825A | B | 7.4e-08 | 3.7e-07 | 1.3e-07 | 1.913E-07 | 1.572E-07 | 7.400E-08 | 3.700E-07 |
| 16482 | 825 | S825R | B | 4.4e-07 | 2.5e-07 | 2.8e-07 | 3.233E-07 | 1.021E-07 | 2.500E-07 | 4.400E-07 |
| 16483 | 825 | S825N | B | 2e-07 | 1.9e-07 | 5.2e-07 | 3.033E-07 | 1.877E-07 | 1.900E-07 | 5.200E-07 |
| 16484 | 825 | S825D | B | 3.7e-07 | 3.6e-07 | 2.4e-07 | 3.233E-07 | 7.234E-08 | 2.400E-07 | 3.700E-07 |
| 16485 | 825 | S825C | B | 1.7e-07 | 2e-07 | 2.7e-07 | 2.133E-07 | 5.132E-08 | 1.700E-07 | 2.700E-07 |
| 16486 | 825 | S825E | B | 2e-07 | 2.3e-07 | 2.1e-07 | 2.133E-07 | 1.528E-08 | 2.000E-07 | 2.300E-07 |
| 16487 | 825 | S825Q | B | 2.5e-07 | 2.1e-07 | 4.6e-07 | 3.067E-07 | 1.343E-07 | 2.100E-07 | 4.600E-07 |
| 16488 | 825 | S825G | B | 1.4e-07 | 3e-07 | 3e-07 | 2.467E-07 | 9.238E-08 | 1.400E-07 | 3.000E-07 |
| 16489 | 825 | S825H | B | 9.8e-08 | 4e-07 | 2.2e-07 | 2.393E-07 | 1.519E-07 | 9.800E-08 | 4.000E-07 |

| | | | | | | | | | | |
|---|---|---|---|---|---|---|---|---|---|---|
| 16490 | 825 | S825I | B | 2.8e-07 | 1.9e-07 | 3.3e-07 | 2.667E-07 | 7.095E-08 | 1.900E-07 | 3.300E-07 |
| 16491 | 825 | S825L | B | 1.7e-07 | 3.6e-07 | 2.8e-07 | 2.700E-07 | 9.539E-08 | 1.700E-07 | 3.600E-07 |
| 16492 | 825 | S825K | B | 2.5e-07 | 3e-07 | 4.5e-07 | 3.333E-07 | 1.041E-07 | 2.500E-07 | 4.500E-07 |
| 16493 | 825 | S825M | B | 4.1e-07 | 9.7e-08 | 2.3e-07 | 2.457E-07 | 1.571E-07 | 9.700E-08 | 4.100E-07 |
| 16494 | 825 | S825F | B | 2.8e-07 | 2.9e-07 | 9.1e-08 | 2.203E-07 | 1.121E-07 | 9.100E-08 | 2.900E-07 |
| 16495 | 825 | S825P | B | 2.9e-07 | 4.7e-07 | 2.3e-07 | 3.300E-07 | 1.249E-07 | 2.300E-07 | 4.700E-07 |
| 16496 | 825 | S825S | B | 3.9e-07 | 1.6e-07 | 2.5e-07 | 2.667E-07 | 1.159E-07 | 1.600E-07 | 3.900E-07 |
| 16497 | 825 | S825T | B | 3.5e-07 | 2.7e-07 | 2.8e-07 | 3.000E-07 | 4.359E-08 | 2.700E-07 | 3.500E-07 |
| 16498 | 825 | S825W | B | 2.6e-07 | 4.1e-07 | 1.5e-07 | 2.733E-07 | 1.305E-07 | 1.500E-07 | 4.100E-07 |
| 16499 | 825 | S825Y | B | 1.9e-07 | 1.5e-07 | 3e-07 | 2.133E-07 | 7.767E-08 | 1.500E-07 | 3.000E-07 |
| 16500 | 825 | S825V | B | 2.6e-07 | 3.7e-07 | 3.6e-07 | 3.300E-07 | 6.083E-08 | 2.600E-07 | 3.700E-07 |
| 16501 | 826 | P826A | B | 3.2e-07 | 3.3e-07 | 3.2e-07 | 3.233E-07 | 5.774E-09 | 3.200E-07 | 3.300E-07 |
| 16502 | 826 | P826R | B | 2.3e-07 | 4.5e-07 | 2.6e-07 | 3.133E-07 | 1.193E-07 | 2.300E-07 | 4.500E-07 |
| 16503 | 826 | P826N | B | 1.1e-07 | 3e-07 | 1.1e-07 | 1.733E-07 | 1.097E-07 | 1.100E-07 | 3.000E-07 |
| 16504 | 826 | P826D | B | 2.9e-07 | 5.4e-07 | 1.7e-07 | 3.333E-07 | 1.888E-07 | 1.700E-07 | 5.400E-07 |
| 16505 | 826 | P826C | B | 3.4e-07 | 3.6e-07 | 3e-07 | 3.333E-07 | 3.055E-08 | 3.000E-07 | 3.600E-07 |
| 16506 | 826 | P826E | B | 1.7e-07 | 3e-07 | 2.5e-07 | 2.400E-07 | 6.557E-08 | 1.700E-07 | 3.000E-07 |
| 16507 | 826 | P826Q | B | 4.4e-07 | 2e-07 | 5.7e-07 | 4.033E-07 | 1.877E-07 | 2.000E-07 | 5.700E-07 |
| 16508 | 826 | P826G | B | 2.9e-07 | 3.8e-07 | 5.3e-07 | 4.000E-07 | 1.212E-07 | 2.900E-07 | 5.300E-07 |
| 16509 | 826 | P826H | B | 1.6e-07 | 2.2e-07 | 3e-07 | 2.267E-07 | 7.024E-08 | 1.600E-07 | 3.000E-07 |
| 16510 | 826 | P826I | B | 2e-07 | 5.3e-07 | 2.5e-07 | 3.267E-07 | 1.779E-07 | 2.000E-07 | 5.300E-07 |
| 16511 | 826 | P826L | B | 3.1e-07 | 9.3e-08 | 2.5e-07 | 2.177E-07 | 1.121E-07 | 9.300E-08 | 3.100E-07 |
| 16512 | 826 | P826K | B | 1.9e-07 | 2.1e-07 | 1.8e-07 | 1.933E-07 | 1.528E-08 | 1.800E-07 | 2.100E-07 |
| 16513 | 826 | P826M | B | 5.7e-07 | 2.2e-07 | 3.4e-07 | 3.767E-07 | 1.779E-07 | 2.200E-07 | 5.700E-07 |
| 16514 | 826 | P826F | B | 1.2e-07 | 5.1e-07 | 2.6e-07 | 2.967E-07 | 1.976E-07 | 1.200E-07 | 5.100E-07 |
| 16515 | 826 | P826P | B | 3.2e-07 | 7e-08 | 3e-07 | 2.300E-07 | 1.389E-07 | 7.000E-08 | 3.200E-07 |
| 16516 | 826 | P826S | B | 2.4e-07 | 1.7e-07 | 6.8e-07 | 3.633E-07 | 2.765E-07 | 1.700E-07 | 6.800E-07 |
| 16517 | 826 | P826T | B | 2.3e-07 | 2.5e-07 | 9.2e-08 | 1.907E-07 | 8.603E-08 | 9.200E-08 | 2.500E-07 |
| 16518 | 826 | P826W | B | 2.9e-07 | 1.5e-07 | 2.5e-07 | 2.300E-07 | 7.211E-08 | 1.500E-07 | 2.900E-07 |
| 16519 | 826 | P826Y | B | 2.3e-07 | 1.2e-07 | 2.3e-07 | 1.933E-07 | 6.351E-08 | 1.200E-07 | 2.300E-07 |
| 16520 | 826 | P826V | B | 4.7e-07 | 4.3e-07 | 2.3e-07 | 3.767E-07 | 1.286E-07 | 2.300E-07 | 4.700E-07 |
| 16521 | 827 | V827A | B | 4.2e-07 | 2.4e-07 | 1.8e-07 | 2.800E-07 | 1.249E-07 | 1.800E-07 | 4.200E-07 |
| 16522 | 827 | V827R | B | 2.3e-07 | 1.4e-07 | 2.2e-07 | 1.967E-07 | 4.933E-08 | 1.400E-07 | 2.300E-07 |
| 16523 | 827 | V827N | B | 9.8e-08 | 3.5e-07 | 2.5e-07 | 2.327E-07 | 1.269E-07 | 9.800E-08 | 3.500E-07 |
| 16524 | 827 | V827D | B | 2.7e-07 | 2.1e-07 | 2e-07 | 2.267E-07 | 3.786E-08 | 2.000E-07 | 2.700E-07 |
| 16525 | 827 | V827C | B | 2.1e-07 | 4.4e-07 | 1.8e-07 | 2.767E-07 | 1.422E-07 | 1.800E-07 | 4.400E-07 |
| 16526 | 827 | V827E | B | 2.1e-07 | 2.1e-07 | 3.9e-07 | 2.700E-07 | 1.039E-07 | 2.100E-07 | 3.900E-07 |
| 16527 | 827 | V827Q | B | 1.8e-07 | 3.3e-07 | 2.6e-07 | 2.567E-07 | 7.506E-08 | 1.800E-07 | 3.300E-07 |
| 16528 | 827 | V827G | B | 3.5e-07 | 1.6e-07 | 2e-07 | 2.367E-07 | 1.002E-07 | 1.600E-07 | 3.500E-07 |
| 16529 | 827 | V827H | B | 3.8e-07 | 1.6e-07 | 2.4e-07 | 2.600E-07 | 1.114E-07 | 1.600E-07 | 3.800E-07 |
| 16530 | 827 | V827I | B | 1.4e-07 | 3.5e-07 | 2.5e-07 | 2.467E-07 | 1.050E-07 | 1.400E-07 | 3.500E-07 |
| 16531 | 827 | V827L | B | 3.2e-07 | 3.4e-07 | 1.6e-07 | 2.733E-07 | 9.866E-08 | 1.600E-07 | 3.400E-07 |
| 16532 | 827 | V827K | B | 1.3e-07 | 2.9e-07 | 4.6e-07 | 2.933E-07 | 1.650E-07 | 1.300E-07 | 4.600E-07 |
| 16533 | 827 | V827M | B | 1.6e-07 | 2e-07 | 3.8e-07 | 2.467E-07 | 1.172E-07 | 1.600E-07 | 3.800E-07 |
| 16534 | 827 | V827F | B | 1.8e-07 | 2.6e-07 | 2.4e-07 | 2.267E-07 | 4.163E-08 | 1.800E-07 | 2.600E-07 |
| 16535 | 827 | V827P | B | 4.8e-07 | 2e-07 | 1.7e-07 | 2.833E-07 | 1.710E-07 | 1.700E-07 | 4.800E-07 |
| 16536 | 827 | V827S | B | 1.8e-07 | 4.5e-07 | 3.1e-07 | 3.133E-07 | 1.350E-07 | 1.800E-07 | 4.500E-07 |
| 16537 | 827 | V827T | B | 1.2e-07 | 2.5e-07 | 3.6e-07 | 2.433E-07 | 1.201E-07 | 1.200E-07 | 3.600E-07 |
| 16538 | 827 | V827W | B | 2.5e-07 | 1.4e-07 | 2.9e-07 | 2.267E-07 | 7.767E-08 | 1.400E-07 | 2.900E-07 |
| 16539 | 827 | V827Y | B | 2.8e-07 | 1.3e-07 | 1.9e-07 | 2.000E-07 | 7.550E-08 | 1.300E-07 | 2.800E-07 |
| 16540 | 827 | V827V | B | 3e-07 | 2.3e-07 | 2.3e-07 | 2.533E-07 | 4.041E-08 | 2.300E-07 | 3.000E-07 |
| 16541 | 828 | T828A | B | 3e-07 | 4.2e-07 | 2.3e-07 | 3.167E-07 | 9.609E-08 | 2.300E-07 | 4.200E-07 |
| 16542 | 828 | T828R | B | 2.2e-07 | 2.6e-07 | 1e-07 | 1.933E-07 | 8.327E-08 | 1.000E-07 | 2.600E-07 |
| 16543 | 828 | T828N | B | 3.7e-07 | 2.6e-07 | 1.9e-07 | 2.733E-07 | 9.074E-08 | 1.900E-07 | 3.700E-07 |
| 16544 | 828 | T828D | B | 2.2e-07 | 2.2e-07 | 2.4e-07 | 2.267E-07 | 1.155E-08 | 2.200E-07 | 2.400E-07 |
| 16545 | 828 | T828C | B | 2e-07 | 1.4e-07 | 2.4e-07 | 1.933E-07 | 5.033E-08 | 1.400E-07 | 2.400E-07 |
| 16546 | 828 | T828E | B | 2.3e-07 | 2.5e-07 | 3.9e-07 | 2.900E-07 | 8.718E-08 | 2.300E-07 | 3.900E-07 |
| 16547 | 828 | T828Q | B | 1.6e-07 | 2.6e-07 | 2.4e-07 | 2.200E-07 | 5.292E-08 | 1.600E-07 | 2.600E-07 |
| 16548 | 828 | T828G | B | 2.4e-07 | 2.7e-07 | 2.5e-07 | 2.533E-07 | 1.528E-08 | 2.400E-07 | 2.700E-07 |
| 16549 | 828 | T828H | B | 2.5e-07 | 3.4e-07 | 1.6e-07 | 2.500E-07 | 9.000E-08 | 1.600E-07 | 3.400E-07 |
| 16550 | 828 | T828I | B | 4.8e-07 | 1.5e-07 | 4e-07 | 3.433E-07 | 1.721E-07 | 1.500E-07 | 4.800E-07 |
| 16551 | 828 | T828L | B | 4.3e-07 | 2.8e-07 | 3.3e-07 | 3.467E-07 | 7.638E-08 | 2.800E-07 | 4.300E-07 |
| 16552 | 828 | T828K | B | 1.4e-07 | 4.5e-07 | 3e-07 | 2.967E-07 | 1.550E-07 | 1.400E-07 | 4.500E-07 |
| 16553 | 828 | T828M | B | 6.2e-07 | 1.4e-07 | 1.6e-07 | 3.067E-07 | 2.715E-07 | 1.400E-07 | 6.200E-07 |
| 16554 | 828 | T828F | B | 2.7e-07 | 3.3e-07 | 2.3e-07 | 2.767E-07 | 5.033E-08 | 2.300E-07 | 3.300E-07 |
| 16555 | 828 | T828P | B | 1.8e-07 | 5.4e-07 | 2.1e-07 | 3.100E-07 | 1.997E-07 | 1.800E-07 | 5.400E-07 |
| 16556 | 828 | T828S | B | 1.5e-07 | 1.7e-07 | 1.5e-07 | 1.567E-07 | 1.155E-08 | 1.500E-07 | 1.700E-07 |
| 16557 | 828 | T828T | B | 4.1e-07 | 1.7e-07 | 2.3e-07 | 2.700E-07 | 1.249E-07 | 1.700E-07 | 4.100E-07 |
| 16558 | 828 | T828W | B | 1.6e-07 | 8.9e-08 | 2e-07 | 1.497E-07 | 5.622E-08 | 8.900E-08 | 2.000E-07 |
| 16559 | 828 | T828Y | B | 3.7e-07 | 4.5e-07 | 1.2e-07 | 3.133E-07 | 1.721E-07 | 1.200E-07 | 4.500E-07 |
| 16560 | 828 | T828V | B | 1.6e-07 | 5.3e-07 | 2.4e-07 | 3.100E-07 | 1.947E-07 | 1.600E-07 | 5.300E-07 |
| 16561 | 829 | K829A | B | 4.4e-07 | 2e-07 | 1.5e-07 | 2.633E-07 | 1.550E-07 | 1.500E-07 | 4.400E-07 |
| 16562 | 829 | K829R | B | 3.2e-07 | 1.6e-07 | 2.4e-07 | 2.400E-07 | 8.000E-08 | 1.600E-07 | 3.200E-07 |
| 16563 | 829 | K829N | B | 3.9e-07 | 1.3e-07 | 1.6e-07 | 2.267E-07 | 1.422E-07 | 1.300E-07 | 3.900E-07 |
| 16564 | 829 | K829D | B | 1.5e-07 | 1.8e-07 | 4.1e-07 | 2.467E-07 | 1.422E-07 | 1.500E-07 | 4.100E-07 |
| 16565 | 829 | K829C | B | 2.3e-07 | 9.3e-08 | 1.3e-07 | 1.510E-07 | 7.087E-08 | 9.300E-08 | 2.300E-07 |
| 16566 | 829 | K829E | B | 1.3e-07 | 2.1e-07 | 2.1e-07 | 1.833E-07 | 4.619E-08 | 1.300E-07 | 2.100E-07 |
| 16567 | 829 | K829Q | B | 4.4e-07 | 1.8e-07 | 3.2e-07 | 3.133E-07 | 1.301E-07 | 1.800E-07 | 4.400E-07 |
| 16568 | 829 | K829G | B | 3.6e-07 | 3.2e-07 | 3.3e-07 | 3.367E-07 | 2.082E-08 | 3.200E-07 | 3.600E-07 |
| 16569 | 829 | K829H | B | 2.3e-07 | 2.2e-07 | 2.3e-07 | 2.267E-07 | 5.774E-09 | 2.200E-07 | 2.300E-07 |
| 16570 | 829 | K829I | B | 1.3e-07 | 2.9e-07 | 4.7e-07 | 2.967E-07 | 1.701E-07 | 1.300E-07 | 4.700E-07 |
| 16571 | 829 | K829L | B | 2.9e-07 | 2.1e-07 | 3.9e-07 | 2.967E-07 | 9.018E-08 | 2.100E-07 | 3.900E-07 |
| 16572 | 829 | K829K | B | 6.1e-08 | 1.4e-07 | 2.2e-07 | 1.403E-07 | 7.950E-08 | 6.100E-08 | 2.200E-07 |
| 16573 | 829 | K829M | B | 2.4e-07 | 1.8e-07 | 3.1e-07 | 2.433E-07 | 6.506E-08 | 1.800E-07 | 3.100E-07 |
| 16574 | 829 | K829F | B | 2.6e-07 | 8e-08 | 5.2e-07 | 2.867E-07 | 2.212E-07 | 8.000E-08 | 5.200E-07 |

| | | | | | | | | | | |
|---|---|---|---|---|---|---|---|---|---|---|
| 16575 | 829 | K829P | B | 1.3e-07 | 3.5e-07 | 2.3e-07 | 2.367E-07 | 1.102E-07 | 1.300E-07 | 3.500E-07 |
| 16576 | 829 | K829S | B | 1.4e-07 | 1.3e-07 | 1.9e-07 | 1.533E-07 | 3.215E-08 | 1.300E-07 | 1.900E-07 |
| 16577 | 829 | K829T | B | 2.2e-07 | 2e-07 | 5.2e-07 | 3.133E-07 | 1.793E-07 | 2.000E-07 | 5.200E-07 |
| 16578 | 829 | K829W | B | 3.5e-07 | 1.8e-07 | 2.7e-07 | 2.667E-07 | 8.505E-08 | 1.800E-07 | 3.500E-07 |
| 16579 | 829 | K829Y | B | 9e-08 | 1.7e-07 | 2.9e-07 | 1.833E-07 | 1.007E-07 | 9.000E-08 | 2.900E-07 |
| 16580 | 829 | K829V | B | 1.4e-07 | 2.3e-07 | 2.4e-07 | 2.033E-07 | 5.508E-08 | 1.400E-07 | 2.400E-07 |
| 16581 | 830 | S830A | B | 7.7e-07 | 2.8e-07 | 3.7e-07 | 4.733E-07 | 2.608E-07 | 2.800E-07 | 7.700E-07 |
| 16582 | 830 | S830R | B | 1.3e-07 | 2.4e-07 | 1.3e-07 | 1.667E-07 | 6.351E-08 | 1.300E-07 | 2.400E-07 |
| 16583 | 830 | S830N | B | 1.6e-07 | 4.2e-07 | 1.4e-07 | 2.400E-07 | 1.562E-07 | 1.400E-07 | 4.200E-07 |
| 16584 | 830 | S830D | B | 2.8e-07 | 3.2e-07 | 2.4e-07 | 2.800E-07 | 4.000E-08 | 2.400E-07 | 3.200E-07 |
| 16585 | 830 | S830C | B | 1.5e-07 | 4e-07 | 1.8e-07 | 2.433E-07 | 1.365E-07 | 1.500E-07 | 4.000E-07 |
| 16586 | 830 | S830E | B | 2.7e-07 | 2.2e-07 | 1.3e-07 | 2.067E-07 | 7.095E-08 | 1.300E-07 | 2.700E-07 |
| 16587 | 830 | S830Q | B | 5.1e-07 | 2.2e-07 | 1.5e-07 | 2.933E-07 | 1.909E-07 | 1.500E-07 | 5.100E-07 |
| 16588 | 830 | S830G | B | 2.6e-07 | 1.5e-07 | 3.2e-07 | 2.433E-07 | 8.622E-08 | 1.500E-07 | 3.200E-07 |
| 16589 | 830 | S830H | B | 2.1e-07 | 2e-07 | 1.8e-07 | 1.967E-07 | 1.528E-08 | 1.800E-07 | 2.100E-07 |
| 16590 | 830 | S830I | B | 6.1e-07 | 3.3e-07 | 1.3e-07 | 3.567E-07 | 2.411E-07 | 1.300E-07 | 6.100E-07 |
| 16591 | 830 | S830L | B | 4.3e-07 | 2.3e-07 | 4.3e-07 | 3.633E-07 | 1.155E-07 | 2.300E-07 | 4.300E-07 |
| 16592 | 830 | S830K | B | 2.3e-07 | 4e-07 | 1.9e-07 | 2.733E-07 | 1.115E-07 | 1.900E-07 | 4.000E-07 |
| 16593 | 830 | S830M | B | 3.5e-07 | 4.5e-07 | 4.1e-07 | 4.033E-07 | 5.033E-08 | 3.500E-07 | 4.500E-07 |
| 16594 | 830 | S830F | B | 2.2e-07 | 4.9e-07 | 3.3e-07 | 3.467E-07 | 1.358E-07 | 2.200E-07 | 4.900E-07 |
| 16595 | 830 | S830P | B | 1.6e-07 | 4.9e-07 | 1.8e-07 | 2.767E-07 | 1.850E-07 | 1.600E-07 | 4.900E-07 |
| 16596 | 830 | S830S | B | 2.8e-07 | 1.3e-07 | 2.8e-07 | 2.300E-07 | 8.660E-08 | 1.300E-07 | 2.800E-07 |
| 16597 | 830 | S830T | B | 2.6e-07 | 3.4e-07 | 2.2e-07 | 2.733E-07 | 6.110E-08 | 2.200E-07 | 3.400E-07 |
| 16598 | 830 | S830W | B | 1.3e-07 | 1.9e-07 | 3.2e-07 | 2.133E-07 | 9.713E-08 | 1.300E-07 | 3.200E-07 |
| 16599 | 830 | S830Y | B | 3.8e-07 | 3.1e-07 | 2.4e-07 | 3.100E-07 | 7.000E-08 | 2.400E-07 | 3.800E-07 |
| 16600 | 830 | S830V | B | 1.6e-07 | 1.4e-07 | 1.9e-07 | 1.633E-07 | 2.517E-08 | 1.400E-07 | 1.900E-07 |
| 16601 | 831 | F831A | B | 1.2e-07 | 5.3e-07 | 3.1e-07 | 3.200E-07 | 2.052E-07 | 1.200E-07 | 5.300E-07 |
| 16602 | 831 | F831R | B | 1.7e-07 | 1.4e-07 | 1.9e-07 | 1.667E-07 | 2.517E-08 | 1.400E-07 | 1.900E-07 |
| 16603 | 831 | F831N | B | 1.7e-07 | 2e-07 | 3.3e-07 | 2.333E-07 | 8.505E-08 | 1.700E-07 | 3.300E-07 |
| 16604 | 831 | F831D | B | 3.5e-07 | 4.2e-07 | 6.3e-07 | 4.667E-07 | 1.457E-07 | 3.500E-07 | 6.300E-07 |
| 16605 | 831 | F831C | B | 1.6e-07 | 2.6e-07 | 3.6e-07 | 2.600E-07 | 1.000E-07 | 1.600E-07 | 3.600E-07 |
| 16606 | 831 | F831E | B | 1.4e-07 | 3.3e-07 | 3e-07 | 2.567E-07 | 1.021E-07 | 1.400E-07 | 3.300E-07 |
| 16607 | 831 | F831Q | B | 3.2e-07 | 3.4e-07 | 1.9e-07 | 2.833E-07 | 8.145E-08 | 1.900E-07 | 3.400E-07 |
| 16608 | 831 | F831G | B | 3.5e-07 | 2.6e-07 | 2e-07 | 2.700E-07 | 7.550E-08 | 2.000E-07 | 3.500E-07 |
| 16609 | 831 | F831H | B | 2.5e-07 | 3.5e-07 | 2.8e-07 | 2.933E-07 | 5.132E-08 | 2.500E-07 | 3.500E-07 |
| 16610 | 831 | F831I | B | 2.3e-07 | 1.3e-07 | 3.7e-07 | 2.433E-07 | 1.206E-07 | 1.300E-07 | 3.700E-07 |
| 16611 | 831 | F831L | B | 3e-07 | 3.2e-07 | 1.4e-07 | 2.533E-07 | 9.866E-08 | 1.400E-07 | 3.200E-07 |
| 16612 | 831 | F831K | B | 1.8e-07 | 2e-07 | 1.2e-07 | 1.667E-07 | 4.163E-08 | 1.200E-07 | 2.000E-07 |
| 16613 | 831 | F831M | B | 3.8e-07 | 2.9e-07 | 2e-07 | 2.900E-07 | 9.000E-08 | 2.000E-07 | 3.800E-07 |
| 16614 | 831 | F831F | B | 2.2e-07 | 2.3e-07 | 4.1e-07 | 2.867E-07 | 1.069E-07 | 2.200E-07 | 4.100E-07 |
| 16615 | 831 | F831P | B | 1.8e-07 | 1.1e-07 | 2.1e-07 | 1.667E-07 | 5.132E-08 | 1.100E-07 | 2.100E-07 |
| 16616 | 831 | F831S | B | 3.3e-07 | 1.4e-07 | 2.2e-07 | 2.300E-07 | 9.539E-08 | 1.400E-07 | 3.300E-07 |
| 16617 | 831 | F831T | B | 1.4e-07 | 1.8e-07 | 2.2e-07 | 1.800E-07 | 4.000E-08 | 1.400E-07 | 2.200E-07 |
| 16618 | 831 | F831W | B | 1.3e-07 | 2.7e-07 | 9.4e-08 | 1.647E-07 | 9.298E-08 | 9.400E-08 | 2.700E-07 |
| 16619 | 831 | F831Y | B | 8.9e-08 | 6e-07 | 3.4e-07 | 3.430E-07 | 2.555E-07 | 8.900E-08 | 6.000E-07 |
| 16620 | 831 | F831V | B | 2e-07 | 4.8e-07 | 4.1e-07 | 3.633E-07 | 1.457E-07 | 2.000E-07 | 4.800E-07 |
| 16621 | 832 | N832A | B | 3e-07 | 3.8e-07 | 2.1e-07 | 2.967E-07 | 8.505E-08 | 2.100E-07 | 3.800E-07 |
| 16622 | 832 | N832R | B | 1.1e-07 | 5.4e-07 | 1.3e-07 | 2.600E-07 | 2.427E-07 | 1.100E-07 | 5.400E-07 |
| 16623 | 832 | N832N | B | 1.4e-07 | 3.4e-07 | 2.9e-07 | 2.567E-07 | 1.041E-07 | 1.400E-07 | 3.400E-07 |
| 16624 | 832 | N832D | B | 2.9e-07 | 4.8e-07 | 3.6e-07 | 3.767E-07 | 9.609E-08 | 2.900E-07 | 4.800E-07 |
| 16625 | 832 | N832C | B | 3e-07 | 1.1e-07 | 3.7e-07 | 2.600E-07 | 1.345E-07 | 1.100E-07 | 3.700E-07 |
| 16626 | 832 | N832E | B | 5.2e-07 | 2.2e-07 | 2.5e-07 | 3.300E-07 | 1.652E-07 | 2.200E-07 | 5.200E-07 |
| 16627 | 832 | N832Q | B | 3.7e-07 | 3.2e-07 | 1.7e-07 | 2.867E-07 | 1.041E-07 | 1.700E-07 | 3.700E-07 |
| 16628 | 832 | N832G | B | 1.5e-07 | 2.2e-07 | 4.2e-07 | 2.633E-07 | 1.401E-07 | 1.500E-07 | 4.200E-07 |
| 16629 | 832 | N832H | B | 2.7e-07 | 1.7e-07 | 4.1e-07 | 2.833E-07 | 1.206E-07 | 1.700E-07 | 4.100E-07 |
| 16630 | 832 | N832I | B | 1.3e-07 | 2.6e-07 | 3.8e-07 | 2.567E-07 | 1.250E-07 | 1.300E-07 | 3.800E-07 |
| 16631 | 832 | N832L | B | 4.5e-07 | 1.4e-07 | 1.3e-07 | 2.400E-07 | 1.819E-07 | 1.300E-07 | 4.500E-07 |
| 16632 | 832 | N832K | B | 2.8e-07 | 4.2e-07 | 1.6e-07 | 2.867E-07 | 1.301E-07 | 1.600E-07 | 4.200E-07 |
| 16633 | 832 | N832M | B | 1.9e-07 | 3.5e-07 | 1.2e-07 | 2.200E-07 | 1.179E-07 | 1.200E-07 | 3.500E-07 |
| 16634 | 832 | N832F | B | 1.6e-07 | 1.3e-07 | 3.3e-07 | 2.067E-07 | 1.079E-07 | 1.300E-07 | 3.300E-07 |
| 16635 | 832 | N832P | B | 2.3e-07 | 3.4e-07 | 1.4e-07 | 2.367E-07 | 1.002E-07 | 1.400E-07 | 3.400E-07 |
| 16636 | 832 | N832S | B | 2.3e-07 | 1.9e-07 | 3.1e-07 | 2.433E-07 | 6.110E-08 | 1.900E-07 | 3.100E-07 |
| 16637 | 832 | N832T | B | 3.1e-07 | 1.7e-07 | 7e-07 | 3.933E-07 | 2.747E-07 | 1.700E-07 | 7.000E-07 |
| 16638 | 832 | N832W | B | 2.7e-07 | 2.4e-07 | 2.7e-07 | 2.600E-07 | 1.732E-08 | 2.400E-07 | 2.700E-07 |
| 16639 | 832 | N832Y | B | 3.4e-07 | 2e-07 | 1e-07 | 2.133E-07 | 1.206E-07 | 1.000E-07 | 3.400E-07 |
| 16640 | 832 | N832V | B | 2.1e-07 | 1.5e-07 | 2.8e-07 | 2.133E-07 | 6.506E-08 | 1.500E-07 | 2.800E-07 |
| 16641 | 833 | R833A | B | 3.6e-07 | 1.3e-07 | 2.5e-07 | 2.467E-07 | 1.150E-07 | 1.300E-07 | 3.600E-07 |
| 16642 | 833 | R833R | B | 1.3e-07 | 1.6e-07 | 2.1e-07 | 1.667E-07 | 4.041E-08 | 1.300E-07 | 2.100E-07 |
| 16643 | 833 | R833N | B | 2.9e-07 | 4.1e-07 | 2.4e-07 | 3.133E-07 | 8.737E-08 | 2.400E-07 | 4.100E-07 |
| 16644 | 833 | R833D | B | 3.6e-07 | 1.9e-07 | 3.1e-07 | 2.867E-07 | 8.737E-08 | 1.900E-07 | 3.600E-07 |
| 16645 | 833 | R833C | B | 7.2e-07 | 3.7e-07 | 2.4e-07 | 4.433E-07 | 2.483E-07 | 2.400E-07 | 7.200E-07 |
| 16646 | 833 | R833E | B | 2.3e-07 | 2.1e-07 | 2.3e-07 | 2.233E-07 | 1.155E-08 | 2.100E-07 | 2.300E-07 |
| 16647 | 833 | R833Q | B | 3e-07 | 1.4e-07 | 4.4e-07 | 2.933E-07 | 1.501E-07 | 1.400E-07 | 4.400E-07 |
| 16648 | 833 | R833G | B | 1.7e-07 | 3.7e-07 | 2.4e-07 | 2.600E-07 | 1.015E-07 | 1.700E-07 | 3.700E-07 |
| 16649 | 833 | R833H | B | 3.4e-07 | 2.9e-07 | 1.6e-07 | 2.633E-07 | 9.292E-08 | 1.600E-07 | 3.400E-07 |
| 16650 | 833 | R833I | B | 2.3e-07 | 1.9e-07 | 2.2e-07 | 2.133E-07 | 2.082E-08 | 1.900E-07 | 2.300E-07 |
| 16651 | 833 | R833L | B | 1.5e-07 | 2.7e-07 | 4.3e-07 | 2.833E-07 | 1.405E-07 | 1.500E-07 | 4.300E-07 |
| 16652 | 833 | R833K | B | 4.1e-07 | 2.4e-07 | 1.2e-07 | 2.567E-07 | 1.457E-07 | 1.200E-07 | 4.100E-07 |
| 16653 | 833 | R833M | B | 2.5e-07 | 3.9e-07 | 1.5e-07 | 2.633E-07 | 1.206E-07 | 1.500E-07 | 3.900E-07 |
| 16654 | 833 | R833F | B | 1.8e-07 | 7.4e-07 | 1.6e-07 | 3.600E-07 | 3.292E-07 | 1.600E-07 | 7.400E-07 |
| 16655 | 833 | R833P | B | 4.4e-07 | 2.3e-07 | 4e-07 | 3.567E-07 | 1.115E-07 | 2.300E-07 | 4.400E-07 |
| 16656 | 833 | R833S | B | 2.2e-07 | 2.7e-07 | 1.5e-07 | 2.133E-07 | 6.028E-08 | 1.500E-07 | 2.700E-07 |
| 16657 | 833 | R833T | B | 1.5e-07 | 8.2e-08 | 3e-07 | 1.773E-07 | 1.115E-07 | 8.200E-08 | 3.000E-07 |
| 16658 | 833 | R833W | B | 1.1e-07 | 1.3e-07 | 2.6e-07 | 1.667E-07 | 8.145E-08 | 1.100E-07 | 2.600E-07 |
| 16659 | 833 | R833Y | B | 2.8e-07 | 1.9e-07 | 2.9e-07 | 2.533E-07 | 5.508E-08 | 1.900E-07 | 2.900E-07 |

| | | | | | | | | | | |
|---|---|---|---|---|---|---|---|---|---|---|
| 16660 | 833 | R833V | B | 2.5e-07 | 3.1e-07 | 3.9e-07 | 3.167E-07 | 7.024E-08 | 2.500E-07 | 3.900E-07 |
| 16661 | 834 | G834A | B | 1.7e-07 | 1.5e-07 | 5.6e-07 | 2.933E-07 | 2.312E-07 | 1.500E-07 | 5.600E-07 |
| 16662 | 834 | G834R | B | 2.5e-07 | 1.5e-07 | 2.8e-07 | 2.267E-07 | 6.807E-08 | 1.500E-07 | 2.800E-07 |
| 16663 | 834 | G834N | B | 1.5e-07 | 1.3e-07 | 2.7e-07 | 1.833E-07 | 7.572E-08 | 1.300E-07 | 2.700E-07 |
| 16664 | 834 | G834D | B | 1.6e-07 | 3.9e-07 | 2.9e-07 | 2.800E-07 | 1.153E-07 | 1.600E-07 | 3.900E-07 |
| 16665 | 834 | G834C | B | 2.9e-07 | 3e-07 | 4.5e-07 | 3.467E-07 | 8.963E-08 | 2.900E-07 | 4.500E-07 |
| 16666 | 834 | G834E | B | 3.9e-07 | 3e-07 | 2e-07 | 2.967E-07 | 9.504E-08 | 2.000E-07 | 3.900E-07 |
| 16667 | 834 | G834Q | B | 3e-07 | 1.4e-07 | 9.8e-08 | 1.793E-07 | 1.066E-07 | 9.800E-08 | 3.000E-07 |
| 16668 | 834 | G834G | B | 3.4e-07 | 3.1e-07 | 2.5e-07 | 3.000E-07 | 4.583E-08 | 2.500E-07 | 3.400E-07 |
| 16669 | 834 | G834H | B | 1.4e-07 | 5.1e-07 | 1.4e-07 | 2.633E-07 | 2.136E-07 | 1.400E-07 | 5.100E-07 |
| 16670 | 834 | G834I | B | 1.5e-07 | 8.8e-08 | 3.2e-07 | 1.860E-07 | 1.201E-07 | 8.800E-08 | 3.200E-07 |
| 16671 | 834 | G834L | B | 3.2e-07 | 2.1e-07 | 1.2e-07 | 2.167E-07 | 1.002E-07 | 1.200E-07 | 3.200E-07 |
| 16672 | 834 | G834K | B | 1.5e-07 | 3.5e-07 | 3.1e-07 | 2.700E-07 | 1.058E-07 | 1.500E-07 | 3.500E-07 |
| 16673 | 834 | G834M | B | 4e-07 | 1.8e-07 | 3.1e-07 | 2.967E-07 | 1.106E-07 | 1.800E-07 | 4.000E-07 |
| 16674 | 834 | G834F | B | 3.1e-07 | 1.8e-07 | 3.5e-07 | 2.800E-07 | 8.888E-08 | 1.800E-07 | 3.500E-07 |
| 16675 | 834 | G834P | B | 1.5e-07 | 2.5e-07 | 4.2e-07 | 2.733E-07 | 1.365E-07 | 1.500E-07 | 4.200E-07 |
| 16676 | 834 | G834S | B | 2.1e-07 | 4e-07 | 4.6e-07 | 3.567E-07 | 1.305E-07 | 2.100E-07 | 4.600E-07 |
| 16677 | 834 | G834T | B | 6.1e-07 | 4.1e-07 | 2e-07 | 4.067E-07 | 2.050E-07 | 2.000E-07 | 6.100E-07 |
| 16678 | 834 | G834W | B | 5.5e-07 | 1.1e-07 | 1.1e-07 | 2.567E-07 | 2.540E-07 | 1.100E-07 | 5.500E-07 |
| 16679 | 834 | G834Y | B | 1.8e-07 | 2.4e-07 | 4e-07 | 2.733E-07 | 1.137E-07 | 1.800E-07 | 4.000E-07 |
| 16680 | 834 | G834V | B | 1.8e-07 | 2.3e-07 | 1e-07 | 1.700E-07 | 6.557E-08 | 1.000E-07 | 2.300E-07 |
| 16681 | 835 | E835A | B | 2.2e-07 | 2.3e-07 | 1.6e-07 | 2.033E-07 | 3.786E-08 | 1.600E-07 | 2.300E-07 |
| 16682 | 835 | E835R | B | 2.4e-07 | 1.6e-07 | 2e-07 | 2.000E-07 | 4.000E-08 | 1.600E-07 | 2.400E-07 |
| 16683 | 835 | E835N | B | 4.8e-07 | 4.7e-07 | 1e-06 | 6.500E-07 | 3.032E-07 | 4.700E-07 | 1.000E-06 |
| 16684 | 835 | E835D | B | 2.2e-07 | 4.1e-07 | 3.2e-07 | 3.167E-07 | 9.504E-08 | 2.200E-07 | 4.100E-07 |
| 16685 | 835 | E835C | B | 3.2e-07 | 3e-07 | 2.8e-07 | 3.000E-07 | 2.000E-08 | 2.800E-07 | 3.200E-07 |
| 16686 | 835 | E835E | B | 2.3e-07 | 1.7e-07 | 1.7e-07 | 1.900E-07 | 3.464E-08 | 1.700E-07 | 2.300E-07 |
| 16687 | 835 | E835Q | B | 2.3e-07 | 2.6e-07 | 2.9e-07 | 2.600E-07 | 3.000E-08 | 2.300E-07 | 2.900E-07 |
| 16688 | 835 | E835G | B | 2.1e-07 | 2.8e-07 | 4e-07 | 2.967E-07 | 9.609E-08 | 2.100E-07 | 4.000E-07 |
| 16689 | 835 | E835H | B | 1.2e-07 | 1.6e-07 | 2.7e-07 | 1.833E-07 | 7.767E-08 | 1.200E-07 | 2.700E-07 |
| 16690 | 835 | E835I | B | 2.4e-07 | 2.1e-07 | 2.2e-07 | 2.233E-07 | 1.528E-08 | 2.100E-07 | 2.400E-07 |
| 16691 | 835 | E835L | B | 2e-07 | 3.1e-07 | 2.5e-07 | 2.533E-07 | 5.508E-08 | 2.000E-07 | 3.100E-07 |
| 16692 | 835 | E835K | B | 1.1e-07 | 1.6e-07 | 2.8e-07 | 1.833E-07 | 8.737E-08 | 1.100E-07 | 2.800E-07 |
| 16693 | 835 | E835M | B | 1.7e-07 | 2.3e-07 | 4.4e-07 | 2.800E-07 | 1.418E-07 | 1.700E-07 | 4.400E-07 |
| 16694 | 835 | E835F | B | 1.6e-07 | 2.5e-07 | 2.3e-07 | 2.133E-07 | 4.726E-08 | 1.600E-07 | 2.500E-07 |
| 16695 | 835 | E835P | B | 5e-07 | 2.3e-07 | 1.5e-07 | 2.933E-07 | 1.834E-07 | 1.500E-07 | 5.000E-07 |
| 16696 | 835 | E835S | B | 2.6e-07 | 1.3e-07 | 9.7e-08 | 1.623E-07 | 8.618E-08 | 9.700E-08 | 2.600E-07 |
| 16697 | 835 | E835T | B | 2.9e-07 | 2.2e-07 | 1.6e-07 | 2.233E-07 | 6.506E-08 | 1.600E-07 | 2.900E-07 |
| 16698 | 835 | E835W | B | 3.6e-07 | 1.1e-07 | 3.1e-07 | 2.600E-07 | 1.323E-07 | 1.100E-07 | 3.600E-07 |
| 16699 | 835 | E835Y | B | 1.9e-07 | 1.5e-07 | 3.1e-07 | 2.167E-07 | 8.327E-08 | 1.500E-07 | 3.100E-07 |
| 16700 | 835 | E835V | B | 1.6e-07 | 4.1e-07 | 1.7e-07 | 2.467E-07 | 1.415E-07 | 1.600E-07 | 4.100E-07 |
| 16701 | 836 | C836A | B | 1.5e-07 | 2.6e-07 | 1.6e-07 | 1.900E-07 | 6.083E-08 | 1.500E-07 | 2.600E-07 |
| 16702 | 836 | C836R | B | 1.5e-07 | 2.5e-07 | 1.6e-07 | 1.867E-07 | 5.508E-08 | 1.500E-07 | 2.500E-07 |
| 16703 | 836 | C836N | B | 1.6e-07 | 1.6e-07 | 1.8e-07 | 1.667E-07 | 1.155E-08 | 1.600E-07 | 1.800E-07 |
| 16704 | 836 | C836D | B | 3e-07 | 2.2e-07 | 4.5e-07 | 3.233E-07 | 1.168E-07 | 2.200E-07 | 4.500E-07 |
| 16705 | 836 | C836C | B | 1.5e-07 | 3.9e-07 | 2.3e-07 | 2.567E-07 | 1.222E-07 | 1.500E-07 | 3.900E-07 |
| 16706 | 836 | C836E | B | 2.5e-07 | 2.4e-07 | 3.3e-07 | 2.733E-07 | 4.933E-08 | 2.400E-07 | 3.300E-07 |
| 16707 | 836 | C836Q | B | 4.4e-07 | 2.8e-07 | 7.8e-07 | 5.000E-07 | 2.553E-07 | 2.800E-07 | 7.800E-07 |
| 16708 | 836 | C836G | B | 5.5e-07 | 1.4e-07 | 3.1e-07 | 3.333E-07 | 2.060E-07 | 1.400E-07 | 5.500E-07 |
| 16709 | 836 | C836H | B | 4.8e-07 | 1.9e-07 | 2.8e-07 | 3.167E-07 | 1.484E-07 | 1.900E-07 | 4.800E-07 |
| 16710 | 836 | C836I | B | 5.1e-07 | 2.4e-07 | 2.6e-07 | 3.367E-07 | 1.504E-07 | 2.400E-07 | 5.100E-07 |
| 16711 | 836 | C836L | B | 2.5e-07 | 3e-07 | 1.7e-07 | 2.400E-07 | 6.557E-08 | 1.700E-07 | 3.000E-07 |
| 16712 | 836 | C836K | B | 2e-07 | 3.6e-07 | 2.8e-07 | 2.800E-07 | 8.000E-08 | 2.000E-07 | 3.600E-07 |
| 16713 | 836 | C836M | B | 1.7e-07 | 9.6e-08 | 2.1e-07 | 1.587E-07 | 5.784E-08 | 9.600E-08 | 2.100E-07 |
| 16714 | 836 | C836F | B | 5.4e-07 | 1.6e-07 | 4.1e-07 | 3.700E-07 | 1.931E-07 | 1.600E-07 | 5.400E-07 |
| 16715 | 836 | C836P | B | 2.5e-07 | 2.5e-07 | 1.6e-07 | 2.200E-07 | 5.196E-08 | 1.600E-07 | 2.500E-07 |
| 16716 | 836 | C836S | B | 2.2e-07 | 1.9e-07 | 4.3e-07 | 2.800E-07 | 1.308E-07 | 1.900E-07 | 4.300E-07 |
| 16717 | 836 | C836T | B | 3.5e-07 | 2.5e-07 | 2.5e-07 | 2.833E-07 | 5.774E-08 | 2.500E-07 | 3.500E-07 |
| 16718 | 836 | C836W | B | 2.4e-07 | 4.7e-07 | 3.6e-07 | 3.567E-07 | 1.150E-07 | 2.400E-07 | 4.700E-07 |
| 16719 | 836 | C836Y | B | 2.6e-07 | 1.3e-07 | 6.2e-07 | 3.367E-07 | 2.538E-07 | 1.300E-07 | 6.200E-07 |
| 16720 | 836 | C836V | B | 4e-07 | 3.3e-07 | 1.5e-07 | 2.933E-07 | 1.290E-07 | 1.500E-07 | 4.000E-07 |
| 16721 | 837 | E837A | C | 3e-07 | 4.2e-07 | 2e-07 | 3.067E-07 | 1.102E-07 | 2.000E-07 | 4.200E-07 |
| 16722 | 837 | E837R | C | 4e-07 | 2.3e-07 | 1.8e-07 | 2.700E-07 | 1.153E-07 | 1.800E-07 | 4.000E-07 |
| 16723 | 837 | E837N | C | 3.6e-07 | 2.3e-07 | 3.5e-07 | 3.133E-07 | 7.234E-08 | 2.300E-07 | 3.600E-07 |
| 16724 | 837 | E837D | C | 3.1e-07 | 1.1e-07 | 4.8e-07 | 3.000E-07 | 1.852E-07 | 1.100E-07 | 4.800E-07 |
| 16725 | 837 | E837C | C | 3.7e-07 | 2.6e-07 | 1.5e-07 | 2.600E-07 | 1.100E-07 | 1.500E-07 | 3.700E-07 |
| 16726 | 837 | E837E | C | 3e-07 | 2.5e-07 | 2.9e-07 | 2.800E-07 | 2.646E-08 | 2.500E-07 | 3.000E-07 |
| 16727 | 837 | E837Q | C | 1.2e-07 | 2.1e-07 | 3.4e-07 | 2.233E-07 | 1.106E-07 | 1.200E-07 | 3.400E-07 |
| 16728 | 837 | E837G | C | 1.9e-07 | 3.5e-07 | 3.3e-07 | 2.900E-07 | 8.718E-08 | 1.900E-07 | 3.500E-07 |
| 16729 | 837 | E837H | C | 2.6e-07 | 2.2e-07 | 2.5e-07 | 2.433E-07 | 2.082E-08 | 2.200E-07 | 2.600E-07 |
| 16730 | 837 | E837I | C | 4e-07 | 3.4e-07 | 3.5e-07 | 3.633E-07 | 3.215E-08 | 3.400E-07 | 4.000E-07 |
| 16731 | 837 | E837L | C | 1.6e-07 | 7.9e-07 | 2.6e-07 | 4.033E-07 | 3.386E-07 | 1.600E-07 | 7.900E-07 |
| 16732 | 837 | E837K | C | 2.2e-07 | 3.2e-07 | 2.6e-07 | 2.667E-07 | 5.033E-08 | 2.200E-07 | 3.200E-07 |
| 16733 | 837 | E837M | C | 3.5e-07 | 3e-07 | 3.5e-07 | 3.333E-07 | 2.887E-08 | 3.000E-07 | 3.500E-07 |
| 16734 | 837 | E837F | C | 2.5e-07 | 3.8e-07 | 5.3e-07 | 3.867E-07 | 1.401E-07 | 2.500E-07 | 5.300E-07 |
| 16735 | 837 | E837P | C | 3.3e-07 | 2.3e-07 | 1.9e-07 | 2.500E-07 | 7.211E-08 | 1.900E-07 | 3.300E-07 |
| 16736 | 837 | E837S | C | 2.6e-07 | 1.6e-07 | 4.7e-07 | 2.967E-07 | 1.582E-07 | 1.600E-07 | 4.700E-07 |
| 16737 | 837 | E837T | C | 2e-07 | 1.5e-07 | 2e-07 | 1.833E-07 | 2.887E-08 | 1.500E-07 | 2.000E-07 |
| 16738 | 837 | E837W | C | 1.5e-07 | 1.9e-07 | 2.4e-07 | 1.933E-07 | 4.509E-08 | 1.500E-07 | 2.400E-07 |
| 16739 | 837 | E837Y | C | 1e-07 | 1.4e-07 | 3.5e-07 | 1.967E-07 | 1.343E-07 | 1.000E-07 | 3.500E-07 |
| 16740 | 837 | E837V | C | 3.4e-07 | 6.1e-07 | 2.9e-07 | 4.133E-07 | 1.721E-07 | 2.900E-07 | 6.100E-07 |
| 16741 | 838 | V838A | C | 3.2e-07 | 8e-08 | 1.5e-07 | 1.833E-07 | 1.234E-07 | 8.000E-08 | 3.200E-07 |
| 16742 | 838 | V838R | C | 2.6e-07 | 2.2e-07 | 3.7e-07 | 2.833E-07 | 7.767E-08 | 2.200E-07 | 3.700E-07 |
| 16743 | 838 | V838N | C | 4.7e-07 | 6.4e-07 | 7.1e-08 | 3.937E-07 | 2.921E-07 | 7.100E-08 | 6.400E-07 |
| 16744 | 838 | V838D | C | 3.3e-07 | 3.2e-07 | 4e-07 | 3.500E-07 | 4.359E-08 | 3.200E-07 | 4.000E-07 |

| | | | | | | | | | | |
|---|---|---|---|---|---|---|---|---|---|---|
| 16745 | 838 | V838C | C | 4.1e-07 | 3.4e-07 | 3.9e-07 | 3.800E-07 | 3.606E-08 | 3.400E-07 | 4.100E-07 |
| 16746 | 838 | V838E | C | 1.6e-07 | 1.6e-07 | 2.6e-07 | 1.933E-07 | 5.774E-08 | 1.600E-07 | 2.600E-07 |
| 16747 | 838 | V838Q | C | 3.6e-07 | 2.3e-07 | 3.2e-07 | 3.033E-07 | 6.658E-08 | 2.300E-07 | 3.600E-07 |
| 16748 | 838 | V838G | C | 1.4e-07 | 5e-07 | 1.5e-07 | 2.633E-07 | 2.050E-07 | 1.400E-07 | 5.000E-07 |
| 16749 | 838 | V838H | C | 1.6e-07 | 1.9e-07 | 3e-07 | 2.167E-07 | 7.371E-08 | 1.600E-07 | 3.000E-07 |
| 16750 | 838 | V838I | C | 1.8e-07 | 3.3e-07 | 1.2e-07 | 2.100E-07 | 1.082E-07 | 1.200E-07 | 3.300E-07 |
| 16751 | 838 | V838L | C | 4.8e-07 | 3.1e-07 | 8.9e-08 | 2.930E-07 | 1.961E-07 | 8.900E-08 | 4.800E-07 |
| 16752 | 838 | V838K | C | 4.8e-07 | 1.3e-07 | 4.3e-07 | 3.467E-07 | 1.893E-07 | 1.300E-07 | 4.800E-07 |
| 16753 | 838 | V838M | C | 2.1e-07 | 4.8e-07 | 1.5e-07 | 2.800E-07 | 1.758E-07 | 1.500E-07 | 4.800E-07 |
| 16754 | 838 | V838F | C | 1.3e-07 | 1.7e-07 | 2.3e-07 | 1.767E-07 | 5.033E-08 | 1.300E-07 | 2.300E-07 |
| 16755 | 838 | V838P | C | 6.7e-07 | 2.3e-07 | 1.8e-07 | 3.600E-07 | 2.696E-07 | 1.800E-07 | 6.700E-07 |
| 16756 | 838 | V838S | C | 2.9e-07 | 1.9e-07 | 1.6e-07 | 2.133E-07 | 6.807E-08 | 1.600E-07 | 2.900E-07 |
| 16757 | 838 | V838T | C | 1.7e-07 | 3.4e-07 | 2.5e-07 | 2.533E-07 | 8.505E-08 | 1.700E-07 | 3.400E-07 |
| 16758 | 838 | V838W | C | 4.1e-07 | 1.5e-07 | 1.6e-07 | 2.400E-07 | 1.473E-07 | 1.500E-07 | 4.100E-07 |
| 16759 | 838 | V838Y | C | 1.6e-07 | 2e-07 | 3.2e-07 | 2.267E-07 | 8.327E-08 | 1.600E-07 | 3.200E-07 |
| 16760 | 838 | V838V | C | 4.4e-07 | 7.6e-07 | 2.9e-07 | 4.967E-07 | 2.401E-07 | 2.900E-07 | 7.600E-07 |
| 16761 | 839 | Q839A | C | 4e-07 | 4.7e-07 | 1.9e-07 | 3.533E-07 | 1.457E-07 | 1.900E-07 | 4.700E-07 |
| 16762 | 839 | Q839R | C | 2.2e-07 | 2.2e-07 | 1.5e-07 | 1.967E-07 | 4.041E-08 | 1.500E-07 | 2.200E-07 |
| 16763 | 839 | Q839N | C | 2.1e-07 | 2e-07 | 1.2e-07 | 1.767E-07 | 4.933E-08 | 1.200E-07 | 2.100E-07 |
| 16764 | 839 | Q839D | C | 3.1e-07 | 4.7e-07 | 3.6e-07 | 3.800E-07 | 8.185E-08 | 3.100E-07 | 4.700E-07 |
| 16765 | 839 | Q839C | C | 1.2e-07 | 2.5e-07 | 3.5e-07 | 2.400E-07 | 1.153E-07 | 1.200E-07 | 3.500E-07 |
| 16766 | 839 | Q839E | C | 3.9e-07 | 1.5e-07 | 2.3e-07 | 2.567E-07 | 1.222E-07 | 1.500E-07 | 3.900E-07 |
| 16767 | 839 | Q839Q | C | 1.4e-07 | 2.9e-07 | 1.8e-07 | 2.033E-07 | 7.767E-08 | 1.400E-07 | 2.900E-07 |
| 16768 | 839 | Q839G | C | 3.1e-07 | 8.5e-08 | 2.4e-07 | 2.117E-07 | 1.151E-07 | 8.500E-08 | 3.100E-07 |
| 16769 | 839 | Q839H | C | 3.9e-07 | 3e-07 | 1.6e-07 | 2.833E-07 | 1.159E-07 | 1.600E-07 | 3.900E-07 |
| 16770 | 839 | Q839I | C | 2.1e-07 | 1.7e-07 | 1.3e-07 | 1.700E-07 | 4.000E-08 | 1.300E-07 | 2.100E-07 |
| 16771 | 839 | Q839L | C | 3.2e-07 | 2.5e-07 | 2.1e-07 | 2.600E-07 | 5.568E-08 | 2.100E-07 | 3.200E-07 |
| 16772 | 839 | Q839K | C | 2.9e-07 | 2e-07 | 1.7e-07 | 2.200E-07 | 6.245E-08 | 1.700E-07 | 2.900E-07 |
| 16773 | 839 | Q839M | C | 1.8e-07 | 2.1e-07 | 3.5e-07 | 2.467E-07 | 9.074E-08 | 1.800E-07 | 3.500E-07 |
| 16774 | 839 | Q839F | C | 1.8e-07 | 1.6e-07 | 4.9e-07 | 2.767E-07 | 1.850E-07 | 1.600E-07 | 4.900E-07 |
| 16775 | 839 | Q839P | C | 2.9e-07 | 1.6e-07 | 2.5e-07 | 2.333E-07 | 6.658E-08 | 1.600E-07 | 2.900E-07 |
| 16776 | 839 | Q839S | C | 2.1e-07 | 4.5e-07 | 1.5e-07 | 2.700E-07 | 1.587E-07 | 1.500E-07 | 4.500E-07 |
| 16777 | 839 | Q839T | C | 3e-07 | 1.3e-07 | 3.3e-07 | 2.533E-07 | 1.079E-07 | 1.300E-07 | 3.300E-07 |
| 16778 | 839 | Q839W | C | 1.8e-07 | 1.8e-07 | 1.9e-07 | 1.833E-07 | 5.774E-09 | 1.800E-07 | 1.900E-07 |
| 16779 | 839 | Q839Y | C | 2.7e-07 | 1.9e-07 | 4.1e-07 | 2.900E-07 | 1.114E-07 | 1.900E-07 | 4.100E-07 |
| 16780 | 839 | Q839V | C | 4.3e-07 | 6e-07 | 3.6e-07 | 4.633E-07 | 1.234E-07 | 3.600E-07 | 6.000E-07 |
| 16781 | 840 | L840A | C | 4.3e-07 | 1.3e-07 | 4.2e-07 | 3.267E-07 | 1.704E-07 | 1.300E-07 | 4.300E-07 |
| 16782 | 840 | L840R | C | 3e-07 | 2.2e-07 | 1.8e-07 | 2.333E-07 | 6.110E-08 | 1.800E-07 | 3.000E-07 |
| 16783 | 840 | L840N | C | 1.4e-07 | 2.7e-07 | 1.4e-07 | 1.833E-07 | 7.506E-08 | 1.400E-07 | 2.700E-07 |
| 16784 | 840 | L840D | C | 1.6e-07 | 1.9e-07 | 5.3e-07 | 2.933E-07 | 2.055E-07 | 1.600E-07 | 5.300E-07 |
| 16785 | 840 | L840C | C | 2.9e-07 | 3.3e-07 | 2.2e-07 | 2.800E-07 | 5.568E-08 | 2.200E-07 | 3.300E-07 |
| 16786 | 840 | L840E | C | 2.8e-07 | 3.5e-07 | 1.2e-07 | 2.500E-07 | 1.179E-07 | 1.200E-07 | 3.500E-07 |
| 16787 | 840 | L840Q | C | 2.6e-07 | 1.7e-07 | 1.9e-07 | 2.067E-07 | 4.726E-08 | 1.700E-07 | 2.600E-07 |
| 16788 | 840 | L840G | C | 4.2e-07 | 3.8e-07 | 1.4e-07 | 3.133E-07 | 1.514E-07 | 1.400E-07 | 4.200E-07 |
| 16789 | 840 | L840H | C | 4.2e-07 | 1.6e-07 | 2.3e-07 | 2.700E-07 | 1.345E-07 | 1.600E-07 | 4.200E-07 |
| 16790 | 840 | L840I | C | 2e-07 | 2e-07 | 1.7e-07 | 1.900E-07 | 1.732E-08 | 1.700E-07 | 2.000E-07 |
| 16791 | 840 | L840L | C | 3.1e-07 | 3e-07 | 3e-07 | 3.033E-07 | 5.774E-09 | 3.000E-07 | 3.100E-07 |
| 16792 | 840 | L840K | C | 1.6e-07 | 2.3e-07 | 1.8e-07 | 1.900E-07 | 3.606E-08 | 1.600E-07 | 2.300E-07 |
| 16793 | 840 | L840M | C | 7.9e-07 | 3.9e-07 | 3.8e-07 | 5.200E-07 | 2.339E-07 | 3.800E-07 | 7.900E-07 |
| 16794 | 840 | L840F | C | 2.7e-07 | 9.7e-08 | 3.6e-07 | 2.423E-07 | 1.337E-07 | 9.700E-08 | 3.600E-07 |
| 16795 | 840 | L840P | C | 3.8e-07 | 3.2e-07 | 2e-07 | 3.000E-07 | 9.165E-08 | 2.000E-07 | 3.800E-07 |
| 16796 | 840 | L840S | C | 1.5e-07 | 4.2e-07 | 1.7e-07 | 2.467E-07 | 1.504E-07 | 1.500E-07 | 4.200E-07 |
| 16797 | 840 | L840T | C | 2.3e-07 | 1e-07 | 2.9e-07 | 2.067E-07 | 9.713E-08 | 1.000E-07 | 2.900E-07 |
| 16798 | 840 | L840W | C | 1.9e-07 | 3.4e-07 | 4.6e-07 | 3.300E-07 | 1.353E-07 | 1.900E-07 | 4.600E-07 |
| 16799 | 840 | L840Y | C | 2.1e-07 | 5.7e-07 | 2e-07 | 3.267E-07 | 2.108E-07 | 2.000E-07 | 5.700E-07 |
| 16800 | 840 | L840V | C | 1.3e-07 | 2.6e-07 | 5.5e-07 | 3.133E-07 | 2.150E-07 | 1.300E-07 | 5.500E-07 |
| 16801 | 841 | V841A | C | 2.6e-07 | 3e-07 | 2.5e-07 | 2.700E-07 | 2.646E-08 | 2.500E-07 | 3.000E-07 |
| 16802 | 841 | V841R | C | 2.9e-07 | 3.3e-07 | 3.4e-07 | 3.200E-07 | 2.646E-08 | 2.900E-07 | 3.400E-07 |
| 16803 | 841 | V841N | C | 1.9e-07 | 4e-07 | 1.7e-07 | 2.533E-07 | 1.274E-07 | 1.700E-07 | 4.000E-07 |
| 16804 | 841 | V841D | C | 1.9e-07 | 2.4e-07 | 1.1e-07 | 1.800E-07 | 6.557E-08 | 1.100E-07 | 2.400E-07 |
| 16805 | 841 | V841C | C | 2.9e-07 | 1.9e-07 | 2.2e-07 | 2.333E-07 | 5.132E-08 | 1.900E-07 | 2.900E-07 |
| 16806 | 841 | V841E | C | 1.7e-07 | 1.8e-07 | 2.8e-07 | 2.100E-07 | 6.083E-08 | 1.700E-07 | 2.800E-07 |
| 16807 | 841 | V841Q | C | 1.8e-07 | 3.2e-07 | 2.9e-07 | 2.633E-07 | 7.371E-08 | 1.800E-07 | 3.200E-07 |
| 16808 | 841 | V841G | C | 1.6e-07 | 2.2e-07 | 2.1e-07 | 1.967E-07 | 3.215E-08 | 1.600E-07 | 2.200E-07 |
| 16809 | 841 | V841H | C | 7.4e-08 | 3.2e-07 | 1.2e-07 | 1.713E-07 | 1.308E-07 | 7.400E-08 | 3.200E-07 |
| 16810 | 841 | V841I | C | 1.4e-07 | 3.6e-07 | 2.4e-07 | 2.467E-07 | 1.102E-07 | 1.400E-07 | 3.600E-07 |
| 16811 | 841 | V841L | C | 1.2e-07 | 4.1e-07 | 2e-07 | 2.433E-07 | 1.498E-07 | 1.200E-07 | 4.100E-07 |
| 16812 | 841 | V841K | C | 1.5e-07 | 2.5e-07 | 2.7e-07 | 2.233E-07 | 6.429E-08 | 1.500E-07 | 2.700E-07 |
| 16813 | 841 | V841M | C | 2.6e-07 | 2e-07 | 1.8e-07 | 2.133E-07 | 4.163E-08 | 1.800E-07 | 2.600E-07 |
| 16814 | 841 | V841F | C | 2.4e-07 | 2.5e-07 | 5.4e-07 | 3.433E-07 | 1.704E-07 | 2.400E-07 | 5.400E-07 |
| 16815 | 841 | V841P | C | 8.5e-07 | 1.9e-07 | 3.3e-07 | 4.567E-07 | 3.478E-07 | 1.900E-07 | 8.500E-07 |
| 16816 | 841 | V841S | C | 2.1e-07 | 2.1e-07 | 1.8e-07 | 2.000E-07 | 1.732E-08 | 1.800E-07 | 2.100E-07 |
| 16817 | 841 | V841T | C | 3.2e-07 | 3.2e-07 | 2e-07 | 2.800E-07 | 6.928E-08 | 2.000E-07 | 3.200E-07 |
| 16818 | 841 | V841W | C | 1.7e-07 | 2.6e-07 | 3e-07 | 2.433E-07 | 6.658E-08 | 1.700E-07 | 3.000E-07 |
| 16819 | 841 | V841Y | C | 2.4e-07 | 1.5e-07 | 3.2e-07 | 2.367E-07 | 8.505E-08 | 1.500E-07 | 3.200E-07 |
| 16820 | 841 | V841V | C | 3.3e-07 | 2.4e-07 | 3.5e-07 | 3.067E-07 | 5.859E-08 | 2.400E-07 | 3.500E-07 |
| 16821 | 842 | E842A | C | 3.9e-07 | 4e-07 | 3e-07 | 3.633E-07 | 5.508E-08 | 3.000E-07 | 4.000E-07 |
| 16822 | 842 | E842R | C | 2.9e-07 | 3.9e-07 | 1.7e-07 | 2.833E-07 | 1.102E-07 | 1.700E-07 | 3.900E-07 |
| 16823 | 842 | E842N | C | 1.6e-07 | 2.1e-07 | 1.6e-07 | 1.767E-07 | 2.887E-08 | 1.600E-07 | 2.100E-07 |
| 16824 | 842 | E842D | C | 2.8e-07 | 2.7e-07 | 2.5e-07 | 2.667E-07 | 1.528E-08 | 2.500E-07 | 2.800E-07 |
| 16825 | 842 | E842C | C | 3.5e-07 | 2.1e-07 | 4.6e-07 | 3.400E-07 | 1.253E-07 | 2.100E-07 | 4.600E-07 |
| 16826 | 842 | E842E | C | 2.5e-07 | 2e-07 | 6e-07 | 3.500E-07 | 2.179E-07 | 2.000E-07 | 6.000E-07 |
| 16827 | 842 | E842Q | C | 3e-07 | 4.8e-07 | 1.7e-07 | 3.167E-07 | 1.557E-07 | 1.700E-07 | 4.800E-07 |
| 16828 | 842 | E842G | C | 3.8e-07 | 3.2e-07 | 2.1e-07 | 3.033E-07 | 8.622E-08 | 2.100E-07 | 3.800E-07 |
| 16829 | 842 | E842H | C | 2.7e-07 | 2.1e-07 | 1.4e-07 | 2.067E-07 | 6.506E-08 | 1.400E-07 | 2.700E-07 |

| | | | | | | | | | | |
|---|---|---|---|---|---|---|---|---|---|---|
| 16830 | 842 | E842I | C | 8.6e-08 | 1.6e-07 | 2.5e-07 | 1.653E-07 | 8.213E-08 | 8.600E-08 | 2.500E-07 |
| 16831 | 842 | E842L | C | 1.7e-07 | 2.5e-07 | 2.4e-07 | 2.200E-07 | 4.359E-08 | 1.700E-07 | 2.500E-07 |
| 16832 | 842 | E842K | C | 2.9e-07 | 1.4e-07 | 4.2e-07 | 2.833E-07 | 1.401E-07 | 1.400E-07 | 4.200E-07 |
| 16833 | 842 | E842M | C | 3.9e-07 | 7.8e-07 | 1.1e-07 | 4.267E-07 | 3.365E-07 | 1.100E-07 | 7.800E-07 |
| 16834 | 842 | E842F | C | 2.5e-07 | 1.9e-07 | 4e-07 | 2.800E-07 | 1.082E-07 | 1.900E-07 | 4.000E-07 |
| 16835 | 842 | E842P | C | 3.5e-07 | 4.2e-07 | 2.4e-07 | 3.367E-07 | 9.074E-08 | 2.400E-07 | 4.200E-07 |
| 16836 | 842 | E842S | C | 2e-07 | 3.2e-07 | 2.1e-07 | 2.433E-07 | 6.658E-08 | 2.000E-07 | 3.200E-07 |
| 16837 | 842 | E842T | C | 4.2e-07 | 2.6e-07 | 2.4e-07 | 3.067E-07 | 9.866E-08 | 2.400E-07 | 4.200E-07 |
| 16838 | 842 | E842W | C | 1.8e-07 | 1.6e-07 | 2.2e-07 | 1.867E-07 | 3.055E-08 | 1.600E-07 | 2.200E-07 |
| 16839 | 842 | E842Y | C | 2e-07 | 1.2e-07 | 1.5e-07 | 1.567E-07 | 4.041E-08 | 1.200E-07 | 2.000E-07 |
| 16840 | 842 | E842V | C | 8.5e-08 | 3.5e-07 | 3.2e-07 | 2.517E-07 | 1.451E-07 | 8.500E-08 | 3.500E-07 |
| 16841 | 843 | S843A | C | 2.4e-07 | 2.9e-07 | 2.9e-07 | 2.733E-07 | 2.887E-08 | 2.400E-07 | 2.900E-07 |
| 16842 | 843 | S843R | C | 3e-07 | 2.2e-07 | 2.2e-07 | 2.467E-07 | 4.619E-08 | 2.200E-07 | 3.000E-07 |
| 16843 | 843 | S843N | C | 1.1e-07 | 3e-07 | 3.3e-07 | 2.467E-07 | 1.193E-07 | 1.100E-07 | 3.300E-07 |
| 16844 | 843 | S843D | C | 2.2e-07 | 2.2e-07 | 2.2e-07 | 2.200E-07 | 0.000E+00 | 2.200E-07 | 2.200E-07 |
| 16845 | 843 | S843C | C | 1.6e-07 | 2e-07 | 2.2e-07 | 1.933E-07 | 3.055E-08 | 1.600E-07 | 2.200E-07 |
| 16846 | 843 | S843E | C | 2.6e-07 | 2.4e-07 | 2.4e-07 | 2.467E-07 | 1.155E-08 | 2.400E-07 | 2.600E-07 |
| 16847 | 843 | S843Q | C | 2.8e-07 | 3.3e-07 | 3.3e-07 | 3.133E-07 | 2.887E-08 | 2.800E-07 | 3.300E-07 |
| 16848 | 843 | S843G | C | 2.6e-07 | 1.2e-07 | 2.7e-07 | 2.167E-07 | 8.386E-08 | 1.200E-07 | 2.700E-07 |
| 16849 | 843 | S843H | C | 2.8e-07 | 6e-08 | 2.3e-07 | 1.900E-07 | 1.153E-07 | 6.000E-08 | 2.800E-07 |
| 16850 | 843 | S843I | C | 2.6e-07 | 5.7e-07 | 2.9e-07 | 3.733E-07 | 1.710E-07 | 2.600E-07 | 5.700E-07 |
| 16851 | 843 | S843L | C | 2.7e-07 | 2e-07 | 3.3e-07 | 2.667E-07 | 6.506E-08 | 2.000E-07 | 3.300E-07 |
| 16852 | 843 | S843K | C | 2.7e-07 | 2e-07 | 2.7e-07 | 2.467E-07 | 4.041E-08 | 2.000E-07 | 2.700E-07 |
| 16853 | 843 | S843M | C | 6.9e-07 | 4.2e-07 | 2e-07 | 4.367E-07 | 2.454E-07 | 2.000E-07 | 6.900E-07 |
| 16854 | 843 | S843F | C | 2.1e-07 | 2.3e-07 | 1.4e-07 | 1.933E-07 | 4.726E-08 | 1.400E-07 | 2.300E-07 |
| 16855 | 843 | S843P | C | 2.7e-07 | 1.5e-07 | 1.7e-07 | 1.967E-07 | 6.429E-08 | 1.500E-07 | 2.700E-07 |
| 16856 | 843 | S843S | C | 3.1e-07 | 2.4e-07 | 1.8e-07 | 2.433E-07 | 6.506E-08 | 1.800E-07 | 3.100E-07 |
| 16857 | 843 | S843T | C | 1.9e-07 | 1.9e-07 | 2e-07 | 1.933E-07 | 5.774E-09 | 1.900E-07 | 2.000E-07 |
| 16858 | 843 | S843W | C | 2.7e-07 | 4e-07 | 2.5e-07 | 3.067E-07 | 8.145E-08 | 2.500E-07 | 4.000E-07 |
| 16859 | 843 | S843Y | C | 4e-07 | 2e-07 | 1.7e-07 | 2.567E-07 | 1.250E-07 | 1.700E-07 | 4.000E-07 |
| 16860 | 843 | S843V | C | 3.9e-07 | 3.9e-07 | 3.9e-07 | 3.900E-07 | 0.000E+00 | 3.900E-07 | 3.900E-07 |
| 16861 | 844 | G844A | C | 8.7e-08 | 3.6e-07 | 3.5e-07 | 2.657E-07 | 1.548E-07 | 8.700E-08 | 3.600E-07 |
| 16862 | 844 | G844R | C | 3.1e-07 | 2.3e-07 | 2.7e-07 | 2.700E-07 | 4.000E-08 | 2.300E-07 | 3.100E-07 |
| 16863 | 844 | G844N | C | 1.2e-07 | 7.8e-07 | 1.9e-07 | 3.633E-07 | 3.625E-07 | 1.200E-07 | 7.800E-07 |
| 16864 | 844 | G844D | C | 1.9e-07 | 1.1e-07 | 2.8e-07 | 1.933E-07 | 8.505E-08 | 1.100E-07 | 2.800E-07 |
| 16865 | 844 | G844C | C | 2e-07 | 2.1e-07 | 1.5e-07 | 1.867E-07 | 3.215E-08 | 1.500E-07 | 2.100E-07 |
| 16866 | 844 | G844E | C | 1.3e-07 | 1.2e-07 | 1.5e-07 | 1.333E-07 | 1.528E-08 | 1.200E-07 | 1.500E-07 |
| 16867 | 844 | G844Q | C | 2e-07 | 3.3e-07 | 1.3e-07 | 2.200E-07 | 1.015E-07 | 1.300E-07 | 3.300E-07 |
| 16868 | 844 | G844G | C | 1.8e-07 | 2.9e-07 | 1.6e-07 | 2.100E-07 | 7.000E-08 | 1.600E-07 | 2.900E-07 |
| 16869 | 844 | G844H | C | 1.6e-07 | 2.9e-07 | 2.8e-07 | 2.433E-07 | 7.234E-08 | 1.600E-07 | 2.900E-07 |
| 16870 | 844 | G844I | C | 3.1e-07 | 1.2e-07 | 2.8e-07 | 2.367E-07 | 1.021E-07 | 1.200E-07 | 3.100E-07 |
| 16871 | 844 | G844L | C | 1.6e-07 | 2.2e-07 | 5.8e-07 | 3.200E-07 | 2.272E-07 | 1.600E-07 | 5.800E-07 |
| 16872 | 844 | G844K | C | 1.8e-07 | 2e-07 | 3.3e-07 | 2.367E-07 | 8.145E-08 | 1.800E-07 | 3.300E-07 |
| 16873 | 844 | G844M | C | 4.2e-07 | 2.4e-07 | 2.1e-07 | 2.900E-07 | 1.136E-07 | 2.100E-07 | 4.200E-07 |
| 16874 | 844 | G844F | C | 2.5e-07 | 1.1e-07 | 4.1e-07 | 2.567E-07 | 1.501E-07 | 1.100E-07 | 4.100E-07 |
| 16875 | 844 | G844P | C | 2.9e-07 | 2.2e-07 | 3.1e-07 | 2.733E-07 | 4.726E-08 | 2.200E-07 | 3.100E-07 |
| 16876 | 844 | G844S | C | 3.8e-07 | 2.7e-07 | 2.9e-07 | 3.133E-07 | 5.859E-08 | 2.700E-07 | 3.800E-07 |
| 16877 | 844 | G844T | C | 1.9e-07 | 2.7e-07 | 2.3e-07 | 2.300E-07 | 4.000E-08 | 1.900E-07 | 2.700E-07 |
| 16878 | 844 | G844W | C | 3.8e-07 | 4.2e-07 | 1.6e-07 | 3.200E-07 | 1.400E-07 | 1.600E-07 | 4.200E-07 |
| 16879 | 844 | G844Y | C | 3.3e-07 | 3.4e-07 | 4.4e-07 | 3.700E-07 | 6.083E-08 | 3.300E-07 | 4.400E-07 |
| 16880 | 844 | G844V | C | 2.6e-07 | 2.3e-07 | 2.3e-07 | 2.400E-07 | 1.732E-08 | 2.300E-07 | 2.600E-07 |
| 16881 | 845 | G845A | C | 1.8e-07 | 4.2e-07 | 2.2e-07 | 2.733E-07 | 1.286E-07 | 1.800E-07 | 4.200E-07 |
| 16882 | 845 | G845R | C | 1.3e-07 | 2.5e-07 | 2.1e-07 | 1.967E-07 | 6.110E-08 | 1.300E-07 | 2.500E-07 |
| 16883 | 845 | G845N | C | 2.7e-07 | 2.5e-07 | 1.3e-07 | 2.167E-07 | 7.572E-08 | 1.300E-07 | 2.700E-07 |
| 16884 | 845 | G845D | C | 3e-07 | 2.2e-07 | 1.3e-07 | 2.167E-07 | 8.505E-08 | 1.300E-07 | 3.000E-07 |
| 16885 | 845 | G845C | C | 2.5e-07 | 2.8e-07 | 8.4e-07 | 4.567E-07 | 3.323E-07 | 2.500E-07 | 8.400E-07 |
| 16886 | 845 | G845E | C | 3.8e-07 | 2.4e-07 | 2.9e-07 | 3.033E-07 | 7.095E-08 | 2.400E-07 | 3.800E-07 |
| 16887 | 845 | G845Q | C | 2.1e-07 | 3.4e-07 | 1.4e-07 | 2.300E-07 | 1.015E-07 | 1.400E-07 | 3.400E-07 |
| 16888 | 845 | G845G | C | 1.1e-07 | 3e-07 | 8.6e-08 | 1.653E-07 | 1.172E-07 | 8.600E-08 | 3.000E-07 |
| 16889 | 845 | G845H | C | 2.1e-07 | 1.7e-07 | 1.6e-07 | 1.800E-07 | 2.646E-08 | 1.600E-07 | 2.100E-07 |
| 16890 | 845 | G845I | C | 3.6e-07 | 1.4e-07 | 2.4e-07 | 2.467E-07 | 1.102E-07 | 1.400E-07 | 3.600E-07 |
| 16891 | 845 | G845L | C | 2.3e-07 | 2.3e-07 | 4.6e-07 | 3.067E-07 | 1.328E-07 | 2.300E-07 | 4.600E-07 |
| 16892 | 845 | G845K | C | 1.4e-07 | 1.8e-07 | 2.9e-07 | 2.033E-07 | 7.767E-08 | 1.400E-07 | 2.900E-07 |
| 16893 | 845 | G845M | C | 3.2e-07 | 1.9e-07 | 2e-07 | 2.367E-07 | 7.234E-08 | 1.900E-07 | 3.200E-07 |
| 16894 | 845 | G845F | C | 2e-07 | 8.8e-08 | 2.5e-07 | 1.793E-07 | 8.295E-08 | 8.800E-08 | 2.500E-07 |
| 16895 | 845 | G845P | C | 2.2e-07 | 1.3e-07 | 4.7e-07 | 2.733E-07 | 1.762E-07 | 1.300E-07 | 4.700E-07 |
| 16896 | 845 | G845S | C | 2.7e-07 | 4.2e-07 | 5.2e-07 | 4.033E-07 | 1.258E-07 | 2.700E-07 | 5.200E-07 |
| 16897 | 845 | G845T | C | 3.2e-07 | 2.1e-07 | 3.1e-07 | 2.800E-07 | 6.083E-08 | 2.100E-07 | 3.200E-07 |
| 16898 | 845 | G845W | C | 1.7e-07 | 1.6e-07 | 3.5e-07 | 2.267E-07 | 1.069E-07 | 1.600E-07 | 3.500E-07 |
| 16899 | 845 | G845Y | C | 3.4e-07 | 3.4e-07 | 3.1e-07 | 3.300E-07 | 1.732E-08 | 3.100E-07 | 3.400E-07 |
| 16900 | 845 | G845V | C | 3.6e-07 | 4.1e-07 | 2.2e-07 | 3.300E-07 | 9.849E-08 | 2.200E-07 | 4.100E-07 |
| 16901 | 846 | G846A | C | 2.5e-07 | 2.4e-07 | 7.8e-07 | 4.233E-07 | 3.089E-07 | 2.400E-07 | 7.800E-07 |
| 16902 | 846 | G846R | C | 2e-07 | 2.7e-07 | 3.3e-07 | 2.667E-07 | 6.506E-08 | 2.000E-07 | 3.300E-07 |
| 16903 | 846 | G846N | C | 2.6e-07 | 2.7e-07 | 2.3e-07 | 2.533E-07 | 2.082E-08 | 2.300E-07 | 2.700E-07 |
| 16904 | 846 | G846D | C | 2.4e-07 | 1.9e-07 | 2.7e-07 | 2.333E-07 | 4.041E-08 | 1.900E-07 | 2.700E-07 |
| 16905 | 846 | G846C | C | 5.1e-07 | 3.2e-07 | 2.2e-07 | 3.500E-07 | 1.473E-07 | 2.200E-07 | 5.100E-07 |
| 16906 | 846 | G846E | C | 1.4e-07 | 2e-07 | 4.3e-07 | 2.567E-07 | 1.531E-07 | 1.400E-07 | 4.300E-07 |
| 16907 | 846 | G846Q | C | 2.5e-07 | 2.3e-07 | 2.7e-07 | 2.500E-07 | 2.000E-08 | 2.300E-07 | 2.700E-07 |
| 16908 | 846 | G846G | C | 2.3e-07 | 3.5e-07 | 3.1e-07 | 2.967E-07 | 6.110E-08 | 2.300E-07 | 3.500E-07 |
| 16909 | 846 | G846H | C | 3.5e-07 | 2.3e-07 | 1.9e-07 | 2.567E-07 | 8.327E-08 | 1.900E-07 | 3.500E-07 |
| 16910 | 846 | G846I | C | 5.1e-07 | 2.7e-07 | 2.5e-07 | 3.433E-07 | 1.447E-07 | 2.500E-07 | 5.100E-07 |
| 16911 | 846 | G846L | C | 1.5e-07 | 7.6e-07 | 1.3e-07 | 3.467E-07 | 3.581E-07 | 1.300E-07 | 7.600E-07 |
| 16912 | 846 | G846K | C | 2.7e-07 | 2.1e-07 | 3e-07 | 2.600E-07 | 4.583E-08 | 2.100E-07 | 3.000E-07 |
| 16913 | 846 | G846M | C | 2.4e-07 | 2.6e-07 | 1.6e-07 | 2.200E-07 | 5.292E-08 | 1.600E-07 | 2.600E-07 |
| 16914 | 846 | G846F | C | 4.7e-07 | 1.5e-07 | 2.3e-07 | 2.833E-07 | 1.665E-07 | 1.500E-07 | 4.700E-07 |

| | | | | | | | | | | |
|---|---|---|---|---|---|---|---|---|---|---|
| 16915 | 846 | G846P | C | 2.2e-07 | 2.8e-07 | 1.4e-07 | 2.133E-07 | 7.024E-08 | 1.400E-07 | 2.800E-07 |
| 16916 | 846 | G846S | C | 1.4e-07 | 1.6e-07 | 4.2e-07 | 2.400E-07 | 1.562E-07 | 1.400E-07 | 4.200E-07 |
| 16917 | 846 | G846T | C | 1.7e-07 | 1.3e-07 | 2.7e-07 | 1.900E-07 | 7.211E-08 | 1.300E-07 | 2.700E-07 |
| 16918 | 846 | G846W | C | 2.8e-07 | 2.4e-07 | 3.8e-07 | 3.000E-07 | 7.211E-08 | 2.400E-07 | 3.800E-07 |
| 16919 | 846 | G846Y | C | 3.4e-07 | 1.8e-07 | 2.1e-07 | 2.433E-07 | 8.505E-08 | 1.800E-07 | 3.400E-07 |
| 16920 | 846 | G846V | C | 3e-07 | 1.5e-07 | 2.3e-07 | 2.267E-07 | 7.506E-08 | 1.500E-07 | 3.000E-07 |
| 16921 | 847 | L847A | C | 1.9e-07 | 3.1e-07 | 2.4e-07 | 2.467E-07 | 6.028E-08 | 1.900E-07 | 3.100E-07 |
| 16922 | 847 | L847R | C | 2.7e-07 | 4.1e-07 | 2.2e-07 | 3.000E-07 | 9.849E-08 | 2.200E-07 | 4.100E-07 |
| 16923 | 847 | L847N | C | 2.9e-07 | 3.3e-07 | 1.3e-07 | 2.500E-07 | 1.058E-07 | 1.300E-07 | 3.300E-07 |
| 16924 | 847 | L847D | C | 1.7e-07 | 1.9e-07 | 1.4e-07 | 1.667E-07 | 2.517E-08 | 1.400E-07 | 1.900E-07 |
| 16925 | 847 | L847C | C | 4.4e-07 | 9.7e-08 | 3.5e-07 | 2.957E-07 | 1.778E-07 | 9.700E-08 | 4.400E-07 |
| 16926 | 847 | L847E | C | 1e-07 | 3.8e-07 | 1.8e-07 | 2.200E-07 | 1.442E-07 | 1.000E-07 | 3.800E-07 |
| 16927 | 847 | L847Q | C | 1.5e-07 | 3e-07 | 1.6e-07 | 2.033E-07 | 8.386E-08 | 1.500E-07 | 3.000E-07 |
| 16928 | 847 | L847G | C | 3.7e-07 | 2.6e-07 | 4.6e-07 | 3.633E-07 | 1.002E-07 | 2.600E-07 | 4.600E-07 |
| 16929 | 847 | L847H | C | 2.4e-07 | 1.2e-07 | 4.2e-07 | 2.600E-07 | 1.510E-07 | 1.200E-07 | 4.200E-07 |
| 16930 | 847 | L847I | C | 5.7e-07 | 2.4e-07 | 2.9e-07 | 3.667E-07 | 1.779E-07 | 2.400E-07 | 5.700E-07 |
| 16931 | 847 | L847L | C | 1.1e-07 | 3.3e-07 | 3.2e-07 | 2.533E-07 | 1.242E-07 | 1.100E-07 | 3.300E-07 |
| 16932 | 847 | L847K | C | 1.5e-07 | 3.1e-07 | 4.9e-07 | 3.167E-07 | 1.701E-07 | 1.500E-07 | 4.900E-07 |
| 16933 | 847 | L847M | C | 4.2e-07 | 3.3e-07 | 4e-07 | 3.833E-07 | 4.726E-08 | 3.300E-07 | 4.200E-07 |
| 16934 | 847 | L847F | C | 3e-07 | 5.4e-07 | 3.3e-07 | 3.900E-07 | 1.308E-07 | 3.000E-07 | 5.400E-07 |
| 16935 | 847 | L847P | C | 1.4e-07 | 3.3e-07 | 2.6e-07 | 2.433E-07 | 9.609E-08 | 1.400E-07 | 3.300E-07 |
| 16936 | 847 | L847S | C | 3e-07 | 4.4e-07 | 3e-07 | 3.467E-07 | 8.083E-08 | 3.000E-07 | 4.400E-07 |
| 16937 | 847 | L847T | C | 5.2e-07 | 3.8e-07 | 2.8e-07 | 3.933E-07 | 1.206E-07 | 2.800E-07 | 5.200E-07 |
| 16938 | 847 | L847W | C | 2.3e-07 | 3.8e-07 | 3.6e-07 | 3.233E-07 | 8.145E-08 | 2.300E-07 | 3.800E-07 |
| 16939 | 847 | L847Y | C | 1.3e-07 | 3.3e-07 | 3.6e-07 | 2.733E-07 | 1.250E-07 | 1.300E-07 | 3.600E-07 |
| 16940 | 847 | L847V | C | 2.8e-07 | 1.8e-07 | 5.4e-07 | 3.333E-07 | 1.858E-07 | 1.800E-07 | 5.400E-07 |
| 16941 | 848 | V848A | C | 1.1e-07 | 5.8e-07 | 1.6e-07 | 2.833E-07 | 2.581E-07 | 1.100E-07 | 5.800E-07 |
| 16942 | 848 | V848R | C | 2.3e-07 | 1.4e-07 | 2.4e-07 | 2.033E-07 | 5.508E-08 | 1.400E-07 | 2.400E-07 |
| 16943 | 848 | V848N | C | 2.7e-07 | 8.6e-08 | 2e-07 | 1.853E-07 | 9.287E-08 | 8.600E-08 | 2.700E-07 |
| 16944 | 848 | V848D | C | 5.7e-07 | 3.2e-07 | 3.3e-07 | 4.067E-07 | 1.415E-07 | 3.200E-07 | 5.700E-07 |
| 16945 | 848 | V848C | C | 3.3e-07 | 3e-07 | 1.6e-07 | 2.633E-07 | 9.074E-08 | 1.600E-07 | 3.300E-07 |
| 16946 | 848 | V848E | C | 3e-07 | 4.1e-07 | 3.6e-07 | 3.567E-07 | 5.508E-08 | 3.000E-07 | 4.100E-07 |
| 16947 | 848 | V848Q | C | 2.5e-07 | 3.4e-07 | 2.6e-07 | 2.833E-07 | 4.933E-08 | 2.500E-07 | 3.400E-07 |
| 16948 | 848 | V848G | C | 2e-07 | 2.4e-07 | 2.7e-07 | 2.367E-07 | 3.512E-08 | 2.000E-07 | 2.700E-07 |
| 16949 | 848 | V848H | C | 5.9e-08 | 2.7e-07 | 3e-07 | 2.097E-07 | 1.313E-07 | 5.900E-08 | 3.000E-07 |
| 16950 | 848 | V848I | C | 2.1e-07 | 2.4e-07 | 3.5e-07 | 2.667E-07 | 7.371E-08 | 2.100E-07 | 3.500E-07 |
| 16951 | 848 | V848L | C | 2.7e-07 | 3.1e-07 | 2e-07 | 2.600E-07 | 5.568E-08 | 2.000E-07 | 3.100E-07 |
| 16952 | 848 | V848K | C | 2.3e-07 | 1.9e-07 | 1.8e-07 | 2.000E-07 | 2.646E-08 | 1.800E-07 | 2.300E-07 |
| 16953 | 848 | V848M | C | 2.8e-07 | 4.8e-07 | 1.8e-07 | 3.133E-07 | 1.528E-07 | 1.800E-07 | 4.800E-07 |
| 16954 | 848 | V848F | C | 1.8e-07 | 4.4e-07 | 3e-07 | 3.067E-07 | 1.301E-07 | 1.800E-07 | 4.400E-07 |
| 16955 | 848 | V848P | C | 7.7e-07 | 2.5e-07 | 2.9e-07 | 4.367E-07 | 2.894E-07 | 2.500E-07 | 7.700E-07 |
| 16956 | 848 | V848S | C | 1.9e-07 | 3e-07 | 2.6e-07 | 2.500E-07 | 5.568E-08 | 1.900E-07 | 3.000E-07 |
| 16957 | 848 | V848T | C | 8.7e-07 | 2.7e-07 | 3e-07 | 4.800E-07 | 3.381E-07 | 2.700E-07 | 8.700E-07 |
| 16958 | 848 | V848W | C | 1.5e-07 | 3.5e-07 | 2.7e-07 | 2.567E-07 | 1.007E-07 | 1.500E-07 | 3.500E-07 |
| 16959 | 848 | V848Y | C | 4e-07 | 2.3e-07 | 6.9e-07 | 4.400E-07 | 2.326E-07 | 2.300E-07 | 6.900E-07 |
| 16960 | 848 | V848V | C | 1.6e-07 | 2.2e-07 | 2.4e-07 | 2.067E-07 | 4.163E-08 | 1.600E-07 | 2.400E-07 |
| 16961 | 849 | Q849A | C | 4e-07 | 3.5e-07 | 3.1e-07 | 3.533E-07 | 4.509E-08 | 3.100E-07 | 4.000E-07 |
| 16962 | 849 | Q849R | C | 1.3e-07 | 3.5e-07 | 1.6e-07 | 2.133E-07 | 1.193E-07 | 1.300E-07 | 3.500E-07 |
| 16963 | 849 | Q849N | C | 4.4e-08 | 1.4e-07 | 7.6e-08 | 8.667E-08 | 4.888E-08 | 4.400E-08 | 1.400E-07 |
| 16964 | 849 | Q849D | C | 3.2e-07 | 2.9e-07 | 4.1e-07 | 3.400E-07 | 6.245E-08 | 2.900E-07 | 4.100E-07 |
| 16965 | 849 | Q849C | C | 3e-07 | 4.1e-07 | 1.6e-07 | 2.900E-07 | 1.253E-07 | 1.600E-07 | 4.100E-07 |
| 16966 | 849 | Q849E | C | 3e-07 | 4.7e-07 | 2.6e-07 | 3.433E-07 | 1.115E-07 | 2.600E-07 | 4.700E-07 |
| 16967 | 849 | Q849Q | C | 1.9e-07 | 1.9e-07 | 2.3e-07 | 2.033E-07 | 2.309E-08 | 1.900E-07 | 2.300E-07 |
| 16968 | 849 | Q849G | C | 3.4e-07 | 3.9e-07 | 3.9e-07 | 3.733E-07 | 2.887E-08 | 3.400E-07 | 3.900E-07 |
| 16969 | 849 | Q849H | C | 1.4e-07 | 3.5e-07 | 1.1e-07 | 2.000E-07 | 1.308E-07 | 1.100E-07 | 3.500E-07 |
| 16970 | 849 | Q849I | C | 5.8e-07 | 2.9e-07 | 3.4e-07 | 4.033E-07 | 1.550E-07 | 2.900E-07 | 5.800E-07 |
| 16971 | 849 | Q849L | C | 2.3e-07 | 5.7e-07 | 2.7e-07 | 3.567E-07 | 1.858E-07 | 2.300E-07 | 5.700E-07 |
| 16972 | 849 | Q849K | C | 1.6e-07 | 2.7e-07 | 4e-07 | 2.767E-07 | 1.201E-07 | 1.600E-07 | 4.000E-07 |
| 16973 | 849 | Q849M | C | 1.5e-07 | 3.8e-07 | 4.1e-07 | 3.133E-07 | 1.422E-07 | 1.500E-07 | 4.100E-07 |
| 16974 | 849 | Q849F | C | 1.5e-07 | 1.4e-07 | 4.2e-07 | 2.367E-07 | 1.589E-07 | 1.400E-07 | 4.200E-07 |
| 16975 | 849 | Q849P | C | 1.9e-07 | 1.9e-07 | 2.8e-07 | 2.200E-07 | 5.196E-08 | 1.900E-07 | 2.800E-07 |
| 16976 | 849 | Q849S | C | 2.3e-07 | 2.3e-07 | 2.3e-07 | 2.300E-07 | 0.000E+00 | 2.300E-07 | 2.300E-07 |
| 16977 | 849 | Q849T | C | 1.3e-07 | 1.6e-07 | 3e-07 | 1.967E-07 | 9.074E-08 | 1.300E-07 | 3.000E-07 |
| 16978 | 849 | Q849W | C | 2.3e-07 | 2.2e-07 | 1.9e-07 | 2.133E-07 | 2.082E-08 | 1.900E-07 | 2.300E-07 |
| 16979 | 849 | Q849Y | C | 3.3e-07 | 3.9e-07 | 2.3e-07 | 3.167E-07 | 8.083E-08 | 2.300E-07 | 3.900E-07 |
| 16980 | 849 | Q849V | C | 3.3e-07 | 2.1e-07 | 1.1e-07 | 2.167E-07 | 1.102E-07 | 1.100E-07 | 3.300E-07 |
| 16981 | 850 | P850A | C | 1.1e-07 | 1.5e-07 | 1.5e-07 | 1.367E-07 | 2.309E-08 | 1.100E-07 | 1.500E-07 |
| 16982 | 850 | P850R | C | 1.6e-07 | 1.3e-07 | 3.6e-07 | 2.167E-07 | 1.250E-07 | 1.300E-07 | 3.600E-07 |
| 16983 | 850 | P850N | C | 2.7e-07 | 1.8e-07 | 4.1e-07 | 2.867E-07 | 1.159E-07 | 1.800E-07 | 4.100E-07 |
| 16984 | 850 | P850D | C | 3.9e-07 | 2.4e-07 | 3.8e-07 | 3.367E-07 | 8.386E-08 | 2.400E-07 | 3.900E-07 |
| 16985 | 850 | P850C | C | 2.1e-07 | 8.6e-08 | 9.5e-08 | 1.303E-07 | 6.914E-08 | 8.600E-08 | 2.100E-07 |
| 16986 | 850 | P850E | C | 9.5e-08 | 3.5e-07 | 4.3e-07 | 2.917E-07 | 1.750E-07 | 9.500E-08 | 4.300E-07 |
| 16987 | 850 | P850Q | C | 1.4e-07 | 2e-07 | 1.7e-07 | 1.700E-07 | 3.000E-08 | 1.400E-07 | 2.000E-07 |
| 16988 | 850 | P850G | C | 2.1e-07 | 2.6e-07 | 2.9e-07 | 2.533E-07 | 4.041E-08 | 2.100E-07 | 2.900E-07 |
| 16989 | 850 | P850H | C | 4.6e-07 | 1.2e-07 | 3.1e-07 | 2.967E-07 | 1.704E-07 | 1.200E-07 | 4.600E-07 |
| 16990 | 850 | P850I | C | 3.1e-07 | 2.7e-07 | 2.9e-07 | 2.900E-07 | 2.000E-08 | 2.700E-07 | 3.100E-07 |
| 16991 | 850 | P850L | C | 2.4e-07 | 2.8e-07 | 5.9e-07 | 3.700E-07 | 1.916E-07 | 2.400E-07 | 5.900E-07 |
| 16992 | 850 | P850K | C | 1.1e-07 | 3e-07 | 2e-07 | 2.033E-07 | 9.504E-08 | 1.100E-07 | 3.000E-07 |
| 16993 | 850 | P850M | C | 4.7e-07 | 2.4e-07 | 1.7e-07 | 2.933E-07 | 1.570E-07 | 1.700E-07 | 4.700E-07 |
| 16994 | 850 | P850F | C | 3.1e-07 | 2.2e-07 | 2.8e-07 | 2.700E-07 | 4.583E-08 | 2.200E-07 | 3.100E-07 |
| 16995 | 850 | P850P | C | 1e-07 | 5.3e-07 | 2.6e-07 | 2.967E-07 | 2.173E-07 | 1.000E-07 | 5.300E-07 |
| 16996 | 850 | P850S | C | 1.6e-07 | 4e-07 | 1.3e-07 | 2.300E-07 | 1.480E-07 | 1.300E-07 | 4.000E-07 |
| 16997 | 850 | P850T | C | 2.1e-07 | 1.9e-07 | 3.4e-07 | 2.467E-07 | 8.145E-08 | 1.900E-07 | 3.400E-07 |
| 16998 | 850 | P850W | C | 1.5e-07 | 5.5e-07 | 2.7e-07 | 3.233E-07 | 2.053E-07 | 1.500E-07 | 5.500E-07 |
| 16999 | 850 | P850Y | C | 7.2e-07 | 1.7e-07 | 2e-07 | 3.633E-07 | 3.092E-07 | 1.700E-07 | 7.200E-07 |

| | | | | | | | | | | |
|---|---|---|---|---|---|---|---|---|---|---|
| 17000 | 850 | P850V | C | 3.6e-07 | 2.8e-07 | 2.6e-07 | 3.000E-07 | 5.292E-08 | 2.600E-07 | 3.600E-07 |
| 17001 | 851 | G851A | C | 9.1e-08 | 2.7e-07 | 3.1e-07 | 2.237E-07 | 1.166E-07 | 9.100E-08 | 3.100E-07 |
| 17002 | 851 | G851R | C | 7.8e-07 | 1.8e-07 | 1.9e-07 | 3.833E-07 | 3.436E-07 | 1.800E-07 | 7.800E-07 |
| 17003 | 851 | G851N | C | 2.4e-07 | 3.8e-07 | 1.1e-07 | 2.433E-07 | 1.350E-07 | 1.100E-07 | 3.800E-07 |
| 17004 | 851 | G851D | C | 1.6e-07 | 3.6e-07 | 3.9e-07 | 3.033E-07 | 1.250E-07 | 1.600E-07 | 3.900E-07 |
| 17005 | 851 | G851C | C | 2e-07 | 2.3e-07 | 1.4e-07 | 1.900E-07 | 4.583E-08 | 1.400E-07 | 2.300E-07 |
| 17006 | 851 | G851E | C | 4.5e-07 | 4.7e-07 | 1.8e-07 | 3.667E-07 | 1.620E-07 | 1.800E-07 | 4.700E-07 |
| 17007 | 851 | G851Q | C | 2.4e-07 | 2.4e-07 | 3.5e-07 | 2.767E-07 | 6.351E-08 | 2.400E-07 | 3.500E-07 |
| 17008 | 851 | G851G | C | 1.7e-07 | 2.5e-07 | 2.8e-07 | 2.333E-07 | 5.686E-08 | 1.700E-07 | 2.800E-07 |
| 17009 | 851 | G851H | C | 2.2e-07 | 1.9e-07 | 3.7e-07 | 2.600E-07 | 9.644E-08 | 1.900E-07 | 3.700E-07 |
| 17010 | 851 | G851I | C | 2.9e-07 | 4.2e-07 | 1.2e-07 | 2.767E-07 | 1.504E-07 | 1.200E-07 | 4.200E-07 |
| 17011 | 851 | G851L | C | 3.7e-07 | 2.1e-07 | 2.4e-07 | 2.733E-07 | 8.505E-08 | 2.100E-07 | 3.700E-07 |
| 17012 | 851 | G851K | C | 4.5e-07 | 1.5e-07 | 2.6e-07 | 2.867E-07 | 1.518E-07 | 1.500E-07 | 4.500E-07 |
| 17013 | 851 | G851M | C | 3e-07 | 4e-07 | 4.7e-07 | 3.900E-07 | 8.544E-08 | 3.000E-07 | 4.700E-07 |
| 17014 | 851 | G851F | C | 2.1e-07 | 1.7e-07 | 5.5e-07 | 3.100E-07 | 2.088E-07 | 1.700E-07 | 5.500E-07 |
| 17015 | 851 | G851P | C | 4.2e-07 | 4.8e-07 | 2.5e-07 | 3.833E-07 | 1.193E-07 | 2.500E-07 | 4.800E-07 |
| 17016 | 851 | G851S | C | 3e-07 | 1.9e-07 | 2.1e-07 | 2.333E-07 | 5.859E-08 | 1.900E-07 | 3.000E-07 |
| 17017 | 851 | G851T | C | 2e-07 | 3e-07 | 1.6e-07 | 2.200E-07 | 7.211E-08 | 1.600E-07 | 3.000E-07 |
| 17018 | 851 | G851W | C | 1.5e-07 | 2.8e-07 | 2e-07 | 2.100E-07 | 6.557E-08 | 1.500E-07 | 2.800E-07 |
| 17019 | 851 | G851Y | C | 2.3e-07 | 2e-07 | 1.4e-07 | 1.900E-07 | 4.583E-08 | 1.400E-07 | 2.300E-07 |
| 17020 | 851 | G851V | C | 1.8e-07 | 4e-07 | 3e-07 | 2.933E-07 | 1.102E-07 | 1.800E-07 | 4.000E-07 |
| 17021 | 852 | G852A | C | 3.8e-07 | 4e-07 | 4.6e-07 | 4.133E-07 | 4.163E-08 | 3.800E-07 | 4.600E-07 |
| 17022 | 852 | G852R | C | 3.9e-07 | 2.4e-07 | 1.6e-07 | 2.633E-07 | 1.168E-07 | 1.600E-07 | 3.900E-07 |
| 17023 | 852 | G852N | C | 1.1e-07 | 2.6e-07 | 1e-07 | 1.567E-07 | 8.963E-08 | 1.000E-07 | 2.600E-07 |
| 17024 | 852 | G852D | C | 2.1e-07 | 2.3e-07 | 2.1e-07 | 2.167E-07 | 1.155E-08 | 2.100E-07 | 2.300E-07 |
| 17025 | 852 | G852C | C | 4.3e-07 | 1.6e-07 | 3.1e-07 | 3.000E-07 | 1.353E-07 | 1.600E-07 | 4.300E-07 |
| 17026 | 852 | G852E | C | 2.1e-07 | 6.2e-07 | 2.5e-07 | 3.600E-07 | 2.261E-07 | 2.100E-07 | 6.200E-07 |
| 17027 | 852 | G852Q | C | 3.1e-07 | 3.2e-07 | 4.4e-07 | 3.567E-07 | 7.234E-08 | 3.100E-07 | 4.400E-07 |
| 17028 | 852 | G852G | C | 2.3e-07 | 2.4e-07 | 3.5e-07 | 2.733E-07 | 6.658E-08 | 2.300E-07 | 3.500E-07 |
| 17029 | 852 | G852H | C | 5e-07 | 4.4e-07 | 2.1e-07 | 3.833E-07 | 1.531E-07 | 2.100E-07 | 5.000E-07 |
| 17030 | 852 | G852I | C | 1.2e-07 | 3.7e-07 | 1.3e-07 | 2.067E-07 | 1.415E-07 | 1.200E-07 | 3.700E-07 |
| 17031 | 852 | G852L | C | 5.4e-07 | 3.6e-07 | 2.3e-07 | 3.767E-07 | 1.557E-07 | 2.300E-07 | 5.400E-07 |
| 17032 | 852 | G852K | C | 3.2e-07 | 4.4e-07 | 2.9e-07 | 3.500E-07 | 7.937E-08 | 2.900E-07 | 4.400E-07 |
| 17033 | 852 | G852M | C | 2.3e-07 | 6.1e-07 | 2.1e-07 | 3.500E-07 | 2.254E-07 | 2.100E-07 | 6.100E-07 |
| 17034 | 852 | G852F | C | 1.7e-07 | 1.9e-07 | 2.1e-07 | 1.900E-07 | 2.000E-08 | 1.700E-07 | 2.100E-07 |
| 17035 | 852 | G852P | C | 2.2e-07 | 3.4e-07 | 2.3e-07 | 2.633E-07 | 6.658E-08 | 2.200E-07 | 3.400E-07 |
| 17036 | 852 | G852S | C | 8.4e-08 | 2.5e-07 | 3.5e-07 | 2.280E-07 | 1.344E-07 | 8.400E-08 | 3.500E-07 |
| 17037 | 852 | G852T | C | 2.9e-07 | 3.9e-07 | 4.4e-07 | 3.733E-07 | 7.638E-08 | 2.900E-07 | 4.400E-07 |
| 17038 | 852 | G852W | C | 1.4e-07 | 9.4e-08 | 2.6e-07 | 1.647E-07 | 8.570E-08 | 9.400E-08 | 2.600E-07 |
| 17039 | 852 | G852Y | C | 1.5e-07 | 1.3e-07 | 1.2e-07 | 1.333E-07 | 1.528E-08 | 1.200E-07 | 1.500E-07 |
| 17040 | 852 | G852V | C | 3.8e-07 | 2.8e-07 | 1.5e-07 | 2.700E-07 | 1.153E-07 | 1.500E-07 | 3.800E-07 |
| 17041 | 853 | S853A | C | 8.1e-07 | 2.4e-07 | 2.3e-07 | 4.267E-07 | 3.320E-07 | 2.300E-07 | 8.100E-07 |
| 17042 | 853 | S853R | C | 2.4e-07 | 3e-07 | 3.4e-07 | 2.933E-07 | 5.033E-08 | 2.400E-07 | 3.400E-07 |
| 17043 | 853 | S853N | C | 3.6e-07 | 1.7e-07 | 2.2e-07 | 2.500E-07 | 9.849E-08 | 1.700E-07 | 3.600E-07 |
| 17044 | 853 | S853D | C | 3e-07 | 4.2e-07 | 2e-07 | 3.067E-07 | 1.102E-07 | 2.000E-07 | 4.200E-07 |
| 17045 | 853 | S853C | C | 4.3e-07 | 5.5e-07 | 2.8e-07 | 4.200E-07 | 1.353E-07 | 2.800E-07 | 5.500E-07 |
| 17046 | 853 | S853E | C | 4.5e-07 | 1.6e-07 | 1.5e-07 | 2.533E-07 | 1.704E-07 | 1.500E-07 | 4.500E-07 |
| 17047 | 853 | S853Q | C | 3.6e-07 | 6e-07 | 3.8e-07 | 4.467E-07 | 1.332E-07 | 3.600E-07 | 6.000E-07 |
| 17048 | 853 | S853G | C | 1.1e-07 | 3.5e-07 | 2.6e-07 | 2.400E-07 | 1.212E-07 | 1.100E-07 | 3.500E-07 |
| 17049 | 853 | S853H | C | 2.9e-07 | 1.8e-07 | 2e-07 | 2.233E-07 | 5.859E-08 | 1.800E-07 | 2.900E-07 |
| 17050 | 853 | S853I | C | 3.4e-07 | 2e-07 | 3.1e-07 | 2.833E-07 | 7.371E-08 | 2.000E-07 | 3.400E-07 |
| 17051 | 853 | S853L | C | 4.3e-07 | 1.9e-07 | 1.7e-07 | 2.633E-07 | 1.447E-07 | 1.700E-07 | 4.300E-07 |
| 17052 | 853 | S853K | C | 2.2e-07 | 2.9e-07 | 3.2e-07 | 2.767E-07 | 5.132E-08 | 2.200E-07 | 3.200E-07 |
| 17053 | 853 | S853M | C | 3.3e-07 | 2.2e-07 | 2.6e-07 | 2.700E-07 | 5.568E-08 | 2.200E-07 | 3.300E-07 |
| 17054 | 853 | S853F | C | 1.6e-07 | 2.1e-07 | 4.5e-07 | 2.733E-07 | 1.550E-07 | 1.600E-07 | 4.500E-07 |
| 17055 | 853 | S853P | C | 3e-07 | 2.3e-07 | 4.3e-07 | 3.200E-07 | 1.015E-07 | 2.300E-07 | 4.300E-07 |
| 17056 | 853 | S853S | C | 3.3e-07 | 3.1e-07 | 3.4e-07 | 3.267E-07 | 1.528E-08 | 3.100E-07 | 3.400E-07 |
| 17057 | 853 | S853T | C | 4.3e-07 | 2.1e-07 | 3.3e-07 | 3.233E-07 | 1.102E-07 | 2.100E-07 | 4.300E-07 |
| 17058 | 853 | S853W | C | 4.2e-07 | 1.9e-07 | 1.5e-07 | 2.533E-07 | 1.457E-07 | 1.500E-07 | 4.200E-07 |
| 17059 | 853 | S853Y | C | 1.5e-07 | 1.9e-07 | 2e-07 | 1.800E-07 | 2.646E-08 | 1.500E-07 | 2.000E-07 |
| 17060 | 853 | S853V | C | 2.3e-07 | 1.6e-07 | 2.4e-07 | 2.100E-07 | 4.359E-08 | 1.600E-07 | 2.400E-07 |
| 17061 | 854 | L854A | C | 1.3e-07 | 6.6e-08 | 1.7e-07 | 1.220E-07 | 5.246E-08 | 6.600E-08 | 1.700E-07 |
| 17062 | 854 | L854R | C | 3.9e-07 | 3.1e-07 | 2.7e-07 | 3.233E-07 | 6.110E-08 | 2.700E-07 | 3.900E-07 |
| 17063 | 854 | L854N | C | 1.6e-07 | 2.4e-07 | 2e-07 | 2.000E-07 | 4.000E-08 | 1.600E-07 | 2.400E-07 |
| 17064 | 854 | L854D | C | 4e-07 | 2.4e-07 | 3.9e-07 | 3.433E-07 | 8.963E-08 | 2.400E-07 | 4.000E-07 |
| 17065 | 854 | L854C | C | 3.7e-07 | 5.6e-07 | 4.4e-07 | 4.567E-07 | 9.609E-08 | 3.700E-07 | 5.600E-07 |
| 17066 | 854 | L854E | C | 4.2e-07 | 1.4e-07 | 1.8e-07 | 2.467E-07 | 1.514E-07 | 1.400E-07 | 4.200E-07 |
| 17067 | 854 | L854Q | C | 1.9e-07 | 1.4e-07 | 3.4e-07 | 2.233E-07 | 1.041E-07 | 1.400E-07 | 3.400E-07 |
| 17068 | 854 | L854G | C | 2.1e-07 | 9.5e-08 | 1.8e-07 | 1.617E-07 | 5.965E-08 | 9.500E-08 | 2.100E-07 |
| 17069 | 854 | L854H | C | 2.5e-07 | 3e-07 | 1.8e-07 | 2.433E-07 | 6.028E-08 | 1.800E-07 | 3.000E-07 |
| 17070 | 854 | L854I | C | 4.9e-07 | 2.9e-07 | 3.5e-07 | 3.767E-07 | 1.026E-07 | 2.900E-07 | 4.900E-07 |
| 17071 | 854 | L854L | C | 4e-07 | 2.5e-07 | 2.8e-07 | 3.100E-07 | 7.937E-08 | 2.500E-07 | 4.000E-07 |
| 17072 | 854 | L854K | C | 1.9e-07 | 2.7e-07 | 2.4e-07 | 2.333E-07 | 4.041E-08 | 1.900E-07 | 2.700E-07 |
| 17073 | 854 | L854M | C | 2.5e-07 | 2.2e-07 | 1.3e-07 | 2.000E-07 | 6.245E-08 | 1.300E-07 | 2.500E-07 |
| 17074 | 854 | L854F | C | 3e-07 | 2.7e-07 | 3.5e-07 | 3.067E-07 | 4.041E-08 | 2.700E-07 | 3.500E-07 |
| 17075 | 854 | L854P | C | 1.4e-07 | 3.8e-07 | 3.5e-07 | 2.900E-07 | 1.308E-07 | 1.400E-07 | 3.800E-07 |
| 17076 | 854 | L854S | C | 2.7e-07 | 3.6e-07 | 4.8e-07 | 3.700E-07 | 1.054E-07 | 2.700E-07 | 4.800E-07 |
| 17077 | 854 | L854T | C | 5.2e-07 | 3.7e-07 | 1.1e-07 | 3.333E-07 | 2.074E-07 | 1.100E-07 | 5.200E-07 |
| 17078 | 854 | L854W | C | 9.1e-08 | 3e-07 | 2.6e-07 | 2.170E-07 | 1.109E-07 | 9.100E-08 | 3.000E-07 |
| 17079 | 854 | L854Y | C | 2.7e-07 | 4e-07 | 1.5e-07 | 2.733E-07 | 1.250E-07 | 1.500E-07 | 4.000E-07 |
| 17080 | 854 | L854V | C | 2.1e-07 | 3.6e-07 | 4.1e-07 | 3.267E-07 | 1.041E-07 | 2.100E-07 | 4.100E-07 |
| 17081 | 855 | R855A | C | 2.9e-07 | 3.6e-07 | 3.2e-07 | 3.233E-07 | 3.512E-08 | 2.900E-07 | 3.600E-07 |
| 17082 | 855 | R855R | C | 2.6e-07 | 2.4e-07 | 3.1e-07 | 2.700E-07 | 3.606E-08 | 2.400E-07 | 3.100E-07 |
| 17083 | 855 | R855N | C | 1.8e-07 | 3.3e-07 | 2.3e-07 | 2.467E-07 | 7.638E-08 | 1.800E-07 | 3.300E-07 |
| 17084 | 855 | R855D | C | 2e-07 | 2.7e-07 | 2.5e-07 | 2.400E-07 | 3.606E-08 | 2.000E-07 | 2.700E-07 |

| | | | | | | | | | | |
|---|---|---|---|---|---|---|---|---|---|---|
| 17085 | 855 | R855C | C | 4.1e-07 | 2.4e-07 | 1.9e-07 | 2.800E-07 | 1.153E-07 | 1.900E-07 | 4.100E-07 |
| 17086 | 855 | R855E | C | 2e-07 | 2.7e-07 | 4.6e-07 | 3.100E-07 | 1.345E-07 | 2.000E-07 | 4.600E-07 |
| 17087 | 855 | R855Q | C | 3.3e-07 | 5.4e-07 | 2.5e-07 | 3.733E-07 | 1.498E-07 | 2.500E-07 | 5.400E-07 |
| 17088 | 855 | R855G | C | 1.2e-07 | 1.9e-07 | 2.3e-07 | 1.800E-07 | 5.568E-08 | 1.200E-07 | 2.300E-07 |
| 17089 | 855 | R855H | C | 3.6e-07 | 2.5e-07 | 2.1e-07 | 2.733E-07 | 7.767E-08 | 2.100E-07 | 3.600E-07 |
| 17090 | 855 | R855I | C | 7.8e-08 | 1.9e-07 | 1.8e-07 | 1.493E-07 | 6.198E-08 | 7.800E-08 | 1.900E-07 |
| 17091 | 855 | R855L | C | 3e-07 | 3e-07 | 1.7e-07 | 2.567E-07 | 7.506E-08 | 1.700E-07 | 3.000E-07 |
| 17092 | 855 | R855K | C | 5.1e-07 | 3.2e-07 | 4.2e-07 | 4.167E-07 | 9.504E-08 | 3.200E-07 | 5.100E-07 |
| 17093 | 855 | R855M | C | 4.7e-07 | 2.6e-07 | 3.3e-07 | 3.533E-07 | 1.069E-07 | 2.600E-07 | 4.700E-07 |
| 17094 | 855 | R855F | C | 3.6e-07 | 1.2e-07 | 9.4e-08 | 1.913E-07 | 1.466E-07 | 9.400E-08 | 3.600E-07 |
| 17095 | 855 | R855P | C | 1.6e-07 | 3.7e-07 | 1.3e-07 | 2.200E-07 | 1.308E-07 | 1.300E-07 | 3.700E-07 |
| 17096 | 855 | R855S | C | 2.7e-07 | 2.4e-07 | 3.7e-07 | 2.933E-07 | 6.807E-08 | 2.400E-07 | 3.700E-07 |
| 17097 | 855 | R855T | C | 1e-07 | 2.5e-07 | 1.9e-07 | 1.800E-07 | 7.550E-08 | 1.000E-07 | 2.500E-07 |
| 17098 | 855 | R855W | C | 1.7e-07 | 2.2e-07 | 3.5e-07 | 2.467E-07 | 9.292E-08 | 1.700E-07 | 3.500E-07 |
| 17099 | 855 | R855Y | C | 1.6e-07 | 1.3e-07 | 6.9e-08 | 1.197E-07 | 4.637E-08 | 6.900E-08 | 1.600E-07 |
| 17100 | 855 | R855V | C | 5.6e-07 | 5.5e-07 | 1.8e-07 | 4.300E-07 | 2.166E-07 | 1.800E-07 | 5.600E-07 |
| 17101 | 856 | L856A | C | 1.9e-07 | 1e-06 | 4.4e-07 | 5.433E-07 | 4.148E-07 | 1.900E-07 | 1.000E-06 |
| 17102 | 856 | L856R | C | 2.8e-07 | 1.6e-07 | 1.1e-07 | 1.833E-07 | 8.737E-08 | 1.100E-07 | 2.800E-07 |
| 17103 | 856 | L856N | C | 3.9e-07 | 3.5e-07 | 2.7e-07 | 3.367E-07 | 6.110E-08 | 2.700E-07 | 3.900E-07 |
| 17104 | 856 | L856D | C | 4.3e-07 | 3.2e-07 | 1.2e-07 | 2.900E-07 | 1.572E-07 | 1.200E-07 | 4.300E-07 |
| 17105 | 856 | L856C | C | 1.8e-07 | 1.1e-07 | 1.4e-07 | 1.433E-07 | 3.512E-08 | 1.100E-07 | 1.800E-07 |
| 17106 | 856 | L856E | C | 1.5e-07 | 5.2e-07 | 2.4e-07 | 3.033E-07 | 1.930E-07 | 1.500E-07 | 5.200E-07 |
| 17107 | 856 | L856Q | C | 1.8e-07 | 2.5e-07 | 3.7e-07 | 2.667E-07 | 9.609E-08 | 1.800E-07 | 3.700E-07 |
| 17108 | 856 | L856G | C | 3.9e-07 | 2.8e-07 | 4.4e-07 | 3.700E-07 | 8.185E-08 | 2.800E-07 | 4.400E-07 |
| 17109 | 856 | L856H | C | 3.2e-07 | 1.8e-07 | 3e-07 | 2.667E-07 | 7.572E-08 | 1.800E-07 | 3.200E-07 |
| 17110 | 856 | L856I | C | 1.7e-07 | 1.5e-07 | 3.9e-07 | 2.367E-07 | 1.332E-07 | 1.500E-07 | 3.900E-07 |
| 17111 | 856 | L856L | C | 2.9e-07 | 2.1e-07 | 5.4e-07 | 3.467E-07 | 1.721E-07 | 2.100E-07 | 5.400E-07 |
| 17112 | 856 | L856K | C | 1.7e-07 | 1.7e-07 | 3.7e-07 | 2.367E-07 | 1.155E-07 | 1.700E-07 | 3.700E-07 |
| 17113 | 856 | L856M | C | 3.1e-07 | 5.9e-07 | 1.9e-07 | 3.633E-07 | 2.053E-07 | 1.900E-07 | 5.900E-07 |
| 17114 | 856 | L856F | C | 2.9e-07 | 1.1e-07 | 2.9e-07 | 2.300E-07 | 1.039E-07 | 1.100E-07 | 2.900E-07 |
| 17115 | 856 | L856P | C | 3.5e-07 | 1.5e-07 | 3.1e-07 | 2.700E-07 | 1.058E-07 | 1.500E-07 | 3.500E-07 |
| 17116 | 856 | L856S | C | 1e-07 | 2e-07 | 2.1e-07 | 1.700E-07 | 6.083E-08 | 1.000E-07 | 2.100E-07 |
| 17117 | 856 | L856T | C | 3e-07 | 3.9e-07 | 4.6e-07 | 3.833E-07 | 8.021E-08 | 3.000E-07 | 4.600E-07 |
| 17118 | 856 | L856W | C | 1.7e-07 | 1.6e-07 | 1.7e-07 | 1.667E-07 | 5.774E-09 | 1.600E-07 | 1.700E-07 |
| 17119 | 856 | L856Y | C | 2.3e-07 | 1.8e-07 | 3e-07 | 2.367E-07 | 6.028E-08 | 1.800E-07 | 3.000E-07 |
| 17120 | 856 | L856V | C | 2e-07 | 2.2e-07 | 1.9e-07 | 2.033E-07 | 1.528E-08 | 1.900E-07 | 2.200E-07 |
| 17121 | 857 | S857A | C | 3.2e-07 | 1.5e-07 | 3.1e-07 | 2.600E-07 | 9.539E-08 | 1.500E-07 | 3.200E-07 |
| 17122 | 857 | S857R | C | 2.3e-07 | 2.6e-07 | 2.4e-07 | 2.433E-07 | 1.528E-08 | 2.300E-07 | 2.600E-07 |
| 17123 | 857 | S857N | C | 3.1e-07 | 2.9e-07 | 3.7e-07 | 3.233E-07 | 4.163E-08 | 2.900E-07 | 3.700E-07 |
| 17124 | 857 | S857D | C | 1.7e-07 | 1.3e-07 | 3.1e-07 | 2.033E-07 | 9.452E-08 | 1.300E-07 | 3.100E-07 |
| 17125 | 857 | S857C | C | 4e-07 | 2e-07 | 2.8e-07 | 2.933E-07 | 1.007E-07 | 2.000E-07 | 4.000E-07 |
| 17126 | 857 | S857E | C | 3.1e-07 | 3.9e-07 | 1.8e-07 | 2.933E-07 | 1.060E-07 | 1.800E-07 | 3.900E-07 |
| 17127 | 857 | S857Q | C | 2e-07 | 1.5e-07 | 1.4e-07 | 1.633E-07 | 3.215E-08 | 1.400E-07 | 2.000E-07 |
| 17128 | 857 | S857G | C | 2.3e-07 | 2.4e-07 | 3.1e-07 | 2.600E-07 | 4.359E-08 | 2.300E-07 | 3.100E-07 |
| 17129 | 857 | S857H | C | 2.7e-07 | 1e-07 | 3.2e-07 | 2.300E-07 | 1.153E-07 | 1.000E-07 | 3.200E-07 |
| 17130 | 857 | S857I | C | 2e-07 | 4e-07 | 1.9e-07 | 2.633E-07 | 1.185E-07 | 1.900E-07 | 4.000E-07 |
| 17131 | 857 | S857L | C | 5.4e-07 | 9.6e-08 | 4.9e-07 | 3.753E-07 | 2.432E-07 | 9.600E-08 | 5.400E-07 |
| 17132 | 857 | S857K | C | 3.1e-07 | 3.1e-07 | 3.1e-07 | 3.100E-07 | 0.000E+00 | 3.100E-07 | 3.100E-07 |
| 17133 | 857 | S857M | C | 3.8e-07 | 1.1e-07 | 2e-07 | 2.300E-07 | 1.375E-07 | 1.100E-07 | 3.800E-07 |
| 17134 | 857 | S857F | C | 2.6e-07 | 1.4e-07 | 2.1e-07 | 2.033E-07 | 6.028E-08 | 1.400E-07 | 2.600E-07 |
| 17135 | 857 | S857P | C | 3.5e-07 | 1.7e-07 | 2.3e-07 | 2.500E-07 | 9.165E-08 | 1.700E-07 | 3.500E-07 |
| 17136 | 857 | S857S | C | 2.1e-07 | 2e-07 | 2.2e-07 | 2.100E-07 | 1.000E-08 | 2.000E-07 | 2.200E-07 |
| 17137 | 857 | S857T | C | 8.4e-08 | 1.9e-07 | 2.9e-07 | 1.880E-07 | 1.030E-07 | 8.400E-08 | 2.900E-07 |
| 17138 | 857 | S857W | C | 2.6e-07 | 3.8e-07 | 4.4e-07 | 3.600E-07 | 9.165E-08 | 2.600E-07 | 4.400E-07 |
| 17139 | 857 | S857Y | C | 2e-07 | 1.8e-07 | 7.3e-07 | 3.700E-07 | 3.119E-07 | 1.800E-07 | 7.300E-07 |
| 17140 | 857 | S857V | C | 1.6e-07 | 2.6e-07 | 2.4e-07 | 2.200E-07 | 5.292E-08 | 1.600E-07 | 2.600E-07 |
| 17141 | 858 | C858A | C | 2.2e-07 | 4.9e-07 | 1.7e-07 | 2.933E-07 | 1.721E-07 | 1.700E-07 | 4.900E-07 |
| 17142 | 858 | C858R | C | 5.2e-07 | 2.4e-07 | 4.5e-07 | 4.033E-07 | 1.457E-07 | 2.400E-07 | 5.200E-07 |
| 17143 | 858 | C858N | C | 4.4e-07 | 2e-07 | 2e-07 | 2.800E-07 | 1.386E-07 | 2.000E-07 | 4.400E-07 |
| 17144 | 858 | C858D | C | 3.6e-07 | 3.7e-07 | 1.4e-07 | 2.900E-07 | 1.300E-07 | 1.400E-07 | 3.700E-07 |
| 17145 | 858 | C858C | C | 2e-07 | 3.2e-07 | 2.9e-07 | 2.700E-07 | 6.245E-08 | 2.000E-07 | 3.200E-07 |
| 17146 | 858 | C858E | C | 1.9e-07 | 3.1e-07 | 3.5e-07 | 2.833E-07 | 8.327E-08 | 1.900E-07 | 3.500E-07 |
| 17147 | 858 | C858Q | C | 1.8e-07 | 1.9e-07 | 2.7e-07 | 2.133E-07 | 4.933E-08 | 1.800E-07 | 2.700E-07 |
| 17148 | 858 | C858G | C | 1.1e-07 | 3.1e-07 | 2e-07 | 2.067E-07 | 1.002E-07 | 1.100E-07 | 3.100E-07 |
| 17149 | 858 | C858H | C | 2.9e-07 | 3e-07 | 1.7e-07 | 2.533E-07 | 7.234E-08 | 1.700E-07 | 3.000E-07 |
| 17150 | 858 | C858I | C | 4.2e-07 | 4.7e-07 | 7.7e-08 | 3.223E-07 | 2.139E-07 | 7.700E-08 | 4.700E-07 |
| 17151 | 858 | C858L | C | 3.2e-07 | 1.9e-07 | 8.2e-08 | 1.973E-07 | 1.192E-07 | 8.200E-08 | 3.200E-07 |
| 17152 | 858 | C858K | C | 2.9e-07 | 2.3e-07 | 1.3e-07 | 2.167E-07 | 8.083E-08 | 1.300E-07 | 2.900E-07 |
| 17153 | 858 | C858M | C | 3.9e-07 | 2.8e-07 | 3e-07 | 3.233E-07 | 5.859E-08 | 2.800E-07 | 3.900E-07 |
| 17154 | 858 | C858F | C | 3.6e-07 | 3e-07 | 3.9e-07 | 3.500E-07 | 4.583E-08 | 3.000E-07 | 3.900E-07 |
| 17155 | 858 | C858P | C | 2.8e-07 | 2.8e-07 | 2.2e-07 | 2.600E-07 | 3.464E-08 | 2.200E-07 | 2.800E-07 |
| 17156 | 858 | C858S | C | 3.1e-07 | 3.5e-07 | 2.5e-07 | 3.033E-07 | 5.033E-08 | 2.500E-07 | 3.500E-07 |
| 17157 | 858 | C858T | C | 3.4e-07 | 1.1e-07 | 3.3e-07 | 2.600E-07 | 1.300E-07 | 1.100E-07 | 3.400E-07 |
| 17158 | 858 | C858W | C | 1.5e-07 | 3.1e-07 | 2.8e-07 | 2.467E-07 | 8.505E-08 | 1.500E-07 | 3.100E-07 |
| 17159 | 858 | C858Y | C | 3.4e-07 | 2.5e-07 | 2.1e-07 | 2.667E-07 | 6.658E-08 | 2.100E-07 | 3.400E-07 |
| 17160 | 858 | C858V | C | 2.4e-07 | 2.7e-07 | 2.2e-07 | 2.433E-07 | 2.517E-08 | 2.200E-07 | 2.700E-07 |
| 17161 | 859 | A859A | C | 1.9e-07 | 1.6e-07 | 2.3e-07 | 1.933E-07 | 3.512E-08 | 1.600E-07 | 2.300E-07 |
| 17162 | 859 | A859R | C | 1.3e-07 | 4.7e-07 | 3.2e-07 | 3.067E-07 | 1.704E-07 | 1.300E-07 | 4.700E-07 |
| 17163 | 859 | A859N | C | 3.2e-07 | 4.2e-07 | 3.1e-07 | 3.500E-07 | 6.083E-08 | 3.100E-07 | 4.200E-07 |
| 17164 | 859 | A859D | C | 3.9e-07 | 2.5e-07 | 6.5e-07 | 4.300E-07 | 2.030E-07 | 2.500E-07 | 6.500E-07 |
| 17165 | 859 | A859C | C | 3.6e-07 | 3.2e-07 | 4.6e-07 | 3.800E-07 | 7.211E-08 | 3.200E-07 | 4.600E-07 |
| 17166 | 859 | A859E | C | 1.2e-07 | 3.4e-07 | 2.1e-07 | 2.233E-07 | 1.106E-07 | 1.200E-07 | 3.400E-07 |
| 17167 | 859 | A859Q | C | 2.2e-07 | 2.3e-07 | 2.1e-07 | 2.200E-07 | 1.000E-08 | 2.100E-07 | 2.300E-07 |
| 17168 | 859 | A859G | C | 2.4e-07 | 2.7e-07 | 2.1e-07 | 2.400E-07 | 3.000E-08 | 2.100E-07 | 2.700E-07 |
| 17169 | 859 | A859H | C | 5.3e-07 | 5.8e-07 | 3.8e-07 | 4.967E-07 | 1.041E-07 | 3.800E-07 | 5.800E-07 |

| | | | | | | | | | | |
|---|---|---|---|---|---|---|---|---|---|---|
| 17170 | 859 | A859I | C | 2.9e-07 | 3.7e-07 | 2.6e-07 | 3.067E-07 | 5.686E-08 | 2.600E-07 | 3.700E-07 |
| 17171 | 859 | A859L | C | 1.6e-07 | 1.7e-07 | 3.9e-07 | 2.400E-07 | 1.300E-07 | 1.600E-07 | 3.900E-07 |
| 17172 | 859 | A859K | C | 3e-07 | 3e-07 | 1.6e-07 | 2.533E-07 | 8.083E-08 | 1.600E-07 | 3.000E-07 |
| 17173 | 859 | A859M | C | 3e-07 | 6.8e-08 | 1.8e-07 | 1.827E-07 | 1.160E-07 | 6.800E-08 | 3.000E-07 |
| 17174 | 859 | A859F | C | 3.5e-07 | 1.3e-07 | 2.4e-07 | 2.400E-07 | 1.100E-07 | 1.300E-07 | 3.500E-07 |
| 17175 | 859 | A859P | C | 1.7e-07 | 2.2e-07 | 3.4e-07 | 2.433E-07 | 8.737E-08 | 1.700E-07 | 3.400E-07 |
| 17176 | 859 | A859S | C | 1.2e-07 | 2.2e-07 | 2e-07 | 1.800E-07 | 5.292E-08 | 1.200E-07 | 2.200E-07 |
| 17177 | 859 | A859T | C | 1.5e-07 | 3.3e-07 | 3.7e-07 | 2.833E-07 | 1.172E-07 | 1.500E-07 | 3.700E-07 |
| 17178 | 859 | A859W | C | 2.6e-07 | 2.3e-07 | 3.8e-07 | 2.900E-07 | 7.937E-08 | 2.300E-07 | 3.800E-07 |
| 17179 | 859 | A859Y | C | 3.1e-07 | 1.6e-07 | 6.3e-07 | 3.667E-07 | 2.401E-07 | 1.600E-07 | 6.300E-07 |
| 17180 | 859 | A859V | C | 4.7e-07 | 3.4e-07 | 1.7e-07 | 3.267E-07 | 1.504E-07 | 1.700E-07 | 4.700E-07 |
| 17181 | 860 | A860A | C | 2.7e-07 | 4.5e-07 | 5e-07 | 4.067E-07 | 1.210E-07 | 2.700E-07 | 5.000E-07 |
| 17182 | 860 | A860R | C | 2.8e-07 | 2.1e-07 | 2.6e-07 | 2.500E-07 | 3.606E-08 | 2.100E-07 | 2.800E-07 |
| 17183 | 860 | A860N | C | 4.6e-07 | 3.8e-07 | 2.1e-07 | 3.500E-07 | 1.277E-07 | 2.100E-07 | 4.600E-07 |
| 17184 | 860 | A860D | C | 1.9e-07 | 1.9e-07 | 4.4e-07 | 2.733E-07 | 1.443E-07 | 1.900E-07 | 4.400E-07 |
| 17185 | 860 | A860C | C | 2.1e-07 | 2.4e-07 | 3.2e-07 | 2.567E-07 | 5.686E-08 | 2.100E-07 | 3.200E-07 |
| 17186 | 860 | A860E | C | 2e-07 | 2.1e-07 | 3.8e-07 | 2.633E-07 | 1.012E-07 | 2.000E-07 | 3.800E-07 |
| 17187 | 860 | A860Q | C | 2.8e-07 | 2.6e-07 | 2.6e-07 | 2.667E-07 | 1.155E-08 | 2.600E-07 | 2.800E-07 |
| 17188 | 860 | A860G | C | 4.8e-07 | 1.9e-07 | 2.2e-07 | 2.967E-07 | 1.595E-07 | 1.900E-07 | 4.800E-07 |
| 17189 | 860 | A860H | C | 1.8e-07 | 2e-07 | 2.8e-07 | 2.200E-07 | 5.292E-08 | 1.800E-07 | 2.800E-07 |
| 17190 | 860 | A860I | C | 3.5e-07 | 2.7e-07 | 1.5e-07 | 2.567E-07 | 1.007E-07 | 1.500E-07 | 3.500E-07 |
| 17191 | 860 | A860L | C | 2.3e-07 | 5.9e-07 | 5.2e-07 | 4.467E-07 | 1.909E-07 | 2.300E-07 | 5.900E-07 |
| 17192 | 860 | A860K | C | 3.8e-07 | 3.5e-07 | 2.1e-07 | 3.133E-07 | 9.074E-08 | 2.100E-07 | 3.800E-07 |
| 17193 | 860 | A860M | C | 4e-07 | 1.3e-07 | 2.8e-07 | 2.700E-07 | 1.353E-07 | 1.300E-07 | 4.000E-07 |
| 17194 | 860 | A860F | C | 3.4e-07 | 2.3e-07 | 5.4e-07 | 3.700E-07 | 1.572E-07 | 2.300E-07 | 5.400E-07 |
| 17195 | 860 | A860P | C | 1e-07 | 1.5e-07 | 1.4e-07 | 1.300E-07 | 2.646E-08 | 1.000E-07 | 1.500E-07 |
| 17196 | 860 | A860S | C | 1.4e-07 | 1.6e-07 | 4.7e-07 | 2.567E-07 | 1.850E-07 | 1.400E-07 | 4.700E-07 |
| 17197 | 860 | A860T | C | 1.7e-07 | 1.5e-07 | 1.2e-07 | 1.467E-07 | 2.517E-08 | 1.200E-07 | 1.700E-07 |
| 17198 | 860 | A860W | C | 2.3e-07 | 2.1e-07 | 1.1e-07 | 1.833E-07 | 6.429E-08 | 1.100E-07 | 2.300E-07 |
| 17199 | 860 | A860Y | C | 4.9e-07 | 3.1e-07 | 4.6e-07 | 4.200E-07 | 9.644E-08 | 3.100E-07 | 4.900E-07 |
| 17200 | 860 | A860V | C | 3.4e-07 | 3.1e-07 | 3.3e-07 | 3.267E-07 | 1.528E-08 | 3.100E-07 | 3.400E-07 |
| 17201 | 861 | S861A | C | 4e-07 | 1.7e-07 | 1.1e-07 | 2.267E-07 | 1.531E-07 | 1.100E-07 | 4.000E-07 |
| 17202 | 861 | S861R | C | 3.2e-07 | 1.6e-07 | 2.7e-07 | 2.500E-07 | 8.185E-08 | 1.600E-07 | 3.200E-07 |
| 17203 | 861 | S861N | C | 2e-07 | 2e-07 | 2.3e-07 | 2.100E-07 | 1.732E-08 | 2.000E-07 | 2.300E-07 |
| 17204 | 861 | S861D | C | 4.2e-07 | 1.1e-07 | 1.2e-07 | 2.167E-07 | 1.762E-07 | 1.100E-07 | 4.200E-07 |
| 17205 | 861 | S861C | C | 1.4e-07 | 3.8e-07 | 1.9e-07 | 2.367E-07 | 1.266E-07 | 1.400E-07 | 3.800E-07 |
| 17206 | 861 | S861E | C | 2.4e-07 | 2.3e-07 | 2e-07 | 2.233E-07 | 2.082E-08 | 2.000E-07 | 2.400E-07 |
| 17207 | 861 | S861Q | C | 2.2e-07 | 3.8e-07 | 4.4e-07 | 3.467E-07 | 1.137E-07 | 2.200E-07 | 4.400E-07 |
| 17208 | 861 | S861G | C | 3.9e-07 | 1.7e-07 | 2.7e-07 | 2.767E-07 | 1.102E-07 | 1.700E-07 | 3.900E-07 |
| 17209 | 861 | S861H | C | 3.7e-07 | 3.1e-07 | 1.6e-07 | 2.800E-07 | 1.082E-07 | 1.600E-07 | 3.700E-07 |
| 17210 | 861 | S861I | C | 2.6e-07 | 2.7e-07 | 7.8e-08 | 2.027E-07 | 1.081E-07 | 7.800E-08 | 2.700E-07 |
| 17211 | 861 | S861L | C | 2.3e-07 | 3.9e-07 | 5.9e-07 | 4.033E-07 | 1.804E-07 | 2.300E-07 | 5.900E-07 |
| 17212 | 861 | S861K | C | 3.4e-07 | 1.4e-07 | 3.6e-07 | 2.800E-07 | 1.217E-07 | 1.400E-07 | 3.600E-07 |
| 17213 | 861 | S861M | C | 1.4e-07 | 1.7e-07 | 2.1e-07 | 1.733E-07 | 3.512E-08 | 1.400E-07 | 2.100E-07 |
| 17214 | 861 | S861F | C | 5e-07 | 2e-07 | 2e-07 | 3.000E-07 | 1.732E-07 | 2.000E-07 | 5.000E-07 |
| 17215 | 861 | S861P | C | 7.2e-07 | 3.9e-07 | 2.2e-07 | 4.433E-07 | 2.542E-07 | 2.200E-07 | 7.200E-07 |
| 17216 | 861 | S861S | C | 1.6e-07 | 1.2e-07 | 2.5e-07 | 1.767E-07 | 6.658E-08 | 1.200E-07 | 2.500E-07 |
| 17217 | 861 | S861T | C | 2.2e-07 | 1.5e-07 | 2.6e-07 | 2.100E-07 | 5.568E-08 | 1.500E-07 | 2.600E-07 |
| 17218 | 861 | S861W | C | 3e-07 | 2.5e-07 | 1.1e-07 | 2.200E-07 | 9.849E-08 | 1.100E-07 | 3.000E-07 |
| 17219 | 861 | S861Y | C | 2.6e-07 | 2.1e-07 | 1.3e-07 | 2.000E-07 | 6.557E-08 | 1.300E-07 | 2.600E-07 |
| 17220 | 861 | S861V | C | 5.3e-07 | 1.5e-07 | 1.6e-07 | 2.800E-07 | 2.166E-07 | 1.500E-07 | 5.300E-07 |
| 17221 | 862 | G862A | C | 1.2e-07 | 1.1e-07 | 2.3e-07 | 1.533E-07 | 6.658E-08 | 1.100E-07 | 2.300E-07 |
| 17222 | 862 | G862R | C | 2.7e-07 | 3.3e-07 | 1.4e-07 | 2.467E-07 | 9.713E-08 | 1.400E-07 | 3.300E-07 |
| 17223 | 862 | G862N | C | 8.4e-08 | 1.9e-07 | 3.2e-07 | 1.980E-07 | 1.182E-07 | 8.400E-08 | 3.200E-07 |
| 17224 | 862 | G862D | C | 1.4e-07 | 1.6e-07 | 2.2e-07 | 1.733E-07 | 4.163E-08 | 1.400E-07 | 2.200E-07 |
| 17225 | 862 | G862C | C | 2.8e-07 | 1.7e-07 | 2.5e-07 | 2.333E-07 | 5.686E-08 | 1.700E-07 | 2.800E-07 |
| 17226 | 862 | G862E | C | 7.2e-07 | 2.7e-07 | 1e-07 | 3.633E-07 | 3.204E-07 | 1.000E-07 | 7.200E-07 |
| 17227 | 862 | G862Q | C | 2.8e-07 | 7.4e-07 | 2e-07 | 4.067E-07 | 2.914E-07 | 2.000E-07 | 7.400E-07 |
| 17228 | 862 | G862G | C | 2.1e-07 | 1.3e-07 | 2.7e-07 | 2.033E-07 | 7.024E-08 | 1.300E-07 | 2.700E-07 |
| 17229 | 862 | G862H | C | 3.8e-07 | 7e-07 | 2.7e-07 | 4.500E-07 | 2.234E-07 | 2.700E-07 | 7.000E-07 |
| 17230 | 862 | G862I | C | 1.7e-07 | 3.1e-07 | 2.2e-07 | 2.333E-07 | 7.095E-08 | 1.700E-07 | 3.100E-07 |
| 17231 | 862 | G862L | C | 3.3e-07 | 2.1e-07 | 3.7e-07 | 3.033E-07 | 8.327E-08 | 2.100E-07 | 3.700E-07 |
| 17232 | 862 | G862K | C | 3.1e-07 | 1e-07 | 1.2e-07 | 1.767E-07 | 1.159E-07 | 1.000E-07 | 3.100E-07 |
| 17233 | 862 | G862M | C | 4e-07 | 2e-07 | 4.3e-07 | 3.433E-07 | 1.250E-07 | 2.000E-07 | 4.300E-07 |
| 17234 | 862 | G862F | C | 3e-07 | 2.3e-07 | 3.6e-07 | 2.967E-07 | 6.506E-08 | 2.300E-07 | 3.600E-07 |
| 17235 | 862 | G862P | C | 1.9e-07 | 2.1e-07 | 3.8e-07 | 2.600E-07 | 1.044E-07 | 1.900E-07 | 3.800E-07 |
| 17236 | 862 | G862S | C | 2.3e-07 | 2.1e-07 | 4.3e-07 | 2.900E-07 | 1.217E-07 | 2.100E-07 | 4.300E-07 |
| 17237 | 862 | G862T | C | 2.1e-07 | 2.5e-07 | 2e-07 | 2.200E-07 | 2.646E-08 | 2.000E-07 | 2.500E-07 |
| 17238 | 862 | G862W | C | 1.8e-07 | 2.6e-07 | 1.1e-07 | 1.833E-07 | 7.506E-08 | 1.100E-07 | 2.600E-07 |
| 17239 | 862 | G862Y | C | 3.1e-07 | 1.1e-07 | 2.2e-07 | 2.133E-07 | 1.002E-07 | 1.100E-07 | 3.100E-07 |
| 17240 | 862 | G862V | C | 1.4e-07 | 2.4e-07 | 2.1e-07 | 1.967E-07 | 5.132E-08 | 1.400E-07 | 2.400E-07 |
| 17241 | 863 | F863A | C | 2.9e-07 | 2.8e-07 | 2e-07 | 2.567E-07 | 4.933E-08 | 2.000E-07 | 2.900E-07 |
| 17242 | 863 | F863R | C | 1.4e-07 | 4.8e-07 | 3.5e-07 | 3.233E-07 | 1.716E-07 | 1.400E-07 | 4.800E-07 |
| 17243 | 863 | F863N | C | 3.9e-07 | 3.6e-07 | 8.9e-07 | 5.467E-07 | 2.977E-07 | 3.600E-07 | 8.900E-07 |
| 17244 | 863 | F863D | C | 3.5e-07 | 2.5e-07 | 1.7e-07 | 2.567E-07 | 9.018E-08 | 1.700E-07 | 3.500E-07 |
| 17245 | 863 | F863C | C | 1e-07 | 4.8e-07 | 1.7e-07 | 2.500E-07 | 2.022E-07 | 1.000E-07 | 4.800E-07 |
| 17246 | 863 | F863E | C | 2.3e-07 | 2.5e-07 | 4.1e-07 | 2.967E-07 | 9.866E-08 | 2.300E-07 | 4.100E-07 |
| 17247 | 863 | F863Q | C | 1.9e-07 | 2.4e-07 | 3.3e-07 | 2.533E-07 | 7.095E-08 | 1.900E-07 | 3.300E-07 |
| 17248 | 863 | F863G | C | 1e-07 | 1.5e-07 | 2.2e-07 | 1.567E-07 | 6.028E-08 | 1.000E-07 | 2.200E-07 |
| 17249 | 863 | F863H | C | 3.8e-07 | 6.7e-08 | 1.5e-07 | 1.990E-07 | 1.622E-07 | 6.700E-08 | 3.800E-07 |
| 17250 | 863 | F863I | C | 3.3e-07 | 3.5e-07 | 2.1e-07 | 2.967E-07 | 7.572E-08 | 2.100E-07 | 3.500E-07 |
| 17251 | 863 | F863L | C | 3e-07 | 1.6e-07 | 2.2e-07 | 2.267E-07 | 7.024E-08 | 1.600E-07 | 3.000E-07 |
| 17252 | 863 | F863K | C | 2.5e-07 | 2.2e-07 | 1.7e-07 | 2.133E-07 | 4.041E-08 | 1.700E-07 | 2.500E-07 |
| 17253 | 863 | F863M | C | 4.7e-07 | 1.4e-07 | 1.4e-07 | 2.500E-07 | 1.905E-07 | 1.400E-07 | 4.700E-07 |
| 17254 | 863 | F863F | C | 2.4e-07 | 4.1e-07 | 1.6e-07 | 2.700E-07 | 1.277E-07 | 1.600E-07 | 4.100E-07 |

| | | | | | | | | | | |
|---|---|---|---|---|---|---|---|---|---|---|
| 17255 | 863 | F863P | C | 2.1e-07 | 1.9e-07 | 2.9e-07 | 2.300E-07 | 5.292E-08 | 1.900E-07 | 2.900E-07 |
| 17256 | 863 | F863S | C | 1.4e-07 | 1.5e-07 | 2.1e-07 | 1.667E-07 | 3.786E-08 | 1.400E-07 | 2.100E-07 |
| 17257 | 863 | F863T | C | 2.2e-07 | 1.9e-07 | 1.8e-07 | 1.967E-07 | 2.082E-08 | 1.800E-07 | 2.200E-07 |
| 17258 | 863 | F863W | C | 4.4e-07 | 4.5e-07 | 1.1e-07 | 3.333E-07 | 1.935E-07 | 1.100E-07 | 4.500E-07 |
| 17259 | 863 | F863Y | C | 1.8e-07 | 1.3e-07 | 1.5e-07 | 1.533E-07 | 2.517E-08 | 1.300E-07 | 1.800E-07 |
| 17260 | 863 | F863V | C | 1.4e-07 | 3.4e-07 | 5.6e-07 | 3.467E-07 | 2.101E-07 | 1.400E-07 | 5.600E-07 |
| 17261 | 864 | T864A | C | 3.1e-07 | 1.7e-07 | 4.1e-07 | 2.967E-07 | 1.206E-07 | 1.700E-07 | 4.100E-07 |
| 17262 | 864 | T864R | C | 2.4e-07 | 2.4e-07 | 3.7e-07 | 2.833E-07 | 7.506E-08 | 2.400E-07 | 3.700E-07 |
| 17263 | 864 | T864N | C | 1.4e-07 | 2.6e-07 | 1.3e-07 | 1.767E-07 | 7.234E-08 | 1.300E-07 | 2.600E-07 |
| 17264 | 864 | T864D | C | 2.6e-07 | 5.5e-07 | 1.6e-07 | 3.233E-07 | 2.026E-07 | 1.600E-07 | 5.500E-07 |
| 17265 | 864 | T864C | C | 5.1e-07 | 1.7e-07 | 2e-07 | 2.933E-07 | 1.882E-07 | 1.700E-07 | 5.100E-07 |
| 17266 | 864 | T864E | C | 3.2e-07 | 1.5e-07 | 3.1e-07 | 2.600E-07 | 9.539E-08 | 1.500E-07 | 3.200E-07 |
| 17267 | 864 | T864Q | C | 1.8e-07 | 1.8e-07 | 5.5e-07 | 3.033E-07 | 2.136E-07 | 1.800E-07 | 5.500E-07 |
| 17268 | 864 | T864G | C | 3.3e-07 | 1.1e-07 | 1.8e-07 | 2.067E-07 | 1.124E-07 | 1.100E-07 | 3.300E-07 |
| 17269 | 864 | T864H | C | 9.6e-08 | 2.6e-07 | 1.5e-07 | 1.687E-07 | 8.358E-08 | 9.600E-08 | 2.600E-07 |
| 17270 | 864 | T864I | C | 2.8e-07 | 2e-07 | 3.4e-07 | 2.733E-07 | 7.024E-08 | 2.000E-07 | 3.400E-07 |
| 17271 | 864 | T864L | C | 3.3e-07 | 2.6e-07 | 3.6e-07 | 3.167E-07 | 5.132E-08 | 2.600E-07 | 3.600E-07 |
| 17272 | 864 | T864K | C | 2.6e-07 | 1.4e-07 | 3.1e-07 | 2.367E-07 | 8.737E-08 | 1.400E-07 | 3.100E-07 |
| 17273 | 864 | T864M | C | 1.4e-07 | 4e-07 | 5.5e-07 | 3.633E-07 | 2.074E-07 | 1.400E-07 | 5.500E-07 |
| 17274 | 864 | T864F | C | 2e-07 | 1.6e-07 | 2.2e-07 | 1.933E-07 | 3.055E-08 | 1.600E-07 | 2.200E-07 |
| 17275 | 864 | T864P | C | 2e-07 | 3.8e-07 | 4.1e-07 | 3.300E-07 | 1.136E-07 | 2.000E-07 | 4.100E-07 |
| 17276 | 864 | T864S | C | 1.8e-07 | 1.8e-07 | 3.6e-07 | 2.400E-07 | 1.039E-07 | 1.800E-07 | 3.600E-07 |
| 17277 | 864 | T864T | C | 2.2e-07 | 6.1e-07 | 3e-07 | 3.767E-07 | 2.060E-07 | 2.200E-07 | 6.100E-07 |
| 17278 | 864 | T864W | C | 2.7e-07 | 1.3e-07 | 3e-07 | 2.333E-07 | 9.074E-08 | 1.300E-07 | 3.000E-07 |
| 17279 | 864 | T864Y | C | 2.1e-07 | 3.7e-07 | 3.1e-07 | 2.967E-07 | 8.083E-08 | 2.100E-07 | 3.700E-07 |
| 17280 | 864 | T864V | C | 2.2e-07 | 2.3e-07 | 2.2e-07 | 2.233E-07 | 5.774E-09 | 2.200E-07 | 2.300E-07 |
| 17281 | 865 | F865A | C | 4.8e-07 | 4.4e-07 | 1.9e-07 | 3.700E-07 | 1.572E-07 | 1.900E-07 | 4.800E-07 |
| 17282 | 865 | F865R | C | 5.9e-07 | 3.2e-07 | 3.2e-07 | 4.100E-07 | 1.559E-07 | 3.200E-07 | 5.900E-07 |
| 17283 | 865 | F865N | C | 2.2e-07 | 1.6e-07 | 3.4e-07 | 2.400E-07 | 9.165E-08 | 1.600E-07 | 3.400E-07 |
| 17284 | 865 | F865D | C | 1.3e-07 | 2.5e-07 | 5.5e-07 | 3.100E-07 | 2.163E-07 | 1.300E-07 | 5.500E-07 |
| 17285 | 865 | F865C | C | 2.8e-07 | 2.5e-07 | 3.7e-07 | 3.000E-07 | 6.245E-08 | 2.500E-07 | 3.700E-07 |
| 17286 | 865 | F865E | C | 1.5e-07 | 1.5e-07 | 2.5e-07 | 1.833E-07 | 5.774E-08 | 1.500E-07 | 2.500E-07 |
| 17287 | 865 | F865Q | C | 3.4e-07 | 2.1e-07 | 2.9e-07 | 2.800E-07 | 6.557E-08 | 2.100E-07 | 3.400E-07 |
| 17288 | 865 | F865G | C | 3.9e-07 | 3.4e-07 | 2.4e-07 | 3.233E-07 | 7.638E-08 | 2.400E-07 | 3.900E-07 |
| 17289 | 865 | F865H | C | 7.8e-07 | 1.2e-07 | 3.5e-07 | 4.167E-07 | 3.350E-07 | 1.200E-07 | 7.800E-07 |
| 17290 | 865 | F865I | C | 5.4e-07 | 2.7e-07 | 1.2e-07 | 3.100E-07 | 2.128E-07 | 1.200E-07 | 5.400E-07 |
| 17291 | 865 | F865L | C | 3.3e-07 | 2.4e-07 | 4e-07 | 3.233E-07 | 8.021E-08 | 2.400E-07 | 4.000E-07 |
| 17292 | 865 | F865K | C | 2.8e-07 | 4.8e-07 | 3.4e-07 | 3.667E-07 | 1.026E-07 | 2.800E-07 | 4.800E-07 |
| 17293 | 865 | F865M | C | 1.7e-07 | 3.1e-07 | 2.6e-07 | 2.467E-07 | 7.095E-08 | 1.700E-07 | 3.100E-07 |
| 17294 | 865 | F865F | C | 2.3e-07 | 2.2e-07 | 2.7e-07 | 2.400E-07 | 2.646E-08 | 2.200E-07 | 2.700E-07 |
| 17295 | 865 | F865P | C | 2.3e-07 | 2.2e-07 | 1.9e-07 | 2.133E-07 | 2.082E-08 | 1.900E-07 | 2.300E-07 |
| 17296 | 865 | F865S | C | 4.6e-07 | 9.8e-07 | 1.9e-07 | 5.433E-07 | 4.015E-07 | 1.900E-07 | 9.800E-07 |
| 17297 | 865 | F865T | C | 2.7e-07 | 3.2e-07 | 2.4e-07 | 2.767E-07 | 4.041E-08 | 2.400E-07 | 3.200E-07 |
| 17298 | 865 | F865W | C | 2.3e-07 | 1.9e-07 | 2.4e-07 | 2.200E-07 | 2.646E-08 | 1.900E-07 | 2.400E-07 |
| 17299 | 865 | F865Y | C | 1.1e-07 | 3.2e-07 | 2.9e-07 | 2.400E-07 | 1.136E-07 | 1.100E-07 | 3.200E-07 |
| 17300 | 865 | F865V | C | 2.8e-07 | 2.3e-07 | 3.2e-07 | 2.767E-07 | 4.509E-08 | 2.300E-07 | 3.200E-07 |
| 17301 | 866 | T866A | C | 2.4e-07 | 1.9e-07 | 2.4e-07 | 2.233E-07 | 2.887E-08 | 1.900E-07 | 2.400E-07 |
| 17302 | 866 | T866R | C | 2.3e-07 | 1.1e-07 | 3.9e-07 | 2.433E-07 | 1.405E-07 | 1.100E-07 | 3.900E-07 |
| 17303 | 866 | T866N | C | 2.2e-07 | 3.6e-07 | 2.3e-07 | 2.700E-07 | 7.810E-08 | 2.200E-07 | 3.600E-07 |
| 17304 | 866 | T866D | C | 9.5e-08 | 2.7e-07 | 4.5e-07 | 2.717E-07 | 1.775E-07 | 9.500E-08 | 4.500E-07 |
| 17305 | 866 | T866C | C | 1.5e-07 | 2.4e-07 | 1e-07 | 1.633E-07 | 7.095E-08 | 1.000E-07 | 2.400E-07 |
| 17306 | 866 | T866E | C | 1.5e-07 | 1.7e-07 | 3.3e-07 | 2.167E-07 | 9.866E-08 | 1.500E-07 | 3.300E-07 |
| 17307 | 866 | T866Q | C | 2.8e-07 | 2.9e-07 | 3e-07 | 2.900E-07 | 1.000E-08 | 2.800E-07 | 3.000E-07 |
| 17308 | 866 | T866G | C | 3.6e-07 | 1.9e-07 | 2.4e-07 | 2.633E-07 | 8.737E-08 | 1.900E-07 | 3.600E-07 |
| 17309 | 866 | T866H | C | 1.8e-07 | 1.5e-07 | 4e-07 | 2.433E-07 | 1.365E-07 | 1.500E-07 | 4.000E-07 |
| 17310 | 866 | T866I | C | 2.8e-07 | 2.9e-07 | 2.4e-07 | 2.700E-07 | 2.646E-08 | 2.400E-07 | 2.900E-07 |
| 17311 | 866 | T866L | C | 4.5e-07 | 1.5e-07 | 1.9e-07 | 2.633E-07 | 1.629E-07 | 1.500E-07 | 4.500E-07 |
| 17312 | 866 | T866K | C | 1.9e-07 | 2e-07 | 2.2e-07 | 2.033E-07 | 1.528E-08 | 1.900E-07 | 2.200E-07 |
| 17313 | 866 | T866M | C | 2.9e-07 | 2.2e-07 | 3.7e-07 | 2.933E-07 | 7.506E-08 | 2.200E-07 | 3.700E-07 |
| 17314 | 866 | T866F | C | 2.4e-07 | 2.8e-07 | 1.2e-07 | 2.133E-07 | 8.327E-08 | 1.200E-07 | 2.800E-07 |
| 17315 | 866 | T866P | C | 3.4e-07 | 1e-07 | 3.7e-07 | 2.700E-07 | 1.480E-07 | 1.000E-07 | 3.700E-07 |
| 17316 | 866 | T866S | C | 2.9e-07 | 3.4e-07 | 3.6e-07 | 3.300E-07 | 3.606E-08 | 2.900E-07 | 3.600E-07 |
| 17317 | 866 | T866T | C | 4.3e-07 | 2.6e-07 | 2.7e-07 | 3.200E-07 | 9.539E-08 | 2.600E-07 | 4.300E-07 |
| 17318 | 866 | T866W | C | 2e-07 | 1.9e-07 | 1.7e-07 | 1.867E-07 | 1.528E-08 | 1.700E-07 | 2.000E-07 |
| 17319 | 866 | T866Y | C | 1.8e-07 | 2.2e-07 | 4.2e-07 | 2.733E-07 | 1.286E-07 | 1.800E-07 | 4.200E-07 |
| 17320 | 866 | T866V | C | 1.5e-07 | 2.2e-07 | 3.1e-07 | 2.267E-07 | 8.021E-08 | 1.500E-07 | 3.100E-07 |
| 17321 | 867 | D867A | C | 3.5e-07 | 1.5e-07 | 1.7e-07 | 2.233E-07 | 1.102E-07 | 1.500E-07 | 3.500E-07 |
| 17322 | 867 | D867R | C | 4.8e-07 | 2.3e-07 | 2e-07 | 3.033E-07 | 1.537E-07 | 2.000E-07 | 4.800E-07 |
| 17323 | 867 | D867N | C | 1.4e-07 | 5.1e-07 | 2.3e-07 | 2.933E-07 | 1.930E-07 | 1.400E-07 | 5.100E-07 |
| 17324 | 867 | D867D | C | 1.7e-07 | 4.2e-07 | 3.3e-07 | 3.067E-07 | 1.266E-07 | 1.700E-07 | 4.200E-07 |
| 17325 | 867 | D867C | C | 4.1e-07 | 3.2e-07 | 2.1e-07 | 3.133E-07 | 1.002E-07 | 2.100E-07 | 4.100E-07 |
| 17326 | 867 | D867E | C | 3.4e-07 | 4e-07 | 2.4e-07 | 3.267E-07 | 8.083E-08 | 2.400E-07 | 4.000E-07 |
| 17327 | 867 | D867Q | C | 3.1e-07 | 3.1e-07 | 1.5e-07 | 2.567E-07 | 9.238E-08 | 1.500E-07 | 3.100E-07 |
| 17328 | 867 | D867G | C | 2.4e-07 | 1.9e-07 | 3.2e-07 | 2.500E-07 | 6.557E-08 | 1.900E-07 | 3.200E-07 |
| 17329 | 867 | D867H | C | 4.4e-07 | 2.8e-07 | 2e-07 | 3.067E-07 | 1.222E-07 | 2.000E-07 | 4.400E-07 |
| 17330 | 867 | D867I | C | 1.4e-07 | 2.5e-07 | 1.9e-07 | 1.933E-07 | 5.508E-08 | 1.400E-07 | 2.500E-07 |
| 17331 | 867 | D867L | C | 1.1e-07 | 3.7e-07 | 2.4e-07 | 2.400E-07 | 1.300E-07 | 1.100E-07 | 3.700E-07 |
| 17332 | 867 | D867K | C | 1.5e-07 | 1.7e-07 | 3.1e-07 | 2.100E-07 | 8.718E-08 | 1.500E-07 | 3.100E-07 |
| 17333 | 867 | D867M | C | 1.8e-07 | 2e-07 | 3e-07 | 2.267E-07 | 6.429E-08 | 1.800E-07 | 3.000E-07 |
| 17334 | 867 | D867F | C | 1.8e-07 | 7.1e-07 | 1.6e-07 | 3.500E-07 | 3.119E-07 | 1.600E-07 | 7.100E-07 |
| 17335 | 867 | D867P | C | 1.5e-07 | 2.8e-07 | 1.8e-07 | 2.033E-07 | 6.807E-08 | 1.500E-07 | 2.800E-07 |
| 17336 | 867 | D867S | C | 2.6e-07 | 1.8e-07 | 1.7e-07 | 2.033E-07 | 4.933E-08 | 1.700E-07 | 2.600E-07 |
| 17337 | 867 | D867T | C | 2.6e-07 | 1.5e-07 | 2.4e-07 | 2.167E-07 | 5.859E-08 | 1.500E-07 | 2.600E-07 |
| 17338 | 867 | D867W | C | 3e-07 | 2.3e-07 | 6.1e-07 | 3.800E-07 | 2.022E-07 | 2.300E-07 | 6.100E-07 |
| 17339 | 867 | D867Y | C | 3.2e-07 | 1.6e-07 | 1.3e-07 | 2.033E-07 | 1.021E-07 | 1.300E-07 | 3.200E-07 |

| | | | | | | | | | | |
|---|---|---|---|---|---|---|---|---|---|---|
| 17340 | 867 | D867V | C | 2.3e-07 | 1.8e-07 | 3.2e-07 | 2.433E-07 | 7.095E-08 | 1.800E-07 | 3.200E-07 |
| 17341 | 868 | Y868A | C | 1.9e-07 | 1.5e-07 | 1.4e-07 | 1.600E-07 | 2.646E-08 | 1.400E-07 | 1.900E-07 |
| 17342 | 868 | Y868R | C | 7.2e-07 | 3.8e-07 | 1.7e-07 | 4.233E-07 | 2.775E-07 | 1.700E-07 | 7.200E-07 |
| 17343 | 868 | Y868N | C | 3.9e-07 | 1.5e-07 | 2.4e-07 | 2.600E-07 | 1.212E-07 | 1.500E-07 | 3.900E-07 |
| 17344 | 868 | Y868D | C | 2.2e-07 | 2.4e-07 | 2.5e-07 | 2.367E-07 | 1.528E-08 | 2.200E-07 | 2.500E-07 |
| 17345 | 868 | Y868C | C | 2.1e-07 | 2.9e-07 | 3e-07 | 2.667E-07 | 4.933E-08 | 2.100E-07 | 3.000E-07 |
| 17346 | 868 | Y868E | C | 1.8e-07 | 1.6e-07 | 3.3e-07 | 2.233E-07 | 9.292E-08 | 1.600E-07 | 3.300E-07 |
| 17347 | 868 | Y868Q | C | 2.6e-07 | 4.4e-07 | 1.3e-07 | 2.767E-07 | 1.557E-07 | 1.300E-07 | 4.400E-07 |
| 17348 | 868 | Y868G | C | 2.5e-07 | 6.1e-07 | 3.7e-07 | 4.100E-07 | 1.833E-07 | 2.500E-07 | 6.100E-07 |
| 17349 | 868 | Y868H | C | 1.1e-07 | 2.3e-07 | 3.3e-07 | 2.233E-07 | 1.102E-07 | 1.100E-07 | 3.300E-07 |
| 17350 | 868 | Y868I | C | 1.4e-07 | 3.9e-07 | 3.5e-07 | 2.933E-07 | 1.343E-07 | 1.400E-07 | 3.900E-07 |
| 17351 | 868 | Y868L | C | 4.1e-07 | 3.7e-07 | 1.5e-07 | 3.100E-07 | 1.400E-07 | 1.500E-07 | 4.100E-07 |
| 17352 | 868 | Y868K | C | 1.8e-07 | 2e-07 | 3.6e-07 | 2.467E-07 | 9.866E-08 | 1.800E-07 | 3.600E-07 |
| 17353 | 868 | Y868M | C | 3e-07 | 3.3e-07 | 1.2e-07 | 2.500E-07 | 1.136E-07 | 1.200E-07 | 3.300E-07 |
| 17354 | 868 | Y868F | C | 3.2e-07 | 3.4e-07 | 3.4e-07 | 3.333E-07 | 1.155E-08 | 3.200E-07 | 3.400E-07 |
| 17355 | 868 | Y868P | C | 3.2e-07 | 2e-07 | 5.4e-07 | 3.533E-07 | 1.724E-07 | 2.000E-07 | 5.400E-07 |
| 17356 | 868 | Y868S | C | 3.2e-07 | 3.1e-07 | 2.7e-07 | 3.000E-07 | 2.646E-08 | 2.700E-07 | 3.200E-07 |
| 17357 | 868 | Y868T | C | 3.5e-07 | 1.7e-07 | 3.7e-07 | 2.967E-07 | 1.102E-07 | 1.700E-07 | 3.700E-07 |
| 17358 | 868 | Y868W | C | 4.5e-07 | 3.5e-07 | 9.2e-08 | 2.973E-07 | 1.847E-07 | 9.200E-08 | 4.500E-07 |
| 17359 | 868 | Y868Y | C | 3.7e-07 | 2.1e-07 | 2.9e-07 | 2.900E-07 | 8.000E-08 | 2.100E-07 | 3.700E-07 |
| 17360 | 868 | Y868V | C | 2.1e-07 | 1.6e-07 | 1.5e-07 | 1.733E-07 | 3.215E-08 | 1.500E-07 | 2.100E-07 |
| 17361 | 869 | T869A | C | 1.4e-07 | 7.6e-08 | 2.5e-07 | 1.553E-07 | 8.801E-08 | 7.600E-08 | 2.500E-07 |
| 17362 | 869 | T869R | C | 2.5e-07 | 3.1e-07 | 5.9e-07 | 3.833E-07 | 1.815E-07 | 2.500E-07 | 5.900E-07 |
| 17363 | 869 | T869N | C | 3.6e-07 | 2.1e-07 | 2.7e-07 | 2.800E-07 | 7.550E-08 | 2.100E-07 | 3.600E-07 |
| 17364 | 869 | T869D | C | 3.1e-07 | 2.5e-07 | 3.4e-07 | 3.000E-07 | 4.583E-08 | 2.500E-07 | 3.400E-07 |
| 17365 | 869 | T869C | C | 2.7e-07 | 1.6e-07 | 9.2e-08 | 1.740E-07 | 8.982E-08 | 9.200E-08 | 2.700E-07 |
| 17366 | 869 | T869E | C | 2.8e-07 | 3.1e-07 | 2.7e-07 | 2.867E-07 | 2.082E-08 | 2.700E-07 | 3.100E-07 |
| 17367 | 869 | T869Q | C | 1.7e-07 | 2.5e-07 | 1.8e-07 | 2.000E-07 | 4.359E-08 | 1.700E-07 | 2.500E-07 |
| 17368 | 869 | T869G | C | 2.2e-07 | 2.4e-07 | 1.4e-07 | 2.000E-07 | 5.292E-08 | 1.400E-07 | 2.400E-07 |
| 17369 | 869 | T869H | C | 2.4e-07 | 1.5e-07 | 1.6e-07 | 1.833E-07 | 4.933E-08 | 1.500E-07 | 2.400E-07 |
| 17370 | 869 | T869I | C | 3.1e-07 | 2.5e-07 | 3e-07 | 2.867E-07 | 3.215E-08 | 2.500E-07 | 3.100E-07 |
| 17371 | 869 | T869L | C | 3.8e-07 | 2.3e-07 | 3.1e-07 | 3.067E-07 | 7.506E-08 | 2.300E-07 | 3.800E-07 |
| 17372 | 869 | T869K | C | 2.8e-07 | 2.9e-07 | 1.7e-07 | 2.467E-07 | 6.658E-08 | 1.700E-07 | 2.900E-07 |
| 17373 | 869 | T869M | C | 6.1e-07 | 4.3e-07 | 3.6e-07 | 4.667E-07 | 1.290E-07 | 3.600E-07 | 6.100E-07 |
| 17374 | 869 | T869F | C | 2.1e-07 | 3.2e-07 | 1.6e-07 | 2.300E-07 | 8.185E-08 | 1.600E-07 | 3.200E-07 |
| 17375 | 869 | T869P | C | 3e-07 | 3.8e-07 | 4.3e-07 | 3.700E-07 | 6.557E-08 | 3.000E-07 | 4.300E-07 |
| 17376 | 869 | T869S | C | 4.2e-07 | 2.5e-07 | 2.4e-07 | 3.033E-07 | 1.012E-07 | 2.400E-07 | 4.200E-07 |
| 17377 | 869 | T869T | C | 5.1e-07 | 6.2e-08 | 5.2e-07 | 3.640E-07 | 2.616E-07 | 6.200E-08 | 5.200E-07 |
| 17378 | 869 | T869W | C | 2.1e-07 | 5.5e-07 | 4.9e-07 | 4.167E-07 | 1.815E-07 | 2.100E-07 | 5.500E-07 |
| 17379 | 869 | T869Y | C | 5.1e-07 | 3.9e-07 | 2.2e-07 | 3.733E-07 | 1.457E-07 | 2.200E-07 | 5.100E-07 |
| 17380 | 869 | T869V | C | 4.3e-07 | 1.1e-07 | 2.8e-07 | 2.733E-07 | 1.601E-07 | 1.100E-07 | 4.300E-07 |
| 17381 | 870 | M870A | C | 1.6e-07 | 1.1e-07 | 1.9e-07 | 1.533E-07 | 4.041E-08 | 1.100E-07 | 1.900E-07 |
| 17382 | 870 | M870R | C | 1.6e-07 | 1.7e-07 | 1.5e-07 | 1.600E-07 | 1.000E-08 | 1.500E-07 | 1.700E-07 |
| 17383 | 870 | M870N | C | 2.4e-07 | 1.8e-07 | 5e-07 | 3.067E-07 | 1.701E-07 | 1.800E-07 | 5.000E-07 |
| 17384 | 870 | M870D | C | 2.6e-07 | 5e-07 | 3.3e-07 | 3.633E-07 | 1.234E-07 | 2.600E-07 | 5.000E-07 |
| 17385 | 870 | M870C | C | 2.8e-07 | 2e-07 | 1.7e-07 | 2.167E-07 | 5.686E-08 | 1.700E-07 | 2.800E-07 |
| 17386 | 870 | M870E | C | 2.1e-07 | 2e-07 | 1.6e-07 | 1.900E-07 | 2.646E-08 | 1.600E-07 | 2.100E-07 |
| 17387 | 870 | M870Q | C | 1.8e-07 | 1.7e-07 | 8.3e-08 | 1.443E-07 | 5.335E-08 | 8.300E-08 | 1.800E-07 |
| 17388 | 870 | M870G | C | 2.3e-07 | 2.8e-07 | 3.7e-07 | 2.933E-07 | 7.095E-08 | 2.300E-07 | 3.700E-07 |
| 17389 | 870 | M870H | C | 3.3e-07 | 1.9e-07 | 9.4e-08 | 2.047E-07 | 1.187E-07 | 9.400E-08 | 3.300E-07 |
| 17390 | 870 | M870I | C | 2.5e-07 | 1.8e-07 | 2.9e-07 | 2.400E-07 | 5.568E-08 | 1.800E-07 | 2.900E-07 |
| 17391 | 870 | M870L | C | 1.6e-07 | 4.2e-07 | 4.4e-07 | 3.400E-07 | 1.562E-07 | 1.600E-07 | 4.400E-07 |
| 17392 | 870 | M870K | C | 3.8e-07 | 3.3e-07 | 2.1e-07 | 3.067E-07 | 8.737E-08 | 2.100E-07 | 3.800E-07 |
| 17393 | 870 | M870M | C | 2.9e-07 | 1.5e-07 | 2.3e-07 | 2.233E-07 | 7.024E-08 | 1.500E-07 | 2.900E-07 |
| 17394 | 870 | M870F | C | 1.5e-07 | 2.3e-07 | 2.4e-07 | 2.067E-07 | 4.933E-08 | 1.500E-07 | 2.400E-07 |
| 17395 | 870 | M870P | C | 2.5e-07 | 2.7e-07 | 2.4e-07 | 2.533E-07 | 1.528E-08 | 2.400E-07 | 2.700E-07 |
| 17396 | 870 | M870S | C | 2.1e-07 | 1.9e-07 | 2.8e-07 | 2.267E-07 | 4.726E-08 | 1.900E-07 | 2.800E-07 |
| 17397 | 870 | M870T | C | 1.2e-07 | 2.9e-07 | 1.7e-07 | 1.933E-07 | 8.737E-08 | 1.200E-07 | 2.900E-07 |
| 17398 | 870 | M870W | C | 1.6e-07 | 4e-07 | 3.1e-07 | 2.900E-07 | 1.212E-07 | 1.600E-07 | 4.000E-07 |
| 17399 | 870 | M870Y | C | 3.3e-07 | 3.7e-07 | 3e-07 | 3.333E-07 | 3.512E-08 | 3.000E-07 | 3.700E-07 |
| 17400 | 870 | M870V | C | 3.1e-07 | 1.9e-07 | 4.7e-07 | 3.233E-07 | 1.405E-07 | 1.900E-07 | 4.700E-07 |
| 17401 | 871 | D871A | C | 1.6e-07 | 2.2e-07 | 2.3e-07 | 2.033E-07 | 3.786E-08 | 1.600E-07 | 2.300E-07 |
| 17402 | 871 | D871R | C | 2.4e-07 | 1.6e-07 | 4e-07 | 2.667E-07 | 1.222E-07 | 1.600E-07 | 4.000E-07 |
| 17403 | 871 | D871N | C | 1.4e-07 | 3.1e-07 | 2.4e-07 | 2.300E-07 | 8.544E-08 | 1.400E-07 | 3.100E-07 |
| 17404 | 871 | D871D | C | 1.7e-07 | 2.4e-07 | 2.7e-07 | 2.267E-07 | 5.132E-08 | 1.700E-07 | 2.700E-07 |
| 17405 | 871 | D871C | C | 3.8e-07 | 1.1e-07 | 1.6e-07 | 2.167E-07 | 1.436E-07 | 1.100E-07 | 3.800E-07 |
| 17406 | 871 | D871E | C | 2.2e-07 | 2.6e-07 | 1e-07 | 1.933E-07 | 8.327E-08 | 1.000E-07 | 2.600E-07 |
| 17407 | 871 | D871Q | C | 2.9e-07 | 3.2e-07 | 2.6e-07 | 2.900E-07 | 3.000E-08 | 2.600E-07 | 3.200E-07 |
| 17408 | 871 | D871G | C | 5.1e-07 | 2.9e-07 | 1.9e-07 | 3.300E-07 | 1.637E-07 | 1.900E-07 | 5.100E-07 |
| 17409 | 871 | D871H | C | 3.6e-07 | 2.3e-07 | 2.7e-07 | 2.867E-07 | 6.658E-08 | 2.300E-07 | 3.600E-07 |
| 17410 | 871 | D871I | C | 2.8e-07 | 6e-07 | 3e-07 | 3.933E-07 | 1.793E-07 | 2.800E-07 | 6.000E-07 |
| 17411 | 871 | D871L | C | 2.2e-07 | 1.7e-07 | 3.2e-07 | 2.367E-07 | 7.638E-08 | 1.700E-07 | 3.200E-07 |
| 17412 | 871 | D871K | C | 3.9e-07 | 3.8e-07 | 1.5e-07 | 3.067E-07 | 1.358E-07 | 1.500E-07 | 3.900E-07 |
| 17413 | 871 | D871M | C | 2.1e-07 | 1.6e-07 | 1.3e-07 | 1.667E-07 | 4.041E-08 | 1.300E-07 | 2.100E-07 |
| 17414 | 871 | D871F | C | 3.6e-07 | 2.2e-07 | 2.4e-07 | 2.733E-07 | 7.572E-08 | 2.200E-07 | 3.600E-07 |
| 17415 | 871 | D871P | C | 2.9e-07 | 2.8e-07 | 1.4e-07 | 2.367E-07 | 8.386E-08 | 1.400E-07 | 2.900E-07 |
| 17416 | 871 | D871S | C | 1.3e-07 | 1.6e-07 | 6.3e-07 | 3.067E-07 | 2.804E-07 | 1.300E-07 | 6.300E-07 |
| 17417 | 871 | D871T | C | 3.7e-07 | 4.3e-07 | 1.9e-07 | 3.300E-07 | 1.249E-07 | 1.900E-07 | 4.300E-07 |
| 17418 | 871 | D871W | C | 2.2e-07 | 2.1e-07 | 3.9e-07 | 2.733E-07 | 1.012E-07 | 2.100E-07 | 3.900E-07 |
| 17419 | 871 | D871Y | C | 2e-07 | 1.6e-07 | 4e-07 | 2.533E-07 | 1.286E-07 | 1.600E-07 | 4.000E-07 |
| 17420 | 871 | D871V | C | 1.6e-07 | 2.4e-07 | 2.1e-07 | 2.033E-07 | 4.041E-08 | 1.600E-07 | 2.400E-07 |
| 17421 | 872 | W872A | C | 1.2e-07 | 3.5e-07 | 1.8e-07 | 2.167E-07 | 1.193E-07 | 1.200E-07 | 3.500E-07 |
| 17422 | 872 | W872R | C | 5.3e-07 | 1.4e-07 | 1.4e-07 | 2.700E-07 | 2.252E-07 | 1.400E-07 | 5.300E-07 |
| 17423 | 872 | W872N | C | 4e-07 | 6.2e-07 | 2.3e-07 | 4.167E-07 | 1.955E-07 | 2.300E-07 | 6.200E-07 |
| 17424 | 872 | W872D | C | 2.2e-07 | 2.6e-07 | 1.5e-07 | 2.100E-07 | 5.568E-08 | 1.500E-07 | 2.600E-07 |

| | | | | | | | | | | |
|---|---|---|---|---|---|---|---|---|---|---|
| 17425 | 872 | W872C | C | 1.7e-07 | 2.1e-07 | 1.1e-07 | 1.633E-07 | 5.033E-08 | 1.100E-07 | 2.100E-07 |
| 17426 | 872 | W872E | C | 2.8e-07 | 2.6e-07 | 2.6e-07 | 2.667E-07 | 1.155E-08 | 2.600E-07 | 2.800E-07 |
| 17427 | 872 | W872Q | C | 1.5e-07 | 2.5e-07 | 1.8e-07 | 1.933E-07 | 5.132E-08 | 1.500E-07 | 2.500E-07 |
| 17428 | 872 | W872G | C | 1.4e-07 | 1.9e-07 | 3.2e-07 | 2.167E-07 | 9.292E-08 | 1.400E-07 | 3.200E-07 |
| 17429 | 872 | W872H | C | 3.1e-07 | 1.6e-07 | 3.9e-07 | 2.867E-07 | 1.168E-07 | 1.600E-07 | 3.900E-07 |
| 17430 | 872 | W872I | C | 4.1e-07 | 5.2e-07 | 3.4e-07 | 4.233E-07 | 9.074E-08 | 3.400E-07 | 5.200E-07 |
| 17431 | 872 | W872L | C | 1.1e-07 | 2.1e-07 | 2.4e-07 | 1.867E-07 | 6.807E-08 | 1.100E-07 | 2.400E-07 |
| 17432 | 872 | W872K | C | 2.6e-07 | 1.7e-07 | 2.3e-07 | 2.200E-07 | 4.583E-08 | 1.700E-07 | 2.600E-07 |
| 17433 | 872 | W872M | C | 2.3e-07 | 2e-07 | 1.9e-07 | 2.067E-07 | 2.082E-08 | 1.900E-07 | 2.300E-07 |
| 17434 | 872 | W872F | C | 1.6e-07 | 1.8e-07 | 3.1e-07 | 2.167E-07 | 8.145E-08 | 1.600E-07 | 3.100E-07 |
| 17435 | 872 | W872P | C | 4.6e-07 | 1e-07 | 1.5e-07 | 2.367E-07 | 1.950E-07 | 1.000E-07 | 4.600E-07 |
| 17436 | 872 | W872S | C | 8.5e-07 | 1.1e-07 | 2e-07 | 3.867E-07 | 4.038E-07 | 1.100E-07 | 8.500E-07 |
| 17437 | 872 | W872T | C | 2.6e-07 | 1.6e-07 | 3.8e-07 | 2.667E-07 | 1.102E-07 | 1.600E-07 | 3.800E-07 |
| 17438 | 872 | W872W | C | 3e-07 | 1.8e-07 | 2.6e-07 | 2.467E-07 | 6.110E-08 | 1.800E-07 | 3.000E-07 |
| 17439 | 872 | W872Y | C | 1.7e-07 | 2.4e-07 | 1.6e-07 | 1.900E-07 | 4.359E-08 | 1.600E-07 | 2.400E-07 |
| 17440 | 872 | W872V | C | 1.1e-07 | 1.9e-07 | 2.5e-07 | 1.833E-07 | 7.024E-08 | 1.100E-07 | 2.500E-07 |
| 17441 | 873 | V873A | C | 2.1e-07 | 2.4e-07 | 2.5e-07 | 2.333E-07 | 2.082E-08 | 2.100E-07 | 2.500E-07 |
| 17442 | 873 | V873R | C | 1.9e-07 | 2.9e-07 | 2e-07 | 2.267E-07 | 5.508E-08 | 1.900E-07 | 2.900E-07 |
| 17443 | 873 | V873N | C | 2.9e-07 | 2.8e-07 | 2.5e-07 | 2.733E-07 | 2.082E-08 | 2.500E-07 | 2.900E-07 |
| 17444 | 873 | V873D | C | 1.4e-07 | 2.5e-07 | 5e-07 | 2.967E-07 | 1.845E-07 | 1.400E-07 | 5.000E-07 |
| 17445 | 873 | V873C | C | 1.4e-07 | 2.4e-07 | 2.1e-07 | 1.967E-07 | 5.132E-08 | 1.400E-07 | 2.400E-07 |
| 17446 | 873 | V873E | C | 5e-07 | 1.7e-07 | 1.8e-07 | 2.833E-07 | 1.877E-07 | 1.700E-07 | 5.000E-07 |
| 17447 | 873 | V873Q | C | 1.5e-07 | 1.9e-07 | 3.1e-07 | 2.167E-07 | 8.327E-08 | 1.500E-07 | 3.100E-07 |
| 17448 | 873 | V873G | C | 2.9e-07 | 2.8e-07 | 2.4e-07 | 2.700E-07 | 2.646E-08 | 2.400E-07 | 2.900E-07 |
| 17449 | 873 | V873H | C | 2.8e-07 | 1.4e-07 | 2.2e-07 | 2.133E-07 | 7.024E-08 | 1.400E-07 | 2.800E-07 |
| 17450 | 873 | V873I | C | 5.7e-07 | 3.4e-07 | 1.8e-07 | 3.633E-07 | 1.960E-07 | 1.800E-07 | 5.700E-07 |
| 17451 | 873 | V873L | C | 2.5e-07 | 3e-07 | 1.4e-07 | 2.300E-07 | 8.185E-08 | 1.400E-07 | 3.000E-07 |
| 17452 | 873 | V873K | C | 1.9e-07 | 1.7e-07 | 1.2e-07 | 1.600E-07 | 3.606E-08 | 1.200E-07 | 1.900E-07 |
| 17453 | 873 | V873M | C | 3.2e-07 | 1.7e-07 | 4.1e-07 | 3.000E-07 | 1.212E-07 | 1.700E-07 | 4.100E-07 |
| 17454 | 873 | V873F | C | 2.1e-07 | 1.4e-07 | 2e-07 | 1.833E-07 | 3.786E-08 | 1.400E-07 | 2.100E-07 |
| 17455 | 873 | V873P | C | 2e-07 | 2.3e-07 | 7.6e-08 | 1.687E-07 | 8.164E-08 | 7.600E-08 | 2.300E-07 |
| 17456 | 873 | V873S | C | 1.9e-07 | 1.4e-07 | 2.1e-07 | 1.800E-07 | 3.606E-08 | 1.400E-07 | 2.100E-07 |
| 17457 | 873 | V873T | C | 3.7e-07 | 1.3e-07 | 1.4e-07 | 2.133E-07 | 1.358E-07 | 1.300E-07 | 3.700E-07 |
| 17458 | 873 | V873W | C | 2.8e-07 | 1.4e-07 | 2.7e-07 | 2.300E-07 | 7.810E-08 | 1.400E-07 | 2.800E-07 |
| 17459 | 873 | V873Y | C | 1.3e-07 | 3.4e-07 | 3e-07 | 2.567E-07 | 1.115E-07 | 1.300E-07 | 3.400E-07 |
| 17460 | 873 | V873V | C | 1.9e-07 | 2.8e-07 | 1.6e-07 | 2.100E-07 | 6.245E-08 | 1.600E-07 | 2.800E-07 |
| 17461 | 874 | R874A | C | 1.3e-07 | 2.7e-07 | 2.8e-07 | 2.267E-07 | 8.386E-08 | 1.300E-07 | 2.800E-07 |
| 17462 | 874 | R874R | C | 1.9e-07 | 5.1e-07 | 1.7e-07 | 2.900E-07 | 1.908E-07 | 1.700E-07 | 5.100E-07 |
| 17463 | 874 | R874N | C | 3.7e-07 | 1.5e-07 | 2e-07 | 2.400E-07 | 1.153E-07 | 1.500E-07 | 3.700E-07 |
| 17464 | 874 | R874D | C | 1.6e-07 | 2.6e-07 | 8.4e-07 | 4.200E-07 | 3.672E-07 | 1.600E-07 | 8.400E-07 |
| 17465 | 874 | R874C | C | 4.1e-07 | 4e-07 | 1.8e-07 | 3.300E-07 | 1.300E-07 | 1.800E-07 | 4.100E-07 |
| 17466 | 874 | R874E | C | 2.4e-07 | 3e-07 | 2.1e-07 | 2.500E-07 | 4.583E-08 | 2.100E-07 | 3.000E-07 |
| 17467 | 874 | R874Q | C | 1.2e-07 | 1.1e-07 | 2.6e-07 | 1.633E-07 | 8.386E-08 | 1.100E-07 | 2.600E-07 |
| 17468 | 874 | R874G | C | 1.6e-07 | 2.8e-07 | 1.5e-07 | 1.967E-07 | 7.234E-08 | 1.500E-07 | 2.800E-07 |
| 17469 | 874 | R874H | C | 3.3e-07 | 3.2e-07 | 9.9e-07 | 5.467E-07 | 3.840E-07 | 3.200E-07 | 9.900E-07 |
| 17470 | 874 | R874I | C | 4.4e-07 | 1.6e-07 | 3.4e-07 | 3.133E-07 | 1.419E-07 | 1.600E-07 | 4.400E-07 |
| 17471 | 874 | R874L | C | 3.4e-07 | 2.1e-07 | 1.9e-07 | 2.467E-07 | 8.145E-08 | 1.900E-07 | 3.400E-07 |
| 17472 | 874 | R874K | C | 3.3e-07 | 2e-07 | 1.9e-07 | 2.400E-07 | 7.810E-08 | 1.900E-07 | 3.300E-07 |
| 17473 | 874 | R874M | C | 1.2e-07 | 1.6e-07 | 1.9e-07 | 1.567E-07 | 3.512E-08 | 1.200E-07 | 1.900E-07 |
| 17474 | 874 | R874F | C | 3e-07 | 2.7e-07 | 5.9e-07 | 3.867E-07 | 1.767E-07 | 2.700E-07 | 5.900E-07 |
| 17475 | 874 | R874P | C | 2.9e-07 | 1.5e-07 | 3.2e-07 | 2.533E-07 | 9.074E-08 | 1.500E-07 | 3.200E-07 |
| 17476 | 874 | R874S | C | 3.8e-07 | 3.5e-07 | 5.4e-07 | 4.233E-07 | 1.021E-07 | 3.500E-07 | 5.400E-07 |
| 17477 | 874 | R874T | C | 3.8e-07 | 3.8e-07 | 3e-07 | 3.533E-07 | 4.619E-08 | 3.000E-07 | 3.800E-07 |
| 17478 | 874 | R874W | C | 1.6e-07 | 2.3e-07 | 1.9e-07 | 1.933E-07 | 3.512E-08 | 1.600E-07 | 2.300E-07 |
| 17479 | 874 | R874Y | C | 3.6e-07 | 7.8e-08 | 4.1e-07 | 2.827E-07 | 1.790E-07 | 7.800E-08 | 4.100E-07 |
| 17480 | 874 | R874V | C | 2.3e-07 | 2.4e-07 | 4.4e-07 | 3.033E-07 | 1.185E-07 | 2.300E-07 | 4.400E-07 |
| 17481 | 875 | Q875A | C | 3.3e-07 | 2.7e-07 | 1.1e-07 | 2.367E-07 | 1.137E-07 | 1.100E-07 | 3.300E-07 |
| 17482 | 875 | Q875R | C | 2.6e-07 | 3.2e-07 | 2.3e-07 | 2.700E-07 | 4.583E-08 | 2.300E-07 | 3.200E-07 |
| 17483 | 875 | Q875N | C | 2.9e-07 | 2.7e-07 | 2.1e-07 | 2.567E-07 | 4.163E-08 | 2.100E-07 | 2.900E-07 |
| 17484 | 875 | Q875D | C | 2e-07 | 2.9e-07 | 3.6e-07 | 2.833E-07 | 8.021E-08 | 2.000E-07 | 3.600E-07 |
| 17485 | 875 | Q875C | C | 2.2e-07 | 1.7e-07 | 2.2e-07 | 2.033E-07 | 2.887E-08 | 1.700E-07 | 2.200E-07 |
| 17486 | 875 | Q875E | C | 3.2e-07 | 2.4e-07 | 3e-07 | 2.867E-07 | 4.163E-08 | 2.400E-07 | 3.200E-07 |
| 17487 | 875 | Q875Q | C | 2.1e-07 | 1.7e-07 | 1.5e-07 | 1.767E-07 | 3.055E-08 | 1.500E-07 | 2.100E-07 |
| 17488 | 875 | Q875G | C | 6.3e-07 | 3.2e-07 | 3.6e-07 | 4.367E-07 | 1.686E-07 | 3.200E-07 | 6.300E-07 |
| 17489 | 875 | Q875H | C | 1.5e-07 | 2.4e-07 | 5.9e-07 | 3.267E-07 | 2.325E-07 | 1.500E-07 | 5.900E-07 |
| 17490 | 875 | Q875I | C | 2.6e-07 | 7.9e-07 | 1.4e-07 | 3.967E-07 | 3.459E-07 | 1.400E-07 | 7.900E-07 |
| 17491 | 875 | Q875L | C | 2.4e-07 | 3.7e-07 | 3.6e-07 | 3.233E-07 | 7.234E-08 | 2.400E-07 | 3.700E-07 |
| 17492 | 875 | Q875K | C | 1.6e-07 | 3.4e-07 | 8e-08 | 1.933E-07 | 1.332E-07 | 8.000E-08 | 3.400E-07 |
| 17493 | 875 | Q875M | C | 2.2e-07 | 2.5e-07 | 2.4e-07 | 2.367E-07 | 1.528E-08 | 2.200E-07 | 2.500E-07 |
| 17494 | 875 | Q875F | C | 2e-07 | 4.6e-07 | 1.1e-07 | 2.567E-07 | 1.818E-07 | 1.100E-07 | 4.600E-07 |
| 17495 | 875 | Q875P | C | 5.8e-07 | 1.4e-07 | 2.9e-07 | 3.367E-07 | 2.237E-07 | 1.400E-07 | 5.800E-07 |
| 17496 | 875 | Q875S | C | 1.2e-07 | 2.8e-07 | 1.9e-07 | 1.967E-07 | 8.021E-08 | 1.200E-07 | 2.800E-07 |
| 17497 | 875 | Q875T | C | 2.8e-07 | 3.2e-07 | 2.4e-07 | 2.800E-07 | 4.000E-08 | 2.400E-07 | 3.200E-07 |
| 17498 | 875 | Q875W | C | 3.4e-07 | 2.1e-07 | 2.1e-07 | 2.533E-07 | 7.506E-08 | 2.100E-07 | 3.400E-07 |
| 17499 | 875 | Q875Y | C | 3.8e-07 | 1.8e-07 | 6.6e-08 | 2.087E-07 | 1.590E-07 | 6.600E-08 | 3.800E-07 |
| 17500 | 875 | Q875V | C | 2.1e-07 | 5e-07 | 4.1e-07 | 3.733E-07 | 1.484E-07 | 2.100E-07 | 5.000E-07 |
| 17501 | 876 | A876A | C | 6.3e-07 | 2.7e-07 | 2.7e-07 | 3.900E-07 | 2.078E-07 | 2.700E-07 | 6.300E-07 |
| 17502 | 876 | A876R | C | 2.3e-07 | 2.3e-07 | 2.4e-07 | 2.333E-07 | 5.774E-09 | 2.300E-07 | 2.400E-07 |
| 17503 | 876 | A876N | C | 2.2e-07 | 1.8e-07 | 4.8e-07 | 2.933E-07 | 1.629E-07 | 1.800E-07 | 4.800E-07 |
| 17504 | 876 | A876D | C | 2e-07 | 2.9e-07 | 2.1e-07 | 2.333E-07 | 4.933E-08 | 2.000E-07 | 2.900E-07 |
| 17505 | 876 | A876C | C | 4.7e-07 | 2e-07 | 2.3e-07 | 3.000E-07 | 1.480E-07 | 2.000E-07 | 4.700E-07 |
| 17506 | 876 | A876E | C | 2.4e-07 | 1.5e-07 | 3.3e-07 | 2.400E-07 | 9.000E-08 | 1.500E-07 | 3.300E-07 |
| 17507 | 876 | A876Q | C | 2.2e-07 | 5.4e-07 | 1.2e-07 | 2.933E-07 | 2.194E-07 | 1.200E-07 | 5.400E-07 |
| 17508 | 876 | A876G | C | 9.5e-08 | 3.9e-07 | 3.1e-07 | 2.650E-07 | 1.526E-07 | 9.500E-08 | 3.900E-07 |
| 17509 | 876 | A876H | C | 3e-07 | 1.2e-07 | 3.3e-07 | 2.500E-07 | 1.136E-07 | 1.200E-07 | 3.300E-07 |

| | | | | | | | | | | |
|---|---|---|---|---|---|---|---|---|---|---|
| 17510 | 876 | A876I | C | 2.1e-07 | 4.9e-07 | 3.3e-07 | 3.433E-07 | 1.405E-07 | 2.100E-07 | 4.900E-07 |
| 17511 | 876 | A876L | C | 3.8e-07 | 5.1e-07 | 3.4e-07 | 4.100E-07 | 8.888E-08 | 3.400E-07 | 5.100E-07 |
| 17512 | 876 | A876K | C | 4.9e-07 | 3e-07 | 4.7e-07 | 4.200E-07 | 1.044E-07 | 3.000E-07 | 4.900E-07 |
| 17513 | 876 | A876M | C | 2.1e-07 | 2.5e-07 | 3.3e-07 | 2.633E-07 | 6.110E-08 | 2.100E-07 | 3.300E-07 |
| 17514 | 876 | A876F | C | 1.1e-07 | 3.1e-07 | 4.5e-07 | 2.900E-07 | 1.709E-07 | 1.100E-07 | 4.500E-07 |
| 17515 | 876 | A876P | C | 2.8e-07 | 2.3e-07 | 3.3e-07 | 2.800E-07 | 5.000E-08 | 2.300E-07 | 3.300E-07 |
| 17516 | 876 | A876S | C | 8.3e-08 | 2.4e-07 | 1.8e-07 | 1.677E-07 | 7.922E-08 | 8.300E-08 | 2.400E-07 |
| 17517 | 876 | A876T | C | 1.9e-07 | 3.5e-07 | 1.4e-07 | 2.267E-07 | 1.097E-07 | 1.400E-07 | 3.500E-07 |
| 17518 | 876 | A876W | C | 2e-07 | 1e-07 | 3.3e-07 | 2.100E-07 | 1.153E-07 | 1.000E-07 | 3.300E-07 |
| 17519 | 876 | A876Y | C | 2.3e-07 | 1.2e-07 | 3.4e-07 | 2.300E-07 | 1.100E-07 | 1.200E-07 | 3.400E-07 |
| 17520 | 876 | A876V | C | 2.7e-07 | 1.8e-07 | 2.3e-07 | 2.267E-07 | 4.509E-08 | 1.800E-07 | 2.700E-07 |
| 17521 | 877 | P877A | C | 4.9e-07 | 7.2e-07 | 1e-07 | 4.367E-07 | 3.134E-07 | 1.000E-07 | 7.200E-07 |
| 17522 | 877 | P877R | C | 6.9e-07 | 5.3e-07 | 2.7e-07 | 4.967E-07 | 2.120E-07 | 2.700E-07 | 6.900E-07 |
| 17523 | 877 | P877N | C | 2.4e-07 | 3.8e-07 | 1.5e-07 | 2.567E-07 | 1.159E-07 | 1.500E-07 | 3.800E-07 |
| 17524 | 877 | P877D | C | 3.8e-07 | 2.5e-07 | 5.8e-07 | 4.033E-07 | 1.662E-07 | 2.500E-07 | 5.800E-07 |
| 17525 | 877 | P877C | C | 3.2e-07 | 2.6e-07 | 2.3e-07 | 2.700E-07 | 4.583E-08 | 2.300E-07 | 3.200E-07 |
| 17526 | 877 | P877E | C | 3.6e-07 | 3.4e-07 | 1.8e-07 | 2.933E-07 | 9.866E-08 | 1.800E-07 | 3.600E-07 |
| 17527 | 877 | P877Q | C | 7.9e-08 | 2.5e-07 | 2.2e-07 | 1.830E-07 | 9.131E-08 | 7.900E-08 | 2.500E-07 |
| 17528 | 877 | P877G | C | 1.9e-07 | 1.8e-07 | 3.6e-07 | 2.433E-07 | 1.012E-07 | 1.800E-07 | 3.600E-07 |
| 17529 | 877 | P877H | C | 2.2e-07 | 3.2e-07 | 2.4e-07 | 2.600E-07 | 5.292E-08 | 2.200E-07 | 3.200E-07 |
| 17530 | 877 | P877I | C | 9.2e-08 | 1.3e-07 | 3.8e-07 | 2.007E-07 | 1.565E-07 | 9.200E-08 | 3.800E-07 |
| 17531 | 877 | P877L | C | 3.1e-07 | 3.8e-07 | 2.7e-07 | 3.200E-07 | 5.568E-08 | 2.700E-07 | 3.800E-07 |
| 17532 | 877 | P877K | C | 5e-07 | 1.1e-07 | 3e-07 | 3.033E-07 | 1.950E-07 | 1.100E-07 | 5.000E-07 |
| 17533 | 877 | P877M | C | 4.8e-07 | 2.6e-07 | 2.9e-07 | 3.433E-07 | 1.193E-07 | 2.600E-07 | 4.800E-07 |
| 17534 | 877 | P877F | C | 3.4e-07 | 2.8e-07 | 2.9e-07 | 3.033E-07 | 3.215E-08 | 2.800E-07 | 3.400E-07 |
| 17535 | 877 | P877P | C | 2.8e-07 | 1.2e-07 | 1.3e-07 | 1.767E-07 | 8.963E-08 | 1.200E-07 | 2.800E-07 |
| 17536 | 877 | P877S | C | 2.7e-07 | 3.9e-07 | 1.5e-07 | 2.700E-07 | 1.200E-07 | 1.500E-07 | 3.900E-07 |
| 17537 | 877 | P877T | C | 3.8e-07 | 2.1e-07 | 1.9e-07 | 2.600E-07 | 1.044E-07 | 1.900E-07 | 3.800E-07 |
| 17538 | 877 | P877W | C | 1.1e-07 | 1.3e-07 | 1.5e-07 | 1.300E-07 | 2.000E-08 | 1.100E-07 | 1.500E-07 |
| 17539 | 877 | P877Y | C | 5.1e-07 | 2.6e-07 | 1.5e-07 | 3.067E-07 | 1.845E-07 | 1.500E-07 | 5.100E-07 |
| 17540 | 877 | P877V | C | 4.1e-07 | 2.5e-07 | 2.6e-07 | 3.067E-07 | 8.963E-08 | 2.500E-07 | 4.100E-07 |
| 17541 | 878 | G878A | C | 2.5e-07 | 3.9e-07 | 2.8e-07 | 3.067E-07 | 7.371E-08 | 2.500E-07 | 3.900E-07 |
| 17542 | 878 | G878R | C | 7e-07 | 1.7e-07 | 2.9e-07 | 3.867E-07 | 2.779E-07 | 1.700E-07 | 7.000E-07 |
| 17543 | 878 | G878N | C | 1.3e-07 | 2.3e-07 | 3.7e-07 | 2.433E-07 | 1.206E-07 | 1.300E-07 | 3.700E-07 |
| 17544 | 878 | G878D | C | 3.4e-07 | 2.8e-07 | 2.6e-07 | 2.933E-07 | 4.163E-08 | 2.600E-07 | 3.400E-07 |
| 17545 | 878 | G878C | C | 1.7e-07 | 1.7e-07 | 2.4e-07 | 1.933E-07 | 4.041E-08 | 1.700E-07 | 2.400E-07 |
| 17546 | 878 | G878E | C | 2.3e-07 | 2.8e-07 | 3.4e-07 | 2.833E-07 | 5.508E-08 | 2.300E-07 | 3.400E-07 |
| 17547 | 878 | G878Q | C | 3.6e-07 | 1.1e-07 | 2.9e-07 | 2.533E-07 | 1.290E-07 | 1.100E-07 | 3.600E-07 |
| 17548 | 878 | G878G | C | 2.2e-07 | 3.2e-07 | 3e-07 | 2.800E-07 | 5.292E-08 | 2.200E-07 | 3.200E-07 |
| 17549 | 878 | G878H | C | 1.9e-07 | 1.2e-07 | 2.6e-07 | 1.900E-07 | 7.000E-08 | 1.200E-07 | 2.600E-07 |
| 17550 | 878 | G878I | C | 9.3e-08 | 2.2e-07 | 2.8e-07 | 1.977E-07 | 9.548E-08 | 9.300E-08 | 2.800E-07 |
| 17551 | 878 | G878L | C | 1.9e-07 | 2.1e-07 | 2.7e-07 | 2.233E-07 | 4.163E-08 | 1.900E-07 | 2.700E-07 |
| 17552 | 878 | G878K | C | 1.7e-07 | 2.2e-07 | 8.3e-08 | 1.577E-07 | 6.933E-08 | 8.300E-08 | 2.200E-07 |
| 17553 | 878 | G878M | C | 2.9e-07 | 6.1e-07 | 4.1e-07 | 4.367E-07 | 1.617E-07 | 2.900E-07 | 6.100E-07 |
| 17554 | 878 | G878F | C | 1.6e-07 | 1.1e-07 | 2.2e-07 | 1.633E-07 | 5.508E-08 | 1.100E-07 | 2.200E-07 |
| 17555 | 878 | G878P | C | 1.8e-07 | 3e-07 | 1.7e-07 | 2.167E-07 | 7.234E-08 | 1.700E-07 | 3.000E-07 |
| 17556 | 878 | G878S | C | 1.2e-07 | 1.5e-07 | 5.5e-07 | 2.733E-07 | 2.401E-07 | 1.200E-07 | 5.500E-07 |
| 17557 | 878 | G878T | C | 3.3e-07 | 3.2e-07 | 3.9e-07 | 3.467E-07 | 3.786E-08 | 3.200E-07 | 3.900E-07 |
| 17558 | 878 | G878W | C | 1.7e-07 | 2.3e-07 | 3.8e-07 | 2.600E-07 | 1.082E-07 | 1.700E-07 | 3.800E-07 |
| 17559 | 878 | G878Y | C | 2.5e-07 | 7.1e-07 | 2.7e-07 | 4.100E-07 | 2.600E-07 | 2.500E-07 | 7.100E-07 |
| 17560 | 878 | G878V | C | 5.8e-07 | 3.2e-07 | 2.1e-07 | 3.700E-07 | 1.900E-07 | 2.100E-07 | 5.800E-07 |
| 17561 | 879 | K879A | C | 3e-07 | 5e-07 | 2.5e-07 | 3.500E-07 | 1.323E-07 | 2.500E-07 | 5.000E-07 |
| 17562 | 879 | K879R | C | 5.2e-07 | 1.6e-07 | 2.2e-07 | 3.000E-07 | 1.929E-07 | 1.600E-07 | 5.200E-07 |
| 17563 | 879 | K879N | C | 2.5e-07 | 2e-07 | 1.8e-07 | 2.100E-07 | 3.606E-08 | 1.800E-07 | 2.500E-07 |
| 17564 | 879 | K879D | C | 2.3e-07 | 4.8e-07 | 2.5e-07 | 3.200E-07 | 1.389E-07 | 2.300E-07 | 4.800E-07 |
| 17565 | 879 | K879C | C | 1.4e-07 | 1.7e-07 | 2.7e-07 | 1.933E-07 | 6.807E-08 | 1.400E-07 | 2.700E-07 |
| 17566 | 879 | K879E | C | 4e-07 | 2.7e-07 | 2.4e-07 | 3.033E-07 | 8.505E-08 | 2.400E-07 | 4.000E-07 |
| 17567 | 879 | K879Q | C | 4.1e-07 | 6.8e-07 | 2.3e-07 | 4.400E-07 | 2.265E-07 | 2.300E-07 | 6.800E-07 |
| 17568 | 879 | K879G | C | 2.1e-07 | 5.1e-07 | 3.2e-07 | 3.467E-07 | 1.518E-07 | 2.100E-07 | 5.100E-07 |
| 17569 | 879 | K879H | C | 2.3e-07 | 2.1e-07 | 2.4e-07 | 2.267E-07 | 1.528E-08 | 2.100E-07 | 2.400E-07 |
| 17570 | 879 | K879I | C | 3.4e-07 | 2.3e-07 | 3.4e-07 | 3.033E-07 | 6.351E-08 | 2.300E-07 | 3.400E-07 |
| 17571 | 879 | K879L | C | 2.2e-07 | 1.7e-07 | 3e-07 | 2.300E-07 | 6.557E-08 | 1.700E-07 | 3.000E-07 |
| 17572 | 879 | K879K | C | 4.7e-07 | 2.4e-07 | 5.3e-07 | 4.133E-07 | 1.531E-07 | 2.400E-07 | 5.300E-07 |
| 17573 | 879 | K879M | C | 1.6e-07 | 2e-07 | 9.2e-08 | 1.507E-07 | 5.460E-08 | 9.200E-08 | 2.000E-07 |
| 17574 | 879 | K879F | C | 2e-07 | 2.2e-07 | 3e-07 | 2.400E-07 | 5.292E-08 | 2.000E-07 | 3.000E-07 |
| 17575 | 879 | K879P | C | 1.5e-07 | 2e-07 | 2.2e-07 | 1.900E-07 | 3.606E-08 | 1.500E-07 | 2.200E-07 |
| 17576 | 879 | K879S | C | 1.8e-07 | 3.4e-07 | 3.5e-07 | 2.900E-07 | 9.539E-08 | 1.800E-07 | 3.500E-07 |
| 17577 | 879 | K879T | C | 3e-07 | 2.5e-07 | 2.1e-07 | 2.533E-07 | 4.509E-08 | 2.100E-07 | 3.000E-07 |
| 17578 | 879 | K879W | C | 3.5e-07 | 2.2e-07 | 1.7e-07 | 2.467E-07 | 9.292E-08 | 1.700E-07 | 3.500E-07 |
| 17579 | 879 | K879Y | C | 3.9e-07 | 2e-07 | 2.5e-07 | 2.800E-07 | 9.849E-08 | 2.000E-07 | 3.900E-07 |
| 17580 | 879 | K879V | C | 2.2e-07 | 2.8e-07 | 1.9e-07 | 2.300E-07 | 4.583E-08 | 1.900E-07 | 2.800E-07 |
| 17581 | 880 | G880A | C | 1.9e-07 | 2.1e-07 | 6.4e-07 | 3.467E-07 | 2.542E-07 | 1.900E-07 | 6.400E-07 |
| 17582 | 880 | G880R | C | 3.8e-07 | 1.8e-07 | 4.2e-07 | 3.267E-07 | 1.286E-07 | 1.800E-07 | 4.200E-07 |
| 17583 | 880 | G880N | C | 2.1e-07 | 2.5e-07 | 2.5e-07 | 2.367E-07 | 2.309E-08 | 2.100E-07 | 2.500E-07 |
| 17584 | 880 | G880D | C | 3.7e-07 | 1.8e-07 | 2.9e-07 | 2.800E-07 | 9.539E-08 | 1.800E-07 | 3.700E-07 |
| 17585 | 880 | G880C | C | 2.8e-07 | 2.1e-07 | 4.2e-07 | 3.033E-07 | 1.069E-07 | 2.100E-07 | 4.200E-07 |
| 17586 | 880 | G880E | C | 1.4e-07 | 1e-07 | 1.1e-07 | 1.167E-07 | 2.082E-08 | 1.000E-07 | 1.400E-07 |
| 17587 | 880 | G880Q | C | 3.7e-07 | 3e-07 | 6.2e-07 | 4.300E-07 | 1.682E-07 | 3.000E-07 | 6.200E-07 |
| 17588 | 880 | G880G | C | 2.2e-07 | 1.4e-07 | 1.8e-07 | 1.800E-07 | 4.000E-08 | 1.400E-07 | 2.200E-07 |
| 17589 | 880 | G880H | C | 3.6e-07 | 3.1e-07 | 1.1e-07 | 2.600E-07 | 1.323E-07 | 1.100E-07 | 3.600E-07 |
| 17590 | 880 | G880I | C | 3.6e-07 | 7.2e-08 | 2.7e-07 | 2.340E-07 | 1.473E-07 | 7.200E-08 | 3.600E-07 |
| 17591 | 880 | G880L | C | 1.3e-07 | 1.3e-07 | 2.8e-07 | 1.800E-07 | 8.660E-08 | 1.300E-07 | 2.800E-07 |
| 17592 | 880 | G880K | C | 1.6e-07 | 3.5e-07 | 2.6e-07 | 2.567E-07 | 9.504E-08 | 1.600E-07 | 3.500E-07 |
| 17593 | 880 | G880M | C | 4.3e-07 | 2.9e-07 | 1.7e-07 | 2.967E-07 | 1.301E-07 | 1.700E-07 | 4.300E-07 |
| 17594 | 880 | G880F | C | 1.5e-07 | 1.1e-07 | 1.8e-07 | 1.467E-07 | 3.512E-08 | 1.100E-07 | 1.800E-07 |

| | | | | | | | | | | |
|---|---|---|---|---|---|---|---|---|---|---|
| 17595 | 880 | G880P | C | 6.9e-07 | 3.2e-07 | 2.2e-07 | 4.100E-07 | 2.476E-07 | 2.200E-07 | 6.900E-07 |
| 17596 | 880 | G880S | C | 2.7e-07 | 2.6e-07 | 2.9e-07 | 2.733E-07 | 1.528E-08 | 2.600E-07 | 2.900E-07 |
| 17597 | 880 | G880T | C | 2.4e-07 | 1.4e-07 | 2.4e-07 | 2.067E-07 | 5.774E-08 | 1.400E-07 | 2.400E-07 |
| 17598 | 880 | G880W | C | 2e-07 | 2.9e-07 | 2.6e-07 | 2.500E-07 | 4.583E-08 | 2.000E-07 | 2.900E-07 |
| 17599 | 880 | G880Y | C | 2e-07 | 1.2e-07 | 3.6e-07 | 2.267E-07 | 1.222E-07 | 1.200E-07 | 3.600E-07 |
| 17600 | 880 | G880V | C | 4.1e-07 | 1.6e-07 | 1.7e-07 | 2.467E-07 | 1.415E-07 | 1.600E-07 | 4.100E-07 |
| 17601 | 881 | L881A | C | 2.4e-07 | 4.2e-07 | 3.1e-07 | 3.233E-07 | 9.074E-08 | 2.400E-07 | 4.200E-07 |
| 17602 | 881 | L881R | C | 2e-07 | 6.1e-07 | 1.4e-07 | 3.167E-07 | 2.558E-07 | 1.400E-07 | 6.100E-07 |
| 17603 | 881 | L881N | C | 2.5e-07 | 1.2e-07 | 2.7e-07 | 2.133E-07 | 8.145E-08 | 1.200E-07 | 2.700E-07 |
| 17604 | 881 | L881D | C | 2.2e-07 | 3.3e-07 | 2.2e-07 | 2.567E-07 | 6.351E-08 | 2.200E-07 | 3.300E-07 |
| 17605 | 881 | L881C | C | 4.6e-07 | 2.8e-07 | 1.6e-07 | 3.000E-07 | 1.510E-07 | 1.600E-07 | 4.600E-07 |
| 17606 | 881 | L881E | C | 2.6e-07 | 3.2e-07 | 2.4e-07 | 2.733E-07 | 4.163E-08 | 2.400E-07 | 3.200E-07 |
| 17607 | 881 | L881Q | C | 2.2e-07 | 2.8e-07 | 2.1e-07 | 2.367E-07 | 3.786E-08 | 2.100E-07 | 2.800E-07 |
| 17608 | 881 | L881G | C | 1.5e-07 | 2.6e-07 | 3.4e-07 | 2.500E-07 | 9.539E-08 | 1.500E-07 | 3.400E-07 |
| 17609 | 881 | L881H | C | 3.1e-07 | 7.8e-08 | 3.8e-07 | 2.560E-07 | 1.581E-07 | 7.800E-08 | 3.800E-07 |
| 17610 | 881 | L881I | C | 3.7e-07 | 3.2e-07 | 2.7e-07 | 3.200E-07 | 5.000E-08 | 2.700E-07 | 3.700E-07 |
| 17611 | 881 | L881L | C | 3.9e-07 | 2.6e-07 | 1.9e-07 | 2.800E-07 | 1.015E-07 | 1.900E-07 | 3.900E-07 |
| 17612 | 881 | L881K | C | 1.8e-07 | 2.1e-07 | 2.5e-07 | 2.133E-07 | 3.512E-08 | 1.800E-07 | 2.500E-07 |
| 17613 | 881 | L881M | C | 1.8e-07 | 3.9e-07 | 2.8e-07 | 2.833E-07 | 1.050E-07 | 1.800E-07 | 3.900E-07 |
| 17614 | 881 | L881F | C | 2.6e-07 | 2.6e-07 | 4e-07 | 3.067E-07 | 8.083E-08 | 2.600E-07 | 4.000E-07 |
| 17615 | 881 | L881P | C | 2.9e-07 | 2.5e-07 | 1.8e-07 | 2.400E-07 | 5.568E-08 | 1.800E-07 | 2.900E-07 |
| 17616 | 881 | L881S | C | 1.9e-07 | 1.6e-07 | 5.1e-07 | 2.867E-07 | 1.940E-07 | 1.600E-07 | 5.100E-07 |
| 17617 | 881 | L881T | C | 2.3e-07 | 3.5e-07 | 2e-07 | 2.600E-07 | 7.937E-08 | 2.000E-07 | 3.500E-07 |
| 17618 | 881 | L881W | C | 5.7e-07 | 2.8e-07 | 2e-07 | 3.500E-07 | 1.947E-07 | 2.000E-07 | 5.700E-07 |
| 17619 | 881 | L881Y | C | 2.7e-07 | 1.7e-07 | 2.8e-07 | 2.400E-07 | 6.083E-08 | 1.700E-07 | 2.800E-07 |
| 17620 | 881 | L881V | C | 2.7e-07 | 3.4e-07 | 4.5e-07 | 3.533E-07 | 9.074E-08 | 2.700E-07 | 4.500E-07 |
| 17621 | 882 | E882A | C | 3e-07 | 3.4e-07 | 3.7e-07 | 3.367E-07 | 3.512E-08 | 3.000E-07 | 3.700E-07 |
| 17622 | 882 | E882R | C | 1.6e-07 | 1.7e-07 | 3.8e-07 | 2.367E-07 | 1.242E-07 | 1.600E-07 | 3.800E-07 |
| 17623 | 882 | E882N | C | 2.1e-07 | 2.7e-07 | 1.6e-07 | 2.133E-07 | 5.508E-08 | 1.600E-07 | 2.700E-07 |
| 17624 | 882 | E882D | C | 1.4e-07 | 5.1e-07 | 2.2e-07 | 2.900E-07 | 1.947E-07 | 1.400E-07 | 5.100E-07 |
| 17625 | 882 | E882C | C | 5.8e-08 | 1.5e-07 | 2.2e-07 | 1.427E-07 | 8.125E-08 | 5.800E-08 | 2.200E-07 |
| 17626 | 882 | E882E | C | 1.4e-07 | 2.5e-07 | 2.4e-07 | 2.100E-07 | 6.083E-08 | 1.400E-07 | 2.500E-07 |
| 17627 | 882 | E882Q | C | 5e-07 | 1.2e-07 | 1.6e-07 | 2.600E-07 | 2.088E-07 | 1.200E-07 | 5.000E-07 |
| 17628 | 882 | E882G | C | 4.5e-07 | 2.1e-07 | 2.5e-07 | 3.033E-07 | 1.286E-07 | 2.100E-07 | 4.500E-07 |
| 17629 | 882 | E882H | C | 2.3e-07 | 5.8e-07 | 2.3e-07 | 3.467E-07 | 2.021E-07 | 2.300E-07 | 5.800E-07 |
| 17630 | 882 | E882I | C | 2.6e-07 | 3.9e-07 | 3.5e-07 | 3.333E-07 | 6.658E-08 | 2.600E-07 | 3.900E-07 |
| 17631 | 882 | E882L | C | 2.2e-07 | 1.9e-07 | 6.3e-07 | 3.467E-07 | 2.458E-07 | 1.900E-07 | 6.300E-07 |
| 17632 | 882 | E882K | C | 2.2e-07 | 2.1e-07 | 2.5e-07 | 2.267E-07 | 2.082E-08 | 2.100E-07 | 2.500E-07 |
| 17633 | 882 | E882M | C | 2.7e-07 | 1.1e-06 | 1.8e-07 | 5.167E-07 | 5.072E-07 | 1.800E-07 | 1.100E-06 |
| 17634 | 882 | E882F | C | 2.5e-07 | 5.1e-07 | 2.7e-07 | 3.433E-07 | 1.447E-07 | 2.500E-07 | 5.100E-07 |
| 17635 | 882 | E882P | C | 2.5e-07 | 2e-07 | 3.1e-07 | 2.533E-07 | 5.508E-08 | 2.000E-07 | 3.100E-07 |
| 17636 | 882 | E882S | C | 2.3e-07 | 2.1e-07 | 2.2e-07 | 2.200E-07 | 1.000E-08 | 2.100E-07 | 2.300E-07 |
| 17637 | 882 | E882T | C | 1.8e-07 | 3e-07 | 4.4e-07 | 3.067E-07 | 1.301E-07 | 1.800E-07 | 4.400E-07 |
| 17638 | 882 | E882W | C | 3.9e-07 | 2.6e-07 | 2.4e-07 | 2.967E-07 | 8.145E-08 | 2.400E-07 | 3.900E-07 |
| 17639 | 882 | E882Y | C | 3.4e-07 | 2.5e-07 | 3.9e-07 | 3.267E-07 | 7.095E-08 | 2.500E-07 | 3.900E-07 |
| 17640 | 882 | E882V | C | 3.3e-07 | 2.9e-07 | 6.5e-07 | 4.233E-07 | 1.973E-07 | 2.900E-07 | 6.500E-07 |
| 17641 | 883 | W883A | C | 3.5e-07 | 1.7e-07 | 3.4e-07 | 2.867E-07 | 1.012E-07 | 1.700E-07 | 3.500E-07 |
| 17642 | 883 | W883R | C | 2.3e-07 | 1.2e-07 | 1.7e-07 | 1.733E-07 | 5.508E-08 | 1.200E-07 | 2.300E-07 |
| 17643 | 883 | W883N | C | 4.7e-07 | 1.9e-07 | 5.3e-07 | 3.967E-07 | 1.815E-07 | 1.900E-07 | 5.300E-07 |
| 17644 | 883 | W883D | C | 2.8e-07 | 1.8e-07 | 1.8e-07 | 2.133E-07 | 5.774E-08 | 1.800E-07 | 2.800E-07 |
| 17645 | 883 | W883C | C | 1.4e-07 | 4e-07 | 4.3e-07 | 3.233E-07 | 1.595E-07 | 1.400E-07 | 4.300E-07 |
| 17646 | 883 | W883E | C | 2.4e-07 | 1.6e-07 | 4.3e-07 | 2.767E-07 | 1.387E-07 | 1.600E-07 | 4.300E-07 |
| 17647 | 883 | W883Q | C | 1.5e-07 | 2.3e-07 | 1.2e-07 | 1.667E-07 | 5.686E-08 | 1.200E-07 | 2.300E-07 |
| 17648 | 883 | W883G | C | 1.6e-07 | 1e-07 | 1.7e-07 | 1.433E-07 | 3.786E-08 | 1.000E-07 | 1.700E-07 |
| 17649 | 883 | W883H | C | 1.6e-07 | 2.3e-07 | 3.8e-07 | 2.567E-07 | 1.124E-07 | 1.600E-07 | 3.800E-07 |
| 17650 | 883 | W883I | C | 3.4e-07 | 2.7e-07 | 2e-07 | 2.700E-07 | 7.000E-08 | 2.000E-07 | 3.400E-07 |
| 17651 | 883 | W883L | C | 4.7e-07 | 1.8e-07 | 1.4e-07 | 2.633E-07 | 1.801E-07 | 1.400E-07 | 4.700E-07 |
| 17652 | 883 | W883K | C | 2.5e-07 | 2.9e-07 | 2.8e-07 | 2.733E-07 | 2.082E-08 | 2.500E-07 | 2.900E-07 |
| 17653 | 883 | W883M | C | 1.9e-07 | 2.2e-07 | 2.5e-07 | 2.200E-07 | 3.000E-08 | 1.900E-07 | 2.500E-07 |
| 17654 | 883 | W883F | C | 1.6e-07 | 8.2e-08 | 2.3e-07 | 1.573E-07 | 7.404E-08 | 8.200E-08 | 2.300E-07 |
| 17655 | 883 | W883P | C | 2.1e-07 | 1.2e-07 | 3.2e-07 | 2.167E-07 | 1.002E-07 | 1.200E-07 | 3.200E-07 |
| 17656 | 883 | W883S | C | 4.3e-07 | 1.8e-07 | 2.1e-07 | 2.733E-07 | 1.365E-07 | 1.800E-07 | 4.300E-07 |
| 17657 | 883 | W883T | C | 3.1e-07 | 4.9e-07 | 2.4e-07 | 3.467E-07 | 1.290E-07 | 2.400E-07 | 4.900E-07 |
| 17658 | 883 | W883W | C | 1.9e-07 | 2.3e-07 | 2.8e-07 | 2.333E-07 | 4.509E-08 | 1.900E-07 | 2.800E-07 |
| 17659 | 883 | W883Y | C | 5.5e-07 | 3.4e-07 | 2.6e-07 | 3.833E-07 | 1.498E-07 | 2.600E-07 | 5.500E-07 |
| 17660 | 883 | W883V | C | 3.3e-07 | 2.5e-07 | 7.3e-07 | 4.367E-07 | 2.572E-07 | 2.500E-07 | 7.300E-07 |
| 17661 | 884 | V884A | C | 5.7e-08 | 3e-07 | 1.9e-07 | 1.823E-07 | 1.217E-07 | 5.700E-08 | 3.000E-07 |
| 17662 | 884 | V884R | C | 2.5e-07 | 3.5e-07 | 3.2e-07 | 3.067E-07 | 5.132E-08 | 2.500E-07 | 3.500E-07 |
| 17663 | 884 | V884N | C | 3.5e-07 | 3.4e-07 | 2.7e-07 | 3.200E-07 | 4.359E-08 | 2.700E-07 | 3.500E-07 |
| 17664 | 884 | V884D | C | 2.2e-07 | 1.9e-07 | 4.7e-07 | 2.933E-07 | 1.537E-07 | 1.900E-07 | 4.700E-07 |
| 17665 | 884 | V884C | C | 2.9e-07 | 1.3e-07 | 5.7e-07 | 3.300E-07 | 2.227E-07 | 1.300E-07 | 5.700E-07 |
| 17666 | 884 | V884E | C | 2.9e-07 | 3.7e-07 | 2e-07 | 2.867E-07 | 8.505E-08 | 2.000E-07 | 3.700E-07 |
| 17667 | 884 | V884Q | C | 2.3e-07 | 2.2e-07 | 3e-07 | 2.500E-07 | 4.359E-08 | 2.200E-07 | 3.000E-07 |
| 17668 | 884 | V884G | C | 3.1e-07 | 1.7e-07 | 3.5e-07 | 2.767E-07 | 9.452E-08 | 1.700E-07 | 3.500E-07 |
| 17669 | 884 | V884H | C | 1.2e-07 | 2e-07 | 1.1e-07 | 1.433E-07 | 4.933E-08 | 1.100E-07 | 2.000E-07 |
| 17670 | 884 | V884I | C | 2.3e-07 | 1.4e-07 | 2.3e-07 | 2.000E-07 | 5.196E-08 | 1.400E-07 | 2.300E-07 |
| 17671 | 884 | V884L | C | 2.4e-07 | 2.3e-07 | 2.1e-07 | 2.267E-07 | 1.528E-08 | 2.100E-07 | 2.400E-07 |
| 17672 | 884 | V884K | C | 5.3e-07 | 2.5e-07 | 2.5e-07 | 3.433E-07 | 1.617E-07 | 2.500E-07 | 5.300E-07 |
| 17673 | 884 | V884M | C | 2.4e-07 | 2e-07 | 2.2e-07 | 2.200E-07 | 2.000E-08 | 2.000E-07 | 2.400E-07 |
| 17674 | 884 | V884F | C | 2.3e-07 | 4.4e-07 | 1.4e-07 | 2.700E-07 | 1.539E-07 | 1.400E-07 | 4.400E-07 |
| 17675 | 884 | V884P | C | 2.2e-07 | 2.5e-07 | 1.6e-07 | 2.100E-07 | 4.583E-08 | 1.600E-07 | 2.500E-07 |
| 17676 | 884 | V884S | C | 3.9e-07 | 2.9e-07 | 2.7e-07 | 3.167E-07 | 6.429E-08 | 2.700E-07 | 3.900E-07 |
| 17677 | 884 | V884T | C | 2.3e-07 | 2e-07 | 1.3e-07 | 1.867E-07 | 5.132E-08 | 1.300E-07 | 2.300E-07 |
| 17678 | 884 | V884W | C | 5.2e-07 | 2.3e-07 | 1.7e-07 | 3.067E-07 | 1.872E-07 | 1.700E-07 | 5.200E-07 |
| 17679 | 884 | V884Y | C | 2.1e-07 | 1.5e-07 | 4.3e-07 | 2.633E-07 | 1.474E-07 | 1.500E-07 | 4.300E-07 |

| | | | | | | | | | | |
|---|---|---|---|---|---|---|---|---|---|---|
| 17680 | 884 | V884V | C | 1.6e-07 | 1.4e-07 | 2.5e-07 | 1.833E-07 | 5.859E-08 | 1.400E-07 | 2.500E-07 |
| 17681 | 885 | A885A | C | 5.9e-07 | 2e-07 | 6e-07 | 4.633E-07 | 2.281E-07 | 2.000E-07 | 6.000E-07 |
| 17682 | 885 | A885R | C | 1.2e-07 | 2.3e-07 | 1.4e-07 | 1.633E-07 | 5.859E-08 | 1.200E-07 | 2.300E-07 |
| 17683 | 885 | A885N | C | 3.3e-07 | 6.1e-07 | 2.9e-07 | 4.100E-07 | 1.744E-07 | 2.900E-07 | 6.100E-07 |
| 17684 | 885 | A885D | C | 2.5e-07 | 2.9e-07 | 2.4e-07 | 2.600E-07 | 2.646E-08 | 2.400E-07 | 2.900E-07 |
| 17685 | 885 | A885C | C | 3.8e-07 | 2.8e-07 | 6.8e-07 | 4.467E-07 | 2.082E-07 | 2.800E-07 | 6.800E-07 |
| 17686 | 885 | A885E | C | 2e-07 | 2.5e-07 | 3e-07 | 2.500E-07 | 5.000E-08 | 2.000E-07 | 3.000E-07 |
| 17687 | 885 | A885Q | C | 5.7e-07 | 2.3e-07 | 2.4e-07 | 3.467E-07 | 1.935E-07 | 2.300E-07 | 5.700E-07 |
| 17688 | 885 | A885G | C | 1.8e-07 | 2.5e-07 | 3.8e-07 | 2.700E-07 | 1.015E-07 | 1.800E-07 | 3.800E-07 |
| 17689 | 885 | A885H | C | 3.2e-07 | 2.3e-07 | 2.5e-07 | 2.667E-07 | 4.726E-08 | 2.300E-07 | 3.200E-07 |
| 17690 | 885 | A885I | C | 3e-07 | 4.7e-07 | 2.8e-07 | 3.500E-07 | 1.044E-07 | 2.800E-07 | 4.700E-07 |
| 17691 | 885 | A885L | C | 2.9e-07 | 3.3e-07 | 2.1e-07 | 2.767E-07 | 6.110E-08 | 2.100E-07 | 3.300E-07 |
| 17692 | 885 | A885K | C | 1.4e-07 | 1.3e-07 | 1.8e-07 | 1.500E-07 | 2.646E-08 | 1.300E-07 | 1.800E-07 |
| 17693 | 885 | A885M | C | 2.5e-07 | 1.4e-07 | 1.6e-07 | 1.833E-07 | 5.859E-08 | 1.400E-07 | 2.500E-07 |
| 17694 | 885 | A885F | C | 3.9e-07 | 2e-07 | 9.8e-08 | 2.293E-07 | 1.482E-07 | 9.800E-08 | 3.900E-07 |
| 17695 | 885 | A885P | C | 1.2e-07 | 6.6e-07 | 2.8e-07 | 3.533E-07 | 2.774E-07 | 1.200E-07 | 6.600E-07 |
| 17696 | 885 | A885S | C | 1.7e-07 | 2.3e-07 | 3.9e-07 | 2.633E-07 | 1.137E-07 | 1.700E-07 | 3.900E-07 |
| 17697 | 885 | A885T | C | 2.5e-07 | 2.7e-07 | 4.1e-07 | 3.100E-07 | 8.718E-08 | 2.500E-07 | 4.100E-07 |
| 17698 | 885 | A885W | C | 3.2e-07 | 2.4e-07 | 2e-07 | 2.533E-07 | 6.110E-08 | 2.000E-07 | 3.200E-07 |
| 17699 | 885 | A885Y | C | 3.5e-07 | 1.6e-07 | 1.7e-07 | 2.267E-07 | 1.069E-07 | 1.600E-07 | 3.500E-07 |
| 17700 | 885 | A885V | C | 1.5e-07 | 8.7e-08 | 8.6e-08 | 1.077E-07 | 3.667E-08 | 8.600E-08 | 1.500E-07 |
| 17701 | 886 | D886A | C | 2e-07 | 2.7e-07 | 1.6e-07 | 2.100E-07 | 5.568E-08 | 1.600E-07 | 2.700E-07 |
| 17702 | 886 | D886R | C | 3.9e-07 | 3e-07 | 3.7e-07 | 3.533E-07 | 4.726E-08 | 3.000E-07 | 3.900E-07 |
| 17703 | 886 | D886N | C | 3.5e-07 | 1.4e-07 | 3.1e-07 | 2.667E-07 | 1.115E-07 | 1.400E-07 | 3.500E-07 |
| 17704 | 886 | D886D | C | 4e-07 | 2e-07 | 1.9e-07 | 2.633E-07 | 1.185E-07 | 1.900E-07 | 4.000E-07 |
| 17705 | 886 | D886C | C | 2.8e-07 | 1.9e-07 | 2.1e-07 | 2.267E-07 | 4.726E-08 | 1.900E-07 | 2.800E-07 |
| 17706 | 886 | D886E | C | 3.8e-07 | 3.4e-07 | 1.4e-07 | 2.867E-07 | 1.286E-07 | 1.400E-07 | 3.800E-07 |
| 17707 | 886 | D886Q | C | 2e-07 | 3.5e-07 | 2.3e-07 | 2.600E-07 | 7.937E-08 | 2.000E-07 | 3.500E-07 |
| 17708 | 886 | D886G | C | 2.6e-07 | 2.5e-07 | 3.4e-07 | 2.833E-07 | 4.933E-08 | 2.500E-07 | 3.400E-07 |
| 17709 | 886 | D886H | C | 4.5e-07 | 2.9e-07 | 1.8e-07 | 3.067E-07 | 1.358E-07 | 1.800E-07 | 4.500E-07 |
| 17710 | 886 | D886I | C | 1.4e-07 | 3.8e-07 | 2.7e-07 | 2.633E-07 | 1.201E-07 | 1.400E-07 | 3.800E-07 |
| 17711 | 886 | D886L | C | 1.7e-07 | 4.3e-07 | 2.4e-07 | 2.800E-07 | 1.345E-07 | 1.700E-07 | 4.300E-07 |
| 17712 | 886 | D886K | C | 2e-07 | 1.5e-07 | 2.8e-07 | 2.100E-07 | 6.557E-08 | 1.500E-07 | 2.800E-07 |
| 17713 | 886 | D886M | C | 3.8e-07 | 3.5e-07 | 1.8e-07 | 3.033E-07 | 1.079E-07 | 1.800E-07 | 3.800E-07 |
| 17714 | 886 | D886F | C | 1.4e-07 | 3.8e-07 | 2.2e-07 | 2.467E-07 | 1.222E-07 | 1.400E-07 | 3.800E-07 |
| 17715 | 886 | D886P | C | 1.2e-07 | 3.3e-07 | 1.6e-07 | 2.033E-07 | 1.115E-07 | 1.200E-07 | 3.300E-07 |
| 17716 | 886 | D886S | C | 1.7e-07 | 2.7e-07 | 3.9e-07 | 2.767E-07 | 1.102E-07 | 1.700E-07 | 3.900E-07 |
| 17717 | 886 | D886T | C | 4.5e-07 | 1.8e-07 | 4.8e-07 | 3.700E-07 | 1.652E-07 | 1.800E-07 | 4.800E-07 |
| 17718 | 886 | D886W | C | 2.4e-07 | 4.8e-07 | 2.4e-07 | 3.200E-07 | 1.386E-07 | 2.400E-07 | 4.800E-07 |
| 17719 | 886 | D886Y | C | 5.2e-07 | 9.7e-08 | 1.3e-07 | 2.490E-07 | 2.353E-07 | 9.700E-08 | 5.200E-07 |
| 17720 | 886 | D886V | C | 2.1e-07 | 1.5e-07 | 4.8e-07 | 2.800E-07 | 1.758E-07 | 1.500E-07 | 4.800E-07 |
| 17721 | 887 | V887A | C | 1.8e-07 | 2.2e-07 | 2.2e-07 | 2.067E-07 | 2.309E-08 | 1.800E-07 | 2.200E-07 |
| 17722 | 887 | V887R | C | 1.9e-07 | 4.5e-07 | 2.3e-07 | 2.900E-07 | 1.400E-07 | 1.900E-07 | 4.500E-07 |
| 17723 | 887 | V887N | C | 4.3e-07 | 2.6e-07 | 4.3e-07 | 3.733E-07 | 9.815E-08 | 2.600E-07 | 4.300E-07 |
| 17724 | 887 | V887D | C | 1.5e-07 | 1.8e-07 | 1.5e-07 | 1.600E-07 | 1.732E-08 | 1.500E-07 | 1.800E-07 |
| 17725 | 887 | V887C | C | 2.4e-07 | 2e-07 | 7e-08 | 1.700E-07 | 8.888E-08 | 7.000E-08 | 2.400E-07 |
| 17726 | 887 | V887E | C | 2.1e-07 | 1.6e-07 | 1.1e-07 | 1.600E-07 | 5.000E-08 | 1.100E-07 | 2.100E-07 |
| 17727 | 887 | V887Q | C | 2.3e-07 | 1.5e-07 | 1.4e-07 | 1.733E-07 | 4.933E-08 | 1.400E-07 | 2.300E-07 |
| 17728 | 887 | V887G | C | 1.7e-07 | 2.7e-07 | 1.7e-07 | 2.033E-07 | 5.774E-08 | 1.700E-07 | 2.700E-07 |
| 17729 | 887 | V887H | C | 1.4e-07 | 1.8e-07 | 2.7e-07 | 1.967E-07 | 6.658E-08 | 1.400E-07 | 2.700E-07 |
| 17730 | 887 | V887I | C | 9.9e-08 | 3.6e-07 | 2.8e-07 | 2.463E-07 | 1.337E-07 | 9.900E-08 | 3.600E-07 |
| 17731 | 887 | V887L | C | 1.7e-07 | 2.7e-07 | 3.5e-07 | 2.633E-07 | 9.018E-08 | 1.700E-07 | 3.500E-07 |
| 17732 | 887 | V887K | C | 2.4e-07 | 3.1e-07 | 2.4e-07 | 2.633E-07 | 4.041E-08 | 2.400E-07 | 3.100E-07 |
| 17733 | 887 | V887M | C | 1.5e-07 | 2.2e-07 | 5.4e-07 | 3.033E-07 | 2.079E-07 | 1.500E-07 | 5.400E-07 |
| 17734 | 887 | V887F | C | 3.4e-07 | 2e-07 | 2.3e-07 | 2.567E-07 | 7.371E-08 | 2.000E-07 | 3.400E-07 |
| 17735 | 887 | V887P | C | 3.1e-07 | 2.9e-07 | 3.1e-07 | 3.033E-07 | 1.155E-08 | 2.900E-07 | 3.100E-07 |
| 17736 | 887 | V887S | C | 3e-07 | 2.1e-07 | 3.7e-07 | 2.933E-07 | 8.021E-08 | 2.100E-07 | 3.700E-07 |
| 17737 | 887 | V887T | C | 6e-08 | 2e-07 | 3.7e-07 | 2.100E-07 | 1.552E-07 | 6.000E-08 | 3.700E-07 |
| 17738 | 887 | V887W | C | 1.6e-07 | 1.4e-07 | 2.5e-07 | 1.833E-07 | 5.859E-08 | 1.400E-07 | 2.500E-07 |
| 17739 | 887 | V887Y | C | 1.3e-07 | 3.5e-07 | 5.4e-07 | 3.400E-07 | 2.052E-07 | 1.300E-07 | 5.400E-07 |
| 17740 | 887 | V887V | C | 2.2e-07 | 3.2e-07 | 1.2e-07 | 2.200E-07 | 1.000E-07 | 1.200E-07 | 3.200E-07 |
| 17741 | 888 | N888A | C | 1.8e-07 | 4e-07 | 3.1e-07 | 2.967E-07 | 1.106E-07 | 1.800E-07 | 4.000E-07 |
| 17742 | 888 | N888R | C | 4.9e-07 | 2.7e-07 | 1.6e-07 | 3.067E-07 | 1.680E-07 | 1.600E-07 | 4.900E-07 |
| 17743 | 888 | N888N | C | 2.7e-07 | 5.4e-07 | 2.1e-07 | 3.400E-07 | 1.758E-07 | 2.100E-07 | 5.400E-07 |
| 17744 | 888 | N888D | C | 2.5e-07 | 1.3e-07 | 1.5e-07 | 1.767E-07 | 6.429E-08 | 1.300E-07 | 2.500E-07 |
| 17745 | 888 | N888C | C | 4.8e-07 | 1.5e-07 | 3.4e-07 | 3.233E-07 | 1.656E-07 | 1.500E-07 | 4.800E-07 |
| 17746 | 888 | N888E | C | 2e-07 | 2e-07 | 1.9e-07 | 1.967E-07 | 5.774E-09 | 1.900E-07 | 2.000E-07 |
| 17747 | 888 | N888Q | C | 1.9e-07 | 4.4e-07 | 1.3e-07 | 2.533E-07 | 1.644E-07 | 1.300E-07 | 4.400E-07 |
| 17748 | 888 | N888G | C | 2.1e-07 | 3.6e-07 | 3.3e-07 | 3.000E-07 | 7.937E-08 | 2.100E-07 | 3.600E-07 |
| 17749 | 888 | N888H | C | 2e-07 | 2.6e-07 | 1.9e-07 | 2.167E-07 | 3.786E-08 | 1.900E-07 | 2.600E-07 |
| 17750 | 888 | N888I | C | 2.1e-07 | 4.1e-07 | 2.1e-07 | 2.767E-07 | 1.155E-07 | 2.100E-07 | 4.100E-07 |
| 17751 | 888 | N888L | C | 3.6e-07 | 3e-07 | 5.6e-07 | 4.067E-07 | 1.361E-07 | 3.000E-07 | 5.600E-07 |
| 17752 | 888 | N888K | C | 2.9e-07 | 4.8e-07 | 7.1e-08 | 2.803E-07 | 2.047E-07 | 7.100E-08 | 4.800E-07 |
| 17753 | 888 | N888M | C | 3.6e-07 | 5.4e-07 | 8.3e-07 | 5.767E-07 | 2.371E-07 | 3.600E-07 | 8.300E-07 |
| 17754 | 888 | N888F | C | 3.6e-07 | 2.1e-07 | 2.7e-07 | 2.800E-07 | 7.550E-08 | 2.100E-07 | 3.600E-07 |
| 17755 | 888 | N888P | C | 1.9e-07 | 1.9e-07 | 3.3e-07 | 2.367E-07 | 8.083E-08 | 1.900E-07 | 3.300E-07 |
| 17756 | 888 | N888S | C | 2.8e-07 | 2.2e-07 | 2.1e-07 | 2.367E-07 | 3.786E-08 | 2.100E-07 | 2.800E-07 |
| 17757 | 888 | N888T | C | 3.5e-07 | 6.4e-07 | 6.1e-07 | 5.333E-07 | 1.595E-07 | 3.500E-07 | 6.400E-07 |
| 17758 | 888 | N888W | C | 1.3e-07 | 5.2e-07 | 1.5e-07 | 2.667E-07 | 2.196E-07 | 1.300E-07 | 5.200E-07 |
| 17759 | 888 | N888Y | C | 3.4e-07 | 1.6e-07 | 9.6e-08 | 1.987E-07 | 1.265E-07 | 9.600E-08 | 3.400E-07 |
| 17760 | 888 | N888V | C | 5.5e-07 | 1.9e-07 | 3.3e-07 | 3.567E-07 | 1.815E-07 | 1.900E-07 | 5.500E-07 |
| 17761 | 889 | P889A | C | 3.6e-07 | 5.6e-07 | 2.4e-07 | 3.867E-07 | 1.617E-07 | 2.400E-07 | 5.600E-07 |
| 17762 | 889 | P889R | C | 9.3e-08 | 3.6e-07 | 5.4e-07 | 3.310E-07 | 2.249E-07 | 9.300E-08 | 5.400E-07 |
| 17763 | 889 | P889N | C | 1.4e-07 | 1.4e-07 | 1.7e-07 | 1.500E-07 | 1.732E-08 | 1.400E-07 | 1.700E-07 |
| 17764 | 889 | P889D | C | 2.7e-07 | 1.8e-07 | 4.8e-07 | 3.100E-07 | 1.539E-07 | 1.800E-07 | 4.800E-07 |

| | | | | | | | | | | |
|---|---|---|---|---|---|---|---|---|---|---|
| 17765 | 889 | P889C | C | 9.7e-08 | 1.2e-07 | 4.6e-07 | 2.257E-07 | 2.033E-07 | 9.700E-08 | 4.600E-07 |
| 17766 | 889 | P889E | C | 2.6e-07 | 1.9e-07 | 3.4e-07 | 2.633E-07 | 7.506E-08 | 1.900E-07 | 3.400E-07 |
| 17767 | 889 | P889Q | C | 3e-07 | 3.9e-07 | 2.7e-07 | 3.200E-07 | 6.245E-08 | 2.700E-07 | 3.900E-07 |
| 17768 | 889 | P889G | C | 2.5e-07 | 9.5e-08 | 2.7e-07 | 2.050E-07 | 9.579E-08 | 9.500E-08 | 2.700E-07 |
| 17769 | 889 | P889H | C | 2.6e-07 | 2.1e-07 | 1.3e-07 | 2.000E-07 | 6.557E-08 | 1.300E-07 | 2.600E-07 |
| 17770 | 889 | P889I | C | 2.1e-07 | 3.4e-07 | 3.5e-07 | 3.000E-07 | 7.810E-08 | 2.100E-07 | 3.500E-07 |
| 17771 | 889 | P889L | C | 2.7e-07 | 3.4e-07 | 2.9e-07 | 3.000E-07 | 3.606E-08 | 2.700E-07 | 3.400E-07 |
| 17772 | 889 | P889K | C | 6.6e-07 | 2.2e-07 | 1.2e-07 | 3.333E-07 | 2.873E-07 | 1.200E-07 | 6.600E-07 |
| 17773 | 889 | P889M | C | 2.7e-07 | 2.5e-07 | 5.4e-07 | 3.533E-07 | 1.620E-07 | 2.500E-07 | 5.400E-07 |
| 17774 | 889 | P889F | C | 2.1e-07 | 3.4e-07 | 2.2e-07 | 2.567E-07 | 7.234E-08 | 2.100E-07 | 3.400E-07 |
| 17775 | 889 | P889P | C | 1.7e-07 | 1.7e-07 | 2e-07 | 1.800E-07 | 1.732E-08 | 1.700E-07 | 2.000E-07 |
| 17776 | 889 | P889S | C | 3.8e-07 | 2.7e-07 | 1e-07 | 2.500E-07 | 1.411E-07 | 1.000E-07 | 3.800E-07 |
| 17777 | 889 | P889T | C | 2.5e-07 | 2.7e-07 | 2.9e-07 | 2.700E-07 | 2.000E-08 | 2.500E-07 | 2.900E-07 |
| 17778 | 889 | P889W | C | 1.4e-07 | 3e-07 | 8.2e-07 | 4.200E-07 | 3.555E-07 | 1.400E-07 | 8.200E-07 |
| 17779 | 889 | P889Y | C | 1.9e-07 | 1.9e-07 | 2.5e-07 | 2.100E-07 | 3.464E-08 | 1.900E-07 | 2.500E-07 |
| 17780 | 889 | P889V | C | 4.5e-07 | 4.3e-07 | 4.6e-07 | 4.467E-07 | 1.528E-08 | 4.300E-07 | 4.600E-07 |
| 17781 | 890 | N890A | C | 1.8e-07 | 2.1e-07 | 1.9e-07 | 1.933E-07 | 1.528E-08 | 1.800E-07 | 2.100E-07 |
| 17782 | 890 | N890R | C | 2.9e-07 | 2.1e-07 | 8.2e-07 | 4.400E-07 | 3.315E-07 | 2.100E-07 | 8.200E-07 |
| 17783 | 890 | N890N | C | 3.6e-07 | 3.7e-07 | 2.6e-07 | 3.300E-07 | 6.083E-08 | 2.600E-07 | 3.700E-07 |
| 17784 | 890 | N890D | C | 3.9e-07 | 2.6e-07 | 3.1e-07 | 3.200E-07 | 6.557E-08 | 2.600E-07 | 3.900E-07 |
| 17785 | 890 | N890C | C | 1.4e-07 | 1.4e-07 | 2.7e-07 | 1.833E-07 | 7.506E-08 | 1.400E-07 | 2.700E-07 |
| 17786 | 890 | N890E | C | 3.1e-07 | 1.6e-07 | 2.8e-07 | 2.500E-07 | 7.937E-08 | 1.600E-07 | 3.100E-07 |
| 17787 | 890 | N890Q | C | 4e-07 | 5.5e-07 | 1.5e-07 | 3.667E-07 | 2.021E-07 | 1.500E-07 | 5.500E-07 |
| 17788 | 890 | N890G | C | 1.5e-07 | 2.9e-07 | 3.7e-07 | 2.700E-07 | 1.114E-07 | 1.500E-07 | 3.700E-07 |
| 17789 | 890 | N890H | C | 3.6e-07 | 2.3e-07 | 3.2e-07 | 3.033E-07 | 6.658E-08 | 2.300E-07 | 3.600E-07 |
| 17790 | 890 | N890I | C | 1.9e-07 | 4.5e-07 | 1.6e-07 | 2.667E-07 | 1.595E-07 | 1.600E-07 | 4.500E-07 |
| 17791 | 890 | N890L | C | 1.8e-07 | 2.8e-07 | 1.4e-07 | 2.000E-07 | 7.211E-08 | 1.400E-07 | 2.800E-07 |
| 17792 | 890 | N890K | C | 1.3e-07 | 2.5e-07 | 2.4e-07 | 2.067E-07 | 6.658E-08 | 1.300E-07 | 2.500E-07 |
| 17793 | 890 | N890M | C | 2.7e-07 | 2.6e-07 | 1.8e-07 | 2.367E-07 | 4.933E-08 | 1.800E-07 | 2.700E-07 |
| 17794 | 890 | N890F | C | 2.2e-07 | 2.8e-07 | 4.3e-07 | 3.100E-07 | 1.082E-07 | 2.200E-07 | 4.300E-07 |
| 17795 | 890 | N890P | C | 2.9e-07 | 9.1e-08 | 3.2e-07 | 2.337E-07 | 1.245E-07 | 9.100E-08 | 3.200E-07 |
| 17796 | 890 | N890S | C | 1.9e-07 | 2.6e-07 | 2.7e-07 | 2.400E-07 | 4.359E-08 | 1.900E-07 | 2.700E-07 |
| 17797 | 890 | N890T | C | 2.4e-07 | 1.9e-07 | 5.4e-07 | 3.233E-07 | 1.893E-07 | 1.900E-07 | 5.400E-07 |
| 17798 | 890 | N890W | C | 1e-07 | 2.4e-07 | 2.4e-07 | 1.933E-07 | 8.083E-08 | 1.000E-07 | 2.400E-07 |
| 17799 | 890 | N890Y | C | 4.2e-07 | 1.4e-07 | 4.8e-07 | 3.467E-07 | 1.815E-07 | 1.400E-07 | 4.800E-07 |
| 17800 | 890 | N890V | C | 2.3e-07 | 2.3e-07 | 2.6e-07 | 2.400E-07 | 1.732E-08 | 2.300E-07 | 2.600E-07 |
| 17801 | 891 | S891A | C | 6.5e-07 | 1.7e-07 | 5.3e-07 | 4.500E-07 | 2.498E-07 | 1.700E-07 | 6.500E-07 |
| 17802 | 891 | S891R | C | 3e-07 | 3.1e-07 | 2e-07 | 2.700E-07 | 6.083E-08 | 2.000E-07 | 3.100E-07 |
| 17803 | 891 | S891N | C | 1.2e-07 | 1.3e-07 | 2.9e-07 | 1.800E-07 | 9.539E-08 | 1.200E-07 | 2.900E-07 |
| 17804 | 891 | S891D | C | 4.5e-07 | 2.4e-07 | 1.2e-07 | 2.700E-07 | 1.670E-07 | 1.200E-07 | 4.500E-07 |
| 17805 | 891 | S891C | C | 2e-07 | 3.3e-07 | 2.8e-07 | 2.700E-07 | 6.557E-08 | 2.000E-07 | 3.300E-07 |
| 17806 | 891 | S891E | C | 1.9e-07 | 4e-07 | 3.1e-07 | 3.000E-07 | 1.054E-07 | 1.900E-07 | 4.000E-07 |
| 17807 | 891 | S891Q | C | 1.6e-07 | 4.3e-07 | 2.1e-07 | 2.667E-07 | 1.436E-07 | 1.600E-07 | 4.300E-07 |
| 17808 | 891 | S891G | C | 4.3e-07 | 2.5e-07 | 3.3e-07 | 3.367E-07 | 9.018E-08 | 2.500E-07 | 4.300E-07 |
| 17809 | 891 | S891H | C | 1.6e-07 | 3.6e-07 | 3.3e-07 | 2.833E-07 | 1.079E-07 | 1.600E-07 | 3.600E-07 |
| 17810 | 891 | S891I | C | 3.7e-07 | 4.6e-07 | 1.4e-07 | 3.233E-07 | 1.650E-07 | 1.400E-07 | 4.600E-07 |
| 17811 | 891 | S891L | C | 1.3e-07 | 1.7e-07 | 2.5e-07 | 1.833E-07 | 6.110E-08 | 1.300E-07 | 2.500E-07 |
| 17812 | 891 | S891K | C | 2.3e-07 | 4.1e-07 | 1.4e-07 | 2.600E-07 | 1.375E-07 | 1.400E-07 | 4.100E-07 |
| 17813 | 891 | S891M | C | 3e-07 | 3.9e-07 | 2.4e-07 | 3.100E-07 | 7.550E-08 | 2.400E-07 | 3.900E-07 |
| 17814 | 891 | S891F | C | 2.7e-07 | 2e-07 | 2.1e-07 | 2.267E-07 | 3.786E-08 | 2.000E-07 | 2.700E-07 |
| 17815 | 891 | S891P | C | 3.2e-07 | 1.3e-07 | 2.8e-07 | 2.433E-07 | 1.002E-07 | 1.300E-07 | 3.200E-07 |
| 17816 | 891 | S891S | C | 2.1e-07 | 1.3e-07 | 2.8e-07 | 2.067E-07 | 7.506E-08 | 1.300E-07 | 2.800E-07 |
| 17817 | 891 | S891T | C | 3.4e-07 | 3.5e-07 | 1.9e-07 | 2.933E-07 | 8.963E-08 | 1.900E-07 | 3.500E-07 |
| 17818 | 891 | S891W | C | 4.7e-07 | 2.1e-07 | 2.5e-07 | 3.100E-07 | 1.400E-07 | 2.100E-07 | 4.700E-07 |
| 17819 | 891 | S891Y | C | 2.2e-07 | 1.2e-07 | 1.9e-07 | 1.767E-07 | 5.132E-08 | 1.200E-07 | 2.200E-07 |
| 17820 | 891 | S891V | C | 7.4e-08 | 1.3e-07 | 1.2e-07 | 1.080E-07 | 2.987E-08 | 7.400E-08 | 1.300E-07 |
| 17821 | 892 | G892A | C | 2.1e-07 | 2.8e-07 | 1.2e-07 | 2.033E-07 | 8.021E-08 | 1.200E-07 | 2.800E-07 |
| 17822 | 892 | G892R | C | 2.6e-07 | 4e-07 | 5.1e-07 | 3.900E-07 | 1.253E-07 | 2.600E-07 | 5.100E-07 |
| 17823 | 892 | G892N | C | 1e-07 | 4.1e-07 | 1.3e-07 | 2.133E-07 | 1.710E-07 | 1.000E-07 | 4.100E-07 |
| 17824 | 892 | G892D | C | 2e-07 | 2.7e-07 | 2.1e-07 | 2.267E-07 | 3.786E-08 | 2.000E-07 | 2.700E-07 |
| 17825 | 892 | G892C | C | 3.3e-07 | 1.6e-07 | 2.8e-07 | 2.567E-07 | 8.737E-08 | 1.600E-07 | 3.300E-07 |
| 17826 | 892 | G892E | C | 3.2e-07 | 3.1e-07 | 2.9e-07 | 3.067E-07 | 1.528E-08 | 2.900E-07 | 3.200E-07 |
| 17827 | 892 | G892Q | C | 1.7e-07 | 2.5e-07 | 4.8e-07 | 3.000E-07 | 1.609E-07 | 1.700E-07 | 4.800E-07 |
| 17828 | 892 | G892G | C | 3e-07 | 3.1e-07 | 1.8e-07 | 2.633E-07 | 7.234E-08 | 1.800E-07 | 3.100E-07 |
| 17829 | 892 | G892H | C | 3.1e-07 | 1.8e-07 | 2e-07 | 2.300E-07 | 7.000E-08 | 1.800E-07 | 3.100E-07 |
| 17830 | 892 | G892I | C | 3e-07 | 6.8e-08 | 1.8e-07 | 1.827E-07 | 1.160E-07 | 6.800E-08 | 3.000E-07 |
| 17831 | 892 | G892L | C | 3.6e-07 | 3.1e-07 | 1.5e-07 | 2.733E-07 | 1.097E-07 | 1.500E-07 | 3.600E-07 |
| 17832 | 892 | G892K | C | 4.6e-07 | 2e-07 | 4.8e-07 | 3.800E-07 | 1.562E-07 | 2.000E-07 | 4.800E-07 |
| 17833 | 892 | G892M | C | 3.1e-07 | 3.3e-07 | 3.4e-07 | 3.267E-07 | 1.528E-08 | 3.100E-07 | 3.400E-07 |
| 17834 | 892 | G892F | C | 1.4e-07 | 1.9e-07 | 1.7e-07 | 1.667E-07 | 2.517E-08 | 1.400E-07 | 1.900E-07 |
| 17835 | 892 | G892P | C | 2e-07 | 2.9e-07 | 3.7e-07 | 2.867E-07 | 8.505E-08 | 2.000E-07 | 3.700E-07 |
| 17836 | 892 | G892S | C | 3.6e-07 | 2.2e-07 | 1.4e-07 | 2.400E-07 | 1.114E-07 | 1.400E-07 | 3.600E-07 |
| 17837 | 892 | G892T | C | 3.1e-07 | 2.8e-07 | 1.2e-07 | 2.367E-07 | 1.021E-07 | 1.200E-07 | 3.100E-07 |
| 17838 | 892 | G892W | C | 2.1e-07 | 1.6e-07 | 2.7e-07 | 2.133E-07 | 5.508E-08 | 1.600E-07 | 2.700E-07 |
| 17839 | 892 | G892Y | C | 2.4e-07 | 3.8e-07 | 1.7e-07 | 2.633E-07 | 1.069E-07 | 1.700E-07 | 3.800E-07 |
| 17840 | 892 | G892V | C | 2.2e-07 | 3.2e-07 | 2.4e-07 | 2.600E-07 | 5.292E-08 | 2.200E-07 | 3.200E-07 |
| 17841 | 893 | G893A | C | 3.6e-07 | 3.5e-07 | 1.3e-07 | 2.800E-07 | 1.300E-07 | 1.300E-07 | 3.600E-07 |
| 17842 | 893 | G893R | C | 3.2e-07 | 2.3e-07 | 2.1e-07 | 2.533E-07 | 5.859E-08 | 2.100E-07 | 3.200E-07 |
| 17843 | 893 | G893N | C | 5.1e-07 | 2.9e-07 | 2.3e-07 | 3.433E-07 | 1.474E-07 | 2.300E-07 | 5.100E-07 |
| 17844 | 893 | G893D | C | 1.5e-07 | 3.5e-07 | 2.3e-07 | 2.433E-07 | 1.007E-07 | 1.500E-07 | 3.500E-07 |
| 17845 | 893 | G893C | C | 2.2e-07 | 5.1e-07 | 1.7e-07 | 3.000E-07 | 1.836E-07 | 1.700E-07 | 5.100E-07 |
| 17846 | 893 | G893E | C | 3.1e-07 | 2.5e-07 | 2e-07 | 2.533E-07 | 5.508E-08 | 2.000E-07 | 3.100E-07 |
| 17847 | 893 | G893Q | C | 1.3e-07 | 2.5e-07 | 3.3e-07 | 2.367E-07 | 1.007E-07 | 1.300E-07 | 3.300E-07 |
| 17848 | 893 | G893G | C | 4.4e-07 | 2e-07 | 3.5e-07 | 3.300E-07 | 1.212E-07 | 2.000E-07 | 4.400E-07 |
| 17849 | 893 | G893H | C | 3e-07 | 2.5e-07 | 1.8e-07 | 2.433E-07 | 6.028E-08 | 1.800E-07 | 3.000E-07 |

| | | | | | | | | | | |
|---|---|---|---|---|---|---|---|---|---|---|
| 17850 | 893 | G893I | C | 3.5e-07 | 3.1e-07 | 2.3e-07 | 2.967E-07 | 6.110E-08 | 2.300E-07 | 3.500E-07 |
| 17851 | 893 | G893L | C | 3e-07 | 5.6e-07 | 3.5e-07 | 4.033E-07 | 1.380E-07 | 3.000E-07 | 5.600E-07 |
| 17852 | 893 | G893K | C | 2.8e-07 | 1.8e-07 | 2e-07 | 2.200E-07 | 5.292E-08 | 1.800E-07 | 2.800E-07 |
| 17853 | 893 | G893M | C | 6.3e-07 | 4e-07 | 3.2e-07 | 4.500E-07 | 1.609E-07 | 3.200E-07 | 6.300E-07 |
| 17854 | 893 | G893F | C | 4e-07 | 3e-07 | 2.2e-07 | 3.067E-07 | 9.018E-08 | 2.200E-07 | 4.000E-07 |
| 17855 | 893 | G893P | C | 2.9e-07 | 6e-07 | 1.4e-07 | 3.433E-07 | 2.346E-07 | 1.400E-07 | 6.000E-07 |
| 17856 | 893 | G893S | C | 2.3e-07 | 2.4e-07 | 3.1e-07 | 2.600E-07 | 4.359E-08 | 2.300E-07 | 3.100E-07 |
| 17857 | 893 | G893T | C | 1.6e-07 | 2.1e-07 | 2.3e-07 | 2.000E-07 | 3.606E-08 | 1.600E-07 | 2.300E-07 |
| 17858 | 893 | G893W | C | 1.6e-07 | 1.4e-07 | 1.7e-07 | 1.567E-07 | 1.528E-08 | 1.400E-07 | 1.700E-07 |
| 17859 | 893 | G893Y | C | 2.7e-07 | 2.5e-07 | 2.8e-07 | 2.667E-07 | 1.528E-08 | 2.500E-07 | 2.800E-07 |
| 17860 | 893 | G893V | C | 3.3e-07 | 3.5e-07 | 2.3e-07 | 3.033E-07 | 6.429E-08 | 2.300E-07 | 3.500E-07 |
| 17861 | 894 | S894A | C | 3.6e-07 | 2.9e-07 | 4.2e-07 | 3.567E-07 | 6.506E-08 | 2.900E-07 | 4.200E-07 |
| 17862 | 894 | S894R | C | 1.6e-07 | 2.1e-07 | 2.6e-07 | 2.100E-07 | 5.000E-08 | 1.600E-07 | 2.600E-07 |
| 17863 | 894 | S894N | C | 2.5e-07 | 2.6e-07 | 1.6e-07 | 2.233E-07 | 5.508E-08 | 1.600E-07 | 2.600E-07 |
| 17864 | 894 | S894D | C | 2.8e-07 | 6.7e-08 | 2.7e-07 | 2.057E-07 | 1.202E-07 | 6.700E-08 | 2.800E-07 |
| 17865 | 894 | S894C | C | 2.6e-07 | 2.7e-07 | 2.3e-07 | 2.533E-07 | 2.082E-08 | 2.300E-07 | 2.700E-07 |
| 17866 | 894 | S894E | C | 1.6e-07 | 2.4e-07 | 2.3e-07 | 2.100E-07 | 4.359E-08 | 1.600E-07 | 2.400E-07 |
| 17867 | 894 | S894Q | C | 2.4e-07 | 2.8e-07 | 5.2e-07 | 3.467E-07 | 1.514E-07 | 2.400E-07 | 5.200E-07 |
| 17868 | 894 | S894G | C | 3.9e-07 | 2.7e-07 | 2.8e-07 | 3.133E-07 | 6.658E-08 | 2.700E-07 | 3.900E-07 |
| 17869 | 894 | S894H | C | 2.3e-07 | 2.1e-07 | 1.9e-07 | 2.100E-07 | 2.000E-08 | 1.900E-07 | 2.300E-07 |
| 17870 | 894 | S894I | C | 4.6e-07 | 3.4e-07 | 1.5e-07 | 3.167E-07 | 1.563E-07 | 1.500E-07 | 4.600E-07 |
| 17871 | 894 | S894L | C | 2.5e-07 | 1.3e-07 | 1.8e-07 | 1.867E-07 | 6.028E-08 | 1.300E-07 | 2.500E-07 |
| 17872 | 894 | S894K | C | 3.5e-07 | 3.2e-07 | 2.2e-07 | 2.967E-07 | 6.807E-08 | 2.200E-07 | 3.500E-07 |
| 17873 | 894 | S894M | C | 2.4e-07 | 2.5e-07 | 1.9e-07 | 2.267E-07 | 3.215E-08 | 1.900E-07 | 2.500E-07 |
| 17874 | 894 | S894F | C | 1.4e-07 | 3.8e-07 | 2.3e-07 | 2.500E-07 | 1.212E-07 | 1.400E-07 | 3.800E-07 |
| 17875 | 894 | S894P | C | 2.4e-07 | 2.3e-07 | 2.3e-07 | 2.333E-07 | 5.774E-09 | 2.300E-07 | 2.400E-07 |
| 17876 | 894 | S894S | C | 1.8e-07 | 2.2e-07 | 2.7e-07 | 2.233E-07 | 4.509E-08 | 1.800E-07 | 2.700E-07 |
| 17877 | 894 | S894T | C | 2.4e-07 | 2.8e-07 | 3.4e-07 | 2.867E-07 | 5.033E-08 | 2.400E-07 | 3.400E-07 |
| 17878 | 894 | S894W | C | 2.1e-07 | 6.3e-07 | 1.5e-07 | 3.300E-07 | 2.615E-07 | 1.500E-07 | 6.300E-07 |
| 17879 | 894 | S894Y | C | 1.4e-07 | 1.8e-07 | 1.6e-07 | 1.600E-07 | 2.000E-08 | 1.400E-07 | 1.800E-07 |
| 17880 | 894 | S894V | C | 2.2e-07 | 2.6e-07 | 2.4e-07 | 2.400E-07 | 2.000E-08 | 2.200E-07 | 2.600E-07 |
| 17881 | 895 | I895A | C | 2e-07 | 3.4e-07 | 2.2e-07 | 2.533E-07 | 7.572E-08 | 2.000E-07 | 3.400E-07 |
| 17882 | 895 | I895R | C | 1e-07 | 2.9e-07 | 2.6e-07 | 2.167E-07 | 1.021E-07 | 1.000E-07 | 2.900E-07 |
| 17883 | 895 | I895N | C | 3.9e-07 | 2.2e-07 | 2.9e-07 | 3.000E-07 | 8.544E-08 | 2.200E-07 | 3.900E-07 |
| 17884 | 895 | I895D | C | 4.2e-07 | 2e-07 | 3.9e-07 | 3.367E-07 | 1.193E-07 | 2.000E-07 | 4.200E-07 |
| 17885 | 895 | I895C | C | 1.5e-07 | 2.2e-07 | 2.9e-07 | 2.200E-07 | 7.000E-08 | 1.500E-07 | 2.900E-07 |
| 17886 | 895 | I895E | C | 6.8e-07 | 3.2e-07 | 4e-07 | 4.667E-07 | 1.890E-07 | 3.200E-07 | 6.800E-07 |
| 17887 | 895 | I895Q | C | 3.2e-07 | 1.4e-07 | 2.8e-07 | 2.467E-07 | 9.452E-08 | 1.400E-07 | 3.200E-07 |
| 17888 | 895 | I895G | C | 3e-07 | 1.4e-07 | 3.3e-07 | 2.567E-07 | 1.021E-07 | 1.400E-07 | 3.300E-07 |
| 17889 | 895 | I895H | C | 3e-07 | 1.5e-07 | 1.9e-07 | 2.133E-07 | 7.767E-08 | 1.500E-07 | 3.000E-07 |
| 17890 | 895 | I895I | C | 5e-07 | 2e-07 | 4.3e-07 | 3.767E-07 | 1.570E-07 | 2.000E-07 | 5.000E-07 |
| 17891 | 895 | I895L | C | 4.7e-07 | 2.5e-07 | 3.4e-07 | 3.533E-07 | 1.106E-07 | 2.500E-07 | 4.700E-07 |
| 17892 | 895 | I895K | C | 2.5e-07 | 3.3e-07 | 3.9e-07 | 3.233E-07 | 7.024E-08 | 2.500E-07 | 3.900E-07 |
| 17893 | 895 | I895M | C | 3.1e-07 | 2.3e-07 | 1.1e-07 | 2.167E-07 | 1.007E-07 | 1.100E-07 | 3.100E-07 |
| 17894 | 895 | I895F | C | 3.5e-07 | 3.8e-07 | 2.5e-07 | 3.267E-07 | 6.807E-08 | 2.500E-07 | 3.800E-07 |
| 17895 | 895 | I895P | C | 2e-07 | 4.2e-07 | 1.5e-07 | 2.567E-07 | 1.436E-07 | 1.500E-07 | 4.200E-07 |
| 17896 | 895 | I895S | C | 1.5e-07 | 1.8e-07 | 5.2e-07 | 2.833E-07 | 2.055E-07 | 1.500E-07 | 5.200E-07 |
| 17897 | 895 | I895T | C | 3.6e-07 | 3.9e-07 | 1.3e-07 | 2.933E-07 | 1.422E-07 | 1.300E-07 | 3.900E-07 |
| 17898 | 895 | I895W | C | 2.5e-07 | 5e-07 | 2.6e-07 | 3.367E-07 | 1.415E-07 | 2.500E-07 | 5.000E-07 |
| 17899 | 895 | I895Y | C | 2.3e-07 | 3.2e-07 | 1.9e-07 | 2.467E-07 | 6.658E-08 | 1.900E-07 | 3.200E-07 |
| 17900 | 895 | I895V | C | 2.3e-07 | 4.1e-07 | 2e-07 | 2.800E-07 | 1.136E-07 | 2.000E-07 | 4.100E-07 |
| 17901 | 896 | Y896A | C | 6.4e-08 | 3e-07 | 3.5e-07 | 2.380E-07 | 1.527E-07 | 6.400E-08 | 3.500E-07 |
| 17902 | 896 | Y896R | C | 1.6e-07 | 2.6e-07 | 3.1e-07 | 2.433E-07 | 7.638E-08 | 1.600E-07 | 3.100E-07 |
| 17903 | 896 | Y896N | C | 1.3e-07 | 4e-07 | 2.7e-07 | 2.667E-07 | 1.350E-07 | 1.300E-07 | 4.000E-07 |
| 17904 | 896 | Y896D | C | 2.6e-07 | 2e-07 | 8.6e-08 | 1.820E-07 | 8.839E-08 | 8.600E-08 | 2.600E-07 |
| 17905 | 896 | Y896C | C | 2.6e-07 | 2.5e-07 | 5.4e-07 | 3.500E-07 | 1.646E-07 | 2.500E-07 | 5.400E-07 |
| 17906 | 896 | Y896E | C | 1.7e-07 | 6.7e-07 | 2.1e-07 | 3.500E-07 | 2.778E-07 | 1.700E-07 | 6.700E-07 |
| 17907 | 896 | Y896Q | C | 2.5e-07 | 2.6e-07 | 1.4e-07 | 2.167E-07 | 6.658E-08 | 1.400E-07 | 2.600E-07 |
| 17908 | 896 | Y896G | C | 2.5e-07 | 4.6e-07 | 2.3e-07 | 3.133E-07 | 1.274E-07 | 2.300E-07 | 4.600E-07 |
| 17909 | 896 | Y896H | C | 2.1e-07 | 1.5e-07 | 3.3e-07 | 2.300E-07 | 9.165E-08 | 1.500E-07 | 3.300E-07 |
| 17910 | 896 | Y896I | C | 1.9e-07 | 4.3e-07 | 2.3e-07 | 2.833E-07 | 1.286E-07 | 1.900E-07 | 4.300E-07 |
| 17911 | 896 | Y896L | C | 3.3e-07 | 4.4e-07 | 3.6e-07 | 3.767E-07 | 5.686E-08 | 3.300E-07 | 4.400E-07 |
| 17912 | 896 | Y896K | C | 2.4e-07 | 2.8e-07 | 1.7e-07 | 2.300E-07 | 5.568E-08 | 1.700E-07 | 2.800E-07 |
| 17913 | 896 | Y896M | C | 4.9e-07 | 3.3e-07 | 2.6e-07 | 3.600E-07 | 1.179E-07 | 2.600E-07 | 4.900E-07 |
| 17914 | 896 | Y896F | C | 3.8e-07 | 4.1e-07 | 3e-07 | 3.633E-07 | 5.686E-08 | 3.000E-07 | 4.100E-07 |
| 17915 | 896 | Y896P | C | 1.6e-07 | 2.8e-07 | 6.1e-07 | 3.500E-07 | 2.330E-07 | 1.600E-07 | 6.100E-07 |
| 17916 | 896 | Y896S | C | 5.5e-07 | 3.8e-07 | 2.7e-07 | 4.000E-07 | 1.411E-07 | 2.700E-07 | 5.500E-07 |
| 17917 | 896 | Y896T | C | 3e-07 | 2.6e-07 | 2.3e-07 | 2.633E-07 | 3.512E-08 | 2.300E-07 | 3.000E-07 |
| 17918 | 896 | Y896W | C | 2e-07 | 3.2e-07 | 2.2e-07 | 2.467E-07 | 6.429E-08 | 2.000E-07 | 3.200E-07 |
| 17919 | 896 | Y896Y | C | 3.4e-07 | 9.3e-08 | 3.2e-07 | 2.510E-07 | 1.372E-07 | 9.300E-08 | 3.400E-07 |
| 17920 | 896 | Y896V | C | 7.5e-08 | 3.7e-07 | 1.6e-07 | 2.017E-07 | 1.518E-07 | 7.500E-08 | 3.700E-07 |
| 17921 | 897 | N897A | C | 3e-07 | 5.4e-07 | 2.4e-07 | 3.600E-07 | 1.587E-07 | 2.400E-07 | 5.400E-07 |
| 17922 | 897 | N897R | C | 1.7e-07 | 1.6e-07 | 2.3e-07 | 1.867E-07 | 3.786E-08 | 1.600E-07 | 2.300E-07 |
| 17923 | 897 | N897N | C | 4.2e-07 | 1.9e-07 | 2.2e-07 | 2.767E-07 | 1.250E-07 | 1.900E-07 | 4.200E-07 |
| 17924 | 897 | N897D | C | 2.8e-07 | 2.7e-07 | 1.4e-07 | 2.300E-07 | 7.810E-08 | 1.400E-07 | 2.800E-07 |
| 17925 | 897 | N897C | C | 2.3e-07 | 1.9e-07 | 9e-08 | 1.700E-07 | 7.211E-08 | 9.000E-08 | 2.300E-07 |
| 17926 | 897 | N897E | C | 1.6e-07 | 1.4e-07 | 3.9e-07 | 2.300E-07 | 1.389E-07 | 1.400E-07 | 3.900E-07 |
| 17927 | 897 | N897Q | C | 1.2e-07 | 2.1e-07 | 2e-07 | 1.767E-07 | 4.933E-08 | 1.200E-07 | 2.100E-07 |
| 17928 | 897 | N897G | C | 2.7e-07 | 1.7e-07 | 5.9e-07 | 3.433E-07 | 2.194E-07 | 1.700E-07 | 5.900E-07 |
| 17929 | 897 | N897H | C | 3.1e-07 | 1.5e-07 | 2.1e-07 | 2.233E-07 | 8.083E-08 | 1.500E-07 | 3.100E-07 |
| 17930 | 897 | N897I | C | 3.5e-07 | 1.7e-07 | 3.6e-07 | 2.933E-07 | 1.069E-07 | 1.700E-07 | 3.600E-07 |
| 17931 | 897 | N897L | C | 2.5e-07 | 1.9e-07 | 1.1e-07 | 1.833E-07 | 7.024E-08 | 1.100E-07 | 2.500E-07 |
| 17932 | 897 | N897K | C | 2.7e-07 | 1.8e-07 | 3.1e-07 | 2.533E-07 | 6.658E-08 | 1.800E-07 | 3.100E-07 |
| 17933 | 897 | N897M | C | 2.7e-07 | 4.6e-07 | 3.8e-07 | 3.700E-07 | 9.539E-08 | 2.700E-07 | 4.600E-07 |
| 17934 | 897 | N897F | C | 5.3e-07 | 1.7e-07 | 5e-07 | 4.000E-07 | 1.997E-07 | 1.700E-07 | 5.300E-07 |

| | | | | | | | | | | |
|---|---|---|---|---|---|---|---|---|---|---|
| 17935 | 897 | N897P | C | 1.6e-07 | 3.9e-07 | 5.7e-07 | 3.733E-07 | 2.055E-07 | 1.600E-07 | 5.700E-07 |
| 17936 | 897 | N897S | C | 3.1e-07 | 2.5e-07 | 1.8e-07 | 2.467E-07 | 6.506E-08 | 1.800E-07 | 3.100E-07 |
| 17937 | 897 | N897T | C | 3e-07 | 1.6e-07 | 1.5e-07 | 2.033E-07 | 8.386E-08 | 1.500E-07 | 3.000E-07 |
| 17938 | 897 | N897W | C | 3.4e-07 | 2e-07 | 3.3e-07 | 2.900E-07 | 7.810E-08 | 2.000E-07 | 3.400E-07 |
| 17939 | 897 | N897Y | C | 2.5e-07 | 2.9e-07 | 2.4e-07 | 2.600E-07 | 2.646E-08 | 2.400E-07 | 2.900E-07 |
| 17940 | 897 | N897V | C | 2e-07 | 1.4e-07 | 1.6e-07 | 1.667E-07 | 3.055E-08 | 1.400E-07 | 2.000E-07 |
| 17941 | 898 | Q898A | C | 8.6e-08 | 1.9e-07 | 4.4e-07 | 2.387E-07 | 1.819E-07 | 8.600E-08 | 4.400E-07 |
| 17942 | 898 | Q898R | C | 1.7e-07 | 2e-07 | 2.6e-07 | 2.100E-07 | 4.583E-08 | 1.700E-07 | 2.600E-07 |
| 17943 | 898 | Q898N | C | 2e-07 | 2.1e-07 | 2.5e-07 | 2.200E-07 | 2.646E-08 | 2.000E-07 | 2.500E-07 |
| 17944 | 898 | Q898D | C | 1.3e-07 | 2.5e-07 | 2.4e-07 | 2.067E-07 | 6.658E-08 | 1.300E-07 | 2.500E-07 |
| 17945 | 898 | Q898C | C | 2.2e-07 | 2.1e-07 | 1.3e-07 | 1.867E-07 | 4.933E-08 | 1.300E-07 | 2.200E-07 |
| 17946 | 898 | Q898E | C | 3.2e-07 | 2.3e-07 | 1.7e-07 | 2.400E-07 | 7.550E-08 | 1.700E-07 | 3.200E-07 |
| 17947 | 898 | Q898Q | C | 2.8e-07 | 5.4e-07 | 1.5e-07 | 3.233E-07 | 1.986E-07 | 1.500E-07 | 5.400E-07 |
| 17948 | 898 | Q898G | C | 2.2e-07 | 3.1e-07 | 1.6e-07 | 2.300E-07 | 7.550E-08 | 1.600E-07 | 3.100E-07 |
| 17949 | 898 | Q898H | C | 2.8e-07 | 3.2e-07 | 1.1e-07 | 2.367E-07 | 1.115E-07 | 1.100E-07 | 3.200E-07 |
| 17950 | 898 | Q898I | C | 2.5e-07 | 1.4e-07 | 4e-07 | 2.633E-07 | 1.305E-07 | 1.400E-07 | 4.000E-07 |
| 17951 | 898 | Q898L | C | 1.8e-07 | 3.2e-07 | 2.8e-07 | 2.600E-07 | 7.211E-08 | 1.800E-07 | 3.200E-07 |
| 17952 | 898 | Q898K | C | 1.3e-07 | 2.4e-07 | 2.8e-07 | 2.167E-07 | 7.767E-08 | 1.300E-07 | 2.800E-07 |
| 17953 | 898 | Q898M | C | 1.7e-07 | 3.7e-07 | 2.6e-07 | 2.667E-07 | 1.002E-07 | 1.700E-07 | 3.700E-07 |
| 17954 | 898 | Q898F | C | 4e-07 | 3e-07 | 1.9e-07 | 2.967E-07 | 1.050E-07 | 1.900E-07 | 4.000E-07 |
| 17955 | 898 | Q898P | C | 1.1e-06 | 3.1e-07 | 4.3e-07 | 6.133E-07 | 4.257E-07 | 3.100E-07 | 1.100E-06 |
| 17956 | 898 | Q898S | C | 2.8e-07 | 2.4e-07 | 2.7e-07 | 2.633E-07 | 2.082E-08 | 2.400E-07 | 2.800E-07 |
| 17957 | 898 | Q898T | C | 5.2e-07 | 2.5e-07 | 2.2e-07 | 3.300E-07 | 1.652E-07 | 2.200E-07 | 5.200E-07 |
| 17958 | 898 | Q898W | C | 2.5e-07 | 3.4e-07 | 1.8e-07 | 2.567E-07 | 8.021E-08 | 1.800E-07 | 3.400E-07 |
| 17959 | 898 | Q898Y | C | 3.8e-07 | 2e-07 | 4.5e-07 | 3.433E-07 | 1.290E-07 | 2.000E-07 | 4.500E-07 |
| 17960 | 898 | Q898V | C | 2.3e-07 | 2.5e-07 | 2.6e-07 | 2.467E-07 | 1.528E-08 | 2.300E-07 | 2.600E-07 |
| 17961 | 899 | R899A | C | 1.4e-07 | 2e-07 | 2.4e-07 | 1.933E-07 | 5.033E-08 | 1.400E-07 | 2.400E-07 |
| 17962 | 899 | R899R | C | 1.7e-07 | 2.6e-07 | 2.4e-07 | 2.233E-07 | 4.726E-08 | 1.700E-07 | 2.600E-07 |
| 17963 | 899 | R899N | C | 6.4e-07 | 3.4e-07 | 4.5e-07 | 4.767E-07 | 1.518E-07 | 3.400E-07 | 6.400E-07 |
| 17964 | 899 | R899D | C | 2.2e-07 | 3.2e-07 | 2.3e-07 | 2.567E-07 | 5.508E-08 | 2.200E-07 | 3.200E-07 |
| 17965 | 899 | R899C | C | 2.3e-07 | 3.5e-07 | 3e-07 | 2.933E-07 | 6.028E-08 | 2.300E-07 | 3.500E-07 |
| 17966 | 899 | R899E | C | 4e-07 | 3.6e-07 | 2.4e-07 | 3.333E-07 | 8.327E-08 | 2.400E-07 | 4.000E-07 |
| 17967 | 899 | R899Q | C | 1.7e-07 | 4.8e-07 | 2e-07 | 2.833E-07 | 1.710E-07 | 1.700E-07 | 4.800E-07 |
| 17968 | 899 | R899G | C | 4.4e-07 | 4e-07 | 1.5e-07 | 3.300E-07 | 1.572E-07 | 1.500E-07 | 4.400E-07 |
| 17969 | 899 | R899H | C | 5.6e-07 | 2.4e-07 | 3.1e-07 | 3.700E-07 | 1.682E-07 | 2.400E-07 | 5.600E-07 |
| 17970 | 899 | R899I | C | 4e-07 | 1.2e-07 | 1.9e-07 | 2.367E-07 | 1.457E-07 | 1.200E-07 | 4.000E-07 |
| 17971 | 899 | R899L | C | 2.4e-07 | 2.8e-07 | 1.9e-07 | 2.367E-07 | 4.509E-08 | 1.900E-07 | 2.800E-07 |
| 17972 | 899 | R899K | C | 1.6e-07 | 2.4e-07 | 1.9e-07 | 1.967E-07 | 4.041E-08 | 1.600E-07 | 2.400E-07 |
| 17973 | 899 | R899M | C | 2.3e-07 | 1.5e-07 | 1.1e-07 | 1.633E-07 | 6.110E-08 | 1.100E-07 | 2.300E-07 |
| 17974 | 899 | R899F | C | 3.1e-07 | 2.4e-07 | 2.3e-07 | 2.600E-07 | 4.359E-08 | 2.300E-07 | 3.100E-07 |
| 17975 | 899 | R899P | C | 2.6e-07 | 1.5e-07 | 2.2e-07 | 2.100E-07 | 5.568E-08 | 1.500E-07 | 2.600E-07 |
| 17976 | 899 | R899S | C | 2.9e-07 | 1.6e-07 | 1.2e-07 | 1.900E-07 | 8.888E-08 | 1.200E-07 | 2.900E-07 |
| 17977 | 899 | R899T | C | 1.6e-07 | 2.1e-07 | 1.6e-07 | 1.767E-07 | 2.887E-08 | 1.600E-07 | 2.100E-07 |
| 17978 | 899 | R899W | C | 2.2e-07 | 3.1e-07 | 1.3e-07 | 2.200E-07 | 9.000E-08 | 1.300E-07 | 3.100E-07 |
| 17979 | 899 | R899Y | C | 9.6e-08 | 2.4e-07 | 3.3e-07 | 2.220E-07 | 1.180E-07 | 9.600E-08 | 3.300E-07 |
| 17980 | 899 | R899V | C | 3e-07 | 1.7e-07 | 2.4e-07 | 2.367E-07 | 6.506E-08 | 1.700E-07 | 3.000E-07 |
| 17981 | 900 | F900A | C | 2.6e-07 | 2.6e-07 | 4.2e-07 | 3.133E-07 | 9.238E-08 | 2.600E-07 | 4.200E-07 |
| 17982 | 900 | F900R | C | 1.6e-07 | 3.5e-07 | 2e-07 | 2.367E-07 | 1.002E-07 | 1.600E-07 | 3.500E-07 |
| 17983 | 900 | F900N | C | 2.3e-07 | 1.9e-07 | 2.1e-07 | 2.100E-07 | 2.000E-08 | 1.900E-07 | 2.300E-07 |
| 17984 | 900 | F900D | C | 1.5e-07 | 2e-07 | 6.5e-07 | 3.333E-07 | 2.754E-07 | 1.500E-07 | 6.500E-07 |
| 17985 | 900 | F900C | C | 1.7e-07 | 2.9e-07 | 1.2e-07 | 1.933E-07 | 8.737E-08 | 1.200E-07 | 2.900E-07 |
| 17986 | 900 | F900E | C | 2.4e-07 | 3e-07 | 1.1e-07 | 2.167E-07 | 9.713E-08 | 1.100E-07 | 3.000E-07 |
| 17987 | 900 | F900Q | C | 3.9e-07 | 2.4e-07 | 3.1e-07 | 3.133E-07 | 7.506E-08 | 2.400E-07 | 3.900E-07 |
| 17988 | 900 | F900G | C | 1.4e-07 | 3.7e-07 | 5.2e-07 | 3.433E-07 | 1.914E-07 | 1.400E-07 | 5.200E-07 |
| 17989 | 900 | F900H | C | 4e-07 | 4.6e-07 | 1.6e-07 | 3.400E-07 | 1.587E-07 | 1.600E-07 | 4.600E-07 |
| 17990 | 900 | F900I | C | 2e-07 | 3.2e-07 | 3.4e-07 | 2.867E-07 | 7.572E-08 | 2.000E-07 | 3.400E-07 |
| 17991 | 900 | F900L | C | 6.9e-07 | 1.4e-07 | 2.2e-07 | 3.500E-07 | 2.972E-07 | 1.400E-07 | 6.900E-07 |
| 17992 | 900 | F900K | C | 3.3e-07 | 3.1e-07 | 2.5e-07 | 2.967E-07 | 4.163E-08 | 2.500E-07 | 3.300E-07 |
| 17993 | 900 | F900M | C | 1.1e-07 | 1.4e-07 | 1.5e-07 | 1.333E-07 | 2.082E-08 | 1.100E-07 | 1.500E-07 |
| 17994 | 900 | F900F | C | 1.4e-07 | 3.4e-07 | 1.7e-07 | 2.167E-07 | 1.079E-07 | 1.400E-07 | 3.400E-07 |
| 17995 | 900 | F900P | C | 1.7e-07 | 2.7e-07 | 2.5e-07 | 2.300E-07 | 5.292E-08 | 1.700E-07 | 2.700E-07 |
| 17996 | 900 | F900S | C | 2.5e-07 | 1.8e-07 | 1.8e-07 | 2.033E-07 | 4.041E-08 | 1.800E-07 | 2.500E-07 |
| 17997 | 900 | F900T | C | 1.4e-07 | 3.9e-07 | 2.6e-07 | 2.633E-07 | 1.250E-07 | 1.400E-07 | 3.900E-07 |
| 17998 | 900 | F900W | C | 3.9e-07 | 2e-07 | 1.9e-07 | 2.600E-07 | 1.127E-07 | 1.900E-07 | 3.900E-07 |
| 17999 | 900 | F900Y | C | 6.2e-07 | 2.3e-07 | 2.7e-07 | 3.733E-07 | 2.146E-07 | 2.300E-07 | 6.200E-07 |
| 18000 | 900 | F900V | C | 1e-07 | 2.2e-07 | 3.5e-07 | 2.233E-07 | 1.250E-07 | 1.000E-07 | 3.500E-07 |
| 18001 | 901 | K901A | C | 2.5e-07 | 4e-07 | 2.5e-07 | 3.000E-07 | 8.660E-08 | 2.500E-07 | 4.000E-07 |
| 18002 | 901 | K901R | C | 1.5e-07 | 2.6e-07 | 3.9e-07 | 2.667E-07 | 1.201E-07 | 1.500E-07 | 3.900E-07 |
| 18003 | 901 | K901N | C | 1.8e-07 | 1.7e-07 | 2.5e-07 | 2.000E-07 | 4.359E-08 | 1.700E-07 | 2.500E-07 |
| 18004 | 901 | K901D | C | 4.4e-07 | 2.2e-07 | 2.2e-07 | 2.933E-07 | 1.270E-07 | 2.200E-07 | 4.400E-07 |
| 18005 | 901 | K901C | C | 3.9e-07 | 5.7e-07 | 1.9e-07 | 3.833E-07 | 1.901E-07 | 1.900E-07 | 5.700E-07 |
| 18006 | 901 | K901E | C | 2.8e-07 | 2.6e-07 | 1.7e-07 | 2.367E-07 | 5.859E-08 | 1.700E-07 | 2.800E-07 |
| 18007 | 901 | K901Q | C | 1.4e-07 | 3.5e-07 | 2.2e-07 | 2.367E-07 | 1.060E-07 | 1.400E-07 | 3.500E-07 |
| 18008 | 901 | K901G | C | 3.3e-07 | 3.8e-07 | 3.4e-07 | 3.500E-07 | 2.646E-08 | 3.300E-07 | 3.800E-07 |
| 18009 | 901 | K901H | C | 1.3e-07 | 2.7e-07 | 1.9e-07 | 1.967E-07 | 7.024E-08 | 1.300E-07 | 2.700E-07 |
| 18010 | 901 | K901I | C | 2.9e-07 | 2.4e-07 | 3.6e-07 | 2.967E-07 | 6.028E-08 | 2.400E-07 | 3.600E-07 |
| 18011 | 901 | K901L | C | 2.2e-07 | 3e-07 | 2.3e-07 | 2.500E-07 | 4.359E-08 | 2.200E-07 | 3.000E-07 |
| 18012 | 901 | K901K | C | 2.3e-07 | 2.9e-07 | 2e-07 | 2.400E-07 | 4.583E-08 | 2.000E-07 | 2.900E-07 |
| 18013 | 901 | K901M | C | 3.5e-07 | 2.4e-07 | 3.6e-07 | 3.167E-07 | 6.658E-08 | 2.400E-07 | 3.600E-07 |
| 18014 | 901 | K901F | C | 1.9e-07 | 3.6e-07 | 2.2e-07 | 2.567E-07 | 9.074E-08 | 1.900E-07 | 3.600E-07 |
| 18015 | 901 | K901P | C | 1.3e-07 | 2.4e-07 | 8.9e-08 | 1.530E-07 | 7.808E-08 | 8.900E-08 | 2.400E-07 |
| 18016 | 901 | K901S | C | 1.6e-07 | 2.9e-07 | 3.9e-07 | 2.800E-07 | 1.153E-07 | 1.600E-07 | 3.900E-07 |
| 18017 | 901 | K901T | C | 2e-07 | 1.9e-07 | 2.4e-07 | 2.100E-07 | 2.646E-08 | 1.900E-07 | 2.400E-07 |
| 18018 | 901 | K901W | C | 1.6e-07 | 1.6e-07 | 3.4e-07 | 2.200E-07 | 1.039E-07 | 1.600E-07 | 3.400E-07 |
| 18019 | 901 | K901Y | C | 2.8e-07 | 1.2e-07 | 2.6e-07 | 2.200E-07 | 8.718E-08 | 1.200E-07 | 2.800E-07 |

| | | | | | | | | | | |
|---|---|---|---|---|---|---|---|---|---|---|
| 18020 | 901 | K901V | C | 2e-07 | 4.3e-07 | 2e-07 | 2.767E-07 | 1.328E-07 | 2.000E-07 | 4.300E-07 |
| 18021 | 902 | G902A | C | 4.6e-07 | 3.1e-07 | 6.7e-07 | 4.800E-07 | 1.808E-07 | 3.100E-07 | 6.700E-07 |
| 18022 | 902 | G902R | C | 2.4e-07 | 3.1e-07 | 1.6e-07 | 2.367E-07 | 7.506E-08 | 1.600E-07 | 3.100E-07 |
| 18023 | 902 | G902N | C | 1.2e-07 | 2.9e-07 | 4.4e-07 | 2.833E-07 | 1.601E-07 | 1.200E-07 | 4.400E-07 |
| 18024 | 902 | G902D | C | 1.8e-07 | 1.9e-07 | 2.3e-07 | 2.000E-07 | 2.646E-08 | 1.800E-07 | 2.300E-07 |
| 18025 | 902 | G902C | C | 3.4e-07 | 2e-07 | 1.9e-07 | 2.433E-07 | 8.386E-08 | 1.900E-07 | 3.400E-07 |
| 18026 | 902 | G902E | C | 5.7e-07 | 8.6e-08 | 3e-07 | 3.187E-07 | 2.425E-07 | 8.600E-08 | 5.700E-07 |
| 18027 | 902 | G902Q | C | 1.8e-07 | 4.1e-07 | 2.3e-07 | 2.733E-07 | 1.210E-07 | 1.800E-07 | 4.100E-07 |
| 18028 | 902 | G902G | C | 1.5e-07 | 2.2e-07 | 5.9e-07 | 3.200E-07 | 2.364E-07 | 1.500E-07 | 5.900E-07 |
| 18029 | 902 | G902H | C | 1.4e-07 | 6e-07 | 1.9e-07 | 3.100E-07 | 2.524E-07 | 1.400E-07 | 6.000E-07 |
| 18030 | 902 | G902I | C | 3.8e-07 | 1.7e-07 | 2.7e-07 | 2.733E-07 | 1.050E-07 | 1.700E-07 | 3.800E-07 |
| 18031 | 902 | G902L | C | 6.9e-07 | 2.7e-07 | 1.2e-07 | 3.600E-07 | 2.955E-07 | 1.200E-07 | 6.900E-07 |
| 18032 | 902 | G902K | C | 2.2e-07 | 2.4e-07 | 3.9e-07 | 2.833E-07 | 9.292E-08 | 2.200E-07 | 3.900E-07 |
| 18033 | 902 | G902M | C | 2.2e-07 | 2.4e-07 | 2.6e-07 | 2.400E-07 | 2.000E-08 | 2.200E-07 | 2.600E-07 |
| 18034 | 902 | G902F | C | 5.4e-07 | 1.8e-07 | 8.9e-08 | 2.697E-07 | 2.385E-07 | 8.900E-08 | 5.400E-07 |
| 18035 | 902 | G902P | C | 2.6e-07 | 2.1e-07 | 1.1e-07 | 1.933E-07 | 7.638E-08 | 1.100E-07 | 2.600E-07 |
| 18036 | 902 | G902S | C | 4.5e-07 | 1.6e-07 | 2.1e-07 | 2.733E-07 | 1.550E-07 | 1.600E-07 | 4.500E-07 |
| 18037 | 902 | G902T | C | 4.1e-07 | 4.9e-07 | 5.2e-07 | 4.733E-07 | 5.686E-08 | 4.100E-07 | 5.200E-07 |
| 18038 | 902 | G902W | C | 1.8e-07 | 3.9e-07 | 2.6e-07 | 2.767E-07 | 1.060E-07 | 1.800E-07 | 3.900E-07 |
| 18039 | 902 | G902Y | C | 2e-07 | 1.1e-07 | 1.9e-07 | 1.667E-07 | 4.933E-08 | 1.100E-07 | 2.000E-07 |
| 18040 | 902 | G902V | C | 2.2e-07 | 8.8e-08 | 1.9e-07 | 1.660E-07 | 6.920E-08 | 8.800E-08 | 2.200E-07 |
| 18041 | 903 | R903A | C | 3.5e-07 | 3.2e-07 | 1.9e-07 | 2.867E-07 | 8.505E-08 | 1.900E-07 | 3.500E-07 |
| 18042 | 903 | R903R | C | 2e-07 | 1.6e-07 | 2.4e-07 | 2.000E-07 | 4.000E-08 | 1.600E-07 | 2.400E-07 |
| 18043 | 903 | R903N | C | 2.3e-07 | 4.7e-07 | 1.9e-07 | 2.967E-07 | 1.514E-07 | 1.900E-07 | 4.700E-07 |
| 18044 | 903 | R903D | C | 2.4e-07 | 2.2e-07 | 1.1e-07 | 1.900E-07 | 7.000E-08 | 1.100E-07 | 2.400E-07 |
| 18045 | 903 | R903C | C | 1.2e-07 | 4.9e-07 | 2.3e-07 | 2.800E-07 | 1.900E-07 | 1.200E-07 | 4.900E-07 |
| 18046 | 903 | R903E | C | 2.9e-07 | 4.2e-07 | 2.7e-07 | 3.267E-07 | 8.145E-08 | 2.700E-07 | 4.200E-07 |
| 18047 | 903 | R903Q | C | 2e-07 | 4e-07 | 1.8e-07 | 2.600E-07 | 1.217E-07 | 1.800E-07 | 4.000E-07 |
| 18048 | 903 | R903G | C | 2e-07 | 3.2e-07 | 2.4e-07 | 2.533E-07 | 6.110E-08 | 2.000E-07 | 3.200E-07 |
| 18049 | 903 | R903H | C | 2.6e-07 | 2.4e-07 | 2.3e-07 | 2.433E-07 | 1.528E-08 | 2.300E-07 | 2.600E-07 |
| 18050 | 903 | R903I | C | 2e-07 | 2.1e-07 | 1.5e-07 | 1.867E-07 | 3.215E-08 | 1.500E-07 | 2.100E-07 |
| 18051 | 903 | R903L | C | 2.5e-07 | 1.9e-07 | 3e-07 | 2.467E-07 | 5.508E-08 | 1.900E-07 | 3.000E-07 |
| 18052 | 903 | R903K | C | 2.4e-07 | 4.2e-07 | 4.9e-07 | 3.833E-07 | 1.290E-07 | 2.400E-07 | 4.900E-07 |
| 18053 | 903 | R903M | C | 4.2e-07 | 2.6e-07 | 2.7e-07 | 3.167E-07 | 8.963E-08 | 2.600E-07 | 4.200E-07 |
| 18054 | 903 | R903F | C | 2.1e-07 | 2.4e-07 | 1.7e-07 | 2.067E-07 | 3.512E-08 | 1.700E-07 | 2.400E-07 |
| 18055 | 903 | R903P | C | 2.1e-07 | 4.8e-07 | 3.5e-07 | 3.467E-07 | 1.350E-07 | 2.100E-07 | 4.800E-07 |
| 18056 | 903 | R903S | C | 3e-07 | 3.9e-07 | 6.3e-07 | 4.400E-07 | 1.706E-07 | 3.000E-07 | 6.300E-07 |
| 18057 | 903 | R903T | C | 6.2e-07 | 2.2e-07 | 5.8e-07 | 4.733E-07 | 2.203E-07 | 2.200E-07 | 6.200E-07 |
| 18058 | 903 | R903W | C | 1.8e-07 | 1.7e-07 | 1.3e-07 | 1.600E-07 | 2.646E-08 | 1.300E-07 | 1.800E-07 |
| 18059 | 903 | R903Y | C | 1.9e-07 | 3.9e-07 | 3.9e-07 | 3.233E-07 | 1.155E-07 | 1.900E-07 | 3.900E-07 |
| 18060 | 903 | R903V | C | 2.5e-07 | 2.8e-07 | 1.9e-07 | 2.400E-07 | 4.583E-08 | 1.900E-07 | 2.800E-07 |
| 18061 | 904 | F904A | C | 3.7e-07 | 3.7e-07 | 4.7e-07 | 4.033E-07 | 5.774E-08 | 3.700E-07 | 4.700E-07 |
| 18062 | 904 | F904R | C | 2.4e-07 | 1.8e-07 | 2.4e-07 | 2.200E-07 | 3.464E-08 | 1.800E-07 | 2.400E-07 |
| 18063 | 904 | F904N | C | 3.1e-07 | 1.9e-07 | 1.9e-07 | 2.300E-07 | 6.928E-08 | 1.900E-07 | 3.100E-07 |
| 18064 | 904 | F904D | C | 1.4e-07 | 2.4e-07 | 2.8e-07 | 2.200E-07 | 7.211E-08 | 1.400E-07 | 2.800E-07 |
| 18065 | 904 | F904C | C | 2.5e-07 | 2e-07 | 1.7e-07 | 2.067E-07 | 4.041E-08 | 1.700E-07 | 2.500E-07 |
| 18066 | 904 | F904E | C | 3.1e-07 | 2.2e-07 | 4.8e-07 | 3.367E-07 | 1.320E-07 | 2.200E-07 | 4.800E-07 |
| 18067 | 904 | F904Q | C | 2.6e-07 | 2.2e-07 | 1.8e-07 | 2.200E-07 | 4.000E-08 | 1.800E-07 | 2.600E-07 |
| 18068 | 904 | F904G | C | 2.4e-07 | 2.5e-07 | 1.6e-07 | 2.167E-07 | 4.933E-08 | 1.600E-07 | 2.500E-07 |
| 18069 | 904 | F904H | C | 1.6e-07 | 2.2e-07 | 2.2e-07 | 2.000E-07 | 3.464E-08 | 1.600E-07 | 2.200E-07 |
| 18070 | 904 | F904I | C | 2.2e-07 | 1.2e-07 | 2.2e-07 | 1.867E-07 | 5.774E-08 | 1.200E-07 | 2.200E-07 |
| 18071 | 904 | F904L | C | 3.5e-07 | 1.9e-07 | 3.3e-07 | 2.900E-07 | 8.718E-08 | 1.900E-07 | 3.500E-07 |
| 18072 | 904 | F904K | C | 1.7e-07 | 7.8e-07 | 3.2e-07 | 4.233E-07 | 3.179E-07 | 1.700E-07 | 7.800E-07 |
| 18073 | 904 | F904M | C | 3.4e-07 | 2.8e-07 | 2.4e-07 | 2.867E-07 | 5.033E-08 | 2.400E-07 | 3.400E-07 |
| 18074 | 904 | F904F | C | 2.9e-07 | 2.6e-07 | 1.8e-07 | 2.433E-07 | 5.686E-08 | 1.800E-07 | 2.900E-07 |
| 18075 | 904 | F904P | C | 3.3e-07 | 2.2e-07 | 2.1e-07 | 2.533E-07 | 6.658E-08 | 2.100E-07 | 3.300E-07 |
| 18076 | 904 | F904S | C | 1.6e-07 | 3.6e-07 | 2.5e-07 | 2.567E-07 | 1.002E-07 | 1.600E-07 | 3.600E-07 |
| 18077 | 904 | F904T | C | 1.5e-07 | 2.7e-07 | 4e-07 | 2.733E-07 | 1.250E-07 | 1.500E-07 | 4.000E-07 |
| 18078 | 904 | F904W | C | 3.2e-07 | 2.9e-07 | 3.1e-07 | 3.067E-07 | 1.528E-08 | 2.900E-07 | 3.200E-07 |
| 18079 | 904 | F904Y | C | 2.9e-07 | 5e-07 | 2.4e-07 | 3.433E-07 | 1.380E-07 | 2.400E-07 | 5.000E-07 |
| 18080 | 904 | F904V | C | 5.1e-07 | 3e-07 | 5.7e-07 | 4.600E-07 | 1.418E-07 | 3.000E-07 | 5.700E-07 |
| 18081 | 905 | T905A | C | 2.5e-07 | 1.4e-07 | 3.4e-07 | 2.433E-07 | 1.002E-07 | 1.400E-07 | 3.400E-07 |
| 18082 | 905 | T905R | C | 2.4e-07 | 1.6e-07 | 2.2e-07 | 2.067E-07 | 4.163E-08 | 1.600E-07 | 2.400E-07 |
| 18083 | 905 | T905N | C | 2e-07 | 1.5e-07 | 1.8e-07 | 1.767E-07 | 2.517E-08 | 1.500E-07 | 2.000E-07 |
| 18084 | 905 | T905D | C | 7.7e-07 | 3.5e-07 | 2.6e-07 | 4.600E-07 | 2.722E-07 | 2.600E-07 | 7.700E-07 |
| 18085 | 905 | T905C | C | 2.9e-07 | 5.5e-07 | 2.7e-07 | 3.700E-07 | 1.562E-07 | 2.700E-07 | 5.500E-07 |
| 18086 | 905 | T905E | C | 3.9e-07 | 4.1e-07 | 9.6e-08 | 2.987E-07 | 1.758E-07 | 9.600E-08 | 4.100E-07 |
| 18087 | 905 | T905Q | C | 8.9e-08 | 5.1e-07 | 2.9e-07 | 2.963E-07 | 2.106E-07 | 8.900E-08 | 5.100E-07 |
| 18088 | 905 | T905G | C | 1.6e-07 | 1.2e-07 | 1.7e-07 | 1.500E-07 | 2.646E-08 | 1.200E-07 | 1.700E-07 |
| 18089 | 905 | T905H | C | 1.4e-07 | 1.9e-07 | 4.7e-07 | 2.667E-07 | 1.779E-07 | 1.400E-07 | 4.700E-07 |
| 18090 | 905 | T905I | C | 3.3e-07 | 3.7e-07 | 1.9e-07 | 2.967E-07 | 9.452E-08 | 1.900E-07 | 3.700E-07 |
| 18091 | 905 | T905L | C | 2.5e-07 | 1.9e-07 | 3.1e-07 | 2.500E-07 | 6.000E-08 | 1.900E-07 | 3.100E-07 |
| 18092 | 905 | T905K | C | 3.1e-07 | 2.4e-07 | 2.5e-07 | 2.667E-07 | 3.786E-08 | 2.400E-07 | 3.100E-07 |
| 18093 | 905 | T905M | C | 3.5e-07 | 1.7e-07 | 3e-07 | 2.733E-07 | 9.292E-08 | 1.700E-07 | 3.500E-07 |
| 18094 | 905 | T905F | C | 2e-07 | 2.8e-07 | 2.5e-07 | 2.433E-07 | 4.041E-08 | 2.000E-07 | 2.800E-07 |
| 18095 | 905 | T905P | C | 4.2e-07 | 1.6e-07 | 3.2e-07 | 3.000E-07 | 1.311E-07 | 1.600E-07 | 4.200E-07 |
| 18096 | 905 | T905S | C | 9.2e-07 | 1.6e-07 | 2.6e-07 | 4.467E-07 | 4.130E-07 | 1.600E-07 | 9.200E-07 |
| 18097 | 905 | T905T | C | 3.3e-07 | 4e-07 | 3.5e-07 | 3.600E-07 | 3.606E-08 | 3.300E-07 | 4.000E-07 |
| 18098 | 905 | T905W | C | 1.9e-07 | 1.9e-07 | 5.3e-07 | 3.033E-07 | 1.963E-07 | 1.900E-07 | 5.300E-07 |
| 18099 | 905 | T905Y | C | 8.8e-08 | 4.9e-07 | 1.7e-07 | 2.493E-07 | 2.124E-07 | 8.800E-08 | 4.900E-07 |
| 18100 | 905 | T905V | C | 3.3e-07 | 3.4e-07 | 1.6e-07 | 2.767E-07 | 1.012E-07 | 1.600E-07 | 3.400E-07 |
| 18101 | 906 | L906A | C | 3.5e-07 | 2.5e-07 | 4.9e-07 | 3.633E-07 | 1.206E-07 | 2.500E-07 | 4.900E-07 |
| 18102 | 906 | L906R | C | 4.4e-07 | 3.8e-07 | 3.3e-07 | 3.833E-07 | 5.508E-08 | 3.300E-07 | 4.400E-07 |
| 18103 | 906 | L906N | C | 2.1e-07 | 1e-07 | 1.1e-07 | 1.400E-07 | 6.083E-08 | 1.000E-07 | 2.100E-07 |
| 18104 | 906 | L906D | C | 1.9e-07 | 5.5e-07 | 5.7e-07 | 4.367E-07 | 2.139E-07 | 1.900E-07 | 5.700E-07 |

| | | | | | | | | | | |
|---|---|---|---|---|---|---|---|---|---|---|
| 18105 | 906 | L906C | C | 1.8e-07 | 8.8e-08 | 1.7e-07 | 1.460E-07 | 5.048E-08 | 8.800E-08 | 1.800E-07 |
| 18106 | 906 | L906E | C | 1.8e-07 | 2.1e-07 | 2e-07 | 1.967E-07 | 1.528E-08 | 1.800E-07 | 2.100E-07 |
| 18107 | 906 | L906Q | C | 2.3e-07 | 5.3e-07 | 1.3e-07 | 2.967E-07 | 2.082E-07 | 1.300E-07 | 5.300E-07 |
| 18108 | 906 | L906G | C | 4.1e-07 | 1.7e-07 | 1.5e-07 | 2.433E-07 | 1.447E-07 | 1.500E-07 | 4.100E-07 |
| 18109 | 906 | L906H | C | 1.1e-07 | 2.3e-07 | 2.2e-07 | 1.867E-07 | 6.658E-08 | 1.100E-07 | 2.300E-07 |
| 18110 | 906 | L906I | C | 2.8e-07 | 2e-07 | 2.2e-07 | 2.333E-07 | 4.163E-08 | 2.000E-07 | 2.800E-07 |
| 18111 | 906 | L906L | C | 1.8e-07 | 3.4e-07 | 1.9e-07 | 2.367E-07 | 8.963E-08 | 1.800E-07 | 3.400E-07 |
| 18112 | 906 | L906K | C | 2.5e-07 | 2.5e-07 | 3.6e-07 | 2.867E-07 | 6.351E-08 | 2.500E-07 | 3.600E-07 |
| 18113 | 906 | L906M | C | 2.4e-07 | 3.9e-07 | 1.1e-07 | 2.467E-07 | 1.401E-07 | 1.100E-07 | 3.900E-07 |
| 18114 | 906 | L906F | C | 3.6e-07 | 1.1e-07 | 2.7e-07 | 2.467E-07 | 1.266E-07 | 1.100E-07 | 3.600E-07 |
| 18115 | 906 | L906P | C | 1.8e-07 | 2.2e-07 | 2.6e-07 | 2.200E-07 | 4.000E-08 | 1.800E-07 | 2.600E-07 |
| 18116 | 906 | L906S | C | 2.5e-07 | 1.9e-07 | 1.7e-07 | 2.033E-07 | 4.163E-08 | 1.700E-07 | 2.500E-07 |
| 18117 | 906 | L906T | C | 3.7e-07 | 4.9e-07 | 2.5e-07 | 3.700E-07 | 1.200E-07 | 2.500E-07 | 4.900E-07 |
| 18118 | 906 | L906W | C | 6.4e-07 | 1.3e-07 | 1.6e-07 | 3.100E-07 | 2.862E-07 | 1.300E-07 | 6.400E-07 |
| 18119 | 906 | L906Y | C | 2.3e-07 | 3.1e-07 | 3.6e-07 | 3.000E-07 | 6.557E-08 | 2.300E-07 | 3.600E-07 |
| 18120 | 906 | L906V | C | 3.4e-07 | 3e-07 | 2.9e-07 | 3.100E-07 | 2.646E-08 | 2.900E-07 | 3.400E-07 |
| 18121 | 907 | S907A | C | 2.8e-07 | 2.5e-07 | 1.8e-07 | 2.367E-07 | 5.132E-08 | 1.800E-07 | 2.800E-07 |
| 18122 | 907 | S907R | C | 4.5e-07 | 7.8e-08 | 2.4e-07 | 2.560E-07 | 1.865E-07 | 7.800E-08 | 4.500E-07 |
| 18123 | 907 | S907N | C | 2.2e-07 | 3.5e-07 | 2.5e-07 | 2.733E-07 | 6.807E-08 | 2.200E-07 | 3.500E-07 |
| 18124 | 907 | S907D | C | 2e-07 | 1.9e-07 | 2.4e-07 | 2.100E-07 | 2.646E-08 | 1.900E-07 | 2.400E-07 |
| 18125 | 907 | S907C | C | 1.5e-07 | 1.8e-07 | 1.6e-07 | 1.633E-07 | 1.528E-08 | 1.500E-07 | 1.800E-07 |
| 18126 | 907 | S907E | C | 2.1e-07 | 2.4e-07 | 2.6e-07 | 2.367E-07 | 2.517E-08 | 2.100E-07 | 2.600E-07 |
| 18127 | 907 | S907Q | C | 2.4e-07 | 3.1e-07 | 3.6e-07 | 3.033E-07 | 6.028E-08 | 2.400E-07 | 3.600E-07 |
| 18128 | 907 | S907G | C | 4.2e-07 | 4.6e-07 | 3e-07 | 3.933E-07 | 8.327E-08 | 3.000E-07 | 4.600E-07 |
| 18129 | 907 | S907H | C | 2.5e-07 | 4.2e-07 | 3.7e-07 | 3.467E-07 | 8.737E-08 | 2.500E-07 | 4.200E-07 |
| 18130 | 907 | S907I | C | 1.2e-07 | 2.7e-07 | 1.1e-07 | 1.667E-07 | 8.963E-08 | 1.100E-07 | 2.700E-07 |
| 18131 | 907 | S907L | C | 2e-07 | 3.3e-07 | 4.1e-07 | 3.133E-07 | 1.060E-07 | 2.000E-07 | 4.100E-07 |
| 18132 | 907 | S907K | C | 3.3e-07 | 2.4e-07 | 2.1e-07 | 2.600E-07 | 6.245E-08 | 2.100E-07 | 3.300E-07 |
| 18133 | 907 | S907M | C | 4.1e-07 | 1.4e-07 | 3e-07 | 2.833E-07 | 1.358E-07 | 1.400E-07 | 4.100E-07 |
| 18134 | 907 | S907F | C | 2.5e-07 | 2.4e-07 | 2.2e-07 | 2.367E-07 | 1.528E-08 | 2.200E-07 | 2.500E-07 |
| 18135 | 907 | S907P | C | 2.7e-07 | 2.3e-07 | 4.5e-07 | 3.167E-07 | 1.172E-07 | 2.300E-07 | 4.500E-07 |
| 18136 | 907 | S907S | C | 1.4e-07 | 2e-07 | 2.8e-07 | 2.067E-07 | 7.024E-08 | 1.400E-07 | 2.800E-07 |
| 18137 | 907 | S907T | C | 2.8e-07 | 2e-07 | 1.6e-07 | 2.133E-07 | 6.110E-08 | 1.600E-07 | 2.800E-07 |
| 18138 | 907 | S907W | C | 3.2e-07 | 3.2e-07 | 1.4e-07 | 2.600E-07 | 1.039E-07 | 1.400E-07 | 3.200E-07 |
| 18139 | 907 | S907Y | C | 3.1e-07 | 1e-07 | 2.5e-07 | 2.200E-07 | 1.082E-07 | 1.000E-07 | 3.100E-07 |
| 18140 | 907 | S907V | C | 3.3e-07 | 4.1e-07 | 4.5e-07 | 3.967E-07 | 6.110E-08 | 3.300E-07 | 4.500E-07 |
| 18141 | 908 | V908A | C | 3.5e-07 | 1.4e-07 | 1.8e-07 | 2.233E-07 | 1.115E-07 | 1.400E-07 | 3.500E-07 |
| 18142 | 908 | V908R | C | 2.2e-07 | 1.5e-07 | 2.3e-07 | 2.000E-07 | 4.359E-08 | 1.500E-07 | 2.300E-07 |
| 18143 | 908 | V908N | C | 3.5e-07 | 3.9e-07 | 2e-07 | 3.133E-07 | 1.002E-07 | 2.000E-07 | 3.900E-07 |
| 18144 | 908 | V908D | C | 4.1e-07 | 3.1e-07 | 3.2e-07 | 3.467E-07 | 5.508E-08 | 3.100E-07 | 4.100E-07 |
| 18145 | 908 | V908C | C | 2.7e-07 | 1.5e-07 | 9.2e-08 | 1.707E-07 | 9.078E-08 | 9.200E-08 | 2.700E-07 |
| 18146 | 908 | V908E | C | 1.5e-07 | 1.5e-07 | 3e-07 | 2.000E-07 | 8.660E-08 | 1.500E-07 | 3.000E-07 |
| 18147 | 908 | V908Q | C | 2.4e-07 | 3.5e-07 | 1.9e-07 | 2.600E-07 | 8.185E-08 | 1.900E-07 | 3.500E-07 |
| 18148 | 908 | V908G | C | 4.4e-07 | 3.7e-07 | 1.6e-07 | 3.233E-07 | 1.457E-07 | 1.600E-07 | 4.400E-07 |
| 18149 | 908 | V908H | C | 4.3e-07 | 5.9e-07 | 1.5e-07 | 3.900E-07 | 2.227E-07 | 1.500E-07 | 5.900E-07 |
| 18150 | 908 | V908I | C | 1.2e-07 | 5.4e-07 | 2.2e-07 | 2.933E-07 | 2.194E-07 | 1.200E-07 | 5.400E-07 |
| 18151 | 908 | V908L | C | 2e-07 | 3e-07 | 3.3e-07 | 2.767E-07 | 6.807E-08 | 2.000E-07 | 3.300E-07 |
| 18152 | 908 | V908K | C | 3.1e-07 | 2.8e-07 | 2.5e-07 | 2.800E-07 | 3.000E-08 | 2.500E-07 | 3.100E-07 |
| 18153 | 908 | V908M | C | 1.4e-07 | 5e-07 | 3.8e-07 | 3.400E-07 | 1.833E-07 | 1.400E-07 | 5.000E-07 |
| 18154 | 908 | V908F | C | 5.7e-08 | 2.1e-07 | 5.1e-07 | 2.590E-07 | 2.304E-07 | 5.700E-08 | 5.100E-07 |
| 18155 | 908 | V908P | C | 2.7e-07 | 2.8e-07 | 1.6e-07 | 2.367E-07 | 6.658E-08 | 1.600E-07 | 2.800E-07 |
| 18156 | 908 | V908S | C | 3.8e-07 | 3.8e-07 | 1.2e-07 | 2.933E-07 | 1.501E-07 | 1.200E-07 | 3.800E-07 |
| 18157 | 908 | V908T | C | 2.8e-07 | 2e-07 | 1.5e-07 | 2.100E-07 | 6.557E-08 | 1.500E-07 | 2.800E-07 |
| 18158 | 908 | V908W | C | 1.7e-07 | 4.3e-07 | 2.4e-07 | 2.800E-07 | 1.345E-07 | 1.700E-07 | 4.300E-07 |
| 18159 | 908 | V908Y | C | 1.8e-07 | 2.2e-07 | 4.5e-07 | 2.833E-07 | 1.457E-07 | 1.800E-07 | 4.500E-07 |
| 18160 | 908 | V908V | C | 1.6e-07 | 3.7e-07 | 1.5e-07 | 2.267E-07 | 1.242E-07 | 1.500E-07 | 3.700E-07 |
| 18161 | 909 | D909A | C | 2.5e-07 | 4.1e-07 | 2.3e-07 | 2.967E-07 | 9.866E-08 | 2.300E-07 | 4.100E-07 |
| 18162 | 909 | D909R | C | 4e-07 | 1.9e-07 | 3.4e-07 | 3.100E-07 | 1.082E-07 | 1.900E-07 | 4.000E-07 |
| 18163 | 909 | D909N | C | 2.8e-07 | 1.2e-07 | 2.9e-07 | 2.300E-07 | 9.539E-08 | 1.200E-07 | 2.900E-07 |
| 18164 | 909 | D909D | C | 1.1e-07 | 1.2e-07 | 3e-07 | 1.767E-07 | 1.069E-07 | 1.100E-07 | 3.000E-07 |
| 18165 | 909 | D909C | C | 2.2e-07 | 2.5e-07 | 2.3e-07 | 2.333E-07 | 1.528E-08 | 2.200E-07 | 2.500E-07 |
| 18166 | 909 | D909E | C | 3.8e-07 | 1.6e-07 | 2.1e-07 | 2.500E-07 | 1.153E-07 | 1.600E-07 | 3.800E-07 |
| 18167 | 909 | D909Q | C | 2.7e-07 | 2.5e-07 | 1.7e-07 | 2.300E-07 | 5.292E-08 | 1.700E-07 | 2.700E-07 |
| 18168 | 909 | D909G | C | 1.7e-07 | 2.4e-07 | 3e-07 | 2.367E-07 | 6.506E-08 | 1.700E-07 | 3.000E-07 |
| 18169 | 909 | D909H | C | 2.9e-07 | 2e-07 | 1.4e-07 | 2.100E-07 | 7.550E-08 | 1.400E-07 | 2.900E-07 |
| 18170 | 909 | D909I | C | 1.8e-07 | 2.1e-07 | 2.2e-07 | 2.033E-07 | 2.082E-08 | 1.800E-07 | 2.200E-07 |
| 18171 | 909 | D909L | C | 3e-07 | 2.3e-07 | 1.7e-07 | 2.333E-07 | 6.506E-08 | 1.700E-07 | 3.000E-07 |
| 18172 | 909 | D909K | C | 8e-08 | 2.3e-07 | 3.8e-07 | 2.300E-07 | 1.500E-07 | 8.000E-08 | 3.800E-07 |
| 18173 | 909 | D909M | C | 2.1e-07 | 3.4e-07 | 3.5e-07 | 3.000E-07 | 7.810E-08 | 2.100E-07 | 3.500E-07 |
| 18174 | 909 | D909F | C | 2.5e-07 | 1.7e-07 | 2.2e-07 | 2.133E-07 | 4.041E-08 | 1.700E-07 | 2.500E-07 |
| 18175 | 909 | D909P | C | 3e-07 | 2.5e-07 | 1.8e-07 | 2.433E-07 | 6.028E-08 | 1.800E-07 | 3.000E-07 |
| 18176 | 909 | D909S | C | 2.2e-07 | 2.7e-07 | 3.2e-07 | 2.700E-07 | 5.000E-08 | 2.200E-07 | 3.200E-07 |
| 18177 | 909 | D909T | C | 1.8e-07 | 2.7e-07 | 1.5e-07 | 2.000E-07 | 6.245E-08 | 1.500E-07 | 2.700E-07 |
| 18178 | 909 | D909W | C | 1.8e-07 | 2.3e-07 | 4.1e-07 | 2.733E-07 | 1.210E-07 | 1.800E-07 | 4.100E-07 |
| 18179 | 909 | D909Y | C | 2.2e-07 | 2.5e-07 | 3e-07 | 2.567E-07 | 4.041E-08 | 2.200E-07 | 3.000E-07 |
| 18180 | 909 | D909V | C | 8.1e-07 | 2.1e-07 | 5.2e-07 | 5.133E-07 | 3.001E-07 | 2.100E-07 | 8.100E-07 |
| 18181 | 910 | R910A | C | 2.7e-07 | 1.4e-07 | 2.1e-07 | 2.067E-07 | 6.506E-08 | 1.400E-07 | 2.700E-07 |
| 18182 | 910 | R910R | C | 1.9e-07 | 1.3e-07 | 4.2e-07 | 2.467E-07 | 1.531E-07 | 1.300E-07 | 4.200E-07 |
| 18183 | 910 | R910N | C | 1.6e-07 | 3.2e-07 | 2.5e-07 | 2.433E-07 | 8.021E-08 | 1.600E-07 | 3.200E-07 |
| 18184 | 910 | R910D | C | 1.2e-07 | 2.7e-07 | 1.4e-07 | 1.767E-07 | 8.145E-08 | 1.200E-07 | 2.700E-07 |
| 18185 | 910 | R910C | C | 2.1e-07 | 6.9e-08 | 2.8e-07 | 1.863E-07 | 1.075E-07 | 6.900E-08 | 2.800E-07 |
| 18186 | 910 | R910E | C | 1e-07 | 2.6e-07 | 5.1e-07 | 2.900E-07 | 2.066E-07 | 1.000E-07 | 5.100E-07 |
| 18187 | 910 | R910Q | C | 2e-07 | 1.8e-07 | 2.7e-07 | 2.167E-07 | 4.726E-08 | 1.800E-07 | 2.700E-07 |
| 18188 | 910 | R910G | C | 2.3e-07 | 2.5e-07 | 1.4e-07 | 2.067E-07 | 5.859E-08 | 1.400E-07 | 2.500E-07 |
| 18189 | 910 | R910H | C | 2.6e-07 | 2.8e-07 | 2.3e-07 | 2.567E-07 | 2.517E-08 | 2.300E-07 | 2.800E-07 |

| | | | | | | | | | | |
|---|---|---|---|---|---|---|---|---|---|---|
| 18190 | 910 | R910I | C | 1.1e-07 | 2.5e-07 | 6.5e-07 | 3.367E-07 | 2.802E-07 | 1.100E-07 | 6.500E-07 |
| 18191 | 910 | R910L | C | 4.6e-07 | 4.8e-07 | 1.8e-07 | 3.733E-07 | 1.677E-07 | 1.800E-07 | 4.800E-07 |
| 18192 | 910 | R910K | C | 5.8e-07 | 2.2e-07 | 7e-08 | 2.900E-07 | 2.621E-07 | 7.000E-08 | 5.800E-07 |
| 18193 | 910 | R910M | C | 2.6e-07 | 4.9e-07 | 2.4e-07 | 3.300E-07 | 1.389E-07 | 2.400E-07 | 4.900E-07 |
| 18194 | 910 | R910F | C | 2.2e-07 | 4.1e-07 | 2e-07 | 2.767E-07 | 1.159E-07 | 2.000E-07 | 4.100E-07 |
| 18195 | 910 | R910P | C | 2.4e-07 | 1.2e-07 | 2.3e-07 | 1.967E-07 | 6.658E-08 | 1.200E-07 | 2.400E-07 |
| 18196 | 910 | R910S | C | 6.8e-08 | 1.6e-07 | 4.1e-07 | 2.127E-07 | 1.770E-07 | 6.800E-08 | 4.100E-07 |
| 18197 | 910 | R910T | C | 1.5e-07 | 2.6e-07 | 2e-07 | 2.033E-07 | 5.508E-08 | 1.500E-07 | 2.600E-07 |
| 18198 | 910 | R910W | C | 1.4e-07 | 3e-07 | 1.2e-07 | 1.867E-07 | 9.866E-08 | 1.200E-07 | 3.000E-07 |
| 18199 | 910 | R910Y | C | 1.7e-07 | 2e-07 | 4.3e-07 | 2.667E-07 | 1.422E-07 | 1.700E-07 | 4.300E-07 |
| 18200 | 910 | R910V | C | 2.3e-07 | 2.2e-07 | 4.4e-07 | 2.967E-07 | 1.242E-07 | 2.200E-07 | 4.400E-07 |
| 18201 | 911 | S911A | C | 3.3e-07 | 3.9e-07 | 2.2e-07 | 3.133E-07 | 8.622E-08 | 2.200E-07 | 3.900E-07 |
| 18202 | 911 | S911R | C | 1.9e-07 | 3e-07 | 2.1e-07 | 2.333E-07 | 5.859E-08 | 1.900E-07 | 3.000E-07 |
| 18203 | 911 | S911N | C | 4.4e-07 | 4.2e-07 | 2.3e-07 | 3.633E-07 | 1.159E-07 | 2.300E-07 | 4.400E-07 |
| 18204 | 911 | S911D | C | 4.3e-07 | 3.6e-07 | 6.4e-07 | 4.767E-07 | 1.457E-07 | 3.600E-07 | 6.400E-07 |
| 18205 | 911 | S911C | C | 2.8e-07 | 6.9e-07 | 3.4e-07 | 4.367E-07 | 2.214E-07 | 2.800E-07 | 6.900E-07 |
| 18206 | 911 | S911E | C | 4.7e-07 | 1.6e-07 | 3.1e-07 | 3.133E-07 | 1.550E-07 | 1.600E-07 | 4.700E-07 |
| 18207 | 911 | S911Q | C | 1.2e-07 | 1.2e-07 | 2.5e-07 | 1.633E-07 | 7.506E-08 | 1.200E-07 | 2.500E-07 |
| 18208 | 911 | S911G | C | 4.3e-07 | 1.6e-07 | 1.7e-07 | 2.533E-07 | 1.531E-07 | 1.600E-07 | 4.300E-07 |
| 18209 | 911 | S911H | C | 1.6e-07 | 2.9e-07 | 1.6e-07 | 2.033E-07 | 7.506E-08 | 1.600E-07 | 2.900E-07 |
| 18210 | 911 | S911I | C | 4.1e-07 | 2.5e-07 | 5.4e-07 | 4.000E-07 | 1.453E-07 | 2.500E-07 | 5.400E-07 |
| 18211 | 911 | S911L | C | 4.3e-07 | 4.2e-07 | 1.7e-07 | 3.400E-07 | 1.473E-07 | 1.700E-07 | 4.300E-07 |
| 18212 | 911 | S911K | C | 4.3e-07 | 3e-07 | 1.7e-07 | 3.000E-07 | 1.300E-07 | 1.700E-07 | 4.300E-07 |
| 18213 | 911 | S911M | C | 2.2e-07 | 3.2e-07 | 1.1e-07 | 2.167E-07 | 1.050E-07 | 1.100E-07 | 3.200E-07 |
| 18214 | 911 | S911F | C | 3.2e-07 | 2.3e-07 | 2.7e-07 | 2.733E-07 | 4.509E-08 | 2.300E-07 | 3.200E-07 |
| 18215 | 911 | S911P | C | 3.1e-07 | 1.3e-07 | 2.5e-07 | 2.300E-07 | 9.165E-08 | 1.300E-07 | 3.100E-07 |
| 18216 | 911 | S911S | C | 2.1e-07 | 6.2e-07 | 3.4e-07 | 3.900E-07 | 2.095E-07 | 2.100E-07 | 6.200E-07 |
| 18217 | 911 | S911T | C | 3.8e-07 | 4e-07 | 3.8e-07 | 3.867E-07 | 1.155E-08 | 3.800E-07 | 4.000E-07 |
| 18218 | 911 | S911W | C | 2.8e-07 | 1.8e-07 | 1.5e-07 | 2.033E-07 | 6.807E-08 | 1.500E-07 | 2.800E-07 |
| 18219 | 911 | S911Y | C | 2e-07 | 9.9e-08 | 4.5e-07 | 2.497E-07 | 1.807E-07 | 9.900E-08 | 4.500E-07 |
| 18220 | 911 | S911V | C | 4.2e-07 | 2.1e-07 | 3.5e-07 | 3.267E-07 | 1.069E-07 | 2.100E-07 | 4.200E-07 |
| 18221 | 912 | K912A | C | 1.4e-07 | 2.8e-07 | 2e-07 | 2.067E-07 | 7.024E-08 | 1.400E-07 | 2.800E-07 |
| 18222 | 912 | K912R | C | 4e-07 | 3e-07 | 2.9e-07 | 3.300E-07 | 6.083E-08 | 2.900E-07 | 4.000E-07 |
| 18223 | 912 | K912N | C | 8.9e-08 | 5.5e-07 | 1.3e-07 | 2.563E-07 | 2.551E-07 | 8.900E-08 | 5.500E-07 |
| 18224 | 912 | K912D | C | 1.7e-07 | 3.3e-07 | 4.2e-07 | 3.067E-07 | 1.266E-07 | 1.700E-07 | 4.200E-07 |
| 18225 | 912 | K912C | C | 2.9e-07 | 3e-07 | 2e-07 | 2.633E-07 | 5.508E-08 | 2.000E-07 | 3.000E-07 |
| 18226 | 912 | K912E | C | 1e-07 | 2.7e-07 | 2.5e-07 | 2.067E-07 | 9.292E-08 | 1.000E-07 | 2.700E-07 |
| 18227 | 912 | K912Q | C | 3.4e-07 | 2.1e-07 | 2.9e-07 | 2.800E-07 | 6.557E-08 | 2.100E-07 | 3.400E-07 |
| 18228 | 912 | K912G | C | 3.1e-07 | 2.1e-07 | 3.8e-07 | 3.000E-07 | 8.544E-08 | 2.100E-07 | 3.800E-07 |
| 18229 | 912 | K912H | C | 1.7e-07 | 1.1e-07 | 4.7e-07 | 2.500E-07 | 1.929E-07 | 1.100E-07 | 4.700E-07 |
| 18230 | 912 | K912I | C | 9.7e-08 | 5.8e-07 | 1.9e-07 | 2.890E-07 | 2.563E-07 | 9.700E-08 | 5.800E-07 |
| 18231 | 912 | K912L | C | 2.8e-07 | 2.5e-07 | 2.4e-07 | 2.567E-07 | 2.082E-08 | 2.400E-07 | 2.800E-07 |
| 18232 | 912 | K912K | C | 2.7e-07 | 4.3e-07 | 3.1e-07 | 3.367E-07 | 8.327E-08 | 2.700E-07 | 4.300E-07 |
| 18233 | 912 | K912M | C | 2.3e-07 | 2.8e-07 | 2e-07 | 2.367E-07 | 4.041E-08 | 2.000E-07 | 2.800E-07 |
| 18234 | 912 | K912F | C | 1.2e-07 | 3e-07 | 1.7e-07 | 1.967E-07 | 9.292E-08 | 1.200E-07 | 3.000E-07 |
| 18235 | 912 | K912P | C | 2.7e-07 | 1.6e-07 | 2.8e-07 | 2.367E-07 | 6.658E-08 | 1.600E-07 | 2.800E-07 |
| 18236 | 912 | K912S | C | 1.7e-07 | 2e-07 | 2.2e-07 | 1.967E-07 | 2.517E-08 | 1.700E-07 | 2.200E-07 |
| 18237 | 912 | K912T | C | 4.5e-07 | 3.7e-07 | 1.5e-07 | 3.233E-07 | 1.553E-07 | 1.500E-07 | 4.500E-07 |
| 18238 | 912 | K912W | C | 2.6e-07 | 3.4e-07 | 2.4e-07 | 2.800E-07 | 5.292E-08 | 2.400E-07 | 3.400E-07 |
| 18239 | 912 | K912Y | C | 2.1e-07 | 5.4e-07 | 4.8e-07 | 4.100E-07 | 1.758E-07 | 2.100E-07 | 5.400E-07 |
| 18240 | 912 | K912V | C | 2.2e-07 | 2.2e-07 | 1.3e-07 | 1.900E-07 | 5.196E-08 | 1.300E-07 | 2.200E-07 |
| 18241 | 913 | N913A | C | 1.5e-07 | 4.1e-07 | 5.2e-07 | 3.600E-07 | 1.900E-07 | 1.500E-07 | 5.200E-07 |
| 18242 | 913 | N913R | C | 2.1e-07 | 1.7e-07 | 3.8e-07 | 2.533E-07 | 1.115E-07 | 1.700E-07 | 3.800E-07 |
| 18243 | 913 | N913N | C | 3.3e-07 | 4.2e-07 | 1.4e-07 | 2.967E-07 | 1.429E-07 | 1.400E-07 | 4.200E-07 |
| 18244 | 913 | N913D | C | 5.4e-07 | 2.6e-07 | 2e-07 | 3.333E-07 | 1.815E-07 | 2.000E-07 | 5.400E-07 |
| 18245 | 913 | N913C | C | 3.4e-07 | 3.1e-07 | 2.7e-07 | 3.067E-07 | 3.512E-08 | 2.700E-07 | 3.400E-07 |
| 18246 | 913 | N913E | C | 1.8e-07 | 2.7e-07 | 3.5e-07 | 2.667E-07 | 8.505E-08 | 1.800E-07 | 3.500E-07 |
| 18247 | 913 | N913Q | C | 4.2e-07 | 1.5e-07 | 1.5e-07 | 2.400E-07 | 1.559E-07 | 1.500E-07 | 4.200E-07 |
| 18248 | 913 | N913G | C | 3.2e-07 | 3.3e-07 | 1.8e-07 | 2.767E-07 | 8.386E-08 | 1.800E-07 | 3.300E-07 |
| 18249 | 913 | N913H | C | 2.3e-07 | 2.3e-07 | 1.5e-07 | 2.033E-07 | 4.619E-08 | 1.500E-07 | 2.300E-07 |
| 18250 | 913 | N913I | C | 1.8e-07 | 1.7e-07 | 2.1e-07 | 1.867E-07 | 2.082E-08 | 1.700E-07 | 2.100E-07 |
| 18251 | 913 | N913L | C | 6.3e-07 | 2e-07 | 4.7e-07 | 4.333E-07 | 2.173E-07 | 2.000E-07 | 6.300E-07 |
| 18252 | 913 | N913K | C | 2.8e-07 | 3.9e-07 | 1.3e-07 | 2.667E-07 | 1.305E-07 | 1.300E-07 | 3.900E-07 |
| 18253 | 913 | N913M | C | 2.1e-07 | 2.6e-07 | 1.7e-07 | 2.133E-07 | 4.509E-08 | 1.700E-07 | 2.600E-07 |
| 18254 | 913 | N913F | C | 1.5e-07 | 1.8e-07 | 2e-07 | 1.767E-07 | 2.517E-08 | 1.500E-07 | 2.000E-07 |
| 18255 | 913 | N913P | C | 1.4e-07 | 1.6e-07 | 4.1e-07 | 2.367E-07 | 1.504E-07 | 1.400E-07 | 4.100E-07 |
| 18256 | 913 | N913S | C | 2.5e-07 | 3.5e-07 | 1.3e-07 | 2.433E-07 | 1.102E-07 | 1.300E-07 | 3.500E-07 |
| 18257 | 913 | N913T | C | 1.8e-07 | 2.8e-07 | 2.2e-07 | 2.267E-07 | 5.033E-08 | 1.800E-07 | 2.800E-07 |
| 18258 | 913 | N913W | C | 3.6e-07 | 3.7e-07 | 2.7e-07 | 3.333E-07 | 5.508E-08 | 2.700E-07 | 3.700E-07 |
| 18259 | 913 | N913Y | C | 3.3e-07 | 3.5e-07 | 2.9e-07 | 3.233E-07 | 3.055E-08 | 2.900E-07 | 3.500E-07 |
| 18260 | 913 | N913V | C | 2.6e-07 | 2.8e-07 | 1.8e-07 | 2.400E-07 | 5.292E-08 | 1.800E-07 | 2.800E-07 |
| 18261 | 914 | T914A | C | 3.2e-07 | 4.4e-07 | 2.7e-07 | 3.433E-07 | 8.737E-08 | 2.700E-07 | 4.400E-07 |
| 18262 | 914 | T914R | C | 4.3e-07 | 2.3e-07 | 2.7e-07 | 3.100E-07 | 1.058E-07 | 2.300E-07 | 4.300E-07 |
| 18263 | 914 | T914N | C | 2.3e-07 | 8.9e-08 | 3.4e-07 | 2.197E-07 | 1.258E-07 | 8.900E-08 | 3.400E-07 |
| 18264 | 914 | T914D | C | 1.9e-07 | 1.5e-07 | 4e-07 | 2.467E-07 | 1.343E-07 | 1.500E-07 | 4.000E-07 |
| 18265 | 914 | T914C | C | 2e-07 | 3.1e-07 | 4.4e-07 | 3.167E-07 | 1.201E-07 | 2.000E-07 | 4.400E-07 |
| 18266 | 914 | T914E | C | 1.7e-07 | 2.8e-07 | 1.1e-07 | 1.867E-07 | 8.622E-08 | 1.100E-07 | 2.800E-07 |
| 18267 | 914 | T914Q | C | 4.7e-07 | 2.9e-07 | 2.7e-07 | 3.433E-07 | 1.102E-07 | 2.700E-07 | 4.700E-07 |
| 18268 | 914 | T914G | C | 1.9e-07 | 3.4e-07 | 1.2e-07 | 2.167E-07 | 1.124E-07 | 1.200E-07 | 3.400E-07 |
| 18269 | 914 | T914H | C | 4.2e-07 | 2.4e-07 | 2.6e-07 | 3.067E-07 | 9.866E-08 | 2.400E-07 | 4.200E-07 |
| 18270 | 914 | T914I | C | 2.9e-07 | 1.7e-07 | 1.3e-07 | 1.967E-07 | 8.327E-08 | 1.300E-07 | 2.900E-07 |
| 18271 | 914 | T914L | C | 5.5e-07 | 4.5e-07 | 1.8e-07 | 3.933E-07 | 1.914E-07 | 1.800E-07 | 5.500E-07 |
| 18272 | 914 | T914K | C | 1.8e-07 | 2.7e-07 | 1.6e-07 | 2.033E-07 | 5.859E-08 | 1.600E-07 | 2.700E-07 |
| 18273 | 914 | T914M | C | 3e-07 | 4.4e-07 | 1.5e-07 | 2.967E-07 | 1.450E-07 | 1.500E-07 | 4.400E-07 |
| 18274 | 914 | T914F | C | 1.1e-07 | 4.1e-07 | 1.6e-07 | 2.267E-07 | 1.607E-07 | 1.100E-07 | 4.100E-07 |

| | | | | | | | | | | |
|---|---|---|---|---|---|---|---|---|---|---|
| 18275 | 914 | T914P | C | 1.4e-07 | 2.8e-07 | 1.9e-07 | 2.033E-07 | 7.095E-08 | 1.400E-07 | 2.800E-07 |
| 18276 | 914 | T914S | C | 2.2e-07 | 3e-07 | 2.1e-07 | 2.433E-07 | 4.933E-08 | 2.100E-07 | 3.000E-07 |
| 18277 | 914 | T914T | C | 3.4e-07 | 4.2e-07 | 3.4e-07 | 3.667E-07 | 4.619E-08 | 3.400E-07 | 4.200E-07 |
| 18278 | 914 | T914W | C | 3.8e-07 | 1.9e-07 | 2.2e-07 | 2.633E-07 | 1.021E-07 | 1.900E-07 | 3.800E-07 |
| 18279 | 914 | T914Y | C | 4.7e-07 | 2e-07 | 2.2e-07 | 2.967E-07 | 1.504E-07 | 2.000E-07 | 4.700E-07 |
| 18280 | 914 | T914V | C | 2.1e-07 | 2.8e-07 | 1.9e-07 | 2.267E-07 | 4.726E-08 | 1.900E-07 | 2.800E-07 |
| 18281 | 915 | L915A | C | 2.8e-07 | 6e-07 | 6.6e-07 | 5.133E-07 | 2.043E-07 | 2.800E-07 | 6.600E-07 |
| 18282 | 915 | L915R | C | 2.8e-07 | 1.9e-07 | 5.2e-07 | 3.300E-07 | 1.706E-07 | 1.900E-07 | 5.200E-07 |
| 18283 | 915 | L915N | C | 9.9e-08 | 4.4e-07 | 3.9e-07 | 3.097E-07 | 1.841E-07 | 9.900E-08 | 4.400E-07 |
| 18284 | 915 | L915D | C | 2.6e-07 | 5.2e-07 | 2.1e-07 | 3.300E-07 | 1.664E-07 | 2.100E-07 | 5.200E-07 |
| 18285 | 915 | L915C | C | 2.5e-07 | 1.2e-07 | 2.8e-07 | 2.167E-07 | 8.505E-08 | 1.200E-07 | 2.800E-07 |
| 18286 | 915 | L915E | C | 4e-07 | 2.2e-07 | 1.9e-07 | 2.700E-07 | 1.136E-07 | 1.900E-07 | 4.000E-07 |
| 18287 | 915 | L915Q | C | 5.6e-07 | 1.8e-07 | 1.5e-07 | 2.967E-07 | 2.285E-07 | 1.500E-07 | 5.600E-07 |
| 18288 | 915 | L915G | C | 1.5e-07 | 3.5e-07 | 4.1e-07 | 3.033E-07 | 1.361E-07 | 1.500E-07 | 4.100E-07 |
| 18289 | 915 | L915H | C | 2.2e-07 | 1.3e-07 | 2.8e-07 | 2.100E-07 | 7.550E-08 | 1.300E-07 | 2.800E-07 |
| 18290 | 915 | L915I | C | 2.2e-07 | 4.3e-07 | 5.8e-07 | 4.100E-07 | 1.808E-07 | 2.200E-07 | 5.800E-07 |
| 18291 | 915 | L915L | C | 2.6e-07 | 2.4e-07 | 1.7e-07 | 2.233E-07 | 4.726E-08 | 1.700E-07 | 2.600E-07 |
| 18292 | 915 | L915K | C | 1.5e-07 | 1.2e-07 | 1.2e-07 | 1.300E-07 | 1.732E-08 | 1.200E-07 | 1.500E-07 |
| 18293 | 915 | L915M | C | 2.9e-07 | 2.8e-07 | 3e-07 | 2.900E-07 | 1.000E-08 | 2.800E-07 | 3.000E-07 |
| 18294 | 915 | L915F | C | 7.3e-07 | 5e-07 | 1.9e-07 | 4.733E-07 | 2.710E-07 | 1.900E-07 | 7.300E-07 |
| 18295 | 915 | L915P | C | 2.5e-07 | 5.9e-07 | 2.5e-07 | 3.633E-07 | 1.963E-07 | 2.500E-07 | 5.900E-07 |
| 18296 | 915 | L915S | C | 1.6e-07 | 1.8e-07 | 3.8e-07 | 2.400E-07 | 1.217E-07 | 1.600E-07 | 3.800E-07 |
| 18297 | 915 | L915T | C | 3e-07 | 2.8e-07 | 1.3e-07 | 2.367E-07 | 9.292E-08 | 1.300E-07 | 3.000E-07 |
| 18298 | 915 | L915W | C | 1.1e-07 | 3.4e-07 | 2.5e-07 | 2.333E-07 | 1.159E-07 | 1.100E-07 | 3.400E-07 |
| 18299 | 915 | L915Y | C | 4e-07 | 1.5e-07 | 1.1e-07 | 2.200E-07 | 1.572E-07 | 1.100E-07 | 4.000E-07 |
| 18300 | 915 | L915V | C | 1.1e-06 | 1.8e-07 | 4.2e-07 | 5.667E-07 | 4.772E-07 | 1.800E-07 | 1.100E-06 |
| 18301 | 916 | Y916A | C | 2.2e-07 | 1.5e-07 | 1.6e-07 | 1.767E-07 | 3.786E-08 | 1.500E-07 | 2.200E-07 |
| 18302 | 916 | Y916R | C | 1.8e-07 | 2.8e-07 | 1.1e-07 | 1.900E-07 | 8.544E-08 | 1.100E-07 | 2.800E-07 |
| 18303 | 916 | Y916N | C | 3.6e-07 | 3.9e-07 | 1.9e-07 | 3.133E-07 | 1.079E-07 | 1.900E-07 | 3.900E-07 |
| 18304 | 916 | Y916D | C | 2.4e-07 | 2.8e-07 | 2e-07 | 2.400E-07 | 4.000E-08 | 2.000E-07 | 2.800E-07 |
| 18305 | 916 | Y916C | C | 2.3e-07 | 5e-07 | 2.6e-07 | 3.300E-07 | 1.480E-07 | 2.300E-07 | 5.000E-07 |
| 18306 | 916 | Y916E | C | 2.3e-07 | 3.1e-07 | 1.7e-07 | 2.367E-07 | 7.024E-08 | 1.700E-07 | 3.100E-07 |
| 18307 | 916 | Y916Q | C | 1.6e-07 | 9.2e-08 | 2.3e-07 | 1.607E-07 | 6.900E-08 | 9.200E-08 | 2.300E-07 |
| 18308 | 916 | Y916G | C | 4.8e-07 | 2.4e-07 | 3.9e-07 | 3.700E-07 | 1.212E-07 | 2.400E-07 | 4.800E-07 |
| 18309 | 916 | Y916H | C | 1.6e-07 | 3.2e-07 | 2.8e-07 | 2.533E-07 | 8.327E-08 | 1.600E-07 | 3.200E-07 |
| 18310 | 916 | Y916I | C | 2.4e-07 | 2.2e-07 | 4.2e-07 | 2.933E-07 | 1.102E-07 | 2.200E-07 | 4.200E-07 |
| 18311 | 916 | Y916L | C | 2.3e-07 | 4e-07 | 1.4e-07 | 2.567E-07 | 1.320E-07 | 1.400E-07 | 4.000E-07 |
| 18312 | 916 | Y916K | C | 1.1e-07 | 3.9e-07 | 1.8e-07 | 2.267E-07 | 1.457E-07 | 1.100E-07 | 3.900E-07 |
| 18313 | 916 | Y916M | C | 1.6e-07 | 1.4e-07 | 2.6e-07 | 1.867E-07 | 6.429E-08 | 1.400E-07 | 2.600E-07 |
| 18314 | 916 | Y916F | C | 5.1e-07 | 2.1e-07 | 3.1e-07 | 3.433E-07 | 1.528E-07 | 2.100E-07 | 5.100E-07 |
| 18315 | 916 | Y916P | C | 3e-07 | 2.7e-07 | 3.6e-07 | 3.100E-07 | 4.583E-08 | 2.700E-07 | 3.600E-07 |
| 18316 | 916 | Y916S | C | 1.8e-07 | 1.4e-07 | 2e-07 | 1.733E-07 | 3.055E-08 | 1.400E-07 | 2.000E-07 |
| 18317 | 916 | Y916T | C | 3.7e-07 | 3.8e-07 | 2.4e-07 | 3.300E-07 | 7.810E-08 | 2.400E-07 | 3.800E-07 |
| 18318 | 916 | Y916W | C | 1.7e-07 | 2.1e-07 | 2.8e-07 | 2.200E-07 | 5.568E-08 | 1.700E-07 | 2.800E-07 |
| 18319 | 916 | Y916Y | C | 3.2e-07 | 2.1e-07 | 2.3e-07 | 2.533E-07 | 5.859E-08 | 2.100E-07 | 3.200E-07 |
| 18320 | 916 | Y916V | C | 2.7e-07 | 2.9e-07 | 2.3e-07 | 2.633E-07 | 3.055E-08 | 2.300E-07 | 2.900E-07 |
| 18321 | 917 | L917A | C | 2.5e-07 | 2.7e-07 | 1.8e-07 | 2.333E-07 | 4.726E-08 | 1.800E-07 | 2.700E-07 |
| 18322 | 917 | L917R | C | 2.1e-07 | 3.9e-07 | 1.9e-07 | 2.633E-07 | 1.102E-07 | 1.900E-07 | 3.900E-07 |
| 18323 | 917 | L917N | C | 2e-07 | 1.7e-07 | 2.7e-07 | 2.133E-07 | 5.132E-08 | 1.700E-07 | 2.700E-07 |
| 18324 | 917 | L917D | C | 4.4e-07 | 2e-07 | 2.8e-07 | 3.067E-07 | 1.222E-07 | 2.000E-07 | 4.400E-07 |
| 18325 | 917 | L917C | C | 3.6e-07 | 5.2e-07 | 2.3e-07 | 3.700E-07 | 1.453E-07 | 2.300E-07 | 5.200E-07 |
| 18326 | 917 | L917E | C | 2.7e-07 | 2.1e-07 | 2.5e-07 | 2.433E-07 | 3.055E-08 | 2.100E-07 | 2.700E-07 |
| 18327 | 917 | L917Q | C | 1.6e-07 | 1.9e-07 | 2.3e-07 | 1.933E-07 | 3.512E-08 | 1.600E-07 | 2.300E-07 |
| 18328 | 917 | L917G | C | 2.2e-07 | 5.6e-07 | 1.6e-07 | 3.133E-07 | 2.157E-07 | 1.600E-07 | 5.600E-07 |
| 18329 | 917 | L917H | C | 2.5e-07 | 2.1e-07 | 1.9e-07 | 2.167E-07 | 3.055E-08 | 1.900E-07 | 2.500E-07 |
| 18330 | 917 | L917I | C | 2.4e-07 | 1.4e-07 | 3.1e-07 | 2.300E-07 | 8.544E-08 | 1.400E-07 | 3.100E-07 |
| 18331 | 917 | L917L | C | 1.4e-07 | 2.9e-07 | 2.6e-07 | 2.300E-07 | 7.937E-08 | 1.400E-07 | 2.900E-07 |
| 18332 | 917 | L917K | C | 1.8e-07 | 3.3e-07 | 2.1e-07 | 2.400E-07 | 7.937E-08 | 1.800E-07 | 3.300E-07 |
| 18333 | 917 | L917M | C | 1.6e-07 | 3.3e-07 | 1.5e-07 | 2.133E-07 | 1.012E-07 | 1.500E-07 | 3.300E-07 |
| 18334 | 917 | L917F | C | 2.8e-07 | 3.7e-07 | 1.3e-07 | 2.600E-07 | 1.212E-07 | 1.300E-07 | 3.700E-07 |
| 18335 | 917 | L917P | C | 4.4e-07 | 2.6e-07 | 1.7e-07 | 2.900E-07 | 1.375E-07 | 1.700E-07 | 4.400E-07 |
| 18336 | 917 | L917S | C | 4.9e-07 | 4.2e-07 | 3.6e-07 | 4.233E-07 | 6.506E-08 | 3.600E-07 | 4.900E-07 |
| 18337 | 917 | L917T | C | 2.7e-07 | 4.2e-07 | 2.5e-07 | 3.133E-07 | 9.292E-08 | 2.500E-07 | 4.200E-07 |
| 18338 | 917 | L917W | C | 3.5e-07 | 2.2e-07 | 9.4e-08 | 2.213E-07 | 1.280E-07 | 9.400E-08 | 3.500E-07 |
| 18339 | 917 | L917Y | C | 1.5e-07 | 1.2e-07 | 3.2e-07 | 1.967E-07 | 1.079E-07 | 1.200E-07 | 3.200E-07 |
| 18340 | 917 | L917V | C | 5.6e-07 | 1.9e-07 | 3.2e-07 | 3.567E-07 | 1.877E-07 | 1.900E-07 | 5.600E-07 |
| 18341 | 918 | Q918A | C | 2.2e-07 | 1.8e-07 | 2.8e-07 | 2.267E-07 | 5.033E-08 | 1.800E-07 | 2.800E-07 |
| 18342 | 918 | Q918R | C | 2.2e-07 | 3.2e-07 | 2.9e-07 | 2.767E-07 | 5.132E-08 | 2.200E-07 | 3.200E-07 |
| 18343 | 918 | Q918N | C | 2.1e-07 | 7e-08 | 2.2e-07 | 1.667E-07 | 8.386E-08 | 7.000E-08 | 2.200E-07 |
| 18344 | 918 | Q918D | C | 2.2e-07 | 2e-07 | 1.3e-07 | 1.833E-07 | 4.726E-08 | 1.300E-07 | 2.200E-07 |
| 18345 | 918 | Q918C | C | 1.5e-07 | 1.9e-07 | 2.3e-07 | 1.900E-07 | 4.000E-08 | 1.500E-07 | 2.300E-07 |
| 18346 | 918 | Q918E | C | 1.3e-07 | 2.4e-07 | 3.5e-07 | 2.400E-07 | 1.100E-07 | 1.300E-07 | 3.500E-07 |
| 18347 | 918 | Q918Q | C | 9.5e-08 | 3.3e-07 | 2e-07 | 2.083E-07 | 1.177E-07 | 9.500E-08 | 3.300E-07 |
| 18348 | 918 | Q918G | C | 1.6e-07 | 1.6e-07 | 3.9e-07 | 2.367E-07 | 1.328E-07 | 1.600E-07 | 3.900E-07 |
| 18349 | 918 | Q918H | C | 2.6e-07 | 2.3e-07 | 2.1e-07 | 2.333E-07 | 2.517E-08 | 2.100E-07 | 2.600E-07 |
| 18350 | 918 | Q918I | C | 4.4e-07 | 2.3e-07 | 3.2e-07 | 3.300E-07 | 1.054E-07 | 2.300E-07 | 4.400E-07 |
| 18351 | 918 | Q918L | C | 2.2e-07 | 1.4e-07 | 4e-07 | 2.533E-07 | 1.332E-07 | 1.400E-07 | 4.000E-07 |
| 18352 | 918 | Q918K | C | 3.3e-07 | 1.9e-07 | 1.4e-07 | 2.200E-07 | 9.849E-08 | 1.400E-07 | 3.300E-07 |
| 18353 | 918 | Q918M | C | 3e-07 | 1.6e-07 | 9.8e-08 | 1.860E-07 | 1.035E-07 | 9.800E-08 | 3.000E-07 |
| 18354 | 918 | Q918F | C | 2.8e-07 | 2e-07 | 8.9e-07 | 4.567E-07 | 3.774E-07 | 2.000E-07 | 8.900E-07 |
| 18355 | 918 | Q918P | C | 2.4e-07 | 1.7e-07 | 2.9e-07 | 2.333E-07 | 6.028E-08 | 1.700E-07 | 2.900E-07 |
| 18356 | 918 | Q918S | C | 2.5e-07 | 1.9e-07 | 3.6e-07 | 2.667E-07 | 8.622E-08 | 1.900E-07 | 3.600E-07 |
| 18357 | 918 | Q918T | C | 3.6e-07 | 1.7e-07 | 3.2e-07 | 2.833E-07 | 1.002E-07 | 1.700E-07 | 3.600E-07 |
| 18358 | 918 | Q918W | C | 2.5e-07 | 3.2e-07 | 2.2e-07 | 2.633E-07 | 5.132E-08 | 2.200E-07 | 3.200E-07 |
| 18359 | 918 | Q918Y | C | 2.5e-07 | 5.2e-07 | 2.3e-07 | 3.333E-07 | 1.620E-07 | 2.300E-07 | 5.200E-07 |

| | | | | | | | | | | |
|---|---|---|---|---|---|---|---|---|---|---|
| 18360 | 918 | Q918V | C | 4.6e-07 | 2.6e-07 | 3.1e-07 | 3.433E-07 | 1.041E-07 | 2.600E-07 | 4.600E-07 |
| 18361 | 919 | M919A | C | 2.3e-07 | 2.2e-07 | 4.8e-07 | 3.100E-07 | 1.473E-07 | 2.200E-07 | 4.800E-07 |
| 18362 | 919 | M919R | C | 1.8e-07 | 1.7e-07 | 2.6e-07 | 2.033E-07 | 4.933E-08 | 1.700E-07 | 2.600E-07 |
| 18363 | 919 | M919N | C | 2.2e-07 | 4.7e-07 | 2.1e-07 | 3.000E-07 | 1.473E-07 | 2.100E-07 | 4.700E-07 |
| 18364 | 919 | M919D | C | 3.6e-07 | 1.7e-07 | 2e-07 | 2.433E-07 | 1.021E-07 | 1.700E-07 | 3.600E-07 |
| 18365 | 919 | M919C | C | 3.4e-07 | 2.3e-07 | 2.5e-07 | 2.733E-07 | 5.859E-08 | 2.300E-07 | 3.400E-07 |
| 18366 | 919 | M919E | C | 2.4e-07 | 2.7e-07 | 2.8e-07 | 2.633E-07 | 2.082E-08 | 2.400E-07 | 2.800E-07 |
| 18367 | 919 | M919Q | C | 2.1e-07 | 2.4e-07 | 1.3e-07 | 1.933E-07 | 5.686E-08 | 1.300E-07 | 2.400E-07 |
| 18368 | 919 | M919G | C | 1.8e-07 | 5.6e-07 | 5.1e-07 | 4.167E-07 | 2.065E-07 | 1.800E-07 | 5.600E-07 |
| 18369 | 919 | M919H | C | 3.9e-07 | 1.5e-07 | 2.6e-07 | 2.667E-07 | 1.201E-07 | 1.500E-07 | 3.900E-07 |
| 18370 | 919 | M919I | C | 1.2e-07 | 3e-07 | 5e-07 | 3.067E-07 | 1.901E-07 | 1.200E-07 | 5.000E-07 |
| 18371 | 919 | M919L | C | 2.1e-07 | 1.6e-07 | 4.8e-07 | 2.833E-07 | 1.721E-07 | 1.600E-07 | 4.800E-07 |
| 18372 | 919 | M919K | C | 4.4e-07 | 1.3e-07 | 2.1e-07 | 2.600E-07 | 1.609E-07 | 1.300E-07 | 4.400E-07 |
| 18373 | 919 | M919M | C | 4.6e-07 | 2.1e-07 | 3.4e-07 | 3.367E-07 | 1.250E-07 | 2.100E-07 | 4.600E-07 |
| 18374 | 919 | M919F | C | 1e-07 | 2.5e-07 | 1.7e-07 | 1.733E-07 | 7.506E-08 | 1.000E-07 | 2.500E-07 |
| 18375 | 919 | M919P | C | 4.6e-07 | 2.7e-07 | 1.3e-07 | 2.867E-07 | 1.656E-07 | 1.300E-07 | 4.600E-07 |
| 18376 | 919 | M919S | C | 2.4e-07 | 2.7e-07 | 1.6e-07 | 2.233E-07 | 5.686E-08 | 1.600E-07 | 2.700E-07 |
| 18377 | 919 | M919T | C | 2.9e-07 | 2.8e-07 | 1.5e-07 | 2.400E-07 | 7.810E-08 | 1.500E-07 | 2.900E-07 |
| 18378 | 919 | M919W | C | 3.1e-07 | 2.3e-07 | 2.2e-07 | 2.533E-07 | 4.933E-08 | 2.200E-07 | 3.100E-07 |
| 18379 | 919 | M919Y | C | 2.6e-07 | 2.4e-07 | 2.5e-07 | 2.500E-07 | 1.000E-08 | 2.400E-07 | 2.600E-07 |
| 18380 | 919 | M919V | C | 1e-07 | 1.9e-07 | 2.2e-07 | 1.700E-07 | 6.245E-08 | 1.000E-07 | 2.200E-07 |
| 18381 | 920 | N920A | C | 1.4e-07 | 1.6e-07 | 1.6e-07 | 1.533E-07 | 1.155E-08 | 1.400E-07 | 1.600E-07 |
| 18382 | 920 | N920R | C | 2.5e-07 | 2e-07 | 2.7e-07 | 2.400E-07 | 3.606E-08 | 2.000E-07 | 2.700E-07 |
| 18383 | 920 | N920N | C | 1.6e-07 | 2.6e-07 | 2.1e-07 | 2.100E-07 | 5.000E-08 | 1.600E-07 | 2.600E-07 |
| 18384 | 920 | N920D | C | 2.9e-07 | 1.9e-07 | 4.7e-07 | 3.167E-07 | 1.419E-07 | 1.900E-07 | 4.700E-07 |
| 18385 | 920 | N920C | C | 7.8e-07 | 4.6e-07 | 2.8e-07 | 5.067E-07 | 2.532E-07 | 2.800E-07 | 7.800E-07 |
| 18386 | 920 | N920E | C | 1.3e-07 | 2.7e-07 | 2.2e-07 | 2.067E-07 | 7.095E-08 | 1.300E-07 | 2.700E-07 |
| 18387 | 920 | N920Q | C | 5.8e-07 | 2.7e-07 | 2e-07 | 3.500E-07 | 2.022E-07 | 2.000E-07 | 5.800E-07 |
| 18388 | 920 | N920G | C | 2.4e-07 | 3.9e-07 | 2.1e-07 | 2.800E-07 | 9.644E-08 | 2.100E-07 | 3.900E-07 |
| 18389 | 920 | N920H | C | 1.5e-07 | 1.6e-07 | 2e-07 | 1.700E-07 | 2.646E-08 | 1.500E-07 | 2.000E-07 |
| 18390 | 920 | N920I | C | 2.5e-07 | 4.1e-07 | 3.7e-07 | 3.433E-07 | 8.327E-08 | 2.500E-07 | 4.100E-07 |
| 18391 | 920 | N920L | C | 2e-07 | 1.2e-07 | 2.3e-07 | 1.833E-07 | 5.686E-08 | 1.200E-07 | 2.300E-07 |
| 18392 | 920 | N920K | C | 6.8e-07 | 3.9e-07 | 2.7e-07 | 4.467E-07 | 2.108E-07 | 2.700E-07 | 6.800E-07 |
| 18393 | 920 | N920M | C | 1.9e-07 | 3.7e-07 | 3e-07 | 2.867E-07 | 9.074E-08 | 1.900E-07 | 3.700E-07 |
| 18394 | 920 | N920F | C | 2.8e-07 | 3.8e-07 | 2.3e-07 | 2.967E-07 | 7.638E-08 | 2.300E-07 | 3.800E-07 |
| 18395 | 920 | N920P | C | 2.6e-07 | 6.2e-07 | 2.4e-07 | 3.733E-07 | 2.139E-07 | 2.400E-07 | 6.200E-07 |
| 18396 | 920 | N920S | C | 1.8e-07 | 2.4e-07 | 3e-07 | 2.400E-07 | 6.000E-08 | 1.800E-07 | 3.000E-07 |
| 18397 | 920 | N920T | C | 3.9e-07 | 3e-07 | 3.5e-07 | 3.467E-07 | 4.509E-08 | 3.000E-07 | 3.900E-07 |
| 18398 | 920 | N920W | C | 3.8e-07 | 1.1e-07 | 2.9e-07 | 2.600E-07 | 1.375E-07 | 1.100E-07 | 3.800E-07 |
| 18399 | 920 | N920Y | C | 3.7e-07 | 4.2e-07 | 2.6e-07 | 3.500E-07 | 8.185E-08 | 2.600E-07 | 4.200E-07 |
| 18400 | 920 | N920V | C | 1.3e-07 | 2.3e-07 | 4.5e-07 | 2.700E-07 | 1.637E-07 | 1.300E-07 | 4.500E-07 |
| 18401 | 921 | S921A | C | 2.9e-07 | 3.4e-07 | 2.6e-07 | 2.967E-07 | 4.041E-08 | 2.600E-07 | 3.400E-07 |
| 18402 | 921 | S921R | C | 3.9e-07 | 3.6e-07 | 1.7e-07 | 3.067E-07 | 1.193E-07 | 1.700E-07 | 3.900E-07 |
| 18403 | 921 | S921N | C | 2.4e-07 | 3.7e-07 | 1.3e-07 | 2.467E-07 | 1.201E-07 | 1.300E-07 | 3.700E-07 |
| 18404 | 921 | S921D | C | 1e-07 | 2.9e-07 | 3.6e-07 | 2.500E-07 | 1.345E-07 | 1.000E-07 | 3.600E-07 |
| 18405 | 921 | S921C | C | 1e-07 | 2.7e-07 | 8.4e-08 | 1.513E-07 | 1.031E-07 | 8.400E-08 | 2.700E-07 |
| 18406 | 921 | S921E | C | 3.9e-07 | 1.8e-07 | 3e-07 | 2.900E-07 | 1.054E-07 | 1.800E-07 | 3.900E-07 |
| 18407 | 921 | S921Q | C | 3.8e-07 | 2.3e-07 | 2.3e-07 | 2.800E-07 | 8.660E-08 | 2.300E-07 | 3.800E-07 |
| 18408 | 921 | S921G | C | 1.3e-07 | 3.4e-07 | 2.1e-07 | 2.267E-07 | 1.060E-07 | 1.300E-07 | 3.400E-07 |
| 18409 | 921 | S921H | C | 1.8e-07 | 1.7e-07 | 3e-07 | 2.167E-07 | 7.234E-08 | 1.700E-07 | 3.000E-07 |
| 18410 | 921 | S921I | C | 2.8e-07 | 2e-07 | 2.2e-07 | 2.333E-07 | 4.163E-08 | 2.000E-07 | 2.800E-07 |
| 18411 | 921 | S921L | C | 4e-07 | 2.1e-07 | 1.7e-07 | 2.600E-07 | 1.229E-07 | 1.700E-07 | 4.000E-07 |
| 18412 | 921 | S921K | C | 2.4e-07 | 9e-07 | 1.9e-07 | 4.433E-07 | 3.963E-07 | 1.900E-07 | 9.000E-07 |
| 18413 | 921 | S921M | C | 2.2e-07 | 5e-07 | 8.6e-08 | 2.687E-07 | 2.112E-07 | 8.600E-08 | 5.000E-07 |
| 18414 | 921 | S921F | C | 3e-07 | 2e-07 | 3.6e-07 | 2.867E-07 | 8.083E-08 | 2.000E-07 | 3.600E-07 |
| 18415 | 921 | S921P | C | 3.1e-07 | 2.6e-07 | 2.4e-07 | 2.700E-07 | 3.606E-08 | 2.400E-07 | 3.100E-07 |
| 18416 | 921 | S921S | C | 2.7e-07 | 1.7e-07 | 2.4e-07 | 2.267E-07 | 5.132E-08 | 1.700E-07 | 2.700E-07 |
| 18417 | 921 | S921T | C | 2.4e-07 | 2.3e-07 | 4e-07 | 2.900E-07 | 9.539E-08 | 2.300E-07 | 4.000E-07 |
| 18418 | 921 | S921W | C | 3e-07 | 1.5e-07 | 3.4e-07 | 2.633E-07 | 1.002E-07 | 1.500E-07 | 3.400E-07 |
| 18419 | 921 | S921Y | C | 2.4e-07 | 2.2e-07 | 1.8e-07 | 2.133E-07 | 3.055E-08 | 1.800E-07 | 2.400E-07 |
| 18420 | 921 | S921V | C | 3.5e-07 | 4.3e-07 | 3.2e-07 | 3.667E-07 | 5.686E-08 | 3.200E-07 | 4.300E-07 |
| 18421 | 922 | L922A | C | 1.5e-07 | 2.8e-07 | 1.8e-07 | 2.033E-07 | 6.807E-08 | 1.500E-07 | 2.800E-07 |
| 18422 | 922 | L922R | C | 1.1e-07 | 2.5e-07 | 3.1e-07 | 2.233E-07 | 1.026E-07 | 1.100E-07 | 3.100E-07 |
| 18423 | 922 | L922N | C | 3e-07 | 2.2e-07 | 2.7e-07 | 2.633E-07 | 4.041E-08 | 2.200E-07 | 3.000E-07 |
| 18424 | 922 | L922D | C | 6.2e-07 | 2.5e-07 | 1.9e-07 | 3.533E-07 | 2.329E-07 | 1.900E-07 | 6.200E-07 |
| 18425 | 922 | L922C | C | 1.9e-07 | 3e-07 | 2.3e-07 | 2.400E-07 | 5.568E-08 | 1.900E-07 | 3.000E-07 |
| 18426 | 922 | L922E | C | 6.8e-08 | 1.5e-07 | 2.3e-07 | 1.493E-07 | 8.100E-08 | 6.800E-08 | 2.300E-07 |
| 18427 | 922 | L922Q | C | 2.2e-07 | 2e-07 | 1.2e-07 | 1.800E-07 | 5.292E-08 | 1.200E-07 | 2.200E-07 |
| 18428 | 922 | L922G | C | 1.1e-07 | 3.4e-07 | 2.8e-07 | 2.433E-07 | 1.193E-07 | 1.100E-07 | 3.400E-07 |
| 18429 | 922 | L922H | C | 2.6e-07 | 2.6e-07 | 4.2e-07 | 3.133E-07 | 9.238E-08 | 2.600E-07 | 4.200E-07 |
| 18430 | 922 | L922I | C | 2.8e-07 | 3e-07 | 2.4e-07 | 2.733E-07 | 3.055E-08 | 2.400E-07 | 3.000E-07 |
| 18431 | 922 | L922L | C | 1.9e-07 | 1.3e-07 | 1.2e-06 | 5.067E-07 | 6.012E-07 | 1.300E-07 | 1.200E-06 |
| 18432 | 922 | L922K | C | 1.9e-07 | 3.1e-07 | 2.6e-07 | 2.533E-07 | 6.028E-08 | 1.900E-07 | 3.100E-07 |
| 18433 | 922 | L922M | C | 1.2e-07 | 5.5e-07 | 2.1e-07 | 2.933E-07 | 2.268E-07 | 1.200E-07 | 5.500E-07 |
| 18434 | 922 | L922F | C | 1.7e-07 | 2.5e-07 | 3.7e-07 | 2.633E-07 | 1.007E-07 | 1.700E-07 | 3.700E-07 |
| 18435 | 922 | L922P | C | 5.3e-07 | 3.8e-07 | 2.4e-07 | 3.833E-07 | 1.450E-07 | 2.400E-07 | 5.300E-07 |
| 18436 | 922 | L922S | C | 7.8e-08 | 3.5e-07 | 3.4e-07 | 2.560E-07 | 1.542E-07 | 7.800E-08 | 3.500E-07 |
| 18437 | 922 | L922T | C | 2.1e-07 | 2e-07 | 4.6e-07 | 2.900E-07 | 1.473E-07 | 2.000E-07 | 4.600E-07 |
| 18438 | 922 | L922W | C | 3.1e-07 | 1.8e-07 | 1.7e-07 | 2.200E-07 | 7.810E-08 | 1.700E-07 | 3.100E-07 |
| 18439 | 922 | L922Y | C | 2e-07 | 2.2e-07 | 1.7e-07 | 1.967E-07 | 2.517E-08 | 1.700E-07 | 2.200E-07 |
| 18440 | 922 | L922V | C | 2.5e-07 | 1.3e-07 | 1.6e-07 | 1.800E-07 | 6.245E-08 | 1.300E-07 | 2.500E-07 |
| 18441 | 923 | R923A | C | 2.1e-07 | 1.8e-07 | 2.1e-07 | 2.000E-07 | 1.732E-08 | 1.800E-07 | 2.100E-07 |
| 18442 | 923 | R923R | C | 2.1e-07 | 4.1e-07 | 1.6e-07 | 2.600E-07 | 1.323E-07 | 1.600E-07 | 4.100E-07 |
| 18443 | 923 | R923N | C | 2.9e-07 | 3.1e-07 | 7.9e-08 | 2.263E-07 | 1.280E-07 | 7.900E-08 | 3.100E-07 |
| 18444 | 923 | R923D | C | 4.6e-07 | 2.3e-07 | 1.9e-07 | 2.933E-07 | 1.457E-07 | 1.900E-07 | 4.600E-07 |

| | | | | | | | | | | |
|---|---|---|---|---|---|---|---|---|---|---|
| 18445 | 923 | R923C | C | 3e-07 | 2.5e-07 | 9.9e-08 | 2.163E-07 | 1.046E-07 | 9.900E-08 | 3.000E-07 |
| 18446 | 923 | R923E | C | 2.1e-07 | 3.8e-07 | 2e-07 | 2.633E-07 | 1.012E-07 | 2.000E-07 | 3.800E-07 |
| 18447 | 923 | R923Q | C | 4.2e-07 | 3.2e-07 | 3.5e-07 | 3.633E-07 | 5.132E-08 | 3.200E-07 | 4.200E-07 |
| 18448 | 923 | R923G | C | 6.5e-07 | 2.3e-07 | 2.9e-07 | 3.900E-07 | 2.272E-07 | 2.300E-07 | 6.500E-07 |
| 18449 | 923 | R923H | C | 2.4e-07 | 7.1e-07 | 2.8e-07 | 4.100E-07 | 2.606E-07 | 2.400E-07 | 7.100E-07 |
| 18450 | 923 | R923I | C | 2e-07 | 1.9e-07 | 1.4e-07 | 1.767E-07 | 3.215E-08 | 1.400E-07 | 2.000E-07 |
| 18451 | 923 | R923L | C | 2.6e-07 | 3.9e-07 | 1e-07 | 2.500E-07 | 1.453E-07 | 1.000E-07 | 3.900E-07 |
| 18452 | 923 | R923K | C | 3.7e-07 | 2e-07 | 2e-07 | 2.567E-07 | 9.815E-08 | 2.000E-07 | 3.700E-07 |
| 18453 | 923 | R923M | C | 2.6e-07 | 1.2e-07 | 2.4e-07 | 2.067E-07 | 7.572E-08 | 1.200E-07 | 2.600E-07 |
| 18454 | 923 | R923F | C | 1.9e-07 | 3.6e-07 | 4.5e-07 | 3.333E-07 | 1.320E-07 | 1.900E-07 | 4.500E-07 |
| 18455 | 923 | R923P | C | 1.6e-07 | 2.7e-07 | 1.6e-07 | 1.967E-07 | 6.351E-08 | 1.600E-07 | 2.700E-07 |
| 18456 | 923 | R923S | C | 1.7e-07 | 1.6e-07 | 1.7e-07 | 1.667E-07 | 5.774E-09 | 1.600E-07 | 1.700E-07 |
| 18457 | 923 | R923T | C | 1.1e-07 | 4.2e-07 | 3.1e-07 | 2.800E-07 | 1.572E-07 | 1.100E-07 | 4.200E-07 |
| 18458 | 923 | R923W | C | 1.8e-07 | 1.6e-07 | 1.8e-07 | 1.733E-07 | 1.155E-08 | 1.600E-07 | 1.800E-07 |
| 18459 | 923 | R923Y | C | 2e-07 | 3.2e-07 | 3.8e-07 | 3.000E-07 | 9.165E-08 | 2.000E-07 | 3.800E-07 |
| 18460 | 923 | R923V | C | 3.7e-07 | 3.7e-07 | 2e-07 | 3.133E-07 | 9.815E-08 | 2.000E-07 | 3.700E-07 |
| 18461 | 924 | A924A | C | 4.8e-07 | 1.3e-07 | 3.7e-07 | 3.267E-07 | 1.790E-07 | 1.300E-07 | 4.800E-07 |
| 18462 | 924 | A924R | C | 1.3e-07 | 2.7e-07 | 2.2e-07 | 2.067E-07 | 7.095E-08 | 1.300E-07 | 2.700E-07 |
| 18463 | 924 | A924N | C | 8.3e-08 | 8e-08 | 4.2e-07 | 1.943E-07 | 1.954E-07 | 8.000E-08 | 4.200E-07 |
| 18464 | 924 | A924D | C | 5.1e-07 | 3.5e-07 | 2.7e-07 | 3.767E-07 | 1.222E-07 | 2.700E-07 | 5.100E-07 |
| 18465 | 924 | A924C | C | 1.9e-07 | 3.2e-07 | 2.9e-07 | 2.667E-07 | 6.807E-08 | 1.900E-07 | 3.200E-07 |
| 18466 | 924 | A924E | C | 4.7e-07 | 1.5e-07 | 1.9e-07 | 2.700E-07 | 1.744E-07 | 1.500E-07 | 4.700E-07 |
| 18467 | 924 | A924Q | C | 2.7e-07 | 1.1e-07 | 2.7e-07 | 2.167E-07 | 9.238E-08 | 1.100E-07 | 2.700E-07 |
| 18468 | 924 | A924G | C | 2.4e-07 | 3.5e-07 | 2.9e-07 | 2.933E-07 | 5.508E-08 | 2.400E-07 | 3.500E-07 |
| 18469 | 924 | A924H | C | 2.7e-07 | 3.6e-07 | 3.3e-07 | 3.200E-07 | 4.583E-08 | 2.700E-07 | 3.600E-07 |
| 18470 | 924 | A924I | C | 2.5e-07 | 2.7e-07 | 3.7e-07 | 2.967E-07 | 6.429E-08 | 2.500E-07 | 3.700E-07 |
| 18471 | 924 | A924L | C | 2.4e-07 | 2.3e-07 | 2.3e-07 | 2.333E-07 | 5.774E-09 | 2.300E-07 | 2.400E-07 |
| 18472 | 924 | A924K | C | 3e-07 | 1.5e-07 | 2.9e-07 | 2.467E-07 | 8.386E-08 | 1.500E-07 | 3.000E-07 |
| 18473 | 924 | A924M | C | 6.2e-08 | 7.1e-07 | 2.9e-07 | 3.540E-07 | 3.287E-07 | 6.200E-08 | 7.100E-07 |
| 18474 | 924 | A924F | C | 4.4e-07 | 3.4e-07 | 3e-07 | 3.600E-07 | 7.211E-08 | 3.000E-07 | 4.400E-07 |
| 18475 | 924 | A924P | C | 5.2e-07 | 5.6e-07 | 2.7e-07 | 4.500E-07 | 1.572E-07 | 2.700E-07 | 5.600E-07 |
| 18476 | 924 | A924S | C | 1.5e-07 | 1.7e-07 | 9.4e-08 | 1.380E-07 | 3.940E-08 | 9.400E-08 | 1.700E-07 |
| 18477 | 924 | A924T | C | 2.4e-07 | 2.8e-07 | 5e-07 | 3.400E-07 | 1.400E-07 | 2.400E-07 | 5.000E-07 |
| 18478 | 924 | A924W | C | 3.3e-07 | 3.6e-07 | 3.2e-07 | 3.367E-07 | 2.082E-08 | 3.200E-07 | 3.600E-07 |
| 18479 | 924 | A924Y | C | 1.8e-07 | 2e-07 | 3.9e-07 | 2.567E-07 | 1.159E-07 | 1.800E-07 | 3.900E-07 |
| 18480 | 924 | A924V | C | 3.6e-07 | 2.6e-07 | 1.2e-07 | 2.467E-07 | 1.206E-07 | 1.200E-07 | 3.600E-07 |
| 18481 | 925 | E925A | C | 2.9e-07 | 1.8e-07 | 3.2e-07 | 2.633E-07 | 7.371E-08 | 1.800E-07 | 3.200E-07 |
| 18482 | 925 | E925R | C | 1.3e-07 | 1.8e-07 | 1.8e-07 | 1.633E-07 | 2.887E-08 | 1.300E-07 | 1.800E-07 |
| 18483 | 925 | E925N | C | 2.4e-07 | 2.1e-07 | 2.2e-07 | 2.233E-07 | 1.528E-08 | 2.100E-07 | 2.400E-07 |
| 18484 | 925 | E925D | C | 5.7e-07 | 3.1e-07 | 1.9e-07 | 3.567E-07 | 1.943E-07 | 1.900E-07 | 5.700E-07 |
| 18485 | 925 | E925C | C | 3.6e-07 | 2.3e-07 | 3.9e-07 | 3.267E-07 | 8.505E-08 | 2.300E-07 | 3.900E-07 |
| 18486 | 925 | E925E | C | 3.5e-07 | 2.1e-07 | 3e-07 | 2.867E-07 | 7.095E-08 | 2.100E-07 | 3.500E-07 |
| 18487 | 925 | E925Q | C | 4.6e-07 | 3.2e-07 | 1.8e-07 | 3.200E-07 | 1.400E-07 | 1.800E-07 | 4.600E-07 |
| 18488 | 925 | E925G | C | 1.8e-07 | 4.5e-07 | 1.6e-07 | 2.633E-07 | 1.620E-07 | 1.600E-07 | 4.500E-07 |
| 18489 | 925 | E925H | C | 3.9e-07 | 2.4e-07 | 2.1e-07 | 2.800E-07 | 9.644E-08 | 2.100E-07 | 3.900E-07 |
| 18490 | 925 | E925I | C | 1.7e-07 | 6.1e-08 | 3e-07 | 1.770E-07 | 1.197E-07 | 6.100E-08 | 3.000E-07 |
| 18491 | 925 | E925L | C | 2.1e-07 | 7.2e-07 | 1.6e-07 | 3.633E-07 | 3.099E-07 | 1.600E-07 | 7.200E-07 |
| 18492 | 925 | E925K | C | 2.4e-07 | 1e-07 | 2.6e-07 | 2.000E-07 | 8.718E-08 | 1.000E-07 | 2.600E-07 |
| 18493 | 925 | E925M | C | 9.3e-08 | 2e-07 | 1.8e-07 | 1.577E-07 | 5.689E-08 | 9.300E-08 | 2.000E-07 |
| 18494 | 925 | E925F | C | 2.1e-07 | 2.8e-07 | 2.5e-07 | 2.467E-07 | 3.512E-08 | 2.100E-07 | 2.800E-07 |
| 18495 | 925 | E925P | C | 2.7e-07 | 5.1e-07 | 1.7e-07 | 3.167E-07 | 1.747E-07 | 1.700E-07 | 5.100E-07 |
| 18496 | 925 | E925S | C | 1.6e-07 | 2.4e-07 | 1.1e-07 | 1.700E-07 | 6.557E-08 | 1.100E-07 | 2.400E-07 |
| 18497 | 925 | E925T | C | 1.6e-07 | 1.9e-07 | 3.1e-07 | 2.200E-07 | 7.937E-08 | 1.600E-07 | 3.100E-07 |
| 18498 | 925 | E925W | C | 1.8e-07 | 1.2e-07 | 1.3e-07 | 1.433E-07 | 3.215E-08 | 1.200E-07 | 1.800E-07 |
| 18499 | 925 | E925Y | C | 3.9e-07 | 4.5e-07 | 5.4e-07 | 4.600E-07 | 7.550E-08 | 3.900E-07 | 5.400E-07 |
| 18500 | 925 | E925V | C | 3.5e-07 | 2.5e-07 | 2.2e-07 | 2.733E-07 | 6.807E-08 | 2.200E-07 | 3.500E-07 |
| 18501 | 926 | D926A | C | 2.2e-07 | 1.4e-07 | 2.4e-07 | 2.000E-07 | 5.292E-08 | 1.400E-07 | 2.400E-07 |
| 18502 | 926 | D926R | C | 1.8e-07 | 1.8e-07 | 2.5e-07 | 2.033E-07 | 4.041E-08 | 1.800E-07 | 2.500E-07 |
| 18503 | 926 | D926N | C | 9.8e-08 | 3e-07 | 3.1e-07 | 2.360E-07 | 1.196E-07 | 9.800E-08 | 3.100E-07 |
| 18504 | 926 | D926D | C | 2.5e-07 | 2.3e-07 | 2.9e-07 | 2.567E-07 | 3.055E-08 | 2.300E-07 | 2.900E-07 |
| 18505 | 926 | D926C | C | 1.1e-07 | 3.9e-07 | 4.2e-07 | 3.067E-07 | 1.710E-07 | 1.100E-07 | 4.200E-07 |
| 18506 | 926 | D926E | C | 2.2e-07 | 4.6e-07 | 2.3e-07 | 3.033E-07 | 1.358E-07 | 2.200E-07 | 4.600E-07 |
| 18507 | 926 | D926Q | C | 3.4e-07 | 1.8e-07 | 2.9e-07 | 2.700E-07 | 8.185E-08 | 1.800E-07 | 3.400E-07 |
| 18508 | 926 | D926G | C | 2.2e-07 | 3.8e-07 | 3.2e-07 | 3.067E-07 | 8.083E-08 | 2.200E-07 | 3.800E-07 |
| 18509 | 926 | D926H | C | 1.5e-07 | 2.2e-07 | 3.5e-07 | 2.400E-07 | 1.015E-07 | 1.500E-07 | 3.500E-07 |
| 18510 | 926 | D926I | C | 1e-07 | 2.1e-07 | 5.3e-07 | 2.800E-07 | 2.234E-07 | 1.000E-07 | 5.300E-07 |
| 18511 | 926 | D926L | C | 2.7e-07 | 2.2e-07 | 1.8e-07 | 2.233E-07 | 4.509E-08 | 1.800E-07 | 2.700E-07 |
| 18512 | 926 | D926K | C | 1.7e-07 | 3e-07 | 1.1e-07 | 1.933E-07 | 9.713E-08 | 1.100E-07 | 3.000E-07 |
| 18513 | 926 | D926M | C | 1.8e-07 | 1.2e-07 | 1.4e-07 | 1.467E-07 | 3.055E-08 | 1.200E-07 | 1.800E-07 |
| 18514 | 926 | D926F | C | 2.6e-07 | 2.4e-07 | 4.2e-07 | 3.067E-07 | 9.866E-08 | 2.400E-07 | 4.200E-07 |
| 18515 | 926 | D926P | C | 2.8e-07 | 2.4e-07 | 2.3e-07 | 2.500E-07 | 2.646E-08 | 2.300E-07 | 2.800E-07 |
| 18516 | 926 | D926S | C | 2.8e-07 | 5.6e-07 | 1.3e-07 | 3.233E-07 | 2.183E-07 | 1.300E-07 | 5.600E-07 |
| 18517 | 926 | D926T | C | 4.1e-07 | 3.3e-07 | 2.9e-07 | 3.433E-07 | 6.110E-08 | 2.900E-07 | 4.100E-07 |
| 18518 | 926 | D926W | C | 1.5e-07 | 1.1e-07 | 1.2e-07 | 1.267E-07 | 2.082E-08 | 1.100E-07 | 1.500E-07 |
| 18519 | 926 | D926Y | C | 2.3e-07 | 8.6e-07 | 1.3e-07 | 4.067E-07 | 3.958E-07 | 1.300E-07 | 8.600E-07 |
| 18520 | 926 | D926V | C | 5.1e-07 | 2.1e-07 | 1.7e-07 | 2.967E-07 | 1.858E-07 | 1.700E-07 | 5.100E-07 |
| 18521 | 927 | T927A | C | 3.2e-07 | 4.7e-07 | 3.3e-07 | 3.733E-07 | 8.386E-08 | 3.200E-07 | 4.700E-07 |
| 18522 | 927 | T927R | C | 2.4e-07 | 2.7e-07 | 1.8e-07 | 2.300E-07 | 4.583E-08 | 1.800E-07 | 2.700E-07 |
| 18523 | 927 | T927N | C | 1.6e-07 | 3e-07 | 3.9e-07 | 2.833E-07 | 1.159E-07 | 1.600E-07 | 3.900E-07 |
| 18524 | 927 | T927D | C | 2.5e-07 | 5.3e-07 | 6.5e-07 | 4.767E-07 | 2.053E-07 | 2.500E-07 | 6.500E-07 |
| 18525 | 927 | T927C | C | 3.6e-07 | 3.8e-07 | 2e-07 | 3.133E-07 | 9.866E-08 | 2.000E-07 | 3.800E-07 |
| 18526 | 927 | T927E | C | 8.3e-08 | 2.8e-07 | 1.6e-07 | 1.743E-07 | 9.928E-08 | 8.300E-08 | 2.800E-07 |
| 18527 | 927 | T927Q | C | 4.7e-07 | 2e-07 | 1.3e-07 | 2.667E-07 | 1.795E-07 | 1.300E-07 | 4.700E-07 |
| 18528 | 927 | T927G | C | 2e-07 | 2.1e-07 | 3.6e-07 | 2.567E-07 | 8.963E-08 | 2.000E-07 | 3.600E-07 |
| 18529 | 927 | T927H | C | 2e-07 | 3.2e-07 | 1.4e-07 | 2.200E-07 | 9.165E-08 | 1.400E-07 | 3.200E-07 |

| | | | | | | | | | | |
|---|---|---|---|---|---|---|---|---|---|---|
| 18530 | 927 | T927I | C | 1.8e-07 | 4.2e-07 | 4.7e-07 | 3.567E-07 | 1.550E-07 | 1.800E-07 | 4.700E-07 |
| 18531 | 927 | T927L | C | 3.2e-07 | 2.2e-07 | 1.3e-07 | 2.233E-07 | 9.504E-08 | 1.300E-07 | 3.200E-07 |
| 18532 | 927 | T927K | C | 1.4e-07 | 6.3e-08 | 2.3e-07 | 1.443E-07 | 8.358E-08 | 6.300E-08 | 2.300E-07 |
| 18533 | 927 | T927M | C | 2.3e-07 | 4.3e-07 | 5.8e-07 | 4.133E-07 | 1.756E-07 | 2.300E-07 | 5.800E-07 |
| 18534 | 927 | T927F | C | 4.8e-07 | 2e-07 | 1.4e-07 | 2.733E-07 | 1.815E-07 | 1.400E-07 | 4.800E-07 |
| 18535 | 927 | T927P | C | 2.7e-07 | 1.7e-07 | 2.8e-07 | 2.400E-07 | 6.083E-08 | 1.700E-07 | 2.800E-07 |
| 18536 | 927 | T927S | C | 2.7e-07 | 2.2e-07 | 3.2e-07 | 2.700E-07 | 5.000E-08 | 2.200E-07 | 3.200E-07 |
| 18537 | 927 | T927T | C | 5.1e-07 | 1.2e-07 | 3e-07 | 3.100E-07 | 1.952E-07 | 1.200E-07 | 5.100E-07 |
| 18538 | 927 | T927W | C | 2.7e-07 | 2.1e-07 | 2.2e-07 | 2.333E-07 | 3.215E-08 | 2.100E-07 | 2.700E-07 |
| 18539 | 927 | T927Y | C | 4.9e-07 | 2.3e-07 | 2.5e-07 | 3.233E-07 | 1.447E-07 | 2.300E-07 | 4.900E-07 |
| 18540 | 927 | T927V | C | 1.5e-07 | 2.2e-07 | 1.5e-07 | 1.733E-07 | 4.041E-08 | 1.500E-07 | 2.200E-07 |
| 18541 | 928 | A928A | C | 1.1e-07 | 2.1e-07 | 1.5e-07 | 1.567E-07 | 5.033E-08 | 1.100E-07 | 2.100E-07 |
| 18542 | 928 | A928R | C | 2.2e-07 | 4.8e-07 | 1.6e-07 | 2.867E-07 | 1.701E-07 | 1.600E-07 | 4.800E-07 |
| 18543 | 928 | A928N | C | 3e-07 | 2.4e-07 | 2.9e-07 | 2.767E-07 | 3.215E-08 | 2.400E-07 | 3.000E-07 |
| 18544 | 928 | A928D | C | 2.5e-07 | 1.3e-07 | 2.9e-07 | 2.233E-07 | 8.327E-08 | 1.300E-07 | 2.900E-07 |
| 18545 | 928 | A928C | C | 2.4e-07 | 2.2e-07 | 2.8e-07 | 2.467E-07 | 3.055E-08 | 2.200E-07 | 2.800E-07 |
| 18546 | 928 | A928E | C | 5e-07 | 2.8e-07 | 1.2e-07 | 3.000E-07 | 1.908E-07 | 1.200E-07 | 5.000E-07 |
| 18547 | 928 | A928Q | C | 9.8e-08 | 6.3e-08 | 9.7e-08 | 8.600E-08 | 1.992E-08 | 6.300E-08 | 9.800E-08 |
| 18548 | 928 | A928G | C | 2.1e-07 | 2.3e-07 | 3.3e-07 | 2.567E-07 | 6.429E-08 | 2.100E-07 | 3.300E-07 |
| 18549 | 928 | A928H | C | 3.6e-07 | 2e-07 | 6e-07 | 3.867E-07 | 2.013E-07 | 2.000E-07 | 6.000E-07 |
| 18550 | 928 | A928I | C | 1.1e-07 | 1.7e-07 | 3.4e-07 | 2.067E-07 | 1.193E-07 | 1.100E-07 | 3.400E-07 |
| 18551 | 928 | A928L | C | 2.8e-07 | 1.5e-07 | 3e-07 | 2.433E-07 | 8.145E-08 | 1.500E-07 | 3.000E-07 |
| 18552 | 928 | A928K | C | 3.1e-07 | 1.4e-07 | 1.7e-07 | 2.067E-07 | 9.074E-08 | 1.400E-07 | 3.100E-07 |
| 18553 | 928 | A928M | C | 3.5e-07 | 2.8e-07 | 1.5e-07 | 2.600E-07 | 1.015E-07 | 1.500E-07 | 3.500E-07 |
| 18554 | 928 | A928F | C | 1.5e-07 | 5.4e-07 | 2.2e-07 | 3.033E-07 | 2.079E-07 | 1.500E-07 | 5.400E-07 |
| 18555 | 928 | A928P | C | 1.3e-07 | 2.9e-07 | 2.7e-07 | 2.300E-07 | 8.718E-08 | 1.300E-07 | 2.900E-07 |
| 18556 | 928 | A928S | C | 1.9e-07 | 3e-07 | 1.3e-07 | 2.067E-07 | 8.622E-08 | 1.300E-07 | 3.000E-07 |
| 18557 | 928 | A928T | C | 2.9e-07 | 2.1e-07 | 2.1e-07 | 2.367E-07 | 4.619E-08 | 2.100E-07 | 2.900E-07 |
| 18558 | 928 | A928W | C | 3.4e-07 | 5.8e-07 | 2.1e-07 | 3.767E-07 | 1.877E-07 | 2.100E-07 | 5.800E-07 |
| 18559 | 928 | A928Y | C | 4e-07 | 3.1e-07 | 5.7e-07 | 4.267E-07 | 1.320E-07 | 3.100E-07 | 5.700E-07 |
| 18560 | 928 | A928V | C | 1.8e-07 | 1.1e-07 | 2.6e-07 | 1.833E-07 | 7.506E-08 | 1.100E-07 | 2.600E-07 |
| 18561 | 929 | V929A | C | 2.9e-07 | 2.1e-07 | 1.2e-07 | 2.067E-07 | 8.505E-08 | 1.200E-07 | 2.900E-07 |
| 18562 | 929 | V929R | C | 4.2e-07 | 4.7e-07 | 1.3e-07 | 3.400E-07 | 1.836E-07 | 1.300E-07 | 4.700E-07 |
| 18563 | 929 | V929N | C | 2e-07 | 1.4e-07 | 2e-07 | 1.800E-07 | 3.464E-08 | 1.400E-07 | 2.000E-07 |
| 18564 | 929 | V929D | C | 1.4e-07 | 3.8e-07 | 2e-07 | 2.400E-07 | 1.249E-07 | 1.400E-07 | 3.800E-07 |
| 18565 | 929 | V929C | C | 2.3e-07 | 2.5e-07 | 2.9e-07 | 2.567E-07 | 3.055E-08 | 2.300E-07 | 2.900E-07 |
| 18566 | 929 | V929E | C | 1.7e-07 | 2.6e-07 | 4.4e-07 | 2.900E-07 | 1.375E-07 | 1.700E-07 | 4.400E-07 |
| 18567 | 929 | V929Q | C | 1.1e-07 | 1.8e-07 | 2e-07 | 1.633E-07 | 4.726E-08 | 1.100E-07 | 2.000E-07 |
| 18568 | 929 | V929G | C | 1.8e-07 | 2.3e-07 | 2.9e-07 | 2.333E-07 | 5.508E-08 | 1.800E-07 | 2.900E-07 |
| 18569 | 929 | V929H | C | 2.1e-07 | 2e-07 | 2.3e-07 | 2.133E-07 | 1.528E-08 | 2.000E-07 | 2.300E-07 |
| 18570 | 929 | V929I | C | 3.6e-07 | 1.7e-07 | 1.7e-07 | 2.333E-07 | 1.097E-07 | 1.700E-07 | 3.600E-07 |
| 18571 | 929 | V929L | C | 2.7e-07 | 1.9e-07 | 6.5e-07 | 3.700E-07 | 2.458E-07 | 1.900E-07 | 6.500E-07 |
| 18572 | 929 | V929K | C | 1.2e-07 | 3e-07 | 4.6e-07 | 2.933E-07 | 1.701E-07 | 1.200E-07 | 4.600E-07 |
| 18573 | 929 | V929M | C | 4e-07 | 4.2e-07 | 4.9e-07 | 4.367E-07 | 4.726E-08 | 4.000E-07 | 4.900E-07 |
| 18574 | 929 | V929F | C | 2.3e-07 | 4e-07 | 3.3e-07 | 3.200E-07 | 8.544E-08 | 2.300E-07 | 4.000E-07 |
| 18575 | 929 | V929P | C | 2e-07 | 2e-07 | 3.9e-07 | 2.633E-07 | 1.097E-07 | 2.000E-07 | 3.900E-07 |
| 18576 | 929 | V929S | C | 6.7e-08 | 3e-07 | 2.7e-07 | 2.123E-07 | 1.268E-07 | 6.700E-08 | 3.000E-07 |
| 18577 | 929 | V929T | C | 4.6e-07 | 2.4e-07 | 4e-07 | 3.667E-07 | 1.137E-07 | 2.400E-07 | 4.600E-07 |
| 18578 | 929 | V929W | C | 3.7e-07 | 2.3e-07 | 1.2e-07 | 2.400E-07 | 1.253E-07 | 1.200E-07 | 3.700E-07 |
| 18579 | 929 | V929Y | C | 2.7e-07 | 2.5e-07 | 2.6e-07 | 2.600E-07 | 1.000E-08 | 2.500E-07 | 2.700E-07 |
| 18580 | 929 | V929V | C | 2.4e-07 | 1.6e-07 | 3.8e-07 | 2.600E-07 | 1.114E-07 | 1.600E-07 | 3.800E-07 |
| 18581 | 930 | Y930A | C | 1.6e-07 | 1.6e-07 | 2.2e-07 | 1.800E-07 | 3.464E-08 | 1.600E-07 | 2.200E-07 |
| 18582 | 930 | Y930R | C | 4.4e-07 | 2e-07 | 3.1e-07 | 3.167E-07 | 1.201E-07 | 2.000E-07 | 4.400E-07 |
| 18583 | 930 | Y930N | C | 2.9e-07 | 6.5e-07 | 1.7e-07 | 3.700E-07 | 2.498E-07 | 1.700E-07 | 6.500E-07 |
| 18584 | 930 | Y930D | C | 3e-07 | 1.6e-07 | 4.4e-07 | 3.000E-07 | 1.400E-07 | 1.600E-07 | 4.400E-07 |
| 18585 | 930 | Y930C | C | 2.4e-07 | 1.4e-07 | 6.6e-07 | 3.467E-07 | 2.759E-07 | 1.400E-07 | 6.600E-07 |
| 18586 | 930 | Y930E | C | 2.2e-07 | 2e-07 | 6.1e-07 | 3.433E-07 | 2.312E-07 | 2.000E-07 | 6.100E-07 |
| 18587 | 930 | Y930Q | C | 3.2e-07 | 3.6e-07 | 2e-07 | 2.933E-07 | 8.327E-08 | 2.000E-07 | 3.600E-07 |
| 18588 | 930 | Y930G | C | 3e-07 | 2.2e-07 | 6e-07 | 3.733E-07 | 2.003E-07 | 2.200E-07 | 6.000E-07 |
| 18589 | 930 | Y930H | C | 3.3e-07 | 1.9e-07 | 3.2e-07 | 2.800E-07 | 7.810E-08 | 1.900E-07 | 3.300E-07 |
| 18590 | 930 | Y930I | C | 2.2e-07 | 1.5e-07 | 5.3e-07 | 3.000E-07 | 2.022E-07 | 1.500E-07 | 5.300E-07 |
| 18591 | 930 | Y930L | C | 1.4e-07 | 2.5e-07 | 1.2e-07 | 1.700E-07 | 7.000E-08 | 1.200E-07 | 2.500E-07 |
| 18592 | 930 | Y930K | C | 3.1e-07 | 2e-07 | 2.9e-07 | 2.667E-07 | 5.859E-08 | 2.000E-07 | 3.100E-07 |
| 18593 | 930 | Y930M | C | 3.7e-07 | 1.6e-07 | 4e-07 | 3.100E-07 | 1.308E-07 | 1.600E-07 | 4.000E-07 |
| 18594 | 930 | Y930F | C | 5.5e-07 | 1.5e-07 | 1.8e-07 | 2.933E-07 | 2.228E-07 | 1.500E-07 | 5.500E-07 |
| 18595 | 930 | Y930P | C | 3.6e-07 | 2e-07 | 4.4e-07 | 3.333E-07 | 1.222E-07 | 2.000E-07 | 4.400E-07 |
| 18596 | 930 | Y930S | C | 2e-07 | 1.2e-07 | 3e-07 | 2.067E-07 | 9.018E-08 | 1.200E-07 | 3.000E-07 |
| 18597 | 930 | Y930T | C | 1.9e-07 | 1.9e-07 | 1.4e-07 | 1.733E-07 | 2.887E-08 | 1.400E-07 | 1.900E-07 |
| 18598 | 930 | Y930W | C | 1.4e-07 | 4.7e-07 | 3.1e-07 | 3.067E-07 | 1.650E-07 | 1.400E-07 | 4.700E-07 |
| 18599 | 930 | Y930Y | C | 2.9e-07 | 1.9e-07 | 3.9e-07 | 2.900E-07 | 1.000E-07 | 1.900E-07 | 3.900E-07 |
| 18600 | 930 | Y930V | C | 1.7e-07 | 4.1e-07 | 3.5e-07 | 3.100E-07 | 1.249E-07 | 1.700E-07 | 4.100E-07 |
| 18601 | 931 | Y931A | C | 3.8e-07 | 2.8e-07 | 3.1e-07 | 3.233E-07 | 5.132E-08 | 2.800E-07 | 3.800E-07 |
| 18602 | 931 | Y931R | C | 3.2e-07 | 2.5e-07 | 3.6e-07 | 3.100E-07 | 5.568E-08 | 2.500E-07 | 3.600E-07 |
| 18603 | 931 | Y931N | C | 3.4e-07 | 3.5e-07 | 2.5e-07 | 3.133E-07 | 5.508E-08 | 2.500E-07 | 3.500E-07 |
| 18604 | 931 | Y931D | C | 3.4e-07 | 3.3e-07 | 3.1e-07 | 3.267E-07 | 1.528E-08 | 3.100E-07 | 3.400E-07 |
| 18605 | 931 | Y931C | C | 2.7e-07 | 8.7e-08 | 3.3e-07 | 2.290E-07 | 1.266E-07 | 8.700E-08 | 3.300E-07 |
| 18606 | 931 | Y931E | C | 5.2e-07 | 2.8e-07 | 3e-07 | 3.667E-07 | 1.332E-07 | 2.800E-07 | 5.200E-07 |
| 18607 | 931 | Y931Q | C | 2.6e-07 | 2.4e-07 | 2e-07 | 2.333E-07 | 3.055E-08 | 2.000E-07 | 2.600E-07 |
| 18608 | 931 | Y931G | C | 3.7e-07 | 1.9e-07 | 2.4e-07 | 2.667E-07 | 9.292E-08 | 1.900E-07 | 3.700E-07 |
| 18609 | 931 | Y931H | C | 4.2e-07 | 2.1e-07 | 2.7e-07 | 3.000E-07 | 1.082E-07 | 2.100E-07 | 4.200E-07 |
| 18610 | 931 | Y931I | C | 1.8e-07 | 2.8e-07 | 1.9e-07 | 2.167E-07 | 5.508E-08 | 1.800E-07 | 2.800E-07 |
| 18611 | 931 | Y931L | C | 1.3e-07 | 2.2e-07 | 2e-07 | 1.833E-07 | 4.726E-08 | 1.300E-07 | 2.200E-07 |
| 18612 | 931 | Y931K | C | 3.5e-07 | 3.4e-07 | 5.4e-07 | 4.100E-07 | 1.127E-07 | 3.400E-07 | 5.400E-07 |
| 18613 | 931 | Y931M | C | 1e-07 | 2.8e-07 | 8.4e-08 | 1.547E-07 | 1.088E-07 | 8.400E-08 | 2.800E-07 |
| 18614 | 931 | Y931F | C | 1.6e-07 | 2.2e-07 | 3.1e-07 | 2.300E-07 | 7.550E-08 | 1.600E-07 | 3.100E-07 |

| | | | | | | | | | | |
|---|---|---|---|---|---|---|---|---|---|---|
| 18615 | 931 | Y931P | C | 3.2e-07 | 2.5e-07 | 4.7e-07 | 3.467E-07 | 1.124E-07 | 2.500E-07 | 4.700E-07 |
| 18616 | 931 | Y931S | C | 2.9e-07 | 4.1e-07 | 2e-07 | 3.000E-07 | 1.054E-07 | 2.000E-07 | 4.100E-07 |
| 18617 | 931 | Y931T | C | 4.8e-07 | 2.4e-07 | 2.7e-07 | 3.300E-07 | 1.308E-07 | 2.400E-07 | 4.800E-07 |
| 18618 | 931 | Y931W | C | 2.2e-07 | 2.2e-07 | 2.2e-07 | 2.200E-07 | 0.000E+00 | 2.200E-07 | 2.200E-07 |
| 18619 | 931 | Y931Y | C | 3.1e-07 | 1.3e-07 | 2.7e-07 | 2.367E-07 | 9.452E-08 | 1.300E-07 | 3.100E-07 |
| 18620 | 931 | Y931V | C | 2e-07 | 2e-07 | 2.3e-07 | 2.100E-07 | 1.732E-08 | 2.000E-07 | 2.300E-07 |
| 18621 | 932 | C932A | C | 4.1e-07 | 2.7e-07 | 4.6e-07 | 3.800E-07 | 9.849E-08 | 2.700E-07 | 4.600E-07 |
| 18622 | 932 | C932R | C | 4.3e-07 | 2.2e-07 | 2.1e-07 | 2.867E-07 | 1.242E-07 | 2.100E-07 | 4.300E-07 |
| 18623 | 932 | C932N | C | 2.2e-07 | 1.7e-07 | 2.9e-07 | 2.267E-07 | 6.028E-08 | 1.700E-07 | 2.900E-07 |
| 18624 | 932 | C932D | C | 2e-07 | 2.6e-07 | 2.3e-07 | 2.300E-07 | 3.000E-08 | 2.000E-07 | 2.600E-07 |
| 18625 | 932 | C932C | C | 1.9e-07 | 1.8e-07 | 2.9e-07 | 2.200E-07 | 6.083E-08 | 1.800E-07 | 2.900E-07 |
| 18626 | 932 | C932E | C | 1.1e-07 | 2e-07 | 1.8e-07 | 1.633E-07 | 4.726E-08 | 1.100E-07 | 2.000E-07 |
| 18627 | 932 | C932Q | C | 7.8e-08 | 1.8e-07 | 3.4e-07 | 1.993E-07 | 1.321E-07 | 7.800E-08 | 3.400E-07 |
| 18628 | 932 | C932G | C | 1.4e-07 | 2.2e-07 | 4e-07 | 2.533E-07 | 1.332E-07 | 1.400E-07 | 4.000E-07 |
| 18629 | 932 | C932H | C | 3.6e-07 | 2e-07 | 4.5e-07 | 3.367E-07 | 1.266E-07 | 2.000E-07 | 4.500E-07 |
| 18630 | 932 | C932I | C | 4.8e-07 | 4e-07 | 4.1e-07 | 4.300E-07 | 4.359E-08 | 4.000E-07 | 4.800E-07 |
| 18631 | 932 | C932L | C | 2.1e-07 | 1.2e-07 | 2.2e-07 | 1.833E-07 | 5.508E-08 | 1.200E-07 | 2.200E-07 |
| 18632 | 932 | C932K | C | 3e-07 | 4.2e-07 | 2.6e-07 | 3.267E-07 | 8.327E-08 | 2.600E-07 | 4.200E-07 |
| 18633 | 932 | C932M | C | 2.9e-07 | 3.7e-07 | 3.4e-07 | 3.333E-07 | 4.041E-08 | 2.900E-07 | 3.700E-07 |
| 18634 | 932 | C932F | C | 3.7e-07 | 7.7e-08 | 2.4e-07 | 2.290E-07 | 1.468E-07 | 7.700E-08 | 3.700E-07 |
| 18635 | 932 | C932P | C | 2.7e-07 | 2.8e-07 | 1.7e-07 | 2.400E-07 | 6.083E-08 | 1.700E-07 | 2.800E-07 |
| 18636 | 932 | C932S | C | 1.5e-07 | 4.4e-07 | 2.7e-07 | 2.867E-07 | 1.457E-07 | 1.500E-07 | 4.400E-07 |
| 18637 | 932 | C932T | C | 2.7e-07 | 6.8e-08 | 1.3e-07 | 1.560E-07 | 1.035E-07 | 6.800E-08 | 2.700E-07 |
| 18638 | 932 | C932W | C | 9.5e-08 | 2e-07 | 3.3e-07 | 2.083E-07 | 1.177E-07 | 9.500E-08 | 3.300E-07 |
| 18639 | 932 | C932Y | C | 3.3e-07 | 2.3e-07 | 1.5e-07 | 2.367E-07 | 9.018E-08 | 1.500E-07 | 3.300E-07 |
| 18640 | 932 | C932V | C | 1.4e-07 | 5.7e-07 | 2.1e-07 | 3.067E-07 | 2.307E-07 | 1.400E-07 | 5.700E-07 |
| 18641 | 933 | A933A | C | 1.5e-07 | 2.7e-07 | 5.8e-07 | 3.333E-07 | 2.219E-07 | 1.500E-07 | 5.800E-07 |
| 18642 | 933 | A933R | C | 2.5e-07 | 2.4e-07 | 3.8e-07 | 2.900E-07 | 7.810E-08 | 2.400E-07 | 3.800E-07 |
| 18643 | 933 | A933N | C | 2.2e-07 | 3.1e-07 | 3.4e-07 | 2.900E-07 | 6.245E-08 | 2.200E-07 | 3.400E-07 |
| 18644 | 933 | A933D | C | 3.4e-07 | 4.7e-07 | 2e-07 | 3.367E-07 | 1.350E-07 | 2.000E-07 | 4.700E-07 |
| 18645 | 933 | A933C | C | 1.5e-07 | 1.2e-07 | 4.4e-07 | 2.367E-07 | 1.767E-07 | 1.200E-07 | 4.400E-07 |
| 18646 | 933 | A933E | C | 1.5e-07 | 2.5e-07 | 3.5e-07 | 2.500E-07 | 1.000E-07 | 1.500E-07 | 3.500E-07 |
| 18647 | 933 | A933Q | C | 2.6e-07 | 2.4e-07 | 2.2e-07 | 2.400E-07 | 2.000E-08 | 2.200E-07 | 2.600E-07 |
| 18648 | 933 | A933G | C | 1.8e-07 | 1.8e-07 | 4.9e-07 | 2.833E-07 | 1.790E-07 | 1.800E-07 | 4.900E-07 |
| 18649 | 933 | A933H | C | 1.7e-07 | 1.6e-07 | 3e-07 | 2.100E-07 | 7.810E-08 | 1.600E-07 | 3.000E-07 |
| 18650 | 933 | A933I | C | 1.4e-07 | 2.4e-07 | 1.8e-07 | 1.867E-07 | 5.033E-08 | 1.400E-07 | 2.400E-07 |
| 18651 | 933 | A933L | C | 3.9e-07 | 7.9e-08 | 1.5e-07 | 2.063E-07 | 1.630E-07 | 7.900E-08 | 3.900E-07 |
| 18652 | 933 | A933K | C | 2.6e-07 | 2.1e-07 | 4e-07 | 2.900E-07 | 9.849E-08 | 2.100E-07 | 4.000E-07 |
| 18653 | 933 | A933M | C | 3.1e-07 | 3.9e-07 | 2.7e-07 | 3.233E-07 | 6.110E-08 | 2.700E-07 | 3.900E-07 |
| 18654 | 933 | A933F | C | 1.1e-07 | 1.2e-07 | 2.8e-07 | 1.700E-07 | 9.539E-08 | 1.100E-07 | 2.800E-07 |
| 18655 | 933 | A933P | C | 3.3e-07 | 1.6e-07 | 2.3e-07 | 2.400E-07 | 8.544E-08 | 1.600E-07 | 3.300E-07 |
| 18656 | 933 | A933S | C | 2.9e-07 | 1.4e-07 | 3.9e-07 | 2.733E-07 | 1.258E-07 | 1.400E-07 | 3.900E-07 |
| 18657 | 933 | A933T | C | 3.5e-07 | 1.9e-07 | 1e-07 | 2.133E-07 | 1.266E-07 | 1.000E-07 | 3.500E-07 |
| 18658 | 933 | A933W | C | 3.7e-07 | 2.1e-07 | 2.6e-07 | 2.800E-07 | 8.185E-08 | 2.100E-07 | 3.700E-07 |
| 18659 | 933 | A933Y | C | 2.3e-07 | 1.9e-07 | 2.5e-07 | 2.233E-07 | 3.055E-08 | 1.900E-07 | 2.500E-07 |
| 18660 | 933 | A933V | C | 3.6e-07 | 2.9e-07 | 2.2e-07 | 2.900E-07 | 7.000E-08 | 2.200E-07 | 3.600E-07 |
| 18661 | 934 | R934A | C | 2.8e-07 | 1.5e-07 | 3.3e-07 | 2.533E-07 | 9.292E-08 | 1.500E-07 | 3.300E-07 |
| 18662 | 934 | R934R | C | 1.2e-07 | 3.3e-07 | 3.1e-07 | 2.533E-07 | 1.159E-07 | 1.200E-07 | 3.300E-07 |
| 18663 | 934 | R934N | C | 2.5e-07 | 5e-07 | 2.4e-07 | 3.300E-07 | 1.473E-07 | 2.400E-07 | 5.000E-07 |
| 18664 | 934 | R934D | C | 1.7e-07 | 5.5e-07 | 5.5e-07 | 4.233E-07 | 2.194E-07 | 1.700E-07 | 5.500E-07 |
| 18665 | 934 | R934C | C | 5.6e-07 | 1.7e-07 | 5.4e-07 | 4.233E-07 | 2.196E-07 | 1.700E-07 | 5.600E-07 |
| 18666 | 934 | R934E | C | 3.6e-07 | 2.8e-07 | 2.9e-07 | 3.100E-07 | 4.359E-08 | 2.800E-07 | 3.600E-07 |
| 18667 | 934 | R934Q | C | 4.5e-07 | 5.6e-07 | 1.6e-07 | 3.900E-07 | 2.066E-07 | 1.600E-07 | 5.600E-07 |
| 18668 | 934 | R934G | C | 2.3e-07 | 4.8e-07 | 2e-07 | 3.033E-07 | 1.537E-07 | 2.000E-07 | 4.800E-07 |
| 18669 | 934 | R934H | C | 1.2e-07 | 5.2e-07 | 4e-07 | 3.467E-07 | 2.053E-07 | 1.200E-07 | 5.200E-07 |
| 18670 | 934 | R934I | C | 3.9e-07 | 4e-07 | 3.1e-07 | 3.667E-07 | 4.933E-08 | 3.100E-07 | 4.000E-07 |
| 18671 | 934 | R934L | C | 1.8e-07 | 2.2e-07 | 4e-07 | 2.667E-07 | 1.172E-07 | 1.800E-07 | 4.000E-07 |
| 18672 | 934 | R934K | C | 3e-07 | 1.4e-07 | 3.5e-07 | 2.633E-07 | 1.097E-07 | 1.400E-07 | 3.500E-07 |
| 18673 | 934 | R934M | C | 2.5e-07 | 3.4e-07 | 5.3e-07 | 3.733E-07 | 1.429E-07 | 2.500E-07 | 5.300E-07 |
| 18674 | 934 | R934F | C | 2.9e-07 | 4e-07 | 4.4e-07 | 3.767E-07 | 7.767E-08 | 2.900E-07 | 4.400E-07 |
| 18675 | 934 | R934P | C | 3.2e-07 | 1.9e-07 | 3.7e-07 | 2.933E-07 | 9.292E-08 | 1.900E-07 | 3.700E-07 |
| 18676 | 934 | R934S | C | 4.6e-07 | 3.1e-07 | 1e-07 | 2.900E-07 | 1.808E-07 | 1.000E-07 | 4.600E-07 |
| 18677 | 934 | R934T | C | 2.2e-07 | 1.4e-07 | 1.8e-07 | 1.800E-07 | 4.000E-08 | 1.400E-07 | 2.200E-07 |
| 18678 | 934 | R934W | C | 1.8e-07 | 3.2e-07 | 3.9e-07 | 2.967E-07 | 1.069E-07 | 1.800E-07 | 3.900E-07 |
| 18679 | 934 | R934Y | C | 2.7e-07 | 6.8e-07 | 1.6e-07 | 3.700E-07 | 2.740E-07 | 1.600E-07 | 6.800E-07 |
| 18680 | 934 | R934V | C | 4e-07 | 2e-07 | 2.3e-07 | 2.767E-07 | 1.079E-07 | 2.000E-07 | 4.000E-07 |
| 18681 | 935 | N935A | C | 2.2e-07 | 2.5e-07 | 2.6e-07 | 2.433E-07 | 2.082E-08 | 2.200E-07 | 2.600E-07 |
| 18682 | 935 | N935R | C | 2.6e-07 | 1.4e-07 | 1.8e-07 | 1.933E-07 | 6.110E-08 | 1.400E-07 | 2.600E-07 |
| 18683 | 935 | N935N | C | 4.2e-07 | 1.6e-07 | 2.8e-07 | 2.867E-07 | 1.301E-07 | 1.600E-07 | 4.200E-07 |
| 18684 | 935 | N935D | C | 1.6e-07 | 5.8e-07 | 1.3e-07 | 2.900E-07 | 2.516E-07 | 1.300E-07 | 5.800E-07 |
| 18685 | 935 | N935C | C | 4.5e-07 | 1.9e-07 | 2e-07 | 2.800E-07 | 1.473E-07 | 1.900E-07 | 4.500E-07 |
| 18686 | 935 | N935E | C | 1.3e-07 | 4.3e-07 | 2.2e-07 | 2.600E-07 | 1.539E-07 | 1.300E-07 | 4.300E-07 |
| 18687 | 935 | N935Q | C | 2.5e-07 | 8.2e-07 | 3.1e-07 | 4.600E-07 | 3.132E-07 | 2.500E-07 | 8.200E-07 |
| 18688 | 935 | N935G | C | 2.4e-07 | 2.3e-07 | 5.3e-07 | 3.333E-07 | 1.704E-07 | 2.300E-07 | 5.300E-07 |
| 18689 | 935 | N935H | C | 1.9e-07 | 3.7e-07 | 1.6e-07 | 2.400E-07 | 1.136E-07 | 1.600E-07 | 3.700E-07 |
| 18690 | 935 | N935I | C | 3.6e-07 | 2.1e-07 | 2.3e-07 | 2.667E-07 | 8.145E-08 | 2.100E-07 | 3.600E-07 |
| 18691 | 935 | N935L | C | 1.7e-07 | 2.1e-07 | 3.1e-07 | 2.300E-07 | 7.211E-08 | 1.700E-07 | 3.100E-07 |
| 18692 | 935 | N935K | C | 1.5e-07 | 2.4e-07 | 2.4e-07 | 2.100E-07 | 5.196E-08 | 1.500E-07 | 2.400E-07 |
| 18693 | 935 | N935M | C | 2e-07 | 2e-07 | 8.1e-07 | 4.033E-07 | 3.522E-07 | 2.000E-07 | 8.100E-07 |
| 18694 | 935 | N935F | C | 3.1e-07 | 4e-07 | 2.5e-07 | 3.200E-07 | 7.550E-08 | 2.500E-07 | 4.000E-07 |
| 18695 | 935 | N935P | C | 3.2e-07 | 2.1e-07 | 2.2e-07 | 2.500E-07 | 6.083E-08 | 2.100E-07 | 3.200E-07 |
| 18696 | 935 | N935S | C | 3.2e-07 | 1.6e-07 | 1.8e-07 | 2.200E-07 | 8.718E-08 | 1.600E-07 | 3.200E-07 |
| 18697 | 935 | N935T | C | 2.8e-07 | 4.2e-07 | 2.4e-07 | 3.133E-07 | 9.452E-08 | 2.400E-07 | 4.200E-07 |
| 18698 | 935 | N935W | C | 3.4e-07 | 2.4e-07 | 4.1e-07 | 3.300E-07 | 8.544E-08 | 2.400E-07 | 4.100E-07 |
| 18699 | 935 | N935Y | C | 3.2e-07 | 2.5e-07 | 4.1e-07 | 3.267E-07 | 8.021E-08 | 2.500E-07 | 4.100E-07 |

| | | | | | | | | | | |
|---|---|---|---|---|---|---|---|---|---|---|
| 18700 | 935 | N935V | C | 4.3e-07 | 2e-07 | 2.3e-07 | 2.867E-07 | 1.250E-07 | 2.000E-07 | 4.300E-07 |
| 18701 | 936 | L936A | C | 1.4e-07 | 2e-07 | 2.3e-07 | 1.900E-07 | 4.583E-08 | 1.400E-07 | 2.300E-07 |
| 18702 | 936 | L936R | C | 2.3e-07 | 3.6e-07 | 2.3e-07 | 2.733E-07 | 7.506E-08 | 2.300E-07 | 3.600E-07 |
| 18703 | 936 | L936N | C | 9.5e-08 | 2.1e-07 | 1.7e-07 | 1.583E-07 | 5.838E-08 | 9.500E-08 | 2.100E-07 |
| 18704 | 936 | L936D | C | 2.3e-07 | 2.5e-07 | 2.8e-07 | 2.533E-07 | 2.517E-08 | 2.300E-07 | 2.800E-07 |
| 18705 | 936 | L936C | C | 2.8e-07 | 2.8e-07 | 1.8e-07 | 2.467E-07 | 5.774E-08 | 1.800E-07 | 2.800E-07 |
| 18706 | 936 | L936E | C | 4.1e-07 | 3.8e-07 | 1.4e-07 | 3.100E-07 | 1.480E-07 | 1.400E-07 | 4.100E-07 |
| 18707 | 936 | L936Q | C | 3.3e-07 | 3.2e-07 | 3.2e-07 | 3.233E-07 | 5.774E-09 | 3.200E-07 | 3.300E-07 |
| 18708 | 936 | L936G | C | 1.9e-07 | 2.5e-07 | 1.8e-07 | 2.067E-07 | 3.786E-08 | 1.800E-07 | 2.500E-07 |
| 18709 | 936 | L936H | C | 3.1e-07 | 1.5e-07 | 3.2e-07 | 2.600E-07 | 9.539E-08 | 1.500E-07 | 3.200E-07 |
| 18710 | 936 | L936I | C | 2.4e-07 | 3.4e-07 | 2.9e-07 | 2.900E-07 | 5.000E-08 | 2.400E-07 | 3.400E-07 |
| 18711 | 936 | L936L | C | 4.2e-07 | 2.3e-07 | 1.1e-07 | 2.533E-07 | 1.563E-07 | 1.100E-07 | 4.200E-07 |
| 18712 | 936 | L936K | C | 6e-07 | 5.2e-07 | 2.5e-07 | 4.567E-07 | 1.834E-07 | 2.500E-07 | 6.000E-07 |
| 18713 | 936 | L936M | C | 5.2e-07 | 2.7e-07 | 1.3e-07 | 3.067E-07 | 1.976E-07 | 1.300E-07 | 5.200E-07 |
| 18714 | 936 | L936F | C | 7.1e-08 | 3.6e-07 | 2.4e-07 | 2.237E-07 | 1.452E-07 | 7.100E-08 | 3.600E-07 |
| 18715 | 936 | L936P | C | 6.8e-07 | 2.4e-07 | 1.3e-07 | 3.500E-07 | 2.910E-07 | 1.300E-07 | 6.800E-07 |
| 18716 | 936 | L936S | C | 1.6e-07 | 3.1e-07 | 2.4e-07 | 2.367E-07 | 7.506E-08 | 1.600E-07 | 3.100E-07 |
| 18717 | 936 | L936T | C | 1.7e-07 | 2.4e-07 | 2.1e-07 | 2.067E-07 | 3.512E-08 | 1.700E-07 | 2.400E-07 |
| 18718 | 936 | L936W | C | 2.5e-07 | 3.2e-07 | 4.1e-07 | 3.267E-07 | 8.021E-08 | 2.500E-07 | 4.100E-07 |
| 18719 | 936 | L936Y | C | 1.9e-07 | 1.5e-07 | 2.1e-07 | 1.833E-07 | 3.055E-08 | 1.500E-07 | 2.100E-07 |
| 18720 | 936 | L936V | C | 2.7e-07 | 2e-07 | 3.1e-07 | 2.600E-07 | 5.568E-08 | 2.000E-07 | 3.100E-07 |
| 18721 | 937 | G937A | C | 3.6e-07 | 2e-07 | 2.1e-07 | 2.567E-07 | 8.963E-08 | 2.000E-07 | 3.600E-07 |
| 18722 | 937 | G937R | C | 1.8e-07 | 4.2e-07 | 2.4e-07 | 2.800E-07 | 1.249E-07 | 1.800E-07 | 4.200E-07 |
| 18723 | 937 | G937N | C | 2.1e-07 | 3.8e-07 | 2.8e-07 | 2.900E-07 | 8.544E-08 | 2.100E-07 | 3.800E-07 |
| 18724 | 937 | G937D | C | 1.9e-07 | 1.1e-07 | 2e-07 | 1.667E-07 | 4.933E-08 | 1.100E-07 | 2.000E-07 |
| 18725 | 937 | G937C | C | 2.3e-07 | 9.8e-08 | 1e-07 | 1.427E-07 | 7.564E-08 | 9.800E-08 | 2.300E-07 |
| 18726 | 937 | G937E | C | 1.4e-07 | 2.3e-07 | 1.5e-07 | 1.733E-07 | 4.933E-08 | 1.400E-07 | 2.300E-07 |
| 18727 | 937 | G937Q | C | 2.1e-07 | 2.1e-07 | 2.1e-07 | 2.100E-07 | 0.000E+00 | 2.100E-07 | 2.100E-07 |
| 18728 | 937 | G937G | C | 1.7e-07 | 2.5e-07 | 4.3e-07 | 2.833E-07 | 1.332E-07 | 1.700E-07 | 4.300E-07 |
| 18729 | 937 | G937H | C | 2.2e-07 | 2.5e-07 | 2.8e-07 | 2.500E-07 | 3.000E-08 | 2.200E-07 | 2.800E-07 |
| 18730 | 937 | G937I | C | 1.7e-07 | 1.6e-07 | 2.7e-07 | 2.000E-07 | 6.083E-08 | 1.600E-07 | 2.700E-07 |
| 18731 | 937 | G937L | C | 2.1e-07 | 4.2e-07 | 4.2e-07 | 3.500E-07 | 1.212E-07 | 2.100E-07 | 4.200E-07 |
| 18732 | 937 | G937K | C | 4.1e-07 | 2.9e-07 | 3.5e-07 | 3.500E-07 | 6.000E-08 | 2.900E-07 | 4.100E-07 |
| 18733 | 937 | G937M | C | 3e-07 | 3.4e-07 | 1.3e-07 | 2.567E-07 | 1.115E-07 | 1.300E-07 | 3.400E-07 |
| 18734 | 937 | G937F | C | 4.8e-07 | 3.6e-07 | 4.4e-07 | 4.267E-07 | 6.110E-08 | 3.600E-07 | 4.800E-07 |
| 18735 | 937 | G937P | C | 2.6e-07 | 3e-07 | 4e-07 | 3.200E-07 | 7.211E-08 | 2.600E-07 | 4.000E-07 |
| 18736 | 937 | G937S | C | 1.5e-07 | 1.7e-07 | 2e-07 | 1.733E-07 | 2.517E-08 | 1.500E-07 | 2.000E-07 |
| 18737 | 937 | G937T | C | 2.7e-07 | 8.2e-08 | 4.2e-07 | 2.573E-07 | 1.694E-07 | 8.200E-08 | 4.200E-07 |
| 18738 | 937 | G937W | C | 1.7e-07 | 2.4e-07 | 1.5e-07 | 1.867E-07 | 4.726E-08 | 1.500E-07 | 2.400E-07 |
| 18739 | 937 | G937Y | C | 1.8e-07 | 2.9e-07 | 1.5e-07 | 2.067E-07 | 7.371E-08 | 1.500E-07 | 2.900E-07 |
| 18740 | 937 | G937V | C | 2.5e-07 | 6.4e-07 | 2.1e-07 | 3.667E-07 | 2.376E-07 | 2.100E-07 | 6.400E-07 |
| 18741 | 938 | P938A | C | 5.5e-07 | 2.7e-07 | 1.8e-07 | 3.333E-07 | 1.930E-07 | 1.800E-07 | 5.500E-07 |
| 18742 | 938 | P938R | C | 3e-07 | 2.7e-07 | 5e-07 | 3.567E-07 | 1.250E-07 | 2.700E-07 | 5.000E-07 |
| 18743 | 938 | P938N | C | 2.9e-07 | 1.5e-07 | 1.2e-07 | 1.867E-07 | 9.074E-08 | 1.200E-07 | 2.900E-07 |
| 18744 | 938 | P938D | C | 1.6e-07 | 4.5e-07 | 2.1e-07 | 2.733E-07 | 1.550E-07 | 1.600E-07 | 4.500E-07 |
| 18745 | 938 | P938C | C | 2e-07 | 2.2e-07 | 1.6e-07 | 1.933E-07 | 3.055E-08 | 1.600E-07 | 2.200E-07 |
| 18746 | 938 | P938E | C | 2e-07 | 2.9e-07 | 2.1e-07 | 2.333E-07 | 4.933E-08 | 2.000E-07 | 2.900E-07 |
| 18747 | 938 | P938Q | C | 4.9e-08 | 1.1e-07 | 3e-07 | 1.530E-07 | 1.309E-07 | 4.900E-08 | 3.000E-07 |
| 18748 | 938 | P938G | C | 3.5e-07 | 2.8e-07 | 5e-07 | 3.767E-07 | 1.124E-07 | 2.800E-07 | 5.000E-07 |
| 18749 | 938 | P938H | C | 1.3e-07 | 1.8e-07 | 2e-07 | 1.700E-07 | 3.606E-08 | 1.300E-07 | 2.000E-07 |
| 18750 | 938 | P938I | C | 2.5e-07 | 2.4e-07 | 3.1e-07 | 2.667E-07 | 3.786E-08 | 2.400E-07 | 3.100E-07 |
| 18751 | 938 | P938L | C | 2.1e-07 | 2.5e-07 | 1.4e-07 | 2.000E-07 | 5.568E-08 | 1.400E-07 | 2.500E-07 |
| 18752 | 938 | P938K | C | 2.4e-07 | 1.9e-07 | 1.6e-07 | 1.967E-07 | 4.041E-08 | 1.600E-07 | 2.400E-07 |
| 18753 | 938 | P938M | C | 1.9e-07 | 2.7e-07 | 1.5e-07 | 2.033E-07 | 6.110E-08 | 1.500E-07 | 2.700E-07 |
| 18754 | 938 | P938F | C | 1.4e-07 | 2.2e-07 | 2.9e-07 | 2.167E-07 | 7.506E-08 | 1.400E-07 | 2.900E-07 |
| 18755 | 938 | P938P | C | 1.1e-07 | 3.5e-07 | 1.7e-07 | 2.100E-07 | 1.249E-07 | 1.100E-07 | 3.500E-07 |
| 18756 | 938 | P938S | C | 3.6e-07 | 9.7e-08 | 2.1e-07 | 2.223E-07 | 1.319E-07 | 9.700E-08 | 3.600E-07 |
| 18757 | 938 | P938T | C | 5.6e-07 | 1.7e-07 | 1.5e-07 | 2.933E-07 | 2.312E-07 | 1.500E-07 | 5.600E-07 |
| 18758 | 938 | P938W | C | 2.3e-07 | 8.9e-08 | 1.8e-07 | 1.663E-07 | 7.149E-08 | 8.900E-08 | 2.300E-07 |
| 18759 | 938 | P938Y | C | 2.7e-07 | 3.9e-07 | 7.1e-08 | 2.437E-07 | 1.611E-07 | 7.100E-08 | 3.900E-07 |
| 18760 | 938 | P938V | C | 4.8e-07 | 5.6e-07 | 3.3e-07 | 4.567E-07 | 1.168E-07 | 3.300E-07 | 5.600E-07 |
| 18761 | 939 | S939A | C | 2.6e-07 | 2.3e-07 | 1.8e-07 | 2.233E-07 | 4.041E-08 | 1.800E-07 | 2.600E-07 |
| 18762 | 939 | S939R | C | 1.5e-07 | 2.4e-07 | 5.1e-07 | 3.000E-07 | 1.873E-07 | 1.500E-07 | 5.100E-07 |
| 18763 | 939 | S939N | C | 1.1e-07 | 2.8e-07 | 1.3e-07 | 1.733E-07 | 9.292E-08 | 1.100E-07 | 2.800E-07 |
| 18764 | 939 | S939D | C | 1.5e-07 | 1.2e-07 | 2.3e-07 | 1.667E-07 | 5.686E-08 | 1.200E-07 | 2.300E-07 |
| 18765 | 939 | S939C | C | 3.5e-07 | 3.2e-07 | 2.6e-07 | 3.100E-07 | 4.583E-08 | 2.600E-07 | 3.500E-07 |
| 18766 | 939 | S939E | C | 2.4e-07 | 2.9e-07 | 1.9e-07 | 2.400E-07 | 5.000E-08 | 1.900E-07 | 2.900E-07 |
| 18767 | 939 | S939Q | C | 3e-07 | 2.8e-07 | 3.6e-07 | 3.133E-07 | 4.163E-08 | 2.800E-07 | 3.600E-07 |
| 18768 | 939 | S939G | C | 2.9e-07 | 4.7e-07 | 9.6e-08 | 2.853E-07 | 1.870E-07 | 9.600E-08 | 4.700E-07 |
| 18769 | 939 | S939H | C | 3.6e-07 | 2.6e-07 | 2.7e-07 | 2.967E-07 | 5.508E-08 | 2.600E-07 | 3.600E-07 |
| 18770 | 939 | S939I | C | 2e-07 | 2.8e-07 | 2.7e-07 | 2.500E-07 | 4.359E-08 | 2.000E-07 | 2.800E-07 |
| 18771 | 939 | S939L | C | 3.4e-07 | 3.2e-07 | 9.5e-08 | 2.517E-07 | 1.360E-07 | 9.500E-08 | 3.400E-07 |
| 18772 | 939 | S939K | C | 2.4e-07 | 2.8e-07 | 1.9e-07 | 2.367E-07 | 4.509E-08 | 1.900E-07 | 2.800E-07 |
| 18773 | 939 | S939M | C | 3.3e-07 | 4.4e-07 | 1.3e-07 | 3.000E-07 | 1.572E-07 | 1.300E-07 | 4.400E-07 |
| 18774 | 939 | S939F | C | 2.3e-07 | 2e-07 | 3.4e-07 | 2.567E-07 | 7.371E-08 | 2.000E-07 | 3.400E-07 |
| 18775 | 939 | S939P | C | 5.1e-07 | 1.9e-07 | 2.3e-07 | 3.100E-07 | 1.744E-07 | 1.900E-07 | 5.100E-07 |
| 18776 | 939 | S939S | C | 4.6e-07 | 2.3e-07 | 2.8e-07 | 3.233E-07 | 1.210E-07 | 2.300E-07 | 4.600E-07 |
| 18777 | 939 | S939T | C | 7.5e-07 | 3e-07 | 2.9e-07 | 4.467E-07 | 2.627E-07 | 2.900E-07 | 7.500E-07 |
| 18778 | 939 | S939W | C | 1.1e-07 | 4.4e-07 | 3.6e-07 | 3.033E-07 | 1.721E-07 | 1.100E-07 | 4.400E-07 |
| 18779 | 939 | S939Y | C | 3.5e-07 | 3.3e-07 | 5.2e-07 | 4.000E-07 | 1.044E-07 | 3.300E-07 | 5.200E-07 |
| 18780 | 939 | S939V | C | 2e-07 | 2.6e-07 | 2.8e-07 | 2.467E-07 | 4.163E-08 | 2.000E-07 | 2.800E-07 |
| 18781 | 940 | F940A | C | 2.6e-07 | 2.5e-07 | 1.3e-07 | 2.133E-07 | 7.234E-08 | 1.300E-07 | 2.600E-07 |
| 18782 | 940 | F940R | C | 1.4e-07 | 2.6e-07 | 3.4e-07 | 2.467E-07 | 1.007E-07 | 1.400E-07 | 3.400E-07 |
| 18783 | 940 | F940N | C | 2.9e-07 | 3e-07 | 9.7e-08 | 2.290E-07 | 1.144E-07 | 9.700E-08 | 3.000E-07 |
| 18784 | 940 | F940D | C | 2.6e-07 | 2.7e-07 | 3e-07 | 2.767E-07 | 2.082E-08 | 2.600E-07 | 3.000E-07 |

| | | | | | | | | | | |
|---|---|---|---|---|---|---|---|---|---|---|
| 18785 | 940 | F940C | C | 2e-07 | 2.1e-07 | 9.4e-08 | 1.680E-07 | 6.428E-08 | 9.400E-08 | 2.100E-07 |
| 18786 | 940 | F940E | C | 3.5e-07 | 3.2e-07 | 1.6e-07 | 2.767E-07 | 1.021E-07 | 1.600E-07 | 3.500E-07 |
| 18787 | 940 | F940Q | C | 3.6e-07 | 1.7e-07 | 2.1e-07 | 2.467E-07 | 1.002E-07 | 1.700E-07 | 3.600E-07 |
| 18788 | 940 | F940G | C | 3.6e-07 | 2.3e-07 | 2.3e-07 | 2.733E-07 | 7.506E-08 | 2.300E-07 | 3.600E-07 |
| 18789 | 940 | F940H | C | 1.5e-07 | 2.9e-07 | 5.5e-07 | 3.300E-07 | 2.030E-07 | 1.500E-07 | 5.500E-07 |
| 18790 | 940 | F940I | C | 2.6e-07 | 3.2e-07 | 2.2e-07 | 2.667E-07 | 5.033E-08 | 2.200E-07 | 3.200E-07 |
| 18791 | 940 | F940L | C | 5.2e-07 | 1.4e-07 | 6.9e-07 | 4.500E-07 | 2.816E-07 | 1.400E-07 | 6.900E-07 |
| 18792 | 940 | F940K | C | 2.8e-07 | 3.3e-07 | 3.9e-07 | 3.333E-07 | 5.508E-08 | 2.800E-07 | 3.900E-07 |
| 18793 | 940 | F940M | C | 3.3e-07 | 2e-07 | 3.5e-07 | 2.933E-07 | 8.145E-08 | 2.000E-07 | 3.500E-07 |
| 18794 | 940 | F940F | C | 2.4e-07 | 2.2e-07 | 2.6e-07 | 2.400E-07 | 2.000E-08 | 2.200E-07 | 2.600E-07 |
| 18795 | 940 | F940P | C | 7e-08 | 4.8e-07 | 2.1e-07 | 2.533E-07 | 2.084E-07 | 7.000E-08 | 4.800E-07 |
| 18796 | 940 | F940S | C | 3.4e-07 | 1.8e-07 | 1.9e-07 | 2.367E-07 | 8.963E-08 | 1.800E-07 | 3.400E-07 |
| 18797 | 940 | F940T | C | 1.8e-07 | 3.1e-07 | 1.2e-07 | 2.033E-07 | 9.713E-08 | 1.200E-07 | 3.100E-07 |
| 18798 | 940 | F940W | C | 1.1e-07 | 2.2e-07 | 1.5e-07 | 1.600E-07 | 5.568E-08 | 1.100E-07 | 2.200E-07 |
| 18799 | 940 | F940Y | C | 1.6e-07 | 2.8e-07 | 2.3e-07 | 2.233E-07 | 6.028E-08 | 1.600E-07 | 2.800E-07 |
| 18800 | 940 | F940V | C | 1.5e-07 | 3.3e-07 | 3.1e-07 | 2.633E-07 | 9.866E-08 | 1.500E-07 | 3.300E-07 |
| 18801 | 941 | Y941A | C | 3.8e-07 | 5.5e-07 | 1.9e-07 | 3.733E-07 | 1.801E-07 | 1.900E-07 | 5.500E-07 |
| 18802 | 941 | Y941R | C | 4.4e-07 | 3.9e-07 | 1.3e-07 | 3.200E-07 | 1.664E-07 | 1.300E-07 | 4.400E-07 |
| 18803 | 941 | Y941N | C | 3.4e-07 | 2.4e-07 | 1.3e-07 | 2.367E-07 | 1.050E-07 | 1.300E-07 | 3.400E-07 |
| 18804 | 941 | Y941D | C | 1.7e-07 | 1.9e-07 | 2.5e-07 | 2.033E-07 | 4.163E-08 | 1.700E-07 | 2.500E-07 |
| 18805 | 941 | Y941C | C | 4.8e-07 | 1.7e-07 | 3.4e-07 | 3.300E-07 | 1.552E-07 | 1.700E-07 | 4.800E-07 |
| 18806 | 941 | Y941E | C | 3.1e-07 | 1.8e-07 | 4.6e-07 | 3.167E-07 | 1.401E-07 | 1.800E-07 | 4.600E-07 |
| 18807 | 941 | Y941Q | C | 1.5e-07 | 5.2e-07 | 1.1e-07 | 2.600E-07 | 2.261E-07 | 1.100E-07 | 5.200E-07 |
| 18808 | 941 | Y941G | C | 2e-07 | 1.4e-07 | 2.2e-07 | 1.867E-07 | 4.163E-08 | 1.400E-07 | 2.200E-07 |
| 18809 | 941 | Y941H | C | 3.7e-07 | 1.9e-07 | 1.7e-07 | 2.433E-07 | 1.102E-07 | 1.700E-07 | 3.700E-07 |
| 18810 | 941 | Y941I | C | 6e-07 | 2e-07 | 2.6e-07 | 3.533E-07 | 2.157E-07 | 2.000E-07 | 6.000E-07 |
| 18811 | 941 | Y941L | C | 5.2e-07 | 5.4e-07 | 2e-07 | 4.200E-07 | 1.908E-07 | 2.000E-07 | 5.400E-07 |
| 18812 | 941 | Y941K | C | 1.6e-07 | 1.3e-07 | 1.3e-07 | 1.400E-07 | 1.732E-08 | 1.300E-07 | 1.600E-07 |
| 18813 | 941 | Y941M | C | 2.2e-07 | 4.6e-07 | 1.1e-07 | 2.633E-07 | 1.790E-07 | 1.100E-07 | 4.600E-07 |
| 18814 | 941 | Y941F | C | 3.5e-07 | 3.9e-07 | 2.7e-07 | 3.367E-07 | 6.110E-08 | 2.700E-07 | 3.900E-07 |
| 18815 | 941 | Y941P | C | 1.9e-07 | 1.5e-07 | 3.1e-07 | 2.167E-07 | 8.327E-08 | 1.500E-07 | 3.100E-07 |
| 18816 | 941 | Y941S | C | 1.6e-07 | 2.5e-07 | 5.2e-07 | 3.100E-07 | 1.873E-07 | 1.600E-07 | 5.200E-07 |
| 18817 | 941 | Y941T | C | 2.4e-07 | 2.6e-07 | 1.4e-07 | 2.133E-07 | 6.429E-08 | 1.400E-07 | 2.600E-07 |
| 18818 | 941 | Y941W | C | 2.7e-07 | 3e-07 | 2.6e-07 | 2.767E-07 | 2.082E-08 | 2.600E-07 | 3.000E-07 |
| 18819 | 941 | Y941Y | C | 3.6e-07 | 2.5e-07 | 2.6e-07 | 2.900E-07 | 6.083E-08 | 2.500E-07 | 3.600E-07 |
| 18820 | 941 | Y941V | C | 2.4e-07 | 2e-07 | 3.6e-07 | 2.667E-07 | 8.327E-08 | 2.000E-07 | 3.600E-07 |
| 18821 | 942 | F942A | C | 3.9e-07 | 1.4e-07 | 3.6e-07 | 2.967E-07 | 1.365E-07 | 1.400E-07 | 3.900E-07 |
| 18822 | 942 | F942R | C | 3.1e-07 | 6.8e-07 | 2.7e-07 | 4.200E-07 | 2.261E-07 | 2.700E-07 | 6.800E-07 |
| 18823 | 942 | F942N | C | 1.6e-07 | 2.5e-07 | 2.1e-07 | 2.067E-07 | 4.509E-08 | 1.600E-07 | 2.500E-07 |
| 18824 | 942 | F942D | C | 2.4e-07 | 3e-07 | 5.6e-07 | 3.667E-07 | 1.701E-07 | 2.400E-07 | 5.600E-07 |
| 18825 | 942 | F942C | C | 2.2e-07 | 2e-07 | 2.6e-07 | 2.267E-07 | 3.055E-08 | 2.000E-07 | 2.600E-07 |
| 18826 | 942 | F942E | C | 5.2e-07 | 2.7e-07 | 2.7e-07 | 3.533E-07 | 1.443E-07 | 2.700E-07 | 5.200E-07 |
| 18827 | 942 | F942Q | C | 3.5e-07 | 3.6e-07 | 5.2e-07 | 4.100E-07 | 9.539E-08 | 3.500E-07 | 5.200E-07 |
| 18828 | 942 | F942G | C | 2.1e-07 | 1.3e-07 | 4.5e-07 | 2.633E-07 | 1.665E-07 | 1.300E-07 | 4.500E-07 |
| 18829 | 942 | F942H | C | 2.4e-07 | 2.2e-07 | 9.2e-08 | 1.840E-07 | 8.030E-08 | 9.200E-08 | 2.400E-07 |
| 18830 | 942 | F942I | C | 2.5e-07 | 4.2e-07 | 2.6e-07 | 3.100E-07 | 9.539E-08 | 2.500E-07 | 4.200E-07 |
| 18831 | 942 | F942L | C | 3.3e-07 | 1.9e-07 | 1.8e-07 | 2.333E-07 | 8.386E-08 | 1.800E-07 | 3.300E-07 |
| 18832 | 942 | F942K | C | 2.3e-07 | 1.8e-07 | 1.8e-07 | 1.967E-07 | 2.887E-08 | 1.800E-07 | 2.300E-07 |
| 18833 | 942 | F942M | C | 2.5e-07 | 1.7e-07 | 4.9e-07 | 3.033E-07 | 1.665E-07 | 1.700E-07 | 4.900E-07 |
| 18834 | 942 | F942F | C | 1.7e-07 | 9.4e-08 | 3.2e-07 | 1.947E-07 | 1.150E-07 | 9.400E-08 | 3.200E-07 |
| 18835 | 942 | F942P | C | 3.5e-07 | 2.7e-07 | 2.2e-07 | 2.800E-07 | 6.557E-08 | 2.200E-07 | 3.500E-07 |
| 18836 | 942 | F942S | C | 1.3e-07 | 1.4e-07 | 2.8e-07 | 1.833E-07 | 8.386E-08 | 1.300E-07 | 2.800E-07 |
| 18837 | 942 | F942T | C | 2.5e-07 | 1.1e-07 | 3.5e-07 | 2.367E-07 | 1.206E-07 | 1.100E-07 | 3.500E-07 |
| 18838 | 942 | F942W | C | 1.6e-07 | 1.4e-07 | 1.7e-07 | 1.567E-07 | 1.528E-08 | 1.400E-07 | 1.700E-07 |
| 18839 | 942 | F942Y | C | 1.4e-07 | 2.7e-07 | 1.6e-07 | 1.900E-07 | 7.000E-08 | 1.400E-07 | 2.700E-07 |
| 18840 | 942 | F942V | C | 3.9e-07 | 2.1e-07 | 2.4e-07 | 2.800E-07 | 9.644E-08 | 2.100E-07 | 3.900E-07 |
| 18841 | 943 | D943A | C | 3.7e-07 | 3.1e-07 | 3e-07 | 3.267E-07 | 3.786E-08 | 3.000E-07 | 3.700E-07 |
| 18842 | 943 | D943R | C | 2.7e-07 | 4.7e-07 | 2e-07 | 3.133E-07 | 1.401E-07 | 2.000E-07 | 4.700E-07 |
| 18843 | 943 | D943N | C | 3.8e-07 | 3.8e-07 | 2.9e-07 | 3.500E-07 | 5.196E-08 | 2.900E-07 | 3.800E-07 |
| 18844 | 943 | D943D | C | 1.7e-07 | 1.4e-07 | 1.9e-07 | 1.667E-07 | 2.517E-08 | 1.400E-07 | 1.900E-07 |
| 18845 | 943 | D943C | C | 3.2e-07 | 2e-07 | 3.5e-07 | 2.900E-07 | 7.937E-08 | 2.000E-07 | 3.500E-07 |
| 18846 | 943 | D943E | C | 2.7e-07 | 1.6e-07 | 3.6e-07 | 2.633E-07 | 1.002E-07 | 1.600E-07 | 3.600E-07 |
| 18847 | 943 | D943Q | C | 2.1e-07 | 2.6e-07 | 2.2e-07 | 2.300E-07 | 2.646E-08 | 2.100E-07 | 2.600E-07 |
| 18848 | 943 | D943G | C | 5.3e-07 | 3.2e-07 | 3.3e-07 | 3.933E-07 | 1.185E-07 | 3.200E-07 | 5.300E-07 |
| 18849 | 943 | D943H | C | 2.6e-07 | 2.3e-07 | 1.9e-07 | 2.267E-07 | 3.512E-08 | 1.900E-07 | 2.600E-07 |
| 18850 | 943 | D943I | C | 2.9e-07 | 1.8e-07 | 4.9e-07 | 3.200E-07 | 1.572E-07 | 1.800E-07 | 4.900E-07 |
| 18851 | 943 | D943L | C | 2.7e-07 | 2.5e-07 | 8.6e-08 | 2.020E-07 | 1.010E-07 | 8.600E-08 | 2.700E-07 |
| 18852 | 943 | D943K | C | 2.1e-07 | 1.5e-07 | 1.2e-07 | 1.600E-07 | 4.583E-08 | 1.200E-07 | 2.100E-07 |
| 18853 | 943 | D943M | C | 2.9e-07 | 5e-07 | 2.3e-07 | 3.400E-07 | 1.418E-07 | 2.300E-07 | 5.000E-07 |
| 18854 | 943 | D943F | C | 2.5e-07 | 1.8e-07 | 1.6e-07 | 1.967E-07 | 4.726E-08 | 1.600E-07 | 2.500E-07 |
| 18855 | 943 | D943P | C | 1.8e-07 | 1.1e-07 | 2.7e-07 | 1.867E-07 | 8.021E-08 | 1.100E-07 | 2.700E-07 |
| 18856 | 943 | D943S | C | 1.4e-07 | 2.3e-07 | 2.2e-07 | 1.967E-07 | 4.933E-08 | 1.400E-07 | 2.300E-07 |
| 18857 | 943 | D943T | C | 3e-07 | 4.8e-07 | 1.5e-07 | 3.100E-07 | 1.652E-07 | 1.500E-07 | 4.800E-07 |
| 18858 | 943 | D943W | C | 3.2e-07 | 1.5e-07 | 1.9e-07 | 2.200E-07 | 8.888E-08 | 1.500E-07 | 3.200E-07 |
| 18859 | 943 | D943Y | C | 3.4e-07 | 2.1e-07 | 2.5e-07 | 2.667E-07 | 6.658E-08 | 2.100E-07 | 3.400E-07 |
| 18860 | 943 | D943V | C | 2.9e-07 | 3.5e-07 | 2.5e-07 | 2.967E-07 | 5.033E-08 | 2.500E-07 | 3.500E-07 |
| 18861 | 944 | Y944A | C | 2.2e-07 | 1.9e-07 | 1.9e-07 | 2.000E-07 | 1.732E-08 | 1.900E-07 | 2.200E-07 |
| 18862 | 944 | Y944R | C | 1.2e-07 | 5.2e-07 | 3.9e-07 | 3.433E-07 | 2.040E-07 | 1.200E-07 | 5.200E-07 |
| 18863 | 944 | Y944N | C | 1.7e-07 | 4.1e-07 | 2.1e-07 | 2.633E-07 | 1.286E-07 | 1.700E-07 | 4.100E-07 |
| 18864 | 944 | Y944D | C | 3e-07 | 2.5e-07 | 1.2e-07 | 2.233E-07 | 9.292E-08 | 1.200E-07 | 3.000E-07 |
| 18865 | 944 | Y944C | C | 1.5e-07 | 1.1e-07 | 2.7e-07 | 1.767E-07 | 8.327E-08 | 1.100E-07 | 2.700E-07 |
| 18866 | 944 | Y944E | C | 2.3e-07 | 3.3e-07 | 2.5e-07 | 2.700E-07 | 5.292E-08 | 2.300E-07 | 3.300E-07 |
| 18867 | 944 | Y944Q | C | 3.2e-07 | 3.3e-07 | 2.1e-07 | 2.867E-07 | 6.658E-08 | 2.100E-07 | 3.300E-07 |
| 18868 | 944 | Y944G | C | 1.2e-07 | 1.9e-07 | 2.2e-07 | 1.767E-07 | 5.132E-08 | 1.200E-07 | 2.200E-07 |
| 18869 | 944 | Y944H | C | 1.1e-07 | 6e-07 | 2.2e-07 | 3.100E-07 | 2.571E-07 | 1.100E-07 | 6.000E-07 |

| | | | | | | | | | | |
|---|---|---|---|---|---|---|---|---|---|---|
| 18870 | 944 | Y944I | C | 5.6e-07 | 2e-07 | 4.8e-07 | 4.133E-07 | 1.890E-07 | 2.000E-07 | 5.600E-07 |
| 18871 | 944 | Y944L | C | 1.5e-07 | 3.8e-07 | 2.5e-07 | 2.600E-07 | 1.153E-07 | 1.500E-07 | 3.800E-07 |
| 18872 | 944 | Y944K | C | 2.4e-07 | 5.3e-07 | 1.6e-07 | 3.100E-07 | 1.947E-07 | 1.600E-07 | 5.300E-07 |
| 18873 | 944 | Y944M | C | 2.6e-07 | 9.8e-08 | 2.8e-07 | 2.127E-07 | 9.981E-08 | 9.800E-08 | 2.800E-07 |
| 18874 | 944 | Y944F | C | 2.8e-07 | 2.2e-07 | 2e-07 | 2.333E-07 | 4.163E-08 | 2.000E-07 | 2.800E-07 |
| 18875 | 944 | Y944P | C | 1.7e-07 | 2.7e-07 | 3.3e-07 | 2.567E-07 | 8.083E-08 | 1.700E-07 | 3.300E-07 |
| 18876 | 944 | Y944S | C | 2.7e-07 | 1.2e-07 | 2.2e-07 | 2.033E-07 | 7.638E-08 | 1.200E-07 | 2.700E-07 |
| 18877 | 944 | Y944T | C | 3.3e-07 | 2.3e-07 | 3.8e-07 | 3.133E-07 | 7.638E-08 | 2.300E-07 | 3.800E-07 |
| 18878 | 944 | Y944W | C | 2.5e-07 | 2.1e-07 | 8.2e-08 | 1.807E-07 | 8.776E-08 | 8.200E-08 | 2.500E-07 |
| 18879 | 944 | Y944Y | C | 5.9e-07 | 1.7e-07 | 3.7e-07 | 3.767E-07 | 2.101E-07 | 1.700E-07 | 5.900E-07 |
| 18880 | 944 | Y944V | C | 3.1e-07 | 5.3e-07 | 3.4e-07 | 3.933E-07 | 1.193E-07 | 3.100E-07 | 5.300E-07 |
| 18881 | 945 | W945A | C | 2.2e-07 | 3.6e-07 | 3.2e-07 | 3.000E-07 | 7.211E-08 | 2.200E-07 | 3.600E-07 |
| 18882 | 945 | W945R | C | 1.7e-07 | 5.2e-07 | 4.1e-07 | 3.667E-07 | 1.790E-07 | 1.700E-07 | 5.200E-07 |
| 18883 | 945 | W945N | C | 2.7e-07 | 3.2e-07 | 1.7e-07 | 2.533E-07 | 7.638E-08 | 1.700E-07 | 3.200E-07 |
| 18884 | 945 | W945D | C | 1.8e-07 | 2.8e-07 | 2.3e-07 | 2.300E-07 | 5.000E-08 | 1.800E-07 | 2.800E-07 |
| 18885 | 945 | W945C | C | 2.6e-07 | 7.9e-07 | 3.2e-07 | 4.567E-07 | 2.902E-07 | 2.600E-07 | 7.900E-07 |
| 18886 | 945 | W945E | C | 5.3e-07 | 5.8e-07 | 2.4e-07 | 4.500E-07 | 1.836E-07 | 2.400E-07 | 5.800E-07 |
| 18887 | 945 | W945Q | C | 2.6e-07 | 2.2e-07 | 9.8e-08 | 1.927E-07 | 8.439E-08 | 9.800E-08 | 2.600E-07 |
| 18888 | 945 | W945G | C | 1.6e-07 | 3.6e-07 | 1.9e-07 | 2.367E-07 | 1.079E-07 | 1.600E-07 | 3.600E-07 |
| 18889 | 945 | W945H | C | 3.1e-07 | 1.7e-07 | 2.3e-07 | 2.367E-07 | 7.024E-08 | 1.700E-07 | 3.100E-07 |
| 18890 | 945 | W945I | C | 1.9e-07 | 2.9e-07 | 2.3e-07 | 2.367E-07 | 5.033E-08 | 1.900E-07 | 2.900E-07 |
| 18891 | 945 | W945L | C | 1.2e-07 | 3.9e-07 | 3.6e-07 | 2.900E-07 | 1.480E-07 | 1.200E-07 | 3.900E-07 |
| 18892 | 945 | W945K | C | 1.9e-07 | 1.2e-07 | 1.4e-07 | 1.500E-07 | 3.606E-08 | 1.200E-07 | 1.900E-07 |
| 18893 | 945 | W945M | C | 3.3e-07 | 2.8e-07 | 1.9e-07 | 2.667E-07 | 7.095E-08 | 1.900E-07 | 3.300E-07 |
| 18894 | 945 | W945F | C | 1.6e-07 | 2.2e-07 | 5.2e-07 | 3.000E-07 | 1.929E-07 | 1.600E-07 | 5.200E-07 |
| 18895 | 945 | W945P | C | 1.8e-07 | 5.8e-07 | 4.9e-07 | 4.167E-07 | 2.098E-07 | 1.800E-07 | 5.800E-07 |
| 18896 | 945 | W945S | C | 3.1e-07 | 4.2e-07 | 2.9e-07 | 3.400E-07 | 7.000E-08 | 2.900E-07 | 4.200E-07 |
| 18897 | 945 | W945T | C | 2.9e-07 | 2.6e-07 | 1.9e-07 | 2.467E-07 | 5.132E-08 | 1.900E-07 | 2.900E-07 |
| 18898 | 945 | W945W | C | 1.5e-07 | 3.5e-07 | 2.2e-07 | 2.400E-07 | 1.015E-07 | 1.500E-07 | 3.500E-07 |
| 18899 | 945 | W945Y | C | 3.7e-07 | 2e-07 | 4.4e-07 | 3.367E-07 | 1.234E-07 | 2.000E-07 | 4.400E-07 |
| 18900 | 945 | W945V | C | 3.4e-07 | 2.9e-07 | 4.3e-07 | 3.533E-07 | 7.095E-08 | 2.900E-07 | 4.300E-07 |
| 18901 | 946 | G946A | C | 2.6e-07 | 1.1e-07 | 9.3e-08 | 1.543E-07 | 9.190E-08 | 9.300E-08 | 2.600E-07 |
| 18902 | 946 | G946R | C | 1.6e-07 | 1.7e-07 | 1.8e-07 | 1.700E-07 | 1.000E-08 | 1.600E-07 | 1.800E-07 |
| 18903 | 946 | G946N | C | 3.8e-07 | 3.2e-07 | 1.8e-07 | 2.933E-07 | 1.026E-07 | 1.800E-07 | 3.800E-07 |
| 18904 | 946 | G946D | C | 3.1e-07 | 2.3e-07 | 1.3e-07 | 2.233E-07 | 9.018E-08 | 1.300E-07 | 3.100E-07 |
| 18905 | 946 | G946C | C | 1.5e-07 | 2.4e-07 | 2.5e-07 | 2.133E-07 | 5.508E-08 | 1.500E-07 | 2.500E-07 |
| 18906 | 946 | G946E | C | 4.4e-07 | 2.1e-07 | 1.9e-07 | 2.800E-07 | 1.389E-07 | 1.900E-07 | 4.400E-07 |
| 18907 | 946 | G946Q | C | 1.1e-07 | 2.5e-07 | 3e-07 | 2.200E-07 | 9.849E-08 | 1.100E-07 | 3.000E-07 |
| 18908 | 946 | G946G | C | 2.1e-07 | 2.3e-07 | 1.6e-07 | 2.000E-07 | 3.606E-08 | 1.600E-07 | 2.300E-07 |
| 18909 | 946 | G946H | C | 2.9e-07 | 4.4e-07 | 3.4e-07 | 3.567E-07 | 7.638E-08 | 2.900E-07 | 4.400E-07 |
| 18910 | 946 | G946I | C | 2.1e-07 | 2.1e-07 | 2.9e-07 | 2.367E-07 | 4.619E-08 | 2.100E-07 | 2.900E-07 |
| 18911 | 946 | G946L | C | 1.9e-07 | 2.6e-07 | 3e-07 | 2.500E-07 | 5.568E-08 | 1.900E-07 | 3.000E-07 |
| 18912 | 946 | G946K | C | 2.1e-07 | 1.9e-07 | 3.7e-07 | 2.567E-07 | 9.866E-08 | 1.900E-07 | 3.700E-07 |
| 18913 | 946 | G946M | C | 3e-07 | 3.2e-07 | 2.6e-07 | 2.933E-07 | 3.055E-08 | 2.600E-07 | 3.200E-07 |
| 18914 | 946 | G946F | C | 2.1e-07 | 2.7e-07 | 1.9e-07 | 2.233E-07 | 4.163E-08 | 1.900E-07 | 2.700E-07 |
| 18915 | 946 | G946P | C | 2.3e-07 | 1.8e-07 | 1.2e-07 | 1.767E-07 | 5.508E-08 | 1.200E-07 | 2.300E-07 |
| 18916 | 946 | G946S | C | 3.7e-07 | 2.3e-07 | 1.3e-07 | 2.433E-07 | 1.206E-07 | 1.300E-07 | 3.700E-07 |
| 18917 | 946 | G946T | C | 3.4e-07 | 2.5e-07 | 2.1e-07 | 2.667E-07 | 6.658E-08 | 2.100E-07 | 3.400E-07 |
| 18918 | 946 | G946W | C | 1.7e-07 | 4.9e-07 | 1.4e-07 | 2.667E-07 | 1.940E-07 | 1.400E-07 | 4.900E-07 |
| 18919 | 946 | G946Y | C | 2e-07 | 3.1e-07 | 2.3e-07 | 2.467E-07 | 5.686E-08 | 2.000E-07 | 3.100E-07 |
| 18920 | 946 | G946V | C | 2e-07 | 2.2e-07 | 2.7e-07 | 2.300E-07 | 3.606E-08 | 2.000E-07 | 2.700E-07 |
| 18921 | 947 | Q947A | C | 3e-07 | 3e-07 | 1.9e-07 | 2.633E-07 | 6.351E-08 | 1.900E-07 | 3.000E-07 |
| 18922 | 947 | Q947R | C | 3.1e-07 | 2.6e-07 | 2.3e-07 | 2.667E-07 | 4.041E-08 | 2.300E-07 | 3.100E-07 |
| 18923 | 947 | Q947N | C | 2.3e-07 | 2.9e-07 | 5.6e-07 | 3.600E-07 | 1.758E-07 | 2.300E-07 | 5.600E-07 |
| 18924 | 947 | Q947D | C | 1.8e-07 | 1.8e-07 | 1.7e-07 | 1.767E-07 | 5.774E-09 | 1.700E-07 | 1.800E-07 |
| 18925 | 947 | Q947C | C | 4.8e-07 | 2.4e-07 | 2.6e-07 | 3.267E-07 | 1.332E-07 | 2.400E-07 | 4.800E-07 |
| 18926 | 947 | Q947E | C | 1.5e-07 | 1.7e-07 | 4.3e-07 | 2.500E-07 | 1.562E-07 | 1.500E-07 | 4.300E-07 |
| 18927 | 947 | Q947Q | C | 2.9e-07 | 1.4e-07 | 5.4e-07 | 3.233E-07 | 2.021E-07 | 1.400E-07 | 5.400E-07 |
| 18928 | 947 | Q947G | C | 1.7e-07 | 2.3e-07 | 3.2e-07 | 2.400E-07 | 7.550E-08 | 1.700E-07 | 3.200E-07 |
| 18929 | 947 | Q947H | C | 2.2e-07 | 1.8e-07 | 3.6e-07 | 2.533E-07 | 9.452E-08 | 1.800E-07 | 3.600E-07 |
| 18930 | 947 | Q947I | C | 5.1e-07 | 1.9e-07 | 1.4e-07 | 2.800E-07 | 2.007E-07 | 1.400E-07 | 5.100E-07 |
| 18931 | 947 | Q947L | C | 4.1e-07 | 7.1e-07 | 3.6e-07 | 4.933E-07 | 1.893E-07 | 3.600E-07 | 7.100E-07 |
| 18932 | 947 | Q947K | C | 1.5e-07 | 1.8e-07 | 2e-07 | 1.767E-07 | 2.517E-08 | 1.500E-07 | 2.000E-07 |
| 18933 | 947 | Q947M | C | 2.7e-07 | 4e-07 | 6.7e-07 | 4.467E-07 | 2.040E-07 | 2.700E-07 | 6.700E-07 |
| 18934 | 947 | Q947F | C | 2.6e-07 | 3.7e-07 | 2.8e-07 | 3.033E-07 | 5.859E-08 | 2.600E-07 | 3.700E-07 |
| 18935 | 947 | Q947P | C | 2.4e-07 | 2.1e-07 | 2.2e-07 | 2.233E-07 | 1.528E-08 | 2.100E-07 | 2.400E-07 |
| 18936 | 947 | Q947S | C | 3.3e-07 | 1.9e-07 | 2.1e-07 | 2.433E-07 | 7.572E-08 | 1.900E-07 | 3.300E-07 |
| 18937 | 947 | Q947T | C | 2.1e-07 | 3e-07 | 3.6e-07 | 2.900E-07 | 7.550E-08 | 2.100E-07 | 3.600E-07 |
| 18938 | 947 | Q947W | C | 2.3e-07 | 3.3e-07 | 2.2e-07 | 2.600E-07 | 6.083E-08 | 2.200E-07 | 3.300E-07 |
| 18939 | 947 | Q947Y | C | 2.4e-07 | 1.9e-07 | 2e-07 | 2.100E-07 | 2.646E-08 | 1.900E-07 | 2.400E-07 |
| 18940 | 947 | Q947V | C | 3.6e-07 | 3.3e-07 | 2.9e-07 | 3.267E-07 | 3.512E-08 | 2.900E-07 | 3.600E-07 |
| 18941 | 948 | G948A | C | 2.3e-07 | 1.6e-07 | 5.1e-07 | 3.000E-07 | 1.852E-07 | 1.600E-07 | 5.100E-07 |
| 18942 | 948 | G948R | C | 2.1e-07 | 1.8e-07 | 2.6e-07 | 2.167E-07 | 4.041E-08 | 1.800E-07 | 2.600E-07 |
| 18943 | 948 | G948N | C | 3.3e-07 | 1.5e-07 | 1.9e-07 | 2.233E-07 | 9.452E-08 | 1.500E-07 | 3.300E-07 |
| 18944 | 948 | G948D | C | 7.2e-07 | 2.5e-07 | 1.2e-07 | 3.633E-07 | 3.156E-07 | 1.200E-07 | 7.200E-07 |
| 18945 | 948 | G948C | C | 2.3e-07 | 2.1e-07 | 1.8e-07 | 2.067E-07 | 2.517E-08 | 1.800E-07 | 2.300E-07 |
| 18946 | 948 | G948E | C | 1.9e-07 | 2.3e-07 | 3.4e-07 | 2.533E-07 | 7.767E-08 | 1.900E-07 | 3.400E-07 |
| 18947 | 948 | G948Q | C | 4e-07 | 1.1e-06 | 1.7e-07 | 5.567E-07 | 4.844E-07 | 1.700E-07 | 1.100E-06 |
| 18948 | 948 | G948G | C | 3.9e-07 | 1.8e-07 | 1.8e-07 | 2.500E-07 | 1.212E-07 | 1.800E-07 | 3.900E-07 |
| 18949 | 948 | G948H | C | 4.7e-07 | 2.8e-07 | 1.7e-07 | 3.067E-07 | 1.518E-07 | 1.700E-07 | 4.700E-07 |
| 18950 | 948 | G948I | C | 2.5e-07 | 2.5e-07 | 1.5e-07 | 2.167E-07 | 5.774E-08 | 1.500E-07 | 2.500E-07 |
| 18951 | 948 | G948L | C | 3.9e-07 | 2.6e-07 | 3.4e-07 | 3.300E-07 | 6.557E-08 | 2.600E-07 | 3.900E-07 |
| 18952 | 948 | G948K | C | 1.8e-07 | 1.1e-07 | 9.3e-08 | 1.277E-07 | 4.611E-08 | 9.300E-08 | 1.800E-07 |
| 18953 | 948 | G948M | C | 2.1e-07 | 2.3e-07 | 5.9e-07 | 3.433E-07 | 2.139E-07 | 2.100E-07 | 5.900E-07 |
| 18954 | 948 | G948F | C | 1.1e-07 | 4.1e-07 | 1.5e-07 | 2.233E-07 | 1.629E-07 | 1.100E-07 | 4.100E-07 |

| | | | | | | | | | | |
|---|---|---|---|---|---|---|---|---|---|---|
| 18955 | 948 | G948P | C | 2.4e-07 | 1.8e-07 | 2.9e-07 | 2.367E-07 | 5.508E-08 | 1.800E-07 | 2.900E-07 |
| 18956 | 948 | G948S | C | 2.6e-07 | 1.3e-07 | 3.6e-07 | 2.500E-07 | 1.153E-07 | 1.300E-07 | 3.600E-07 |
| 18957 | 948 | G948T | C | 1.1e-07 | 2.4e-07 | 4.9e-07 | 2.800E-07 | 1.931E-07 | 1.100E-07 | 4.900E-07 |
| 18958 | 948 | G948W | C | 2.5e-07 | 8.5e-08 | 3.1e-07 | 2.150E-07 | 1.165E-07 | 8.500E-08 | 3.100E-07 |
| 18959 | 948 | G948Y | C | 7.5e-07 | 2.7e-07 | 3.4e-07 | 4.533E-07 | 2.593E-07 | 2.700E-07 | 7.500E-07 |
| 18960 | 948 | G948V | C | 2.2e-07 | 2.5e-07 | 3.3e-07 | 2.667E-07 | 5.686E-08 | 2.200E-07 | 3.300E-07 |
| 18961 | 949 | T949A | C | 4.1e-07 | 2e-07 | 2.7e-07 | 2.933E-07 | 1.069E-07 | 2.000E-07 | 4.100E-07 |
| 18962 | 949 | T949R | C | 1.5e-07 | 2.4e-07 | 2.7e-07 | 2.200E-07 | 6.245E-08 | 1.500E-07 | 2.700E-07 |
| 18963 | 949 | T949N | C | 1.8e-07 | 2.8e-07 | 2.3e-07 | 2.300E-07 | 5.000E-08 | 1.800E-07 | 2.800E-07 |
| 18964 | 949 | T949D | C | 2.3e-07 | 3.7e-07 | 4e-07 | 3.333E-07 | 9.074E-08 | 2.300E-07 | 4.000E-07 |
| 18965 | 949 | T949C | C | 1.8e-07 | 1.7e-07 | 1.7e-07 | 1.733E-07 | 5.774E-09 | 1.700E-07 | 1.800E-07 |
| 18966 | 949 | T949E | C | 3.1e-07 | 1.9e-07 | 2.5e-07 | 2.500E-07 | 6.000E-08 | 1.900E-07 | 3.100E-07 |
| 18967 | 949 | T949Q | C | 4.5e-07 | 5.4e-07 | 2e-07 | 3.967E-07 | 1.762E-07 | 2.000E-07 | 5.400E-07 |
| 18968 | 949 | T949G | C | 2.8e-07 | 1.3e-07 | 2.9e-07 | 2.333E-07 | 8.963E-08 | 1.300E-07 | 2.900E-07 |
| 18969 | 949 | T949H | C | 2.8e-07 | 2.7e-07 | 1.7e-07 | 2.400E-07 | 6.083E-08 | 1.700E-07 | 2.800E-07 |
| 18970 | 949 | T949I | C | 2.6e-07 | 2e-07 | 2.6e-07 | 2.400E-07 | 3.464E-08 | 2.000E-07 | 2.600E-07 |
| 18971 | 949 | T949L | C | 3.1e-07 | 2.5e-07 | 1.4e-07 | 2.333E-07 | 8.622E-08 | 1.400E-07 | 3.100E-07 |
| 18972 | 949 | T949K | C | 2.7e-07 | 2.6e-07 | 4.4e-07 | 3.233E-07 | 1.012E-07 | 2.600E-07 | 4.400E-07 |
| 18973 | 949 | T949M | C | 3.1e-07 | 2.6e-07 | 2.5e-07 | 2.733E-07 | 3.215E-08 | 2.500E-07 | 3.100E-07 |
| 18974 | 949 | T949F | C | 3.2e-07 | 1.6e-07 | 5.3e-07 | 3.367E-07 | 1.856E-07 | 1.600E-07 | 5.300E-07 |
| 18975 | 949 | T949P | C | 6e-07 | 3e-07 | 2.9e-07 | 3.967E-07 | 1.762E-07 | 2.900E-07 | 6.000E-07 |
| 18976 | 949 | T949S | C | 2.6e-07 | 1.6e-07 | 3.9e-07 | 2.700E-07 | 1.153E-07 | 1.600E-07 | 3.900E-07 |
| 18977 | 949 | T949T | C | 2e-07 | 1.9e-07 | 1.8e-07 | 1.900E-07 | 1.000E-08 | 1.800E-07 | 2.000E-07 |
| 18978 | 949 | T949W | C | 2.8e-07 | 1.9e-07 | 1.3e-07 | 2.000E-07 | 7.550E-08 | 1.300E-07 | 2.800E-07 |
| 18979 | 949 | T949Y | C | 2.6e-07 | 2.3e-07 | 6.6e-07 | 3.833E-07 | 2.401E-07 | 2.300E-07 | 6.600E-07 |
| 18980 | 949 | T949V | C | 2.3e-07 | 2.7e-07 | 6.4e-07 | 3.800E-07 | 2.261E-07 | 2.300E-07 | 6.400E-07 |
| 18981 | 950 | L950A | C | 1.8e-07 | 2.2e-07 | 2.5e-07 | 2.167E-07 | 3.512E-08 | 1.800E-07 | 2.500E-07 |
| 18982 | 950 | L950R | C | 1.6e-07 | 3.2e-07 | 2.1e-07 | 2.300E-07 | 8.185E-08 | 1.600E-07 | 3.200E-07 |
| 18983 | 950 | L950N | C | 2.6e-07 | 2.9e-07 | 2e-07 | 2.500E-07 | 4.583E-08 | 2.000E-07 | 2.900E-07 |
| 18984 | 950 | L950D | C | 2.9e-07 | 2.5e-07 | 4.3e-07 | 3.233E-07 | 9.452E-08 | 2.500E-07 | 4.300E-07 |
| 18985 | 950 | L950C | C | 1.3e-07 | 3.4e-07 | 3.8e-07 | 2.833E-07 | 1.343E-07 | 1.300E-07 | 3.800E-07 |
| 18986 | 950 | L950E | C | 4e-07 | 3.5e-07 | 1.3e-07 | 2.933E-07 | 1.436E-07 | 1.300E-07 | 4.000E-07 |
| 18987 | 950 | L950Q | C | 2.6e-07 | 2.5e-07 | 1.7e-07 | 2.267E-07 | 4.933E-08 | 1.700E-07 | 2.600E-07 |
| 18988 | 950 | L950G | C | 4.1e-07 | 1.2e-07 | 2.3e-07 | 2.533E-07 | 1.464E-07 | 1.200E-07 | 4.100E-07 |
| 18989 | 950 | L950H | C | 2.4e-07 | 3.6e-07 | 1.4e-07 | 2.467E-07 | 1.102E-07 | 1.400E-07 | 3.600E-07 |
| 18990 | 950 | L950I | C | 1.4e-07 | 1.6e-07 | 1.3e-07 | 1.433E-07 | 1.528E-08 | 1.300E-07 | 1.600E-07 |
| 18991 | 950 | L950L | C | 5e-07 | 1.7e-07 | 2.6e-07 | 3.100E-07 | 1.706E-07 | 1.700E-07 | 5.000E-07 |
| 18992 | 950 | L950K | C | 6.5e-07 | 2.6e-07 | 5.7e-07 | 4.933E-07 | 2.060E-07 | 2.600E-07 | 6.500E-07 |
| 18993 | 950 | L950M | C | 1.3e-07 | 2.5e-07 | 1.6e-07 | 1.800E-07 | 6.245E-08 | 1.300E-07 | 2.500E-07 |
| 18994 | 950 | L950F | C | 6.1e-07 | 6.7e-07 | 2.4e-07 | 5.067E-07 | 2.329E-07 | 2.400E-07 | 6.700E-07 |
| 18995 | 950 | L950P | C | 3.9e-07 | 1.6e-07 | 3.1e-07 | 2.867E-07 | 1.168E-07 | 1.600E-07 | 3.900E-07 |
| 18996 | 950 | L950S | C | 4.8e-07 | 2.3e-07 | 3.4e-07 | 3.500E-07 | 1.253E-07 | 2.300E-07 | 4.800E-07 |
| 18997 | 950 | L950T | C | 2.4e-07 | 1.7e-07 | 1.5e-07 | 1.867E-07 | 4.726E-08 | 1.500E-07 | 2.400E-07 |
| 18998 | 950 | L950W | C | 2.2e-07 | 2.9e-07 | 1.7e-07 | 2.267E-07 | 6.028E-08 | 1.700E-07 | 2.900E-07 |
| 18999 | 950 | L950Y | C | 2.5e-07 | 1.6e-07 | 4.6e-07 | 2.900E-07 | 1.539E-07 | 1.600E-07 | 4.600E-07 |
| 19000 | 950 | L950V | C | 1.7e-07 | 4.1e-07 | 6.1e-07 | 3.967E-07 | 2.203E-07 | 1.700E-07 | 6.100E-07 |
| 19001 | 951 | V951A | C | 1.8e-07 | 4.5e-07 | 4.4e-07 | 3.567E-07 | 1.531E-07 | 1.800E-07 | 4.500E-07 |
| 19002 | 951 | V951R | C | 1.2e-07 | 4.9e-07 | 4e-07 | 3.367E-07 | 1.930E-07 | 1.200E-07 | 4.900E-07 |
| 19003 | 951 | V951N | C | 3.4e-07 | 2.9e-07 | 2.7e-07 | 3.000E-07 | 3.606E-08 | 2.700E-07 | 3.400E-07 |
| 19004 | 951 | V951D | C | 4e-07 | 4e-07 | 2.8e-07 | 3.600E-07 | 6.928E-08 | 2.800E-07 | 4.000E-07 |
| 19005 | 951 | V951C | C | 3.3e-07 | 3.9e-07 | 1.5e-07 | 2.900E-07 | 1.249E-07 | 1.500E-07 | 3.900E-07 |
| 19006 | 951 | V951E | C | 3.2e-07 | 1.6e-07 | 2.9e-07 | 2.567E-07 | 8.505E-08 | 1.600E-07 | 3.200E-07 |
| 19007 | 951 | V951Q | C | 1e-07 | 1.7e-07 | 1.3e-07 | 1.333E-07 | 3.512E-08 | 1.000E-07 | 1.700E-07 |
| 19008 | 951 | V951G | C | 2.1e-07 | 1.6e-07 | 1.6e-07 | 1.767E-07 | 2.887E-08 | 1.600E-07 | 2.100E-07 |
| 19009 | 951 | V951H | C | 2.9e-07 | 4e-07 | 3.1e-07 | 3.333E-07 | 5.859E-08 | 2.900E-07 | 4.000E-07 |
| 19010 | 951 | V951I | C | 5.8e-07 | 2.9e-07 | 2.8e-07 | 3.833E-07 | 1.704E-07 | 2.800E-07 | 5.800E-07 |
| 19011 | 951 | V951L | C | 4e-07 | 4.2e-07 | 2e-07 | 3.400E-07 | 1.217E-07 | 2.000E-07 | 4.200E-07 |
| 19012 | 951 | V951K | C | 1.7e-07 | 4e-07 | 1.5e-07 | 2.400E-07 | 1.389E-07 | 1.500E-07 | 4.000E-07 |
| 19013 | 951 | V951M | C | 2.4e-07 | 4.1e-07 | 3.9e-07 | 3.467E-07 | 9.292E-08 | 2.400E-07 | 4.100E-07 |
| 19014 | 951 | V951F | C | 2e-07 | 2.2e-07 | 5.5e-07 | 3.233E-07 | 1.966E-07 | 2.000E-07 | 5.500E-07 |
| 19015 | 951 | V951P | C | 2.8e-07 | 1.9e-07 | 4.1e-07 | 2.933E-07 | 1.106E-07 | 1.900E-07 | 4.100E-07 |
| 19016 | 951 | V951S | C | 1.8e-07 | 2.8e-07 | 3.8e-07 | 2.800E-07 | 1.000E-07 | 1.800E-07 | 3.800E-07 |
| 19017 | 951 | V951T | C | 2.8e-07 | 2e-07 | 1.2e-07 | 2.000E-07 | 8.000E-08 | 1.200E-07 | 2.800E-07 |
| 19018 | 951 | V951W | C | 2.5e-07 | 3e-07 | 4.6e-07 | 3.367E-07 | 1.097E-07 | 2.500E-07 | 4.600E-07 |
| 19019 | 951 | V951Y | C | 2.3e-07 | 2.1e-07 | 4.2e-07 | 2.867E-07 | 1.159E-07 | 2.100E-07 | 4.200E-07 |
| 19020 | 951 | V951V | C | 2.9e-07 | 2e-07 | 2.5e-07 | 2.467E-07 | 4.509E-08 | 2.000E-07 | 2.900E-07 |
| 19021 | 952 | T952A | C | 2.9e-07 | 2.5e-07 | 5.4e-07 | 3.600E-07 | 1.572E-07 | 2.500E-07 | 5.400E-07 |
| 19022 | 952 | T952R | C | 4.8e-07 | 3.3e-07 | 1.1e-07 | 3.067E-07 | 1.861E-07 | 1.100E-07 | 4.800E-07 |
| 19023 | 952 | T952N | C | 3.3e-07 | 2.5e-07 | 3.7e-07 | 3.167E-07 | 6.110E-08 | 2.500E-07 | 3.700E-07 |
| 19024 | 952 | T952D | C | 2.9e-07 | 2.7e-07 | 2.9e-07 | 2.833E-07 | 1.155E-08 | 2.700E-07 | 2.900E-07 |
| 19025 | 952 | T952C | C | 3.4e-07 | 2.9e-07 | 2.9e-07 | 3.067E-07 | 2.887E-08 | 2.900E-07 | 3.400E-07 |
| 19026 | 952 | T952E | C | 3.8e-07 | 9.3e-07 | 1.6e-07 | 4.900E-07 | 3.966E-07 | 1.600E-07 | 9.300E-07 |
| 19027 | 952 | T952Q | C | 2.8e-07 | 1.8e-07 | 2.1e-07 | 2.233E-07 | 5.132E-08 | 1.800E-07 | 2.800E-07 |
| 19028 | 952 | T952G | C | 3e-07 | 2.1e-07 | 2.5e-07 | 2.533E-07 | 4.509E-08 | 2.100E-07 | 3.000E-07 |
| 19029 | 952 | T952H | C | 1.5e-07 | 5.3e-07 | 2.7e-07 | 3.167E-07 | 1.943E-07 | 1.500E-07 | 5.300E-07 |
| 19030 | 952 | T952I | C | 2.6e-07 | 2.8e-07 | 2.9e-07 | 2.767E-07 | 1.528E-08 | 2.600E-07 | 2.900E-07 |
| 19031 | 952 | T952L | C | 2e-07 | 4.4e-07 | 2.9e-07 | 3.100E-07 | 1.212E-07 | 2.000E-07 | 4.400E-07 |
| 19032 | 952 | T952K | C | 3.2e-07 | 2.5e-07 | 5.8e-07 | 3.833E-07 | 1.739E-07 | 2.500E-07 | 5.800E-07 |
| 19033 | 952 | T952M | C | 1.2e-07 | 1.5e-07 | 2.4e-07 | 1.700E-07 | 6.245E-08 | 1.200E-07 | 2.400E-07 |
| 19034 | 952 | T952F | C | 3.1e-07 | 1.5e-07 | 2.5e-07 | 2.367E-07 | 8.083E-08 | 1.500E-07 | 3.100E-07 |
| 19035 | 952 | T952P | C | 2e-07 | 2.4e-07 | 2.2e-07 | 2.200E-07 | 2.000E-08 | 2.000E-07 | 2.400E-07 |
| 19036 | 952 | T952S | C | 3.8e-07 | 1.4e-07 | 3.9e-07 | 3.033E-07 | 1.415E-07 | 1.400E-07 | 3.900E-07 |
| 19037 | 952 | T952T | C | 3.2e-07 | 3e-07 | 5.6e-07 | 3.933E-07 | 1.447E-07 | 3.000E-07 | 5.600E-07 |
| 19038 | 952 | T952W | C | 1.8e-07 | 2.9e-07 | 1.4e-07 | 2.033E-07 | 7.767E-08 | 1.400E-07 | 2.900E-07 |
| 19039 | 952 | T952Y | C | 3.4e-07 | 2.4e-07 | 5.1e-07 | 3.633E-07 | 1.365E-07 | 2.400E-07 | 5.100E-07 |

| | | | | | | | | | | |
|---|---|---|---|---|---|---|---|---|---|---|
| 19040 | 952 | T952V | C | 1.9e-07 | 1.7e-07 | 4.5e-07 | 2.700E-07 | 1.562E-07 | 1.700E-07 | 4.500E-07 |
| 19041 | 953 | V953A | C | 2.4e-07 | 2.3e-07 | 1.9e-07 | 2.200E-07 | 2.646E-08 | 1.900E-07 | 2.400E-07 |
| 19042 | 953 | V953R | C | 2.2e-07 | 1.9e-07 | 4.5e-07 | 2.867E-07 | 1.422E-07 | 1.900E-07 | 4.500E-07 |
| 19043 | 953 | V953N | C | 2.1e-07 | 1.5e-07 | 4.6e-07 | 2.733E-07 | 1.644E-07 | 1.500E-07 | 4.600E-07 |
| 19044 | 953 | V953D | C | 1.7e-07 | 4.9e-07 | 4.1e-07 | 3.567E-07 | 1.665E-07 | 1.700E-07 | 4.900E-07 |
| 19045 | 953 | V953C | C | 2.1e-07 | 2.3e-07 | 2.4e-07 | 2.267E-07 | 1.528E-08 | 2.100E-07 | 2.400E-07 |
| 19046 | 953 | V953E | C | 3.6e-07 | 2.1e-07 | 2.6e-07 | 2.767E-07 | 7.638E-08 | 2.100E-07 | 3.600E-07 |
| 19047 | 953 | V953Q | C | 2.3e-07 | 3.4e-07 | 2.3e-07 | 2.667E-07 | 6.351E-08 | 2.300E-07 | 3.400E-07 |
| 19048 | 953 | V953G | C | 2.4e-07 | 4.5e-07 | 2.2e-07 | 3.033E-07 | 1.274E-07 | 2.200E-07 | 4.500E-07 |
| 19049 | 953 | V953H | C | 5.2e-07 | 3.2e-07 | 2.3e-07 | 3.567E-07 | 1.484E-07 | 2.300E-07 | 5.200E-07 |
| 19050 | 953 | V953I | C | 2.1e-07 | 3.8e-07 | 2.6e-07 | 2.833E-07 | 8.737E-08 | 2.100E-07 | 3.800E-07 |
| 19051 | 953 | V953L | C | 9.4e-08 | 2.1e-07 | 4.9e-07 | 2.647E-07 | 2.036E-07 | 9.400E-08 | 4.900E-07 |
| 19052 | 953 | V953K | C | 1.2e-07 | 1.2e-07 | 1.8e-07 | 1.400E-07 | 3.464E-08 | 1.200E-07 | 1.800E-07 |
| 19053 | 953 | V953M | C | 2.3e-07 | 3.9e-07 | 4.4e-07 | 3.533E-07 | 1.097E-07 | 2.300E-07 | 4.400E-07 |
| 19054 | 953 | V953F | C | 1.6e-07 | 2.7e-07 | 2.4e-07 | 2.233E-07 | 5.686E-08 | 1.600E-07 | 2.700E-07 |
| 19055 | 953 | V953P | C | 1.9e-07 | 9e-08 | 2e-07 | 1.600E-07 | 6.083E-08 | 9.000E-08 | 2.000E-07 |
| 19056 | 953 | V953S | C | 2e-07 | 1.5e-07 | 2.4e-07 | 1.967E-07 | 4.509E-08 | 1.500E-07 | 2.400E-07 |
| 19057 | 953 | V953T | C | 3e-07 | 2.5e-07 | 1.7e-07 | 2.400E-07 | 6.557E-08 | 1.700E-07 | 3.000E-07 |
| 19058 | 953 | V953W | C | 3.8e-07 | 2.6e-07 | 2.3e-07 | 2.900E-07 | 7.937E-08 | 2.300E-07 | 3.800E-07 |
| 19059 | 953 | V953Y | C | 2e-07 | 1.6e-07 | 3.5e-07 | 2.367E-07 | 1.002E-07 | 1.600E-07 | 3.500E-07 |
| 19060 | 953 | V953V | C | 3.8e-07 | 4.4e-07 | 3.4e-07 | 3.867E-07 | 5.033E-08 | 3.400E-07 | 4.400E-07 |
| 19061 | 954 | S954A | C | 1.9e-07 | 4.3e-07 | 1.8e-07 | 2.667E-07 | 1.415E-07 | 1.800E-07 | 4.300E-07 |
| 19062 | 954 | S954R | C | 2.4e-07 | 2.5e-07 | 4.5e-07 | 3.133E-07 | 1.185E-07 | 2.400E-07 | 4.500E-07 |
| 19063 | 954 | S954N | C | 2.1e-07 | 1.7e-07 | 1.8e-07 | 1.867E-07 | 2.082E-08 | 1.700E-07 | 2.100E-07 |
| 19064 | 954 | S954D | C | 2.5e-07 | 3e-07 | 3.3e-07 | 2.933E-07 | 4.041E-08 | 2.500E-07 | 3.300E-07 |
| 19065 | 954 | S954C | C | 2e-07 | 2.6e-07 | 3.3e-07 | 2.633E-07 | 6.506E-08 | 2.000E-07 | 3.300E-07 |
| 19066 | 954 | S954E | C | 3.2e-07 | 2.9e-07 | 2.1e-07 | 2.733E-07 | 5.686E-08 | 2.100E-07 | 3.200E-07 |
| 19067 | 954 | S954Q | C | 3.5e-07 | 3.4e-07 | 3.1e-07 | 3.333E-07 | 2.082E-08 | 3.100E-07 | 3.500E-07 |
| 19068 | 954 | S954G | C | 3.1e-07 | 2.6e-07 | 3e-07 | 2.900E-07 | 2.646E-08 | 2.600E-07 | 3.100E-07 |
| 19069 | 954 | S954H | C | 1.7e-07 | 2.7e-07 | 1.2e-07 | 1.867E-07 | 7.638E-08 | 1.200E-07 | 2.700E-07 |
| 19070 | 954 | S954I | C | 1.6e-07 | 2.4e-07 | 3.6e-07 | 2.533E-07 | 1.007E-07 | 1.600E-07 | 3.600E-07 |
| 19071 | 954 | S954L | C | 3e-07 | 3.2e-07 | 1e-07 | 2.400E-07 | 1.217E-07 | 1.000E-07 | 3.200E-07 |
| 19072 | 954 | S954K | C | 3.6e-07 | 2.1e-07 | 1.8e-07 | 2.500E-07 | 9.644E-08 | 1.800E-07 | 3.600E-07 |
| 19073 | 954 | S954M | C | 4.5e-07 | 4.7e-07 | 2.4e-07 | 3.867E-07 | 1.274E-07 | 2.400E-07 | 4.700E-07 |
| 19074 | 954 | S954F | C | 2e-07 | 1.4e-07 | 4.6e-07 | 2.667E-07 | 1.701E-07 | 1.400E-07 | 4.600E-07 |
| 19075 | 954 | S954P | C | 3.5e-07 | 1.7e-07 | 6.2e-07 | 3.800E-07 | 2.265E-07 | 1.700E-07 | 6.200E-07 |
| 19076 | 954 | S954S | C | 2.9e-07 | 3e-07 | 2.4e-07 | 2.767E-07 | 3.215E-08 | 2.400E-07 | 3.000E-07 |
| 19077 | 954 | S954T | C | 1.6e-07 | 1.6e-07 | 3.9e-07 | 2.367E-07 | 1.328E-07 | 1.600E-07 | 3.900E-07 |
| 19078 | 954 | S954W | C | 2.6e-07 | 1.4e-07 | 4.6e-07 | 2.867E-07 | 1.617E-07 | 1.400E-07 | 4.600E-07 |
| 19079 | 954 | S954Y | C | 2.2e-07 | 3.5e-07 | 4.2e-07 | 3.300E-07 | 1.015E-07 | 2.200E-07 | 4.200E-07 |
| 19080 | 954 | S954V | C | 2.1e-07 | 5.2e-07 | 4.6e-07 | 3.967E-07 | 1.644E-07 | 2.100E-07 | 5.200E-07 |
| 19081 | 955 | S955A | C | 7.5e-08 | 2.9e-07 | 1.9e-07 | 1.850E-07 | 1.076E-07 | 7.500E-08 | 2.900E-07 |
| 19082 | 955 | S955R | C | 1.2e-07 | 4e-07 | 2.8e-07 | 2.667E-07 | 1.405E-07 | 1.200E-07 | 4.000E-07 |
| 19083 | 955 | S955N | C | 5.2e-07 | 2.3e-07 | 2.6e-07 | 3.367E-07 | 1.595E-07 | 2.300E-07 | 5.200E-07 |
| 19084 | 955 | S955D | C | 1.7e-07 | 5.4e-07 | 1.5e-07 | 2.867E-07 | 2.196E-07 | 1.500E-07 | 5.400E-07 |
| 19085 | 955 | S955C | C | 1.5e-07 | 1.5e-07 | 2.3e-07 | 1.767E-07 | 4.619E-08 | 1.500E-07 | 2.300E-07 |
| 19086 | 955 | S955E | C | 8.8e-08 | 3e-07 | 2.3e-07 | 2.060E-07 | 1.080E-07 | 8.800E-08 | 3.000E-07 |
| 19087 | 955 | S955Q | C | 2.2e-07 | 9.3e-08 | 4.9e-07 | 2.677E-07 | 2.027E-07 | 9.300E-08 | 4.900E-07 |
| 19088 | 955 | S955G | C | 2.4e-07 | 1.9e-07 | 2.9e-07 | 2.400E-07 | 5.000E-08 | 1.900E-07 | 2.900E-07 |
| 19089 | 955 | S955H | C | 2.4e-07 | 4.6e-07 | 2.8e-07 | 3.267E-07 | 1.172E-07 | 2.400E-07 | 4.600E-07 |
| 19090 | 955 | S955I | C | 2.2e-07 | 2.5e-07 | 3.8e-07 | 2.833E-07 | 8.505E-08 | 2.200E-07 | 3.800E-07 |
| 19091 | 955 | S955L | C | 1.2e-07 | 5.9e-07 | 2.9e-07 | 3.333E-07 | 2.380E-07 | 1.200E-07 | 5.900E-07 |
| 19092 | 955 | S955K | C | 2.5e-07 | 3.4e-07 | 1.9e-07 | 2.600E-07 | 7.550E-08 | 1.900E-07 | 3.400E-07 |
| 19093 | 955 | S955M | C | 1.6e-07 | 3.1e-07 | 6.5e-08 | 1.783E-07 | 1.235E-07 | 6.500E-08 | 3.100E-07 |
| 19094 | 955 | S955F | C | 1.1e-07 | 3.9e-07 | 4.6e-07 | 3.200E-07 | 1.852E-07 | 1.100E-07 | 4.600E-07 |
| 19095 | 955 | S955P | C | 2.4e-07 | 1.7e-07 | 2.6e-07 | 2.233E-07 | 4.726E-08 | 1.700E-07 | 2.600E-07 |
| 19096 | 955 | S955S | C | 4e-07 | 2e-07 | 2e-07 | 2.667E-07 | 1.155E-07 | 2.000E-07 | 4.000E-07 |
| 19097 | 955 | S955T | C | 1.4e-07 | 2.3e-07 | 2.3e-07 | 2.000E-07 | 5.196E-08 | 1.400E-07 | 2.300E-07 |
| 19098 | 955 | S955W | C | 1.6e-07 | 2.4e-07 | 4e-07 | 2.667E-07 | 1.222E-07 | 1.600E-07 | 4.000E-07 |
| 19099 | 955 | S955Y | C | 2.9e-07 | 2.8e-07 | 3.6e-07 | 3.100E-07 | 4.359E-08 | 2.800E-07 | 3.600E-07 |
| 19100 | 955 | S955V | C | 1.6e-07 | 4.7e-07 | 4.7e-07 | 3.667E-07 | 1.790E-07 | 1.600E-07 | 4.700E-07 |
| 19101 | 956 | A956A | C | 1.6e-07 | 2e-07 | 2.3e-07 | 1.967E-07 | 3.512E-08 | 1.600E-07 | 2.300E-07 |
| 19102 | 956 | A956R | C | 2e-07 | 1.8e-07 | 2.4e-07 | 2.067E-07 | 3.055E-08 | 1.800E-07 | 2.400E-07 |
| 19103 | 956 | A956N | C | 1e-07 | 2e-07 | 1.3e-07 | 1.433E-07 | 5.132E-08 | 1.000E-07 | 2.000E-07 |
| 19104 | 956 | A956D | C | 2.4e-07 | 2.7e-07 | 1.1e-07 | 2.067E-07 | 8.505E-08 | 1.100E-07 | 2.700E-07 |
| 19105 | 956 | A956C | C | 3e-07 | 1.7e-07 | 3.1e-07 | 2.600E-07 | 7.810E-08 | 1.700E-07 | 3.100E-07 |
| 19106 | 956 | A956E | C | 1.4e-07 | 2e-07 | 6.9e-08 | 1.363E-07 | 6.558E-08 | 6.900E-08 | 2.000E-07 |
| 19107 | 956 | A956Q | C | 2.3e-07 | 2.4e-07 | 5.4e-07 | 3.367E-07 | 1.762E-07 | 2.300E-07 | 5.400E-07 |
| 19108 | 956 | A956G | C | 2.3e-07 | 3.8e-07 | 3.4e-07 | 3.167E-07 | 7.767E-08 | 2.300E-07 | 3.800E-07 |
| 19109 | 956 | A956H | C | 3.2e-07 | 8.4e-08 | 1e-07 | 1.680E-07 | 1.319E-07 | 8.400E-08 | 3.200E-07 |
| 19110 | 956 | A956I | C | 3.3e-07 | 3.9e-07 | 2.1e-07 | 3.100E-07 | 9.165E-08 | 2.100E-07 | 3.900E-07 |
| 19111 | 956 | A956L | C | 4.2e-07 | 2.4e-07 | 2.5e-07 | 3.033E-07 | 1.012E-07 | 2.400E-07 | 4.200E-07 |
| 19112 | 956 | A956K | C | 5.4e-07 | 3.2e-07 | 3e-07 | 3.867E-07 | 1.332E-07 | 3.000E-07 | 5.400E-07 |
| 19113 | 956 | A956M | C | 1.2e-07 | 3.2e-07 | 2.3e-07 | 2.233E-07 | 1.002E-07 | 1.200E-07 | 3.200E-07 |
| 19114 | 956 | A956F | C | 3.3e-07 | 2.7e-07 | 2.5e-07 | 2.833E-07 | 4.163E-08 | 2.500E-07 | 3.300E-07 |
| 19115 | 956 | A956P | C | 2.5e-07 | 1.5e-07 | 5.3e-07 | 3.100E-07 | 1.970E-07 | 1.500E-07 | 5.300E-07 |
| 19116 | 956 | A956S | C | 2.6e-07 | 2.6e-07 | 4.1e-07 | 3.100E-07 | 8.660E-08 | 2.600E-07 | 4.100E-07 |
| 19117 | 956 | A956T | C | 3.3e-07 | 1.2e-07 | 9.8e-08 | 1.827E-07 | 1.281E-07 | 9.800E-08 | 3.300E-07 |
| 19118 | 956 | A956W | C | 1.6e-07 | 4.1e-07 | 2.6e-07 | 2.767E-07 | 1.258E-07 | 1.600E-07 | 4.100E-07 |
| 19119 | 956 | A956Y | C | 3.7e-07 | 3.2e-07 | 2.4e-07 | 3.100E-07 | 6.557E-08 | 2.400E-07 | 3.700E-07 |
| 19120 | 956 | A956V | C | 2.2e-07 | 2.1e-07 | 1.7e-07 | 2.000E-07 | 2.646E-08 | 1.700E-07 | 2.200E-07 |
| 19121 | 957 | S957A | C | 2.7e-07 | 4.4e-07 | 5.2e-07 | 4.100E-07 | 1.277E-07 | 2.700E-07 | 5.200E-07 |
| 19122 | 957 | S957R | C | 9.6e-08 | 3.2e-07 | 1.5e-07 | 1.887E-07 | 1.169E-07 | 9.600E-08 | 3.200E-07 |
| 19123 | 957 | S957N | C | 3.9e-07 | 3.6e-07 | 2e-07 | 3.167E-07 | 1.021E-07 | 2.000E-07 | 3.900E-07 |
| 19124 | 957 | S957D | C | 4e-07 | 4.3e-07 | 4e-07 | 4.100E-07 | 1.732E-08 | 4.000E-07 | 4.300E-07 |

| | | | | | | | | | | |
|---|---|---|---|---|---|---|---|---|---|---|
| 19125 | 957 | S957C | C | 2e-07 | 3.2e-07 | 1.3e-07 | 2.167E-07 | 9.609E-08 | 1.300E-07 | 3.200E-07 |
| 19126 | 957 | S957E | C | 2e-07 | 4.7e-07 | 2.4e-07 | 3.033E-07 | 1.457E-07 | 2.000E-07 | 4.700E-07 |
| 19127 | 957 | S957Q | C | 2.3e-07 | 1.6e-07 | 1.9e-07 | 1.933E-07 | 3.512E-08 | 1.600E-07 | 2.300E-07 |
| 19128 | 957 | S957G | C | 2.1e-07 | 3.6e-07 | 1.6e-07 | 2.433E-07 | 1.041E-07 | 1.600E-07 | 3.600E-07 |
| 19129 | 957 | S957H | C | 2.9e-07 | 5.6e-07 | 1.8e-07 | 3.433E-07 | 1.955E-07 | 1.800E-07 | 5.600E-07 |
| 19130 | 957 | S957I | C | 2.3e-07 | 3.4e-07 | 2.8e-07 | 2.833E-07 | 5.508E-08 | 2.300E-07 | 3.400E-07 |
| 19131 | 957 | S957L | C | 2.6e-07 | 2.9e-07 | 1.9e-07 | 2.467E-07 | 5.132E-08 | 1.900E-07 | 2.900E-07 |
| 19132 | 957 | S957K | C | 4.2e-07 | 4.7e-07 | 3e-07 | 3.967E-07 | 8.737E-08 | 3.000E-07 | 4.700E-07 |
| 19133 | 957 | S957M | C | 4.1e-07 | 9.5e-07 | 1.9e-07 | 5.167E-07 | 3.911E-07 | 1.900E-07 | 9.500E-07 |
| 19134 | 957 | S957F | C | 4.7e-07 | 3.8e-07 | 1.9e-07 | 3.467E-07 | 1.429E-07 | 1.900E-07 | 4.700E-07 |
| 19135 | 957 | S957P | C | 1.5e-07 | 2.8e-07 | 2.9e-07 | 2.400E-07 | 7.810E-08 | 1.500E-07 | 2.900E-07 |
| 19136 | 957 | S957S | C | 4.2e-07 | 1.9e-07 | 3e-07 | 3.033E-07 | 1.150E-07 | 1.900E-07 | 4.200E-07 |
| 19137 | 957 | S957T | C | 5.2e-07 | 2e-07 | 2.6e-07 | 3.267E-07 | 1.701E-07 | 2.000E-07 | 5.200E-07 |
| 19138 | 957 | S957W | C | 1.5e-07 | 1.3e-07 | 1.6e-07 | 1.467E-07 | 1.528E-08 | 1.300E-07 | 1.600E-07 |
| 19139 | 957 | S957Y | C | 2.6e-07 | 2.1e-07 | 1.7e-07 | 2.133E-07 | 4.509E-08 | 1.700E-07 | 2.600E-07 |
| 19140 | 957 | S957V | C | 2.1e-07 | 4.3e-07 | 3e-07 | 3.133E-07 | 1.106E-07 | 2.100E-07 | 4.300E-07 |
| 19141 | 958 | T958A | C | 1.8e-07 | 1.6e-07 | 1.9e-07 | 1.767E-07 | 1.528E-08 | 1.600E-07 | 1.900E-07 |
| 19142 | 958 | T958R | C | 1.6e-07 | 4.2e-07 | 3.7e-07 | 3.167E-07 | 1.380E-07 | 1.600E-07 | 4.200E-07 |
| 19143 | 958 | T958N | C | 2.2e-07 | 2.7e-07 | 2.2e-07 | 2.367E-07 | 2.887E-08 | 2.200E-07 | 2.700E-07 |
| 19144 | 958 | T958D | C | 4.5e-07 | 3.2e-07 | 2.2e-07 | 3.300E-07 | 1.153E-07 | 2.200E-07 | 4.500E-07 |
| 19145 | 958 | T958C | C | 3.3e-07 | 2.6e-07 | 1.9e-07 | 2.600E-07 | 7.000E-08 | 1.900E-07 | 3.300E-07 |
| 19146 | 958 | T958E | C | 5.4e-07 | 3.5e-07 | 2e-07 | 3.633E-07 | 1.704E-07 | 2.000E-07 | 5.400E-07 |
| 19147 | 958 | T958Q | C | 5.1e-07 | 4.6e-07 | 1.8e-07 | 3.833E-07 | 1.779E-07 | 1.800E-07 | 5.100E-07 |
| 19148 | 958 | T958G | C | 1.9e-07 | 3.1e-07 | 1.3e-07 | 2.100E-07 | 9.165E-08 | 1.300E-07 | 3.100E-07 |
| 19149 | 958 | T958H | C | 3.5e-07 | 4.1e-07 | 4.3e-07 | 3.967E-07 | 4.163E-08 | 3.500E-07 | 4.300E-07 |
| 19150 | 958 | T958I | C | 3.5e-07 | 2.6e-07 | 2.4e-07 | 2.833E-07 | 5.859E-08 | 2.400E-07 | 3.500E-07 |
| 19151 | 958 | T958L | C | 3.8e-07 | 3.2e-07 | 1.5e-07 | 2.833E-07 | 1.193E-07 | 1.500E-07 | 3.800E-07 |
| 19152 | 958 | T958K | C | 2.1e-07 | 5.8e-07 | 1.2e-07 | 3.033E-07 | 2.438E-07 | 1.200E-07 | 5.800E-07 |
| 19153 | 958 | T958M | C | 2.4e-07 | 1.6e-07 | 1.8e-07 | 1.933E-07 | 4.163E-08 | 1.600E-07 | 2.400E-07 |
| 19154 | 958 | T958F | C | 3.3e-07 | 2.9e-07 | 2.1e-07 | 2.767E-07 | 6.110E-08 | 2.100E-07 | 3.300E-07 |
| 19155 | 958 | T958P | C | 2.5e-07 | 1.6e-07 | 1.7e-07 | 1.933E-07 | 4.933E-08 | 1.600E-07 | 2.500E-07 |
| 19156 | 958 | T958S | C | 1.9e-07 | 2e-07 | 1.7e-07 | 1.867E-07 | 1.528E-08 | 1.700E-07 | 2.000E-07 |
| 19157 | 958 | T958T | C | 3.4e-07 | 1.7e-07 | 1.6e-07 | 2.233E-07 | 1.012E-07 | 1.600E-07 | 3.400E-07 |
| 19158 | 958 | T958W | C | 3.2e-07 | 1.7e-07 | 2.4e-07 | 2.433E-07 | 7.506E-08 | 1.700E-07 | 3.200E-07 |
| 19159 | 958 | T958Y | C | 2.6e-07 | 2.2e-07 | 2.5e-07 | 2.433E-07 | 2.082E-08 | 2.200E-07 | 2.600E-07 |
| 19160 | 958 | T958V | C | 3.8e-07 | 1.5e-07 | 1e-07 | 2.100E-07 | 1.493E-07 | 1.000E-07 | 3.800E-07 |
| 19161 | 959 | K959A | C | 2.1e-07 | 3.4e-07 | 1.4e-07 | 2.300E-07 | 1.015E-07 | 1.400E-07 | 3.400E-07 |
| 19162 | 959 | K959R | C | 4.9e-07 | 3.4e-07 | 2.2e-07 | 3.500E-07 | 1.353E-07 | 2.200E-07 | 4.900E-07 |
| 19163 | 959 | K959N | C | 5.2e-07 | 2.4e-07 | 2.3e-07 | 3.300E-07 | 1.646E-07 | 2.300E-07 | 5.200E-07 |
| 19164 | 959 | K959D | C | 2.5e-07 | 2.7e-07 | 1.5e-07 | 2.233E-07 | 6.429E-08 | 1.500E-07 | 2.700E-07 |
| 19165 | 959 | K959C | C | 2.9e-07 | 2e-07 | 4.6e-07 | 3.167E-07 | 1.320E-07 | 2.000E-07 | 4.600E-07 |
| 19166 | 959 | K959E | C | 7.7e-07 | 2.5e-07 | 1.5e-07 | 3.900E-07 | 3.329E-07 | 1.500E-07 | 7.700E-07 |
| 19167 | 959 | K959Q | C | 3e-07 | 1.9e-07 | 1.8e-07 | 2.233E-07 | 6.658E-08 | 1.800E-07 | 3.000E-07 |
| 19168 | 959 | K959G | C | 3.6e-07 | 1.6e-07 | 2.4e-07 | 2.533E-07 | 1.007E-07 | 1.600E-07 | 3.600E-07 |
| 19169 | 959 | K959H | C | 1e-07 | 1.2e-07 | 3.1e-07 | 1.767E-07 | 1.159E-07 | 1.000E-07 | 3.100E-07 |
| 19170 | 959 | K959I | C | 1.2e-07 | 1.6e-07 | 2.2e-07 | 1.667E-07 | 5.033E-08 | 1.200E-07 | 2.200E-07 |
| 19171 | 959 | K959L | C | 3.5e-07 | 1.5e-07 | 2.2e-07 | 2.400E-07 | 1.015E-07 | 1.500E-07 | 3.500E-07 |
| 19172 | 959 | K959K | C | 1.4e-07 | 2.6e-07 | 2.3e-07 | 2.100E-07 | 6.245E-08 | 1.400E-07 | 2.600E-07 |
| 19173 | 959 | K959M | C | 1.1e-07 | 4.3e-07 | 7.8e-07 | 4.400E-07 | 3.351E-07 | 1.100E-07 | 7.800E-07 |
| 19174 | 959 | K959F | C | 1.6e-07 | 1.8e-07 | 2.3e-07 | 1.900E-07 | 3.606E-08 | 1.600E-07 | 2.300E-07 |
| 19175 | 959 | K959P | C | 3.1e-07 | 2.9e-07 | 2.6e-07 | 2.867E-07 | 2.517E-08 | 2.600E-07 | 3.100E-07 |
| 19176 | 959 | K959S | C | 1.9e-07 | 1.9e-07 | 4e-07 | 2.600E-07 | 1.212E-07 | 1.900E-07 | 4.000E-07 |
| 19177 | 959 | K959T | C | 2.3e-07 | 3.5e-07 | 1.8e-07 | 2.533E-07 | 8.737E-08 | 1.800E-07 | 3.500E-07 |
| 19178 | 959 | K959W | C | 5e-07 | 2.1e-07 | 1.9e-07 | 3.000E-07 | 1.735E-07 | 1.900E-07 | 5.000E-07 |
| 19179 | 959 | K959Y | C | 7.1e-07 | 2.8e-07 | 1.6e-07 | 3.833E-07 | 2.892E-07 | 1.600E-07 | 7.100E-07 |
| 19180 | 959 | K959V | C | 3.7e-07 | 1.6e-07 | 2.5e-07 | 2.600E-07 | 1.054E-07 | 1.600E-07 | 3.700E-07 |
| 19181 | 960 | G960A | C | 2.3e-07 | 2.9e-07 | 4.3e-07 | 3.167E-07 | 1.026E-07 | 2.300E-07 | 4.300E-07 |
| 19182 | 960 | G960R | C | 1.2e-07 | 3.4e-07 | 1.7e-07 | 2.100E-07 | 1.153E-07 | 1.200E-07 | 3.400E-07 |
| 19183 | 960 | G960N | C | 3.6e-07 | 4.8e-07 | 3.5e-07 | 3.967E-07 | 7.234E-08 | 3.500E-07 | 4.800E-07 |
| 19184 | 960 | G960D | C | 1.9e-07 | 2e-07 | 4.6e-07 | 2.833E-07 | 1.531E-07 | 1.900E-07 | 4.600E-07 |
| 19185 | 960 | G960C | C | 2.8e-07 | 1.2e-07 | 3e-07 | 2.333E-07 | 9.866E-08 | 1.200E-07 | 3.000E-07 |
| 19186 | 960 | G960E | C | 6.7e-07 | 2.2e-07 | 2.1e-07 | 3.667E-07 | 2.627E-07 | 2.100E-07 | 6.700E-07 |
| 19187 | 960 | G960Q | C | 2.9e-07 | 4.2e-07 | 3.4e-07 | 3.500E-07 | 6.557E-08 | 2.900E-07 | 4.200E-07 |
| 19188 | 960 | G960G | C | 7.4e-08 | 2.4e-07 | 1.6e-07 | 1.580E-07 | 8.302E-08 | 7.400E-08 | 2.400E-07 |
| 19189 | 960 | G960H | C | 1.7e-07 | 2.6e-07 | 3.5e-07 | 2.600E-07 | 9.000E-08 | 1.700E-07 | 3.500E-07 |
| 19190 | 960 | G960I | C | 1.8e-07 | 2.2e-07 | 3e-07 | 2.333E-07 | 6.110E-08 | 1.800E-07 | 3.000E-07 |
| 19191 | 960 | G960L | C | 2.9e-07 | 1.1e-07 | 2.9e-07 | 2.300E-07 | 1.039E-07 | 1.100E-07 | 2.900E-07 |
| 19192 | 960 | G960K | C | 3.5e-07 | 2.3e-07 | 1.8e-07 | 2.533E-07 | 8.737E-08 | 1.800E-07 | 3.500E-07 |
| 19193 | 960 | G960M | C | 5.2e-07 | 1.5e-07 | 4.1e-07 | 3.600E-07 | 1.900E-07 | 1.500E-07 | 5.200E-07 |
| 19194 | 960 | G960F | C | 6.4e-07 | 9.8e-08 | 6.7e-07 | 4.693E-07 | 3.219E-07 | 9.800E-08 | 6.700E-07 |
| 19195 | 960 | G960P | C | 3.1e-07 | 2.6e-07 | 1.6e-07 | 2.433E-07 | 7.638E-08 | 1.600E-07 | 3.100E-07 |
| 19196 | 960 | G960S | C | 3.1e-07 | 2e-07 | 3.7e-07 | 2.933E-07 | 8.622E-08 | 2.000E-07 | 3.700E-07 |
| 19197 | 960 | G960T | C | 1.3e-07 | 1.9e-07 | 3.2e-07 | 2.133E-07 | 9.713E-08 | 1.300E-07 | 3.200E-07 |
| 19198 | 960 | G960W | C | 3.3e-07 | 2.6e-07 | 4.5e-07 | 3.467E-07 | 9.609E-08 | 2.600E-07 | 4.500E-07 |
| 19199 | 960 | G960Y | C | 1.4e-07 | 3.2e-07 | 2.2e-07 | 2.267E-07 | 9.018E-08 | 1.400E-07 | 3.200E-07 |
| 19200 | 960 | G960V | C | 2e-07 | 2.3e-07 | 2.5e-07 | 2.267E-07 | 2.517E-08 | 2.000E-07 | 2.500E-07 |
| 19201 | 961 | P961A | C | 3.3e-07 | 2.4e-07 | 2.6e-07 | 2.767E-07 | 4.726E-08 | 2.400E-07 | 3.300E-07 |
| 19202 | 961 | P961R | C | 1.4e-07 | 6.4e-07 | 1.4e-07 | 3.067E-07 | 2.887E-07 | 1.400E-07 | 6.400E-07 |
| 19203 | 961 | P961N | C | 3.9e-07 | 1.8e-07 | 4.5e-07 | 3.400E-07 | 1.418E-07 | 1.800E-07 | 4.500E-07 |
| 19204 | 961 | P961D | C | 2.4e-07 | 3.5e-07 | 2.5e-07 | 2.800E-07 | 6.083E-08 | 2.400E-07 | 3.500E-07 |
| 19205 | 961 | P961C | C | 2.6e-07 | 4.4e-07 | 5.4e-07 | 4.133E-07 | 1.419E-07 | 2.600E-07 | 5.400E-07 |
| 19206 | 961 | P961E | C | 3.3e-07 | 3e-07 | 9.6e-08 | 2.420E-07 | 1.273E-07 | 9.600E-08 | 3.300E-07 |
| 19207 | 961 | P961Q | C | 2.3e-07 | 2.6e-07 | 2.9e-07 | 2.600E-07 | 3.000E-08 | 2.300E-07 | 2.900E-07 |
| 19208 | 961 | P961G | C | 2.2e-07 | 3.1e-07 | 1.3e-07 | 2.200E-07 | 9.000E-08 | 1.300E-07 | 3.100E-07 |
| 19209 | 961 | P961H | C | 9.6e-08 | 6.5e-07 | 2.9e-07 | 3.453E-07 | 2.811E-07 | 9.600E-08 | 6.500E-07 |

| | | | | | | | | | | |
|---|---|---|---|---|---|---|---|---|---|---|
| 19210 | 961 | P961I | C | 2.3e-07 | 2.9e-07 | 1.8e-07 | 2.333E-07 | 5.508E-08 | 1.800E-07 | 2.900E-07 |
| 19211 | 961 | P961L | C | 2.3e-07 | 6.8e-07 | 2.3e-07 | 3.800E-07 | 2.598E-07 | 2.300E-07 | 6.800E-07 |
| 19212 | 961 | P961K | C | 2.7e-07 | 3.6e-07 | 2.9e-07 | 3.067E-07 | 4.726E-08 | 2.700E-07 | 3.600E-07 |
| 19213 | 961 | P961M | C | 3e-07 | 3.3e-07 | 2.7e-07 | 3.000E-07 | 3.000E-08 | 2.700E-07 | 3.300E-07 |
| 19214 | 961 | P961F | C | 1.9e-07 | 3.5e-07 | 2.8e-07 | 2.733E-07 | 8.021E-08 | 1.900E-07 | 3.500E-07 |
| 19215 | 961 | P961P | C | 2.1e-07 | 5.6e-07 | 2.2e-07 | 3.300E-07 | 1.992E-07 | 2.100E-07 | 5.600E-07 |
| 19216 | 961 | P961S | C | 7.1e-07 | 1.9e-07 | 2.8e-07 | 3.933E-07 | 2.779E-07 | 1.900E-07 | 7.100E-07 |
| 19217 | 961 | P961T | C | 4.6e-07 | 2.3e-07 | 2.5e-07 | 3.133E-07 | 1.274E-07 | 2.300E-07 | 4.600E-07 |
| 19218 | 961 | P961W | C | 2e-07 | 3.8e-07 | 2.7e-07 | 2.833E-07 | 9.074E-08 | 2.000E-07 | 3.800E-07 |
| 19219 | 961 | P961Y | C | 2.7e-07 | 1.1e-07 | 2e-07 | 1.933E-07 | 8.021E-08 | 1.100E-07 | 2.700E-07 |
| 19220 | 961 | P961V | C | 5.3e-08 | 4.9e-07 | 1.7e-07 | 2.377E-07 | 2.262E-07 | 5.300E-08 | 4.900E-07 |
| 19221 | 962 | S962A | C | 3.4e-07 | 2.1e-07 | 3.6e-07 | 3.033E-07 | 8.145E-08 | 2.100E-07 | 3.600E-07 |
| 19222 | 962 | S962R | C | 2.9e-07 | 1e-07 | 2.6e-07 | 2.167E-07 | 1.021E-07 | 1.000E-07 | 2.900E-07 |
| 19223 | 962 | S962N | C | 3.3e-07 | 2.4e-07 | 1.8e-07 | 2.500E-07 | 7.550E-08 | 1.800E-07 | 3.300E-07 |
| 19224 | 962 | S962D | C | 5.9e-07 | 2.9e-07 | 4.8e-07 | 4.533E-07 | 1.518E-07 | 2.900E-07 | 5.900E-07 |
| 19225 | 962 | S962C | C | 1.6e-07 | 1.5e-07 | 1.2e-07 | 1.433E-07 | 2.082E-08 | 1.200E-07 | 1.600E-07 |
| 19226 | 962 | S962E | C | 1.9e-07 | 3.5e-07 | 2.7e-07 | 2.700E-07 | 8.000E-08 | 1.900E-07 | 3.500E-07 |
| 19227 | 962 | S962Q | C | 1.2e-07 | 4.4e-07 | 2.5e-07 | 2.700E-07 | 1.609E-07 | 1.200E-07 | 4.400E-07 |
| 19228 | 962 | S962G | C | 4.5e-07 | 2.3e-07 | 4.1e-07 | 3.633E-07 | 1.172E-07 | 2.300E-07 | 4.500E-07 |
| 19229 | 962 | S962H | C | 2.7e-07 | 1.9e-07 | 2.3e-07 | 2.300E-07 | 4.000E-08 | 1.900E-07 | 2.700E-07 |
| 19230 | 962 | S962I | C | 2.4e-07 | 7.5e-07 | 1.3e-07 | 3.733E-07 | 3.308E-07 | 1.300E-07 | 7.500E-07 |
| 19231 | 962 | S962L | C | 3.4e-07 | 2e-07 | 2.5e-07 | 2.633E-07 | 7.095E-08 | 2.000E-07 | 3.400E-07 |
| 19232 | 962 | S962K | C | 2.1e-07 | 1.6e-07 | 1.6e-07 | 1.767E-07 | 2.887E-08 | 1.600E-07 | 2.100E-07 |
| 19233 | 962 | S962M | C | 6.3e-07 | 1.1e-07 | 2.9e-07 | 3.433E-07 | 2.641E-07 | 1.100E-07 | 6.300E-07 |
| 19234 | 962 | S962F | C | 2.1e-07 | 1.9e-07 | 3.5e-07 | 2.500E-07 | 8.718E-08 | 1.900E-07 | 3.500E-07 |
| 19235 | 962 | S962P | C | 3.5e-07 | 1.4e-07 | 1.6e-07 | 2.167E-07 | 1.159E-07 | 1.400E-07 | 3.500E-07 |
| 19236 | 962 | S962S | C | 2.5e-07 | 1.3e-07 | 3.6e-07 | 2.467E-07 | 1.150E-07 | 1.300E-07 | 3.600E-07 |
| 19237 | 962 | S962T | C | 3.7e-07 | 3e-07 | 2.8e-07 | 3.167E-07 | 4.726E-08 | 2.800E-07 | 3.700E-07 |
| 19238 | 962 | S962W | C | 1.5e-07 | 3.1e-07 | 2e-07 | 2.200E-07 | 8.185E-08 | 1.500E-07 | 3.100E-07 |
| 19239 | 962 | S962Y | C | 7.9e-08 | 1.4e-07 | 5.3e-07 | 2.497E-07 | 2.447E-07 | 7.900E-08 | 5.300E-07 |
| 19240 | 962 | S962V | C | 1.5e-07 | 1.9e-07 | 2.8e-07 | 2.067E-07 | 6.658E-08 | 1.500E-07 | 2.800E-07 |
| 19241 | 963 | V963A | C | 2.1e-07 | 1.5e-07 | 4.4e-07 | 2.667E-07 | 1.531E-07 | 1.500E-07 | 4.400E-07 |
| 19242 | 963 | V963R | C | 1.6e-07 | 3.5e-07 | 1.2e-07 | 2.100E-07 | 1.229E-07 | 1.200E-07 | 3.500E-07 |
| 19243 | 963 | V963N | C | 2.7e-07 | 2.4e-07 | 3.4e-07 | 2.833E-07 | 5.132E-08 | 2.400E-07 | 3.400E-07 |
| 19244 | 963 | V963D | C | 2.4e-07 | 5.7e-07 | 3.4e-07 | 3.833E-07 | 1.692E-07 | 2.400E-07 | 5.700E-07 |
| 19245 | 963 | V963C | C | 1.5e-07 | 3.6e-07 | 4.5e-07 | 3.200E-07 | 1.539E-07 | 1.500E-07 | 4.500E-07 |
| 19246 | 963 | V963E | C | 1.8e-07 | 3.1e-07 | 1.8e-07 | 2.233E-07 | 7.506E-08 | 1.800E-07 | 3.100E-07 |
| 19247 | 963 | V963Q | C | 3.1e-07 | 1.3e-07 | 2.6e-07 | 2.333E-07 | 9.292E-08 | 1.300E-07 | 3.100E-07 |
| 19248 | 963 | V963G | C | 1.2e-07 | 3.8e-07 | 4.1e-07 | 3.033E-07 | 1.595E-07 | 1.200E-07 | 4.100E-07 |
| 19249 | 963 | V963H | C | 3e-07 | 2e-07 | 1.9e-07 | 2.300E-07 | 6.083E-08 | 1.900E-07 | 3.000E-07 |
| 19250 | 963 | V963I | C | 4.8e-07 | 1.8e-07 | 2.3e-07 | 2.967E-07 | 1.607E-07 | 1.800E-07 | 4.800E-07 |
| 19251 | 963 | V963L | C | 2.9e-07 | 1.6e-07 | 2.7e-07 | 2.400E-07 | 7.000E-08 | 1.600E-07 | 2.900E-07 |
| 19252 | 963 | V963K | C | 1.4e-07 | 2.4e-07 | 1.3e-07 | 1.700E-07 | 6.083E-08 | 1.300E-07 | 2.400E-07 |
| 19253 | 963 | V963M | C | 2.6e-07 | 2.4e-07 | 1.7e-07 | 2.233E-07 | 4.726E-08 | 1.700E-07 | 2.600E-07 |
| 19254 | 963 | V963F | C | 3.9e-07 | 5.4e-07 | 1.4e-07 | 3.567E-07 | 2.021E-07 | 1.400E-07 | 5.400E-07 |
| 19255 | 963 | V963P | C | 3.9e-07 | 3e-07 | 1.4e-07 | 2.767E-07 | 1.266E-07 | 1.400E-07 | 3.900E-07 |
| 19256 | 963 | V963S | C | 2.2e-07 | 4.4e-07 | 4.7e-07 | 3.767E-07 | 1.365E-07 | 2.200E-07 | 4.700E-07 |
| 19257 | 963 | V963T | C | 3.2e-07 | 1.8e-07 | 3.5e-07 | 2.833E-07 | 9.074E-08 | 1.800E-07 | 3.500E-07 |
| 19258 | 963 | V963W | C | 3.1e-07 | 1.7e-07 | 5.3e-07 | 3.367E-07 | 1.815E-07 | 1.700E-07 | 5.300E-07 |
| 19259 | 963 | V963Y | C | 2.1e-07 | 2.4e-07 | 2.5e-07 | 2.333E-07 | 2.082E-08 | 2.100E-07 | 2.500E-07 |
| 19260 | 963 | V963V | C | 2.5e-07 | 3.7e-07 | 1.7e-07 | 2.633E-07 | 1.007E-07 | 1.700E-07 | 3.700E-07 |
| 19261 | 964 | F964A | C | 3e-07 | 4.7e-07 | 4.1e-07 | 3.933E-07 | 8.622E-08 | 3.000E-07 | 4.700E-07 |
| 19262 | 964 | F964R | C | 5.4e-07 | 1.2e-07 | 5e-07 | 3.867E-07 | 2.318E-07 | 1.200E-07 | 5.400E-07 |
| 19263 | 964 | F964N | C | 9.6e-08 | 2.8e-07 | 3.3e-07 | 2.353E-07 | 1.232E-07 | 9.600E-08 | 3.300E-07 |
| 19264 | 964 | F964D | C | 1.4e-07 | 2.6e-07 | 2.8e-07 | 2.267E-07 | 7.572E-08 | 1.400E-07 | 2.800E-07 |
| 19265 | 964 | F964C | C | 3.2e-07 | 2.6e-07 | 5.5e-07 | 3.767E-07 | 1.531E-07 | 2.600E-07 | 5.500E-07 |
| 19266 | 964 | F964E | C | 4e-07 | 2.1e-07 | 3.3e-07 | 3.133E-07 | 9.609E-08 | 2.100E-07 | 4.000E-07 |
| 19267 | 964 | F964Q | C | 2.1e-07 | 3.2e-07 | 1.9e-07 | 2.400E-07 | 7.000E-08 | 1.900E-07 | 3.200E-07 |
| 19268 | 964 | F964G | C | 3.3e-07 | 2.3e-07 | 2.5e-07 | 2.700E-07 | 5.292E-08 | 2.300E-07 | 3.300E-07 |
| 19269 | 964 | F964H | C | 4.5e-07 | 2.7e-07 | 1e-07 | 2.733E-07 | 1.750E-07 | 1.000E-07 | 4.500E-07 |
| 19270 | 964 | F964I | C | 2.4e-07 | 5.2e-07 | 2.2e-07 | 3.267E-07 | 1.677E-07 | 2.200E-07 | 5.200E-07 |
| 19271 | 964 | F964L | C | 3e-07 | 2.8e-07 | 1.9e-07 | 2.567E-07 | 5.859E-08 | 1.900E-07 | 3.000E-07 |
| 19272 | 964 | F964K | C | 1.7e-07 | 4.1e-07 | 6.3e-07 | 4.033E-07 | 2.301E-07 | 1.700E-07 | 6.300E-07 |
| 19273 | 964 | F964M | C | 3.9e-07 | 2.4e-07 | 6.1e-07 | 4.133E-07 | 1.861E-07 | 2.400E-07 | 6.100E-07 |
| 19274 | 964 | F964F | C | 2.8e-07 | 4.9e-07 | 1.8e-07 | 3.167E-07 | 1.582E-07 | 1.800E-07 | 4.900E-07 |
| 19275 | 964 | F964P | C | 3.8e-07 | 3.2e-07 | 1.8e-07 | 2.933E-07 | 1.026E-07 | 1.800E-07 | 3.800E-07 |
| 19276 | 964 | F964S | C | 2.1e-07 | 2.6e-07 | 1.8e-07 | 2.167E-07 | 4.041E-08 | 1.800E-07 | 2.600E-07 |
| 19277 | 964 | F964T | C | 3.8e-07 | 3.6e-07 | 2.1e-07 | 3.167E-07 | 9.292E-08 | 2.100E-07 | 3.800E-07 |
| 19278 | 964 | F964W | C | 4e-07 | 1.3e-07 | 4.5e-07 | 3.267E-07 | 1.721E-07 | 1.300E-07 | 4.500E-07 |
| 19279 | 964 | F964Y | C | 2.8e-07 | 2.9e-07 | 3e-07 | 2.900E-07 | 1.000E-08 | 2.800E-07 | 3.000E-07 |
| 19280 | 964 | F964V | C | 1.9e-07 | 3.8e-07 | 3.2e-07 | 2.967E-07 | 9.713E-08 | 1.900E-07 | 3.800E-07 |
| 19281 | 965 | P965A | C | 2.6e-07 | 2.1e-07 | 1.8e-07 | 2.167E-07 | 4.041E-08 | 1.800E-07 | 2.600E-07 |
| 19282 | 965 | P965R | C | 3.1e-07 | 9.1e-08 | 9.1e-08 | 1.640E-07 | 1.264E-07 | 9.100E-08 | 3.100E-07 |
| 19283 | 965 | P965N | C | 2.3e-07 | 1.6e-07 | 2e-07 | 1.967E-07 | 3.512E-08 | 1.600E-07 | 2.300E-07 |
| 19284 | 965 | P965D | C | 1.4e-07 | 2.3e-07 | 1.4e-07 | 1.700E-07 | 5.196E-08 | 1.400E-07 | 2.300E-07 |
| 19285 | 965 | P965C | C | 2.8e-07 | 3.4e-07 | 3.3e-07 | 3.167E-07 | 3.215E-08 | 2.800E-07 | 3.400E-07 |
| 19286 | 965 | P965E | C | 3.1e-07 | 3.8e-07 | 3.1e-07 | 3.333E-07 | 4.041E-08 | 3.100E-07 | 3.800E-07 |
| 19287 | 965 | P965Q | C | 3e-07 | 1.5e-07 | 2.3e-07 | 2.267E-07 | 7.506E-08 | 1.500E-07 | 3.000E-07 |
| 19288 | 965 | P965G | C | 7.6e-08 | 2.9e-07 | 4e-07 | 2.553E-07 | 1.648E-07 | 7.600E-08 | 4.000E-07 |
| 19289 | 965 | P965H | C | 5.7e-07 | 2.3e-07 | 1.9e-07 | 3.300E-07 | 2.088E-07 | 1.900E-07 | 5.700E-07 |
| 19290 | 965 | P965I | C | 2.1e-07 | 3.2e-07 | 1.8e-07 | 2.367E-07 | 7.371E-08 | 1.800E-07 | 3.200E-07 |
| 19291 | 965 | P965L | C | 4.4e-07 | 2.4e-07 | 1.5e-07 | 2.767E-07 | 1.484E-07 | 1.500E-07 | 4.400E-07 |
| 19292 | 965 | P965K | C | 2.1e-07 | 1.1e-07 | 3.5e-07 | 2.233E-07 | 1.206E-07 | 1.100E-07 | 3.500E-07 |
| 19293 | 965 | P965M | C | 6.6e-07 | 1.2e-07 | 2.9e-07 | 3.567E-07 | 2.761E-07 | 1.200E-07 | 6.600E-07 |
| 19294 | 965 | P965F | C | 2.2e-07 | 8.8e-08 | 2e-07 | 1.693E-07 | 7.114E-08 | 8.800E-08 | 2.200E-07 |

| | | | | | | | | | | |
|---|---|---|---|---|---|---|---|---|---|---|
| 19295 | 965 | P965P | C | 1.4e-07 | 1.8e-07 | 2.1e-07 | 1.767E-07 | 3.512E-08 | 1.400E-07 | 2.100E-07 |
| 19296 | 965 | P965S | C | 4e-07 | 2.9e-07 | 1.4e-07 | 2.767E-07 | 1.305E-07 | 1.400E-07 | 4.000E-07 |
| 19297 | 965 | P965T | C | 1.9e-07 | 1.8e-07 | 1.3e-07 | 1.667E-07 | 3.215E-08 | 1.300E-07 | 1.900E-07 |
| 19298 | 965 | P965W | C | 1.8e-07 | 3e-07 | 4.7e-07 | 3.167E-07 | 1.457E-07 | 1.800E-07 | 4.700E-07 |
| 19299 | 965 | P965Y | C | 2.7e-07 | 1.7e-07 | 1.5e-07 | 1.967E-07 | 6.429E-08 | 1.500E-07 | 2.700E-07 |
| 19300 | 965 | P965V | C | 1.7e-07 | 1.6e-07 | 4.1e-07 | 2.467E-07 | 1.415E-07 | 1.600E-07 | 4.100E-07 |
| 19301 | 966 | L966A | C | 4.1e-07 | 2.3e-07 | 4.9e-07 | 3.767E-07 | 1.332E-07 | 2.300E-07 | 4.900E-07 |
| 19302 | 966 | L966R | C | 7.5e-07 | 1.8e-07 | 1.4e-07 | 3.567E-07 | 3.412E-07 | 1.400E-07 | 7.500E-07 |
| 19303 | 966 | L966N | C | 2.8e-07 | 5.4e-07 | 4.2e-07 | 4.133E-07 | 1.301E-07 | 2.800E-07 | 5.400E-07 |
| 19304 | 966 | L966D | C | 2.6e-07 | 1.6e-07 | 2.3e-07 | 2.167E-07 | 5.132E-08 | 1.600E-07 | 2.600E-07 |
| 19305 | 966 | L966C | C | 2.7e-07 | 1.4e-07 | 2.8e-07 | 2.300E-07 | 7.810E-08 | 1.400E-07 | 2.800E-07 |
| 19306 | 966 | L966E | C | 2.3e-07 | 4.2e-07 | 1.3e-07 | 2.600E-07 | 1.473E-07 | 1.300E-07 | 4.200E-07 |
| 19307 | 966 | L966Q | C | 1.6e-07 | 2.9e-07 | 2.7e-07 | 2.400E-07 | 7.000E-08 | 1.600E-07 | 2.900E-07 |
| 19308 | 966 | L966G | C | 3.4e-07 | 2.2e-07 | 4.4e-07 | 3.333E-07 | 1.102E-07 | 2.200E-07 | 4.400E-07 |
| 19309 | 966 | L966H | C | 2e-07 | 3.1e-07 | 2.8e-07 | 2.633E-07 | 5.686E-08 | 2.000E-07 | 3.100E-07 |
| 19310 | 966 | L966I | C | 3.5e-07 | 2.8e-07 | 2.4e-07 | 2.900E-07 | 5.568E-08 | 2.400E-07 | 3.500E-07 |
| 19311 | 966 | L966L | C | 1.2e-07 | 1.8e-07 | 1.5e-07 | 1.500E-07 | 3.000E-08 | 1.200E-07 | 1.800E-07 |
| 19312 | 966 | L966K | C | 1.7e-07 | 2.6e-07 | 2.3e-07 | 2.200E-07 | 4.583E-08 | 1.700E-07 | 2.600E-07 |
| 19313 | 966 | L966M | C | 2.9e-07 | 5e-07 | 1.2e-07 | 3.033E-07 | 1.904E-07 | 1.200E-07 | 5.000E-07 |
| 19314 | 966 | L966F | C | 3.1e-07 | 2.2e-07 | 1.9e-07 | 2.400E-07 | 6.245E-08 | 1.900E-07 | 3.100E-07 |
| 19315 | 966 | L966P | C | 2e-07 | 2.2e-07 | 1.4e-07 | 1.867E-07 | 4.163E-08 | 1.400E-07 | 2.200E-07 |
| 19316 | 966 | L966S | C | 2.2e-07 | 2.4e-07 | 2.5e-07 | 2.367E-07 | 1.528E-08 | 2.200E-07 | 2.500E-07 |
| 19317 | 966 | L966T | C | 4.6e-07 | 1.4e-07 | 1.9e-07 | 2.633E-07 | 1.721E-07 | 1.400E-07 | 4.600E-07 |
| 19318 | 966 | L966W | C | 2.4e-07 | 2.2e-07 | 4e-07 | 2.867E-07 | 9.866E-08 | 2.200E-07 | 4.000E-07 |
| 19319 | 966 | L966Y | C | 2.4e-07 | 1.5e-07 | 3.8e-07 | 2.567E-07 | 1.159E-07 | 1.500E-07 | 3.800E-07 |
| 19320 | 966 | L966V | C | 6.8e-08 | 9.4e-08 | 2.7e-07 | 1.440E-07 | 1.099E-07 | 6.800E-08 | 2.700E-07 |
| 19321 | 967 | A967A | C | 8.8e-07 | 3.3e-07 | 2.8e-07 | 4.967E-07 | 3.329E-07 | 2.800E-07 | 8.800E-07 |
| 19322 | 967 | A967R | C | 1.6e-07 | 1.9e-07 | 3.1e-07 | 2.200E-07 | 7.937E-08 | 1.600E-07 | 3.100E-07 |
| 19323 | 967 | A967N | C | 2.4e-07 | 3.7e-07 | 5.4e-07 | 3.833E-07 | 1.504E-07 | 2.400E-07 | 5.400E-07 |
| 19324 | 967 | A967D | C | 2.2e-07 | 2.4e-07 | 3.7e-07 | 2.767E-07 | 8.145E-08 | 2.200E-07 | 3.700E-07 |
| 19325 | 967 | A967C | C | 3.1e-07 | 2.5e-07 | 2.4e-07 | 2.667E-07 | 3.786E-08 | 2.400E-07 | 3.100E-07 |
| 19326 | 967 | A967E | C | 3.2e-07 | 3.4e-07 | 1.5e-07 | 2.700E-07 | 1.044E-07 | 1.500E-07 | 3.400E-07 |
| 19327 | 967 | A967Q | C | 1.7e-07 | 5.7e-07 | 2.2e-07 | 3.200E-07 | 2.179E-07 | 1.700E-07 | 5.700E-07 |
| 19328 | 967 | A967G | C | 2.7e-07 | 3.6e-07 | 2.3e-07 | 2.867E-07 | 6.658E-08 | 2.300E-07 | 3.600E-07 |
| 19329 | 967 | A967H | C | 2.8e-07 | 2e-07 | 1.1e-07 | 1.967E-07 | 8.505E-08 | 1.100E-07 | 2.800E-07 |
| 19330 | 967 | A967I | C | 2.2e-07 | 1.6e-07 | 1.2e-07 | 1.667E-07 | 5.033E-08 | 1.200E-07 | 2.200E-07 |
| 19331 | 967 | A967L | C | 1.6e-07 | 5.6e-07 | 4.5e-07 | 3.900E-07 | 2.066E-07 | 1.600E-07 | 5.600E-07 |
| 19332 | 967 | A967K | C | 1.1e-07 | 5.5e-07 | 1.4e-07 | 2.667E-07 | 2.458E-07 | 1.100E-07 | 5.500E-07 |
| 19333 | 967 | A967M | C | 1.9e-07 | 4.4e-07 | 1.5e-07 | 2.600E-07 | 1.572E-07 | 1.500E-07 | 4.400E-07 |
| 19334 | 967 | A967F | C | 2e-07 | 1.1e-07 | 3e-07 | 2.033E-07 | 9.504E-08 | 1.100E-07 | 3.000E-07 |
| 19335 | 967 | A967P | C | 1.4e-07 | 2e-07 | 2.2e-07 | 1.867E-07 | 4.163E-08 | 1.400E-07 | 2.200E-07 |
| 19336 | 967 | A967S | C | 2.8e-07 | 4.5e-07 | 2.3e-07 | 3.200E-07 | 1.153E-07 | 2.300E-07 | 4.500E-07 |
| 19337 | 967 | A967T | C | 1.4e-07 | 2.4e-07 | 1.5e-07 | 1.767E-07 | 5.508E-08 | 1.400E-07 | 2.400E-07 |
| 19338 | 967 | A967W | C | 3.5e-07 | 3.7e-07 | 2.9e-07 | 3.367E-07 | 4.163E-08 | 2.900E-07 | 3.700E-07 |
| 19339 | 967 | A967Y | C | 2.4e-07 | 1.6e-07 | 2.3e-07 | 2.100E-07 | 4.359E-08 | 1.600E-07 | 2.400E-07 |
| 19340 | 967 | A967V | C | 2.4e-07 | 1.6e-07 | 3.7e-07 | 2.567E-07 | 1.060E-07 | 1.600E-07 | 3.700E-07 |
| 19341 | 968 | P968A | C | 3.5e-07 | 1.8e-07 | 1.7e-07 | 2.333E-07 | 1.012E-07 | 1.700E-07 | 3.500E-07 |
| 19342 | 968 | P968R | C | 4.5e-07 | 2.2e-07 | 2.2e-07 | 2.967E-07 | 1.328E-07 | 2.200E-07 | 4.500E-07 |
| 19343 | 968 | P968N | C | 3.3e-07 | 6e-07 | 8.4e-08 | 3.380E-07 | 2.581E-07 | 8.400E-08 | 6.000E-07 |
| 19344 | 968 | P968D | C | 2.2e-07 | 3e-07 | 4e-07 | 3.067E-07 | 9.018E-08 | 2.200E-07 | 4.000E-07 |
| 19345 | 968 | P968C | C | 2.6e-07 | 1.7e-07 | 2.8e-07 | 2.367E-07 | 5.859E-08 | 1.700E-07 | 2.800E-07 |
| 19346 | 968 | P968E | C | 2.1e-07 | 3.5e-07 | 2.2e-07 | 2.600E-07 | 7.810E-08 | 2.100E-07 | 3.500E-07 |
| 19347 | 968 | P968Q | C | 1.9e-07 | 5.4e-07 | 4e-07 | 3.767E-07 | 1.762E-07 | 1.900E-07 | 5.400E-07 |
| 19348 | 968 | P968G | C | 2.4e-07 | 2.1e-07 | 2.8e-07 | 2.433E-07 | 3.512E-08 | 2.100E-07 | 2.800E-07 |
| 19349 | 968 | P968H | C | 2.6e-07 | 3.4e-07 | 6e-07 | 4.000E-07 | 1.778E-07 | 2.600E-07 | 6.000E-07 |
| 19350 | 968 | P968I | C | 1.3e-07 | 3.4e-07 | 1.4e-07 | 2.033E-07 | 1.185E-07 | 1.300E-07 | 3.400E-07 |
| 19351 | 968 | P968L | C | 3.6e-07 | 3.3e-07 | 1.1e-07 | 2.667E-07 | 1.365E-07 | 1.100E-07 | 3.600E-07 |
| 19352 | 968 | P968K | C | 2.4e-07 | 2.1e-07 | 1.4e-07 | 1.967E-07 | 5.132E-08 | 1.400E-07 | 2.400E-07 |
| 19353 | 968 | P968M | C | 2.2e-07 | 2.3e-07 | 2e-07 | 2.167E-07 | 1.528E-08 | 2.000E-07 | 2.300E-07 |
| 19354 | 968 | P968F | C | 2.9e-07 | 1.1e-07 | 2.6e-07 | 2.200E-07 | 9.644E-08 | 1.100E-07 | 2.900E-07 |
| 19355 | 968 | P968P | C | 1.7e-07 | 4.9e-07 | 2.5e-07 | 3.033E-07 | 1.665E-07 | 1.700E-07 | 4.900E-07 |
| 19356 | 968 | P968S | C | 3.7e-07 | 3e-07 | 1.9e-07 | 2.867E-07 | 9.074E-08 | 1.900E-07 | 3.700E-07 |
| 19357 | 968 | P968T | C | 8.3e-08 | 7.7e-08 | 8.4e-08 | 8.133E-08 | 3.786E-09 | 7.700E-08 | 8.400E-08 |
| 19358 | 968 | P968W | C | 1.2e-07 | 3.2e-07 | 2.1e-07 | 2.167E-07 | 1.002E-07 | 1.200E-07 | 3.200E-07 |
| 19359 | 968 | P968Y | C | 1.2e-07 | 3.4e-07 | 2.3e-07 | 2.300E-07 | 1.100E-07 | 1.200E-07 | 3.400E-07 |
| 19360 | 968 | P968V | C | 3.1e-07 | 2.1e-07 | 1.4e-07 | 2.200E-07 | 8.544E-08 | 1.400E-07 | 3.100E-07 |
| 19361 | 969 | S969A | C | 3e-07 | 6.8e-07 | 2.5e-07 | 4.100E-07 | 2.352E-07 | 2.500E-07 | 6.800E-07 |
| 19362 | 969 | S969R | C | 3.5e-07 | 1.5e-07 | 1.8e-07 | 2.267E-07 | 1.079E-07 | 1.500E-07 | 3.500E-07 |
| 19363 | 969 | S969N | C | 4.5e-07 | 1.5e-07 | 3.4e-07 | 3.133E-07 | 1.518E-07 | 1.500E-07 | 4.500E-07 |
| 19364 | 969 | S969D | C | 1.7e-07 | 2.4e-07 | 4.3e-07 | 2.800E-07 | 1.345E-07 | 1.700E-07 | 4.300E-07 |
| 19365 | 969 | S969C | C | 2.5e-07 | 1.8e-07 | 3.8e-07 | 2.700E-07 | 1.015E-07 | 1.800E-07 | 3.800E-07 |
| 19366 | 969 | S969E | C | 1.3e-07 | 1e-07 | 2.9e-07 | 1.733E-07 | 1.021E-07 | 1.000E-07 | 2.900E-07 |
| 19367 | 969 | S969Q | C | 5.6e-08 | 1.2e-07 | 1.9e-07 | 1.220E-07 | 6.702E-08 | 5.600E-08 | 1.900E-07 |
| 19368 | 969 | S969G | C | 2.1e-07 | 2.1e-07 | 1.6e-07 | 1.933E-07 | 2.887E-08 | 1.600E-07 | 2.100E-07 |
| 19369 | 969 | S969H | C | 1.7e-07 | 8.9e-08 | 4.2e-07 | 2.263E-07 | 1.725E-07 | 8.900E-08 | 4.200E-07 |
| 19370 | 969 | S969I | C | 2e-07 | 4.1e-07 | 2.2e-07 | 2.767E-07 | 1.159E-07 | 2.000E-07 | 4.100E-07 |
| 19371 | 969 | S969L | C | 1.9e-07 | 1.5e-07 | 1.6e-07 | 1.667E-07 | 2.082E-08 | 1.500E-07 | 1.900E-07 |
| 19372 | 969 | S969K | C | 4.5e-07 | 2.2e-07 | 1.2e-07 | 2.633E-07 | 1.692E-07 | 1.200E-07 | 4.500E-07 |
| 19373 | 969 | S969M | C | 3.5e-07 | 1.7e-07 | 4.3e-07 | 3.167E-07 | 1.332E-07 | 1.700E-07 | 4.300E-07 |
| 19374 | 969 | S969F | C | 3.5e-07 | 3.5e-07 | 2.5e-07 | 3.167E-07 | 5.774E-08 | 2.500E-07 | 3.500E-07 |
| 19375 | 969 | S969P | C | 2.3e-07 | 9.2e-08 | 1.1e-07 | 1.440E-07 | 7.502E-08 | 9.200E-08 | 2.300E-07 |
| 19376 | 969 | S969S | C | 2.9e-07 | 2e-07 | 3.4e-07 | 2.767E-07 | 7.095E-08 | 2.000E-07 | 3.400E-07 |
| 19377 | 969 | S969T | C | 4.9e-07 | 7.5e-07 | 1.4e-07 | 4.600E-07 | 3.061E-07 | 1.400E-07 | 7.500E-07 |
| 19378 | 969 | S969W | C | 2.1e-07 | 1.9e-07 | 3.1e-07 | 2.367E-07 | 6.429E-08 | 1.900E-07 | 3.100E-07 |
| 19379 | 969 | S969Y | C | 1.3e-07 | 2.6e-07 | 4.7e-07 | 2.867E-07 | 1.716E-07 | 1.300E-07 | 4.700E-07 |

| | | | | | | | | | | |
|---|---|---|---|---|---|---|---|---|---|---|
| 19380 | 969 | S969V | C | 3e-07 | 1.3e-07 | 4.3e-07 | 2.867E-07 | 1.504E-07 | 1.300E-07 | 4.300E-07 |
| 19381 | 970 | S970A | C | 3.9e-07 | 2e-07 | 2.5e-07 | 2.800E-07 | 9.849E-08 | 2.000E-07 | 3.900E-07 |
| 19382 | 970 | S970R | C | 5.5e-07 | 3.2e-07 | 2.7e-07 | 3.800E-07 | 1.493E-07 | 2.700E-07 | 5.500E-07 |
| 19383 | 970 | S970N | C | 4.7e-07 | 1.9e-07 | 2.1e-07 | 2.900E-07 | 1.562E-07 | 1.900E-07 | 4.700E-07 |
| 19384 | 970 | S970D | C | 4.4e-07 | 1.2e-07 | 1.3e-07 | 2.300E-07 | 1.819E-07 | 1.200E-07 | 4.400E-07 |
| 19385 | 970 | S970C | C | 1.6e-07 | 2.6e-07 | 2.6e-07 | 2.267E-07 | 5.774E-08 | 1.600E-07 | 2.600E-07 |
| 19386 | 970 | S970E | C | 3.1e-07 | 1.9e-07 | 2e-07 | 2.333E-07 | 6.658E-08 | 1.900E-07 | 3.100E-07 |
| 19387 | 970 | S970Q | C | 4.3e-07 | 3.6e-07 | 4.9e-07 | 4.267E-07 | 6.506E-08 | 3.600E-07 | 4.900E-07 |
| 19388 | 970 | S970G | C | 1.7e-07 | 1.2e-07 | 2.5e-07 | 1.800E-07 | 6.557E-08 | 1.200E-07 | 2.500E-07 |
| 19389 | 970 | S970H | C | 1.8e-07 | 1.4e-07 | 2.7e-07 | 1.967E-07 | 6.658E-08 | 1.400E-07 | 2.700E-07 |
| 19390 | 970 | S970I | C | 2e-07 | 3.5e-07 | 1.6e-07 | 2.367E-07 | 1.002E-07 | 1.600E-07 | 3.500E-07 |
| 19391 | 970 | S970L | C | 1.9e-07 | 6e-07 | 7.1e-07 | 5.000E-07 | 2.740E-07 | 1.900E-07 | 7.100E-07 |
| 19392 | 970 | S970K | C | 2.4e-07 | 2.9e-07 | 1.3e-07 | 2.200E-07 | 8.185E-08 | 1.300E-07 | 2.900E-07 |
| 19393 | 970 | S970M | C | 1.8e-07 | 2.5e-07 | 3.2e-07 | 2.500E-07 | 7.000E-08 | 1.800E-07 | 3.200E-07 |
| 19394 | 970 | S970F | C | 3.9e-07 | 5.4e-07 | 2.7e-07 | 4.000E-07 | 1.353E-07 | 2.700E-07 | 5.400E-07 |
| 19395 | 970 | S970P | C | 1.1e-07 | 1.9e-07 | 4.2e-07 | 2.400E-07 | 1.609E-07 | 1.100E-07 | 4.200E-07 |
| 19396 | 970 | S970S | C | 2.9e-07 | 3e-07 | 2.1e-07 | 2.667E-07 | 4.933E-08 | 2.100E-07 | 3.000E-07 |
| 19397 | 970 | S970T | C | 1.2e-07 | 1.4e-07 | 2.2e-07 | 1.600E-07 | 5.292E-08 | 1.200E-07 | 2.200E-07 |
| 19398 | 970 | S970W | C | 7.4e-07 | 1.9e-07 | 1.3e-07 | 3.533E-07 | 3.362E-07 | 1.300E-07 | 7.400E-07 |
| 19399 | 970 | S970Y | C | 2.4e-07 | 2.8e-07 | 7e-08 | 1.967E-07 | 1.115E-07 | 7.000E-08 | 2.800E-07 |
| 19400 | 970 | S970V | C | 1.6e-07 | 2.3e-07 | 1.3e-07 | 1.733E-07 | 5.132E-08 | 1.300E-07 | 2.300E-07 |
| 19401 | 971 | K971A | C | 4.4e-07 | 2.3e-07 | 1.9e-07 | 2.867E-07 | 1.343E-07 | 1.900E-07 | 4.400E-07 |
| 19402 | 971 | K971R | C | 1.3e-07 | 3e-07 | 4e-07 | 2.767E-07 | 1.365E-07 | 1.300E-07 | 4.000E-07 |
| 19403 | 971 | K971N | C | 2e-07 | 2.6e-07 | 1.7e-07 | 2.100E-07 | 4.583E-08 | 1.700E-07 | 2.600E-07 |
| 19404 | 971 | K971D | C | 1.5e-07 | 2.7e-07 | 1.9e-07 | 2.033E-07 | 6.110E-08 | 1.500E-07 | 2.700E-07 |
| 19405 | 971 | K971C | C | 5.3e-07 | 1.6e-07 | 3.3e-07 | 3.400E-07 | 1.852E-07 | 1.600E-07 | 5.300E-07 |
| 19406 | 971 | K971E | C | 2.5e-07 | 1e-07 | 2.5e-07 | 2.000E-07 | 8.660E-08 | 1.000E-07 | 2.500E-07 |
| 19407 | 971 | K971Q | C | 2.4e-07 | 2.2e-07 | 1.7e-07 | 2.100E-07 | 3.606E-08 | 1.700E-07 | 2.400E-07 |
| 19408 | 971 | K971G | C | 3.1e-07 | 4.6e-07 | 3.1e-07 | 3.600E-07 | 8.660E-08 | 3.100E-07 | 4.600E-07 |
| 19409 | 971 | K971H | C | 2.8e-07 | 2.2e-07 | 1.9e-07 | 2.300E-07 | 4.583E-08 | 1.900E-07 | 2.800E-07 |
| 19410 | 971 | K971I | C | 1.9e-07 | 1.5e-07 | 1.1e-07 | 1.500E-07 | 4.000E-08 | 1.100E-07 | 1.900E-07 |
| 19411 | 971 | K971L | C | 8.7e-07 | 4e-07 | 3.7e-07 | 5.467E-07 | 2.804E-07 | 3.700E-07 | 8.700E-07 |
| 19412 | 971 | K971K | C | 2.9e-07 | 1.7e-07 | 1.8e-07 | 2.133E-07 | 6.658E-08 | 1.700E-07 | 2.900E-07 |
| 19413 | 971 | K971M | C | 3.2e-07 | 2.2e-07 | 1.3e-07 | 2.233E-07 | 9.504E-08 | 1.300E-07 | 3.200E-07 |
| 19414 | 971 | K971F | C | 3.3e-07 | 2.5e-07 | 1.3e-07 | 2.367E-07 | 1.007E-07 | 1.300E-07 | 3.300E-07 |
| 19415 | 971 | K971P | C | 3.8e-07 | 3.1e-07 | 1.1e-07 | 2.667E-07 | 1.401E-07 | 1.100E-07 | 3.800E-07 |
| 19416 | 971 | K971S | C | 2.7e-07 | 1.8e-07 | 1e-07 | 1.833E-07 | 8.505E-08 | 1.000E-07 | 2.700E-07 |
| 19417 | 971 | K971T | C | 2e-07 | 2.3e-07 | 5.8e-07 | 3.367E-07 | 2.113E-07 | 2.000E-07 | 5.800E-07 |
| 19418 | 971 | K971W | C | 2.8e-07 | 1.8e-07 | 2.1e-07 | 2.233E-07 | 5.132E-08 | 1.800E-07 | 2.800E-07 |
| 19419 | 971 | K971Y | C | 3.9e-07 | 3.6e-07 | 2.7e-07 | 3.400E-07 | 6.245E-08 | 2.700E-07 | 3.900E-07 |
| 19420 | 971 | K971V | C | 6.5e-07 | 2.6e-07 | 1.9e-07 | 3.667E-07 | 2.479E-07 | 1.900E-07 | 6.500E-07 |
| 19421 | 972 | S972A | C | 4.2e-07 | 1.4e-07 | 2.4e-07 | 2.667E-07 | 1.419E-07 | 1.400E-07 | 4.200E-07 |
| 19422 | 972 | S972R | C | 2.1e-07 | 1.1e-07 | 2e-07 | 1.733E-07 | 5.508E-08 | 1.100E-07 | 2.100E-07 |
| 19423 | 972 | S972N | C | 1.6e-07 | 2.8e-07 | 3.1e-07 | 2.500E-07 | 7.937E-08 | 1.600E-07 | 3.100E-07 |
| 19424 | 972 | S972D | C | 2.6e-07 | 2e-07 | 3.2e-07 | 2.600E-07 | 6.000E-08 | 2.000E-07 | 3.200E-07 |
| 19425 | 972 | S972C | C | 2e-07 | 7.8e-07 | 3e-07 | 4.267E-07 | 3.101E-07 | 2.000E-07 | 7.800E-07 |
| 19426 | 972 | S972E | C | 1.4e-07 | 2.6e-07 | 4e-07 | 2.667E-07 | 1.301E-07 | 1.400E-07 | 4.000E-07 |
| 19427 | 972 | S972Q | C | 1.5e-07 | 1.9e-07 | 1.9e-07 | 1.767E-07 | 2.309E-08 | 1.500E-07 | 1.900E-07 |
| 19428 | 972 | S972G | C | 2.4e-07 | 3e-07 | 2.8e-07 | 2.733E-07 | 3.055E-08 | 2.400E-07 | 3.000E-07 |
| 19429 | 972 | S972H | C | 2.1e-07 | 2.1e-07 | 2.3e-07 | 2.167E-07 | 1.155E-08 | 2.100E-07 | 2.300E-07 |
| 19430 | 972 | S972I | C | 1.6e-07 | 1.8e-07 | 2.4e-07 | 1.933E-07 | 4.163E-08 | 1.600E-07 | 2.400E-07 |
| 19431 | 972 | S972L | C | 1.2e-07 | 2.1e-07 | 1.2e-07 | 1.500E-07 | 5.196E-08 | 1.200E-07 | 2.100E-07 |
| 19432 | 972 | S972K | C | 1.7e-07 | 1.9e-07 | 2.2e-07 | 1.933E-07 | 2.517E-08 | 1.700E-07 | 2.200E-07 |
| 19433 | 972 | S972M | C | 6.6e-07 | 3.1e-07 | 3.4e-07 | 4.367E-07 | 1.940E-07 | 3.100E-07 | 6.600E-07 |
| 19434 | 972 | S972F | C | 1.9e-07 | 3e-07 | 5.2e-07 | 3.367E-07 | 1.680E-07 | 1.900E-07 | 5.200E-07 |
| 19435 | 972 | S972P | C | 2.7e-07 | 2.8e-07 | 3e-07 | 2.833E-07 | 1.528E-08 | 2.700E-07 | 3.000E-07 |
| 19436 | 972 | S972S | C | 4.7e-07 | 2.7e-07 | 2.1e-07 | 3.167E-07 | 1.361E-07 | 2.100E-07 | 4.700E-07 |
| 19437 | 972 | S972T | C | 2.7e-07 | 4.3e-07 | 1.7e-07 | 2.900E-07 | 1.311E-07 | 1.700E-07 | 4.300E-07 |
| 19438 | 972 | S972W | C | 3.7e-07 | 5.8e-07 | 3.4e-07 | 4.300E-07 | 1.308E-07 | 3.400E-07 | 5.800E-07 |
| 19439 | 972 | S972Y | C | 2.1e-07 | 1.9e-07 | 2.7e-07 | 2.233E-07 | 4.163E-08 | 1.900E-07 | 2.700E-07 |
| 19440 | 972 | S972V | C | 4e-07 | 2.6e-07 | 1.9e-07 | 2.833E-07 | 1.069E-07 | 1.900E-07 | 4.000E-07 |
| 19441 | 973 | T973A | C | 1.7e-07 | 3.2e-07 | 3e-07 | 2.633E-07 | 8.145E-08 | 1.700E-07 | 3.200E-07 |
| 19442 | 973 | T973R | C | 1.3e-07 | 2.4e-07 | 3.5e-07 | 2.400E-07 | 1.100E-07 | 1.300E-07 | 3.500E-07 |
| 19443 | 973 | T973N | C | 1.6e-07 | 1.6e-07 | 2e-07 | 1.733E-07 | 2.309E-08 | 1.600E-07 | 2.000E-07 |
| 19444 | 973 | T973D | C | 3.3e-07 | 4.3e-07 | 1.8e-07 | 3.133E-07 | 1.258E-07 | 1.800E-07 | 4.300E-07 |
| 19445 | 973 | T973C | C | 6.8e-07 | 2.6e-07 | 2.8e-07 | 4.067E-07 | 2.369E-07 | 2.600E-07 | 6.800E-07 |
| 19446 | 973 | T973E | C | 2.5e-07 | 4.9e-07 | 1.4e-07 | 2.933E-07 | 1.790E-07 | 1.400E-07 | 4.900E-07 |
| 19447 | 973 | T973Q | C | 5.4e-07 | 2.4e-07 | 2.6e-07 | 3.467E-07 | 1.677E-07 | 2.400E-07 | 5.400E-07 |
| 19448 | 973 | T973G | C | 1.3e-07 | 2.6e-07 | 2.9e-07 | 2.267E-07 | 8.505E-08 | 1.300E-07 | 2.900E-07 |
| 19449 | 973 | T973H | C | 3.8e-07 | 1.9e-07 | 1e-07 | 2.233E-07 | 1.429E-07 | 1.000E-07 | 3.800E-07 |
| 19450 | 973 | T973I | C | 1.1e-07 | 1.1e-07 | 2.4e-07 | 1.533E-07 | 7.506E-08 | 1.100E-07 | 2.400E-07 |
| 19451 | 973 | T973L | C | 2.8e-07 | 4.3e-07 | 2.9e-07 | 3.333E-07 | 8.386E-08 | 2.800E-07 | 4.300E-07 |
| 19452 | 973 | T973K | C | 3.7e-07 | 5.2e-07 | 3.1e-07 | 4.000E-07 | 1.082E-07 | 3.100E-07 | 5.200E-07 |
| 19453 | 973 | T973M | C | 2.6e-07 | 2.7e-07 | 1.8e-07 | 2.367E-07 | 4.933E-08 | 1.800E-07 | 2.700E-07 |
| 19454 | 973 | T973F | C | 5e-07 | 2.4e-07 | 3e-07 | 3.467E-07 | 1.361E-07 | 2.400E-07 | 5.000E-07 |
| 19455 | 973 | T973P | C | 1.7e-07 | 1.7e-07 | 1.9e-07 | 1.767E-07 | 1.155E-08 | 1.700E-07 | 1.900E-07 |
| 19456 | 973 | T973S | C | 7.5e-07 | 1.7e-07 | 5.5e-07 | 4.900E-07 | 2.946E-07 | 1.700E-07 | 7.500E-07 |
| 19457 | 973 | T973T | C | 2.2e-07 | 1.8e-07 | 1.8e-07 | 1.933E-07 | 2.309E-08 | 1.800E-07 | 2.200E-07 |
| 19458 | 973 | T973W | C | 2.8e-07 | 2.7e-07 | 1.5e-07 | 2.333E-07 | 7.234E-08 | 1.500E-07 | 2.800E-07 |
| 19459 | 973 | T973Y | C | 2.3e-07 | 1.3e-07 | 1.1e-07 | 1.567E-07 | 6.429E-08 | 1.100E-07 | 2.300E-07 |
| 19460 | 973 | T973V | C | 5.5e-07 | 2.7e-07 | 3.4e-07 | 3.867E-07 | 1.457E-07 | 2.700E-07 | 5.500E-07 |
| 19461 | 974 | S974A | C | 1.9e-07 | 3.8e-07 | 1.6e-07 | 2.433E-07 | 1.193E-07 | 1.600E-07 | 3.800E-07 |
| 19462 | 974 | S974R | C | 1.8e-07 | 2.7e-07 | 4.2e-07 | 2.900E-07 | 1.212E-07 | 1.800E-07 | 4.200E-07 |
| 19463 | 974 | S974N | C | 3.9e-07 | 2.4e-07 | 1.8e-07 | 2.700E-07 | 1.082E-07 | 1.800E-07 | 3.900E-07 |
| 19464 | 974 | S974D | C | 2.3e-07 | 4.4e-07 | 3.5e-07 | 3.400E-07 | 1.054E-07 | 2.300E-07 | 4.400E-07 |

| | | | | | | | | | | |
|---|---|---|---|---|---|---|---|---|---|---|
| 19465 | 974 | S974C | C | 2.4e-07 | 1.2e-07 | 3.4e-07 | 2.333E-07 | 1.102E-07 | 1.200E-07 | 3.400E-07 |
| 19466 | 974 | S974E | C | 3e-07 | 5.6e-07 | 2e-07 | 3.533E-07 | 1.858E-07 | 2.000E-07 | 5.600E-07 |
| 19467 | 974 | S974Q | C | 2.4e-07 | 2.8e-07 | 2.3e-07 | 2.500E-07 | 2.646E-08 | 2.300E-07 | 2.800E-07 |
| 19468 | 974 | S974G | C | 3.4e-07 | 1.3e-07 | 2.9e-07 | 2.533E-07 | 1.097E-07 | 1.300E-07 | 3.400E-07 |
| 19469 | 974 | S974H | C | 1.5e-07 | 3.2e-07 | 2.8e-07 | 2.500E-07 | 8.888E-08 | 1.500E-07 | 3.200E-07 |
| 19470 | 974 | S974I | C | 7.8e-08 | 2e-07 | 4.2e-07 | 2.327E-07 | 1.733E-07 | 7.800E-08 | 4.200E-07 |
| 19471 | 974 | S974L | C | 2.9e-07 | 1.5e-07 | 2.9e-07 | 2.433E-07 | 8.083E-08 | 1.500E-07 | 2.900E-07 |
| 19472 | 974 | S974K | C | 4.1e-07 | 2.3e-07 | 3.8e-07 | 3.400E-07 | 9.644E-08 | 2.300E-07 | 4.100E-07 |
| 19473 | 974 | S974M | C | 9.3e-08 | 4.3e-07 | 3.9e-07 | 3.043E-07 | 1.841E-07 | 9.300E-08 | 4.300E-07 |
| 19474 | 974 | S974F | C | 9.2e-07 | 5.1e-07 | 3.6e-07 | 5.967E-07 | 2.899E-07 | 3.600E-07 | 9.200E-07 |
| 19475 | 974 | S974P | C | 4.2e-07 | 8.4e-08 | 2.8e-07 | 2.613E-07 | 1.688E-07 | 8.400E-08 | 4.200E-07 |
| 19476 | 974 | S974S | C | 1.9e-07 | 4.8e-07 | 2.5e-07 | 3.067E-07 | 1.531E-07 | 1.900E-07 | 4.800E-07 |
| 19477 | 974 | S974T | C | 6.6e-08 | 2.6e-07 | 1.4e-07 | 1.553E-07 | 9.790E-08 | 6.600E-08 | 2.600E-07 |
| 19478 | 974 | S974W | C | 4.1e-07 | 3.7e-07 | 1.3e-07 | 3.033E-07 | 1.514E-07 | 1.300E-07 | 4.100E-07 |
| 19479 | 974 | S974Y | C | 3.2e-07 | 1.9e-07 | 2.1e-07 | 2.400E-07 | 7.000E-08 | 1.900E-07 | 3.200E-07 |
| 19480 | 974 | S974V | C | 2.3e-07 | 2.4e-07 | 4.5e-07 | 3.067E-07 | 1.242E-07 | 2.300E-07 | 4.500E-07 |
| 19481 | 975 | G975A | C | 3.2e-07 | 2.4e-07 | 2.5e-07 | 2.700E-07 | 4.359E-08 | 2.400E-07 | 3.200E-07 |
| 19482 | 975 | G975R | C | 4e-07 | 1e-07 | 6.9e-07 | 3.967E-07 | 2.950E-07 | 1.000E-07 | 6.900E-07 |
| 19483 | 975 | G975N | C | 1.7e-07 | 3.1e-07 | 2.4e-07 | 2.400E-07 | 7.000E-08 | 1.700E-07 | 3.100E-07 |
| 19484 | 975 | G975D | C | 3.4e-07 | 1.3e-07 | 4.5e-07 | 3.067E-07 | 1.626E-07 | 1.300E-07 | 4.500E-07 |
| 19485 | 975 | G975C | C | 2.2e-07 | 1.2e-07 | 2.6e-07 | 2.000E-07 | 7.211E-08 | 1.200E-07 | 2.600E-07 |
| 19486 | 975 | G975E | C | 2.4e-07 | 1.4e-07 | 2.8e-07 | 2.200E-07 | 7.211E-08 | 1.400E-07 | 2.800E-07 |
| 19487 | 975 | G975Q | C | 3.1e-07 | 3.3e-07 | 2.8e-07 | 3.067E-07 | 2.517E-08 | 2.800E-07 | 3.300E-07 |
| 19488 | 975 | G975G | C | 3.2e-07 | 1.7e-07 | 1.8e-07 | 2.233E-07 | 8.386E-08 | 1.700E-07 | 3.200E-07 |
| 19489 | 975 | G975H | C | 2e-07 | 9.7e-08 | 3e-07 | 1.990E-07 | 1.015E-07 | 9.700E-08 | 3.000E-07 |
| 19490 | 975 | G975I | C | 3.4e-07 | 2.2e-07 | 3.8e-07 | 3.133E-07 | 8.327E-08 | 2.200E-07 | 3.800E-07 |
| 19491 | 975 | G975L | C | 1.8e-07 | 2.5e-07 | 4.2e-07 | 2.833E-07 | 1.234E-07 | 1.800E-07 | 4.200E-07 |
| 19492 | 975 | G975K | C | 5.4e-07 | 3.1e-07 | 2.7e-07 | 3.733E-07 | 1.457E-07 | 2.700E-07 | 5.400E-07 |
| 19493 | 975 | G975M | C | 2.5e-07 | 3.5e-07 | 2.8e-07 | 2.933E-07 | 5.132E-08 | 2.500E-07 | 3.500E-07 |
| 19494 | 975 | G975F | C | 3.2e-07 | 1.9e-07 | 1.3e-07 | 2.133E-07 | 9.713E-08 | 1.300E-07 | 3.200E-07 |
| 19495 | 975 | G975P | C | 3.6e-07 | 3e-07 | 3.8e-07 | 3.467E-07 | 4.163E-08 | 3.000E-07 | 3.800E-07 |
| 19496 | 975 | G975S | C | 5.7e-07 | 1.4e-07 | 2.4e-07 | 3.167E-07 | 2.250E-07 | 1.400E-07 | 5.700E-07 |
| 19497 | 975 | G975T | C | 3.8e-07 | 3.9e-07 | 1.4e-07 | 3.033E-07 | 1.415E-07 | 1.400E-07 | 3.900E-07 |
| 19498 | 975 | G975W | C | 4.5e-07 | 2.2e-07 | 3e-07 | 3.233E-07 | 1.168E-07 | 2.200E-07 | 4.500E-07 |
| 19499 | 975 | G975Y | C | 5.5e-07 | 2e-07 | 2.1e-07 | 3.200E-07 | 1.992E-07 | 2.000E-07 | 5.500E-07 |
| 19500 | 975 | G975V | C | 2.9e-07 | 2e-07 | 5.6e-07 | 3.500E-07 | 1.873E-07 | 2.000E-07 | 5.600E-07 |
| 19501 | 976 | G976A | C | 2.6e-07 | 8e-08 | 1.3e-07 | 1.567E-07 | 9.292E-08 | 8.000E-08 | 2.600E-07 |
| 19502 | 976 | G976R | C | 9.9e-08 | 3.4e-07 | 1.3e-07 | 1.897E-07 | 1.311E-07 | 9.900E-08 | 3.400E-07 |
| 19503 | 976 | G976N | C | 1.8e-07 | 3e-07 | 3.7e-07 | 2.833E-07 | 9.609E-08 | 1.800E-07 | 3.700E-07 |
| 19504 | 976 | G976D | C | 2.5e-07 | 2.2e-07 | 3.1e-07 | 2.600E-07 | 4.583E-08 | 2.200E-07 | 3.100E-07 |
| 19505 | 976 | G976C | C | 1.7e-07 | 3.9e-07 | 3.3e-07 | 2.967E-07 | 1.137E-07 | 1.700E-07 | 3.900E-07 |
| 19506 | 976 | G976E | C | 4.3e-07 | 2.5e-07 | 2.8e-07 | 3.200E-07 | 9.644E-08 | 2.500E-07 | 4.300E-07 |
| 19507 | 976 | G976Q | C | 2.4e-07 | 1.3e-07 | 2.1e-07 | 1.933E-07 | 5.686E-08 | 1.300E-07 | 2.400E-07 |
| 19508 | 976 | G976G | C | 3.1e-07 | 1.2e-07 | 2.3e-07 | 2.200E-07 | 9.539E-08 | 1.200E-07 | 3.100E-07 |
| 19509 | 976 | G976H | C | 3.6e-07 | 1.8e-07 | 2.5e-07 | 2.633E-07 | 9.074E-08 | 1.800E-07 | 3.600E-07 |
| 19510 | 976 | G976I | C | 4e-07 | 2.5e-07 | 2.1e-07 | 2.867E-07 | 1.002E-07 | 2.100E-07 | 4.000E-07 |
| 19511 | 976 | G976L | C | 1.5e-07 | 2.2e-07 | 3.1e-07 | 2.267E-07 | 8.021E-08 | 1.500E-07 | 3.100E-07 |
| 19512 | 976 | G976K | C | 4.9e-07 | 3.1e-07 | 3.6e-07 | 3.867E-07 | 9.292E-08 | 3.100E-07 | 4.900E-07 |
| 19513 | 976 | G976M | C | 2.7e-07 | 1.4e-07 | 1.1e-07 | 1.733E-07 | 8.505E-08 | 1.100E-07 | 2.700E-07 |
| 19514 | 976 | G976F | C | 9.9e-08 | 5.3e-07 | 3e-07 | 3.097E-07 | 2.157E-07 | 9.900E-08 | 5.300E-07 |
| 19515 | 976 | G976P | C | 1.8e-07 | 2.1e-07 | 2.4e-07 | 2.100E-07 | 3.000E-08 | 1.800E-07 | 2.400E-07 |
| 19516 | 976 | G976S | C | 1.3e-07 | 7.8e-07 | 2e-07 | 3.700E-07 | 3.568E-07 | 1.300E-07 | 7.800E-07 |
| 19517 | 976 | G976T | C | 3.5e-07 | 2.7e-07 | 1.5e-07 | 2.567E-07 | 1.007E-07 | 1.500E-07 | 3.500E-07 |
| 19518 | 976 | G976W | C | 2.6e-07 | 1.7e-07 | 2.2e-07 | 2.167E-07 | 4.509E-08 | 1.700E-07 | 2.600E-07 |
| 19519 | 976 | G976Y | C | 2.6e-07 | 3.4e-07 | 2.5e-07 | 2.833E-07 | 4.933E-08 | 2.500E-07 | 3.400E-07 |
| 19520 | 976 | G976V | C | 1.2e-07 | 5.8e-07 | 1.4e-07 | 2.800E-07 | 2.600E-07 | 1.200E-07 | 5.800E-07 |
| 19521 | 977 | T977A | C | 3.2e-07 | 7.5e-07 | 2.7e-07 | 4.467E-07 | 2.639E-07 | 2.700E-07 | 7.500E-07 |
| 19522 | 977 | T977R | C | 2.8e-07 | 1.8e-07 | 2.1e-07 | 2.233E-07 | 5.132E-08 | 1.800E-07 | 2.800E-07 |
| 19523 | 977 | T977N | C | 2.9e-07 | 3.6e-07 | 2.5e-07 | 3.000E-07 | 5.568E-08 | 2.500E-07 | 3.600E-07 |
| 19524 | 977 | T977D | C | 2.4e-07 | 4.8e-07 | 2e-07 | 3.067E-07 | 1.514E-07 | 2.000E-07 | 4.800E-07 |
| 19525 | 977 | T977C | C | 3.3e-07 | 2.7e-07 | 3e-07 | 3.000E-07 | 3.000E-08 | 2.700E-07 | 3.300E-07 |
| 19526 | 977 | T977E | C | 3.1e-07 | 3.2e-07 | 3.3e-07 | 3.200E-07 | 1.000E-08 | 3.100E-07 | 3.300E-07 |
| 19527 | 977 | T977Q | C | 3e-07 | 7.8e-08 | 1.4e-07 | 1.727E-07 | 1.145E-07 | 7.800E-08 | 3.000E-07 |
| 19528 | 977 | T977G | C | 4e-07 | 3.9e-07 | 1.8e-07 | 3.233E-07 | 1.242E-07 | 1.800E-07 | 4.000E-07 |
| 19529 | 977 | T977H | C | 3.2e-07 | 1.9e-07 | 3e-07 | 2.700E-07 | 7.000E-08 | 1.900E-07 | 3.200E-07 |
| 19530 | 977 | T977I | C | 1.4e-07 | 2.7e-07 | 4.8e-07 | 2.967E-07 | 1.716E-07 | 1.400E-07 | 4.800E-07 |
| 19531 | 977 | T977L | C | 2.5e-07 | 1.5e-07 | 2.4e-07 | 2.133E-07 | 5.508E-08 | 1.500E-07 | 2.500E-07 |
| 19532 | 977 | T977K | C | 4.8e-07 | 3.9e-07 | 4.1e-07 | 4.267E-07 | 4.726E-08 | 3.900E-07 | 4.800E-07 |
| 19533 | 977 | T977M | C | 2e-07 | 1.6e-07 | 9.8e-08 | 1.527E-07 | 5.139E-08 | 9.800E-08 | 2.000E-07 |
| 19534 | 977 | T977F | C | 2.5e-07 | 1.3e-07 | 2e-07 | 1.933E-07 | 6.028E-08 | 1.300E-07 | 2.500E-07 |
| 19535 | 977 | T977P | C | 2.6e-07 | 4e-07 | 2.2e-07 | 2.933E-07 | 9.452E-08 | 2.200E-07 | 4.000E-07 |
| 19536 | 977 | T977S | C | 2.7e-07 | 3.2e-07 | 3.6e-07 | 3.167E-07 | 4.509E-08 | 2.700E-07 | 3.600E-07 |
| 19537 | 977 | T977T | C | 2.8e-07 | 3e-07 | 2.4e-07 | 2.733E-07 | 3.055E-08 | 2.400E-07 | 3.000E-07 |
| 19538 | 977 | T977W | C | 4.1e-07 | 9.1e-08 | 4.4e-07 | 3.137E-07 | 1.934E-07 | 9.100E-08 | 4.400E-07 |
| 19539 | 977 | T977Y | C | 3.1e-07 | 1.5e-07 | 4e-07 | 2.867E-07 | 1.266E-07 | 1.500E-07 | 4.000E-07 |
| 19540 | 977 | T977V | C | 2.8e-07 | 1.9e-07 | 3.7e-07 | 2.800E-07 | 9.000E-08 | 1.900E-07 | 3.700E-07 |
| 19541 | 978 | A978A | C | 3e-07 | 2.4e-07 | 2.5e-07 | 2.633E-07 | 3.215E-08 | 2.400E-07 | 3.000E-07 |
| 19542 | 978 | A978R | C | 3e-07 | 1e-07 | 3.1e-07 | 2.367E-07 | 1.185E-07 | 1.000E-07 | 3.100E-07 |
| 19543 | 978 | A978N | C | 3.4e-07 | 5.4e-07 | 1.5e-07 | 3.433E-07 | 1.950E-07 | 1.500E-07 | 5.400E-07 |
| 19544 | 978 | A978D | C | 4e-07 | 2.9e-07 | 2.2e-07 | 3.033E-07 | 9.074E-08 | 2.200E-07 | 4.000E-07 |
| 19545 | 978 | A978C | C | 2.7e-07 | 1.7e-07 | 1.8e-07 | 2.067E-07 | 5.508E-08 | 1.700E-07 | 2.700E-07 |
| 19546 | 978 | A978E | C | 2.9e-07 | 2.6e-07 | 2.3e-07 | 2.600E-07 | 3.000E-08 | 2.300E-07 | 2.900E-07 |
| 19547 | 978 | A978Q | C | 2.9e-07 | 3.6e-07 | 4.8e-07 | 3.767E-07 | 9.609E-08 | 2.900E-07 | 4.800E-07 |
| 19548 | 978 | A978G | C | 2e-07 | 3e-07 | 4.1e-07 | 3.033E-07 | 1.050E-07 | 2.000E-07 | 4.100E-07 |
| 19549 | 978 | A978H | C | 2.1e-07 | 2.9e-07 | 3e-07 | 2.667E-07 | 4.933E-08 | 2.100E-07 | 3.000E-07 |

| | | | | | | | | | | |
|---|---|---|---|---|---|---|---|---|---|---|
| 19550 | 978 | A978I | C | 1.6e-07 | 2.4e-07 | 2.1e-07 | 2.033E-07 | 4.041E-08 | 1.600E-07 | 2.400E-07 |
| 19551 | 978 | A978L | C | 3.9e-07 | 2.1e-07 | 2.3e-07 | 2.767E-07 | 9.866E-08 | 2.100E-07 | 3.900E-07 |
| 19552 | 978 | A978K | C | 2e-07 | 1.8e-07 | 3.3e-07 | 2.367E-07 | 8.145E-08 | 1.800E-07 | 3.300E-07 |
| 19553 | 978 | A978M | C | 1.8e-07 | 3.8e-07 | 2.7e-07 | 2.767E-07 | 1.002E-07 | 1.800E-07 | 3.800E-07 |
| 19554 | 978 | A978F | C | 5.5e-07 | 2.2e-07 | 1.1e-07 | 2.933E-07 | 2.290E-07 | 1.100E-07 | 5.500E-07 |
| 19555 | 978 | A978P | C | 1.5e-07 | 1.8e-07 | 3.3e-07 | 2.200E-07 | 9.644E-08 | 1.500E-07 | 3.300E-07 |
| 19556 | 978 | A978S | C | 3.2e-07 | 5.1e-07 | 2.2e-07 | 3.500E-07 | 1.473E-07 | 2.200E-07 | 5.100E-07 |
| 19557 | 978 | A978T | C | 2e-07 | 1.8e-07 | 1.3e-07 | 1.700E-07 | 3.606E-08 | 1.300E-07 | 2.000E-07 |
| 19558 | 978 | A978W | C | 3.7e-07 | 1.2e-07 | 2e-07 | 2.300E-07 | 1.277E-07 | 1.200E-07 | 3.700E-07 |
| 19559 | 978 | A978Y | C | 1.3e-07 | 1.8e-07 | 3.1e-07 | 2.067E-07 | 9.292E-08 | 1.300E-07 | 3.100E-07 |
| 19560 | 978 | A978V | C | 2e-07 | 3.6e-07 | 3.7e-07 | 3.100E-07 | 9.539E-08 | 2.000E-07 | 3.700E-07 |
| 19561 | 979 | A979A | C | 4.1e-07 | 1.5e-07 | 2.8e-07 | 2.800E-07 | 1.300E-07 | 1.500E-07 | 4.100E-07 |
| 19562 | 979 | A979R | C | 2.4e-07 | 2.1e-07 | 1.9e-07 | 2.133E-07 | 2.517E-08 | 1.900E-07 | 2.400E-07 |
| 19563 | 979 | A979N | C | 2.4e-07 | 3.9e-07 | 2.7e-07 | 3.000E-07 | 7.937E-08 | 2.400E-07 | 3.900E-07 |
| 19564 | 979 | A979D | C | 4.5e-07 | 3.3e-07 | 3.7e-07 | 3.833E-07 | 6.110E-08 | 3.300E-07 | 4.500E-07 |
| 19565 | 979 | A979C | C | 3.8e-07 | 1.5e-07 | 3.2e-07 | 2.833E-07 | 1.193E-07 | 1.500E-07 | 3.800E-07 |
| 19566 | 979 | A979E | C | 3.5e-07 | 1.7e-07 | 2.7e-07 | 2.633E-07 | 9.018E-08 | 1.700E-07 | 3.500E-07 |
| 19567 | 979 | A979Q | C | 1.6e-07 | 1.6e-07 | 4.2e-07 | 2.467E-07 | 1.501E-07 | 1.600E-07 | 4.200E-07 |
| 19568 | 979 | A979G | C | 1.8e-07 | 1.2e-07 | 1.4e-07 | 1.467E-07 | 3.055E-08 | 1.200E-07 | 1.800E-07 |
| 19569 | 979 | A979H | C | 3.4e-07 | 1.9e-07 | 2.7e-07 | 2.667E-07 | 7.506E-08 | 1.900E-07 | 3.400E-07 |
| 19570 | 979 | A979I | C | 3e-07 | 1.1e-07 | 2.1e-07 | 2.067E-07 | 9.504E-08 | 1.100E-07 | 3.000E-07 |
| 19571 | 979 | A979L | C | 4.2e-07 | 1.9e-07 | 1.8e-07 | 2.633E-07 | 1.358E-07 | 1.800E-07 | 4.200E-07 |
| 19572 | 979 | A979K | C | 1.6e-07 | 1.3e-07 | 2.7e-07 | 1.867E-07 | 7.371E-08 | 1.300E-07 | 2.700E-07 |
| 19573 | 979 | A979M | C | 3.4e-07 | 3e-07 | 1e-07 | 2.467E-07 | 1.286E-07 | 1.000E-07 | 3.400E-07 |
| 19574 | 979 | A979F | C | 1.8e-07 | 4.8e-07 | 3.9e-07 | 3.500E-07 | 1.539E-07 | 1.800E-07 | 4.800E-07 |
| 19575 | 979 | A979P | C | 7.5e-07 | 2.7e-07 | 2.4e-07 | 4.200E-07 | 2.862E-07 | 2.400E-07 | 7.500E-07 |
| 19576 | 979 | A979S | C | 1.7e-07 | 6.8e-08 | 3.1e-07 | 1.827E-07 | 1.215E-07 | 6.800E-08 | 3.100E-07 |
| 19577 | 979 | A979T | C | 3.8e-07 | 2.4e-07 | 1.8e-07 | 2.667E-07 | 1.026E-07 | 1.800E-07 | 3.800E-07 |
| 19578 | 979 | A979W | C | 1.9e-07 | 3.1e-07 | 2.7e-07 | 2.567E-07 | 6.110E-08 | 1.900E-07 | 3.100E-07 |
| 19579 | 979 | A979Y | C | 1.7e-07 | 3.2e-07 | 2.9e-07 | 2.600E-07 | 7.937E-08 | 1.700E-07 | 3.200E-07 |
| 19580 | 979 | A979V | C | 2e-07 | 2.7e-07 | 3e-07 | 2.567E-07 | 5.132E-08 | 2.000E-07 | 3.000E-07 |
| 19581 | 980 | L980A | C | 2.6e-07 | 2.1e-07 | 2.7e-07 | 2.467E-07 | 3.215E-08 | 2.100E-07 | 2.700E-07 |
| 19582 | 980 | L980R | C | 3.8e-07 | 1.8e-07 | 2.2e-07 | 2.600E-07 | 1.058E-07 | 1.800E-07 | 3.800E-07 |
| 19583 | 980 | L980N | C | 2.6e-07 | 3.2e-07 | 3.2e-07 | 3.000E-07 | 3.464E-08 | 2.600E-07 | 3.200E-07 |
| 19584 | 980 | L980D | C | 1.1e-07 | 2.7e-07 | 2.4e-07 | 2.067E-07 | 8.505E-08 | 1.100E-07 | 2.700E-07 |
| 19585 | 980 | L980C | C | 3.1e-07 | 3.2e-07 | 2.4e-07 | 2.900E-07 | 4.359E-08 | 2.400E-07 | 3.200E-07 |
| 19586 | 980 | L980E | C | 2.1e-07 | 2.4e-07 | 4.5e-07 | 3.000E-07 | 1.308E-07 | 2.100E-07 | 4.500E-07 |
| 19587 | 980 | L980Q | C | 5.5e-07 | 2.5e-07 | 3.1e-07 | 3.700E-07 | 1.587E-07 | 2.500E-07 | 5.500E-07 |
| 19588 | 980 | L980G | C | 4.8e-07 | 2.6e-07 | 2.4e-07 | 3.267E-07 | 1.332E-07 | 2.400E-07 | 4.800E-07 |
| 19589 | 980 | L980H | C | 2.1e-07 | 2.3e-07 | 3.2e-07 | 2.533E-07 | 5.859E-08 | 2.100E-07 | 3.200E-07 |
| 19590 | 980 | L980I | C | 1.4e-07 | 2.2e-07 | 2.8e-07 | 2.133E-07 | 7.024E-08 | 1.400E-07 | 2.800E-07 |
| 19591 | 980 | L980L | C | 2.7e-07 | 2.7e-07 | 3.1e-07 | 2.833E-07 | 2.309E-08 | 2.700E-07 | 3.100E-07 |
| 19592 | 980 | L980K | C | 2.9e-07 | 3.4e-07 | 3.8e-07 | 3.367E-07 | 4.509E-08 | 2.900E-07 | 3.800E-07 |
| 19593 | 980 | L980M | C | 2.5e-07 | 9.9e-08 | 3.8e-07 | 2.430E-07 | 1.406E-07 | 9.900E-08 | 3.800E-07 |
| 19594 | 980 | L980F | C | 6.1e-08 | 1.3e-07 | 5.9e-07 | 2.603E-07 | 2.876E-07 | 6.100E-08 | 5.900E-07 |
| 19595 | 980 | L980P | C | 1.6e-07 | 2.3e-07 | 2.2e-07 | 2.033E-07 | 3.786E-08 | 1.600E-07 | 2.300E-07 |
| 19596 | 980 | L980S | C | 3.2e-07 | 2.7e-07 | 3.3e-07 | 3.067E-07 | 3.215E-08 | 2.700E-07 | 3.300E-07 |
| 19597 | 980 | L980T | C | 5e-07 | 3e-07 | 2.3e-07 | 3.433E-07 | 1.401E-07 | 2.300E-07 | 5.000E-07 |
| 19598 | 980 | L980W | C | 2.1e-07 | 3.3e-07 | 3.4e-07 | 2.933E-07 | 7.234E-08 | 2.100E-07 | 3.400E-07 |
| 19599 | 980 | L980Y | C | 1.4e-07 | 2.1e-07 | 2.9e-07 | 2.133E-07 | 7.506E-08 | 1.400E-07 | 2.900E-07 |
| 19600 | 980 | L980V | C | 4.2e-07 | 1.5e-07 | 2.1e-07 | 2.600E-07 | 1.418E-07 | 1.500E-07 | 4.200E-07 |
| 19601 | 981 | G981A | C | 2.9e-07 | 3.3e-07 | 3.6e-07 | 3.267E-07 | 3.512E-08 | 2.900E-07 | 3.600E-07 |
| 19602 | 981 | G981R | C | 2.3e-07 | 3.4e-07 | 3.6e-07 | 3.100E-07 | 7.000E-08 | 2.300E-07 | 3.600E-07 |
| 19603 | 981 | G981N | C | 2.7e-07 | 2.7e-07 | 1.2e-07 | 2.200E-07 | 8.660E-08 | 1.200E-07 | 2.700E-07 |
| 19604 | 981 | G981D | C | 1.6e-07 | 2.1e-07 | 2.9e-07 | 2.200E-07 | 6.557E-08 | 1.600E-07 | 2.900E-07 |
| 19605 | 981 | G981C | C | 3.9e-07 | 2.8e-07 | 1.4e-07 | 2.700E-07 | 1.253E-07 | 1.400E-07 | 3.900E-07 |
| 19606 | 981 | G981E | C | 6.1e-07 | 1.2e-07 | 2.7e-07 | 3.333E-07 | 2.511E-07 | 1.200E-07 | 6.100E-07 |
| 19607 | 981 | G981Q | C | 2.2e-07 | 3.3e-07 | 4.1e-07 | 3.200E-07 | 9.539E-08 | 2.200E-07 | 4.100E-07 |
| 19608 | 981 | G981G | C | 2.9e-07 | 4.1e-07 | 2.3e-07 | 3.100E-07 | 9.165E-08 | 2.300E-07 | 4.100E-07 |
| 19609 | 981 | G981H | C | 1.7e-07 | 2.8e-07 | 6.3e-07 | 3.600E-07 | 2.402E-07 | 1.700E-07 | 6.300E-07 |
| 19610 | 981 | G981I | C | 4.5e-07 | 1.5e-07 | 2.2e-07 | 2.733E-07 | 1.570E-07 | 1.500E-07 | 4.500E-07 |
| 19611 | 981 | G981L | C | 2.5e-07 | 3.3e-07 | 3.9e-07 | 3.233E-07 | 7.024E-08 | 2.500E-07 | 3.900E-07 |
| 19612 | 981 | G981K | C | 9.5e-08 | 5.3e-07 | 1.8e-07 | 2.683E-07 | 2.306E-07 | 9.500E-08 | 5.300E-07 |
| 19613 | 981 | G981M | C | 2.9e-07 | 1.1e-07 | 2.4e-07 | 2.133E-07 | 9.292E-08 | 1.100E-07 | 2.900E-07 |
| 19614 | 981 | G981F | C | 1.6e-07 | 1.7e-07 | 3.6e-07 | 2.300E-07 | 1.127E-07 | 1.600E-07 | 3.600E-07 |
| 19615 | 981 | G981P | C | 1.7e-07 | 1.6e-07 | 3.9e-07 | 2.400E-07 | 1.300E-07 | 1.600E-07 | 3.900E-07 |
| 19616 | 981 | G981S | C | 2.7e-07 | 2.7e-07 | 2e-07 | 2.467E-07 | 4.041E-08 | 2.000E-07 | 2.700E-07 |
| 19617 | 981 | G981T | C | 2.5e-07 | 2.1e-07 | 2.6e-07 | 2.400E-07 | 2.646E-08 | 2.100E-07 | 2.600E-07 |
| 19618 | 981 | G981W | C | 1.6e-07 | 3.6e-07 | 5.4e-07 | 3.533E-07 | 1.901E-07 | 1.600E-07 | 5.400E-07 |
| 19619 | 981 | G981Y | C | 1.8e-07 | 4.1e-07 | 3.4e-07 | 3.100E-07 | 1.179E-07 | 1.800E-07 | 4.100E-07 |
| 19620 | 981 | G981V | C | 1.5e-07 | 2.6e-07 | 2e-07 | 2.033E-07 | 5.508E-08 | 1.500E-07 | 2.600E-07 |
| 19621 | 982 | C982A | C | 2.1e-07 | 3.3e-07 | 3.4e-07 | 2.933E-07 | 7.234E-08 | 2.100E-07 | 3.400E-07 |
| 19622 | 982 | C982R | C | 3.3e-07 | 2.4e-07 | 3.2e-07 | 2.967E-07 | 4.933E-08 | 2.400E-07 | 3.300E-07 |
| 19623 | 982 | C982N | C | 1.3e-07 | 2.6e-07 | 3e-07 | 2.300E-07 | 8.888E-08 | 1.300E-07 | 3.000E-07 |
| 19624 | 982 | C982D | C | 6.6e-07 | 1.6e-07 | 1.3e-07 | 3.167E-07 | 2.977E-07 | 1.300E-07 | 6.600E-07 |
| 19625 | 982 | C982C | C | 3.8e-07 | 1.3e-07 | 2.3e-07 | 2.467E-07 | 1.258E-07 | 1.300E-07 | 3.800E-07 |
| 19626 | 982 | C982E | C | 2e-07 | 3.5e-07 | 3.4e-07 | 2.967E-07 | 8.386E-08 | 2.000E-07 | 3.500E-07 |
| 19627 | 982 | C982Q | C | 1.8e-07 | 3.4e-07 | 1.7e-07 | 2.300E-07 | 9.539E-08 | 1.700E-07 | 3.400E-07 |
| 19628 | 982 | C982G | C | 2.4e-07 | 3.4e-07 | 2.1e-07 | 2.633E-07 | 6.807E-08 | 2.100E-07 | 3.400E-07 |
| 19629 | 982 | C982H | C | 1.5e-07 | 2.7e-07 | 2.3e-07 | 2.167E-07 | 6.110E-08 | 1.500E-07 | 2.700E-07 |
| 19630 | 982 | C982I | C | 2.3e-07 | 2.5e-07 | 1.9e-07 | 2.233E-07 | 3.055E-08 | 1.900E-07 | 2.500E-07 |
| 19631 | 982 | C982L | C | 1.7e-07 | 2.3e-07 | 1.4e-07 | 1.800E-07 | 4.583E-08 | 1.400E-07 | 2.300E-07 |
| 19632 | 982 | C982K | C | 1.3e-07 | 8.8e-08 | 2.6e-07 | 1.593E-07 | 8.967E-08 | 8.800E-08 | 2.600E-07 |
| 19633 | 982 | C982M | C | 5.1e-07 | 2e-07 | 2.4e-07 | 3.167E-07 | 1.686E-07 | 2.000E-07 | 5.100E-07 |
| 19634 | 982 | C982F | C | 1.6e-07 | 2.3e-07 | 1.7e-07 | 1.867E-07 | 3.786E-08 | 1.600E-07 | 2.300E-07 |

| | | | | | | | | | | |
|---|---|---|---|---|---|---|---|---|---|---|
| 19635 | 982 | C982P | C | 2.8e-07 | 2.1e-07 | 3.9e-07 | 2.933E-07 | 9.074E-08 | 2.100E-07 | 3.900E-07 |
| 19636 | 982 | C982S | C | 2.4e-07 | 4.1e-07 | 2e-07 | 2.833E-07 | 1.115E-07 | 2.000E-07 | 4.100E-07 |
| 19637 | 982 | C982T | C | 2.7e-07 | 2.5e-07 | 2.4e-07 | 2.533E-07 | 1.528E-08 | 2.400E-07 | 2.700E-07 |
| 19638 | 982 | C982W | C | 2.4e-07 | 1.3e-07 | 4.6e-07 | 2.767E-07 | 1.680E-07 | 1.300E-07 | 4.600E-07 |
| 19639 | 982 | C982Y | C | 2.3e-07 | 2.7e-07 | 2.8e-07 | 2.600E-07 | 2.646E-08 | 2.300E-07 | 2.800E-07 |
| 19640 | 982 | C982V | C | 4.7e-07 | 3.1e-07 | 3.6e-07 | 3.800E-07 | 8.185E-08 | 3.100E-07 | 4.700E-07 |
| 19641 | 983 | L983A | C | 3.3e-07 | 3.6e-07 | 2e-07 | 2.967E-07 | 8.505E-08 | 2.000E-07 | 3.600E-07 |
| 19642 | 983 | L983R | C | 3.5e-07 | 3.2e-07 | 1.7e-07 | 2.800E-07 | 9.644E-08 | 1.700E-07 | 3.500E-07 |
| 19643 | 983 | L983N | C | 3.2e-07 | 1.6e-07 | 3.9e-07 | 2.900E-07 | 1.179E-07 | 1.600E-07 | 3.900E-07 |
| 19644 | 983 | L983D | C | 3.8e-07 | 2.1e-07 | 3.1e-07 | 3.000E-07 | 8.544E-08 | 2.100E-07 | 3.800E-07 |
| 19645 | 983 | L983C | C | 1.9e-07 | 2.5e-07 | 5.6e-07 | 3.333E-07 | 1.986E-07 | 1.900E-07 | 5.600E-07 |
| 19646 | 983 | L983E | C | 4.6e-07 | 1.2e-07 | 8e-08 | 2.200E-07 | 2.088E-07 | 8.000E-08 | 4.600E-07 |
| 19647 | 983 | L983Q | C | 2.1e-07 | 1.4e-07 | 2.5e-07 | 2.000E-07 | 5.568E-08 | 1.400E-07 | 2.500E-07 |
| 19648 | 983 | L983G | C | 3.2e-07 | 3.9e-07 | 1.8e-07 | 2.967E-07 | 1.069E-07 | 1.800E-07 | 3.900E-07 |
| 19649 | 983 | L983H | C | 1.2e-07 | 2.5e-07 | 1.9e-07 | 1.867E-07 | 6.506E-08 | 1.200E-07 | 2.500E-07 |
| 19650 | 983 | L983I | C | 1.8e-07 | 3.8e-07 | 3.5e-07 | 3.033E-07 | 1.079E-07 | 1.800E-07 | 3.800E-07 |
| 19651 | 983 | L983L | C | 1.4e-07 | 5e-07 | 1.5e-07 | 2.633E-07 | 2.050E-07 | 1.400E-07 | 5.000E-07 |
| 19652 | 983 | L983K | C | 3.9e-07 | 3.6e-07 | 3.1e-07 | 3.533E-07 | 4.041E-08 | 3.100E-07 | 3.900E-07 |
| 19653 | 983 | L983M | C | 6.9e-07 | 1.3e-07 | 7.1e-07 | 5.100E-07 | 3.292E-07 | 1.300E-07 | 7.100E-07 |
| 19654 | 983 | L983F | C | 3e-07 | 4.3e-07 | 2e-07 | 3.100E-07 | 1.153E-07 | 2.000E-07 | 4.300E-07 |
| 19655 | 983 | L983P | C | 3.1e-07 | 1.6e-07 | 1.6e-07 | 2.100E-07 | 8.660E-08 | 1.600E-07 | 3.100E-07 |
| 19656 | 983 | L983S | C | 2.7e-07 | 1.8e-07 | 4.3e-07 | 2.933E-07 | 1.266E-07 | 1.800E-07 | 4.300E-07 |
| 19657 | 983 | L983T | C | 2.9e-07 | 1.6e-07 | 2.6e-07 | 2.367E-07 | 6.807E-08 | 1.600E-07 | 2.900E-07 |
| 19658 | 983 | L983W | C | 2.3e-07 | 5.5e-07 | 4.2e-07 | 4.000E-07 | 1.609E-07 | 2.300E-07 | 5.500E-07 |
| 19659 | 983 | L983Y | C | 4.4e-07 | 1.6e-07 | 2e-07 | 2.667E-07 | 1.514E-07 | 1.600E-07 | 4.400E-07 |
| 19660 | 983 | L983V | C | 1.3e-07 | 3e-07 | 4.7e-07 | 3.000E-07 | 1.700E-07 | 1.300E-07 | 4.700E-07 |
| 19661 | 984 | V984A | C | 5.7e-07 | 3.8e-07 | 2.4e-07 | 3.967E-07 | 1.656E-07 | 2.400E-07 | 5.700E-07 |
| 19662 | 984 | V984R | C | 1.6e-07 | 2.5e-07 | 2.8e-07 | 2.300E-07 | 6.245E-08 | 1.600E-07 | 2.800E-07 |
| 19663 | 984 | V984N | C | 3.8e-07 | 2.2e-07 | 1.4e-07 | 2.467E-07 | 1.222E-07 | 1.400E-07 | 3.800E-07 |
| 19664 | 984 | V984D | C | 2.6e-07 | 1.5e-07 | 2.3e-07 | 2.133E-07 | 5.686E-08 | 1.500E-07 | 2.600E-07 |
| 19665 | 984 | V984C | C | 1e-07 | 2.3e-07 | 1.8e-07 | 1.700E-07 | 6.557E-08 | 1.000E-07 | 2.300E-07 |
| 19666 | 984 | V984E | C | 1.8e-07 | 6.4e-07 | 2.3e-07 | 3.500E-07 | 2.524E-07 | 1.800E-07 | 6.400E-07 |
| 19667 | 984 | V984Q | C | 3e-07 | 2.2e-07 | 5.7e-07 | 3.633E-07 | 1.834E-07 | 2.200E-07 | 5.700E-07 |
| 19668 | 984 | V984G | C | 2.7e-07 | 2.1e-07 | 4e-07 | 2.933E-07 | 9.713E-08 | 2.100E-07 | 4.000E-07 |
| 19669 | 984 | V984H | C | 2.7e-07 | 1.4e-07 | 2.6e-07 | 2.233E-07 | 7.234E-08 | 1.400E-07 | 2.700E-07 |
| 19670 | 984 | V984I | C | 1.5e-07 | 2.1e-07 | 3.1e-07 | 2.233E-07 | 8.083E-08 | 1.500E-07 | 3.100E-07 |
| 19671 | 984 | V984L | C | 5.1e-07 | 1.3e-07 | 1.9e-07 | 2.767E-07 | 2.043E-07 | 1.300E-07 | 5.100E-07 |
| 19672 | 984 | V984K | C | 1.5e-07 | 3.2e-07 | 2.8e-07 | 2.500E-07 | 8.888E-08 | 1.500E-07 | 3.200E-07 |
| 19673 | 984 | V984M | C | 1.5e-07 | 2.9e-07 | 3e-07 | 2.467E-07 | 8.386E-08 | 1.500E-07 | 3.000E-07 |
| 19674 | 984 | V984F | C | 2.9e-07 | 1.4e-07 | 2.8e-07 | 2.367E-07 | 8.386E-08 | 1.400E-07 | 2.900E-07 |
| 19675 | 984 | V984P | C | 3.2e-07 | 6.4e-08 | 2.7e-07 | 2.180E-07 | 1.357E-07 | 6.400E-08 | 3.200E-07 |
| 19676 | 984 | V984S | C | 2.6e-07 | 1.7e-07 | 9.8e-08 | 1.760E-07 | 8.117E-08 | 9.800E-08 | 2.600E-07 |
| 19677 | 984 | V984T | C | 3.8e-07 | 5.3e-07 | 3.1e-07 | 4.067E-07 | 1.124E-07 | 3.100E-07 | 5.300E-07 |
| 19678 | 984 | V984W | C | 1.6e-07 | 2.4e-07 | 1.7e-07 | 1.900E-07 | 4.359E-08 | 1.600E-07 | 2.400E-07 |
| 19679 | 984 | V984Y | C | 3.4e-07 | 1.9e-07 | 2.3e-07 | 2.533E-07 | 7.767E-08 | 1.900E-07 | 3.400E-07 |
| 19680 | 984 | V984V | C | 3.6e-07 | 4.6e-07 | 1.2e-07 | 3.133E-07 | 1.747E-07 | 1.200E-07 | 4.600E-07 |
| 19681 | 985 | K985A | C | 2.4e-07 | 2e-07 | 1.7e-07 | 2.033E-07 | 3.512E-08 | 1.700E-07 | 2.400E-07 |
| 19682 | 985 | K985R | C | 3.5e-07 | 1.4e-07 | 2.4e-07 | 2.433E-07 | 1.050E-07 | 1.400E-07 | 3.500E-07 |
| 19683 | 985 | K985N | C | 1.3e-07 | 3.1e-07 | 2.8e-07 | 2.400E-07 | 9.644E-08 | 1.300E-07 | 3.100E-07 |
| 19684 | 985 | K985D | C | 2.4e-07 | 5.7e-07 | 1.8e-07 | 3.300E-07 | 2.100E-07 | 1.800E-07 | 5.700E-07 |
| 19685 | 985 | K985C | C | 1.3e-07 | 4.3e-07 | 3e-07 | 2.867E-07 | 1.504E-07 | 1.300E-07 | 4.300E-07 |
| 19686 | 985 | K985E | C | 3.8e-07 | 3.7e-07 | 3.7e-07 | 3.733E-07 | 5.774E-09 | 3.700E-07 | 3.800E-07 |
| 19687 | 985 | K985Q | C | 3.6e-07 | 1.4e-07 | 1.6e-07 | 2.200E-07 | 1.217E-07 | 1.400E-07 | 3.600E-07 |
| 19688 | 985 | K985G | C | 3.5e-07 | 3.5e-07 | 2.2e-07 | 3.067E-07 | 7.506E-08 | 2.200E-07 | 3.500E-07 |
| 19689 | 985 | K985H | C | 2.3e-07 | 3.1e-07 | 3.9e-07 | 3.100E-07 | 8.000E-08 | 2.300E-07 | 3.900E-07 |
| 19690 | 985 | K985I | C | 3.8e-07 | 3.1e-07 | 2.4e-07 | 3.100E-07 | 7.000E-08 | 2.400E-07 | 3.800E-07 |
| 19691 | 985 | K985L | C | 2.2e-07 | 2.1e-07 | 2.5e-07 | 2.267E-07 | 2.082E-08 | 2.100E-07 | 2.500E-07 |
| 19692 | 985 | K985K | C | 4.8e-07 | 1.8e-07 | 1.3e-07 | 2.633E-07 | 1.893E-07 | 1.300E-07 | 4.800E-07 |
| 19693 | 985 | K985M | C | 2.2e-07 | 2.9e-07 | 5.5e-07 | 3.533E-07 | 1.739E-07 | 2.200E-07 | 5.500E-07 |
| 19694 | 985 | K985F | C | 4.5e-07 | 3.4e-07 | 1.7e-07 | 3.200E-07 | 1.411E-07 | 1.700E-07 | 4.500E-07 |
| 19695 | 985 | K985P | C | 4.8e-07 | 3.4e-07 | 2.1e-07 | 3.433E-07 | 1.350E-07 | 2.100E-07 | 4.800E-07 |
| 19696 | 985 | K985S | C | 1.3e-07 | 3e-07 | 4e-07 | 2.767E-07 | 1.365E-07 | 1.300E-07 | 4.000E-07 |
| 19697 | 985 | K985T | C | 1.7e-07 | 4.8e-07 | 8.5e-08 | 2.450E-07 | 2.079E-07 | 8.500E-08 | 4.800E-07 |
| 19698 | 985 | K985W | C | 4.6e-07 | 1.7e-07 | 1.1e-07 | 2.467E-07 | 1.872E-07 | 1.100E-07 | 4.600E-07 |
| 19699 | 985 | K985Y | C | 2.3e-07 | 4.9e-07 | 2.4e-07 | 3.200E-07 | 1.473E-07 | 2.300E-07 | 4.900E-07 |
| 19700 | 985 | K985V | C | 2.3e-07 | 1.7e-07 | 1.9e-07 | 1.967E-07 | 3.055E-08 | 1.700E-07 | 2.300E-07 |
| 19701 | 986 | D986A | C | 3.4e-07 | 4.1e-07 | 1.4e-07 | 2.967E-07 | 1.401E-07 | 1.400E-07 | 4.100E-07 |
| 19702 | 986 | D986R | C | 1.4e-07 | 3.1e-07 | 1.2e-07 | 1.900E-07 | 1.044E-07 | 1.200E-07 | 3.100E-07 |
| 19703 | 986 | D986N | C | 1.8e-07 | 4.5e-07 | 2.1e-07 | 2.800E-07 | 1.480E-07 | 1.800E-07 | 4.500E-07 |
| 19704 | 986 | D986D | C | 1.7e-07 | 3.7e-07 | 4.1e-07 | 3.167E-07 | 1.286E-07 | 1.700E-07 | 4.100E-07 |
| 19705 | 986 | D986C | C | 2e-07 | 2.8e-07 | 2.7e-07 | 2.500E-07 | 4.359E-08 | 2.000E-07 | 2.800E-07 |
| 19706 | 986 | D986E | C | 1.4e-07 | 2.9e-07 | 2.2e-07 | 2.167E-07 | 7.506E-08 | 1.400E-07 | 2.900E-07 |
| 19707 | 986 | D986Q | C | 1.2e-07 | 4.4e-07 | 3.4e-07 | 3.000E-07 | 1.637E-07 | 1.200E-07 | 4.400E-07 |
| 19708 | 986 | D986G | C | 2.6e-07 | 9.8e-08 | 1.6e-07 | 1.727E-07 | 8.174E-08 | 9.800E-08 | 2.600E-07 |
| 19709 | 986 | D986H | C | 1.9e-07 | 2.3e-07 | 2.2e-07 | 2.133E-07 | 2.082E-08 | 1.900E-07 | 2.300E-07 |
| 19710 | 986 | D986I | C | 2.1e-07 | 1.5e-07 | 2.5e-07 | 2.033E-07 | 5.033E-08 | 1.500E-07 | 2.500E-07 |
| 19711 | 986 | D986L | C | 2.7e-07 | 2.8e-07 | 3.9e-07 | 3.133E-07 | 6.658E-08 | 2.700E-07 | 3.900E-07 |
| 19712 | 986 | D986K | C | 2.2e-07 | 4.6e-07 | 2.5e-07 | 3.100E-07 | 1.308E-07 | 2.200E-07 | 4.600E-07 |
| 19713 | 986 | D986M | C | 4.4e-07 | 2.3e-07 | 2.6e-07 | 3.100E-07 | 1.136E-07 | 2.300E-07 | 4.400E-07 |
| 19714 | 986 | D986F | C | 2.5e-07 | 3.1e-07 | 4.6e-07 | 3.400E-07 | 1.082E-07 | 2.500E-07 | 4.600E-07 |
| 19715 | 986 | D986P | C | 1.7e-07 | 2.2e-07 | 3.2e-07 | 2.367E-07 | 7.638E-08 | 1.700E-07 | 3.200E-07 |
| 19716 | 986 | D986S | C | 1.9e-07 | 3.2e-07 | 2.1e-07 | 2.400E-07 | 7.000E-08 | 1.900E-07 | 3.200E-07 |
| 19717 | 986 | D986T | C | 1.6e-07 | 4.5e-07 | 3.4e-07 | 3.167E-07 | 1.464E-07 | 1.600E-07 | 4.500E-07 |
| 19718 | 986 | D986W | C | 3.5e-07 | 3.1e-07 | 3.6e-07 | 3.400E-07 | 2.646E-08 | 3.100E-07 | 3.600E-07 |
| 19719 | 986 | D986Y | C | 2.4e-07 | 1.9e-07 | 4.2e-07 | 2.833E-07 | 1.210E-07 | 1.900E-07 | 4.200E-07 |

| | | | | | | | | | | |
|---|---|---|---|---|---|---|---|---|---|---|
| 19720 | 986 | D986V | C | 3.3e-07 | 2.4e-07 | 2.3e-07 | 2.667E-07 | 5.508E-08 | 2.300E-07 | 3.300E-07 |
| 19721 | 987 | Y987A | C | 1.9e-07 | 3.4e-07 | 1.1e-07 | 2.133E-07 | 1.168E-07 | 1.100E-07 | 3.400E-07 |
| 19722 | 987 | Y987R | C | 1.7e-07 | 2.1e-07 | 1.8e-07 | 1.867E-07 | 2.082E-08 | 1.700E-07 | 2.100E-07 |
| 19723 | 987 | Y987N | C | 4.6e-07 | 4.6e-07 | 3.7e-07 | 4.300E-07 | 5.196E-08 | 3.700E-07 | 4.600E-07 |
| 19724 | 987 | Y987D | C | 1.8e-07 | 2.6e-07 | 3.4e-07 | 2.600E-07 | 8.000E-08 | 1.800E-07 | 3.400E-07 |
| 19725 | 987 | Y987C | C | 2.4e-07 | 1.5e-07 | 3e-07 | 2.300E-07 | 7.550E-08 | 1.500E-07 | 3.000E-07 |
| 19726 | 987 | Y987E | C | 4.6e-07 | 4.7e-07 | 3.9e-07 | 4.400E-07 | 4.359E-08 | 3.900E-07 | 4.700E-07 |
| 19727 | 987 | Y987Q | C | 4.7e-07 | 4.9e-07 | 3.7e-07 | 4.433E-07 | 6.429E-08 | 3.700E-07 | 4.900E-07 |
| 19728 | 987 | Y987G | C | 2.1e-07 | 1.9e-07 | 2.4e-07 | 2.133E-07 | 2.517E-08 | 1.900E-07 | 2.400E-07 |
| 19729 | 987 | Y987H | C | 3.8e-07 | 3.4e-07 | 1.6e-07 | 2.933E-07 | 1.172E-07 | 1.600E-07 | 3.800E-07 |
| 19730 | 987 | Y987I | C | 1.9e-07 | 3.9e-07 | 1.6e-07 | 2.467E-07 | 1.250E-07 | 1.600E-07 | 3.900E-07 |
| 19731 | 987 | Y987L | C | 2.8e-07 | 2.6e-07 | 2.6e-07 | 2.667E-07 | 1.155E-08 | 2.600E-07 | 2.800E-07 |
| 19732 | 987 | Y987K | C | 2.5e-07 | 2.9e-07 | 1.5e-07 | 2.300E-07 | 7.211E-08 | 1.500E-07 | 2.900E-07 |
| 19733 | 987 | Y987M | C | 1.5e-07 | 5.5e-07 | 3.3e-07 | 3.433E-07 | 2.003E-07 | 1.500E-07 | 5.500E-07 |
| 19734 | 987 | Y987F | C | 1.5e-07 | 3.7e-07 | 2e-07 | 2.400E-07 | 1.153E-07 | 1.500E-07 | 3.700E-07 |
| 19735 | 987 | Y987P | C | 3.3e-07 | 1.8e-07 | 3.7e-07 | 2.933E-07 | 1.002E-07 | 1.800E-07 | 3.700E-07 |
| 19736 | 987 | Y987S | C | 2e-07 | 2.5e-07 | 1.7e-07 | 2.067E-07 | 4.041E-08 | 1.700E-07 | 2.500E-07 |
| 19737 | 987 | Y987T | C | 2.1e-07 | 2.5e-07 | 2.7e-07 | 2.433E-07 | 3.055E-08 | 2.100E-07 | 2.700E-07 |
| 19738 | 987 | Y987W | C | 2.9e-07 | 2.3e-07 | 3.4e-07 | 2.867E-07 | 5.508E-08 | 2.300E-07 | 3.400E-07 |
| 19739 | 987 | Y987Y | C | 3.5e-07 | 3.1e-07 | 1.3e-07 | 2.633E-07 | 1.172E-07 | 1.300E-07 | 3.500E-07 |
| 19740 | 987 | Y987V | C | 4.6e-07 | 2.8e-07 | 1.5e-07 | 2.967E-07 | 1.557E-07 | 1.500E-07 | 4.600E-07 |
| 19741 | 988 | F988A | C | 2.4e-07 | 2.2e-07 | 2.3e-07 | 2.300E-07 | 1.000E-08 | 2.200E-07 | 2.400E-07 |
| 19742 | 988 | F988R | C | 2.2e-07 | 1.3e-07 | 2.2e-07 | 1.900E-07 | 5.196E-08 | 1.300E-07 | 2.200E-07 |
| 19743 | 988 | F988N | C | 3.8e-07 | 2.7e-07 | 3.2e-07 | 3.233E-07 | 5.508E-08 | 2.700E-07 | 3.800E-07 |
| 19744 | 988 | F988D | C | 1.1e-07 | 4.4e-07 | 4.6e-07 | 3.367E-07 | 1.966E-07 | 1.100E-07 | 4.600E-07 |
| 19745 | 988 | F988C | C | 2.5e-07 | 4.4e-07 | 2.2e-07 | 3.033E-07 | 1.193E-07 | 2.200E-07 | 4.400E-07 |
| 19746 | 988 | F988E | C | 2.9e-07 | 8.8e-08 | 1.6e-07 | 1.793E-07 | 1.024E-07 | 8.800E-08 | 2.900E-07 |
| 19747 | 988 | F988Q | C | 1e-07 | 3.7e-07 | 4.3e-07 | 3.000E-07 | 1.758E-07 | 1.000E-07 | 4.300E-07 |
| 19748 | 988 | F988G | C | 2.1e-07 | 1.4e-07 | 5.3e-07 | 2.933E-07 | 2.079E-07 | 1.400E-07 | 5.300E-07 |
| 19749 | 988 | F988H | C | 2.6e-07 | 1.4e-07 | 3.4e-07 | 2.467E-07 | 1.007E-07 | 1.400E-07 | 3.400E-07 |
| 19750 | 988 | F988I | C | 3.6e-07 | 2.7e-07 | 2.1e-07 | 2.800E-07 | 7.550E-08 | 2.100E-07 | 3.600E-07 |
| 19751 | 988 | F988L | C | 3.7e-07 | 2.9e-07 | 2.3e-07 | 2.967E-07 | 7.024E-08 | 2.300E-07 | 3.700E-07 |
| 19752 | 988 | F988K | C | 3e-07 | 2.6e-07 | 1.1e-07 | 2.233E-07 | 1.002E-07 | 1.100E-07 | 3.000E-07 |
| 19753 | 988 | F988M | C | 5.2e-07 | 1.5e-07 | 1.2e-07 | 2.633E-07 | 2.228E-07 | 1.200E-07 | 5.200E-07 |
| 19754 | 988 | F988F | C | 2.2e-07 | 1.8e-07 | 2.5e-07 | 2.167E-07 | 3.512E-08 | 1.800E-07 | 2.500E-07 |
| 19755 | 988 | F988P | C | 2.7e-07 | 1.6e-07 | 3.3e-07 | 2.533E-07 | 8.622E-08 | 1.600E-07 | 3.300E-07 |
| 19756 | 988 | F988S | C | 2.1e-07 | 3.2e-07 | 3.5e-07 | 2.933E-07 | 7.371E-08 | 2.100E-07 | 3.500E-07 |
| 19757 | 988 | F988T | C | 1.2e-07 | 2.5e-07 | 2.9e-07 | 2.200E-07 | 8.888E-08 | 1.200E-07 | 2.900E-07 |
| 19758 | 988 | F988W | C | 2.2e-07 | 4e-07 | 1.7e-07 | 2.633E-07 | 1.210E-07 | 1.700E-07 | 4.000E-07 |
| 19759 | 988 | F988Y | C | 2.6e-07 | 1.8e-07 | 1.5e-07 | 1.967E-07 | 5.686E-08 | 1.500E-07 | 2.600E-07 |
| 19760 | 988 | F988V | C | 1.9e-07 | 1.5e-07 | 1.9e-07 | 1.767E-07 | 2.309E-08 | 1.500E-07 | 1.900E-07 |
| 19761 | 989 | P989A | C | 2.7e-07 | 2e-07 | 2.2e-07 | 2.300E-07 | 3.606E-08 | 2.000E-07 | 2.700E-07 |
| 19762 | 989 | P989R | C | 4.3e-07 | 5e-07 | 2.1e-07 | 3.800E-07 | 1.513E-07 | 2.100E-07 | 5.000E-07 |
| 19763 | 989 | P989N | C | 3.5e-07 | 2.7e-07 | 1.9e-07 | 2.700E-07 | 8.000E-08 | 1.900E-07 | 3.500E-07 |
| 19764 | 989 | P989D | C | 3.7e-07 | 2.3e-07 | 1.6e-07 | 2.533E-07 | 1.069E-07 | 1.600E-07 | 3.700E-07 |
| 19765 | 989 | P989C | C | 2e-07 | 1.8e-07 | 2.4e-07 | 2.067E-07 | 3.055E-08 | 1.800E-07 | 2.400E-07 |
| 19766 | 989 | P989E | C | 2.9e-07 | 3.2e-07 | 6.7e-07 | 4.267E-07 | 2.113E-07 | 2.900E-07 | 6.700E-07 |
| 19767 | 989 | P989Q | C | 1.5e-07 | 1.6e-07 | 3.8e-07 | 2.300E-07 | 1.300E-07 | 1.500E-07 | 3.800E-07 |
| 19768 | 989 | P989G | C | 5.7e-07 | 4e-07 | 1.4e-07 | 3.700E-07 | 2.166E-07 | 1.400E-07 | 5.700E-07 |
| 19769 | 989 | P989H | C | 1.6e-07 | 1.3e-07 | 1.6e-07 | 1.500E-07 | 1.732E-08 | 1.300E-07 | 1.600E-07 |
| 19770 | 989 | P989I | C | 3.3e-07 | 3.1e-07 | 1.8e-07 | 2.733E-07 | 8.145E-08 | 1.800E-07 | 3.300E-07 |
| 19771 | 989 | P989L | C | 1.8e-07 | 2.6e-07 | 1.8e-07 | 2.067E-07 | 4.619E-08 | 1.800E-07 | 2.600E-07 |
| 19772 | 989 | P989K | C | 3.7e-07 | 3e-07 | 1.7e-07 | 2.800E-07 | 1.015E-07 | 1.700E-07 | 3.700E-07 |
| 19773 | 989 | P989M | C | 1.1e-07 | 2.3e-07 | 3.9e-07 | 2.433E-07 | 1.405E-07 | 1.100E-07 | 3.900E-07 |
| 19774 | 989 | P989F | C | 2.5e-07 | 1.2e-07 | 4.5e-07 | 2.733E-07 | 1.662E-07 | 1.200E-07 | 4.500E-07 |
| 19775 | 989 | P989P | C | 3.6e-07 | 1.4e-07 | 2.5e-07 | 2.500E-07 | 1.100E-07 | 1.400E-07 | 3.600E-07 |
| 19776 | 989 | P989S | C | 2.3e-07 | 4.6e-07 | 2.8e-07 | 3.233E-07 | 1.210E-07 | 2.300E-07 | 4.600E-07 |
| 19777 | 989 | P989T | C | 2.1e-07 | 4.4e-07 | 1.2e-07 | 2.567E-07 | 1.650E-07 | 1.200E-07 | 4.400E-07 |
| 19778 | 989 | P989W | C | 2.4e-07 | 2.7e-07 | 4.3e-07 | 3.133E-07 | 1.021E-07 | 2.400E-07 | 4.300E-07 |
| 19779 | 989 | P989Y | C | 2.5e-07 | 1.8e-07 | 4.8e-07 | 3.033E-07 | 1.570E-07 | 1.800E-07 | 4.800E-07 |
| 19780 | 989 | P989V | C | 2e-07 | 2.8e-07 | 4e-07 | 2.933E-07 | 1.007E-07 | 2.000E-07 | 4.000E-07 |
| 19781 | 990 | E990A | C | 2e-07 | 2.4e-07 | 1.6e-07 | 2.000E-07 | 4.000E-08 | 1.600E-07 | 2.400E-07 |
| 19782 | 990 | E990R | C | 2.4e-07 | 4.2e-07 | 3.4e-07 | 3.333E-07 | 9.018E-08 | 2.400E-07 | 4.200E-07 |
| 19783 | 990 | E990N | C | 1e-07 | 3.4e-07 | 1.8e-07 | 2.067E-07 | 1.222E-07 | 1.000E-07 | 3.400E-07 |
| 19784 | 990 | E990D | C | 2.2e-07 | 2.5e-07 | 2.9e-07 | 2.533E-07 | 3.512E-08 | 2.200E-07 | 2.900E-07 |
| 19785 | 990 | E990C | C | 3.5e-07 | 1.2e-07 | 4.9e-07 | 3.200E-07 | 1.868E-07 | 1.200E-07 | 4.900E-07 |
| 19786 | 990 | E990E | C | 4.6e-07 | 1.9e-07 | 3.6e-07 | 3.367E-07 | 1.365E-07 | 1.900E-07 | 4.600E-07 |
| 19787 | 990 | E990Q | C | 2.2e-07 | 2.5e-07 | 1.8e-07 | 2.167E-07 | 3.512E-08 | 1.800E-07 | 2.500E-07 |
| 19788 | 990 | E990G | C | 7.8e-08 | 2.5e-07 | 2e-07 | 1.760E-07 | 8.848E-08 | 7.800E-08 | 2.500E-07 |
| 19789 | 990 | E990H | C | 5.3e-07 | 1e-07 | 1.7e-07 | 2.667E-07 | 2.307E-07 | 1.000E-07 | 5.300E-07 |
| 19790 | 990 | E990I | C | 1.6e-07 | 3.1e-07 | 2.9e-07 | 2.533E-07 | 8.145E-08 | 1.600E-07 | 3.100E-07 |
| 19791 | 990 | E990L | C | 3.8e-07 | 3.2e-07 | 3.2e-07 | 3.400E-07 | 3.464E-08 | 3.200E-07 | 3.800E-07 |
| 19792 | 990 | E990K | C | 2.3e-07 | 1.6e-07 | 2.1e-07 | 2.000E-07 | 3.606E-08 | 1.600E-07 | 2.300E-07 |
| 19793 | 990 | E990M | C | 2.1e-07 | 2e-07 | 2.4e-07 | 2.167E-07 | 2.082E-08 | 2.000E-07 | 2.400E-07 |
| 19794 | 990 | E990F | C | 3.8e-07 | 3.6e-07 | 2.2e-07 | 3.200E-07 | 8.718E-08 | 2.200E-07 | 3.800E-07 |
| 19795 | 990 | E990P | C | 1.4e-07 | 4.3e-07 | 1.9e-07 | 2.533E-07 | 1.550E-07 | 1.400E-07 | 4.300E-07 |
| 19796 | 990 | E990S | C | 2.6e-07 | 3.5e-07 | 3.6e-07 | 3.233E-07 | 5.508E-08 | 2.600E-07 | 3.600E-07 |
| 19797 | 990 | E990T | C | 1.7e-07 | 1.3e-07 | 1.8e-07 | 1.600E-07 | 2.646E-08 | 1.300E-07 | 1.800E-07 |
| 19798 | 990 | E990W | C | 3.3e-07 | 1.9e-07 | 2.2e-07 | 2.467E-07 | 7.371E-08 | 1.900E-07 | 3.300E-07 |
| 19799 | 990 | E990Y | C | 1.7e-07 | 2.8e-07 | 1.4e-07 | 1.967E-07 | 7.371E-08 | 1.400E-07 | 2.800E-07 |
| 19800 | 990 | E990V | C | 2.8e-07 | 3.2e-07 | 2.6e-07 | 2.867E-07 | 3.055E-08 | 2.600E-07 | 3.200E-07 |
| 19801 | 991 | P991A | C | 1.6e-07 | 2.8e-07 | 2.4e-07 | 2.267E-07 | 6.110E-08 | 1.600E-07 | 2.800E-07 |
| 19802 | 991 | P991R | C | 3e-07 | 2.2e-07 | 3.9e-07 | 3.033E-07 | 8.505E-08 | 2.200E-07 | 3.900E-07 |
| 19803 | 991 | P991N | C | 2.1e-07 | 3e-07 | 1.6e-07 | 2.233E-07 | 7.095E-08 | 1.600E-07 | 3.000E-07 |
| 19804 | 991 | P991D | C | 2.6e-07 | 3.6e-07 | 5.5e-07 | 3.900E-07 | 1.473E-07 | 2.600E-07 | 5.500E-07 |

| | | | | | | | | | | |
|---|---|---|---|---|---|---|---|---|---|---|
| 19805 | 991 | P991C | C | 1.9e-07 | 1.7e-07 | 2e-07 | 1.867E-07 | 1.528E-08 | 1.700E-07 | 2.000E-07 |
| 19806 | 991 | P991E | C | 2.2e-07 | 3e-07 | 3.6e-07 | 2.933E-07 | 7.024E-08 | 2.200E-07 | 3.600E-07 |
| 19807 | 991 | P991Q | C | 2.5e-07 | 2.8e-07 | 2.1e-07 | 2.467E-07 | 3.512E-08 | 2.100E-07 | 2.800E-07 |
| 19808 | 991 | P991G | C | 2.9e-07 | 2.5e-07 | 3.2e-07 | 2.867E-07 | 3.512E-08 | 2.500E-07 | 3.200E-07 |
| 19809 | 991 | P991H | C | 3.8e-07 | 1.3e-07 | 1.6e-07 | 2.233E-07 | 1.365E-07 | 1.300E-07 | 3.800E-07 |
| 19810 | 991 | P991I | C | 8.2e-08 | 2.5e-07 | 3.8e-07 | 2.373E-07 | 1.494E-07 | 8.200E-08 | 3.800E-07 |
| 19811 | 991 | P991L | C | 2.5e-07 | 4.2e-07 | 2.2e-07 | 2.967E-07 | 1.079E-07 | 2.200E-07 | 4.200E-07 |
| 19812 | 991 | P991K | C | 2.9e-07 | 1.9e-07 | 2.2e-07 | 2.333E-07 | 5.132E-08 | 1.900E-07 | 2.900E-07 |
| 19813 | 991 | P991M | C | 6.9e-07 | 1.9e-07 | 3.1e-07 | 3.967E-07 | 2.610E-07 | 1.900E-07 | 6.900E-07 |
| 19814 | 991 | P991F | C | 1.6e-07 | 3.3e-07 | 1.6e-07 | 2.167E-07 | 9.815E-08 | 1.600E-07 | 3.300E-07 |
| 19815 | 991 | P991P | C | 3e-07 | 2.2e-07 | 1.6e-07 | 2.267E-07 | 7.024E-08 | 1.600E-07 | 3.000E-07 |
| 19816 | 991 | P991S | C | 1.7e-07 | 1.7e-07 | 2.5e-07 | 1.967E-07 | 4.619E-08 | 1.700E-07 | 2.500E-07 |
| 19817 | 991 | P991T | C | 2.3e-07 | 2.3e-07 | 4.9e-07 | 3.167E-07 | 1.501E-07 | 2.300E-07 | 4.900E-07 |
| 19818 | 991 | P991W | C | 2.6e-07 | 1.3e-07 | 1.9e-07 | 1.933E-07 | 6.506E-08 | 1.300E-07 | 2.600E-07 |
| 19819 | 991 | P991Y | C | 1.8e-07 | 2.3e-07 | 2e-07 | 2.033E-07 | 2.517E-08 | 1.800E-07 | 2.300E-07 |
| 19820 | 991 | P991V | C | 2.2e-07 | 3.4e-07 | 2.6e-07 | 2.733E-07 | 6.110E-08 | 2.200E-07 | 3.400E-07 |
| 19821 | 992 | V992A | C | 3.1e-07 | 4.6e-07 | 4.1e-07 | 3.933E-07 | 7.638E-08 | 3.100E-07 | 4.600E-07 |
| 19822 | 992 | V992R | C | 9.1e-08 | 2.4e-07 | 1.6e-07 | 1.637E-07 | 7.457E-08 | 9.100E-08 | 2.400E-07 |
| 19823 | 992 | V992N | C | 2e-07 | 1.4e-07 | 1.6e-07 | 1.667E-07 | 3.055E-08 | 1.400E-07 | 2.000E-07 |
| 19824 | 992 | V992D | C | 3.7e-07 | 3.2e-07 | 1.6e-07 | 2.833E-07 | 1.097E-07 | 1.600E-07 | 3.700E-07 |
| 19825 | 992 | V992C | C | 3.4e-07 | 2.2e-07 | 2.1e-07 | 2.567E-07 | 7.234E-08 | 2.100E-07 | 3.400E-07 |
| 19826 | 992 | V992E | C | 4.3e-07 | 3e-07 | 9.4e-08 | 2.747E-07 | 1.694E-07 | 9.400E-08 | 4.300E-07 |
| 19827 | 992 | V992Q | C | 1.2e-07 | 4.7e-07 | 2.2e-07 | 2.700E-07 | 1.803E-07 | 1.200E-07 | 4.700E-07 |
| 19828 | 992 | V992G | C | 1.5e-07 | 3.3e-07 | 3.7e-07 | 2.833E-07 | 1.172E-07 | 1.500E-07 | 3.700E-07 |
| 19829 | 992 | V992H | C | 2.7e-07 | 1.4e-07 | 2.1e-07 | 2.067E-07 | 6.506E-08 | 1.400E-07 | 2.700E-07 |
| 19830 | 992 | V992I | C | 4.9e-07 | 1.3e-07 | 3.9e-07 | 3.367E-07 | 1.858E-07 | 1.300E-07 | 4.900E-07 |
| 19831 | 992 | V992L | C | 2.7e-07 | 2.6e-07 | 3.1e-07 | 2.800E-07 | 2.646E-08 | 2.600E-07 | 3.100E-07 |
| 19832 | 992 | V992K | C | 2.4e-07 | 2e-07 | 3e-07 | 2.467E-07 | 5.033E-08 | 2.000E-07 | 3.000E-07 |
| 19833 | 992 | V992M | C | 3.2e-07 | 1.8e-07 | 3.7e-07 | 2.900E-07 | 9.849E-08 | 1.800E-07 | 3.700E-07 |
| 19834 | 992 | V992F | C | 3.7e-07 | 3e-07 | 2.7e-07 | 3.133E-07 | 5.132E-08 | 2.700E-07 | 3.700E-07 |
| 19835 | 992 | V992P | C | 3.2e-07 | 4.9e-07 | 1.4e-07 | 3.167E-07 | 1.750E-07 | 1.400E-07 | 4.900E-07 |
| 19836 | 992 | V992S | C | 2.1e-07 | 3e-07 | 2.3e-07 | 2.467E-07 | 4.726E-08 | 2.100E-07 | 3.000E-07 |
| 19837 | 992 | V992T | C | 2.1e-07 | 2.3e-07 | 1.8e-07 | 2.067E-07 | 2.517E-08 | 1.800E-07 | 2.300E-07 |
| 19838 | 992 | V992W | C | 6.3e-07 | 2.5e-07 | 2.3e-07 | 3.700E-07 | 2.254E-07 | 2.300E-07 | 6.300E-07 |
| 19839 | 992 | V992Y | C | 2.2e-07 | 3.1e-07 | 2.3e-07 | 2.533E-07 | 4.933E-08 | 2.200E-07 | 3.100E-07 |
| 19840 | 992 | V992V | C | 2.7e-07 | 2e-07 | 2.7e-07 | 2.467E-07 | 4.041E-08 | 2.000E-07 | 2.700E-07 |
| 19841 | 993 | T993A | C | 1.7e-07 | 3e-07 | 2.1e-07 | 2.267E-07 | 6.658E-08 | 1.700E-07 | 3.000E-07 |
| 19842 | 993 | T993R | C | 3e-07 | 2.1e-07 | 1.8e-07 | 2.300E-07 | 6.245E-08 | 1.800E-07 | 3.000E-07 |
| 19843 | 993 | T993N | C | 2.4e-07 | 1.7e-07 | 2.5e-07 | 2.200E-07 | 4.359E-08 | 1.700E-07 | 2.500E-07 |
| 19844 | 993 | T993D | C | 1.8e-07 | 3.7e-07 | 1.5e-07 | 2.333E-07 | 1.193E-07 | 1.500E-07 | 3.700E-07 |
| 19845 | 993 | T993C | C | 1.8e-07 | 1.5e-07 | 2.4e-07 | 1.900E-07 | 4.583E-08 | 1.500E-07 | 2.400E-07 |
| 19846 | 993 | T993E | C | 2.9e-07 | 2.3e-07 | 4e-07 | 3.067E-07 | 8.622E-08 | 2.300E-07 | 4.000E-07 |
| 19847 | 993 | T993Q | C | 2.3e-07 | 2.2e-07 | 1.4e-07 | 1.967E-07 | 4.933E-08 | 1.400E-07 | 2.300E-07 |
| 19848 | 993 | T993G | C | 2e-07 | 2e-07 | 2.7e-07 | 2.233E-07 | 4.041E-08 | 2.000E-07 | 2.700E-07 |
| 19849 | 993 | T993H | C | 1.6e-07 | 2.3e-07 | 2e-07 | 1.967E-07 | 3.512E-08 | 1.600E-07 | 2.300E-07 |
| 19850 | 993 | T993I | C | 2.8e-07 | 1.4e-07 | 2.6e-07 | 2.267E-07 | 7.572E-08 | 1.400E-07 | 2.800E-07 |
| 19851 | 993 | T993L | C | 3.1e-07 | 2.5e-07 | 4.9e-07 | 3.500E-07 | 1.249E-07 | 2.500E-07 | 4.900E-07 |
| 19852 | 993 | T993K | C | 3.2e-07 | 2.8e-07 | 2.1e-07 | 2.700E-07 | 5.568E-08 | 2.100E-07 | 3.200E-07 |
| 19853 | 993 | T993M | C | 3.3e-07 | 2.9e-07 | 1.7e-07 | 2.633E-07 | 8.327E-08 | 1.700E-07 | 3.300E-07 |
| 19854 | 993 | T993F | C | 3.1e-07 | 1.1e-07 | 3.5e-07 | 2.567E-07 | 1.286E-07 | 1.100E-07 | 3.500E-07 |
| 19855 | 993 | T993P | C | 3.1e-07 | 3.1e-07 | 2.7e-07 | 2.967E-07 | 2.309E-08 | 2.700E-07 | 3.100E-07 |
| 19856 | 993 | T993S | C | 1.1e-07 | 2.9e-07 | 1.3e-07 | 1.767E-07 | 9.866E-08 | 1.100E-07 | 2.900E-07 |
| 19857 | 993 | T993T | C | 2.5e-07 | 2.9e-07 | 2e-07 | 2.467E-07 | 4.509E-08 | 2.000E-07 | 2.900E-07 |
| 19858 | 993 | T993W | C | 3.1e-07 | 3.9e-07 | 1.6e-07 | 2.867E-07 | 1.168E-07 | 1.600E-07 | 3.900E-07 |
| 19859 | 993 | T993Y | C | 2.2e-07 | 2.5e-07 | 2.5e-07 | 2.400E-07 | 1.732E-08 | 2.200E-07 | 2.500E-07 |
| 19860 | 993 | T993V | C | 2.2e-07 | 3.3e-07 | 1.6e-07 | 2.367E-07 | 8.622E-08 | 1.600E-07 | 3.300E-07 |
| 19861 | 994 | V994A | C | 1.9e-07 | 2.5e-07 | 1.9e-07 | 2.100E-07 | 3.464E-08 | 1.900E-07 | 2.500E-07 |
| 19862 | 994 | V994R | C | 4.9e-07 | 2.1e-07 | 2.8e-07 | 3.267E-07 | 1.457E-07 | 2.100E-07 | 4.900E-07 |
| 19863 | 994 | V994N | C | 1.5e-07 | 2.4e-07 | 1.5e-07 | 1.800E-07 | 5.196E-08 | 1.500E-07 | 2.400E-07 |
| 19864 | 994 | V994D | C | 3.4e-07 | 1.3e-07 | 1.3e-07 | 2.000E-07 | 1.212E-07 | 1.300E-07 | 3.400E-07 |
| 19865 | 994 | V994C | C | 9.3e-08 | 2.6e-07 | 5.2e-07 | 2.910E-07 | 2.152E-07 | 9.300E-08 | 5.200E-07 |
| 19866 | 994 | V994E | C | 1.6e-07 | 2.4e-07 | 2.2e-07 | 2.067E-07 | 4.163E-08 | 1.600E-07 | 2.400E-07 |
| 19867 | 994 | V994Q | C | 4.1e-07 | 1.8e-07 | 3.5e-07 | 3.133E-07 | 1.193E-07 | 1.800E-07 | 4.100E-07 |
| 19868 | 994 | V994G | C | 2.1e-07 | 5.6e-07 | 1.8e-07 | 3.167E-07 | 2.113E-07 | 1.800E-07 | 5.600E-07 |
| 19869 | 994 | V994H | C | 2.1e-07 | 2.4e-07 | 5.6e-07 | 3.367E-07 | 1.940E-07 | 2.100E-07 | 5.600E-07 |
| 19870 | 994 | V994I | C | 2.6e-07 | 2.5e-07 | 2.1e-07 | 2.400E-07 | 2.646E-08 | 2.100E-07 | 2.600E-07 |
| 19871 | 994 | V994L | C | 1.8e-07 | 1.8e-07 | 1.6e-07 | 1.733E-07 | 1.155E-08 | 1.600E-07 | 1.800E-07 |
| 19872 | 994 | V994K | C | 2e-07 | 2e-07 | 1.7e-07 | 1.900E-07 | 1.732E-08 | 1.700E-07 | 2.000E-07 |
| 19873 | 994 | V994M | C | 1.8e-07 | 2.3e-07 | 2.8e-07 | 2.300E-07 | 5.000E-08 | 1.800E-07 | 2.800E-07 |
| 19874 | 994 | V994F | C | 2.5e-07 | 2.9e-07 | 2.1e-07 | 2.500E-07 | 4.000E-08 | 2.100E-07 | 2.900E-07 |
| 19875 | 994 | V994P | C | 1.5e-07 | 2e-07 | 2.2e-07 | 1.900E-07 | 3.606E-08 | 1.500E-07 | 2.200E-07 |
| 19876 | 994 | V994S | C | 1.6e-07 | 2.9e-07 | 4.8e-07 | 3.100E-07 | 1.609E-07 | 1.600E-07 | 4.800E-07 |
| 19877 | 994 | V994T | C | 2e-07 | 7.7e-08 | 2.3e-07 | 1.690E-07 | 8.107E-08 | 7.700E-08 | 2.300E-07 |
| 19878 | 994 | V994W | C | 3.2e-07 | 2.9e-07 | 2.3e-07 | 2.800E-07 | 4.583E-08 | 2.300E-07 | 3.200E-07 |
| 19879 | 994 | V994Y | C | 2.9e-07 | 2.3e-07 | 3e-07 | 2.733E-07 | 3.786E-08 | 2.300E-07 | 3.000E-07 |
| 19880 | 994 | V994V | C | 4.3e-07 | 2.1e-07 | 3.5e-07 | 3.300E-07 | 1.114E-07 | 2.100E-07 | 4.300E-07 |
| 19881 | 995 | S995A | C | 2.9e-07 | 3.2e-07 | 2e-07 | 2.700E-07 | 6.245E-08 | 2.000E-07 | 3.200E-07 |
| 19882 | 995 | S995R | C | 2.1e-07 | 2.7e-07 | 1.6e-07 | 2.133E-07 | 5.508E-08 | 1.600E-07 | 2.700E-07 |
| 19883 | 995 | S995N | C | 2.2e-07 | 1.8e-07 | 3e-07 | 2.333E-07 | 6.110E-08 | 1.800E-07 | 3.000E-07 |
| 19884 | 995 | S995D | C | 1.7e-07 | 1.9e-07 | 1.9e-07 | 1.833E-07 | 1.155E-08 | 1.700E-07 | 1.900E-07 |
| 19885 | 995 | S995C | C | 2e-07 | 3.2e-07 | 4.7e-07 | 3.300E-07 | 1.353E-07 | 2.000E-07 | 4.700E-07 |
| 19886 | 995 | S995E | C | 1.9e-07 | 3.1e-07 | 1.4e-07 | 2.133E-07 | 8.737E-08 | 1.400E-07 | 3.100E-07 |
| 19887 | 995 | S995Q | C | 1.2e-07 | 1.8e-07 | 2.1e-07 | 1.700E-07 | 4.583E-08 | 1.200E-07 | 2.100E-07 |
| 19888 | 995 | S995G | C | 3e-07 | 4.3e-07 | 2.8e-07 | 3.367E-07 | 8.145E-08 | 2.800E-07 | 4.300E-07 |
| 19889 | 995 | S995H | C | 1.1e-07 | 2.2e-07 | 2.4e-07 | 1.900E-07 | 7.000E-08 | 1.100E-07 | 2.400E-07 |

| | | | | | | | | | | |
|---|---|---|---|---|---|---|---|---|---|---|
| 19890 | 995 | S995I | C | 2.8e-07 | 1.3e-07 | 2.2e-07 | 2.100E-07 | 7.550E-08 | 1.300E-07 | 2.800E-07 |
| 19891 | 995 | S995L | C | 3.9e-07 | 3.7e-07 | 3.1e-07 | 3.567E-07 | 4.163E-08 | 3.100E-07 | 3.900E-07 |
| 19892 | 995 | S995K | C | 1.9e-07 | 1.7e-07 | 1.5e-07 | 1.700E-07 | 2.000E-08 | 1.500E-07 | 1.900E-07 |
| 19893 | 995 | S995M | C | 2e-07 | 2.9e-07 | 1.1e-07 | 2.000E-07 | 9.000E-08 | 1.100E-07 | 2.900E-07 |
| 19894 | 995 | S995F | C | 3.4e-07 | 2.2e-07 | 4e-07 | 3.200E-07 | 9.165E-08 | 2.200E-07 | 4.000E-07 |
| 19895 | 995 | S995P | C | 2e-07 | 3.5e-07 | 3.2e-07 | 2.900E-07 | 7.937E-08 | 2.000E-07 | 3.500E-07 |
| 19896 | 995 | S995S | C | 2.5e-07 | 4.1e-07 | 1.8e-07 | 2.800E-07 | 1.179E-07 | 1.800E-07 | 4.100E-07 |
| 19897 | 995 | S995T | C | 1e-07 | 1.4e-07 | 5e-07 | 2.467E-07 | 2.203E-07 | 1.000E-07 | 5.000E-07 |
| 19898 | 995 | S995W | C | 1.2e-07 | 2.7e-07 | 2.4e-07 | 2.100E-07 | 7.937E-08 | 1.200E-07 | 2.700E-07 |
| 19899 | 995 | S995Y | C | 3.7e-07 | 2.6e-07 | 2.6e-07 | 2.967E-07 | 6.351E-08 | 2.600E-07 | 3.700E-07 |
| 19900 | 995 | S995V | C | 1.6e-07 | 2.1e-07 | 2.5e-07 | 2.067E-07 | 4.509E-08 | 1.600E-07 | 2.500E-07 |
| 19901 | 996 | W996A | C | 2.9e-07 | 1.7e-07 | 3.2e-07 | 2.600E-07 | 7.937E-08 | 1.700E-07 | 3.200E-07 |
| 19902 | 996 | W996R | C | 1.5e-07 | 2.3e-07 | 4.2e-07 | 2.667E-07 | 1.387E-07 | 1.500E-07 | 4.200E-07 |
| 19903 | 996 | W996N | C | 1.8e-07 | 1.1e-07 | 3.1e-07 | 2.000E-07 | 1.015E-07 | 1.100E-07 | 3.100E-07 |
| 19904 | 996 | W996D | C | 3.1e-07 | 1.6e-07 | 2e-07 | 2.233E-07 | 7.767E-08 | 1.600E-07 | 3.100E-07 |
| 19905 | 996 | W996C | C | 5e-07 | 4.6e-07 | 3.9e-07 | 4.500E-07 | 5.568E-08 | 3.900E-07 | 5.000E-07 |
| 19906 | 996 | W996E | C | 1.7e-07 | 3.5e-07 | 5.1e-07 | 3.433E-07 | 1.701E-07 | 1.700E-07 | 5.100E-07 |
| 19907 | 996 | W996Q | C | 1.7e-07 | 2.4e-07 | 3.8e-07 | 2.633E-07 | 1.069E-07 | 1.700E-07 | 3.800E-07 |
| 19908 | 996 | W996G | C | 2.2e-07 | 2.2e-07 | 2.2e-07 | 2.200E-07 | 0.000E+00 | 2.200E-07 | 2.200E-07 |
| 19909 | 996 | W996H | C | 1.8e-07 | 3.4e-07 | 3.3e-07 | 2.833E-07 | 8.963E-08 | 1.800E-07 | 3.400E-07 |
| 19910 | 996 | W996I | C | 1.7e-07 | 1.8e-07 | 1.1e-07 | 1.533E-07 | 3.786E-08 | 1.100E-07 | 1.800E-07 |
| 19911 | 996 | W996L | C | 1.3e-07 | 1e-07 | 2.9e-07 | 1.733E-07 | 1.021E-07 | 1.000E-07 | 2.900E-07 |
| 19912 | 996 | W996K | C | 3.9e-07 | 1.7e-07 | 1.7e-07 | 2.433E-07 | 1.270E-07 | 1.700E-07 | 3.900E-07 |
| 19913 | 996 | W996M | C | 4.3e-07 | 2.9e-07 | 3.3e-07 | 3.500E-07 | 7.211E-08 | 2.900E-07 | 4.300E-07 |
| 19914 | 996 | W996F | C | 1.8e-07 | 1.8e-07 | 4e-07 | 2.533E-07 | 1.270E-07 | 1.800E-07 | 4.000E-07 |
| 19915 | 996 | W996P | C | 1.5e-07 | 3.4e-07 | 5.7e-07 | 3.533E-07 | 2.103E-07 | 1.500E-07 | 5.700E-07 |
| 19916 | 996 | W996S | C | 4.5e-07 | 3.2e-07 | 1.8e-07 | 3.167E-07 | 1.350E-07 | 1.800E-07 | 4.500E-07 |
| 19917 | 996 | W996T | C | 2.9e-07 | 2.2e-07 | 2e-07 | 2.367E-07 | 4.726E-08 | 2.000E-07 | 2.900E-07 |
| 19918 | 996 | W996W | C | 3.4e-07 | 2.5e-07 | 2e-07 | 2.633E-07 | 7.095E-08 | 2.000E-07 | 3.400E-07 |
| 19919 | 996 | W996Y | C | 1.9e-07 | 4.1e-07 | 3.5e-07 | 3.167E-07 | 1.137E-07 | 1.900E-07 | 4.100E-07 |
| 19920 | 996 | W996V | C | 2.2e-07 | 2.4e-07 | 3.3e-07 | 2.633E-07 | 5.859E-08 | 2.200E-07 | 3.300E-07 |
| 19921 | 997 | N997A | C | 4.1e-07 | 3.4e-07 | 7.4e-08 | 2.747E-07 | 1.773E-07 | 7.400E-08 | 4.100E-07 |
| 19922 | 997 | N997R | C | 1.1e-07 | 1.7e-07 | 2.6e-07 | 1.800E-07 | 7.550E-08 | 1.100E-07 | 2.600E-07 |
| 19923 | 997 | N997N | C | 2e-07 | 2.9e-07 | 2.3e-07 | 2.400E-07 | 4.583E-08 | 2.000E-07 | 2.900E-07 |
| 19924 | 997 | N997D | C | 1e-07 | 1.9e-07 | 2.7e-07 | 1.867E-07 | 8.505E-08 | 1.000E-07 | 2.700E-07 |
| 19925 | 997 | N997C | C | 2.2e-07 | 3.7e-07 | 3e-07 | 2.967E-07 | 7.506E-08 | 2.200E-07 | 3.700E-07 |
| 19926 | 997 | N997E | C | 1.9e-07 | 2.2e-07 | 2.5e-07 | 2.200E-07 | 3.000E-08 | 1.900E-07 | 2.500E-07 |
| 19927 | 997 | N997Q | C | 2.3e-07 | 9.5e-08 | 1.7e-07 | 1.650E-07 | 6.764E-08 | 9.500E-08 | 2.300E-07 |
| 19928 | 997 | N997G | C | 1.2e-07 | 2.8e-07 | 1.7e-07 | 1.900E-07 | 8.185E-08 | 1.200E-07 | 2.800E-07 |
| 19929 | 997 | N997H | C | 3.5e-07 | 2.4e-07 | 3.1e-07 | 3.000E-07 | 5.568E-08 | 2.400E-07 | 3.500E-07 |
| 19930 | 997 | N997I | C | 3.6e-07 | 3.6e-07 | 1.6e-07 | 2.933E-07 | 1.155E-07 | 1.600E-07 | 3.600E-07 |
| 19931 | 997 | N997L | C | 6.7e-08 | 3.2e-07 | 5.5e-07 | 3.123E-07 | 2.416E-07 | 6.700E-08 | 5.500E-07 |
| 19932 | 997 | N997K | C | 1.8e-07 | 2.8e-07 | 2.7e-07 | 2.433E-07 | 5.508E-08 | 1.800E-07 | 2.800E-07 |
| 19933 | 997 | N997M | C | 4.8e-07 | 2.5e-07 | 2e-07 | 3.100E-07 | 1.493E-07 | 2.000E-07 | 4.800E-07 |
| 19934 | 997 | N997F | C | 3e-07 | 1.6e-07 | 5.3e-07 | 3.300E-07 | 1.868E-07 | 1.600E-07 | 5.300E-07 |
| 19935 | 997 | N997P | C | 4.2e-07 | 2.3e-07 | 1.6e-07 | 2.700E-07 | 1.345E-07 | 1.600E-07 | 4.200E-07 |
| 19936 | 997 | N997S | C | 2e-07 | 2e-07 | 4.3e-07 | 2.767E-07 | 1.328E-07 | 2.000E-07 | 4.300E-07 |
| 19937 | 997 | N997T | C | 1.7e-07 | 2.4e-07 | 3.1e-07 | 2.400E-07 | 7.000E-08 | 1.700E-07 | 3.100E-07 |
| 19938 | 997 | N997W | C | 3.5e-07 | 2.6e-07 | 3.2e-07 | 3.100E-07 | 4.583E-08 | 2.600E-07 | 3.500E-07 |
| 19939 | 997 | N997Y | C | 2.7e-07 | 1.5e-07 | 4.5e-07 | 2.900E-07 | 1.510E-07 | 1.500E-07 | 4.500E-07 |
| 19940 | 997 | N997V | C | 1.2e-07 | 2.9e-07 | 4.2e-07 | 2.767E-07 | 1.504E-07 | 1.200E-07 | 4.200E-07 |
| 19941 | 998 | S998A | C | 2e-07 | 2.5e-07 | 7.3e-08 | 1.743E-07 | 9.125E-08 | 7.300E-08 | 2.500E-07 |
| 19942 | 998 | S998R | C | 3.5e-07 | 2.5e-07 | 3.4e-07 | 3.133E-07 | 5.508E-08 | 2.500E-07 | 3.500E-07 |
| 19943 | 998 | S998N | C | 4.2e-07 | 1e-07 | 3.6e-07 | 2.933E-07 | 1.701E-07 | 1.000E-07 | 4.200E-07 |
| 19944 | 998 | S998D | C | 5.5e-07 | 2.4e-07 | 4e-07 | 3.967E-07 | 1.550E-07 | 2.400E-07 | 5.500E-07 |
| 19945 | 998 | S998C | C | 2.5e-07 | 6.4e-08 | 2.3e-07 | 1.813E-07 | 1.021E-07 | 6.400E-08 | 2.500E-07 |
| 19946 | 998 | S998E | C | 8.3e-08 | 1.6e-07 | 3.7e-07 | 2.043E-07 | 1.485E-07 | 8.300E-08 | 3.700E-07 |
| 19947 | 998 | S998Q | C | 2.2e-07 | 2.3e-07 | 2.1e-07 | 2.200E-07 | 1.000E-08 | 2.100E-07 | 2.300E-07 |
| 19948 | 998 | S998G | C | 4.6e-07 | 1.5e-07 | 1.3e-07 | 2.467E-07 | 1.850E-07 | 1.300E-07 | 4.600E-07 |
| 19949 | 998 | S998H | C | 2.6e-07 | 9.8e-08 | 5.2e-07 | 2.927E-07 | 2.129E-07 | 9.800E-08 | 5.200E-07 |
| 19950 | 998 | S998I | C | 4.7e-07 | 3.6e-07 | 2.6e-07 | 3.633E-07 | 1.050E-07 | 2.600E-07 | 4.700E-07 |
| 19951 | 998 | S998L | C | 2.3e-07 | 3e-07 | 1.6e-07 | 2.300E-07 | 7.000E-08 | 1.600E-07 | 3.000E-07 |
| 19952 | 998 | S998K | C | 2.1e-07 | 1.5e-07 | 1e-07 | 1.533E-07 | 5.508E-08 | 1.000E-07 | 2.100E-07 |
| 19953 | 998 | S998M | C | 2.5e-07 | 2.2e-07 | 2.7e-07 | 2.467E-07 | 2.517E-08 | 2.200E-07 | 2.700E-07 |
| 19954 | 998 | S998F | C | 2.8e-07 | 2.6e-07 | 2.7e-07 | 2.700E-07 | 1.000E-08 | 2.600E-07 | 2.800E-07 |
| 19955 | 998 | S998P | C | 1.3e-07 | 2.3e-07 | 1.8e-07 | 1.800E-07 | 5.000E-08 | 1.300E-07 | 2.300E-07 |
| 19956 | 998 | S998S | C | 1.6e-07 | 1.8e-07 | 3e-07 | 2.133E-07 | 7.572E-08 | 1.600E-07 | 3.000E-07 |
| 19957 | 998 | S998T | C | 3e-07 | 3.2e-07 | 2.4e-07 | 2.867E-07 | 4.163E-08 | 2.400E-07 | 3.200E-07 |
| 19958 | 998 | S998W | C | 2.2e-07 | 7.1e-07 | 1.8e-07 | 3.700E-07 | 2.951E-07 | 1.800E-07 | 7.100E-07 |
| 19959 | 998 | S998Y | C | 2.9e-07 | 2.8e-07 | 3.7e-07 | 3.133E-07 | 4.933E-08 | 2.800E-07 | 3.700E-07 |
| 19960 | 998 | S998V | C | 4.6e-07 | 1.6e-07 | 3.1e-07 | 3.100E-07 | 1.500E-07 | 1.600E-07 | 4.600E-07 |
| 19961 | 999 | G999A | C | 2.2e-07 | 4.1e-07 | 1.8e-07 | 2.700E-07 | 1.229E-07 | 1.800E-07 | 4.100E-07 |
| 19962 | 999 | G999R | C | 4e-07 | 2.4e-07 | 2.4e-07 | 2.933E-07 | 9.238E-08 | 2.400E-07 | 4.000E-07 |
| 19963 | 999 | G999N | C | 2.7e-07 | 2.2e-07 | 2.5e-07 | 2.467E-07 | 2.517E-08 | 2.200E-07 | 2.700E-07 |
| 19964 | 999 | G999D | C | 1.8e-07 | 2.7e-07 | 2e-07 | 2.167E-07 | 4.726E-08 | 1.800E-07 | 2.700E-07 |
| 19965 | 999 | G999C | C | 4.3e-07 | 1.3e-07 | 2e-07 | 2.533E-07 | 1.570E-07 | 1.300E-07 | 4.300E-07 |
| 19966 | 999 | G999E | C | 2.1e-07 | 2.5e-07 | 6.2e-07 | 3.600E-07 | 2.261E-07 | 2.100E-07 | 6.200E-07 |
| 19967 | 999 | G999Q | C | 2.1e-07 | 1.8e-07 | 7.3e-07 | 3.733E-07 | 3.092E-07 | 1.800E-07 | 7.300E-07 |
| 19968 | 999 | G999G | C | 4e-07 | 4.2e-07 | 2.8e-07 | 3.667E-07 | 7.572E-08 | 2.800E-07 | 4.200E-07 |
| 19969 | 999 | G999H | C | 1.5e-07 | 2.5e-07 | 2.8e-07 | 2.267E-07 | 6.807E-08 | 1.500E-07 | 2.800E-07 |
| 19970 | 999 | G999I | C | 2e-07 | 4.6e-07 | 9e-08 | 2.500E-07 | 1.900E-07 | 9.000E-08 | 4.600E-07 |
| 19971 | 999 | G999L | C | 2.7e-07 | 2e-07 | 2.5e-07 | 2.400E-07 | 3.606E-08 | 2.000E-07 | 2.700E-07 |
| 19972 | 999 | G999K | C | 2.6e-07 | 1.6e-07 | 1.5e-07 | 1.900E-07 | 6.083E-08 | 1.500E-07 | 2.600E-07 |
| 19973 | 999 | G999M | C | 3.2e-07 | 2.3e-07 | 1.5e-07 | 2.333E-07 | 8.505E-08 | 1.500E-07 | 3.200E-07 |
| 19974 | 999 | G999F | C | 2.2e-07 | 1.8e-07 | 1.1e-07 | 1.700E-07 | 5.568E-08 | 1.100E-07 | 2.200E-07 |

| | | | | | | | | | | |
|---|---|---|---|---|---|---|---|---|---|---|
| 19975 | 999 | G999P | C | 2.1e-07 | 1.3e-07 | 3e-07 | 2.133E-07 | 8.505E-08 | 1.300E-07 | 3.000E-07 |
| 19976 | 999 | G999S | C | 3.9e-07 | 2.2e-07 | 8.7e-08 | 2.323E-07 | 1.519E-07 | 8.700E-08 | 3.900E-07 |
| 19977 | 999 | G999T | C | 4.2e-07 | 1.8e-07 | 4.1e-07 | 3.367E-07 | 1.358E-07 | 1.800E-07 | 4.200E-07 |
| 19978 | 999 | G999W | C | 3.1e-07 | 3.3e-07 | 2.3e-07 | 2.900E-07 | 5.292E-08 | 2.300E-07 | 3.300E-07 |
| 19979 | 999 | G999Y | C | 2.7e-07 | 4.4e-07 | 3.2e-07 | 3.433E-07 | 8.737E-08 | 2.700E-07 | 4.400E-07 |
| 19980 | 999 | G999V | C | 4.4e-07 | 3.5e-07 | 4.3e-07 | 4.067E-07 | 4.933E-08 | 3.500E-07 | 4.400E-07 |
| 19981 | 1000 | A1000A | C | 2.4e-07 | 2.7e-07 | 2.3e-07 | 2.467E-07 | 2.082E-08 | 2.300E-07 | 2.700E-07 |
| 19982 | 1000 | A1000R | C | 3.5e-07 | 3.1e-07 | 4.3e-07 | 3.633E-07 | 6.110E-08 | 3.100E-07 | 4.300E-07 |
| 19983 | 1000 | A1000N | C | 2.3e-07 | 2.1e-07 | 4.6e-07 | 3.000E-07 | 1.389E-07 | 2.100E-07 | 4.600E-07 |
| 19984 | 1000 | A1000D | C | 3e-07 | 6.9e-07 | 3.1e-07 | 4.333E-07 | 2.223E-07 | 3.000E-07 | 6.900E-07 |
| 19985 | 1000 | A1000C | C | 2.3e-07 | 2.7e-07 | 4.2e-07 | 3.067E-07 | 1.002E-07 | 2.300E-07 | 4.200E-07 |
| 19986 | 1000 | A1000E | C | 4.1e-07 | 8.9e-08 | 3.4e-07 | 2.797E-07 | 1.688E-07 | 8.900E-08 | 4.100E-07 |
| 19987 | 1000 | A1000Q | C | 4.3e-07 | 1.6e-07 | 2.4e-07 | 2.767E-07 | 1.387E-07 | 1.600E-07 | 4.300E-07 |
| 19988 | 1000 | A1000G | C | 1e-06 | 2.4e-07 | 2.1e-07 | 4.833E-07 | 4.477E-07 | 2.100E-07 | 1.000E-06 |
| 19989 | 1000 | A1000H | C | 2.8e-07 | 4.4e-07 | 4.5e-07 | 3.900E-07 | 9.539E-08 | 2.800E-07 | 4.500E-07 |
| 19990 | 1000 | A1000I | C | 3.6e-07 | 1.5e-07 | 2e-07 | 2.367E-07 | 1.097E-07 | 1.500E-07 | 3.600E-07 |
| 19991 | 1000 | A1000L | C | 3.3e-07 | 5.9e-07 | 1.9e-07 | 3.700E-07 | 2.030E-07 | 1.900E-07 | 5.900E-07 |
| 19992 | 1000 | A1000K | C | 1.4e-07 | 3e-07 | 2.3e-07 | 2.233E-07 | 8.021E-08 | 1.400E-07 | 3.000E-07 |
| 19993 | 1000 | A1000M | C | 1.4e-07 | 2.7e-07 | 1.8e-07 | 1.967E-07 | 6.658E-08 | 1.400E-07 | 2.700E-07 |
| 19994 | 1000 | A1000F | C | 5.1e-07 | 2.5e-07 | 1.4e-07 | 3.000E-07 | 1.900E-07 | 1.400E-07 | 5.100E-07 |
| 19995 | 1000 | A1000P | C | 3e-07 | 2.1e-07 | 6.2e-07 | 3.767E-07 | 2.155E-07 | 2.100E-07 | 6.200E-07 |
| 19996 | 1000 | A1000S | C | 2.2e-07 | 5e-07 | 1.7e-07 | 2.967E-07 | 1.779E-07 | 1.700E-07 | 5.000E-07 |
| 19997 | 1000 | A1000T | C | 3e-07 | 2.1e-07 | 2.8e-07 | 2.633E-07 | 4.726E-08 | 2.100E-07 | 3.000E-07 |
| 19998 | 1000 | A1000W | C | 2.2e-07 | 2.2e-07 | 2.5e-07 | 2.300E-07 | 1.732E-08 | 2.200E-07 | 2.500E-07 |
| 19999 | 1000 | A1000Y | C | 1.8e-07 | 2.4e-07 | 2.1e-07 | 2.100E-07 | 3.000E-08 | 1.800E-07 | 2.400E-07 |
| 20000 | 1000 | A1000V | C | 2.5e-07 | 1.3e-07 | 1.4e-07 | 1.733E-07 | 6.658E-08 | 1.300E-07 | 2.500E-07 |
| 20001 | 1001 | L1001A | C | 1.5e-07 | 1.3e-07 | 3e-07 | 1.933E-07 | 9.292E-08 | 1.300E-07 | 3.000E-07 |
| 20002 | 1001 | L1001R | C | 2.9e-07 | 2.6e-07 | 1.8e-07 | 2.433E-07 | 5.686E-08 | 1.800E-07 | 2.900E-07 |
| 20003 | 1001 | L1001N | C | 2.3e-07 | 2e-07 | 2.5e-07 | 2.267E-07 | 2.517E-08 | 2.000E-07 | 2.500E-07 |
| 20004 | 1001 | L1001D | C | 2e-07 | 7e-08 | 2.9e-07 | 1.867E-07 | 1.106E-07 | 7.000E-08 | 2.900E-07 |
| 20005 | 1001 | L1001C | C | 3.5e-07 | 2.2e-07 | 2.3e-07 | 2.667E-07 | 7.234E-08 | 2.200E-07 | 3.500E-07 |
| 20006 | 1001 | L1001E | C | 7.3e-08 | 1.9e-07 | 3.6e-07 | 2.077E-07 | 1.443E-07 | 7.300E-08 | 3.600E-07 |
| 20007 | 1001 | L1001Q | C | 4e-07 | 3e-07 | 2.9e-07 | 3.300E-07 | 6.083E-08 | 2.900E-07 | 4.000E-07 |
| 20008 | 1001 | L1001G | C | 9.2e-07 | 9.9e-08 | 2.1e-07 | 4.097E-07 | 4.454E-07 | 9.900E-08 | 9.200E-07 |
| 20009 | 1001 | L1001H | C | 3.6e-07 | 1.1e-07 | 1.9e-07 | 2.200E-07 | 1.277E-07 | 1.100E-07 | 3.600E-07 |
| 20010 | 1001 | L1001I | C | 3.8e-07 | 4e-07 | 1.1e-07 | 2.967E-07 | 1.620E-07 | 1.100E-07 | 4.000E-07 |
| 20011 | 1001 | L1001L | C | 1.6e-07 | 1.5e-07 | 2e-07 | 1.700E-07 | 2.646E-08 | 1.500E-07 | 2.000E-07 |
| 20012 | 1001 | L1001K | C | 2e-07 | 1.8e-07 | 2.9e-07 | 2.233E-07 | 5.859E-08 | 1.800E-07 | 2.900E-07 |
| 20013 | 1001 | L1001M | C | 1.6e-07 | 2.2e-07 | 2.8e-07 | 2.200E-07 | 6.000E-08 | 1.600E-07 | 2.800E-07 |
| 20014 | 1001 | L1001F | C | 3.1e-07 | 2.5e-07 | 2e-07 | 2.533E-07 | 5.508E-08 | 2.000E-07 | 3.100E-07 |
| 20015 | 1001 | L1001P | C | 3.7e-07 | 2.9e-07 | 2.8e-07 | 3.133E-07 | 4.933E-08 | 2.800E-07 | 3.700E-07 |
| 20016 | 1001 | L1001S | C | 4.5e-07 | 3.5e-07 | 2e-07 | 3.333E-07 | 1.258E-07 | 2.000E-07 | 4.500E-07 |
| 20017 | 1001 | L1001T | C | 1.2e-07 | 2.5e-07 | 3e-07 | 2.233E-07 | 9.292E-08 | 1.200E-07 | 3.000E-07 |
| 20018 | 1001 | L1001W | C | 2.7e-07 | 1.3e-07 | 1.2e-07 | 1.733E-07 | 8.386E-08 | 1.200E-07 | 2.700E-07 |
| 20019 | 1001 | L1001Y | C | 3.2e-07 | 2.6e-07 | 1.9e-07 | 2.567E-07 | 6.506E-08 | 1.900E-07 | 3.200E-07 |
| 20020 | 1001 | L1001V | C | 2.5e-07 | 4e-07 | 2.4e-07 | 2.967E-07 | 8.963E-08 | 2.400E-07 | 4.000E-07 |
| 20021 | 1002 | T1002A | C | 2.9e-07 | 9.5e-08 | 2.4e-07 | 2.083E-07 | 1.013E-07 | 9.500E-08 | 2.900E-07 |
| 20022 | 1002 | T1002R | C | 2.3e-07 | 2.6e-07 | 1.9e-07 | 2.267E-07 | 3.512E-08 | 1.900E-07 | 2.600E-07 |
| 20023 | 1002 | T1002N | C | 2.4e-07 | 4.8e-07 | 2.6e-07 | 3.267E-07 | 1.332E-07 | 2.400E-07 | 4.800E-07 |
| 20024 | 1002 | T1002D | C | 1.8e-07 | 3e-07 | 3.6e-07 | 2.800E-07 | 9.165E-08 | 1.800E-07 | 3.600E-07 |
| 20025 | 1002 | T1002C | C | 2.4e-07 | 1.7e-07 | 6.9e-07 | 3.667E-07 | 2.822E-07 | 1.700E-07 | 6.900E-07 |
| 20026 | 1002 | T1002E | C | 1.9e-07 | 1.9e-07 | 2.6e-07 | 2.133E-07 | 4.041E-08 | 1.900E-07 | 2.600E-07 |
| 20027 | 1002 | T1002Q | C | 2.9e-07 | 1.9e-07 | 3.9e-07 | 2.900E-07 | 1.000E-07 | 1.900E-07 | 3.900E-07 |
| 20028 | 1002 | T1002G | C | 1.5e-07 | 3.1e-07 | 1.3e-07 | 1.967E-07 | 9.866E-08 | 1.300E-07 | 3.100E-07 |
| 20029 | 1002 | T1002H | C | 3.4e-07 | 2.4e-07 | 4.7e-07 | 3.500E-07 | 1.153E-07 | 2.400E-07 | 4.700E-07 |
| 20030 | 1002 | T1002I | C | 2.7e-07 | 5.5e-07 | 4.7e-07 | 4.300E-07 | 1.442E-07 | 2.700E-07 | 5.500E-07 |
| 20031 | 1002 | T1002L | C | 3.8e-07 | 1.8e-07 | 2.7e-07 | 2.767E-07 | 1.002E-07 | 1.800E-07 | 3.800E-07 |
| 20032 | 1002 | T1002K | C | 1.5e-07 | 3.5e-07 | 2e-07 | 2.333E-07 | 1.041E-07 | 1.500E-07 | 3.500E-07 |
| 20033 | 1002 | T1002M | C | 3.4e-07 | 2.8e-07 | 2.5e-07 | 2.900E-07 | 4.583E-08 | 2.500E-07 | 3.400E-07 |
| 20034 | 1002 | T1002F | C | 3.5e-07 | 2e-07 | 4.3e-07 | 3.267E-07 | 1.168E-07 | 2.000E-07 | 4.300E-07 |
| 20035 | 1002 | T1002P | C | 4.7e-07 | 3.8e-07 | 2.4e-07 | 3.633E-07 | 1.159E-07 | 2.400E-07 | 4.700E-07 |
| 20036 | 1002 | T1002S | C | 1.1e-07 | 3.1e-07 | 2.2e-07 | 2.133E-07 | 1.002E-07 | 1.100E-07 | 3.100E-07 |
| 20037 | 1002 | T1002T | C | 2e-07 | 2.2e-07 | 2.2e-07 | 2.133E-07 | 1.155E-08 | 2.000E-07 | 2.200E-07 |
| 20038 | 1002 | T1002W | C | 4.7e-07 | 1.1e-07 | 2.3e-07 | 2.700E-07 | 1.833E-07 | 1.100E-07 | 4.700E-07 |
| 20039 | 1002 | T1002Y | C | 6.8e-07 | 4.9e-07 | 3.7e-07 | 5.133E-07 | 1.563E-07 | 3.700E-07 | 6.800E-07 |
| 20040 | 1002 | T1002V | C | 3.2e-07 | 2.4e-07 | 1.7e-07 | 2.433E-07 | 7.506E-08 | 1.700E-07 | 3.200E-07 |
| 20041 | 1003 | S1003A | C | 1.9e-07 | 2.3e-07 | 2.9e-07 | 2.367E-07 | 5.033E-08 | 1.900E-07 | 2.900E-07 |
| 20042 | 1003 | S1003R | C | 6.5e-07 | 3.3e-07 | 1.4e-07 | 3.733E-07 | 2.577E-07 | 1.400E-07 | 6.500E-07 |
| 20043 | 1003 | S1003N | C | 1.9e-07 | 1.8e-07 | 3.3e-07 | 2.333E-07 | 8.386E-08 | 1.800E-07 | 3.300E-07 |
| 20044 | 1003 | S1003D | C | 6.9e-07 | 1.5e-07 | 1.8e-07 | 3.400E-07 | 3.035E-07 | 1.500E-07 | 6.900E-07 |
| 20045 | 1003 | S1003C | C | 2.2e-07 | 2.4e-07 | 2.7e-07 | 2.433E-07 | 2.517E-08 | 2.200E-07 | 2.700E-07 |
| 20046 | 1003 | S1003E | C | 3.1e-07 | 1.2e-07 | 7.1e-07 | 3.800E-07 | 3.012E-07 | 1.200E-07 | 7.100E-07 |
| 20047 | 1003 | S1003Q | C | 2.1e-07 | 5.4e-07 | 1.8e-07 | 3.100E-07 | 1.997E-07 | 1.800E-07 | 5.400E-07 |
| 20048 | 1003 | S1003G | C | 4e-07 | 3e-07 | 4.2e-07 | 3.733E-07 | 6.429E-08 | 3.000E-07 | 4.200E-07 |
| 20049 | 1003 | S1003H | C | 2.8e-07 | 1.8e-07 | 2.5e-07 | 2.367E-07 | 5.132E-08 | 1.800E-07 | 2.800E-07 |
| 20050 | 1003 | S1003I | C | 3.1e-07 | 3.1e-07 | 4.9e-07 | 3.700E-07 | 1.039E-07 | 3.100E-07 | 4.900E-07 |
| 20051 | 1003 | S1003L | C | 2.2e-07 | 2.2e-07 | 3.3e-07 | 2.567E-07 | 6.351E-08 | 2.200E-07 | 3.300E-07 |
| 20052 | 1003 | S1003K | C | 1.2e-07 | 2.2e-07 | 1.5e-07 | 1.633E-07 | 5.132E-08 | 1.200E-07 | 2.200E-07 |
| 20053 | 1003 | S1003M | C | 3.6e-07 | 2.9e-07 | 5.4e-07 | 3.967E-07 | 1.290E-07 | 2.900E-07 | 5.400E-07 |
| 20054 | 1003 | S1003F | C | 2.4e-07 | 2.1e-07 | 3.5e-07 | 2.667E-07 | 7.371E-08 | 2.100E-07 | 3.500E-07 |
| 20055 | 1003 | S1003P | C | 5.4e-07 | 7.6e-08 | 2.9e-07 | 3.020E-07 | 2.322E-07 | 7.600E-08 | 5.400E-07 |
| 20056 | 1003 | S1003S | C | 2.2e-07 | 3.4e-07 | 3.1e-07 | 2.900E-07 | 6.245E-08 | 2.200E-07 | 3.400E-07 |
| 20057 | 1003 | S1003T | C | 3.7e-07 | 2.1e-07 | 9.4e-08 | 2.247E-07 | 1.386E-07 | 9.400E-08 | 3.700E-07 |
| 20058 | 1003 | S1003W | C | 1.7e-07 | 2.6e-07 | 3.5e-07 | 2.600E-07 | 9.000E-08 | 1.700E-07 | 3.500E-07 |
| 20059 | 1003 | S1003Y | C | 1.5e-07 | 3.9e-07 | 4.6e-07 | 3.333E-07 | 1.626E-07 | 1.500E-07 | 4.600E-07 |

| | | | | | | | | | | |
|---|---|---|---|---|---|---|---|---|---|---|
| 20060 | 1003 | S1003V | C | 3.6e-07 | 2.2e-07 | 3.5e-07 | 3.100E-07 | 7.810E-08 | 2.200E-07 | 3.600E-07 |
| 20061 | 1004 | G1004A | C | 3.3e-07 | 1.7e-07 | 1.2e-07 | 2.067E-07 | 1.097E-07 | 1.200E-07 | 3.300E-07 |
| 20062 | 1004 | G1004R | C | 7.7e-08 | 2.9e-07 | 1.7e-07 | 1.790E-07 | 1.068E-07 | 7.700E-08 | 2.900E-07 |
| 20063 | 1004 | G1004N | C | 1.5e-07 | 3.1e-07 | 2.2e-07 | 2.267E-07 | 8.021E-08 | 1.500E-07 | 3.100E-07 |
| 20064 | 1004 | G1004D | C | 2.8e-07 | 1.6e-07 | 3.1e-07 | 2.500E-07 | 7.937E-08 | 1.600E-07 | 3.100E-07 |
| 20065 | 1004 | G1004C | C | 3.9e-07 | 1.9e-07 | 2.7e-07 | 2.833E-07 | 1.007E-07 | 1.900E-07 | 3.900E-07 |
| 20066 | 1004 | G1004E | C | 3.7e-07 | 2.2e-07 | 2.4e-07 | 2.767E-07 | 8.145E-08 | 2.200E-07 | 3.700E-07 |
| 20067 | 1004 | G1004Q | C | 2.6e-07 | 1.2e-07 | 2.3e-07 | 2.033E-07 | 7.371E-08 | 1.200E-07 | 2.600E-07 |
| 20068 | 1004 | G1004G | C | 3.1e-07 | 1.4e-07 | 1.6e-07 | 2.033E-07 | 9.292E-08 | 1.400E-07 | 3.100E-07 |
| 20069 | 1004 | G1004H | C | 1.7e-07 | 9.5e-08 | 1.9e-07 | 1.517E-07 | 5.008E-08 | 9.500E-08 | 1.900E-07 |
| 20070 | 1004 | G1004I | C | 1.2e-07 | 4.1e-07 | 5.6e-07 | 3.633E-07 | 2.237E-07 | 1.200E-07 | 5.600E-07 |
| 20071 | 1004 | G1004L | C | 1.7e-07 | 2.3e-07 | 1.5e-07 | 1.833E-07 | 4.163E-08 | 1.500E-07 | 2.300E-07 |
| 20072 | 1004 | G1004K | C | 4e-07 | 1.4e-07 | 5e-07 | 3.467E-07 | 1.858E-07 | 1.400E-07 | 5.000E-07 |
| 20073 | 1004 | G1004M | C | 2.8e-07 | 3.1e-07 | 1.5e-07 | 2.467E-07 | 8.505E-08 | 1.500E-07 | 3.100E-07 |
| 20074 | 1004 | G1004F | C | 1.5e-07 | 2.1e-07 | 2e-07 | 1.867E-07 | 3.215E-08 | 1.500E-07 | 2.100E-07 |
| 20075 | 1004 | G1004P | C | 2.3e-07 | 2.2e-07 | 8.1e-07 | 4.200E-07 | 3.378E-07 | 2.200E-07 | 8.100E-07 |
| 20076 | 1004 | G1004S | C | 1.6e-07 | 1.8e-07 | 2.4e-07 | 1.933E-07 | 4.163E-08 | 1.600E-07 | 2.400E-07 |
| 20077 | 1004 | G1004T | C | 3.2e-07 | 1.7e-07 | 8.4e-08 | 1.913E-07 | 1.194E-07 | 8.400E-08 | 3.200E-07 |
| 20078 | 1004 | G1004W | C | 2e-07 | 3.4e-07 | 2.3e-07 | 2.567E-07 | 7.371E-08 | 2.000E-07 | 3.400E-07 |
| 20079 | 1004 | G1004Y | C | 2.9e-07 | 3.8e-07 | 2.1e-07 | 2.933E-07 | 8.505E-08 | 2.100E-07 | 3.800E-07 |
| 20080 | 1004 | G1004V | C | 1.3e-07 | 2.4e-07 | 3.1e-07 | 2.267E-07 | 9.074E-08 | 1.300E-07 | 3.100E-07 |
| 20081 | 1005 | V1005A | C | 3.1e-07 | 2e-07 | 2.6e-07 | 2.567E-07 | 5.508E-08 | 2.000E-07 | 3.100E-07 |
| 20082 | 1005 | V1005R | C | 1.7e-07 | 3.7e-07 | 3.8e-07 | 3.067E-07 | 1.185E-07 | 1.700E-07 | 3.800E-07 |
| 20083 | 1005 | V1005N | C | 2.1e-07 | 9.3e-08 | 2.9e-07 | 1.977E-07 | 9.908E-08 | 9.300E-08 | 2.900E-07 |
| 20084 | 1005 | V1005D | C | 3.9e-07 | 1.7e-07 | 1.5e-07 | 2.367E-07 | 1.332E-07 | 1.500E-07 | 3.900E-07 |
| 20085 | 1005 | V1005C | C | 1.3e-07 | 1.7e-07 | 2.1e-07 | 1.700E-07 | 4.000E-08 | 1.300E-07 | 2.100E-07 |
| 20086 | 1005 | V1005E | C | 5.9e-07 | 1.9e-07 | 2.4e-07 | 3.400E-07 | 2.179E-07 | 1.900E-07 | 5.900E-07 |
| 20087 | 1005 | V1005Q | C | 2.7e-07 | 2e-07 | 1.6e-07 | 2.100E-07 | 5.568E-08 | 1.600E-07 | 2.700E-07 |
| 20088 | 1005 | V1005G | C | 1.5e-07 | 1.8e-07 | 4.5e-07 | 2.600E-07 | 1.652E-07 | 1.500E-07 | 4.500E-07 |
| 20089 | 1005 | V1005H | C | 2.1e-07 | 1.8e-07 | 2e-07 | 1.967E-07 | 1.528E-08 | 1.800E-07 | 2.100E-07 |
| 20090 | 1005 | V1005I | C | 2.2e-07 | 2.6e-07 | 3.8e-07 | 2.867E-07 | 8.327E-08 | 2.200E-07 | 3.800E-07 |
| 20091 | 1005 | V1005L | C | 2.9e-07 | 2.1e-07 | 3.3e-07 | 2.767E-07 | 6.110E-08 | 2.100E-07 | 3.300E-07 |
| 20092 | 1005 | V1005K | C | 1.6e-07 | 1.7e-07 | 6.6e-07 | 3.300E-07 | 2.858E-07 | 1.600E-07 | 6.600E-07 |
| 20093 | 1005 | V1005M | C | 1.8e-07 | 1.7e-07 | 2.6e-07 | 2.033E-07 | 4.933E-08 | 1.700E-07 | 2.600E-07 |
| 20094 | 1005 | V1005F | C | 1.3e-07 | 3.4e-07 | 2.7e-07 | 2.467E-07 | 1.069E-07 | 1.300E-07 | 3.400E-07 |
| 20095 | 1005 | V1005P | C | 4.1e-07 | 2.9e-07 | 2.1e-07 | 3.033E-07 | 1.007E-07 | 2.100E-07 | 4.100E-07 |
| 20096 | 1005 | V1005S | C | 3.3e-07 | 2e-07 | 2.2e-07 | 2.500E-07 | 7.000E-08 | 2.000E-07 | 3.300E-07 |
| 20097 | 1005 | V1005T | C | 4.4e-07 | 1.8e-07 | 2.4e-07 | 2.867E-07 | 1.361E-07 | 1.800E-07 | 4.400E-07 |
| 20098 | 1005 | V1005W | C | 1.7e-07 | 2.4e-07 | 2.3e-07 | 2.133E-07 | 3.786E-08 | 1.700E-07 | 2.400E-07 |
| 20099 | 1005 | V1005Y | C | 3.4e-07 | 5e-07 | 2.6e-07 | 3.667E-07 | 1.222E-07 | 2.600E-07 | 5.000E-07 |
| 20100 | 1005 | V1005V | C | 3.8e-07 | 6.5e-07 | 2.9e-07 | 4.400E-07 | 1.873E-07 | 2.900E-07 | 6.500E-07 |
| 20101 | 1006 | H1006A | C | 3.5e-07 | 3.7e-07 | 4e-07 | 3.733E-07 | 2.517E-08 | 3.500E-07 | 4.000E-07 |
| 20102 | 1006 | H1006R | C | 1.4e-07 | 2.9e-07 | 3e-07 | 2.433E-07 | 8.963E-08 | 1.400E-07 | 3.000E-07 |
| 20103 | 1006 | H1006N | C | 4.9e-08 | 2.7e-07 | 1.4e-07 | 1.530E-07 | 1.111E-07 | 4.900E-08 | 2.700E-07 |
| 20104 | 1006 | H1006D | C | 3.2e-07 | 1.8e-07 | 1.5e-07 | 2.167E-07 | 9.074E-08 | 1.500E-07 | 3.200E-07 |
| 20105 | 1006 | H1006C | C | 1.5e-07 | 6e-07 | 5.7e-08 | 2.690E-07 | 2.904E-07 | 5.700E-08 | 6.000E-07 |
| 20106 | 1006 | H1006E | C | 2.7e-07 | 2.2e-07 | 2.1e-07 | 2.333E-07 | 3.215E-08 | 2.100E-07 | 2.700E-07 |
| 20107 | 1006 | H1006Q | C | 3.8e-07 | 1.2e-07 | 4e-07 | 3.000E-07 | 1.562E-07 | 1.200E-07 | 4.000E-07 |
| 20108 | 1006 | H1006G | C | 3.3e-07 | 2.1e-07 | 2e-07 | 2.467E-07 | 7.234E-08 | 2.000E-07 | 3.300E-07 |
| 20109 | 1006 | H1006H | C | 5.3e-07 | 6.7e-08 | 2.5e-07 | 2.823E-07 | 2.332E-07 | 6.700E-08 | 5.300E-07 |
| 20110 | 1006 | H1006I | C | 2.6e-07 | 2.2e-07 | 4.8e-07 | 3.200E-07 | 1.400E-07 | 2.200E-07 | 4.800E-07 |
| 20111 | 1006 | H1006L | C | 3.6e-07 | 2.5e-07 | 2.9e-07 | 3.000E-07 | 5.568E-08 | 2.500E-07 | 3.600E-07 |
| 20112 | 1006 | H1006K | C | 2.9e-07 | 2.9e-07 | 2.4e-07 | 2.733E-07 | 2.887E-08 | 2.400E-07 | 2.900E-07 |
| 20113 | 1006 | H1006M | C | 2.9e-07 | 1.9e-07 | 2e-07 | 2.267E-07 | 5.508E-08 | 1.900E-07 | 2.900E-07 |
| 20114 | 1006 | H1006F | C | 2.4e-07 | 3.2e-07 | 2.4e-07 | 2.667E-07 | 4.619E-08 | 2.400E-07 | 3.200E-07 |
| 20115 | 1006 | H1006P | C | 1.2e-07 | 3.3e-07 | 3.1e-07 | 2.533E-07 | 1.159E-07 | 1.200E-07 | 3.300E-07 |
| 20116 | 1006 | H1006S | C | 3.1e-07 | 2.7e-07 | 4.7e-07 | 3.500E-07 | 1.058E-07 | 2.700E-07 | 4.700E-07 |
| 20117 | 1006 | H1006T | C | 3.1e-07 | 2.2e-07 | 1.4e-07 | 2.233E-07 | 8.505E-08 | 1.400E-07 | 3.100E-07 |
| 20118 | 1006 | H1006W | C | 3e-07 | 4.4e-07 | 2.5e-07 | 3.300E-07 | 9.849E-08 | 2.500E-07 | 4.400E-07 |
| 20119 | 1006 | H1006Y | C | 1.7e-07 | 1.4e-07 | 1.8e-07 | 1.633E-07 | 2.082E-08 | 1.400E-07 | 1.800E-07 |
| 20120 | 1006 | H1006V | C | 2.1e-07 | 3.5e-07 | 2.1e-07 | 2.567E-07 | 8.083E-08 | 2.100E-07 | 3.500E-07 |
| 20121 | 1007 | T1007A | C | 3.2e-07 | 1.1e-07 | 1.8e-07 | 2.033E-07 | 1.069E-07 | 1.100E-07 | 3.200E-07 |
| 20122 | 1007 | T1007R | C | 3.9e-07 | 2e-07 | 3.9e-07 | 3.267E-07 | 1.097E-07 | 2.000E-07 | 3.900E-07 |
| 20123 | 1007 | T1007N | C | 2.2e-07 | 1.6e-07 | 4.4e-07 | 2.733E-07 | 1.474E-07 | 1.600E-07 | 4.400E-07 |
| 20124 | 1007 | T1007D | C | 4.3e-07 | 2.9e-07 | 3.1e-07 | 3.433E-07 | 7.572E-08 | 2.900E-07 | 4.300E-07 |
| 20125 | 1007 | T1007C | C | 1.2e-07 | 1.9e-07 | 3.3e-07 | 2.133E-07 | 1.069E-07 | 1.200E-07 | 3.300E-07 |
| 20126 | 1007 | T1007E | C | 4e-07 | 3.8e-07 | 2.7e-07 | 3.500E-07 | 7.000E-08 | 2.700E-07 | 4.000E-07 |
| 20127 | 1007 | T1007Q | C | 2.6e-07 | 1.8e-07 | 3.1e-07 | 2.500E-07 | 6.557E-08 | 1.800E-07 | 3.100E-07 |
| 20128 | 1007 | T1007G | C | 1.7e-07 | 1.3e-07 | 6.1e-07 | 3.033E-07 | 2.663E-07 | 1.300E-07 | 6.100E-07 |
| 20129 | 1007 | T1007H | C | 1.5e-07 | 1.9e-07 | 3e-07 | 2.133E-07 | 7.767E-08 | 1.500E-07 | 3.000E-07 |
| 20130 | 1007 | T1007I | C | 3e-07 | 2.4e-07 | 2.2e-07 | 2.533E-07 | 4.163E-08 | 2.200E-07 | 3.000E-07 |
| 20131 | 1007 | T1007L | C | 3.5e-07 | 6.2e-08 | 2.7e-07 | 2.273E-07 | 1.487E-07 | 6.200E-08 | 3.500E-07 |
| 20132 | 1007 | T1007K | C | 2.7e-07 | 2.7e-07 | 3.2e-07 | 2.867E-07 | 2.887E-08 | 2.700E-07 | 3.200E-07 |
| 20133 | 1007 | T1007M | C | 1.8e-07 | 4.3e-07 | 1.9e-07 | 2.667E-07 | 1.415E-07 | 1.800E-07 | 4.300E-07 |
| 20134 | 1007 | T1007F | C | 2.4e-07 | 2.8e-07 | 8.9e-08 | 2.030E-07 | 1.007E-07 | 8.900E-08 | 2.800E-07 |
| 20135 | 1007 | T1007P | C | 4.7e-07 | 3.7e-07 | 6.5e-07 | 4.967E-07 | 1.419E-07 | 3.700E-07 | 6.500E-07 |
| 20136 | 1007 | T1007S | C | 2.3e-07 | 2.4e-07 | 2.7e-07 | 2.467E-07 | 2.082E-08 | 2.300E-07 | 2.700E-07 |
| 20137 | 1007 | T1007T | C | 2.4e-07 | 2.3e-07 | 1.5e-07 | 2.067E-07 | 4.933E-08 | 1.500E-07 | 2.400E-07 |
| 20138 | 1007 | T1007W | C | 5.1e-07 | 2.1e-07 | 3.4e-07 | 3.533E-07 | 1.504E-07 | 2.100E-07 | 5.100E-07 |
| 20139 | 1007 | T1007Y | C | 1.9e-07 | 1.6e-07 | 2.9e-07 | 2.133E-07 | 6.807E-08 | 1.600E-07 | 2.900E-07 |
| 20140 | 1007 | T1007V | C | 1.8e-07 | 3.9e-07 | 4e-07 | 3.233E-07 | 1.242E-07 | 1.800E-07 | 4.000E-07 |
| 20141 | 1008 | F1008A | C | 2.8e-07 | 5e-07 | 2.6e-07 | 3.467E-07 | 1.332E-07 | 2.600E-07 | 5.000E-07 |
| 20142 | 1008 | F1008R | C | 3.8e-07 | 2.4e-07 | 2.9e-07 | 3.033E-07 | 7.095E-08 | 2.400E-07 | 3.800E-07 |
| 20143 | 1008 | F1008N | C | 2.9e-07 | 4.7e-07 | 1.3e-07 | 2.967E-07 | 1.701E-07 | 1.300E-07 | 4.700E-07 |
| 20144 | 1008 | F1008D | C | 2.8e-07 | 2e-07 | 3.3e-07 | 2.700E-07 | 6.557E-08 | 2.000E-07 | 3.300E-07 |

| | | | | | | | | | | |
|---|---|---|---|---|---|---|---|---|---|---|
| 20145 | 1008 | F1008C | C | 1.7e-07 | 2.4e-07 | 4.3e-07 | 2.800E-07 | 1.345E-07 | 1.700E-07 | 4.300E-07 |
| 20146 | 1008 | F1008E | C | 3.3e-07 | 2.3e-07 | 1.9e-07 | 2.500E-07 | 7.211E-08 | 1.900E-07 | 3.300E-07 |
| 20147 | 1008 | F1008Q | C | 1.9e-07 | 2.9e-07 | 5.5e-07 | 3.433E-07 | 1.858E-07 | 1.900E-07 | 5.500E-07 |
| 20148 | 1008 | F1008G | C | 1.7e-07 | 3e-07 | 2.7e-07 | 2.467E-07 | 6.807E-08 | 1.700E-07 | 3.000E-07 |
| 20149 | 1008 | F1008H | C | 9.9e-08 | 2e-07 | 2e-07 | 1.663E-07 | 5.831E-08 | 9.900E-08 | 2.000E-07 |
| 20150 | 1008 | F1008I | C | 2.9e-07 | 1.9e-07 | 2.1e-07 | 2.300E-07 | 5.292E-08 | 1.900E-07 | 2.900E-07 |
| 20151 | 1008 | F1008L | C | 1.1e-07 | 3.1e-07 | 1.9e-07 | 2.033E-07 | 1.007E-07 | 1.100E-07 | 3.100E-07 |
| 20152 | 1008 | F1008K | C | 1.7e-07 | 2.4e-07 | 3.3e-07 | 2.467E-07 | 8.021E-08 | 1.700E-07 | 3.300E-07 |
| 20153 | 1008 | F1008M | C | 3.5e-07 | 3.1e-07 | 2.3e-07 | 2.967E-07 | 6.110E-08 | 2.300E-07 | 3.500E-07 |
| 20154 | 1008 | F1008F | C | 1.2e-07 | 2e-07 | 1.1e-07 | 1.433E-07 | 4.933E-08 | 1.100E-07 | 2.000E-07 |
| 20155 | 1008 | F1008P | C | 1.2e-07 | 2.3e-07 | 2.5e-07 | 2.000E-07 | 7.000E-08 | 1.200E-07 | 2.500E-07 |
| 20156 | 1008 | F1008S | C | 3.8e-07 | 2e-07 | 2.7e-07 | 2.833E-07 | 9.074E-08 | 2.000E-07 | 3.800E-07 |
| 20157 | 1008 | F1008T | C | 1.9e-07 | 6.8e-07 | 1.9e-07 | 3.533E-07 | 2.829E-07 | 1.900E-07 | 6.800E-07 |
| 20158 | 1008 | F1008W | C | 3.7e-07 | 1.8e-07 | 3.1e-07 | 2.867E-07 | 9.713E-08 | 1.800E-07 | 3.700E-07 |
| 20159 | 1008 | F1008Y | C | 1.3e-07 | 2.4e-07 | 1.6e-07 | 1.767E-07 | 5.686E-08 | 1.300E-07 | 2.400E-07 |
| 20160 | 1008 | F1008V | C | 2.8e-07 | 3.2e-07 | 2.6e-07 | 2.867E-07 | 3.055E-08 | 2.600E-07 | 3.200E-07 |
| 20161 | 1009 | P1009A | C | 3.6e-07 | 2.3e-07 | 2.9e-07 | 2.933E-07 | 6.506E-08 | 2.300E-07 | 3.600E-07 |
| 20162 | 1009 | P1009R | C | 2.4e-07 | 3.8e-07 | 5.2e-07 | 3.800E-07 | 1.400E-07 | 2.400E-07 | 5.200E-07 |
| 20163 | 1009 | P1009N | C | 2e-07 | 1.3e-07 | 7.9e-07 | 3.733E-07 | 3.625E-07 | 1.300E-07 | 7.900E-07 |
| 20164 | 1009 | P1009D | C | 3.9e-07 | 2.7e-07 | 1.2e-07 | 2.600E-07 | 1.353E-07 | 1.200E-07 | 3.900E-07 |
| 20165 | 1009 | P1009C | C | 1e-07 | 2.4e-07 | 1.9e-07 | 1.767E-07 | 7.095E-08 | 1.000E-07 | 2.400E-07 |
| 20166 | 1009 | P1009E | C | 2.1e-07 | 1.9e-07 | 2.7e-07 | 2.233E-07 | 4.163E-08 | 1.900E-07 | 2.700E-07 |
| 20167 | 1009 | P1009Q | C | 2.2e-07 | 3.1e-07 | 4.1e-07 | 3.133E-07 | 9.504E-08 | 2.200E-07 | 4.100E-07 |
| 20168 | 1009 | P1009G | C | 3.2e-07 | 1.3e-07 | 2.1e-07 | 2.200E-07 | 9.539E-08 | 1.300E-07 | 3.200E-07 |
| 20169 | 1009 | P1009H | C | 2.7e-07 | 1.6e-07 | 3.5e-07 | 2.600E-07 | 9.539E-08 | 1.600E-07 | 3.500E-07 |
| 20170 | 1009 | P1009I | C | 9e-08 | 2.8e-07 | 1.1e-07 | 1.600E-07 | 1.044E-07 | 9.000E-08 | 2.800E-07 |
| 20171 | 1009 | P1009L | C | 2.1e-07 | 1.6e-07 | 3e-07 | 2.233E-07 | 7.095E-08 | 1.600E-07 | 3.000E-07 |
| 20172 | 1009 | P1009K | C | 2.8e-07 | 2.8e-07 | 1.5e-07 | 2.367E-07 | 7.506E-08 | 1.500E-07 | 2.800E-07 |
| 20173 | 1009 | P1009M | C | 2e-07 | 1.9e-07 | 2.8e-07 | 2.233E-07 | 4.933E-08 | 1.900E-07 | 2.800E-07 |
| 20174 | 1009 | P1009F | C | 2.9e-07 | 3.5e-07 | 4.6e-07 | 3.667E-07 | 8.622E-08 | 2.900E-07 | 4.600E-07 |
| 20175 | 1009 | P1009P | C | 1.9e-07 | 2.6e-07 | 4e-07 | 2.833E-07 | 1.069E-07 | 1.900E-07 | 4.000E-07 |
| 20176 | 1009 | P1009S | C | 1.8e-07 | 3.8e-07 | 3.7e-07 | 3.100E-07 | 1.127E-07 | 1.800E-07 | 3.800E-07 |
| 20177 | 1009 | P1009T | C | 3.7e-07 | 2.5e-07 | 2.7e-07 | 2.967E-07 | 6.429E-08 | 2.500E-07 | 3.700E-07 |
| 20178 | 1009 | P1009W | C | 2.6e-07 | 2.4e-07 | 2e-07 | 2.333E-07 | 3.055E-08 | 2.000E-07 | 2.600E-07 |
| 20179 | 1009 | P1009Y | C | 1.4e-07 | 2.5e-07 | 2e-07 | 1.967E-07 | 5.508E-08 | 1.400E-07 | 2.500E-07 |
| 20180 | 1009 | P1009V | C | 2.6e-07 | 2.7e-07 | 1.4e-07 | 2.233E-07 | 7.234E-08 | 1.400E-07 | 2.700E-07 |
| 20181 | 1010 | A1010A | C | 2.2e-07 | 2.1e-07 | 3e-07 | 2.433E-07 | 4.933E-08 | 2.100E-07 | 3.000E-07 |
| 20182 | 1010 | A1010R | C | 2.2e-07 | 3.8e-07 | 3.2e-07 | 3.067E-07 | 8.083E-08 | 2.200E-07 | 3.800E-07 |
| 20183 | 1010 | A1010N | C | 2.1e-07 | 1.5e-07 | 3.4e-07 | 2.333E-07 | 9.713E-08 | 1.500E-07 | 3.400E-07 |
| 20184 | 1010 | A1010D | C | 3.2e-07 | 4.1e-07 | 1e-07 | 2.767E-07 | 1.595E-07 | 1.000E-07 | 4.100E-07 |
| 20185 | 1010 | A1010C | C | 4.8e-07 | 2.3e-07 | 2.4e-07 | 3.167E-07 | 1.415E-07 | 2.300E-07 | 4.800E-07 |
| 20186 | 1010 | A1010E | C | 4.1e-07 | 3.1e-07 | 1.5e-07 | 2.900E-07 | 1.311E-07 | 1.500E-07 | 4.100E-07 |
| 20187 | 1010 | A1010Q | C | 3.2e-07 | 2.2e-07 | 3.6e-07 | 3.000E-07 | 7.211E-08 | 2.200E-07 | 3.600E-07 |
| 20188 | 1010 | A1010G | C | 1.8e-07 | 1.5e-07 | 3.1e-07 | 2.133E-07 | 8.505E-08 | 1.500E-07 | 3.100E-07 |
| 20189 | 1010 | A1010H | C | 1.9e-07 | 1.5e-07 | 2.5e-07 | 1.967E-07 | 5.033E-08 | 1.500E-07 | 2.500E-07 |
| 20190 | 1010 | A1010I | C | 3.8e-07 | 2.3e-07 | 3.4e-07 | 3.167E-07 | 7.767E-08 | 2.300E-07 | 3.800E-07 |
| 20191 | 1010 | A1010L | C | 5.5e-07 | 1.9e-07 | 1.6e-07 | 3.000E-07 | 2.170E-07 | 1.600E-07 | 5.500E-07 |
| 20192 | 1010 | A1010K | C | 2.3e-07 | 3.2e-07 | 2.7e-07 | 2.733E-07 | 4.509E-08 | 2.300E-07 | 3.200E-07 |
| 20193 | 1010 | A1010M | C | 1.5e-07 | 3.3e-07 | 3.8e-07 | 2.867E-07 | 1.210E-07 | 1.500E-07 | 3.800E-07 |
| 20194 | 1010 | A1010F | C | 2.1e-07 | 1.3e-07 | 2.6e-07 | 2.000E-07 | 6.557E-08 | 1.300E-07 | 2.600E-07 |
| 20195 | 1010 | A1010P | C | 3.5e-07 | 3.1e-07 | 2.3e-07 | 2.967E-07 | 6.110E-08 | 2.300E-07 | 3.500E-07 |
| 20196 | 1010 | A1010S | C | 2.5e-07 | 3.3e-07 | 5.5e-07 | 3.767E-07 | 1.553E-07 | 2.500E-07 | 5.500E-07 |
| 20197 | 1010 | A1010T | C | 2.4e-07 | 3e-07 | 9.2e-07 | 4.867E-07 | 3.765E-07 | 2.400E-07 | 9.200E-07 |
| 20198 | 1010 | A1010W | C | 2e-07 | 2.6e-07 | 4.6e-07 | 3.067E-07 | 1.361E-07 | 2.000E-07 | 4.600E-07 |
| 20199 | 1010 | A1010Y | C | 1.4e-07 | 1.1e-07 | 3.1e-07 | 1.867E-07 | 1.079E-07 | 1.100E-07 | 3.100E-07 |
| 20200 | 1010 | A1010V | C | 4.2e-07 | 4.2e-07 | 2.2e-07 | 3.533E-07 | 1.155E-07 | 2.200E-07 | 4.200E-07 |
| 20201 | 1011 | V1011A | C | 1.9e-07 | 1.4e-07 | 4.3e-07 | 2.533E-07 | 1.550E-07 | 1.400E-07 | 4.300E-07 |
| 20202 | 1011 | V1011R | C | 1.6e-07 | 4e-07 | 2.5e-07 | 2.700E-07 | 1.212E-07 | 1.600E-07 | 4.000E-07 |
| 20203 | 1011 | V1011N | C | 2.2e-07 | 3.3e-07 | 2.5e-07 | 2.667E-07 | 5.686E-08 | 2.200E-07 | 3.300E-07 |
| 20204 | 1011 | V1011D | C | 1.5e-07 | 2.9e-07 | 2.9e-07 | 2.433E-07 | 8.083E-08 | 1.500E-07 | 2.900E-07 |
| 20205 | 1011 | V1011C | C | 5.1e-07 | 3.1e-07 | 1.1e-07 | 3.100E-07 | 2.000E-07 | 1.100E-07 | 5.100E-07 |
| 20206 | 1011 | V1011E | C | 3.3e-07 | 1.4e-07 | 4e-07 | 2.900E-07 | 1.345E-07 | 1.400E-07 | 4.000E-07 |
| 20207 | 1011 | V1011Q | C | 3.8e-07 | 5.9e-07 | 4.5e-07 | 4.733E-07 | 1.069E-07 | 3.800E-07 | 5.900E-07 |
| 20208 | 1011 | V1011G | C | 2.5e-07 | 4.3e-07 | 3e-07 | 3.267E-07 | 9.292E-08 | 2.500E-07 | 4.300E-07 |
| 20209 | 1011 | V1011H | C | 2.8e-07 | 3.7e-07 | 9.1e-08 | 2.470E-07 | 1.424E-07 | 9.100E-08 | 3.700E-07 |
| 20210 | 1011 | V1011I | C | 2.2e-07 | 2.9e-07 | 1.2e-07 | 2.100E-07 | 8.544E-08 | 1.200E-07 | 2.900E-07 |
| 20211 | 1011 | V1011L | C | 1.7e-07 | 3.4e-07 | 5.9e-07 | 3.667E-07 | 2.113E-07 | 1.700E-07 | 5.900E-07 |
| 20212 | 1011 | V1011K | C | 1.8e-07 | 9.7e-08 | 2.6e-07 | 1.790E-07 | 8.150E-08 | 9.700E-08 | 2.600E-07 |
| 20213 | 1011 | V1011M | C | 2.3e-07 | 3.6e-07 | 6.6e-08 | 2.187E-07 | 1.473E-07 | 6.600E-08 | 3.600E-07 |
| 20214 | 1011 | V1011F | C | 1.8e-07 | 6.7e-07 | 4e-07 | 4.167E-07 | 2.454E-07 | 1.800E-07 | 6.700E-07 |
| 20215 | 1011 | V1011P | C | 2.4e-07 | 2.7e-07 | 1.4e-07 | 2.167E-07 | 6.807E-08 | 1.400E-07 | 2.700E-07 |
| 20216 | 1011 | V1011S | C | 1.5e-07 | 2.5e-07 | 3.1e-07 | 2.367E-07 | 8.083E-08 | 1.500E-07 | 3.100E-07 |
| 20217 | 1011 | V1011T | C | 2.1e-07 | 2.5e-07 | 1.1e-07 | 1.900E-07 | 7.211E-08 | 1.100E-07 | 2.500E-07 |
| 20218 | 1011 | V1011W | C | 3e-07 | 2.7e-07 | 3.7e-07 | 3.133E-07 | 5.132E-08 | 2.700E-07 | 3.700E-07 |
| 20219 | 1011 | V1011Y | C | 1.2e-07 | 4.7e-07 | 3.6e-07 | 3.167E-07 | 1.790E-07 | 1.200E-07 | 4.700E-07 |
| 20220 | 1011 | V1011V | C | 1.7e-07 | 1.5e-07 | 2.4e-07 | 1.867E-07 | 4.726E-08 | 1.500E-07 | 2.400E-07 |
| 20221 | 1012 | L1012A | C | 4.6e-07 | 3e-07 | 1.9e-07 | 3.167E-07 | 1.358E-07 | 1.900E-07 | 4.600E-07 |
| 20222 | 1012 | L1012R | C | 2.2e-07 | 2.8e-07 | 2.4e-07 | 2.467E-07 | 3.055E-08 | 2.200E-07 | 2.800E-07 |
| 20223 | 1012 | L1012N | C | 2.1e-07 | 2.8e-07 | 4e-07 | 2.967E-07 | 9.609E-08 | 2.100E-07 | 4.000E-07 |
| 20224 | 1012 | L1012D | C | 3.5e-07 | 3.6e-07 | 2.7e-07 | 3.267E-07 | 4.933E-08 | 2.700E-07 | 3.600E-07 |
| 20225 | 1012 | L1012C | C | 4.6e-07 | 2.8e-07 | 2.3e-07 | 3.233E-07 | 1.210E-07 | 2.300E-07 | 4.600E-07 |
| 20226 | 1012 | L1012E | C | 3.4e-07 | 4.6e-07 | 2.1e-07 | 3.367E-07 | 1.250E-07 | 2.100E-07 | 4.600E-07 |
| 20227 | 1012 | L1012Q | C | 1.6e-07 | 3.9e-07 | 3.4e-07 | 2.967E-07 | 1.210E-07 | 1.600E-07 | 3.900E-07 |
| 20228 | 1012 | L1012G | C | 3.1e-07 | 3.1e-07 | 4.6e-07 | 3.600E-07 | 8.660E-08 | 3.100E-07 | 4.600E-07 |
| 20229 | 1012 | L1012H | C | 1.7e-07 | 3.9e-07 | 3.1e-07 | 2.900E-07 | 1.114E-07 | 1.700E-07 | 3.900E-07 |

| | | | | | | | | | | |
|---|---|---|---|---|---|---|---|---|---|---|
| 20230 | 1012 | L1012I | C | 2.7e-07 | 2.4e-07 | 1.4e-07 | 2.167E-07 | 6.807E-08 | 1.400E-07 | 2.700E-07 |
| 20231 | 1012 | L1012L | C | 4.6e-07 | 1.6e-07 | 4e-07 | 3.400E-07 | 1.587E-07 | 1.600E-07 | 4.600E-07 |
| 20232 | 1012 | L1012K | C | 1.3e-07 | 3.1e-07 | 1.1e-07 | 1.833E-07 | 1.102E-07 | 1.100E-07 | 3.100E-07 |
| 20233 | 1012 | L1012M | C | 3.1e-07 | 2.7e-07 | 3.9e-07 | 3.233E-07 | 6.110E-08 | 2.700E-07 | 3.900E-07 |
| 20234 | 1012 | L1012F | C | 1.5e-07 | 1.7e-07 | 1.4e-07 | 1.533E-07 | 1.528E-08 | 1.400E-07 | 1.700E-07 |
| 20235 | 1012 | L1012P | C | 2.1e-07 | 9.9e-08 | 2.9e-07 | 1.997E-07 | 9.592E-08 | 9.900E-08 | 2.900E-07 |
| 20236 | 1012 | L1012S | C | 3.5e-07 | 2.9e-07 | 1.8e-07 | 2.733E-07 | 8.622E-08 | 1.800E-07 | 3.500E-07 |
| 20237 | 1012 | L1012T | C | 2.3e-07 | 2.6e-07 | 1.7e-07 | 2.200E-07 | 4.583E-08 | 1.700E-07 | 2.600E-07 |
| 20238 | 1012 | L1012W | C | 2.9e-07 | 2.6e-07 | 2.3e-07 | 2.600E-07 | 3.000E-08 | 2.300E-07 | 2.900E-07 |
| 20239 | 1012 | L1012Y | C | 6.4e-07 | 1.5e-07 | 2.4e-07 | 3.433E-07 | 2.608E-07 | 1.500E-07 | 6.400E-07 |
| 20240 | 1012 | L1012V | C | 2.9e-07 | 2.1e-07 | 2.3e-07 | 2.433E-07 | 4.163E-08 | 2.100E-07 | 2.900E-07 |
| 20241 | 1013 | Q1013A | C | 1.2e-07 | 5.4e-07 | 2.4e-07 | 3.000E-07 | 2.163E-07 | 1.200E-07 | 5.400E-07 |
| 20242 | 1013 | Q1013R | C | 1.9e-07 | 1.5e-07 | 1.8e-07 | 1.733E-07 | 2.082E-08 | 1.500E-07 | 1.900E-07 |
| 20243 | 1013 | Q1013N | C | 2.8e-07 | 3.8e-07 | 3.5e-07 | 3.367E-07 | 5.132E-08 | 2.800E-07 | 3.800E-07 |
| 20244 | 1013 | Q1013D | C | 3.4e-07 | 3.2e-07 | 2.2e-07 | 2.933E-07 | 6.429E-08 | 2.200E-07 | 3.400E-07 |
| 20245 | 1013 | Q1013C | C | 2.1e-07 | 2.4e-07 | 2e-07 | 2.167E-07 | 2.082E-08 | 2.000E-07 | 2.400E-07 |
| 20246 | 1013 | Q1013E | C | 1.9e-07 | 2.1e-07 | 1.7e-07 | 1.900E-07 | 2.000E-08 | 1.700E-07 | 2.100E-07 |
| 20247 | 1013 | Q1013Q | C | 7.3e-08 | 2.6e-07 | 1.6e-07 | 1.643E-07 | 9.358E-08 | 7.300E-08 | 2.600E-07 |
| 20248 | 1013 | Q1013G | C | 3.3e-07 | 3.7e-07 | 2.8e-07 | 3.267E-07 | 4.509E-08 | 2.800E-07 | 3.700E-07 |
| 20249 | 1013 | Q1013H | C | 2.2e-07 | 1.7e-07 | 1.3e-07 | 1.733E-07 | 4.509E-08 | 1.300E-07 | 2.200E-07 |
| 20250 | 1013 | Q1013I | C | 4.3e-07 | 1.6e-07 | 1.6e-07 | 2.500E-07 | 1.559E-07 | 1.600E-07 | 4.300E-07 |
| 20251 | 1013 | Q1013L | C | 1.3e-07 | 1.8e-07 | 1.9e-07 | 1.667E-07 | 3.215E-08 | 1.300E-07 | 1.900E-07 |
| 20252 | 1013 | Q1013K | C | 2.4e-07 | 2.2e-07 | 4.1e-07 | 2.900E-07 | 1.044E-07 | 2.200E-07 | 4.100E-07 |
| 20253 | 1013 | Q1013M | C | 2.2e-07 | 2.7e-07 | 2.3e-07 | 2.400E-07 | 2.646E-08 | 2.200E-07 | 2.700E-07 |
| 20254 | 1013 | Q1013F | C | 2.9e-07 | 1.4e-07 | 2.6e-07 | 2.300E-07 | 7.937E-08 | 1.400E-07 | 2.900E-07 |
| 20255 | 1013 | Q1013P | C | 1.3e-07 | 4.6e-07 | 3e-07 | 2.967E-07 | 1.650E-07 | 1.300E-07 | 4.600E-07 |
| 20256 | 1013 | Q1013S | C | 2.7e-07 | 2.3e-07 | 4e-07 | 3.000E-07 | 8.888E-08 | 2.300E-07 | 4.000E-07 |
| 20257 | 1013 | Q1013T | C | 5.1e-07 | 1.7e-07 | 2.3e-07 | 3.033E-07 | 1.815E-07 | 1.700E-07 | 5.100E-07 |
| 20258 | 1013 | Q1013W | C | 2e-07 | 2.6e-07 | 2.3e-07 | 2.300E-07 | 3.000E-08 | 2.000E-07 | 2.600E-07 |
| 20259 | 1013 | Q1013Y | C | 3.6e-07 | 4.7e-07 | 1.2e-07 | 3.167E-07 | 1.790E-07 | 1.200E-07 | 4.700E-07 |
| 20260 | 1013 | Q1013V | C | 2.4e-07 | 2e-07 | 4e-07 | 2.800E-07 | 1.058E-07 | 2.000E-07 | 4.000E-07 |
| 20261 | 1014 | S1014A | C | 4.4e-07 | 1.6e-07 | 2.9e-07 | 2.967E-07 | 1.401E-07 | 1.600E-07 | 4.400E-07 |
| 20262 | 1014 | S1014R | C | 1.7e-07 | 1.3e-07 | 4.1e-07 | 2.367E-07 | 1.514E-07 | 1.300E-07 | 4.100E-07 |
| 20263 | 1014 | S1014N | C | 1.1e-06 | 3.3e-07 | 3e-07 | 5.767E-07 | 4.535E-07 | 3.000E-07 | 1.100E-06 |
| 20264 | 1014 | S1014D | C | 3.3e-07 | 2.1e-07 | 3.1e-07 | 2.833E-07 | 6.429E-08 | 2.100E-07 | 3.300E-07 |
| 20265 | 1014 | S1014C | C | 2.2e-07 | 2.3e-07 | 1.8e-07 | 2.100E-07 | 2.646E-08 | 1.800E-07 | 2.300E-07 |
| 20266 | 1014 | S1014E | C | 2.1e-07 | 1.5e-07 | 2.9e-07 | 2.167E-07 | 7.024E-08 | 1.500E-07 | 2.900E-07 |
| 20267 | 1014 | S1014Q | C | 7.5e-08 | 1.5e-07 | 1.5e-07 | 1.250E-07 | 4.330E-08 | 7.500E-08 | 1.500E-07 |
| 20268 | 1014 | S1014G | C | 3e-07 | 1.3e-07 | 1.9e-07 | 2.067E-07 | 8.622E-08 | 1.300E-07 | 3.000E-07 |
| 20269 | 1014 | S1014H | C | 2.5e-07 | 2.8e-07 | 2.6e-07 | 2.633E-07 | 1.528E-08 | 2.500E-07 | 2.800E-07 |
| 20270 | 1014 | S1014I | C | 1.9e-07 | 1.8e-07 | 2.6e-07 | 2.100E-07 | 4.359E-08 | 1.800E-07 | 2.600E-07 |
| 20271 | 1014 | S1014L | C | 3.5e-07 | 6.3e-07 | 2.7e-07 | 4.167E-07 | 1.890E-07 | 2.700E-07 | 6.300E-07 |
| 20272 | 1014 | S1014K | C | 2.2e-07 | 2.7e-07 | 1.9e-07 | 2.267E-07 | 4.041E-08 | 1.900E-07 | 2.700E-07 |
| 20273 | 1014 | S1014M | C | 2.1e-07 | 4e-07 | 3.7e-07 | 3.267E-07 | 1.021E-07 | 2.100E-07 | 4.000E-07 |
| 20274 | 1014 | S1014F | C | 1.7e-07 | 4.9e-07 | 3.2e-07 | 3.267E-07 | 1.601E-07 | 1.700E-07 | 4.900E-07 |
| 20275 | 1014 | S1014P | C | 3.4e-07 | 1.8e-07 | 4e-07 | 3.067E-07 | 1.137E-07 | 1.800E-07 | 4.000E-07 |
| 20276 | 1014 | S1014S | C | 2.5e-07 | 2.2e-07 | 1.8e-07 | 2.167E-07 | 3.512E-08 | 1.800E-07 | 2.500E-07 |
| 20277 | 1014 | S1014T | C | 1.7e-07 | 2.8e-07 | 4.1e-07 | 2.867E-07 | 1.201E-07 | 1.700E-07 | 4.100E-07 |
| 20278 | 1014 | S1014W | C | 2.5e-07 | 2.4e-07 | 1.8e-07 | 2.233E-07 | 3.786E-08 | 1.800E-07 | 2.500E-07 |
| 20279 | 1014 | S1014Y | C | 2.7e-07 | 4.1e-07 | 2.2e-07 | 3.000E-07 | 9.849E-08 | 2.200E-07 | 4.100E-07 |
| 20280 | 1014 | S1014V | C | 2.4e-07 | 2.7e-07 | 4.2e-07 | 3.100E-07 | 9.644E-08 | 2.400E-07 | 4.200E-07 |
| 20281 | 1015 | S1015A | C | 2.6e-07 | 2.3e-07 | 2.3e-07 | 2.400E-07 | 1.732E-08 | 2.300E-07 | 2.600E-07 |
| 20282 | 1015 | S1015R | C | 1.8e-07 | 2.4e-07 | 3.3e-07 | 2.500E-07 | 7.550E-08 | 1.800E-07 | 3.300E-07 |
| 20283 | 1015 | S1015N | C | 3.4e-07 | 7.6e-07 | 4.4e-07 | 5.133E-07 | 2.194E-07 | 3.400E-07 | 7.600E-07 |
| 20284 | 1015 | S1015D | C | 2.4e-07 | 8.9e-08 | 1.5e-07 | 1.597E-07 | 7.596E-08 | 8.900E-08 | 2.400E-07 |
| 20285 | 1015 | S1015C | C | 3.5e-07 | 1.5e-07 | 2.5e-07 | 2.500E-07 | 1.000E-07 | 1.500E-07 | 3.500E-07 |
| 20286 | 1015 | S1015E | C | 3.5e-07 | 1.6e-07 | 2.6e-07 | 2.567E-07 | 9.504E-08 | 1.600E-07 | 3.500E-07 |
| 20287 | 1015 | S1015Q | C | 1.7e-07 | 2.9e-07 | 3.8e-07 | 2.800E-07 | 1.054E-07 | 1.700E-07 | 3.800E-07 |
| 20288 | 1015 | S1015G | C | 2.6e-07 | 2.1e-07 | 3.8e-07 | 2.833E-07 | 8.737E-08 | 2.100E-07 | 3.800E-07 |
| 20289 | 1015 | S1015H | C | 1.5e-07 | 2.5e-07 | 2e-07 | 2.000E-07 | 5.000E-08 | 1.500E-07 | 2.500E-07 |
| 20290 | 1015 | S1015I | C | 1.8e-07 | 3.3e-07 | 3.9e-07 | 3.000E-07 | 1.082E-07 | 1.800E-07 | 3.900E-07 |
| 20291 | 1015 | S1015L | C | 1.2e-07 | 1.5e-07 | 2.3e-07 | 1.667E-07 | 5.686E-08 | 1.200E-07 | 2.300E-07 |
| 20292 | 1015 | S1015K | C | 2.2e-07 | 1.3e-07 | 2e-07 | 1.833E-07 | 4.726E-08 | 1.300E-07 | 2.200E-07 |
| 20293 | 1015 | S1015M | C | 2.4e-07 | 2.5e-07 | 2e-07 | 2.300E-07 | 2.646E-08 | 2.000E-07 | 2.500E-07 |
| 20294 | 1015 | S1015F | C | 1.5e-07 | 1.3e-07 | 1.7e-07 | 1.500E-07 | 2.000E-08 | 1.300E-07 | 1.700E-07 |
| 20295 | 1015 | S1015P | C | 2.1e-07 | 1.6e-07 | 2.8e-07 | 2.167E-07 | 6.028E-08 | 1.600E-07 | 2.800E-07 |
| 20296 | 1015 | S1015S | C | 2e-07 | 2.4e-07 | 2.3e-07 | 2.233E-07 | 2.082E-08 | 2.000E-07 | 2.400E-07 |
| 20297 | 1015 | S1015T | C | 2e-07 | 1.7e-07 | 1.5e-07 | 1.733E-07 | 2.517E-08 | 1.500E-07 | 2.000E-07 |
| 20298 | 1015 | S1015W | C | 3.4e-07 | 4.3e-07 | 4.6e-07 | 4.100E-07 | 6.245E-08 | 3.400E-07 | 4.600E-07 |
| 20299 | 1015 | S1015Y | C | 2.4e-07 | 3.3e-07 | 5e-07 | 3.567E-07 | 1.320E-07 | 2.400E-07 | 5.000E-07 |
| 20300 | 1015 | S1015V | C | 1.5e-07 | 1.6e-07 | 3.4e-07 | 2.167E-07 | 1.069E-07 | 1.500E-07 | 3.400E-07 |
| 20301 | 1016 | G1016A | C | 2.8e-07 | 2.3e-07 | 1.9e-07 | 2.333E-07 | 4.509E-08 | 1.900E-07 | 2.800E-07 |
| 20302 | 1016 | G1016R | C | 3.3e-07 | 2e-07 | 4e-07 | 3.100E-07 | 1.015E-07 | 2.000E-07 | 4.000E-07 |
| 20303 | 1016 | G1016N | C | 7.6e-07 | 2.2e-07 | 1.7e-07 | 3.833E-07 | 3.272E-07 | 1.700E-07 | 7.600E-07 |
| 20304 | 1016 | G1016D | C | 1.7e-07 | 2.8e-07 | 3.8e-07 | 2.767E-07 | 1.050E-07 | 1.700E-07 | 3.800E-07 |
| 20305 | 1016 | G1016C | C | 2.9e-07 | 1.8e-07 | 2.4e-07 | 2.367E-07 | 5.508E-08 | 1.800E-07 | 2.900E-07 |
| 20306 | 1016 | G1016E | C | 1.7e-07 | 3.2e-07 | 2.8e-07 | 2.567E-07 | 7.767E-08 | 1.700E-07 | 3.200E-07 |
| 20307 | 1016 | G1016Q | C | 2.8e-07 | 1.3e-07 | 2e-07 | 2.033E-07 | 7.506E-08 | 1.300E-07 | 2.800E-07 |
| 20308 | 1016 | G1016G | C | 2.6e-07 | 2e-07 | 2.3e-07 | 2.300E-07 | 3.000E-08 | 2.000E-07 | 2.600E-07 |
| 20309 | 1016 | G1016H | C | 3e-07 | 5.3e-07 | 2.9e-07 | 3.733E-07 | 1.358E-07 | 2.900E-07 | 5.300E-07 |
| 20310 | 1016 | G1016I | C | 2e-07 | 1.5e-07 | 2.4e-07 | 1.967E-07 | 4.509E-08 | 1.500E-07 | 2.400E-07 |
| 20311 | 1016 | G1016L | C | 2.8e-07 | 2.1e-07 | 2.4e-07 | 2.433E-07 | 3.512E-08 | 2.100E-07 | 2.800E-07 |
| 20312 | 1016 | G1016K | C | 4.5e-07 | 1.5e-07 | 2.3e-07 | 2.767E-07 | 1.553E-07 | 1.500E-07 | 4.500E-07 |
| 20313 | 1016 | G1016M | C | 2.7e-07 | 2.3e-07 | 1.2e-07 | 2.067E-07 | 7.767E-08 | 1.200E-07 | 2.700E-07 |
| 20314 | 1016 | G1016F | C | 1.9e-07 | 4.9e-07 | 4.8e-07 | 3.867E-07 | 1.704E-07 | 1.900E-07 | 4.900E-07 |

| | | | | | | | | | | |
|---|---|---|---|---|---|---|---|---|---|---|
| 20315 | 1016 | G1016P | C | 2.3e-07 | 1.3e-07 | 2.5e-07 | 2.033E-07 | 6.429E-08 | 1.300E-07 | 2.500E-07 |
| 20316 | 1016 | G1016S | C | 3.7e-07 | 3.6e-07 | 2.2e-07 | 3.167E-07 | 8.386E-08 | 2.200E-07 | 3.700E-07 |
| 20317 | 1016 | G1016T | C | 3.2e-07 | 4.3e-07 | 2e-07 | 3.167E-07 | 1.150E-07 | 2.000E-07 | 4.300E-07 |
| 20318 | 1016 | G1016W | C | 2.9e-07 | 3.8e-07 | 2.5e-07 | 3.067E-07 | 6.658E-08 | 2.500E-07 | 3.800E-07 |
| 20319 | 1016 | G1016Y | C | 2.4e-07 | 3.2e-07 | 2e-07 | 2.533E-07 | 6.110E-08 | 2.000E-07 | 3.200E-07 |
| 20320 | 1016 | G1016V | C | 3.7e-07 | 1.9e-07 | 3.2e-07 | 2.933E-07 | 9.292E-08 | 1.900E-07 | 3.700E-07 |
| 20321 | 1017 | L1017A | C | 4.4e-07 | 3.4e-07 | 2.5e-07 | 3.433E-07 | 9.504E-08 | 2.500E-07 | 4.400E-07 |
| 20322 | 1017 | L1017R | C | 5.8e-07 | 5.4e-07 | 3.9e-07 | 5.033E-07 | 1.002E-07 | 3.900E-07 | 5.800E-07 |
| 20323 | 1017 | L1017N | C | 1.9e-07 | 1.5e-07 | 1.8e-07 | 1.733E-07 | 2.082E-08 | 1.500E-07 | 1.900E-07 |
| 20324 | 1017 | L1017D | C | 3.2e-07 | 2.4e-07 | 2.8e-07 | 2.800E-07 | 4.000E-08 | 2.400E-07 | 3.200E-07 |
| 20325 | 1017 | L1017C | C | 1.2e-07 | 2.3e-07 | 2.9e-07 | 2.133E-07 | 8.622E-08 | 1.200E-07 | 2.900E-07 |
| 20326 | 1017 | L1017E | C | 2.8e-07 | 6.1e-07 | 2.2e-07 | 3.700E-07 | 2.100E-07 | 2.200E-07 | 6.100E-07 |
| 20327 | 1017 | L1017Q | C | 3.5e-07 | 2.3e-07 | 1.7e-07 | 2.500E-07 | 9.165E-08 | 1.700E-07 | 3.500E-07 |
| 20328 | 1017 | L1017G | C | 2.8e-07 | 1.4e-07 | 2.3e-07 | 2.167E-07 | 7.095E-08 | 1.400E-07 | 2.800E-07 |
| 20329 | 1017 | L1017H | C | 3.1e-07 | 2.2e-07 | 3.3e-07 | 2.867E-07 | 5.859E-08 | 2.200E-07 | 3.300E-07 |
| 20330 | 1017 | L1017I | C | 2.4e-07 | 2e-07 | 6.8e-07 | 3.733E-07 | 2.663E-07 | 2.000E-07 | 6.800E-07 |
| 20331 | 1017 | L1017L | C | 1.7e-07 | 2.1e-07 | 3.5e-07 | 2.433E-07 | 9.452E-08 | 1.700E-07 | 3.500E-07 |
| 20332 | 1017 | L1017K | C | 3.2e-07 | 2.5e-07 | 1.1e-07 | 2.267E-07 | 1.069E-07 | 1.100E-07 | 3.200E-07 |
| 20333 | 1017 | L1017M | C | 2.7e-07 | 2.2e-07 | 2.3e-07 | 2.400E-07 | 2.646E-08 | 2.200E-07 | 2.700E-07 |
| 20334 | 1017 | L1017F | C | 4.5e-07 | 4e-07 | 2.1e-07 | 3.533E-07 | 1.266E-07 | 2.100E-07 | 4.500E-07 |
| 20335 | 1017 | L1017P | C | 1.9e-07 | 1.5e-07 | 2.6e-07 | 2.000E-07 | 5.568E-08 | 1.500E-07 | 2.600E-07 |
| 20336 | 1017 | L1017S | C | 1.3e-07 | 2.1e-07 | 2.7e-07 | 2.033E-07 | 7.024E-08 | 1.300E-07 | 2.700E-07 |
| 20337 | 1017 | L1017T | C | 1.2e-07 | 3.1e-07 | 2e-07 | 2.100E-07 | 9.539E-08 | 1.200E-07 | 3.100E-07 |
| 20338 | 1017 | L1017W | C | 1.7e-07 | 2.4e-07 | 2.1e-07 | 2.067E-07 | 3.512E-08 | 1.700E-07 | 2.400E-07 |
| 20339 | 1017 | L1017Y | C | 1.8e-07 | 2.4e-07 | 3.3e-07 | 2.500E-07 | 7.550E-08 | 1.800E-07 | 3.300E-07 |
| 20340 | 1017 | L1017V | C | 1.9e-07 | 3e-07 | 3.4e-07 | 2.767E-07 | 7.767E-08 | 1.900E-07 | 3.400E-07 |
| 20341 | 1018 | Y1018A | C | 1.7e-07 | 4.3e-07 | 2.1e-07 | 2.700E-07 | 1.400E-07 | 1.700E-07 | 4.300E-07 |
| 20342 | 1018 | Y1018R | C | 2.2e-07 | 7.7e-07 | 1.9e-07 | 3.933E-07 | 3.265E-07 | 1.900E-07 | 7.700E-07 |
| 20343 | 1018 | Y1018N | C | 2.2e-07 | 1.6e-07 | 2.8e-07 | 2.200E-07 | 6.000E-08 | 1.600E-07 | 2.800E-07 |
| 20344 | 1018 | Y1018D | C | 2.5e-07 | 6.8e-07 | 1.3e-07 | 3.533E-07 | 2.892E-07 | 1.300E-07 | 6.800E-07 |
| 20345 | 1018 | Y1018C | C | 3.2e-07 | 1.6e-07 | 1.7e-07 | 2.167E-07 | 8.963E-08 | 1.600E-07 | 3.200E-07 |
| 20346 | 1018 | Y1018E | C | 4.1e-07 | 4.3e-07 | 3.5e-07 | 3.967E-07 | 4.163E-08 | 3.500E-07 | 4.300E-07 |
| 20347 | 1018 | Y1018Q | C | 1.9e-07 | 2.4e-07 | 1.9e-07 | 2.067E-07 | 2.887E-08 | 1.900E-07 | 2.400E-07 |
| 20348 | 1018 | Y1018G | C | 2e-07 | 2.3e-07 | 2.8e-07 | 2.367E-07 | 4.041E-08 | 2.000E-07 | 2.800E-07 |
| 20349 | 1018 | Y1018H | C | 2.2e-07 | 9.6e-08 | 2.7e-07 | 1.953E-07 | 8.958E-08 | 9.600E-08 | 2.700E-07 |
| 20350 | 1018 | Y1018I | C | 2.4e-07 | 3.4e-07 | 2.6e-07 | 2.800E-07 | 5.292E-08 | 2.400E-07 | 3.400E-07 |
| 20351 | 1018 | Y1018L | C | 2.7e-07 | 2.8e-07 | 2.6e-07 | 2.700E-07 | 1.000E-08 | 2.600E-07 | 2.800E-07 |
| 20352 | 1018 | Y1018K | C | 2.1e-07 | 1.5e-07 | 2.2e-07 | 1.933E-07 | 3.786E-08 | 1.500E-07 | 2.200E-07 |
| 20353 | 1018 | Y1018M | C | 1.8e-07 | 2e-07 | 1.6e-07 | 1.800E-07 | 2.000E-08 | 1.600E-07 | 2.000E-07 |
| 20354 | 1018 | Y1018F | C | 1.4e-07 | 1.8e-07 | 1e-07 | 1.400E-07 | 4.000E-08 | 1.000E-07 | 1.800E-07 |
| 20355 | 1018 | Y1018P | C | 2.3e-07 | 4.1e-07 | 4e-07 | 3.467E-07 | 1.012E-07 | 2.300E-07 | 4.100E-07 |
| 20356 | 1018 | Y1018S | C | 4.7e-07 | 4.2e-07 | 2.8e-07 | 3.900E-07 | 9.849E-08 | 2.800E-07 | 4.700E-07 |
| 20357 | 1018 | Y1018T | C | 2.8e-07 | 1.6e-07 | 1.6e-07 | 2.000E-07 | 6.928E-08 | 1.600E-07 | 2.800E-07 |
| 20358 | 1018 | Y1018W | C | 3.8e-07 | 1e-07 | 2.2e-07 | 2.333E-07 | 1.405E-07 | 1.000E-07 | 3.800E-07 |
| 20359 | 1018 | Y1018Y | C | 2.9e-07 | 2.8e-07 | 3.5e-07 | 3.067E-07 | 3.786E-08 | 2.800E-07 | 3.500E-07 |
| 20360 | 1018 | Y1018V | C | 2.4e-07 | 3.3e-07 | 1.4e-07 | 2.367E-07 | 9.504E-08 | 1.400E-07 | 3.300E-07 |
| 20361 | 1019 | S1019A | C | 4e-07 | 3.7e-07 | 3.9e-07 | 3.867E-07 | 1.528E-08 | 3.700E-07 | 4.000E-07 |
| 20362 | 1019 | S1019R | C | 2.7e-07 | 6.9e-07 | 4.1e-07 | 4.567E-07 | 2.139E-07 | 2.700E-07 | 6.900E-07 |
| 20363 | 1019 | S1019N | C | 4.7e-07 | 2.9e-07 | 3.9e-07 | 3.833E-07 | 9.018E-08 | 2.900E-07 | 4.700E-07 |
| 20364 | 1019 | S1019D | C | 1.6e-07 | 2.7e-07 | 2.2e-07 | 2.167E-07 | 5.508E-08 | 1.600E-07 | 2.700E-07 |
| 20365 | 1019 | S1019C | C | 1.7e-07 | 1.4e-07 | 3.1e-07 | 2.067E-07 | 9.074E-08 | 1.400E-07 | 3.100E-07 |
| 20366 | 1019 | S1019E | C | 2.1e-07 | 2.1e-07 | 3.8e-07 | 2.667E-07 | 9.815E-08 | 2.100E-07 | 3.800E-07 |
| 20367 | 1019 | S1019Q | C | 3.4e-07 | 3.7e-07 | 3.3e-07 | 3.467E-07 | 2.082E-08 | 3.300E-07 | 3.700E-07 |
| 20368 | 1019 | S1019G | C | 2.9e-07 | 3.2e-07 | 3e-07 | 3.033E-07 | 1.528E-08 | 2.900E-07 | 3.200E-07 |
| 20369 | 1019 | S1019H | C | 2.4e-07 | 5.2e-07 | 1.4e-07 | 3.000E-07 | 1.970E-07 | 1.400E-07 | 5.200E-07 |
| 20370 | 1019 | S1019I | C | 1.9e-07 | 3.6e-07 | 4.8e-07 | 3.433E-07 | 1.457E-07 | 1.900E-07 | 4.800E-07 |
| 20371 | 1019 | S1019L | C | 3.5e-07 | 4.1e-07 | 4.9e-07 | 4.167E-07 | 7.024E-08 | 3.500E-07 | 4.900E-07 |
| 20372 | 1019 | S1019K | C | 2.9e-07 | 2.7e-07 | 2.1e-07 | 2.567E-07 | 4.163E-08 | 2.100E-07 | 2.900E-07 |
| 20373 | 1019 | S1019M | C | 4.9e-07 | 2.1e-07 | 2.1e-07 | 3.033E-07 | 1.617E-07 | 2.100E-07 | 4.900E-07 |
| 20374 | 1019 | S1019F | C | 3.3e-07 | 2.1e-07 | 3e-07 | 2.800E-07 | 6.245E-08 | 2.100E-07 | 3.300E-07 |
| 20375 | 1019 | S1019P | C | 9.2e-08 | 1.9e-07 | 8.7e-07 | 3.840E-07 | 4.237E-07 | 9.200E-08 | 8.700E-07 |
| 20376 | 1019 | S1019S | C | 1.8e-07 | 2.2e-07 | 1.2e-07 | 1.733E-07 | 5.033E-08 | 1.200E-07 | 2.200E-07 |
| 20377 | 1019 | S1019T | C | 4.6e-07 | 9.1e-08 | 1.5e-07 | 2.337E-07 | 1.982E-07 | 9.100E-08 | 4.600E-07 |
| 20378 | 1019 | S1019W | C | 1.6e-07 | 3e-07 | 2.3e-07 | 2.300E-07 | 7.000E-08 | 1.600E-07 | 3.000E-07 |
| 20379 | 1019 | S1019Y | C | 2.6e-07 | 2.4e-07 | 3.6e-07 | 2.867E-07 | 6.429E-08 | 2.400E-07 | 3.600E-07 |
| 20380 | 1019 | S1019V | C | 4.3e-07 | 3.1e-07 | 1.6e-07 | 3.000E-07 | 1.353E-07 | 1.600E-07 | 4.300E-07 |
| 20381 | 1020 | L1020A | C | 1.9e-07 | 2e-07 | 3.8e-07 | 2.567E-07 | 1.069E-07 | 1.900E-07 | 3.800E-07 |
| 20382 | 1020 | L1020R | C | 5.7e-08 | 1.9e-07 | 1.8e-07 | 1.423E-07 | 7.407E-08 | 5.700E-08 | 1.900E-07 |
| 20383 | 1020 | L1020N | C | 5.5e-07 | 4.8e-07 | 2.1e-07 | 4.133E-07 | 1.795E-07 | 2.100E-07 | 5.500E-07 |
| 20384 | 1020 | L1020D | C | 3e-07 | 1.7e-07 | 2.7e-07 | 2.467E-07 | 6.807E-08 | 1.700E-07 | 3.000E-07 |
| 20385 | 1020 | L1020C | C | 6.1e-07 | 1.6e-07 | 3e-07 | 3.567E-07 | 2.303E-07 | 1.600E-07 | 6.100E-07 |
| 20386 | 1020 | L1020E | C | 1.7e-07 | 2.1e-07 | 4.6e-07 | 2.800E-07 | 1.572E-07 | 1.700E-07 | 4.600E-07 |
| 20387 | 1020 | L1020Q | C | 2.4e-07 | 2e-07 | 3.1e-07 | 2.500E-07 | 5.568E-08 | 2.000E-07 | 3.100E-07 |
| 20388 | 1020 | L1020G | C | 1.9e-07 | 2.5e-07 | 2.4e-07 | 2.267E-07 | 3.215E-08 | 1.900E-07 | 2.500E-07 |
| 20389 | 1020 | L1020H | C | 3.8e-07 | 1.7e-07 | 3.4e-07 | 2.967E-07 | 1.115E-07 | 1.700E-07 | 3.800E-07 |
| 20390 | 1020 | L1020I | C | 1.5e-07 | 2e-07 | 1.6e-07 | 1.700E-07 | 2.646E-08 | 1.500E-07 | 2.000E-07 |
| 20391 | 1020 | L1020L | C | 2.7e-07 | 1.2e-07 | 1.6e-07 | 1.833E-07 | 7.767E-08 | 1.200E-07 | 2.700E-07 |
| 20392 | 1020 | L1020K | C | 8.4e-08 | 1.6e-07 | 6.8e-07 | 3.080E-07 | 3.244E-07 | 8.400E-08 | 6.800E-07 |
| 20393 | 1020 | L1020M | C | 4e-07 | 2.7e-07 | 1.8e-07 | 2.833E-07 | 1.106E-07 | 1.800E-07 | 4.000E-07 |
| 20394 | 1020 | L1020F | C | 3.6e-07 | 3e-07 | 3.4e-07 | 3.333E-07 | 3.055E-08 | 3.000E-07 | 3.600E-07 |
| 20395 | 1020 | L1020P | C | 3.4e-07 | 3.5e-07 | 1.8e-07 | 2.900E-07 | 9.539E-08 | 1.800E-07 | 3.500E-07 |
| 20396 | 1020 | L1020S | C | 2.5e-07 | 2.3e-07 | 2.9e-07 | 2.567E-07 | 3.055E-08 | 2.300E-07 | 2.900E-07 |
| 20397 | 1020 | L1020T | C | 1.8e-07 | 2.7e-07 | 3e-07 | 2.500E-07 | 6.245E-08 | 1.800E-07 | 3.000E-07 |
| 20398 | 1020 | L1020W | C | 4.4e-07 | 2.3e-07 | 1.9e-07 | 2.867E-07 | 1.343E-07 | 1.900E-07 | 4.400E-07 |
| 20399 | 1020 | L1020Y | C | 1.2e-07 | 2.4e-07 | 1.2e-07 | 1.600E-07 | 6.928E-08 | 1.200E-07 | 2.400E-07 |

| | | | | | | | | | | |
|---|---|---|---|---|---|---|---|---|---|---|
| 20400 | 1020 | L1020V | C | 3.4e-07 | 1.8e-07 | 2.7e-07 | 2.633E-07 | 8.021E-08 | 1.800E-07 | 3.400E-07 |
| 20401 | 1021 | S1021A | C | 4e-07 | 2.4e-07 | 2.1e-07 | 2.833E-07 | 1.021E-07 | 2.100E-07 | 4.000E-07 |
| 20402 | 1021 | S1021R | C | 2.5e-07 | 7.7e-07 | 1.7e-07 | 3.967E-07 | 3.258E-07 | 1.700E-07 | 7.700E-07 |
| 20403 | 1021 | S1021N | C | 2.6e-07 | 2.7e-07 | 3.2e-07 | 2.833E-07 | 3.215E-08 | 2.600E-07 | 3.200E-07 |
| 20404 | 1021 | S1021D | C | 3.7e-07 | 1e-07 | 1.7e-07 | 2.133E-07 | 1.401E-07 | 1.000E-07 | 3.700E-07 |
| 20405 | 1021 | S1021C | C | 1.6e-07 | 4.5e-07 | 4.1e-07 | 3.400E-07 | 1.572E-07 | 1.600E-07 | 4.500E-07 |
| 20406 | 1021 | S1021E | C | 1.8e-07 | 4e-07 | 2.2e-07 | 2.667E-07 | 1.172E-07 | 1.800E-07 | 4.000E-07 |
| 20407 | 1021 | S1021Q | C | 1.9e-07 | 1.6e-07 | 4.9e-07 | 2.800E-07 | 1.825E-07 | 1.600E-07 | 4.900E-07 |
| 20408 | 1021 | S1021G | C | 3.5e-07 | 1.8e-07 | 4e-07 | 3.100E-07 | 1.153E-07 | 1.800E-07 | 4.000E-07 |
| 20409 | 1021 | S1021H | C | 2.4e-07 | 1.6e-07 | 1.7e-07 | 1.900E-07 | 4.359E-08 | 1.600E-07 | 2.400E-07 |
| 20410 | 1021 | S1021I | C | 1.7e-07 | 5.7e-07 | 3.6e-07 | 3.667E-07 | 2.001E-07 | 1.700E-07 | 5.700E-07 |
| 20411 | 1021 | S1021L | C | 1.7e-07 | 4.2e-07 | 2.4e-07 | 2.767E-07 | 1.290E-07 | 1.700E-07 | 4.200E-07 |
| 20412 | 1021 | S1021K | C | 3.4e-07 | 1.3e-07 | 2.9e-07 | 2.533E-07 | 1.097E-07 | 1.300E-07 | 3.400E-07 |
| 20413 | 1021 | S1021M | C | 5.8e-07 | 2.6e-07 | 2.7e-07 | 3.700E-07 | 1.819E-07 | 2.600E-07 | 5.800E-07 |
| 20414 | 1021 | S1021F | C | 1.9e-07 | 3.3e-07 | 2.5e-07 | 2.567E-07 | 7.024E-08 | 1.900E-07 | 3.300E-07 |
| 20415 | 1021 | S1021P | C | 1.5e-07 | 1.3e-07 | 3.7e-07 | 2.167E-07 | 1.332E-07 | 1.300E-07 | 3.700E-07 |
| 20416 | 1021 | S1021S | C | 2.4e-07 | 2.5e-07 | 6.4e-08 | 1.847E-07 | 1.046E-07 | 6.400E-08 | 2.500E-07 |
| 20417 | 1021 | S1021T | C | 2.5e-07 | 2.4e-07 | 3.5e-07 | 2.800E-07 | 6.083E-08 | 2.400E-07 | 3.500E-07 |
| 20418 | 1021 | S1021W | C | 1.7e-07 | 2.5e-07 | 3.5e-07 | 2.567E-07 | 9.018E-08 | 1.700E-07 | 3.500E-07 |
| 20419 | 1021 | S1021Y | C | 1.3e-07 | 2.2e-07 | 5.3e-07 | 2.933E-07 | 2.098E-07 | 1.300E-07 | 5.300E-07 |
| 20420 | 1021 | S1021V | C | 1.7e-07 | 3e-07 | 4.3e-07 | 3.000E-07 | 1.300E-07 | 1.700E-07 | 4.300E-07 |
| 20421 | 1022 | S1022A | C | 2.7e-07 | 2.9e-07 | 2.5e-07 | 2.700E-07 | 2.000E-08 | 2.500E-07 | 2.900E-07 |
| 20422 | 1022 | S1022R | C | 1e-07 | 2.3e-07 | 2.2e-07 | 1.833E-07 | 7.234E-08 | 1.000E-07 | 2.300E-07 |
| 20423 | 1022 | S1022N | C | 2.2e-07 | 1.3e-07 | 1.3e-07 | 1.600E-07 | 5.196E-08 | 1.300E-07 | 2.200E-07 |
| 20424 | 1022 | S1022D | C | 3.1e-07 | 2.7e-07 | 2.4e-07 | 2.733E-07 | 3.512E-08 | 2.400E-07 | 3.100E-07 |
| 20425 | 1022 | S1022C | C | 2.8e-07 | 2.9e-07 | 1.3e-07 | 2.333E-07 | 8.963E-08 | 1.300E-07 | 2.900E-07 |
| 20426 | 1022 | S1022E | C | 1.1e-06 | 2.7e-07 | 1.9e-07 | 5.200E-07 | 5.039E-07 | 1.900E-07 | 1.100E-06 |
| 20427 | 1022 | S1022Q | C | 1.9e-07 | 4e-07 | 3.9e-07 | 3.267E-07 | 1.185E-07 | 1.900E-07 | 4.000E-07 |
| 20428 | 1022 | S1022G | C | 2.4e-07 | 4.7e-07 | 2e-07 | 3.033E-07 | 1.457E-07 | 2.000E-07 | 4.700E-07 |
| 20429 | 1022 | S1022H | C | 1.2e-07 | 2.3e-07 | 2.2e-07 | 1.900E-07 | 6.083E-08 | 1.200E-07 | 2.300E-07 |
| 20430 | 1022 | S1022I | C | 2.5e-07 | 1e-07 | 1.6e-07 | 1.700E-07 | 7.550E-08 | 1.000E-07 | 2.500E-07 |
| 20431 | 1022 | S1022L | C | 2.6e-07 | 2.2e-07 | 1.6e-07 | 2.133E-07 | 5.033E-08 | 1.600E-07 | 2.600E-07 |
| 20432 | 1022 | S1022K | C | 2.1e-07 | 1.2e-07 | 5.5e-07 | 2.933E-07 | 2.268E-07 | 1.200E-07 | 5.500E-07 |
| 20433 | 1022 | S1022M | C | 7.1e-07 | 1.9e-07 | 1.4e-07 | 3.467E-07 | 3.156E-07 | 1.400E-07 | 7.100E-07 |
| 20434 | 1022 | S1022F | C | 5.2e-07 | 2.5e-07 | 2.4e-07 | 3.367E-07 | 1.589E-07 | 2.400E-07 | 5.200E-07 |
| 20435 | 1022 | S1022P | C | 1.4e-07 | 1.7e-07 | 2.7e-07 | 1.933E-07 | 6.807E-08 | 1.400E-07 | 2.700E-07 |
| 20436 | 1022 | S1022S | C | 7.6e-07 | 1.4e-07 | 2.9e-07 | 3.967E-07 | 3.235E-07 | 1.400E-07 | 7.600E-07 |
| 20437 | 1022 | S1022T | C | 5.3e-07 | 1.8e-07 | 4.7e-07 | 3.933E-07 | 1.872E-07 | 1.800E-07 | 5.300E-07 |
| 20438 | 1022 | S1022W | C | 4.4e-07 | 1.5e-07 | 2.7e-07 | 2.867E-07 | 1.457E-07 | 1.500E-07 | 4.400E-07 |
| 20439 | 1022 | S1022Y | C | 1.5e-07 | 1.2e-07 | 2.9e-07 | 1.867E-07 | 9.074E-08 | 1.200E-07 | 2.900E-07 |
| 20440 | 1022 | S1022V | C | 2.7e-07 | 2.8e-07 | 4.2e-07 | 3.233E-07 | 8.386E-08 | 2.700E-07 | 4.200E-07 |
| 20441 | 1023 | V1023A | C | 2.8e-07 | 2.1e-07 | 2.1e-07 | 2.333E-07 | 4.041E-08 | 2.100E-07 | 2.800E-07 |
| 20442 | 1023 | V1023R | C | 5.3e-07 | 5.3e-07 | 2.9e-07 | 4.500E-07 | 1.386E-07 | 2.900E-07 | 5.300E-07 |
| 20443 | 1023 | V1023N | C | 1.9e-07 | 4.8e-07 | 4.1e-07 | 3.600E-07 | 1.513E-07 | 1.900E-07 | 4.800E-07 |
| 20444 | 1023 | V1023D | C | 2e-07 | 1.9e-07 | 2.3e-07 | 2.067E-07 | 2.082E-08 | 1.900E-07 | 2.300E-07 |
| 20445 | 1023 | V1023C | C | 2.5e-07 | 2.1e-07 | 1.9e-07 | 2.167E-07 | 3.055E-08 | 1.900E-07 | 2.500E-07 |
| 20446 | 1023 | V1023E | C | 1.9e-07 | 2.2e-07 | 4.6e-07 | 2.900E-07 | 1.480E-07 | 1.900E-07 | 4.600E-07 |
| 20447 | 1023 | V1023Q | C | 2.1e-07 | 1.2e-07 | 2.4e-07 | 1.900E-07 | 6.245E-08 | 1.200E-07 | 2.400E-07 |
| 20448 | 1023 | V1023G | C | 2.4e-07 | 3.1e-07 | 2.2e-07 | 2.567E-07 | 4.726E-08 | 2.200E-07 | 3.100E-07 |
| 20449 | 1023 | V1023H | C | 3.4e-07 | 3.2e-07 | 2.8e-07 | 3.133E-07 | 3.055E-08 | 2.800E-07 | 3.400E-07 |
| 20450 | 1023 | V1023I | C | 5.3e-07 | 2.2e-07 | 4e-07 | 3.833E-07 | 1.557E-07 | 2.200E-07 | 5.300E-07 |
| 20451 | 1023 | V1023L | C | 1.6e-07 | 1e-07 | 4.2e-07 | 2.267E-07 | 1.701E-07 | 1.000E-07 | 4.200E-07 |
| 20452 | 1023 | V1023K | C | 2.2e-07 | 3.9e-07 | 1.7e-07 | 2.600E-07 | 1.153E-07 | 1.700E-07 | 3.900E-07 |
| 20453 | 1023 | V1023M | C | 2.8e-07 | 3.2e-07 | 3.5e-07 | 3.167E-07 | 3.512E-08 | 2.800E-07 | 3.500E-07 |
| 20454 | 1023 | V1023F | C | 2e-07 | 2.9e-07 | 1.8e-07 | 2.233E-07 | 5.859E-08 | 1.800E-07 | 2.900E-07 |
| 20455 | 1023 | V1023P | C | 1.4e-07 | 2.6e-07 | 2.1e-07 | 2.033E-07 | 6.028E-08 | 1.400E-07 | 2.600E-07 |
| 20456 | 1023 | V1023S | C | 3.6e-07 | 2.1e-07 | 1.7e-07 | 2.467E-07 | 1.002E-07 | 1.700E-07 | 3.600E-07 |
| 20457 | 1023 | V1023T | C | 3.6e-07 | 2.9e-07 | 1.2e-07 | 2.567E-07 | 1.234E-07 | 1.200E-07 | 3.600E-07 |
| 20458 | 1023 | V1023W | C | 2.2e-07 | 1.6e-07 | 3.1e-07 | 2.300E-07 | 7.550E-08 | 1.600E-07 | 3.100E-07 |
| 20459 | 1023 | V1023Y | C | 2.4e-07 | 1.4e-07 | 1.8e-07 | 1.867E-07 | 5.033E-08 | 1.400E-07 | 2.400E-07 |
| 20460 | 1023 | V1023V | C | 5e-07 | 1.3e-07 | 2.3e-07 | 2.867E-07 | 1.914E-07 | 1.300E-07 | 5.000E-07 |
| 20461 | 1024 | V1024A | C | 1.3e-07 | 2e-07 | 2.3e-07 | 1.867E-07 | 5.132E-08 | 1.300E-07 | 2.300E-07 |
| 20462 | 1024 | V1024R | C | 2.9e-07 | 4.3e-07 | 1.1e-07 | 2.767E-07 | 1.604E-07 | 1.100E-07 | 4.300E-07 |
| 20463 | 1024 | V1024N | C | 5.6e-07 | 3.1e-07 | 2.5e-07 | 3.733E-07 | 1.644E-07 | 2.500E-07 | 5.600E-07 |
| 20464 | 1024 | V1024D | C | 1.1e-07 | 4.6e-07 | 1.5e-07 | 2.400E-07 | 1.916E-07 | 1.100E-07 | 4.600E-07 |
| 20465 | 1024 | V1024C | C | 2.1e-07 | 9.6e-08 | 3.3e-07 | 2.120E-07 | 1.170E-07 | 9.600E-08 | 3.300E-07 |
| 20466 | 1024 | V1024E | C | 2.7e-07 | 2.5e-07 | 2.9e-07 | 2.700E-07 | 2.000E-08 | 2.500E-07 | 2.900E-07 |
| 20467 | 1024 | V1024Q | C | 3e-07 | 2.5e-07 | 2e-07 | 2.500E-07 | 5.000E-08 | 2.000E-07 | 3.000E-07 |
| 20468 | 1024 | V1024G | C | 1.1e-07 | 2.6e-07 | 5.4e-07 | 3.033E-07 | 2.183E-07 | 1.100E-07 | 5.400E-07 |
| 20469 | 1024 | V1024H | C | 2.6e-07 | 1.6e-07 | 5.7e-07 | 3.300E-07 | 2.138E-07 | 1.600E-07 | 5.700E-07 |
| 20470 | 1024 | V1024I | C | 1.3e-07 | 2.6e-07 | 2.1e-07 | 2.000E-07 | 6.557E-08 | 1.300E-07 | 2.600E-07 |
| 20471 | 1024 | V1024L | C | 1.7e-07 | 2.5e-07 | 1.7e-07 | 1.967E-07 | 4.619E-08 | 1.700E-07 | 2.500E-07 |
| 20472 | 1024 | V1024K | C | 3.1e-07 | 1.3e-07 | 5.9e-07 | 3.433E-07 | 2.318E-07 | 1.300E-07 | 5.900E-07 |
| 20473 | 1024 | V1024M | C | 2.8e-07 | 3.9e-07 | 2.3e-07 | 3.000E-07 | 8.185E-08 | 2.300E-07 | 3.900E-07 |
| 20474 | 1024 | V1024F | C | 2.4e-07 | 1.1e-07 | 1.9e-07 | 1.800E-07 | 6.557E-08 | 1.100E-07 | 2.400E-07 |
| 20475 | 1024 | V1024P | C | 1.5e-07 | 1.8e-07 | 2.7e-07 | 2.000E-07 | 6.245E-08 | 1.500E-07 | 2.700E-07 |
| 20476 | 1024 | V1024S | C | 2.5e-07 | 2.6e-07 | 3.8e-07 | 2.967E-07 | 7.234E-08 | 2.500E-07 | 3.800E-07 |
| 20477 | 1024 | V1024T | C | 3.4e-07 | 2.8e-07 | 2.2e-07 | 2.800E-07 | 6.000E-08 | 2.200E-07 | 3.400E-07 |
| 20478 | 1024 | V1024W | C | 2.8e-07 | 1.9e-07 | 2.9e-07 | 2.533E-07 | 5.508E-08 | 1.900E-07 | 2.900E-07 |
| 20479 | 1024 | V1024Y | C | 2.4e-07 | 3.9e-07 | 1e-07 | 2.433E-07 | 1.450E-07 | 1.000E-07 | 3.900E-07 |
| 20480 | 1024 | V1024V | C | 1.4e-07 | 2.7e-07 | 3.9e-07 | 2.667E-07 | 1.250E-07 | 1.400E-07 | 3.900E-07 |
| 20481 | 1025 | T1025A | C | 1.7e-07 | 3.4e-07 | 2.3e-07 | 2.467E-07 | 8.622E-08 | 1.700E-07 | 3.400E-07 |
| 20482 | 1025 | T1025R | C | 2.7e-07 | 4.2e-07 | 2.7e-07 | 3.200E-07 | 8.660E-08 | 2.700E-07 | 4.200E-07 |
| 20483 | 1025 | T1025N | C | 2.3e-07 | 3.6e-07 | 2.4e-07 | 2.767E-07 | 7.234E-08 | 2.300E-07 | 3.600E-07 |
| 20484 | 1025 | T1025D | C | 1.6e-07 | 1.8e-07 | 2.2e-07 | 1.867E-07 | 3.055E-08 | 1.600E-07 | 2.200E-07 |

| | | | | | | | | | | |
|---|---|---|---|---|---|---|---|---|---|---|
| 20485 | 1025 | T1025C | C | 3.4e-07 | 1.5e-07 | 2.5e-07 | 2.467E-07 | 9.504E-08 | 1.500E-07 | 3.400E-07 |
| 20486 | 1025 | T1025E | C | 2.6e-07 | 8.9e-08 | 3.4e-07 | 2.297E-07 | 1.282E-07 | 8.900E-08 | 3.400E-07 |
| 20487 | 1025 | T1025Q | C | 2.5e-07 | 2.1e-07 | 3.6e-07 | 2.733E-07 | 7.767E-08 | 2.100E-07 | 3.600E-07 |
| 20488 | 1025 | T1025G | C | 1.6e-07 | 3.8e-07 | 3e-07 | 2.800E-07 | 1.114E-07 | 1.600E-07 | 3.800E-07 |
| 20489 | 1025 | T1025H | C | 3e-07 | 2.6e-07 | 3e-07 | 2.867E-07 | 2.309E-08 | 2.600E-07 | 3.000E-07 |
| 20490 | 1025 | T1025I | C | 1.7e-07 | 5.4e-07 | 2.5e-07 | 3.200E-07 | 1.947E-07 | 1.700E-07 | 5.400E-07 |
| 20491 | 1025 | T1025L | C | 3.6e-07 | 1.7e-07 | 2.6e-07 | 2.633E-07 | 9.504E-08 | 1.700E-07 | 3.600E-07 |
| 20492 | 1025 | T1025K | C | 1.2e-07 | 6.2e-07 | 1.7e-07 | 3.033E-07 | 2.754E-07 | 1.200E-07 | 6.200E-07 |
| 20493 | 1025 | T1025M | C | 3e-07 | 2.6e-07 | 2.7e-07 | 2.767E-07 | 2.082E-08 | 2.600E-07 | 3.000E-07 |
| 20494 | 1025 | T1025F | C | 1.7e-07 | 2.4e-07 | 2.9e-07 | 2.333E-07 | 6.028E-08 | 1.700E-07 | 2.900E-07 |
| 20495 | 1025 | T1025P | C | 1.3e-07 | 1.7e-07 | 2.1e-07 | 1.700E-07 | 4.000E-08 | 1.300E-07 | 2.100E-07 |
| 20496 | 1025 | T1025S | C | 2.3e-07 | 1.7e-07 | 1.9e-07 | 1.967E-07 | 3.055E-08 | 1.700E-07 | 2.300E-07 |
| 20497 | 1025 | T1025T | C | 1.8e-07 | 4.3e-07 | 3.2e-07 | 3.100E-07 | 1.253E-07 | 1.800E-07 | 4.300E-07 |
| 20498 | 1025 | T1025W | C | 2.4e-07 | 1.9e-07 | 2.3e-07 | 2.200E-07 | 2.646E-08 | 1.900E-07 | 2.400E-07 |
| 20499 | 1025 | T1025Y | C | 2.4e-07 | 3.7e-07 | 2.6e-07 | 2.900E-07 | 7.000E-08 | 2.400E-07 | 3.700E-07 |
| 20500 | 1025 | T1025V | C | 1.6e-07 | 1.2e-07 | 1.7e-07 | 1.500E-07 | 2.646E-08 | 1.200E-07 | 1.700E-07 |
| 20501 | 1026 | V1026A | C | 1.5e-07 | 3.3e-07 | 1.4e-07 | 2.067E-07 | 1.069E-07 | 1.400E-07 | 3.300E-07 |
| 20502 | 1026 | V1026R | C | 2.9e-07 | 3.5e-07 | 2.8e-07 | 3.067E-07 | 3.786E-08 | 2.800E-07 | 3.500E-07 |
| 20503 | 1026 | V1026N | C | 3.3e-07 | 1.1e-07 | 1.2e-07 | 1.867E-07 | 1.242E-07 | 1.100E-07 | 3.300E-07 |
| 20504 | 1026 | V1026D | C | 7.4e-07 | 4e-07 | 4.8e-07 | 5.400E-07 | 1.778E-07 | 4.000E-07 | 7.400E-07 |
| 20505 | 1026 | V1026C | C | 2.2e-07 | 2.6e-07 | 1.9e-07 | 2.233E-07 | 3.512E-08 | 1.900E-07 | 2.600E-07 |
| 20506 | 1026 | V1026E | C | 5.3e-07 | 3.4e-07 | 4.1e-07 | 4.267E-07 | 9.609E-08 | 3.400E-07 | 5.300E-07 |
| 20507 | 1026 | V1026Q | C | 1.8e-07 | 3.8e-07 | 2.4e-07 | 2.667E-07 | 1.026E-07 | 1.800E-07 | 3.800E-07 |
| 20508 | 1026 | V1026G | C | 2.6e-07 | 3.6e-07 | 2.8e-07 | 3.000E-07 | 5.292E-08 | 2.600E-07 | 3.600E-07 |
| 20509 | 1026 | V1026H | C | 2.7e-07 | 2.4e-07 | 1.2e-07 | 2.100E-07 | 7.937E-08 | 1.200E-07 | 2.700E-07 |
| 20510 | 1026 | V1026I | C | 1.5e-07 | 2e-07 | 2e-07 | 1.833E-07 | 2.887E-08 | 1.500E-07 | 2.000E-07 |
| 20511 | 1026 | V1026L | C | 1.7e-07 | 1.7e-07 | 3.3e-07 | 2.233E-07 | 9.238E-08 | 1.700E-07 | 3.300E-07 |
| 20512 | 1026 | V1026K | C | 4.9e-07 | 2.8e-07 | 2.2e-07 | 3.300E-07 | 1.418E-07 | 2.200E-07 | 4.900E-07 |
| 20513 | 1026 | V1026M | C | 3.3e-07 | 1.7e-07 | 2.9e-07 | 2.633E-07 | 8.327E-08 | 1.700E-07 | 3.300E-07 |
| 20514 | 1026 | V1026F | C | 1.7e-07 | 4.5e-07 | 2.8e-07 | 3.000E-07 | 1.411E-07 | 1.700E-07 | 4.500E-07 |
| 20515 | 1026 | V1026P | C | 3.6e-07 | 1.8e-07 | 2.9e-07 | 2.767E-07 | 9.074E-08 | 1.800E-07 | 3.600E-07 |
| 20516 | 1026 | V1026S | C | 2.1e-07 | 1.4e-07 | 3.1e-07 | 2.200E-07 | 8.544E-08 | 1.400E-07 | 3.100E-07 |
| 20517 | 1026 | V1026T | C | 1.7e-07 | 2e-07 | 2.8e-07 | 2.167E-07 | 5.686E-08 | 1.700E-07 | 2.800E-07 |
| 20518 | 1026 | V1026W | C | 2.1e-07 | 8.1e-08 | 1.8e-07 | 1.570E-07 | 6.751E-08 | 8.100E-08 | 2.100E-07 |
| 20519 | 1026 | V1026Y | C | 3.1e-07 | 3e-07 | 2.1e-07 | 2.733E-07 | 5.508E-08 | 2.100E-07 | 3.100E-07 |
| 20520 | 1026 | V1026V | C | 5.7e-07 | 1.9e-07 | 2.8e-07 | 3.467E-07 | 1.986E-07 | 1.900E-07 | 5.700E-07 |
| 20521 | 1027 | P1027A | C | 3.2e-07 | 4.3e-07 | 4e-07 | 3.833E-07 | 5.686E-08 | 3.200E-07 | 4.300E-07 |
| 20522 | 1027 | P1027R | C | 2.7e-07 | 2.5e-07 | 2.7e-07 | 2.633E-07 | 1.155E-08 | 2.500E-07 | 2.700E-07 |
| 20523 | 1027 | P1027N | C | 2.1e-07 | 3.8e-07 | 2e-07 | 2.633E-07 | 1.012E-07 | 2.000E-07 | 3.800E-07 |
| 20524 | 1027 | P1027D | C | 1.9e-07 | 1.1e-07 | 6.4e-08 | 1.213E-07 | 6.376E-08 | 6.400E-08 | 1.900E-07 |
| 20525 | 1027 | P1027C | C | 4.3e-07 | 1.8e-07 | 4.5e-07 | 3.533E-07 | 1.504E-07 | 1.800E-07 | 4.500E-07 |
| 20526 | 1027 | P1027E | C | 2.3e-07 | 2.4e-07 | 3e-07 | 2.567E-07 | 3.786E-08 | 2.300E-07 | 3.000E-07 |
| 20527 | 1027 | P1027Q | C | 1.6e-07 | 2.8e-07 | 3.5e-07 | 2.633E-07 | 9.609E-08 | 1.600E-07 | 3.500E-07 |
| 20528 | 1027 | P1027G | C | 2.3e-07 | 2.3e-07 | 4.1e-07 | 2.900E-07 | 1.039E-07 | 2.300E-07 | 4.100E-07 |
| 20529 | 1027 | P1027H | C | 2.3e-07 | 2.9e-07 | 1.5e-07 | 2.233E-07 | 7.024E-08 | 1.500E-07 | 2.900E-07 |
| 20530 | 1027 | P1027I | C | 3.2e-07 | 2.1e-07 | 3.5e-07 | 2.933E-07 | 7.371E-08 | 2.100E-07 | 3.500E-07 |
| 20531 | 1027 | P1027L | C | 3e-07 | 2.1e-07 | 3.4e-07 | 2.833E-07 | 6.658E-08 | 2.100E-07 | 3.400E-07 |
| 20532 | 1027 | P1027K | C | 1.7e-07 | 2.2e-07 | 1e-06 | 4.633E-07 | 4.654E-07 | 1.700E-07 | 1.000E-06 |
| 20533 | 1027 | P1027M | C | 1.1e-07 | 2.7e-07 | 3.5e-07 | 2.433E-07 | 1.222E-07 | 1.100E-07 | 3.500E-07 |
| 20534 | 1027 | P1027F | C | 2.8e-07 | 1.9e-07 | 1.8e-07 | 2.167E-07 | 5.508E-08 | 1.800E-07 | 2.800E-07 |
| 20535 | 1027 | P1027P | C | 3.2e-07 | 2.4e-07 | 3.2e-07 | 2.933E-07 | 4.619E-08 | 2.400E-07 | 3.200E-07 |
| 20536 | 1027 | P1027S | C | 3.6e-07 | 1.7e-07 | 2.2e-07 | 2.500E-07 | 9.849E-08 | 1.700E-07 | 3.600E-07 |
| 20537 | 1027 | P1027T | C | 3.9e-07 | 2.2e-07 | 4.1e-07 | 3.400E-07 | 1.044E-07 | 2.200E-07 | 4.100E-07 |
| 20538 | 1027 | P1027W | C | 2.9e-07 | 3.1e-07 | 2.6e-07 | 2.867E-07 | 2.517E-08 | 2.600E-07 | 3.100E-07 |
| 20539 | 1027 | P1027Y | C | 1.9e-07 | 2e-07 | 1.8e-07 | 1.900E-07 | 1.000E-08 | 1.800E-07 | 2.000E-07 |
| 20540 | 1027 | P1027V | C | 1.5e-07 | 4.7e-07 | 3e-07 | 3.067E-07 | 1.601E-07 | 1.500E-07 | 4.700E-07 |
| 20541 | 1028 | S1028A | C | 3.6e-07 | 1.7e-07 | 1.6e-07 | 2.300E-07 | 1.127E-07 | 1.600E-07 | 3.600E-07 |
| 20542 | 1028 | S1028R | C | 2.1e-07 | 3.1e-07 | 4.1e-07 | 3.100E-07 | 1.000E-07 | 2.100E-07 | 4.100E-07 |
| 20543 | 1028 | S1028N | C | 1.1e-07 | 4.4e-07 | 3.4e-07 | 2.967E-07 | 1.692E-07 | 1.100E-07 | 4.400E-07 |
| 20544 | 1028 | S1028D | C | 3.4e-07 | 3.9e-07 | 2.4e-07 | 3.233E-07 | 7.638E-08 | 2.400E-07 | 3.900E-07 |
| 20545 | 1028 | S1028C | C | 3.5e-07 | 2.5e-07 | 2.3e-07 | 2.767E-07 | 6.429E-08 | 2.300E-07 | 3.500E-07 |
| 20546 | 1028 | S1028E | C | 4.8e-07 | 1.5e-07 | 2.6e-07 | 2.967E-07 | 1.680E-07 | 1.500E-07 | 4.800E-07 |
| 20547 | 1028 | S1028Q | C | 1.6e-07 | 3.4e-07 | 6.7e-08 | 1.890E-07 | 1.388E-07 | 6.700E-08 | 3.400E-07 |
| 20548 | 1028 | S1028G | C | 2.9e-07 | 3.1e-07 | 1.9e-07 | 2.633E-07 | 6.429E-08 | 1.900E-07 | 3.100E-07 |
| 20549 | 1028 | S1028H | C | 3.1e-07 | 2.2e-07 | 3.3e-07 | 2.867E-07 | 5.859E-08 | 2.200E-07 | 3.300E-07 |
| 20550 | 1028 | S1028I | C | 2.9e-07 | 1.4e-07 | 4.2e-07 | 2.833E-07 | 1.401E-07 | 1.400E-07 | 4.200E-07 |
| 20551 | 1028 | S1028L | C | 3.7e-07 | 1.5e-07 | 3e-07 | 2.733E-07 | 1.124E-07 | 1.500E-07 | 3.700E-07 |
| 20552 | 1028 | S1028K | C | 3.4e-07 | 2.6e-07 | 3.8e-07 | 3.267E-07 | 6.110E-08 | 2.600E-07 | 3.800E-07 |
| 20553 | 1028 | S1028M | C | 3.3e-07 | 1.8e-07 | 3.6e-07 | 2.900E-07 | 9.644E-08 | 1.800E-07 | 3.600E-07 |
| 20554 | 1028 | S1028F | C | 2.6e-07 | 3.7e-07 | 3.5e-07 | 3.267E-07 | 5.859E-08 | 2.600E-07 | 3.700E-07 |
| 20555 | 1028 | S1028P | C | 2.3e-07 | 3.5e-07 | 4.1e-07 | 3.300E-07 | 9.165E-08 | 2.300E-07 | 4.100E-07 |
| 20556 | 1028 | S1028S | C | 2.1e-07 | 1.6e-07 | 2.1e-07 | 1.933E-07 | 2.887E-08 | 1.600E-07 | 2.100E-07 |
| 20557 | 1028 | S1028T | C | 2.4e-07 | 1.6e-07 | 3.4e-07 | 2.467E-07 | 9.018E-08 | 1.600E-07 | 3.400E-07 |
| 20558 | 1028 | S1028W | C | 2e-07 | 2.1e-07 | 5.5e-07 | 3.200E-07 | 1.992E-07 | 2.000E-07 | 5.500E-07 |
| 20559 | 1028 | S1028Y | C | 2.2e-07 | 4.9e-07 | 4.4e-07 | 3.833E-07 | 1.436E-07 | 2.200E-07 | 4.900E-07 |
| 20560 | 1028 | S1028V | C | 3.7e-07 | 2.2e-07 | 1.5e-07 | 2.467E-07 | 1.124E-07 | 1.500E-07 | 3.700E-07 |
| 20561 | 1029 | S1029A | C | 2.4e-07 | 4.9e-07 | 2.8e-07 | 3.367E-07 | 1.343E-07 | 2.400E-07 | 4.900E-07 |
| 20562 | 1029 | S1029R | C | 2.5e-07 | 3e-07 | 2.6e-07 | 2.700E-07 | 2.646E-08 | 2.500E-07 | 3.000E-07 |
| 20563 | 1029 | S1029N | C | 2.3e-07 | 4.4e-07 | 1.2e-07 | 2.633E-07 | 1.626E-07 | 1.200E-07 | 4.400E-07 |
| 20564 | 1029 | S1029D | C | 2.1e-07 | 1.8e-07 | 1.7e-07 | 1.867E-07 | 2.082E-08 | 1.700E-07 | 2.100E-07 |
| 20565 | 1029 | S1029C | C | 1.4e-07 | 3.6e-07 | 1.6e-07 | 2.200E-07 | 1.217E-07 | 1.400E-07 | 3.600E-07 |
| 20566 | 1029 | S1029E | C | 1.6e-07 | 1.9e-07 | 2.1e-07 | 1.867E-07 | 2.517E-08 | 1.600E-07 | 2.100E-07 |
| 20567 | 1029 | S1029Q | C | 3.8e-07 | 5.1e-07 | 2.7e-07 | 3.867E-07 | 1.201E-07 | 2.700E-07 | 5.100E-07 |
| 20568 | 1029 | S1029G | C | 2.9e-07 | 3.1e-07 | 2.6e-07 | 2.867E-07 | 2.517E-08 | 2.600E-07 | 3.100E-07 |
| 20569 | 1029 | S1029H | C | 4.1e-07 | 5.1e-07 | 4.1e-07 | 4.433E-07 | 5.774E-08 | 4.100E-07 | 5.100E-07 |

| | | | | | | | | | | |
|---|---|---|---|---|---|---|---|---|---|---|
| 20570 | 1029 | S1029I | C | 2.6e-07 | 6.2e-07 | 1.3e-07 | 3.367E-07 | 2.538E-07 | 1.300E-07 | 6.200E-07 |
| 20571 | 1029 | S1029L | C | 2.3e-07 | 5.5e-07 | 2.6e-07 | 3.467E-07 | 1.767E-07 | 2.300E-07 | 5.500E-07 |
| 20572 | 1029 | S1029K | C | 1.9e-07 | 1.9e-07 | 1.7e-07 | 1.833E-07 | 1.155E-08 | 1.700E-07 | 1.900E-07 |
| 20573 | 1029 | S1029M | C | 5.5e-07 | 2.6e-07 | 2.1e-07 | 3.400E-07 | 1.836E-07 | 2.100E-07 | 5.500E-07 |
| 20574 | 1029 | S1029F | C | 2.5e-07 | 2.4e-07 | 6.3e-07 | 3.733E-07 | 2.223E-07 | 2.400E-07 | 6.300E-07 |
| 20575 | 1029 | S1029P | C | 2.5e-07 | 3e-07 | 3.3e-07 | 2.933E-07 | 4.041E-08 | 2.500E-07 | 3.300E-07 |
| 20576 | 1029 | S1029S | C | 2.9e-07 | 3.3e-07 | 3.9e-07 | 3.367E-07 | 5.033E-08 | 2.900E-07 | 3.900E-07 |
| 20577 | 1029 | S1029T | C | 1.5e-07 | 5e-07 | 3.3e-07 | 3.267E-07 | 1.750E-07 | 1.500E-07 | 5.000E-07 |
| 20578 | 1029 | S1029W | C | 2.4e-07 | 2.3e-07 | 2.2e-07 | 2.300E-07 | 1.000E-08 | 2.200E-07 | 2.400E-07 |
| 20579 | 1029 | S1029Y | C | 2.8e-07 | 1.1e-07 | 3e-07 | 2.300E-07 | 1.044E-07 | 1.100E-07 | 3.000E-07 |
| 20580 | 1029 | S1029V | C | 2.2e-07 | 1.8e-07 | 2.6e-07 | 2.200E-07 | 4.000E-08 | 1.800E-07 | 2.600E-07 |
| 20581 | 1030 | S1030A | C | 2.6e-07 | 3.7e-07 | 2e-07 | 2.767E-07 | 8.622E-08 | 2.000E-07 | 3.700E-07 |
| 20582 | 1030 | S1030R | C | 2.1e-07 | 1.7e-07 | 3.3e-07 | 2.367E-07 | 8.327E-08 | 1.700E-07 | 3.300E-07 |
| 20583 | 1030 | S1030N | C | 2.7e-07 | 2.4e-07 | 1.8e-07 | 2.300E-07 | 4.583E-08 | 1.800E-07 | 2.700E-07 |
| 20584 | 1030 | S1030D | C | 3.7e-07 | 2.2e-07 | 1.7e-07 | 2.533E-07 | 1.041E-07 | 1.700E-07 | 3.700E-07 |
| 20585 | 1030 | S1030C | C | 3.8e-07 | 3.6e-07 | 2.2e-07 | 3.200E-07 | 8.718E-08 | 2.200E-07 | 3.800E-07 |
| 20586 | 1030 | S1030E | C | 2.5e-07 | 2.8e-07 | 3.1e-07 | 2.800E-07 | 3.000E-08 | 2.500E-07 | 3.100E-07 |
| 20587 | 1030 | S1030Q | C | 2e-07 | 3.3e-07 | 2e-07 | 2.433E-07 | 7.506E-08 | 2.000E-07 | 3.300E-07 |
| 20588 | 1030 | S1030G | C | 1.2e-07 | 3e-07 | 2.1e-07 | 2.100E-07 | 9.000E-08 | 1.200E-07 | 3.000E-07 |
| 20589 | 1030 | S1030H | C | 2.4e-07 | 3.7e-07 | 3.3e-07 | 3.133E-07 | 6.658E-08 | 2.400E-07 | 3.700E-07 |
| 20590 | 1030 | S1030I | C | 1.8e-07 | 2.8e-07 | 2.9e-07 | 2.500E-07 | 6.083E-08 | 1.800E-07 | 2.900E-07 |
| 20591 | 1030 | S1030L | C | 1.9e-07 | 2.9e-07 | 3.1e-07 | 2.633E-07 | 6.429E-08 | 1.900E-07 | 3.100E-07 |
| 20592 | 1030 | S1030K | C | 2.1e-07 | 2e-07 | 2.2e-07 | 2.100E-07 | 1.000E-08 | 2.000E-07 | 2.200E-07 |
| 20593 | 1030 | S1030M | C | 3.6e-07 | 4e-07 | 4.2e-07 | 3.933E-07 | 3.055E-08 | 3.600E-07 | 4.200E-07 |
| 20594 | 1030 | S1030F | C | 2.2e-07 | 3.5e-07 | 3.9e-07 | 3.200E-07 | 8.888E-08 | 2.200E-07 | 3.900E-07 |
| 20595 | 1030 | S1030P | C | 3.3e-07 | 3.7e-07 | 2.6e-07 | 3.200E-07 | 5.568E-08 | 2.600E-07 | 3.700E-07 |
| 20596 | 1030 | S1030S | C | 1.7e-07 | 2.7e-07 | 8e-08 | 1.733E-07 | 9.504E-08 | 8.000E-08 | 2.700E-07 |
| 20597 | 1030 | S1030T | C | 1.2e-07 | 2.5e-07 | 5.5e-07 | 3.067E-07 | 2.205E-07 | 1.200E-07 | 5.500E-07 |
| 20598 | 1030 | S1030W | C | 5.2e-07 | 3e-07 | 2.1e-07 | 3.433E-07 | 1.595E-07 | 2.100E-07 | 5.200E-07 |
| 20599 | 1030 | S1030Y | C | 2.5e-07 | 3.9e-07 | 3.5e-07 | 3.300E-07 | 7.211E-08 | 2.500E-07 | 3.900E-07 |
| 20600 | 1030 | S1030V | C | 3.9e-07 | 3.3e-07 | 2.9e-07 | 3.367E-07 | 5.033E-08 | 2.900E-07 | 3.900E-07 |
| 20601 | 1031 | L1031A | C | 5.5e-07 | 3.3e-07 | 1.6e-07 | 3.467E-07 | 1.955E-07 | 1.600E-07 | 5.500E-07 |
| 20602 | 1031 | L1031R | C | 2.1e-07 | 2.5e-07 | 1.7e-07 | 2.100E-07 | 4.000E-08 | 1.700E-07 | 2.500E-07 |
| 20603 | 1031 | L1031N | C | 2.7e-07 | 2.2e-07 | 4.5e-07 | 3.133E-07 | 1.210E-07 | 2.200E-07 | 4.500E-07 |
| 20604 | 1031 | L1031D | C | 1.7e-07 | 1.5e-07 | 2.9e-07 | 2.033E-07 | 7.572E-08 | 1.500E-07 | 2.900E-07 |
| 20605 | 1031 | L1031C | C | 2.2e-07 | 2.6e-07 | 2.1e-07 | 2.300E-07 | 2.646E-08 | 2.100E-07 | 2.600E-07 |
| 20606 | 1031 | L1031E | C | 1.6e-07 | 1.2e-07 | 1.8e-07 | 1.533E-07 | 3.055E-08 | 1.200E-07 | 1.800E-07 |
| 20607 | 1031 | L1031Q | C | 1.9e-07 | 3e-07 | 1.7e-07 | 2.200E-07 | 7.000E-08 | 1.700E-07 | 3.000E-07 |
| 20608 | 1031 | L1031G | C | 1.7e-07 | 1e-07 | 1.7e-07 | 1.467E-07 | 4.041E-08 | 1.000E-07 | 1.700E-07 |
| 20609 | 1031 | L1031H | C | 9.9e-08 | 2.8e-07 | 2.4e-07 | 2.063E-07 | 9.508E-08 | 9.900E-08 | 2.800E-07 |
| 20610 | 1031 | L1031I | C | 4.4e-07 | 2.3e-07 | 3e-07 | 3.233E-07 | 1.069E-07 | 2.300E-07 | 4.400E-07 |
| 20611 | 1031 | L1031L | C | 1.4e-07 | 2e-07 | 2.2e-07 | 1.867E-07 | 4.163E-08 | 1.400E-07 | 2.200E-07 |
| 20612 | 1031 | L1031K | C | 2e-07 | 1e-07 | 2.9e-07 | 1.967E-07 | 9.504E-08 | 1.000E-07 | 2.900E-07 |
| 20613 | 1031 | L1031M | C | 2.4e-07 | 1.5e-07 | 4.6e-07 | 2.833E-07 | 1.595E-07 | 1.500E-07 | 4.600E-07 |
| 20614 | 1031 | L1031F | C | 9.4e-08 | 1.8e-07 | 2.2e-07 | 1.647E-07 | 6.438E-08 | 9.400E-08 | 2.200E-07 |
| 20615 | 1031 | L1031P | C | 2.2e-07 | 1.7e-07 | 8.7e-07 | 4.200E-07 | 3.905E-07 | 1.700E-07 | 8.700E-07 |
| 20616 | 1031 | L1031S | C | 1.5e-07 | 2.3e-07 | 3.2e-07 | 2.333E-07 | 8.505E-08 | 1.500E-07 | 3.200E-07 |
| 20617 | 1031 | L1031T | C | 2.1e-07 | 2.2e-07 | 1.7e-07 | 2.000E-07 | 2.646E-08 | 1.700E-07 | 2.200E-07 |
| 20618 | 1031 | L1031W | C | 2.9e-07 | 1.2e-07 | 1.9e-07 | 2.000E-07 | 8.544E-08 | 1.200E-07 | 2.900E-07 |
| 20619 | 1031 | L1031Y | C | 3.7e-07 | 2.5e-07 | 2.9e-07 | 3.033E-07 | 6.110E-08 | 2.500E-07 | 3.700E-07 |
| 20620 | 1031 | L1031V | C | 3e-07 | 1.5e-07 | 2.2e-07 | 2.233E-07 | 7.506E-08 | 1.500E-07 | 3.000E-07 |
| 20621 | 1032 | G1032A | C | 1.7e-07 | 3.2e-07 | 9.4e-07 | 4.767E-07 | 4.082E-07 | 1.700E-07 | 9.400E-07 |
| 20622 | 1032 | G1032R | C | 2.9e-07 | 2.5e-07 | 2.3e-07 | 2.567E-07 | 3.055E-08 | 2.300E-07 | 2.900E-07 |
| 20623 | 1032 | G1032N | C | 1.4e-07 | 3.7e-07 | 3.1e-07 | 2.733E-07 | 1.193E-07 | 1.400E-07 | 3.700E-07 |
| 20624 | 1032 | G1032D | C | 3.2e-07 | 2.3e-07 | 5.5e-07 | 3.667E-07 | 1.650E-07 | 2.300E-07 | 5.500E-07 |
| 20625 | 1032 | G1032C | C | 2.2e-07 | 1.9e-07 | 2.1e-07 | 2.067E-07 | 1.528E-08 | 1.900E-07 | 2.200E-07 |
| 20626 | 1032 | G1032E | C | 1.9e-07 | 2.3e-07 | 2.7e-07 | 2.300E-07 | 4.000E-08 | 1.900E-07 | 2.700E-07 |
| 20627 | 1032 | G1032Q | C | 1.9e-07 | 4.7e-07 | 1.7e-07 | 2.767E-07 | 1.677E-07 | 1.700E-07 | 4.700E-07 |
| 20628 | 1032 | G1032G | C | 1.8e-07 | 3.9e-07 | 2e-07 | 2.567E-07 | 1.159E-07 | 1.800E-07 | 3.900E-07 |
| 20629 | 1032 | G1032H | C | 2.8e-07 | 2e-07 | 9.7e-08 | 1.923E-07 | 9.174E-08 | 9.700E-08 | 2.800E-07 |
| 20630 | 1032 | G1032I | C | 3.9e-07 | 1.3e-07 | 2.1e-07 | 2.433E-07 | 1.332E-07 | 1.300E-07 | 3.900E-07 |
| 20631 | 1032 | G1032L | C | 1.7e-07 | 1.4e-07 | 2.5e-07 | 1.867E-07 | 5.686E-08 | 1.400E-07 | 2.500E-07 |
| 20632 | 1032 | G1032K | C | 1.1e-07 | 1.6e-07 | 2.6e-07 | 1.767E-07 | 7.638E-08 | 1.100E-07 | 2.600E-07 |
| 20633 | 1032 | G1032M | C | 2e-07 | 1.4e-07 | 1.8e-07 | 1.733E-07 | 3.055E-08 | 1.400E-07 | 2.000E-07 |
| 20634 | 1032 | G1032F | C | 2.3e-07 | 3.5e-07 | 3.6e-07 | 3.133E-07 | 7.234E-08 | 2.300E-07 | 3.600E-07 |
| 20635 | 1032 | G1032P | C | 1.4e-07 | 1.5e-07 | 4.5e-07 | 2.467E-07 | 1.762E-07 | 1.400E-07 | 4.500E-07 |
| 20636 | 1032 | G1032S | C | 3e-07 | 2.5e-07 | 2e-07 | 2.500E-07 | 5.000E-08 | 2.000E-07 | 3.000E-07 |
| 20637 | 1032 | G1032T | C | 1.5e-07 | 3.2e-07 | 3.8e-07 | 2.833E-07 | 1.193E-07 | 1.500E-07 | 3.800E-07 |
| 20638 | 1032 | G1032W | C | 2e-07 | 1.5e-07 | 1.2e-07 | 1.567E-07 | 4.041E-08 | 1.200E-07 | 2.000E-07 |
| 20639 | 1032 | G1032Y | C | 1.3e-07 | 2.4e-07 | 3.4e-07 | 2.367E-07 | 1.050E-07 | 1.300E-07 | 3.400E-07 |
| 20640 | 1032 | G1032V | C | 3.1e-07 | 1.2e-07 | 1.3e-07 | 1.867E-07 | 1.069E-07 | 1.200E-07 | 3.100E-07 |
| 20641 | 1033 | T1033A | C | 2.8e-07 | 1.5e-07 | 3.3e-07 | 2.533E-07 | 9.292E-08 | 1.500E-07 | 3.300E-07 |
| 20642 | 1033 | T1033R | C | 1.7e-07 | 9.8e-08 | 1.7e-07 | 1.460E-07 | 4.157E-08 | 9.800E-08 | 1.700E-07 |
| 20643 | 1033 | T1033N | C | 1.2e-07 | 1.7e-07 | 2.1e-07 | 1.667E-07 | 4.509E-08 | 1.200E-07 | 2.100E-07 |
| 20644 | 1033 | T1033D | C | 1.8e-07 | 2.2e-07 | 5.1e-07 | 3.033E-07 | 1.801E-07 | 1.800E-07 | 5.100E-07 |
| 20645 | 1033 | T1033C | C | 4e-07 | 1.7e-07 | 2.8e-07 | 2.833E-07 | 1.150E-07 | 1.700E-07 | 4.000E-07 |
| 20646 | 1033 | T1033E | C | 2.1e-07 | 2.2e-07 | 2.4e-07 | 2.233E-07 | 1.528E-08 | 2.100E-07 | 2.400E-07 |
| 20647 | 1033 | T1033Q | C | 1.2e-07 | 2e-07 | 3.3e-07 | 2.167E-07 | 1.060E-07 | 1.200E-07 | 3.300E-07 |
| 20648 | 1033 | T1033G | C | 4e-07 | 3.3e-07 | 2.9e-07 | 3.400E-07 | 5.568E-08 | 2.900E-07 | 4.000E-07 |
| 20649 | 1033 | T1033H | C | 5.2e-07 | 2.6e-07 | 5.3e-07 | 4.367E-07 | 1.531E-07 | 2.600E-07 | 5.300E-07 |
| 20650 | 1033 | T1033I | C | 7.9e-07 | 1.2e-07 | 2.5e-07 | 3.867E-07 | 3.553E-07 | 1.200E-07 | 7.900E-07 |
| 20651 | 1033 | T1033L | C | 4e-07 | 3.3e-07 | 2.6e-07 | 3.300E-07 | 7.000E-08 | 2.600E-07 | 4.000E-07 |
| 20652 | 1033 | T1033K | C | 1.4e-07 | 2.5e-07 | 7.5e-08 | 1.550E-07 | 8.846E-08 | 7.500E-08 | 2.500E-07 |
| 20653 | 1033 | T1033M | C | 2.1e-07 | 3.5e-07 | 2.4e-07 | 2.667E-07 | 7.371E-08 | 2.100E-07 | 3.500E-07 |
| 20654 | 1033 | T1033F | C | 2.8e-07 | 2.9e-07 | 1.4e-07 | 2.367E-07 | 8.386E-08 | 1.400E-07 | 2.900E-07 |

| | | | | | | | | | | |
|---|---|---|---|---|---|---|---|---|---|---|
| 20655 | 1033 | T1033P | C | 2.8e-07 | 8.9e-07 | 2.1e-07 | 4.600E-07 | 3.740E-07 | 2.100E-07 | 8.900E-07 |
| 20656 | 1033 | T1033S | C | 2.8e-07 | 9.1e-07 | 2.6e-07 | 4.833E-07 | 3.696E-07 | 2.600E-07 | 9.100E-07 |
| 20657 | 1033 | T1033T | C | 3.4e-07 | 1.9e-07 | 2.3e-07 | 2.533E-07 | 7.767E-08 | 1.900E-07 | 3.400E-07 |
| 20658 | 1033 | T1033W | C | 2.8e-07 | 3e-07 | 1.9e-07 | 2.567E-07 | 5.859E-08 | 1.900E-07 | 3.000E-07 |
| 20659 | 1033 | T1033Y | C | 2.4e-07 | 2.2e-07 | 1.7e-07 | 2.100E-07 | 3.606E-08 | 1.700E-07 | 2.400E-07 |
| 20660 | 1033 | T1033V | C | 2.2e-07 | 3.2e-07 | 2e-07 | 2.467E-07 | 6.429E-08 | 2.000E-07 | 3.200E-07 |
| 20661 | 1034 | Q1034A | C | 3.8e-07 | 2.8e-07 | 1.5e-07 | 2.700E-07 | 1.153E-07 | 1.500E-07 | 3.800E-07 |
| 20662 | 1034 | Q1034R | C | 7.2e-07 | 2.4e-07 | 1.7e-07 | 3.767E-07 | 2.994E-07 | 1.700E-07 | 7.200E-07 |
| 20663 | 1034 | Q1034N | C | 4.2e-07 | 1.1e-07 | 2.1e-07 | 2.467E-07 | 1.582E-07 | 1.100E-07 | 4.200E-07 |
| 20664 | 1034 | Q1034D | C | 2e-07 | 2.7e-07 | 5.4e-07 | 3.367E-07 | 1.795E-07 | 2.000E-07 | 5.400E-07 |
| 20665 | 1034 | Q1034C | C | 2.7e-07 | 3.6e-07 | 1.5e-07 | 2.600E-07 | 1.054E-07 | 1.500E-07 | 3.600E-07 |
| 20666 | 1034 | Q1034E | C | 2.4e-07 | 4.2e-07 | 2.3e-07 | 2.967E-07 | 1.069E-07 | 2.300E-07 | 4.200E-07 |
| 20667 | 1034 | Q1034Q | C | 1.7e-07 | 2.5e-07 | 2.8e-07 | 2.333E-07 | 5.686E-08 | 1.700E-07 | 2.800E-07 |
| 20668 | 1034 | Q1034G | C | 3.2e-07 | 3.9e-07 | 3.9e-07 | 3.667E-07 | 4.041E-08 | 3.200E-07 | 3.900E-07 |
| 20669 | 1034 | Q1034H | C | 1.5e-07 | 2e-07 | 2.5e-07 | 2.000E-07 | 5.000E-08 | 1.500E-07 | 2.500E-07 |
| 20670 | 1034 | Q1034I | C | 3.5e-07 | 6.3e-07 | 3.9e-07 | 4.567E-07 | 1.514E-07 | 3.500E-07 | 6.300E-07 |
| 20671 | 1034 | Q1034L | C | 1.5e-07 | 2.4e-07 | 5.5e-07 | 3.133E-07 | 2.098E-07 | 1.500E-07 | 5.500E-07 |
| 20672 | 1034 | Q1034K | C | 3.3e-07 | 1.8e-07 | 1.6e-07 | 2.233E-07 | 9.292E-08 | 1.600E-07 | 3.300E-07 |
| 20673 | 1034 | Q1034M | C | 2.1e-07 | 3.8e-07 | 3.5e-07 | 3.133E-07 | 9.074E-08 | 2.100E-07 | 3.800E-07 |
| 20674 | 1034 | Q1034F | C | 2.9e-07 | 7.8e-07 | 3.5e-07 | 4.733E-07 | 2.673E-07 | 2.900E-07 | 7.800E-07 |
| 20675 | 1034 | Q1034P | C | 9.6e-08 | 2.1e-07 | 3.5e-07 | 2.187E-07 | 1.272E-07 | 9.600E-08 | 3.500E-07 |
| 20676 | 1034 | Q1034S | C | 1.9e-07 | 1.8e-07 | 3.1e-07 | 2.267E-07 | 7.234E-08 | 1.800E-07 | 3.100E-07 |
| 20677 | 1034 | Q1034T | C | 1.4e-07 | 5.6e-07 | 2.6e-07 | 3.200E-07 | 2.163E-07 | 1.400E-07 | 5.600E-07 |
| 20678 | 1034 | Q1034W | C | 3.6e-07 | 2.4e-07 | 1.7e-07 | 2.567E-07 | 9.609E-08 | 1.700E-07 | 3.600E-07 |
| 20679 | 1034 | Q1034Y | C | 7.7e-07 | 3.5e-07 | 1.8e-07 | 4.333E-07 | 3.037E-07 | 1.800E-07 | 7.700E-07 |
| 20680 | 1034 | Q1034V | C | 2.8e-07 | 1.5e-07 | 3.2e-07 | 2.500E-07 | 8.888E-08 | 1.500E-07 | 3.200E-07 |
| 20681 | 1035 | T1035A | C | 2.4e-07 | 4.2e-07 | 1.7e-07 | 2.767E-07 | 1.290E-07 | 1.700E-07 | 4.200E-07 |
| 20682 | 1035 | T1035R | C | 2e-07 | 4.9e-07 | 2.8e-07 | 3.233E-07 | 1.498E-07 | 2.000E-07 | 4.900E-07 |
| 20683 | 1035 | T1035N | C | 1.1e-07 | 1.9e-07 | 2.9e-07 | 1.967E-07 | 9.018E-08 | 1.100E-07 | 2.900E-07 |
| 20684 | 1035 | T1035D | C | 2.8e-07 | 3.4e-07 | 3.3e-07 | 3.167E-07 | 3.215E-08 | 2.800E-07 | 3.400E-07 |
| 20685 | 1035 | T1035C | C | 4.9e-07 | 4e-07 | 5.8e-07 | 4.900E-07 | 9.000E-08 | 4.000E-07 | 5.800E-07 |
| 20686 | 1035 | T1035E | C | 3.1e-07 | 3.6e-07 | 1.6e-07 | 2.767E-07 | 1.041E-07 | 1.600E-07 | 3.600E-07 |
| 20687 | 1035 | T1035Q | C | 1.7e-07 | 3.6e-07 | 3.5e-07 | 2.933E-07 | 1.069E-07 | 1.700E-07 | 3.600E-07 |
| 20688 | 1035 | T1035G | C | 1.7e-07 | 2.5e-07 | 2e-07 | 2.067E-07 | 4.041E-08 | 1.700E-07 | 2.500E-07 |
| 20689 | 1035 | T1035H | C | 2.5e-07 | 1.9e-07 | 2.5e-07 | 2.300E-07 | 3.464E-08 | 1.900E-07 | 2.500E-07 |
| 20690 | 1035 | T1035I | C | 5.5e-07 | 2e-07 | 3.7e-07 | 3.733E-07 | 1.750E-07 | 2.000E-07 | 5.500E-07 |
| 20691 | 1035 | T1035L | C | 2e-07 | 1.8e-07 | 3e-07 | 2.267E-07 | 6.429E-08 | 1.800E-07 | 3.000E-07 |
| 20692 | 1035 | T1035K | C | 2.3e-07 | 1.6e-07 | 2e-07 | 1.967E-07 | 3.512E-08 | 1.600E-07 | 2.300E-07 |
| 20693 | 1035 | T1035M | C | 1.3e-07 | 2.8e-07 | 2.3e-07 | 2.133E-07 | 7.638E-08 | 1.300E-07 | 2.800E-07 |
| 20694 | 1035 | T1035F | C | 1.4e-07 | 2.7e-07 | 2.1e-07 | 2.067E-07 | 6.506E-08 | 1.400E-07 | 2.700E-07 |
| 20695 | 1035 | T1035P | C | 5.6e-07 | 2.4e-07 | 2.5e-07 | 3.500E-07 | 1.819E-07 | 2.400E-07 | 5.600E-07 |
| 20696 | 1035 | T1035S | C | 1.2e-07 | 2.1e-07 | 2.3e-07 | 1.867E-07 | 5.859E-08 | 1.200E-07 | 2.300E-07 |
| 20697 | 1035 | T1035T | C | 1.1e-06 | 2.1e-07 | 1.7e-07 | 4.933E-07 | 5.258E-07 | 1.700E-07 | 1.100E-06 |
| 20698 | 1035 | T1035W | C | 2e-07 | 1.7e-07 | 3e-07 | 2.233E-07 | 6.807E-08 | 1.700E-07 | 3.000E-07 |
| 20699 | 1035 | T1035Y | C | 2.1e-07 | 1.9e-07 | 2.2e-07 | 2.067E-07 | 1.528E-08 | 1.900E-07 | 2.200E-07 |
| 20700 | 1035 | T1035V | C | 1.8e-07 | 2.2e-07 | 3.5e-07 | 2.500E-07 | 8.888E-08 | 1.800E-07 | 3.500E-07 |
| 20701 | 1036 | Y1036A | C | 2.4e-07 | 4.3e-07 | 2.8e-07 | 3.167E-07 | 1.002E-07 | 2.400E-07 | 4.300E-07 |
| 20702 | 1036 | Y1036R | C | 6.5e-07 | 1.7e-07 | 2.6e-07 | 3.600E-07 | 2.551E-07 | 1.700E-07 | 6.500E-07 |
| 20703 | 1036 | Y1036N | C | 2.8e-07 | 1.4e-07 | 2.9e-07 | 2.367E-07 | 8.386E-08 | 1.400E-07 | 2.900E-07 |
| 20704 | 1036 | Y1036D | C | 1.4e-07 | 2.4e-07 | 2e-07 | 1.933E-07 | 5.033E-08 | 1.400E-07 | 2.400E-07 |
| 20705 | 1036 | Y1036C | C | 2.8e-07 | 2.1e-07 | 3e-07 | 2.633E-07 | 4.726E-08 | 2.100E-07 | 3.000E-07 |
| 20706 | 1036 | Y1036E | C | 2.4e-07 | 2.9e-07 | 1.4e-07 | 2.233E-07 | 7.638E-08 | 1.400E-07 | 2.900E-07 |
| 20707 | 1036 | Y1036Q | C | 4.4e-07 | 4e-07 | 2.6e-07 | 3.667E-07 | 9.452E-08 | 2.600E-07 | 4.400E-07 |
| 20708 | 1036 | Y1036G | C | 3.6e-07 | 2.3e-07 | 2.7e-07 | 2.867E-07 | 6.658E-08 | 2.300E-07 | 3.600E-07 |
| 20709 | 1036 | Y1036H | C | 2.7e-07 | 1.8e-07 | 3.5e-07 | 2.667E-07 | 8.505E-08 | 1.800E-07 | 3.500E-07 |
| 20710 | 1036 | Y1036I | C | 3.1e-07 | 4.8e-07 | 2.3e-07 | 3.400E-07 | 1.277E-07 | 2.300E-07 | 4.800E-07 |
| 20711 | 1036 | Y1036L | C | 1.3e-07 | 1.9e-07 | 1.5e-07 | 1.567E-07 | 3.055E-08 | 1.300E-07 | 1.900E-07 |
| 20712 | 1036 | Y1036K | C | 4.1e-07 | 2.8e-07 | 2.1e-07 | 3.000E-07 | 1.015E-07 | 2.100E-07 | 4.100E-07 |
| 20713 | 1036 | Y1036M | C | 1.7e-07 | 2.1e-07 | 2.2e-07 | 2.000E-07 | 2.646E-08 | 1.700E-07 | 2.200E-07 |
| 20714 | 1036 | Y1036F | C | 1.4e-07 | 1.4e-07 | 2.4e-07 | 1.733E-07 | 5.774E-08 | 1.400E-07 | 2.400E-07 |
| 20715 | 1036 | Y1036P | C | 2.8e-07 | 2.5e-07 | 2.5e-07 | 2.600E-07 | 1.732E-08 | 2.500E-07 | 2.800E-07 |
| 20716 | 1036 | Y1036S | C | 1.6e-07 | 3.5e-07 | 1.3e-07 | 2.133E-07 | 1.193E-07 | 1.300E-07 | 3.500E-07 |
| 20717 | 1036 | Y1036T | C | 2.8e-07 | 3.5e-07 | 1.5e-07 | 2.600E-07 | 1.015E-07 | 1.500E-07 | 3.500E-07 |
| 20718 | 1036 | Y1036W | C | 2.1e-07 | 1.5e-07 | 2.8e-07 | 2.133E-07 | 6.506E-08 | 1.500E-07 | 2.800E-07 |
| 20719 | 1036 | Y1036Y | C | 5.2e-07 | 1.3e-07 | 4.6e-07 | 3.700E-07 | 2.100E-07 | 1.300E-07 | 5.200E-07 |
| 20720 | 1036 | Y1036V | C | 2e-07 | 3.4e-07 | 2.2e-07 | 2.533E-07 | 7.572E-08 | 2.000E-07 | 3.400E-07 |
| 20721 | 1037 | I1037A | C | 1.9e-07 | 2.6e-07 | 2.2e-07 | 2.233E-07 | 3.512E-08 | 1.900E-07 | 2.600E-07 |
| 20722 | 1037 | I1037R | C | 2.9e-07 | 1.9e-07 | 1.2e-07 | 2.000E-07 | 8.544E-08 | 1.200E-07 | 2.900E-07 |
| 20723 | 1037 | I1037N | C | 2.4e-07 | 1.9e-07 | 1.8e-07 | 2.033E-07 | 3.215E-08 | 1.800E-07 | 2.400E-07 |
| 20724 | 1037 | I1037D | C | 3e-07 | 3.5e-07 | 2.5e-07 | 3.000E-07 | 5.000E-08 | 2.500E-07 | 3.500E-07 |
| 20725 | 1037 | I1037C | C | 1.8e-07 | 2.5e-07 | 4.6e-07 | 2.967E-07 | 1.457E-07 | 1.800E-07 | 4.600E-07 |
| 20726 | 1037 | I1037E | C | 4.3e-07 | 2e-07 | 2.1e-07 | 2.800E-07 | 1.300E-07 | 2.000E-07 | 4.300E-07 |
| 20727 | 1037 | I1037Q | C | 3.4e-07 | 4e-07 | 2.6e-07 | 3.333E-07 | 7.024E-08 | 2.600E-07 | 4.000E-07 |
| 20728 | 1037 | I1037G | C | 4.9e-07 | 3e-07 | 3.1e-07 | 3.667E-07 | 1.069E-07 | 3.000E-07 | 4.900E-07 |
| 20729 | 1037 | I1037H | C | 3.5e-07 | 1.5e-07 | 3.1e-07 | 2.700E-07 | 1.058E-07 | 1.500E-07 | 3.500E-07 |
| 20730 | 1037 | I1037I | C | 2e-07 | 2.4e-07 | 2.1e-07 | 2.167E-07 | 2.082E-08 | 2.000E-07 | 2.400E-07 |
| 20731 | 1037 | I1037L | C | 1.7e-07 | 1.7e-07 | 2.5e-07 | 1.967E-07 | 4.619E-08 | 1.700E-07 | 2.500E-07 |
| 20732 | 1037 | I1037K | C | 3.7e-07 | 3.1e-07 | 2.8e-07 | 3.200E-07 | 4.583E-08 | 2.800E-07 | 3.700E-07 |
| 20733 | 1037 | I1037M | C | 1.9e-07 | 7.3e-08 | 4e-07 | 2.210E-07 | 1.657E-07 | 7.300E-08 | 4.000E-07 |
| 20734 | 1037 | I1037F | C | 2.9e-07 | 4.3e-07 | 1.8e-07 | 3.000E-07 | 1.253E-07 | 1.800E-07 | 4.300E-07 |
| 20735 | 1037 | I1037P | C | 1.8e-07 | 2.9e-07 | 1.3e-07 | 2.000E-07 | 8.185E-08 | 1.300E-07 | 2.900E-07 |
| 20736 | 1037 | I1037S | C | 1.7e-07 | 1.2e-07 | 3.3e-07 | 2.067E-07 | 1.097E-07 | 1.200E-07 | 3.300E-07 |
| 20737 | 1037 | I1037T | C | 2.4e-07 | 2.3e-07 | 2.9e-07 | 2.533E-07 | 3.215E-08 | 2.300E-07 | 2.900E-07 |
| 20738 | 1037 | I1037W | C | 1.9e-07 | 2.2e-07 | 9.1e-08 | 1.670E-07 | 6.751E-08 | 9.100E-08 | 2.200E-07 |
| 20739 | 1037 | I1037Y | C | 3.2e-07 | 4.6e-07 | 1.7e-07 | 3.167E-07 | 1.450E-07 | 1.700E-07 | 4.600E-07 |

| | | | | | | | | | | |
|---|---|---|---|---|---|---|---|---|---|---|
| 20740 | 1037 | I1037V | C | 2.6e-07 | 1.9e-07 | 4.4e-07 | 2.967E-07 | 1.290E-07 | 1.900E-07 | 4.400E-07 |
| 20741 | 1038 | C1038A | C | 1.5e-07 | 2.9e-07 | 3.7e-07 | 2.700E-07 | 1.114E-07 | 1.500E-07 | 3.700E-07 |
| 20742 | 1038 | C1038R | C | 1.5e-07 | 2.5e-07 | 3.6e-07 | 2.533E-07 | 1.050E-07 | 1.500E-07 | 3.600E-07 |
| 20743 | 1038 | C1038N | C | 1.4e-07 | 5.8e-07 | 2.1e-07 | 3.100E-07 | 2.364E-07 | 1.400E-07 | 5.800E-07 |
| 20744 | 1038 | C1038D | C | 1.4e-07 | 3.4e-07 | 1.1e-07 | 1.967E-07 | 1.250E-07 | 1.100E-07 | 3.400E-07 |
| 20745 | 1038 | C1038C | C | 1.4e-07 | 1.2e-07 | 2.8e-07 | 1.800E-07 | 8.718E-08 | 1.200E-07 | 2.800E-07 |
| 20746 | 1038 | C1038E | C | 1.3e-07 | 3.4e-07 | 1.2e-07 | 1.967E-07 | 1.242E-07 | 1.200E-07 | 3.400E-07 |
| 20747 | 1038 | C1038Q | C | 2.1e-07 | 3.1e-07 | 2.4e-07 | 2.533E-07 | 5.132E-08 | 2.100E-07 | 3.100E-07 |
| 20748 | 1038 | C1038G | C | 2.5e-07 | 4.1e-07 | 1.9e-07 | 2.833E-07 | 1.137E-07 | 1.900E-07 | 4.100E-07 |
| 20749 | 1038 | C1038H | C | 2.5e-07 | 4.7e-07 | 1.3e-07 | 2.833E-07 | 1.724E-07 | 1.300E-07 | 4.700E-07 |
| 20750 | 1038 | C1038I | C | 2.6e-07 | 3.6e-07 | 2.1e-07 | 2.767E-07 | 7.638E-08 | 2.100E-07 | 3.600E-07 |
| 20751 | 1038 | C1038L | C | 3.1e-07 | 2.6e-07 | 3.1e-07 | 2.933E-07 | 2.887E-08 | 2.600E-07 | 3.100E-07 |
| 20752 | 1038 | C1038K | C | 1.6e-07 | 2.2e-07 | 1.3e-07 | 1.700E-07 | 4.583E-08 | 1.300E-07 | 2.200E-07 |
| 20753 | 1038 | C1038M | C | 1.9e-07 | 2.5e-07 | 3e-07 | 2.467E-07 | 5.508E-08 | 1.900E-07 | 3.000E-07 |
| 20754 | 1038 | C1038F | C | 4.1e-07 | 3.4e-07 | 3.5e-07 | 3.667E-07 | 3.786E-08 | 3.400E-07 | 4.100E-07 |
| 20755 | 1038 | C1038P | C | 3.7e-07 | 3.1e-07 | 3.3e-07 | 3.367E-07 | 3.055E-08 | 3.100E-07 | 3.700E-07 |
| 20756 | 1038 | C1038S | C | 2.5e-07 | 2.9e-07 | 2.3e-07 | 2.567E-07 | 3.055E-08 | 2.300E-07 | 2.900E-07 |
| 20757 | 1038 | C1038T | C | 2.4e-07 | 3.3e-07 | 2.7e-07 | 2.800E-07 | 4.583E-08 | 2.400E-07 | 3.300E-07 |
| 20758 | 1038 | C1038W | C | 3e-07 | 1.8e-07 | 1.8e-07 | 2.200E-07 | 6.928E-08 | 1.800E-07 | 3.000E-07 |
| 20759 | 1038 | C1038Y | C | 3e-07 | 1.4e-07 | 3.3e-07 | 2.567E-07 | 1.021E-07 | 1.400E-07 | 3.300E-07 |
| 20760 | 1038 | C1038V | C | 3.4e-07 | 3.2e-07 | 2.9e-07 | 3.167E-07 | 2.517E-08 | 2.900E-07 | 3.400E-07 |
| 20761 | 1039 | N1039A | C | 2.5e-07 | 1.2e-07 | 1.3e-07 | 1.667E-07 | 7.234E-08 | 1.200E-07 | 2.500E-07 |
| 20762 | 1039 | N1039R | C | 3e-07 | 1.4e-07 | 1.7e-07 | 2.033E-07 | 8.505E-08 | 1.400E-07 | 3.000E-07 |
| 20763 | 1039 | N1039N | C | 2.2e-07 | 4e-07 | 1.7e-07 | 2.633E-07 | 1.210E-07 | 1.700E-07 | 4.000E-07 |
| 20764 | 1039 | N1039D | C | 1.8e-07 | 4.2e-07 | 3.8e-07 | 3.267E-07 | 1.286E-07 | 1.800E-07 | 4.200E-07 |
| 20765 | 1039 | N1039C | C | 3.6e-07 | 3e-07 | 2.7e-07 | 3.100E-07 | 4.583E-08 | 2.700E-07 | 3.600E-07 |
| 20766 | 1039 | N1039E | C | 1.9e-07 | 2.6e-07 | 2.5e-07 | 2.333E-07 | 3.786E-08 | 1.900E-07 | 2.600E-07 |
| 20767 | 1039 | N1039Q | C | 1.5e-07 | 4.3e-07 | 3.2e-07 | 3.000E-07 | 1.411E-07 | 1.500E-07 | 4.300E-07 |
| 20768 | 1039 | N1039G | C | 2.6e-07 | 1.4e-07 | 1.7e-07 | 1.900E-07 | 6.245E-08 | 1.400E-07 | 2.600E-07 |
| 20769 | 1039 | N1039H | C | 1.6e-07 | 3.8e-07 | 2.1e-07 | 2.500E-07 | 1.153E-07 | 1.600E-07 | 3.800E-07 |
| 20770 | 1039 | N1039I | C | 1.8e-07 | 2.6e-07 | 3.4e-07 | 2.600E-07 | 8.000E-08 | 1.800E-07 | 3.400E-07 |
| 20771 | 1039 | N1039L | C | 3.6e-07 | 1.6e-07 | 2.9e-07 | 2.700E-07 | 1.015E-07 | 1.600E-07 | 3.600E-07 |
| 20772 | 1039 | N1039K | C | 3.1e-07 | 9.2e-08 | 2.5e-07 | 2.173E-07 | 1.126E-07 | 9.200E-08 | 3.100E-07 |
| 20773 | 1039 | N1039M | C | 5.3e-07 | 2.3e-07 | 2.5e-07 | 3.367E-07 | 1.677E-07 | 2.300E-07 | 5.300E-07 |
| 20774 | 1039 | N1039F | C | 1.5e-07 | 5.9e-07 | 2.5e-07 | 3.300E-07 | 2.307E-07 | 1.500E-07 | 5.900E-07 |
| 20775 | 1039 | N1039P | C | 3.3e-07 | 4.7e-07 | 1.1e-07 | 3.033E-07 | 1.815E-07 | 1.100E-07 | 4.700E-07 |
| 20776 | 1039 | N1039S | C | 2e-07 | 1.4e-07 | 1.1e-07 | 1.500E-07 | 4.583E-08 | 1.100E-07 | 2.000E-07 |
| 20777 | 1039 | N1039T | C | 2.7e-07 | 2.6e-07 | 3.4e-07 | 2.900E-07 | 4.359E-08 | 2.600E-07 | 3.400E-07 |
| 20778 | 1039 | N1039W | C | 1.7e-07 | 1.8e-07 | 2e-07 | 1.833E-07 | 1.528E-08 | 1.700E-07 | 2.000E-07 |
| 20779 | 1039 | N1039Y | C | 2.8e-07 | 4e-07 | 2e-07 | 2.933E-07 | 1.007E-07 | 2.000E-07 | 4.000E-07 |
| 20780 | 1039 | N1039V | C | 2.1e-07 | 1.2e-07 | 2.7e-07 | 2.000E-07 | 7.550E-08 | 1.200E-07 | 2.700E-07 |
| 20781 | 1040 | V1040A | C | 2.1e-07 | 1.7e-07 | 2.3e-07 | 2.033E-07 | 3.055E-08 | 1.700E-07 | 2.300E-07 |
| 20782 | 1040 | V1040R | C | 1.6e-07 | 3e-07 | 2.9e-07 | 2.500E-07 | 7.810E-08 | 1.600E-07 | 3.000E-07 |
| 20783 | 1040 | V1040N | C | 2e-07 | 3.1e-07 | 1.8e-07 | 2.300E-07 | 7.000E-08 | 1.800E-07 | 3.100E-07 |
| 20784 | 1040 | V1040D | C | 2.7e-07 | 2.8e-07 | 9.5e-08 | 2.150E-07 | 1.040E-07 | 9.500E-08 | 2.800E-07 |
| 20785 | 1040 | V1040C | C | 1.2e-07 | 6.2e-07 | 1.7e-07 | 3.033E-07 | 2.754E-07 | 1.200E-07 | 6.200E-07 |
| 20786 | 1040 | V1040E | C | 2.2e-07 | 2.7e-07 | 2.3e-07 | 2.400E-07 | 2.646E-08 | 2.200E-07 | 2.700E-07 |
| 20787 | 1040 | V1040Q | C | 1.3e-07 | 7.4e-08 | 1.5e-07 | 1.180E-07 | 3.940E-08 | 7.400E-08 | 1.500E-07 |
| 20788 | 1040 | V1040G | C | 7.8e-07 | 3.8e-07 | 1.6e-07 | 4.400E-07 | 3.143E-07 | 1.600E-07 | 7.800E-07 |
| 20789 | 1040 | V1040H | C | 2.6e-07 | 2.3e-07 | 3.3e-07 | 2.733E-07 | 5.132E-08 | 2.300E-07 | 3.300E-07 |
| 20790 | 1040 | V1040I | C | 3.5e-07 | 4.5e-07 | 1.8e-07 | 3.267E-07 | 1.365E-07 | 1.800E-07 | 4.500E-07 |
| 20791 | 1040 | V1040L | C | 2e-07 | 3.7e-07 | 1.9e-07 | 2.533E-07 | 1.012E-07 | 1.900E-07 | 3.700E-07 |
| 20792 | 1040 | V1040K | C | 3.8e-07 | 3.7e-07 | 4e-07 | 3.833E-07 | 1.528E-08 | 3.700E-07 | 4.000E-07 |
| 20793 | 1040 | V1040M | C | 3.4e-07 | 2.8e-07 | 2.9e-07 | 3.033E-07 | 3.215E-08 | 2.800E-07 | 3.400E-07 |
| 20794 | 1040 | V1040F | C | 4.8e-07 | 3.6e-07 | 4.5e-07 | 4.300E-07 | 6.245E-08 | 3.600E-07 | 4.800E-07 |
| 20795 | 1040 | V1040P | C | 1.8e-07 | 1.9e-07 | 2.7e-07 | 2.133E-07 | 4.933E-08 | 1.800E-07 | 2.700E-07 |
| 20796 | 1040 | V1040S | C | 2.2e-07 | 3.4e-07 | 2.4e-07 | 2.667E-07 | 6.429E-08 | 2.200E-07 | 3.400E-07 |
| 20797 | 1040 | V1040T | C | 2.6e-07 | 1.4e-07 | 2.5e-07 | 2.167E-07 | 6.658E-08 | 1.400E-07 | 2.600E-07 |
| 20798 | 1040 | V1040W | C | 2.9e-07 | 1e-07 | 2.9e-07 | 2.267E-07 | 1.097E-07 | 1.000E-07 | 2.900E-07 |
| 20799 | 1040 | V1040Y | C | 4.7e-07 | 3.1e-07 | 2.8e-07 | 3.533E-07 | 1.021E-07 | 2.800E-07 | 4.700E-07 |
| 20800 | 1040 | V1040V | C | 6.5e-07 | 2.7e-07 | 1.1e-07 | 3.433E-07 | 2.774E-07 | 1.100E-07 | 6.500E-07 |
| 20801 | 1041 | N1041A | C | 2.4e-07 | 1.4e-07 | 2.2e-07 | 2.000E-07 | 5.292E-08 | 1.400E-07 | 2.400E-07 |
| 20802 | 1041 | N1041R | C | 8.2e-07 | 3.1e-07 | 2.6e-07 | 4.633E-07 | 3.099E-07 | 2.600E-07 | 8.200E-07 |
| 20803 | 1041 | N1041N | C | 1.5e-07 | 2.2e-07 | 3.5e-07 | 2.400E-07 | 1.015E-07 | 1.500E-07 | 3.500E-07 |
| 20804 | 1041 | N1041D | C | 2.3e-07 | 3.9e-07 | 3.4e-07 | 3.200E-07 | 8.185E-08 | 2.300E-07 | 3.900E-07 |
| 20805 | 1041 | N1041C | C | 2.2e-07 | 2.6e-07 | 2.6e-07 | 2.467E-07 | 2.309E-08 | 2.200E-07 | 2.600E-07 |
| 20806 | 1041 | N1041E | C | 2.1e-07 | 1.5e-07 | 3.7e-07 | 2.433E-07 | 1.137E-07 | 1.500E-07 | 3.700E-07 |
| 20807 | 1041 | N1041Q | C | 1.4e-07 | 2.1e-07 | 4e-07 | 2.500E-07 | 1.345E-07 | 1.400E-07 | 4.000E-07 |
| 20808 | 1041 | N1041G | C | 2.8e-07 | 2.3e-07 | 2.9e-07 | 2.667E-07 | 3.215E-08 | 2.300E-07 | 2.900E-07 |
| 20809 | 1041 | N1041H | C | 1.4e-07 | 2.5e-07 | 2e-07 | 1.967E-07 | 5.508E-08 | 1.400E-07 | 2.500E-07 |
| 20810 | 1041 | N1041I | C | 1.7e-07 | 3.6e-07 | 1.8e-07 | 2.367E-07 | 1.069E-07 | 1.700E-07 | 3.600E-07 |
| 20811 | 1041 | N1041L | C | 1.2e-07 | 2.7e-07 | 3e-07 | 2.300E-07 | 9.644E-08 | 1.200E-07 | 3.000E-07 |
| 20812 | 1041 | N1041K | C | 3.3e-07 | 1.7e-07 | 3.5e-07 | 2.833E-07 | 9.866E-08 | 1.700E-07 | 3.500E-07 |
| 20813 | 1041 | N1041M | C | 6.3e-07 | 1.5e-07 | 3.3e-07 | 3.700E-07 | 2.425E-07 | 1.500E-07 | 6.300E-07 |
| 20814 | 1041 | N1041F | C | 1.5e-07 | 2.4e-07 | 3.6e-07 | 2.500E-07 | 1.054E-07 | 1.500E-07 | 3.600E-07 |
| 20815 | 1041 | N1041P | C | 1.8e-07 | 3.4e-07 | 1.6e-07 | 2.267E-07 | 9.866E-08 | 1.600E-07 | 3.400E-07 |
| 20816 | 1041 | N1041S | C | 1.3e-07 | 1.8e-07 | 9.7e-08 | 1.357E-07 | 4.179E-08 | 9.700E-08 | 1.800E-07 |
| 20817 | 1041 | N1041T | C | 3.5e-07 | 2.8e-07 | 1.9e-07 | 2.733E-07 | 8.021E-08 | 1.900E-07 | 3.500E-07 |
| 20818 | 1041 | N1041W | C | 2e-07 | 4.7e-07 | 1.6e-07 | 2.767E-07 | 1.686E-07 | 1.600E-07 | 4.700E-07 |
| 20819 | 1041 | N1041Y | C | 2.1e-07 | 1.5e-07 | 8.5e-08 | 1.483E-07 | 6.252E-08 | 8.500E-08 | 2.100E-07 |
| 20820 | 1041 | N1041V | C | 2.8e-07 | 4.8e-07 | 1.6e-07 | 3.067E-07 | 1.617E-07 | 1.600E-07 | 4.800E-07 |
| 20821 | 1042 | H1042A | C | 3.5e-07 | 3.1e-07 | 2.8e-07 | 3.133E-07 | 3.512E-08 | 2.800E-07 | 3.500E-07 |
| 20822 | 1042 | H1042R | C | 1e-06 | 9.5e-07 | 3.9e-07 | 7.800E-07 | 3.387E-07 | 3.900E-07 | 1.000E-06 |
| 20823 | 1042 | H1042N | C | 3.2e-07 | 5e-07 | 3.6e-07 | 3.933E-07 | 9.452E-08 | 3.200E-07 | 5.000E-07 |
| 20824 | 1042 | H1042D | C | 1.3e-07 | 2.2e-07 | 9.2e-08 | 1.473E-07 | 6.574E-08 | 9.200E-08 | 2.200E-07 |

| | | | | | | | | | | |
|---|---|---|---|---|---|---|---|---|---|---|
| 20825 | 1042 | H1042C | C | 2e-07 | 1.8e-07 | 1.3e-07 | 1.700E-07 | 3.606E-08 | 1.300E-07 | 2.000E-07 |
| 20826 | 1042 | H1042E | C | 2.1e-07 | 2.7e-07 | 8.8e-08 | 1.893E-07 | 9.274E-08 | 8.800E-08 | 2.700E-07 |
| 20827 | 1042 | H1042Q | C | 1.7e-07 | 4.8e-07 | 2.4e-07 | 2.967E-07 | 1.626E-07 | 1.700E-07 | 4.800E-07 |
| 20828 | 1042 | H1042G | C | 3.2e-07 | 2.4e-07 | 4e-07 | 3.200E-07 | 8.000E-08 | 2.400E-07 | 4.000E-07 |
| 20829 | 1042 | H1042H | C | 5e-07 | 4.9e-07 | 2.7e-07 | 4.200E-07 | 1.300E-07 | 2.700E-07 | 5.000E-07 |
| 20830 | 1042 | H1042I | C | 2.3e-07 | 2e-07 | 2.1e-07 | 2.133E-07 | 1.528E-08 | 2.000E-07 | 2.300E-07 |
| 20831 | 1042 | H1042L | C | 1.4e-07 | 2.6e-07 | 2.6e-07 | 2.200E-07 | 6.928E-08 | 1.400E-07 | 2.600E-07 |
| 20832 | 1042 | H1042K | C | 1.4e-07 | 3.6e-07 | 2.3e-07 | 2.433E-07 | 1.106E-07 | 1.400E-07 | 3.600E-07 |
| 20833 | 1042 | H1042M | C | 1.4e-07 | 4.2e-07 | 3e-07 | 2.867E-07 | 1.405E-07 | 1.400E-07 | 4.200E-07 |
| 20834 | 1042 | H1042F | C | 2.8e-07 | 2e-07 | 1.3e-07 | 2.033E-07 | 7.506E-08 | 1.300E-07 | 2.800E-07 |
| 20835 | 1042 | H1042P | C | 3.9e-07 | 2.7e-07 | 1.6e-07 | 2.733E-07 | 1.150E-07 | 1.600E-07 | 3.900E-07 |
| 20836 | 1042 | H1042S | C | 2.3e-07 | 4.1e-07 | 2.3e-07 | 2.900E-07 | 1.039E-07 | 2.300E-07 | 4.100E-07 |
| 20837 | 1042 | H1042T | C | 1.9e-07 | 3.6e-07 | 4.6e-07 | 3.367E-07 | 1.365E-07 | 1.900E-07 | 4.600E-07 |
| 20838 | 1042 | H1042W | C | 2.2e-07 | 2.2e-07 | 1.7e-07 | 2.033E-07 | 2.887E-08 | 1.700E-07 | 2.200E-07 |
| 20839 | 1042 | H1042Y | C | 3.2e-07 | 1.5e-07 | 5.8e-07 | 3.500E-07 | 2.166E-07 | 1.500E-07 | 5.800E-07 |
| 20840 | 1042 | H1042V | C | 3.2e-07 | 3.6e-07 | 1.4e-07 | 2.733E-07 | 1.172E-07 | 1.400E-07 | 3.600E-07 |
| 20841 | 1043 | K1043A | C | 1.8e-07 | 1.3e-07 | 4.2e-07 | 2.433E-07 | 1.550E-07 | 1.300E-07 | 4.200E-07 |
| 20842 | 1043 | K1043R | C | 1.5e-07 | 2.2e-07 | 2.5e-07 | 2.067E-07 | 5.132E-08 | 1.500E-07 | 2.500E-07 |
| 20843 | 1043 | K1043N | C | 2.2e-07 | 8.4e-08 | 2.4e-07 | 1.813E-07 | 8.488E-08 | 8.400E-08 | 2.400E-07 |
| 20844 | 1043 | K1043D | C | 2.1e-07 | 1.6e-07 | 3.6e-07 | 2.433E-07 | 1.041E-07 | 1.600E-07 | 3.600E-07 |
| 20845 | 1043 | K1043C | C | 4.1e-07 | 2e-07 | 3.3e-07 | 3.133E-07 | 1.060E-07 | 2.000E-07 | 4.100E-07 |
| 20846 | 1043 | K1043E | C | 3.4e-07 | 2.7e-07 | 3.4e-07 | 3.167E-07 | 4.041E-08 | 2.700E-07 | 3.400E-07 |
| 20847 | 1043 | K1043Q | C | 2.9e-07 | 2.9e-07 | 2.2e-07 | 2.667E-07 | 4.041E-08 | 2.200E-07 | 2.900E-07 |
| 20848 | 1043 | K1043G | C | 8.9e-08 | 1.9e-07 | 3.3e-07 | 2.030E-07 | 1.210E-07 | 8.900E-08 | 3.300E-07 |
| 20849 | 1043 | K1043H | C | 2.7e-07 | 2e-07 | 2.7e-07 | 2.467E-07 | 4.041E-08 | 2.000E-07 | 2.700E-07 |
| 20850 | 1043 | K1043I | C | 1.7e-07 | 7.6e-07 | 2.8e-07 | 4.033E-07 | 3.137E-07 | 1.700E-07 | 7.600E-07 |
| 20851 | 1043 | K1043L | C | 1.3e-07 | 2.9e-07 | 2.1e-07 | 2.100E-07 | 8.000E-08 | 1.300E-07 | 2.900E-07 |
| 20852 | 1043 | K1043K | C | 1.3e-07 | 1.9e-07 | 1.7e-07 | 1.633E-07 | 3.055E-08 | 1.300E-07 | 1.900E-07 |
| 20853 | 1043 | K1043M | C | 2.9e-07 | 2e-07 | 2.2e-07 | 2.367E-07 | 4.726E-08 | 2.000E-07 | 2.900E-07 |
| 20854 | 1043 | K1043F | C | 1.1e-07 | 2.3e-07 | 1.4e-07 | 1.600E-07 | 6.245E-08 | 1.100E-07 | 2.300E-07 |
| 20855 | 1043 | K1043P | C | 1.4e-07 | 1.8e-07 | 1.4e-07 | 1.533E-07 | 2.309E-08 | 1.400E-07 | 1.800E-07 |
| 20856 | 1043 | K1043S | C | 2.1e-07 | 2.7e-07 | 1.4e-07 | 2.067E-07 | 6.506E-08 | 1.400E-07 | 2.700E-07 |
| 20857 | 1043 | K1043T | C | 1.8e-07 | 4.8e-07 | 1.5e-07 | 2.700E-07 | 1.825E-07 | 1.500E-07 | 4.800E-07 |
| 20858 | 1043 | K1043W | C | 2.3e-07 | 2.7e-07 | 1.7e-07 | 2.233E-07 | 5.033E-08 | 1.700E-07 | 2.700E-07 |
| 20859 | 1043 | K1043Y | C | 1.1e-07 | 2.4e-07 | 1.6e-07 | 1.700E-07 | 6.557E-08 | 1.100E-07 | 2.400E-07 |
| 20860 | 1043 | K1043V | C | 3.1e-07 | 2.1e-07 | 2.9e-07 | 2.700E-07 | 5.292E-08 | 2.100E-07 | 3.100E-07 |
| 20861 | 1044 | P1044A | C | 4.4e-07 | 4.4e-07 | 7.7e-07 | 5.500E-07 | 1.905E-07 | 4.400E-07 | 7.700E-07 |
| 20862 | 1044 | P1044R | C | 2e-07 | 3.9e-07 | 2.2e-07 | 2.700E-07 | 1.044E-07 | 2.000E-07 | 3.900E-07 |
| 20863 | 1044 | P1044N | C | 2e-07 | 2.6e-07 | 2.1e-07 | 2.233E-07 | 3.215E-08 | 2.000E-07 | 2.600E-07 |
| 20864 | 1044 | P1044D | C | 2.5e-07 | 2.7e-07 | 2.8e-07 | 2.667E-07 | 1.528E-08 | 2.500E-07 | 2.800E-07 |
| 20865 | 1044 | P1044C | C | 2.8e-07 | 4.4e-07 | 2.5e-07 | 3.233E-07 | 1.021E-07 | 2.500E-07 | 4.400E-07 |
| 20866 | 1044 | P1044E | C | 2.5e-07 | 2.8e-07 | 1.1e-07 | 2.133E-07 | 9.074E-08 | 1.100E-07 | 2.800E-07 |
| 20867 | 1044 | P1044Q | C | 3.7e-07 | 3.1e-07 | 3.4e-07 | 3.400E-07 | 3.000E-08 | 3.100E-07 | 3.700E-07 |
| 20868 | 1044 | P1044G | C | 3.9e-07 | 2e-07 | 2.2e-07 | 2.700E-07 | 1.044E-07 | 2.000E-07 | 3.900E-07 |
| 20869 | 1044 | P1044H | C | 3.2e-07 | 2.3e-07 | 1.9e-07 | 2.467E-07 | 6.658E-08 | 1.900E-07 | 3.200E-07 |
| 20870 | 1044 | P1044I | C | 1.4e-07 | 2.3e-07 | 2.7e-07 | 2.133E-07 | 6.658E-08 | 1.400E-07 | 2.700E-07 |
| 20871 | 1044 | P1044L | C | 1.6e-07 | 5e-07 | 1.6e-07 | 2.733E-07 | 1.963E-07 | 1.600E-07 | 5.000E-07 |
| 20872 | 1044 | P1044K | C | 2.1e-07 | 3e-07 | 1.9e-07 | 2.333E-07 | 5.859E-08 | 1.900E-07 | 3.000E-07 |
| 20873 | 1044 | P1044M | C | 3.5e-07 | 3.9e-07 | 2.7e-07 | 3.367E-07 | 6.110E-08 | 2.700E-07 | 3.900E-07 |
| 20874 | 1044 | P1044F | C | 1e-07 | 1.1e-07 | 4.7e-07 | 2.267E-07 | 2.108E-07 | 1.000E-07 | 4.700E-07 |
| 20875 | 1044 | P1044P | C | 2.1e-07 | 1.3e-07 | 3.7e-07 | 2.367E-07 | 1.222E-07 | 1.300E-07 | 3.700E-07 |
| 20876 | 1044 | P1044S | C | 2.9e-07 | 2.2e-07 | 3.1e-07 | 2.733E-07 | 4.726E-08 | 2.200E-07 | 3.100E-07 |
| 20877 | 1044 | P1044T | C | 2.6e-07 | 2.4e-07 | 2.2e-07 | 2.400E-07 | 2.000E-08 | 2.200E-07 | 2.600E-07 |
| 20878 | 1044 | P1044W | C | 1.6e-07 | 1e-07 | 1.7e-07 | 1.433E-07 | 3.786E-08 | 1.000E-07 | 1.700E-07 |
| 20879 | 1044 | P1044Y | C | 3.5e-07 | 1.8e-07 | 1e-07 | 2.100E-07 | 1.277E-07 | 1.000E-07 | 3.500E-07 |
| 20880 | 1044 | P1044V | C | 2.5e-07 | 1.6e-07 | 2.5e-07 | 2.200E-07 | 5.196E-08 | 1.600E-07 | 2.500E-07 |
| 20881 | 1045 | S1045A | C | 1.7e-07 | 2.5e-07 | 2.8e-07 | 2.333E-07 | 5.686E-08 | 1.700E-07 | 2.800E-07 |
| 20882 | 1045 | S1045R | C | 3.1e-07 | 2.5e-07 | 4.7e-07 | 3.433E-07 | 1.137E-07 | 2.500E-07 | 4.700E-07 |
| 20883 | 1045 | S1045N | C | 7.1e-07 | 1.4e-07 | 2.4e-07 | 3.633E-07 | 3.044E-07 | 1.400E-07 | 7.100E-07 |
| 20884 | 1045 | S1045D | C | 2.7e-07 | 2e-07 | 1.6e-07 | 2.100E-07 | 5.568E-08 | 1.600E-07 | 2.700E-07 |
| 20885 | 1045 | S1045C | C | 1.1e-07 | 5.5e-07 | 1.4e-07 | 2.667E-07 | 2.458E-07 | 1.100E-07 | 5.500E-07 |
| 20886 | 1045 | S1045E | C | 3.2e-07 | 2.5e-07 | 2.1e-07 | 2.600E-07 | 5.568E-08 | 2.100E-07 | 3.200E-07 |
| 20887 | 1045 | S1045Q | C | 3.1e-07 | 1.6e-07 | 1.3e-07 | 2.000E-07 | 9.644E-08 | 1.300E-07 | 3.100E-07 |
| 20888 | 1045 | S1045G | C | 3.6e-07 | 2.4e-07 | 3.1e-07 | 3.033E-07 | 6.028E-08 | 2.400E-07 | 3.600E-07 |
| 20889 | 1045 | S1045H | C | 2.6e-07 | 2.3e-07 | 3.2e-07 | 2.700E-07 | 4.583E-08 | 2.300E-07 | 3.200E-07 |
| 20890 | 1045 | S1045I | C | 4e-07 | 5e-07 | 2.4e-07 | 3.800E-07 | 1.311E-07 | 2.400E-07 | 5.000E-07 |
| 20891 | 1045 | S1045L | C | 3.3e-07 | 3.8e-07 | 3.7e-07 | 3.600E-07 | 2.646E-08 | 3.300E-07 | 3.800E-07 |
| 20892 | 1045 | S1045K | C | 1.3e-07 | 3e-07 | 2.6e-07 | 2.300E-07 | 8.888E-08 | 1.300E-07 | 3.000E-07 |
| 20893 | 1045 | S1045M | C | 2.7e-07 | 2.7e-07 | 1.4e-07 | 2.267E-07 | 7.506E-08 | 1.400E-07 | 2.700E-07 |
| 20894 | 1045 | S1045F | C | 4.8e-07 | 1.6e-07 | 2.3e-07 | 2.900E-07 | 1.682E-07 | 1.600E-07 | 4.800E-07 |
| 20895 | 1045 | S1045P | C | 1.5e-07 | 3.3e-07 | 1.6e-07 | 2.133E-07 | 1.012E-07 | 1.500E-07 | 3.300E-07 |
| 20896 | 1045 | S1045S | C | 3.5e-07 | 4.7e-07 | 1.7e-07 | 3.300E-07 | 1.510E-07 | 1.700E-07 | 4.700E-07 |
| 20897 | 1045 | S1045T | C | 5.8e-07 | 4e-07 | 1.1e-07 | 3.633E-07 | 2.371E-07 | 1.100E-07 | 5.800E-07 |
| 20898 | 1045 | S1045W | C | 1.9e-07 | 5.7e-07 | 1.5e-07 | 3.033E-07 | 2.318E-07 | 1.500E-07 | 5.700E-07 |
| 20899 | 1045 | S1045Y | C | 3.4e-07 | 2.6e-07 | 7.9e-07 | 4.633E-07 | 2.857E-07 | 2.600E-07 | 7.900E-07 |
| 20900 | 1045 | S1045V | C | 3.9e-07 | 2e-07 | 1.9e-07 | 2.600E-07 | 1.127E-07 | 1.900E-07 | 3.900E-07 |
| 20901 | 1046 | N1046A | C | 2.9e-07 | 5.2e-07 | 1.7e-07 | 3.267E-07 | 1.779E-07 | 1.700E-07 | 5.200E-07 |
| 20902 | 1046 | N1046R | C | 6.3e-07 | 3e-07 | 1.7e-07 | 3.667E-07 | 2.371E-07 | 1.700E-07 | 6.300E-07 |
| 20903 | 1046 | N1046N | C | 2.8e-07 | 1.4e-07 | 2.3e-07 | 2.167E-07 | 7.095E-08 | 1.400E-07 | 2.800E-07 |
| 20904 | 1046 | N1046D | C | 2.8e-07 | 2.7e-07 | 1.7e-07 | 2.400E-07 | 6.083E-08 | 1.700E-07 | 2.800E-07 |
| 20905 | 1046 | N1046C | C | 1.8e-07 | 2.8e-07 | 3e-07 | 2.533E-07 | 6.429E-08 | 1.800E-07 | 3.000E-07 |
| 20906 | 1046 | N1046E | C | 2.4e-07 | 5.4e-07 | 3.5e-07 | 3.767E-07 | 1.518E-07 | 2.400E-07 | 5.400E-07 |
| 20907 | 1046 | N1046Q | C | 1.4e-07 | 3e-07 | 1.4e-07 | 1.933E-07 | 9.238E-08 | 1.400E-07 | 3.000E-07 |
| 20908 | 1046 | N1046G | C | 2.4e-07 | 5e-07 | 2.9e-07 | 3.433E-07 | 1.380E-07 | 2.400E-07 | 5.000E-07 |
| 20909 | 1046 | N1046H | C | 2e-07 | 4.2e-07 | 2.5e-07 | 2.900E-07 | 1.153E-07 | 2.000E-07 | 4.200E-07 |

| | | | | | | | | | | |
|---|---|---|---|---|---|---|---|---|---|---|
| 20910 | 1046 | N1046I | C | 1.6e-07 | 1.7e-07 | 1.5e-07 | 1.600E-07 | 1.000E-08 | 1.500E-07 | 1.700E-07 |
| 20911 | 1046 | N1046L | C | 7.5e-07 | 3e-07 | 2.5e-07 | 4.333E-07 | 2.754E-07 | 2.500E-07 | 7.500E-07 |
| 20912 | 1046 | N1046K | C | 2e-07 | 4.4e-07 | 1.7e-07 | 2.700E-07 | 1.480E-07 | 1.700E-07 | 4.400E-07 |
| 20913 | 1046 | N1046M | C | 1.6e-07 | 1.8e-07 | 5.7e-07 | 3.033E-07 | 2.312E-07 | 1.600E-07 | 5.700E-07 |
| 20914 | 1046 | N1046F | C | 2.8e-07 | 3.7e-07 | 2.2e-07 | 2.900E-07 | 7.550E-08 | 2.200E-07 | 3.700E-07 |
| 20915 | 1046 | N1046P | C | 1.9e-07 | 2.6e-07 | 2.5e-07 | 2.333E-07 | 3.786E-08 | 1.900E-07 | 2.600E-07 |
| 20916 | 1046 | N1046S | C | 2.3e-07 | 2.1e-07 | 1.5e-07 | 1.967E-07 | 4.163E-08 | 1.500E-07 | 2.300E-07 |
| 20917 | 1046 | N1046T | C | 1.1e-07 | 2.6e-07 | 3.9e-08 | 1.363E-07 | 1.128E-07 | 3.900E-08 | 2.600E-07 |
| 20918 | 1046 | N1046W | C | 3.5e-07 | 9.7e-08 | 2.9e-07 | 2.457E-07 | 1.322E-07 | 9.700E-08 | 3.500E-07 |
| 20919 | 1046 | N1046Y | C | 2.3e-07 | 2.9e-07 | 2.1e-07 | 2.433E-07 | 4.163E-08 | 2.100E-07 | 2.900E-07 |
| 20920 | 1046 | N1046V | C | 3.9e-07 | 5.6e-07 | 2.1e-07 | 3.867E-07 | 1.750E-07 | 2.100E-07 | 5.600E-07 |
| 20921 | 1047 | T1047A | C | 1.7e-07 | 4.2e-07 | 3.4e-07 | 3.100E-07 | 1.277E-07 | 1.700E-07 | 4.200E-07 |
| 20922 | 1047 | T1047R | C | 3.7e-07 | 2.2e-07 | 1.5e-07 | 2.467E-07 | 1.124E-07 | 1.500E-07 | 3.700E-07 |
| 20923 | 1047 | T1047N | C | 1.4e-07 | 3.1e-07 | 2.7e-07 | 2.400E-07 | 8.888E-08 | 1.400E-07 | 3.100E-07 |
| 20924 | 1047 | T1047D | C | 3e-07 | 3.3e-07 | 2.2e-07 | 2.833E-07 | 5.686E-08 | 2.200E-07 | 3.300E-07 |
| 20925 | 1047 | T1047C | C | 2.8e-07 | 3.3e-07 | 1.3e-07 | 2.467E-07 | 1.041E-07 | 1.300E-07 | 3.300E-07 |
| 20926 | 1047 | T1047E | C | 3.6e-07 | 3.1e-07 | 5e-07 | 3.900E-07 | 9.849E-08 | 3.100E-07 | 5.000E-07 |
| 20927 | 1047 | T1047Q | C | 2.3e-07 | 4.7e-07 | 2.5e-07 | 3.167E-07 | 1.332E-07 | 2.300E-07 | 4.700E-07 |
| 20928 | 1047 | T1047G | C | 1.3e-07 | 8.2e-08 | 1.4e-07 | 1.173E-07 | 3.101E-08 | 8.200E-08 | 1.400E-07 |
| 20929 | 1047 | T1047H | C | 2e-07 | 7.4e-08 | 1.7e-07 | 1.480E-07 | 6.582E-08 | 7.400E-08 | 2.000E-07 |
| 20930 | 1047 | T1047I | C | 1.5e-07 | 1.9e-07 | 2.2e-07 | 1.867E-07 | 3.512E-08 | 1.500E-07 | 2.200E-07 |
| 20931 | 1047 | T1047L | C | 3.2e-07 | 1.8e-07 | 3e-07 | 2.667E-07 | 7.572E-08 | 1.800E-07 | 3.200E-07 |
| 20932 | 1047 | T1047K | C | 9.7e-08 | 2.2e-07 | 3.5e-07 | 2.223E-07 | 1.265E-07 | 9.700E-08 | 3.500E-07 |
| 20933 | 1047 | T1047M | C | 2.6e-07 | 5.2e-07 | 4e-07 | 3.933E-07 | 1.301E-07 | 2.600E-07 | 5.200E-07 |
| 20934 | 1047 | T1047F | C | 2.1e-07 | 2.9e-07 | 2e-07 | 2.333E-07 | 4.933E-08 | 2.000E-07 | 2.900E-07 |
| 20935 | 1047 | T1047P | C | 3e-07 | 3.1e-07 | 3.4e-07 | 3.167E-07 | 2.082E-08 | 3.000E-07 | 3.400E-07 |
| 20936 | 1047 | T1047S | C | 4.7e-07 | 5.8e-07 | 3.6e-07 | 4.700E-07 | 1.100E-07 | 3.600E-07 | 5.800E-07 |
| 20937 | 1047 | T1047T | C | 1e-07 | 2.3e-07 | 1.2e-07 | 1.500E-07 | 7.000E-08 | 1.000E-07 | 2.300E-07 |
| 20938 | 1047 | T1047W | C | 1.7e-07 | 2.3e-07 | 2.9e-07 | 2.300E-07 | 6.000E-08 | 1.700E-07 | 2.900E-07 |
| 20939 | 1047 | T1047Y | C | 1.8e-07 | 1.2e-07 | 2.4e-07 | 1.800E-07 | 6.000E-08 | 1.200E-07 | 2.400E-07 |
| 20940 | 1047 | T1047V | C | 1.9e-07 | 1.9e-07 | 8.4e-08 | 1.547E-07 | 6.120E-08 | 8.400E-08 | 1.900E-07 |
| 20941 | 1048 | K1048A | C | 3.3e-07 | 4.1e-07 | 2.8e-07 | 3.400E-07 | 6.557E-08 | 2.800E-07 | 4.100E-07 |
| 20942 | 1048 | K1048R | C | 2e-07 | 3.5e-07 | 2.6e-07 | 2.700E-07 | 7.550E-08 | 2.000E-07 | 3.500E-07 |
| 20943 | 1048 | K1048N | C | 3e-07 | 2.6e-07 | 5.1e-07 | 3.567E-07 | 1.343E-07 | 2.600E-07 | 5.100E-07 |
| 20944 | 1048 | K1048D | C | 1.6e-07 | 3.4e-07 | 1.8e-07 | 2.267E-07 | 9.866E-08 | 1.600E-07 | 3.400E-07 |
| 20945 | 1048 | K1048C | C | 2.5e-07 | 1.8e-07 | 2.9e-07 | 2.400E-07 | 5.568E-08 | 1.800E-07 | 2.900E-07 |
| 20946 | 1048 | K1048E | C | 3.6e-07 | 2.1e-07 | 2.8e-07 | 2.833E-07 | 7.506E-08 | 2.100E-07 | 3.600E-07 |
| 20947 | 1048 | K1048Q | C | 1.8e-07 | 2.3e-07 | 4.2e-07 | 2.767E-07 | 1.266E-07 | 1.800E-07 | 4.200E-07 |
| 20948 | 1048 | K1048G | C | 2.1e-07 | 2.9e-07 | 2.7e-07 | 2.567E-07 | 4.163E-08 | 2.100E-07 | 2.900E-07 |
| 20949 | 1048 | K1048H | C | 1.9e-07 | 1.6e-07 | 1.9e-07 | 1.800E-07 | 1.732E-08 | 1.600E-07 | 1.900E-07 |
| 20950 | 1048 | K1048I | C | 5.5e-07 | 1.9e-07 | 2e-07 | 3.133E-07 | 2.050E-07 | 1.900E-07 | 5.500E-07 |
| 20951 | 1048 | K1048L | C | 1.5e-07 | 3.1e-07 | 1.8e-07 | 2.133E-07 | 8.505E-08 | 1.500E-07 | 3.100E-07 |
| 20952 | 1048 | K1048K | C | 6e-07 | 2e-07 | 3.5e-07 | 3.833E-07 | 2.021E-07 | 2.000E-07 | 6.000E-07 |
| 20953 | 1048 | K1048M | C | 4.1e-07 | 1.5e-07 | 1.6e-07 | 2.400E-07 | 1.473E-07 | 1.500E-07 | 4.100E-07 |
| 20954 | 1048 | K1048F | C | 1.2e-07 | 3e-07 | 3.4e-07 | 2.533E-07 | 1.172E-07 | 1.200E-07 | 3.400E-07 |
| 20955 | 1048 | K1048P | C | 6.8e-07 | 2.6e-07 | 2.3e-07 | 3.900E-07 | 2.516E-07 | 2.300E-07 | 6.800E-07 |
| 20956 | 1048 | K1048S | C | 1.6e-07 | 2.7e-07 | 3.5e-07 | 2.600E-07 | 9.539E-08 | 1.600E-07 | 3.500E-07 |
| 20957 | 1048 | K1048T | C | 2.7e-07 | 2e-07 | 9.4e-07 | 4.700E-07 | 4.085E-07 | 2.000E-07 | 9.400E-07 |
| 20958 | 1048 | K1048W | C | 1e-07 | 2e-07 | 2.2e-07 | 1.733E-07 | 6.429E-08 | 1.000E-07 | 2.200E-07 |
| 20959 | 1048 | K1048Y | C | 3.1e-07 | 2.7e-07 | 2.5e-07 | 2.767E-07 | 3.055E-08 | 2.500E-07 | 3.100E-07 |
| 20960 | 1048 | K1048V | C | 1.9e-07 | 2.2e-07 | 2.8e-07 | 2.300E-07 | 4.583E-08 | 1.900E-07 | 2.800E-07 |
| 20961 | 1049 | V1049A | C | 1.6e-07 | 2.2e-07 | 3e-07 | 2.267E-07 | 7.024E-08 | 1.600E-07 | 3.000E-07 |
| 20962 | 1049 | V1049R | C | 5.7e-07 | 2.7e-07 | 3.6e-07 | 4.000E-07 | 1.539E-07 | 2.700E-07 | 5.700E-07 |
| 20963 | 1049 | V1049N | C | 1.9e-07 | 6.6e-08 | 2.7e-07 | 1.753E-07 | 1.028E-07 | 6.600E-08 | 2.700E-07 |
| 20964 | 1049 | V1049D | C | 2.4e-07 | 1.1e-07 | 4.3e-07 | 2.600E-07 | 1.609E-07 | 1.100E-07 | 4.300E-07 |
| 20965 | 1049 | V1049C | C | 2.7e-07 | 2.1e-07 | 2.2e-07 | 2.333E-07 | 3.215E-08 | 2.100E-07 | 2.700E-07 |
| 20966 | 1049 | V1049E | C | 2.7e-07 | 1.6e-07 | 4e-07 | 2.767E-07 | 1.201E-07 | 1.600E-07 | 4.000E-07 |
| 20967 | 1049 | V1049Q | C | 1.6e-07 | 2e-07 | 1.8e-07 | 1.800E-07 | 2.000E-08 | 1.600E-07 | 2.000E-07 |
| 20968 | 1049 | V1049G | C | 5.1e-07 | 5.8e-07 | 9.1e-08 | 3.937E-07 | 2.644E-07 | 9.100E-08 | 5.800E-07 |
| 20969 | 1049 | V1049H | C | 4.5e-07 | 1.7e-07 | 3.4e-07 | 3.200E-07 | 1.411E-07 | 1.700E-07 | 4.500E-07 |
| 20970 | 1049 | V1049I | C | 1.6e-07 | 3.4e-07 | 1.1e-07 | 2.033E-07 | 1.210E-07 | 1.100E-07 | 3.400E-07 |
| 20971 | 1049 | V1049L | C | 3.3e-07 | 5.3e-07 | 1.6e-07 | 3.400E-07 | 1.852E-07 | 1.600E-07 | 5.300E-07 |
| 20972 | 1049 | V1049K | C | 3.4e-07 | 1.9e-07 | 1.4e-07 | 2.233E-07 | 1.041E-07 | 1.400E-07 | 3.400E-07 |
| 20973 | 1049 | V1049M | C | 2.7e-07 | 1.5e-07 | 3.9e-07 | 2.700E-07 | 1.200E-07 | 1.500E-07 | 3.900E-07 |
| 20974 | 1049 | V1049F | C | 2.3e-07 | 1.4e-07 | 4.1e-07 | 2.600E-07 | 1.375E-07 | 1.400E-07 | 4.100E-07 |
| 20975 | 1049 | V1049P | C | 1.6e-07 | 2.9e-07 | 2.2e-07 | 2.233E-07 | 6.506E-08 | 1.600E-07 | 2.900E-07 |
| 20976 | 1049 | V1049S | C | 3.5e-07 | 3.5e-07 | 2.1e-07 | 3.033E-07 | 8.083E-08 | 2.100E-07 | 3.500E-07 |
| 20977 | 1049 | V1049T | C | 1.2e-07 | 1.8e-07 | 2.8e-07 | 1.933E-07 | 8.083E-08 | 1.200E-07 | 2.800E-07 |
| 20978 | 1049 | V1049W | C | 1.2e-07 | 3e-07 | 1e-07 | 1.733E-07 | 1.102E-07 | 1.000E-07 | 3.000E-07 |
| 20979 | 1049 | V1049Y | C | 2.5e-07 | 1.2e-07 | 2.1e-07 | 1.933E-07 | 6.658E-08 | 1.200E-07 | 2.500E-07 |
| 20980 | 1049 | V1049V | C | 2.7e-07 | 1.7e-07 | 7.3e-08 | 1.710E-07 | 9.850E-08 | 7.300E-08 | 2.700E-07 |
| 20981 | 1050 | D1050A | C | 4.7e-07 | 1.3e-07 | 2e-07 | 2.667E-07 | 1.795E-07 | 1.300E-07 | 4.700E-07 |
| 20982 | 1050 | D1050R | C | 3.4e-07 | 3.3e-07 | 2.2e-07 | 2.967E-07 | 6.658E-08 | 2.200E-07 | 3.400E-07 |
| 20983 | 1050 | D1050N | C | 1.1e-07 | 2.9e-07 | 3.5e-07 | 2.500E-07 | 1.249E-07 | 1.100E-07 | 3.500E-07 |
| 20984 | 1050 | D1050D | C | 4.4e-07 | 5.6e-07 | 2.5e-07 | 4.167E-07 | 1.563E-07 | 2.500E-07 | 5.600E-07 |
| 20985 | 1050 | D1050C | C | 4.8e-07 | 2.6e-07 | 4.1e-07 | 3.833E-07 | 1.124E-07 | 2.600E-07 | 4.800E-07 |
| 20986 | 1050 | D1050E | C | 1.2e-07 | 1.3e-07 | 1.8e-07 | 1.433E-07 | 3.215E-08 | 1.200E-07 | 1.800E-07 |
| 20987 | 1050 | D1050Q | C | 2.9e-07 | 2.6e-07 | 5.1e-07 | 3.533E-07 | 1.365E-07 | 2.600E-07 | 5.100E-07 |
| 20988 | 1050 | D1050G | C | 2e-07 | 2.1e-07 | 2.6e-07 | 2.233E-07 | 3.215E-08 | 2.000E-07 | 2.600E-07 |
| 20989 | 1050 | D1050H | C | 3.6e-07 | 2.2e-07 | 2.3e-07 | 2.700E-07 | 7.810E-08 | 2.200E-07 | 3.600E-07 |
| 20990 | 1050 | D1050I | C | 2.6e-07 | 2.2e-07 | 4e-07 | 2.933E-07 | 9.452E-08 | 2.200E-07 | 4.000E-07 |
| 20991 | 1050 | D1050L | C | 2.6e-07 | 1.9e-07 | 1.6e-07 | 2.033E-07 | 5.132E-08 | 1.600E-07 | 2.600E-07 |
| 20992 | 1050 | D1050K | C | 2.1e-07 | 1.6e-07 | 2.1e-07 | 1.933E-07 | 2.887E-08 | 1.600E-07 | 2.100E-07 |
| 20993 | 1050 | D1050M | C | 3.8e-07 | 1.2e-07 | 2.4e-07 | 2.467E-07 | 1.301E-07 | 1.200E-07 | 3.800E-07 |
| 20994 | 1050 | D1050F | C | 1.7e-07 | 2e-07 | 3.9e-07 | 2.533E-07 | 1.193E-07 | 1.700E-07 | 3.900E-07 |

| | | | | | | | | | | |
|---|---|---|---|---|---|---|---|---|---|---|
| 20995 | 1050 | D1050P | C | 5e-07 | 2e-07 | 3.5e-07 | 3.500E-07 | 1.500E-07 | 2.000E-07 | 5.000E-07 |
| 20996 | 1050 | D1050S | C | 2e-07 | 2e-07 | 2.5e-07 | 2.167E-07 | 2.887E-08 | 2.000E-07 | 2.500E-07 |
| 20997 | 1050 | D1050T | C | 4.3e-07 | 2.3e-07 | 3.7e-07 | 3.433E-07 | 1.026E-07 | 2.300E-07 | 4.300E-07 |
| 20998 | 1050 | D1050W | C | 7.2e-08 | 3.2e-07 | 4.3e-07 | 2.740E-07 | 1.834E-07 | 7.200E-08 | 4.300E-07 |
| 20999 | 1050 | D1050Y | C | 1.4e-07 | 2.9e-07 | 3.5e-07 | 2.600E-07 | 1.082E-07 | 1.400E-07 | 3.500E-07 |
| 21000 | 1050 | D1050V | C | 3.7e-07 | 1.7e-07 | 3.1e-07 | 2.833E-07 | 1.026E-07 | 1.700E-07 | 3.700E-07 |
| 21001 | 1051 | K1051A | C | 1.8e-07 | 1.9e-07 | 1.9e-07 | 1.867E-07 | 5.774E-09 | 1.800E-07 | 1.900E-07 |
| 21002 | 1051 | K1051R | C | 3.1e-07 | 3.7e-07 | 4.3e-07 | 3.700E-07 | 6.000E-08 | 3.100E-07 | 4.300E-07 |
| 21003 | 1051 | K1051N | C | 3.2e-07 | 1.9e-07 | 3.2e-07 | 2.767E-07 | 7.506E-08 | 1.900E-07 | 3.200E-07 |
| 21004 | 1051 | K1051D | C | 6.3e-07 | 3.3e-07 | 1.8e-07 | 3.800E-07 | 2.291E-07 | 1.800E-07 | 6.300E-07 |
| 21005 | 1051 | K1051C | C | 9.9e-08 | 3.6e-07 | 5.7e-07 | 3.430E-07 | 2.360E-07 | 9.900E-08 | 5.700E-07 |
| 21006 | 1051 | K1051E | C | 1.9e-07 | 4.7e-07 | 1.2e-07 | 2.600E-07 | 1.852E-07 | 1.200E-07 | 4.700E-07 |
| 21007 | 1051 | K1051Q | C | 1.7e-07 | 1.9e-07 | 1.3e-07 | 1.633E-07 | 3.055E-08 | 1.300E-07 | 1.900E-07 |
| 21008 | 1051 | K1051G | C | 2.4e-07 | 3e-07 | 2.1e-07 | 2.500E-07 | 4.583E-08 | 2.100E-07 | 3.000E-07 |
| 21009 | 1051 | K1051H | C | 2.7e-07 | 1e-07 | 1.9e-07 | 1.867E-07 | 8.505E-08 | 1.000E-07 | 2.700E-07 |
| 21010 | 1051 | K1051I | C | 3.3e-07 | 3e-07 | 4.8e-07 | 3.700E-07 | 9.644E-08 | 3.000E-07 | 4.800E-07 |
| 21011 | 1051 | K1051L | C | 5.1e-07 | 2.7e-07 | 2.9e-07 | 3.567E-07 | 1.332E-07 | 2.700E-07 | 5.100E-07 |
| 21012 | 1051 | K1051K | C | 2.6e-07 | 4.6e-07 | 2.2e-07 | 3.133E-07 | 1.286E-07 | 2.200E-07 | 4.600E-07 |
| 21013 | 1051 | K1051M | C | 1.4e-07 | 1.9e-07 | 2.1e-07 | 1.800E-07 | 3.606E-08 | 1.400E-07 | 2.100E-07 |
| 21014 | 1051 | K1051F | C | 1.8e-07 | 1.4e-07 | 1.1e-07 | 1.433E-07 | 3.512E-08 | 1.100E-07 | 1.800E-07 |
| 21015 | 1051 | K1051P | C | 1.7e-07 | 3.9e-07 | 2.7e-07 | 2.767E-07 | 1.102E-07 | 1.700E-07 | 3.900E-07 |
| 21016 | 1051 | K1051S | C | 1.4e-07 | 4.1e-07 | 2.5e-07 | 2.667E-07 | 1.358E-07 | 1.400E-07 | 4.100E-07 |
| 21017 | 1051 | K1051T | C | 5.8e-07 | 1.2e-07 | 1.1e-07 | 2.700E-07 | 2.685E-07 | 1.100E-07 | 5.800E-07 |
| 21018 | 1051 | K1051W | C | 1.6e-07 | 3.7e-07 | 2.1e-07 | 2.467E-07 | 1.097E-07 | 1.600E-07 | 3.700E-07 |
| 21019 | 1051 | K1051Y | C | 2.1e-07 | 3.4e-07 | 3.4e-07 | 2.967E-07 | 7.506E-08 | 2.100E-07 | 3.400E-07 |
| 21020 | 1051 | K1051V | C | 2.5e-07 | 1.9e-07 | 2.8e-07 | 2.400E-07 | 4.583E-08 | 1.900E-07 | 2.800E-07 |
| 21021 | 1052 | K1052A | C | 2.8e-07 | 7.2e-07 | 2.3e-07 | 4.100E-07 | 2.696E-07 | 2.300E-07 | 7.200E-07 |
| 21022 | 1052 | K1052R | C | 3.2e-07 | 1.7e-07 | 3.9e-07 | 2.933E-07 | 1.124E-07 | 1.700E-07 | 3.900E-07 |
| 21023 | 1052 | K1052N | C | 3.3e-07 | 2.3e-07 | 8.5e-08 | 2.150E-07 | 1.232E-07 | 8.500E-08 | 3.300E-07 |
| 21024 | 1052 | K1052D | C | 1.7e-07 | 2e-07 | 3.9e-07 | 2.533E-07 | 1.193E-07 | 1.700E-07 | 3.900E-07 |
| 21025 | 1052 | K1052C | C | 2.6e-07 | 3.4e-07 | 4.7e-07 | 3.567E-07 | 1.060E-07 | 2.600E-07 | 4.700E-07 |
| 21026 | 1052 | K1052E | C | 5.4e-07 | 2e-07 | 1.9e-07 | 3.100E-07 | 1.992E-07 | 1.900E-07 | 5.400E-07 |
| 21027 | 1052 | K1052Q | C | 7.3e-07 | 2.2e-07 | 1.1e-07 | 3.533E-07 | 3.308E-07 | 1.100E-07 | 7.300E-07 |
| 21028 | 1052 | K1052G | C | 1.7e-07 | 2e-07 | 2.8e-07 | 2.167E-07 | 5.686E-08 | 1.700E-07 | 2.800E-07 |
| 21029 | 1052 | K1052H | C | 1.7e-07 | 2.5e-07 | 1.7e-07 | 1.967E-07 | 4.619E-08 | 1.700E-07 | 2.500E-07 |
| 21030 | 1052 | K1052I | C | 1e-07 | 9.8e-08 | 4.6e-07 | 2.193E-07 | 2.084E-07 | 9.800E-08 | 4.600E-07 |
| 21031 | 1052 | K1052L | C | 3.6e-07 | 4.2e-07 | 2e-07 | 3.267E-07 | 1.137E-07 | 2.000E-07 | 4.200E-07 |
| 21032 | 1052 | K1052K | C | 1.9e-07 | 3.2e-07 | 3.1e-07 | 2.733E-07 | 7.234E-08 | 1.900E-07 | 3.200E-07 |
| 21033 | 1052 | K1052M | C | 1.6e-07 | 5.7e-07 | 5.4e-07 | 4.233E-07 | 2.285E-07 | 1.600E-07 | 5.700E-07 |
| 21034 | 1052 | K1052F | C | 1.7e-07 | 2.5e-07 | 1.8e-07 | 2.000E-07 | 4.359E-08 | 1.700E-07 | 2.500E-07 |
| 21035 | 1052 | K1052P | C | 2.9e-07 | 2.4e-07 | 2.5e-07 | 2.600E-07 | 2.646E-08 | 2.400E-07 | 2.900E-07 |
| 21036 | 1052 | K1052S | C | 3.9e-07 | 3.3e-07 | 2.3e-07 | 3.167E-07 | 8.083E-08 | 2.300E-07 | 3.900E-07 |
| 21037 | 1052 | K1052T | C | 2.5e-07 | 4.2e-07 | 2.6e-07 | 3.100E-07 | 9.539E-08 | 2.500E-07 | 4.200E-07 |
| 21038 | 1052 | K1052W | C | 2.3e-07 | 1.8e-07 | 2.5e-07 | 2.200E-07 | 3.606E-08 | 1.800E-07 | 2.500E-07 |
| 21039 | 1052 | K1052Y | C | 2e-07 | 1.5e-07 | 2.5e-07 | 2.000E-07 | 5.000E-08 | 1.500E-07 | 2.500E-07 |
| 21040 | 1052 | K1052V | C | 2.5e-07 | 1.2e-07 | 3.5e-07 | 2.400E-07 | 1.153E-07 | 1.200E-07 | 3.500E-07 |
| 21041 | 1053 | V1053A | C | 1.8e-07 | 3.2e-07 | 3.1e-07 | 2.700E-07 | 7.810E-08 | 1.800E-07 | 3.200E-07 |
| 21042 | 1053 | V1053R | C | 1.6e-07 | 2.8e-07 | 4.5e-07 | 2.967E-07 | 1.457E-07 | 1.600E-07 | 4.500E-07 |
| 21043 | 1053 | V1053N | C | 4.8e-07 | 2e-07 | 2.3e-07 | 3.033E-07 | 1.537E-07 | 2.000E-07 | 4.800E-07 |
| 21044 | 1053 | V1053D | C | 3.2e-07 | 2.5e-07 | 1.2e-07 | 2.300E-07 | 1.015E-07 | 1.200E-07 | 3.200E-07 |
| 21045 | 1053 | V1053C | C | 3.8e-07 | 1.4e-07 | 2.7e-07 | 2.633E-07 | 1.201E-07 | 1.400E-07 | 3.800E-07 |
| 21046 | 1053 | V1053E | C | 2.9e-07 | 9.5e-08 | 2.3e-07 | 2.050E-07 | 9.987E-08 | 9.500E-08 | 2.900E-07 |
| 21047 | 1053 | V1053Q | C | 1.5e-07 | 1.3e-07 | 2.8e-07 | 1.867E-07 | 8.145E-08 | 1.300E-07 | 2.800E-07 |
| 21048 | 1053 | V1053G | C | 2.5e-07 | 3.6e-07 | 5.4e-07 | 3.833E-07 | 1.464E-07 | 2.500E-07 | 5.400E-07 |
| 21049 | 1053 | V1053H | C | 3.2e-07 | 1.8e-07 | 2.1e-07 | 2.367E-07 | 7.371E-08 | 1.800E-07 | 3.200E-07 |
| 21050 | 1053 | V1053I | C | 2e-07 | 3.5e-07 | 7.5e-07 | 4.333E-07 | 2.843E-07 | 2.000E-07 | 7.500E-07 |
| 21051 | 1053 | V1053L | C | 4.5e-07 | 1.2e-07 | 4.1e-07 | 3.267E-07 | 1.801E-07 | 1.200E-07 | 4.500E-07 |
| 21052 | 1053 | V1053K | C | 1.3e-07 | 2e-07 | 1.6e-07 | 1.633E-07 | 3.512E-08 | 1.300E-07 | 2.000E-07 |
| 21053 | 1053 | V1053M | C | 1.7e-07 | 5.9e-07 | 1.4e-07 | 3.000E-07 | 2.516E-07 | 1.400E-07 | 5.900E-07 |
| 21054 | 1053 | V1053F | C | 2.5e-07 | 2.5e-07 | 1.3e-07 | 2.100E-07 | 6.928E-08 | 1.300E-07 | 2.500E-07 |
| 21055 | 1053 | V1053P | C | 1.1e-06 | 2.9e-07 | 2.5e-07 | 5.467E-07 | 4.796E-07 | 2.500E-07 | 1.100E-06 |
| 21056 | 1053 | V1053S | C | 2.9e-07 | 1.6e-07 | 1.9e-07 | 2.133E-07 | 6.807E-08 | 1.600E-07 | 2.900E-07 |
| 21057 | 1053 | V1053T | C | 3.2e-07 | 5e-07 | 2.2e-07 | 3.467E-07 | 1.419E-07 | 2.200E-07 | 5.000E-07 |
| 21058 | 1053 | V1053W | C | 1.5e-07 | 2.4e-07 | 1.4e-07 | 1.767E-07 | 5.508E-08 | 1.400E-07 | 2.400E-07 |
| 21059 | 1053 | V1053Y | C | 2.1e-07 | 1.4e-07 | 3.1e-07 | 2.200E-07 | 8.544E-08 | 1.400E-07 | 3.100E-07 |
| 21060 | 1053 | V1053V | C | 6e-07 | 5.7e-07 | 2.3e-07 | 4.667E-07 | 2.055E-07 | 2.300E-07 | 6.000E-07 |
| 21061 | 1054 | E1054A | C | 4e-07 | 2.7e-07 | 2.9e-07 | 3.200E-07 | 7.000E-08 | 2.700E-07 | 4.000E-07 |
| 21062 | 1054 | E1054R | C | 1.9e-07 | 2.7e-07 | 3.3e-07 | 2.633E-07 | 7.024E-08 | 1.900E-07 | 3.300E-07 |
| 21063 | 1054 | E1054N | C | 2.5e-07 | 2.5e-07 | 1.7e-07 | 2.233E-07 | 4.619E-08 | 1.700E-07 | 2.500E-07 |
| 21064 | 1054 | E1054D | C | 5e-07 | 9.8e-08 | 3.9e-07 | 3.293E-07 | 2.078E-07 | 9.800E-08 | 5.000E-07 |
| 21065 | 1054 | E1054C | C | 1.9e-07 | 2.1e-07 | 1.5e-07 | 1.833E-07 | 3.055E-08 | 1.500E-07 | 2.100E-07 |
| 21066 | 1054 | E1054E | C | 3e-07 | 2.5e-07 | 3.3e-07 | 2.933E-07 | 4.041E-08 | 2.500E-07 | 3.300E-07 |
| 21067 | 1054 | E1054Q | C | 2.3e-07 | 1.6e-07 | 5.4e-07 | 3.100E-07 | 2.022E-07 | 1.600E-07 | 5.400E-07 |
| 21068 | 1054 | E1054G | C | 1.2e-07 | 2.7e-07 | 3.9e-07 | 2.600E-07 | 1.353E-07 | 1.200E-07 | 3.900E-07 |
| 21069 | 1054 | E1054H | C | 3.6e-07 | 2.9e-07 | 1.8e-07 | 2.767E-07 | 9.074E-08 | 1.800E-07 | 3.600E-07 |
| 21070 | 1054 | E1054I | C | 2.1e-07 | 3.2e-07 | 2.9e-07 | 2.733E-07 | 5.686E-08 | 2.100E-07 | 3.200E-07 |
| 21071 | 1054 | E1054L | C | 2.4e-07 | 2.1e-07 | 4e-07 | 2.833E-07 | 1.021E-07 | 2.100E-07 | 4.000E-07 |
| 21072 | 1054 | E1054K | C | 5.3e-07 | 1.8e-07 | 1.6e-07 | 2.900E-07 | 2.081E-07 | 1.600E-07 | 5.300E-07 |
| 21073 | 1054 | E1054M | C | 2.8e-07 | 1.7e-07 | 2.9e-07 | 2.467E-07 | 6.658E-08 | 1.700E-07 | 2.900E-07 |
| 21074 | 1054 | E1054F | C | 2.9e-07 | 1.7e-07 | 3.9e-07 | 2.833E-07 | 1.102E-07 | 1.700E-07 | 3.900E-07 |
| 21075 | 1054 | E1054P | C | 3.6e-07 | 3.2e-07 | 1.4e-07 | 2.733E-07 | 1.172E-07 | 1.400E-07 | 3.600E-07 |
| 21076 | 1054 | E1054S | C | 3.7e-07 | 4.2e-07 | 2.6e-07 | 3.500E-07 | 8.185E-08 | 2.600E-07 | 4.200E-07 |
| 21077 | 1054 | E1054T | C | 2.7e-07 | 4e-07 | 3e-07 | 3.233E-07 | 6.807E-08 | 2.700E-07 | 4.000E-07 |
| 21078 | 1054 | E1054W | C | 4.7e-07 | 6.3e-07 | 1.8e-07 | 4.267E-07 | 2.281E-07 | 1.800E-07 | 6.300E-07 |
| 21079 | 1054 | E1054Y | C | 4.2e-07 | 9.1e-08 | 1.7e-07 | 2.270E-07 | 1.717E-07 | 9.100E-08 | 4.200E-07 |

| | | | | | | | | | | |
|---|---|---|---|---|---|---|---|---|---|---|
| 21080 | 1054 | E1054V | C | 1.7e-07 | 2.1e-07 | 1.6e-07 | 1.800E-07 | 2.646E-08 | 1.600E-07 | 2.100E-07 |
| 21081 | 1055 | P1055A | C | 1.6e-07 | 1.3e-07 | 4.6e-07 | 2.500E-07 | 1.825E-07 | 1.300E-07 | 4.600E-07 |
| 21082 | 1055 | P1055R | C | 2.7e-07 | 9.1e-08 | 1.6e-07 | 1.737E-07 | 9.028E-08 | 9.100E-08 | 2.700E-07 |
| 21083 | 1055 | P1055N | C | 1.9e-07 | 2.1e-07 | 6e-08 | 1.533E-07 | 8.145E-08 | 6.000E-08 | 2.100E-07 |
| 21084 | 1055 | P1055D | C | 1.6e-07 | 3.3e-07 | 3.1e-07 | 2.667E-07 | 9.292E-08 | 1.600E-07 | 3.300E-07 |
| 21085 | 1055 | P1055C | C | 1.9e-07 | 2.4e-07 | 4.1e-07 | 2.800E-07 | 1.153E-07 | 1.900E-07 | 4.100E-07 |
| 21086 | 1055 | P1055E | C | 1.2e-07 | 2e-07 | 3.1e-07 | 2.100E-07 | 9.539E-08 | 1.200E-07 | 3.100E-07 |
| 21087 | 1055 | P1055Q | C | 4.5e-07 | 3.5e-07 | 1.5e-07 | 3.167E-07 | 1.528E-07 | 1.500E-07 | 4.500E-07 |
| 21088 | 1055 | P1055G | C | 2e-07 | 3.3e-07 | 2.1e-07 | 2.467E-07 | 7.234E-08 | 2.000E-07 | 3.300E-07 |
| 21089 | 1055 | P1055H | C | 2.5e-07 | 5e-07 | 1.9e-07 | 3.133E-07 | 1.644E-07 | 1.900E-07 | 5.000E-07 |
| 21090 | 1055 | P1055I | C | 2.7e-07 | 1.7e-07 | 2.9e-07 | 2.433E-07 | 6.429E-08 | 1.700E-07 | 2.900E-07 |
| 21091 | 1055 | P1055L | C | 1.4e-07 | 1.9e-07 | 2.5e-07 | 1.933E-07 | 5.508E-08 | 1.400E-07 | 2.500E-07 |
| 21092 | 1055 | P1055K | C | 3.1e-07 | 3.5e-07 | 2.8e-07 | 3.133E-07 | 3.512E-08 | 2.800E-07 | 3.500E-07 |
| 21093 | 1055 | P1055M | C | 1.1e-07 | 3.1e-07 | 7.8e-07 | 4.000E-07 | 3.439E-07 | 1.100E-07 | 7.800E-07 |
| 21094 | 1055 | P1055F | C | 1.9e-07 | 3.9e-07 | 1.9e-07 | 2.567E-07 | 1.155E-07 | 1.900E-07 | 3.900E-07 |
| 21095 | 1055 | P1055P | C | 1.2e-07 | 2.3e-07 | 3.3e-07 | 2.267E-07 | 1.050E-07 | 1.200E-07 | 3.300E-07 |
| 21096 | 1055 | P1055S | C | 2.4e-07 | 2.3e-07 | 3.7e-07 | 2.800E-07 | 7.810E-08 | 2.300E-07 | 3.700E-07 |
| 21097 | 1055 | P1055T | C | 3.2e-07 | 3.3e-07 | 4.8e-07 | 3.767E-07 | 8.963E-08 | 3.200E-07 | 4.800E-07 |
| 21098 | 1055 | P1055W | C | 1.9e-07 | 1.2e-07 | 2.7e-07 | 1.933E-07 | 7.506E-08 | 1.200E-07 | 2.700E-07 |
| 21099 | 1055 | P1055Y | C | 2.1e-07 | 2.3e-07 | 2.6e-07 | 2.333E-07 | 2.517E-08 | 2.100E-07 | 2.600E-07 |
| 21100 | 1055 | P1055V | C | 3.3e-07 | 2.5e-07 | 2.2e-07 | 2.667E-07 | 5.686E-08 | 2.200E-07 | 3.300E-07 |
| 21101 | 1056 | K1056A | C | 3.3e-07 | 1.8e-07 | 1.8e-07 | 2.300E-07 | 8.660E-08 | 1.800E-07 | 3.300E-07 |
| 21102 | 1056 | K1056R | C | 3.4e-07 | 3e-07 | 2e-07 | 2.800E-07 | 7.211E-08 | 2.000E-07 | 3.400E-07 |
| 21103 | 1056 | K1056N | C | 1.7e-07 | 2e-07 | 2.5e-07 | 2.067E-07 | 4.041E-08 | 1.700E-07 | 2.500E-07 |
| 21104 | 1056 | K1056D | C | 1.5e-07 | 1.5e-07 | 2.5e-07 | 1.833E-07 | 5.774E-08 | 1.500E-07 | 2.500E-07 |
| 21105 | 1056 | K1056C | C | 1.4e-07 | 3e-07 | 2.2e-07 | 2.200E-07 | 8.000E-08 | 1.400E-07 | 3.000E-07 |
| 21106 | 1056 | K1056E | C | 5.4e-07 | 1.1e-07 | 9.8e-08 | 2.493E-07 | 2.518E-07 | 9.800E-08 | 5.400E-07 |
| 21107 | 1056 | K1056Q | C | 3.5e-07 | 3e-07 | 3.8e-07 | 3.433E-07 | 4.041E-08 | 3.000E-07 | 3.800E-07 |
| 21108 | 1056 | K1056G | C | 2.8e-07 | 2.4e-07 | 2.4e-07 | 2.533E-07 | 2.309E-08 | 2.400E-07 | 2.800E-07 |
| 21109 | 1056 | K1056H | C | 2.4e-07 | 3.4e-07 | 2.3e-07 | 2.700E-07 | 6.083E-08 | 2.300E-07 | 3.400E-07 |
| 21110 | 1056 | K1056I | C | 3.9e-07 | 2.2e-07 | 2.8e-07 | 2.967E-07 | 8.622E-08 | 2.200E-07 | 3.900E-07 |
| 21111 | 1056 | K1056L | C | 2e-07 | 1.2e-07 | 1.5e-07 | 1.567E-07 | 4.041E-08 | 1.200E-07 | 2.000E-07 |
| 21112 | 1056 | K1056K | C | 2e-07 | 1.1e-07 | 1.8e-07 | 1.633E-07 | 4.726E-08 | 1.100E-07 | 2.000E-07 |
| 21113 | 1056 | K1056M | C | 1.5e-07 | 2e-07 | 1.3e-07 | 1.600E-07 | 3.606E-08 | 1.300E-07 | 2.000E-07 |
| 21114 | 1056 | K1056F | C | 2.9e-07 | 2.7e-07 | 1.6e-07 | 2.400E-07 | 7.000E-08 | 1.600E-07 | 2.900E-07 |
| 21115 | 1056 | K1056P | C | 4.2e-07 | 1.8e-07 | 3.4e-07 | 3.133E-07 | 1.222E-07 | 1.800E-07 | 4.200E-07 |
| 21116 | 1056 | K1056S | C | 4.3e-07 | 2.1e-07 | 2.3e-07 | 2.900E-07 | 1.217E-07 | 2.100E-07 | 4.300E-07 |
| 21117 | 1056 | K1056T | C | 2.2e-07 | 2e-07 | 1.5e-07 | 1.900E-07 | 3.606E-08 | 1.500E-07 | 2.200E-07 |
| 21118 | 1056 | K1056W | C | 1.2e-07 | 2.9e-07 | 6.4e-07 | 3.500E-07 | 2.651E-07 | 1.200E-07 | 6.400E-07 |
| 21119 | 1056 | K1056Y | C | 2.8e-07 | 2.1e-07 | 1.8e-07 | 2.233E-07 | 5.132E-08 | 1.800E-07 | 2.800E-07 |
| 21120 | 1056 | K1056V | C | 1.8e-07 | 1.5e-07 | 2.3e-07 | 1.867E-07 | 4.041E-08 | 1.500E-07 | 2.300E-07 |
| 21121 | 1057 | S1057A | C | 2.1e-07 | 2.2e-07 | 4e-07 | 2.767E-07 | 1.069E-07 | 2.100E-07 | 4.000E-07 |
| 21122 | 1057 | S1057R | C | 2.8e-07 | 3e-07 | 2.7e-07 | 2.833E-07 | 1.528E-08 | 2.700E-07 | 3.000E-07 |
| 21123 | 1057 | S1057N | C | 2.1e-07 | 2.1e-07 | 2.4e-07 | 2.200E-07 | 1.732E-08 | 2.100E-07 | 2.400E-07 |
| 21124 | 1057 | S1057D | C | 2.2e-07 | 1.6e-07 | 4.9e-07 | 2.900E-07 | 1.758E-07 | 1.600E-07 | 4.900E-07 |
| 21125 | 1057 | S1057C | C | 1.7e-07 | 5.4e-07 | 1.1e-07 | 2.733E-07 | 2.329E-07 | 1.100E-07 | 5.400E-07 |
| 21126 | 1057 | S1057E | C | 3.2e-07 | 2.4e-07 | 2.1e-07 | 2.567E-07 | 5.686E-08 | 2.100E-07 | 3.200E-07 |
| 21127 | 1057 | S1057Q | C | 3.8e-07 | 2.2e-07 | 2.3e-07 | 2.767E-07 | 8.963E-08 | 2.200E-07 | 3.800E-07 |
| 21128 | 1057 | S1057G | C | 3.1e-07 | 9e-08 | 3.4e-07 | 2.467E-07 | 1.365E-07 | 9.000E-08 | 3.400E-07 |
| 21129 | 1057 | S1057H | C | 4e-07 | 2.3e-07 | 4.5e-07 | 3.600E-07 | 1.153E-07 | 2.300E-07 | 4.500E-07 |
| 21130 | 1057 | S1057I | C | 1.4e-07 | 3.2e-07 | 2.7e-07 | 2.433E-07 | 9.292E-08 | 1.400E-07 | 3.200E-07 |
| 21131 | 1057 | S1057L | C | 1e-07 | 2.3e-07 | 3.7e-07 | 2.333E-07 | 1.350E-07 | 1.000E-07 | 3.700E-07 |
| 21132 | 1057 | S1057K | C | 2.8e-07 | 2.6e-07 | 3e-07 | 2.800E-07 | 2.000E-08 | 2.600E-07 | 3.000E-07 |
| 21133 | 1057 | S1057M | C | 3.2e-07 | 1.7e-07 | 3.3e-07 | 2.733E-07 | 8.963E-08 | 1.700E-07 | 3.300E-07 |
| 21134 | 1057 | S1057F | C | 3.6e-07 | 1.7e-07 | 3.6e-07 | 2.967E-07 | 1.097E-07 | 1.700E-07 | 3.600E-07 |
| 21135 | 1057 | S1057P | C | 1.3e-07 | 4e-07 | 6.9e-07 | 4.067E-07 | 2.801E-07 | 1.300E-07 | 6.900E-07 |
| 21136 | 1057 | S1057S | C | 1.2e-07 | 3.6e-07 | 3.1e-07 | 2.633E-07 | 1.266E-07 | 1.200E-07 | 3.600E-07 |
| 21137 | 1057 | S1057T | C | 1.8e-07 | 1.9e-07 | 2e-07 | 1.900E-07 | 1.000E-08 | 1.800E-07 | 2.000E-07 |
| 21138 | 1057 | S1057W | C | 1.7e-07 | 1e-07 | 4.4e-07 | 2.367E-07 | 1.795E-07 | 1.000E-07 | 4.400E-07 |
| 21139 | 1057 | S1057Y | C | 7.3e-08 | 8.2e-07 | 1.5e-07 | 3.477E-07 | 4.109E-07 | 7.300E-08 | 8.200E-07 |
| 21140 | 1057 | S1057V | C | 1.3e-07 | 9.8e-08 | 4.4e-07 | 2.227E-07 | 1.889E-07 | 9.800E-08 | 4.400E-07 |
| 21141 | 1058 | C1058A | C | 3.7e-07 | 2.9e-07 | 2.5e-07 | 3.033E-07 | 6.110E-08 | 2.500E-07 | 3.700E-07 |
| 21142 | 1058 | C1058R | C | 2.1e-07 | 4.5e-07 | 3.5e-07 | 3.367E-07 | 1.206E-07 | 2.100E-07 | 4.500E-07 |
| 21143 | 1058 | C1058N | C | 4.4e-07 | 4e-07 | 2.1e-07 | 3.500E-07 | 1.229E-07 | 2.100E-07 | 4.400E-07 |
| 21144 | 1058 | C1058D | C | 1.3e-07 | 2.1e-07 | 3e-07 | 2.133E-07 | 8.505E-08 | 1.300E-07 | 3.000E-07 |
| 21145 | 1058 | C1058C | C | 1.9e-07 | 3.6e-07 | 1.9e-07 | 2.467E-07 | 9.815E-08 | 1.900E-07 | 3.600E-07 |
| 21146 | 1058 | C1058E | C | 3.5e-07 | 1.2e-07 | 4.7e-07 | 3.133E-07 | 1.779E-07 | 1.200E-07 | 4.700E-07 |
| 21147 | 1058 | C1058Q | C | 2.4e-07 | 4.1e-07 | 3.3e-07 | 3.267E-07 | 8.505E-08 | 2.400E-07 | 4.100E-07 |
| 21148 | 1058 | C1058G | C | 2e-07 | 3e-07 | 3.9e-07 | 2.967E-07 | 9.504E-08 | 2.000E-07 | 3.900E-07 |
| 21149 | 1058 | C1058H | C | 1.6e-07 | 2.7e-07 | 3.6e-07 | 2.633E-07 | 1.002E-07 | 1.600E-07 | 3.600E-07 |
| 21150 | 1058 | C1058I | C | 6.9e-07 | 3.2e-07 | 2.8e-07 | 4.300E-07 | 2.261E-07 | 2.800E-07 | 6.900E-07 |
| 21151 | 1058 | C1058L | C | 2.9e-07 | 2.8e-07 | 3.5e-07 | 3.067E-07 | 3.786E-08 | 2.800E-07 | 3.500E-07 |
| 21152 | 1058 | C1058K | C | 2.4e-07 | 2.2e-07 | 5.6e-07 | 3.400E-07 | 1.908E-07 | 2.200E-07 | 5.600E-07 |
| 21153 | 1058 | C1058M | C | 4.7e-07 | 6.6e-07 | 3.6e-07 | 4.967E-07 | 1.518E-07 | 3.600E-07 | 6.600E-07 |
| 21154 | 1058 | C1058F | C | 3e-07 | 1.3e-07 | 1.8e-07 | 2.033E-07 | 8.737E-08 | 1.300E-07 | 3.000E-07 |
| 21155 | 1058 | C1058P | C | 2.2e-07 | 6.7e-08 | 5.3e-07 | 2.723E-07 | 2.359E-07 | 6.700E-08 | 5.300E-07 |
| 21156 | 1058 | C1058S | C | 1.9e-07 | 2.4e-07 | 7.7e-08 | 1.690E-07 | 8.350E-08 | 7.700E-08 | 2.400E-07 |
| 21157 | 1058 | C1058T | C | 3.8e-07 | 2e-07 | 4.3e-07 | 3.367E-07 | 1.210E-07 | 2.000E-07 | 4.300E-07 |
| 21158 | 1058 | C1058W | C | 2.3e-07 | 4.2e-07 | 2.8e-07 | 3.100E-07 | 9.849E-08 | 2.300E-07 | 4.200E-07 |
| 21159 | 1058 | C1058Y | C | 2.1e-07 | 3.1e-07 | 2.6e-07 | 2.600E-07 | 5.000E-08 | 2.100E-07 | 3.100E-07 |
| 21160 | 1058 | C1058V | C | 2.1e-07 | 2.1e-07 | 3.8e-07 | 2.667E-07 | 9.815E-08 | 2.100E-07 | 3.800E-07 |
| 21161 | 1059 | D1059A | D | 1.6e-07 | 3.9e-07 | 2e-07 | 2.500E-07 | 1.229E-07 | 1.600E-07 | 3.900E-07 |
| 21162 | 1059 | D1059R | D | 2.1e-07 | 2.5e-07 | 2.5e-07 | 2.367E-07 | 2.309E-08 | 2.100E-07 | 2.500E-07 |
| 21163 | 1059 | D1059N | D | 2.1e-07 | 2.4e-07 | 1.2e-07 | 1.900E-07 | 6.245E-08 | 1.200E-07 | 2.400E-07 |
| 21164 | 1059 | D1059D | D | 1.6e-07 | 2.8e-07 | 5.9e-07 | 3.433E-07 | 2.219E-07 | 1.600E-07 | 5.900E-07 |

| | | | | | | | | | | |
|---|---|---|---|---|---|---|---|---|---|---|
| 21165 | 1059 | D1059C | D | 2.2e-07 | 1.3e-07 | 1.8e-07 | 1.767E-07 | 4.509E-08 | 1.300E-07 | 2.200E-07 |
| 21166 | 1059 | D1059E | D | 1.9e-07 | 1.6e-07 | 4.4e-07 | 2.633E-07 | 1.537E-07 | 1.600E-07 | 4.400E-07 |
| 21167 | 1059 | D1059Q | D | 2.4e-07 | 3.2e-07 | 2.2e-07 | 2.600E-07 | 5.292E-08 | 2.200E-07 | 3.200E-07 |
| 21168 | 1059 | D1059G | D | 2.7e-07 | 4.7e-07 | 1.6e-07 | 3.000E-07 | 1.572E-07 | 1.600E-07 | 4.700E-07 |
| 21169 | 1059 | D1059H | D | 2e-07 | 4.7e-07 | 4e-07 | 3.567E-07 | 1.401E-07 | 2.000E-07 | 4.700E-07 |
| 21170 | 1059 | D1059I | D | 5.4e-07 | 2.8e-07 | 1.9e-07 | 3.367E-07 | 1.818E-07 | 1.900E-07 | 5.400E-07 |
| 21171 | 1059 | D1059L | D | 2.7e-07 | 2.6e-07 | 2.1e-07 | 2.467E-07 | 3.215E-08 | 2.100E-07 | 2.700E-07 |
| 21172 | 1059 | D1059K | D | 1.1e-07 | 2.8e-07 | 2.4e-07 | 2.100E-07 | 8.888E-08 | 1.100E-07 | 2.800E-07 |
| 21173 | 1059 | D1059M | D | 1.9e-07 | 3.3e-07 | 2.6e-07 | 2.600E-07 | 7.000E-08 | 1.900E-07 | 3.300E-07 |
| 21174 | 1059 | D1059F | D | 2.4e-07 | 1.3e-07 | 3.9e-07 | 2.533E-07 | 1.305E-07 | 1.300E-07 | 3.900E-07 |
| 21175 | 1059 | D1059P | D | 1.8e-07 | 3.1e-07 | 2.6e-07 | 2.500E-07 | 6.557E-08 | 1.800E-07 | 3.100E-07 |
| 21176 | 1059 | D1059S | D | 3.6e-07 | 3.1e-07 | 2.7e-07 | 3.133E-07 | 4.509E-08 | 2.700E-07 | 3.600E-07 |
| 21177 | 1059 | D1059T | D | 6.7e-07 | 3.2e-07 | 2.3e-07 | 4.067E-07 | 2.325E-07 | 2.300E-07 | 6.700E-07 |
| 21178 | 1059 | D1059W | D | 2.5e-07 | 3e-07 | 3.3e-07 | 2.933E-07 | 4.041E-08 | 2.500E-07 | 3.300E-07 |
| 21179 | 1059 | D1059Y | D | 1.9e-07 | 3.5e-07 | 2.7e-07 | 2.700E-07 | 8.000E-08 | 1.900E-07 | 3.500E-07 |
| 21180 | 1059 | D1059V | D | 2.1e-07 | 1.9e-07 | 3.9e-07 | 2.633E-07 | 1.102E-07 | 1.900E-07 | 3.900E-07 |
| 21181 | 1060 | I1060A | D | 6.4e-07 | 1.2e-07 | 2e-07 | 3.200E-07 | 2.800E-07 | 1.200E-07 | 6.400E-07 |
| 21182 | 1060 | I1060R | D | 2.9e-07 | 2.9e-07 | 3.4e-07 | 3.067E-07 | 2.887E-08 | 2.900E-07 | 3.400E-07 |
| 21183 | 1060 | I1060N | D | 1.8e-07 | 3e-07 | 2e-07 | 2.267E-07 | 6.429E-08 | 1.800E-07 | 3.000E-07 |
| 21184 | 1060 | I1060D | D | 2.8e-07 | 9e-08 | 3e-07 | 2.233E-07 | 1.159E-07 | 9.000E-08 | 3.000E-07 |
| 21185 | 1060 | I1060C | D | 2.4e-07 | 3.7e-07 | 2.8e-07 | 2.967E-07 | 6.658E-08 | 2.400E-07 | 3.700E-07 |
| 21186 | 1060 | I1060E | D | 2.8e-07 | 1.8e-07 | 1.9e-07 | 2.167E-07 | 5.508E-08 | 1.800E-07 | 2.800E-07 |
| 21187 | 1060 | I1060Q | D | 1.2e-07 | 1.9e-07 | 1.1e-07 | 1.400E-07 | 4.359E-08 | 1.100E-07 | 1.900E-07 |
| 21188 | 1060 | I1060G | D | 3.5e-07 | 2e-07 | 2.4e-07 | 2.633E-07 | 7.767E-08 | 2.000E-07 | 3.500E-07 |
| 21189 | 1060 | I1060H | D | 3e-07 | 3.3e-07 | 1.9e-07 | 2.733E-07 | 7.371E-08 | 1.900E-07 | 3.300E-07 |
| 21190 | 1060 | I1060I | D | 5.9e-07 | 1.8e-07 | 2.7e-07 | 3.467E-07 | 2.155E-07 | 1.800E-07 | 5.900E-07 |
| 21191 | 1060 | I1060L | D | 3.8e-07 | 1.5e-07 | 3.6e-07 | 2.967E-07 | 1.274E-07 | 1.500E-07 | 3.800E-07 |
| 21192 | 1060 | I1060K | D | 2e-07 | 2.3e-07 | 2.2e-07 | 2.167E-07 | 1.528E-08 | 2.000E-07 | 2.300E-07 |
| 21193 | 1060 | I1060M | D | 1.2e-07 | 3.7e-07 | 1.5e-07 | 2.133E-07 | 1.365E-07 | 1.200E-07 | 3.700E-07 |
| 21194 | 1060 | I1060F | D | 3.9e-07 | 1.8e-07 | 3.4e-07 | 3.033E-07 | 1.097E-07 | 1.800E-07 | 3.900E-07 |
| 21195 | 1060 | I1060P | D | 2.6e-07 | 4.8e-07 | 2.5e-07 | 3.300E-07 | 1.300E-07 | 2.500E-07 | 4.800E-07 |
| 21196 | 1060 | I1060S | D | 1.7e-07 | 5e-07 | 3.3e-07 | 3.333E-07 | 1.650E-07 | 1.700E-07 | 5.000E-07 |
| 21197 | 1060 | I1060T | D | 6.2e-07 | 1.5e-07 | 3.2e-07 | 3.633E-07 | 2.380E-07 | 1.500E-07 | 6.200E-07 |
| 21198 | 1060 | I1060W | D | 1.8e-07 | 5.9e-07 | 3.8e-07 | 3.833E-07 | 2.050E-07 | 1.800E-07 | 5.900E-07 |
| 21199 | 1060 | I1060Y | D | 2.9e-07 | 2.3e-07 | 1.8e-07 | 2.333E-07 | 5.508E-08 | 1.800E-07 | 2.900E-07 |
| 21200 | 1060 | I1060V | D | 3.7e-07 | 2.7e-07 | 2.2e-07 | 2.867E-07 | 7.638E-08 | 2.200E-07 | 3.700E-07 |
| 21201 | 1061 | Q1061A | D | 2.1e-07 | 4.4e-07 | 2.8e-07 | 3.100E-07 | 1.179E-07 | 2.100E-07 | 4.400E-07 |
| 21202 | 1061 | Q1061R | D | 2.5e-07 | 1.3e-07 | 4.6e-07 | 2.800E-07 | 1.670E-07 | 1.300E-07 | 4.600E-07 |
| 21203 | 1061 | Q1061N | D | 3.5e-07 | 1.4e-07 | 2.2e-07 | 2.367E-07 | 1.060E-07 | 1.400E-07 | 3.500E-07 |
| 21204 | 1061 | Q1061D | D | 2.4e-07 | 4.6e-07 | 3.1e-07 | 3.367E-07 | 1.124E-07 | 2.400E-07 | 4.600E-07 |
| 21205 | 1061 | Q1061C | D | 3.7e-07 | 1.8e-07 | 2.6e-07 | 2.700E-07 | 9.539E-08 | 1.800E-07 | 3.700E-07 |
| 21206 | 1061 | Q1061E | D | 1.6e-07 | 3.2e-07 | 2.9e-07 | 2.567E-07 | 8.505E-08 | 1.600E-07 | 3.200E-07 |
| 21207 | 1061 | Q1061Q | D | 3.5e-07 | 2.2e-07 | 2e-07 | 2.567E-07 | 8.145E-08 | 2.000E-07 | 3.500E-07 |
| 21208 | 1061 | Q1061G | D | 2.4e-07 | 1.3e-07 | 2e-07 | 1.900E-07 | 5.568E-08 | 1.300E-07 | 2.400E-07 |
| 21209 | 1061 | Q1061H | D | 4.3e-07 | 1.8e-07 | 2.3e-07 | 2.800E-07 | 1.323E-07 | 1.800E-07 | 4.300E-07 |
| 21210 | 1061 | Q1061I | D | 2.6e-07 | 1.8e-07 | 3.7e-07 | 2.700E-07 | 9.539E-08 | 1.800E-07 | 3.700E-07 |
| 21211 | 1061 | Q1061L | D | 1.4e-07 | 1.7e-07 | 3.3e-07 | 2.133E-07 | 1.021E-07 | 1.400E-07 | 3.300E-07 |
| 21212 | 1061 | Q1061K | D | 7.4e-08 | 2.7e-07 | 3.2e-07 | 2.213E-07 | 1.300E-07 | 7.400E-08 | 3.200E-07 |
| 21213 | 1061 | Q1061M | D | 1.6e-07 | 3.9e-07 | 3.6e-07 | 3.033E-07 | 1.250E-07 | 1.600E-07 | 3.900E-07 |
| 21214 | 1061 | Q1061F | D | 1.4e-07 | 2.5e-07 | 2.8e-07 | 2.233E-07 | 7.371E-08 | 1.400E-07 | 2.800E-07 |
| 21215 | 1061 | Q1061P | D | 1.9e-07 | 2.2e-07 | 4.4e-07 | 2.833E-07 | 1.365E-07 | 1.900E-07 | 4.400E-07 |
| 21216 | 1061 | Q1061S | D | 3.5e-07 | 3.4e-07 | 2.5e-07 | 3.133E-07 | 5.508E-08 | 2.500E-07 | 3.500E-07 |
| 21217 | 1061 | Q1061T | D | 2.9e-07 | 2.4e-07 | 5.1e-07 | 3.467E-07 | 1.436E-07 | 2.400E-07 | 5.100E-07 |
| 21218 | 1061 | Q1061W | D | 3.3e-07 | 4.8e-07 | 1.2e-07 | 3.100E-07 | 1.808E-07 | 1.200E-07 | 4.800E-07 |
| 21219 | 1061 | Q1061Y | D | 3.6e-07 | 4.4e-07 | 1.2e-07 | 3.067E-07 | 1.665E-07 | 1.200E-07 | 4.400E-07 |
| 21220 | 1061 | Q1061V | D | 2.7e-07 | 1.1e-07 | 6.8e-07 | 3.533E-07 | 2.940E-07 | 1.100E-07 | 6.800E-07 |
| 21221 | 1062 | M1062A | D | 2.5e-07 | 2.8e-07 | 2.7e-07 | 2.667E-07 | 1.528E-08 | 2.500E-07 | 2.800E-07 |
| 21222 | 1062 | M1062R | D | 1.6e-07 | 2.1e-07 | 1.8e-07 | 1.833E-07 | 2.517E-08 | 1.600E-07 | 2.100E-07 |
| 21223 | 1062 | M1062N | D | 4.4e-07 | 2.9e-07 | 5.4e-07 | 4.233E-07 | 1.258E-07 | 2.900E-07 | 5.400E-07 |
| 21224 | 1062 | M1062D | D | 2.7e-07 | 3.4e-07 | 1.7e-07 | 2.600E-07 | 8.544E-08 | 1.700E-07 | 3.400E-07 |
| 21225 | 1062 | M1062C | D | 2.3e-07 | 1.1e-07 | 1.7e-07 | 1.700E-07 | 6.000E-08 | 1.100E-07 | 2.300E-07 |
| 21226 | 1062 | M1062E | D | 2.4e-07 | 4.1e-07 | 1.2e-07 | 2.567E-07 | 1.457E-07 | 1.200E-07 | 4.100E-07 |
| 21227 | 1062 | M1062Q | D | 2.7e-07 | 1.3e-07 | 3.6e-07 | 2.533E-07 | 1.159E-07 | 1.300E-07 | 3.600E-07 |
| 21228 | 1062 | M1062G | D | 3.3e-07 | 1.3e-07 | 1.6e-07 | 2.067E-07 | 1.079E-07 | 1.300E-07 | 3.300E-07 |
| 21229 | 1062 | M1062H | D | 1.1e-07 | 2.9e-07 | 2.4e-07 | 2.133E-07 | 9.292E-08 | 1.100E-07 | 2.900E-07 |
| 21230 | 1062 | M1062I | D | 3.6e-07 | 2.8e-07 | 3.7e-07 | 3.367E-07 | 4.933E-08 | 2.800E-07 | 3.700E-07 |
| 21231 | 1062 | M1062L | D | 1.9e-07 | 2.3e-07 | 1e-07 | 1.733E-07 | 6.658E-08 | 1.000E-07 | 2.300E-07 |
| 21232 | 1062 | M1062K | D | 2.4e-07 | 3.1e-07 | 3.3e-07 | 2.933E-07 | 4.726E-08 | 2.400E-07 | 3.300E-07 |
| 21233 | 1062 | M1062M | D | 4.2e-07 | 3.1e-07 | 1.1e-07 | 2.800E-07 | 1.572E-07 | 1.100E-07 | 4.200E-07 |
| 21234 | 1062 | M1062F | D | 1e-07 | 2.5e-07 | 5.6e-07 | 3.033E-07 | 2.346E-07 | 1.000E-07 | 5.600E-07 |
| 21235 | 1062 | M1062P | D | 5.7e-07 | 2.6e-07 | 1.8e-07 | 3.367E-07 | 2.060E-07 | 1.800E-07 | 5.700E-07 |
| 21236 | 1062 | M1062S | D | 2.7e-07 | 2.1e-07 | 2.1e-07 | 2.300E-07 | 3.464E-08 | 2.100E-07 | 2.700E-07 |
| 21237 | 1062 | M1062T | D | 1.2e-07 | 1.8e-07 | 2.9e-07 | 1.967E-07 | 8.622E-08 | 1.200E-07 | 2.900E-07 |
| 21238 | 1062 | M1062W | D | 1.4e-07 | 1.8e-07 | 3e-07 | 2.067E-07 | 8.327E-08 | 1.400E-07 | 3.000E-07 |
| 21239 | 1062 | M1062Y | D | 1.7e-07 | 2.8e-07 | 2.6e-07 | 2.367E-07 | 5.859E-08 | 1.700E-07 | 2.800E-07 |
| 21240 | 1062 | M1062V | D | 3.9e-07 | 2.3e-07 | 3e-07 | 3.067E-07 | 8.021E-08 | 2.300E-07 | 3.900E-07 |
| 21241 | 1063 | T1063A | D | 3.1e-07 | 1.5e-07 | 2.8e-07 | 2.467E-07 | 8.505E-08 | 1.500E-07 | 3.100E-07 |
| 21242 | 1063 | T1063R | D | 2e-07 | 3.6e-07 | 4.4e-07 | 3.333E-07 | 1.222E-07 | 2.000E-07 | 4.400E-07 |
| 21243 | 1063 | T1063N | D | 5.1e-07 | 2e-07 | 1.6e-07 | 2.900E-07 | 1.916E-07 | 1.600E-07 | 5.100E-07 |
| 21244 | 1063 | T1063D | D | 1.9e-07 | 1.9e-07 | 1.9e-07 | 1.900E-07 | 0.000E+00 | 1.900E-07 | 1.900E-07 |
| 21245 | 1063 | T1063C | D | 1.3e-07 | 2e-07 | 2.4e-07 | 1.900E-07 | 5.568E-08 | 1.300E-07 | 2.400E-07 |
| 21246 | 1063 | T1063E | D | 7.4e-07 | 2.2e-07 | 1.5e-07 | 3.700E-07 | 3.223E-07 | 1.500E-07 | 7.400E-07 |
| 21247 | 1063 | T1063Q | D | 4.2e-07 | 3.1e-07 | 2.9e-07 | 3.400E-07 | 7.000E-08 | 2.900E-07 | 4.200E-07 |
| 21248 | 1063 | T1063G | D | 3.2e-07 | 2.8e-07 | 4.8e-07 | 3.600E-07 | 1.058E-07 | 2.800E-07 | 4.800E-07 |
| 21249 | 1063 | T1063H | D | 1.3e-07 | 2.6e-07 | 2.3e-07 | 2.067E-07 | 6.807E-08 | 1.300E-07 | 2.600E-07 |

| | | | | | | | | | | |
|---|---|---|---|---|---|---|---|---|---|---|
| 21250 | 1063 | T1063I | D | 2.1e-07 | 2e-07 | 9.9e-08 | 1.697E-07 | 6.140E-08 | 9.900E-08 | 2.100E-07 |
| 21251 | 1063 | T1063L | D | 3.9e-07 | 1.2e-07 | 4.8e-07 | 3.300E-07 | 1.873E-07 | 1.200E-07 | 4.800E-07 |
| 21252 | 1063 | T1063K | D | 2.2e-07 | 1.2e-07 | 3.5e-07 | 2.300E-07 | 1.153E-07 | 1.200E-07 | 3.500E-07 |
| 21253 | 1063 | T1063M | D | 3.1e-07 | 5.6e-07 | 2.4e-07 | 3.700E-07 | 1.682E-07 | 2.400E-07 | 5.600E-07 |
| 21254 | 1063 | T1063F | D | 5.7e-07 | 1.6e-07 | 1.8e-07 | 3.033E-07 | 2.312E-07 | 1.600E-07 | 5.700E-07 |
| 21255 | 1063 | T1063P | D | 4.9e-07 | 1.1e-07 | 2.3e-07 | 2.767E-07 | 1.943E-07 | 1.100E-07 | 4.900E-07 |
| 21256 | 1063 | T1063S | D | 3.4e-07 | 1.6e-07 | 2.6e-07 | 2.533E-07 | 9.018E-08 | 1.600E-07 | 3.400E-07 |
| 21257 | 1063 | T1063T | D | 1.5e-07 | 3.4e-07 | 3.4e-07 | 2.767E-07 | 1.097E-07 | 1.500E-07 | 3.400E-07 |
| 21258 | 1063 | T1063W | D | 2.2e-07 | 3.4e-07 | 1.4e-07 | 2.333E-07 | 1.007E-07 | 1.400E-07 | 3.400E-07 |
| 21259 | 1063 | T1063Y | D | 2.8e-07 | 2.7e-07 | 1.5e-07 | 2.333E-07 | 7.234E-08 | 1.500E-07 | 2.800E-07 |
| 21260 | 1063 | T1063V | D | 3.8e-07 | 2.2e-07 | 2.8e-07 | 2.933E-07 | 8.083E-08 | 2.200E-07 | 3.800E-07 |
| 21261 | 1064 | Q1064A | D | 3.3e-07 | 2.8e-07 | 3.1e-07 | 3.067E-07 | 2.517E-08 | 2.800E-07 | 3.300E-07 |
| 21262 | 1064 | Q1064R | D | 1.9e-07 | 4.8e-07 | 2.9e-07 | 3.200E-07 | 1.473E-07 | 1.900E-07 | 4.800E-07 |
| 21263 | 1064 | Q1064N | D | 1.9e-07 | 1.7e-07 | 1.9e-07 | 1.833E-07 | 1.155E-08 | 1.700E-07 | 1.900E-07 |
| 21264 | 1064 | Q1064D | D | 3.1e-07 | 3.6e-07 | 2e-07 | 2.900E-07 | 8.185E-08 | 2.000E-07 | 3.600E-07 |
| 21265 | 1064 | Q1064C | D | 5.4e-07 | 2.1e-07 | 1.2e-07 | 2.900E-07 | 2.211E-07 | 1.200E-07 | 5.400E-07 |
| 21266 | 1064 | Q1064E | D | 2.6e-07 | 2.3e-07 | 2.7e-07 | 2.533E-07 | 2.082E-08 | 2.300E-07 | 2.700E-07 |
| 21267 | 1064 | Q1064Q | D | 3.1e-07 | 3.1e-07 | 2.8e-07 | 3.000E-07 | 1.732E-08 | 2.800E-07 | 3.100E-07 |
| 21268 | 1064 | Q1064G | D | 3.4e-07 | 1.8e-07 | 2.7e-07 | 2.633E-07 | 8.021E-08 | 1.800E-07 | 3.400E-07 |
| 21269 | 1064 | Q1064H | D | 2.7e-07 | 1.7e-07 | 1.4e-07 | 1.933E-07 | 6.807E-08 | 1.400E-07 | 2.700E-07 |
| 21270 | 1064 | Q1064I | D | 3.6e-07 | 2.8e-07 | 2.7e-07 | 3.033E-07 | 4.933E-08 | 2.700E-07 | 3.600E-07 |
| 21271 | 1064 | Q1064L | D | 2.3e-07 | 3.1e-07 | 3e-07 | 2.800E-07 | 4.359E-08 | 2.300E-07 | 3.100E-07 |
| 21272 | 1064 | Q1064K | D | 3.9e-07 | 2.4e-07 | 2.6e-07 | 2.967E-07 | 8.145E-08 | 2.400E-07 | 3.900E-07 |
| 21273 | 1064 | Q1064M | D | 1.8e-07 | 3.2e-07 | 1.7e-07 | 2.233E-07 | 8.386E-08 | 1.700E-07 | 3.200E-07 |
| 21274 | 1064 | Q1064F | D | 3.5e-07 | 1.6e-07 | 3.8e-07 | 2.967E-07 | 1.193E-07 | 1.600E-07 | 3.800E-07 |
| 21275 | 1064 | Q1064P | D | 2.8e-07 | 3.2e-07 | 2e-07 | 2.667E-07 | 6.110E-08 | 2.000E-07 | 3.200E-07 |
| 21276 | 1064 | Q1064S | D | 1.1e-07 | 1.1e-07 | 4e-07 | 2.067E-07 | 1.674E-07 | 1.100E-07 | 4.000E-07 |
| 21277 | 1064 | Q1064T | D | 4.8e-07 | 3.7e-07 | 2.2e-07 | 3.567E-07 | 1.305E-07 | 2.200E-07 | 4.800E-07 |
| 21278 | 1064 | Q1064W | D | 3.7e-07 | 2.3e-07 | 4e-07 | 3.333E-07 | 9.074E-08 | 2.300E-07 | 4.000E-07 |
| 21279 | 1064 | Q1064Y | D | 1.7e-07 | 2.1e-07 | 2.4e-07 | 2.067E-07 | 3.512E-08 | 1.700E-07 | 2.400E-07 |
| 21280 | 1064 | Q1064V | D | 2.2e-07 | 4.4e-07 | 4.2e-07 | 3.600E-07 | 1.217E-07 | 2.200E-07 | 4.400E-07 |
| 21281 | 1065 | S1065A | D | 3.6e-07 | 1.8e-07 | 1.3e-07 | 2.233E-07 | 1.210E-07 | 1.300E-07 | 3.600E-07 |
| 21282 | 1065 | S1065R | D | 3.6e-07 | 1e-07 | 2.5e-07 | 2.367E-07 | 1.305E-07 | 1.000E-07 | 3.600E-07 |
| 21283 | 1065 | S1065N | D | 1.5e-07 | 1.8e-07 | 3.6e-07 | 2.300E-07 | 1.136E-07 | 1.500E-07 | 3.600E-07 |
| 21284 | 1065 | S1065D | D | 1.4e-07 | 2.1e-07 | 4.9e-07 | 2.800E-07 | 1.852E-07 | 1.400E-07 | 4.900E-07 |
| 21285 | 1065 | S1065C | D | 2.5e-07 | 7.7e-07 | 2e-07 | 4.067E-07 | 3.156E-07 | 2.000E-07 | 7.700E-07 |
| 21286 | 1065 | S1065E | D | 5.9e-07 | 2e-07 | 1.2e-07 | 3.033E-07 | 2.515E-07 | 1.200E-07 | 5.900E-07 |
| 21287 | 1065 | S1065Q | D | 1.8e-07 | 3.2e-07 | 2.6e-07 | 2.533E-07 | 7.024E-08 | 1.800E-07 | 3.200E-07 |
| 21288 | 1065 | S1065G | D | 6.6e-08 | 1.9e-07 | 4e-07 | 2.187E-07 | 1.688E-07 | 6.600E-08 | 4.000E-07 |
| 21289 | 1065 | S1065H | D | 1.8e-07 | 2.3e-07 | 3.1e-07 | 2.400E-07 | 6.557E-08 | 1.800E-07 | 3.100E-07 |
| 21290 | 1065 | S1065I | D | 3.7e-07 | 1.8e-07 | 3.2e-07 | 2.900E-07 | 9.849E-08 | 1.800E-07 | 3.700E-07 |
| 21291 | 1065 | S1065L | D | 4e-07 | 4e-07 | 2e-07 | 3.333E-07 | 1.155E-07 | 2.000E-07 | 4.000E-07 |
| 21292 | 1065 | S1065K | D | 2e-07 | 1.7e-07 | 4.5e-07 | 2.733E-07 | 1.537E-07 | 1.700E-07 | 4.500E-07 |
| 21293 | 1065 | S1065M | D | 2.2e-07 | 4.2e-07 | 1.7e-07 | 2.700E-07 | 1.323E-07 | 1.700E-07 | 4.200E-07 |
| 21294 | 1065 | S1065F | D | 4e-07 | 4e-07 | 2.3e-07 | 3.433E-07 | 9.815E-08 | 2.300E-07 | 4.000E-07 |
| 21295 | 1065 | S1065P | D | 9.3e-08 | 3.2e-07 | 9.2e-07 | 4.443E-07 | 4.273E-07 | 9.300E-08 | 9.200E-07 |
| 21296 | 1065 | S1065S | D | 2e-07 | 4.9e-07 | 1.9e-07 | 2.933E-07 | 1.704E-07 | 1.900E-07 | 4.900E-07 |
| 21297 | 1065 | S1065T | D | 2.3e-07 | 7.1e-08 | 1.4e-07 | 1.470E-07 | 7.973E-08 | 7.100E-08 | 2.300E-07 |
| 21298 | 1065 | S1065W | D | 2.2e-07 | 2.6e-07 | 2.9e-07 | 2.567E-07 | 3.512E-08 | 2.200E-07 | 2.900E-07 |
| 21299 | 1065 | S1065Y | D | 1.6e-07 | 4.4e-07 | 1.4e-07 | 2.467E-07 | 1.677E-07 | 1.400E-07 | 4.400E-07 |
| 21300 | 1065 | S1065V | D | 5.1e-07 | 2.4e-07 | 2.2e-07 | 3.233E-07 | 1.620E-07 | 2.200E-07 | 5.100E-07 |
| 21301 | 1066 | P1066A | D | 1.8e-07 | 2.3e-07 | 2.8e-07 | 2.300E-07 | 5.000E-08 | 1.800E-07 | 2.800E-07 |
| 21302 | 1066 | P1066R | D | 3.6e-07 | 1.7e-07 | 3.4e-07 | 2.900E-07 | 1.044E-07 | 1.700E-07 | 3.600E-07 |
| 21303 | 1066 | P1066N | D | 3e-07 | 1.3e-07 | 1.9e-07 | 2.067E-07 | 8.622E-08 | 1.300E-07 | 3.000E-07 |
| 21304 | 1066 | P1066D | D | 5.1e-07 | 3.8e-07 | 3.7e-07 | 4.200E-07 | 7.810E-08 | 3.700E-07 | 5.100E-07 |
| 21305 | 1066 | P1066C | D | 2.5e-07 | 1.9e-07 | 1.6e-07 | 2.000E-07 | 4.583E-08 | 1.600E-07 | 2.500E-07 |
| 21306 | 1066 | P1066E | D | 3.8e-07 | 2.8e-07 | 4.7e-07 | 3.767E-07 | 9.504E-08 | 2.800E-07 | 4.700E-07 |
| 21307 | 1066 | P1066Q | D | 1.3e-07 | 1.7e-07 | 2.7e-07 | 1.900E-07 | 7.211E-08 | 1.300E-07 | 2.700E-07 |
| 21308 | 1066 | P1066G | D | 5.8e-07 | 4.1e-07 | 2.5e-07 | 4.133E-07 | 1.650E-07 | 2.500E-07 | 5.800E-07 |
| 21309 | 1066 | P1066H | D | 2.2e-07 | 1.2e-07 | 2.6e-07 | 2.000E-07 | 7.211E-08 | 1.200E-07 | 2.600E-07 |
| 21310 | 1066 | P1066I | D | 1.9e-07 | 1.7e-07 | 3.8e-07 | 2.467E-07 | 1.159E-07 | 1.700E-07 | 3.800E-07 |
| 21311 | 1066 | P1066L | D | 3.3e-07 | 3.9e-07 | 3.3e-07 | 3.500E-07 | 3.464E-08 | 3.300E-07 | 3.900E-07 |
| 21312 | 1066 | P1066K | D | 8e-08 | 1.9e-07 | 1.7e-07 | 1.467E-07 | 5.859E-08 | 8.000E-08 | 1.900E-07 |
| 21313 | 1066 | P1066M | D | 2.8e-07 | 2.5e-07 | 6.4e-07 | 3.900E-07 | 2.170E-07 | 2.500E-07 | 6.400E-07 |
| 21314 | 1066 | P1066F | D | 1.8e-07 | 1.9e-07 | 3.4e-07 | 2.367E-07 | 8.963E-08 | 1.800E-07 | 3.400E-07 |
| 21315 | 1066 | P1066P | D | 2.4e-07 | 2.6e-07 | 2.6e-07 | 2.533E-07 | 1.155E-08 | 2.400E-07 | 2.600E-07 |
| 21316 | 1066 | P1066S | D | 3e-07 | 2.4e-07 | 5.4e-07 | 3.600E-07 | 1.587E-07 | 2.400E-07 | 5.400E-07 |
| 21317 | 1066 | P1066T | D | 6.3e-07 | 6.4e-08 | 2.8e-07 | 3.247E-07 | 2.856E-07 | 6.400E-08 | 6.300E-07 |
| 21318 | 1066 | P1066W | D | 4.6e-07 | 2.7e-07 | 3.1e-07 | 3.467E-07 | 1.002E-07 | 2.700E-07 | 4.600E-07 |
| 21319 | 1066 | P1066Y | D | 4.7e-07 | 4.5e-07 | 3.1e-07 | 4.100E-07 | 8.718E-08 | 3.100E-07 | 4.700E-07 |
| 21320 | 1066 | P1066V | D | 2.6e-07 | 1.9e-07 | 6.2e-07 | 3.567E-07 | 2.307E-07 | 1.900E-07 | 6.200E-07 |
| 21321 | 1067 | S1067A | D | 2.2e-07 | 3.9e-07 | 5.5e-07 | 3.867E-07 | 1.650E-07 | 2.200E-07 | 5.500E-07 |
| 21322 | 1067 | S1067R | D | 4.3e-07 | 3.3e-07 | 2.1e-07 | 3.233E-07 | 1.102E-07 | 2.100E-07 | 4.300E-07 |
| 21323 | 1067 | S1067N | D | 1.1e-07 | 1.6e-07 | 1.3e-07 | 1.333E-07 | 2.517E-08 | 1.100E-07 | 1.600E-07 |
| 21324 | 1067 | S1067D | D | 3.8e-07 | 2.8e-07 | 2.8e-07 | 3.133E-07 | 5.774E-08 | 2.800E-07 | 3.800E-07 |
| 21325 | 1067 | S1067C | D | 3.9e-07 | 3.5e-07 | 2.2e-07 | 3.200E-07 | 8.888E-08 | 2.200E-07 | 3.900E-07 |
| 21326 | 1067 | S1067E | D | 5.6e-07 | 1.9e-07 | 1.8e-07 | 3.100E-07 | 2.166E-07 | 1.800E-07 | 5.600E-07 |
| 21327 | 1067 | S1067Q | D | 3.1e-07 | 1.6e-07 | 2.1e-07 | 2.267E-07 | 7.638E-08 | 1.600E-07 | 3.100E-07 |
| 21328 | 1067 | S1067G | D | 1.2e-07 | 2.9e-07 | 1.8e-07 | 1.967E-07 | 8.622E-08 | 1.200E-07 | 2.900E-07 |
| 21329 | 1067 | S1067H | D | 1.8e-07 | 2e-07 | 3.3e-07 | 2.367E-07 | 8.145E-08 | 1.800E-07 | 3.300E-07 |
| 21330 | 1067 | S1067I | D | 2.1e-07 | 2.4e-07 | 1.7e-07 | 2.067E-07 | 3.512E-08 | 1.700E-07 | 2.400E-07 |
| 21331 | 1067 | S1067L | D | 9.4e-08 | 5.9e-08 | 1.7e-07 | 1.077E-07 | 5.675E-08 | 5.900E-08 | 1.700E-07 |
| 21332 | 1067 | S1067K | D | 1.5e-07 | 1.9e-07 | 1.2e-07 | 1.533E-07 | 3.512E-08 | 1.200E-07 | 1.900E-07 |
| 21333 | 1067 | S1067M | D | 1.6e-07 | 1.3e-07 | 5.1e-07 | 2.667E-07 | 2.113E-07 | 1.300E-07 | 5.100E-07 |
| 21334 | 1067 | S1067F | D | 3.6e-07 | 3.2e-07 | 1.4e-07 | 2.733E-07 | 1.172E-07 | 1.400E-07 | 3.600E-07 |

| | | | | | | | | | | |
|---|---|---|---|---|---|---|---|---|---|---|
| 21335 | 1067 | S1067P | D | 1.3e-07 | 3.3e-07 | 1.7e-07 | 2.100E-07 | 1.058E-07 | 1.300E-07 | 3.300E-07 |
| 21336 | 1067 | S1067S | D | 1.7e-07 | 2.2e-07 | 3.2e-07 | 2.367E-07 | 7.638E-08 | 1.700E-07 | 3.200E-07 |
| 21337 | 1067 | S1067T | D | 4.3e-07 | 3.4e-07 | 2.5e-07 | 3.400E-07 | 9.000E-08 | 2.500E-07 | 4.300E-07 |
| 21338 | 1067 | S1067W | D | 4e-07 | 4.1e-07 | 6.7e-07 | 4.933E-07 | 1.531E-07 | 4.000E-07 | 6.700E-07 |
| 21339 | 1067 | S1067Y | D | 3.7e-07 | 1.9e-07 | 3e-07 | 2.867E-07 | 9.074E-08 | 1.900E-07 | 3.700E-07 |
| 21340 | 1067 | S1067V | D | 3.8e-07 | 2e-07 | 2.4e-07 | 2.733E-07 | 9.452E-08 | 2.000E-07 | 3.800E-07 |
| 21341 | 1068 | S1068A | D | 2.8e-07 | 1.3e-07 | 1.9e-07 | 2.000E-07 | 7.550E-08 | 1.300E-07 | 2.800E-07 |
| 21342 | 1068 | S1068R | D | 6.6e-07 | 2.5e-07 | 9.8e-08 | 3.360E-07 | 2.907E-07 | 9.800E-08 | 6.600E-07 |
| 21343 | 1068 | S1068N | D | 3.3e-07 | 1.4e-07 | 2.5e-07 | 2.400E-07 | 9.539E-08 | 1.400E-07 | 3.300E-07 |
| 21344 | 1068 | S1068D | D | 2.7e-07 | 4.3e-07 | 3.5e-07 | 3.500E-07 | 8.000E-08 | 2.700E-07 | 4.300E-07 |
| 21345 | 1068 | S1068C | D | 2e-07 | 1.4e-07 | 3e-07 | 2.133E-07 | 8.083E-08 | 1.400E-07 | 3.000E-07 |
| 21346 | 1068 | S1068E | D | 3.3e-07 | 5.6e-07 | 4.6e-07 | 4.500E-07 | 1.153E-07 | 3.300E-07 | 5.600E-07 |
| 21347 | 1068 | S1068Q | D | 2e-07 | 1.4e-07 | 2.8e-07 | 2.067E-07 | 7.024E-08 | 1.400E-07 | 2.800E-07 |
| 21348 | 1068 | S1068G | D | 3.7e-07 | 4e-07 | 2.3e-07 | 3.333E-07 | 9.074E-08 | 2.300E-07 | 4.000E-07 |
| 21349 | 1068 | S1068H | D | 4.6e-07 | 3e-07 | 1.8e-07 | 3.133E-07 | 1.405E-07 | 1.800E-07 | 4.600E-07 |
| 21350 | 1068 | S1068I | D | 4.2e-07 | 1.3e-07 | 2.6e-07 | 2.700E-07 | 1.453E-07 | 1.300E-07 | 4.200E-07 |
| 21351 | 1068 | S1068L | D | 1.6e-07 | 2.2e-07 | 1.9e-07 | 1.900E-07 | 3.000E-08 | 1.600E-07 | 2.200E-07 |
| 21352 | 1068 | S1068K | D | 3.8e-07 | 1.9e-07 | 9.7e-07 | 5.133E-07 | 4.067E-07 | 1.900E-07 | 9.700E-07 |
| 21353 | 1068 | S1068M | D | 3.5e-07 | 1.7e-07 | 1.4e-07 | 2.200E-07 | 1.136E-07 | 1.400E-07 | 3.500E-07 |
| 21354 | 1068 | S1068F | D | 2.5e-07 | 3.3e-07 | 3.8e-07 | 3.200E-07 | 6.557E-08 | 2.500E-07 | 3.800E-07 |
| 21355 | 1068 | S1068P | D | 2.7e-07 | 5e-07 | 2.5e-07 | 3.400E-07 | 1.389E-07 | 2.500E-07 | 5.000E-07 |
| 21356 | 1068 | S1068S | D | 2.3e-07 | 1.5e-07 | 4.6e-07 | 2.800E-07 | 1.609E-07 | 1.500E-07 | 4.600E-07 |
| 21357 | 1068 | S1068T | D | 2.4e-07 | 2.4e-07 | 2.9e-07 | 2.567E-07 | 2.887E-08 | 2.400E-07 | 2.900E-07 |
| 21358 | 1068 | S1068W | D | 5.3e-07 | 2.6e-07 | 6.1e-07 | 4.667E-07 | 1.834E-07 | 2.600E-07 | 6.100E-07 |
| 21359 | 1068 | S1068Y | D | 3.2e-07 | 2.6e-07 | 1.6e-07 | 2.467E-07 | 8.083E-08 | 1.600E-07 | 3.200E-07 |
| 21360 | 1068 | S1068V | D | 2.1e-07 | 1.4e-07 | 2.3e-07 | 1.933E-07 | 4.726E-08 | 1.400E-07 | 2.300E-07 |
| 21361 | 1069 | L1069A | D | 2.3e-07 | 2.1e-07 | 2.6e-07 | 2.333E-07 | 2.517E-08 | 2.100E-07 | 2.600E-07 |
| 21362 | 1069 | L1069R | D | 1.9e-07 | 1.4e-07 | 2.8e-07 | 2.033E-07 | 7.095E-08 | 1.400E-07 | 2.800E-07 |
| 21363 | 1069 | L1069N | D | 1.9e-07 | 1.9e-07 | 1.9e-07 | 1.900E-07 | 0.000E+00 | 1.900E-07 | 1.900E-07 |
| 21364 | 1069 | L1069D | D | 2.9e-07 | 1.7e-07 | 2.5e-07 | 2.367E-07 | 6.110E-08 | 1.700E-07 | 2.900E-07 |
| 21365 | 1069 | L1069C | D | 4.9e-07 | 2.2e-07 | 1.1e-07 | 2.733E-07 | 1.955E-07 | 1.100E-07 | 4.900E-07 |
| 21366 | 1069 | L1069E | D | 2.2e-07 | 2.6e-07 | 2.3e-07 | 2.367E-07 | 2.082E-08 | 2.200E-07 | 2.600E-07 |
| 21367 | 1069 | L1069Q | D | 3.8e-07 | 1.9e-07 | 2.6e-07 | 2.767E-07 | 9.609E-08 | 1.900E-07 | 3.800E-07 |
| 21368 | 1069 | L1069G | D | 1.6e-07 | 2.2e-07 | 7.4e-07 | 3.733E-07 | 3.190E-07 | 1.600E-07 | 7.400E-07 |
| 21369 | 1069 | L1069H | D | 1.6e-07 | 1.7e-07 | 9.9e-08 | 1.430E-07 | 3.843E-08 | 9.900E-08 | 1.700E-07 |
| 21370 | 1069 | L1069I | D | 2.6e-07 | 1.9e-07 | 1.5e-07 | 2.000E-07 | 5.568E-08 | 1.500E-07 | 2.600E-07 |
| 21371 | 1069 | L1069L | D | 1.9e-07 | 3.3e-07 | 3.3e-07 | 2.833E-07 | 8.083E-08 | 1.900E-07 | 3.300E-07 |
| 21372 | 1069 | L1069K | D | 4.4e-07 | 3.9e-07 | 2.7e-07 | 3.667E-07 | 8.737E-08 | 2.700E-07 | 4.400E-07 |
| 21373 | 1069 | L1069M | D | 1.7e-07 | 3.2e-07 | 2.2e-07 | 2.367E-07 | 7.638E-08 | 1.700E-07 | 3.200E-07 |
| 21374 | 1069 | L1069F | D | 9.3e-08 | 3.1e-07 | 4.1e-07 | 2.710E-07 | 1.621E-07 | 9.300E-08 | 4.100E-07 |
| 21375 | 1069 | L1069P | D | 3.5e-07 | 2.3e-07 | 1.6e-07 | 2.467E-07 | 9.609E-08 | 1.600E-07 | 3.500E-07 |
| 21376 | 1069 | L1069S | D | 2e-07 | 2e-07 | 4e-07 | 2.667E-07 | 1.155E-07 | 2.000E-07 | 4.000E-07 |
| 21377 | 1069 | L1069T | D | 1.9e-07 | 2.4e-07 | 9.2e-08 | 1.740E-07 | 7.529E-08 | 9.200E-08 | 2.400E-07 |
| 21378 | 1069 | L1069W | D | 3.9e-07 | 1.7e-07 | 3.3e-07 | 2.967E-07 | 1.137E-07 | 1.700E-07 | 3.900E-07 |
| 21379 | 1069 | L1069Y | D | 3.4e-07 | 2.1e-07 | 1.4e-07 | 2.300E-07 | 1.015E-07 | 1.400E-07 | 3.400E-07 |
| 21380 | 1069 | L1069V | D | 6.1e-07 | 2e-07 | 2e-07 | 3.367E-07 | 2.367E-07 | 2.000E-07 | 6.100E-07 |
| 21381 | 1070 | S1070A | D | 2.2e-07 | 1.7e-07 | 2.8e-07 | 2.233E-07 | 5.508E-08 | 1.700E-07 | 2.800E-07 |
| 21382 | 1070 | S1070R | D | 5.3e-07 | 3.9e-07 | 3.8e-07 | 4.333E-07 | 8.386E-08 | 3.800E-07 | 5.300E-07 |
| 21383 | 1070 | S1070N | D | 2.6e-07 | 2.7e-07 | 2.5e-07 | 2.600E-07 | 1.000E-08 | 2.500E-07 | 2.700E-07 |
| 21384 | 1070 | S1070D | D | 2.9e-07 | 3.3e-07 | 2.5e-07 | 2.900E-07 | 4.000E-08 | 2.500E-07 | 3.300E-07 |
| 21385 | 1070 | S1070C | D | 3.7e-07 | 1.4e-07 | 3.4e-07 | 2.833E-07 | 1.250E-07 | 1.400E-07 | 3.700E-07 |
| 21386 | 1070 | S1070E | D | 2.2e-07 | 8.4e-07 | 3.4e-07 | 4.667E-07 | 3.288E-07 | 2.200E-07 | 8.400E-07 |
| 21387 | 1070 | S1070Q | D | 2.1e-07 | 2.7e-07 | 2.3e-07 | 2.367E-07 | 3.055E-08 | 2.100E-07 | 2.700E-07 |
| 21388 | 1070 | S1070G | D | 2e-07 | 1.2e-07 | 4.1e-07 | 2.433E-07 | 1.498E-07 | 1.200E-07 | 4.100E-07 |
| 21389 | 1070 | S1070H | D | 2.3e-07 | 2.1e-07 | 2.3e-07 | 2.233E-07 | 1.155E-08 | 2.100E-07 | 2.300E-07 |
| 21390 | 1070 | S1070I | D | 1.5e-07 | 7.5e-08 | 2.8e-07 | 1.683E-07 | 1.037E-07 | 7.500E-08 | 2.800E-07 |
| 21391 | 1070 | S1070L | D | 8.3e-07 | 1.3e-07 | 9.8e-08 | 3.527E-07 | 4.137E-07 | 9.800E-08 | 8.300E-07 |
| 21392 | 1070 | S1070K | D | 3.7e-07 | 1.2e-07 | 6.6e-07 | 3.833E-07 | 2.702E-07 | 1.200E-07 | 6.600E-07 |
| 21393 | 1070 | S1070M | D | 3.2e-07 | 2.3e-07 | 2.1e-07 | 2.533E-07 | 5.859E-08 | 2.100E-07 | 3.200E-07 |
| 21394 | 1070 | S1070F | D | 2.7e-07 | 2.7e-07 | 4.2e-07 | 3.200E-07 | 8.660E-08 | 2.700E-07 | 4.200E-07 |
| 21395 | 1070 | S1070P | D | 1.5e-07 | 1.3e-07 | 4.6e-07 | 2.467E-07 | 1.850E-07 | 1.300E-07 | 4.600E-07 |
| 21396 | 1070 | S1070S | D | 3.1e-07 | 2e-07 | 2.1e-07 | 2.400E-07 | 6.083E-08 | 2.000E-07 | 3.100E-07 |
| 21397 | 1070 | S1070T | D | 3.3e-07 | 4.7e-07 | 3.5e-07 | 3.833E-07 | 7.572E-08 | 3.300E-07 | 4.700E-07 |
| 21398 | 1070 | S1070W | D | 3.5e-07 | 1.7e-07 | 1.5e-07 | 2.233E-07 | 1.102E-07 | 1.500E-07 | 3.500E-07 |
| 21399 | 1070 | S1070Y | D | 3.1e-07 | 3.7e-07 | 1.6e-07 | 2.800E-07 | 1.082E-07 | 1.600E-07 | 3.700E-07 |
| 21400 | 1070 | S1070V | D | 1.8e-07 | 4.3e-07 | 1.9e-07 | 2.667E-07 | 1.415E-07 | 1.800E-07 | 4.300E-07 |
| 21401 | 1071 | A1071A | D | 1e-07 | 1.4e-07 | 3e-07 | 1.800E-07 | 1.058E-07 | 1.000E-07 | 3.000E-07 |
| 21402 | 1071 | A1071R | D | 2.9e-07 | 3.2e-07 | 1.4e-07 | 2.500E-07 | 9.644E-08 | 1.400E-07 | 3.200E-07 |
| 21403 | 1071 | A1071N | D | 2.6e-07 | 3.4e-07 | 1.3e-07 | 2.433E-07 | 1.060E-07 | 1.300E-07 | 3.400E-07 |
| 21404 | 1071 | A1071D | D | 1.5e-07 | 2.3e-07 | 6.5e-07 | 3.433E-07 | 2.686E-07 | 1.500E-07 | 6.500E-07 |
| 21405 | 1071 | A1071C | D | 3.2e-07 | 4.1e-07 | 2.6e-07 | 3.300E-07 | 7.550E-08 | 2.600E-07 | 4.100E-07 |
| 21406 | 1071 | A1071E | D | 1.9e-07 | 2e-07 | 4.1e-07 | 2.667E-07 | 1.242E-07 | 1.900E-07 | 4.100E-07 |
| 21407 | 1071 | A1071Q | D | 3e-07 | 2.3e-07 | 2.3e-07 | 2.533E-07 | 4.041E-08 | 2.300E-07 | 3.000E-07 |
| 21408 | 1071 | A1071G | D | 3.2e-07 | 1e-07 | 2.4e-07 | 2.200E-07 | 1.114E-07 | 1.000E-07 | 3.200E-07 |
| 21409 | 1071 | A1071H | D | 1.8e-07 | 1.9e-07 | 1.5e-07 | 1.733E-07 | 2.082E-08 | 1.500E-07 | 1.900E-07 |
| 21410 | 1071 | A1071I | D | 1.7e-07 | 3.5e-07 | 3e-07 | 2.733E-07 | 9.292E-08 | 1.700E-07 | 3.500E-07 |
| 21411 | 1071 | A1071L | D | 3.4e-07 | 3.5e-07 | 2.9e-07 | 3.267E-07 | 3.215E-08 | 2.900E-07 | 3.500E-07 |
| 21412 | 1071 | A1071K | D | 1.9e-07 | 1.6e-07 | 1.4e-07 | 1.633E-07 | 2.517E-08 | 1.400E-07 | 1.900E-07 |
| 21413 | 1071 | A1071M | D | 1.5e-07 | 5.4e-07 | 5.1e-07 | 4.000E-07 | 2.170E-07 | 1.500E-07 | 5.400E-07 |
| 21414 | 1071 | A1071F | D | 2.4e-07 | 1.8e-07 | 2.3e-07 | 2.167E-07 | 3.215E-08 | 1.800E-07 | 2.400E-07 |
| 21415 | 1071 | A1071P | D | 2.1e-07 | 1.5e-07 | 2.7e-07 | 2.100E-07 | 6.000E-08 | 1.500E-07 | 2.700E-07 |
| 21416 | 1071 | A1071S | D | 5.2e-07 | 2.1e-07 | 3e-07 | 3.433E-07 | 1.595E-07 | 2.100E-07 | 5.200E-07 |
| 21417 | 1071 | A1071T | D | 2.9e-07 | 6.4e-07 | 2.4e-07 | 3.900E-07 | 2.179E-07 | 2.400E-07 | 6.400E-07 |
| 21418 | 1071 | A1071W | D | 1.5e-07 | 2.5e-07 | 1.7e-07 | 1.900E-07 | 5.292E-08 | 1.500E-07 | 2.500E-07 |
| 21419 | 1071 | A1071Y | D | 2.6e-07 | 2.2e-07 | 2.5e-07 | 2.433E-07 | 2.082E-08 | 2.200E-07 | 2.600E-07 |

| | | | | | | | | | | |
|---|---|---|---|---|---|---|---|---|---|---|
| 21420 | 1071 | A1071V | D | 3.6e-07 | 2.9e-07 | 1e-07 | 2.500E-07 | 1.345E-07 | 1.000E-07 | 3.600E-07 |
| 21421 | 1072 | S1072A | D | 2e-07 | 3.2e-07 | 3.3e-07 | 2.833E-07 | 7.234E-08 | 2.000E-07 | 3.300E-07 |
| 21422 | 1072 | S1072R | D | 1.4e-07 | 1.5e-07 | 2.5e-07 | 1.800E-07 | 6.083E-08 | 1.400E-07 | 2.500E-07 |
| 21423 | 1072 | S1072N | D | 1.9e-07 | 3.3e-07 | 1.7e-07 | 2.300E-07 | 8.718E-08 | 1.700E-07 | 3.300E-07 |
| 21424 | 1072 | S1072D | D | 2.1e-07 | 2.7e-07 | 1.5e-07 | 2.100E-07 | 6.000E-08 | 1.500E-07 | 2.700E-07 |
| 21425 | 1072 | S1072C | D | 2.9e-07 | 1.7e-07 | 1.5e-07 | 2.033E-07 | 7.572E-08 | 1.500E-07 | 2.900E-07 |
| 21426 | 1072 | S1072E | D | 2.6e-07 | 3.3e-07 | 4e-07 | 3.300E-07 | 7.000E-08 | 2.600E-07 | 4.000E-07 |
| 21427 | 1072 | S1072Q | D | 2.6e-07 | 1.1e-07 | 1.8e-07 | 1.833E-07 | 7.506E-08 | 1.100E-07 | 2.600E-07 |
| 21428 | 1072 | S1072G | D | 4.1e-07 | 2.5e-07 | 3.8e-07 | 3.467E-07 | 8.505E-08 | 2.500E-07 | 4.100E-07 |
| 21429 | 1072 | S1072H | D | 4e-07 | 3.3e-07 | 3.2e-07 | 3.500E-07 | 4.359E-08 | 3.200E-07 | 4.000E-07 |
| 21430 | 1072 | S1072I | D | 9.8e-08 | 1.9e-07 | 2.6e-07 | 1.827E-07 | 8.125E-08 | 9.800E-08 | 2.600E-07 |
| 21431 | 1072 | S1072L | D | 2.1e-07 | 2.4e-07 | 3.2e-07 | 2.567E-07 | 5.686E-08 | 2.100E-07 | 3.200E-07 |
| 21432 | 1072 | S1072K | D | 2.3e-07 | 2.5e-07 | 1.9e-07 | 2.233E-07 | 3.055E-08 | 1.900E-07 | 2.500E-07 |
| 21433 | 1072 | S1072M | D | 3e-07 | 2e-07 | 2.7e-07 | 2.567E-07 | 5.132E-08 | 2.000E-07 | 3.000E-07 |
| 21434 | 1072 | S1072F | D | 5.5e-07 | 2.6e-07 | 1.9e-07 | 3.333E-07 | 1.909E-07 | 1.900E-07 | 5.500E-07 |
| 21435 | 1072 | S1072P | D | 2.5e-07 | 3.5e-07 | 1.9e-07 | 2.633E-07 | 8.083E-08 | 1.900E-07 | 3.500E-07 |
| 21436 | 1072 | S1072S | D | 3.6e-07 | 2e-07 | 1.5e-07 | 2.367E-07 | 1.097E-07 | 1.500E-07 | 3.600E-07 |
| 21437 | 1072 | S1072T | D | 2.3e-07 | 2.7e-07 | 1.8e-07 | 2.267E-07 | 4.509E-08 | 1.800E-07 | 2.700E-07 |
| 21438 | 1072 | S1072W | D | 6e-07 | 3.1e-07 | 3.3e-07 | 4.133E-07 | 1.620E-07 | 3.100E-07 | 6.000E-07 |
| 21439 | 1072 | S1072Y | D | 2.8e-07 | 2.2e-07 | 3.8e-07 | 2.933E-07 | 8.083E-08 | 2.200E-07 | 3.800E-07 |
| 21440 | 1072 | S1072V | D | 1.7e-07 | 2.9e-07 | 3.1e-07 | 2.567E-07 | 7.572E-08 | 1.700E-07 | 3.100E-07 |
| 21441 | 1073 | V1073A | D | 5.5e-07 | 1.9e-07 | 2.3e-07 | 3.233E-07 | 1.973E-07 | 1.900E-07 | 5.500E-07 |
| 21442 | 1073 | V1073R | D | 3e-07 | 4.7e-07 | 1.8e-07 | 3.167E-07 | 1.457E-07 | 1.800E-07 | 4.700E-07 |
| 21443 | 1073 | V1073N | D | 1.6e-07 | 2.7e-07 | 1e-07 | 1.767E-07 | 8.622E-08 | 1.000E-07 | 2.700E-07 |
| 21444 | 1073 | V1073D | D | 2.5e-07 | 3e-07 | 3.4e-07 | 2.967E-07 | 4.509E-08 | 2.500E-07 | 3.400E-07 |
| 21445 | 1073 | V1073C | D | 1.4e-07 | 7e-07 | 2.5e-07 | 3.633E-07 | 2.967E-07 | 1.400E-07 | 7.000E-07 |
| 21446 | 1073 | V1073E | D | 1.8e-07 | 2.8e-07 | 1.8e-07 | 2.133E-07 | 5.774E-08 | 1.800E-07 | 2.800E-07 |
| 21447 | 1073 | V1073Q | D | 1.3e-07 | 3e-07 | 2.9e-07 | 2.400E-07 | 9.539E-08 | 1.300E-07 | 3.000E-07 |
| 21448 | 1073 | V1073G | D | 2e-07 | 3.7e-07 | 3.1e-07 | 2.933E-07 | 8.622E-08 | 2.000E-07 | 3.700E-07 |
| 21449 | 1073 | V1073H | D | 3.1e-07 | 3e-07 | 1.9e-07 | 2.667E-07 | 6.658E-08 | 1.900E-07 | 3.100E-07 |
| 21450 | 1073 | V1073I | D | 3.8e-07 | 2.8e-07 | 2.2e-07 | 2.933E-07 | 8.083E-08 | 2.200E-07 | 3.800E-07 |
| 21451 | 1073 | V1073L | D | 2.7e-07 | 1.6e-07 | 2.3e-07 | 2.200E-07 | 5.568E-08 | 1.600E-07 | 2.700E-07 |
| 21452 | 1073 | V1073K | D | 2.8e-07 | 1.6e-07 | 2.1e-07 | 2.167E-07 | 6.028E-08 | 1.600E-07 | 2.800E-07 |
| 21453 | 1073 | V1073M | D | 2.9e-07 | 2.1e-07 | 2.6e-07 | 2.533E-07 | 4.041E-08 | 2.100E-07 | 2.900E-07 |
| 21454 | 1073 | V1073F | D | 2.1e-07 | 1.8e-07 | 3.2e-07 | 2.367E-07 | 7.371E-08 | 1.800E-07 | 3.200E-07 |
| 21455 | 1073 | V1073P | D | 1.2e-07 | 3.4e-07 | 3.7e-07 | 2.767E-07 | 1.365E-07 | 1.200E-07 | 3.700E-07 |
| 21456 | 1073 | V1073S | D | 2.5e-07 | 4.5e-07 | 6e-07 | 4.333E-07 | 1.756E-07 | 2.500E-07 | 6.000E-07 |
| 21457 | 1073 | V1073T | D | 1.2e-07 | 2.1e-07 | 2.6e-07 | 1.967E-07 | 7.095E-08 | 1.200E-07 | 2.600E-07 |
| 21458 | 1073 | V1073W | D | 1.3e-07 | 3.1e-07 | 2.2e-07 | 2.200E-07 | 9.000E-08 | 1.300E-07 | 3.100E-07 |
| 21459 | 1073 | V1073Y | D | 3.8e-07 | 1.8e-07 | 8.9e-08 | 2.163E-07 | 1.489E-07 | 8.900E-08 | 3.800E-07 |
| 21460 | 1073 | V1073V | D | 3.9e-07 | 2e-07 | 2.5e-07 | 2.800E-07 | 9.849E-08 | 2.000E-07 | 3.900E-07 |
| 21461 | 1074 | G1074A | D | 4.7e-07 | 2e-07 | 1.3e-07 | 2.667E-07 | 1.795E-07 | 1.300E-07 | 4.700E-07 |
| 21462 | 1074 | G1074R | D | 2.1e-07 | 5.7e-07 | 2e-07 | 3.267E-07 | 2.108E-07 | 2.000E-07 | 5.700E-07 |
| 21463 | 1074 | G1074N | D | 3e-07 | 2.6e-07 | 2.1e-07 | 2.567E-07 | 4.509E-08 | 2.100E-07 | 3.000E-07 |
| 21464 | 1074 | G1074D | D | 2.3e-07 | 3.5e-07 | 2.4e-07 | 2.733E-07 | 6.658E-08 | 2.300E-07 | 3.500E-07 |
| 21465 | 1074 | G1074C | D | 8.1e-07 | 2.4e-07 | 1.5e-07 | 4.000E-07 | 3.579E-07 | 1.500E-07 | 8.100E-07 |
| 21466 | 1074 | G1074E | D | 2.8e-07 | 3.3e-07 | 3.6e-07 | 3.233E-07 | 4.041E-08 | 2.800E-07 | 3.600E-07 |
| 21467 | 1074 | G1074Q | D | 2.5e-07 | 1.4e-07 | 1.6e-07 | 1.833E-07 | 5.859E-08 | 1.400E-07 | 2.500E-07 |
| 21468 | 1074 | G1074G | D | 1.4e-07 | 3.1e-07 | 2.5e-07 | 2.333E-07 | 8.622E-08 | 1.400E-07 | 3.100E-07 |
| 21469 | 1074 | G1074H | D | 2.8e-07 | 4.5e-07 | 2e-07 | 3.100E-07 | 1.277E-07 | 2.000E-07 | 4.500E-07 |
| 21470 | 1074 | G1074I | D | 2.4e-07 | 3.5e-07 | 3.9e-07 | 3.267E-07 | 7.767E-08 | 2.400E-07 | 3.900E-07 |
| 21471 | 1074 | G1074L | D | 4.6e-07 | 3e-07 | 2.3e-07 | 3.300E-07 | 1.179E-07 | 2.300E-07 | 4.600E-07 |
| 21472 | 1074 | G1074K | D | 1.7e-07 | 2.2e-07 | 2.1e-07 | 2.000E-07 | 2.646E-08 | 1.700E-07 | 2.200E-07 |
| 21473 | 1074 | G1074M | D | 1.5e-07 | 2.8e-07 | 4.3e-07 | 2.867E-07 | 1.401E-07 | 1.500E-07 | 4.300E-07 |
| 21474 | 1074 | G1074F | D | 1.3e-07 | 1.6e-07 | 3.7e-07 | 2.200E-07 | 1.308E-07 | 1.300E-07 | 3.700E-07 |
| 21475 | 1074 | G1074P | D | 2e-07 | 2.2e-07 | 1.9e-07 | 2.033E-07 | 1.528E-08 | 1.900E-07 | 2.200E-07 |
| 21476 | 1074 | G1074S | D | 6.5e-07 | 1.5e-07 | 2.2e-07 | 3.400E-07 | 2.707E-07 | 1.500E-07 | 6.500E-07 |
| 21477 | 1074 | G1074T | D | 2.6e-07 | 1.8e-07 | 2.1e-07 | 2.167E-07 | 4.041E-08 | 1.800E-07 | 2.600E-07 |
| 21478 | 1074 | G1074W | D | 2.6e-07 | 2.6e-07 | 2.7e-07 | 2.633E-07 | 5.774E-09 | 2.600E-07 | 2.700E-07 |
| 21479 | 1074 | G1074Y | D | 2.1e-07 | 2.4e-07 | 4.1e-07 | 2.867E-07 | 1.079E-07 | 2.100E-07 | 4.100E-07 |
| 21480 | 1074 | G1074V | D | 2.4e-07 | 2.7e-07 | 3.3e-07 | 2.800E-07 | 4.583E-08 | 2.400E-07 | 3.300E-07 |
| 21481 | 1075 | D1075A | D | 3e-07 | 1.1e-07 | 1.7e-07 | 1.933E-07 | 9.713E-08 | 1.100E-07 | 3.000E-07 |
| 21482 | 1075 | D1075R | D | 3.3e-07 | 1.7e-07 | 1.5e-07 | 2.167E-07 | 9.866E-08 | 1.500E-07 | 3.300E-07 |
| 21483 | 1075 | D1075N | D | 4.1e-07 | 9.4e-07 | 2.4e-07 | 5.300E-07 | 3.651E-07 | 2.400E-07 | 9.400E-07 |
| 21484 | 1075 | D1075D | D | 3.6e-07 | 5.3e-07 | 1.9e-07 | 3.600E-07 | 1.700E-07 | 1.900E-07 | 5.300E-07 |
| 21485 | 1075 | D1075C | D | 2.3e-07 | 6.2e-08 | 4.6e-07 | 2.507E-07 | 1.998E-07 | 6.200E-08 | 4.600E-07 |
| 21486 | 1075 | D1075E | D | 2.3e-07 | 2.3e-07 | 3.9e-07 | 2.833E-07 | 9.238E-08 | 2.300E-07 | 3.900E-07 |
| 21487 | 1075 | D1075Q | D | 3e-07 | 1.9e-07 | 2.7e-07 | 2.533E-07 | 5.686E-08 | 1.900E-07 | 3.000E-07 |
| 21488 | 1075 | D1075G | D | 2e-07 | 3.9e-07 | 2.2e-07 | 2.700E-07 | 1.044E-07 | 2.000E-07 | 3.900E-07 |
| 21489 | 1075 | D1075H | D | 2.9e-07 | 2.3e-07 | 2.1e-07 | 2.433E-07 | 4.163E-08 | 2.100E-07 | 2.900E-07 |
| 21490 | 1075 | D1075I | D | 1.9e-07 | 2.5e-07 | 6.6e-08 | 1.687E-07 | 9.384E-08 | 6.600E-08 | 2.500E-07 |
| 21491 | 1075 | D1075L | D | 1.6e-07 | 3e-07 | 2.5e-07 | 2.367E-07 | 7.095E-08 | 1.600E-07 | 3.000E-07 |
| 21492 | 1075 | D1075K | D | 2.3e-07 | 1.8e-07 | 1.8e-07 | 1.967E-07 | 2.887E-08 | 1.800E-07 | 2.300E-07 |
| 21493 | 1075 | D1075M | D | 2e-07 | 2.2e-07 | 1.5e-07 | 1.900E-07 | 3.606E-08 | 1.500E-07 | 2.200E-07 |
| 21494 | 1075 | D1075F | D | 1.3e-07 | 2.5e-07 | 3.3e-07 | 2.367E-07 | 1.007E-07 | 1.300E-07 | 3.300E-07 |
| 21495 | 1075 | D1075P | D | 1.9e-07 | 3.3e-07 | 1.2e-07 | 2.133E-07 | 1.069E-07 | 1.200E-07 | 3.300E-07 |
| 21496 | 1075 | D1075S | D | 1.9e-07 | 2.7e-07 | 2.5e-07 | 2.367E-07 | 4.163E-08 | 1.900E-07 | 2.700E-07 |
| 21497 | 1075 | D1075T | D | 1.2e-07 | 3.8e-07 | 4e-07 | 3.000E-07 | 1.562E-07 | 1.200E-07 | 4.000E-07 |
| 21498 | 1075 | D1075W | D | 1.4e-07 | 1.9e-07 | 8.6e-08 | 1.387E-07 | 5.201E-08 | 8.600E-08 | 1.900E-07 |
| 21499 | 1075 | D1075Y | D | 1.6e-07 | 1.7e-07 | 2.9e-07 | 2.067E-07 | 7.234E-08 | 1.600E-07 | 2.900E-07 |
| 21500 | 1075 | D1075V | D | 2.7e-07 | 4.3e-07 | 3.1e-07 | 3.367E-07 | 8.327E-08 | 2.700E-07 | 4.300E-07 |
| 21501 | 1076 | R1076A | D | 1.3e-07 | 2.1e-07 | 3e-07 | 2.133E-07 | 8.505E-08 | 1.300E-07 | 3.000E-07 |
| 21502 | 1076 | R1076R | D | 1.3e-07 | 2.2e-07 | 2e-07 | 1.833E-07 | 4.726E-08 | 1.300E-07 | 2.200E-07 |
| 21503 | 1076 | R1076N | D | 2.2e-07 | 9.8e-08 | 3.7e-07 | 2.293E-07 | 1.362E-07 | 9.800E-08 | 3.700E-07 |
| 21504 | 1076 | R1076D | D | 9.4e-08 | 2.4e-07 | 3.3e-07 | 2.213E-07 | 1.191E-07 | 9.400E-08 | 3.300E-07 |

| | | | | | | | | | | |
|---|---|---|---|---|---|---|---|---|---|---|
| 21505 | 1076 | R1076C | D | 1.7e-07 | 3.9e-07 | 2.8e-07 | 2.800E-07 | 1.100E-07 | 1.700E-07 | 3.900E-07 |
| 21506 | 1076 | R1076E | D | 1.5e-07 | 2.5e-07 | 1.4e-07 | 1.800E-07 | 6.083E-08 | 1.400E-07 | 2.500E-07 |
| 21507 | 1076 | R1076Q | D | 9.9e-08 | 1.9e-07 | 2.3e-07 | 1.730E-07 | 6.713E-08 | 9.900E-08 | 2.300E-07 |
| 21508 | 1076 | R1076G | D | 1.9e-07 | 3.4e-07 | 4e-07 | 3.100E-07 | 1.082E-07 | 1.900E-07 | 4.000E-07 |
| 21509 | 1076 | R1076H | D | 1.1e-07 | 1.8e-07 | 3.4e-07 | 2.100E-07 | 1.179E-07 | 1.100E-07 | 3.400E-07 |
| 21510 | 1076 | R1076I | D | 2e-07 | 2.1e-07 | 2.2e-07 | 2.100E-07 | 1.000E-08 | 2.000E-07 | 2.200E-07 |
| 21511 | 1076 | R1076L | D | 1.9e-07 | 3.1e-07 | 2.2e-07 | 2.400E-07 | 6.245E-08 | 1.900E-07 | 3.100E-07 |
| 21512 | 1076 | R1076K | D | 2.6e-07 | 2.4e-07 | 2.6e-07 | 2.533E-07 | 1.155E-08 | 2.400E-07 | 2.600E-07 |
| 21513 | 1076 | R1076M | D | 3.9e-07 | 2.7e-07 | 4e-07 | 3.533E-07 | 7.234E-08 | 2.700E-07 | 4.000E-07 |
| 21514 | 1076 | R1076F | D | 4.2e-07 | 1.3e-07 | 2e-07 | 2.500E-07 | 1.513E-07 | 1.300E-07 | 4.200E-07 |
| 21515 | 1076 | R1076P | D | 2.6e-07 | 2e-07 | 3.3e-07 | 2.633E-07 | 6.506E-08 | 2.000E-07 | 3.300E-07 |
| 21516 | 1076 | R1076S | D | 3.8e-07 | 2.9e-07 | 4.3e-07 | 3.667E-07 | 7.095E-08 | 2.900E-07 | 4.300E-07 |
| 21517 | 1076 | R1076T | D | 2.2e-07 | 2.4e-07 | 3.7e-07 | 2.767E-07 | 8.145E-08 | 2.200E-07 | 3.700E-07 |
| 21518 | 1076 | R1076W | D | 2e-07 | 3.5e-07 | 2.2e-07 | 2.567E-07 | 8.145E-08 | 2.000E-07 | 3.500E-07 |
| 21519 | 1076 | R1076Y | D | 2.4e-07 | 2.2e-07 | 3.6e-07 | 2.733E-07 | 7.572E-08 | 2.200E-07 | 3.600E-07 |
| 21520 | 1076 | R1076V | D | 2.6e-07 | 3.2e-07 | 2e-07 | 2.600E-07 | 6.000E-08 | 2.000E-07 | 3.200E-07 |
| 21521 | 1077 | V1077A | D | 1.6e-07 | 2.1e-07 | 2.5e-07 | 2.067E-07 | 4.509E-08 | 1.600E-07 | 2.500E-07 |
| 21522 | 1077 | V1077R | D | 3.6e-07 | 1e-07 | 1.7e-07 | 2.100E-07 | 1.345E-07 | 1.000E-07 | 3.600E-07 |
| 21523 | 1077 | V1077N | D | 1.3e-07 | 3.3e-07 | 4.8e-07 | 3.133E-07 | 1.756E-07 | 1.300E-07 | 4.800E-07 |
| 21524 | 1077 | V1077D | D | 1.3e-07 | 3.3e-07 | 1.5e-07 | 2.033E-07 | 1.102E-07 | 1.300E-07 | 3.300E-07 |
| 21525 | 1077 | V1077C | D | 1.6e-07 | 3.2e-07 | 1.7e-07 | 2.167E-07 | 8.963E-08 | 1.600E-07 | 3.200E-07 |
| 21526 | 1077 | V1077E | D | 2.2e-07 | 2.6e-07 | 2.8e-07 | 2.533E-07 | 3.055E-08 | 2.200E-07 | 2.800E-07 |
| 21527 | 1077 | V1077Q | D | 1.8e-07 | 2.4e-07 | 1.5e-07 | 1.900E-07 | 4.583E-08 | 1.500E-07 | 2.400E-07 |
| 21528 | 1077 | V1077G | D | 1.3e-07 | 2.3e-07 | 2.2e-07 | 1.933E-07 | 5.508E-08 | 1.300E-07 | 2.300E-07 |
| 21529 | 1077 | V1077H | D | 2.5e-07 | 4.7e-07 | 1.9e-07 | 3.033E-07 | 1.474E-07 | 1.900E-07 | 4.700E-07 |
| 21530 | 1077 | V1077I | D | 2.1e-07 | 2e-07 | 2.4e-07 | 2.167E-07 | 2.082E-08 | 2.000E-07 | 2.400E-07 |
| 21531 | 1077 | V1077L | D | 3.6e-07 | 4.5e-07 | 2.7e-07 | 3.600E-07 | 9.000E-08 | 2.700E-07 | 4.500E-07 |
| 21532 | 1077 | V1077K | D | 4.2e-07 | 3e-07 | 4.1e-07 | 3.767E-07 | 6.658E-08 | 3.000E-07 | 4.200E-07 |
| 21533 | 1077 | V1077M | D | 2.5e-07 | 4.9e-07 | 2.4e-07 | 3.267E-07 | 1.415E-07 | 2.400E-07 | 4.900E-07 |
| 21534 | 1077 | V1077F | D | 4.4e-07 | 2.1e-07 | 2.9e-07 | 3.133E-07 | 1.168E-07 | 2.100E-07 | 4.400E-07 |
| 21535 | 1077 | V1077P | D | 5e-07 | 2.9e-07 | 2.2e-07 | 3.367E-07 | 1.457E-07 | 2.200E-07 | 5.000E-07 |
| 21536 | 1077 | V1077S | D | 1.8e-07 | 1.4e-07 | 1e-07 | 1.400E-07 | 4.000E-08 | 1.000E-07 | 1.800E-07 |
| 21537 | 1077 | V1077T | D | 2.9e-07 | 4.2e-07 | 1.6e-07 | 2.900E-07 | 1.300E-07 | 1.600E-07 | 4.200E-07 |
| 21538 | 1077 | V1077W | D | 9.6e-08 | 1.2e-07 | 1.9e-07 | 1.353E-07 | 4.884E-08 | 9.600E-08 | 1.900E-07 |
| 21539 | 1077 | V1077Y | D | 2.2e-07 | 3e-07 | 2e-07 | 2.400E-07 | 5.292E-08 | 2.000E-07 | 3.000E-07 |
| 21540 | 1077 | V1077V | D | 2.1e-07 | 5.2e-07 | 2.7e-07 | 3.333E-07 | 1.644E-07 | 2.100E-07 | 5.200E-07 |
| 21541 | 1078 | T1078A | D | 2.6e-07 | 4.2e-07 | 2.3e-07 | 3.033E-07 | 1.021E-07 | 2.300E-07 | 4.200E-07 |
| 21542 | 1078 | T1078R | D | 5.2e-07 | 2.1e-07 | 1.3e-07 | 2.867E-07 | 2.060E-07 | 1.300E-07 | 5.200E-07 |
| 21543 | 1078 | T1078N | D | 3.4e-07 | 3.2e-07 | 3e-07 | 3.200E-07 | 2.000E-08 | 3.000E-07 | 3.400E-07 |
| 21544 | 1078 | T1078D | D | 4.5e-07 | 2.6e-07 | 5.8e-07 | 4.300E-07 | 1.609E-07 | 2.600E-07 | 5.800E-07 |
| 21545 | 1078 | T1078C | D | 2.1e-07 | 1.3e-07 | 4.8e-07 | 2.733E-07 | 1.834E-07 | 1.300E-07 | 4.800E-07 |
| 21546 | 1078 | T1078E | D | 2.3e-07 | 1.8e-07 | 2.4e-07 | 2.167E-07 | 3.215E-08 | 1.800E-07 | 2.400E-07 |
| 21547 | 1078 | T1078Q | D | 2.1e-07 | 1.5e-07 | 2.7e-07 | 2.100E-07 | 6.000E-08 | 1.500E-07 | 2.700E-07 |
| 21548 | 1078 | T1078G | D | 4.5e-07 | 2.3e-07 | 3.5e-07 | 3.433E-07 | 1.102E-07 | 2.300E-07 | 4.500E-07 |
| 21549 | 1078 | T1078H | D | 2.8e-07 | 1.9e-07 | 2.3e-07 | 2.333E-07 | 4.509E-08 | 1.900E-07 | 2.800E-07 |
| 21550 | 1078 | T1078I | D | 2.2e-07 | 3.3e-07 | 8.5e-08 | 2.117E-07 | 1.227E-07 | 8.500E-08 | 3.300E-07 |
| 21551 | 1078 | T1078L | D | 2.4e-07 | 1.9e-07 | 2.7e-07 | 2.333E-07 | 4.041E-08 | 1.900E-07 | 2.700E-07 |
| 21552 | 1078 | T1078K | D | 1.2e-07 | 3.4e-07 | 1.4e-07 | 2.000E-07 | 1.217E-07 | 1.200E-07 | 3.400E-07 |
| 21553 | 1078 | T1078M | D | 3.3e-07 | 3.4e-07 | 2.2e-07 | 2.967E-07 | 6.658E-08 | 2.200E-07 | 3.400E-07 |
| 21554 | 1078 | T1078F | D | 1.1e-07 | 4.5e-07 | 2.9e-07 | 2.833E-07 | 1.701E-07 | 1.100E-07 | 4.500E-07 |
| 21555 | 1078 | T1078P | D | 5e-07 | 2.9e-07 | 3.9e-07 | 3.933E-07 | 1.050E-07 | 2.900E-07 | 5.000E-07 |
| 21556 | 1078 | T1078S | D | 1.9e-07 | 4.7e-07 | 2.3e-07 | 2.967E-07 | 1.514E-07 | 1.900E-07 | 4.700E-07 |
| 21557 | 1078 | T1078T | D | 1.5e-07 | 2.3e-07 | 2.8e-07 | 2.200E-07 | 6.557E-08 | 1.500E-07 | 2.800E-07 |
| 21558 | 1078 | T1078W | D | 1.9e-07 | 3.6e-07 | 1e-07 | 2.167E-07 | 1.320E-07 | 1.000E-07 | 3.600E-07 |
| 21559 | 1078 | T1078Y | D | 2.5e-07 | 1.7e-07 | 1.4e-07 | 1.867E-07 | 5.686E-08 | 1.400E-07 | 2.500E-07 |
| 21560 | 1078 | T1078V | D | 3.6e-07 | 8.8e-08 | 3.8e-07 | 2.760E-07 | 1.631E-07 | 8.800E-08 | 3.800E-07 |
| 21561 | 1079 | I1079A | D | 1.8e-07 | 3e-07 | 2.5e-07 | 2.433E-07 | 6.028E-08 | 1.800E-07 | 3.000E-07 |
| 21562 | 1079 | I1079R | D | 1.6e-07 | 4.5e-07 | 2.6e-07 | 2.900E-07 | 1.473E-07 | 1.600E-07 | 4.500E-07 |
| 21563 | 1079 | I1079N | D | 2.6e-07 | 2e-07 | 1.5e-07 | 2.033E-07 | 5.508E-08 | 1.500E-07 | 2.600E-07 |
| 21564 | 1079 | I1079D | D | 1.7e-07 | 8.7e-08 | 1.6e-07 | 1.390E-07 | 4.531E-08 | 8.700E-08 | 1.700E-07 |
| 21565 | 1079 | I1079C | D | 1.6e-07 | 3e-07 | 2.6e-07 | 2.400E-07 | 7.211E-08 | 1.600E-07 | 3.000E-07 |
| 21566 | 1079 | I1079E | D | 5.3e-07 | 4.1e-07 | 2.9e-07 | 4.100E-07 | 1.200E-07 | 2.900E-07 | 5.300E-07 |
| 21567 | 1079 | I1079Q | D | 3.5e-07 | 1.1e-07 | 2.9e-07 | 2.500E-07 | 1.249E-07 | 1.100E-07 | 3.500E-07 |
| 21568 | 1079 | I1079G | D | 2.8e-07 | 3.1e-07 | 1.1e-07 | 2.333E-07 | 1.079E-07 | 1.100E-07 | 3.100E-07 |
| 21569 | 1079 | I1079H | D | 3.4e-07 | 2.5e-07 | 3.8e-07 | 3.233E-07 | 6.658E-08 | 2.500E-07 | 3.800E-07 |
| 21570 | 1079 | I1079I | D | 1.9e-07 | 4.9e-07 | 2.3e-07 | 3.033E-07 | 1.629E-07 | 1.900E-07 | 4.900E-07 |
| 21571 | 1079 | I1079L | D | 5.7e-07 | 3.1e-07 | 2.4e-07 | 3.733E-07 | 1.739E-07 | 2.400E-07 | 5.700E-07 |
| 21572 | 1079 | I1079K | D | 4.9e-07 | 3e-07 | 6.9e-07 | 4.933E-07 | 1.950E-07 | 3.000E-07 | 6.900E-07 |
| 21573 | 1079 | I1079M | D | 1e-07 | 2.3e-07 | 1.2e-07 | 1.500E-07 | 7.000E-08 | 1.000E-07 | 2.300E-07 |
| 21574 | 1079 | I1079F | D | 2.6e-07 | 5e-07 | 2.8e-07 | 3.467E-07 | 1.332E-07 | 2.600E-07 | 5.000E-07 |
| 21575 | 1079 | I1079P | D | 2.7e-07 | 3.3e-07 | 2.2e-07 | 2.733E-07 | 5.508E-08 | 2.200E-07 | 3.300E-07 |
| 21576 | 1079 | I1079S | D | 3.6e-07 | 2.4e-07 | 1.9e-07 | 2.633E-07 | 8.737E-08 | 1.900E-07 | 3.600E-07 |
| 21577 | 1079 | I1079T | D | 2e-07 | 6.3e-07 | 3.5e-07 | 3.933E-07 | 2.183E-07 | 2.000E-07 | 6.300E-07 |
| 21578 | 1079 | I1079W | D | 1.6e-07 | 2.8e-07 | 3e-07 | 2.467E-07 | 7.572E-08 | 1.600E-07 | 3.000E-07 |
| 21579 | 1079 | I1079Y | D | 2.9e-07 | 2.5e-07 | 4.4e-07 | 3.267E-07 | 1.002E-07 | 2.500E-07 | 4.400E-07 |
| 21580 | 1079 | I1079V | D | 2.2e-07 | 1.8e-07 | 1.5e-07 | 1.833E-07 | 3.512E-08 | 1.500E-07 | 2.200E-07 |
| 21581 | 1080 | T1080A | D | 4.3e-07 | 4.5e-07 | 2e-07 | 3.600E-07 | 1.389E-07 | 2.000E-07 | 4.500E-07 |
| 21582 | 1080 | T1080R | D | 1.9e-07 | 2.9e-07 | 3.9e-07 | 2.900E-07 | 1.000E-07 | 1.900E-07 | 3.900E-07 |
| 21583 | 1080 | T1080N | D | 3.8e-07 | 8.3e-07 | 2.5e-07 | 4.867E-07 | 3.044E-07 | 2.500E-07 | 8.300E-07 |
| 21584 | 1080 | T1080D | D | 1.5e-07 | 1.4e-07 | 2.6e-07 | 1.833E-07 | 6.658E-08 | 1.400E-07 | 2.600E-07 |
| 21585 | 1080 | T1080C | D | 3.4e-07 | 5.7e-07 | 3.3e-07 | 4.133E-07 | 1.358E-07 | 3.300E-07 | 5.700E-07 |
| 21586 | 1080 | T1080E | D | 2e-07 | 1.7e-07 | 2.5e-07 | 2.067E-07 | 4.041E-08 | 1.700E-07 | 2.500E-07 |
| 21587 | 1080 | T1080Q | D | 1.7e-07 | 1.2e-07 | 3.6e-07 | 2.167E-07 | 1.266E-07 | 1.200E-07 | 3.600E-07 |
| 21588 | 1080 | T1080G | D | 3.7e-07 | 3.9e-07 | 3.1e-07 | 3.567E-07 | 4.163E-08 | 3.100E-07 | 3.900E-07 |
| 21589 | 1080 | T1080H | D | 3.2e-07 | 2.8e-07 | 2e-07 | 2.667E-07 | 6.110E-08 | 2.000E-07 | 3.200E-07 |

| | | | | | | | | | | |
|---|---|---|---|---|---|---|---|---|---|---|
| 21590 | 1080 | T1080I | D | 1.2e-07 | 2.1e-07 | 2e-07 | 1.767E-07 | 4.933E-08 | 1.200E-07 | 2.100E-07 |
| 21591 | 1080 | T1080L | D | 3.3e-07 | 2e-07 | 2.2e-07 | 2.500E-07 | 7.000E-08 | 2.000E-07 | 3.300E-07 |
| 21592 | 1080 | T1080K | D | 2.3e-07 | 2.8e-07 | 2.1e-07 | 2.400E-07 | 3.606E-08 | 2.100E-07 | 2.800E-07 |
| 21593 | 1080 | T1080M | D | 1.9e-07 | 1.9e-07 | 2.9e-07 | 2.233E-07 | 5.774E-08 | 1.900E-07 | 2.900E-07 |
| 21594 | 1080 | T1080F | D | 3.6e-07 | 1.5e-07 | 2.4e-07 | 2.500E-07 | 1.054E-07 | 1.500E-07 | 3.600E-07 |
| 21595 | 1080 | T1080P | D | 2.3e-07 | 3.6e-07 | 2.3e-07 | 2.733E-07 | 7.506E-08 | 2.300E-07 | 3.600E-07 |
| 21596 | 1080 | T1080S | D | 2.2e-07 | 1.3e-07 | 2e-07 | 1.833E-07 | 4.726E-08 | 1.300E-07 | 2.200E-07 |
| 21597 | 1080 | T1080T | D | 1.2e-07 | 2.7e-07 | 2.4e-07 | 2.100E-07 | 7.937E-08 | 1.200E-07 | 2.700E-07 |
| 21598 | 1080 | T1080W | D | 4.2e-07 | 2.4e-07 | 2.2e-07 | 2.933E-07 | 1.102E-07 | 2.200E-07 | 4.200E-07 |
| 21599 | 1080 | T1080Y | D | 3e-07 | 3.8e-07 | 1.7e-07 | 2.833E-07 | 1.060E-07 | 1.700E-07 | 3.800E-07 |
| 21600 | 1080 | T1080V | D | 1.3e-07 | 3.5e-07 | 5.4e-07 | 3.400E-07 | 2.052E-07 | 1.300E-07 | 5.400E-07 |
| 21601 | 1081 | C1081A | D | 1.9e-07 | 4e-07 | 2.1e-07 | 2.667E-07 | 1.159E-07 | 1.900E-07 | 4.000E-07 |
| 21602 | 1081 | C1081R | D | 2.9e-07 | 1.7e-07 | 2.2e-07 | 2.267E-07 | 6.028E-08 | 1.700E-07 | 2.900E-07 |
| 21603 | 1081 | C1081N | D | 2.2e-07 | 5.4e-07 | 3.6e-07 | 3.733E-07 | 1.604E-07 | 2.200E-07 | 5.400E-07 |
| 21604 | 1081 | C1081D | D | 4.2e-07 | 2.6e-07 | 2.1e-07 | 2.967E-07 | 1.097E-07 | 2.100E-07 | 4.200E-07 |
| 21605 | 1081 | C1081C | D | 2.5e-07 | 2.2e-07 | 1.8e-07 | 2.167E-07 | 3.512E-08 | 1.800E-07 | 2.500E-07 |
| 21606 | 1081 | C1081E | D | 4.4e-07 | 3e-07 | 1.2e-07 | 2.867E-07 | 1.604E-07 | 1.200E-07 | 4.400E-07 |
| 21607 | 1081 | C1081Q | D | 1.6e-07 | 2e-07 | 1.5e-07 | 1.700E-07 | 2.646E-08 | 1.500E-07 | 2.000E-07 |
| 21608 | 1081 | C1081G | D | 1.6e-07 | 2.8e-07 | 2.4e-07 | 2.267E-07 | 6.110E-08 | 1.600E-07 | 2.800E-07 |
| 21609 | 1081 | C1081H | D | 2.4e-07 | 3.9e-07 | 5.5e-07 | 3.933E-07 | 1.550E-07 | 2.400E-07 | 5.500E-07 |
| 21610 | 1081 | C1081I | D | 2.3e-07 | 2.3e-07 | 2.3e-07 | 2.300E-07 | 0.000E+00 | 2.300E-07 | 2.300E-07 |
| 21611 | 1081 | C1081L | D | 2.1e-07 | 3.9e-07 | 3.4e-07 | 3.133E-07 | 9.292E-08 | 2.100E-07 | 3.900E-07 |
| 21612 | 1081 | C1081K | D | 1.7e-07 | 9.2e-08 | 2.9e-07 | 1.840E-07 | 9.974E-08 | 9.200E-08 | 2.900E-07 |
| 21613 | 1081 | C1081M | D | 3.2e-07 | 1.4e-07 | 2.4e-07 | 2.333E-07 | 9.018E-08 | 1.400E-07 | 3.200E-07 |
| 21614 | 1081 | C1081F | D | 3.6e-07 | 1.2e-07 | 1.9e-07 | 2.233E-07 | 1.234E-07 | 1.200E-07 | 3.600E-07 |
| 21615 | 1081 | C1081P | D | 2.9e-07 | 3.4e-07 | 2.9e-07 | 3.067E-07 | 2.887E-08 | 2.900E-07 | 3.400E-07 |
| 21616 | 1081 | C1081S | D | 9.1e-08 | 3.3e-07 | 3.4e-07 | 2.537E-07 | 1.410E-07 | 9.100E-08 | 3.400E-07 |
| 21617 | 1081 | C1081T | D | 3.1e-07 | 2.7e-07 | 1e-07 | 2.267E-07 | 1.115E-07 | 1.000E-07 | 3.100E-07 |
| 21618 | 1081 | C1081W | D | 6.1e-07 | 1.8e-07 | 3.5e-07 | 3.800E-07 | 2.166E-07 | 1.800E-07 | 6.100E-07 |
| 21619 | 1081 | C1081Y | D | 4.1e-07 | 1.7e-07 | 2.7e-07 | 2.833E-07 | 1.206E-07 | 1.700E-07 | 4.100E-07 |
| 21620 | 1081 | C1081V | D | 1.1e-07 | 1.4e-07 | 1.8e-07 | 1.433E-07 | 3.512E-08 | 1.100E-07 | 1.800E-07 |
| 21621 | 1082 | R1082A | D | 3.4e-07 | 3.7e-07 | 2.3e-07 | 3.133E-07 | 7.371E-08 | 2.300E-07 | 3.700E-07 |
| 21622 | 1082 | R1082R | D | 1.8e-07 | 1.6e-07 | 3.9e-07 | 2.433E-07 | 1.274E-07 | 1.600E-07 | 3.900E-07 |
| 21623 | 1082 | R1082N | D | 6.6e-07 | 6.9e-07 | 2.3e-07 | 5.267E-07 | 2.574E-07 | 2.300E-07 | 6.900E-07 |
| 21624 | 1082 | R1082D | D | 4.7e-07 | 2.4e-07 | 4.6e-07 | 3.900E-07 | 1.300E-07 | 2.400E-07 | 4.700E-07 |
| 21625 | 1082 | R1082C | D | 1.3e-07 | 4.7e-07 | 2.9e-07 | 2.967E-07 | 1.701E-07 | 1.300E-07 | 4.700E-07 |
| 21626 | 1082 | R1082E | D | 2.5e-07 | 2.2e-07 | 1.6e-07 | 2.100E-07 | 4.583E-08 | 1.600E-07 | 2.500E-07 |
| 21627 | 1082 | R1082Q | D | 2.8e-07 | 2.9e-07 | 4.1e-07 | 3.267E-07 | 7.234E-08 | 2.800E-07 | 4.100E-07 |
| 21628 | 1082 | R1082G | D | 2.3e-07 | 2.7e-07 | 3.3e-07 | 2.767E-07 | 5.033E-08 | 2.300E-07 | 3.300E-07 |
| 21629 | 1082 | R1082H | D | 2.5e-07 | 2e-07 | 2.4e-07 | 2.300E-07 | 2.646E-08 | 2.000E-07 | 2.500E-07 |
| 21630 | 1082 | R1082I | D | 1.9e-07 | 2.2e-07 | 4.7e-07 | 2.933E-07 | 1.537E-07 | 1.900E-07 | 4.700E-07 |
| 21631 | 1082 | R1082L | D | 3.1e-07 | 3.4e-07 | 4e-07 | 3.500E-07 | 4.583E-08 | 3.100E-07 | 4.000E-07 |
| 21632 | 1082 | R1082K | D | 1.9e-07 | 3.5e-07 | 3.6e-07 | 3.000E-07 | 9.539E-08 | 1.900E-07 | 3.600E-07 |
| 21633 | 1082 | R1082M | D | 1.4e-07 | 2.3e-07 | 9.4e-08 | 1.547E-07 | 6.918E-08 | 9.400E-08 | 2.300E-07 |
| 21634 | 1082 | R1082F | D | 3e-07 | 1.9e-07 | 5.9e-07 | 3.600E-07 | 2.066E-07 | 1.900E-07 | 5.900E-07 |
| 21635 | 1082 | R1082P | D | 9.9e-08 | 2.8e-07 | 7.5e-07 | 3.763E-07 | 3.360E-07 | 9.900E-08 | 7.500E-07 |
| 21636 | 1082 | R1082S | D | 1.8e-07 | 2.8e-07 | 1.6e-07 | 2.067E-07 | 6.429E-08 | 1.600E-07 | 2.800E-07 |
| 21637 | 1082 | R1082T | D | 2.6e-07 | 3.4e-07 | 3e-07 | 3.000E-07 | 4.000E-08 | 2.600E-07 | 3.400E-07 |
| 21638 | 1082 | R1082W | D | 1.5e-07 | 1.2e-07 | 2.1e-07 | 1.600E-07 | 4.583E-08 | 1.200E-07 | 2.100E-07 |
| 21639 | 1082 | R1082Y | D | 2.8e-07 | 4.6e-07 | 1.5e-07 | 2.967E-07 | 1.557E-07 | 1.500E-07 | 4.600E-07 |
| 21640 | 1082 | R1082V | D | 1.8e-07 | 3e-07 | 2.6e-07 | 2.467E-07 | 6.110E-08 | 1.800E-07 | 3.000E-07 |
| 21641 | 1083 | A1083A | D | 2.2e-07 | 5.2e-07 | 4.1e-07 | 3.833E-07 | 1.518E-07 | 2.200E-07 | 5.200E-07 |
| 21642 | 1083 | A1083R | D | 1.6e-07 | 2.1e-07 | 4e-07 | 2.567E-07 | 1.266E-07 | 1.600E-07 | 4.000E-07 |
| 21643 | 1083 | A1083N | D | 1.5e-07 | 2.6e-07 | 5.1e-07 | 3.067E-07 | 1.845E-07 | 1.500E-07 | 5.100E-07 |
| 21644 | 1083 | A1083D | D | 2.1e-07 | 3.4e-07 | 1.3e-07 | 2.267E-07 | 1.060E-07 | 1.300E-07 | 3.400E-07 |
| 21645 | 1083 | A1083C | D | 3.3e-07 | 2.6e-07 | 2.2e-07 | 2.700E-07 | 5.568E-08 | 2.200E-07 | 3.300E-07 |
| 21646 | 1083 | A1083E | D | 1.9e-07 | 2.3e-07 | 1.8e-07 | 2.000E-07 | 2.646E-08 | 1.800E-07 | 2.300E-07 |
| 21647 | 1083 | A1083Q | D | 1.7e-07 | 2.9e-07 | 1.3e-07 | 1.967E-07 | 8.327E-08 | 1.300E-07 | 2.900E-07 |
| 21648 | 1083 | A1083G | D | 1.2e-07 | 1.9e-07 | 2.3e-07 | 1.800E-07 | 5.568E-08 | 1.200E-07 | 2.300E-07 |
| 21649 | 1083 | A1083H | D | 1.6e-07 | 2.6e-07 | 2.6e-07 | 2.267E-07 | 5.774E-08 | 1.600E-07 | 2.600E-07 |
| 21650 | 1083 | A1083I | D | 2.5e-07 | 2.7e-07 | 2.2e-07 | 2.467E-07 | 2.517E-08 | 2.200E-07 | 2.700E-07 |
| 21651 | 1083 | A1083L | D | 3.7e-07 | 3.8e-07 | 3.4e-07 | 3.633E-07 | 2.082E-08 | 3.400E-07 | 3.800E-07 |
| 21652 | 1083 | A1083K | D | 2.3e-07 | 2.2e-07 | 2.9e-07 | 2.467E-07 | 3.786E-08 | 2.200E-07 | 2.900E-07 |
| 21653 | 1083 | A1083M | D | 3.7e-07 | 3e-07 | 2.1e-07 | 2.933E-07 | 8.021E-08 | 2.100E-07 | 3.700E-07 |
| 21654 | 1083 | A1083F | D | 4.6e-07 | 5.2e-07 | 2.9e-07 | 4.233E-07 | 1.193E-07 | 2.900E-07 | 5.200E-07 |
| 21655 | 1083 | A1083P | D | 3.1e-07 | 2.6e-07 | 3.3e-07 | 3.000E-07 | 3.606E-08 | 2.600E-07 | 3.300E-07 |
| 21656 | 1083 | A1083S | D | 3.3e-07 | 1.8e-07 | 2.2e-07 | 2.433E-07 | 7.767E-08 | 1.800E-07 | 3.300E-07 |
| 21657 | 1083 | A1083T | D | 5.2e-07 | 2e-07 | 1.8e-07 | 3.000E-07 | 1.908E-07 | 1.800E-07 | 5.200E-07 |
| 21658 | 1083 | A1083W | D | 1.3e-07 | 3.6e-07 | 4e-07 | 2.967E-07 | 1.457E-07 | 1.300E-07 | 4.000E-07 |
| 21659 | 1083 | A1083Y | D | 1.8e-07 | 2.8e-07 | 1.2e-07 | 1.933E-07 | 8.083E-08 | 1.200E-07 | 2.800E-07 |
| 21660 | 1083 | A1083V | D | 4.1e-07 | 8.4e-08 | 5e-07 | 3.313E-07 | 2.189E-07 | 8.400E-08 | 5.000E-07 |
| 21661 | 1084 | S1084A | D | 3.3e-07 | 3.3e-07 | 4e-07 | 3.533E-07 | 4.041E-08 | 3.300E-07 | 4.000E-07 |
| 21662 | 1084 | S1084R | D | 2.4e-07 | 3.2e-07 | 1.7e-07 | 2.433E-07 | 7.506E-08 | 1.700E-07 | 3.200E-07 |
| 21663 | 1084 | S1084N | D | 5.5e-07 | 2.2e-07 | 2.2e-07 | 3.300E-07 | 1.905E-07 | 2.200E-07 | 5.500E-07 |
| 21664 | 1084 | S1084D | D | 2.7e-07 | 3.1e-07 | 2.9e-07 | 2.900E-07 | 2.000E-08 | 2.700E-07 | 3.100E-07 |
| 21665 | 1084 | S1084C | D | 4.7e-07 | 5e-07 | 2.7e-07 | 4.133E-07 | 1.250E-07 | 2.700E-07 | 5.000E-07 |
| 21666 | 1084 | S1084E | D | 4.8e-07 | 4.4e-07 | 1.8e-07 | 3.667E-07 | 1.629E-07 | 1.800E-07 | 4.800E-07 |
| 21667 | 1084 | S1084Q | D | 2.4e-07 | 1.2e-07 | 3.6e-07 | 2.400E-07 | 1.200E-07 | 1.200E-07 | 3.600E-07 |
| 21668 | 1084 | S1084G | D | 3.7e-07 | 2.8e-07 | 1.7e-07 | 2.733E-07 | 1.002E-07 | 1.700E-07 | 3.700E-07 |
| 21669 | 1084 | S1084H | D | 3.2e-07 | 2.4e-07 | 4e-07 | 3.200E-07 | 8.000E-08 | 2.400E-07 | 4.000E-07 |
| 21670 | 1084 | S1084I | D | 1.9e-07 | 4.4e-07 | 2e-07 | 2.767E-07 | 1.415E-07 | 1.900E-07 | 4.400E-07 |
| 21671 | 1084 | S1084L | D | 1.2e-07 | 2.9e-07 | 2.4e-07 | 2.167E-07 | 8.737E-08 | 1.200E-07 | 2.900E-07 |
| 21672 | 1084 | S1084K | D | 2.7e-07 | 1.4e-07 | 2.7e-07 | 2.267E-07 | 7.506E-08 | 1.400E-07 | 2.700E-07 |
| 21673 | 1084 | S1084M | D | 1.7e-07 | 2.9e-07 | 7.8e-07 | 4.133E-07 | 3.232E-07 | 1.700E-07 | 7.800E-07 |
| 21674 | 1084 | S1084F | D | 8.5e-08 | 7.5e-07 | 1.4e-07 | 3.250E-07 | 3.691E-07 | 8.500E-08 | 7.500E-07 |

| | | | | | | | | | | |
|---|---|---|---|---|---|---|---|---|---|---|
| 21675 | 1084 | S1084P | D | 2.5e-07 | 1.6e-07 | 2.2e-07 | 2.100E-07 | 4.583E-08 | 1.600E-07 | 2.500E-07 |
| 21676 | 1084 | S1084S | D | 9.7e-07 | 2e-07 | 2.4e-07 | 4.700E-07 | 4.335E-07 | 2.000E-07 | 9.700E-07 |
| 21677 | 1084 | S1084T | D | 3.1e-07 | 1.2e-07 | 5.2e-07 | 3.167E-07 | 2.001E-07 | 1.200E-07 | 5.200E-07 |
| 21678 | 1084 | S1084W | D | 1.3e-07 | 3.8e-07 | 2e-07 | 2.367E-07 | 1.290E-07 | 1.300E-07 | 3.800E-07 |
| 21679 | 1084 | S1084Y | D | 4.6e-07 | 2.2e-07 | 2.1e-07 | 2.967E-07 | 1.415E-07 | 2.100E-07 | 4.600E-07 |
| 21680 | 1084 | S1084V | D | 2.8e-07 | 2.2e-07 | 2.5e-07 | 2.500E-07 | 3.000E-08 | 2.200E-07 | 2.800E-07 |
| 21681 | 1085 | Q1085A | D | 1.9e-07 | 3.1e-07 | 1.7e-07 | 2.233E-07 | 7.572E-08 | 1.700E-07 | 3.100E-07 |
| 21682 | 1085 | Q1085R | D | 7.6e-08 | 3.3e-07 | 1.5e-07 | 1.853E-07 | 1.306E-07 | 7.600E-08 | 3.300E-07 |
| 21683 | 1085 | Q1085N | D | 4.8e-07 | 4.2e-07 | 1.5e-07 | 3.500E-07 | 1.758E-07 | 1.500E-07 | 4.800E-07 |
| 21684 | 1085 | Q1085D | D | 4.5e-07 | 4.8e-07 | 2.7e-07 | 4.000E-07 | 1.136E-07 | 2.700E-07 | 4.800E-07 |
| 21685 | 1085 | Q1085C | D | 3.4e-07 | 3.3e-07 | 3.3e-07 | 3.333E-07 | 5.774E-09 | 3.300E-07 | 3.400E-07 |
| 21686 | 1085 | Q1085E | D | 4.1e-07 | 1.8e-07 | 4.7e-07 | 3.533E-07 | 1.531E-07 | 1.800E-07 | 4.700E-07 |
| 21687 | 1085 | Q1085Q | D | 4.5e-07 | 1.6e-07 | 3.3e-07 | 3.133E-07 | 1.457E-07 | 1.600E-07 | 4.500E-07 |
| 21688 | 1085 | Q1085G | D | 2.1e-07 | 2.6e-07 | 1.7e-07 | 2.133E-07 | 4.509E-08 | 1.700E-07 | 2.600E-07 |
| 21689 | 1085 | Q1085H | D | 1.7e-07 | 4.7e-07 | 2.5e-07 | 2.967E-07 | 1.553E-07 | 1.700E-07 | 4.700E-07 |
| 21690 | 1085 | Q1085I | D | 2.6e-07 | 7.5e-07 | 1.6e-07 | 3.900E-07 | 3.158E-07 | 1.600E-07 | 7.500E-07 |
| 21691 | 1085 | Q1085L | D | 2e-07 | 2.2e-07 | 1.5e-07 | 1.900E-07 | 3.606E-08 | 1.500E-07 | 2.200E-07 |
| 21692 | 1085 | Q1085K | D | 3.2e-07 | 1.2e-07 | 2.1e-07 | 2.167E-07 | 1.002E-07 | 1.200E-07 | 3.200E-07 |
| 21693 | 1085 | Q1085M | D | 2.1e-07 | 2.8e-07 | 4.3e-07 | 3.067E-07 | 1.124E-07 | 2.100E-07 | 4.300E-07 |
| 21694 | 1085 | Q1085F | D | 2.3e-07 | 2.9e-07 | 2.1e-07 | 2.433E-07 | 4.163E-08 | 2.100E-07 | 2.900E-07 |
| 21695 | 1085 | Q1085P | D | 3.4e-07 | 4.5e-07 | 3.1e-07 | 3.667E-07 | 7.371E-08 | 3.100E-07 | 4.500E-07 |
| 21696 | 1085 | Q1085S | D | 4.1e-07 | 3.5e-07 | 2.2e-07 | 3.267E-07 | 9.713E-08 | 2.200E-07 | 4.100E-07 |
| 21697 | 1085 | Q1085T | D | 1.8e-07 | 3.8e-07 | 2.5e-07 | 2.700E-07 | 1.015E-07 | 1.800E-07 | 3.800E-07 |
| 21698 | 1085 | Q1085W | D | 1.9e-07 | 1.8e-07 | 5.4e-07 | 3.033E-07 | 2.050E-07 | 1.800E-07 | 5.400E-07 |
| 21699 | 1085 | Q1085Y | D | 2.5e-07 | 1.5e-07 | 2.9e-07 | 2.300E-07 | 7.211E-08 | 1.500E-07 | 2.900E-07 |
| 21700 | 1085 | Q1085V | D | 2.7e-07 | 4.3e-07 | 3.3e-07 | 3.433E-07 | 8.083E-08 | 2.700E-07 | 4.300E-07 |
| 21701 | 1086 | D1086A | D | 2.9e-07 | 1.8e-07 | 9.8e-08 | 1.893E-07 | 9.634E-08 | 9.800E-08 | 2.900E-07 |
| 21702 | 1086 | D1086R | D | 2.6e-07 | 4e-07 | 1.3e-07 | 2.633E-07 | 1.350E-07 | 1.300E-07 | 4.000E-07 |
| 21703 | 1086 | D1086N | D | 2.5e-07 | 3.4e-07 | 2.9e-07 | 2.933E-07 | 4.509E-08 | 2.500E-07 | 3.400E-07 |
| 21704 | 1086 | D1086D | D | 1.8e-07 | 1.9e-07 | 2.4e-07 | 2.033E-07 | 3.215E-08 | 1.800E-07 | 2.400E-07 |
| 21705 | 1086 | D1086C | D | 2.9e-07 | 1.6e-07 | 3e-07 | 2.500E-07 | 7.810E-08 | 1.600E-07 | 3.000E-07 |
| 21706 | 1086 | D1086E | D | 1.5e-07 | 1.5e-07 | 1.9e-07 | 1.633E-07 | 2.309E-08 | 1.500E-07 | 1.900E-07 |
| 21707 | 1086 | D1086Q | D | 2.6e-07 | 1.7e-07 | 2.3e-07 | 2.200E-07 | 4.583E-08 | 1.700E-07 | 2.600E-07 |
| 21708 | 1086 | D1086G | D | 3.2e-07 | 2.9e-07 | 2e-07 | 2.700E-07 | 6.245E-08 | 2.000E-07 | 3.200E-07 |
| 21709 | 1086 | D1086H | D | 1.9e-07 | 3.5e-07 | 3.5e-07 | 2.967E-07 | 9.238E-08 | 1.900E-07 | 3.500E-07 |
| 21710 | 1086 | D1086I | D | 4.8e-07 | 2.6e-07 | 2.7e-07 | 3.367E-07 | 1.242E-07 | 2.600E-07 | 4.800E-07 |
| 21711 | 1086 | D1086L | D | 5e-07 | 2.7e-07 | 3e-07 | 3.567E-07 | 1.250E-07 | 2.700E-07 | 5.000E-07 |
| 21712 | 1086 | D1086K | D | 2e-07 | 1.8e-07 | 1.4e-07 | 1.733E-07 | 3.055E-08 | 1.400E-07 | 2.000E-07 |
| 21713 | 1086 | D1086M | D | 3.2e-07 | 1.7e-07 | 3.1e-07 | 2.667E-07 | 8.386E-08 | 1.700E-07 | 3.200E-07 |
| 21714 | 1086 | D1086F | D | 4.5e-07 | 2.7e-07 | 1.7e-07 | 2.967E-07 | 1.419E-07 | 1.700E-07 | 4.500E-07 |
| 21715 | 1086 | D1086P | D | 1.8e-07 | 4.1e-07 | 2.4e-07 | 2.767E-07 | 1.193E-07 | 1.800E-07 | 4.100E-07 |
| 21716 | 1086 | D1086S | D | 2.3e-07 | 3.2e-07 | 1.2e-07 | 2.233E-07 | 1.002E-07 | 1.200E-07 | 3.200E-07 |
| 21717 | 1086 | D1086T | D | 1.8e-07 | 1e-07 | 3.1e-07 | 1.967E-07 | 1.060E-07 | 1.000E-07 | 3.100E-07 |
| 21718 | 1086 | D1086W | D | 1.7e-07 | 2.7e-07 | 2.7e-07 | 2.367E-07 | 5.774E-08 | 1.700E-07 | 2.700E-07 |
| 21719 | 1086 | D1086Y | D | 4.8e-07 | 2e-07 | 2.5e-07 | 3.100E-07 | 1.493E-07 | 2.000E-07 | 4.800E-07 |
| 21720 | 1086 | D1086V | D | 3.5e-07 | 4.2e-07 | 2.3e-07 | 3.333E-07 | 9.609E-08 | 2.300E-07 | 4.200E-07 |
| 21721 | 1087 | V1087A | D | 9.4e-08 | 5.3e-07 | 2.5e-07 | 2.913E-07 | 2.209E-07 | 9.400E-08 | 5.300E-07 |
| 21722 | 1087 | V1087R | D | 1.5e-07 | 3.2e-07 | 2e-07 | 2.233E-07 | 8.737E-08 | 1.500E-07 | 3.200E-07 |
| 21723 | 1087 | V1087N | D | 1.3e-07 | 2.4e-07 | 2.3e-07 | 2.000E-07 | 6.083E-08 | 1.300E-07 | 2.400E-07 |
| 21724 | 1087 | V1087D | D | 4.3e-07 | 5.9e-08 | 3.4e-07 | 2.763E-07 | 1.935E-07 | 5.900E-08 | 4.300E-07 |
| 21725 | 1087 | V1087C | D | 2.2e-07 | 1.1e-07 | 2.7e-07 | 2.000E-07 | 8.185E-08 | 1.100E-07 | 2.700E-07 |
| 21726 | 1087 | V1087E | D | 1.9e-07 | 2.8e-07 | 2.4e-07 | 2.367E-07 | 4.509E-08 | 1.900E-07 | 2.800E-07 |
| 21727 | 1087 | V1087Q | D | 1.9e-07 | 2.2e-07 | 1.8e-07 | 1.967E-07 | 2.082E-08 | 1.800E-07 | 2.200E-07 |
| 21728 | 1087 | V1087G | D | 1.7e-07 | 1.1e-07 | 2.7e-07 | 1.833E-07 | 8.083E-08 | 1.100E-07 | 2.700E-07 |
| 21729 | 1087 | V1087H | D | 3.4e-07 | 1.3e-07 | 1.6e-07 | 2.100E-07 | 1.136E-07 | 1.300E-07 | 3.400E-07 |
| 21730 | 1087 | V1087I | D | 1.9e-07 | 2.8e-07 | 3.4e-07 | 2.700E-07 | 7.550E-08 | 1.900E-07 | 3.400E-07 |
| 21731 | 1087 | V1087L | D | 1.7e-07 | 1.6e-07 | 4.3e-07 | 2.533E-07 | 1.531E-07 | 1.600E-07 | 4.300E-07 |
| 21732 | 1087 | V1087K | D | 1.1e-07 | 2.2e-07 | 4.6e-07 | 2.633E-07 | 1.790E-07 | 1.100E-07 | 4.600E-07 |
| 21733 | 1087 | V1087M | D | 5.4e-07 | 1.9e-07 | 3.4e-07 | 3.567E-07 | 1.756E-07 | 1.900E-07 | 5.400E-07 |
| 21734 | 1087 | V1087F | D | 2.7e-07 | 3.7e-07 | 1.7e-07 | 2.700E-07 | 1.000E-07 | 1.700E-07 | 3.700E-07 |
| 21735 | 1087 | V1087P | D | 3.3e-07 | 1.6e-07 | 1.8e-07 | 2.233E-07 | 9.292E-08 | 1.600E-07 | 3.300E-07 |
| 21736 | 1087 | V1087S | D | 2.7e-07 | 1.5e-07 | 1.9e-07 | 2.033E-07 | 6.110E-08 | 1.500E-07 | 2.700E-07 |
| 21737 | 1087 | V1087T | D | 1.6e-07 | 1e-07 | 2.1e-07 | 1.567E-07 | 5.508E-08 | 1.000E-07 | 2.100E-07 |
| 21738 | 1087 | V1087W | D | 2.4e-07 | 3.3e-07 | 2.1e-07 | 2.600E-07 | 6.245E-08 | 2.100E-07 | 3.300E-07 |
| 21739 | 1087 | V1087Y | D | 2.9e-07 | 3.5e-07 | 1.6e-07 | 2.667E-07 | 9.713E-08 | 1.600E-07 | 3.500E-07 |
| 21740 | 1087 | V1087V | D | 3.2e-07 | 3.4e-07 | 1.1e-07 | 2.567E-07 | 1.274E-07 | 1.100E-07 | 3.400E-07 |
| 21741 | 1088 | N1088A | D | 6.8e-07 | 2.7e-07 | 1.5e-07 | 3.667E-07 | 2.779E-07 | 1.500E-07 | 6.800E-07 |
| 21742 | 1088 | N1088R | D | 4e-07 | 3.4e-07 | 3.5e-07 | 3.633E-07 | 3.215E-08 | 3.400E-07 | 4.000E-07 |
| 21743 | 1088 | N1088N | D | 3.3e-07 | 1.8e-07 | 1.9e-07 | 2.333E-07 | 8.386E-08 | 1.800E-07 | 3.300E-07 |
| 21744 | 1088 | N1088D | D | 1.7e-07 | 1.5e-07 | 1.4e-07 | 1.533E-07 | 1.528E-08 | 1.400E-07 | 1.700E-07 |
| 21745 | 1088 | N1088C | D | 3e-07 | 3e-07 | 1.5e-07 | 2.500E-07 | 8.660E-08 | 1.500E-07 | 3.000E-07 |
| 21746 | 1088 | N1088E | D | 3.6e-07 | 1.7e-07 | 1.8e-07 | 2.367E-07 | 1.069E-07 | 1.700E-07 | 3.600E-07 |
| 21747 | 1088 | N1088Q | D | 3.6e-07 | 2.1e-07 | 3.5e-07 | 3.067E-07 | 8.386E-08 | 2.100E-07 | 3.600E-07 |
| 21748 | 1088 | N1088G | D | 4.3e-07 | 2e-07 | 1.2e-06 | 6.100E-07 | 5.237E-07 | 2.000E-07 | 1.200E-06 |
| 21749 | 1088 | N1088H | D | 2.4e-07 | 3e-07 | 2.3e-07 | 2.567E-07 | 3.786E-08 | 2.300E-07 | 3.000E-07 |
| 21750 | 1088 | N1088I | D | 2.6e-07 | 1.7e-07 | 3.5e-07 | 2.600E-07 | 9.000E-08 | 1.700E-07 | 3.500E-07 |
| 21751 | 1088 | N1088L | D | 1.7e-07 | 2.9e-07 | 3.3e-07 | 2.633E-07 | 8.327E-08 | 1.700E-07 | 3.300E-07 |
| 21752 | 1088 | N1088K | D | 2.1e-07 | 1.9e-07 | 3.4e-07 | 2.467E-07 | 8.145E-08 | 1.900E-07 | 3.400E-07 |
| 21753 | 1088 | N1088M | D | 1.8e-07 | 2.7e-07 | 2.5e-07 | 2.333E-07 | 4.726E-08 | 1.800E-07 | 2.700E-07 |
| 21754 | 1088 | N1088F | D | 2.9e-07 | 1.8e-07 | 2e-07 | 2.233E-07 | 5.859E-08 | 1.800E-07 | 2.900E-07 |
| 21755 | 1088 | N1088P | D | 5.1e-07 | 2.1e-07 | 3.1e-07 | 3.433E-07 | 1.528E-07 | 2.100E-07 | 5.100E-07 |
| 21756 | 1088 | N1088S | D | 5.9e-07 | 5.2e-07 | 1.6e-07 | 4.233E-07 | 2.307E-07 | 1.600E-07 | 5.900E-07 |
| 21757 | 1088 | N1088T | D | 2.9e-07 | 2.9e-07 | 5.7e-07 | 3.833E-07 | 1.617E-07 | 2.900E-07 | 5.700E-07 |
| 21758 | 1088 | N1088W | D | 9.3e-08 | 3.2e-07 | 2.5e-07 | 2.210E-07 | 1.162E-07 | 9.300E-08 | 3.200E-07 |
| 21759 | 1088 | N1088Y | D | 7.5e-07 | 5.2e-07 | 3.4e-07 | 5.367E-07 | 2.055E-07 | 3.400E-07 | 7.500E-07 |

| | | | | | | | | | | |
|---|---|---|---|---|---|---|---|---|---|---|
| 21760 | 1088 | N1088V | D | 5.1e-07 | 4.8e-07 | 2.3e-07 | 4.067E-07 | 1.537E-07 | 2.300E-07 | 5.100E-07 |
| 21761 | 1089 | T1089A | D | 1.9e-07 | 2.9e-07 | 2.4e-07 | 2.400E-07 | 5.000E-08 | 1.900E-07 | 2.900E-07 |
| 21762 | 1089 | T1089R | D | 1.9e-07 | 1.7e-07 | 3.2e-07 | 2.267E-07 | 8.145E-08 | 1.700E-07 | 3.200E-07 |
| 21763 | 1089 | T1089N | D | 1.2e-07 | 3e-07 | 2.8e-07 | 2.333E-07 | 9.866E-08 | 1.200E-07 | 3.000E-07 |
| 21764 | 1089 | T1089D | D | 2.2e-07 | 3e-07 | 5.1e-07 | 3.433E-07 | 1.498E-07 | 2.200E-07 | 5.100E-07 |
| 21765 | 1089 | T1089C | D | 1.9e-07 | 1.8e-07 | 1.3e-07 | 1.667E-07 | 3.215E-08 | 1.300E-07 | 1.900E-07 |
| 21766 | 1089 | T1089E | D | 1.7e-07 | 2.5e-07 | 2.5e-07 | 2.233E-07 | 4.619E-08 | 1.700E-07 | 2.500E-07 |
| 21767 | 1089 | T1089Q | D | 3.4e-07 | 2.1e-07 | 1.8e-07 | 2.433E-07 | 8.505E-08 | 1.800E-07 | 3.400E-07 |
| 21768 | 1089 | T1089G | D | 5.1e-07 | 3e-07 | 1.5e-07 | 3.200E-07 | 1.808E-07 | 1.500E-07 | 5.100E-07 |
| 21769 | 1089 | T1089H | D | 2.1e-07 | 8.5e-08 | 1e-07 | 1.317E-07 | 6.825E-08 | 8.500E-08 | 2.100E-07 |
| 21770 | 1089 | T1089I | D | 1.4e-07 | 8.4e-08 | 2.8e-07 | 1.680E-07 | 1.010E-07 | 8.400E-08 | 2.800E-07 |
| 21771 | 1089 | T1089L | D | 1.9e-07 | 1.3e-07 | 1e-07 | 1.400E-07 | 4.583E-08 | 1.000E-07 | 1.900E-07 |
| 21772 | 1089 | T1089K | D | 1.3e-07 | 3.4e-07 | 3.3e-07 | 2.667E-07 | 1.185E-07 | 1.300E-07 | 3.400E-07 |
| 21773 | 1089 | T1089M | D | 1.8e-07 | 1.1e-07 | 1.6e-07 | 1.500E-07 | 3.606E-08 | 1.100E-07 | 1.800E-07 |
| 21774 | 1089 | T1089F | D | 3.7e-07 | 1.7e-07 | 1.2e-07 | 2.200E-07 | 1.323E-07 | 1.200E-07 | 3.700E-07 |
| 21775 | 1089 | T1089P | D | 2.5e-07 | 1.9e-07 | 2.1e-07 | 2.167E-07 | 3.055E-08 | 1.900E-07 | 2.500E-07 |
| 21776 | 1089 | T1089S | D | 7.7e-07 | 1.7e-07 | 2.8e-07 | 4.067E-07 | 3.194E-07 | 1.700E-07 | 7.700E-07 |
| 21777 | 1089 | T1089T | D | 2.1e-07 | 2.5e-07 | 1.1e-07 | 1.900E-07 | 7.211E-08 | 1.100E-07 | 2.500E-07 |
| 21778 | 1089 | T1089W | D | 1.9e-07 | 2.7e-07 | 4.5e-07 | 3.033E-07 | 1.332E-07 | 1.900E-07 | 4.500E-07 |
| 21779 | 1089 | T1089Y | D | 2.5e-07 | 2.3e-07 | 2.2e-07 | 2.333E-07 | 1.528E-08 | 2.200E-07 | 2.500E-07 |
| 21780 | 1089 | T1089V | D | 4.2e-07 | 2.1e-07 | 1.6e-07 | 2.633E-07 | 1.380E-07 | 1.600E-07 | 4.200E-07 |
| 21781 | 1090 | A1090A | D | 5.7e-07 | 1.4e-07 | 1.8e-07 | 2.967E-07 | 2.376E-07 | 1.400E-07 | 5.700E-07 |
| 21782 | 1090 | A1090R | D | 9.9e-08 | 3.7e-08 | 5.7e-08 | 6.433E-08 | 3.164E-08 | 3.700E-08 | 9.900E-08 |
| 21783 | 1090 | A1090N | D | 8.5e-08 | 1.8e-07 | 6.1e-08 | 1.087E-07 | 6.293E-08 | 6.100E-08 | 1.800E-07 |
| 21784 | 1090 | A1090D | D | 1.2e-07 | 1.1e-07 | 2.3e-07 | 1.533E-07 | 6.658E-08 | 1.100E-07 | 2.300E-07 |
| 21785 | 1090 | A1090C | D | 2.5e-07 | 2e-07 | 3.4e-07 | 2.633E-07 | 7.095E-08 | 2.000E-07 | 3.400E-07 |
| 21786 | 1090 | A1090E | D | 1.3e-07 | 1.2e-07 | 2.9e-07 | 1.800E-07 | 9.539E-08 | 1.200E-07 | 2.900E-07 |
| 21787 | 1090 | A1090Q | D | 3.5e-07 | 2.4e-07 | 1.4e-07 | 2.433E-07 | 1.050E-07 | 1.400E-07 | 3.500E-07 |
| 21788 | 1090 | A1090G | D | 1.8e-07 | 2.8e-07 | 2.8e-07 | 2.467E-07 | 5.774E-08 | 1.800E-07 | 2.800E-07 |
| 21789 | 1090 | A1090H | D | 5.1e-08 | 1.1e-07 | 5.5e-08 | 7.200E-08 | 3.297E-08 | 5.100E-08 | 1.100E-07 |
| 21790 | 1090 | A1090I | D | 1.3e-07 | 1.9e-07 | 9.1e-08 | 1.370E-07 | 4.987E-08 | 9.100E-08 | 1.900E-07 |
| 21791 | 1090 | A1090L | D | 9.3e-08 | 1.7e-07 | 6.1e-08 | 1.080E-07 | 5.603E-08 | 6.100E-08 | 1.700E-07 |
| 21792 | 1090 | A1090K | D | 2.1e-07 | 1e-07 | 1.1e-07 | 1.400E-07 | 6.083E-08 | 1.000E-07 | 2.100E-07 |
| 21793 | 1090 | A1090M | D | 1.1e-07 | 1e-07 | 1.8e-07 | 1.300E-07 | 4.359E-08 | 1.000E-07 | 1.800E-07 |
| 21794 | 1090 | A1090F | D | 8.8e-08 | 1.5e-07 | 1.6e-07 | 1.327E-07 | 3.900E-08 | 8.800E-08 | 1.600E-07 |
| 21795 | 1090 | A1090P | D | 7.4e-07 | 1.4e-07 | 2.4e-07 | 3.733E-07 | 3.215E-07 | 1.400E-07 | 7.400E-07 |
| 21796 | 1090 | A1090S | D | 1.3e-07 | 7e-08 | 1.3e-07 | 1.100E-07 | 3.464E-08 | 7.000E-08 | 1.300E-07 |
| 21797 | 1090 | A1090T | D | 1.7e-07 | 3.1e-07 | 1.2e-07 | 2.000E-07 | 9.849E-08 | 1.200E-07 | 3.100E-07 |
| 21798 | 1090 | A1090W | D | 7.5e-08 | 1.7e-07 | 1.5e-07 | 1.317E-07 | 5.008E-08 | 7.500E-08 | 1.700E-07 |
| 21799 | 1090 | A1090Y | D | 8.8e-08 | 1e-07 | 2.1e-07 | 1.327E-07 | 6.724E-08 | 8.800E-08 | 2.100E-07 |
| 21800 | 1090 | A1090V | D | 2.1e-07 | 1.5e-07 | 7.9e-08 | 1.463E-07 | 6.558E-08 | 7.900E-08 | 2.100E-07 |
| 21801 | 1091 | V1091A | D | 2.5e-07 | 3.4e-07 | 1.6e-07 | 2.500E-07 | 9.000E-08 | 1.600E-07 | 3.400E-07 |
| 21802 | 1091 | V1091R | D | 1.9e-07 | 3.3e-07 | 2.4e-07 | 2.533E-07 | 7.095E-08 | 1.900E-07 | 3.300E-07 |
| 21803 | 1091 | V1091N | D | 2.9e-07 | 2.2e-07 | 2.8e-07 | 2.633E-07 | 3.786E-08 | 2.200E-07 | 2.900E-07 |
| 21804 | 1091 | V1091D | D | 2e-07 | 3.2e-07 | 2.1e-07 | 2.433E-07 | 6.658E-08 | 2.000E-07 | 3.200E-07 |
| 21805 | 1091 | V1091C | D | 3.3e-07 | 1.9e-07 | 1.6e-07 | 2.267E-07 | 9.074E-08 | 1.600E-07 | 3.300E-07 |
| 21806 | 1091 | V1091E | D | 1.1e-07 | 2.1e-07 | 2.8e-07 | 2.000E-07 | 8.544E-08 | 1.100E-07 | 2.800E-07 |
| 21807 | 1091 | V1091Q | D | 1.2e-07 | 1.9e-07 | 4.2e-07 | 2.433E-07 | 1.570E-07 | 1.200E-07 | 4.200E-07 |
| 21808 | 1091 | V1091G | D | 2.7e-07 | 3e-07 | 1.5e-07 | 2.400E-07 | 7.937E-08 | 1.500E-07 | 3.000E-07 |
| 21809 | 1091 | V1091H | D | 1.8e-07 | 2.8e-07 | 2.7e-07 | 2.433E-07 | 5.508E-08 | 1.800E-07 | 2.800E-07 |
| 21810 | 1091 | V1091I | D | 2.7e-07 | 1.7e-07 | 2.7e-07 | 2.367E-07 | 5.774E-08 | 1.700E-07 | 2.700E-07 |
| 21811 | 1091 | V1091L | D | 3.3e-07 | 1.7e-07 | 1.9e-07 | 2.300E-07 | 8.718E-08 | 1.700E-07 | 3.300E-07 |
| 21812 | 1091 | V1091K | D | 2e-07 | 1.2e-07 | 1.3e-07 | 1.500E-07 | 4.359E-08 | 1.200E-07 | 2.000E-07 |
| 21813 | 1091 | V1091M | D | 2e-07 | 2.1e-07 | 5e-07 | 3.033E-07 | 1.704E-07 | 2.000E-07 | 5.000E-07 |
| 21814 | 1091 | V1091F | D | 4.1e-07 | 1.8e-07 | 4e-07 | 3.300E-07 | 1.300E-07 | 1.800E-07 | 4.100E-07 |
| 21815 | 1091 | V1091P | D | 2.6e-07 | 4.1e-07 | 2.7e-07 | 3.133E-07 | 8.386E-08 | 2.600E-07 | 4.100E-07 |
| 21816 | 1091 | V1091S | D | 4.2e-07 | 1.8e-07 | 3.2e-07 | 3.067E-07 | 1.206E-07 | 1.800E-07 | 4.200E-07 |
| 21817 | 1091 | V1091T | D | 2.7e-07 | 2.2e-07 | 2.3e-07 | 2.400E-07 | 2.646E-08 | 2.200E-07 | 2.700E-07 |
| 21818 | 1091 | V1091W | D | 2.4e-07 | 2.3e-07 | 5.7e-07 | 3.467E-07 | 1.935E-07 | 2.300E-07 | 5.700E-07 |
| 21819 | 1091 | V1091Y | D | 2.4e-07 | 1.5e-07 | 2.9e-07 | 2.267E-07 | 7.095E-08 | 1.500E-07 | 2.900E-07 |
| 21820 | 1091 | V1091V | D | 3.1e-07 | 8.5e-08 | 2e-07 | 1.983E-07 | 1.125E-07 | 8.500E-08 | 3.100E-07 |
| 21821 | 1092 | A1092A | D | 1.6e-07 | 1.9e-07 | 1.9e-07 | 1.800E-07 | 1.732E-08 | 1.600E-07 | 1.900E-07 |
| 21822 | 1092 | A1092R | D | 2.3e-07 | 1.5e-07 | 4e-07 | 2.600E-07 | 1.277E-07 | 1.500E-07 | 4.000E-07 |
| 21823 | 1092 | A1092N | D | 3.3e-07 | 1.4e-07 | 2.8e-07 | 2.500E-07 | 9.849E-08 | 1.400E-07 | 3.300E-07 |
| 21824 | 1092 | A1092D | D | 1.8e-07 | 4.1e-07 | 3.9e-07 | 3.267E-07 | 1.274E-07 | 1.800E-07 | 4.100E-07 |
| 21825 | 1092 | A1092C | D | 3.1e-07 | 3.2e-07 | 3.4e-07 | 3.233E-07 | 1.528E-08 | 3.100E-07 | 3.400E-07 |
| 21826 | 1092 | A1092E | D | 3.7e-07 | 2.1e-07 | 2.5e-07 | 2.767E-07 | 8.327E-08 | 2.100E-07 | 3.700E-07 |
| 21827 | 1092 | A1092Q | D | 2.4e-07 | 2e-07 | 1.9e-07 | 2.100E-07 | 2.646E-08 | 1.900E-07 | 2.400E-07 |
| 21828 | 1092 | A1092G | D | 2e-07 | 2.8e-07 | 1.1e-07 | 1.967E-07 | 8.505E-08 | 1.100E-07 | 2.800E-07 |
| 21829 | 1092 | A1092H | D | 3e-07 | 2.3e-07 | 2.4e-07 | 2.567E-07 | 3.786E-08 | 2.300E-07 | 3.000E-07 |
| 21830 | 1092 | A1092I | D | 1.8e-07 | 1.6e-07 | 1.5e-07 | 1.633E-07 | 1.528E-08 | 1.500E-07 | 1.800E-07 |
| 21831 | 1092 | A1092L | D | 1.5e-07 | 1.6e-07 | 2e-07 | 1.700E-07 | 2.646E-08 | 1.500E-07 | 2.000E-07 |
| 21832 | 1092 | A1092K | D | 2e-07 | 3.3e-07 | 2.6e-07 | 2.633E-07 | 6.506E-08 | 2.000E-07 | 3.300E-07 |
| 21833 | 1092 | A1092M | D | 2.8e-07 | 8.4e-08 | 3.5e-07 | 2.380E-07 | 1.379E-07 | 8.400E-08 | 3.500E-07 |
| 21834 | 1092 | A1092F | D | 3.6e-07 | 2.4e-07 | 1.9e-07 | 2.633E-07 | 8.737E-08 | 1.900E-07 | 3.600E-07 |
| 21835 | 1092 | A1092P | D | 1.9e-07 | 2.6e-07 | 2.4e-07 | 2.300E-07 | 3.606E-08 | 1.900E-07 | 2.600E-07 |
| 21836 | 1092 | A1092S | D | 2.6e-07 | 2.1e-07 | 1e-06 | 4.900E-07 | 4.424E-07 | 2.100E-07 | 1.000E-06 |
| 21837 | 1092 | A1092T | D | 3.7e-07 | 1.9e-07 | 2.3e-07 | 2.633E-07 | 9.452E-08 | 1.900E-07 | 3.700E-07 |
| 21838 | 1092 | A1092W | D | 2.7e-07 | 1.4e-07 | 2.3e-07 | 2.133E-07 | 6.658E-08 | 1.400E-07 | 2.700E-07 |
| 21839 | 1092 | A1092Y | D | 2.7e-07 | 3.9e-07 | 2.9e-07 | 3.167E-07 | 6.429E-08 | 2.700E-07 | 3.900E-07 |
| 21840 | 1092 | A1092V | D | 2e-07 | 2.4e-07 | 2.9e-07 | 2.433E-07 | 4.509E-08 | 2.000E-07 | 2.900E-07 |
| 21841 | 1093 | W1093A | D | 2.1e-07 | 4e-07 | 4.2e-07 | 3.433E-07 | 1.159E-07 | 2.100E-07 | 4.200E-07 |
| 21842 | 1093 | W1093R | D | 3.4e-07 | 4.7e-07 | 4.1e-07 | 4.067E-07 | 6.506E-08 | 3.400E-07 | 4.700E-07 |
| 21843 | 1093 | W1093N | D | 2.2e-07 | 4.1e-07 | 1.4e-07 | 2.567E-07 | 1.387E-07 | 1.400E-07 | 4.100E-07 |
| 21844 | 1093 | W1093D | D | 1.8e-07 | 2e-07 | 2.6e-07 | 2.133E-07 | 4.163E-08 | 1.800E-07 | 2.600E-07 |

| | | | | | | | | | | |
|---|---|---|---|---|---|---|---|---|---|---|
| 21845 | 1093 | W1093C | D | 2.2e-07 | 3.2e-07 | 5.8e-07 | 3.733E-07 | 1.858E-07 | 2.200E-07 | 5.800E-07 |
| 21846 | 1093 | W1093E | D | 1.7e-07 | 2.6e-07 | 3.2e-07 | 2.500E-07 | 7.550E-08 | 1.700E-07 | 3.200E-07 |
| 21847 | 1093 | W1093Q | D | 2.2e-07 | 2.6e-07 | 1.5e-07 | 2.100E-07 | 5.568E-08 | 1.500E-07 | 2.600E-07 |
| 21848 | 1093 | W1093G | D | 2.8e-07 | 3.8e-07 | 2.7e-07 | 3.100E-07 | 6.083E-08 | 2.700E-07 | 3.800E-07 |
| 21849 | 1093 | W1093H | D | 3.3e-07 | 2.4e-07 | 6.3e-07 | 4.000E-07 | 2.042E-07 | 2.400E-07 | 6.300E-07 |
| 21850 | 1093 | W1093I | D | 2.5e-07 | 2e-07 | 3e-07 | 2.500E-07 | 5.000E-08 | 2.000E-07 | 3.000E-07 |
| 21851 | 1093 | W1093L | D | 2.4e-07 | 1e-07 | 1.1e-07 | 1.500E-07 | 7.810E-08 | 1.000E-07 | 2.400E-07 |
| 21852 | 1093 | W1093K | D | 2.6e-07 | 4e-07 | 1.7e-07 | 2.767E-07 | 1.159E-07 | 1.700E-07 | 4.000E-07 |
| 21853 | 1093 | W1093M | D | 3e-07 | 2.2e-07 | 4.1e-07 | 3.100E-07 | 9.539E-08 | 2.200E-07 | 4.100E-07 |
| 21854 | 1093 | W1093F | D | 2.7e-07 | 1e-07 | 3.4e-07 | 2.367E-07 | 1.234E-07 | 1.000E-07 | 3.400E-07 |
| 21855 | 1093 | W1093P | D | 2e-07 | 2.4e-07 | 1.9e-07 | 2.100E-07 | 2.646E-08 | 1.900E-07 | 2.400E-07 |
| 21856 | 1093 | W1093S | D | 2.9e-07 | 2.5e-07 | 3.1e-07 | 2.833E-07 | 3.055E-08 | 2.500E-07 | 3.100E-07 |
| 21857 | 1093 | W1093T | D | 3.3e-07 | 2.4e-07 | 1.7e-07 | 2.467E-07 | 8.021E-08 | 1.700E-07 | 3.300E-07 |
| 21858 | 1093 | W1093W | D | 8.7e-08 | 1.7e-07 | 1.8e-07 | 1.457E-07 | 5.105E-08 | 8.700E-08 | 1.800E-07 |
| 21859 | 1093 | W1093Y | D | 3.6e-07 | 2.1e-07 | 4e-07 | 3.233E-07 | 1.002E-07 | 2.100E-07 | 4.000E-07 |
| 21860 | 1093 | W1093V | D | 3.9e-07 | 1.3e-07 | 3.5e-07 | 2.900E-07 | 1.400E-07 | 1.300E-07 | 3.900E-07 |
| 21861 | 1094 | Y1094A | D | 5.9e-07 | 2e-07 | 2.6e-07 | 3.500E-07 | 2.100E-07 | 2.000E-07 | 5.900E-07 |
| 21862 | 1094 | Y1094R | D | 2.3e-07 | 2.2e-07 | 3e-07 | 2.500E-07 | 4.359E-08 | 2.200E-07 | 3.000E-07 |
| 21863 | 1094 | Y1094N | D | 2.2e-07 | 3.8e-07 | 1.7e-07 | 2.567E-07 | 1.097E-07 | 1.700E-07 | 3.800E-07 |
| 21864 | 1094 | Y1094D | D | 2.7e-07 | 3.6e-07 | 1.1e-07 | 2.467E-07 | 1.266E-07 | 1.100E-07 | 3.600E-07 |
| 21865 | 1094 | Y1094C | D | 3.3e-07 | 1.7e-07 | 1.2e-07 | 2.067E-07 | 1.097E-07 | 1.200E-07 | 3.300E-07 |
| 21866 | 1094 | Y1094E | D | 1.9e-07 | 1.8e-07 | 1.8e-07 | 1.833E-07 | 5.774E-09 | 1.800E-07 | 1.900E-07 |
| 21867 | 1094 | Y1094Q | D | 1.6e-07 | 4e-07 | 4.5e-07 | 3.367E-07 | 1.550E-07 | 1.600E-07 | 4.500E-07 |
| 21868 | 1094 | Y1094G | D | 1.3e-07 | 3.5e-07 | 6.5e-08 | 1.817E-07 | 1.494E-07 | 6.500E-08 | 3.500E-07 |
| 21869 | 1094 | Y1094H | D | 2.4e-07 | 2.1e-07 | 2e-07 | 2.167E-07 | 2.082E-08 | 2.000E-07 | 2.400E-07 |
| 21870 | 1094 | Y1094I | D | 2.9e-07 | 2.3e-07 | 3.9e-07 | 3.033E-07 | 8.083E-08 | 2.300E-07 | 3.900E-07 |
| 21871 | 1094 | Y1094L | D | 2.8e-07 | 2.4e-07 | 2.3e-07 | 2.500E-07 | 2.646E-08 | 2.300E-07 | 2.800E-07 |
| 21872 | 1094 | Y1094K | D | 4.1e-07 | 2.5e-07 | 3.1e-07 | 3.233E-07 | 8.083E-08 | 2.500E-07 | 4.100E-07 |
| 21873 | 1094 | Y1094M | D | 1.9e-07 | 5e-07 | 1.6e-07 | 2.833E-07 | 1.882E-07 | 1.600E-07 | 5.000E-07 |
| 21874 | 1094 | Y1094F | D | 2.6e-07 | 1.2e-07 | 2.3e-07 | 2.033E-07 | 7.371E-08 | 1.200E-07 | 2.600E-07 |
| 21875 | 1094 | Y1094P | D | 2e-07 | 1.8e-07 | 1.5e-07 | 1.767E-07 | 2.517E-08 | 1.500E-07 | 2.000E-07 |
| 21876 | 1094 | Y1094S | D | 2.5e-07 | 2.9e-07 | 8.3e-07 | 4.567E-07 | 3.239E-07 | 2.500E-07 | 8.300E-07 |
| 21877 | 1094 | Y1094T | D | 1.9e-07 | 1.4e-07 | 1.1e-07 | 1.467E-07 | 4.041E-08 | 1.100E-07 | 1.900E-07 |
| 21878 | 1094 | Y1094W | D | 2.9e-07 | 7.4e-08 | 4.8e-07 | 2.813E-07 | 2.031E-07 | 7.400E-08 | 4.800E-07 |
| 21879 | 1094 | Y1094Y | D | 2.1e-07 | 8.3e-08 | 2.4e-07 | 1.777E-07 | 8.334E-08 | 8.300E-08 | 2.400E-07 |
| 21880 | 1094 | Y1094V | D | 2.7e-07 | 1.8e-07 | 2.6e-07 | 2.367E-07 | 4.933E-08 | 1.800E-07 | 2.700E-07 |
| 21881 | 1095 | Q1095A | D | 2.3e-07 | 1.5e-07 | 2.3e-07 | 2.033E-07 | 4.619E-08 | 1.500E-07 | 2.300E-07 |
| 21882 | 1095 | Q1095R | D | 1.6e-07 | 4.3e-07 | 2.8e-07 | 2.900E-07 | 1.353E-07 | 1.600E-07 | 4.300E-07 |
| 21883 | 1095 | Q1095N | D | 2.5e-07 | 1.5e-07 | 1.5e-07 | 1.833E-07 | 5.774E-08 | 1.500E-07 | 2.500E-07 |
| 21884 | 1095 | Q1095D | D | 3e-07 | 1.8e-07 | 1.8e-07 | 2.200E-07 | 6.928E-08 | 1.800E-07 | 3.000E-07 |
| 21885 | 1095 | Q1095C | D | 3.8e-07 | 3.5e-07 | 1.7e-07 | 3.000E-07 | 1.136E-07 | 1.700E-07 | 3.800E-07 |
| 21886 | 1095 | Q1095E | D | 4.6e-07 | 2.3e-07 | 3.3e-07 | 3.400E-07 | 1.153E-07 | 2.300E-07 | 4.600E-07 |
| 21887 | 1095 | Q1095Q | D | 4.8e-07 | 2.2e-07 | 2e-07 | 3.000E-07 | 1.562E-07 | 2.000E-07 | 4.800E-07 |
| 21888 | 1095 | Q1095G | D | 2.3e-07 | 3.3e-07 | 2.6e-07 | 2.733E-07 | 5.132E-08 | 2.300E-07 | 3.300E-07 |
| 21889 | 1095 | Q1095H | D | 1.6e-07 | 5.2e-07 | 2.8e-07 | 3.200E-07 | 1.833E-07 | 1.600E-07 | 5.200E-07 |
| 21890 | 1095 | Q1095I | D | 3.2e-07 | 2.8e-07 | 2.3e-07 | 2.767E-07 | 4.509E-08 | 2.300E-07 | 3.200E-07 |
| 21891 | 1095 | Q1095L | D | 3e-07 | 2.9e-07 | 1.4e-07 | 2.433E-07 | 8.963E-08 | 1.400E-07 | 3.000E-07 |
| 21892 | 1095 | Q1095K | D | 2.7e-07 | 2e-07 | 1.6e-07 | 2.100E-07 | 5.568E-08 | 1.600E-07 | 2.700E-07 |
| 21893 | 1095 | Q1095M | D | 1.8e-07 | 2.6e-07 | 2.4e-07 | 2.267E-07 | 4.163E-08 | 1.800E-07 | 2.600E-07 |
| 21894 | 1095 | Q1095F | D | 2.7e-07 | 1.7e-07 | 4.4e-07 | 2.933E-07 | 1.365E-07 | 1.700E-07 | 4.400E-07 |
| 21895 | 1095 | Q1095P | D | 3.3e-07 | 1.6e-07 | 4e-07 | 2.967E-07 | 1.234E-07 | 1.600E-07 | 4.000E-07 |
| 21896 | 1095 | Q1095S | D | 1.8e-07 | 3.1e-07 | 7.5e-07 | 4.133E-07 | 2.987E-07 | 1.800E-07 | 7.500E-07 |
| 21897 | 1095 | Q1095T | D | 2.1e-07 | 1.2e-07 | 1.8e-07 | 1.700E-07 | 4.583E-08 | 1.200E-07 | 2.100E-07 |
| 21898 | 1095 | Q1095W | D | 1.7e-07 | 3.3e-07 | 3e-07 | 2.667E-07 | 8.505E-08 | 1.700E-07 | 3.300E-07 |
| 21899 | 1095 | Q1095Y | D | 3e-07 | 2e-07 | 2.1e-07 | 2.367E-07 | 5.508E-08 | 2.000E-07 | 3.000E-07 |
| 21900 | 1095 | Q1095V | D | 3.3e-07 | 2.5e-07 | 2.2e-07 | 2.667E-07 | 5.686E-08 | 2.200E-07 | 3.300E-07 |
| 21901 | 1096 | Q1096A | D | 9e-08 | 2.8e-07 | 6.2e-07 | 3.300E-07 | 2.685E-07 | 9.000E-08 | 6.200E-07 |
| 21902 | 1096 | Q1096R | D | 2.9e-07 | 4e-07 | 3.5e-07 | 3.467E-07 | 5.508E-08 | 2.900E-07 | 4.000E-07 |
| 21903 | 1096 | Q1096N | D | 3e-07 | 7.5e-08 | 1.6e-07 | 1.783E-07 | 1.136E-07 | 7.500E-08 | 3.000E-07 |
| 21904 | 1096 | Q1096D | D | 6.4e-08 | 3.3e-07 | 2.8e-07 | 2.247E-07 | 1.414E-07 | 6.400E-08 | 3.300E-07 |
| 21905 | 1096 | Q1096C | D | 3.6e-07 | 3.2e-07 | 1.1e-07 | 2.633E-07 | 1.343E-07 | 1.100E-07 | 3.600E-07 |
| 21906 | 1096 | Q1096E | D | 1.8e-07 | 2.9e-07 | 2.7e-07 | 2.467E-07 | 5.859E-08 | 1.800E-07 | 2.900E-07 |
| 21907 | 1096 | Q1096Q | D | 1.9e-07 | 2.4e-07 | 1.3e-07 | 1.867E-07 | 5.508E-08 | 1.300E-07 | 2.400E-07 |
| 21908 | 1096 | Q1096G | D | 1.3e-07 | 1.8e-07 | 1.6e-07 | 1.567E-07 | 2.517E-08 | 1.300E-07 | 1.800E-07 |
| 21909 | 1096 | Q1096H | D | 7.8e-08 | 1.4e-07 | 3.2e-07 | 1.793E-07 | 1.257E-07 | 7.800E-08 | 3.200E-07 |
| 21910 | 1096 | Q1096I | D | 2.2e-07 | 1.1e-07 | 4.1e-07 | 2.467E-07 | 1.518E-07 | 1.100E-07 | 4.100E-07 |
| 21911 | 1096 | Q1096L | D | 4.4e-07 | 5.3e-07 | 2.1e-07 | 3.933E-07 | 1.650E-07 | 2.100E-07 | 5.300E-07 |
| 21912 | 1096 | Q1096K | D | 4.3e-07 | 3e-07 | 1.5e-07 | 2.933E-07 | 1.401E-07 | 1.500E-07 | 4.300E-07 |
| 21913 | 1096 | Q1096M | D | 7.1e-07 | 4.4e-07 | 4.4e-07 | 5.300E-07 | 1.559E-07 | 4.400E-07 | 7.100E-07 |
| 21914 | 1096 | Q1096F | D | 2.9e-07 | 3e-07 | 2.2e-07 | 2.700E-07 | 4.359E-08 | 2.200E-07 | 3.000E-07 |
| 21915 | 1096 | Q1096P | D | 2.8e-07 | 1.8e-07 | 2.6e-07 | 2.400E-07 | 5.292E-08 | 1.800E-07 | 2.800E-07 |
| 21916 | 1096 | Q1096S | D | 2.2e-07 | 3.8e-07 | 2e-07 | 2.667E-07 | 9.866E-08 | 2.000E-07 | 3.800E-07 |
| 21917 | 1096 | Q1096T | D | 2.4e-07 | 9.6e-08 | 5.7e-07 | 3.020E-07 | 2.430E-07 | 9.600E-08 | 5.700E-07 |
| 21918 | 1096 | Q1096W | D | 2.2e-07 | 2.3e-07 | 6.6e-08 | 1.720E-07 | 9.193E-08 | 6.600E-08 | 2.300E-07 |
| 21919 | 1096 | Q1096Y | D | 1.7e-07 | 3.5e-07 | 1.5e-07 | 2.233E-07 | 1.102E-07 | 1.500E-07 | 3.500E-07 |
| 21920 | 1096 | Q1096V | D | 4e-07 | 2e-07 | 5.1e-08 | 2.170E-07 | 1.751E-07 | 5.100E-08 | 4.000E-07 |
| 21921 | 1097 | K1097A | D | 1.9e-07 | 4.3e-07 | 1.5e-07 | 2.567E-07 | 1.514E-07 | 1.500E-07 | 4.300E-07 |
| 21922 | 1097 | K1097R | D | 4.6e-07 | 6e-07 | 4.8e-07 | 5.133E-07 | 7.572E-08 | 4.600E-07 | 6.000E-07 |
| 21923 | 1097 | K1097N | D | 3.6e-07 | 2.6e-07 | 1.5e-07 | 2.567E-07 | 1.050E-07 | 1.500E-07 | 3.600E-07 |
| 21924 | 1097 | K1097D | D | 3.2e-07 | 2.9e-07 | 1.8e-07 | 2.633E-07 | 7.371E-08 | 1.800E-07 | 3.200E-07 |
| 21925 | 1097 | K1097C | D | 2e-07 | 2.3e-07 | 2.8e-07 | 2.367E-07 | 4.041E-08 | 2.000E-07 | 2.800E-07 |
| 21926 | 1097 | K1097E | D | 3.2e-07 | 2.4e-07 | 1.5e-07 | 2.367E-07 | 8.505E-08 | 1.500E-07 | 3.200E-07 |
| 21927 | 1097 | K1097Q | D | 2.9e-07 | 4e-07 | 2.6e-07 | 3.167E-07 | 7.371E-08 | 2.600E-07 | 4.000E-07 |
| 21928 | 1097 | K1097G | D | 1.7e-07 | 6.7e-07 | 2.4e-07 | 3.600E-07 | 2.707E-07 | 1.700E-07 | 6.700E-07 |
| 21929 | 1097 | K1097H | D | 2.4e-07 | 4.3e-07 | 3.6e-07 | 3.433E-07 | 9.609E-08 | 2.400E-07 | 4.300E-07 |

| | | | | | | | | | | |
|---|---|---|---|---|---|---|---|---|---|---|
| 21930 | 1097 | K1097I | D | 5.3e-07 | 2.1e-07 | 1.8e-07 | 3.067E-07 | 1.940E-07 | 1.800E-07 | 5.300E-07 |
| 21931 | 1097 | K1097L | D | 1.9e-07 | 4.4e-07 | 1.3e-07 | 2.533E-07 | 1.644E-07 | 1.300E-07 | 4.400E-07 |
| 21932 | 1097 | K1097K | D | 2.7e-07 | 1.5e-07 | 2.4e-07 | 2.200E-07 | 6.245E-08 | 1.500E-07 | 2.700E-07 |
| 21933 | 1097 | K1097M | D | 1.1e-07 | 3.1e-07 | 3.2e-07 | 2.467E-07 | 1.185E-07 | 1.100E-07 | 3.200E-07 |
| 21934 | 1097 | K1097F | D | 3.3e-07 | 3.9e-07 | 2e-07 | 3.067E-07 | 9.713E-08 | 2.000E-07 | 3.900E-07 |
| 21935 | 1097 | K1097P | D | 1e-07 | 2.8e-07 | 2.5e-07 | 2.100E-07 | 9.644E-08 | 1.000E-07 | 2.800E-07 |
| 21936 | 1097 | K1097S | D | 6.5e-07 | 2.8e-07 | 5e-07 | 4.767E-07 | 1.861E-07 | 2.800E-07 | 6.500E-07 |
| 21937 | 1097 | K1097T | D | 2.2e-07 | 8.1e-07 | 1.4e-07 | 3.900E-07 | 3.659E-07 | 1.400E-07 | 8.100E-07 |
| 21938 | 1097 | K1097W | D | 2.5e-07 | 1.6e-07 | 2.7e-07 | 2.267E-07 | 5.859E-08 | 1.600E-07 | 2.700E-07 |
| 21939 | 1097 | K1097Y | D | 3e-07 | 2.2e-07 | 1.4e-07 | 2.200E-07 | 8.000E-08 | 1.400E-07 | 3.000E-07 |
| 21940 | 1097 | K1097V | D | 3e-07 | 3.3e-07 | 1.7e-07 | 2.667E-07 | 8.505E-08 | 1.700E-07 | 3.300E-07 |
| 21941 | 1098 | P1098A | D | 3.8e-07 | 3.9e-07 | 2.6e-07 | 3.433E-07 | 7.234E-08 | 2.600E-07 | 3.900E-07 |
| 21942 | 1098 | P1098R | D | 3.3e-07 | 3.8e-07 | 2.1e-07 | 3.067E-07 | 8.737E-08 | 2.100E-07 | 3.800E-07 |
| 21943 | 1098 | P1098N | D | 1.9e-07 | 1.6e-07 | 2.1e-07 | 1.867E-07 | 2.517E-08 | 1.600E-07 | 2.100E-07 |
| 21944 | 1098 | P1098D | D | 4.3e-07 | 3.8e-07 | 1.7e-07 | 3.267E-07 | 1.380E-07 | 1.700E-07 | 4.300E-07 |
| 21945 | 1098 | P1098C | D | 3.7e-07 | 1.8e-07 | 2.8e-07 | 2.767E-07 | 9.504E-08 | 1.800E-07 | 3.700E-07 |
| 21946 | 1098 | P1098E | D | 1.2e-07 | 3.5e-07 | 3.5e-07 | 2.733E-07 | 1.328E-07 | 1.200E-07 | 3.500E-07 |
| 21947 | 1098 | P1098Q | D | 2.3e-07 | 2.7e-07 | 3.5e-07 | 2.833E-07 | 6.110E-08 | 2.300E-07 | 3.500E-07 |
| 21948 | 1098 | P1098G | D | 2.3e-07 | 7.3e-07 | 2e-07 | 3.867E-07 | 2.977E-07 | 2.000E-07 | 7.300E-07 |
| 21949 | 1098 | P1098H | D | 1.6e-07 | 1.5e-07 | 5.5e-07 | 2.867E-07 | 2.281E-07 | 1.500E-07 | 5.500E-07 |
| 21950 | 1098 | P1098I | D | 1.3e-07 | 3.4e-07 | 4.8e-07 | 3.167E-07 | 1.762E-07 | 1.300E-07 | 4.800E-07 |
| 21951 | 1098 | P1098L | D | 2.7e-07 | 2.1e-07 | 1.6e-07 | 2.133E-07 | 5.508E-08 | 1.600E-07 | 2.700E-07 |
| 21952 | 1098 | P1098K | D | 6.7e-07 | 1.7e-07 | 2.9e-07 | 3.767E-07 | 2.610E-07 | 1.700E-07 | 6.700E-07 |
| 21953 | 1098 | P1098M | D | 2.2e-07 | 3.1e-07 | 1.5e-07 | 2.267E-07 | 8.021E-08 | 1.500E-07 | 3.100E-07 |
| 21954 | 1098 | P1098F | D | 1.9e-07 | 2.7e-07 | 2.8e-07 | 2.467E-07 | 4.933E-08 | 1.900E-07 | 2.800E-07 |
| 21955 | 1098 | P1098P | D | 3e-07 | 2e-07 | 3.6e-07 | 2.867E-07 | 8.083E-08 | 2.000E-07 | 3.600E-07 |
| 21956 | 1098 | P1098S | D | 4.6e-07 | 4e-07 | 2.3e-07 | 3.633E-07 | 1.193E-07 | 2.300E-07 | 4.600E-07 |
| 21957 | 1098 | P1098T | D | 1.2e-07 | 2.5e-07 | 2.2e-07 | 1.967E-07 | 6.807E-08 | 1.200E-07 | 2.500E-07 |
| 21958 | 1098 | P1098W | D | 2e-07 | 2.1e-07 | 2.7e-07 | 2.267E-07 | 3.786E-08 | 2.000E-07 | 2.700E-07 |
| 21959 | 1098 | P1098Y | D | 1.9e-07 | 7.7e-07 | 1.5e-07 | 3.700E-07 | 3.470E-07 | 1.500E-07 | 7.700E-07 |
| 21960 | 1098 | P1098V | D | 3.5e-07 | 5e-07 | 3.1e-07 | 3.867E-07 | 1.002E-07 | 3.100E-07 | 5.000E-07 |
| 21961 | 1099 | G1099A | D | 4.9e-07 | 2.1e-07 | 3e-07 | 3.333E-07 | 1.429E-07 | 2.100E-07 | 4.900E-07 |
| 21962 | 1099 | G1099R | D | 3.9e-07 | 1.1e-07 | 4.2e-07 | 3.067E-07 | 1.710E-07 | 1.100E-07 | 4.200E-07 |
| 21963 | 1099 | G1099N | D | 1.8e-07 | 6.1e-07 | 2.9e-07 | 3.600E-07 | 2.234E-07 | 1.800E-07 | 6.100E-07 |
| 21964 | 1099 | G1099D | D | 2.2e-07 | 3.6e-07 | 2e-07 | 2.600E-07 | 8.718E-08 | 2.000E-07 | 3.600E-07 |
| 21965 | 1099 | G1099C | D | 4.1e-07 | 2.2e-07 | 1.7e-07 | 2.667E-07 | 1.266E-07 | 1.700E-07 | 4.100E-07 |
| 21966 | 1099 | G1099E | D | 3.7e-07 | 1e-07 | 2.7e-07 | 2.467E-07 | 1.365E-07 | 1.000E-07 | 3.700E-07 |
| 21967 | 1099 | G1099Q | D | 3.5e-07 | 1.8e-07 | 6.7e-07 | 4.000E-07 | 2.488E-07 | 1.800E-07 | 6.700E-07 |
| 21968 | 1099 | G1099G | D | 2.4e-07 | 4.8e-07 | 3e-07 | 3.400E-07 | 1.249E-07 | 2.400E-07 | 4.800E-07 |
| 21969 | 1099 | G1099H | D | 4.8e-07 | 4.9e-07 | 2e-07 | 3.900E-07 | 1.646E-07 | 2.000E-07 | 4.900E-07 |
| 21970 | 1099 | G1099I | D | 2.5e-07 | 6.7e-07 | 2.2e-07 | 3.800E-07 | 2.516E-07 | 2.200E-07 | 6.700E-07 |
| 21971 | 1099 | G1099L | D | 1.6e-07 | 3.2e-07 | 4e-07 | 2.933E-07 | 1.222E-07 | 1.600E-07 | 4.000E-07 |
| 21972 | 1099 | G1099K | D | 2.7e-07 | 1.8e-07 | 1.8e-07 | 2.100E-07 | 5.196E-08 | 1.800E-07 | 2.700E-07 |
| 21973 | 1099 | G1099M | D | 1.5e-07 | 1.4e-07 | 3.3e-07 | 2.067E-07 | 1.069E-07 | 1.400E-07 | 3.300E-07 |
| 21974 | 1099 | G1099F | D | 3e-07 | 2.4e-07 | 2.8e-07 | 2.733E-07 | 3.055E-08 | 2.400E-07 | 3.000E-07 |
| 21975 | 1099 | G1099P | D | 9.7e-08 | 4e-07 | 2.9e-07 | 2.623E-07 | 1.534E-07 | 9.700E-08 | 4.000E-07 |
| 21976 | 1099 | G1099S | D | 3e-07 | 2.7e-07 | 1.5e-07 | 2.400E-07 | 7.937E-08 | 1.500E-07 | 3.000E-07 |
| 21977 | 1099 | G1099T | D | 1.2e-07 | 4.4e-07 | 2.1e-07 | 2.567E-07 | 1.650E-07 | 1.200E-07 | 4.400E-07 |
| 21978 | 1099 | G1099W | D | 1.7e-07 | 1.9e-07 | 3.6e-07 | 2.400E-07 | 1.044E-07 | 1.700E-07 | 3.600E-07 |
| 21979 | 1099 | G1099Y | D | 2.2e-07 | 2.1e-07 | 2.3e-07 | 2.200E-07 | 1.000E-08 | 2.100E-07 | 2.300E-07 |
| 21980 | 1099 | G1099V | D | 5.4e-07 | 2.1e-07 | 1.7e-07 | 3.067E-07 | 2.031E-07 | 1.700E-07 | 5.400E-07 |
| 21981 | 1100 | K1100A | D | 5.2e-07 | 2.7e-07 | 2e-07 | 3.300E-07 | 1.682E-07 | 2.000E-07 | 5.200E-07 |
| 21982 | 1100 | K1100R | D | 2.3e-07 | 1.9e-07 | 3.4e-07 | 2.533E-07 | 7.767E-08 | 1.900E-07 | 3.400E-07 |
| 21983 | 1100 | K1100N | D | 1.8e-07 | 1.1e-07 | 2e-07 | 1.633E-07 | 4.726E-08 | 1.100E-07 | 2.000E-07 |
| 21984 | 1100 | K1100D | D | 7e-07 | 1.5e-07 | 1.3e-07 | 3.267E-07 | 3.235E-07 | 1.300E-07 | 7.000E-07 |
| 21985 | 1100 | K1100C | D | 2.2e-07 | 1.8e-07 | 2.3e-07 | 2.100E-07 | 2.646E-08 | 1.800E-07 | 2.300E-07 |
| 21986 | 1100 | K1100E | D | 2e-07 | 2e-07 | 3.4e-07 | 2.467E-07 | 8.083E-08 | 2.000E-07 | 3.400E-07 |
| 21987 | 1100 | K1100Q | D | 1.8e-07 | 5e-07 | 3.7e-07 | 3.500E-07 | 1.609E-07 | 1.800E-07 | 5.000E-07 |
| 21988 | 1100 | K1100G | D | 1.4e-07 | 1.5e-07 | 1.1e-07 | 1.333E-07 | 2.082E-08 | 1.100E-07 | 1.500E-07 |
| 21989 | 1100 | K1100H | D | 1.4e-07 | 2.2e-07 | 4.4e-07 | 2.667E-07 | 1.553E-07 | 1.400E-07 | 4.400E-07 |
| 21990 | 1100 | K1100I | D | 3.4e-07 | 1.4e-07 | 1.2e-07 | 2.000E-07 | 1.217E-07 | 1.200E-07 | 3.400E-07 |
| 21991 | 1100 | K1100L | D | 2.3e-07 | 2.1e-07 | 3.7e-07 | 2.700E-07 | 8.718E-08 | 2.100E-07 | 3.700E-07 |
| 21992 | 1100 | K1100K | D | 2.2e-07 | 2.8e-07 | 2.5e-07 | 2.500E-07 | 3.000E-08 | 2.200E-07 | 2.800E-07 |
| 21993 | 1100 | K1100M | D | 1e-07 | 1.5e-07 | 3.7e-07 | 2.067E-07 | 1.436E-07 | 1.000E-07 | 3.700E-07 |
| 21994 | 1100 | K1100F | D | 1.4e-07 | 1.8e-07 | 2.2e-07 | 1.800E-07 | 4.000E-08 | 1.400E-07 | 2.200E-07 |
| 21995 | 1100 | K1100P | D | 2.5e-07 | 1.6e-07 | 1.7e-07 | 1.933E-07 | 4.933E-08 | 1.600E-07 | 2.500E-07 |
| 21996 | 1100 | K1100S | D | 3e-07 | 1.2e-07 | 3.5e-07 | 2.567E-07 | 1.210E-07 | 1.200E-07 | 3.500E-07 |
| 21997 | 1100 | K1100T | D | 2.2e-07 | 2.1e-07 | 2.5e-07 | 2.267E-07 | 2.082E-08 | 2.100E-07 | 2.500E-07 |
| 21998 | 1100 | K1100W | D | 1.8e-07 | 3.7e-07 | 3.6e-07 | 3.033E-07 | 1.069E-07 | 1.800E-07 | 3.700E-07 |
| 21999 | 1100 | K1100Y | D | 4e-07 | 2e-07 | 5.6e-07 | 3.867E-07 | 1.804E-07 | 2.000E-07 | 5.600E-07 |
| 22000 | 1100 | K1100V | D | 2.8e-07 | 1.2e-07 | 4e-07 | 2.667E-07 | 1.405E-07 | 1.200E-07 | 4.000E-07 |
| 22001 | 1101 | A1101A | D | 1.1e-07 | 4.8e-07 | 2.8e-07 | 2.900E-07 | 1.852E-07 | 1.100E-07 | 4.800E-07 |
| 22002 | 1101 | A1101R | D | 4.1e-07 | 4.5e-07 | 2.6e-07 | 3.733E-07 | 1.002E-07 | 2.600E-07 | 4.500E-07 |
| 22003 | 1101 | A1101N | D | 1e-07 | 2e-07 | 1.8e-07 | 1.600E-07 | 5.292E-08 | 1.000E-07 | 2.000E-07 |
| 22004 | 1101 | A1101D | D | 1.9e-07 | 3.6e-07 | 1.7e-07 | 2.400E-07 | 1.044E-07 | 1.700E-07 | 3.600E-07 |
| 22005 | 1101 | A1101C | D | 3.7e-07 | 1.9e-07 | 2.4e-07 | 2.667E-07 | 9.292E-08 | 1.900E-07 | 3.700E-07 |
| 22006 | 1101 | A1101E | D | 2.1e-07 | 1.7e-07 | 3.7e-07 | 2.500E-07 | 1.058E-07 | 1.700E-07 | 3.700E-07 |
| 22007 | 1101 | A1101Q | D | 3.8e-07 | 3e-07 | 3.1e-07 | 3.300E-07 | 4.359E-08 | 3.000E-07 | 3.800E-07 |
| 22008 | 1101 | A1101G | D | 3.9e-07 | 2.9e-07 | 2e-07 | 2.933E-07 | 9.504E-08 | 2.000E-07 | 3.900E-07 |
| 22009 | 1101 | A1101H | D | 3.4e-07 | 9.3e-08 | 4.5e-07 | 2.943E-07 | 1.828E-07 | 9.300E-08 | 4.500E-07 |
| 22010 | 1101 | A1101I | D | 3.1e-07 | 1.8e-07 | 1.7e-07 | 2.200E-07 | 7.810E-08 | 1.700E-07 | 3.100E-07 |
| 22011 | 1101 | A1101L | D | 1.4e-07 | 2.1e-07 | 3.3e-07 | 2.267E-07 | 9.609E-08 | 1.400E-07 | 3.300E-07 |
| 22012 | 1101 | A1101K | D | 2.9e-07 | 1.9e-07 | 6.2e-07 | 3.667E-07 | 2.250E-07 | 1.900E-07 | 6.200E-07 |
| 22013 | 1101 | A1101M | D | 1.3e-07 | 2.2e-07 | 1.6e-07 | 1.700E-07 | 4.583E-08 | 1.300E-07 | 2.200E-07 |
| 22014 | 1101 | A1101F | D | 2.4e-07 | 3.2e-07 | 2e-07 | 2.533E-07 | 6.110E-08 | 2.000E-07 | 3.200E-07 |

| | | | | | | | | | | |
|---|---|---|---|---|---|---|---|---|---|---|
| 22015 | 1101 | A1101P | D | 3.5e-07 | 2.2e-07 | 8.1e-07 | 4.600E-07 | 3.100E-07 | 2.200E-07 | 8.100E-07 |
| 22016 | 1101 | A1101S | D | 1.8e-07 | 1.1e-07 | 2.5e-07 | 1.800E-07 | 7.000E-08 | 1.100E-07 | 2.500E-07 |
| 22017 | 1101 | A1101T | D | 2.8e-07 | 3.3e-07 | 2.9e-07 | 3.000E-07 | 2.646E-08 | 2.800E-07 | 3.300E-07 |
| 22018 | 1101 | A1101W | D | 2.3e-07 | 3.2e-07 | 2.7e-07 | 2.733E-07 | 4.509E-08 | 2.300E-07 | 3.200E-07 |
| 22019 | 1101 | A1101Y | D | 2.1e-07 | 5e-07 | 3.2e-07 | 3.433E-07 | 1.464E-07 | 2.100E-07 | 5.000E-07 |
| 22020 | 1101 | A1101V | D | 2.1e-07 | 6.8e-07 | 1.9e-07 | 3.600E-07 | 2.773E-07 | 1.900E-07 | 6.800E-07 |
| 22021 | 1102 | P1102A | D | 2.5e-07 | 1.8e-07 | 1.7e-07 | 2.000E-07 | 4.359E-08 | 1.700E-07 | 2.500E-07 |
| 22022 | 1102 | P1102R | D | 6.7e-07 | 2.4e-07 | 2.2e-07 | 3.767E-07 | 2.542E-07 | 2.200E-07 | 6.700E-07 |
| 22023 | 1102 | P1102N | D | 1.2e-07 | 2.2e-07 | 2.7e-07 | 2.033E-07 | 7.638E-08 | 1.200E-07 | 2.700E-07 |
| 22024 | 1102 | P1102D | D | 1.9e-07 | 1.2e-07 | 2.6e-07 | 1.900E-07 | 7.000E-08 | 1.200E-07 | 2.600E-07 |
| 22025 | 1102 | P1102C | D | 2.5e-07 | 6.2e-07 | 3.3e-07 | 4.000E-07 | 1.947E-07 | 2.500E-07 | 6.200E-07 |
| 22026 | 1102 | P1102E | D | 3e-07 | 1.9e-07 | 1.4e-07 | 2.100E-07 | 8.185E-08 | 1.400E-07 | 3.000E-07 |
| 22027 | 1102 | P1102Q | D | 4.9e-07 | 1.4e-07 | 2.7e-07 | 3.000E-07 | 1.769E-07 | 1.400E-07 | 4.900E-07 |
| 22028 | 1102 | P1102G | D | 2.9e-07 | 2.3e-07 | 2.9e-07 | 2.700E-07 | 3.464E-08 | 2.300E-07 | 2.900E-07 |
| 22029 | 1102 | P1102H | D | 1.6e-07 | 2.7e-07 | 5.8e-07 | 3.367E-07 | 2.178E-07 | 1.600E-07 | 5.800E-07 |
| 22030 | 1102 | P1102I | D | 2.3e-07 | 4.4e-07 | 2.7e-07 | 3.133E-07 | 1.115E-07 | 2.300E-07 | 4.400E-07 |
| 22031 | 1102 | P1102L | D | 2e-07 | 2.4e-07 | 2.3e-07 | 2.233E-07 | 2.082E-08 | 2.000E-07 | 2.400E-07 |
| 22032 | 1102 | P1102K | D | 4.4e-07 | 3.7e-07 | 2e-07 | 3.367E-07 | 1.234E-07 | 2.000E-07 | 4.400E-07 |
| 22033 | 1102 | P1102M | D | 4.6e-07 | 2.6e-07 | 2.6e-07 | 3.267E-07 | 1.155E-07 | 2.600E-07 | 4.600E-07 |
| 22034 | 1102 | P1102F | D | 1.4e-07 | 2.5e-07 | 2.6e-07 | 2.167E-07 | 6.658E-08 | 1.400E-07 | 2.600E-07 |
| 22035 | 1102 | P1102P | D | 3.7e-07 | 1.9e-07 | 7.9e-08 | 2.130E-07 | 1.469E-07 | 7.900E-08 | 3.700E-07 |
| 22036 | 1102 | P1102S | D | 1.8e-07 | 1.9e-07 | 1.8e-07 | 1.833E-07 | 5.774E-09 | 1.800E-07 | 1.900E-07 |
| 22037 | 1102 | P1102T | D | 3.4e-07 | 1.5e-07 | 1.6e-07 | 2.167E-07 | 1.069E-07 | 1.500E-07 | 3.400E-07 |
| 22038 | 1102 | P1102W | D | 3.6e-07 | 4.2e-07 | 3e-07 | 3.600E-07 | 6.000E-08 | 3.000E-07 | 4.200E-07 |
| 22039 | 1102 | P1102Y | D | 2.5e-07 | 4.3e-07 | 2e-07 | 2.933E-07 | 1.210E-07 | 2.000E-07 | 4.300E-07 |
| 22040 | 1102 | P1102V | D | 1.6e-07 | 1e-07 | 3.8e-08 | 9.933E-08 | 6.100E-08 | 3.800E-08 | 1.600E-07 |
| 22041 | 1103 | K1103A | D | 3.3e-07 | 3.2e-07 | 2.3e-07 | 2.933E-07 | 5.508E-08 | 2.300E-07 | 3.300E-07 |
| 22042 | 1103 | K1103R | D | 2.1e-07 | 1.3e-07 | 2.1e-07 | 1.833E-07 | 4.619E-08 | 1.300E-07 | 2.100E-07 |
| 22043 | 1103 | K1103N | D | 3.5e-07 | 2.7e-07 | 4.1e-07 | 3.433E-07 | 7.024E-08 | 2.700E-07 | 4.100E-07 |
| 22044 | 1103 | K1103D | D | 3.1e-07 | 6.1e-08 | 3.8e-07 | 2.503E-07 | 1.677E-07 | 6.100E-08 | 3.800E-07 |
| 22045 | 1103 | K1103C | D | 1.4e-07 | 2.1e-07 | 1.4e-07 | 1.633E-07 | 4.041E-08 | 1.400E-07 | 2.100E-07 |
| 22046 | 1103 | K1103E | D | 1.3e-07 | 2e-07 | 1.8e-07 | 1.700E-07 | 3.606E-08 | 1.300E-07 | 2.000E-07 |
| 22047 | 1103 | K1103Q | D | 2.2e-07 | 3.3e-07 | 3.1e-07 | 2.867E-07 | 5.859E-08 | 2.200E-07 | 3.300E-07 |
| 22048 | 1103 | K1103G | D | 1.4e-07 | 1.7e-07 | 1.5e-07 | 1.533E-07 | 1.528E-08 | 1.400E-07 | 1.700E-07 |
| 22049 | 1103 | K1103H | D | 3.4e-07 | 2.7e-07 | 1.6e-07 | 2.567E-07 | 9.074E-08 | 1.600E-07 | 3.400E-07 |
| 22050 | 1103 | K1103I | D | 4.2e-07 | 2.4e-07 | 2.3e-07 | 2.967E-07 | 1.069E-07 | 2.300E-07 | 4.200E-07 |
| 22051 | 1103 | K1103L | D | 1.8e-07 | 2e-07 | 4.5e-07 | 2.767E-07 | 1.504E-07 | 1.800E-07 | 4.500E-07 |
| 22052 | 1103 | K1103K | D | 2.7e-07 | 3.6e-07 | 1.3e-07 | 2.533E-07 | 1.159E-07 | 1.300E-07 | 3.600E-07 |
| 22053 | 1103 | K1103M | D | 4.7e-07 | 2.7e-07 | 2.5e-07 | 3.300E-07 | 1.217E-07 | 2.500E-07 | 4.700E-07 |
| 22054 | 1103 | K1103F | D | 1.6e-07 | 1.8e-07 | 4e-07 | 2.467E-07 | 1.332E-07 | 1.600E-07 | 4.000E-07 |
| 22055 | 1103 | K1103P | D | 2.1e-07 | 2.5e-07 | 3.9e-07 | 2.833E-07 | 9.452E-08 | 2.100E-07 | 3.900E-07 |
| 22056 | 1103 | K1103S | D | 4.1e-07 | 1.8e-07 | 2.7e-07 | 2.867E-07 | 1.159E-07 | 1.800E-07 | 4.100E-07 |
| 22057 | 1103 | K1103T | D | 1.9e-07 | 1.4e-07 | 2.7e-07 | 2.000E-07 | 6.557E-08 | 1.400E-07 | 2.700E-07 |
| 22058 | 1103 | K1103W | D | 6.1e-07 | 1.6e-07 | 4.8e-07 | 4.167E-07 | 2.316E-07 | 1.600E-07 | 6.100E-07 |
| 22059 | 1103 | K1103Y | D | 3.8e-07 | 4.5e-07 | 6.4e-08 | 2.980E-07 | 2.057E-07 | 6.400E-08 | 4.500E-07 |
| 22060 | 1103 | K1103V | D | 1.4e-07 | 2.3e-07 | 2.3e-07 | 2.000E-07 | 5.196E-08 | 1.400E-07 | 2.300E-07 |
| 22061 | 1104 | L1104A | D | 2e-07 | 1.8e-07 | 4e-07 | 2.600E-07 | 1.217E-07 | 1.800E-07 | 4.000E-07 |
| 22062 | 1104 | L1104R | D | 2.3e-07 | 5.5e-07 | 2.3e-07 | 3.367E-07 | 1.848E-07 | 2.300E-07 | 5.500E-07 |
| 22063 | 1104 | L1104N | D | 2.2e-07 | 2.5e-07 | 1.9e-07 | 2.200E-07 | 3.000E-08 | 1.900E-07 | 2.500E-07 |
| 22064 | 1104 | L1104D | D | 9.1e-07 | 1.7e-07 | 1.3e-07 | 4.033E-07 | 4.392E-07 | 1.300E-07 | 9.100E-07 |
| 22065 | 1104 | L1104C | D | 3.8e-07 | 2.6e-07 | 3.5e-07 | 3.300E-07 | 6.245E-08 | 2.600E-07 | 3.800E-07 |
| 22066 | 1104 | L1104E | D | 2.9e-07 | 3.7e-07 | 1.2e-07 | 2.600E-07 | 1.277E-07 | 1.200E-07 | 3.700E-07 |
| 22067 | 1104 | L1104Q | D | 2.3e-07 | 1.5e-07 | 1.9e-07 | 1.900E-07 | 4.000E-08 | 1.500E-07 | 2.300E-07 |
| 22068 | 1104 | L1104G | D | 4.3e-07 | 3.5e-07 | 1.3e-07 | 3.033E-07 | 1.553E-07 | 1.300E-07 | 4.300E-07 |
| 22069 | 1104 | L1104H | D | 1.9e-07 | 4.5e-07 | 2.7e-07 | 3.033E-07 | 1.332E-07 | 1.900E-07 | 4.500E-07 |
| 22070 | 1104 | L1104I | D | 3.4e-07 | 2.4e-07 | 2.4e-07 | 2.733E-07 | 5.774E-08 | 2.400E-07 | 3.400E-07 |
| 22071 | 1104 | L1104L | D | 4.2e-07 | 2.8e-07 | 5.4e-08 | 2.513E-07 | 1.847E-07 | 5.400E-08 | 4.200E-07 |
| 22072 | 1104 | L1104K | D | 3.7e-07 | 2e-07 | 2.6e-07 | 2.767E-07 | 8.622E-08 | 2.000E-07 | 3.700E-07 |
| 22073 | 1104 | L1104M | D | 1.3e-07 | 1.6e-07 | 2e-07 | 1.633E-07 | 3.512E-08 | 1.300E-07 | 2.000E-07 |
| 22074 | 1104 | L1104F | D | 2.6e-07 | 2.8e-07 | 3.2e-07 | 2.867E-07 | 3.055E-08 | 2.600E-07 | 3.200E-07 |
| 22075 | 1104 | L1104P | D | 1.7e-07 | 6.8e-07 | 3.3e-07 | 3.933E-07 | 2.608E-07 | 1.700E-07 | 6.800E-07 |
| 22076 | 1104 | L1104S | D | 2.8e-07 | 4.4e-07 | 2.5e-07 | 3.233E-07 | 1.021E-07 | 2.500E-07 | 4.400E-07 |
| 22077 | 1104 | L1104T | D | 4.5e-07 | 1.5e-07 | 6e-08 | 2.200E-07 | 2.042E-07 | 6.000E-08 | 4.500E-07 |
| 22078 | 1104 | L1104W | D | 2.4e-07 | 4.4e-07 | 2e-07 | 2.933E-07 | 1.286E-07 | 2.000E-07 | 4.400E-07 |
| 22079 | 1104 | L1104Y | D | 1.5e-07 | 4.8e-07 | 2.5e-07 | 2.933E-07 | 1.692E-07 | 1.500E-07 | 4.800E-07 |
| 22080 | 1104 | L1104V | D | 1.2e-07 | 2.2e-07 | 1.8e-07 | 1.733E-07 | 5.033E-08 | 1.200E-07 | 2.200E-07 |
| 22081 | 1105 | L1105A | D | 2e-07 | 6e-07 | 1.6e-07 | 3.200E-07 | 2.433E-07 | 1.600E-07 | 6.000E-07 |
| 22082 | 1105 | L1105R | D | 2.1e-07 | 5.1e-07 | 2.9e-07 | 3.367E-07 | 1.553E-07 | 2.100E-07 | 5.100E-07 |
| 22083 | 1105 | L1105N | D | 6.1e-08 | 2.3e-07 | 3.2e-07 | 2.037E-07 | 1.315E-07 | 6.100E-08 | 3.200E-07 |
| 22084 | 1105 | L1105D | D | 3.2e-07 | 9.2e-08 | 1.9e-07 | 2.007E-07 | 1.144E-07 | 9.200E-08 | 3.200E-07 |
| 22085 | 1105 | L1105C | D | 6e-07 | 4.5e-07 | 1.4e-07 | 3.967E-07 | 2.346E-07 | 1.400E-07 | 6.000E-07 |
| 22086 | 1105 | L1105E | D | 3e-07 | 1.2e-07 | 6.7e-07 | 3.633E-07 | 2.804E-07 | 1.200E-07 | 6.700E-07 |
| 22087 | 1105 | L1105Q | D | 1.3e-07 | 2e-07 | 1.3e-07 | 1.533E-07 | 4.041E-08 | 1.300E-07 | 2.000E-07 |
| 22088 | 1105 | L1105G | D | 3e-07 | 5.3e-07 | 2.9e-07 | 3.733E-07 | 1.358E-07 | 2.900E-07 | 5.300E-07 |
| 22089 | 1105 | L1105H | D | 2.1e-07 | 2.1e-07 | 2.2e-07 | 2.133E-07 | 5.774E-09 | 2.100E-07 | 2.200E-07 |
| 22090 | 1105 | L1105I | D | 6.4e-07 | 1.7e-07 | 1.6e-07 | 3.233E-07 | 2.743E-07 | 1.600E-07 | 6.400E-07 |
| 22091 | 1105 | L1105L | D | 5.2e-07 | 1.6e-07 | 2.4e-07 | 3.067E-07 | 1.890E-07 | 1.600E-07 | 5.200E-07 |
| 22092 | 1105 | L1105K | D | 1.9e-07 | 4.8e-07 | 2.2e-07 | 2.967E-07 | 1.595E-07 | 1.900E-07 | 4.800E-07 |
| 22093 | 1105 | L1105M | D | 7.9e-07 | 1.9e-07 | 3.6e-07 | 4.467E-07 | 3.092E-07 | 1.900E-07 | 7.900E-07 |
| 22094 | 1105 | L1105F | D | 1.8e-07 | 2.1e-07 | 1.7e-07 | 1.867E-07 | 2.082E-08 | 1.700E-07 | 2.100E-07 |
| 22095 | 1105 | L1105P | D | 4.6e-07 | 3.2e-07 | 2.4e-07 | 3.400E-07 | 1.114E-07 | 2.400E-07 | 4.600E-07 |
| 22096 | 1105 | L1105S | D | 1.9e-07 | 1.8e-07 | 3.7e-07 | 2.467E-07 | 1.069E-07 | 1.800E-07 | 3.700E-07 |
| 22097 | 1105 | L1105T | D | 2.8e-07 | 2.9e-07 | 5.4e-07 | 3.700E-07 | 1.473E-07 | 2.800E-07 | 5.400E-07 |
| 22098 | 1105 | L1105W | D | 2.5e-07 | 2.9e-07 | 2.4e-07 | 2.600E-07 | 2.646E-08 | 2.400E-07 | 2.900E-07 |
| 22099 | 1105 | L1105Y | D | 2e-07 | 2.8e-07 | 2e-07 | 2.267E-07 | 4.619E-08 | 2.000E-07 | 2.800E-07 |

| | | | | | | | | | | |
|---|---|---|---|---|---|---|---|---|---|---|
| 22100 | 1105 | L1105V | D | 2.1e-07 | 2.9e-07 | 1.1e-06 | 5.333E-07 | 4.924E-07 | 2.100E-07 | 1.100E-06 |
| 22101 | 1106 | I1106A | D | 7.2e-07 | 2.4e-07 | 1e-07 | 3.533E-07 | 3.252E-07 | 1.000E-07 | 7.200E-07 |
| 22102 | 1106 | I1106R | D | 8.8e-07 | 3.4e-07 | 2.4e-07 | 4.867E-07 | 3.443E-07 | 2.400E-07 | 8.800E-07 |
| 22103 | 1106 | I1106N | D | 2.8e-07 | 5.1e-07 | 9.3e-08 | 2.943E-07 | 2.089E-07 | 9.300E-08 | 5.100E-07 |
| 22104 | 1106 | I1106D | D | 1.5e-07 | 1.9e-07 | 3.7e-07 | 2.367E-07 | 1.172E-07 | 1.500E-07 | 3.700E-07 |
| 22105 | 1106 | I1106C | D | 3.3e-07 | 4.5e-07 | 2.2e-07 | 3.333E-07 | 1.150E-07 | 2.200E-07 | 4.500E-07 |
| 22106 | 1106 | I1106E | D | 1.2e-07 | 3.1e-07 | 3.3e-07 | 2.533E-07 | 1.159E-07 | 1.200E-07 | 3.300E-07 |
| 22107 | 1106 | I1106Q | D | 3.6e-07 | 1.3e-07 | 2.1e-07 | 2.333E-07 | 1.168E-07 | 1.300E-07 | 3.600E-07 |
| 22108 | 1106 | I1106G | D | 2.7e-07 | 1.8e-07 | 5e-07 | 3.167E-07 | 1.650E-07 | 1.800E-07 | 5.000E-07 |
| 22109 | 1106 | I1106H | D | 4.5e-07 | 3e-07 | 4.6e-07 | 4.033E-07 | 8.963E-08 | 3.000E-07 | 4.600E-07 |
| 22110 | 1106 | I1106I | D | 5.7e-07 | 2.7e-07 | 4.5e-07 | 4.300E-07 | 1.510E-07 | 2.700E-07 | 5.700E-07 |
| 22111 | 1106 | I1106L | D | 2.1e-07 | 3.3e-07 | 6e-07 | 3.800E-07 | 1.997E-07 | 2.100E-07 | 6.000E-07 |
| 22112 | 1106 | I1106K | D | 3.9e-07 | 1.8e-07 | 2.1e-07 | 2.600E-07 | 1.136E-07 | 1.800E-07 | 3.900E-07 |
| 22113 | 1106 | I1106M | D | 8.4e-08 | 2.7e-07 | 2.2e-07 | 1.913E-07 | 9.626E-08 | 8.400E-08 | 2.700E-07 |
| 22114 | 1106 | I1106F | D | 2.8e-07 | 1.4e-07 | 1.6e-07 | 1.933E-07 | 7.572E-08 | 1.400E-07 | 2.800E-07 |
| 22115 | 1106 | I1106P | D | 1.7e-07 | 3e-07 | 1.9e-07 | 2.200E-07 | 7.000E-08 | 1.700E-07 | 3.000E-07 |
| 22116 | 1106 | I1106S | D | 9.4e-08 | 3.7e-07 | 2.1e-07 | 2.247E-07 | 1.386E-07 | 9.400E-08 | 3.700E-07 |
| 22117 | 1106 | I1106T | D | 1.5e-07 | 4.3e-07 | 3e-07 | 2.933E-07 | 1.401E-07 | 1.500E-07 | 4.300E-07 |
| 22118 | 1106 | I1106W | D | 4.4e-07 | 1.9e-07 | 1.1e-07 | 2.467E-07 | 1.721E-07 | 1.100E-07 | 4.400E-07 |
| 22119 | 1106 | I1106Y | D | 1.5e-07 | 1.9e-07 | 2.8e-07 | 2.067E-07 | 6.658E-08 | 1.500E-07 | 2.800E-07 |
| 22120 | 1106 | I1106V | D | 3.4e-07 | 1.5e-07 | 3.4e-07 | 2.767E-07 | 1.097E-07 | 1.500E-07 | 3.400E-07 |
| 22121 | 1107 | Y1107A | D | 9.2e-08 | 2.1e-07 | 2.4e-07 | 1.807E-07 | 7.824E-08 | 9.200E-08 | 2.400E-07 |
| 22122 | 1107 | Y1107R | D | 4.4e-07 | 1.6e-07 | 3.7e-07 | 3.233E-07 | 1.457E-07 | 1.600E-07 | 4.400E-07 |
| 22123 | 1107 | Y1107N | D | 4.5e-07 | 3.2e-07 | 1.8e-07 | 3.167E-07 | 1.350E-07 | 1.800E-07 | 4.500E-07 |
| 22124 | 1107 | Y1107D | D | 3e-07 | 3.2e-07 | 3.5e-07 | 3.233E-07 | 2.517E-08 | 3.000E-07 | 3.500E-07 |
| 22125 | 1107 | Y1107C | D | 4.4e-07 | 8.7e-08 | 1.8e-07 | 2.357E-07 | 1.830E-07 | 8.700E-08 | 4.400E-07 |
| 22126 | 1107 | Y1107E | D | 2.6e-07 | 4.8e-07 | 3.2e-07 | 3.533E-07 | 1.137E-07 | 2.600E-07 | 4.800E-07 |
| 22127 | 1107 | Y1107Q | D | 1.3e-07 | 1.8e-07 | 2.4e-07 | 1.833E-07 | 5.508E-08 | 1.300E-07 | 2.400E-07 |
| 22128 | 1107 | Y1107G | D | 4.8e-07 | 2.1e-07 | 1.8e-07 | 2.900E-07 | 1.652E-07 | 1.800E-07 | 4.800E-07 |
| 22129 | 1107 | Y1107H | D | 2e-07 | 2.3e-07 | 3.2e-07 | 2.500E-07 | 6.245E-08 | 2.000E-07 | 3.200E-07 |
| 22130 | 1107 | Y1107I | D | 2.9e-07 | 2e-07 | 1.3e-07 | 2.067E-07 | 8.021E-08 | 1.300E-07 | 2.900E-07 |
| 22131 | 1107 | Y1107L | D | 1.8e-07 | 1.4e-07 | 2.7e-07 | 1.967E-07 | 6.658E-08 | 1.400E-07 | 2.700E-07 |
| 22132 | 1107 | Y1107K | D | 1e-07 | 1.6e-07 | 1.4e-07 | 1.333E-07 | 3.055E-08 | 1.000E-07 | 1.600E-07 |
| 22133 | 1107 | Y1107M | D | 2.3e-07 | 1.5e-07 | 3e-07 | 2.267E-07 | 7.506E-08 | 1.500E-07 | 3.000E-07 |
| 22134 | 1107 | Y1107F | D | 6.6e-07 | 2.5e-07 | 5.2e-07 | 4.767E-07 | 2.084E-07 | 2.500E-07 | 6.600E-07 |
| 22135 | 1107 | Y1107P | D | 4.9e-07 | 6.9e-07 | 2.6e-07 | 4.800E-07 | 2.152E-07 | 2.600E-07 | 6.900E-07 |
| 22136 | 1107 | Y1107S | D | 1.7e-07 | 9.6e-08 | 2.4e-07 | 1.687E-07 | 7.201E-08 | 9.600E-08 | 2.400E-07 |
| 22137 | 1107 | Y1107T | D | 6.6e-07 | 2.1e-07 | 2.4e-07 | 3.700E-07 | 2.516E-07 | 2.100E-07 | 6.600E-07 |
| 22138 | 1107 | Y1107W | D | 2.9e-07 | 3.4e-07 | 2.1e-07 | 2.800E-07 | 6.557E-08 | 2.100E-07 | 3.400E-07 |
| 22139 | 1107 | Y1107Y | D | 1.3e-07 | 3.8e-07 | 2.1e-07 | 2.400E-07 | 1.277E-07 | 1.300E-07 | 3.800E-07 |
| 22140 | 1107 | Y1107V | D | 2.5e-07 | 2.8e-07 | 1.5e-07 | 2.267E-07 | 6.807E-08 | 1.500E-07 | 2.800E-07 |
| 22141 | 1108 | S1108A | D | 5.7e-07 | 1.3e-07 | 9.8e-08 | 2.660E-07 | 2.638E-07 | 9.800E-08 | 5.700E-07 |
| 22142 | 1108 | S1108R | D | 2.1e-07 | 1.5e-07 | 1.4e-07 | 1.667E-07 | 3.786E-08 | 1.400E-07 | 2.100E-07 |
| 22143 | 1108 | S1108N | D | 1.7e-07 | 1.9e-07 | 1.6e-07 | 1.733E-07 | 1.528E-08 | 1.600E-07 | 1.900E-07 |
| 22144 | 1108 | S1108D | D | 4.3e-07 | 2.6e-07 | 2.1e-07 | 3.000E-07 | 1.153E-07 | 2.100E-07 | 4.300E-07 |
| 22145 | 1108 | S1108C | D | 1.6e-07 | 9.6e-08 | 1.1e-07 | 1.220E-07 | 3.365E-08 | 9.600E-08 | 1.600E-07 |
| 22146 | 1108 | S1108E | D | 3.5e-07 | 9.9e-08 | 1.4e-07 | 1.963E-07 | 1.346E-07 | 9.900E-08 | 3.500E-07 |
| 22147 | 1108 | S1108Q | D | 1.6e-07 | 1.6e-07 | 1.3e-07 | 1.500E-07 | 1.732E-08 | 1.300E-07 | 1.600E-07 |
| 22148 | 1108 | S1108G | D | 8.3e-08 | 1.3e-07 | 1.7e-07 | 1.277E-07 | 4.355E-08 | 8.300E-08 | 1.700E-07 |
| 22149 | 1108 | S1108H | D | 1.3e-07 | 1.2e-07 | 2.2e-07 | 1.567E-07 | 5.508E-08 | 1.200E-07 | 2.200E-07 |
| 22150 | 1108 | S1108I | D | 1.8e-07 | 1.4e-07 | 1.8e-07 | 1.667E-07 | 2.309E-08 | 1.400E-07 | 1.800E-07 |
| 22151 | 1108 | S1108L | D | 1.3e-07 | 1.9e-07 | 2.3e-07 | 1.833E-07 | 5.033E-08 | 1.300E-07 | 2.300E-07 |
| 22152 | 1108 | S1108K | D | 1.4e-07 | 1.3e-07 | 2.9e-07 | 1.867E-07 | 8.963E-08 | 1.300E-07 | 2.900E-07 |
| 22153 | 1108 | S1108M | D | 3.2e-07 | 2.3e-07 | 1.5e-07 | 2.333E-07 | 8.505E-08 | 1.500E-07 | 3.200E-07 |
| 22154 | 1108 | S1108F | D | 2.8e-07 | 1.3e-07 | 7.9e-08 | 1.630E-07 | 1.045E-07 | 7.900E-08 | 2.800E-07 |
| 22155 | 1108 | S1108P | D | 1.2e-07 | 1.3e-07 | 5.7e-07 | 2.733E-07 | 2.570E-07 | 1.200E-07 | 5.700E-07 |
| 22156 | 1108 | S1108S | D | 2.2e-07 | 3.4e-07 | 3.3e-07 | 2.967E-07 | 6.658E-08 | 2.200E-07 | 3.400E-07 |
| 22157 | 1108 | S1108T | D | 1.1e-07 | 2.5e-07 | 4.9e-07 | 2.833E-07 | 1.922E-07 | 1.100E-07 | 4.900E-07 |
| 22158 | 1108 | S1108W | D | 1.7e-07 | 1.4e-07 | 2.8e-07 | 1.967E-07 | 7.371E-08 | 1.400E-07 | 2.800E-07 |
| 22159 | 1108 | S1108Y | D | 1.7e-07 | 2e-07 | 2.8e-07 | 2.167E-07 | 5.686E-08 | 1.700E-07 | 2.800E-07 |
| 22160 | 1108 | S1108V | D | 3e-07 | 2.2e-07 | 2.1e-07 | 2.433E-07 | 4.933E-08 | 2.100E-07 | 3.000E-07 |
| 22161 | 1109 | A1109A | D | 2.9e-07 | 2.2e-07 | 3e-07 | 2.700E-07 | 4.359E-08 | 2.200E-07 | 3.000E-07 |
| 22162 | 1109 | A1109R | D | 3.9e-07 | 1.5e-07 | 1.5e-07 | 2.300E-07 | 1.386E-07 | 1.500E-07 | 3.900E-07 |
| 22163 | 1109 | A1109N | D | 5.1e-07 | 6.7e-07 | 2.4e-07 | 4.733E-07 | 2.173E-07 | 2.400E-07 | 6.700E-07 |
| 22164 | 1109 | A1109D | D | 1.9e-07 | 1.4e-07 | 1.5e-07 | 1.600E-07 | 2.646E-08 | 1.400E-07 | 1.900E-07 |
| 22165 | 1109 | A1109C | D | 3.8e-07 | 2.9e-07 | 1.3e-07 | 2.667E-07 | 1.266E-07 | 1.300E-07 | 3.800E-07 |
| 22166 | 1109 | A1109E | D | 2.3e-07 | 2.9e-07 | 1.8e-07 | 2.333E-07 | 5.508E-08 | 1.800E-07 | 2.900E-07 |
| 22167 | 1109 | A1109Q | D | 1e-07 | 1.1e-07 | 2.4e-07 | 1.500E-07 | 7.810E-08 | 1.000E-07 | 2.400E-07 |
| 22168 | 1109 | A1109G | D | 1.7e-07 | 4.1e-07 | 3.4e-07 | 3.067E-07 | 1.234E-07 | 1.700E-07 | 4.100E-07 |
| 22169 | 1109 | A1109H | D | 1.7e-07 | 1.6e-07 | 2.6e-07 | 1.967E-07 | 5.508E-08 | 1.600E-07 | 2.600E-07 |
| 22170 | 1109 | A1109I | D | 2.6e-07 | 5.4e-07 | 2.2e-07 | 3.400E-07 | 1.744E-07 | 2.200E-07 | 5.400E-07 |
| 22171 | 1109 | A1109L | D | 7.9e-08 | 2.7e-07 | 1.1e-07 | 1.530E-07 | 1.025E-07 | 7.900E-08 | 2.700E-07 |
| 22172 | 1109 | A1109K | D | 5.4e-08 | 1.6e-07 | 1.2e-07 | 1.113E-07 | 5.353E-08 | 5.400E-08 | 1.600E-07 |
| 22173 | 1109 | A1109M | D | 1.9e-07 | 4.1e-07 | 3.2e-07 | 3.067E-07 | 1.106E-07 | 1.900E-07 | 4.100E-07 |
| 22174 | 1109 | A1109F | D | 9.3e-08 | 1.7e-07 | 2.1e-07 | 1.577E-07 | 5.947E-08 | 9.300E-08 | 2.100E-07 |
| 22175 | 1109 | A1109P | D | 2.2e-07 | 1.7e-07 | 3.6e-07 | 2.500E-07 | 9.849E-08 | 1.700E-07 | 3.600E-07 |
| 22176 | 1109 | A1109S | D | 4e-07 | 2.6e-07 | 3e-07 | 3.200E-07 | 7.211E-08 | 2.600E-07 | 4.000E-07 |
| 22177 | 1109 | A1109T | D | 2.8e-07 | 2.7e-07 | 3.3e-07 | 2.933E-07 | 3.215E-08 | 2.700E-07 | 3.300E-07 |
| 22178 | 1109 | A1109W | D | 2e-07 | 2.6e-07 | 1.1e-07 | 1.900E-07 | 7.550E-08 | 1.100E-07 | 2.600E-07 |
| 22179 | 1109 | A1109Y | D | 2.6e-07 | 2.3e-07 | 1.6e-07 | 2.167E-07 | 5.132E-08 | 1.600E-07 | 2.600E-07 |
| 22180 | 1109 | A1109V | D | 2.2e-07 | 2.7e-07 | 2.8e-07 | 2.567E-07 | 3.215E-08 | 2.200E-07 | 2.800E-07 |
| 22181 | 1110 | S1110A | D | 3.1e-07 | 2.1e-07 | 1.4e-07 | 2.200E-07 | 8.544E-08 | 1.400E-07 | 3.100E-07 |
| 22182 | 1110 | S1110R | D | 3.3e-07 | 1.5e-07 | 4.8e-07 | 3.200E-07 | 1.652E-07 | 1.500E-07 | 4.800E-07 |
| 22183 | 1110 | S1110N | D | 1.8e-07 | 5.1e-07 | 2e-07 | 2.967E-07 | 1.850E-07 | 1.800E-07 | 5.100E-07 |
| 22184 | 1110 | S1110D | D | 4.7e-07 | 1.7e-07 | 2.4e-07 | 2.933E-07 | 1.570E-07 | 1.700E-07 | 4.700E-07 |

| | | | | | | | | | | |
|---|---|---|---|---|---|---|---|---|---|---|
| 22185 | 1110 | S1110C | D | 4.9e-07 | 1.6e-07 | 3.2e-07 | 3.233E-07 | 1.650E-07 | 1.600E-07 | 4.900E-07 |
| 22186 | 1110 | S1110E | D | 2.1e-07 | 1.2e-07 | 3.8e-07 | 2.367E-07 | 1.320E-07 | 1.200E-07 | 3.800E-07 |
| 22187 | 1110 | S1110Q | D | 3e-07 | 1.1e-07 | 2.7e-07 | 2.267E-07 | 1.021E-07 | 1.100E-07 | 3.000E-07 |
| 22188 | 1110 | S1110G | D | 3.2e-07 | 1e-07 | 3.2e-07 | 2.467E-07 | 1.270E-07 | 1.000E-07 | 3.200E-07 |
| 22189 | 1110 | S1110H | D | 3.3e-07 | 3.4e-07 | 1.1e-07 | 2.600E-07 | 1.300E-07 | 1.100E-07 | 3.400E-07 |
| 22190 | 1110 | S1110I | D | 3.7e-07 | 5.4e-07 | 2.3e-07 | 3.800E-07 | 1.552E-07 | 2.300E-07 | 5.400E-07 |
| 22191 | 1110 | S1110L | D | 1.9e-07 | 6.1e-07 | 1.4e-07 | 3.133E-07 | 2.581E-07 | 1.400E-07 | 6.100E-07 |
| 22192 | 1110 | S1110K | D | 1.9e-07 | 2.5e-07 | 1.9e-07 | 2.100E-07 | 3.464E-08 | 1.900E-07 | 2.500E-07 |
| 22193 | 1110 | S1110M | D | 3.8e-07 | 3.2e-07 | 1e-07 | 2.667E-07 | 1.474E-07 | 1.000E-07 | 3.800E-07 |
| 22194 | 1110 | S1110F | D | 1.9e-07 | 2.6e-07 | 2.5e-07 | 2.333E-07 | 3.786E-08 | 1.900E-07 | 2.600E-07 |
| 22195 | 1110 | S1110P | D | 3.1e-07 | 2.3e-07 | 3.8e-07 | 3.067E-07 | 7.506E-08 | 2.300E-07 | 3.800E-07 |
| 22196 | 1110 | S1110S | D | 2.2e-07 | 2.5e-07 | 3.9e-07 | 2.867E-07 | 9.074E-08 | 2.200E-07 | 3.900E-07 |
| 22197 | 1110 | S1110T | D | 1.8e-07 | 3.2e-07 | 5.3e-07 | 3.433E-07 | 1.762E-07 | 1.800E-07 | 5.300E-07 |
| 22198 | 1110 | S1110W | D | 2.5e-07 | 2.9e-07 | 1.4e-07 | 2.267E-07 | 7.767E-08 | 1.400E-07 | 2.900E-07 |
| 22199 | 1110 | S1110Y | D | 1.2e-07 | 1.1e-07 | 2.4e-07 | 1.567E-07 | 7.234E-08 | 1.100E-07 | 2.400E-07 |
| 22200 | 1110 | S1110V | D | 4.1e-07 | 3.1e-07 | 2.6e-07 | 3.267E-07 | 7.638E-08 | 2.600E-07 | 4.100E-07 |
| 22201 | 1111 | F1111A | D | 1.7e-07 | 1.3e-07 | 1.7e-07 | 1.567E-07 | 2.309E-08 | 1.300E-07 | 1.700E-07 |
| 22202 | 1111 | F1111R | D | 1.4e-07 | 5.8e-08 | 1.8e-07 | 1.260E-07 | 6.219E-08 | 5.800E-08 | 1.800E-07 |
| 22203 | 1111 | F1111N | D | 9.6e-08 | 8.5e-08 | 1.5e-07 | 1.103E-07 | 3.479E-08 | 8.500E-08 | 1.500E-07 |
| 22204 | 1111 | F1111D | D | 1.4e-07 | 7.8e-08 | 1.4e-07 | 1.193E-07 | 3.580E-08 | 7.800E-08 | 1.400E-07 |
| 22205 | 1111 | F1111C | D | 1.7e-07 | 2.2e-07 | 3.1e-07 | 2.333E-07 | 7.095E-08 | 1.700E-07 | 3.100E-07 |
| 22206 | 1111 | F1111E | D | 1.4e-07 | 1.1e-07 | 1.5e-07 | 1.333E-07 | 2.082E-08 | 1.100E-07 | 1.500E-07 |
| 22207 | 1111 | F1111Q | D | 7.6e-08 | 1.5e-07 | 2.7e-07 | 1.653E-07 | 9.790E-08 | 7.600E-08 | 2.700E-07 |
| 22208 | 1111 | F1111G | D | 8.8e-08 | 1.8e-07 | 1.2e-07 | 1.293E-07 | 4.670E-08 | 8.800E-08 | 1.800E-07 |
| 22209 | 1111 | F1111H | D | 2.3e-07 | 6.5e-08 | 2e-07 | 1.650E-07 | 8.789E-08 | 6.500E-08 | 2.300E-07 |
| 22210 | 1111 | F1111I | D | 1.5e-07 | 3.3e-07 | 1.8e-07 | 2.200E-07 | 9.644E-08 | 1.500E-07 | 3.300E-07 |
| 22211 | 1111 | F1111L | D | 4.2e-07 | 1.4e-07 | 2.2e-07 | 2.600E-07 | 1.442E-07 | 1.400E-07 | 4.200E-07 |
| 22212 | 1111 | F1111K | D | 2.5e-07 | 1.4e-07 | 1.6e-07 | 1.833E-07 | 5.859E-08 | 1.400E-07 | 2.500E-07 |
| 22213 | 1111 | F1111M | D | 3.9e-07 | 1.8e-07 | 1.6e-07 | 2.433E-07 | 1.274E-07 | 1.600E-07 | 3.900E-07 |
| 22214 | 1111 | F1111F | D | 3.1e-07 | 2.8e-07 | 3.6e-07 | 3.167E-07 | 4.041E-08 | 2.800E-07 | 3.600E-07 |
| 22215 | 1111 | F1111P | D | 2.4e-07 | 3.2e-07 | 2.2e-07 | 2.600E-07 | 5.292E-08 | 2.200E-07 | 3.200E-07 |
| 22216 | 1111 | F1111S | D | 2.5e-07 | 1.5e-07 | 2.8e-07 | 2.267E-07 | 6.807E-08 | 1.500E-07 | 2.800E-07 |
| 22217 | 1111 | F1111T | D | 2.2e-07 | 5.4e-07 | 1.8e-07 | 3.133E-07 | 1.973E-07 | 1.800E-07 | 5.400E-07 |
| 22218 | 1111 | F1111W | D | 1.5e-07 | 1.6e-07 | 3.5e-07 | 2.200E-07 | 1.127E-07 | 1.500E-07 | 3.500E-07 |
| 22219 | 1111 | F1111Y | D | 5.6e-07 | 1.5e-07 | 3.5e-07 | 3.533E-07 | 2.050E-07 | 1.500E-07 | 5.600E-07 |
| 22220 | 1111 | F1111V | D | 1.1e-07 | 2.8e-07 | 2.5e-07 | 2.133E-07 | 9.074E-08 | 1.100E-07 | 2.800E-07 |
| 22221 | 1112 | L1112A | D | 2e-07 | 3.9e-07 | 1.5e-07 | 2.467E-07 | 1.266E-07 | 1.500E-07 | 3.900E-07 |
| 22222 | 1112 | L1112R | D | 2.8e-07 | 3.7e-07 | 1.4e-07 | 2.633E-07 | 1.159E-07 | 1.400E-07 | 3.700E-07 |
| 22223 | 1112 | L1112N | D | 3.3e-07 | 2.8e-07 | 2.3e-07 | 2.800E-07 | 5.000E-08 | 2.300E-07 | 3.300E-07 |
| 22224 | 1112 | L1112D | D | 3.5e-07 | 1.6e-07 | 2e-07 | 2.367E-07 | 1.002E-07 | 1.600E-07 | 3.500E-07 |
| 22225 | 1112 | L1112C | D | 1.1e-07 | 1.1e-07 | 2.7e-07 | 1.633E-07 | 9.238E-08 | 1.100E-07 | 2.700E-07 |
| 22226 | 1112 | L1112E | D | 2.9e-07 | 3.3e-07 | 2.3e-07 | 2.833E-07 | 5.033E-08 | 2.300E-07 | 3.300E-07 |
| 22227 | 1112 | L1112Q | D | 3.3e-07 | 2.9e-07 | 3.4e-07 | 3.200E-07 | 2.646E-08 | 2.900E-07 | 3.400E-07 |
| 22228 | 1112 | L1112G | D | 2.8e-07 | 5.7e-07 | 3.3e-07 | 3.933E-07 | 1.550E-07 | 2.800E-07 | 5.700E-07 |
| 22229 | 1112 | L1112H | D | 2.1e-07 | 2.8e-07 | 1.5e-07 | 2.133E-07 | 6.506E-08 | 1.500E-07 | 2.800E-07 |
| 22230 | 1112 | L1112I | D | 3.8e-07 | 3.1e-07 | 1.3e-07 | 2.733E-07 | 1.290E-07 | 1.300E-07 | 3.800E-07 |
| 22231 | 1112 | L1112L | D | 2.7e-07 | 1.9e-07 | 3.8e-07 | 2.800E-07 | 9.539E-08 | 1.900E-07 | 3.800E-07 |
| 22232 | 1112 | L1112K | D | 3.2e-07 | 3.2e-07 | 3.2e-07 | 3.200E-07 | 6.484E-23 | 3.200E-07 | 3.200E-07 |
| 22233 | 1112 | L1112M | D | 1.7e-07 | 3.8e-07 | 3.9e-07 | 3.133E-07 | 1.242E-07 | 1.700E-07 | 3.900E-07 |
| 22234 | 1112 | L1112F | D | 5.1e-07 | 5.8e-07 | 3.8e-07 | 4.900E-07 | 1.015E-07 | 3.800E-07 | 5.800E-07 |
| 22235 | 1112 | L1112P | D | 2.7e-07 | 1.6e-07 | 2e-07 | 2.100E-07 | 5.568E-08 | 1.600E-07 | 2.700E-07 |
| 22236 | 1112 | L1112S | D | 2e-07 | 2.9e-07 | 1.9e-07 | 2.267E-07 | 5.508E-08 | 1.900E-07 | 2.900E-07 |
| 22237 | 1112 | L1112T | D | 4.1e-07 | 2.6e-07 | 3.1e-07 | 3.267E-07 | 7.638E-08 | 2.600E-07 | 4.100E-07 |
| 22238 | 1112 | L1112W | D | 1.2e-07 | 2.4e-07 | 2.2e-07 | 1.933E-07 | 6.429E-08 | 1.200E-07 | 2.400E-07 |
| 22239 | 1112 | L1112Y | D | 2.2e-07 | 1.5e-07 | 4.7e-07 | 2.800E-07 | 1.682E-07 | 1.500E-07 | 4.700E-07 |
| 22240 | 1112 | L1112V | D | 3.1e-07 | 3.2e-07 | 3e-07 | 3.100E-07 | 1.000E-08 | 3.000E-07 | 3.200E-07 |
| 22241 | 1113 | Y1113A | D | 2e-07 | 2e-07 | 2.2e-07 | 2.067E-07 | 1.155E-08 | 2.000E-07 | 2.200E-07 |
| 22242 | 1113 | Y1113R | D | 1.2e-07 | 2.5e-07 | 1.8e-07 | 1.833E-07 | 6.506E-08 | 1.200E-07 | 2.500E-07 |
| 22243 | 1113 | Y1113N | D | 1.9e-07 | 3.6e-07 | 2.2e-07 | 2.567E-07 | 9.074E-08 | 1.900E-07 | 3.600E-07 |
| 22244 | 1113 | Y1113D | D | 3.7e-07 | 1.8e-07 | 1.7e-07 | 2.400E-07 | 1.127E-07 | 1.700E-07 | 3.700E-07 |
| 22245 | 1113 | Y1113C | D | 1.7e-07 | 3.1e-07 | 8.8e-08 | 1.893E-07 | 1.123E-07 | 8.800E-08 | 3.100E-07 |
| 22246 | 1113 | Y1113E | D | 4.2e-07 | 1.7e-07 | 1.9e-07 | 2.600E-07 | 1.389E-07 | 1.700E-07 | 4.200E-07 |
| 22247 | 1113 | Y1113Q | D | 1.9e-07 | 2.8e-07 | 1.3e-07 | 2.000E-07 | 7.550E-08 | 1.300E-07 | 2.800E-07 |
| 22248 | 1113 | Y1113G | D | 3.3e-07 | 1.5e-07 | 3.2e-07 | 2.667E-07 | 1.012E-07 | 1.500E-07 | 3.300E-07 |
| 22249 | 1113 | Y1113H | D | 3.2e-07 | 3.1e-07 | 7.8e-08 | 2.360E-07 | 1.369E-07 | 7.800E-08 | 3.200E-07 |
| 22250 | 1113 | Y1113I | D | 2.9e-07 | 2.2e-07 | 3.1e-07 | 2.733E-07 | 4.726E-08 | 2.200E-07 | 3.100E-07 |
| 22251 | 1113 | Y1113L | D | 2.8e-07 | 2.2e-07 | 2.5e-07 | 2.500E-07 | 3.000E-08 | 2.200E-07 | 2.800E-07 |
| 22252 | 1113 | Y1113K | D | 2.3e-07 | 3.4e-07 | 4.8e-07 | 3.500E-07 | 1.253E-07 | 2.300E-07 | 4.800E-07 |
| 22253 | 1113 | Y1113M | D | 1.4e-07 | 9.1e-07 | 3.1e-07 | 4.533E-07 | 4.045E-07 | 1.400E-07 | 9.100E-07 |
| 22254 | 1113 | Y1113F | D | 1.7e-07 | 3.2e-07 | 4.1e-07 | 3.000E-07 | 1.212E-07 | 1.700E-07 | 4.100E-07 |
| 22255 | 1113 | Y1113P | D | 3.8e-07 | 1.8e-07 | 1.5e-07 | 2.367E-07 | 1.250E-07 | 1.500E-07 | 3.800E-07 |
| 22256 | 1113 | Y1113S | D | 2.3e-07 | 2.2e-07 | 4.1e-07 | 2.867E-07 | 1.069E-07 | 2.200E-07 | 4.100E-07 |
| 22257 | 1113 | Y1113T | D | 6.1e-07 | 1.2e-07 | 2.9e-07 | 3.400E-07 | 2.488E-07 | 1.200E-07 | 6.100E-07 |
| 22258 | 1113 | Y1113W | D | 1.2e-07 | 2.4e-07 | 3.1e-07 | 2.233E-07 | 9.609E-08 | 1.200E-07 | 3.100E-07 |
| 22259 | 1113 | Y1113Y | D | 3.5e-07 | 2.2e-07 | 2.4e-07 | 2.700E-07 | 7.000E-08 | 2.200E-07 | 3.500E-07 |
| 22260 | 1113 | Y1113V | D | 2.9e-07 | 2.8e-07 | 8.7e-07 | 4.800E-07 | 3.378E-07 | 2.800E-07 | 8.700E-07 |
| 22261 | 1114 | S1114A | D | 3.2e-07 | 4e-07 | 2e-07 | 3.067E-07 | 1.007E-07 | 2.000E-07 | 4.000E-07 |
| 22262 | 1114 | S1114R | D | 1.6e-07 | 3.5e-07 | 3.8e-07 | 2.967E-07 | 1.193E-07 | 1.600E-07 | 3.800E-07 |
| 22263 | 1114 | S1114N | D | 3.2e-07 | 2.7e-07 | 4.5e-07 | 3.467E-07 | 9.292E-08 | 2.700E-07 | 4.500E-07 |
| 22264 | 1114 | S1114D | D | 3.7e-07 | 2.3e-07 | 3.9e-07 | 3.300E-07 | 8.718E-08 | 2.300E-07 | 3.900E-07 |
| 22265 | 1114 | S1114C | D | 5.3e-07 | 2.7e-07 | 7.3e-07 | 5.100E-07 | 2.307E-07 | 2.700E-07 | 7.300E-07 |
| 22266 | 1114 | S1114E | D | 1.8e-07 | 3.5e-07 | 3.5e-07 | 2.933E-07 | 9.815E-08 | 1.800E-07 | 3.500E-07 |
| 22267 | 1114 | S1114Q | D | 3.1e-07 | 1.1e-07 | 6.1e-07 | 3.433E-07 | 2.517E-07 | 1.100E-07 | 6.100E-07 |
| 22268 | 1114 | S1114G | D | 3.7e-07 | 3.2e-07 | 4.5e-07 | 3.800E-07 | 6.557E-08 | 3.200E-07 | 4.500E-07 |
| 22269 | 1114 | S1114H | D | 2.2e-07 | 3.7e-07 | 2.2e-07 | 2.700E-07 | 8.660E-08 | 2.200E-07 | 3.700E-07 |

| | | | | | | | | | | |
|---|---|---|---|---|---|---|---|---|---|---|
| 22270 | 1114 | S1114I | D | 2.5e-07 | 5.1e-07 | 4.3e-07 | 3.967E-07 | 1.332E-07 | 2.500E-07 | 5.100E-07 |
| 22271 | 1114 | S1114L | D | 3.5e-07 | 3.4e-07 | 3.7e-07 | 3.533E-07 | 1.528E-08 | 3.400E-07 | 3.700E-07 |
| 22272 | 1114 | S1114K | D | 2.6e-07 | 2.6e-07 | 4e-07 | 3.067E-07 | 8.083E-08 | 2.600E-07 | 4.000E-07 |
| 22273 | 1114 | S1114M | D | 2e-07 | 1.6e-07 | 6e-07 | 3.200E-07 | 2.433E-07 | 1.600E-07 | 6.000E-07 |
| 22274 | 1114 | S1114F | D | 1.4e-07 | 1.9e-07 | 4.7e-07 | 2.667E-07 | 1.779E-07 | 1.400E-07 | 4.700E-07 |
| 22275 | 1114 | S1114P | D | 3.1e-07 | 5.6e-07 | 3e-07 | 3.900E-07 | 1.473E-07 | 3.000E-07 | 5.600E-07 |
| 22276 | 1114 | S1114S | D | 1.6e-07 | 2.8e-07 | 3.1e-07 | 2.500E-07 | 7.937E-08 | 1.600E-07 | 3.100E-07 |
| 22277 | 1114 | S1114T | D | 2.1e-07 | 1.8e-07 | 7.4e-08 | 1.547E-07 | 7.145E-08 | 7.400E-08 | 2.100E-07 |
| 22278 | 1114 | S1114W | D | 6.9e-07 | 2.6e-07 | 2.6e-07 | 4.033E-07 | 2.483E-07 | 2.600E-07 | 6.900E-07 |
| 22279 | 1114 | S1114Y | D | 2.8e-07 | 1.4e-07 | 4.5e-07 | 2.900E-07 | 1.552E-07 | 1.400E-07 | 4.500E-07 |
| 22280 | 1114 | S1114V | D | 2.1e-07 | 2.2e-07 | 1.6e-07 | 1.967E-07 | 3.215E-08 | 1.600E-07 | 2.200E-07 |
| 22281 | 1115 | G1115A | D | 3.2e-07 | 3.2e-07 | 2.1e-07 | 2.833E-07 | 6.351E-08 | 2.100E-07 | 3.200E-07 |
| 22282 | 1115 | G1115R | D | 2.2e-07 | 2.6e-07 | 1.7e-07 | 2.167E-07 | 4.509E-08 | 1.700E-07 | 2.600E-07 |
| 22283 | 1115 | G1115N | D | 1.4e-07 | 7.3e-08 | 1.7e-07 | 1.277E-07 | 4.966E-08 | 7.300E-08 | 1.700E-07 |
| 22284 | 1115 | G1115D | D | 2.9e-07 | 5.2e-07 | 2.6e-07 | 3.567E-07 | 1.422E-07 | 2.600E-07 | 5.200E-07 |
| 22285 | 1115 | G1115C | D | 1.8e-07 | 2.3e-07 | 2.5e-07 | 2.200E-07 | 3.606E-08 | 1.800E-07 | 2.500E-07 |
| 22286 | 1115 | G1115E | D | 1.7e-07 | 3e-07 | 3e-07 | 2.567E-07 | 7.506E-08 | 1.700E-07 | 3.000E-07 |
| 22287 | 1115 | G1115Q | D | 3e-07 | 2.3e-07 | 1.2e-07 | 2.167E-07 | 9.074E-08 | 1.200E-07 | 3.000E-07 |
| 22288 | 1115 | G1115G | D | 7.1e-07 | 1.7e-07 | 5.3e-07 | 4.700E-07 | 2.750E-07 | 1.700E-07 | 7.100E-07 |
| 22289 | 1115 | G1115H | D | 2.1e-07 | 3e-07 | 3e-07 | 2.700E-07 | 5.196E-08 | 2.100E-07 | 3.000E-07 |
| 22290 | 1115 | G1115I | D | 1.4e-07 | 3.3e-07 | 2.9e-07 | 2.533E-07 | 1.002E-07 | 1.400E-07 | 3.300E-07 |
| 22291 | 1115 | G1115L | D | 1.3e-07 | 5.8e-07 | 4.4e-07 | 3.833E-07 | 2.303E-07 | 1.300E-07 | 5.800E-07 |
| 22292 | 1115 | G1115K | D | 6.8e-07 | 1e-07 | 1.7e-07 | 3.167E-07 | 3.166E-07 | 1.000E-07 | 6.800E-07 |
| 22293 | 1115 | G1115M | D | 2e-07 | 2.6e-07 | 1.8e-07 | 2.133E-07 | 4.163E-08 | 1.800E-07 | 2.600E-07 |
| 22294 | 1115 | G1115F | D | 1.2e-07 | 6e-08 | 3.4e-07 | 1.733E-07 | 1.474E-07 | 6.000E-08 | 3.400E-07 |
| 22295 | 1115 | G1115P | D | 2.4e-07 | 4e-07 | 2.3e-07 | 2.900E-07 | 9.539E-08 | 2.300E-07 | 4.000E-07 |
| 22296 | 1115 | G1115S | D | 3.5e-07 | 3.9e-07 | 1.8e-07 | 3.067E-07 | 1.115E-07 | 1.800E-07 | 3.900E-07 |
| 22297 | 1115 | G1115T | D | 1.7e-07 | 1.9e-07 | 2.5e-07 | 2.033E-07 | 4.163E-08 | 1.700E-07 | 2.500E-07 |
| 22298 | 1115 | G1115W | D | 2.2e-07 | 4.6e-07 | 9.4e-08 | 2.580E-07 | 1.859E-07 | 9.400E-08 | 4.600E-07 |
| 22299 | 1115 | G1115Y | D | 2.3e-07 | 2.8e-07 | 3.1e-07 | 2.733E-07 | 4.041E-08 | 2.300E-07 | 3.100E-07 |
| 22300 | 1115 | G1115V | D | 1.5e-07 | 4.3e-07 | 2.1e-07 | 2.633E-07 | 1.474E-07 | 1.500E-07 | 4.300E-07 |
| 22301 | 1116 | V1116A | D | 5.5e-07 | 3.7e-07 | 2.5e-07 | 3.900E-07 | 1.510E-07 | 2.500E-07 | 5.500E-07 |
| 22302 | 1116 | V1116R | D | 2.8e-07 | 2.4e-07 | 3.1e-07 | 2.767E-07 | 3.512E-08 | 2.400E-07 | 3.100E-07 |
| 22303 | 1116 | V1116N | D | 2e-07 | 3.7e-07 | 1.6e-07 | 2.433E-07 | 1.115E-07 | 1.600E-07 | 3.700E-07 |
| 22304 | 1116 | V1116D | D | 1.8e-07 | 3.4e-07 | 1.6e-07 | 2.267E-07 | 9.866E-08 | 1.600E-07 | 3.400E-07 |
| 22305 | 1116 | V1116C | D | 1.4e-07 | 2e-07 | 3.4e-07 | 2.267E-07 | 1.026E-07 | 1.400E-07 | 3.400E-07 |
| 22306 | 1116 | V1116E | D | 1.7e-07 | 3e-07 | 4.2e-07 | 2.967E-07 | 1.250E-07 | 1.700E-07 | 4.200E-07 |
| 22307 | 1116 | V1116Q | D | 2.4e-07 | 3.8e-07 | 2.2e-07 | 2.800E-07 | 8.718E-08 | 2.200E-07 | 3.800E-07 |
| 22308 | 1116 | V1116G | D | 4.4e-07 | 2.2e-07 | 8.4e-07 | 5.000E-07 | 3.143E-07 | 2.200E-07 | 8.400E-07 |
| 22309 | 1116 | V1116H | D | 2.5e-07 | 2.3e-07 | 4e-07 | 2.933E-07 | 9.292E-08 | 2.300E-07 | 4.000E-07 |
| 22310 | 1116 | V1116I | D | 3.4e-07 | 3.3e-07 | 2.8e-07 | 3.167E-07 | 3.215E-08 | 2.800E-07 | 3.400E-07 |
| 22311 | 1116 | V1116L | D | 2.2e-07 | 2.3e-07 | 1.6e-07 | 2.033E-07 | 3.786E-08 | 1.600E-07 | 2.300E-07 |
| 22312 | 1116 | V1116K | D | 1.8e-07 | 2.5e-07 | 3.5e-07 | 2.600E-07 | 8.544E-08 | 1.800E-07 | 3.500E-07 |
| 22313 | 1116 | V1116M | D | 2.6e-07 | 1.2e-07 | 2.2e-07 | 2.000E-07 | 7.211E-08 | 1.200E-07 | 2.600E-07 |
| 22314 | 1116 | V1116F | D | 1.2e-07 | 3.6e-07 | 2.1e-07 | 2.300E-07 | 1.212E-07 | 1.200E-07 | 3.600E-07 |
| 22315 | 1116 | V1116P | D | 2.1e-07 | 7.1e-08 | 3.7e-07 | 2.170E-07 | 1.496E-07 | 7.100E-08 | 3.700E-07 |
| 22316 | 1116 | V1116S | D | 1.9e-07 | 2.3e-07 | 2.3e-07 | 2.167E-07 | 2.309E-08 | 1.900E-07 | 2.300E-07 |
| 22317 | 1116 | V1116T | D | 3.5e-07 | 2.6e-07 | 3.2e-07 | 3.100E-07 | 4.583E-08 | 2.600E-07 | 3.500E-07 |
| 22318 | 1116 | V1116W | D | 2.9e-07 | 2.5e-07 | 1.7e-07 | 2.367E-07 | 6.110E-08 | 1.700E-07 | 2.900E-07 |
| 22319 | 1116 | V1116Y | D | 2.8e-07 | 2.1e-07 | 1.5e-07 | 2.133E-07 | 6.506E-08 | 1.500E-07 | 2.800E-07 |
| 22320 | 1116 | V1116V | D | 3.9e-07 | 1.7e-07 | 2.9e-07 | 2.833E-07 | 1.102E-07 | 1.700E-07 | 3.900E-07 |
| 22321 | 1117 | P1117A | D | 3.3e-07 | 3.4e-07 | 1.9e-07 | 2.867E-07 | 8.386E-08 | 1.900E-07 | 3.400E-07 |
| 22322 | 1117 | P1117R | D | 8.2e-08 | 3.1e-07 | 4.9e-07 | 2.940E-07 | 2.045E-07 | 8.200E-08 | 4.900E-07 |
| 22323 | 1117 | P1117N | D | 1.5e-07 | 2.8e-07 | 2.9e-07 | 2.400E-07 | 7.810E-08 | 1.500E-07 | 2.900E-07 |
| 22324 | 1117 | P1117D | D | 2.5e-07 | 2e-07 | 1.9e-07 | 2.133E-07 | 3.215E-08 | 1.900E-07 | 2.500E-07 |
| 22325 | 1117 | P1117C | D | 2.5e-07 | 4.2e-07 | 3.9e-07 | 3.533E-07 | 9.074E-08 | 2.500E-07 | 4.200E-07 |
| 22326 | 1117 | P1117E | D | 1.2e-07 | 1.6e-07 | 1.5e-07 | 1.433E-07 | 2.082E-08 | 1.200E-07 | 1.600E-07 |
| 22327 | 1117 | P1117Q | D | 4.2e-07 | 2.6e-07 | 1.6e-07 | 2.800E-07 | 1.311E-07 | 1.600E-07 | 4.200E-07 |
| 22328 | 1117 | P1117G | D | 2.9e-07 | 2.9e-07 | 3e-07 | 2.933E-07 | 5.774E-09 | 2.900E-07 | 3.000E-07 |
| 22329 | 1117 | P1117H | D | 2.9e-07 | 1.5e-07 | 2.3e-07 | 2.233E-07 | 7.024E-08 | 1.500E-07 | 2.900E-07 |
| 22330 | 1117 | P1117I | D | 1.6e-07 | 2.3e-07 | 1.5e-07 | 1.800E-07 | 4.359E-08 | 1.500E-07 | 2.300E-07 |
| 22331 | 1117 | P1117L | D | 1e-07 | 4e-07 | 1.8e-07 | 2.267E-07 | 1.553E-07 | 1.000E-07 | 4.000E-07 |
| 22332 | 1117 | P1117K | D | 1.9e-07 | 2.6e-07 | 1.9e-07 | 2.133E-07 | 4.041E-08 | 1.900E-07 | 2.600E-07 |
| 22333 | 1117 | P1117M | D | 3.4e-07 | 8.7e-08 | 2.8e-07 | 2.357E-07 | 1.322E-07 | 8.700E-08 | 3.400E-07 |
| 22334 | 1117 | P1117F | D | 2.1e-07 | 1.2e-07 | 2.9e-07 | 2.067E-07 | 8.505E-08 | 1.200E-07 | 2.900E-07 |
| 22335 | 1117 | P1117P | D | 4.2e-07 | 1.9e-07 | 6.9e-07 | 4.333E-07 | 2.503E-07 | 1.900E-07 | 6.900E-07 |
| 22336 | 1117 | P1117S | D | 2.6e-07 | 3.5e-07 | 4.4e-07 | 3.500E-07 | 9.000E-08 | 2.600E-07 | 4.400E-07 |
| 22337 | 1117 | P1117T | D | 1.4e-07 | 1.9e-07 | 7.4e-08 | 1.347E-07 | 5.818E-08 | 7.400E-08 | 1.900E-07 |
| 22338 | 1117 | P1117W | D | 2.2e-07 | 2.5e-07 | 3.8e-07 | 2.833E-07 | 8.505E-08 | 2.200E-07 | 3.800E-07 |
| 22339 | 1117 | P1117Y | D | 1.9e-07 | 9e-08 | 5.3e-07 | 2.700E-07 | 2.307E-07 | 9.000E-08 | 5.300E-07 |
| 22340 | 1117 | P1117V | D | 3.7e-07 | 5.3e-07 | 3.5e-07 | 4.167E-07 | 9.866E-08 | 3.500E-07 | 5.300E-07 |
| 22341 | 1118 | S1118A | D | 1.7e-07 | 2.2e-07 | 3.7e-07 | 2.533E-07 | 1.041E-07 | 1.700E-07 | 3.700E-07 |
| 22342 | 1118 | S1118R | D | 1.1e-07 | 3.3e-07 | 2.9e-07 | 2.433E-07 | 1.172E-07 | 1.100E-07 | 3.300E-07 |
| 22343 | 1118 | S1118N | D | 2.1e-07 | 2e-07 | 3e-07 | 2.367E-07 | 5.508E-08 | 2.000E-07 | 3.000E-07 |
| 22344 | 1118 | S1118D | D | 1.5e-07 | 2.5e-07 | 1.8e-07 | 1.933E-07 | 5.132E-08 | 1.500E-07 | 2.500E-07 |
| 22345 | 1118 | S1118C | D | 2e-07 | 1e-07 | 1.3e-07 | 1.433E-07 | 5.132E-08 | 1.000E-07 | 2.000E-07 |
| 22346 | 1118 | S1118E | D | 2.3e-07 | 4e-07 | 2.4e-07 | 2.900E-07 | 9.539E-08 | 2.300E-07 | 4.000E-07 |
| 22347 | 1118 | S1118Q | D | 2.2e-07 | 4.1e-07 | 2.6e-07 | 2.967E-07 | 1.002E-07 | 2.200E-07 | 4.100E-07 |
| 22348 | 1118 | S1118G | D | 3.4e-07 | 1.4e-07 | 2.2e-07 | 2.333E-07 | 1.007E-07 | 1.400E-07 | 3.400E-07 |
| 22349 | 1118 | S1118H | D | 3.3e-07 | 1.7e-07 | 2.9e-07 | 2.633E-07 | 8.327E-08 | 1.700E-07 | 3.300E-07 |
| 22350 | 1118 | S1118I | D | 3.7e-07 | 3.3e-07 | 3.8e-07 | 3.600E-07 | 2.646E-08 | 3.300E-07 | 3.800E-07 |
| 22351 | 1118 | S1118L | D | 1.8e-07 | 1.6e-07 | 3.3e-07 | 2.233E-07 | 9.292E-08 | 1.600E-07 | 3.300E-07 |
| 22352 | 1118 | S1118K | D | 3.2e-07 | 2.1e-07 | 2.2e-07 | 2.500E-07 | 6.083E-08 | 2.100E-07 | 3.200E-07 |
| 22353 | 1118 | S1118M | D | 7.3e-08 | 1.8e-07 | 1.6e-07 | 1.377E-07 | 5.689E-08 | 7.300E-08 | 1.800E-07 |
| 22354 | 1118 | S1118F | D | 3.2e-07 | 4.5e-07 | 1.8e-07 | 3.167E-07 | 1.350E-07 | 1.800E-07 | 4.500E-07 |

| | | | | | | | | | | |
|---|---|---|---|---|---|---|---|---|---|---|
| 22355 | 1118 | S1118P | D | 2e-07 | 4.5e-07 | 2.3e-07 | 2.933E-07 | 1.365E-07 | 2.000E-07 | 4.500E-07 |
| 22356 | 1118 | S1118S | D | 1.7e-07 | 1.2e-07 | 3.3e-07 | 2.067E-07 | 1.097E-07 | 1.200E-07 | 3.300E-07 |
| 22357 | 1118 | S1118T | D | 2.2e-07 | 2.6e-07 | 2.8e-07 | 2.533E-07 | 3.055E-08 | 2.200E-07 | 2.800E-07 |
| 22358 | 1118 | S1118W | D | 8.5e-07 | 2.7e-07 | 1.5e-07 | 4.233E-07 | 3.743E-07 | 1.500E-07 | 8.500E-07 |
| 22359 | 1118 | S1118Y | D | 2.7e-07 | 3.6e-07 | 2.9e-07 | 3.067E-07 | 4.726E-08 | 2.700E-07 | 3.600E-07 |
| 22360 | 1118 | S1118V | D | 2.3e-07 | 3e-07 | 3.3e-07 | 2.867E-07 | 5.132E-08 | 2.300E-07 | 3.300E-07 |
| 22361 | 1119 | R1119A | D | 4e-07 | 1.4e-07 | 2.4e-07 | 2.600E-07 | 1.311E-07 | 1.400E-07 | 4.000E-07 |
| 22362 | 1119 | R1119R | D | 4e-07 | 1.8e-07 | 1.8e-07 | 2.533E-07 | 1.270E-07 | 1.800E-07 | 4.000E-07 |
| 22363 | 1119 | R1119N | D | 1.6e-07 | 1.9e-07 | 2.5e-07 | 2.000E-07 | 4.583E-08 | 1.600E-07 | 2.500E-07 |
| 22364 | 1119 | R1119D | D | 2.5e-07 | 3.5e-07 | 2.4e-07 | 2.800E-07 | 6.083E-08 | 2.400E-07 | 3.500E-07 |
| 22365 | 1119 | R1119C | D | 1.9e-07 | 1.4e-07 | 3.7e-07 | 2.333E-07 | 1.210E-07 | 1.400E-07 | 3.700E-07 |
| 22366 | 1119 | R1119E | D | 4.7e-07 | 3.3e-07 | 2.4e-07 | 3.467E-07 | 1.159E-07 | 2.400E-07 | 4.700E-07 |
| 22367 | 1119 | R1119Q | D | 3.2e-07 | 5.3e-07 | 3.3e-07 | 3.933E-07 | 1.185E-07 | 3.200E-07 | 5.300E-07 |
| 22368 | 1119 | R1119G | D | 3.9e-07 | 3.3e-07 | 2.9e-07 | 3.367E-07 | 5.033E-08 | 2.900E-07 | 3.900E-07 |
| 22369 | 1119 | R1119H | D | 2.7e-07 | 2.4e-07 | 9.4e-08 | 2.013E-07 | 9.416E-08 | 9.400E-08 | 2.700E-07 |
| 22370 | 1119 | R1119I | D | 5.5e-07 | 6.1e-07 | 1.2e-07 | 4.267E-07 | 2.673E-07 | 1.200E-07 | 6.100E-07 |
| 22371 | 1119 | R1119L | D | 2.2e-07 | 2.6e-07 | 4.9e-07 | 3.233E-07 | 1.457E-07 | 2.200E-07 | 4.900E-07 |
| 22372 | 1119 | R1119K | D | 2.3e-07 | 3e-07 | 3.5e-07 | 2.933E-07 | 6.028E-08 | 2.300E-07 | 3.500E-07 |
| 22373 | 1119 | R1119M | D | 2.1e-07 | 1.1e-07 | 2.3e-07 | 1.833E-07 | 6.429E-08 | 1.100E-07 | 2.300E-07 |
| 22374 | 1119 | R1119F | D | 1.8e-07 | 2.6e-07 | 2.9e-07 | 2.433E-07 | 5.686E-08 | 1.800E-07 | 2.900E-07 |
| 22375 | 1119 | R1119P | D | 1.3e-07 | 2.9e-07 | 2.3e-07 | 2.167E-07 | 8.083E-08 | 1.300E-07 | 2.900E-07 |
| 22376 | 1119 | R1119S | D | 4e-07 | 3.8e-07 | 2.2e-07 | 3.333E-07 | 9.866E-08 | 2.200E-07 | 4.000E-07 |
| 22377 | 1119 | R1119T | D | 4.6e-07 | 2.8e-07 | 2.5e-07 | 3.300E-07 | 1.136E-07 | 2.500E-07 | 4.600E-07 |
| 22378 | 1119 | R1119W | D | 1.5e-07 | 3.8e-07 | 3.6e-07 | 2.967E-07 | 1.274E-07 | 1.500E-07 | 3.800E-07 |
| 22379 | 1119 | R1119Y | D | 1.2e-07 | 2.4e-07 | 2.1e-07 | 1.900E-07 | 6.245E-08 | 1.200E-07 | 2.400E-07 |
| 22380 | 1119 | R1119V | D | 2.3e-07 | 3.5e-07 | 3.3e-07 | 3.033E-07 | 6.429E-08 | 2.300E-07 | 3.500E-07 |
| 22381 | 1120 | F1120A | D | 2.2e-07 | 2.8e-07 | 1.6e-07 | 2.200E-07 | 6.000E-08 | 1.600E-07 | 2.800E-07 |
| 22382 | 1120 | F1120R | D | 5.2e-07 | 2.3e-07 | 2.8e-07 | 3.433E-07 | 1.550E-07 | 2.300E-07 | 5.200E-07 |
| 22383 | 1120 | F1120N | D | 2.9e-07 | 6.5e-07 | 2.4e-07 | 3.933E-07 | 2.237E-07 | 2.400E-07 | 6.500E-07 |
| 22384 | 1120 | F1120D | D | 2.4e-07 | 2.4e-07 | 2e-07 | 2.267E-07 | 2.309E-08 | 2.000E-07 | 2.400E-07 |
| 22385 | 1120 | F1120C | D | 1.2e-07 | 5.7e-07 | 3.7e-07 | 3.533E-07 | 2.255E-07 | 1.200E-07 | 5.700E-07 |
| 22386 | 1120 | F1120E | D | 1.4e-07 | 2.1e-07 | 1.7e-07 | 1.733E-07 | 3.512E-08 | 1.400E-07 | 2.100E-07 |
| 22387 | 1120 | F1120Q | D | 3.2e-07 | 2.5e-07 | 2.5e-07 | 2.733E-07 | 4.041E-08 | 2.500E-07 | 3.200E-07 |
| 22388 | 1120 | F1120G | D | 2.8e-07 | 4.9e-07 | 2.9e-07 | 3.533E-07 | 1.185E-07 | 2.800E-07 | 4.900E-07 |
| 22389 | 1120 | F1120H | D | 9.9e-08 | 2.9e-07 | 2.8e-07 | 2.230E-07 | 1.075E-07 | 9.900E-08 | 2.900E-07 |
| 22390 | 1120 | F1120I | D | 2e-07 | 1.9e-07 | 4.1e-07 | 2.667E-07 | 1.242E-07 | 1.900E-07 | 4.100E-07 |
| 22391 | 1120 | F1120L | D | 2.1e-07 | 2.8e-07 | 3.7e-07 | 2.867E-07 | 8.021E-08 | 2.100E-07 | 3.700E-07 |
| 22392 | 1120 | F1120K | D | 1.4e-07 | 1e-07 | 4.8e-07 | 2.400E-07 | 2.088E-07 | 1.000E-07 | 4.800E-07 |
| 22393 | 1120 | F1120M | D | 1.4e-07 | 2.3e-07 | 1.8e-07 | 1.833E-07 | 4.509E-08 | 1.400E-07 | 2.300E-07 |
| 22394 | 1120 | F1120F | D | 2.7e-07 | 3.2e-07 | 1.1e-07 | 2.333E-07 | 1.097E-07 | 1.100E-07 | 3.200E-07 |
| 22395 | 1120 | F1120P | D | 1.8e-07 | 2.8e-07 | 4.6e-07 | 3.067E-07 | 1.419E-07 | 1.800E-07 | 4.600E-07 |
| 22396 | 1120 | F1120S | D | 2.6e-07 | 2.6e-07 | 2.9e-07 | 2.700E-07 | 1.732E-08 | 2.600E-07 | 2.900E-07 |
| 22397 | 1120 | F1120T | D | 3.6e-07 | 2e-07 | 1.4e-07 | 2.333E-07 | 1.137E-07 | 1.400E-07 | 3.600E-07 |
| 22398 | 1120 | F1120W | D | 2.1e-07 | 2.4e-07 | 6.9e-08 | 1.730E-07 | 9.131E-08 | 6.900E-08 | 2.400E-07 |
| 22399 | 1120 | F1120Y | D | 3.5e-07 | 3.7e-07 | 2.5e-07 | 3.233E-07 | 6.429E-08 | 2.500E-07 | 3.700E-07 |
| 22400 | 1120 | F1120V | D | 2.2e-07 | 1.6e-07 | 1.5e-07 | 1.767E-07 | 3.786E-08 | 1.500E-07 | 2.200E-07 |
| 22401 | 1121 | S1121A | D | 3.2e-07 | 1.9e-07 | 6.4e-07 | 3.833E-07 | 2.316E-07 | 1.900E-07 | 6.400E-07 |
| 22402 | 1121 | S1121R | D | 3.9e-07 | 2.9e-07 | 2e-07 | 2.933E-07 | 9.504E-08 | 2.000E-07 | 3.900E-07 |
| 22403 | 1121 | S1121N | D | 3.7e-07 | 2.1e-07 | 1e-07 | 2.267E-07 | 1.358E-07 | 1.000E-07 | 3.700E-07 |
| 22404 | 1121 | S1121D | D | 1.3e-07 | 1.4e-07 | 3.1e-07 | 1.933E-07 | 1.012E-07 | 1.300E-07 | 3.100E-07 |
| 22405 | 1121 | S1121C | D | 1.7e-07 | 2.2e-07 | 1.7e-07 | 1.867E-07 | 2.887E-08 | 1.700E-07 | 2.200E-07 |
| 22406 | 1121 | S1121E | D | 3e-07 | 2.4e-07 | 2.4e-07 | 2.600E-07 | 3.464E-08 | 2.400E-07 | 3.000E-07 |
| 22407 | 1121 | S1121Q | D | 1.4e-07 | 4.4e-07 | 3.1e-07 | 2.967E-07 | 1.504E-07 | 1.400E-07 | 4.400E-07 |
| 22408 | 1121 | S1121G | D | 1.6e-07 | 2.7e-07 | 1.1e-07 | 1.800E-07 | 8.185E-08 | 1.100E-07 | 2.700E-07 |
| 22409 | 1121 | S1121H | D | 1.5e-07 | 3e-07 | 1.6e-07 | 2.033E-07 | 8.386E-08 | 1.500E-07 | 3.000E-07 |
| 22410 | 1121 | S1121I | D | 2.5e-07 | 1.5e-07 | 1.8e-07 | 1.933E-07 | 5.132E-08 | 1.500E-07 | 2.500E-07 |
| 22411 | 1121 | S1121L | D | 1.9e-07 | 2.3e-07 | 2.6e-07 | 2.267E-07 | 3.512E-08 | 1.900E-07 | 2.600E-07 |
| 22412 | 1121 | S1121K | D | 1.8e-07 | 3.7e-07 | 3.4e-07 | 2.967E-07 | 1.021E-07 | 1.800E-07 | 3.700E-07 |
| 22413 | 1121 | S1121M | D | 1.6e-07 | 3.3e-07 | 5.2e-07 | 3.367E-07 | 1.801E-07 | 1.600E-07 | 5.200E-07 |
| 22414 | 1121 | S1121F | D | 2.1e-07 | 1.9e-07 | 4e-07 | 2.667E-07 | 1.159E-07 | 1.900E-07 | 4.000E-07 |
| 22415 | 1121 | S1121P | D | 5.3e-07 | 2.9e-07 | 1.3e-07 | 3.167E-07 | 2.013E-07 | 1.300E-07 | 5.300E-07 |
| 22416 | 1121 | S1121S | D | 2.5e-07 | 4.8e-07 | 3.7e-07 | 3.667E-07 | 1.150E-07 | 2.500E-07 | 4.800E-07 |
| 22417 | 1121 | S1121T | D | 2.5e-07 | 2.1e-07 | 2.6e-07 | 2.400E-07 | 2.646E-08 | 2.100E-07 | 2.600E-07 |
| 22418 | 1121 | S1121W | D | 1.8e-07 | 4.7e-07 | 3.4e-07 | 3.300E-07 | 1.453E-07 | 1.800E-07 | 4.700E-07 |
| 22419 | 1121 | S1121Y | D | 2.5e-07 | 1.2e-07 | 3.4e-07 | 2.367E-07 | 1.106E-07 | 1.200E-07 | 3.400E-07 |
| 22420 | 1121 | S1121V | D | 3.3e-07 | 2.7e-07 | 6.6e-07 | 4.200E-07 | 2.100E-07 | 2.700E-07 | 6.600E-07 |
| 22421 | 1122 | G1122A | D | 2.3e-07 | 2.7e-07 | 3.4e-07 | 2.800E-07 | 5.568E-08 | 2.300E-07 | 3.400E-07 |
| 22422 | 1122 | G1122R | D | 2.1e-07 | 3.4e-07 | 3.9e-07 | 3.133E-07 | 9.292E-08 | 2.100E-07 | 3.900E-07 |
| 22423 | 1122 | G1122N | D | 2.1e-07 | 2.1e-07 | 2.4e-07 | 2.200E-07 | 1.732E-08 | 2.100E-07 | 2.400E-07 |
| 22424 | 1122 | G1122D | D | 1.7e-07 | 2.1e-07 | 1.3e-07 | 1.700E-07 | 4.000E-08 | 1.300E-07 | 2.100E-07 |
| 22425 | 1122 | G1122C | D | 4.2e-07 | 3.9e-07 | 3.5e-07 | 3.867E-07 | 3.512E-08 | 3.500E-07 | 4.200E-07 |
| 22426 | 1122 | G1122E | D | 2.4e-07 | 5.5e-07 | 6e-07 | 4.633E-07 | 1.950E-07 | 2.400E-07 | 6.000E-07 |
| 22427 | 1122 | G1122Q | D | 3.6e-07 | 2.1e-07 | 3.1e-07 | 2.933E-07 | 7.638E-08 | 2.100E-07 | 3.600E-07 |
| 22428 | 1122 | G1122G | D | 2.4e-07 | 3.1e-07 | 1.6e-07 | 2.367E-07 | 7.506E-08 | 1.600E-07 | 3.100E-07 |
| 22429 | 1122 | G1122H | D | 1.3e-07 | 2.6e-07 | 1.7e-07 | 1.867E-07 | 6.658E-08 | 1.300E-07 | 2.600E-07 |
| 22430 | 1122 | G1122I | D | 1.5e-07 | 1.9e-07 | 4.8e-07 | 2.733E-07 | 1.801E-07 | 1.500E-07 | 4.800E-07 |
| 22431 | 1122 | G1122L | D | 3.4e-07 | 3.3e-07 | 2.8e-07 | 3.167E-07 | 3.215E-08 | 2.800E-07 | 3.400E-07 |
| 22432 | 1122 | G1122K | D | 3e-07 | 1.7e-07 | 2.2e-07 | 2.300E-07 | 6.557E-08 | 1.700E-07 | 3.000E-07 |
| 22433 | 1122 | G1122M | D | 3.3e-07 | 1.4e-06 | 1.5e-07 | 6.267E-07 | 6.757E-07 | 1.500E-07 | 1.400E-06 |
| 22434 | 1122 | G1122F | D | 2.6e-07 | 3.2e-07 | 1.8e-07 | 2.533E-07 | 7.024E-08 | 1.800E-07 | 3.200E-07 |
| 22435 | 1122 | G1122P | D | 1.8e-07 | 2.2e-07 | 2.8e-07 | 2.267E-07 | 5.033E-08 | 1.800E-07 | 2.800E-07 |
| 22436 | 1122 | G1122S | D | 1.8e-07 | 1.6e-07 | 3.2e-07 | 2.200E-07 | 8.718E-08 | 1.600E-07 | 3.200E-07 |
| 22437 | 1122 | G1122T | D | 2.6e-07 | 2.2e-07 | 3.1e-07 | 2.633E-07 | 4.509E-08 | 2.200E-07 | 3.100E-07 |
| 22438 | 1122 | G1122W | D | 3e-07 | 1.8e-07 | 2.3e-07 | 2.367E-07 | 6.028E-08 | 1.800E-07 | 3.000E-07 |
| 22439 | 1122 | G1122Y | D | 3.5e-07 | 4.2e-07 | 2.6e-07 | 3.433E-07 | 8.021E-08 | 2.600E-07 | 4.200E-07 |

| | | | | | | | | | | |
|---|---|---|---|---|---|---|---|---|---|---|
| 22440 | 1122 | G1122V | D | 1.6e-07 | 2.8e-07 | 5.9e-07 | 3.433E-07 | 2.219E-07 | 1.600E-07 | 5.900E-07 |
| 22441 | 1123 | S1123A | D | 4.3e-07 | 2.1e-07 | 1.7e-07 | 2.700E-07 | 1.400E-07 | 1.700E-07 | 4.300E-07 |
| 22442 | 1123 | S1123R | D | 2.6e-07 | 1.4e-07 | 3.4e-07 | 2.467E-07 | 1.007E-07 | 1.400E-07 | 3.400E-07 |
| 22443 | 1123 | S1123N | D | 4e-07 | 3.1e-07 | 3e-07 | 3.367E-07 | 5.508E-08 | 3.000E-07 | 4.000E-07 |
| 22444 | 1123 | S1123D | D | 2e-07 | 1.7e-07 | 2.2e-07 | 1.967E-07 | 2.517E-08 | 1.700E-07 | 2.200E-07 |
| 22445 | 1123 | S1123C | D | 2.8e-07 | 4.5e-07 | 3.5e-07 | 3.600E-07 | 8.544E-08 | 2.800E-07 | 4.500E-07 |
| 22446 | 1123 | S1123E | D | 2.2e-07 | 2.1e-07 | 2.8e-07 | 2.367E-07 | 3.786E-08 | 2.100E-07 | 2.800E-07 |
| 22447 | 1123 | S1123Q | D | 1.9e-07 | 1.9e-07 | 2.5e-07 | 2.100E-07 | 3.464E-08 | 1.900E-07 | 2.500E-07 |
| 22448 | 1123 | S1123G | D | 1.8e-07 | 5.6e-07 | 4.9e-07 | 4.100E-07 | 2.022E-07 | 1.800E-07 | 5.600E-07 |
| 22449 | 1123 | S1123H | D | 4.5e-07 | 3.5e-07 | 1.6e-07 | 3.200E-07 | 1.473E-07 | 1.600E-07 | 4.500E-07 |
| 22450 | 1123 | S1123I | D | 1.8e-07 | 1.6e-07 | 1.1e-07 | 1.500E-07 | 3.606E-08 | 1.100E-07 | 1.800E-07 |
| 22451 | 1123 | S1123L | D | 2.2e-07 | 3.8e-07 | 1.8e-07 | 2.600E-07 | 1.058E-07 | 1.800E-07 | 3.800E-07 |
| 22452 | 1123 | S1123K | D | 2.5e-07 | 2.8e-07 | 1.6e-07 | 2.300E-07 | 6.245E-08 | 1.600E-07 | 2.800E-07 |
| 22453 | 1123 | S1123M | D | 5.1e-07 | 2.2e-07 | 1.9e-07 | 3.067E-07 | 1.767E-07 | 1.900E-07 | 5.100E-07 |
| 22454 | 1123 | S1123F | D | 1.8e-07 | 2.6e-07 | 1.9e-07 | 2.100E-07 | 4.359E-08 | 1.800E-07 | 2.600E-07 |
| 22455 | 1123 | S1123P | D | 2.6e-07 | 2.1e-07 | 4.5e-07 | 3.067E-07 | 1.266E-07 | 2.100E-07 | 4.500E-07 |
| 22456 | 1123 | S1123S | D | 1.6e-07 | 4.2e-07 | 2e-07 | 2.600E-07 | 1.400E-07 | 1.600E-07 | 4.200E-07 |
| 22457 | 1123 | S1123T | D | 2.2e-07 | 1.5e-07 | 2.5e-07 | 2.067E-07 | 5.132E-08 | 1.500E-07 | 2.500E-07 |
| 22458 | 1123 | S1123W | D | 4.4e-07 | 1.9e-07 | 3.6e-07 | 3.300E-07 | 1.277E-07 | 1.900E-07 | 4.400E-07 |
| 22459 | 1123 | S1123Y | D | 2e-07 | 1.7e-07 | 2.8e-07 | 2.167E-07 | 5.686E-08 | 1.700E-07 | 2.800E-07 |
| 22460 | 1123 | S1123V | D | 3.4e-07 | 2.2e-07 | 3.9e-07 | 3.167E-07 | 8.737E-08 | 2.200E-07 | 3.900E-07 |
| 22461 | 1124 | R1124A | D | 3e-07 | 2.1e-07 | 2.1e-07 | 2.400E-07 | 5.196E-08 | 2.100E-07 | 3.000E-07 |
| 22462 | 1124 | R1124R | D | 3.2e-07 | 3.6e-07 | 2.1e-07 | 2.967E-07 | 7.767E-08 | 2.100E-07 | 3.600E-07 |
| 22463 | 1124 | R1124N | D | 5e-07 | 1.7e-07 | 1.3e-07 | 2.667E-07 | 2.031E-07 | 1.300E-07 | 5.000E-07 |
| 22464 | 1124 | R1124D | D | 1.4e-07 | 3.4e-07 | 3e-07 | 2.600E-07 | 1.058E-07 | 1.400E-07 | 3.400E-07 |
| 22465 | 1124 | R1124C | D | 1.4e-07 | 3.1e-07 | 2.7e-07 | 2.400E-07 | 8.888E-08 | 1.400E-07 | 3.100E-07 |
| 22466 | 1124 | R1124E | D | 6.8e-07 | 5.4e-07 | 1.9e-07 | 4.700E-07 | 2.524E-07 | 1.900E-07 | 6.800E-07 |
| 22467 | 1124 | R1124Q | D | 1.9e-07 | 8.7e-07 | 2.3e-07 | 4.300E-07 | 3.816E-07 | 1.900E-07 | 8.700E-07 |
| 22468 | 1124 | R1124G | D | 2.8e-07 | 4.1e-07 | 2.9e-07 | 3.267E-07 | 7.234E-08 | 2.800E-07 | 4.100E-07 |
| 22469 | 1124 | R1124H | D | 5e-07 | 2.4e-07 | 1.5e-07 | 2.967E-07 | 1.818E-07 | 1.500E-07 | 5.000E-07 |
| 22470 | 1124 | R1124I | D | 2e-07 | 5.9e-07 | 2.3e-07 | 3.400E-07 | 2.170E-07 | 2.000E-07 | 5.900E-07 |
| 22471 | 1124 | R1124L | D | 3e-07 | 1.4e-07 | 2.8e-07 | 2.400E-07 | 8.718E-08 | 1.400E-07 | 3.000E-07 |
| 22472 | 1124 | R1124K | D | 1.5e-07 | 2.9e-07 | 2e-07 | 2.133E-07 | 7.095E-08 | 1.500E-07 | 2.900E-07 |
| 22473 | 1124 | R1124M | D | 4.4e-07 | 1.8e-07 | 2.2e-07 | 2.800E-07 | 1.400E-07 | 1.800E-07 | 4.400E-07 |
| 22474 | 1124 | R1124F | D | 3.1e-07 | 1.3e-07 | 1.5e-07 | 1.967E-07 | 9.866E-08 | 1.300E-07 | 3.100E-07 |
| 22475 | 1124 | R1124P | D | 3e-07 | 2.7e-07 | 3.9e-07 | 3.200E-07 | 6.245E-08 | 2.700E-07 | 3.900E-07 |
| 22476 | 1124 | R1124S | D | 2.3e-07 | 2.2e-07 | 1.8e-07 | 2.100E-07 | 2.646E-08 | 1.800E-07 | 2.300E-07 |
| 22477 | 1124 | R1124T | D | 2.1e-07 | 1.9e-07 | 2.5e-07 | 2.167E-07 | 3.055E-08 | 1.900E-07 | 2.500E-07 |
| 22478 | 1124 | R1124W | D | 2.1e-07 | 3.6e-07 | 3e-07 | 2.900E-07 | 7.550E-08 | 2.100E-07 | 3.600E-07 |
| 22479 | 1124 | R1124Y | D | 2.4e-07 | 2.9e-07 | 3.6e-07 | 2.967E-07 | 6.028E-08 | 2.400E-07 | 3.600E-07 |
| 22480 | 1124 | R1124V | D | 2e-07 | 3.2e-07 | 3.5e-07 | 2.900E-07 | 7.937E-08 | 2.000E-07 | 3.500E-07 |
| 22481 | 1125 | S1125A | D | 2.6e-07 | 3.3e-07 | 2.9e-07 | 2.933E-07 | 3.512E-08 | 2.600E-07 | 3.300E-07 |
| 22482 | 1125 | S1125R | D | 2.5e-07 | 4.3e-07 | 3.8e-07 | 3.533E-07 | 9.292E-08 | 2.500E-07 | 4.300E-07 |
| 22483 | 1125 | S1125N | D | 4e-07 | 1.6e-07 | 2.3e-07 | 2.633E-07 | 1.234E-07 | 1.600E-07 | 4.000E-07 |
| 22484 | 1125 | S1125D | D | 1.6e-07 | 3.2e-07 | 2.9e-07 | 2.567E-07 | 8.505E-08 | 1.600E-07 | 3.200E-07 |
| 22485 | 1125 | S1125C | D | 3.6e-07 | 2.1e-07 | 2.3e-07 | 2.667E-07 | 8.145E-08 | 2.100E-07 | 3.600E-07 |
| 22486 | 1125 | S1125E | D | 5.5e-07 | 1.5e-07 | 3.3e-07 | 3.433E-07 | 2.003E-07 | 1.500E-07 | 5.500E-07 |
| 22487 | 1125 | S1125Q | D | 3.1e-07 | 4.4e-07 | 1.5e-07 | 3.000E-07 | 1.453E-07 | 1.500E-07 | 4.400E-07 |
| 22488 | 1125 | S1125G | D | 3.6e-07 | 1.7e-07 | 2.1e-07 | 2.467E-07 | 1.002E-07 | 1.700E-07 | 3.600E-07 |
| 22489 | 1125 | S1125H | D | 2.6e-07 | 1.9e-07 | 2.4e-07 | 2.300E-07 | 3.606E-08 | 1.900E-07 | 2.600E-07 |
| 22490 | 1125 | S1125I | D | 2.8e-07 | 4.3e-07 | 9e-08 | 2.667E-07 | 1.704E-07 | 9.000E-08 | 4.300E-07 |
| 22491 | 1125 | S1125L | D | 3e-07 | 4.1e-07 | 1.7e-07 | 2.933E-07 | 1.201E-07 | 1.700E-07 | 4.100E-07 |
| 22492 | 1125 | S1125K | D | 1.8e-07 | 1.3e-07 | 2.3e-07 | 1.800E-07 | 5.000E-08 | 1.300E-07 | 2.300E-07 |
| 22493 | 1125 | S1125M | D | 2.3e-07 | 2.4e-07 | 3.6e-07 | 2.767E-07 | 7.234E-08 | 2.300E-07 | 3.600E-07 |
| 22494 | 1125 | S1125F | D | 2.3e-07 | 1.7e-07 | 5.6e-07 | 3.200E-07 | 2.100E-07 | 1.700E-07 | 5.600E-07 |
| 22495 | 1125 | S1125P | D | 1.6e-07 | 4e-07 | 3.7e-07 | 3.100E-07 | 1.308E-07 | 1.600E-07 | 4.000E-07 |
| 22496 | 1125 | S1125S | D | 3.2e-07 | 3.8e-07 | 3.3e-07 | 3.433E-07 | 3.215E-08 | 3.200E-07 | 3.800E-07 |
| 22497 | 1125 | S1125T | D | 4.6e-07 | 3.7e-07 | 2.1e-07 | 3.467E-07 | 1.266E-07 | 2.100E-07 | 4.600E-07 |
| 22498 | 1125 | S1125W | D | 1.8e-07 | 2e-07 | 1.4e-07 | 1.733E-07 | 3.055E-08 | 1.400E-07 | 2.000E-07 |
| 22499 | 1125 | S1125Y | D | 3.8e-07 | 2.5e-07 | 2.9e-07 | 3.067E-07 | 6.658E-08 | 2.500E-07 | 3.800E-07 |
| 22500 | 1125 | S1125V | D | 2.4e-07 | 1.1e-07 | 2.1e-07 | 1.867E-07 | 6.807E-08 | 1.100E-07 | 2.400E-07 |
| 22501 | 1126 | G1126A | D | 3.2e-07 | 1.8e-07 | 3.4e-07 | 2.800E-07 | 8.718E-08 | 1.800E-07 | 3.400E-07 |
| 22502 | 1126 | G1126R | D | 1.2e-07 | 4.7e-07 | 3.7e-07 | 3.200E-07 | 1.803E-07 | 1.200E-07 | 4.700E-07 |
| 22503 | 1126 | G1126N | D | 2.1e-07 | 2.4e-07 | 3e-07 | 2.500E-07 | 4.583E-08 | 2.100E-07 | 3.000E-07 |
| 22504 | 1126 | G1126D | D | 2.2e-07 | 2.8e-07 | 6.4e-08 | 1.880E-07 | 1.115E-07 | 6.400E-08 | 2.800E-07 |
| 22505 | 1126 | G1126C | D | 5e-07 | 3.2e-07 | 3.9e-07 | 4.033E-07 | 9.074E-08 | 3.200E-07 | 5.000E-07 |
| 22506 | 1126 | G1126E | D | 1.9e-07 | 2.5e-07 | 1.8e-07 | 2.067E-07 | 3.786E-08 | 1.800E-07 | 2.500E-07 |
| 22507 | 1126 | G1126Q | D | 2e-07 | 3.2e-07 | 1.8e-07 | 2.333E-07 | 7.572E-08 | 1.800E-07 | 3.200E-07 |
| 22508 | 1126 | G1126G | D | 4e-07 | 4.2e-07 | 3.1e-07 | 3.767E-07 | 5.859E-08 | 3.100E-07 | 4.200E-07 |
| 22509 | 1126 | G1126H | D | 1.6e-07 | 2.9e-07 | 1.7e-07 | 2.067E-07 | 7.234E-08 | 1.600E-07 | 2.900E-07 |
| 22510 | 1126 | G1126I | D | 2.1e-07 | 2.9e-07 | 3.8e-07 | 2.933E-07 | 8.505E-08 | 2.100E-07 | 3.800E-07 |
| 22511 | 1126 | G1126L | D | 3.7e-07 | 1.5e-07 | 3.1e-07 | 2.767E-07 | 1.137E-07 | 1.500E-07 | 3.700E-07 |
| 22512 | 1126 | G1126K | D | 1.4e-07 | 3.4e-07 | 9.9e-08 | 1.930E-07 | 1.289E-07 | 9.900E-08 | 3.400E-07 |
| 22513 | 1126 | G1126M | D | 4.3e-07 | 1.3e-07 | 3.5e-07 | 3.033E-07 | 1.553E-07 | 1.300E-07 | 4.300E-07 |
| 22514 | 1126 | G1126F | D | 2.6e-07 | 1.4e-07 | 2.5e-07 | 2.167E-07 | 6.658E-08 | 1.400E-07 | 2.600E-07 |
| 22515 | 1126 | G1126P | D | 3.7e-07 | 2.4e-07 | 1.7e-07 | 2.600E-07 | 1.015E-07 | 1.700E-07 | 3.700E-07 |
| 22516 | 1126 | G1126S | D | 1.1e-07 | 1.2e-07 | 1.8e-07 | 1.367E-07 | 3.786E-08 | 1.100E-07 | 1.800E-07 |
| 22517 | 1126 | G1126T | D | 3.4e-07 | 3.1e-07 | 2e-07 | 2.833E-07 | 7.371E-08 | 2.000E-07 | 3.400E-07 |
| 22518 | 1126 | G1126W | D | 1.4e-07 | 2.2e-07 | 2.7e-07 | 2.100E-07 | 6.557E-08 | 1.400E-07 | 2.700E-07 |
| 22519 | 1126 | G1126Y | D | 2.9e-07 | 4.3e-07 | 2.7e-07 | 3.300E-07 | 8.718E-08 | 2.700E-07 | 4.300E-07 |
| 22520 | 1126 | G1126V | D | 2.1e-07 | 1.9e-07 | 3.1e-07 | 2.367E-07 | 6.429E-08 | 1.900E-07 | 3.100E-07 |
| 22521 | 1127 | T1127A | D | 1.6e-07 | 2.6e-07 | 2.1e-07 | 2.100E-07 | 5.000E-08 | 1.600E-07 | 2.600E-07 |
| 22522 | 1127 | T1127R | D | 2.3e-07 | 2.9e-07 | 4e-07 | 3.067E-07 | 8.622E-08 | 2.300E-07 | 4.000E-07 |
| 22523 | 1127 | T1127N | D | 3.6e-07 | 1.5e-07 | 2.3e-07 | 2.467E-07 | 1.060E-07 | 1.500E-07 | 3.600E-07 |
| 22524 | 1127 | T1127D | D | 3.6e-07 | 2.6e-07 | 2.7e-07 | 2.967E-07 | 5.508E-08 | 2.600E-07 | 3.600E-07 |

| | | | | | | | | | | |
|---|---|---|---|---|---|---|---|---|---|---|
| 22525 | 1127 | T1127C | D | 2.6e-07 | 1.1e-07 | 1.9e-07 | 1.867E-07 | 7.506E-08 | 1.100E-07 | 2.600E-07 |
| 22526 | 1127 | T1127E | D | 2e-07 | 3.7e-07 | 3.1e-07 | 2.933E-07 | 8.622E-08 | 2.000E-07 | 3.700E-07 |
| 22527 | 1127 | T1127Q | D | 3e-07 | 2e-07 | 3.7e-07 | 2.900E-07 | 8.544E-08 | 2.000E-07 | 3.700E-07 |
| 22528 | 1127 | T1127G | D | 3.3e-07 | 1.8e-07 | 5.9e-07 | 3.667E-07 | 2.074E-07 | 1.800E-07 | 5.900E-07 |
| 22529 | 1127 | T1127H | D | 9.5e-08 | 4.2e-07 | 1.8e-07 | 2.317E-07 | 1.685E-07 | 9.500E-08 | 4.200E-07 |
| 22530 | 1127 | T1127I | D | 4.4e-07 | 2.7e-07 | 1.5e-07 | 2.867E-07 | 1.457E-07 | 1.500E-07 | 4.400E-07 |
| 22531 | 1127 | T1127L | D | 3.4e-07 | 1.6e-07 | 3.7e-07 | 2.900E-07 | 1.136E-07 | 1.600E-07 | 3.700E-07 |
| 22532 | 1127 | T1127K | D | 1.6e-07 | 3.5e-07 | 1.5e-07 | 2.200E-07 | 1.127E-07 | 1.500E-07 | 3.500E-07 |
| 22533 | 1127 | T1127M | D | 4.4e-07 | 2.3e-07 | 3e-07 | 3.233E-07 | 1.069E-07 | 2.300E-07 | 4.400E-07 |
| 22534 | 1127 | T1127F | D | 7.1e-07 | 2.4e-07 | 2.8e-07 | 4.100E-07 | 2.606E-07 | 2.400E-07 | 7.100E-07 |
| 22535 | 1127 | T1127P | D | 3e-07 | 2e-07 | 3.6e-07 | 2.867E-07 | 8.083E-08 | 2.000E-07 | 3.600E-07 |
| 22536 | 1127 | T1127S | D | 1.5e-07 | 1.4e-07 | 2.2e-07 | 1.700E-07 | 4.359E-08 | 1.400E-07 | 2.200E-07 |
| 22537 | 1127 | T1127T | D | 3.1e-07 | 3.9e-07 | 2.6e-07 | 3.200E-07 | 6.557E-08 | 2.600E-07 | 3.900E-07 |
| 22538 | 1127 | T1127W | D | 3.2e-07 | 4.7e-07 | 1.4e-07 | 3.100E-07 | 1.652E-07 | 1.400E-07 | 4.700E-07 |
| 22539 | 1127 | T1127Y | D | 1.6e-07 | 2.4e-07 | 3.4e-07 | 2.467E-07 | 9.018E-08 | 1.600E-07 | 3.400E-07 |
| 22540 | 1127 | T1127V | D | 2.3e-07 | 2.6e-07 | 2.5e-07 | 2.467E-07 | 1.528E-08 | 2.300E-07 | 2.600E-07 |
| 22541 | 1128 | D1128A | D | 2.4e-07 | 2.8e-07 | 2.7e-07 | 2.633E-07 | 2.082E-08 | 2.400E-07 | 2.800E-07 |
| 22542 | 1128 | D1128R | D | 1.3e-07 | 3.6e-07 | 2.7e-07 | 2.533E-07 | 1.159E-07 | 1.300E-07 | 3.600E-07 |
| 22543 | 1128 | D1128N | D | 2.6e-07 | 3.9e-07 | 2e-07 | 2.833E-07 | 9.713E-08 | 2.000E-07 | 3.900E-07 |
| 22544 | 1128 | D1128D | D | 5.2e-07 | 1.2e-07 | 1.8e-07 | 2.733E-07 | 2.157E-07 | 1.200E-07 | 5.200E-07 |
| 22545 | 1128 | D1128C | D | 3e-07 | 3.6e-07 | 4.2e-07 | 3.600E-07 | 6.000E-08 | 3.000E-07 | 4.200E-07 |
| 22546 | 1128 | D1128E | D | 4.4e-07 | 2e-07 | 2.2e-07 | 2.867E-07 | 1.332E-07 | 2.000E-07 | 4.400E-07 |
| 22547 | 1128 | D1128Q | D | 1.7e-07 | 2.6e-07 | 1.1e-07 | 1.800E-07 | 7.550E-08 | 1.100E-07 | 2.600E-07 |
| 22548 | 1128 | D1128G | D | 3.4e-07 | 1.7e-07 | 7.6e-07 | 4.233E-07 | 3.037E-07 | 1.700E-07 | 7.600E-07 |
| 22549 | 1128 | D1128H | D | 1.4e-07 | 2.3e-07 | 3.1e-07 | 2.267E-07 | 8.505E-08 | 1.400E-07 | 3.100E-07 |
| 22550 | 1128 | D1128I | D | 1.9e-07 | 3.3e-07 | 2.7e-07 | 2.633E-07 | 7.024E-08 | 1.900E-07 | 3.300E-07 |
| 22551 | 1128 | D1128L | D | 4.2e-07 | 2.2e-07 | 3e-07 | 3.133E-07 | 1.007E-07 | 2.200E-07 | 4.200E-07 |
| 22552 | 1128 | D1128K | D | 2.8e-07 | 1.1e-07 | 2.9e-07 | 2.267E-07 | 1.012E-07 | 1.100E-07 | 2.900E-07 |
| 22553 | 1128 | D1128M | D | 3.3e-07 | 3.1e-07 | 3e-07 | 3.133E-07 | 1.528E-08 | 3.000E-07 | 3.300E-07 |
| 22554 | 1128 | D1128F | D | 1.4e-07 | 4.6e-07 | 7.4e-08 | 2.247E-07 | 2.065E-07 | 7.400E-08 | 4.600E-07 |
| 22555 | 1128 | D1128P | D | 2.1e-07 | 2.5e-07 | 2.1e-07 | 2.233E-07 | 2.309E-08 | 2.100E-07 | 2.500E-07 |
| 22556 | 1128 | D1128S | D | 3.6e-07 | 1.2e-07 | 2.4e-07 | 2.400E-07 | 1.200E-07 | 1.200E-07 | 3.600E-07 |
| 22557 | 1128 | D1128T | D | 3e-07 | 2.8e-07 | 2.1e-07 | 2.633E-07 | 4.726E-08 | 2.100E-07 | 3.000E-07 |
| 22558 | 1128 | D1128W | D | 1.9e-07 | 2.2e-07 | 2e-07 | 2.033E-07 | 1.528E-08 | 1.900E-07 | 2.200E-07 |
| 22559 | 1128 | D1128Y | D | 3.2e-07 | 2.3e-07 | 1.3e-07 | 2.267E-07 | 9.504E-08 | 1.300E-07 | 3.200E-07 |
| 22560 | 1128 | D1128V | D | 2.7e-07 | 2.2e-07 | 2.6e-07 | 2.500E-07 | 2.646E-08 | 2.200E-07 | 2.700E-07 |
| 22561 | 1129 | F1129A | D | 1e-07 | 4.3e-07 | 3e-07 | 2.767E-07 | 1.662E-07 | 1.000E-07 | 4.300E-07 |
| 22562 | 1129 | F1129R | D | 2.2e-07 | 1.5e-07 | 2.4e-07 | 2.033E-07 | 4.726E-08 | 1.500E-07 | 2.400E-07 |
| 22563 | 1129 | F1129N | D | 1.2e-07 | 4e-07 | 1.5e-07 | 2.233E-07 | 1.537E-07 | 1.200E-07 | 4.000E-07 |
| 22564 | 1129 | F1129D | D | 1.7e-07 | 3.4e-07 | 2.6e-07 | 2.567E-07 | 8.505E-08 | 1.700E-07 | 3.400E-07 |
| 22565 | 1129 | F1129C | D | 5.7e-07 | 1.8e-07 | 2.8e-07 | 3.433E-07 | 2.026E-07 | 1.800E-07 | 5.700E-07 |
| 22566 | 1129 | F1129E | D | 3.8e-07 | 2e-07 | 3e-07 | 2.933E-07 | 9.018E-08 | 2.000E-07 | 3.800E-07 |
| 22567 | 1129 | F1129Q | D | 1.8e-07 | 2.1e-07 | 3.3e-07 | 2.400E-07 | 7.937E-08 | 1.800E-07 | 3.300E-07 |
| 22568 | 1129 | F1129G | D | 2.2e-07 | 2.6e-07 | 3.6e-07 | 2.800E-07 | 7.211E-08 | 2.200E-07 | 3.600E-07 |
| 22569 | 1129 | F1129H | D | 1.6e-07 | 6.2e-07 | 2.8e-07 | 3.533E-07 | 2.386E-07 | 1.600E-07 | 6.200E-07 |
| 22570 | 1129 | F1129I | D | 6.5e-07 | 3.5e-07 | 3.5e-07 | 4.500E-07 | 1.732E-07 | 3.500E-07 | 6.500E-07 |
| 22571 | 1129 | F1129L | D | 2.2e-07 | 2e-07 | 2.5e-07 | 2.233E-07 | 2.517E-08 | 2.000E-07 | 2.500E-07 |
| 22572 | 1129 | F1129K | D | 2e-07 | 2.2e-07 | 3.7e-07 | 2.633E-07 | 9.292E-08 | 2.000E-07 | 3.700E-07 |
| 22573 | 1129 | F1129M | D | 1.2e-07 | 1.4e-07 | 2.3e-07 | 1.633E-07 | 5.859E-08 | 1.200E-07 | 2.300E-07 |
| 22574 | 1129 | F1129F | D | 2.4e-07 | 3.5e-07 | 1.8e-07 | 2.567E-07 | 8.622E-08 | 1.800E-07 | 3.500E-07 |
| 22575 | 1129 | F1129P | D | 3.9e-07 | 1.7e-07 | 3.3e-07 | 2.967E-07 | 1.137E-07 | 1.700E-07 | 3.900E-07 |
| 22576 | 1129 | F1129S | D | 4.7e-07 | 3.2e-07 | 1.2e-07 | 3.033E-07 | 1.756E-07 | 1.200E-07 | 4.700E-07 |
| 22577 | 1129 | F1129T | D | 2.2e-07 | 3.1e-07 | 3.3e-07 | 2.867E-07 | 5.859E-08 | 2.200E-07 | 3.300E-07 |
| 22578 | 1129 | F1129W | D | 1.4e-07 | 1.7e-07 | 3.8e-07 | 2.300E-07 | 1.308E-07 | 1.400E-07 | 3.800E-07 |
| 22579 | 1129 | F1129Y | D | 2e-07 | 1.7e-07 | 1.8e-07 | 1.833E-07 | 1.528E-08 | 1.700E-07 | 2.000E-07 |
| 22580 | 1129 | F1129V | D | 2.4e-07 | 2.4e-07 | 2.3e-07 | 2.367E-07 | 5.774E-09 | 2.300E-07 | 2.400E-07 |
| 22581 | 1130 | T1130A | D | 2.8e-07 | 2.3e-07 | 1.5e-07 | 2.200E-07 | 6.557E-08 | 1.500E-07 | 2.800E-07 |
| 22582 | 1130 | T1130R | D | 1.5e-07 | 4.4e-07 | 2.1e-07 | 2.667E-07 | 1.531E-07 | 1.500E-07 | 4.400E-07 |
| 22583 | 1130 | T1130N | D | 2.6e-07 | 2.3e-07 | 5.2e-07 | 3.367E-07 | 1.595E-07 | 2.300E-07 | 5.200E-07 |
| 22584 | 1130 | T1130D | D | 2.2e-07 | 3.2e-07 | 1.3e-07 | 2.233E-07 | 9.504E-08 | 1.300E-07 | 3.200E-07 |
| 22585 | 1130 | T1130C | D | 2e-07 | 3.3e-07 | 2.9e-07 | 2.733E-07 | 6.658E-08 | 2.000E-07 | 3.300E-07 |
| 22586 | 1130 | T1130E | D | 1.3e-07 | 2.4e-07 | 2.3e-07 | 2.000E-07 | 6.083E-08 | 1.300E-07 | 2.400E-07 |
| 22587 | 1130 | T1130Q | D | 3.6e-07 | 3e-07 | 1.9e-07 | 2.833E-07 | 8.622E-08 | 1.900E-07 | 3.600E-07 |
| 22588 | 1130 | T1130G | D | 1.8e-07 | 3.5e-07 | 4.8e-07 | 3.367E-07 | 1.504E-07 | 1.800E-07 | 4.800E-07 |
| 22589 | 1130 | T1130H | D | 3.3e-07 | 2.1e-07 | 3.7e-07 | 3.033E-07 | 8.327E-08 | 2.100E-07 | 3.700E-07 |
| 22590 | 1130 | T1130I | D | 2.3e-07 | 1e-07 | 2.5e-07 | 1.933E-07 | 8.145E-08 | 1.000E-07 | 2.500E-07 |
| 22591 | 1130 | T1130L | D | 2e-07 | 2.4e-07 | 1.5e-07 | 1.967E-07 | 4.509E-08 | 1.500E-07 | 2.400E-07 |
| 22592 | 1130 | T1130K | D | 3.4e-07 | 2.7e-07 | 2.9e-07 | 3.000E-07 | 3.606E-08 | 2.700E-07 | 3.400E-07 |
| 22593 | 1130 | T1130M | D | 1.7e-07 | 1.7e-07 | 1.5e-07 | 1.633E-07 | 1.155E-08 | 1.500E-07 | 1.700E-07 |
| 22594 | 1130 | T1130F | D | 3e-07 | 1.2e-07 | 3.4e-07 | 2.533E-07 | 1.172E-07 | 1.200E-07 | 3.400E-07 |
| 22595 | 1130 | T1130P | D | 4.2e-07 | 2e-07 | 2.6e-07 | 2.933E-07 | 1.137E-07 | 2.000E-07 | 4.200E-07 |
| 22596 | 1130 | T1130S | D | 2e-07 | 1.1e-07 | 6.3e-07 | 3.133E-07 | 2.779E-07 | 1.100E-07 | 6.300E-07 |
| 22597 | 1130 | T1130T | D | 3.7e-07 | 1.2e-07 | 4.5e-07 | 3.133E-07 | 1.721E-07 | 1.200E-07 | 4.500E-07 |
| 22598 | 1130 | T1130W | D | 3.7e-07 | 2.5e-07 | 1.6e-07 | 2.600E-07 | 1.054E-07 | 1.600E-07 | 3.700E-07 |
| 22599 | 1130 | T1130Y | D | 3.3e-07 | 4e-07 | 2.6e-07 | 3.300E-07 | 7.000E-08 | 2.600E-07 | 4.000E-07 |
| 22600 | 1130 | T1130V | D | 4.7e-07 | 3.5e-07 | 1.9e-07 | 3.367E-07 | 1.405E-07 | 1.900E-07 | 4.700E-07 |
| 22601 | 1131 | L1131A | D | 1.4e-07 | 1.4e-07 | 4.3e-07 | 2.367E-07 | 1.674E-07 | 1.400E-07 | 4.300E-07 |
| 22602 | 1131 | L1131R | D | 2.6e-07 | 5e-07 | 6.1e-08 | 2.737E-07 | 2.198E-07 | 6.100E-08 | 5.000E-07 |
| 22603 | 1131 | L1131N | D | 1.4e-07 | 2.6e-07 | 2.9e-07 | 2.300E-07 | 7.937E-08 | 1.400E-07 | 2.900E-07 |
| 22604 | 1131 | L1131D | D | 2.6e-07 | 2.3e-07 | 4e-07 | 2.967E-07 | 9.074E-08 | 2.300E-07 | 4.000E-07 |
| 22605 | 1131 | L1131C | D | 3.6e-07 | 2.1e-07 | 3.5e-07 | 3.067E-07 | 8.386E-08 | 2.100E-07 | 3.600E-07 |
| 22606 | 1131 | L1131E | D | 1.7e-07 | 4.6e-07 | 2.9e-07 | 3.067E-07 | 1.457E-07 | 1.700E-07 | 4.600E-07 |
| 22607 | 1131 | L1131Q | D | 3.7e-07 | 2.6e-07 | 2.7e-07 | 3.000E-07 | 6.083E-08 | 2.600E-07 | 3.700E-07 |
| 22608 | 1131 | L1131G | D | 4e-07 | 1.1e-07 | 2e-07 | 2.367E-07 | 1.484E-07 | 1.100E-07 | 4.000E-07 |
| 22609 | 1131 | L1131H | D | 4.3e-07 | 2.3e-07 | 2.2e-07 | 2.933E-07 | 1.185E-07 | 2.200E-07 | 4.300E-07 |

| | | | | | | | | | | |
|---|---|---|---|---|---|---|---|---|---|---|
| 22610 | 1131 | L1131I | D | 2.1e-07 | 5.6e-07 | 4.9e-07 | 4.200E-07 | 1.852E-07 | 2.100E-07 | 5.600E-07 |
| 22611 | 1131 | L1131L | D | 1.5e-07 | 1.4e-07 | 3.2e-07 | 2.033E-07 | 1.012E-07 | 1.400E-07 | 3.200E-07 |
| 22612 | 1131 | L1131K | D | 3.9e-07 | 3.1e-07 | 2.2e-07 | 3.067E-07 | 8.505E-08 | 2.200E-07 | 3.900E-07 |
| 22613 | 1131 | L1131M | D | 1.3e-07 | 1.7e-07 | 3.7e-07 | 2.233E-07 | 1.286E-07 | 1.300E-07 | 3.700E-07 |
| 22614 | 1131 | L1131F | D | 2.8e-07 | 2.7e-07 | 3e-07 | 2.833E-07 | 1.528E-08 | 2.700E-07 | 3.000E-07 |
| 22615 | 1131 | L1131P | D | 2.8e-07 | 1.7e-07 | 3.3e-07 | 2.600E-07 | 8.185E-08 | 1.700E-07 | 3.300E-07 |
| 22616 | 1131 | L1131S | D | 1.3e-07 | 1.8e-07 | 2.9e-07 | 2.000E-07 | 8.185E-08 | 1.300E-07 | 2.900E-07 |
| 22617 | 1131 | L1131T | D | 2.8e-07 | 3.7e-07 | 4.7e-07 | 3.733E-07 | 9.504E-08 | 2.800E-07 | 4.700E-07 |
| 22618 | 1131 | L1131W | D | 1.8e-07 | 3.6e-07 | 1.5e-07 | 2.300E-07 | 1.136E-07 | 1.500E-07 | 3.600E-07 |
| 22619 | 1131 | L1131Y | D | 2.6e-07 | 1.1e-07 | 2.3e-07 | 2.000E-07 | 7.937E-08 | 1.100E-07 | 2.600E-07 |
| 22620 | 1131 | L1131V | D | 1.7e-07 | 1.9e-07 | 1.3e-07 | 1.633E-07 | 3.055E-08 | 1.300E-07 | 1.900E-07 |
| 22621 | 1132 | T1132A | D | 3.8e-07 | 4.2e-07 | 1.1e-07 | 3.033E-07 | 1.686E-07 | 1.100E-07 | 4.200E-07 |
| 22622 | 1132 | T1132R | D | 4.1e-07 | 3.2e-07 | 3.6e-07 | 3.633E-07 | 4.509E-08 | 3.200E-07 | 4.100E-07 |
| 22623 | 1132 | T1132N | D | 2.7e-07 | 1.3e-07 | 3.2e-07 | 2.400E-07 | 9.849E-08 | 1.300E-07 | 3.200E-07 |
| 22624 | 1132 | T1132D | D | 2.6e-07 | 2.8e-07 | 3e-07 | 2.800E-07 | 2.000E-08 | 2.600E-07 | 3.000E-07 |
| 22625 | 1132 | T1132C | D | 3.6e-07 | 2.1e-07 | 4.5e-07 | 3.400E-07 | 1.212E-07 | 2.100E-07 | 4.500E-07 |
| 22626 | 1132 | T1132E | D | 3.9e-07 | 2.5e-07 | 2.6e-07 | 3.000E-07 | 7.810E-08 | 2.500E-07 | 3.900E-07 |
| 22627 | 1132 | T1132Q | D | 4.1e-07 | 1.7e-07 | 1.7e-07 | 2.500E-07 | 1.386E-07 | 1.700E-07 | 4.100E-07 |
| 22628 | 1132 | T1132G | D | 3.3e-07 | 2.3e-07 | 2.7e-07 | 2.767E-07 | 5.033E-08 | 2.300E-07 | 3.300E-07 |
| 22629 | 1132 | T1132H | D | 2.1e-07 | 1.8e-07 | 1.5e-07 | 1.800E-07 | 3.000E-08 | 1.500E-07 | 2.100E-07 |
| 22630 | 1132 | T1132I | D | 2.2e-07 | 3.4e-07 | 3.8e-07 | 3.133E-07 | 8.327E-08 | 2.200E-07 | 3.800E-07 |
| 22631 | 1132 | T1132L | D | 2.6e-07 | 1.6e-07 | 2e-07 | 2.067E-07 | 5.033E-08 | 1.600E-07 | 2.600E-07 |
| 22632 | 1132 | T1132K | D | 2.4e-07 | 2.6e-07 | 3.1e-07 | 2.700E-07 | 3.606E-08 | 2.400E-07 | 3.100E-07 |
| 22633 | 1132 | T1132M | D | 3.5e-07 | 2.1e-07 | 1.7e-07 | 2.433E-07 | 9.452E-08 | 1.700E-07 | 3.500E-07 |
| 22634 | 1132 | T1132F | D | 9.9e-08 | 2.3e-07 | 3.6e-07 | 2.297E-07 | 1.305E-07 | 9.900E-08 | 3.600E-07 |
| 22635 | 1132 | T1132P | D | 1.3e-07 | 2.2e-07 | 1.9e-07 | 1.800E-07 | 4.583E-08 | 1.300E-07 | 2.200E-07 |
| 22636 | 1132 | T1132S | D | 1.6e-07 | 2.3e-07 | 1.8e-07 | 1.900E-07 | 3.606E-08 | 1.600E-07 | 2.300E-07 |
| 22637 | 1132 | T1132T | D | 3.4e-07 | 2.8e-07 | 2.8e-07 | 3.000E-07 | 3.464E-08 | 2.800E-07 | 3.400E-07 |
| 22638 | 1132 | T1132W | D | 3e-07 | 3.4e-07 | 1.7e-07 | 2.700E-07 | 8.888E-08 | 1.700E-07 | 3.400E-07 |
| 22639 | 1132 | T1132Y | D | 3.6e-07 | 8.1e-08 | 2.6e-07 | 2.337E-07 | 1.414E-07 | 8.100E-08 | 3.600E-07 |
| 22640 | 1132 | T1132V | D | 2.5e-07 | 2.2e-07 | 5.3e-07 | 3.333E-07 | 1.710E-07 | 2.200E-07 | 5.300E-07 |
| 22641 | 1133 | I1133A | D | 1.7e-07 | 1.7e-07 | 3.1e-07 | 2.167E-07 | 8.083E-08 | 1.700E-07 | 3.100E-07 |
| 22642 | 1133 | I1133R | D | 1.1e-07 | 1.3e-07 | 1.7e-07 | 1.367E-07 | 3.055E-08 | 1.100E-07 | 1.700E-07 |
| 22643 | 1133 | I1133N | D | 1.5e-07 | 5.3e-07 | 1e-07 | 2.600E-07 | 2.352E-07 | 1.000E-07 | 5.300E-07 |
| 22644 | 1133 | I1133D | D | 1.9e-07 | 4.8e-07 | 3e-07 | 3.233E-07 | 1.464E-07 | 1.900E-07 | 4.800E-07 |
| 22645 | 1133 | I1133C | D | 6.9e-07 | 3.4e-07 | 3.4e-07 | 4.567E-07 | 2.021E-07 | 3.400E-07 | 6.900E-07 |
| 22646 | 1133 | I1133E | D | 1.3e-07 | 2.5e-07 | 2e-07 | 1.933E-07 | 6.028E-08 | 1.300E-07 | 2.500E-07 |
| 22647 | 1133 | I1133Q | D | 1.1e-07 | 2.5e-07 | 1.4e-07 | 1.667E-07 | 7.371E-08 | 1.100E-07 | 2.500E-07 |
| 22648 | 1133 | I1133G | D | 2.3e-07 | 2.3e-07 | 3.9e-07 | 2.833E-07 | 9.238E-08 | 2.300E-07 | 3.900E-07 |
| 22649 | 1133 | I1133H | D | 1.3e-07 | 2.4e-07 | 1.6e-07 | 1.767E-07 | 5.686E-08 | 1.300E-07 | 2.400E-07 |
| 22650 | 1133 | I1133I | D | 1e-07 | 2.9e-07 | 1.8e-07 | 1.900E-07 | 9.539E-08 | 1.000E-07 | 2.900E-07 |
| 22651 | 1133 | I1133L | D | 3.1e-07 | 1.8e-07 | 1.9e-07 | 2.267E-07 | 7.234E-08 | 1.800E-07 | 3.100E-07 |
| 22652 | 1133 | I1133K | D | 1.9e-07 | 3.3e-07 | 2.5e-07 | 2.567E-07 | 7.024E-08 | 1.900E-07 | 3.300E-07 |
| 22653 | 1133 | I1133M | D | 1.9e-07 | 2.7e-07 | 1.9e-07 | 2.167E-07 | 4.619E-08 | 1.900E-07 | 2.700E-07 |
| 22654 | 1133 | I1133F | D | 2.7e-07 | 9.2e-08 | 1.3e-07 | 1.640E-07 | 9.374E-08 | 9.200E-08 | 2.700E-07 |
| 22655 | 1133 | I1133P | D | 1.9e-07 | 1.6e-07 | 2.2e-07 | 1.900E-07 | 3.000E-08 | 1.600E-07 | 2.200E-07 |
| 22656 | 1133 | I1133S | D | 2.5e-07 | 1.2e-07 | 2e-07 | 1.900E-07 | 6.557E-08 | 1.200E-07 | 2.500E-07 |
| 22657 | 1133 | I1133T | D | 3.2e-07 | 3.4e-07 | 1.7e-07 | 2.767E-07 | 9.292E-08 | 1.700E-07 | 3.400E-07 |
| 22658 | 1133 | I1133W | D | 1.8e-07 | 9.9e-08 | 4.3e-07 | 2.363E-07 | 1.725E-07 | 9.900E-08 | 4.300E-07 |
| 22659 | 1133 | I1133Y | D | 3.1e-07 | 2.2e-07 | 2.4e-07 | 2.567E-07 | 4.726E-08 | 2.200E-07 | 3.100E-07 |
| 22660 | 1133 | I1133V | D | 2.2e-07 | 4.2e-07 | 2.5e-07 | 2.967E-07 | 1.079E-07 | 2.200E-07 | 4.200E-07 |
| 22661 | 1134 | S1134A | D | 3e-07 | 2.5e-07 | 2.2e-07 | 2.567E-07 | 4.041E-08 | 2.200E-07 | 3.000E-07 |
| 22662 | 1134 | S1134R | D | 2.3e-07 | 1.6e-07 | 3.7e-07 | 2.533E-07 | 1.069E-07 | 1.600E-07 | 3.700E-07 |
| 22663 | 1134 | S1134N | D | 2.2e-07 | 4e-07 | 1.5e-07 | 2.567E-07 | 1.290E-07 | 1.500E-07 | 4.000E-07 |
| 22664 | 1134 | S1134D | D | 2.9e-07 | 1.8e-07 | 6.4e-07 | 3.700E-07 | 2.402E-07 | 1.800E-07 | 6.400E-07 |
| 22665 | 1134 | S1134C | D | 2.6e-07 | 4.3e-07 | 4.9e-07 | 3.933E-07 | 1.193E-07 | 2.600E-07 | 4.900E-07 |
| 22666 | 1134 | S1134E | D | 3.9e-07 | 1.6e-07 | 2.4e-07 | 2.633E-07 | 1.168E-07 | 1.600E-07 | 3.900E-07 |
| 22667 | 1134 | S1134Q | D | 2.8e-07 | 1.9e-07 | 3.5e-07 | 2.733E-07 | 8.021E-08 | 1.900E-07 | 3.500E-07 |
| 22668 | 1134 | S1134G | D | 3.4e-07 | 3.4e-07 | 2.4e-07 | 3.067E-07 | 5.774E-08 | 2.400E-07 | 3.400E-07 |
| 22669 | 1134 | S1134H | D | 3.5e-07 | 2.6e-07 | 8.1e-07 | 4.733E-07 | 2.950E-07 | 2.600E-07 | 8.100E-07 |
| 22670 | 1134 | S1134I | D | 2.2e-07 | 7e-07 | 1.5e-07 | 3.567E-07 | 2.994E-07 | 1.500E-07 | 7.000E-07 |
| 22671 | 1134 | S1134L | D | 4.7e-07 | 1.7e-07 | 1.5e-07 | 2.633E-07 | 1.793E-07 | 1.500E-07 | 4.700E-07 |
| 22672 | 1134 | S1134K | D | 1.6e-07 | 7.7e-07 | 2.4e-07 | 3.900E-07 | 3.315E-07 | 1.600E-07 | 7.700E-07 |
| 22673 | 1134 | S1134M | D | 3.8e-07 | 1.9e-07 | 3.4e-07 | 3.033E-07 | 1.002E-07 | 1.900E-07 | 3.800E-07 |
| 22674 | 1134 | S1134F | D | 3.1e-07 | 3.5e-07 | 6.5e-08 | 2.417E-07 | 1.543E-07 | 6.500E-08 | 3.500E-07 |
| 22675 | 1134 | S1134P | D | 3e-07 | 2e-07 | 9.3e-08 | 1.977E-07 | 1.035E-07 | 9.300E-08 | 3.000E-07 |
| 22676 | 1134 | S1134S | D | 2.2e-07 | 2.6e-07 | 3.7e-07 | 2.833E-07 | 7.767E-08 | 2.200E-07 | 3.700E-07 |
| 22677 | 1134 | S1134T | D | 2.8e-07 | 8.5e-08 | 2.5e-07 | 2.050E-07 | 1.050E-07 | 8.500E-08 | 2.800E-07 |
| 22678 | 1134 | S1134W | D | 6.3e-07 | 2.3e-07 | 2.8e-07 | 3.800E-07 | 2.179E-07 | 2.300E-07 | 6.300E-07 |
| 22679 | 1134 | S1134Y | D | 2.6e-07 | 3.2e-07 | 1.3e-07 | 2.367E-07 | 9.713E-08 | 1.300E-07 | 3.200E-07 |
| 22680 | 1134 | S1134V | D | 2.2e-07 | 1.7e-07 | 2.2e-07 | 2.033E-07 | 2.887E-08 | 1.700E-07 | 2.200E-07 |
| 22681 | 1135 | S1135A | D | 1.2e-07 | 2.2e-07 | 2.9e-07 | 2.100E-07 | 8.544E-08 | 1.200E-07 | 2.900E-07 |
| 22682 | 1135 | S1135R | D | 2.7e-07 | 1.7e-07 | 3.3e-07 | 2.567E-07 | 8.083E-08 | 1.700E-07 | 3.300E-07 |
| 22683 | 1135 | S1135N | D | 1.9e-07 | 2.8e-07 | 6.2e-07 | 3.633E-07 | 2.268E-07 | 1.900E-07 | 6.200E-07 |
| 22684 | 1135 | S1135D | D | 4e-07 | 1.5e-07 | 6.1e-07 | 3.867E-07 | 2.303E-07 | 1.500E-07 | 6.100E-07 |
| 22685 | 1135 | S1135C | D | 2.7e-07 | 1.7e-07 | 4e-07 | 2.800E-07 | 1.153E-07 | 1.700E-07 | 4.000E-07 |
| 22686 | 1135 | S1135E | D | 1.9e-07 | 1.2e-07 | 2.4e-07 | 1.833E-07 | 6.028E-08 | 1.200E-07 | 2.400E-07 |
| 22687 | 1135 | S1135Q | D | 2.7e-07 | 1.8e-07 | 2.6e-07 | 2.367E-07 | 4.933E-08 | 1.800E-07 | 2.700E-07 |
| 22688 | 1135 | S1135G | D | 2.6e-07 | 5.7e-07 | 1.5e-07 | 3.267E-07 | 2.178E-07 | 1.500E-07 | 5.700E-07 |
| 22689 | 1135 | S1135H | D | 2.9e-07 | 2.8e-07 | 1.6e-07 | 2.433E-07 | 7.234E-08 | 1.600E-07 | 2.900E-07 |
| 22690 | 1135 | S1135I | D | 2.3e-07 | 4.4e-07 | 1.2e-07 | 2.633E-07 | 1.626E-07 | 1.200E-07 | 4.400E-07 |
| 22691 | 1135 | S1135L | D | 1.2e-07 | 3.9e-07 | 4.3e-07 | 3.133E-07 | 1.686E-07 | 1.200E-07 | 4.300E-07 |
| 22692 | 1135 | S1135K | D | 3.4e-07 | 2.8e-07 | 1.8e-07 | 2.667E-07 | 8.083E-08 | 1.800E-07 | 3.400E-07 |
| 22693 | 1135 | S1135M | D | 3e-07 | 1.7e-07 | 3.8e-07 | 2.833E-07 | 1.060E-07 | 1.700E-07 | 3.800E-07 |
| 22694 | 1135 | S1135F | D | 3.1e-07 | 3.9e-07 | 1.4e-07 | 2.800E-07 | 1.277E-07 | 1.400E-07 | 3.900E-07 |

| | | | | | | | | | | |
|---|---|---|---|---|---|---|---|---|---|---|
| 22695 | 1135 | S1135P | D | 1.8e-07 | 1.8e-07 | 2e-07 | 1.867E-07 | 1.155E-08 | 1.800E-07 | 2.000E-07 |
| 22696 | 1135 | S1135S | D | 2.3e-07 | 1.3e-07 | 6.6e-08 | 1.420E-07 | 8.266E-08 | 6.600E-08 | 2.300E-07 |
| 22697 | 1135 | S1135T | D | 2.9e-07 | 2.5e-07 | 3.9e-07 | 3.100E-07 | 7.211E-08 | 2.500E-07 | 3.900E-07 |
| 22698 | 1135 | S1135W | D | 3.5e-07 | 1e-07 | 1.2e-07 | 1.900E-07 | 1.389E-07 | 1.000E-07 | 3.500E-07 |
| 22699 | 1135 | S1135Y | D | 1.7e-07 | 3.2e-07 | 2.5e-07 | 2.467E-07 | 7.506E-08 | 1.700E-07 | 3.200E-07 |
| 22700 | 1135 | S1135V | D | 3.6e-07 | 1.2e-07 | 8.3e-08 | 1.877E-07 | 1.504E-07 | 8.300E-08 | 3.600E-07 |
| 22701 | 1136 | L1136A | D | 2e-07 | 3.3e-07 | 3.6e-07 | 2.967E-07 | 8.505E-08 | 2.000E-07 | 3.600E-07 |
| 22702 | 1136 | L1136R | D | 2.9e-07 | 2.6e-07 | 3e-07 | 2.833E-07 | 2.082E-08 | 2.600E-07 | 3.000E-07 |
| 22703 | 1136 | L1136N | D | 6.3e-07 | 3.1e-07 | 3e-07 | 4.133E-07 | 1.877E-07 | 3.000E-07 | 6.300E-07 |
| 22704 | 1136 | L1136D | D | 3.9e-07 | 2.2e-07 | 3.8e-07 | 3.300E-07 | 9.539E-08 | 2.200E-07 | 3.900E-07 |
| 22705 | 1136 | L1136C | D | 2.8e-07 | 1.3e-07 | 3.5e-07 | 2.533E-07 | 1.124E-07 | 1.300E-07 | 3.500E-07 |
| 22706 | 1136 | L1136E | D | 3.6e-07 | 2.1e-07 | 3.5e-07 | 3.067E-07 | 8.386E-08 | 2.100E-07 | 3.600E-07 |
| 22707 | 1136 | L1136Q | D | 1e-07 | 2e-07 | 2.7e-07 | 1.900E-07 | 8.544E-08 | 1.000E-07 | 2.700E-07 |
| 22708 | 1136 | L1136G | D | 1.8e-07 | 2.3e-07 | 1.7e-07 | 1.933E-07 | 3.215E-08 | 1.700E-07 | 2.300E-07 |
| 22709 | 1136 | L1136H | D | 6.1e-07 | 3.3e-07 | 1.8e-07 | 3.733E-07 | 2.183E-07 | 1.800E-07 | 6.100E-07 |
| 22710 | 1136 | L1136I | D | 5.2e-07 | 2.2e-07 | 4e-07 | 3.800E-07 | 1.510E-07 | 2.200E-07 | 5.200E-07 |
| 22711 | 1136 | L1136L | D | 1.8e-07 | 2.5e-07 | 2.5e-07 | 2.267E-07 | 4.041E-08 | 1.800E-07 | 2.500E-07 |
| 22712 | 1136 | L1136K | D | 4.3e-07 | 2.1e-07 | 3.1e-07 | 3.167E-07 | 1.102E-07 | 2.100E-07 | 4.300E-07 |
| 22713 | 1136 | L1136M | D | 2.6e-07 | 2.6e-07 | 2.1e-07 | 2.433E-07 | 2.887E-08 | 2.100E-07 | 2.600E-07 |
| 22714 | 1136 | L1136F | D | 1.5e-07 | 2.6e-07 | 2.3e-07 | 2.133E-07 | 5.686E-08 | 1.500E-07 | 2.600E-07 |
| 22715 | 1136 | L1136P | D | 5.9e-07 | 2e-07 | 3.4e-07 | 3.767E-07 | 1.976E-07 | 2.000E-07 | 5.900E-07 |
| 22716 | 1136 | L1136S | D | 2.8e-07 | 2.6e-07 | 8.4e-08 | 2.080E-07 | 1.079E-07 | 8.400E-08 | 2.800E-07 |
| 22717 | 1136 | L1136T | D | 1.8e-07 | 2.7e-07 | 2.1e-07 | 2.200E-07 | 4.583E-08 | 1.800E-07 | 2.700E-07 |
| 22718 | 1136 | L1136W | D | 3e-07 | 2.3e-07 | 1.8e-07 | 2.367E-07 | 6.028E-08 | 1.800E-07 | 3.000E-07 |
| 22719 | 1136 | L1136Y | D | 1.3e-07 | 1.7e-07 | 3.5e-07 | 2.167E-07 | 1.172E-07 | 1.300E-07 | 3.500E-07 |
| 22720 | 1136 | L1136V | D | 1.4e-07 | 1.3e-07 | 1.8e-07 | 1.500E-07 | 2.646E-08 | 1.300E-07 | 1.800E-07 |
| 22721 | 1137 | Q1137A | D | 1.1e-07 | 3.7e-07 | 2e-07 | 2.267E-07 | 1.320E-07 | 1.100E-07 | 3.700E-07 |
| 22722 | 1137 | Q1137R | D | 1.8e-07 | 1.4e-07 | 5.8e-07 | 3.000E-07 | 2.433E-07 | 1.400E-07 | 5.800E-07 |
| 22723 | 1137 | Q1137N | D | 2e-07 | 2.7e-07 | 2.6e-07 | 2.433E-07 | 3.786E-08 | 2.000E-07 | 2.700E-07 |
| 22724 | 1137 | Q1137D | D | 4.2e-07 | 3.6e-07 | 4e-07 | 3.933E-07 | 3.055E-08 | 3.600E-07 | 4.200E-07 |
| 22725 | 1137 | Q1137C | D | 2.4e-07 | 1.5e-07 | 3.4e-07 | 2.433E-07 | 9.504E-08 | 1.500E-07 | 3.400E-07 |
| 22726 | 1137 | Q1137E | D | 4e-07 | 4.1e-07 | 3.6e-07 | 3.900E-07 | 2.646E-08 | 3.600E-07 | 4.100E-07 |
| 22727 | 1137 | Q1137Q | D | 2.2e-07 | 3.5e-07 | 3.3e-07 | 3.000E-07 | 7.000E-08 | 2.200E-07 | 3.500E-07 |
| 22728 | 1137 | Q1137G | D | 1.7e-07 | 1.1e-07 | 2.6e-07 | 1.800E-07 | 7.550E-08 | 1.100E-07 | 2.600E-07 |
| 22729 | 1137 | Q1137H | D | 3.5e-07 | 2.3e-07 | 1.5e-07 | 2.433E-07 | 1.007E-07 | 1.500E-07 | 3.500E-07 |
| 22730 | 1137 | Q1137I | D | 5.8e-07 | 2e-07 | 4.1e-07 | 3.967E-07 | 1.904E-07 | 2.000E-07 | 5.800E-07 |
| 22731 | 1137 | Q1137L | D | 4.5e-07 | 1.2e-07 | 3e-07 | 2.900E-07 | 1.652E-07 | 1.200E-07 | 4.500E-07 |
| 22732 | 1137 | Q1137K | D | 1.9e-07 | 2.6e-07 | 1.2e-07 | 1.900E-07 | 7.000E-08 | 1.200E-07 | 2.600E-07 |
| 22733 | 1137 | Q1137M | D | 2.6e-07 | 2.2e-07 | 2.1e-07 | 2.300E-07 | 2.646E-08 | 2.100E-07 | 2.600E-07 |
| 22734 | 1137 | Q1137F | D | 1.9e-07 | 2.5e-07 | 1.7e-07 | 2.033E-07 | 4.163E-08 | 1.700E-07 | 2.500E-07 |
| 22735 | 1137 | Q1137P | D | 1.6e-07 | 1.1e-07 | 3.5e-07 | 2.067E-07 | 1.266E-07 | 1.100E-07 | 3.500E-07 |
| 22736 | 1137 | Q1137S | D | 2.5e-07 | 2.6e-07 | 3.2e-07 | 2.767E-07 | 3.786E-08 | 2.500E-07 | 3.200E-07 |
| 22737 | 1137 | Q1137T | D | 4.3e-07 | 2.3e-07 | 9.3e-08 | 2.510E-07 | 1.695E-07 | 9.300E-08 | 4.300E-07 |
| 22738 | 1137 | Q1137W | D | 1.4e-07 | 2.8e-07 | 4.1e-07 | 2.767E-07 | 1.350E-07 | 1.400E-07 | 4.100E-07 |
| 22739 | 1137 | Q1137Y | D | 2.4e-07 | 1.5e-07 | 2.1e-07 | 2.000E-07 | 4.583E-08 | 1.500E-07 | 2.400E-07 |
| 22740 | 1137 | Q1137V | D | 3.4e-07 | 3.2e-07 | 3.1e-07 | 3.233E-07 | 1.528E-08 | 3.100E-07 | 3.400E-07 |
| 22741 | 1138 | P1138A | D | 2.6e-07 | 1.7e-07 | 2.8e-07 | 2.367E-07 | 5.859E-08 | 1.700E-07 | 2.800E-07 |
| 22742 | 1138 | P1138R | D | 1.7e-07 | 4.5e-07 | 2.4e-07 | 2.867E-07 | 1.457E-07 | 1.700E-07 | 4.500E-07 |
| 22743 | 1138 | P1138N | D | 2.4e-07 | 2.4e-07 | 1.8e-07 | 2.200E-07 | 3.464E-08 | 1.800E-07 | 2.400E-07 |
| 22744 | 1138 | P1138D | D | 3.2e-07 | 3.8e-07 | 1.6e-07 | 2.867E-07 | 1.137E-07 | 1.600E-07 | 3.800E-07 |
| 22745 | 1138 | P1138C | D | 3.8e-07 | 1e-07 | 1.6e-07 | 2.133E-07 | 1.474E-07 | 1.000E-07 | 3.800E-07 |
| 22746 | 1138 | P1138E | D | 2.3e-07 | 2.2e-07 | 2e-07 | 2.167E-07 | 1.528E-08 | 2.000E-07 | 2.300E-07 |
| 22747 | 1138 | P1138Q | D | 1.7e-07 | 4.7e-07 | 1.3e-07 | 2.567E-07 | 1.858E-07 | 1.300E-07 | 4.700E-07 |
| 22748 | 1138 | P1138G | D | 1.9e-07 | 2.3e-07 | 5.1e-07 | 3.100E-07 | 1.744E-07 | 1.900E-07 | 5.100E-07 |
| 22749 | 1138 | P1138H | D | 2.3e-07 | 1.8e-07 | 2.1e-07 | 2.067E-07 | 2.517E-08 | 1.800E-07 | 2.300E-07 |
| 22750 | 1138 | P1138I | D | 1.8e-07 | 3.3e-07 | 3.4e-07 | 2.833E-07 | 8.963E-08 | 1.800E-07 | 3.400E-07 |
| 22751 | 1138 | P1138L | D | 1.9e-07 | 4.1e-07 | 2.4e-07 | 2.800E-07 | 1.153E-07 | 1.900E-07 | 4.100E-07 |
| 22752 | 1138 | P1138K | D | 1.5e-07 | 3.1e-07 | 4e-07 | 2.867E-07 | 1.266E-07 | 1.500E-07 | 4.000E-07 |
| 22753 | 1138 | P1138M | D | 3.9e-07 | 2.5e-07 | 1.5e-07 | 2.633E-07 | 1.206E-07 | 1.500E-07 | 3.900E-07 |
| 22754 | 1138 | P1138F | D | 2.1e-07 | 2e-07 | 2e-07 | 2.033E-07 | 5.774E-09 | 2.000E-07 | 2.100E-07 |
| 22755 | 1138 | P1138P | D | 2e-07 | 2.1e-07 | 2e-07 | 2.033E-07 | 5.774E-09 | 2.000E-07 | 2.100E-07 |
| 22756 | 1138 | P1138S | D | 3e-07 | 3.5e-07 | 2e-07 | 2.833E-07 | 7.638E-08 | 2.000E-07 | 3.500E-07 |
| 22757 | 1138 | P1138T | D | 2.1e-07 | 3.7e-07 | 4e-07 | 3.267E-07 | 1.021E-07 | 2.100E-07 | 4.000E-07 |
| 22758 | 1138 | P1138W | D | 2.7e-07 | 3.4e-07 | 1.6e-07 | 2.567E-07 | 9.074E-08 | 1.600E-07 | 3.400E-07 |
| 22759 | 1138 | P1138Y | D | 1.6e-07 | 4.2e-07 | 2.2e-07 | 2.667E-07 | 1.361E-07 | 1.600E-07 | 4.200E-07 |
| 22760 | 1138 | P1138V | D | 1.8e-07 | 3.6e-07 | 1.6e-07 | 2.333E-07 | 1.102E-07 | 1.600E-07 | 3.600E-07 |
| 22761 | 1139 | E1139A | D | 2e-07 | 2.8e-07 | 4e-07 | 2.933E-07 | 1.007E-07 | 2.000E-07 | 4.000E-07 |
| 22762 | 1139 | E1139R | D | 2.9e-07 | 2e-07 | 6.1e-07 | 3.667E-07 | 2.155E-07 | 2.000E-07 | 6.100E-07 |
| 22763 | 1139 | E1139N | D | 2.7e-07 | 1.2e-07 | 1.8e-07 | 1.900E-07 | 7.550E-08 | 1.200E-07 | 2.700E-07 |
| 22764 | 1139 | E1139D | D | 2.3e-07 | 3.8e-07 | 3.1e-07 | 3.067E-07 | 7.506E-08 | 2.300E-07 | 3.800E-07 |
| 22765 | 1139 | E1139C | D | 3e-07 | 2.1e-07 | 5e-07 | 3.367E-07 | 1.484E-07 | 2.100E-07 | 5.000E-07 |
| 22766 | 1139 | E1139E | D | 3.9e-07 | 2.3e-07 | 4.5e-07 | 3.567E-07 | 1.137E-07 | 2.300E-07 | 4.500E-07 |
| 22767 | 1139 | E1139Q | D | 1.8e-07 | 2.4e-07 | 1.1e-07 | 1.767E-07 | 6.506E-08 | 1.100E-07 | 2.400E-07 |
| 22768 | 1139 | E1139G | D | 4.6e-07 | 6e-07 | 2.7e-07 | 4.433E-07 | 1.656E-07 | 2.700E-07 | 6.000E-07 |
| 22769 | 1139 | E1139H | D | 3.8e-07 | 2.6e-07 | 1.4e-07 | 2.600E-07 | 1.200E-07 | 1.400E-07 | 3.800E-07 |
| 22770 | 1139 | E1139I | D | 3e-07 | 5.9e-07 | 2.5e-07 | 3.800E-07 | 1.836E-07 | 2.500E-07 | 5.900E-07 |
| 22771 | 1139 | E1139L | D | 6.3e-07 | 4.7e-07 | 4e-07 | 5.000E-07 | 1.179E-07 | 4.000E-07 | 6.300E-07 |
| 22772 | 1139 | E1139K | D | 2e-07 | 1.2e-07 | 3.4e-07 | 2.200E-07 | 1.114E-07 | 1.200E-07 | 3.400E-07 |
| 22773 | 1139 | E1139M | D | 3.6e-07 | 2.1e-07 | 3.1e-07 | 2.933E-07 | 7.638E-08 | 2.100E-07 | 3.600E-07 |
| 22774 | 1139 | E1139F | D | 3.3e-07 | 2.2e-07 | 2.1e-07 | 2.533E-07 | 6.658E-08 | 2.100E-07 | 3.300E-07 |
| 22775 | 1139 | E1139P | D | 4e-07 | 1.6e-07 | 3.2e-07 | 2.933E-07 | 1.222E-07 | 1.600E-07 | 4.000E-07 |
| 22776 | 1139 | E1139S | D | 2.9e-07 | 3.5e-07 | 3.2e-07 | 3.200E-07 | 3.000E-08 | 2.900E-07 | 3.500E-07 |
| 22777 | 1139 | E1139T | D | 1.3e-07 | 2.8e-07 | 1.9e-07 | 2.000E-07 | 7.550E-08 | 1.300E-07 | 2.800E-07 |
| 22778 | 1139 | E1139W | D | 3.1e-07 | 1.3e-07 | 4.7e-07 | 3.033E-07 | 1.701E-07 | 1.300E-07 | 4.700E-07 |
| 22779 | 1139 | E1139Y | D | 1.3e-07 | 1.8e-07 | 5.8e-07 | 2.967E-07 | 2.466E-07 | 1.300E-07 | 5.800E-07 |

| | | | | | | | | | | |
|---|---|---|---|---|---|---|---|---|---|---|
| 22780 | 1139 | E1139V | D | 2.7e-07 | 3.2e-07 | 2.4e-07 | 2.767E-07 | 4.041E-08 | 2.400E-07 | 3.200E-07 |
| 22781 | 1140 | D1140A | D | 2.5e-07 | 3.1e-07 | 2.5e-07 | 2.700E-07 | 3.464E-08 | 2.500E-07 | 3.100E-07 |
| 22782 | 1140 | D1140R | D | 2.5e-07 | 2.2e-07 | 1.6e-07 | 2.100E-07 | 4.583E-08 | 1.600E-07 | 2.500E-07 |
| 22783 | 1140 | D1140N | D | 2.2e-07 | 2.1e-07 | 2.4e-07 | 2.233E-07 | 1.528E-08 | 2.100E-07 | 2.400E-07 |
| 22784 | 1140 | D1140D | D | 4e-07 | 3.3e-07 | 2.3e-07 | 3.200E-07 | 8.544E-08 | 2.300E-07 | 4.000E-07 |
| 22785 | 1140 | D1140C | D | 4.3e-07 | 6.9e-07 | 2.2e-07 | 4.467E-07 | 2.354E-07 | 2.200E-07 | 6.900E-07 |
| 22786 | 1140 | D1140E | D | 3.4e-07 | 1.8e-07 | 1.8e-07 | 2.333E-07 | 9.238E-08 | 1.800E-07 | 3.400E-07 |
| 22787 | 1140 | D1140Q | D | 3.6e-07 | 3.2e-07 | 1.9e-07 | 2.900E-07 | 8.888E-08 | 1.900E-07 | 3.600E-07 |
| 22788 | 1140 | D1140G | D | 9.7e-08 | 2.4e-07 | 2.8e-07 | 2.057E-07 | 9.621E-08 | 9.700E-08 | 2.800E-07 |
| 22789 | 1140 | D1140H | D | 1.7e-07 | 1.9e-07 | 2.4e-07 | 2.000E-07 | 3.606E-08 | 1.700E-07 | 2.400E-07 |
| 22790 | 1140 | D1140I | D | 1.9e-07 | 4.3e-07 | 2.1e-07 | 2.767E-07 | 1.332E-07 | 1.900E-07 | 4.300E-07 |
| 22791 | 1140 | D1140L | D | 4.8e-07 | 2.2e-07 | 2.7e-07 | 3.233E-07 | 1.380E-07 | 2.200E-07 | 4.800E-07 |
| 22792 | 1140 | D1140K | D | 4.6e-07 | 2e-07 | 3.1e-07 | 3.233E-07 | 1.305E-07 | 2.000E-07 | 4.600E-07 |
| 22793 | 1140 | D1140M | D | 3.8e-07 | 2.2e-07 | 1.4e-07 | 2.467E-07 | 1.222E-07 | 1.400E-07 | 3.800E-07 |
| 22794 | 1140 | D1140F | D | 5e-07 | 2.6e-07 | 1.4e-07 | 3.000E-07 | 1.833E-07 | 1.400E-07 | 5.000E-07 |
| 22795 | 1140 | D1140P | D | 2.3e-07 | 4.3e-07 | 2.4e-07 | 3.000E-07 | 1.127E-07 | 2.300E-07 | 4.300E-07 |
| 22796 | 1140 | D1140S | D | 2.5e-07 | 1.7e-07 | 6.7e-07 | 3.633E-07 | 2.686E-07 | 1.700E-07 | 6.700E-07 |
| 22797 | 1140 | D1140T | D | 2.4e-07 | 3.7e-07 | 3.3e-07 | 3.133E-07 | 6.658E-08 | 2.400E-07 | 3.700E-07 |
| 22798 | 1140 | D1140W | D | 1.5e-07 | 1.1e-07 | 1.2e-07 | 1.267E-07 | 2.082E-08 | 1.100E-07 | 1.500E-07 |
| 22799 | 1140 | D1140Y | D | 4.2e-07 | 1.7e-07 | 2.6e-07 | 2.833E-07 | 1.266E-07 | 1.700E-07 | 4.200E-07 |
| 22800 | 1140 | D1140V | D | 2.8e-07 | 2.3e-07 | 2.9e-07 | 2.667E-07 | 3.215E-08 | 2.300E-07 | 2.900E-07 |
| 22801 | 1141 | F1141A | D | 3.9e-07 | 2.7e-07 | 1.9e-07 | 2.833E-07 | 1.007E-07 | 1.900E-07 | 3.900E-07 |
| 22802 | 1141 | F1141R | D | 2.3e-07 | 1.9e-07 | 4.5e-07 | 2.900E-07 | 1.400E-07 | 1.900E-07 | 4.500E-07 |
| 22803 | 1141 | F1141N | D | 2.5e-07 | 1.6e-07 | 2.4e-07 | 2.167E-07 | 4.933E-08 | 1.600E-07 | 2.500E-07 |
| 22804 | 1141 | F1141D | D | 1.9e-07 | 2.2e-07 | 3.6e-07 | 2.567E-07 | 9.074E-08 | 1.900E-07 | 3.600E-07 |
| 22805 | 1141 | F1141C | D | 2.7e-07 | 2.4e-07 | 3.1e-07 | 2.733E-07 | 3.512E-08 | 2.400E-07 | 3.100E-07 |
| 22806 | 1141 | F1141E | D | 2.2e-07 | 2.2e-07 | 2.4e-07 | 2.267E-07 | 1.155E-08 | 2.200E-07 | 2.400E-07 |
| 22807 | 1141 | F1141Q | D | 1.6e-07 | 6.3e-07 | 2.1e-07 | 3.333E-07 | 2.581E-07 | 1.600E-07 | 6.300E-07 |
| 22808 | 1141 | F1141G | D | 1.3e-07 | 2.5e-07 | 1.7e-07 | 1.833E-07 | 6.110E-08 | 1.300E-07 | 2.500E-07 |
| 22809 | 1141 | F1141H | D | 2.6e-07 | 4.2e-07 | 2.5e-07 | 3.100E-07 | 9.539E-08 | 2.500E-07 | 4.200E-07 |
| 22810 | 1141 | F1141I | D | 3.4e-07 | 2.4e-07 | 2e-07 | 2.600E-07 | 7.211E-08 | 2.000E-07 | 3.400E-07 |
| 22811 | 1141 | F1141L | D | 1.7e-07 | 2.7e-07 | 4.8e-07 | 3.067E-07 | 1.582E-07 | 1.700E-07 | 4.800E-07 |
| 22812 | 1141 | F1141K | D | 2.8e-07 | 3.9e-07 | 2.2e-07 | 2.967E-07 | 8.622E-08 | 2.200E-07 | 3.900E-07 |
| 22813 | 1141 | F1141M | D | 4.3e-07 | 2.2e-07 | 1.8e-07 | 2.767E-07 | 1.343E-07 | 1.800E-07 | 4.300E-07 |
| 22814 | 1141 | F1141F | D | 5.1e-07 | 3.3e-07 | 1.5e-07 | 3.300E-07 | 1.800E-07 | 1.500E-07 | 5.100E-07 |
| 22815 | 1141 | F1141P | D | 3.8e-07 | 2.2e-07 | 2.8e-07 | 2.933E-07 | 8.083E-08 | 2.200E-07 | 3.800E-07 |
| 22816 | 1141 | F1141S | D | 2.3e-07 | 5.2e-07 | 2.2e-07 | 3.233E-07 | 1.704E-07 | 2.200E-07 | 5.200E-07 |
| 22817 | 1141 | F1141T | D | 2.8e-07 | 1.1e-07 | 3.5e-07 | 2.467E-07 | 1.234E-07 | 1.100E-07 | 3.500E-07 |
| 22818 | 1141 | F1141W | D | 3.5e-07 | 1.6e-07 | 1.2e-07 | 2.100E-07 | 1.229E-07 | 1.200E-07 | 3.500E-07 |
| 22819 | 1141 | F1141Y | D | 3.2e-07 | 5.2e-07 | 3.3e-07 | 3.900E-07 | 1.127E-07 | 3.200E-07 | 5.200E-07 |
| 22820 | 1141 | F1141V | D | 2.7e-07 | 2.4e-07 | 3.2e-07 | 2.767E-07 | 4.041E-08 | 2.400E-07 | 3.200E-07 |
| 22821 | 1142 | A1142A | D | 3.2e-07 | 2.6e-07 | 1.9e-07 | 2.567E-07 | 6.506E-08 | 1.900E-07 | 3.200E-07 |
| 22822 | 1142 | A1142R | D | 2.6e-07 | 5.6e-07 | 2.3e-07 | 3.500E-07 | 1.825E-07 | 2.300E-07 | 5.600E-07 |
| 22823 | 1142 | A1142N | D | 3.6e-07 | 1.1e-07 | 1e-07 | 1.900E-07 | 1.473E-07 | 1.000E-07 | 3.600E-07 |
| 22824 | 1142 | A1142D | D | 2.1e-07 | 1e-07 | 2e-07 | 1.700E-07 | 6.083E-08 | 1.000E-07 | 2.100E-07 |
| 22825 | 1142 | A1142C | D | 2.8e-07 | 2.9e-07 | 2.8e-07 | 2.833E-07 | 5.774E-09 | 2.800E-07 | 2.900E-07 |
| 22826 | 1142 | A1142E | D | 1.7e-07 | 2.3e-07 | 3.9e-07 | 2.633E-07 | 1.137E-07 | 1.700E-07 | 3.900E-07 |
| 22827 | 1142 | A1142Q | D | 1.9e-07 | 2.4e-07 | 2.9e-07 | 2.400E-07 | 5.000E-08 | 1.900E-07 | 2.900E-07 |
| 22828 | 1142 | A1142G | D | 2.7e-07 | 1e-07 | 2.5e-07 | 2.067E-07 | 9.292E-08 | 1.000E-07 | 2.700E-07 |
| 22829 | 1142 | A1142H | D | 3e-07 | 1.8e-07 | 3.3e-07 | 2.700E-07 | 7.937E-08 | 1.800E-07 | 3.300E-07 |
| 22830 | 1142 | A1142I | D | 1.3e-07 | 2e-07 | 1.5e-07 | 1.600E-07 | 3.606E-08 | 1.300E-07 | 2.000E-07 |
| 22831 | 1142 | A1142L | D | 2.8e-07 | 1.3e-07 | 3.5e-07 | 2.533E-07 | 1.124E-07 | 1.300E-07 | 3.500E-07 |
| 22832 | 1142 | A1142K | D | 1.2e-07 | 2.5e-07 | 2.5e-07 | 2.067E-07 | 7.506E-08 | 1.200E-07 | 2.500E-07 |
| 22833 | 1142 | A1142M | D | 4.8e-07 | 2.2e-07 | 3e-07 | 3.333E-07 | 1.332E-07 | 2.200E-07 | 4.800E-07 |
| 22834 | 1142 | A1142F | D | 2.6e-07 | 3.4e-07 | 1.9e-07 | 2.633E-07 | 7.506E-08 | 1.900E-07 | 3.400E-07 |
| 22835 | 1142 | A1142P | D | 2.1e-07 | 3.5e-07 | 2.7e-07 | 2.767E-07 | 7.024E-08 | 2.100E-07 | 3.500E-07 |
| 22836 | 1142 | A1142S | D | 9.2e-08 | 1.8e-07 | 4.6e-07 | 2.440E-07 | 1.922E-07 | 9.200E-08 | 4.600E-07 |
| 22837 | 1142 | A1142T | D | 1.6e-07 | 1.9e-07 | 5.2e-07 | 2.900E-07 | 1.997E-07 | 1.600E-07 | 5.200E-07 |
| 22838 | 1142 | A1142W | D | 1.4e-07 | 3.5e-07 | 4.5e-07 | 3.133E-07 | 1.582E-07 | 1.400E-07 | 4.500E-07 |
| 22839 | 1142 | A1142Y | D | 3.7e-07 | 8.7e-07 | 2.4e-07 | 4.933E-07 | 3.326E-07 | 2.400E-07 | 8.700E-07 |
| 22840 | 1142 | A1142V | D | 2.1e-07 | 2.8e-07 | 2.7e-07 | 2.533E-07 | 3.786E-08 | 2.100E-07 | 2.800E-07 |
| 22841 | 1143 | T1143A | D | 2.4e-07 | 3.7e-07 | 1.7e-07 | 2.600E-07 | 1.015E-07 | 1.700E-07 | 3.700E-07 |
| 22842 | 1143 | T1143R | D | 2.9e-07 | 9.7e-08 | 1.8e-07 | 1.890E-07 | 9.681E-08 | 9.700E-08 | 2.900E-07 |
| 22843 | 1143 | T1143N | D | 3.2e-07 | 2.8e-07 | 3e-07 | 3.000E-07 | 2.000E-08 | 2.800E-07 | 3.200E-07 |
| 22844 | 1143 | T1143D | D | 3e-07 | 1.5e-07 | 3.7e-07 | 2.733E-07 | 1.124E-07 | 1.500E-07 | 3.700E-07 |
| 22845 | 1143 | T1143C | D | 3.1e-07 | 6.3e-07 | 3.1e-07 | 4.167E-07 | 1.848E-07 | 3.100E-07 | 6.300E-07 |
| 22846 | 1143 | T1143E | D | 2.7e-07 | 4.5e-07 | 2.3e-07 | 3.167E-07 | 1.172E-07 | 2.300E-07 | 4.500E-07 |
| 22847 | 1143 | T1143Q | D | 2.6e-07 | 1.2e-07 | 2.6e-07 | 2.133E-07 | 8.083E-08 | 1.200E-07 | 2.600E-07 |
| 22848 | 1143 | T1143G | D | 2.8e-07 | 3.1e-07 | 1.4e-07 | 2.433E-07 | 9.074E-08 | 1.400E-07 | 3.100E-07 |
| 22849 | 1143 | T1143H | D | 3.4e-07 | 5.8e-07 | 3.1e-07 | 4.100E-07 | 1.480E-07 | 3.100E-07 | 5.800E-07 |
| 22850 | 1143 | T1143I | D | 1.3e-07 | 5.2e-07 | 2.3e-07 | 2.933E-07 | 2.026E-07 | 1.300E-07 | 5.200E-07 |
| 22851 | 1143 | T1143L | D | 1.5e-07 | 2.5e-07 | 5.7e-07 | 3.233E-07 | 2.194E-07 | 1.500E-07 | 5.700E-07 |
| 22852 | 1143 | T1143K | D | 2.8e-07 | 2.7e-07 | 2.8e-07 | 2.767E-07 | 5.774E-09 | 2.700E-07 | 2.800E-07 |
| 22853 | 1143 | T1143M | D | 5.6e-07 | 1.8e-07 | 2e-07 | 3.133E-07 | 2.139E-07 | 1.800E-07 | 5.600E-07 |
| 22854 | 1143 | T1143F | D | 5.2e-07 | 3.7e-07 | 3.9e-07 | 4.267E-07 | 8.145E-08 | 3.700E-07 | 5.200E-07 |
| 22855 | 1143 | T1143P | D | 7e-07 | 2.8e-07 | 2.8e-07 | 4.200E-07 | 2.425E-07 | 2.800E-07 | 7.000E-07 |
| 22856 | 1143 | T1143S | D | 7.6e-07 | 2.8e-07 | 4.8e-07 | 5.067E-07 | 2.411E-07 | 2.800E-07 | 7.600E-07 |
| 22857 | 1143 | T1143T | D | 2e-07 | 4.7e-07 | 2.8e-07 | 3.167E-07 | 1.387E-07 | 2.000E-07 | 4.700E-07 |
| 22858 | 1143 | T1143W | D | 3e-07 | 7.1e-07 | 1.1e-07 | 3.733E-07 | 3.066E-07 | 1.100E-07 | 7.100E-07 |
| 22859 | 1143 | T1143Y | D | 1.9e-07 | 1.7e-07 | 2.9e-07 | 2.167E-07 | 6.429E-08 | 1.700E-07 | 2.900E-07 |
| 22860 | 1143 | T1143V | D | 1.4e-07 | 1.4e-07 | 2e-07 | 1.600E-07 | 3.464E-08 | 1.400E-07 | 2.000E-07 |
| 22861 | 1144 | Y1144A | D | 3.2e-07 | 5.8e-07 | 3.2e-07 | 4.067E-07 | 1.501E-07 | 3.200E-07 | 5.800E-07 |
| 22862 | 1144 | Y1144R | D | 1.9e-07 | 3.4e-07 | 1.2e-07 | 2.167E-07 | 1.124E-07 | 1.200E-07 | 3.400E-07 |
| 22863 | 1144 | Y1144N | D | 2.6e-07 | 2.8e-07 | 2.1e-07 | 2.500E-07 | 3.606E-08 | 2.100E-07 | 2.800E-07 |
| 22864 | 1144 | Y1144D | D | 2.9e-07 | 3.1e-07 | 3.7e-07 | 3.233E-07 | 4.163E-08 | 2.900E-07 | 3.700E-07 |

| | | | | | | | | | | |
|---|---|---|---|---|---|---|---|---|---|---|
| 22865 | 1144 | Y1144C | D | 2.1e-07 | 2.8e-07 | 1.1e-07 | 2.000E-07 | 8.544E-08 | 1.100E-07 | 2.800E-07 |
| 22866 | 1144 | Y1144E | D | 3.2e-07 | 1.8e-07 | 4.9e-07 | 3.300E-07 | 1.552E-07 | 1.800E-07 | 4.900E-07 |
| 22867 | 1144 | Y1144Q | D | 4.7e-07 | 5.4e-07 | 9.3e-07 | 6.467E-07 | 2.479E-07 | 4.700E-07 | 9.300E-07 |
| 22868 | 1144 | Y1144G | D | 1.6e-07 | 2.8e-07 | 2.9e-07 | 2.433E-07 | 7.234E-08 | 1.600E-07 | 2.900E-07 |
| 22869 | 1144 | Y1144H | D | 2.8e-07 | 2.4e-07 | 1.2e-07 | 2.133E-07 | 8.327E-08 | 1.200E-07 | 2.800E-07 |
| 22870 | 1144 | Y1144I | D | 3.4e-07 | 2.8e-07 | 2.1e-07 | 2.767E-07 | 6.506E-08 | 2.100E-07 | 3.400E-07 |
| 22871 | 1144 | Y1144L | D | 4.1e-07 | 1.5e-07 | 1e-07 | 2.200E-07 | 1.664E-07 | 1.000E-07 | 4.100E-07 |
| 22872 | 1144 | Y1144K | D | 1.6e-07 | 4.6e-07 | 4e-07 | 3.400E-07 | 1.587E-07 | 1.600E-07 | 4.600E-07 |
| 22873 | 1144 | Y1144M | D | 2e-07 | 1.6e-07 | 2.7e-07 | 2.100E-07 | 5.568E-08 | 1.600E-07 | 2.700E-07 |
| 22874 | 1144 | Y1144F | D | 1.9e-07 | 2.4e-07 | 5.7e-07 | 3.333E-07 | 2.065E-07 | 1.900E-07 | 5.700E-07 |
| 22875 | 1144 | Y1144P | D | 2.7e-07 | 5.7e-07 | 2.5e-07 | 3.633E-07 | 1.793E-07 | 2.500E-07 | 5.700E-07 |
| 22876 | 1144 | Y1144S | D | 4.1e-07 | 3.5e-07 | 1.5e-07 | 3.033E-07 | 1.361E-07 | 1.500E-07 | 4.100E-07 |
| 22877 | 1144 | Y1144T | D | 2.8e-07 | 2e-07 | 2.6e-07 | 2.467E-07 | 4.163E-08 | 2.000E-07 | 2.800E-07 |
| 22878 | 1144 | Y1144W | D | 1.9e-07 | 1.1e-07 | 3.9e-07 | 2.300E-07 | 1.442E-07 | 1.100E-07 | 3.900E-07 |
| 22879 | 1144 | Y1144Y | D | 1.6e-07 | 2.2e-07 | 9.1e-08 | 1.570E-07 | 6.455E-08 | 9.100E-08 | 2.200E-07 |
| 22880 | 1144 | Y1144V | D | 1.7e-07 | 5.2e-07 | 1.9e-07 | 2.933E-07 | 1.966E-07 | 1.700E-07 | 5.200E-07 |
| 22881 | 1145 | Y1145A | D | 2.2e-07 | 4.3e-07 | 2.8e-07 | 3.100E-07 | 1.082E-07 | 2.200E-07 | 4.300E-07 |
| 22882 | 1145 | Y1145R | D | 1.9e-07 | 1.2e-07 | 1.8e-07 | 1.633E-07 | 3.786E-08 | 1.200E-07 | 1.900E-07 |
| 22883 | 1145 | Y1145N | D | 2e-07 | 1.5e-07 | 2.9e-07 | 2.133E-07 | 7.095E-08 | 1.500E-07 | 2.900E-07 |
| 22884 | 1145 | Y1145D | D | 6e-07 | 2.4e-07 | 3.3e-07 | 3.900E-07 | 1.873E-07 | 2.400E-07 | 6.000E-07 |
| 22885 | 1145 | Y1145C | D | 2.8e-07 | 3.7e-07 | 2.5e-07 | 3.000E-07 | 6.245E-08 | 2.500E-07 | 3.700E-07 |
| 22886 | 1145 | Y1145E | D | 1.9e-07 | 4e-07 | 2.3e-07 | 2.733E-07 | 1.115E-07 | 1.900E-07 | 4.000E-07 |
| 22887 | 1145 | Y1145Q | D | 2.2e-07 | 3e-07 | 1.8e-07 | 2.333E-07 | 6.110E-08 | 1.800E-07 | 3.000E-07 |
| 22888 | 1145 | Y1145G | D | 2.7e-07 | 1.1e-07 | 2.4e-07 | 2.067E-07 | 8.505E-08 | 1.100E-07 | 2.700E-07 |
| 22889 | 1145 | Y1145H | D | 1.3e-07 | 2.2e-07 | 3.3e-07 | 2.267E-07 | 1.002E-07 | 1.300E-07 | 3.300E-07 |
| 22890 | 1145 | Y1145I | D | 4.2e-07 | 3.6e-07 | 2.7e-07 | 3.500E-07 | 7.550E-08 | 2.700E-07 | 4.200E-07 |
| 22891 | 1145 | Y1145L | D | 1.9e-07 | 2.1e-07 | 2.7e-07 | 2.233E-07 | 4.163E-08 | 1.900E-07 | 2.700E-07 |
| 22892 | 1145 | Y1145K | D | 2.2e-07 | 1.7e-07 | 3.1e-07 | 2.333E-07 | 7.095E-08 | 1.700E-07 | 3.100E-07 |
| 22893 | 1145 | Y1145M | D | 3.4e-07 | 1.9e-07 | 2.3e-07 | 2.533E-07 | 7.767E-08 | 1.900E-07 | 3.400E-07 |
| 22894 | 1145 | Y1145F | D | 3e-07 | 2.1e-07 | 1.7e-07 | 2.267E-07 | 6.658E-08 | 1.700E-07 | 3.000E-07 |
| 22895 | 1145 | Y1145P | D | 2.7e-07 | 3.4e-07 | 3.9e-07 | 3.333E-07 | 6.028E-08 | 2.700E-07 | 3.900E-07 |
| 22896 | 1145 | Y1145S | D | 2e-07 | 6.8e-07 | 2.1e-07 | 3.633E-07 | 2.743E-07 | 2.000E-07 | 6.800E-07 |
| 22897 | 1145 | Y1145T | D | 5.8e-07 | 2.7e-07 | 3.1e-07 | 3.867E-07 | 1.686E-07 | 2.700E-07 | 5.800E-07 |
| 22898 | 1145 | Y1145W | D | 1.3e-07 | 2.4e-07 | 2.6e-07 | 2.100E-07 | 7.000E-08 | 1.300E-07 | 2.600E-07 |
| 22899 | 1145 | Y1145Y | D | 1.5e-07 | 1.7e-07 | 1.5e-07 | 1.567E-07 | 1.155E-08 | 1.500E-07 | 1.700E-07 |
| 22900 | 1145 | Y1145V | D | 3.2e-07 | 5.6e-07 | 3.2e-07 | 4.000E-07 | 1.386E-07 | 3.200E-07 | 5.600E-07 |
| 22901 | 1146 | C1146A | D | 3.5e-07 | 1.1e-07 | 4.1e-07 | 2.900E-07 | 1.587E-07 | 1.100E-07 | 4.100E-07 |
| 22902 | 1146 | C1146R | D | 2.3e-07 | 2.1e-07 | 1.5e-07 | 1.967E-07 | 4.163E-08 | 1.500E-07 | 2.300E-07 |
| 22903 | 1146 | C1146N | D | 2.5e-07 | 5.5e-07 | 6.8e-07 | 4.933E-07 | 2.205E-07 | 2.500E-07 | 6.800E-07 |
| 22904 | 1146 | C1146D | D | 1.5e-07 | 2.4e-07 | 3.3e-07 | 2.400E-07 | 9.000E-08 | 1.500E-07 | 3.300E-07 |
| 22905 | 1146 | C1146C | D | 3.7e-07 | 1.8e-07 | 2.5e-07 | 2.667E-07 | 9.609E-08 | 1.800E-07 | 3.700E-07 |
| 22906 | 1146 | C1146E | D | 3.4e-07 | 2.4e-07 | 2.6e-07 | 2.800E-07 | 5.292E-08 | 2.400E-07 | 3.400E-07 |
| 22907 | 1146 | C1146Q | D | 1.7e-07 | 1.3e-07 | 3.9e-07 | 2.300E-07 | 1.400E-07 | 1.300E-07 | 3.900E-07 |
| 22908 | 1146 | C1146G | D | 4.3e-07 | 2.7e-07 | 4.6e-07 | 3.867E-07 | 1.021E-07 | 2.700E-07 | 4.600E-07 |
| 22909 | 1146 | C1146H | D | 3.9e-07 | 1.6e-07 | 2.6e-07 | 2.700E-07 | 1.153E-07 | 1.600E-07 | 3.900E-07 |
| 22910 | 1146 | C1146I | D | 2.2e-07 | 2.7e-07 | 3.9e-07 | 2.933E-07 | 8.737E-08 | 2.200E-07 | 3.900E-07 |
| 22911 | 1146 | C1146L | D | 2.6e-07 | 3.1e-07 | 5.1e-07 | 3.600E-07 | 1.323E-07 | 2.600E-07 | 5.100E-07 |
| 22912 | 1146 | C1146K | D | 7.5e-08 | 1.8e-07 | 4.5e-07 | 2.350E-07 | 1.935E-07 | 7.500E-08 | 4.500E-07 |
| 22913 | 1146 | C1146M | D | 1.4e-07 | 2.2e-07 | 2.2e-07 | 1.933E-07 | 4.619E-08 | 1.400E-07 | 2.200E-07 |
| 22914 | 1146 | C1146F | D | 2.8e-07 | 2.1e-07 | 5.7e-07 | 3.533E-07 | 1.909E-07 | 2.100E-07 | 5.700E-07 |
| 22915 | 1146 | C1146P | D | 1.1e-07 | 1.3e-07 | 2e-07 | 1.467E-07 | 4.726E-08 | 1.100E-07 | 2.000E-07 |
| 22916 | 1146 | C1146S | D | 2.8e-07 | 3.2e-07 | 2.9e-07 | 2.967E-07 | 2.082E-08 | 2.800E-07 | 3.200E-07 |
| 22917 | 1146 | C1146T | D | 1.8e-07 | 2.2e-07 | 4.3e-07 | 2.767E-07 | 1.343E-07 | 1.800E-07 | 4.300E-07 |
| 22918 | 1146 | C1146W | D | 5.1e-07 | 1.2e-07 | 3.2e-07 | 3.167E-07 | 1.950E-07 | 1.200E-07 | 5.100E-07 |
| 22919 | 1146 | C1146Y | D | 2.5e-07 | 1.9e-07 | 2.6e-07 | 2.333E-07 | 3.786E-08 | 1.900E-07 | 2.600E-07 |
| 22920 | 1146 | C1146V | D | 3.3e-07 | 6.1e-07 | 1.4e-07 | 3.600E-07 | 2.364E-07 | 1.400E-07 | 6.100E-07 |
| 22921 | 1147 | Q1147A | D | 2e-07 | 1.6e-07 | 2.8e-07 | 2.133E-07 | 6.110E-08 | 1.600E-07 | 2.800E-07 |
| 22922 | 1147 | Q1147R | D | 1.4e-07 | 5.2e-07 | 4.3e-07 | 3.633E-07 | 1.986E-07 | 1.400E-07 | 5.200E-07 |
| 22923 | 1147 | Q1147N | D | 3.7e-07 | 1.9e-07 | 2.6e-07 | 2.733E-07 | 9.074E-08 | 1.900E-07 | 3.700E-07 |
| 22924 | 1147 | Q1147D | D | 2.8e-07 | 8.4e-07 | 3.4e-07 | 4.867E-07 | 3.075E-07 | 2.800E-07 | 8.400E-07 |
| 22925 | 1147 | Q1147C | D | 5.3e-07 | 6.1e-07 | 1.9e-07 | 4.433E-07 | 2.230E-07 | 1.900E-07 | 6.100E-07 |
| 22926 | 1147 | Q1147E | D | 3.2e-07 | 1.9e-07 | 1e-07 | 2.033E-07 | 1.106E-07 | 1.000E-07 | 3.200E-07 |
| 22927 | 1147 | Q1147Q | D | 2.1e-07 | 1.4e-07 | 9.1e-08 | 1.470E-07 | 5.981E-08 | 9.100E-08 | 2.100E-07 |
| 22928 | 1147 | Q1147G | D | 5.2e-07 | 3e-07 | 2.7e-07 | 3.633E-07 | 1.365E-07 | 2.700E-07 | 5.200E-07 |
| 22929 | 1147 | Q1147H | D | 1.1e-07 | 2.5e-07 | 1.2e-07 | 1.600E-07 | 7.810E-08 | 1.100E-07 | 2.500E-07 |
| 22930 | 1147 | Q1147I | D | 1.4e-07 | 2.1e-07 | 1.6e-07 | 1.700E-07 | 3.606E-08 | 1.400E-07 | 2.100E-07 |
| 22931 | 1147 | Q1147L | D | 2.6e-07 | 3.5e-07 | 1.3e-07 | 2.467E-07 | 1.106E-07 | 1.300E-07 | 3.500E-07 |
| 22932 | 1147 | Q1147K | D | 4.4e-07 | 2.2e-07 | 4.3e-07 | 3.633E-07 | 1.242E-07 | 2.200E-07 | 4.400E-07 |
| 22933 | 1147 | Q1147M | D | 6.7e-07 | 1.6e-07 | 1.6e-07 | 3.300E-07 | 2.944E-07 | 1.600E-07 | 6.700E-07 |
| 22934 | 1147 | Q1147F | D | 3.9e-07 | 1.9e-07 | 1.9e-07 | 2.567E-07 | 1.155E-07 | 1.900E-07 | 3.900E-07 |
| 22935 | 1147 | Q1147P | D | 3.5e-07 | 2e-07 | 4e-07 | 3.167E-07 | 1.041E-07 | 2.000E-07 | 4.000E-07 |
| 22936 | 1147 | Q1147S | D | 2.8e-07 | 1.2e-07 | 4.5e-07 | 2.833E-07 | 1.650E-07 | 1.200E-07 | 4.500E-07 |
| 22937 | 1147 | Q1147T | D | 2.7e-07 | 8.2e-08 | 1.5e-07 | 1.673E-07 | 9.519E-08 | 8.200E-08 | 2.700E-07 |
| 22938 | 1147 | Q1147W | D | 3.4e-07 | 2.4e-07 | 6.7e-08 | 2.157E-07 | 1.381E-07 | 6.700E-08 | 3.400E-07 |
| 22939 | 1147 | Q1147Y | D | 8.9e-08 | 3.9e-07 | 2.5e-07 | 2.430E-07 | 1.506E-07 | 8.900E-08 | 3.900E-07 |
| 22940 | 1147 | Q1147V | D | 2.2e-07 | 1.7e-07 | 9.3e-08 | 1.610E-07 | 6.398E-08 | 9.300E-08 | 2.200E-07 |
| 22941 | 1148 | Q1148A | D | 2e-07 | 1.9e-07 | 1.8e-07 | 1.900E-07 | 1.000E-08 | 1.800E-07 | 2.000E-07 |
| 22942 | 1148 | Q1148R | D | 2.9e-07 | 2e-07 | 2.9e-07 | 2.600E-07 | 5.196E-08 | 2.000E-07 | 2.900E-07 |
| 22943 | 1148 | Q1148N | D | 2e-07 | 1e-07 | 2.3e-07 | 1.767E-07 | 6.807E-08 | 1.000E-07 | 2.300E-07 |
| 22944 | 1148 | Q1148D | D | 1.7e-07 | 2.8e-07 | 2.2e-07 | 2.233E-07 | 5.508E-08 | 1.700E-07 | 2.800E-07 |
| 22945 | 1148 | Q1148C | D | 1.9e-07 | 9.7e-08 | 4.9e-07 | 2.590E-07 | 2.054E-07 | 9.700E-08 | 4.900E-07 |
| 22946 | 1148 | Q1148E | D | 2.1e-07 | 1.5e-07 | 1.6e-07 | 1.733E-07 | 3.215E-08 | 1.500E-07 | 2.100E-07 |
| 22947 | 1148 | Q1148Q | D | 1.2e-07 | 2.4e-07 | 4.8e-07 | 2.800E-07 | 1.833E-07 | 1.200E-07 | 4.800E-07 |
| 22948 | 1148 | Q1148G | D | 2.8e-07 | 2.6e-07 | 1.1e-07 | 2.167E-07 | 9.292E-08 | 1.100E-07 | 2.800E-07 |
| 22949 | 1148 | Q1148H | D | 2.6e-07 | 3.2e-07 | 1.7e-07 | 2.500E-07 | 7.550E-08 | 1.700E-07 | 3.200E-07 |

| | | | | | | | | | | |
|---|---|---|---|---|---|---|---|---|---|---|
| 22950 | 1148 | Q1148I | D | 6.8e-08 | 1.8e-07 | 2.1e-07 | 1.527E-07 | 7.484E-08 | 6.800E-08 | 2.100E-07 |
| 22951 | 1148 | Q1148L | D | 5.8e-07 | 2e-07 | 1.6e-07 | 3.133E-07 | 2.318E-07 | 1.600E-07 | 5.800E-07 |
| 22952 | 1148 | Q1148K | D | 2.5e-07 | 3.9e-07 | 1.7e-07 | 2.700E-07 | 1.114E-07 | 1.700E-07 | 3.900E-07 |
| 22953 | 1148 | Q1148M | D | 4.6e-07 | 4.3e-07 | 2e-07 | 3.633E-07 | 1.422E-07 | 2.000E-07 | 4.600E-07 |
| 22954 | 1148 | Q1148F | D | 1.4e-07 | 3.9e-07 | 4.6e-07 | 3.300E-07 | 1.682E-07 | 1.400E-07 | 4.600E-07 |
| 22955 | 1148 | Q1148P | D | 8.3e-08 | 3.4e-07 | 2.5e-07 | 2.243E-07 | 1.304E-07 | 8.300E-08 | 3.400E-07 |
| 22956 | 1148 | Q1148S | D | 1.2e-07 | 1.6e-07 | 3.2e-07 | 2.000E-07 | 1.058E-07 | 1.200E-07 | 3.200E-07 |
| 22957 | 1148 | Q1148T | D | 4.3e-07 | 1.7e-07 | 1.5e-07 | 2.500E-07 | 1.562E-07 | 1.500E-07 | 4.300E-07 |
| 22958 | 1148 | Q1148W | D | 5.3e-07 | 4.7e-07 | 3e-07 | 4.333E-07 | 1.193E-07 | 3.000E-07 | 5.300E-07 |
| 22959 | 1148 | Q1148Y | D | 3.8e-07 | 3.1e-07 | 3.2e-07 | 3.367E-07 | 3.786E-08 | 3.100E-07 | 3.800E-07 |
| 22960 | 1148 | Q1148V | D | 3.3e-07 | 1.7e-07 | 9.3e-08 | 1.977E-07 | 1.209E-07 | 9.300E-08 | 3.300E-07 |
| 22961 | 1149 | H1149A | D | 2.9e-07 | 3.5e-07 | 7e-07 | 4.467E-07 | 2.214E-07 | 2.900E-07 | 7.000E-07 |
| 22962 | 1149 | H1149R | D | 2.3e-07 | 1.7e-07 | 1.5e-07 | 1.833E-07 | 4.163E-08 | 1.500E-07 | 2.300E-07 |
| 22963 | 1149 | H1149N | D | 1.3e-07 | 2e-07 | 1.3e-07 | 1.533E-07 | 4.041E-08 | 1.300E-07 | 2.000E-07 |
| 22964 | 1149 | H1149D | D | 3.3e-07 | 2e-07 | 2.1e-07 | 2.467E-07 | 7.234E-08 | 2.000E-07 | 3.300E-07 |
| 22965 | 1149 | H1149C | D | 1.3e-07 | 2.5e-07 | 2.3e-07 | 2.033E-07 | 6.429E-08 | 1.300E-07 | 2.500E-07 |
| 22966 | 1149 | H1149E | D | 1.8e-07 | 2.3e-07 | 1.5e-07 | 1.867E-07 | 4.041E-08 | 1.500E-07 | 2.300E-07 |
| 22967 | 1149 | H1149Q | D | 1.8e-07 | 9.6e-08 | 3e-07 | 1.920E-07 | 1.025E-07 | 9.600E-08 | 3.000E-07 |
| 22968 | 1149 | H1149G | D | 5.5e-07 | 3.6e-07 | 6.1e-07 | 5.067E-07 | 1.305E-07 | 3.600E-07 | 6.100E-07 |
| 22969 | 1149 | H1149H | D | 3.4e-07 | 2.7e-07 | 6.5e-07 | 4.200E-07 | 2.022E-07 | 2.700E-07 | 6.500E-07 |
| 22970 | 1149 | H1149I | D | 2.3e-07 | 8e-07 | 4.4e-07 | 4.900E-07 | 2.883E-07 | 2.300E-07 | 8.000E-07 |
| 22971 | 1149 | H1149L | D | 3.2e-07 | 3.4e-07 | 3.4e-07 | 3.333E-07 | 1.155E-08 | 3.200E-07 | 3.400E-07 |
| 22972 | 1149 | H1149K | D | 5e-07 | 4e-07 | 4.1e-07 | 4.367E-07 | 5.508E-08 | 4.000E-07 | 5.000E-07 |
| 22973 | 1149 | H1149M | D | 7.1e-07 | 2.3e-07 | 5e-07 | 4.800E-07 | 2.406E-07 | 2.300E-07 | 7.100E-07 |
| 22974 | 1149 | H1149F | D | 5.7e-07 | 1.7e-07 | 8.3e-07 | 5.233E-07 | 3.325E-07 | 1.700E-07 | 8.300E-07 |
| 22975 | 1149 | H1149P | D | 5.8e-07 | 9.2e-07 | 4e-07 | 6.333E-07 | 2.641E-07 | 4.000E-07 | 9.200E-07 |
| 22976 | 1149 | H1149S | D | 1.3e-07 | 1.7e-07 | 1.5e-07 | 1.500E-07 | 2.000E-08 | 1.300E-07 | 1.700E-07 |
| 22977 | 1149 | H1149T | D | 1.5e-07 | 1.9e-07 | 1.6e-07 | 1.667E-07 | 2.082E-08 | 1.500E-07 | 1.900E-07 |
| 22978 | 1149 | H1149W | D | 4.7e-07 | 1.8e-07 | 4.8e-07 | 3.767E-07 | 1.704E-07 | 1.800E-07 | 4.800E-07 |
| 22979 | 1149 | H1149Y | D | 4.5e-07 | 2.9e-07 | 4.5e-07 | 3.967E-07 | 9.238E-08 | 2.900E-07 | 4.500E-07 |
| 22980 | 1149 | H1149V | D | 3e-07 | 5e-07 | 4.1e-07 | 4.033E-07 | 1.002E-07 | 3.000E-07 | 5.000E-07 |
| 22981 | 1150 | Y1150A | D | 2.3e-07 | 2.9e-07 | 3.6e-07 | 2.933E-07 | 6.506E-08 | 2.300E-07 | 3.600E-07 |
| 22982 | 1150 | Y1150R | D | 2.3e-07 | 1.5e-07 | 3.9e-07 | 2.567E-07 | 1.222E-07 | 1.500E-07 | 3.900E-07 |
| 22983 | 1150 | Y1150N | D | 1.7e-07 | 2.4e-07 | 1.4e-07 | 1.833E-07 | 5.132E-08 | 1.400E-07 | 2.400E-07 |
| 22984 | 1150 | Y1150D | D | 2.4e-07 | 2.3e-07 | 1.8e-07 | 2.167E-07 | 3.215E-08 | 1.800E-07 | 2.400E-07 |
| 22985 | 1150 | Y1150C | D | 2.5e-07 | 4.9e-07 | 3.2e-07 | 3.533E-07 | 1.234E-07 | 2.500E-07 | 4.900E-07 |
| 22986 | 1150 | Y1150E | D | 1.9e-07 | 1.9e-07 | 1.3e-07 | 1.700E-07 | 3.464E-08 | 1.300E-07 | 1.900E-07 |
| 22987 | 1150 | Y1150Q | D | 1.4e-07 | 3.3e-07 | 1.9e-07 | 2.200E-07 | 9.849E-08 | 1.400E-07 | 3.300E-07 |
| 22988 | 1150 | Y1150G | D | 4.5e-07 | 5e-07 | 1.7e-07 | 3.733E-07 | 1.779E-07 | 1.700E-07 | 5.000E-07 |
| 22989 | 1150 | Y1150H | D | 2.5e-07 | 3.2e-07 | 2e-07 | 2.567E-07 | 6.028E-08 | 2.000E-07 | 3.200E-07 |
| 22990 | 1150 | Y1150I | D | 5.6e-07 | 2.3e-07 | 4.6e-07 | 4.167E-07 | 1.692E-07 | 2.300E-07 | 5.600E-07 |
| 22991 | 1150 | Y1150L | D | 1.7e-07 | 3.5e-07 | 4e-07 | 3.067E-07 | 1.210E-07 | 1.700E-07 | 4.000E-07 |
| 22992 | 1150 | Y1150K | D | 1.9e-07 | 2.7e-07 | 1.9e-07 | 2.167E-07 | 4.619E-08 | 1.900E-07 | 2.700E-07 |
| 22993 | 1150 | Y1150M | D | 3.1e-07 | 2.7e-07 | 2.2e-07 | 2.667E-07 | 4.509E-08 | 2.200E-07 | 3.100E-07 |
| 22994 | 1150 | Y1150F | D | 3.4e-07 | 3.9e-07 | 2.1e-07 | 3.133E-07 | 9.292E-08 | 2.100E-07 | 3.900E-07 |
| 22995 | 1150 | Y1150P | D | 1.8e-07 | 3.4e-07 | 5.1e-07 | 3.433E-07 | 1.650E-07 | 1.800E-07 | 5.100E-07 |
| 22996 | 1150 | Y1150S | D | 1.2e-07 | 1.5e-07 | 1.3e-07 | 1.333E-07 | 1.528E-08 | 1.200E-07 | 1.500E-07 |
| 22997 | 1150 | Y1150T | D | 9.7e-08 | 2.5e-07 | 2.6e-07 | 2.023E-07 | 9.136E-08 | 9.700E-08 | 2.600E-07 |
| 22998 | 1150 | Y1150W | D | 2.5e-07 | 2e-07 | 1.6e-07 | 2.033E-07 | 4.509E-08 | 1.600E-07 | 2.500E-07 |
| 22999 | 1150 | Y1150Y | D | 8.9e-08 | 3.9e-07 | 1.8e-07 | 2.197E-07 | 1.544E-07 | 8.900E-08 | 3.900E-07 |
| 23000 | 1150 | Y1150V | D | 4.3e-07 | 1.7e-07 | 1.5e-07 | 2.500E-07 | 1.562E-07 | 1.500E-07 | 4.300E-07 |
| 23001 | 1151 | T1151A | D | 1.5e-07 | 2.1e-07 | 4.9e-07 | 2.833E-07 | 1.815E-07 | 1.500E-07 | 4.900E-07 |
| 23002 | 1151 | T1151R | D | 1.8e-07 | 2.8e-07 | 1.4e-07 | 2.000E-07 | 7.211E-08 | 1.400E-07 | 2.800E-07 |
| 23003 | 1151 | T1151N | D | 9.9e-08 | 1.4e-07 | 1.9e-07 | 1.430E-07 | 4.557E-08 | 9.900E-08 | 1.900E-07 |
| 23004 | 1151 | T1151D | D | 2.2e-07 | 1.8e-07 | 3.2e-07 | 2.400E-07 | 7.211E-08 | 1.800E-07 | 3.200E-07 |
| 23005 | 1151 | T1151C | D | 2.6e-07 | 4.4e-07 | 2.2e-07 | 3.067E-07 | 1.172E-07 | 2.200E-07 | 4.400E-07 |
| 23006 | 1151 | T1151E | D | 4.4e-07 | 1.5e-07 | 2e-07 | 2.633E-07 | 1.550E-07 | 1.500E-07 | 4.400E-07 |
| 23007 | 1151 | T1151Q | D | 1.7e-07 | 1.3e-07 | 2.4e-07 | 1.800E-07 | 5.568E-08 | 1.300E-07 | 2.400E-07 |
| 23008 | 1151 | T1151G | D | 2.1e-07 | 1.8e-07 | 4.7e-07 | 2.867E-07 | 1.595E-07 | 1.800E-07 | 4.700E-07 |
| 23009 | 1151 | T1151H | D | 1.6e-07 | 2e-07 | 1.9e-07 | 1.833E-07 | 2.082E-08 | 1.600E-07 | 2.000E-07 |
| 23010 | 1151 | T1151I | D | 4.2e-07 | 4.3e-07 | 2e-07 | 3.500E-07 | 1.300E-07 | 2.000E-07 | 4.300E-07 |
| 23011 | 1151 | T1151L | D | 2.1e-07 | 3e-07 | 1.6e-07 | 2.233E-07 | 7.095E-08 | 1.600E-07 | 3.000E-07 |
| 23012 | 1151 | T1151K | D | 6.1e-07 | 2.7e-07 | 3.5e-07 | 4.100E-07 | 1.778E-07 | 2.700E-07 | 6.100E-07 |
| 23013 | 1151 | T1151M | D | 2.1e-07 | 1.6e-07 | 1.8e-07 | 1.833E-07 | 2.517E-08 | 1.600E-07 | 2.100E-07 |
| 23014 | 1151 | T1151F | D | 1.8e-07 | 2.1e-07 | 1.6e-07 | 1.833E-07 | 2.517E-08 | 1.600E-07 | 2.100E-07 |
| 23015 | 1151 | T1151P | D | 5.9e-07 | 4.2e-07 | 2.9e-07 | 4.333E-07 | 1.504E-07 | 2.900E-07 | 5.900E-07 |
| 23016 | 1151 | T1151S | D | 2.3e-07 | 1.6e-07 | 2.2e-07 | 2.033E-07 | 3.786E-08 | 1.600E-07 | 2.300E-07 |
| 23017 | 1151 | T1151T | D | 5.5e-07 | 3.2e-07 | 1.7e-07 | 3.467E-07 | 1.914E-07 | 1.700E-07 | 5.500E-07 |
| 23018 | 1151 | T1151W | D | 2e-07 | 2.8e-07 | 2e-07 | 2.267E-07 | 4.619E-08 | 2.000E-07 | 2.800E-07 |
| 23019 | 1151 | T1151Y | D | 4.5e-07 | 5.1e-07 | 3.4e-07 | 4.333E-07 | 8.622E-08 | 3.400E-07 | 5.100E-07 |
| 23020 | 1151 | T1151V | D | 2.2e-07 | 2e-07 | 2.2e-07 | 2.133E-07 | 1.155E-08 | 2.000E-07 | 2.200E-07 |
| 23021 | 1152 | T1152A | D | 3.3e-07 | 3.3e-07 | 7.5e-07 | 4.700E-07 | 2.425E-07 | 3.300E-07 | 7.500E-07 |
| 23022 | 1152 | T1152R | D | 3.2e-07 | 8.4e-08 | 3.3e-07 | 2.447E-07 | 1.392E-07 | 8.400E-08 | 3.300E-07 |
| 23023 | 1152 | T1152N | D | 2.7e-07 | 2e-07 | 2.9e-07 | 2.533E-07 | 4.726E-08 | 2.000E-07 | 2.900E-07 |
| 23024 | 1152 | T1152D | D | 1.6e-07 | 1.7e-07 | 1.6e-07 | 1.633E-07 | 5.774E-09 | 1.600E-07 | 1.700E-07 |
| 23025 | 1152 | T1152C | D | 5.5e-07 | 5.1e-07 | 3.1e-07 | 4.567E-07 | 1.286E-07 | 3.100E-07 | 5.500E-07 |
| 23026 | 1152 | T1152E | D | 2.5e-07 | 2.1e-07 | 1.9e-07 | 2.167E-07 | 3.055E-08 | 1.900E-07 | 2.500E-07 |
| 23027 | 1152 | T1152Q | D | 2.8e-07 | 9e-08 | 8.9e-08 | 1.530E-07 | 1.100E-07 | 8.900E-08 | 2.800E-07 |
| 23028 | 1152 | T1152G | D | 3.9e-07 | 3.8e-07 | 3.8e-07 | 3.833E-07 | 5.774E-09 | 3.800E-07 | 3.900E-07 |
| 23029 | 1152 | T1152H | D | 1.1e-07 | 1.7e-07 | 3.2e-07 | 2.000E-07 | 1.082E-07 | 1.100E-07 | 3.200E-07 |
| 23030 | 1152 | T1152I | D | 1.3e-07 | 2.5e-07 | 2.5e-07 | 2.100E-07 | 6.928E-08 | 1.300E-07 | 2.500E-07 |
| 23031 | 1152 | T1152L | D | 2.9e-07 | 2.1e-07 | 3.7e-07 | 2.900E-07 | 8.000E-08 | 2.100E-07 | 3.700E-07 |
| 23032 | 1152 | T1152K | D | 1.3e-07 | 1.9e-07 | 2.6e-07 | 1.933E-07 | 6.506E-08 | 1.300E-07 | 2.600E-07 |
| 23033 | 1152 | T1152M | D | 2.2e-07 | 1.9e-07 | 1.3e-07 | 1.800E-07 | 4.583E-08 | 1.300E-07 | 2.200E-07 |
| 23034 | 1152 | T1152F | D | 2.4e-07 | 3.6e-07 | 1.8e-07 | 2.600E-07 | 9.165E-08 | 1.800E-07 | 3.600E-07 |

| | | | | | | | | | | |
|---|---|---|---|---|---|---|---|---|---|---|
| 23035 | 1152 | T1152P | D | 1.8e-07 | 1.8e-07 | 4.5e-07 | 2.700E-07 | 1.559E-07 | 1.800E-07 | 4.500E-07 |
| 23036 | 1152 | T1152S | D | 2.2e-07 | 1.3e-07 | 2.8e-07 | 2.100E-07 | 7.550E-08 | 1.300E-07 | 2.800E-07 |
| 23037 | 1152 | T1152T | D | 1.4e-07 | 3.9e-07 | 5.1e-07 | 3.467E-07 | 1.888E-07 | 1.400E-07 | 5.100E-07 |
| 23038 | 1152 | T1152W | D | 2.9e-07 | 2.1e-07 | 2e-07 | 2.333E-07 | 4.933E-08 | 2.000E-07 | 2.900E-07 |
| 23039 | 1152 | T1152Y | D | 1.6e-07 | 2.6e-07 | 2.7e-07 | 2.300E-07 | 6.083E-08 | 1.600E-07 | 2.700E-07 |
| 23040 | 1152 | T1152V | D | 1.4e-07 | 2e-07 | 1.9e-07 | 1.767E-07 | 3.215E-08 | 1.400E-07 | 2.000E-07 |
| 23041 | 1153 | P1153A | D | 4.8e-07 | 9.8e-07 | 4.1e-07 | 6.233E-07 | 3.109E-07 | 4.100E-07 | 9.800E-07 |
| 23042 | 1153 | P1153R | D | 4.1e-07 | 1.6e-07 | 1.8e-07 | 2.500E-07 | 1.389E-07 | 1.600E-07 | 4.100E-07 |
| 23043 | 1153 | P1153N | D | 2.9e-07 | 4.6e-07 | 2.5e-07 | 3.333E-07 | 1.115E-07 | 2.500E-07 | 4.600E-07 |
| 23044 | 1153 | P1153D | D | 2e-07 | 2.5e-07 | 3.7e-07 | 2.733E-07 | 8.737E-08 | 2.000E-07 | 3.700E-07 |
| 23045 | 1153 | P1153C | D | 1.1e-07 | 1.7e-07 | 1.7e-07 | 1.500E-07 | 3.464E-08 | 1.100E-07 | 1.700E-07 |
| 23046 | 1153 | P1153E | D | 2.8e-07 | 3e-07 | 1.9e-07 | 2.567E-07 | 5.859E-08 | 1.900E-07 | 3.000E-07 |
| 23047 | 1153 | P1153Q | D | 2.6e-07 | 1.8e-07 | 2.2e-07 | 2.200E-07 | 4.000E-08 | 1.800E-07 | 2.600E-07 |
| 23048 | 1153 | P1153G | D | 1.6e-07 | 1.9e-07 | 3.7e-07 | 2.400E-07 | 1.136E-07 | 1.600E-07 | 3.700E-07 |
| 23049 | 1153 | P1153H | D | 2.2e-07 | 7.5e-07 | 2.5e-07 | 4.067E-07 | 2.977E-07 | 2.200E-07 | 7.500E-07 |
| 23050 | 1153 | P1153I | D | 3.3e-07 | 2.6e-07 | 2.3e-07 | 2.733E-07 | 5.132E-08 | 2.300E-07 | 3.300E-07 |
| 23051 | 1153 | P1153L | D | 1.4e-07 | 1.9e-07 | 1e-07 | 1.433E-07 | 4.509E-08 | 1.000E-07 | 1.900E-07 |
| 23052 | 1153 | P1153K | D | 5.9e-07 | 3.2e-07 | 2.5e-07 | 3.867E-07 | 1.795E-07 | 2.500E-07 | 5.900E-07 |
| 23053 | 1153 | P1153M | D | 3.6e-07 | 2.2e-07 | 3.6e-07 | 3.133E-07 | 8.083E-08 | 2.200E-07 | 3.600E-07 |
| 23054 | 1153 | P1153F | D | 1.4e-07 | 2.8e-07 | 2.5e-07 | 2.233E-07 | 7.371E-08 | 1.400E-07 | 2.800E-07 |
| 23055 | 1153 | P1153P | D | 3.5e-07 | 3e-07 | 1.7e-07 | 2.733E-07 | 9.292E-08 | 1.700E-07 | 3.500E-07 |
| 23056 | 1153 | P1153S | D | 2.9e-07 | 2.3e-07 | 2.1e-07 | 2.433E-07 | 4.163E-08 | 2.100E-07 | 2.900E-07 |
| 23057 | 1153 | P1153T | D | 1.3e-07 | 2.5e-07 | 1e-07 | 1.600E-07 | 7.937E-08 | 1.000E-07 | 2.500E-07 |
| 23058 | 1153 | P1153W | D | 2.3e-07 | 2e-07 | 4.1e-07 | 2.800E-07 | 1.136E-07 | 2.000E-07 | 4.100E-07 |
| 23059 | 1153 | P1153Y | D | 3.2e-07 | 3.1e-07 | 1.7e-07 | 2.667E-07 | 8.386E-08 | 1.700E-07 | 3.200E-07 |
| 23060 | 1153 | P1153V | D | 5.8e-08 | 4.1e-07 | 3.3e-07 | 2.660E-07 | 1.845E-07 | 5.800E-08 | 4.100E-07 |
| 23061 | 1154 | P1154A | D | 2.4e-07 | 8.9e-08 | 2.2e-07 | 1.830E-07 | 8.202E-08 | 8.900E-08 | 2.400E-07 |
| 23062 | 1154 | P1154R | D | 7.2e-08 | 2e-07 | 2e-07 | 1.573E-07 | 7.390E-08 | 7.200E-08 | 2.000E-07 |
| 23063 | 1154 | P1154N | D | 2.3e-07 | 4.1e-07 | 2.2e-07 | 2.867E-07 | 1.069E-07 | 2.200E-07 | 4.100E-07 |
| 23064 | 1154 | P1154D | D | 2.6e-07 | 2.3e-07 | 2.8e-07 | 2.567E-07 | 2.517E-08 | 2.300E-07 | 2.800E-07 |
| 23065 | 1154 | P1154C | D | 5.1e-07 | 2.4e-07 | 1.7e-07 | 3.067E-07 | 1.795E-07 | 1.700E-07 | 5.100E-07 |
| 23066 | 1154 | P1154E | D | 3.9e-07 | 2.4e-07 | 5.7e-07 | 4.000E-07 | 1.652E-07 | 2.400E-07 | 5.700E-07 |
| 23067 | 1154 | P1154Q | D | 3.7e-07 | 7.3e-08 | 4.4e-07 | 2.943E-07 | 1.948E-07 | 7.300E-08 | 4.400E-07 |
| 23068 | 1154 | P1154G | D | 1.7e-07 | 3.2e-07 | 1.7e-07 | 2.200E-07 | 8.660E-08 | 1.700E-07 | 3.200E-07 |
| 23069 | 1154 | P1154H | D | 1.6e-07 | 1.1e-07 | 3.5e-07 | 2.067E-07 | 1.266E-07 | 1.100E-07 | 3.500E-07 |
| 23070 | 1154 | P1154I | D | 2.8e-07 | 4.3e-07 | 2.1e-07 | 3.067E-07 | 1.124E-07 | 2.100E-07 | 4.300E-07 |
| 23071 | 1154 | P1154L | D | 2.4e-07 | 5e-07 | 3.1e-07 | 3.500E-07 | 1.345E-07 | 2.400E-07 | 5.000E-07 |
| 23072 | 1154 | P1154K | D | 2.5e-07 | 1.3e-07 | 8.8e-08 | 1.560E-07 | 8.407E-08 | 8.800E-08 | 2.500E-07 |
| 23073 | 1154 | P1154M | D | 1.6e-07 | 1.8e-07 | 2.5e-07 | 1.967E-07 | 4.726E-08 | 1.600E-07 | 2.500E-07 |
| 23074 | 1154 | P1154F | D | 4.1e-07 | 3.4e-07 | 1.7e-07 | 3.067E-07 | 1.234E-07 | 1.700E-07 | 4.100E-07 |
| 23075 | 1154 | P1154P | D | 4.1e-07 | 2.8e-07 | 2.8e-07 | 3.233E-07 | 7.506E-08 | 2.800E-07 | 4.100E-07 |
| 23076 | 1154 | P1154S | D | 4e-07 | 1.4e-07 | 3.7e-07 | 3.033E-07 | 1.422E-07 | 1.400E-07 | 4.000E-07 |
| 23077 | 1154 | P1154T | D | 3.4e-07 | 1.9e-07 | 1.7e-07 | 2.333E-07 | 9.292E-08 | 1.700E-07 | 3.400E-07 |
| 23078 | 1154 | P1154W | D | 3.4e-07 | 2.9e-07 | 4.6e-07 | 3.633E-07 | 8.737E-08 | 2.900E-07 | 4.600E-07 |
| 23079 | 1154 | P1154Y | D | 2e-07 | 3.7e-07 | 1.4e-07 | 2.367E-07 | 1.193E-07 | 1.400E-07 | 3.700E-07 |
| 23080 | 1154 | P1154V | D | 4.4e-07 | 2.1e-07 | 1.5e-07 | 2.667E-07 | 1.531E-07 | 1.500E-07 | 4.400E-07 |
| 23081 | 1155 | T1155A | D | 2.2e-07 | 4.4e-07 | 3.3e-07 | 3.300E-07 | 1.100E-07 | 2.200E-07 | 4.400E-07 |
| 23082 | 1155 | T1155R | D | 2.3e-07 | 1.9e-07 | 1.6e-07 | 1.933E-07 | 3.512E-08 | 1.600E-07 | 2.300E-07 |
| 23083 | 1155 | T1155N | D | 1.7e-07 | 4.5e-07 | 1.2e-07 | 2.467E-07 | 1.779E-07 | 1.200E-07 | 4.500E-07 |
| 23084 | 1155 | T1155D | D | 2.9e-07 | 1.4e-07 | 3.3e-07 | 2.533E-07 | 1.002E-07 | 1.400E-07 | 3.300E-07 |
| 23085 | 1155 | T1155C | D | 1.4e-07 | 1.7e-07 | 1.7e-07 | 1.600E-07 | 1.732E-08 | 1.400E-07 | 1.700E-07 |
| 23086 | 1155 | T1155E | D | 2.2e-07 | 1.6e-07 | 1.3e-07 | 1.700E-07 | 4.583E-08 | 1.300E-07 | 2.200E-07 |
| 23087 | 1155 | T1155Q | D | 3.5e-07 | 2.3e-07 | 3.7e-07 | 3.167E-07 | 7.572E-08 | 2.300E-07 | 3.700E-07 |
| 23088 | 1155 | T1155G | D | 3.1e-07 | 4.2e-07 | 2.6e-07 | 3.300E-07 | 8.185E-08 | 2.600E-07 | 4.200E-07 |
| 23089 | 1155 | T1155H | D | 2.4e-07 | 2.9e-07 | 2.9e-07 | 2.733E-07 | 2.887E-08 | 2.400E-07 | 2.900E-07 |
| 23090 | 1155 | T1155I | D | 1.9e-07 | 3.5e-07 | 5.7e-07 | 3.700E-07 | 1.908E-07 | 1.900E-07 | 5.700E-07 |
| 23091 | 1155 | T1155L | D | 2.9e-07 | 3.5e-07 | 1.5e-07 | 2.633E-07 | 1.026E-07 | 1.500E-07 | 3.500E-07 |
| 23092 | 1155 | T1155K | D | 2.6e-07 | 1.9e-07 | 3.7e-07 | 2.733E-07 | 9.074E-08 | 1.900E-07 | 3.700E-07 |
| 23093 | 1155 | T1155M | D | 2.4e-07 | 4.5e-07 | 3.1e-07 | 3.333E-07 | 1.069E-07 | 2.400E-07 | 4.500E-07 |
| 23094 | 1155 | T1155F | D | 4e-07 | 1.6e-07 | 4.7e-07 | 3.433E-07 | 1.626E-07 | 1.600E-07 | 4.700E-07 |
| 23095 | 1155 | T1155P | D | 2.4e-07 | 1.5e-07 | 3.9e-07 | 2.600E-07 | 1.212E-07 | 1.500E-07 | 3.900E-07 |
| 23096 | 1155 | T1155S | D | 2.2e-07 | 1.8e-07 | 1.7e-07 | 1.900E-07 | 2.646E-08 | 1.700E-07 | 2.200E-07 |
| 23097 | 1155 | T1155T | D | 3.9e-07 | 2.5e-07 | 1.7e-07 | 2.700E-07 | 1.114E-07 | 1.700E-07 | 3.900E-07 |
| 23098 | 1155 | T1155W | D | 4.4e-07 | 2.4e-07 | 1.4e-07 | 2.733E-07 | 1.528E-07 | 1.400E-07 | 4.400E-07 |
| 23099 | 1155 | T1155Y | D | 4.4e-07 | 2.7e-07 | 1e-07 | 2.700E-07 | 1.700E-07 | 1.000E-07 | 4.400E-07 |
| 23100 | 1155 | T1155V | D | 5.4e-07 | 1.4e-07 | 2.4e-07 | 3.067E-07 | 2.082E-07 | 1.400E-07 | 5.400E-07 |
| 23101 | 1156 | F1156A | D | 5.8e-07 | 2.5e-07 | 2.9e-07 | 3.733E-07 | 1.801E-07 | 2.500E-07 | 5.800E-07 |
| 23102 | 1156 | F1156R | D | 4.7e-07 | 2.5e-07 | 1.1e-07 | 2.767E-07 | 1.815E-07 | 1.100E-07 | 4.700E-07 |
| 23103 | 1156 | F1156N | D | 1.7e-07 | 2e-07 | 3.2e-07 | 2.300E-07 | 7.937E-08 | 1.700E-07 | 3.200E-07 |
| 23104 | 1156 | F1156D | D | 2e-07 | 6e-07 | 1.6e-07 | 3.200E-07 | 2.433E-07 | 1.600E-07 | 6.000E-07 |
| 23105 | 1156 | F1156C | D | 2.1e-07 | 5e-07 | 1.9e-07 | 3.000E-07 | 1.735E-07 | 1.900E-07 | 5.000E-07 |
| 23106 | 1156 | F1156E | D | 4.4e-07 | 2.4e-07 | 2.2e-07 | 3.000E-07 | 1.217E-07 | 2.200E-07 | 4.400E-07 |
| 23107 | 1156 | F1156Q | D | 2.7e-07 | 1.5e-07 | 2.5e-07 | 2.233E-07 | 6.429E-08 | 1.500E-07 | 2.700E-07 |
| 23108 | 1156 | F1156G | D | 1.6e-07 | 1.7e-07 | 1.9e-07 | 1.733E-07 | 1.528E-08 | 1.600E-07 | 1.900E-07 |
| 23109 | 1156 | F1156H | D | 3.7e-07 | 1.7e-07 | 9.5e-08 | 2.117E-07 | 1.422E-07 | 9.500E-08 | 3.700E-07 |
| 23110 | 1156 | F1156I | D | 4.7e-07 | 3.1e-07 | 2.4e-07 | 3.400E-07 | 1.179E-07 | 2.400E-07 | 4.700E-07 |
| 23111 | 1156 | F1156L | D | 2e-07 | 1.5e-07 | 1.2e-07 | 1.567E-07 | 4.041E-08 | 1.200E-07 | 2.000E-07 |
| 23112 | 1156 | F1156K | D | 1.3e-07 | 1.9e-07 | 1.5e-07 | 1.567E-07 | 3.055E-08 | 1.300E-07 | 1.900E-07 |
| 23113 | 1156 | F1156M | D | 3.5e-07 | 6.1e-07 | 2.8e-07 | 4.133E-07 | 1.739E-07 | 2.800E-07 | 6.100E-07 |
| 23114 | 1156 | F1156F | D | 4.4e-07 | 7.3e-07 | 1.8e-07 | 4.500E-07 | 2.751E-07 | 1.800E-07 | 7.300E-07 |
| 23115 | 1156 | F1156P | D | 3.9e-07 | 3.9e-07 | 2.3e-07 | 3.367E-07 | 9.238E-08 | 2.300E-07 | 3.900E-07 |
| 23116 | 1156 | F1156S | D | 3.2e-07 | 1.9e-07 | 3.6e-07 | 2.900E-07 | 8.888E-08 | 1.900E-07 | 3.600E-07 |
| 23117 | 1156 | F1156T | D | 2.1e-07 | 2.8e-07 | 2.6e-07 | 2.500E-07 | 3.606E-08 | 2.100E-07 | 2.800E-07 |
| 23118 | 1156 | F1156W | D | 2.9e-07 | 2.7e-07 | 3.6e-07 | 3.067E-07 | 4.726E-08 | 2.700E-07 | 3.600E-07 |
| 23119 | 1156 | F1156Y | D | 5.5e-07 | 1.7e-07 | 5e-07 | 4.067E-07 | 2.065E-07 | 1.700E-07 | 5.500E-07 |

| | | | | | | | | | | |
|---|---|---|---|---|---|---|---|---|---|---|
| 23120 | 1156 | F1156V | D | 2.1e-07 | 2.4e-07 | 2.4e-07 | 2.300E-07 | 1.732E-08 | 2.100E-07 | 2.400E-07 |
| 23121 | 1157 | G1157A | D | 2.8e-07 | 2.5e-07 | 2.5e-07 | 2.600E-07 | 1.732E-08 | 2.500E-07 | 2.800E-07 |
| 23122 | 1157 | G1157R | D | 3.6e-07 | 1.3e-07 | 2.3e-07 | 2.400E-07 | 1.153E-07 | 1.300E-07 | 3.600E-07 |
| 23123 | 1157 | G1157N | D | 2.2e-07 | 2.5e-07 | 2.3e-07 | 2.333E-07 | 1.528E-08 | 2.200E-07 | 2.500E-07 |
| 23124 | 1157 | G1157D | D | 2.7e-07 | 2.7e-07 | 1.7e-07 | 2.367E-07 | 5.774E-08 | 1.700E-07 | 2.700E-07 |
| 23125 | 1157 | G1157C | D | 1.6e-07 | 4.8e-07 | 2.3e-07 | 2.900E-07 | 1.682E-07 | 1.600E-07 | 4.800E-07 |
| 23126 | 1157 | G1157E | D | 1.1e-07 | 4.2e-07 | 1.8e-07 | 2.367E-07 | 1.626E-07 | 1.100E-07 | 4.200E-07 |
| 23127 | 1157 | G1157Q | D | 3e-07 | 2.9e-07 | 3.9e-07 | 3.267E-07 | 5.508E-08 | 2.900E-07 | 3.900E-07 |
| 23128 | 1157 | G1157G | D | 4e-07 | 2.9e-07 | 6.3e-07 | 4.400E-07 | 1.735E-07 | 2.900E-07 | 6.300E-07 |
| 23129 | 1157 | G1157H | D | 3.1e-07 | 2e-07 | 2.5e-07 | 2.533E-07 | 5.508E-08 | 2.000E-07 | 3.100E-07 |
| 23130 | 1157 | G1157I | D | 2.7e-07 | 2.1e-07 | 2.6e-07 | 2.467E-07 | 3.215E-08 | 2.100E-07 | 2.700E-07 |
| 23131 | 1157 | G1157L | D | 2.1e-07 | 3.8e-07 | 2.6e-07 | 2.833E-07 | 8.737E-08 | 2.100E-07 | 3.800E-07 |
| 23132 | 1157 | G1157K | D | 1.5e-07 | 2.3e-07 | 8.5e-08 | 1.550E-07 | 7.263E-08 | 8.500E-08 | 2.300E-07 |
| 23133 | 1157 | G1157M | D | 1.1e-07 | 4.6e-07 | 3.9e-07 | 3.200E-07 | 1.852E-07 | 1.100E-07 | 4.600E-07 |
| 23134 | 1157 | G1157F | D | 1.2e-07 | 3.3e-07 | 1.8e-07 | 2.100E-07 | 1.082E-07 | 1.200E-07 | 3.300E-07 |
| 23135 | 1157 | G1157P | D | 2.7e-07 | 1.5e-07 | 3.1e-07 | 2.433E-07 | 8.327E-08 | 1.500E-07 | 3.100E-07 |
| 23136 | 1157 | G1157S | D | 3.8e-07 | 2.2e-07 | 1.1e-07 | 2.367E-07 | 1.358E-07 | 1.100E-07 | 3.800E-07 |
| 23137 | 1157 | G1157T | D | 3.6e-07 | 2.8e-07 | 3.4e-07 | 3.267E-07 | 4.163E-08 | 2.800E-07 | 3.600E-07 |
| 23138 | 1157 | G1157W | D | 3.4e-07 | 3.9e-07 | 7.2e-07 | 4.833E-07 | 2.065E-07 | 3.400E-07 | 7.200E-07 |
| 23139 | 1157 | G1157Y | D | 1.9e-07 | 1.3e-07 | 3.5e-07 | 2.233E-07 | 1.137E-07 | 1.300E-07 | 3.500E-07 |
| 23140 | 1157 | G1157V | D | 2.4e-07 | 4.8e-07 | 4.3e-07 | 3.833E-07 | 1.266E-07 | 2.400E-07 | 4.800E-07 |
| 23141 | 1158 | Q1158A | D | 2.8e-07 | 8e-07 | 2.6e-07 | 4.467E-07 | 3.062E-07 | 2.600E-07 | 8.000E-07 |
| 23142 | 1158 | Q1158R | D | 2.4e-07 | 1.3e-07 | 1.2e-07 | 1.633E-07 | 6.658E-08 | 1.200E-07 | 2.400E-07 |
| 23143 | 1158 | Q1158N | D | 2.2e-07 | 1.5e-07 | 2e-07 | 1.900E-07 | 3.606E-08 | 1.500E-07 | 2.200E-07 |
| 23144 | 1158 | Q1158D | D | 1.5e-07 | 2.1e-07 | 4.9e-07 | 2.833E-07 | 1.815E-07 | 1.500E-07 | 4.900E-07 |
| 23145 | 1158 | Q1158C | D | 4e-07 | 2.4e-07 | 1.3e-07 | 2.567E-07 | 1.358E-07 | 1.300E-07 | 4.000E-07 |
| 23146 | 1158 | Q1158E | D | 2.5e-07 | 1.2e-07 | 3.6e-07 | 2.433E-07 | 1.201E-07 | 1.200E-07 | 3.600E-07 |
| 23147 | 1158 | Q1158Q | D | 1.8e-07 | 1.7e-07 | 2.2e-07 | 1.900E-07 | 2.646E-08 | 1.700E-07 | 2.200E-07 |
| 23148 | 1158 | Q1158G | D | 4.7e-07 | 2.3e-07 | 2.6e-07 | 3.200E-07 | 1.308E-07 | 2.300E-07 | 4.700E-07 |
| 23149 | 1158 | Q1158H | D | 1.6e-07 | 1.3e-07 | 5.4e-07 | 2.767E-07 | 2.285E-07 | 1.300E-07 | 5.400E-07 |
| 23150 | 1158 | Q1158I | D | 4.1e-07 | 2.7e-07 | 1.8e-07 | 2.867E-07 | 1.159E-07 | 1.800E-07 | 4.100E-07 |
| 23151 | 1158 | Q1158L | D | 1.7e-07 | 2.5e-07 | 1.6e-07 | 1.933E-07 | 4.933E-08 | 1.600E-07 | 2.500E-07 |
| 23152 | 1158 | Q1158K | D | 1.5e-07 | 2.6e-07 | 1.4e-07 | 1.833E-07 | 6.658E-08 | 1.400E-07 | 2.600E-07 |
| 23153 | 1158 | Q1158M | D | 2.3e-07 | 1.4e-07 | 3.2e-07 | 2.300E-07 | 9.000E-08 | 1.400E-07 | 3.200E-07 |
| 23154 | 1158 | Q1158F | D | 3.3e-07 | 2.2e-07 | 3.7e-07 | 3.067E-07 | 7.767E-08 | 2.200E-07 | 3.700E-07 |
| 23155 | 1158 | Q1158P | D | 3.6e-07 | 3.7e-07 | 1.2e-07 | 2.833E-07 | 1.415E-07 | 1.200E-07 | 3.700E-07 |
| 23156 | 1158 | Q1158S | D | 2.3e-07 | 1.9e-07 | 2.6e-07 | 2.267E-07 | 3.512E-08 | 1.900E-07 | 2.600E-07 |
| 23157 | 1158 | Q1158T | D | 2e-07 | 8.5e-08 | 3.3e-07 | 2.050E-07 | 1.226E-07 | 8.500E-08 | 3.300E-07 |
| 23158 | 1158 | Q1158W | D | 3.5e-07 | 1.4e-07 | 1.9e-07 | 2.267E-07 | 1.097E-07 | 1.400E-07 | 3.500E-07 |
| 23159 | 1158 | Q1158Y | D | 3e-07 | 1.6e-07 | 1.8e-07 | 2.133E-07 | 7.572E-08 | 1.600E-07 | 3.000E-07 |
| 23160 | 1158 | Q1158V | D | 1.6e-07 | 1.6e-07 | 3.8e-07 | 2.333E-07 | 1.270E-07 | 1.600E-07 | 3.800E-07 |
| 23161 | 1159 | G1159A | D | 3.5e-07 | 1.7e-07 | 4.3e-07 | 3.167E-07 | 1.332E-07 | 1.700E-07 | 4.300E-07 |
| 23162 | 1159 | G1159R | D | 1.3e-07 | 3.1e-07 | 1.7e-07 | 2.033E-07 | 9.452E-08 | 1.300E-07 | 3.100E-07 |
| 23163 | 1159 | G1159N | D | 1.2e-07 | 1.3e-07 | 3.7e-07 | 2.067E-07 | 1.415E-07 | 1.200E-07 | 3.700E-07 |
| 23164 | 1159 | G1159D | D | 2.3e-07 | 3.5e-07 | 9.7e-08 | 2.257E-07 | 1.266E-07 | 9.700E-08 | 3.500E-07 |
| 23165 | 1159 | G1159C | D | 2.8e-07 | 2.5e-07 | 4.4e-07 | 3.233E-07 | 1.021E-07 | 2.500E-07 | 4.400E-07 |
| 23166 | 1159 | G1159E | D | 2.8e-07 | 2.7e-07 | 6.6e-07 | 4.033E-07 | 2.223E-07 | 2.700E-07 | 6.600E-07 |
| 23167 | 1159 | G1159Q | D | 3.6e-07 | 1.8e-07 | 3.6e-07 | 3.000E-07 | 1.039E-07 | 1.800E-07 | 3.600E-07 |
| 23168 | 1159 | G1159G | D | 6.7e-07 | 2.2e-07 | 3.5e-07 | 4.133E-07 | 2.316E-07 | 2.200E-07 | 6.700E-07 |
| 23169 | 1159 | G1159H | D | 2.8e-07 | 2.2e-07 | 2.7e-07 | 2.567E-07 | 3.215E-08 | 2.200E-07 | 2.800E-07 |
| 23170 | 1159 | G1159I | D | 4.2e-07 | 1.7e-07 | 3.9e-07 | 3.267E-07 | 1.365E-07 | 1.700E-07 | 4.200E-07 |
| 23171 | 1159 | G1159L | D | 2.2e-07 | 3.6e-07 | 2.7e-07 | 2.833E-07 | 7.095E-08 | 2.200E-07 | 3.600E-07 |
| 23172 | 1159 | G1159K | D | 2.1e-07 | 1.5e-07 | 3.4e-07 | 2.333E-07 | 9.713E-08 | 1.500E-07 | 3.400E-07 |
| 23173 | 1159 | G1159M | D | 1.6e-07 | 1.5e-07 | 4.7e-07 | 2.600E-07 | 1.819E-07 | 1.500E-07 | 4.700E-07 |
| 23174 | 1159 | G1159F | D | 3.5e-07 | 2.1e-07 | 1.4e-07 | 2.333E-07 | 1.069E-07 | 1.400E-07 | 3.500E-07 |
| 23175 | 1159 | G1159P | D | 1.9e-07 | 2.5e-07 | 2.8e-07 | 2.400E-07 | 4.583E-08 | 1.900E-07 | 2.800E-07 |
| 23176 | 1159 | G1159S | D | 1.3e-07 | 2.4e-07 | 2.1e-07 | 1.933E-07 | 5.686E-08 | 1.300E-07 | 2.400E-07 |
| 23177 | 1159 | G1159T | D | 3e-07 | 2.7e-07 | 2.5e-07 | 2.733E-07 | 2.517E-08 | 2.500E-07 | 3.000E-07 |
| 23178 | 1159 | G1159W | D | 1.9e-07 | 3.7e-07 | 1.8e-07 | 2.467E-07 | 1.069E-07 | 1.800E-07 | 3.700E-07 |
| 23179 | 1159 | G1159Y | D | 2.9e-07 | 2e-07 | 1.9e-07 | 2.267E-07 | 5.508E-08 | 1.900E-07 | 2.900E-07 |
| 23180 | 1159 | G1159V | D | 2.8e-07 | 1.5e-07 | 2.8e-07 | 2.367E-07 | 7.506E-08 | 1.500E-07 | 2.800E-07 |
| 23181 | 1160 | T1160A | D | 1.2e-07 | 4.8e-07 | 2.3e-07 | 2.767E-07 | 1.845E-07 | 1.200E-07 | 4.800E-07 |
| 23182 | 1160 | T1160R | D | 3.1e-07 | 3.6e-07 | 3.7e-07 | 3.467E-07 | 3.215E-08 | 3.100E-07 | 3.700E-07 |
| 23183 | 1160 | T1160N | D | 2e-07 | 5.4e-07 | 3.6e-07 | 3.667E-07 | 1.701E-07 | 2.000E-07 | 5.400E-07 |
| 23184 | 1160 | T1160D | D | 4.2e-07 | 5.9e-07 | 4.6e-07 | 4.900E-07 | 8.888E-08 | 4.200E-07 | 5.900E-07 |
| 23185 | 1160 | T1160C | D | 2.3e-07 | 4.5e-07 | 2.2e-07 | 3.000E-07 | 1.300E-07 | 2.200E-07 | 4.500E-07 |
| 23186 | 1160 | T1160E | D | 2e-07 | 2.8e-07 | 2e-07 | 2.267E-07 | 4.619E-08 | 2.000E-07 | 2.800E-07 |
| 23187 | 1160 | T1160Q | D | 4.1e-07 | 4.3e-07 | 5.5e-07 | 4.633E-07 | 7.572E-08 | 4.100E-07 | 5.500E-07 |
| 23188 | 1160 | T1160G | D | 8.3e-07 | 4.9e-07 | 2.1e-07 | 5.100E-07 | 3.105E-07 | 2.100E-07 | 8.300E-07 |
| 23189 | 1160 | T1160H | D | 2.5e-07 | 9.8e-08 | 2.8e-07 | 2.093E-07 | 9.758E-08 | 9.800E-08 | 2.800E-07 |
| 23190 | 1160 | T1160I | D | 1.7e-07 | 2.5e-07 | 1.4e-07 | 1.867E-07 | 5.686E-08 | 1.400E-07 | 2.500E-07 |
| 23191 | 1160 | T1160L | D | 3.7e-07 | 2.7e-07 | 2.6e-07 | 3.000E-07 | 6.083E-08 | 2.600E-07 | 3.700E-07 |
| 23192 | 1160 | T1160K | D | 3e-07 | 3.9e-07 | 1.8e-07 | 2.900E-07 | 1.054E-07 | 1.800E-07 | 3.900E-07 |
| 23193 | 1160 | T1160M | D | 1.9e-07 | 7.6e-07 | 1.3e-07 | 3.600E-07 | 3.477E-07 | 1.300E-07 | 7.600E-07 |
| 23194 | 1160 | T1160F | D | 3e-07 | 2e-07 | 1.3e-07 | 2.100E-07 | 8.544E-08 | 1.300E-07 | 3.000E-07 |
| 23195 | 1160 | T1160P | D | 1.2e-07 | 2.4e-07 | 2.9e-07 | 2.167E-07 | 8.737E-08 | 1.200E-07 | 2.900E-07 |
| 23196 | 1160 | T1160S | D | 2.7e-07 | 2.6e-07 | 2.5e-07 | 2.600E-07 | 1.000E-08 | 2.500E-07 | 2.700E-07 |
| 23197 | 1160 | T1160T | D | 1.8e-07 | 1.8e-07 | 5e-07 | 2.867E-07 | 1.848E-07 | 1.800E-07 | 5.000E-07 |
| 23198 | 1160 | T1160W | D | 1.7e-07 | 3.3e-07 | 2e-07 | 2.333E-07 | 8.505E-08 | 1.700E-07 | 3.300E-07 |
| 23199 | 1160 | T1160Y | D | 2.3e-07 | 1.2e-07 | 4.2e-07 | 2.567E-07 | 1.518E-07 | 1.200E-07 | 4.200E-07 |
| 23200 | 1160 | T1160V | D | 1.3e-07 | 1.8e-07 | 2.6e-07 | 1.900E-07 | 6.557E-08 | 1.300E-07 | 2.600E-07 |
| 23201 | 1161 | K1161A | D | 1.3e-07 | 3.9e-07 | 4.1e-07 | 3.100E-07 | 1.562E-07 | 1.300E-07 | 4.100E-07 |
| 23202 | 1161 | K1161R | D | 2.2e-07 | 2.5e-07 | 1.6e-07 | 2.100E-07 | 4.583E-08 | 1.600E-07 | 2.500E-07 |
| 23203 | 1161 | K1161N | D | 1.7e-07 | 1.3e-07 | 2.8e-07 | 1.933E-07 | 7.767E-08 | 1.300E-07 | 2.800E-07 |
| 23204 | 1161 | K1161D | D | 2e-07 | 2.6e-07 | 2.9e-07 | 2.500E-07 | 4.583E-08 | 2.000E-07 | 2.900E-07 |

| | | | | | | | | | | |
|---|---|---|---|---|---|---|---|---|---|---|
| 23205 | 1161 | K1161C | D | 1.6e-07 | 4e-07 | 9e-08 | 2.167E-07 | 1.626E-07 | 9.000E-08 | 4.000E-07 |
| 23206 | 1161 | K1161E | D | 3e-07 | 2.7e-07 | 5.5e-07 | 3.733E-07 | 1.537E-07 | 2.700E-07 | 5.500E-07 |
| 23207 | 1161 | K1161Q | D | 2.3e-07 | 1.3e-07 | 5.1e-07 | 2.900E-07 | 1.970E-07 | 1.300E-07 | 5.100E-07 |
| 23208 | 1161 | K1161G | D | 4.1e-07 | 2.6e-07 | 2.9e-07 | 3.200E-07 | 7.937E-08 | 2.600E-07 | 4.100E-07 |
| 23209 | 1161 | K1161H | D | 7.1e-08 | 1.5e-07 | 5.1e-07 | 2.437E-07 | 2.340E-07 | 7.100E-08 | 5.100E-07 |
| 23210 | 1161 | K1161I | D | 2.4e-07 | 2.8e-07 | 2.8e-07 | 2.667E-07 | 2.309E-08 | 2.400E-07 | 2.800E-07 |
| 23211 | 1161 | K1161L | D | 2.3e-07 | 2.2e-07 | 1.5e-07 | 2.000E-07 | 4.359E-08 | 1.500E-07 | 2.300E-07 |
| 23212 | 1161 | K1161K | D | 3e-07 | 6e-07 | 1e-07 | 3.333E-07 | 2.517E-07 | 1.000E-07 | 6.000E-07 |
| 23213 | 1161 | K1161M | D | 5.5e-07 | 3.5e-07 | 2.6e-07 | 3.867E-07 | 1.484E-07 | 2.600E-07 | 5.500E-07 |
| 23214 | 1161 | K1161F | D | 4.3e-07 | 4.1e-07 | 2.1e-07 | 3.500E-07 | 1.217E-07 | 2.100E-07 | 4.300E-07 |
| 23215 | 1161 | K1161P | D | 2.1e-07 | 1.7e-07 | 3e-07 | 2.267E-07 | 6.658E-08 | 1.700E-07 | 3.000E-07 |
| 23216 | 1161 | K1161S | D | 3.2e-07 | 1.9e-07 | 1.9e-07 | 2.333E-07 | 7.506E-08 | 1.900E-07 | 3.200E-07 |
| 23217 | 1161 | K1161T | D | 2.3e-07 | 1.9e-07 | 3.4e-07 | 2.533E-07 | 7.767E-08 | 1.900E-07 | 3.400E-07 |
| 23218 | 1161 | K1161W | D | 2.1e-07 | 2.2e-07 | 3.1e-07 | 2.467E-07 | 5.508E-08 | 2.100E-07 | 3.100E-07 |
| 23219 | 1161 | K1161Y | D | 4e-07 | 6.3e-07 | 1e-07 | 3.767E-07 | 2.658E-07 | 1.000E-07 | 6.300E-07 |
| 23220 | 1161 | K1161V | D | 5.5e-07 | 1.5e-07 | 1.2e-07 | 2.733E-07 | 2.401E-07 | 1.200E-07 | 5.500E-07 |
| 23221 | 1162 | V1162A | D | 3e-07 | 2.7e-07 | 2.1e-07 | 2.600E-07 | 4.583E-08 | 2.100E-07 | 3.000E-07 |
| 23222 | 1162 | V1162R | D | 2.3e-07 | 1.9e-07 | 1.8e-07 | 2.000E-07 | 2.646E-08 | 1.800E-07 | 2.300E-07 |
| 23223 | 1162 | V1162N | D | 2.3e-07 | 4.2e-07 | 2.2e-07 | 2.900E-07 | 1.127E-07 | 2.200E-07 | 4.200E-07 |
| 23224 | 1162 | V1162D | D | 3e-07 | 3.3e-07 | 4e-07 | 3.433E-07 | 5.132E-08 | 3.000E-07 | 4.000E-07 |
| 23225 | 1162 | V1162C | D | 4.2e-07 | 3.5e-07 | 1.4e-07 | 3.033E-07 | 1.457E-07 | 1.400E-07 | 4.200E-07 |
| 23226 | 1162 | V1162E | D | 1.4e-07 | 2.9e-07 | 3.4e-07 | 2.567E-07 | 1.041E-07 | 1.400E-07 | 3.400E-07 |
| 23227 | 1162 | V1162Q | D | 2.1e-07 | 1.7e-07 | 1.1e-07 | 1.633E-07 | 5.033E-08 | 1.100E-07 | 2.100E-07 |
| 23228 | 1162 | V1162G | D | 2.9e-07 | 3.1e-07 | 1e-07 | 2.333E-07 | 1.159E-07 | 1.000E-07 | 3.100E-07 |
| 23229 | 1162 | V1162H | D | 1.8e-07 | 1.7e-07 | 1.9e-07 | 1.800E-07 | 1.000E-08 | 1.700E-07 | 1.900E-07 |
| 23230 | 1162 | V1162I | D | 6.8e-07 | 2.2e-07 | 3.4e-07 | 4.133E-07 | 2.386E-07 | 2.200E-07 | 6.800E-07 |
| 23231 | 1162 | V1162L | D | 2.1e-07 | 3.1e-07 | 4.5e-07 | 3.233E-07 | 1.206E-07 | 2.100E-07 | 4.500E-07 |
| 23232 | 1162 | V1162K | D | 1.8e-07 | 1.4e-07 | 2.4e-07 | 1.867E-07 | 5.033E-08 | 1.400E-07 | 2.400E-07 |
| 23233 | 1162 | V1162M | D | 6.5e-07 | 3.6e-07 | 1.7e-07 | 3.933E-07 | 2.417E-07 | 1.700E-07 | 6.500E-07 |
| 23234 | 1162 | V1162F | D | 3e-07 | 2.2e-07 | 1.8e-07 | 2.333E-07 | 6.110E-08 | 1.800E-07 | 3.000E-07 |
| 23235 | 1162 | V1162P | D | 2.6e-07 | 1.5e-07 | 2.1e-07 | 2.067E-07 | 5.508E-08 | 1.500E-07 | 2.600E-07 |
| 23236 | 1162 | V1162S | D | 2.5e-07 | 3.7e-07 | 1.8e-07 | 2.667E-07 | 9.609E-08 | 1.800E-07 | 3.700E-07 |
| 23237 | 1162 | V1162T | D | 1.2e-07 | 2.4e-07 | 8.4e-07 | 4.000E-07 | 3.857E-07 | 1.200E-07 | 8.400E-07 |
| 23238 | 1162 | V1162W | D | 2.7e-07 | 1.8e-07 | 5.9e-07 | 3.467E-07 | 2.155E-07 | 1.800E-07 | 5.900E-07 |
| 23239 | 1162 | V1162Y | D | 3.7e-07 | 1.4e-07 | 5.2e-07 | 3.433E-07 | 1.914E-07 | 1.400E-07 | 5.200E-07 |
| 23240 | 1162 | V1162V | D | 3.1e-07 | 1.4e-07 | 1.2e-07 | 1.900E-07 | 1.044E-07 | 1.200E-07 | 3.100E-07 |
| 23241 | 1163 | E1163A | D | 1.3e-07 | 1.2e-07 | 2.1e-07 | 1.533E-07 | 4.933E-08 | 1.200E-07 | 2.100E-07 |
| 23242 | 1163 | E1163R | D | 1.6e-07 | 2.8e-07 | 3e-07 | 2.467E-07 | 7.572E-08 | 1.600E-07 | 3.000E-07 |
| 23243 | 1163 | E1163N | D | 2.2e-07 | 4.8e-07 | 1.6e-07 | 2.867E-07 | 1.701E-07 | 1.600E-07 | 4.800E-07 |
| 23244 | 1163 | E1163D | D | 2.3e-07 | 2.7e-07 | 3.2e-07 | 2.733E-07 | 4.509E-08 | 2.300E-07 | 3.200E-07 |
| 23245 | 1163 | E1163C | D | 2.1e-07 | 2.9e-07 | 1.6e-07 | 2.200E-07 | 6.557E-08 | 1.600E-07 | 2.900E-07 |
| 23246 | 1163 | E1163E | D | 2e-07 | 2.9e-07 | 2.7e-07 | 2.533E-07 | 4.726E-08 | 2.000E-07 | 2.900E-07 |
| 23247 | 1163 | E1163Q | D | 2.4e-07 | 3.2e-07 | 1.2e-07 | 2.267E-07 | 1.007E-07 | 1.200E-07 | 3.200E-07 |
| 23248 | 1163 | E1163G | D | 1.8e-07 | 2.3e-07 | 1.9e-07 | 2.000E-07 | 2.646E-08 | 1.800E-07 | 2.300E-07 |
| 23249 | 1163 | E1163H | D | 2.1e-07 | 3.2e-07 | 1.5e-07 | 2.267E-07 | 8.622E-08 | 1.500E-07 | 3.200E-07 |
| 23250 | 1163 | E1163I | D | 2.1e-07 | 3.4e-07 | 4.5e-07 | 3.333E-07 | 1.201E-07 | 2.100E-07 | 4.500E-07 |
| 23251 | 1163 | E1163L | D | 2.9e-07 | 3.4e-07 | 2.7e-07 | 3.000E-07 | 3.606E-08 | 2.700E-07 | 3.400E-07 |
| 23252 | 1163 | E1163K | D | 2.6e-07 | 1.1e-07 | 2.6e-07 | 2.100E-07 | 8.660E-08 | 1.100E-07 | 2.600E-07 |
| 23253 | 1163 | E1163M | D | 2.6e-07 | 1.7e-07 | 3.1e-07 | 2.467E-07 | 7.095E-08 | 1.700E-07 | 3.100E-07 |
| 23254 | 1163 | E1163F | D | 3.7e-07 | 4.4e-07 | 3.4e-07 | 3.833E-07 | 5.132E-08 | 3.400E-07 | 4.400E-07 |
| 23255 | 1163 | E1163P | D | 2.4e-07 | 1.3e-07 | 1.5e-07 | 1.733E-07 | 5.859E-08 | 1.300E-07 | 2.400E-07 |
| 23256 | 1163 | E1163S | D | 2.6e-07 | 2e-07 | 1.2e-07 | 1.933E-07 | 7.024E-08 | 1.200E-07 | 2.600E-07 |
| 23257 | 1163 | E1163T | D | 1.8e-07 | 4.2e-07 | 3.4e-07 | 3.133E-07 | 1.222E-07 | 1.800E-07 | 4.200E-07 |
| 23258 | 1163 | E1163W | D | 1.3e-07 | 1e-07 | 1.9e-07 | 1.400E-07 | 4.583E-08 | 1.000E-07 | 1.900E-07 |
| 23259 | 1163 | E1163Y | D | 2.6e-07 | 3.9e-07 | 1.6e-07 | 2.700E-07 | 1.153E-07 | 1.600E-07 | 3.900E-07 |
| 23260 | 1163 | E1163V | D | 1.1e-07 | 1.3e-07 | 3.2e-07 | 1.867E-07 | 1.159E-07 | 1.100E-07 | 3.200E-07 |
| 23261 | 1164 | I1164A | D | 1.4e-07 | 2.1e-07 | 1.2e-07 | 1.567E-07 | 4.726E-08 | 1.200E-07 | 2.100E-07 |
| 23262 | 1164 | I1164R | D | 1.2e-07 | 2.1e-07 | 1.5e-07 | 1.600E-07 | 4.583E-08 | 1.200E-07 | 2.100E-07 |
| 23263 | 1164 | I1164N | D | 3.8e-07 | 2.6e-07 | 4.8e-07 | 3.733E-07 | 1.102E-07 | 2.600E-07 | 4.800E-07 |
| 23264 | 1164 | I1164D | D | 2e-07 | 3.9e-07 | 1.9e-07 | 2.600E-07 | 1.127E-07 | 1.900E-07 | 3.900E-07 |
| 23265 | 1164 | I1164C | D | 3.1e-07 | 3.9e-07 | 2.8e-07 | 3.267E-07 | 5.686E-08 | 2.800E-07 | 3.900E-07 |
| 23266 | 1164 | I1164E | D | 1.5e-07 | 2.1e-07 | 3e-07 | 2.200E-07 | 7.550E-08 | 1.500E-07 | 3.000E-07 |
| 23267 | 1164 | I1164Q | D | 1.4e-07 | 5.8e-07 | 1.7e-07 | 2.967E-07 | 2.458E-07 | 1.400E-07 | 5.800E-07 |
| 23268 | 1164 | I1164G | D | 5.7e-07 | 1.6e-07 | 5e-07 | 4.100E-07 | 2.193E-07 | 1.600E-07 | 5.700E-07 |
| 23269 | 1164 | I1164H | D | 2.6e-07 | 2.1e-07 | 3e-07 | 2.567E-07 | 4.509E-08 | 2.100E-07 | 3.000E-07 |
| 23270 | 1164 | I1164I | D | 2.8e-07 | 1.3e-07 | 2.2e-07 | 2.100E-07 | 7.550E-08 | 1.300E-07 | 2.800E-07 |
| 23271 | 1164 | I1164L | D | 2.8e-07 | 3.8e-07 | 3.2e-07 | 3.267E-07 | 5.033E-08 | 2.800E-07 | 3.800E-07 |
| 23272 | 1164 | I1164K | D | 1.4e-07 | 2.5e-07 | 3.1e-07 | 2.333E-07 | 8.622E-08 | 1.400E-07 | 3.100E-07 |
| 23273 | 1164 | I1164M | D | 3.2e-07 | 1.6e-07 | 1.5e-07 | 2.100E-07 | 9.539E-08 | 1.500E-07 | 3.200E-07 |
| 23274 | 1164 | I1164F | D | 5.2e-07 | 2e-07 | 2.6e-07 | 3.267E-07 | 1.701E-07 | 2.000E-07 | 5.200E-07 |
| 23275 | 1164 | I1164P | D | 3.7e-07 | 2.2e-07 | 5.5e-07 | 3.800E-07 | 1.652E-07 | 2.200E-07 | 5.500E-07 |
| 23276 | 1164 | I1164S | D | 1.3e-07 | 2.6e-07 | 1.4e-07 | 1.767E-07 | 7.234E-08 | 1.300E-07 | 2.600E-07 |
| 23277 | 1164 | I1164T | D | 4.1e-07 | 3e-07 | 2.1e-07 | 3.067E-07 | 1.002E-07 | 2.100E-07 | 4.100E-07 |
| 23278 | 1164 | I1164W | D | 3.3e-07 | 1.9e-07 | 1.9e-07 | 2.367E-07 | 8.083E-08 | 1.900E-07 | 3.300E-07 |
| 23279 | 1164 | I1164Y | D | 1.2e-07 | 1.9e-07 | 1.2e-07 | 1.433E-07 | 4.041E-08 | 1.200E-07 | 1.900E-07 |
| 23280 | 1164 | I1164V | D | 2.9e-07 | 1.8e-07 | 2.2e-07 | 2.300E-07 | 5.568E-08 | 1.800E-07 | 2.900E-07 |
| 23281 | 1165 | K1165A | D | 1.5e-07 | 3.8e-07 | 4.9e-07 | 3.400E-07 | 1.735E-07 | 1.500E-07 | 4.900E-07 |
| 23282 | 1165 | K1165R | D | 1.8e-07 | 3.8e-07 | 3.6e-07 | 3.067E-07 | 1.102E-07 | 1.800E-07 | 3.800E-07 |
| 23283 | 1165 | K1165N | D | 1e-07 | 2.1e-07 | 2.3e-07 | 1.800E-07 | 7.000E-08 | 1.000E-07 | 2.300E-07 |
| 23284 | 1165 | K1165D | D | 2.4e-07 | 3.2e-07 | 2e-07 | 2.533E-07 | 6.110E-08 | 2.000E-07 | 3.200E-07 |
| 23285 | 1165 | K1165C | D | 2e-07 | 1.7e-07 | 2.4e-07 | 2.033E-07 | 3.512E-08 | 1.700E-07 | 2.400E-07 |
| 23286 | 1165 | K1165E | D | 1.5e-07 | 1.7e-07 | 2e-07 | 1.733E-07 | 2.517E-08 | 1.500E-07 | 2.000E-07 |
| 23287 | 1165 | K1165Q | D | 2.3e-07 | 1.3e-07 | 2e-07 | 1.867E-07 | 5.132E-08 | 1.300E-07 | 2.300E-07 |
| 23288 | 1165 | K1165G | D | 1.3e-07 | 2.9e-07 | 5.5e-07 | 3.233E-07 | 2.120E-07 | 1.300E-07 | 5.500E-07 |
| 23289 | 1165 | K1165H | D | 2.4e-07 | 2.6e-07 | 3.4e-07 | 2.800E-07 | 5.292E-08 | 2.400E-07 | 3.400E-07 |

| | | | | | | | | | | |
|---|---|---|---|---|---|---|---|---|---|---|
| 23290 | 1165 | K1165I | D | 2.4e-07 | 1.7e-07 | 4.8e-07 | 2.967E-07 | 1.626E-07 | 1.700E-07 | 4.800E-07 |
| 23291 | 1165 | K1165L | D | 3.3e-07 | 4.1e-07 | 1.5e-07 | 2.967E-07 | 1.332E-07 | 1.500E-07 | 4.100E-07 |
| 23292 | 1165 | K1165K | D | 4.1e-07 | 3.1e-07 | 1.8e-07 | 3.000E-07 | 1.153E-07 | 1.800E-07 | 4.100E-07 |
| 23293 | 1165 | K1165M | D | 2.6e-07 | 4.1e-07 | 1.8e-07 | 2.833E-07 | 1.168E-07 | 1.800E-07 | 4.100E-07 |
| 23294 | 1165 | K1165F | D | 3.2e-07 | 2.7e-07 | 2.9e-07 | 2.933E-07 | 2.517E-08 | 2.700E-07 | 3.200E-07 |
| 23295 | 1165 | K1165P | D | 1.8e-07 | 3.2e-07 | 2.5e-07 | 2.500E-07 | 7.000E-08 | 1.800E-07 | 3.200E-07 |
| 23296 | 1165 | K1165S | D | 2.8e-07 | 2.1e-07 | 2.8e-07 | 2.567E-07 | 4.041E-08 | 2.100E-07 | 2.800E-07 |
| 23297 | 1165 | K1165T | D | 1.1e-07 | 3.4e-07 | 3e-07 | 2.500E-07 | 1.229E-07 | 1.100E-07 | 3.400E-07 |
| 23298 | 1165 | K1165W | D | 2.3e-07 | 2.3e-07 | 1.5e-07 | 2.033E-07 | 4.619E-08 | 1.500E-07 | 2.300E-07 |
| 23299 | 1165 | K1165Y | D | 2.7e-07 | 2.5e-07 | 2.3e-07 | 2.500E-07 | 2.000E-08 | 2.300E-07 | 2.700E-07 |
| 23300 | 1165 | K1165V | D | 1e-07 | 2.1e-07 | 5.3e-07 | 2.800E-07 | 2.234E-07 | 1.000E-07 | 5.300E-07 |
| 23301 | 1166 | R1166A | D | 3.4e-07 | 2e-07 | 2.8e-07 | 2.733E-07 | 7.024E-08 | 2.000E-07 | 3.400E-07 |
| 23302 | 1166 | R1166R | D | 2e-07 | 2.4e-07 | 1.3e-07 | 1.900E-07 | 5.568E-08 | 1.300E-07 | 2.400E-07 |
| 23303 | 1166 | R1166N | D | 2.8e-07 | 1.4e-07 | 2.7e-07 | 2.300E-07 | 7.810E-08 | 1.400E-07 | 2.800E-07 |
| 23304 | 1166 | R1166D | D | 2.2e-07 | 2.5e-07 | 1.4e-07 | 2.033E-07 | 5.686E-08 | 1.400E-07 | 2.500E-07 |
| 23305 | 1166 | R1166C | D | 2.9e-07 | 2.5e-07 | 2.3e-07 | 2.567E-07 | 3.055E-08 | 2.300E-07 | 2.900E-07 |
| 23306 | 1166 | R1166E | D | 1.5e-07 | 3.3e-07 | 4e-07 | 2.933E-07 | 1.290E-07 | 1.500E-07 | 4.000E-07 |
| 23307 | 1166 | R1166Q | D | 3.3e-07 | 3.7e-07 | 3.8e-07 | 3.600E-07 | 2.646E-08 | 3.300E-07 | 3.800E-07 |
| 23308 | 1166 | R1166G | D | 2.4e-07 | 3.1e-07 | 3.1e-07 | 2.867E-07 | 4.041E-08 | 2.400E-07 | 3.100E-07 |
| 23309 | 1166 | R1166H | D | 1.5e-07 | 2e-07 | 3.3e-07 | 2.267E-07 | 9.292E-08 | 1.500E-07 | 3.300E-07 |
| 23310 | 1166 | R1166I | D | 4e-07 | 2.1e-07 | 3e-07 | 3.033E-07 | 9.504E-08 | 2.100E-07 | 4.000E-07 |
| 23311 | 1166 | R1166L | D | 2.6e-07 | 6.2e-07 | 3.2e-07 | 4.000E-07 | 1.929E-07 | 2.600E-07 | 6.200E-07 |
| 23312 | 1166 | R1166K | D | 1.6e-07 | 1.7e-07 | 6.6e-07 | 3.300E-07 | 2.858E-07 | 1.600E-07 | 6.600E-07 |
| 23313 | 1166 | R1166M | D | 1.4e-07 | 2.7e-07 | 3e-07 | 2.367E-07 | 8.505E-08 | 1.400E-07 | 3.000E-07 |
| 23314 | 1166 | R1166F | D | 2.9e-07 | 1.7e-07 | 1.2e-07 | 1.933E-07 | 8.737E-08 | 1.200E-07 | 2.900E-07 |
| 23315 | 1166 | R1166P | D | 2e-07 | 2e-07 | 1.4e-07 | 1.800E-07 | 3.464E-08 | 1.400E-07 | 2.000E-07 |
| 23316 | 1166 | R1166S | D | 1.3e-07 | 1e-07 | 3.4e-07 | 1.900E-07 | 1.308E-07 | 1.000E-07 | 3.400E-07 |
| 23317 | 1166 | R1166T | D | 1.8e-07 | 4.6e-07 | 3.5e-07 | 3.300E-07 | 1.411E-07 | 1.800E-07 | 4.600E-07 |
| 23318 | 1166 | R1166W | D | 1.9e-07 | 2.7e-07 | 3.7e-07 | 2.767E-07 | 9.018E-08 | 1.900E-07 | 3.700E-07 |
| 23319 | 1166 | R1166Y | D | 1.7e-07 | 2.1e-07 | 2.7e-07 | 2.167E-07 | 5.033E-08 | 1.700E-07 | 2.700E-07 |
| 23320 | 1166 | R1166V | D | 2.6e-07 | 4.2e-07 | 2.5e-07 | 3.100E-07 | 9.539E-08 | 2.500E-07 | 4.200E-07 |
| 23321 | 1167 | T1167A | D | 3.1e-07 | 1.2e-07 | 2.8e-07 | 2.367E-07 | 1.021E-07 | 1.200E-07 | 3.100E-07 |
| 23322 | 1167 | T1167R | D | 2.9e-07 | 2.6e-07 | 4.6e-07 | 3.367E-07 | 1.079E-07 | 2.600E-07 | 4.600E-07 |
| 23323 | 1167 | T1167N | D | 1.9e-07 | 1e-07 | 2e-07 | 1.633E-07 | 5.508E-08 | 1.000E-07 | 2.000E-07 |
| 23324 | 1167 | T1167D | D | 3.1e-07 | 2.9e-07 | 2e-07 | 2.667E-07 | 5.859E-08 | 2.000E-07 | 3.100E-07 |
| 23325 | 1167 | T1167C | D | 2.6e-07 | 3.3e-07 | 1.8e-07 | 2.567E-07 | 7.506E-08 | 1.800E-07 | 3.300E-07 |
| 23326 | 1167 | T1167E | D | 2.1e-07 | 2.4e-07 | 2e-07 | 2.167E-07 | 2.082E-08 | 2.000E-07 | 2.400E-07 |
| 23327 | 1167 | T1167Q | D | 1.2e-07 | 2.8e-07 | 2.6e-07 | 2.200E-07 | 8.718E-08 | 1.200E-07 | 2.800E-07 |
| 23328 | 1167 | T1167G | D | 2.2e-07 | 3.1e-07 | 1.8e-07 | 2.367E-07 | 6.658E-08 | 1.800E-07 | 3.100E-07 |
| 23329 | 1167 | T1167H | D | 3.7e-07 | 3.6e-07 | 1.4e-07 | 2.900E-07 | 1.300E-07 | 1.400E-07 | 3.700E-07 |
| 23330 | 1167 | T1167I | D | 2.5e-07 | 2.8e-07 | 1.7e-07 | 2.333E-07 | 5.686E-08 | 1.700E-07 | 2.800E-07 |
| 23331 | 1167 | T1167L | D | 2e-07 | 1.7e-07 | 2.3e-07 | 2.000E-07 | 3.000E-08 | 1.700E-07 | 2.300E-07 |
| 23332 | 1167 | T1167K | D | 3.9e-07 | 2.3e-07 | 1.4e-07 | 2.533E-07 | 1.266E-07 | 1.400E-07 | 3.900E-07 |
| 23333 | 1167 | T1167M | D | 2.2e-07 | 2.3e-07 | 5e-07 | 3.167E-07 | 1.589E-07 | 2.200E-07 | 5.000E-07 |
| 23334 | 1167 | T1167F | D | 1.9e-07 | 3e-07 | 2.4e-07 | 2.433E-07 | 5.508E-08 | 1.900E-07 | 3.000E-07 |
| 23335 | 1167 | T1167P | D | 2.7e-07 | 4.3e-07 | 2.7e-07 | 3.233E-07 | 9.238E-08 | 2.700E-07 | 4.300E-07 |
| 23336 | 1167 | T1167S | D | 1.4e-07 | 3.6e-07 | 1.9e-07 | 2.300E-07 | 1.153E-07 | 1.400E-07 | 3.600E-07 |
| 23337 | 1167 | T1167T | D | 2.7e-07 | 2.4e-07 | 3.6e-07 | 2.900E-07 | 6.245E-08 | 2.400E-07 | 3.600E-07 |
| 23338 | 1167 | T1167W | D | 4.8e-07 | 2.2e-07 | 2e-07 | 3.000E-07 | 1.562E-07 | 2.000E-07 | 4.800E-07 |
| 23339 | 1167 | T1167Y | D | 2.8e-07 | 8e-08 | 2.6e-07 | 2.067E-07 | 1.102E-07 | 8.000E-08 | 2.800E-07 |
| 23340 | 1167 | T1167V | D | 4.2e-07 | 2.4e-07 | 3.6e-07 | 3.400E-07 | 9.165E-08 | 2.400E-07 | 4.200E-07 |
| 23341 | 1168 | V1168A | D | 2.4e-07 | 4e-07 | 3.5e-07 | 3.300E-07 | 8.185E-08 | 2.400E-07 | 4.000E-07 |
| 23342 | 1168 | V1168R | D | 2.4e-07 | 1.7e-07 | 5.4e-07 | 3.167E-07 | 1.966E-07 | 1.700E-07 | 5.400E-07 |
| 23343 | 1168 | V1168N | D | 4e-07 | 2.8e-07 | 2.7e-07 | 3.167E-07 | 7.234E-08 | 2.700E-07 | 4.000E-07 |
| 23344 | 1168 | V1168D | D | 3.2e-07 | 2.1e-07 | 8.5e-08 | 2.050E-07 | 1.176E-07 | 8.500E-08 | 3.200E-07 |
| 23345 | 1168 | V1168C | D | 1.1e-07 | 2.1e-07 | 2.4e-07 | 1.867E-07 | 6.807E-08 | 1.100E-07 | 2.400E-07 |
| 23346 | 1168 | V1168E | D | 4.5e-07 | 2.4e-07 | 4.3e-07 | 3.733E-07 | 1.159E-07 | 2.400E-07 | 4.500E-07 |
| 23347 | 1168 | V1168Q | D | 3.2e-07 | 1.4e-07 | 2.2e-07 | 2.267E-07 | 9.018E-08 | 1.400E-07 | 3.200E-07 |
| 23348 | 1168 | V1168G | D | 4.2e-07 | 1.7e-07 | 2.4e-07 | 2.767E-07 | 1.290E-07 | 1.700E-07 | 4.200E-07 |
| 23349 | 1168 | V1168H | D | 3.3e-07 | 1.3e-07 | 4e-07 | 2.867E-07 | 1.401E-07 | 1.300E-07 | 4.000E-07 |
| 23350 | 1168 | V1168I | D | 2.7e-07 | 6.5e-08 | 3.5e-07 | 2.283E-07 | 1.470E-07 | 6.500E-08 | 3.500E-07 |
| 23351 | 1168 | V1168L | D | 3e-07 | 2.5e-07 | 2.8e-07 | 2.767E-07 | 2.517E-08 | 2.500E-07 | 3.000E-07 |
| 23352 | 1168 | V1168K | D | 2.4e-07 | 1.6e-07 | 4.7e-07 | 2.900E-07 | 1.609E-07 | 1.600E-07 | 4.700E-07 |
| 23353 | 1168 | V1168M | D | 2.6e-07 | 2.8e-07 | 3.2e-07 | 2.867E-07 | 3.055E-08 | 2.600E-07 | 3.200E-07 |
| 23354 | 1168 | V1168F | D | 3.2e-07 | 1.5e-07 | 3.1e-07 | 2.600E-07 | 9.539E-08 | 1.500E-07 | 3.200E-07 |
| 23355 | 1168 | V1168P | D | 5.4e-07 | 3.6e-07 | 2.2e-07 | 3.733E-07 | 1.604E-07 | 2.200E-07 | 5.400E-07 |
| 23356 | 1168 | V1168S | D | 4.6e-07 | 2.9e-07 | 1.3e-07 | 2.933E-07 | 1.650E-07 | 1.300E-07 | 4.600E-07 |
| 23357 | 1168 | V1168T | D | 2.6e-07 | 2.9e-07 | 3.1e-07 | 2.867E-07 | 2.517E-08 | 2.600E-07 | 3.100E-07 |
| 23358 | 1168 | V1168W | D | 2.1e-07 | 5.5e-07 | 3e-07 | 3.533E-07 | 1.762E-07 | 2.100E-07 | 5.500E-07 |
| 23359 | 1168 | V1168Y | D | 4.8e-07 | 2.6e-07 | 2.1e-07 | 3.167E-07 | 1.436E-07 | 2.100E-07 | 4.800E-07 |
| 23360 | 1168 | V1168V | D | 3.6e-07 | 7.1e-07 | 2.1e-07 | 4.267E-07 | 2.566E-07 | 2.100E-07 | 7.100E-07 |
| 23361 | 1169 | A1169A | D | 2.8e-07 | 4.4e-07 | 4.5e-07 | 3.900E-07 | 9.539E-08 | 2.800E-07 | 4.500E-07 |
| 23362 | 1169 | A1169R | D | 1.6e-07 | 1.8e-07 | 2e-07 | 1.800E-07 | 2.000E-08 | 1.600E-07 | 2.000E-07 |
| 23363 | 1169 | A1169N | D | 5.7e-08 | 1.6e-07 | 1.5e-07 | 1.223E-07 | 5.680E-08 | 5.700E-08 | 1.600E-07 |
| 23364 | 1169 | A1169D | D | 2.4e-07 | 3e-07 | 5.3e-07 | 3.567E-07 | 1.531E-07 | 2.400E-07 | 5.300E-07 |
| 23365 | 1169 | A1169C | D | 2.8e-07 | 1.4e-07 | 3.6e-07 | 2.600E-07 | 1.114E-07 | 1.400E-07 | 3.600E-07 |
| 23366 | 1169 | A1169E | D | 2.7e-07 | 1.8e-07 | 2.9e-07 | 2.467E-07 | 5.859E-08 | 1.800E-07 | 2.900E-07 |
| 23367 | 1169 | A1169Q | D | 2e-07 | 1.8e-07 | 3.4e-07 | 2.400E-07 | 8.718E-08 | 1.800E-07 | 3.400E-07 |
| 23368 | 1169 | A1169G | D | 2.5e-07 | 2.2e-07 | 2.4e-07 | 2.367E-07 | 1.528E-08 | 2.200E-07 | 2.500E-07 |
| 23369 | 1169 | A1169H | D | 2.6e-07 | 1.7e-07 | 2.8e-07 | 2.367E-07 | 5.859E-08 | 1.700E-07 | 2.800E-07 |
| 23370 | 1169 | A1169I | D | 1.3e-07 | 1.8e-07 | 4.2e-07 | 2.433E-07 | 1.550E-07 | 1.300E-07 | 4.200E-07 |
| 23371 | 1169 | A1169L | D | 4.1e-07 | 1.6e-07 | 1.8e-07 | 2.500E-07 | 1.389E-07 | 1.600E-07 | 4.100E-07 |
| 23372 | 1169 | A1169K | D | 1.9e-07 | 2e-07 | 3.2e-07 | 2.367E-07 | 7.234E-08 | 1.900E-07 | 3.200E-07 |
| 23373 | 1169 | A1169M | D | 3.7e-07 | 3e-07 | 1.4e-07 | 2.700E-07 | 1.179E-07 | 1.400E-07 | 3.700E-07 |
| 23374 | 1169 | A1169F | D | 1.5e-07 | 3e-07 | 2.9e-07 | 2.467E-07 | 8.386E-08 | 1.500E-07 | 3.000E-07 |

| | | | | | | | | | | |
|---|---|---|---|---|---|---|---|---|---|---|
| 23375 | 1169 | A1169P | D | 2e-07 | 3.4e-07 | 3.8e-07 | 3.067E-07 | 9.452E-08 | 2.000E-07 | 3.800E-07 |
| 23376 | 1169 | A1169S | D | 3e-07 | 2.1e-07 | 1.3e-07 | 2.133E-07 | 8.505E-08 | 1.300E-07 | 3.000E-07 |
| 23377 | 1169 | A1169T | D | 2.6e-07 | 2.5e-07 | 2e-07 | 2.367E-07 | 3.215E-08 | 2.000E-07 | 2.600E-07 |
| 23378 | 1169 | A1169W | D | 2.4e-07 | 2.1e-07 | 1.7e-07 | 2.067E-07 | 3.512E-08 | 1.700E-07 | 2.400E-07 |
| 23379 | 1169 | A1169Y | D | 2.3e-07 | 1.1e-07 | 3.1e-07 | 2.167E-07 | 1.007E-07 | 1.100E-07 | 3.100E-07 |
| 23380 | 1169 | A1169V | D | 2.2e-07 | 2.9e-07 | 3.5e-07 | 2.867E-07 | 6.506E-08 | 2.200E-07 | 3.500E-07 |
| 23381 | 1170 | A1170A | D | 3.5e-07 | 1.2e-07 | 4.2e-07 | 2.967E-07 | 1.570E-07 | 1.200E-07 | 4.200E-07 |
| 23382 | 1170 | A1170R | D | 2.2e-07 | 1.7e-07 | 2.6e-07 | 2.167E-07 | 4.509E-08 | 1.700E-07 | 2.600E-07 |
| 23383 | 1170 | A1170N | D | 2.8e-07 | 3.5e-07 | 2.3e-07 | 2.867E-07 | 6.028E-08 | 2.300E-07 | 3.500E-07 |
| 23384 | 1170 | A1170D | D | 1.8e-07 | 2.6e-07 | 2.9e-07 | 2.433E-07 | 5.686E-08 | 1.800E-07 | 2.900E-07 |
| 23385 | 1170 | A1170C | D | 2.8e-07 | 4.3e-07 | 3.1e-07 | 3.400E-07 | 7.937E-08 | 2.800E-07 | 4.300E-07 |
| 23386 | 1170 | A1170E | D | 2.9e-07 | 2.1e-07 | 1.7e-07 | 2.233E-07 | 6.110E-08 | 1.700E-07 | 2.900E-07 |
| 23387 | 1170 | A1170Q | D | 1.6e-07 | 2.2e-07 | 2e-07 | 1.933E-07 | 3.055E-08 | 1.600E-07 | 2.200E-07 |
| 23388 | 1170 | A1170G | D | 2.1e-07 | 1.7e-07 | 1.4e-07 | 1.733E-07 | 3.512E-08 | 1.400E-07 | 2.100E-07 |
| 23389 | 1170 | A1170H | D | 1.7e-07 | 3.3e-07 | 4e-07 | 3.000E-07 | 1.179E-07 | 1.700E-07 | 4.000E-07 |
| 23390 | 1170 | A1170I | D | 2.6e-07 | 1.5e-07 | 3.1e-07 | 2.400E-07 | 8.185E-08 | 1.500E-07 | 3.100E-07 |
| 23391 | 1170 | A1170L | D | 2.6e-07 | 7.8e-08 | 2.3e-07 | 1.893E-07 | 9.758E-08 | 7.800E-08 | 2.600E-07 |
| 23392 | 1170 | A1170K | D | 2.1e-07 | 1.6e-07 | 2.5e-07 | 2.067E-07 | 4.509E-08 | 1.600E-07 | 2.500E-07 |
| 23393 | 1170 | A1170M | D | 5e-07 | 2.5e-07 | 1.4e-07 | 2.967E-07 | 1.845E-07 | 1.400E-07 | 5.000E-07 |
| 23394 | 1170 | A1170F | D | 2.2e-07 | 1.4e-07 | 2.6e-07 | 2.067E-07 | 6.110E-08 | 1.400E-07 | 2.600E-07 |
| 23395 | 1170 | A1170P | D | 3.5e-07 | 2.4e-07 | 5.5e-07 | 3.800E-07 | 1.572E-07 | 2.400E-07 | 5.500E-07 |
| 23396 | 1170 | A1170S | D | 1.8e-07 | 5.4e-07 | 3.5e-07 | 3.567E-07 | 1.801E-07 | 1.800E-07 | 5.400E-07 |
| 23397 | 1170 | A1170T | D | 3.9e-07 | 1.1e-07 | 2.4e-07 | 2.467E-07 | 1.401E-07 | 1.100E-07 | 3.900E-07 |
| 23398 | 1170 | A1170W | D | 1.3e-07 | 1.3e-07 | 1.2e-07 | 1.267E-07 | 5.774E-09 | 1.200E-07 | 1.300E-07 |
| 23399 | 1170 | A1170Y | D | 2.5e-07 | 2e-07 | 2.7e-07 | 2.400E-07 | 3.606E-08 | 2.000E-07 | 2.700E-07 |
| 23400 | 1170 | A1170V | D | 2.5e-07 | 1.6e-07 | 4.4e-07 | 2.833E-07 | 1.429E-07 | 1.600E-07 | 4.400E-07 |
| 23401 | 1171 | P1171A | D | 2.3e-07 | 4.1e-07 | 1.9e-07 | 2.767E-07 | 1.172E-07 | 1.900E-07 | 4.100E-07 |
| 23402 | 1171 | P1171R | D | 9.9e-08 | 2.1e-07 | 1.5e-07 | 1.530E-07 | 5.556E-08 | 9.900E-08 | 2.100E-07 |
| 23403 | 1171 | P1171N | D | 3.6e-07 | 2.1e-07 | 5.3e-07 | 3.667E-07 | 1.601E-07 | 2.100E-07 | 5.300E-07 |
| 23404 | 1171 | P1171D | D | 2.8e-07 | 2.5e-07 | 3.9e-07 | 3.067E-07 | 7.371E-08 | 2.500E-07 | 3.900E-07 |
| 23405 | 1171 | P1171C | D | 3.4e-07 | 1.4e-07 | 2.2e-07 | 2.333E-07 | 1.007E-07 | 1.400E-07 | 3.400E-07 |
| 23406 | 1171 | P1171E | D | 2.6e-07 | 2.3e-07 | 2.6e-07 | 2.500E-07 | 1.732E-08 | 2.300E-07 | 2.600E-07 |
| 23407 | 1171 | P1171Q | D | 2.5e-07 | 1.2e-07 | 1.6e-07 | 1.767E-07 | 6.658E-08 | 1.200E-07 | 2.500E-07 |
| 23408 | 1171 | P1171G | D | 2.8e-07 | 1.9e-07 | 2.2e-07 | 2.300E-07 | 4.583E-08 | 1.900E-07 | 2.800E-07 |
| 23409 | 1171 | P1171H | D | 1.7e-07 | 1.2e-07 | 3.6e-07 | 2.167E-07 | 1.266E-07 | 1.200E-07 | 3.600E-07 |
| 23410 | 1171 | P1171I | D | 3.4e-07 | 4.7e-07 | 2.9e-07 | 3.667E-07 | 9.292E-08 | 2.900E-07 | 4.700E-07 |
| 23411 | 1171 | P1171L | D | 2.8e-07 | 2.7e-07 | 4.1e-07 | 3.200E-07 | 7.810E-08 | 2.700E-07 | 4.100E-07 |
| 23412 | 1171 | P1171K | D | 2.9e-07 | 5.1e-07 | 1.4e-07 | 3.133E-07 | 1.861E-07 | 1.400E-07 | 5.100E-07 |
| 23413 | 1171 | P1171M | D | 2.9e-07 | 1.6e-07 | 2.7e-07 | 2.400E-07 | 7.000E-08 | 1.600E-07 | 2.900E-07 |
| 23414 | 1171 | P1171F | D | 1.7e-07 | 2e-07 | 1.2e-07 | 1.633E-07 | 4.041E-08 | 1.200E-07 | 2.000E-07 |
| 23415 | 1171 | P1171P | D | 3.5e-07 | 1.7e-07 | 2.9e-07 | 2.700E-07 | 9.165E-08 | 1.700E-07 | 3.500E-07 |
| 23416 | 1171 | P1171S | D | 1.6e-07 | 2.2e-07 | 2.7e-07 | 2.167E-07 | 5.508E-08 | 1.600E-07 | 2.700E-07 |
| 23417 | 1171 | P1171T | D | 2.2e-07 | 2.6e-07 | 1.9e-07 | 2.233E-07 | 3.512E-08 | 1.900E-07 | 2.600E-07 |
| 23418 | 1171 | P1171W | D | 4.4e-07 | 1.5e-07 | 3e-07 | 2.967E-07 | 1.450E-07 | 1.500E-07 | 4.400E-07 |
| 23419 | 1171 | P1171Y | D | 1.5e-07 | 2.5e-07 | 1.5e-07 | 1.833E-07 | 5.774E-08 | 1.500E-07 | 2.500E-07 |
| 23420 | 1171 | P1171V | D | 2.7e-07 | 2.7e-07 | 4.3e-07 | 3.233E-07 | 9.238E-08 | 2.700E-07 | 4.300E-07 |
| 23421 | 1172 | S1172A | D | 1.9e-07 | 2.5e-07 | 2.1e-07 | 2.167E-07 | 3.055E-08 | 1.900E-07 | 2.500E-07 |
| 23422 | 1172 | S1172R | D | 8.7e-08 | 3.1e-07 | 3.3e-07 | 2.423E-07 | 1.349E-07 | 8.700E-08 | 3.300E-07 |
| 23423 | 1172 | S1172N | D | 3.3e-07 | 2.4e-07 | 2.8e-07 | 2.833E-07 | 4.509E-08 | 2.400E-07 | 3.300E-07 |
| 23424 | 1172 | S1172D | D | 4.3e-07 | 3.3e-07 | 2.2e-07 | 3.267E-07 | 1.050E-07 | 2.200E-07 | 4.300E-07 |
| 23425 | 1172 | S1172C | D | 4.4e-07 | 3.4e-07 | 1.4e-07 | 3.067E-07 | 1.528E-07 | 1.400E-07 | 4.400E-07 |
| 23426 | 1172 | S1172E | D | 3.1e-07 | 2.1e-07 | 4.4e-07 | 3.200E-07 | 1.153E-07 | 2.100E-07 | 4.400E-07 |
| 23427 | 1172 | S1172Q | D | 1.1e-07 | 2e-07 | 3.2e-07 | 2.100E-07 | 1.054E-07 | 1.100E-07 | 3.200E-07 |
| 23428 | 1172 | S1172G | D | 1.7e-07 | 1.1e-07 | 2.4e-07 | 1.733E-07 | 6.506E-08 | 1.100E-07 | 2.400E-07 |
| 23429 | 1172 | S1172H | D | 1.7e-07 | 2.2e-07 | 3e-07 | 2.300E-07 | 6.557E-08 | 1.700E-07 | 3.000E-07 |
| 23430 | 1172 | S1172I | D | 9.8e-08 | 4.2e-07 | 2.9e-07 | 2.693E-07 | 1.620E-07 | 9.800E-08 | 4.200E-07 |
| 23431 | 1172 | S1172L | D | 2e-07 | 2.2e-07 | 9.8e-08 | 1.727E-07 | 6.543E-08 | 9.800E-08 | 2.200E-07 |
| 23432 | 1172 | S1172K | D | 2.7e-07 | 1.6e-07 | 2.9e-07 | 2.400E-07 | 7.000E-08 | 1.600E-07 | 2.900E-07 |
| 23433 | 1172 | S1172M | D | 3.1e-07 | 9.6e-08 | 1.9e-07 | 1.987E-07 | 1.073E-07 | 9.600E-08 | 3.100E-07 |
| 23434 | 1172 | S1172F | D | 1.2e-07 | 3.5e-07 | 1.9e-07 | 2.200E-07 | 1.179E-07 | 1.200E-07 | 3.500E-07 |
| 23435 | 1172 | S1172P | D | 1.3e-07 | 4.3e-07 | 2.2e-07 | 2.600E-07 | 1.539E-07 | 1.300E-07 | 4.300E-07 |
| 23436 | 1172 | S1172S | D | 7.6e-08 | 1.9e-07 | 5.2e-07 | 2.620E-07 | 2.306E-07 | 7.600E-08 | 5.200E-07 |
| 23437 | 1172 | S1172T | D | 2.7e-07 | 1.3e-07 | 4.2e-07 | 2.733E-07 | 1.450E-07 | 1.300E-07 | 4.200E-07 |
| 23438 | 1172 | S1172W | D | 3.2e-07 | 9.7e-08 | 6.4e-07 | 3.523E-07 | 2.729E-07 | 9.700E-08 | 6.400E-07 |
| 23439 | 1172 | S1172Y | D | 2.6e-07 | 1.1e-07 | 2.5e-07 | 2.067E-07 | 8.386E-08 | 1.100E-07 | 2.600E-07 |
| 23440 | 1172 | S1172V | D | 3.2e-07 | 2.7e-07 | 2.2e-07 | 2.700E-07 | 5.000E-08 | 2.200E-07 | 3.200E-07 |
| 23441 | 1173 | V1173A | D | 2.1e-07 | 3.4e-07 | 1.8e-07 | 2.433E-07 | 8.505E-08 | 1.800E-07 | 3.400E-07 |
| 23442 | 1173 | V1173R | D | 2.4e-07 | 2.7e-07 | 2.4e-07 | 2.500E-07 | 1.732E-08 | 2.400E-07 | 2.700E-07 |
| 23443 | 1173 | V1173N | D | 5.4e-07 | 2.1e-07 | 2.5e-07 | 3.333E-07 | 1.801E-07 | 2.100E-07 | 5.400E-07 |
| 23444 | 1173 | V1173D | D | 2.1e-07 | 3.1e-07 | 5.1e-07 | 3.433E-07 | 1.528E-07 | 2.100E-07 | 5.100E-07 |
| 23445 | 1173 | V1173C | D | 3.6e-07 | 1.9e-07 | 2.9e-07 | 2.800E-07 | 8.544E-08 | 1.900E-07 | 3.600E-07 |
| 23446 | 1173 | V1173E | D | 2.5e-07 | 2e-07 | 5e-07 | 3.167E-07 | 1.607E-07 | 2.000E-07 | 5.000E-07 |
| 23447 | 1173 | V1173Q | D | 4.4e-07 | 2e-07 | 2.7e-07 | 3.033E-07 | 1.234E-07 | 2.000E-07 | 4.400E-07 |
| 23448 | 1173 | V1173G | D | 3.2e-07 | 1.7e-07 | 1.6e-07 | 2.167E-07 | 8.963E-08 | 1.600E-07 | 3.200E-07 |
| 23449 | 1173 | V1173H | D | 1.3e-07 | 3.3e-07 | 2.4e-07 | 2.333E-07 | 1.002E-07 | 1.300E-07 | 3.300E-07 |
| 23450 | 1173 | V1173I | D | 2.2e-07 | 3.1e-07 | 3.7e-07 | 3.000E-07 | 7.550E-08 | 2.200E-07 | 3.700E-07 |
| 23451 | 1173 | V1173L | D | 2.9e-07 | 2.4e-07 | 2e-07 | 2.433E-07 | 4.509E-08 | 2.000E-07 | 2.900E-07 |
| 23452 | 1173 | V1173K | D | 3.7e-07 | 1.6e-07 | 2.9e-07 | 2.733E-07 | 1.060E-07 | 1.600E-07 | 3.700E-07 |
| 23453 | 1173 | V1173M | D | 1.7e-07 | 1.9e-07 | 3.2e-07 | 2.267E-07 | 8.145E-08 | 1.700E-07 | 3.200E-07 |
| 23454 | 1173 | V1173F | D | 1.2e-07 | 3.3e-07 | 3.7e-07 | 2.733E-07 | 1.343E-07 | 1.200E-07 | 3.700E-07 |
| 23455 | 1173 | V1173P | D | 3.1e-07 | 9.4e-08 | 2.9e-07 | 2.313E-07 | 1.194E-07 | 9.400E-08 | 3.100E-07 |
| 23456 | 1173 | V1173S | D | 3.1e-07 | 2.6e-07 | 1.2e-07 | 2.300E-07 | 9.849E-08 | 1.200E-07 | 3.100E-07 |
| 23457 | 1173 | V1173T | D | 3.3e-07 | 2.2e-07 | 1.9e-07 | 2.467E-07 | 7.371E-08 | 1.900E-07 | 3.300E-07 |
| 23458 | 1173 | V1173W | D | 4e-07 | 2.9e-07 | 2.2e-07 | 3.033E-07 | 9.074E-08 | 2.200E-07 | 4.000E-07 |
| 23459 | 1173 | V1173Y | D | 4e-07 | 2.2e-07 | 2e-07 | 2.733E-07 | 1.102E-07 | 2.000E-07 | 4.000E-07 |

| | | | | | | | | | | |
|---|---|---|---|---|---|---|---|---|---|---|
| 23460 | 1173 | V1173V | D | 6.4e-07 | 2.5e-07 | 2.3e-07 | 3.733E-07 | 2.312E-07 | 2.300E-07 | 6.400E-07 |
| 23461 | 1174 | F1174A | D | 1.4e-07 | 1.5e-07 | 2.4e-07 | 1.767E-07 | 5.508E-08 | 1.400E-07 | 2.400E-07 |
| 23462 | 1174 | F1174R | D | 2.8e-07 | 2.6e-07 | 2.4e-07 | 2.600E-07 | 2.000E-08 | 2.400E-07 | 2.800E-07 |
| 23463 | 1174 | F1174N | D | 2.5e-07 | 1.7e-07 | 4.1e-07 | 2.767E-07 | 1.222E-07 | 1.700E-07 | 4.100E-07 |
| 23464 | 1174 | F1174D | D | 3e-07 | 2.8e-07 | 2.1e-07 | 2.633E-07 | 4.726E-08 | 2.100E-07 | 3.000E-07 |
| 23465 | 1174 | F1174C | D | 3.3e-07 | 1.7e-07 | 2.1e-07 | 2.367E-07 | 8.327E-08 | 1.700E-07 | 3.300E-07 |
| 23466 | 1174 | F1174E | D | 3.3e-07 | 3.1e-07 | 5.4e-07 | 3.933E-07 | 1.274E-07 | 3.100E-07 | 5.400E-07 |
| 23467 | 1174 | F1174Q | D | 2.1e-07 | 1.8e-07 | 2.2e-07 | 2.033E-07 | 2.082E-08 | 1.800E-07 | 2.200E-07 |
| 23468 | 1174 | F1174G | D | 9.4e-08 | 3.8e-07 | 1.8e-07 | 2.180E-07 | 1.467E-07 | 9.400E-08 | 3.800E-07 |
| 23469 | 1174 | F1174H | D | 2.6e-07 | 2.7e-07 | 1.4e-07 | 2.233E-07 | 7.234E-08 | 1.400E-07 | 2.700E-07 |
| 23470 | 1174 | F1174I | D | 2.2e-07 | 3.1e-07 | 2.4e-07 | 2.567E-07 | 4.726E-08 | 2.200E-07 | 3.100E-07 |
| 23471 | 1174 | F1174L | D | 2.6e-07 | 1.5e-07 | 3.8e-07 | 2.633E-07 | 1.150E-07 | 1.500E-07 | 3.800E-07 |
| 23472 | 1174 | F1174K | D | 2.2e-07 | 3e-07 | 2.2e-07 | 2.467E-07 | 4.619E-08 | 2.200E-07 | 3.000E-07 |
| 23473 | 1174 | F1174M | D | 2.2e-07 | 3.1e-07 | 2.2e-07 | 2.500E-07 | 5.196E-08 | 2.200E-07 | 3.100E-07 |
| 23474 | 1174 | F1174F | D | 1.2e-07 | 2.2e-07 | 4.9e-07 | 2.767E-07 | 1.914E-07 | 1.200E-07 | 4.900E-07 |
| 23475 | 1174 | F1174P | D | 1.2e-07 | 5.8e-07 | 2.1e-07 | 3.033E-07 | 2.438E-07 | 1.200E-07 | 5.800E-07 |
| 23476 | 1174 | F1174S | D | 2e-07 | 2e-07 | 4.2e-07 | 2.733E-07 | 1.270E-07 | 2.000E-07 | 4.200E-07 |
| 23477 | 1174 | F1174T | D | 3.2e-07 | 2.6e-07 | 3.2e-07 | 3.000E-07 | 3.464E-08 | 2.600E-07 | 3.200E-07 |
| 23478 | 1174 | F1174W | D | 6e-07 | 2.8e-07 | 5.7e-07 | 4.833E-07 | 1.767E-07 | 2.800E-07 | 6.000E-07 |
| 23479 | 1174 | F1174Y | D | 4.7e-07 | 1.4e-07 | 1.6e-07 | 2.567E-07 | 1.850E-07 | 1.400E-07 | 4.700E-07 |
| 23480 | 1174 | F1174V | D | 2.8e-07 | 3.2e-07 | 2e-07 | 2.667E-07 | 6.110E-08 | 2.000E-07 | 3.200E-07 |
| 23481 | 1175 | I1175A | D | 1.3e-07 | 2.8e-07 | 2.7e-07 | 2.267E-07 | 8.386E-08 | 1.300E-07 | 2.800E-07 |
| 23482 | 1175 | I1175R | D | 1.3e-07 | 2.7e-07 | 2.5e-07 | 2.167E-07 | 7.572E-08 | 1.300E-07 | 2.700E-07 |
| 23483 | 1175 | I1175N | D | 3.9e-07 | 3.7e-07 | 8.2e-08 | 2.807E-07 | 1.723E-07 | 8.200E-08 | 3.900E-07 |
| 23484 | 1175 | I1175D | D | 1.9e-07 | 2.1e-07 | 1.8e-07 | 1.933E-07 | 1.528E-08 | 1.800E-07 | 2.100E-07 |
| 23485 | 1175 | I1175C | D | 2.5e-07 | 3.5e-07 | 2.9e-07 | 2.967E-07 | 5.033E-08 | 2.500E-07 | 3.500E-07 |
| 23486 | 1175 | I1175E | D | 3.2e-07 | 4.5e-07 | 2.3e-07 | 3.333E-07 | 1.106E-07 | 2.300E-07 | 4.500E-07 |
| 23487 | 1175 | I1175Q | D | 1.7e-07 | 2.5e-07 | 9.5e-08 | 1.717E-07 | 7.751E-08 | 9.500E-08 | 2.500E-07 |
| 23488 | 1175 | I1175G | D | 2.7e-07 | 2.3e-07 | 2e-07 | 2.333E-07 | 3.512E-08 | 2.000E-07 | 2.700E-07 |
| 23489 | 1175 | I1175H | D | 4.6e-07 | 2.5e-07 | 1.6e-07 | 2.900E-07 | 1.539E-07 | 1.600E-07 | 4.600E-07 |
| 23490 | 1175 | I1175I | D | 3.5e-07 | 2.9e-07 | 3.3e-07 | 3.233E-07 | 3.055E-08 | 2.900E-07 | 3.500E-07 |
| 23491 | 1175 | I1175L | D | 1.8e-07 | 4.4e-07 | 2.2e-07 | 2.800E-07 | 1.400E-07 | 1.800E-07 | 4.400E-07 |
| 23492 | 1175 | I1175K | D | 2.9e-07 | 1.3e-07 | 1.4e-07 | 1.867E-07 | 8.963E-08 | 1.300E-07 | 2.900E-07 |
| 23493 | 1175 | I1175M | D | 1e-07 | 4.8e-07 | 2.4e-07 | 2.733E-07 | 1.922E-07 | 1.000E-07 | 4.800E-07 |
| 23494 | 1175 | I1175F | D | 1.5e-07 | 1.6e-07 | 1.2e-07 | 1.433E-07 | 2.082E-08 | 1.200E-07 | 1.600E-07 |
| 23495 | 1175 | I1175P | D | 2.5e-07 | 8.3e-07 | 1.7e-07 | 4.167E-07 | 3.602E-07 | 1.700E-07 | 8.300E-07 |
| 23496 | 1175 | I1175S | D | 1.4e-07 | 2.5e-07 | 3.7e-07 | 2.533E-07 | 1.150E-07 | 1.400E-07 | 3.700E-07 |
| 23497 | 1175 | I1175T | D | 4e-07 | 2.9e-07 | 3.2e-07 | 3.367E-07 | 5.686E-08 | 2.900E-07 | 4.000E-07 |
| 23498 | 1175 | I1175W | D | 1.5e-07 | 2e-07 | 2.8e-07 | 2.100E-07 | 6.557E-08 | 1.500E-07 | 2.800E-07 |
| 23499 | 1175 | I1175Y | D | 2.7e-07 | 2.2e-07 | 3.1e-07 | 2.667E-07 | 4.509E-08 | 2.200E-07 | 3.100E-07 |
| 23500 | 1175 | I1175V | D | 2.1e-07 | 5.5e-07 | 1.8e-07 | 3.133E-07 | 2.055E-07 | 1.800E-07 | 5.500E-07 |
| 23501 | 1176 | F1176A | D | 3.6e-07 | 3.4e-07 | 2.9e-07 | 3.300E-07 | 3.606E-08 | 2.900E-07 | 3.600E-07 |
| 23502 | 1176 | F1176R | D | 3.5e-07 | 3e-07 | 4.2e-07 | 3.567E-07 | 6.028E-08 | 3.000E-07 | 4.200E-07 |
| 23503 | 1176 | F1176N | D | 1.9e-07 | 1.6e-07 | 1.4e-07 | 1.633E-07 | 2.517E-08 | 1.400E-07 | 1.900E-07 |
| 23504 | 1176 | F1176D | D | 7e-08 | 4e-07 | 1.6e-07 | 2.100E-07 | 1.706E-07 | 7.000E-08 | 4.000E-07 |
| 23505 | 1176 | F1176C | D | 6.2e-07 | 2.7e-07 | 2.3e-07 | 3.733E-07 | 2.146E-07 | 2.300E-07 | 6.200E-07 |
| 23506 | 1176 | F1176E | D | 3.4e-07 | 3.3e-07 | 4.6e-07 | 3.767E-07 | 7.234E-08 | 3.300E-07 | 4.600E-07 |
| 23507 | 1176 | F1176Q | D | 5.3e-07 | 1.2e-07 | 1.8e-07 | 2.767E-07 | 2.214E-07 | 1.200E-07 | 5.300E-07 |
| 23508 | 1176 | F1176G | D | 5e-07 | 2.6e-07 | 2.4e-07 | 3.333E-07 | 1.447E-07 | 2.400E-07 | 5.000E-07 |
| 23509 | 1176 | F1176H | D | 2.6e-07 | 1.9e-07 | 3.2e-07 | 2.567E-07 | 6.506E-08 | 1.900E-07 | 3.200E-07 |
| 23510 | 1176 | F1176I | D | 5.6e-07 | 2.7e-07 | 2.7e-07 | 3.667E-07 | 1.674E-07 | 2.700E-07 | 5.600E-07 |
| 23511 | 1176 | F1176L | D | 2.8e-07 | 2.1e-07 | 3.1e-07 | 2.667E-07 | 5.132E-08 | 2.100E-07 | 3.100E-07 |
| 23512 | 1176 | F1176K | D | 1.6e-07 | 4.3e-07 | 1.8e-07 | 2.567E-07 | 1.504E-07 | 1.600E-07 | 4.300E-07 |
| 23513 | 1176 | F1176M | D | 2.7e-07 | 2e-07 | 3.3e-07 | 2.667E-07 | 6.506E-08 | 2.000E-07 | 3.300E-07 |
| 23514 | 1176 | F1176F | D | 2.6e-07 | 1.4e-07 | 2.1e-07 | 2.033E-07 | 6.028E-08 | 1.400E-07 | 2.600E-07 |
| 23515 | 1176 | F1176P | D | 3.6e-07 | 2.2e-07 | 2.3e-07 | 2.700E-07 | 7.810E-08 | 2.200E-07 | 3.600E-07 |
| 23516 | 1176 | F1176S | D | 1.2e-07 | 2.7e-07 | 3.4e-07 | 2.433E-07 | 1.124E-07 | 1.200E-07 | 3.400E-07 |
| 23517 | 1176 | F1176T | D | 2.7e-07 | 3.2e-07 | 3.5e-07 | 3.133E-07 | 4.041E-08 | 2.700E-07 | 3.500E-07 |
| 23518 | 1176 | F1176W | D | 3.6e-07 | 2.3e-07 | 5.2e-07 | 3.700E-07 | 1.453E-07 | 2.300E-07 | 5.200E-07 |
| 23519 | 1176 | F1176Y | D | 4.7e-07 | 1.6e-07 | 3.3e-07 | 3.200E-07 | 1.552E-07 | 1.600E-07 | 4.700E-07 |
| 23520 | 1176 | F1176V | D | 6.5e-07 | 2.1e-07 | 1.1e-07 | 3.233E-07 | 2.873E-07 | 1.100E-07 | 6.500E-07 |
| 23521 | 1177 | P1177A | D | 1.6e-07 | 4.7e-07 | 3.4e-07 | 3.233E-07 | 1.557E-07 | 1.600E-07 | 4.700E-07 |
| 23522 | 1177 | P1177R | D | 2.4e-07 | 1.7e-07 | 1.7e-07 | 1.933E-07 | 4.041E-08 | 1.700E-07 | 2.400E-07 |
| 23523 | 1177 | P1177N | D | 3.7e-07 | 2.1e-07 | 3.1e-07 | 2.967E-07 | 8.083E-08 | 2.100E-07 | 3.700E-07 |
| 23524 | 1177 | P1177D | D | 2.9e-07 | 3.7e-07 | 4e-07 | 3.533E-07 | 5.686E-08 | 2.900E-07 | 4.000E-07 |
| 23525 | 1177 | P1177C | D | 2.2e-07 | 3.8e-07 | 3.5e-07 | 3.167E-07 | 8.505E-08 | 2.200E-07 | 3.800E-07 |
| 23526 | 1177 | P1177E | D | 1.8e-07 | 1.6e-07 | 1.8e-07 | 1.733E-07 | 1.155E-08 | 1.600E-07 | 1.800E-07 |
| 23527 | 1177 | P1177Q | D | 1.2e-07 | 2.5e-07 | 1.4e-07 | 1.700E-07 | 7.000E-08 | 1.200E-07 | 2.500E-07 |
| 23528 | 1177 | P1177G | D | 1.9e-07 | 1.4e-07 | 1.3e-07 | 1.533E-07 | 3.215E-08 | 1.300E-07 | 1.900E-07 |
| 23529 | 1177 | P1177H | D | 1.7e-07 | 2.7e-07 | 2.8e-07 | 2.400E-07 | 6.083E-08 | 1.700E-07 | 2.800E-07 |
| 23530 | 1177 | P1177I | D | 1.9e-07 | 3.5e-07 | 2.4e-07 | 2.600E-07 | 8.185E-08 | 1.900E-07 | 3.500E-07 |
| 23531 | 1177 | P1177L | D | 2.4e-07 | 2.6e-07 | 1.5e-07 | 2.167E-07 | 5.859E-08 | 1.500E-07 | 2.600E-07 |
| 23532 | 1177 | P1177K | D | 2.3e-07 | 2.1e-07 | 2.7e-07 | 2.367E-07 | 3.055E-08 | 2.100E-07 | 2.700E-07 |
| 23533 | 1177 | P1177M | D | 3.8e-07 | 2.8e-07 | 3.1e-07 | 3.233E-07 | 5.132E-08 | 2.800E-07 | 3.800E-07 |
| 23534 | 1177 | P1177F | D | 1e-07 | 2.5e-07 | 2.3e-07 | 1.933E-07 | 8.145E-08 | 1.000E-07 | 2.500E-07 |
| 23535 | 1177 | P1177P | D | 5.3e-07 | 6.5e-07 | 3e-07 | 4.933E-07 | 1.779E-07 | 3.000E-07 | 6.500E-07 |
| 23536 | 1177 | P1177S | D | 3.5e-07 | 2.4e-07 | 1.9e-07 | 2.600E-07 | 8.185E-08 | 1.900E-07 | 3.500E-07 |
| 23537 | 1177 | P1177T | D | 4.6e-07 | 5.6e-07 | 3.3e-07 | 4.500E-07 | 1.153E-07 | 3.300E-07 | 5.600E-07 |
| 23538 | 1177 | P1177W | D | 2.7e-07 | 3.1e-07 | 2.5e-07 | 2.767E-07 | 3.055E-08 | 2.500E-07 | 3.100E-07 |
| 23539 | 1177 | P1177Y | D | 3.7e-07 | 1.4e-07 | 1.6e-07 | 2.233E-07 | 1.274E-07 | 1.400E-07 | 3.700E-07 |
| 23540 | 1177 | P1177V | D | 2.1e-07 | 2e-07 | 2.5e-07 | 2.200E-07 | 2.646E-08 | 2.000E-07 | 2.500E-07 |
| 23541 | 1178 | P1178A | D | 1.9e-07 | 3.1e-07 | 4e-07 | 3.000E-07 | 1.054E-07 | 1.900E-07 | 4.000E-07 |
| 23542 | 1178 | P1178R | D | 2.8e-07 | 2.4e-07 | 2e-07 | 2.400E-07 | 4.000E-08 | 2.000E-07 | 2.800E-07 |
| 23543 | 1178 | P1178N | D | 2.2e-07 | 1.5e-07 | 3.7e-07 | 2.467E-07 | 1.124E-07 | 1.500E-07 | 3.700E-07 |
| 23544 | 1178 | P1178D | D | 3.3e-07 | 4.9e-07 | 2.8e-07 | 3.667E-07 | 1.097E-07 | 2.800E-07 | 4.900E-07 |

| | | | | | | | | | | |
|---|---|---|---|---|---|---|---|---|---|---|
| 23545 | 1178 | P1178C | D | 2.9e-07 | 2.2e-07 | 2.9e-07 | 2.667E-07 | 4.041E-08 | 2.200E-07 | 2.900E-07 |
| 23546 | 1178 | P1178E | D | 1.1e-07 | 3.3e-07 | 2.5e-07 | 2.300E-07 | 1.114E-07 | 1.100E-07 | 3.300E-07 |
| 23547 | 1178 | P1178Q | D | 3.2e-07 | 2.7e-07 | 1.9e-07 | 2.600E-07 | 6.557E-08 | 1.900E-07 | 3.200E-07 |
| 23548 | 1178 | P1178G | D | 1.6e-07 | 2.3e-07 | 2.5e-07 | 2.133E-07 | 4.726E-08 | 1.600E-07 | 2.500E-07 |
| 23549 | 1178 | P1178H | D | 3.3e-07 | 2.8e-07 | 3.3e-07 | 3.133E-07 | 2.887E-08 | 2.800E-07 | 3.300E-07 |
| 23550 | 1178 | P1178I | D | 1.6e-07 | 2.8e-07 | 1.9e-07 | 2.100E-07 | 6.245E-08 | 1.600E-07 | 2.800E-07 |
| 23551 | 1178 | P1178L | D | 1.6e-07 | 3.7e-07 | 2.2e-07 | 2.500E-07 | 1.082E-07 | 1.600E-07 | 3.700E-07 |
| 23552 | 1178 | P1178K | D | 1.1e-07 | 3.7e-07 | 2.1e-07 | 2.300E-07 | 1.311E-07 | 1.100E-07 | 3.700E-07 |
| 23553 | 1178 | P1178M | D | 2.3e-07 | 3.5e-07 | 2e-07 | 2.600E-07 | 7.937E-08 | 2.000E-07 | 3.500E-07 |
| 23554 | 1178 | P1178F | D | 2.3e-07 | 1.9e-07 | 1.7e-07 | 1.967E-07 | 3.055E-08 | 1.700E-07 | 2.300E-07 |
| 23555 | 1178 | P1178P | D | 3.2e-07 | 2.4e-07 | 3.6e-07 | 3.067E-07 | 6.110E-08 | 2.400E-07 | 3.600E-07 |
| 23556 | 1178 | P1178S | D | 3.6e-07 | 1.3e-07 | 1.4e-07 | 2.100E-07 | 1.300E-07 | 1.300E-07 | 3.600E-07 |
| 23557 | 1178 | P1178T | D | 1.8e-07 | 2.3e-07 | 2.8e-07 | 2.300E-07 | 5.000E-08 | 1.800E-07 | 2.800E-07 |
| 23558 | 1178 | P1178W | D | 2.5e-07 | 3.2e-07 | 3.7e-07 | 3.133E-07 | 6.028E-08 | 2.500E-07 | 3.700E-07 |
| 23559 | 1178 | P1178Y | D | 6.7e-07 | 2.7e-07 | 2.8e-07 | 4.067E-07 | 2.281E-07 | 2.700E-07 | 6.700E-07 |
| 23560 | 1178 | P1178V | D | 5.1e-07 | 3.4e-07 | 1.4e-07 | 3.300E-07 | 1.852E-07 | 1.400E-07 | 5.100E-07 |
| 23561 | 1179 | S1179A | D | 5e-07 | 2.5e-07 | 7.5e-08 | 2.750E-07 | 2.136E-07 | 7.500E-08 | 5.000E-07 |
| 23562 | 1179 | S1179R | D | 2.1e-07 | 2.9e-07 | 2.6e-07 | 2.533E-07 | 4.041E-08 | 2.100E-07 | 2.900E-07 |
| 23563 | 1179 | S1179N | D | 1.1e-07 | 1.7e-07 | 3e-07 | 1.933E-07 | 9.713E-08 | 1.100E-07 | 3.000E-07 |
| 23564 | 1179 | S1179D | D | 7.3e-07 | 2.2e-07 | 3e-07 | 4.167E-07 | 2.743E-07 | 2.200E-07 | 7.300E-07 |
| 23565 | 1179 | S1179C | D | 3e-07 | 2.5e-07 | 1.4e-07 | 2.300E-07 | 8.185E-08 | 1.400E-07 | 3.000E-07 |
| 23566 | 1179 | S1179E | D | 3.9e-07 | 2e-07 | 4.7e-07 | 3.533E-07 | 1.387E-07 | 2.000E-07 | 4.700E-07 |
| 23567 | 1179 | S1179Q | D | 2.7e-07 | 2.4e-07 | 4.2e-07 | 3.100E-07 | 9.644E-08 | 2.400E-07 | 4.200E-07 |
| 23568 | 1179 | S1179G | D | 3.5e-07 | 1.9e-07 | 3.4e-07 | 2.933E-07 | 8.963E-08 | 1.900E-07 | 3.500E-07 |
| 23569 | 1179 | S1179H | D | 2.7e-07 | 2.4e-07 | 3e-07 | 2.700E-07 | 3.000E-08 | 2.400E-07 | 3.000E-07 |
| 23570 | 1179 | S1179I | D | 2.8e-07 | 1.9e-07 | 1.4e-07 | 2.033E-07 | 7.095E-08 | 1.400E-07 | 2.800E-07 |
| 23571 | 1179 | S1179L | D | 2.8e-07 | 3.6e-07 | 1.8e-07 | 2.733E-07 | 9.018E-08 | 1.800E-07 | 3.600E-07 |
| 23572 | 1179 | S1179K | D | 3.4e-07 | 1.1e-07 | 2.1e-07 | 2.200E-07 | 1.153E-07 | 1.100E-07 | 3.400E-07 |
| 23573 | 1179 | S1179M | D | 2.3e-07 | 1.8e-07 | 1.9e-07 | 2.000E-07 | 2.646E-08 | 1.800E-07 | 2.300E-07 |
| 23574 | 1179 | S1179F | D | 2.3e-07 | 1.4e-07 | 2.4e-07 | 2.033E-07 | 5.508E-08 | 1.400E-07 | 2.400E-07 |
| 23575 | 1179 | S1179P | D | 5.9e-07 | 2.1e-07 | 2.4e-07 | 3.467E-07 | 2.113E-07 | 2.100E-07 | 5.900E-07 |
| 23576 | 1179 | S1179S | D | 1.8e-07 | 3.3e-07 | 2.7e-07 | 2.600E-07 | 7.550E-08 | 1.800E-07 | 3.300E-07 |
| 23577 | 1179 | S1179T | D | 3.8e-07 | 2.7e-07 | 2.1e-07 | 2.867E-07 | 8.622E-08 | 2.100E-07 | 3.800E-07 |
| 23578 | 1179 | S1179W | D | 3.3e-07 | 2.7e-07 | 2.6e-07 | 2.867E-07 | 3.786E-08 | 2.600E-07 | 3.300E-07 |
| 23579 | 1179 | S1179Y | D | 1.8e-07 | 1.6e-07 | 1.2e-07 | 1.533E-07 | 3.055E-08 | 1.200E-07 | 1.800E-07 |
| 23580 | 1179 | S1179V | D | 1.2e-07 | 2.1e-07 | 2.4e-07 | 1.900E-07 | 6.245E-08 | 1.200E-07 | 2.400E-07 |
| 23581 | 1180 | D1180A | D | 7.9e-08 | 1.3e-07 | 2.2e-07 | 1.430E-07 | 7.139E-08 | 7.900E-08 | 2.200E-07 |
| 23582 | 1180 | D1180R | D | 2.3e-07 | 1.9e-07 | 1.8e-07 | 2.000E-07 | 2.646E-08 | 1.800E-07 | 2.300E-07 |
| 23583 | 1180 | D1180N | D | 2.8e-07 | 6e-07 | 2.3e-07 | 3.700E-07 | 2.007E-07 | 2.300E-07 | 6.000E-07 |
| 23584 | 1180 | D1180D | D | 2.1e-07 | 3.2e-07 | 2.7e-07 | 2.667E-07 | 5.508E-08 | 2.100E-07 | 3.200E-07 |
| 23585 | 1180 | D1180C | D | 1.9e-07 | 2e-07 | 2.8e-07 | 2.233E-07 | 4.933E-08 | 1.900E-07 | 2.800E-07 |
| 23586 | 1180 | D1180E | D | 2.7e-07 | 3.1e-07 | 2.3e-07 | 2.700E-07 | 4.000E-08 | 2.300E-07 | 3.100E-07 |
| 23587 | 1180 | D1180Q | D | 1.8e-07 | 3.4e-07 | 2.1e-07 | 2.433E-07 | 8.505E-08 | 1.800E-07 | 3.400E-07 |
| 23588 | 1180 | D1180G | D | 3.4e-07 | 1.2e-07 | 1.8e-07 | 2.133E-07 | 1.137E-07 | 1.200E-07 | 3.400E-07 |
| 23589 | 1180 | D1180H | D | 2.3e-07 | 2.7e-07 | 1.3e-07 | 2.100E-07 | 7.211E-08 | 1.300E-07 | 2.700E-07 |
| 23590 | 1180 | D1180I | D | 2.1e-07 | 2e-07 | 2.3e-07 | 2.133E-07 | 1.528E-08 | 2.000E-07 | 2.300E-07 |
| 23591 | 1180 | D1180L | D | 2.1e-07 | 6.7e-07 | 2.6e-07 | 3.800E-07 | 2.524E-07 | 2.100E-07 | 6.700E-07 |
| 23592 | 1180 | D1180K | D | 1.7e-07 | 1.9e-07 | 1.6e-07 | 1.733E-07 | 1.528E-08 | 1.600E-07 | 1.900E-07 |
| 23593 | 1180 | D1180M | D | 3.4e-07 | 3.3e-07 | 5.1e-07 | 3.933E-07 | 1.012E-07 | 3.300E-07 | 5.100E-07 |
| 23594 | 1180 | D1180F | D | 2.3e-07 | 1.7e-07 | 3.3e-07 | 2.433E-07 | 8.083E-08 | 1.700E-07 | 3.300E-07 |
| 23595 | 1180 | D1180P | D | 1.8e-07 | 1.7e-07 | 2.5e-07 | 2.000E-07 | 4.359E-08 | 1.700E-07 | 2.500E-07 |
| 23596 | 1180 | D1180S | D | 2e-07 | 1.7e-07 | 4.1e-07 | 2.600E-07 | 1.308E-07 | 1.700E-07 | 4.100E-07 |
| 23597 | 1180 | D1180T | D | 3e-07 | 2.5e-07 | 3.2e-07 | 2.900E-07 | 3.606E-08 | 2.500E-07 | 3.200E-07 |
| 23598 | 1180 | D1180W | D | 1.2e-07 | 3.9e-07 | 1.6e-07 | 2.233E-07 | 1.457E-07 | 1.200E-07 | 3.900E-07 |
| 23599 | 1180 | D1180Y | D | 1.7e-07 | 2.2e-07 | 3.6e-07 | 2.500E-07 | 9.849E-08 | 1.700E-07 | 3.600E-07 |
| 23600 | 1180 | D1180V | D | 4.4e-07 | 3.2e-07 | 2.1e-07 | 3.233E-07 | 1.150E-07 | 2.100E-07 | 4.400E-07 |
| 23601 | 1181 | E1181A | D | 5.1e-07 | 2.4e-07 | 1.8e-07 | 3.100E-07 | 1.758E-07 | 1.800E-07 | 5.100E-07 |
| 23602 | 1181 | E1181R | D | 2.5e-07 | 2.6e-07 | 4.5e-07 | 3.200E-07 | 1.127E-07 | 2.500E-07 | 4.500E-07 |
| 23603 | 1181 | E1181N | D | 3.2e-07 | 5.5e-07 | 2.2e-07 | 3.633E-07 | 1.692E-07 | 2.200E-07 | 5.500E-07 |
| 23604 | 1181 | E1181D | D | 1.9e-07 | 1.4e-07 | 2e-07 | 1.767E-07 | 3.215E-08 | 1.400E-07 | 2.000E-07 |
| 23605 | 1181 | E1181C | D | 2.2e-07 | 3.3e-07 | 8.1e-08 | 2.103E-07 | 1.248E-07 | 8.100E-08 | 3.300E-07 |
| 23606 | 1181 | E1181E | D | 1.9e-07 | 2.2e-07 | 3e-07 | 2.367E-07 | 5.686E-08 | 1.900E-07 | 3.000E-07 |
| 23607 | 1181 | E1181Q | D | 1.5e-07 | 4.8e-07 | 2.3e-07 | 2.867E-07 | 1.721E-07 | 1.500E-07 | 4.800E-07 |
| 23608 | 1181 | E1181G | D | 1.9e-07 | 2.2e-07 | 2.8e-07 | 2.300E-07 | 4.583E-08 | 1.900E-07 | 2.800E-07 |
| 23609 | 1181 | E1181H | D | 2e-07 | 1.1e-07 | 2.7e-07 | 1.933E-07 | 8.021E-08 | 1.100E-07 | 2.700E-07 |
| 23610 | 1181 | E1181I | D | 1.8e-07 | 2e-07 | 1.8e-07 | 1.867E-07 | 1.155E-08 | 1.800E-07 | 2.000E-07 |
| 23611 | 1181 | E1181L | D | 2.1e-07 | 2.6e-07 | 1.7e-07 | 2.133E-07 | 4.509E-08 | 1.700E-07 | 2.600E-07 |
| 23612 | 1181 | E1181K | D | 1.2e-07 | 1.5e-07 | 2.4e-07 | 1.700E-07 | 6.245E-08 | 1.200E-07 | 2.400E-07 |
| 23613 | 1181 | E1181M | D | 2.3e-07 | 2.1e-07 | 2.4e-07 | 2.267E-07 | 1.528E-08 | 2.100E-07 | 2.400E-07 |
| 23614 | 1181 | E1181F | D | 3.9e-07 | 2.8e-07 | 5e-07 | 3.900E-07 | 1.100E-07 | 2.800E-07 | 5.000E-07 |
| 23615 | 1181 | E1181P | D | 1.2e-07 | 1.9e-07 | 2.3e-07 | 1.800E-07 | 5.568E-08 | 1.200E-07 | 2.300E-07 |
| 23616 | 1181 | E1181S | D | 1.5e-07 | 3.2e-07 | 1.4e-07 | 2.033E-07 | 1.012E-07 | 1.400E-07 | 3.200E-07 |
| 23617 | 1181 | E1181T | D | 4e-07 | 2.1e-07 | 1.5e-07 | 2.533E-07 | 1.305E-07 | 1.500E-07 | 4.000E-07 |
| 23618 | 1181 | E1181W | D | 1.7e-07 | 4e-07 | 1.6e-07 | 2.433E-07 | 1.358E-07 | 1.600E-07 | 4.000E-07 |
| 23619 | 1181 | E1181Y | D | 3.3e-07 | 1.6e-07 | 3.7e-07 | 2.867E-07 | 1.115E-07 | 1.600E-07 | 3.700E-07 |
| 23620 | 1181 | E1181V | D | 1.4e-07 | 2.1e-07 | 3.4e-07 | 2.300E-07 | 1.015E-07 | 1.400E-07 | 3.400E-07 |
| 23621 | 1182 | Q1182A | D | 4e-07 | 4.5e-07 | 5e-07 | 4.500E-07 | 5.000E-08 | 4.000E-07 | 5.000E-07 |
| 23622 | 1182 | Q1182R | D | 3.8e-07 | 4.7e-07 | 7.8e-08 | 3.093E-07 | 2.053E-07 | 7.800E-08 | 4.700E-07 |
| 23623 | 1182 | Q1182N | D | 2e-07 | 1.8e-07 | 1.5e-07 | 1.767E-07 | 2.517E-08 | 1.500E-07 | 2.000E-07 |
| 23624 | 1182 | Q1182D | D | 2.8e-07 | 1.7e-07 | 4.7e-07 | 3.067E-07 | 1.518E-07 | 1.700E-07 | 4.700E-07 |
| 23625 | 1182 | Q1182C | D | 3.2e-07 | 3.4e-07 | 4.2e-07 | 3.600E-07 | 5.292E-08 | 3.200E-07 | 4.200E-07 |
| 23626 | 1182 | Q1182E | D | 2.7e-07 | 1.6e-07 | 2.8e-07 | 2.367E-07 | 6.658E-08 | 1.600E-07 | 2.800E-07 |
| 23627 | 1182 | Q1182Q | D | 3e-07 | 1.3e-07 | 2.6e-07 | 2.300E-07 | 8.888E-08 | 1.300E-07 | 3.000E-07 |
| 23628 | 1182 | Q1182G | D | 1.3e-07 | 1.5e-07 | 3.3e-07 | 2.033E-07 | 1.102E-07 | 1.300E-07 | 3.300E-07 |
| 23629 | 1182 | Q1182H | D | 2.3e-07 | 2.2e-07 | 8.9e-08 | 1.797E-07 | 7.868E-08 | 8.900E-08 | 2.300E-07 |

| | | | | | | | | | | |
|---|---|---|---|---|---|---|---|---|---|---|
| 23630 | 1182 | Q1182I | D | 1.6e-07 | 3.7e-07 | 5.1e-07 | 3.467E-07 | 1.762E-07 | 1.600E-07 | 5.100E-07 |
| 23631 | 1182 | Q1182L | D | 2.3e-07 | 2.9e-07 | 2.3e-07 | 2.500E-07 | 3.464E-08 | 2.300E-07 | 2.900E-07 |
| 23632 | 1182 | Q1182K | D | 3.5e-07 | 2.9e-07 | 2.2e-07 | 2.867E-07 | 6.506E-08 | 2.200E-07 | 3.500E-07 |
| 23633 | 1182 | Q1182M | D | 3.6e-07 | 1e-07 | 2.3e-07 | 2.300E-07 | 1.300E-07 | 1.000E-07 | 3.600E-07 |
| 23634 | 1182 | Q1182F | D | 3.5e-07 | 1.3e-07 | 1.9e-07 | 2.233E-07 | 1.137E-07 | 1.300E-07 | 3.500E-07 |
| 23635 | 1182 | Q1182P | D | 2.2e-07 | 1.4e-07 | 2.5e-07 | 2.033E-07 | 5.686E-08 | 1.400E-07 | 2.500E-07 |
| 23636 | 1182 | Q1182S | D | 2.5e-07 | 3.5e-07 | 2.3e-07 | 2.767E-07 | 6.429E-08 | 2.300E-07 | 3.500E-07 |
| 23637 | 1182 | Q1182T | D | 3.4e-07 | 3e-07 | 2.9e-07 | 3.100E-07 | 2.646E-08 | 2.900E-07 | 3.400E-07 |
| 23638 | 1182 | Q1182W | D | 2e-07 | 3.8e-07 | 4.6e-07 | 3.467E-07 | 1.332E-07 | 2.000E-07 | 4.600E-07 |
| 23639 | 1182 | Q1182Y | D | 2e-07 | 2.6e-07 | 2.7e-07 | 2.433E-07 | 3.786E-08 | 2.000E-07 | 2.700E-07 |
| 23640 | 1182 | Q1182V | D | 3.1e-07 | 2.1e-07 | 1.7e-07 | 2.300E-07 | 7.211E-08 | 1.700E-07 | 3.100E-07 |
| 23641 | 1183 | L1183A | D | 1.2e-07 | 3.2e-07 | 1.2e-07 | 1.867E-07 | 1.155E-07 | 1.200E-07 | 3.200E-07 |
| 23642 | 1183 | L1183R | D | 4.3e-07 | 1.1e-07 | 1.4e-07 | 2.267E-07 | 1.767E-07 | 1.100E-07 | 4.300E-07 |
| 23643 | 1183 | L1183N | D | 1.5e-07 | 2.5e-07 | 2.3e-07 | 2.100E-07 | 5.292E-08 | 1.500E-07 | 2.500E-07 |
| 23644 | 1183 | L1183D | D | 5.6e-07 | 3.9e-07 | 3e-07 | 4.167E-07 | 1.320E-07 | 3.000E-07 | 5.600E-07 |
| 23645 | 1183 | L1183C | D | 1.3e-07 | 2.2e-07 | 1.6e-07 | 1.700E-07 | 4.583E-08 | 1.300E-07 | 2.200E-07 |
| 23646 | 1183 | L1183E | D | 1.3e-07 | 2.6e-07 | 3.5e-07 | 2.467E-07 | 1.106E-07 | 1.300E-07 | 3.500E-07 |
| 23647 | 1183 | L1183Q | D | 7.4e-07 | 2.6e-07 | 2.2e-07 | 4.067E-07 | 2.894E-07 | 2.200E-07 | 7.400E-07 |
| 23648 | 1183 | L1183G | D | 3.7e-07 | 6.6e-07 | 1e-07 | 3.767E-07 | 2.801E-07 | 1.000E-07 | 6.600E-07 |
| 23649 | 1183 | L1183H | D | 1.5e-07 | 4.2e-07 | 3.7e-07 | 3.133E-07 | 1.436E-07 | 1.500E-07 | 4.200E-07 |
| 23650 | 1183 | L1183I | D | 4.3e-07 | 3.3e-07 | 3.3e-07 | 3.633E-07 | 5.774E-08 | 3.300E-07 | 4.300E-07 |
| 23651 | 1183 | L1183L | D | 1.8e-07 | 3.7e-07 | 4.1e-07 | 3.200E-07 | 1.229E-07 | 1.800E-07 | 4.100E-07 |
| 23652 | 1183 | L1183K | D | 2.2e-07 | 2.6e-07 | 1.6e-07 | 2.133E-07 | 5.033E-08 | 1.600E-07 | 2.600E-07 |
| 23653 | 1183 | L1183M | D | 2.2e-07 | 2e-07 | 3.5e-07 | 2.567E-07 | 8.145E-08 | 2.000E-07 | 3.500E-07 |
| 23654 | 1183 | L1183F | D | 1.9e-07 | 3.5e-07 | 1.8e-07 | 2.400E-07 | 9.539E-08 | 1.800E-07 | 3.500E-07 |
| 23655 | 1183 | L1183P | D | 4.2e-07 | 4.2e-07 | 3.4e-07 | 3.933E-07 | 4.619E-08 | 3.400E-07 | 4.200E-07 |
| 23656 | 1183 | L1183S | D | 2.7e-07 | 1.7e-07 | 2.2e-07 | 2.200E-07 | 5.000E-08 | 1.700E-07 | 2.700E-07 |
| 23657 | 1183 | L1183T | D | 2.1e-07 | 2.1e-07 | 3.3e-07 | 2.500E-07 | 6.928E-08 | 2.100E-07 | 3.300E-07 |
| 23658 | 1183 | L1183W | D | 2.2e-07 | 3.3e-07 | 2.6e-07 | 2.700E-07 | 5.568E-08 | 2.200E-07 | 3.300E-07 |
| 23659 | 1183 | L1183Y | D | 2.5e-07 | 1.7e-07 | 2.2e-07 | 2.133E-07 | 4.041E-08 | 1.700E-07 | 2.500E-07 |
| 23660 | 1183 | L1183V | D | 1.2e-07 | 1.8e-07 | 2.1e-07 | 1.700E-07 | 4.583E-08 | 1.200E-07 | 2.100E-07 |
| 23661 | 1184 | K1184A | D | 1.2e-07 | 3.1e-07 | 2.2e-07 | 2.167E-07 | 9.504E-08 | 1.200E-07 | 3.100E-07 |
| 23662 | 1184 | K1184R | D | 2.3e-07 | 2.5e-07 | 5.4e-07 | 3.400E-07 | 1.735E-07 | 2.300E-07 | 5.400E-07 |
| 23663 | 1184 | K1184N | D | 1.9e-07 | 1.2e-07 | 1.5e-07 | 1.533E-07 | 3.512E-08 | 1.200E-07 | 1.900E-07 |
| 23664 | 1184 | K1184D | D | 1.8e-07 | 2.7e-07 | 1.3e-07 | 1.933E-07 | 7.095E-08 | 1.300E-07 | 2.700E-07 |
| 23665 | 1184 | K1184C | D | 2.6e-07 | 2.2e-07 | 1.8e-07 | 2.200E-07 | 4.000E-08 | 1.800E-07 | 2.600E-07 |
| 23666 | 1184 | K1184E | D | 2.7e-07 | 2.9e-07 | 1.1e-07 | 2.233E-07 | 9.866E-08 | 1.100E-07 | 2.900E-07 |
| 23667 | 1184 | K1184Q | D | 2.2e-07 | 1.6e-07 | 1.5e-07 | 1.767E-07 | 3.786E-08 | 1.500E-07 | 2.200E-07 |
| 23668 | 1184 | K1184G | D | 1.2e-07 | 5.7e-07 | 3.7e-07 | 3.533E-07 | 2.255E-07 | 1.200E-07 | 5.700E-07 |
| 23669 | 1184 | K1184H | D | 1.1e-07 | 1.9e-07 | 3e-07 | 2.000E-07 | 9.539E-08 | 1.100E-07 | 3.000E-07 |
| 23670 | 1184 | K1184I | D | 2.7e-07 | 6.8e-07 | 1.8e-07 | 3.767E-07 | 2.665E-07 | 1.800E-07 | 6.800E-07 |
| 23671 | 1184 | K1184L | D | 2.3e-07 | 2.8e-07 | 4.5e-07 | 3.200E-07 | 1.153E-07 | 2.300E-07 | 4.500E-07 |
| 23672 | 1184 | K1184K | D | 2e-07 | 1.7e-07 | 3.2e-07 | 2.300E-07 | 7.937E-08 | 1.700E-07 | 3.200E-07 |
| 23673 | 1184 | K1184M | D | 2.3e-07 | 1.1e-07 | 1.8e-07 | 1.733E-07 | 6.028E-08 | 1.100E-07 | 2.300E-07 |
| 23674 | 1184 | K1184F | D | 2.4e-07 | 2.3e-07 | 2.4e-07 | 2.367E-07 | 5.774E-09 | 2.300E-07 | 2.400E-07 |
| 23675 | 1184 | K1184P | D | 4.5e-07 | 2.4e-07 | 2e-07 | 2.967E-07 | 1.343E-07 | 2.000E-07 | 4.500E-07 |
| 23676 | 1184 | K1184S | D | 3.4e-07 | 2e-07 | 1.3e-07 | 2.233E-07 | 1.069E-07 | 1.300E-07 | 3.400E-07 |
| 23677 | 1184 | K1184T | D | 2.8e-07 | 3.3e-07 | 3.2e-07 | 3.100E-07 | 2.646E-08 | 2.800E-07 | 3.300E-07 |
| 23678 | 1184 | K1184W | D | 4.2e-07 | 3.1e-07 | 1.6e-07 | 2.967E-07 | 1.305E-07 | 1.600E-07 | 4.200E-07 |
| 23679 | 1184 | K1184Y | D | 3.7e-07 | 1.2e-07 | 6.4e-07 | 3.767E-07 | 2.601E-07 | 1.200E-07 | 6.400E-07 |
| 23680 | 1184 | K1184V | D | 1.6e-07 | 1.7e-07 | 2.2e-07 | 1.833E-07 | 3.215E-08 | 1.600E-07 | 2.200E-07 |
| 23681 | 1185 | S1185A | D | 3.7e-07 | 1.6e-07 | 4e-07 | 3.100E-07 | 1.308E-07 | 1.600E-07 | 4.000E-07 |
| 23682 | 1185 | S1185R | D | 3.4e-07 | 4e-07 | 1.4e-07 | 2.933E-07 | 1.361E-07 | 1.400E-07 | 4.000E-07 |
| 23683 | 1185 | S1185N | D | 4.4e-07 | 1.3e-07 | 5.9e-07 | 3.867E-07 | 2.346E-07 | 1.300E-07 | 5.900E-07 |
| 23684 | 1185 | S1185D | D | 2.1e-07 | 1.6e-07 | 2.4e-07 | 2.033E-07 | 4.041E-08 | 1.600E-07 | 2.400E-07 |
| 23685 | 1185 | S1185C | D | 1.5e-07 | 7.1e-07 | 2.3e-07 | 3.633E-07 | 3.029E-07 | 1.500E-07 | 7.100E-07 |
| 23686 | 1185 | S1185E | D | 2.4e-07 | 3.2e-07 | 2.3e-07 | 2.633E-07 | 4.933E-08 | 2.300E-07 | 3.200E-07 |
| 23687 | 1185 | S1185Q | D | 2e-07 | 2.4e-07 | 1.8e-07 | 2.067E-07 | 3.055E-08 | 1.800E-07 | 2.400E-07 |
| 23688 | 1185 | S1185G | D | 2.9e-07 | 2.2e-07 | 1.1e-07 | 2.067E-07 | 9.074E-08 | 1.100E-07 | 2.900E-07 |
| 23689 | 1185 | S1185H | D | 1.1e-07 | 2.4e-07 | 2e-07 | 1.833E-07 | 6.658E-08 | 1.100E-07 | 2.400E-07 |
| 23690 | 1185 | S1185I | D | 2.5e-07 | 1.3e-07 | 3e-07 | 2.267E-07 | 8.737E-08 | 1.300E-07 | 3.000E-07 |
| 23691 | 1185 | S1185L | D | 3.2e-07 | 2.3e-07 | 3.7e-07 | 3.067E-07 | 7.095E-08 | 2.300E-07 | 3.700E-07 |
| 23692 | 1185 | S1185K | D | 2e-07 | 4e-07 | 1.4e-07 | 2.467E-07 | 1.361E-07 | 1.400E-07 | 4.000E-07 |
| 23693 | 1185 | S1185M | D | 1.2e-07 | 4.2e-07 | 1.4e-07 | 2.267E-07 | 1.677E-07 | 1.200E-07 | 4.200E-07 |
| 23694 | 1185 | S1185F | D | 1.7e-07 | 2e-07 | 1.6e-07 | 1.767E-07 | 2.082E-08 | 1.600E-07 | 2.000E-07 |
| 23695 | 1185 | S1185P | D | 3.4e-07 | 3.6e-07 | 2.7e-07 | 3.233E-07 | 4.726E-08 | 2.700E-07 | 3.600E-07 |
| 23696 | 1185 | S1185S | D | 1.4e-07 | 3.9e-07 | 3.7e-07 | 3.000E-07 | 1.389E-07 | 1.400E-07 | 3.900E-07 |
| 23697 | 1185 | S1185T | D | 5.2e-07 | 1e-07 | 1.1e-07 | 2.433E-07 | 2.397E-07 | 1.000E-07 | 5.200E-07 |
| 23698 | 1185 | S1185W | D | 3.1e-07 | 3.7e-07 | 2.7e-07 | 3.167E-07 | 5.033E-08 | 2.700E-07 | 3.700E-07 |
| 23699 | 1185 | S1185Y | D | 3.2e-07 | 2.3e-07 | 3e-07 | 2.833E-07 | 4.726E-08 | 2.300E-07 | 3.200E-07 |
| 23700 | 1185 | S1185V | D | 2.7e-07 | 1.8e-07 | 4.2e-07 | 2.900E-07 | 1.212E-07 | 1.800E-07 | 4.200E-07 |
| 23701 | 1186 | G1186A | D | 3.1e-07 | 1.9e-07 | 4.5e-07 | 3.167E-07 | 1.301E-07 | 1.900E-07 | 4.500E-07 |
| 23702 | 1186 | G1186R | D | 1.8e-07 | 2.8e-07 | 1.9e-07 | 2.167E-07 | 5.508E-08 | 1.800E-07 | 2.800E-07 |
| 23703 | 1186 | G1186N | D | 2.9e-07 | 4e-07 | 2.7e-07 | 3.200E-07 | 7.000E-08 | 2.700E-07 | 4.000E-07 |
| 23704 | 1186 | G1186D | D | 2.8e-07 | 1.8e-07 | 2.5e-07 | 2.367E-07 | 5.132E-08 | 1.800E-07 | 2.800E-07 |
| 23705 | 1186 | G1186C | D | 1.8e-07 | 2.4e-07 | 1.7e-07 | 1.967E-07 | 3.786E-08 | 1.700E-07 | 2.400E-07 |
| 23706 | 1186 | G1186E | D | 2.4e-07 | 3.2e-07 | 3e-07 | 2.867E-07 | 4.163E-08 | 2.400E-07 | 3.200E-07 |
| 23707 | 1186 | G1186Q | D | 2.2e-07 | 1.2e-07 | 2.6e-07 | 2.000E-07 | 7.211E-08 | 1.200E-07 | 2.600E-07 |
| 23708 | 1186 | G1186G | D | 1.8e-07 | 1.6e-07 | 1.6e-07 | 1.667E-07 | 1.155E-08 | 1.600E-07 | 1.800E-07 |
| 23709 | 1186 | G1186H | D | 2.4e-07 | 4.8e-07 | 4.4e-07 | 3.867E-07 | 1.286E-07 | 2.400E-07 | 4.800E-07 |
| 23710 | 1186 | G1186I | D | 2.4e-07 | 2.3e-07 | 3.1e-07 | 2.600E-07 | 4.359E-08 | 2.300E-07 | 3.100E-07 |
| 23711 | 1186 | G1186L | D | 3.1e-07 | 2e-07 | 3.8e-07 | 2.967E-07 | 9.074E-08 | 2.000E-07 | 3.800E-07 |
| 23712 | 1186 | G1186K | D | 3.8e-07 | 2.2e-07 | 2.3e-07 | 2.767E-07 | 8.963E-08 | 2.200E-07 | 3.800E-07 |
| 23713 | 1186 | G1186M | D | 1.8e-07 | 2.6e-07 | 1.7e-07 | 2.033E-07 | 4.933E-08 | 1.700E-07 | 2.600E-07 |
| 23714 | 1186 | G1186F | D | 4.7e-07 | 2.6e-07 | 2.1e-07 | 3.133E-07 | 1.380E-07 | 2.100E-07 | 4.700E-07 |

| | | | | | | | | | | |
|---|---|---|---|---|---|---|---|---|---|---|
| 23715 | 1186 | G1186P | D | 1.7e-07 | 1.8e-07 | 2.8e-07 | 2.100E-07 | 6.083E-08 | 1.700E-07 | 2.800E-07 |
| 23716 | 1186 | G1186S | D | 2.1e-07 | 2e-07 | 2.7e-07 | 2.267E-07 | 3.786E-08 | 2.000E-07 | 2.700E-07 |
| 23717 | 1186 | G1186T | D | 5.2e-07 | 1.5e-07 | 3.7e-07 | 3.467E-07 | 1.861E-07 | 1.500E-07 | 5.200E-07 |
| 23718 | 1186 | G1186W | D | 2.7e-07 | 2.9e-07 | 1.5e-07 | 2.367E-07 | 7.572E-08 | 1.500E-07 | 2.900E-07 |
| 23719 | 1186 | G1186Y | D | 2.5e-07 | 1.7e-07 | 1.7e-07 | 1.967E-07 | 4.619E-08 | 1.700E-07 | 2.500E-07 |
| 23720 | 1186 | G1186V | D | 3.7e-07 | 1.2e-07 | 2.6e-07 | 2.500E-07 | 1.253E-07 | 1.200E-07 | 3.700E-07 |
| 23721 | 1187 | T1187A | D | 6.4e-07 | 2.6e-07 | 7.2e-08 | 3.240E-07 | 2.894E-07 | 7.200E-08 | 6.400E-07 |
| 23722 | 1187 | T1187R | D | 2.2e-07 | 3.7e-07 | 1.2e-07 | 2.367E-07 | 1.258E-07 | 1.200E-07 | 3.700E-07 |
| 23723 | 1187 | T1187N | D | 2.6e-07 | 2.5e-07 | 4.6e-07 | 3.233E-07 | 1.185E-07 | 2.500E-07 | 4.600E-07 |
| 23724 | 1187 | T1187D | D | 1.3e-07 | 2.7e-07 | 3.9e-07 | 2.633E-07 | 1.301E-07 | 1.300E-07 | 3.900E-07 |
| 23725 | 1187 | T1187C | D | 1.2e-07 | 2.2e-07 | 2e-07 | 1.800E-07 | 5.292E-08 | 1.200E-07 | 2.200E-07 |
| 23726 | 1187 | T1187E | D | 2.7e-07 | 1.4e-07 | 1.7e-07 | 1.933E-07 | 6.807E-08 | 1.400E-07 | 2.700E-07 |
| 23727 | 1187 | T1187Q | D | 2.1e-07 | 3.8e-07 | 1.2e-07 | 2.367E-07 | 1.320E-07 | 1.200E-07 | 3.800E-07 |
| 23728 | 1187 | T1187G | D | 2.9e-07 | 2.1e-07 | 4.3e-07 | 3.100E-07 | 1.114E-07 | 2.100E-07 | 4.300E-07 |
| 23729 | 1187 | T1187H | D | 2.9e-07 | 1.3e-07 | 6.4e-07 | 3.533E-07 | 2.608E-07 | 1.300E-07 | 6.400E-07 |
| 23730 | 1187 | T1187I | D | 1.1e-07 | 2.7e-07 | 2.1e-07 | 1.967E-07 | 8.083E-08 | 1.100E-07 | 2.700E-07 |
| 23731 | 1187 | T1187L | D | 3.7e-07 | 8.5e-08 | 2.4e-07 | 2.317E-07 | 1.427E-07 | 8.500E-08 | 3.700E-07 |
| 23732 | 1187 | T1187K | D | 2.7e-07 | 3.6e-07 | 3.1e-07 | 3.133E-07 | 4.509E-08 | 2.700E-07 | 3.600E-07 |
| 23733 | 1187 | T1187M | D | 3.4e-07 | 3.8e-07 | 2.5e-07 | 3.233E-07 | 6.658E-08 | 2.500E-07 | 3.800E-07 |
| 23734 | 1187 | T1187F | D | 2.4e-07 | 3.7e-07 | 3e-07 | 3.033E-07 | 6.506E-08 | 2.400E-07 | 3.700E-07 |
| 23735 | 1187 | T1187P | D | 2e-07 | 3.2e-07 | 3e-07 | 2.733E-07 | 6.429E-08 | 2.000E-07 | 3.200E-07 |
| 23736 | 1187 | T1187S | D | 3.6e-07 | 1.7e-07 | 1.7e-07 | 2.333E-07 | 1.097E-07 | 1.700E-07 | 3.600E-07 |
| 23737 | 1187 | T1187T | D | 2.5e-07 | 1.8e-07 | 5.1e-07 | 3.133E-07 | 1.739E-07 | 1.800E-07 | 5.100E-07 |
| 23738 | 1187 | T1187W | D | 4.1e-07 | 2.4e-07 | 3.3e-07 | 3.267E-07 | 8.505E-08 | 2.400E-07 | 4.100E-07 |
| 23739 | 1187 | T1187Y | D | 2.9e-07 | 1.6e-07 | 1.4e-07 | 1.967E-07 | 8.145E-08 | 1.400E-07 | 2.900E-07 |
| 23740 | 1187 | T1187V | D | 1.1e-07 | 2.7e-07 | 1.8e-07 | 1.867E-07 | 8.021E-08 | 1.100E-07 | 2.700E-07 |
| 23741 | 1188 | A1188A | D | 2.8e-07 | 2.4e-07 | 2.1e-07 | 2.433E-07 | 3.512E-08 | 2.100E-07 | 2.800E-07 |
| 23742 | 1188 | A1188R | D | 3.5e-07 | 4.2e-07 | 1.6e-07 | 3.100E-07 | 1.345E-07 | 1.600E-07 | 4.200E-07 |
| 23743 | 1188 | A1188N | D | 2.7e-07 | 2.9e-07 | 1.7e-07 | 2.433E-07 | 6.429E-08 | 1.700E-07 | 2.900E-07 |
| 23744 | 1188 | A1188D | D | 3.5e-07 | 2.7e-07 | 2.1e-07 | 2.767E-07 | 7.024E-08 | 2.100E-07 | 3.500E-07 |
| 23745 | 1188 | A1188C | D | 1.9e-07 | 1.7e-07 | 2.5e-07 | 2.033E-07 | 4.163E-08 | 1.700E-07 | 2.500E-07 |
| 23746 | 1188 | A1188E | D | 4.7e-07 | 2.2e-07 | 2.2e-07 | 3.033E-07 | 1.443E-07 | 2.200E-07 | 4.700E-07 |
| 23747 | 1188 | A1188Q | D | 1.7e-07 | 4.2e-07 | 2.6e-07 | 2.833E-07 | 1.266E-07 | 1.700E-07 | 4.200E-07 |
| 23748 | 1188 | A1188G | D | 3e-07 | 3.4e-07 | 2.1e-07 | 2.833E-07 | 6.658E-08 | 2.100E-07 | 3.400E-07 |
| 23749 | 1188 | A1188H | D | 4.6e-07 | 2.3e-07 | 2.5e-07 | 3.133E-07 | 1.274E-07 | 2.300E-07 | 4.600E-07 |
| 23750 | 1188 | A1188I | D | 4.7e-07 | 4.1e-07 | 2.6e-07 | 3.800E-07 | 1.082E-07 | 2.600E-07 | 4.700E-07 |
| 23751 | 1188 | A1188L | D | 1.1e-07 | 3.3e-07 | 1.9e-07 | 2.100E-07 | 1.114E-07 | 1.100E-07 | 3.300E-07 |
| 23752 | 1188 | A1188K | D | 1.6e-07 | 2.1e-07 | 2.5e-07 | 2.067E-07 | 4.509E-08 | 1.600E-07 | 2.500E-07 |
| 23753 | 1188 | A1188M | D | 3.9e-07 | 3.5e-07 | 2.7e-07 | 3.367E-07 | 6.110E-08 | 2.700E-07 | 3.900E-07 |
| 23754 | 1188 | A1188F | D | 5.6e-07 | 1.8e-07 | 1.4e-07 | 2.933E-07 | 2.318E-07 | 1.400E-07 | 5.600E-07 |
| 23755 | 1188 | A1188P | D | 2.2e-07 | 2.4e-07 | 2e-07 | 2.200E-07 | 2.000E-08 | 2.000E-07 | 2.400E-07 |
| 23756 | 1188 | A1188S | D | 1.7e-07 | 4e-07 | 6.8e-07 | 4.167E-07 | 2.554E-07 | 1.700E-07 | 6.800E-07 |
| 23757 | 1188 | A1188T | D | 3.1e-07 | 3.1e-07 | 3.5e-07 | 3.233E-07 | 2.309E-08 | 3.100E-07 | 3.500E-07 |
| 23758 | 1188 | A1188W | D | 1.9e-07 | 1.4e-07 | 1.6e-07 | 1.633E-07 | 2.517E-08 | 1.400E-07 | 1.900E-07 |
| 23759 | 1188 | A1188Y | D | 2e-07 | 1.8e-07 | 2e-07 | 1.933E-07 | 1.155E-08 | 1.800E-07 | 2.000E-07 |
| 23760 | 1188 | A1188V | D | 4e-07 | 1.6e-07 | 2.8e-07 | 2.800E-07 | 1.200E-07 | 1.600E-07 | 4.000E-07 |
| 23761 | 1189 | S1189A | D | 3.6e-07 | 1.3e-07 | 3.2e-07 | 2.700E-07 | 1.229E-07 | 1.300E-07 | 3.600E-07 |
| 23762 | 1189 | S1189R | D | 2.4e-07 | 1.6e-07 | 3.3e-07 | 2.433E-07 | 8.505E-08 | 1.600E-07 | 3.300E-07 |
| 23763 | 1189 | S1189N | D | 2.4e-07 | 2.1e-07 | 3.2e-07 | 2.567E-07 | 5.686E-08 | 2.100E-07 | 3.200E-07 |
| 23764 | 1189 | S1189D | D | 3e-07 | 5.1e-07 | 2.3e-07 | 3.467E-07 | 1.457E-07 | 2.300E-07 | 5.100E-07 |
| 23765 | 1189 | S1189C | D | 5.6e-07 | 5.1e-07 | 3.7e-07 | 4.800E-07 | 9.849E-08 | 3.700E-07 | 5.600E-07 |
| 23766 | 1189 | S1189E | D | 3e-07 | 2e-07 | 2.6e-07 | 2.533E-07 | 5.033E-08 | 2.000E-07 | 3.000E-07 |
| 23767 | 1189 | S1189Q | D | 3.1e-07 | 3.6e-07 | 2.2e-07 | 2.967E-07 | 7.095E-08 | 2.200E-07 | 3.600E-07 |
| 23768 | 1189 | S1189G | D | 3.3e-07 | 2.9e-07 | 2.3e-07 | 2.833E-07 | 5.033E-08 | 2.300E-07 | 3.300E-07 |
| 23769 | 1189 | S1189H | D | 2.2e-07 | 1.8e-07 | 2.9e-07 | 2.300E-07 | 5.568E-08 | 1.800E-07 | 2.900E-07 |
| 23770 | 1189 | S1189I | D | 3.8e-07 | 3.2e-07 | 3.2e-07 | 3.400E-07 | 3.464E-08 | 3.200E-07 | 3.800E-07 |
| 23771 | 1189 | S1189L | D | 9.7e-08 | 1.8e-07 | 2.9e-07 | 1.890E-07 | 9.681E-08 | 9.700E-08 | 2.900E-07 |
| 23772 | 1189 | S1189K | D | 2.3e-07 | 2.3e-07 | 2.8e-07 | 2.467E-07 | 2.887E-08 | 2.300E-07 | 2.800E-07 |
| 23773 | 1189 | S1189M | D | 1.6e-07 | 2.9e-07 | 1.9e-07 | 2.133E-07 | 6.807E-08 | 1.600E-07 | 2.900E-07 |
| 23774 | 1189 | S1189F | D | 5.4e-07 | 1.9e-07 | 2e-07 | 3.100E-07 | 1.992E-07 | 1.900E-07 | 5.400E-07 |
| 23775 | 1189 | S1189P | D | 2.7e-07 | 1.9e-07 | 1.6e-07 | 2.067E-07 | 5.686E-08 | 1.600E-07 | 2.700E-07 |
| 23776 | 1189 | S1189S | D | 6.5e-07 | 1.6e-07 | 3.3e-07 | 3.800E-07 | 2.488E-07 | 1.600E-07 | 6.500E-07 |
| 23777 | 1189 | S1189T | D | 5e-08 | 2.1e-07 | 2.3e-07 | 1.633E-07 | 9.866E-08 | 5.000E-08 | 2.300E-07 |
| 23778 | 1189 | S1189W | D | 2.1e-07 | 5e-07 | 4.1e-07 | 3.733E-07 | 1.484E-07 | 2.100E-07 | 5.000E-07 |
| 23779 | 1189 | S1189Y | D | 2.9e-07 | 3.5e-07 | 1.7e-07 | 2.700E-07 | 9.165E-08 | 1.700E-07 | 3.500E-07 |
| 23780 | 1189 | S1189V | D | 3.4e-07 | 2.8e-07 | 1.4e-07 | 2.533E-07 | 1.026E-07 | 1.400E-07 | 3.400E-07 |
| 23781 | 1190 | V1190A | D | 3.1e-07 | 1.2e-07 | 4.7e-07 | 3.000E-07 | 1.752E-07 | 1.200E-07 | 4.700E-07 |
| 23782 | 1190 | V1190R | D | 7.2e-08 | 3e-07 | 2.4e-07 | 2.040E-07 | 1.182E-07 | 7.200E-08 | 3.000E-07 |
| 23783 | 1190 | V1190N | D | 5.3e-07 | 1.8e-07 | 4e-07 | 3.700E-07 | 1.769E-07 | 1.800E-07 | 5.300E-07 |
| 23784 | 1190 | V1190D | D | 3.7e-07 | 2.4e-07 | 2.8e-07 | 2.967E-07 | 6.658E-08 | 2.400E-07 | 3.700E-07 |
| 23785 | 1190 | V1190C | D | 2.7e-07 | 3.5e-07 | 3.7e-07 | 3.300E-07 | 5.292E-08 | 2.700E-07 | 3.700E-07 |
| 23786 | 1190 | V1190E | D | 3.3e-07 | 2.2e-07 | 4.5e-07 | 3.333E-07 | 1.150E-07 | 2.200E-07 | 4.500E-07 |
| 23787 | 1190 | V1190Q | D | 1.3e-07 | 2e-07 | 2.1e-07 | 1.800E-07 | 4.359E-08 | 1.300E-07 | 2.100E-07 |
| 23788 | 1190 | V1190G | D | 3.3e-07 | 1.3e-07 | 1.1e-07 | 1.900E-07 | 1.217E-07 | 1.100E-07 | 3.300E-07 |
| 23789 | 1190 | V1190H | D | 2.4e-07 | 1.9e-07 | 2.5e-07 | 2.267E-07 | 3.215E-08 | 1.900E-07 | 2.500E-07 |
| 23790 | 1190 | V1190I | D | 4.1e-07 | 2.3e-07 | 3.7e-07 | 3.367E-07 | 9.452E-08 | 2.300E-07 | 4.100E-07 |
| 23791 | 1190 | V1190L | D | 2.5e-07 | 2.6e-07 | 2.2e-07 | 2.433E-07 | 2.082E-08 | 2.200E-07 | 2.600E-07 |
| 23792 | 1190 | V1190K | D | 2.1e-07 | 2.3e-07 | 2.3e-07 | 2.233E-07 | 1.155E-08 | 2.100E-07 | 2.300E-07 |
| 23793 | 1190 | V1190M | D | 3.8e-07 | 2.1e-07 | 1.7e-07 | 2.533E-07 | 1.115E-07 | 1.700E-07 | 3.800E-07 |
| 23794 | 1190 | V1190F | D | 2.3e-07 | 1.4e-07 | 1.3e-07 | 1.667E-07 | 5.508E-08 | 1.300E-07 | 2.300E-07 |
| 23795 | 1190 | V1190P | D | 5.9e-07 | 2.1e-07 | 3.3e-07 | 3.767E-07 | 1.943E-07 | 2.100E-07 | 5.900E-07 |
| 23796 | 1190 | V1190S | D | 4.4e-07 | 1.5e-07 | 4.8e-07 | 3.567E-07 | 1.801E-07 | 1.500E-07 | 4.800E-07 |
| 23797 | 1190 | V1190T | D | 4.4e-07 | 1.9e-07 | 3.5e-07 | 3.267E-07 | 1.266E-07 | 1.900E-07 | 4.400E-07 |
| 23798 | 1190 | V1190W | D | 3.1e-07 | 2e-07 | 3.1e-07 | 2.733E-07 | 6.351E-08 | 2.000E-07 | 3.100E-07 |
| 23799 | 1190 | V1190Y | D | 2.5e-07 | 2.4e-07 | 2.4e-07 | 2.433E-07 | 5.774E-09 | 2.400E-07 | 2.500E-07 |

| | | | | | | | | | | |
|---|---|---|---|---|---|---|---|---|---|---|
| 23800 | 1190 | V1190V | D | 2.6e-07 | 1.6e-07 | 9.8e-08 | 1.727E-07 | 8.174E-08 | 9.800E-08 | 2.600E-07 |
| 23801 | 1191 | V1191A | D | 2.9e-07 | 2.2e-07 | 2.9e-07 | 2.667E-07 | 4.041E-08 | 2.200E-07 | 2.900E-07 |
| 23802 | 1191 | V1191R | D | 1.6e-07 | 2.6e-07 | 1.8e-07 | 2.000E-07 | 5.292E-08 | 1.600E-07 | 2.600E-07 |
| 23803 | 1191 | V1191N | D | 2.8e-07 | 2.8e-07 | 2.7e-07 | 2.767E-07 | 5.774E-09 | 2.700E-07 | 2.800E-07 |
| 23804 | 1191 | V1191D | D | 1.8e-07 | 1.3e-07 | 5.1e-07 | 2.733E-07 | 2.065E-07 | 1.300E-07 | 5.100E-07 |
| 23805 | 1191 | V1191C | D | 3.1e-07 | 2.5e-07 | 1.7e-07 | 2.433E-07 | 7.024E-08 | 1.700E-07 | 3.100E-07 |
| 23806 | 1191 | V1191E | D | 1.3e-07 | 1.1e-07 | 1.8e-07 | 1.400E-07 | 3.606E-08 | 1.100E-07 | 1.800E-07 |
| 23807 | 1191 | V1191Q | D | 2.4e-07 | 1.7e-07 | 2.6e-07 | 2.233E-07 | 4.726E-08 | 1.700E-07 | 2.600E-07 |
| 23808 | 1191 | V1191G | D | 3.1e-07 | 2.5e-07 | 1.8e-07 | 2.467E-07 | 6.506E-08 | 1.800E-07 | 3.100E-07 |
| 23809 | 1191 | V1191H | D | 1.1e-07 | 9.6e-08 | 2e-07 | 1.353E-07 | 5.644E-08 | 9.600E-08 | 2.000E-07 |
| 23810 | 1191 | V1191I | D | 1.8e-07 | 2.8e-07 | 1.1e-07 | 1.900E-07 | 8.544E-08 | 1.100E-07 | 2.800E-07 |
| 23811 | 1191 | V1191L | D | 1.9e-07 | 4.8e-07 | 5.4e-07 | 4.033E-07 | 1.872E-07 | 1.900E-07 | 5.400E-07 |
| 23812 | 1191 | V1191K | D | 1.4e-07 | 2.3e-07 | 3.9e-07 | 2.533E-07 | 1.266E-07 | 1.400E-07 | 3.900E-07 |
| 23813 | 1191 | V1191M | D | 2.6e-07 | 2.7e-07 | 5.6e-07 | 3.633E-07 | 1.704E-07 | 2.600E-07 | 5.600E-07 |
| 23814 | 1191 | V1191F | D | 1.9e-07 | 2e-07 | 3.3e-07 | 2.400E-07 | 7.810E-08 | 1.900E-07 | 3.300E-07 |
| 23815 | 1191 | V1191P | D | 2.3e-07 | 2e-07 | 4.5e-07 | 2.933E-07 | 1.365E-07 | 2.000E-07 | 4.500E-07 |
| 23816 | 1191 | V1191S | D | 4.1e-07 | 2.4e-07 | 2.7e-07 | 3.067E-07 | 9.074E-08 | 2.400E-07 | 4.100E-07 |
| 23817 | 1191 | V1191T | D | 1.6e-07 | 2.8e-07 | 3.4e-07 | 2.600E-07 | 9.165E-08 | 1.600E-07 | 3.400E-07 |
| 23818 | 1191 | V1191W | D | 2e-07 | 2e-07 | 2.3e-07 | 2.100E-07 | 1.732E-08 | 2.000E-07 | 2.300E-07 |
| 23819 | 1191 | V1191Y | D | 1.3e-07 | 2.3e-07 | 2.1e-07 | 1.900E-07 | 5.292E-08 | 1.300E-07 | 2.300E-07 |
| 23820 | 1191 | V1191V | D | 3.1e-07 | 3.8e-07 | 3e-07 | 3.300E-07 | 4.359E-08 | 3.000E-07 | 3.800E-07 |
| 23821 | 1192 | C1192A | D | 1.6e-07 | 3.4e-07 | 2.9e-07 | 2.633E-07 | 9.292E-08 | 1.600E-07 | 3.400E-07 |
| 23822 | 1192 | C1192R | D | 1.6e-07 | 1.5e-07 | 2.5e-07 | 1.867E-07 | 5.508E-08 | 1.500E-07 | 2.500E-07 |
| 23823 | 1192 | C1192N | D | 1.5e-07 | 2.7e-07 | 5.2e-07 | 3.133E-07 | 1.888E-07 | 1.500E-07 | 5.200E-07 |
| 23824 | 1192 | C1192D | D | 3.2e-07 | 1.7e-07 | 9.6e-07 | 4.833E-07 | 4.196E-07 | 1.700E-07 | 9.600E-07 |
| 23825 | 1192 | C1192C | D | 3.7e-07 | 3.4e-07 | 2.4e-07 | 3.167E-07 | 6.807E-08 | 2.400E-07 | 3.700E-07 |
| 23826 | 1192 | C1192E | D | 3.6e-07 | 2.1e-07 | 3.1e-07 | 2.933E-07 | 7.638E-08 | 2.100E-07 | 3.600E-07 |
| 23827 | 1192 | C1192Q | D | 1.6e-07 | 5.1e-07 | 7e-08 | 2.467E-07 | 2.325E-07 | 7.000E-08 | 5.100E-07 |
| 23828 | 1192 | C1192G | D | 2.9e-07 | 5.2e-07 | 1.5e-07 | 3.200E-07 | 1.868E-07 | 1.500E-07 | 5.200E-07 |
| 23829 | 1192 | C1192H | D | 3.4e-07 | 4.4e-07 | 3.6e-07 | 3.800E-07 | 5.292E-08 | 3.400E-07 | 4.400E-07 |
| 23830 | 1192 | C1192I | D | 3e-07 | 1.3e-07 | 1.2e-07 | 1.833E-07 | 1.012E-07 | 1.200E-07 | 3.000E-07 |
| 23831 | 1192 | C1192L | D | 1.3e-07 | 2.3e-07 | 1.6e-07 | 1.733E-07 | 5.132E-08 | 1.300E-07 | 2.300E-07 |
| 23832 | 1192 | C1192K | D | 2.3e-07 | 1.4e-07 | 2.7e-07 | 2.133E-07 | 6.658E-08 | 1.400E-07 | 2.700E-07 |
| 23833 | 1192 | C1192M | D | 2.7e-07 | 2.2e-07 | 2e-07 | 2.300E-07 | 3.606E-08 | 2.000E-07 | 2.700E-07 |
| 23834 | 1192 | C1192F | D | 2.2e-07 | 1.8e-07 | 2.2e-07 | 2.067E-07 | 2.309E-08 | 1.800E-07 | 2.200E-07 |
| 23835 | 1192 | C1192P | D | 4.3e-07 | 2.7e-07 | 2.2e-07 | 3.067E-07 | 1.097E-07 | 2.200E-07 | 4.300E-07 |
| 23836 | 1192 | C1192S | D | 3e-07 | 3.3e-07 | 1e-06 | 5.433E-07 | 3.958E-07 | 3.000E-07 | 1.000E-06 |
| 23837 | 1192 | C1192T | D | 1.8e-07 | 1.8e-07 | 8.4e-07 | 4.000E-07 | 3.811E-07 | 1.800E-07 | 8.400E-07 |
| 23838 | 1192 | C1192W | D | 1.9e-07 | 4.4e-07 | 2.5e-07 | 2.933E-07 | 1.305E-07 | 1.900E-07 | 4.400E-07 |
| 23839 | 1192 | C1192Y | D | 2.4e-07 | 1.1e-07 | 2.4e-07 | 1.967E-07 | 7.506E-08 | 1.100E-07 | 2.400E-07 |
| 23840 | 1192 | C1192V | D | 4.3e-07 | 2.4e-07 | 6.8e-07 | 4.500E-07 | 2.207E-07 | 2.400E-07 | 6.800E-07 |
| 23841 | 1193 | L1193A | D | 2.3e-07 | 2.1e-07 | 1.3e-07 | 1.900E-07 | 5.292E-08 | 1.300E-07 | 2.300E-07 |
| 23842 | 1193 | L1193R | D | 2.3e-07 | 1.7e-07 | 3.7e-07 | 2.567E-07 | 1.026E-07 | 1.700E-07 | 3.700E-07 |
| 23843 | 1193 | L1193N | D | 4.1e-07 | 1.3e-07 | 3.5e-07 | 2.967E-07 | 1.474E-07 | 1.300E-07 | 4.100E-07 |
| 23844 | 1193 | L1193D | D | 3.7e-07 | 3.3e-07 | 1.7e-07 | 2.900E-07 | 1.058E-07 | 1.700E-07 | 3.700E-07 |
| 23845 | 1193 | L1193C | D | 1.5e-07 | 1.8e-07 | 1.9e-07 | 1.733E-07 | 2.082E-08 | 1.500E-07 | 1.900E-07 |
| 23846 | 1193 | L1193E | D | 5e-07 | 2.2e-07 | 3.2e-07 | 3.467E-07 | 1.419E-07 | 2.200E-07 | 5.000E-07 |
| 23847 | 1193 | L1193Q | D | 3.9e-07 | 2e-07 | 1.8e-07 | 2.567E-07 | 1.159E-07 | 1.800E-07 | 3.900E-07 |
| 23848 | 1193 | L1193G | D | 2.7e-07 | 4.6e-07 | 3.6e-07 | 3.633E-07 | 9.504E-08 | 2.700E-07 | 4.600E-07 |
| 23849 | 1193 | L1193H | D | 2.3e-07 | 1.5e-07 | 2.7e-07 | 2.167E-07 | 6.110E-08 | 1.500E-07 | 2.700E-07 |
| 23850 | 1193 | L1193I | D | 4.6e-07 | 2.1e-07 | 3.7e-07 | 3.467E-07 | 1.266E-07 | 2.100E-07 | 4.600E-07 |
| 23851 | 1193 | L1193L | D | 4.7e-07 | 1.6e-07 | 4.9e-07 | 3.733E-07 | 1.850E-07 | 1.600E-07 | 4.900E-07 |
| 23852 | 1193 | L1193K | D | 3.1e-07 | 1.4e-07 | 2.9e-07 | 2.467E-07 | 9.292E-08 | 1.400E-07 | 3.100E-07 |
| 23853 | 1193 | L1193M | D | 5.9e-07 | 1.3e-07 | 2.5e-07 | 3.233E-07 | 2.386E-07 | 1.300E-07 | 5.900E-07 |
| 23854 | 1193 | L1193F | D | 1.9e-07 | 3.6e-07 | 2.3e-07 | 2.600E-07 | 8.888E-08 | 1.900E-07 | 3.600E-07 |
| 23855 | 1193 | L1193P | D | 2.5e-07 | 3.9e-07 | 2.9e-07 | 3.100E-07 | 7.211E-08 | 2.500E-07 | 3.900E-07 |
| 23856 | 1193 | L1193S | D | 1.9e-07 | 2.4e-07 | 2.6e-07 | 2.300E-07 | 3.606E-08 | 1.900E-07 | 2.600E-07 |
| 23857 | 1193 | L1193T | D | 2.5e-07 | 4.8e-07 | 3e-07 | 3.433E-07 | 1.210E-07 | 2.500E-07 | 4.800E-07 |
| 23858 | 1193 | L1193W | D | 5.6e-07 | 3e-07 | 1.5e-07 | 3.367E-07 | 2.074E-07 | 1.500E-07 | 5.600E-07 |
| 23859 | 1193 | L1193Y | D | 7.4e-07 | 1.9e-07 | 1.7e-07 | 3.667E-07 | 3.235E-07 | 1.700E-07 | 7.400E-07 |
| 23860 | 1193 | L1193V | D | 2.8e-07 | 2.2e-07 | 1.3e-07 | 2.100E-07 | 7.550E-08 | 1.300E-07 | 2.800E-07 |
| 23861 | 1194 | L1194A | D | 3.2e-07 | 1.6e-07 | 2.9e-07 | 2.567E-07 | 8.505E-08 | 1.600E-07 | 3.200E-07 |
| 23862 | 1194 | L1194R | D | 2.3e-07 | 2.3e-07 | 6.5e-07 | 3.700E-07 | 2.425E-07 | 2.300E-07 | 6.500E-07 |
| 23863 | 1194 | L1194N | D | 2.1e-07 | 4.6e-07 | 3.7e-07 | 3.467E-07 | 1.266E-07 | 2.100E-07 | 4.600E-07 |
| 23864 | 1194 | L1194D | D | 2.6e-07 | 4.3e-07 | 2.6e-07 | 3.167E-07 | 9.815E-08 | 2.600E-07 | 4.300E-07 |
| 23865 | 1194 | L1194C | D | 3.3e-07 | 1.6e-07 | 1.7e-07 | 2.200E-07 | 9.539E-08 | 1.600E-07 | 3.300E-07 |
| 23866 | 1194 | L1194E | D | 3.2e-07 | 2.2e-07 | 1.6e-07 | 2.333E-07 | 8.083E-08 | 1.600E-07 | 3.200E-07 |
| 23867 | 1194 | L1194Q | D | 1.8e-07 | 2.7e-07 | 1.7e-07 | 2.067E-07 | 5.508E-08 | 1.700E-07 | 2.700E-07 |
| 23868 | 1194 | L1194G | D | 2.4e-07 | 1.9e-07 | 1.2e-07 | 1.833E-07 | 6.028E-08 | 1.200E-07 | 2.400E-07 |
| 23869 | 1194 | L1194H | D | 2.1e-07 | 2.8e-07 | 3.6e-07 | 2.833E-07 | 7.506E-08 | 2.100E-07 | 3.600E-07 |
| 23870 | 1194 | L1194I | D | 3.6e-07 | 3.1e-07 | 4.6e-07 | 3.767E-07 | 7.638E-08 | 3.100E-07 | 4.600E-07 |
| 23871 | 1194 | L1194L | D | 1.8e-07 | 1.9e-07 | 2.3e-07 | 2.000E-07 | 2.646E-08 | 1.800E-07 | 2.300E-07 |
| 23872 | 1194 | L1194K | D | 3e-07 | 1.4e-07 | 2.5e-07 | 2.300E-07 | 8.185E-08 | 1.400E-07 | 3.000E-07 |
| 23873 | 1194 | L1194M | D | 2.5e-07 | 3.1e-07 | 3.7e-07 | 3.100E-07 | 6.000E-08 | 2.500E-07 | 3.700E-07 |
| 23874 | 1194 | L1194F | D | 3e-07 | 8.9e-08 | 2.9e-07 | 2.263E-07 | 1.190E-07 | 8.900E-08 | 3.000E-07 |
| 23875 | 1194 | L1194P | D | 1.7e-07 | 2e-07 | 2.7e-07 | 2.133E-07 | 5.132E-08 | 1.700E-07 | 2.700E-07 |
| 23876 | 1194 | L1194S | D | 1.9e-07 | 4.3e-07 | 1.8e-07 | 2.667E-07 | 1.415E-07 | 1.800E-07 | 4.300E-07 |
| 23877 | 1194 | L1194T | D | 2.1e-07 | 2.2e-07 | 3.2e-07 | 2.500E-07 | 6.083E-08 | 2.100E-07 | 3.200E-07 |
| 23878 | 1194 | L1194W | D | 2.4e-07 | 5.5e-07 | 1.8e-07 | 3.233E-07 | 1.986E-07 | 1.800E-07 | 5.500E-07 |
| 23879 | 1194 | L1194Y | D | 2e-07 | 2.3e-07 | 1.3e-07 | 1.867E-07 | 5.132E-08 | 1.300E-07 | 2.300E-07 |
| 23880 | 1194 | L1194V | D | 2.8e-07 | 4.1e-07 | 2.3e-07 | 3.067E-07 | 9.292E-08 | 2.300E-07 | 4.100E-07 |
| 23881 | 1195 | N1195A | D | 1.5e-07 | 3.3e-07 | 3.3e-07 | 2.700E-07 | 1.039E-07 | 1.500E-07 | 3.300E-07 |
| 23882 | 1195 | N1195R | D | 3.5e-07 | 2.6e-07 | 2.2e-07 | 2.767E-07 | 6.658E-08 | 2.200E-07 | 3.500E-07 |
| 23883 | 1195 | N1195N | D | 2.8e-07 | 3e-07 | 2.8e-07 | 2.867E-07 | 1.155E-08 | 2.800E-07 | 3.000E-07 |
| 23884 | 1195 | N1195D | D | 2.7e-07 | 7.6e-07 | 2.4e-07 | 4.233E-07 | 2.919E-07 | 2.400E-07 | 7.600E-07 |

| | | | | | | | | | | |
|---|---|---|---|---|---|---|---|---|---|---|
| 23885 | 1195 | N1195C | D | 2.9e-07 | 2.6e-07 | 4.4e-07 | 3.300E-07 | 9.644E-08 | 2.600E-07 | 4.400E-07 |
| 23886 | 1195 | N1195E | D | 2.3e-07 | 2.3e-07 | 1.1e-07 | 1.900E-07 | 6.928E-08 | 1.100E-07 | 2.300E-07 |
| 23887 | 1195 | N1195Q | D | 2.1e-07 | 1.6e-07 | 1.8e-07 | 1.833E-07 | 2.517E-08 | 1.600E-07 | 2.100E-07 |
| 23888 | 1195 | N1195G | D | 2.1e-07 | 2.2e-07 | 2.4e-07 | 2.233E-07 | 1.528E-08 | 2.100E-07 | 2.400E-07 |
| 23889 | 1195 | N1195H | D | 2.1e-07 | 1.3e-07 | 5.4e-07 | 2.933E-07 | 2.173E-07 | 1.300E-07 | 5.400E-07 |
| 23890 | 1195 | N1195I | D | 3.5e-07 | 2.3e-07 | 3.5e-07 | 3.100E-07 | 6.928E-08 | 2.300E-07 | 3.500E-07 |
| 23891 | 1195 | N1195L | D | 2.3e-07 | 2.5e-07 | 2.3e-07 | 2.367E-07 | 1.155E-08 | 2.300E-07 | 2.500E-07 |
| 23892 | 1195 | N1195K | D | 4.6e-07 | 1.9e-07 | 1.8e-07 | 2.767E-07 | 1.589E-07 | 1.800E-07 | 4.600E-07 |
| 23893 | 1195 | N1195M | D | 3.2e-07 | 3.2e-07 | 6.6e-07 | 4.333E-07 | 1.963E-07 | 3.200E-07 | 6.600E-07 |
| 23894 | 1195 | N1195F | D | 2.5e-07 | 1.8e-07 | 7.5e-07 | 3.933E-07 | 3.109E-07 | 1.800E-07 | 7.500E-07 |
| 23895 | 1195 | N1195P | D | 5.4e-07 | 1.1e-07 | 2e-07 | 2.833E-07 | 2.268E-07 | 1.100E-07 | 5.400E-07 |
| 23896 | 1195 | N1195S | D | 2.7e-07 | 3.2e-07 | 2.6e-07 | 2.833E-07 | 3.215E-08 | 2.600E-07 | 3.200E-07 |
| 23897 | 1195 | N1195T | D | 1.8e-07 | 1.4e-07 | 1.8e-07 | 1.667E-07 | 2.309E-08 | 1.400E-07 | 1.800E-07 |
| 23898 | 1195 | N1195W | D | 1.5e-07 | 2e-07 | 1.2e-07 | 1.567E-07 | 4.041E-08 | 1.200E-07 | 2.000E-07 |
| 23899 | 1195 | N1195Y | D | 1.7e-07 | 1.7e-07 | 4.7e-07 | 2.700E-07 | 1.732E-07 | 1.700E-07 | 4.700E-07 |
| 23900 | 1195 | N1195V | D | 2.3e-07 | 2.9e-07 | 1e-07 | 2.067E-07 | 9.713E-08 | 1.000E-07 | 2.900E-07 |
| 23901 | 1196 | N1196A | D | 2.3e-07 | 2e-07 | 1.1e-07 | 1.800E-07 | 6.245E-08 | 1.100E-07 | 2.300E-07 |
| 23902 | 1196 | N1196R | D | 1.3e-07 | 2.1e-07 | 5e-07 | 2.800E-07 | 1.947E-07 | 1.300E-07 | 5.000E-07 |
| 23903 | 1196 | N1196N | D | 2.3e-07 | 3.8e-07 | 2.1e-07 | 2.733E-07 | 9.292E-08 | 2.100E-07 | 3.800E-07 |
| 23904 | 1196 | N1196D | D | 1.1e-07 | 2.3e-07 | 1.2e-07 | 1.533E-07 | 6.658E-08 | 1.100E-07 | 2.300E-07 |
| 23905 | 1196 | N1196C | D | 1.4e-07 | 1.4e-07 | 2.3e-07 | 1.700E-07 | 5.196E-08 | 1.400E-07 | 2.300E-07 |
| 23906 | 1196 | N1196E | D | 2.7e-07 | 2.6e-07 | 1.6e-07 | 2.300E-07 | 6.083E-08 | 1.600E-07 | 2.700E-07 |
| 23907 | 1196 | N1196Q | D | 2.3e-07 | 3.3e-07 | 2.4e-07 | 2.667E-07 | 5.508E-08 | 2.300E-07 | 3.300E-07 |
| 23908 | 1196 | N1196G | D | 6.8e-08 | 1.2e-07 | 1.7e-07 | 1.193E-07 | 5.100E-08 | 6.800E-08 | 1.700E-07 |
| 23909 | 1196 | N1196H | D | 3.4e-07 | 1.6e-07 | 1.8e-07 | 2.267E-07 | 9.866E-08 | 1.600E-07 | 3.400E-07 |
| 23910 | 1196 | N1196I | D | 3.5e-07 | 4e-07 | 2.5e-07 | 3.333E-07 | 7.638E-08 | 2.500E-07 | 4.000E-07 |
| 23911 | 1196 | N1196L | D | 1.9e-07 | 2.5e-07 | 2.7e-07 | 2.367E-07 | 4.163E-08 | 1.900E-07 | 2.700E-07 |
| 23912 | 1196 | N1196K | D | 3.6e-07 | 1.7e-07 | 1.5e-07 | 2.267E-07 | 1.159E-07 | 1.500E-07 | 3.600E-07 |
| 23913 | 1196 | N1196M | D | 3.5e-07 | 2.2e-07 | 3e-07 | 2.900E-07 | 6.557E-08 | 2.200E-07 | 3.500E-07 |
| 23914 | 1196 | N1196F | D | 2.1e-07 | 2.4e-07 | 2.6e-07 | 2.367E-07 | 2.517E-08 | 2.100E-07 | 2.600E-07 |
| 23915 | 1196 | N1196P | D | 7.7e-08 | 2.1e-07 | 1.9e-07 | 1.590E-07 | 7.171E-08 | 7.700E-08 | 2.100E-07 |
| 23916 | 1196 | N1196S | D | 2.1e-07 | 3.3e-07 | 1.4e-07 | 2.267E-07 | 9.609E-08 | 1.400E-07 | 3.300E-07 |
| 23917 | 1196 | N1196T | D | 3.3e-07 | 4.4e-07 | 1.6e-07 | 3.100E-07 | 1.411E-07 | 1.600E-07 | 4.400E-07 |
| 23918 | 1196 | N1196W | D | 2.8e-07 | 1.7e-07 | 1.7e-07 | 2.067E-07 | 6.351E-08 | 1.700E-07 | 2.800E-07 |
| 23919 | 1196 | N1196Y | D | 3.1e-07 | 2.4e-07 | 1.1e-07 | 2.200E-07 | 1.015E-07 | 1.100E-07 | 3.100E-07 |
| 23920 | 1196 | N1196V | D | 1.7e-07 | 2.9e-07 | 2.1e-07 | 2.233E-07 | 6.110E-08 | 1.700E-07 | 2.900E-07 |
| 23921 | 1197 | F1197A | D | 1.3e-07 | 2.5e-07 | 3.6e-07 | 2.467E-07 | 1.150E-07 | 1.300E-07 | 3.600E-07 |
| 23922 | 1197 | F1197R | D | 1.3e-07 | 5.1e-07 | 3.6e-07 | 3.333E-07 | 1.914E-07 | 1.300E-07 | 5.100E-07 |
| 23923 | 1197 | F1197N | D | 2e-07 | 1.6e-07 | 1.9e-07 | 1.833E-07 | 2.082E-08 | 1.600E-07 | 2.000E-07 |
| 23924 | 1197 | F1197D | D | 1.9e-07 | 2.9e-07 | 4.1e-07 | 2.967E-07 | 1.102E-07 | 1.900E-07 | 4.100E-07 |
| 23925 | 1197 | F1197C | D | 2.9e-07 | 3.4e-07 | 2e-07 | 2.767E-07 | 7.095E-08 | 2.000E-07 | 3.400E-07 |
| 23926 | 1197 | F1197E | D | 3e-07 | 3.2e-07 | 4.1e-07 | 3.433E-07 | 5.859E-08 | 3.000E-07 | 4.100E-07 |
| 23927 | 1197 | F1197Q | D | 1.1e-07 | 4.7e-07 | 2.4e-07 | 2.733E-07 | 1.823E-07 | 1.100E-07 | 4.700E-07 |
| 23928 | 1197 | F1197G | D | 3.7e-07 | 3.6e-07 | 1.7e-07 | 3.000E-07 | 1.127E-07 | 1.700E-07 | 3.700E-07 |
| 23929 | 1197 | F1197H | D | 3.1e-07 | 3.6e-07 | 5e-07 | 3.900E-07 | 9.849E-08 | 3.100E-07 | 5.000E-07 |
| 23930 | 1197 | F1197I | D | 2.1e-07 | 1.4e-07 | 3.9e-07 | 2.467E-07 | 1.290E-07 | 1.400E-07 | 3.900E-07 |
| 23931 | 1197 | F1197L | D | 1.7e-07 | 1.2e-07 | 1.3e-07 | 1.400E-07 | 2.646E-08 | 1.200E-07 | 1.700E-07 |
| 23932 | 1197 | F1197K | D | 1.7e-07 | 7e-08 | 1.4e-07 | 1.267E-07 | 5.132E-08 | 7.000E-08 | 1.700E-07 |
| 23933 | 1197 | F1197M | D | 8.3e-07 | 3.3e-07 | 1.3e-07 | 4.300E-07 | 3.606E-07 | 1.300E-07 | 8.300E-07 |
| 23934 | 1197 | F1197F | D | 2e-07 | 3.9e-07 | 3.7e-07 | 3.200E-07 | 1.044E-07 | 2.000E-07 | 3.900E-07 |
| 23935 | 1197 | F1197P | D | 3.1e-07 | 2.5e-07 | 1e-07 | 2.200E-07 | 1.082E-07 | 1.000E-07 | 3.100E-07 |
| 23936 | 1197 | F1197S | D | 1.4e-07 | 1.8e-07 | 2.4e-07 | 1.867E-07 | 5.033E-08 | 1.400E-07 | 2.400E-07 |
| 23937 | 1197 | F1197T | D | 2.8e-07 | 1.8e-07 | 2e-07 | 2.200E-07 | 5.292E-08 | 1.800E-07 | 2.800E-07 |
| 23938 | 1197 | F1197W | D | 2.2e-07 | 4.3e-07 | 3e-07 | 3.167E-07 | 1.060E-07 | 2.200E-07 | 4.300E-07 |
| 23939 | 1197 | F1197Y | D | 6.1e-07 | 7.1e-07 | 8.3e-08 | 4.677E-07 | 3.369E-07 | 8.300E-08 | 7.100E-07 |
| 23940 | 1197 | F1197V | D | 1.3e-07 | 3.3e-07 | 1.5e-07 | 2.033E-07 | 1.102E-07 | 1.300E-07 | 3.300E-07 |
| 23941 | 1198 | Y1198A | D | 2e-07 | 5.4e-07 | 3.4e-07 | 3.600E-07 | 1.709E-07 | 2.000E-07 | 5.400E-07 |
| 23942 | 1198 | Y1198R | D | 2.7e-07 | 1.8e-07 | 3.7e-07 | 2.733E-07 | 9.504E-08 | 1.800E-07 | 3.700E-07 |
| 23943 | 1198 | Y1198N | D | 3.1e-07 | 1.8e-07 | 3.8e-07 | 2.900E-07 | 1.015E-07 | 1.800E-07 | 3.800E-07 |
| 23944 | 1198 | Y1198D | D | 2e-07 | 1.9e-07 | 2.6e-07 | 2.167E-07 | 3.786E-08 | 1.900E-07 | 2.600E-07 |
| 23945 | 1198 | Y1198C | D | 4e-07 | 3.3e-07 | 2.7e-07 | 3.333E-07 | 6.506E-08 | 2.700E-07 | 4.000E-07 |
| 23946 | 1198 | Y1198E | D | 2.7e-07 | 2.5e-07 | 2.7e-07 | 2.633E-07 | 1.155E-08 | 2.500E-07 | 2.700E-07 |
| 23947 | 1198 | Y1198Q | D | 1.7e-07 | 2.9e-07 | 6.9e-08 | 1.763E-07 | 1.106E-07 | 6.900E-08 | 2.900E-07 |
| 23948 | 1198 | Y1198G | D | 1.4e-07 | 1.7e-07 | 2e-07 | 1.700E-07 | 3.000E-08 | 1.400E-07 | 2.000E-07 |
| 23949 | 1198 | Y1198H | D | 5e-07 | 2.3e-07 | 2.6e-07 | 3.300E-07 | 1.480E-07 | 2.300E-07 | 5.000E-07 |
| 23950 | 1198 | Y1198I | D | 1e-07 | 2.8e-07 | 5e-07 | 2.933E-07 | 2.003E-07 | 1.000E-07 | 5.000E-07 |
| 23951 | 1198 | Y1198L | D | 3.3e-07 | 1.3e-07 | 1.7e-07 | 2.100E-07 | 1.058E-07 | 1.300E-07 | 3.300E-07 |
| 23952 | 1198 | Y1198K | D | 3.1e-07 | 2.9e-07 | 1e-07 | 2.333E-07 | 1.159E-07 | 1.000E-07 | 3.100E-07 |
| 23953 | 1198 | Y1198M | D | 3.6e-07 | 1.2e-07 | 3.7e-07 | 2.833E-07 | 1.415E-07 | 1.200E-07 | 3.700E-07 |
| 23954 | 1198 | Y1198F | D | 2.8e-07 | 3.7e-07 | 2.2e-07 | 2.900E-07 | 7.550E-08 | 2.200E-07 | 3.700E-07 |
| 23955 | 1198 | Y1198P | D | 1.9e-07 | 1.8e-07 | 2.9e-07 | 2.200E-07 | 6.083E-08 | 1.800E-07 | 2.900E-07 |
| 23956 | 1198 | Y1198S | D | 4.7e-07 | 2.6e-07 | 1.2e-07 | 2.833E-07 | 1.762E-07 | 1.200E-07 | 4.700E-07 |
| 23957 | 1198 | Y1198T | D | 2e-07 | 3.5e-07 | 1.7e-07 | 2.400E-07 | 9.644E-08 | 1.700E-07 | 3.500E-07 |
| 23958 | 1198 | Y1198W | D | 1e-07 | 2.4e-07 | 2.6e-07 | 2.000E-07 | 8.718E-08 | 1.000E-07 | 2.600E-07 |
| 23959 | 1198 | Y1198Y | D | 6.3e-07 | 2.5e-07 | 3.5e-07 | 4.100E-07 | 1.970E-07 | 2.500E-07 | 6.300E-07 |
| 23960 | 1198 | Y1198V | D | 2.6e-07 | 2.6e-07 | 2e-07 | 2.400E-07 | 3.464E-08 | 2.000E-07 | 2.600E-07 |
| 23961 | 1199 | P1199A | D | 3.6e-07 | 3.1e-07 | 1.6e-07 | 2.767E-07 | 1.041E-07 | 1.600E-07 | 3.600E-07 |
| 23962 | 1199 | P1199R | D | 1.7e-07 | 1.7e-07 | 2.3e-07 | 1.900E-07 | 3.464E-08 | 1.700E-07 | 2.300E-07 |
| 23963 | 1199 | P1199N | D | 3.3e-07 | 2.6e-07 | 2e-07 | 2.633E-07 | 6.506E-08 | 2.000E-07 | 3.300E-07 |
| 23964 | 1199 | P1199D | D | 3.3e-07 | 3.8e-07 | 1.8e-07 | 2.967E-07 | 1.041E-07 | 1.800E-07 | 3.800E-07 |
| 23965 | 1199 | P1199C | D | 6.6e-08 | 1.3e-07 | 3.3e-07 | 1.753E-07 | 1.377E-07 | 6.600E-08 | 3.300E-07 |
| 23966 | 1199 | P1199E | D | 3.3e-07 | 4.7e-07 | 5.5e-07 | 4.500E-07 | 1.114E-07 | 3.300E-07 | 5.500E-07 |
| 23967 | 1199 | P1199Q | D | 2.4e-07 | 1.9e-07 | 5.6e-07 | 3.300E-07 | 2.007E-07 | 1.900E-07 | 5.600E-07 |
| 23968 | 1199 | P1199G | D | 2.3e-07 | 2.6e-07 | 4.2e-07 | 3.033E-07 | 1.021E-07 | 2.300E-07 | 4.200E-07 |
| 23969 | 1199 | P1199H | D | 3.1e-07 | 1.3e-07 | 1.4e-07 | 1.933E-07 | 1.012E-07 | 1.300E-07 | 3.100E-07 |

| | | | | | | | | | | |
|---|---|---|---|---|---|---|---|---|---|---|
| 23970 | 1199 | P1199I | D | 2.5e-07 | 2.9e-07 | 2.3e-07 | 2.567E-07 | 3.055E-08 | 2.300E-07 | 2.900E-07 |
| 23971 | 1199 | P1199L | D | 3.8e-07 | 2.5e-07 | 1.4e-07 | 2.567E-07 | 1.201E-07 | 1.400E-07 | 3.800E-07 |
| 23972 | 1199 | P1199K | D | 2.2e-07 | 2.4e-07 | 1.3e-07 | 1.967E-07 | 5.859E-08 | 1.300E-07 | 2.400E-07 |
| 23973 | 1199 | P1199M | D | 3.4e-07 | 3.3e-07 | 1.6e-07 | 2.767E-07 | 1.012E-07 | 1.600E-07 | 3.400E-07 |
| 23974 | 1199 | P1199F | D | 3.5e-07 | 7.4e-08 | 1.3e-07 | 1.847E-07 | 1.459E-07 | 7.400E-08 | 3.500E-07 |
| 23975 | 1199 | P1199P | D | 2.4e-07 | 5.9e-07 | 2.7e-07 | 3.667E-07 | 1.940E-07 | 2.400E-07 | 5.900E-07 |
| 23976 | 1199 | P1199S | D | 2.2e-07 | 3e-07 | 3.5e-07 | 2.900E-07 | 6.557E-08 | 2.200E-07 | 3.500E-07 |
| 23977 | 1199 | P1199T | D | 3.1e-07 | 1.4e-07 | 3.2e-07 | 2.567E-07 | 1.012E-07 | 1.400E-07 | 3.200E-07 |
| 23978 | 1199 | P1199W | D | 2.6e-07 | 6.3e-07 | 1.8e-07 | 3.567E-07 | 2.401E-07 | 1.800E-07 | 6.300E-07 |
| 23979 | 1199 | P1199Y | D | 1e-07 | 2e-07 | 1.5e-07 | 1.500E-07 | 5.000E-08 | 1.000E-07 | 2.000E-07 |
| 23980 | 1199 | P1199V | D | 3.5e-07 | 1.8e-07 | 7.4e-08 | 2.013E-07 | 1.392E-07 | 7.400E-08 | 3.500E-07 |
| 23981 | 1200 | R1200A | D | 1.3e-07 | 4.7e-07 | 2.4e-07 | 2.800E-07 | 1.735E-07 | 1.300E-07 | 4.700E-07 |
| 23982 | 1200 | R1200R | D | 1.8e-07 | 2.2e-07 | 1.8e-07 | 1.933E-07 | 2.309E-08 | 1.800E-07 | 2.200E-07 |
| 23983 | 1200 | R1200N | D | 1.8e-07 | 2.1e-07 | 2.6e-07 | 2.167E-07 | 4.041E-08 | 1.800E-07 | 2.600E-07 |
| 23984 | 1200 | R1200D | D | 8.6e-07 | 4e-07 | 3.6e-07 | 5.400E-07 | 2.778E-07 | 3.600E-07 | 8.600E-07 |
| 23985 | 1200 | R1200C | D | 3.8e-07 | 6e-08 | 2e-07 | 2.133E-07 | 1.604E-07 | 6.000E-08 | 3.800E-07 |
| 23986 | 1200 | R1200E | D | 1.8e-07 | 1.6e-07 | 6.8e-07 | 3.400E-07 | 2.946E-07 | 1.600E-07 | 6.800E-07 |
| 23987 | 1200 | R1200Q | D | 1.3e-07 | 3.6e-07 | 2e-07 | 2.300E-07 | 1.179E-07 | 1.300E-07 | 3.600E-07 |
| 23988 | 1200 | R1200G | D | 1.8e-07 | 2.1e-07 | 2.5e-07 | 2.133E-07 | 3.512E-08 | 1.800E-07 | 2.500E-07 |
| 23989 | 1200 | R1200H | D | 1.9e-07 | 2.5e-07 | 3.2e-07 | 2.533E-07 | 6.506E-08 | 1.900E-07 | 3.200E-07 |
| 23990 | 1200 | R1200I | D | 2.5e-07 | 2.6e-07 | 2.7e-07 | 2.600E-07 | 1.000E-08 | 2.500E-07 | 2.700E-07 |
| 23991 | 1200 | R1200L | D | 3.5e-07 | 4.2e-07 | 9.5e-08 | 2.883E-07 | 1.711E-07 | 9.500E-08 | 4.200E-07 |
| 23992 | 1200 | R1200K | D | 2.2e-07 | 1.5e-07 | 2.4e-07 | 2.033E-07 | 4.726E-08 | 1.500E-07 | 2.400E-07 |
| 23993 | 1200 | R1200M | D | 2.4e-07 | 2.6e-07 | 1.3e-07 | 2.100E-07 | 7.000E-08 | 1.300E-07 | 2.600E-07 |
| 23994 | 1200 | R1200F | D | 2.7e-07 | 2.9e-07 | 1.6e-07 | 2.400E-07 | 7.000E-08 | 1.600E-07 | 2.900E-07 |
| 23995 | 1200 | R1200P | D | 2.9e-07 | 3.1e-07 | 2.2e-07 | 2.733E-07 | 4.726E-08 | 2.200E-07 | 3.100E-07 |
| 23996 | 1200 | R1200S | D | 2.1e-07 | 2.1e-07 | 5.9e-07 | 3.367E-07 | 2.194E-07 | 2.100E-07 | 5.900E-07 |
| 23997 | 1200 | R1200T | D | 3.8e-07 | 2.4e-07 | 3.7e-07 | 3.300E-07 | 7.810E-08 | 2.400E-07 | 3.800E-07 |
| 23998 | 1200 | R1200W | D | 1.9e-07 | 1.5e-07 | 1.5e-07 | 1.633E-07 | 2.309E-08 | 1.500E-07 | 1.900E-07 |
| 23999 | 1200 | R1200Y | D | 6.3e-07 | 2e-07 | 5e-07 | 4.433E-07 | 2.205E-07 | 2.000E-07 | 6.300E-07 |
| 24000 | 1200 | R1200V | D | 5.2e-07 | 5e-07 | 2.4e-07 | 4.200E-07 | 1.562E-07 | 2.400E-07 | 5.200E-07 |
| 24001 | 1201 | E1201A | D | 2.4e-07 | 2.5e-07 | 2.2e-07 | 2.367E-07 | 1.528E-08 | 2.200E-07 | 2.500E-07 |
| 24002 | 1201 | E1201R | D | 2.7e-07 | 3.4e-07 | 2.7e-07 | 2.933E-07 | 4.041E-08 | 2.700E-07 | 3.400E-07 |
| 24003 | 1201 | E1201N | D | 1.2e-07 | 4.8e-07 | 2.2e-07 | 2.733E-07 | 1.858E-07 | 1.200E-07 | 4.800E-07 |
| 24004 | 1201 | E1201D | D | 3.6e-07 | 1.6e-07 | 2.2e-07 | 2.467E-07 | 1.026E-07 | 1.600E-07 | 3.600E-07 |
| 24005 | 1201 | E1201C | D | 6.4e-07 | 8.5e-08 | 1.8e-07 | 3.017E-07 | 2.968E-07 | 8.500E-08 | 6.400E-07 |
| 24006 | 1201 | E1201E | D | 3.5e-07 | 1.8e-07 | 3.3e-07 | 2.867E-07 | 9.292E-08 | 1.800E-07 | 3.500E-07 |
| 24007 | 1201 | E1201Q | D | 2.6e-07 | 3.5e-07 | 3.6e-07 | 3.233E-07 | 5.508E-08 | 2.600E-07 | 3.600E-07 |
| 24008 | 1201 | E1201G | D | 1.7e-07 | 2.4e-07 | 1.5e-07 | 1.867E-07 | 4.726E-08 | 1.500E-07 | 2.400E-07 |
| 24009 | 1201 | E1201H | D | 4.9e-07 | 3.3e-07 | 3.1e-07 | 3.767E-07 | 9.866E-08 | 3.100E-07 | 4.900E-07 |
| 24010 | 1201 | E1201I | D | 2.8e-07 | 4e-07 | 2.5e-07 | 3.100E-07 | 7.937E-08 | 2.500E-07 | 4.000E-07 |
| 24011 | 1201 | E1201L | D | 2.1e-07 | 2.4e-07 | 2.5e-07 | 2.333E-07 | 2.082E-08 | 2.100E-07 | 2.500E-07 |
| 24012 | 1201 | E1201K | D | 4e-07 | 2.5e-07 | 2.8e-07 | 3.100E-07 | 7.937E-08 | 2.500E-07 | 4.000E-07 |
| 24013 | 1201 | E1201M | D | 1.4e-07 | 1.6e-07 | 1.9e-07 | 1.633E-07 | 2.517E-08 | 1.400E-07 | 1.900E-07 |
| 24014 | 1201 | E1201F | D | 3.2e-07 | 1.4e-07 | 4e-07 | 2.867E-07 | 1.332E-07 | 1.400E-07 | 4.000E-07 |
| 24015 | 1201 | E1201P | D | 2.2e-07 | 1.3e-07 | 3.5e-07 | 2.333E-07 | 1.106E-07 | 1.300E-07 | 3.500E-07 |
| 24016 | 1201 | E1201S | D | 5.4e-07 | 2.7e-07 | 2.5e-07 | 3.533E-07 | 1.620E-07 | 2.500E-07 | 5.400E-07 |
| 24017 | 1201 | E1201T | D | 3.9e-07 | 2.9e-07 | 1.5e-07 | 2.767E-07 | 1.206E-07 | 1.500E-07 | 3.900E-07 |
| 24018 | 1201 | E1201W | D | 2.2e-07 | 2.6e-07 | 1.9e-07 | 2.233E-07 | 3.512E-08 | 1.900E-07 | 2.600E-07 |
| 24019 | 1201 | E1201Y | D | 4.2e-07 | 5.3e-07 | 2e-07 | 3.833E-07 | 1.680E-07 | 2.000E-07 | 5.300E-07 |
| 24020 | 1201 | E1201V | D | 3.1e-07 | 5.2e-07 | 2.4e-07 | 3.567E-07 | 1.457E-07 | 2.400E-07 | 5.200E-07 |
| 24021 | 1202 | A1202A | D | 2.9e-07 | 2.5e-07 | 1e-07 | 2.133E-07 | 1.002E-07 | 1.000E-07 | 2.900E-07 |
| 24022 | 1202 | A1202R | D | 1.7e-07 | 1.3e-07 | 1.6e-07 | 1.533E-07 | 2.082E-08 | 1.300E-07 | 1.700E-07 |
| 24023 | 1202 | A1202N | D | 2.3e-07 | 4.2e-07 | 3.3e-07 | 3.267E-07 | 9.504E-08 | 2.300E-07 | 4.200E-07 |
| 24024 | 1202 | A1202D | D | 3.7e-07 | 4.7e-07 | 2.4e-07 | 3.600E-07 | 1.153E-07 | 2.400E-07 | 4.700E-07 |
| 24025 | 1202 | A1202C | D | 1.9e-07 | 3.2e-07 | 5e-07 | 3.367E-07 | 1.557E-07 | 1.900E-07 | 5.000E-07 |
| 24026 | 1202 | A1202E | D | 2e-07 | 2.5e-07 | 4e-07 | 2.833E-07 | 1.041E-07 | 2.000E-07 | 4.000E-07 |
| 24027 | 1202 | A1202Q | D | 1.8e-07 | 3.3e-07 | 2.5e-07 | 2.533E-07 | 7.506E-08 | 1.800E-07 | 3.300E-07 |
| 24028 | 1202 | A1202G | D | 1.4e-07 | 6.8e-07 | 1.9e-07 | 3.367E-07 | 2.984E-07 | 1.400E-07 | 6.800E-07 |
| 24029 | 1202 | A1202H | D | 1.8e-07 | 1.5e-07 | 3.2e-07 | 2.167E-07 | 9.074E-08 | 1.500E-07 | 3.200E-07 |
| 24030 | 1202 | A1202I | D | 4.1e-07 | 2e-07 | 3.6e-07 | 3.233E-07 | 1.097E-07 | 2.000E-07 | 4.100E-07 |
| 24031 | 1202 | A1202L | D | 3e-07 | 4.3e-07 | 2.1e-07 | 3.133E-07 | 1.106E-07 | 2.100E-07 | 4.300E-07 |
| 24032 | 1202 | A1202K | D | 3.4e-07 | 4.6e-07 | 2.8e-07 | 3.600E-07 | 9.165E-08 | 2.800E-07 | 4.600E-07 |
| 24033 | 1202 | A1202M | D | 7.7e-07 | 1.7e-07 | 1.6e-07 | 3.667E-07 | 3.493E-07 | 1.600E-07 | 7.700E-07 |
| 24034 | 1202 | A1202F | D | 2.8e-07 | 4.4e-07 | 1.1e-07 | 2.767E-07 | 1.650E-07 | 1.100E-07 | 4.400E-07 |
| 24035 | 1202 | A1202P | D | 4.9e-07 | 2.9e-07 | 2.9e-07 | 3.567E-07 | 1.155E-07 | 2.900E-07 | 4.900E-07 |
| 24036 | 1202 | A1202S | D | 3.3e-07 | 1.3e-07 | 1.8e-07 | 2.133E-07 | 1.041E-07 | 1.300E-07 | 3.300E-07 |
| 24037 | 1202 | A1202T | D | 2.3e-07 | 2.4e-07 | 2.7e-07 | 2.467E-07 | 2.082E-08 | 2.300E-07 | 2.700E-07 |
| 24038 | 1202 | A1202W | D | 1.8e-07 | 2.7e-07 | 1.5e-07 | 2.000E-07 | 6.245E-08 | 1.500E-07 | 2.700E-07 |
| 24039 | 1202 | A1202Y | D | 3.2e-07 | 2.1e-07 | 1.8e-07 | 2.367E-07 | 7.371E-08 | 1.800E-07 | 3.200E-07 |
| 24040 | 1202 | A1202V | D | 3.9e-07 | 3.9e-07 | 2.2e-07 | 3.333E-07 | 9.815E-08 | 2.200E-07 | 3.900E-07 |
| 24041 | 1203 | K1203A | D | 3.4e-07 | 3.3e-07 | 2.8e-07 | 3.167E-07 | 3.215E-08 | 2.800E-07 | 3.400E-07 |
| 24042 | 1203 | K1203R | D | 1.6e-07 | 2.8e-07 | 1.4e-07 | 1.933E-07 | 7.572E-08 | 1.400E-07 | 2.800E-07 |
| 24043 | 1203 | K1203N | D | 3.5e-07 | 2.1e-07 | 1.2e-07 | 2.267E-07 | 1.159E-07 | 1.200E-07 | 3.500E-07 |
| 24044 | 1203 | K1203D | D | 2.6e-07 | 3.9e-07 | 3.5e-07 | 3.333E-07 | 6.658E-08 | 2.600E-07 | 3.900E-07 |
| 24045 | 1203 | K1203C | D | 4.4e-07 | 1.4e-07 | 1.9e-07 | 2.567E-07 | 1.607E-07 | 1.400E-07 | 4.400E-07 |
| 24046 | 1203 | K1203E | D | 7.7e-07 | 2.9e-07 | 2.7e-07 | 4.433E-07 | 2.831E-07 | 2.700E-07 | 7.700E-07 |
| 24047 | 1203 | K1203Q | D | 1.2e-07 | 1.2e-07 | 1.8e-07 | 1.400E-07 | 3.464E-08 | 1.200E-07 | 1.800E-07 |
| 24048 | 1203 | K1203G | D | 2e-07 | 3.3e-07 | 2.7e-07 | 2.667E-07 | 6.506E-08 | 2.000E-07 | 3.300E-07 |
| 24049 | 1203 | K1203H | D | 3.9e-07 | 3.2e-07 | 1.2e-07 | 2.767E-07 | 1.401E-07 | 1.200E-07 | 3.900E-07 |
| 24050 | 1203 | K1203I | D | 1.1e-07 | 2.3e-07 | 3.2e-07 | 2.200E-07 | 1.054E-07 | 1.100E-07 | 3.200E-07 |
| 24051 | 1203 | K1203L | D | 1.6e-07 | 3e-07 | 2e-07 | 2.200E-07 | 7.211E-08 | 1.600E-07 | 3.000E-07 |
| 24052 | 1203 | K1203K | D | 1.4e-07 | 3.4e-07 | 1.6e-07 | 2.133E-07 | 1.102E-07 | 1.400E-07 | 3.400E-07 |
| 24053 | 1203 | K1203M | D | 2.8e-07 | 3e-07 | 2.8e-07 | 2.867E-07 | 1.155E-08 | 2.800E-07 | 3.000E-07 |
| 24054 | 1203 | K1203F | D | 2.7e-07 | 1.6e-07 | 4.5e-07 | 2.933E-07 | 1.464E-07 | 1.600E-07 | 4.500E-07 |

| | | | | | | | | | | |
|---|---|---|---|---|---|---|---|---|---|---|
| 24055 | 1203 | K1203P | D | 3.3e-07 | 3.5e-07 | 2.7e-07 | 3.167E-07 | 4.163E-08 | 2.700E-07 | 3.500E-07 |
| 24056 | 1203 | K1203S | D | 3.2e-07 | 4.9e-07 | 3e-07 | 3.700E-07 | 1.044E-07 | 3.000E-07 | 4.900E-07 |
| 24057 | 1203 | K1203T | D | 3.4e-07 | 1.5e-07 | 2.5e-07 | 2.467E-07 | 9.504E-08 | 1.500E-07 | 3.400E-07 |
| 24058 | 1203 | K1203W | D | 1e-07 | 9.9e-08 | 2.4e-07 | 1.463E-07 | 8.112E-08 | 9.900E-08 | 2.400E-07 |
| 24059 | 1203 | K1203Y | D | 2.4e-07 | 1.3e-07 | 1.5e-07 | 1.733E-07 | 5.859E-08 | 1.300E-07 | 2.400E-07 |
| 24060 | 1203 | K1203V | D | 2.9e-07 | 4.2e-07 | 1.9e-07 | 3.000E-07 | 1.153E-07 | 1.900E-07 | 4.200E-07 |
| 24061 | 1204 | V1204A | D | 1.5e-07 | 1.3e-07 | 5.6e-07 | 2.800E-07 | 2.427E-07 | 1.300E-07 | 5.600E-07 |
| 24062 | 1204 | V1204R | D | 2.7e-07 | 2.2e-07 | 1.2e-07 | 2.033E-07 | 7.638E-08 | 1.200E-07 | 2.700E-07 |
| 24063 | 1204 | V1204N | D | 1.9e-07 | 4.2e-07 | 2.5e-07 | 2.867E-07 | 1.193E-07 | 1.900E-07 | 4.200E-07 |
| 24064 | 1204 | V1204D | D | 2e-07 | 3.5e-07 | 4.4e-07 | 3.300E-07 | 1.212E-07 | 2.000E-07 | 4.400E-07 |
| 24065 | 1204 | V1204C | D | 2.6e-07 | 1.3e-07 | 2.2e-07 | 2.033E-07 | 6.658E-08 | 1.300E-07 | 2.600E-07 |
| 24066 | 1204 | V1204E | D | 4.5e-07 | 1.8e-07 | 1.3e-07 | 2.533E-07 | 1.721E-07 | 1.300E-07 | 4.500E-07 |
| 24067 | 1204 | V1204Q | D | 2.7e-07 | 2.9e-07 | 2.9e-07 | 2.833E-07 | 1.155E-08 | 2.700E-07 | 2.900E-07 |
| 24068 | 1204 | V1204G | D | 1.9e-07 | 5.6e-07 | 1.8e-07 | 3.100E-07 | 2.166E-07 | 1.800E-07 | 5.600E-07 |
| 24069 | 1204 | V1204H | D | 2.6e-07 | 4.8e-07 | 2.4e-07 | 3.267E-07 | 1.332E-07 | 2.400E-07 | 4.800E-07 |
| 24070 | 1204 | V1204I | D | 1.9e-07 | 4.8e-07 | 3.6e-07 | 3.433E-07 | 1.457E-07 | 1.900E-07 | 4.800E-07 |
| 24071 | 1204 | V1204L | D | 2.3e-07 | 1.9e-07 | 2.7e-07 | 2.300E-07 | 4.000E-08 | 1.900E-07 | 2.700E-07 |
| 24072 | 1204 | V1204K | D | 1.6e-07 | 1.5e-07 | 3.7e-07 | 2.267E-07 | 1.242E-07 | 1.500E-07 | 3.700E-07 |
| 24073 | 1204 | V1204M | D | 2.3e-07 | 3.4e-07 | 5.3e-07 | 3.667E-07 | 1.518E-07 | 2.300E-07 | 5.300E-07 |
| 24074 | 1204 | V1204F | D | 2.9e-07 | 5.1e-07 | 3.6e-07 | 3.867E-07 | 1.124E-07 | 2.900E-07 | 5.100E-07 |
| 24075 | 1204 | V1204P | D | 2.9e-07 | 1.4e-07 | 4.1e-07 | 2.800E-07 | 1.353E-07 | 1.400E-07 | 4.100E-07 |
| 24076 | 1204 | V1204S | D | 1.4e-07 | 2.8e-07 | 2.5e-07 | 2.233E-07 | 7.371E-08 | 1.400E-07 | 2.800E-07 |
| 24077 | 1204 | V1204T | D | 1.6e-07 | 2.7e-07 | 1.8e-07 | 2.033E-07 | 5.859E-08 | 1.600E-07 | 2.700E-07 |
| 24078 | 1204 | V1204W | D | 1.8e-07 | 3.7e-07 | 3.6e-07 | 3.033E-07 | 1.069E-07 | 1.800E-07 | 3.700E-07 |
| 24079 | 1204 | V1204Y | D | 1.6e-07 | 1.2e-07 | 1.4e-07 | 1.400E-07 | 2.000E-08 | 1.200E-07 | 1.600E-07 |
| 24080 | 1204 | V1204V | D | 1e-07 | 5.9e-07 | 4.9e-07 | 3.933E-07 | 2.589E-07 | 1.000E-07 | 5.900E-07 |
| 24081 | 1205 | Q1205A | D | 2.6e-07 | 3.2e-07 | 1.5e-07 | 2.433E-07 | 8.622E-08 | 1.500E-07 | 3.200E-07 |
| 24082 | 1205 | Q1205R | D | 1.8e-07 | 1.9e-07 | 2.9e-07 | 2.200E-07 | 6.083E-08 | 1.800E-07 | 2.900E-07 |
| 24083 | 1205 | Q1205N | D | 1.9e-07 | 3.3e-07 | 2.5e-07 | 2.567E-07 | 7.024E-08 | 1.900E-07 | 3.300E-07 |
| 24084 | 1205 | Q1205D | D | 1.9e-07 | 1.5e-07 | 2.2e-07 | 1.867E-07 | 3.512E-08 | 1.500E-07 | 2.200E-07 |
| 24085 | 1205 | Q1205C | D | 2.5e-07 | 1.8e-07 | 2.3e-07 | 2.200E-07 | 3.606E-08 | 1.800E-07 | 2.500E-07 |
| 24086 | 1205 | Q1205E | D | 3.9e-07 | 3.9e-07 | 1.8e-07 | 3.200E-07 | 1.212E-07 | 1.800E-07 | 3.900E-07 |
| 24087 | 1205 | Q1205Q | D | 3.6e-07 | 1.8e-07 | 5.3e-07 | 3.567E-07 | 1.750E-07 | 1.800E-07 | 5.300E-07 |
| 24088 | 1205 | Q1205G | D | 2.9e-07 | 3.4e-07 | 2.8e-07 | 3.033E-07 | 3.215E-08 | 2.800E-07 | 3.400E-07 |
| 24089 | 1205 | Q1205H | D | 2.5e-07 | 1.8e-07 | 2e-07 | 2.100E-07 | 3.606E-08 | 1.800E-07 | 2.500E-07 |
| 24090 | 1205 | Q1205I | D | 4.4e-07 | 1.3e-07 | 1.4e-07 | 2.367E-07 | 1.762E-07 | 1.300E-07 | 4.400E-07 |
| 24091 | 1205 | Q1205L | D | 3e-07 | 3.8e-07 | 2.7e-07 | 3.167E-07 | 5.686E-08 | 2.700E-07 | 3.800E-07 |
| 24092 | 1205 | Q1205K | D | 3.4e-07 | 2.2e-07 | 2.2e-07 | 2.600E-07 | 6.928E-08 | 2.200E-07 | 3.400E-07 |
| 24093 | 1205 | Q1205M | D | 2.3e-07 | 2e-07 | 2.7e-07 | 2.333E-07 | 3.512E-08 | 2.000E-07 | 2.700E-07 |
| 24094 | 1205 | Q1205F | D | 4.8e-07 | 2.3e-07 | 1.6e-07 | 2.900E-07 | 1.682E-07 | 1.600E-07 | 4.800E-07 |
| 24095 | 1205 | Q1205P | D | 2e-07 | 5.3e-07 | 2.4e-07 | 3.233E-07 | 1.801E-07 | 2.000E-07 | 5.300E-07 |
| 24096 | 1205 | Q1205S | D | 2.9e-07 | 1.9e-07 | 2.6e-07 | 2.467E-07 | 5.132E-08 | 1.900E-07 | 2.900E-07 |
| 24097 | 1205 | Q1205T | D | 2.7e-07 | 1.1e-07 | 1.1e-07 | 1.633E-07 | 9.238E-08 | 1.100E-07 | 2.700E-07 |
| 24098 | 1205 | Q1205W | D | 1.1e-07 | 2.6e-07 | 2.4e-07 | 2.033E-07 | 8.145E-08 | 1.100E-07 | 2.600E-07 |
| 24099 | 1205 | Q1205Y | D | 3.4e-07 | 2.5e-07 | 2.9e-07 | 2.933E-07 | 4.509E-08 | 2.500E-07 | 3.400E-07 |
| 24100 | 1205 | Q1205V | D | 1.7e-07 | 2e-07 | 3.5e-07 | 2.400E-07 | 9.644E-08 | 1.700E-07 | 3.500E-07 |
| 24101 | 1206 | W1206A | D | 3.9e-07 | 1.4e-07 | 2.6e-07 | 2.633E-07 | 1.250E-07 | 1.400E-07 | 3.900E-07 |
| 24102 | 1206 | W1206R | D | 1.7e-07 | 3.2e-07 | 2.4e-07 | 2.433E-07 | 7.506E-08 | 1.700E-07 | 3.200E-07 |
| 24103 | 1206 | W1206N | D | 4.6e-07 | 2.1e-07 | 2.3e-07 | 3.000E-07 | 1.389E-07 | 2.100E-07 | 4.600E-07 |
| 24104 | 1206 | W1206D | D | 2.4e-07 | 2.4e-07 | 2.9e-07 | 2.567E-07 | 2.887E-08 | 2.400E-07 | 2.900E-07 |
| 24105 | 1206 | W1206C | D | 2.4e-07 | 6e-07 | 3.6e-07 | 4.000E-07 | 1.833E-07 | 2.400E-07 | 6.000E-07 |
| 24106 | 1206 | W1206E | D | 3.5e-07 | 1.8e-07 | 3.1e-07 | 2.800E-07 | 8.888E-08 | 1.800E-07 | 3.500E-07 |
| 24107 | 1206 | W1206Q | D | 2.5e-07 | 1.8e-07 | 1.7e-07 | 2.000E-07 | 4.359E-08 | 1.700E-07 | 2.500E-07 |
| 24108 | 1206 | W1206G | D | 3.8e-07 | 3.7e-07 | 2.1e-07 | 3.200E-07 | 9.539E-08 | 2.100E-07 | 3.800E-07 |
| 24109 | 1206 | W1206H | D | 1.6e-07 | 4.3e-07 | 1.7e-07 | 2.533E-07 | 1.531E-07 | 1.600E-07 | 4.300E-07 |
| 24110 | 1206 | W1206I | D | 1.8e-07 | 3.1e-07 | 2.3e-07 | 2.400E-07 | 6.557E-08 | 1.800E-07 | 3.100E-07 |
| 24111 | 1206 | W1206L | D | 2.9e-07 | 1.4e-07 | 2.4e-07 | 2.233E-07 | 7.638E-08 | 1.400E-07 | 2.900E-07 |
| 24112 | 1206 | W1206K | D | 2e-07 | 2.7e-07 | 2.2e-07 | 2.300E-07 | 3.606E-08 | 2.000E-07 | 2.700E-07 |
| 24113 | 1206 | W1206M | D | 7.1e-07 | 1.4e-07 | 3.4e-07 | 3.967E-07 | 2.892E-07 | 1.400E-07 | 7.100E-07 |
| 24114 | 1206 | W1206F | D | 2.7e-07 | 2.3e-07 | 3.2e-07 | 2.733E-07 | 4.509E-08 | 2.300E-07 | 3.200E-07 |
| 24115 | 1206 | W1206P | D | 1.4e-07 | 4.2e-07 | 3.1e-07 | 2.900E-07 | 1.411E-07 | 1.400E-07 | 4.200E-07 |
| 24116 | 1206 | W1206S | D | 3e-07 | 9.1e-08 | 4.4e-07 | 2.770E-07 | 1.756E-07 | 9.100E-08 | 4.400E-07 |
| 24117 | 1206 | W1206T | D | 2.7e-07 | 3.1e-07 | 1.9e-07 | 2.567E-07 | 6.110E-08 | 1.900E-07 | 3.100E-07 |
| 24118 | 1206 | W1206W | D | 1.7e-07 | 2.4e-07 | 3.8e-07 | 2.633E-07 | 1.069E-07 | 1.700E-07 | 3.800E-07 |
| 24119 | 1206 | W1206Y | D | 2.2e-07 | 3.3e-07 | 1.1e-07 | 2.200E-07 | 1.100E-07 | 1.100E-07 | 3.300E-07 |
| 24120 | 1206 | W1206V | D | 2.7e-07 | 3e-07 | 2.1e-07 | 2.600E-07 | 4.583E-08 | 2.100E-07 | 3.000E-07 |
| 24121 | 1207 | K1207A | D | 2.6e-07 | 2.6e-07 | 9.6e-08 | 2.053E-07 | 9.469E-08 | 9.600E-08 | 2.600E-07 |
| 24122 | 1207 | K1207R | D | 2.5e-07 | 1.5e-07 | 3.1e-07 | 2.367E-07 | 8.083E-08 | 1.500E-07 | 3.100E-07 |
| 24123 | 1207 | K1207N | D | 1.5e-07 | 1.9e-07 | 2.8e-07 | 2.067E-07 | 6.658E-08 | 1.500E-07 | 2.800E-07 |
| 24124 | 1207 | K1207D | D | 3.3e-07 | 3e-07 | 4.3e-07 | 3.533E-07 | 6.807E-08 | 3.000E-07 | 4.300E-07 |
| 24125 | 1207 | K1207C | D | 2.2e-07 | 2.5e-07 | 3.5e-07 | 2.733E-07 | 6.807E-08 | 2.200E-07 | 3.500E-07 |
| 24126 | 1207 | K1207E | D | 3.8e-07 | 4.8e-07 | 2.8e-07 | 3.800E-07 | 1.000E-07 | 2.800E-07 | 4.800E-07 |
| 24127 | 1207 | K1207Q | D | 3.7e-07 | 4.8e-07 | 1.4e-07 | 3.300E-07 | 1.735E-07 | 1.400E-07 | 4.800E-07 |
| 24128 | 1207 | K1207G | D | 2.4e-07 | 2.4e-07 | 1.4e-07 | 2.067E-07 | 5.774E-08 | 1.400E-07 | 2.400E-07 |
| 24129 | 1207 | K1207H | D | 1.4e-07 | 1.7e-07 | 2.9e-07 | 2.000E-07 | 7.937E-08 | 1.400E-07 | 2.900E-07 |
| 24130 | 1207 | K1207I | D | 3.1e-07 | 1.3e-07 | 1.1e-07 | 1.833E-07 | 1.102E-07 | 1.100E-07 | 3.100E-07 |
| 24131 | 1207 | K1207L | D | 7.1e-07 | 3e-07 | 2.7e-07 | 4.267E-07 | 2.458E-07 | 2.700E-07 | 7.100E-07 |
| 24132 | 1207 | K1207K | D | 1.3e-07 | 2.2e-07 | 1.9e-07 | 1.800E-07 | 4.583E-08 | 1.300E-07 | 2.200E-07 |
| 24133 | 1207 | K1207M | D | 1.8e-07 | 1.6e-07 | 1.4e-07 | 1.600E-07 | 2.000E-08 | 1.400E-07 | 1.800E-07 |
| 24134 | 1207 | K1207F | D | 1.5e-07 | 3e-07 | 2.8e-07 | 2.433E-07 | 8.145E-08 | 1.500E-07 | 3.000E-07 |
| 24135 | 1207 | K1207P | D | 2.3e-07 | 1.2e-07 | 2.2e-07 | 1.900E-07 | 6.083E-08 | 1.200E-07 | 2.300E-07 |
| 24136 | 1207 | K1207S | D | 1.7e-07 | 2.3e-07 | 2.5e-07 | 2.167E-07 | 4.163E-08 | 1.700E-07 | 2.500E-07 |
| 24137 | 1207 | K1207T | D | 1.8e-07 | 2.4e-07 | 2.3e-07 | 2.167E-07 | 3.215E-08 | 1.800E-07 | 2.400E-07 |
| 24138 | 1207 | K1207W | D | 1.6e-07 | 2.1e-07 | 2.5e-07 | 2.067E-07 | 4.509E-08 | 1.600E-07 | 2.500E-07 |
| 24139 | 1207 | K1207Y | D | 4e-07 | 1.9e-07 | 1.3e-07 | 2.400E-07 | 1.418E-07 | 1.300E-07 | 4.000E-07 |

| | | | | | | | | | | |
|---|---|---|---|---|---|---|---|---|---|---|
| 24140 | 1207 | K1207V | D | 2.2e-07 | 2.6e-07 | 3.1e-07 | 2.633E-07 | 4.509E-08 | 2.200E-07 | 3.100E-07 |
| 24141 | 1208 | V1208A | D | 1.5e-07 | 2.3e-07 | 3.5e-07 | 2.433E-07 | 1.007E-07 | 1.500E-07 | 3.500E-07 |
| 24142 | 1208 | V1208R | D | 2e-07 | 1.9e-07 | 1.6e-07 | 1.833E-07 | 2.082E-08 | 1.600E-07 | 2.000E-07 |
| 24143 | 1208 | V1208N | D | 2.9e-07 | 3.8e-07 | 2.7e-07 | 3.133E-07 | 5.859E-08 | 2.700E-07 | 3.800E-07 |
| 24144 | 1208 | V1208D | D | 5.7e-07 | 1.9e-07 | 3.1e-07 | 3.567E-07 | 1.943E-07 | 1.900E-07 | 5.700E-07 |
| 24145 | 1208 | V1208C | D | 4.3e-07 | 1.8e-07 | 1.8e-07 | 2.633E-07 | 1.443E-07 | 1.800E-07 | 4.300E-07 |
| 24146 | 1208 | V1208E | D | 3.8e-07 | 2.3e-07 | 2e-07 | 2.700E-07 | 9.644E-08 | 2.000E-07 | 3.800E-07 |
| 24147 | 1208 | V1208Q | D | 3.5e-07 | 2.1e-07 | 5.2e-07 | 3.600E-07 | 1.552E-07 | 2.100E-07 | 5.200E-07 |
| 24148 | 1208 | V1208G | D | 3e-07 | 1.4e-07 | 2e-07 | 2.133E-07 | 8.083E-08 | 1.400E-07 | 3.000E-07 |
| 24149 | 1208 | V1208H | D | 2.7e-07 | 4e-07 | 3.2e-07 | 3.300E-07 | 6.557E-08 | 2.700E-07 | 4.000E-07 |
| 24150 | 1208 | V1208I | D | 3e-07 | 5.3e-07 | 3.1e-07 | 3.800E-07 | 1.300E-07 | 3.000E-07 | 5.300E-07 |
| 24151 | 1208 | V1208L | D | 4.8e-07 | 2.1e-07 | 2.2e-07 | 3.033E-07 | 1.531E-07 | 2.100E-07 | 4.800E-07 |
| 24152 | 1208 | V1208K | D | 2.9e-07 | 3.5e-07 | 1.1e-06 | 5.800E-07 | 4.513E-07 | 2.900E-07 | 1.100E-06 |
| 24153 | 1208 | V1208M | D | 2.9e-07 | 2.1e-07 | 2.9e-07 | 2.633E-07 | 4.619E-08 | 2.100E-07 | 2.900E-07 |
| 24154 | 1208 | V1208F | D | 1.6e-07 | 1.2e-07 | 2e-07 | 1.600E-07 | 4.000E-08 | 1.200E-07 | 2.000E-07 |
| 24155 | 1208 | V1208P | D | 1.3e-07 | 1.8e-07 | 1.2e-07 | 1.433E-07 | 3.215E-08 | 1.200E-07 | 1.800E-07 |
| 24156 | 1208 | V1208S | D | 2.8e-07 | 3.2e-07 | 3.7e-07 | 3.233E-07 | 4.509E-08 | 2.800E-07 | 3.700E-07 |
| 24157 | 1208 | V1208T | D | 1.6e-07 | 1.7e-07 | 1.9e-07 | 1.733E-07 | 1.528E-08 | 1.600E-07 | 1.900E-07 |
| 24158 | 1208 | V1208W | D | 1.2e-07 | 2.5e-07 | 9.9e-08 | 1.563E-07 | 8.179E-08 | 9.900E-08 | 2.500E-07 |
| 24159 | 1208 | V1208Y | D | 4.2e-07 | 2.5e-07 | 2.9e-07 | 3.200E-07 | 8.888E-08 | 2.500E-07 | 4.200E-07 |
| 24160 | 1208 | V1208V | D | 2.4e-07 | 2.6e-07 | 1.8e-07 | 2.267E-07 | 4.163E-08 | 1.800E-07 | 2.600E-07 |
| 24161 | 1209 | D1209A | D | 2.6e-07 | 1.8e-07 | 1.7e-07 | 2.033E-07 | 4.933E-08 | 1.700E-07 | 2.600E-07 |
| 24162 | 1209 | D1209R | D | 2.7e-07 | 2.3e-07 | 2.4e-07 | 2.467E-07 | 2.082E-08 | 2.300E-07 | 2.700E-07 |
| 24163 | 1209 | D1209N | D | 3.3e-07 | 1.8e-07 | 1.8e-07 | 2.300E-07 | 8.660E-08 | 1.800E-07 | 3.300E-07 |
| 24164 | 1209 | D1209D | D | 3.3e-07 | 7.8e-07 | 2.6e-07 | 4.567E-07 | 2.822E-07 | 2.600E-07 | 7.800E-07 |
| 24165 | 1209 | D1209C | D | 4e-07 | 2.7e-07 | 3.7e-07 | 3.467E-07 | 6.807E-08 | 2.700E-07 | 4.000E-07 |
| 24166 | 1209 | D1209E | D | 3.8e-07 | 2.4e-07 | 2.7e-07 | 2.967E-07 | 7.371E-08 | 2.400E-07 | 3.800E-07 |
| 24167 | 1209 | D1209Q | D | 3.5e-07 | 3.8e-07 | 4.5e-07 | 3.933E-07 | 5.132E-08 | 3.500E-07 | 4.500E-07 |
| 24168 | 1209 | D1209G | D | 2.4e-07 | 1.2e-07 | 1.6e-07 | 1.733E-07 | 6.110E-08 | 1.200E-07 | 2.400E-07 |
| 24169 | 1209 | D1209H | D | 8.7e-08 | 1.7e-07 | 2.2e-07 | 1.590E-07 | 6.718E-08 | 8.700E-08 | 2.200E-07 |
| 24170 | 1209 | D1209I | D | 4.4e-07 | 1.4e-07 | 2.1e-07 | 2.633E-07 | 1.570E-07 | 1.400E-07 | 4.400E-07 |
| 24171 | 1209 | D1209L | D | 3.4e-07 | 3.3e-07 | 2.5e-07 | 3.067E-07 | 4.933E-08 | 2.500E-07 | 3.400E-07 |
| 24172 | 1209 | D1209K | D | 1.3e-07 | 2.4e-07 | 1.3e-07 | 1.667E-07 | 6.351E-08 | 1.300E-07 | 2.400E-07 |
| 24173 | 1209 | D1209M | D | 1.3e-07 | 4.4e-07 | 1.8e-07 | 2.500E-07 | 1.664E-07 | 1.300E-07 | 4.400E-07 |
| 24174 | 1209 | D1209F | D | 2.1e-07 | 1.7e-07 | 3.9e-07 | 2.567E-07 | 1.172E-07 | 1.700E-07 | 3.900E-07 |
| 24175 | 1209 | D1209P | D | 1.6e-07 | 1.6e-07 | 5.7e-07 | 2.967E-07 | 2.367E-07 | 1.600E-07 | 5.700E-07 |
| 24176 | 1209 | D1209S | D | 1.6e-07 | 3e-07 | 8e-08 | 1.800E-07 | 1.114E-07 | 8.000E-08 | 3.000E-07 |
| 24177 | 1209 | D1209T | D | 4.1e-07 | 1.3e-07 | 2.7e-07 | 2.700E-07 | 1.400E-07 | 1.300E-07 | 4.100E-07 |
| 24178 | 1209 | D1209W | D | 3.5e-07 | 3.2e-07 | 1.2e-07 | 2.633E-07 | 1.250E-07 | 1.200E-07 | 3.500E-07 |
| 24179 | 1209 | D1209Y | D | 2.3e-07 | 2.3e-07 | 1.7e-07 | 2.100E-07 | 3.464E-08 | 1.700E-07 | 2.300E-07 |
| 24180 | 1209 | D1209V | D | 2.8e-07 | 1.9e-07 | 2.9e-07 | 2.533E-07 | 5.508E-08 | 1.900E-07 | 2.900E-07 |
| 24181 | 1210 | N1210A | D | 2.1e-07 | 1.6e-07 | 5.6e-07 | 3.100E-07 | 2.179E-07 | 1.600E-07 | 5.600E-07 |
| 24182 | 1210 | N1210R | D | 1.4e-07 | 1.8e-07 | 3.6e-07 | 2.267E-07 | 1.172E-07 | 1.400E-07 | 3.600E-07 |
| 24183 | 1210 | N1210N | D | 1.4e-07 | 2.7e-07 | 5.1e-07 | 3.067E-07 | 1.877E-07 | 1.400E-07 | 5.100E-07 |
| 24184 | 1210 | N1210D | D | 2.2e-07 | 1.2e-07 | 2e-07 | 1.800E-07 | 5.292E-08 | 1.200E-07 | 2.200E-07 |
| 24185 | 1210 | N1210C | D | 4e-07 | 3.3e-07 | 2.7e-07 | 3.333E-07 | 6.506E-08 | 2.700E-07 | 4.000E-07 |
| 24186 | 1210 | N1210E | D | 1.2e-07 | 1.6e-07 | 2.1e-07 | 1.633E-07 | 4.509E-08 | 1.200E-07 | 2.100E-07 |
| 24187 | 1210 | N1210Q | D | 3.6e-07 | 1.8e-07 | 4e-07 | 3.133E-07 | 1.172E-07 | 1.800E-07 | 4.000E-07 |
| 24188 | 1210 | N1210G | D | 3.3e-07 | 2.5e-07 | 8.5e-08 | 2.217E-07 | 1.249E-07 | 8.500E-08 | 3.300E-07 |
| 24189 | 1210 | N1210H | D | 3.9e-07 | 4.2e-07 | 1.5e-07 | 3.200E-07 | 1.480E-07 | 1.500E-07 | 4.200E-07 |
| 24190 | 1210 | N1210I | D | 3.3e-07 | 4e-07 | 2.2e-07 | 3.167E-07 | 9.074E-08 | 2.200E-07 | 4.000E-07 |
| 24191 | 1210 | N1210L | D | 1.7e-07 | 2.7e-07 | 4e-07 | 2.800E-07 | 1.153E-07 | 1.700E-07 | 4.000E-07 |
| 24192 | 1210 | N1210K | D | 3.5e-07 | 2.7e-07 | 4.4e-07 | 3.533E-07 | 8.505E-08 | 2.700E-07 | 4.400E-07 |
| 24193 | 1210 | N1210M | D | 2.9e-07 | 7.5e-07 | 3.4e-07 | 4.600E-07 | 2.524E-07 | 2.900E-07 | 7.500E-07 |
| 24194 | 1210 | N1210F | D | 2.1e-07 | 1.5e-07 | 4.3e-07 | 2.633E-07 | 1.474E-07 | 1.500E-07 | 4.300E-07 |
| 24195 | 1210 | N1210P | D | 2.3e-07 | 1.2e-07 | 2.6e-07 | 2.033E-07 | 7.371E-08 | 1.200E-07 | 2.600E-07 |
| 24196 | 1210 | N1210S | D | 2.1e-07 | 3.4e-07 | 1.9e-07 | 2.467E-07 | 8.145E-08 | 1.900E-07 | 3.400E-07 |
| 24197 | 1210 | N1210T | D | 2.9e-07 | 2.2e-07 | 2.7e-07 | 2.600E-07 | 3.606E-08 | 2.200E-07 | 2.900E-07 |
| 24198 | 1210 | N1210W | D | 3.5e-07 | 3e-07 | 2e-07 | 2.833E-07 | 7.638E-08 | 2.000E-07 | 3.500E-07 |
| 24199 | 1210 | N1210Y | D | 2.9e-07 | 1.8e-07 | 5.1e-07 | 3.267E-07 | 1.680E-07 | 1.800E-07 | 5.100E-07 |
| 24200 | 1210 | N1210V | D | 3.1e-07 | 4.1e-07 | 3.2e-07 | 3.467E-07 | 5.508E-08 | 3.100E-07 | 4.100E-07 |
| 24201 | 1211 | A1211A | D | 1.9e-07 | 4.5e-07 | 3.1e-07 | 3.167E-07 | 1.301E-07 | 1.900E-07 | 4.500E-07 |
| 24202 | 1211 | A1211R | D | 1.1e-07 | 2.4e-07 | 2.5e-07 | 2.000E-07 | 7.810E-08 | 1.100E-07 | 2.500E-07 |
| 24203 | 1211 | A1211N | D | 1.7e-07 | 2.9e-07 | 1.7e-07 | 2.100E-07 | 6.928E-08 | 1.700E-07 | 2.900E-07 |
| 24204 | 1211 | A1211D | D | 3.6e-07 | 2.2e-07 | 4.8e-07 | 3.533E-07 | 1.301E-07 | 2.200E-07 | 4.800E-07 |
| 24205 | 1211 | A1211C | D | 3.2e-07 | 2e-07 | 3e-07 | 2.733E-07 | 6.429E-08 | 2.000E-07 | 3.200E-07 |
| 24206 | 1211 | A1211E | D | 2.9e-07 | 1.7e-07 | 2.4e-07 | 2.333E-07 | 6.028E-08 | 1.700E-07 | 2.900E-07 |
| 24207 | 1211 | A1211Q | D | 2.7e-07 | 4.4e-07 | 3.1e-07 | 3.400E-07 | 8.888E-08 | 2.700E-07 | 4.400E-07 |
| 24208 | 1211 | A1211G | D | 1.3e-07 | 3.1e-07 | 5.8e-07 | 3.400E-07 | 2.265E-07 | 1.300E-07 | 5.800E-07 |
| 24209 | 1211 | A1211H | D | 2.5e-07 | 3.7e-07 | 2.8e-07 | 3.000E-07 | 6.245E-08 | 2.500E-07 | 3.700E-07 |
| 24210 | 1211 | A1211I | D | 2.1e-07 | 2e-07 | 2.8e-07 | 2.300E-07 | 4.359E-08 | 2.000E-07 | 2.800E-07 |
| 24211 | 1211 | A1211L | D | 3.8e-07 | 1.1e-07 | 3.7e-07 | 2.867E-07 | 1.531E-07 | 1.100E-07 | 3.800E-07 |
| 24212 | 1211 | A1211K | D | 4.7e-07 | 2.6e-07 | 2.3e-07 | 3.200E-07 | 1.308E-07 | 2.300E-07 | 4.700E-07 |
| 24213 | 1211 | A1211M | D | 2.5e-07 | 3.4e-07 | 2.3e-07 | 2.733E-07 | 5.859E-08 | 2.300E-07 | 3.400E-07 |
| 24214 | 1211 | A1211F | D | 9.8e-08 | 2.7e-07 | 1.8e-07 | 1.827E-07 | 8.603E-08 | 9.800E-08 | 2.700E-07 |
| 24215 | 1211 | A1211P | D | 2.2e-07 | 2.8e-07 | 3.5e-07 | 2.833E-07 | 6.506E-08 | 2.200E-07 | 3.500E-07 |
| 24216 | 1211 | A1211S | D | 1.8e-07 | 4.4e-07 | 4.8e-07 | 3.667E-07 | 1.629E-07 | 1.800E-07 | 4.800E-07 |
| 24217 | 1211 | A1211T | D | 1.3e-07 | 2e-07 | 2e-07 | 1.767E-07 | 4.041E-08 | 1.300E-07 | 2.000E-07 |
| 24218 | 1211 | A1211W | D | 1.4e-07 | 1.9e-07 | 1.9e-07 | 1.733E-07 | 2.887E-08 | 1.400E-07 | 1.900E-07 |
| 24219 | 1211 | A1211Y | D | 2.7e-07 | 1.4e-07 | 1.1e-07 | 1.733E-07 | 8.505E-08 | 1.100E-07 | 2.700E-07 |
| 24220 | 1211 | A1211V | D | 1.2e-07 | 8.8e-08 | 1.6e-07 | 1.227E-07 | 3.607E-08 | 8.800E-08 | 1.600E-07 |
| 24221 | 1212 | L1212A | D | 3.7e-07 | 2e-07 | 1.6e-07 | 2.433E-07 | 1.115E-07 | 1.600E-07 | 3.700E-07 |
| 24222 | 1212 | L1212R | D | 3e-07 | 1.2e-07 | 1.5e-07 | 1.900E-07 | 9.644E-08 | 1.200E-07 | 3.000E-07 |
| 24223 | 1212 | L1212N | D | 2.1e-07 | 3.1e-07 | 4e-07 | 3.067E-07 | 9.504E-08 | 2.100E-07 | 4.000E-07 |
| 24224 | 1212 | L1212D | D | 2.1e-07 | 2.3e-07 | 4.6e-07 | 3.000E-07 | 1.389E-07 | 2.100E-07 | 4.600E-07 |

| | | | | | | | | | | |
|---|---|---|---|---|---|---|---|---|---|---|
| 24225 | 1212 | L1212C | D | 2e-07 | 2.6e-07 | 3.7e-07 | 2.767E-07 | 8.622E-08 | 2.000E-07 | 3.700E-07 |
| 24226 | 1212 | L1212E | D | 2e-07 | 2.8e-07 | 1.7e-07 | 2.167E-07 | 5.686E-08 | 1.700E-07 | 2.800E-07 |
| 24227 | 1212 | L1212Q | D | 1.2e-07 | 2e-07 | 2.9e-07 | 2.033E-07 | 8.505E-08 | 1.200E-07 | 2.900E-07 |
| 24228 | 1212 | L1212G | D | 1.9e-07 | 3.7e-07 | 2.6e-07 | 2.733E-07 | 9.074E-08 | 1.900E-07 | 3.700E-07 |
| 24229 | 1212 | L1212H | D | 3.5e-07 | 7.7e-07 | 3.2e-07 | 4.800E-07 | 2.516E-07 | 3.200E-07 | 7.700E-07 |
| 24230 | 1212 | L1212I | D | 2.2e-07 | 3e-07 | 1.9e-07 | 2.367E-07 | 5.686E-08 | 1.900E-07 | 3.000E-07 |
| 24231 | 1212 | L1212L | D | 2.4e-07 | 1.7e-07 | 2.2e-07 | 2.100E-07 | 3.606E-08 | 1.700E-07 | 2.400E-07 |
| 24232 | 1212 | L1212K | D | 2.9e-07 | 3.4e-07 | 1.3e-07 | 2.533E-07 | 1.097E-07 | 1.300E-07 | 3.400E-07 |
| 24233 | 1212 | L1212M | D | 1.5e-07 | 1.6e-07 | 2.6e-07 | 1.900E-07 | 6.083E-08 | 1.500E-07 | 2.600E-07 |
| 24234 | 1212 | L1212F | D | 2.6e-07 | 2.2e-07 | 1.4e-07 | 2.067E-07 | 6.110E-08 | 1.400E-07 | 2.600E-07 |
| 24235 | 1212 | L1212P | D | 2.2e-07 | 2.1e-07 | 2e-07 | 2.100E-07 | 1.000E-08 | 2.000E-07 | 2.200E-07 |
| 24236 | 1212 | L1212S | D | 2.2e-07 | 4.3e-07 | 2.3e-07 | 2.933E-07 | 1.185E-07 | 2.200E-07 | 4.300E-07 |
| 24237 | 1212 | L1212T | D | 2.8e-07 | 3e-07 | 3e-07 | 2.933E-07 | 1.155E-08 | 2.800E-07 | 3.000E-07 |
| 24238 | 1212 | L1212W | D | 8.3e-08 | 3.5e-07 | 2.9e-07 | 2.410E-07 | 1.401E-07 | 8.300E-08 | 3.500E-07 |
| 24239 | 1212 | L1212Y | D | 1.6e-07 | 2.6e-07 | 1.7e-07 | 1.967E-07 | 5.508E-08 | 1.600E-07 | 2.600E-07 |
| 24240 | 1212 | L1212V | D | 2.7e-07 | 5.4e-07 | 3.4e-07 | 3.833E-07 | 1.401E-07 | 2.700E-07 | 5.400E-07 |
| 24241 | 1213 | Q1213A | D | 4e-07 | 1.7e-07 | 4.2e-07 | 3.300E-07 | 1.389E-07 | 1.700E-07 | 4.200E-07 |
| 24242 | 1213 | Q1213R | D | 2e-07 | 2.7e-07 | 1.4e-07 | 2.033E-07 | 6.506E-08 | 1.400E-07 | 2.700E-07 |
| 24243 | 1213 | Q1213N | D | 1.7e-07 | 4.8e-07 | 1.6e-07 | 2.700E-07 | 1.819E-07 | 1.600E-07 | 4.800E-07 |
| 24244 | 1213 | Q1213D | D | 2.1e-07 | 2.8e-07 | 2.6e-07 | 2.500E-07 | 3.606E-08 | 2.100E-07 | 2.800E-07 |
| 24245 | 1213 | Q1213C | D | 1.2e-07 | 3.1e-07 | 2.7e-07 | 2.333E-07 | 1.002E-07 | 1.200E-07 | 3.100E-07 |
| 24246 | 1213 | Q1213E | D | 2.3e-07 | 2e-07 | 4.8e-07 | 3.033E-07 | 1.537E-07 | 2.000E-07 | 4.800E-07 |
| 24247 | 1213 | Q1213Q | D | 3.5e-07 | 3e-07 | 2.8e-07 | 3.100E-07 | 3.606E-08 | 2.800E-07 | 3.500E-07 |
| 24248 | 1213 | Q1213G | D | 2.2e-07 | 1.8e-07 | 7.5e-07 | 3.833E-07 | 3.182E-07 | 1.800E-07 | 7.500E-07 |
| 24249 | 1213 | Q1213H | D | 2.2e-07 | 2.7e-07 | 2.6e-07 | 2.500E-07 | 2.646E-08 | 2.200E-07 | 2.700E-07 |
| 24250 | 1213 | Q1213I | D | 3.3e-07 | 3e-07 | 1.3e-07 | 2.533E-07 | 1.079E-07 | 1.300E-07 | 3.300E-07 |
| 24251 | 1213 | Q1213L | D | 4.4e-07 | 1.9e-07 | 5.1e-07 | 3.800E-07 | 1.682E-07 | 1.900E-07 | 5.100E-07 |
| 24252 | 1213 | Q1213K | D | 8.4e-08 | 2.4e-07 | 3.6e-07 | 2.280E-07 | 1.384E-07 | 8.400E-08 | 3.600E-07 |
| 24253 | 1213 | Q1213M | D | 2.5e-07 | 3.7e-07 | 2.9e-07 | 3.033E-07 | 6.110E-08 | 2.500E-07 | 3.700E-07 |
| 24254 | 1213 | Q1213F | D | 3.2e-07 | 2.3e-07 | 1.6e-07 | 2.367E-07 | 8.021E-08 | 1.600E-07 | 3.200E-07 |
| 24255 | 1213 | Q1213P | D | 1.9e-07 | 4.3e-07 | 2.4e-07 | 2.867E-07 | 1.266E-07 | 1.900E-07 | 4.300E-07 |
| 24256 | 1213 | Q1213S | D | 1.6e-07 | 4.8e-07 | 2.6e-07 | 3.000E-07 | 1.637E-07 | 1.600E-07 | 4.800E-07 |
| 24257 | 1213 | Q1213T | D | 2.5e-07 | 2.6e-07 | 2.2e-07 | 2.433E-07 | 2.082E-08 | 2.200E-07 | 2.600E-07 |
| 24258 | 1213 | Q1213W | D | 1.6e-07 | 3.5e-07 | 1.7e-07 | 2.267E-07 | 1.069E-07 | 1.600E-07 | 3.500E-07 |
| 24259 | 1213 | Q1213Y | D | 2e-07 | 3.2e-07 | 2.4e-07 | 2.533E-07 | 6.110E-08 | 2.000E-07 | 3.200E-07 |
| 24260 | 1213 | Q1213V | D | 2.1e-07 | 4.7e-07 | 4.1e-07 | 3.633E-07 | 1.361E-07 | 2.100E-07 | 4.700E-07 |
| 24261 | 1214 | S1214A | D | 7.5e-08 | 7.1e-07 | 3.5e-07 | 3.783E-07 | 3.184E-07 | 7.500E-08 | 7.100E-07 |
| 24262 | 1214 | S1214R | D | 2.3e-07 | 2.5e-07 | 1.9e-07 | 2.233E-07 | 3.055E-08 | 1.900E-07 | 2.500E-07 |
| 24263 | 1214 | S1214N | D | 1.1e-07 | 3.2e-07 | 3.4e-07 | 2.567E-07 | 1.274E-07 | 1.100E-07 | 3.400E-07 |
| 24264 | 1214 | S1214D | D | 4.8e-07 | 1.9e-07 | 3e-07 | 3.233E-07 | 1.464E-07 | 1.900E-07 | 4.800E-07 |
| 24265 | 1214 | S1214C | D | 1.9e-07 | 3.1e-07 | 3e-07 | 2.667E-07 | 6.658E-08 | 1.900E-07 | 3.100E-07 |
| 24266 | 1214 | S1214E | D | 3.1e-07 | 2.3e-07 | 3.9e-07 | 3.100E-07 | 8.000E-08 | 2.300E-07 | 3.900E-07 |
| 24267 | 1214 | S1214Q | D | 2e-07 | 2.8e-07 | 3.5e-07 | 2.767E-07 | 7.506E-08 | 2.000E-07 | 3.500E-07 |
| 24268 | 1214 | S1214G | D | 2.8e-07 | 1.7e-07 | 7.9e-08 | 1.763E-07 | 1.006E-07 | 7.900E-08 | 2.800E-07 |
| 24269 | 1214 | S1214H | D | 2.9e-07 | 2.6e-07 | 2.4e-07 | 2.633E-07 | 2.517E-08 | 2.400E-07 | 2.900E-07 |
| 24270 | 1214 | S1214I | D | 2.5e-07 | 3.1e-07 | 2.1e-07 | 2.567E-07 | 5.033E-08 | 2.100E-07 | 3.100E-07 |
| 24271 | 1214 | S1214L | D | 2.1e-07 | 3.1e-07 | 5e-07 | 3.400E-07 | 1.473E-07 | 2.100E-07 | 5.000E-07 |
| 24272 | 1214 | S1214K | D | 3.5e-07 | 3.6e-07 | 1.3e-07 | 2.800E-07 | 1.300E-07 | 1.300E-07 | 3.600E-07 |
| 24273 | 1214 | S1214M | D | 2.6e-07 | 4.5e-07 | 3.3e-07 | 3.467E-07 | 9.609E-08 | 2.600E-07 | 4.500E-07 |
| 24274 | 1214 | S1214F | D | 4.6e-07 | 1.7e-07 | 3.7e-07 | 3.333E-07 | 1.484E-07 | 1.700E-07 | 4.600E-07 |
| 24275 | 1214 | S1214P | D | 1.4e-07 | 1.6e-07 | 1.7e-07 | 1.567E-07 | 1.528E-08 | 1.400E-07 | 1.700E-07 |
| 24276 | 1214 | S1214S | D | 1.1e-07 | 3.2e-07 | 2.2e-07 | 2.167E-07 | 1.050E-07 | 1.100E-07 | 3.200E-07 |
| 24277 | 1214 | S1214T | D | 4.7e-07 | 2.7e-07 | 2.5e-07 | 3.300E-07 | 1.217E-07 | 2.500E-07 | 4.700E-07 |
| 24278 | 1214 | S1214W | D | 6.1e-07 | 1.8e-07 | 4.5e-07 | 4.133E-07 | 2.173E-07 | 1.800E-07 | 6.100E-07 |
| 24279 | 1214 | S1214Y | D | 1.8e-07 | 1.6e-07 | 2.7e-07 | 2.033E-07 | 5.859E-08 | 1.600E-07 | 2.700E-07 |
| 24280 | 1214 | S1214V | D | 2.9e-07 | 3.1e-07 | 1.6e-07 | 2.533E-07 | 8.145E-08 | 1.600E-07 | 3.100E-07 |
| 24281 | 1215 | G1215A | D | 1e-07 | 1.6e-07 | 3.1e-07 | 1.900E-07 | 1.082E-07 | 1.000E-07 | 3.100E-07 |
| 24282 | 1215 | G1215R | D | 4.6e-07 | 2.5e-07 | 1.4e-07 | 2.833E-07 | 1.626E-07 | 1.400E-07 | 4.600E-07 |
| 24283 | 1215 | G1215N | D | 1.3e-07 | 2.6e-07 | 1.2e-07 | 1.700E-07 | 7.810E-08 | 1.200E-07 | 2.600E-07 |
| 24284 | 1215 | G1215D | D | 3.7e-07 | 1.2e-07 | 1.3e-07 | 2.067E-07 | 1.415E-07 | 1.200E-07 | 3.700E-07 |
| 24285 | 1215 | G1215C | D | 1.7e-07 | 4.1e-07 | 2.8e-07 | 2.867E-07 | 1.201E-07 | 1.700E-07 | 4.100E-07 |
| 24286 | 1215 | G1215E | D | 4.3e-07 | 1.2e-07 | 3e-07 | 2.833E-07 | 1.557E-07 | 1.200E-07 | 4.300E-07 |
| 24287 | 1215 | G1215Q | D | 2.6e-07 | 2.6e-07 | 4.2e-07 | 3.133E-07 | 9.238E-08 | 2.600E-07 | 4.200E-07 |
| 24288 | 1215 | G1215G | D | 2.8e-07 | 1.8e-07 | 2e-07 | 2.200E-07 | 5.292E-08 | 1.800E-07 | 2.800E-07 |
| 24289 | 1215 | G1215H | D | 2.4e-07 | 3.7e-07 | 2.2e-07 | 2.767E-07 | 8.145E-08 | 2.200E-07 | 3.700E-07 |
| 24290 | 1215 | G1215I | D | 2.9e-07 | 2.8e-07 | 3.2e-07 | 2.967E-07 | 2.082E-08 | 2.800E-07 | 3.200E-07 |
| 24291 | 1215 | G1215L | D | 1.5e-07 | 1.8e-07 | 4.6e-07 | 2.633E-07 | 1.710E-07 | 1.500E-07 | 4.600E-07 |
| 24292 | 1215 | G1215K | D | 4.5e-07 | 2.3e-07 | 1.5e-07 | 2.767E-07 | 1.553E-07 | 1.500E-07 | 4.500E-07 |
| 24293 | 1215 | G1215M | D | 2e-07 | 4e-07 | 1.2e-07 | 2.400E-07 | 1.442E-07 | 1.200E-07 | 4.000E-07 |
| 24294 | 1215 | G1215F | D | 1.9e-07 | 1e-07 | 2e-07 | 1.633E-07 | 5.508E-08 | 1.000E-07 | 2.000E-07 |
| 24295 | 1215 | G1215P | D | 2e-07 | 3.8e-07 | 2.1e-07 | 2.633E-07 | 1.012E-07 | 2.000E-07 | 3.800E-07 |
| 24296 | 1215 | G1215S | D | 1.5e-07 | 2.2e-07 | 2.2e-07 | 1.967E-07 | 4.041E-08 | 1.500E-07 | 2.200E-07 |
| 24297 | 1215 | G1215T | D | 2.8e-07 | 2.2e-07 | 1.8e-07 | 2.267E-07 | 5.033E-08 | 1.800E-07 | 2.800E-07 |
| 24298 | 1215 | G1215W | D | 3.6e-07 | 1.9e-07 | 1.4e-07 | 2.300E-07 | 1.153E-07 | 1.400E-07 | 3.600E-07 |
| 24299 | 1215 | G1215Y | D | 1.6e-07 | 2.7e-07 | 1.7e-07 | 2.000E-07 | 6.083E-08 | 1.600E-07 | 2.700E-07 |
| 24300 | 1215 | G1215V | D | 3.1e-07 | 1.1e-07 | 2.1e-07 | 2.100E-07 | 1.000E-07 | 1.100E-07 | 3.100E-07 |
| 24301 | 1216 | N1216A | D | 2.7e-07 | 2.3e-07 | 3.7e-07 | 2.900E-07 | 7.211E-08 | 2.300E-07 | 3.700E-07 |
| 24302 | 1216 | N1216R | D | 9.3e-08 | 8.6e-08 | 2.2e-07 | 1.330E-07 | 7.543E-08 | 8.600E-08 | 2.200E-07 |
| 24303 | 1216 | N1216N | D | 2.2e-07 | 2.3e-07 | 2.5e-07 | 2.333E-07 | 1.528E-08 | 2.200E-07 | 2.500E-07 |
| 24304 | 1216 | N1216D | D | 3.3e-07 | 2.3e-07 | 5.7e-08 | 2.057E-07 | 1.381E-07 | 5.700E-08 | 3.300E-07 |
| 24305 | 1216 | N1216C | D | 4e-07 | 3.3e-07 | 2.9e-07 | 3.400E-07 | 5.568E-08 | 2.900E-07 | 4.000E-07 |
| 24306 | 1216 | N1216E | D | 2.3e-07 | 2.2e-07 | 1.8e-07 | 2.100E-07 | 2.646E-08 | 1.800E-07 | 2.300E-07 |
| 24307 | 1216 | N1216Q | D | 3.4e-07 | 3.7e-07 | 2.3e-07 | 3.133E-07 | 7.371E-08 | 2.300E-07 | 3.700E-07 |
| 24308 | 1216 | N1216G | D | 1.1e-07 | 3e-07 | 2.8e-07 | 2.300E-07 | 1.044E-07 | 1.100E-07 | 3.000E-07 |
| 24309 | 1216 | N1216H | D | 2.1e-07 | 2.1e-07 | 2.3e-07 | 2.167E-07 | 1.155E-08 | 2.100E-07 | 2.300E-07 |

| | | | | | | | | | | |
|---|---|---|---|---|---|---|---|---|---|---|
| 24310 | 1216 | N1216I | D | 2.8e-07 | 3.7e-07 | 1.6e-07 | 2.700E-07 | 1.054E-07 | 1.600E-07 | 3.700E-07 |
| 24311 | 1216 | N1216L | D | 2.1e-07 | 1.3e-07 | 4.6e-07 | 2.667E-07 | 1.721E-07 | 1.300E-07 | 4.600E-07 |
| 24312 | 1216 | N1216K | D | 3.8e-07 | 2.1e-07 | 2.3e-07 | 2.733E-07 | 9.292E-08 | 2.100E-07 | 3.800E-07 |
| 24313 | 1216 | N1216M | D | 1.8e-07 | 1e-07 | 1.9e-07 | 1.567E-07 | 4.933E-08 | 1.000E-07 | 1.900E-07 |
| 24314 | 1216 | N1216F | D | 3.5e-07 | 2e-07 | 1.4e-07 | 2.300E-07 | 1.082E-07 | 1.400E-07 | 3.500E-07 |
| 24315 | 1216 | N1216P | D | 1.9e-07 | 1.9e-07 | 1.4e-07 | 1.733E-07 | 2.887E-08 | 1.400E-07 | 1.900E-07 |
| 24316 | 1216 | N1216S | D | 3.9e-07 | 1.7e-07 | 2e-07 | 2.533E-07 | 1.193E-07 | 1.700E-07 | 3.900E-07 |
| 24317 | 1216 | N1216T | D | 1.5e-07 | 1.9e-07 | 2.9e-07 | 2.100E-07 | 7.211E-08 | 1.500E-07 | 2.900E-07 |
| 24318 | 1216 | N1216W | D | 2.7e-07 | 2e-07 | 1.8e-07 | 2.167E-07 | 4.726E-08 | 1.800E-07 | 2.700E-07 |
| 24319 | 1216 | N1216Y | D | 3.7e-07 | 2.1e-07 | 3.5e-07 | 3.100E-07 | 8.718E-08 | 2.100E-07 | 3.700E-07 |
| 24320 | 1216 | N1216V | D | 3.4e-07 | 2.4e-07 | 3.7e-07 | 3.167E-07 | 6.807E-08 | 2.400E-07 | 3.700E-07 |
| 24321 | 1217 | S1217A | D | 1.4e-07 | 4.4e-07 | 2e-07 | 2.600E-07 | 1.587E-07 | 1.400E-07 | 4.400E-07 |
| 24322 | 1217 | S1217R | D | 2.1e-07 | 3.2e-07 | 1.1e-07 | 2.133E-07 | 1.050E-07 | 1.100E-07 | 3.200E-07 |
| 24323 | 1217 | S1217N | D | 3.5e-07 | 2.1e-07 | 1.3e-07 | 2.300E-07 | 1.114E-07 | 1.300E-07 | 3.500E-07 |
| 24324 | 1217 | S1217D | D | 2.6e-07 | 3.2e-07 | 1.6e-07 | 2.467E-07 | 8.083E-08 | 1.600E-07 | 3.200E-07 |
| 24325 | 1217 | S1217C | D | 2.8e-07 | 2e-07 | 1.7e-07 | 2.167E-07 | 5.686E-08 | 1.700E-07 | 2.800E-07 |
| 24326 | 1217 | S1217E | D | 4e-07 | 3.4e-07 | 1.8e-07 | 3.067E-07 | 1.137E-07 | 1.800E-07 | 4.000E-07 |
| 24327 | 1217 | S1217Q | D | 2.1e-07 | 2e-07 | 5.5e-07 | 3.200E-07 | 1.992E-07 | 2.000E-07 | 5.500E-07 |
| 24328 | 1217 | S1217G | D | 3.2e-07 | 4.2e-07 | 1.5e-07 | 2.967E-07 | 1.365E-07 | 1.500E-07 | 4.200E-07 |
| 24329 | 1217 | S1217H | D | 3.6e-07 | 2.2e-07 | 2.1e-07 | 2.633E-07 | 8.386E-08 | 2.100E-07 | 3.600E-07 |
| 24330 | 1217 | S1217I | D | 1.6e-07 | 1.6e-07 | 1.1e-07 | 1.433E-07 | 2.887E-08 | 1.100E-07 | 1.600E-07 |
| 24331 | 1217 | S1217L | D | 2.1e-07 | 3e-07 | 7.6e-07 | 4.233E-07 | 2.950E-07 | 2.100E-07 | 7.600E-07 |
| 24332 | 1217 | S1217K | D | 5.2e-07 | 2.9e-07 | 1.7e-07 | 3.267E-07 | 1.779E-07 | 1.700E-07 | 5.200E-07 |
| 24333 | 1217 | S1217M | D | 1.4e-07 | 1.5e-07 | 4.1e-07 | 2.333E-07 | 1.531E-07 | 1.400E-07 | 4.100E-07 |
| 24334 | 1217 | S1217F | D | 2.1e-07 | 2.9e-07 | 4.9e-07 | 3.300E-07 | 1.442E-07 | 2.100E-07 | 4.900E-07 |
| 24335 | 1217 | S1217P | D | 1.2e-07 | 4.6e-07 | 1.1e-07 | 2.300E-07 | 1.992E-07 | 1.100E-07 | 4.600E-07 |
| 24336 | 1217 | S1217S | D | 5.4e-07 | 1.4e-07 | 2.5e-07 | 3.100E-07 | 2.066E-07 | 1.400E-07 | 5.400E-07 |
| 24337 | 1217 | S1217T | D | 3.2e-07 | 2.7e-07 | 1.7e-07 | 2.533E-07 | 7.638E-08 | 1.700E-07 | 3.200E-07 |
| 24338 | 1217 | S1217W | D | 6.6e-07 | 1.7e-07 | 3.3e-07 | 3.867E-07 | 2.499E-07 | 1.700E-07 | 6.600E-07 |
| 24339 | 1217 | S1217Y | D | 2.8e-07 | 2.3e-07 | 3e-07 | 2.700E-07 | 3.606E-08 | 2.300E-07 | 3.000E-07 |
| 24340 | 1217 | S1217V | D | 4.6e-07 | 3.1e-07 | 2e-07 | 3.233E-07 | 1.305E-07 | 2.000E-07 | 4.600E-07 |
| 24341 | 1218 | Q1218A | D | 3.8e-07 | 2.8e-07 | 1.5e-07 | 2.700E-07 | 1.153E-07 | 1.500E-07 | 3.800E-07 |
| 24342 | 1218 | Q1218R | D | 2.4e-07 | 2.6e-07 | 3.3e-07 | 2.767E-07 | 4.726E-08 | 2.400E-07 | 3.300E-07 |
| 24343 | 1218 | Q1218N | D | 1.3e-07 | 2.3e-07 | 2.6e-07 | 2.067E-07 | 6.807E-08 | 1.300E-07 | 2.600E-07 |
| 24344 | 1218 | Q1218D | D | 3.5e-07 | 1.7e-07 | 3.4e-07 | 2.867E-07 | 1.012E-07 | 1.700E-07 | 3.500E-07 |
| 24345 | 1218 | Q1218C | D | 4.7e-07 | 1.7e-07 | 4.6e-07 | 3.667E-07 | 1.704E-07 | 1.700E-07 | 4.700E-07 |
| 24346 | 1218 | Q1218E | D | 2.9e-07 | 3.7e-07 | 4e-07 | 3.533E-07 | 5.686E-08 | 2.900E-07 | 4.000E-07 |
| 24347 | 1218 | Q1218Q | D | 2.1e-07 | 2.7e-07 | 3e-07 | 2.600E-07 | 4.583E-08 | 2.100E-07 | 3.000E-07 |
| 24348 | 1218 | Q1218G | D | 1.4e-07 | 1.9e-07 | 2.6e-07 | 1.967E-07 | 6.028E-08 | 1.400E-07 | 2.600E-07 |
| 24349 | 1218 | Q1218H | D | 2.4e-07 | 3.2e-07 | 3.4e-07 | 3.000E-07 | 5.292E-08 | 2.400E-07 | 3.400E-07 |
| 24350 | 1218 | Q1218I | D | 1.6e-07 | 2.4e-07 | 4e-07 | 2.667E-07 | 1.222E-07 | 1.600E-07 | 4.000E-07 |
| 24351 | 1218 | Q1218L | D | 1.9e-07 | 3.4e-07 | 2.1e-07 | 2.467E-07 | 8.145E-08 | 1.900E-07 | 3.400E-07 |
| 24352 | 1218 | Q1218K | D | 1.6e-07 | 1.8e-07 | 2.1e-07 | 1.833E-07 | 2.517E-08 | 1.600E-07 | 2.100E-07 |
| 24353 | 1218 | Q1218M | D | 1.4e-07 | 2.5e-07 | 2.1e-07 | 2.000E-07 | 5.568E-08 | 1.400E-07 | 2.500E-07 |
| 24354 | 1218 | Q1218F | D | 1.6e-07 | 3.8e-07 | 2.8e-07 | 2.733E-07 | 1.102E-07 | 1.600E-07 | 3.800E-07 |
| 24355 | 1218 | Q1218P | D | 2.6e-07 | 1.7e-07 | 3.9e-07 | 2.733E-07 | 1.106E-07 | 1.700E-07 | 3.900E-07 |
| 24356 | 1218 | Q1218S | D | 9.1e-08 | 2.8e-07 | 4.6e-07 | 2.770E-07 | 1.845E-07 | 9.100E-08 | 4.600E-07 |
| 24357 | 1218 | Q1218T | D | 2.5e-07 | 1.3e-07 | 2.7e-07 | 2.167E-07 | 7.572E-08 | 1.300E-07 | 2.700E-07 |
| 24358 | 1218 | Q1218W | D | 3.1e-07 | 3.1e-07 | 3.8e-07 | 3.333E-07 | 4.041E-08 | 3.100E-07 | 3.800E-07 |
| 24359 | 1218 | Q1218Y | D | 2.1e-07 | 1.7e-07 | 3.5e-07 | 2.433E-07 | 9.452E-08 | 1.700E-07 | 3.500E-07 |
| 24360 | 1218 | Q1218V | D | 1.8e-07 | 1.7e-07 | 2.2e-07 | 1.900E-07 | 2.646E-08 | 1.700E-07 | 2.200E-07 |
| 24361 | 1219 | E1219A | D | 3.4e-07 | 2.5e-07 | 1.6e-07 | 2.500E-07 | 9.000E-08 | 1.600E-07 | 3.400E-07 |
| 24362 | 1219 | E1219R | D | 2.2e-07 | 1.8e-07 | 2.5e-07 | 2.167E-07 | 3.512E-08 | 1.800E-07 | 2.500E-07 |
| 24363 | 1219 | E1219N | D | 2.5e-07 | 1.8e-07 | 2.3e-07 | 2.200E-07 | 3.606E-08 | 1.800E-07 | 2.500E-07 |
| 24364 | 1219 | E1219D | D | 9.7e-08 | 3.8e-07 | 1.5e-07 | 2.090E-07 | 1.504E-07 | 9.700E-08 | 3.800E-07 |
| 24365 | 1219 | E1219C | D | 3.8e-07 | 4.1e-07 | 3e-07 | 3.633E-07 | 5.686E-08 | 3.000E-07 | 4.100E-07 |
| 24366 | 1219 | E1219E | D | 3.1e-07 | 1.6e-07 | 1.8e-07 | 2.167E-07 | 8.145E-08 | 1.600E-07 | 3.100E-07 |
| 24367 | 1219 | E1219Q | D | 2.3e-07 | 1.4e-07 | 2e-07 | 1.900E-07 | 4.583E-08 | 1.400E-07 | 2.300E-07 |
| 24368 | 1219 | E1219G | D | 4.8e-07 | 5.8e-07 | 2.8e-07 | 4.467E-07 | 1.528E-07 | 2.800E-07 | 5.800E-07 |
| 24369 | 1219 | E1219H | D | 1.8e-07 | 1.9e-07 | 2.8e-07 | 2.167E-07 | 5.508E-08 | 1.800E-07 | 2.800E-07 |
| 24370 | 1219 | E1219I | D | 2e-07 | 3.4e-07 | 2e-07 | 2.467E-07 | 8.083E-08 | 2.000E-07 | 3.400E-07 |
| 24371 | 1219 | E1219L | D | 1.4e-07 | 3.8e-07 | 1.4e-07 | 2.200E-07 | 1.386E-07 | 1.400E-07 | 3.800E-07 |
| 24372 | 1219 | E1219K | D | 9.8e-08 | 2.4e-07 | 7.5e-08 | 1.377E-07 | 8.937E-08 | 7.500E-08 | 2.400E-07 |
| 24373 | 1219 | E1219M | D | 8.3e-07 | 3.8e-07 | 2.6e-07 | 4.900E-07 | 3.005E-07 | 2.600E-07 | 8.300E-07 |
| 24374 | 1219 | E1219F | D | 2.1e-07 | 2.7e-07 | 1.9e-07 | 2.233E-07 | 4.163E-08 | 1.900E-07 | 2.700E-07 |
| 24375 | 1219 | E1219P | D | 3.3e-07 | 2.7e-07 | 1.7e-07 | 2.567E-07 | 8.083E-08 | 1.700E-07 | 3.300E-07 |
| 24376 | 1219 | E1219S | D | 3.3e-07 | 1.9e-07 | 1.8e-07 | 2.333E-07 | 8.386E-08 | 1.800E-07 | 3.300E-07 |
| 24377 | 1219 | E1219T | D | 2.8e-07 | 1.8e-07 | 2.9e-07 | 2.500E-07 | 6.083E-08 | 1.800E-07 | 2.900E-07 |
| 24378 | 1219 | E1219W | D | 2.9e-07 | 4.8e-07 | 2.5e-07 | 3.400E-07 | 1.229E-07 | 2.500E-07 | 4.800E-07 |
| 24379 | 1219 | E1219Y | D | 1.3e-07 | 5.2e-07 | 2.3e-07 | 2.933E-07 | 2.026E-07 | 1.300E-07 | 5.200E-07 |
| 24380 | 1219 | E1219V | D | 1.8e-07 | 1.5e-07 | 6.5e-08 | 1.317E-07 | 5.965E-08 | 6.500E-08 | 1.800E-07 |
| 24381 | 1220 | S1220A | D | 2.1e-07 | 5.3e-07 | 1.5e-07 | 2.967E-07 | 2.043E-07 | 1.500E-07 | 5.300E-07 |
| 24382 | 1220 | S1220R | D | 5.6e-07 | 3.4e-07 | 2.2e-07 | 3.733E-07 | 1.724E-07 | 2.200E-07 | 5.600E-07 |
| 24383 | 1220 | S1220N | D | 4.3e-07 | 1.7e-07 | 1.6e-07 | 2.533E-07 | 1.531E-07 | 1.600E-07 | 4.300E-07 |
| 24384 | 1220 | S1220D | D | 2.7e-07 | 1.2e-07 | 4.8e-07 | 2.900E-07 | 1.808E-07 | 1.200E-07 | 4.800E-07 |
| 24385 | 1220 | S1220C | D | 2.3e-07 | 3.2e-07 | 1.5e-07 | 2.333E-07 | 8.505E-08 | 1.500E-07 | 3.200E-07 |
| 24386 | 1220 | S1220E | D | 1.1e-07 | 3.6e-07 | 1.2e-07 | 1.967E-07 | 1.415E-07 | 1.100E-07 | 3.600E-07 |
| 24387 | 1220 | S1220Q | D | 1.3e-07 | 2e-07 | 4.5e-07 | 2.600E-07 | 1.682E-07 | 1.300E-07 | 4.500E-07 |
| 24388 | 1220 | S1220G | D | 3.6e-07 | 1.5e-07 | 2.9e-07 | 2.667E-07 | 1.069E-07 | 1.500E-07 | 3.600E-07 |
| 24389 | 1220 | S1220H | D | 2.7e-07 | 2.2e-07 | 2.2e-07 | 2.367E-07 | 2.887E-08 | 2.200E-07 | 2.700E-07 |
| 24390 | 1220 | S1220I | D | 2.3e-07 | 2.9e-07 | 2.3e-07 | 2.500E-07 | 3.464E-08 | 2.300E-07 | 2.900E-07 |
| 24391 | 1220 | S1220L | D | 2.8e-07 | 1.7e-07 | 1.8e-07 | 2.100E-07 | 6.083E-08 | 1.700E-07 | 2.800E-07 |
| 24392 | 1220 | S1220K | D | 3.3e-07 | 1.2e-07 | 3.2e-07 | 2.567E-07 | 1.185E-07 | 1.200E-07 | 3.300E-07 |
| 24393 | 1220 | S1220M | D | 2.1e-07 | 2.4e-07 | 1.1e-07 | 1.867E-07 | 6.807E-08 | 1.100E-07 | 2.400E-07 |
| 24394 | 1220 | S1220F | D | 2.5e-07 | 2.7e-07 | 4.2e-07 | 3.133E-07 | 9.292E-08 | 2.500E-07 | 4.200E-07 |

| | | | | | | | | | | |
|---|---|---|---|---|---|---|---|---|---|---|
| 24395 | 1220 | S1220P | D | 1.8e-07 | 2.7e-07 | 1.7e-07 | 2.067E-07 | 5.508E-08 | 1.700E-07 | 2.700E-07 |
| 24396 | 1220 | S1220S | D | 1.5e-07 | 1.8e-07 | 3.8e-07 | 2.367E-07 | 1.250E-07 | 1.500E-07 | 3.800E-07 |
| 24397 | 1220 | S1220T | D | 1.6e-07 | 3.6e-07 | 2.2e-07 | 2.467E-07 | 1.026E-07 | 1.600E-07 | 3.600E-07 |
| 24398 | 1220 | S1220W | D | 2.7e-07 | 3e-07 | 3.7e-07 | 3.133E-07 | 5.132E-08 | 2.700E-07 | 3.700E-07 |
| 24399 | 1220 | S1220Y | D | 2.6e-07 | 1.2e-07 | 2.4e-07 | 2.067E-07 | 7.572E-08 | 1.200E-07 | 2.600E-07 |
| 24400 | 1220 | S1220V | D | 1.5e-07 | 2.2e-07 | 2.4e-07 | 2.033E-07 | 4.726E-08 | 1.500E-07 | 2.400E-07 |
| 24401 | 1221 | V1221A | D | 1.1e-07 | 3e-07 | 2e-07 | 2.033E-07 | 9.504E-08 | 1.100E-07 | 3.000E-07 |
| 24402 | 1221 | V1221R | D | 4.8e-07 | 1.1e-07 | 2.8e-07 | 2.900E-07 | 1.852E-07 | 1.100E-07 | 4.800E-07 |
| 24403 | 1221 | V1221N | D | 4.1e-07 | 4.7e-07 | 2.1e-07 | 3.633E-07 | 1.361E-07 | 2.100E-07 | 4.700E-07 |
| 24404 | 1221 | V1221D | D | 1.7e-07 | 1.4e-07 | 1.5e-07 | 1.533E-07 | 1.528E-08 | 1.400E-07 | 1.700E-07 |
| 24405 | 1221 | V1221C | D | 2.2e-07 | 1.8e-07 | 2.8e-07 | 2.267E-07 | 5.033E-08 | 1.800E-07 | 2.800E-07 |
| 24406 | 1221 | V1221E | D | 1.5e-07 | 1.8e-07 | 3.2e-07 | 2.167E-07 | 9.074E-08 | 1.500E-07 | 3.200E-07 |
| 24407 | 1221 | V1221Q | D | 4.4e-07 | 3.2e-07 | 3e-07 | 3.533E-07 | 7.572E-08 | 3.000E-07 | 4.400E-07 |
| 24408 | 1221 | V1221G | D | 1.3e-07 | 2.6e-07 | 1.5e-07 | 1.800E-07 | 7.000E-08 | 1.300E-07 | 2.600E-07 |
| 24409 | 1221 | V1221H | D | 4.2e-07 | 3.4e-07 | 2.6e-07 | 3.400E-07 | 8.000E-08 | 2.600E-07 | 4.200E-07 |
| 24410 | 1221 | V1221I | D | 2.1e-07 | 2e-07 | 2.7e-07 | 2.267E-07 | 3.786E-08 | 2.000E-07 | 2.700E-07 |
| 24411 | 1221 | V1221L | D | 2.5e-07 | 1.7e-07 | 1.1e-07 | 1.767E-07 | 7.024E-08 | 1.100E-07 | 2.500E-07 |
| 24412 | 1221 | V1221K | D | 3.3e-07 | 5.4e-07 | 2e-07 | 3.567E-07 | 1.716E-07 | 2.000E-07 | 5.400E-07 |
| 24413 | 1221 | V1221M | D | 2.7e-07 | 3.6e-07 | 5.1e-07 | 3.800E-07 | 1.212E-07 | 2.700E-07 | 5.100E-07 |
| 24414 | 1221 | V1221F | D | 3.4e-07 | 1.6e-07 | 3.2e-07 | 2.733E-07 | 9.866E-08 | 1.600E-07 | 3.400E-07 |
| 24415 | 1221 | V1221P | D | 2.2e-07 | 1.7e-07 | 3.1e-07 | 2.333E-07 | 7.095E-08 | 1.700E-07 | 3.100E-07 |
| 24416 | 1221 | V1221S | D | 1.8e-07 | 3.8e-07 | 2.6e-07 | 2.733E-07 | 1.007E-07 | 1.800E-07 | 3.800E-07 |
| 24417 | 1221 | V1221T | D | 7.4e-07 | 2.6e-07 | 1.6e-07 | 3.867E-07 | 3.101E-07 | 1.600E-07 | 7.400E-07 |
| 24418 | 1221 | V1221W | D | 2.1e-07 | 6.5e-08 | 3.9e-07 | 2.217E-07 | 1.628E-07 | 6.500E-08 | 3.900E-07 |
| 24419 | 1221 | V1221Y | D | 2.2e-07 | 3.1e-07 | 2.4e-07 | 2.567E-07 | 4.726E-08 | 2.200E-07 | 3.100E-07 |
| 24420 | 1221 | V1221V | D | 1.4e-07 | 2.2e-07 | 1.8e-07 | 1.800E-07 | 4.000E-08 | 1.400E-07 | 2.200E-07 |
| 24421 | 1222 | T1222A | D | 1.8e-07 | 2e-07 | 2.3e-07 | 2.033E-07 | 2.517E-08 | 1.800E-07 | 2.300E-07 |
| 24422 | 1222 | T1222R | D | 2.4e-07 | 3.1e-07 | 4.7e-07 | 3.400E-07 | 1.179E-07 | 2.400E-07 | 4.700E-07 |
| 24423 | 1222 | T1222N | D | 4.3e-07 | 1.9e-07 | 3.2e-07 | 3.133E-07 | 1.201E-07 | 1.900E-07 | 4.300E-07 |
| 24424 | 1222 | T1222D | D | 1.3e-07 | 3.4e-07 | 2.1e-07 | 2.267E-07 | 1.060E-07 | 1.300E-07 | 3.400E-07 |
| 24425 | 1222 | T1222C | D | 3.4e-07 | 2.5e-07 | 1.2e-07 | 2.367E-07 | 1.106E-07 | 1.200E-07 | 3.400E-07 |
| 24426 | 1222 | T1222E | D | 3.7e-07 | 3e-07 | 4.8e-07 | 3.833E-07 | 9.074E-08 | 3.000E-07 | 4.800E-07 |
| 24427 | 1222 | T1222Q | D | 1e-07 | 4.6e-07 | 3.7e-07 | 3.100E-07 | 1.873E-07 | 1.000E-07 | 4.600E-07 |
| 24428 | 1222 | T1222G | D | 2e-07 | 4.5e-07 | 2.4e-07 | 2.967E-07 | 1.343E-07 | 2.000E-07 | 4.500E-07 |
| 24429 | 1222 | T1222H | D | 2.3e-07 | 2.8e-07 | 1.7e-07 | 2.267E-07 | 5.508E-08 | 1.700E-07 | 2.800E-07 |
| 24430 | 1222 | T1222I | D | 3.1e-07 | 1.9e-07 | 4.7e-07 | 3.233E-07 | 1.405E-07 | 1.900E-07 | 4.700E-07 |
| 24431 | 1222 | T1222L | D | 2.7e-07 | 2.8e-07 | 6.4e-08 | 2.047E-07 | 1.219E-07 | 6.400E-08 | 2.800E-07 |
| 24432 | 1222 | T1222K | D | 3.7e-07 | 2.8e-07 | 1.5e-07 | 2.667E-07 | 1.106E-07 | 1.500E-07 | 3.700E-07 |
| 24433 | 1222 | T1222M | D | 2.7e-07 | 2.4e-07 | 1.8e-07 | 2.300E-07 | 4.583E-08 | 1.800E-07 | 2.700E-07 |
| 24434 | 1222 | T1222F | D | 3.9e-07 | 3.3e-07 | 4.3e-07 | 3.833E-07 | 5.033E-08 | 3.300E-07 | 4.300E-07 |
| 24435 | 1222 | T1222P | D | 2.3e-07 | 3.1e-07 | 2.2e-07 | 2.533E-07 | 4.933E-08 | 2.200E-07 | 3.100E-07 |
| 24436 | 1222 | T1222S | D | 2.4e-07 | 2.5e-07 | 1.8e-07 | 2.233E-07 | 3.786E-08 | 1.800E-07 | 2.500E-07 |
| 24437 | 1222 | T1222T | D | 3e-07 | 2.2e-07 | 1.7e-07 | 2.300E-07 | 6.557E-08 | 1.700E-07 | 3.000E-07 |
| 24438 | 1222 | T1222W | D | 2.4e-07 | 4.9e-07 | 2.1e-07 | 3.133E-07 | 1.537E-07 | 2.100E-07 | 4.900E-07 |
| 24439 | 1222 | T1222Y | D | 1.6e-07 | 7.3e-07 | 1.5e-07 | 3.467E-07 | 3.320E-07 | 1.500E-07 | 7.300E-07 |
| 24440 | 1222 | T1222V | D | 2.4e-07 | 1.4e-07 | 2.9e-07 | 2.233E-07 | 7.638E-08 | 1.400E-07 | 2.900E-07 |
| 24441 | 1223 | E1223A | D | 1.4e-07 | 2.4e-07 | 3.1e-07 | 2.300E-07 | 8.544E-08 | 1.400E-07 | 3.100E-07 |
| 24442 | 1223 | E1223R | D | 3.9e-07 | 2.4e-07 | 3.4e-07 | 3.233E-07 | 7.638E-08 | 2.400E-07 | 3.900E-07 |
| 24443 | 1223 | E1223N | D | 2.3e-07 | 2.8e-07 | 1.1e-07 | 2.067E-07 | 8.737E-08 | 1.100E-07 | 2.800E-07 |
| 24444 | 1223 | E1223D | D | 3.8e-07 | 3.8e-07 | 5.1e-07 | 4.233E-07 | 7.506E-08 | 3.800E-07 | 5.100E-07 |
| 24445 | 1223 | E1223C | D | 2.3e-07 | 2.2e-07 | 4e-07 | 2.833E-07 | 1.012E-07 | 2.200E-07 | 4.000E-07 |
| 24446 | 1223 | E1223E | D | 2.7e-07 | 2e-07 | 2.3e-07 | 2.333E-07 | 3.512E-08 | 2.000E-07 | 2.700E-07 |
| 24447 | 1223 | E1223Q | D | 1.3e-07 | 1.7e-07 | 5.5e-07 | 2.833E-07 | 2.318E-07 | 1.300E-07 | 5.500E-07 |
| 24448 | 1223 | E1223G | D | 3.3e-07 | 3.2e-07 | 2.4e-07 | 2.967E-07 | 4.933E-08 | 2.400E-07 | 3.300E-07 |
| 24449 | 1223 | E1223H | D | 2.6e-07 | 1.6e-07 | 2.6e-07 | 2.267E-07 | 5.774E-08 | 1.600E-07 | 2.600E-07 |
| 24450 | 1223 | E1223I | D | 2e-07 | 5e-07 | 2.3e-07 | 3.100E-07 | 1.652E-07 | 2.000E-07 | 5.000E-07 |
| 24451 | 1223 | E1223L | D | 3.4e-07 | 2.8e-07 | 4.1e-07 | 3.433E-07 | 6.506E-08 | 2.800E-07 | 4.100E-07 |
| 24452 | 1223 | E1223K | D | 2e-07 | 3.1e-07 | 1.5e-07 | 2.200E-07 | 8.185E-08 | 1.500E-07 | 3.100E-07 |
| 24453 | 1223 | E1223M | D | 3.1e-07 | 3.4e-07 | 4.2e-07 | 3.567E-07 | 5.686E-08 | 3.100E-07 | 4.200E-07 |
| 24454 | 1223 | E1223F | D | 1.4e-07 | 2e-07 | 3.4e-07 | 2.267E-07 | 1.026E-07 | 1.400E-07 | 3.400E-07 |
| 24455 | 1223 | E1223P | D | 2e-07 | 2.9e-07 | 2.4e-07 | 2.433E-07 | 4.509E-08 | 2.000E-07 | 2.900E-07 |
| 24456 | 1223 | E1223S | D | 2.9e-07 | 2.1e-07 | 1.5e-07 | 2.167E-07 | 7.024E-08 | 1.500E-07 | 2.900E-07 |
| 24457 | 1223 | E1223T | D | 1.8e-07 | 1.1e-07 | 1.1e-07 | 1.333E-07 | 4.041E-08 | 1.100E-07 | 1.800E-07 |
| 24458 | 1223 | E1223W | D | 5.4e-07 | 4e-07 | 2.4e-07 | 3.933E-07 | 1.501E-07 | 2.400E-07 | 5.400E-07 |
| 24459 | 1223 | E1223Y | D | 3.2e-07 | 3e-07 | 4.2e-07 | 3.467E-07 | 6.429E-08 | 3.000E-07 | 4.200E-07 |
| 24460 | 1223 | E1223V | D | 4.8e-07 | 2.2e-07 | 2.4e-07 | 3.133E-07 | 1.447E-07 | 2.200E-07 | 4.800E-07 |
| 24461 | 1224 | Q1224A | D | 3.1e-07 | 2.1e-07 | 3.1e-07 | 2.767E-07 | 5.774E-08 | 2.100E-07 | 3.100E-07 |
| 24462 | 1224 | Q1224R | D | 2.4e-07 | 2.7e-07 | 4.6e-07 | 3.233E-07 | 1.193E-07 | 2.400E-07 | 4.600E-07 |
| 24463 | 1224 | Q1224N | D | 2.6e-07 | 1.5e-07 | 2e-07 | 2.033E-07 | 5.508E-08 | 1.500E-07 | 2.600E-07 |
| 24464 | 1224 | Q1224D | D | 2.8e-07 | 3.3e-07 | 1.4e-07 | 2.500E-07 | 9.849E-08 | 1.400E-07 | 3.300E-07 |
| 24465 | 1224 | Q1224C | D | 5e-07 | 1.9e-07 | 1.9e-07 | 2.933E-07 | 1.790E-07 | 1.900E-07 | 5.000E-07 |
| 24466 | 1224 | Q1224E | D | 1.9e-07 | 6.7e-07 | 3.1e-07 | 3.900E-07 | 2.498E-07 | 1.900E-07 | 6.700E-07 |
| 24467 | 1224 | Q1224Q | D | 3e-07 | 3.4e-07 | 4.5e-07 | 3.633E-07 | 7.767E-08 | 3.000E-07 | 4.500E-07 |
| 24468 | 1224 | Q1224G | D | 1.9e-07 | 1.3e-07 | 3.9e-07 | 2.367E-07 | 1.361E-07 | 1.300E-07 | 3.900E-07 |
| 24469 | 1224 | Q1224H | D | 4.5e-07 | 5.1e-07 | 2.1e-07 | 3.900E-07 | 1.587E-07 | 2.100E-07 | 5.100E-07 |
| 24470 | 1224 | Q1224I | D | 2.8e-07 | 3.1e-07 | 4.1e-07 | 3.333E-07 | 6.807E-08 | 2.800E-07 | 4.100E-07 |
| 24471 | 1224 | Q1224L | D | 3.5e-07 | 2.2e-07 | 2.6e-07 | 2.767E-07 | 6.658E-08 | 2.200E-07 | 3.500E-07 |
| 24472 | 1224 | Q1224K | D | 3.9e-07 | 3e-07 | 1.8e-07 | 2.900E-07 | 1.054E-07 | 1.800E-07 | 3.900E-07 |
| 24473 | 1224 | Q1224M | D | 1.8e-07 | 1.8e-07 | 2.7e-07 | 2.100E-07 | 5.196E-08 | 1.800E-07 | 2.700E-07 |
| 24474 | 1224 | Q1224F | D | 2e-07 | 1.3e-07 | 3.5e-07 | 2.267E-07 | 1.124E-07 | 1.300E-07 | 3.500E-07 |
| 24475 | 1224 | Q1224P | D | 6.9e-07 | 3.1e-07 | 1.5e-07 | 3.833E-07 | 2.774E-07 | 1.500E-07 | 6.900E-07 |
| 24476 | 1224 | Q1224S | D | 1.9e-07 | 3.7e-07 | 1.9e-07 | 2.500E-07 | 1.039E-07 | 1.900E-07 | 3.700E-07 |
| 24477 | 1224 | Q1224T | D | 2e-07 | 2.9e-07 | 1.7e-07 | 2.200E-07 | 6.245E-08 | 1.700E-07 | 2.900E-07 |
| 24478 | 1224 | Q1224W | D | 2.9e-07 | 1.5e-07 | 1.7e-07 | 2.033E-07 | 7.572E-08 | 1.500E-07 | 2.900E-07 |
| 24479 | 1224 | Q1224Y | D | 3.1e-07 | 2.6e-07 | 2.3e-07 | 2.667E-07 | 4.041E-08 | 2.300E-07 | 3.100E-07 |

| | | | | | | | | | | |
|---|---|---|---|---|---|---|---|---|---|---|
| 24480 | 1224 | Q1224V | D | 5.5e-07 | 4.3e-07 | 1.5e-07 | 3.767E-07 | 2.053E-07 | 1.500E-07 | 5.500E-07 |
| 24481 | 1225 | D1225A | D | 2.7e-07 | 1.3e-07 | 1.7e-07 | 1.900E-07 | 7.211E-08 | 1.300E-07 | 2.700E-07 |
| 24482 | 1225 | D1225R | D | 2e-07 | 1.9e-07 | 3e-07 | 2.300E-07 | 6.083E-08 | 1.900E-07 | 3.000E-07 |
| 24483 | 1225 | D1225N | D | 5.4e-07 | 2e-07 | 1.4e-07 | 2.933E-07 | 2.157E-07 | 1.400E-07 | 5.400E-07 |
| 24484 | 1225 | D1225D | D | 5.4e-07 | 2.5e-07 | 2.8e-07 | 3.567E-07 | 1.595E-07 | 2.500E-07 | 5.400E-07 |
| 24485 | 1225 | D1225C | D | 2.1e-07 | 1.7e-07 | 3.7e-07 | 2.500E-07 | 1.058E-07 | 1.700E-07 | 3.700E-07 |
| 24486 | 1225 | D1225E | D | 2.8e-07 | 2.3e-07 | 1.8e-07 | 2.300E-07 | 5.000E-08 | 1.800E-07 | 2.800E-07 |
| 24487 | 1225 | D1225Q | D | 4.2e-07 | 2.5e-07 | 1.9e-07 | 2.867E-07 | 1.193E-07 | 1.900E-07 | 4.200E-07 |
| 24488 | 1225 | D1225G | D | 2.4e-07 | 1.9e-07 | 6.2e-08 | 1.640E-07 | 9.180E-08 | 6.200E-08 | 2.400E-07 |
| 24489 | 1225 | D1225H | D | 2.8e-07 | 2.2e-07 | 2.5e-07 | 2.500E-07 | 3.000E-08 | 2.200E-07 | 2.800E-07 |
| 24490 | 1225 | D1225I | D | 1.9e-07 | 2.5e-07 | 3.8e-07 | 2.733E-07 | 9.713E-08 | 1.900E-07 | 3.800E-07 |
| 24491 | 1225 | D1225L | D | 1.8e-07 | 2.3e-07 | 2.8e-07 | 2.300E-07 | 5.000E-08 | 1.800E-07 | 2.800E-07 |
| 24492 | 1225 | D1225K | D | 4.4e-07 | 1.8e-07 | 3.9e-07 | 3.367E-07 | 1.380E-07 | 1.800E-07 | 4.400E-07 |
| 24493 | 1225 | D1225M | D | 4e-07 | 2.1e-07 | 3e-07 | 3.033E-07 | 9.504E-08 | 2.100E-07 | 4.000E-07 |
| 24494 | 1225 | D1225F | D | 3.4e-07 | 2.7e-07 | 2.1e-07 | 2.733E-07 | 6.506E-08 | 2.100E-07 | 3.400E-07 |
| 24495 | 1225 | D1225P | D | 2.1e-07 | 2.7e-07 | 1.9e-07 | 2.233E-07 | 4.163E-08 | 1.900E-07 | 2.700E-07 |
| 24496 | 1225 | D1225S | D | 2.4e-07 | 2.6e-07 | 2.3e-07 | 2.433E-07 | 1.528E-08 | 2.300E-07 | 2.600E-07 |
| 24497 | 1225 | D1225T | D | 2.6e-07 | 3e-07 | 2.3e-07 | 2.633E-07 | 3.512E-08 | 2.300E-07 | 3.000E-07 |
| 24498 | 1225 | D1225W | D | 3.5e-07 | 1.9e-07 | 2.5e-07 | 2.633E-07 | 8.083E-08 | 1.900E-07 | 3.500E-07 |
| 24499 | 1225 | D1225Y | D | 4.3e-07 | 9.8e-08 | 2.4e-07 | 2.560E-07 | 1.666E-07 | 9.800E-08 | 4.300E-07 |
| 24500 | 1225 | D1225V | D | 3.6e-07 | 2.2e-07 | 1.5e-07 | 2.433E-07 | 1.069E-07 | 1.500E-07 | 3.600E-07 |
| 24501 | 1226 | S1226A | D | 2e-07 | 4e-07 | 1.9e-07 | 2.633E-07 | 1.185E-07 | 1.900E-07 | 4.000E-07 |
| 24502 | 1226 | S1226R | D | 2.3e-07 | 2e-07 | 2.7e-07 | 2.333E-07 | 3.512E-08 | 2.000E-07 | 2.700E-07 |
| 24503 | 1226 | S1226N | D | 1.5e-07 | 2.1e-07 | 3e-07 | 2.200E-07 | 7.550E-08 | 1.500E-07 | 3.000E-07 |
| 24504 | 1226 | S1226D | D | 1.9e-07 | 2.3e-07 | 3.5e-07 | 2.567E-07 | 8.327E-08 | 1.900E-07 | 3.500E-07 |
| 24505 | 1226 | S1226C | D | 1.4e-07 | 2.1e-07 | 2.2e-07 | 1.900E-07 | 4.359E-08 | 1.400E-07 | 2.200E-07 |
| 24506 | 1226 | S1226E | D | 3.6e-07 | 2.6e-07 | 3.2e-07 | 3.133E-07 | 5.033E-08 | 2.600E-07 | 3.600E-07 |
| 24507 | 1226 | S1226Q | D | 1.3e-07 | 2.2e-07 | 1.3e-07 | 1.600E-07 | 5.196E-08 | 1.300E-07 | 2.200E-07 |
| 24508 | 1226 | S1226G | D | 3.6e-07 | 2e-07 | 1.7e-07 | 2.433E-07 | 1.021E-07 | 1.700E-07 | 3.600E-07 |
| 24509 | 1226 | S1226H | D | 3.1e-07 | 1.5e-07 | 6.2e-07 | 3.600E-07 | 2.390E-07 | 1.500E-07 | 6.200E-07 |
| 24510 | 1226 | S1226I | D | 1.7e-07 | 4.7e-07 | 2.7e-07 | 3.033E-07 | 1.528E-07 | 1.700E-07 | 4.700E-07 |
| 24511 | 1226 | S1226L | D | 2.7e-07 | 3.4e-07 | 2.6e-07 | 2.900E-07 | 4.359E-08 | 2.600E-07 | 3.400E-07 |
| 24512 | 1226 | S1226K | D | 1.8e-07 | 5.5e-07 | 1.9e-07 | 3.067E-07 | 2.108E-07 | 1.800E-07 | 5.500E-07 |
| 24513 | 1226 | S1226M | D | 4.3e-07 | 3.1e-07 | 1.8e-07 | 3.067E-07 | 1.250E-07 | 1.800E-07 | 4.300E-07 |
| 24514 | 1226 | S1226F | D | 2.5e-07 | 3.1e-07 | 3.3e-07 | 2.967E-07 | 4.163E-08 | 2.500E-07 | 3.300E-07 |
| 24515 | 1226 | S1226P | D | 2.2e-07 | 3.1e-07 | 2.5e-07 | 2.600E-07 | 4.583E-08 | 2.200E-07 | 3.100E-07 |
| 24516 | 1226 | S1226S | D | 1.9e-07 | 3.8e-07 | 1.3e-07 | 2.333E-07 | 1.305E-07 | 1.300E-07 | 3.800E-07 |
| 24517 | 1226 | S1226T | D | 4.9e-07 | 2.2e-07 | 1.5e-07 | 2.867E-07 | 1.795E-07 | 1.500E-07 | 4.900E-07 |
| 24518 | 1226 | S1226W | D | 4e-07 | 1.6e-07 | 2.7e-07 | 2.767E-07 | 1.201E-07 | 1.600E-07 | 4.000E-07 |
| 24519 | 1226 | S1226Y | D | 1.5e-07 | 3.1e-07 | 2.6e-07 | 2.400E-07 | 8.185E-08 | 1.500E-07 | 3.100E-07 |
| 24520 | 1226 | S1226V | D | 2.5e-07 | 1.6e-07 | 3.3e-07 | 2.467E-07 | 8.505E-08 | 1.600E-07 | 3.300E-07 |
| 24521 | 1227 | K1227A | D | 1.5e-07 | 5.2e-07 | 2.6e-07 | 3.100E-07 | 1.900E-07 | 1.500E-07 | 5.200E-07 |
| 24522 | 1227 | K1227R | D | 2e-07 | 1.7e-07 | 3.1e-07 | 2.267E-07 | 7.371E-08 | 1.700E-07 | 3.100E-07 |
| 24523 | 1227 | K1227N | D | 2.2e-07 | 1.3e-07 | 1.9e-07 | 1.800E-07 | 4.583E-08 | 1.300E-07 | 2.200E-07 |
| 24524 | 1227 | K1227D | D | 2.3e-07 | 3.3e-07 | 2.7e-07 | 2.767E-07 | 5.033E-08 | 2.300E-07 | 3.300E-07 |
| 24525 | 1227 | K1227C | D | 2.1e-07 | 3.1e-07 | 1.7e-07 | 2.300E-07 | 7.211E-08 | 1.700E-07 | 3.100E-07 |
| 24526 | 1227 | K1227E | D | 2.5e-07 | 3.6e-07 | 3.1e-07 | 3.067E-07 | 5.508E-08 | 2.500E-07 | 3.600E-07 |
| 24527 | 1227 | K1227Q | D | 1.9e-07 | 1.6e-07 | 1.1e-07 | 1.533E-07 | 4.041E-08 | 1.100E-07 | 1.900E-07 |
| 24528 | 1227 | K1227G | D | 1e-07 | 1.1e-07 | 2.9e-07 | 1.667E-07 | 1.069E-07 | 1.000E-07 | 2.900E-07 |
| 24529 | 1227 | K1227H | D | 2.5e-07 | 2.8e-07 | 6e-07 | 3.767E-07 | 1.940E-07 | 2.500E-07 | 6.000E-07 |
| 24530 | 1227 | K1227I | D | 1.2e-07 | 2.6e-07 | 2.7e-07 | 2.167E-07 | 8.386E-08 | 1.200E-07 | 2.700E-07 |
| 24531 | 1227 | K1227L | D | 5.5e-07 | 1.7e-07 | 4.2e-07 | 3.800E-07 | 1.931E-07 | 1.700E-07 | 5.500E-07 |
| 24532 | 1227 | K1227K | D | 4.1e-07 | 2.1e-07 | 2e-07 | 2.733E-07 | 1.185E-07 | 2.000E-07 | 4.100E-07 |
| 24533 | 1227 | K1227M | D | 1.9e-07 | 3.7e-07 | 3.1e-07 | 2.900E-07 | 9.165E-08 | 1.900E-07 | 3.700E-07 |
| 24534 | 1227 | K1227F | D | 1.6e-07 | 3.9e-07 | 1.5e-07 | 2.333E-07 | 1.358E-07 | 1.500E-07 | 3.900E-07 |
| 24535 | 1227 | K1227P | D | 4.9e-07 | 3.1e-07 | 3.3e-07 | 3.767E-07 | 9.866E-08 | 3.100E-07 | 4.900E-07 |
| 24536 | 1227 | K1227S | D | 3.1e-07 | 3e-07 | 3.1e-07 | 3.067E-07 | 5.774E-09 | 3.000E-07 | 3.100E-07 |
| 24537 | 1227 | K1227T | D | 2.7e-07 | 4.8e-07 | 3.6e-07 | 3.700E-07 | 1.054E-07 | 2.700E-07 | 4.800E-07 |
| 24538 | 1227 | K1227W | D | 1.6e-07 | 2.5e-07 | 2.8e-07 | 2.300E-07 | 6.245E-08 | 1.600E-07 | 2.800E-07 |
| 24539 | 1227 | K1227Y | D | 3.4e-07 | 2.4e-07 | 2.1e-07 | 2.633E-07 | 6.807E-08 | 2.100E-07 | 3.400E-07 |
| 24540 | 1227 | K1227V | D | 2.3e-07 | 3.9e-07 | 2.3e-07 | 2.833E-07 | 9.238E-08 | 2.300E-07 | 3.900E-07 |
| 24541 | 1228 | D1228A | D | 3.1e-07 | 4.5e-07 | 2e-07 | 3.200E-07 | 1.253E-07 | 2.000E-07 | 4.500E-07 |
| 24542 | 1228 | D1228R | D | 2.3e-07 | 1.8e-07 | 3.8e-07 | 2.633E-07 | 1.041E-07 | 1.800E-07 | 3.800E-07 |
| 24543 | 1228 | D1228N | D | 2.6e-07 | 1.2e-07 | 4.7e-07 | 2.833E-07 | 1.762E-07 | 1.200E-07 | 4.700E-07 |
| 24544 | 1228 | D1228D | D | 2.8e-07 | 3.2e-07 | 2.8e-07 | 2.933E-07 | 2.309E-08 | 2.800E-07 | 3.200E-07 |
| 24545 | 1228 | D1228C | D | 2.8e-07 | 2e-07 | 3e-07 | 2.600E-07 | 5.292E-08 | 2.000E-07 | 3.000E-07 |
| 24546 | 1228 | D1228E | D | 2.2e-07 | 6e-07 | 2.2e-07 | 3.467E-07 | 2.194E-07 | 2.200E-07 | 6.000E-07 |
| 24547 | 1228 | D1228Q | D | 4.1e-07 | 3.1e-07 | 5.8e-07 | 4.333E-07 | 1.365E-07 | 3.100E-07 | 5.800E-07 |
| 24548 | 1228 | D1228G | D | 1e-07 | 4.1e-07 | 2.9e-07 | 2.667E-07 | 1.563E-07 | 1.000E-07 | 4.100E-07 |
| 24549 | 1228 | D1228H | D | 3.4e-07 | 3.1e-07 | 2.4e-07 | 2.967E-07 | 5.132E-08 | 2.400E-07 | 3.400E-07 |
| 24550 | 1228 | D1228I | D | 1.3e-07 | 3.2e-07 | 6.7e-07 | 3.733E-07 | 2.739E-07 | 1.300E-07 | 6.700E-07 |
| 24551 | 1228 | D1228L | D | 2.2e-07 | 3.5e-07 | 1.6e-07 | 2.433E-07 | 9.713E-08 | 1.600E-07 | 3.500E-07 |
| 24552 | 1228 | D1228K | D | 3.1e-07 | 3.1e-07 | 1.5e-07 | 2.567E-07 | 9.238E-08 | 1.500E-07 | 3.100E-07 |
| 24553 | 1228 | D1228M | D | 3.5e-07 | 1.3e-07 | 2e-07 | 2.267E-07 | 1.124E-07 | 1.300E-07 | 3.500E-07 |
| 24554 | 1228 | D1228F | D | 1.6e-07 | 2.7e-07 | 1.2e-07 | 1.833E-07 | 7.767E-08 | 1.200E-07 | 2.700E-07 |
| 24555 | 1228 | D1228P | D | 4.7e-07 | 4.3e-07 | 3.3e-07 | 4.100E-07 | 7.211E-08 | 3.300E-07 | 4.700E-07 |
| 24556 | 1228 | D1228S | D | 1.1e-07 | 1e-07 | 9.7e-08 | 1.023E-07 | 6.807E-09 | 9.700E-08 | 1.100E-07 |
| 24557 | 1228 | D1228T | D | 1.6e-07 | 2.5e-07 | 2.3e-07 | 2.133E-07 | 4.726E-08 | 1.600E-07 | 2.500E-07 |
| 24558 | 1228 | D1228W | D | 3.7e-07 | 3.6e-07 | 2.6e-07 | 3.300E-07 | 6.083E-08 | 2.600E-07 | 3.700E-07 |
| 24559 | 1228 | D1228Y | D | 3e-07 | 4.6e-07 | 3.1e-07 | 3.567E-07 | 8.963E-08 | 3.000E-07 | 4.600E-07 |
| 24560 | 1228 | D1228V | D | 3.4e-07 | 2.6e-07 | 2.5e-07 | 2.833E-07 | 4.933E-08 | 2.500E-07 | 3.400E-07 |
| 24561 | 1229 | S1229A | D | 2.6e-07 | 1.9e-07 | 1.2e-07 | 1.900E-07 | 7.000E-08 | 1.200E-07 | 2.600E-07 |
| 24562 | 1229 | S1229R | D | 2.2e-07 | 1.9e-07 | 3.1e-07 | 2.400E-07 | 6.245E-08 | 1.900E-07 | 3.100E-07 |
| 24563 | 1229 | S1229N | D | 3.3e-07 | 2.2e-07 | 1.6e-07 | 2.367E-07 | 8.622E-08 | 1.600E-07 | 3.300E-07 |
| 24564 | 1229 | S1229D | D | 2.9e-07 | 2.1e-07 | 2.5e-07 | 2.500E-07 | 4.000E-08 | 2.100E-07 | 2.900E-07 |

| | | | | | | | | | | |
|---|---|---|---|---|---|---|---|---|---|---|
| 24565 | 1229 | S1229C | D | 3.2e-07 | 1.7e-07 | 2.2e-07 | 2.367E-07 | 7.638E-08 | 1.700E-07 | 3.200E-07 |
| 24566 | 1229 | S1229E | D | 2.4e-07 | 1.2e-07 | 2.1e-07 | 1.900E-07 | 6.245E-08 | 1.200E-07 | 2.400E-07 |
| 24567 | 1229 | S1229Q | D | 2.2e-07 | 4.3e-07 | 2e-07 | 2.833E-07 | 1.274E-07 | 2.000E-07 | 4.300E-07 |
| 24568 | 1229 | S1229G | D | 2.1e-07 | 3.7e-07 | 2.3e-07 | 2.700E-07 | 8.718E-08 | 2.100E-07 | 3.700E-07 |
| 24569 | 1229 | S1229H | D | 6.1e-07 | 2.2e-07 | 3.5e-07 | 3.933E-07 | 1.986E-07 | 2.200E-07 | 6.100E-07 |
| 24570 | 1229 | S1229I | D | 1.5e-07 | 1.5e-07 | 1.1e-07 | 1.367E-07 | 2.309E-08 | 1.100E-07 | 1.500E-07 |
| 24571 | 1229 | S1229L | D | 2.1e-07 | 1.4e-07 | 3e-07 | 2.167E-07 | 8.021E-08 | 1.400E-07 | 3.000E-07 |
| 24572 | 1229 | S1229K | D | 3e-07 | 1.7e-07 | 2.9e-07 | 2.533E-07 | 7.234E-08 | 1.700E-07 | 3.000E-07 |
| 24573 | 1229 | S1229M | D | 5.6e-07 | 3.5e-07 | 3.9e-07 | 4.333E-07 | 1.115E-07 | 3.500E-07 | 5.600E-07 |
| 24574 | 1229 | S1229F | D | 2.8e-07 | 3.5e-07 | 2.6e-07 | 2.967E-07 | 4.726E-08 | 2.600E-07 | 3.500E-07 |
| 24575 | 1229 | S1229P | D | 1.7e-07 | 2.2e-07 | 1.7e-07 | 1.867E-07 | 2.887E-08 | 1.700E-07 | 2.200E-07 |
| 24576 | 1229 | S1229S | D | 7.3e-07 | 2.4e-07 | 5.2e-07 | 4.967E-07 | 2.458E-07 | 2.400E-07 | 7.300E-07 |
| 24577 | 1229 | S1229T | D | 3.7e-07 | 7.2e-08 | 2.4e-07 | 2.273E-07 | 1.494E-07 | 7.200E-08 | 3.700E-07 |
| 24578 | 1229 | S1229W | D | 1.9e-07 | 3.4e-07 | 2.3e-07 | 2.533E-07 | 7.767E-08 | 1.900E-07 | 3.400E-07 |
| 24579 | 1229 | S1229Y | D | 2.2e-07 | 1.9e-07 | 2.1e-07 | 2.067E-07 | 1.528E-08 | 1.900E-07 | 2.200E-07 |
| 24580 | 1229 | S1229V | D | 3.9e-07 | 3.6e-07 | 4.3e-07 | 3.933E-07 | 3.512E-08 | 3.600E-07 | 4.300E-07 |
| 24581 | 1230 | T1230A | D | 4.6e-07 | 3.2e-07 | 2.2e-07 | 3.333E-07 | 1.206E-07 | 2.200E-07 | 4.600E-07 |
| 24582 | 1230 | T1230R | D | 3.1e-07 | 3.5e-07 | 2.4e-07 | 3.000E-07 | 5.568E-08 | 2.400E-07 | 3.500E-07 |
| 24583 | 1230 | T1230N | D | 1.1e-07 | 2.7e-07 | 2.6e-07 | 2.133E-07 | 8.963E-08 | 1.100E-07 | 2.700E-07 |
| 24584 | 1230 | T1230D | D | 2.3e-07 | 1.9e-07 | 1.5e-07 | 1.900E-07 | 4.000E-08 | 1.500E-07 | 2.300E-07 |
| 24585 | 1230 | T1230C | D | 5.6e-07 | 4.9e-07 | 4.8e-07 | 5.100E-07 | 4.359E-08 | 4.800E-07 | 5.600E-07 |
| 24586 | 1230 | T1230E | D | 1.9e-07 | 3.1e-07 | 1.9e-07 | 2.300E-07 | 6.928E-08 | 1.900E-07 | 3.100E-07 |
| 24587 | 1230 | T1230Q | D | 7.6e-08 | 2.8e-07 | 2.7e-07 | 2.087E-07 | 1.150E-07 | 7.600E-08 | 2.800E-07 |
| 24588 | 1230 | T1230G | D | 1.8e-07 | 1.5e-07 | 4.3e-07 | 2.533E-07 | 1.537E-07 | 1.500E-07 | 4.300E-07 |
| 24589 | 1230 | T1230H | D | 3e-07 | 3.4e-07 | 2.9e-07 | 3.100E-07 | 2.646E-08 | 2.900E-07 | 3.400E-07 |
| 24590 | 1230 | T1230I | D | 3.3e-07 | 3.4e-07 | 2.6e-07 | 3.100E-07 | 4.359E-08 | 2.600E-07 | 3.400E-07 |
| 24591 | 1230 | T1230L | D | 4.4e-07 | 2.4e-07 | 2.2e-07 | 3.000E-07 | 1.217E-07 | 2.200E-07 | 4.400E-07 |
| 24592 | 1230 | T1230K | D | 3.2e-07 | 1.3e-07 | 1.8e-07 | 2.100E-07 | 9.849E-08 | 1.300E-07 | 3.200E-07 |
| 24593 | 1230 | T1230M | D | 2.3e-07 | 1.7e-07 | 2e-07 | 2.000E-07 | 3.000E-08 | 1.700E-07 | 2.300E-07 |
| 24594 | 1230 | T1230F | D | 2.4e-07 | 1.4e-07 | 2.6e-07 | 2.133E-07 | 6.429E-08 | 1.400E-07 | 2.600E-07 |
| 24595 | 1230 | T1230P | D | 2e-07 | 3.1e-07 | 1.8e-07 | 2.300E-07 | 7.000E-08 | 1.800E-07 | 3.100E-07 |
| 24596 | 1230 | T1230S | D | 4e-07 | 1.8e-07 | 1.3e-07 | 2.367E-07 | 1.436E-07 | 1.300E-07 | 4.000E-07 |
| 24597 | 1230 | T1230T | D | 3.5e-07 | 2.9e-07 | 1.5e-07 | 2.633E-07 | 1.026E-07 | 1.500E-07 | 3.500E-07 |
| 24598 | 1230 | T1230W | D | 1.8e-07 | 2.8e-07 | 4.2e-07 | 2.933E-07 | 1.206E-07 | 1.800E-07 | 4.200E-07 |
| 24599 | 1230 | T1230Y | D | 3.9e-07 | 1.3e-07 | 2.1e-07 | 2.433E-07 | 1.332E-07 | 1.300E-07 | 3.900E-07 |
| 24600 | 1230 | T1230V | D | 3.4e-07 | 1.5e-07 | 4e-07 | 2.967E-07 | 1.305E-07 | 1.500E-07 | 4.000E-07 |
| 24601 | 1231 | Y1231A | D | 4e-07 | 2.8e-07 | 3.1e-07 | 3.300E-07 | 6.245E-08 | 2.800E-07 | 4.000E-07 |
| 24602 | 1231 | Y1231R | D | 2.1e-07 | 3.3e-07 | 2e-07 | 2.467E-07 | 7.234E-08 | 2.000E-07 | 3.300E-07 |
| 24603 | 1231 | Y1231N | D | 1.7e-07 | 2.8e-07 | 5.6e-07 | 3.367E-07 | 2.011E-07 | 1.700E-07 | 5.600E-07 |
| 24604 | 1231 | Y1231D | D | 3.5e-07 | 1.6e-07 | 2e-07 | 2.367E-07 | 1.002E-07 | 1.600E-07 | 3.500E-07 |
| 24605 | 1231 | Y1231C | D | 3.7e-07 | 1.8e-07 | 3.6e-07 | 3.033E-07 | 1.069E-07 | 1.800E-07 | 3.700E-07 |
| 24606 | 1231 | Y1231E | D | 4.9e-07 | 1.3e-07 | 1.5e-07 | 2.567E-07 | 2.023E-07 | 1.300E-07 | 4.900E-07 |
| 24607 | 1231 | Y1231Q | D | 2.9e-07 | 2.9e-07 | 1.5e-07 | 2.433E-07 | 8.083E-08 | 1.500E-07 | 2.900E-07 |
| 24608 | 1231 | Y1231G | D | 3.7e-07 | 1.2e-07 | 2.6e-07 | 2.500E-07 | 1.253E-07 | 1.200E-07 | 3.700E-07 |
| 24609 | 1231 | Y1231H | D | 4e-07 | 2.9e-07 | 1.5e-07 | 2.800E-07 | 1.253E-07 | 1.500E-07 | 4.000E-07 |
| 24610 | 1231 | Y1231I | D | 2.3e-07 | 1.4e-07 | 1.6e-07 | 1.767E-07 | 4.726E-08 | 1.400E-07 | 2.300E-07 |
| 24611 | 1231 | Y1231L | D | 1.6e-07 | 4.2e-07 | 2.6e-07 | 2.800E-07 | 1.311E-07 | 1.600E-07 | 4.200E-07 |
| 24612 | 1231 | Y1231K | D | 3.2e-07 | 3.1e-07 | 2.3e-07 | 2.867E-07 | 4.933E-08 | 2.300E-07 | 3.200E-07 |
| 24613 | 1231 | Y1231M | D | 3.6e-07 | 2.8e-07 | 4.6e-07 | 3.667E-07 | 9.018E-08 | 2.800E-07 | 4.600E-07 |
| 24614 | 1231 | Y1231F | D | 5.4e-07 | 1.6e-07 | 1.7e-07 | 2.900E-07 | 2.166E-07 | 1.600E-07 | 5.400E-07 |
| 24615 | 1231 | Y1231P | D | 1.7e-07 | 2.4e-07 | 5.7e-07 | 3.267E-07 | 2.136E-07 | 1.700E-07 | 5.700E-07 |
| 24616 | 1231 | Y1231S | D | 1.6e-07 | 1.5e-07 | 2.4e-07 | 1.833E-07 | 4.933E-08 | 1.500E-07 | 2.400E-07 |
| 24617 | 1231 | Y1231T | D | 1.1e-07 | 2.1e-07 | 3.1e-07 | 2.100E-07 | 1.000E-07 | 1.100E-07 | 3.100E-07 |
| 24618 | 1231 | Y1231W | D | 3.4e-07 | 2.2e-07 | 2.4e-07 | 2.667E-07 | 6.429E-08 | 2.200E-07 | 3.400E-07 |
| 24619 | 1231 | Y1231Y | D | 4.3e-07 | 2.3e-07 | 2.5e-07 | 3.033E-07 | 1.102E-07 | 2.300E-07 | 4.300E-07 |
| 24620 | 1231 | Y1231V | D | 3.8e-07 | 3e-07 | 2.4e-07 | 3.067E-07 | 7.024E-08 | 2.400E-07 | 3.800E-07 |
| 24621 | 1232 | S1232A | D | 3.9e-07 | 2.8e-07 | 1.5e-07 | 2.733E-07 | 1.201E-07 | 1.500E-07 | 3.900E-07 |
| 24622 | 1232 | S1232R | D | 1.6e-07 | 1.5e-07 | 2.5e-07 | 1.867E-07 | 5.508E-08 | 1.500E-07 | 2.500E-07 |
| 24623 | 1232 | S1232N | D | 1.8e-07 | 1.9e-07 | 1.2e-07 | 1.633E-07 | 3.786E-08 | 1.200E-07 | 1.900E-07 |
| 24624 | 1232 | S1232D | D | 3.6e-07 | 3.8e-07 | 2.5e-07 | 3.300E-07 | 7.000E-08 | 2.500E-07 | 3.800E-07 |
| 24625 | 1232 | S1232C | D | 1.6e-07 | 3e-07 | 1.6e-07 | 2.067E-07 | 8.083E-08 | 1.600E-07 | 3.000E-07 |
| 24626 | 1232 | S1232E | D | 3.3e-07 | 1.8e-07 | 1.3e-07 | 2.133E-07 | 1.041E-07 | 1.300E-07 | 3.300E-07 |
| 24627 | 1232 | S1232Q | D | 3.1e-07 | 1.3e-07 | 2.5e-07 | 2.300E-07 | 9.165E-08 | 1.300E-07 | 3.100E-07 |
| 24628 | 1232 | S1232G | D | 2.1e-07 | 1.8e-07 | 1.7e-07 | 1.867E-07 | 2.082E-08 | 1.700E-07 | 2.100E-07 |
| 24629 | 1232 | S1232H | D | 4e-07 | 4e-07 | 2.8e-07 | 3.600E-07 | 6.928E-08 | 2.800E-07 | 4.000E-07 |
| 24630 | 1232 | S1232I | D | 1.9e-07 | 2.2e-07 | 5.2e-07 | 3.100E-07 | 1.825E-07 | 1.900E-07 | 5.200E-07 |
| 24631 | 1232 | S1232L | D | 3.7e-07 | 2.6e-07 | 2.6e-07 | 2.967E-07 | 6.351E-08 | 2.600E-07 | 3.700E-07 |
| 24632 | 1232 | S1232K | D | 3.4e-07 | 2.5e-07 | 3.1e-07 | 3.000E-07 | 4.583E-08 | 2.500E-07 | 3.400E-07 |
| 24633 | 1232 | S1232M | D | 2e-07 | 4.4e-07 | 1.4e-07 | 2.600E-07 | 1.587E-07 | 1.400E-07 | 4.400E-07 |
| 24634 | 1232 | S1232F | D | 1.9e-07 | 2.3e-07 | 3.4e-07 | 2.533E-07 | 7.767E-08 | 1.900E-07 | 3.400E-07 |
| 24635 | 1232 | S1232P | D | 2.6e-07 | 8.7e-08 | 3.2e-07 | 2.223E-07 | 1.210E-07 | 8.700E-08 | 3.200E-07 |
| 24636 | 1232 | S1232S | D | 4.5e-07 | 1.9e-07 | 1.9e-07 | 2.767E-07 | 1.501E-07 | 1.900E-07 | 4.500E-07 |
| 24637 | 1232 | S1232T | D | 3.3e-07 | 1.9e-07 | 1.4e-07 | 2.200E-07 | 9.849E-08 | 1.400E-07 | 3.300E-07 |
| 24638 | 1232 | S1232W | D | 4.3e-07 | 1.6e-07 | 2.8e-07 | 2.900E-07 | 1.353E-07 | 1.600E-07 | 4.300E-07 |
| 24639 | 1232 | S1232Y | D | 2.4e-07 | 1.9e-07 | 1.5e-07 | 1.933E-07 | 4.509E-08 | 1.500E-07 | 2.400E-07 |
| 24640 | 1232 | S1232V | D | 2.6e-07 | 2.7e-07 | 3.1e-07 | 2.800E-07 | 2.646E-08 | 2.600E-07 | 3.100E-07 |
| 24641 | 1233 | L1233A | D | 1.5e-07 | 3.6e-07 | 1.5e-07 | 2.200E-07 | 1.212E-07 | 1.500E-07 | 3.600E-07 |
| 24642 | 1233 | L1233R | D | 1.8e-07 | 3.6e-07 | 6.4e-07 | 3.933E-07 | 2.318E-07 | 1.800E-07 | 6.400E-07 |
| 24643 | 1233 | L1233N | D | 1.1e-07 | 2e-07 | 3e-07 | 2.033E-07 | 9.504E-08 | 1.100E-07 | 3.000E-07 |
| 24644 | 1233 | L1233D | D | 2.2e-07 | 1.9e-07 | 3.3e-07 | 2.467E-07 | 7.371E-08 | 1.900E-07 | 3.300E-07 |
| 24645 | 1233 | L1233C | D | 1.4e-07 | 2.3e-07 | 1.2e-07 | 1.633E-07 | 5.859E-08 | 1.200E-07 | 2.300E-07 |
| 24646 | 1233 | L1233E | D | 2e-07 | 5.6e-07 | 5.5e-07 | 4.367E-07 | 2.050E-07 | 2.000E-07 | 5.600E-07 |
| 24647 | 1233 | L1233Q | D | 3.7e-07 | 2.9e-07 | 3.3e-07 | 3.300E-07 | 4.000E-08 | 2.900E-07 | 3.700E-07 |
| 24648 | 1233 | L1233G | D | 3.3e-07 | 2.2e-07 | 2.7e-07 | 2.733E-07 | 5.508E-08 | 2.200E-07 | 3.300E-07 |
| 24649 | 1233 | L1233H | D | 1.4e-07 | 1.2e-07 | 1.1e-07 | 1.233E-07 | 1.528E-08 | 1.100E-07 | 1.400E-07 |

| | | | | | | | | | | |
|---|---|---|---|---|---|---|---|---|---|---|
| 24650 | 1233 | L1233I | D | 1.2e-07 | 4.5e-07 | 4.7e-07 | 3.467E-07 | 1.966E-07 | 1.200E-07 | 4.700E-07 |
| 24651 | 1233 | L1233L | D | 1.2e-07 | 6.5e-07 | 3.9e-07 | 3.867E-07 | 2.650E-07 | 1.200E-07 | 6.500E-07 |
| 24652 | 1233 | L1233K | D | 4e-07 | 2.3e-07 | 2.6e-07 | 2.967E-07 | 9.074E-08 | 2.300E-07 | 4.000E-07 |
| 24653 | 1233 | L1233M | D | 3e-07 | 1.6e-07 | 3.5e-07 | 2.700E-07 | 9.849E-08 | 1.600E-07 | 3.500E-07 |
| 24654 | 1233 | L1233F | D | 3e-07 | 1.5e-07 | 2.2e-07 | 2.233E-07 | 7.506E-08 | 1.500E-07 | 3.000E-07 |
| 24655 | 1233 | L1233P | D | 2.8e-07 | 3.2e-07 | 1.7e-07 | 2.567E-07 | 7.767E-08 | 1.700E-07 | 3.200E-07 |
| 24656 | 1233 | L1233S | D | 1.1e-07 | 1.9e-07 | 2.8e-07 | 1.933E-07 | 8.505E-08 | 1.100E-07 | 2.800E-07 |
| 24657 | 1233 | L1233T | D | 3.4e-07 | 1.9e-07 | 3.6e-07 | 2.967E-07 | 9.292E-08 | 1.900E-07 | 3.600E-07 |
| 24658 | 1233 | L1233W | D | 2.5e-07 | 2.2e-07 | 2.3e-07 | 2.333E-07 | 1.528E-08 | 2.200E-07 | 2.500E-07 |
| 24659 | 1233 | L1233Y | D | 3.8e-07 | 2.9e-07 | 3.5e-07 | 3.400E-07 | 4.583E-08 | 2.900E-07 | 3.800E-07 |
| 24660 | 1233 | L1233V | D | 2.3e-07 | 3.1e-07 | 8.2e-07 | 4.533E-07 | 3.201E-07 | 2.300E-07 | 8.200E-07 |
| 24661 | 1234 | S1234A | D | 5.8e-07 | 5.1e-07 | 1.6e-07 | 4.167E-07 | 2.250E-07 | 1.600E-07 | 5.800E-07 |
| 24662 | 1234 | S1234R | D | 4e-07 | 1.6e-07 | 3.1e-07 | 2.900E-07 | 1.212E-07 | 1.600E-07 | 4.000E-07 |
| 24663 | 1234 | S1234N | D | 3.5e-07 | 2.4e-07 | 2.5e-07 | 2.800E-07 | 6.083E-08 | 2.400E-07 | 3.500E-07 |
| 24664 | 1234 | S1234D | D | 2.8e-07 | 2.5e-07 | 3.6e-07 | 2.967E-07 | 5.686E-08 | 2.500E-07 | 3.600E-07 |
| 24665 | 1234 | S1234C | D | 1.6e-07 | 4.1e-07 | 2.2e-07 | 2.633E-07 | 1.305E-07 | 1.600E-07 | 4.100E-07 |
| 24666 | 1234 | S1234E | D | 3.3e-07 | 1.4e-07 | 5.3e-07 | 3.333E-07 | 1.950E-07 | 1.400E-07 | 5.300E-07 |
| 24667 | 1234 | S1234Q | D | 3.4e-07 | 4.1e-07 | 2e-07 | 3.167E-07 | 1.069E-07 | 2.000E-07 | 4.100E-07 |
| 24668 | 1234 | S1234G | D | 1.8e-07 | 3e-07 | 2.5e-07 | 2.433E-07 | 6.028E-08 | 1.800E-07 | 3.000E-07 |
| 24669 | 1234 | S1234H | D | 1.2e-07 | 4.4e-07 | 8.7e-07 | 4.767E-07 | 3.763E-07 | 1.200E-07 | 8.700E-07 |
| 24670 | 1234 | S1234I | D | 8.9e-08 | 2.7e-07 | 3.3e-07 | 2.297E-07 | 1.255E-07 | 8.900E-08 | 3.300E-07 |
| 24671 | 1234 | S1234L | D | 2.3e-07 | 2.9e-07 | 2e-07 | 2.400E-07 | 4.583E-08 | 2.000E-07 | 2.900E-07 |
| 24672 | 1234 | S1234K | D | 2.6e-07 | 2.3e-07 | 3.1e-07 | 2.667E-07 | 4.041E-08 | 2.300E-07 | 3.100E-07 |
| 24673 | 1234 | S1234M | D | 1.4e-07 | 8.8e-08 | 2.2e-07 | 1.493E-07 | 6.649E-08 | 8.800E-08 | 2.200E-07 |
| 24674 | 1234 | S1234F | D | 4.5e-07 | 1.7e-07 | 2.8e-07 | 3.000E-07 | 1.411E-07 | 1.700E-07 | 4.500E-07 |
| 24675 | 1234 | S1234P | D | 2.5e-07 | 2.8e-07 | 4.7e-07 | 3.333E-07 | 1.193E-07 | 2.500E-07 | 4.700E-07 |
| 24676 | 1234 | S1234S | D | 3.3e-07 | 2.7e-07 | 1.3e-07 | 2.433E-07 | 1.026E-07 | 1.300E-07 | 3.300E-07 |
| 24677 | 1234 | S1234T | D | 2.4e-07 | 3.9e-07 | 2.1e-07 | 2.800E-07 | 9.644E-08 | 2.100E-07 | 3.900E-07 |
| 24678 | 1234 | S1234W | D | 2e-07 | 2.7e-07 | 2.7e-07 | 2.467E-07 | 4.041E-08 | 2.000E-07 | 2.700E-07 |
| 24679 | 1234 | S1234Y | D | 1.6e-07 | 2.9e-07 | 3.2e-07 | 2.567E-07 | 8.505E-08 | 1.600E-07 | 3.200E-07 |
| 24680 | 1234 | S1234V | D | 2.3e-07 | 4.6e-07 | 3.5e-07 | 3.467E-07 | 1.150E-07 | 2.300E-07 | 4.600E-07 |
| 24681 | 1235 | S1235A | D | 4.3e-07 | 1.4e-07 | 2.5e-07 | 2.733E-07 | 1.464E-07 | 1.400E-07 | 4.300E-07 |
| 24682 | 1235 | S1235R | D | 3.2e-07 | 2.8e-07 | 1.4e-07 | 2.467E-07 | 9.452E-08 | 1.400E-07 | 3.200E-07 |
| 24683 | 1235 | S1235N | D | 3.8e-07 | 2.9e-07 | 1.7e-07 | 2.800E-07 | 1.054E-07 | 1.700E-07 | 3.800E-07 |
| 24684 | 1235 | S1235D | D | 2.9e-07 | 3.4e-07 | 1.5e-07 | 2.600E-07 | 9.849E-08 | 1.500E-07 | 3.400E-07 |
| 24685 | 1235 | S1235C | D | 4.9e-07 | 1.9e-07 | 2.5e-07 | 3.100E-07 | 1.587E-07 | 1.900E-07 | 4.900E-07 |
| 24686 | 1235 | S1235E | D | 3.2e-07 | 1.1e-07 | 8.2e-07 | 4.167E-07 | 3.647E-07 | 1.100E-07 | 8.200E-07 |
| 24687 | 1235 | S1235Q | D | 3.1e-07 | 2.3e-07 | 2.9e-07 | 2.767E-07 | 4.163E-08 | 2.300E-07 | 3.100E-07 |
| 24688 | 1235 | S1235G | D | 1.4e-07 | 3.3e-07 | 3.7e-07 | 2.800E-07 | 1.229E-07 | 1.400E-07 | 3.700E-07 |
| 24689 | 1235 | S1235H | D | 1.5e-07 | 3.7e-07 | 2.1e-07 | 2.433E-07 | 1.137E-07 | 1.500E-07 | 3.700E-07 |
| 24690 | 1235 | S1235I | D | 5.1e-07 | 3.3e-07 | 4.1e-07 | 4.167E-07 | 9.018E-08 | 3.300E-07 | 5.100E-07 |
| 24691 | 1235 | S1235L | D | 2.4e-07 | 1.4e-07 | 1.4e-07 | 1.733E-07 | 5.774E-08 | 1.400E-07 | 2.400E-07 |
| 24692 | 1235 | S1235K | D | 2.4e-07 | 4e-07 | 2.2e-07 | 2.867E-07 | 9.866E-08 | 2.200E-07 | 4.000E-07 |
| 24693 | 1235 | S1235M | D | 1.4e-07 | 3.1e-07 | 3.3e-07 | 2.600E-07 | 1.044E-07 | 1.400E-07 | 3.300E-07 |
| 24694 | 1235 | S1235F | D | 2.5e-07 | 1.9e-07 | 2e-07 | 2.133E-07 | 3.215E-08 | 1.900E-07 | 2.500E-07 |
| 24695 | 1235 | S1235P | D | 2.5e-07 | 2.7e-07 | 2.4e-07 | 2.533E-07 | 1.528E-08 | 2.400E-07 | 2.700E-07 |
| 24696 | 1235 | S1235S | D | 3.4e-07 | 2.6e-07 | 6.7e-07 | 4.233E-07 | 2.173E-07 | 2.600E-07 | 6.700E-07 |
| 24697 | 1235 | S1235T | D | 3e-07 | 1.1e-06 | 4.1e-07 | 6.033E-07 | 4.336E-07 | 3.000E-07 | 1.100E-06 |
| 24698 | 1235 | S1235W | D | 1.6e-07 | 3.4e-07 | 3.2e-07 | 2.733E-07 | 9.866E-08 | 1.600E-07 | 3.400E-07 |
| 24699 | 1235 | S1235Y | D | 5e-07 | 2.2e-07 | 2.9e-07 | 3.367E-07 | 1.457E-07 | 2.200E-07 | 5.000E-07 |
| 24700 | 1235 | S1235V | D | 1.9e-07 | 6.2e-07 | 2.2e-07 | 3.433E-07 | 2.401E-07 | 1.900E-07 | 6.200E-07 |
| 24701 | 1236 | T1236A | D | 3.2e-07 | 1.7e-07 | 2.2e-07 | 2.367E-07 | 7.638E-08 | 1.700E-07 | 3.200E-07 |
| 24702 | 1236 | T1236R | D | 1.8e-07 | 1.8e-07 | 3e-07 | 2.200E-07 | 6.928E-08 | 1.800E-07 | 3.000E-07 |
| 24703 | 1236 | T1236N | D | 1.1e-07 | 2.8e-07 | 2.6e-07 | 2.167E-07 | 9.292E-08 | 1.100E-07 | 2.800E-07 |
| 24704 | 1236 | T1236D | D | 6.5e-07 | 1.6e-07 | 2.9e-07 | 3.667E-07 | 2.538E-07 | 1.600E-07 | 6.500E-07 |
| 24705 | 1236 | T1236C | D | 2.1e-07 | 2e-07 | 3.2e-07 | 2.433E-07 | 6.658E-08 | 2.000E-07 | 3.200E-07 |
| 24706 | 1236 | T1236E | D | 1.8e-07 | 2.5e-07 | 2.6e-07 | 2.300E-07 | 4.359E-08 | 1.800E-07 | 2.600E-07 |
| 24707 | 1236 | T1236Q | D | 3.3e-07 | 4.9e-07 | 1.7e-07 | 3.300E-07 | 1.600E-07 | 1.700E-07 | 4.900E-07 |
| 24708 | 1236 | T1236G | D | 3.4e-07 | 3.6e-07 | 1.7e-07 | 2.900E-07 | 1.044E-07 | 1.700E-07 | 3.600E-07 |
| 24709 | 1236 | T1236H | D | 2.8e-07 | 1.4e-07 | 1.5e-07 | 1.900E-07 | 7.810E-08 | 1.400E-07 | 2.800E-07 |
| 24710 | 1236 | T1236I | D | 2.8e-07 | 3.5e-07 | 2.5e-07 | 2.933E-07 | 5.132E-08 | 2.500E-07 | 3.500E-07 |
| 24711 | 1236 | T1236L | D | 3.6e-07 | 1.6e-07 | 2.9e-07 | 2.700E-07 | 1.015E-07 | 1.600E-07 | 3.600E-07 |
| 24712 | 1236 | T1236K | D | 2.5e-07 | 2.6e-07 | 1.7e-07 | 2.267E-07 | 4.933E-08 | 1.700E-07 | 2.600E-07 |
| 24713 | 1236 | T1236M | D | 4.3e-07 | 3.4e-07 | 1.6e-07 | 3.100E-07 | 1.375E-07 | 1.600E-07 | 4.300E-07 |
| 24714 | 1236 | T1236F | D | 2.7e-07 | 2.2e-07 | 4.2e-07 | 3.033E-07 | 1.041E-07 | 2.200E-07 | 4.200E-07 |
| 24715 | 1236 | T1236P | D | 3.9e-07 | 3.3e-07 | 2.4e-07 | 3.200E-07 | 7.550E-08 | 2.400E-07 | 3.900E-07 |
| 24716 | 1236 | T1236S | D | 1.9e-07 | 2.5e-07 | 5e-07 | 3.133E-07 | 1.644E-07 | 1.900E-07 | 5.000E-07 |
| 24717 | 1236 | T1236T | D | 2.2e-07 | 1.4e-07 | 5.9e-07 | 3.167E-07 | 2.401E-07 | 1.400E-07 | 5.900E-07 |
| 24718 | 1236 | T1236W | D | 3.3e-07 | 3.2e-07 | 1.3e-07 | 2.600E-07 | 1.127E-07 | 1.300E-07 | 3.300E-07 |
| 24719 | 1236 | T1236Y | D | 2e-07 | 1.9e-07 | 1.1e-07 | 1.667E-07 | 4.933E-08 | 1.100E-07 | 2.000E-07 |
| 24720 | 1236 | T1236V | D | 3.5e-07 | 2.5e-07 | 3.7e-07 | 3.233E-07 | 6.429E-08 | 2.500E-07 | 3.700E-07 |
| 24721 | 1237 | L1237A | D | 3.1e-07 | 4.9e-07 | 2.3e-07 | 3.433E-07 | 1.332E-07 | 2.300E-07 | 4.900E-07 |
| 24722 | 1237 | L1237R | D | 3.5e-07 | 1.6e-07 | 4.8e-07 | 3.300E-07 | 1.609E-07 | 1.600E-07 | 4.800E-07 |
| 24723 | 1237 | L1237N | D | 6.8e-07 | 2.4e-07 | 2.7e-07 | 3.967E-07 | 2.458E-07 | 2.400E-07 | 6.800E-07 |
| 24724 | 1237 | L1237D | D | 1.6e-07 | 1.3e-07 | 2.6e-07 | 1.833E-07 | 6.807E-08 | 1.300E-07 | 2.600E-07 |
| 24725 | 1237 | L1237C | D | 1.5e-07 | 3.1e-07 | 2.3e-07 | 2.300E-07 | 8.000E-08 | 1.500E-07 | 3.100E-07 |
| 24726 | 1237 | L1237E | D | 3e-07 | 4.8e-07 | 2.5e-07 | 3.433E-07 | 1.210E-07 | 2.500E-07 | 4.800E-07 |
| 24727 | 1237 | L1237Q | D | 1.9e-07 | 3e-07 | 2.3e-07 | 2.400E-07 | 5.568E-08 | 1.900E-07 | 3.000E-07 |
| 24728 | 1237 | L1237G | D | 3.3e-07 | 3.3e-07 | 3e-07 | 3.200E-07 | 1.732E-08 | 3.000E-07 | 3.300E-07 |
| 24729 | 1237 | L1237H | D | 1.1e-07 | 5.1e-07 | 2.1e-07 | 2.767E-07 | 2.082E-07 | 1.100E-07 | 5.100E-07 |
| 24730 | 1237 | L1237I | D | 2.9e-07 | 2e-07 | 2.9e-07 | 2.600E-07 | 5.196E-08 | 2.000E-07 | 2.900E-07 |
| 24731 | 1237 | L1237L | D | 2.5e-07 | 4e-07 | 3e-07 | 3.167E-07 | 7.638E-08 | 2.500E-07 | 4.000E-07 |
| 24732 | 1237 | L1237K | D | 2.2e-07 | 2.7e-07 | 2.2e-07 | 2.367E-07 | 2.887E-08 | 2.200E-07 | 2.700E-07 |
| 24733 | 1237 | L1237M | D | 3.7e-07 | 3.9e-07 | 2.4e-07 | 3.333E-07 | 8.145E-08 | 2.400E-07 | 3.900E-07 |
| 24734 | 1237 | L1237F | D | 3.1e-07 | 1.3e-07 | 2.1e-07 | 2.167E-07 | 9.018E-08 | 1.300E-07 | 3.100E-07 |

| | | | | | | | | | | |
|---|---|---|---|---|---|---|---|---|---|---|
| 24735 | 1237 | L1237P | D | 2.5e-07 | 2.1e-07 | 2.9e-07 | 2.500E-07 | 4.000E-08 | 2.100E-07 | 2.900E-07 |
| 24736 | 1237 | L1237S | D | 1.6e-07 | 3.1e-07 | 2.1e-07 | 2.267E-07 | 7.638E-08 | 1.600E-07 | 3.100E-07 |
| 24737 | 1237 | L1237T | D | 5e-07 | 4.1e-07 | 3.5e-07 | 4.200E-07 | 7.550E-08 | 3.500E-07 | 5.000E-07 |
| 24738 | 1237 | L1237W | D | 3.6e-07 | 2.8e-07 | 3.2e-07 | 3.200E-07 | 4.000E-08 | 2.800E-07 | 3.600E-07 |
| 24739 | 1237 | L1237Y | D | 5e-07 | 5.9e-07 | 2.7e-07 | 4.533E-07 | 1.650E-07 | 2.700E-07 | 5.900E-07 |
| 24740 | 1237 | L1237V | D | 2.8e-07 | 2.5e-07 | 1.9e-07 | 2.400E-07 | 4.583E-08 | 1.900E-07 | 2.800E-07 |
| 24741 | 1238 | T1238A | D | 2.8e-07 | 7.8e-07 | 1.7e-07 | 4.100E-07 | 3.251E-07 | 1.700E-07 | 7.800E-07 |
| 24742 | 1238 | T1238R | D | 1.5e-07 | 3.5e-07 | 4.6e-07 | 3.200E-07 | 1.572E-07 | 1.500E-07 | 4.600E-07 |
| 24743 | 1238 | T1238N | D | 3e-07 | 5.2e-07 | 6e-07 | 4.733E-07 | 1.553E-07 | 3.000E-07 | 6.000E-07 |
| 24744 | 1238 | T1238D | D | 4.1e-07 | 1.4e-07 | 2.6e-07 | 2.700E-07 | 1.353E-07 | 1.400E-07 | 4.100E-07 |
| 24745 | 1238 | T1238C | D | 3.8e-07 | 2.1e-07 | 3.5e-07 | 3.133E-07 | 9.074E-08 | 2.100E-07 | 3.800E-07 |
| 24746 | 1238 | T1238E | D | 3e-07 | 2.9e-07 | 4e-07 | 3.300E-07 | 6.083E-08 | 2.900E-07 | 4.000E-07 |
| 24747 | 1238 | T1238Q | D | 3.4e-07 | 5e-07 | 5e-07 | 4.467E-07 | 9.238E-08 | 3.400E-07 | 5.000E-07 |
| 24748 | 1238 | T1238G | D | 3.1e-07 | 1.5e-07 | 1.4e-07 | 2.000E-07 | 9.539E-08 | 1.400E-07 | 3.100E-07 |
| 24749 | 1238 | T1238H | D | 1.3e-07 | 2.1e-07 | 1.9e-07 | 1.767E-07 | 4.163E-08 | 1.300E-07 | 2.100E-07 |
| 24750 | 1238 | T1238I | D | 3.6e-07 | 1.8e-07 | 4e-07 | 3.133E-07 | 1.172E-07 | 1.800E-07 | 4.000E-07 |
| 24751 | 1238 | T1238L | D | 2.8e-07 | 3e-07 | 3.5e-07 | 3.100E-07 | 3.606E-08 | 2.800E-07 | 3.500E-07 |
| 24752 | 1238 | T1238K | D | 2.9e-07 | 1e-07 | 1.7e-07 | 1.867E-07 | 9.609E-08 | 1.000E-07 | 2.900E-07 |
| 24753 | 1238 | T1238M | D | 3.3e-07 | 2.5e-07 | 4e-07 | 3.267E-07 | 7.506E-08 | 2.500E-07 | 4.000E-07 |
| 24754 | 1238 | T1238F | D | 1.5e-07 | 6.4e-07 | 1.9e-07 | 3.267E-07 | 2.721E-07 | 1.500E-07 | 6.400E-07 |
| 24755 | 1238 | T1238P | D | 2.9e-07 | 4.2e-07 | 3.1e-07 | 3.400E-07 | 7.000E-08 | 2.900E-07 | 4.200E-07 |
| 24756 | 1238 | T1238S | D | 1.9e-07 | 2.7e-07 | 2.4e-07 | 2.333E-07 | 4.041E-08 | 1.900E-07 | 2.700E-07 |
| 24757 | 1238 | T1238T | D | 4.4e-07 | 3.5e-07 | 1.4e-07 | 3.100E-07 | 1.539E-07 | 1.400E-07 | 4.400E-07 |
| 24758 | 1238 | T1238W | D | 2.2e-07 | 1.5e-07 | 1.3e-07 | 1.667E-07 | 4.726E-08 | 1.300E-07 | 2.200E-07 |
| 24759 | 1238 | T1238Y | D | 2.1e-07 | 5.2e-07 | 2.6e-07 | 3.300E-07 | 1.664E-07 | 2.100E-07 | 5.200E-07 |
| 24760 | 1238 | T1238V | D | 1.6e-07 | 2e-07 | 1.8e-07 | 1.800E-07 | 2.000E-08 | 1.600E-07 | 2.000E-07 |
| 24761 | 1239 | L1239A | D | 3.8e-07 | 2.5e-07 | 2.2e-07 | 2.833E-07 | 8.505E-08 | 2.200E-07 | 3.800E-07 |
| 24762 | 1239 | L1239R | D | 3.8e-07 | 4.1e-07 | 3.8e-07 | 3.900E-07 | 1.732E-08 | 3.800E-07 | 4.100E-07 |
| 24763 | 1239 | L1239N | D | 1.8e-07 | 2.3e-07 | 6.3e-07 | 3.467E-07 | 2.466E-07 | 1.800E-07 | 6.300E-07 |
| 24764 | 1239 | L1239D | D | 3.8e-07 | 1.8e-07 | 3e-07 | 2.867E-07 | 1.007E-07 | 1.800E-07 | 3.800E-07 |
| 24765 | 1239 | L1239C | D | 4.4e-07 | 2.4e-07 | 3.3e-07 | 3.367E-07 | 1.002E-07 | 2.400E-07 | 4.400E-07 |
| 24766 | 1239 | L1239E | D | 1.8e-07 | 4.4e-07 | 2.5e-07 | 2.900E-07 | 1.345E-07 | 1.800E-07 | 4.400E-07 |
| 24767 | 1239 | L1239Q | D | 1.9e-07 | 2e-07 | 2e-07 | 1.967E-07 | 5.774E-09 | 1.900E-07 | 2.000E-07 |
| 24768 | 1239 | L1239G | D | 7.1e-07 | 4e-07 | 7.8e-07 | 6.300E-07 | 2.022E-07 | 4.000E-07 | 7.800E-07 |
| 24769 | 1239 | L1239H | D | 2.8e-07 | 3.5e-07 | 2.7e-07 | 3.000E-07 | 4.359E-08 | 2.700E-07 | 3.500E-07 |
| 24770 | 1239 | L1239I | D | 2.1e-07 | 6e-07 | 2.4e-07 | 3.500E-07 | 2.170E-07 | 2.100E-07 | 6.000E-07 |
| 24771 | 1239 | L1239L | D | 2.2e-07 | 1.5e-07 | 2.7e-07 | 2.133E-07 | 6.028E-08 | 1.500E-07 | 2.700E-07 |
| 24772 | 1239 | L1239K | D | 3.1e-07 | 1e-07 | 1.3e-07 | 1.800E-07 | 1.136E-07 | 1.000E-07 | 3.100E-07 |
| 24773 | 1239 | L1239M | D | 2.8e-07 | 2e-07 | 1.9e-07 | 2.233E-07 | 4.933E-08 | 1.900E-07 | 2.800E-07 |
| 24774 | 1239 | L1239F | D | 2.3e-07 | 4.4e-07 | 2.2e-07 | 2.967E-07 | 1.242E-07 | 2.200E-07 | 4.400E-07 |
| 24775 | 1239 | L1239P | D | 1.7e-07 | 3.5e-07 | 3e-07 | 2.733E-07 | 9.292E-08 | 1.700E-07 | 3.500E-07 |
| 24776 | 1239 | L1239S | D | 4.7e-07 | 2.5e-07 | 2.6e-07 | 3.267E-07 | 1.242E-07 | 2.500E-07 | 4.700E-07 |
| 24777 | 1239 | L1239T | D | 2e-07 | 2e-07 | 2.9e-07 | 2.300E-07 | 5.196E-08 | 2.000E-07 | 2.900E-07 |
| 24778 | 1239 | L1239W | D | 2e-07 | 2.1e-07 | 2.6e-07 | 2.233E-07 | 3.215E-08 | 2.000E-07 | 2.600E-07 |
| 24779 | 1239 | L1239Y | D | 7.7e-08 | 3.3e-07 | 4.2e-07 | 2.757E-07 | 1.778E-07 | 7.700E-08 | 4.200E-07 |
| 24780 | 1239 | L1239V | D | 2.3e-07 | 2.5e-07 | 1.8e-07 | 2.200E-07 | 3.606E-08 | 1.800E-07 | 2.500E-07 |
| 24781 | 1240 | S1240A | D | 2.1e-07 | 1.4e-07 | 1.9e-07 | 1.800E-07 | 3.606E-08 | 1.400E-07 | 2.100E-07 |
| 24782 | 1240 | S1240R | D | 2.5e-07 | 5.5e-07 | 2.4e-07 | 3.467E-07 | 1.762E-07 | 2.400E-07 | 5.500E-07 |
| 24783 | 1240 | S1240N | D | 1.6e-07 | 1.4e-07 | 1e-07 | 1.333E-07 | 3.055E-08 | 1.000E-07 | 1.600E-07 |
| 24784 | 1240 | S1240D | D | 3.1e-07 | 3.6e-07 | 2.7e-07 | 3.133E-07 | 4.509E-08 | 2.700E-07 | 3.600E-07 |
| 24785 | 1240 | S1240C | D | 2.8e-07 | 3.5e-07 | 3.3e-07 | 3.200E-07 | 3.606E-08 | 2.800E-07 | 3.500E-07 |
| 24786 | 1240 | S1240E | D | 4.2e-07 | 3.5e-07 | 2.6e-07 | 3.433E-07 | 8.021E-08 | 2.600E-07 | 4.200E-07 |
| 24787 | 1240 | S1240Q | D | 2.5e-07 | 1.7e-07 | 2.2e-07 | 2.133E-07 | 4.041E-08 | 1.700E-07 | 2.500E-07 |
| 24788 | 1240 | S1240G | D | 1.6e-07 | 2e-07 | 3.1e-07 | 2.233E-07 | 7.767E-08 | 1.600E-07 | 3.100E-07 |
| 24789 | 1240 | S1240H | D | 2.4e-07 | 4.2e-07 | 2.3e-07 | 2.967E-07 | 1.069E-07 | 2.300E-07 | 4.200E-07 |
| 24790 | 1240 | S1240I | D | 2.5e-07 | 4e-07 | 2.4e-07 | 2.967E-07 | 8.963E-08 | 2.400E-07 | 4.000E-07 |
| 24791 | 1240 | S1240L | D | 4.2e-07 | 3.1e-07 | 4.3e-07 | 3.867E-07 | 6.658E-08 | 3.100E-07 | 4.300E-07 |
| 24792 | 1240 | S1240K | D | 3.3e-07 | 2.7e-07 | 1.6e-07 | 2.533E-07 | 8.622E-08 | 1.600E-07 | 3.300E-07 |
| 24793 | 1240 | S1240M | D | 2.8e-07 | 3.1e-07 | 1.5e-07 | 2.467E-07 | 8.505E-08 | 1.500E-07 | 3.100E-07 |
| 24794 | 1240 | S1240F | D | 2.5e-07 | 2.3e-07 | 2.2e-07 | 2.333E-07 | 1.528E-08 | 2.200E-07 | 2.500E-07 |
| 24795 | 1240 | S1240P | D | 2.6e-07 | 1.6e-07 | 2.6e-07 | 2.267E-07 | 5.774E-08 | 1.600E-07 | 2.600E-07 |
| 24796 | 1240 | S1240S | D | 1.1e-07 | 3.1e-07 | 4e-07 | 2.733E-07 | 1.484E-07 | 1.100E-07 | 4.000E-07 |
| 24797 | 1240 | S1240T | D | 1.6e-07 | 1.5e-07 | 2e-07 | 1.700E-07 | 2.646E-08 | 1.500E-07 | 2.000E-07 |
| 24798 | 1240 | S1240W | D | 2e-07 | 2.7e-07 | 2.8e-07 | 2.500E-07 | 4.359E-08 | 2.000E-07 | 2.800E-07 |
| 24799 | 1240 | S1240Y | D | 1.7e-07 | 1.8e-07 | 2.5e-07 | 2.000E-07 | 4.359E-08 | 1.700E-07 | 2.500E-07 |
| 24800 | 1240 | S1240V | D | 2.4e-07 | 6.2e-07 | 3.9e-07 | 4.167E-07 | 1.914E-07 | 2.400E-07 | 6.200E-07 |
| 24801 | 1241 | K1241A | D | 3.7e-07 | 2.3e-07 | 1.9e-07 | 2.633E-07 | 9.452E-08 | 1.900E-07 | 3.700E-07 |
| 24802 | 1241 | K1241R | D | 1.1e-07 | 1.5e-07 | 1.9e-07 | 1.500E-07 | 4.000E-08 | 1.100E-07 | 1.900E-07 |
| 24803 | 1241 | K1241N | D | 5.3e-07 | 2.1e-07 | 1e-07 | 2.800E-07 | 2.234E-07 | 1.000E-07 | 5.300E-07 |
| 24804 | 1241 | K1241D | D | 3e-07 | 1.4e-07 | 4.3e-07 | 2.900E-07 | 1.453E-07 | 1.400E-07 | 4.300E-07 |
| 24805 | 1241 | K1241C | D | 2.8e-07 | 2.9e-07 | 2.6e-07 | 2.767E-07 | 1.528E-08 | 2.600E-07 | 2.900E-07 |
| 24806 | 1241 | K1241E | D | 3.3e-07 | 3.1e-07 | 3.8e-07 | 3.400E-07 | 3.606E-08 | 3.100E-07 | 3.800E-07 |
| 24807 | 1241 | K1241Q | D | 1.7e-07 | 2.6e-07 | 1.9e-07 | 2.067E-07 | 4.726E-08 | 1.700E-07 | 2.600E-07 |
| 24808 | 1241 | K1241G | D | 2.9e-07 | 3e-07 | 2.3e-07 | 2.733E-07 | 3.786E-08 | 2.300E-07 | 3.000E-07 |
| 24809 | 1241 | K1241H | D | 1.7e-07 | 2.4e-07 | 2.9e-07 | 2.333E-07 | 6.028E-08 | 1.700E-07 | 2.900E-07 |
| 24810 | 1241 | K1241I | D | 3.8e-07 | 2e-07 | 3.2e-07 | 3.000E-07 | 9.165E-08 | 2.000E-07 | 3.800E-07 |
| 24811 | 1241 | K1241L | D | 2.2e-07 | 2.6e-07 | 4e-07 | 2.933E-07 | 9.452E-08 | 2.200E-07 | 4.000E-07 |
| 24812 | 1241 | K1241K | D | 2.3e-07 | 5.4e-07 | 1.6e-07 | 3.100E-07 | 2.022E-07 | 1.600E-07 | 5.400E-07 |
| 24813 | 1241 | K1241M | D | 1.6e-07 | 1.6e-07 | 2.3e-07 | 1.833E-07 | 4.041E-08 | 1.600E-07 | 2.300E-07 |
| 24814 | 1241 | K1241F | D | 2.3e-07 | 1.1e-07 | 1.3e-07 | 1.567E-07 | 6.429E-08 | 1.100E-07 | 2.300E-07 |
| 24815 | 1241 | K1241P | D | 1.8e-07 | 1.9e-07 | 3e-07 | 2.233E-07 | 6.658E-08 | 1.800E-07 | 3.000E-07 |
| 24816 | 1241 | K1241S | D | 3.3e-07 | 2.3e-07 | 1.9e-07 | 2.500E-07 | 7.211E-08 | 1.900E-07 | 3.300E-07 |
| 24817 | 1241 | K1241T | D | 5.7e-07 | 5.4e-07 | 1.7e-07 | 4.267E-07 | 2.228E-07 | 1.700E-07 | 5.700E-07 |
| 24818 | 1241 | K1241W | D | 3.1e-07 | 1.7e-07 | 2.8e-07 | 2.533E-07 | 7.371E-08 | 1.700E-07 | 3.100E-07 |
| 24819 | 1241 | K1241Y | D | 2.1e-07 | 2.1e-07 | 1.3e-07 | 1.833E-07 | 4.619E-08 | 1.300E-07 | 2.100E-07 |

| | | | | | | | | | | |
|---|---|---|---|---|---|---|---|---|---|---|
| 24820 | 1241 | K1241V | D | 2.3e-07 | 2.2e-07 | 3.1e-07 | 2.533E-07 | 4.933E-08 | 2.200E-07 | 3.100E-07 |
| 24821 | 1242 | A1242A | D | 1.6e-07 | 4.2e-07 | 2.3e-07 | 2.700E-07 | 1.345E-07 | 1.600E-07 | 4.200E-07 |
| 24822 | 1242 | A1242R | D | 2.6e-07 | 2e-07 | 2.2e-07 | 2.267E-07 | 3.055E-08 | 2.000E-07 | 2.600E-07 |
| 24823 | 1242 | A1242N | D | 2.1e-07 | 3.6e-07 | 1.1e-07 | 2.267E-07 | 1.258E-07 | 1.100E-07 | 3.600E-07 |
| 24824 | 1242 | A1242D | D | 2.2e-07 | 1.7e-07 | 2e-07 | 1.967E-07 | 2.517E-08 | 1.700E-07 | 2.200E-07 |
| 24825 | 1242 | A1242C | D | 1e-07 | 3.4e-07 | 5.2e-07 | 3.200E-07 | 2.107E-07 | 1.000E-07 | 5.200E-07 |
| 24826 | 1242 | A1242E | D | 3.1e-07 | 4.4e-07 | 3.2e-07 | 3.567E-07 | 7.234E-08 | 3.100E-07 | 4.400E-07 |
| 24827 | 1242 | A1242Q | D | 2e-07 | 1.5e-07 | 1.9e-07 | 1.800E-07 | 2.646E-08 | 1.500E-07 | 2.000E-07 |
| 24828 | 1242 | A1242G | D | 1.3e-07 | 5.9e-07 | 1.5e-07 | 2.900E-07 | 2.600E-07 | 1.300E-07 | 5.900E-07 |
| 24829 | 1242 | A1242H | D | 2.2e-07 | 7.6e-08 | 3.3e-07 | 2.087E-07 | 1.274E-07 | 7.600E-08 | 3.300E-07 |
| 24830 | 1242 | A1242I | D | 7.4e-07 | 3e-07 | 4.7e-07 | 5.033E-07 | 2.219E-07 | 3.000E-07 | 7.400E-07 |
| 24831 | 1242 | A1242L | D | 3.2e-07 | 1.8e-07 | 2.2e-07 | 2.400E-07 | 7.211E-08 | 1.800E-07 | 3.200E-07 |
| 24832 | 1242 | A1242K | D | 2.2e-07 | 5.7e-07 | 1.9e-07 | 3.267E-07 | 2.113E-07 | 1.900E-07 | 5.700E-07 |
| 24833 | 1242 | A1242M | D | 2e-07 | 1.5e-07 | 2.4e-07 | 1.967E-07 | 4.509E-08 | 1.500E-07 | 2.400E-07 |
| 24834 | 1242 | A1242F | D | 1.5e-07 | 2.7e-07 | 1.7e-07 | 1.967E-07 | 6.429E-08 | 1.500E-07 | 2.700E-07 |
| 24835 | 1242 | A1242P | D | 2.5e-07 | 3.6e-07 | 3.3e-07 | 3.133E-07 | 5.686E-08 | 2.500E-07 | 3.600E-07 |
| 24836 | 1242 | A1242S | D | 4.6e-07 | 2.1e-07 | 3e-07 | 3.233E-07 | 1.266E-07 | 2.100E-07 | 4.600E-07 |
| 24837 | 1242 | A1242T | D | 5.9e-08 | 2.6e-07 | 2.7e-07 | 1.963E-07 | 1.190E-07 | 5.900E-08 | 2.700E-07 |
| 24838 | 1242 | A1242W | D | 5.7e-07 | 3.2e-07 | 3.3e-07 | 4.067E-07 | 1.415E-07 | 3.200E-07 | 5.700E-07 |
| 24839 | 1242 | A1242Y | D | 1.2e-07 | 3.2e-07 | 4.8e-07 | 3.067E-07 | 1.804E-07 | 1.200E-07 | 4.800E-07 |
| 24840 | 1242 | A1242V | D | 4.1e-07 | 5.8e-07 | 4e-07 | 4.633E-07 | 1.012E-07 | 4.000E-07 | 5.800E-07 |
| 24841 | 1243 | D1243A | D | 3.1e-07 | 1.6e-07 | 2.7e-07 | 2.467E-07 | 7.767E-08 | 1.600E-07 | 3.100E-07 |
| 24842 | 1243 | D1243R | D | 1.8e-07 | 2.6e-07 | 1.5e-07 | 1.967E-07 | 5.686E-08 | 1.500E-07 | 2.600E-07 |
| 24843 | 1243 | D1243N | D | 1.8e-07 | 6.1e-07 | 3.1e-07 | 3.667E-07 | 2.205E-07 | 1.800E-07 | 6.100E-07 |
| 24844 | 1243 | D1243D | D | 1.7e-07 | 2.3e-07 | 1.4e-07 | 1.800E-07 | 4.583E-08 | 1.400E-07 | 2.300E-07 |
| 24845 | 1243 | D1243C | D | 2.3e-07 | 2.9e-07 | 4.2e-07 | 3.133E-07 | 9.713E-08 | 2.300E-07 | 4.200E-07 |
| 24846 | 1243 | D1243E | D | 1.3e-07 | 1.9e-07 | 1.9e-07 | 1.700E-07 | 3.464E-08 | 1.300E-07 | 1.900E-07 |
| 24847 | 1243 | D1243Q | D | 2e-07 | 2.8e-07 | 2.3e-07 | 2.367E-07 | 4.041E-08 | 2.000E-07 | 2.800E-07 |
| 24848 | 1243 | D1243G | D | 3.1e-07 | 1.2e-07 | 3.7e-07 | 2.667E-07 | 1.305E-07 | 1.200E-07 | 3.700E-07 |
| 24849 | 1243 | D1243H | D | 1.3e-07 | 2.8e-07 | 1.8e-07 | 1.967E-07 | 7.638E-08 | 1.300E-07 | 2.800E-07 |
| 24850 | 1243 | D1243I | D | 2.1e-07 | 1.6e-07 | 1.4e-07 | 1.700E-07 | 3.606E-08 | 1.400E-07 | 2.100E-07 |
| 24851 | 1243 | D1243L | D | 2e-07 | 3.6e-07 | 3.2e-07 | 2.933E-07 | 8.327E-08 | 2.000E-07 | 3.600E-07 |
| 24852 | 1243 | D1243K | D | 1.7e-07 | 2.2e-07 | 3.1e-07 | 2.333E-07 | 7.095E-08 | 1.700E-07 | 3.100E-07 |
| 24853 | 1243 | D1243M | D | 4e-07 | 3.1e-07 | 2.2e-07 | 3.100E-07 | 9.000E-08 | 2.200E-07 | 4.000E-07 |
| 24854 | 1243 | D1243F | D | 3.5e-07 | 2.3e-07 | 6.1e-07 | 3.967E-07 | 1.943E-07 | 2.300E-07 | 6.100E-07 |
| 24855 | 1243 | D1243P | D | 1.8e-07 | 2e-07 | 3.9e-07 | 2.567E-07 | 1.159E-07 | 1.800E-07 | 3.900E-07 |
| 24856 | 1243 | D1243S | D | 6.2e-07 | 1.1e-07 | 3.1e-07 | 3.467E-07 | 2.570E-07 | 1.100E-07 | 6.200E-07 |
| 24857 | 1243 | D1243T | D | 1.6e-07 | 4.2e-07 | 3.8e-07 | 3.200E-07 | 1.400E-07 | 1.600E-07 | 4.200E-07 |
| 24858 | 1243 | D1243W | D | 1.3e-07 | 2.7e-07 | 3.2e-07 | 2.400E-07 | 9.849E-08 | 1.300E-07 | 3.200E-07 |
| 24859 | 1243 | D1243Y | D | 3.6e-07 | 2e-07 | 1.8e-07 | 2.467E-07 | 9.866E-08 | 1.800E-07 | 3.600E-07 |
| 24860 | 1243 | D1243V | D | 1.5e-07 | 1.7e-07 | 1.9e-07 | 1.700E-07 | 2.000E-08 | 1.500E-07 | 1.900E-07 |
| 24861 | 1244 | Y1244A | D | 1.8e-07 | 1.8e-07 | 6.5e-07 | 3.367E-07 | 2.714E-07 | 1.800E-07 | 6.500E-07 |
| 24862 | 1244 | Y1244R | D | 2.4e-07 | 1.4e-07 | 3.7e-07 | 2.500E-07 | 1.153E-07 | 1.400E-07 | 3.700E-07 |
| 24863 | 1244 | Y1244N | D | 4e-07 | 3.5e-07 | 3.8e-07 | 3.767E-07 | 2.517E-08 | 3.500E-07 | 4.000E-07 |
| 24864 | 1244 | Y1244D | D | 1.1e-07 | 2.7e-07 | 2.4e-07 | 2.067E-07 | 8.505E-08 | 1.100E-07 | 2.700E-07 |
| 24865 | 1244 | Y1244C | D | 2.4e-07 | 1.9e-07 | 1.4e-07 | 1.900E-07 | 5.000E-08 | 1.400E-07 | 2.400E-07 |
| 24866 | 1244 | Y1244E | D | 1.2e-07 | 2.9e-07 | 3.5e-07 | 2.533E-07 | 1.193E-07 | 1.200E-07 | 3.500E-07 |
| 24867 | 1244 | Y1244Q | D | 2.2e-07 | 2.9e-07 | 5e-07 | 3.367E-07 | 1.457E-07 | 2.200E-07 | 5.000E-07 |
| 24868 | 1244 | Y1244G | D | 3.4e-07 | 1.8e-07 | 4.4e-07 | 3.200E-07 | 1.311E-07 | 1.800E-07 | 4.400E-07 |
| 24869 | 1244 | Y1244H | D | 5e-07 | 3.1e-07 | 2.2e-07 | 3.433E-07 | 1.429E-07 | 2.200E-07 | 5.000E-07 |
| 24870 | 1244 | Y1244I | D | 1.4e-07 | 7e-07 | 2.2e-07 | 3.533E-07 | 3.029E-07 | 1.400E-07 | 7.000E-07 |
| 24871 | 1244 | Y1244L | D | 1.1e-07 | 1.7e-07 | 2e-07 | 1.600E-07 | 4.583E-08 | 1.100E-07 | 2.000E-07 |
| 24872 | 1244 | Y1244K | D | 5.2e-07 | 2.3e-07 | 1.9e-07 | 3.133E-07 | 1.801E-07 | 1.900E-07 | 5.200E-07 |
| 24873 | 1244 | Y1244M | D | 2.9e-07 | 2.3e-07 | 3e-07 | 2.733E-07 | 3.786E-08 | 2.300E-07 | 3.000E-07 |
| 24874 | 1244 | Y1244F | D | 3.5e-07 | 1.6e-07 | 2.4e-07 | 2.500E-07 | 9.539E-08 | 1.600E-07 | 3.500E-07 |
| 24875 | 1244 | Y1244P | D | 1.4e-07 | 2.2e-07 | 3.1e-07 | 2.233E-07 | 8.505E-08 | 1.400E-07 | 3.100E-07 |
| 24876 | 1244 | Y1244S | D | 1.9e-07 | 6.3e-07 | 1.3e-07 | 3.167E-07 | 2.730E-07 | 1.300E-07 | 6.300E-07 |
| 24877 | 1244 | Y1244T | D | 4.3e-07 | 3.5e-07 | 2.8e-07 | 3.533E-07 | 7.506E-08 | 2.800E-07 | 4.300E-07 |
| 24878 | 1244 | Y1244W | D | 2.4e-07 | 1.4e-07 | 3.4e-07 | 2.400E-07 | 1.000E-07 | 1.400E-07 | 3.400E-07 |
| 24879 | 1244 | Y1244Y | D | 2.3e-07 | 2.9e-07 | 1.9e-07 | 2.367E-07 | 5.033E-08 | 1.900E-07 | 2.900E-07 |
| 24880 | 1244 | Y1244V | D | 3.3e-07 | 1.2e-07 | 2.9e-07 | 2.467E-07 | 1.115E-07 | 1.200E-07 | 3.300E-07 |
| 24881 | 1245 | E1245A | D | 2.8e-07 | 1.1e-07 | 2.3e-07 | 2.067E-07 | 8.737E-08 | 1.100E-07 | 2.800E-07 |
| 24882 | 1245 | E1245R | D | 1.5e-07 | 9.9e-08 | 3.7e-07 | 2.063E-07 | 1.440E-07 | 9.900E-08 | 3.700E-07 |
| 24883 | 1245 | E1245N | D | 2.5e-07 | 1.5e-07 | 4e-07 | 2.667E-07 | 1.258E-07 | 1.500E-07 | 4.000E-07 |
| 24884 | 1245 | E1245D | D | 3e-07 | 1.8e-07 | 3.6e-07 | 2.800E-07 | 9.165E-08 | 1.800E-07 | 3.600E-07 |
| 24885 | 1245 | E1245C | D | 2.3e-07 | 1.2e-07 | 2.2e-07 | 1.900E-07 | 6.083E-08 | 1.200E-07 | 2.300E-07 |
| 24886 | 1245 | E1245E | D | 1.8e-07 | 2.1e-07 | 2e-07 | 1.967E-07 | 1.528E-08 | 1.800E-07 | 2.100E-07 |
| 24887 | 1245 | E1245Q | D | 3.5e-07 | 2.7e-07 | 1.6e-07 | 2.600E-07 | 9.539E-08 | 1.600E-07 | 3.500E-07 |
| 24888 | 1245 | E1245G | D | 3.1e-07 | 2.4e-07 | 2.9e-07 | 2.800E-07 | 3.606E-08 | 2.400E-07 | 3.100E-07 |
| 24889 | 1245 | E1245H | D | 5e-07 | 1.6e-07 | 3.3e-07 | 3.300E-07 | 1.700E-07 | 1.600E-07 | 5.000E-07 |
| 24890 | 1245 | E1245I | D | 2.2e-07 | 1.6e-07 | 2.4e-07 | 2.067E-07 | 4.163E-08 | 1.600E-07 | 2.400E-07 |
| 24891 | 1245 | E1245L | D | 5e-07 | 1.2e-07 | 2.8e-07 | 3.000E-07 | 1.908E-07 | 1.200E-07 | 5.000E-07 |
| 24892 | 1245 | E1245K | D | 4.6e-07 | 1.8e-07 | 3.5e-07 | 3.300E-07 | 1.411E-07 | 1.800E-07 | 4.600E-07 |
| 24893 | 1245 | E1245M | D | 3.5e-07 | 4.2e-07 | 3.2e-07 | 3.633E-07 | 5.132E-08 | 3.200E-07 | 4.200E-07 |
| 24894 | 1245 | E1245F | D | 4e-07 | 7.4e-08 | 3.9e-07 | 2.880E-07 | 1.854E-07 | 7.400E-08 | 4.000E-07 |
| 24895 | 1245 | E1245P | D | 1.2e-07 | 1.1e-07 | 3.5e-07 | 1.933E-07 | 1.358E-07 | 1.100E-07 | 3.500E-07 |
| 24896 | 1245 | E1245S | D | 3.5e-07 | 4.1e-07 | 2.3e-07 | 3.300E-07 | 9.165E-08 | 2.300E-07 | 4.100E-07 |
| 24897 | 1245 | E1245T | D | 2.5e-07 | 2.9e-07 | 4.4e-07 | 3.267E-07 | 1.002E-07 | 2.500E-07 | 4.400E-07 |
| 24898 | 1245 | E1245W | D | 1.7e-07 | 2.6e-07 | 3.5e-07 | 2.600E-07 | 9.000E-08 | 1.700E-07 | 3.500E-07 |
| 24899 | 1245 | E1245Y | D | 9.1e-08 | 5.6e-07 | 2.2e-07 | 2.903E-07 | 2.423E-07 | 9.100E-08 | 5.600E-07 |
| 24900 | 1245 | E1245V | D | 1.2e-07 | 3.8e-07 | 4e-07 | 3.000E-07 | 1.562E-07 | 1.200E-07 | 4.000E-07 |
| 24901 | 1246 | K1246A | D | 1.5e-07 | 1.8e-07 | 2e-07 | 1.767E-07 | 2.517E-08 | 1.500E-07 | 2.000E-07 |
| 24902 | 1246 | K1246R | D | 2.4e-07 | 3.1e-07 | 2.8e-07 | 2.767E-07 | 3.512E-08 | 2.400E-07 | 3.100E-07 |
| 24903 | 1246 | K1246N | D | 1.8e-07 | 2.4e-07 | 1.2e-07 | 1.800E-07 | 6.000E-08 | 1.200E-07 | 2.400E-07 |
| 24904 | 1246 | K1246D | D | 1.9e-07 | 3.8e-07 | 4.5e-07 | 3.400E-07 | 1.345E-07 | 1.900E-07 | 4.500E-07 |

| | | | | | | | | | | |
|---|---|---|---|---|---|---|---|---|---|---|
| 24905 | 1246 | K1246C | D | 2.2e-07 | 2e-07 | 4.7e-07 | 2.967E-07 | 1.504E-07 | 2.000E-07 | 4.700E-07 |
| 24906 | 1246 | K1246E | D | 3.1e-07 | 1.5e-07 | 1.2e-07 | 1.933E-07 | 1.021E-07 | 1.200E-07 | 3.100E-07 |
| 24907 | 1246 | K1246Q | D | 4.6e-07 | 3.3e-07 | 4.7e-07 | 4.200E-07 | 7.810E-08 | 3.300E-07 | 4.700E-07 |
| 24908 | 1246 | K1246G | D | 1.2e-07 | 2e-07 | 2.5e-07 | 1.900E-07 | 6.557E-08 | 1.200E-07 | 2.500E-07 |
| 24909 | 1246 | K1246H | D | 2e-07 | 1.5e-07 | 1.8e-07 | 1.767E-07 | 2.517E-08 | 1.500E-07 | 2.000E-07 |
| 24910 | 1246 | K1246I | D | 2e-07 | 3.4e-07 | 3.1e-07 | 2.833E-07 | 7.371E-08 | 2.000E-07 | 3.400E-07 |
| 24911 | 1246 | K1246L | D | 2.7e-07 | 3.7e-07 | 2.2e-07 | 2.867E-07 | 7.638E-08 | 2.200E-07 | 3.700E-07 |
| 24912 | 1246 | K1246K | D | 2.6e-07 | 5.8e-07 | 2.1e-07 | 3.500E-07 | 2.007E-07 | 2.100E-07 | 5.800E-07 |
| 24913 | 1246 | K1246M | D | 2.2e-07 | 2.5e-07 | 2.7e-07 | 2.467E-07 | 2.517E-08 | 2.200E-07 | 2.700E-07 |
| 24914 | 1246 | K1246F | D | 3.1e-07 | 3.6e-07 | 4.5e-07 | 3.733E-07 | 7.095E-08 | 3.100E-07 | 4.500E-07 |
| 24915 | 1246 | K1246P | D | 3.4e-07 | 1.7e-07 | 1.8e-07 | 2.300E-07 | 9.539E-08 | 1.700E-07 | 3.400E-07 |
| 24916 | 1246 | K1246S | D | 1.7e-07 | 2.8e-07 | 1.9e-07 | 2.133E-07 | 5.859E-08 | 1.700E-07 | 2.800E-07 |
| 24917 | 1246 | K1246T | D | 3.1e-07 | 1.7e-07 | 3.1e-07 | 2.633E-07 | 8.083E-08 | 1.700E-07 | 3.100E-07 |
| 24918 | 1246 | K1246W | D | 3.7e-07 | 2.7e-07 | 2e-07 | 2.800E-07 | 8.544E-08 | 2.000E-07 | 3.700E-07 |
| 24919 | 1246 | K1246Y | D | 1.8e-07 | 2e-07 | 3.5e-07 | 2.433E-07 | 9.292E-08 | 1.800E-07 | 3.500E-07 |
| 24920 | 1246 | K1246V | D | 2.4e-07 | 5.2e-07 | 1.4e-07 | 3.000E-07 | 1.970E-07 | 1.400E-07 | 5.200E-07 |
| 24921 | 1247 | H1247A | D | 2.8e-07 | 1.4e-07 | 1.1e-07 | 1.767E-07 | 9.074E-08 | 1.100E-07 | 2.800E-07 |
| 24922 | 1247 | H1247R | D | 2.5e-07 | 3.2e-07 | 1.5e-07 | 2.400E-07 | 8.544E-08 | 1.500E-07 | 3.200E-07 |
| 24923 | 1247 | H1247N | D | 3.9e-07 | 2e-07 | 2.5e-07 | 2.800E-07 | 9.849E-08 | 2.000E-07 | 3.900E-07 |
| 24924 | 1247 | H1247D | D | 1.9e-07 | 3.5e-07 | 2.6e-07 | 2.667E-07 | 8.021E-08 | 1.900E-07 | 3.500E-07 |
| 24925 | 1247 | H1247C | D | 1.9e-07 | 2.2e-07 | 2.2e-07 | 2.100E-07 | 1.732E-08 | 1.900E-07 | 2.200E-07 |
| 24926 | 1247 | H1247E | D | 4.1e-07 | 3.1e-07 | 1.7e-07 | 2.967E-07 | 1.206E-07 | 1.700E-07 | 4.100E-07 |
| 24927 | 1247 | H1247Q | D | 1.9e-07 | 1.9e-07 | 1.4e-07 | 1.733E-07 | 2.887E-08 | 1.400E-07 | 1.900E-07 |
| 24928 | 1247 | H1247G | D | 4.6e-07 | 6.5e-07 | 2e-07 | 4.367E-07 | 2.259E-07 | 2.000E-07 | 6.500E-07 |
| 24929 | 1247 | H1247H | D | 1.3e-07 | 1e-07 | 1.9e-07 | 1.400E-07 | 4.583E-08 | 1.000E-07 | 1.900E-07 |
| 24930 | 1247 | H1247I | D | 2.9e-07 | 3e-07 | 1.8e-07 | 2.567E-07 | 6.658E-08 | 1.800E-07 | 3.000E-07 |
| 24931 | 1247 | H1247L | D | 3e-07 | 3.3e-07 | 1.3e-07 | 2.533E-07 | 1.079E-07 | 1.300E-07 | 3.300E-07 |
| 24932 | 1247 | H1247K | D | 1.8e-07 | 1.2e-07 | 1.8e-07 | 1.600E-07 | 3.464E-08 | 1.200E-07 | 1.800E-07 |
| 24933 | 1247 | H1247M | D | 6.4e-07 | 2.1e-07 | 3.8e-07 | 4.100E-07 | 2.166E-07 | 2.100E-07 | 6.400E-07 |
| 24934 | 1247 | H1247F | D | 1.3e-07 | 2.4e-07 | 5.3e-07 | 3.000E-07 | 2.066E-07 | 1.300E-07 | 5.300E-07 |
| 24935 | 1247 | H1247P | D | 1.2e-07 | 2.1e-07 | 2.3e-07 | 1.867E-07 | 5.859E-08 | 1.200E-07 | 2.300E-07 |
| 24936 | 1247 | H1247S | D | 4.1e-07 | 6.4e-07 | 2.4e-07 | 4.300E-07 | 2.007E-07 | 2.400E-07 | 6.400E-07 |
| 24937 | 1247 | H1247T | D | 2.8e-07 | 1.5e-07 | 1.2e-07 | 1.833E-07 | 8.505E-08 | 1.200E-07 | 2.800E-07 |
| 24938 | 1247 | H1247W | D | 2.1e-07 | 4.7e-07 | 2.1e-07 | 2.967E-07 | 1.501E-07 | 2.100E-07 | 4.700E-07 |
| 24939 | 1247 | H1247Y | D | 1.4e-07 | 8e-08 | 1.8e-07 | 1.333E-07 | 5.033E-08 | 8.000E-08 | 1.800E-07 |
| 24940 | 1247 | H1247V | D | 2.8e-07 | 2e-07 | 1.2e-07 | 2.000E-07 | 8.000E-08 | 1.200E-07 | 2.800E-07 |
| 24941 | 1248 | K1248A | D | 1.7e-07 | 4.5e-07 | 2.4e-07 | 2.867E-07 | 1.457E-07 | 1.700E-07 | 4.500E-07 |
| 24942 | 1248 | K1248R | D | 2.1e-07 | 4.5e-07 | 3e-07 | 3.200E-07 | 1.212E-07 | 2.100E-07 | 4.500E-07 |
| 24943 | 1248 | K1248N | D | 2.6e-07 | 1.3e-07 | 2.7e-07 | 2.200E-07 | 7.810E-08 | 1.300E-07 | 2.700E-07 |
| 24944 | 1248 | K1248D | D | 1.9e-07 | 1.4e-07 | 2.2e-07 | 1.833E-07 | 4.041E-08 | 1.400E-07 | 2.200E-07 |
| 24945 | 1248 | K1248C | D | 2.6e-07 | 4e-07 | 5.1e-07 | 3.900E-07 | 1.253E-07 | 2.600E-07 | 5.100E-07 |
| 24946 | 1248 | K1248E | D | 4.2e-07 | 4e-07 | 1.7e-07 | 3.300E-07 | 1.389E-07 | 1.700E-07 | 4.200E-07 |
| 24947 | 1248 | K1248Q | D | 2.5e-07 | 2.1e-07 | 3.4e-07 | 2.667E-07 | 6.658E-08 | 2.100E-07 | 3.400E-07 |
| 24948 | 1248 | K1248G | D | 3.7e-07 | 2.9e-07 | 2.8e-07 | 3.133E-07 | 4.933E-08 | 2.800E-07 | 3.700E-07 |
| 24949 | 1248 | K1248H | D | 2.7e-07 | 1.7e-07 | 8.2e-08 | 1.740E-07 | 9.406E-08 | 8.200E-08 | 2.700E-07 |
| 24950 | 1248 | K1248I | D | 3.3e-07 | 2.9e-07 | 2.8e-07 | 3.000E-07 | 2.646E-08 | 2.800E-07 | 3.300E-07 |
| 24951 | 1248 | K1248L | D | 1.5e-07 | 3.8e-07 | 2.9e-07 | 2.733E-07 | 1.159E-07 | 1.500E-07 | 3.800E-07 |
| 24952 | 1248 | K1248K | D | 4.5e-07 | 2.2e-07 | 1.9e-07 | 2.867E-07 | 1.422E-07 | 1.900E-07 | 4.500E-07 |
| 24953 | 1248 | K1248M | D | 3.4e-07 | 9e-08 | 4.5e-07 | 2.933E-07 | 1.845E-07 | 9.000E-08 | 4.500E-07 |
| 24954 | 1248 | K1248F | D | 2e-07 | 3.1e-07 | 1.6e-07 | 2.233E-07 | 7.767E-08 | 1.600E-07 | 3.100E-07 |
| 24955 | 1248 | K1248P | D | 1.2e-07 | 3.1e-07 | 2.9e-07 | 2.400E-07 | 1.044E-07 | 1.200E-07 | 3.100E-07 |
| 24956 | 1248 | K1248S | D | 1.4e-07 | 2.1e-07 | 2.6e-07 | 2.033E-07 | 6.028E-08 | 1.400E-07 | 2.600E-07 |
| 24957 | 1248 | K1248T | D | 2.8e-07 | 2.2e-07 | 2.6e-07 | 2.533E-07 | 3.055E-08 | 2.200E-07 | 2.800E-07 |
| 24958 | 1248 | K1248W | D | 1.9e-07 | 4.4e-07 | 5.5e-07 | 3.933E-07 | 1.845E-07 | 1.900E-07 | 5.500E-07 |
| 24959 | 1248 | K1248Y | D | 4.4e-07 | 5.1e-07 | 3e-07 | 4.167E-07 | 1.069E-07 | 3.000E-07 | 5.100E-07 |
| 24960 | 1248 | K1248V | D | 4.2e-07 | 3.2e-07 | 2e-07 | 3.133E-07 | 1.102E-07 | 2.000E-07 | 4.200E-07 |
| 24961 | 1249 | V1249A | D | 1.9e-07 | 3.5e-07 | 2e-07 | 2.467E-07 | 8.963E-08 | 1.900E-07 | 3.500E-07 |
| 24962 | 1249 | V1249R | D | 1.6e-07 | 2.8e-07 | 3.9e-07 | 2.767E-07 | 1.150E-07 | 1.600E-07 | 3.900E-07 |
| 24963 | 1249 | V1249N | D | 1.7e-07 | 4e-07 | 1.2e-07 | 2.300E-07 | 1.493E-07 | 1.200E-07 | 4.000E-07 |
| 24964 | 1249 | V1249D | D | 1.6e-07 | 1.4e-07 | 1.8e-07 | 1.600E-07 | 2.000E-08 | 1.400E-07 | 1.800E-07 |
| 24965 | 1249 | V1249C | D | 2.7e-07 | 1.6e-07 | 2.4e-07 | 2.233E-07 | 5.686E-08 | 1.600E-07 | 2.700E-07 |
| 24966 | 1249 | V1249E | D | 1.8e-07 | 3.2e-07 | 2.7e-07 | 2.567E-07 | 7.095E-08 | 1.800E-07 | 3.200E-07 |
| 24967 | 1249 | V1249Q | D | 1.8e-07 | 1.6e-07 | 3.4e-07 | 2.267E-07 | 9.866E-08 | 1.600E-07 | 3.400E-07 |
| 24968 | 1249 | V1249G | D | 3.2e-07 | 1.3e-07 | 1e-07 | 1.833E-07 | 1.193E-07 | 1.000E-07 | 3.200E-07 |
| 24969 | 1249 | V1249H | D | 1.8e-07 | 2.1e-07 | 2.7e-07 | 2.200E-07 | 4.583E-08 | 1.800E-07 | 2.700E-07 |
| 24970 | 1249 | V1249I | D | 1.7e-07 | 2.2e-07 | 5.7e-08 | 1.490E-07 | 8.350E-08 | 5.700E-08 | 2.200E-07 |
| 24971 | 1249 | V1249L | D | 2.5e-07 | 5e-07 | 1.1e-07 | 2.867E-07 | 1.976E-07 | 1.100E-07 | 5.000E-07 |
| 24972 | 1249 | V1249K | D | 3.1e-07 | 3.3e-07 | 1.3e-07 | 2.567E-07 | 1.102E-07 | 1.300E-07 | 3.300E-07 |
| 24973 | 1249 | V1249M | D | 4.1e-07 | 1.7e-07 | 4.1e-07 | 3.300E-07 | 1.386E-07 | 1.700E-07 | 4.100E-07 |
| 24974 | 1249 | V1249F | D | 3e-07 | 3.2e-07 | 1.7e-07 | 2.633E-07 | 8.145E-08 | 1.700E-07 | 3.200E-07 |
| 24975 | 1249 | V1249P | D | 4.9e-07 | 7.6e-07 | 3.5e-07 | 5.333E-07 | 2.084E-07 | 3.500E-07 | 7.600E-07 |
| 24976 | 1249 | V1249S | D | 1.9e-07 | 1.3e-07 | 2.9e-07 | 2.033E-07 | 8.083E-08 | 1.300E-07 | 2.900E-07 |
| 24977 | 1249 | V1249T | D | 5.4e-07 | 1e-07 | 1.8e-07 | 2.733E-07 | 2.344E-07 | 1.000E-07 | 5.400E-07 |
| 24978 | 1249 | V1249W | D | 1.6e-07 | 3e-07 | 1.5e-07 | 2.033E-07 | 8.386E-08 | 1.500E-07 | 3.000E-07 |
| 24979 | 1249 | V1249Y | D | 1.2e-07 | 2.5e-07 | 1.2e-07 | 1.633E-07 | 7.506E-08 | 1.200E-07 | 2.500E-07 |
| 24980 | 1249 | V1249V | D | 1.7e-07 | 3.9e-07 | 1.4e-07 | 2.333E-07 | 1.365E-07 | 1.400E-07 | 3.900E-07 |
| 24981 | 1250 | Y1250A | D | 2e-07 | 1.9e-07 | 2.8e-07 | 2.233E-07 | 4.933E-08 | 1.900E-07 | 2.800E-07 |
| 24982 | 1250 | Y1250R | D | 1.2e-07 | 2.7e-07 | 1.4e-07 | 1.767E-07 | 8.145E-08 | 1.200E-07 | 2.700E-07 |
| 24983 | 1250 | Y1250N | D | 3.9e-07 | 1.4e-07 | 3.2e-07 | 2.833E-07 | 1.290E-07 | 1.400E-07 | 3.900E-07 |
| 24984 | 1250 | Y1250D | D | 2.2e-07 | 2e-07 | 4.7e-07 | 2.967E-07 | 1.504E-07 | 2.000E-07 | 4.700E-07 |
| 24985 | 1250 | Y1250C | D | 4.3e-07 | 1.9e-07 | 3.1e-07 | 3.100E-07 | 1.200E-07 | 1.900E-07 | 4.300E-07 |
| 24986 | 1250 | Y1250E | D | 1.8e-07 | 2.4e-07 | 3.5e-07 | 2.567E-07 | 8.622E-08 | 1.800E-07 | 3.500E-07 |
| 24987 | 1250 | Y1250Q | D | 2e-07 | 1.2e-07 | 2.1e-07 | 1.767E-07 | 4.933E-08 | 1.200E-07 | 2.100E-07 |
| 24988 | 1250 | Y1250G | D | 3.8e-07 | 2.3e-07 | 1.9e-07 | 2.667E-07 | 1.002E-07 | 1.900E-07 | 3.800E-07 |
| 24989 | 1250 | Y1250H | D | 5e-07 | 1.4e-07 | 2e-07 | 2.800E-07 | 1.929E-07 | 1.400E-07 | 5.000E-07 |

| | | | | | | | | | | |
|---|---|---|---|---|---|---|---|---|---|---|
| 24990 | 1250 | Y1250I | D | 1.7e-07 | 1.5e-07 | 4.5e-07 | 2.567E-07 | 1.677E-07 | 1.500E-07 | 4.500E-07 |
| 24991 | 1250 | Y1250L | D | 2.2e-07 | 3.6e-07 | 1.3e-07 | 2.367E-07 | 1.159E-07 | 1.300E-07 | 3.600E-07 |
| 24992 | 1250 | Y1250K | D | 2.1e-07 | 2.3e-07 | 2.6e-07 | 2.333E-07 | 2.517E-08 | 2.100E-07 | 2.600E-07 |
| 24993 | 1250 | Y1250M | D | 2.3e-07 | 3.1e-07 | 2.8e-07 | 2.733E-07 | 4.041E-08 | 2.300E-07 | 3.100E-07 |
| 24994 | 1250 | Y1250F | D | 2.7e-07 | 2.2e-07 | 3.8e-07 | 2.900E-07 | 8.185E-08 | 2.200E-07 | 3.800E-07 |
| 24995 | 1250 | Y1250P | D | 1.4e-07 | 1.4e-07 | 3.3e-07 | 2.033E-07 | 1.097E-07 | 1.400E-07 | 3.300E-07 |
| 24996 | 1250 | Y1250S | D | 5e-07 | 1.7e-07 | 2.4e-07 | 3.033E-07 | 1.739E-07 | 1.700E-07 | 5.000E-07 |
| 24997 | 1250 | Y1250T | D | 2.5e-07 | 3.2e-07 | 1.5e-07 | 2.400E-07 | 8.544E-08 | 1.500E-07 | 3.200E-07 |
| 24998 | 1250 | Y1250W | D | 3.1e-07 | 2.6e-07 | 1.7e-07 | 2.467E-07 | 7.095E-08 | 1.700E-07 | 3.100E-07 |
| 24999 | 1250 | Y1250Y | D | 3.3e-07 | 3.3e-07 | 2.2e-07 | 2.933E-07 | 6.351E-08 | 2.200E-07 | 3.300E-07 |
| 25000 | 1250 | Y1250V | D | 2.3e-07 | 1.2e-07 | 4.5e-07 | 2.667E-07 | 1.680E-07 | 1.200E-07 | 4.500E-07 |
| 25001 | 1251 | A1251A | D | 1.8e-07 | 3.6e-07 | 2.4e-07 | 2.600E-07 | 9.165E-08 | 1.800E-07 | 3.600E-07 |
| 25002 | 1251 | A1251R | D | 3.8e-07 | 3.8e-07 | 8.3e-07 | 5.300E-07 | 2.598E-07 | 3.800E-07 | 8.300E-07 |
| 25003 | 1251 | A1251N | D | 3.3e-07 | 4.2e-07 | 1.6e-07 | 3.033E-07 | 1.320E-07 | 1.600E-07 | 4.200E-07 |
| 25004 | 1251 | A1251D | D | 1.5e-07 | 2.1e-07 | 3.7e-07 | 2.433E-07 | 1.137E-07 | 1.500E-07 | 3.700E-07 |
| 25005 | 1251 | A1251C | D | 1.9e-07 | 2.2e-07 | 2.1e-07 | 2.067E-07 | 1.528E-08 | 1.900E-07 | 2.200E-07 |
| 25006 | 1251 | A1251E | D | 2.7e-07 | 2.1e-07 | 2.7e-07 | 2.500E-07 | 3.464E-08 | 2.100E-07 | 2.700E-07 |
| 25007 | 1251 | A1251Q | D | 1.5e-07 | 2.3e-07 | 2.7e-07 | 2.167E-07 | 6.110E-08 | 1.500E-07 | 2.700E-07 |
| 25008 | 1251 | A1251G | D | 3.2e-07 | 8.2e-07 | 2.1e-07 | 4.500E-07 | 3.251E-07 | 2.100E-07 | 8.200E-07 |
| 25009 | 1251 | A1251H | D | 1.7e-07 | 2.8e-07 | 4.7e-07 | 3.067E-07 | 1.518E-07 | 1.700E-07 | 4.700E-07 |
| 25010 | 1251 | A1251I | D | 3.2e-07 | 6.1e-07 | 2.6e-07 | 3.967E-07 | 1.872E-07 | 2.600E-07 | 6.100E-07 |
| 25011 | 1251 | A1251L | D | 3.8e-07 | 2.2e-07 | 3.5e-07 | 3.167E-07 | 8.505E-08 | 2.200E-07 | 3.800E-07 |
| 25012 | 1251 | A1251K | D | 1.9e-07 | 8e-08 | 2.7e-07 | 1.800E-07 | 9.539E-08 | 8.000E-08 | 2.700E-07 |
| 25013 | 1251 | A1251M | D | 1e-07 | 2.5e-07 | 2.2e-07 | 1.900E-07 | 7.937E-08 | 1.000E-07 | 2.500E-07 |
| 25014 | 1251 | A1251F | D | 2.6e-07 | 2.6e-07 | 4e-07 | 3.067E-07 | 8.083E-08 | 2.600E-07 | 4.000E-07 |
| 25015 | 1251 | A1251P | D | 6.4e-08 | 4e-07 | 9.6e-08 | 1.867E-07 | 1.854E-07 | 6.400E-08 | 4.000E-07 |
| 25016 | 1251 | A1251S | D | 2.7e-07 | 3.1e-07 | 1.5e-07 | 2.433E-07 | 8.327E-08 | 1.500E-07 | 3.100E-07 |
| 25017 | 1251 | A1251T | D | 1.5e-07 | 3.1e-07 | 2.4e-07 | 2.333E-07 | 8.021E-08 | 1.500E-07 | 3.100E-07 |
| 25018 | 1251 | A1251W | D | 1.6e-07 | 2.5e-07 | 2.2e-07 | 2.100E-07 | 4.583E-08 | 1.600E-07 | 2.500E-07 |
| 25019 | 1251 | A1251Y | D | 4.5e-07 | 3.8e-07 | 2.2e-07 | 3.500E-07 | 1.179E-07 | 2.200E-07 | 4.500E-07 |
| 25020 | 1251 | A1251V | D | 1.1e-07 | 1.9e-07 | 3.2e-07 | 2.067E-07 | 1.060E-07 | 1.100E-07 | 3.200E-07 |
| 25021 | 1252 | C1252A | D | 2.6e-07 | 1.3e-07 | 2.1e-07 | 2.000E-07 | 6.557E-08 | 1.300E-07 | 2.600E-07 |
| 25022 | 1252 | C1252R | D | 2.4e-07 | 2.8e-07 | 4.1e-07 | 3.100E-07 | 8.888E-08 | 2.400E-07 | 4.100E-07 |
| 25023 | 1252 | C1252N | D | 2.3e-07 | 1.7e-07 | 2.3e-07 | 2.100E-07 | 3.464E-08 | 1.700E-07 | 2.300E-07 |
| 25024 | 1252 | C1252D | D | 2e-07 | 2.7e-07 | 3e-07 | 2.567E-07 | 5.132E-08 | 2.000E-07 | 3.000E-07 |
| 25025 | 1252 | C1252C | D | 3.1e-07 | 2.5e-07 | 1.2e-07 | 2.267E-07 | 9.713E-08 | 1.200E-07 | 3.100E-07 |
| 25026 | 1252 | C1252E | D | 5.5e-07 | 3.2e-07 | 2.5e-07 | 3.733E-07 | 1.570E-07 | 2.500E-07 | 5.500E-07 |
| 25027 | 1252 | C1252Q | D | 2.2e-07 | 3.4e-07 | 1.7e-07 | 2.433E-07 | 8.737E-08 | 1.700E-07 | 3.400E-07 |
| 25028 | 1252 | C1252G | D | 2e-07 | 2.5e-07 | 1.4e-07 | 1.967E-07 | 5.508E-08 | 1.400E-07 | 2.500E-07 |
| 25029 | 1252 | C1252H | D | 2.3e-07 | 2.4e-07 | 3.4e-07 | 2.700E-07 | 6.083E-08 | 2.300E-07 | 3.400E-07 |
| 25030 | 1252 | C1252I | D | 3.1e-07 | 2.5e-07 | 2e-07 | 2.533E-07 | 5.508E-08 | 2.000E-07 | 3.100E-07 |
| 25031 | 1252 | C1252L | D | 2.8e-07 | 2.2e-07 | 1.8e-07 | 2.267E-07 | 5.033E-08 | 1.800E-07 | 2.800E-07 |
| 25032 | 1252 | C1252K | D | 3e-07 | 5e-07 | 4.3e-07 | 4.100E-07 | 1.015E-07 | 3.000E-07 | 5.000E-07 |
| 25033 | 1252 | C1252M | D | 3.2e-07 | 1.5e-07 | 1.4e-07 | 2.033E-07 | 1.012E-07 | 1.400E-07 | 3.200E-07 |
| 25034 | 1252 | C1252F | D | 7.5e-08 | 1.7e-07 | 4.7e-07 | 2.383E-07 | 2.062E-07 | 7.500E-08 | 4.700E-07 |
| 25035 | 1252 | C1252P | D | 3.1e-07 | 2.3e-07 | 2.3e-07 | 2.567E-07 | 4.619E-08 | 2.300E-07 | 3.100E-07 |
| 25036 | 1252 | C1252S | D | 2.5e-07 | 3.1e-07 | 2.7e-07 | 2.767E-07 | 3.055E-08 | 2.500E-07 | 3.100E-07 |
| 25037 | 1252 | C1252T | D | 4.1e-07 | 4.9e-07 | 2e-07 | 3.667E-07 | 1.498E-07 | 2.000E-07 | 4.900E-07 |
| 25038 | 1252 | C1252W | D | 1.2e-07 | 4.4e-07 | 1.3e-07 | 2.300E-07 | 1.819E-07 | 1.200E-07 | 4.400E-07 |
| 25039 | 1252 | C1252Y | D | 2.4e-07 | 2.6e-07 | 2.1e-07 | 2.367E-07 | 2.517E-08 | 2.100E-07 | 2.600E-07 |
| 25040 | 1252 | C1252V | D | 3.9e-07 | 4.6e-07 | 2.9e-07 | 3.800E-07 | 8.544E-08 | 2.900E-07 | 4.600E-07 |
| 25041 | 1253 | E1253A | D | 1.7e-07 | 1.9e-07 | 1.9e-07 | 1.833E-07 | 1.155E-08 | 1.700E-07 | 1.900E-07 |
| 25042 | 1253 | E1253R | D | 6.5e-07 | 2.9e-07 | 4e-07 | 4.467E-07 | 1.845E-07 | 2.900E-07 | 6.500E-07 |
| 25043 | 1253 | E1253N | D | 2.2e-07 | 1.1e-07 | 1.7e-07 | 1.667E-07 | 5.508E-08 | 1.100E-07 | 2.200E-07 |
| 25044 | 1253 | E1253D | D | 2.9e-07 | 1e-07 | 2.1e-07 | 2.000E-07 | 9.539E-08 | 1.000E-07 | 2.900E-07 |
| 25045 | 1253 | E1253C | D | 3.9e-07 | 2.1e-07 | 2.3e-07 | 2.767E-07 | 9.866E-08 | 2.100E-07 | 3.900E-07 |
| 25046 | 1253 | E1253E | D | 2.8e-07 | 3.7e-07 | 1.3e-07 | 2.600E-07 | 1.212E-07 | 1.300E-07 | 3.700E-07 |
| 25047 | 1253 | E1253Q | D | 4.4e-07 | 2.9e-07 | 2.1e-07 | 3.133E-07 | 1.168E-07 | 2.100E-07 | 4.400E-07 |
| 25048 | 1253 | E1253G | D | 5.4e-07 | 1.3e-07 | 3.1e-07 | 3.267E-07 | 2.055E-07 | 1.300E-07 | 5.400E-07 |
| 25049 | 1253 | E1253H | D | 2.9e-07 | 2.1e-07 | 3.5e-07 | 2.833E-07 | 7.024E-08 | 2.100E-07 | 3.500E-07 |
| 25050 | 1253 | E1253I | D | 3.5e-07 | 3.1e-07 | 1.5e-07 | 2.700E-07 | 1.058E-07 | 1.500E-07 | 3.500E-07 |
| 25051 | 1253 | E1253L | D | 2.3e-07 | 2.2e-07 | 3e-07 | 2.500E-07 | 4.359E-08 | 2.200E-07 | 3.000E-07 |
| 25052 | 1253 | E1253K | D | 3.7e-07 | 4e-07 | 2.8e-07 | 3.500E-07 | 6.245E-08 | 2.800E-07 | 4.000E-07 |
| 25053 | 1253 | E1253M | D | 1e-07 | 1.9e-07 | 1.9e-07 | 1.600E-07 | 5.196E-08 | 1.000E-07 | 1.900E-07 |
| 25054 | 1253 | E1253F | D | 1.2e-07 | 3.2e-07 | 2.4e-07 | 2.267E-07 | 1.007E-07 | 1.200E-07 | 3.200E-07 |
| 25055 | 1253 | E1253P | D | 2.2e-07 | 3.2e-07 | 3.1e-07 | 2.833E-07 | 5.508E-08 | 2.200E-07 | 3.200E-07 |
| 25056 | 1253 | E1253S | D | 3.4e-07 | 1e-07 | 3.2e-07 | 2.533E-07 | 1.332E-07 | 1.000E-07 | 3.400E-07 |
| 25057 | 1253 | E1253T | D | 2.1e-07 | 2.2e-07 | 1.3e-07 | 1.867E-07 | 4.933E-08 | 1.300E-07 | 2.200E-07 |
| 25058 | 1253 | E1253W | D | 4.5e-07 | 1.2e-07 | 2.1e-07 | 2.600E-07 | 1.706E-07 | 1.200E-07 | 4.500E-07 |
| 25059 | 1253 | E1253Y | D | 5e-07 | 1.8e-07 | 1.9e-07 | 2.900E-07 | 1.819E-07 | 1.800E-07 | 5.000E-07 |
| 25060 | 1253 | E1253V | D | 2.4e-07 | 2.6e-07 | 3.4e-07 | 2.800E-07 | 5.292E-08 | 2.400E-07 | 3.400E-07 |
| 25061 | 1254 | V1254A | D | 1.2e-07 | 4.7e-07 | 1.5e-07 | 2.467E-07 | 1.940E-07 | 1.200E-07 | 4.700E-07 |
| 25062 | 1254 | V1254R | D | 4e-07 | 1.2e-07 | 3.8e-07 | 3.000E-07 | 1.562E-07 | 1.200E-07 | 4.000E-07 |
| 25063 | 1254 | V1254N | D | 3.2e-07 | 1.8e-07 | 1.8e-07 | 2.267E-07 | 8.083E-08 | 1.800E-07 | 3.200E-07 |
| 25064 | 1254 | V1254D | D | 2.3e-07 | 3.5e-07 | 1.3e-07 | 2.367E-07 | 1.102E-07 | 1.300E-07 | 3.500E-07 |
| 25065 | 1254 | V1254C | D | 3.8e-07 | 1.8e-07 | 2.8e-07 | 2.800E-07 | 1.000E-07 | 1.800E-07 | 3.800E-07 |
| 25066 | 1254 | V1254E | D | 1.7e-07 | 2.6e-07 | 2.5e-07 | 2.267E-07 | 4.933E-08 | 1.700E-07 | 2.600E-07 |
| 25067 | 1254 | V1254Q | D | 3.4e-07 | 3.1e-07 | 9.9e-08 | 2.497E-07 | 1.313E-07 | 9.900E-08 | 3.400E-07 |
| 25068 | 1254 | V1254G | D | 3.8e-07 | 4.1e-07 | 1.8e-07 | 3.233E-07 | 1.250E-07 | 1.800E-07 | 4.100E-07 |
| 25069 | 1254 | V1254H | D | 4.8e-07 | 1.3e-07 | 1.9e-07 | 2.667E-07 | 1.872E-07 | 1.300E-07 | 4.800E-07 |
| 25070 | 1254 | V1254I | D | 2.6e-07 | 2.1e-07 | 2.4e-07 | 2.367E-07 | 2.517E-08 | 2.100E-07 | 2.600E-07 |
| 25071 | 1254 | V1254L | D | 2.6e-07 | 2.6e-07 | 1.7e-07 | 2.300E-07 | 5.196E-08 | 1.700E-07 | 2.600E-07 |
| 25072 | 1254 | V1254K | D | 1.3e-07 | 1.1e-07 | 2.3e-07 | 1.567E-07 | 6.429E-08 | 1.100E-07 | 2.300E-07 |
| 25073 | 1254 | V1254M | D | 1.7e-07 | 2.4e-07 | 2.6e-07 | 2.233E-07 | 4.726E-08 | 1.700E-07 | 2.600E-07 |
| 25074 | 1254 | V1254F | D | 1e-07 | 1.8e-07 | 6.9e-07 | 3.233E-07 | 3.201E-07 | 1.000E-07 | 6.900E-07 |

| | | | | | | | | | | |
|---|---|---|---|---|---|---|---|---|---|---|
| 25075 | 1254 | V1254P | D | 1.5e-07 | 2.4e-07 | 3.7e-07 | 2.533E-07 | 1.106E-07 | 1.500E-07 | 3.700E-07 |
| 25076 | 1254 | V1254S | D | 2e-07 | 1.4e-07 | 2.3e-07 | 1.900E-07 | 4.583E-08 | 1.400E-07 | 2.300E-07 |
| 25077 | 1254 | V1254T | D | 2.4e-07 | 1.2e-07 | 1.5e-07 | 1.700E-07 | 6.245E-08 | 1.200E-07 | 2.400E-07 |
| 25078 | 1254 | V1254W | D | 1.8e-07 | 5.4e-07 | 4.2e-07 | 3.800E-07 | 1.833E-07 | 1.800E-07 | 5.400E-07 |
| 25079 | 1254 | V1254Y | D | 1.6e-07 | 1.3e-07 | 2.6e-07 | 1.833E-07 | 6.807E-08 | 1.300E-07 | 2.600E-07 |
| 25080 | 1254 | V1254V | D | 7.8e-08 | 2.1e-07 | 2.2e-07 | 1.693E-07 | 7.925E-08 | 7.800E-08 | 2.200E-07 |
| 25081 | 1255 | T1255A | D | 3.1e-07 | 2.1e-07 | 4e-07 | 3.067E-07 | 9.504E-08 | 2.100E-07 | 4.000E-07 |
| 25082 | 1255 | T1255R | D | 2.5e-07 | 1.6e-07 | 3.5e-07 | 2.533E-07 | 9.504E-08 | 1.600E-07 | 3.500E-07 |
| 25083 | 1255 | T1255N | D | 3e-07 | 3.3e-07 | 3.7e-07 | 3.333E-07 | 3.512E-08 | 3.000E-07 | 3.700E-07 |
| 25084 | 1255 | T1255D | D | 2e-07 | 3.3e-07 | 3.9e-07 | 3.067E-07 | 9.713E-08 | 2.000E-07 | 3.900E-07 |
| 25085 | 1255 | T1255C | D | 3.9e-07 | 4.5e-07 | 2.6e-07 | 3.667E-07 | 9.713E-08 | 2.600E-07 | 4.500E-07 |
| 25086 | 1255 | T1255E | D | 1.5e-07 | 3.4e-07 | 4.5e-07 | 3.133E-07 | 1.518E-07 | 1.500E-07 | 4.500E-07 |
| 25087 | 1255 | T1255Q | D | 3e-07 | 3.3e-07 | 1.8e-07 | 2.700E-07 | 7.937E-08 | 1.800E-07 | 3.300E-07 |
| 25088 | 1255 | T1255G | D | 2.8e-07 | 2e-07 | 2.5e-07 | 2.433E-07 | 4.041E-08 | 2.000E-07 | 2.800E-07 |
| 25089 | 1255 | T1255H | D | 3.9e-07 | 4.2e-07 | 1.1e-07 | 3.067E-07 | 1.710E-07 | 1.100E-07 | 4.200E-07 |
| 25090 | 1255 | T1255I | D | 2.7e-07 | 1.3e-07 | 2.2e-07 | 2.067E-07 | 7.095E-08 | 1.300E-07 | 2.700E-07 |
| 25091 | 1255 | T1255L | D | 4.1e-07 | 5e-07 | 9.2e-07 | 6.100E-07 | 2.722E-07 | 4.100E-07 | 9.200E-07 |
| 25092 | 1255 | T1255K | D | 1.7e-07 | 1.1e-07 | 1.7e-07 | 1.500E-07 | 3.464E-08 | 1.100E-07 | 1.700E-07 |
| 25093 | 1255 | T1255M | D | 1.8e-07 | 3.4e-07 | 1.6e-07 | 2.267E-07 | 9.866E-08 | 1.600E-07 | 3.400E-07 |
| 25094 | 1255 | T1255F | D | 3.4e-07 | 3e-07 | 2.9e-07 | 3.100E-07 | 2.646E-08 | 2.900E-07 | 3.400E-07 |
| 25095 | 1255 | T1255P | D | 1.1e-07 | 1.7e-07 | 1.4e-07 | 1.400E-07 | 3.000E-08 | 1.100E-07 | 1.700E-07 |
| 25096 | 1255 | T1255S | D | 1.6e-07 | 1.7e-07 | 2.7e-07 | 2.000E-07 | 6.083E-08 | 1.600E-07 | 2.700E-07 |
| 25097 | 1255 | T1255T | D | 2.7e-07 | 1.4e-07 | 2.7e-07 | 2.267E-07 | 7.506E-08 | 1.400E-07 | 2.700E-07 |
| 25098 | 1255 | T1255W | D | 3.4e-07 | 4.7e-07 | 3.3e-07 | 3.800E-07 | 7.810E-08 | 3.300E-07 | 4.700E-07 |
| 25099 | 1255 | T1255Y | D | 3.3e-07 | 1.2e-07 | 1.3e-07 | 1.933E-07 | 1.185E-07 | 1.200E-07 | 3.300E-07 |
| 25100 | 1255 | T1255V | D | 2.4e-07 | 1.9e-07 | 3.1e-07 | 2.467E-07 | 6.028E-08 | 1.900E-07 | 3.100E-07 |
| 25101 | 1256 | H1256A | D | 3.5e-07 | 3.5e-07 | 2e-07 | 3.000E-07 | 8.660E-08 | 2.000E-07 | 3.500E-07 |
| 25102 | 1256 | H1256R | D | 3.2e-07 | 2.3e-07 | 2.1e-07 | 2.533E-07 | 5.859E-08 | 2.100E-07 | 3.200E-07 |
| 25103 | 1256 | H1256N | D | 2e-07 | 1.7e-07 | 2e-07 | 1.900E-07 | 1.732E-08 | 1.700E-07 | 2.000E-07 |
| 25104 | 1256 | H1256D | D | 3.3e-07 | 1.7e-07 | 2e-07 | 2.333E-07 | 8.505E-08 | 1.700E-07 | 3.300E-07 |
| 25105 | 1256 | H1256C | D | 3.5e-07 | 2.6e-07 | 4.2e-07 | 3.433E-07 | 8.021E-08 | 2.600E-07 | 4.200E-07 |
| 25106 | 1256 | H1256E | D | 1.5e-07 | 3.3e-07 | 2.9e-07 | 2.567E-07 | 9.452E-08 | 1.500E-07 | 3.300E-07 |
| 25107 | 1256 | H1256Q | D | 1.5e-07 | 4.6e-07 | 3.3e-07 | 3.133E-07 | 1.557E-07 | 1.500E-07 | 4.600E-07 |
| 25108 | 1256 | H1256G | D | 2.3e-07 | 3.3e-07 | 3.4e-07 | 3.000E-07 | 6.083E-08 | 2.300E-07 | 3.400E-07 |
| 25109 | 1256 | H1256H | D | 1.6e-07 | 9.8e-08 | 4.1e-07 | 2.227E-07 | 1.652E-07 | 9.800E-08 | 4.100E-07 |
| 25110 | 1256 | H1256I | D | 2e-07 | 1.5e-07 | 1.7e-07 | 1.733E-07 | 2.517E-08 | 1.500E-07 | 2.000E-07 |
| 25111 | 1256 | H1256L | D | 2.4e-07 | 2.2e-07 | 2.5e-07 | 2.367E-07 | 1.528E-08 | 2.200E-07 | 2.500E-07 |
| 25112 | 1256 | H1256K | D | 1.4e-07 | 2e-07 | 2.2e-07 | 1.867E-07 | 4.163E-08 | 1.400E-07 | 2.200E-07 |
| 25113 | 1256 | H1256M | D | 1.8e-07 | 1.8e-07 | 2.8e-07 | 2.133E-07 | 5.774E-08 | 1.800E-07 | 2.800E-07 |
| 25114 | 1256 | H1256F | D | 3.5e-07 | 2.5e-07 | 2.2e-07 | 2.733E-07 | 6.807E-08 | 2.200E-07 | 3.500E-07 |
| 25115 | 1256 | H1256P | D | 3.6e-07 | 3.2e-07 | 5.9e-07 | 4.233E-07 | 1.457E-07 | 3.200E-07 | 5.900E-07 |
| 25116 | 1256 | H1256S | D | 1.5e-07 | 2.2e-07 | 2.6e-07 | 2.100E-07 | 5.568E-08 | 1.500E-07 | 2.600E-07 |
| 25117 | 1256 | H1256T | D | 2.8e-07 | 1.5e-07 | 2.2e-07 | 2.167E-07 | 6.506E-08 | 1.500E-07 | 2.800E-07 |
| 25118 | 1256 | H1256W | D | 1.7e-07 | 2.5e-07 | 2.5e-07 | 2.233E-07 | 4.619E-08 | 1.700E-07 | 2.500E-07 |
| 25119 | 1256 | H1256Y | D | 1e-06 | 1.3e-07 | 3.6e-07 | 4.967E-07 | 4.508E-07 | 1.300E-07 | 1.000E-06 |
| 25120 | 1256 | H1256V | D | 2.2e-07 | 2.7e-07 | 2e-07 | 2.300E-07 | 3.606E-08 | 2.000E-07 | 2.700E-07 |
| 25121 | 1257 | Q1257A | D | 5e-07 | 5.5e-07 | 3.6e-07 | 4.700E-07 | 9.849E-08 | 3.600E-07 | 5.500E-07 |
| 25122 | 1257 | Q1257R | D | 3.1e-07 | 4.7e-07 | 1.5e-07 | 3.100E-07 | 1.600E-07 | 1.500E-07 | 4.700E-07 |
| 25123 | 1257 | Q1257N | D | 1.4e-07 | 3.4e-07 | 2.5e-07 | 2.433E-07 | 1.002E-07 | 1.400E-07 | 3.400E-07 |
| 25124 | 1257 | Q1257D | D | 2e-07 | 2.8e-07 | 2.3e-07 | 2.367E-07 | 4.041E-08 | 2.000E-07 | 2.800E-07 |
| 25125 | 1257 | Q1257C | D | 1.9e-07 | 1.7e-07 | 2.2e-07 | 1.933E-07 | 2.517E-08 | 1.700E-07 | 2.200E-07 |
| 25126 | 1257 | Q1257E | D | 3.1e-07 | 2.5e-07 | 1.7e-07 | 2.433E-07 | 7.024E-08 | 1.700E-07 | 3.100E-07 |
| 25127 | 1257 | Q1257Q | D | 4.3e-07 | 2e-07 | 3.2e-07 | 3.167E-07 | 1.150E-07 | 2.000E-07 | 4.300E-07 |
| 25128 | 1257 | Q1257G | D | 3.8e-07 | 4.1e-07 | 2e-07 | 3.300E-07 | 1.136E-07 | 2.000E-07 | 4.100E-07 |
| 25129 | 1257 | Q1257H | D | 2.6e-07 | 1.5e-07 | 1.7e-07 | 1.933E-07 | 5.859E-08 | 1.500E-07 | 2.600E-07 |
| 25130 | 1257 | Q1257I | D | 2.1e-07 | 2.4e-07 | 3.9e-07 | 2.800E-07 | 9.644E-08 | 2.100E-07 | 3.900E-07 |
| 25131 | 1257 | Q1257L | D | 1.7e-07 | 1.2e-07 | 2.1e-07 | 1.667E-07 | 4.509E-08 | 1.200E-07 | 2.100E-07 |
| 25132 | 1257 | Q1257K | D | 1.9e-07 | 2.5e-07 | 2.4e-07 | 2.267E-07 | 3.215E-08 | 1.900E-07 | 2.500E-07 |
| 25133 | 1257 | Q1257M | D | 3.3e-07 | 2e-07 | 2.8e-07 | 2.700E-07 | 6.557E-08 | 2.000E-07 | 3.300E-07 |
| 25134 | 1257 | Q1257F | D | 5.4e-07 | 3.4e-07 | 1.2e-07 | 3.333E-07 | 2.101E-07 | 1.200E-07 | 5.400E-07 |
| 25135 | 1257 | Q1257P | D | 3.8e-07 | 3e-07 | 1e-07 | 2.600E-07 | 1.442E-07 | 1.000E-07 | 3.800E-07 |
| 25136 | 1257 | Q1257S | D | 5e-07 | 1.6e-07 | 1.8e-07 | 2.800E-07 | 1.908E-07 | 1.600E-07 | 5.000E-07 |
| 25137 | 1257 | Q1257T | D | 2.1e-07 | 2.9e-07 | 2.3e-07 | 2.433E-07 | 4.163E-08 | 2.100E-07 | 2.900E-07 |
| 25138 | 1257 | Q1257W | D | 5.4e-07 | 1e-07 | 2.5e-07 | 2.967E-07 | 2.237E-07 | 1.000E-07 | 5.400E-07 |
| 25139 | 1257 | Q1257Y | D | 1.5e-07 | 1.4e-07 | 2.3e-07 | 1.733E-07 | 4.933E-08 | 1.400E-07 | 2.300E-07 |
| 25140 | 1257 | Q1257V | D | 6.7e-07 | 2.8e-07 | 3.2e-07 | 4.233E-07 | 2.146E-07 | 2.800E-07 | 6.700E-07 |
| 25141 | 1258 | G1258A | D | 2.5e-07 | 1.3e-07 | 2.1e-07 | 1.967E-07 | 6.110E-08 | 1.300E-07 | 2.500E-07 |
| 25142 | 1258 | G1258R | D | 2.3e-07 | 1.8e-07 | 1.9e-07 | 2.000E-07 | 2.646E-08 | 1.800E-07 | 2.300E-07 |
| 25143 | 1258 | G1258N | D | 1.9e-07 | 1.9e-07 | 2e-07 | 1.933E-07 | 5.774E-09 | 1.900E-07 | 2.000E-07 |
| 25144 | 1258 | G1258D | D | 2e-07 | 2.9e-07 | 2e-07 | 2.300E-07 | 5.196E-08 | 2.000E-07 | 2.900E-07 |
| 25145 | 1258 | G1258C | D | 4.3e-07 | 3.6e-07 | 3e-07 | 3.633E-07 | 6.506E-08 | 3.000E-07 | 4.300E-07 |
| 25146 | 1258 | G1258E | D | 6e-07 | 1.5e-07 | 4e-07 | 3.833E-07 | 2.255E-07 | 1.500E-07 | 6.000E-07 |
| 25147 | 1258 | G1258Q | D | 2.8e-07 | 3.7e-07 | 2e-07 | 2.833E-07 | 8.505E-08 | 2.000E-07 | 3.700E-07 |
| 25148 | 1258 | G1258G | D | 1.1e-07 | 2.7e-07 | 5.5e-07 | 3.100E-07 | 2.227E-07 | 1.100E-07 | 5.500E-07 |
| 25149 | 1258 | G1258H | D | 1.1e-07 | 2e-07 | 3.9e-07 | 2.333E-07 | 1.429E-07 | 1.100E-07 | 3.900E-07 |
| 25150 | 1258 | G1258I | D | 1.3e-07 | 2.3e-07 | 3.7e-07 | 2.433E-07 | 1.206E-07 | 1.300E-07 | 3.700E-07 |
| 25151 | 1258 | G1258L | D | 3.5e-07 | 2.2e-07 | 1.8e-07 | 2.500E-07 | 8.888E-08 | 1.800E-07 | 3.500E-07 |
| 25152 | 1258 | G1258K | D | 3.9e-07 | 1.9e-07 | 2.4e-07 | 2.733E-07 | 1.041E-07 | 1.900E-07 | 3.900E-07 |
| 25153 | 1258 | G1258M | D | 4.2e-07 | 2.9e-07 | 2.5e-07 | 3.200E-07 | 8.888E-08 | 2.500E-07 | 4.200E-07 |
| 25154 | 1258 | G1258F | D | 2.7e-07 | 1.5e-07 | 1.2e-07 | 1.800E-07 | 7.937E-08 | 1.200E-07 | 2.700E-07 |
| 25155 | 1258 | G1258P | D | 3.8e-07 | 1.9e-07 | 2.6e-07 | 2.767E-07 | 9.609E-08 | 1.900E-07 | 3.800E-07 |
| 25156 | 1258 | G1258S | D | 2.3e-07 | 2.6e-07 | 1.2e-07 | 2.033E-07 | 7.371E-08 | 1.200E-07 | 2.600E-07 |
| 25157 | 1258 | G1258T | D | 1.1e-07 | 2.9e-07 | 3.2e-07 | 2.400E-07 | 1.136E-07 | 1.100E-07 | 3.200E-07 |
| 25158 | 1258 | G1258W | D | 1.7e-07 | 5e-07 | 2.8e-07 | 3.167E-07 | 1.680E-07 | 1.700E-07 | 5.000E-07 |
| 25159 | 1258 | G1258Y | D | 2.5e-07 | 3.2e-07 | 1.5e-07 | 2.400E-07 | 8.544E-08 | 1.500E-07 | 3.200E-07 |

| | | | | | | | | | | |
|---|---|---|---|---|---|---|---|---|---|---|
| 25160 | 1258 | G1258V | D | 4.9e-07 | 3.2e-07 | 1.7e-07 | 3.267E-07 | 1.601E-07 | 1.700E-07 | 4.900E-07 |
| 25161 | 1259 | L1259A | D | 1.9e-07 | 2.2e-07 | 4e-07 | 2.700E-07 | 1.136E-07 | 1.900E-07 | 4.000E-07 |
| 25162 | 1259 | L1259R | D | 2.2e-07 | 3.3e-07 | 2.9e-07 | 2.800E-07 | 5.568E-08 | 2.200E-07 | 3.300E-07 |
| 25163 | 1259 | L1259N | D | 2.3e-07 | 1.9e-07 | 3e-07 | 2.400E-07 | 5.568E-08 | 1.900E-07 | 3.000E-07 |
| 25164 | 1259 | L1259D | D | 9.8e-08 | 2.3e-07 | 2.6e-07 | 1.960E-07 | 8.619E-08 | 9.800E-08 | 2.600E-07 |
| 25165 | 1259 | L1259C | D | 1.7e-07 | 3.3e-07 | 4.7e-07 | 3.233E-07 | 1.501E-07 | 1.700E-07 | 4.700E-07 |
| 25166 | 1259 | L1259E | D | 1.5e-07 | 5.5e-07 | 1.4e-07 | 2.800E-07 | 2.339E-07 | 1.400E-07 | 5.500E-07 |
| 25167 | 1259 | L1259Q | D | 1.5e-07 | 2e-07 | 2.1e-07 | 1.867E-07 | 3.215E-08 | 1.500E-07 | 2.100E-07 |
| 25168 | 1259 | L1259G | D | 2.7e-07 | 3.9e-07 | 1.8e-07 | 2.800E-07 | 1.054E-07 | 1.800E-07 | 3.900E-07 |
| 25169 | 1259 | L1259H | D | 1.6e-07 | 1.3e-07 | 1.6e-07 | 1.500E-07 | 1.732E-08 | 1.300E-07 | 1.600E-07 |
| 25170 | 1259 | L1259I | D | 5.4e-07 | 3.2e-07 | 2.8e-07 | 3.800E-07 | 1.400E-07 | 2.800E-07 | 5.400E-07 |
| 25171 | 1259 | L1259L | D | 4.5e-07 | 3.5e-07 | 4.6e-07 | 4.200E-07 | 6.083E-08 | 3.500E-07 | 4.600E-07 |
| 25172 | 1259 | L1259K | D | 2.6e-07 | 3e-07 | 1.9e-07 | 2.500E-07 | 5.568E-08 | 1.900E-07 | 3.000E-07 |
| 25173 | 1259 | L1259M | D | 2.8e-07 | 1.2e-07 | 1.8e-07 | 1.933E-07 | 8.083E-08 | 1.200E-07 | 2.800E-07 |
| 25174 | 1259 | L1259F | D | 1.7e-07 | 3.3e-07 | 2.1e-07 | 2.367E-07 | 8.327E-08 | 1.700E-07 | 3.300E-07 |
| 25175 | 1259 | L1259P | D | 2e-07 | 2.8e-07 | 1.7e-07 | 2.167E-07 | 5.686E-08 | 1.700E-07 | 2.800E-07 |
| 25176 | 1259 | L1259S | D | 5.9e-07 | 1.4e-07 | 2e-07 | 3.100E-07 | 2.443E-07 | 1.400E-07 | 5.900E-07 |
| 25177 | 1259 | L1259T | D | 1e-07 | 3e-07 | 3e-07 | 2.333E-07 | 1.155E-07 | 1.000E-07 | 3.000E-07 |
| 25178 | 1259 | L1259W | D | 2.1e-07 | 1.3e-07 | 2.2e-07 | 1.867E-07 | 4.933E-08 | 1.300E-07 | 2.200E-07 |
| 25179 | 1259 | L1259Y | D | 2.9e-07 | 1.4e-07 | 3.8e-07 | 2.700E-07 | 1.212E-07 | 1.400E-07 | 3.800E-07 |
| 25180 | 1259 | L1259V | D | 5.7e-07 | 3.7e-07 | 2.3e-07 | 3.900E-07 | 1.709E-07 | 2.300E-07 | 5.700E-07 |
| 25181 | 1260 | S1260A | D | 1.7e-07 | 2.1e-07 | 2.6e-07 | 2.133E-07 | 4.509E-08 | 1.700E-07 | 2.600E-07 |
| 25182 | 1260 | S1260R | D | 2.3e-07 | 1.9e-07 | 1.3e-07 | 1.833E-07 | 5.033E-08 | 1.300E-07 | 2.300E-07 |
| 25183 | 1260 | S1260N | D | 1.9e-07 | 1.6e-07 | 2.8e-07 | 2.100E-07 | 6.245E-08 | 1.600E-07 | 2.800E-07 |
| 25184 | 1260 | S1260D | D | 1.6e-07 | 1.6e-07 | 2e-07 | 1.733E-07 | 2.309E-08 | 1.600E-07 | 2.000E-07 |
| 25185 | 1260 | S1260C | D | 1.6e-07 | 1e-07 | 1.8e-07 | 1.467E-07 | 4.163E-08 | 1.000E-07 | 1.800E-07 |
| 25186 | 1260 | S1260E | D | 2.5e-07 | 5.6e-07 | 2.2e-07 | 3.433E-07 | 1.882E-07 | 2.200E-07 | 5.600E-07 |
| 25187 | 1260 | S1260Q | D | 2.8e-07 | 1.7e-07 | 1.2e-07 | 1.900E-07 | 8.185E-08 | 1.200E-07 | 2.800E-07 |
| 25188 | 1260 | S1260G | D | 1.7e-07 | 2e-07 | 1.8e-07 | 1.833E-07 | 1.528E-08 | 1.700E-07 | 2.000E-07 |
| 25189 | 1260 | S1260H | D | 2e-07 | 2.4e-07 | 1.5e-07 | 1.967E-07 | 4.509E-08 | 1.500E-07 | 2.400E-07 |
| 25190 | 1260 | S1260I | D | 2.5e-07 | 2.2e-07 | 1.7e-07 | 2.133E-07 | 4.041E-08 | 1.700E-07 | 2.500E-07 |
| 25191 | 1260 | S1260L | D | 3.7e-07 | 3.4e-07 | 2.9e-07 | 3.333E-07 | 4.041E-08 | 2.900E-07 | 3.700E-07 |
| 25192 | 1260 | S1260K | D | 2.9e-07 | 3.3e-07 | 2.9e-07 | 3.033E-07 | 2.309E-08 | 2.900E-07 | 3.300E-07 |
| 25193 | 1260 | S1260M | D | 1.7e-07 | 1.6e-07 | 1e-07 | 1.433E-07 | 3.786E-08 | 1.000E-07 | 1.700E-07 |
| 25194 | 1260 | S1260F | D | 2.9e-07 | 1.8e-07 | 2.2e-07 | 2.300E-07 | 5.568E-08 | 1.800E-07 | 2.900E-07 |
| 25195 | 1260 | S1260P | D | 2.1e-07 | 2.7e-07 | 4.7e-07 | 3.167E-07 | 1.361E-07 | 2.100E-07 | 4.700E-07 |
| 25196 | 1260 | S1260S | D | 5.2e-07 | 2.7e-07 | 5.4e-07 | 4.433E-07 | 1.504E-07 | 2.700E-07 | 5.400E-07 |
| 25197 | 1260 | S1260T | D | 2.1e-07 | 1.9e-07 | 1.5e-07 | 1.833E-07 | 3.055E-08 | 1.500E-07 | 2.100E-07 |
| 25198 | 1260 | S1260W | D | 3.6e-07 | 3.4e-07 | 4.7e-07 | 3.900E-07 | 7.000E-08 | 3.400E-07 | 4.700E-07 |
| 25199 | 1260 | S1260Y | D | 2.4e-07 | 1.7e-07 | 2e-07 | 2.033E-07 | 3.512E-08 | 1.700E-07 | 2.400E-07 |
| 25200 | 1260 | S1260V | D | 1.4e-07 | 1.9e-07 | 5.1e-07 | 2.800E-07 | 2.007E-07 | 1.400E-07 | 5.100E-07 |
| 25201 | 1261 | S1261A | D | 4.5e-07 | 2.7e-07 | 2.5e-07 | 3.233E-07 | 1.102E-07 | 2.500E-07 | 4.500E-07 |
| 25202 | 1261 | S1261R | D | 5.7e-07 | 2.6e-07 | 4.1e-07 | 4.133E-07 | 1.550E-07 | 2.600E-07 | 5.700E-07 |
| 25203 | 1261 | S1261N | D | 1.6e-07 | 1.9e-07 | 1.5e-07 | 1.667E-07 | 2.082E-08 | 1.500E-07 | 1.900E-07 |
| 25204 | 1261 | S1261D | D | 2e-07 | 1.8e-07 | 2.3e-07 | 2.033E-07 | 2.517E-08 | 1.800E-07 | 2.300E-07 |
| 25205 | 1261 | S1261C | D | 3e-07 | 4.9e-07 | 3.2e-07 | 3.700E-07 | 1.044E-07 | 3.000E-07 | 4.900E-07 |
| 25206 | 1261 | S1261E | D | 2.6e-07 | 1.9e-07 | 2.6e-07 | 2.367E-07 | 4.041E-08 | 1.900E-07 | 2.600E-07 |
| 25207 | 1261 | S1261Q | D | 3.2e-07 | 2.6e-07 | 1.9e-07 | 2.567E-07 | 6.506E-08 | 1.900E-07 | 3.200E-07 |
| 25208 | 1261 | S1261G | D | 4e-07 | 2.4e-07 | 2.3e-07 | 2.900E-07 | 9.539E-08 | 2.300E-07 | 4.000E-07 |
| 25209 | 1261 | S1261H | D | 9e-08 | 3.1e-07 | 3e-07 | 2.333E-07 | 1.242E-07 | 9.000E-08 | 3.100E-07 |
| 25210 | 1261 | S1261I | D | 3.1e-07 | 1.4e-07 | 3.2e-07 | 2.567E-07 | 1.012E-07 | 1.400E-07 | 3.200E-07 |
| 25211 | 1261 | S1261L | D | 2.4e-07 | 4.4e-07 | 3.6e-07 | 3.467E-07 | 1.007E-07 | 2.400E-07 | 4.400E-07 |
| 25212 | 1261 | S1261K | D | 2.9e-07 | 3.7e-07 | 9.2e-08 | 2.507E-07 | 1.431E-07 | 9.200E-08 | 3.700E-07 |
| 25213 | 1261 | S1261M | D | 3.7e-07 | 2.5e-07 | 2.4e-07 | 2.867E-07 | 7.234E-08 | 2.400E-07 | 3.700E-07 |
| 25214 | 1261 | S1261F | D | 3e-07 | 5.2e-07 | 2.5e-07 | 3.567E-07 | 1.436E-07 | 2.500E-07 | 5.200E-07 |
| 25215 | 1261 | S1261P | D | 4.3e-07 | 4e-07 | 4.1e-07 | 4.133E-07 | 1.528E-08 | 4.000E-07 | 4.300E-07 |
| 25216 | 1261 | S1261S | D | 9e-08 | 5.5e-07 | 2.8e-07 | 3.067E-07 | 2.312E-07 | 9.000E-08 | 5.500E-07 |
| 25217 | 1261 | S1261T | D | 2.2e-07 | 1.7e-07 | 2e-07 | 1.967E-07 | 2.517E-08 | 1.700E-07 | 2.200E-07 |
| 25218 | 1261 | S1261W | D | 2.8e-07 | 4.4e-07 | 1.2e-07 | 2.800E-07 | 1.600E-07 | 1.200E-07 | 4.400E-07 |
| 25219 | 1261 | S1261Y | D | 1.3e-07 | 2.7e-07 | 3e-07 | 2.333E-07 | 9.074E-08 | 1.300E-07 | 3.000E-07 |
| 25220 | 1261 | S1261V | D | 3.1e-07 | 2.3e-07 | 4.3e-07 | 3.233E-07 | 1.007E-07 | 2.300E-07 | 4.300E-07 |
| 25221 | 1262 | P1262A | D | 1.3e-07 | 2.7e-07 | 2e-07 | 2.000E-07 | 7.000E-08 | 1.300E-07 | 2.700E-07 |
| 25222 | 1262 | P1262R | D | 3.8e-07 | 1.6e-07 | 1.4e-07 | 2.267E-07 | 1.332E-07 | 1.400E-07 | 3.800E-07 |
| 25223 | 1262 | P1262N | D | 6e-07 | 1.9e-07 | 4.4e-07 | 4.100E-07 | 2.066E-07 | 1.900E-07 | 6.000E-07 |
| 25224 | 1262 | P1262D | D | 2.4e-07 | 1.9e-07 | 5.3e-07 | 3.200E-07 | 1.836E-07 | 1.900E-07 | 5.300E-07 |
| 25225 | 1262 | P1262C | D | 1.9e-07 | 2.6e-07 | 2.7e-07 | 2.400E-07 | 4.359E-08 | 1.900E-07 | 2.700E-07 |
| 25226 | 1262 | P1262E | D | 3.2e-07 | 6e-07 | 1.5e-07 | 3.567E-07 | 2.272E-07 | 1.500E-07 | 6.000E-07 |
| 25227 | 1262 | P1262Q | D | 2.6e-07 | 2e-07 | 1.3e-07 | 1.967E-07 | 6.506E-08 | 1.300E-07 | 2.600E-07 |
| 25228 | 1262 | P1262G | D | 7.8e-07 | 3.4e-07 | 4.7e-07 | 5.300E-07 | 2.261E-07 | 3.400E-07 | 7.800E-07 |
| 25229 | 1262 | P1262H | D | 4.3e-07 | 1.4e-07 | 1.2e-07 | 2.300E-07 | 1.735E-07 | 1.200E-07 | 4.300E-07 |
| 25230 | 1262 | P1262I | D | 3.8e-07 | 1.7e-07 | 3.1e-07 | 2.867E-07 | 1.069E-07 | 1.700E-07 | 3.800E-07 |
| 25231 | 1262 | P1262L | D | 1.4e-07 | 1.9e-07 | 2.6e-07 | 1.967E-07 | 6.028E-08 | 1.400E-07 | 2.600E-07 |
| 25232 | 1262 | P1262K | D | 1.5e-07 | 5.6e-07 | 1.5e-07 | 2.867E-07 | 2.367E-07 | 1.500E-07 | 5.600E-07 |
| 25233 | 1262 | P1262M | D | 4.2e-07 | 3.2e-07 | 2.3e-07 | 3.233E-07 | 9.504E-08 | 2.300E-07 | 4.200E-07 |
| 25234 | 1262 | P1262F | D | 2.8e-07 | 2.7e-07 | 2.2e-07 | 2.567E-07 | 3.215E-08 | 2.200E-07 | 2.800E-07 |
| 25235 | 1262 | P1262P | D | 1.5e-07 | 2.1e-07 | 4e-07 | 2.533E-07 | 1.305E-07 | 1.500E-07 | 4.000E-07 |
| 25236 | 1262 | P1262S | D | 2.6e-07 | 4.3e-07 | 1.4e-07 | 2.767E-07 | 1.457E-07 | 1.400E-07 | 4.300E-07 |
| 25237 | 1262 | P1262T | D | 3.1e-07 | 4.1e-07 | 1.9e-07 | 3.033E-07 | 1.102E-07 | 1.900E-07 | 4.100E-07 |
| 25238 | 1262 | P1262W | D | 3.2e-07 | 4.3e-07 | 3.8e-07 | 3.767E-07 | 5.508E-08 | 3.200E-07 | 4.300E-07 |
| 25239 | 1262 | P1262Y | D | 1.4e-07 | 1.8e-07 | 3.7e-07 | 2.300E-07 | 1.229E-07 | 1.400E-07 | 3.700E-07 |
| 25240 | 1262 | P1262V | D | 3.1e-07 | 1.6e-07 | 2.6e-07 | 2.433E-07 | 7.638E-08 | 1.600E-07 | 3.100E-07 |
| 25241 | 1263 | V1263A | D | 1.7e-07 | 1.9e-07 | 3.6e-07 | 2.400E-07 | 1.044E-07 | 1.700E-07 | 3.600E-07 |
| 25242 | 1263 | V1263R | D | 3.6e-07 | 2.7e-07 | 1.3e-07 | 2.533E-07 | 1.159E-07 | 1.300E-07 | 3.600E-07 |
| 25243 | 1263 | V1263N | D | 2.4e-07 | 1.4e-07 | 2.1e-07 | 1.967E-07 | 5.132E-08 | 1.400E-07 | 2.400E-07 |
| 25244 | 1263 | V1263D | D | 2.3e-07 | 1.4e-07 | 1.4e-07 | 1.700E-07 | 5.196E-08 | 1.400E-07 | 2.300E-07 |

| | | | | | | | | | | |
|---|---|---|---|---|---|---|---|---|---|---|
| 25245 | 1263 | V1263C | D | 1.8e-07 | 3.1e-07 | 2e-07 | 2.300E-07 | 7.000E-08 | 1.800E-07 | 3.100E-07 |
| 25246 | 1263 | V1263E | D | 3.4e-07 | 2.3e-07 | 2.7e-07 | 2.800E-07 | 5.568E-08 | 2.300E-07 | 3.400E-07 |
| 25247 | 1263 | V1263Q | D | 1.6e-07 | 2.3e-07 | 2.9e-07 | 2.267E-07 | 6.506E-08 | 1.600E-07 | 2.900E-07 |
| 25248 | 1263 | V1263G | D | 2.6e-07 | 2e-07 | 1.5e-07 | 2.033E-07 | 5.508E-08 | 1.500E-07 | 2.600E-07 |
| 25249 | 1263 | V1263H | D | 2.9e-07 | 8.8e-07 | 3.5e-07 | 5.067E-07 | 3.247E-07 | 2.900E-07 | 8.800E-07 |
| 25250 | 1263 | V1263I | D | 2.3e-07 | 3.1e-07 | 2.2e-07 | 2.533E-07 | 4.933E-08 | 2.200E-07 | 3.100E-07 |
| 25251 | 1263 | V1263L | D | 1.9e-07 | 1.8e-07 | 3.1e-07 | 2.267E-07 | 7.234E-08 | 1.800E-07 | 3.100E-07 |
| 25252 | 1263 | V1263K | D | 4.7e-07 | 3.8e-07 | 2.6e-07 | 3.700E-07 | 1.054E-07 | 2.600E-07 | 4.700E-07 |
| 25253 | 1263 | V1263M | D | 1.9e-07 | 1.2e-07 | 1.7e-07 | 1.600E-07 | 3.606E-08 | 1.200E-07 | 1.900E-07 |
| 25254 | 1263 | V1263F | D | 2.7e-07 | 1.6e-07 | 3.3e-07 | 2.533E-07 | 8.622E-08 | 1.600E-07 | 3.300E-07 |
| 25255 | 1263 | V1263P | D | 1.9e-07 | 1.9e-07 | 2.6e-07 | 2.133E-07 | 4.041E-08 | 1.900E-07 | 2.600E-07 |
| 25256 | 1263 | V1263S | D | 6.1e-07 | 3.2e-07 | 2.4e-07 | 3.900E-07 | 1.947E-07 | 2.400E-07 | 6.100E-07 |
| 25257 | 1263 | V1263T | D | 1.7e-07 | 2.1e-07 | 3.2e-07 | 2.333E-07 | 7.767E-08 | 1.700E-07 | 3.200E-07 |
| 25258 | 1263 | V1263W | D | 4.8e-07 | 2.1e-07 | 2e-07 | 2.967E-07 | 1.589E-07 | 2.000E-07 | 4.800E-07 |
| 25259 | 1263 | V1263Y | D | 3.6e-07 | 3.1e-07 | 1.6e-07 | 2.767E-07 | 1.041E-07 | 1.600E-07 | 3.600E-07 |
| 25260 | 1263 | V1263V | D | 1.7e-07 | 3.7e-07 | 1.9e-07 | 2.433E-07 | 1.102E-07 | 1.700E-07 | 3.700E-07 |
| 25261 | 1264 | T1264A | D | 5.3e-07 | 4.4e-07 | 4e-07 | 4.567E-07 | 6.658E-08 | 4.000E-07 | 5.300E-07 |
| 25262 | 1264 | T1264R | D | 2.5e-07 | 3.9e-07 | 2.3e-07 | 2.900E-07 | 8.718E-08 | 2.300E-07 | 3.900E-07 |
| 25263 | 1264 | T1264N | D | 2.2e-07 | 2.1e-07 | 1.3e-07 | 1.867E-07 | 4.933E-08 | 1.300E-07 | 2.200E-07 |
| 25264 | 1264 | T1264D | D | 3.1e-07 | 1.4e-07 | 4.4e-07 | 2.967E-07 | 1.504E-07 | 1.400E-07 | 4.400E-07 |
| 25265 | 1264 | T1264C | D | 6.6e-08 | 4.7e-07 | 4.6e-07 | 3.320E-07 | 2.304E-07 | 6.600E-08 | 4.700E-07 |
| 25266 | 1264 | T1264E | D | 2e-07 | 1.6e-07 | 3.6e-07 | 2.400E-07 | 1.058E-07 | 1.600E-07 | 3.600E-07 |
| 25267 | 1264 | T1264Q | D | 1.5e-07 | 5.4e-07 | 2.1e-07 | 3.000E-07 | 2.100E-07 | 1.500E-07 | 5.400E-07 |
| 25268 | 1264 | T1264G | D | 2.5e-07 | 2.3e-07 | 2.4e-07 | 2.400E-07 | 1.000E-08 | 2.300E-07 | 2.500E-07 |
| 25269 | 1264 | T1264H | D | 1.6e-07 | 2.3e-07 | 3.4e-07 | 2.433E-07 | 9.074E-08 | 1.600E-07 | 3.400E-07 |
| 25270 | 1264 | T1264I | D | 3.2e-07 | 2.5e-07 | 2e-07 | 2.567E-07 | 6.028E-08 | 2.000E-07 | 3.200E-07 |
| 25271 | 1264 | T1264L | D | 3.4e-07 | 2.7e-07 | 2e-07 | 2.700E-07 | 7.000E-08 | 2.000E-07 | 3.400E-07 |
| 25272 | 1264 | T1264K | D | 2.1e-07 | 4e-07 | 1.9e-07 | 2.667E-07 | 1.159E-07 | 1.900E-07 | 4.000E-07 |
| 25273 | 1264 | T1264M | D | 1.9e-07 | 3.2e-07 | 3.4e-07 | 2.833E-07 | 8.145E-08 | 1.900E-07 | 3.400E-07 |
| 25274 | 1264 | T1264F | D | 1.7e-07 | 2.5e-07 | 4.2e-07 | 2.800E-07 | 1.277E-07 | 1.700E-07 | 4.200E-07 |
| 25275 | 1264 | T1264P | D | 3e-07 | 1.9e-07 | 3.5e-07 | 2.800E-07 | 8.185E-08 | 1.900E-07 | 3.500E-07 |
| 25276 | 1264 | T1264S | D | 3.4e-07 | 3.4e-07 | 3.6e-07 | 3.467E-07 | 1.155E-08 | 3.400E-07 | 3.600E-07 |
| 25277 | 1264 | T1264T | D | 4.6e-07 | 4.5e-07 | 2.8e-07 | 3.967E-07 | 1.012E-07 | 2.800E-07 | 4.600E-07 |
| 25278 | 1264 | T1264W | D | 7.6e-08 | 2.1e-07 | 5.6e-07 | 2.820E-07 | 2.499E-07 | 7.600E-08 | 5.600E-07 |
| 25279 | 1264 | T1264Y | D | 2.9e-07 | 3e-07 | 2.8e-07 | 2.900E-07 | 1.000E-08 | 2.800E-07 | 3.000E-07 |
| 25280 | 1264 | T1264V | D | 1.4e-07 | 1.5e-07 | 2.4e-07 | 1.767E-07 | 5.508E-08 | 1.400E-07 | 2.400E-07 |
| 25281 | 1265 | K1265A | D | 1.2e-07 | 2.2e-07 | 2e-07 | 1.800E-07 | 5.292E-08 | 1.200E-07 | 2.200E-07 |
| 25282 | 1265 | K1265R | D | 2.4e-07 | 2.4e-07 | 2.3e-07 | 2.367E-07 | 5.774E-09 | 2.300E-07 | 2.400E-07 |
| 25283 | 1265 | K1265N | D | 2.7e-07 | 1.1e-07 | 5e-07 | 2.933E-07 | 1.960E-07 | 1.100E-07 | 5.000E-07 |
| 25284 | 1265 | K1265D | D | 4.5e-07 | 2.2e-07 | 2.4e-07 | 3.033E-07 | 1.274E-07 | 2.200E-07 | 4.500E-07 |
| 25285 | 1265 | K1265C | D | 4.6e-07 | 3.7e-07 | 3.7e-07 | 4.000E-07 | 5.196E-08 | 3.700E-07 | 4.600E-07 |
| 25286 | 1265 | K1265E | D | 2.3e-07 | 1.9e-07 | 3.5e-07 | 2.567E-07 | 8.327E-08 | 1.900E-07 | 3.500E-07 |
| 25287 | 1265 | K1265Q | D | 4.4e-07 | 1.5e-07 | 2.1e-07 | 2.667E-07 | 1.531E-07 | 1.500E-07 | 4.400E-07 |
| 25288 | 1265 | K1265G | D | 3.5e-07 | 1.6e-07 | 2.9e-07 | 2.667E-07 | 9.713E-08 | 1.600E-07 | 3.500E-07 |
| 25289 | 1265 | K1265H | D | 2.6e-07 | 2.2e-07 | 2.7e-07 | 2.500E-07 | 2.646E-08 | 2.200E-07 | 2.700E-07 |
| 25290 | 1265 | K1265I | D | 4.3e-07 | 1.5e-07 | 1.5e-07 | 2.433E-07 | 1.617E-07 | 1.500E-07 | 4.300E-07 |
| 25291 | 1265 | K1265L | D | 1.8e-07 | 4.6e-07 | 4.9e-07 | 3.767E-07 | 1.710E-07 | 1.800E-07 | 4.900E-07 |
| 25292 | 1265 | K1265K | D | 3.3e-07 | 2e-07 | 1e-07 | 2.100E-07 | 1.153E-07 | 1.000E-07 | 3.300E-07 |
| 25293 | 1265 | K1265M | D | 2e-07 | 3.5e-07 | 3e-07 | 2.833E-07 | 7.638E-08 | 2.000E-07 | 3.500E-07 |
| 25294 | 1265 | K1265F | D | 3.5e-07 | 2.1e-07 | 3.3e-07 | 2.967E-07 | 7.572E-08 | 2.100E-07 | 3.500E-07 |
| 25295 | 1265 | K1265P | D | 8.2e-08 | 2.5e-07 | 9.4e-08 | 1.420E-07 | 9.372E-08 | 8.200E-08 | 2.500E-07 |
| 25296 | 1265 | K1265S | D | 2.4e-07 | 1.9e-07 | 6.6e-07 | 3.633E-07 | 2.581E-07 | 1.900E-07 | 6.600E-07 |
| 25297 | 1265 | K1265T | D | 2.9e-07 | 5.1e-07 | 2e-07 | 3.333E-07 | 1.595E-07 | 2.000E-07 | 5.100E-07 |
| 25298 | 1265 | K1265W | D | 4.5e-07 | 3.1e-07 | 3.2e-07 | 3.600E-07 | 7.810E-08 | 3.100E-07 | 4.500E-07 |
| 25299 | 1265 | K1265Y | D | 2.5e-07 | 3.3e-07 | 4.8e-07 | 3.533E-07 | 1.168E-07 | 2.500E-07 | 4.800E-07 |
| 25300 | 1265 | K1265V | D | 3.4e-07 | 3.2e-07 | 1.6e-07 | 2.733E-07 | 9.866E-08 | 1.600E-07 | 3.400E-07 |
| 25301 | 1266 | S1266A | D | 2.9e-07 | 3.7e-07 | 9.9e-08 | 2.530E-07 | 1.392E-07 | 9.900E-08 | 3.700E-07 |
| 25302 | 1266 | S1266R | D | 2e-07 | 2.7e-07 | 2.7e-07 | 2.467E-07 | 4.041E-08 | 2.000E-07 | 2.700E-07 |
| 25303 | 1266 | S1266N | D | 3.4e-07 | 2.1e-07 | 1.5e-07 | 2.333E-07 | 9.713E-08 | 1.500E-07 | 3.400E-07 |
| 25304 | 1266 | S1266D | D | 1.6e-07 | 2.7e-07 | 2.5e-07 | 2.267E-07 | 5.859E-08 | 1.600E-07 | 2.700E-07 |
| 25305 | 1266 | S1266C | D | 3.4e-07 | 1.4e-07 | 2.7e-07 | 2.500E-07 | 1.015E-07 | 1.400E-07 | 3.400E-07 |
| 25306 | 1266 | S1266E | D | 2.4e-07 | 1.6e-07 | 2e-07 | 2.000E-07 | 4.000E-08 | 1.600E-07 | 2.400E-07 |
| 25307 | 1266 | S1266Q | D | 2.7e-07 | 2.1e-07 | 2.1e-07 | 2.300E-07 | 3.464E-08 | 2.100E-07 | 2.700E-07 |
| 25308 | 1266 | S1266G | D | 2.4e-07 | 6e-07 | 3.2e-07 | 3.867E-07 | 1.890E-07 | 2.400E-07 | 6.000E-07 |
| 25309 | 1266 | S1266H | D | 2.8e-07 | 1.9e-07 | 1.7e-07 | 2.133E-07 | 5.859E-08 | 1.700E-07 | 2.800E-07 |
| 25310 | 1266 | S1266I | D | 7.1e-07 | 1.9e-07 | 6.1e-07 | 5.033E-07 | 2.759E-07 | 1.900E-07 | 7.100E-07 |
| 25311 | 1266 | S1266L | D | 2.3e-07 | 2.6e-07 | 4e-07 | 2.967E-07 | 9.074E-08 | 2.300E-07 | 4.000E-07 |
| 25312 | 1266 | S1266K | D | 3.4e-07 | 3.6e-07 | 1.2e-07 | 2.733E-07 | 1.332E-07 | 1.200E-07 | 3.600E-07 |
| 25313 | 1266 | S1266M | D | 3e-07 | 3.2e-07 | 4.2e-07 | 3.467E-07 | 6.429E-08 | 3.000E-07 | 4.200E-07 |
| 25314 | 1266 | S1266F | D | 1.7e-07 | 4e-07 | 1.8e-07 | 2.500E-07 | 1.300E-07 | 1.700E-07 | 4.000E-07 |
| 25315 | 1266 | S1266P | D | 3.5e-07 | 2e-07 | 2.1e-07 | 2.533E-07 | 8.386E-08 | 2.000E-07 | 3.500E-07 |
| 25316 | 1266 | S1266S | D | 1.8e-07 | 1.5e-07 | 4.2e-07 | 2.500E-07 | 1.480E-07 | 1.500E-07 | 4.200E-07 |
| 25317 | 1266 | S1266T | D | 2.7e-07 | 2.8e-07 | 1.1e-07 | 2.200E-07 | 9.539E-08 | 1.100E-07 | 2.800E-07 |
| 25318 | 1266 | S1266W | D | 1.5e-07 | 2.9e-07 | 2.4e-07 | 2.267E-07 | 7.095E-08 | 1.500E-07 | 2.900E-07 |
| 25319 | 1266 | S1266Y | D | 2e-07 | 9.8e-08 | 2.1e-07 | 1.693E-07 | 6.198E-08 | 9.800E-08 | 2.100E-07 |
| 25320 | 1266 | S1266V | D | 1.2e-07 | 5.8e-07 | 2.7e-07 | 3.233E-07 | 2.346E-07 | 1.200E-07 | 5.800E-07 |
| 25321 | 1267 | F1267A | D | 2.8e-07 | 2e-07 | 2.8e-07 | 2.533E-07 | 4.619E-08 | 2.000E-07 | 2.800E-07 |
| 25322 | 1267 | F1267R | D | 6.3e-07 | 4.5e-07 | 1.7e-07 | 4.167E-07 | 2.318E-07 | 1.700E-07 | 6.300E-07 |
| 25323 | 1267 | F1267N | D | 2.1e-07 | 3.3e-07 | 3.2e-07 | 2.867E-07 | 6.658E-08 | 2.100E-07 | 3.300E-07 |
| 25324 | 1267 | F1267D | D | 2.2e-07 | 2.8e-07 | 2.8e-07 | 2.600E-07 | 3.464E-08 | 2.200E-07 | 2.800E-07 |
| 25325 | 1267 | F1267C | D | 1.6e-07 | 2.2e-07 | 2e-07 | 1.933E-07 | 3.055E-08 | 1.600E-07 | 2.200E-07 |
| 25326 | 1267 | F1267E | D | 2e-07 | 6.6e-07 | 3.9e-07 | 4.167E-07 | 2.312E-07 | 2.000E-07 | 6.600E-07 |
| 25327 | 1267 | F1267Q | D | 1.4e-07 | 7e-07 | 4.8e-07 | 4.400E-07 | 2.821E-07 | 1.400E-07 | 7.000E-07 |
| 25328 | 1267 | F1267G | D | 1.1e-07 | 3.9e-07 | 2.3e-07 | 2.433E-07 | 1.405E-07 | 1.100E-07 | 3.900E-07 |
| 25329 | 1267 | F1267H | D | 2.2e-07 | 1.9e-07 | 2e-07 | 2.033E-07 | 1.528E-08 | 1.900E-07 | 2.200E-07 |

| | | | | | | | | | | |
|---|---|---|---|---|---|---|---|---|---|---|
| 25330 | 1267 | F1267I | D | 2.8e-07 | 1.2e-07 | 2.8e-07 | 2.267E-07 | 9.238E-08 | 1.200E-07 | 2.800E-07 |
| 25331 | 1267 | F1267L | D | 2.8e-07 | 3.2e-07 | 2e-07 | 2.667E-07 | 6.110E-08 | 2.000E-07 | 3.200E-07 |
| 25332 | 1267 | F1267K | D | 1.7e-07 | 2.3e-07 | 3.7e-07 | 2.567E-07 | 1.026E-07 | 1.700E-07 | 3.700E-07 |
| 25333 | 1267 | F1267M | D | 8.2e-08 | 1.8e-07 | 2.8e-07 | 1.807E-07 | 9.900E-08 | 8.200E-08 | 2.800E-07 |
| 25334 | 1267 | F1267F | D | 2.4e-07 | 3.1e-07 | 4e-07 | 3.167E-07 | 8.021E-08 | 2.400E-07 | 4.000E-07 |
| 25335 | 1267 | F1267P | D | 1.3e-07 | 2.3e-07 | 5.6e-07 | 3.067E-07 | 2.250E-07 | 1.300E-07 | 5.600E-07 |
| 25336 | 1267 | F1267S | D | 2.2e-07 | 2.9e-07 | 7.2e-07 | 4.100E-07 | 2.707E-07 | 2.200E-07 | 7.200E-07 |
| 25337 | 1267 | F1267T | D | 4.3e-07 | 3.2e-07 | 5.8e-07 | 4.433E-07 | 1.305E-07 | 3.200E-07 | 5.800E-07 |
| 25338 | 1267 | F1267W | D | 3.5e-07 | 4.3e-07 | 2e-07 | 3.267E-07 | 1.168E-07 | 2.000E-07 | 4.300E-07 |
| 25339 | 1267 | F1267Y | D | 1.7e-07 | 1.5e-07 | 4.7e-07 | 2.633E-07 | 1.793E-07 | 1.500E-07 | 4.700E-07 |
| 25340 | 1267 | F1267V | D | 4.4e-07 | 2.5e-07 | 2.8e-07 | 3.233E-07 | 1.021E-07 | 2.500E-07 | 4.400E-07 |
| 25341 | 1268 | N1268A | D | 4.3e-07 | 1.5e-07 | 2.6e-07 | 2.800E-07 | 1.411E-07 | 1.500E-07 | 4.300E-07 |
| 25342 | 1268 | N1268R | D | 1.7e-07 | 1.7e-07 | 4.5e-07 | 2.633E-07 | 1.617E-07 | 1.700E-07 | 4.500E-07 |
| 25343 | 1268 | N1268N | D | 1.1e-07 | 1.8e-07 | 3e-07 | 1.967E-07 | 9.609E-08 | 1.100E-07 | 3.000E-07 |
| 25344 | 1268 | N1268D | D | 2.8e-07 | 1.9e-07 | 2.8e-07 | 2.500E-07 | 5.196E-08 | 1.900E-07 | 2.800E-07 |
| 25345 | 1268 | N1268C | D | 5e-07 | 2.9e-07 | 2.1e-07 | 3.333E-07 | 1.498E-07 | 2.100E-07 | 5.000E-07 |
| 25346 | 1268 | N1268E | D | 2.8e-07 | 3.4e-07 | 1.4e-07 | 2.533E-07 | 1.026E-07 | 1.400E-07 | 3.400E-07 |
| 25347 | 1268 | N1268Q | D | 5.1e-07 | 1.2e-07 | 2.5e-07 | 2.933E-07 | 1.986E-07 | 1.200E-07 | 5.100E-07 |
| 25348 | 1268 | N1268G | D | 1.9e-07 | 1.5e-07 | 2.8e-07 | 2.067E-07 | 6.658E-08 | 1.500E-07 | 2.800E-07 |
| 25349 | 1268 | N1268H | D | 2e-07 | 1.6e-07 | 3.3e-07 | 2.300E-07 | 8.888E-08 | 1.600E-07 | 3.300E-07 |
| 25350 | 1268 | N1268I | D | 9.3e-08 | 3.9e-07 | 2.6e-07 | 2.477E-07 | 1.489E-07 | 9.300E-08 | 3.900E-07 |
| 25351 | 1268 | N1268L | D | 3.3e-07 | 3.6e-07 | 9.1e-08 | 2.603E-07 | 1.474E-07 | 9.100E-08 | 3.600E-07 |
| 25352 | 1268 | N1268K | D | 2.9e-07 | 2.9e-07 | 1.6e-07 | 2.467E-07 | 7.506E-08 | 1.600E-07 | 2.900E-07 |
| 25353 | 1268 | N1268M | D | 3.1e-07 | 2.7e-07 | 2.3e-07 | 2.700E-07 | 4.000E-08 | 2.300E-07 | 3.100E-07 |
| 25354 | 1268 | N1268F | D | 3.8e-07 | 3.4e-07 | 2.3e-07 | 3.167E-07 | 7.767E-08 | 2.300E-07 | 3.800E-07 |
| 25355 | 1268 | N1268P | D | 1.2e-07 | 3.2e-07 | 2.1e-07 | 2.167E-07 | 1.002E-07 | 1.200E-07 | 3.200E-07 |
| 25356 | 1268 | N1268S | D | 1.8e-07 | 4e-07 | 2e-07 | 2.600E-07 | 1.217E-07 | 1.800E-07 | 4.000E-07 |
| 25357 | 1268 | N1268T | D | 1.3e-07 | 2.7e-07 | 2.2e-07 | 2.067E-07 | 7.095E-08 | 1.300E-07 | 2.700E-07 |
| 25358 | 1268 | N1268W | D | 3.4e-07 | 1.6e-07 | 5e-07 | 3.333E-07 | 1.701E-07 | 1.600E-07 | 5.000E-07 |
| 25359 | 1268 | N1268Y | D | 1.7e-07 | 8.4e-07 | 1.6e-07 | 3.900E-07 | 3.897E-07 | 1.600E-07 | 8.400E-07 |
| 25360 | 1268 | N1268V | D | 2.7e-07 | 2.3e-07 | 3e-07 | 2.667E-07 | 3.512E-08 | 2.300E-07 | 3.000E-07 |
| 25361 | 1269 | R1269A | D | 3.2e-07 | 3.1e-07 | 2.6e-07 | 2.967E-07 | 3.215E-08 | 2.600E-07 | 3.200E-07 |
| 25362 | 1269 | R1269R | D | 5.5e-07 | 1.7e-07 | 2.8e-07 | 3.333E-07 | 1.955E-07 | 1.700E-07 | 5.500E-07 |
| 25363 | 1269 | R1269N | D | 3.8e-07 | 8.2e-08 | 2.7e-07 | 2.440E-07 | 1.507E-07 | 8.200E-08 | 3.800E-07 |
| 25364 | 1269 | R1269D | D | 2.2e-07 | 2.8e-07 | 1.4e-07 | 2.133E-07 | 7.024E-08 | 1.400E-07 | 2.800E-07 |
| 25365 | 1269 | R1269C | D | 2e-07 | 4.5e-07 | 1.3e-07 | 2.600E-07 | 1.682E-07 | 1.300E-07 | 4.500E-07 |
| 25366 | 1269 | R1269E | D | 1.6e-07 | 3.6e-07 | 3.5e-07 | 2.900E-07 | 1.127E-07 | 1.600E-07 | 3.600E-07 |
| 25367 | 1269 | R1269Q | D | 1.2e-07 | 4.8e-07 | 1.7e-07 | 2.567E-07 | 1.950E-07 | 1.200E-07 | 4.800E-07 |
| 25368 | 1269 | R1269G | D | 2.5e-07 | 2.6e-07 | 2.1e-07 | 2.400E-07 | 2.646E-08 | 2.100E-07 | 2.600E-07 |
| 25369 | 1269 | R1269H | D | 4.7e-07 | 2.9e-07 | 3e-07 | 3.533E-07 | 1.012E-07 | 2.900E-07 | 4.700E-07 |
| 25370 | 1269 | R1269I | D | 2.3e-07 | 1e-07 | 1.6e-07 | 1.633E-07 | 6.506E-08 | 1.000E-07 | 2.300E-07 |
| 25371 | 1269 | R1269L | D | 4.6e-07 | 3.1e-07 | 3.3e-07 | 3.667E-07 | 8.145E-08 | 3.100E-07 | 4.600E-07 |
| 25372 | 1269 | R1269K | D | 8.3e-08 | 2.3e-07 | 3.8e-07 | 2.310E-07 | 1.485E-07 | 8.300E-08 | 3.800E-07 |
| 25373 | 1269 | R1269M | D | 2e-07 | 2.7e-07 | 2.4e-07 | 2.367E-07 | 3.512E-08 | 2.000E-07 | 2.700E-07 |
| 25374 | 1269 | R1269F | D | 1.4e-07 | 2.9e-07 | 3.1e-07 | 2.467E-07 | 9.292E-08 | 1.400E-07 | 3.100E-07 |
| 25375 | 1269 | R1269P | D | 4e-07 | 2.4e-07 | 2.2e-07 | 2.867E-07 | 9.866E-08 | 2.200E-07 | 4.000E-07 |
| 25376 | 1269 | R1269S | D | 3.2e-07 | 5.1e-07 | 2.2e-07 | 3.500E-07 | 1.473E-07 | 2.200E-07 | 5.100E-07 |
| 25377 | 1269 | R1269T | D | 4e-07 | 3.5e-07 | 3.9e-07 | 3.800E-07 | 2.646E-08 | 3.500E-07 | 4.000E-07 |
| 25378 | 1269 | R1269W | D | 1.3e-07 | 3.1e-07 | 1.7e-07 | 2.033E-07 | 9.452E-08 | 1.300E-07 | 3.100E-07 |
| 25379 | 1269 | R1269Y | D | 2.5e-07 | 2.6e-07 | 2.2e-07 | 2.433E-07 | 2.082E-08 | 2.200E-07 | 2.600E-07 |
| 25380 | 1269 | R1269V | D | 3.2e-07 | 1e-07 | 1.9e-07 | 2.033E-07 | 1.106E-07 | 1.000E-07 | 3.200E-07 |
| 25381 | 1270 | G1270A | D | 1e-07 | 1.9e-07 | 3.9e-07 | 2.267E-07 | 1.484E-07 | 1.000E-07 | 3.900E-07 |
| 25382 | 1270 | G1270R | D | 2e-07 | 2.7e-07 | 2.8e-07 | 2.500E-07 | 4.359E-08 | 2.000E-07 | 2.800E-07 |
| 25383 | 1270 | G1270N | D | 2e-07 | 9.9e-07 | 3e-07 | 4.967E-07 | 4.302E-07 | 2.000E-07 | 9.900E-07 |
| 25384 | 1270 | G1270D | D | 2e-07 | 3.4e-07 | 1.3e-07 | 2.233E-07 | 1.069E-07 | 1.300E-07 | 3.400E-07 |
| 25385 | 1270 | G1270C | D | 2.8e-07 | 5.3e-07 | 2.1e-07 | 3.400E-07 | 1.682E-07 | 2.100E-07 | 5.300E-07 |
| 25386 | 1270 | G1270E | D | 2.4e-07 | 2.6e-07 | 2.2e-07 | 2.400E-07 | 2.000E-08 | 2.200E-07 | 2.600E-07 |
| 25387 | 1270 | G1270Q | D | 3.5e-07 | 3.3e-07 | 1e-07 | 2.600E-07 | 1.389E-07 | 1.000E-07 | 3.500E-07 |
| 25388 | 1270 | G1270G | D | 3.2e-07 | 2.9e-07 | 4.3e-07 | 3.467E-07 | 7.371E-08 | 2.900E-07 | 4.300E-07 |
| 25389 | 1270 | G1270H | D | 2.4e-07 | 1.2e-07 | 2.2e-07 | 1.933E-07 | 6.429E-08 | 1.200E-07 | 2.400E-07 |
| 25390 | 1270 | G1270I | D | 2.7e-07 | 2.4e-07 | 2.9e-07 | 2.667E-07 | 2.517E-08 | 2.400E-07 | 2.900E-07 |
| 25391 | 1270 | G1270L | D | 2.3e-07 | 3.5e-07 | 1.9e-07 | 2.567E-07 | 8.327E-08 | 1.900E-07 | 3.500E-07 |
| 25392 | 1270 | G1270K | D | 1.5e-07 | 1e-07 | 1.9e-07 | 1.467E-07 | 4.509E-08 | 1.000E-07 | 1.900E-07 |
| 25393 | 1270 | G1270M | D | 3.2e-07 | 2.5e-07 | 1.4e-07 | 2.367E-07 | 9.074E-08 | 1.400E-07 | 3.200E-07 |
| 25394 | 1270 | G1270F | D | 3.6e-07 | 2.3e-07 | 2.9e-07 | 2.933E-07 | 6.506E-08 | 2.300E-07 | 3.600E-07 |
| 25395 | 1270 | G1270P | D | 2.6e-07 | 1.9e-07 | 1.3e-07 | 1.933E-07 | 6.506E-08 | 1.300E-07 | 2.600E-07 |
| 25396 | 1270 | G1270S | D | 2.5e-07 | 2.1e-07 | 2.6e-07 | 2.400E-07 | 2.646E-08 | 2.100E-07 | 2.600E-07 |
| 25397 | 1270 | G1270T | D | 2e-07 | 2.8e-07 | 2e-07 | 2.267E-07 | 4.619E-08 | 2.000E-07 | 2.800E-07 |
| 25398 | 1270 | G1270W | D | 3.2e-07 | 4.4e-07 | 2.3e-07 | 3.300E-07 | 1.054E-07 | 2.300E-07 | 4.400E-07 |
| 25399 | 1270 | G1270Y | D | 1.2e-07 | 3.7e-07 | 2.9e-07 | 2.600E-07 | 1.277E-07 | 1.200E-07 | 3.700E-07 |
| 25400 | 1270 | G1270V | D | 7.2e-07 | 3.2e-07 | 3e-07 | 4.467E-07 | 2.369E-07 | 3.000E-07 | 7.200E-07 |
| 25401 | 1271 | E1271A | D | 3.6e-07 | 3.1e-07 | 2.4e-07 | 3.033E-07 | 6.028E-08 | 2.400E-07 | 3.600E-07 |
| 25402 | 1271 | E1271R | D | 2.4e-07 | 4.5e-07 | 2.5e-07 | 3.133E-07 | 1.185E-07 | 2.400E-07 | 4.500E-07 |
| 25403 | 1271 | E1271N | D | 3.2e-07 | 2.3e-07 | 3.1e-07 | 2.867E-07 | 4.933E-08 | 2.300E-07 | 3.200E-07 |
| 25404 | 1271 | E1271D | D | 3.2e-07 | 2.3e-07 | 3e-07 | 2.833E-07 | 4.726E-08 | 2.300E-07 | 3.200E-07 |
| 25405 | 1271 | E1271C | D | 1.6e-07 | 2.1e-07 | 2e-07 | 1.900E-07 | 2.646E-08 | 1.600E-07 | 2.100E-07 |
| 25406 | 1271 | E1271E | D | 1.1e-07 | 6.4e-07 | 3.3e-07 | 3.600E-07 | 2.663E-07 | 1.100E-07 | 6.400E-07 |
| 25407 | 1271 | E1271Q | D | 9.1e-08 | 4.8e-07 | 2.6e-07 | 2.770E-07 | 1.951E-07 | 9.100E-08 | 4.800E-07 |
| 25408 | 1271 | E1271G | D | 1.7e-07 | 3.1e-07 | 5.8e-07 | 3.533E-07 | 2.084E-07 | 1.700E-07 | 5.800E-07 |
| 25409 | 1271 | E1271H | D | 2.2e-07 | 2.7e-07 | 4.9e-07 | 3.267E-07 | 1.436E-07 | 2.200E-07 | 4.900E-07 |
| 25410 | 1271 | E1271I | D | 2.2e-07 | 2.4e-07 | 4e-07 | 2.867E-07 | 9.866E-08 | 2.200E-07 | 4.000E-07 |
| 25411 | 1271 | E1271L | D | 1.2e-07 | 3e-07 | 1.7e-07 | 1.967E-07 | 9.292E-08 | 1.200E-07 | 3.000E-07 |
| 25412 | 1271 | E1271K | D | 1.2e-07 | 2.6e-07 | 2.3e-07 | 2.033E-07 | 7.371E-08 | 1.200E-07 | 2.600E-07 |
| 25413 | 1271 | E1271M | D | 1.6e-07 | 1.8e-07 | 1.5e-07 | 1.633E-07 | 1.528E-08 | 1.500E-07 | 1.800E-07 |
| 25414 | 1271 | E1271F | D | 6.3e-07 | 5.4e-07 | 2.4e-07 | 4.700E-07 | 2.042E-07 | 2.400E-07 | 6.300E-07 |

| | | | | | | | | | | |
|---|---|---|---|---|---|---|---|---|---|---|
| 25415 | 1271 | E1271P | D | 1.8e-07 | 1.5e-07 | 3.2e-07 | 2.167E-07 | 9.074E-08 | 1.500E-07 | 3.200E-07 |
| 25416 | 1271 | E1271S | D | 2.7e-07 | 2.7e-07 | 1.9e-07 | 2.433E-07 | 4.619E-08 | 1.900E-07 | 2.700E-07 |
| 25417 | 1271 | E1271T | D | 1.9e-07 | 2.7e-07 | 2.2e-07 | 2.267E-07 | 4.041E-08 | 1.900E-07 | 2.700E-07 |
| 25418 | 1271 | E1271W | D | 2.2e-07 | 4.1e-07 | 2.5e-07 | 2.933E-07 | 1.021E-07 | 2.200E-07 | 4.100E-07 |
| 25419 | 1271 | E1271Y | D | 2.1e-07 | 1.9e-07 | 2.6e-07 | 2.200E-07 | 3.606E-08 | 1.900E-07 | 2.600E-07 |
| 25420 | 1271 | E1271V | D | 2.8e-07 | 1.4e-07 | 3.1e-07 | 2.433E-07 | 9.074E-08 | 1.400E-07 | 3.100E-07 |
| 25421 | 1272 | C1272A | D | 1.6e-07 | 2.6e-07 | 1.4e-07 | 1.867E-07 | 6.429E-08 | 1.400E-07 | 2.600E-07 |
| 25422 | 1272 | C1272R | D | 2.6e-07 | 2.5e-07 | 1.3e-07 | 2.133E-07 | 7.234E-08 | 1.300E-07 | 2.600E-07 |
| 25423 | 1272 | C1272N | D | 3.4e-07 | 2.6e-07 | 1.4e-07 | 2.467E-07 | 1.007E-07 | 1.400E-07 | 3.400E-07 |
| 25424 | 1272 | C1272D | D | 1.3e-07 | 2e-07 | 6.9e-07 | 3.400E-07 | 3.051E-07 | 1.300E-07 | 6.900E-07 |
| 25425 | 1272 | C1272C | D | 1.3e-07 | 2.6e-07 | 1.9e-07 | 1.933E-07 | 6.506E-08 | 1.300E-07 | 2.600E-07 |
| 25426 | 1272 | C1272E | D | 1e-07 | 1.7e-07 | 4e-07 | 2.233E-07 | 1.570E-07 | 1.000E-07 | 4.000E-07 |
| 25427 | 1272 | C1272Q | D | 1.3e-07 | 3.3e-07 | 1.5e-07 | 2.033E-07 | 1.102E-07 | 1.300E-07 | 3.300E-07 |
| 25428 | 1272 | C1272G | D | 1.5e-07 | 2.9e-07 | 1.6e-07 | 2.000E-07 | 7.810E-08 | 1.500E-07 | 2.900E-07 |
| 25429 | 1272 | C1272H | D | 3.4e-07 | 1.8e-07 | 1.3e-07 | 2.167E-07 | 1.097E-07 | 1.300E-07 | 3.400E-07 |
| 25430 | 1272 | C1272I | D | 1.3e-07 | 1.9e-07 | 2.4e-07 | 1.867E-07 | 5.508E-08 | 1.300E-07 | 2.400E-07 |
| 25431 | 1272 | C1272L | D | 2.1e-07 | 2.7e-07 | 2.7e-07 | 2.500E-07 | 3.464E-08 | 2.100E-07 | 2.700E-07 |
| 25432 | 1272 | C1272K | D | 1.6e-07 | 2.4e-07 | 3.3e-07 | 2.433E-07 | 8.505E-08 | 1.600E-07 | 3.300E-07 |
| 25433 | 1272 | C1272M | D | 1.1e-07 | 2.7e-07 | 3.1e-07 | 2.300E-07 | 1.058E-07 | 1.100E-07 | 3.100E-07 |
| 25434 | 1272 | C1272F | D | 2.9e-07 | 6e-07 | 2.5e-07 | 3.800E-07 | 1.916E-07 | 2.500E-07 | 6.000E-07 |
| 25435 | 1272 | C1272P | D | 9.1e-07 | 1.7e-07 | 2.8e-07 | 4.533E-07 | 3.993E-07 | 1.700E-07 | 9.100E-07 |
| 25436 | 1272 | C1272S | D | 3.5e-07 | 1.7e-07 | 3.1e-07 | 2.767E-07 | 9.452E-08 | 1.700E-07 | 3.500E-07 |
| 25437 | 1272 | C1272T | D | 2.1e-07 | 7.4e-08 | 2e-07 | 1.613E-07 | 7.580E-08 | 7.400E-08 | 2.100E-07 |
| 25438 | 1272 | C1272W | D | 7.4e-07 | 1.4e-07 | 2.8e-07 | 3.867E-07 | 3.139E-07 | 1.400E-07 | 7.400E-07 |
| 25439 | 1272 | C1272Y | D | 1.8e-07 | 2.1e-07 | 2e-07 | 1.967E-07 | 1.528E-08 | 1.800E-07 | 2.100E-07 |
| 25440 | 1272 | C1272V | D | 2.9e-07 | 3.3e-07 | 7.6e-07 | 4.600E-07 | 2.606E-07 | 2.900E-07 | 7.600E-07 |
| 25441 | 1273 | E1273A | E | 2.5e-07 | 1.6e-07 | 2.8e-07 | 2.300E-07 | 6.245E-08 | 1.600E-07 | 2.800E-07 |
| 25442 | 1273 | E1273R | E | 2.8e-07 | 2.3e-07 | 4.3e-07 | 3.133E-07 | 1.041E-07 | 2.300E-07 | 4.300E-07 |
| 25443 | 1273 | E1273N | E | 1.6e-07 | 1.2e-07 | 2e-07 | 1.600E-07 | 4.000E-08 | 1.200E-07 | 2.000E-07 |
| 25444 | 1273 | E1273D | E | 3.1e-07 | 5.2e-07 | 1.7e-07 | 3.333E-07 | 1.762E-07 | 1.700E-07 | 5.200E-07 |
| 25445 | 1273 | E1273C | E | 2.2e-07 | 2.2e-07 | 2.3e-07 | 2.233E-07 | 5.774E-09 | 2.200E-07 | 2.300E-07 |
| 25446 | 1273 | E1273E | E | 1.7e-07 | 2e-07 | 4.1e-07 | 2.600E-07 | 1.308E-07 | 1.700E-07 | 4.100E-07 |
| 25447 | 1273 | E1273Q | E | 1.6e-07 | 1.5e-07 | 3.1e-07 | 2.067E-07 | 8.963E-08 | 1.500E-07 | 3.100E-07 |
| 25448 | 1273 | E1273G | E | 1.8e-07 | 3e-07 | 3.3e-07 | 2.700E-07 | 7.937E-08 | 1.800E-07 | 3.300E-07 |
| 25449 | 1273 | E1273H | E | 2.9e-07 | 2.4e-07 | 1.9e-07 | 2.400E-07 | 5.000E-08 | 1.900E-07 | 2.900E-07 |
| 25450 | 1273 | E1273I | E | 4.9e-07 | 4.6e-07 | 3.2e-07 | 4.233E-07 | 9.074E-08 | 3.200E-07 | 4.900E-07 |
| 25451 | 1273 | E1273L | E | 5e-07 | 2.5e-07 | 2.5e-07 | 3.333E-07 | 1.443E-07 | 2.500E-07 | 5.000E-07 |
| 25452 | 1273 | E1273K | E | 2.8e-07 | 3.4e-07 | 2.6e-07 | 2.933E-07 | 4.163E-08 | 2.600E-07 | 3.400E-07 |
| 25453 | 1273 | E1273M | E | 2.4e-07 | 1.6e-07 | 1.5e-07 | 1.833E-07 | 4.933E-08 | 1.500E-07 | 2.400E-07 |
| 25454 | 1273 | E1273F | E | 1.9e-07 | 1.7e-07 | 1.6e-07 | 1.733E-07 | 1.528E-08 | 1.600E-07 | 1.900E-07 |
| 25455 | 1273 | E1273P | E | 2.5e-07 | 3.6e-07 | 6.8e-07 | 4.300E-07 | 2.234E-07 | 2.500E-07 | 6.800E-07 |
| 25456 | 1273 | E1273S | E | 1.1e-07 | 2.3e-07 | 1.6e-07 | 1.667E-07 | 6.028E-08 | 1.100E-07 | 2.300E-07 |
| 25457 | 1273 | E1273T | E | 3.4e-07 | 2.9e-07 | 2e-07 | 2.767E-07 | 7.095E-08 | 2.000E-07 | 3.400E-07 |
| 25458 | 1273 | E1273W | E | 2.4e-07 | 2.2e-07 | 3.8e-07 | 2.800E-07 | 8.718E-08 | 2.200E-07 | 3.800E-07 |
| 25459 | 1273 | E1273Y | E | 1.6e-07 | 2.3e-07 | 8.1e-08 | 1.570E-07 | 7.455E-08 | 8.100E-08 | 2.300E-07 |
| 25460 | 1273 | E1273V | E | 5.8e-07 | 4.9e-07 | 2.7e-07 | 4.467E-07 | 1.595E-07 | 2.700E-07 | 5.800E-07 |
| 25461 | 1274 | V1274A | E | 3e-07 | 1.4e-07 | 2.4e-07 | 2.267E-07 | 8.083E-08 | 1.400E-07 | 3.000E-07 |
| 25462 | 1274 | V1274R | E | 2.5e-07 | 3.2e-07 | 2.2e-07 | 2.633E-07 | 5.132E-08 | 2.200E-07 | 3.200E-07 |
| 25463 | 1274 | V1274N | E | 1.6e-07 | 1.4e-07 | 1.4e-07 | 1.467E-07 | 1.155E-08 | 1.400E-07 | 1.600E-07 |
| 25464 | 1274 | V1274D | E | 1.6e-07 | 1.1e-07 | 1.8e-07 | 1.500E-07 | 3.606E-08 | 1.100E-07 | 1.800E-07 |
| 25465 | 1274 | V1274C | E | 2.5e-07 | 3.9e-07 | 1.2e-07 | 2.533E-07 | 1.350E-07 | 1.200E-07 | 3.900E-07 |
| 25466 | 1274 | V1274E | E | 1.3e-07 | 1e-07 | 1.2e-07 | 1.167E-07 | 1.528E-08 | 1.000E-07 | 1.300E-07 |
| 25467 | 1274 | V1274Q | E | 3e-07 | 1.8e-07 | 9e-08 | 1.900E-07 | 1.054E-07 | 9.000E-08 | 3.000E-07 |
| 25468 | 1274 | V1274G | E | 3.4e-07 | 7.9e-07 | 2.2e-07 | 4.500E-07 | 3.005E-07 | 2.200E-07 | 7.900E-07 |
| 25469 | 1274 | V1274H | E | 2.4e-07 | 2.6e-07 | 2.1e-07 | 2.367E-07 | 2.517E-08 | 2.100E-07 | 2.600E-07 |
| 25470 | 1274 | V1274I | E | 3.2e-07 | 2.8e-07 | 1.5e-07 | 2.500E-07 | 8.888E-08 | 1.500E-07 | 3.200E-07 |
| 25471 | 1274 | V1274L | E | 3.5e-07 | 3.1e-07 | 1.4e-07 | 2.667E-07 | 1.115E-07 | 1.400E-07 | 3.500E-07 |
| 25472 | 1274 | V1274K | E | 1.4e-07 | 2e-07 | 2.1e-07 | 1.833E-07 | 3.786E-08 | 1.400E-07 | 2.100E-07 |
| 25473 | 1274 | V1274M | E | 3.3e-07 | 1.4e-07 | 3.9e-07 | 2.867E-07 | 1.305E-07 | 1.400E-07 | 3.900E-07 |
| 25474 | 1274 | V1274F | E | 1.4e-07 | 2.7e-07 | 3.3e-07 | 2.467E-07 | 9.713E-08 | 1.400E-07 | 3.300E-07 |
| 25475 | 1274 | V1274P | E | 3e-07 | 4.2e-07 | 3e-07 | 3.400E-07 | 6.928E-08 | 3.000E-07 | 4.200E-07 |
| 25476 | 1274 | V1274S | E | 2e-07 | 7.3e-08 | 2.1e-07 | 1.610E-07 | 7.637E-08 | 7.300E-08 | 2.100E-07 |
| 25477 | 1274 | V1274T | E | 2.5e-07 | 3.3e-07 | 1.4e-07 | 2.400E-07 | 9.539E-08 | 1.400E-07 | 3.300E-07 |
| 25478 | 1274 | V1274W | E | 4.5e-07 | 2.9e-07 | 1.8e-07 | 3.067E-07 | 1.358E-07 | 1.800E-07 | 4.500E-07 |
| 25479 | 1274 | V1274Y | E | 2.8e-07 | 1.5e-07 | 2.2e-07 | 2.167E-07 | 6.506E-08 | 1.500E-07 | 2.800E-07 |
| 25480 | 1274 | V1274V | E | 4.3e-07 | 3.8e-07 | 2.1e-07 | 3.400E-07 | 1.153E-07 | 2.100E-07 | 4.300E-07 |
| 25481 | 1275 | Q1275A | E | 3.5e-07 | 1.2e-07 | 3.4e-07 | 2.700E-07 | 1.300E-07 | 1.200E-07 | 3.500E-07 |
| 25482 | 1275 | Q1275R | E | 2.7e-07 | 2.9e-07 | 3.5e-07 | 3.033E-07 | 4.163E-08 | 2.700E-07 | 3.500E-07 |
| 25483 | 1275 | Q1275N | E | 1.2e-07 | 6.5e-07 | 1e-07 | 2.900E-07 | 3.119E-07 | 1.000E-07 | 6.500E-07 |
| 25484 | 1275 | Q1275D | E | 6.3e-07 | 4.6e-07 | 2e-07 | 4.300E-07 | 2.166E-07 | 2.000E-07 | 6.300E-07 |
| 25485 | 1275 | Q1275C | E | 3.5e-07 | 2.7e-07 | 1.8e-07 | 2.667E-07 | 8.505E-08 | 1.800E-07 | 3.500E-07 |
| 25486 | 1275 | Q1275E | E | 2.5e-07 | 5.1e-07 | 5.5e-07 | 4.367E-07 | 1.629E-07 | 2.500E-07 | 5.500E-07 |
| 25487 | 1275 | Q1275Q | E | 2.2e-07 | 1.6e-07 | 2e-07 | 1.933E-07 | 3.055E-08 | 1.600E-07 | 2.200E-07 |
| 25488 | 1275 | Q1275G | E | 3e-07 | 3e-07 | 2.4e-07 | 2.800E-07 | 3.464E-08 | 2.400E-07 | 3.000E-07 |
| 25489 | 1275 | Q1275H | E | 3e-07 | 7.8e-08 | 4.5e-07 | 2.760E-07 | 1.872E-07 | 7.800E-08 | 4.500E-07 |
| 25490 | 1275 | Q1275I | E | 3.1e-07 | 3.5e-07 | 3.4e-07 | 3.333E-07 | 2.082E-08 | 3.100E-07 | 3.500E-07 |
| 25491 | 1275 | Q1275L | E | 2e-07 | 4.9e-07 | 2.4e-07 | 3.100E-07 | 1.572E-07 | 2.000E-07 | 4.900E-07 |
| 25492 | 1275 | Q1275K | E | 5.9e-07 | 1.1e-07 | 2.2e-07 | 3.067E-07 | 2.515E-07 | 1.100E-07 | 5.900E-07 |
| 25493 | 1275 | Q1275M | E | 5.5e-07 | 3.1e-07 | 1.5e-07 | 3.367E-07 | 2.013E-07 | 1.500E-07 | 5.500E-07 |
| 25494 | 1275 | Q1275F | E | 3.6e-07 | 2.7e-07 | 2.5e-07 | 2.933E-07 | 5.859E-08 | 2.500E-07 | 3.600E-07 |
| 25495 | 1275 | Q1275P | E | 1.2e-07 | 2.8e-07 | 3e-07 | 2.333E-07 | 9.866E-08 | 1.200E-07 | 3.000E-07 |
| 25496 | 1275 | Q1275S | E | 2.1e-07 | 1.9e-07 | 5.4e-07 | 3.133E-07 | 1.966E-07 | 1.900E-07 | 5.400E-07 |
| 25497 | 1275 | Q1275T | E | 2.8e-07 | 2.4e-07 | 3.7e-07 | 2.967E-07 | 6.658E-08 | 2.400E-07 | 3.700E-07 |
| 25498 | 1275 | Q1275W | E | 3.2e-07 | 4.1e-07 | 1.8e-07 | 3.033E-07 | 1.159E-07 | 1.800E-07 | 4.100E-07 |
| 25499 | 1275 | Q1275Y | E | 5.2e-07 | 1.6e-07 | 2.8e-07 | 3.200E-07 | 1.833E-07 | 1.600E-07 | 5.200E-07 |

| | | | | | | | | | | |
|---|---|---|---|---|---|---|---|---|---|---|
| 25500 | 1275 | Q1275V | E | 1.1e-07 | 4.3e-07 | 2e-07 | 2.467E-07 | 1.650E-07 | 1.100E-07 | 4.300E-07 |
| 25501 | 1276 | L1276A | E | 1.8e-07 | 4.4e-07 | 3.6e-07 | 3.267E-07 | 1.332E-07 | 1.800E-07 | 4.400E-07 |
| 25502 | 1276 | L1276R | E | 2.2e-07 | 1.5e-07 | 3.5e-07 | 2.400E-07 | 1.015E-07 | 1.500E-07 | 3.500E-07 |
| 25503 | 1276 | L1276N | E | 3e-07 | 1.7e-07 | 3.7e-07 | 2.800E-07 | 1.015E-07 | 1.700E-07 | 3.700E-07 |
| 25504 | 1276 | L1276D | E | 5.6e-07 | 1.3e-07 | 2.8e-07 | 3.233E-07 | 2.183E-07 | 1.300E-07 | 5.600E-07 |
| 25505 | 1276 | L1276C | E | 1.9e-07 | 3e-07 | 5.3e-07 | 3.400E-07 | 1.735E-07 | 1.900E-07 | 5.300E-07 |
| 25506 | 1276 | L1276E | E | 2e-07 | 4.9e-07 | 2.6e-07 | 3.167E-07 | 1.531E-07 | 2.000E-07 | 4.900E-07 |
| 25507 | 1276 | L1276Q | E | 3.6e-07 | 3.1e-07 | 3.7e-07 | 3.467E-07 | 3.215E-08 | 3.100E-07 | 3.700E-07 |
| 25508 | 1276 | L1276G | E | 2.6e-07 | 2.4e-07 | 3.7e-07 | 2.900E-07 | 7.000E-08 | 2.400E-07 | 3.700E-07 |
| 25509 | 1276 | L1276H | E | 2.1e-07 | 4.7e-07 | 3.9e-07 | 3.567E-07 | 1.332E-07 | 2.100E-07 | 4.700E-07 |
| 25510 | 1276 | L1276I | E | 1.6e-07 | 2.4e-07 | 2.8e-07 | 2.267E-07 | 6.110E-08 | 1.600E-07 | 2.800E-07 |
| 25511 | 1276 | L1276L | E | 2.3e-07 | 1.6e-07 | 7e-07 | 3.633E-07 | 2.937E-07 | 1.600E-07 | 7.000E-07 |
| 25512 | 1276 | L1276K | E | 4.5e-07 | 9e-07 | 1.8e-07 | 5.100E-07 | 3.637E-07 | 1.800E-07 | 9.000E-07 |
| 25513 | 1276 | L1276M | E | 5.1e-07 | 2.8e-07 | 2e-07 | 3.300E-07 | 1.609E-07 | 2.000E-07 | 5.100E-07 |
| 25514 | 1276 | L1276F | E | 2.1e-07 | 1.6e-07 | 1.2e-07 | 1.633E-07 | 4.509E-08 | 1.200E-07 | 2.100E-07 |
| 25515 | 1276 | L1276P | E | 1.4e-07 | 2.2e-07 | 1.7e-07 | 1.767E-07 | 4.041E-08 | 1.400E-07 | 2.200E-07 |
| 25516 | 1276 | L1276S | E | 3.4e-07 | 1.9e-07 | 2.1e-07 | 2.467E-07 | 8.145E-08 | 1.900E-07 | 3.400E-07 |
| 25517 | 1276 | L1276T | E | 2.6e-07 | 1.9e-07 | 4.1e-07 | 2.867E-07 | 1.124E-07 | 1.900E-07 | 4.100E-07 |
| 25518 | 1276 | L1276W | E | 7.2e-07 | 2.3e-07 | 2.1e-07 | 3.867E-07 | 2.888E-07 | 2.100E-07 | 7.200E-07 |
| 25519 | 1276 | L1276Y | E | 2.4e-07 | 2.9e-07 | 2.7e-07 | 2.667E-07 | 2.517E-08 | 2.400E-07 | 2.900E-07 |
| 25520 | 1276 | L1276V | E | 3.3e-07 | 2.7e-07 | 5.3e-07 | 3.767E-07 | 1.361E-07 | 2.700E-07 | 5.300E-07 |
| 25521 | 1277 | V1277A | E | 2.8e-07 | 3.3e-07 | 1.9e-07 | 2.667E-07 | 7.095E-08 | 1.900E-07 | 3.300E-07 |
| 25522 | 1277 | V1277R | E | 3.5e-07 | 3.6e-07 | 1.9e-07 | 3.000E-07 | 9.539E-08 | 1.900E-07 | 3.600E-07 |
| 25523 | 1277 | V1277N | E | 2.6e-07 | 2e-07 | 2.1e-07 | 2.233E-07 | 3.215E-08 | 2.000E-07 | 2.600E-07 |
| 25524 | 1277 | V1277D | E | 2.7e-07 | 2.3e-07 | 2.7e-07 | 2.567E-07 | 2.309E-08 | 2.300E-07 | 2.700E-07 |
| 25525 | 1277 | V1277C | E | 5.7e-07 | 4.4e-07 | 9.2e-08 | 3.673E-07 | 2.471E-07 | 9.200E-08 | 5.700E-07 |
| 25526 | 1277 | V1277E | E | 2.1e-07 | 1.7e-07 | 2.8e-07 | 2.200E-07 | 5.568E-08 | 1.700E-07 | 2.800E-07 |
| 25527 | 1277 | V1277Q | E | 2.3e-07 | 2.6e-07 | 5.4e-07 | 3.433E-07 | 1.710E-07 | 2.300E-07 | 5.400E-07 |
| 25528 | 1277 | V1277G | E | 2.3e-07 | 2.1e-07 | 2.7e-07 | 2.367E-07 | 3.055E-08 | 2.100E-07 | 2.700E-07 |
| 25529 | 1277 | V1277H | E | 3.7e-07 | 2.5e-07 | 2.2e-07 | 2.800E-07 | 7.937E-08 | 2.200E-07 | 3.700E-07 |
| 25530 | 1277 | V1277I | E | 2.7e-07 | 3.1e-07 | 1.4e-07 | 2.400E-07 | 8.888E-08 | 1.400E-07 | 3.100E-07 |
| 25531 | 1277 | V1277L | E | 4.2e-07 | 3.1e-07 | 3.9e-07 | 3.733E-07 | 5.686E-08 | 3.100E-07 | 4.200E-07 |
| 25532 | 1277 | V1277K | E | 8e-07 | 2.4e-07 | 2.9e-07 | 4.433E-07 | 3.099E-07 | 2.400E-07 | 8.000E-07 |
| 25533 | 1277 | V1277M | E | 4.9e-07 | 2e-07 | 5.8e-07 | 4.233E-07 | 1.986E-07 | 2.000E-07 | 5.800E-07 |
| 25534 | 1277 | V1277F | E | 2e-07 | 2.6e-07 | 2e-07 | 2.200E-07 | 3.464E-08 | 2.000E-07 | 2.600E-07 |
| 25535 | 1277 | V1277P | E | 5e-07 | 1.9e-07 | 1.9e-07 | 2.933E-07 | 1.790E-07 | 1.900E-07 | 5.000E-07 |
| 25536 | 1277 | V1277S | E | 3.4e-07 | 1.9e-07 | 1.5e-07 | 2.267E-07 | 1.002E-07 | 1.500E-07 | 3.400E-07 |
| 25537 | 1277 | V1277T | E | 3.1e-07 | 1.5e-07 | 3.4e-07 | 2.667E-07 | 1.021E-07 | 1.500E-07 | 3.400E-07 |
| 25538 | 1277 | V1277W | E | 2.4e-07 | 3.6e-07 | 2.6e-07 | 2.867E-07 | 6.429E-08 | 2.400E-07 | 3.600E-07 |
| 25539 | 1277 | V1277Y | E | 9.3e-08 | 2.5e-07 | 1.7e-07 | 1.710E-07 | 7.850E-08 | 9.300E-08 | 2.500E-07 |
| 25540 | 1277 | V1277V | E | 1.9e-07 | 7.5e-08 | 2.7e-07 | 1.783E-07 | 9.802E-08 | 7.500E-08 | 2.700E-07 |
| 25541 | 1278 | E1278A | E | 1.3e-07 | 3.5e-07 | 2.3e-07 | 2.367E-07 | 1.102E-07 | 1.300E-07 | 3.500E-07 |
| 25542 | 1278 | E1278R | E | 2.4e-07 | 3.3e-07 | 2.8e-07 | 2.833E-07 | 4.509E-08 | 2.400E-07 | 3.300E-07 |
| 25543 | 1278 | E1278N | E | 5.7e-07 | 2.8e-07 | 3.4e-07 | 3.967E-07 | 1.531E-07 | 2.800E-07 | 5.700E-07 |
| 25544 | 1278 | E1278D | E | 1.9e-07 | 2.5e-07 | 2.3e-07 | 2.233E-07 | 3.055E-08 | 1.900E-07 | 2.500E-07 |
| 25545 | 1278 | E1278C | E | 2.3e-07 | 2.9e-07 | 1.7e-07 | 2.300E-07 | 6.000E-08 | 1.700E-07 | 2.900E-07 |
| 25546 | 1278 | E1278E | E | 1.1e-07 | 2.4e-07 | 2.4e-07 | 1.967E-07 | 7.506E-08 | 1.100E-07 | 2.400E-07 |
| 25547 | 1278 | E1278Q | E | 2.8e-07 | 2.7e-07 | 2.3e-07 | 2.600E-07 | 2.646E-08 | 2.300E-07 | 2.800E-07 |
| 25548 | 1278 | E1278G | E | 9.4e-07 | 4.1e-07 | 8.5e-08 | 4.783E-07 | 4.316E-07 | 8.500E-08 | 9.400E-07 |
| 25549 | 1278 | E1278H | E | 5.2e-07 | 3.1e-07 | 1.1e-07 | 3.133E-07 | 2.050E-07 | 1.100E-07 | 5.200E-07 |
| 25550 | 1278 | E1278I | E | 1.2e-07 | 2.8e-07 | 3e-07 | 2.333E-07 | 9.866E-08 | 1.200E-07 | 3.000E-07 |
| 25551 | 1278 | E1278L | E | 2.8e-07 | 3e-07 | 3.2e-07 | 3.000E-07 | 2.000E-08 | 2.800E-07 | 3.200E-07 |
| 25552 | 1278 | E1278K | E | 3.4e-07 | 6e-07 | 3.4e-07 | 4.267E-07 | 1.501E-07 | 3.400E-07 | 6.000E-07 |
| 25553 | 1278 | E1278M | E | 2.3e-07 | 1.2e-07 | 3.2e-07 | 2.233E-07 | 1.002E-07 | 1.200E-07 | 3.200E-07 |
| 25554 | 1278 | E1278F | E | 4.7e-07 | 2.3e-07 | 1.4e-07 | 2.800E-07 | 1.706E-07 | 1.400E-07 | 4.700E-07 |
| 25555 | 1278 | E1278P | E | 2.9e-07 | 1.6e-07 | 2.2e-07 | 2.233E-07 | 6.506E-08 | 1.600E-07 | 2.900E-07 |
| 25556 | 1278 | E1278S | E | 1.3e-07 | 4.1e-07 | 3.8e-07 | 3.067E-07 | 1.537E-07 | 1.300E-07 | 4.100E-07 |
| 25557 | 1278 | E1278T | E | 3.4e-07 | 8.5e-07 | 2.6e-07 | 4.833E-07 | 3.201E-07 | 2.600E-07 | 8.500E-07 |
| 25558 | 1278 | E1278W | E | 1.1e-07 | 5.7e-07 | 5.1e-07 | 3.967E-07 | 2.501E-07 | 1.100E-07 | 5.700E-07 |
| 25559 | 1278 | E1278Y | E | 3.8e-07 | 3.8e-07 | 3.4e-07 | 3.667E-07 | 2.309E-08 | 3.400E-07 | 3.800E-07 |
| 25560 | 1278 | E1278V | E | 2.8e-07 | 1.6e-07 | 2.4e-07 | 2.267E-07 | 6.110E-08 | 1.600E-07 | 2.800E-07 |
| 25561 | 1279 | S1279A | E | 3e-07 | 6.8e-07 | 1.5e-07 | 3.767E-07 | 2.732E-07 | 1.500E-07 | 6.800E-07 |
| 25562 | 1279 | S1279R | E | 1.7e-07 | 2.8e-07 | 3.2e-07 | 2.567E-07 | 7.767E-08 | 1.700E-07 | 3.200E-07 |
| 25563 | 1279 | S1279N | E | 6.5e-07 | 5.1e-07 | 1.7e-07 | 4.433E-07 | 2.468E-07 | 1.700E-07 | 6.500E-07 |
| 25564 | 1279 | S1279D | E | 3.7e-07 | 2.1e-07 | 2.5e-07 | 2.767E-07 | 8.327E-08 | 2.100E-07 | 3.700E-07 |
| 25565 | 1279 | S1279C | E | 1.3e-07 | 1.1e-07 | 2.8e-07 | 1.733E-07 | 9.292E-08 | 1.100E-07 | 2.800E-07 |
| 25566 | 1279 | S1279E | E | 3.1e-07 | 2.6e-07 | 3.3e-07 | 3.000E-07 | 3.606E-08 | 2.600E-07 | 3.300E-07 |
| 25567 | 1279 | S1279Q | E | 2.5e-07 | 1.9e-07 | 2.3e-07 | 2.233E-07 | 3.055E-08 | 1.900E-07 | 2.500E-07 |
| 25568 | 1279 | S1279G | E | 1.6e-07 | 2.9e-07 | 2.2e-07 | 2.233E-07 | 6.506E-08 | 1.600E-07 | 2.900E-07 |
| 25569 | 1279 | S1279H | E | 2.8e-07 | 1.3e-07 | 5e-07 | 3.033E-07 | 1.861E-07 | 1.300E-07 | 5.000E-07 |
| 25570 | 1279 | S1279I | E | 2.4e-07 | 1.9e-07 | 2.4e-07 | 2.233E-07 | 2.887E-08 | 1.900E-07 | 2.400E-07 |
| 25571 | 1279 | S1279L | E | 4.8e-07 | 2.4e-07 | 2.3e-07 | 3.167E-07 | 1.415E-07 | 2.300E-07 | 4.800E-07 |
| 25572 | 1279 | S1279K | E | 2.8e-07 | 2e-07 | 1.3e-07 | 2.033E-07 | 7.506E-08 | 1.300E-07 | 2.800E-07 |
| 25573 | 1279 | S1279M | E | 1.4e-07 | 2.7e-07 | 3.1e-07 | 2.400E-07 | 8.888E-08 | 1.400E-07 | 3.100E-07 |
| 25574 | 1279 | S1279F | E | 4.2e-07 | 6.7e-08 | 1.4e-07 | 2.090E-07 | 1.863E-07 | 6.700E-08 | 4.200E-07 |
| 25575 | 1279 | S1279P | E | 2.7e-07 | 1.2e-07 | 3.9e-07 | 2.600E-07 | 1.353E-07 | 1.200E-07 | 3.900E-07 |
| 25576 | 1279 | S1279S | E | 4.5e-07 | 2.5e-07 | 2.9e-07 | 3.300E-07 | 1.058E-07 | 2.500E-07 | 4.500E-07 |
| 25577 | 1279 | S1279T | E | 2.4e-07 | 2.7e-07 | 4.6e-07 | 3.233E-07 | 1.193E-07 | 2.400E-07 | 4.600E-07 |
| 25578 | 1279 | S1279W | E | 3e-07 | 1.8e-07 | 3e-07 | 2.600E-07 | 6.928E-08 | 1.800E-07 | 3.000E-07 |
| 25579 | 1279 | S1279Y | E | 3.6e-07 | 3.4e-07 | 1.8e-07 | 2.933E-07 | 9.866E-08 | 1.800E-07 | 3.600E-07 |
| 25580 | 1279 | S1279V | E | 3.7e-07 | 3e-07 | 1.7e-07 | 2.800E-07 | 1.015E-07 | 1.700E-07 | 3.700E-07 |
| 25581 | 1280 | G1280A | E | 1.4e-07 | 1.3e-07 | 1.3e-07 | 1.333E-07 | 5.774E-09 | 1.300E-07 | 1.400E-07 |
| 25582 | 1280 | G1280R | E | 7.2e-08 | 3.3e-07 | 2.2e-07 | 2.073E-07 | 1.295E-07 | 7.200E-08 | 3.300E-07 |
| 25583 | 1280 | G1280N | E | 1.6e-07 | 2.9e-07 | 4.1e-07 | 2.867E-07 | 1.250E-07 | 1.600E-07 | 4.100E-07 |
| 25584 | 1280 | G1280D | E | 1.7e-07 | 1.3e-07 | 3.6e-07 | 2.200E-07 | 1.229E-07 | 1.300E-07 | 3.600E-07 |

| | | | | | | | | | | |
|---|---|---|---|---|---|---|---|---|---|---|
| 25585 | 1280 | G1280C | E | 2.8e-07 | 2.7e-07 | 2.5e-07 | 2.667E-07 | 1.528E-08 | 2.500E-07 | 2.800E-07 |
| 25586 | 1280 | G1280E | E | 2.7e-07 | 2.3e-07 | 3.3e-07 | 2.767E-07 | 5.033E-08 | 2.300E-07 | 3.300E-07 |
| 25587 | 1280 | G1280Q | E | 2.7e-07 | 2.9e-07 | 2e-07 | 2.533E-07 | 4.726E-08 | 2.000E-07 | 2.900E-07 |
| 25588 | 1280 | G1280G | E | 4e-07 | 4.2e-07 | 1.6e-07 | 3.267E-07 | 1.447E-07 | 1.600E-07 | 4.200E-07 |
| 25589 | 1280 | G1280H | E | 3.6e-07 | 1e-07 | 2.4e-07 | 2.333E-07 | 1.301E-07 | 1.000E-07 | 3.600E-07 |
| 25590 | 1280 | G1280I | E | 4.2e-07 | 1.9e-07 | 3.7e-07 | 3.267E-07 | 1.210E-07 | 1.900E-07 | 4.200E-07 |
| 25591 | 1280 | G1280L | E | 3.9e-07 | 3.5e-07 | 3.4e-07 | 3.600E-07 | 2.646E-08 | 3.400E-07 | 3.900E-07 |
| 25592 | 1280 | G1280K | E | 3.2e-07 | 1.5e-07 | 2.5e-07 | 2.400E-07 | 8.544E-08 | 1.500E-07 | 3.200E-07 |
| 25593 | 1280 | G1280M | E | 3.6e-07 | 3e-07 | 1.5e-07 | 2.700E-07 | 1.082E-07 | 1.500E-07 | 3.600E-07 |
| 25594 | 1280 | G1280F | E | 4e-07 | 1.4e-07 | 1.5e-07 | 2.300E-07 | 1.473E-07 | 1.400E-07 | 4.000E-07 |
| 25595 | 1280 | G1280P | E | 4.1e-07 | 1.9e-07 | 1.1e-07 | 2.367E-07 | 1.553E-07 | 1.100E-07 | 4.100E-07 |
| 25596 | 1280 | G1280S | E | 1.4e-07 | 2.6e-07 | 2.8e-07 | 2.267E-07 | 7.572E-08 | 1.400E-07 | 2.800E-07 |
| 25597 | 1280 | G1280T | E | 2.3e-07 | 1.9e-07 | 1.8e-07 | 2.000E-07 | 2.646E-08 | 1.800E-07 | 2.300E-07 |
| 25598 | 1280 | G1280W | E | 4.4e-07 | 1.7e-07 | 2.8e-07 | 2.967E-07 | 1.358E-07 | 1.700E-07 | 4.400E-07 |
| 25599 | 1280 | G1280Y | E | 2.1e-07 | 3.6e-07 | 3.7e-07 | 3.133E-07 | 8.963E-08 | 2.100E-07 | 3.700E-07 |
| 25600 | 1280 | G1280V | E | 1.8e-07 | 2.1e-07 | 4e-07 | 2.633E-07 | 1.193E-07 | 1.800E-07 | 4.000E-07 |
| 25601 | 1281 | G1281A | E | 2.3e-07 | 2.3e-07 | 3e-07 | 2.533E-07 | 4.041E-08 | 2.300E-07 | 3.000E-07 |
| 25602 | 1281 | G1281R | E | 2.8e-07 | 2.1e-07 | 1.9e-07 | 2.267E-07 | 4.726E-08 | 1.900E-07 | 2.800E-07 |
| 25603 | 1281 | G1281N | E | 1.5e-07 | 2.6e-07 | 2.7e-07 | 2.267E-07 | 6.658E-08 | 1.500E-07 | 2.700E-07 |
| 25604 | 1281 | G1281D | E | 3.2e-07 | 2.2e-07 | 2e-07 | 2.467E-07 | 6.429E-08 | 2.000E-07 | 3.200E-07 |
| 25605 | 1281 | G1281C | E | 1.9e-07 | 1.8e-07 | 2.2e-07 | 1.967E-07 | 2.082E-08 | 1.800E-07 | 2.200E-07 |
| 25606 | 1281 | G1281E | E | 1.4e-07 | 1.3e-07 | 3.8e-07 | 2.167E-07 | 1.415E-07 | 1.300E-07 | 3.800E-07 |
| 25607 | 1281 | G1281Q | E | 3.8e-07 | 1.4e-07 | 3e-07 | 2.733E-07 | 1.222E-07 | 1.400E-07 | 3.800E-07 |
| 25608 | 1281 | G1281G | E | 1.4e-07 | 4.4e-07 | 3e-07 | 2.933E-07 | 1.501E-07 | 1.400E-07 | 4.400E-07 |
| 25609 | 1281 | G1281H | E | 1.5e-07 | 1.8e-07 | 3.1e-07 | 2.133E-07 | 8.505E-08 | 1.500E-07 | 3.100E-07 |
| 25610 | 1281 | G1281I | E | 1.4e-07 | 2.2e-07 | 2e-07 | 1.867E-07 | 4.163E-08 | 1.400E-07 | 2.200E-07 |
| 25611 | 1281 | G1281L | E | 1.5e-07 | 3.4e-07 | 3.7e-07 | 2.867E-07 | 1.193E-07 | 1.500E-07 | 3.700E-07 |
| 25612 | 1281 | G1281K | E | 3.5e-07 | 3.4e-07 | 3.6e-07 | 3.500E-07 | 1.000E-08 | 3.400E-07 | 3.600E-07 |
| 25613 | 1281 | G1281M | E | 1.9e-07 | 6e-07 | 2.7e-07 | 3.533E-07 | 2.173E-07 | 1.900E-07 | 6.000E-07 |
| 25614 | 1281 | G1281F | E | 4.9e-07 | 3.5e-07 | 1.2e-07 | 3.200E-07 | 1.868E-07 | 1.200E-07 | 4.900E-07 |
| 25615 | 1281 | G1281P | E | 1.6e-07 | 2.7e-07 | 3e-07 | 2.433E-07 | 7.371E-08 | 1.600E-07 | 3.000E-07 |
| 25616 | 1281 | G1281S | E | 1.6e-07 | 1.8e-07 | 2.4e-07 | 1.933E-07 | 4.163E-08 | 1.600E-07 | 2.400E-07 |
| 25617 | 1281 | G1281T | E | 3.2e-07 | 1.5e-07 | 2.4e-07 | 2.367E-07 | 8.505E-08 | 1.500E-07 | 3.200E-07 |
| 25618 | 1281 | G1281W | E | 2.6e-07 | 2.4e-07 | 3.1e-07 | 2.700E-07 | 3.606E-08 | 2.400E-07 | 3.100E-07 |
| 25619 | 1281 | G1281Y | E | 5.6e-07 | 2.9e-07 | 1.8e-07 | 3.433E-07 | 1.955E-07 | 1.800E-07 | 5.600E-07 |
| 25620 | 1281 | G1281V | E | 1.4e-07 | 1.5e-07 | 1.3e-07 | 1.400E-07 | 1.000E-08 | 1.300E-07 | 1.500E-07 |
| 25621 | 1282 | G1282A | E | 2.2e-07 | 3.5e-07 | 2.7e-07 | 2.800E-07 | 6.557E-08 | 2.200E-07 | 3.500E-07 |
| 25622 | 1282 | G1282R | E | 3.2e-07 | 2.8e-07 | 2.7e-07 | 2.900E-07 | 2.646E-08 | 2.700E-07 | 3.200E-07 |
| 25623 | 1282 | G1282N | E | 1.4e-07 | 3.1e-07 | 2.3e-07 | 2.267E-07 | 8.505E-08 | 1.400E-07 | 3.100E-07 |
| 25624 | 1282 | G1282D | E | 3e-07 | 2.6e-07 | 2.2e-07 | 2.600E-07 | 4.000E-08 | 2.200E-07 | 3.000E-07 |
| 25625 | 1282 | G1282C | E | 1.8e-07 | 2e-07 | 1.4e-07 | 1.733E-07 | 3.055E-08 | 1.400E-07 | 2.000E-07 |
| 25626 | 1282 | G1282E | E | 3.3e-07 | 1.1e-07 | 2.3e-07 | 2.233E-07 | 1.102E-07 | 1.100E-07 | 3.300E-07 |
| 25627 | 1282 | G1282Q | E | 2.3e-07 | 2.8e-07 | 1.6e-07 | 2.233E-07 | 6.028E-08 | 1.600E-07 | 2.800E-07 |
| 25628 | 1282 | G1282G | E | 3.5e-07 | 2.2e-07 | 2.2e-07 | 2.633E-07 | 7.506E-08 | 2.200E-07 | 3.500E-07 |
| 25629 | 1282 | G1282H | E | 1.6e-07 | 3.9e-07 | 3.1e-07 | 2.867E-07 | 1.168E-07 | 1.600E-07 | 3.900E-07 |
| 25630 | 1282 | G1282I | E | 1.8e-07 | 3e-07 | 3.4e-07 | 2.733E-07 | 8.327E-08 | 1.800E-07 | 3.400E-07 |
| 25631 | 1282 | G1282L | E | 2.7e-07 | 3.8e-07 | 3.8e-07 | 3.433E-07 | 6.351E-08 | 2.700E-07 | 3.800E-07 |
| 25632 | 1282 | G1282K | E | 1.8e-07 | 4e-07 | 2.9e-07 | 2.900E-07 | 1.100E-07 | 1.800E-07 | 4.000E-07 |
| 25633 | 1282 | G1282M | E | 3.3e-07 | 4.3e-07 | 1.6e-07 | 3.067E-07 | 1.365E-07 | 1.600E-07 | 4.300E-07 |
| 25634 | 1282 | G1282F | E | 3.7e-07 | 2.4e-07 | 3e-07 | 3.033E-07 | 6.506E-08 | 2.400E-07 | 3.700E-07 |
| 25635 | 1282 | G1282P | E | 3.7e-07 | 3.4e-07 | 2.9e-07 | 3.333E-07 | 4.041E-08 | 2.900E-07 | 3.700E-07 |
| 25636 | 1282 | G1282S | E | 4.5e-07 | 2e-07 | 2.5e-07 | 3.000E-07 | 1.323E-07 | 2.000E-07 | 4.500E-07 |
| 25637 | 1282 | G1282T | E | 2.3e-07 | 2e-07 | 3.9e-07 | 2.733E-07 | 1.021E-07 | 2.000E-07 | 3.900E-07 |
| 25638 | 1282 | G1282W | E | 3e-07 | 2.9e-07 | 2.1e-07 | 2.667E-07 | 4.933E-08 | 2.100E-07 | 3.000E-07 |
| 25639 | 1282 | G1282Y | E | 3.1e-07 | 3.3e-07 | 4.8e-07 | 3.733E-07 | 9.292E-08 | 3.100E-07 | 4.800E-07 |
| 25640 | 1282 | G1282V | E | 2.4e-07 | 1.6e-07 | 2.5e-07 | 2.167E-07 | 4.933E-08 | 1.600E-07 | 2.500E-07 |
| 25641 | 1283 | L1283A | E | 2.4e-07 | 2.4e-07 | 1.6e-07 | 2.133E-07 | 4.619E-08 | 1.600E-07 | 2.400E-07 |
| 25642 | 1283 | L1283R | E | 3e-07 | 1.9e-07 | 2.4e-07 | 2.433E-07 | 5.508E-08 | 1.900E-07 | 3.000E-07 |
| 25643 | 1283 | L1283N | E | 2.5e-07 | 2e-07 | 4.7e-07 | 3.067E-07 | 1.436E-07 | 2.000E-07 | 4.700E-07 |
| 25644 | 1283 | L1283D | E | 1.9e-07 | 3.1e-07 | 2.3e-07 | 2.433E-07 | 6.110E-08 | 1.900E-07 | 3.100E-07 |
| 25645 | 1283 | L1283C | E | 5.4e-08 | 1.7e-07 | 3e-07 | 1.747E-07 | 1.231E-07 | 5.400E-08 | 3.000E-07 |
| 25646 | 1283 | L1283E | E | 1.6e-07 | 1.9e-07 | 7.4e-08 | 1.413E-07 | 6.021E-08 | 7.400E-08 | 1.900E-07 |
| 25647 | 1283 | L1283Q | E | 2.8e-07 | 2.3e-07 | 2.2e-07 | 2.433E-07 | 3.215E-08 | 2.200E-07 | 2.800E-07 |
| 25648 | 1283 | L1283G | E | 2.4e-07 | 2e-07 | 5.2e-07 | 3.200E-07 | 1.744E-07 | 2.000E-07 | 5.200E-07 |
| 25649 | 1283 | L1283H | E | 2.9e-07 | 7e-07 | 3.2e-07 | 4.367E-07 | 2.285E-07 | 2.900E-07 | 7.000E-07 |
| 25650 | 1283 | L1283I | E | 4.1e-07 | 1.1e-07 | 4e-07 | 3.067E-07 | 1.704E-07 | 1.100E-07 | 4.100E-07 |
| 25651 | 1283 | L1283L | E | 7.9e-08 | 3.2e-07 | 2.2e-07 | 2.063E-07 | 1.211E-07 | 7.900E-08 | 3.200E-07 |
| 25652 | 1283 | L1283K | E | 3.2e-07 | 2.9e-07 | 2.4e-07 | 2.833E-07 | 4.041E-08 | 2.400E-07 | 3.200E-07 |
| 25653 | 1283 | L1283M | E | 2.1e-07 | 6e-07 | 2.6e-07 | 3.567E-07 | 2.122E-07 | 2.100E-07 | 6.000E-07 |
| 25654 | 1283 | L1283F | E | 1.9e-07 | 2.7e-07 | 2.7e-07 | 2.433E-07 | 4.619E-08 | 1.900E-07 | 2.700E-07 |
| 25655 | 1283 | L1283P | E | 1.3e-07 | 1.7e-07 | 3.1e-07 | 2.033E-07 | 9.452E-08 | 1.300E-07 | 3.100E-07 |
| 25656 | 1283 | L1283S | E | 5.4e-08 | 3.6e-07 | 5.2e-07 | 3.113E-07 | 2.368E-07 | 5.400E-08 | 5.200E-07 |
| 25657 | 1283 | L1283T | E | 1.9e-07 | 2.7e-07 | 1.7e-07 | 2.100E-07 | 5.292E-08 | 1.700E-07 | 2.700E-07 |
| 25658 | 1283 | L1283W | E | 5.2e-07 | 4.3e-07 | 3.6e-07 | 4.367E-07 | 8.021E-08 | 3.600E-07 | 5.200E-07 |
| 25659 | 1283 | L1283Y | E | 6.1e-07 | 1.6e-07 | 1.9e-07 | 3.200E-07 | 2.516E-07 | 1.600E-07 | 6.100E-07 |
| 25660 | 1283 | L1283V | E | 1.3e-07 | 2.2e-07 | 1.2e-07 | 1.567E-07 | 5.508E-08 | 1.200E-07 | 2.200E-07 |
| 25661 | 1284 | V1284A | E | 2.7e-07 | 5.2e-07 | 2e-07 | 3.300E-07 | 1.682E-07 | 2.000E-07 | 5.200E-07 |
| 25662 | 1284 | V1284R | E | 1.3e-07 | 2.7e-07 | 3.5e-07 | 2.500E-07 | 1.114E-07 | 1.300E-07 | 3.500E-07 |
| 25663 | 1284 | V1284N | E | 3.8e-07 | 2.7e-07 | 1.1e-07 | 2.533E-07 | 1.358E-07 | 1.100E-07 | 3.800E-07 |
| 25664 | 1284 | V1284D | E | 2.6e-07 | 3.4e-07 | 3e-07 | 3.000E-07 | 4.000E-08 | 2.600E-07 | 3.400E-07 |
| 25665 | 1284 | V1284C | E | 2.9e-07 | 3.5e-07 | 3.5e-07 | 3.300E-07 | 3.464E-08 | 2.900E-07 | 3.500E-07 |
| 25666 | 1284 | V1284E | E | 5.2e-07 | 2.1e-07 | 2.8e-07 | 3.367E-07 | 1.626E-07 | 2.100E-07 | 5.200E-07 |
| 25667 | 1284 | V1284Q | E | 4.6e-07 | 3.1e-07 | 2.1e-07 | 3.267E-07 | 1.258E-07 | 2.100E-07 | 4.600E-07 |
| 25668 | 1284 | V1284G | E | 1.8e-07 | 2.4e-07 | 3.4e-07 | 2.533E-07 | 8.083E-08 | 1.800E-07 | 3.400E-07 |
| 25669 | 1284 | V1284H | E | 1.2e-07 | 3.3e-07 | 2.7e-07 | 2.400E-07 | 1.082E-07 | 1.200E-07 | 3.300E-07 |

| | | | | | | | | | | |
|---|---|---|---|---|---|---|---|---|---|---|
| 25670 | 1284 | V1284I | E | 4.1e-07 | 3.3e-07 | 2.7e-07 | 3.367E-07 | 7.024E-08 | 2.700E-07 | 4.100E-07 |
| 25671 | 1284 | V1284L | E | 1e-07 | 2.2e-07 | 1.9e-07 | 1.700E-07 | 6.245E-08 | 1.000E-07 | 2.200E-07 |
| 25672 | 1284 | V1284K | E | 2.5e-07 | 3e-07 | 1.2e-07 | 2.233E-07 | 9.292E-08 | 1.200E-07 | 3.000E-07 |
| 25673 | 1284 | V1284M | E | 1.7e-07 | 2.2e-07 | 3e-07 | 2.300E-07 | 6.557E-08 | 1.700E-07 | 3.000E-07 |
| 25674 | 1284 | V1284F | E | 4.1e-08 | 2e-07 | 1.9e-07 | 1.437E-07 | 8.905E-08 | 4.100E-08 | 2.000E-07 |
| 25675 | 1284 | V1284P | E | 3.9e-07 | 2.3e-07 | 2.5e-07 | 2.900E-07 | 8.718E-08 | 2.300E-07 | 3.900E-07 |
| 25676 | 1284 | V1284S | E | 5.1e-08 | 5.4e-07 | 2.4e-07 | 2.770E-07 | 2.466E-07 | 5.100E-08 | 5.400E-07 |
| 25677 | 1284 | V1284T | E | 2.9e-07 | 2.4e-07 | 3.6e-07 | 2.967E-07 | 6.028E-08 | 2.400E-07 | 3.600E-07 |
| 25678 | 1284 | V1284W | E | 2e-07 | 4.1e-07 | 4e-07 | 3.367E-07 | 1.185E-07 | 2.000E-07 | 4.100E-07 |
| 25679 | 1284 | V1284Y | E | 1.7e-07 | 1.2e-07 | 3e-07 | 1.967E-07 | 9.292E-08 | 1.200E-07 | 3.000E-07 |
| 25680 | 1284 | V1284V | E | 2e-07 | 1.6e-07 | 3.8e-07 | 2.467E-07 | 1.172E-07 | 1.600E-07 | 3.800E-07 |
| 25681 | 1285 | Q1285A | E | 1.5e-07 | 2e-07 | 2e-07 | 1.833E-07 | 2.887E-08 | 1.500E-07 | 2.000E-07 |
| 25682 | 1285 | Q1285R | E | 2.8e-07 | 1.8e-07 | 2e-07 | 2.200E-07 | 5.292E-08 | 1.800E-07 | 2.800E-07 |
| 25683 | 1285 | Q1285N | E | 1.3e-07 | 1.2e-07 | 1.4e-07 | 1.300E-07 | 1.000E-08 | 1.200E-07 | 1.400E-07 |
| 25684 | 1285 | Q1285D | E | 2.4e-07 | 2.7e-07 | 1.6e-07 | 2.233E-07 | 5.686E-08 | 1.600E-07 | 2.700E-07 |
| 25685 | 1285 | Q1285C | E | 3.2e-07 | 2.2e-07 | 1.8e-07 | 2.400E-07 | 7.211E-08 | 1.800E-07 | 3.200E-07 |
| 25686 | 1285 | Q1285E | E | 1.8e-07 | 1.8e-07 | 2.4e-07 | 2.000E-07 | 3.464E-08 | 1.800E-07 | 2.400E-07 |
| 25687 | 1285 | Q1285Q | E | 3.5e-07 | 2.3e-07 | 1.8e-07 | 2.533E-07 | 8.737E-08 | 1.800E-07 | 3.500E-07 |
| 25688 | 1285 | Q1285G | E | 1.2e-07 | 2.9e-07 | 1.7e-07 | 1.933E-07 | 8.737E-08 | 1.200E-07 | 2.900E-07 |
| 25689 | 1285 | Q1285H | E | 4.8e-07 | 2.5e-07 | 2.7e-07 | 3.333E-07 | 1.274E-07 | 2.500E-07 | 4.800E-07 |
| 25690 | 1285 | Q1285I | E | 2.4e-07 | 2.2e-07 | 3e-07 | 2.533E-07 | 4.163E-08 | 2.200E-07 | 3.000E-07 |
| 25691 | 1285 | Q1285L | E | 1.5e-07 | 2e-07 | 3.5e-07 | 2.333E-07 | 1.041E-07 | 1.500E-07 | 3.500E-07 |
| 25692 | 1285 | Q1285K | E | 2.4e-07 | 1.7e-07 | 2e-07 | 2.033E-07 | 3.512E-08 | 1.700E-07 | 2.400E-07 |
| 25693 | 1285 | Q1285M | E | 2.1e-07 | 4.1e-07 | 3.2e-07 | 3.133E-07 | 1.002E-07 | 2.100E-07 | 4.100E-07 |
| 25694 | 1285 | Q1285F | E | 1.7e-07 | 1.3e-07 | 1.3e-07 | 1.433E-07 | 2.309E-08 | 1.300E-07 | 1.700E-07 |
| 25695 | 1285 | Q1285P | E | 3.3e-07 | 2.4e-07 | 2.1e-07 | 2.600E-07 | 6.245E-08 | 2.100E-07 | 3.300E-07 |
| 25696 | 1285 | Q1285S | E | 1.4e-07 | 1.6e-07 | 2.7e-07 | 1.900E-07 | 7.000E-08 | 1.400E-07 | 2.700E-07 |
| 25697 | 1285 | Q1285T | E | 2.9e-07 | 5.3e-07 | 3.1e-07 | 3.767E-07 | 1.332E-07 | 2.900E-07 | 5.300E-07 |
| 25698 | 1285 | Q1285W | E | 4e-07 | 3.6e-07 | 4.5e-07 | 4.033E-07 | 4.509E-08 | 3.600E-07 | 4.500E-07 |
| 25699 | 1285 | Q1285Y | E | 2.7e-07 | 2.9e-07 | 3.6e-07 | 3.067E-07 | 4.726E-08 | 2.700E-07 | 3.600E-07 |
| 25700 | 1285 | Q1285V | E | 2.1e-07 | 6.6e-07 | 3.7e-07 | 4.133E-07 | 2.281E-07 | 2.100E-07 | 6.600E-07 |
| 25701 | 1286 | P1286A | E | 1.4e-07 | 4.6e-07 | 3.5e-07 | 3.167E-07 | 1.626E-07 | 1.400E-07 | 4.600E-07 |
| 25702 | 1286 | P1286R | E | 1.9e-07 | 1.2e-07 | 1.7e-07 | 1.600E-07 | 3.606E-08 | 1.200E-07 | 1.900E-07 |
| 25703 | 1286 | P1286N | E | 4.4e-07 | 4.4e-07 | 1.4e-07 | 3.400E-07 | 1.732E-07 | 1.400E-07 | 4.400E-07 |
| 25704 | 1286 | P1286D | E | 5e-07 | 3.5e-07 | 2.4e-07 | 3.633E-07 | 1.305E-07 | 2.400E-07 | 5.000E-07 |
| 25705 | 1286 | P1286C | E | 2e-07 | 1.4e-07 | 1.6e-07 | 1.667E-07 | 3.055E-08 | 1.400E-07 | 2.000E-07 |
| 25706 | 1286 | P1286E | E | 2.7e-07 | 1.8e-07 | 2.6e-07 | 2.367E-07 | 4.933E-08 | 1.800E-07 | 2.700E-07 |
| 25707 | 1286 | P1286Q | E | 2.6e-07 | 3.5e-07 | 1.8e-07 | 2.633E-07 | 8.505E-08 | 1.800E-07 | 3.500E-07 |
| 25708 | 1286 | P1286G | E | 2.8e-07 | 1.2e-07 | 1.7e-07 | 1.900E-07 | 8.185E-08 | 1.200E-07 | 2.800E-07 |
| 25709 | 1286 | P1286H | E | 1.8e-07 | 2.3e-07 | 3.2e-07 | 2.433E-07 | 7.095E-08 | 1.800E-07 | 3.200E-07 |
| 25710 | 1286 | P1286I | E | 1.6e-07 | 7.2e-07 | 4.3e-07 | 4.367E-07 | 2.801E-07 | 1.600E-07 | 7.200E-07 |
| 25711 | 1286 | P1286L | E | 1.5e-07 | 3.7e-07 | 2e-07 | 2.400E-07 | 1.153E-07 | 1.500E-07 | 3.700E-07 |
| 25712 | 1286 | P1286K | E | 2.1e-07 | 1.9e-07 | 3.5e-07 | 2.500E-07 | 8.718E-08 | 1.900E-07 | 3.500E-07 |
| 25713 | 1286 | P1286M | E | 5e-07 | 3.3e-07 | 3.2e-07 | 3.833E-07 | 1.012E-07 | 3.200E-07 | 5.000E-07 |
| 25714 | 1286 | P1286F | E | 2.5e-07 | 2.7e-07 | 2.1e-07 | 2.433E-07 | 3.055E-08 | 2.100E-07 | 2.700E-07 |
| 25715 | 1286 | P1286P | E | 3.3e-07 | 2e-07 | 1.7e-07 | 2.333E-07 | 8.505E-08 | 1.700E-07 | 3.300E-07 |
| 25716 | 1286 | P1286S | E | 3.6e-07 | 1.3e-07 | 2.1e-07 | 2.333E-07 | 1.168E-07 | 1.300E-07 | 3.600E-07 |
| 25717 | 1286 | P1286T | E | 1.3e-07 | 2.5e-07 | 2.4e-07 | 2.067E-07 | 6.658E-08 | 1.300E-07 | 2.500E-07 |
| 25718 | 1286 | P1286W | E | 1.6e-07 | 2.2e-07 | 3.8e-07 | 2.533E-07 | 1.137E-07 | 1.600E-07 | 3.800E-07 |
| 25719 | 1286 | P1286Y | E | 3.2e-07 | 4.8e-07 | 1.8e-07 | 3.267E-07 | 1.501E-07 | 1.800E-07 | 4.800E-07 |
| 25720 | 1286 | P1286V | E | 1.9e-07 | 3.3e-07 | 8.8e-08 | 2.027E-07 | 1.215E-07 | 8.800E-08 | 3.300E-07 |
| 25721 | 1287 | G1287A | E | 3e-07 | 2.5e-07 | 2.1e-07 | 2.533E-07 | 4.509E-08 | 2.100E-07 | 3.000E-07 |
| 25722 | 1287 | G1287R | E | 1.7e-07 | 3e-07 | 2.8e-07 | 2.500E-07 | 7.000E-08 | 1.700E-07 | 3.000E-07 |
| 25723 | 1287 | G1287N | E | 4.2e-07 | 4e-07 | 1.7e-07 | 3.300E-07 | 1.389E-07 | 1.700E-07 | 4.200E-07 |
| 25724 | 1287 | G1287D | E | 1.4e-07 | 2.2e-07 | 4.4e-07 | 2.667E-07 | 1.553E-07 | 1.400E-07 | 4.400E-07 |
| 25725 | 1287 | G1287C | E | 2.4e-07 | 1.5e-07 | 7.6e-07 | 3.833E-07 | 3.293E-07 | 1.500E-07 | 7.600E-07 |
| 25726 | 1287 | G1287E | E | 3.8e-07 | 1.7e-07 | 2.2e-07 | 2.567E-07 | 1.097E-07 | 1.700E-07 | 3.800E-07 |
| 25727 | 1287 | G1287Q | E | 1.9e-07 | 3.2e-07 | 2e-07 | 2.367E-07 | 7.234E-08 | 1.900E-07 | 3.200E-07 |
| 25728 | 1287 | G1287G | E | 2.1e-07 | 1.7e-07 | 3.2e-07 | 2.333E-07 | 7.767E-08 | 1.700E-07 | 3.200E-07 |
| 25729 | 1287 | G1287H | E | 2.3e-07 | 3.4e-07 | 4e-07 | 3.233E-07 | 8.622E-08 | 2.300E-07 | 4.000E-07 |
| 25730 | 1287 | G1287I | E | 2.8e-07 | 1.1e-07 | 3.1e-07 | 2.333E-07 | 1.079E-07 | 1.100E-07 | 3.100E-07 |
| 25731 | 1287 | G1287L | E | 3.5e-07 | 3.6e-07 | 1.1e-07 | 2.733E-07 | 1.415E-07 | 1.100E-07 | 3.600E-07 |
| 25732 | 1287 | G1287K | E | 2.3e-07 | 1.7e-07 | 2.5e-07 | 2.167E-07 | 4.163E-08 | 1.700E-07 | 2.500E-07 |
| 25733 | 1287 | G1287M | E | 2.8e-07 | 2.2e-07 | 1.9e-07 | 2.300E-07 | 4.583E-08 | 1.900E-07 | 2.800E-07 |
| 25734 | 1287 | G1287F | E | 1.6e-07 | 3.4e-07 | 1.7e-07 | 2.233E-07 | 1.012E-07 | 1.600E-07 | 3.400E-07 |
| 25735 | 1287 | G1287P | E | 1.6e-07 | 3e-07 | 2.5e-07 | 2.367E-07 | 7.095E-08 | 1.600E-07 | 3.000E-07 |
| 25736 | 1287 | G1287S | E | 2e-07 | 4.1e-07 | 1.7e-07 | 2.600E-07 | 1.308E-07 | 1.700E-07 | 4.100E-07 |
| 25737 | 1287 | G1287T | E | 2.1e-07 | 1.7e-07 | 2.9e-07 | 2.233E-07 | 6.110E-08 | 1.700E-07 | 2.900E-07 |
| 25738 | 1287 | G1287W | E | 2.1e-07 | 3.8e-07 | 3e-07 | 2.967E-07 | 8.505E-08 | 2.100E-07 | 3.800E-07 |
| 25739 | 1287 | G1287Y | E | 1.3e-07 | 5.7e-07 | 3.8e-07 | 3.600E-07 | 2.207E-07 | 1.300E-07 | 5.700E-07 |
| 25740 | 1287 | G1287V | E | 2e-07 | 3.9e-07 | 2.1e-07 | 2.667E-07 | 1.069E-07 | 2.000E-07 | 3.900E-07 |
| 25741 | 1288 | G1288A | E | 3.9e-07 | 2.2e-07 | 1.3e-07 | 2.467E-07 | 1.320E-07 | 1.300E-07 | 3.900E-07 |
| 25742 | 1288 | G1288R | E | 3.5e-07 | 2.4e-07 | 2e-07 | 2.633E-07 | 7.767E-08 | 2.000E-07 | 3.500E-07 |
| 25743 | 1288 | G1288N | E | 2.2e-07 | 9.1e-08 | 2.6e-07 | 1.903E-07 | 8.832E-08 | 9.100E-08 | 2.600E-07 |
| 25744 | 1288 | G1288D | E | 3e-07 | 3.3e-07 | 1.5e-07 | 2.600E-07 | 9.644E-08 | 1.500E-07 | 3.300E-07 |
| 25745 | 1288 | G1288C | E | 1e-07 | 2.3e-07 | 2e-07 | 1.767E-07 | 6.807E-08 | 1.000E-07 | 2.300E-07 |
| 25746 | 1288 | G1288E | E | 1.5e-07 | 2.1e-07 | 2.3e-07 | 1.967E-07 | 4.163E-08 | 1.500E-07 | 2.300E-07 |
| 25747 | 1288 | G1288Q | E | 3.1e-07 | 1.9e-07 | 1.8e-07 | 2.267E-07 | 7.234E-08 | 1.800E-07 | 3.100E-07 |
| 25748 | 1288 | G1288G | E | 4.2e-07 | 1.7e-07 | 2.3e-07 | 2.733E-07 | 1.305E-07 | 1.700E-07 | 4.200E-07 |
| 25749 | 1288 | G1288H | E | 2.6e-07 | 1.6e-07 | 3.5e-07 | 2.567E-07 | 9.504E-08 | 1.600E-07 | 3.500E-07 |
| 25750 | 1288 | G1288I | E | 2.9e-07 | 6.5e-07 | 2.6e-07 | 4.000E-07 | 2.170E-07 | 2.600E-07 | 6.500E-07 |
| 25751 | 1288 | G1288L | E | 1.8e-07 | 2.9e-07 | 3e-07 | 2.567E-07 | 6.658E-08 | 1.800E-07 | 3.000E-07 |
| 25752 | 1288 | G1288K | E | 1.9e-07 | 2.4e-07 | 2.1e-07 | 2.133E-07 | 2.517E-08 | 1.900E-07 | 2.400E-07 |
| 25753 | 1288 | G1288M | E | 1.2e-07 | 3.5e-07 | 2.3e-07 | 2.333E-07 | 1.150E-07 | 1.200E-07 | 3.500E-07 |
| 25754 | 1288 | G1288F | E | 9.7e-08 | 1.1e-07 | 2.5e-07 | 1.523E-07 | 8.483E-08 | 9.700E-08 | 2.500E-07 |

| | | | | | | | | | | |
|---|---|---|---|---|---|---|---|---|---|---|
| 25755 | 1288 | G1288P | E | 3.9e-07 | 5.8e-07 | 2.1e-07 | 3.933E-07 | 1.850E-07 | 2.100E-07 | 5.800E-07 |
| 25756 | 1288 | G1288S | E | 3.6e-07 | 2.6e-07 | 3.6e-07 | 3.267E-07 | 5.774E-08 | 2.600E-07 | 3.600E-07 |
| 25757 | 1288 | G1288T | E | 2.1e-07 | 2.1e-07 | 5.4e-07 | 3.200E-07 | 1.905E-07 | 2.100E-07 | 5.400E-07 |
| 25758 | 1288 | G1288W | E | 2.7e-07 | 2.7e-07 | 1.5e-07 | 2.300E-07 | 6.928E-08 | 1.500E-07 | 2.700E-07 |
| 25759 | 1288 | G1288Y | E | 1.5e-07 | 3.2e-07 | 3.7e-07 | 2.800E-07 | 1.153E-07 | 1.500E-07 | 3.700E-07 |
| 25760 | 1288 | G1288V | E | 2.4e-07 | 2.8e-07 | 1.5e-07 | 2.233E-07 | 6.658E-08 | 1.500E-07 | 2.800E-07 |
| 25761 | 1289 | S1289A | E | 3.2e-07 | 2.6e-07 | 3.1e-07 | 2.967E-07 | 3.215E-08 | 2.600E-07 | 3.200E-07 |
| 25762 | 1289 | S1289R | E | 3.8e-07 | 4.1e-07 | 8.1e-08 | 2.903E-07 | 1.819E-07 | 8.100E-08 | 4.100E-07 |
| 25763 | 1289 | S1289N | E | 2e-07 | 5.8e-07 | 1.4e-07 | 3.067E-07 | 2.386E-07 | 1.400E-07 | 5.800E-07 |
| 25764 | 1289 | S1289D | E | 2.6e-07 | 3.5e-07 | 3.3e-07 | 3.133E-07 | 4.726E-08 | 2.600E-07 | 3.500E-07 |
| 25765 | 1289 | S1289C | E | 1.6e-07 | 2.5e-07 | 2.2e-07 | 2.100E-07 | 4.583E-08 | 1.600E-07 | 2.500E-07 |
| 25766 | 1289 | S1289E | E | 2.5e-07 | 2.1e-07 | 2.6e-07 | 2.400E-07 | 2.646E-08 | 2.100E-07 | 2.600E-07 |
| 25767 | 1289 | S1289Q | E | 2.7e-07 | 4.7e-07 | 1.3e-07 | 2.900E-07 | 1.709E-07 | 1.300E-07 | 4.700E-07 |
| 25768 | 1289 | S1289G | E | 3.6e-07 | 2.3e-07 | 4.4e-07 | 3.433E-07 | 1.060E-07 | 2.300E-07 | 4.400E-07 |
| 25769 | 1289 | S1289H | E | 1.6e-07 | 2.9e-07 | 1.1e-07 | 1.867E-07 | 9.292E-08 | 1.100E-07 | 2.900E-07 |
| 25770 | 1289 | S1289I | E | 1.7e-07 | 9e-08 | 4.3e-07 | 2.300E-07 | 1.778E-07 | 9.000E-08 | 4.300E-07 |
| 25771 | 1289 | S1289L | E | 2.4e-07 | 1.4e-07 | 2.1e-07 | 1.967E-07 | 5.132E-08 | 1.400E-07 | 2.400E-07 |
| 25772 | 1289 | S1289K | E | 5.7e-07 | 3e-07 | 2.7e-07 | 3.800E-07 | 1.652E-07 | 2.700E-07 | 5.700E-07 |
| 25773 | 1289 | S1289M | E | 3.1e-07 | 3.5e-07 | 3e-07 | 3.200E-07 | 2.646E-08 | 3.000E-07 | 3.500E-07 |
| 25774 | 1289 | S1289F | E | 6.6e-07 | 1.9e-07 | 2.2e-07 | 3.567E-07 | 2.631E-07 | 1.900E-07 | 6.600E-07 |
| 25775 | 1289 | S1289P | E | 2.5e-07 | 1.7e-07 | 1.8e-07 | 2.000E-07 | 4.359E-08 | 1.700E-07 | 2.500E-07 |
| 25776 | 1289 | S1289S | E | 2.3e-07 | 3.3e-07 | 1.2e-06 | 5.867E-07 | 5.335E-07 | 2.300E-07 | 1.200E-06 |
| 25777 | 1289 | S1289T | E | 4e-07 | 4.1e-07 | 1.9e-07 | 3.333E-07 | 1.242E-07 | 1.900E-07 | 4.100E-07 |
| 25778 | 1289 | S1289W | E | 5.4e-07 | 3.4e-07 | 2.8e-07 | 3.867E-07 | 1.361E-07 | 2.800E-07 | 5.400E-07 |
| 25779 | 1289 | S1289Y | E | 2.9e-07 | 2.3e-07 | 1.2e-07 | 2.133E-07 | 8.622E-08 | 1.200E-07 | 2.900E-07 |
| 25780 | 1289 | S1289V | E | 1.7e-07 | 3.4e-07 | 4.2e-07 | 3.100E-07 | 1.277E-07 | 1.700E-07 | 4.200E-07 |
| 25781 | 1290 | L1290A | E | 1.2e-07 | 1.5e-07 | 2.8e-07 | 1.833E-07 | 8.505E-08 | 1.200E-07 | 2.800E-07 |
| 25782 | 1290 | L1290R | E | 3.2e-07 | 1.6e-07 | 2.6e-07 | 2.467E-07 | 8.083E-08 | 1.600E-07 | 3.200E-07 |
| 25783 | 1290 | L1290N | E | 3.4e-07 | 2.7e-07 | 1.8e-07 | 2.633E-07 | 8.021E-08 | 1.800E-07 | 3.400E-07 |
| 25784 | 1290 | L1290D | E | 2.3e-07 | 2e-07 | 2.1e-07 | 2.133E-07 | 1.528E-08 | 2.000E-07 | 2.300E-07 |
| 25785 | 1290 | L1290C | E | 1.3e-07 | 1.4e-07 | 3e-07 | 1.900E-07 | 9.539E-08 | 1.300E-07 | 3.000E-07 |
| 25786 | 1290 | L1290E | E | 2.4e-07 | 3.8e-07 | 2.9e-07 | 3.033E-07 | 7.095E-08 | 2.400E-07 | 3.800E-07 |
| 25787 | 1290 | L1290Q | E | 9.1e-08 | 1.5e-07 | 6.9e-07 | 3.103E-07 | 3.301E-07 | 9.100E-08 | 6.900E-07 |
| 25788 | 1290 | L1290G | E | 7.6e-07 | 3.4e-07 | 2e-07 | 4.333E-07 | 2.914E-07 | 2.000E-07 | 7.600E-07 |
| 25789 | 1290 | L1290H | E | 1.8e-07 | 3.3e-07 | 1.6e-07 | 2.233E-07 | 9.292E-08 | 1.600E-07 | 3.300E-07 |
| 25790 | 1290 | L1290I | E | 2.7e-07 | 4.8e-07 | 1.8e-07 | 3.100E-07 | 1.539E-07 | 1.800E-07 | 4.800E-07 |
| 25791 | 1290 | L1290L | E | 1.4e-07 | 3.1e-07 | 2e-07 | 2.167E-07 | 8.622E-08 | 1.400E-07 | 3.100E-07 |
| 25792 | 1290 | L1290K | E | 2.1e-07 | 3.1e-07 | 2.6e-07 | 2.600E-07 | 5.000E-08 | 2.100E-07 | 3.100E-07 |
| 25793 | 1290 | L1290M | E | 1.8e-07 | 1.7e-07 | 3.4e-07 | 2.300E-07 | 9.539E-08 | 1.700E-07 | 3.400E-07 |
| 25794 | 1290 | L1290F | E | 2e-07 | 3.1e-07 | 1.4e-07 | 2.167E-07 | 8.622E-08 | 1.400E-07 | 3.100E-07 |
| 25795 | 1290 | L1290P | E | 1.4e-07 | 1.6e-07 | 2.6e-07 | 1.867E-07 | 6.429E-08 | 1.400E-07 | 2.600E-07 |
| 25796 | 1290 | L1290S | E | 1.8e-07 | 2.1e-07 | 1.9e-07 | 1.933E-07 | 1.528E-08 | 1.800E-07 | 2.100E-07 |
| 25797 | 1290 | L1290T | E | 8.8e-08 | 1.1e-07 | 1.2e-07 | 1.060E-07 | 1.637E-08 | 8.800E-08 | 1.200E-07 |
| 25798 | 1290 | L1290W | E | 4.1e-07 | 1.8e-07 | 3.1e-07 | 3.000E-07 | 1.153E-07 | 1.800E-07 | 4.100E-07 |
| 25799 | 1290 | L1290Y | E | 3.4e-07 | 3.8e-07 | 5.1e-07 | 4.100E-07 | 8.888E-08 | 3.400E-07 | 5.100E-07 |
| 25800 | 1290 | L1290V | E | 3.1e-07 | 2.6e-07 | 2.6e-07 | 2.767E-07 | 2.887E-08 | 2.600E-07 | 3.100E-07 |
| 25801 | 1291 | R1291A | E | 6.5e-07 | 1e-07 | 1.2e-07 | 2.900E-07 | 3.119E-07 | 1.000E-07 | 6.500E-07 |
| 25802 | 1291 | R1291R | E | 1.7e-07 | 5.8e-07 | 4.6e-07 | 4.033E-07 | 2.108E-07 | 1.700E-07 | 5.800E-07 |
| 25803 | 1291 | R1291N | E | 2.5e-07 | 1.9e-07 | 3e-07 | 2.467E-07 | 5.508E-08 | 1.900E-07 | 3.000E-07 |
| 25804 | 1291 | R1291D | E | 2.6e-07 | 1.4e-07 | 1.2e-07 | 1.733E-07 | 7.572E-08 | 1.200E-07 | 2.600E-07 |
| 25805 | 1291 | R1291C | E | 1.2e-07 | 1.4e-07 | 2.7e-07 | 1.767E-07 | 8.145E-08 | 1.200E-07 | 2.700E-07 |
| 25806 | 1291 | R1291E | E | 2.3e-07 | 3.3e-07 | 2.3e-07 | 2.633E-07 | 5.774E-08 | 2.300E-07 | 3.300E-07 |
| 25807 | 1291 | R1291Q | E | 1.8e-07 | 2.2e-07 | 1.4e-07 | 1.800E-07 | 4.000E-08 | 1.400E-07 | 2.200E-07 |
| 25808 | 1291 | R1291G | E | 3.1e-07 | 1.6e-07 | 1.5e-07 | 2.067E-07 | 8.963E-08 | 1.500E-07 | 3.100E-07 |
| 25809 | 1291 | R1291H | E | 4.6e-07 | 2.6e-07 | 1.6e-07 | 2.933E-07 | 1.528E-07 | 1.600E-07 | 4.600E-07 |
| 25810 | 1291 | R1291I | E | 4.2e-07 | 7.4e-07 | 2.4e-07 | 4.667E-07 | 2.532E-07 | 2.400E-07 | 7.400E-07 |
| 25811 | 1291 | R1291L | E | 2.3e-07 | 2.8e-07 | 1.8e-07 | 2.300E-07 | 5.000E-08 | 1.800E-07 | 2.800E-07 |
| 25812 | 1291 | R1291K | E | 2.7e-07 | 2.3e-07 | 4.9e-07 | 3.300E-07 | 1.400E-07 | 2.300E-07 | 4.900E-07 |
| 25813 | 1291 | R1291M | E | 1.8e-07 | 3.2e-07 | 3.6e-07 | 2.867E-07 | 9.452E-08 | 1.800E-07 | 3.600E-07 |
| 25814 | 1291 | R1291F | E | 2.3e-07 | 2.4e-07 | 2.1e-07 | 2.267E-07 | 1.528E-08 | 2.100E-07 | 2.400E-07 |
| 25815 | 1291 | R1291P | E | 3e-07 | 1.7e-07 | 8e-08 | 1.833E-07 | 1.106E-07 | 8.000E-08 | 3.000E-07 |
| 25816 | 1291 | R1291S | E | 1.4e-07 | 1.4e-07 | 5.4e-07 | 2.733E-07 | 2.309E-07 | 1.400E-07 | 5.400E-07 |
| 25817 | 1291 | R1291T | E | 2.8e-07 | 2.9e-07 | 3.4e-07 | 3.033E-07 | 3.215E-08 | 2.800E-07 | 3.400E-07 |
| 25818 | 1291 | R1291W | E | 1.9e-07 | 2e-07 | 3e-07 | 2.300E-07 | 6.083E-08 | 1.900E-07 | 3.000E-07 |
| 25819 | 1291 | R1291Y | E | 1.3e-07 | 2.1e-07 | 3.4e-07 | 2.267E-07 | 1.060E-07 | 1.300E-07 | 3.400E-07 |
| 25820 | 1291 | R1291V | E | 1.7e-07 | 3.1e-07 | 2.5e-07 | 2.433E-07 | 7.024E-08 | 1.700E-07 | 3.100E-07 |
| 25821 | 1292 | L1292A | E | 1.7e-07 | 2.8e-07 | 1.4e-07 | 1.967E-07 | 7.371E-08 | 1.400E-07 | 2.800E-07 |
| 25822 | 1292 | L1292R | E | 1.8e-07 | 3.4e-07 | 2.6e-07 | 2.600E-07 | 8.000E-08 | 1.800E-07 | 3.400E-07 |
| 25823 | 1292 | L1292N | E | 4.3e-07 | 1.6e-07 | 1.8e-07 | 2.567E-07 | 1.504E-07 | 1.600E-07 | 4.300E-07 |
| 25824 | 1292 | L1292D | E | 2.4e-07 | 1.5e-07 | 2.6e-07 | 2.167E-07 | 5.859E-08 | 1.500E-07 | 2.600E-07 |
| 25825 | 1292 | L1292C | E | 3e-07 | 2.3e-07 | 1.9e-07 | 2.400E-07 | 5.568E-08 | 1.900E-07 | 3.000E-07 |
| 25826 | 1292 | L1292E | E | 2.4e-07 | 5e-07 | 2.2e-07 | 3.200E-07 | 1.562E-07 | 2.200E-07 | 5.000E-07 |
| 25827 | 1292 | L1292Q | E | 5.9e-07 | 2.9e-07 | 2.8e-07 | 3.867E-07 | 1.762E-07 | 2.800E-07 | 5.900E-07 |
| 25828 | 1292 | L1292G | E | 2.5e-07 | 1e-07 | 2.8e-07 | 2.100E-07 | 9.644E-08 | 1.000E-07 | 2.800E-07 |
| 25829 | 1292 | L1292H | E | 1.8e-07 | 1.7e-07 | 2.4e-07 | 1.967E-07 | 3.786E-08 | 1.700E-07 | 2.400E-07 |
| 25830 | 1292 | L1292I | E | 1e-07 | 2e-07 | 7.6e-07 | 3.533E-07 | 3.557E-07 | 1.000E-07 | 7.600E-07 |
| 25831 | 1292 | L1292L | E | 2.2e-07 | 1.1e-07 | 2.8e-07 | 2.033E-07 | 8.622E-08 | 1.100E-07 | 2.800E-07 |
| 25832 | 1292 | L1292K | E | 1.7e-07 | 1.2e-07 | 2.3e-07 | 1.733E-07 | 5.508E-08 | 1.200E-07 | 2.300E-07 |
| 25833 | 1292 | L1292M | E | 1.7e-07 | 3.1e-07 | 3e-07 | 2.600E-07 | 7.810E-08 | 1.700E-07 | 3.100E-07 |
| 25834 | 1292 | L1292F | E | 1.7e-07 | 2.6e-07 | 4.6e-07 | 2.967E-07 | 1.484E-07 | 1.700E-07 | 4.600E-07 |
| 25835 | 1292 | L1292P | E | 2.3e-07 | 2.4e-07 | 2.9e-07 | 2.533E-07 | 3.215E-08 | 2.300E-07 | 2.900E-07 |
| 25836 | 1292 | L1292S | E | 1.8e-07 | 3e-07 | 2.2e-07 | 2.333E-07 | 6.110E-08 | 1.800E-07 | 3.000E-07 |
| 25837 | 1292 | L1292T | E | 4.9e-07 | 3.3e-07 | 3.2e-07 | 3.800E-07 | 9.539E-08 | 3.200E-07 | 4.900E-07 |
| 25838 | 1292 | L1292W | E | 1.9e-07 | 4.2e-07 | 2.3e-07 | 2.800E-07 | 1.229E-07 | 1.900E-07 | 4.200E-07 |
| 25839 | 1292 | L1292Y | E | 2.3e-07 | 1.8e-07 | 1.8e-07 | 1.967E-07 | 2.887E-08 | 1.800E-07 | 2.300E-07 |

| | | | | | | | | | | |
|---|---|---|---|---|---|---|---|---|---|---|
| 25840 | 1292 | L1292V | E | 2.3e-07 | 4.7e-07 | 4.5e-07 | 3.833E-07 | 1.332E-07 | 2.300E-07 | 4.700E-07 |
| 25841 | 1293 | S1293A | E | 2.9e-07 | 3.1e-07 | 1.9e-07 | 2.633E-07 | 6.429E-08 | 1.900E-07 | 3.100E-07 |
| 25842 | 1293 | S1293R | E | 1.3e-07 | 3.4e-07 | 2.5e-07 | 2.400E-07 | 1.054E-07 | 1.300E-07 | 3.400E-07 |
| 25843 | 1293 | S1293N | E | 2.1e-07 | 3.2e-07 | 4.2e-07 | 3.167E-07 | 1.050E-07 | 2.100E-07 | 4.200E-07 |
| 25844 | 1293 | S1293D | E | 2.6e-07 | 2.3e-07 | 9.4e-08 | 1.947E-07 | 8.846E-08 | 9.400E-08 | 2.600E-07 |
| 25845 | 1293 | S1293C | E | 1.2e-07 | 1.4e-07 | 1.1e-07 | 1.233E-07 | 1.528E-08 | 1.100E-07 | 1.400E-07 |
| 25846 | 1293 | S1293E | E | 2e-07 | 1.9e-07 | 6.2e-08 | 1.507E-07 | 7.695E-08 | 6.200E-08 | 2.000E-07 |
| 25847 | 1293 | S1293Q | E | 2.7e-07 | 4.2e-08 | 2.6e-07 | 1.907E-07 | 1.288E-07 | 4.200E-08 | 2.700E-07 |
| 25848 | 1293 | S1293G | E | 7.1e-07 | 1.9e-07 | 1.5e-07 | 3.500E-07 | 3.124E-07 | 1.500E-07 | 7.100E-07 |
| 25849 | 1293 | S1293H | E | 9.1e-08 | 1.6e-07 | 3.5e-07 | 2.003E-07 | 1.341E-07 | 9.100E-08 | 3.500E-07 |
| 25850 | 1293 | S1293I | E | 3.2e-07 | 2.4e-07 | 1.1e-07 | 2.233E-07 | 1.060E-07 | 1.100E-07 | 3.200E-07 |
| 25851 | 1293 | S1293L | E | 1.3e-07 | 2e-07 | 1.6e-07 | 1.633E-07 | 3.512E-08 | 1.300E-07 | 2.000E-07 |
| 25852 | 1293 | S1293K | E | 3e-07 | 1.9e-07 | 2.4e-07 | 2.433E-07 | 5.508E-08 | 1.900E-07 | 3.000E-07 |
| 25853 | 1293 | S1293M | E | 2.5e-07 | 1.4e-07 | 4.4e-07 | 2.767E-07 | 1.518E-07 | 1.400E-07 | 4.400E-07 |
| 25854 | 1293 | S1293F | E | 3.5e-07 | 3.5e-07 | 2.7e-07 | 3.233E-07 | 4.619E-08 | 2.700E-07 | 3.500E-07 |
| 25855 | 1293 | S1293P | E | 1.9e-07 | 2.9e-07 | 1.8e-07 | 2.200E-07 | 6.083E-08 | 1.800E-07 | 2.900E-07 |
| 25856 | 1293 | S1293S | E | 3.6e-07 | 1.5e-07 | 1.8e-07 | 2.300E-07 | 1.136E-07 | 1.500E-07 | 3.600E-07 |
| 25857 | 1293 | S1293T | E | 3.4e-07 | 4.3e-07 | 1.8e-07 | 3.167E-07 | 1.266E-07 | 1.800E-07 | 4.300E-07 |
| 25858 | 1293 | S1293W | E | 4.5e-07 | 2.1e-07 | 1.9e-07 | 2.833E-07 | 1.447E-07 | 1.900E-07 | 4.500E-07 |
| 25859 | 1293 | S1293Y | E | 4.3e-07 | 2.7e-07 | 4.7e-07 | 3.900E-07 | 1.058E-07 | 2.700E-07 | 4.700E-07 |
| 25860 | 1293 | S1293V | E | 2.2e-07 | 4.3e-07 | 1.6e-07 | 2.700E-07 | 1.418E-07 | 1.600E-07 | 4.300E-07 |
| 25861 | 1294 | C1294A | E | 2.5e-07 | 4.2e-07 | 1.3e-07 | 2.667E-07 | 1.457E-07 | 1.300E-07 | 4.200E-07 |
| 25862 | 1294 | C1294R | E | 1.8e-07 | 1.8e-07 | 1.3e-07 | 1.633E-07 | 2.887E-08 | 1.300E-07 | 1.800E-07 |
| 25863 | 1294 | C1294N | E | 2e-07 | 1.7e-07 | 1.5e-07 | 1.733E-07 | 2.517E-08 | 1.500E-07 | 2.000E-07 |
| 25864 | 1294 | C1294D | E | 2.4e-07 | 5e-07 | 2.8e-07 | 3.400E-07 | 1.400E-07 | 2.400E-07 | 5.000E-07 |
| 25865 | 1294 | C1294C | E | 3.4e-07 | 2.3e-07 | 1.5e-07 | 2.400E-07 | 9.539E-08 | 1.500E-07 | 3.400E-07 |
| 25866 | 1294 | C1294E | E | 1.8e-07 | 1.5e-07 | 2.3e-07 | 1.867E-07 | 4.041E-08 | 1.500E-07 | 2.300E-07 |
| 25867 | 1294 | C1294Q | E | 2.8e-07 | 3.2e-07 | 2.1e-07 | 2.700E-07 | 5.568E-08 | 2.100E-07 | 3.200E-07 |
| 25868 | 1294 | C1294G | E | 3.5e-07 | 2.2e-07 | 3.3e-07 | 3.000E-07 | 7.000E-08 | 2.200E-07 | 3.500E-07 |
| 25869 | 1294 | C1294H | E | 4.4e-08 | 1.9e-07 | 3.7e-07 | 2.013E-07 | 1.633E-07 | 4.400E-08 | 3.700E-07 |
| 25870 | 1294 | C1294I | E | 4.3e-07 | 5e-07 | 1.1e-07 | 3.467E-07 | 2.079E-07 | 1.100E-07 | 5.000E-07 |
| 25871 | 1294 | C1294L | E | 1.6e-07 | 2.5e-07 | 1.4e-07 | 1.833E-07 | 5.859E-08 | 1.400E-07 | 2.500E-07 |
| 25872 | 1294 | C1294K | E | 2.5e-07 | 2e-07 | 1.6e-07 | 2.033E-07 | 4.509E-08 | 1.600E-07 | 2.500E-07 |
| 25873 | 1294 | C1294M | E | 1.7e-07 | 3.5e-07 | 4.2e-07 | 3.133E-07 | 1.290E-07 | 1.700E-07 | 4.200E-07 |
| 25874 | 1294 | C1294F | E | 2.6e-07 | 1.3e-07 | 8.5e-08 | 1.583E-07 | 9.088E-08 | 8.500E-08 | 2.600E-07 |
| 25875 | 1294 | C1294P | E | 2e-07 | 3.5e-07 | 1.2e-07 | 2.233E-07 | 1.168E-07 | 1.200E-07 | 3.500E-07 |
| 25876 | 1294 | C1294S | E | 2.1e-07 | 2.7e-07 | 1.7e-07 | 2.167E-07 | 5.033E-08 | 1.700E-07 | 2.700E-07 |
| 25877 | 1294 | C1294T | E | 1.9e-07 | 2.9e-07 | 3.8e-07 | 2.867E-07 | 9.504E-08 | 1.900E-07 | 3.800E-07 |
| 25878 | 1294 | C1294W | E | 2.2e-07 | 2.1e-07 | 2.9e-07 | 2.400E-07 | 4.359E-08 | 2.100E-07 | 2.900E-07 |
| 25879 | 1294 | C1294Y | E | 2.2e-07 | 4.3e-07 | 2.2e-07 | 2.900E-07 | 1.212E-07 | 2.200E-07 | 4.300E-07 |
| 25880 | 1294 | C1294V | E | 4.1e-07 | 7.3e-07 | 2e-07 | 4.467E-07 | 2.669E-07 | 2.000E-07 | 7.300E-07 |
| 25881 | 1295 | A1295A | E | 2.9e-07 | 2.5e-07 | 2.6e-07 | 2.667E-07 | 2.082E-08 | 2.500E-07 | 2.900E-07 |
| 25882 | 1295 | A1295R | E | 1.4e-07 | 1.8e-07 | 2.2e-07 | 1.800E-07 | 4.000E-08 | 1.400E-07 | 2.200E-07 |
| 25883 | 1295 | A1295N | E | 1.8e-07 | 5e-07 | 2.4e-07 | 3.067E-07 | 1.701E-07 | 1.800E-07 | 5.000E-07 |
| 25884 | 1295 | A1295D | E | 4.3e-07 | 2.9e-07 | 1.5e-07 | 2.900E-07 | 1.400E-07 | 1.500E-07 | 4.300E-07 |
| 25885 | 1295 | A1295C | E | 3.7e-07 | 1.5e-07 | 1.9e-07 | 2.367E-07 | 1.172E-07 | 1.500E-07 | 3.700E-07 |
| 25886 | 1295 | A1295E | E | 3.6e-07 | 2e-07 | 2.3e-07 | 2.633E-07 | 8.505E-08 | 2.000E-07 | 3.600E-07 |
| 25887 | 1295 | A1295Q | E | 8.5e-08 | 2.9e-07 | 3.3e-07 | 2.350E-07 | 1.314E-07 | 8.500E-08 | 3.300E-07 |
| 25888 | 1295 | A1295G | E | 5.2e-07 | 3.6e-07 | 1.7e-07 | 3.500E-07 | 1.752E-07 | 1.700E-07 | 5.200E-07 |
| 25889 | 1295 | A1295H | E | 1.1e-06 | 2.2e-07 | 3e-07 | 5.400E-07 | 4.866E-07 | 2.200E-07 | 1.100E-06 |
| 25890 | 1295 | A1295I | E | 5.3e-07 | 2.1e-07 | 2.3e-07 | 3.233E-07 | 1.793E-07 | 2.100E-07 | 5.300E-07 |
| 25891 | 1295 | A1295L | E | 2.8e-07 | 2.1e-07 | 2.3e-07 | 2.400E-07 | 3.606E-08 | 2.100E-07 | 2.800E-07 |
| 25892 | 1295 | A1295K | E | 3.2e-07 | 1.1e-07 | 3.7e-07 | 2.667E-07 | 1.380E-07 | 1.100E-07 | 3.700E-07 |
| 25893 | 1295 | A1295M | E | 3.3e-07 | 4.2e-07 | 2.4e-07 | 3.300E-07 | 9.000E-08 | 2.400E-07 | 4.200E-07 |
| 25894 | 1295 | A1295F | E | 1.3e-07 | 3e-07 | 1.9e-07 | 2.067E-07 | 8.622E-08 | 1.300E-07 | 3.000E-07 |
| 25895 | 1295 | A1295P | E | 7.4e-07 | 2.9e-07 | 2.9e-07 | 4.400E-07 | 2.598E-07 | 2.900E-07 | 7.400E-07 |
| 25896 | 1295 | A1295S | E | 2.9e-07 | 2.4e-07 | 3e-07 | 2.767E-07 | 3.215E-08 | 2.400E-07 | 3.000E-07 |
| 25897 | 1295 | A1295T | E | 4.8e-07 | 2.6e-07 | 2e-07 | 3.133E-07 | 1.474E-07 | 2.000E-07 | 4.800E-07 |
| 25898 | 1295 | A1295W | E | 5.1e-07 | 2.5e-07 | 3.3e-07 | 3.633E-07 | 1.332E-07 | 2.500E-07 | 5.100E-07 |
| 25899 | 1295 | A1295Y | E | 2.7e-07 | 2.3e-07 | 2.5e-07 | 2.500E-07 | 2.000E-08 | 2.300E-07 | 2.700E-07 |
| 25900 | 1295 | A1295V | E | 2.2e-07 | 2.6e-07 | 4.5e-07 | 3.100E-07 | 1.229E-07 | 2.200E-07 | 4.500E-07 |
| 25901 | 1296 | A1296A | E | 3e-07 | 2.4e-07 | 1.7e-07 | 2.367E-07 | 6.506E-08 | 1.700E-07 | 3.000E-07 |
| 25902 | 1296 | A1296R | E | 1.6e-07 | 3.8e-07 | 8.7e-08 | 2.090E-07 | 1.525E-07 | 8.700E-08 | 3.800E-07 |
| 25903 | 1296 | A1296N | E | 1.6e-07 | 1.4e-07 | 1.6e-07 | 1.533E-07 | 1.155E-08 | 1.400E-07 | 1.600E-07 |
| 25904 | 1296 | A1296D | E | 3.1e-07 | 3e-07 | 2.4e-07 | 2.833E-07 | 3.786E-08 | 2.400E-07 | 3.100E-07 |
| 25905 | 1296 | A1296C | E | 2.8e-07 | 2.6e-07 | 1.6e-07 | 2.333E-07 | 6.429E-08 | 1.600E-07 | 2.800E-07 |
| 25906 | 1296 | A1296E | E | 1.3e-07 | 4.1e-07 | 2.5e-07 | 2.633E-07 | 1.405E-07 | 1.300E-07 | 4.100E-07 |
| 25907 | 1296 | A1296Q | E | 1.6e-07 | 1.5e-07 | 3.4e-07 | 2.167E-07 | 1.069E-07 | 1.500E-07 | 3.400E-07 |
| 25908 | 1296 | A1296G | E | 2.5e-07 | 1.4e-07 | 3.7e-07 | 2.533E-07 | 1.150E-07 | 1.400E-07 | 3.700E-07 |
| 25909 | 1296 | A1296H | E | 1.2e-07 | 1.6e-07 | 1.9e-07 | 1.567E-07 | 3.512E-08 | 1.200E-07 | 1.900E-07 |
| 25910 | 1296 | A1296I | E | 3.3e-07 | 3.5e-07 | 4.8e-07 | 3.867E-07 | 8.145E-08 | 3.300E-07 | 4.800E-07 |
| 25911 | 1296 | A1296L | E | 5.6e-07 | 4.5e-07 | 2.4e-07 | 4.167E-07 | 1.626E-07 | 2.400E-07 | 5.600E-07 |
| 25912 | 1296 | A1296K | E | 1.3e-07 | 2.5e-07 | 2.7e-07 | 2.167E-07 | 7.572E-08 | 1.300E-07 | 2.700E-07 |
| 25913 | 1296 | A1296M | E | 1.4e-07 | 3.7e-07 | 2.2e-07 | 2.433E-07 | 1.168E-07 | 1.400E-07 | 3.700E-07 |
| 25914 | 1296 | A1296F | E | 1.1e-07 | 3.6e-07 | 1.1e-07 | 1.933E-07 | 1.443E-07 | 1.100E-07 | 3.600E-07 |
| 25915 | 1296 | A1296P | E | 2.4e-07 | 1.3e-07 | 2e-07 | 1.900E-07 | 5.568E-08 | 1.300E-07 | 2.400E-07 |
| 25916 | 1296 | A1296S | E | 1.9e-07 | 1.2e-07 | 2.9e-07 | 2.000E-07 | 8.544E-08 | 1.200E-07 | 2.900E-07 |
| 25917 | 1296 | A1296T | E | 2.1e-07 | 4.9e-07 | 2.2e-07 | 3.067E-07 | 1.589E-07 | 2.100E-07 | 4.900E-07 |
| 25918 | 1296 | A1296W | E | 2.9e-07 | 1.9e-07 | 2.5e-07 | 2.433E-07 | 5.033E-08 | 1.900E-07 | 2.900E-07 |
| 25919 | 1296 | A1296Y | E | 2.5e-07 | 2.1e-07 | 3.1e-07 | 2.567E-07 | 5.033E-08 | 2.100E-07 | 3.100E-07 |
| 25920 | 1296 | A1296V | E | 2.2e-07 | 2.9e-07 | 2e-07 | 2.367E-07 | 4.726E-08 | 2.000E-07 | 2.900E-07 |
| 25921 | 1297 | S1297A | E | 2.1e-07 | 3.7e-07 | 1.7e-07 | 2.500E-07 | 1.058E-07 | 1.700E-07 | 3.700E-07 |
| 25922 | 1297 | S1297R | E | 3.2e-07 | 4e-07 | 2.7e-07 | 3.300E-07 | 6.557E-08 | 2.700E-07 | 4.000E-07 |
| 25923 | 1297 | S1297N | E | 3.7e-07 | 2.7e-07 | 2.7e-07 | 3.033E-07 | 5.774E-08 | 2.700E-07 | 3.700E-07 |
| 25924 | 1297 | S1297D | E | 2.2e-07 | 1.4e-07 | 4.4e-07 | 2.667E-07 | 1.553E-07 | 1.400E-07 | 4.400E-07 |

| | | | | | | | | | | |
|---|---|---|---|---|---|---|---|---|---|---|
| 25925 | 1297 | S1297C | E | 5e-07 | 2.6e-07 | 2.5e-07 | 3.367E-07 | 1.415E-07 | 2.500E-07 | 5.000E-07 |
| 25926 | 1297 | S1297E | E | 1.5e-07 | 4.3e-07 | 2.4e-07 | 2.733E-07 | 1.429E-07 | 1.500E-07 | 4.300E-07 |
| 25927 | 1297 | S1297Q | E | 1.8e-07 | 1.6e-07 | 1.8e-07 | 1.733E-07 | 1.155E-08 | 1.600E-07 | 1.800E-07 |
| 25928 | 1297 | S1297G | E | 2.5e-07 | 1.7e-07 | 1.5e-07 | 1.900E-07 | 5.292E-08 | 1.500E-07 | 2.500E-07 |
| 25929 | 1297 | S1297H | E | 1.7e-07 | 3.6e-07 | 1.5e-07 | 2.267E-07 | 1.159E-07 | 1.500E-07 | 3.600E-07 |
| 25930 | 1297 | S1297I | E | 1.3e-07 | 2.1e-07 | 2.1e-07 | 1.833E-07 | 4.619E-08 | 1.300E-07 | 2.100E-07 |
| 25931 | 1297 | S1297L | E | 1.7e-07 | 5.2e-07 | 5e-07 | 3.967E-07 | 1.966E-07 | 1.700E-07 | 5.200E-07 |
| 25932 | 1297 | S1297K | E | 2.2e-07 | 2.3e-07 | 1.1e-07 | 1.867E-07 | 6.658E-08 | 1.100E-07 | 2.300E-07 |
| 25933 | 1297 | S1297M | E | 2.5e-07 | 2e-07 | 3.3e-07 | 2.600E-07 | 6.557E-08 | 2.000E-07 | 3.300E-07 |
| 25934 | 1297 | S1297F | E | 3.5e-07 | 3.7e-07 | 4.3e-07 | 3.833E-07 | 4.163E-08 | 3.500E-07 | 4.300E-07 |
| 25935 | 1297 | S1297P | E | 3.7e-07 | 2.3e-07 | 2.4e-07 | 2.800E-07 | 7.810E-08 | 2.300E-07 | 3.700E-07 |
| 25936 | 1297 | S1297S | E | 4.3e-07 | 6.3e-07 | 3.5e-07 | 4.700E-07 | 1.442E-07 | 3.500E-07 | 6.300E-07 |
| 25937 | 1297 | S1297T | E | 2.9e-07 | 2.8e-07 | 2.1e-07 | 2.600E-07 | 4.359E-08 | 2.100E-07 | 2.900E-07 |
| 25938 | 1297 | S1297W | E | 2.9e-07 | 2.6e-07 | 3.1e-07 | 2.867E-07 | 2.517E-08 | 2.600E-07 | 3.100E-07 |
| 25939 | 1297 | S1297Y | E | 1.3e-07 | 2.2e-07 | 2.7e-07 | 2.067E-07 | 7.095E-08 | 1.300E-07 | 2.700E-07 |
| 25940 | 1297 | S1297V | E | 3.8e-07 | 2.1e-07 | 2.2e-07 | 2.700E-07 | 9.539E-08 | 2.100E-07 | 3.800E-07 |
| 25941 | 1298 | G1298A | E | 8.3e-08 | 3.5e-07 | 1.2e-07 | 1.843E-07 | 1.447E-07 | 8.300E-08 | 3.500E-07 |
| 25942 | 1298 | G1298R | E | 3.5e-07 | 1.3e-07 | 2.8e-07 | 2.533E-07 | 1.124E-07 | 1.300E-07 | 3.500E-07 |
| 25943 | 1298 | G1298N | E | 2.9e-07 | 2.8e-07 | 2.2e-07 | 2.633E-07 | 3.786E-08 | 2.200E-07 | 2.900E-07 |
| 25944 | 1298 | G1298D | E | 9.5e-08 | 1.4e-07 | 2.3e-07 | 1.550E-07 | 6.874E-08 | 9.500E-08 | 2.300E-07 |
| 25945 | 1298 | G1298C | E | 1.4e-07 | 1.3e-07 | 2.1e-07 | 1.600E-07 | 4.359E-08 | 1.300E-07 | 2.100E-07 |
| 25946 | 1298 | G1298E | E | 1.5e-07 | 1.5e-07 | 5.3e-07 | 2.767E-07 | 2.194E-07 | 1.500E-07 | 5.300E-07 |
| 25947 | 1298 | G1298Q | E | 3.1e-07 | 3.6e-07 | 4.3e-07 | 3.667E-07 | 6.028E-08 | 3.100E-07 | 4.300E-07 |
| 25948 | 1298 | G1298G | E | 4.9e-07 | 1.8e-07 | 1e-07 | 2.567E-07 | 2.060E-07 | 1.000E-07 | 4.900E-07 |
| 25949 | 1298 | G1298H | E | 2.2e-07 | 2e-07 | 1.4e-07 | 1.867E-07 | 4.163E-08 | 1.400E-07 | 2.200E-07 |
| 25950 | 1298 | G1298I | E | 2.3e-07 | 1.9e-07 | 2.1e-07 | 2.100E-07 | 2.000E-08 | 1.900E-07 | 2.300E-07 |
| 25951 | 1298 | G1298L | E | 1.8e-07 | 2.3e-07 | 3e-07 | 2.367E-07 | 6.028E-08 | 1.800E-07 | 3.000E-07 |
| 25952 | 1298 | G1298K | E | 2.3e-07 | 3e-07 | 4e-07 | 3.100E-07 | 8.544E-08 | 2.300E-07 | 4.000E-07 |
| 25953 | 1298 | G1298M | E | 3e-07 | 2e-07 | 3.3e-07 | 2.767E-07 | 6.807E-08 | 2.000E-07 | 3.300E-07 |
| 25954 | 1298 | G1298F | E | 1.8e-07 | 3.1e-07 | 2.1e-07 | 2.333E-07 | 6.807E-08 | 1.800E-07 | 3.100E-07 |
| 25955 | 1298 | G1298P | E | 2.6e-07 | 3.5e-07 | 2.9e-07 | 3.000E-07 | 4.583E-08 | 2.600E-07 | 3.500E-07 |
| 25956 | 1298 | G1298S | E | 1.8e-07 | 3.7e-07 | 2.3e-07 | 2.600E-07 | 9.849E-08 | 1.800E-07 | 3.700E-07 |
| 25957 | 1298 | G1298T | E | 2.5e-07 | 5.7e-07 | 2.6e-07 | 3.600E-07 | 1.819E-07 | 2.500E-07 | 5.700E-07 |
| 25958 | 1298 | G1298W | E | 5.3e-07 | 5.3e-07 | 1.7e-07 | 4.100E-07 | 2.078E-07 | 1.700E-07 | 5.300E-07 |
| 25959 | 1298 | G1298Y | E | 3.6e-07 | 2.6e-07 | 1.9e-07 | 2.700E-07 | 8.544E-08 | 1.900E-07 | 3.600E-07 |
| 25960 | 1298 | G1298V | E | 1.9e-07 | 1.4e-07 | 1.6e-07 | 1.633E-07 | 2.517E-08 | 1.400E-07 | 1.900E-07 |
| 25961 | 1299 | F1299A | E | 9.2e-08 | 1.7e-07 | 1.8e-07 | 1.473E-07 | 4.818E-08 | 9.200E-08 | 1.800E-07 |
| 25962 | 1299 | F1299R | E | 3.9e-07 | 1.4e-07 | 2.4e-07 | 2.567E-07 | 1.258E-07 | 1.400E-07 | 3.900E-07 |
| 25963 | 1299 | F1299N | E | 3.7e-07 | 3.7e-07 | 3e-07 | 3.467E-07 | 4.041E-08 | 3.000E-07 | 3.700E-07 |
| 25964 | 1299 | F1299D | E | 2.2e-07 | 1.7e-07 | 5.6e-07 | 3.167E-07 | 2.122E-07 | 1.700E-07 | 5.600E-07 |
| 25965 | 1299 | F1299C | E | 1.7e-07 | 3.5e-07 | 2.2e-07 | 2.467E-07 | 9.292E-08 | 1.700E-07 | 3.500E-07 |
| 25966 | 1299 | F1299E | E | 1.8e-07 | 3.2e-07 | 1.5e-07 | 2.167E-07 | 9.074E-08 | 1.500E-07 | 3.200E-07 |
| 25967 | 1299 | F1299Q | E | 1.1e-07 | 2.1e-07 | 4.2e-07 | 2.467E-07 | 1.582E-07 | 1.100E-07 | 4.200E-07 |
| 25968 | 1299 | F1299G | E | 3.8e-07 | 3.4e-07 | 2.9e-07 | 3.367E-07 | 4.509E-08 | 2.900E-07 | 3.800E-07 |
| 25969 | 1299 | F1299H | E | 3.7e-07 | 2.8e-07 | 5.5e-07 | 4.000E-07 | 1.375E-07 | 2.800E-07 | 5.500E-07 |
| 25970 | 1299 | F1299I | E | 3.1e-07 | 4.1e-07 | 3.4e-07 | 3.533E-07 | 5.132E-08 | 3.100E-07 | 4.100E-07 |
| 25971 | 1299 | F1299L | E | 1.5e-07 | 2.4e-07 | 2.3e-07 | 2.067E-07 | 4.933E-08 | 1.500E-07 | 2.400E-07 |
| 25972 | 1299 | F1299K | E | 1.9e-07 | 1.9e-07 | 7.2e-08 | 1.507E-07 | 6.813E-08 | 7.200E-08 | 1.900E-07 |
| 25973 | 1299 | F1299M | E | 2e-07 | 2.4e-07 | 2.9e-07 | 2.433E-07 | 4.509E-08 | 2.000E-07 | 2.900E-07 |
| 25974 | 1299 | F1299F | E | 4.1e-07 | 1.3e-07 | 1.9e-07 | 2.433E-07 | 1.474E-07 | 1.300E-07 | 4.100E-07 |
| 25975 | 1299 | F1299P | E | 4e-07 | 3e-07 | 1.9e-07 | 2.967E-07 | 1.050E-07 | 1.900E-07 | 4.000E-07 |
| 25976 | 1299 | F1299S | E | 6.5e-07 | 7.6e-07 | 8.5e-08 | 4.983E-07 | 3.622E-07 | 8.500E-08 | 7.600E-07 |
| 25977 | 1299 | F1299T | E | 2.4e-07 | 9.3e-08 | 6.4e-07 | 3.243E-07 | 2.831E-07 | 9.300E-08 | 6.400E-07 |
| 25978 | 1299 | F1299W | E | 2.9e-07 | 2.1e-07 | 3.9e-07 | 2.967E-07 | 9.018E-08 | 2.100E-07 | 3.900E-07 |
| 25979 | 1299 | F1299Y | E | 1.2e-07 | 3.7e-07 | 1.8e-07 | 2.233E-07 | 1.305E-07 | 1.200E-07 | 3.700E-07 |
| 25980 | 1299 | F1299V | E | 2.1e-07 | 1.8e-07 | 2.3e-07 | 2.067E-07 | 2.517E-08 | 1.800E-07 | 2.300E-07 |
| 25981 | 1300 | N1300A | E | 2.3e-07 | 2.5e-07 | 1.5e-07 | 2.100E-07 | 5.292E-08 | 1.500E-07 | 2.500E-07 |
| 25982 | 1300 | N1300R | E | 7.6e-08 | 1.3e-07 | 1.9e-07 | 1.320E-07 | 5.703E-08 | 7.600E-08 | 1.900E-07 |
| 25983 | 1300 | N1300N | E | 2.8e-07 | 4.3e-07 | 1.3e-07 | 2.800E-07 | 1.500E-07 | 1.300E-07 | 4.300E-07 |
| 25984 | 1300 | N1300D | E | 2.3e-07 | 2.6e-07 | 1.4e-07 | 2.100E-07 | 6.245E-08 | 1.400E-07 | 2.600E-07 |
| 25985 | 1300 | N1300C | E | 1.9e-07 | 1.6e-07 | 2.3e-07 | 1.933E-07 | 3.512E-08 | 1.600E-07 | 2.300E-07 |
| 25986 | 1300 | N1300E | E | 3.5e-07 | 3.6e-07 | 1.6e-07 | 2.900E-07 | 1.127E-07 | 1.600E-07 | 3.600E-07 |
| 25987 | 1300 | N1300Q | E | 3.8e-07 | 3e-07 | 1.6e-07 | 2.800E-07 | 1.114E-07 | 1.600E-07 | 3.800E-07 |
| 25988 | 1300 | N1300G | E | 1.5e-07 | 3.2e-07 | 2e-07 | 2.233E-07 | 8.737E-08 | 1.500E-07 | 3.200E-07 |
| 25989 | 1300 | N1300H | E | 2.5e-07 | 1.5e-07 | 1.6e-07 | 1.867E-07 | 5.508E-08 | 1.500E-07 | 2.500E-07 |
| 25990 | 1300 | N1300I | E | 2.3e-07 | 2e-07 | 1.4e-07 | 1.900E-07 | 4.583E-08 | 1.400E-07 | 2.300E-07 |
| 25991 | 1300 | N1300L | E | 1.4e-07 | 2.3e-07 | 1.4e-07 | 1.700E-07 | 5.196E-08 | 1.400E-07 | 2.300E-07 |
| 25992 | 1300 | N1300K | E | 1.7e-07 | 2.6e-07 | 2e-07 | 2.100E-07 | 4.583E-08 | 1.700E-07 | 2.600E-07 |
| 25993 | 1300 | N1300M | E | 6.3e-08 | 2.7e-07 | 3.4e-07 | 2.243E-07 | 1.440E-07 | 6.300E-08 | 3.400E-07 |
| 25994 | 1300 | N1300F | E | 2.3e-07 | 1.6e-07 | 2.2e-07 | 2.033E-07 | 3.786E-08 | 1.600E-07 | 2.300E-07 |
| 25995 | 1300 | N1300P | E | 1.3e-07 | 1.6e-07 | 1.6e-07 | 1.500E-07 | 1.732E-08 | 1.300E-07 | 1.600E-07 |
| 25996 | 1300 | N1300S | E | 2.2e-07 | 2e-07 | 2.1e-07 | 2.100E-07 | 1.000E-08 | 2.000E-07 | 2.200E-07 |
| 25997 | 1300 | N1300T | E | 1.5e-07 | 3.3e-07 | 4e-07 | 2.933E-07 | 1.290E-07 | 1.500E-07 | 4.000E-07 |
| 25998 | 1300 | N1300W | E | 2.6e-07 | 1.9e-07 | 3.2e-07 | 2.567E-07 | 6.506E-08 | 1.900E-07 | 3.200E-07 |
| 25999 | 1300 | N1300Y | E | 1.9e-07 | 4.7e-07 | 3.3e-07 | 3.300E-07 | 1.400E-07 | 1.900E-07 | 4.700E-07 |
| 26000 | 1300 | N1300V | E | 1e-07 | 3.2e-07 | 2.4e-07 | 2.200E-07 | 1.114E-07 | 1.000E-07 | 3.200E-07 |
| 26001 | 1301 | I1301A | E | 5.5e-07 | 3.9e-07 | 5.8e-07 | 5.067E-07 | 1.021E-07 | 3.900E-07 | 5.800E-07 |
| 26002 | 1301 | I1301R | E | 2.5e-07 | 3.1e-07 | 1.9e-07 | 2.500E-07 | 6.000E-08 | 1.900E-07 | 3.100E-07 |
| 26003 | 1301 | I1301N | E | 3.9e-07 | 2.2e-07 | 3.2e-07 | 3.100E-07 | 8.544E-08 | 2.200E-07 | 3.900E-07 |
| 26004 | 1301 | I1301D | E | 9.5e-08 | 2.3e-07 | 2.4e-07 | 1.883E-07 | 8.098E-08 | 9.500E-08 | 2.400E-07 |
| 26005 | 1301 | I1301C | E | 2.5e-07 | 1.3e-07 | 2.9e-07 | 2.233E-07 | 8.327E-08 | 1.300E-07 | 2.900E-07 |
| 26006 | 1301 | I1301E | E | 2.7e-07 | 2.4e-07 | 2.9e-07 | 2.667E-07 | 2.517E-08 | 2.400E-07 | 2.900E-07 |
| 26007 | 1301 | I1301Q | E | 4.2e-07 | 2.9e-07 | 4.9e-07 | 4.000E-07 | 1.015E-07 | 2.900E-07 | 4.900E-07 |
| 26008 | 1301 | I1301G | E | 8e-08 | 2.3e-07 | 2.5e-07 | 1.867E-07 | 9.292E-08 | 8.000E-08 | 2.500E-07 |
| 26009 | 1301 | I1301H | E | 2e-07 | 2e-07 | 2.7e-07 | 2.233E-07 | 4.041E-08 | 2.000E-07 | 2.700E-07 |

| | | | | | | | | | | |
|---|---|---|---|---|---|---|---|---|---|---|
| 26010 | 1301 | I1301I | E | 1.8e-07 | 3.6e-07 | 1.5e-07 | 2.300E-07 | 1.136E-07 | 1.500E-07 | 3.600E-07 |
| 26011 | 1301 | I1301L | E | 4.3e-07 | 9.8e-08 | 2.2e-07 | 2.493E-07 | 1.679E-07 | 9.800E-08 | 4.300E-07 |
| 26012 | 1301 | I1301K | E | 4.2e-07 | 1.2e-07 | 1.6e-07 | 2.333E-07 | 1.629E-07 | 1.200E-07 | 4.200E-07 |
| 26013 | 1301 | I1301M | E | 2.1e-07 | 3.5e-07 | 3.2e-07 | 2.933E-07 | 7.371E-08 | 2.100E-07 | 3.500E-07 |
| 26014 | 1301 | I1301F | E | 1.4e-07 | 5.2e-07 | 4e-07 | 3.533E-07 | 1.943E-07 | 1.400E-07 | 5.200E-07 |
| 26015 | 1301 | I1301P | E | 1.8e-07 | 3e-07 | 1.6e-07 | 2.133E-07 | 7.572E-08 | 1.600E-07 | 3.000E-07 |
| 26016 | 1301 | I1301S | E | 2.8e-07 | 2.5e-07 | 2e-07 | 2.433E-07 | 4.041E-08 | 2.000E-07 | 2.800E-07 |
| 26017 | 1301 | I1301T | E | 1.8e-07 | 4.1e-07 | 5e-07 | 3.633E-07 | 1.650E-07 | 1.800E-07 | 5.000E-07 |
| 26018 | 1301 | I1301W | E | 2.8e-07 | 4.5e-07 | 4.7e-07 | 4.000E-07 | 1.044E-07 | 2.800E-07 | 4.700E-07 |
| 26019 | 1301 | I1301Y | E | 2.9e-07 | 4.1e-07 | 4.1e-07 | 3.700E-07 | 6.928E-08 | 2.900E-07 | 4.100E-07 |
| 26020 | 1301 | I1301V | E | 2.5e-07 | 4.9e-07 | 4.7e-07 | 4.033E-07 | 1.332E-07 | 2.500E-07 | 4.900E-07 |
| 26021 | 1302 | K1302A | E | 4.6e-07 | 1.9e-07 | 3.2e-07 | 3.233E-07 | 1.350E-07 | 1.900E-07 | 4.600E-07 |
| 26022 | 1302 | K1302R | E | 4.7e-07 | 2.1e-07 | 4e-07 | 3.600E-07 | 1.345E-07 | 2.100E-07 | 4.700E-07 |
| 26023 | 1302 | K1302N | E | 2.6e-07 | 4.8e-08 | 2.9e-07 | 1.993E-07 | 1.319E-07 | 4.800E-08 | 2.900E-07 |
| 26024 | 1302 | K1302D | E | 2.5e-07 | 3.8e-07 | 3.6e-07 | 3.300E-07 | 7.000E-08 | 2.500E-07 | 3.800E-07 |
| 26025 | 1302 | K1302C | E | 3.3e-07 | 2.5e-07 | 1.9e-07 | 2.567E-07 | 7.024E-08 | 1.900E-07 | 3.300E-07 |
| 26026 | 1302 | K1302E | E | 2.1e-07 | 3.6e-07 | 1.6e-07 | 2.433E-07 | 1.041E-07 | 1.600E-07 | 3.600E-07 |
| 26027 | 1302 | K1302Q | E | 2.9e-07 | 1.9e-07 | 1.9e-07 | 2.233E-07 | 5.774E-08 | 1.900E-07 | 2.900E-07 |
| 26028 | 1302 | K1302G | E | 3.5e-07 | 1.1e-07 | 2.5e-07 | 2.367E-07 | 1.206E-07 | 1.100E-07 | 3.500E-07 |
| 26029 | 1302 | K1302H | E | 3.5e-07 | 2e-07 | 4e-07 | 3.167E-07 | 1.041E-07 | 2.000E-07 | 4.000E-07 |
| 26030 | 1302 | K1302I | E | 1.2e-07 | 2.7e-07 | 1.1e-07 | 1.667E-07 | 8.963E-08 | 1.100E-07 | 2.700E-07 |
| 26031 | 1302 | K1302L | E | 3.1e-07 | 7.3e-07 | 8.4e-08 | 3.747E-07 | 3.278E-07 | 8.400E-08 | 7.300E-07 |
| 26032 | 1302 | K1302K | E | 3.9e-07 | 2.3e-07 | 2.3e-07 | 2.833E-07 | 9.238E-08 | 2.300E-07 | 3.900E-07 |
| 26033 | 1302 | K1302M | E | 3.4e-07 | 1.6e-07 | 3.5e-07 | 2.833E-07 | 1.069E-07 | 1.600E-07 | 3.500E-07 |
| 26034 | 1302 | K1302F | E | 1.7e-07 | 1.8e-07 | 2.9e-07 | 2.133E-07 | 6.658E-08 | 1.700E-07 | 2.900E-07 |
| 26035 | 1302 | K1302P | E | 1.2e-07 | 4.4e-07 | 1.6e-07 | 2.400E-07 | 1.744E-07 | 1.200E-07 | 4.400E-07 |
| 26036 | 1302 | K1302S | E | 2.3e-07 | 3.2e-07 | 1.9e-07 | 2.467E-07 | 6.658E-08 | 1.900E-07 | 3.200E-07 |
| 26037 | 1302 | K1302T | E | 2.5e-07 | 2.5e-07 | 2.4e-07 | 2.467E-07 | 5.774E-09 | 2.400E-07 | 2.500E-07 |
| 26038 | 1302 | K1302W | E | 1.8e-07 | 2.7e-07 | 1.9e-07 | 2.133E-07 | 4.933E-08 | 1.800E-07 | 2.700E-07 |
| 26039 | 1302 | K1302Y | E | 5e-07 | 2.3e-07 | 2.4e-07 | 3.233E-07 | 1.531E-07 | 2.300E-07 | 5.000E-07 |
| 26040 | 1302 | K1302V | E | 2.2e-07 | 1.3e-07 | 3.3e-07 | 2.267E-07 | 1.002E-07 | 1.300E-07 | 3.300E-07 |
| 26041 | 1303 | D1303A | E | 2e-07 | 5.2e-07 | 1.1e-07 | 2.767E-07 | 2.155E-07 | 1.100E-07 | 5.200E-07 |
| 26042 | 1303 | D1303R | E | 1.1e-07 | 2.1e-07 | 2.2e-07 | 1.800E-07 | 6.083E-08 | 1.100E-07 | 2.200E-07 |
| 26043 | 1303 | D1303N | E | 2.2e-07 | 1.8e-07 | 3.2e-07 | 2.400E-07 | 7.211E-08 | 1.800E-07 | 3.200E-07 |
| 26044 | 1303 | D1303D | E | 1.9e-07 | 2.4e-07 | 6.3e-08 | 1.643E-07 | 9.125E-08 | 6.300E-08 | 2.400E-07 |
| 26045 | 1303 | D1303C | E | 1.8e-07 | 2e-07 | 6e-07 | 3.267E-07 | 2.369E-07 | 1.800E-07 | 6.000E-07 |
| 26046 | 1303 | D1303E | E | 1.3e-07 | 1.2e-07 | 2.4e-07 | 1.633E-07 | 6.658E-08 | 1.200E-07 | 2.400E-07 |
| 26047 | 1303 | D1303Q | E | 2.3e-07 | 1.3e-07 | 2.2e-07 | 1.933E-07 | 5.508E-08 | 1.300E-07 | 2.300E-07 |
| 26048 | 1303 | D1303G | E | 3.1e-07 | 2.7e-07 | 3.8e-07 | 3.200E-07 | 5.568E-08 | 2.700E-07 | 3.800E-07 |
| 26049 | 1303 | D1303H | E | 8.4e-08 | 3.2e-07 | 2.2e-07 | 2.080E-07 | 1.185E-07 | 8.400E-08 | 3.200E-07 |
| 26050 | 1303 | D1303I | E | 1.9e-07 | 1.3e-07 | 2.6e-07 | 1.933E-07 | 6.506E-08 | 1.300E-07 | 2.600E-07 |
| 26051 | 1303 | D1303L | E | 2.2e-07 | 9.1e-08 | 1.2e-07 | 1.437E-07 | 6.768E-08 | 9.100E-08 | 2.200E-07 |
| 26052 | 1303 | D1303K | E | 1.5e-07 | 5.2e-07 | 1.1e-07 | 2.600E-07 | 2.261E-07 | 1.100E-07 | 5.200E-07 |
| 26053 | 1303 | D1303M | E | 2.4e-07 | 3e-07 | 1e-07 | 2.133E-07 | 1.026E-07 | 1.000E-07 | 3.000E-07 |
| 26054 | 1303 | D1303F | E | 3.5e-07 | 2.1e-07 | 3.6e-07 | 3.067E-07 | 8.386E-08 | 2.100E-07 | 3.600E-07 |
| 26055 | 1303 | D1303P | E | 2.9e-07 | 1.8e-07 | 2.9e-07 | 2.533E-07 | 6.351E-08 | 1.800E-07 | 2.900E-07 |
| 26056 | 1303 | D1303S | E | 1e-07 | 2.6e-07 | 2.2e-07 | 1.933E-07 | 8.327E-08 | 1.000E-07 | 2.600E-07 |
| 26057 | 1303 | D1303T | E | 3.4e-07 | 3.5e-07 | 1.8e-07 | 2.900E-07 | 9.539E-08 | 1.800E-07 | 3.500E-07 |
| 26058 | 1303 | D1303W | E | 3.2e-07 | 3.6e-07 | 1.9e-07 | 2.900E-07 | 8.888E-08 | 1.900E-07 | 3.600E-07 |
| 26059 | 1303 | D1303Y | E | 3e-07 | 1.5e-07 | 3.3e-07 | 2.600E-07 | 9.644E-08 | 1.500E-07 | 3.300E-07 |
| 26060 | 1303 | D1303V | E | 2.2e-07 | 8e-08 | 2e-07 | 1.667E-07 | 7.572E-08 | 8.000E-08 | 2.200E-07 |
| 26061 | 1304 | T1304A | E | 5.2e-07 | 1.7e-07 | 1.5e-07 | 2.800E-07 | 2.081E-07 | 1.500E-07 | 5.200E-07 |
| 26062 | 1304 | T1304R | E | 2.7e-07 | 2e-07 | 6.7e-08 | 1.790E-07 | 1.031E-07 | 6.700E-08 | 2.700E-07 |
| 26063 | 1304 | T1304N | E | 3.9e-07 | 1.1e-07 | 3e-07 | 2.667E-07 | 1.429E-07 | 1.100E-07 | 3.900E-07 |
| 26064 | 1304 | T1304D | E | 2.4e-07 | 3.2e-07 | 1.3e-07 | 2.300E-07 | 9.539E-08 | 1.300E-07 | 3.200E-07 |
| 26065 | 1304 | T1304C | E | 1.1e-07 | 3.6e-07 | 3.5e-07 | 2.733E-07 | 1.415E-07 | 1.100E-07 | 3.600E-07 |
| 26066 | 1304 | T1304E | E | 2.3e-07 | 3.4e-07 | 1.4e-07 | 2.367E-07 | 1.002E-07 | 1.400E-07 | 3.400E-07 |
| 26067 | 1304 | T1304Q | E | 1.5e-07 | 1.9e-07 | 2.6e-07 | 2.000E-07 | 5.568E-08 | 1.500E-07 | 2.600E-07 |
| 26068 | 1304 | T1304G | E | 4.1e-07 | 2e-07 | 3.4e-07 | 3.167E-07 | 1.069E-07 | 2.000E-07 | 4.100E-07 |
| 26069 | 1304 | T1304H | E | 4e-07 | 2.9e-07 | 2.2e-07 | 3.033E-07 | 9.074E-08 | 2.200E-07 | 4.000E-07 |
| 26070 | 1304 | T1304I | E | 4e-07 | 1.3e-07 | 2.7e-07 | 2.667E-07 | 1.350E-07 | 1.300E-07 | 4.000E-07 |
| 26071 | 1304 | T1304L | E | 2.1e-07 | 3.1e-07 | 2.8e-07 | 2.667E-07 | 5.132E-08 | 2.100E-07 | 3.100E-07 |
| 26072 | 1304 | T1304K | E | 3.5e-07 | 8.2e-08 | 1.3e-07 | 1.873E-07 | 1.429E-07 | 8.200E-08 | 3.500E-07 |
| 26073 | 1304 | T1304M | E | 2.3e-07 | 2.3e-07 | 2.6e-07 | 2.400E-07 | 1.732E-08 | 2.300E-07 | 2.600E-07 |
| 26074 | 1304 | T1304F | E | 1.7e-07 | 3.1e-07 | 2.6e-07 | 2.467E-07 | 7.095E-08 | 1.700E-07 | 3.100E-07 |
| 26075 | 1304 | T1304P | E | 3.3e-07 | 1.1e-07 | 1.7e-07 | 2.033E-07 | 1.137E-07 | 1.100E-07 | 3.300E-07 |
| 26076 | 1304 | T1304S | E | 5.7e-07 | 7.5e-07 | 1.7e-07 | 4.967E-07 | 2.969E-07 | 1.700E-07 | 7.500E-07 |
| 26077 | 1304 | T1304T | E | 3.9e-07 | 2.4e-07 | 1.7e-07 | 2.667E-07 | 1.124E-07 | 1.700E-07 | 3.900E-07 |
| 26078 | 1304 | T1304W | E | 1.9e-07 | 4.3e-07 | 4e-07 | 3.400E-07 | 1.308E-07 | 1.900E-07 | 4.300E-07 |
| 26079 | 1304 | T1304Y | E | 3.4e-07 | 5.2e-07 | 2.4e-07 | 3.667E-07 | 1.419E-07 | 2.400E-07 | 5.200E-07 |
| 26080 | 1304 | T1304V | E | 1e-07 | 1.4e-07 | 6e-08 | 1.000E-07 | 4.000E-08 | 6.000E-08 | 1.400E-07 |
| 26081 | 1305 | Y1305A | E | 3e-07 | 4.6e-07 | 2.4e-07 | 3.333E-07 | 1.137E-07 | 2.400E-07 | 4.600E-07 |
| 26082 | 1305 | Y1305R | E | 2.5e-07 | 1.5e-07 | 7.5e-08 | 1.583E-07 | 8.780E-08 | 7.500E-08 | 2.500E-07 |
| 26083 | 1305 | Y1305N | E | 1.6e-07 | 2.4e-07 | 4.1e-07 | 2.700E-07 | 1.277E-07 | 1.600E-07 | 4.100E-07 |
| 26084 | 1305 | Y1305D | E | 1.7e-07 | 7.3e-08 | 2.7e-07 | 1.710E-07 | 9.850E-08 | 7.300E-08 | 2.700E-07 |
| 26085 | 1305 | Y1305C | E | 4.8e-07 | 9.9e-08 | 2.5e-07 | 2.763E-07 | 1.919E-07 | 9.900E-08 | 4.800E-07 |
| 26086 | 1305 | Y1305E | E | 2.8e-07 | 2.1e-07 | 3.2e-07 | 2.700E-07 | 5.568E-08 | 2.100E-07 | 3.200E-07 |
| 26087 | 1305 | Y1305Q | E | 2.2e-07 | 1.5e-07 | 3.3e-07 | 2.333E-07 | 9.074E-08 | 1.500E-07 | 3.300E-07 |
| 26088 | 1305 | Y1305G | E | 3.9e-07 | 2.6e-07 | 1e-07 | 2.500E-07 | 1.453E-07 | 1.000E-07 | 3.900E-07 |
| 26089 | 1305 | Y1305H | E | 3.9e-07 | 2.6e-07 | 3.5e-07 | 3.333E-07 | 6.658E-08 | 2.600E-07 | 3.900E-07 |
| 26090 | 1305 | Y1305I | E | 4e-07 | 2.5e-07 | 4.7e-07 | 3.733E-07 | 1.124E-07 | 2.500E-07 | 4.700E-07 |
| 26091 | 1305 | Y1305L | E | 2.3e-07 | 2.1e-07 | 3.4e-07 | 2.600E-07 | 7.000E-08 | 2.100E-07 | 3.400E-07 |
| 26092 | 1305 | Y1305K | E | 2e-07 | 1.6e-07 | 1.7e-07 | 1.767E-07 | 2.082E-08 | 1.600E-07 | 2.000E-07 |
| 26093 | 1305 | Y1305M | E | 2.3e-07 | 2.2e-07 | 2.7e-07 | 2.400E-07 | 2.646E-08 | 2.200E-07 | 2.700E-07 |
| 26094 | 1305 | Y1305F | E | 3.7e-07 | 1.8e-07 | 1.9e-07 | 2.467E-07 | 1.069E-07 | 1.800E-07 | 3.700E-07 |

| | | | | | | | | | | |
|---|---|---|---|---|---|---|---|---|---|---|
| 26095 | 1305 | Y1305P | E | 4.6e-07 | 1.5e-07 | 5.5e-07 | 3.867E-07 | 2.098E-07 | 1.500E-07 | 5.500E-07 |
| 26096 | 1305 | Y1305S | E | 2.5e-07 | 2.7e-07 | 2.3e-07 | 2.500E-07 | 2.000E-08 | 2.300E-07 | 2.700E-07 |
| 26097 | 1305 | Y1305T | E | 2e-07 | 4.3e-07 | 2.9e-07 | 3.067E-07 | 1.159E-07 | 2.000E-07 | 4.300E-07 |
| 26098 | 1305 | Y1305W | E | 2.5e-07 | 3.5e-07 | 5.7e-07 | 3.900E-07 | 1.637E-07 | 2.500E-07 | 5.700E-07 |
| 26099 | 1305 | Y1305Y | E | 2.2e-07 | 3.6e-07 | 1.2e-07 | 2.333E-07 | 1.206E-07 | 1.200E-07 | 3.600E-07 |
| 26100 | 1305 | Y1305V | E | 5.1e-07 | 2.1e-07 | 2.1e-07 | 3.100E-07 | 1.732E-07 | 2.100E-07 | 5.100E-07 |
| 26101 | 1306 | I1306A | E | 2.8e-07 | 1.8e-07 | 2.1e-07 | 2.233E-07 | 5.132E-08 | 1.800E-07 | 2.800E-07 |
| 26102 | 1306 | I1306R | E | 4.1e-07 | 1.2e-07 | 1.5e-07 | 2.267E-07 | 1.595E-07 | 1.200E-07 | 4.100E-07 |
| 26103 | 1306 | I1306N | E | 3.5e-07 | 2.8e-07 | 6.5e-07 | 4.267E-07 | 1.966E-07 | 2.800E-07 | 6.500E-07 |
| 26104 | 1306 | I1306D | E | 2e-07 | 2.3e-07 | 2.6e-07 | 2.300E-07 | 3.000E-08 | 2.000E-07 | 2.600E-07 |
| 26105 | 1306 | I1306C | E | 2e-07 | 2e-07 | 2.9e-07 | 2.300E-07 | 5.196E-08 | 2.000E-07 | 2.900E-07 |
| 26106 | 1306 | I1306E | E | 3.8e-07 | 2.1e-07 | 4.4e-07 | 3.433E-07 | 1.193E-07 | 2.100E-07 | 4.400E-07 |
| 26107 | 1306 | I1306Q | E | 2.7e-07 | 2.5e-07 | 1.4e-07 | 2.200E-07 | 7.000E-08 | 1.400E-07 | 2.700E-07 |
| 26108 | 1306 | I1306G | E | 3.3e-07 | 2e-07 | 1.4e-07 | 2.233E-07 | 9.713E-08 | 1.400E-07 | 3.300E-07 |
| 26109 | 1306 | I1306H | E | 1.6e-07 | 1.5e-07 | 3.4e-07 | 2.167E-07 | 1.069E-07 | 1.500E-07 | 3.400E-07 |
| 26110 | 1306 | I1306I | E | 1.6e-07 | 2.3e-07 | 2e-07 | 1.967E-07 | 3.512E-08 | 1.600E-07 | 2.300E-07 |
| 26111 | 1306 | I1306L | E | 1.8e-07 | 2.9e-07 | 3.9e-07 | 2.867E-07 | 1.050E-07 | 1.800E-07 | 3.900E-07 |
| 26112 | 1306 | I1306K | E | 2.8e-07 | 1.6e-07 | 5.8e-07 | 3.400E-07 | 2.163E-07 | 1.600E-07 | 5.800E-07 |
| 26113 | 1306 | I1306M | E | 2e-07 | 3.5e-07 | 3.1e-07 | 2.867E-07 | 7.767E-08 | 2.000E-07 | 3.500E-07 |
| 26114 | 1306 | I1306F | E | 2.8e-07 | 1.6e-07 | 2e-07 | 2.133E-07 | 6.110E-08 | 1.600E-07 | 2.800E-07 |
| 26115 | 1306 | I1306P | E | 3.3e-07 | 4.4e-07 | 2.2e-07 | 3.300E-07 | 1.100E-07 | 2.200E-07 | 4.400E-07 |
| 26116 | 1306 | I1306S | E | 2.2e-07 | 1.1e-07 | 2.5e-07 | 1.933E-07 | 7.371E-08 | 1.100E-07 | 2.500E-07 |
| 26117 | 1306 | I1306T | E | 1.7e-07 | 1.3e-07 | 1.2e-07 | 1.400E-07 | 2.646E-08 | 1.200E-07 | 1.700E-07 |
| 26118 | 1306 | I1306W | E | 6.2e-07 | 1.2e-07 | 2.5e-07 | 3.300E-07 | 2.594E-07 | 1.200E-07 | 6.200E-07 |
| 26119 | 1306 | I1306Y | E | 2.5e-07 | 4.1e-07 | 1.6e-07 | 2.733E-07 | 1.266E-07 | 1.600E-07 | 4.100E-07 |
| 26120 | 1306 | I1306V | E | 2e-07 | 3e-07 | 2.7e-07 | 2.567E-07 | 5.132E-08 | 2.000E-07 | 3.000E-07 |
| 26121 | 1307 | H1307A | E | 2.1e-07 | 3.4e-07 | 2e-07 | 2.500E-07 | 7.810E-08 | 2.000E-07 | 3.400E-07 |
| 26122 | 1307 | H1307R | E | 2.4e-07 | 4.5e-07 | 2e-07 | 2.967E-07 | 1.343E-07 | 2.000E-07 | 4.500E-07 |
| 26123 | 1307 | H1307N | E | 3.2e-07 | 3.6e-07 | 2.8e-07 | 3.200E-07 | 4.000E-08 | 2.800E-07 | 3.600E-07 |
| 26124 | 1307 | H1307D | E | 1.7e-07 | 2.5e-07 | 3.1e-07 | 2.433E-07 | 7.024E-08 | 1.700E-07 | 3.100E-07 |
| 26125 | 1307 | H1307C | E | 1.9e-07 | 2.4e-07 | 3.2e-07 | 2.500E-07 | 6.557E-08 | 1.900E-07 | 3.200E-07 |
| 26126 | 1307 | H1307E | E | 2.4e-07 | 3e-07 | 4.2e-07 | 3.200E-07 | 9.165E-08 | 2.400E-07 | 4.200E-07 |
| 26127 | 1307 | H1307Q | E | 3.1e-07 | 3.4e-07 | 2.5e-07 | 3.000E-07 | 4.583E-08 | 2.500E-07 | 3.400E-07 |
| 26128 | 1307 | H1307G | E | 2.3e-07 | 1.2e-07 | 3.2e-07 | 2.233E-07 | 1.002E-07 | 1.200E-07 | 3.200E-07 |
| 26129 | 1307 | H1307H | E | 2.9e-07 | 3.2e-07 | 4.8e-07 | 3.633E-07 | 1.021E-07 | 2.900E-07 | 4.800E-07 |
| 26130 | 1307 | H1307I | E | 4.8e-07 | 1.3e-07 | 1.4e-07 | 2.500E-07 | 1.992E-07 | 1.300E-07 | 4.800E-07 |
| 26131 | 1307 | H1307L | E | 2.8e-07 | 2.3e-07 | 8.7e-07 | 4.600E-07 | 3.559E-07 | 2.300E-07 | 8.700E-07 |
| 26132 | 1307 | H1307K | E | 1.4e-07 | 1.8e-07 | 2.8e-07 | 2.000E-07 | 7.211E-08 | 1.400E-07 | 2.800E-07 |
| 26133 | 1307 | H1307M | E | 4.2e-07 | 2.6e-07 | 2.4e-07 | 3.067E-07 | 9.866E-08 | 2.400E-07 | 4.200E-07 |
| 26134 | 1307 | H1307F | E | 1.8e-07 | 2.5e-07 | 5.9e-07 | 3.400E-07 | 2.193E-07 | 1.800E-07 | 5.900E-07 |
| 26135 | 1307 | H1307P | E | 2e-07 | 1.7e-07 | 1e-07 | 1.567E-07 | 5.132E-08 | 1.000E-07 | 2.000E-07 |
| 26136 | 1307 | H1307S | E | 4.6e-07 | 3.7e-07 | 1.4e-07 | 3.233E-07 | 1.650E-07 | 1.400E-07 | 4.600E-07 |
| 26137 | 1307 | H1307T | E | 2.4e-07 | 1.8e-07 | 2.9e-07 | 2.367E-07 | 5.508E-08 | 1.800E-07 | 2.900E-07 |
| 26138 | 1307 | H1307W | E | 6.5e-08 | 1.3e-07 | 1e-07 | 9.833E-08 | 3.253E-08 | 6.500E-08 | 1.300E-07 |
| 26139 | 1307 | H1307Y | E | 1.4e-07 | 1.5e-07 | 6.3e-07 | 3.067E-07 | 2.801E-07 | 1.400E-07 | 6.300E-07 |
| 26140 | 1307 | H1307V | E | 2.5e-07 | 1.5e-07 | 3.4e-07 | 2.467E-07 | 9.504E-08 | 1.500E-07 | 3.400E-07 |
| 26141 | 1308 | W1308A | E | 4.9e-07 | 1.4e-07 | 3.6e-07 | 3.300E-07 | 1.769E-07 | 1.400E-07 | 4.900E-07 |
| 26142 | 1308 | W1308R | E | 2.7e-07 | 1.6e-07 | 7.7e-07 | 4.000E-07 | 3.251E-07 | 1.600E-07 | 7.700E-07 |
| 26143 | 1308 | W1308N | E | 1.1e-07 | 3.7e-07 | 4.8e-07 | 3.200E-07 | 1.900E-07 | 1.100E-07 | 4.800E-07 |
| 26144 | 1308 | W1308D | E | 2.7e-07 | 2.6e-07 | 1.6e-07 | 2.300E-07 | 6.083E-08 | 1.600E-07 | 2.700E-07 |
| 26145 | 1308 | W1308C | E | 1.7e-07 | 2.3e-07 | 3.1e-07 | 2.367E-07 | 7.024E-08 | 1.700E-07 | 3.100E-07 |
| 26146 | 1308 | W1308E | E | 1.6e-07 | 2.1e-07 | 1.7e-07 | 1.800E-07 | 2.646E-08 | 1.600E-07 | 2.100E-07 |
| 26147 | 1308 | W1308Q | E | 1e-07 | 4.9e-07 | 4e-07 | 3.300E-07 | 2.042E-07 | 1.000E-07 | 4.900E-07 |
| 26148 | 1308 | W1308G | E | 2.1e-07 | 5.4e-07 | 7e-07 | 4.833E-07 | 2.499E-07 | 2.100E-07 | 7.000E-07 |
| 26149 | 1308 | W1308H | E | 2.4e-07 | 2.3e-07 | 2.9e-07 | 2.533E-07 | 3.215E-08 | 2.300E-07 | 2.900E-07 |
| 26150 | 1308 | W1308I | E | 1.5e-07 | 3.3e-07 | 3.6e-07 | 2.800E-07 | 1.136E-07 | 1.500E-07 | 3.600E-07 |
| 26151 | 1308 | W1308L | E | 1.7e-07 | 2.5e-07 | 2.3e-07 | 2.167E-07 | 4.163E-08 | 1.700E-07 | 2.500E-07 |
| 26152 | 1308 | W1308K | E | 2.2e-07 | 2.7e-07 | 2.2e-07 | 2.367E-07 | 2.887E-08 | 2.200E-07 | 2.700E-07 |
| 26153 | 1308 | W1308M | E | 2.1e-07 | 3.1e-07 | 3.2e-07 | 2.800E-07 | 6.083E-08 | 2.100E-07 | 3.200E-07 |
| 26154 | 1308 | W1308F | E | 1.7e-07 | 2.6e-07 | 2.7e-07 | 2.333E-07 | 5.508E-08 | 1.700E-07 | 2.700E-07 |
| 26155 | 1308 | W1308P | E | 4.8e-07 | 2e-07 | 3.3e-07 | 3.367E-07 | 1.401E-07 | 2.000E-07 | 4.800E-07 |
| 26156 | 1308 | W1308S | E | 2.4e-07 | 1.7e-07 | 3.1e-07 | 2.400E-07 | 7.000E-08 | 1.700E-07 | 3.100E-07 |
| 26157 | 1308 | W1308T | E | 1.2e-07 | 2.6e-07 | 1e-07 | 1.600E-07 | 8.718E-08 | 1.000E-07 | 2.600E-07 |
| 26158 | 1308 | W1308W | E | 8.9e-08 | 4.4e-07 | 2.7e-07 | 2.663E-07 | 1.755E-07 | 8.900E-08 | 4.400E-07 |
| 26159 | 1308 | W1308Y | E | 1.6e-07 | 3e-07 | 2.4e-07 | 2.333E-07 | 7.024E-08 | 1.600E-07 | 3.000E-07 |
| 26160 | 1308 | W1308V | E | 1.3e-07 | 3.5e-07 | 2.8e-07 | 2.533E-07 | 1.124E-07 | 1.300E-07 | 3.500E-07 |
| 26161 | 1309 | V1309A | E | 1.5e-07 | 1.1e-07 | 5e-07 | 2.533E-07 | 2.146E-07 | 1.100E-07 | 5.000E-07 |
| 26162 | 1309 | V1309R | E | 2.3e-07 | 2.2e-07 | 4.9e-07 | 3.133E-07 | 1.531E-07 | 2.200E-07 | 4.900E-07 |
| 26163 | 1309 | V1309N | E | 4.5e-07 | 1.9e-07 | 1.9e-07 | 2.767E-07 | 1.501E-07 | 1.900E-07 | 4.500E-07 |
| 26164 | 1309 | V1309D | E | 1.9e-07 | 3.6e-07 | 4.3e-07 | 3.267E-07 | 1.234E-07 | 1.900E-07 | 4.300E-07 |
| 26165 | 1309 | V1309C | E | 2.2e-07 | 1.9e-07 | 2.7e-07 | 2.267E-07 | 4.041E-08 | 1.900E-07 | 2.700E-07 |
| 26166 | 1309 | V1309E | E | 2.2e-07 | 3.5e-07 | 3.7e-07 | 3.133E-07 | 8.145E-08 | 2.200E-07 | 3.700E-07 |
| 26167 | 1309 | V1309Q | E | 2.8e-07 | 3.4e-07 | 2.2e-07 | 2.800E-07 | 6.000E-08 | 2.200E-07 | 3.400E-07 |
| 26168 | 1309 | V1309G | E | 1.8e-07 | 3.6e-07 | 4.2e-07 | 3.200E-07 | 1.249E-07 | 1.800E-07 | 4.200E-07 |
| 26169 | 1309 | V1309H | E | 5.2e-07 | 2.3e-07 | 3.6e-07 | 3.700E-07 | 1.453E-07 | 2.300E-07 | 5.200E-07 |
| 26170 | 1309 | V1309I | E | 1.1e-07 | 3.2e-07 | 1.4e-07 | 1.900E-07 | 1.136E-07 | 1.100E-07 | 3.200E-07 |
| 26171 | 1309 | V1309L | E | 4.3e-07 | 2.3e-07 | 1.2e-07 | 2.600E-07 | 1.572E-07 | 1.200E-07 | 4.300E-07 |
| 26172 | 1309 | V1309K | E | 1e-07 | 2.3e-07 | 2.3e-07 | 1.867E-07 | 7.506E-08 | 1.000E-07 | 2.300E-07 |
| 26173 | 1309 | V1309M | E | 1.5e-07 | 1.6e-07 | 5.3e-07 | 2.800E-07 | 2.166E-07 | 1.500E-07 | 5.300E-07 |
| 26174 | 1309 | V1309F | E | 2.1e-07 | 1.7e-07 | 1.5e-07 | 1.767E-07 | 3.055E-08 | 1.500E-07 | 2.100E-07 |
| 26175 | 1309 | V1309P | E | 3e-07 | 2.9e-07 | 5e-07 | 3.633E-07 | 1.185E-07 | 2.900E-07 | 5.000E-07 |
| 26176 | 1309 | V1309S | E | 3.4e-07 | 7.5e-07 | 1.6e-07 | 4.167E-07 | 3.024E-07 | 1.600E-07 | 7.500E-07 |
| 26177 | 1309 | V1309T | E | 2.5e-07 | 2.5e-07 | 2.5e-07 | 2.500E-07 | 0.000E+00 | 2.500E-07 | 2.500E-07 |
| 26178 | 1309 | V1309W | E | 1.4e-07 | 2.4e-07 | 4.5e-07 | 2.767E-07 | 1.582E-07 | 1.400E-07 | 4.500E-07 |
| 26179 | 1309 | V1309Y | E | 2.2e-07 | 1.5e-07 | 3e-07 | 2.233E-07 | 7.506E-08 | 1.500E-07 | 3.000E-07 |

| | | | | | | | | | | |
|---|---|---|---|---|---|---|---|---|---|---|
| 26180 | 1309 | V1309V | E | 2.5e-07 | 1.7e-07 | 3e-07 | 2.400E-07 | 6.557E-08 | 1.700E-07 | 3.000E-07 |
| 26181 | 1310 | R1310A | E | 1.6e-07 | 2.8e-07 | 2.5e-07 | 2.300E-07 | 6.245E-08 | 1.600E-07 | 2.800E-07 |
| 26182 | 1310 | R1310R | E | 2.4e-07 | 1.8e-07 | 2.3e-07 | 2.167E-07 | 3.215E-08 | 1.800E-07 | 2.400E-07 |
| 26183 | 1310 | R1310N | E | 1.4e-07 | 3.2e-07 | 2.1e-07 | 2.233E-07 | 9.074E-08 | 1.400E-07 | 3.200E-07 |
| 26184 | 1310 | R1310D | E | 2e-07 | 6e-07 | 3.3e-07 | 3.767E-07 | 2.040E-07 | 2.000E-07 | 6.000E-07 |
| 26185 | 1310 | R1310C | E | 1.8e-07 | 1.3e-07 | 3.9e-07 | 2.333E-07 | 1.380E-07 | 1.300E-07 | 3.900E-07 |
| 26186 | 1310 | R1310E | E | 2.4e-07 | 2.5e-07 | 5.3e-07 | 3.400E-07 | 1.646E-07 | 2.400E-07 | 5.300E-07 |
| 26187 | 1310 | R1310Q | E | 2.8e-07 | 3.1e-07 | 2.5e-07 | 2.800E-07 | 3.000E-08 | 2.500E-07 | 3.100E-07 |
| 26188 | 1310 | R1310G | E | 1.9e-07 | 2.2e-07 | 3.1e-07 | 2.400E-07 | 6.245E-08 | 1.900E-07 | 3.100E-07 |
| 26189 | 1310 | R1310H | E | 2e-07 | 2.3e-07 | 1.9e-07 | 2.067E-07 | 2.082E-08 | 1.900E-07 | 2.300E-07 |
| 26190 | 1310 | R1310I | E | 1.1e-07 | 2.6e-07 | 2.1e-07 | 1.933E-07 | 7.638E-08 | 1.100E-07 | 2.600E-07 |
| 26191 | 1310 | R1310L | E | 3.3e-07 | 5.6e-07 | 2.3e-07 | 3.733E-07 | 1.692E-07 | 2.300E-07 | 5.600E-07 |
| 26192 | 1310 | R1310K | E | 1.1e-07 | 5.4e-07 | 3.5e-07 | 3.333E-07 | 2.155E-07 | 1.100E-07 | 5.400E-07 |
| 26193 | 1310 | R1310M | E | 1.6e-07 | 1.8e-07 | 2.3e-07 | 1.900E-07 | 3.606E-08 | 1.600E-07 | 2.300E-07 |
| 26194 | 1310 | R1310F | E | 1.8e-07 | 2.1e-07 | 3.4e-07 | 2.433E-07 | 8.505E-08 | 1.800E-07 | 3.400E-07 |
| 26195 | 1310 | R1310P | E | 1.7e-07 | 4.7e-07 | 1.7e-07 | 2.700E-07 | 1.732E-07 | 1.700E-07 | 4.700E-07 |
| 26196 | 1310 | R1310S | E | 2.9e-07 | 3.8e-07 | 1.9e-07 | 2.867E-07 | 9.504E-08 | 1.900E-07 | 3.800E-07 |
| 26197 | 1310 | R1310T | E | 2.8e-07 | 5.2e-07 | 3.2e-07 | 3.733E-07 | 1.286E-07 | 2.800E-07 | 5.200E-07 |
| 26198 | 1310 | R1310W | E | 2.7e-07 | 3.8e-07 | 3.2e-07 | 3.233E-07 | 5.508E-08 | 2.700E-07 | 3.800E-07 |
| 26199 | 1310 | R1310Y | E | 1.7e-07 | 2.6e-07 | 1.4e-07 | 1.900E-07 | 6.245E-08 | 1.400E-07 | 2.600E-07 |
| 26200 | 1310 | R1310V | E | 3.1e-07 | 2.8e-07 | 2.6e-07 | 2.833E-07 | 2.517E-08 | 2.600E-07 | 3.100E-07 |
| 26201 | 1311 | Q1311A | E | 1.8e-07 | 3e-07 | 2.5e-07 | 2.433E-07 | 6.028E-08 | 1.800E-07 | 3.000E-07 |
| 26202 | 1311 | Q1311R | E | 2.7e-07 | 2e-07 | 1.4e-07 | 2.033E-07 | 6.506E-08 | 1.400E-07 | 2.700E-07 |
| 26203 | 1311 | Q1311N | E | 2.6e-07 | 2.3e-07 | 5.2e-07 | 3.367E-07 | 1.595E-07 | 2.300E-07 | 5.200E-07 |
| 26204 | 1311 | Q1311D | E | 1.5e-07 | 4.6e-07 | 2.1e-07 | 2.733E-07 | 1.644E-07 | 1.500E-07 | 4.600E-07 |
| 26205 | 1311 | Q1311C | E | 1.6e-07 | 5e-07 | 2.5e-07 | 3.033E-07 | 1.762E-07 | 1.600E-07 | 5.000E-07 |
| 26206 | 1311 | Q1311E | E | 2.1e-07 | 1.8e-07 | 1.6e-07 | 1.833E-07 | 2.517E-08 | 1.600E-07 | 2.100E-07 |
| 26207 | 1311 | Q1311Q | E | 1.9e-07 | 3.4e-07 | 1.1e-07 | 2.133E-07 | 1.168E-07 | 1.100E-07 | 3.400E-07 |
| 26208 | 1311 | Q1311G | E | 2.1e-07 | 2.6e-07 | 1.4e-07 | 2.033E-07 | 6.028E-08 | 1.400E-07 | 2.600E-07 |
| 26209 | 1311 | Q1311H | E | 2.7e-07 | 2.1e-07 | 2.2e-07 | 2.333E-07 | 3.215E-08 | 2.100E-07 | 2.700E-07 |
| 26210 | 1311 | Q1311I | E | 2.4e-07 | 2.4e-07 | 3.7e-07 | 2.833E-07 | 7.506E-08 | 2.400E-07 | 3.700E-07 |
| 26211 | 1311 | Q1311L | E | 3e-07 | 2e-07 | 7.9e-08 | 1.930E-07 | 1.107E-07 | 7.900E-08 | 3.000E-07 |
| 26212 | 1311 | Q1311K | E | 1.7e-07 | 1.7e-07 | 3.5e-07 | 2.300E-07 | 1.039E-07 | 1.700E-07 | 3.500E-07 |
| 26213 | 1311 | Q1311M | E | 3e-07 | 2e-07 | 5.6e-07 | 3.533E-07 | 1.858E-07 | 2.000E-07 | 5.600E-07 |
| 26214 | 1311 | Q1311F | E | 2.9e-07 | 3.7e-07 | 1.4e-07 | 2.667E-07 | 1.168E-07 | 1.400E-07 | 3.700E-07 |
| 26215 | 1311 | Q1311P | E | 2.9e-07 | 4.8e-07 | 2.7e-07 | 3.467E-07 | 1.159E-07 | 2.700E-07 | 4.800E-07 |
| 26216 | 1311 | Q1311S | E | 3.5e-07 | 1.1e-07 | 2e-07 | 2.200E-07 | 1.212E-07 | 1.100E-07 | 3.500E-07 |
| 26217 | 1311 | Q1311T | E | 5.2e-07 | 3.4e-07 | 2.7e-07 | 3.767E-07 | 1.290E-07 | 2.700E-07 | 5.200E-07 |
| 26218 | 1311 | Q1311W | E | 2.3e-07 | 4.7e-07 | 1.9e-07 | 2.967E-07 | 1.514E-07 | 1.900E-07 | 4.700E-07 |
| 26219 | 1311 | Q1311Y | E | 1.7e-07 | 2e-07 | 2.3e-07 | 2.000E-07 | 3.000E-08 | 1.700E-07 | 2.300E-07 |
| 26220 | 1311 | Q1311V | E | 2.4e-07 | 2.3e-07 | 1.9e-07 | 2.200E-07 | 2.646E-08 | 1.900E-07 | 2.400E-07 |
| 26221 | 1312 | A1312A | E | 3.5e-07 | 2.8e-07 | 2e-07 | 2.767E-07 | 7.506E-08 | 2.000E-07 | 3.500E-07 |
| 26222 | 1312 | A1312R | E | 3.3e-07 | 1.5e-07 | 3.1e-07 | 2.633E-07 | 9.866E-08 | 1.500E-07 | 3.300E-07 |
| 26223 | 1312 | A1312N | E | 1.8e-07 | 2.2e-07 | 2.1e-07 | 2.033E-07 | 2.082E-08 | 1.800E-07 | 2.200E-07 |
| 26224 | 1312 | A1312D | E | 1.7e-07 | 3.3e-07 | 2.5e-07 | 2.500E-07 | 8.000E-08 | 1.700E-07 | 3.300E-07 |
| 26225 | 1312 | A1312C | E | 6.3e-08 | 1.3e-07 | 2.7e-07 | 1.543E-07 | 1.056E-07 | 6.300E-08 | 2.700E-07 |
| 26226 | 1312 | A1312E | E | 3e-07 | 4.6e-07 | 1.8e-07 | 3.133E-07 | 1.405E-07 | 1.800E-07 | 4.600E-07 |
| 26227 | 1312 | A1312Q | E | 2.7e-07 | 2.4e-07 | 3.4e-07 | 2.833E-07 | 5.132E-08 | 2.400E-07 | 3.400E-07 |
| 26228 | 1312 | A1312G | E | 1.3e-07 | 2.8e-07 | 2.7e-07 | 2.267E-07 | 8.386E-08 | 1.300E-07 | 2.800E-07 |
| 26229 | 1312 | A1312H | E | 2.2e-07 | 3.1e-07 | 3.5e-07 | 2.933E-07 | 6.658E-08 | 2.200E-07 | 3.500E-07 |
| 26230 | 1312 | A1312I | E | 4.1e-07 | 2.5e-07 | 2.8e-07 | 3.133E-07 | 8.505E-08 | 2.500E-07 | 4.100E-07 |
| 26231 | 1312 | A1312L | E | 4e-07 | 1.6e-07 | 2.6e-07 | 2.733E-07 | 1.206E-07 | 1.600E-07 | 4.000E-07 |
| 26232 | 1312 | A1312K | E | 3e-07 | 3.2e-07 | 4.4e-07 | 3.533E-07 | 7.572E-08 | 3.000E-07 | 4.400E-07 |
| 26233 | 1312 | A1312M | E | 2.7e-07 | 2.5e-07 | 2e-07 | 2.400E-07 | 3.606E-08 | 2.000E-07 | 2.700E-07 |
| 26234 | 1312 | A1312F | E | 2.3e-07 | 2.8e-07 | 1.5e-07 | 2.200E-07 | 6.557E-08 | 1.500E-07 | 2.800E-07 |
| 26235 | 1312 | A1312P | E | 2.6e-07 | 3.5e-07 | 2.3e-07 | 2.800E-07 | 6.245E-08 | 2.300E-07 | 3.500E-07 |
| 26236 | 1312 | A1312S | E | 2.4e-07 | 2.3e-07 | 2.3e-07 | 2.333E-07 | 5.774E-09 | 2.300E-07 | 2.400E-07 |
| 26237 | 1312 | A1312T | E | 2.8e-07 | 2.7e-07 | 1.8e-07 | 2.433E-07 | 5.508E-08 | 1.800E-07 | 2.800E-07 |
| 26238 | 1312 | A1312W | E | 5.9e-07 | 1.8e-07 | 2.4e-07 | 3.367E-07 | 2.214E-07 | 1.800E-07 | 5.900E-07 |
| 26239 | 1312 | A1312Y | E | 1.5e-07 | 7e-07 | 1.9e-07 | 3.467E-07 | 3.066E-07 | 1.500E-07 | 7.000E-07 |
| 26240 | 1312 | A1312V | E | 3.5e-07 | 1.4e-07 | 6e-07 | 3.633E-07 | 2.303E-07 | 1.400E-07 | 6.000E-07 |
| 26241 | 1313 | P1313A | E | 1.6e-07 | 4.7e-07 | 2.8e-07 | 3.033E-07 | 1.563E-07 | 1.600E-07 | 4.700E-07 |
| 26242 | 1313 | P1313R | E | 5.2e-07 | 1.9e-07 | 3.8e-07 | 3.633E-07 | 1.656E-07 | 1.900E-07 | 5.200E-07 |
| 26243 | 1313 | P1313N | E | 4.6e-07 | 1.7e-07 | 3.7e-07 | 3.333E-07 | 1.484E-07 | 1.700E-07 | 4.600E-07 |
| 26244 | 1313 | P1313D | E | 2.1e-07 | 2.6e-07 | 3.1e-07 | 2.600E-07 | 5.000E-08 | 2.100E-07 | 3.100E-07 |
| 26245 | 1313 | P1313C | E | 3.7e-07 | 3.8e-07 | 2.3e-07 | 3.267E-07 | 8.386E-08 | 2.300E-07 | 3.800E-07 |
| 26246 | 1313 | P1313E | E | 3.1e-07 | 2.7e-07 | 1.8e-07 | 2.533E-07 | 6.658E-08 | 1.800E-07 | 3.100E-07 |
| 26247 | 1313 | P1313Q | E | 2.5e-07 | 3.2e-07 | 2.7e-07 | 2.800E-07 | 3.606E-08 | 2.500E-07 | 3.200E-07 |
| 26248 | 1313 | P1313G | E | 1.8e-07 | 2.3e-07 | 2.4e-07 | 2.167E-07 | 3.215E-08 | 1.800E-07 | 2.400E-07 |
| 26249 | 1313 | P1313H | E | 3e-07 | 1.5e-07 | 5.8e-07 | 3.433E-07 | 2.183E-07 | 1.500E-07 | 5.800E-07 |
| 26250 | 1313 | P1313I | E | 3.9e-07 | 3.3e-07 | 7.4e-07 | 4.867E-07 | 2.214E-07 | 3.300E-07 | 7.400E-07 |
| 26251 | 1313 | P1313L | E | 1.7e-07 | 1.8e-07 | 2.9e-07 | 2.133E-07 | 6.658E-08 | 1.700E-07 | 2.900E-07 |
| 26252 | 1313 | P1313K | E | 7.4e-08 | 2.1e-07 | 6.6e-07 | 3.147E-07 | 3.067E-07 | 7.400E-08 | 6.600E-07 |
| 26253 | 1313 | P1313M | E | 3.4e-07 | 4.3e-07 | 1.7e-07 | 3.133E-07 | 1.320E-07 | 1.700E-07 | 4.300E-07 |
| 26254 | 1313 | P1313F | E | 2.6e-07 | 3.2e-07 | 1.5e-07 | 2.433E-07 | 8.622E-08 | 1.500E-07 | 3.200E-07 |
| 26255 | 1313 | P1313P | E | 4.2e-07 | 1.9e-07 | 3.4e-07 | 3.167E-07 | 1.168E-07 | 1.900E-07 | 4.200E-07 |
| 26256 | 1313 | P1313S | E | 4.5e-07 | 1.7e-07 | 3.6e-07 | 3.267E-07 | 1.429E-07 | 1.700E-07 | 4.500E-07 |
| 26257 | 1313 | P1313T | E | 2.7e-07 | 1.2e-07 | 1.8e-07 | 1.900E-07 | 7.550E-08 | 1.200E-07 | 2.700E-07 |
| 26258 | 1313 | P1313W | E | 4.7e-07 | 2.7e-07 | 2.9e-07 | 3.433E-07 | 1.102E-07 | 2.700E-07 | 4.700E-07 |
| 26259 | 1313 | P1313Y | E | 2.1e-07 | 2.5e-07 | 3.8e-07 | 2.800E-07 | 8.888E-08 | 2.100E-07 | 3.800E-07 |
| 26260 | 1313 | P1313V | E | 2e-07 | 3.7e-07 | 3e-07 | 2.900E-07 | 8.544E-08 | 2.000E-07 | 3.700E-07 |
| 26261 | 1314 | G1314A | E | 1.6e-07 | 1.5e-07 | 8.4e-07 | 3.833E-07 | 3.955E-07 | 1.500E-07 | 8.400E-07 |
| 26262 | 1314 | G1314R | E | 1.7e-07 | 2.1e-07 | 2.5e-07 | 2.100E-07 | 4.000E-08 | 1.700E-07 | 2.500E-07 |
| 26263 | 1314 | G1314N | E | 1.8e-07 | 3.3e-07 | 1.8e-07 | 2.300E-07 | 8.660E-08 | 1.800E-07 | 3.300E-07 |
| 26264 | 1314 | G1314D | E | 8.9e-07 | 2.7e-07 | 2.8e-07 | 4.800E-07 | 3.551E-07 | 2.700E-07 | 8.900E-07 |

| | | | | | | | | | | |
|---|---|---|---|---|---|---|---|---|---|---|
| 26265 | 1314 | G1314C | E | 1.9e-07 | 1.5e-07 | 3.1e-07 | 2.167E-07 | 8.327E-08 | 1.500E-07 | 3.100E-07 |
| 26266 | 1314 | G1314E | E | 2.6e-07 | 3.3e-07 | 1.9e-07 | 2.600E-07 | 7.000E-08 | 1.900E-07 | 3.300E-07 |
| 26267 | 1314 | G1314Q | E | 2.9e-07 | 3.2e-07 | 1.2e-07 | 2.433E-07 | 1.079E-07 | 1.200E-07 | 3.200E-07 |
| 26268 | 1314 | G1314G | E | 3.2e-07 | 4.5e-07 | 1.4e-07 | 3.033E-07 | 1.557E-07 | 1.400E-07 | 4.500E-07 |
| 26269 | 1314 | G1314H | E | 2.5e-07 | 3.6e-07 | 2.9e-07 | 3.000E-07 | 5.568E-08 | 2.500E-07 | 3.600E-07 |
| 26270 | 1314 | G1314I | E | 4.5e-07 | 1.2e-07 | 3.2e-07 | 2.967E-07 | 1.662E-07 | 1.200E-07 | 4.500E-07 |
| 26271 | 1314 | G1314L | E | 2.3e-07 | 2.9e-07 | 3.8e-07 | 3.000E-07 | 7.550E-08 | 2.300E-07 | 3.800E-07 |
| 26272 | 1314 | G1314K | E | 2.1e-07 | 2e-07 | 7.3e-07 | 3.800E-07 | 3.032E-07 | 2.000E-07 | 7.300E-07 |
| 26273 | 1314 | G1314M | E | 3.3e-07 | 3.6e-07 | 2.7e-07 | 3.200E-07 | 4.583E-08 | 2.700E-07 | 3.600E-07 |
| 26274 | 1314 | G1314F | E | 4.7e-07 | 1.6e-07 | 1.8e-07 | 2.700E-07 | 1.735E-07 | 1.600E-07 | 4.700E-07 |
| 26275 | 1314 | G1314P | E | 2.5e-07 | 2.4e-07 | 3.2e-07 | 2.700E-07 | 4.359E-08 | 2.400E-07 | 3.200E-07 |
| 26276 | 1314 | G1314S | E | 2.2e-07 | 2.4e-07 | 2.4e-07 | 2.333E-07 | 1.155E-08 | 2.200E-07 | 2.400E-07 |
| 26277 | 1314 | G1314T | E | 1.4e-07 | 3.1e-07 | 2.4e-07 | 2.300E-07 | 8.544E-08 | 1.400E-07 | 3.100E-07 |
| 26278 | 1314 | G1314W | E | 4.6e-07 | 1.2e-07 | 3.9e-07 | 3.233E-07 | 1.795E-07 | 1.200E-07 | 4.600E-07 |
| 26279 | 1314 | G1314Y | E | 1.1e-07 | 2.2e-07 | 3.7e-07 | 2.333E-07 | 1.305E-07 | 1.100E-07 | 3.700E-07 |
| 26280 | 1314 | G1314V | E | 2.8e-07 | 3.2e-07 | 2.8e-07 | 2.933E-07 | 2.309E-08 | 2.800E-07 | 3.200E-07 |
| 26281 | 1315 | K1315A | E | 8.5e-08 | 1.6e-07 | 1.8e-07 | 1.417E-07 | 5.008E-08 | 8.500E-08 | 1.800E-07 |
| 26282 | 1315 | K1315R | E | 4.6e-07 | 3.1e-07 | 2.2e-07 | 3.300E-07 | 1.212E-07 | 2.200E-07 | 4.600E-07 |
| 26283 | 1315 | K1315N | E | 2.1e-07 | 3.1e-07 | 4e-07 | 3.067E-07 | 9.504E-08 | 2.100E-07 | 4.000E-07 |
| 26284 | 1315 | K1315D | E | 4.3e-07 | 2.9e-07 | 2.5e-07 | 3.233E-07 | 9.452E-08 | 2.500E-07 | 4.300E-07 |
| 26285 | 1315 | K1315C | E | 2e-07 | 2.2e-07 | 6e-07 | 3.400E-07 | 2.254E-07 | 2.000E-07 | 6.000E-07 |
| 26286 | 1315 | K1315E | E | 2.1e-07 | 3.6e-07 | 2.5e-07 | 2.733E-07 | 7.767E-08 | 2.100E-07 | 3.600E-07 |
| 26287 | 1315 | K1315Q | E | 5.1e-07 | 2.4e-07 | 2.7e-07 | 3.400E-07 | 1.480E-07 | 2.400E-07 | 5.100E-07 |
| 26288 | 1315 | K1315G | E | 3.2e-07 | 3.3e-07 | 4.3e-07 | 3.600E-07 | 6.083E-08 | 3.200E-07 | 4.300E-07 |
| 26289 | 1315 | K1315H | E | 2.6e-07 | 1.9e-07 | 2.4e-07 | 2.300E-07 | 3.606E-08 | 1.900E-07 | 2.600E-07 |
| 26290 | 1315 | K1315I | E | 2.2e-07 | 2.3e-07 | 3.6e-07 | 2.700E-07 | 7.810E-08 | 2.200E-07 | 3.600E-07 |
| 26291 | 1315 | K1315L | E | 7.8e-07 | 1.7e-07 | 2e-07 | 3.833E-07 | 3.439E-07 | 1.700E-07 | 7.800E-07 |
| 26292 | 1315 | K1315K | E | 8.2e-08 | 9e-08 | 1.5e-07 | 1.073E-07 | 3.717E-08 | 8.200E-08 | 1.500E-07 |
| 26293 | 1315 | K1315M | E | 9.5e-07 | 7.1e-08 | 2.9e-07 | 4.370E-07 | 4.576E-07 | 7.100E-08 | 9.500E-07 |
| 26294 | 1315 | K1315F | E | 3.2e-07 | 1.4e-07 | 3e-07 | 2.533E-07 | 9.866E-08 | 1.400E-07 | 3.200E-07 |
| 26295 | 1315 | K1315P | E | 2.2e-07 | 2.2e-07 | 2.4e-07 | 2.267E-07 | 1.155E-08 | 2.200E-07 | 2.400E-07 |
| 26296 | 1315 | K1315S | E | 4.4e-07 | 3.4e-07 | 2e-07 | 3.267E-07 | 1.206E-07 | 2.000E-07 | 4.400E-07 |
| 26297 | 1315 | K1315T | E | 1.9e-07 | 2.3e-07 | 2e-07 | 2.067E-07 | 2.082E-08 | 1.900E-07 | 2.300E-07 |
| 26298 | 1315 | K1315W | E | 1.6e-07 | 1.5e-07 | 1.3e-07 | 1.467E-07 | 1.528E-08 | 1.300E-07 | 1.600E-07 |
| 26299 | 1315 | K1315Y | E | 1.9e-07 | 3.2e-07 | 1.9e-07 | 2.333E-07 | 7.506E-08 | 1.900E-07 | 3.200E-07 |
| 26300 | 1315 | K1315V | E | 1.5e-07 | 2.6e-07 | 1.1e-07 | 1.733E-07 | 7.767E-08 | 1.100E-07 | 2.600E-07 |
| 26301 | 1316 | G1316A | E | 2.2e-07 | 1.5e-07 | 1.8e-07 | 1.833E-07 | 3.512E-08 | 1.500E-07 | 2.200E-07 |
| 26302 | 1316 | G1316R | E | 2.3e-07 | 1.7e-07 | 2.3e-07 | 2.100E-07 | 3.464E-08 | 1.700E-07 | 2.300E-07 |
| 26303 | 1316 | G1316N | E | 2.6e-07 | 1.7e-07 | 2.7e-07 | 2.333E-07 | 5.508E-08 | 1.700E-07 | 2.700E-07 |
| 26304 | 1316 | G1316D | E | 2.3e-07 | 4e-07 | 2.6e-07 | 2.967E-07 | 9.074E-08 | 2.300E-07 | 4.000E-07 |
| 26305 | 1316 | G1316C | E | 2.9e-07 | 1.2e-07 | 1.6e-07 | 1.900E-07 | 8.888E-08 | 1.200E-07 | 2.900E-07 |
| 26306 | 1316 | G1316E | E | 1.7e-07 | 1.8e-07 | 2.3e-07 | 1.933E-07 | 3.215E-08 | 1.700E-07 | 2.300E-07 |
| 26307 | 1316 | G1316Q | E | 1.9e-07 | 1.2e-07 | 1.6e-07 | 1.567E-07 | 3.512E-08 | 1.200E-07 | 1.900E-07 |
| 26308 | 1316 | G1316G | E | 1.4e-07 | 2.1e-07 | 2.7e-07 | 2.067E-07 | 6.506E-08 | 1.400E-07 | 2.700E-07 |
| 26309 | 1316 | G1316H | E | 5.3e-07 | 1.2e-07 | 2.7e-07 | 3.067E-07 | 2.074E-07 | 1.200E-07 | 5.300E-07 |
| 26310 | 1316 | G1316I | E | 2.8e-07 | 2.2e-07 | 2.3e-07 | 2.433E-07 | 3.215E-08 | 2.200E-07 | 2.800E-07 |
| 26311 | 1316 | G1316L | E | 1.2e-07 | 3.2e-07 | 1.6e-07 | 2.000E-07 | 1.058E-07 | 1.200E-07 | 3.200E-07 |
| 26312 | 1316 | G1316K | E | 2.3e-07 | 2.9e-07 | 2.2e-07 | 2.467E-07 | 3.786E-08 | 2.200E-07 | 2.900E-07 |
| 26313 | 1316 | G1316M | E | 1.2e-07 | 4.4e-07 | 1.5e-07 | 2.367E-07 | 1.767E-07 | 1.200E-07 | 4.400E-07 |
| 26314 | 1316 | G1316F | E | 4e-07 | 1.7e-07 | 2.3e-07 | 2.667E-07 | 1.193E-07 | 1.700E-07 | 4.000E-07 |
| 26315 | 1316 | G1316P | E | 2e-07 | 3.5e-07 | 1.9e-07 | 2.467E-07 | 8.963E-08 | 1.900E-07 | 3.500E-07 |
| 26316 | 1316 | G1316S | E | 2.9e-07 | 2.4e-07 | 6.3e-07 | 3.867E-07 | 2.122E-07 | 2.400E-07 | 6.300E-07 |
| 26317 | 1316 | G1316T | E | 3.5e-07 | 2.1e-07 | 2.9e-07 | 2.833E-07 | 7.024E-08 | 2.100E-07 | 3.500E-07 |
| 26318 | 1316 | G1316W | E | 2.5e-07 | 2.6e-07 | 4.3e-07 | 3.133E-07 | 1.012E-07 | 2.500E-07 | 4.300E-07 |
| 26319 | 1316 | G1316Y | E | 3.9e-07 | 2.1e-07 | 2.4e-07 | 2.800E-07 | 9.644E-08 | 2.100E-07 | 3.900E-07 |
| 26320 | 1316 | G1316V | E | 2.3e-07 | 4.4e-07 | 1.2e-07 | 2.633E-07 | 1.626E-07 | 1.200E-07 | 4.400E-07 |
| 26321 | 1317 | L1317A | E | 2.1e-07 | 1.8e-07 | 3.9e-07 | 2.600E-07 | 1.136E-07 | 1.800E-07 | 3.900E-07 |
| 26322 | 1317 | L1317R | E | 2.9e-07 | 2.6e-07 | 8.1e-08 | 2.103E-07 | 1.130E-07 | 8.100E-08 | 2.900E-07 |
| 26323 | 1317 | L1317N | E | 3.5e-07 | 1.7e-07 | 1.5e-07 | 2.233E-07 | 1.102E-07 | 1.500E-07 | 3.500E-07 |
| 26324 | 1317 | L1317D | E | 1.8e-07 | 2.7e-07 | 2.5e-07 | 2.333E-07 | 4.726E-08 | 1.800E-07 | 2.700E-07 |
| 26325 | 1317 | L1317C | E | 5.9e-07 | 4.6e-07 | 1.8e-07 | 4.100E-07 | 2.095E-07 | 1.800E-07 | 5.900E-07 |
| 26326 | 1317 | L1317E | E | 3.6e-07 | 1.4e-07 | 1.6e-07 | 2.200E-07 | 1.217E-07 | 1.400E-07 | 3.600E-07 |
| 26327 | 1317 | L1317Q | E | 2.6e-07 | 2.3e-07 | 1.9e-07 | 2.267E-07 | 3.512E-08 | 1.900E-07 | 2.600E-07 |
| 26328 | 1317 | L1317G | E | 2.1e-07 | 2.2e-07 | 8.6e-08 | 1.720E-07 | 7.465E-08 | 8.600E-08 | 2.200E-07 |
| 26329 | 1317 | L1317H | E | 1.8e-07 | 3.8e-07 | 1.5e-07 | 2.367E-07 | 1.250E-07 | 1.500E-07 | 3.800E-07 |
| 26330 | 1317 | L1317I | E | 4.1e-07 | 4e-07 | 1.4e-07 | 3.167E-07 | 1.531E-07 | 1.400E-07 | 4.100E-07 |
| 26331 | 1317 | L1317L | E | 3.5e-07 | 2.9e-07 | 5.4e-07 | 3.933E-07 | 1.305E-07 | 2.900E-07 | 5.400E-07 |
| 26332 | 1317 | L1317K | E | 1.6e-07 | 3.3e-07 | 2e-07 | 2.300E-07 | 8.888E-08 | 1.600E-07 | 3.300E-07 |
| 26333 | 1317 | L1317M | E | 2.4e-07 | 1.6e-07 | 2.1e-07 | 2.033E-07 | 4.041E-08 | 1.600E-07 | 2.400E-07 |
| 26334 | 1317 | L1317F | E | 3.9e-07 | 2.7e-07 | 3.9e-07 | 3.500E-07 | 6.928E-08 | 2.700E-07 | 3.900E-07 |
| 26335 | 1317 | L1317P | E | 2.2e-07 | 2e-07 | 3.1e-07 | 2.433E-07 | 5.859E-08 | 2.000E-07 | 3.100E-07 |
| 26336 | 1317 | L1317S | E | 3.1e-07 | 2.1e-07 | 2.7e-07 | 2.633E-07 | 5.033E-08 | 2.100E-07 | 3.100E-07 |
| 26337 | 1317 | L1317T | E | 1.3e-07 | 3.4e-07 | 3.3e-07 | 2.667E-07 | 1.185E-07 | 1.300E-07 | 3.400E-07 |
| 26338 | 1317 | L1317W | E | 3.8e-07 | 2e-07 | 2.1e-07 | 2.633E-07 | 1.012E-07 | 2.000E-07 | 3.800E-07 |
| 26339 | 1317 | L1317Y | E | 6.2e-07 | 2.5e-07 | 1.8e-07 | 3.500E-07 | 2.364E-07 | 1.800E-07 | 6.200E-07 |
| 26340 | 1317 | L1317V | E | 4.8e-07 | 2e-07 | 2.1e-07 | 2.967E-07 | 1.589E-07 | 2.000E-07 | 4.800E-07 |
| 26341 | 1318 | E1318A | E | 3.5e-07 | 1.3e-07 | 2.2e-07 | 2.333E-07 | 1.106E-07 | 1.300E-07 | 3.500E-07 |
| 26342 | 1318 | E1318R | E | 2.5e-07 | 1.9e-07 | 2.8e-07 | 2.400E-07 | 4.583E-08 | 1.900E-07 | 2.800E-07 |
| 26343 | 1318 | E1318N | E | 2.9e-07 | 2e-07 | 1.6e-07 | 2.167E-07 | 6.658E-08 | 1.600E-07 | 2.900E-07 |
| 26344 | 1318 | E1318D | E | 3.1e-07 | 3.3e-07 | 4.5e-07 | 3.633E-07 | 7.572E-08 | 3.100E-07 | 4.500E-07 |
| 26345 | 1318 | E1318C | E | 2.5e-07 | 2.9e-07 | 4.4e-07 | 3.267E-07 | 1.002E-07 | 2.500E-07 | 4.400E-07 |
| 26346 | 1318 | E1318E | E | 2.7e-07 | 1.8e-07 | 1.3e-07 | 1.933E-07 | 7.095E-08 | 1.300E-07 | 2.700E-07 |
| 26347 | 1318 | E1318Q | E | 2.9e-07 | 2.2e-07 | 1.7e-07 | 2.267E-07 | 6.028E-08 | 1.700E-07 | 2.900E-07 |
| 26348 | 1318 | E1318G | E | 2.1e-07 | 3.8e-07 | 3e-07 | 2.967E-07 | 8.505E-08 | 2.100E-07 | 3.800E-07 |
| 26349 | 1318 | E1318H | E | 1.8e-07 | 1.9e-07 | 1.6e-07 | 1.767E-07 | 1.528E-08 | 1.600E-07 | 1.900E-07 |

| | | | | | | | | | | |
|---|---|---|---|---|---|---|---|---|---|---|
| 26350 | 1318 | E1318I | E | 1.8e-07 | 2.3e-07 | 4.1e-07 | 2.733E-07 | 1.210E-07 | 1.800E-07 | 4.100E-07 |
| 26351 | 1318 | E1318L | E | 4.1e-07 | 1.5e-07 | 1.7e-07 | 2.433E-07 | 1.447E-07 | 1.500E-07 | 4.100E-07 |
| 26352 | 1318 | E1318K | E | 4.4e-07 | 1.6e-07 | 3.1e-07 | 3.033E-07 | 1.401E-07 | 1.600E-07 | 4.400E-07 |
| 26353 | 1318 | E1318M | E | 3e-07 | 1.5e-07 | 2.8e-07 | 2.433E-07 | 8.145E-08 | 1.500E-07 | 3.000E-07 |
| 26354 | 1318 | E1318F | E | 1.6e-07 | 2.4e-07 | 3.3e-07 | 2.433E-07 | 8.505E-08 | 1.600E-07 | 3.300E-07 |
| 26355 | 1318 | E1318P | E | 1.2e-07 | 1.8e-07 | 1.4e-07 | 1.467E-07 | 3.055E-08 | 1.200E-07 | 1.800E-07 |
| 26356 | 1318 | E1318S | E | 2e-07 | 3.2e-07 | 1.7e-07 | 2.300E-07 | 7.937E-08 | 1.700E-07 | 3.200E-07 |
| 26357 | 1318 | E1318T | E | 3.5e-07 | 1.9e-07 | 1.3e-07 | 2.233E-07 | 1.137E-07 | 1.300E-07 | 3.500E-07 |
| 26358 | 1318 | E1318W | E | 2.4e-07 | 3.4e-07 | 2.3e-07 | 2.700E-07 | 6.083E-08 | 2.300E-07 | 3.400E-07 |
| 26359 | 1318 | E1318Y | E | 3.4e-07 | 2e-07 | 2.9e-07 | 2.767E-07 | 7.095E-08 | 2.000E-07 | 3.400E-07 |
| 26360 | 1318 | E1318V | E | 1.1e-07 | 1.7e-07 | 1.9e-07 | 1.567E-07 | 4.163E-08 | 1.100E-07 | 1.900E-07 |
| 26361 | 1319 | W1319A | E | 4.3e-07 | 2.1e-07 | 2.4e-07 | 2.933E-07 | 1.193E-07 | 2.100E-07 | 4.300E-07 |
| 26362 | 1319 | W1319R | E | 2.3e-07 | 2.5e-07 | 3.2e-07 | 2.667E-07 | 4.726E-08 | 2.300E-07 | 3.200E-07 |
| 26363 | 1319 | W1319N | E | 1.9e-07 | 2.7e-07 | 2.3e-07 | 2.300E-07 | 4.000E-08 | 1.900E-07 | 2.700E-07 |
| 26364 | 1319 | W1319D | E | 6e-07 | 2.2e-07 | 3.8e-07 | 4.000E-07 | 1.908E-07 | 2.200E-07 | 6.000E-07 |
| 26365 | 1319 | W1319C | E | 1.4e-07 | 1.4e-07 | 1.2e-07 | 1.333E-07 | 1.155E-08 | 1.200E-07 | 1.400E-07 |
| 26366 | 1319 | W1319E | E | 5.8e-07 | 3.4e-07 | 3.3e-07 | 4.167E-07 | 1.415E-07 | 3.300E-07 | 5.800E-07 |
| 26367 | 1319 | W1319Q | E | 2.3e-07 | 9.6e-08 | 2e-07 | 1.753E-07 | 7.032E-08 | 9.600E-08 | 2.300E-07 |
| 26368 | 1319 | W1319G | E | 1.2e-07 | 4.6e-07 | 1.3e-07 | 2.367E-07 | 1.935E-07 | 1.200E-07 | 4.600E-07 |
| 26369 | 1319 | W1319H | E | 3.1e-07 | 1.3e-07 | 2.8e-07 | 2.400E-07 | 9.644E-08 | 1.300E-07 | 3.100E-07 |
| 26370 | 1319 | W1319I | E | 2.6e-07 | 4.7e-07 | 3.5e-07 | 3.600E-07 | 1.054E-07 | 2.600E-07 | 4.700E-07 |
| 26371 | 1319 | W1319L | E | 2e-07 | 2.6e-07 | 2.9e-07 | 2.500E-07 | 4.583E-08 | 2.000E-07 | 2.900E-07 |
| 26372 | 1319 | W1319K | E | 2.7e-07 | 4.3e-07 | 5.5e-07 | 4.167E-07 | 1.405E-07 | 2.700E-07 | 5.500E-07 |
| 26373 | 1319 | W1319M | E | 2.4e-07 | 3.2e-07 | 4e-07 | 3.200E-07 | 8.000E-08 | 2.400E-07 | 4.000E-07 |
| 26374 | 1319 | W1319F | E | 1.6e-07 | 2.3e-07 | 1.8e-07 | 1.900E-07 | 3.606E-08 | 1.600E-07 | 2.300E-07 |
| 26375 | 1319 | W1319P | E | 8.5e-08 | 4.3e-07 | 2.7e-07 | 2.617E-07 | 1.727E-07 | 8.500E-08 | 4.300E-07 |
| 26376 | 1319 | W1319S | E | 1.9e-07 | 2.6e-07 | 3.3e-07 | 2.600E-07 | 7.000E-08 | 1.900E-07 | 3.300E-07 |
| 26377 | 1319 | W1319T | E | 3.4e-07 | 2e-07 | 2.5e-07 | 2.633E-07 | 7.095E-08 | 2.000E-07 | 3.400E-07 |
| 26378 | 1319 | W1319W | E | 1.8e-07 | 3.9e-07 | 1.9e-07 | 2.533E-07 | 1.185E-07 | 1.800E-07 | 3.900E-07 |
| 26379 | 1319 | W1319Y | E | 3.5e-07 | 4.1e-07 | 1.1e-07 | 2.900E-07 | 1.587E-07 | 1.100E-07 | 4.100E-07 |
| 26380 | 1319 | W1319V | E | 4.2e-07 | 6.9e-07 | 3.5e-07 | 4.867E-07 | 1.795E-07 | 3.500E-07 | 6.900E-07 |
| 26381 | 1320 | V1320A | E | 2.9e-07 | 1.5e-07 | 1.5e-07 | 1.967E-07 | 8.083E-08 | 1.500E-07 | 2.900E-07 |
| 26382 | 1320 | V1320R | E | 6.6e-07 | 2.8e-07 | 3.1e-07 | 4.167E-07 | 2.113E-07 | 2.800E-07 | 6.600E-07 |
| 26383 | 1320 | V1320N | E | 3.6e-07 | 2.2e-07 | 1.6e-07 | 2.467E-07 | 1.026E-07 | 1.600E-07 | 3.600E-07 |
| 26384 | 1320 | V1320D | E | 3.6e-07 | 2.2e-07 | 4.2e-07 | 3.333E-07 | 1.026E-07 | 2.200E-07 | 4.200E-07 |
| 26385 | 1320 | V1320C | E | 3e-07 | 2.2e-07 | 1.7e-07 | 2.300E-07 | 6.557E-08 | 1.700E-07 | 3.000E-07 |
| 26386 | 1320 | V1320E | E | 1.8e-07 | 1.6e-07 | 3e-07 | 2.133E-07 | 7.572E-08 | 1.600E-07 | 3.000E-07 |
| 26387 | 1320 | V1320Q | E | 2e-07 | 4.8e-07 | 4.2e-07 | 3.667E-07 | 1.474E-07 | 2.000E-07 | 4.800E-07 |
| 26388 | 1320 | V1320G | E | 9.1e-08 | 2.7e-07 | 3e-07 | 2.203E-07 | 1.130E-07 | 9.100E-08 | 3.000E-07 |
| 26389 | 1320 | V1320H | E | 2.1e-07 | 2.1e-07 | 3.1e-07 | 2.433E-07 | 5.774E-08 | 2.100E-07 | 3.100E-07 |
| 26390 | 1320 | V1320I | E | 5.2e-07 | 3.3e-07 | 1.1e-07 | 3.200E-07 | 2.052E-07 | 1.100E-07 | 5.200E-07 |
| 26391 | 1320 | V1320L | E | 6.6e-07 | 4.9e-07 | 3.1e-07 | 4.867E-07 | 1.750E-07 | 3.100E-07 | 6.600E-07 |
| 26392 | 1320 | V1320K | E | 4.3e-07 | 1.8e-07 | 3.5e-07 | 3.200E-07 | 1.277E-07 | 1.800E-07 | 4.300E-07 |
| 26393 | 1320 | V1320M | E | 2.5e-07 | 2.2e-07 | 2.5e-07 | 2.400E-07 | 1.732E-08 | 2.200E-07 | 2.500E-07 |
| 26394 | 1320 | V1320F | E | 3.2e-07 | 3.5e-07 | 1.7e-07 | 2.800E-07 | 9.644E-08 | 1.700E-07 | 3.500E-07 |
| 26395 | 1320 | V1320P | E | 2.8e-07 | 1.7e-07 | 1.6e-07 | 2.033E-07 | 6.658E-08 | 1.600E-07 | 2.800E-07 |
| 26396 | 1320 | V1320S | E | 2.9e-07 | 2.4e-07 | 2.3e-07 | 2.533E-07 | 3.215E-08 | 2.300E-07 | 2.900E-07 |
| 26397 | 1320 | V1320T | E | 5.6e-07 | 2.6e-07 | 3e-07 | 3.733E-07 | 1.629E-07 | 2.600E-07 | 5.600E-07 |
| 26398 | 1320 | V1320W | E | 1.2e-07 | 6e-07 | 4.1e-07 | 3.767E-07 | 2.417E-07 | 1.200E-07 | 6.000E-07 |
| 26399 | 1320 | V1320Y | E | 2.9e-07 | 1.4e-07 | 2e-07 | 2.100E-07 | 7.550E-08 | 1.400E-07 | 2.900E-07 |
| 26400 | 1320 | V1320V | E | 4.4e-07 | 2.6e-07 | 1.9e-07 | 2.967E-07 | 1.290E-07 | 1.900E-07 | 4.400E-07 |
| 26401 | 1321 | A1321A | E | 1.6e-07 | 1.1e-07 | 3.4e-07 | 2.033E-07 | 1.210E-07 | 1.100E-07 | 3.400E-07 |
| 26402 | 1321 | A1321R | E | 2.2e-07 | 3.4e-07 | 2.1e-07 | 2.567E-07 | 7.234E-08 | 2.100E-07 | 3.400E-07 |
| 26403 | 1321 | A1321N | E | 1.9e-07 | 2.4e-07 | 3.7e-07 | 2.667E-07 | 9.292E-08 | 1.900E-07 | 3.700E-07 |
| 26404 | 1321 | A1321D | E | 1.7e-07 | 2.2e-07 | 2.7e-07 | 2.200E-07 | 5.000E-08 | 1.700E-07 | 2.700E-07 |
| 26405 | 1321 | A1321C | E | 3.2e-07 | 1e-07 | 4.8e-07 | 3.000E-07 | 1.908E-07 | 1.000E-07 | 4.800E-07 |
| 26406 | 1321 | A1321E | E | 2.2e-07 | 3.2e-07 | 1.4e-07 | 2.267E-07 | 9.018E-08 | 1.400E-07 | 3.200E-07 |
| 26407 | 1321 | A1321Q | E | 3.1e-07 | 2.1e-07 | 3e-07 | 2.733E-07 | 5.508E-08 | 2.100E-07 | 3.100E-07 |
| 26408 | 1321 | A1321G | E | 2e-07 | 4.5e-07 | 2.4e-07 | 2.967E-07 | 1.343E-07 | 2.000E-07 | 4.500E-07 |
| 26409 | 1321 | A1321H | E | 4.1e-07 | 1.3e-07 | 2.9e-07 | 2.767E-07 | 1.405E-07 | 1.300E-07 | 4.100E-07 |
| 26410 | 1321 | A1321I | E | 1.6e-07 | 2.8e-07 | 1.5e-07 | 1.967E-07 | 7.234E-08 | 1.500E-07 | 2.800E-07 |
| 26411 | 1321 | A1321L | E | 1.2e-07 | 3.2e-07 | 1.6e-07 | 2.000E-07 | 1.058E-07 | 1.200E-07 | 3.200E-07 |
| 26412 | 1321 | A1321K | E | 2.3e-07 | 3.1e-07 | 3.6e-07 | 3.000E-07 | 6.557E-08 | 2.300E-07 | 3.600E-07 |
| 26413 | 1321 | A1321M | E | 1.9e-07 | 3.8e-07 | 1.7e-07 | 2.467E-07 | 1.159E-07 | 1.700E-07 | 3.800E-07 |
| 26414 | 1321 | A1321F | E | 3.9e-07 | 6.7e-07 | 2.7e-07 | 4.433E-07 | 2.053E-07 | 2.700E-07 | 6.700E-07 |
| 26415 | 1321 | A1321P | E | 2e-07 | 2.7e-07 | 3e-07 | 2.567E-07 | 5.132E-08 | 2.000E-07 | 3.000E-07 |
| 26416 | 1321 | A1321S | E | 4e-07 | 2.7e-07 | 1.7e-07 | 2.800E-07 | 1.153E-07 | 1.700E-07 | 4.000E-07 |
| 26417 | 1321 | A1321T | E | 1.6e-07 | 9.1e-08 | 1.6e-07 | 1.370E-07 | 3.984E-08 | 9.100E-08 | 1.600E-07 |
| 26418 | 1321 | A1321W | E | 1.2e-07 | 1.2e-07 | 2.6e-07 | 1.667E-07 | 8.083E-08 | 1.200E-07 | 2.600E-07 |
| 26419 | 1321 | A1321Y | E | 1.3e-07 | 1.3e-07 | 1.8e-07 | 1.467E-07 | 2.887E-08 | 1.300E-07 | 1.800E-07 |
| 26420 | 1321 | A1321V | E | 1.8e-07 | 6.5e-07 | 2.5e-07 | 3.600E-07 | 2.536E-07 | 1.800E-07 | 6.500E-07 |
| 26421 | 1322 | R1322A | E | 1.8e-07 | 7.2e-07 | 3.4e-07 | 4.133E-07 | 2.774E-07 | 1.800E-07 | 7.200E-07 |
| 26422 | 1322 | R1322R | E | 2.3e-07 | 2.7e-07 | 2.9e-07 | 2.633E-07 | 3.055E-08 | 2.300E-07 | 2.900E-07 |
| 26423 | 1322 | R1322N | E | 4.5e-07 | 2.5e-07 | 7.1e-07 | 4.700E-07 | 2.307E-07 | 2.500E-07 | 7.100E-07 |
| 26424 | 1322 | R1322D | E | 2e-07 | 3.8e-07 | 3e-07 | 2.933E-07 | 9.018E-08 | 2.000E-07 | 3.800E-07 |
| 26425 | 1322 | R1322C | E | 5.1e-07 | 5.9e-07 | 3.4e-07 | 4.800E-07 | 1.277E-07 | 3.400E-07 | 5.900E-07 |
| 26426 | 1322 | R1322E | E | 2.8e-07 | 3.6e-07 | 2.1e-07 | 2.833E-07 | 7.506E-08 | 2.100E-07 | 3.600E-07 |
| 26427 | 1322 | R1322Q | E | 1.3e-07 | 6.9e-07 | 3.3e-07 | 3.833E-07 | 2.838E-07 | 1.300E-07 | 6.900E-07 |
| 26428 | 1322 | R1322G | E | 7.2e-07 | 2.4e-07 | 5e-07 | 4.867E-07 | 2.403E-07 | 2.400E-07 | 7.200E-07 |
| 26429 | 1322 | R1322H | E | 3.8e-07 | 2.1e-07 | 4.4e-07 | 3.433E-07 | 1.193E-07 | 2.100E-07 | 4.400E-07 |
| 26430 | 1322 | R1322I | E | 4.1e-07 | 2.4e-07 | 3.6e-07 | 3.367E-07 | 8.737E-08 | 2.400E-07 | 4.100E-07 |
| 26431 | 1322 | R1322L | E | 3.3e-07 | 2.6e-07 | 2.7e-07 | 2.867E-07 | 3.786E-08 | 2.600E-07 | 3.300E-07 |
| 26432 | 1322 | R1322K | E | 3.7e-07 | 4.5e-07 | 2.7e-07 | 3.633E-07 | 9.018E-08 | 2.700E-07 | 4.500E-07 |
| 26433 | 1322 | R1322M | E | 3.2e-07 | 7.1e-07 | 3.4e-07 | 4.567E-07 | 2.196E-07 | 3.200E-07 | 7.100E-07 |
| 26434 | 1322 | R1322F | E | 4.6e-07 | 2e-07 | 4.4e-07 | 3.667E-07 | 1.447E-07 | 2.000E-07 | 4.600E-07 |

| | | | | | | | | | | |
|---|---|---|---|---|---|---|---|---|---|---|
| 26435 | 1322 | R1322P | E | 4e-07 | 7.6e-07 | 2.9e-07 | 4.833E-07 | 2.458E-07 | 2.900E-07 | 7.600E-07 |
| 26436 | 1322 | R1322S | E | 3.6e-07 | 6.6e-07 | 1.2e-06 | 7.400E-07 | 4.257E-07 | 3.600E-07 | 1.200E-06 |
| 26437 | 1322 | R1322T | E | 3.3e-07 | 3.4e-07 | 3.1e-07 | 3.267E-07 | 1.528E-08 | 3.100E-07 | 3.400E-07 |
| 26438 | 1322 | R1322W | E | 2.7e-07 | 2.6e-07 | 4.7e-07 | 3.333E-07 | 1.185E-07 | 2.600E-07 | 4.700E-07 |
| 26439 | 1322 | R1322Y | E | 5.5e-07 | 3.7e-07 | 1.8e-07 | 3.667E-07 | 1.850E-07 | 1.800E-07 | 5.500E-07 |
| 26440 | 1322 | R1322V | E | 2.5e-07 | 6e-07 | 7.9e-07 | 5.467E-07 | 2.739E-07 | 2.500E-07 | 7.900E-07 |
| 26441 | 1323 | I1323A | E | 3e-07 | 1.7e-07 | 3.6e-07 | 2.767E-07 | 9.713E-08 | 1.700E-07 | 3.600E-07 |
| 26442 | 1323 | I1323R | E | 3.6e-07 | 2.4e-07 | 2.1e-07 | 2.700E-07 | 7.937E-08 | 2.100E-07 | 3.600E-07 |
| 26443 | 1323 | I1323N | E | 2.8e-07 | 4.4e-07 | 3.3e-07 | 3.500E-07 | 8.185E-08 | 2.800E-07 | 4.400E-07 |
| 26444 | 1323 | I1323D | E | 1.6e-07 | 2e-07 | 4.7e-07 | 2.767E-07 | 1.686E-07 | 1.600E-07 | 4.700E-07 |
| 26445 | 1323 | I1323C | E | 1.4e-07 | 2.4e-07 | 9e-08 | 1.567E-07 | 7.638E-08 | 9.000E-08 | 2.400E-07 |
| 26446 | 1323 | I1323E | E | 2.4e-07 | 2.2e-07 | 4.8e-07 | 3.133E-07 | 1.447E-07 | 2.200E-07 | 4.800E-07 |
| 26447 | 1323 | I1323Q | E | 1.6e-07 | 2.2e-07 | 1.8e-07 | 1.867E-07 | 3.055E-08 | 1.600E-07 | 2.200E-07 |
| 26448 | 1323 | I1323G | E | 3.6e-07 | 1.9e-07 | 1.6e-07 | 2.367E-07 | 1.079E-07 | 1.600E-07 | 3.600E-07 |
| 26449 | 1323 | I1323H | E | 2.9e-07 | 1e-07 | 2.7e-07 | 2.200E-07 | 1.044E-07 | 1.000E-07 | 2.900E-07 |
| 26450 | 1323 | I1323I | E | 3.9e-07 | 2.1e-07 | 1.3e-07 | 2.433E-07 | 1.332E-07 | 1.300E-07 | 3.900E-07 |
| 26451 | 1323 | I1323L | E | 3.8e-07 | 5.8e-07 | 1.6e-07 | 3.733E-07 | 2.101E-07 | 1.600E-07 | 5.800E-07 |
| 26452 | 1323 | I1323K | E | 3.1e-07 | 3.9e-07 | 6.2e-07 | 4.400E-07 | 1.609E-07 | 3.100E-07 | 6.200E-07 |
| 26453 | 1323 | I1323M | E | 1.2e-07 | 1.8e-07 | 2.4e-07 | 1.800E-07 | 6.000E-08 | 1.200E-07 | 2.400E-07 |
| 26454 | 1323 | I1323F | E | 1.4e-07 | 3.2e-07 | 2e-07 | 2.200E-07 | 9.165E-08 | 1.400E-07 | 3.200E-07 |
| 26455 | 1323 | I1323P | E | 3e-07 | 1.4e-07 | 2.7e-07 | 2.367E-07 | 8.505E-08 | 1.400E-07 | 3.000E-07 |
| 26456 | 1323 | I1323S | E | 2.5e-07 | 6.3e-07 | 2.6e-07 | 3.800E-07 | 2.166E-07 | 2.500E-07 | 6.300E-07 |
| 26457 | 1323 | I1323T | E | 3.4e-07 | 4.3e-07 | 3.2e-07 | 3.633E-07 | 5.859E-08 | 3.200E-07 | 4.300E-07 |
| 26458 | 1323 | I1323W | E | 4.7e-07 | 2.6e-07 | 1.8e-07 | 3.033E-07 | 1.498E-07 | 1.800E-07 | 4.700E-07 |
| 26459 | 1323 | I1323Y | E | 1.8e-07 | 3.6e-07 | 4.2e-07 | 3.200E-07 | 1.249E-07 | 1.800E-07 | 4.200E-07 |
| 26460 | 1323 | I1323V | E | 3.4e-07 | 3.7e-07 | 6.6e-08 | 2.587E-07 | 1.675E-07 | 6.600E-08 | 3.700E-07 |
| 26461 | 1324 | Y1324A | E | 1.5e-07 | 3.4e-07 | 2.4e-07 | 2.433E-07 | 9.504E-08 | 1.500E-07 | 3.400E-07 |
| 26462 | 1324 | Y1324R | E | 1e-07 | 1.8e-07 | 1.9e-07 | 1.567E-07 | 4.933E-08 | 1.000E-07 | 1.900E-07 |
| 26463 | 1324 | Y1324N | E | 1.9e-07 | 3.8e-07 | 5.9e-07 | 3.867E-07 | 2.001E-07 | 1.900E-07 | 5.900E-07 |
| 26464 | 1324 | Y1324D | E | 2.3e-07 | 3.3e-07 | 4.3e-07 | 3.300E-07 | 1.000E-07 | 2.300E-07 | 4.300E-07 |
| 26465 | 1324 | Y1324C | E | 2.8e-07 | 3.6e-07 | 2.1e-07 | 2.833E-07 | 7.506E-08 | 2.100E-07 | 3.600E-07 |
| 26466 | 1324 | Y1324E | E | 1.5e-07 | 2.5e-07 | 1.3e-07 | 1.767E-07 | 6.429E-08 | 1.300E-07 | 2.500E-07 |
| 26467 | 1324 | Y1324Q | E | 3e-07 | 3e-07 | 2.7e-07 | 2.900E-07 | 1.732E-08 | 2.700E-07 | 3.000E-07 |
| 26468 | 1324 | Y1324G | E | 2.3e-07 | 2.4e-07 | 2.9e-07 | 2.533E-07 | 3.215E-08 | 2.300E-07 | 2.900E-07 |
| 26469 | 1324 | Y1324H | E | 2.1e-07 | 3.9e-07 | 2.3e-07 | 2.767E-07 | 9.866E-08 | 2.100E-07 | 3.900E-07 |
| 26470 | 1324 | Y1324I | E | 4.8e-07 | 4.2e-07 | 3.4e-07 | 4.133E-07 | 7.024E-08 | 3.400E-07 | 4.800E-07 |
| 26471 | 1324 | Y1324L | E | 4.2e-07 | 2e-07 | 2.1e-07 | 2.767E-07 | 1.242E-07 | 2.000E-07 | 4.200E-07 |
| 26472 | 1324 | Y1324K | E | 2.6e-07 | 3e-07 | 1.8e-07 | 2.467E-07 | 6.110E-08 | 1.800E-07 | 3.000E-07 |
| 26473 | 1324 | Y1324M | E | 4.4e-07 | 3e-07 | 1.3e-07 | 2.900E-07 | 1.552E-07 | 1.300E-07 | 4.400E-07 |
| 26474 | 1324 | Y1324F | E | 5.2e-07 | 2.4e-07 | 2.8e-07 | 3.467E-07 | 1.514E-07 | 2.400E-07 | 5.200E-07 |
| 26475 | 1324 | Y1324P | E | 1.9e-07 | 1.8e-07 | 2.1e-07 | 1.933E-07 | 1.528E-08 | 1.800E-07 | 2.100E-07 |
| 26476 | 1324 | Y1324S | E | 1.8e-07 | 2.9e-07 | 2.3e-07 | 2.333E-07 | 5.508E-08 | 1.800E-07 | 2.900E-07 |
| 26477 | 1324 | Y1324T | E | 8.1e-07 | 2.1e-07 | 3.3e-07 | 4.500E-07 | 3.175E-07 | 2.100E-07 | 8.100E-07 |
| 26478 | 1324 | Y1324W | E | 6.2e-07 | 8.7e-08 | 3.3e-07 | 3.457E-07 | 2.668E-07 | 8.700E-08 | 6.200E-07 |
| 26479 | 1324 | Y1324Y | E | 1.5e-07 | 3.1e-07 | 2.5e-07 | 2.367E-07 | 8.083E-08 | 1.500E-07 | 3.100E-07 |
| 26480 | 1324 | Y1324V | E | 1.7e-07 | 9.2e-08 | 1.8e-07 | 1.473E-07 | 4.818E-08 | 9.200E-08 | 1.800E-07 |
| 26481 | 1325 | P1325A | E | 3.7e-07 | 2.2e-07 | 3e-07 | 2.967E-07 | 7.506E-08 | 2.200E-07 | 3.700E-07 |
| 26482 | 1325 | P1325R | E | 1.5e-07 | 1.7e-07 | 2.8e-07 | 2.000E-07 | 7.000E-08 | 1.500E-07 | 2.800E-07 |
| 26483 | 1325 | P1325N | E | 2.4e-07 | 1.5e-07 | 3.6e-07 | 2.500E-07 | 1.054E-07 | 1.500E-07 | 3.600E-07 |
| 26484 | 1325 | P1325D | E | 1.9e-07 | 1.8e-07 | 2.2e-07 | 1.967E-07 | 2.082E-08 | 1.800E-07 | 2.200E-07 |
| 26485 | 1325 | P1325C | E | 2.7e-07 | 1.2e-07 | 1.8e-07 | 1.900E-07 | 7.550E-08 | 1.200E-07 | 2.700E-07 |
| 26486 | 1325 | P1325E | E | 2.2e-07 | 4e-07 | 2.4e-07 | 2.867E-07 | 9.866E-08 | 2.200E-07 | 4.000E-07 |
| 26487 | 1325 | P1325Q | E | 2.1e-07 | 3.9e-07 | 2.8e-07 | 2.933E-07 | 9.074E-08 | 2.100E-07 | 3.900E-07 |
| 26488 | 1325 | P1325G | E | 1.8e-07 | 3.2e-07 | 4.2e-07 | 3.067E-07 | 1.206E-07 | 1.800E-07 | 4.200E-07 |
| 26489 | 1325 | P1325H | E | 3e-07 | 2.2e-07 | 2.9e-07 | 2.700E-07 | 4.359E-08 | 2.200E-07 | 3.000E-07 |
| 26490 | 1325 | P1325I | E | 1.8e-07 | 2.2e-07 | 1.6e-07 | 1.867E-07 | 3.055E-08 | 1.600E-07 | 2.200E-07 |
| 26491 | 1325 | P1325L | E | 1.8e-07 | 3.6e-07 | 3e-07 | 2.800E-07 | 9.165E-08 | 1.800E-07 | 3.600E-07 |
| 26492 | 1325 | P1325K | E | 2e-07 | 4.1e-07 | 2.5e-07 | 2.867E-07 | 1.097E-07 | 2.000E-07 | 4.100E-07 |
| 26493 | 1325 | P1325M | E | 5.8e-07 | 1.6e-07 | 2.2e-07 | 3.200E-07 | 2.272E-07 | 1.600E-07 | 5.800E-07 |
| 26494 | 1325 | P1325F | E | 1.6e-07 | 3.9e-07 | 3.7e-07 | 3.067E-07 | 1.274E-07 | 1.600E-07 | 3.900E-07 |
| 26495 | 1325 | P1325P | E | 5.2e-07 | 2.6e-07 | 3.3e-07 | 3.700E-07 | 1.345E-07 | 2.600E-07 | 5.200E-07 |
| 26496 | 1325 | P1325S | E | 3.5e-07 | 1.5e-07 | 2.7e-07 | 2.567E-07 | 1.007E-07 | 1.500E-07 | 3.500E-07 |
| 26497 | 1325 | P1325T | E | 1.9e-07 | 2.3e-07 | 2.9e-07 | 2.367E-07 | 5.033E-08 | 1.900E-07 | 2.900E-07 |
| 26498 | 1325 | P1325W | E | 2.2e-07 | 1.8e-07 | 2.1e-07 | 2.033E-07 | 2.082E-08 | 1.800E-07 | 2.200E-07 |
| 26499 | 1325 | P1325Y | E | 4.4e-07 | 2.5e-07 | 4.7e-07 | 3.867E-07 | 1.193E-07 | 2.500E-07 | 4.700E-07 |
| 26500 | 1325 | P1325V | E | 2e-07 | 3.3e-07 | 2.3e-07 | 2.533E-07 | 6.807E-08 | 2.000E-07 | 3.300E-07 |
| 26501 | 1326 | T1326A | E | 2.7e-07 | 3.3e-07 | 5.4e-07 | 3.800E-07 | 1.418E-07 | 2.700E-07 | 5.400E-07 |
| 26502 | 1326 | T1326R | E | 2e-07 | 3e-07 | 1.4e-07 | 2.133E-07 | 8.083E-08 | 1.400E-07 | 3.000E-07 |
| 26503 | 1326 | T1326N | E | 3.2e-07 | 2.2e-07 | 2.6e-07 | 2.667E-07 | 5.033E-08 | 2.200E-07 | 3.200E-07 |
| 26504 | 1326 | T1326D | E | 2.5e-07 | 3.1e-07 | 5e-07 | 3.533E-07 | 1.305E-07 | 2.500E-07 | 5.000E-07 |
| 26505 | 1326 | T1326C | E | 2.4e-07 | 4.1e-07 | 3.3e-07 | 3.267E-07 | 8.505E-08 | 2.400E-07 | 4.100E-07 |
| 26506 | 1326 | T1326E | E | 3e-07 | 4.6e-07 | 2e-07 | 3.200E-07 | 1.311E-07 | 2.000E-07 | 4.600E-07 |
| 26507 | 1326 | T1326Q | E | 3.4e-07 | 2.1e-07 | 2.6e-07 | 2.700E-07 | 6.557E-08 | 2.100E-07 | 3.400E-07 |
| 26508 | 1326 | T1326G | E | 5.1e-07 | 3.1e-07 | 1.1e-07 | 3.100E-07 | 2.000E-07 | 1.100E-07 | 5.100E-07 |
| 26509 | 1326 | T1326H | E | 3e-07 | 2.5e-07 | 1.2e-07 | 2.233E-07 | 9.292E-08 | 1.200E-07 | 3.000E-07 |
| 26510 | 1326 | T1326I | E | 1.8e-07 | 1.6e-07 | 2.5e-07 | 1.967E-07 | 4.726E-08 | 1.600E-07 | 2.500E-07 |
| 26511 | 1326 | T1326L | E | 1.4e-07 | 3.3e-07 | 3.1e-07 | 2.600E-07 | 1.044E-07 | 1.400E-07 | 3.300E-07 |
| 26512 | 1326 | T1326K | E | 2e-07 | 2.2e-07 | 4.5e-07 | 2.900E-07 | 1.389E-07 | 2.000E-07 | 4.500E-07 |
| 26513 | 1326 | T1326M | E | 3.4e-07 | 4.4e-07 | 4.3e-07 | 4.033E-07 | 5.508E-08 | 3.400E-07 | 4.400E-07 |
| 26514 | 1326 | T1326F | E | 1.9e-07 | 3.3e-07 | 1.7e-07 | 2.300E-07 | 8.718E-08 | 1.700E-07 | 3.300E-07 |
| 26515 | 1326 | T1326P | E | 4e-07 | 2.8e-07 | 3.4e-07 | 3.400E-07 | 6.000E-08 | 2.800E-07 | 4.000E-07 |
| 26516 | 1326 | T1326S | E | 2.6e-07 | 2.5e-07 | 1.3e-07 | 2.133E-07 | 7.234E-08 | 1.300E-07 | 2.600E-07 |
| 26517 | 1326 | T1326T | E | 2.3e-07 | 1.9e-07 | 4.1e-07 | 2.767E-07 | 1.172E-07 | 1.900E-07 | 4.100E-07 |
| 26518 | 1326 | T1326W | E | 1.1e-07 | 5e-07 | 2.3e-07 | 2.800E-07 | 1.997E-07 | 1.100E-07 | 5.000E-07 |
| 26519 | 1326 | T1326Y | E | 2.9e-07 | 2.3e-07 | 3.9e-07 | 3.033E-07 | 8.083E-08 | 2.300E-07 | 3.900E-07 |

| | | | | | | | | | | |
|---|---|---|---|---|---|---|---|---|---|---|
| 26520 | 1326 | T1326V | E | 2.3e-07 | 1.7e-07 | 4.5e-07 | 2.833E-07 | 1.474E-07 | 1.700E-07 | 4.500E-07 |
| 26521 | 1327 | N1327A | E | 1.9e-07 | 1.7e-07 | 2.1e-07 | 1.900E-07 | 2.000E-08 | 1.700E-07 | 2.100E-07 |
| 26522 | 1327 | N1327R | E | 1.5e-07 | 3.4e-07 | 1.1e-07 | 2.000E-07 | 1.229E-07 | 1.100E-07 | 3.400E-07 |
| 26523 | 1327 | N1327N | E | 2.9e-07 | 6e-07 | 2.8e-07 | 3.900E-07 | 1.819E-07 | 2.800E-07 | 6.000E-07 |
| 26524 | 1327 | N1327D | E | 3.8e-07 | 2.4e-07 | 5.8e-07 | 4.000E-07 | 1.709E-07 | 2.400E-07 | 5.800E-07 |
| 26525 | 1327 | N1327C | E | 1.1e-07 | 1.7e-07 | 1.2e-07 | 1.333E-07 | 3.215E-08 | 1.100E-07 | 1.700E-07 |
| 26526 | 1327 | N1327E | E | 2.5e-07 | 1.6e-07 | 3.4e-07 | 2.500E-07 | 9.000E-08 | 1.600E-07 | 3.400E-07 |
| 26527 | 1327 | N1327Q | E | 2.4e-07 | 3.7e-07 | 1.5e-07 | 2.533E-07 | 1.106E-07 | 1.500E-07 | 3.700E-07 |
| 26528 | 1327 | N1327G | E | 3.6e-07 | 2e-07 | 4.2e-07 | 3.267E-07 | 1.137E-07 | 2.000E-07 | 4.200E-07 |
| 26529 | 1327 | N1327H | E | 2.8e-07 | 2e-07 | 4.3e-07 | 3.033E-07 | 1.168E-07 | 2.000E-07 | 4.300E-07 |
| 26530 | 1327 | N1327I | E | 1.3e-07 | 2.8e-07 | 2.7e-07 | 2.267E-07 | 8.386E-08 | 1.300E-07 | 2.800E-07 |
| 26531 | 1327 | N1327L | E | 5.1e-08 | 3.1e-07 | 2.9e-07 | 2.170E-07 | 1.441E-07 | 5.100E-08 | 3.100E-07 |
| 26532 | 1327 | N1327K | E | 2.3e-07 | 1.7e-07 | 3e-07 | 2.333E-07 | 6.506E-08 | 1.700E-07 | 3.000E-07 |
| 26533 | 1327 | N1327M | E | 2.1e-07 | 6.2e-07 | 2.9e-07 | 3.733E-07 | 2.173E-07 | 2.100E-07 | 6.200E-07 |
| 26534 | 1327 | N1327F | E | 3.1e-07 | 2.7e-07 | 3e-07 | 2.933E-07 | 2.082E-08 | 2.700E-07 | 3.100E-07 |
| 26535 | 1327 | N1327P | E | 8.9e-08 | 2.1e-07 | 2.3e-07 | 1.763E-07 | 7.629E-08 | 8.900E-08 | 2.300E-07 |
| 26536 | 1327 | N1327S | E | 6.1e-07 | 3.5e-07 | 3.3e-07 | 4.300E-07 | 1.562E-07 | 3.300E-07 | 6.100E-07 |
| 26537 | 1327 | N1327T | E | 3.3e-07 | 1.3e-07 | 3.4e-07 | 2.667E-07 | 1.185E-07 | 1.300E-07 | 3.400E-07 |
| 26538 | 1327 | N1327W | E | 6.4e-07 | 2.1e-07 | 1e-07 | 3.167E-07 | 2.854E-07 | 1.000E-07 | 6.400E-07 |
| 26539 | 1327 | N1327Y | E | 2.7e-07 | 3.5e-07 | 3.7e-07 | 3.300E-07 | 5.292E-08 | 2.700E-07 | 3.700E-07 |
| 26540 | 1327 | N1327V | E | 1.9e-07 | 4.1e-07 | 2.1e-07 | 2.700E-07 | 1.217E-07 | 1.900E-07 | 4.100E-07 |
| 26541 | 1328 | G1328A | E | 2.9e-07 | 3.1e-07 | 2.1e-07 | 2.700E-07 | 5.292E-08 | 2.100E-07 | 3.100E-07 |
| 26542 | 1328 | G1328R | E | 2.6e-07 | 3.1e-07 | 4.4e-07 | 3.367E-07 | 9.292E-08 | 2.600E-07 | 4.400E-07 |
| 26543 | 1328 | G1328N | E | 3e-07 | 2.2e-07 | 2.8e-07 | 2.667E-07 | 4.163E-08 | 2.200E-07 | 3.000E-07 |
| 26544 | 1328 | G1328D | E | 2.8e-07 | 2.8e-07 | 3.3e-07 | 2.967E-07 | 2.887E-08 | 2.800E-07 | 3.300E-07 |
| 26545 | 1328 | G1328C | E | 7.2e-07 | 3.6e-07 | 1.8e-07 | 4.200E-07 | 2.750E-07 | 1.800E-07 | 7.200E-07 |
| 26546 | 1328 | G1328E | E | 1.5e-07 | 1.3e-07 | 1e-07 | 1.267E-07 | 2.517E-08 | 1.000E-07 | 1.500E-07 |
| 26547 | 1328 | G1328Q | E | 2.5e-07 | 4.5e-07 | 2.1e-07 | 3.033E-07 | 1.286E-07 | 2.100E-07 | 4.500E-07 |
| 26548 | 1328 | G1328G | E | 2.6e-07 | 1.8e-07 | 1.9e-07 | 2.100E-07 | 4.359E-08 | 1.800E-07 | 2.600E-07 |
| 26549 | 1328 | G1328H | E | 2.5e-07 | 2.5e-07 | 2e-07 | 2.333E-07 | 2.887E-08 | 2.000E-07 | 2.500E-07 |
| 26550 | 1328 | G1328I | E | 1.6e-07 | 4.2e-07 | 1.7e-07 | 2.500E-07 | 1.473E-07 | 1.600E-07 | 4.200E-07 |
| 26551 | 1328 | G1328L | E | 3.7e-07 | 1.9e-07 | 6.8e-07 | 4.133E-07 | 2.479E-07 | 1.900E-07 | 6.800E-07 |
| 26552 | 1328 | G1328K | E | 5.4e-07 | 2.9e-07 | 2.2e-07 | 3.500E-07 | 1.682E-07 | 2.200E-07 | 5.400E-07 |
| 26553 | 1328 | G1328M | E | 1.7e-07 | 2.5e-07 | 2.7e-07 | 2.300E-07 | 5.292E-08 | 1.700E-07 | 2.700E-07 |
| 26554 | 1328 | G1328F | E | 2e-07 | 1.4e-07 | 3.7e-07 | 2.367E-07 | 1.193E-07 | 1.400E-07 | 3.700E-07 |
| 26555 | 1328 | G1328P | E | 3.3e-07 | 2.7e-07 | 1.8e-07 | 2.600E-07 | 7.550E-08 | 1.800E-07 | 3.300E-07 |
| 26556 | 1328 | G1328S | E | 9.1e-08 | 3.5e-07 | 3.3e-07 | 2.570E-07 | 1.441E-07 | 9.100E-08 | 3.500E-07 |
| 26557 | 1328 | G1328T | E | 1.7e-07 | 2.7e-07 | 1.8e-07 | 2.067E-07 | 5.508E-08 | 1.700E-07 | 2.700E-07 |
| 26558 | 1328 | G1328W | E | 2.4e-07 | 2.6e-07 | 3.7e-07 | 2.900E-07 | 7.000E-08 | 2.400E-07 | 3.700E-07 |
| 26559 | 1328 | G1328Y | E | 2.3e-07 | 1.3e-07 | 2.8e-07 | 2.133E-07 | 7.638E-08 | 1.300E-07 | 2.800E-07 |
| 26560 | 1328 | G1328V | E | 3.8e-07 | 4.2e-07 | 2e-07 | 3.333E-07 | 1.172E-07 | 2.000E-07 | 4.200E-07 |
| 26561 | 1329 | Y1329A | E | 2.1e-07 | 3.1e-07 | 2.2e-07 | 2.467E-07 | 5.508E-08 | 2.100E-07 | 3.100E-07 |
| 26562 | 1329 | Y1329R | E | 1e-07 | 1.6e-07 | 1.6e-07 | 1.400E-07 | 3.464E-08 | 1.000E-07 | 1.600E-07 |
| 26563 | 1329 | Y1329N | E | 4e-07 | 2.8e-07 | 5.7e-08 | 2.457E-07 | 1.741E-07 | 5.700E-08 | 4.000E-07 |
| 26564 | 1329 | Y1329D | E | 3.1e-07 | 1.5e-07 | 4.9e-07 | 3.167E-07 | 1.701E-07 | 1.500E-07 | 4.900E-07 |
| 26565 | 1329 | Y1329C | E | 1.4e-07 | 2.7e-07 | 4.5e-07 | 2.867E-07 | 1.557E-07 | 1.400E-07 | 4.500E-07 |
| 26566 | 1329 | Y1329E | E | 2e-07 | 8.1e-08 | 1.5e-07 | 1.437E-07 | 5.975E-08 | 8.100E-08 | 2.000E-07 |
| 26567 | 1329 | Y1329Q | E | 8.7e-08 | 1.3e-07 | 1.4e-07 | 1.190E-07 | 2.816E-08 | 8.700E-08 | 1.400E-07 |
| 26568 | 1329 | Y1329G | E | 2.8e-07 | 4.8e-07 | 8.4e-08 | 2.813E-07 | 1.980E-07 | 8.400E-08 | 4.800E-07 |
| 26569 | 1329 | Y1329H | E | 1.8e-07 | 1.7e-07 | 2.2e-07 | 1.900E-07 | 2.646E-08 | 1.700E-07 | 2.200E-07 |
| 26570 | 1329 | Y1329I | E | 2.4e-07 | 7.8e-08 | 1.3e-07 | 1.493E-07 | 8.271E-08 | 7.800E-08 | 2.400E-07 |
| 26571 | 1329 | Y1329L | E | 3.6e-07 | 3.6e-07 | 3.5e-07 | 3.567E-07 | 5.774E-09 | 3.500E-07 | 3.600E-07 |
| 26572 | 1329 | Y1329K | E | 1.8e-07 | 2e-07 | 1.4e-07 | 1.733E-07 | 3.055E-08 | 1.400E-07 | 2.000E-07 |
| 26573 | 1329 | Y1329M | E | 2.6e-07 | 3.1e-07 | 1.1e-07 | 2.267E-07 | 1.041E-07 | 1.100E-07 | 3.100E-07 |
| 26574 | 1329 | Y1329F | E | 2.6e-07 | 2.4e-07 | 1.9e-07 | 2.300E-07 | 3.606E-08 | 1.900E-07 | 2.600E-07 |
| 26575 | 1329 | Y1329P | E | 4.8e-07 | 2.2e-07 | 1.1e-07 | 2.700E-07 | 1.900E-07 | 1.100E-07 | 4.800E-07 |
| 26576 | 1329 | Y1329S | E | 1.2e-07 | 1.5e-07 | 1.6e-07 | 1.433E-07 | 2.082E-08 | 1.200E-07 | 1.600E-07 |
| 26577 | 1329 | Y1329T | E | 3.3e-07 | 1.4e-07 | 2.3e-07 | 2.333E-07 | 9.504E-08 | 1.400E-07 | 3.300E-07 |
| 26578 | 1329 | Y1329W | E | 1.6e-07 | 2.4e-07 | 3.4e-07 | 2.467E-07 | 9.018E-08 | 1.600E-07 | 3.400E-07 |
| 26579 | 1329 | Y1329Y | E | 3.7e-07 | 2.3e-07 | 2.6e-07 | 2.867E-07 | 7.371E-08 | 2.300E-07 | 3.700E-07 |
| 26580 | 1329 | Y1329V | E | 2.4e-07 | 2.6e-07 | 2.6e-07 | 2.533E-07 | 1.155E-08 | 2.400E-07 | 2.600E-07 |
| 26581 | 1330 | T1330A | E | 2.3e-07 | 2.9e-07 | 1.9e-07 | 2.367E-07 | 5.033E-08 | 1.900E-07 | 2.900E-07 |
| 26582 | 1330 | T1330R | E | 2.2e-07 | 2e-07 | 2.1e-07 | 2.100E-07 | 1.000E-08 | 2.000E-07 | 2.200E-07 |
| 26583 | 1330 | T1330N | E | 2e-07 | 4.3e-07 | 4.5e-07 | 3.600E-07 | 1.389E-07 | 2.000E-07 | 4.500E-07 |
| 26584 | 1330 | T1330D | E | 2.1e-07 | 1.7e-07 | 1.1e-07 | 1.633E-07 | 5.033E-08 | 1.100E-07 | 2.100E-07 |
| 26585 | 1330 | T1330C | E | 1.2e-07 | 2.4e-07 | 2.3e-07 | 1.967E-07 | 6.658E-08 | 1.200E-07 | 2.400E-07 |
| 26586 | 1330 | T1330E | E | 1.9e-07 | 1.4e-07 | 2.8e-07 | 2.033E-07 | 7.095E-08 | 1.400E-07 | 2.800E-07 |
| 26587 | 1330 | T1330Q | E | 2.1e-07 | 2.6e-07 | 3.2e-07 | 2.633E-07 | 5.508E-08 | 2.100E-07 | 3.200E-07 |
| 26588 | 1330 | T1330G | E | 2.4e-07 | 3e-07 | 2.3e-07 | 2.567E-07 | 3.786E-08 | 2.300E-07 | 3.000E-07 |
| 26589 | 1330 | T1330H | E | 2.8e-07 | 1.7e-07 | 3.1e-07 | 2.533E-07 | 7.371E-08 | 1.700E-07 | 3.100E-07 |
| 26590 | 1330 | T1330I | E | 4e-07 | 1.7e-07 | 2.4e-07 | 2.700E-07 | 1.179E-07 | 1.700E-07 | 4.000E-07 |
| 26591 | 1330 | T1330L | E | 3.2e-07 | 2e-07 | 1.7e-07 | 2.300E-07 | 7.937E-08 | 1.700E-07 | 3.200E-07 |
| 26592 | 1330 | T1330K | E | 2e-07 | 2.8e-07 | 3e-07 | 2.600E-07 | 5.292E-08 | 2.000E-07 | 3.000E-07 |
| 26593 | 1330 | T1330M | E | 1.5e-07 | 1.4e-07 | 2.4e-07 | 1.767E-07 | 5.508E-08 | 1.400E-07 | 2.400E-07 |
| 26594 | 1330 | T1330F | E | 3.3e-07 | 1.7e-07 | 2.1e-07 | 2.367E-07 | 8.327E-08 | 1.700E-07 | 3.300E-07 |
| 26595 | 1330 | T1330P | E | 1.4e-07 | 1.6e-07 | 2.6e-07 | 1.867E-07 | 6.429E-08 | 1.400E-07 | 2.600E-07 |
| 26596 | 1330 | T1330S | E | 3.1e-07 | 7.5e-07 | 3.1e-07 | 4.567E-07 | 2.540E-07 | 3.100E-07 | 7.500E-07 |
| 26597 | 1330 | T1330T | E | 4e-07 | 1.9e-07 | 2.2e-07 | 2.700E-07 | 1.136E-07 | 1.900E-07 | 4.000E-07 |
| 26598 | 1330 | T1330W | E | 3.2e-07 | 1.8e-07 | 2.2e-07 | 2.400E-07 | 7.211E-08 | 1.800E-07 | 3.200E-07 |
| 26599 | 1330 | T1330Y | E | 2e-07 | 1.5e-07 | 3.5e-07 | 2.333E-07 | 1.041E-07 | 1.500E-07 | 3.500E-07 |
| 26600 | 1330 | T1330V | E | 2e-07 | 2.5e-07 | 2.3e-07 | 2.267E-07 | 2.517E-08 | 2.000E-07 | 2.500E-07 |
| 26601 | 1331 | R1331A | E | 2.6e-07 | 2.3e-07 | 2.5e-07 | 2.467E-07 | 1.528E-08 | 2.300E-07 | 2.600E-07 |
| 26602 | 1331 | R1331R | E | 2.1e-07 | 2.3e-07 | 3.2e-07 | 2.533E-07 | 5.859E-08 | 2.100E-07 | 3.200E-07 |
| 26603 | 1331 | R1331N | E | 5.7e-07 | 2.1e-07 | 3.4e-07 | 3.733E-07 | 1.823E-07 | 2.100E-07 | 5.700E-07 |
| 26604 | 1331 | R1331D | E | 5.3e-07 | 2.5e-07 | 4.6e-07 | 4.133E-07 | 1.457E-07 | 2.500E-07 | 5.300E-07 |

| | | | | | | | | | | |
|---|---|---|---|---|---|---|---|---|---|---|
| 26605 | 1331 | R1331C | E | 1.6e-07 | 2.7e-07 | 2.2e-07 | 2.167E-07 | 5.508E-08 | 1.600E-07 | 2.700E-07 |
| 26606 | 1331 | R1331E | E | 6e-07 | 4.9e-07 | 3.4e-07 | 4.767E-07 | 1.305E-07 | 3.400E-07 | 6.000E-07 |
| 26607 | 1331 | R1331Q | E | 4.6e-07 | 3.4e-07 | 2.4e-07 | 3.467E-07 | 1.102E-07 | 2.400E-07 | 4.600E-07 |
| 26608 | 1331 | R1331G | E | 2.6e-07 | 4.6e-07 | 2.9e-07 | 3.367E-07 | 1.079E-07 | 2.600E-07 | 4.600E-07 |
| 26609 | 1331 | R1331H | E | 3.6e-07 | 1.4e-07 | 3.5e-07 | 2.833E-07 | 1.242E-07 | 1.400E-07 | 3.600E-07 |
| 26610 | 1331 | R1331I | E | 3.6e-07 | 3.2e-07 | 1.6e-07 | 2.800E-07 | 1.058E-07 | 1.600E-07 | 3.600E-07 |
| 26611 | 1331 | R1331L | E | 5e-07 | 1.4e-07 | 2.5e-07 | 2.967E-07 | 1.845E-07 | 1.400E-07 | 5.000E-07 |
| 26612 | 1331 | R1331K | E | 3.2e-07 | 2.3e-07 | 3.1e-07 | 2.867E-07 | 4.933E-08 | 2.300E-07 | 3.200E-07 |
| 26613 | 1331 | R1331M | E | 3.4e-07 | 1.4e-07 | 2.3e-07 | 2.367E-07 | 1.002E-07 | 1.400E-07 | 3.400E-07 |
| 26614 | 1331 | R1331F | E | 2.5e-07 | 2.9e-07 | 1.9e-07 | 2.433E-07 | 5.033E-08 | 1.900E-07 | 2.900E-07 |
| 26615 | 1331 | R1331P | E | 4e-07 | 2.3e-07 | 3.8e-07 | 3.367E-07 | 9.292E-08 | 2.300E-07 | 4.000E-07 |
| 26616 | 1331 | R1331S | E | 7.7e-07 | 2.9e-07 | 5e-07 | 5.200E-07 | 2.406E-07 | 2.900E-07 | 7.700E-07 |
| 26617 | 1331 | R1331T | E | 2.1e-07 | 6.5e-07 | 1.2e-06 | 6.867E-07 | 4.960E-07 | 2.100E-07 | 1.200E-06 |
| 26618 | 1331 | R1331W | E | 1.6e-07 | 5.6e-07 | 5.3e-07 | 4.167E-07 | 2.228E-07 | 1.600E-07 | 5.600E-07 |
| 26619 | 1331 | R1331Y | E | 8e-08 | 2.6e-07 | 3.1e-07 | 2.167E-07 | 1.210E-07 | 8.000E-08 | 3.100E-07 |
| 26620 | 1331 | R1331V | E | 1.5e-07 | 3e-07 | 2.8e-07 | 2.433E-07 | 8.145E-08 | 1.500E-07 | 3.000E-07 |
| 26621 | 1332 | Y1332A | E | 1.5e-07 | 1.6e-07 | 3e-07 | 2.033E-07 | 8.386E-08 | 1.500E-07 | 3.000E-07 |
| 26622 | 1332 | Y1332R | E | 4.7e-07 | 1.6e-07 | 1.6e-07 | 2.633E-07 | 1.790E-07 | 1.600E-07 | 4.700E-07 |
| 26623 | 1332 | Y1332N | E | 2.4e-07 | 1.2e-07 | 6e-07 | 3.200E-07 | 2.498E-07 | 1.200E-07 | 6.000E-07 |
| 26624 | 1332 | Y1332D | E | 2.4e-07 | 2.8e-07 | 2.1e-07 | 2.433E-07 | 3.512E-08 | 2.100E-07 | 2.800E-07 |
| 26625 | 1332 | Y1332C | E | 1.9e-07 | 3.9e-07 | 2e-07 | 2.600E-07 | 1.127E-07 | 1.900E-07 | 3.900E-07 |
| 26626 | 1332 | Y1332E | E | 2.4e-07 | 3.5e-07 | 3e-07 | 2.967E-07 | 5.508E-08 | 2.400E-07 | 3.500E-07 |
| 26627 | 1332 | Y1332Q | E | 1.7e-07 | 2.6e-07 | 3.8e-07 | 2.700E-07 | 1.054E-07 | 1.700E-07 | 3.800E-07 |
| 26628 | 1332 | Y1332G | E | 5.1e-07 | 2.2e-07 | 1.2e-07 | 2.833E-07 | 2.026E-07 | 1.200E-07 | 5.100E-07 |
| 26629 | 1332 | Y1332H | E | 2e-07 | 2.9e-07 | 1.6e-07 | 2.167E-07 | 6.658E-08 | 1.600E-07 | 2.900E-07 |
| 26630 | 1332 | Y1332I | E | 2.4e-07 | 3e-07 | 3e-07 | 2.800E-07 | 3.464E-08 | 2.400E-07 | 3.000E-07 |
| 26631 | 1332 | Y1332L | E | 9.9e-08 | 2.1e-07 | 3e-07 | 2.030E-07 | 1.007E-07 | 9.900E-08 | 3.000E-07 |
| 26632 | 1332 | Y1332K | E | 6.5e-07 | 3.3e-07 | 2.9e-07 | 4.233E-07 | 1.973E-07 | 2.900E-07 | 6.500E-07 |
| 26633 | 1332 | Y1332M | E | 1.8e-07 | 1.9e-07 | 3.3e-07 | 2.333E-07 | 8.386E-08 | 1.800E-07 | 3.300E-07 |
| 26634 | 1332 | Y1332F | E | 1.6e-07 | 4.8e-07 | 8.9e-08 | 2.430E-07 | 2.083E-07 | 8.900E-08 | 4.800E-07 |
| 26635 | 1332 | Y1332P | E | 4e-07 | 3e-07 | 5.2e-07 | 4.067E-07 | 1.102E-07 | 3.000E-07 | 5.200E-07 |
| 26636 | 1332 | Y1332S | E | 3.4e-07 | 2.9e-07 | 1.4e-07 | 2.567E-07 | 1.041E-07 | 1.400E-07 | 3.400E-07 |
| 26637 | 1332 | Y1332T | E | 2.8e-07 | 1.6e-07 | 3.9e-07 | 2.767E-07 | 1.150E-07 | 1.600E-07 | 3.900E-07 |
| 26638 | 1332 | Y1332W | E | 4.1e-07 | 6.2e-07 | 1.7e-07 | 4.000E-07 | 2.252E-07 | 1.700E-07 | 6.200E-07 |
| 26639 | 1332 | Y1332Y | E | 2.1e-07 | 5.9e-07 | 3.4e-07 | 3.800E-07 | 1.931E-07 | 2.100E-07 | 5.900E-07 |
| 26640 | 1332 | Y1332V | E | 2.9e-07 | 1.6e-07 | 1.3e-07 | 1.933E-07 | 8.505E-08 | 1.300E-07 | 2.900E-07 |
| 26641 | 1333 | A1333A | E | 3.2e-07 | 1.3e-07 | 3.3e-07 | 2.600E-07 | 1.127E-07 | 1.300E-07 | 3.300E-07 |
| 26642 | 1333 | A1333R | E | 1.7e-07 | 2e-07 | 1.8e-07 | 1.833E-07 | 1.528E-08 | 1.700E-07 | 2.000E-07 |
| 26643 | 1333 | A1333N | E | 2.6e-07 | 2.2e-07 | 4.1e-07 | 2.967E-07 | 1.002E-07 | 2.200E-07 | 4.100E-07 |
| 26644 | 1333 | A1333D | E | 1.9e-07 | 9.7e-08 | 3.4e-07 | 2.090E-07 | 1.226E-07 | 9.700E-08 | 3.400E-07 |
| 26645 | 1333 | A1333C | E | 2e-07 | 2.2e-07 | 1.9e-07 | 2.033E-07 | 1.528E-08 | 1.900E-07 | 2.200E-07 |
| 26646 | 1333 | A1333E | E | 6.6e-07 | 4.7e-07 | 2.4e-07 | 4.567E-07 | 2.103E-07 | 2.400E-07 | 6.600E-07 |
| 26647 | 1333 | A1333Q | E | 2.6e-07 | 9.6e-08 | 1.5e-07 | 1.687E-07 | 8.358E-08 | 9.600E-08 | 2.600E-07 |
| 26648 | 1333 | A1333G | E | 7.4e-07 | 1.6e-07 | 2.5e-07 | 3.833E-07 | 3.121E-07 | 1.600E-07 | 7.400E-07 |
| 26649 | 1333 | A1333H | E | 1.2e-07 | 2.2e-07 | 2.2e-07 | 1.867E-07 | 5.774E-08 | 1.200E-07 | 2.200E-07 |
| 26650 | 1333 | A1333I | E | 1.8e-07 | 1.4e-07 | 4.8e-07 | 2.667E-07 | 1.858E-07 | 1.400E-07 | 4.800E-07 |
| 26651 | 1333 | A1333L | E | 2.2e-07 | 2e-07 | 2.1e-07 | 2.100E-07 | 1.000E-08 | 2.000E-07 | 2.200E-07 |
| 26652 | 1333 | A1333K | E | 4.6e-07 | 2.3e-07 | 2.1e-07 | 3.000E-07 | 1.389E-07 | 2.100E-07 | 4.600E-07 |
| 26653 | 1333 | A1333M | E | 3.6e-07 | 1.3e-07 | 3e-07 | 2.633E-07 | 1.193E-07 | 1.300E-07 | 3.600E-07 |
| 26654 | 1333 | A1333F | E | 2.5e-07 | 1.9e-07 | 5.1e-07 | 3.167E-07 | 1.701E-07 | 1.900E-07 | 5.100E-07 |
| 26655 | 1333 | A1333P | E | 2.2e-07 | 2.7e-07 | 2.8e-07 | 2.567E-07 | 3.215E-08 | 2.200E-07 | 2.800E-07 |
| 26656 | 1333 | A1333S | E | 1.4e-07 | 1.9e-07 | 2.5e-07 | 1.933E-07 | 5.508E-08 | 1.400E-07 | 2.500E-07 |
| 26657 | 1333 | A1333T | E | 1.8e-07 | 3.3e-07 | 1.8e-07 | 2.300E-07 | 8.660E-08 | 1.800E-07 | 3.300E-07 |
| 26658 | 1333 | A1333W | E | 2e-07 | 1.5e-07 | 1.8e-07 | 1.767E-07 | 2.517E-08 | 1.500E-07 | 2.000E-07 |
| 26659 | 1333 | A1333Y | E | 2.1e-07 | 1.8e-07 | 4.4e-07 | 2.767E-07 | 1.422E-07 | 1.800E-07 | 4.400E-07 |
| 26660 | 1333 | A1333V | E | 6.8e-07 | 1.7e-07 | 2.7e-07 | 3.733E-07 | 2.702E-07 | 1.700E-07 | 6.800E-07 |
| 26661 | 1334 | D1334A | E | 1.7e-07 | 2.2e-07 | 3.2e-07 | 2.367E-07 | 7.638E-08 | 1.700E-07 | 3.200E-07 |
| 26662 | 1334 | D1334R | E | 3.1e-07 | 1.7e-07 | 2.4e-07 | 2.400E-07 | 7.000E-08 | 1.700E-07 | 3.100E-07 |
| 26663 | 1334 | D1334N | E | 2.9e-07 | 2.8e-07 | 1.3e-07 | 2.333E-07 | 8.963E-08 | 1.300E-07 | 2.900E-07 |
| 26664 | 1334 | D1334D | E | 2.4e-07 | 3.2e-07 | 3e-07 | 2.867E-07 | 4.163E-08 | 2.400E-07 | 3.200E-07 |
| 26665 | 1334 | D1334C | E | 1.8e-07 | 4.2e-07 | 1.8e-07 | 2.600E-07 | 1.386E-07 | 1.800E-07 | 4.200E-07 |
| 26666 | 1334 | D1334E | E | 2.8e-07 | 1.5e-07 | 3.2e-07 | 2.500E-07 | 8.888E-08 | 1.500E-07 | 3.200E-07 |
| 26667 | 1334 | D1334Q | E | 1.2e-07 | 2.2e-07 | 1.7e-07 | 1.700E-07 | 5.000E-08 | 1.200E-07 | 2.200E-07 |
| 26668 | 1334 | D1334G | E | 1.7e-07 | 2.8e-07 | 1.4e-07 | 1.967E-07 | 7.371E-08 | 1.400E-07 | 2.800E-07 |
| 26669 | 1334 | D1334H | E | 4.1e-07 | 2.8e-07 | 2.2e-07 | 3.033E-07 | 9.713E-08 | 2.200E-07 | 4.100E-07 |
| 26670 | 1334 | D1334I | E | 4.6e-07 | 1.7e-07 | 1.3e-07 | 2.533E-07 | 1.801E-07 | 1.300E-07 | 4.600E-07 |
| 26671 | 1334 | D1334L | E | 1.4e-07 | 2.6e-07 | 3.9e-07 | 2.633E-07 | 1.250E-07 | 1.400E-07 | 3.900E-07 |
| 26672 | 1334 | D1334K | E | 2e-07 | 4e-07 | 3.5e-07 | 3.167E-07 | 1.041E-07 | 2.000E-07 | 4.000E-07 |
| 26673 | 1334 | D1334M | E | 3.3e-07 | 5.1e-07 | 2.9e-07 | 3.767E-07 | 1.172E-07 | 2.900E-07 | 5.100E-07 |
| 26674 | 1334 | D1334F | E | 3.4e-07 | 3e-07 | 1.6e-07 | 2.667E-07 | 9.452E-08 | 1.600E-07 | 3.400E-07 |
| 26675 | 1334 | D1334P | E | 2.7e-07 | 2e-07 | 2.7e-07 | 2.467E-07 | 4.041E-08 | 2.000E-07 | 2.700E-07 |
| 26676 | 1334 | D1334S | E | 1.1e-07 | 1.9e-07 | 9.9e-08 | 1.330E-07 | 4.967E-08 | 9.900E-08 | 1.900E-07 |
| 26677 | 1334 | D1334T | E | 3.8e-07 | 3.7e-07 | 2.6e-07 | 3.367E-07 | 6.658E-08 | 2.600E-07 | 3.800E-07 |
| 26678 | 1334 | D1334W | E | 6.6e-07 | 3.3e-07 | 2.1e-07 | 4.000E-07 | 2.330E-07 | 2.100E-07 | 6.600E-07 |
| 26679 | 1334 | D1334Y | E | 1.6e-07 | 2.7e-07 | 3.1e-07 | 2.467E-07 | 7.767E-08 | 1.600E-07 | 3.100E-07 |
| 26680 | 1334 | D1334V | E | 4.2e-07 | 1.8e-07 | 2.2e-07 | 2.733E-07 | 1.286E-07 | 1.800E-07 | 4.200E-07 |
| 26681 | 1335 | S1335A | E | 2.6e-07 | 3.1e-07 | 1.8e-07 | 2.500E-07 | 6.557E-08 | 1.800E-07 | 3.100E-07 |
| 26682 | 1335 | S1335R | E | 3.2e-07 | 1.7e-07 | 1.1e-07 | 2.000E-07 | 1.082E-07 | 1.100E-07 | 3.200E-07 |
| 26683 | 1335 | S1335N | E | 1.4e-07 | 1.6e-07 | 2.3e-07 | 1.767E-07 | 4.726E-08 | 1.400E-07 | 2.300E-07 |
| 26684 | 1335 | S1335D | E | 2.4e-07 | 4.9e-07 | 2.7e-07 | 3.333E-07 | 1.365E-07 | 2.400E-07 | 4.900E-07 |
| 26685 | 1335 | S1335C | E | 2e-07 | 1.7e-07 | 5.3e-07 | 3.000E-07 | 1.997E-07 | 1.700E-07 | 5.300E-07 |
| 26686 | 1335 | S1335E | E | 1.1e-07 | 2e-07 | 2.1e-07 | 1.733E-07 | 5.508E-08 | 1.100E-07 | 2.100E-07 |
| 26687 | 1335 | S1335Q | E | 2.3e-07 | 2.1e-07 | 6.1e-07 | 3.500E-07 | 2.254E-07 | 2.100E-07 | 6.100E-07 |
| 26688 | 1335 | S1335G | E | 1.1e-07 | 2.3e-07 | 1.3e-07 | 1.567E-07 | 6.429E-08 | 1.100E-07 | 2.300E-07 |
| 26689 | 1335 | S1335H | E | 1.1e-07 | 4.9e-07 | 2.4e-07 | 2.800E-07 | 1.931E-07 | 1.100E-07 | 4.900E-07 |

| | | | | | | | | | | |
|---|---|---|---|---|---|---|---|---|---|---|
| 26690 | 1335 | S1335I | E | 2.3e-07 | 1.4e-07 | 5e-07 | 2.900E-07 | 1.873E-07 | 1.400E-07 | 5.000E-07 |
| 26691 | 1335 | S1335L | E | 4.7e-07 | 1.8e-07 | 1.3e-07 | 2.600E-07 | 1.836E-07 | 1.300E-07 | 4.700E-07 |
| 26692 | 1335 | S1335K | E | 3.4e-07 | 3.4e-07 | 7.5e-07 | 4.767E-07 | 2.367E-07 | 3.400E-07 | 7.500E-07 |
| 26693 | 1335 | S1335M | E | 2.1e-07 | 3.6e-07 | 4e-07 | 3.233E-07 | 1.002E-07 | 2.100E-07 | 4.000E-07 |
| 26694 | 1335 | S1335F | E | 2.9e-07 | 2.3e-07 | 3.1e-07 | 2.767E-07 | 4.163E-08 | 2.300E-07 | 3.100E-07 |
| 26695 | 1335 | S1335P | E | 3.2e-07 | 1.9e-07 | 2.8e-07 | 2.633E-07 | 6.658E-08 | 1.900E-07 | 3.200E-07 |
| 26696 | 1335 | S1335S | E | 2.1e-07 | 3.1e-07 | 2.5e-07 | 2.567E-07 | 5.033E-08 | 2.100E-07 | 3.100E-07 |
| 26697 | 1335 | S1335T | E | 1.6e-07 | 1.1e-07 | 1.3e-07 | 1.333E-07 | 2.517E-08 | 1.100E-07 | 1.600E-07 |
| 26698 | 1335 | S1335W | E | 2.5e-07 | 3e-07 | 1.2e-07 | 2.233E-07 | 9.292E-08 | 1.200E-07 | 3.000E-07 |
| 26699 | 1335 | S1335Y | E | 2.8e-07 | 9.4e-07 | 3e-07 | 5.067E-07 | 3.754E-07 | 2.800E-07 | 9.400E-07 |
| 26700 | 1335 | S1335V | E | 1.9e-07 | 2.7e-07 | 8.2e-08 | 1.807E-07 | 9.435E-08 | 8.200E-08 | 2.700E-07 |
| 26701 | 1336 | V1336A | E | 9.7e-08 | 2.7e-07 | 1.7e-07 | 1.790E-07 | 8.685E-08 | 9.700E-08 | 2.700E-07 |
| 26702 | 1336 | V1336R | E | 2e-07 | 2.8e-07 | 5.6e-07 | 3.467E-07 | 1.890E-07 | 2.000E-07 | 5.600E-07 |
| 26703 | 1336 | V1336N | E | 9.9e-08 | 3.1e-07 | 1.5e-07 | 1.863E-07 | 1.101E-07 | 9.900E-08 | 3.100E-07 |
| 26704 | 1336 | V1336D | E | 3.1e-07 | 2.5e-07 | 3.2e-07 | 2.933E-07 | 3.786E-08 | 2.500E-07 | 3.200E-07 |
| 26705 | 1336 | V1336C | E | 2.8e-07 | 1.8e-07 | 1.8e-07 | 2.133E-07 | 5.774E-08 | 1.800E-07 | 2.800E-07 |
| 26706 | 1336 | V1336E | E | 2.9e-07 | 3.4e-07 | 2.2e-07 | 2.833E-07 | 6.028E-08 | 2.200E-07 | 3.400E-07 |
| 26707 | 1336 | V1336Q | E | 1.7e-07 | 1.3e-07 | 3.1e-07 | 2.033E-07 | 9.452E-08 | 1.300E-07 | 3.100E-07 |
| 26708 | 1336 | V1336G | E | 1.9e-07 | 2.4e-07 | 2e-07 | 2.100E-07 | 2.646E-08 | 1.900E-07 | 2.400E-07 |
| 26709 | 1336 | V1336H | E | 2.6e-07 | 1.4e-07 | 2.4e-07 | 2.133E-07 | 6.429E-08 | 1.400E-07 | 2.600E-07 |
| 26710 | 1336 | V1336I | E | 2.2e-07 | 3.3e-07 | 1.8e-07 | 2.433E-07 | 7.767E-08 | 1.800E-07 | 3.300E-07 |
| 26711 | 1336 | V1336L | E | 2e-07 | 3.1e-07 | 2e-07 | 2.367E-07 | 6.351E-08 | 2.000E-07 | 3.100E-07 |
| 26712 | 1336 | V1336K | E | 3e-07 | 2e-07 | 4.3e-07 | 3.100E-07 | 1.153E-07 | 2.000E-07 | 4.300E-07 |
| 26713 | 1336 | V1336M | E | 3.2e-07 | 1.8e-07 | 3.1e-07 | 2.700E-07 | 7.810E-08 | 1.800E-07 | 3.200E-07 |
| 26714 | 1336 | V1336F | E | 1.7e-07 | 2e-07 | 1.7e-07 | 1.800E-07 | 1.732E-08 | 1.700E-07 | 2.000E-07 |
| 26715 | 1336 | V1336P | E | 2e-07 | 1.7e-07 | 4.8e-07 | 2.833E-07 | 1.710E-07 | 1.700E-07 | 4.800E-07 |
| 26716 | 1336 | V1336S | E | 1.6e-07 | 2e-07 | 3.5e-07 | 2.367E-07 | 1.002E-07 | 1.600E-07 | 3.500E-07 |
| 26717 | 1336 | V1336T | E | 2.3e-07 | 1.9e-07 | 2.3e-07 | 2.167E-07 | 2.309E-08 | 1.900E-07 | 2.300E-07 |
| 26718 | 1336 | V1336W | E | 4e-07 | 1.4e-07 | 2.9e-07 | 2.767E-07 | 1.305E-07 | 1.400E-07 | 4.000E-07 |
| 26719 | 1336 | V1336Y | E | 1.7e-07 | 1.3e-07 | 1.8e-07 | 1.600E-07 | 2.646E-08 | 1.300E-07 | 1.800E-07 |
| 26720 | 1336 | V1336V | E | 2.9e-07 | 2e-07 | 4.5e-07 | 3.133E-07 | 1.266E-07 | 2.000E-07 | 4.500E-07 |
| 26721 | 1337 | K1337A | E | 2.2e-07 | 3.5e-07 | 1.5e-07 | 2.400E-07 | 1.015E-07 | 1.500E-07 | 3.500E-07 |
| 26722 | 1337 | K1337R | E | 1.9e-07 | 2.8e-07 | 2e-07 | 2.233E-07 | 4.933E-08 | 1.900E-07 | 2.800E-07 |
| 26723 | 1337 | K1337N | E | 1.8e-07 | 1.5e-07 | 3.5e-07 | 2.267E-07 | 1.079E-07 | 1.500E-07 | 3.500E-07 |
| 26724 | 1337 | K1337D | E | 1.2e-07 | 1.8e-07 | 3.3e-07 | 2.100E-07 | 1.082E-07 | 1.200E-07 | 3.300E-07 |
| 26725 | 1337 | K1337C | E | 3.6e-07 | 2.4e-07 | 4.2e-07 | 3.400E-07 | 9.165E-08 | 2.400E-07 | 4.200E-07 |
| 26726 | 1337 | K1337E | E | 1.8e-07 | 1.9e-07 | 2.3e-07 | 2.000E-07 | 2.646E-08 | 1.800E-07 | 2.300E-07 |
| 26727 | 1337 | K1337Q | E | 2e-07 | 3e-07 | 3.9e-07 | 2.967E-07 | 9.504E-08 | 2.000E-07 | 3.900E-07 |
| 26728 | 1337 | K1337G | E | 1.8e-07 | 2.4e-07 | 4.3e-07 | 2.833E-07 | 1.305E-07 | 1.800E-07 | 4.300E-07 |
| 26729 | 1337 | K1337H | E | 1.7e-07 | 3.6e-07 | 1.6e-07 | 2.300E-07 | 1.127E-07 | 1.600E-07 | 3.600E-07 |
| 26730 | 1337 | K1337I | E | 1.5e-07 | 2.2e-07 | 4.2e-07 | 2.633E-07 | 1.401E-07 | 1.500E-07 | 4.200E-07 |
| 26731 | 1337 | K1337L | E | 3.5e-07 | 9.9e-08 | 3.4e-07 | 2.630E-07 | 1.421E-07 | 9.900E-08 | 3.500E-07 |
| 26732 | 1337 | K1337K | E | 2.2e-07 | 1.8e-07 | 4.8e-07 | 2.933E-07 | 1.629E-07 | 1.800E-07 | 4.800E-07 |
| 26733 | 1337 | K1337M | E | 3e-07 | 1.7e-07 | 2.7e-07 | 2.467E-07 | 6.807E-08 | 1.700E-07 | 3.000E-07 |
| 26734 | 1337 | K1337F | E | 1.5e-07 | 3.2e-07 | 3.5e-07 | 2.733E-07 | 1.079E-07 | 1.500E-07 | 3.500E-07 |
| 26735 | 1337 | K1337P | E | 5.2e-07 | 2.8e-07 | 2e-07 | 3.333E-07 | 1.665E-07 | 2.000E-07 | 5.200E-07 |
| 26736 | 1337 | K1337S | E | 2.2e-07 | 1.7e-07 | 3e-07 | 2.300E-07 | 6.557E-08 | 1.700E-07 | 3.000E-07 |
| 26737 | 1337 | K1337T | E | 4.7e-07 | 1.4e-07 | 2.3e-07 | 2.800E-07 | 1.706E-07 | 1.400E-07 | 4.700E-07 |
| 26738 | 1337 | K1337W | E | 1.7e-07 | 3.9e-07 | 4.2e-07 | 3.267E-07 | 1.365E-07 | 1.700E-07 | 4.200E-07 |
| 26739 | 1337 | K1337Y | E | 2.3e-07 | 9.3e-08 | 2.4e-07 | 1.877E-07 | 8.214E-08 | 9.300E-08 | 2.400E-07 |
| 26740 | 1337 | K1337V | E | 1.2e-07 | 2.7e-07 | 1.9e-07 | 1.933E-07 | 7.506E-08 | 1.200E-07 | 2.700E-07 |
| 26741 | 1338 | G1338A | E | 1.4e-07 | 2.2e-07 | 5.3e-07 | 2.967E-07 | 2.060E-07 | 1.400E-07 | 5.300E-07 |
| 26742 | 1338 | G1338R | E | 1.5e-07 | 2.3e-07 | 2.9e-07 | 2.233E-07 | 7.024E-08 | 1.500E-07 | 2.900E-07 |
| 26743 | 1338 | G1338N | E | 1.8e-07 | 1.9e-07 | 1.6e-07 | 1.767E-07 | 1.528E-08 | 1.600E-07 | 1.900E-07 |
| 26744 | 1338 | G1338D | E | 1.4e-07 | 1.7e-07 | 2.2e-07 | 1.767E-07 | 4.041E-08 | 1.400E-07 | 2.200E-07 |
| 26745 | 1338 | G1338C | E | 1.8e-07 | 3.2e-07 | 1.3e-07 | 2.100E-07 | 9.849E-08 | 1.300E-07 | 3.200E-07 |
| 26746 | 1338 | G1338E | E | 1.9e-07 | 6e-07 | 1.3e-07 | 3.067E-07 | 2.558E-07 | 1.300E-07 | 6.000E-07 |
| 26747 | 1338 | G1338Q | E | 3.9e-07 | 2.2e-07 | 2.3e-07 | 2.800E-07 | 9.539E-08 | 2.200E-07 | 3.900E-07 |
| 26748 | 1338 | G1338G | E | 2.8e-07 | 1.6e-07 | 1.5e-07 | 1.967E-07 | 7.234E-08 | 1.500E-07 | 2.800E-07 |
| 26749 | 1338 | G1338H | E | 2.9e-07 | 2.8e-07 | 2.2e-07 | 2.633E-07 | 3.786E-08 | 2.200E-07 | 2.900E-07 |
| 26750 | 1338 | G1338I | E | 1.6e-07 | 1.7e-07 | 3e-07 | 2.100E-07 | 7.810E-08 | 1.600E-07 | 3.000E-07 |
| 26751 | 1338 | G1338L | E | 2.9e-07 | 2.8e-07 | 3e-07 | 2.900E-07 | 1.000E-08 | 2.800E-07 | 3.000E-07 |
| 26752 | 1338 | G1338K | E | 1.9e-07 | 2.2e-07 | 1.8e-07 | 1.967E-07 | 2.082E-08 | 1.800E-07 | 2.200E-07 |
| 26753 | 1338 | G1338M | E | 9.2e-08 | 1.2e-07 | 4.5e-07 | 2.207E-07 | 1.991E-07 | 9.200E-08 | 4.500E-07 |
| 26754 | 1338 | G1338F | E | 4.7e-07 | 2.7e-07 | 2.2e-07 | 3.200E-07 | 1.323E-07 | 2.200E-07 | 4.700E-07 |
| 26755 | 1338 | G1338P | E | 1.2e-07 | 1.4e-07 | 2.6e-07 | 1.733E-07 | 7.572E-08 | 1.200E-07 | 2.600E-07 |
| 26756 | 1338 | G1338S | E | 2.4e-07 | 1.9e-07 | 2.4e-07 | 2.233E-07 | 2.887E-08 | 1.900E-07 | 2.400E-07 |
| 26757 | 1338 | G1338T | E | 2.6e-07 | 2e-07 | 1.7e-07 | 2.100E-07 | 4.583E-08 | 1.700E-07 | 2.600E-07 |
| 26758 | 1338 | G1338W | E | 2.7e-07 | 2e-07 | 4.7e-07 | 3.133E-07 | 1.401E-07 | 2.000E-07 | 4.700E-07 |
| 26759 | 1338 | G1338Y | E | 1.9e-07 | 2.7e-07 | 2.4e-07 | 2.333E-07 | 4.041E-08 | 1.900E-07 | 2.700E-07 |
| 26760 | 1338 | G1338V | E | 3.3e-07 | 5.4e-07 | 2.1e-07 | 3.600E-07 | 1.670E-07 | 2.100E-07 | 5.400E-07 |
| 26761 | 1339 | R1339A | E | 3.1e-07 | 2.4e-07 | 2.1e-07 | 2.533E-07 | 5.132E-08 | 2.100E-07 | 3.100E-07 |
| 26762 | 1339 | R1339R | E | 2.6e-07 | 2.3e-07 | 2.2e-07 | 2.367E-07 | 2.082E-08 | 2.200E-07 | 2.600E-07 |
| 26763 | 1339 | R1339N | E | 2.7e-07 | 1.1e-07 | 3e-07 | 2.267E-07 | 1.021E-07 | 1.100E-07 | 3.000E-07 |
| 26764 | 1339 | R1339D | E | 9.7e-08 | 3.8e-07 | 1.6e-07 | 2.123E-07 | 1.486E-07 | 9.700E-08 | 3.800E-07 |
| 26765 | 1339 | R1339C | E | 1.8e-07 | 3.4e-07 | 3.9e-07 | 3.033E-07 | 1.097E-07 | 1.800E-07 | 3.900E-07 |
| 26766 | 1339 | R1339E | E | 8e-08 | 1.4e-07 | 2e-07 | 1.400E-07 | 6.000E-08 | 8.000E-08 | 2.000E-07 |
| 26767 | 1339 | R1339Q | E | 2.3e-07 | 2.8e-07 | 3e-07 | 2.700E-07 | 3.606E-08 | 2.300E-07 | 3.000E-07 |
| 26768 | 1339 | R1339G | E | 1.4e-07 | 4.4e-07 | 2.7e-07 | 2.833E-07 | 1.504E-07 | 1.400E-07 | 4.400E-07 |
| 26769 | 1339 | R1339H | E | 1.7e-07 | 1.9e-07 | 1.7e-07 | 1.767E-07 | 1.155E-08 | 1.700E-07 | 1.900E-07 |
| 26770 | 1339 | R1339I | E | 2e-07 | 2.8e-07 | 3.3e-07 | 2.700E-07 | 6.557E-08 | 2.000E-07 | 3.300E-07 |
| 26771 | 1339 | R1339L | E | 3.1e-07 | 1.1e-07 | 2.9e-07 | 2.367E-07 | 1.102E-07 | 1.100E-07 | 3.100E-07 |
| 26772 | 1339 | R1339K | E | 1.7e-07 | 2.4e-07 | 2e-07 | 2.033E-07 | 3.512E-08 | 1.700E-07 | 2.400E-07 |
| 26773 | 1339 | R1339M | E | 5.8e-07 | 3.1e-07 | 1.3e-07 | 3.400E-07 | 2.265E-07 | 1.300E-07 | 5.800E-07 |
| 26774 | 1339 | R1339F | E | 1.1e-07 | 5e-07 | 2.4e-07 | 2.833E-07 | 1.986E-07 | 1.100E-07 | 5.000E-07 |

| | | | | | | | | | | |
|---|---|---|---|---|---|---|---|---|---|---|
| 26775 | 1339 | R1339P | E | 1.5e-07 | 2.1e-07 | 2.8e-07 | 2.133E-07 | 6.506E-08 | 1.500E-07 | 2.800E-07 |
| 26776 | 1339 | R1339S | E | 2.4e-07 | 3.4e-07 | 2.5e-07 | 2.767E-07 | 5.508E-08 | 2.400E-07 | 3.400E-07 |
| 26777 | 1339 | R1339T | E | 2.6e-07 | 1.8e-07 | 2e-07 | 2.133E-07 | 4.163E-08 | 1.800E-07 | 2.600E-07 |
| 26778 | 1339 | R1339W | E | 2.8e-07 | 2.5e-07 | 1.5e-07 | 2.267E-07 | 6.807E-08 | 1.500E-07 | 2.800E-07 |
| 26779 | 1339 | R1339Y | E | 2.8e-07 | 2.2e-07 | 4.6e-07 | 3.200E-07 | 1.249E-07 | 2.200E-07 | 4.600E-07 |
| 26780 | 1339 | R1339V | E | 2.1e-07 | 2.3e-07 | 2e-07 | 2.133E-07 | 1.528E-08 | 2.000E-07 | 2.300E-07 |
| 26781 | 1340 | F1340A | E | 3e-07 | 3.3e-07 | 4e-07 | 3.433E-07 | 5.132E-08 | 3.000E-07 | 4.000E-07 |
| 26782 | 1340 | F1340R | E | 2.7e-07 | 4.8e-07 | 7.9e-08 | 2.763E-07 | 2.006E-07 | 7.900E-08 | 4.800E-07 |
| 26783 | 1340 | F1340N | E | 3e-07 | 2.2e-07 | 3.3e-07 | 2.833E-07 | 5.686E-08 | 2.200E-07 | 3.300E-07 |
| 26784 | 1340 | F1340D | E | 2.8e-07 | 2.9e-07 | 6.4e-07 | 4.033E-07 | 2.050E-07 | 2.800E-07 | 6.400E-07 |
| 26785 | 1340 | F1340C | E | 1.1e-07 | 3.2e-07 | 9.8e-08 | 1.760E-07 | 1.249E-07 | 9.800E-08 | 3.200E-07 |
| 26786 | 1340 | F1340E | E | 2.3e-07 | 3.3e-07 | 3.1e-07 | 2.900E-07 | 5.292E-08 | 2.300E-07 | 3.300E-07 |
| 26787 | 1340 | F1340Q | E | 3e-07 | 2e-07 | 1.5e-07 | 2.167E-07 | 7.638E-08 | 1.500E-07 | 3.000E-07 |
| 26788 | 1340 | F1340G | E | 7.2e-08 | 3.6e-07 | 2.7e-07 | 2.340E-07 | 1.473E-07 | 7.200E-08 | 3.600E-07 |
| 26789 | 1340 | F1340H | E | 2.8e-07 | 3.4e-07 | 2.1e-07 | 2.767E-07 | 6.506E-08 | 2.100E-07 | 3.400E-07 |
| 26790 | 1340 | F1340I | E | 1.1e-07 | 3.6e-07 | 6.7e-07 | 3.800E-07 | 2.805E-07 | 1.100E-07 | 6.700E-07 |
| 26791 | 1340 | F1340L | E | 1.9e-07 | 2.1e-07 | 2.6e-07 | 2.200E-07 | 3.606E-08 | 1.900E-07 | 2.600E-07 |
| 26792 | 1340 | F1340K | E | 2.9e-07 | 3.3e-07 | 2.1e-07 | 2.767E-07 | 6.110E-08 | 2.100E-07 | 3.300E-07 |
| 26793 | 1340 | F1340M | E | 2.7e-07 | 2e-07 | 1.6e-07 | 2.100E-07 | 5.568E-08 | 1.600E-07 | 2.700E-07 |
| 26794 | 1340 | F1340F | E | 2.8e-07 | 5.6e-07 | 2.4e-07 | 3.600E-07 | 1.744E-07 | 2.400E-07 | 5.600E-07 |
| 26795 | 1340 | F1340P | E | 2.4e-07 | 1.8e-07 | 1.7e-07 | 1.967E-07 | 3.786E-08 | 1.700E-07 | 2.400E-07 |
| 26796 | 1340 | F1340S | E | 2.2e-07 | 1.8e-07 | 2.2e-07 | 2.067E-07 | 2.309E-08 | 1.800E-07 | 2.200E-07 |
| 26797 | 1340 | F1340T | E | 4.7e-07 | 2.1e-07 | 3.9e-07 | 3.567E-07 | 1.332E-07 | 2.100E-07 | 4.700E-07 |
| 26798 | 1340 | F1340W | E | 1.7e-07 | 4.3e-07 | 2.7e-07 | 2.900E-07 | 1.311E-07 | 1.700E-07 | 4.300E-07 |
| 26799 | 1340 | F1340Y | E | 1.8e-07 | 4e-07 | 2.9e-07 | 2.900E-07 | 1.100E-07 | 1.800E-07 | 4.000E-07 |
| 26800 | 1340 | F1340V | E | 1.6e-07 | 1.9e-07 | 4e-07 | 2.500E-07 | 1.308E-07 | 1.600E-07 | 4.000E-07 |
| 26801 | 1341 | T1341A | E | 3.2e-07 | 2.3e-07 | 3.8e-07 | 3.100E-07 | 7.550E-08 | 2.300E-07 | 3.800E-07 |
| 26802 | 1341 | T1341R | E | 3.4e-07 | 2.5e-07 | 1.8e-07 | 2.567E-07 | 8.021E-08 | 1.800E-07 | 3.400E-07 |
| 26803 | 1341 | T1341N | E | 3.3e-07 | 5.7e-08 | 1.5e-07 | 1.790E-07 | 1.388E-07 | 5.700E-08 | 3.300E-07 |
| 26804 | 1341 | T1341D | E | 2e-07 | 3.4e-07 | 1.1e-07 | 2.167E-07 | 1.159E-07 | 1.100E-07 | 3.400E-07 |
| 26805 | 1341 | T1341C | E | 1.6e-07 | 3.2e-07 | 3e-07 | 2.600E-07 | 8.718E-08 | 1.600E-07 | 3.200E-07 |
| 26806 | 1341 | T1341E | E | 4.5e-07 | 1.6e-07 | 2.1e-07 | 2.733E-07 | 1.550E-07 | 1.600E-07 | 4.500E-07 |
| 26807 | 1341 | T1341Q | E | 2e-07 | 5.6e-07 | 1.6e-07 | 3.067E-07 | 2.203E-07 | 1.600E-07 | 5.600E-07 |
| 26808 | 1341 | T1341G | E | 2.4e-07 | 9.3e-08 | 3.4e-07 | 2.243E-07 | 1.242E-07 | 9.300E-08 | 3.400E-07 |
| 26809 | 1341 | T1341H | E | 5.1e-07 | 3.5e-07 | 4e-07 | 4.200E-07 | 8.185E-08 | 3.500E-07 | 5.100E-07 |
| 26810 | 1341 | T1341I | E | 2.6e-07 | 3e-07 | 1.9e-07 | 2.500E-07 | 5.568E-08 | 1.900E-07 | 3.000E-07 |
| 26811 | 1341 | T1341L | E | 3.5e-07 | 4.1e-07 | 3e-07 | 3.533E-07 | 5.508E-08 | 3.000E-07 | 4.100E-07 |
| 26812 | 1341 | T1341K | E | 5.6e-07 | 1.8e-07 | 3.5e-07 | 3.633E-07 | 1.904E-07 | 1.800E-07 | 5.600E-07 |
| 26813 | 1341 | T1341M | E | 9.4e-08 | 5.7e-07 | 1.7e-07 | 2.780E-07 | 2.557E-07 | 9.400E-08 | 5.700E-07 |
| 26814 | 1341 | T1341F | E | 2.4e-07 | 2.3e-07 | 2.2e-07 | 2.300E-07 | 1.000E-08 | 2.200E-07 | 2.400E-07 |
| 26815 | 1341 | T1341P | E | 2.8e-07 | 2.3e-07 | 8.9e-08 | 1.997E-07 | 9.905E-08 | 8.900E-08 | 2.800E-07 |
| 26816 | 1341 | T1341S | E | 1.3e-07 | 4.5e-07 | 1.2e-07 | 2.333E-07 | 1.877E-07 | 1.200E-07 | 4.500E-07 |
| 26817 | 1341 | T1341T | E | 1.9e-07 | 2e-07 | 1.5e-07 | 1.800E-07 | 2.646E-08 | 1.500E-07 | 2.000E-07 |
| 26818 | 1341 | T1341W | E | 1.6e-07 | 1.9e-07 | 1.5e-07 | 1.667E-07 | 2.082E-08 | 1.500E-07 | 1.900E-07 |
| 26819 | 1341 | T1341Y | E | 2.4e-07 | 1.9e-07 | 3.1e-07 | 2.467E-07 | 6.028E-08 | 1.900E-07 | 3.100E-07 |
| 26820 | 1341 | T1341V | E | 3.5e-07 | 5e-07 | 3.6e-07 | 4.033E-07 | 8.386E-08 | 3.500E-07 | 5.000E-07 |
| 26821 | 1342 | I1342A | E | 1.7e-07 | 2.4e-07 | 2e-07 | 2.033E-07 | 3.512E-08 | 1.700E-07 | 2.400E-07 |
| 26822 | 1342 | I1342R | E | 2.7e-07 | 1.5e-07 | 2.4e-07 | 2.200E-07 | 6.245E-08 | 1.500E-07 | 2.700E-07 |
| 26823 | 1342 | I1342N | E | 2e-07 | 1.4e-07 | 2.1e-07 | 1.833E-07 | 3.786E-08 | 1.400E-07 | 2.100E-07 |
| 26824 | 1342 | I1342D | E | 1.5e-07 | 2.7e-07 | 3e-07 | 2.400E-07 | 7.937E-08 | 1.500E-07 | 3.000E-07 |
| 26825 | 1342 | I1342C | E | 2.5e-07 | 3.7e-07 | 3.9e-07 | 3.367E-07 | 7.572E-08 | 2.500E-07 | 3.900E-07 |
| 26826 | 1342 | I1342E | E | 1.5e-07 | 2.8e-07 | 5.8e-07 | 3.367E-07 | 2.205E-07 | 1.500E-07 | 5.800E-07 |
| 26827 | 1342 | I1342Q | E | 2.2e-07 | 3.9e-07 | 1.5e-07 | 2.533E-07 | 1.234E-07 | 1.500E-07 | 3.900E-07 |
| 26828 | 1342 | I1342G | E | 4.7e-07 | 3.9e-07 | 1.3e-07 | 3.300E-07 | 1.778E-07 | 1.300E-07 | 4.700E-07 |
| 26829 | 1342 | I1342H | E | 1.7e-07 | 8.2e-08 | 1.3e-07 | 1.273E-07 | 4.406E-08 | 8.200E-08 | 1.700E-07 |
| 26830 | 1342 | I1342I | E | 3.1e-07 | 1.2e-07 | 1.5e-07 | 1.933E-07 | 1.021E-07 | 1.200E-07 | 3.100E-07 |
| 26831 | 1342 | I1342L | E | 1.5e-07 | 3.1e-07 | 1.7e-07 | 2.100E-07 | 8.718E-08 | 1.500E-07 | 3.100E-07 |
| 26832 | 1342 | I1342K | E | 2.1e-07 | 1.7e-07 | 2.4e-07 | 2.067E-07 | 3.512E-08 | 1.700E-07 | 2.400E-07 |
| 26833 | 1342 | I1342M | E | 1.6e-07 | 6.2e-07 | 1.9e-07 | 3.233E-07 | 2.574E-07 | 1.600E-07 | 6.200E-07 |
| 26834 | 1342 | I1342F | E | 3.6e-07 | 5.9e-07 | 1.7e-07 | 3.733E-07 | 2.103E-07 | 1.700E-07 | 5.900E-07 |
| 26835 | 1342 | I1342P | E | 3.7e-07 | 3e-07 | 7.2e-08 | 2.473E-07 | 1.558E-07 | 7.200E-08 | 3.700E-07 |
| 26836 | 1342 | I1342S | E | 1.7e-07 | 1.7e-07 | 2.2e-07 | 1.867E-07 | 2.887E-08 | 1.700E-07 | 2.200E-07 |
| 26837 | 1342 | I1342T | E | 1.6e-07 | 2.2e-07 | 1.2e-07 | 1.667E-07 | 5.033E-08 | 1.200E-07 | 2.200E-07 |
| 26838 | 1342 | I1342W | E | 1.2e-07 | 1e-07 | 3.5e-07 | 1.900E-07 | 1.389E-07 | 1.000E-07 | 3.500E-07 |
| 26839 | 1342 | I1342Y | E | 1.6e-07 | 2.9e-07 | 1.9e-07 | 2.133E-07 | 6.807E-08 | 1.600E-07 | 2.900E-07 |
| 26840 | 1342 | I1342V | E | 2.9e-07 | 1.7e-07 | 3.8e-07 | 2.800E-07 | 1.054E-07 | 1.700E-07 | 3.800E-07 |
| 26841 | 1343 | S1343A | E | 1.8e-07 | 3e-07 | 4.3e-07 | 3.033E-07 | 1.250E-07 | 1.800E-07 | 4.300E-07 |
| 26842 | 1343 | S1343R | E | 2.5e-07 | 2.9e-07 | 4.6e-07 | 3.333E-07 | 1.115E-07 | 2.500E-07 | 4.600E-07 |
| 26843 | 1343 | S1343N | E | 3.4e-07 | 3e-07 | 1.5e-07 | 2.633E-07 | 1.002E-07 | 1.500E-07 | 3.400E-07 |
| 26844 | 1343 | S1343D | E | 3.6e-07 | 1.8e-07 | 8.7e-07 | 4.700E-07 | 3.579E-07 | 1.800E-07 | 8.700E-07 |
| 26845 | 1343 | S1343C | E | 2.6e-07 | 1.2e-07 | 1.9e-07 | 1.900E-07 | 7.000E-08 | 1.200E-07 | 2.600E-07 |
| 26846 | 1343 | S1343E | E | 2.8e-07 | 2.2e-07 | 7.4e-07 | 4.133E-07 | 2.845E-07 | 2.200E-07 | 7.400E-07 |
| 26847 | 1343 | S1343Q | E | 3.5e-07 | 5e-07 | 3.7e-07 | 4.067E-07 | 8.145E-08 | 3.500E-07 | 5.000E-07 |
| 26848 | 1343 | S1343G | E | 2.8e-07 | 3.1e-07 | 1.2e-07 | 2.367E-07 | 1.021E-07 | 1.200E-07 | 3.100E-07 |
| 26849 | 1343 | S1343H | E | 2e-07 | 1.8e-07 | 2.9e-07 | 2.233E-07 | 5.859E-08 | 1.800E-07 | 2.900E-07 |
| 26850 | 1343 | S1343I | E | 1.9e-07 | 3.4e-07 | 5.3e-07 | 3.533E-07 | 1.704E-07 | 1.900E-07 | 5.300E-07 |
| 26851 | 1343 | S1343L | E | 7.4e-07 | 7.9e-07 | 3.8e-07 | 6.367E-07 | 2.237E-07 | 3.800E-07 | 7.900E-07 |
| 26852 | 1343 | S1343K | E | 3e-07 | 2.9e-07 | 2.5e-07 | 2.800E-07 | 2.646E-08 | 2.500E-07 | 3.000E-07 |
| 26853 | 1343 | S1343M | E | 2.1e-07 | 2.1e-07 | 2.5e-07 | 2.233E-07 | 2.309E-08 | 2.100E-07 | 2.500E-07 |
| 26854 | 1343 | S1343F | E | 2.3e-07 | 1.6e-07 | 2.5e-07 | 2.133E-07 | 4.726E-08 | 1.600E-07 | 2.500E-07 |
| 26855 | 1343 | S1343P | E | 1.7e-07 | 6e-07 | 1.8e-07 | 3.167E-07 | 2.454E-07 | 1.700E-07 | 6.000E-07 |
| 26856 | 1343 | S1343S | E | 2.9e-07 | 2e-07 | 1.8e-07 | 2.233E-07 | 5.859E-08 | 1.800E-07 | 2.900E-07 |
| 26857 | 1343 | S1343T | E | 4.3e-07 | 2.8e-07 | 4.2e-07 | 3.767E-07 | 8.386E-08 | 2.800E-07 | 4.300E-07 |
| 26858 | 1343 | S1343W | E | 1.5e-07 | 3.2e-07 | 3.4e-07 | 2.700E-07 | 1.044E-07 | 1.500E-07 | 3.400E-07 |
| 26859 | 1343 | S1343Y | E | 2.5e-07 | 2.2e-07 | 1.7e-07 | 2.133E-07 | 4.041E-08 | 1.700E-07 | 2.500E-07 |

| | | | | | | | | | | |
|---|---|---|---|---|---|---|---|---|---|---|
| 26860 | 1343 | S1343V | E | 3.6e-07 | 2.1e-07 | 2.1e-07 | 2.600E-07 | 8.660E-08 | 2.100E-07 | 3.600E-07 |
| 26861 | 1344 | A1344A | E | 2.4e-07 | 1.6e-07 | 1.7e-07 | 1.900E-07 | 4.359E-08 | 1.600E-07 | 2.400E-07 |
| 26862 | 1344 | A1344R | E | 1.9e-07 | 3.8e-07 | 4.6e-07 | 3.433E-07 | 1.387E-07 | 1.900E-07 | 4.600E-07 |
| 26863 | 1344 | A1344N | E | 3.8e-07 | 2.2e-07 | 1.2e-07 | 2.400E-07 | 1.311E-07 | 1.200E-07 | 3.800E-07 |
| 26864 | 1344 | A1344D | E | 2.4e-07 | 2.9e-07 | 2.7e-07 | 2.667E-07 | 2.517E-08 | 2.400E-07 | 2.900E-07 |
| 26865 | 1344 | A1344C | E | 2.6e-07 | 9.8e-08 | 2.4e-07 | 1.993E-07 | 8.833E-08 | 9.800E-08 | 2.600E-07 |
| 26866 | 1344 | A1344E | E | 1.4e-07 | 2.1e-07 | 1.9e-07 | 1.800E-07 | 3.606E-08 | 1.400E-07 | 2.100E-07 |
| 26867 | 1344 | A1344Q | E | 2.6e-07 | 2.4e-07 | 1.6e-07 | 2.200E-07 | 5.292E-08 | 1.600E-07 | 2.600E-07 |
| 26868 | 1344 | A1344G | E | 2e-07 | 2.4e-07 | 3e-07 | 2.467E-07 | 5.033E-08 | 2.000E-07 | 3.000E-07 |
| 26869 | 1344 | A1344H | E | 2.2e-07 | 2.3e-07 | 3e-07 | 2.500E-07 | 4.359E-08 | 2.200E-07 | 3.000E-07 |
| 26870 | 1344 | A1344I | E | 2.5e-07 | 1.7e-07 | 4.9e-07 | 3.033E-07 | 1.665E-07 | 1.700E-07 | 4.900E-07 |
| 26871 | 1344 | A1344L | E | 2.6e-07 | 6.7e-07 | 3.3e-07 | 4.200E-07 | 2.193E-07 | 2.600E-07 | 6.700E-07 |
| 26872 | 1344 | A1344K | E | 2e-07 | 2.8e-07 | 1.8e-07 | 2.200E-07 | 5.292E-08 | 1.800E-07 | 2.800E-07 |
| 26873 | 1344 | A1344M | E | 1.8e-07 | 2.3e-07 | 2.2e-07 | 2.100E-07 | 2.646E-08 | 1.800E-07 | 2.300E-07 |
| 26874 | 1344 | A1344F | E | 1e-07 | 2.5e-07 | 1.4e-07 | 1.633E-07 | 7.767E-08 | 1.000E-07 | 2.500E-07 |
| 26875 | 1344 | A1344P | E | 2.2e-07 | 2.5e-07 | 3.3e-07 | 2.667E-07 | 5.686E-08 | 2.200E-07 | 3.300E-07 |
| 26876 | 1344 | A1344S | E | 2.4e-07 | 1.8e-07 | 2.5e-07 | 2.233E-07 | 3.786E-08 | 1.800E-07 | 2.500E-07 |
| 26877 | 1344 | A1344T | E | 7.2e-08 | 2.6e-07 | 1.1e-07 | 1.473E-07 | 9.940E-08 | 7.200E-08 | 2.600E-07 |
| 26878 | 1344 | A1344W | E | 1.8e-07 | 2.4e-07 | 2.2e-07 | 2.133E-07 | 3.055E-08 | 1.800E-07 | 2.400E-07 |
| 26879 | 1344 | A1344Y | E | 3.9e-07 | 1.6e-07 | 2.6e-07 | 2.700E-07 | 1.153E-07 | 1.600E-07 | 3.900E-07 |
| 26880 | 1344 | A1344V | E | 2e-07 | 1.5e-07 | 3.2e-07 | 2.233E-07 | 8.737E-08 | 1.500E-07 | 3.200E-07 |
| 26881 | 1345 | D1345A | E | 1e-07 | 4.1e-07 | 1.5e-07 | 2.200E-07 | 1.664E-07 | 1.000E-07 | 4.100E-07 |
| 26882 | 1345 | D1345R | E | 3e-07 | 2e-07 | 2.5e-07 | 2.500E-07 | 5.000E-08 | 2.000E-07 | 3.000E-07 |
| 26883 | 1345 | D1345N | E | 2.8e-07 | 2.5e-07 | 2e-07 | 2.433E-07 | 4.041E-08 | 2.000E-07 | 2.800E-07 |
| 26884 | 1345 | D1345D | E | 1.7e-07 | 2e-07 | 1.4e-07 | 1.700E-07 | 3.000E-08 | 1.400E-07 | 2.000E-07 |
| 26885 | 1345 | D1345C | E | 3.9e-07 | 1.7e-07 | 1.7e-07 | 2.433E-07 | 1.270E-07 | 1.700E-07 | 3.900E-07 |
| 26886 | 1345 | D1345E | E | 4e-07 | 8.1e-08 | 2.1e-07 | 2.303E-07 | 1.605E-07 | 8.100E-08 | 4.000E-07 |
| 26887 | 1345 | D1345Q | E | 2.8e-07 | 1.8e-07 | 2.7e-07 | 2.433E-07 | 5.508E-08 | 1.800E-07 | 2.800E-07 |
| 26888 | 1345 | D1345G | E | 3.4e-07 | 3.6e-07 | 3.3e-07 | 3.433E-07 | 1.528E-08 | 3.300E-07 | 3.600E-07 |
| 26889 | 1345 | D1345H | E | 2.3e-07 | 1.7e-07 | 1.9e-07 | 1.967E-07 | 3.055E-08 | 1.700E-07 | 2.300E-07 |
| 26890 | 1345 | D1345I | E | 2.2e-07 | 2.7e-07 | 2.4e-07 | 2.433E-07 | 2.517E-08 | 2.200E-07 | 2.700E-07 |
| 26891 | 1345 | D1345L | E | 3.9e-07 | 2.8e-07 | 3.2e-07 | 3.300E-07 | 5.568E-08 | 2.800E-07 | 3.900E-07 |
| 26892 | 1345 | D1345K | E | 2.4e-07 | 2.3e-07 | 2.2e-07 | 2.300E-07 | 1.000E-08 | 2.200E-07 | 2.400E-07 |
| 26893 | 1345 | D1345M | E | 2.4e-07 | 2.3e-07 | 1.9e-07 | 2.200E-07 | 2.646E-08 | 1.900E-07 | 2.400E-07 |
| 26894 | 1345 | D1345F | E | 1.4e-07 | 1.4e-07 | 4.1e-07 | 2.300E-07 | 1.559E-07 | 1.400E-07 | 4.100E-07 |
| 26895 | 1345 | D1345P | E | 5.1e-07 | 2.4e-07 | 1.9e-07 | 3.133E-07 | 1.721E-07 | 1.900E-07 | 5.100E-07 |
| 26896 | 1345 | D1345S | E | 3.2e-07 | 3.5e-07 | 2.2e-07 | 2.967E-07 | 6.807E-08 | 2.200E-07 | 3.500E-07 |
| 26897 | 1345 | D1345T | E | 3.9e-07 | 1.9e-07 | 1.7e-07 | 2.500E-07 | 1.217E-07 | 1.700E-07 | 3.900E-07 |
| 26898 | 1345 | D1345W | E | 3e-07 | 1.9e-07 | 1.9e-07 | 2.267E-07 | 6.351E-08 | 1.900E-07 | 3.000E-07 |
| 26899 | 1345 | D1345Y | E | 1.3e-07 | 2.3e-07 | 1.9e-07 | 1.833E-07 | 5.033E-08 | 1.300E-07 | 2.300E-07 |
| 26900 | 1345 | D1345V | E | 2e-07 | 3.2e-07 | 1.9e-07 | 2.367E-07 | 7.234E-08 | 1.900E-07 | 3.200E-07 |
| 26901 | 1346 | T1346A | E | 3.3e-07 | 2.9e-07 | 2.8e-07 | 3.000E-07 | 2.646E-08 | 2.800E-07 | 3.300E-07 |
| 26902 | 1346 | T1346R | E | 1.9e-07 | 2.5e-07 | 3.5e-07 | 2.633E-07 | 8.083E-08 | 1.900E-07 | 3.500E-07 |
| 26903 | 1346 | T1346N | E | 1.6e-07 | 2.6e-07 | 2.1e-07 | 2.100E-07 | 5.000E-08 | 1.600E-07 | 2.600E-07 |
| 26904 | 1346 | T1346D | E | 2.7e-07 | 4.5e-07 | 1.9e-07 | 3.033E-07 | 1.332E-07 | 1.900E-07 | 4.500E-07 |
| 26905 | 1346 | T1346C | E | 7.8e-08 | 2.3e-07 | 1.1e-07 | 1.393E-07 | 8.013E-08 | 7.800E-08 | 2.300E-07 |
| 26906 | 1346 | T1346E | E | 3.6e-07 | 2.5e-07 | 2.6e-07 | 2.900E-07 | 6.083E-08 | 2.500E-07 | 3.600E-07 |
| 26907 | 1346 | T1346Q | E | 3.9e-07 | 1.8e-07 | 1.5e-07 | 2.400E-07 | 1.308E-07 | 1.500E-07 | 3.900E-07 |
| 26908 | 1346 | T1346G | E | 2.5e-07 | 2.8e-07 | 2.8e-07 | 2.700E-07 | 1.732E-08 | 2.500E-07 | 2.800E-07 |
| 26909 | 1346 | T1346H | E | 3e-07 | 2.4e-07 | 2e-07 | 2.467E-07 | 5.033E-08 | 2.000E-07 | 3.000E-07 |
| 26910 | 1346 | T1346I | E | 5.2e-07 | 2e-07 | 1.4e-07 | 2.867E-07 | 2.043E-07 | 1.400E-07 | 5.200E-07 |
| 26911 | 1346 | T1346L | E | 1.8e-07 | 2.5e-07 | 3.7e-07 | 2.667E-07 | 9.609E-08 | 1.800E-07 | 3.700E-07 |
| 26912 | 1346 | T1346K | E | 2.6e-07 | 1.2e-07 | 3.3e-07 | 2.367E-07 | 1.069E-07 | 1.200E-07 | 3.300E-07 |
| 26913 | 1346 | T1346M | E | 5.3e-07 | 4e-07 | 2.1e-07 | 3.800E-07 | 1.609E-07 | 2.100E-07 | 5.300E-07 |
| 26914 | 1346 | T1346F | E | 2.3e-07 | 2.6e-07 | 1.4e-07 | 2.100E-07 | 6.245E-08 | 1.400E-07 | 2.600E-07 |
| 26915 | 1346 | T1346P | E | 2.1e-07 | 1.9e-07 | 1e-07 | 1.667E-07 | 5.859E-08 | 1.000E-07 | 2.100E-07 |
| 26916 | 1346 | T1346S | E | 3.2e-07 | 1.6e-07 | 4e-07 | 2.933E-07 | 1.222E-07 | 1.600E-07 | 4.000E-07 |
| 26917 | 1346 | T1346T | E | 2.1e-07 | 1.2e-07 | 2e-07 | 1.767E-07 | 4.933E-08 | 1.200E-07 | 2.100E-07 |
| 26918 | 1346 | T1346W | E | 2.9e-07 | 4.4e-07 | 4.6e-07 | 3.967E-07 | 9.292E-08 | 2.900E-07 | 4.600E-07 |
| 26919 | 1346 | T1346Y | E | 4.3e-07 | 2.7e-07 | 4.2e-07 | 3.733E-07 | 8.963E-08 | 2.700E-07 | 4.300E-07 |
| 26920 | 1346 | T1346V | E | 5.6e-07 | 2.2e-07 | 7.5e-07 | 5.100E-07 | 2.685E-07 | 2.200E-07 | 7.500E-07 |
| 26921 | 1347 | S1347A | E | 2.5e-07 | 2.1e-07 | 4.6e-07 | 3.067E-07 | 1.343E-07 | 2.100E-07 | 4.600E-07 |
| 26922 | 1347 | S1347R | E | 1.5e-07 | 4e-07 | 2e-07 | 2.500E-07 | 1.323E-07 | 1.500E-07 | 4.000E-07 |
| 26923 | 1347 | S1347N | E | 2.2e-07 | 1.9e-07 | 4.4e-07 | 2.833E-07 | 1.365E-07 | 1.900E-07 | 4.400E-07 |
| 26924 | 1347 | S1347D | E | 4.5e-07 | 2e-07 | 1.4e-07 | 2.633E-07 | 1.644E-07 | 1.400E-07 | 4.500E-07 |
| 26925 | 1347 | S1347C | E | 4.2e-07 | 3.1e-07 | 2.1e-07 | 3.133E-07 | 1.050E-07 | 2.100E-07 | 4.200E-07 |
| 26926 | 1347 | S1347E | E | 2.3e-07 | 1.5e-07 | 1.7e-07 | 1.833E-07 | 4.163E-08 | 1.500E-07 | 2.300E-07 |
| 26927 | 1347 | S1347Q | E | 3e-07 | 1.3e-07 | 1.4e-07 | 1.900E-07 | 9.539E-08 | 1.300E-07 | 3.000E-07 |
| 26928 | 1347 | S1347G | E | 2.1e-07 | 4.8e-07 | 2.3e-07 | 3.067E-07 | 1.504E-07 | 2.100E-07 | 4.800E-07 |
| 26929 | 1347 | S1347H | E | 2.8e-07 | 2e-07 | 3e-07 | 2.600E-07 | 5.292E-08 | 2.000E-07 | 3.000E-07 |
| 26930 | 1347 | S1347I | E | 4.1e-07 | 2.1e-07 | 1.4e-07 | 2.533E-07 | 1.401E-07 | 1.400E-07 | 4.100E-07 |
| 26931 | 1347 | S1347L | E | 2.2e-07 | 1.6e-07 | 3.1e-07 | 2.300E-07 | 7.550E-08 | 1.600E-07 | 3.100E-07 |
| 26932 | 1347 | S1347K | E | 3.5e-07 | 1.5e-07 | 4.4e-07 | 3.133E-07 | 1.484E-07 | 1.500E-07 | 4.400E-07 |
| 26933 | 1347 | S1347M | E | 2e-07 | 2.9e-07 | 3.4e-07 | 2.767E-07 | 7.095E-08 | 2.000E-07 | 3.400E-07 |
| 26934 | 1347 | S1347F | E | 2.6e-07 | 1.7e-07 | 3e-07 | 2.433E-07 | 6.658E-08 | 1.700E-07 | 3.000E-07 |
| 26935 | 1347 | S1347P | E | 2.8e-07 | 6.4e-07 | 1.6e-07 | 3.600E-07 | 2.498E-07 | 1.600E-07 | 6.400E-07 |
| 26936 | 1347 | S1347S | E | 2.9e-07 | 3e-07 | 3.2e-07 | 3.033E-07 | 1.528E-08 | 2.900E-07 | 3.200E-07 |
| 26937 | 1347 | S1347T | E | 1.3e-07 | 2.2e-07 | 1.8e-07 | 1.767E-07 | 4.509E-08 | 1.300E-07 | 2.200E-07 |
| 26938 | 1347 | S1347W | E | 5e-07 | 1.6e-07 | 3.5e-07 | 3.367E-07 | 1.704E-07 | 1.600E-07 | 5.000E-07 |
| 26939 | 1347 | S1347Y | E | 5.6e-07 | 1.1e-07 | 4e-07 | 3.567E-07 | 2.281E-07 | 1.100E-07 | 5.600E-07 |
| 26940 | 1347 | S1347V | E | 1.4e-07 | 9.2e-08 | 2e-07 | 1.440E-07 | 5.411E-08 | 9.200E-08 | 2.000E-07 |
| 26941 | 1348 | K1348A | E | 3e-07 | 3.6e-07 | 3.2e-07 | 3.267E-07 | 3.055E-08 | 3.000E-07 | 3.600E-07 |
| 26942 | 1348 | K1348R | E | 1.7e-07 | 4.9e-07 | 4.4e-07 | 3.667E-07 | 1.721E-07 | 1.700E-07 | 4.900E-07 |
| 26943 | 1348 | K1348N | E | 2e-07 | 1.8e-07 | 1.4e-07 | 1.733E-07 | 3.055E-08 | 1.400E-07 | 2.000E-07 |
| 26944 | 1348 | K1348D | E | 2.8e-07 | 3.2e-07 | 2.2e-07 | 2.733E-07 | 5.033E-08 | 2.200E-07 | 3.200E-07 |

| | | | | | | | | | | |
|---|---|---|---|---|---|---|---|---|---|---|
| 26945 | 1348 | K1348C | E | 2.1e-07 | 1.4e-07 | 2.4e-07 | 1.967E-07 | 5.132E-08 | 1.400E-07 | 2.400E-07 |
| 26946 | 1348 | K1348E | E | 2.8e-07 | 3.5e-07 | 5e-07 | 3.767E-07 | 1.124E-07 | 2.800E-07 | 5.000E-07 |
| 26947 | 1348 | K1348Q | E | 2.7e-07 | 3e-07 | 2.6e-07 | 2.767E-07 | 2.082E-08 | 2.600E-07 | 3.000E-07 |
| 26948 | 1348 | K1348G | E | 5.3e-07 | 3.3e-07 | 3e-07 | 3.867E-07 | 1.250E-07 | 3.000E-07 | 5.300E-07 |
| 26949 | 1348 | K1348H | E | 1.7e-07 | 1.6e-07 | 2.5e-07 | 1.933E-07 | 4.933E-08 | 1.600E-07 | 2.500E-07 |
| 26950 | 1348 | K1348I | E | 1.1e-07 | 1e-07 | 1.8e-07 | 1.300E-07 | 4.359E-08 | 1.000E-07 | 1.800E-07 |
| 26951 | 1348 | K1348L | E | 2.7e-07 | 1.9e-07 | 4.5e-07 | 3.033E-07 | 1.332E-07 | 1.900E-07 | 4.500E-07 |
| 26952 | 1348 | K1348K | E | 3e-07 | 4.6e-07 | 2.6e-07 | 3.400E-07 | 1.058E-07 | 2.600E-07 | 4.600E-07 |
| 26953 | 1348 | K1348M | E | 6.1e-07 | 2.2e-07 | 5.8e-08 | 2.960E-07 | 2.837E-07 | 5.800E-08 | 6.100E-07 |
| 26954 | 1348 | K1348F | E | 1.8e-07 | 3.6e-07 | 1.8e-07 | 2.400E-07 | 1.039E-07 | 1.800E-07 | 3.600E-07 |
| 26955 | 1348 | K1348P | E | 5.3e-07 | 3e-07 | 1.7e-07 | 3.333E-07 | 1.823E-07 | 1.700E-07 | 5.300E-07 |
| 26956 | 1348 | K1348S | E | 2.3e-07 | 3.9e-07 | 1.9e-07 | 2.700E-07 | 1.058E-07 | 1.900E-07 | 3.900E-07 |
| 26957 | 1348 | K1348T | E | 2.4e-07 | 3.3e-07 | 2.6e-07 | 2.767E-07 | 4.726E-08 | 2.400E-07 | 3.300E-07 |
| 26958 | 1348 | K1348W | E | 1.9e-07 | 4.4e-07 | 7e-07 | 4.433E-07 | 2.550E-07 | 1.900E-07 | 7.000E-07 |
| 26959 | 1348 | K1348Y | E | 3.2e-07 | 3.5e-07 | 1.3e-07 | 2.667E-07 | 1.193E-07 | 1.300E-07 | 3.500E-07 |
| 26960 | 1348 | K1348V | E | 3.6e-07 | 5.9e-07 | 4.1e-07 | 4.533E-07 | 1.210E-07 | 3.600E-07 | 5.900E-07 |
| 26961 | 1349 | N1349A | E | 2e-07 | 9.9e-08 | 1.8e-07 | 1.597E-07 | 5.348E-08 | 9.900E-08 | 2.000E-07 |
| 26962 | 1349 | N1349R | E | 2.7e-07 | 5.1e-07 | 2e-07 | 3.267E-07 | 1.626E-07 | 2.000E-07 | 5.100E-07 |
| 26963 | 1349 | N1349N | E | 2.3e-07 | 1.4e-07 | 1.4e-07 | 1.700E-07 | 5.196E-08 | 1.400E-07 | 2.300E-07 |
| 26964 | 1349 | N1349D | E | 1.9e-07 | 3.4e-07 | 1.2e-07 | 2.167E-07 | 1.124E-07 | 1.200E-07 | 3.400E-07 |
| 26965 | 1349 | N1349C | E | 1.6e-07 | 1.5e-07 | 3.2e-07 | 2.100E-07 | 9.539E-08 | 1.500E-07 | 3.200E-07 |
| 26966 | 1349 | N1349E | E | 1.9e-07 | 2e-07 | 2.5e-07 | 2.133E-07 | 3.215E-08 | 1.900E-07 | 2.500E-07 |
| 26967 | 1349 | N1349Q | E | 2.6e-07 | 1.9e-07 | 4.1e-07 | 2.867E-07 | 1.124E-07 | 1.900E-07 | 4.100E-07 |
| 26968 | 1349 | N1349G | E | 3.9e-07 | 4.1e-07 | 8e-08 | 2.933E-07 | 1.850E-07 | 8.000E-08 | 4.100E-07 |
| 26969 | 1349 | N1349H | E | 2.5e-07 | 3.7e-07 | 2.1e-07 | 2.767E-07 | 8.327E-08 | 2.100E-07 | 3.700E-07 |
| 26970 | 1349 | N1349I | E | 2.3e-07 | 3.3e-07 | 2.4e-07 | 2.667E-07 | 5.508E-08 | 2.300E-07 | 3.300E-07 |
| 26971 | 1349 | N1349L | E | 3.5e-07 | 1.9e-07 | 3.5e-07 | 2.967E-07 | 9.238E-08 | 1.900E-07 | 3.500E-07 |
| 26972 | 1349 | N1349K | E | 1.8e-07 | 2.3e-07 | 2.8e-07 | 2.300E-07 | 5.000E-08 | 1.800E-07 | 2.800E-07 |
| 26973 | 1349 | N1349M | E | 2.3e-07 | 4.7e-07 | 1.4e-07 | 2.800E-07 | 1.706E-07 | 1.400E-07 | 4.700E-07 |
| 26974 | 1349 | N1349F | E | 3.8e-07 | 2.2e-07 | 9.1e-08 | 2.303E-07 | 1.448E-07 | 9.100E-08 | 3.800E-07 |
| 26975 | 1349 | N1349P | E | 3.5e-07 | 2.2e-07 | 2.5e-07 | 2.733E-07 | 6.807E-08 | 2.200E-07 | 3.500E-07 |
| 26976 | 1349 | N1349S | E | 2.6e-07 | 1.7e-07 | 3.4e-07 | 2.567E-07 | 8.505E-08 | 1.700E-07 | 3.400E-07 |
| 26977 | 1349 | N1349T | E | 2.3e-07 | 4.8e-07 | 2.5e-07 | 3.200E-07 | 1.389E-07 | 2.300E-07 | 4.800E-07 |
| 26978 | 1349 | N1349W | E | 4e-07 | 3.7e-07 | 2.6e-07 | 3.433E-07 | 7.371E-08 | 2.600E-07 | 4.000E-07 |
| 26979 | 1349 | N1349Y | E | 1.5e-07 | 5.1e-07 | 3.5e-07 | 3.367E-07 | 1.804E-07 | 1.500E-07 | 5.100E-07 |
| 26980 | 1349 | N1349V | E | 2.6e-07 | 7.8e-08 | 1.8e-07 | 1.727E-07 | 9.122E-08 | 7.800E-08 | 2.600E-07 |
| 26981 | 1350 | T1350A | E | 1.5e-07 | 2e-07 | 3.3e-07 | 2.267E-07 | 9.292E-08 | 1.500E-07 | 3.300E-07 |
| 26982 | 1350 | T1350R | E | 1.7e-07 | 1.2e-07 | 2.1e-07 | 1.667E-07 | 4.509E-08 | 1.200E-07 | 2.100E-07 |
| 26983 | 1350 | T1350N | E | 3.7e-07 | 1.5e-07 | 2.5e-07 | 2.567E-07 | 1.102E-07 | 1.500E-07 | 3.700E-07 |
| 26984 | 1350 | T1350D | E | 1.7e-07 | 2.1e-07 | 2e-07 | 1.933E-07 | 2.082E-08 | 1.700E-07 | 2.100E-07 |
| 26985 | 1350 | T1350C | E | 5e-07 | 1.5e-07 | 2.3e-07 | 2.933E-07 | 1.834E-07 | 1.500E-07 | 5.000E-07 |
| 26986 | 1350 | T1350E | E | 2.1e-07 | 1.8e-07 | 3.1e-07 | 2.333E-07 | 6.807E-08 | 1.800E-07 | 3.100E-07 |
| 26987 | 1350 | T1350Q | E | 1.7e-07 | 2.1e-07 | 2.2e-07 | 2.000E-07 | 2.646E-08 | 1.700E-07 | 2.200E-07 |
| 26988 | 1350 | T1350G | E | 2.7e-07 | 1.8e-07 | 1.6e-07 | 2.033E-07 | 5.859E-08 | 1.600E-07 | 2.700E-07 |
| 26989 | 1350 | T1350H | E | 1.8e-07 | 2.9e-07 | 1.9e-07 | 2.200E-07 | 6.083E-08 | 1.800E-07 | 2.900E-07 |
| 26990 | 1350 | T1350I | E | 9.1e-08 | 1.1e-07 | 2.4e-07 | 1.470E-07 | 8.110E-08 | 9.100E-08 | 2.400E-07 |
| 26991 | 1350 | T1350L | E | 2e-07 | 2.4e-07 | 2.4e-07 | 2.267E-07 | 2.309E-08 | 2.000E-07 | 2.400E-07 |
| 26992 | 1350 | T1350K | E | 2.4e-07 | 1.9e-07 | 2.1e-07 | 2.133E-07 | 2.517E-08 | 1.900E-07 | 2.400E-07 |
| 26993 | 1350 | T1350M | E | 2.5e-07 | 3.5e-07 | 2.7e-07 | 2.900E-07 | 5.292E-08 | 2.500E-07 | 3.500E-07 |
| 26994 | 1350 | T1350F | E | 1.9e-07 | 1.6e-07 | 4.3e-07 | 2.600E-07 | 1.480E-07 | 1.600E-07 | 4.300E-07 |
| 26995 | 1350 | T1350P | E | 2.4e-07 | 3.9e-07 | 1.7e-07 | 2.667E-07 | 1.124E-07 | 1.700E-07 | 3.900E-07 |
| 26996 | 1350 | T1350S | E | 5.5e-07 | 2.3e-07 | 1.1e-07 | 2.967E-07 | 2.274E-07 | 1.100E-07 | 5.500E-07 |
| 26997 | 1350 | T1350T | E | 3.4e-07 | 1.3e-07 | 1.2e-07 | 1.967E-07 | 1.242E-07 | 1.200E-07 | 3.400E-07 |
| 26998 | 1350 | T1350W | E | 2.3e-07 | 3.6e-07 | 3e-07 | 2.967E-07 | 6.506E-08 | 2.300E-07 | 3.600E-07 |
| 26999 | 1350 | T1350Y | E | 1.7e-07 | 3.2e-07 | 3.3e-07 | 2.733E-07 | 8.963E-08 | 1.700E-07 | 3.300E-07 |
| 27000 | 1350 | T1350V | E | 1.9e-07 | 2.6e-07 | 4.4e-07 | 2.967E-07 | 1.290E-07 | 1.900E-07 | 4.400E-07 |
| 27001 | 1351 | A1351A | E | 2.3e-07 | 2.6e-07 | 6.5e-07 | 3.800E-07 | 2.343E-07 | 2.300E-07 | 6.500E-07 |
| 27002 | 1351 | A1351R | E | 2.2e-07 | 3.7e-07 | 2.7e-07 | 2.867E-07 | 7.638E-08 | 2.200E-07 | 3.700E-07 |
| 27003 | 1351 | A1351N | E | 1.7e-07 | 2.8e-07 | 2e-07 | 2.167E-07 | 5.686E-08 | 1.700E-07 | 2.800E-07 |
| 27004 | 1351 | A1351D | E | 3e-07 | 4.9e-07 | 3.8e-07 | 3.900E-07 | 9.539E-08 | 3.000E-07 | 4.900E-07 |
| 27005 | 1351 | A1351C | E | 2.2e-07 | 1.6e-07 | 2.7e-07 | 2.167E-07 | 5.508E-08 | 1.600E-07 | 2.700E-07 |
| 27006 | 1351 | A1351E | E | 3.3e-07 | 1.8e-07 | 7.9e-07 | 4.333E-07 | 3.179E-07 | 1.800E-07 | 7.900E-07 |
| 27007 | 1351 | A1351Q | E | 2.9e-07 | 5.1e-07 | 1.6e-07 | 3.200E-07 | 1.769E-07 | 1.600E-07 | 5.100E-07 |
| 27008 | 1351 | A1351G | E | 7e-07 | 1.6e-07 | 3.8e-07 | 4.133E-07 | 2.715E-07 | 1.600E-07 | 7.000E-07 |
| 27009 | 1351 | A1351H | E | 3.6e-07 | 1.8e-07 | 1.7e-07 | 2.367E-07 | 1.069E-07 | 1.700E-07 | 3.600E-07 |
| 27010 | 1351 | A1351I | E | 2.2e-07 | 3.9e-07 | 2.4e-07 | 2.833E-07 | 9.292E-08 | 2.200E-07 | 3.900E-07 |
| 27011 | 1351 | A1351L | E | 4.1e-07 | 1.4e-07 | 3.5e-07 | 3.000E-07 | 1.418E-07 | 1.400E-07 | 4.100E-07 |
| 27012 | 1351 | A1351K | E | 1.1e-07 | 1.6e-07 | 3e-07 | 1.900E-07 | 9.849E-08 | 1.100E-07 | 3.000E-07 |
| 27013 | 1351 | A1351M | E | 3.7e-07 | 2.4e-07 | 1.9e-07 | 2.667E-07 | 9.292E-08 | 1.900E-07 | 3.700E-07 |
| 27014 | 1351 | A1351F | E | 2.8e-07 | 3.5e-07 | 3.5e-07 | 3.267E-07 | 4.041E-08 | 2.800E-07 | 3.500E-07 |
| 27015 | 1351 | A1351P | E | 2.5e-07 | 2.5e-07 | 2.6e-07 | 2.533E-07 | 5.774E-09 | 2.500E-07 | 2.600E-07 |
| 27016 | 1351 | A1351S | E | 2.3e-07 | 1.9e-07 | 5.1e-07 | 3.100E-07 | 1.744E-07 | 1.900E-07 | 5.100E-07 |
| 27017 | 1351 | A1351T | E | 2.6e-07 | 1.9e-07 | 6.4e-07 | 3.633E-07 | 2.421E-07 | 1.900E-07 | 6.400E-07 |
| 27018 | 1351 | A1351W | E | 2.9e-07 | 2.3e-07 | 1.4e-07 | 2.200E-07 | 7.550E-08 | 1.400E-07 | 2.900E-07 |
| 27019 | 1351 | A1351Y | E | 8.6e-08 | 1.4e-07 | 1.9e-07 | 1.387E-07 | 5.201E-08 | 8.600E-08 | 1.900E-07 |
| 27020 | 1351 | A1351V | E | 1.8e-07 | 3.4e-07 | 3.5e-07 | 2.900E-07 | 9.539E-08 | 1.800E-07 | 3.500E-07 |
| 27021 | 1352 | Y1352A | E | 2.6e-07 | 1.6e-07 | 2.6e-07 | 2.267E-07 | 5.774E-08 | 1.600E-07 | 2.600E-07 |
| 27022 | 1352 | Y1352R | E | 2e-07 | 2.2e-07 | 2e-07 | 2.067E-07 | 1.155E-08 | 2.000E-07 | 2.200E-07 |
| 27023 | 1352 | Y1352N | E | 3.8e-07 | 3.7e-07 | 2.2e-07 | 3.233E-07 | 8.963E-08 | 2.200E-07 | 3.800E-07 |
| 27024 | 1352 | Y1352D | E | 1.3e-07 | 9.9e-08 | 1.6e-07 | 1.297E-07 | 3.050E-08 | 9.900E-08 | 1.600E-07 |
| 27025 | 1352 | Y1352C | E | 2.8e-07 | 2e-07 | 4.5e-07 | 3.100E-07 | 1.277E-07 | 2.000E-07 | 4.500E-07 |
| 27026 | 1352 | Y1352E | E | 1.4e-07 | 1.8e-07 | 2.9e-07 | 2.033E-07 | 7.767E-08 | 1.400E-07 | 2.900E-07 |
| 27027 | 1352 | Y1352Q | E | 4.4e-07 | 4.4e-07 | 2e-07 | 3.600E-07 | 1.386E-07 | 2.000E-07 | 4.400E-07 |
| 27028 | 1352 | Y1352G | E | 1.3e-07 | 2.2e-07 | 2.5e-07 | 2.000E-07 | 6.245E-08 | 1.300E-07 | 2.500E-07 |
| 27029 | 1352 | Y1352H | E | 4.3e-07 | 1.7e-07 | 2.4e-07 | 2.800E-07 | 1.345E-07 | 1.700E-07 | 4.300E-07 |

| | | | | | | | | | | |
|---|---|---|---|---|---|---|---|---|---|---|
| 27030 | 1352 | Y1352I | E | 3.2e-07 | 3e-07 | 1.8e-07 | 2.667E-07 | 7.572E-08 | 1.800E-07 | 3.200E-07 |
| 27031 | 1352 | Y1352L | E | 3.1e-07 | 2.4e-07 | 1.1e-07 | 2.200E-07 | 1.015E-07 | 1.100E-07 | 3.100E-07 |
| 27032 | 1352 | Y1352K | E | 1.3e-07 | 2.5e-07 | 3.1e-07 | 2.300E-07 | 9.165E-08 | 1.300E-07 | 3.100E-07 |
| 27033 | 1352 | Y1352M | E | 2.8e-07 | 2e-07 | 1.8e-07 | 2.200E-07 | 5.292E-08 | 1.800E-07 | 2.800E-07 |
| 27034 | 1352 | Y1352F | E | 4.1e-07 | 1.8e-07 | 4.8e-07 | 3.567E-07 | 1.570E-07 | 1.800E-07 | 4.800E-07 |
| 27035 | 1352 | Y1352P | E | 1.8e-07 | 2.2e-07 | 7.2e-08 | 1.573E-07 | 7.656E-08 | 7.200E-08 | 2.200E-07 |
| 27036 | 1352 | Y1352S | E | 1.8e-07 | 1.4e-07 | 3.1e-07 | 2.100E-07 | 8.888E-08 | 1.400E-07 | 3.100E-07 |
| 27037 | 1352 | Y1352T | E | 4.3e-07 | 2.1e-07 | 2.2e-07 | 2.867E-07 | 1.242E-07 | 2.100E-07 | 4.300E-07 |
| 27038 | 1352 | Y1352W | E | 5.7e-07 | 3.1e-07 | 1.6e-07 | 3.467E-07 | 2.074E-07 | 1.600E-07 | 5.700E-07 |
| 27039 | 1352 | Y1352Y | E | 4.1e-07 | 1.8e-07 | 5.3e-07 | 3.733E-07 | 1.779E-07 | 1.800E-07 | 5.300E-07 |
| 27040 | 1352 | Y1352V | E | 5e-07 | 2e-07 | 1.7e-07 | 2.900E-07 | 1.825E-07 | 1.700E-07 | 5.000E-07 |
| 27041 | 1353 | L1353A | E | 4e-07 | 1.3e-07 | 1.7e-07 | 2.333E-07 | 1.457E-07 | 1.300E-07 | 4.000E-07 |
| 27042 | 1353 | L1353R | E | 5.1e-07 | 3e-07 | 1.9e-07 | 3.333E-07 | 1.626E-07 | 1.900E-07 | 5.100E-07 |
| 27043 | 1353 | L1353N | E | 5e-07 | 1.4e-07 | 3.2e-07 | 3.200E-07 | 1.800E-07 | 1.400E-07 | 5.000E-07 |
| 27044 | 1353 | L1353D | E | 3.3e-07 | 1.7e-07 | 5e-07 | 3.333E-07 | 1.650E-07 | 1.700E-07 | 5.000E-07 |
| 27045 | 1353 | L1353C | E | 1.6e-07 | 2.2e-07 | 1.5e-07 | 1.767E-07 | 3.786E-08 | 1.500E-07 | 2.200E-07 |
| 27046 | 1353 | L1353E | E | 2.1e-07 | 1.5e-07 | 2.7e-07 | 2.100E-07 | 6.000E-08 | 1.500E-07 | 2.700E-07 |
| 27047 | 1353 | L1353Q | E | 4.1e-07 | 2.2e-07 | 1.7e-07 | 2.667E-07 | 1.266E-07 | 1.700E-07 | 4.100E-07 |
| 27048 | 1353 | L1353G | E | 1.9e-07 | 2.3e-07 | 1.1e-07 | 1.767E-07 | 6.110E-08 | 1.100E-07 | 2.300E-07 |
| 27049 | 1353 | L1353H | E | 2.6e-07 | 3.3e-07 | 3.1e-07 | 3.000E-07 | 3.606E-08 | 2.600E-07 | 3.300E-07 |
| 27050 | 1353 | L1353I | E | 4.2e-07 | 2.3e-07 | 2.7e-07 | 3.067E-07 | 1.002E-07 | 2.300E-07 | 4.200E-07 |
| 27051 | 1353 | L1353L | E | 2.7e-07 | 8.4e-08 | 2.7e-07 | 2.080E-07 | 1.074E-07 | 8.400E-08 | 2.700E-07 |
| 27052 | 1353 | L1353K | E | 2.6e-07 | 1.5e-07 | 2.1e-07 | 2.067E-07 | 5.508E-08 | 1.500E-07 | 2.600E-07 |
| 27053 | 1353 | L1353M | E | 3.1e-07 | 2.1e-07 | 1.3e-07 | 2.167E-07 | 9.018E-08 | 1.300E-07 | 3.100E-07 |
| 27054 | 1353 | L1353F | E | 2.6e-07 | 1.4e-07 | 3.8e-07 | 2.600E-07 | 1.200E-07 | 1.400E-07 | 3.800E-07 |
| 27055 | 1353 | L1353P | E | 2e-07 | 2.2e-07 | 1.5e-07 | 1.900E-07 | 3.606E-08 | 1.500E-07 | 2.200E-07 |
| 27056 | 1353 | L1353S | E | 1.8e-07 | 3.3e-07 | 1.7e-07 | 2.267E-07 | 8.963E-08 | 1.700E-07 | 3.300E-07 |
| 27057 | 1353 | L1353T | E | 3.6e-07 | 1.9e-07 | 1.9e-07 | 2.467E-07 | 9.815E-08 | 1.900E-07 | 3.600E-07 |
| 27058 | 1353 | L1353W | E | 2e-07 | 1.4e-07 | 2.7e-07 | 2.033E-07 | 6.506E-08 | 1.400E-07 | 2.700E-07 |
| 27059 | 1353 | L1353Y | E | 4.3e-07 | 1.5e-07 | 3.1e-07 | 2.967E-07 | 1.405E-07 | 1.500E-07 | 4.300E-07 |
| 27060 | 1353 | L1353V | E | 2.2e-07 | 4e-07 | 4e-07 | 3.400E-07 | 1.039E-07 | 2.200E-07 | 4.000E-07 |
| 27061 | 1354 | Q1354A | E | 4.6e-07 | 1e-07 | 1.2e-07 | 2.267E-07 | 2.023E-07 | 1.000E-07 | 4.600E-07 |
| 27062 | 1354 | Q1354R | E | 6.9e-08 | 4.1e-07 | 3.2e-07 | 2.663E-07 | 1.767E-07 | 6.900E-08 | 4.100E-07 |
| 27063 | 1354 | Q1354N | E | 2.6e-07 | 3.2e-07 | 2.5e-07 | 2.767E-07 | 3.786E-08 | 2.500E-07 | 3.200E-07 |
| 27064 | 1354 | Q1354D | E | 5.4e-07 | 1.7e-07 | 1.6e-07 | 2.900E-07 | 2.166E-07 | 1.600E-07 | 5.400E-07 |
| 27065 | 1354 | Q1354C | E | 2.9e-07 | 2e-07 | 1.8e-07 | 2.233E-07 | 5.859E-08 | 1.800E-07 | 2.900E-07 |
| 27066 | 1354 | Q1354E | E | 4.6e-07 | 2.8e-07 | 8.1e-08 | 2.737E-07 | 1.896E-07 | 8.100E-08 | 4.600E-07 |
| 27067 | 1354 | Q1354Q | E | 2.3e-07 | 3.5e-07 | 2.1e-07 | 2.633E-07 | 7.572E-08 | 2.100E-07 | 3.500E-07 |
| 27068 | 1354 | Q1354G | E | 2.3e-07 | 1.4e-07 | 2.7e-07 | 2.133E-07 | 6.658E-08 | 1.400E-07 | 2.700E-07 |
| 27069 | 1354 | Q1354H | E | 2.6e-07 | 2.5e-07 | 1.1e-07 | 2.067E-07 | 8.386E-08 | 1.100E-07 | 2.600E-07 |
| 27070 | 1354 | Q1354I | E | 4.3e-07 | 4.2e-07 | 3.9e-07 | 4.133E-07 | 2.082E-08 | 3.900E-07 | 4.300E-07 |
| 27071 | 1354 | Q1354L | E | 2.8e-07 | 1.4e-07 | 1.2e-07 | 1.800E-07 | 8.718E-08 | 1.200E-07 | 2.800E-07 |
| 27072 | 1354 | Q1354K | E | 2.7e-07 | 1.6e-07 | 1.9e-07 | 2.067E-07 | 5.686E-08 | 1.600E-07 | 2.700E-07 |
| 27073 | 1354 | Q1354M | E | 1.1e-07 | 1.9e-07 | 2.5e-07 | 1.833E-07 | 7.024E-08 | 1.100E-07 | 2.500E-07 |
| 27074 | 1354 | Q1354F | E | 2.6e-07 | 4.6e-07 | 3.5e-07 | 3.567E-07 | 1.002E-07 | 2.600E-07 | 4.600E-07 |
| 27075 | 1354 | Q1354P | E | 2.5e-07 | 4.2e-07 | 3.2e-07 | 3.300E-07 | 8.544E-08 | 2.500E-07 | 4.200E-07 |
| 27076 | 1354 | Q1354S | E | 2.4e-07 | 1.9e-07 | 2e-07 | 2.100E-07 | 2.646E-08 | 1.900E-07 | 2.400E-07 |
| 27077 | 1354 | Q1354T | E | 1.8e-07 | 9.7e-08 | 1.2e-07 | 1.323E-07 | 4.285E-08 | 9.700E-08 | 1.800E-07 |
| 27078 | 1354 | Q1354W | E | 2.8e-07 | 9.7e-08 | 1.9e-07 | 1.890E-07 | 9.150E-08 | 9.700E-08 | 2.800E-07 |
| 27079 | 1354 | Q1354Y | E | 7.5e-08 | 2.4e-07 | 2e-07 | 1.717E-07 | 8.607E-08 | 7.500E-08 | 2.400E-07 |
| 27080 | 1354 | Q1354V | E | 2.1e-07 | 3.2e-07 | 5.2e-07 | 3.500E-07 | 1.572E-07 | 2.100E-07 | 5.200E-07 |
| 27081 | 1355 | M1355A | E | 1.6e-07 | 1.9e-07 | 2.4e-07 | 1.967E-07 | 4.041E-08 | 1.600E-07 | 2.400E-07 |
| 27082 | 1355 | M1355R | E | 2.3e-07 | 3.6e-07 | 8.1e-08 | 2.237E-07 | 1.396E-07 | 8.100E-08 | 3.600E-07 |
| 27083 | 1355 | M1355N | E | 2.6e-07 | 4.7e-07 | 8e-08 | 2.700E-07 | 1.952E-07 | 8.000E-08 | 4.700E-07 |
| 27084 | 1355 | M1355D | E | 2e-07 | 5.9e-07 | 4e-07 | 3.967E-07 | 1.950E-07 | 2.000E-07 | 5.900E-07 |
| 27085 | 1355 | M1355C | E | 6.7e-07 | 2.2e-07 | 3.7e-07 | 4.200E-07 | 2.291E-07 | 2.200E-07 | 6.700E-07 |
| 27086 | 1355 | M1355E | E | 2.3e-07 | 5.1e-07 | 2e-07 | 3.133E-07 | 1.710E-07 | 2.000E-07 | 5.100E-07 |
| 27087 | 1355 | M1355Q | E | 2.6e-07 | 2.1e-07 | 4.2e-07 | 2.967E-07 | 1.097E-07 | 2.100E-07 | 4.200E-07 |
| 27088 | 1355 | M1355G | E | 3.7e-07 | 2.7e-07 | 3.5e-07 | 3.300E-07 | 5.292E-08 | 2.700E-07 | 3.700E-07 |
| 27089 | 1355 | M1355H | E | 1.5e-07 | 5.8e-07 | 1.1e-07 | 2.800E-07 | 2.606E-07 | 1.100E-07 | 5.800E-07 |
| 27090 | 1355 | M1355I | E | 2.1e-07 | 3.2e-07 | 2.8e-07 | 2.700E-07 | 5.568E-08 | 2.100E-07 | 3.200E-07 |
| 27091 | 1355 | M1355L | E | 2.7e-07 | 1.5e-07 | 2.1e-07 | 2.100E-07 | 6.000E-08 | 1.500E-07 | 2.700E-07 |
| 27092 | 1355 | M1355K | E | 2.2e-07 | 3.6e-07 | 2.6e-07 | 2.800E-07 | 7.211E-08 | 2.200E-07 | 3.600E-07 |
| 27093 | 1355 | M1355M | E | 2.8e-07 | 2.3e-07 | 2.8e-07 | 2.633E-07 | 2.887E-08 | 2.300E-07 | 2.800E-07 |
| 27094 | 1355 | M1355F | E | 2.1e-07 | 1.5e-07 | 3.7e-07 | 2.433E-07 | 1.137E-07 | 1.500E-07 | 3.700E-07 |
| 27095 | 1355 | M1355P | E | 2.5e-07 | 4.6e-07 | 3.9e-07 | 3.667E-07 | 1.069E-07 | 2.500E-07 | 4.600E-07 |
| 27096 | 1355 | M1355S | E | 2.8e-07 | 2.4e-07 | 2.4e-07 | 2.533E-07 | 2.309E-08 | 2.400E-07 | 2.800E-07 |
| 27097 | 1355 | M1355T | E | 2.4e-07 | 5.6e-07 | 2.3e-07 | 3.433E-07 | 1.877E-07 | 2.300E-07 | 5.600E-07 |
| 27098 | 1355 | M1355W | E | 3.7e-07 | 2.4e-07 | 2.8e-07 | 2.967E-07 | 6.658E-08 | 2.400E-07 | 3.700E-07 |
| 27099 | 1355 | M1355Y | E | 1.7e-07 | 9.8e-08 | 5e-07 | 2.560E-07 | 2.144E-07 | 9.800E-08 | 5.000E-07 |
| 27100 | 1355 | M1355V | E | 2.2e-07 | 3.9e-07 | 5e-07 | 3.700E-07 | 1.411E-07 | 2.200E-07 | 5.000E-07 |
| 27101 | 1356 | N1356A | E | 2.1e-07 | 7.8e-07 | 2.5e-07 | 4.133E-07 | 3.182E-07 | 2.100E-07 | 7.800E-07 |
| 27102 | 1356 | N1356R | E | 6e-07 | 2.3e-07 | 3e-07 | 3.767E-07 | 1.966E-07 | 2.300E-07 | 6.000E-07 |
| 27103 | 1356 | N1356N | E | 1.5e-07 | 1.7e-07 | 1.3e-07 | 1.500E-07 | 2.000E-08 | 1.300E-07 | 1.700E-07 |
| 27104 | 1356 | N1356D | E | 3e-07 | 3.7e-07 | 3.6e-07 | 3.433E-07 | 3.786E-08 | 3.000E-07 | 3.700E-07 |
| 27105 | 1356 | N1356C | E | 1.1e-07 | 2.2e-07 | 1.8e-07 | 1.700E-07 | 5.568E-08 | 1.100E-07 | 2.200E-07 |
| 27106 | 1356 | N1356E | E | 2.4e-07 | 2.5e-07 | 2.2e-07 | 2.367E-07 | 1.528E-08 | 2.200E-07 | 2.500E-07 |
| 27107 | 1356 | N1356Q | E | 5.2e-07 | 1.2e-07 | 1.8e-07 | 2.733E-07 | 2.157E-07 | 1.200E-07 | 5.200E-07 |
| 27108 | 1356 | N1356G | E | 2.9e-07 | 2.9e-07 | 4e-07 | 3.267E-07 | 6.351E-08 | 2.900E-07 | 4.000E-07 |
| 27109 | 1356 | N1356H | E | 2e-07 | 9.3e-08 | 3.2e-07 | 2.043E-07 | 1.136E-07 | 9.300E-08 | 3.200E-07 |
| 27110 | 1356 | N1356I | E | 2.2e-07 | 5.5e-07 | 3e-07 | 3.567E-07 | 1.721E-07 | 2.200E-07 | 5.500E-07 |
| 27111 | 1356 | N1356L | E | 1.6e-07 | 3e-07 | 1.6e-07 | 2.067E-07 | 8.083E-08 | 1.600E-07 | 3.000E-07 |
| 27112 | 1356 | N1356K | E | 4.7e-07 | 2.9e-07 | 2.2e-07 | 3.267E-07 | 1.290E-07 | 2.200E-07 | 4.700E-07 |
| 27113 | 1356 | N1356M | E | 3.2e-07 | 2.6e-07 | 2.6e-07 | 2.800E-07 | 3.464E-08 | 2.600E-07 | 3.200E-07 |
| 27114 | 1356 | N1356F | E | 2.8e-07 | 3.1e-07 | 3.4e-07 | 3.100E-07 | 3.000E-08 | 2.800E-07 | 3.400E-07 |

| | | | | | | | | | | |
|---|---|---|---|---|---|---|---|---|---|---|
| 27115 | 1356 | N1356P | E | 5e-07 | 1.8e-07 | 3.4e-07 | 3.400E-07 | 1.600E-07 | 1.800E-07 | 5.000E-07 |
| 27116 | 1356 | N1356S | E | 2.2e-07 | 2.9e-07 | 4.9e-07 | 3.333E-07 | 1.401E-07 | 2.200E-07 | 4.900E-07 |
| 27117 | 1356 | N1356T | E | 4.1e-07 | 1.6e-07 | 2.9e-07 | 2.867E-07 | 1.250E-07 | 1.600E-07 | 4.100E-07 |
| 27118 | 1356 | N1356W | E | 2.6e-07 | 3.6e-07 | 2.9e-07 | 3.033E-07 | 5.132E-08 | 2.600E-07 | 3.600E-07 |
| 27119 | 1356 | N1356Y | E | 1.9e-07 | 3.1e-07 | 2.6e-07 | 2.533E-07 | 6.028E-08 | 1.900E-07 | 3.100E-07 |
| 27120 | 1356 | N1356V | E | 1.8e-07 | 1.4e-07 | 5.6e-08 | 1.253E-07 | 6.329E-08 | 5.600E-08 | 1.800E-07 |
| 27121 | 1357 | S1357A | E | 1.9e-07 | 1.9e-07 | 1.9e-07 | 1.900E-07 | 0.000E+00 | 1.900E-07 | 1.900E-07 |
| 27122 | 1357 | S1357R | E | 2.3e-07 | 9.7e-08 | 1.5e-07 | 1.590E-07 | 6.696E-08 | 9.700E-08 | 2.300E-07 |
| 27123 | 1357 | S1357N | E | 2.2e-07 | 2.7e-07 | 2.8e-07 | 2.567E-07 | 3.215E-08 | 2.200E-07 | 2.800E-07 |
| 27124 | 1357 | S1357D | E | 3.3e-07 | 2.4e-07 | 3.2e-07 | 2.967E-07 | 4.933E-08 | 2.400E-07 | 3.300E-07 |
| 27125 | 1357 | S1357C | E | 2.2e-07 | 2.9e-07 | 3.4e-07 | 2.833E-07 | 6.028E-08 | 2.200E-07 | 3.400E-07 |
| 27126 | 1357 | S1357E | E | 3.7e-07 | 2.2e-07 | 3.5e-07 | 3.133E-07 | 8.145E-08 | 2.200E-07 | 3.700E-07 |
| 27127 | 1357 | S1357Q | E | 2.4e-07 | 1.7e-07 | 2e-07 | 2.033E-07 | 3.512E-08 | 1.700E-07 | 2.400E-07 |
| 27128 | 1357 | S1357G | E | 2.9e-07 | 2.8e-07 | 3.2e-07 | 2.967E-07 | 2.082E-08 | 2.800E-07 | 3.200E-07 |
| 27129 | 1357 | S1357H | E | 3.2e-07 | 2.1e-07 | 4.7e-07 | 3.333E-07 | 1.305E-07 | 2.100E-07 | 4.700E-07 |
| 27130 | 1357 | S1357I | E | 4.5e-07 | 2.1e-07 | 1.9e-07 | 2.833E-07 | 1.447E-07 | 1.900E-07 | 4.500E-07 |
| 27131 | 1357 | S1357L | E | 2.9e-07 | 3.9e-07 | 3.7e-07 | 3.500E-07 | 5.292E-08 | 2.900E-07 | 3.900E-07 |
| 27132 | 1357 | S1357K | E | 1.4e-07 | 2.8e-07 | 2.8e-07 | 2.333E-07 | 8.083E-08 | 1.400E-07 | 2.800E-07 |
| 27133 | 1357 | S1357M | E | 1.4e-07 | 2.1e-07 | 2.4e-07 | 1.967E-07 | 5.132E-08 | 1.400E-07 | 2.400E-07 |
| 27134 | 1357 | S1357F | E | 1.7e-07 | 2.4e-07 | 2.1e-07 | 2.067E-07 | 3.512E-08 | 1.700E-07 | 2.400E-07 |
| 27135 | 1357 | S1357P | E | 1.4e-07 | 2.3e-07 | 1.8e-07 | 1.833E-07 | 4.509E-08 | 1.400E-07 | 2.300E-07 |
| 27136 | 1357 | S1357S | E | 2.6e-07 | 2.9e-07 | 5.5e-07 | 3.667E-07 | 1.595E-07 | 2.600E-07 | 5.500E-07 |
| 27137 | 1357 | S1357T | E | 6.3e-07 | 2.5e-07 | 1.6e-07 | 3.467E-07 | 2.495E-07 | 1.600E-07 | 6.300E-07 |
| 27138 | 1357 | S1357W | E | 3.5e-07 | 4.1e-07 | 5.2e-07 | 4.267E-07 | 8.622E-08 | 3.500E-07 | 5.200E-07 |
| 27139 | 1357 | S1357Y | E | 3.2e-07 | 2e-07 | 2.9e-07 | 2.700E-07 | 6.245E-08 | 2.000E-07 | 3.200E-07 |
| 27140 | 1357 | S1357V | E | 2.8e-07 | 1.4e-07 | 3.5e-07 | 2.567E-07 | 1.069E-07 | 1.400E-07 | 3.500E-07 |
| 27141 | 1358 | L1358A | E | 2.4e-07 | 2e-07 | 3.1e-07 | 2.500E-07 | 5.568E-08 | 2.000E-07 | 3.100E-07 |
| 27142 | 1358 | L1358R | E | 1.3e-07 | 2.7e-07 | 2.6e-07 | 2.200E-07 | 7.810E-08 | 1.300E-07 | 2.700E-07 |
| 27143 | 1358 | L1358N | E | 2.9e-07 | 1.5e-07 | 1.7e-07 | 2.033E-07 | 7.572E-08 | 1.500E-07 | 2.900E-07 |
| 27144 | 1358 | L1358D | E | 2.3e-07 | 2.7e-07 | 2.1e-07 | 2.367E-07 | 3.055E-08 | 2.100E-07 | 2.700E-07 |
| 27145 | 1358 | L1358C | E | 1.1e-07 | 1.5e-07 | 2.9e-07 | 1.833E-07 | 9.452E-08 | 1.100E-07 | 2.900E-07 |
| 27146 | 1358 | L1358E | E | 2e-07 | 2.4e-07 | 5.3e-07 | 3.233E-07 | 1.801E-07 | 2.000E-07 | 5.300E-07 |
| 27147 | 1358 | L1358Q | E | 2.6e-07 | 4e-07 | 1.6e-07 | 2.733E-07 | 1.206E-07 | 1.600E-07 | 4.000E-07 |
| 27148 | 1358 | L1358G | E | 1.2e-07 | 4e-07 | 1.9e-07 | 2.367E-07 | 1.457E-07 | 1.200E-07 | 4.000E-07 |
| 27149 | 1358 | L1358H | E | 3.5e-07 | 1.4e-07 | 1.8e-07 | 2.233E-07 | 1.115E-07 | 1.400E-07 | 3.500E-07 |
| 27150 | 1358 | L1358I | E | 4e-07 | 2.6e-07 | 2.1e-07 | 2.900E-07 | 9.849E-08 | 2.100E-07 | 4.000E-07 |
| 27151 | 1358 | L1358L | E | 1.7e-07 | 5e-07 | 3.4e-07 | 3.367E-07 | 1.650E-07 | 1.700E-07 | 5.000E-07 |
| 27152 | 1358 | L1358K | E | 2.3e-07 | 2.5e-07 | 2.8e-07 | 2.533E-07 | 2.517E-08 | 2.300E-07 | 2.800E-07 |
| 27153 | 1358 | L1358M | E | 2e-07 | 1.4e-07 | 1.7e-07 | 1.700E-07 | 3.000E-08 | 1.400E-07 | 2.000E-07 |
| 27154 | 1358 | L1358F | E | 1.1e-07 | 2.3e-07 | 3e-07 | 2.133E-07 | 9.609E-08 | 1.100E-07 | 3.000E-07 |
| 27155 | 1358 | L1358P | E | 4.5e-07 | 3.4e-07 | 7.5e-07 | 5.133E-07 | 2.122E-07 | 3.400E-07 | 7.500E-07 |
| 27156 | 1358 | L1358S | E | 2.1e-07 | 3e-07 | 2.2e-07 | 2.433E-07 | 4.933E-08 | 2.100E-07 | 3.000E-07 |
| 27157 | 1358 | L1358T | E | 2.3e-07 | 1.9e-07 | 9.3e-07 | 4.500E-07 | 4.162E-07 | 1.900E-07 | 9.300E-07 |
| 27158 | 1358 | L1358W | E | 2.6e-07 | 1.7e-07 | 1.4e-07 | 1.900E-07 | 6.245E-08 | 1.400E-07 | 2.600E-07 |
| 27159 | 1358 | L1358Y | E | 2.8e-07 | 2.4e-07 | 4.9e-07 | 3.367E-07 | 1.343E-07 | 2.400E-07 | 4.900E-07 |
| 27160 | 1358 | L1358V | E | 3.8e-07 | 1.6e-07 | 2.5e-07 | 2.633E-07 | 1.106E-07 | 1.600E-07 | 3.800E-07 |
| 27161 | 1359 | R1359A | E | 1e-07 | 2.5e-07 | 4e-07 | 2.500E-07 | 1.500E-07 | 1.000E-07 | 4.000E-07 |
| 27162 | 1359 | R1359R | E | 2.8e-07 | 2.1e-07 | 2.2e-07 | 2.367E-07 | 3.786E-08 | 2.100E-07 | 2.800E-07 |
| 27163 | 1359 | R1359N | E | 2.4e-07 | 3e-07 | 6.8e-07 | 4.067E-07 | 2.386E-07 | 2.400E-07 | 6.800E-07 |
| 27164 | 1359 | R1359D | E | 2.9e-07 | 2.5e-07 | 2.4e-07 | 2.600E-07 | 2.646E-08 | 2.400E-07 | 2.900E-07 |
| 27165 | 1359 | R1359C | E | 2.7e-07 | 2.7e-07 | 2.8e-07 | 2.733E-07 | 5.774E-09 | 2.700E-07 | 2.800E-07 |
| 27166 | 1359 | R1359E | E | 3.5e-07 | 4.1e-07 | 2.6e-07 | 3.400E-07 | 7.550E-08 | 2.600E-07 | 4.100E-07 |
| 27167 | 1359 | R1359Q | E | 4.8e-07 | 4.4e-07 | 2e-07 | 3.733E-07 | 1.514E-07 | 2.000E-07 | 4.800E-07 |
| 27168 | 1359 | R1359G | E | 3.7e-07 | 2.7e-07 | 3.2e-07 | 3.200E-07 | 5.000E-08 | 2.700E-07 | 3.700E-07 |
| 27169 | 1359 | R1359H | E | 2.2e-07 | 2.6e-07 | 1.8e-07 | 2.200E-07 | 4.000E-08 | 1.800E-07 | 2.600E-07 |
| 27170 | 1359 | R1359I | E | 2.1e-07 | 2e-07 | 1.7e-07 | 1.933E-07 | 2.082E-08 | 1.700E-07 | 2.100E-07 |
| 27171 | 1359 | R1359L | E | 3.1e-07 | 1.9e-07 | 2.9e-07 | 2.633E-07 | 6.429E-08 | 1.900E-07 | 3.100E-07 |
| 27172 | 1359 | R1359K | E | 6.7e-07 | 1.7e-07 | 2.8e-07 | 3.733E-07 | 2.627E-07 | 1.700E-07 | 6.700E-07 |
| 27173 | 1359 | R1359M | E | 2.2e-07 | 1.8e-07 | 3.3e-07 | 2.433E-07 | 7.767E-08 | 1.800E-07 | 3.300E-07 |
| 27174 | 1359 | R1359F | E | 2e-07 | 3.3e-07 | 6.7e-07 | 4.000E-07 | 2.427E-07 | 2.000E-07 | 6.700E-07 |
| 27175 | 1359 | R1359P | E | 1.5e-07 | 2.7e-07 | 1.7e-07 | 1.967E-07 | 6.429E-08 | 1.500E-07 | 2.700E-07 |
| 27176 | 1359 | R1359S | E | 6.2e-07 | 6e-07 | 2.3e-07 | 4.833E-07 | 2.196E-07 | 2.300E-07 | 6.200E-07 |
| 27177 | 1359 | R1359T | E | 1.7e-07 | 2.2e-07 | 3.1e-07 | 2.333E-07 | 7.095E-08 | 1.700E-07 | 3.100E-07 |
| 27178 | 1359 | R1359W | E | 2.7e-07 | 1.7e-07 | 3.2e-07 | 2.533E-07 | 7.638E-08 | 1.700E-07 | 3.200E-07 |
| 27179 | 1359 | R1359Y | E | 2.7e-07 | 3.1e-07 | 2e-07 | 2.600E-07 | 5.568E-08 | 2.000E-07 | 3.100E-07 |
| 27180 | 1359 | R1359V | E | 5e-08 | 2.4e-07 | 2.5e-07 | 1.800E-07 | 1.127E-07 | 5.000E-08 | 2.500E-07 |
| 27181 | 1360 | A1360A | E | 2.2e-07 | 3.6e-07 | 1.6e-07 | 2.467E-07 | 1.026E-07 | 1.600E-07 | 3.600E-07 |
| 27182 | 1360 | A1360R | E | 1.6e-07 | 2.5e-07 | 1.9e-07 | 2.000E-07 | 4.583E-08 | 1.600E-07 | 2.500E-07 |
| 27183 | 1360 | A1360N | E | 1e-07 | 2.3e-07 | 3e-07 | 2.100E-07 | 1.015E-07 | 1.000E-07 | 3.000E-07 |
| 27184 | 1360 | A1360D | E | 1.8e-07 | 1.2e-07 | 2.1e-07 | 1.700E-07 | 4.583E-08 | 1.200E-07 | 2.100E-07 |
| 27185 | 1360 | A1360C | E | 1.8e-07 | 2.8e-07 | 4.5e-07 | 3.033E-07 | 1.365E-07 | 1.800E-07 | 4.500E-07 |
| 27186 | 1360 | A1360E | E | 1.9e-07 | 3.1e-07 | 2.9e-07 | 2.633E-07 | 6.429E-08 | 1.900E-07 | 3.100E-07 |
| 27187 | 1360 | A1360Q | E | 4e-07 | 2.3e-07 | 1.5e-07 | 2.600E-07 | 1.277E-07 | 1.500E-07 | 4.000E-07 |
| 27188 | 1360 | A1360G | E | 1.8e-07 | 1.9e-07 | 2.6e-07 | 2.100E-07 | 4.359E-08 | 1.800E-07 | 2.600E-07 |
| 27189 | 1360 | A1360H | E | 1.2e-07 | 3.1e-07 | 3.9e-07 | 2.733E-07 | 1.387E-07 | 1.200E-07 | 3.900E-07 |
| 27190 | 1360 | A1360I | E | 2.3e-07 | 1.5e-07 | 2.1e-07 | 1.967E-07 | 4.163E-08 | 1.500E-07 | 2.300E-07 |
| 27191 | 1360 | A1360L | E | 3.2e-07 | 2.7e-07 | 1.3e-07 | 2.400E-07 | 9.849E-08 | 1.300E-07 | 3.200E-07 |
| 27192 | 1360 | A1360K | E | 2.7e-07 | 3.8e-07 | 2.3e-07 | 2.933E-07 | 7.767E-08 | 2.300E-07 | 3.800E-07 |
| 27193 | 1360 | A1360M | E | 1.7e-07 | 2.7e-07 | 1.8e-07 | 2.067E-07 | 5.508E-08 | 1.700E-07 | 2.700E-07 |
| 27194 | 1360 | A1360F | E | 3.9e-07 | 2.1e-07 | 2.5e-07 | 2.833E-07 | 9.452E-08 | 2.100E-07 | 3.900E-07 |
| 27195 | 1360 | A1360P | E | 2.6e-07 | 2e-07 | 1.7e-07 | 2.100E-07 | 4.583E-08 | 1.700E-07 | 2.600E-07 |
| 27196 | 1360 | A1360S | E | 6.8e-07 | 1.4e-07 | 3.1e-07 | 3.767E-07 | 2.761E-07 | 1.400E-07 | 6.800E-07 |
| 27197 | 1360 | A1360T | E | 4.4e-07 | 2.7e-07 | 1.5e-07 | 2.867E-07 | 1.457E-07 | 1.500E-07 | 4.400E-07 |
| 27198 | 1360 | A1360W | E | 1.6e-07 | 2.1e-07 | 4.3e-07 | 2.667E-07 | 1.436E-07 | 1.600E-07 | 4.300E-07 |
| 27199 | 1360 | A1360Y | E | 2.9e-07 | 1.5e-07 | 2.8e-07 | 2.400E-07 | 7.810E-08 | 1.500E-07 | 2.900E-07 |

| | | | | | | | | | | |
|---|---|---|---|---|---|---|---|---|---|---|
| 27200 | 1360 | A1360V | E | 2.3e-07 | 5.1e-07 | 3.6e-07 | 3.667E-07 | 1.401E-07 | 2.300E-07 | 5.100E-07 |
| 27201 | 1361 | E1361A | E | 1.9e-07 | 3.8e-07 | 1.4e-07 | 2.367E-07 | 1.266E-07 | 1.400E-07 | 3.800E-07 |
| 27202 | 1361 | E1361R | E | 1.3e-07 | 2.6e-07 | 2.3e-07 | 2.067E-07 | 6.807E-08 | 1.300E-07 | 2.600E-07 |
| 27203 | 1361 | E1361N | E | 1.9e-07 | 3.7e-07 | 1.2e-07 | 2.267E-07 | 1.290E-07 | 1.200E-07 | 3.700E-07 |
| 27204 | 1361 | E1361D | E | 1.2e-07 | 3.6e-07 | 3.2e-07 | 2.667E-07 | 1.286E-07 | 1.200E-07 | 3.600E-07 |
| 27205 | 1361 | E1361C | E | 1.8e-07 | 2.8e-07 | 2.6e-07 | 2.400E-07 | 5.292E-08 | 1.800E-07 | 2.800E-07 |
| 27206 | 1361 | E1361E | E | 1.6e-07 | 2.2e-07 | 6.8e-08 | 1.493E-07 | 7.656E-08 | 6.800E-08 | 2.200E-07 |
| 27207 | 1361 | E1361Q | E | 1.4e-07 | 3.8e-07 | 5.3e-07 | 3.500E-07 | 1.967E-07 | 1.400E-07 | 5.300E-07 |
| 27208 | 1361 | E1361G | E | 4.8e-07 | 1.6e-07 | 3.1e-07 | 3.167E-07 | 1.601E-07 | 1.600E-07 | 4.800E-07 |
| 27209 | 1361 | E1361H | E | 6.4e-07 | 1.1e-07 | 3.5e-07 | 3.667E-07 | 2.654E-07 | 1.100E-07 | 6.400E-07 |
| 27210 | 1361 | E1361I | E | 2.7e-07 | 4.3e-07 | 1.5e-07 | 2.833E-07 | 1.405E-07 | 1.500E-07 | 4.300E-07 |
| 27211 | 1361 | E1361L | E | 3.9e-07 | 3.1e-07 | 2.8e-07 | 3.267E-07 | 5.686E-08 | 2.800E-07 | 3.900E-07 |
| 27212 | 1361 | E1361K | E | 3.4e-07 | 4.5e-07 | 1.9e-07 | 3.267E-07 | 1.305E-07 | 1.900E-07 | 4.500E-07 |
| 27213 | 1361 | E1361M | E | 4e-07 | 5.6e-07 | 2.1e-07 | 3.900E-07 | 1.752E-07 | 2.100E-07 | 5.600E-07 |
| 27214 | 1361 | E1361F | E | 2.8e-07 | 2.9e-07 | 1.4e-07 | 2.367E-07 | 8.386E-08 | 1.400E-07 | 2.900E-07 |
| 27215 | 1361 | E1361P | E | 3.7e-07 | 8e-07 | 1.3e-07 | 4.333E-07 | 3.395E-07 | 1.300E-07 | 8.000E-07 |
| 27216 | 1361 | E1361S | E | 1.6e-07 | 1.8e-07 | 1.5e-07 | 1.633E-07 | 1.528E-08 | 1.500E-07 | 1.800E-07 |
| 27217 | 1361 | E1361T | E | 4.3e-07 | 3.2e-07 | 1.3e-07 | 2.933E-07 | 1.518E-07 | 1.300E-07 | 4.300E-07 |
| 27218 | 1361 | E1361W | E | 2.5e-07 | 3.3e-07 | 2.5e-07 | 2.767E-07 | 4.619E-08 | 2.500E-07 | 3.300E-07 |
| 27219 | 1361 | E1361Y | E | 3.3e-07 | 3.2e-07 | 3.5e-07 | 3.333E-07 | 1.528E-08 | 3.200E-07 | 3.500E-07 |
| 27220 | 1361 | E1361V | E | 2.6e-07 | 1.7e-07 | 1.4e-07 | 1.900E-07 | 6.245E-08 | 1.400E-07 | 2.600E-07 |
| 27221 | 1362 | D1362A | E | 3.7e-07 | 3.3e-07 | 2.7e-07 | 3.233E-07 | 5.033E-08 | 2.700E-07 | 3.700E-07 |
| 27222 | 1362 | D1362R | E | 3.6e-07 | 2e-07 | 2.2e-07 | 2.600E-07 | 8.718E-08 | 2.000E-07 | 3.600E-07 |
| 27223 | 1362 | D1362N | E | 9.5e-08 | 3.5e-07 | 3.6e-07 | 2.683E-07 | 1.502E-07 | 9.500E-08 | 3.600E-07 |
| 27224 | 1362 | D1362D | E | 1.4e-07 | 1.7e-07 | 3.5e-07 | 2.200E-07 | 1.136E-07 | 1.400E-07 | 3.500E-07 |
| 27225 | 1362 | D1362C | E | 5.5e-07 | 3.6e-07 | 3.1e-07 | 4.067E-07 | 1.266E-07 | 3.100E-07 | 5.500E-07 |
| 27226 | 1362 | D1362E | E | 4.7e-07 | 4.2e-07 | 2.2e-07 | 3.700E-07 | 1.323E-07 | 2.200E-07 | 4.700E-07 |
| 27227 | 1362 | D1362Q | E | 2.1e-07 | 1.6e-07 | 1.8e-07 | 1.833E-07 | 2.517E-08 | 1.600E-07 | 2.100E-07 |
| 27228 | 1362 | D1362G | E | 4e-07 | 4.7e-07 | 2.2e-07 | 3.633E-07 | 1.290E-07 | 2.200E-07 | 4.700E-07 |
| 27229 | 1362 | D1362H | E | 2.6e-07 | 3.6e-07 | 1.6e-07 | 2.600E-07 | 1.000E-07 | 1.600E-07 | 3.600E-07 |
| 27230 | 1362 | D1362I | E | 1.9e-07 | 1.9e-07 | 2.1e-07 | 1.967E-07 | 1.155E-08 | 1.900E-07 | 2.100E-07 |
| 27231 | 1362 | D1362L | E | 3.8e-07 | 4.3e-07 | 2.6e-07 | 3.567E-07 | 8.737E-08 | 2.600E-07 | 4.300E-07 |
| 27232 | 1362 | D1362K | E | 3.2e-07 | 4e-07 | 1.8e-07 | 3.000E-07 | 1.114E-07 | 1.800E-07 | 4.000E-07 |
| 27233 | 1362 | D1362M | E | 2.8e-07 | 4.1e-07 | 2.4e-07 | 3.100E-07 | 8.888E-08 | 2.400E-07 | 4.100E-07 |
| 27234 | 1362 | D1362F | E | 2.2e-07 | 4.2e-07 | 5.9e-07 | 4.100E-07 | 1.852E-07 | 2.200E-07 | 5.900E-07 |
| 27235 | 1362 | D1362P | E | 1.5e-07 | 3.1e-07 | 2.1e-07 | 2.233E-07 | 8.083E-08 | 1.500E-07 | 3.100E-07 |
| 27236 | 1362 | D1362S | E | 1.4e-07 | 1.8e-07 | 1.7e-07 | 1.633E-07 | 2.082E-08 | 1.400E-07 | 1.800E-07 |
| 27237 | 1362 | D1362T | E | 1.1e-07 | 4.2e-07 | 1.6e-07 | 2.300E-07 | 1.664E-07 | 1.100E-07 | 4.200E-07 |
| 27238 | 1362 | D1362W | E | 9.8e-08 | 2.4e-07 | 2.2e-07 | 1.860E-07 | 7.686E-08 | 9.800E-08 | 2.400E-07 |
| 27239 | 1362 | D1362Y | E | 9.5e-08 | 2.1e-07 | 2.3e-07 | 1.783E-07 | 7.286E-08 | 9.500E-08 | 2.300E-07 |
| 27240 | 1362 | D1362V | E | 1.7e-07 | 3.3e-07 | 3.7e-07 | 2.900E-07 | 1.058E-07 | 1.700E-07 | 3.700E-07 |
| 27241 | 1363 | T1363A | E | 1.7e-07 | 2.8e-07 | 3.8e-07 | 2.767E-07 | 1.050E-07 | 1.700E-07 | 3.800E-07 |
| 27242 | 1363 | T1363R | E | 3e-07 | 4.8e-07 | 1.8e-07 | 3.200E-07 | 1.510E-07 | 1.800E-07 | 4.800E-07 |
| 27243 | 1363 | T1363N | E | 3.3e-07 | 1.4e-07 | 2.6e-07 | 2.433E-07 | 9.609E-08 | 1.400E-07 | 3.300E-07 |
| 27244 | 1363 | T1363D | E | 2.4e-07 | 2.7e-07 | 3.4e-07 | 2.833E-07 | 5.132E-08 | 2.400E-07 | 3.400E-07 |
| 27245 | 1363 | T1363C | E | 2.3e-07 | 1.9e-07 | 1.9e-07 | 2.033E-07 | 2.309E-08 | 1.900E-07 | 2.300E-07 |
| 27246 | 1363 | T1363E | E | 5e-07 | 3.3e-07 | 2.7e-07 | 3.667E-07 | 1.193E-07 | 2.700E-07 | 5.000E-07 |
| 27247 | 1363 | T1363Q | E | 2e-07 | 2.8e-07 | 2.6e-07 | 2.467E-07 | 4.163E-08 | 2.000E-07 | 2.800E-07 |
| 27248 | 1363 | T1363G | E | 1.1e-07 | 2.8e-07 | 1.2e-07 | 1.700E-07 | 9.539E-08 | 1.100E-07 | 2.800E-07 |
| 27249 | 1363 | T1363H | E | 2e-07 | 1.7e-07 | 3.3e-07 | 2.333E-07 | 8.505E-08 | 1.700E-07 | 3.300E-07 |
| 27250 | 1363 | T1363I | E | 1.6e-07 | 1.4e-07 | 1.5e-07 | 1.500E-07 | 1.000E-08 | 1.400E-07 | 1.600E-07 |
| 27251 | 1363 | T1363L | E | 2.7e-07 | 2.1e-07 | 2e-07 | 2.267E-07 | 3.786E-08 | 2.000E-07 | 2.700E-07 |
| 27252 | 1363 | T1363K | E | 1.1e-07 | 1.7e-07 | 3e-07 | 1.933E-07 | 9.713E-08 | 1.100E-07 | 3.000E-07 |
| 27253 | 1363 | T1363M | E | 1.4e-07 | 2.4e-07 | 2.1e-07 | 1.967E-07 | 5.132E-08 | 1.400E-07 | 2.400E-07 |
| 27254 | 1363 | T1363F | E | 2.4e-07 | 3.2e-07 | 2.6e-07 | 2.733E-07 | 4.163E-08 | 2.400E-07 | 3.200E-07 |
| 27255 | 1363 | T1363P | E | 3e-07 | 1.7e-07 | 3.9e-07 | 2.867E-07 | 1.106E-07 | 1.700E-07 | 3.900E-07 |
| 27256 | 1363 | T1363S | E | 3.9e-07 | 4e-07 | 2.9e-07 | 3.600E-07 | 6.083E-08 | 2.900E-07 | 4.000E-07 |
| 27257 | 1363 | T1363T | E | 1.1e-07 | 3.2e-07 | 5e-07 | 3.100E-07 | 1.952E-07 | 1.100E-07 | 5.000E-07 |
| 27258 | 1363 | T1363W | E | 2.5e-07 | 1.6e-07 | 2.8e-07 | 2.300E-07 | 6.245E-08 | 1.600E-07 | 2.800E-07 |
| 27259 | 1363 | T1363Y | E | 2.9e-07 | 2.4e-07 | 2.4e-07 | 2.567E-07 | 2.887E-08 | 2.400E-07 | 2.900E-07 |
| 27260 | 1363 | T1363V | E | 4.8e-07 | 3.1e-07 | 2.7e-07 | 3.533E-07 | 1.115E-07 | 2.700E-07 | 4.800E-07 |
| 27261 | 1364 | A1364A | E | 3.3e-07 | 2.7e-07 | 2.9e-07 | 2.967E-07 | 3.055E-08 | 2.700E-07 | 3.300E-07 |
| 27262 | 1364 | A1364R | E | 3e-07 | 1.7e-07 | 1.7e-07 | 2.133E-07 | 7.506E-08 | 1.700E-07 | 3.000E-07 |
| 27263 | 1364 | A1364N | E | 1.7e-07 | 2.1e-07 | 2.3e-07 | 2.033E-07 | 3.055E-08 | 1.700E-07 | 2.300E-07 |
| 27264 | 1364 | A1364D | E | 1.3e-07 | 1.5e-07 | 1.5e-07 | 1.433E-07 | 1.155E-08 | 1.300E-07 | 1.500E-07 |
| 27265 | 1364 | A1364C | E | 1.7e-07 | 2.5e-07 | 2.2e-07 | 2.133E-07 | 4.041E-08 | 1.700E-07 | 2.500E-07 |
| 27266 | 1364 | A1364E | E | 2.7e-07 | 2.4e-07 | 2.1e-07 | 2.400E-07 | 3.000E-08 | 2.100E-07 | 2.700E-07 |
| 27267 | 1364 | A1364Q | E | 2.9e-07 | 2.1e-07 | 1.9e-07 | 2.300E-07 | 5.292E-08 | 1.900E-07 | 2.900E-07 |
| 27268 | 1364 | A1364G | E | 2.5e-07 | 1.6e-07 | 1e-07 | 1.700E-07 | 7.550E-08 | 1.000E-07 | 2.500E-07 |
| 27269 | 1364 | A1364H | E | 3.8e-07 | 2.7e-07 | 1.5e-07 | 2.667E-07 | 1.150E-07 | 1.500E-07 | 3.800E-07 |
| 27270 | 1364 | A1364I | E | 1.5e-07 | 2.3e-07 | 1.2e-07 | 1.667E-07 | 5.686E-08 | 1.200E-07 | 2.300E-07 |
| 27271 | 1364 | A1364L | E | 5.1e-07 | 1.4e-07 | 2.5e-07 | 3.000E-07 | 1.900E-07 | 1.400E-07 | 5.100E-07 |
| 27272 | 1364 | A1364K | E | 1.7e-07 | 1.9e-07 | 2.6e-07 | 2.067E-07 | 4.726E-08 | 1.700E-07 | 2.600E-07 |
| 27273 | 1364 | A1364M | E | 3e-07 | 2.1e-07 | 1.3e-07 | 2.133E-07 | 8.505E-08 | 1.300E-07 | 3.000E-07 |
| 27274 | 1364 | A1364F | E | 4.4e-07 | 7.6e-08 | 2.4e-07 | 2.520E-07 | 1.823E-07 | 7.600E-08 | 4.400E-07 |
| 27275 | 1364 | A1364P | E | 4.8e-07 | 2.6e-07 | 2.5e-07 | 3.300E-07 | 1.300E-07 | 2.500E-07 | 4.800E-07 |
| 27276 | 1364 | A1364S | E | 2.2e-07 | 2.9e-07 | 2e-07 | 2.367E-07 | 4.726E-08 | 2.000E-07 | 2.900E-07 |
| 27277 | 1364 | A1364T | E | 2.8e-07 | 3.1e-07 | 2.5e-07 | 2.800E-07 | 3.000E-08 | 2.500E-07 | 3.100E-07 |
| 27278 | 1364 | A1364W | E | 1.9e-07 | 3e-07 | 2.9e-07 | 2.600E-07 | 6.083E-08 | 1.900E-07 | 3.000E-07 |
| 27279 | 1364 | A1364Y | E | 3.5e-07 | 1.5e-07 | 2.8e-07 | 2.600E-07 | 1.015E-07 | 1.500E-07 | 3.500E-07 |
| 27280 | 1364 | A1364V | E | 3.8e-07 | 4e-07 | 1.4e-07 | 3.067E-07 | 1.447E-07 | 1.400E-07 | 4.000E-07 |
| 27281 | 1365 | V1365A | E | 2.2e-07 | 1.2e-07 | 4e-07 | 2.467E-07 | 1.419E-07 | 1.200E-07 | 4.000E-07 |
| 27282 | 1365 | V1365R | E | 3.7e-07 | 2.2e-07 | 7.4e-07 | 4.433E-07 | 2.676E-07 | 2.200E-07 | 7.400E-07 |
| 27283 | 1365 | V1365N | E | 1.6e-07 | 1.1e-07 | 1.4e-07 | 1.367E-07 | 2.517E-08 | 1.100E-07 | 1.600E-07 |
| 27284 | 1365 | V1365D | E | 3.4e-07 | 1.9e-07 | 2.6e-07 | 2.633E-07 | 7.506E-08 | 1.900E-07 | 3.400E-07 |

| | | | | | | | | | | |
|---|---|---|---|---|---|---|---|---|---|---|
| 27285 | 1365 | V1365C | E | 1.3e-07 | 4.7e-07 | 1.7e-07 | 2.567E-07 | 1.858E-07 | 1.300E-07 | 4.700E-07 |
| 27286 | 1365 | V1365E | E | 1.4e-07 | 1.9e-07 | 2.1e-07 | 1.800E-07 | 3.606E-08 | 1.400E-07 | 2.100E-07 |
| 27287 | 1365 | V1365Q | E | 1.9e-07 | 1.1e-07 | 3.2e-07 | 2.067E-07 | 1.060E-07 | 1.100E-07 | 3.200E-07 |
| 27288 | 1365 | V1365G | E | 2.6e-07 | 3.5e-07 | 2.1e-07 | 2.733E-07 | 7.095E-08 | 2.100E-07 | 3.500E-07 |
| 27289 | 1365 | V1365H | E | 1.4e-07 | 2.4e-07 | 1.4e-07 | 1.733E-07 | 5.774E-08 | 1.400E-07 | 2.400E-07 |
| 27290 | 1365 | V1365I | E | 7.2e-08 | 2.2e-07 | 2.4e-07 | 1.773E-07 | 9.177E-08 | 7.200E-08 | 2.400E-07 |
| 27291 | 1365 | V1365L | E | 3.6e-07 | 2.7e-07 | 2e-07 | 2.767E-07 | 8.021E-08 | 2.000E-07 | 3.600E-07 |
| 27292 | 1365 | V1365K | E | 4.5e-07 | 6.5e-07 | 2.4e-07 | 4.467E-07 | 2.050E-07 | 2.400E-07 | 6.500E-07 |
| 27293 | 1365 | V1365M | E | 2.4e-07 | 2.3e-07 | 2.1e-07 | 2.267E-07 | 1.528E-08 | 2.100E-07 | 2.400E-07 |
| 27294 | 1365 | V1365F | E | 1.6e-07 | 5.5e-07 | 4.1e-07 | 3.733E-07 | 1.976E-07 | 1.600E-07 | 5.500E-07 |
| 27295 | 1365 | V1365P | E | 3.7e-07 | 3.6e-07 | 2.2e-07 | 3.167E-07 | 8.386E-08 | 2.200E-07 | 3.700E-07 |
| 27296 | 1365 | V1365S | E | 2.7e-07 | 2.7e-07 | 2.4e-07 | 2.600E-07 | 1.732E-08 | 2.400E-07 | 2.700E-07 |
| 27297 | 1365 | V1365T | E | 2.7e-07 | 1.8e-07 | 1.6e-07 | 2.033E-07 | 5.859E-08 | 1.600E-07 | 2.700E-07 |
| 27298 | 1365 | V1365W | E | 1.3e-07 | 1.8e-07 | 2.9e-07 | 2.000E-07 | 8.185E-08 | 1.300E-07 | 2.900E-07 |
| 27299 | 1365 | V1365Y | E | 1.9e-07 | 1.7e-07 | 1.3e-07 | 1.633E-07 | 3.055E-08 | 1.300E-07 | 1.900E-07 |
| 27300 | 1365 | V1365V | E | 2.9e-07 | 2.3e-07 | 3.8e-07 | 3.000E-07 | 7.550E-08 | 2.300E-07 | 3.800E-07 |
| 27301 | 1366 | Y1366A | E | 3.2e-07 | 3.2e-07 | 1.6e-07 | 2.667E-07 | 9.238E-08 | 1.600E-07 | 3.200E-07 |
| 27302 | 1366 | Y1366R | E | 2.2e-07 | 4.2e-07 | 1.7e-07 | 2.700E-07 | 1.323E-07 | 1.700E-07 | 4.200E-07 |
| 27303 | 1366 | Y1366N | E | 1.6e-07 | 2.5e-07 | 1.2e-07 | 1.767E-07 | 6.658E-08 | 1.200E-07 | 2.500E-07 |
| 27304 | 1366 | Y1366D | E | 1.8e-07 | 4.4e-07 | 2.2e-07 | 2.800E-07 | 1.400E-07 | 1.800E-07 | 4.400E-07 |
| 27305 | 1366 | Y1366C | E | 3.1e-07 | 2.9e-07 | 3.6e-07 | 3.200E-07 | 3.606E-08 | 2.900E-07 | 3.600E-07 |
| 27306 | 1366 | Y1366E | E | 2.8e-07 | 3.2e-07 | 2.9e-07 | 2.967E-07 | 2.082E-08 | 2.800E-07 | 3.200E-07 |
| 27307 | 1366 | Y1366Q | E | 2.4e-07 | 5e-07 | 2.2e-07 | 3.200E-07 | 1.562E-07 | 2.200E-07 | 5.000E-07 |
| 27308 | 1366 | Y1366G | E | 4.1e-07 | 2.5e-07 | 3.2e-07 | 3.267E-07 | 8.021E-08 | 2.500E-07 | 4.100E-07 |
| 27309 | 1366 | Y1366H | E | 2.2e-07 | 2.2e-07 | 2.7e-07 | 2.367E-07 | 2.887E-08 | 2.200E-07 | 2.700E-07 |
| 27310 | 1366 | Y1366I | E | 3.6e-07 | 5.9e-07 | 1.6e-07 | 3.700E-07 | 2.152E-07 | 1.600E-07 | 5.900E-07 |
| 27311 | 1366 | Y1366L | E | 2e-07 | 3.3e-07 | 2.1e-07 | 2.467E-07 | 7.234E-08 | 2.000E-07 | 3.300E-07 |
| 27312 | 1366 | Y1366K | E | 2.9e-07 | 1.2e-07 | 3.6e-07 | 2.567E-07 | 1.234E-07 | 1.200E-07 | 3.600E-07 |
| 27313 | 1366 | Y1366M | E | 9.9e-08 | 3.8e-07 | 1.6e-07 | 2.130E-07 | 1.478E-07 | 9.900E-08 | 3.800E-07 |
| 27314 | 1366 | Y1366F | E | 4.6e-07 | 4e-07 | 1.1e-07 | 3.233E-07 | 1.872E-07 | 1.100E-07 | 4.600E-07 |
| 27315 | 1366 | Y1366P | E | 3.5e-07 | 6.5e-07 | 3.5e-07 | 4.500E-07 | 1.732E-07 | 3.500E-07 | 6.500E-07 |
| 27316 | 1366 | Y1366S | E | 2.1e-07 | 2.6e-07 | 3.1e-07 | 2.600E-07 | 5.000E-08 | 2.100E-07 | 3.100E-07 |
| 27317 | 1366 | Y1366T | E | 2.6e-07 | 2.9e-07 | 2.9e-07 | 2.800E-07 | 1.732E-08 | 2.600E-07 | 2.900E-07 |
| 27318 | 1366 | Y1366W | E | 2.1e-07 | 3.2e-07 | 2.5e-07 | 2.600E-07 | 5.568E-08 | 2.100E-07 | 3.200E-07 |
| 27319 | 1366 | Y1366Y | E | 2e-07 | 2e-07 | 4.8e-07 | 2.933E-07 | 1.617E-07 | 2.000E-07 | 4.800E-07 |
| 27320 | 1366 | Y1366V | E | 3.5e-07 | 9.3e-07 | 2.8e-07 | 5.200E-07 | 3.568E-07 | 2.800E-07 | 9.300E-07 |
| 27321 | 1367 | Y1367A | E | 1.2e-07 | 2.3e-07 | 2e-07 | 1.833E-07 | 5.686E-08 | 1.200E-07 | 2.300E-07 |
| 27322 | 1367 | Y1367R | E | 2.7e-07 | 2.8e-07 | 2.6e-07 | 2.700E-07 | 1.000E-08 | 2.600E-07 | 2.800E-07 |
| 27323 | 1367 | Y1367N | E | 2.7e-07 | 1.2e-07 | 3.3e-07 | 2.400E-07 | 1.082E-07 | 1.200E-07 | 3.300E-07 |
| 27324 | 1367 | Y1367D | E | 1.8e-07 | 1.7e-07 | 3.1e-07 | 2.200E-07 | 7.810E-08 | 1.700E-07 | 3.100E-07 |
| 27325 | 1367 | Y1367C | E | 1.4e-07 | 4.3e-07 | 2.7e-07 | 2.800E-07 | 1.453E-07 | 1.400E-07 | 4.300E-07 |
| 27326 | 1367 | Y1367E | E | 3.8e-07 | 3.3e-07 | 2e-07 | 3.033E-07 | 9.292E-08 | 2.000E-07 | 3.800E-07 |
| 27327 | 1367 | Y1367Q | E | 3.4e-07 | 1.3e-07 | 3.1e-07 | 2.600E-07 | 1.136E-07 | 1.300E-07 | 3.400E-07 |
| 27328 | 1367 | Y1367G | E | 2.8e-07 | 1.7e-07 | 2e-07 | 2.167E-07 | 5.686E-08 | 1.700E-07 | 2.800E-07 |
| 27329 | 1367 | Y1367H | E | 2.4e-07 | 2.1e-07 | 6.6e-07 | 3.700E-07 | 2.516E-07 | 2.100E-07 | 6.600E-07 |
| 27330 | 1367 | Y1367I | E | 1.8e-07 | 6.8e-07 | 2.6e-07 | 3.733E-07 | 2.686E-07 | 1.800E-07 | 6.800E-07 |
| 27331 | 1367 | Y1367L | E | 1.4e-07 | 2.9e-07 | 2.8e-07 | 2.367E-07 | 8.386E-08 | 1.400E-07 | 2.900E-07 |
| 27332 | 1367 | Y1367K | E | 1.4e-07 | 4.6e-07 | 2.8e-07 | 2.933E-07 | 1.604E-07 | 1.400E-07 | 4.600E-07 |
| 27333 | 1367 | Y1367M | E | 5.5e-07 | 8.2e-08 | 3.9e-07 | 3.407E-07 | 2.379E-07 | 8.200E-08 | 5.500E-07 |
| 27334 | 1367 | Y1367F | E | 2.2e-07 | 2.8e-07 | 1.7e-07 | 2.233E-07 | 5.508E-08 | 1.700E-07 | 2.800E-07 |
| 27335 | 1367 | Y1367P | E | 1.8e-07 | 2.3e-07 | 1.4e-07 | 1.833E-07 | 4.509E-08 | 1.400E-07 | 2.300E-07 |
| 27336 | 1367 | Y1367S | E | 2.7e-07 | 2.7e-07 | 2.2e-07 | 2.533E-07 | 2.887E-08 | 2.200E-07 | 2.700E-07 |
| 27337 | 1367 | Y1367T | E | 2.5e-07 | 2.2e-07 | 2.3e-07 | 2.333E-07 | 1.528E-08 | 2.200E-07 | 2.500E-07 |
| 27338 | 1367 | Y1367W | E | 2.5e-07 | 2.1e-07 | 2.8e-07 | 2.467E-07 | 3.512E-08 | 2.100E-07 | 2.800E-07 |
| 27339 | 1367 | Y1367Y | E | 4e-07 | 1.8e-07 | 2e-07 | 2.600E-07 | 1.217E-07 | 1.800E-07 | 4.000E-07 |
| 27340 | 1367 | Y1367V | E | 2.1e-07 | 3.5e-07 | 1.8e-07 | 2.467E-07 | 9.074E-08 | 1.800E-07 | 3.500E-07 |
| 27341 | 1368 | C1368A | E | 1.9e-07 | 1.6e-07 | 2.9e-07 | 2.133E-07 | 6.807E-08 | 1.600E-07 | 2.900E-07 |
| 27342 | 1368 | C1368R | E | 1.8e-07 | 1.3e-07 | 1.7e-07 | 1.600E-07 | 2.646E-08 | 1.300E-07 | 1.800E-07 |
| 27343 | 1368 | C1368N | E | 3.1e-07 | 1.9e-07 | 3.2e-07 | 2.733E-07 | 7.234E-08 | 1.900E-07 | 3.200E-07 |
| 27344 | 1368 | C1368D | E | 1.8e-07 | 3.3e-07 | 1.6e-07 | 2.233E-07 | 9.292E-08 | 1.600E-07 | 3.300E-07 |
| 27345 | 1368 | C1368C | E | 1.3e-07 | 2.5e-07 | 1.7e-07 | 1.833E-07 | 6.110E-08 | 1.300E-07 | 2.500E-07 |
| 27346 | 1368 | C1368E | E | 2.7e-07 | 2.5e-07 | 3e-07 | 2.733E-07 | 2.517E-08 | 2.500E-07 | 3.000E-07 |
| 27347 | 1368 | C1368Q | E | 1.8e-07 | 1.9e-07 | 2.6e-07 | 2.100E-07 | 4.359E-08 | 1.800E-07 | 2.600E-07 |
| 27348 | 1368 | C1368G | E | 2.6e-07 | 2e-07 | 1.3e-07 | 1.967E-07 | 6.506E-08 | 1.300E-07 | 2.600E-07 |
| 27349 | 1368 | C1368H | E | 3.9e-07 | 2.9e-07 | 2e-07 | 2.933E-07 | 9.504E-08 | 2.000E-07 | 3.900E-07 |
| 27350 | 1368 | C1368I | E | 1.7e-07 | 2.2e-07 | 3.3e-07 | 2.400E-07 | 8.185E-08 | 1.700E-07 | 3.300E-07 |
| 27351 | 1368 | C1368L | E | 2.1e-07 | 4.4e-07 | 1.7e-07 | 2.733E-07 | 1.457E-07 | 1.700E-07 | 4.400E-07 |
| 27352 | 1368 | C1368K | E | 2.8e-07 | 4.8e-07 | 2.5e-07 | 3.367E-07 | 1.250E-07 | 2.500E-07 | 4.800E-07 |
| 27353 | 1368 | C1368M | E | 2.7e-07 | 3.1e-07 | 2.3e-07 | 2.700E-07 | 4.000E-08 | 2.300E-07 | 3.100E-07 |
| 27354 | 1368 | C1368F | E | 2.2e-07 | 2.5e-07 | 2.3e-07 | 2.333E-07 | 1.528E-08 | 2.200E-07 | 2.500E-07 |
| 27355 | 1368 | C1368P | E | 3.9e-07 | 1.5e-07 | 3.9e-07 | 3.100E-07 | 1.386E-07 | 1.500E-07 | 3.900E-07 |
| 27356 | 1368 | C1368S | E | 1.7e-07 | 4.6e-07 | 2e-07 | 2.767E-07 | 1.595E-07 | 1.700E-07 | 4.600E-07 |
| 27357 | 1368 | C1368T | E | 3.3e-07 | 3.7e-07 | 2.2e-07 | 3.067E-07 | 7.767E-08 | 2.200E-07 | 3.700E-07 |
| 27358 | 1368 | C1368W | E | 2.4e-07 | 4.2e-07 | 2.8e-07 | 3.133E-07 | 9.452E-08 | 2.400E-07 | 4.200E-07 |
| 27359 | 1368 | C1368Y | E | 1.7e-07 | 4.5e-07 | 2.8e-07 | 3.000E-07 | 1.411E-07 | 1.700E-07 | 4.500E-07 |
| 27360 | 1368 | C1368V | E | 1.2e-07 | 2.2e-07 | 2.8e-07 | 2.067E-07 | 8.083E-08 | 1.200E-07 | 2.800E-07 |
| 27361 | 1369 | S1369A | E | 1.7e-07 | 1.8e-07 | 1.8e-07 | 1.767E-07 | 5.774E-09 | 1.700E-07 | 1.800E-07 |
| 27362 | 1369 | S1369R | E | 2.6e-07 | 4.7e-07 | 2.8e-07 | 3.367E-07 | 1.159E-07 | 2.600E-07 | 4.700E-07 |
| 27363 | 1369 | S1369N | E | 4.2e-07 | 2.4e-07 | 3.4e-07 | 3.333E-07 | 9.018E-08 | 2.400E-07 | 4.200E-07 |
| 27364 | 1369 | S1369D | E | 2.2e-07 | 1.3e-07 | 1.8e-07 | 1.767E-07 | 4.509E-08 | 1.300E-07 | 2.200E-07 |
| 27365 | 1369 | S1369C | E | 2.7e-07 | 2.8e-07 | 2.9e-07 | 2.800E-07 | 1.000E-08 | 2.700E-07 | 2.900E-07 |
| 27366 | 1369 | S1369E | E | 4.2e-07 | 1.7e-07 | 2.6e-07 | 2.833E-07 | 1.266E-07 | 1.700E-07 | 4.200E-07 |
| 27367 | 1369 | S1369Q | E | 2.1e-07 | 2.5e-07 | 3.2e-07 | 2.600E-07 | 5.568E-08 | 2.100E-07 | 3.200E-07 |
| 27368 | 1369 | S1369G | E | 5.6e-07 | 2e-07 | 2.2e-07 | 3.267E-07 | 2.023E-07 | 2.000E-07 | 5.600E-07 |
| 27369 | 1369 | S1369H | E | 1.5e-07 | 4.5e-07 | 2.8e-07 | 2.933E-07 | 1.504E-07 | 1.500E-07 | 4.500E-07 |

| | | | | | | | | | | |
|---|---|---|---|---|---|---|---|---|---|---|
| 27370 | 1369 | S1369I | E | 4.7e-07 | 2.8e-07 | 2.3e-07 | 3.267E-07 | 1.266E-07 | 2.300E-07 | 4.700E-07 |
| 27371 | 1369 | S1369L | E | 2.8e-07 | 3.2e-07 | 1.9e-07 | 2.633E-07 | 6.658E-08 | 1.900E-07 | 3.200E-07 |
| 27372 | 1369 | S1369K | E | 1.1e-07 | 1e-07 | 3.3e-07 | 1.800E-07 | 1.300E-07 | 1.000E-07 | 3.300E-07 |
| 27373 | 1369 | S1369M | E | 1.8e-07 | 2.1e-07 | 1.9e-07 | 1.933E-07 | 1.528E-08 | 1.800E-07 | 2.100E-07 |
| 27374 | 1369 | S1369F | E | 2.8e-07 | 3.9e-07 | 1e-07 | 2.567E-07 | 1.464E-07 | 1.000E-07 | 3.900E-07 |
| 27375 | 1369 | S1369P | E | 1.1e-07 | 2.4e-07 | 3.2e-07 | 2.233E-07 | 1.060E-07 | 1.100E-07 | 3.200E-07 |
| 27376 | 1369 | S1369S | E | 1.4e-07 | 2.1e-07 | 9.4e-08 | 1.480E-07 | 5.841E-08 | 9.400E-08 | 2.100E-07 |
| 27377 | 1369 | S1369T | E | 2.9e-07 | 3.2e-07 | 1.6e-07 | 2.567E-07 | 8.505E-08 | 1.600E-07 | 3.200E-07 |
| 27378 | 1369 | S1369W | E | 4.1e-07 | 2e-07 | 1.7e-07 | 2.600E-07 | 1.308E-07 | 1.700E-07 | 4.100E-07 |
| 27379 | 1369 | S1369Y | E | 3.4e-07 | 1.6e-07 | 1.5e-07 | 2.167E-07 | 1.069E-07 | 1.500E-07 | 3.400E-07 |
| 27380 | 1369 | S1369V | E | 3.2e-07 | 2.6e-07 | 3.6e-07 | 3.133E-07 | 5.033E-08 | 2.600E-07 | 3.600E-07 |
| 27381 | 1370 | R1370A | E | 1.9e-07 | 2.9e-07 | 2.7e-07 | 2.500E-07 | 5.292E-08 | 1.900E-07 | 2.900E-07 |
| 27382 | 1370 | R1370R | E | 5e-07 | 4.1e-07 | 4.3e-07 | 4.467E-07 | 4.726E-08 | 4.100E-07 | 5.000E-07 |
| 27383 | 1370 | R1370N | E | 3.2e-07 | 2.6e-07 | 4.7e-07 | 3.500E-07 | 1.082E-07 | 2.600E-07 | 4.700E-07 |
| 27384 | 1370 | R1370D | E | 4.6e-07 | 2.7e-07 | 3.8e-07 | 3.700E-07 | 9.539E-08 | 2.700E-07 | 4.600E-07 |
| 27385 | 1370 | R1370C | E | 1.8e-07 | 3.9e-07 | 7.5e-07 | 4.400E-07 | 2.883E-07 | 1.800E-07 | 7.500E-07 |
| 27386 | 1370 | R1370E | E | 2.8e-07 | 9.3e-08 | 3.2e-07 | 2.310E-07 | 1.212E-07 | 9.300E-08 | 3.200E-07 |
| 27387 | 1370 | R1370Q | E | 2.3e-07 | 1.2e-07 | 2.2e-07 | 1.900E-07 | 6.083E-08 | 1.200E-07 | 2.300E-07 |
| 27388 | 1370 | R1370G | E | 2.1e-07 | 3.6e-07 | 4.9e-07 | 3.533E-07 | 1.401E-07 | 2.100E-07 | 4.900E-07 |
| 27389 | 1370 | R1370H | E | 4.7e-07 | 1.9e-07 | 3.6e-07 | 3.400E-07 | 1.411E-07 | 1.900E-07 | 4.700E-07 |
| 27390 | 1370 | R1370I | E | 2.6e-07 | 2.5e-07 | 4.9e-07 | 3.333E-07 | 1.358E-07 | 2.500E-07 | 4.900E-07 |
| 27391 | 1370 | R1370L | E | 2.9e-07 | 2.7e-07 | 9.4e-07 | 5.000E-07 | 3.812E-07 | 2.700E-07 | 9.400E-07 |
| 27392 | 1370 | R1370K | E | 1.2e-07 | 5.6e-07 | 3.1e-07 | 3.300E-07 | 2.207E-07 | 1.200E-07 | 5.600E-07 |
| 27393 | 1370 | R1370M | E | 1.8e-07 | 2.3e-07 | 3.4e-07 | 2.500E-07 | 8.185E-08 | 1.800E-07 | 3.400E-07 |
| 27394 | 1370 | R1370F | E | 2.1e-07 | 1.3e-07 | 4.3e-07 | 2.567E-07 | 1.553E-07 | 1.300E-07 | 4.300E-07 |
| 27395 | 1370 | R1370P | E | 5.8e-07 | 3.1e-07 | 5.3e-07 | 4.733E-07 | 1.436E-07 | 3.100E-07 | 5.800E-07 |
| 27396 | 1370 | R1370S | E | 3.8e-07 | 2.7e-07 | 3.2e-07 | 3.233E-07 | 5.508E-08 | 2.700E-07 | 3.800E-07 |
| 27397 | 1370 | R1370T | E | 2.5e-07 | 2.3e-07 | 5.9e-07 | 3.567E-07 | 2.023E-07 | 2.300E-07 | 5.900E-07 |
| 27398 | 1370 | R1370W | E | 1.7e-07 | 1.9e-07 | 2.8e-07 | 2.133E-07 | 5.859E-08 | 1.700E-07 | 2.800E-07 |
| 27399 | 1370 | R1370Y | E | 3.4e-07 | 2.3e-07 | 1.9e-07 | 2.533E-07 | 7.767E-08 | 1.900E-07 | 3.400E-07 |
| 27400 | 1370 | R1370V | E | 3.1e-07 | 5.1e-07 | 1.6e-07 | 3.267E-07 | 1.756E-07 | 1.600E-07 | 5.100E-07 |
| 27401 | 1371 | W1371A | E | 2.7e-07 | 5.1e-07 | 2.6e-07 | 3.467E-07 | 1.415E-07 | 2.600E-07 | 5.100E-07 |
| 27402 | 1371 | W1371R | E | 7.6e-08 | 1.7e-07 | 4.8e-07 | 2.420E-07 | 2.114E-07 | 7.600E-08 | 4.800E-07 |
| 27403 | 1371 | W1371N | E | 1.2e-07 | 1e-07 | 2.9e-07 | 1.700E-07 | 1.044E-07 | 1.000E-07 | 2.900E-07 |
| 27404 | 1371 | W1371D | E | 2.3e-07 | 1.6e-07 | 2.9e-07 | 2.267E-07 | 6.506E-08 | 1.600E-07 | 2.900E-07 |
| 27405 | 1371 | W1371C | E | 2.7e-07 | 2.7e-07 | 1.7e-07 | 2.367E-07 | 5.774E-08 | 1.700E-07 | 2.700E-07 |
| 27406 | 1371 | W1371E | E | 1.9e-07 | 1.1e-07 | 9.9e-08 | 1.330E-07 | 4.967E-08 | 9.900E-08 | 1.900E-07 |
| 27407 | 1371 | W1371Q | E | 4.2e-07 | 1.2e-07 | 2.3e-07 | 2.567E-07 | 1.518E-07 | 1.200E-07 | 4.200E-07 |
| 27408 | 1371 | W1371G | E | 2.3e-07 | 2.5e-07 | 3.2e-07 | 2.667E-07 | 4.726E-08 | 2.300E-07 | 3.200E-07 |
| 27409 | 1371 | W1371H | E | 2.6e-07 | 1.2e-07 | 4.6e-07 | 2.800E-07 | 1.709E-07 | 1.200E-07 | 4.600E-07 |
| 27410 | 1371 | W1371I | E | 2.5e-07 | 1.8e-07 | 3e-07 | 2.433E-07 | 6.028E-08 | 1.800E-07 | 3.000E-07 |
| 27411 | 1371 | W1371L | E | 1.3e-07 | 4.2e-07 | 3.5e-07 | 3.000E-07 | 1.513E-07 | 1.300E-07 | 4.200E-07 |
| 27412 | 1371 | W1371K | E | 4.4e-07 | 2.6e-07 | 1.7e-07 | 2.900E-07 | 1.375E-07 | 1.700E-07 | 4.400E-07 |
| 27413 | 1371 | W1371M | E | 2.5e-07 | 4.2e-07 | 2.8e-07 | 3.167E-07 | 9.074E-08 | 2.500E-07 | 4.200E-07 |
| 27414 | 1371 | W1371F | E | 5.1e-07 | 2.8e-07 | 2.7e-07 | 3.533E-07 | 1.358E-07 | 2.700E-07 | 5.100E-07 |
| 27415 | 1371 | W1371P | E | 2.5e-07 | 2.6e-07 | 1.6e-07 | 2.233E-07 | 5.508E-08 | 1.600E-07 | 2.600E-07 |
| 27416 | 1371 | W1371S | E | 2.9e-07 | 2.2e-07 | 3.1e-07 | 2.733E-07 | 4.726E-08 | 2.200E-07 | 3.100E-07 |
| 27417 | 1371 | W1371T | E | 1.6e-07 | 5.9e-07 | 1.2e-07 | 2.900E-07 | 2.606E-07 | 1.200E-07 | 5.900E-07 |
| 27418 | 1371 | W1371W | E | 1.3e-07 | 1.1e-07 | 3.4e-07 | 1.933E-07 | 1.274E-07 | 1.100E-07 | 3.400E-07 |
| 27419 | 1371 | W1371Y | E | 1.7e-07 | 2.4e-07 | 2.8e-07 | 2.300E-07 | 5.568E-08 | 1.700E-07 | 2.800E-07 |
| 27420 | 1371 | W1371V | E | 2.1e-07 | 2.3e-07 | 3e-07 | 2.467E-07 | 4.726E-08 | 2.100E-07 | 3.000E-07 |
| 27421 | 1372 | G1372A | E | 4.1e-07 | 3.6e-07 | 7.6e-08 | 2.820E-07 | 1.801E-07 | 7.600E-08 | 4.100E-07 |
| 27422 | 1372 | G1372R | E | 2.2e-07 | 4.1e-07 | 2.5e-07 | 2.933E-07 | 1.021E-07 | 2.200E-07 | 4.100E-07 |
| 27423 | 1372 | G1372N | E | 2.2e-07 | 1.4e-07 | 2e-07 | 1.867E-07 | 4.163E-08 | 1.400E-07 | 2.200E-07 |
| 27424 | 1372 | G1372D | E | 8.4e-08 | 1.1e-07 | 5.9e-07 | 2.613E-07 | 2.849E-07 | 8.400E-08 | 5.900E-07 |
| 27425 | 1372 | G1372C | E | 1.7e-07 | 4.4e-07 | 3.6e-07 | 3.233E-07 | 1.387E-07 | 1.700E-07 | 4.400E-07 |
| 27426 | 1372 | G1372E | E | 2.7e-07 | 2.4e-07 | 1.4e-07 | 2.167E-07 | 6.807E-08 | 1.400E-07 | 2.700E-07 |
| 27427 | 1372 | G1372Q | E | 1.3e-07 | 1.6e-07 | 1.6e-07 | 1.500E-07 | 1.732E-08 | 1.300E-07 | 1.600E-07 |
| 27428 | 1372 | G1372G | E | 1.4e-07 | 3.1e-07 | 4.5e-07 | 3.000E-07 | 1.552E-07 | 1.400E-07 | 4.500E-07 |
| 27429 | 1372 | G1372H | E | 1.1e-07 | 1.8e-07 | 1.6e-07 | 1.500E-07 | 3.606E-08 | 1.100E-07 | 1.800E-07 |
| 27430 | 1372 | G1372I | E | 1.7e-07 | 4.6e-07 | 1.3e-07 | 2.533E-07 | 1.801E-07 | 1.300E-07 | 4.600E-07 |
| 27431 | 1372 | G1372L | E | 2.1e-07 | 2.9e-07 | 2.3e-07 | 2.433E-07 | 4.163E-08 | 2.100E-07 | 2.900E-07 |
| 27432 | 1372 | G1372K | E | 2.9e-07 | 3.3e-07 | 1.5e-07 | 2.567E-07 | 9.452E-08 | 1.500E-07 | 3.300E-07 |
| 27433 | 1372 | G1372M | E | 2.1e-07 | 2.4e-07 | 4e-07 | 2.833E-07 | 1.021E-07 | 2.100E-07 | 4.000E-07 |
| 27434 | 1372 | G1372F | E | 1.9e-07 | 2.8e-07 | 3.3e-07 | 2.667E-07 | 7.095E-08 | 1.900E-07 | 3.300E-07 |
| 27435 | 1372 | G1372P | E | 2.3e-07 | 1e-07 | 1.5e-07 | 1.600E-07 | 6.557E-08 | 1.000E-07 | 2.300E-07 |
| 27436 | 1372 | G1372S | E | 2e-07 | 1e-07 | 1.9e-07 | 1.633E-07 | 5.508E-08 | 1.000E-07 | 2.000E-07 |
| 27437 | 1372 | G1372T | E | 1.3e-07 | 8.1e-08 | 2e-07 | 1.370E-07 | 5.981E-08 | 8.100E-08 | 2.000E-07 |
| 27438 | 1372 | G1372W | E | 1.1e-07 | 6.4e-07 | 2.6e-07 | 3.367E-07 | 2.732E-07 | 1.100E-07 | 6.400E-07 |
| 27439 | 1372 | G1372Y | E | 1.3e-07 | 1.4e-07 | 1.1e-07 | 1.267E-07 | 1.528E-08 | 1.100E-07 | 1.400E-07 |
| 27440 | 1372 | G1372V | E | 2e-07 | 3.2e-07 | 2.7e-07 | 2.633E-07 | 6.028E-08 | 2.000E-07 | 3.200E-07 |
| 27441 | 1373 | G1373A | E | 5.4e-07 | 2.4e-07 | 5e-07 | 4.267E-07 | 1.629E-07 | 2.400E-07 | 5.400E-07 |
| 27442 | 1373 | G1373R | E | 1.2e-07 | 8.2e-08 | 1.2e-07 | 1.073E-07 | 2.194E-08 | 8.200E-08 | 1.200E-07 |
| 27443 | 1373 | G1373N | E | 8.8e-08 | 4.4e-08 | 6.3e-08 | 6.500E-08 | 2.207E-08 | 4.400E-08 | 8.800E-08 |
| 27444 | 1373 | G1373D | E | 1e-07 | 6.2e-08 | 4.6e-07 | 2.073E-07 | 2.196E-07 | 6.200E-08 | 4.600E-07 |
| 27445 | 1373 | G1373C | E | 2.8e-07 | 2.7e-07 | 2.2e-07 | 2.567E-07 | 3.215E-08 | 2.200E-07 | 2.800E-07 |
| 27446 | 1373 | G1373E | E | 2.9e-07 | 7.7e-08 | 7.4e-08 | 1.470E-07 | 1.239E-07 | 7.400E-08 | 2.900E-07 |
| 27447 | 1373 | G1373Q | E | 7.9e-08 | 2.7e-08 | 2.4e-08 | 4.333E-08 | 3.092E-08 | 2.400E-08 | 7.900E-08 |
| 27448 | 1373 | G1373G | E | 1.5e-07 | 2.8e-07 | 1.8e-07 | 2.033E-07 | 6.807E-08 | 1.500E-07 | 2.800E-07 |
| 27449 | 1373 | G1373H | E | 1.2e-07 | 9.2e-08 | 7.8e-08 | 9.667E-08 | 2.139E-08 | 7.800E-08 | 1.200E-07 |
| 27450 | 1373 | G1373I | E | 3.3e-07 | 6.6e-07 | 2.9e-07 | 4.267E-07 | 2.031E-07 | 2.900E-07 | 6.600E-07 |
| 27451 | 1373 | G1373L | E | 1.6e-07 | 2.2e-07 | 4.2e-07 | 2.667E-07 | 1.361E-07 | 1.600E-07 | 4.200E-07 |
| 27452 | 1373 | G1373K | E | 1.6e-07 | 1.4e-07 | 3.7e-07 | 2.233E-07 | 1.274E-07 | 1.400E-07 | 3.700E-07 |
| 27453 | 1373 | G1373M | E | 2.3e-07 | 5.6e-07 | 2e-07 | 3.300E-07 | 1.997E-07 | 2.000E-07 | 5.600E-07 |
| 27454 | 1373 | G1373F | E | 9.6e-08 | 4.4e-07 | 7e-08 | 2.020E-07 | 2.065E-07 | 7.000E-08 | 4.400E-07 |

| | | | | | | | | | | |
|---|---|---|---|---|---|---|---|---|---|---|
| 27455 | 1373 | G1373P | E | 3.1e-07 | 2.3e-07 | 3.9e-07 | 3.100E-07 | 8.000E-08 | 2.300E-07 | 3.900E-07 |
| 27456 | 1373 | G1373S | E | 8.5e-08 | 9.2e-08 | 7.1e-08 | 8.267E-08 | 1.069E-08 | 7.100E-08 | 9.200E-08 |
| 27457 | 1373 | G1373T | E | 1.1e-07 | 2.3e-08 | 6.5e-08 | 6.600E-08 | 4.351E-08 | 2.300E-08 | 1.100E-07 |
| 27458 | 1373 | G1373W | E | 1.1e-07 | 2.1e-07 | 1.4e-07 | 1.533E-07 | 5.132E-08 | 1.100E-07 | 2.100E-07 |
| 27459 | 1373 | G1373Y | E | 2.8e-07 | 2.1e-07 | 1e-07 | 1.967E-07 | 9.074E-08 | 1.000E-07 | 2.800E-07 |
| 27460 | 1373 | G1373V | E | 1.9e-07 | 1.6e-07 | 1.5e-07 | 1.667E-07 | 2.082E-08 | 1.500E-07 | 1.900E-07 |
| 27461 | 1374 | D1374A | E | 3.3e-07 | 7.4e-07 | 5e-07 | 5.233E-07 | 2.060E-07 | 3.300E-07 | 7.400E-07 |
| 27462 | 1374 | D1374R | E | 9.2e-08 | 1.6e-07 | 9.2e-08 | 1.147E-07 | 3.926E-08 | 9.200E-08 | 1.600E-07 |
| 27463 | 1374 | D1374N | E | 6.1e-07 | 3.5e-07 | 4.1e-07 | 4.567E-07 | 1.361E-07 | 3.500E-07 | 6.100E-07 |
| 27464 | 1374 | D1374D | E | 2.2e-07 | 2.1e-07 | 2.4e-07 | 2.233E-07 | 1.528E-08 | 2.100E-07 | 2.400E-07 |
| 27465 | 1374 | D1374C | E | 3.2e-07 | 1.3e-07 | 2.2e-07 | 2.233E-07 | 9.504E-08 | 1.300E-07 | 3.200E-07 |
| 27466 | 1374 | D1374E | E | 2.3e-07 | 2e-07 | 2.2e-07 | 2.167E-07 | 1.528E-08 | 2.000E-07 | 2.300E-07 |
| 27467 | 1374 | D1374Q | E | 1.1e-07 | 2e-07 | 1.8e-07 | 1.633E-07 | 4.726E-08 | 1.100E-07 | 2.000E-07 |
| 27468 | 1374 | D1374G | E | 2e-07 | 4.1e-07 | 2.4e-07 | 2.833E-07 | 1.115E-07 | 2.000E-07 | 4.100E-07 |
| 27469 | 1374 | D1374H | E | 1.1e-07 | 1.9e-07 | 4.9e-07 | 2.633E-07 | 2.003E-07 | 1.100E-07 | 4.900E-07 |
| 27470 | 1374 | D1374I | E | 3.9e-07 | 3.2e-07 | 4.9e-07 | 4.000E-07 | 8.544E-08 | 3.200E-07 | 4.900E-07 |
| 27471 | 1374 | D1374L | E | 2.7e-07 | 1.9e-07 | 9.8e-07 | 4.800E-07 | 4.349E-07 | 1.900E-07 | 9.800E-07 |
| 27472 | 1374 | D1374K | E | 2.6e-07 | 3.3e-07 | 1.5e-07 | 2.467E-07 | 9.074E-08 | 1.500E-07 | 3.300E-07 |
| 27473 | 1374 | D1374M | E | 1.3e-07 | 3.8e-07 | 4.8e-07 | 3.300E-07 | 1.803E-07 | 1.300E-07 | 4.800E-07 |
| 27474 | 1374 | D1374F | E | 1.2e-07 | 4e-07 | 3.1e-07 | 2.767E-07 | 1.429E-07 | 1.200E-07 | 4.000E-07 |
| 27475 | 1374 | D1374P | E | 2.6e-07 | 5.2e-07 | 3.1e-07 | 3.633E-07 | 1.380E-07 | 2.600E-07 | 5.200E-07 |
| 27476 | 1374 | D1374S | E | 3e-07 | 1.9e-07 | 2.4e-07 | 2.433E-07 | 5.508E-08 | 1.900E-07 | 3.000E-07 |
| 27477 | 1374 | D1374T | E | 3e-07 | 1.9e-07 | 3.3e-07 | 2.733E-07 | 7.371E-08 | 1.900E-07 | 3.300E-07 |
| 27478 | 1374 | D1374W | E | 1.4e-07 | 2.2e-07 | 3.2e-07 | 2.267E-07 | 9.018E-08 | 1.400E-07 | 3.200E-07 |
| 27479 | 1374 | D1374Y | E | 4.7e-07 | 8e-07 | 2.5e-07 | 5.067E-07 | 2.768E-07 | 2.500E-07 | 8.000E-07 |
| 27480 | 1374 | D1374V | E | 2e-07 | 6e-07 | 3.3e-07 | 3.767E-07 | 2.040E-07 | 2.000E-07 | 6.000E-07 |
| 27481 | 1375 | G1375A | E | 1.8e-07 | 2.6e-07 | 2.7e-07 | 2.367E-07 | 4.933E-08 | 1.800E-07 | 2.700E-07 |
| 27482 | 1375 | G1375R | E | 1.8e-07 | 9.6e-08 | 1.8e-07 | 1.520E-07 | 4.850E-08 | 9.600E-08 | 1.800E-07 |
| 27483 | 1375 | G1375N | E | 5.7e-08 | 9.5e-08 | 9.2e-08 | 8.133E-08 | 2.113E-08 | 5.700E-08 | 9.500E-08 |
| 27484 | 1375 | G1375D | E | 2.9e-07 | 1.8e-07 | 1.7e-07 | 2.133E-07 | 6.658E-08 | 1.700E-07 | 2.900E-07 |
| 27485 | 1375 | G1375C | E | 7e-08 | 1.7e-07 | 2.7e-07 | 1.700E-07 | 1.000E-07 | 7.000E-08 | 2.700E-07 |
| 27486 | 1375 | G1375E | E | 1.5e-07 | 1.9e-07 | 1.6e-07 | 1.667E-07 | 2.082E-08 | 1.500E-07 | 1.900E-07 |
| 27487 | 1375 | G1375Q | E | 1e-07 | 2.7e-08 | 1.2e-07 | 8.233E-08 | 4.895E-08 | 2.700E-08 | 1.200E-07 |
| 27488 | 1375 | G1375G | E | 8.5e-08 | 3.4e-07 | 2.2e-07 | 2.150E-07 | 1.276E-07 | 8.500E-08 | 3.400E-07 |
| 27489 | 1375 | G1375H | E | 8.9e-08 | 1.1e-07 | 1.1e-07 | 1.030E-07 | 1.212E-08 | 8.900E-08 | 1.100E-07 |
| 27490 | 1375 | G1375I | E | 1.2e-07 | 2.2e-07 | 3.4e-07 | 2.267E-07 | 1.102E-07 | 1.200E-07 | 3.400E-07 |
| 27491 | 1375 | G1375L | E | 1.7e-07 | 6.7e-07 | 1.3e-07 | 3.233E-07 | 3.009E-07 | 1.300E-07 | 6.700E-07 |
| 27492 | 1375 | G1375K | E | 1.8e-07 | 1.8e-07 | 7.6e-08 | 1.453E-07 | 6.004E-08 | 7.600E-08 | 1.800E-07 |
| 27493 | 1375 | G1375M | E | 2.9e-07 | 4.5e-07 | 1.8e-07 | 3.067E-07 | 1.358E-07 | 1.800E-07 | 4.500E-07 |
| 27494 | 1375 | G1375F | E | 1.9e-07 | 7.5e-08 | 2e-07 | 1.550E-07 | 6.946E-08 | 7.500E-08 | 2.000E-07 |
| 27495 | 1375 | G1375P | E | 4.7e-07 | 3.7e-07 | 3.9e-07 | 4.100E-07 | 5.292E-08 | 3.700E-07 | 4.700E-07 |
| 27496 | 1375 | G1375S | E | 5.8e-08 | 6.9e-08 | 1.1e-07 | 7.900E-08 | 2.740E-08 | 5.800E-08 | 1.100E-07 |
| 27497 | 1375 | G1375T | E | 9e-08 | 1e-07 | 7.5e-08 | 8.833E-08 | 1.258E-08 | 7.500E-08 | 1.000E-07 |
| 27498 | 1375 | G1375W | E | 2.5e-07 | 4e-07 | 2.1e-07 | 2.867E-07 | 1.002E-07 | 2.100E-07 | 4.000E-07 |
| 27499 | 1375 | G1375Y | E | 1.8e-07 | 3.2e-07 | 2e-07 | 2.333E-07 | 7.572E-08 | 1.800E-07 | 3.200E-07 |
| 27500 | 1375 | G1375V | E | 2.2e-07 | 3.4e-07 | 1.9e-07 | 2.500E-07 | 7.937E-08 | 1.900E-07 | 3.400E-07 |
| 27501 | 1376 | F1376A | E | 1.7e-07 | 2.1e-07 | 4.9e-07 | 2.900E-07 | 1.744E-07 | 1.700E-07 | 4.900E-07 |
| 27502 | 1376 | F1376R | E | 1.4e-07 | 6.2e-07 | 1.4e-07 | 3.000E-07 | 2.771E-07 | 1.400E-07 | 6.200E-07 |
| 27503 | 1376 | F1376N | E | 2.9e-07 | 3.2e-07 | 2.5e-07 | 2.867E-07 | 3.512E-08 | 2.500E-07 | 3.200E-07 |
| 27504 | 1376 | F1376D | E | 2.6e-07 | 5.1e-07 | 4.5e-07 | 4.067E-07 | 1.305E-07 | 2.600E-07 | 5.100E-07 |
| 27505 | 1376 | F1376C | E | 2e-07 | 2.8e-07 | 5e-07 | 3.267E-07 | 1.553E-07 | 2.000E-07 | 5.000E-07 |
| 27506 | 1376 | F1376E | E | 3.4e-07 | 2.7e-07 | 1.4e-07 | 2.500E-07 | 1.015E-07 | 1.400E-07 | 3.400E-07 |
| 27507 | 1376 | F1376Q | E | 1.1e-07 | 1e-07 | 7.1e-08 | 9.367E-08 | 2.026E-08 | 7.100E-08 | 1.100E-07 |
| 27508 | 1376 | F1376G | E | 1.9e-07 | 7e-08 | 2.6e-07 | 1.733E-07 | 9.609E-08 | 7.000E-08 | 2.600E-07 |
| 27509 | 1376 | F1376H | E | 3.2e-07 | 2e-07 | 2.6e-07 | 2.600E-07 | 6.000E-08 | 2.000E-07 | 3.200E-07 |
| 27510 | 1376 | F1376I | E | 3.1e-07 | 4.5e-07 | 3e-07 | 3.533E-07 | 8.386E-08 | 3.000E-07 | 4.500E-07 |
| 27511 | 1376 | F1376L | E | 1.2e-07 | 3.4e-07 | 1.8e-07 | 2.133E-07 | 1.137E-07 | 1.200E-07 | 3.400E-07 |
| 27512 | 1376 | F1376K | E | 1.2e-07 | 1.8e-07 | 1.5e-07 | 1.500E-07 | 3.000E-08 | 1.200E-07 | 1.800E-07 |
| 27513 | 1376 | F1376M | E | 5.7e-07 | 5.2e-07 | 1.4e-07 | 4.100E-07 | 2.352E-07 | 1.400E-07 | 5.700E-07 |
| 27514 | 1376 | F1376F | E | 5.8e-07 | 1e-07 | 2.4e-07 | 3.067E-07 | 2.468E-07 | 1.000E-07 | 5.800E-07 |
| 27515 | 1376 | F1376P | E | 4.4e-07 | 3.5e-07 | 2.4e-07 | 3.433E-07 | 1.002E-07 | 2.400E-07 | 4.400E-07 |
| 27516 | 1376 | F1376S | E | 2.4e-07 | 1.6e-07 | 1.9e-07 | 1.967E-07 | 4.041E-08 | 1.600E-07 | 2.400E-07 |
| 27517 | 1376 | F1376T | E | 3.6e-07 | 2.3e-07 | 6e-07 | 3.967E-07 | 1.877E-07 | 2.300E-07 | 6.000E-07 |
| 27518 | 1376 | F1376W | E | 2.2e-07 | 3.2e-07 | 2.7e-07 | 2.700E-07 | 5.000E-08 | 2.200E-07 | 3.200E-07 |
| 27519 | 1376 | F1376Y | E | 1.9e-07 | 3.2e-07 | 3.4e-07 | 2.833E-07 | 8.145E-08 | 1.900E-07 | 3.400E-07 |
| 27520 | 1376 | F1376V | E | 4.7e-07 | 1.7e-07 | 2.3e-07 | 2.900E-07 | 1.587E-07 | 1.700E-07 | 4.700E-07 |
| 27521 | 1377 | Y1377A | E | 2e-07 | 4.2e-07 | 3.5e-07 | 3.233E-07 | 1.124E-07 | 2.000E-07 | 4.200E-07 |
| 27522 | 1377 | Y1377R | E | 1.5e-07 | 8.5e-08 | 2.4e-07 | 1.583E-07 | 7.784E-08 | 8.500E-08 | 2.400E-07 |
| 27523 | 1377 | Y1377N | E | 2.3e-07 | 1.3e-07 | 1.1e-07 | 1.567E-07 | 6.429E-08 | 1.100E-07 | 2.300E-07 |
| 27524 | 1377 | Y1377D | E | 2e-07 | 2e-07 | 1.8e-07 | 1.933E-07 | 1.155E-08 | 1.800E-07 | 2.000E-07 |
| 27525 | 1377 | Y1377C | E | 2e-07 | 1.5e-07 | 1.9e-07 | 1.800E-07 | 2.646E-08 | 1.500E-07 | 2.000E-07 |
| 27526 | 1377 | Y1377E | E | 4.3e-07 | 1.6e-07 | 1.3e-07 | 2.400E-07 | 1.652E-07 | 1.300E-07 | 4.300E-07 |
| 27527 | 1377 | Y1377Q | E | 1.4e-07 | 1.1e-07 | 1.6e-07 | 1.367E-07 | 2.517E-08 | 1.100E-07 | 1.600E-07 |
| 27528 | 1377 | Y1377G | E | 2.7e-07 | 2.3e-07 | 5.3e-07 | 3.433E-07 | 1.629E-07 | 2.300E-07 | 5.300E-07 |
| 27529 | 1377 | Y1377H | E | 1e-07 | 9.9e-08 | 1.3e-07 | 1.097E-07 | 1.762E-08 | 9.900E-08 | 1.300E-07 |
| 27530 | 1377 | Y1377I | E | 1.8e-07 | 1.6e-07 | 1.6e-07 | 1.667E-07 | 1.155E-08 | 1.600E-07 | 1.800E-07 |
| 27531 | 1377 | Y1377L | E | 3.3e-07 | 4.9e-07 | 2.6e-07 | 3.600E-07 | 1.179E-07 | 2.600E-07 | 4.900E-07 |
| 27532 | 1377 | Y1377K | E | 2.6e-07 | 9.4e-08 | 1.2e-07 | 1.580E-07 | 8.929E-08 | 9.400E-08 | 2.600E-07 |
| 27533 | 1377 | Y1377M | E | 2.4e-07 | 2.2e-07 | 3.4e-07 | 2.667E-07 | 6.429E-08 | 2.200E-07 | 3.400E-07 |
| 27534 | 1377 | Y1377F | E | 3.2e-07 | 3e-07 | 2.8e-07 | 3.000E-07 | 2.000E-08 | 2.800E-07 | 3.200E-07 |
| 27535 | 1377 | Y1377P | E | 1.6e-07 | 7.2e-07 | 3.1e-07 | 3.967E-07 | 2.899E-07 | 1.600E-07 | 7.200E-07 |
| 27536 | 1377 | Y1377S | E | 2.4e-07 | 7.1e-08 | 1.6e-07 | 1.570E-07 | 8.454E-08 | 7.100E-08 | 2.400E-07 |
| 27537 | 1377 | Y1377T | E | 4.6e-08 | 1.7e-07 | 1.1e-07 | 1.087E-07 | 6.201E-08 | 4.600E-08 | 1.700E-07 |
| 27538 | 1377 | Y1377W | E | 2.1e-07 | 3.5e-07 | 8.3e-08 | 2.143E-07 | 1.336E-07 | 8.300E-08 | 3.500E-07 |
| 27539 | 1377 | Y1377Y | E | 2.6e-07 | 4.6e-07 | 2.2e-07 | 3.133E-07 | 1.286E-07 | 2.200E-07 | 4.600E-07 |

| | | | | | | | | | | |
|---|---|---|---|---|---|---|---|---|---|---|
| 27540 | 1377 | Y1377V | E | 6.5e-07 | 5.7e-07 | 2.1e-07 | 4.767E-07 | 2.344E-07 | 2.100E-07 | 6.500E-07 |
| 27541 | 1378 | A1378A | E | 1.2e-07 | 6.3e-07 | 2e-07 | 3.167E-07 | 2.743E-07 | 1.200E-07 | 6.300E-07 |
| 27542 | 1378 | A1378R | E | 3.9e-07 | 2.1e-07 | 2.1e-07 | 2.700E-07 | 1.039E-07 | 2.100E-07 | 3.900E-07 |
| 27543 | 1378 | A1378N | E | 2.2e-07 | 2.7e-07 | 4e-07 | 2.967E-07 | 9.292E-08 | 2.200E-07 | 4.000E-07 |
| 27544 | 1378 | A1378D | E | 1.9e-07 | 3.3e-07 | 1.8e-07 | 2.333E-07 | 8.386E-08 | 1.800E-07 | 3.300E-07 |
| 27545 | 1378 | A1378C | E | 1.6e-07 | 2.4e-07 | 2.4e-07 | 2.133E-07 | 4.619E-08 | 1.600E-07 | 2.400E-07 |
| 27546 | 1378 | A1378E | E | 1.8e-07 | 1.9e-07 | 1.1e-07 | 1.600E-07 | 4.359E-08 | 1.100E-07 | 1.900E-07 |
| 27547 | 1378 | A1378Q | E | 4e-07 | 2.8e-07 | 2.1e-07 | 2.967E-07 | 9.609E-08 | 2.100E-07 | 4.000E-07 |
| 27548 | 1378 | A1378G | E | 2.1e-07 | 3.1e-07 | 1.9e-07 | 2.367E-07 | 6.429E-08 | 1.900E-07 | 3.100E-07 |
| 27549 | 1378 | A1378H | E | 2.3e-07 | 2e-07 | 1.3e-07 | 1.867E-07 | 5.132E-08 | 1.300E-07 | 2.300E-07 |
| 27550 | 1378 | A1378I | E | 2.4e-07 | 3.8e-07 | 4.4e-07 | 3.533E-07 | 1.026E-07 | 2.400E-07 | 4.400E-07 |
| 27551 | 1378 | A1378L | E | 3.2e-07 | 2.3e-07 | 2.8e-07 | 2.767E-07 | 4.509E-08 | 2.300E-07 | 3.200E-07 |
| 27552 | 1378 | A1378K | E | 2.6e-07 | 3.9e-07 | 2.3e-07 | 2.933E-07 | 8.505E-08 | 2.300E-07 | 3.900E-07 |
| 27553 | 1378 | A1378M | E | 1.4e-07 | 2.1e-07 | 1.2e-07 | 1.567E-07 | 4.726E-08 | 1.200E-07 | 2.100E-07 |
| 27554 | 1378 | A1378F | E | 2.4e-07 | 2.2e-07 | 2.2e-07 | 2.267E-07 | 1.155E-08 | 2.200E-07 | 2.400E-07 |
| 27555 | 1378 | A1378P | E | 3.5e-07 | 2.8e-07 | 2.3e-07 | 2.867E-07 | 6.028E-08 | 2.300E-07 | 3.500E-07 |
| 27556 | 1378 | A1378S | E | 2.8e-07 | 3.3e-07 | 1.7e-07 | 2.600E-07 | 8.185E-08 | 1.700E-07 | 3.300E-07 |
| 27557 | 1378 | A1378T | E | 1.5e-07 | 3e-07 | 3.2e-07 | 2.567E-07 | 9.292E-08 | 1.500E-07 | 3.200E-07 |
| 27558 | 1378 | A1378W | E | 1.6e-07 | 2.5e-07 | 8.8e-07 | 4.300E-07 | 3.923E-07 | 1.600E-07 | 8.800E-07 |
| 27559 | 1378 | A1378Y | E | 3.8e-07 | 3.5e-07 | 2.1e-07 | 3.133E-07 | 9.074E-08 | 2.100E-07 | 3.800E-07 |
| 27560 | 1378 | A1378V | E | 1.3e-07 | 2.9e-07 | 3.3e-07 | 2.500E-07 | 1.058E-07 | 1.300E-07 | 3.300E-07 |
| 27561 | 1379 | M1379A | E | 3.1e-07 | 2.6e-07 | 2e-07 | 2.567E-07 | 5.508E-08 | 2.000E-07 | 3.100E-07 |
| 27562 | 1379 | M1379R | E | 2.5e-07 | 3.4e-07 | 5.1e-07 | 3.667E-07 | 1.320E-07 | 2.500E-07 | 5.100E-07 |
| 27563 | 1379 | M1379N | E | 4e-07 | 3.1e-07 | 4.2e-07 | 3.767E-07 | 5.859E-08 | 3.100E-07 | 4.200E-07 |
| 27564 | 1379 | M1379D | E | 2.6e-07 | 2.3e-07 | 1.4e-07 | 2.100E-07 | 6.245E-08 | 1.400E-07 | 2.600E-07 |
| 27565 | 1379 | M1379C | E | 2.1e-07 | 2.4e-07 | 2.7e-07 | 2.400E-07 | 3.000E-08 | 2.100E-07 | 2.700E-07 |
| 27566 | 1379 | M1379E | E | 5e-07 | 4.2e-07 | 3.4e-07 | 4.200E-07 | 8.000E-08 | 3.400E-07 | 5.000E-07 |
| 27567 | 1379 | M1379Q | E | 1.6e-07 | 3e-07 | 5.7e-07 | 3.433E-07 | 2.084E-07 | 1.600E-07 | 5.700E-07 |
| 27568 | 1379 | M1379G | E | 2.7e-07 | 4.1e-07 | 1.6e-07 | 2.800E-07 | 1.253E-07 | 1.600E-07 | 4.100E-07 |
| 27569 | 1379 | M1379H | E | 3e-07 | 2.6e-07 | 3.7e-07 | 3.100E-07 | 5.568E-08 | 2.600E-07 | 3.700E-07 |
| 27570 | 1379 | M1379I | E | 1.7e-07 | 5.1e-07 | 3.6e-07 | 3.467E-07 | 1.704E-07 | 1.700E-07 | 5.100E-07 |
| 27571 | 1379 | M1379L | E | 2.8e-07 | 4.4e-07 | 2e-07 | 3.067E-07 | 1.222E-07 | 2.000E-07 | 4.400E-07 |
| 27572 | 1379 | M1379K | E | 2.2e-07 | 5.7e-07 | 3.2e-07 | 3.700E-07 | 1.803E-07 | 2.200E-07 | 5.700E-07 |
| 27573 | 1379 | M1379M | E | 3.4e-07 | 3.4e-07 | 3e-07 | 3.267E-07 | 2.309E-08 | 3.000E-07 | 3.400E-07 |
| 27574 | 1379 | M1379F | E | 3.2e-07 | 6.5e-07 | 3.3e-07 | 4.333E-07 | 1.877E-07 | 3.200E-07 | 6.500E-07 |
| 27575 | 1379 | M1379P | E | 2.5e-07 | 1.3e-07 | 3.4e-07 | 2.400E-07 | 1.054E-07 | 1.300E-07 | 3.400E-07 |
| 27576 | 1379 | M1379S | E | 4.8e-07 | 1.7e-07 | 1.8e-07 | 2.767E-07 | 1.762E-07 | 1.700E-07 | 4.800E-07 |
| 27577 | 1379 | M1379T | E | 2e-07 | 1.1e-07 | 3.1e-07 | 2.067E-07 | 1.002E-07 | 1.100E-07 | 3.100E-07 |
| 27578 | 1379 | M1379W | E | 4.2e-07 | 4.1e-07 | 2.1e-07 | 3.467E-07 | 1.185E-07 | 2.100E-07 | 4.200E-07 |
| 27579 | 1379 | M1379Y | E | 2.5e-07 | 3e-07 | 1.8e-07 | 2.433E-07 | 6.028E-08 | 1.800E-07 | 3.000E-07 |
| 27580 | 1379 | M1379V | E | 1.1e-07 | 4.4e-07 | 6.2e-07 | 3.900E-07 | 2.587E-07 | 1.100E-07 | 6.200E-07 |
| 27581 | 1380 | D1380A | E | 4.3e-07 | 1.8e-07 | 1.5e-07 | 2.533E-07 | 1.537E-07 | 1.500E-07 | 4.300E-07 |
| 27582 | 1380 | D1380R | E | 1.2e-07 | 2.2e-07 | 1.3e-07 | 1.567E-07 | 5.508E-08 | 1.200E-07 | 2.200E-07 |
| 27583 | 1380 | D1380N | E | 3.6e-07 | 1.8e-07 | 3e-07 | 2.800E-07 | 9.165E-08 | 1.800E-07 | 3.600E-07 |
| 27584 | 1380 | D1380D | E | 4e-07 | 2.5e-07 | 2.9e-07 | 3.133E-07 | 7.767E-08 | 2.500E-07 | 4.000E-07 |
| 27585 | 1380 | D1380C | E | 1.9e-07 | 2e-07 | 2.9e-07 | 2.267E-07 | 5.508E-08 | 1.900E-07 | 2.900E-07 |
| 27586 | 1380 | D1380E | E | 1.8e-07 | 4e-07 | 1.8e-07 | 2.533E-07 | 1.270E-07 | 1.800E-07 | 4.000E-07 |
| 27587 | 1380 | D1380Q | E | 1.7e-07 | 2.7e-07 | 2.6e-07 | 2.333E-07 | 5.508E-08 | 1.700E-07 | 2.700E-07 |
| 27588 | 1380 | D1380G | E | 2.1e-07 | 3.3e-07 | 1.8e-07 | 2.400E-07 | 7.937E-08 | 1.800E-07 | 3.300E-07 |
| 27589 | 1380 | D1380H | E | 1.9e-07 | 2.1e-07 | 3.4e-07 | 2.467E-07 | 8.145E-08 | 1.900E-07 | 3.400E-07 |
| 27590 | 1380 | D1380I | E | 7.6e-07 | 3.8e-07 | 1.6e-07 | 4.333E-07 | 3.035E-07 | 1.600E-07 | 7.600E-07 |
| 27591 | 1380 | D1380L | E | 2.7e-07 | 2.2e-07 | 2.8e-07 | 2.567E-07 | 3.215E-08 | 2.200E-07 | 2.800E-07 |
| 27592 | 1380 | D1380K | E | 3.3e-07 | 1.7e-07 | 1.4e-07 | 2.133E-07 | 1.021E-07 | 1.400E-07 | 3.300E-07 |
| 27593 | 1380 | D1380M | E | 1.2e-07 | 2.4e-07 | 2.5e-07 | 2.033E-07 | 7.234E-08 | 1.200E-07 | 2.500E-07 |
| 27594 | 1380 | D1380F | E | 3e-07 | 2.8e-07 | 3.8e-07 | 3.200E-07 | 5.292E-08 | 2.800E-07 | 3.800E-07 |
| 27595 | 1380 | D1380P | E | 2.4e-07 | 2e-07 | 2.7e-07 | 2.367E-07 | 3.512E-08 | 2.000E-07 | 2.700E-07 |
| 27596 | 1380 | D1380S | E | 1.8e-07 | 1.8e-07 | 2.8e-07 | 2.133E-07 | 5.774E-08 | 1.800E-07 | 2.800E-07 |
| 27597 | 1380 | D1380T | E | 2.2e-07 | 2.2e-07 | 3.1e-07 | 2.500E-07 | 5.196E-08 | 2.200E-07 | 3.100E-07 |
| 27598 | 1380 | D1380W | E | 4e-07 | 3.5e-07 | 3.7e-07 | 3.733E-07 | 2.517E-08 | 3.500E-07 | 4.000E-07 |
| 27599 | 1380 | D1380Y | E | 8.8e-08 | 1.2e-07 | 2e-07 | 1.360E-07 | 5.769E-08 | 8.800E-08 | 2.000E-07 |
| 27600 | 1380 | D1380V | E | 2.2e-07 | 2.5e-07 | 2.3e-07 | 2.333E-07 | 1.528E-08 | 2.200E-07 | 2.500E-07 |
| 27601 | 1381 | Y1381A | E | 1.2e-07 | 1.5e-07 | 1.7e-07 | 1.467E-07 | 2.517E-08 | 1.200E-07 | 1.700E-07 |
| 27602 | 1381 | Y1381R | E | 1.3e-07 | 2.1e-07 | 3.4e-07 | 2.267E-07 | 1.060E-07 | 1.300E-07 | 3.400E-07 |
| 27603 | 1381 | Y1381N | E | 1.5e-07 | 1.8e-07 | 2e-07 | 1.767E-07 | 2.517E-08 | 1.500E-07 | 2.000E-07 |
| 27604 | 1381 | Y1381D | E | 2.6e-07 | 2.6e-07 | 4.6e-07 | 3.267E-07 | 1.155E-07 | 2.600E-07 | 4.600E-07 |
| 27605 | 1381 | Y1381C | E | 1.7e-07 | 3.5e-07 | 3.3e-07 | 2.833E-07 | 9.866E-08 | 1.700E-07 | 3.500E-07 |
| 27606 | 1381 | Y1381E | E | 3.1e-07 | 5.8e-07 | 3.3e-07 | 4.067E-07 | 1.504E-07 | 3.100E-07 | 5.800E-07 |
| 27607 | 1381 | Y1381Q | E | 2.8e-07 | 2.4e-07 | 2e-07 | 2.400E-07 | 4.000E-08 | 2.000E-07 | 2.800E-07 |
| 27608 | 1381 | Y1381G | E | 1.6e-07 | 3.4e-07 | 3.5e-07 | 2.833E-07 | 1.069E-07 | 1.600E-07 | 3.500E-07 |
| 27609 | 1381 | Y1381H | E | 2.9e-07 | 2.4e-07 | 2.8e-07 | 2.700E-07 | 2.646E-08 | 2.400E-07 | 2.900E-07 |
| 27610 | 1381 | Y1381I | E | 3.7e-07 | 3.1e-07 | 1.8e-07 | 2.867E-07 | 9.713E-08 | 1.800E-07 | 3.700E-07 |
| 27611 | 1381 | Y1381L | E | 1.4e-07 | 1.2e-07 | 1.3e-07 | 1.300E-07 | 1.000E-08 | 1.200E-07 | 1.400E-07 |
| 27612 | 1381 | Y1381K | E | 1.5e-07 | 2.8e-07 | 1.7e-07 | 2.000E-07 | 7.000E-08 | 1.500E-07 | 2.800E-07 |
| 27613 | 1381 | Y1381M | E | 3e-07 | 5e-07 | 5.4e-07 | 4.467E-07 | 1.286E-07 | 3.000E-07 | 5.400E-07 |
| 27614 | 1381 | Y1381F | E | 1.8e-07 | 5.1e-07 | 1.6e-07 | 2.833E-07 | 1.966E-07 | 1.600E-07 | 5.100E-07 |
| 27615 | 1381 | Y1381P | E | 1.8e-07 | 1.8e-07 | 1.1e-07 | 1.567E-07 | 4.041E-08 | 1.100E-07 | 1.800E-07 |
| 27616 | 1381 | Y1381S | E | 3.2e-07 | 2.8e-07 | 1.4e-07 | 2.467E-07 | 9.452E-08 | 1.400E-07 | 3.200E-07 |
| 27617 | 1381 | Y1381T | E | 3.6e-07 | 2.3e-07 | 2.1e-07 | 2.667E-07 | 8.145E-08 | 2.100E-07 | 3.600E-07 |
| 27618 | 1381 | Y1381W | E | 1.8e-07 | 3.1e-07 | 2.8e-07 | 2.567E-07 | 6.807E-08 | 1.800E-07 | 3.100E-07 |
| 27619 | 1381 | Y1381Y | E | 1.8e-07 | 1.9e-07 | 2.1e-07 | 1.933E-07 | 1.528E-08 | 1.800E-07 | 2.100E-07 |
| 27620 | 1381 | Y1381V | E | 6e-07 | 2.3e-07 | 2.9e-07 | 3.733E-07 | 1.986E-07 | 2.300E-07 | 6.000E-07 |
| 27621 | 1382 | W1382A | E | 2.2e-07 | 2.1e-07 | 1.6e-07 | 1.967E-07 | 3.215E-08 | 1.600E-07 | 2.200E-07 |
| 27622 | 1382 | W1382R | E | 1.9e-07 | 2.2e-07 | 2.2e-07 | 2.100E-07 | 1.732E-08 | 1.900E-07 | 2.200E-07 |
| 27623 | 1382 | W1382N | E | 2.7e-07 | 2.8e-07 | 2.5e-07 | 2.667E-07 | 1.528E-08 | 2.500E-07 | 2.800E-07 |
| 27624 | 1382 | W1382D | E | 2.2e-07 | 4.6e-07 | 2.4e-07 | 3.067E-07 | 1.332E-07 | 2.200E-07 | 4.600E-07 |

| | | | | | | | | | | |
|---|---|---|---|---|---|---|---|---|---|---|
| 27625 | 1382 | W1382C | E | 8.2e-08 | 3.1e-07 | 3.4e-07 | 2.440E-07 | 1.411E-07 | 8.200E-08 | 3.400E-07 |
| 27626 | 1382 | W1382E | E | 2.2e-07 | 2.1e-07 | 2.6e-07 | 2.300E-07 | 2.646E-08 | 2.100E-07 | 2.600E-07 |
| 27627 | 1382 | W1382Q | E | 1.3e-07 | 1.8e-07 | 1e-07 | 1.367E-07 | 4.041E-08 | 1.000E-07 | 1.800E-07 |
| 27628 | 1382 | W1382G | E | 6.6e-07 | 2.1e-07 | 2.2e-07 | 3.633E-07 | 2.570E-07 | 2.100E-07 | 6.600E-07 |
| 27629 | 1382 | W1382H | E | 3.4e-07 | 1.8e-07 | 2.9e-07 | 2.700E-07 | 8.185E-08 | 1.800E-07 | 3.400E-07 |
| 27630 | 1382 | W1382I | E | 2.5e-07 | 1.2e-07 | 3.6e-07 | 2.433E-07 | 1.201E-07 | 1.200E-07 | 3.600E-07 |
| 27631 | 1382 | W1382L | E | 1.7e-07 | 1.4e-07 | 3.4e-07 | 2.167E-07 | 1.079E-07 | 1.400E-07 | 3.400E-07 |
| 27632 | 1382 | W1382K | E | 1.8e-07 | 1.5e-07 | 1.3e-07 | 1.533E-07 | 2.517E-08 | 1.300E-07 | 1.800E-07 |
| 27633 | 1382 | W1382M | E | 1.8e-07 | 4.7e-07 | 1.9e-07 | 2.800E-07 | 1.646E-07 | 1.800E-07 | 4.700E-07 |
| 27634 | 1382 | W1382F | E | 2.6e-07 | 3.6e-07 | 1.7e-07 | 2.633E-07 | 9.504E-08 | 1.700E-07 | 3.600E-07 |
| 27635 | 1382 | W1382P | E | 2.3e-07 | 2.8e-07 | 2.8e-07 | 2.633E-07 | 2.887E-08 | 2.300E-07 | 2.800E-07 |
| 27636 | 1382 | W1382S | E | 4.5e-07 | 3.4e-07 | 3e-07 | 3.633E-07 | 7.767E-08 | 3.000E-07 | 4.500E-07 |
| 27637 | 1382 | W1382T | E | 3.3e-07 | 1.2e-07 | 2e-07 | 2.167E-07 | 1.060E-07 | 1.200E-07 | 3.300E-07 |
| 27638 | 1382 | W1382W | E | 2.6e-07 | 1.7e-07 | 1.9e-07 | 2.067E-07 | 4.726E-08 | 1.700E-07 | 2.600E-07 |
| 27639 | 1382 | W1382Y | E | 1.6e-07 | 4.2e-07 | 2e-07 | 2.600E-07 | 1.400E-07 | 1.600E-07 | 4.200E-07 |
| 27640 | 1382 | W1382V | E | 3.9e-07 | 2e-07 | 3.6e-07 | 3.167E-07 | 1.021E-07 | 2.000E-07 | 3.900E-07 |
| 27641 | 1383 | G1383A | E | 1.7e-07 | 1.1e-07 | 1.9e-07 | 1.567E-07 | 4.163E-08 | 1.100E-07 | 1.900E-07 |
| 27642 | 1383 | G1383R | E | 1.9e-07 | 1.6e-07 | 1.5e-07 | 1.667E-07 | 2.082E-08 | 1.500E-07 | 1.900E-07 |
| 27643 | 1383 | G1383N | E | 2.3e-07 | 2.8e-07 | 2.7e-07 | 2.600E-07 | 2.646E-08 | 2.300E-07 | 2.800E-07 |
| 27644 | 1383 | G1383D | E | 2.3e-07 | 2.5e-07 | 4.7e-07 | 3.167E-07 | 1.332E-07 | 2.300E-07 | 4.700E-07 |
| 27645 | 1383 | G1383C | E | 2.4e-07 | 2.2e-07 | 2e-07 | 2.200E-07 | 2.000E-08 | 2.000E-07 | 2.400E-07 |
| 27646 | 1383 | G1383E | E | 2.9e-07 | 1.9e-07 | 3e-07 | 2.600E-07 | 6.083E-08 | 1.900E-07 | 3.000E-07 |
| 27647 | 1383 | G1383Q | E | 2.9e-07 | 2.7e-07 | 4.3e-07 | 3.300E-07 | 8.718E-08 | 2.700E-07 | 4.300E-07 |
| 27648 | 1383 | G1383G | E | 2.5e-07 | 2e-07 | 2.2e-07 | 2.233E-07 | 2.517E-08 | 2.000E-07 | 2.500E-07 |
| 27649 | 1383 | G1383H | E | 1.9e-07 | 5.4e-07 | 1.8e-07 | 3.033E-07 | 2.050E-07 | 1.800E-07 | 5.400E-07 |
| 27650 | 1383 | G1383I | E | 5.8e-08 | 6e-07 | 2.4e-07 | 2.993E-07 | 2.758E-07 | 5.800E-08 | 6.000E-07 |
| 27651 | 1383 | G1383L | E | 3.5e-07 | 1.3e-07 | 2.3e-07 | 2.367E-07 | 1.102E-07 | 1.300E-07 | 3.500E-07 |
| 27652 | 1383 | G1383K | E | 2e-07 | 1.9e-07 | 2.1e-07 | 2.000E-07 | 1.000E-08 | 1.900E-07 | 2.100E-07 |
| 27653 | 1383 | G1383M | E | 3e-07 | 2.9e-07 | 3e-07 | 2.967E-07 | 5.774E-09 | 2.900E-07 | 3.000E-07 |
| 27654 | 1383 | G1383F | E | 2e-07 | 2.1e-07 | 3.7e-07 | 2.600E-07 | 9.539E-08 | 2.000E-07 | 3.700E-07 |
| 27655 | 1383 | G1383P | E | 1.2e-07 | 1.5e-07 | 2.9e-07 | 1.867E-07 | 9.074E-08 | 1.200E-07 | 2.900E-07 |
| 27656 | 1383 | G1383S | E | 2.6e-07 | 1.4e-07 | 2.3e-07 | 2.100E-07 | 6.245E-08 | 1.400E-07 | 2.600E-07 |
| 27657 | 1383 | G1383T | E | 3.8e-07 | 4.2e-07 | 2e-07 | 3.333E-07 | 1.172E-07 | 2.000E-07 | 4.200E-07 |
| 27658 | 1383 | G1383W | E | 1.4e-07 | 2.8e-07 | 1.9e-07 | 2.033E-07 | 7.095E-08 | 1.400E-07 | 2.800E-07 |
| 27659 | 1383 | G1383Y | E | 4.1e-07 | 3e-07 | 3.5e-07 | 3.533E-07 | 5.508E-08 | 3.000E-07 | 4.100E-07 |
| 27660 | 1383 | G1383V | E | 2.4e-07 | 2.5e-07 | 4.2e-07 | 3.033E-07 | 1.012E-07 | 2.400E-07 | 4.200E-07 |
| 27661 | 1384 | Q1384A | E | 2.2e-07 | 3.2e-07 | 1.9e-07 | 2.433E-07 | 6.807E-08 | 1.900E-07 | 3.200E-07 |
| 27662 | 1384 | Q1384R | E | 4e-07 | 1.2e-07 | 2.2e-07 | 2.467E-07 | 1.419E-07 | 1.200E-07 | 4.000E-07 |
| 27663 | 1384 | Q1384N | E | 1.2e-07 | 1.5e-07 | 5.7e-07 | 2.800E-07 | 2.516E-07 | 1.200E-07 | 5.700E-07 |
| 27664 | 1384 | Q1384D | E | 2.4e-07 | 2.8e-07 | 2.3e-07 | 2.500E-07 | 2.646E-08 | 2.300E-07 | 2.800E-07 |
| 27665 | 1384 | Q1384C | E | 3e-07 | 4e-07 | 2.1e-07 | 3.033E-07 | 9.504E-08 | 2.100E-07 | 4.000E-07 |
| 27666 | 1384 | Q1384E | E | 2.8e-07 | 1.1e-07 | 3.4e-07 | 2.433E-07 | 1.193E-07 | 1.100E-07 | 3.400E-07 |
| 27667 | 1384 | Q1384Q | E | 3.3e-07 | 2.1e-07 | 2.9e-07 | 2.767E-07 | 6.110E-08 | 2.100E-07 | 3.300E-07 |
| 27668 | 1384 | Q1384G | E | 2.6e-07 | 2.9e-07 | 2.5e-07 | 2.667E-07 | 2.082E-08 | 2.500E-07 | 2.900E-07 |
| 27669 | 1384 | Q1384H | E | 2.3e-07 | 4e-07 | 5.4e-07 | 3.900E-07 | 1.552E-07 | 2.300E-07 | 5.400E-07 |
| 27670 | 1384 | Q1384I | E | 4e-07 | 2.7e-07 | 2.6e-07 | 3.100E-07 | 7.810E-08 | 2.600E-07 | 4.000E-07 |
| 27671 | 1384 | Q1384L | E | 6.9e-07 | 3.1e-07 | 2.3e-07 | 4.100E-07 | 2.458E-07 | 2.300E-07 | 6.900E-07 |
| 27672 | 1384 | Q1384K | E | 2.8e-07 | 4.2e-07 | 3.3e-07 | 3.433E-07 | 7.095E-08 | 2.800E-07 | 4.200E-07 |
| 27673 | 1384 | Q1384M | E | 2.8e-07 | 1.2e-07 | 2.2e-07 | 2.067E-07 | 8.083E-08 | 1.200E-07 | 2.800E-07 |
| 27674 | 1384 | Q1384F | E | 2.6e-07 | 1.8e-07 | 3.5e-07 | 2.633E-07 | 8.505E-08 | 1.800E-07 | 3.500E-07 |
| 27675 | 1384 | Q1384P | E | 1.3e-07 | 2.4e-07 | 2.7e-07 | 2.133E-07 | 7.371E-08 | 1.300E-07 | 2.700E-07 |
| 27676 | 1384 | Q1384S | E | 2.2e-07 | 2.6e-07 | 4.7e-07 | 3.167E-07 | 1.343E-07 | 2.200E-07 | 4.700E-07 |
| 27677 | 1384 | Q1384T | E | 1.8e-07 | 7.3e-08 | 2.1e-07 | 1.543E-07 | 7.202E-08 | 7.300E-08 | 2.100E-07 |
| 27678 | 1384 | Q1384W | E | 3.4e-07 | 3.7e-07 | 1.8e-07 | 2.967E-07 | 1.021E-07 | 1.800E-07 | 3.700E-07 |
| 27679 | 1384 | Q1384Y | E | 3.9e-07 | 2.8e-07 | 1.3e-07 | 2.667E-07 | 1.305E-07 | 1.300E-07 | 3.900E-07 |
| 27680 | 1384 | Q1384V | E | 6.3e-07 | 3.1e-07 | 1.7e-07 | 3.700E-07 | 2.358E-07 | 1.700E-07 | 6.300E-07 |
| 27681 | 1385 | G1385A | E | 1.9e-07 | 2e-07 | 6.5e-07 | 3.467E-07 | 2.627E-07 | 1.900E-07 | 6.500E-07 |
| 27682 | 1385 | G1385R | E | 2.9e-07 | 3.4e-07 | 4.7e-07 | 3.667E-07 | 9.292E-08 | 2.900E-07 | 4.700E-07 |
| 27683 | 1385 | G1385N | E | 3.4e-07 | 2.3e-07 | 1.7e-07 | 2.467E-07 | 8.622E-08 | 1.700E-07 | 3.400E-07 |
| 27684 | 1385 | G1385D | E | 1.4e-07 | 1.6e-07 | 1.7e-07 | 1.567E-07 | 1.528E-08 | 1.400E-07 | 1.700E-07 |
| 27685 | 1385 | G1385C | E | 3e-07 | 1.2e-07 | 3.4e-07 | 2.533E-07 | 1.172E-07 | 1.200E-07 | 3.400E-07 |
| 27686 | 1385 | G1385E | E | 2.9e-07 | 2.9e-07 | 1.8e-07 | 2.533E-07 | 6.351E-08 | 1.800E-07 | 2.900E-07 |
| 27687 | 1385 | G1385Q | E | 2.1e-07 | 1.5e-07 | 2.2e-07 | 1.933E-07 | 3.786E-08 | 1.500E-07 | 2.200E-07 |
| 27688 | 1385 | G1385G | E | 2e-07 | 5.5e-07 | 4.3e-07 | 3.933E-07 | 1.779E-07 | 2.000E-07 | 5.500E-07 |
| 27689 | 1385 | G1385H | E | 2e-07 | 2.3e-07 | 1.3e-07 | 1.867E-07 | 5.132E-08 | 1.300E-07 | 2.300E-07 |
| 27690 | 1385 | G1385I | E | 4.4e-07 | 4.4e-07 | 1.7e-07 | 3.500E-07 | 1.559E-07 | 1.700E-07 | 4.400E-07 |
| 27691 | 1385 | G1385L | E | 3.9e-07 | 3.5e-07 | 2.4e-07 | 3.267E-07 | 7.767E-08 | 2.400E-07 | 3.900E-07 |
| 27692 | 1385 | G1385K | E | 3.6e-07 | 1.9e-07 | 4.3e-07 | 3.267E-07 | 1.234E-07 | 1.900E-07 | 4.300E-07 |
| 27693 | 1385 | G1385M | E | 2.5e-07 | 4.3e-07 | 2e-07 | 2.933E-07 | 1.210E-07 | 2.000E-07 | 4.300E-07 |
| 27694 | 1385 | G1385F | E | 1.7e-07 | 1.9e-07 | 2.9e-07 | 2.167E-07 | 6.429E-08 | 1.700E-07 | 2.900E-07 |
| 27695 | 1385 | G1385P | E | 4.3e-07 | 1.7e-07 | 1.3e-07 | 2.433E-07 | 1.629E-07 | 1.300E-07 | 4.300E-07 |
| 27696 | 1385 | G1385S | E | 3.5e-07 | 3.6e-07 | 1.6e-07 | 2.900E-07 | 1.127E-07 | 1.600E-07 | 3.600E-07 |
| 27697 | 1385 | G1385T | E | 2.5e-07 | 2.6e-07 | 4.5e-07 | 3.200E-07 | 1.127E-07 | 2.500E-07 | 4.500E-07 |
| 27698 | 1385 | G1385W | E | 2e-07 | 3.9e-07 | 1.8e-07 | 2.567E-07 | 1.159E-07 | 1.800E-07 | 3.900E-07 |
| 27699 | 1385 | G1385Y | E | 2.3e-07 | 3.3e-07 | 4.8e-07 | 3.467E-07 | 1.258E-07 | 2.300E-07 | 4.800E-07 |
| 27700 | 1385 | G1385V | E | 3.3e-07 | 9.6e-08 | 3.1e-07 | 2.453E-07 | 1.297E-07 | 9.600E-08 | 3.300E-07 |
| 27701 | 1386 | T1386A | E | 5.3e-07 | 8.8e-08 | 2.7e-07 | 2.960E-07 | 2.221E-07 | 8.800E-08 | 5.300E-07 |
| 27702 | 1386 | T1386R | E | 3.5e-07 | 3.9e-07 | 1.9e-07 | 3.100E-07 | 1.058E-07 | 1.900E-07 | 3.900E-07 |
| 27703 | 1386 | T1386N | E | 2.6e-07 | 9.1e-08 | 3e-07 | 2.170E-07 | 1.109E-07 | 9.100E-08 | 3.000E-07 |
| 27704 | 1386 | T1386D | E | 2.1e-07 | 2.8e-07 | 2.2e-07 | 2.367E-07 | 3.786E-08 | 2.100E-07 | 2.800E-07 |
| 27705 | 1386 | T1386C | E | 1.8e-07 | 2.6e-07 | 3.2e-07 | 2.533E-07 | 7.024E-08 | 1.800E-07 | 3.200E-07 |
| 27706 | 1386 | T1386E | E | 2.7e-07 | 8.3e-08 | 3.2e-07 | 2.243E-07 | 1.249E-07 | 8.300E-08 | 3.200E-07 |
| 27707 | 1386 | T1386Q | E | 3.1e-07 | 3.6e-07 | 5.2e-07 | 3.967E-07 | 1.097E-07 | 3.100E-07 | 5.200E-07 |
| 27708 | 1386 | T1386G | E | 4.3e-07 | 4e-07 | 1.4e-07 | 3.233E-07 | 1.595E-07 | 1.400E-07 | 4.300E-07 |
| 27709 | 1386 | T1386H | E | 4.7e-07 | 3.1e-07 | 3.1e-07 | 3.633E-07 | 9.238E-08 | 3.100E-07 | 4.700E-07 |

| | | | | | | | | | | |
|---|---|---|---|---|---|---|---|---|---|---|
| 27710 | 1386 | T1386I | E | 1.6e-07 | 2.2e-07 | 2.2e-07 | 2.000E-07 | 3.464E-08 | 1.600E-07 | 2.200E-07 |
| 27711 | 1386 | T1386L | E | 3.9e-07 | 4.2e-07 | 2.6e-07 | 3.567E-07 | 8.505E-08 | 2.600E-07 | 4.200E-07 |
| 27712 | 1386 | T1386K | E | 2.3e-07 | 2.7e-07 | 2.1e-07 | 2.367E-07 | 3.055E-08 | 2.100E-07 | 2.700E-07 |
| 27713 | 1386 | T1386M | E | 2.9e-07 | 3.2e-07 | 2.1e-07 | 2.733E-07 | 5.686E-08 | 2.100E-07 | 3.200E-07 |
| 27714 | 1386 | T1386F | E | 2.6e-07 | 4.7e-07 | 1.7e-07 | 3.000E-07 | 1.539E-07 | 1.700E-07 | 4.700E-07 |
| 27715 | 1386 | T1386P | E | 1.1e-07 | 1.7e-07 | 1.5e-07 | 1.433E-07 | 3.055E-08 | 1.100E-07 | 1.700E-07 |
| 27716 | 1386 | T1386S | E | 3.3e-07 | 1.7e-07 | 2.3e-07 | 2.433E-07 | 8.083E-08 | 1.700E-07 | 3.300E-07 |
| 27717 | 1386 | T1386T | E | 3.3e-07 | 1.3e-07 | 2.4e-07 | 2.333E-07 | 1.002E-07 | 1.300E-07 | 3.300E-07 |
| 27718 | 1386 | T1386W | E | 2.4e-07 | 5.9e-07 | 2e-07 | 3.433E-07 | 2.146E-07 | 2.000E-07 | 5.900E-07 |
| 27719 | 1386 | T1386Y | E | 1.7e-07 | 2.5e-07 | 1.8e-07 | 2.000E-07 | 4.359E-08 | 1.700E-07 | 2.500E-07 |
| 27720 | 1386 | T1386V | E | 3e-07 | 2.7e-07 | 1.9e-07 | 2.533E-07 | 5.686E-08 | 1.900E-07 | 3.000E-07 |
| 27721 | 1387 | L1387A | E | 2.7e-07 | 3.3e-07 | 3.2e-07 | 3.067E-07 | 3.215E-08 | 2.700E-07 | 3.300E-07 |
| 27722 | 1387 | L1387R | E | 1.9e-07 | 8.9e-08 | 2.2e-07 | 1.663E-07 | 6.863E-08 | 8.900E-08 | 2.200E-07 |
| 27723 | 1387 | L1387N | E | 1.8e-07 | 6.3e-07 | 3.1e-07 | 3.733E-07 | 2.316E-07 | 1.800E-07 | 6.300E-07 |
| 27724 | 1387 | L1387D | E | 3.9e-07 | 1.8e-07 | 6.3e-07 | 4.000E-07 | 2.252E-07 | 1.800E-07 | 6.300E-07 |
| 27725 | 1387 | L1387C | E | 1.9e-07 | 2.1e-07 | 2.1e-07 | 2.033E-07 | 1.155E-08 | 1.900E-07 | 2.100E-07 |
| 27726 | 1387 | L1387E | E | 2.3e-07 | 2.7e-07 | 2.8e-07 | 2.600E-07 | 2.646E-08 | 2.300E-07 | 2.800E-07 |
| 27727 | 1387 | L1387Q | E | 2.6e-07 | 2.7e-07 | 1.9e-07 | 2.400E-07 | 4.359E-08 | 1.900E-07 | 2.700E-07 |
| 27728 | 1387 | L1387G | E | 4.2e-07 | 2.7e-07 | 1.7e-07 | 2.867E-07 | 1.258E-07 | 1.700E-07 | 4.200E-07 |
| 27729 | 1387 | L1387H | E | 2.6e-07 | 2.4e-07 | 1.8e-07 | 2.267E-07 | 4.163E-08 | 1.800E-07 | 2.600E-07 |
| 27730 | 1387 | L1387I | E | 2.5e-07 | 2.9e-07 | 1.2e-07 | 2.200E-07 | 8.888E-08 | 1.200E-07 | 2.900E-07 |
| 27731 | 1387 | L1387L | E | 1.7e-07 | 1.4e-07 | 3.1e-07 | 2.067E-07 | 9.074E-08 | 1.400E-07 | 3.100E-07 |
| 27732 | 1387 | L1387K | E | 3.8e-07 | 2.5e-07 | 1.2e-07 | 2.500E-07 | 1.300E-07 | 1.200E-07 | 3.800E-07 |
| 27733 | 1387 | L1387M | E | 2.1e-07 | 3.8e-07 | 9.1e-07 | 5.000E-07 | 3.651E-07 | 2.100E-07 | 9.100E-07 |
| 27734 | 1387 | L1387F | E | 1.6e-07 | 3.1e-07 | 1.7e-07 | 2.133E-07 | 8.386E-08 | 1.600E-07 | 3.100E-07 |
| 27735 | 1387 | L1387P | E | 2.3e-07 | 2.2e-07 | 1.9e-07 | 2.133E-07 | 2.082E-08 | 1.900E-07 | 2.300E-07 |
| 27736 | 1387 | L1387S | E | 1.4e-07 | 2.5e-07 | 2.7e-07 | 2.200E-07 | 7.000E-08 | 1.400E-07 | 2.700E-07 |
| 27737 | 1387 | L1387T | E | 1.9e-07 | 1.8e-07 | 1.9e-07 | 1.867E-07 | 5.774E-09 | 1.800E-07 | 1.900E-07 |
| 27738 | 1387 | L1387W | E | 1.2e-07 | 3.8e-07 | 2.4e-07 | 2.467E-07 | 1.301E-07 | 1.200E-07 | 3.800E-07 |
| 27739 | 1387 | L1387Y | E | 1.9e-07 | 1.3e-07 | 2.5e-07 | 1.900E-07 | 6.000E-08 | 1.300E-07 | 2.500E-07 |
| 27740 | 1387 | L1387V | E | 4.5e-07 | 3.5e-07 | 1.7e-07 | 3.233E-07 | 1.419E-07 | 1.700E-07 | 4.500E-07 |
| 27741 | 1388 | V1388A | E | 3.7e-07 | 2.7e-07 | 1.4e-07 | 2.600E-07 | 1.153E-07 | 1.400E-07 | 3.700E-07 |
| 27742 | 1388 | V1388R | E | 2.5e-07 | 3.2e-07 | 2.4e-07 | 2.700E-07 | 4.359E-08 | 2.400E-07 | 3.200E-07 |
| 27743 | 1388 | V1388N | E | 1.8e-07 | 2.8e-07 | 2.6e-07 | 2.400E-07 | 5.292E-08 | 1.800E-07 | 2.800E-07 |
| 27744 | 1388 | V1388D | E | 4e-07 | 1.2e-07 | 1.2e-07 | 2.133E-07 | 1.617E-07 | 1.200E-07 | 4.000E-07 |
| 27745 | 1388 | V1388C | E | 1.7e-07 | 1.9e-07 | 1.8e-07 | 1.800E-07 | 1.000E-08 | 1.700E-07 | 1.900E-07 |
| 27746 | 1388 | V1388E | E | 2.7e-07 | 2.2e-07 | 1.8e-07 | 2.233E-07 | 4.509E-08 | 1.800E-07 | 2.700E-07 |
| 27747 | 1388 | V1388Q | E | 2.9e-07 | 3.5e-07 | 2.5e-07 | 2.967E-07 | 5.033E-08 | 2.500E-07 | 3.500E-07 |
| 27748 | 1388 | V1388G | E | 1.7e-07 | 2.3e-07 | 2.9e-07 | 2.300E-07 | 6.000E-08 | 1.700E-07 | 2.900E-07 |
| 27749 | 1388 | V1388H | E | 2.2e-07 | 1.5e-07 | 5.6e-07 | 3.100E-07 | 2.193E-07 | 1.500E-07 | 5.600E-07 |
| 27750 | 1388 | V1388I | E | 3e-07 | 4.1e-07 | 1.5e-07 | 2.867E-07 | 1.305E-07 | 1.500E-07 | 4.100E-07 |
| 27751 | 1388 | V1388L | E | 4.4e-07 | 4.3e-07 | 1.2e-07 | 3.300E-07 | 1.819E-07 | 1.200E-07 | 4.400E-07 |
| 27752 | 1388 | V1388K | E | 3.3e-07 | 1.6e-07 | 3.6e-07 | 2.833E-07 | 1.079E-07 | 1.600E-07 | 3.600E-07 |
| 27753 | 1388 | V1388M | E | 1.7e-07 | 1.3e-07 | 2e-07 | 1.667E-07 | 3.512E-08 | 1.300E-07 | 2.000E-07 |
| 27754 | 1388 | V1388F | E | 2.2e-07 | 2.3e-07 | 2.5e-07 | 2.333E-07 | 1.528E-08 | 2.200E-07 | 2.500E-07 |
| 27755 | 1388 | V1388P | E | 1.3e-07 | 1.8e-07 | 3.3e-07 | 2.133E-07 | 1.041E-07 | 1.300E-07 | 3.300E-07 |
| 27756 | 1388 | V1388S | E | 1.8e-07 | 2.4e-07 | 1.9e-07 | 2.033E-07 | 3.215E-08 | 1.800E-07 | 2.400E-07 |
| 27757 | 1388 | V1388T | E | 3.4e-07 | 1.5e-07 | 2e-07 | 2.300E-07 | 9.849E-08 | 1.500E-07 | 3.400E-07 |
| 27758 | 1388 | V1388W | E | 3.8e-07 | 2.3e-07 | 2.7e-07 | 2.933E-07 | 7.767E-08 | 2.300E-07 | 3.800E-07 |
| 27759 | 1388 | V1388Y | E | 4.3e-07 | 7.6e-08 | 2.5e-07 | 2.520E-07 | 1.770E-07 | 7.600E-08 | 4.300E-07 |
| 27760 | 1388 | V1388V | E | 3e-07 | 2e-07 | 2.1e-07 | 2.367E-07 | 5.508E-08 | 2.000E-07 | 3.000E-07 |
| 27761 | 1389 | T1389A | E | 1.6e-07 | 3.5e-07 | 2e-07 | 2.367E-07 | 1.002E-07 | 1.600E-07 | 3.500E-07 |
| 27762 | 1389 | T1389R | E | 3.6e-07 | 1.3e-07 | 1.6e-07 | 2.167E-07 | 1.250E-07 | 1.300E-07 | 3.600E-07 |
| 27763 | 1389 | T1389N | E | 2.5e-07 | 1.9e-07 | 1.5e-07 | 1.967E-07 | 5.033E-08 | 1.500E-07 | 2.500E-07 |
| 27764 | 1389 | T1389D | E | 3.8e-07 | 2e-07 | 3e-07 | 2.933E-07 | 9.018E-08 | 2.000E-07 | 3.800E-07 |
| 27765 | 1389 | T1389C | E | 2.4e-07 | 2.8e-07 | 2.7e-07 | 2.633E-07 | 2.082E-08 | 2.400E-07 | 2.800E-07 |
| 27766 | 1389 | T1389E | E | 2.6e-07 | 2.1e-07 | 2.2e-07 | 2.300E-07 | 2.646E-08 | 2.100E-07 | 2.600E-07 |
| 27767 | 1389 | T1389Q | E | 3.2e-07 | 2.4e-07 | 2.2e-07 | 2.600E-07 | 5.292E-08 | 2.200E-07 | 3.200E-07 |
| 27768 | 1389 | T1389G | E | 1.6e-07 | 3.7e-07 | 2.1e-07 | 2.467E-07 | 1.097E-07 | 1.600E-07 | 3.700E-07 |
| 27769 | 1389 | T1389H | E | 2.6e-07 | 1.7e-07 | 2.9e-07 | 2.400E-07 | 6.245E-08 | 1.700E-07 | 2.900E-07 |
| 27770 | 1389 | T1389I | E | 1.8e-07 | 2.6e-07 | 5.3e-07 | 3.233E-07 | 1.834E-07 | 1.800E-07 | 5.300E-07 |
| 27771 | 1389 | T1389L | E | 3.5e-07 | 1.6e-07 | 1.9e-07 | 2.333E-07 | 1.021E-07 | 1.600E-07 | 3.500E-07 |
| 27772 | 1389 | T1389K | E | 2.7e-07 | 1.1e-07 | 2.2e-07 | 2.000E-07 | 8.185E-08 | 1.100E-07 | 2.700E-07 |
| 27773 | 1389 | T1389M | E | 2.1e-07 | 2.8e-07 | 6.4e-07 | 3.767E-07 | 2.307E-07 | 2.100E-07 | 6.400E-07 |
| 27774 | 1389 | T1389F | E | 1.7e-07 | 2.5e-07 | 1.1e-07 | 1.767E-07 | 7.024E-08 | 1.100E-07 | 2.500E-07 |
| 27775 | 1389 | T1389P | E | 2.1e-07 | 1.4e-07 | 2.2e-07 | 1.900E-07 | 4.359E-08 | 1.400E-07 | 2.200E-07 |
| 27776 | 1389 | T1389S | E | 2.7e-07 | 1.5e-07 | 9.8e-08 | 1.727E-07 | 8.821E-08 | 9.800E-08 | 2.700E-07 |
| 27777 | 1389 | T1389T | E | 3e-07 | 1.7e-07 | 3.1e-07 | 2.600E-07 | 7.810E-08 | 1.700E-07 | 3.100E-07 |
| 27778 | 1389 | T1389W | E | 1.5e-07 | 2.3e-07 | 2.2e-07 | 2.000E-07 | 4.359E-08 | 1.500E-07 | 2.300E-07 |
| 27779 | 1389 | T1389Y | E | 1.3e-07 | 3.2e-07 | 5.2e-07 | 3.233E-07 | 1.950E-07 | 1.300E-07 | 5.200E-07 |
| 27780 | 1389 | T1389V | E | 2.2e-07 | 3.3e-07 | 3.2e-07 | 2.900E-07 | 6.083E-08 | 2.200E-07 | 3.300E-07 |
| 27781 | 1390 | V1390A | E | 1.9e-07 | 5.3e-07 | 2.7e-07 | 3.300E-07 | 1.778E-07 | 1.900E-07 | 5.300E-07 |
| 27782 | 1390 | V1390R | E | 1.8e-07 | 2.4e-07 | 1.9e-07 | 2.033E-07 | 3.215E-08 | 1.800E-07 | 2.400E-07 |
| 27783 | 1390 | V1390N | E | 7.5e-08 | 3.9e-07 | 2.2e-07 | 2.283E-07 | 1.577E-07 | 7.500E-08 | 3.900E-07 |
| 27784 | 1390 | V1390D | E | 2.3e-07 | 1.7e-07 | 3.3e-07 | 2.433E-07 | 8.083E-08 | 1.700E-07 | 3.300E-07 |
| 27785 | 1390 | V1390C | E | 2.1e-07 | 1.7e-07 | 4.9e-07 | 2.900E-07 | 1.744E-07 | 1.700E-07 | 4.900E-07 |
| 27786 | 1390 | V1390E | E | 1.9e-07 | 1.6e-07 | 2.9e-07 | 2.133E-07 | 6.807E-08 | 1.600E-07 | 2.900E-07 |
| 27787 | 1390 | V1390Q | E | 2.4e-07 | 3.3e-07 | 2.5e-07 | 2.733E-07 | 4.933E-08 | 2.400E-07 | 3.300E-07 |
| 27788 | 1390 | V1390G | E | 3e-07 | 1.7e-07 | 1.3e-07 | 2.000E-07 | 8.888E-08 | 1.300E-07 | 3.000E-07 |
| 27789 | 1390 | V1390H | E | 3.7e-07 | 1.5e-07 | 2.1e-07 | 2.433E-07 | 1.137E-07 | 1.500E-07 | 3.700E-07 |
| 27790 | 1390 | V1390I | E | 2.1e-07 | 3.1e-07 | 2.5e-07 | 2.567E-07 | 5.033E-08 | 2.100E-07 | 3.100E-07 |
| 27791 | 1390 | V1390L | E | 3.4e-07 | 5e-07 | 4.8e-07 | 4.400E-07 | 8.718E-08 | 3.400E-07 | 5.000E-07 |
| 27792 | 1390 | V1390K | E | 3.9e-07 | 5.1e-07 | 1.9e-07 | 3.633E-07 | 1.617E-07 | 1.900E-07 | 5.100E-07 |
| 27793 | 1390 | V1390M | E | 3.2e-07 | 3.8e-07 | 1.6e-07 | 2.867E-07 | 1.137E-07 | 1.600E-07 | 3.800E-07 |
| 27794 | 1390 | V1390F | E | 1.9e-07 | 1.7e-07 | 3.9e-07 | 2.500E-07 | 1.217E-07 | 1.700E-07 | 3.900E-07 |

| | | | | | | | | | | |
|---|---|---|---|---|---|---|---|---|---|---|
| 27795 | 1390 | V1390P | E | 2.7e-07 | 1.5e-07 | 2.7e-07 | 2.300E-07 | 6.928E-08 | 1.500E-07 | 2.700E-07 |
| 27796 | 1390 | V1390S | E | 1.3e-07 | 3.5e-07 | 1.8e-07 | 2.200E-07 | 1.153E-07 | 1.300E-07 | 3.500E-07 |
| 27797 | 1390 | V1390T | E | 1.1e-07 | 1.9e-07 | 1.3e-07 | 1.433E-07 | 4.163E-08 | 1.100E-07 | 1.900E-07 |
| 27798 | 1390 | V1390W | E | 2.2e-07 | 2.3e-07 | 3.3e-07 | 2.600E-07 | 6.083E-08 | 2.200E-07 | 3.300E-07 |
| 27799 | 1390 | V1390Y | E | 4.5e-07 | 2.9e-07 | 2.7e-07 | 3.367E-07 | 9.866E-08 | 2.700E-07 | 4.500E-07 |
| 27800 | 1390 | V1390V | E | 6.4e-08 | 1.5e-07 | 2.9e-07 | 1.680E-07 | 1.141E-07 | 6.400E-08 | 2.900E-07 |
| 27801 | 1391 | S1391A | E | 3.3e-07 | 1.5e-07 | 1.9e-07 | 2.233E-07 | 9.452E-08 | 1.500E-07 | 3.300E-07 |
| 27802 | 1391 | S1391R | E | 1.5e-07 | 2.6e-07 | 2.9e-07 | 2.333E-07 | 7.371E-08 | 1.500E-07 | 2.900E-07 |
| 27803 | 1391 | S1391N | E | 3.1e-07 | 1.3e-07 | 2.6e-07 | 2.333E-07 | 9.292E-08 | 1.300E-07 | 3.100E-07 |
| 27804 | 1391 | S1391D | E | 3e-07 | 3.4e-07 | 3.4e-07 | 3.267E-07 | 2.309E-08 | 3.000E-07 | 3.400E-07 |
| 27805 | 1391 | S1391C | E | 4.4e-07 | 4e-07 | 1.7e-07 | 3.367E-07 | 1.457E-07 | 1.700E-07 | 4.400E-07 |
| 27806 | 1391 | S1391E | E | 3.3e-07 | 1.4e-07 | 3.6e-07 | 2.767E-07 | 1.193E-07 | 1.400E-07 | 3.600E-07 |
| 27807 | 1391 | S1391Q | E | 1.4e-07 | 2.7e-07 | 1.7e-07 | 1.933E-07 | 6.807E-08 | 1.400E-07 | 2.700E-07 |
| 27808 | 1391 | S1391G | E | 1.4e-07 | 3e-07 | 2.5e-07 | 2.300E-07 | 8.185E-08 | 1.400E-07 | 3.000E-07 |
| 27809 | 1391 | S1391H | E | 4e-07 | 2.9e-07 | 3.2e-07 | 3.367E-07 | 5.686E-08 | 2.900E-07 | 4.000E-07 |
| 27810 | 1391 | S1391I | E | 2.6e-07 | 3.4e-07 | 3.2e-07 | 3.067E-07 | 4.163E-08 | 2.600E-07 | 3.400E-07 |
| 27811 | 1391 | S1391L | E | 1.8e-07 | 1.5e-07 | 4e-07 | 2.433E-07 | 1.365E-07 | 1.500E-07 | 4.000E-07 |
| 27812 | 1391 | S1391K | E | 2.7e-07 | 2.5e-07 | 3.1e-07 | 2.767E-07 | 3.055E-08 | 2.500E-07 | 3.100E-07 |
| 27813 | 1391 | S1391M | E | 1.8e-07 | 3e-07 | 5.8e-07 | 3.533E-07 | 2.053E-07 | 1.800E-07 | 5.800E-07 |
| 27814 | 1391 | S1391F | E | 1.8e-07 | 2.7e-07 | 4.2e-07 | 2.900E-07 | 1.212E-07 | 1.800E-07 | 4.200E-07 |
| 27815 | 1391 | S1391P | E | 2.2e-07 | 3.4e-07 | 1.5e-07 | 2.367E-07 | 9.609E-08 | 1.500E-07 | 3.400E-07 |
| 27816 | 1391 | S1391S | E | 1.7e-07 | 1.8e-07 | 2.9e-07 | 2.133E-07 | 6.658E-08 | 1.700E-07 | 2.900E-07 |
| 27817 | 1391 | S1391T | E | 3.4e-07 | 2e-07 | 2.9e-07 | 2.767E-07 | 7.095E-08 | 2.000E-07 | 3.400E-07 |
| 27818 | 1391 | S1391W | E | 2.3e-07 | 1.5e-07 | 3.2e-07 | 2.333E-07 | 8.505E-08 | 1.500E-07 | 3.200E-07 |
| 27819 | 1391 | S1391Y | E | 2.2e-07 | 1.1e-07 | 7.2e-07 | 3.500E-07 | 3.251E-07 | 1.100E-07 | 7.200E-07 |
| 27820 | 1391 | S1391V | E | 2.8e-07 | 3.5e-07 | 2.5e-07 | 2.933E-07 | 5.132E-08 | 2.500E-07 | 3.500E-07 |
| 27821 | 1392 | S1392A | E | 3.3e-07 | 1.7e-07 | 1.8e-07 | 2.267E-07 | 8.963E-08 | 1.700E-07 | 3.300E-07 |
| 27822 | 1392 | S1392R | E | 5.1e-07 | 2.7e-07 | 9.1e-08 | 2.903E-07 | 2.102E-07 | 9.100E-08 | 5.100E-07 |
| 27823 | 1392 | S1392N | E | 4.9e-07 | 2.2e-07 | 3.4e-07 | 3.500E-07 | 1.353E-07 | 2.200E-07 | 4.900E-07 |
| 27824 | 1392 | S1392D | E | 2.9e-07 | 3.5e-07 | 3.1e-07 | 3.167E-07 | 3.055E-08 | 2.900E-07 | 3.500E-07 |
| 27825 | 1392 | S1392C | E | 1.4e-07 | 1.6e-07 | 1.3e-07 | 1.433E-07 | 1.528E-08 | 1.300E-07 | 1.600E-07 |
| 27826 | 1392 | S1392E | E | 1.7e-07 | 2.6e-07 | 1.5e-07 | 1.933E-07 | 5.859E-08 | 1.500E-07 | 2.600E-07 |
| 27827 | 1392 | S1392Q | E | 2.2e-07 | 1.6e-07 | 2.7e-07 | 2.167E-07 | 5.508E-08 | 1.600E-07 | 2.700E-07 |
| 27828 | 1392 | S1392G | E | 1.8e-07 | 2.7e-07 | 2.7e-07 | 2.400E-07 | 5.196E-08 | 1.800E-07 | 2.700E-07 |
| 27829 | 1392 | S1392H | E | 3.1e-07 | 1.3e-07 | 1.1e-07 | 1.833E-07 | 1.102E-07 | 1.100E-07 | 3.100E-07 |
| 27830 | 1392 | S1392I | E | 1.9e-07 | 5.2e-07 | 3.4e-07 | 3.500E-07 | 1.652E-07 | 1.900E-07 | 5.200E-07 |
| 27831 | 1392 | S1392L | E | 1.5e-07 | 1.8e-07 | 2.3e-07 | 1.867E-07 | 4.041E-08 | 1.500E-07 | 2.300E-07 |
| 27832 | 1392 | S1392K | E | 3.1e-07 | 2.5e-07 | 3.3e-07 | 2.967E-07 | 4.163E-08 | 2.500E-07 | 3.300E-07 |
| 27833 | 1392 | S1392M | E | 2.4e-07 | 1.7e-07 | 2.3e-07 | 2.133E-07 | 3.786E-08 | 1.700E-07 | 2.400E-07 |
| 27834 | 1392 | S1392F | E | 2.5e-07 | 2.1e-07 | 7.2e-07 | 3.933E-07 | 2.836E-07 | 2.100E-07 | 7.200E-07 |
| 27835 | 1392 | S1392P | E | 1.6e-07 | 1.8e-07 | 2.6e-07 | 2.000E-07 | 5.292E-08 | 1.600E-07 | 2.600E-07 |
| 27836 | 1392 | S1392S | E | 2.3e-07 | 1.8e-07 | 2.4e-07 | 2.167E-07 | 3.215E-08 | 1.800E-07 | 2.400E-07 |
| 27837 | 1392 | S1392T | E | 2.5e-07 | 2.3e-07 | 1.1e-07 | 1.967E-07 | 7.572E-08 | 1.100E-07 | 2.500E-07 |
| 27838 | 1392 | S1392W | E | 3.7e-07 | 2.8e-07 | 1.3e-07 | 2.600E-07 | 1.212E-07 | 1.300E-07 | 3.700E-07 |
| 27839 | 1392 | S1392Y | E | 1.4e-07 | 1.8e-07 | 4e-07 | 2.400E-07 | 1.400E-07 | 1.400E-07 | 4.000E-07 |
| 27840 | 1392 | S1392V | E | 1.7e-07 | 3.7e-07 | 2.5e-07 | 2.633E-07 | 1.007E-07 | 1.700E-07 | 3.700E-07 |
| 27841 | 1393 | A1393A | E | 3.2e-07 | 1.1e-07 | 2.6e-07 | 2.300E-07 | 1.082E-07 | 1.100E-07 | 3.200E-07 |
| 27842 | 1393 | A1393R | E | 2e-07 | 1.8e-07 | 1.5e-07 | 1.767E-07 | 2.517E-08 | 1.500E-07 | 2.000E-07 |
| 27843 | 1393 | A1393N | E | 2e-07 | 1.9e-07 | 2.4e-07 | 2.100E-07 | 2.646E-08 | 1.900E-07 | 2.400E-07 |
| 27844 | 1393 | A1393D | E | 1.9e-07 | 5.2e-07 | 2.1e-07 | 3.067E-07 | 1.850E-07 | 1.900E-07 | 5.200E-07 |
| 27845 | 1393 | A1393C | E | 4.4e-07 | 4.6e-07 | 2e-07 | 3.667E-07 | 1.447E-07 | 2.000E-07 | 4.600E-07 |
| 27846 | 1393 | A1393E | E | 4.8e-07 | 1.4e-06 | 1.1e-07 | 6.633E-07 | 6.643E-07 | 1.100E-07 | 1.400E-06 |
| 27847 | 1393 | A1393Q | E | 1.8e-07 | 2.1e-07 | 3.8e-07 | 2.567E-07 | 1.079E-07 | 1.800E-07 | 3.800E-07 |
| 27848 | 1393 | A1393G | E | 3.6e-07 | 2.5e-07 | 4.8e-07 | 3.633E-07 | 1.150E-07 | 2.500E-07 | 4.800E-07 |
| 27849 | 1393 | A1393H | E | 1.4e-07 | 5.6e-07 | 1.4e-07 | 2.800E-07 | 2.425E-07 | 1.400E-07 | 5.600E-07 |
| 27850 | 1393 | A1393I | E | 3.4e-07 | 2.8e-07 | 1.1e-07 | 2.433E-07 | 1.193E-07 | 1.100E-07 | 3.400E-07 |
| 27851 | 1393 | A1393L | E | 1.2e-07 | 1.6e-07 | 2.4e-07 | 1.733E-07 | 6.110E-08 | 1.200E-07 | 2.400E-07 |
| 27852 | 1393 | A1393K | E | 2.9e-07 | 4.2e-07 | 4.7e-07 | 3.933E-07 | 9.292E-08 | 2.900E-07 | 4.700E-07 |
| 27853 | 1393 | A1393M | E | 1.6e-07 | 2e-07 | 1.5e-07 | 1.700E-07 | 2.646E-08 | 1.500E-07 | 2.000E-07 |
| 27854 | 1393 | A1393F | E | 2.5e-07 | 1.4e-07 | 2.5e-07 | 2.133E-07 | 6.351E-08 | 1.400E-07 | 2.500E-07 |
| 27855 | 1393 | A1393P | E | 1.3e-07 | 1.3e-07 | 3.6e-07 | 2.067E-07 | 1.328E-07 | 1.300E-07 | 3.600E-07 |
| 27856 | 1393 | A1393S | E | 3.7e-07 | 2e-07 | 4e-07 | 3.233E-07 | 1.079E-07 | 2.000E-07 | 4.000E-07 |
| 27857 | 1393 | A1393T | E | 2.3e-07 | 5.1e-07 | 3.3e-07 | 3.567E-07 | 1.419E-07 | 2.300E-07 | 5.100E-07 |
| 27858 | 1393 | A1393W | E | 2.8e-07 | 1.5e-07 | 3e-07 | 2.433E-07 | 8.145E-08 | 1.500E-07 | 3.000E-07 |
| 27859 | 1393 | A1393Y | E | 3.9e-07 | 2.3e-07 | 3.3e-07 | 3.167E-07 | 8.083E-08 | 2.300E-07 | 3.900E-07 |
| 27860 | 1393 | A1393V | E | 2.6e-07 | 2.5e-07 | 2.4e-07 | 2.500E-07 | 1.000E-08 | 2.400E-07 | 2.600E-07 |
| 27861 | 1394 | S1394A | E | 2.6e-07 | 4.7e-07 | 2.9e-07 | 3.400E-07 | 1.136E-07 | 2.600E-07 | 4.700E-07 |
| 27862 | 1394 | S1394R | E | 2.1e-07 | 9.6e-08 | 2.6e-07 | 1.887E-07 | 8.406E-08 | 9.600E-08 | 2.600E-07 |
| 27863 | 1394 | S1394N | E | 1.3e-07 | 3.2e-07 | 6.4e-07 | 3.633E-07 | 2.577E-07 | 1.300E-07 | 6.400E-07 |
| 27864 | 1394 | S1394D | E | 1.6e-07 | 9.3e-08 | 2.4e-07 | 1.643E-07 | 7.360E-08 | 9.300E-08 | 2.400E-07 |
| 27865 | 1394 | S1394C | E | 2.9e-07 | 2.2e-07 | 1.5e-07 | 2.200E-07 | 7.000E-08 | 1.500E-07 | 2.900E-07 |
| 27866 | 1394 | S1394E | E | 1.4e-07 | 2.7e-07 | 2.4e-07 | 2.167E-07 | 6.807E-08 | 1.400E-07 | 2.700E-07 |
| 27867 | 1394 | S1394Q | E | 1.4e-07 | 2.1e-07 | 2.6e-07 | 2.033E-07 | 6.028E-08 | 1.400E-07 | 2.600E-07 |
| 27868 | 1394 | S1394G | E | 5.6e-07 | 1.6e-07 | 2.7e-07 | 3.300E-07 | 2.066E-07 | 1.600E-07 | 5.600E-07 |
| 27869 | 1394 | S1394H | E | 3.3e-07 | 3.5e-07 | 3.3e-07 | 3.367E-07 | 1.155E-08 | 3.300E-07 | 3.500E-07 |
| 27870 | 1394 | S1394I | E | 2.6e-07 | 2.8e-07 | 9.7e-08 | 2.123E-07 | 1.004E-07 | 9.700E-08 | 2.800E-07 |
| 27871 | 1394 | S1394L | E | 1.5e-07 | 1.6e-07 | 4e-07 | 2.367E-07 | 1.415E-07 | 1.500E-07 | 4.000E-07 |
| 27872 | 1394 | S1394K | E | 2.9e-07 | 2.7e-07 | 2.8e-07 | 2.800E-07 | 1.000E-08 | 2.700E-07 | 2.900E-07 |
| 27873 | 1394 | S1394M | E | 1.7e-07 | 2.7e-07 | 7.7e-08 | 1.723E-07 | 9.652E-08 | 7.700E-08 | 2.700E-07 |
| 27874 | 1394 | S1394F | E | 2e-07 | 1.9e-07 | 2.7e-07 | 2.200E-07 | 4.359E-08 | 1.900E-07 | 2.700E-07 |
| 27875 | 1394 | S1394P | E | 2e-07 | 3e-07 | 3.1e-07 | 2.700E-07 | 6.083E-08 | 2.000E-07 | 3.100E-07 |
| 27876 | 1394 | S1394S | E | 1.9e-07 | 2.7e-07 | 4.2e-07 | 2.933E-07 | 1.168E-07 | 1.900E-07 | 4.200E-07 |
| 27877 | 1394 | S1394T | E | 2.9e-07 | 1.9e-07 | 1.3e-07 | 2.033E-07 | 8.083E-08 | 1.300E-07 | 2.900E-07 |
| 27878 | 1394 | S1394W | E | 3.1e-07 | 1.6e-07 | 2.5e-07 | 2.400E-07 | 7.550E-08 | 1.600E-07 | 3.100E-07 |
| 27879 | 1394 | S1394Y | E | 1.1e-07 | 2e-07 | 2.5e-07 | 1.867E-07 | 7.095E-08 | 1.100E-07 | 2.500E-07 |

| | | | | | | | | | | |
|---|---|---|---|---|---|---|---|---|---|---|
| 27880 | 1394 | S1394V | E | 2.3e-07 | 1.9e-07 | 2.7e-07 | 2.300E-07 | 4.000E-08 | 1.900E-07 | 2.700E-07 |
| 27881 | 1395 | T1395A | E | 7e-07 | 3e-07 | 1.6e-07 | 3.867E-07 | 2.802E-07 | 1.600E-07 | 7.000E-07 |
| 27882 | 1395 | T1395R | E | 2.5e-07 | 4.6e-07 | 3.6e-07 | 3.567E-07 | 1.050E-07 | 2.500E-07 | 4.600E-07 |
| 27883 | 1395 | T1395N | E | 4.7e-07 | 2e-07 | 3.4e-07 | 3.367E-07 | 1.350E-07 | 2.000E-07 | 4.700E-07 |
| 27884 | 1395 | T1395D | E | 2.1e-07 | 5e-07 | 3.1e-07 | 3.400E-07 | 1.473E-07 | 2.100E-07 | 5.000E-07 |
| 27885 | 1395 | T1395C | E | 1.4e-07 | 2e-07 | 2.2e-07 | 1.867E-07 | 4.163E-08 | 1.400E-07 | 2.200E-07 |
| 27886 | 1395 | T1395E | E | 2.6e-07 | 2.2e-07 | 1.9e-07 | 2.233E-07 | 3.512E-08 | 1.900E-07 | 2.600E-07 |
| 27887 | 1395 | T1395Q | E | 1e-07 | 3.2e-07 | 1.9e-07 | 2.033E-07 | 1.106E-07 | 1.000E-07 | 3.200E-07 |
| 27888 | 1395 | T1395G | E | 1.5e-07 | 8.1e-08 | 2e-07 | 1.437E-07 | 5.975E-08 | 8.100E-08 | 2.000E-07 |
| 27889 | 1395 | T1395H | E | 3.7e-07 | 2.3e-07 | 4e-07 | 3.333E-07 | 9.074E-08 | 2.300E-07 | 4.000E-07 |
| 27890 | 1395 | T1395I | E | 1.4e-07 | 2.6e-07 | 3.4e-07 | 2.467E-07 | 1.007E-07 | 1.400E-07 | 3.400E-07 |
| 27891 | 1395 | T1395L | E | 3.3e-07 | 3.5e-07 | 2.4e-07 | 3.067E-07 | 5.859E-08 | 2.400E-07 | 3.500E-07 |
| 27892 | 1395 | T1395K | E | 1.6e-07 | 3.5e-07 | 5.5e-07 | 3.533E-07 | 1.950E-07 | 1.600E-07 | 5.500E-07 |
| 27893 | 1395 | T1395M | E | 1.6e-07 | 2.9e-07 | 5.2e-07 | 3.233E-07 | 1.823E-07 | 1.600E-07 | 5.200E-07 |
| 27894 | 1395 | T1395F | E | 2.1e-07 | 1.9e-07 | 2.3e-07 | 2.100E-07 | 2.000E-08 | 1.900E-07 | 2.300E-07 |
| 27895 | 1395 | T1395P | E | 5.4e-07 | 2.5e-07 | 4.5e-07 | 4.133E-07 | 1.484E-07 | 2.500E-07 | 5.400E-07 |
| 27896 | 1395 | T1395S | E | 2.3e-07 | 3e-07 | 3.2e-07 | 2.833E-07 | 4.726E-08 | 2.300E-07 | 3.200E-07 |
| 27897 | 1395 | T1395T | E | 3.3e-07 | 4.8e-07 | 4.5e-08 | 2.850E-07 | 2.210E-07 | 4.500E-08 | 4.800E-07 |
| 27898 | 1395 | T1395W | E | 1.1e-07 | 1.7e-07 | 3.8e-07 | 2.200E-07 | 1.418E-07 | 1.100E-07 | 3.800E-07 |
| 27899 | 1395 | T1395Y | E | 4.4e-07 | 4.9e-07 | 4.2e-07 | 4.500E-07 | 3.606E-08 | 4.200E-07 | 4.900E-07 |
| 27900 | 1395 | T1395V | E | 1.4e-07 | 2.1e-07 | 1.2e-07 | 1.567E-07 | 4.726E-08 | 1.200E-07 | 2.100E-07 |
| 27901 | 1396 | K1396A | E | 2.3e-07 | 3.1e-07 | 1.5e-07 | 2.300E-07 | 8.000E-08 | 1.500E-07 | 3.100E-07 |
| 27902 | 1396 | K1396R | E | 4.7e-07 | 3.3e-07 | 2.3e-07 | 3.433E-07 | 1.206E-07 | 2.300E-07 | 4.700E-07 |
| 27903 | 1396 | K1396N | E | 3.5e-07 | 2.4e-07 | 1.5e-07 | 2.467E-07 | 1.002E-07 | 1.500E-07 | 3.500E-07 |
| 27904 | 1396 | K1396D | E | 1.6e-07 | 1.9e-07 | 6.4e-07 | 3.300E-07 | 2.689E-07 | 1.600E-07 | 6.400E-07 |
| 27905 | 1396 | K1396C | E | 3.8e-07 | 1.8e-07 | 2.4e-07 | 2.667E-07 | 1.026E-07 | 1.800E-07 | 3.800E-07 |
| 27906 | 1396 | K1396E | E | 6.5e-07 | 2.2e-07 | 1.6e-07 | 3.433E-07 | 2.673E-07 | 1.600E-07 | 6.500E-07 |
| 27907 | 1396 | K1396Q | E | 2.7e-07 | 1.9e-07 | 3.9e-07 | 2.833E-07 | 1.007E-07 | 1.900E-07 | 3.900E-07 |
| 27908 | 1396 | K1396G | E | 1.5e-07 | 1.9e-07 | 3.3e-07 | 2.233E-07 | 9.452E-08 | 1.500E-07 | 3.300E-07 |
| 27909 | 1396 | K1396H | E | 3.1e-07 | 3.7e-07 | 1.3e-07 | 2.700E-07 | 1.249E-07 | 1.300E-07 | 3.700E-07 |
| 27910 | 1396 | K1396I | E | 6.6e-07 | 1.6e-07 | 2.3e-07 | 3.500E-07 | 2.707E-07 | 1.600E-07 | 6.600E-07 |
| 27911 | 1396 | K1396L | E | 2.6e-07 | 6.7e-07 | 2.6e-07 | 3.967E-07 | 2.367E-07 | 2.600E-07 | 6.700E-07 |
| 27912 | 1396 | K1396K | E | 2.1e-07 | 1.5e-07 | 2.2e-07 | 1.933E-07 | 3.786E-08 | 1.500E-07 | 2.200E-07 |
| 27913 | 1396 | K1396M | E | 4.6e-07 | 2.3e-07 | 2.7e-07 | 3.200E-07 | 1.229E-07 | 2.300E-07 | 4.600E-07 |
| 27914 | 1396 | K1396F | E | 2.5e-07 | 1.8e-07 | 2.7e-07 | 2.333E-07 | 4.726E-08 | 1.800E-07 | 2.700E-07 |
| 27915 | 1396 | K1396P | E | 2.6e-07 | 4e-07 | 3.3e-07 | 3.300E-07 | 7.000E-08 | 2.600E-07 | 4.000E-07 |
| 27916 | 1396 | K1396S | E | 2.2e-07 | 2.4e-07 | 1.9e-07 | 2.167E-07 | 2.517E-08 | 1.900E-07 | 2.400E-07 |
| 27917 | 1396 | K1396T | E | 1.4e-07 | 2.7e-07 | 3.3e-07 | 2.467E-07 | 9.713E-08 | 1.400E-07 | 3.300E-07 |
| 27918 | 1396 | K1396W | E | 1.9e-07 | 1.6e-07 | 2.2e-07 | 1.900E-07 | 3.000E-08 | 1.600E-07 | 2.200E-07 |
| 27919 | 1396 | K1396Y | E | 2.1e-07 | 7.3e-07 | 2.4e-07 | 3.933E-07 | 2.919E-07 | 2.100E-07 | 7.300E-07 |
| 27920 | 1396 | K1396V | E | 1.2e-07 | 1.8e-07 | 1.4e-07 | 1.467E-07 | 3.055E-08 | 1.200E-07 | 1.800E-07 |
| 27921 | 1397 | G1397A | E | 2e-07 | 2.4e-07 | 3.6e-07 | 2.667E-07 | 8.327E-08 | 2.000E-07 | 3.600E-07 |
| 27922 | 1397 | G1397R | E | 2.8e-07 | 5.3e-07 | 2.1e-07 | 3.400E-07 | 1.682E-07 | 2.100E-07 | 5.300E-07 |
| 27923 | 1397 | G1397N | E | 1.9e-07 | 3.3e-07 | 2.3e-07 | 2.500E-07 | 7.211E-08 | 1.900E-07 | 3.300E-07 |
| 27924 | 1397 | G1397D | E | 5e-07 | 2.5e-07 | 2.5e-07 | 3.333E-07 | 1.443E-07 | 2.500E-07 | 5.000E-07 |
| 27925 | 1397 | G1397C | E | 3.3e-07 | 2.7e-07 | 4.1e-07 | 3.367E-07 | 7.024E-08 | 2.700E-07 | 4.100E-07 |
| 27926 | 1397 | G1397E | E | 2.1e-07 | 6.3e-08 | 2e-07 | 1.577E-07 | 8.214E-08 | 6.300E-08 | 2.100E-07 |
| 27927 | 1397 | G1397Q | E | 2.8e-07 | 2.1e-07 | 3.4e-07 | 2.767E-07 | 6.506E-08 | 2.100E-07 | 3.400E-07 |
| 27928 | 1397 | G1397G | E | 1.8e-07 | 2e-07 | 4.7e-07 | 2.833E-07 | 1.620E-07 | 1.800E-07 | 4.700E-07 |
| 27929 | 1397 | G1397H | E | 2.5e-07 | 3.1e-07 | 2.5e-07 | 2.700E-07 | 3.464E-08 | 2.500E-07 | 3.100E-07 |
| 27930 | 1397 | G1397I | E | 3.4e-07 | 2.4e-07 | 3e-07 | 2.933E-07 | 5.033E-08 | 2.400E-07 | 3.400E-07 |
| 27931 | 1397 | G1397L | E | 5.2e-07 | 2.3e-07 | 2.2e-07 | 3.233E-07 | 1.704E-07 | 2.200E-07 | 5.200E-07 |
| 27932 | 1397 | G1397K | E | 8.8e-08 | 2.2e-07 | 2.4e-07 | 1.827E-07 | 8.259E-08 | 8.800E-08 | 2.400E-07 |
| 27933 | 1397 | G1397M | E | 1.7e-07 | 2.1e-07 | 2.2e-07 | 2.000E-07 | 2.646E-08 | 1.700E-07 | 2.200E-07 |
| 27934 | 1397 | G1397F | E | 2.5e-07 | 4.5e-07 | 2.7e-07 | 3.233E-07 | 1.102E-07 | 2.500E-07 | 4.500E-07 |
| 27935 | 1397 | G1397P | E | 1.1e-07 | 1.9e-07 | 1.3e-07 | 1.433E-07 | 4.163E-08 | 1.100E-07 | 1.900E-07 |
| 27936 | 1397 | G1397S | E | 1.7e-07 | 1.4e-07 | 2.2e-07 | 1.767E-07 | 4.041E-08 | 1.400E-07 | 2.200E-07 |
| 27937 | 1397 | G1397T | E | 1.4e-07 | 1.6e-07 | 2.4e-07 | 1.800E-07 | 5.292E-08 | 1.400E-07 | 2.400E-07 |
| 27938 | 1397 | G1397W | E | 2.5e-07 | 3.7e-07 | 1.4e-07 | 2.533E-07 | 1.150E-07 | 1.400E-07 | 3.700E-07 |
| 27939 | 1397 | G1397Y | E | 3.3e-07 | 2.9e-07 | 7.6e-07 | 4.600E-07 | 2.606E-07 | 2.900E-07 | 7.600E-07 |
| 27940 | 1397 | G1397V | E | 1.8e-07 | 2.7e-07 | 6.4e-07 | 3.633E-07 | 2.438E-07 | 1.800E-07 | 6.400E-07 |
| 27941 | 1398 | P1398A | E | 1.5e-07 | 2.6e-07 | 1.8e-07 | 1.967E-07 | 5.686E-08 | 1.500E-07 | 2.600E-07 |
| 27942 | 1398 | P1398R | E | 1.2e-07 | 1.2e-07 | 4.5e-07 | 2.300E-07 | 1.905E-07 | 1.200E-07 | 4.500E-07 |
| 27943 | 1398 | P1398N | E | 3.7e-07 | 2.6e-07 | 4.1e-07 | 3.467E-07 | 7.767E-08 | 2.600E-07 | 4.100E-07 |
| 27944 | 1398 | P1398D | E | 4.6e-07 | 2.6e-07 | 2.9e-07 | 3.367E-07 | 1.079E-07 | 2.600E-07 | 4.600E-07 |
| 27945 | 1398 | P1398C | E | 5.5e-07 | 8.5e-08 | 4.5e-07 | 3.617E-07 | 2.448E-07 | 8.500E-08 | 5.500E-07 |
| 27946 | 1398 | P1398E | E | 2.4e-07 | 2.2e-07 | 2.1e-07 | 2.233E-07 | 1.528E-08 | 2.100E-07 | 2.400E-07 |
| 27947 | 1398 | P1398Q | E | 1.7e-07 | 2e-07 | 5.3e-07 | 3.000E-07 | 1.997E-07 | 1.700E-07 | 5.300E-07 |
| 27948 | 1398 | P1398G | E | 2.1e-07 | 2.4e-07 | 3.1e-07 | 2.533E-07 | 5.132E-08 | 2.100E-07 | 3.100E-07 |
| 27949 | 1398 | P1398H | E | 1.9e-07 | 3.2e-07 | 2.1e-07 | 2.400E-07 | 7.000E-08 | 1.900E-07 | 3.200E-07 |
| 27950 | 1398 | P1398I | E | 2.3e-07 | 3.1e-07 | 2.1e-07 | 2.500E-07 | 5.292E-08 | 2.100E-07 | 3.100E-07 |
| 27951 | 1398 | P1398L | E | 4e-07 | 1.7e-07 | 3.7e-07 | 3.133E-07 | 1.250E-07 | 1.700E-07 | 4.000E-07 |
| 27952 | 1398 | P1398K | E | 2.1e-07 | 4.5e-07 | 2.2e-07 | 2.933E-07 | 1.358E-07 | 2.100E-07 | 4.500E-07 |
| 27953 | 1398 | P1398M | E | 2.9e-07 | 1.9e-07 | 1.6e-07 | 2.133E-07 | 6.807E-08 | 1.600E-07 | 2.900E-07 |
| 27954 | 1398 | P1398F | E | 3.7e-07 | 2.1e-07 | 2.2e-07 | 2.667E-07 | 8.963E-08 | 2.100E-07 | 3.700E-07 |
| 27955 | 1398 | P1398P | E | 3.4e-07 | 2.3e-07 | 2.8e-07 | 2.833E-07 | 5.508E-08 | 2.300E-07 | 3.400E-07 |
| 27956 | 1398 | P1398S | E | 3.1e-07 | 1.6e-07 | 2.4e-07 | 2.367E-07 | 7.506E-08 | 1.600E-07 | 3.100E-07 |
| 27957 | 1398 | P1398T | E | 1.7e-07 | 4.5e-07 | 2.4e-07 | 2.867E-07 | 1.457E-07 | 1.700E-07 | 4.500E-07 |
| 27958 | 1398 | P1398W | E | 3.9e-07 | 1.5e-07 | 3.1e-07 | 2.833E-07 | 1.222E-07 | 1.500E-07 | 3.900E-07 |
| 27959 | 1398 | P1398Y | E | 2.5e-07 | 2e-07 | 9.8e-08 | 1.827E-07 | 7.747E-08 | 9.800E-08 | 2.500E-07 |
| 27960 | 1398 | P1398V | E | 3.1e-07 | 2.2e-07 | 2.4e-07 | 2.567E-07 | 4.726E-08 | 2.200E-07 | 3.100E-07 |
| 27961 | 1399 | S1399A | E | 3.1e-07 | 1.8e-07 | 5.8e-07 | 3.567E-07 | 2.040E-07 | 1.800E-07 | 5.800E-07 |
| 27962 | 1399 | S1399R | E | 2.9e-07 | 2.9e-07 | 1.9e-07 | 2.567E-07 | 5.774E-08 | 1.900E-07 | 2.900E-07 |
| 27963 | 1399 | S1399N | E | 5.8e-07 | 1.9e-07 | 3.3e-07 | 3.667E-07 | 1.976E-07 | 1.900E-07 | 5.800E-07 |
| 27964 | 1399 | S1399D | E | 3e-07 | 3.7e-07 | 1.5e-07 | 2.733E-07 | 1.124E-07 | 1.500E-07 | 3.700E-07 |

| | | | | | | | | | | |
|---|---|---|---|---|---|---|---|---|---|---|
| 27965 | 1399 | S1399C | E | 2.6e-07 | 2.6e-07 | 2.2e-07 | 2.467E-07 | 2.309E-08 | 2.200E-07 | 2.600E-07 |
| 27966 | 1399 | S1399E | E | 2.1e-07 | 4.6e-07 | 3.2e-07 | 3.300E-07 | 1.253E-07 | 2.100E-07 | 4.600E-07 |
| 27967 | 1399 | S1399Q | E | 1.9e-07 | 3.1e-07 | 3.1e-07 | 2.700E-07 | 6.928E-08 | 1.900E-07 | 3.100E-07 |
| 27968 | 1399 | S1399G | E | 1.7e-07 | 1.9e-07 | 3.1e-07 | 2.233E-07 | 7.572E-08 | 1.700E-07 | 3.100E-07 |
| 27969 | 1399 | S1399H | E | 1.2e-07 | 1.6e-07 | 3e-07 | 1.933E-07 | 9.452E-08 | 1.200E-07 | 3.000E-07 |
| 27970 | 1399 | S1399I | E | 1.3e-07 | 7.3e-07 | 2.3e-07 | 3.633E-07 | 3.215E-07 | 1.300E-07 | 7.300E-07 |
| 27971 | 1399 | S1399L | E | 4.6e-07 | 1.1e-07 | 2e-07 | 2.567E-07 | 1.818E-07 | 1.100E-07 | 4.600E-07 |
| 27972 | 1399 | S1399K | E | 2.8e-07 | 1.6e-07 | 3.5e-07 | 2.633E-07 | 9.609E-08 | 1.600E-07 | 3.500E-07 |
| 27973 | 1399 | S1399M | E | 3.1e-07 | 4.6e-07 | 2.5e-07 | 3.400E-07 | 1.082E-07 | 2.500E-07 | 4.600E-07 |
| 27974 | 1399 | S1399F | E | 2.2e-07 | 1.6e-07 | 4.1e-07 | 2.633E-07 | 1.305E-07 | 1.600E-07 | 4.100E-07 |
| 27975 | 1399 | S1399P | E | 2.4e-07 | 2.4e-07 | 1.9e-07 | 2.233E-07 | 2.887E-08 | 1.900E-07 | 2.400E-07 |
| 27976 | 1399 | S1399S | E | 3.4e-07 | 1.3e-07 | 1.5e-07 | 2.067E-07 | 1.159E-07 | 1.300E-07 | 3.400E-07 |
| 27977 | 1399 | S1399T | E | 3.3e-07 | 2.6e-07 | 1.9e-07 | 2.600E-07 | 7.000E-08 | 1.900E-07 | 3.300E-07 |
| 27978 | 1399 | S1399W | E | 2.1e-07 | 1.6e-07 | 2.5e-07 | 2.067E-07 | 4.509E-08 | 1.600E-07 | 2.500E-07 |
| 27979 | 1399 | S1399Y | E | 5.7e-07 | 2.6e-07 | 2.3e-07 | 3.533E-07 | 1.882E-07 | 2.300E-07 | 5.700E-07 |
| 27980 | 1399 | S1399V | E | 1.7e-07 | 3.3e-07 | 6e-07 | 3.667E-07 | 2.173E-07 | 1.700E-07 | 6.000E-07 |
| 27981 | 1400 | V1400A | E | 3.2e-07 | 1.7e-07 | 1.4e-07 | 2.100E-07 | 9.644E-08 | 1.400E-07 | 3.200E-07 |
| 27982 | 1400 | V1400R | E | 2.5e-07 | 8.7e-08 | 3.3e-07 | 2.223E-07 | 1.238E-07 | 8.700E-08 | 3.300E-07 |
| 27983 | 1400 | V1400N | E | 2.1e-07 | 2.9e-07 | 2.5e-07 | 2.500E-07 | 4.000E-08 | 2.100E-07 | 2.900E-07 |
| 27984 | 1400 | V1400D | E | 2.2e-07 | 3.6e-07 | 7.3e-08 | 2.177E-07 | 1.435E-07 | 7.300E-08 | 3.600E-07 |
| 27985 | 1400 | V1400C | E | 2e-07 | 1e-07 | 5.2e-07 | 2.733E-07 | 2.194E-07 | 1.000E-07 | 5.200E-07 |
| 27986 | 1400 | V1400E | E | 3.6e-07 | 2.3e-07 | 3.1e-07 | 3.000E-07 | 6.557E-08 | 2.300E-07 | 3.600E-07 |
| 27987 | 1400 | V1400Q | E | 1.9e-07 | 2e-07 | 2.3e-07 | 2.067E-07 | 2.082E-08 | 1.900E-07 | 2.300E-07 |
| 27988 | 1400 | V1400G | E | 2.9e-07 | 3.1e-07 | 2.2e-07 | 2.733E-07 | 4.726E-08 | 2.200E-07 | 3.100E-07 |
| 27989 | 1400 | V1400H | E | 5.6e-07 | 1.9e-07 | 2.4e-07 | 3.300E-07 | 2.007E-07 | 1.900E-07 | 5.600E-07 |
| 27990 | 1400 | V1400I | E | 4.1e-07 | 1.9e-07 | 2.4e-07 | 2.800E-07 | 1.153E-07 | 1.900E-07 | 4.100E-07 |
| 27991 | 1400 | V1400L | E | 5.4e-07 | 3.9e-07 | 2.2e-07 | 3.833E-07 | 1.601E-07 | 2.200E-07 | 5.400E-07 |
| 27992 | 1400 | V1400K | E | 2.5e-07 | 1.8e-07 | 3.3e-07 | 2.533E-07 | 7.506E-08 | 1.800E-07 | 3.300E-07 |
| 27993 | 1400 | V1400M | E | 9.2e-08 | 3.8e-07 | 1.6e-07 | 2.107E-07 | 1.505E-07 | 9.200E-08 | 3.800E-07 |
| 27994 | 1400 | V1400F | E | 1.8e-07 | 1.7e-07 | 2.7e-07 | 2.067E-07 | 5.508E-08 | 1.700E-07 | 2.700E-07 |
| 27995 | 1400 | V1400P | E | 2.5e-07 | 2.2e-07 | 3.5e-07 | 2.733E-07 | 6.807E-08 | 2.200E-07 | 3.500E-07 |
| 27996 | 1400 | V1400S | E | 1.7e-07 | 4.7e-07 | 3.1e-07 | 3.167E-07 | 1.501E-07 | 1.700E-07 | 4.700E-07 |
| 27997 | 1400 | V1400T | E | 3.1e-07 | 2.3e-07 | 1.2e-07 | 2.200E-07 | 9.539E-08 | 1.200E-07 | 3.100E-07 |
| 27998 | 1400 | V1400W | E | 1e-07 | 1.1e-07 | 3.3e-07 | 1.800E-07 | 1.300E-07 | 1.000E-07 | 3.300E-07 |
| 27999 | 1400 | V1400Y | E | 1.4e-07 | 1.9e-07 | 2.8e-07 | 2.033E-07 | 7.095E-08 | 1.400E-07 | 2.800E-07 |
| 28000 | 1400 | V1400V | E | 2.5e-07 | 9e-08 | 3e-07 | 2.133E-07 | 1.097E-07 | 9.000E-08 | 3.000E-07 |
| 28001 | 1401 | F1401A | E | 1.2e-07 | 2.3e-07 | 2.5e-07 | 2.000E-07 | 7.000E-08 | 1.200E-07 | 2.500E-07 |
| 28002 | 1401 | F1401R | E | 2.7e-07 | 3.2e-07 | 2.1e-07 | 2.667E-07 | 5.508E-08 | 2.100E-07 | 3.200E-07 |
| 28003 | 1401 | F1401N | E | 3.9e-07 | 2.1e-07 | 4.2e-07 | 3.400E-07 | 1.136E-07 | 2.100E-07 | 4.200E-07 |
| 28004 | 1401 | F1401D | E | 5.5e-07 | 2.4e-07 | 5.1e-07 | 4.333E-07 | 1.686E-07 | 2.400E-07 | 5.500E-07 |
| 28005 | 1401 | F1401C | E | 1.6e-07 | 2e-07 | 1.9e-07 | 1.833E-07 | 2.082E-08 | 1.600E-07 | 2.000E-07 |
| 28006 | 1401 | F1401E | E | 1.9e-07 | 4.3e-07 | 1.2e-07 | 2.467E-07 | 1.626E-07 | 1.200E-07 | 4.300E-07 |
| 28007 | 1401 | F1401Q | E | 2.5e-07 | 3.3e-07 | 1.9e-07 | 2.567E-07 | 7.024E-08 | 1.900E-07 | 3.300E-07 |
| 28008 | 1401 | F1401G | E | 1.2e-07 | 1.9e-07 | 3.5e-07 | 2.200E-07 | 1.179E-07 | 1.200E-07 | 3.500E-07 |
| 28009 | 1401 | F1401H | E | 1.6e-07 | 2.6e-07 | 3.3e-07 | 2.500E-07 | 8.544E-08 | 1.600E-07 | 3.300E-07 |
| 28010 | 1401 | F1401I | E | 5.5e-07 | 2.4e-07 | 1.4e-07 | 3.100E-07 | 2.138E-07 | 1.400E-07 | 5.500E-07 |
| 28011 | 1401 | F1401L | E | 2.2e-07 | 1.7e-07 | 1.9e-07 | 1.933E-07 | 2.517E-08 | 1.700E-07 | 2.200E-07 |
| 28012 | 1401 | F1401K | E | 4.7e-07 | 3e-07 | 2.1e-07 | 3.267E-07 | 1.320E-07 | 2.100E-07 | 4.700E-07 |
| 28013 | 1401 | F1401M | E | 3.3e-07 | 1.4e-07 | 1.2e-07 | 1.967E-07 | 1.159E-07 | 1.200E-07 | 3.300E-07 |
| 28014 | 1401 | F1401F | E | 6.8e-07 | 3.3e-07 | 1.6e-07 | 3.900E-07 | 2.651E-07 | 1.600E-07 | 6.800E-07 |
| 28015 | 1401 | F1401P | E | 1.9e-07 | 2.5e-07 | 4e-07 | 2.800E-07 | 1.082E-07 | 1.900E-07 | 4.000E-07 |
| 28016 | 1401 | F1401S | E | 2.4e-07 | 3.1e-07 | 5.5e-07 | 3.667E-07 | 1.626E-07 | 2.400E-07 | 5.500E-07 |
| 28017 | 1401 | F1401T | E | 1.8e-07 | 1.9e-07 | 2.1e-07 | 1.933E-07 | 1.528E-08 | 1.800E-07 | 2.100E-07 |
| 28018 | 1401 | F1401W | E | 1.8e-07 | 1e-07 | 2e-07 | 1.600E-07 | 5.292E-08 | 1.000E-07 | 2.000E-07 |
| 28019 | 1401 | F1401Y | E | 2.4e-07 | 3.3e-07 | 3.9e-07 | 3.200E-07 | 7.550E-08 | 2.400E-07 | 3.900E-07 |
| 28020 | 1401 | F1401V | E | 3e-07 | 4e-07 | 3.4e-07 | 3.467E-07 | 5.033E-08 | 3.000E-07 | 4.000E-07 |
| 28021 | 1402 | P1402A | E | 1.5e-07 | 2.1e-07 | 2.2e-07 | 1.933E-07 | 3.786E-08 | 1.500E-07 | 2.200E-07 |
| 28022 | 1402 | P1402R | E | 2.3e-07 | 2.3e-07 | 6.3e-07 | 3.633E-07 | 2.309E-07 | 2.300E-07 | 6.300E-07 |
| 28023 | 1402 | P1402N | E | 3.8e-07 | 2.2e-07 | 5.3e-07 | 3.767E-07 | 1.550E-07 | 2.200E-07 | 5.300E-07 |
| 28024 | 1402 | P1402D | E | 2.2e-07 | 3e-07 | 2.6e-07 | 2.600E-07 | 4.000E-08 | 2.200E-07 | 3.000E-07 |
| 28025 | 1402 | P1402C | E | 1.8e-07 | 2.6e-07 | 3.3e-07 | 2.567E-07 | 7.506E-08 | 1.800E-07 | 3.300E-07 |
| 28026 | 1402 | P1402E | E | 2.2e-07 | 2.9e-07 | 3.3e-07 | 2.800E-07 | 5.568E-08 | 2.200E-07 | 3.300E-07 |
| 28027 | 1402 | P1402Q | E | 2.1e-07 | 2.7e-07 | 2.5e-07 | 2.433E-07 | 3.055E-08 | 2.100E-07 | 2.700E-07 |
| 28028 | 1402 | P1402G | E | 3.1e-07 | 3.5e-07 | 1.7e-07 | 2.767E-07 | 9.452E-08 | 1.700E-07 | 3.500E-07 |
| 28029 | 1402 | P1402H | E | 1.9e-07 | 3.3e-07 | 2.8e-07 | 2.667E-07 | 7.095E-08 | 1.900E-07 | 3.300E-07 |
| 28030 | 1402 | P1402I | E | 4.4e-07 | 4e-07 | 3.2e-07 | 3.867E-07 | 6.110E-08 | 3.200E-07 | 4.400E-07 |
| 28031 | 1402 | P1402L | E | 1.1e-07 | 1.1e-07 | 2.3e-07 | 1.500E-07 | 6.928E-08 | 1.100E-07 | 2.300E-07 |
| 28032 | 1402 | P1402K | E | 4.2e-07 | 1.1e-07 | 2.4e-07 | 2.567E-07 | 1.557E-07 | 1.100E-07 | 4.200E-07 |
| 28033 | 1402 | P1402M | E | 4.2e-07 | 3.1e-07 | 1.7e-07 | 3.000E-07 | 1.253E-07 | 1.700E-07 | 4.200E-07 |
| 28034 | 1402 | P1402F | E | 9.2e-08 | 1.8e-07 | 1.2e-07 | 1.307E-07 | 4.496E-08 | 9.200E-08 | 1.800E-07 |
| 28035 | 1402 | P1402P | E | 3.1e-07 | 4.2e-07 | 3.6e-07 | 3.633E-07 | 5.508E-08 | 3.100E-07 | 4.200E-07 |
| 28036 | 1402 | P1402S | E | 3.4e-07 | 1.9e-07 | 1.9e-07 | 2.400E-07 | 8.660E-08 | 1.900E-07 | 3.400E-07 |
| 28037 | 1402 | P1402T | E | 1.7e-07 | 7.4e-07 | 3.2e-07 | 4.100E-07 | 2.955E-07 | 1.700E-07 | 7.400E-07 |
| 28038 | 1402 | P1402W | E | 1.1e-06 | 1.5e-07 | 4e-07 | 5.500E-07 | 4.924E-07 | 1.500E-07 | 1.100E-06 |
| 28039 | 1402 | P1402Y | E | 2e-07 | 3.8e-07 | 1.9e-07 | 2.567E-07 | 1.069E-07 | 1.900E-07 | 3.800E-07 |
| 28040 | 1402 | P1402V | E | 1.8e-07 | 3.1e-07 | 2.4e-07 | 2.433E-07 | 6.506E-08 | 1.800E-07 | 3.100E-07 |
| 28041 | 1403 | L1403A | E | 3.4e-07 | 3.9e-07 | 2.4e-07 | 3.233E-07 | 7.638E-08 | 2.400E-07 | 3.900E-07 |
| 28042 | 1403 | L1403R | E | 2.3e-07 | 1.9e-07 | 2.9e-07 | 2.367E-07 | 5.033E-08 | 1.900E-07 | 2.900E-07 |
| 28043 | 1403 | L1403N | E | 3e-07 | 1.6e-07 | 3.4e-07 | 2.667E-07 | 9.452E-08 | 1.600E-07 | 3.400E-07 |
| 28044 | 1403 | L1403D | E | 1.5e-07 | 1.7e-07 | 3.1e-07 | 2.100E-07 | 8.718E-08 | 1.500E-07 | 3.100E-07 |
| 28045 | 1403 | L1403C | E | 2.7e-07 | 2.3e-07 | 4e-07 | 3.000E-07 | 8.888E-08 | 2.300E-07 | 4.000E-07 |
| 28046 | 1403 | L1403E | E | 2.9e-07 | 1.5e-07 | 1.6e-07 | 2.000E-07 | 7.810E-08 | 1.500E-07 | 2.900E-07 |
| 28047 | 1403 | L1403Q | E | 1.9e-07 | 2.8e-07 | 1.9e-07 | 2.200E-07 | 5.196E-08 | 1.900E-07 | 2.800E-07 |
| 28048 | 1403 | L1403G | E | 3.5e-07 | 3.3e-07 | 1.7e-07 | 2.833E-07 | 9.866E-08 | 1.700E-07 | 3.500E-07 |
| 28049 | 1403 | L1403H | E | 4.9e-07 | 8e-07 | 3.4e-07 | 5.433E-07 | 2.346E-07 | 3.400E-07 | 8.000E-07 |

| | | | | | | | | | | |
|---|---|---|---|---|---|---|---|---|---|---|
| 28050 | 1403 | L1403I | E | 1.6e-07 | 2.1e-07 | 3.3e-07 | 2.333E-07 | 8.737E-08 | 1.600E-07 | 3.300E-07 |
| 28051 | 1403 | L1403L | E | 7.5e-07 | 3.3e-07 | 2.4e-07 | 4.400E-07 | 2.722E-07 | 2.400E-07 | 7.500E-07 |
| 28052 | 1403 | L1403K | E | 1.3e-07 | 5e-07 | 6.6e-07 | 4.300E-07 | 2.718E-07 | 1.300E-07 | 6.600E-07 |
| 28053 | 1403 | L1403M | E | 3.8e-07 | 2.3e-07 | 3.1e-07 | 3.067E-07 | 7.506E-08 | 2.300E-07 | 3.800E-07 |
| 28054 | 1403 | L1403F | E | 3.2e-07 | 1.8e-07 | 3e-07 | 2.667E-07 | 7.572E-08 | 1.800E-07 | 3.200E-07 |
| 28055 | 1403 | L1403P | E | 1.4e-07 | 3.5e-07 | 2.4e-07 | 2.433E-07 | 1.050E-07 | 1.400E-07 | 3.500E-07 |
| 28056 | 1403 | L1403S | E | 1.5e-07 | 2.5e-07 | 2.7e-07 | 2.233E-07 | 6.429E-08 | 1.500E-07 | 2.700E-07 |
| 28057 | 1403 | L1403T | E | 3.6e-07 | 2.7e-07 | 1.7e-07 | 2.667E-07 | 9.504E-08 | 1.700E-07 | 3.600E-07 |
| 28058 | 1403 | L1403W | E | 4.5e-07 | 5.6e-07 | 4.4e-07 | 4.833E-07 | 6.658E-08 | 4.400E-07 | 5.600E-07 |
| 28059 | 1403 | L1403Y | E | 1.9e-07 | 2.3e-07 | 3.2e-07 | 2.467E-07 | 6.658E-08 | 1.900E-07 | 3.200E-07 |
| 28060 | 1403 | L1403V | E | 3.3e-07 | 3.1e-07 | 1.9e-07 | 2.767E-07 | 7.572E-08 | 1.900E-07 | 3.300E-07 |
| 28061 | 1404 | A1404A | E | 2.2e-07 | 2.5e-07 | 1.2e-07 | 1.967E-07 | 6.807E-08 | 1.200E-07 | 2.500E-07 |
| 28062 | 1404 | A1404R | E | 1.4e-07 | 2.1e-07 | 4.6e-07 | 2.700E-07 | 1.682E-07 | 1.400E-07 | 4.600E-07 |
| 28063 | 1404 | A1404N | E | 1.4e-07 | 3.8e-07 | 1.4e-07 | 2.200E-07 | 1.386E-07 | 1.400E-07 | 3.800E-07 |
| 28064 | 1404 | A1404D | E | 4.6e-07 | 3.4e-07 | 6e-07 | 4.667E-07 | 1.301E-07 | 3.400E-07 | 6.000E-07 |
| 28065 | 1404 | A1404C | E | 3.8e-07 | 4.7e-07 | 2.9e-07 | 3.800E-07 | 9.000E-08 | 2.900E-07 | 4.700E-07 |
| 28066 | 1404 | A1404E | E | 4.3e-07 | 1.9e-07 | 1.9e-07 | 2.700E-07 | 1.386E-07 | 1.900E-07 | 4.300E-07 |
| 28067 | 1404 | A1404Q | E | 1.8e-07 | 2.5e-07 | 2.7e-07 | 2.333E-07 | 4.726E-08 | 1.800E-07 | 2.700E-07 |
| 28068 | 1404 | A1404G | E | 1.9e-07 | 3.6e-07 | 1.2e-07 | 2.233E-07 | 1.234E-07 | 1.200E-07 | 3.600E-07 |
| 28069 | 1404 | A1404H | E | 1.5e-07 | 1.6e-07 | 3.3e-07 | 2.133E-07 | 1.012E-07 | 1.500E-07 | 3.300E-07 |
| 28070 | 1404 | A1404I | E | 1.8e-07 | 1.8e-07 | 3.2e-07 | 2.267E-07 | 8.083E-08 | 1.800E-07 | 3.200E-07 |
| 28071 | 1404 | A1404L | E | 3.8e-07 | 1.5e-07 | 4.1e-07 | 3.133E-07 | 1.422E-07 | 1.500E-07 | 4.100E-07 |
| 28072 | 1404 | A1404K | E | 2.5e-07 | 3.6e-07 | 2.3e-07 | 2.800E-07 | 7.000E-08 | 2.300E-07 | 3.600E-07 |
| 28073 | 1404 | A1404M | E | 4.3e-07 | 2.7e-07 | 3.2e-07 | 3.400E-07 | 8.185E-08 | 2.700E-07 | 4.300E-07 |
| 28074 | 1404 | A1404F | E | 1.9e-07 | 1.7e-07 | 3.5e-07 | 2.367E-07 | 9.866E-08 | 1.700E-07 | 3.500E-07 |
| 28075 | 1404 | A1404P | E | 4.4e-07 | 4.3e-07 | 2.2e-07 | 3.633E-07 | 1.242E-07 | 2.200E-07 | 4.400E-07 |
| 28076 | 1404 | A1404S | E | 3.2e-07 | 2.6e-07 | 2.3e-07 | 2.700E-07 | 4.583E-08 | 2.300E-07 | 3.200E-07 |
| 28077 | 1404 | A1404T | E | 1.2e-07 | 2.3e-07 | 4.3e-07 | 2.600E-07 | 1.572E-07 | 1.200E-07 | 4.300E-07 |
| 28078 | 1404 | A1404W | E | 1.5e-07 | 2.2e-07 | 1.8e-07 | 1.833E-07 | 3.512E-08 | 1.500E-07 | 2.200E-07 |
| 28079 | 1404 | A1404Y | E | 1.5e-07 | 1.5e-07 | 4.7e-07 | 2.567E-07 | 1.848E-07 | 1.500E-07 | 4.700E-07 |
| 28080 | 1404 | A1404V | E | 3.8e-07 | 3.1e-07 | 1.5e-07 | 2.800E-07 | 1.179E-07 | 1.500E-07 | 3.800E-07 |
| 28081 | 1405 | P1405A | E | 4e-07 | 2.8e-07 | 2.1e-07 | 2.967E-07 | 9.609E-08 | 2.100E-07 | 4.000E-07 |
| 28082 | 1405 | P1405R | E | 2.2e-07 | 4.8e-07 | 2e-07 | 3.000E-07 | 1.562E-07 | 2.000E-07 | 4.800E-07 |
| 28083 | 1405 | P1405N | E | 1.1e-07 | 2.2e-07 | 4.8e-07 | 2.700E-07 | 1.900E-07 | 1.100E-07 | 4.800E-07 |
| 28084 | 1405 | P1405D | E | 3e-07 | 1.4e-07 | 4.7e-07 | 3.033E-07 | 1.650E-07 | 1.400E-07 | 4.700E-07 |
| 28085 | 1405 | P1405C | E | 1.4e-07 | 6.7e-07 | 2.9e-07 | 3.667E-07 | 2.732E-07 | 1.400E-07 | 6.700E-07 |
| 28086 | 1405 | P1405E | E | 4.4e-07 | 3.3e-07 | 2.3e-07 | 3.333E-07 | 1.050E-07 | 2.300E-07 | 4.400E-07 |
| 28087 | 1405 | P1405Q | E | 1.2e-07 | 2.5e-07 | 3.2e-07 | 2.300E-07 | 1.015E-07 | 1.200E-07 | 3.200E-07 |
| 28088 | 1405 | P1405G | E | 8e-07 | 1.6e-07 | 2e-07 | 3.867E-07 | 3.585E-07 | 1.600E-07 | 8.000E-07 |
| 28089 | 1405 | P1405H | E | 3e-07 | 1.2e-07 | 1.8e-07 | 2.000E-07 | 9.165E-08 | 1.200E-07 | 3.000E-07 |
| 28090 | 1405 | P1405I | E | 1.5e-07 | 1.6e-07 | 1.7e-07 | 1.600E-07 | 1.000E-08 | 1.500E-07 | 1.700E-07 |
| 28091 | 1405 | P1405L | E | 2.4e-07 | 3.4e-07 | 2.5e-07 | 2.767E-07 | 5.508E-08 | 2.400E-07 | 3.400E-07 |
| 28092 | 1405 | P1405K | E | 1.3e-07 | 1.7e-07 | 1.3e-07 | 1.433E-07 | 2.309E-08 | 1.300E-07 | 1.700E-07 |
| 28093 | 1405 | P1405M | E | 3.6e-07 | 3.4e-07 | 2.6e-07 | 3.200E-07 | 5.292E-08 | 2.600E-07 | 3.600E-07 |
| 28094 | 1405 | P1405F | E | 2.2e-07 | 3.2e-07 | 2.2e-07 | 2.533E-07 | 5.774E-08 | 2.200E-07 | 3.200E-07 |
| 28095 | 1405 | P1405P | E | 1.4e-07 | 1.9e-07 | 1.6e-07 | 1.633E-07 | 2.517E-08 | 1.400E-07 | 1.900E-07 |
| 28096 | 1405 | P1405S | E | 2.7e-07 | 7e-08 | 2.3e-07 | 1.900E-07 | 1.058E-07 | 7.000E-08 | 2.700E-07 |
| 28097 | 1405 | P1405T | E | 2.3e-07 | 2.4e-07 | 3.2e-07 | 2.633E-07 | 4.933E-08 | 2.300E-07 | 3.200E-07 |
| 28098 | 1405 | P1405W | E | 1.8e-07 | 2.9e-07 | 2.5e-07 | 2.400E-07 | 5.568E-08 | 1.800E-07 | 2.900E-07 |
| 28099 | 1405 | P1405Y | E | 4.3e-07 | 1.4e-07 | 1.7e-07 | 2.467E-07 | 1.595E-07 | 1.400E-07 | 4.300E-07 |
| 28100 | 1405 | P1405V | E | 2.2e-07 | 6e-07 | 1.6e-07 | 3.267E-07 | 2.386E-07 | 1.600E-07 | 6.000E-07 |
| 28101 | 1406 | S1406A | E | 2.4e-07 | 1.8e-07 | 2.2e-07 | 2.133E-07 | 3.055E-08 | 1.800E-07 | 2.400E-07 |
| 28102 | 1406 | S1406R | E | 2e-07 | 2.4e-07 | 1.8e-07 | 2.067E-07 | 3.055E-08 | 1.800E-07 | 2.400E-07 |
| 28103 | 1406 | S1406N | E | 1.7e-07 | 4.6e-07 | 2.7e-07 | 3.000E-07 | 1.473E-07 | 1.700E-07 | 4.600E-07 |
| 28104 | 1406 | S1406D | E | 3.5e-07 | 4.9e-07 | 2.8e-07 | 3.733E-07 | 1.069E-07 | 2.800E-07 | 4.900E-07 |
| 28105 | 1406 | S1406C | E | 3.1e-07 | 1.7e-07 | 2e-07 | 2.267E-07 | 7.371E-08 | 1.700E-07 | 3.100E-07 |
| 28106 | 1406 | S1406E | E | 2.4e-07 | 3.6e-07 | 2.1e-07 | 2.700E-07 | 7.937E-08 | 2.100E-07 | 3.600E-07 |
| 28107 | 1406 | S1406Q | E | 4.5e-07 | 2.9e-07 | 2.6e-07 | 3.333E-07 | 1.021E-07 | 2.600E-07 | 4.500E-07 |
| 28108 | 1406 | S1406G | E | 5.1e-07 | 1.8e-07 | 3.4e-07 | 3.433E-07 | 1.650E-07 | 1.800E-07 | 5.100E-07 |
| 28109 | 1406 | S1406H | E | 3.6e-07 | 2.9e-07 | 1.8e-07 | 2.767E-07 | 9.074E-08 | 1.800E-07 | 3.600E-07 |
| 28110 | 1406 | S1406I | E | 3.2e-07 | 2.7e-07 | 3.3e-07 | 3.067E-07 | 3.215E-08 | 2.700E-07 | 3.300E-07 |
| 28111 | 1406 | S1406L | E | 2.1e-07 | 2.7e-07 | 1.9e-07 | 2.233E-07 | 4.163E-08 | 1.900E-07 | 2.700E-07 |
| 28112 | 1406 | S1406K | E | 8.1e-07 | 1.1e-07 | 3.9e-07 | 4.367E-07 | 3.523E-07 | 1.100E-07 | 8.100E-07 |
| 28113 | 1406 | S1406M | E | 2.2e-07 | 1.7e-07 | 1.8e-07 | 1.900E-07 | 2.646E-08 | 1.700E-07 | 2.200E-07 |
| 28114 | 1406 | S1406F | E | 1.8e-07 | 2.9e-07 | 1.7e-07 | 2.133E-07 | 6.658E-08 | 1.700E-07 | 2.900E-07 |
| 28115 | 1406 | S1406P | E | 8.3e-08 | 3.6e-07 | 2.7e-07 | 2.377E-07 | 1.413E-07 | 8.300E-08 | 3.600E-07 |
| 28116 | 1406 | S1406S | E | 1.2e-07 | 3e-07 | 1.2e-07 | 1.800E-07 | 1.039E-07 | 1.200E-07 | 3.000E-07 |
| 28117 | 1406 | S1406T | E | 1.3e-07 | 1.5e-07 | 4.2e-07 | 2.333E-07 | 1.620E-07 | 1.300E-07 | 4.200E-07 |
| 28118 | 1406 | S1406W | E | 2.4e-07 | 9.6e-08 | 2.5e-07 | 1.953E-07 | 8.617E-08 | 9.600E-08 | 2.500E-07 |
| 28119 | 1406 | S1406Y | E | 2.1e-07 | 2.8e-07 | 1.5e-07 | 2.133E-07 | 6.506E-08 | 1.500E-07 | 2.800E-07 |
| 28120 | 1406 | S1406V | E | 2.8e-07 | 2.3e-07 | 2.5e-07 | 2.533E-07 | 2.517E-08 | 2.300E-07 | 2.800E-07 |
| 28121 | 1407 | S1407A | E | 1.6e-07 | 1.8e-07 | 1.9e-07 | 1.767E-07 | 1.528E-08 | 1.600E-07 | 1.900E-07 |
| 28122 | 1407 | S1407R | E | 3.2e-07 | 1.5e-07 | 4.2e-07 | 2.967E-07 | 1.365E-07 | 1.500E-07 | 4.200E-07 |
| 28123 | 1407 | S1407N | E | 3.2e-07 | 4.8e-07 | 2.4e-07 | 3.467E-07 | 1.222E-07 | 2.400E-07 | 4.800E-07 |
| 28124 | 1407 | S1407D | E | 1.4e-07 | 5.2e-07 | 2.5e-07 | 3.033E-07 | 1.955E-07 | 1.400E-07 | 5.200E-07 |
| 28125 | 1407 | S1407C | E | 1.8e-07 | 2.2e-07 | 2.9e-07 | 2.300E-07 | 5.568E-08 | 1.800E-07 | 2.900E-07 |
| 28126 | 1407 | S1407E | E | 3.4e-07 | 1.2e-07 | 2.9e-07 | 2.500E-07 | 1.153E-07 | 1.200E-07 | 3.400E-07 |
| 28127 | 1407 | S1407Q | E | 2.3e-07 | 1.9e-07 | 2.5e-07 | 2.233E-07 | 3.055E-08 | 1.900E-07 | 2.500E-07 |
| 28128 | 1407 | S1407G | E | 2.7e-07 | 3e-07 | 2.9e-07 | 2.867E-07 | 1.528E-08 | 2.700E-07 | 3.000E-07 |
| 28129 | 1407 | S1407H | E | 2.6e-07 | 2.7e-07 | 1.3e-07 | 2.200E-07 | 7.810E-08 | 1.300E-07 | 2.700E-07 |
| 28130 | 1407 | S1407I | E | 2e-07 | 3.6e-07 | 1.6e-07 | 2.400E-07 | 1.058E-07 | 1.600E-07 | 3.600E-07 |
| 28131 | 1407 | S1407L | E | 2.3e-07 | 1.6e-07 | 2.7e-07 | 2.200E-07 | 5.568E-08 | 1.600E-07 | 2.700E-07 |
| 28132 | 1407 | S1407K | E | 2.6e-07 | 4.8e-07 | 2.6e-07 | 3.333E-07 | 1.270E-07 | 2.600E-07 | 4.800E-07 |
| 28133 | 1407 | S1407M | E | 3e-07 | 5.9e-07 | 2.4e-07 | 3.767E-07 | 1.872E-07 | 2.400E-07 | 5.900E-07 |
| 28134 | 1407 | S1407F | E | 3.3e-07 | 1.1e-07 | 3.2e-07 | 2.533E-07 | 1.242E-07 | 1.100E-07 | 3.300E-07 |

| | | | | | | | | | | |
|---|---|---|---|---|---|---|---|---|---|---|
| 28135 | 1407 | S1407P | E | 2.2e-07 | 1.8e-07 | 1.5e-07 | 1.833E-07 | 3.512E-08 | 1.500E-07 | 2.200E-07 |
| 28136 | 1407 | S1407S | E | 4.8e-07 | 4.7e-07 | 2.4e-07 | 3.967E-07 | 1.358E-07 | 2.400E-07 | 4.800E-07 |
| 28137 | 1407 | S1407T | E | 3.6e-07 | 2.8e-07 | 4.1e-07 | 3.500E-07 | 6.557E-08 | 2.800E-07 | 4.100E-07 |
| 28138 | 1407 | S1407W | E | 1.4e-07 | 2.9e-07 | 1.6e-07 | 1.967E-07 | 8.145E-08 | 1.400E-07 | 2.900E-07 |
| 28139 | 1407 | S1407Y | E | 3.6e-07 | 2.2e-07 | 3.6e-07 | 3.133E-07 | 8.083E-08 | 2.200E-07 | 3.600E-07 |
| 28140 | 1407 | S1407V | E | 1.5e-07 | 3.3e-07 | 2.6e-07 | 2.467E-07 | 9.074E-08 | 1.500E-07 | 3.300E-07 |
| 28141 | 1408 | K1408A | E | 1.6e-07 | 1.1e-07 | 1.8e-07 | 1.500E-07 | 3.606E-08 | 1.100E-07 | 1.800E-07 |
| 28142 | 1408 | K1408R | E | 2.7e-07 | 1.9e-07 | 3.2e-07 | 2.600E-07 | 6.557E-08 | 1.900E-07 | 3.200E-07 |
| 28143 | 1408 | K1408N | E | 1.5e-07 | 2e-07 | 1.6e-07 | 1.700E-07 | 2.646E-08 | 1.500E-07 | 2.000E-07 |
| 28144 | 1408 | K1408D | E | 3.2e-07 | 2.7e-07 | 2.7e-07 | 2.867E-07 | 2.887E-08 | 2.700E-07 | 3.200E-07 |
| 28145 | 1408 | K1408C | E | 4.9e-07 | 1.6e-07 | 2e-07 | 2.833E-07 | 1.801E-07 | 1.600E-07 | 4.900E-07 |
| 28146 | 1408 | K1408E | E | 4.3e-07 | 5.9e-07 | 1.1e-07 | 3.767E-07 | 2.444E-07 | 1.100E-07 | 5.900E-07 |
| 28147 | 1408 | K1408Q | E | 2.9e-07 | 2.1e-07 | 6.4e-07 | 3.800E-07 | 2.287E-07 | 2.100E-07 | 6.400E-07 |
| 28148 | 1408 | K1408G | E | 1.4e-07 | 2.4e-07 | 3.6e-07 | 2.467E-07 | 1.102E-07 | 1.400E-07 | 3.600E-07 |
| 28149 | 1408 | K1408H | E | 4e-07 | 3.3e-07 | 3.5e-07 | 3.600E-07 | 3.606E-08 | 3.300E-07 | 4.000E-07 |
| 28150 | 1408 | K1408I | E | 4.4e-07 | 2.1e-07 | 2.2e-07 | 2.900E-07 | 1.300E-07 | 2.100E-07 | 4.400E-07 |
| 28151 | 1408 | K1408L | E | 2.6e-07 | 1.8e-07 | 2.4e-07 | 2.267E-07 | 4.163E-08 | 1.800E-07 | 2.600E-07 |
| 28152 | 1408 | K1408K | E | 2.3e-07 | 1.6e-07 | 2.1e-07 | 2.000E-07 | 3.606E-08 | 1.600E-07 | 2.300E-07 |
| 28153 | 1408 | K1408M | E | 1.3e-07 | 2e-07 | 1.6e-07 | 1.633E-07 | 3.512E-08 | 1.300E-07 | 2.000E-07 |
| 28154 | 1408 | K1408F | E | 3e-07 | 3e-07 | 2.3e-07 | 2.767E-07 | 4.041E-08 | 2.300E-07 | 3.000E-07 |
| 28155 | 1408 | K1408P | E | 2.7e-07 | 1.4e-07 | 4.7e-07 | 2.933E-07 | 1.662E-07 | 1.400E-07 | 4.700E-07 |
| 28156 | 1408 | K1408S | E | 2e-07 | 1.3e-07 | 3.7e-07 | 2.333E-07 | 1.234E-07 | 1.300E-07 | 3.700E-07 |
| 28157 | 1408 | K1408T | E | 2.4e-07 | 4.6e-07 | 1.8e-07 | 2.933E-07 | 1.474E-07 | 1.800E-07 | 4.600E-07 |
| 28158 | 1408 | K1408W | E | 3.4e-07 | 1.3e-07 | 4.1e-07 | 2.933E-07 | 1.457E-07 | 1.300E-07 | 4.100E-07 |
| 28159 | 1408 | K1408Y | E | 1.6e-07 | 1.5e-07 | 3.3e-07 | 2.133E-07 | 1.012E-07 | 1.500E-07 | 3.300E-07 |
| 28160 | 1408 | K1408V | E | 1.6e-07 | 2.8e-07 | 4e-07 | 2.800E-07 | 1.200E-07 | 1.600E-07 | 4.000E-07 |
| 28161 | 1409 | S1409A | E | 4.5e-07 | 1.9e-07 | 2e-07 | 2.800E-07 | 1.473E-07 | 1.900E-07 | 4.500E-07 |
| 28162 | 1409 | S1409R | E | 6e-07 | 2.8e-07 | 2.7e-07 | 3.833E-07 | 1.877E-07 | 2.700E-07 | 6.000E-07 |
| 28163 | 1409 | S1409N | E | 2e-07 | 2.5e-07 | 2.1e-07 | 2.200E-07 | 2.646E-08 | 2.000E-07 | 2.500E-07 |
| 28164 | 1409 | S1409D | E | 1.9e-07 | 2.2e-07 | 2e-07 | 2.033E-07 | 1.528E-08 | 1.900E-07 | 2.200E-07 |
| 28165 | 1409 | S1409C | E | 3e-07 | 2e-07 | 1.2e-07 | 2.067E-07 | 9.018E-08 | 1.200E-07 | 3.000E-07 |
| 28166 | 1409 | S1409E | E | 3e-07 | 3.2e-07 | 4e-07 | 3.400E-07 | 5.292E-08 | 3.000E-07 | 4.000E-07 |
| 28167 | 1409 | S1409Q | E | 1.4e-07 | 2.6e-07 | 1.5e-07 | 1.833E-07 | 6.658E-08 | 1.400E-07 | 2.600E-07 |
| 28168 | 1409 | S1409G | E | 2.7e-07 | 1.7e-07 | 1.9e-07 | 2.100E-07 | 5.292E-08 | 1.700E-07 | 2.700E-07 |
| 28169 | 1409 | S1409H | E | 2.7e-07 | 5.1e-07 | 2.1e-07 | 3.300E-07 | 1.587E-07 | 2.100E-07 | 5.100E-07 |
| 28170 | 1409 | S1409I | E | 1.6e-07 | 2.2e-07 | 2e-07 | 1.933E-07 | 3.055E-08 | 1.600E-07 | 2.200E-07 |
| 28171 | 1409 | S1409L | E | 3.2e-07 | 2.7e-07 | 1.2e-07 | 2.367E-07 | 1.041E-07 | 1.200E-07 | 3.200E-07 |
| 28172 | 1409 | S1409K | E | 2.9e-07 | 2.4e-07 | 4.1e-07 | 3.133E-07 | 8.737E-08 | 2.400E-07 | 4.100E-07 |
| 28173 | 1409 | S1409M | E | 3.9e-07 | 2.5e-07 | 2.1e-07 | 2.833E-07 | 9.452E-08 | 2.100E-07 | 3.900E-07 |
| 28174 | 1409 | S1409F | E | 9.7e-08 | 2.3e-07 | 9.9e-08 | 1.420E-07 | 7.622E-08 | 9.700E-08 | 2.300E-07 |
| 28175 | 1409 | S1409P | E | 3.4e-07 | 2.5e-07 | 2.9e-07 | 2.933E-07 | 4.509E-08 | 2.500E-07 | 3.400E-07 |
| 28176 | 1409 | S1409S | E | 2.3e-07 | 2.3e-07 | 1.8e-07 | 2.133E-07 | 2.887E-08 | 1.800E-07 | 2.300E-07 |
| 28177 | 1409 | S1409T | E | 2.7e-07 | 1.8e-07 | 2.1e-07 | 2.200E-07 | 4.583E-08 | 1.800E-07 | 2.700E-07 |
| 28178 | 1409 | S1409W | E | 4.4e-07 | 3.1e-07 | 3.9e-07 | 3.800E-07 | 6.557E-08 | 3.100E-07 | 4.400E-07 |
| 28179 | 1409 | S1409Y | E | 2.4e-07 | 1.5e-07 | 9.4e-08 | 1.613E-07 | 7.366E-08 | 9.400E-08 | 2.400E-07 |
| 28180 | 1409 | S1409V | E | 9.7e-08 | 1.9e-07 | 1.7e-07 | 1.523E-07 | 4.895E-08 | 9.700E-08 | 1.900E-07 |
| 28181 | 1410 | T1410A | E | 2.7e-07 | 1.4e-07 | 4.8e-07 | 2.967E-07 | 1.716E-07 | 1.400E-07 | 4.800E-07 |
| 28182 | 1410 | T1410R | E | 2.4e-07 | 1.6e-07 | 3.8e-07 | 2.600E-07 | 1.114E-07 | 1.600E-07 | 3.800E-07 |
| 28183 | 1410 | T1410N | E | 3.2e-07 | 1.3e-07 | 3.5e-07 | 2.667E-07 | 1.193E-07 | 1.300E-07 | 3.500E-07 |
| 28184 | 1410 | T1410D | E | 3.7e-07 | 3.9e-07 | 5e-07 | 4.200E-07 | 7.000E-08 | 3.700E-07 | 5.000E-07 |
| 28185 | 1410 | T1410C | E | 7e-07 | 3.4e-07 | 1.9e-07 | 4.100E-07 | 2.621E-07 | 1.900E-07 | 7.000E-07 |
| 28186 | 1410 | T1410E | E | 2e-07 | 4.4e-07 | 3.8e-07 | 3.400E-07 | 1.249E-07 | 2.000E-07 | 4.400E-07 |
| 28187 | 1410 | T1410Q | E | 4.7e-07 | 3.1e-07 | 3.6e-07 | 3.800E-07 | 8.185E-08 | 3.100E-07 | 4.700E-07 |
| 28188 | 1410 | T1410G | E | 2.1e-07 | 1.6e-07 | 1.9e-07 | 1.867E-07 | 2.517E-08 | 1.600E-07 | 2.100E-07 |
| 28189 | 1410 | T1410H | E | 3.5e-07 | 1.8e-07 | 2.8e-07 | 2.700E-07 | 8.544E-08 | 1.800E-07 | 3.500E-07 |
| 28190 | 1410 | T1410I | E | 6.2e-07 | 3.1e-07 | 2.6e-07 | 3.967E-07 | 1.950E-07 | 2.600E-07 | 6.200E-07 |
| 28191 | 1410 | T1410L | E | 2e-07 | 4e-07 | 2e-07 | 2.667E-07 | 1.155E-07 | 2.000E-07 | 4.000E-07 |
| 28192 | 1410 | T1410K | E | 2e-07 | 3.6e-07 | 3.9e-07 | 3.167E-07 | 1.021E-07 | 2.000E-07 | 3.900E-07 |
| 28193 | 1410 | T1410M | E | 2.3e-07 | 1.1e-07 | 3.6e-07 | 2.333E-07 | 1.250E-07 | 1.100E-07 | 3.600E-07 |
| 28194 | 1410 | T1410F | E | 2.3e-07 | 2.1e-07 | 2.4e-07 | 2.267E-07 | 1.528E-08 | 2.100E-07 | 2.400E-07 |
| 28195 | 1410 | T1410P | E | 3.5e-07 | 2.8e-07 | 1.7e-07 | 2.667E-07 | 9.074E-08 | 1.700E-07 | 3.500E-07 |
| 28196 | 1410 | T1410S | E | 3.4e-07 | 1.9e-07 | 4e-07 | 3.100E-07 | 1.082E-07 | 1.900E-07 | 4.000E-07 |
| 28197 | 1410 | T1410T | E | 3.7e-07 | 3.3e-07 | 1.8e-07 | 2.933E-07 | 1.002E-07 | 1.800E-07 | 3.700E-07 |
| 28198 | 1410 | T1410W | E | 2.3e-07 | 1.8e-07 | 9e-08 | 1.667E-07 | 7.095E-08 | 9.000E-08 | 2.300E-07 |
| 28199 | 1410 | T1410Y | E | 1.8e-07 | 2.9e-07 | 3.8e-07 | 2.833E-07 | 1.002E-07 | 1.800E-07 | 3.800E-07 |
| 28200 | 1410 | T1410V | E | 2e-07 | 4.6e-07 | 1.9e-07 | 2.833E-07 | 1.531E-07 | 1.900E-07 | 4.600E-07 |
| 28201 | 1411 | S1411A | E | 3.5e-07 | 3.1e-07 | 3.3e-07 | 3.300E-07 | 2.000E-08 | 3.100E-07 | 3.500E-07 |
| 28202 | 1411 | S1411R | E | 2.1e-07 | 3.8e-07 | 2.3e-07 | 2.733E-07 | 9.292E-08 | 2.100E-07 | 3.800E-07 |
| 28203 | 1411 | S1411N | E | 1.1e-07 | 6.9e-07 | 2.2e-07 | 3.400E-07 | 3.081E-07 | 1.100E-07 | 6.900E-07 |
| 28204 | 1411 | S1411D | E | 3.1e-07 | 1.1e-07 | 1.6e-07 | 1.933E-07 | 1.041E-07 | 1.100E-07 | 3.100E-07 |
| 28205 | 1411 | S1411C | E | 3e-07 | 1.8e-07 | 2.6e-07 | 2.467E-07 | 6.110E-08 | 1.800E-07 | 3.000E-07 |
| 28206 | 1411 | S1411E | E | 4.6e-07 | 1.5e-07 | 4.5e-07 | 3.533E-07 | 1.762E-07 | 1.500E-07 | 4.600E-07 |
| 28207 | 1411 | S1411Q | E | 3.2e-07 | 1.6e-07 | 1.3e-07 | 2.033E-07 | 1.021E-07 | 1.300E-07 | 3.200E-07 |
| 28208 | 1411 | S1411G | E | 2e-07 | 2.2e-07 | 3.7e-07 | 2.633E-07 | 9.292E-08 | 2.000E-07 | 3.700E-07 |
| 28209 | 1411 | S1411H | E | 3.8e-07 | 5e-07 | 2.9e-07 | 3.900E-07 | 1.054E-07 | 2.900E-07 | 5.000E-07 |
| 28210 | 1411 | S1411I | E | 2.2e-07 | 2.6e-07 | 1.4e-07 | 2.067E-07 | 6.110E-08 | 1.400E-07 | 2.600E-07 |
| 28211 | 1411 | S1411L | E | 2.4e-07 | 2.2e-07 | 1.9e-07 | 2.167E-07 | 2.517E-08 | 1.900E-07 | 2.400E-07 |
| 28212 | 1411 | S1411K | E | 5e-07 | 8.5e-07 | 2.4e-07 | 5.300E-07 | 3.061E-07 | 2.400E-07 | 8.500E-07 |
| 28213 | 1411 | S1411M | E | 4.8e-07 | 1.6e-07 | 2.1e-07 | 2.833E-07 | 1.721E-07 | 1.600E-07 | 4.800E-07 |
| 28214 | 1411 | S1411F | E | 4.7e-07 | 4e-07 | 4.4e-07 | 4.367E-07 | 3.512E-08 | 4.000E-07 | 4.700E-07 |
| 28215 | 1411 | S1411P | E | 4.9e-07 | 1.2e-07 | 1.8e-07 | 2.633E-07 | 1.986E-07 | 1.200E-07 | 4.900E-07 |
| 28216 | 1411 | S1411S | E | 2e-07 | 3.8e-07 | 3.6e-07 | 3.133E-07 | 9.866E-08 | 2.000E-07 | 3.800E-07 |
| 28217 | 1411 | S1411T | E | 1.7e-07 | 3.4e-07 | 9e-08 | 2.000E-07 | 1.277E-07 | 9.000E-08 | 3.400E-07 |
| 28218 | 1411 | S1411W | E | 2e-07 | 1.5e-07 | 4.3e-07 | 2.600E-07 | 1.493E-07 | 1.500E-07 | 4.300E-07 |
| 28219 | 1411 | S1411Y | E | 1.5e-07 | 4.7e-07 | 2.7e-07 | 2.967E-07 | 1.617E-07 | 1.500E-07 | 4.700E-07 |

| | | | | | | | | | | |
|---|---|---|---|---|---|---|---|---|---|---|
| 28220 | 1411 | S1411V | E | 2.8e-07 | 2.9e-07 | 2.7e-07 | 2.800E-07 | 1.000E-08 | 2.700E-07 | 2.900E-07 |
| 28221 | 1412 | G1412A | E | 1.7e-07 | 3.4e-07 | 4.7e-07 | 3.267E-07 | 1.504E-07 | 1.700E-07 | 4.700E-07 |
| 28222 | 1412 | G1412R | E | 2e-07 | 5.4e-07 | 1.7e-07 | 3.033E-07 | 2.055E-07 | 1.700E-07 | 5.400E-07 |
| 28223 | 1412 | G1412N | E | 1e-07 | 5.6e-07 | 2.9e-07 | 3.167E-07 | 2.312E-07 | 1.000E-07 | 5.600E-07 |
| 28224 | 1412 | G1412D | E | 1.3e-07 | 3.6e-07 | 2e-07 | 2.300E-07 | 1.179E-07 | 1.300E-07 | 3.600E-07 |
| 28225 | 1412 | G1412C | E | 3.6e-07 | 4.8e-07 | 2.1e-07 | 3.500E-07 | 1.353E-07 | 2.100E-07 | 4.800E-07 |
| 28226 | 1412 | G1412E | E | 2.7e-07 | 1.5e-07 | 2.1e-07 | 2.100E-07 | 6.000E-08 | 1.500E-07 | 2.700E-07 |
| 28227 | 1412 | G1412Q | E | 3.2e-07 | 3.2e-07 | 2.6e-07 | 3.000E-07 | 3.464E-08 | 2.600E-07 | 3.200E-07 |
| 28228 | 1412 | G1412G | E | 2.3e-07 | 1.4e-07 | 3e-07 | 2.233E-07 | 8.021E-08 | 1.400E-07 | 3.000E-07 |
| 28229 | 1412 | G1412H | E | 1.3e-07 | 2.3e-07 | 1.9e-07 | 1.833E-07 | 5.033E-08 | 1.300E-07 | 2.300E-07 |
| 28230 | 1412 | G1412I | E | 7.1e-08 | 2.7e-07 | 2.6e-07 | 2.003E-07 | 1.121E-07 | 7.100E-08 | 2.700E-07 |
| 28231 | 1412 | G1412L | E | 3.4e-07 | 1.7e-07 | 4.5e-07 | 3.200E-07 | 1.411E-07 | 1.700E-07 | 4.500E-07 |
| 28232 | 1412 | G1412K | E | 3e-07 | 3.3e-07 | 1.6e-07 | 2.633E-07 | 9.074E-08 | 1.600E-07 | 3.300E-07 |
| 28233 | 1412 | G1412M | E | 2.9e-07 | 2.2e-07 | 1.3e-07 | 2.133E-07 | 8.021E-08 | 1.300E-07 | 2.900E-07 |
| 28234 | 1412 | G1412F | E | 1.4e-07 | 2.3e-07 | 2.1e-07 | 1.933E-07 | 4.726E-08 | 1.400E-07 | 2.300E-07 |
| 28235 | 1412 | G1412P | E | 1.6e-07 | 2.2e-07 | 1.7e-07 | 1.833E-07 | 3.215E-08 | 1.600E-07 | 2.200E-07 |
| 28236 | 1412 | G1412S | E | 2.6e-07 | 1.9e-07 | 2.6e-07 | 2.367E-07 | 4.041E-08 | 1.900E-07 | 2.600E-07 |
| 28237 | 1412 | G1412T | E | 1.7e-07 | 2.5e-07 | 1.4e-07 | 1.867E-07 | 5.686E-08 | 1.400E-07 | 2.500E-07 |
| 28238 | 1412 | G1412W | E | 1.9e-07 | 6.5e-07 | 1.7e-07 | 3.367E-07 | 2.715E-07 | 1.700E-07 | 6.500E-07 |
| 28239 | 1412 | G1412Y | E | 8.3e-08 | 5e-07 | 4.2e-07 | 3.343E-07 | 2.213E-07 | 8.300E-08 | 5.000E-07 |
| 28240 | 1412 | G1412V | E | 1.1e-07 | 1.5e-07 | 2.6e-07 | 1.733E-07 | 7.767E-08 | 1.100E-07 | 2.600E-07 |
| 28241 | 1413 | G1413A | E | 2.6e-07 | 3.4e-07 | 2.4e-07 | 2.800E-07 | 5.292E-08 | 2.400E-07 | 3.400E-07 |
| 28242 | 1413 | G1413R | E | 5.4e-07 | 2.1e-07 | 5e-07 | 4.167E-07 | 1.801E-07 | 2.100E-07 | 5.400E-07 |
| 28243 | 1413 | G1413N | E | 2.5e-07 | 3.5e-07 | 4.1e-07 | 3.367E-07 | 8.083E-08 | 2.500E-07 | 4.100E-07 |
| 28244 | 1413 | G1413D | E | 4.8e-07 | 5.6e-07 | 3.4e-07 | 4.600E-07 | 1.114E-07 | 3.400E-07 | 5.600E-07 |
| 28245 | 1413 | G1413C | E | 2.8e-07 | 1.4e-07 | 2.8e-07 | 2.333E-07 | 8.083E-08 | 1.400E-07 | 2.800E-07 |
| 28246 | 1413 | G1413E | E | 5.3e-07 | 2.1e-07 | 1.6e-07 | 3.000E-07 | 2.007E-07 | 1.600E-07 | 5.300E-07 |
| 28247 | 1413 | G1413Q | E | 2.7e-07 | 8.3e-08 | 2e-07 | 1.843E-07 | 9.448E-08 | 8.300E-08 | 2.700E-07 |
| 28248 | 1413 | G1413G | E | 6e-08 | 3.2e-07 | 3.9e-07 | 2.567E-07 | 1.739E-07 | 6.000E-08 | 3.900E-07 |
| 28249 | 1413 | G1413H | E | 1.5e-07 | 1.8e-07 | 1.7e-07 | 1.667E-07 | 1.528E-08 | 1.500E-07 | 1.800E-07 |
| 28250 | 1413 | G1413I | E | 2e-07 | 1.5e-07 | 2.1e-07 | 1.867E-07 | 3.215E-08 | 1.500E-07 | 2.100E-07 |
| 28251 | 1413 | G1413L | E | 7.1e-07 | 1.2e-07 | 2e-07 | 3.433E-07 | 3.201E-07 | 1.200E-07 | 7.100E-07 |
| 28252 | 1413 | G1413K | E | 1.6e-07 | 1.4e-07 | 1.6e-07 | 1.533E-07 | 1.155E-08 | 1.400E-07 | 1.600E-07 |
| 28253 | 1413 | G1413M | E | 2.5e-07 | 1.5e-07 | 4.9e-07 | 2.967E-07 | 1.747E-07 | 1.500E-07 | 4.900E-07 |
| 28254 | 1413 | G1413F | E | 1.7e-07 | 4.4e-07 | 1.5e-07 | 2.533E-07 | 1.620E-07 | 1.500E-07 | 4.400E-07 |
| 28255 | 1413 | G1413P | E | 1.8e-07 | 3.1e-07 | 1.6e-07 | 2.167E-07 | 8.145E-08 | 1.600E-07 | 3.100E-07 |
| 28256 | 1413 | G1413S | E | 1.3e-07 | 1.4e-07 | 2e-07 | 1.567E-07 | 3.786E-08 | 1.300E-07 | 2.000E-07 |
| 28257 | 1413 | G1413T | E | 3.4e-07 | 5.1e-07 | 1.8e-07 | 3.433E-07 | 1.650E-07 | 1.800E-07 | 5.100E-07 |
| 28258 | 1413 | G1413W | E | 1.6e-07 | 2.9e-07 | 1.8e-07 | 2.100E-07 | 7.000E-08 | 1.600E-07 | 2.900E-07 |
| 28259 | 1413 | G1413Y | E | 2.9e-07 | 3.5e-07 | 2.2e-07 | 2.867E-07 | 6.506E-08 | 2.200E-07 | 3.500E-07 |
| 28260 | 1413 | G1413V | E | 1.7e-07 | 2.8e-07 | 1.4e-07 | 1.967E-07 | 7.371E-08 | 1.400E-07 | 2.800E-07 |
| 28261 | 1414 | T1414A | E | 2.5e-07 | 1.3e-07 | 1.4e-07 | 1.733E-07 | 6.658E-08 | 1.300E-07 | 2.500E-07 |
| 28262 | 1414 | T1414R | E | 2.3e-07 | 3.9e-07 | 2.2e-07 | 2.800E-07 | 9.539E-08 | 2.200E-07 | 3.900E-07 |
| 28263 | 1414 | T1414N | E | 2.3e-07 | 2.4e-07 | 3e-07 | 2.567E-07 | 3.786E-08 | 2.300E-07 | 3.000E-07 |
| 28264 | 1414 | T1414D | E | 3.1e-07 | 2.6e-07 | 4.8e-07 | 3.500E-07 | 1.153E-07 | 2.600E-07 | 4.800E-07 |
| 28265 | 1414 | T1414C | E | 3.4e-07 | 2.1e-07 | 4.3e-07 | 3.267E-07 | 1.106E-07 | 2.100E-07 | 4.300E-07 |
| 28266 | 1414 | T1414E | E | 2.4e-07 | 3.2e-07 | 2.4e-07 | 2.667E-07 | 4.619E-08 | 2.400E-07 | 3.200E-07 |
| 28267 | 1414 | T1414Q | E | 2e-07 | 2.2e-07 | 3.1e-07 | 2.433E-07 | 5.859E-08 | 2.000E-07 | 3.100E-07 |
| 28268 | 1414 | T1414G | E | 2.4e-07 | 2.7e-07 | 1.3e-07 | 2.133E-07 | 7.371E-08 | 1.300E-07 | 2.700E-07 |
| 28269 | 1414 | T1414H | E | 3.3e-07 | 3.2e-07 | 4.1e-07 | 3.533E-07 | 4.933E-08 | 3.200E-07 | 4.100E-07 |
| 28270 | 1414 | T1414I | E | 5.4e-07 | 2e-07 | 3e-07 | 3.467E-07 | 1.747E-07 | 2.000E-07 | 5.400E-07 |
| 28271 | 1414 | T1414L | E | 2.1e-07 | 1.8e-07 | 4e-07 | 2.633E-07 | 1.193E-07 | 1.800E-07 | 4.000E-07 |
| 28272 | 1414 | T1414K | E | 5.1e-07 | 2.9e-07 | 5.6e-07 | 4.533E-07 | 1.436E-07 | 2.900E-07 | 5.600E-07 |
| 28273 | 1414 | T1414M | E | 1.2e-07 | 3.4e-07 | 2.6e-07 | 2.400E-07 | 1.114E-07 | 1.200E-07 | 3.400E-07 |
| 28274 | 1414 | T1414F | E | 4.4e-07 | 6.5e-07 | 2.8e-07 | 4.567E-07 | 1.856E-07 | 2.800E-07 | 6.500E-07 |
| 28275 | 1414 | T1414P | E | 1.8e-07 | 7.4e-07 | 2.1e-07 | 3.767E-07 | 3.150E-07 | 1.800E-07 | 7.400E-07 |
| 28276 | 1414 | T1414S | E | 1.4e-07 | 1.3e-07 | 2.9e-07 | 1.867E-07 | 8.963E-08 | 1.300E-07 | 2.900E-07 |
| 28277 | 1414 | T1414T | E | 1.9e-07 | 4.2e-07 | 2.2e-07 | 2.767E-07 | 1.250E-07 | 1.900E-07 | 4.200E-07 |
| 28278 | 1414 | T1414W | E | 2.9e-07 | 1.4e-07 | 2.6e-07 | 2.300E-07 | 7.937E-08 | 1.400E-07 | 2.900E-07 |
| 28279 | 1414 | T1414Y | E | 3.5e-07 | 2.5e-07 | 1.9e-07 | 2.633E-07 | 8.083E-08 | 1.900E-07 | 3.500E-07 |
| 28280 | 1414 | T1414V | E | 5.1e-07 | 2.4e-07 | 3.5e-07 | 3.667E-07 | 1.358E-07 | 2.400E-07 | 5.100E-07 |
| 28281 | 1415 | A1415A | E | 4.3e-07 | 7.4e-08 | 4.8e-07 | 3.280E-07 | 2.214E-07 | 7.400E-08 | 4.800E-07 |
| 28282 | 1415 | A1415R | E | 1.9e-07 | 3.3e-07 | 1.2e-07 | 2.133E-07 | 1.069E-07 | 1.200E-07 | 3.300E-07 |
| 28283 | 1415 | A1415N | E | 2.4e-07 | 1.4e-07 | 3.6e-07 | 2.467E-07 | 1.102E-07 | 1.400E-07 | 3.600E-07 |
| 28284 | 1415 | A1415D | E | 1.7e-07 | 1.9e-07 | 1.9e-07 | 1.833E-07 | 1.155E-08 | 1.700E-07 | 1.900E-07 |
| 28285 | 1415 | A1415C | E | 3.1e-07 | 4.8e-07 | 2.1e-07 | 3.333E-07 | 1.365E-07 | 2.100E-07 | 4.800E-07 |
| 28286 | 1415 | A1415E | E | 1.2e-07 | 2.6e-07 | 1.2e-07 | 1.667E-07 | 8.083E-08 | 1.200E-07 | 2.600E-07 |
| 28287 | 1415 | A1415Q | E | 1.9e-07 | 1.3e-07 | 2.9e-07 | 2.033E-07 | 8.083E-08 | 1.300E-07 | 2.900E-07 |
| 28288 | 1415 | A1415G | E | 2.4e-07 | 1.6e-07 | 2.5e-07 | 2.167E-07 | 4.933E-08 | 1.600E-07 | 2.500E-07 |
| 28289 | 1415 | A1415H | E | 3e-07 | 2.1e-07 | 1.6e-07 | 2.233E-07 | 7.095E-08 | 1.600E-07 | 3.000E-07 |
| 28290 | 1415 | A1415I | E | 1.8e-07 | 3e-07 | 2.9e-07 | 2.567E-07 | 6.658E-08 | 1.800E-07 | 3.000E-07 |
| 28291 | 1415 | A1415L | E | 1.7e-07 | 3.9e-07 | 1.3e-07 | 2.300E-07 | 1.400E-07 | 1.300E-07 | 3.900E-07 |
| 28292 | 1415 | A1415K | E | 1.5e-07 | 4.2e-07 | 1.9e-07 | 2.533E-07 | 1.457E-07 | 1.500E-07 | 4.200E-07 |
| 28293 | 1415 | A1415M | E | 2.2e-07 | 3.3e-07 | 4.9e-07 | 3.467E-07 | 1.358E-07 | 2.200E-07 | 4.900E-07 |
| 28294 | 1415 | A1415F | E | 4.8e-07 | 2.1e-07 | 2e-07 | 2.967E-07 | 1.589E-07 | 2.000E-07 | 4.800E-07 |
| 28295 | 1415 | A1415P | E | 2.1e-07 | 1e-07 | 2e-07 | 1.700E-07 | 6.083E-08 | 1.000E-07 | 2.100E-07 |
| 28296 | 1415 | A1415S | E | 1.4e-07 | 3.4e-07 | 2.7e-07 | 2.500E-07 | 1.015E-07 | 1.400E-07 | 3.400E-07 |
| 28297 | 1415 | A1415T | E | 7e-08 | 2.3e-07 | 3.3e-07 | 2.100E-07 | 1.311E-07 | 7.000E-08 | 3.300E-07 |
| 28298 | 1415 | A1415W | E | 3.8e-07 | 2.6e-07 | 2e-07 | 2.800E-07 | 9.165E-08 | 2.000E-07 | 3.800E-07 |
| 28299 | 1415 | A1415Y | E | 3.5e-07 | 2.1e-07 | 5e-07 | 3.533E-07 | 1.450E-07 | 2.100E-07 | 5.000E-07 |
| 28300 | 1415 | A1415V | E | 3e-07 | 2.3e-07 | 1.2e-07 | 2.167E-07 | 9.074E-08 | 1.200E-07 | 3.000E-07 |
| 28301 | 1416 | A1416A | E | 4e-07 | 2.9e-07 | 3.1e-07 | 3.333E-07 | 5.859E-08 | 2.900E-07 | 4.000E-07 |
| 28302 | 1416 | A1416R | E | 5.3e-07 | 1.5e-07 | 1.7e-07 | 2.833E-07 | 2.139E-07 | 1.500E-07 | 5.300E-07 |
| 28303 | 1416 | A1416N | E | 2.7e-07 | 4.3e-07 | 3.7e-07 | 3.567E-07 | 8.083E-08 | 2.700E-07 | 4.300E-07 |
| 28304 | 1416 | A1416D | E | 1.8e-07 | 3.4e-07 | 3.7e-07 | 2.967E-07 | 1.021E-07 | 1.800E-07 | 3.700E-07 |

| | | | | | | | | | | |
|---|---|---|---|---|---|---|---|---|---|---|
| 28305 | 1416 | A1416C | E | 2.5e-07 | 2.8e-07 | 2e-07 | 2.433E-07 | 4.041E-08 | 2.000E-07 | 2.800E-07 |
| 28306 | 1416 | A1416E | E | 5.1e-07 | 1.9e-07 | 2e-07 | 3.000E-07 | 1.819E-07 | 1.900E-07 | 5.100E-07 |
| 28307 | 1416 | A1416Q | E | 1.4e-07 | 2.5e-07 | 2.1e-07 | 2.000E-07 | 5.568E-08 | 1.400E-07 | 2.500E-07 |
| 28308 | 1416 | A1416G | E | 2.9e-07 | 2.4e-07 | 4.7e-07 | 3.333E-07 | 1.210E-07 | 2.400E-07 | 4.700E-07 |
| 28309 | 1416 | A1416H | E | 3.2e-07 | 2.2e-07 | 1.7e-07 | 2.367E-07 | 7.638E-08 | 1.700E-07 | 3.200E-07 |
| 28310 | 1416 | A1416I | E | 1e-07 | 7.5e-08 | 2.9e-07 | 1.550E-07 | 1.176E-07 | 7.500E-08 | 2.900E-07 |
| 28311 | 1416 | A1416L | E | 2.5e-07 | 3.9e-07 | 1.6e-07 | 2.667E-07 | 1.159E-07 | 1.600E-07 | 3.900E-07 |
| 28312 | 1416 | A1416K | E | 7.1e-08 | 1.3e-07 | 2.5e-07 | 1.503E-07 | 9.122E-08 | 7.100E-08 | 2.500E-07 |
| 28313 | 1416 | A1416M | E | 3.1e-07 | 2.1e-07 | 2.3e-07 | 2.500E-07 | 5.292E-08 | 2.100E-07 | 3.100E-07 |
| 28314 | 1416 | A1416F | E | 2.5e-07 | 2.5e-07 | 1.7e-07 | 2.233E-07 | 4.619E-08 | 1.700E-07 | 2.500E-07 |
| 28315 | 1416 | A1416P | E | 1e-07 | 2.2e-07 | 3.2e-07 | 2.133E-07 | 1.102E-07 | 1.000E-07 | 3.200E-07 |
| 28316 | 1416 | A1416S | E | 9.5e-08 | 7.2e-07 | 2.8e-07 | 3.650E-07 | 3.211E-07 | 9.500E-08 | 7.200E-07 |
| 28317 | 1416 | A1416T | E | 2.7e-07 | 2.1e-07 | 3.1e-07 | 2.633E-07 | 5.033E-08 | 2.100E-07 | 3.100E-07 |
| 28318 | 1416 | A1416W | E | 1.2e-07 | 5.3e-07 | 1.2e-07 | 2.567E-07 | 2.367E-07 | 1.200E-07 | 5.300E-07 |
| 28319 | 1416 | A1416Y | E | 2.1e-07 | 1.9e-07 | 1.7e-07 | 1.900E-07 | 2.000E-08 | 1.700E-07 | 2.100E-07 |
| 28320 | 1416 | A1416V | E | 2e-07 | 2.1e-07 | 3.6e-07 | 2.567E-07 | 8.963E-08 | 2.000E-07 | 3.600E-07 |
| 28321 | 1417 | L1417A | E | 5e-07 | 1.4e-07 | 2.1e-07 | 2.833E-07 | 1.909E-07 | 1.400E-07 | 5.000E-07 |
| 28322 | 1417 | L1417R | E | 2.1e-07 | 3e-07 | 2.3e-07 | 2.467E-07 | 4.726E-08 | 2.100E-07 | 3.000E-07 |
| 28323 | 1417 | L1417N | E | 2.1e-07 | 1.5e-07 | 1.1e-07 | 1.567E-07 | 5.033E-08 | 1.100E-07 | 2.100E-07 |
| 28324 | 1417 | L1417D | E | 1.5e-07 | 5.7e-07 | 2.4e-07 | 3.200E-07 | 2.211E-07 | 1.500E-07 | 5.700E-07 |
| 28325 | 1417 | L1417C | E | 1.8e-07 | 3.1e-07 | 2.6e-07 | 2.500E-07 | 6.557E-08 | 1.800E-07 | 3.100E-07 |
| 28326 | 1417 | L1417E | E | 2.5e-07 | 5.3e-07 | 2.3e-07 | 3.367E-07 | 1.677E-07 | 2.300E-07 | 5.300E-07 |
| 28327 | 1417 | L1417Q | E | 3.4e-07 | 8e-07 | 7.1e-08 | 4.037E-07 | 3.686E-07 | 7.100E-08 | 8.000E-07 |
| 28328 | 1417 | L1417G | E | 2e-07 | 2.3e-07 | 2.7e-07 | 2.333E-07 | 3.512E-08 | 2.000E-07 | 2.700E-07 |
| 28329 | 1417 | L1417H | E | 4.2e-07 | 4.5e-07 | 6.6e-08 | 3.120E-07 | 2.136E-07 | 6.600E-08 | 4.500E-07 |
| 28330 | 1417 | L1417I | E | 4e-07 | 8.2e-08 | 4.8e-07 | 3.207E-07 | 2.105E-07 | 8.200E-08 | 4.800E-07 |
| 28331 | 1417 | L1417L | E | 5.3e-07 | 1.4e-07 | 2.4e-07 | 3.033E-07 | 2.026E-07 | 1.400E-07 | 5.300E-07 |
| 28332 | 1417 | L1417K | E | 4.1e-07 | 1.6e-07 | 1.2e-07 | 2.300E-07 | 1.572E-07 | 1.200E-07 | 4.100E-07 |
| 28333 | 1417 | L1417M | E | 1.5e-07 | 3.5e-07 | 1.3e-07 | 2.100E-07 | 1.217E-07 | 1.300E-07 | 3.500E-07 |
| 28334 | 1417 | L1417F | E | 3.3e-07 | 1.9e-07 | 2.8e-07 | 2.667E-07 | 7.095E-08 | 1.900E-07 | 3.300E-07 |
| 28335 | 1417 | L1417P | E | 1.7e-07 | 1.6e-07 | 1.9e-07 | 1.733E-07 | 1.528E-08 | 1.600E-07 | 1.900E-07 |
| 28336 | 1417 | L1417S | E | 1.5e-07 | 2.3e-07 | 1.2e-07 | 1.667E-07 | 5.686E-08 | 1.200E-07 | 2.300E-07 |
| 28337 | 1417 | L1417T | E | 1.9e-07 | 2.6e-07 | 2.5e-07 | 2.333E-07 | 3.786E-08 | 1.900E-07 | 2.600E-07 |
| 28338 | 1417 | L1417W | E | 2.7e-07 | 3.1e-07 | 2e-07 | 2.600E-07 | 5.568E-08 | 2.000E-07 | 3.100E-07 |
| 28339 | 1417 | L1417Y | E | 1.5e-07 | 1.5e-07 | 4.3e-07 | 2.433E-07 | 1.617E-07 | 1.500E-07 | 4.300E-07 |
| 28340 | 1417 | L1417V | E | 6e-07 | 1.9e-07 | 2.4e-07 | 3.433E-07 | 2.237E-07 | 1.900E-07 | 6.000E-07 |
| 28341 | 1418 | G1418A | E | 4.3e-07 | 1.7e-07 | 3.5e-07 | 3.167E-07 | 1.332E-07 | 1.700E-07 | 4.300E-07 |
| 28342 | 1418 | G1418R | E | 1.9e-07 | 1.9e-07 | 2.8e-07 | 2.200E-07 | 5.196E-08 | 1.900E-07 | 2.800E-07 |
| 28343 | 1418 | G1418N | E | 2.4e-07 | 1.9e-07 | 2.6e-07 | 2.300E-07 | 3.606E-08 | 1.900E-07 | 2.600E-07 |
| 28344 | 1418 | G1418D | E | 2e-07 | 2.3e-07 | 2.4e-07 | 2.233E-07 | 2.082E-08 | 2.000E-07 | 2.400E-07 |
| 28345 | 1418 | G1418C | E | 1.9e-07 | 1.7e-07 | 2.5e-07 | 2.033E-07 | 4.163E-08 | 1.700E-07 | 2.500E-07 |
| 28346 | 1418 | G1418E | E | 2.4e-07 | 2.3e-07 | 3.7e-07 | 2.800E-07 | 7.810E-08 | 2.300E-07 | 3.700E-07 |
| 28347 | 1418 | G1418Q | E | 2.3e-07 | 2.4e-07 | 5.9e-07 | 3.533E-07 | 2.050E-07 | 2.300E-07 | 5.900E-07 |
| 28348 | 1418 | G1418G | E | 2e-07 | 2e-07 | 2.4e-07 | 2.133E-07 | 2.309E-08 | 2.000E-07 | 2.400E-07 |
| 28349 | 1418 | G1418H | E | 2.8e-07 | 3e-07 | 2.1e-07 | 2.633E-07 | 4.726E-08 | 2.100E-07 | 3.000E-07 |
| 28350 | 1418 | G1418I | E | 1.6e-07 | 2.2e-07 | 3.1e-07 | 2.300E-07 | 7.550E-08 | 1.600E-07 | 3.100E-07 |
| 28351 | 1418 | G1418L | E | 2.9e-07 | 4.2e-07 | 2.3e-07 | 3.133E-07 | 9.713E-08 | 2.300E-07 | 4.200E-07 |
| 28352 | 1418 | G1418K | E | 3.4e-07 | 2.8e-07 | 1.8e-07 | 2.667E-07 | 8.083E-08 | 1.800E-07 | 3.400E-07 |
| 28353 | 1418 | G1418M | E | 2.2e-07 | 3.3e-07 | 2.2e-07 | 2.567E-07 | 6.351E-08 | 2.200E-07 | 3.300E-07 |
| 28354 | 1418 | G1418F | E | 3.5e-07 | 1.9e-07 | 2.3e-07 | 2.567E-07 | 8.327E-08 | 1.900E-07 | 3.500E-07 |
| 28355 | 1418 | G1418P | E | 2.2e-07 | 2.4e-07 | 1.6e-07 | 2.067E-07 | 4.163E-08 | 1.600E-07 | 2.400E-07 |
| 28356 | 1418 | G1418S | E | 1.8e-07 | 1.8e-07 | 2.1e-07 | 1.900E-07 | 1.732E-08 | 1.800E-07 | 2.100E-07 |
| 28357 | 1418 | G1418T | E | 5.3e-07 | 8.8e-08 | 1e-07 | 2.393E-07 | 2.518E-07 | 8.800E-08 | 5.300E-07 |
| 28358 | 1418 | G1418W | E | 1.3e-07 | 2.8e-07 | 3.8e-07 | 2.633E-07 | 1.258E-07 | 1.300E-07 | 3.800E-07 |
| 28359 | 1418 | G1418Y | E | 2.4e-07 | 4.8e-07 | 1.5e-07 | 2.900E-07 | 1.706E-07 | 1.500E-07 | 4.800E-07 |
| 28360 | 1418 | G1418V | E | 5.6e-07 | 2.8e-07 | 1.9e-07 | 3.433E-07 | 1.930E-07 | 1.900E-07 | 5.600E-07 |
| 28361 | 1419 | C1419A | E | 1.4e-07 | 2.9e-07 | 2.1e-07 | 2.133E-07 | 7.506E-08 | 1.400E-07 | 2.900E-07 |
| 28362 | 1419 | C1419R | E | 2.2e-07 | 3.4e-07 | 6.7e-07 | 4.100E-07 | 2.330E-07 | 2.200E-07 | 6.700E-07 |
| 28363 | 1419 | C1419N | E | 3.3e-07 | 2.4e-07 | 3.1e-07 | 2.933E-07 | 4.726E-08 | 2.400E-07 | 3.300E-07 |
| 28364 | 1419 | C1419D | E | 3.1e-07 | 3.4e-07 | 1.4e-07 | 2.633E-07 | 1.079E-07 | 1.400E-07 | 3.400E-07 |
| 28365 | 1419 | C1419C | E | 2.8e-07 | 9.6e-08 | 2.3e-07 | 2.020E-07 | 9.514E-08 | 9.600E-08 | 2.800E-07 |
| 28366 | 1419 | C1419E | E | 2.9e-07 | 4.8e-07 | 4.3e-07 | 4.000E-07 | 9.849E-08 | 2.900E-07 | 4.800E-07 |
| 28367 | 1419 | C1419Q | E | 3.3e-07 | 2.9e-07 | 1.3e-07 | 2.500E-07 | 1.058E-07 | 1.300E-07 | 3.300E-07 |
| 28368 | 1419 | C1419G | E | 1.5e-07 | 1.8e-07 | 6.7e-07 | 3.333E-07 | 2.919E-07 | 1.500E-07 | 6.700E-07 |
| 28369 | 1419 | C1419H | E | 3.7e-07 | 6.3e-07 | 2.2e-07 | 4.067E-07 | 2.074E-07 | 2.200E-07 | 6.300E-07 |
| 28370 | 1419 | C1419I | E | 3e-07 | 5e-07 | 3.4e-07 | 3.800E-07 | 1.058E-07 | 3.000E-07 | 5.000E-07 |
| 28371 | 1419 | C1419L | E | 2.6e-07 | 1.6e-07 | 2.5e-07 | 2.233E-07 | 5.508E-08 | 1.600E-07 | 2.600E-07 |
| 28372 | 1419 | C1419K | E | 2e-07 | 1.1e-07 | 1.4e-07 | 1.500E-07 | 4.583E-08 | 1.100E-07 | 2.000E-07 |
| 28373 | 1419 | C1419M | E | 8.4e-07 | 2.9e-07 | 1.8e-07 | 4.367E-07 | 3.536E-07 | 1.800E-07 | 8.400E-07 |
| 28374 | 1419 | C1419F | E | 3.5e-07 | 1.5e-07 | 2.9e-07 | 2.633E-07 | 1.026E-07 | 1.500E-07 | 3.500E-07 |
| 28375 | 1419 | C1419P | E | 3.4e-07 | 3.3e-07 | 2.5e-07 | 3.067E-07 | 4.933E-08 | 2.500E-07 | 3.400E-07 |
| 28376 | 1419 | C1419S | E | 1.7e-07 | 2.2e-07 | 1.6e-07 | 1.833E-07 | 3.215E-08 | 1.600E-07 | 2.200E-07 |
| 28377 | 1419 | C1419T | E | 4.1e-07 | 3.4e-07 | 2e-07 | 3.167E-07 | 1.069E-07 | 2.000E-07 | 4.100E-07 |
| 28378 | 1419 | C1419W | E | 2.7e-07 | 1.8e-07 | 1.4e-07 | 1.967E-07 | 6.658E-08 | 1.400E-07 | 2.700E-07 |
| 28379 | 1419 | C1419Y | E | 1.5e-07 | 2.1e-07 | 1.4e-07 | 1.667E-07 | 3.786E-08 | 1.400E-07 | 2.100E-07 |
| 28380 | 1419 | C1419V | E | 1.8e-07 | 1.3e-07 | 2.5e-07 | 1.867E-07 | 6.028E-08 | 1.300E-07 | 2.500E-07 |
| 28381 | 1420 | L1420A | E | 1.1e-07 | 1.7e-07 | 4.3e-07 | 2.367E-07 | 1.701E-07 | 1.100E-07 | 4.300E-07 |
| 28382 | 1420 | L1420R | E | 3e-07 | 2.7e-07 | 2.8e-07 | 2.833E-07 | 1.528E-08 | 2.700E-07 | 3.000E-07 |
| 28383 | 1420 | L1420N | E | 9.4e-08 | 2.8e-07 | 4.1e-07 | 2.613E-07 | 1.588E-07 | 9.400E-08 | 4.100E-07 |
| 28384 | 1420 | L1420D | E | 2.7e-07 | 1.8e-07 | 2.3e-07 | 2.267E-07 | 4.509E-08 | 1.800E-07 | 2.700E-07 |
| 28385 | 1420 | L1420C | E | 1.7e-07 | 1.7e-07 | 1.8e-07 | 1.733E-07 | 5.774E-09 | 1.700E-07 | 1.800E-07 |
| 28386 | 1420 | L1420E | E | 3.2e-07 | 1.5e-07 | 1.4e-07 | 2.033E-07 | 1.012E-07 | 1.400E-07 | 3.200E-07 |
| 28387 | 1420 | L1420Q | E | 2e-07 | 4.7e-07 | 1.4e-07 | 2.700E-07 | 1.758E-07 | 1.400E-07 | 4.700E-07 |
| 28388 | 1420 | L1420G | E | 1.7e-07 | 1.9e-07 | 2.2e-07 | 1.933E-07 | 2.517E-08 | 1.700E-07 | 2.200E-07 |
| 28389 | 1420 | L1420H | E | 2.9e-07 | 3.8e-07 | 2.1e-07 | 2.933E-07 | 8.505E-08 | 2.100E-07 | 3.800E-07 |

| | | | | | | | | | | |
|---|---|---|---|---|---|---|---|---|---|---|
| 28390 | 1420 | L1420I | E | 1.1e-07 | 3.3e-07 | 4.3e-07 | 2.900E-07 | 1.637E-07 | 1.100E-07 | 4.300E-07 |
| 28391 | 1420 | L1420L | E | 3.7e-07 | 2e-07 | 1.9e-07 | 2.533E-07 | 1.012E-07 | 1.900E-07 | 3.700E-07 |
| 28392 | 1420 | L1420K | E | 1.9e-07 | 5.1e-07 | 2.6e-07 | 3.200E-07 | 1.682E-07 | 1.900E-07 | 5.100E-07 |
| 28393 | 1420 | L1420M | E | 4.2e-07 | 2.8e-07 | 1.7e-07 | 2.900E-07 | 1.253E-07 | 1.700E-07 | 4.200E-07 |
| 28394 | 1420 | L1420F | E | 1.9e-07 | 2e-07 | 2.3e-07 | 2.067E-07 | 2.082E-08 | 1.900E-07 | 2.300E-07 |
| 28395 | 1420 | L1420P | E | 3.1e-07 | 2.3e-07 | 4.1e-07 | 3.167E-07 | 9.018E-08 | 2.300E-07 | 4.100E-07 |
| 28396 | 1420 | L1420S | E | 2.2e-07 | 3.4e-07 | 3.1e-07 | 2.900E-07 | 6.245E-08 | 2.200E-07 | 3.400E-07 |
| 28397 | 1420 | L1420T | E | 3.9e-07 | 1.5e-07 | 1.9e-07 | 2.433E-07 | 1.286E-07 | 1.500E-07 | 3.900E-07 |
| 28398 | 1420 | L1420W | E | 3.3e-07 | 2.3e-07 | 3e-07 | 2.867E-07 | 5.132E-08 | 2.300E-07 | 3.300E-07 |
| 28399 | 1420 | L1420Y | E | 3.4e-07 | 7e-07 | 3.9e-07 | 4.767E-07 | 1.950E-07 | 3.400E-07 | 7.000E-07 |
| 28400 | 1420 | L1420V | E | 5.4e-07 | 1.6e-07 | 2e-07 | 3.000E-07 | 2.088E-07 | 1.600E-07 | 5.400E-07 |
| 28401 | 1421 | V1421A | E | 3.3e-07 | 1.9e-07 | 2.3e-07 | 2.500E-07 | 7.211E-08 | 1.900E-07 | 3.300E-07 |
| 28402 | 1421 | V1421R | E | 1.8e-07 | 3.2e-07 | 2.6e-07 | 2.533E-07 | 7.024E-08 | 1.800E-07 | 3.200E-07 |
| 28403 | 1421 | V1421N | E | 1.9e-07 | 2.7e-07 | 3.1e-07 | 2.567E-07 | 6.110E-08 | 1.900E-07 | 3.100E-07 |
| 28404 | 1421 | V1421D | E | 4.3e-07 | 3.2e-07 | 2.6e-07 | 3.367E-07 | 8.622E-08 | 2.600E-07 | 4.300E-07 |
| 28405 | 1421 | V1421C | E | 3.2e-07 | 1.3e-07 | 1.2e-07 | 1.900E-07 | 1.127E-07 | 1.200E-07 | 3.200E-07 |
| 28406 | 1421 | V1421E | E | 2.3e-07 | 3.5e-07 | 3.6e-07 | 3.133E-07 | 7.234E-08 | 2.300E-07 | 3.600E-07 |
| 28407 | 1421 | V1421Q | E | 4e-07 | 2.3e-07 | 3.2e-07 | 3.167E-07 | 8.505E-08 | 2.300E-07 | 4.000E-07 |
| 28408 | 1421 | V1421G | E | 2.1e-07 | 1.2e-07 | 3.9e-07 | 2.400E-07 | 1.375E-07 | 1.200E-07 | 3.900E-07 |
| 28409 | 1421 | V1421H | E | 4.3e-07 | 9.6e-08 | 3.7e-07 | 2.987E-07 | 1.781E-07 | 9.600E-08 | 4.300E-07 |
| 28410 | 1421 | V1421I | E | 2.2e-07 | 2.4e-07 | 1e-07 | 1.867E-07 | 7.572E-08 | 1.000E-07 | 2.400E-07 |
| 28411 | 1421 | V1421L | E | 2.3e-07 | 2.1e-07 | 3e-07 | 2.467E-07 | 4.726E-08 | 2.100E-07 | 3.000E-07 |
| 28412 | 1421 | V1421K | E | 3.9e-07 | 1.9e-07 | 2.1e-07 | 2.633E-07 | 1.102E-07 | 1.900E-07 | 3.900E-07 |
| 28413 | 1421 | V1421M | E | 2.6e-07 | 1.8e-07 | 5.2e-07 | 3.200E-07 | 1.778E-07 | 1.800E-07 | 5.200E-07 |
| 28414 | 1421 | V1421F | E | 6.5e-07 | 2.7e-07 | 1.6e-07 | 3.600E-07 | 2.571E-07 | 1.600E-07 | 6.500E-07 |
| 28415 | 1421 | V1421P | E | 4.1e-07 | 1.7e-07 | 1.3e-07 | 2.367E-07 | 1.514E-07 | 1.300E-07 | 4.100E-07 |
| 28416 | 1421 | V1421S | E | 1.2e-07 | 1.7e-07 | 2.8e-07 | 1.900E-07 | 8.185E-08 | 1.200E-07 | 2.800E-07 |
| 28417 | 1421 | V1421T | E | 1.3e-07 | 2.2e-07 | 2.1e-07 | 1.867E-07 | 4.933E-08 | 1.300E-07 | 2.200E-07 |
| 28418 | 1421 | V1421W | E | 2.9e-07 | 2e-07 | 7.2e-08 | 1.873E-07 | 1.096E-07 | 7.200E-08 | 2.900E-07 |
| 28419 | 1421 | V1421Y | E | 3e-07 | 2.3e-07 | 2.4e-07 | 2.567E-07 | 3.786E-08 | 2.300E-07 | 3.000E-07 |
| 28420 | 1421 | V1421V | E | 3.5e-07 | 2.6e-07 | 3.7e-07 | 3.267E-07 | 5.859E-08 | 2.600E-07 | 3.700E-07 |
| 28421 | 1422 | K1422A | E | 2.5e-07 | 2.3e-07 | 1.8e-07 | 2.200E-07 | 3.606E-08 | 1.800E-07 | 2.500E-07 |
| 28422 | 1422 | K1422R | E | 2.1e-07 | 2.4e-07 | 3.1e-07 | 2.533E-07 | 5.132E-08 | 2.100E-07 | 3.100E-07 |
| 28423 | 1422 | K1422N | E | 2.1e-07 | 1.2e-07 | 3e-07 | 2.100E-07 | 9.000E-08 | 1.200E-07 | 3.000E-07 |
| 28424 | 1422 | K1422D | E | 2.3e-07 | 2.5e-07 | 3.8e-07 | 2.867E-07 | 8.145E-08 | 2.300E-07 | 3.800E-07 |
| 28425 | 1422 | K1422C | E | 1.8e-07 | 6.3e-07 | 2.9e-07 | 3.667E-07 | 2.346E-07 | 1.800E-07 | 6.300E-07 |
| 28426 | 1422 | K1422E | E | 2.7e-07 | 3.2e-07 | 1.8e-07 | 2.567E-07 | 7.095E-08 | 1.800E-07 | 3.200E-07 |
| 28427 | 1422 | K1422Q | E | 2.4e-07 | 2.6e-07 | 3.7e-07 | 2.900E-07 | 7.000E-08 | 2.400E-07 | 3.700E-07 |
| 28428 | 1422 | K1422G | E | 2.4e-07 | 2.3e-07 | 2.9e-07 | 2.533E-07 | 3.215E-08 | 2.300E-07 | 2.900E-07 |
| 28429 | 1422 | K1422H | E | 1.3e-07 | 3.4e-07 | 4.2e-07 | 2.967E-07 | 1.498E-07 | 1.300E-07 | 4.200E-07 |
| 28430 | 1422 | K1422I | E | 2.6e-07 | 3.7e-07 | 2.4e-07 | 2.900E-07 | 7.000E-08 | 2.400E-07 | 3.700E-07 |
| 28431 | 1422 | K1422L | E | 1.9e-07 | 1.9e-07 | 2.7e-07 | 2.167E-07 | 4.619E-08 | 1.900E-07 | 2.700E-07 |
| 28432 | 1422 | K1422K | E | 2.2e-07 | 1.8e-07 | 3.4e-07 | 2.467E-07 | 8.327E-08 | 1.800E-07 | 3.400E-07 |
| 28433 | 1422 | K1422M | E | 3.8e-07 | 2.2e-07 | 5e-07 | 3.667E-07 | 1.405E-07 | 2.200E-07 | 5.000E-07 |
| 28434 | 1422 | K1422F | E | 1.8e-07 | 2.3e-07 | 1.2e-07 | 1.767E-07 | 5.508E-08 | 1.200E-07 | 2.300E-07 |
| 28435 | 1422 | K1422P | E | 8.3e-08 | 1.6e-07 | 2e-07 | 1.477E-07 | 5.947E-08 | 8.300E-08 | 2.000E-07 |
| 28436 | 1422 | K1422S | E | 2.8e-07 | 1.2e-07 | 1.6e-07 | 1.867E-07 | 8.327E-08 | 1.200E-07 | 2.800E-07 |
| 28437 | 1422 | K1422T | E | 5.6e-07 | 3.6e-07 | 1.2e-07 | 3.467E-07 | 2.203E-07 | 1.200E-07 | 5.600E-07 |
| 28438 | 1422 | K1422W | E | 3.3e-07 | 2.5e-07 | 2.2e-07 | 2.667E-07 | 5.686E-08 | 2.200E-07 | 3.300E-07 |
| 28439 | 1422 | K1422Y | E | 2.6e-07 | 2.2e-07 | 1.6e-07 | 2.133E-07 | 5.033E-08 | 1.600E-07 | 2.600E-07 |
| 28440 | 1422 | K1422V | E | 4.6e-07 | 2.1e-07 | 1.8e-07 | 2.833E-07 | 1.537E-07 | 1.800E-07 | 4.600E-07 |
| 28441 | 1423 | D1423A | E | 1.8e-07 | 2e-07 | 2.7e-07 | 2.167E-07 | 4.726E-08 | 1.800E-07 | 2.700E-07 |
| 28442 | 1423 | D1423R | E | 2.3e-07 | 2e-07 | 3.7e-07 | 2.667E-07 | 9.074E-08 | 2.000E-07 | 3.700E-07 |
| 28443 | 1423 | D1423N | E | 3.3e-07 | 3.5e-07 | 1.8e-07 | 2.867E-07 | 9.292E-08 | 1.800E-07 | 3.500E-07 |
| 28444 | 1423 | D1423D | E | 1.2e-07 | 3.1e-07 | 3.2e-07 | 2.500E-07 | 1.127E-07 | 1.200E-07 | 3.200E-07 |
| 28445 | 1423 | D1423C | E | 1.9e-07 | 1.3e-07 | 1.4e-07 | 1.533E-07 | 3.215E-08 | 1.300E-07 | 1.900E-07 |
| 28446 | 1423 | D1423E | E | 1.1e-07 | 1.5e-07 | 1.8e-07 | 1.467E-07 | 3.512E-08 | 1.100E-07 | 1.800E-07 |
| 28447 | 1423 | D1423Q | E | 1.6e-07 | 3.1e-07 | 1.9e-07 | 2.200E-07 | 7.937E-08 | 1.600E-07 | 3.100E-07 |
| 28448 | 1423 | D1423G | E | 1.9e-07 | 2.7e-07 | 2.9e-07 | 2.500E-07 | 5.292E-08 | 1.900E-07 | 2.900E-07 |
| 28449 | 1423 | D1423H | E | 3.6e-07 | 3.2e-07 | 2.3e-07 | 3.033E-07 | 6.658E-08 | 2.300E-07 | 3.600E-07 |
| 28450 | 1423 | D1423I | E | 2.7e-07 | 2.1e-07 | 3.2e-07 | 2.667E-07 | 5.508E-08 | 2.100E-07 | 3.200E-07 |
| 28451 | 1423 | D1423L | E | 1.1e-07 | 3.1e-07 | 1.8e-07 | 2.000E-07 | 1.015E-07 | 1.100E-07 | 3.100E-07 |
| 28452 | 1423 | D1423K | E | 3.4e-07 | 3.3e-07 | 2e-07 | 2.900E-07 | 7.810E-08 | 2.000E-07 | 3.400E-07 |
| 28453 | 1423 | D1423M | E | 4e-07 | 5.1e-07 | 4e-07 | 4.367E-07 | 6.351E-08 | 4.000E-07 | 5.100E-07 |
| 28454 | 1423 | D1423F | E | 2.7e-07 | 1.6e-07 | 1.8e-07 | 2.033E-07 | 5.859E-08 | 1.600E-07 | 2.700E-07 |
| 28455 | 1423 | D1423P | E | 2.2e-07 | 2.7e-07 | 1.9e-07 | 2.267E-07 | 4.041E-08 | 1.900E-07 | 2.700E-07 |
| 28456 | 1423 | D1423S | E | 3e-07 | 1.1e-07 | 2.9e-07 | 2.333E-07 | 1.069E-07 | 1.100E-07 | 3.000E-07 |
| 28457 | 1423 | D1423T | E | 1.9e-07 | 5.1e-07 | 2.6e-07 | 3.200E-07 | 1.682E-07 | 1.900E-07 | 5.100E-07 |
| 28458 | 1423 | D1423W | E | 5.1e-07 | 1.2e-07 | 5.8e-08 | 2.293E-07 | 2.450E-07 | 5.800E-08 | 5.100E-07 |
| 28459 | 1423 | D1423Y | E | 3.9e-07 | 1.3e-07 | 3.5e-07 | 2.900E-07 | 1.400E-07 | 1.300E-07 | 3.900E-07 |
| 28460 | 1423 | D1423V | E | 2.6e-07 | 3.7e-07 | 1.8e-07 | 2.700E-07 | 9.539E-08 | 1.800E-07 | 3.700E-07 |
| 28461 | 1424 | Y1424A | E | 1.4e-07 | 2.1e-07 | 1.6e-07 | 1.700E-07 | 3.606E-08 | 1.400E-07 | 2.100E-07 |
| 28462 | 1424 | Y1424R | E | 4.4e-07 | 1.8e-07 | 2.7e-07 | 2.967E-07 | 1.320E-07 | 1.800E-07 | 4.400E-07 |
| 28463 | 1424 | Y1424N | E | 2.8e-07 | 6e-07 | 5.7e-07 | 4.833E-07 | 1.767E-07 | 2.800E-07 | 6.000E-07 |
| 28464 | 1424 | Y1424D | E | 2.5e-07 | 2e-07 | 1.2e-07 | 1.900E-07 | 6.557E-08 | 1.200E-07 | 2.500E-07 |
| 28465 | 1424 | Y1424C | E | 1.1e-07 | 1.5e-07 | 5.4e-07 | 2.667E-07 | 2.376E-07 | 1.100E-07 | 5.400E-07 |
| 28466 | 1424 | Y1424E | E | 1.8e-07 | 2.2e-07 | 1.5e-07 | 1.833E-07 | 3.512E-08 | 1.500E-07 | 2.200E-07 |
| 28467 | 1424 | Y1424Q | E | 3.5e-07 | 2.2e-07 | 1.2e-07 | 2.300E-07 | 1.153E-07 | 1.200E-07 | 3.500E-07 |
| 28468 | 1424 | Y1424G | E | 1.7e-07 | 3.6e-07 | 1.9e-07 | 2.400E-07 | 1.044E-07 | 1.700E-07 | 3.600E-07 |
| 28469 | 1424 | Y1424H | E | 3.7e-07 | 2.9e-07 | 3.3e-07 | 3.300E-07 | 4.000E-08 | 2.900E-07 | 3.700E-07 |
| 28470 | 1424 | Y1424I | E | 2.1e-07 | 3.9e-07 | 1.4e-07 | 2.467E-07 | 1.290E-07 | 1.400E-07 | 3.900E-07 |
| 28471 | 1424 | Y1424L | E | 2.9e-07 | 4.7e-07 | 1.9e-07 | 3.167E-07 | 1.419E-07 | 1.900E-07 | 4.700E-07 |
| 28472 | 1424 | Y1424K | E | 1.5e-07 | 4.4e-07 | 2.9e-07 | 2.933E-07 | 1.450E-07 | 1.500E-07 | 4.400E-07 |
| 28473 | 1424 | Y1424M | E | 1.6e-07 | 1.5e-07 | 2.3e-07 | 1.800E-07 | 4.359E-08 | 1.500E-07 | 2.300E-07 |
| 28474 | 1424 | Y1424F | E | 3.2e-07 | 2.5e-07 | 8.5e-08 | 2.183E-07 | 1.207E-07 | 8.500E-08 | 3.200E-07 |

| | | | | | | | | | | |
|---|---|---|---|---|---|---|---|---|---|---|
| 28475 | 1424 | Y1424P | E | 3.8e-07 | 1.1e-07 | 1.9e-07 | 2.267E-07 | 1.387E-07 | 1.100E-07 | 3.800E-07 |
| 28476 | 1424 | Y1424S | E | 4.7e-07 | 2.2e-07 | 2.1e-07 | 3.000E-07 | 1.473E-07 | 2.100E-07 | 4.700E-07 |
| 28477 | 1424 | Y1424T | E | 8.6e-07 | 2.4e-07 | 1.6e-07 | 4.200E-07 | 3.831E-07 | 1.600E-07 | 8.600E-07 |
| 28478 | 1424 | Y1424W | E | 1.6e-07 | 1.4e-07 | 3.1e-07 | 2.033E-07 | 9.292E-08 | 1.400E-07 | 3.100E-07 |
| 28479 | 1424 | Y1424Y | E | 2.3e-07 | 1.7e-07 | 7.5e-07 | 3.833E-07 | 3.190E-07 | 1.700E-07 | 7.500E-07 |
| 28480 | 1424 | Y1424V | E | 3e-07 | 2e-07 | 3.3e-07 | 2.767E-07 | 6.807E-08 | 2.000E-07 | 3.300E-07 |
| 28481 | 1425 | F1425A | E | 2.3e-07 | 2.1e-07 | 1.6e-07 | 2.000E-07 | 3.606E-08 | 1.600E-07 | 2.300E-07 |
| 28482 | 1425 | F1425R | E | 8.5e-08 | 2.3e-07 | 3.9e-07 | 2.350E-07 | 1.526E-07 | 8.500E-08 | 3.900E-07 |
| 28483 | 1425 | F1425N | E | 2.4e-07 | 5.1e-07 | 2.9e-07 | 3.467E-07 | 1.436E-07 | 2.400E-07 | 5.100E-07 |
| 28484 | 1425 | F1425D | E | 2.7e-07 | 2.6e-07 | 2.4e-07 | 2.567E-07 | 1.528E-08 | 2.400E-07 | 2.700E-07 |
| 28485 | 1425 | F1425C | E | 1.5e-07 | 5.2e-07 | 3.1e-07 | 3.267E-07 | 1.856E-07 | 1.500E-07 | 5.200E-07 |
| 28486 | 1425 | F1425E | E | 2.5e-07 | 1.9e-07 | 5.6e-07 | 3.333E-07 | 1.986E-07 | 1.900E-07 | 5.600E-07 |
| 28487 | 1425 | F1425Q | E | 4.6e-07 | 3.9e-07 | 1.6e-07 | 3.367E-07 | 1.570E-07 | 1.600E-07 | 4.600E-07 |
| 28488 | 1425 | F1425G | E | 3.7e-07 | 1.7e-07 | 1.5e-07 | 2.300E-07 | 1.217E-07 | 1.500E-07 | 3.700E-07 |
| 28489 | 1425 | F1425H | E | 2.6e-07 | 3.3e-07 | 3.1e-07 | 3.000E-07 | 3.606E-08 | 2.600E-07 | 3.300E-07 |
| 28490 | 1425 | F1425I | E | 3.1e-07 | 1.3e-07 | 1.8e-07 | 2.067E-07 | 9.292E-08 | 1.300E-07 | 3.100E-07 |
| 28491 | 1425 | F1425L | E | 3.2e-07 | 1.5e-07 | 2.5e-07 | 2.400E-07 | 8.544E-08 | 1.500E-07 | 3.200E-07 |
| 28492 | 1425 | F1425K | E | 4.9e-07 | 2.8e-07 | 3.2e-07 | 3.633E-07 | 1.115E-07 | 2.800E-07 | 4.900E-07 |
| 28493 | 1425 | F1425M | E | 3.6e-07 | 1.7e-07 | 1.6e-07 | 2.300E-07 | 1.127E-07 | 1.600E-07 | 3.600E-07 |
| 28494 | 1425 | F1425F | E | 3.7e-07 | 2e-07 | 3.9e-07 | 3.200E-07 | 1.044E-07 | 2.000E-07 | 3.900E-07 |
| 28495 | 1425 | F1425P | E | 5.6e-07 | 2.4e-07 | 2.2e-07 | 3.400E-07 | 1.908E-07 | 2.200E-07 | 5.600E-07 |
| 28496 | 1425 | F1425S | E | 1.5e-07 | 2.5e-07 | 3.5e-07 | 2.500E-07 | 1.000E-07 | 1.500E-07 | 3.500E-07 |
| 28497 | 1425 | F1425T | E | 8.2e-08 | 2.1e-07 | 3e-07 | 1.973E-07 | 1.096E-07 | 8.200E-08 | 3.000E-07 |
| 28498 | 1425 | F1425W | E | 1e-07 | 2.5e-07 | 2.5e-07 | 2.000E-07 | 8.660E-08 | 1.000E-07 | 2.500E-07 |
| 28499 | 1425 | F1425Y | E | 3e-07 | 1.7e-07 | 1.5e-07 | 2.067E-07 | 8.145E-08 | 1.500E-07 | 3.000E-07 |
| 28500 | 1425 | F1425V | E | 5.8e-07 | 2.9e-07 | 1.2e-07 | 3.300E-07 | 2.326E-07 | 1.200E-07 | 5.800E-07 |
| 28501 | 1426 | P1426A | E | 4e-07 | 7e-07 | 2.6e-07 | 4.533E-07 | 2.248E-07 | 2.600E-07 | 7.000E-07 |
| 28502 | 1426 | P1426R | E | 3.3e-07 | 9.9e-08 | 2.2e-07 | 2.163E-07 | 1.155E-07 | 9.900E-08 | 3.300E-07 |
| 28503 | 1426 | P1426N | E | 1.5e-07 | 2.9e-07 | 2.4e-07 | 2.267E-07 | 7.095E-08 | 1.500E-07 | 2.900E-07 |
| 28504 | 1426 | P1426D | E | 2.3e-07 | 2.7e-07 | 2.7e-07 | 2.567E-07 | 2.309E-08 | 2.300E-07 | 2.700E-07 |
| 28505 | 1426 | P1426C | E | 2.4e-07 | 3.7e-07 | 2.2e-07 | 2.767E-07 | 8.145E-08 | 2.200E-07 | 3.700E-07 |
| 28506 | 1426 | P1426E | E | 2.6e-07 | 3.6e-07 | 3.7e-07 | 3.300E-07 | 6.083E-08 | 2.600E-07 | 3.700E-07 |
| 28507 | 1426 | P1426Q | E | 1.8e-07 | 6.8e-07 | 2e-07 | 3.533E-07 | 2.831E-07 | 1.800E-07 | 6.800E-07 |
| 28508 | 1426 | P1426G | E | 7.5e-08 | 3.2e-07 | 2e-07 | 1.983E-07 | 1.225E-07 | 7.500E-08 | 3.200E-07 |
| 28509 | 1426 | P1426H | E | 6.9e-08 | 2.7e-07 | 2e-07 | 1.797E-07 | 1.020E-07 | 6.900E-08 | 2.700E-07 |
| 28510 | 1426 | P1426I | E | 3.3e-07 | 1.6e-07 | 1.9e-07 | 2.267E-07 | 9.074E-08 | 1.600E-07 | 3.300E-07 |
| 28511 | 1426 | P1426L | E | 3.5e-07 | 1.8e-07 | 1.9e-07 | 2.400E-07 | 9.539E-08 | 1.800E-07 | 3.500E-07 |
| 28512 | 1426 | P1426K | E | 3.2e-07 | 6e-07 | 2.9e-07 | 4.033E-07 | 1.710E-07 | 2.900E-07 | 6.000E-07 |
| 28513 | 1426 | P1426M | E | 2.8e-07 | 1.2e-07 | 1.6e-07 | 1.867E-07 | 8.327E-08 | 1.200E-07 | 2.800E-07 |
| 28514 | 1426 | P1426F | E | 2.7e-07 | 2.9e-07 | 2.4e-07 | 2.667E-07 | 2.517E-08 | 2.400E-07 | 2.900E-07 |
| 28515 | 1426 | P1426P | E | 2.1e-07 | 2.7e-07 | 1.4e-07 | 2.067E-07 | 6.506E-08 | 1.400E-07 | 2.700E-07 |
| 28516 | 1426 | P1426S | E | 2e-07 | 1.7e-07 | 1.9e-07 | 1.867E-07 | 1.528E-08 | 1.700E-07 | 2.000E-07 |
| 28517 | 1426 | P1426T | E | 4.2e-07 | 1.3e-07 | 4e-07 | 3.167E-07 | 1.620E-07 | 1.300E-07 | 4.200E-07 |
| 28518 | 1426 | P1426W | E | 2.6e-07 | 3.1e-07 | 5e-07 | 3.567E-07 | 1.266E-07 | 2.600E-07 | 5.000E-07 |
| 28519 | 1426 | P1426Y | E | 1.7e-07 | 2e-07 | 2.9e-07 | 2.200E-07 | 6.245E-08 | 1.700E-07 | 2.900E-07 |
| 28520 | 1426 | P1426V | E | 1.4e-07 | 2e-07 | 4.5e-07 | 2.633E-07 | 1.644E-07 | 1.400E-07 | 4.500E-07 |
| 28521 | 1427 | E1427A | E | 1.4e-07 | 1.4e-07 | 3e-07 | 1.933E-07 | 9.238E-08 | 1.400E-07 | 3.000E-07 |
| 28522 | 1427 | E1427R | E | 1.3e-07 | 2.3e-07 | 4.7e-07 | 2.767E-07 | 1.747E-07 | 1.300E-07 | 4.700E-07 |
| 28523 | 1427 | E1427N | E | 1.5e-07 | 3.3e-07 | 2.3e-07 | 2.367E-07 | 9.018E-08 | 1.500E-07 | 3.300E-07 |
| 28524 | 1427 | E1427D | E | 3.1e-07 | 2.7e-07 | 4.9e-07 | 3.567E-07 | 1.172E-07 | 2.700E-07 | 4.900E-07 |
| 28525 | 1427 | E1427C | E | 4e-07 | 4.1e-07 | 8.7e-07 | 5.600E-07 | 2.685E-07 | 4.000E-07 | 8.700E-07 |
| 28526 | 1427 | E1427E | E | 2.1e-07 | 2.2e-07 | 2.7e-07 | 2.333E-07 | 3.215E-08 | 2.100E-07 | 2.700E-07 |
| 28527 | 1427 | E1427Q | E | 2.1e-07 | 3.5e-07 | 3.6e-07 | 3.067E-07 | 8.386E-08 | 2.100E-07 | 3.600E-07 |
| 28528 | 1427 | E1427G | E | 1.6e-07 | 2.2e-07 | 3.4e-07 | 2.400E-07 | 9.165E-08 | 1.600E-07 | 3.400E-07 |
| 28529 | 1427 | E1427H | E | 3.1e-07 | 2.7e-07 | 3.2e-07 | 3.000E-07 | 2.646E-08 | 2.700E-07 | 3.200E-07 |
| 28530 | 1427 | E1427I | E | 3.2e-07 | 1.5e-07 | 4e-07 | 2.900E-07 | 1.277E-07 | 1.500E-07 | 4.000E-07 |
| 28531 | 1427 | E1427L | E | 1.1e-07 | 2e-07 | 1.5e-07 | 1.533E-07 | 4.509E-08 | 1.100E-07 | 2.000E-07 |
| 28532 | 1427 | E1427K | E | 2.1e-07 | 2.3e-07 | 1.5e-07 | 1.967E-07 | 4.163E-08 | 1.500E-07 | 2.300E-07 |
| 28533 | 1427 | E1427M | E | 3.3e-07 | 1.7e-07 | 2.9e-07 | 2.633E-07 | 8.327E-08 | 1.700E-07 | 3.300E-07 |
| 28534 | 1427 | E1427F | E | 1.6e-07 | 2.3e-07 | 4.5e-07 | 2.800E-07 | 1.513E-07 | 1.600E-07 | 4.500E-07 |
| 28535 | 1427 | E1427P | E | 5.1e-07 | 2e-07 | 2.6e-07 | 3.233E-07 | 1.644E-07 | 2.000E-07 | 5.100E-07 |
| 28536 | 1427 | E1427S | E | 2.9e-07 | 2.2e-07 | 2.6e-07 | 2.567E-07 | 3.512E-08 | 2.200E-07 | 2.900E-07 |
| 28537 | 1427 | E1427T | E | 1e-07 | 1.3e-07 | 2.2e-07 | 1.500E-07 | 6.245E-08 | 1.000E-07 | 2.200E-07 |
| 28538 | 1427 | E1427W | E | 1.4e-07 | 2.1e-07 | 3.3e-07 | 2.267E-07 | 9.609E-08 | 1.400E-07 | 3.300E-07 |
| 28539 | 1427 | E1427Y | E | 1.8e-07 | 1.3e-07 | 2.6e-07 | 1.900E-07 | 6.557E-08 | 1.300E-07 | 2.600E-07 |
| 28540 | 1427 | E1427V | E | 1.2e-07 | 2.9e-07 | 5.2e-07 | 3.100E-07 | 2.007E-07 | 1.200E-07 | 5.200E-07 |
| 28541 | 1428 | P1428A | E | 1.5e-07 | 5.8e-08 | 1.8e-07 | 1.293E-07 | 6.357E-08 | 5.800E-08 | 1.800E-07 |
| 28542 | 1428 | P1428R | E | 4.3e-07 | 1.8e-07 | 4.5e-07 | 3.533E-07 | 1.504E-07 | 1.800E-07 | 4.500E-07 |
| 28543 | 1428 | P1428N | E | 3.2e-07 | 5.1e-07 | 2e-07 | 3.433E-07 | 1.563E-07 | 2.000E-07 | 5.100E-07 |
| 28544 | 1428 | P1428D | E | 4e-07 | 1.4e-07 | 3.2e-07 | 2.867E-07 | 1.332E-07 | 1.400E-07 | 4.000E-07 |
| 28545 | 1428 | P1428C | E | 2.8e-07 | 1.7e-07 | 2.7e-07 | 2.400E-07 | 6.083E-08 | 1.700E-07 | 2.800E-07 |
| 28546 | 1428 | P1428E | E | 2.3e-07 | 1.6e-07 | 2.4e-07 | 2.100E-07 | 4.359E-08 | 1.600E-07 | 2.400E-07 |
| 28547 | 1428 | P1428Q | E | 5.4e-07 | 1.3e-07 | 3.1e-07 | 3.267E-07 | 2.055E-07 | 1.300E-07 | 5.400E-07 |
| 28548 | 1428 | P1428G | E | 2e-07 | 1.7e-07 | 2e-07 | 1.900E-07 | 1.732E-08 | 1.700E-07 | 2.000E-07 |
| 28549 | 1428 | P1428H | E | 3.4e-07 | 2.3e-07 | 2.2e-07 | 2.633E-07 | 6.658E-08 | 2.200E-07 | 3.400E-07 |
| 28550 | 1428 | P1428I | E | 1.8e-07 | 2.3e-07 | 2.5e-07 | 2.200E-07 | 3.606E-08 | 1.800E-07 | 2.500E-07 |
| 28551 | 1428 | P1428L | E | 2.5e-07 | 3.6e-07 | 1.9e-07 | 2.667E-07 | 8.622E-08 | 1.900E-07 | 3.600E-07 |
| 28552 | 1428 | P1428K | E | 1.9e-07 | 4.2e-07 | 3.5e-07 | 3.200E-07 | 1.179E-07 | 1.900E-07 | 4.200E-07 |
| 28553 | 1428 | P1428M | E | 2.4e-07 | 3.6e-07 | 2.8e-07 | 2.933E-07 | 6.110E-08 | 2.400E-07 | 3.600E-07 |
| 28554 | 1428 | P1428F | E | 3.9e-07 | 4.6e-07 | 2.2e-07 | 3.567E-07 | 1.234E-07 | 2.200E-07 | 4.600E-07 |
| 28555 | 1428 | P1428P | E | 1.7e-07 | 3.3e-07 | 4.5e-07 | 3.167E-07 | 1.405E-07 | 1.700E-07 | 4.500E-07 |
| 28556 | 1428 | P1428S | E | 2.6e-07 | 1.9e-07 | 1.7e-07 | 2.067E-07 | 4.726E-08 | 1.700E-07 | 2.600E-07 |
| 28557 | 1428 | P1428T | E | 3.6e-07 | 2e-07 | 2.1e-07 | 2.567E-07 | 8.963E-08 | 2.000E-07 | 3.600E-07 |
| 28558 | 1428 | P1428W | E | 2.5e-07 | 3.3e-07 | 2.5e-07 | 2.767E-07 | 4.619E-08 | 2.500E-07 | 3.300E-07 |
| 28559 | 1428 | P1428Y | E | 2.3e-07 | 2.6e-07 | 2.2e-07 | 2.367E-07 | 2.082E-08 | 2.200E-07 | 2.600E-07 |

| | | | | | | | | | | |
|---|---|---|---|---|---|---|---|---|---|---|
| 28560 | 1428 | P1428V | E | 3e-07 | 2.7e-07 | 3.4e-07 | 3.033E-07 | 3.512E-08 | 2.700E-07 | 3.400E-07 |
| 28561 | 1429 | V1429A | E | 2.9e-07 | 3.5e-07 | 9.4e-08 | 2.447E-07 | 1.339E-07 | 9.400E-08 | 3.500E-07 |
| 28562 | 1429 | V1429R | E | 2.8e-07 | 2e-07 | 3.8e-07 | 2.867E-07 | 9.018E-08 | 2.000E-07 | 3.800E-07 |
| 28563 | 1429 | V1429N | E | 2.7e-07 | 2.9e-07 | 2.4e-07 | 2.667E-07 | 2.517E-08 | 2.400E-07 | 2.900E-07 |
| 28564 | 1429 | V1429D | E | 3.1e-07 | 4.9e-07 | 1.9e-07 | 3.300E-07 | 1.510E-07 | 1.900E-07 | 4.900E-07 |
| 28565 | 1429 | V1429C | E | 1.6e-07 | 6.6e-08 | 7.2e-08 | 9.933E-08 | 5.262E-08 | 6.600E-08 | 1.600E-07 |
| 28566 | 1429 | V1429E | E | 9e-08 | 3.6e-07 | 2.7e-07 | 2.400E-07 | 1.375E-07 | 9.000E-08 | 3.600E-07 |
| 28567 | 1429 | V1429Q | E | 1.6e-07 | 3.3e-07 | 1.6e-07 | 2.167E-07 | 9.815E-08 | 1.600E-07 | 3.300E-07 |
| 28568 | 1429 | V1429G | E | 2.5e-07 | 2.6e-07 | 7.7e-08 | 1.957E-07 | 1.029E-07 | 7.700E-08 | 2.600E-07 |
| 28569 | 1429 | V1429H | E | 1.5e-07 | 4.4e-07 | 3.1e-07 | 3.000E-07 | 1.453E-07 | 1.500E-07 | 4.400E-07 |
| 28570 | 1429 | V1429I | E | 2.1e-07 | 1.4e-07 | 6.3e-07 | 3.267E-07 | 2.650E-07 | 1.400E-07 | 6.300E-07 |
| 28571 | 1429 | V1429L | E | 1.8e-07 | 1.4e-07 | 3.3e-07 | 2.167E-07 | 1.002E-07 | 1.400E-07 | 3.300E-07 |
| 28572 | 1429 | V1429K | E | 3.3e-07 | 1.3e-07 | 3.3e-07 | 2.633E-07 | 1.155E-07 | 1.300E-07 | 3.300E-07 |
| 28573 | 1429 | V1429M | E | 2.5e-07 | 1.6e-07 | 1.9e-07 | 2.000E-07 | 4.583E-08 | 1.600E-07 | 2.500E-07 |
| 28574 | 1429 | V1429F | E | 2e-07 | 1.9e-07 | 5.7e-07 | 3.200E-07 | 2.166E-07 | 1.900E-07 | 5.700E-07 |
| 28575 | 1429 | V1429P | E | 9.3e-08 | 2.5e-07 | 1.2e-07 | 1.543E-07 | 8.394E-08 | 9.300E-08 | 2.500E-07 |
| 28576 | 1429 | V1429S | E | 3.6e-07 | 2.9e-07 | 1.8e-07 | 2.767E-07 | 9.074E-08 | 1.800E-07 | 3.600E-07 |
| 28577 | 1429 | V1429T | E | 2.4e-07 | 3.5e-07 | 1.7e-07 | 2.533E-07 | 9.074E-08 | 1.700E-07 | 3.500E-07 |
| 28578 | 1429 | V1429W | E | 2.5e-07 | 1.6e-07 | 2.7e-07 | 2.267E-07 | 5.859E-08 | 1.600E-07 | 2.700E-07 |
| 28579 | 1429 | V1429Y | E | 3.7e-07 | 2.5e-07 | 3.1e-07 | 3.100E-07 | 6.000E-08 | 2.500E-07 | 3.700E-07 |
| 28580 | 1429 | V1429V | E | 1.6e-07 | 1.9e-07 | 3.7e-07 | 2.400E-07 | 1.136E-07 | 1.600E-07 | 3.700E-07 |
| 28581 | 1430 | T1430A | E | 2.7e-07 | 1.4e-07 | 2.6e-07 | 2.233E-07 | 7.234E-08 | 1.400E-07 | 2.700E-07 |
| 28582 | 1430 | T1430R | E | 1.8e-07 | 1.8e-07 | 6.6e-07 | 3.400E-07 | 2.771E-07 | 1.800E-07 | 6.600E-07 |
| 28583 | 1430 | T1430N | E | 2.7e-07 | 2.3e-07 | 2.7e-07 | 2.567E-07 | 2.309E-08 | 2.300E-07 | 2.700E-07 |
| 28584 | 1430 | T1430D | E | 2.8e-07 | 2.4e-07 | 2.8e-07 | 2.667E-07 | 2.309E-08 | 2.400E-07 | 2.800E-07 |
| 28585 | 1430 | T1430C | E | 2.2e-07 | 2.4e-07 | 2.4e-07 | 2.333E-07 | 1.155E-08 | 2.200E-07 | 2.400E-07 |
| 28586 | 1430 | T1430E | E | 1.7e-07 | 2e-07 | 2.1e-07 | 1.933E-07 | 2.082E-08 | 1.700E-07 | 2.100E-07 |
| 28587 | 1430 | T1430Q | E | 1.3e-07 | 4.4e-07 | 3.8e-07 | 3.167E-07 | 1.644E-07 | 1.300E-07 | 4.400E-07 |
| 28588 | 1430 | T1430G | E | 2.3e-07 | 2e-07 | 1.8e-07 | 2.033E-07 | 2.517E-08 | 1.800E-07 | 2.300E-07 |
| 28589 | 1430 | T1430H | E | 2.8e-07 | 3.6e-07 | 1e-07 | 2.467E-07 | 1.332E-07 | 1.000E-07 | 3.600E-07 |
| 28590 | 1430 | T1430I | E | 2.1e-07 | 3e-07 | 1.1e-07 | 2.067E-07 | 9.504E-08 | 1.100E-07 | 3.000E-07 |
| 28591 | 1430 | T1430L | E | 3.5e-07 | 2e-07 | 1.3e-07 | 2.267E-07 | 1.124E-07 | 1.300E-07 | 3.500E-07 |
| 28592 | 1430 | T1430K | E | 1.9e-07 | 3.2e-07 | 3.7e-07 | 2.933E-07 | 9.292E-08 | 1.900E-07 | 3.700E-07 |
| 28593 | 1430 | T1430M | E | 2.5e-07 | 2.8e-07 | 2.1e-07 | 2.467E-07 | 3.512E-08 | 2.100E-07 | 2.800E-07 |
| 28594 | 1430 | T1430F | E | 1.9e-07 | 5e-07 | 3e-07 | 3.300E-07 | 1.572E-07 | 1.900E-07 | 5.000E-07 |
| 28595 | 1430 | T1430P | E | 3.3e-07 | 1.5e-07 | 3.6e-07 | 2.800E-07 | 1.136E-07 | 1.500E-07 | 3.600E-07 |
| 28596 | 1430 | T1430S | E | 1.4e-07 | 1.5e-07 | 1.5e-07 | 1.467E-07 | 5.774E-09 | 1.400E-07 | 1.500E-07 |
| 28597 | 1430 | T1430T | E | 2.5e-07 | 3.1e-07 | 2.2e-07 | 2.600E-07 | 4.583E-08 | 2.200E-07 | 3.100E-07 |
| 28598 | 1430 | T1430W | E | 6.2e-07 | 1.8e-07 | 2e-07 | 3.333E-07 | 2.485E-07 | 1.800E-07 | 6.200E-07 |
| 28599 | 1430 | T1430Y | E | 1.9e-07 | 1.1e-07 | 2e-07 | 1.667E-07 | 4.933E-08 | 1.100E-07 | 2.000E-07 |
| 28600 | 1430 | T1430V | E | 2.5e-07 | 1.7e-07 | 2.5e-07 | 2.233E-07 | 4.619E-08 | 1.700E-07 | 2.500E-07 |
| 28601 | 1431 | V1431A | E | 2e-07 | 2.9e-07 | 2e-07 | 2.300E-07 | 5.196E-08 | 2.000E-07 | 2.900E-07 |
| 28602 | 1431 | V1431R | E | 3e-07 | 1.5e-07 | 2.6e-07 | 2.367E-07 | 7.767E-08 | 1.500E-07 | 3.000E-07 |
| 28603 | 1431 | V1431N | E | 2.4e-07 | 5.7e-08 | 2.4e-07 | 1.790E-07 | 1.057E-07 | 5.700E-08 | 2.400E-07 |
| 28604 | 1431 | V1431D | E | 1.6e-07 | 3.2e-07 | 2.4e-07 | 2.400E-07 | 8.000E-08 | 1.600E-07 | 3.200E-07 |
| 28605 | 1431 | V1431C | E | 7e-07 | 1.9e-07 | 2.6e-07 | 3.833E-07 | 2.765E-07 | 1.900E-07 | 7.000E-07 |
| 28606 | 1431 | V1431E | E | 1.6e-07 | 1.6e-07 | 1.7e-07 | 1.633E-07 | 5.774E-09 | 1.600E-07 | 1.700E-07 |
| 28607 | 1431 | V1431Q | E | 2.1e-07 | 2e-07 | 4.7e-07 | 2.933E-07 | 1.531E-07 | 2.000E-07 | 4.700E-07 |
| 28608 | 1431 | V1431G | E | 1.7e-07 | 1.5e-07 | 2.7e-07 | 1.967E-07 | 6.429E-08 | 1.500E-07 | 2.700E-07 |
| 28609 | 1431 | V1431H | E | 3.3e-07 | 1.4e-07 | 1.4e-07 | 2.033E-07 | 1.097E-07 | 1.400E-07 | 3.300E-07 |
| 28610 | 1431 | V1431I | E | 2.1e-07 | 3.6e-07 | 1.6e-07 | 2.433E-07 | 1.041E-07 | 1.600E-07 | 3.600E-07 |
| 28611 | 1431 | V1431L | E | 2.4e-07 | 3e-07 | 1.3e-07 | 2.233E-07 | 8.622E-08 | 1.300E-07 | 3.000E-07 |
| 28612 | 1431 | V1431K | E | 3.1e-07 | 5.9e-07 | 3.4e-07 | 4.133E-07 | 1.537E-07 | 3.100E-07 | 5.900E-07 |
| 28613 | 1431 | V1431M | E | 2.7e-07 | 1.6e-07 | 1.9e-07 | 2.067E-07 | 5.686E-08 | 1.600E-07 | 2.700E-07 |
| 28614 | 1431 | V1431F | E | 2.3e-07 | 2.1e-07 | 1.3e-07 | 1.900E-07 | 5.292E-08 | 1.300E-07 | 2.300E-07 |
| 28615 | 1431 | V1431P | E | 3.2e-07 | 2.2e-07 | 4.5e-07 | 3.300E-07 | 1.153E-07 | 2.200E-07 | 4.500E-07 |
| 28616 | 1431 | V1431S | E | 3.6e-07 | 1.6e-07 | 2.8e-07 | 2.667E-07 | 1.007E-07 | 1.600E-07 | 3.600E-07 |
| 28617 | 1431 | V1431T | E | 4.8e-07 | 2.2e-07 | 5.9e-07 | 4.300E-07 | 1.900E-07 | 2.200E-07 | 5.900E-07 |
| 28618 | 1431 | V1431W | E | 1.7e-07 | 1.8e-07 | 3.2e-07 | 2.233E-07 | 8.386E-08 | 1.700E-07 | 3.200E-07 |
| 28619 | 1431 | V1431Y | E | 2.7e-07 | 2.3e-07 | 3e-07 | 2.667E-07 | 3.512E-08 | 2.300E-07 | 3.000E-07 |
| 28620 | 1431 | V1431V | E | 3.4e-07 | 3.8e-07 | 2.5e-07 | 3.233E-07 | 6.658E-08 | 2.500E-07 | 3.800E-07 |
| 28621 | 1432 | S1432A | E | 2.4e-07 | 2.4e-07 | 1.9e-07 | 2.233E-07 | 2.887E-08 | 1.900E-07 | 2.400E-07 |
| 28622 | 1432 | S1432R | E | 9e-08 | 2.3e-07 | 1.4e-07 | 1.533E-07 | 7.095E-08 | 9.000E-08 | 2.300E-07 |
| 28623 | 1432 | S1432N | E | 2e-07 | 1.8e-07 | 2.9e-07 | 2.233E-07 | 5.859E-08 | 1.800E-07 | 2.900E-07 |
| 28624 | 1432 | S1432D | E | 1.7e-07 | 8.7e-08 | 1.6e-07 | 1.390E-07 | 4.531E-08 | 8.700E-08 | 1.700E-07 |
| 28625 | 1432 | S1432C | E | 2.7e-07 | 2.9e-07 | 2.1e-07 | 2.567E-07 | 4.163E-08 | 2.100E-07 | 2.900E-07 |
| 28626 | 1432 | S1432E | E | 1.9e-07 | 2e-07 | 2.3e-07 | 2.067E-07 | 2.082E-08 | 1.900E-07 | 2.300E-07 |
| 28627 | 1432 | S1432Q | E | 2.2e-07 | 2.1e-07 | 1.2e-07 | 1.833E-07 | 5.508E-08 | 1.200E-07 | 2.200E-07 |
| 28628 | 1432 | S1432G | E | 2e-07 | 1.2e-07 | 1.8e-07 | 1.667E-07 | 4.163E-08 | 1.200E-07 | 2.000E-07 |
| 28629 | 1432 | S1432H | E | 2.2e-07 | 3.1e-07 | 1.1e-07 | 2.133E-07 | 1.002E-07 | 1.100E-07 | 3.100E-07 |
| 28630 | 1432 | S1432I | E | 2.2e-07 | 4.9e-07 | 2.3e-07 | 3.133E-07 | 1.531E-07 | 2.200E-07 | 4.900E-07 |
| 28631 | 1432 | S1432L | E | 2.3e-07 | 1.5e-07 | 2.7e-07 | 2.167E-07 | 6.110E-08 | 1.500E-07 | 2.700E-07 |
| 28632 | 1432 | S1432K | E | 3.5e-07 | 5e-07 | 3.3e-07 | 3.933E-07 | 9.292E-08 | 3.300E-07 | 5.000E-07 |
| 28633 | 1432 | S1432M | E | 3e-07 | 2.8e-07 | 2.8e-07 | 2.867E-07 | 1.155E-08 | 2.800E-07 | 3.000E-07 |
| 28634 | 1432 | S1432F | E | 3.7e-07 | 2.7e-07 | 4.2e-07 | 3.533E-07 | 7.638E-08 | 2.700E-07 | 4.200E-07 |
| 28635 | 1432 | S1432P | E | 1.1e-07 | 2.4e-07 | 3.4e-07 | 2.300E-07 | 1.153E-07 | 1.100E-07 | 3.400E-07 |
| 28636 | 1432 | S1432S | E | 2.7e-07 | 1.3e-07 | 5.4e-07 | 3.133E-07 | 2.084E-07 | 1.300E-07 | 5.400E-07 |
| 28637 | 1432 | S1432T | E | 2.2e-07 | 2.1e-07 | 1.7e-07 | 2.000E-07 | 2.646E-08 | 1.700E-07 | 2.200E-07 |
| 28638 | 1432 | S1432W | E | 1.2e-07 | 1.8e-07 | 1.4e-07 | 1.467E-07 | 3.055E-08 | 1.200E-07 | 1.800E-07 |
| 28639 | 1432 | S1432Y | E | 2.9e-07 | 3.3e-07 | 4.8e-07 | 3.667E-07 | 1.002E-07 | 2.900E-07 | 4.800E-07 |
| 28640 | 1432 | S1432V | E | 1.4e-07 | 1.5e-07 | 3.4e-07 | 2.100E-07 | 1.127E-07 | 1.400E-07 | 3.400E-07 |
| 28641 | 1433 | W1433A | E | 2e-07 | 2.2e-07 | 3.1e-07 | 2.433E-07 | 5.859E-08 | 2.000E-07 | 3.100E-07 |
| 28642 | 1433 | W1433R | E | 2.1e-07 | 3.6e-07 | 1.5e-07 | 2.400E-07 | 1.082E-07 | 1.500E-07 | 3.600E-07 |
| 28643 | 1433 | W1433N | E | 2e-07 | 2.1e-07 | 3.6e-07 | 2.567E-07 | 8.963E-08 | 2.000E-07 | 3.600E-07 |
| 28644 | 1433 | W1433D | E | 2.3e-07 | 2.4e-07 | 2.2e-07 | 2.300E-07 | 1.000E-08 | 2.200E-07 | 2.400E-07 |

| | | | | | | | | | | |
|---|---|---|---|---|---|---|---|---|---|---|
| 28645 | 1433 | W1433C | E | 1.5e-07 | 2.4e-07 | 4.5e-07 | 2.800E-07 | 1.539E-07 | 1.500E-07 | 4.500E-07 |
| 28646 | 1433 | W1433E | E | 6.5e-07 | 2.2e-07 | 1.8e-07 | 3.500E-07 | 2.606E-07 | 1.800E-07 | 6.500E-07 |
| 28647 | 1433 | W1433Q | E | 4e-07 | 3.3e-07 | 2.2e-07 | 3.167E-07 | 9.074E-08 | 2.200E-07 | 4.000E-07 |
| 28648 | 1433 | W1433G | E | 1.1e-07 | 2.9e-07 | 2.2e-07 | 2.067E-07 | 9.074E-08 | 1.100E-07 | 2.900E-07 |
| 28649 | 1433 | W1433H | E | 2e-07 | 2.7e-07 | 1.5e-07 | 2.067E-07 | 6.028E-08 | 1.500E-07 | 2.700E-07 |
| 28650 | 1433 | W1433I | E | 3.5e-07 | 2.5e-07 | 3.1e-07 | 3.033E-07 | 5.033E-08 | 2.500E-07 | 3.500E-07 |
| 28651 | 1433 | W1433L | E | 2e-07 | 5.3e-07 | 2.5e-07 | 3.267E-07 | 1.779E-07 | 2.000E-07 | 5.300E-07 |
| 28652 | 1433 | W1433K | E | 6.8e-07 | 1.2e-07 | 2.3e-07 | 3.433E-07 | 2.967E-07 | 1.200E-07 | 6.800E-07 |
| 28653 | 1433 | W1433M | E | 4e-07 | 4.7e-07 | 4.1e-07 | 4.267E-07 | 3.786E-08 | 4.000E-07 | 4.700E-07 |
| 28654 | 1433 | W1433F | E | 3.5e-07 | 3.6e-07 | 2.2e-07 | 3.100E-07 | 7.810E-08 | 2.200E-07 | 3.600E-07 |
| 28655 | 1433 | W1433P | E | 2.2e-07 | 2.4e-07 | 3.6e-07 | 2.733E-07 | 7.572E-08 | 2.200E-07 | 3.600E-07 |
| 28656 | 1433 | W1433S | E | 1.8e-07 | 8e-07 | 1e-07 | 3.600E-07 | 3.831E-07 | 1.000E-07 | 8.000E-07 |
| 28657 | 1433 | W1433T | E | 3.9e-07 | 1.5e-07 | 5.6e-07 | 3.667E-07 | 2.060E-07 | 1.500E-07 | 5.600E-07 |
| 28658 | 1433 | W1433W | E | 3.1e-07 | 2.9e-07 | 2.4e-07 | 2.800E-07 | 3.606E-08 | 2.400E-07 | 3.100E-07 |
| 28659 | 1433 | W1433Y | E | 2.2e-07 | 4e-07 | 2e-07 | 2.733E-07 | 1.102E-07 | 2.000E-07 | 4.000E-07 |
| 28660 | 1433 | W1433V | E | 2.7e-07 | 1.9e-07 | 2.9e-07 | 2.500E-07 | 5.292E-08 | 1.900E-07 | 2.900E-07 |
| 28661 | 1434 | N1434A | E | 4.4e-07 | 1.9e-07 | 2.5e-07 | 2.933E-07 | 1.305E-07 | 1.900E-07 | 4.400E-07 |
| 28662 | 1434 | N1434R | E | 2.9e-07 | 1.1e-07 | 6.7e-08 | 1.557E-07 | 1.183E-07 | 6.700E-08 | 2.900E-07 |
| 28663 | 1434 | N1434N | E | 4e-07 | 2.2e-07 | 1.5e-07 | 2.567E-07 | 1.290E-07 | 1.500E-07 | 4.000E-07 |
| 28664 | 1434 | N1434D | E | 2.6e-07 | 2e-07 | 2.2e-07 | 2.267E-07 | 3.055E-08 | 2.000E-07 | 2.600E-07 |
| 28665 | 1434 | N1434C | E | 2.5e-07 | 1.4e-07 | 1.7e-07 | 1.867E-07 | 5.686E-08 | 1.400E-07 | 2.500E-07 |
| 28666 | 1434 | N1434E | E | 9.4e-08 | 3.4e-07 | 1.4e-07 | 1.913E-07 | 1.308E-07 | 9.400E-08 | 3.400E-07 |
| 28667 | 1434 | N1434Q | E | 2.8e-07 | 1.6e-07 | 3.3e-07 | 2.567E-07 | 8.737E-08 | 1.600E-07 | 3.300E-07 |
| 28668 | 1434 | N1434G | E | 2.7e-07 | 1.7e-07 | 1.6e-07 | 2.000E-07 | 6.083E-08 | 1.600E-07 | 2.700E-07 |
| 28669 | 1434 | N1434H | E | 2.5e-07 | 3.7e-07 | 2.4e-07 | 2.867E-07 | 7.234E-08 | 2.400E-07 | 3.700E-07 |
| 28670 | 1434 | N1434I | E | 3.1e-07 | 3.7e-07 | 2.1e-07 | 2.967E-07 | 8.083E-08 | 2.100E-07 | 3.700E-07 |
| 28671 | 1434 | N1434L | E | 2.4e-07 | 1.9e-07 | 2.2e-07 | 2.167E-07 | 2.517E-08 | 1.900E-07 | 2.400E-07 |
| 28672 | 1434 | N1434K | E | 3.6e-07 | 3.1e-07 | 1.1e-07 | 2.600E-07 | 1.323E-07 | 1.100E-07 | 3.600E-07 |
| 28673 | 1434 | N1434M | E | 1.9e-07 | 1.1e-07 | 4e-07 | 2.333E-07 | 1.498E-07 | 1.100E-07 | 4.000E-07 |
| 28674 | 1434 | N1434F | E | 2.8e-07 | 3.3e-07 | 1.5e-07 | 2.533E-07 | 9.292E-08 | 1.500E-07 | 3.300E-07 |
| 28675 | 1434 | N1434P | E | 3.5e-07 | 4.5e-07 | 3.6e-07 | 3.867E-07 | 5.508E-08 | 3.500E-07 | 4.500E-07 |
| 28676 | 1434 | N1434S | E | 2.3e-07 | 2.1e-07 | 5e-07 | 3.133E-07 | 1.620E-07 | 2.100E-07 | 5.000E-07 |
| 28677 | 1434 | N1434T | E | 2.9e-07 | 1.9e-07 | 5.9e-07 | 3.567E-07 | 2.082E-07 | 1.900E-07 | 5.900E-07 |
| 28678 | 1434 | N1434W | E | 1.8e-07 | 5.6e-07 | 2.5e-07 | 3.300E-07 | 2.022E-07 | 1.800E-07 | 5.600E-07 |
| 28679 | 1434 | N1434Y | E | 1.7e-07 | 3.6e-07 | 2.9e-07 | 2.733E-07 | 9.609E-08 | 1.700E-07 | 3.600E-07 |
| 28680 | 1434 | N1434V | E | 3.4e-07 | 1.3e-07 | 1.6e-07 | 2.100E-07 | 1.136E-07 | 1.300E-07 | 3.400E-07 |
| 28681 | 1435 | S1435A | E | 3e-07 | 1.6e-07 | 2.2e-07 | 2.267E-07 | 7.024E-08 | 1.600E-07 | 3.000E-07 |
| 28682 | 1435 | S1435R | E | 1.6e-07 | 2e-07 | 1.7e-07 | 1.767E-07 | 2.082E-08 | 1.600E-07 | 2.000E-07 |
| 28683 | 1435 | S1435N | E | 5.5e-07 | 1.8e-07 | 1.5e-07 | 2.933E-07 | 2.228E-07 | 1.500E-07 | 5.500E-07 |
| 28684 | 1435 | S1435D | E | 4.5e-07 | 1.7e-07 | 3.9e-07 | 3.367E-07 | 1.474E-07 | 1.700E-07 | 4.500E-07 |
| 28685 | 1435 | S1435C | E | 3.7e-07 | 3.5e-07 | 5.6e-07 | 4.267E-07 | 1.159E-07 | 3.500E-07 | 5.600E-07 |
| 28686 | 1435 | S1435E | E | 3.2e-07 | 2.6e-07 | 1.6e-07 | 2.467E-07 | 8.083E-08 | 1.600E-07 | 3.200E-07 |
| 28687 | 1435 | S1435Q | E | 1.3e-07 | 2e-07 | 2.6e-07 | 1.967E-07 | 6.506E-08 | 1.300E-07 | 2.600E-07 |
| 28688 | 1435 | S1435G | E | 2.7e-07 | 2.1e-07 | 5.1e-07 | 3.300E-07 | 1.587E-07 | 2.100E-07 | 5.100E-07 |
| 28689 | 1435 | S1435H | E | 4.1e-07 | 1.7e-07 | 3.2e-07 | 3.000E-07 | 1.212E-07 | 1.700E-07 | 4.100E-07 |
| 28690 | 1435 | S1435I | E | 1.6e-07 | 8.6e-08 | 3.6e-07 | 2.020E-07 | 1.417E-07 | 8.600E-08 | 3.600E-07 |
| 28691 | 1435 | S1435L | E | 1.9e-07 | 3.6e-07 | 4.6e-07 | 3.367E-07 | 1.365E-07 | 1.900E-07 | 4.600E-07 |
| 28692 | 1435 | S1435K | E | 2e-07 | 2.1e-07 | 1e-07 | 1.700E-07 | 6.083E-08 | 1.000E-07 | 2.100E-07 |
| 28693 | 1435 | S1435M | E | 3.8e-07 | 4.6e-07 | 5.2e-07 | 4.533E-07 | 7.024E-08 | 3.800E-07 | 5.200E-07 |
| 28694 | 1435 | S1435F | E | 3.3e-07 | 1.2e-07 | 2.1e-07 | 2.200E-07 | 1.054E-07 | 1.200E-07 | 3.300E-07 |
| 28695 | 1435 | S1435P | E | 3.3e-07 | 3.2e-07 | 2e-07 | 2.833E-07 | 7.234E-08 | 2.000E-07 | 3.300E-07 |
| 28696 | 1435 | S1435S | E | 1.9e-07 | 5.5e-07 | 1.5e-07 | 2.967E-07 | 2.203E-07 | 1.500E-07 | 5.500E-07 |
| 28697 | 1435 | S1435T | E | 3.7e-07 | 1.4e-07 | 5.3e-07 | 3.467E-07 | 1.960E-07 | 1.400E-07 | 5.300E-07 |
| 28698 | 1435 | S1435W | E | 2.2e-07 | 2.7e-07 | 1.5e-07 | 2.133E-07 | 6.028E-08 | 1.500E-07 | 2.700E-07 |
| 28699 | 1435 | S1435Y | E | 2.8e-07 | 1.2e-07 | 2.6e-07 | 2.200E-07 | 8.718E-08 | 1.200E-07 | 2.800E-07 |
| 28700 | 1435 | S1435V | E | 1.7e-07 | 2.9e-07 | 3.5e-07 | 2.700E-07 | 9.165E-08 | 1.700E-07 | 3.500E-07 |
| 28701 | 1436 | G1436A | E | 2.5e-07 | 2.9e-07 | 2.3e-07 | 2.567E-07 | 3.055E-08 | 2.300E-07 | 2.900E-07 |
| 28702 | 1436 | G1436R | E | 9.4e-08 | 3.2e-07 | 2.9e-07 | 2.347E-07 | 1.227E-07 | 9.400E-08 | 3.200E-07 |
| 28703 | 1436 | G1436N | E | 2.2e-07 | 4.1e-07 | 3.5e-07 | 3.267E-07 | 9.713E-08 | 2.200E-07 | 4.100E-07 |
| 28704 | 1436 | G1436D | E | 2.1e-07 | 4.3e-07 | 1.5e-07 | 2.633E-07 | 1.474E-07 | 1.500E-07 | 4.300E-07 |
| 28705 | 1436 | G1436C | E | 2.9e-07 | 4e-07 | 2.9e-07 | 3.267E-07 | 6.351E-08 | 2.900E-07 | 4.000E-07 |
| 28706 | 1436 | G1436E | E | 2.3e-07 | 3.9e-07 | 5.9e-07 | 4.033E-07 | 1.804E-07 | 2.300E-07 | 5.900E-07 |
| 28707 | 1436 | G1436Q | E | 3.9e-07 | 2.7e-07 | 5.3e-07 | 3.967E-07 | 1.301E-07 | 2.700E-07 | 5.300E-07 |
| 28708 | 1436 | G1436G | E | 3.7e-07 | 3.2e-07 | 1.9e-07 | 2.933E-07 | 9.292E-08 | 1.900E-07 | 3.700E-07 |
| 28709 | 1436 | G1436H | E | 2.1e-07 | 4.6e-07 | 2.4e-07 | 3.033E-07 | 1.365E-07 | 2.100E-07 | 4.600E-07 |
| 28710 | 1436 | G1436I | E | 2.5e-07 | 2e-07 | 1.7e-07 | 2.067E-07 | 4.041E-08 | 1.700E-07 | 2.500E-07 |
| 28711 | 1436 | G1436L | E | 4e-07 | 3.3e-07 | 1.5e-07 | 2.933E-07 | 1.290E-07 | 1.500E-07 | 4.000E-07 |
| 28712 | 1436 | G1436K | E | 4.4e-08 | 1.1e-07 | 1.6e-07 | 1.047E-07 | 5.818E-08 | 4.400E-08 | 1.600E-07 |
| 28713 | 1436 | G1436M | E | 4.3e-07 | 3.9e-07 | 3.1e-07 | 3.767E-07 | 6.110E-08 | 3.100E-07 | 4.300E-07 |
| 28714 | 1436 | G1436F | E | 2.7e-07 | 2.2e-07 | 2.4e-07 | 2.433E-07 | 2.517E-08 | 2.200E-07 | 2.700E-07 |
| 28715 | 1436 | G1436P | E | 3.4e-07 | 2.9e-07 | 1.2e-07 | 2.500E-07 | 1.153E-07 | 1.200E-07 | 3.400E-07 |
| 28716 | 1436 | G1436S | E | 3.8e-07 | 3.6e-07 | 2.7e-07 | 3.367E-07 | 5.859E-08 | 2.700E-07 | 3.800E-07 |
| 28717 | 1436 | G1436T | E | 4.7e-07 | 1.7e-07 | 3.4e-07 | 3.267E-07 | 1.504E-07 | 1.700E-07 | 4.700E-07 |
| 28718 | 1436 | G1436W | E | 2.2e-07 | 4.2e-07 | 1.9e-07 | 2.767E-07 | 1.250E-07 | 1.900E-07 | 4.200E-07 |
| 28719 | 1436 | G1436Y | E | 1.6e-07 | 1.3e-07 | 1.4e-07 | 1.433E-07 | 1.528E-08 | 1.300E-07 | 1.600E-07 |
| 28720 | 1436 | G1436V | E | 2.1e-07 | 3.9e-07 | 3.9e-07 | 3.300E-07 | 1.039E-07 | 2.100E-07 | 3.900E-07 |
| 28721 | 1437 | A1437A | E | 6e-07 | 4.7e-07 | 1.4e-07 | 4.033E-07 | 2.371E-07 | 1.400E-07 | 6.000E-07 |
| 28722 | 1437 | A1437R | E | 2.6e-07 | 3.2e-07 | 1.5e-07 | 2.433E-07 | 8.622E-08 | 1.500E-07 | 3.200E-07 |
| 28723 | 1437 | A1437N | E | 1.5e-07 | 3.3e-07 | 2.4e-07 | 2.400E-07 | 9.000E-08 | 1.500E-07 | 3.300E-07 |
| 28724 | 1437 | A1437D | E | 1.9e-07 | 3.4e-07 | 2.5e-07 | 2.600E-07 | 7.550E-08 | 1.900E-07 | 3.400E-07 |
| 28725 | 1437 | A1437C | E | 1.5e-07 | 2.2e-07 | 1.9e-07 | 1.867E-07 | 3.512E-08 | 1.500E-07 | 2.200E-07 |
| 28726 | 1437 | A1437E | E | 1.5e-07 | 1.7e-07 | 2.2e-07 | 1.800E-07 | 3.606E-08 | 1.500E-07 | 2.200E-07 |
| 28727 | 1437 | A1437Q | E | 2.9e-07 | 1.6e-07 | 2.7e-07 | 2.400E-07 | 7.000E-08 | 1.600E-07 | 2.900E-07 |
| 28728 | 1437 | A1437G | E | 2.6e-07 | 3.7e-07 | 2.6e-07 | 2.967E-07 | 6.351E-08 | 2.600E-07 | 3.700E-07 |
| 28729 | 1437 | A1437H | E | 2e-07 | 2.6e-07 | 1.4e-07 | 2.000E-07 | 6.000E-08 | 1.400E-07 | 2.600E-07 |

| | | | | | | | | | | |
|---|---|---|---|---|---|---|---|---|---|---|
| 28730 | 1437 | A1437I | E | 2.7e-07 | 2e-07 | 3e-07 | 2.567E-07 | 5.132E-08 | 2.000E-07 | 3.000E-07 |
| 28731 | 1437 | A1437L | E | 2e-07 | 3e-07 | 2.2e-07 | 2.400E-07 | 5.292E-08 | 2.000E-07 | 3.000E-07 |
| 28732 | 1437 | A1437K | E | 1.7e-07 | 3.3e-07 | 4.9e-07 | 3.300E-07 | 1.600E-07 | 1.700E-07 | 4.900E-07 |
| 28733 | 1437 | A1437M | E | 1.1e-07 | 1.7e-07 | 2.5e-07 | 1.767E-07 | 7.024E-08 | 1.100E-07 | 2.500E-07 |
| 28734 | 1437 | A1437F | E | 1.1e-07 | 2.2e-07 | 3.2e-07 | 2.167E-07 | 1.050E-07 | 1.100E-07 | 3.200E-07 |
| 28735 | 1437 | A1437P | E | 2.4e-07 | 5.9e-07 | 1.7e-07 | 3.333E-07 | 2.250E-07 | 1.700E-07 | 5.900E-07 |
| 28736 | 1437 | A1437S | E | 5.6e-08 | 4.6e-07 | 2.3e-07 | 2.487E-07 | 2.026E-07 | 5.600E-08 | 4.600E-07 |
| 28737 | 1437 | A1437T | E | 2.4e-07 | 3.2e-07 | 4e-07 | 3.200E-07 | 8.000E-08 | 2.400E-07 | 4.000E-07 |
| 28738 | 1437 | A1437W | E | 9e-08 | 4.3e-07 | 1.7e-07 | 2.300E-07 | 1.778E-07 | 9.000E-08 | 4.300E-07 |
| 28739 | 1437 | A1437Y | E | 1.4e-07 | 2.8e-07 | 4.8e-07 | 3.000E-07 | 1.709E-07 | 1.400E-07 | 4.800E-07 |
| 28740 | 1437 | A1437V | E | 3.5e-07 | 2e-07 | 1.3e-07 | 2.267E-07 | 1.124E-07 | 1.300E-07 | 3.500E-07 |
| 28741 | 1438 | L1438A | E | 1.5e-07 | 1.6e-07 | 2.1e-07 | 1.733E-07 | 3.215E-08 | 1.500E-07 | 2.100E-07 |
| 28742 | 1438 | L1438R | E | 1.1e-07 | 1.3e-07 | 1.9e-07 | 1.433E-07 | 4.163E-08 | 1.100E-07 | 1.900E-07 |
| 28743 | 1438 | L1438N | E | 2.6e-07 | 3.8e-07 | 1.8e-07 | 2.733E-07 | 1.007E-07 | 1.800E-07 | 3.800E-07 |
| 28744 | 1438 | L1438D | E | 2.1e-07 | 3e-07 | 3.1e-07 | 2.733E-07 | 5.508E-08 | 2.100E-07 | 3.100E-07 |
| 28745 | 1438 | L1438C | E | 3e-07 | 5.8e-07 | 6.2e-07 | 5.000E-07 | 1.744E-07 | 3.000E-07 | 6.200E-07 |
| 28746 | 1438 | L1438E | E | 4.9e-07 | 1.5e-07 | 3.1e-07 | 3.167E-07 | 1.701E-07 | 1.500E-07 | 4.900E-07 |
| 28747 | 1438 | L1438Q | E | 1.9e-07 | 3.2e-07 | 2.1e-07 | 2.400E-07 | 7.000E-08 | 1.900E-07 | 3.200E-07 |
| 28748 | 1438 | L1438G | E | 2e-07 | 1.4e-07 | 3.8e-07 | 2.400E-07 | 1.249E-07 | 1.400E-07 | 3.800E-07 |
| 28749 | 1438 | L1438H | E | 2.7e-07 | 1.7e-07 | 3.1e-07 | 2.500E-07 | 7.211E-08 | 1.700E-07 | 3.100E-07 |
| 28750 | 1438 | L1438I | E | 2.5e-07 | 3.9e-07 | 2.2e-07 | 2.867E-07 | 9.074E-08 | 2.200E-07 | 3.900E-07 |
| 28751 | 1438 | L1438L | E | 1.3e-07 | 2.7e-07 | 2.2e-07 | 2.067E-07 | 7.095E-08 | 1.300E-07 | 2.700E-07 |
| 28752 | 1438 | L1438K | E | 2.7e-07 | 8.4e-07 | 2.8e-07 | 4.633E-07 | 3.262E-07 | 2.700E-07 | 8.400E-07 |
| 28753 | 1438 | L1438M | E | 1.4e-07 | 2e-07 | 2.5e-07 | 1.967E-07 | 5.508E-08 | 1.400E-07 | 2.500E-07 |
| 28754 | 1438 | L1438F | E | 2.9e-07 | 2.7e-07 | 3.4e-07 | 3.000E-07 | 3.606E-08 | 2.700E-07 | 3.400E-07 |
| 28755 | 1438 | L1438P | E | 1.4e-07 | 1.4e-07 | 1.6e-07 | 1.467E-07 | 1.155E-08 | 1.400E-07 | 1.600E-07 |
| 28756 | 1438 | L1438S | E | 3.8e-07 | 8.7e-07 | 3.1e-07 | 5.200E-07 | 3.051E-07 | 3.100E-07 | 8.700E-07 |
| 28757 | 1438 | L1438T | E | 3e-07 | 3.4e-07 | 3e-07 | 3.133E-07 | 2.309E-08 | 3.000E-07 | 3.400E-07 |
| 28758 | 1438 | L1438W | E | 2.4e-07 | 3.2e-07 | 1.8e-07 | 2.467E-07 | 7.024E-08 | 1.800E-07 | 3.200E-07 |
| 28759 | 1438 | L1438Y | E | 3.7e-07 | 2.2e-07 | 1.2e-07 | 2.367E-07 | 1.258E-07 | 1.200E-07 | 3.700E-07 |
| 28760 | 1438 | L1438V | E | 1.3e-07 | 2.9e-07 | 5.4e-07 | 3.200E-07 | 2.066E-07 | 1.300E-07 | 5.400E-07 |
| 28761 | 1439 | T1439A | E | 4.4e-07 | 2e-07 | 2.2e-07 | 2.867E-07 | 1.332E-07 | 2.000E-07 | 4.400E-07 |
| 28762 | 1439 | T1439R | E | 3.6e-07 | 1.4e-07 | 1.8e-07 | 2.267E-07 | 1.172E-07 | 1.400E-07 | 3.600E-07 |
| 28763 | 1439 | T1439N | E | 2.2e-07 | 2.9e-07 | 2.7e-07 | 2.600E-07 | 3.606E-08 | 2.200E-07 | 2.900E-07 |
| 28764 | 1439 | T1439D | E | 1e-07 | 2.7e-07 | 3.4e-07 | 2.367E-07 | 1.234E-07 | 1.000E-07 | 3.400E-07 |
| 28765 | 1439 | T1439C | E | 3.1e-07 | 4.7e-07 | 3.2e-07 | 3.667E-07 | 8.963E-08 | 3.100E-07 | 4.700E-07 |
| 28766 | 1439 | T1439E | E | 3.3e-07 | 1.7e-07 | 2.8e-07 | 2.600E-07 | 8.185E-08 | 1.700E-07 | 3.300E-07 |
| 28767 | 1439 | T1439Q | E | 1.9e-07 | 9.5e-08 | 1.6e-07 | 1.483E-07 | 4.856E-08 | 9.500E-08 | 1.900E-07 |
| 28768 | 1439 | T1439G | E | 2.6e-07 | 2.7e-07 | 4.9e-07 | 3.400E-07 | 1.300E-07 | 2.600E-07 | 4.900E-07 |
| 28769 | 1439 | T1439H | E | 3.1e-07 | 8.6e-08 | 2e-07 | 1.987E-07 | 1.120E-07 | 8.600E-08 | 3.100E-07 |
| 28770 | 1439 | T1439I | E | 1.8e-07 | 1.7e-07 | 1.4e-07 | 1.633E-07 | 2.082E-08 | 1.400E-07 | 1.800E-07 |
| 28771 | 1439 | T1439L | E | 8.1e-08 | 2.2e-07 | 3.6e-07 | 2.203E-07 | 1.395E-07 | 8.100E-08 | 3.600E-07 |
| 28772 | 1439 | T1439K | E | 2.9e-07 | 1.4e-07 | 3.9e-07 | 2.733E-07 | 1.258E-07 | 1.400E-07 | 3.900E-07 |
| 28773 | 1439 | T1439M | E | 3e-07 | 9.7e-08 | 1.7e-07 | 1.890E-07 | 1.028E-07 | 9.700E-08 | 3.000E-07 |
| 28774 | 1439 | T1439F | E | 1.4e-07 | 4.5e-07 | 2.8e-07 | 2.900E-07 | 1.552E-07 | 1.400E-07 | 4.500E-07 |
| 28775 | 1439 | T1439P | E | 3.3e-07 | 2e-07 | 3.1e-07 | 2.800E-07 | 7.000E-08 | 2.000E-07 | 3.300E-07 |
| 28776 | 1439 | T1439S | E | 2.9e-07 | 2.4e-07 | 4.6e-07 | 3.300E-07 | 1.153E-07 | 2.400E-07 | 4.600E-07 |
| 28777 | 1439 | T1439T | E | 4.6e-07 | 2.6e-07 | 3.2e-07 | 3.467E-07 | 1.026E-07 | 2.600E-07 | 4.600E-07 |
| 28778 | 1439 | T1439W | E | 2.8e-07 | 2e-07 | 1.4e-07 | 2.067E-07 | 7.024E-08 | 1.400E-07 | 2.800E-07 |
| 28779 | 1439 | T1439Y | E | 2.3e-07 | 2.4e-07 | 1.7e-07 | 2.133E-07 | 3.786E-08 | 1.700E-07 | 2.400E-07 |
| 28780 | 1439 | T1439V | E | 3.8e-07 | 1.7e-07 | 2.7e-07 | 2.733E-07 | 1.050E-07 | 1.700E-07 | 3.800E-07 |
| 28781 | 1440 | S1440A | E | 1.8e-07 | 4e-07 | 5e-07 | 3.600E-07 | 1.637E-07 | 1.800E-07 | 5.000E-07 |
| 28782 | 1440 | S1440R | E | 1.4e-07 | 3e-07 | 2.7e-07 | 2.367E-07 | 8.505E-08 | 1.400E-07 | 3.000E-07 |
| 28783 | 1440 | S1440N | E | 3.8e-07 | 2.4e-07 | 2e-07 | 2.733E-07 | 9.452E-08 | 2.000E-07 | 3.800E-07 |
| 28784 | 1440 | S1440D | E | 2.4e-07 | 3.8e-07 | 2.9e-07 | 3.033E-07 | 7.095E-08 | 2.400E-07 | 3.800E-07 |
| 28785 | 1440 | S1440C | E | 2.4e-07 | 3.9e-07 | 3.8e-07 | 3.367E-07 | 8.386E-08 | 2.400E-07 | 3.900E-07 |
| 28786 | 1440 | S1440E | E | 3.3e-07 | 2.1e-07 | 1.8e-07 | 2.400E-07 | 7.937E-08 | 1.800E-07 | 3.300E-07 |
| 28787 | 1440 | S1440Q | E | 1.5e-07 | 5.5e-07 | 1.9e-07 | 2.967E-07 | 2.203E-07 | 1.500E-07 | 5.500E-07 |
| 28788 | 1440 | S1440G | E | 3e-07 | 2.8e-07 | 6.1e-07 | 3.967E-07 | 1.850E-07 | 2.800E-07 | 6.100E-07 |
| 28789 | 1440 | S1440H | E | 2.1e-07 | 2.5e-07 | 3.3e-07 | 2.633E-07 | 6.110E-08 | 2.100E-07 | 3.300E-07 |
| 28790 | 1440 | S1440I | E | 1.9e-07 | 3.1e-07 | 5e-07 | 3.333E-07 | 1.563E-07 | 1.900E-07 | 5.000E-07 |
| 28791 | 1440 | S1440L | E | 3.1e-07 | 3.9e-07 | 4e-07 | 3.667E-07 | 4.933E-08 | 3.100E-07 | 4.000E-07 |
| 28792 | 1440 | S1440K | E | 3.2e-07 | 2.5e-07 | 3.5e-07 | 3.067E-07 | 5.132E-08 | 2.500E-07 | 3.500E-07 |
| 28793 | 1440 | S1440M | E | 2.8e-07 | 2.4e-07 | 3.8e-07 | 3.000E-07 | 7.211E-08 | 2.400E-07 | 3.800E-07 |
| 28794 | 1440 | S1440F | E | 1.6e-07 | 2.1e-07 | 1.7e-07 | 1.800E-07 | 2.646E-08 | 1.600E-07 | 2.100E-07 |
| 28795 | 1440 | S1440P | E | 4e-07 | 2.1e-07 | 1.7e-07 | 2.600E-07 | 1.229E-07 | 1.700E-07 | 4.000E-07 |
| 28796 | 1440 | S1440S | E | 5.2e-07 | 3.3e-07 | 1.4e-07 | 3.300E-07 | 1.900E-07 | 1.400E-07 | 5.200E-07 |
| 28797 | 1440 | S1440T | E | 2.4e-07 | 1.6e-07 | 2.6e-07 | 2.200E-07 | 5.292E-08 | 1.600E-07 | 2.600E-07 |
| 28798 | 1440 | S1440W | E | 1.8e-07 | 2.8e-07 | 1.2e-07 | 1.933E-07 | 8.083E-08 | 1.200E-07 | 2.800E-07 |
| 28799 | 1440 | S1440Y | E | 1.5e-07 | 4.6e-07 | 2.9e-07 | 3.000E-07 | 1.552E-07 | 1.500E-07 | 4.600E-07 |
| 28800 | 1440 | S1440V | E | 3.8e-07 | 5.2e-07 | 1.7e-07 | 3.567E-07 | 1.762E-07 | 1.700E-07 | 5.200E-07 |
| 28801 | 1441 | G1441A | E | 1.1e-07 | 2.7e-07 | 2.7e-07 | 2.167E-07 | 9.238E-08 | 1.100E-07 | 2.700E-07 |
| 28802 | 1441 | G1441R | E | 1.9e-07 | 4.5e-07 | 2.1e-07 | 2.833E-07 | 1.447E-07 | 1.900E-07 | 4.500E-07 |
| 28803 | 1441 | G1441N | E | 3.8e-07 | 2.4e-07 | 1.3e-07 | 2.500E-07 | 1.253E-07 | 1.300E-07 | 3.800E-07 |
| 28804 | 1441 | G1441D | E | 1.8e-07 | 2.7e-07 | 3.8e-07 | 2.767E-07 | 1.002E-07 | 1.800E-07 | 3.800E-07 |
| 28805 | 1441 | G1441C | E | 2.6e-07 | 3.3e-07 | 4e-07 | 3.300E-07 | 7.000E-08 | 2.600E-07 | 4.000E-07 |
| 28806 | 1441 | G1441E | E | 2.5e-07 | 1.3e-07 | 2e-07 | 1.933E-07 | 6.028E-08 | 1.300E-07 | 2.500E-07 |
| 28807 | 1441 | G1441Q | E | 2.3e-07 | 4.3e-07 | 3e-07 | 3.200E-07 | 1.015E-07 | 2.300E-07 | 4.300E-07 |
| 28808 | 1441 | G1441G | E | 1.5e-07 | 2e-07 | 4e-07 | 2.500E-07 | 1.323E-07 | 1.500E-07 | 4.000E-07 |
| 28809 | 1441 | G1441H | E | 1.7e-07 | 3.2e-07 | 2.1e-07 | 2.333E-07 | 7.767E-08 | 1.700E-07 | 3.200E-07 |
| 28810 | 1441 | G1441I | E | 3.4e-07 | 2.9e-07 | 2.2e-07 | 2.833E-07 | 6.028E-08 | 2.200E-07 | 3.400E-07 |
| 28811 | 1441 | G1441L | E | 1.7e-07 | 3.2e-07 | 4.5e-07 | 3.133E-07 | 1.401E-07 | 1.700E-07 | 4.500E-07 |
| 28812 | 1441 | G1441K | E | 3.3e-07 | 2.6e-07 | 1.9e-07 | 2.600E-07 | 7.000E-08 | 1.900E-07 | 3.300E-07 |
| 28813 | 1441 | G1441M | E | 4.5e-07 | 2.9e-07 | 6.1e-07 | 4.500E-07 | 1.600E-07 | 2.900E-07 | 6.100E-07 |
| 28814 | 1441 | G1441F | E | 1.9e-07 | 4.9e-07 | 1.4e-07 | 2.733E-07 | 1.893E-07 | 1.400E-07 | 4.900E-07 |

| | | | | | | | | | | |
|---|---|---|---|---|---|---|---|---|---|---|
| 28815 | 1441 | G1441P | E | 4.5e-07 | 2.4e-07 | 8.5e-08 | 2.583E-07 | 1.832E-07 | 8.500E-08 | 4.500E-07 |
| 28816 | 1441 | G1441S | E | 1.9e-07 | 2.5e-07 | 2.3e-07 | 2.233E-07 | 3.055E-08 | 1.900E-07 | 2.500E-07 |
| 28817 | 1441 | G1441T | E | 1.1e-07 | 3.9e-07 | 3.1e-07 | 2.700E-07 | 1.442E-07 | 1.100E-07 | 3.900E-07 |
| 28818 | 1441 | G1441W | E | 2.2e-07 | 1.5e-07 | 1.3e-07 | 1.667E-07 | 4.726E-08 | 1.300E-07 | 2.200E-07 |
| 28819 | 1441 | G1441Y | E | 2.6e-07 | 1.6e-07 | 2.2e-07 | 2.133E-07 | 5.033E-08 | 1.600E-07 | 2.600E-07 |
| 28820 | 1441 | G1441V | E | 1.9e-07 | 2.9e-07 | 3e-07 | 2.600E-07 | 6.083E-08 | 1.900E-07 | 3.000E-07 |
| 28821 | 1442 | V1442A | E | 2.2e-07 | 2.6e-07 | 1.8e-07 | 2.200E-07 | 4.000E-08 | 1.800E-07 | 2.600E-07 |
| 28822 | 1442 | V1442R | E | 3.1e-07 | 2.5e-07 | 3.9e-07 | 3.167E-07 | 7.024E-08 | 2.500E-07 | 3.900E-07 |
| 28823 | 1442 | V1442N | E | 3.3e-07 | 1.9e-07 | 4.9e-07 | 3.367E-07 | 1.501E-07 | 1.900E-07 | 4.900E-07 |
| 28824 | 1442 | V1442D | E | 1.4e-07 | 5.3e-07 | 3.9e-07 | 3.533E-07 | 1.976E-07 | 1.400E-07 | 5.300E-07 |
| 28825 | 1442 | V1442C | E | 2.6e-07 | 2.2e-07 | 1.1e-07 | 1.967E-07 | 7.767E-08 | 1.100E-07 | 2.600E-07 |
| 28826 | 1442 | V1442E | E | 4.2e-07 | 2e-07 | 3e-07 | 3.067E-07 | 1.102E-07 | 2.000E-07 | 4.200E-07 |
| 28827 | 1442 | V1442Q | E | 1.3e-07 | 2.1e-07 | 5.8e-07 | 3.067E-07 | 2.401E-07 | 1.300E-07 | 5.800E-07 |
| 28828 | 1442 | V1442G | E | 2.9e-07 | 3.1e-07 | 1.9e-07 | 2.633E-07 | 6.429E-08 | 1.900E-07 | 3.100E-07 |
| 28829 | 1442 | V1442H | E | 9.5e-08 | 2.2e-07 | 1.5e-07 | 1.550E-07 | 6.265E-08 | 9.500E-08 | 2.200E-07 |
| 28830 | 1442 | V1442I | E | 2.9e-07 | 2.3e-07 | 4.8e-07 | 3.333E-07 | 1.305E-07 | 2.300E-07 | 4.800E-07 |
| 28831 | 1442 | V1442L | E | 2.3e-07 | 8e-08 | 3.9e-07 | 2.333E-07 | 1.550E-07 | 8.000E-08 | 3.900E-07 |
| 28832 | 1442 | V1442K | E | 2.7e-07 | 2.9e-07 | 2.8e-07 | 2.800E-07 | 1.000E-08 | 2.700E-07 | 2.900E-07 |
| 28833 | 1442 | V1442M | E | 1e-07 | 3e-07 | 2.6e-07 | 2.200E-07 | 1.058E-07 | 1.000E-07 | 3.000E-07 |
| 28834 | 1442 | V1442F | E | 1.5e-07 | 1.5e-07 | 4.3e-07 | 2.433E-07 | 1.617E-07 | 1.500E-07 | 4.300E-07 |
| 28835 | 1442 | V1442P | E | 1.9e-07 | 2.7e-07 | 3.7e-07 | 2.767E-07 | 9.018E-08 | 1.900E-07 | 3.700E-07 |
| 28836 | 1442 | V1442S | E | 1.2e-07 | 2.6e-07 | 3.5e-07 | 2.433E-07 | 1.159E-07 | 1.200E-07 | 3.500E-07 |
| 28837 | 1442 | V1442T | E | 2.9e-07 | 3.9e-07 | 2.7e-07 | 3.167E-07 | 6.429E-08 | 2.700E-07 | 3.900E-07 |
| 28838 | 1442 | V1442W | E | 2.5e-07 | 1.1e-07 | 1.6e-07 | 1.733E-07 | 7.095E-08 | 1.100E-07 | 2.500E-07 |
| 28839 | 1442 | V1442Y | E | 1.7e-07 | 3.5e-07 | 4.2e-07 | 3.133E-07 | 1.290E-07 | 1.700E-07 | 4.200E-07 |
| 28840 | 1442 | V1442V | E | 1.8e-07 | 3.2e-07 | 2.7e-07 | 2.567E-07 | 7.095E-08 | 1.800E-07 | 3.200E-07 |
| 28841 | 1443 | H1443A | E | 3e-07 | 2.2e-07 | 4.5e-07 | 3.233E-07 | 1.168E-07 | 2.200E-07 | 4.500E-07 |
| 28842 | 1443 | H1443R | E | 5.3e-07 | 1.7e-07 | 1.4e-07 | 2.800E-07 | 2.170E-07 | 1.400E-07 | 5.300E-07 |
| 28843 | 1443 | H1443N | E | 3.3e-07 | 1.2e-07 | 1.2e-07 | 1.900E-07 | 1.212E-07 | 1.200E-07 | 3.300E-07 |
| 28844 | 1443 | H1443D | E | 3.9e-07 | 1.3e-07 | 2.1e-07 | 2.433E-07 | 1.332E-07 | 1.300E-07 | 3.900E-07 |
| 28845 | 1443 | H1443C | E | 3.2e-07 | 1.4e-07 | 7.1e-07 | 3.900E-07 | 2.914E-07 | 1.400E-07 | 7.100E-07 |
| 28846 | 1443 | H1443E | E | 6.6e-07 | 2.3e-07 | 2.8e-07 | 3.900E-07 | 2.352E-07 | 2.300E-07 | 6.600E-07 |
| 28847 | 1443 | H1443Q | E | 3.4e-07 | 2.4e-07 | 2.7e-07 | 2.833E-07 | 5.132E-08 | 2.400E-07 | 3.400E-07 |
| 28848 | 1443 | H1443G | E | 4e-07 | 2.4e-07 | 2.5e-07 | 2.967E-07 | 8.963E-08 | 2.400E-07 | 4.000E-07 |
| 28849 | 1443 | H1443H | E | 1.7e-07 | 2.2e-07 | 1.9e-07 | 1.933E-07 | 2.517E-08 | 1.700E-07 | 2.200E-07 |
| 28850 | 1443 | H1443I | E | 3e-07 | 1.9e-07 | 2.4e-07 | 2.433E-07 | 5.508E-08 | 1.900E-07 | 3.000E-07 |
| 28851 | 1443 | H1443L | E | 1.4e-07 | 1.7e-07 | 1.5e-07 | 1.533E-07 | 1.528E-08 | 1.400E-07 | 1.700E-07 |
| 28852 | 1443 | H1443K | E | 4e-07 | 1e-07 | 4.5e-07 | 3.167E-07 | 1.893E-07 | 1.000E-07 | 4.500E-07 |
| 28853 | 1443 | H1443M | E | 4e-07 | 3.4e-07 | 2.6e-07 | 3.333E-07 | 7.024E-08 | 2.600E-07 | 4.000E-07 |
| 28854 | 1443 | H1443F | E | 4.1e-07 | 3.1e-07 | 3.1e-07 | 3.433E-07 | 5.774E-08 | 3.100E-07 | 4.100E-07 |
| 28855 | 1443 | H1443P | E | 3.5e-07 | 2.2e-07 | 5e-07 | 3.567E-07 | 1.401E-07 | 2.200E-07 | 5.000E-07 |
| 28856 | 1443 | H1443S | E | 2e-07 | 1.2e-07 | 2e-07 | 1.733E-07 | 4.619E-08 | 1.200E-07 | 2.000E-07 |
| 28857 | 1443 | H1443T | E | 3.5e-07 | 5e-08 | 2.3e-07 | 2.100E-07 | 1.510E-07 | 5.000E-08 | 3.500E-07 |
| 28858 | 1443 | H1443W | E | 3e-07 | 2.2e-07 | 5e-07 | 3.400E-07 | 1.442E-07 | 2.200E-07 | 5.000E-07 |
| 28859 | 1443 | H1443Y | E | 2.4e-07 | 2.4e-07 | 1.4e-07 | 2.067E-07 | 5.774E-08 | 1.400E-07 | 2.400E-07 |
| 28860 | 1443 | H1443V | E | 2.9e-07 | 1.2e-07 | 2.6e-07 | 2.233E-07 | 9.074E-08 | 1.200E-07 | 2.900E-07 |
| 28861 | 1444 | T1444A | E | 3.5e-07 | 4.2e-07 | 2.3e-07 | 3.333E-07 | 9.609E-08 | 2.300E-07 | 4.200E-07 |
| 28862 | 1444 | T1444R | E | 4.3e-07 | 4.8e-07 | 2.2e-07 | 3.767E-07 | 1.380E-07 | 2.200E-07 | 4.800E-07 |
| 28863 | 1444 | T1444N | E | 2.4e-07 | 1.9e-07 | 2.7e-07 | 2.333E-07 | 4.041E-08 | 1.900E-07 | 2.700E-07 |
| 28864 | 1444 | T1444D | E | 3.8e-07 | 2.6e-07 | 3.1e-07 | 3.167E-07 | 6.028E-08 | 2.600E-07 | 3.800E-07 |
| 28865 | 1444 | T1444C | E | 2.7e-07 | 1.5e-07 | 3e-07 | 2.400E-07 | 7.937E-08 | 1.500E-07 | 3.000E-07 |
| 28866 | 1444 | T1444E | E | 2.6e-07 | 2.7e-07 | 2.9e-07 | 2.733E-07 | 1.528E-08 | 2.600E-07 | 2.900E-07 |
| 28867 | 1444 | T1444Q | E | 9.3e-08 | 1.2e-07 | 2.3e-07 | 1.477E-07 | 7.257E-08 | 9.300E-08 | 2.300E-07 |
| 28868 | 1444 | T1444G | E | 3.7e-07 | 2.8e-07 | 4.7e-07 | 3.733E-07 | 9.504E-08 | 2.800E-07 | 4.700E-07 |
| 28869 | 1444 | T1444H | E | 2.2e-07 | 3.1e-07 | 2.3e-07 | 2.533E-07 | 4.933E-08 | 2.200E-07 | 3.100E-07 |
| 28870 | 1444 | T1444I | E | 2.6e-07 | 2.2e-07 | 1.3e-07 | 2.033E-07 | 6.658E-08 | 1.300E-07 | 2.600E-07 |
| 28871 | 1444 | T1444L | E | 1.7e-07 | 3.2e-07 | 4.7e-07 | 3.200E-07 | 1.500E-07 | 1.700E-07 | 4.700E-07 |
| 28872 | 1444 | T1444K | E | 7.2e-08 | 8.7e-08 | 5.8e-07 | 2.463E-07 | 2.891E-07 | 7.200E-08 | 5.800E-07 |
| 28873 | 1444 | T1444M | E | 4.1e-07 | 2.9e-07 | 3.3e-07 | 3.433E-07 | 6.110E-08 | 2.900E-07 | 4.100E-07 |
| 28874 | 1444 | T1444F | E | 3.4e-07 | 1.9e-07 | 1.8e-07 | 2.367E-07 | 8.963E-08 | 1.800E-07 | 3.400E-07 |
| 28875 | 1444 | T1444P | E | 7.4e-07 | 1.8e-07 | 3.3e-07 | 4.167E-07 | 2.899E-07 | 1.800E-07 | 7.400E-07 |
| 28876 | 1444 | T1444S | E | 2.1e-07 | 2.2e-07 | 5.5e-07 | 3.267E-07 | 1.935E-07 | 2.100E-07 | 5.500E-07 |
| 28877 | 1444 | T1444T | E | 1.7e-07 | 3.1e-07 | 2.4e-07 | 2.400E-07 | 7.000E-08 | 1.700E-07 | 3.100E-07 |
| 28878 | 1444 | T1444W | E | 2.8e-07 | 2.1e-07 | 3.5e-07 | 2.800E-07 | 7.000E-08 | 2.100E-07 | 3.500E-07 |
| 28879 | 1444 | T1444Y | E | 3.6e-07 | 3e-07 | 3.8e-07 | 3.467E-07 | 4.163E-08 | 3.000E-07 | 3.800E-07 |
| 28880 | 1444 | T1444V | E | 3.1e-07 | 1.7e-07 | 2.4e-07 | 2.400E-07 | 7.000E-08 | 1.700E-07 | 3.100E-07 |
| 28881 | 1445 | F1445A | E | 3.7e-07 | 4.9e-07 | 4.3e-07 | 4.300E-07 | 6.000E-08 | 3.700E-07 | 4.900E-07 |
| 28882 | 1445 | F1445R | E | 2.8e-07 | 1.3e-07 | 3.7e-07 | 2.600E-07 | 1.212E-07 | 1.300E-07 | 3.700E-07 |
| 28883 | 1445 | F1445N | E | 1.8e-07 | 2.9e-07 | 2.9e-07 | 2.533E-07 | 6.351E-08 | 1.800E-07 | 2.900E-07 |
| 28884 | 1445 | F1445D | E | 2.7e-07 | 2e-07 | 4.9e-07 | 3.200E-07 | 1.513E-07 | 2.000E-07 | 4.900E-07 |
| 28885 | 1445 | F1445C | E | 3e-07 | 2.2e-07 | 1.8e-07 | 2.333E-07 | 6.110E-08 | 1.800E-07 | 3.000E-07 |
| 28886 | 1445 | F1445E | E | 1.3e-07 | 1.5e-07 | 3.1e-07 | 1.967E-07 | 9.866E-08 | 1.300E-07 | 3.100E-07 |
| 28887 | 1445 | F1445Q | E | 1.4e-07 | 2.7e-07 | 2.8e-07 | 2.300E-07 | 7.810E-08 | 1.400E-07 | 2.800E-07 |
| 28888 | 1445 | F1445G | E | 1.3e-07 | 1.9e-07 | 3.8e-07 | 2.333E-07 | 1.305E-07 | 1.300E-07 | 3.800E-07 |
| 28889 | 1445 | F1445H | E | 2.6e-07 | 3.6e-07 | 4.2e-07 | 3.467E-07 | 8.083E-08 | 2.600E-07 | 4.200E-07 |
| 28890 | 1445 | F1445I | E | 6.3e-08 | 2.2e-07 | 2.2e-07 | 1.677E-07 | 9.064E-08 | 6.300E-08 | 2.200E-07 |
| 28891 | 1445 | F1445L | E | 2.3e-07 | 4.6e-07 | 2.1e-07 | 3.000E-07 | 1.389E-07 | 2.100E-07 | 4.600E-07 |
| 28892 | 1445 | F1445K | E | 1.9e-07 | 3.6e-07 | 2.7e-07 | 2.733E-07 | 8.505E-08 | 1.900E-07 | 3.600E-07 |
| 28893 | 1445 | F1445M | E | 1e-07 | 2.7e-07 | 5.2e-07 | 2.967E-07 | 2.113E-07 | 1.000E-07 | 5.200E-07 |
| 28894 | 1445 | F1445F | E | 1.8e-07 | 1.3e-07 | 1.8e-07 | 1.633E-07 | 2.887E-08 | 1.300E-07 | 1.800E-07 |
| 28895 | 1445 | F1445P | E | 1.2e-07 | 3.3e-07 | 3.2e-07 | 2.567E-07 | 1.185E-07 | 1.200E-07 | 3.300E-07 |
| 28896 | 1445 | F1445S | E | 2.7e-07 | 2.1e-07 | 1.9e-07 | 2.233E-07 | 4.163E-08 | 1.900E-07 | 2.700E-07 |
| 28897 | 1445 | F1445T | E | 2.6e-07 | 1.8e-07 | 2.1e-07 | 2.167E-07 | 4.041E-08 | 1.800E-07 | 2.600E-07 |
| 28898 | 1445 | F1445W | E | 1.2e-07 | 2.4e-07 | 3.2e-07 | 2.267E-07 | 1.007E-07 | 1.200E-07 | 3.200E-07 |
| 28899 | 1445 | F1445Y | E | 4.3e-07 | 8.5e-07 | 2e-07 | 4.933E-07 | 3.296E-07 | 2.000E-07 | 8.500E-07 |

| | | | | | | | | | | |
|---|---|---|---|---|---|---|---|---|---|---|
| 28900 | 1445 | F1445V | E | 2.9e-07 | 5.1e-07 | 2.8e-07 | 3.600E-07 | 1.300E-07 | 2.800E-07 | 5.100E-07 |
| 28901 | 1446 | P1446A | E | 3.1e-07 | 1.3e-07 | 2.5e-07 | 2.300E-07 | 9.165E-08 | 1.300E-07 | 3.100E-07 |
| 28902 | 1446 | P1446R | E | 2.9e-07 | 1.4e-07 | 3.6e-07 | 2.633E-07 | 1.124E-07 | 1.400E-07 | 3.600E-07 |
| 28903 | 1446 | P1446N | E | 3.2e-07 | 2e-07 | 1.8e-07 | 2.333E-07 | 7.572E-08 | 1.800E-07 | 3.200E-07 |
| 28904 | 1446 | P1446D | E | 2.8e-07 | 2e-07 | 3e-07 | 2.600E-07 | 5.292E-08 | 2.000E-07 | 3.000E-07 |
| 28905 | 1446 | P1446C | E | 2.8e-07 | 2.2e-07 | 1.4e-07 | 2.133E-07 | 7.024E-08 | 1.400E-07 | 2.800E-07 |
| 28906 | 1446 | P1446E | E | 2.3e-07 | 4.1e-07 | 1.6e-07 | 2.667E-07 | 1.290E-07 | 1.600E-07 | 4.100E-07 |
| 28907 | 1446 | P1446Q | E | 1.6e-07 | 2e-07 | 1.5e-07 | 1.700E-07 | 2.646E-08 | 1.500E-07 | 2.000E-07 |
| 28908 | 1446 | P1446G | E | 3e-07 | 3.3e-07 | 4.8e-07 | 3.700E-07 | 9.644E-08 | 3.000E-07 | 4.800E-07 |
| 28909 | 1446 | P1446H | E | 9.6e-08 | 3.2e-07 | 2.7e-07 | 2.287E-07 | 1.176E-07 | 9.600E-08 | 3.200E-07 |
| 28910 | 1446 | P1446I | E | 2.2e-07 | 3.2e-07 | 2.4e-07 | 2.600E-07 | 5.292E-08 | 2.200E-07 | 3.200E-07 |
| 28911 | 1446 | P1446L | E | 1.5e-07 | 3.2e-07 | 3.3e-07 | 2.667E-07 | 1.012E-07 | 1.500E-07 | 3.300E-07 |
| 28912 | 1446 | P1446K | E | 1.4e-07 | 2.3e-07 | 3.3e-07 | 2.333E-07 | 9.504E-08 | 1.400E-07 | 3.300E-07 |
| 28913 | 1446 | P1446M | E | 3.1e-07 | 2.1e-07 | 2.3e-07 | 2.500E-07 | 5.292E-08 | 2.100E-07 | 3.100E-07 |
| 28914 | 1446 | P1446F | E | 5e-07 | 2.4e-07 | 2.4e-07 | 3.267E-07 | 1.501E-07 | 2.400E-07 | 5.000E-07 |
| 28915 | 1446 | P1446P | E | 1.5e-07 | 1.9e-07 | 3.4e-07 | 2.267E-07 | 1.002E-07 | 1.500E-07 | 3.400E-07 |
| 28916 | 1446 | P1446S | E | 2.5e-07 | 2.9e-07 | 1.4e-07 | 2.267E-07 | 7.767E-08 | 1.400E-07 | 2.900E-07 |
| 28917 | 1446 | P1446T | E | 1.2e-07 | 3.7e-07 | 1.9e-07 | 2.267E-07 | 1.290E-07 | 1.200E-07 | 3.700E-07 |
| 28918 | 1446 | P1446W | E | 1.6e-07 | 4.5e-07 | 1.2e-07 | 2.433E-07 | 1.801E-07 | 1.200E-07 | 4.500E-07 |
| 28919 | 1446 | P1446Y | E | 4.6e-07 | 1.7e-07 | 1.2e-07 | 2.500E-07 | 1.836E-07 | 1.200E-07 | 4.600E-07 |
| 28920 | 1446 | P1446V | E | 3.3e-07 | 2e-07 | 1.4e-07 | 2.233E-07 | 9.713E-08 | 1.400E-07 | 3.300E-07 |
| 28921 | 1447 | A1447A | E | 4e-07 | 3.5e-07 | 2.2e-07 | 3.233E-07 | 9.292E-08 | 2.200E-07 | 4.000E-07 |
| 28922 | 1447 | A1447R | E | 2e-07 | 2.7e-07 | 2.3e-07 | 2.333E-07 | 3.512E-08 | 2.000E-07 | 2.700E-07 |
| 28923 | 1447 | A1447N | E | 2.1e-07 | 1.8e-07 | 3e-07 | 2.300E-07 | 6.245E-08 | 1.800E-07 | 3.000E-07 |
| 28924 | 1447 | A1447D | E | 2.4e-07 | 1.7e-07 | 5.6e-07 | 3.233E-07 | 2.079E-07 | 1.700E-07 | 5.600E-07 |
| 28925 | 1447 | A1447C | E | 3.4e-07 | 3e-07 | 3e-07 | 3.133E-07 | 2.309E-08 | 3.000E-07 | 3.400E-07 |
| 28926 | 1447 | A1447E | E | 3.1e-07 | 2.1e-07 | 1.4e-07 | 2.200E-07 | 8.544E-08 | 1.400E-07 | 3.100E-07 |
| 28927 | 1447 | A1447Q | E | 3.7e-07 | 8.3e-07 | 2e-07 | 4.667E-07 | 3.259E-07 | 2.000E-07 | 8.300E-07 |
| 28928 | 1447 | A1447G | E | 4.1e-07 | 3.5e-07 | 3.1e-07 | 3.567E-07 | 5.033E-08 | 3.100E-07 | 4.100E-07 |
| 28929 | 1447 | A1447H | E | 2e-07 | 2.5e-07 | 2.3e-07 | 2.267E-07 | 2.517E-08 | 2.000E-07 | 2.500E-07 |
| 28930 | 1447 | A1447I | E | 3.5e-07 | 1.6e-07 | 3.3e-07 | 2.800E-07 | 1.044E-07 | 1.600E-07 | 3.500E-07 |
| 28931 | 1447 | A1447L | E | 5e-07 | 3.1e-07 | 4.6e-07 | 4.233E-07 | 1.002E-07 | 3.100E-07 | 5.000E-07 |
| 28932 | 1447 | A1447K | E | 2.3e-07 | 2.2e-07 | 2.2e-07 | 2.233E-07 | 5.774E-09 | 2.200E-07 | 2.300E-07 |
| 28933 | 1447 | A1447M | E | 2e-07 | 2.9e-07 | 3.8e-07 | 2.900E-07 | 9.000E-08 | 2.000E-07 | 3.800E-07 |
| 28934 | 1447 | A1447F | E | 1.2e-07 | 2.4e-07 | 1.9e-07 | 1.833E-07 | 6.028E-08 | 1.200E-07 | 2.400E-07 |
| 28935 | 1447 | A1447P | E | 1.7e-07 | 2.5e-07 | 2.4e-07 | 2.200E-07 | 4.359E-08 | 1.700E-07 | 2.500E-07 |
| 28936 | 1447 | A1447S | E | 1.9e-07 | 4.1e-07 | 3.9e-07 | 3.300E-07 | 1.217E-07 | 1.900E-07 | 4.100E-07 |
| 28937 | 1447 | A1447T | E | 5.4e-07 | 5.5e-07 | 2.2e-07 | 4.367E-07 | 1.877E-07 | 2.200E-07 | 5.500E-07 |
| 28938 | 1447 | A1447W | E | 2.2e-07 | 4.1e-07 | 3.4e-07 | 3.233E-07 | 9.609E-08 | 2.200E-07 | 4.100E-07 |
| 28939 | 1447 | A1447Y | E | 8.2e-07 | 3.5e-07 | 1.3e-07 | 4.333E-07 | 3.525E-07 | 1.300E-07 | 8.200E-07 |
| 28940 | 1447 | A1447V | E | 2.6e-07 | 2e-07 | 2.9e-07 | 2.500E-07 | 4.583E-08 | 2.000E-07 | 2.900E-07 |
| 28941 | 1448 | V1448A | E | 2.1e-07 | 2.7e-07 | 2.7e-07 | 2.500E-07 | 3.464E-08 | 2.100E-07 | 2.700E-07 |
| 28942 | 1448 | V1448R | E | 3.6e-07 | 2.1e-07 | 1.5e-07 | 2.400E-07 | 1.082E-07 | 1.500E-07 | 3.600E-07 |
| 28943 | 1448 | V1448N | E | 2.1e-07 | 2.5e-07 | 2.4e-07 | 2.333E-07 | 2.082E-08 | 2.100E-07 | 2.500E-07 |
| 28944 | 1448 | V1448D | E | 1.9e-07 | 8.9e-07 | 1.7e-07 | 4.167E-07 | 4.100E-07 | 1.700E-07 | 8.900E-07 |
| 28945 | 1448 | V1448C | E | 2.8e-07 | 3.2e-07 | 5.4e-07 | 3.800E-07 | 1.400E-07 | 2.800E-07 | 5.400E-07 |
| 28946 | 1448 | V1448E | E | 3.4e-07 | 1.8e-07 | 4.3e-07 | 3.167E-07 | 1.266E-07 | 1.800E-07 | 4.300E-07 |
| 28947 | 1448 | V1448Q | E | 1.4e-07 | 2.5e-07 | 5.5e-07 | 3.133E-07 | 2.122E-07 | 1.400E-07 | 5.500E-07 |
| 28948 | 1448 | V1448G | E | 1.6e-07 | 4.2e-07 | 1.5e-07 | 2.433E-07 | 1.531E-07 | 1.500E-07 | 4.200E-07 |
| 28949 | 1448 | V1448H | E | 3.3e-07 | 1.7e-07 | 2.5e-07 | 2.500E-07 | 8.000E-08 | 1.700E-07 | 3.300E-07 |
| 28950 | 1448 | V1448I | E | 1.9e-07 | 2.5e-07 | 1.4e-07 | 1.933E-07 | 5.508E-08 | 1.400E-07 | 2.500E-07 |
| 28951 | 1448 | V1448L | E | 1e-07 | 1.7e-07 | 2.1e-07 | 1.600E-07 | 5.568E-08 | 1.000E-07 | 2.100E-07 |
| 28952 | 1448 | V1448K | E | 2.1e-07 | 4.7e-07 | 1.1e-07 | 2.633E-07 | 1.858E-07 | 1.100E-07 | 4.700E-07 |
| 28953 | 1448 | V1448M | E | 2.2e-07 | 4.4e-07 | 2.9e-07 | 3.167E-07 | 1.124E-07 | 2.200E-07 | 4.400E-07 |
| 28954 | 1448 | V1448F | E | 3.9e-07 | 1.9e-07 | 1.9e-07 | 2.567E-07 | 1.155E-07 | 1.900E-07 | 3.900E-07 |
| 28955 | 1448 | V1448P | E | 2.1e-07 | 4.1e-07 | 1.8e-07 | 2.667E-07 | 1.250E-07 | 1.800E-07 | 4.100E-07 |
| 28956 | 1448 | V1448S | E | 1.3e-07 | 2.7e-07 | 3.1e-07 | 2.367E-07 | 9.452E-08 | 1.300E-07 | 3.100E-07 |
| 28957 | 1448 | V1448T | E | 3.5e-07 | 2.3e-07 | 3.2e-07 | 3.000E-07 | 6.245E-08 | 2.300E-07 | 3.500E-07 |
| 28958 | 1448 | V1448W | E | 3.8e-07 | 1.7e-07 | 2.2e-07 | 2.567E-07 | 1.097E-07 | 1.700E-07 | 3.800E-07 |
| 28959 | 1448 | V1448Y | E | 3.2e-07 | 1.8e-07 | 2.1e-07 | 2.367E-07 | 7.371E-08 | 1.800E-07 | 3.200E-07 |
| 28960 | 1448 | V1448V | E | 1.2e-07 | 3.4e-07 | 1.8e-07 | 2.133E-07 | 1.137E-07 | 1.200E-07 | 3.400E-07 |
| 28961 | 1449 | L1449A | E | 1.4e-07 | 1.8e-07 | 3.6e-07 | 2.267E-07 | 1.172E-07 | 1.400E-07 | 3.600E-07 |
| 28962 | 1449 | L1449R | E | 2.4e-07 | 2.4e-07 | 2.5e-07 | 2.433E-07 | 5.774E-09 | 2.400E-07 | 2.500E-07 |
| 28963 | 1449 | L1449N | E | 5.2e-07 | 1.9e-07 | 1.8e-07 | 2.967E-07 | 1.935E-07 | 1.800E-07 | 5.200E-07 |
| 28964 | 1449 | L1449D | E | 1.8e-07 | 2.2e-07 | 1.6e-07 | 1.867E-07 | 3.055E-08 | 1.600E-07 | 2.200E-07 |
| 28965 | 1449 | L1449C | E | 4.4e-07 | 2e-07 | 1.6e-07 | 2.667E-07 | 1.514E-07 | 1.600E-07 | 4.400E-07 |
| 28966 | 1449 | L1449E | E | 1.7e-07 | 2.8e-07 | 3.4e-07 | 2.633E-07 | 8.622E-08 | 1.700E-07 | 3.400E-07 |
| 28967 | 1449 | L1449Q | E | 3.2e-07 | 1.3e-07 | 2.1e-07 | 2.200E-07 | 9.539E-08 | 1.300E-07 | 3.200E-07 |
| 28968 | 1449 | L1449G | E | 3.6e-07 | 2.4e-07 | 2.7e-07 | 2.900E-07 | 6.245E-08 | 2.400E-07 | 3.600E-07 |
| 28969 | 1449 | L1449H | E | 2.2e-07 | 1.8e-07 | 3.7e-07 | 2.567E-07 | 1.002E-07 | 1.800E-07 | 3.700E-07 |
| 28970 | 1449 | L1449I | E | 3.2e-07 | 2.8e-07 | 2.6e-07 | 2.867E-07 | 3.055E-08 | 2.600E-07 | 3.200E-07 |
| 28971 | 1449 | L1449L | E | 1.8e-07 | 3.4e-07 | 4.6e-07 | 3.267E-07 | 1.405E-07 | 1.800E-07 | 4.600E-07 |
| 28972 | 1449 | L1449K | E | 3.3e-07 | 1.9e-07 | 2.9e-07 | 2.700E-07 | 7.211E-08 | 1.900E-07 | 3.300E-07 |
| 28973 | 1449 | L1449M | E | 6.2e-07 | 1.9e-07 | 2e-07 | 3.367E-07 | 2.454E-07 | 1.900E-07 | 6.200E-07 |
| 28974 | 1449 | L1449F | E | 2.4e-07 | 1.4e-07 | 2.8e-07 | 2.200E-07 | 7.211E-08 | 1.400E-07 | 2.800E-07 |
| 28975 | 1449 | L1449P | E | 1.8e-07 | 2.8e-07 | 3.7e-07 | 2.767E-07 | 9.504E-08 | 1.800E-07 | 3.700E-07 |
| 28976 | 1449 | L1449S | E | 2.4e-07 | 5.7e-07 | 2.6e-07 | 3.567E-07 | 1.850E-07 | 2.400E-07 | 5.700E-07 |
| 28977 | 1449 | L1449T | E | 2.7e-07 | 1.9e-07 | 1.7e-07 | 2.100E-07 | 5.292E-08 | 1.700E-07 | 2.700E-07 |
| 28978 | 1449 | L1449W | E | 2.9e-07 | 3.3e-07 | 2.2e-07 | 2.800E-07 | 5.568E-08 | 2.200E-07 | 3.300E-07 |
| 28979 | 1449 | L1449Y | E | 2.3e-07 | 2.4e-07 | 1.9e-07 | 2.200E-07 | 2.646E-08 | 1.900E-07 | 2.400E-07 |
| 28980 | 1449 | L1449V | E | 2.6e-07 | 2.5e-07 | 1.8e-07 | 2.300E-07 | 4.359E-08 | 1.800E-07 | 2.600E-07 |
| 28981 | 1450 | Q1450A | E | 7.5e-07 | 2e-07 | 2.9e-07 | 4.133E-07 | 2.950E-07 | 2.000E-07 | 7.500E-07 |
| 28982 | 1450 | Q1450R | E | 1.4e-07 | 4.6e-07 | 2e-07 | 2.667E-07 | 1.701E-07 | 1.400E-07 | 4.600E-07 |
| 28983 | 1450 | Q1450N | E | 1.8e-07 | 1.8e-07 | 4.2e-07 | 2.600E-07 | 1.386E-07 | 1.800E-07 | 4.200E-07 |
| 28984 | 1450 | Q1450D | E | 3.6e-07 | 2.9e-07 | 1.3e-07 | 2.600E-07 | 1.179E-07 | 1.300E-07 | 3.600E-07 |

| | | | | | | | | | | |
|---|---|---|---|---|---|---|---|---|---|---|
| 28985 | 1450 | Q1450C | E | 1.6e-07 | 2.1e-07 | 2.4e-07 | 2.033E-07 | 4.041E-08 | 1.600E-07 | 2.400E-07 |
| 28986 | 1450 | Q1450E | E | 1.3e-07 | 1.1e-07 | 2.5e-07 | 1.633E-07 | 7.572E-08 | 1.100E-07 | 2.500E-07 |
| 28987 | 1450 | Q1450Q | E | 3.3e-07 | 3.8e-07 | 3.7e-07 | 3.600E-07 | 2.646E-08 | 3.300E-07 | 3.800E-07 |
| 28988 | 1450 | Q1450G | E | 3.1e-07 | 3.5e-07 | 2.1e-07 | 2.900E-07 | 7.211E-08 | 2.100E-07 | 3.500E-07 |
| 28989 | 1450 | Q1450H | E | 2e-07 | 3.1e-07 | 1.9e-07 | 2.333E-07 | 6.658E-08 | 1.900E-07 | 3.100E-07 |
| 28990 | 1450 | Q1450I | E | 2.6e-07 | 3.2e-07 | 2.8e-07 | 2.867E-07 | 3.055E-08 | 2.600E-07 | 3.200E-07 |
| 28991 | 1450 | Q1450L | E | 3.3e-07 | 2.8e-07 | 1.5e-07 | 2.533E-07 | 9.292E-08 | 1.500E-07 | 3.300E-07 |
| 28992 | 1450 | Q1450K | E | 2e-07 | 1.6e-07 | 5.5e-07 | 3.033E-07 | 2.146E-07 | 1.600E-07 | 5.500E-07 |
| 28993 | 1450 | Q1450M | E | 3.4e-07 | 2.5e-07 | 1.7e-07 | 2.533E-07 | 8.505E-08 | 1.700E-07 | 3.400E-07 |
| 28994 | 1450 | Q1450F | E | 5.2e-07 | 3.2e-07 | 4.6e-07 | 4.333E-07 | 1.026E-07 | 3.200E-07 | 5.200E-07 |
| 28995 | 1450 | Q1450P | E | 7.3e-07 | 2.6e-07 | 1.7e-07 | 3.867E-07 | 3.007E-07 | 1.700E-07 | 7.300E-07 |
| 28996 | 1450 | Q1450S | E | 2e-07 | 3.1e-07 | 2.1e-07 | 2.400E-07 | 6.083E-08 | 2.000E-07 | 3.100E-07 |
| 28997 | 1450 | Q1450T | E | 2.1e-07 | 1.2e-07 | 1.5e-07 | 1.600E-07 | 4.583E-08 | 1.200E-07 | 2.100E-07 |
| 28998 | 1450 | Q1450W | E | 2e-07 | 1.3e-07 | 3.7e-07 | 2.333E-07 | 1.234E-07 | 1.300E-07 | 3.700E-07 |
| 28999 | 1450 | Q1450Y | E | 3.1e-07 | 2.7e-07 | 2.4e-07 | 2.733E-07 | 3.512E-08 | 2.400E-07 | 3.100E-07 |
| 29000 | 1450 | Q1450V | E | 2.6e-07 | 2.9e-07 | 2.8e-07 | 2.767E-07 | 1.528E-08 | 2.600E-07 | 2.900E-07 |
| 29001 | 1451 | S1451A | E | 1.8e-07 | 8.3e-07 | 1.4e-07 | 3.833E-07 | 3.873E-07 | 1.400E-07 | 8.300E-07 |
| 29002 | 1451 | S1451R | E | 1.8e-07 | 2.8e-07 | 2.8e-07 | 2.467E-07 | 5.774E-08 | 1.800E-07 | 2.800E-07 |
| 29003 | 1451 | S1451N | E | 3.4e-07 | 2.9e-07 | 1.3e-07 | 2.533E-07 | 1.097E-07 | 1.300E-07 | 3.400E-07 |
| 29004 | 1451 | S1451D | E | 3.2e-07 | 2.4e-07 | 2.1e-07 | 2.567E-07 | 5.686E-08 | 2.100E-07 | 3.200E-07 |
| 29005 | 1451 | S1451C | E | 2.5e-07 | 3e-07 | 5.4e-07 | 3.633E-07 | 1.550E-07 | 2.500E-07 | 5.400E-07 |
| 29006 | 1451 | S1451E | E | 4.2e-07 | 3.3e-07 | 3.3e-07 | 3.600E-07 | 5.196E-08 | 3.300E-07 | 4.200E-07 |
| 29007 | 1451 | S1451Q | E | 2.8e-07 | 3e-07 | 6e-07 | 3.933E-07 | 1.793E-07 | 2.800E-07 | 6.000E-07 |
| 29008 | 1451 | S1451G | E | 1.3e-07 | 3.5e-07 | 3.3e-07 | 2.700E-07 | 1.217E-07 | 1.300E-07 | 3.500E-07 |
| 29009 | 1451 | S1451H | E | 1.7e-07 | 1.9e-07 | 1.9e-07 | 1.833E-07 | 1.155E-08 | 1.700E-07 | 1.900E-07 |
| 29010 | 1451 | S1451I | E | 6.2e-07 | 2.5e-07 | 5.6e-07 | 4.767E-07 | 1.986E-07 | 2.500E-07 | 6.200E-07 |
| 29011 | 1451 | S1451L | E | 2.3e-07 | 4e-07 | 1.6e-07 | 2.633E-07 | 1.234E-07 | 1.600E-07 | 4.000E-07 |
| 29012 | 1451 | S1451K | E | 1.5e-07 | 1.5e-07 | 1.6e-07 | 1.533E-07 | 5.774E-09 | 1.500E-07 | 1.600E-07 |
| 29013 | 1451 | S1451M | E | 1.6e-07 | 1.7e-07 | 3.1e-07 | 2.133E-07 | 8.386E-08 | 1.600E-07 | 3.100E-07 |
| 29014 | 1451 | S1451F | E | 2e-07 | 4.2e-07 | 1.6e-07 | 2.600E-07 | 1.400E-07 | 1.600E-07 | 4.200E-07 |
| 29015 | 1451 | S1451P | E | 2e-07 | 4.2e-07 | 1.5e-07 | 2.567E-07 | 1.436E-07 | 1.500E-07 | 4.200E-07 |
| 29016 | 1451 | S1451S | E | 6.1e-07 | 4e-07 | 1.6e-07 | 3.900E-07 | 2.252E-07 | 1.600E-07 | 6.100E-07 |
| 29017 | 1451 | S1451T | E | 1.4e-07 | 2.7e-07 | 2.5e-07 | 2.200E-07 | 7.000E-08 | 1.400E-07 | 2.700E-07 |
| 29018 | 1451 | S1451W | E | 4.4e-07 | 1.3e-07 | 3.8e-07 | 3.167E-07 | 1.644E-07 | 1.300E-07 | 4.400E-07 |
| 29019 | 1451 | S1451Y | E | 2.2e-07 | 2.3e-07 | 4.3e-07 | 2.933E-07 | 1.185E-07 | 2.200E-07 | 4.300E-07 |
| 29020 | 1451 | S1451V | E | 3.8e-07 | 1.6e-07 | 5.9e-07 | 3.767E-07 | 2.150E-07 | 1.600E-07 | 5.900E-07 |
| 29021 | 1452 | S1452A | E | 5.2e-07 | 2.2e-07 | 3e-07 | 3.467E-07 | 1.553E-07 | 2.200E-07 | 5.200E-07 |
| 29022 | 1452 | S1452R | E | 2e-07 | 4.3e-07 | 2.2e-07 | 2.833E-07 | 1.274E-07 | 2.000E-07 | 4.300E-07 |
| 29023 | 1452 | S1452N | E | 2.7e-07 | 2.6e-07 | 2.9e-07 | 2.733E-07 | 1.528E-08 | 2.600E-07 | 2.900E-07 |
| 29024 | 1452 | S1452D | E | 2e-07 | 3.3e-07 | 2.2e-07 | 2.500E-07 | 7.000E-08 | 2.000E-07 | 3.300E-07 |
| 29025 | 1452 | S1452C | E | 2.6e-07 | 3.4e-07 | 1.6e-07 | 2.533E-07 | 9.018E-08 | 1.600E-07 | 3.400E-07 |
| 29026 | 1452 | S1452E | E | 1.4e-07 | 3.1e-07 | 2.9e-07 | 2.467E-07 | 9.292E-08 | 1.400E-07 | 3.100E-07 |
| 29027 | 1452 | S1452Q | E | 2.5e-07 | 3.9e-07 | 2.2e-07 | 2.867E-07 | 9.074E-08 | 2.200E-07 | 3.900E-07 |
| 29028 | 1452 | S1452G | E | 3.2e-07 | 4.6e-07 | 2.7e-07 | 3.500E-07 | 9.849E-08 | 2.700E-07 | 4.600E-07 |
| 29029 | 1452 | S1452H | E | 1.6e-07 | 2.1e-07 | 3.5e-07 | 2.400E-07 | 9.849E-08 | 1.600E-07 | 3.500E-07 |
| 29030 | 1452 | S1452I | E | 2.8e-07 | 2.6e-07 | 2.9e-07 | 2.767E-07 | 1.528E-08 | 2.600E-07 | 2.900E-07 |
| 29031 | 1452 | S1452L | E | 2.2e-07 | 1.9e-07 | 1.4e-07 | 1.833E-07 | 4.041E-08 | 1.400E-07 | 2.200E-07 |
| 29032 | 1452 | S1452K | E | 2.9e-07 | 2.4e-07 | 2.7e-07 | 2.667E-07 | 2.517E-08 | 2.400E-07 | 2.900E-07 |
| 29033 | 1452 | S1452M | E | 3e-07 | 1.2e-07 | 2.2e-07 | 2.133E-07 | 9.018E-08 | 1.200E-07 | 3.000E-07 |
| 29034 | 1452 | S1452F | E | 3.6e-07 | 3.1e-07 | 1.9e-07 | 2.867E-07 | 8.737E-08 | 1.900E-07 | 3.600E-07 |
| 29035 | 1452 | S1452P | E | 1.6e-07 | 4.1e-07 | 2.8e-07 | 2.833E-07 | 1.250E-07 | 1.600E-07 | 4.100E-07 |
| 29036 | 1452 | S1452S | E | 1.2e-07 | 3.2e-07 | 1.4e-07 | 1.933E-07 | 1.102E-07 | 1.200E-07 | 3.200E-07 |
| 29037 | 1452 | S1452T | E | 2.2e-07 | 3.4e-07 | 1.8e-07 | 2.467E-07 | 8.327E-08 | 1.800E-07 | 3.400E-07 |
| 29038 | 1452 | S1452W | E | 2.5e-07 | 2.9e-07 | 2.6e-07 | 2.667E-07 | 2.082E-08 | 2.500E-07 | 2.900E-07 |
| 29039 | 1452 | S1452Y | E | 3.7e-07 | 3e-07 | 2e-07 | 2.900E-07 | 8.544E-08 | 2.000E-07 | 3.700E-07 |
| 29040 | 1452 | S1452V | E | 2.4e-07 | 1.7e-07 | 5.9e-07 | 3.333E-07 | 2.250E-07 | 1.700E-07 | 5.900E-07 |
| 29041 | 1453 | G1453A | E | 2.7e-07 | 3.2e-07 | 3.2e-07 | 3.033E-07 | 2.887E-08 | 2.700E-07 | 3.200E-07 |
| 29042 | 1453 | G1453R | E | 2.7e-07 | 2.8e-07 | 4e-07 | 3.167E-07 | 7.234E-08 | 2.700E-07 | 4.000E-07 |
| 29043 | 1453 | G1453N | E | 1.9e-07 | 2.2e-07 | 1.4e-07 | 1.833E-07 | 4.041E-08 | 1.400E-07 | 2.200E-07 |
| 29044 | 1453 | G1453D | E | 3.2e-07 | 1.2e-07 | 2.6e-07 | 2.333E-07 | 1.026E-07 | 1.200E-07 | 3.200E-07 |
| 29045 | 1453 | G1453C | E | 4.3e-07 | 4e-07 | 1.9e-07 | 3.400E-07 | 1.308E-07 | 1.900E-07 | 4.300E-07 |
| 29046 | 1453 | G1453E | E | 2.2e-07 | 2.9e-07 | 1.6e-07 | 2.233E-07 | 6.506E-08 | 1.600E-07 | 2.900E-07 |
| 29047 | 1453 | G1453Q | E | 3.2e-07 | 1.7e-07 | 4.9e-07 | 3.267E-07 | 1.601E-07 | 1.700E-07 | 4.900E-07 |
| 29048 | 1453 | G1453G | E | 1.8e-07 | 2.3e-07 | 1.9e-07 | 2.000E-07 | 2.646E-08 | 1.800E-07 | 2.300E-07 |
| 29049 | 1453 | G1453H | E | 3.6e-07 | 3.5e-07 | 8.9e-08 | 2.663E-07 | 1.537E-07 | 8.900E-08 | 3.600E-07 |
| 29050 | 1453 | G1453I | E | 2.5e-07 | 2e-07 | 7.9e-07 | 4.133E-07 | 3.272E-07 | 2.000E-07 | 7.900E-07 |
| 29051 | 1453 | G1453L | E | 2e-07 | 2.1e-07 | 4.7e-07 | 2.933E-07 | 1.531E-07 | 2.000E-07 | 4.700E-07 |
| 29052 | 1453 | G1453K | E | 3e-07 | 2.3e-07 | 4e-07 | 3.100E-07 | 8.544E-08 | 2.300E-07 | 4.000E-07 |
| 29053 | 1453 | G1453M | E | 3.8e-07 | 7.1e-07 | 1.2e-07 | 4.033E-07 | 2.957E-07 | 1.200E-07 | 7.100E-07 |
| 29054 | 1453 | G1453F | E | 3e-07 | 3.7e-07 | 3.9e-07 | 3.533E-07 | 4.726E-08 | 3.000E-07 | 3.900E-07 |
| 29055 | 1453 | G1453P | E | 2.8e-07 | 3e-07 | 1.5e-07 | 2.433E-07 | 8.145E-08 | 1.500E-07 | 3.000E-07 |
| 29056 | 1453 | G1453S | E | 1.1e-07 | 2.4e-07 | 4.4e-07 | 2.633E-07 | 1.662E-07 | 1.100E-07 | 4.400E-07 |
| 29057 | 1453 | G1453T | E | 1.4e-07 | 4.3e-07 | 1.9e-07 | 2.533E-07 | 1.550E-07 | 1.400E-07 | 4.300E-07 |
| 29058 | 1453 | G1453W | E | 3e-07 | 5.3e-07 | 2.1e-07 | 3.467E-07 | 1.650E-07 | 2.100E-07 | 5.300E-07 |
| 29059 | 1453 | G1453Y | E | 1.5e-07 | 2.4e-07 | 2.4e-07 | 2.100E-07 | 5.196E-08 | 1.500E-07 | 2.400E-07 |
| 29060 | 1453 | G1453V | E | 3.2e-07 | 1.9e-07 | 2.7e-07 | 2.600E-07 | 6.557E-08 | 1.900E-07 | 3.200E-07 |
| 29061 | 1454 | L1454A | E | 1.9e-07 | 1.8e-07 | 4e-07 | 2.567E-07 | 1.242E-07 | 1.800E-07 | 4.000E-07 |
| 29062 | 1454 | L1454R | E | 1.7e-07 | 3.1e-07 | 2.5e-07 | 2.433E-07 | 7.024E-08 | 1.700E-07 | 3.100E-07 |
| 29063 | 1454 | L1454N | E | 2.4e-07 | 1.9e-07 | 3.7e-07 | 2.667E-07 | 9.292E-08 | 1.900E-07 | 3.700E-07 |
| 29064 | 1454 | L1454D | E | 2.4e-07 | 4.5e-07 | 2.7e-07 | 3.200E-07 | 1.136E-07 | 2.400E-07 | 4.500E-07 |
| 29065 | 1454 | L1454C | E | 4.8e-07 | 2.9e-07 | 4.9e-07 | 4.200E-07 | 1.127E-07 | 2.900E-07 | 4.900E-07 |
| 29066 | 1454 | L1454E | E | 4.3e-07 | 2.5e-07 | 1.6e-07 | 2.800E-07 | 1.375E-07 | 1.600E-07 | 4.300E-07 |
| 29067 | 1454 | L1454Q | E | 3.6e-07 | 1.1e-07 | 3e-07 | 2.567E-07 | 1.305E-07 | 1.100E-07 | 3.600E-07 |
| 29068 | 1454 | L1454G | E | 2.5e-07 | 2.3e-07 | 7.7e-07 | 4.167E-07 | 3.062E-07 | 2.300E-07 | 7.700E-07 |
| 29069 | 1454 | L1454H | E | 2.3e-07 | 1.8e-07 | 1.3e-07 | 1.800E-07 | 5.000E-08 | 1.300E-07 | 2.300E-07 |

| | | | | | | | | | | |
|---|---|---|---|---|---|---|---|---|---|---|
| 29070 | 1454 | L1454I | E | 1.6e-07 | 1.6e-07 | 1.2e-07 | 1.467E-07 | 2.309E-08 | 1.200E-07 | 1.600E-07 |
| 29071 | 1454 | L1454L | E | 1.7e-07 | 2e-07 | 5.2e-07 | 2.967E-07 | 1.940E-07 | 1.700E-07 | 5.200E-07 |
| 29072 | 1454 | L1454K | E | 1.5e-07 | 2.3e-07 | 2.2e-07 | 2.000E-07 | 4.359E-08 | 1.500E-07 | 2.300E-07 |
| 29073 | 1454 | L1454M | E | 1.5e-07 | 7.6e-08 | 7.3e-07 | 3.187E-07 | 3.581E-07 | 7.600E-08 | 7.300E-07 |
| 29074 | 1454 | L1454F | E | 1.8e-07 | 2.2e-07 | 4.3e-07 | 2.767E-07 | 1.343E-07 | 1.800E-07 | 4.300E-07 |
| 29075 | 1454 | L1454P | E | 3.9e-07 | 2.5e-07 | 2.1e-07 | 2.833E-07 | 9.452E-08 | 2.100E-07 | 3.900E-07 |
| 29076 | 1454 | L1454S | E | 2.3e-07 | 3.6e-07 | 1.5e-07 | 2.467E-07 | 1.060E-07 | 1.500E-07 | 3.600E-07 |
| 29077 | 1454 | L1454T | E | 1.7e-07 | 1.9e-07 | 1.7e-07 | 1.767E-07 | 1.155E-08 | 1.700E-07 | 1.900E-07 |
| 29078 | 1454 | L1454W | E | 4.2e-07 | 1.4e-07 | 1.5e-07 | 2.367E-07 | 1.589E-07 | 1.400E-07 | 4.200E-07 |
| 29079 | 1454 | L1454Y | E | 2.8e-07 | 2.8e-07 | 2.7e-07 | 2.767E-07 | 5.774E-09 | 2.700E-07 | 2.800E-07 |
| 29080 | 1454 | L1454V | E | 1.6e-07 | 2.3e-07 | 4.1e-07 | 2.667E-07 | 1.290E-07 | 1.600E-07 | 4.100E-07 |
| 29081 | 1455 | Y1455A | E | 9.3e-08 | 2.6e-07 | 2.7e-07 | 2.077E-07 | 9.943E-08 | 9.300E-08 | 2.700E-07 |
| 29082 | 1455 | Y1455R | E | 1.5e-07 | 4.1e-07 | 3.3e-07 | 2.967E-07 | 1.332E-07 | 1.500E-07 | 4.100E-07 |
| 29083 | 1455 | Y1455N | E | 3e-07 | 4.1e-07 | 1.2e-07 | 2.767E-07 | 1.464E-07 | 1.200E-07 | 4.100E-07 |
| 29084 | 1455 | Y1455D | E | 3.5e-07 | 4e-07 | 3e-07 | 3.500E-07 | 5.000E-08 | 3.000E-07 | 4.000E-07 |
| 29085 | 1455 | Y1455C | E | 1.6e-07 | 1.9e-07 | 2.2e-07 | 1.900E-07 | 3.000E-08 | 1.600E-07 | 2.200E-07 |
| 29086 | 1455 | Y1455E | E | 2.5e-07 | 1.3e-07 | 3.2e-07 | 2.333E-07 | 9.609E-08 | 1.300E-07 | 3.200E-07 |
| 29087 | 1455 | Y1455Q | E | 3.7e-07 | 1.8e-07 | 1.9e-07 | 2.467E-07 | 1.069E-07 | 1.800E-07 | 3.700E-07 |
| 29088 | 1455 | Y1455G | E | 1.1e-07 | 2.4e-07 | 2.1e-07 | 1.867E-07 | 6.807E-08 | 1.100E-07 | 2.400E-07 |
| 29089 | 1455 | Y1455H | E | 3.1e-07 | 1.8e-07 | 2.5e-07 | 2.467E-07 | 6.506E-08 | 1.800E-07 | 3.100E-07 |
| 29090 | 1455 | Y1455I | E | 4e-07 | 2.3e-07 | 3.4e-07 | 3.233E-07 | 8.622E-08 | 2.300E-07 | 4.000E-07 |
| 29091 | 1455 | Y1455L | E | 3e-07 | 2e-07 | 2.5e-07 | 2.500E-07 | 5.000E-08 | 2.000E-07 | 3.000E-07 |
| 29092 | 1455 | Y1455K | E | 1.9e-07 | 2.6e-07 | 1.9e-07 | 2.133E-07 | 4.041E-08 | 1.900E-07 | 2.600E-07 |
| 29093 | 1455 | Y1455M | E | 2.4e-07 | 3.2e-07 | 2.9e-07 | 2.833E-07 | 4.041E-08 | 2.400E-07 | 3.200E-07 |
| 29094 | 1455 | Y1455F | E | 9e-08 | 3.4e-07 | 2e-07 | 2.100E-07 | 1.253E-07 | 9.000E-08 | 3.400E-07 |
| 29095 | 1455 | Y1455P | E | 1.7e-07 | 8.8e-08 | 1.4e-07 | 1.327E-07 | 4.149E-08 | 8.800E-08 | 1.700E-07 |
| 29096 | 1455 | Y1455S | E | 3.3e-07 | 4.8e-07 | 2.3e-07 | 3.467E-07 | 1.258E-07 | 2.300E-07 | 4.800E-07 |
| 29097 | 1455 | Y1455T | E | 4.4e-07 | 2.1e-07 | 2.1e-07 | 2.867E-07 | 1.328E-07 | 2.100E-07 | 4.400E-07 |
| 29098 | 1455 | Y1455W | E | 1.5e-07 | 3.7e-07 | 1.9e-07 | 2.367E-07 | 1.172E-07 | 1.500E-07 | 3.700E-07 |
| 29099 | 1455 | Y1455Y | E | 1.9e-07 | 1.9e-07 | 3.4e-07 | 2.400E-07 | 8.660E-08 | 1.900E-07 | 3.400E-07 |
| 29100 | 1455 | Y1455V | E | 6e-07 | 4.2e-07 | 1.4e-07 | 3.867E-07 | 2.318E-07 | 1.400E-07 | 6.000E-07 |
| 29101 | 1456 | S1456A | E | 3e-07 | 5.5e-07 | 1.9e-07 | 3.467E-07 | 1.845E-07 | 1.900E-07 | 5.500E-07 |
| 29102 | 1456 | S1456R | E | 4.3e-07 | 1.5e-07 | 2.7e-07 | 2.833E-07 | 1.405E-07 | 1.500E-07 | 4.300E-07 |
| 29103 | 1456 | S1456N | E | 2.1e-07 | 2.6e-07 | 2.2e-07 | 2.300E-07 | 2.646E-08 | 2.100E-07 | 2.600E-07 |
| 29104 | 1456 | S1456D | E | 1.1e-07 | 1.3e-07 | 3.6e-07 | 2.000E-07 | 1.389E-07 | 1.100E-07 | 3.600E-07 |
| 29105 | 1456 | S1456C | E | 2e-07 | 3.4e-07 | 2.1e-07 | 2.500E-07 | 7.810E-08 | 2.000E-07 | 3.400E-07 |
| 29106 | 1456 | S1456E | E | 2.5e-07 | 3.2e-07 | 3.7e-07 | 3.133E-07 | 6.028E-08 | 2.500E-07 | 3.700E-07 |
| 29107 | 1456 | S1456Q | E | 2.4e-07 | 3.4e-07 | 3.5e-07 | 3.100E-07 | 6.083E-08 | 2.400E-07 | 3.500E-07 |
| 29108 | 1456 | S1456G | E | 2e-07 | 1.6e-07 | 3.4e-07 | 2.333E-07 | 9.452E-08 | 1.600E-07 | 3.400E-07 |
| 29109 | 1456 | S1456H | E | 1.3e-07 | 3.6e-07 | 2.5e-07 | 2.467E-07 | 1.150E-07 | 1.300E-07 | 3.600E-07 |
| 29110 | 1456 | S1456I | E | 1.7e-07 | 3.3e-07 | 2.3e-07 | 2.433E-07 | 8.083E-08 | 1.700E-07 | 3.300E-07 |
| 29111 | 1456 | S1456L | E | 2.5e-07 | 4.8e-07 | 2.4e-07 | 3.233E-07 | 1.358E-07 | 2.400E-07 | 4.800E-07 |
| 29112 | 1456 | S1456K | E | 3e-07 | 1.9e-07 | 2.7e-07 | 2.533E-07 | 5.686E-08 | 1.900E-07 | 3.000E-07 |
| 29113 | 1456 | S1456M | E | 4.2e-07 | 1.9e-07 | 2.4e-07 | 2.833E-07 | 1.210E-07 | 1.900E-07 | 4.200E-07 |
| 29114 | 1456 | S1456F | E | 3.1e-07 | 6.2e-08 | 1.5e-07 | 1.740E-07 | 1.257E-07 | 6.200E-08 | 3.100E-07 |
| 29115 | 1456 | S1456P | E | 2.7e-07 | 5.1e-07 | 3.5e-07 | 3.767E-07 | 1.222E-07 | 2.700E-07 | 5.100E-07 |
| 29116 | 1456 | S1456S | E | 4e-07 | 3e-07 | 2.3e-07 | 3.100E-07 | 8.544E-08 | 2.300E-07 | 4.000E-07 |
| 29117 | 1456 | S1456T | E | 4.2e-07 | 3.2e-07 | 2.2e-07 | 3.200E-07 | 1.000E-07 | 2.200E-07 | 4.200E-07 |
| 29118 | 1456 | S1456W | E | 4.3e-07 | 1.9e-07 | 2.4e-07 | 2.867E-07 | 1.266E-07 | 1.900E-07 | 4.300E-07 |
| 29119 | 1456 | S1456Y | E | 1.8e-07 | 1.9e-07 | 2.7e-07 | 2.133E-07 | 4.933E-08 | 1.800E-07 | 2.700E-07 |
| 29120 | 1456 | S1456V | E | 3e-07 | 3.9e-07 | 3.1e-07 | 3.333E-07 | 4.933E-08 | 3.000E-07 | 3.900E-07 |
| 29121 | 1457 | L1457A | E | 2.9e-07 | 2.3e-07 | 5.3e-07 | 3.500E-07 | 1.587E-07 | 2.300E-07 | 5.300E-07 |
| 29122 | 1457 | L1457R | E | 2.3e-07 | 4.2e-07 | 2.1e-07 | 2.867E-07 | 1.159E-07 | 2.100E-07 | 4.200E-07 |
| 29123 | 1457 | L1457N | E | 1.7e-07 | 4.2e-07 | 3e-07 | 2.967E-07 | 1.250E-07 | 1.700E-07 | 4.200E-07 |
| 29124 | 1457 | L1457D | E | 1.4e-07 | 2.3e-07 | 2.9e-07 | 2.200E-07 | 7.550E-08 | 1.400E-07 | 2.900E-07 |
| 29125 | 1457 | L1457C | E | 1.9e-07 | 3.3e-07 | 1.9e-07 | 2.367E-07 | 8.083E-08 | 1.900E-07 | 3.300E-07 |
| 29126 | 1457 | L1457E | E | 8e-08 | 3.1e-07 | 2.1e-07 | 2.000E-07 | 1.153E-07 | 8.000E-08 | 3.100E-07 |
| 29127 | 1457 | L1457Q | E | 1.9e-07 | 2.8e-07 | 2.4e-07 | 2.367E-07 | 4.509E-08 | 1.900E-07 | 2.800E-07 |
| 29128 | 1457 | L1457G | E | 2.3e-07 | 1.9e-07 | 2.5e-07 | 2.233E-07 | 3.055E-08 | 1.900E-07 | 2.500E-07 |
| 29129 | 1457 | L1457H | E | 9.2e-08 | 2.8e-07 | 2.3e-07 | 2.007E-07 | 9.737E-08 | 9.200E-08 | 2.800E-07 |
| 29130 | 1457 | L1457I | E | 3.6e-07 | 2.3e-07 | 2e-07 | 2.633E-07 | 8.505E-08 | 2.000E-07 | 3.600E-07 |
| 29131 | 1457 | L1457L | E | 4.5e-07 | 2.4e-07 | 3.7e-07 | 3.533E-07 | 1.060E-07 | 2.400E-07 | 4.500E-07 |
| 29132 | 1457 | L1457K | E | 3.8e-07 | 3.1e-07 | 2.3e-07 | 3.067E-07 | 7.506E-08 | 2.300E-07 | 3.800E-07 |
| 29133 | 1457 | L1457M | E | 5.7e-07 | 9.1e-08 | 2.2e-07 | 2.937E-07 | 2.479E-07 | 9.100E-08 | 5.700E-07 |
| 29134 | 1457 | L1457F | E | 2.4e-07 | 2.8e-07 | 2.7e-07 | 2.633E-07 | 2.082E-08 | 2.400E-07 | 2.800E-07 |
| 29135 | 1457 | L1457P | E | 1.9e-07 | 2.3e-07 | 2.7e-07 | 2.300E-07 | 4.000E-08 | 1.900E-07 | 2.700E-07 |
| 29136 | 1457 | L1457S | E | 2.6e-07 | 4.1e-07 | 4.5e-07 | 3.733E-07 | 1.002E-07 | 2.600E-07 | 4.500E-07 |
| 29137 | 1457 | L1457T | E | 3.1e-07 | 2.4e-07 | 2.7e-07 | 2.733E-07 | 3.512E-08 | 2.400E-07 | 3.100E-07 |
| 29138 | 1457 | L1457W | E | 3.6e-07 | 1.6e-07 | 4.1e-07 | 3.100E-07 | 1.323E-07 | 1.600E-07 | 4.100E-07 |
| 29139 | 1457 | L1457Y | E | 1.1e-07 | 1.8e-07 | 1.7e-07 | 1.533E-07 | 3.786E-08 | 1.100E-07 | 1.800E-07 |
| 29140 | 1457 | L1457V | E | 3.9e-07 | 4.5e-07 | 5.4e-07 | 4.600E-07 | 7.550E-08 | 3.900E-07 | 5.400E-07 |
| 29141 | 1458 | S1458A | E | 2.2e-07 | 2e-07 | 1.4e-07 | 1.867E-07 | 4.163E-08 | 1.400E-07 | 2.200E-07 |
| 29142 | 1458 | S1458R | E | 2.4e-07 | 5.6e-07 | 4.3e-07 | 4.100E-07 | 1.609E-07 | 2.400E-07 | 5.600E-07 |
| 29143 | 1458 | S1458N | E | 3.9e-07 | 1.3e-07 | 2.8e-07 | 2.667E-07 | 1.305E-07 | 1.300E-07 | 3.900E-07 |
| 29144 | 1458 | S1458D | E | 2.2e-07 | 2.8e-07 | 1.5e-07 | 2.167E-07 | 6.506E-08 | 1.500E-07 | 2.800E-07 |
| 29145 | 1458 | S1458C | E | 1.1e-07 | 2.5e-07 | 1.8e-07 | 1.800E-07 | 7.000E-08 | 1.100E-07 | 2.500E-07 |
| 29146 | 1458 | S1458E | E | 3.2e-07 | 3e-07 | 3.8e-07 | 3.333E-07 | 4.163E-08 | 3.000E-07 | 3.800E-07 |
| 29147 | 1458 | S1458Q | E | 1.1e-07 | 5.7e-07 | 1.2e-07 | 2.667E-07 | 2.627E-07 | 1.100E-07 | 5.700E-07 |
| 29148 | 1458 | S1458G | E | 4.1e-07 | 2.4e-07 | 3.5e-07 | 3.333E-07 | 8.622E-08 | 2.400E-07 | 4.100E-07 |
| 29149 | 1458 | S1458H | E | 2.1e-07 | 2.2e-07 | 2.6e-07 | 2.300E-07 | 2.646E-08 | 2.100E-07 | 2.600E-07 |
| 29150 | 1458 | S1458I | E | 2.3e-07 | 2.2e-07 | 3.2e-07 | 2.567E-07 | 5.508E-08 | 2.200E-07 | 3.200E-07 |
| 29151 | 1458 | S1458L | E | 5.7e-07 | 5.6e-07 | 5.8e-07 | 5.700E-07 | 1.000E-08 | 5.600E-07 | 5.800E-07 |
| 29152 | 1458 | S1458K | E | 4.3e-07 | 2.9e-07 | 4.2e-07 | 3.800E-07 | 7.810E-08 | 2.900E-07 | 4.300E-07 |
| 29153 | 1458 | S1458M | E | 2.5e-07 | 2.5e-07 | 2.2e-07 | 2.400E-07 | 1.732E-08 | 2.200E-07 | 2.500E-07 |
| 29154 | 1458 | S1458F | E | 2.5e-07 | 6.2e-07 | 3e-07 | 3.900E-07 | 2.007E-07 | 2.500E-07 | 6.200E-07 |

| | | | | | | | | | | |
|---|---|---|---|---|---|---|---|---|---|---|
| 29155 | 1458 | S1458P | E | 4.8e-07 | 3e-07 | 1.5e-07 | 3.100E-07 | 1.652E-07 | 1.500E-07 | 4.800E-07 |
| 29156 | 1458 | S1458S | E | 3e-07 | 8.7e-07 | 3.1e-07 | 4.933E-07 | 3.262E-07 | 3.000E-07 | 8.700E-07 |
| 29157 | 1458 | S1458T | E | 2.1e-07 | 5.3e-07 | 1.5e-07 | 2.967E-07 | 2.043E-07 | 1.500E-07 | 5.300E-07 |
| 29158 | 1458 | S1458W | E | 1.5e-07 | 1.9e-07 | 3.8e-07 | 2.400E-07 | 1.229E-07 | 1.500E-07 | 3.800E-07 |
| 29159 | 1458 | S1458Y | E | 3.9e-07 | 3.8e-07 | 3.3e-07 | 3.667E-07 | 3.215E-08 | 3.300E-07 | 3.900E-07 |
| 29160 | 1458 | S1458V | E | 1.4e-07 | 2.7e-07 | 2.9e-07 | 2.333E-07 | 8.145E-08 | 1.400E-07 | 2.900E-07 |
| 29161 | 1459 | S1459A | E | 4.4e-07 | 2.5e-07 | 1.7e-07 | 2.867E-07 | 1.387E-07 | 1.700E-07 | 4.400E-07 |
| 29162 | 1459 | S1459R | E | 3.4e-07 | 2.1e-07 | 3.1e-07 | 2.867E-07 | 6.807E-08 | 2.100E-07 | 3.400E-07 |
| 29163 | 1459 | S1459N | E | 2e-07 | 2.1e-07 | 2.4e-07 | 2.167E-07 | 2.082E-08 | 2.000E-07 | 2.400E-07 |
| 29164 | 1459 | S1459D | E | 9.2e-08 | 1.3e-07 | 3.4e-07 | 1.873E-07 | 1.336E-07 | 9.200E-08 | 3.400E-07 |
| 29165 | 1459 | S1459C | E | 4.4e-07 | 4.1e-07 | 2.1e-07 | 3.533E-07 | 1.250E-07 | 2.100E-07 | 4.400E-07 |
| 29166 | 1459 | S1459E | E | 3e-07 | 2.1e-07 | 2.1e-07 | 2.400E-07 | 5.196E-08 | 2.100E-07 | 3.000E-07 |
| 29167 | 1459 | S1459Q | E | 2.2e-07 | 3e-07 | 4.2e-07 | 3.133E-07 | 1.007E-07 | 2.200E-07 | 4.200E-07 |
| 29168 | 1459 | S1459G | E | 1.5e-07 | 2.3e-07 | 2.2e-07 | 2.000E-07 | 4.359E-08 | 1.500E-07 | 2.300E-07 |
| 29169 | 1459 | S1459H | E | 3.9e-07 | 4.5e-07 | 8.8e-07 | 5.733E-07 | 2.673E-07 | 3.900E-07 | 8.800E-07 |
| 29170 | 1459 | S1459I | E | 3.3e-07 | 2.6e-07 | 3.1e-07 | 3.000E-07 | 3.606E-08 | 2.600E-07 | 3.300E-07 |
| 29171 | 1459 | S1459L | E | 3.1e-07 | 2.7e-07 | 1.6e-07 | 2.467E-07 | 7.767E-08 | 1.600E-07 | 3.100E-07 |
| 29172 | 1459 | S1459K | E | 2.3e-07 | 2.1e-07 | 1.5e-07 | 1.967E-07 | 4.163E-08 | 1.500E-07 | 2.300E-07 |
| 29173 | 1459 | S1459M | E | 2.4e-07 | 1.8e-07 | 1.7e-07 | 1.967E-07 | 3.786E-08 | 1.700E-07 | 2.400E-07 |
| 29174 | 1459 | S1459F | E | 1.3e-07 | 1.3e-07 | 7.6e-07 | 3.400E-07 | 3.637E-07 | 1.300E-07 | 7.600E-07 |
| 29175 | 1459 | S1459P | E | 1.4e-07 | 1.9e-07 | 1.7e-07 | 1.667E-07 | 2.517E-08 | 1.400E-07 | 1.900E-07 |
| 29176 | 1459 | S1459S | E | 6.6e-07 | 3.4e-07 | 1.1e-07 | 3.700E-07 | 2.762E-07 | 1.100E-07 | 6.600E-07 |
| 29177 | 1459 | S1459T | E | 4.1e-07 | 3.4e-07 | 1.6e-07 | 3.033E-07 | 1.290E-07 | 1.600E-07 | 4.100E-07 |
| 29178 | 1459 | S1459W | E | 5.8e-08 | 3.3e-07 | 1.1e-07 | 1.660E-07 | 1.444E-07 | 5.800E-08 | 3.300E-07 |
| 29179 | 1459 | S1459Y | E | 3.9e-07 | 2e-07 | 1.2e-07 | 2.367E-07 | 1.387E-07 | 1.200E-07 | 3.900E-07 |
| 29180 | 1459 | S1459V | E | 2.7e-07 | 3.1e-07 | 3.3e-07 | 3.033E-07 | 3.055E-08 | 2.700E-07 | 3.300E-07 |
| 29181 | 1460 | V1460A | E | 4.8e-07 | 1.4e-07 | 3.5e-07 | 3.233E-07 | 1.716E-07 | 1.400E-07 | 4.800E-07 |
| 29182 | 1460 | V1460R | E | 3.8e-07 | 2.1e-07 | 2.1e-07 | 2.667E-07 | 9.815E-08 | 2.100E-07 | 3.800E-07 |
| 29183 | 1460 | V1460N | E | 2.3e-07 | 2.6e-07 | 3e-07 | 2.633E-07 | 3.512E-08 | 2.300E-07 | 3.000E-07 |
| 29184 | 1460 | V1460D | E | 7.9e-07 | 4.2e-07 | 4e-07 | 5.367E-07 | 2.196E-07 | 4.000E-07 | 7.900E-07 |
| 29185 | 1460 | V1460C | E | 2.8e-07 | 3.3e-07 | 9e-08 | 2.333E-07 | 1.266E-07 | 9.000E-08 | 3.300E-07 |
| 29186 | 1460 | V1460E | E | 2.7e-07 | 1.9e-07 | 9.5e-08 | 1.850E-07 | 8.761E-08 | 9.500E-08 | 2.700E-07 |
| 29187 | 1460 | V1460Q | E | 1.3e-07 | 3e-07 | 1e-07 | 1.767E-07 | 1.079E-07 | 1.000E-07 | 3.000E-07 |
| 29188 | 1460 | V1460G | E | 1.3e-07 | 2.3e-07 | 2.6e-07 | 2.067E-07 | 6.807E-08 | 1.300E-07 | 2.600E-07 |
| 29189 | 1460 | V1460H | E | 2.1e-07 | 3.2e-07 | 3.3e-07 | 2.867E-07 | 6.658E-08 | 2.100E-07 | 3.300E-07 |
| 29190 | 1460 | V1460I | E | 2.6e-07 | 2.1e-07 | 1.9e-07 | 2.200E-07 | 3.606E-08 | 1.900E-07 | 2.600E-07 |
| 29191 | 1460 | V1460L | E | 1.7e-07 | 2.4e-07 | 1.6e-07 | 1.900E-07 | 4.359E-08 | 1.600E-07 | 2.400E-07 |
| 29192 | 1460 | V1460K | E | 1.6e-07 | 2.4e-07 | 1.9e-07 | 1.967E-07 | 4.041E-08 | 1.600E-07 | 2.400E-07 |
| 29193 | 1460 | V1460M | E | 2.1e-07 | 1.6e-07 | 2.4e-07 | 2.033E-07 | 4.041E-08 | 1.600E-07 | 2.400E-07 |
| 29194 | 1460 | V1460F | E | 4.2e-07 | 8e-08 | 2.7e-07 | 2.567E-07 | 1.704E-07 | 8.000E-08 | 4.200E-07 |
| 29195 | 1460 | V1460P | E | 2.1e-07 | 3.2e-07 | 1.9e-07 | 2.400E-07 | 7.000E-08 | 1.900E-07 | 3.200E-07 |
| 29196 | 1460 | V1460S | E | 4.3e-07 | 1.8e-07 | 3.2e-07 | 3.100E-07 | 1.253E-07 | 1.800E-07 | 4.300E-07 |
| 29197 | 1460 | V1460T | E | 1.8e-07 | 1.1e-07 | 1.5e-07 | 1.467E-07 | 3.512E-08 | 1.100E-07 | 1.800E-07 |
| 29198 | 1460 | V1460W | E | 2.1e-07 | 2.9e-07 | 5.3e-07 | 3.433E-07 | 1.665E-07 | 2.100E-07 | 5.300E-07 |
| 29199 | 1460 | V1460Y | E | 1.8e-07 | 5.5e-07 | 1.6e-07 | 2.967E-07 | 2.196E-07 | 1.600E-07 | 5.500E-07 |
| 29200 | 1460 | V1460V | E | 2.4e-07 | 1.8e-07 | 2.7e-07 | 2.300E-07 | 4.583E-08 | 1.800E-07 | 2.700E-07 |
| 29201 | 1461 | V1461A | E | 2e-07 | 2.4e-07 | 1.6e-07 | 2.000E-07 | 4.000E-08 | 1.600E-07 | 2.400E-07 |
| 29202 | 1461 | V1461R | E | 2e-07 | 5.4e-07 | 1.9e-07 | 3.100E-07 | 1.992E-07 | 1.900E-07 | 5.400E-07 |
| 29203 | 1461 | V1461N | E | 2.2e-07 | 1.1e-06 | 2.8e-07 | 5.333E-07 | 4.917E-07 | 2.200E-07 | 1.100E-06 |
| 29204 | 1461 | V1461D | E | 2e-07 | 2.9e-07 | 2.5e-07 | 2.467E-07 | 4.509E-08 | 2.000E-07 | 2.900E-07 |
| 29205 | 1461 | V1461C | E | 2.6e-07 | 4.5e-07 | 3.7e-07 | 3.600E-07 | 9.539E-08 | 2.600E-07 | 4.500E-07 |
| 29206 | 1461 | V1461E | E | 3.6e-07 | 2.7e-07 | 2.5e-07 | 2.933E-07 | 5.859E-08 | 2.500E-07 | 3.600E-07 |
| 29207 | 1461 | V1461Q | E | 2.3e-07 | 5.3e-07 | 3.2e-07 | 3.600E-07 | 1.539E-07 | 2.300E-07 | 5.300E-07 |
| 29208 | 1461 | V1461G | E | 5.5e-07 | 1.6e-07 | 1.3e-07 | 2.800E-07 | 2.343E-07 | 1.300E-07 | 5.500E-07 |
| 29209 | 1461 | V1461H | E | 2.5e-07 | 1.5e-07 | 3.3e-07 | 2.433E-07 | 9.018E-08 | 1.500E-07 | 3.300E-07 |
| 29210 | 1461 | V1461I | E | 1.3e-07 | 1.1e-07 | 5.5e-07 | 2.633E-07 | 2.485E-07 | 1.100E-07 | 5.500E-07 |
| 29211 | 1461 | V1461L | E | 2.1e-07 | 1.4e-07 | 1.6e-07 | 1.700E-07 | 3.606E-08 | 1.400E-07 | 2.100E-07 |
| 29212 | 1461 | V1461K | E | 2.8e-07 | 2e-07 | 1.6e-07 | 2.133E-07 | 6.110E-08 | 1.600E-07 | 2.800E-07 |
| 29213 | 1461 | V1461M | E | 3e-07 | 2.1e-07 | 5.6e-07 | 3.567E-07 | 1.818E-07 | 2.100E-07 | 5.600E-07 |
| 29214 | 1461 | V1461F | E | 2.2e-07 | 2.5e-07 | 1.5e-07 | 2.067E-07 | 5.132E-08 | 1.500E-07 | 2.500E-07 |
| 29215 | 1461 | V1461P | E | 3.8e-07 | 2.6e-07 | 1.5e-07 | 2.633E-07 | 1.150E-07 | 1.500E-07 | 3.800E-07 |
| 29216 | 1461 | V1461S | E | 1.7e-07 | 2.6e-07 | 4.8e-07 | 3.033E-07 | 1.595E-07 | 1.700E-07 | 4.800E-07 |
| 29217 | 1461 | V1461T | E | 1.9e-07 | 1.5e-07 | 2.2e-07 | 1.867E-07 | 3.512E-08 | 1.500E-07 | 2.200E-07 |
| 29218 | 1461 | V1461W | E | 1.9e-07 | 4.8e-07 | 2e-07 | 2.900E-07 | 1.646E-07 | 1.900E-07 | 4.800E-07 |
| 29219 | 1461 | V1461Y | E | 2e-07 | 2.4e-07 | 1.2e-07 | 1.867E-07 | 6.110E-08 | 1.200E-07 | 2.400E-07 |
| 29220 | 1461 | V1461V | E | 2.8e-07 | 1.7e-07 | 4.1e-07 | 2.867E-07 | 1.201E-07 | 1.700E-07 | 4.100E-07 |
| 29221 | 1462 | T1462A | E | 1.2e-06 | 3.7e-07 | 3.4e-07 | 6.367E-07 | 4.881E-07 | 3.400E-07 | 1.200E-06 |
| 29222 | 1462 | T1462R | E | 3.9e-07 | 2.4e-07 | 2.4e-07 | 2.900E-07 | 8.660E-08 | 2.400E-07 | 3.900E-07 |
| 29223 | 1462 | T1462N | E | 6e-07 | 2.4e-07 | 1.8e-07 | 3.400E-07 | 2.272E-07 | 1.800E-07 | 6.000E-07 |
| 29224 | 1462 | T1462D | E | 6.4e-07 | 1.4e-07 | 1.9e-07 | 3.233E-07 | 2.754E-07 | 1.400E-07 | 6.400E-07 |
| 29225 | 1462 | T1462C | E | 1.2e-07 | 2.7e-07 | 1.3e-07 | 1.733E-07 | 8.386E-08 | 1.200E-07 | 2.700E-07 |
| 29226 | 1462 | T1462E | E | 2.3e-07 | 2.4e-07 | 3.8e-07 | 2.833E-07 | 8.386E-08 | 2.300E-07 | 3.800E-07 |
| 29227 | 1462 | T1462Q | E | 2.3e-07 | 1.7e-07 | 3.5e-07 | 2.500E-07 | 9.165E-08 | 1.700E-07 | 3.500E-07 |
| 29228 | 1462 | T1462G | E | 1.8e-07 | 6.2e-07 | 5.3e-07 | 4.433E-07 | 2.325E-07 | 1.800E-07 | 6.200E-07 |
| 29229 | 1462 | T1462H | E | 1.5e-07 | 1.6e-07 | 2.9e-07 | 2.000E-07 | 7.810E-08 | 1.500E-07 | 2.900E-07 |
| 29230 | 1462 | T1462I | E | 2.8e-07 | 2e-07 | 3.4e-07 | 2.733E-07 | 7.024E-08 | 2.000E-07 | 3.400E-07 |
| 29231 | 1462 | T1462L | E | 4.1e-07 | 7.2e-07 | 3.4e-07 | 4.900E-07 | 2.022E-07 | 3.400E-07 | 7.200E-07 |
| 29232 | 1462 | T1462K | E | 1.1e-07 | 4.1e-07 | 1.8e-07 | 2.333E-07 | 1.570E-07 | 1.100E-07 | 4.100E-07 |
| 29233 | 1462 | T1462M | E | 2.2e-07 | 4e-07 | 2.4e-07 | 2.867E-07 | 9.866E-08 | 2.200E-07 | 4.000E-07 |
| 29234 | 1462 | T1462F | E | 5.1e-07 | 5.6e-07 | 2.7e-07 | 4.467E-07 | 1.550E-07 | 2.700E-07 | 5.600E-07 |
| 29235 | 1462 | T1462P | E | 3.6e-07 | 1.6e-07 | 1.4e-07 | 2.200E-07 | 1.217E-07 | 1.400E-07 | 3.600E-07 |
| 29236 | 1462 | T1462S | E | 2.2e-07 | 3.9e-07 | 2.5e-07 | 2.867E-07 | 9.074E-08 | 2.200E-07 | 3.900E-07 |
| 29237 | 1462 | T1462T | E | 4.6e-08 | 1.6e-07 | 1.7e-07 | 1.253E-07 | 6.889E-08 | 4.600E-08 | 1.700E-07 |
| 29238 | 1462 | T1462W | E | 2.2e-07 | 7.5e-08 | 5.2e-07 | 2.717E-07 | 2.270E-07 | 7.500E-08 | 5.200E-07 |
| 29239 | 1462 | T1462Y | E | 2.3e-07 | 1.8e-07 | 2.5e-07 | 2.200E-07 | 3.606E-08 | 1.800E-07 | 2.500E-07 |

| | | | | | | | | | | |
|---|---|---|---|---|---|---|---|---|---|---|
| 29240 | 1462 | T1462V | E | 1.6e-07 | 2e-07 | 3.9e-07 | 2.500E-07 | 1.229E-07 | 1.600E-07 | 3.900E-07 |
| 29241 | 1463 | V1463A | E | 2.2e-07 | 3.7e-07 | 2.3e-07 | 2.733E-07 | 8.386E-08 | 2.200E-07 | 3.700E-07 |
| 29242 | 1463 | V1463R | E | 2e-07 | 3.3e-07 | 2.6e-07 | 2.633E-07 | 6.506E-08 | 2.000E-07 | 3.300E-07 |
| 29243 | 1463 | V1463N | E | 1.9e-07 | 2.6e-07 | 4.1e-07 | 2.867E-07 | 1.124E-07 | 1.900E-07 | 4.100E-07 |
| 29244 | 1463 | V1463D | E | 2.3e-07 | 3.3e-07 | 2.6e-07 | 2.733E-07 | 5.132E-08 | 2.300E-07 | 3.300E-07 |
| 29245 | 1463 | V1463C | E | 4.1e-07 | 1.6e-07 | 2.8e-07 | 2.833E-07 | 1.250E-07 | 1.600E-07 | 4.100E-07 |
| 29246 | 1463 | V1463E | E | 4.1e-07 | 1.6e-07 | 2.6e-07 | 2.767E-07 | 1.258E-07 | 1.600E-07 | 4.100E-07 |
| 29247 | 1463 | V1463Q | E | 4.6e-07 | 4.1e-07 | 2.2e-07 | 3.633E-07 | 1.266E-07 | 2.200E-07 | 4.600E-07 |
| 29248 | 1463 | V1463G | E | 3e-07 | 6.2e-08 | 2.4e-07 | 2.007E-07 | 1.238E-07 | 6.200E-08 | 3.000E-07 |
| 29249 | 1463 | V1463H | E | 2.3e-07 | 2.5e-07 | 3.7e-07 | 2.833E-07 | 7.572E-08 | 2.300E-07 | 3.700E-07 |
| 29250 | 1463 | V1463I | E | 1.9e-07 | 1.8e-07 | 7.6e-08 | 1.487E-07 | 6.313E-08 | 7.600E-08 | 1.900E-07 |
| 29251 | 1463 | V1463L | E | 2.6e-07 | 5.4e-07 | 2.1e-07 | 3.367E-07 | 1.779E-07 | 2.100E-07 | 5.400E-07 |
| 29252 | 1463 | V1463K | E | 1.9e-07 | 2.2e-07 | 3.3e-07 | 2.467E-07 | 7.371E-08 | 1.900E-07 | 3.300E-07 |
| 29253 | 1463 | V1463M | E | 1.7e-07 | 4.1e-07 | 2.2e-07 | 2.667E-07 | 1.266E-07 | 1.700E-07 | 4.100E-07 |
| 29254 | 1463 | V1463F | E | 7.9e-08 | 2.6e-07 | 4.4e-07 | 2.597E-07 | 1.805E-07 | 7.900E-08 | 4.400E-07 |
| 29255 | 1463 | V1463P | E | 5.3e-07 | 6.6e-07 | 1.4e-07 | 4.433E-07 | 2.706E-07 | 1.400E-07 | 6.600E-07 |
| 29256 | 1463 | V1463S | E | 1.2e-07 | 6.1e-07 | 2.6e-07 | 3.300E-07 | 2.524E-07 | 1.200E-07 | 6.100E-07 |
| 29257 | 1463 | V1463T | E | 1.8e-07 | 3.5e-07 | 2.7e-07 | 2.667E-07 | 8.505E-08 | 1.800E-07 | 3.500E-07 |
| 29258 | 1463 | V1463W | E | 1.9e-07 | 1.9e-07 | 3.2e-07 | 2.333E-07 | 7.506E-08 | 1.900E-07 | 3.200E-07 |
| 29259 | 1463 | V1463Y | E | 2.4e-07 | 1.5e-07 | 1.9e-07 | 1.933E-07 | 4.509E-08 | 1.500E-07 | 2.400E-07 |
| 29260 | 1463 | V1463V | E | 2.1e-07 | 3.7e-07 | 3.9e-07 | 3.233E-07 | 9.866E-08 | 2.100E-07 | 3.900E-07 |
| 29261 | 1464 | P1464A | E | 3e-07 | 1.4e-07 | 1.7e-07 | 2.033E-07 | 8.505E-08 | 1.400E-07 | 3.000E-07 |
| 29262 | 1464 | P1464R | E | 5.3e-07 | 2.9e-07 | 1.6e-07 | 3.267E-07 | 1.877E-07 | 1.600E-07 | 5.300E-07 |
| 29263 | 1464 | P1464N | E | 4.4e-07 | 3.7e-07 | 2e-07 | 3.367E-07 | 1.234E-07 | 2.000E-07 | 4.400E-07 |
| 29264 | 1464 | P1464D | E | 1.7e-07 | 3.3e-07 | 2.6e-07 | 2.533E-07 | 8.021E-08 | 1.700E-07 | 3.300E-07 |
| 29265 | 1464 | P1464C | E | 1.4e-07 | 2.4e-07 | 1.5e-07 | 1.767E-07 | 5.508E-08 | 1.400E-07 | 2.400E-07 |
| 29266 | 1464 | P1464E | E | 1.9e-07 | 3.4e-07 | 2.2e-07 | 2.500E-07 | 7.937E-08 | 1.900E-07 | 3.400E-07 |
| 29267 | 1464 | P1464Q | E | 3.7e-07 | 4.9e-07 | 2.5e-07 | 3.700E-07 | 1.200E-07 | 2.500E-07 | 4.900E-07 |
| 29268 | 1464 | P1464G | E | 1.8e-07 | 2e-07 | 2.9e-07 | 2.233E-07 | 5.859E-08 | 1.800E-07 | 2.900E-07 |
| 29269 | 1464 | P1464H | E | 3.8e-07 | 2.7e-07 | 3e-07 | 3.167E-07 | 5.686E-08 | 2.700E-07 | 3.800E-07 |
| 29270 | 1464 | P1464I | E | 1.9e-07 | 1.9e-07 | 8.9e-08 | 1.563E-07 | 5.831E-08 | 8.900E-08 | 1.900E-07 |
| 29271 | 1464 | P1464L | E | 2.1e-07 | 2.8e-07 | 2.7e-07 | 2.533E-07 | 3.786E-08 | 2.100E-07 | 2.800E-07 |
| 29272 | 1464 | P1464K | E | 2.6e-07 | 5.9e-07 | 1.4e-07 | 3.300E-07 | 2.330E-07 | 1.400E-07 | 5.900E-07 |
| 29273 | 1464 | P1464M | E | 1.5e-07 | 4.8e-07 | 2.8e-07 | 3.033E-07 | 1.662E-07 | 1.500E-07 | 4.800E-07 |
| 29274 | 1464 | P1464F | E | 1.4e-07 | 2.1e-07 | 4.3e-07 | 2.600E-07 | 1.513E-07 | 1.400E-07 | 4.300E-07 |
| 29275 | 1464 | P1464P | E | 4.2e-07 | 4.4e-07 | 6.1e-07 | 4.900E-07 | 1.044E-07 | 4.200E-07 | 6.100E-07 |
| 29276 | 1464 | P1464S | E | 1.4e-07 | 1.5e-07 | 2.5e-07 | 1.800E-07 | 6.083E-08 | 1.400E-07 | 2.500E-07 |
| 29277 | 1464 | P1464T | E | 1.7e-07 | 2.6e-07 | 1.6e-07 | 1.967E-07 | 5.508E-08 | 1.600E-07 | 2.600E-07 |
| 29278 | 1464 | P1464W | E | 1.6e-07 | 1.4e-07 | 1.8e-07 | 1.600E-07 | 2.000E-08 | 1.400E-07 | 1.800E-07 |
| 29279 | 1464 | P1464Y | E | 2.9e-07 | 1.2e-07 | 3.1e-07 | 2.400E-07 | 1.044E-07 | 1.200E-07 | 3.100E-07 |
| 29280 | 1464 | P1464V | E | 2.2e-07 | 2.8e-07 | 2.5e-07 | 2.500E-07 | 3.000E-08 | 2.200E-07 | 2.800E-07 |
| 29281 | 1465 | S1465A | E | 2.2e-07 | 1.6e-07 | 1.2e-07 | 1.667E-07 | 5.033E-08 | 1.200E-07 | 2.200E-07 |
| 29282 | 1465 | S1465R | E | 2.9e-07 | 3.5e-07 | 4e-07 | 3.467E-07 | 5.508E-08 | 2.900E-07 | 4.000E-07 |
| 29283 | 1465 | S1465N | E | 4.1e-07 | 3.5e-07 | 5.4e-07 | 4.333E-07 | 9.713E-08 | 3.500E-07 | 5.400E-07 |
| 29284 | 1465 | S1465D | E | 1.7e-07 | 1.8e-07 | 1.9e-07 | 1.800E-07 | 1.000E-08 | 1.700E-07 | 1.900E-07 |
| 29285 | 1465 | S1465C | E | 2.7e-07 | 1.7e-07 | 2.1e-07 | 2.167E-07 | 5.033E-08 | 1.700E-07 | 2.700E-07 |
| 29286 | 1465 | S1465E | E | 3.3e-07 | 1.4e-07 | 2.2e-07 | 2.300E-07 | 9.539E-08 | 1.400E-07 | 3.300E-07 |
| 29287 | 1465 | S1465Q | E | 3.4e-07 | 3.1e-07 | 2.2e-07 | 2.900E-07 | 6.245E-08 | 2.200E-07 | 3.400E-07 |
| 29288 | 1465 | S1465G | E | 1.8e-07 | 2.1e-07 | 3.9e-07 | 2.600E-07 | 1.136E-07 | 1.800E-07 | 3.900E-07 |
| 29289 | 1465 | S1465H | E | 2.3e-07 | 3.2e-07 | 5.5e-07 | 3.667E-07 | 1.650E-07 | 2.300E-07 | 5.500E-07 |
| 29290 | 1465 | S1465I | E | 4.3e-07 | 3.2e-07 | 3e-07 | 3.500E-07 | 7.000E-08 | 3.000E-07 | 4.300E-07 |
| 29291 | 1465 | S1465L | E | 3.6e-07 | 2.7e-07 | 3.2e-07 | 3.167E-07 | 4.509E-08 | 2.700E-07 | 3.600E-07 |
| 29292 | 1465 | S1465K | E | 1.8e-07 | 2.2e-07 | 3.4e-07 | 2.467E-07 | 8.327E-08 | 1.800E-07 | 3.400E-07 |
| 29293 | 1465 | S1465M | E | 2.9e-07 | 1.2e-07 | 2.4e-07 | 2.167E-07 | 8.737E-08 | 1.200E-07 | 2.900E-07 |
| 29294 | 1465 | S1465F | E | 4.5e-07 | 1.1e-07 | 2.4e-07 | 2.667E-07 | 1.716E-07 | 1.100E-07 | 4.500E-07 |
| 29295 | 1465 | S1465P | E | 3.2e-07 | 2.9e-07 | 2e-07 | 2.700E-07 | 6.245E-08 | 2.000E-07 | 3.200E-07 |
| 29296 | 1465 | S1465S | E | 3.2e-07 | 2.8e-07 | 2.7e-07 | 2.900E-07 | 2.646E-08 | 2.700E-07 | 3.200E-07 |
| 29297 | 1465 | S1465T | E | 4.4e-07 | 6.6e-07 | 4.3e-07 | 5.100E-07 | 1.300E-07 | 4.300E-07 | 6.600E-07 |
| 29298 | 1465 | S1465W | E | 3.2e-07 | 2.2e-07 | 3.2e-07 | 2.867E-07 | 5.774E-08 | 2.200E-07 | 3.200E-07 |
| 29299 | 1465 | S1465Y | E | 1.3e-07 | 1.2e-07 | 5.3e-07 | 2.600E-07 | 2.339E-07 | 1.200E-07 | 5.300E-07 |
| 29300 | 1465 | S1465V | E | 1.3e-07 | 3.6e-07 | 1.8e-07 | 2.233E-07 | 1.210E-07 | 1.300E-07 | 3.600E-07 |
| 29301 | 1466 | S1466A | E | 3.9e-07 | 2.6e-07 | 1.3e-07 | 2.600E-07 | 1.300E-07 | 1.300E-07 | 3.900E-07 |
| 29302 | 1466 | S1466R | E | 1.4e-07 | 2e-07 | 1.9e-07 | 1.767E-07 | 3.215E-08 | 1.400E-07 | 2.000E-07 |
| 29303 | 1466 | S1466N | E | 3.2e-07 | 2.5e-07 | 4.2e-07 | 3.300E-07 | 8.544E-08 | 2.500E-07 | 4.200E-07 |
| 29304 | 1466 | S1466D | E | 1.6e-07 | 1.8e-07 | 2.6e-07 | 2.000E-07 | 5.292E-08 | 1.600E-07 | 2.600E-07 |
| 29305 | 1466 | S1466C | E | 3.2e-07 | 2.6e-07 | 2e-07 | 2.600E-07 | 6.000E-08 | 2.000E-07 | 3.200E-07 |
| 29306 | 1466 | S1466E | E | 2.7e-07 | 2.6e-07 | 2.4e-07 | 2.567E-07 | 1.528E-08 | 2.400E-07 | 2.700E-07 |
| 29307 | 1466 | S1466Q | E | 3.2e-07 | 2.2e-07 | 6.5e-08 | 2.017E-07 | 1.285E-07 | 6.500E-08 | 3.200E-07 |
| 29308 | 1466 | S1466G | E | 1.6e-07 | 2.2e-07 | 1.8e-07 | 1.867E-07 | 3.055E-08 | 1.600E-07 | 2.200E-07 |
| 29309 | 1466 | S1466H | E | 7.9e-07 | 2.1e-07 | 1.9e-07 | 3.967E-07 | 3.408E-07 | 1.900E-07 | 7.900E-07 |
| 29310 | 1466 | S1466I | E | 1.3e-07 | 2.1e-07 | 1.8e-07 | 1.733E-07 | 4.041E-08 | 1.300E-07 | 2.100E-07 |
| 29311 | 1466 | S1466L | E | 3.3e-07 | 2.3e-07 | 1.5e-07 | 2.367E-07 | 9.018E-08 | 1.500E-07 | 3.300E-07 |
| 29312 | 1466 | S1466K | E | 3.8e-07 | 7e-07 | 7.7e-08 | 3.857E-07 | 3.115E-07 | 7.700E-08 | 7.000E-07 |
| 29313 | 1466 | S1466M | E | 2.5e-07 | 1.9e-07 | 6.2e-07 | 3.533E-07 | 2.329E-07 | 1.900E-07 | 6.200E-07 |
| 29314 | 1466 | S1466F | E | 2.3e-07 | 2.5e-07 | 4.3e-07 | 3.033E-07 | 1.102E-07 | 2.300E-07 | 4.300E-07 |
| 29315 | 1466 | S1466P | E | 2.2e-07 | 2.2e-07 | 2e-07 | 2.133E-07 | 1.155E-08 | 2.000E-07 | 2.200E-07 |
| 29316 | 1466 | S1466S | E | 2.5e-07 | 2e-07 | 1.5e-07 | 2.000E-07 | 5.000E-08 | 1.500E-07 | 2.500E-07 |
| 29317 | 1466 | S1466T | E | 1.7e-07 | 1.3e-07 | 2.3e-07 | 1.767E-07 | 5.033E-08 | 1.300E-07 | 2.300E-07 |
| 29318 | 1466 | S1466W | E | 2.3e-07 | 2.6e-07 | 2.6e-07 | 2.500E-07 | 1.732E-08 | 2.300E-07 | 2.600E-07 |
| 29319 | 1466 | S1466Y | E | 2.1e-07 | 2.5e-07 | 2.2e-07 | 2.267E-07 | 2.082E-08 | 2.100E-07 | 2.500E-07 |
| 29320 | 1466 | S1466V | E | 9.3e-08 | 2.4e-07 | 3.1e-07 | 2.143E-07 | 1.108E-07 | 9.300E-08 | 3.100E-07 |
| 29321 | 1467 | S1467A | E | 5e-07 | 9.5e-08 | 2.5e-07 | 2.817E-07 | 2.043E-07 | 9.500E-08 | 5.000E-07 |
| 29322 | 1467 | S1467R | E | 1.4e-07 | 1.5e-07 | 3.3e-07 | 2.067E-07 | 1.069E-07 | 1.400E-07 | 3.300E-07 |
| 29323 | 1467 | S1467N | E | 3.4e-07 | 2.6e-07 | 4.5e-07 | 3.500E-07 | 9.539E-08 | 2.600E-07 | 4.500E-07 |
| 29324 | 1467 | S1467D | E | 4.4e-07 | 1.6e-07 | 2.3e-07 | 2.767E-07 | 1.457E-07 | 1.600E-07 | 4.400E-07 |

| | | | | | | | | | | |
|---|---|---|---|---|---|---|---|---|---|---|
| 29325 | 1467 | S1467C | E | 2e-07 | 5.4e-07 | 2.2e-07 | 3.200E-07 | 1.908E-07 | 2.000E-07 | 5.400E-07 |
| 29326 | 1467 | S1467E | E | 1.5e-07 | 1.9e-07 | 1.5e-07 | 1.633E-07 | 2.309E-08 | 1.500E-07 | 1.900E-07 |
| 29327 | 1467 | S1467Q | E | 3.8e-07 | 5.5e-07 | 1.8e-07 | 3.700E-07 | 1.852E-07 | 1.800E-07 | 5.500E-07 |
| 29328 | 1467 | S1467G | E | 3.8e-07 | 1.7e-07 | 3.5e-07 | 3.000E-07 | 1.136E-07 | 1.700E-07 | 3.800E-07 |
| 29329 | 1467 | S1467H | E | 2e-07 | 2.8e-07 | 1.2e-07 | 2.000E-07 | 8.000E-08 | 1.200E-07 | 2.800E-07 |
| 29330 | 1467 | S1467I | E | 3.5e-07 | 1.6e-07 | 1.1e-07 | 2.067E-07 | 1.266E-07 | 1.100E-07 | 3.500E-07 |
| 29331 | 1467 | S1467L | E | 1.5e-07 | 3.1e-07 | 2.3e-07 | 2.300E-07 | 8.000E-08 | 1.500E-07 | 3.100E-07 |
| 29332 | 1467 | S1467K | E | 1.8e-07 | 1.9e-07 | 3.3e-07 | 2.333E-07 | 8.386E-08 | 1.800E-07 | 3.300E-07 |
| 29333 | 1467 | S1467M | E | 2.7e-07 | 2.1e-07 | 2.1e-07 | 2.300E-07 | 3.464E-08 | 2.100E-07 | 2.700E-07 |
| 29334 | 1467 | S1467F | E | 2e-07 | 2.4e-07 | 3.4e-07 | 2.600E-07 | 7.211E-08 | 2.000E-07 | 3.400E-07 |
| 29335 | 1467 | S1467P | E | 6.4e-07 | 4.3e-07 | 1.7e-07 | 4.133E-07 | 2.354E-07 | 1.700E-07 | 6.400E-07 |
| 29336 | 1467 | S1467S | E | 2.9e-07 | 4.7e-07 | 2.7e-07 | 3.433E-07 | 1.102E-07 | 2.700E-07 | 4.700E-07 |
| 29337 | 1467 | S1467T | E | 3.1e-07 | 2.4e-07 | 2.1e-07 | 2.533E-07 | 5.132E-08 | 2.100E-07 | 3.100E-07 |
| 29338 | 1467 | S1467W | E | 2.5e-07 | 1e-06 | 3e-07 | 5.167E-07 | 4.193E-07 | 2.500E-07 | 1.000E-06 |
| 29339 | 1467 | S1467Y | E | 5.9e-07 | 1.6e-07 | 1.5e-07 | 3.000E-07 | 2.512E-07 | 1.500E-07 | 5.900E-07 |
| 29340 | 1467 | S1467V | E | 2.6e-07 | 2.6e-07 | 3.6e-07 | 2.933E-07 | 5.774E-08 | 2.600E-07 | 3.600E-07 |
| 29341 | 1468 | L1468A | E | 2.7e-07 | 3.1e-07 | 2.7e-07 | 2.833E-07 | 2.309E-08 | 2.700E-07 | 3.100E-07 |
| 29342 | 1468 | L1468R | E | 4.2e-07 | 2e-07 | 2.5e-07 | 2.900E-07 | 1.153E-07 | 2.000E-07 | 4.200E-07 |
| 29343 | 1468 | L1468N | E | 4.6e-07 | 3.7e-07 | 2.6e-07 | 3.633E-07 | 1.002E-07 | 2.600E-07 | 4.600E-07 |
| 29344 | 1468 | L1468D | E | 3.2e-07 | 4e-07 | 5.9e-07 | 4.367E-07 | 1.387E-07 | 3.200E-07 | 5.900E-07 |
| 29345 | 1468 | L1468C | E | 2.3e-07 | 3.3e-07 | 2.1e-07 | 2.567E-07 | 6.429E-08 | 2.100E-07 | 3.300E-07 |
| 29346 | 1468 | L1468E | E | 4.3e-07 | 1.8e-07 | 3.7e-07 | 3.267E-07 | 1.305E-07 | 1.800E-07 | 4.300E-07 |
| 29347 | 1468 | L1468Q | E | 1.5e-07 | 6e-07 | 1.4e-07 | 2.967E-07 | 2.627E-07 | 1.400E-07 | 6.000E-07 |
| 29348 | 1468 | L1468G | E | 2.2e-07 | 2.5e-07 | 5.5e-07 | 3.400E-07 | 1.825E-07 | 2.200E-07 | 5.500E-07 |
| 29349 | 1468 | L1468H | E | 2.5e-07 | 1.8e-07 | 4.3e-07 | 2.867E-07 | 1.290E-07 | 1.800E-07 | 4.300E-07 |
| 29350 | 1468 | L1468I | E | 5.7e-07 | 2.5e-07 | 2.6e-07 | 3.600E-07 | 1.819E-07 | 2.500E-07 | 5.700E-07 |
| 29351 | 1468 | L1468L | E | 4.8e-07 | 2.7e-07 | 3e-07 | 3.500E-07 | 1.136E-07 | 2.700E-07 | 4.800E-07 |
| 29352 | 1468 | L1468K | E | 8.2e-07 | 5.1e-07 | 3.3e-07 | 5.533E-07 | 2.479E-07 | 3.300E-07 | 8.200E-07 |
| 29353 | 1468 | L1468M | E | 3.3e-07 | 1.8e-07 | 4.5e-07 | 3.200E-07 | 1.353E-07 | 1.800E-07 | 4.500E-07 |
| 29354 | 1468 | L1468F | E | 4.5e-07 | 2.3e-07 | 1.5e-07 | 2.767E-07 | 1.553E-07 | 1.500E-07 | 4.500E-07 |
| 29355 | 1468 | L1468P | E | 1.3e-07 | 4.7e-07 | 2.4e-07 | 2.800E-07 | 1.735E-07 | 1.300E-07 | 4.700E-07 |
| 29356 | 1468 | L1468S | E | 2.2e-07 | 3e-07 | 2.9e-07 | 2.700E-07 | 4.359E-08 | 2.200E-07 | 3.000E-07 |
| 29357 | 1468 | L1468T | E | 2.5e-07 | 2.5e-07 | 1.7e-07 | 2.233E-07 | 4.619E-08 | 1.700E-07 | 2.500E-07 |
| 29358 | 1468 | L1468W | E | 4.9e-07 | 5.7e-07 | 2.3e-07 | 4.300E-07 | 1.778E-07 | 2.300E-07 | 5.700E-07 |
| 29359 | 1468 | L1468Y | E | 4.1e-07 | 3.3e-07 | 1.7e-07 | 3.033E-07 | 1.222E-07 | 1.700E-07 | 4.100E-07 |
| 29360 | 1468 | L1468V | E | 1.8e-07 | 2.1e-07 | 2.4e-07 | 2.100E-07 | 3.000E-08 | 1.800E-07 | 2.400E-07 |
| 29361 | 1469 | G1469A | E | 1.8e-07 | 2.7e-07 | 3.1e-07 | 2.533E-07 | 6.658E-08 | 1.800E-07 | 3.100E-07 |
| 29362 | 1469 | G1469R | E | 2e-07 | 3.1e-07 | 5.5e-07 | 3.533E-07 | 1.790E-07 | 2.000E-07 | 5.500E-07 |
| 29363 | 1469 | G1469N | E | 1.8e-07 | 2e-07 | 1.7e-07 | 1.833E-07 | 1.528E-08 | 1.700E-07 | 2.000E-07 |
| 29364 | 1469 | G1469D | E | 1.2e-07 | 4.5e-07 | 2.1e-07 | 2.600E-07 | 1.706E-07 | 1.200E-07 | 4.500E-07 |
| 29365 | 1469 | G1469C | E | 2.6e-07 | 2.1e-07 | 2.5e-07 | 2.400E-07 | 2.646E-08 | 2.100E-07 | 2.600E-07 |
| 29366 | 1469 | G1469E | E | 5.8e-07 | 3.7e-07 | 3.2e-07 | 4.233E-07 | 1.380E-07 | 3.200E-07 | 5.800E-07 |
| 29367 | 1469 | G1469Q | E | 1.9e-07 | 3.3e-07 | 1.6e-07 | 2.267E-07 | 9.074E-08 | 1.600E-07 | 3.300E-07 |
| 29368 | 1469 | G1469G | E | 3.9e-07 | 3.3e-07 | 4.1e-07 | 3.767E-07 | 4.163E-08 | 3.300E-07 | 4.100E-07 |
| 29369 | 1469 | G1469H | E | 2.5e-07 | 3.1e-07 | 2.1e-07 | 2.567E-07 | 5.033E-08 | 2.100E-07 | 3.100E-07 |
| 29370 | 1469 | G1469I | E | 2.3e-07 | 2.8e-07 | 1.7e-07 | 2.267E-07 | 5.508E-08 | 1.700E-07 | 2.800E-07 |
| 29371 | 1469 | G1469L | E | 3.7e-07 | 2.9e-07 | 2.1e-07 | 2.900E-07 | 8.000E-08 | 2.100E-07 | 3.700E-07 |
| 29372 | 1469 | G1469K | E | 1.2e-07 | 1.2e-07 | 1.8e-07 | 1.400E-07 | 3.464E-08 | 1.200E-07 | 1.800E-07 |
| 29373 | 1469 | G1469M | E | 2.9e-07 | 1.6e-07 | 2.7e-07 | 2.400E-07 | 7.000E-08 | 1.600E-07 | 2.900E-07 |
| 29374 | 1469 | G1469F | E | 5.4e-07 | 4e-07 | 2.7e-07 | 4.033E-07 | 1.350E-07 | 2.700E-07 | 5.400E-07 |
| 29375 | 1469 | G1469P | E | 1.9e-07 | 1.8e-07 | 4.1e-07 | 2.600E-07 | 1.300E-07 | 1.800E-07 | 4.100E-07 |
| 29376 | 1469 | G1469S | E | 4.4e-07 | 5.1e-07 | 2.2e-07 | 3.900E-07 | 1.513E-07 | 2.200E-07 | 5.100E-07 |
| 29377 | 1469 | G1469T | E | 3.5e-07 | 2.4e-07 | 3e-07 | 2.967E-07 | 5.508E-08 | 2.400E-07 | 3.500E-07 |
| 29378 | 1469 | G1469W | E | 3.9e-07 | 2.4e-07 | 1.6e-07 | 2.633E-07 | 1.168E-07 | 1.600E-07 | 3.900E-07 |
| 29379 | 1469 | G1469Y | E | 1e-07 | 4e-07 | 1.4e-07 | 2.133E-07 | 1.629E-07 | 1.000E-07 | 4.000E-07 |
| 29380 | 1469 | G1469V | E | 3.6e-07 | 1.3e-07 | 2.3e-07 | 2.400E-07 | 1.153E-07 | 1.300E-07 | 3.600E-07 |
| 29381 | 1470 | T1470A | E | 1.7e-07 | 4.2e-07 | 1.2e-07 | 2.367E-07 | 1.607E-07 | 1.200E-07 | 4.200E-07 |
| 29382 | 1470 | T1470R | E | 2.4e-07 | 3.2e-07 | 2.8e-07 | 2.800E-07 | 4.000E-08 | 2.400E-07 | 3.200E-07 |
| 29383 | 1470 | T1470N | E | 1.8e-07 | 1.1e-07 | 1.7e-07 | 1.533E-07 | 3.786E-08 | 1.100E-07 | 1.800E-07 |
| 29384 | 1470 | T1470D | E | 1.2e-07 | 2.8e-07 | 3.1e-07 | 2.367E-07 | 1.021E-07 | 1.200E-07 | 3.100E-07 |
| 29385 | 1470 | T1470C | E | 2e-07 | 3.2e-07 | 4.1e-07 | 3.100E-07 | 1.054E-07 | 2.000E-07 | 4.100E-07 |
| 29386 | 1470 | T1470E | E | 2e-07 | 2.4e-07 | 1e-07 | 1.800E-07 | 7.211E-08 | 1.000E-07 | 2.400E-07 |
| 29387 | 1470 | T1470Q | E | 3.9e-07 | 2.6e-07 | 4.2e-07 | 3.567E-07 | 8.505E-08 | 2.600E-07 | 4.200E-07 |
| 29388 | 1470 | T1470G | E | 1.7e-07 | 2.8e-07 | 1.4e-07 | 1.967E-07 | 7.371E-08 | 1.400E-07 | 2.800E-07 |
| 29389 | 1470 | T1470H | E | 2.8e-07 | 1.8e-07 | 1.9e-07 | 2.167E-07 | 5.508E-08 | 1.800E-07 | 2.800E-07 |
| 29390 | 1470 | T1470I | E | 1.8e-07 | 2.7e-07 | 1.3e-07 | 1.933E-07 | 7.095E-08 | 1.300E-07 | 2.700E-07 |
| 29391 | 1470 | T1470L | E | 2.2e-07 | 4e-07 | 3.7e-07 | 3.300E-07 | 9.644E-08 | 2.200E-07 | 4.000E-07 |
| 29392 | 1470 | T1470K | E | 1.9e-07 | 2e-07 | 5.2e-07 | 3.033E-07 | 1.877E-07 | 1.900E-07 | 5.200E-07 |
| 29393 | 1470 | T1470M | E | 4.2e-07 | 1.6e-07 | 2.5e-07 | 2.767E-07 | 1.320E-07 | 1.600E-07 | 4.200E-07 |
| 29394 | 1470 | T1470F | E | 5.8e-07 | 1.8e-07 | 2.3e-07 | 3.300E-07 | 2.179E-07 | 1.800E-07 | 5.800E-07 |
| 29395 | 1470 | T1470P | E | 3.2e-07 | 2.3e-07 | 2e-07 | 2.500E-07 | 6.245E-08 | 2.000E-07 | 3.200E-07 |
| 29396 | 1470 | T1470S | E | 1.8e-07 | 1.2e-07 | 2.8e-07 | 1.933E-07 | 8.083E-08 | 1.200E-07 | 2.800E-07 |
| 29397 | 1470 | T1470T | E | 2.1e-07 | 3.3e-07 | 3.2e-07 | 2.867E-07 | 6.658E-08 | 2.100E-07 | 3.300E-07 |
| 29398 | 1470 | T1470W | E | 1.1e-07 | 1.4e-07 | 3.3e-07 | 1.933E-07 | 1.193E-07 | 1.100E-07 | 3.300E-07 |
| 29399 | 1470 | T1470Y | E | 3.3e-07 | 3e-07 | 2.3e-07 | 2.867E-07 | 5.132E-08 | 2.300E-07 | 3.300E-07 |
| 29400 | 1470 | T1470V | E | 1.9e-07 | 2.4e-07 | 1.7e-07 | 2.000E-07 | 3.606E-08 | 1.700E-07 | 2.400E-07 |
| 29401 | 1471 | Q1471A | E | 3e-07 | 1.3e-07 | 2.1e-07 | 2.133E-07 | 8.505E-08 | 1.300E-07 | 3.000E-07 |
| 29402 | 1471 | Q1471R | E | 2.9e-07 | 2.7e-07 | 2.7e-07 | 2.767E-07 | 1.155E-08 | 2.700E-07 | 2.900E-07 |
| 29403 | 1471 | Q1471N | E | 1.6e-07 | 2.7e-07 | 4.7e-07 | 3.000E-07 | 1.572E-07 | 1.600E-07 | 4.700E-07 |
| 29404 | 1471 | Q1471D | E | 2.9e-07 | 3.4e-07 | 2.2e-07 | 2.833E-07 | 6.028E-08 | 2.200E-07 | 3.400E-07 |
| 29405 | 1471 | Q1471C | E | 4.2e-07 | 2.9e-07 | 2.4e-07 | 3.167E-07 | 9.292E-08 | 2.400E-07 | 4.200E-07 |
| 29406 | 1471 | Q1471E | E | 3.3e-07 | 4e-07 | 1.8e-07 | 3.033E-07 | 1.124E-07 | 1.800E-07 | 4.000E-07 |
| 29407 | 1471 | Q1471Q | E | 1.9e-07 | 2.5e-07 | 2e-07 | 2.133E-07 | 3.215E-08 | 1.900E-07 | 2.500E-07 |
| 29408 | 1471 | Q1471G | E | 2.3e-07 | 2.4e-07 | 3.3e-07 | 2.667E-07 | 5.508E-08 | 2.300E-07 | 3.300E-07 |
| 29409 | 1471 | Q1471H | E | 4.1e-07 | 1.5e-07 | 2e-07 | 2.533E-07 | 1.380E-07 | 1.500E-07 | 4.100E-07 |

| | | | | | | | | | | |
|---|---|---|---|---|---|---|---|---|---|---|
| 29410 | 1471 | Q1471I | E | 4.8e-07 | 2.9e-07 | 2.8e-07 | 3.500E-07 | 1.127E-07 | 2.800E-07 | 4.800E-07 |
| 29411 | 1471 | Q1471L | E | 2.9e-07 | 3.8e-07 | 2.6e-07 | 3.100E-07 | 6.245E-08 | 2.600E-07 | 3.800E-07 |
| 29412 | 1471 | Q1471K | E | 3e-07 | 3.1e-07 | 2.9e-07 | 3.000E-07 | 1.000E-08 | 2.900E-07 | 3.100E-07 |
| 29413 | 1471 | Q1471M | E | 7.1e-07 | 2.9e-07 | 6.7e-08 | 3.557E-07 | 3.265E-07 | 6.700E-08 | 7.100E-07 |
| 29414 | 1471 | Q1471F | E | 1.4e-07 | 2.2e-07 | 2.4e-07 | 2.000E-07 | 5.292E-08 | 1.400E-07 | 2.400E-07 |
| 29415 | 1471 | Q1471P | E | 3.4e-07 | 2.9e-07 | 1.2e-07 | 2.500E-07 | 1.153E-07 | 1.200E-07 | 3.400E-07 |
| 29416 | 1471 | Q1471S | E | 4.8e-07 | 6.5e-07 | 1.3e-07 | 4.200E-07 | 2.651E-07 | 1.300E-07 | 6.500E-07 |
| 29417 | 1471 | Q1471T | E | 2.1e-07 | 3.5e-07 | 2.4e-07 | 2.667E-07 | 7.371E-08 | 2.100E-07 | 3.500E-07 |
| 29418 | 1471 | Q1471W | E | 1.5e-07 | 4e-07 | 3.3e-07 | 2.933E-07 | 1.290E-07 | 1.500E-07 | 4.000E-07 |
| 29419 | 1471 | Q1471Y | E | 2.7e-07 | 8.5e-08 | 2.7e-07 | 2.083E-07 | 1.068E-07 | 8.500E-08 | 2.700E-07 |
| 29420 | 1471 | Q1471V | E | 1.4e-07 | 2.1e-07 | 5.6e-07 | 3.033E-07 | 2.250E-07 | 1.400E-07 | 5.600E-07 |
| 29421 | 1472 | T1472A | E | 1.7e-07 | 1.6e-07 | 3.5e-07 | 2.267E-07 | 1.069E-07 | 1.600E-07 | 3.500E-07 |
| 29422 | 1472 | T1472R | E | 3.5e-07 | 2.9e-07 | 3.2e-07 | 3.200E-07 | 3.000E-08 | 2.900E-07 | 3.500E-07 |
| 29423 | 1472 | T1472N | E | 1.2e-07 | 3.1e-07 | 1e-07 | 1.767E-07 | 1.159E-07 | 1.000E-07 | 3.100E-07 |
| 29424 | 1472 | T1472D | E | 2.4e-07 | 2.7e-07 | 1.9e-07 | 2.333E-07 | 4.041E-08 | 1.900E-07 | 2.700E-07 |
| 29425 | 1472 | T1472C | E | 1.1e-07 | 1.7e-07 | 3.2e-07 | 2.000E-07 | 1.082E-07 | 1.100E-07 | 3.200E-07 |
| 29426 | 1472 | T1472E | E | 4.8e-07 | 3.1e-07 | 3.4e-07 | 3.767E-07 | 9.074E-08 | 3.100E-07 | 4.800E-07 |
| 29427 | 1472 | T1472Q | E | 2.4e-07 | 2.2e-07 | 4.7e-07 | 3.100E-07 | 1.389E-07 | 2.200E-07 | 4.700E-07 |
| 29428 | 1472 | T1472G | E | 2.6e-07 | 4.2e-07 | 4e-07 | 3.600E-07 | 8.718E-08 | 2.600E-07 | 4.200E-07 |
| 29429 | 1472 | T1472H | E | 1.7e-07 | 2.4e-07 | 2e-07 | 2.033E-07 | 3.512E-08 | 1.700E-07 | 2.400E-07 |
| 29430 | 1472 | T1472I | E | 4e-07 | 2.4e-07 | 5.9e-07 | 4.100E-07 | 1.752E-07 | 2.400E-07 | 5.900E-07 |
| 29431 | 1472 | T1472L | E | 1.3e-07 | 2.2e-07 | 2.9e-07 | 2.133E-07 | 8.021E-08 | 1.300E-07 | 2.900E-07 |
| 29432 | 1472 | T1472K | E | 1.9e-07 | 2.2e-07 | 4.2e-07 | 2.767E-07 | 1.250E-07 | 1.900E-07 | 4.200E-07 |
| 29433 | 1472 | T1472M | E | 2.7e-07 | 5.7e-07 | 3.1e-07 | 3.833E-07 | 1.629E-07 | 2.700E-07 | 5.700E-07 |
| 29434 | 1472 | T1472F | E | 4e-07 | 1.9e-07 | 1.3e-07 | 2.400E-07 | 1.418E-07 | 1.300E-07 | 4.000E-07 |
| 29435 | 1472 | T1472P | E | 3.3e-07 | 2.6e-07 | 1e-07 | 2.300E-07 | 1.179E-07 | 1.000E-07 | 3.300E-07 |
| 29436 | 1472 | T1472S | E | 3.1e-07 | 3.1e-07 | 2.8e-07 | 3.000E-07 | 1.732E-08 | 2.800E-07 | 3.100E-07 |
| 29437 | 1472 | T1472T | E | 5.7e-07 | 1.1e-07 | 4.7e-07 | 3.833E-07 | 2.419E-07 | 1.100E-07 | 5.700E-07 |
| 29438 | 1472 | T1472W | E | 3.8e-07 | 2.9e-07 | 1.3e-07 | 2.667E-07 | 1.266E-07 | 1.300E-07 | 3.800E-07 |
| 29439 | 1472 | T1472Y | E | 2.9e-07 | 3e-07 | 3.7e-07 | 3.200E-07 | 4.359E-08 | 2.900E-07 | 3.700E-07 |
| 29440 | 1472 | T1472V | E | 6.2e-07 | 4.1e-07 | 2.3e-07 | 4.200E-07 | 1.952E-07 | 2.300E-07 | 6.200E-07 |
| 29441 | 1473 | Y1473A | E | 4.5e-07 | 2.6e-07 | 3.4e-07 | 3.500E-07 | 9.539E-08 | 2.600E-07 | 4.500E-07 |
| 29442 | 1473 | Y1473R | E | 4.5e-07 | 1.7e-07 | 2.1e-07 | 2.767E-07 | 1.514E-07 | 1.700E-07 | 4.500E-07 |
| 29443 | 1473 | Y1473N | E | 2.6e-07 | 1.5e-07 | 2.4e-07 | 2.167E-07 | 5.859E-08 | 1.500E-07 | 2.600E-07 |
| 29444 | 1473 | Y1473D | E | 1.9e-07 | 2.3e-07 | 1.6e-07 | 1.933E-07 | 3.512E-08 | 1.600E-07 | 2.300E-07 |
| 29445 | 1473 | Y1473C | E | 3.1e-07 | 2.8e-07 | 1.8e-07 | 2.567E-07 | 6.807E-08 | 1.800E-07 | 3.100E-07 |
| 29446 | 1473 | Y1473E | E | 1.6e-07 | 2.6e-07 | 3.6e-07 | 2.600E-07 | 1.000E-07 | 1.600E-07 | 3.600E-07 |
| 29447 | 1473 | Y1473Q | E | 1.9e-07 | 1.9e-07 | 2.1e-07 | 1.967E-07 | 1.155E-08 | 1.900E-07 | 2.100E-07 |
| 29448 | 1473 | Y1473G | E | 3.9e-07 | 2.4e-07 | 2e-07 | 2.767E-07 | 1.002E-07 | 2.000E-07 | 3.900E-07 |
| 29449 | 1473 | Y1473H | E | 6.7e-07 | 3.1e-07 | 3.4e-07 | 4.400E-07 | 1.997E-07 | 3.100E-07 | 6.700E-07 |
| 29450 | 1473 | Y1473I | E | 1.3e-07 | 2.2e-07 | 2.2e-07 | 1.900E-07 | 5.196E-08 | 1.300E-07 | 2.200E-07 |
| 29451 | 1473 | Y1473L | E | 1.8e-07 | 1.7e-07 | 5.7e-07 | 3.067E-07 | 2.281E-07 | 1.700E-07 | 5.700E-07 |
| 29452 | 1473 | Y1473K | E | 2.9e-07 | 4.2e-07 | 2.1e-07 | 3.067E-07 | 1.060E-07 | 2.100E-07 | 4.200E-07 |
| 29453 | 1473 | Y1473M | E | 5.8e-07 | 2.5e-07 | 3.8e-07 | 4.033E-07 | 1.662E-07 | 2.500E-07 | 5.800E-07 |
| 29454 | 1473 | Y1473F | E | 1.7e-07 | 2.5e-07 | 1.3e-07 | 1.833E-07 | 6.110E-08 | 1.300E-07 | 2.500E-07 |
| 29455 | 1473 | Y1473P | E | 3.1e-07 | 2.4e-07 | 2.7e-07 | 2.733E-07 | 3.512E-08 | 2.400E-07 | 3.100E-07 |
| 29456 | 1473 | Y1473S | E | 2.9e-07 | 1.8e-07 | 2.2e-07 | 2.300E-07 | 5.568E-08 | 1.800E-07 | 2.900E-07 |
| 29457 | 1473 | Y1473T | E | 2e-07 | 3.5e-07 | 1.8e-07 | 2.433E-07 | 9.292E-08 | 1.800E-07 | 3.500E-07 |
| 29458 | 1473 | Y1473W | E | 2e-07 | 4e-07 | 2.3e-07 | 2.767E-07 | 1.079E-07 | 2.000E-07 | 4.000E-07 |
| 29459 | 1473 | Y1473Y | E | 4e-07 | 3.8e-07 | 3.1e-07 | 3.633E-07 | 4.726E-08 | 3.100E-07 | 4.000E-07 |
| 29460 | 1473 | Y1473V | E | 9.9e-08 | 4e-07 | 2.2e-07 | 2.397E-07 | 1.515E-07 | 9.900E-08 | 4.000E-07 |
| 29461 | 1474 | I1474A | E | 2.9e-07 | 3.2e-07 | 1.5e-07 | 2.533E-07 | 9.074E-08 | 1.500E-07 | 3.200E-07 |
| 29462 | 1474 | I1474R | E | 1.3e-07 | 1.4e-07 | 2.9e-07 | 1.867E-07 | 8.963E-08 | 1.300E-07 | 2.900E-07 |
| 29463 | 1474 | I1474N | E | 2.7e-07 | 3.9e-07 | 2.4e-07 | 3.000E-07 | 7.937E-08 | 2.400E-07 | 3.900E-07 |
| 29464 | 1474 | I1474D | E | 1.8e-07 | 2e-07 | 2.2e-07 | 2.000E-07 | 2.000E-08 | 1.800E-07 | 2.200E-07 |
| 29465 | 1474 | I1474C | E | 1.8e-07 | 2.8e-07 | 1.3e-07 | 1.967E-07 | 7.638E-08 | 1.300E-07 | 2.800E-07 |
| 29466 | 1474 | I1474E | E | 1.5e-07 | 4.2e-07 | 1.1e-07 | 2.267E-07 | 1.686E-07 | 1.100E-07 | 4.200E-07 |
| 29467 | 1474 | I1474Q | E | 1.7e-07 | 2.2e-07 | 3.1e-07 | 2.333E-07 | 7.095E-08 | 1.700E-07 | 3.100E-07 |
| 29468 | 1474 | I1474G | E | 1.5e-07 | 3e-07 | 1.9e-07 | 2.133E-07 | 7.767E-08 | 1.500E-07 | 3.000E-07 |
| 29469 | 1474 | I1474H | E | 2.4e-07 | 2.1e-07 | 4.9e-07 | 3.133E-07 | 1.537E-07 | 2.100E-07 | 4.900E-07 |
| 29470 | 1474 | I1474I | E | 4.3e-07 | 3.1e-07 | 1.6e-07 | 3.000E-07 | 1.353E-07 | 1.600E-07 | 4.300E-07 |
| 29471 | 1474 | I1474L | E | 2.8e-07 | 4.9e-07 | 1.5e-07 | 3.067E-07 | 1.716E-07 | 1.500E-07 | 4.900E-07 |
| 29472 | 1474 | I1474K | E | 1.8e-07 | 3.6e-07 | 2e-07 | 2.467E-07 | 9.866E-08 | 1.800E-07 | 3.600E-07 |
| 29473 | 1474 | I1474M | E | 2e-07 | 2e-07 | 3.6e-07 | 2.533E-07 | 9.238E-08 | 2.000E-07 | 3.600E-07 |
| 29474 | 1474 | I1474F | E | 1.2e-07 | 2.6e-07 | 1.3e-07 | 1.700E-07 | 7.810E-08 | 1.200E-07 | 2.600E-07 |
| 29475 | 1474 | I1474P | E | 4.4e-07 | 1.7e-07 | 3.7e-07 | 3.267E-07 | 1.401E-07 | 1.700E-07 | 4.400E-07 |
| 29476 | 1474 | I1474S | E | 4.7e-07 | 2.1e-07 | 3.2e-07 | 3.333E-07 | 1.305E-07 | 2.100E-07 | 4.700E-07 |
| 29477 | 1474 | I1474T | E | 1.8e-07 | 2.6e-07 | 2.3e-07 | 2.233E-07 | 4.041E-08 | 1.800E-07 | 2.600E-07 |
| 29478 | 1474 | I1474W | E | 1.5e-07 | 2.4e-07 | 2.6e-07 | 2.167E-07 | 5.859E-08 | 1.500E-07 | 2.600E-07 |
| 29479 | 1474 | I1474Y | E | 2.8e-07 | 1.2e-07 | 8.4e-07 | 4.133E-07 | 3.781E-07 | 1.200E-07 | 8.400E-07 |
| 29480 | 1474 | I1474V | E | 5.2e-07 | 3.3e-07 | 8.2e-08 | 3.107E-07 | 2.196E-07 | 8.200E-08 | 5.200E-07 |
| 29481 | 1475 | C1475A | E | 1.4e-07 | 3.9e-07 | 2.8e-07 | 2.700E-07 | 1.253E-07 | 1.400E-07 | 3.900E-07 |
| 29482 | 1475 | C1475R | E | 2.3e-07 | 3.8e-07 | 2.3e-07 | 2.800E-07 | 8.660E-08 | 2.300E-07 | 3.800E-07 |
| 29483 | 1475 | C1475N | E | 9.7e-08 | 1.8e-07 | 1.3e-07 | 1.357E-07 | 4.179E-08 | 9.700E-08 | 1.800E-07 |
| 29484 | 1475 | C1475D | E | 2.5e-07 | 1.3e-07 | 2.3e-07 | 2.033E-07 | 6.429E-08 | 1.300E-07 | 2.500E-07 |
| 29485 | 1475 | C1475C | E | 2.1e-07 | 2.1e-07 | 2.9e-07 | 2.367E-07 | 4.619E-08 | 2.100E-07 | 2.900E-07 |
| 29486 | 1475 | C1475E | E | 4.4e-07 | 3.1e-07 | 3.1e-07 | 3.533E-07 | 7.506E-08 | 3.100E-07 | 4.400E-07 |
| 29487 | 1475 | C1475Q | E | 4.4e-07 | 3.1e-07 | 8.6e-08 | 2.787E-07 | 1.791E-07 | 8.600E-08 | 4.400E-07 |
| 29488 | 1475 | C1475G | E | 2.8e-07 | 2.4e-07 | 1.1e-07 | 2.100E-07 | 8.888E-08 | 1.100E-07 | 2.800E-07 |
| 29489 | 1475 | C1475H | E | 1.4e-07 | 7.2e-08 | 1.4e-07 | 1.173E-07 | 3.926E-08 | 7.200E-08 | 1.400E-07 |
| 29490 | 1475 | C1475I | E | 3.8e-07 | 2.4e-07 | 5.5e-07 | 3.900E-07 | 1.552E-07 | 2.400E-07 | 5.500E-07 |
| 29491 | 1475 | C1475L | E | 1.3e-07 | 2.4e-07 | 1.6e-07 | 1.767E-07 | 5.686E-08 | 1.300E-07 | 2.400E-07 |
| 29492 | 1475 | C1475K | E | 2.6e-07 | 2.7e-07 | 1.5e-07 | 2.267E-07 | 6.658E-08 | 1.500E-07 | 2.700E-07 |
| 29493 | 1475 | C1475M | E | 2.5e-07 | 4.5e-07 | 3.2e-07 | 3.400E-07 | 1.015E-07 | 2.500E-07 | 4.500E-07 |
| 29494 | 1475 | C1475F | E | 9.9e-08 | 2e-07 | 2.5e-07 | 1.830E-07 | 7.692E-08 | 9.900E-08 | 2.500E-07 |

| | | | | | | | | | | |
|---|---|---|---|---|---|---|---|---|---|---|
| 29495 | 1475 | C1475P | E | 3.3e-07 | 3.7e-07 | 2.8e-07 | 3.267E-07 | 4.509E-08 | 2.800E-07 | 3.700E-07 |
| 29496 | 1475 | C1475S | E | 1.7e-07 | 2.9e-07 | 4e-07 | 2.867E-07 | 1.150E-07 | 1.700E-07 | 4.000E-07 |
| 29497 | 1475 | C1475T | E | 4.3e-07 | 3.3e-07 | 1.8e-07 | 3.133E-07 | 1.258E-07 | 1.800E-07 | 4.300E-07 |
| 29498 | 1475 | C1475W | E | 1.7e-07 | 3.5e-07 | 3.3e-07 | 2.833E-07 | 9.866E-08 | 1.700E-07 | 3.500E-07 |
| 29499 | 1475 | C1475Y | E | 2.4e-07 | 2.5e-07 | 4.7e-07 | 3.200E-07 | 1.300E-07 | 2.400E-07 | 4.700E-07 |
| 29500 | 1475 | C1475V | E | 3.4e-07 | 3.4e-07 | 4.1e-07 | 3.633E-07 | 4.041E-08 | 3.400E-07 | 4.100E-07 |
| 29501 | 1476 | N1476A | E | 2.6e-07 | 3.6e-07 | 3.3e-07 | 3.167E-07 | 5.132E-08 | 2.600E-07 | 3.600E-07 |
| 29502 | 1476 | N1476R | E | 3.8e-07 | 1.9e-07 | 4.2e-07 | 3.300E-07 | 1.229E-07 | 1.900E-07 | 4.200E-07 |
| 29503 | 1476 | N1476N | E | 2.3e-07 | 3e-07 | 2.6e-07 | 2.633E-07 | 3.512E-08 | 2.300E-07 | 3.000E-07 |
| 29504 | 1476 | N1476D | E | 1.1e-07 | 4e-07 | 1.6e-07 | 2.233E-07 | 1.550E-07 | 1.100E-07 | 4.000E-07 |
| 29505 | 1476 | N1476C | E | 5.2e-07 | 3.2e-07 | 3e-07 | 3.800E-07 | 1.217E-07 | 3.000E-07 | 5.200E-07 |
| 29506 | 1476 | N1476E | E | 1.9e-07 | 4.4e-07 | 2.3e-07 | 2.867E-07 | 1.343E-07 | 1.900E-07 | 4.400E-07 |
| 29507 | 1476 | N1476Q | E | 3.1e-07 | 2.4e-07 | 1.2e-07 | 2.233E-07 | 9.609E-08 | 1.200E-07 | 3.100E-07 |
| 29508 | 1476 | N1476G | E | 2.7e-07 | 3.4e-07 | 1.6e-07 | 2.567E-07 | 9.074E-08 | 1.600E-07 | 3.400E-07 |
| 29509 | 1476 | N1476H | E | 6.2e-07 | 1.5e-07 | 2.8e-07 | 3.500E-07 | 2.427E-07 | 1.500E-07 | 6.200E-07 |
| 29510 | 1476 | N1476I | E | 3.9e-07 | 1.4e-07 | 1.8e-07 | 2.367E-07 | 1.343E-07 | 1.400E-07 | 3.900E-07 |
| 29511 | 1476 | N1476L | E | 2.1e-07 | 2.4e-07 | 4.4e-07 | 2.967E-07 | 1.250E-07 | 2.100E-07 | 4.400E-07 |
| 29512 | 1476 | N1476K | E | 4e-07 | 2e-07 | 2.8e-07 | 2.933E-07 | 1.007E-07 | 2.000E-07 | 4.000E-07 |
| 29513 | 1476 | N1476M | E | 3.8e-07 | 1.5e-07 | 6.3e-07 | 3.867E-07 | 2.401E-07 | 1.500E-07 | 6.300E-07 |
| 29514 | 1476 | N1476F | E | 3.6e-07 | 2.1e-07 | 3e-07 | 2.900E-07 | 7.550E-08 | 2.100E-07 | 3.600E-07 |
| 29515 | 1476 | N1476P | E | 2.6e-07 | 3.1e-07 | 2.3e-07 | 2.667E-07 | 4.041E-08 | 2.300E-07 | 3.100E-07 |
| 29516 | 1476 | N1476S | E | 3.2e-07 | 2.2e-07 | 1.8e-07 | 2.400E-07 | 7.211E-08 | 1.800E-07 | 3.200E-07 |
| 29517 | 1476 | N1476T | E | 4.8e-07 | 3.5e-07 | 2.3e-07 | 3.533E-07 | 1.250E-07 | 2.300E-07 | 4.800E-07 |
| 29518 | 1476 | N1476W | E | 1.8e-07 | 3.6e-07 | 1.8e-07 | 2.400E-07 | 1.039E-07 | 1.800E-07 | 3.600E-07 |
| 29519 | 1476 | N1476Y | E | 8.3e-07 | 2.4e-07 | 1.4e-07 | 4.033E-07 | 3.729E-07 | 1.400E-07 | 8.300E-07 |
| 29520 | 1476 | N1476V | E | 2e-07 | 5.7e-07 | 9.7e-08 | 2.890E-07 | 2.487E-07 | 9.700E-08 | 5.700E-07 |
| 29521 | 1477 | V1477A | E | 1.7e-07 | 5.7e-07 | 4.1e-07 | 3.833E-07 | 2.013E-07 | 1.700E-07 | 5.700E-07 |
| 29522 | 1477 | V1477R | E | 2.2e-07 | 2.1e-07 | 3.5e-07 | 2.600E-07 | 7.810E-08 | 2.100E-07 | 3.500E-07 |
| 29523 | 1477 | V1477N | E | 2e-07 | 1.4e-07 | 3e-07 | 2.133E-07 | 8.083E-08 | 1.400E-07 | 3.000E-07 |
| 29524 | 1477 | V1477D | E | 6.7e-07 | 3.5e-07 | 1.2e-07 | 3.800E-07 | 2.762E-07 | 1.200E-07 | 6.700E-07 |
| 29525 | 1477 | V1477C | E | 3.1e-07 | 2.4e-07 | 2e-07 | 2.500E-07 | 5.568E-08 | 2.000E-07 | 3.100E-07 |
| 29526 | 1477 | V1477E | E | 1.2e-07 | 1.7e-07 | 2.5e-07 | 1.800E-07 | 6.557E-08 | 1.200E-07 | 2.500E-07 |
| 29527 | 1477 | V1477Q | E | 1.6e-07 | 2.2e-07 | 4.2e-07 | 2.667E-07 | 1.361E-07 | 1.600E-07 | 4.200E-07 |
| 29528 | 1477 | V1477G | E | 1.7e-07 | 1.4e-07 | 4.1e-07 | 2.400E-07 | 1.480E-07 | 1.400E-07 | 4.100E-07 |
| 29529 | 1477 | V1477H | E | 1.6e-07 | 3.5e-07 | 1.7e-07 | 2.267E-07 | 1.069E-07 | 1.600E-07 | 3.500E-07 |
| 29530 | 1477 | V1477I | E | 4.4e-07 | 3.8e-07 | 2.2e-07 | 3.467E-07 | 1.137E-07 | 2.200E-07 | 4.400E-07 |
| 29531 | 1477 | V1477L | E | 1.9e-07 | 3.3e-07 | 2.8e-07 | 2.667E-07 | 7.095E-08 | 1.900E-07 | 3.300E-07 |
| 29532 | 1477 | V1477K | E | 3.3e-07 | 1.8e-07 | 2e-07 | 2.367E-07 | 8.145E-08 | 1.800E-07 | 3.300E-07 |
| 29533 | 1477 | V1477M | E | 2.2e-07 | 3e-07 | 3.9e-07 | 3.033E-07 | 8.505E-08 | 2.200E-07 | 3.900E-07 |
| 29534 | 1477 | V1477F | E | 1.7e-07 | 1.9e-07 | 3.4e-07 | 2.333E-07 | 9.292E-08 | 1.700E-07 | 3.400E-07 |
| 29535 | 1477 | V1477P | E | 2.6e-07 | 3.6e-07 | 2.2e-07 | 2.800E-07 | 7.211E-08 | 2.200E-07 | 3.600E-07 |
| 29536 | 1477 | V1477S | E | 8.2e-07 | 5.1e-07 | 2.3e-07 | 5.200E-07 | 2.951E-07 | 2.300E-07 | 8.200E-07 |
| 29537 | 1477 | V1477T | E | 1.5e-07 | 5.2e-07 | 4.9e-07 | 3.867E-07 | 2.055E-07 | 1.500E-07 | 5.200E-07 |
| 29538 | 1477 | V1477W | E | 2.4e-07 | 2.7e-07 | 3.2e-07 | 2.767E-07 | 4.041E-08 | 2.400E-07 | 3.200E-07 |
| 29539 | 1477 | V1477Y | E | 2.3e-07 | 2.4e-07 | 3e-07 | 2.567E-07 | 3.786E-08 | 2.300E-07 | 3.000E-07 |
| 29540 | 1477 | V1477V | E | 2.6e-07 | 1.3e-07 | 3.1e-07 | 2.333E-07 | 9.292E-08 | 1.300E-07 | 3.100E-07 |
| 29541 | 1478 | N1478A | E | 1.6e-07 | 3.6e-07 | 9.2e-07 | 4.800E-07 | 3.940E-07 | 1.600E-07 | 9.200E-07 |
| 29542 | 1478 | N1478R | E | 1.6e-07 | 2.5e-07 | 2.3e-07 | 2.133E-07 | 4.726E-08 | 1.600E-07 | 2.500E-07 |
| 29543 | 1478 | N1478N | E | 3.1e-07 | 2.1e-07 | 3.7e-07 | 2.967E-07 | 8.083E-08 | 2.100E-07 | 3.700E-07 |
| 29544 | 1478 | N1478D | E | 1.7e-07 | 1.9e-07 | 2e-07 | 1.867E-07 | 1.528E-08 | 1.700E-07 | 2.000E-07 |
| 29545 | 1478 | N1478C | E | 3.6e-07 | 2.8e-07 | 3.9e-07 | 3.433E-07 | 5.686E-08 | 2.800E-07 | 3.900E-07 |
| 29546 | 1478 | N1478E | E | 2e-07 | 2.1e-07 | 2.2e-07 | 2.100E-07 | 1.000E-08 | 2.000E-07 | 2.200E-07 |
| 29547 | 1478 | N1478Q | E | 2.4e-07 | 2.4e-07 | 2.2e-07 | 2.333E-07 | 1.155E-08 | 2.200E-07 | 2.400E-07 |
| 29548 | 1478 | N1478G | E | 1.9e-07 | 1.9e-07 | 2.1e-07 | 1.967E-07 | 1.155E-08 | 1.900E-07 | 2.100E-07 |
| 29549 | 1478 | N1478H | E | 2.3e-07 | 3.1e-07 | 1.6e-07 | 2.333E-07 | 7.506E-08 | 1.600E-07 | 3.100E-07 |
| 29550 | 1478 | N1478I | E | 2.6e-07 | 1.7e-07 | 2.7e-07 | 2.333E-07 | 5.508E-08 | 1.700E-07 | 2.700E-07 |
| 29551 | 1478 | N1478L | E | 4.9e-07 | 2.9e-07 | 4.5e-07 | 4.100E-07 | 1.058E-07 | 2.900E-07 | 4.900E-07 |
| 29552 | 1478 | N1478K | E | 1.8e-07 | 3e-07 | 4.4e-07 | 3.067E-07 | 1.301E-07 | 1.800E-07 | 4.400E-07 |
| 29553 | 1478 | N1478M | E | 1.6e-07 | 3.9e-07 | 4.6e-07 | 3.367E-07 | 1.570E-07 | 1.600E-07 | 4.600E-07 |
| 29554 | 1478 | N1478F | E | 4.5e-07 | 1.3e-07 | 2.9e-07 | 2.900E-07 | 1.600E-07 | 1.300E-07 | 4.500E-07 |
| 29555 | 1478 | N1478P | E | 1.4e-07 | 3.5e-07 | 1.8e-07 | 2.233E-07 | 1.115E-07 | 1.400E-07 | 3.500E-07 |
| 29556 | 1478 | N1478S | E | 4.8e-07 | 2.9e-07 | 3.4e-07 | 3.700E-07 | 9.849E-08 | 2.900E-07 | 4.800E-07 |
| 29557 | 1478 | N1478T | E | 3.8e-07 | 1.2e-07 | 2.2e-07 | 2.400E-07 | 1.311E-07 | 1.200E-07 | 3.800E-07 |
| 29558 | 1478 | N1478W | E | 4.4e-07 | 4e-07 | 1.3e-07 | 3.233E-07 | 1.686E-07 | 1.300E-07 | 4.400E-07 |
| 29559 | 1478 | N1478Y | E | 7.7e-07 | 2.7e-07 | 1.7e-07 | 4.033E-07 | 3.215E-07 | 1.700E-07 | 7.700E-07 |
| 29560 | 1478 | N1478V | E | 4.9e-07 | 4.4e-07 | 1.6e-07 | 3.633E-07 | 1.779E-07 | 1.600E-07 | 4.900E-07 |
| 29561 | 1479 | H1479A | E | 3.7e-07 | 1.7e-07 | 1.1e-07 | 2.167E-07 | 1.361E-07 | 1.100E-07 | 3.700E-07 |
| 29562 | 1479 | H1479R | E | 3.8e-07 | 4.9e-07 | 6.4e-07 | 5.033E-07 | 1.305E-07 | 3.800E-07 | 6.400E-07 |
| 29563 | 1479 | H1479N | E | 2.3e-07 | 3.5e-07 | 2.2e-07 | 2.667E-07 | 7.234E-08 | 2.200E-07 | 3.500E-07 |
| 29564 | 1479 | H1479D | E | 1.5e-07 | 1.9e-07 | 3.3e-07 | 2.233E-07 | 9.452E-08 | 1.500E-07 | 3.300E-07 |
| 29565 | 1479 | H1479C | E | 1.6e-07 | 1.2e-07 | 1.7e-07 | 1.500E-07 | 2.646E-08 | 1.200E-07 | 1.700E-07 |
| 29566 | 1479 | H1479E | E | 1.4e-07 | 2e-07 | 1.8e-07 | 1.733E-07 | 3.055E-08 | 1.400E-07 | 2.000E-07 |
| 29567 | 1479 | H1479Q | E | 1.6e-07 | 2.1e-07 | 1.7e-07 | 1.800E-07 | 2.646E-08 | 1.600E-07 | 2.100E-07 |
| 29568 | 1479 | H1479G | E | 3.6e-07 | 1.6e-07 | 2.5e-07 | 2.567E-07 | 1.002E-07 | 1.600E-07 | 3.600E-07 |
| 29569 | 1479 | H1479H | E | 1.5e-07 | 1.8e-07 | 2.3e-07 | 1.867E-07 | 4.041E-08 | 1.500E-07 | 2.300E-07 |
| 29570 | 1479 | H1479I | E | 5.9e-08 | 2.1e-07 | 1.6e-07 | 1.430E-07 | 7.692E-08 | 5.900E-08 | 2.100E-07 |
| 29571 | 1479 | H1479L | E | 3.7e-07 | 1.4e-07 | 1.9e-07 | 2.333E-07 | 1.210E-07 | 1.400E-07 | 3.700E-07 |
| 29572 | 1479 | H1479K | E | 1.8e-07 | 8.8e-08 | 3.6e-07 | 2.093E-07 | 1.384E-07 | 8.800E-08 | 3.600E-07 |
| 29573 | 1479 | H1479M | E | 2.4e-07 | 3.4e-07 | 3.2e-07 | 3.000E-07 | 5.292E-08 | 2.400E-07 | 3.400E-07 |
| 29574 | 1479 | H1479F | E | 2.6e-07 | 3.2e-07 | 7.7e-08 | 2.190E-07 | 1.266E-07 | 7.700E-08 | 3.200E-07 |
| 29575 | 1479 | H1479P | E | 2.6e-07 | 3.1e-07 | 8.2e-08 | 2.173E-07 | 1.198E-07 | 8.200E-08 | 3.100E-07 |
| 29576 | 1479 | H1479S | E | 2e-07 | 2.9e-07 | 2e-07 | 2.300E-07 | 5.196E-08 | 2.000E-07 | 2.900E-07 |
| 29577 | 1479 | H1479T | E | 2.9e-07 | 2e-07 | 1.8e-07 | 2.233E-07 | 5.859E-08 | 1.800E-07 | 2.900E-07 |
| 29578 | 1479 | H1479W | E | 1.7e-07 | 1.3e-07 | 2.5e-07 | 1.833E-07 | 6.110E-08 | 1.300E-07 | 2.500E-07 |
| 29579 | 1479 | H1479Y | E | 2.8e-07 | 1e-07 | 2.6e-07 | 2.133E-07 | 9.866E-08 | 1.000E-07 | 2.800E-07 |

| | | | | | | | | | | |
|---|---|---|---|---|---|---|---|---|---|---|
| 29580 | 1479 | H1479V | E | 3.2e-07 | 2.8e-07 | 4.7e-07 | 3.567E-07 | 1.002E-07 | 2.800E-07 | 4.700E-07 |
| 29581 | 1480 | K1480A | E | 2.7e-07 | 2.1e-07 | 4.2e-07 | 3.000E-07 | 1.082E-07 | 2.100E-07 | 4.200E-07 |
| 29582 | 1480 | K1480R | E | 3.1e-07 | 2.8e-07 | 2.7e-07 | 2.867E-07 | 2.082E-08 | 2.700E-07 | 3.100E-07 |
| 29583 | 1480 | K1480N | E | 3.5e-07 | 2.3e-07 | 2.1e-07 | 2.633E-07 | 7.572E-08 | 2.100E-07 | 3.500E-07 |
| 29584 | 1480 | K1480D | E | 1.8e-07 | 2.5e-07 | 3.5e-07 | 2.600E-07 | 8.544E-08 | 1.800E-07 | 3.500E-07 |
| 29585 | 1480 | K1480C | E | 4.3e-07 | 3.7e-07 | 3.1e-07 | 3.700E-07 | 6.000E-08 | 3.100E-07 | 4.300E-07 |
| 29586 | 1480 | K1480E | E | 1.9e-07 | 4.6e-07 | 3.7e-07 | 3.400E-07 | 1.375E-07 | 1.900E-07 | 4.600E-07 |
| 29587 | 1480 | K1480Q | E | 3.2e-07 | 1.9e-07 | 4.8e-07 | 3.300E-07 | 1.453E-07 | 1.900E-07 | 4.800E-07 |
| 29588 | 1480 | K1480G | E | 2.3e-07 | 3e-07 | 2.5e-07 | 2.600E-07 | 3.606E-08 | 2.300E-07 | 3.000E-07 |
| 29589 | 1480 | K1480H | E | 3e-07 | 3.1e-07 | 1.8e-07 | 2.633E-07 | 7.234E-08 | 1.800E-07 | 3.100E-07 |
| 29590 | 1480 | K1480I | E | 2.1e-07 | 3.2e-07 | 2.3e-07 | 2.533E-07 | 5.859E-08 | 2.100E-07 | 3.200E-07 |
| 29591 | 1480 | K1480L | E | 2.7e-07 | 4.2e-07 | 1.4e-07 | 2.767E-07 | 1.401E-07 | 1.400E-07 | 4.200E-07 |
| 29592 | 1480 | K1480K | E | 3e-07 | 2.5e-07 | 2.5e-07 | 2.667E-07 | 2.887E-08 | 2.500E-07 | 3.000E-07 |
| 29593 | 1480 | K1480M | E | 5.4e-07 | 1.4e-07 | 1.9e-07 | 2.900E-07 | 2.179E-07 | 1.400E-07 | 5.400E-07 |
| 29594 | 1480 | K1480F | E | 2.1e-07 | 2.4e-07 | 1.6e-07 | 2.033E-07 | 4.041E-08 | 1.600E-07 | 2.400E-07 |
| 29595 | 1480 | K1480P | E | 3e-07 | 3e-07 | 9.3e-08 | 2.310E-07 | 1.195E-07 | 9.300E-08 | 3.000E-07 |
| 29596 | 1480 | K1480S | E | 2.2e-07 | 4.4e-07 | 1.6e-07 | 2.733E-07 | 1.474E-07 | 1.600E-07 | 4.400E-07 |
| 29597 | 1480 | K1480T | E | 1.8e-07 | 2.7e-07 | 2e-07 | 2.167E-07 | 4.726E-08 | 1.800E-07 | 2.700E-07 |
| 29598 | 1480 | K1480W | E | 4.2e-07 | 3.3e-07 | 2.3e-07 | 3.267E-07 | 9.504E-08 | 2.300E-07 | 4.200E-07 |
| 29599 | 1480 | K1480Y | E | 2.3e-07 | 2.4e-07 | 2.2e-07 | 2.300E-07 | 1.000E-08 | 2.200E-07 | 2.400E-07 |
| 29600 | 1480 | K1480V | E | 2.1e-07 | 2.4e-07 | 2.2e-07 | 2.233E-07 | 1.528E-08 | 2.100E-07 | 2.400E-07 |
| 29601 | 1481 | P1481A | E | 3.7e-07 | 1.8e-07 | 1.6e-07 | 2.367E-07 | 1.159E-07 | 1.600E-07 | 3.700E-07 |
| 29602 | 1481 | P1481R | E | 7.6e-07 | 2.7e-07 | 1.6e-07 | 3.967E-07 | 3.194E-07 | 1.600E-07 | 7.600E-07 |
| 29603 | 1481 | P1481N | E | 2.3e-07 | 1.7e-07 | 2.1e-07 | 2.033E-07 | 3.055E-08 | 1.700E-07 | 2.300E-07 |
| 29604 | 1481 | P1481D | E | 3e-07 | 3.5e-07 | 2.4e-07 | 2.967E-07 | 5.508E-08 | 2.400E-07 | 3.500E-07 |
| 29605 | 1481 | P1481C | E | 1.8e-07 | 3.2e-07 | 3.4e-07 | 2.800E-07 | 8.718E-08 | 1.800E-07 | 3.400E-07 |
| 29606 | 1481 | P1481E | E | 3.1e-07 | 3.3e-07 | 2.8e-07 | 3.067E-07 | 2.517E-08 | 2.800E-07 | 3.300E-07 |
| 29607 | 1481 | P1481Q | E | 2e-07 | 4e-07 | 5.2e-07 | 3.733E-07 | 1.617E-07 | 2.000E-07 | 5.200E-07 |
| 29608 | 1481 | P1481G | E | 2.7e-07 | 3.4e-07 | 4.7e-07 | 3.600E-07 | 1.015E-07 | 2.700E-07 | 4.700E-07 |
| 29609 | 1481 | P1481H | E | 2.3e-07 | 1.8e-07 | 7.7e-08 | 1.623E-07 | 7.801E-08 | 7.700E-08 | 2.300E-07 |
| 29610 | 1481 | P1481I | E | 2.8e-07 | 3.4e-07 | 1.4e-07 | 2.533E-07 | 1.026E-07 | 1.400E-07 | 3.400E-07 |
| 29611 | 1481 | P1481L | E | 2.8e-07 | 4.2e-07 | 2.1e-07 | 3.033E-07 | 1.069E-07 | 2.100E-07 | 4.200E-07 |
| 29612 | 1481 | P1481K | E | 2.3e-07 | 2.3e-07 | 2.5e-07 | 2.367E-07 | 1.155E-08 | 2.300E-07 | 2.500E-07 |
| 29613 | 1481 | P1481M | E | 2.3e-07 | 2.1e-07 | 1.9e-07 | 2.100E-07 | 2.000E-08 | 1.900E-07 | 2.300E-07 |
| 29614 | 1481 | P1481F | E | 4e-07 | 1.2e-07 | 1.6e-07 | 2.267E-07 | 1.514E-07 | 1.200E-07 | 4.000E-07 |
| 29615 | 1481 | P1481P | E | 3e-07 | 1.4e-07 | 5.7e-07 | 3.367E-07 | 2.173E-07 | 1.400E-07 | 5.700E-07 |
| 29616 | 1481 | P1481S | E | 2.3e-07 | 2.5e-07 | 1.5e-07 | 2.100E-07 | 5.292E-08 | 1.500E-07 | 2.500E-07 |
| 29617 | 1481 | P1481T | E | 2.6e-07 | 2e-07 | 3.3e-07 | 2.633E-07 | 6.506E-08 | 2.000E-07 | 3.300E-07 |
| 29618 | 1481 | P1481W | E | 1.7e-07 | 3.7e-07 | 2e-07 | 2.467E-07 | 1.079E-07 | 1.700E-07 | 3.700E-07 |
| 29619 | 1481 | P1481Y | E | 2.9e-07 | 2.1e-07 | 2.9e-07 | 2.633E-07 | 4.619E-08 | 2.100E-07 | 2.900E-07 |
| 29620 | 1481 | P1481V | E | 4e-07 | 2.2e-07 | 4.6e-07 | 3.600E-07 | 1.249E-07 | 2.200E-07 | 4.600E-07 |
| 29621 | 1482 | S1482A | E | 1.8e-07 | 2e-07 | 2.2e-07 | 2.000E-07 | 2.000E-08 | 1.800E-07 | 2.200E-07 |
| 29622 | 1482 | S1482R | E | 3.1e-07 | 3.9e-07 | 2e-07 | 3.000E-07 | 9.539E-08 | 2.000E-07 | 3.900E-07 |
| 29623 | 1482 | S1482N | E | 3.9e-07 | 3.5e-07 | 5.8e-07 | 4.400E-07 | 1.229E-07 | 3.500E-07 | 5.800E-07 |
| 29624 | 1482 | S1482D | E | 3.2e-07 | 1.7e-07 | 2.3e-07 | 2.400E-07 | 7.550E-08 | 1.700E-07 | 3.200E-07 |
| 29625 | 1482 | S1482C | E | 2.7e-07 | 3.8e-07 | 1.5e-07 | 2.667E-07 | 1.150E-07 | 1.500E-07 | 3.800E-07 |
| 29626 | 1482 | S1482E | E | 4e-07 | 3.4e-07 | 6.8e-07 | 4.733E-07 | 1.815E-07 | 3.400E-07 | 6.800E-07 |
| 29627 | 1482 | S1482Q | E | 2e-07 | 1.7e-07 | 1.2e-07 | 1.633E-07 | 4.041E-08 | 1.200E-07 | 2.000E-07 |
| 29628 | 1482 | S1482G | E | 3.3e-07 | 2.2e-07 | 1.3e-07 | 2.267E-07 | 1.002E-07 | 1.300E-07 | 3.300E-07 |
| 29629 | 1482 | S1482H | E | 6.7e-07 | 3.9e-07 | 2.5e-07 | 4.367E-07 | 2.139E-07 | 2.500E-07 | 6.700E-07 |
| 29630 | 1482 | S1482I | E | 2.4e-07 | 2.7e-07 | 4.3e-07 | 3.133E-07 | 1.021E-07 | 2.400E-07 | 4.300E-07 |
| 29631 | 1482 | S1482L | E | 2.2e-07 | 1.1e-07 | 2.3e-07 | 1.867E-07 | 6.658E-08 | 1.100E-07 | 2.300E-07 |
| 29632 | 1482 | S1482K | E | 2.5e-07 | 6e-07 | 2.4e-07 | 3.633E-07 | 2.050E-07 | 2.400E-07 | 6.000E-07 |
| 29633 | 1482 | S1482M | E | 1.2e-06 | 1.7e-07 | 1e-07 | 4.900E-07 | 6.159E-07 | 1.000E-07 | 1.200E-06 |
| 29634 | 1482 | S1482F | E | 3.5e-07 | 5.5e-07 | 4.3e-07 | 4.433E-07 | 1.007E-07 | 3.500E-07 | 5.500E-07 |
| 29635 | 1482 | S1482P | E | 3.2e-07 | 2.8e-07 | 2.9e-07 | 2.967E-07 | 2.082E-08 | 2.800E-07 | 3.200E-07 |
| 29636 | 1482 | S1482S | E | 1.8e-07 | 2.7e-07 | 3.3e-07 | 2.600E-07 | 7.550E-08 | 1.800E-07 | 3.300E-07 |
| 29637 | 1482 | S1482T | E | 2.6e-07 | 4.6e-07 | 1.7e-07 | 2.967E-07 | 1.484E-07 | 1.700E-07 | 4.600E-07 |
| 29638 | 1482 | S1482W | E | 3.4e-07 | 1e-07 | 1.8e-07 | 2.067E-07 | 1.222E-07 | 1.000E-07 | 3.400E-07 |
| 29639 | 1482 | S1482Y | E | 2.3e-07 | 3.4e-07 | 2.4e-07 | 2.700E-07 | 6.083E-08 | 2.300E-07 | 3.400E-07 |
| 29640 | 1482 | S1482V | E | 1.2e-07 | 3.1e-07 | 3e-07 | 2.433E-07 | 1.069E-07 | 1.200E-07 | 3.100E-07 |
| 29641 | 1483 | N1483A | E | 1.9e-07 | 4.6e-07 | 1.4e-07 | 2.633E-07 | 1.721E-07 | 1.400E-07 | 4.600E-07 |
| 29642 | 1483 | N1483R | E | 2.8e-07 | 2.1e-07 | 1.4e-07 | 2.100E-07 | 7.000E-08 | 1.400E-07 | 2.800E-07 |
| 29643 | 1483 | N1483N | E | 7.6e-07 | 1.6e-07 | 1.7e-07 | 3.633E-07 | 3.436E-07 | 1.600E-07 | 7.600E-07 |
| 29644 | 1483 | N1483D | E | 1.9e-07 | 2.7e-07 | 7e-08 | 1.767E-07 | 1.007E-07 | 7.000E-08 | 2.700E-07 |
| 29645 | 1483 | N1483C | E | 4.9e-07 | 2.2e-07 | 2.7e-07 | 3.267E-07 | 1.436E-07 | 2.200E-07 | 4.900E-07 |
| 29646 | 1483 | N1483E | E | 3.1e-07 | 2.7e-07 | 3.6e-07 | 3.133E-07 | 4.509E-08 | 2.700E-07 | 3.600E-07 |
| 29647 | 1483 | N1483Q | E | 2.6e-07 | 3.5e-07 | 5.5e-07 | 3.867E-07 | 1.484E-07 | 2.600E-07 | 5.500E-07 |
| 29648 | 1483 | N1483G | E | 3.3e-07 | 4.8e-07 | 6.9e-07 | 5.000E-07 | 1.808E-07 | 3.300E-07 | 6.900E-07 |
| 29649 | 1483 | N1483H | E | 1.7e-07 | 2.4e-07 | 2.6e-07 | 2.233E-07 | 4.726E-08 | 1.700E-07 | 2.600E-07 |
| 29650 | 1483 | N1483I | E | 1.6e-07 | 1.5e-07 | 1.4e-07 | 1.500E-07 | 1.000E-08 | 1.400E-07 | 1.600E-07 |
| 29651 | 1483 | N1483L | E | 2.4e-07 | 1.6e-07 | 2.8e-07 | 2.267E-07 | 6.110E-08 | 1.600E-07 | 2.800E-07 |
| 29652 | 1483 | N1483K | E | 2.8e-07 | 2.6e-07 | 3.1e-07 | 2.833E-07 | 2.517E-08 | 2.600E-07 | 3.100E-07 |
| 29653 | 1483 | N1483M | E | 2.4e-07 | 2.4e-07 | 3.1e-07 | 2.633E-07 | 4.041E-08 | 2.400E-07 | 3.100E-07 |
| 29654 | 1483 | N1483F | E | 1.2e-07 | 2.1e-07 | 3.8e-07 | 2.367E-07 | 1.320E-07 | 1.200E-07 | 3.800E-07 |
| 29655 | 1483 | N1483P | E | 4.5e-07 | 3.2e-07 | 1.6e-07 | 3.100E-07 | 1.453E-07 | 1.600E-07 | 4.500E-07 |
| 29656 | 1483 | N1483S | E | 1.8e-07 | 2.1e-07 | 2e-07 | 1.967E-07 | 1.528E-08 | 1.800E-07 | 2.100E-07 |
| 29657 | 1483 | N1483T | E | 2.5e-07 | 2.1e-07 | 1.2e-07 | 1.933E-07 | 6.658E-08 | 1.200E-07 | 2.500E-07 |
| 29658 | 1483 | N1483W | E | 2.3e-07 | 2.2e-07 | 9.3e-07 | 4.600E-07 | 4.071E-07 | 2.200E-07 | 9.300E-07 |
| 29659 | 1483 | N1483Y | E | 4.7e-07 | 3.5e-07 | 5.2e-07 | 4.467E-07 | 8.737E-08 | 3.500E-07 | 5.200E-07 |
| 29660 | 1483 | N1483V | E | 2.3e-07 | 2.2e-07 | 1.1e-07 | 1.867E-07 | 6.658E-08 | 1.100E-07 | 2.300E-07 |
| 29661 | 1484 | T1484A | E | 1.6e-07 | 2.1e-07 | 2e-07 | 1.900E-07 | 2.646E-08 | 1.600E-07 | 2.100E-07 |
| 29662 | 1484 | T1484R | E | 5.5e-07 | 2.3e-07 | 1.3e-07 | 3.033E-07 | 2.194E-07 | 1.300E-07 | 5.500E-07 |
| 29663 | 1484 | T1484N | E | 6.1e-07 | 1.1e-07 | 2e-07 | 3.067E-07 | 2.665E-07 | 1.100E-07 | 6.100E-07 |
| 29664 | 1484 | T1484D | E | 2.6e-07 | 1.8e-07 | 1.8e-07 | 2.067E-07 | 4.619E-08 | 1.800E-07 | 2.600E-07 |

| | | | | | | | | | | |
|---|---|---|---|---|---|---|---|---|---|---|
| 29665 | 1484 | T1484C | E | 1.5e-07 | 4.5e-07 | 2.6e-07 | 2.867E-07 | 1.518E-07 | 1.500E-07 | 4.500E-07 |
| 29666 | 1484 | T1484E | E | 1.8e-07 | 1.6e-07 | 2.6e-07 | 2.000E-07 | 5.292E-08 | 1.600E-07 | 2.600E-07 |
| 29667 | 1484 | T1484Q | E | 2.8e-07 | 4.3e-07 | 1.8e-07 | 2.967E-07 | 1.258E-07 | 1.800E-07 | 4.300E-07 |
| 29668 | 1484 | T1484G | E | 4.4e-07 | 2.5e-07 | 2.9e-07 | 3.267E-07 | 1.002E-07 | 2.500E-07 | 4.400E-07 |
| 29669 | 1484 | T1484H | E | 4.8e-07 | 7.1e-07 | 2.3e-07 | 4.733E-07 | 2.401E-07 | 2.300E-07 | 7.100E-07 |
| 29670 | 1484 | T1484I | E | 3.9e-07 | 3e-07 | 2.4e-07 | 3.100E-07 | 7.550E-08 | 2.400E-07 | 3.900E-07 |
| 29671 | 1484 | T1484L | E | 2.3e-07 | 2.4e-07 | 1.5e-07 | 2.067E-07 | 4.933E-08 | 1.500E-07 | 2.400E-07 |
| 29672 | 1484 | T1484K | E | 1.7e-07 | 3.3e-07 | 3.5e-07 | 2.833E-07 | 9.866E-08 | 1.700E-07 | 3.500E-07 |
| 29673 | 1484 | T1484M | E | 2.7e-07 | 3e-07 | 1.7e-07 | 2.467E-07 | 6.807E-08 | 1.700E-07 | 3.000E-07 |
| 29674 | 1484 | T1484F | E | 3.9e-07 | 3.4e-07 | 4.7e-07 | 4.000E-07 | 6.557E-08 | 3.400E-07 | 4.700E-07 |
| 29675 | 1484 | T1484P | E | 3.4e-07 | 2.2e-07 | 3.6e-07 | 3.067E-07 | 7.572E-08 | 2.200E-07 | 3.600E-07 |
| 29676 | 1484 | T1484S | E | 1.6e-07 | 2.8e-07 | 5.6e-07 | 3.333E-07 | 2.053E-07 | 1.600E-07 | 5.600E-07 |
| 29677 | 1484 | T1484T | E | 2.8e-07 | 2.8e-07 | 3.2e-07 | 2.933E-07 | 2.309E-08 | 2.800E-07 | 3.200E-07 |
| 29678 | 1484 | T1484W | E | 1.1e-07 | 2.6e-07 | 3.7e-07 | 2.467E-07 | 1.305E-07 | 1.100E-07 | 3.700E-07 |
| 29679 | 1484 | T1484Y | E | 5.3e-07 | 3.1e-07 | 5e-07 | 4.467E-07 | 1.193E-07 | 3.100E-07 | 5.300E-07 |
| 29680 | 1484 | T1484V | E | 2e-07 | 4.4e-07 | 1.2e-07 | 2.533E-07 | 1.665E-07 | 1.200E-07 | 4.400E-07 |
| 29681 | 1485 | K1485A | E | 2.9e-07 | 1.3e-07 | 3.3e-07 | 2.500E-07 | 1.058E-07 | 1.300E-07 | 3.300E-07 |
| 29682 | 1485 | K1485R | E | 1.8e-07 | 4.9e-07 | 2.3e-07 | 3.000E-07 | 1.664E-07 | 1.800E-07 | 4.900E-07 |
| 29683 | 1485 | K1485N | E | 2.7e-07 | 3.8e-07 | 1.9e-07 | 2.800E-07 | 9.539E-08 | 1.900E-07 | 3.800E-07 |
| 29684 | 1485 | K1485D | E | 6.1e-07 | 1.4e-07 | 2.3e-07 | 3.267E-07 | 2.495E-07 | 1.400E-07 | 6.100E-07 |
| 29685 | 1485 | K1485C | E | 1.6e-07 | 2.7e-07 | 2.1e-07 | 2.133E-07 | 5.508E-08 | 1.600E-07 | 2.700E-07 |
| 29686 | 1485 | K1485E | E | 4.7e-07 | 2.4e-07 | 2.6e-07 | 3.233E-07 | 1.274E-07 | 2.400E-07 | 4.700E-07 |
| 29687 | 1485 | K1485Q | E | 3.8e-07 | 4.4e-07 | 1.8e-07 | 3.333E-07 | 1.361E-07 | 1.800E-07 | 4.400E-07 |
| 29688 | 1485 | K1485G | E | 2.3e-07 | 3.5e-07 | 2.3e-07 | 2.700E-07 | 6.928E-08 | 2.300E-07 | 3.500E-07 |
| 29689 | 1485 | K1485H | E | 2e-07 | 3.4e-07 | 2.4e-07 | 2.600E-07 | 7.211E-08 | 2.000E-07 | 3.400E-07 |
| 29690 | 1485 | K1485I | E | 6.8e-07 | 2.1e-07 | 2.1e-07 | 3.667E-07 | 2.714E-07 | 2.100E-07 | 6.800E-07 |
| 29691 | 1485 | K1485L | E | 4e-07 | 1.9e-07 | 4.2e-07 | 3.367E-07 | 1.274E-07 | 1.900E-07 | 4.200E-07 |
| 29692 | 1485 | K1485K | E | 4e-07 | 1.2e-07 | 2.2e-07 | 2.467E-07 | 1.419E-07 | 1.200E-07 | 4.000E-07 |
| 29693 | 1485 | K1485M | E | 3.4e-07 | 1.4e-07 | 2.4e-07 | 2.400E-07 | 1.000E-07 | 1.400E-07 | 3.400E-07 |
| 29694 | 1485 | K1485F | E | 1.9e-07 | 2.4e-07 | 2.7e-07 | 2.333E-07 | 4.041E-08 | 1.900E-07 | 2.700E-07 |
| 29695 | 1485 | K1485P | E | 3.1e-07 | 2e-07 | 1.2e-07 | 2.100E-07 | 9.539E-08 | 1.200E-07 | 3.100E-07 |
| 29696 | 1485 | K1485S | E | 1.6e-07 | 2.2e-07 | 4.4e-07 | 2.733E-07 | 1.474E-07 | 1.600E-07 | 4.400E-07 |
| 29697 | 1485 | K1485T | E | 5e-07 | 3.1e-07 | 2.7e-07 | 3.600E-07 | 1.229E-07 | 2.700E-07 | 5.000E-07 |
| 29698 | 1485 | K1485W | E | 1.9e-07 | 1.8e-07 | 1.4e-07 | 1.700E-07 | 2.646E-08 | 1.400E-07 | 1.900E-07 |
| 29699 | 1485 | K1485Y | E | 1.1e-07 | 1.8e-07 | 3e-07 | 1.967E-07 | 9.609E-08 | 1.100E-07 | 3.000E-07 |
| 29700 | 1485 | K1485V | E | 2.1e-07 | 2.7e-07 | 5.7e-07 | 3.500E-07 | 1.929E-07 | 2.100E-07 | 5.700E-07 |
| 29701 | 1486 | V1486A | E | 9.8e-08 | 1.3e-07 | 3.1e-07 | 1.793E-07 | 1.143E-07 | 9.800E-08 | 3.100E-07 |
| 29702 | 1486 | V1486R | E | 1.9e-07 | 3.5e-07 | 1.9e-07 | 2.433E-07 | 9.238E-08 | 1.900E-07 | 3.500E-07 |
| 29703 | 1486 | V1486N | E | 1.2e-07 | 3.8e-07 | 2.7e-07 | 2.567E-07 | 1.305E-07 | 1.200E-07 | 3.800E-07 |
| 29704 | 1486 | V1486D | E | 2e-07 | 4.8e-07 | 3.4e-07 | 3.400E-07 | 1.400E-07 | 2.000E-07 | 4.800E-07 |
| 29705 | 1486 | V1486C | E | 4.3e-07 | 2.5e-07 | 2e-07 | 2.933E-07 | 1.210E-07 | 2.000E-07 | 4.300E-07 |
| 29706 | 1486 | V1486E | E | 2.8e-07 | 2.9e-07 | 2.3e-07 | 2.667E-07 | 3.215E-08 | 2.300E-07 | 2.900E-07 |
| 29707 | 1486 | V1486Q | E | 2.4e-07 | 3e-07 | 1.4e-07 | 2.267E-07 | 8.083E-08 | 1.400E-07 | 3.000E-07 |
| 29708 | 1486 | V1486G | E | 3.1e-07 | 3.5e-07 | 2.9e-07 | 3.167E-07 | 3.055E-08 | 2.900E-07 | 3.500E-07 |
| 29709 | 1486 | V1486H | E | 2.3e-07 | 3.2e-07 | 1.8e-07 | 2.433E-07 | 7.095E-08 | 1.800E-07 | 3.200E-07 |
| 29710 | 1486 | V1486I | E | 1.9e-07 | 2.1e-07 | 4e-07 | 2.667E-07 | 1.159E-07 | 1.900E-07 | 4.000E-07 |
| 29711 | 1486 | V1486L | E | 2.5e-07 | 2.3e-07 | 2.5e-07 | 2.433E-07 | 1.155E-08 | 2.300E-07 | 2.500E-07 |
| 29712 | 1486 | V1486K | E | 2.3e-07 | 3e-07 | 3.6e-07 | 2.967E-07 | 6.506E-08 | 2.300E-07 | 3.600E-07 |
| 29713 | 1486 | V1486M | E | 2.4e-07 | 2.8e-07 | 2.7e-07 | 2.633E-07 | 2.082E-08 | 2.400E-07 | 2.800E-07 |
| 29714 | 1486 | V1486F | E | 6e-07 | 2.8e-07 | 3.2e-07 | 4.000E-07 | 1.744E-07 | 2.800E-07 | 6.000E-07 |
| 29715 | 1486 | V1486P | E | 2.9e-07 | 8.5e-07 | 3.4e-07 | 4.933E-07 | 3.099E-07 | 2.900E-07 | 8.500E-07 |
| 29716 | 1486 | V1486S | E | 2.2e-07 | 2.7e-07 | 2e-07 | 2.300E-07 | 3.606E-08 | 2.000E-07 | 2.700E-07 |
| 29717 | 1486 | V1486T | E | 3.2e-07 | 3.9e-07 | 3.3e-07 | 3.467E-07 | 3.786E-08 | 3.200E-07 | 3.900E-07 |
| 29718 | 1486 | V1486W | E | 2.2e-07 | 1.8e-07 | 4.3e-07 | 2.767E-07 | 1.343E-07 | 1.800E-07 | 4.300E-07 |
| 29719 | 1486 | V1486Y | E | 5.8e-07 | 5.5e-07 | 1.5e-07 | 4.267E-07 | 2.401E-07 | 1.500E-07 | 5.800E-07 |
| 29720 | 1486 | V1486V | E | 4.4e-07 | 2e-07 | 2.2e-07 | 2.867E-07 | 1.332E-07 | 2.000E-07 | 4.400E-07 |
| 29721 | 1487 | D1487A | E | 2.2e-07 | 2.5e-07 | 1.4e-07 | 2.033E-07 | 5.686E-08 | 1.400E-07 | 2.500E-07 |
| 29722 | 1487 | D1487R | E | 3.1e-07 | 1.6e-07 | 9.8e-08 | 1.893E-07 | 1.090E-07 | 9.800E-08 | 3.100E-07 |
| 29723 | 1487 | D1487N | E | 3.3e-07 | 3.8e-07 | 4.9e-07 | 4.000E-07 | 8.185E-08 | 3.300E-07 | 4.900E-07 |
| 29724 | 1487 | D1487D | E | 6.7e-07 | 2.4e-07 | 2.2e-07 | 3.767E-07 | 2.542E-07 | 2.200E-07 | 6.700E-07 |
| 29725 | 1487 | D1487C | E | 2e-07 | 3.2e-07 | 2.1e-07 | 2.433E-07 | 6.658E-08 | 2.000E-07 | 3.200E-07 |
| 29726 | 1487 | D1487E | E | 1.2e-07 | 2.9e-07 | 3.7e-07 | 2.600E-07 | 1.277E-07 | 1.200E-07 | 3.700E-07 |
| 29727 | 1487 | D1487Q | E | 4e-07 | 2.5e-07 | 1.4e-07 | 2.633E-07 | 1.305E-07 | 1.400E-07 | 4.000E-07 |
| 29728 | 1487 | D1487G | E | 3.7e-07 | 1.9e-07 | 1.3e-07 | 2.300E-07 | 1.249E-07 | 1.300E-07 | 3.700E-07 |
| 29729 | 1487 | D1487H | E | 3.1e-07 | 3.2e-07 | 3.6e-07 | 3.300E-07 | 2.646E-08 | 3.100E-07 | 3.600E-07 |
| 29730 | 1487 | D1487I | E | 2.7e-07 | 3e-07 | 3.1e-07 | 2.933E-07 | 2.082E-08 | 2.700E-07 | 3.100E-07 |
| 29731 | 1487 | D1487L | E | 2.2e-07 | 2e-07 | 4e-07 | 2.733E-07 | 1.102E-07 | 2.000E-07 | 4.000E-07 |
| 29732 | 1487 | D1487K | E | 2.6e-07 | 3.2e-07 | 2.9e-07 | 2.900E-07 | 3.000E-08 | 2.600E-07 | 3.200E-07 |
| 29733 | 1487 | D1487M | E | 4.6e-07 | 3.4e-07 | 9.7e-08 | 2.990E-07 | 1.849E-07 | 9.700E-08 | 4.600E-07 |
| 29734 | 1487 | D1487F | E | 2e-07 | 4.5e-07 | 2.3e-07 | 2.933E-07 | 1.365E-07 | 2.000E-07 | 4.500E-07 |
| 29735 | 1487 | D1487P | E | 2.4e-07 | 2.7e-07 | 2.4e-07 | 2.500E-07 | 1.732E-08 | 2.400E-07 | 2.700E-07 |
| 29736 | 1487 | D1487S | E | 1.8e-07 | 6.1e-07 | 3.8e-07 | 3.900E-07 | 2.152E-07 | 1.800E-07 | 6.100E-07 |
| 29737 | 1487 | D1487T | E | 1.4e-07 | 2.4e-07 | 1.9e-07 | 1.900E-07 | 5.000E-08 | 1.400E-07 | 2.400E-07 |
| 29738 | 1487 | D1487W | E | 3.1e-07 | 9.4e-08 | 3.8e-07 | 2.613E-07 | 1.491E-07 | 9.400E-08 | 3.800E-07 |
| 29739 | 1487 | D1487Y | E | 5.5e-07 | 4.2e-07 | 3e-07 | 4.233E-07 | 1.250E-07 | 3.000E-07 | 5.500E-07 |
| 29740 | 1487 | D1487V | E | 2.3e-07 | 1.5e-07 | 2.3e-07 | 2.033E-07 | 4.619E-08 | 1.500E-07 | 2.300E-07 |
| 29741 | 1488 | K1488A | E | 2.4e-07 | 4.6e-07 | 2e-07 | 3.000E-07 | 1.400E-07 | 2.000E-07 | 4.600E-07 |
| 29742 | 1488 | K1488R | E | 1.6e-07 | 6e-07 | 1.4e-07 | 3.000E-07 | 2.600E-07 | 1.400E-07 | 6.000E-07 |
| 29743 | 1488 | K1488N | E | 3.2e-07 | 3e-07 | 1.4e-07 | 2.533E-07 | 9.866E-08 | 1.400E-07 | 3.200E-07 |
| 29744 | 1488 | K1488D | E | 2.6e-07 | 4.1e-07 | 2.9e-07 | 3.200E-07 | 7.937E-08 | 2.600E-07 | 4.100E-07 |
| 29745 | 1488 | K1488C | E | 2.5e-07 | 2.5e-07 | 2.5e-07 | 2.500E-07 | 0.000E+00 | 2.500E-07 | 2.500E-07 |
| 29746 | 1488 | K1488E | E | 3.2e-07 | 2.2e-07 | 4.1e-07 | 3.167E-07 | 9.504E-08 | 2.200E-07 | 4.100E-07 |
| 29747 | 1488 | K1488Q | E | 3.7e-07 | 2.4e-07 | 1.2e-07 | 2.433E-07 | 1.250E-07 | 1.200E-07 | 3.700E-07 |
| 29748 | 1488 | K1488G | E | 2e-07 | 3.8e-07 | 2.9e-07 | 2.900E-07 | 9.000E-08 | 2.000E-07 | 3.800E-07 |
| 29749 | 1488 | K1488H | E | 1.3e-07 | 1.3e-07 | 2.3e-07 | 1.633E-07 | 5.774E-08 | 1.300E-07 | 2.300E-07 |

| | | | | | | | | | | |
|---|---|---|---|---|---|---|---|---|---|---|
| 29750 | 1488 | K1488I | E | 1.8e-07 | 5.7e-07 | 4.2e-07 | 3.900E-07 | 1.967E-07 | 1.800E-07 | 5.700E-07 |
| 29751 | 1488 | K1488L | E | 1.7e-07 | 8e-07 | 2.3e-07 | 4.000E-07 | 3.477E-07 | 1.700E-07 | 8.000E-07 |
| 29752 | 1488 | K1488K | E | 2.8e-07 | 1.2e-07 | 1.8e-07 | 1.933E-07 | 8.083E-08 | 1.200E-07 | 2.800E-07 |
| 29753 | 1488 | K1488M | E | 9.6e-08 | 2.2e-07 | 3.1e-07 | 2.087E-07 | 1.074E-07 | 9.600E-08 | 3.100E-07 |
| 29754 | 1488 | K1488F | E | 1.2e-07 | 3.7e-07 | 2.6e-07 | 2.500E-07 | 1.253E-07 | 1.200E-07 | 3.700E-07 |
| 29755 | 1488 | K1488P | E | 2.8e-07 | 1.9e-07 | 2.3e-07 | 2.333E-07 | 4.509E-08 | 1.900E-07 | 2.800E-07 |
| 29756 | 1488 | K1488S | E | 3.7e-07 | 2.7e-07 | 2.1e-07 | 2.833E-07 | 8.083E-08 | 2.100E-07 | 3.700E-07 |
| 29757 | 1488 | K1488T | E | 2.2e-07 | 1.2e-07 | 1.6e-07 | 1.667E-07 | 5.033E-08 | 1.200E-07 | 2.200E-07 |
| 29758 | 1488 | K1488W | E | 1.4e-07 | 2.7e-07 | 4e-07 | 2.700E-07 | 1.300E-07 | 1.400E-07 | 4.000E-07 |
| 29759 | 1488 | K1488Y | E | 2.1e-07 | 3e-07 | 2.3e-07 | 2.467E-07 | 4.726E-08 | 2.100E-07 | 3.000E-07 |
| 29760 | 1488 | K1488V | E | 3.9e-07 | 1e-07 | 2.4e-07 | 2.433E-07 | 1.450E-07 | 1.000E-07 | 3.900E-07 |
| 29761 | 1489 | K1489A | E | 3e-07 | 2.8e-07 | 2.1e-07 | 2.633E-07 | 4.726E-08 | 2.100E-07 | 3.000E-07 |
| 29762 | 1489 | K1489R | E | 2.2e-07 | 2.6e-07 | 1.7e-07 | 2.167E-07 | 4.509E-08 | 1.700E-07 | 2.600E-07 |
| 29763 | 1489 | K1489N | E | 1.5e-07 | 1.5e-07 | 1.3e-07 | 1.433E-07 | 1.155E-08 | 1.300E-07 | 1.500E-07 |
| 29764 | 1489 | K1489D | E | 2.8e-07 | 3.1e-07 | 2.5e-07 | 2.800E-07 | 3.000E-08 | 2.500E-07 | 3.100E-07 |
| 29765 | 1489 | K1489C | E | 2e-07 | 1.9e-07 | 2.9e-07 | 2.267E-07 | 5.508E-08 | 1.900E-07 | 2.900E-07 |
| 29766 | 1489 | K1489E | E | 1.7e-07 | 1.6e-07 | 2e-07 | 1.767E-07 | 2.082E-08 | 1.600E-07 | 2.000E-07 |
| 29767 | 1489 | K1489Q | E | 3e-07 | 2.6e-07 | 3.5e-07 | 3.033E-07 | 4.509E-08 | 2.600E-07 | 3.500E-07 |
| 29768 | 1489 | K1489G | E | 2.8e-07 | 5.6e-07 | 2.5e-07 | 3.633E-07 | 1.710E-07 | 2.500E-07 | 5.600E-07 |
| 29769 | 1489 | K1489H | E | 2.4e-07 | 2.5e-07 | 2.8e-07 | 2.567E-07 | 2.082E-08 | 2.400E-07 | 2.800E-07 |
| 29770 | 1489 | K1489I | E | 3.2e-07 | 4e-07 | 4.1e-07 | 3.767E-07 | 4.933E-08 | 3.200E-07 | 4.100E-07 |
| 29771 | 1489 | K1489L | E | 1.6e-07 | 5.1e-07 | 2.2e-07 | 2.967E-07 | 1.872E-07 | 1.600E-07 | 5.100E-07 |
| 29772 | 1489 | K1489K | E | 2.7e-07 | 4.2e-07 | 2.7e-07 | 3.200E-07 | 8.660E-08 | 2.700E-07 | 4.200E-07 |
| 29773 | 1489 | K1489M | E | 2.2e-07 | 8.6e-08 | 2.5e-07 | 1.853E-07 | 8.732E-08 | 8.600E-08 | 2.500E-07 |
| 29774 | 1489 | K1489F | E | 1.6e-07 | 2e-07 | 4.2e-07 | 2.600E-07 | 1.400E-07 | 1.600E-07 | 4.200E-07 |
| 29775 | 1489 | K1489P | E | 3.6e-07 | 2.9e-07 | 1.9e-07 | 2.800E-07 | 8.544E-08 | 1.900E-07 | 3.600E-07 |
| 29776 | 1489 | K1489S | E | 2.6e-07 | 1.8e-07 | 3.2e-07 | 2.533E-07 | 7.024E-08 | 1.800E-07 | 3.200E-07 |
| 29777 | 1489 | K1489T | E | 2.9e-07 | 2.2e-07 | 1.7e-07 | 2.267E-07 | 6.028E-08 | 1.700E-07 | 2.900E-07 |
| 29778 | 1489 | K1489W | E | 1.6e-07 | 3.8e-07 | 4e-07 | 3.133E-07 | 1.332E-07 | 1.600E-07 | 4.000E-07 |
| 29779 | 1489 | K1489Y | E | 3e-07 | 2.2e-07 | 3.1e-07 | 2.767E-07 | 4.933E-08 | 2.200E-07 | 3.100E-07 |
| 29780 | 1489 | K1489V | E | 2.3e-07 | 1.6e-07 | 2e-07 | 1.967E-07 | 3.512E-08 | 1.600E-07 | 2.300E-07 |
| 29781 | 1490 | V1490A | E | 3.9e-07 | 2.5e-07 | 2.2e-07 | 2.867E-07 | 9.074E-08 | 2.200E-07 | 3.900E-07 |
| 29782 | 1490 | V1490R | E | 2e-07 | 1.9e-07 | 2.4e-07 | 2.100E-07 | 2.646E-08 | 1.900E-07 | 2.400E-07 |
| 29783 | 1490 | V1490N | E | 1.9e-07 | 9.9e-08 | 3e-07 | 1.963E-07 | 1.006E-07 | 9.900E-08 | 3.000E-07 |
| 29784 | 1490 | V1490D | E | 2.4e-07 | 2.4e-07 | 3.4e-07 | 2.733E-07 | 5.774E-08 | 2.400E-07 | 3.400E-07 |
| 29785 | 1490 | V1490C | E | 3.1e-07 | 1.7e-07 | 4.2e-07 | 3.000E-07 | 1.253E-07 | 1.700E-07 | 4.200E-07 |
| 29786 | 1490 | V1490E | E | 4.1e-07 | 2e-07 | 1.9e-07 | 2.667E-07 | 1.242E-07 | 1.900E-07 | 4.100E-07 |
| 29787 | 1490 | V1490Q | E | 3.2e-07 | 2.2e-07 | 1.9e-07 | 2.433E-07 | 6.807E-08 | 1.900E-07 | 3.200E-07 |
| 29788 | 1490 | V1490G | E | 2.3e-07 | 2.6e-07 | 3.4e-07 | 2.767E-07 | 5.686E-08 | 2.300E-07 | 3.400E-07 |
| 29789 | 1490 | V1490H | E | 2.9e-07 | 4.2e-07 | 2e-07 | 3.033E-07 | 1.106E-07 | 2.000E-07 | 4.200E-07 |
| 29790 | 1490 | V1490I | E | 2.4e-07 | 5.4e-07 | 2.8e-07 | 3.533E-07 | 1.629E-07 | 2.400E-07 | 5.400E-07 |
| 29791 | 1490 | V1490L | E | 2.8e-07 | 2.5e-07 | 5.5e-07 | 3.600E-07 | 1.652E-07 | 2.500E-07 | 5.500E-07 |
| 29792 | 1490 | V1490K | E | 1.9e-07 | 2e-07 | 3.4e-07 | 2.433E-07 | 8.386E-08 | 1.900E-07 | 3.400E-07 |
| 29793 | 1490 | V1490M | E | 1.4e-07 | 1.6e-07 | 1.3e-07 | 1.433E-07 | 1.528E-08 | 1.300E-07 | 1.600E-07 |
| 29794 | 1490 | V1490F | E | 1.9e-07 | 2.4e-07 | 2e-07 | 2.100E-07 | 2.646E-08 | 1.900E-07 | 2.400E-07 |
| 29795 | 1490 | V1490P | E | 1.8e-07 | 2.1e-07 | 4.6e-07 | 2.833E-07 | 1.537E-07 | 1.800E-07 | 4.600E-07 |
| 29796 | 1490 | V1490S | E | 7.4e-08 | 2.1e-07 | 3.4e-07 | 2.080E-07 | 1.330E-07 | 7.400E-08 | 3.400E-07 |
| 29797 | 1490 | V1490T | E | 2.2e-07 | 2.6e-07 | 2.8e-07 | 2.533E-07 | 3.055E-08 | 2.200E-07 | 2.800E-07 |
| 29798 | 1490 | V1490W | E | 1.9e-07 | 2.8e-07 | 2.9e-07 | 2.533E-07 | 5.508E-08 | 1.900E-07 | 2.900E-07 |
| 29799 | 1490 | V1490Y | E | 1.9e-07 | 4e-07 | 4.4e-07 | 3.433E-07 | 1.343E-07 | 1.900E-07 | 4.400E-07 |
| 29800 | 1490 | V1490V | E | 2.6e-07 | 5.5e-07 | 2.5e-07 | 3.533E-07 | 1.704E-07 | 2.500E-07 | 5.500E-07 |
| 29801 | 1491 | E1491A | E | 2.8e-07 | 4.8e-07 | 2.9e-07 | 3.500E-07 | 1.127E-07 | 2.800E-07 | 4.800E-07 |
| 29802 | 1491 | E1491R | E | 1.8e-07 | 2.9e-07 | 1.9e-07 | 2.200E-07 | 6.083E-08 | 1.800E-07 | 2.900E-07 |
| 29803 | 1491 | E1491N | E | 1.9e-07 | 1.5e-07 | 2.7e-07 | 2.033E-07 | 6.110E-08 | 1.500E-07 | 2.700E-07 |
| 29804 | 1491 | E1491D | E | 4.1e-07 | 2.4e-07 | 1.4e-07 | 2.633E-07 | 1.365E-07 | 1.400E-07 | 4.100E-07 |
| 29805 | 1491 | E1491C | E | 1.4e-07 | 2.5e-07 | 1.9e-07 | 1.933E-07 | 5.508E-08 | 1.400E-07 | 2.500E-07 |
| 29806 | 1491 | E1491E | E | 2.4e-07 | 2.6e-07 | 1.8e-07 | 2.267E-07 | 4.163E-08 | 1.800E-07 | 2.600E-07 |
| 29807 | 1491 | E1491Q | E | 2.8e-07 | 2.5e-07 | 2.1e-07 | 2.467E-07 | 3.512E-08 | 2.100E-07 | 2.800E-07 |
| 29808 | 1491 | E1491G | E | 2.8e-07 | 1.8e-07 | 4.9e-07 | 3.167E-07 | 1.582E-07 | 1.800E-07 | 4.900E-07 |
| 29809 | 1491 | E1491H | E | 2.4e-07 | 1.9e-07 | 2.3e-07 | 2.200E-07 | 2.646E-08 | 1.900E-07 | 2.400E-07 |
| 29810 | 1491 | E1491I | E | 2.7e-07 | 1.7e-07 | 2.7e-07 | 2.367E-07 | 5.774E-08 | 1.700E-07 | 2.700E-07 |
| 29811 | 1491 | E1491L | E | 1.9e-07 | 3.4e-07 | 2.6e-07 | 2.633E-07 | 7.506E-08 | 1.900E-07 | 3.400E-07 |
| 29812 | 1491 | E1491K | E | 1.3e-07 | 2e-07 | 4.5e-07 | 2.600E-07 | 1.682E-07 | 1.300E-07 | 4.500E-07 |
| 29813 | 1491 | E1491M | E | 2.4e-07 | 1e-07 | 3.1e-07 | 2.167E-07 | 1.069E-07 | 1.000E-07 | 3.100E-07 |
| 29814 | 1491 | E1491F | E | 2.2e-07 | 2.3e-07 | 3.8e-07 | 2.767E-07 | 8.963E-08 | 2.200E-07 | 3.800E-07 |
| 29815 | 1491 | E1491P | E | 1.8e-07 | 1.5e-07 | 2.7e-07 | 2.000E-07 | 6.245E-08 | 1.500E-07 | 2.700E-07 |
| 29816 | 1491 | E1491S | E | 3.6e-07 | 2.4e-07 | 1.3e-07 | 2.433E-07 | 1.150E-07 | 1.300E-07 | 3.600E-07 |
| 29817 | 1491 | E1491T | E | 3.5e-07 | 2.5e-07 | 3.7e-07 | 3.233E-07 | 6.429E-08 | 2.500E-07 | 3.700E-07 |
| 29818 | 1491 | E1491W | E | 2.4e-07 | 3.2e-07 | 2.9e-07 | 2.833E-07 | 4.041E-08 | 2.400E-07 | 3.200E-07 |
| 29819 | 1491 | E1491Y | E | 2.1e-07 | 2.2e-07 | 3.4e-07 | 2.567E-07 | 7.234E-08 | 2.100E-07 | 3.400E-07 |
| 29820 | 1491 | E1491V | E | 3.4e-07 | 1.6e-07 | 3.8e-07 | 2.933E-07 | 1.172E-07 | 1.600E-07 | 3.800E-07 |
| 29821 | 1492 | P1492A | E | 4e-07 | 3.1e-07 | 2.1e-07 | 3.067E-07 | 9.504E-08 | 2.100E-07 | 4.000E-07 |
| 29822 | 1492 | P1492R | E | 4.2e-07 | 2.4e-07 | 3.1e-07 | 3.233E-07 | 9.074E-08 | 2.400E-07 | 4.200E-07 |
| 29823 | 1492 | P1492N | E | 1.5e-07 | 1.6e-07 | 3e-07 | 2.033E-07 | 8.386E-08 | 1.500E-07 | 3.000E-07 |
| 29824 | 1492 | P1492D | E | 2e-07 | 2.9e-07 | 1.6e-07 | 2.167E-07 | 6.658E-08 | 1.600E-07 | 2.900E-07 |
| 29825 | 1492 | P1492C | E | 4.4e-07 | 2.7e-07 | 1.8e-07 | 2.967E-07 | 1.320E-07 | 1.800E-07 | 4.400E-07 |
| 29826 | 1492 | P1492E | E | 5.4e-07 | 1.7e-07 | 2.2e-07 | 3.100E-07 | 2.007E-07 | 1.700E-07 | 5.400E-07 |
| 29827 | 1492 | P1492Q | E | 3.8e-07 | 1.5e-07 | 1.9e-07 | 2.400E-07 | 1.229E-07 | 1.500E-07 | 3.800E-07 |
| 29828 | 1492 | P1492G | E | 3.2e-07 | 1.8e-07 | 1.8e-07 | 2.267E-07 | 8.083E-08 | 1.800E-07 | 3.200E-07 |
| 29829 | 1492 | P1492H | E | 2.4e-07 | 1.8e-07 | 1.1e-07 | 1.767E-07 | 6.506E-08 | 1.100E-07 | 2.400E-07 |
| 29830 | 1492 | P1492I | E | 3.2e-07 | 1.6e-07 | 1.4e-07 | 2.067E-07 | 9.866E-08 | 1.400E-07 | 3.200E-07 |
| 29831 | 1492 | P1492L | E | 2e-07 | 4e-07 | 2.3e-07 | 2.767E-07 | 1.079E-07 | 2.000E-07 | 4.000E-07 |
| 29832 | 1492 | P1492K | E | 2.3e-07 | 2e-07 | 8.6e-08 | 1.720E-07 | 7.597E-08 | 8.600E-08 | 2.300E-07 |
| 29833 | 1492 | P1492M | E | 2.6e-07 | 2.3e-07 | 3e-07 | 2.633E-07 | 3.512E-08 | 2.300E-07 | 3.000E-07 |
| 29834 | 1492 | P1492F | E | 1.7e-07 | 1.5e-07 | 4.2e-07 | 2.467E-07 | 1.504E-07 | 1.500E-07 | 4.200E-07 |

| | | | | | | | | | | |
|---|---|---|---|---|---|---|---|---|---|---|
| 29835 | 1492 | P1492P | E | 2.8e-07 | 4.4e-07 | 3.6e-07 | 3.600E-07 | 8.000E-08 | 2.800E-07 | 4.400E-07 |
| 29836 | 1492 | P1492S | E | 9.9e-08 | 1.7e-07 | 1.2e-07 | 1.297E-07 | 3.647E-08 | 9.900E-08 | 1.700E-07 |
| 29837 | 1492 | P1492T | E | 1.4e-07 | 2.8e-07 | 3e-07 | 2.400E-07 | 8.718E-08 | 1.400E-07 | 3.000E-07 |
| 29838 | 1492 | P1492W | E | 2.1e-07 | 1.5e-07 | 4.2e-07 | 2.600E-07 | 1.418E-07 | 1.500E-07 | 4.200E-07 |
| 29839 | 1492 | P1492Y | E | 2.9e-07 | 1.8e-07 | 2.9e-07 | 2.533E-07 | 6.351E-08 | 1.800E-07 | 2.900E-07 |
| 29840 | 1492 | P1492V | E | 7.5e-07 | 2.8e-07 | 3.9e-07 | 4.733E-07 | 2.458E-07 | 2.800E-07 | 7.500E-07 |

Table 8: A scalable Her2-Trastuzumab-Pertuzumab binding affinity landscape. In this table, **ResID** represents the amino acid residue number of PDB entry 6OGE, while **Muta** represents the site-specific missense mutation introduced into the 1492 amino acid residues in PDB entry 6OGE, **Min**, **Max**, **Mean** and **Std** represent the minimum, the maximum, the average and the standard deviation of the $K_d$ values calculated using Prodigy, respectively.

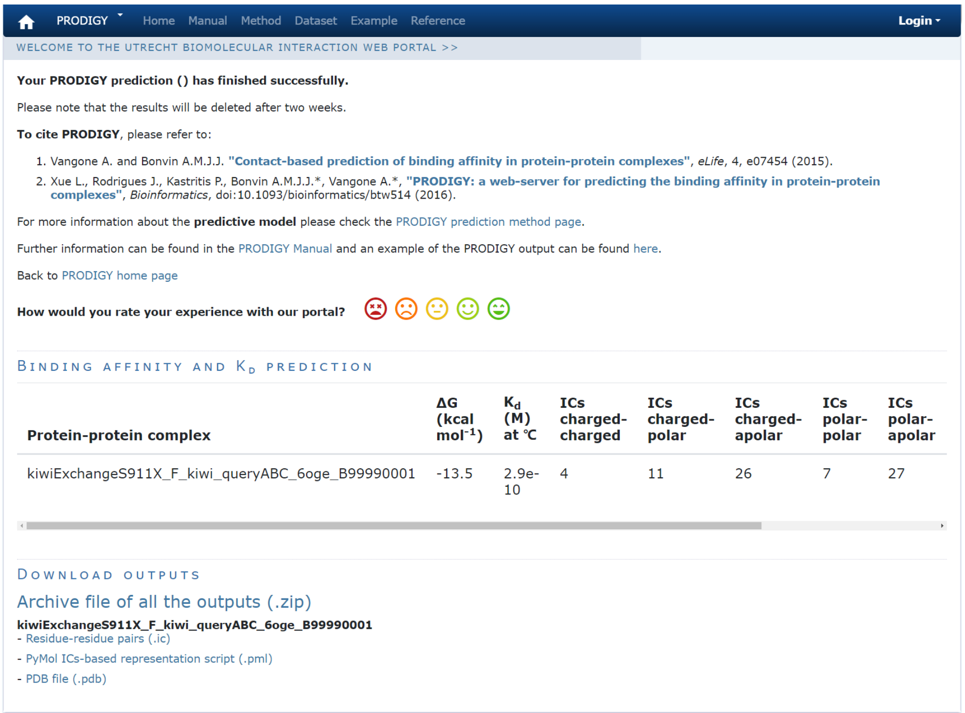

PRODIGY Home Manual Method Dataset Example Reference Login

WELCOME TO THE UTRECHT BIOMOLECULAR INTERACTION WEB PORTAL >>

**Your PRODIGY prediction () has finished successfully.**

Please note that the results will be deleted after two weeks.

**To cite PRODIGY**, please refer to:

1. Vangone A. and Bonvin A.M.J.J. **"Contact-based prediction of binding affinity in protein-protein complexes"**, *eLife*, 4, e07454 (2015).
2. Xue L., Rodrigues J., Kastritis P., Bonvin A.M.J.J.*, Vangone A.*, **"PRODIGY: a web-server for predicting the binding affinity in protein-protein complexes"**, *Bioinformatics*, doi:10.1093/bioinformatics/btw514 (2016).

For more information about the **predictive model** please check the PRODIGY prediction method page.

Further information can be found in the PRODIGY Manual and an example of the PRODIGY output can be found here.

Back to PRODIGY home page

**How would you rate your experience with our portal?**

BINDING AFFINITY AND $K_D$ PREDICTION

| Protein-protein complex | ΔG (kcal mol$^{-1}$) | $K_d$ (M) at ℃ | ICs charged-charged | ICs charged-polar | ICs charged-apolar | ICs polar-polar | ICs polar-apolar |
|---|---|---|---|---|---|---|---|
| kiwiExchangeS911X_F_kiwi_queryABC_6oge_B99990001 | -13.5 | 2.9e-10 | 4 | 11 | 26 | 7 | 27 |

DOWNLOAD OUTPUTS

Archive file of all the outputs (.zip)

**kiwiExchangeS911X_F_kiwi_queryABC_6oge_B99990001**
- Residue-residue pairs (.ic)
- PyMol ICs-based representation script (.pml)
- PDB file (.pdb)

Figure 7: An example (supplementary file fin.pdb) of the calculation by the Prodigy server of the $K_d$ between chain A (Her2) and chain BC (Pertuzumab FAB, Table 1) of PDB entry 6OGE with one site-specific missense mutation S911F in chain C of the heavy chain of Pertuzumab.

**Acknowledgment**

The author is grateful to the communities of structural biology, biophysics, medicinal and computational chemistry and algorithm design, for the continued accumulation of knowledge and data for drug discovery & design, and for the continued development of tools (hardware, software and algorithm) for drug discovery & design.

**Statement of Usage of Artificial Intelligence**

During the preparation of this work, the author used OpenAI's ChatGPT in order to improve the readability of the manuscript, and to make it as concise and short as possible. After using this tool, the author reviewed and edited the content as needed and takes full responsibility for the content of the publication.

**Data Availability**

All research data, including PDB access codes and PDB files of structural models, have been provided. In the generation of a scalable antigen-antibody binding affinity landscape using Trastuzumab as an example, i.e., I have used a set of tools, which are included in a series of in-house python scripts, along with the raw data files. For readers to reproduce the work reported in the manuscript, all files will be provided upon request.

**Author Contributions**

Conceptualization, W.L.; methodology, W.L.; software, W.L.; validation, W.L.; formal analysis, W.L.; investigation, W.L.; resources, W.L.; data duration, W.L.; writing–original draft preparation, W.L.; writing–review and editing, W.L.; visualization, W.L.; supervision, W.L.; project administration, W.L.;

**Conflict of interest**

None.

**Ethical statement**

No ethical approval is required.

**Funding Information**

This research received no external funding.

**ORCID**

Wei Li - `https://orcid.org/0000-0001-6060-7937`
Wei Li - `https://orcid.org/0000-0002-0945-2666`
